## Supplementary Table S1 for "Genome-wide Selection Scan in an Arabian Peninsula Population Identifies a *TNKS* haplotype Linked to Metabolic Traits and Hypertension"

Table S1. The selection tests scores and the allele frequency of all the variants.

| CHR | POS | SNP | iHS | XP-EHH | | | PBS | | | LLRS | Alternate Allele Frequency | | | | | |
|---|---|---|---|---|---|---|---|---|---|---|---|---|---|---|---|---|
| | | | | KWT vs CEU | KWT vs YRI | KWT vs CHB | KWT vs CEU | KWT vs YRI | KWT vs CHB | | KWT | AFR | AMR | EAS | EUR | SAS |
| 1 | 924898 | rs6665000 | 0.699 | -0.651 | 0.816 | -0.322 | -0.057 | -0.064 | -0.064 | 4.808 | 0.87 | 0.71 | 0.94 | 0.95 | 0.96 | 0.98 |
| 1 | 927309 | rs2341362 | 0.693 | -0.651 | 0.816 | -0.322 | -0.057 | -0.064 | -0.064 | 0 | 0.87 | 0.71 | 0.94 | 0.95 | 0.96 | 0.98 |
| 1 | 928836 | rs9777703 | 0.705 | -0.65 | 0.823 | -0.322 | -0.058 | -0.065 | -0.065 | 0 | 0.87 | 0.7 | 0.93 | 0.95 | 0.96 | 0.98 |
| 1 | 947034 | rs2465126 | 0.849 | -0.658 | 1.372 | -0.449 | -0.073 | -0.067 | -0.067 | 1.589 | 0.85 | 0.48 | 0.92 | 0.94 | 0.96 | 0.97 |
| 1 | 948692 | rs2341365 | 1.157 | -0.734 | 1.47 | -0.453 | -0.085 | -0.063 | -0.063 | 0 | 0.83 | 0.46 | 0.91 | 0.92 | 0.96 | 0.97 |
| 1 | 948921 | rs15842 | 0.764 | -0.685 | 1.466 | -0.635 | -0.057 | -0.085 | -0.085 | 0 | 0.86 | 0.72 | 0.93 | 1 | 0.96 | 0.98 |
| 1 | 959842 | rs2710888 | -1.295 | -0.399 | 0.79 | -0.747 | -0.015 | 0.049 | 0.049 | 0 | 0.34 | 0.18 | 0.25 | 0.19 | 0.37 | 0.36 |
| 1 | 962210 | rs3128126 | -1.205 | -0.271 | 1.011 | -0.775 | -0.007 | 0.015 | 0.015 | 0 | 0.32 | 0.3 | 0.26 | 0.19 | 0.39 | 0.37 |
| 1 | 977780 | rs2710875 | 1.581 | -0.687 | 1.418 | -0.712 | -0.049 | -0.093 | -0.093 | 0.096 | 0.84 | 0.43 | 0.9 | 0.97 | 0.92 | 0.96 |
| 1 | 994391 | rs2488991 | 2.193 | -0.516 | 1.927 | -0.761 | -0.021 | -0.103 | -0.103 | 0 | 0.82 | 0.36 | 0.88 | 0.99 | 0.87 | 0.93 |
| 1 | 998395 | rs7526076 | 1.062 | -0.397 | 1.535 | -0.71 | -0.005 | 0.003 | 0.003 | 0.014 | 0.83 | 0.79 | 0.82 | 0.92 | 0.76 | 0.83 |
| 1 | 1005806 | rs3934834 | -0.966 | -0.525 | 1.887 | -0.653 | -0.048 | -0.039 | -0.039 | 0.126 | 0.23 | 0.39 | 0.11 | 0.13 | 0.17 | 0.22 |
| 1 | 1017587 | rs3766191 | -0.593 | -0.609 | 1.872 | -0.851 | -0.004 | -0.003 | -0.003 | 0 | 0.22 | 0.2 | 0.11 | 0.13 | 0.17 | 0.22 |
| 1 | 1021346 | rs10907177 | -0.961 | -0.589 | 1.464 | -0.649 | -0.002 | -0.001 | -0.001 | 0 | 0.25 | 0.24 | 0.13 | 0.12 | 0.18 | 0.19 |
| 1 | 1021415 | rs3737728 | 1.068 | -0.521 | 1.576 | -0.68 | -0.007 | 0.01 | 0.01 | 0 | 0.8 | 0.81 | 0.81 | 0.92 | 0.74 | 0.79 |
| 1 | 1021583 | rs10907178 | -1.154 | -0.706 | 1.428 | -0.386 | -0.002 | -0.001 | -0.001 | 0 | 0.24 | 0.23 | 0.12 | 0.12 | 0.18 | 0.19 |
| 1 | 1025301 | rs9442400 | 0.849 | -0.373 | 1.046 | 0.766 | -0.001 | 0.04 | 0.04 | 0 | 0.93 | 0.93 | 0.9 | 0.74 | 0.92 | 0.87 |
| 1 | 1026707 | rs4074137 | 1.641 | -0.864 | 1 | -0.19 | 0.015 | 0.026 | 0.026 | 0 | 0.55 | 0.35 | 0.69 | 0.92 | 0.6 | 0.68 |
| 1 | 1030565 | rs6687776 | -2.003 | -1.098 | 1.172 | 0.007 | -0.03 | -0.004 | -0.004 | 0 | 0.24 | 0.6 | 0.19 | 0.19 | 0.17 | 0.26 |
| 1 | 1036959 | rs11579015 | -1.216 | -1.099 | 1.272 | 0.33 | 0.001 | 0 | 0 | 1.605 | 0.15 | 0.13 | 0.11 | 0.22 | 0.11 | 0.21 |
| 1 | 1040026 | rs6671356 | -2.343 | -1.1 | 1.457 | 0.367 | -0.056 | 0.03 | 0.03 | 0.33 | 0.24 | 0.54 | 0.17 | 0.22 | 0.13 | 0.24 |
| 1 | 1060174 | rs7548798 | -0.953 | -0.605 | 1.155 | 0.2 | 0.004 | -0.059 | -0.059 | 0 | 0.23 | 0.29 | 0.16 | 0.01 | 0.29 | 0.16 |
| 1 | 1061262 | rs12757754 | -1.517 | -0.653 | 0.748 | 0.311 | 0.024 | -0.03 | -0.03 | 0.094 | 0.18 | 0.08 | 0.11 | 0.22 | 0.1 | 0.2 |
| 1 | 1062638 | rs9442373 | 1.7 | -0.58 | 0.792 | 0.596 | -0.002 | 0.016 | 0.016 | 0 | 0.6 | 0.49 | 0.7 | 0.55 | 0.57 | 0.63 |
| 1 | 1064979 | rs2298217 | -1.081 | -0.288 | 0.365 | 1.654 | -0.004 | -0.027 | -0.027 | 1.38 | 0.12 | 0.05 | 0.11 | 0.41 | 0.12 | 0.15 |
| 1 | 1066403 | rs10907182 | 1.102 | -0.875 | 0.163 | 1.133 | 0.025 | -0.016 | -0.016 | 0 | 0.58 | 0.32 | 0.7 | 0.55 | 0.6 | 0.58 |
| 1 | 1087683 | rs9442380 | 1.482 | -0.922 | 0.45 | 2.021 | -0.001 | 0.064 | 0.064 | 0 | 0.92 | 0.91 | 0.91 | 0.6 | 0.94 | 0.84 |
| 1 | 1097335 | rs9442385 | 1.836 | -0.949 | 0.284 | 2.27 | -0.015 | 0.05 | 0.05 | 0 | 0.88 | 0.82 | 0.9 | 0.63 | 0.96 | 0.89 |
| 1 | 1099342 | rs9660710 | 1.546 | -0.633 | 0.409 | 2.403 | -0.011 | 0.147 | 0.147 | 0 | 0.92 | 0.83 | 0.89 | 0.6 | 0.95 | 0.89 |
| 1 | 1100217 | rs1891905 | 1.534 | -0.804 | 0.293 | 1.968 | -0.01 | 0.17 | 0.17 | 0 | 0.92 | 0.79 | 0.9 | 0.6 | 0.95 | 0.89 |
| 1 | 1106784 | rs4442317 | -0.482 | -1.546 | -0.245 | 0.681 | 0.012 | 0.012 | 0.012 | 0 | 0.35 | 0.31 | 0.22 | 0.18 | 0.21 | 0.43 |
| 1 | 1108637 | rs4970421 | -0.242 | -1.908 | 0.144 | 0.411 | -0.038 | -0.006 | -0.006 | 0 | 0.12 | 0.44 | 0.11 | 0.04 | 0.05 | 0.16 |
| 1 | 1110019 | rs11260542 | -0.601 | -1.202 | -0.353 | 0.622 | 0.082 | 0.116 | 0.116 | 0.272 | 0.07 | 0.28 | 0.06 | 0.13 | 0.09 | 0.1 |
| 1 | 1110586 | rs9442387 | -0.902 | -1.123 | -1.283 | 0.328 | 0.03 | 0.25 | 0.25 | 0.149 | 0.57 | 0.96 | 0.65 | 0.91 | 0.36 | 0.79 |
| 1 | 1113121 | rs12092254 | -0.967 | -0.42 | -0.085 | 1.182 | 0.038 | 0.061 | 0.061 | 0.4 | 0.07 | 0.2 | 0.05 | 0.13 | 0.09 | 0.1 |
| 1 | 1121014 | rs3813204 | -0.749 | -0.92 | -0.89 | 1.509 | -0.011 | -0.172 | -0.172 | 0 | 0.21 | 0.01 | 0.36 | 0.6 | 0.13 | 0.26 |
| 1 | 1121794 | rs11260549 | -0.798 | -0.939 | -0.846 | 1.503 | 0.002 | -0.156 | -0.156 | 0 | 0.21 | 0.04 | 0.36 | 0.6 | 0.13 | 0.26 |
| 1 | 1130727 | rs10907175 | -0.7 | -0.85 | 0.089 | 1.545 | 0.006 | 0.067 | 0.067 | 0 | 0.07 | 0.11 | 0.08 | 0.13 | 0.08 | 0.1 |
| 1 | 1138913 | rs3819001 | -1.67 | -1.177 | 1.058 | 1.399 | -0.02 | 0.111 | 0.111 | 1.016 | 0.11 | 0.49 | 0.09 | 0.15 | 0.06 | 0.07 |
| 1 | 1153113 | rs12036216 | -1.103 | -0.842 | 0.82 | 1.601 | 0.009 | 0.101 | 0.101 | 0 | 0.07 | 0.11 | 0.09 | 0.25 | 0.1 | 0.1 |
| 1 | 1158277 | rs3813199 | -1.127 | -0.868 | 1.014 | 1.162 | 0.022 | 0.105 | 0.105 | 0 | 0.07 | 0.16 | 0.09 | 0.16 | 0.09 | 0.1 |
| 1 | 1163804 | rs7515488 | -2.578 | -1.074 | 1.336 | 1.221 | -0.004 | 0.012 | 0.012 | 0 | 0.16 | 0.24 | 0.16 | 0.18 | 0.15 | 0.18 |
| 1 | 1165310 | rs11260562 | -2.151 | -1.07 | 1.355 | 1.201 | -0.017 | -0.023 | -0.023 | 0 | 0.1 | 0.21 | 0.09 | 0.02 | 0.06 | 0.08 |
| 1 | 1173611 | rs6697886 | -2.624 | -0.976 | 1.657 | 1.28 | 0.006 | 0.104 | 0.104 | 0.377 | 0.14 | 0.42 | 0.12 | 0.17 | 0.13 | 0.17 |
| 1 | 1176597 | rs6675798 | -2.743 | -0.928 | 1.683 | 1.454 | 0.017 | 0.172 | 0.172 | 1.14 | 0.1 | 0.33 | 0.09 | 0.17 | 0.1 | 0.15 |
| 1 | 1178245 | rs11260566 | -2.813 | -0.927 | 1.755 | 1.466 | 0.012 | 0.162 | 0.162 | 0.584 | 0.1 | 0.33 | 0.09 | 0.16 | 0.09 | 0.14 |
| 1 | 1178925 | rs12093154 | -2.235 | -0.865 | 1.441 | 1.501 | -0.002 | 0.081 | 0.081 | 0.443 | 0.08 | 0.15 | 0.07 | 0.17 | 0.09 | 0.15 |
| 1 | 1231656 | rs1749951 | -1.071 | -1.637 | 2.039 | 1.779 | 0.021 | 0 | 0 | 0 | 0.08 | 0.02 | 0.02 | 0.09 | 0.04 | 0.08 |
| 1 | 1239339 | rs11260584 | 2.71 | -1.834 | 2.319 | 1.736 | -0.045 | -0.013 | -0.013 | 0 | 0.82 | 0.31 | 0.86 | 0.83 | 0.92 | 0.85 |
| 1 | 1239935 | rs12410087 | -0.887 | -1.882 | 1.932 | 1.548 | -0.007 | -0.045 | -0.045 | 0 | 0.11 | 0.05 | 0.26 | 0.21 | 0.1 | 0.25 |
| 1 | 1239953 | rs10399878 | 1.463 | -1.804 | 2.41 | 1.872 | -0.018 | 0.002 | 0.002 | 0.713 | 0.88 | 0.84 | 0.91 | 0.84 | 0.95 | 0.89 |
| 1 | 1257593 | rs1810745 | 2.038 | -1.573 | 2.847 | 1.286 | -0.021 | 0.005 | 0.005 | 0 | 0.89 | 0.84 | 0.91 | 0.84 | 0.94 | 0.84 |
| 1 | 1258246 | rs3845293 | 3.1 | -1.546 | 2.884 | 1.276 | -0.042 | -0.005 | -0.005 | 0 | 0.83 | 0.31 | 0.86 | 0.84 | 0.92 | 0.83 |
| 1 | 1264539 | rs307354 | 2.057 | -1.482 | 2.748 | 1.332 | -0.024 | 0.013 | 0.013 | 0 | 0.89 | 0.82 | 0.91 | 0.83 | 0.94 | 0.84 |
| 1 | 1269986 | rs35424002 | -2.242 | -1.479 | 2.924 | 1.271 | 0.005 | 0.031 | 0.031 | 0 | 0.07 | 0.28 | 0.09 | 0.08 | 0.04 | 0.08 |
| 1 | 1273116 | rs307371 | 2.589 | -1.492 | 3.002 | 1.268 | -0.03 | 0.036 | 0.036 | 0 | 0.86 | 0.5 | 0.88 | 0.83 | 0.92 | 0.84 |
| 1 | 1287127 | rs307380 | 2.355 | -1.516 | 2.571 | 1.257 | -0.025 | 0.01 | 0.01 | 0 | 0.87 | 0.7 | 0.89 | 0.83 | 0.92 | 0.83 |
| 1 | 1295323 | rs34389364 | 2.962 | -1.514 | 2.589 | 1.258 | -0.036 | -0.005 | -0.005 | 0 | 0.84 | 0.29 | 0.85 | 0.83 | 0.92 | 0.83 |
| 1 | 1297422 | rs2765021 | 2.307 | -1.4 | 2.079 | 1.307 | -0.022 | 0.007 | 0.007 | 0 | 0.87 | 0.74 | 0.9 | 0.83 | 0.92 | 0.83 |
| 1 | 1310924 | rs2765033 | 0.992 | -1.402 | 1.816 | 1.174 | 0.012 | -0.006 | -0.006 | 0 | 0.94 | 0.98 | 0.98 | 0.91 | 0.97 | 0.92 |
| 1 | 1314015 | rs2649588 | 2.56 | -1.559 | 2.094 | 1.085 | -0.045 | -0.034 | -0.034 | 0 | 0.84 | 0.62 | 0.86 | 0.88 | 0.91 | 0.83 |
| 1 | 1314172 | rs2477777 | 2.56 | -1.559 | 2.094 | 1.085 | -0.045 | -0.034 | -0.034 | 0 | 0.84 | 0.62 | 0.86 | 0.88 | 0.91 | 0.83 |
| 1 | 1335218 | rs2291889 | -1.162 | -1.538 | 1.87 | 1.124 | 0.011 | -0.008 | -0.008 | 0 | 0.05 | 0.02 | 0.02 | 0.08 | 0.03 | 0.08 |
| 1 | 1335302 | rs1240707 | -2.6 | -1.585 | 2.11 | 1.099 | -0.049 | -0.028 | -0.028 | 0 | 0.16 | 0.41 | 0.15 | 0.11 | 0.08 | 0.18 |
| 1 | 1458954 | rs3737714 | -1.905 | -1.48 | 2.214 | 2.411 | 0.034 | 0.435 | 0.435 | 0 | 0.05 | 0.31 | 0.14 | 0.36 | 0.01 | 0.2 |
| 1 | 1472047 | rs3118505 | -1.048 | -1.503 | 1.597 | 2.397 | 0.004 | -0.007 | -0.007 | 0.077 | 0.08 | 0.04 | 0.11 | 0.12 | 0.07 | 0.08 |

| | | | | | | | | | | | | | | | | |
|---|---|---|---|---|---|---|---|---|---|---|---|---|---|---|---|---|
| 1 | 1489670 | rs7531530 | -0.739 | -1.61 | 1.082 | 2.343 | 0.038 | 0.039 | 0.039 | 0 | 0.41 | 0.71 | 0.36 | 0.78 | 0.29 | 0.65 |
| 1 | 1493727 | rs880051 | -0.851 | -1.67 | 0.788 | 2.33 | 0.038 | 0.028 | 0.028 | 0 | 0.38 | 0.69 | 0.34 | 0.78 | 0.24 | 0.66 |
| 1 | 1497824 | rs2296716 | -0.994 | -0.865 | 0.412 | 2.588 | -0.027 | -0.083 | -0.083 | 0 | 0.12 | 0 | 0.1 | 0.4 | 0.12 | 0.29 |
| 1 | 1519068 | rs6687029 | -1.285 | -0.417 | 0.644 | 2.045 | 0.016 | 0 | 0 | 0 | 0.21 | 0.26 | 0.34 | 0.18 | 0.31 | 0.31 |
| 1 | 1521595 | rs6604983 | -1.966 | -0.818 | 0.166 | 2.254 | 0.011 | -0.051 | -0.051 | 0 | 0.11 | 0.03 | 0.25 | 0.17 | 0.12 | 0.19 |
| 1 | 1559971 | rs12748433 | 1.381 | -0.856 | 0.952 | 2.226 | 0.011 | 0.152 | 0.152 | 0 | 0.93 | 0.73 | 0.85 | 0.71 | 0.96 | 0.86 |
| 1 | 1560103 | rs28635343 | -1.24 | -1.397 | 0.364 | 2.412 | 0.01 | -0.012 | -0.012 | 1.271 | 0.22 | 0.17 | 0.3 | 0.29 | 0.17 | 0.31 |
| 1 | 1566160 | rs28464684 | 0.802 | -1.348 | 0.406 | 2.417 | -0.004 | 0.041 | 0.041 | 0 | 0.94 | 0.91 | 0.93 | 0.72 | 0.96 | 0.86 |
| 1 | 1567206 | rs28456011 | -1.375 | -1.42 | 0.21 | 1.732 | -0.02 | 0.002 | 0.002 | 0 | 0.31 | 0.3 | 0.17 | 0.28 | 0.16 | 0.35 |
| 1 | 1619541 | rs28487995 | -1.131 | -0.433 | 0.284 | 1.838 | -0.018 | 0.048 | 0.048 | 0.303 | 0.25 | 0.5 | 0.32 | 0.37 | 0.2 | 0.21 |
| 1 | 1646574 | rs760925 | 1.283 | -0.415 | 0.7 | 2.022 | -0.052 | 0.072 | 0.072 | 0.166 | 0.85 | 0.5 | 0.88 | 0.78 | 0.92 | 0.78 |
| 1 | 1647686 | rs909823 | 1.399 | -0.449 | 0.684 | 2 | -0.06 | 0.059 | 0.059 | 0.424 | 0.84 | 0.51 | 0.88 | 0.78 | 0.92 | 0.78 |
| 1 | 1664019 | rs2294488 | 0.941 | -0.499 | 0.236 | 2.241 | -0.049 | 0.031 | 0.031 | 0.635 | 0.82 | 0.39 | 0.84 | 0.77 | 0.91 | 0.68 |
| 1 | 1664124 | rs2294489 | 1.379 | -0.43 | -0.431 | 2.801 | 0.082 | -0.08 | -0.08 | 1.636 | 0.78 | 0.12 | 0.82 | 0.65 | 0.91 | 0.67 |
| 1 | 1684472 | rs7407 | -0.216 | 0.947 | -1.119 | 2.448 | -0.173 | -0.06 | -0.06 | 0 | 0.32 | 0.06 | 0.56 | 0.35 | 0.51 | 0.39 |
| 1 | 1691050 | rs35672141 | 0.036 | 2.193 | -1.1 | 2.717 | -0.174 | -0.136 | -0.136 | 0 | 0.29 | 0.05 | 0.56 | 0.43 | 0.48 | 0.35 |
| 1 | 1706136 | rs6603811 | 0.157 | 1.573 | -0.099 | 2.468 | -0.074 | -0.036 | -0.036 | 0 | 0.84 | 0.46 | 0.87 | 0.9 | 0.93 | 0.7 |
| 1 | 1706160 | rs7531583 | -0.403 | 0.982 | -0.569 | 2.247 | 0.052 | 0.042 | 0.042 | 1.388 | 0.53 | 0.37 | 0.76 | 0.68 | 0.74 | 0.49 |
| 1 | 1707740 | rs12041925 | 0.786 | 1.141 | -0.675 | 2.438 | 0.002 | -0.004 | -0.004 | 0.548 | 0.22 | 0.24 | 0.18 | 0.19 | 0.23 | 0.12 |
| 1 | 1712230 | rs10907194 | 0.866 | 1.151 | -0.581 | 2.452 | -0.001 | 0 | 0 | 2.969 | 0.22 | 0.19 | 0.18 | 0.21 | 0.23 | 0.12 |
| 1 | 1715011 | rs742359 | 0.879 | 1.149 | -0.582 | 2.501 | -0.001 | 0.001 | 0.001 | 0.004 | 0.22 | 0.19 | 0.18 | 0.14 | 0.23 | 0.12 |
| 1 | 1721479 | rs2272908 | 0.614 | 1.171 | -0.592 | 2.522 | -0.177 | -0.173 | -0.173 | 0 | 0.29 | 0.05 | 0.57 | 0.47 | 0.51 | 0.36 |
| 1 | 1762014 | rs7553640 | -0.464 | 1.409 | -0.126 | 2.822 | -0.076 | -0.031 | -0.031 | 0 | 0.84 | 0.55 | 0.88 | 0.89 | 0.94 | 0.7 |
| 1 | 1776269 | rs4648727 | 0.398 | 1.545 | -0.35 | 2.851 | 0.001 | 0.021 | 0.021 | 0 | 0.54 | 0.37 | 0.3 | 0.36 | 0.44 | 0.33 |
| 1 | 1781220 | rs6681938 | 1.575 | 1.943 | -0.611 | 2.557 | -0.106 | -0.045 | -0.045 | 0 | 0.16 | 0.01 | 0.2 | 0.18 | 0.3 | 0.18 |
| 1 | 1793111 | rs10907192 | -0.437 | 0.913 | -0.245 | 2.114 | -0.081 | -0.034 | -0.034 | 0 | 0.84 | 0.51 | 0.87 | 0.9 | 0.94 | 0.7 |
| 1 | 1801034 | rs4648592 | 0.918 | 0.872 | -0.265 | 2.124 | -0.001 | -0.001 | -0.001 | 0 | 0.21 | 0.19 | 0.18 | 0.1 | 0.23 | 0.12 |
| 1 | 1804302 | rs3855951 | -0.422 | 0.874 | -0.347 | 2.136 | -0.083 | 0.04 | 0.04 | 1.254 | 0.84 | 0.5 | 0.87 | 0.83 | 0.94 | 0.69 |
| 1 | 1810090 | rs7525092 | 0.74 | 0.919 | -0.547 | 2.054 | 0.004 | -0.005 | -0.005 | 0.324 | 0.22 | 0.26 | 0.18 | 0.16 | 0.23 | 0.12 |
| 1 | 1812521 | rs6603802 | 0.504 | 1.265 | -1.098 | 2.165 | -0.125 | -0.121 | -0.121 | 1.902 | 0.29 | 0.07 | 0.57 | 0.51 | 0.5 | 0.36 |
| 1 | 1818129 | rs16824526 | 1.222 | 1.061 | -1.139 | 1.819 | 0.035 | 0.038 | 0.038 | 0 | 0.3 | 0.07 | 0.11 | 0.22 | 0.2 | 0.21 |
| 1 | 1829482 | rs745910 | 1.198 | 1.045 | -1.26 | 1.789 | 0.043 | 0.065 | 0.065 | 0 | 0.31 | 0.07 | 0.1 | 0.17 | 0.2 | 0.21 |
| 1 | 1844046 | rs2474460 | 0.458 | 1.516 | -1.43 | 2.27 | -0.194 | -0.127 | -0.127 | 0 | 0.29 | 0.03 | 0.35 | 0.38 | 0.49 | 0.34 |
| 1 | 1846582 | rs2748987 | -1.528 | 1.309 | -1.471 | 1.983 | 0.029 | 0.013 | 0.013 | 0 | 0.71 | 0.95 | 0.84 | 0.81 | 0.79 | 0.79 |
| 1 | 1846765 | rs2247560 | 0.611 | 1.271 | -1.384 | 2.228 | -0.149 | -0.13 | -0.13 | 0 | 0.29 | 0.06 | 0.51 | 0.48 | 0.49 | 0.39 |
| 1 | 1847531 | rs28710181 | 0.276 | 1.092 | -0.336 | 2.263 | -0.073 | 0.03 | 0.03 | 0.015 | 0.14 | 0.43 | 0.11 | 0.16 | 0.04 | 0.27 |
| 1 | 1852484 | rs28508199 | -0.953 | 2.11 | 0.001 | 2.506 | 0.023 | 0.032 | 0.032 | 0.013 | 0.39 | 0.77 | 0.32 | 0.32 | 0.3 | 0.39 |
| 1 | 1853394 | rs2144687 | 0.273 | 0.498 | -0.145 | 1.675 | -0.076 | 0.02 | 0.02 | 0 | 0.14 | 0.41 | 0.1 | 0.15 | 0.04 | 0.26 |
| 1 | 1854321 | rs2250833 | -1.401 | 1.038 | -0.985 | 1.348 | 0.029 | 0.016 | 0.016 | 0 | 0.7 | 0.95 | 0.83 | 0.81 | 0.78 | 0.79 |
| 1 | 1865298 | rs2803316 | 0.044 | 1.083 | -0.861 | 1.37 | -0.003 | 0.003 | 0.003 | 0 | 0.44 | 0.56 | 0.62 | 0.64 | 0.54 | 0.66 |
| 1 | 1874581 | rs2803329 | -0.874 | 1.367 | -1.765 | 1.532 | 0.038 | 0.021 | 0.021 | 0 | 0.7 | 0.93 | 0.83 | 0.81 | 0.78 | 0.79 |
| 1 | 1875267 | rs2262190 | -0.854 | 0.998 | -1.865 | 1.086 | 0.042 | 0.04 | 0.04 | 0 | 0.69 | 0.95 | 0.83 | 0.82 | 0.79 | 0.8 |
| 1 | 1888193 | rs3820011 | -0.455 | 0.657 | -1.532 | 0.753 | -0.016 | -0.008 | -0.008 | 0.41 | 0.24 | 0.11 | 0.23 | 0.23 | 0.25 | 0.16 |
| 1 | 1900232 | rs16824588 | -0.19 | 0.131 | -1.29 | 0.696 | 0.005 | 0.018 | 0.018 | 0 | 0.53 | 0.43 | 0.44 | 0.31 | 0.43 | 0.33 |
| 1 | 1916587 | rs3795283 | -0.69 | 0.315 | -1.633 | 1.083 | -0.012 | -0.019 | -0.019 | 0.068 | 0.29 | 0.21 | 0.24 | 0.32 | 0.31 | 0.29 |
| 1 | 1926087 | rs6661597 | -0.369 | -0.021 | -1.957 | 1.113 | -0.034 | -0.024 | -0.024 | 8.572 | 0.57 | 0.92 | 0.4 | 0.66 | 0.39 | 0.68 |
| 1 | 1930754 | rs6605081 | 0.079 | -0.185 | -1.227 | 0.562 | 0 | -0.006 | -0.006 | 0 | 0.41 | 0.57 | 0.21 | 0.2 | 0.28 | 0.29 |
| 1 | 1941929 | rs6656398 | 0.724 | 0.272 | -0.826 | 1.286 | 0.001 | 0.007 | 0.007 | 0 | 0.74 | 0.66 | 0.51 | 0.3 | 0.72 | 0.53 |
| 1 | 1946591 | rs13303016 | 0.489 | 0.056 | -0.655 | 1.158 | -0.004 | 0.013 | 0.013 | 0 | 0.84 | 0.76 | 0.73 | 0.82 | 0.83 | 0.77 |
| 1 | 1950621 | rs28603108 | 0.527 | 0.045 | -0.646 | 1.153 | -0.004 | 0.015 | 0.015 | 0 | 0.84 | 0.75 | 0.73 | 0.82 | 0.82 | 0.72 |
| 1 | 1967954 | rs2376803 | 0.667 | -0.274 | -0.927 | 0.929 | 0.001 | 0.001 | 0.001 | 0 | 0.7 | 0.67 | 0.67 | 0.75 | 0.73 | 0.65 |
| 1 | 1991014 | rs1878745 | -0.906 | -0.307 | -1.601 | 2.092 | 0.022 | -0.105 | -0.105 | 0 | 0.24 | 0.07 | 0.2 | 0.48 | 0.19 | 0.29 |
| 1 | 1992748 | rs6605074 | -0.896 | -0.312 | -1.604 | 2.49 | 0.021 | -0.026 | -0.026 | 1.699 | 0.23 | 0.06 | 0.2 | 0.25 | 0.19 | 0.25 |
| 1 | 2000592 | rs2678939 | -0.817 | -2.071 | -2.208 | 1.496 | 0.077 | 0.014 | 0.014 | 0.466 | 0.36 | 0.78 | 0.2 | 0.7 | 0.18 | 0.38 |
| 1 | 2004098 | rs6663158 | -0.824 | -2.224 | -1.669 | 1.353 | 0.014 | 0.008 | 0.008 | 0 | 0.28 | 0.25 | 0.25 | 0.77 | 0.19 | 0.46 |
| 1 | 2006108 | rs2254669 | -1.524 | -2.376 | -1.808 | 1.034 | 0.165 | 0.33 | 0.33 | 1.337 | 0.52 | 0.9 | 0.39 | 0.95 | 0.33 | 0.64 |
| 1 | 2017297 | rs2803310 | -1.327 | -1.456 | -0.655 | 1.35 | 0.041 | 0.043 | 0.043 | 0 | 0.48 | 0.55 | 0.35 | 0.83 | 0.31 | 0.61 |
| 1 | 2024064 | rs2459994 | -0.787 | -1.739 | -0.255 | 1.433 | 0.031 | -0.009 | -0.009 | 0 | 0.3 | 0.22 | 0.27 | 0.64 | 0.21 | 0.47 |
| 1 | 2026361 | rs12755035 | 0.437 | -1.931 | -0.053 | 1.341 | 0.015 | 0.01 | 0.01 | 0 | 0.16 | 0.13 | 0.07 | 0.1 | 0.12 | 0.15 |
| 1 | 2026403 | rs3128326 | -1.873 | -0.761 | -0.262 | 1.124 | 0.035 | 0.257 | 0.257 | 1.086 | 0.55 | 0.89 | 0.58 | 0.93 | 0.47 | 0.7 |
| 1 | 2026749 | rs884080 | -1.852 | -0.767 | -0.036 | 1.109 | 0.041 | 0.18 | 0.18 | 0.976 | 0.55 | 0.81 | 0.57 | 0.94 | 0.47 | 0.7 |
| 1 | 2027901 | rs7513222 | -0.694 | -0.532 | -0.169 | 1.351 | -0.009 | -0.042 | -0.042 | 0 | 0.31 | 0.08 | 0.45 | 0.64 | 0.35 | 0.52 |
| 1 | 2033256 | rs908742 | -0.706 | -0.089 | -0.227 | 1.29 | -0.03 | -0.042 | -0.042 | 0.003 | 0.31 | 0.08 | 0.39 | 0.64 | 0.35 | 0.52 |
| 1 | 2035684 | rs10910030 | 0.883 | 0.277 | 0.512 | 1.597 | -0.003 | 0.001 | 0.001 | 0 | 0.65 | 0.42 | 0.55 | 0.68 | 0.61 | 0.6 |
| 1 | 2038732 | rs12410859 | -0.287 | -0.331 | 0.105 | 1.221 | -0.006 | 0.012 | 0.012 | 0 | 0.24 | 0.07 | 0.36 | 0.19 | 0.29 | 0.31 |
| 1 | 2040898 | rs4648807 | 1.103 | -0.426 | 0.358 | 1.437 | 0.01 | -0.018 | -0.018 | 0 | 0.61 | 0.32 | 0.54 | 0.56 | 0.61 | 0.57 |
| 1 | 2040936 | rs4648808 | -0.485 | -0.415 | 0.409 | 1.465 | -0.012 | 0.081 | 0.081 | 0 | 0.85 | 0.56 | 0.92 | 0.81 | 0.89 | 0.89 |
| 1 | 2044037 | rs925905 | 0.782 | -0.482 | 0.545 | 1.441 | 0.001 | -0.003 | -0.003 | 0.675 | 0.66 | 0.75 | 0.59 | 0.68 | 0.61 | 0.59 |
| 1 | 2051513 | rs3107151 | 0.558 | -0.335 | 0.573 | 1.429 | -0.006 | -0.009 | -0.009 | 0 | 0.67 | 0.8 | 0.59 | 0.68 | 0.61 | 0.58 |

| | | | | | | | | | | | | | | | | |
|---|---|---|---|---|---|---|---|---|---|---|---|---|---|---|---|---|
| 1 | 2058023 | rs3128291 | -0.967 | -0.29 | 0.381 | 1.249 | 0.004 | 0.008 | 0.008 | 0.022 | 0.88 | 0.95 | 0.95 | 0.94 | 0.89 | 0.9 |
| 1 | 2068906 | rs3128296 | -0.232 | -0.304 | 0.113 | 0.984 | -0.022 | -0.001 | -0.001 | 0 | 0.91 | 0.97 | 0.96 | 0.94 | 0.88 | 0.91 |
| 1 | 2069681 | rs3753242 | 0.708 | -0.287 | 0.065 | 1.311 | 0.009 | 0.02 | 0.02 | 0.121 | 0.07 | 0.02 | 0.13 | 0.6 | 0.06 | 0.17 |
| 1 | 2070100 | rs3128297 | 0.225 | -0.288 | 0.135 | 1.294 | -0.004 | -0.001 | -0.001 | 0 | 0.08 | 0.06 | 0.04 | 0.05 | 0.1 | 0.11 |
| 1 | 2071340 | rs424079 | 0.579 | -0.344 | 0.87 | 1.124 | 0.009 | -0.004 | -0.004 | 1.169 | 0.69 | 0.63 | 0.57 | 0.74 | 0.61 | 0.58 |
| 1 | 2074104 | rs12026349 | 0.263 | -0.341 | 1.049 | 1.124 | 0.005 | 0 | 0 | 0 | 0.08 | 0.12 | 0.05 | 0.06 | 0.1 | 0.11 |
| 1 | 2075570 | rs377283 | -0.751 | -0.351 | 1.116 | 1.12 | -0.013 | -0.001 | -0.001 | 0 | 0.29 | 0.2 | 0.41 | 0.26 | 0.39 | 0.42 |
| 1 | 2082505 | rs9442619 | 0.302 | -0.346 | 1.096 | 1.112 | 0.003 | 0 | 0 | 0 | 0.08 | 0.1 | 0.04 | 0.06 | 0.1 | 0.11 |
| 1 | 2082566 | rs2257182 | -1.323 | -0.389 | 1.236 | 1.096 | -0.004 | -0.006 | -0.006 | 4.836 | 0.35 | 0.7 | 0.45 | 0.34 | 0.39 | 0.44 |
| 1 | 2117558 | rs3123595 | -0.291 | -0.211 | 1.15 | 1.078 | 0.031 | 0.017 | 0.017 | 0 | 0.1 | 0.31 | 0.07 | 0.06 | 0.1 | 0.11 |
| 1 | 2118339 | rs262685 | -0.762 | -0.435 | 0.721 | 1.033 | 0.019 | -0.002 | -0.002 | 0 | 0.31 | 0.41 | 0.43 | 0.27 | 0.39 | 0.42 |
| 1 | 2119833 | rs2460002 | -1.166 | -0.74 | -0.757 | -0.161 | -0.067 | 0.176 | 0.176 | 0.057 | 0.68 | 0.96 | 0.8 | 0.94 | 0.67 | 0.86 |
| 1 | 2121118 | rs2503701 | -0.363 | -0.691 | -0.615 | -0.244 | 0.004 | 0.004 | 0.004 | 0.905 | 0.57 | 0.58 | 0.65 | 0.86 | 0.61 | 0.84 |
| 1 | 2146966 | rs7512482 | 0.673 | -1.386 | -1.116 | 0.098 | -0.031 | -0.041 | -0.041 | 0.745 | 0.26 | 0.35 | 0.13 | 0.1 | 0.17 | 0.2 |
| 1 | 2156362 | rs2460000 | -0.881 | -1.114 | -1.272 | 0.405 | -0.009 | 0.036 | 0.036 | 0.423 | 0.65 | 0.77 | 0.76 | 0.8 | 0.64 | 0.87 |
| 1 | 2163568 | rs263533 | 0.132 | 0.155 | -0.956 | 1.441 | -0.023 | -0.026 | -0.026 | 0 | 0.28 | 0.26 | 0.56 | 0.62 | 0.4 | 0.58 |
| 1 | 2173504 | rs263526 | 0.898 | 0.285 | -0.915 | 1.383 | 0.004 | -0.001 | -0.001 | 0 | 0.2 | 0.23 | 0.46 | 0.23 | 0.37 | 0.32 |
| 1 | 2180524 | rs260513 | -0.286 | 0.218 | -0.914 | 1.27 | -0.003 | -0.002 | -0.002 | 0 | 0.32 | 0.33 | 0.56 | 0.41 | 0.45 | 0.41 |
| 1 | 2189580 | rs12084736 | 0.734 | 0.249 | -1.249 | 1.016 | -0.043 | 0.121 | 0.121 | 2.656 | 0.11 | 0.21 | 0.15 | 0.45 | 0.04 | 0.27 |
| 1 | 2192433 | rs10797417 | 0.65 | 0.259 | -1.267 | 1.022 | -0.047 | 0.129 | 0.129 | 3.106 | 0.12 | 0.23 | 0.15 | 0.45 | 0.04 | 0.3 |
| 1 | 2195117 | rs7547453 | 0.599 | 1.199 | -1.59 | 0.557 | -0.041 | -0.038 | -0.038 | 1.155 | 0.2 | 0.12 | 0.5 | 0.39 | 0.38 | 0.33 |
| 1 | 2199057 | rs11588312 | 0.203 | 0.441 | -0.897 | 0.113 | -0.002 | -0.013 | -0.013 | 0 | 0.11 | 0.1 | 0.06 | 0.02 | 0.07 | 0.08 |
| 1 | 2211849 | rs10910047 | -0.194 | 1.079 | -1.138 | 0.43 | 0.012 | -0.003 | -0.003 | 0 | 0.69 | 0.62 | 0.57 | 0.69 | 0.55 | 0.73 |
| 1 | 2218100 | rs12119470 | -0.692 | 0.544 | -1.209 | 0.329 | 0.005 | 0.006 | 0.006 | 0 | 0.17 | 0.12 | 0.1 | 0.07 | 0.12 | 0.18 |
| 1 | 2220649 | rs2017143 | -0.537 | 0.858 | -1.656 | 0.363 | -0.002 | 0.012 | 0.012 | 1.243 | 0.46 | 0.78 | 0.54 | 0.4 | 0.44 | 0.48 |
| 1 | 2221222 | rs2643885 | -0.244 | 1.36 | -0.947 | 0.769 | 0.017 | 0.037 | 0.037 | 0 | 0.14 | 0.19 | 0.38 | 0.32 | 0.23 | 0.34 |
| 1 | 2221534 | rs903924 | -1.372 | 0.844 | -1.309 | 0.47 | -0.026 | -0.04 | -0.04 | 0 | 0.36 | 0.77 | 0.53 | 0.42 | 0.42 | 0.46 |
| 1 | 2222583 | rs903919 | -1.195 | 0.506 | -2.095 | 0.137 | 0.025 | 0.038 | 0.038 | 0.159 | 0.8 | 0.95 | 0.88 | 0.85 | 0.85 | 0.8 |
| 1 | 2223866 | rs884940 | 1.13 | 0.545 | -2.073 | 0.134 | -0.013 | 0.002 | 0.002 | 0.121 | 0.43 | 0.17 | 0.35 | 0.44 | 0.42 | 0.32 |
| 1 | 2224836 | rs12743493 | 1.061 | 0.98 | -1.24 | -0.075 | -0.005 | -0.002 | -0.002 | 0 | 0.43 | 0.47 | 0.36 | 0.43 | 0.37 | 0.31 |
| 1 | 2224876 | rs10910050 | -0.584 | 2.025 | -1.612 | 0.095 | -0.031 | 0.028 | 0.028 | 0 | 0.8 | 0.93 | 0.89 | 0.9 | 0.75 | 0.86 |
| 1 | 2229478 | rs12562937 | 1.196 | 1.623 | -2.011 | 0.141 | -0.044 | -0.09 | -0.09 | 0.004 | 0.14 | 0.01 | 0.23 | 0.34 | 0.21 | 0.2 |
| 1 | 2231463 | rs903916 | 1.571 | 2.153 | -1.88 | -0.258 | -0.046 | 0.149 | 0.149 | 0.954 | 0.31 | 0.09 | 0.17 | 0.07 | 0.3 | 0.12 |
| 1 | 2234632 | rs2279702 | 1.092 | 1.535 | -1.554 | 0.307 | -0.045 | -0.022 | -0.022 | 0 | 0.15 | 0.02 | 0.11 | 0.14 | 0.19 | 0.13 |
| 1 | 2235672 | rs2843159 | -1.057 | 0.764 | -0.401 | 0.36 | 0.008 | 0.115 | 0.115 | 0 | 0.16 | 0.55 | 0.33 | 0.29 | 0.14 | 0.36 |
| 1 | 2240006 | rs2173049 | 0.131 | 0.57 | -0.649 | 0.254 | -0.019 | -0.012 | -0.012 | 0.153 | 0.77 | 0.75 | 0.89 | 0.82 | 0.85 | 0.78 |
| 1 | 2251160 | rs12082516 | -0.75 | 1.065 | -0.793 | 0.042 | -0.014 | 0.015 | 0.015 | 0.172 | 0.29 | 0.37 | 0.25 | 0.4 | 0.24 | 0.5 |
| 1 | 2254910 | rs2645075 | 0.434 | 1.335 | -1.575 | -0.202 | -0.001 | 0 | 0 | 0 | 0.85 | 0.85 | 0.69 | 0.85 | 0.78 | 0.86 |
| 1 | 2261222 | rs2645065 | 1.329 | 0.243 | -2.301 | -0.378 | 0.026 | 0.03 | 0.03 | 5.369 | 0.22 | 0.04 | 0.09 | 0.12 | 0.18 | 0.05 |
| 1 | 2263888 | rs903904 | -0.076 | 0.736 | -2.504 | -0.629 | -0.047 | 0.021 | 0.021 | 0.938 | 0.2 | 0.06 | 0.11 | 0.16 | 0.25 | 0.25 |
| 1 | 2265969 | rs2843146 | -1.116 | 0.566 | -2.537 | -0.548 | -0.091 | 0.062 | 0.062 | 0 | 0.75 | 0.94 | 0.64 | 0.9 | 0.69 | 0.79 |
| 1 | 2269948 | rs2843143 | 1.165 | 0.684 | -2.03 | -0.269 | 0.011 | 0.022 | 0.022 | 2.997 | 0.24 | 0.17 | 0.1 | 0.13 | 0.21 | 0.07 |
| 1 | 2270098 | rs2843142 | -0.493 | 0.246 | -2.214 | 0.095 | -0.036 | -0.054 | -0.054 | 0 | 0.56 | 0.81 | 0.51 | 0.56 | 0.56 | 0.57 |
| 1 | 2280661 | rs2055204 | 0.076 | 0.478 | -1.893 | -0.03 | -0.01 | 0.086 | 0.086 | 0.892 | 0.48 | 0.16 | 0.65 | 0.61 | 0.52 | 0.63 |
| 1 | 2281726 | rs7527871 | 1.709 | 0.44 | -1.998 | 0.01 | 0.012 | -0.045 | -0.045 | 0.26 | 0.17 | 0.08 | 0.18 | 0.36 | 0.15 | 0.27 |
| 1 | 2282135 | rs10910053 | 0.231 | 0.425 | -1.735 | -0.18 | 0.009 | -0.021 | -0.021 | 6.216 | 0.24 | 0.53 | 0.21 | 0.23 | 0.26 | 0.29 |
| 1 | 2283117 | rs4648831 | 1.535 | 0.747 | -2.861 | -0.185 | -0.014 | 0.121 | 0.121 | 0 | 0.22 | 0.04 | 0.09 | 0.02 | 0.2 | 0.13 |
| 1 | 2283161 | rs2840525 | -0.816 | 1.385 | -2.111 | 0.521 | 0.026 | -0.003 | -0.003 | 0.359 | 0.15 | 0.7 | 0.31 | 0.16 | 0.14 | 0.21 |
| 1 | 2284003 | rs2645082 | -0.967 | 1.302 | -2.766 | 0.485 | -0.001 | 0.051 | 0.051 | 4.541 | 0.74 | 0.95 | 0.88 | 0.84 | 0.77 | 0.93 |
| 1 | 2284195 | rs3795272 | 1.126 | 1.588 | -2.668 | 0.61 | 0.027 | -0.034 | -0.034 | 0 | 0.4 | 0.17 | 0.27 | 0.4 | 0.35 | 0.4 |
| 1 | 2286408 | rs2840531 | -0.229 | 2.735 | -2.403 | 0.918 | -0.029 | -0.026 | -0.026 | 2.826 | 0.08 | 0.03 | 0.38 | 0.21 | 0.16 | 0.23 |
| 1 | 2289321 | rs903914 | -0.382 | 1.376 | -2.074 | 0.191 | -0.031 | 0.009 | 0.009 | 0 | 0.17 | 0.06 | 0.16 | 0.15 | 0.25 | 0.14 |
| 1 | 2299651 | rs2840538 | 0.126 | 0.611 | -2.156 | 0.071 | 0.002 | 0.081 | 0.081 | 4.355 | 0.74 | 0.94 | 0.88 | 0.88 | 0.78 | 0.93 |
| 1 | 2301691 | rs6665546 | -1.064 | 0.337 | -2.164 | 0.65 | 0.015 | -0.024 | -0.024 | 0 | 0.27 | 0.11 | 0.14 | 0.31 | 0.21 | 0.23 |
| 1 | 2303566 | rs2840534 | -0.226 | 0.434 | -1.573 | 1.07 | -0.008 | -0.014 | -0.014 | 0 | 0.16 | 0.04 | 0.09 | 0.14 | 0.15 | 0.11 |
| 1 | 2306152 | rs12031614 | -0.003 | 0.567 | -1.63 | 0.825 | 0.037 | -0.02 | -0.02 | 0 | 0.45 | 0.26 | 0.45 | 0.49 | 0.35 | 0.48 |
| 1 | 2306695 | rs2279703 | -0.132 | 0.528 | -1.595 | 0.806 | 0.038 | -0.035 | -0.035 | 0.292 | 0.45 | 0.2 | 0.44 | 0.49 | 0.35 | 0.48 |
| 1 | 2306972 | rs7519807 | 0.358 | 0.371 | -1.832 | 0.761 | -0.008 | -0.158 | -0.158 | 0 | 0.37 | 0.04 | 0.33 | 0.45 | 0.3 | 0.27 |
| 1 | 2307481 | rs12033324 | 0.264 | 0.933 | -1.408 | 0.477 | -0.001 | -0.005 | -0.005 | 0 | 0.06 | 0.06 | 0.09 | 0.04 | 0.03 | 0.18 |
| 1 | 2308029 | rs12070746 | 0.01 | 0.861 | -1.611 | 0.524 | -0.001 | -0.013 | -0.013 | 0 | 0.09 | 0.05 | 0.11 | 0.14 | 0.08 | 0.04 |
| 1 | 2309082 | rs2843160 | -0.59 | 2.717 | -0.857 | 0.891 | -0.014 | 0.046 | 0.046 | 0 | 0.6 | 0.26 | 0.51 | 0.71 | 0.46 | 0.6 |
| 1 | 2310193 | rs903901 | -0.114 | 1.356 | -0.998 | 0.347 | 0 | -0.002 | -0.002 | 0 | 0.07 | 0.07 | 0.11 | 0.07 | 0.05 | 0.3 |
| 1 | 2314319 | rs2643901 | -1.132 | 1.532 | -1.084 | 0.268 | 0 | 0 | 0 | 0 | 0.94 | 0.93 | 0.89 | 0.87 | 0.89 | 0.92 |
| 1 | 2315717 | rs2843127 | -0.858 | 1.348 | -0.95 | 0.35 | 0.025 | -0.078 | -0.078 | 0 | 0.66 | 0.53 | 0.58 | 0.85 | 0.55 | 0.76 |
| 1 | 2319428 | rs903903 | -0.332 | 1.064 | -1.868 | 0.573 | -0.043 | 0.089 | 0.089 | 0 | 0.55 | 0.11 | 0.43 | 0.68 | 0.47 | 0.4 |
| 1 | 2325371 | rs16825102 | 0.354 | 0.875 | -1.447 | 0.831 | -0.016 | -0.041 | -0.041 | 0.065 | 0.08 | 0 | 0.11 | 0.16 | 0.06 | 0.04 |
| 1 | 2326009 | rs1123571 | 0.1 | 0.955 | -1.144 | 0.428 | -0.043 | 0.032 | 0.032 | 0 | 0.35 | 0.24 | 0.46 | 0.32 | 0.45 | 0.34 |
| 1 | 2327075 | rs3736330 | 1.026 | 0.255 | -1.498 | 0.576 | 0.004 | -0.098 | -0.098 | 0 | 0.28 | 0.08 | 0.24 | 0.41 | 0.27 | 0.23 |
| 1 | 2327815 | rs2840532 | -0.132 | -0.181 | -0.754 | 0.515 | -0.005 | -0.002 | -0.002 | 0 | 0.57 | 0.44 | 0.57 | 0.48 | 0.62 | 0.56 |

| | | | | | | | | | | | | | | | | |
|---|---|---|---|---|---|---|---|---|---|---|---|---|---|---|---|---|
| 1 | 2328714 | rs2645081 | -0.439 | -0.152 | -0.259 | 0.474 | -0.007 | 0.002 | 0.002 | 0 | 0.87 | 0.67 | 0.82 | 0.9 | 0.89 | 0.79 |
| 1 | 2329808 | rs3001344 | -1.006 | -0.179 | -0.309 | 0.485 | 0.002 | -0.002 | -0.002 | 0.037 | 0.91 | 0.91 | 0.85 | 0.89 | 0.92 | 0.82 |
| 1 | 2330190 | rs6693447 | 0.863 | -0.151 | -0.483 | 0.45 | -0.033 | 0.04 | 0.04 | 0 | 0.36 | 0.29 | 0.46 | 0.25 | 0.46 | 0.34 |
| 1 | 2333031 | rs3001336 | -0.313 | 0.036 | -0.476 | 0.607 | -0.008 | 0.043 | 0.043 | 0 | 0.86 | 0.59 | 0.82 | 0.83 | 0.89 | 0.79 |
| 1 | 2336210 | rs1129332 | 0.871 | -0.01 | -0.584 | 0.57 | 0.008 | -0.08 | -0.08 | 0 | 0.3 | 0.09 | 0.24 | 0.41 | 0.28 | 0.24 |
| 1 | 2338569 | rs2494427 | 0.171 | 0.213 | 0.127 | 0.686 | 0.002 | 0.041 | 0.041 | 0 | 0.86 | 0.6 | 0.81 | 0.83 | 0.88 | 0.79 |
| 1 | 2338879 | rs6671420 | 0.856 | 0.149 | -0.003 | 0.657 | -0.031 | 0.001 | 0.001 | 0 | 0.69 | 0.5 | 0.73 | 0.71 | 0.75 | 0.69 |
| 1 | 2339074 | rs2494428 | 0.581 | 0.771 | 0.193 | 0.746 | 0.006 | 0.053 | 0.053 | 0.001 | 0.86 | 0.53 | 0.81 | 0.83 | 0.88 | 0.79 |
| 1 | 2351699 | rs3935659 | -0.419 | 0.7 | -0.754 | 0.851 | 0.002 | -0.005 | -0.005 | 0 | 0.34 | 0.31 | 0.28 | 0.52 | 0.28 | 0.44 |
| 1 | 2352146 | rs12731309 | -0.039 | 1.838 | -0.861 | 1.007 | 0.003 | 0 | 0 | 3.509 | 0.12 | 0.14 | 0.35 | 0.16 | 0.22 | 0.22 |
| 1 | 2358944 | rs10797423 | -0.081 | 1.783 | -0.827 | 0.475 | 0.01 | 0.009 | 0.009 | 1.929 | 0.29 | 0.46 | 0.32 | 0.33 | 0.29 | 0.36 |
| 1 | 2359280 | rs2494435 | -0.295 | 1.858 | -0.565 | 0.645 | 0.053 | 0.006 | 0.006 | 0.166 | 0.07 | 0.21 | 0.17 | 0.1 | 0.12 | 0.11 |
| 1 | 2359917 | rs2494436 | -0.902 | 1.435 | -1.264 | 0.664 | 0.018 | 0.018 | 0.018 | 0.74 | 0.65 | 0.75 | 0.68 | 0.76 | 0.71 | 0.78 |
| 1 | 2359940 | rs7515934 | 0.332 | 1.644 | -1.183 | 1.286 | -0.031 | -0.126 | -0.126 | 0.039 | 0.22 | 0.03 | 0.13 | 0.33 | 0.19 | 0.29 |
| 1 | 2359981 | rs12022929 | 0.482 | 1.676 | -1.055 | 1.366 | -0.007 | -0.009 | -0.009 | 0 | 0.07 | 0.04 | 0.11 | 0.24 | 0.1 | 0.19 |
| 1 | 2362020 | rs4531246 | -0.662 | 1.375 | -0.886 | 1.134 | -0.006 | -0.018 | -0.018 | 0.282 | 0.3 | 0.24 | 0.29 | 0.42 | 0.31 | 0.4 |
| 1 | 2362827 | rs2494641 | -0.556 | 1.371 | -0.878 | 0.941 | -0.003 | -0.006 | -0.006 | 0 | 0.3 | 0.27 | 0.25 | 0.38 | 0.3 | 0.39 |
| 1 | 2366316 | rs4648843 | -0.909 | 1.663 | -0.521 | 0.63 | 0.01 | -0.007 | -0.007 | 0 | 0.08 | 0.11 | 0.18 | 0.03 | 0.16 | 0.14 |
| 1 | 2369498 | rs4592207 | -0.912 | 0.931 | -0.074 | 0.606 | -0.001 | 0.006 | 0.006 | 0 | 0.38 | 0.42 | 0.55 | 0.73 | 0.39 | 0.6 |
| 1 | 2373089 | rs6659405 | -0.413 | 0.854 | 0.16 | 1.269 | -0.014 | -0.029 | -0.029 | 0.163 | 0.24 | 0.19 | 0.4 | 0.48 | 0.3 | 0.37 |
| 1 | 2376733 | rs10449222 | -0.013 | 0.707 | 0.246 | 1.268 | -0.016 | 0.033 | 0.033 | 0 | 0.12 | 0.19 | 0.15 | 0.24 | 0.08 | 0.19 |
| 1 | 2377400 | rs4443836 | 0.543 | 0.728 | 0.237 | 0.996 | -0.003 | 0.003 | 0.003 | 0.077 | 0.11 | 0.16 | 0.09 | 0.14 | 0.11 | 0.09 |
| 1 | 2379248 | rs11581548 | -0.15 | 0.828 | 0.112 | 0.82 | -0.028 | -0.053 | -0.053 | 0 | 0.23 | 0.15 | 0.39 | 0.44 | 0.28 | 0.34 |
| 1 | 2387101 | rs4648845 | -1.225 | 0.774 | 0.15 | -0.254 | 0 | -0.023 | -0.023 | 0 | 0.47 | 0.53 | 0.64 | 0.83 | 0.48 | 0.67 |
| 1 | 2390588 | rs10910078 | -1.319 | 0.879 | -0.039 | -0.265 | -0.004 | 0.087 | 0.087 | 0 | 0.49 | 0.68 | 0.67 | 0.87 | 0.5 | 0.67 |
| 1 | 2394226 | rs4648636 | -0.743 | 1.375 | -0.238 | 0.407 | 0 | 0 | 0 | 0 | 0.24 | 0.26 | 0.41 | 0.24 | 0.28 | 0.31 |
| 1 | 2404256 | rs2494626 | -0.766 | 1.122 | -0.417 | 0.188 | -0.004 | -0.026 | -0.026 | 0.065 | 0.35 | 0.51 | 0.33 | 0.19 | 0.31 | 0.25 |
| 1 | 2406887 | rs13376356 | -1.505 | 1.632 | -0.497 | 0.66 | 0.019 | 0.034 | 0.034 | 0.04 | 0.08 | 0.15 | 0.15 | 0.14 | 0.12 | 0.1 |
| 1 | 2409276 | rs12049628 | 0.509 | 1.861 | -1.116 | -0.586 | -0.002 | 0.001 | 0.001 | 0 | 0.63 | 0.57 | 0.66 | 0.79 | 0.61 | 0.7 |
| 1 | 2411451 | rs10797428 | 0.534 | 1.668 | -1.03 | -0.657 | 0.001 | -0.002 | -0.002 | 0 | 0.63 | 0.57 | 0.66 | 0.78 | 0.62 | 0.69 |
| 1 | 2418625 | rs7512269 | -0.68 | 2.087 | -0.971 | -0.635 | 0.015 | 0.028 | 0.028 | 0 | 0.06 | 0.08 | 0.15 | 0.16 | 0.11 | 0.11 |
| 1 | 2418935 | rs10910082 | 0.863 | 1.831 | -0.919 | -0.782 | 0.01 | -0.017 | -0.017 | 1.212 | 0.62 | 0.5 | 0.64 | 0.77 | 0.61 | 0.66 |
| 1 | 2420913 | rs6668720 | 0.683 | 1.908 | -0.893 | -0.847 | 0.003 | -0.005 | -0.005 | 0.05 | 0.66 | 0.6 | 0.67 | 0.79 | 0.65 | 0.68 |
| 1 | 2425992 | rs2477703 | 0.321 | 1.963 | -1.2 | -1.037 | -0.012 | 0.103 | 0.103 | 0 | 0.72 | 0.88 | 0.8 | 0.95 | 0.72 | 0.79 |
| 1 | 2427712 | rs3762444 | 0.502 | 1.487 | -2.218 | -0.787 | -0.1 | -0.11 | -0.11 | 0 | 0.36 | 0.09 | 0.41 | 0.46 | 0.46 | 0.31 |
| 1 | 2430617 | rs2296442 | 0.238 | 1.435 | -2.356 | -0.915 | -0.05 | -0.037 | -0.037 | 0 | 0.37 | 0.17 | 0.45 | 0.48 | 0.48 | 0.33 |
| 1 | 2431057 | rs2477707 | 1.274 | 1.379 | -1.837 | -0.845 | -0.046 | 0.031 | 0.031 | 0.006 | 0.29 | 0.54 | 0.32 | 0.44 | 0.19 | 0.4 |
| 1 | 2439191 | rs1107685 | 1.422 | 1.339 | -1.781 | -0.806 | -0.048 | 0.05 | 0.05 | 0 | 0.28 | 0.5 | 0.31 | 0.45 | 0.2 | 0.41 |
| 1 | 2444414 | rs7535528 | 0.984 | 1.441 | -1.987 | -0.747 | -0.116 | -0.066 | -0.066 | 0.895 | 0.32 | 0.01 | 0.3 | 0.3 | 0.39 | 0.21 |
| 1 | 2445406 | rs2247308 | 0.978 | 0.902 | -1.924 | -0.784 | 0.024 | -0.018 | -0.018 | 0 | 0.32 | 0.72 | 0.35 | 0.47 | 0.21 | 0.45 |
| 1 | 2454709 | rs2494625 | -0.068 | 1.055 | -2.041 | -0.929 | -0.012 | 0.075 | 0.075 | 0 | 0.71 | 0.83 | 0.79 | 0.95 | 0.71 | 0.79 |
| 1 | 2456203 | rs3122922 | -0.546 | 0.865 | -2.094 | -0.995 | -0.023 | 0.046 | 0.046 | 0 | 0.82 | 0.91 | 0.84 | 0.97 | 0.78 | 0.85 |
| 1 | 2465329 | rs11584295 | 0.523 | 0.938 | -2.17 | -0.74 | -0.092 | -0.102 | -0.102 | 0 | 0.36 | 0.1 | 0.44 | 0.47 | 0.5 | 0.33 |
| 1 | 2466633 | rs2494606 | 0.132 | 0.576 | -1.941 | -1.033 | -0.019 | 0.094 | 0.094 | 0.653 | 0.75 | 0.91 | 0.83 | 0.94 | 0.74 | 0.76 |
| 1 | 2488608 | rs1886730 | -0.713 | 0.297 | -1.154 | -0.886 | -0.041 | -0.037 | -0.037 | 0.067 | 0.59 | 0.82 | 0.56 | 0.58 | 0.48 | 0.71 |
| 1 | 2494330 | rs2234167 | 0.226 | 0.015 | -1.148 | -0.943 | 0.063 | 0.091 | 0.091 | 0.079 | 0.25 | 0.1 | 0.09 | 0.06 | 0.15 | 0.17 |
| 1 | 2502780 | rs6667605 | -0.633 | 0.209 | -1.012 | -0.934 | -0.003 | -0.004 | -0.004 | 0 | 0.56 | 0.63 | 0.54 | 0.58 | 0.49 | 0.69 |
| 1 | 2513216 | rs734999 | -0.826 | 0.284 | -1.094 | -0.93 | -0.038 | -0.054 | -0.054 | 0 | 0.57 | 0.86 | 0.54 | 0.57 | 0.47 | 0.69 |
| 1 | 2521430 | rs10910097 | 0.018 | 0.383 | -1.022 | -0.926 | 0.03 | 0.072 | 0.072 | 0 | 0.28 | 0.16 | 0.11 | 0.08 | 0.21 | 0.22 |
| 1 | 2522392 | rs2296443 | 1.509 | 0.53 | -1.28 | -0.891 | -0.118 | -0.084 | -0.084 | 0 | 0.28 | 0.01 | 0.23 | 0.29 | 0.33 | 0.19 |
| 1 | 2524915 | rs10752747 | 0.38 | 0.612 | -0.831 | -0.707 | 0 | 0.016 | 0.016 | 0.301 | 0.31 | 0.61 | 0.45 | 0.53 | 0.31 | 0.54 |
| 1 | 2532416 | rs2985859 | -0.636 | 0.617 | -0.818 | -0.704 | 0 | 0.012 | 0.012 | 0.341 | 0.68 | 0.38 | 0.55 | 0.47 | 0.69 | 0.47 |
| 1 | 2538209 | rs10910105 | 0.41 | 0.617 | -0.796 | -0.694 | 0.002 | 0.047 | 0.047 | 0.183 | 0.31 | 0.54 | 0.45 | 0.53 | 0.31 | 0.53 |
| 1 | 2544640 | rs10752748 | 0.397 | 0.704 | -0.823 | -0.723 | 0.003 | 0.049 | 0.049 | 0.122 | 0.31 | 0.53 | 0.45 | 0.53 | 0.31 | 0.53 |
| 1 | 2546229 | rs6684865 | 0.363 | 0.704 | -0.823 | -0.723 | 0.002 | 0.047 | 0.047 | 0.165 | 0.31 | 0.54 | 0.45 | 0.53 | 0.31 | 0.53 |
| 1 | 2551785 | rs12117343 | 0.098 | 0.697 | -0.9 | -0.796 | 0 | 0.111 | 0.111 | 0 | 0.28 | 0.06 | 0.11 | 0.07 | 0.22 | 0.24 |
| 1 | 2551900 | rs11590198 | 1.901 | 0.529 | -0.065 | -0.76 | 0 | 0.009 | 0.009 | 0.234 | 0.19 | 0.14 | 0.29 | 0.49 | 0.18 | 0.43 |
| 1 | 2553624 | rs3890745 | -0.042 | 0.297 | -0.068 | -0.758 | 0 | 0.039 | 0.039 | 0.335 | 0.32 | 0.58 | 0.45 | 0.53 | 0.31 | 0.51 |
| 1 | 2564183 | rs6698817 | -0.284 | 0.335 | -0.052 | -0.742 | 0.002 | 0.052 | 0.052 | 0.273 | 0.69 | 0.44 | 0.55 | 0.47 | 0.69 | 0.47 |
| 1 | 2700372 | rs4648390 | 1.571 | -0.074 | -0.669 | -0.405 | 0.009 | -0.021 | -0.021 | 0 | 0.18 | 0.09 | 0.29 | 0.48 | 0.16 | 0.43 |
| 1 | 2713327 | rs2377041 | 0.165 | 0.256 | -0.613 | 0.066 | -0.045 | -0.019 | -0.019 | 0 | 0.39 | 0.14 | 0.39 | 0.36 | 0.44 | 0.24 |
| 1 | 2727804 | rs10909880 | 0.109 | 0.269 | -0.756 | 0.263 | -0.07 | -0.031 | -0.031 | 0 | 0.43 | 0.07 | 0.4 | 0.35 | 0.46 | 0.25 |
| 1 | 2733587 | rs10909890 | -1.364 | 0.263 | 0.527 | 0.362 | 0.022 | 0.077 | 0.077 | 0.111 | 0.08 | 0.36 | 0.04 | 0.1 | 0.02 | 0.05 |
| 1 | 2742491 | rs12071313 | -1.151 | 0.298 | 0.238 | 0.399 | -0.016 | 0.019 | 0.019 | 0 | 0.07 | 0.17 | 0.03 | 0.1 | 0.02 | 0.05 |
| 1 | 2749921 | rs4648482 | 0.359 | 0.421 | 0.365 | 0.542 | 0.013 | 0.017 | 0.017 | 0 | 0.27 | 0.37 | 0.26 | 0.3 | 0.29 | 0.16 |
| 1 | 2750437 | rs1563471 | -0.147 | 0.419 | 0.398 | 0.648 | 0.002 | -0.004 | -0.004 | 0 | 0.45 | 0.45 | 0.52 | 0.53 | 0.53 | 0.7 |
| 1 | 2756397 | rs4648383 | 0.014 | 1.051 | 0.36 | 0.955 | 0 | -0.001 | -0.001 | 0 | 0.53 | 0.41 | 0.47 | 0.5 | 0.54 | 0.31 |
| 1 | 2756621 | rs4648384 | 0.142 | 0.919 | 0.245 | 0.968 | 0 | 0.001 | 0.001 | 0.004 | 0.52 | 0.38 | 0.45 | 0.51 | 0.49 | 0.28 |
| 1 | 2761375 | rs12049543 | 0.264 | 0.71 | -0.137 | 0.856 | 0.027 | 0.005 | 0.005 | 0.413 | 0.47 | 0.23 | 0.38 | 0.45 | 0.42 | 0.27 |

| | | | | | | | | | | | | | | | |
|---|---|---|---|---|---|---|---|---|---|---|---|---|---|---|---|
| 1 | 2761863 | rs1456460 | 0.264 | 0.71 | -0.137 | 0.856 | 0.027 | 0.005 | 0.005 | 0.413 | 0.47 | 0.23 | 0.38 | 0.45 | 0.42 | 0.27 |
| 1 | 2784092 | rs897636 | -0.4 | 0.487 | 0.192 | 0.801 | 0.002 | 0.013 | 0.013 | 0 | 0.39 | 0.53 | 0.57 | 0.53 | 0.41 | 0.55 |
| 1 | 2785671 | rs6683974 | -0.893 | 0.429 | 0.541 | 0.864 | 0.026 | -0.005 | -0.005 | 0.656 | 0.06 | 0.29 | 0.03 | 0 | 0.03 | 0.05 |
| 1 | 2794470 | rs6676289 | -1.236 | 0.267 | 0.661 | 1.14 | 0.047 | 0.043 | 0.043 | 1.349 | 0.05 | 0.36 | 0.04 | 0 | 0 | 0.03 |
| 1 | 2794551 | rs10909793 | 0.09 | 0.195 | -0.709 | 0.835 | -0.029 | 0.055 | 0.055 | 0.703 | 0.47 | 0.08 | 0.49 | 0.37 | 0.58 | 0.49 |
| 1 | 2797856 | rs10797342 | -0.171 | 0.48 | -0.156 | 0.792 | -0.001 | 0.002 | 0.002 | 0 | 0.51 | 0.41 | 0.59 | 0.54 | 0.52 | 0.51 |
| 1 | 2817421 | rs2842910 | -1.091 | 0.354 | 0.317 | 1.331 | -0.014 | 0.118 | 0.118 | 0 | 0.2 | 0.44 | 0.28 | 0.36 | 0.18 | 0.35 |
| 1 | 2824989 | rs880725 | -0.608 | 0.582 | -0.185 | 1.252 | -0.027 | -0.04 | -0.04 | 0 | 0.17 | 0.06 | 0.33 | 0.27 | 0.25 | 0.33 |
| 1 | 2833332 | rs7340016 | -0.361 | 0.48 | -0.081 | 1.105 | 0.003 | 0.017 | 0.017 | 0 | 0.22 | 0.32 | 0.24 | 0.35 | 0.23 | 0.2 |
| 1 | 2835692 | rs897634 | -1.117 | 0.293 | -0.521 | 1.037 | 0.002 | -0.076 | -0.076 | 0 | 0.3 | 0.88 | 0.32 | 0.47 | 0.28 | 0.31 |
| 1 | 2842270 | rs2045331 | -1.198 | 0.047 | 0.233 | 1.14 | -0.003 | 0.028 | 0.028 | 0.468 | 0.26 | 0.62 | 0.39 | 0.52 | 0.26 | 0.4 |
| 1 | 2842319 | rs2045332 | -0.968 | 0.058 | 0.568 | 1.145 | 0 | 0.104 | 0.104 | 0.448 | 0.24 | 0.53 | 0.38 | 0.52 | 0.26 | 0.4 |
| 1 | 2847488 | rs2606411 | -1.54 | -0.234 | -0.143 | 0.699 | -0.059 | 0.03 | 0.03 | 0 | 0.55 | 0.91 | 0.65 | 0.69 | 0.53 | 0.62 |
| 1 | 2849190 | rs12403214 | -0.71 | -0.347 | 0.036 | 0.561 | 0 | -0.001 | -0.001 | 0.388 | 0.3 | 0.27 | 0.4 | 0.29 | 0.3 | 0.39 |
| 1 | 2850181 | rs4648441 | -0.244 | -0.373 | 0.018 | 0.544 | -0.002 | 0.002 | 0.002 | 0 | 0.25 | 0.19 | 0.37 | 0.23 | 0.27 | 0.32 |
| 1 | 2853759 | rs16823228 | -0.73 | -0.432 | 0.363 | 0.227 | 0.001 | 0.048 | 0.048 | 0 | 0.33 | 0.44 | 0.42 | 0.57 | 0.33 | 0.44 |
| 1 | 2859724 | rs2842909 | 0.496 | -0.49 | 0.344 | 0.112 | 0.006 | 0.017 | 0.017 | 0.006 | 0.19 | 0.14 | 0.2 | 0.12 | 0.21 | 0.16 |
| 1 | 2862320 | rs7539511 | -0.846 | -0.343 | 0.644 | 0.543 | 0.004 | 0.039 | 0.039 | 0 | 0.31 | 0.38 | 0.41 | 0.51 | 0.32 | 0.36 |
| 1 | 2867961 | rs2842933 | -0.902 | -0.451 | 0.663 | 0.623 | -0.01 | 0.04 | 0.04 | 0 | 0.31 | 0.43 | 0.4 | 0.52 | 0.28 | 0.35 |
| 1 | 2870850 | rs2606418 | -1.124 | -0.424 | 0.408 | 0.634 | -0.017 | 0.047 | 0.047 | 0 | 0.33 | 0.53 | 0.41 | 0.52 | 0.29 | 0.35 |
| 1 | 2876525 | rs10909845 | 0.157 | -0.093 | -0.035 | 0.647 | -0.001 | -0.002 | -0.002 | 0 | 0.76 | 0.75 | 0.62 | 0.45 | 0.72 | 0.64 |
| 1 | 2878366 | rs11583804 | 0.481 | 0.066 | -0.56 | 0.259 | 0.007 | 0.007 | 0.007 | 0.096 | 0.57 | 0.26 | 0.48 | 0.37 | 0.55 | 0.48 |
| 1 | 2879640 | rs878201 | -0.412 | -0.365 | -0.611 | -0.497 | -0.016 | -0.096 | -0.096 | 0 | 0.43 | 0.47 | 0.34 | 0.04 | 0.29 | 0.28 |
| 1 | 2883932 | rs2485945 | -0.597 | 0.348 | -0.521 | 0.7 | 0.002 | 0.04 | 0.04 | 0.442 | 0.25 | 0.55 | 0.22 | 0.31 | 0.18 | 0.31 |
| 1 | 2886988 | rs4072769 | 0.449 | 0.884 | -1.567 | 0.774 | -0.086 | -0.042 | -0.042 | 0 | 0.26 | 0.05 | 0.27 | 0.27 | 0.38 | 0.15 |
| 1 | 2889577 | rs12044030 | -0.211 | -0.545 | -0.696 | 0.724 | -0.057 | -0.012 | -0.012 | 0 | 0.66 | 0.41 | 0.81 | 0.69 | 0.84 | 0.62 |
| 1 | 2891713 | rs12046158 | 0.147 | 0.008 | -1.332 | 1.01 | 0.101 | -0.032 | -0.032 | 0.529 | 0.49 | 0.12 | 0.66 | 0.54 | 0.71 | 0.53 |
| 1 | 2898960 | rs6681741 | -1.007 | 0.775 | -0.153 | 1.356 | 0.084 | 0.197 | 0.197 | 0 | 0.08 | 0.31 | 0.21 | 0.21 | 0.11 | 0.04 |
| 1 | 2902128 | rs1572657 | -0.088 | 0.047 | -0.172 | 0.655 | 0.036 | 0.074 | 0.074 | 0 | 0.67 | 0.84 | 0.9 | 0.92 | 0.82 | 0.68 |
| 1 | 2904056 | rs10797373 | 0.799 | -0.368 | 0.624 | 0.973 | 0.038 | -0.048 | -0.048 | 1.2 | 0.54 | 0.17 | 0.64 | 0.54 | 0.71 | 0.56 |
| 1 | 2905605 | rs10909852 | -1.25 | -0.063 | 1.276 | 1.177 | 0.112 | 0.245 | 0.245 | 0 | 0.09 | 0.41 | 0.22 | 0.21 | 0.11 | 0.04 |
| 1 | 2917377 | rs12025260 | -1.763 | -0.078 | 1.854 | 1.205 | 0.078 | 0.363 | 0.363 | 0.928 | 0.12 | 0.43 | 0.25 | 0.38 | 0.11 | 0.12 |
| 1 | 2919942 | rs12022429 | -0.591 | -0.047 | 1.561 | 1.25 | 0.045 | 0.128 | 0.128 | 0 | 0.07 | 0.18 | 0.21 | 0.2 | 0.11 | 0.04 |
| 1 | 2928421 | rs7534897 | -0.146 | -0.123 | 0.296 | 0.698 | 0.011 | 0.053 | 0.053 | 0 | 0.3 | 0.08 | 0.11 | 0.2 | 0.18 | 0.25 |
| 1 | 2938265 | rs4576609 | -0.755 | 0.414 | 0.537 | 1.055 | 0.036 | 0.127 | 0.127 | 0 | 0.1 | 0.13 | 0.24 | 0.4 | 0.16 | 0.11 |
| 1 | 2938989 | rs3795263 | 0.941 | 0.714 | -0.511 | 0.202 | -0.053 | 0.016 | 0.016 | 0.545 | 0.2 | 0.01 | 0.12 | 0.16 | 0.26 | 0.09 |
| 1 | 2943183 | rs7412983 | -0.488 | 0.875 | 0.238 | 0.446 | 0.095 | 0.241 | 0.241 | 0 | 0.1 | 0.22 | 0.26 | 0.39 | 0.16 | 0.1 |
| 1 | 2956666 | rs2993510 | -1.251 | 0.592 | 0.241 | 0.312 | 0.048 | 0.165 | 0.165 | 0 | 0.15 | 0.65 | 0.31 | 0.39 | 0.17 | 0.1 |
| 1 | 2970464 | rs717795 | -1.081 | 0.605 | 0.375 | 0.362 | 0.117 | 0.356 | 0.356 | 0 | 0.14 | 0.41 | 0.29 | 0.39 | 0.18 | 0.1 |
| 1 | 2973322 | rs2993480 | -1.094 | 0.604 | -0.033 | 0.36 | 0.066 | 0.227 | 0.227 | 0 | 0.14 | 0.53 | 0.3 | 0.39 | 0.17 | 0.1 |
| 1 | 2974742 | rs1125017 | 0.705 | 0.26 | -1.382 | -0.006 | -0.079 | -0.056 | -0.056 | 0 | 0.33 | 0.1 | 0.48 | 0.39 | 0.42 | 0.5 |
| 1 | 2983963 | rs2297828 | 0.656 | -0.042 | -1.771 | -0.336 | -0.068 | 0.107 | 0.107 | 0 | 0.36 | 0.02 | 0.18 | 0.2 | 0.35 | 0.33 |
| 1 | 2984087 | rs2297829 | 0.141 | 0.14 | -1.367 | -0.233 | -0.001 | -0.061 | -0.061 | 0 | 0.33 | 0.13 | 0.38 | 0.41 | 0.35 | 0.37 |
| 1 | 2996602 | rs1569419 | -0.282 | 0.493 | -0.624 | -0.583 | -0.024 | -0.076 | -0.076 | 0 | 0.69 | 0.67 | 0.83 | 0.91 | 0.76 | 0.7 |
| 1 | 3000465 | rs760567 | -0.415 | 1.176 | -1.128 | -0.25 | -0.03 | 0.005 | 0.005 | 0 | 0.94 | 0.99 | 0.9 | 0.97 | 0.85 | 0.97 |
| 1 | 3000924 | rs6695131 | 0.02 | 1.042 | -0.845 | 0.096 | 0 | 0.005 | 0.005 | 1.732 | 0.58 | 0.4 | 0.58 | 0.66 | 0.61 | 0.67 |
| 1 | 3006283 | rs926244 | -0.255 | 0.938 | -0.682 | -0.175 | 0.003 | -0.041 | -0.041 | 0 | 0.3 | 0.35 | 0.24 | 0.13 | 0.32 | 0.14 |
| 1 | 3009969 | rs7543737 | 0.09 | -0.823 | -1.006 | 0.527 | 0.052 | -0.002 | -0.002 | 2.872 | 0.48 | 0.58 | 0.3 | 0.51 | 0.3 | 0.48 |
| 1 | 3013690 | rs2817168 | -0.086 | -0.307 | -1.221 | 0.716 | -0.015 | -0.025 | -0.025 | 0.085 | 0.67 | 0.83 | 0.53 | 0.62 | 0.44 | 0.79 |
| 1 | 3020246 | rs2993493 | -0.273 | -1.128 | -0.926 | 0.577 | -0.034 | 0.019 | 0.019 | 2.157 | 0.22 | 0.28 | 0.23 | 0.3 | 0.15 | 0.22 |
| 1 | 3024265 | rs1890336 | 0.522 | -1.035 | -1.408 | 0.217 | 0.037 | 0.078 | 0.078 | 0.056 | 0.57 | 0.77 | 0.54 | 0.81 | 0.41 | 0.71 |
| 1 | 3025712 | rs4648453 | -0.696 | -0.753 | -1.266 | 0.495 | 0.014 | 0.002 | 0.002 | 0 | 0.17 | 0.06 | 0.11 | 0.11 | 0.13 | 0.17 |
| 1 | 3035227 | rs6690373 | -0.701 | -1.328 | -0.481 | 0.537 | -0.041 | 0.119 | 0.119 | 0 | 0.16 | 0.45 | 0.2 | 0.3 | 0.1 | 0.2 |
| 1 | 3037613 | rs876740 | 0.093 | -0.848 | -1.552 | 0.357 | -0.036 | -0.041 | -0.041 | 0 | 0.63 | 0.91 | 0.49 | 0.61 | 0.42 | 0.61 |
| 1 | 3044181 | rs2817174 | -0.027 | -0.691 | -1.558 | 0.39 | -0.076 | -0.088 | -0.088 | 0.016 | 0.67 | 0.93 | 0.49 | 0.57 | 0.43 | 0.64 |
| 1 | 3047252 | rs2817178 | 0.072 | -0.657 | -1.524 | 0.346 | -0.072 | -0.087 | -0.087 | 0 | 0.65 | 0.93 | 0.51 | 0.55 | 0.44 | 0.57 |
| 1 | 3049713 | rs10909882 | -0.647 | -1.193 | -0.491 | 0.742 | -0.013 | 0.187 | 0.187 | 0 | 0.06 | 0.25 | 0.02 | 0.22 | 0.01 | 0.13 |
| 1 | 3054106 | rs2235176 | -0.746 | -1.187 | -0.36 | 0.265 | -0.03 | -0.006 | -0.006 | 0 | 0.33 | 0.58 | 0.39 | 0.69 | 0.3 | 0.59 |
| 1 | 3055876 | rs10797380 | 0.082 | -0.891 | -0.338 | -0.591 | -0.005 | 0.006 | 0.006 | 0.168 | 0.66 | 0.73 | 0.72 | 0.78 | 0.47 | 0.73 |
| 1 | 3057661 | rs7538096 | 0.264 | -0.992 | -0.447 | -0.625 | -0.006 | -0.008 | -0.008 | 0 | 0.49 | 0.47 | 0.55 | 0.69 | 0.33 | 0.57 |
| 1 | 3062361 | rs2817185 | -1.123 | -0.221 | -0.378 | -0.696 | -0.003 | -0.085 | -0.085 | 0 | 0.29 | 0.32 | 0.16 | 0 | 0.24 | 0.19 |
| 1 | 3072692 | rs731031 | -1.473 | -0.902 | -0.188 | 0.526 | -0.028 | 0.023 | 0.023 | 2.286 | 0.3 | 0.63 | 0.27 | 0.57 | 0.23 | 0.42 |
| 1 | 3074306 | rs2651902 | -0.889 | -1.068 | 0.263 | 0.603 | -0.001 | -0.002 | -0.002 | 0.425 | 0.16 | 0.16 | 0.15 | 0.04 | 0.16 | 0.23 |
| 1 | 3080019 | rs2742673 | -1.887 | -1.255 | -0.372 | 0.069 | -0.022 | 0.058 | 0.058 | 0.632 | 0.59 | 0.82 | 0.56 | 0.71 | 0.48 | 0.69 |
| 1 | 3080855 | rs1798246 | -1.595 | -1.29 | -0.088 | 0.944 | 0 | -0.007 | -0.007 | 0 | 0.55 | 0.4 | 0.51 | 0.48 | 0.47 | 0.69 |
| 1 | 3083712 | rs2651899 | -1.308 | -0.912 | 0.112 | 0.747 | 0.021 | 0.027 | 0.027 | 0.136 | 0.49 | 0.73 | 0.49 | 0.41 | 0.43 | 0.51 |
| 1 | 3086464 | rs7518255 | 0.062 | -0.862 | -0.085 | 0.76 | 0.006 | 0.035 | 0.035 | 0.777 | 0.3 | 0.2 | 0.29 | 0.15 | 0.26 | 0.27 |
| 1 | 3086496 | rs2651935 | -0.907 | -0.338 | 0.305 | 1.034 | -0.009 | 0.041 | 0.041 | 0.217 | 0.15 | 0.34 | 0.11 | 0.23 | 0.12 | 0.18 |
| 1 | 3101762 | rs2651927 | -0.525 | -0.504 | -0.338 | 0.997 | -0.011 | 0.006 | 0.006 | 0 | 0.28 | 0.59 | 0.26 | 0.29 | 0.24 | 0.2 |

| | | | | | | | | | | | | | | | | |
|---|---|---|---|---|---|---|---|---|---|---|---|---|---|---|---|---|
| 1 | 3103826 | rs4233024 | 0.806 | -0.732 | -0.375 | 1.423 | -0.01 | 0.053 | 0.053 | 0 | 0.84 | 0.74 | 0.78 | 0.72 | 0.85 | 0.76 |
| 1 | 3104630 | rs10752733 | 1.188 | -0.917 | -0.392 | 1.484 | -0.033 | 0.019 | 0.019 | 0 | 0.81 | 0.64 | 0.8 | 0.82 | 0.87 | 0.78 |
| 1 | 3107386 | rs12139196 | -0.196 | -1.039 | -0.297 | 1.195 | -0.026 | 0.022 | 0.022 | 0 | 0.26 | 0.51 | 0.22 | 0.29 | 0.19 | 0.2 |
| 1 | 3108496 | rs10737190 | 0.579 | -1.07 | -0.875 | 1.166 | -0.01 | -0.004 | -0.004 | 0 | 0.89 | 0.84 | 0.92 | 0.91 | 0.92 | 0.83 |
| 1 | 3109151 | rs7539775 | 0.75 | -0.927 | -1.28 | 1.215 | 0.056 | -0.011 | -0.011 | 0 | 0.61 | 0.21 | 0.69 | 0.62 | 0.72 | 0.63 |
| 1 | 3110735 | rs2651912 | -0.549 | -0.185 | -1.777 | 1.236 | -0.081 | -0.095 | -0.095 | 1.251 | 0.74 | 0.94 | 0.64 | 0.62 | 0.69 | 0.66 |
| 1 | 3111669 | rs12094678 | -0.898 | -1.646 | -1.124 | 1.23 | 0.096 | -0.165 | -0.165 | 0.619 | 0.36 | 0.94 | 0.27 | 0.45 | 0.18 | 0.47 |
| 1 | 3113680 | rs10909895 | -0.786 | -1.381 | -0.521 | 1.137 | 0.014 | -0.006 | -0.006 | 0.138 | 0.36 | 0.65 | 0.27 | 0.4 | 0.22 | 0.34 |
| 1 | 3119729 | rs4308918 | -1.523 | -0.773 | 0.486 | 0.861 | 0.035 | -0.029 | -0.029 | 0.193 | 0.1 | 0.43 | 0.1 | 0.02 | 0.09 | 0.07 |
| 1 | 3131235 | rs10909901 | -1.736 | -1.814 | -0.713 | 1.188 | 0.074 | -0.093 | -0.093 | 0.318 | 0.44 | 0.93 | 0.34 | 0.42 | 0.26 | 0.6 |
| 1 | 3133439 | rs2817146 | -0.585 | -2.205 | 0.196 | 1.3 | -0.014 | 0.01 | 0.01 | 0 | 0.2 | 0.3 | 0.14 | 0.25 | 0.12 | 0.19 |
| 1 | 3134331 | rs12124147 | 1.116 | -1.764 | -0.428 | 1.494 | -0.147 | -0.079 | -0.079 | 0 | 0.21 | 0.02 | 0.27 | 0.25 | 0.4 | 0.16 |
| 1 | 3140464 | rs16823542 | -0.937 | -2.748 | 1.695 | 1.73 | 0.016 | -0.027 | -0.027 | 0 | 0.07 | 0.17 | 0.07 | 0.01 | 0.08 | 0.03 |
| 1 | 3144745 | rs1553288 | -0.584 | -2.749 | 1.259 | 1.809 | 0.001 | 0.001 | 0.001 | 0 | 0.1 | 0.08 | 0.12 | 0.06 | 0.08 | 0.25 |
| 1 | 3146497 | rs10157420 | -0.737 | -2.847 | 1.183 | 1.875 | -0.023 | 0.012 | 0.012 | 0.111 | 0.18 | 0.31 | 0.12 | 0.23 | 0.1 | 0.19 |
| 1 | 3147030 | rs6424069 | -2.309 | -2.793 | 0.926 | 1.82 | 0.093 | -0.058 | -0.058 | 0 | 0.41 | 0.92 | 0.33 | 0.44 | 0.23 | 0.59 |
| 1 | 3147899 | rs10909905 | -1.821 | -2.422 | 1.456 | 1.975 | -0.026 | 0.006 | 0.006 | 0.827 | 0.29 | 0.37 | 0.24 | 0.31 | 0.18 | 0.44 |
| 1 | 3149145 | rs2817158 | -1.606 | -2.527 | 1.098 | 2.016 | -0.063 | 0.017 | 0.017 | 0.935 | 0.26 | 0.52 | 0.21 | 0.29 | 0.14 | 0.42 |
| 1 | 3150345 | rs12036784 | -1.068 | -1.976 | 0.436 | 2.124 | 0.014 | -0.015 | -0.015 | 1.515 | 0.13 | 0.06 | 0.14 | 0.21 | 0.1 | 0.29 |
| 1 | 3153237 | rs11807062 | -1.466 | -2.065 | 0.65 | 2.419 | 0.009 | -0.013 | -0.013 | 0 | 0.3 | 0.67 | 0.24 | 0.54 | 0.16 | 0.42 |
| 1 | 3155102 | rs2249336 | -1.349 | -2.168 | 0.848 | 1.899 | -0.029 | 0.004 | 0.004 | 0.414 | 0.31 | 0.57 | 0.24 | 0.33 | 0.17 | 0.42 |
| 1 | 3157614 | rs905138 | -1.278 | -2.063 | 0.756 | 1.638 | -0.071 | -0.038 | -0.038 | 0.392 | 0.3 | 0.5 | 0.24 | 0.18 | 0.16 | 0.4 |
| 1 | 3162209 | rs2455118 | -1.317 | -2.03 | -0.16 | 1.598 | 0.072 | -0.004 | -0.004 | 0 | 0.32 | 0.76 | 0.27 | 0.32 | 0.17 | 0.42 |
| 1 | 3168280 | rs10797386 | 0.975 | -1.563 | 0.641 | 1.725 | -0.014 | 0.173 | 0.173 | 0.074 | 0.89 | 0.63 | 0.95 | 0.81 | 0.96 | 0.88 |
| 1 | 3169088 | rs7539489 | 0.144 | -1.363 | -0.9 | 0.991 | -0.057 | -0.088 | -0.088 | 1.436 | 0.38 | 0.15 | 0.35 | 0.6 | 0.55 | 0.45 |
| 1 | 3177480 | rs3002685 | -0.856 | -1.926 | -0.097 | 0.146 | -0.038 | -0.128 | -0.128 | 0 | 0.28 | 0.44 | 0.32 | 0.01 | 0.18 | 0.15 |
| 1 | 3178111 | rs3002686 | -0.812 | -2.062 | -0.12 | 0.53 | -0.049 | 0 | 0 | 3.032 | 0.24 | 0.4 | 0.32 | 0.18 | 0.14 | 0.2 |
| 1 | 3180158 | rs2455107 | -1.63 | -2.121 | 0.343 | 0.496 | -0.028 | -0.011 | -0.011 | 0 | 0.29 | 0.57 | 0.38 | 0.19 | 0.18 | 0.22 |
| 1 | 3180633 | rs2455102 | -2.027 | -2.309 | -0.051 | 0.346 | 0.087 | 0.04 | 0.04 | 0.38 | 0.35 | 0.89 | 0.41 | 0.19 | 0.2 | 0.26 |
| 1 | 3189006 | rs2455125 | -0.96 | -2.183 | 0.261 | 0.276 | 0.021 | 0.001 | 0.001 | 0.872 | 0.25 | 0.73 | 0.32 | 0 | 0.14 | 0.14 |
| 1 | 3196680 | rs16823027 | -0.982 | -2.158 | 0.747 | 0.599 | -0.022 | 0.092 | 0.092 | 0.313 | 0.16 | 0.4 | 0.31 | 0.21 | 0.12 | 0.23 |
| 1 | 3197747 | rs10492940 | -0.76 | -1.661 | 0.169 | 0.387 | -0.007 | 0.112 | 0.112 | 0 | 0.18 | 0 | 0.09 | 0 | 0.12 | 0.11 |
| 1 | 3208375 | rs10909918 | 0.805 | -0.411 | -0.037 | 0.967 | -0.053 | -0.011 | -0.011 | 0 | 0.39 | 0.15 | 0.54 | 0.6 | 0.52 | 0.46 |
| 1 | 3209923 | rs1553291 | -1.075 | -0.799 | 0.122 | 0.941 | 0.007 | -0.038 | -0.038 | 0 | 0.68 | 0.81 | 0.69 | 0.44 | 0.7 | 0.58 |
| 1 | 3211458 | rs4648377 | -0.663 | -1.238 | 0.284 | 0.505 | -0.002 | -0.066 | -0.066 | 0 | 0.62 | 0.36 | 0.8 | 0.99 | 0.78 | 0.74 |
| 1 | 3214732 | rs2455144 | 0.336 | -0.849 | 0.293 | 0.601 | 0.004 | 0.023 | 0.023 | 0 | 0.42 | 0.28 | 0.69 | 0.68 | 0.56 | 0.52 |
| 1 | 3217151 | rs2455137 | 0.528 | -0.837 | 0.338 | 0.699 | -0.004 | -0.002 | -0.002 | 0 | 0.41 | 0.23 | 0.55 | 0.42 | 0.55 | 0.44 |
| 1 | 3219901 | rs2455135 | -0.954 | -0.932 | 0.433 | 0.596 | 0.124 | 0.056 | 0.056 | 0.072 | 0.62 | 0.89 | 0.71 | 0.73 | 0.75 | 0.62 |
| 1 | 3220101 | rs6673393 | 0.014 | -0.873 | 0.688 | 0.618 | 0.071 | 0.067 | 0.067 | 4.015 | 0.06 | 0.5 | 0.04 | 0 | 0 | 0 |
| 1 | 3221371 | rs7517281 | -0.565 | -0.685 | 0.464 | 0.736 | -0.008 | -0.006 | -0.006 | 0.016 | 0.15 | 0.14 | 0.11 | 0.31 | 0.19 | 0.17 |
| 1 | 3225940 | rs2500278 | 0.27 | -0.692 | 0.519 | 0.731 | 0.011 | 0.007 | 0.007 | 0 | 0.43 | 0.29 | 0.57 | 0.43 | 0.58 | 0.45 |
| 1 | 3227188 | rs2483260 | -0.514 | -0.703 | 0.349 | 0.727 | 0.012 | 0.006 | 0.006 | 0 | 0.58 | 0.75 | 0.43 | 0.57 | 0.42 | 0.55 |
| 1 | 3229185 | rs12743017 | -0.006 | -0.593 | 0.545 | 0.656 | -0.008 | -0.05 | -0.05 | 0.752 | 0.11 | 0.17 | 0.06 | 0 | 0.08 | 0.07 |
| 1 | 3236041 | rs2483266 | -0.13 | -0.409 | -0.005 | 0.784 | 0.058 | 0.021 | 0.021 | 0 | 0.48 | 0.73 | 0.38 | 0.55 | 0.37 | 0.5 |
| 1 | 3240199 | rs870465 | 0.517 | 0.946 | -0.37 | 0.602 | -0.041 | -0.032 | -0.032 | 0 | 0.08 | 0.02 | 0.1 | 0.28 | 0.17 | 0.25 |
| 1 | 3241135 | rs1108600 | 1.323 | 0.156 | -0.894 | -0.325 | -0.027 | -0.013 | -0.013 | 0.936 | 0.24 | 0.02 | 0.15 | 0.2 | 0.24 | 0.29 |
| 1 | 3241829 | rs2500271 | 0.539 | 0.824 | 0.16 | -0.16 | 0.031 | -0.012 | -0.012 | 0 | 0.49 | 0.59 | 0.28 | 0.01 | 0.35 | 0.37 |
| 1 | 3243017 | rs10909930 | -0.099 | 0.792 | 0.034 | -0.142 | 0.015 | 0.026 | 0.026 | 0 | 0.11 | 0.24 | 0.11 | 0.2 | 0.15 | 0.18 |
| 1 | 3248098 | rs2483274 | 0.001 | 0.657 | 0 | 0.071 | -0.035 | 0.02 | 0.02 | 0.26 | 0.61 | 0.83 | 0.51 | 0.35 | 0.51 | 0.62 |
| 1 | 3251159 | rs1110307 | -0.746 | 0.941 | 0.429 | 0.27 | 0.024 | 0.111 | 0.111 | 3.222 | 0.1 | 0.52 | 0.22 | 0.14 | 0.06 | 0.12 |
| 1 | 3252104 | rs2483278 | -0.594 | 1.341 | 0.259 | 0.404 | -0.001 | -0.002 | -0.002 | 0 | 0.73 | 0.78 | 0.67 | 0.86 | 0.7 | 0.8 |
| 1 | 3252169 | rs2483279 | 1.237 | 1.406 | -0.232 | 0.454 | -0.027 | 0.118 | 0.118 | 0.107 | 0.39 | 0.08 | 0.21 | 0.2 | 0.39 | 0.42 |
| 1 | 3253889 | rs7516150 | -0.618 | 1.446 | -0.188 | 0.673 | 0.007 | 0.001 | 0.001 | 0 | 0.31 | 0.48 | 0.44 | 0.65 | 0.31 | 0.38 |
| 1 | 3253941 | rs7552331 | -0.613 | 1.467 | -0.441 | 0.595 | 0.005 | 0.031 | 0.031 | 0 | 0.3 | 0.42 | 0.31 | 0.51 | 0.31 | 0.27 |
| 1 | 3259113 | rs882430 | 0.928 | 1.564 | -1.296 | 0.221 | 0.009 | 0.004 | 0.004 | 0 | 0.42 | 0.37 | 0.35 | 0.13 | 0.31 | 0.46 |
| 1 | 3261682 | rs4415513 | 0.336 | 1.938 | -1.295 | 0.814 | 0.009 | 0.011 | 0.011 | 0 | 0.66 | 0.41 | 0.49 | 0.58 | 0.61 | 0.61 |
| 1 | 3261925 | rs4648380 | 0.029 | 1.2 | -2.252 | 0.144 | -0.038 | 0.008 | 0.008 | 1.279 | 0.11 | 0.02 | 0.12 | 0.11 | 0.14 | 0.11 |
| 1 | 3264190 | rs2483289 | 0.215 | 0.63 | -1.934 | 0.175 | 0.001 | 0.05 | 0.05 | 0 | 0.73 | 0.84 | 0.68 | 0.82 | 0.72 | 0.82 |
| 1 | 3269828 | rs2483256 | -0.564 | 0.29 | -2.131 | 0.484 | 0.05 | 0.013 | 0.013 | 0.24 | 0.43 | 0.79 | 0.33 | 0.42 | 0.34 | 0.38 |
| 1 | 3273063 | rs946758 | -0.105 | 0.295 | -2.634 | 0.324 | 0.014 | 0.054 | 0.054 | 0 | 0.53 | 0.86 | 0.4 | 0.33 | 0.41 | 0.55 |
| 1 | 3275149 | rs12748963 | 0.226 | -0.523 | -3.089 | 0.215 | 0.034 | -0.049 | -0.049 | 0 | 0.24 | 0.03 | 0.27 | 0.34 | 0.17 | 0.4 |
| 1 | 3276138 | rs2500286 | -0.199 | -0.511 | -2.285 | -0.874 | -0.015 | -0.088 | -0.088 | 0 | 0.26 | 0.61 | 0.21 | 0.03 | 0.2 | 0.2 |
| 1 | 3276374 | rs12073172 | 0.65 | -0.02 | -2.335 | -0.482 | 0.056 | 0.027 | 0.027 | 0.174 | 0.51 | 0.3 | 0.58 | 0.63 | 0.42 | 0.58 |
| 1 | 3278959 | rs17399569 | 1.347 | -0.098 | -1.868 | -0.692 | -0.002 | -0.001 | -0.001 | 0 | 0.05 | 0.13 | 0.03 | 0.02 | 0.05 | 0.1 |
| 1 | 3279324 | rs7524171 | 0.311 | 0.129 | -2.224 | -0.734 | -0.015 | 0.005 | 0.005 | 0.005 | 0.63 | 0.32 | 0.69 | 0.69 | 0.55 | 0.63 |
| 1 | 3280021 | rs2500262 | -0.909 | 0.174 | -1.372 | -0.635 | 0.04 | 0.009 | 0.009 | 0 | 0.78 | 0.72 | 0.78 | 0.72 | 0.66 | 0.78 |
| 1 | 3285304 | rs4648487 | 0.972 | 0.012 | -1.246 | -1.044 | 0.034 | 0.1 | 0.1 | 2.701 | 0.47 | 0.24 | 0.57 | 0.72 | 0.42 | 0.58 |
| 1 | 3289524 | rs12135987 | 1.678 | -0.359 | -1.382 | -1.108 | 0.013 | -0.003 | -0.003 | 1.899 | 0.35 | 0.15 | 0.53 | 0.68 | 0.33 | 0.56 |
| 1 | 3289729 | rs1333190 | -0.626 | 0.013 | -1.196 | -0.998 | -0.065 | 0.024 | 0.024 | 0 | 0.7 | 0.9 | 0.76 | 0.72 | 0.57 | 0.76 |

| | | | | | | | | | | | | | | | | |
|---|---|---|---|---|---|---|---|---|---|---|---|---|---|---|---|---|
| 1 | 3289971 | rs1333189 | 0.821 | -0.148 | -1.013 | -0.859 | 0.068 | 0.147 | 0.147 | 3.011 | 0.48 | 0.86 | 0.62 | 0.72 | 0.39 | 0.66 |
| 1 | 3291139 | rs2249860 | 0.538 | 0.372 | -0.767 | -0.725 | 0.025 | 0.108 | 0.108 | 0.248 | 0.55 | 0.87 | 0.65 | 0.73 | 0.43 | 0.69 |
| 1 | 3293665 | rs4648489 | 0.214 | 0.892 | -0.778 | -0.341 | -0.033 | 0.007 | 0.007 | 0 | 0.66 | 0.74 | 0.72 | 0.7 | 0.53 | 0.69 |
| 1 | 3294972 | rs1538693 | 0.507 | 1.368 | -0.588 | -0.312 | 0.01 | 0.077 | 0.077 | 0 | 0.77 | 0.65 | 0.67 | 0.53 | 0.7 | 0.74 |
| 1 | 3297638 | rs12085231 | 0.023 | 1.303 | -1.106 | -0.8 | -0.002 | -0.008 | -0.008 | 0 | 0.14 | 0.11 | 0.09 | 0 | 0.13 | 0.07 |
| 1 | 3300807 | rs868688 | 1.364 | 1.342 | -0.876 | -0.941 | -0.003 | -0.003 | -0.003 | 0 | 0.56 | 0.66 | 0.54 | 0.58 | 0.47 | 0.61 |
| 1 | 3304336 | rs10492938 | 0.439 | 1.325 | -1.037 | -1.184 | 0.003 | -0.025 | -0.025 | 3.673 | 0.07 | 0.09 | 0.09 | 0 | 0.13 | 0.08 |
| 1 | 3305016 | rs17399998 | 0.934 | 1.458 | -1.559 | -1.781 | -0.033 | 0.065 | 0.065 | 0 | 0.21 | 0.08 | 0.14 | 0.01 | 0.23 | 0.07 |
| 1 | 3307746 | rs2493278 | -0.616 | 1.537 | -1.647 | -1.673 | -0.016 | -0.045 | -0.045 | 0 | 0.82 | 0.91 | 0.7 | 0.64 | 0.75 | 0.78 |
| 1 | 3308034 | rs2493275 | 0.851 | 1.481 | -1.688 | -1.755 | 0.002 | -0.014 | -0.014 | 0 | 0.13 | 0.1 | 0.05 | 0 | 0.08 | 0.06 |
| 1 | 3312256 | rs17822575 | 1.03 | 1.655 | -1.659 | -1.614 | -0.052 | 0.092 | 0.092 | 0 | 0.21 | 0.04 | 0.14 | 0.01 | 0.23 | 0.08 |
| 1 | 3312914 | rs871822 | 1.282 | 1.256 | -1.486 | -1.569 | -0.008 | 0.004 | 0.004 | 0 | 0.37 | 0.4 | 0.43 | 0.63 | 0.29 | 0.56 |
| 1 | 3315818 | rs7542793 | 1.263 | 1.289 | -2.465 | -1.515 | 0.005 | -0.098 | -0.098 | 0 | 0.36 | 0.08 | 0.35 | 0.62 | 0.3 | 0.52 |
| 1 | 3317780 | rs6424074 | 0.492 | 1.251 | -2.45 | -1.535 | 0.002 | -0.055 | -0.055 | 0 | 0.27 | 0.28 | 0.19 | 0.01 | 0.33 | 0.12 |
| 1 | 3318532 | rs11578011 | 0.3 | 1.166 | -2.165 | -1.192 | 0.001 | -0.005 | -0.005 | 0 | 0.65 | 0.38 | 0.61 | 0.63 | 0.66 | 0.67 |
| 1 | 3320799 | rs12024847 | 0.98 | 1.204 | -2.551 | -0.941 | -0.066 | -0.098 | -0.098 | 0.525 | 0.35 | 0.02 | 0.34 | 0.6 | 0.34 | 0.65 |
| 1 | 3325037 | rs2483225 | -0.855 | 1.497 | -2.013 | -0.906 | -0.043 | 0.123 | 0.123 | 0 | 0.49 | 0.86 | 0.54 | 0.33 | 0.5 | 0.32 |
| 1 | 3328358 | rs870124 | 0.031 | 1.743 | -1.393 | -0.473 | -0.051 | 0.053 | 0.053 | 0 | 0.91 | 0.99 | 0.9 | 1 | 0.84 | 0.96 |
| 1 | 3328659 | rs2493292 | -0.656 | 1.01 | -0.815 | -0.205 | 0.004 | 0.002 | 0.002 | 0 | 0.19 | 0.16 | 0.12 | 0.01 | 0.17 | 0.06 |
| 1 | 3333595 | rs2493285 | -1.379 | 1.156 | -0.391 | 0.475 | 0.049 | 0.158 | 0.158 | 0.166 | 0.07 | 0.16 | 0.18 | 0.31 | 0.09 | 0.08 |
| 1 | 3336200 | rs1984069 | 0.706 | 0.591 | -0.313 | -0.335 | -0.03 | -0.161 | -0.161 | 0 | 0.72 | 0.6 | 0.81 | 0.99 | 0.78 | 0.85 |
| 1 | 3340855 | rs4648500 | 1.199 | 0.674 | -1.263 | 0.235 | 0.013 | -0.054 | -0.054 | 0 | 0.56 | 0.12 | 0.57 | 0.5 | 0.64 | 0.65 |
| 1 | 3342530 | rs2244013 | -1.017 | 0.465 | -0.988 | 0.117 | 0.007 | 0.027 | 0.027 | 0 | 0.3 | 0.77 | 0.36 | 0.2 | 0.26 | 0.16 |
| 1 | 3342804 | rs870171 | -0.922 | 0.818 | -0.258 | 0.61 | 0.017 | 0.07 | 0.07 | 0 | 0.26 | 0.44 | 0.34 | 0.52 | 0.28 | 0.28 |
| 1 | 3349513 | rs2493272 | -1.081 | 0.511 | -0.711 | 0.709 | 0.016 | -0.07 | -0.07 | 0 | 0.3 | 0.8 | 0.35 | 0.57 | 0.25 | 0.29 |
| 1 | 3352872 | rs2236518 | -1.253 | 0.339 | -0.3 | 1.081 | 0.005 | -0.016 | -0.016 | 0 | 0.29 | 0.69 | 0.36 | 0.55 | 0.24 | 0.37 |
| 1 | 3356405 | rs2483250 | 1.504 | 0.696 | 1.054 | 1.212 | 0.051 | 0.169 | 0.169 | 1.132 | 0.89 | 0.59 | 0.82 | 0.76 | 0.92 | 0.86 |
| 1 | 3362256 | rs867814 | 2.688 | 0.736 | 1.55 | 1.376 | 0.017 | 0.149 | 0.149 | 0.315 | 0.82 | 0.52 | 0.75 | 0.59 | 0.84 | 0.83 |
| 1 | 3368143 | rs2487670 | -2.676 | 0.756 | 1.775 | 1.169 | 0.037 | 0.159 | 0.159 | 0 | 0.15 | 0.4 | 0.23 | 0.36 | 0.13 | 0.14 |
| 1 | 3377343 | rs2487681 | -2.681 | 0.768 | 1.511 | 1.173 | 0.002 | 0.072 | 0.072 | 0.145 | 0.16 | 0.58 | 0.25 | 0.36 | 0.13 | 0.14 |
| 1 | 3381698 | rs12562167 | -1.425 | 0.787 | 1.36 | 0.94 | 0.007 | 0.055 | 0.055 | 0 | 0.07 | 0.19 | 0.06 | 0.13 | 0.04 | 0.04 |
| 1 | 3383038 | rs947351 | 0.314 | 1.137 | -0.056 | 0.577 | -0.117 | 0.109 | 0.109 | 5.029 | 0.31 | 0.02 | 0.26 | 0.1 | 0.41 | 0.35 |
| 1 | 3389727 | rs2185639 | 2.68 | 1.092 | 1.351 | 1.484 | 0.008 | 0.079 | 0.079 | 0.271 | 0.84 | 0.39 | 0.76 | 0.67 | 0.86 | 0.87 |
| 1 | 3393037 | rs12124163 | 2.646 | 1.13 | 1.047 | 1.494 | 0.006 | 0.061 | 0.061 | 0.395 | 0.84 | 0.36 | 0.76 | 0.68 | 0.86 | 0.86 |
| 1 | 3400294 | rs2493314 | -1.645 | 1.24 | 0.406 | 1.589 | 0.006 | 0.057 | 0.057 | 0 | 0.09 | 0.15 | 0.18 | 0.31 | 0.1 | 0.14 |
| 1 | 3405296 | rs4648505 | -2.367 | 1.377 | 0.494 | 1.269 | 0.006 | 0.021 | 0.021 | 1.337 | 0.16 | 0.72 | 0.29 | 0.28 | 0.16 | 0.13 |
| 1 | 3406791 | rs2821040 | -2.733 | 1.215 | 0.625 | 1.052 | 0.004 | -0.004 | -0.004 | 0 | 0.15 | 0.84 | 0.27 | 0.19 | 0.1 | 0.06 |
| 1 | 3407007 | rs2821039 | -2.373 | 1.294 | 1.305 | 1.09 | 0.024 | 0.053 | 0.053 | 0.345 | 0.12 | 0.62 | 0.24 | 0.19 | 0.09 | 0.06 |
| 1 | 3421897 | rs2821008 | -1.652 | 1.54 | 1.607 | 1.099 | 0.073 | 0.063 | 0.063 | 0.021 | 0.14 | 0.43 | 0.23 | 0.23 | 0.18 | 0.17 |
| 1 | 3423467 | rs947344 | -1.566 | 1.537 | 1.746 | 1.592 | 0.031 | 0.314 | 0.314 | 0 | 0.14 | 0.37 | 0.2 | 0.43 | 0.14 | 0.14 |
| 1 | 3441264 | rs12093117 | -1.078 | 1.522 | 0.851 | 1.421 | 0.021 | 0.001 | 0.001 | 0 | 0.05 | 0.12 | 0.05 | 0.04 | 0.07 | 0.02 |
| 1 | 3446813 | rs4648392 | 0.507 | 1.372 | 0.327 | 1.415 | 0.013 | -0.018 | -0.018 | 0.031 | 0.06 | 0 | 0.05 | 0.09 | 0.03 | 0.09 |
| 1 | 3450637 | rs10797400 | 0.338 | 1.553 | -0.525 | 0.395 | -0.028 | 0.032 | 0.032 | 0 | 0.3 | 0.1 | 0.28 | 0.22 | 0.34 | 0.2 |
| 1 | 3451005 | rs12119711 | 0.85 | 1.292 | -0.448 | 0.183 | 0.009 | 0.048 | 0.048 | 0 | 0.36 | 0.26 | 0.29 | 0.12 | 0.35 | 0.33 |
| 1 | 3456592 | rs4648524 | -0.279 | 1.292 | -0.541 | 0.753 | -0.032 | -0.007 | -0.007 | 0 | 0.24 | 0.11 | 0.24 | 0.23 | 0.32 | 0.18 |
| 1 | 3456871 | rs10737192 | 0.069 | 0.752 | -0.405 | 0.986 | -0.002 | -0.005 | -0.005 | 0 | 0.25 | 0.1 | 0.23 | 0.22 | 0.25 | 0.18 |
| 1 | 3464451 | rs6685801 | -1.94 | 0.165 | 0.772 | 1.248 | -0.024 | 0.043 | 0.043 | 0 | 0.15 | 0.68 | 0.08 | 0.28 | 0.07 | 0.24 |
| 1 | 3472393 | rs7516641 | -2.498 | 0.201 | 0.707 | 0.136 | 0.015 | 0.017 | 0.017 | 0 | 0.21 | 0.76 | 0.11 | 0.04 | 0.12 | 0.27 |
| 1 | 3472397 | rs7516644 | -1.082 | 0.07 | 0.977 | 0.168 | -0.001 | -0.012 | -0.012 | 0 | 0.06 | 0.1 | 0.02 | 0 | 0.02 | 0.03 |
| 1 | 3472928 | rs878063 | -2.836 | 0.046 | 0.969 | 0.155 | 0.022 | 0.025 | 0.025 | 0.049 | 0.21 | 0.8 | 0.1 | 0.04 | 0.1 | 0.26 |
| 1 | 3473345 | rs9628616 | -2.168 | 0.327 | 2.092 | 0.064 | 0 | -0.108 | -0.108 | 1.696 | 0.17 | 0.44 | 0.1 | 0.02 | 0.1 | 0.21 |
| 1 | 3474626 | rs2821063 | -2.333 | 0.532 | 2.398 | 0.017 | 0.027 | -0.094 | -0.094 | 0.001 | 0.28 | 0.47 | 0.27 | 0.1 | 0.28 | 0.34 |
| 1 | 3477902 | rs2821060 | -1.826 | 0.539 | 2.193 | 0.016 | 0.009 | -0.036 | -0.036 | 0 | 0.22 | 0.24 | 0.26 | 0.05 | 0.27 | 0.19 |
| 1 | 3479742 | rs10909968 | -2.74 | 0.487 | 2.217 | 0.021 | -0.002 | 0.001 | 0.001 | 0.355 | 0.32 | 0.65 | 0.29 | 0.11 | 0.3 | 0.4 |
| 1 | 3483361 | rs880801 | -2.072 | 1.043 | 1.711 | 0.05 | 0.028 | -0.072 | -0.072 | 0 | 0.23 | 0.31 | 0.27 | 0.05 | 0.28 | 0.2 |
| 1 | 3487126 | rs947354 | -0.966 | 0.889 | 1.484 | 0.462 | 0.046 | -0.026 | -0.026 | 0 | 0.1 | 0.27 | 0.18 | 0.04 | 0.12 | 0.13 |
| 1 | 3493850 | rs2821054 | 1.014 | -0.066 | 1.514 | 1.183 | -0.013 | 0.048 | 0.048 | 0 | 0.7 | 0.47 | 0.74 | 0.6 | 0.75 | 0.58 |
| 1 | 3494079 | rs12080677 | -1.106 | -0.103 | 2.045 | 1.132 | 0.02 | -0.08 | -0.08 | 0.448 | 0.17 | 0.39 | 0.14 | 0.04 | 0.14 | 0.18 |
| 1 | 3495672 | rs10909973 | -2.449 | -0.231 | 2.702 | 1.569 | 0.036 | 0.139 | 0.139 | 0 | 0.06 | 0.43 | 0.05 | 0.07 | 0.01 | 0.14 |
| 1 | 3495981 | rs2794341 | 0.48 | -0.357 | 2.438 | 1.919 | -0.002 | 0.028 | 0.028 | 0 | 0.91 | 0.91 | 0.9 | 0.73 | 0.91 | 0.9 |
| 1 | 3496479 | rs2794340 | 0.649 | -0.395 | 2.52 | 1.941 | -0.01 | 0.063 | 0.063 | 0 | 0.9 | 0.87 | 0.89 | 0.71 | 0.92 | 0.89 |
| 1 | 3497109 | rs2821050 | 2.125 | -0.553 | 2.624 | 2.019 | -0.019 | 0.041 | 0.041 | 0 | 0.72 | 0.33 | 0.73 | 0.59 | 0.79 | 0.57 |
| 1 | 3499067 | rs10909974 | -1.304 | -0.485 | 2.248 | 2.382 | -0.011 | 0.096 | 0.096 | 0 | 0.18 | 0.31 | 0.24 | 0.39 | 0.16 | 0.26 |
| 1 | 3504073 | rs12139206 | 1.514 | -0.11 | 1.708 | 2.203 | 0.001 | -0.026 | -0.026 | 0 | 0.74 | 0.21 | 0.69 | 0.61 | 0.78 | 0.61 |
| 1 | 3506497 | rs2794320 | 1.394 | -0.441 | 1.891 | 2.349 | -0.008 | 0.011 | 0.011 | 0 | 0.7 | 0.46 | 0.63 | 0.38 | 0.72 | 0.58 |
| 1 | 3507861 | rs4648527 | 0.815 | -0.484 | 1.908 | 2.359 | -0.023 | 0.074 | 0.074 | 0 | 0.88 | 0.81 | 0.87 | 0.69 | 0.9 | 0.87 |
| 1 | 3510633 | rs6697749 | -0.923 | -0.417 | 2.159 | 2.284 | 0.023 | 0.102 | 0.102 | 0 | 0.19 | 0.35 | 0.24 | 0.44 | 0.17 | 0.3 |
| 1 | 3511295 | rs7544357 | -1.97 | -0.543 | 2.104 | 2.564 | -0.005 | 0 | 0 | 0 | 0.32 | 0.64 | 0.37 | 0.59 | 0.28 | 0.52 |
| 1 | 3512341 | rs2794327 | 0.618 | -1.187 | 1.752 | 1.496 | -0.017 | 0 | 0 | 2.886 | 0.59 | 0.55 | 0.67 | 0.51 | 0.65 | 0.64 |

| | | | | | | | | | | | | | | | | |
|---|---|---|---|---|---|---|---|---|---|---|---|---|---|---|---|---|
| 1 | 3512516 | rs2821025 | 1.254 | -1.298 | 1.545 | 1.175 | -0.001 | -0.023 | -0.023 | 0.036 | 0.91 | 0.85 | 0.94 | 0.98 | 0.91 | 0.89 |
| 1 | 3512800 | rs2821023 | 1.768 | -1.435 | 1.544 | 1.203 | -0.026 | -0.049 | -0.049 | 0 | 0.84 | 0.5 | 0.84 | 0.91 | 0.88 | 0.82 |
| 1 | 3520871 | rs2821017 | -1.056 | -1.227 | 0.963 | 1.288 | 0.004 | 0.001 | 0.001 | 1.328 | 0.44 | 0.48 | 0.47 | 0.55 | 0.35 | 0.46 |
| 1 | 3522561 | rs35701577 | 0.104 | -1.052 | 0.129 | 1.418 | 0.036 | -0.086 | -0.086 | 0.282 | 0.27 | 0.06 | 0.17 | 0.33 | 0.21 | 0.29 |
| 1 | 3531595 | rs4648398 | 1.283 | -0.991 | 0.88 | 1.17 | -0.03 | 0.024 | 0.024 | 0 | 0.8 | 0.45 | 0.67 | 0.8 | 0.83 | 0.79 |
| 1 | 3538501 | rs4364818 | 0.513 | -0.549 | 0.339 | 0.63 | -0.001 | -0.003 | -0.003 | 0 | 0.49 | 0.52 | 0.5 | 0.65 | 0.49 | 0.61 |
| 1 | 3539082 | rs10909989 | 0.406 | -0.667 | 0.232 | 0.313 | 0.042 | -0.129 | -0.129 | 0 | 0.78 | 0.57 | 0.69 | 0.98 | 0.79 | 0.92 |
| 1 | 3539481 | rs10909990 | 0.284 | -0.68 | 0.394 | 0.306 | 0.037 | -0.105 | -0.105 | 0 | 0.79 | 0.67 | 0.71 | 0.98 | 0.79 | 0.92 |
| 1 | 3540024 | rs12757620 | 1.051 | -0.126 | 0.213 | 0.555 | -0.023 | -0.047 | -0.047 | 0 | 0.42 | 0.18 | 0.4 | 0.48 | 0.43 | 0.45 |
| 1 | 3544346 | rs4276857 | 0.104 | -0.252 | -0.216 | 0.973 | -0.072 | -0.095 | -0.095 | 0 | 0.19 | 0.03 | 0.17 | 0.33 | 0.29 | 0.31 |
| 1 | 3545612 | rs1128474 | 0.114 | -0.39 | -0.213 | 0.974 | -0.054 | -0.095 | -0.095 | 0.827 | 0.19 | 0.03 | 0.16 | 0.33 | 0.25 | 0.3 |
| 1 | 3546935 | rs2821007 | 0.482 | -0.387 | -0.273 | 0.865 | -0.003 | 0.037 | 0.037 | 0 | 0.09 | 0.04 | 0.14 | 0 | 0.05 | 0.05 |
| 1 | 3561008 | rs12408890 | 1.153 | -0.135 | -1.391 | 1.039 | -0.003 | 0.008 | 0.008 | 0 | 0.63 | 0.39 | 0.59 | 0.47 | 0.65 | 0.55 |
| 1 | 3566706 | rs2251098 | 0.711 | -0.071 | -0.583 | 0.79 | -0.034 | 0.009 | 0.009 | 0 | 0.81 | 0.39 | 0.74 | 0.8 | 0.9 | 0.81 |
| 1 | 3575929 | rs10910003 | -0.166 | 0.686 | -1.85 | 1.002 | -0.03 | -0.031 | -0.031 | 0 | 0.14 | 0.03 | 0.22 | 0.19 | 0.17 | 0.24 |
| 1 | 3576060 | rs2224718 | 0.278 | 0.331 | -2.111 | 1.023 | 0.02 | -0.008 | -0.008 | 0.179 | 0.52 | 0.24 | 0.55 | 0.49 | 0.61 | 0.6 |
| 1 | 3577321 | rs4648545 | 0.126 | 0.475 | -1.78 | 1.466 | -0.038 | -0.056 | -0.056 | 0 | 0.81 | 0.93 | 0.77 | 0.7 | 0.82 | 0.78 |
| 1 | 3577790 | rs6703610 | -0.509 | 0.087 | -1.411 | 1.243 | 0.001 | -0.002 | -0.002 | 0 | 0.14 | 0.12 | 0.23 | 0.2 | 0.14 | 0.24 |
| 1 | 3585041 | rs7523732 | -0.902 | 0.358 | -0.974 | 0.89 | 0.027 | -0.026 | -0.026 | 0 | 0.05 | 0.16 | 0.09 | 0 | 0.1 | 0.05 |
| 1 | 3586428 | rs1009345 | 0.737 | 0.38 | -2.143 | 0.767 | -0.073 | 0.054 | 0.054 | 2.444 | 0.47 | 0.07 | 0.26 | 0.36 | 0.45 | 0.45 |
| 1 | 3587377 | rs3765695 | -1.487 | 0.218 | -1.808 | 1.268 | 0.001 | 0.008 | 0.008 | 0 | 0.15 | 0.19 | 0.18 | 0.17 | 0.18 | 0.2 |
| 1 | 3592187 | rs3765702 | -0.913 | -0.513 | -1.065 | 1.33 | -0.009 | 0.009 | 0.009 | 0 | 0.36 | 0.56 | 0.52 | 0.44 | 0.39 | 0.51 |
| 1 | 3592436 | rs3765703 | -0.829 | -0.825 | -0.894 | 0.686 | 0.005 | 0.01 | 0.01 | 0 | 0.41 | 0.39 | 0.53 | 0.29 | 0.41 | 0.45 |
| 1 | 3597017 | rs3765705 | -0.148 | -0.327 | 0.17 | 1.236 | 0.004 | 0.016 | 0.016 | 0.772 | 0.11 | 0.26 | 0.02 | 0.11 | 0.05 | 0.12 |
| 1 | 3600799 | rs3765712 | -0.939 | -0.48 | -0.111 | 1.312 | -0.059 | 0.016 | 0.016 | 0.696 | 0.2 | 0.43 | 0.09 | 0.23 | 0.1 | 0.22 |
| 1 | 3601940 | rs3765716 | -0.718 | 0.219 | -0.585 | 1.591 | -0.009 | -0.021 | -0.021 | 0 | 0.16 | 0.11 | 0.28 | 0.2 | 0.2 | 0.16 |
| 1 | 3605097 | rs1885872 | 0.953 | 0.014 | -0.077 | 1.902 | -0.016 | 0.288 | 0.288 | 0 | 0.91 | 0.78 | 0.77 | 0.55 | 0.95 | 0.81 |
| 1 | 3611892 | rs3765731 | -0.056 | 0.075 | -1.076 | 0.543 | -0.003 | 0.1 | 0.1 | 0 | 0.35 | 0.17 | 0.2 | 0.13 | 0.29 | 0.34 |
| 1 | 3615156 | rs12032272 | 0.04 | -0.151 | -1.058 | 0.171 | -0.006 | -0.015 | -0.015 | 0 | 0.15 | 0.21 | 0.24 | 0.09 | 0.16 | 0.16 |
| 1 | 3616601 | rs3765736 | 0.677 | 0.118 | -1.184 | 0.472 | -0.04 | -0.029 | -0.029 | 0.724 | 0.28 | 0.21 | 0.25 | 0.36 | 0.39 | 0.27 |
| 1 | 3628663 | rs6424089 | -0.036 | 0.311 | -1.393 | -0.185 | -0.006 | 0.014 | 0.014 | 5.863 | 0.47 | 0.32 | 0.42 | 0.48 | 0.54 | 0.4 |
| 1 | 3630476 | rs3765761 | 0.766 | 0.251 | -1.621 | -0.708 | -0.011 | 0.035 | 0.035 | 0.004 | 0.16 | 0.07 | 0.13 | 0.02 | 0.16 | 0.11 |
| 1 | 3632608 | rs3753211 | 0.696 | 0.233 | -1.602 | -0.714 | -0.009 | 0.025 | 0.025 | 0 | 0.16 | 0.07 | 0.13 | 0.02 | 0.17 | 0.11 |
| 1 | 3633380 | rs3819967 | 1.152 | -0.134 | -1.549 | -0.35 | -0.043 | -0.023 | -0.023 | 0 | 0.26 | 0.07 | 0.2 | 0.28 | 0.33 | 0.2 |
| 1 | 3637459 | rs12731705 | -0.486 | -0.891 | -1.855 | -1.146 | 0.006 | 0.248 | 0.248 | 0 | 0.48 | 0.2 | 0.42 | 0.04 | 0.47 | 0.35 |
| 1 | 3637613 | rs2146658 | 0.218 | -1.26 | -1.232 | -0.599 | -0.005 | 0.028 | 0.028 | 0.751 | 0.77 | 0.71 | 0.65 | 0.54 | 0.76 | 0.58 |
| 1 | 3645052 | rs12095248 | -1.171 | -1.749 | -0.883 | -0.751 | -0.031 | 0.005 | 0.005 | 0.089 | 0.22 | 0.68 | 0.11 | 0.28 | 0.11 | 0.22 |
| 1 | 3646137 | rs11589885 | 1.054 | -2.08 | -1.283 | -0.995 | 0.13 | 0.142 | 0.142 | 0 | 0.59 | 0.11 | 0.58 | 0.26 | 0.75 | 0.42 |
| 1 | 3647962 | rs1745813 | 0.365 | -2.239 | -0.792 | -1.02 | -0.018 | 0.009 | 0.009 | 0 | 0.7 | 0.41 | 0.62 | 0.49 | 0.79 | 0.51 |
| 1 | 3648879 | rs4648553 | 1.112 | -2.085 | -0.8 | -0.947 | 0.138 | 0.146 | 0.146 | 0 | 0.58 | 0.11 | 0.58 | 0.26 | 0.75 | 0.42 |
| 1 | 3649562 | rs9662633 | 0.138 | -1.977 | -0.673 | -0.873 | -0.016 | 0.07 | 0.07 | 0 | 0.07 | 0.17 | 0.02 | 0.21 | 0.03 | 0.06 |
| 1 | 3651126 | rs1181868 | 1.171 | -2.056 | -0.817 | -0.935 | 0.137 | 0.144 | 0.144 | 0 | 0.58 | 0.12 | 0.58 | 0.27 | 0.75 | 0.43 |
| 1 | 3651318 | rs1181867 | 0.593 | -2.06 | -0.505 | -0.903 | -0.016 | 0.001 | 0.001 | 0 | 0.64 | 0.43 | 0.62 | 0.49 | 0.78 | 0.48 |
| 1 | 3653426 | rs1181865 | -1.072 | -2.139 | -0.608 | -0.863 | 0.03 | -0.099 | -0.099 | 1.29 | 0.81 | 0.94 | 0.7 | 0.56 | 0.87 | 0.64 |
| 1 | 3654747 | rs7515164 | -0.775 | -1.724 | 0.779 | -1 | -0.048 | -0.014 | -0.014 | 0 | 0.12 | 0.38 | 0.06 | 0.06 | 0.07 | 0.19 |
| 1 | 3657759 | rs12117836 | -1.54 | -1.552 | -0.54 | -1.269 | 0.002 | 0.007 | 0.007 | 0 | 0.4 | 0.4 | 0.41 | 0.28 | 0.39 | 0.44 |
| 1 | 3659657 | rs2298222 | -0.616 | -1.518 | -0.529 | -1.265 | -0.013 | 0.041 | 0.041 | 0 | 0.3 | 0.19 | 0.36 | 0.22 | 0.32 | 0.3 |
| 1 | 3659742 | rs2298223 | 0.789 | -1.502 | -0.625 | -1.247 | 0.013 | -0.024 | -0.024 | 0 | 0.05 | 0 | 0.07 | 0.21 | 0.03 | 0.05 |
| 1 | 3662845 | rs3737589 | -1.591 | -1.484 | -0.599 | -1.249 | 0.004 | 0.011 | 0.011 | 0 | 0.42 | 0.4 | 0.41 | 0.29 | 0.4 | 0.44 |
| 1 | 3668004 | rs4648554 | -1.888 | -1.403 | -1.608 | -1.21 | 0.048 | 0.119 | 0.119 | 0 | 0.47 | 0.84 | 0.45 | 0.29 | 0.4 | 0.45 |
| 1 | 3668753 | rs12128253 | -0.211 | -0.806 | -1.329 | -0.892 | -0.043 | 0.094 | 0.094 | 0 | 0.23 | 0.08 | 0.28 | 0.01 | 0.28 | 0.2 |
| 1 | 3671791 | rs1181888 | 0.006 | -0.549 | -1.358 | -0.834 | 0.005 | 0.013 | 0.013 | 0 | 0.16 | 0.1 | 0.09 | 0.07 | 0.13 | 0.19 |
| 1 | 3677933 | rs1181883 | -1.397 | -0.372 | -1.253 | -0.736 | 0.016 | 0.056 | 0.056 | 0 | 0.47 | 0.69 | 0.47 | 0.28 | 0.44 | 0.43 |
| 1 | 3679775 | rs10910024 | -0.633 | 0.514 | -1.323 | -0.67 | -0.016 | 0.014 | 0.014 | 0 | 0.21 | 0.11 | 0.27 | 0 | 0.24 | 0.17 |
| 1 | 3680643 | rs1181878 | -1.14 | -0.244 | -1.341 | -0.411 | -0.004 | 0.003 | 0.003 | 0 | 0.24 | 0.16 | 0.3 | 0.19 | 0.31 | 0.34 |
| 1 | 3680729 | rs1181877 | -1 | -1.011 | -1.225 | -0.907 | 0.032 | 0.056 | 0.056 | 0 | 0.4 | 0.76 | 0.38 | 0.18 | 0.33 | 0.3 |
| 1 | 3681831 | rs1181875 | -0.833 | -1.416 | -1.046 | -0.108 | 0.015 | -0.02 | -0.02 | 0 | 0.18 | 0.13 | 0.18 | 0.57 | 0.12 | 0.44 |
| 1 | 3682070 | rs1181874 | -1.459 | -1.333 | -1.862 | -0.723 | -0.155 | -0.041 | -0.041 | 0 | 0.66 | 0.94 | 0.49 | 0.68 | 0.55 | 0.67 |
| 1 | 3684185 | rs10910025 | 0.526 | -1.725 | -0.855 | -0.977 | -0.037 | -0.097 | -0.097 | 0 | 0.16 | 0.36 | 0.07 | 0.02 | 0.08 | 0.03 |
| 1 | 3689408 | rs2275819 | 0.348 | -1.714 | -1.08 | -0.788 | 0.022 | -0.035 | -0.035 | 0 | 0.12 | 0.04 | 0.15 | 0.57 | 0.09 | 0.35 |
| 1 | 3691727 | rs1175549 | 0.468 | -1.715 | -0.59 | -1.246 | -0.006 | -0.096 | -0.096 | 0 | 0.19 | 0.6 | 0.16 | 0 | 0.22 | 0.14 |
| 1 | 3696000 | rs2799182 | -1.231 | -1.768 | -0.823 | -1.324 | -0.083 | 0.229 | 0.229 | 0 | 0.55 | 0.91 | 0.36 | 0.11 | 0.48 | 0.34 |
| 1 | 3696890 | rs8379 | -1.388 | -1.75 | -0.672 | -1.315 | -0.059 | 0.226 | 0.226 | 0 | 0.55 | 0.88 | 0.36 | 0.11 | 0.48 | 0.34 |
| 1 | 3697035 | rs17411356 | 0.901 | -1.775 | -0.613 | -1.252 | 0.013 | 0.004 | 0.004 | 0.214 | 0.16 | 0.14 | 0.06 | 0.02 | 0.09 | 0.03 |
| 1 | 3701663 | rs2298227 | 0.249 | -1.789 | -0.641 | -1.268 | 0.022 | 0.019 | 0.019 | 2.149 | 0.2 | 0.15 | 0.1 | 0.11 | 0.12 | 0.05 |
| 1 | 3706538 | rs9424284 | -2.066 | -1.78 | -0.95 | -1.128 | -0.154 | -0.06 | -0.06 | 0 | 0.66 | 0.95 | 0.51 | 0.66 | 0.57 | 0.67 |
| 1 | 3709412 | rs12728911 | -0.179 | -1.795 | -0.729 | -1.265 | -0.005 | -0.123 | -0.123 | 0 | 0.29 | 0.56 | 0.16 | 0.05 | 0.23 | 0.09 |
| 1 | 3709487 | rs7513053 | 0.333 | -1.54 | -0.597 | -0.99 | 0.004 | 0.026 | 0.026 | 0 | 0.2 | 0.13 | 0.17 | 0.09 | 0.24 | 0.17 |
| 1 | 3732596 | rs6662706 | -1.033 | -1.512 | -0.644 | -0.976 | -0.05 | 0.325 | 0.325 | 0 | 0.54 | 0.88 | 0.34 | 0.05 | 0.49 | 0.37 |

| | | | | | | | | | | | | | | | | |
|---|---|---|---|---|---|---|---|---|---|---|---|---|---|---|---|---|
| 1 | 3736319 | rs9424303 | -1.928 | -1.529 | -0.657 | -0.845 | -0.13 | -0.053 | -0.053 | 0 | 0.66 | 0.92 | 0.51 | 0.66 | 0.57 | 0.67 |
| 1 | 3741782 | rs7552969 | -1.478 | -1.538 | -0.472 | -0.836 | -0.023 | -0.02 | -0.02 | 0 | 0.62 | 0.75 | 0.45 | 0.57 | 0.54 | 0.65 |
| 1 | 3742009 | rs9424290 | 1.026 | -1.552 | -0.481 | -0.821 | 0.02 | -0.026 | -0.026 | 0 | 0.09 | 0.03 | 0.16 | 0.61 | 0.06 | 0.29 |
| 1 | 3742257 | rs12759497 | 0.795 | -1.641 | -0.369 | -0.899 | -0.009 | -0.031 | -0.031 | 0 | 0.19 | 0.26 | 0.09 | 0.05 | 0.14 | 0.11 |
| 1 | 3743319 | rs6663840 | -0.223 | -1.109 | -0.795 | -0.411 | -0.008 | 0.002 | 0.002 | 0 | 0.44 | 0.57 | 0.31 | 0.57 | 0.33 | 0.51 |
| 1 | 3744084 | rs9424310 | -0.617 | -1.102 | -0.809 | -0.41 | 0.069 | 0.073 | 0.073 | 0 | 0.49 | 0.8 | 0.37 | 0.66 | 0.36 | 0.52 |
| 1 | 3745924 | rs2275831 | 0.561 | -1.088 | -0.828 | -0.394 | -0.004 | 0.075 | 0.075 | 0 | 0.1 | 0.08 | 0.13 | 0.52 | 0.06 | 0.28 |
| 1 | 3747580 | rs6424058 | 1.21 | -1.09 | -0.83 | -0.411 | -0.025 | -0.004 | -0.004 | 0 | 0.34 | 0.22 | 0.49 | 0.34 | 0.43 | 0.33 |
| 1 | 3748085 | rs7522227 | 0.039 | -1.089 | -0.902 | -0.408 | 0.007 | -0.002 | -0.002 | 0 | 0.55 | 0.38 | 0.69 | 0.43 | 0.67 | 0.49 |
| 1 | 3754596 | rs6683827 | -0.723 | -1.081 | -0.922 | -0.397 | -0.002 | 0.07 | 0.07 | 0 | 0.9 | 0.92 | 0.87 | 0.48 | 0.94 | 0.72 |
| 1 | 3755675 | rs1891941 | -0.723 | -1.081 | -0.924 | -0.397 | -0.004 | 0.075 | 0.075 | 0 | 0.9 | 0.92 | 0.87 | 0.48 | 0.94 | 0.72 |
| 1 | 3757749 | rs12744607 | 0.356 | -0.745 | -0.778 | -0.168 | -0.014 | -0.126 | -0.126 | 0 | 0.35 | 0.52 | 0.18 | 0.05 | 0.27 | 0.22 |
| 1 | 3761479 | rs2275822 | 0.723 | -0.445 | -0.824 | 0.008 | -0.015 | -0.002 | -0.002 | 0 | 0.34 | 0.23 | 0.49 | 0.34 | 0.43 | 0.33 |
| 1 | 3772359 | rs4564078 | -0.137 | -0.328 | -0.452 | -0.182 | 0.024 | 0.075 | 0.075 | 0.237 | 0.4 | 0.71 | 0.23 | 0.15 | 0.3 | 0.28 |
| 1 | 3773089 | rs4648426 | -0.231 | -0.274 | -1.05 | -0.155 | -0.014 | -0.055 | -0.055 | 0 | 0.4 | 0.53 | 0.23 | 0.15 | 0.3 | 0.28 |
| 1 | 3781096 | rs10797348 | 0.209 | -0.289 | -0.646 | 0.592 | -0.031 | 0.068 | 0.068 | 0.003 | 0.72 | 0.61 | 0.7 | 0.5 | 0.7 | 0.6 |
| 1 | 3789852 | rs12756705 | -0.617 | 0.531 | -0.518 | 0.968 | -0.004 | 0.001 | 0.001 | 0 | 0.14 | 0.11 | 0.15 | 0.16 | 0.18 | 0.19 |
| 1 | 3793324 | rs6668103 | -1.323 | -0.203 | 0.132 | 0.607 | 0.004 | -0.001 | -0.001 | 0 | 0.27 | 0.44 | 0.24 | 0.22 | 0.29 | 0.29 |
| 1 | 3794273 | rs12132240 | 0.044 | -0.184 | -0.486 | 1.18 | -0.019 | -0.033 | -0.033 | 0 | 0.14 | 0.06 | 0.16 | 0.29 | 0.18 | 0.19 |
| 1 | 3803755 | rs7367066 | -0.917 | -0.5 | -0.424 | 0.902 | 0.017 | -0.001 | -0.001 | 0 | 0.63 | 0.76 | 0.62 | 0.61 | 0.71 | 0.62 |
| 1 | 3805098 | rs7530189 | -0.115 | -0.921 | -0.193 | 0.192 | -0.018 | -0.084 | -0.084 | 0.553 | 0.2 | 0.3 | 0.13 | 0.01 | 0.18 | 0.13 |
| 1 | 3806335 | rs10797351 | 0.769 | -0.974 | -0.159 | 0.169 | -0.016 | -0.026 | -0.026 | 0 | 0.05 | 0.22 | 0.04 | 0 | 0.02 | 0.06 |
| 1 | 3806358 | rs10909819 | 0.747 | -0.938 | -0.172 | 0.248 | 0.01 | -0.02 | -0.02 | 2.02 | 0.13 | 0.1 | 0.11 | 0.23 | 0.13 | 0.16 |
| 1 | 3806973 | rs12073945 | 0.84 | -0.936 | -0.178 | 0.248 | -0.016 | -0.026 | -0.026 | 0.01 | 0.05 | 0.22 | 0.04 | 0 | 0.02 | 0.06 |
| 1 | 3807388 | rs4131373 | 0.326 | -0.807 | -0.131 | 0.315 | -0.003 | -0.04 | -0.04 | 0.274 | 0.19 | 0.25 | 0.12 | 0.01 | 0.18 | 0.13 |
| 1 | 3808362 | rs4446914 | -1.007 | -0.817 | -0.152 | -0.073 | -0.031 | -0.029 | -0.029 | 0 | 0.35 | 0.52 | 0.26 | 0.29 | 0.32 | 0.29 |
| 1 | 3811810 | rs12082157 | 0.023 | -0.705 | -0.332 | -0.087 | -0.029 | -0.068 | -0.068 | 2.405 | 0.21 | 0.37 | 0.13 | 0.06 | 0.2 | 0.13 |
| 1 | 3821782 | rs11589102 | 0.807 | -0.112 | -0.434 | 0.006 | -0.046 | 0.034 | 0.034 | 0 | 0.24 | 0.15 | 0.22 | 0.1 | 0.3 | 0.2 |
| 1 | 3822703 | rs10909830 | -0.59 | -0.557 | -0.346 | 0.205 | -0.003 | -0.005 | -0.005 | 0 | 0.33 | 0.36 | 0.24 | 0.25 | 0.31 | 0.29 |
| 1 | 3842112 | rs34835780 | -0.245 | -0.189 | -0.416 | 0.23 | 0.024 | -0.025 | -0.025 | 0.096 | 0.63 | 0.53 | 0.55 | 0.79 | 0.54 | 0.68 |
| 1 | 3996946 | rs6695346 | -0.519 | -0.735 | -0.65 | 0.271 | -0.051 | -0.01 | -0.01 | 2.853 | 0.33 | 0.51 | 0.19 | 0.41 | 0.16 | 0.32 |
| 1 | 4002498 | rs7538691 | 0.104 | 0.097 | -0.806 | 0.258 | -0.001 | -0.001 | -0.001 | 0 | 0.64 | 0.61 | 0.47 | 0.52 | 0.55 | 0.62 |
| 1 | 4002832 | rs12724233 | 0.059 | -0.124 | -0.349 | -1.235 | -0.006 | -0.149 | -0.149 | 0 | 0.33 | 0.41 | 0.2 | 0.01 | 0.25 | 0.19 |
| 1 | 4002958 | rs10915568 | -0.715 | -0.21 | 0.423 | -0.215 | 0.004 | 0.136 | 0.136 | 0 | 0.12 | 0.34 | 0.12 | 0.25 | 0.12 | 0.29 |
| 1 | 4004250 | rs11583257 | -1.227 | -0.117 | 0.582 | -0.575 | -0.02 | -0.003 | -0.003 | 0 | 0.24 | 0.4 | 0.23 | 0.27 | 0.22 | 0.36 |
| 1 | 4005693 | rs7524279 | -1.244 | -0.135 | 0.6 | -0.572 | -0.022 | -0.018 | -0.018 | 1.146 | 0.25 | 0.41 | 0.21 | 0.26 | 0.22 | 0.35 |
| 1 | 4006060 | rs7548668 | -1.436 | -0.219 | 0.483 | -0.913 | 0.006 | 0.008 | 0.008 | 0 | 0.44 | 0.54 | 0.32 | 0.34 | 0.37 | 0.47 |
| 1 | 4007008 | rs7519349 | -1.146 | -0.267 | 0.45 | -1.001 | 0.008 | 0.019 | 0.019 | 0 | 0.4 | 0.32 | 0.26 | 0.26 | 0.35 | 0.42 |
| 1 | 4007110 | rs7519458 | -1.529 | -0.06 | -0.01 | -1.061 | 0.03 | 0.092 | 0.092 | 0.011 | 0.49 | 0.69 | 0.35 | 0.33 | 0.42 | 0.49 |
| 1 | 4007558 | rs10737395 | -0.934 | 0.389 | -0.067 | -0.513 | -0.045 | 0.032 | 0.032 | 0 | 0.15 | 0.02 | 0.13 | 0.06 | 0.17 | 0.13 |
| 1 | 4007688 | rs12090228 | -0.739 | 0.336 | 0.219 | -0.425 | -0.005 | 0.001 | 0.001 | 0 | 0.14 | 0.22 | 0.1 | 0.14 | 0.12 | 0.21 |
| 1 | 4009071 | rs10799181 | -1.305 | 0.476 | 0.044 | -0.321 | 0 | -0.028 | -0.028 | 0.021 | 0.16 | 0.29 | 0.16 | 0.11 | 0.17 | 0.17 |
| 1 | 4009694 | rs4654479 | -1.174 | 0.546 | 0.02 | -0.261 | 0.007 | -0.021 | -0.021 | 0 | 0.14 | 0.29 | 0.14 | 0.11 | 0.12 | 0.17 |
| 1 | 4009793 | rs4654480 | -0.951 | 0.716 | -0.502 | -0.466 | 0 | -0.006 | -0.006 | 0 | 0.21 | 0.21 | 0.2 | 0.11 | 0.22 | 0.21 |
| 1 | 4009956 | rs6661168 | -0.978 | 1.059 | -0.649 | -0.145 | -0.007 | 0.008 | 0.008 | 0 | 0.13 | 0.11 | 0.19 | 0.05 | 0.17 | 0.14 |
| 1 | 4010743 | rs4654482 | -0.78 | -0.025 | -0.365 | 0.418 | -0.049 | 0.067 | 0.067 | 0.185 | 0.27 | 0.45 | 0.38 | 0.58 | 0.18 | 0.53 |
| 1 | 4012472 | rs11590912 | -0.159 | 1.251 | -0.798 | -0.073 | -0.042 | 0.024 | 0.024 | 0 | 0.19 | 0.11 | 0.23 | 0.1 | 0.32 | 0.09 |
| 1 | 4013618 | rs10799202 | 0.161 | 0.952 | -0.658 | -0.273 | -0.015 | 0.033 | 0.033 | 0 | 0.76 | 0.86 | 0.87 | 0.97 | 0.74 | 0.78 |
| 1 | 4013954 | rs12119556 | -0.166 | 0.756 | -1.092 | 0.295 | -0.026 | -0.106 | -0.106 | 0.293 | 0.14 | 0.02 | 0.37 | 0.48 | 0.16 | 0.27 |
| 1 | 4015382 | rs6681347 | 0.789 | 0.964 | -0.151 | -0.286 | 0.004 | -0.007 | -0.007 | 1.63 | 0.7 | 0.69 | 0.8 | 0.82 | 0.68 | 0.72 |
| 1 | 4017137 | rs7522140 | 0.635 | 0.242 | 0.054 | -0.15 | -0.008 | -0.036 | -0.036 | 0.453 | 0.68 | 0.44 | 0.77 | 0.8 | 0.72 | 0.64 |
| 1 | 4026098 | rs7550609 | -1.277 | 0.2 | 0.581 | 0.043 | -0.009 | 0.07 | 0.07 | 1.862 | 0.08 | 0.56 | 0.07 | 0.08 | 0.01 | 0.03 |
| 1 | 4027590 | rs6693909 | 1.36 | 0.229 | -0.055 | -0.012 | -0.007 | -0.003 | -0.003 | 0.129 | 0.6 | 0.09 | 0.61 | 0.59 | 0.65 | 0.46 |
| 1 | 4029733 | rs10915495 | -1.147 | 0.293 | 0.152 | -0.097 | -0.04 | 0.048 | 0.048 | 2.459 | 0.09 | 0.63 | 0.07 | 0.1 | 0.01 | 0.05 |
| 1 | 4031327 | rs3935162 | 0.522 | 0.382 | -0.427 | -0.195 | -0.08 | 0.155 | 0.155 | 1.316 | 0.74 | 0.93 | 0.85 | 0.99 | 0.69 | 0.68 |
| 1 | 4034097 | rs12031557 | -0.641 | 0.183 | -0.189 | 0.159 | -0.011 | 0.15 | 0.15 | 0 | 0.06 | 0.18 | 0.25 | 0.31 | 0.04 | 0.18 |
| 1 | 4035129 | rs10799224 | 0.349 | 0.602 | -0.488 | 0.02 | -0.034 | 0.084 | 0.084 | 1.519 | 0.76 | 0.89 | 0.86 | 1 | 0.69 | 0.73 |
| 1 | 4036195 | rs10753378 | 0.278 | 0.584 | -0.615 | 0.029 | -0.033 | 0.13 | 0.13 | 0.182 | 0.74 | 0.92 | 0.86 | 1 | 0.71 | 0.76 |
| 1 | 4039896 | rs11587331 | 1.347 | 0.68 | -0.659 | 0.152 | -0.003 | -0.002 | -0.002 | 0.214 | 0.57 | 0.11 | 0.59 | 0.58 | 0.6 | 0.49 |
| 1 | 4052956 | rs10799236 | -0.437 | 0.814 | -0.486 | 0.395 | 0.003 | 0.047 | 0.047 | 0 | 0.08 | 0.12 | 0.21 | 0.31 | 0.1 | 0.22 |
| 1 | 4054067 | rs6691155 | 1.124 | 0.876 | -0.032 | 0.401 | -0.001 | -0.005 | -0.005 | 0 | 0.6 | 0.39 | 0.6 | 0.54 | 0.59 | 0.49 |
| 1 | 4057488 | rs12135298 | 0.661 | 1.016 | 0.366 | 0.827 | 0.015 | 0.01 | 0.01 | 0 | 0.9 | 0.78 | 0.91 | 0.89 | 0.87 | 0.85 |
| 1 | 4061370 | rs6672877 | -1.21 | 0.682 | 0.517 | 0.823 | -0.009 | 0.024 | 0.024 | 0 | 0.24 | 0.6 | 0.29 | 0.29 | 0.2 | 0.29 |
| 1 | 4066781 | rs4128222 | -0.617 | 0.112 | 0.292 | 0.292 | -0.059 | -0.059 | -0.059 | 0 | 0.22 | 0.4 | 0.2 | 0.08 | 0.14 | 0.15 |
| 1 | 4068052 | rs6675979 | -1.881 | 0.936 | 0.195 | 0.238 | 0 | -0.015 | -0.015 | 0.019 | 0.45 | 0.53 | 0.39 | 0.27 | 0.49 | 0.48 |
| 1 | 4068146 | rs12731145 | -0.503 | 1.046 | -0.197 | 0.255 | -0.06 | 0.012 | 0.012 | 0 | 0.22 | 0.12 | 0.19 | 0.18 | 0.32 | 0.33 |
| 1 | 4078448 | rs4129100 | -1.831 | 0.976 | -0.42 | 0.521 | -0.01 | 0.025 | 0.025 | 0.026 | 0.48 | 0.83 | 0.34 | 0.38 | 0.51 | 0.54 |
| 1 | 4080764 | rs4323679 | -1.661 | 1.04 | -0.431 | 0.509 | -0.001 | 0.004 | 0.004 | 0 | 0.46 | 0.61 | 0.33 | 0.39 | 0.51 | 0.54 |
| 1 | 4085098 | rs6426436 | -0.595 | 1.345 | -1.271 | 0.461 | -0.112 | -0.023 | -0.023 | 2.537 | 0.22 | 0.07 | 0.19 | 0.21 | 0.33 | 0.32 |

| | | | | | | | | | | | | | | | | |
|---|---|---|---|---|---|---|---|---|---|---|---|---|---|---|---|---|
| 1 | 4092949 | rs10915659 | -1.218 | 0.635 | -0.516 | 0.405 | -0.028 | -0.018 | -0.018 | 0 | 0.25 | 0.62 | 0.14 | 0.16 | 0.18 | 0.23 |
| 1 | 4094068 | rs12137595 | -0.391 | 0.966 | -1.398 | 0.398 | 0.006 | 0.091 | 0.091 | 0 | 0.16 | 0 | 0.09 | 0.02 | 0.13 | 0.17 |
| 1 | 4104757 | rs12036953 | -1.084 | 2.021 | -0.822 | 1.407 | 0.043 | 0.068 | 0.068 | 0 | 0.09 | 0.15 | 0.13 | 0.28 | 0.21 | 0.17 |
| 1 | 4108282 | rs4654402 | 0.053 | 0.718 | -1.649 | 1.21 | 0.029 | -0.094 | -0.094 | 0 | 0.22 | 0.08 | 0.36 | 0.5 | 0.2 | 0.3 |
| 1 | 4111187 | rs12749761 | -0.557 | 1.183 | -2.186 | 0.715 | -0.083 | 0.033 | 0.033 | 1.184 | 0.13 | 0.02 | 0.25 | 0.07 | 0.26 | 0.29 |
| 1 | 4111741 | rs4553153 | 0.389 | 0.562 | -2.086 | 1.064 | 0.02 | -0.085 | -0.085 | 0 | 0.22 | 0.08 | 0.35 | 0.54 | 0.21 | 0.31 |
| 1 | 4117238 | rs4654404 | -0.161 | -0.539 | -2.128 | -0.189 | -0.029 | 0.001 | 0.001 | 0.002 | 0.35 | 0.21 | 0.41 | 0.33 | 0.53 | 0.41 |
| 1 | 4118987 | rs10799175 | -0.546 | -1.657 | -1.436 | -0.788 | -0.003 | -0.001 | -0.001 | 0 | 0.52 | 0.53 | 0.51 | 0.42 | 0.63 | 0.52 |
| 1 | 4119662 | rs9792946 | -1.092 | -2.097 | -0.881 | -1.13 | -0.028 | -0.05 | -0.05 | 0 | 0.81 | 0.78 | 0.93 | 0.93 | 0.9 | 0.83 |
| 1 | 4120311 | rs7514530 | 0.647 | -2.283 | -0.79 | -1.149 | 0.013 | -0.012 | -0.012 | 0 | 0.33 | 0.25 | 0.43 | 0.5 | 0.28 | 0.33 |
| 1 | 4120898 | rs4233258 | 0.029 | -2.321 | -0.681 | -1.203 | -0.009 | -0.008 | -0.008 | 0 | 0.46 | 0.54 | 0.48 | 0.4 | 0.57 | 0.48 |
| 1 | 4121105 | rs10915373 | 0.037 | -2.324 | -0.682 | -1.205 | -0.009 | -0.008 | -0.008 | 0 | 0.46 | 0.54 | 0.48 | 0.4 | 0.57 | 0.48 |
| 1 | 4121584 | rs7524800 | 0.826 | -2.266 | -0.52 | -0.892 | 0.018 | -0.028 | -0.028 | 0 | 0.3 | 0.2 | 0.43 | 0.4 | 0.28 | 0.31 |
| 1 | 4123099 | rs6669278 | -0.152 | -1.613 | -0.597 | -0.959 | 0.005 | -0.01 | -0.01 | 0 | 0.24 | 0.25 | 0.23 | 0.34 | 0.18 | 0.21 |
| 1 | 4129266 | rs16840701 | 0.944 | -1.519 | -0.931 | -0.922 | -0.003 | -0.035 | -0.035 | 0.316 | 0.15 | 0.1 | 0.19 | 0.28 | 0.15 | 0.13 |
| 1 | 4130348 | rs4130940 | -0.383 | -1.735 | -0.937 | -1.157 | 0 | 0.008 | 0.008 | 0 | 0.39 | 0.33 | 0.43 | 0.33 | 0.55 | 0.43 |
| 1 | 4130684 | rs4391636 | 0.506 | -1.064 | -0.945 | -1.199 | -0.019 | 0.003 | 0.003 | 0.078 | 0.22 | 0.16 | 0.24 | 0.04 | 0.4 | 0.31 |
| 1 | 4132222 | rs12093951 | 0.199 | -1.752 | -0.557 | -0.594 | -0.03 | -0.015 | -0.015 | 0 | 0.12 | 0.24 | 0.06 | 0.05 | 0.04 | 0.07 |
| 1 | 4140030 | rs4436359 | 0.392 | -1.727 | -1.015 | -0.606 | -0.036 | -0.043 | -0.043 | 0 | 0.65 | 0.42 | 0.84 | 0.72 | 0.79 | 0.64 |
| 1 | 4145615 | rs10915421 | 0.424 | -1.447 | -1.729 | -0.35 | 0.05 | -0.081 | -0.081 | 0 | 0.29 | 0.08 | 0.35 | 0.39 | 0.21 | 0.25 |
| 1 | 4146028 | rs10915423 | 0.201 | -0.617 | -1.035 | -0.715 | 0.021 | 0.034 | 0.034 | 0 | 0.82 | 0.93 | 0.94 | 0.91 | 0.9 | 0.78 |
| 1 | 4152715 | rs4462110 | -0.747 | -0.207 | -0.669 | -0.584 | -0.009 | -0.042 | -0.042 | 0 | 0.16 | 0.26 | 0.09 | 0.08 | 0.11 | 0.19 |
| 1 | 4152928 | rs4481820 | -0.723 | -0.105 | -0.847 | -0.983 | -0.017 | -0.097 | -0.097 | 0 | 0.38 | 0.49 | 0.2 | 0.05 | 0.26 | 0.22 |
| 1 | 4158955 | rs10915428 | -0.434 | 0.6 | -1.235 | -0.169 | -0.014 | -0.01 | -0.01 | 0 | 0.37 | 0.27 | 0.54 | 0.53 | 0.51 | 0.56 |
| 1 | 4168066 | rs7542313 | -1.056 | 0.2 | -1.84 | -0.867 | -0.036 | 0.053 | 0.053 | 0 | 0.82 | 0.92 | 0.86 | 1 | 0.73 | 0.94 |
| 1 | 4170048 | rs12077262 | 0.376 | -0.763 | -1.609 | -1.093 | -0.044 | -0.058 | -0.058 | 0.212 | 0.28 | 0.43 | 0.18 | 0.14 | 0.19 | 0.29 |
| 1 | 4170249 | rs7548756 | -0.531 | -1.525 | -2.202 | -0.911 | 0.153 | -0.031 | -0.031 | 0.205 | 0.53 | 0.84 | 0.44 | 0.56 | 0.27 | 0.57 |
| 1 | 4171786 | rs10492944 | 1.464 | -1.118 | -1.972 | -0.864 | 0.018 | 0.005 | 0.005 | 0 | 0.09 | 0.01 | 0.12 | 0.07 | 0.04 | 0.14 |
| 1 | 4174116 | rs10915434 | 0.348 | -0.894 | -1.721 | -0.803 | -0.037 | -0.001 | -0.001 | 0 | 0.31 | 0.53 | 0.18 | 0.28 | 0.2 | 0.3 |
| 1 | 4176106 | rs6670920 | -1.733 | -0.136 | -1.834 | -0.614 | -0.005 | 0.018 | 0.018 | 0 | 0.87 | 0.92 | 0.91 | 1 | 0.83 | 0.96 |
| 1 | 4180842 | rs12040805 | 0.299 | 0.721 | -1.965 | 0.067 | -0.035 | -0.033 | -0.033 | 0 | 0.25 | 0.12 | 0.37 | 0.31 | 0.33 | 0.39 |
| 1 | 4183258 | rs10915438 | 0.066 | 0.391 | -1.714 | -0.047 | -0.012 | -0.007 | -0.007 | 0 | 0.18 | 0.28 | 0.27 | 0.18 | 0.13 | 0.29 |
| 1 | 4185126 | rs10753368 | -0.924 | -0.226 | -1.197 | -0.304 | -0.001 | -0.008 | -0.008 | 0 | 0.53 | 0.56 | 0.72 | 0.9 | 0.51 | 0.75 |
| 1 | 4186309 | rs6426389 | -0.72 | -0.485 | -1.225 | -0.148 | -0.001 | 0.001 | 0.001 | 0 | 0.48 | 0.48 | 0.65 | 0.53 | 0.49 | 0.67 |
| 1 | 4187952 | rs4654487 | -0.38 | -0.659 | -1.555 | -0.5 | -0.009 | 0.011 | 0.011 | 0 | 0.63 | 0.48 | 0.76 | 0.53 | 0.67 | 0.7 |
| 1 | 4190685 | rs4282783 | 0.107 | -0.996 | -1.671 | -0.538 | -0.006 | 0.031 | 0.031 | 0 | 0.88 | 0.84 | 0.92 | 0.63 | 0.91 | 0.9 |
| 1 | 4194597 | rs4131506 | 1.156 | -0.917 | -1.739 | -0.658 | -0.021 | -0.02 | -0.02 | 0.457 | 0.22 | 0.18 | 0.51 | 0.35 | 0.35 | 0.27 |
| 1 | 4197035 | rs6702633 | -0.87 | -1.207 | -1.811 | -0.848 | 0.004 | 0.005 | 0.005 | 0 | 0.47 | 0.41 | 0.65 | 0.54 | 0.43 | 0.67 |
| 1 | 4197642 | rs10915445 | 0.161 | -1.33 | -1.917 | -0.865 | -0.04 | -0.051 | -0.051 | 0 | 0.32 | 0.2 | 0.55 | 0.47 | 0.41 | 0.48 |
| 1 | 4198701 | rs4654490 | -2.379 | -1.224 | -2.023 | -0.742 | -0.046 | -0.03 | -0.03 | 0 | 0.8 | 0.95 | 0.87 | 0.81 | 0.72 | 0.92 |
| 1 | 4201349 | rs16840838 | 1.645 | -1.59 | -1.053 | -0.693 | 0.006 | -0.055 | -0.055 | 0 | 0.21 | 0.24 | 0.14 | 0 | 0.26 | 0.17 |
| 1 | 4201815 | rs4654492 | 1.663 | -1.616 | -1.421 | -0.349 | -0.129 | -0.134 | -0.134 | 0.291 | 0.21 | 0.02 | 0.53 | 0.37 | 0.35 | 0.27 |
| 1 | 4202818 | rs7542208 | -0.936 | -1.652 | -1.162 | -0.776 | 0.008 | 0.025 | 0.025 | 0 | 0.5 | 0.47 | 0.69 | 0.72 | 0.43 | 0.69 |
| 1 | 4206210 | rs9426471 | 0.764 | -1.304 | -0.967 | -0.524 | 0.008 | 0.001 | 0.001 | 0 | 0.34 | 0.28 | 0.25 | 0.28 | 0.29 | 0.25 |
| 1 | 4208542 | rs7539733 | 0.678 | -0.953 | -1.066 | -0.558 | 0.023 | 0.011 | 0.011 | 0 | 0.36 | 0.16 | 0.28 | 0.27 | 0.36 | 0.19 |
| 1 | 4210402 | rs6683453 | 0.035 | -1.176 | -0.935 | -0.803 | 0.006 | 0.004 | 0.004 | 0 | 0.65 | 0.57 | 0.44 | 0.38 | 0.59 | 0.65 |
| 1 | 4210455 | rs9426473 | 2.112 | -0.931 | -1.221 | -0.778 | -0.002 | -0.015 | -0.015 | 0.087 | 0.23 | 0.08 | 0.16 | 0.22 | 0.28 | 0.08 |
| 1 | 4212493 | rs16840866 | 1.626 | -0.769 | -0.956 | -0.706 | 0.008 | -0.082 | -0.082 | 0.107 | 0.23 | 0.36 | 0.13 | 0.05 | 0.21 | 0.21 |
| 1 | 4216165 | rs9988443 | 0.529 | -0.652 | -1.252 | -0.271 | 0.018 | 0.017 | 0.017 | 0 | 0.37 | 0.16 | 0.29 | 0.27 | 0.37 | 0.21 |
| 1 | 4227320 | rs16840873 | 0.929 | -1.027 | -0.861 | -0.058 | -0.035 | 0.02 | 0.02 | 0.724 | 0.07 | 0.16 | 0.01 | 0.15 | 0 | 0.04 |
| 1 | 4227464 | rs12123330 | 1.755 | -1.267 | -1.232 | -0.175 | -0.006 | -0.015 | -0.015 | 0 | 0.23 | 0.09 | 0.16 | 0.23 | 0.28 | 0.07 |
| 1 | 4227634 | rs4468133 | 0.49 | -1.023 | -1.51 | -0.159 | -0.077 | -0.121 | -0.121 | 0 | 0.24 | 0.06 | 0.54 | 0.46 | 0.34 | 0.33 |
| 1 | 4230463 | rs10915459 | 1.068 | -0.901 | -1.339 | -0.034 | -0.023 | -0.014 | -0.014 | 0 | 0.06 | 0 | 0.05 | 0.11 | 0.08 | 0.08 |
| 1 | 4231757 | rs6688343 | 0.6 | -1.319 | -1.028 | -0.845 | 0.001 | 0.004 | 0.004 | 0.662 | 0.48 | 0.44 | 0.27 | 0.29 | 0.44 | 0.28 |
| 1 | 4232794 | rs11583685 | 1.822 | -1.235 | -1.185 | -0.752 | 0.018 | -0.008 | -0.008 | 0.011 | 0.23 | 0.09 | 0.13 | 0.24 | 0.23 | 0.06 |
| 1 | 4233810 | rs6695203 | 0.486 | -1.419 | -0.796 | -0.836 | 0.004 | -0.08 | -0.08 | 0 | 0.32 | 0.49 | 0.19 | 0.17 | 0.29 | 0.26 |
| 1 | 4234792 | rs12743431 | -0.307 | -1.006 | -0.896 | -0.859 | -0.029 | 0.016 | 0.016 | 0 | 0.38 | 0.21 | 0.64 | 0.65 | 0.42 | 0.59 |
| 1 | 4235683 | rs4654500 | 0.834 | -1.075 | -0.605 | -0.527 | 0.008 | -0.005 | -0.005 | 2.303 | 0.19 | 0.17 | 0.15 | 0.29 | 0.1 | 0.3 |
| 1 | 4236982 | rs4233262 | -1.12 | -0.993 | -0.24 | -0.503 | -0.012 | -0.026 | -0.026 | 0 | 0.7 | 0.62 | 0.81 | 0.78 | 0.73 | 0.9 |
| 1 | 4240178 | rs4233264 | -1.239 | -0.962 | -0.241 | -0.503 | -0.002 | -0.005 | -0.005 | 0 | 0.71 | 0.72 | 0.82 | 0.78 | 0.73 | 0.9 |
| 1 | 4250933 | rs11808641 | 0.593 | -0.221 | 0.128 | 0.151 | 0.03 | 0.064 | 0.064 | 0.035 | 0.15 | 0.22 | 0.47 | 0.33 | 0.27 | 0.28 |
| 1 | 4260418 | rs6667065 | 0.897 | -0.51 | -0.114 | -0.165 | 0.022 | 0.096 | 0.096 | 0 | 0.45 | 0.19 | 0.33 | 0.28 | 0.4 | 0.36 |
| 1 | 4263063 | rs4654510 | 1.884 | -0.577 | -0.154 | -0.186 | -0.001 | -0.001 | -0.001 | 0 | 0.11 | 0.1 | 0.05 | 0.1 | 0.08 | 0.16 |
| 1 | 4263677 | rs6687268 | 1.221 | -0.469 | 0.057 | -0.087 | -0.003 | -0.001 | -0.001 | 1.127 | 0.26 | 0.32 | 0.12 | 0.31 | 0.23 | 0.23 |
| 1 | 4263997 | rs4654512 | 0.54 | -0.199 | -0.618 | 0.027 | 0.009 | 0.086 | 0.086 | 0 | 0.49 | 0.25 | 0.38 | 0.36 | 0.47 | 0.46 |
| 1 | 4267089 | rs16840901 | 1.61 | -0.225 | -0.617 | 0.355 | -0.075 | -0.073 | -0.073 | 0.943 | 0.11 | 0.01 | 0.42 | 0.32 | 0.25 | 0.25 |
| 1 | 4269491 | rs6426402 | -0.06 | -0.526 | -0.241 | 0.185 | 0.001 | -0.002 | -0.002 | 0 | 0.41 | 0.48 | 0.58 | 0.63 | 0.48 | 0.51 |
| 1 | 4271526 | rs12022636 | 0.931 | -0.591 | -0.369 | 0.144 | 0.004 | 0.093 | 0.093 | 0 | 0.45 | 0.18 | 0.31 | 0.3 | 0.42 | 0.33 |
| 1 | 4271915 | rs7553814 | -0.025 | -0.542 | -0.292 | 0.182 | 0.011 | -0.008 | -0.008 | 7.537 | 0.4 | 0.5 | 0.57 | 0.63 | 0.46 | 0.5 |

| | | | | | | | | | | | | | | | | |
|---|---|---|---|---|---|---|---|---|---|---|---|---|---|---|---|---|
| 1 | 4276903 | rs10915478 | 0.254 | -0.425 | -0.207 | 0.371 | -0.041 | 0.087 | 0.087 | 0 | 0.9 | 0.75 | 0.94 | 0.77 | 0.96 | 0.88 |
| 1 | 4281160 | rs10799216 | 0.29 | -0.209 | -0.22 | 0.165 | 0.004 | -0.002 | -0.002 | 0 | 0.48 | 0.42 | 0.35 | 0.54 | 0.45 | 0.47 |
| 1 | 4281458 | rs4654424 | 0.773 | 0.011 | -0.642 | 0.088 | -0.056 | -0.063 | -0.063 | 0 | 0.13 | 0.05 | 0.46 | 0.33 | 0.25 | 0.25 |
| 1 | 4282786 | rs11579829 | 0.232 | -0.398 | -0.321 | -0.614 | -0.008 | 0.055 | 0.055 | 0 | 0.34 | 0.2 | 0.26 | 0.21 | 0.34 | 0.21 |
| 1 | 4283315 | rs2132177 | 0.136 | -0.211 | -0.249 | -0.422 | -0.031 | -0.037 | -0.037 | 0.212 | 0.93 | 0.88 | 0.97 | 0.99 | 0.97 | 0.98 |
| 1 | 4284659 | rs903540 | -0.063 | -0.13 | -0.482 | -0.534 | -0.003 | -0.009 | -0.009 | 0 | 0.61 | 0.64 | 0.73 | 0.78 | 0.68 | 0.83 |
| 1 | 4284919 | rs7528938 | 0.298 | 0.683 | -0.434 | -0.026 | 0.033 | 0.021 | 0.021 | 0.115 | 0.68 | 0.55 | 0.41 | 0.4 | 0.6 | 0.57 |
| 1 | 4288803 | rs10915480 | 0.352 | 0.468 | -0.873 | -0.048 | -0.069 | -0.073 | -0.073 | 0 | 0.38 | 0.22 | 0.56 | 0.44 | 0.47 | 0.58 |
| 1 | 4289045 | rs10753374 | -0.987 | -0.269 | -1.453 | -0.337 | -0.061 | -0.122 | -0.122 | 0 | 0.77 | 0.98 | 0.81 | 0.63 | 0.74 | 0.91 |
| 1 | 4289899 | rs10799218 | -0.56 | -0.225 | -0.3 | -0.541 | -0.001 | 0.003 | 0.003 | 0 | 0.67 | 0.71 | 0.76 | 0.5 | 0.7 | 0.83 |
| 1 | 4292038 | rs7534558 | 0.916 | 0.064 | -0.404 | -0.541 | -0.074 | -0.076 | -0.076 | 0 | 0.34 | 0.13 | 0.53 | 0.39 | 0.46 | 0.52 |
| 1 | 4293892 | rs12071761 | -0.374 | -0.063 | 0.419 | -0.94 | 0.007 | -0.003 | -0.003 | 0.655 | 0.1 | 0.21 | 0.1 | 0.09 | 0.1 | 0.15 |
| 1 | 4294000 | rs4512614 | -0.245 | -0.064 | 0.256 | -0.939 | -0.016 | -0.067 | -0.067 | 0 | 0.22 | 0.23 | 0.11 | 0.01 | 0.13 | 0.14 |
| 1 | 4295304 | rs12132422 | -0.633 | 0.228 | -0.099 | -0.729 | -0.028 | -0.092 | -0.092 | 2.617 | 0.32 | 0.47 | 0.2 | 0.11 | 0.24 | 0.29 |
| 1 | 4299727 | rs7517989 | -0.56 | 0.601 | -0.038 | -0.606 | 0.002 | -0.068 | -0.068 | 1.453 | 0.17 | 0.27 | 0.15 | 0 | 0.15 | 0.1 |
| 1 | 4309953 | rs6679489 | -0.546 | 0.552 | -0.568 | -0.35 | 0.028 | -0.026 | -0.026 | 0.046 | 0.6 | 0.86 | 0.72 | 0.41 | 0.66 | 0.77 |
| 1 | 4312205 | rs1908490 | 0.725 | 0.524 | -0.498 | -0.281 | -0.123 | 0.057 | 0.057 | 0 | 0.37 | 0.12 | 0.52 | 0.25 | 0.48 | 0.52 |
| 1 | 4314177 | rs16841022 | 0.261 | 0.503 | -0.379 | -0.301 | 0.001 | -0.06 | -0.06 | 2.436 | 0.17 | 0.26 | 0.14 | 0 | 0.15 | 0.1 |
| 1 | 4315204 | rs966321 | 0.593 | 0.494 | -0.359 | -0.295 | -0.116 | -0.051 | -0.051 | 0 | 0.38 | 0.14 | 0.54 | 0.4 | 0.51 | 0.62 |
| 1 | 4316784 | rs964715 | -0.749 | 0.439 | 0.178 | -0.243 | 0.035 | -0.039 | -0.039 | 0 | 0.6 | 0.88 | 0.73 | 0.42 | 0.67 | 0.77 |
| 1 | 4327635 | rs2101576 | 0.708 | 0.413 | 0.684 | -0.262 | -0.093 | 0.062 | 0.062 | 0 | 0.37 | 0.17 | 0.52 | 0.25 | 0.47 | 0.52 |
| 1 | 4328074 | rs12087250 | -0.106 | 0.419 | 0.578 | -0.257 | 0.002 | -0.12 | -0.12 | 0 | 0.18 | 0.39 | 0.15 | 0 | 0.15 | 0.1 |
| 1 | 4328476 | rs10915494 | -0.22 | 0.42 | 0.663 | -0.239 | 0.008 | -0.006 | -0.006 | 0 | 0.19 | 0.32 | 0.17 | 0.15 | 0.19 | 0.21 |
| 1 | 4330201 | rs17407762 | 0.65 | 0.431 | 0.273 | -0.315 | -0.031 | 0.067 | 0.067 | 0 | 0.13 | 0.03 | 0.12 | 0 | 0.13 | 0.1 |
| 1 | 4330287 | rs6677984 | 0.276 | 0.467 | 0.457 | -0.132 | -0.011 | 0.016 | 0.016 | 0 | 0.42 | 0.59 | 0.56 | 0.28 | 0.48 | 0.55 |
| 1 | 4335893 | rs903541 | -0.556 | 0.695 | 0.088 | -0.112 | 0.001 | -0.015 | -0.015 | 0 | 0.57 | 0.82 | 0.69 | 0.44 | 0.6 | 0.76 |
| 1 | 4341843 | rs1390136 | -1.365 | 0.487 | 0.306 | 0.424 | 0.02 | -0.024 | -0.024 | 2.615 | 0.35 | 0.79 | 0.22 | 0.42 | 0.26 | 0.34 |
| 1 | 4342604 | rs1495244 | -1.3 | 0.481 | 0.517 | 0.421 | 0.007 | -0.009 | -0.009 | 0.181 | 0.34 | 0.72 | 0.22 | 0.42 | 0.26 | 0.35 |
| 1 | 4351888 | rs16838560 | -0.073 | 0.405 | 0.261 | 0.913 | -0.004 | 0.031 | 0.031 | 3.285 | 0.08 | 0.13 | 0.03 | 0.23 | 0.06 | 0.04 |
| 1 | 4352710 | rs16838561 | -0.922 | 0.431 | 0.06 | 0.706 | -0.023 | 0.142 | 0.142 | 1.792 | 0.15 | 0.63 | 0.09 | 0.41 | 0.1 | 0.21 |
| 1 | 4355593 | rs2029490 | -0.521 | 0.816 | -0.744 | 0.442 | -0.004 | -0.008 | -0.008 | 0 | 0.68 | 0.79 | 0.72 | 0.67 | 0.69 | 0.82 |
| 1 | 4364226 | rs693734 | -0.442 | 1.111 | -1.33 | 0.262 | -0.029 | 0.024 | 0.024 | 0 | 0.73 | 0.87 | 0.66 | 0.78 | 0.69 | 0.6 |
| 1 | 4365471 | rs351625 | -0.661 | 1.127 | -1.022 | 0.383 | 0.002 | 0.001 | 0.001 | 0 | 0.35 | 0.37 | 0.35 | 0.35 | 0.42 | 0.29 |
| 1 | 4366890 | rs351623 | -0.055 | 1.152 | -1.474 | -0.152 | -0.058 | 0.109 | 0.109 | 0 | 0.19 | 0.04 | 0.15 | 0 | 0.24 | 0.09 |
| 1 | 4367323 | rs1490413 | -0.154 | 0.64 | -1.411 | -0.131 | -0.001 | -0.002 | -0.002 | 0 | 0.44 | 0.45 | 0.52 | 0.57 | 0.44 | 0.54 |
| 1 | 4367389 | rs16838591 | 1.016 | 0.16 | -2.277 | -0.15 | -0.036 | -0.062 | -0.062 | 0 | 0.26 | 0.02 | 0.24 | 0.33 | 0.23 | 0.24 |
| 1 | 4367810 | rs17448695 | -0.407 | -0.219 | -2.069 | -0.234 | 0.009 | 0.008 | 0.008 | 0 | 0.17 | 0.1 | 0.05 | 0.06 | 0.09 | 0.09 |
| 1 | 4367922 | rs1817914 | 0.204 | -0.281 | -3.086 | -0.736 | -0.047 | 0.103 | 0.103 | 0.216 | 0.68 | 0.98 | 0.63 | 0.89 | 0.71 | 0.76 |
| 1 | 4367980 | rs2934983 | -0.494 | -1.584 | -2.441 | -0.586 | 0.007 | 0.005 | 0.005 | 0 | 0.64 | 0.69 | 0.7 | 0.67 | 0.7 | 0.75 |
| 1 | 4368026 | rs1817913 | -0.217 | -1.067 | -1.435 | -0.168 | 0.006 | -0.006 | -0.006 | 1.213 | 0.38 | 0.32 | 0.3 | 0.45 | 0.28 | 0.26 |
| 1 | 4368271 | rs10915508 | -0.106 | 0.257 | -1.46 | -0.442 | -0.022 | 0.03 | 0.03 | 0 | 0.17 | 0.11 | 0.2 | 0.04 | 0.25 | 0.25 |
| 1 | 4368473 | rs11581089 | 0.92 | 0.647 | -1.487 | -0.259 | -0.049 | 0.03 | 0.03 | 0 | 0.09 | 0.01 | 0.16 | 0.03 | 0.19 | 0.21 |
| 1 | 4368819 | rs351619 | -0.156 | 0.25 | -1.542 | -0.471 | -0.039 | 0.066 | 0.066 | 0 | 0.65 | 0.78 | 0.59 | 0.76 | 0.53 | 0.64 |
| 1 | 4370074 | rs1495239 | 1.069 | -0.088 | -1.737 | -0.097 | 0.014 | -0.026 | -0.026 | 0.02 | 0.14 | 0.07 | 0.27 | 0.31 | 0.1 | 0.23 |
| 1 | 4370385 | rs351615 | -1.692 | 0.031 | -1.731 | -0.581 | -0.002 | 0.025 | 0.025 | 0 | 0.81 | 0.93 | 0.79 | 0.81 | 0.81 | 0.88 |
| 1 | 4370489 | rs12073612 | 0.44 | 0.095 | -0.677 | -0.412 | -0.041 | 0.01 | 0.01 | 0 | 0.14 | 0.31 | 0.08 | 0.14 | 0.06 | 0.08 |
| 1 | 4371949 | rs12132314 | 1.176 | 0.644 | -1.335 | -0.309 | -0.028 | 0.008 | 0.008 | 0.546 | 0.16 | 0.06 | 0.1 | 0.13 | 0.19 | 0.2 |
| 1 | 4372470 | rs716214 | -1.51 | 0.141 | -0.596 | -0.142 | 0 | -0.004 | -0.004 | 0 | 0.32 | 0.65 | 0.3 | 0.18 | 0.35 | 0.35 |
| 1 | 4372992 | rs1353341 | 0.354 | 0.903 | -0.869 | -0.157 | -0.007 | -0.043 | -0.043 | 0 | 0.09 | 0.16 | 0.06 | 0 | 0.08 | 0.03 |
| 1 | 4375859 | rs10915509 | -0.197 | 1.128 | -1.149 | 0.36 | -0.007 | 0.007 | 0.007 | 0 | 0.1 | 0.09 | 0.17 | 0.05 | 0.21 | 0.23 |
| 1 | 4379687 | rs12408012 | -0.427 | -0.451 | -0.963 | 0.762 | -0.018 | 0.01 | 0.01 | 0 | 0.31 | 0.63 | 0.35 | 0.47 | 0.14 | 0.31 |
| 1 | 4380195 | rs1874269 | -0.412 | 0.181 | -0.816 | 0.623 | -0.009 | 0.033 | 0.033 | 0 | 0.34 | 0.44 | 0.35 | 0.48 | 0.35 | 0.3 |
| 1 | 4380518 | rs7418164 | -0.756 | 0.884 | -1.189 | 0.426 | -0.053 | 0.011 | 0.011 | 0 | 0.09 | 0.01 | 0.13 | 0.05 | 0.17 | 0.13 |
| 1 | 4380629 | rs351599 | -0.46 | 0.023 | -0.992 | 0.852 | 0 | -0.002 | -0.002 | 0 | 0.36 | 0.32 | 0.14 | 0.46 | 0.24 | 0.39 |
| 1 | 4381253 | rs2171985 | -1.099 | 0.885 | -0.534 | 0.828 | 0.002 | -0.007 | -0.007 | 0 | 0.13 | 0.18 | 0.14 | 0.08 | 0.15 | 0.14 |
| 1 | 4382789 | rs351605 | -1.281 | 0.822 | -0.68 | 1.091 | -0.008 | -0.009 | -0.009 | 1.501 | 0.56 | 0.79 | 0.4 | 0.48 | 0.54 | 0.64 |
| 1 | 4384056 | rs351606 | -0.515 | 1.077 | 0.041 | 1.37 | -0.003 | -0.012 | -0.012 | 0 | 0.18 | 0.39 | 0.09 | 0.2 | 0.14 | 0.26 |
| 1 | 4384859 | rs351609 | -1.509 | 1.263 | -0.033 | 1.452 | 0.012 | -0.002 | -0.002 | 0.218 | 0.28 | 0.4 | 0.27 | 0.32 | 0.32 | 0.4 |
| 1 | 4387855 | rs656332 | -0.071 | 1.218 | 0.061 | 1.337 | -0.001 | 0 | 0 | 0 | 0.15 | 0.16 | 0.07 | 0.21 | 0.14 | 0.26 |
| 1 | 4388885 | rs641473 | -1.982 | 1.236 | -0.006 | 1.079 | 0.008 | 0.002 | 0.002 | 0 | 0.54 | 0.79 | 0.45 | 0.46 | 0.55 | 0.6 |
| 1 | 4389262 | rs639739 | -0.528 | 1.075 | 0.597 | 1.268 | 0.007 | -0.08 | -0.08 | 0 | 0.17 | 0.4 | 0.09 | 0.05 | 0.13 | 0.2 |
| 1 | 4390289 | rs625079 | -1.835 | 1.133 | 0.704 | 1.353 | 0.013 | 0.038 | 0.038 | 0 | 0.3 | 0.44 | 0.32 | 0.5 | 0.33 | 0.49 |
| 1 | 4395812 | rs579387 | -1.168 | 1.13 | 0.788 | 1.313 | 0.028 | 0.089 | 0.089 | 0 | 0.21 | 0.44 | 0.17 | 0.41 | 0.23 | 0.33 |
| 1 | 4397703 | rs685494 | -1.149 | 1.114 | 0.711 | 1.321 | 0.022 | 0.08 | 0.08 | 0 | 0.21 | 0.4 | 0.17 | 0.41 | 0.23 | 0.33 |
| 1 | 4398015 | rs589585 | -0.322 | 1.094 | 0.638 | 1.193 | 0.002 | -0.029 | -0.029 | 0 | 0.15 | 0.23 | 0.08 | 0.09 | 0.13 | 0.19 |
| 1 | 4398996 | rs498795 | -0.312 | 1.064 | 0.408 | 1.186 | -0.01 | 0.04 | 0.04 | 0 | 0.14 | 0.04 | 0.06 | 0.09 | 0.12 | 0.18 |
| 1 | 4401450 | rs11587563 | -0.031 | 1.081 | 0.528 | 1.1 | 0.005 | 0.065 | 0.065 | 0 | 0.21 | 0.06 | 0.08 | 0.01 | 0.16 | 0.12 |
| 1 | 4405998 | rs10799228 | -0.984 | 1.223 | 0.892 | 1.48 | 0.083 | 0.342 | 0.342 | 0.154 | 0.07 | 0.36 | 0.1 | 0.32 | 0.1 | 0.14 |
| 1 | 4409496 | rs600933 | -1.886 | 1.518 | 0.838 | 1.476 | 0.068 | 0.108 | 0.108 | 0.037 | 0.26 | 0.46 | 0.32 | 0.48 | 0.34 | 0.45 |

| | | | | | | | | | | | | | | | | |
|---|---|---|---|---|---|---|---|---|---|---|---|---|---|---|---|---|
| 1 | 4414045 | rs528291 | -0.963 | 1.399 | 0.58 | 1.464 | -0.022 | 0.009 | 0.009 | 0 | 0.19 | 0.11 | 0.22 | 0.15 | 0.23 | 0.3 |
| 1 | 4418164 | rs1531328 | -1.204 | 1.247 | 0.663 | 1.575 | 0.02 | 0.149 | 0.149 | 0 | 0.1 | 0.18 | 0.21 | 0.39 | 0.14 | 0.24 |
| 1 | 4418319 | rs16838790 | -0.32 | 1.236 | 0.656 | 1.552 | 0.026 | 0.178 | 0.178 | 0 | 0.06 | 0.13 | 0.09 | 0.33 | 0.1 | 0.16 |
| 1 | 4419820 | rs349393 | -1.305 | 1.046 | 0.534 | 1.364 | 0.006 | 0.006 | 0.006 | 0 | 0.3 | 0.33 | 0.3 | 0.38 | 0.32 | 0.37 |
| 1 | 4419997 | rs17410050 | 2.379 | 0.842 | 0.605 | 1.245 | -0.005 | 0.078 | 0.078 | 0.264 | 0.4 | 0.21 | 0.41 | 0.19 | 0.41 | 0.27 |
| 1 | 4422230 | rs16838800 | 0.22 | 0.726 | 0.605 | 1.495 | -0.004 | 0.017 | 0.017 | 0 | 0.12 | 0.11 | 0.19 | 0.29 | 0.11 | 0.17 |
| 1 | 4423061 | rs6656193 | -0.418 | 0.773 | 0.691 | 1.496 | 0.033 | 0.19 | 0.19 | 0 | 0.07 | 0.17 | 0.09 | 0.31 | 0.1 | 0.15 |
| 1 | 4423272 | rs6666204 | -0.418 | 0.773 | 0.691 | 1.496 | 0.033 | 0.19 | 0.19 | 0 | 0.07 | 0.17 | 0.09 | 0.31 | 0.1 | 0.15 |
| 1 | 4425414 | rs75411 | -0.119 | 0.726 | 0.638 | 1.387 | -0.034 | 0.036 | 0.036 | 0 | 0.17 | 0.12 | 0.1 | 0.09 | 0.21 | 0.19 |
| 1 | 4426085 | rs12404619 | 0.027 | 0.677 | 0.649 | 1.452 | 0 | 0.01 | 0.01 | 0 | 0.14 | 0.11 | 0.22 | 0.29 | 0.2 | 0.17 |
| 1 | 4429462 | rs483344 | -2.09 | 0.587 | 0.783 | 1.498 | -0.009 | 0.014 | 0.014 | 0.086 | 0.41 | 0.58 | 0.51 | 0.8 | 0.48 | 0.59 |
| 1 | 4440749 | rs589870 | -0.009 | 0.741 | 0.676 | 1.424 | -0.045 | 0.042 | 0.042 | 0 | 0.16 | 0.07 | 0.1 | 0.09 | 0.21 | 0.19 |
| 1 | 4441089 | rs1845635 | -1.118 | 0.543 | 0.878 | 1.639 | 0.002 | 0.272 | 0.272 | 0 | 0.09 | 0.25 | 0.18 | 0.41 | 0.06 | 0.24 |
| 1 | 4444706 | rs349413 | 0.383 | 0.387 | 0.526 | 1.333 | 0.028 | -0.017 | -0.017 | 0.251 | 0.69 | 0.91 | 0.73 | 0.69 | 0.71 | 0.71 |
| 1 | 4456277 | rs7521665 | -1.582 | 0.461 | 0.948 | 1.16 | 0.06 | 0.599 | 0.599 | 1.034 | 0.06 | 0.43 | 0.19 | 0.42 | 0.01 | 0.12 |
| 1 | 4459689 | rs6679220 | 1.964 | 0.558 | 0.913 | 1.038 | -0.012 | 0.065 | 0.065 | 0.143 | 0.43 | 0.35 | 0.33 | 0.09 | 0.46 | 0.3 |
| 1 | 4462460 | rs349400 | -1.495 | 0.669 | 0.967 | 1.09 | 0.009 | 0.04 | 0.04 | 0 | 0.27 | 0.57 | 0.4 | 0.61 | 0.25 | 0.4 |
| 1 | 4463585 | rs349401 | -0.616 | 0.749 | 0.891 | 1.083 | 0.032 | 0.191 | 0.191 | 0 | 0.16 | 0.36 | 0.25 | 0.51 | 0.17 | 0.3 |
| 1 | 4468884 | rs547923 | 0.015 | 0.679 | 0.647 | 1.093 | 0.024 | -0.028 | -0.028 | 0 | 0.7 | 0.9 | 0.73 | 0.68 | 0.72 | 0.71 |
| 1 | 4471374 | rs10915528 | -0.361 | 0.783 | 0.615 | 1.103 | -0.014 | -0.019 | -0.019 | 0 | 0.09 | 0.04 | 0.15 | 0.1 | 0.08 | 0.12 |
| 1 | 4473263 | rs2926481 | 0.552 | 0.785 | 0.587 | 1.079 | -0.034 | 0.015 | 0.015 | 0 | 0.11 | 0.05 | 0.07 | 0.09 | 0.15 | 0.15 |
| 1 | 4478442 | rs12120157 | -0.235 | 0.9 | 0.517 | 0.941 | -0.041 | -0.035 | -0.035 | 0 | 0.09 | 0.01 | 0.15 | 0.1 | 0.09 | 0.12 |
| 1 | 4481215 | rs4654432 | 1.714 | 0.781 | 0.711 | 0.91 | -0.006 | 0.088 | 0.088 | 0.396 | 0.44 | 0.33 | 0.32 | 0.09 | 0.46 | 0.32 |
| 1 | 4488979 | rs6658372 | 0.955 | 0.366 | 0.56 | 0.898 | -0.03 | -0.017 | -0.017 | 0 | 0.61 | 0.56 | 0.67 | 0.27 | 0.71 | 0.58 |
| 1 | 4489398 | rs6678792 | 1.511 | 0.164 | 0.045 | 0.709 | 0.014 | 0.036 | 0.036 | 0 | 0.51 | 0.45 | 0.34 | 0.21 | 0.56 | 0.43 |
| 1 | 4491460 | rs10915535 | 0.061 | 0.526 | -0.951 | 0.089 | -0.049 | 0.007 | 0.007 | 0.016 | 0.09 | 0.01 | 0.32 | 0.06 | 0.15 | 0.16 |
| 1 | 4493215 | rs1556691 | 0.44 | 0.35 | -0.859 | 0.119 | 0.037 | 0.001 | 0.001 | 0 | 0.67 | 0.88 | 0.85 | 0.7 | 0.72 | 0.72 |
| 1 | 4495287 | rs7546628 | -0.929 | 0.665 | -0.381 | 0.061 | 0.021 | -0.03 | -0.03 | 0.001 | 0.14 | 0.26 | 0.34 | 0.06 | 0.18 | 0.18 |
| 1 | 4495678 | rs1591426 | 1.685 | 0.425 | -0.274 | -0.021 | 0 | 0 | 0 | 0 | 0.51 | 0.5 | 0.61 | 0.61 | 0.61 | 0.5 |
| 1 | 4496256 | rs1762814 | -1.306 | 0.434 | -0.345 | -0.032 | 0.011 | 0.064 | 0.064 | 0 | 0.4 | 0.22 | 0.43 | 0.33 | 0.35 | 0.48 |
| 1 | 4496659 | rs1674877 | -0.613 | 0.375 | -0.369 | -0.051 | 0.022 | 0.019 | 0.019 | 0 | 0.28 | 0.21 | 0.1 | 0.26 | 0.2 | 0.33 |
| 1 | 4497217 | rs1674879 | 0.412 | 0.306 | -0.408 | -0.085 | 0.023 | 0.001 | 0.001 | 0 | 0.18 | 0.09 | 0.06 | 0.19 | 0.12 | 0.21 |
| 1 | 4500436 | rs12731060 | 2.129 | 0.354 | -0.378 | -0.066 | -0.001 | -0.002 | -0.002 | 0 | 0.48 | 0.47 | 0.51 | 0.55 | 0.55 | 0.44 |
| 1 | 4501911 | rs12128230 | -0.057 | 0.346 | -0.296 | 0.026 | -0.055 | 0.028 | 0.028 | 0 | 0.13 | 0.01 | 0.34 | 0.06 | 0.18 | 0.16 |
| 1 | 4502198 | rs780574 | -0.777 | 0.339 | -0.267 | 0.032 | 0.02 | 0.015 | 0.015 | 0 | 0.28 | 0.23 | 0.1 | 0.27 | 0.2 | 0.33 |
| 1 | 4504094 | rs706144 | -1.753 | 0.266 | -0.164 | 0.122 | 0.008 | 0.034 | 0.034 | 0 | 0.45 | 0.43 | 0.46 | 0.33 | 0.4 | 0.52 |
| 1 | 4505966 | rs780587 | 1.081 | 0.007 | 0.113 | 0.631 | 0.015 | 0.083 | 0.083 | 0 | 0.93 | 0.78 | 0.96 | 0.87 | 0.94 | 0.94 |
| 1 | 4507743 | rs780593 | -0.393 | -0.002 | -0.377 | 0.109 | 0.003 | 0.001 | 0.001 | 0 | 0.19 | 0.16 | 0.06 | 0.18 | 0.1 | 0.21 |
| 1 | 4510840 | rs705685 | -0.38 | -0.003 | -0.387 | 0.088 | 0 | -0.001 | -0.001 | 0 | 0.19 | 0.19 | 0.09 | 0.22 | 0.11 | 0.21 |
| 1 | 4512105 | rs780598 | -0.517 | -0.018 | -0.489 | 0.08 | 0.009 | -0.001 | -0.001 | 0 | 0.2 | 0.19 | 0.09 | 0.22 | 0.11 | 0.22 |
| 1 | 4518202 | rs666152 | 0.062 | 0.329 | -0.747 | -0.394 | -0.002 | 0.022 | 0.022 | 0 | 0.38 | 0.27 | 0.41 | 0.74 | 0.37 | 0.37 |
| 1 | 4529544 | rs10915545 | -1.051 | 0.346 | -0.263 | -0.506 | 0.006 | -0.042 | -0.042 | 0 | 0.28 | 0.37 | 0.35 | 0.18 | 0.28 | 0.37 |
| 1 | 4534321 | rs10047182 | -0.623 | 0.361 | -0.554 | -0.622 | -0.003 | -0.003 | -0.003 | 0.298 | 0.17 | 0.21 | 0.19 | 0.15 | 0.11 | 0.25 |
| 1 | 4534350 | rs12143782 | -0.45 | 0.564 | -0.635 | -0.447 | -0.01 | 0.004 | 0.004 | 0 | 0.12 | 0.07 | 0.16 | 0.03 | 0.17 | 0.12 |
| 1 | 4541390 | rs11799990 | -1.08 | -0.433 | -0.228 | -0.365 | -0.04 | -0.016 | -0.016 | 0.826 | 0.19 | 0.72 | 0.12 | 0.13 | 0.06 | 0.2 |
| 1 | 4541425 | rs6426418 | -1.694 | -0.261 | -0.657 | -0.007 | 0.023 | -0.004 | -0.004 | 0 | 0.36 | 0.76 | 0.42 | 0.42 | 0.29 | 0.42 |
| 1 | 4541444 | rs16839089 | -0.897 | -0.597 | 0.224 | -0.279 | -0.066 | -0.027 | -0.027 | 0 | 0.17 | 0.54 | 0.1 | 0.12 | 0.07 | 0.2 |
| 1 | 4543663 | rs6656245 | -0.17 | -0.362 | 0.095 | -0.331 | -0.015 | -0.007 | -0.007 | 0.061 | 0.09 | 0.16 | 0.03 | 0.04 | 0.05 | 0.07 |
| 1 | 4544178 | rs350170 | -1.444 | 0.044 | -0.202 | -0.511 | 0.003 | -0.018 | -0.018 | 0.066 | 0.18 | 0.16 | 0.18 | 0.06 | 0.2 | 0.16 |
| 1 | 4544793 | rs350171 | -0.164 | 0.092 | -0.341 | -0.459 | -0.048 | 0.033 | 0.033 | 1.959 | 0.09 | 0.01 | 0.1 | 0.02 | 0.16 | 0.09 |
| 1 | 4546183 | rs350173 | 0.407 | -0.619 | -0.9 | -1.038 | 0.033 | 0.068 | 0.068 | 0 | 0.15 | 0.03 | 0.05 | 0 | 0.07 | 0.17 |
| 1 | 4549307 | rs705689 | -0.692 | -0.541 | -0.719 | -1.11 | -0.004 | 0.06 | 0.06 | 0 | 0.24 | 0.12 | 0.15 | 0.02 | 0.23 | 0.26 |
| 1 | 4549380 | rs705690 | -1.287 | -0.016 | -0.507 | -0.544 | 0.018 | 0.097 | 0.097 | 0 | 0.87 | 0.76 | 0.8 | 0.65 | 0.82 | 0.79 |
| 1 | 4551471 | rs780612 | -0.197 | 0.034 | -0.705 | -0.591 | -0.039 | 0.126 | 0.126 | 0 | 0.23 | 0.03 | 0.15 | 0.02 | 0.23 | 0.26 |
| 1 | 4552802 | rs1878052 | 1.197 | -0.298 | -0.602 | -0.891 | -0.002 | -0.002 | -0.002 | 0 | 0.43 | 0.38 | 0.41 | 0.56 | 0.47 | 0.42 |
| 1 | 4554214 | rs350193 | 0.765 | 0.193 | -0.755 | -1.065 | -0.038 | 0.02 | 0.02 | 2.524 | 0.07 | 0.01 | 0.08 | 0.02 | 0.12 | 0.08 |
| 1 | 4558933 | rs609506 | 0.066 | -0.239 | -0.432 | -1.018 | -0.007 | -0.028 | -0.028 | 0.845 | 0.66 | 0.58 | 0.78 | 0.86 | 0.7 | 0.62 |
| 1 | 4561962 | rs582598 | 0.693 | -0.11 | -0.746 | -1.401 | -0.05 | 0.054 | 0.054 | 0.706 | 0.12 | 0.01 | 0.1 | 0.02 | 0.17 | 0.11 |
| 1 | 4566053 | rs10753382 | 1.326 | -0.293 | -0.434 | -1.338 | -0.033 | -0.004 | -0.004 | 0.079 | 0.39 | 0.21 | 0.38 | 0.6 | 0.41 | 0.34 |
| 1 | 4566515 | rs4654441 | 1.272 | -0.295 | -0.22 | -1.339 | -0.016 | 0.001 | 0.001 | 0 | 0.39 | 0.29 | 0.38 | 0.6 | 0.42 | 0.34 |
| 1 | 4567165 | rs1018306 | 0.559 | -0.234 | -0.232 | -1.386 | -0.002 | 0.048 | 0.048 | 0 | 0.49 | 0.57 | 0.66 | 0.86 | 0.46 | 0.51 |
| 1 | 4572778 | rs1762826 | 0.173 | -0.29 | -0.648 | -1.601 | -0.032 | 0.149 | 0.149 | 0 | 0.26 | 0.03 | 0.15 | 0.04 | 0.23 | 0.24 |
| 1 | 4573099 | rs4480324 | -0.099 | -0.514 | -0.264 | -1.624 | -0.014 | -0.055 | -0.055 | 8.476 | 0.7 | 0.42 | 0.8 | 0.87 | 0.72 | 0.71 |
| 1 | 4575001 | rs1710424 | -1.626 | -0.477 | 0.148 | -1.644 | -0.008 | -0.018 | -0.018 | 0.683 | 0.83 | 0.75 | 0.88 | 0.88 | 0.84 | 0.79 |
| 1 | 4576970 | rs670296 | -0.139 | -0.057 | -0.424 | -2.424 | -0.002 | 0.004 | 0.004 | 0.263 | 0.62 | 0.38 | 0.39 | 0.16 | 0.61 | 0.63 |
| 1 | 4578495 | rs6703696 | 0.536 | -0.049 | -0.405 | -2.514 | -0.003 | -0.014 | -0.014 | 0.125 | 0.12 | 0.12 | 0.03 | 0.03 | 0.08 | 0.15 |
| 1 | 4582089 | rs518456 | -0.186 | 0.131 | -0.458 | -2.399 | 0.006 | 0.02 | 0.02 | 0 | 0.51 | 0.41 | 0.56 | 0.25 | 0.55 | 0.46 |
| 1 | 4587177 | rs241239 | 0.833 | -0.109 | -1.115 | -2.456 | -0.065 | 0.191 | 0.191 | 0.531 | 0.28 | 0.03 | 0.16 | 0 | 0.29 | 0.28 |
| 1 | 4590550 | rs241233 | -1.577 | -0.293 | -0.493 | -2.152 | 0.001 | 0.006 | 0.006 | 0 | 0.78 | 0.74 | 0.72 | 0.38 | 0.77 | 0.79 |

| | | | | | | | | | | | | | | | | |
|---|---|---|---|---|---|---|---|---|---|---|---|---|---|---|---|---|
| 1 | 4591622 | rs241231 | 0.281 | -0.352 | -0.602 | -2.486 | 0.015 | 0.094 | 0.094 | 0 | 0.27 | 0.12 | 0.14 | 0.03 | 0.2 | 0.24 |
| 1 | 4592382 | rs1515673 | 0.791 | -0.367 | -0.357 | -2.553 | 0.005 | 0.017 | 0.017 | 0 | 0.3 | 0.19 | 0.16 | 0.02 | 0.32 | 0.29 |
| 1 | 4596700 | rs12080165 | 0.124 | -0.428 | 0.132 | -2.291 | -0.004 | -0.038 | -0.038 | 0 | 0.27 | 0.26 | 0.15 | 0.03 | 0.2 | 0.22 |
| 1 | 4598021 | rs12726510 | -1.511 | -0.57 | 0.024 | -2.33 | -0.003 | -0.017 | -0.017 | 1.137 | 0.76 | 0.78 | 0.73 | 0.33 | 0.78 | 0.68 |
| 1 | 4599243 | rs592726 | 0.222 | -0.553 | 0.154 | -2.308 | 0.003 | 0.031 | 0.031 | 0 | 0.49 | 0.42 | 0.58 | 0.3 | 0.58 | 0.46 |
| 1 | 4606471 | rs12092980 | 1.391 | -0.679 | 0.158 | -2.315 | -0.013 | -0.041 | -0.041 | 0.903 | 0.17 | 0.22 | 0.09 | 0.03 | 0.11 | 0.11 |
| 1 | 4609024 | rs16839257 | 1.532 | -0.523 | 0.266 | -2.32 | 0 | -0.015 | -0.015 | 0 | 0.11 | 0.11 | 0.06 | 0 | 0.09 | 0.11 |
| 1 | 4609050 | rs241221 | 0.787 | -0.641 | 0.241 | -2.417 | -0.017 | 0.036 | 0.036 | 0.291 | 0.41 | 0.25 | 0.57 | 0.28 | 0.58 | 0.33 |
| 1 | 4609122 | rs16839263 | 0.078 | -0.668 | 0.182 | -2.447 | -0.03 | -0.081 | -0.081 | 0 | 0.19 | 0.3 | 0.07 | 0.02 | 0.1 | 0.23 |
| 1 | 4611241 | rs241248 | 0.878 | -0.385 | -0.213 | -2.321 | -0.029 | 0.104 | 0.104 | 0.543 | 0.38 | 0.21 | 0.3 | 0.1 | 0.49 | 0.3 |
| 1 | 4611435 | rs7549444 | 2.364 | -0.412 | -0.154 | -2.25 | -0.003 | -0.007 | -0.007 | 0 | 0.13 | 0.15 | 0.12 | 0.1 | 0.15 | 0.11 |
| 1 | 4615331 | rs4654556 | -0.31 | -0.45 | 0.001 | -2.141 | 0.009 | 0.082 | 0.082 | 3.593 | 0.13 | 0.37 | 0.29 | 0.2 | 0.1 | 0.16 |
| 1 | 4618251 | rs6685798 | 1.003 | -0.429 | -0.213 | -2.312 | -0.038 | 0.114 | 0.114 | 0.532 | 0.37 | 0.18 | 0.3 | 0.13 | 0.49 | 0.29 |
| 1 | 4618279 | rs916408 | 0.471 | -0.423 | -0.422 | -2.445 | 0.003 | -0.012 | -0.012 | 0 | 0.3 | 0.22 | 0.18 | 0.01 | 0.24 | 0.21 |
| 1 | 4619038 | rs241264 | -1.649 | -0.426 | -0.489 | -2.417 | 0 | -0.006 | -0.006 | 0 | 0.81 | 0.79 | 0.78 | 0.33 | 0.83 | 0.67 |
| 1 | 4620923 | rs10915560 | -0.862 | -0.262 | -0.626 | -2.638 | -0.018 | -0.119 | -0.119 | 1.269 | 0.39 | 0.49 | 0.2 | 0.03 | 0.25 | 0.34 |
| 1 | 4620996 | rs12120353 | -0.208 | 0.519 | -1.336 | -2.291 | 0.005 | 0.018 | 0.018 | 0 | 0.3 | 0.18 | 0.18 | 0.01 | 0.24 | 0.22 |
| 1 | 4629408 | rs241278 | -0.293 | 0.868 | -1.728 | -1.911 | -0.044 | 0.185 | 0.185 | 0.066 | 0.44 | 0.17 | 0.29 | 0.04 | 0.5 | 0.3 |
| 1 | 4631266 | rs241212 | 0.249 | 1.418 | -1.059 | -1.262 | -0.016 | 0.007 | 0.007 | 0.016 | 0.24 | 0.15 | 0.18 | 0 | 0.33 | 0.16 |
| 1 | 4632828 | rs171317 | 0.603 | 0.84 | -0.606 | -0.599 | 0.04 | 0.091 | 0.091 | 0 | 0.9 | 0.67 | 0.88 | 0.88 | 0.92 | 0.81 |
| 1 | 4634978 | rs1541318 | 0.158 | 1.211 | -1.745 | -1.075 | -0.06 | 0.16 | 0.16 | 0 | 0.35 | 0.15 | 0.22 | 0 | 0.41 | 0.37 |
| 1 | 4636767 | rs1157688 | 0.234 | 1.47 | -1.334 | 0.589 | -0.024 | -0.017 | -0.017 | 0 | 0.89 | 0.93 | 0.61 | 0.63 | 0.76 | 0.88 |
| 1 | 4640902 | rs1024139 | -0.058 | 0.304 | -1.169 | 0.199 | 0.004 | -0.006 | -0.006 | 0 | 0.33 | 0.36 | 0.2 | 0.57 | 0.2 | 0.24 |
| 1 | 4641742 | rs241225 | 0.423 | 0.224 | -0.803 | 0.774 | -0.013 | 0.133 | 0.133 | 0 | 0.94 | 0.85 | 0.91 | 0.72 | 0.94 | 0.78 |
| 1 | 4643432 | rs2078573 | 0.56 | 0.757 | -1.546 | -0.061 | -0.043 | 0.096 | 0.096 | 0 | 0.18 | 0.04 | 0.14 | 0 | 0.24 | 0.08 |
| 1 | 4646379 | rs619364 | -0.106 | 0.571 | -1.518 | -0.042 | 0.009 | -0.006 | -0.006 | 0.739 | 0.54 | 0.43 | 0.59 | 0.6 | 0.64 | 0.47 |
| 1 | 4647469 | rs644647 | -0.272 | 0.323 | -1.711 | -0.142 | -0.012 | -0.006 | -0.006 | 0 | 0.71 | 0.47 | 0.71 | 0.78 | 0.74 | 0.57 |
| 1 | 4648780 | rs10915566 | 0.854 | 0.5 | -1.388 | -0.173 | -0.03 | -0.002 | -0.002 | 0 | 0.24 | 0.35 | 0.15 | 0.18 | 0.12 | 0.15 |
| 1 | 4649316 | rs1849943 | 1.647 | 0.69 | -1.753 | -0.108 | -0.056 | 0.079 | 0.079 | 0 | 0.14 | 0.01 | 0.12 | 0.01 | 0.18 | 0.15 |
| 1 | 4652284 | rs17349020 | 1.949 | 0.694 | -1.756 | -0.112 | -0.023 | 0.069 | 0.069 | 0 | 0.12 | 0.01 | 0.09 | 0.01 | 0.13 | 0.14 |
| 1 | 4653765 | rs164771 | 0.684 | 0.947 | -1.793 | 0.146 | -0.034 | -0.025 | -0.025 | 1.054 | 0.06 | 0.01 | 0.25 | 0.23 | 0.12 | 0.04 |
| 1 | 4657049 | rs526695 | -0.662 | 0.964 | -1.808 | 0.166 | 0.001 | 0.001 | 0.001 | 0.548 | 0.92 | 0.93 | 0.83 | 0.79 | 0.91 | 0.77 |
| 1 | 4658538 | rs12567277 | 0.398 | 0.972 | -1.86 | -0.08 | 0.001 | -0.032 | -0.032 | 0 | 0.57 | 0.78 | 0.33 | 0.54 | 0.4 | 0.49 |
| 1 | 4662408 | rs537761 | 0.498 | 1.256 | -2.02 | -0.238 | -0.061 | -0.03 | -0.03 | 0 | 0.21 | 0.07 | 0.37 | 0.24 | 0.31 | 0.11 |
| 1 | 4665232 | rs164781 | 0.141 | 0.702 | -2.227 | -0.481 | -0.009 | 0.062 | 0.062 | 0 | 0.42 | 0.16 | 0.52 | 0.31 | 0.55 | 0.49 |
| 1 | 4668423 | rs11590266 | 0.041 | 0.919 | -2.091 | -0.656 | -0.067 | 0.207 | 0.207 | 0.123 | 0.38 | 0.09 | 0.27 | 0.07 | 0.45 | 0.36 |
| 1 | 4668670 | rs3896439 | 0.642 | 0.459 | -1.221 | 0.163 | -0.013 | 0.117 | 0.117 | 0.363 | 0.09 | 0.21 | 0.03 | 0.37 | 0.05 | 0.08 |
| 1 | 4672755 | rs6703429 | 0.001 | 0.49 | -1.279 | 0.072 | 0.001 | -0.001 | -0.001 | 0.005 | 0.28 | 0.27 | 0.21 | 0.38 | 0.28 | 0.17 |
| 1 | 4674026 | rs763219 | -0.009 | 0.703 | -1.025 | -0.278 | -0.034 | 0.083 | 0.083 | 0 | 0.39 | 0.24 | 0.28 | 0.08 | 0.42 | 0.46 |
| 1 | 4675555 | rs4654569 | -1.055 | 0.307 | -0.296 | 0.85 | -0.027 | 0.19 | 0.19 | 0.418 | 0.08 | 0.19 | 0.1 | 0.32 | 0.04 | 0.09 |
| 1 | 4677087 | rs2411887 | 0.469 | 0.162 | -0.328 | 0.706 | 0.012 | 0.006 | 0.006 | 0.338 | 0.49 | 0.44 | 0.53 | 0.44 | 0.41 | 0.26 |
| 1 | 4679731 | rs4654574 | -1.311 | 0.682 | -0.66 | 0.822 | 0.001 | 0.024 | 0.024 | 0 | 0.11 | 0.14 | 0.11 | 0.28 | 0.15 | 0.21 |
| 1 | 4679951 | rs1905304 | 0.854 | 0.709 | -0.343 | 0.78 | 0.013 | 0.008 | 0.008 | 0 | 0.5 | 0.59 | 0.54 | 0.45 | 0.42 | 0.26 |
| 1 | 4682230 | rs12026258 | -0.814 | 0.737 | -0.495 | 0.756 | 0.012 | -0.014 | -0.014 | 0 | 0.09 | 0.05 | 0.07 | 0.26 | 0.08 | 0.13 |
| 1 | 4684090 | rs12029673 | -2.087 | 0.757 | -0.486 | 0.785 | -0.004 | -0.007 | -0.007 | 0 | 0.17 | 0.16 | 0.15 | 0.32 | 0.21 | 0.32 |
| 1 | 4685222 | rs1107467 | 1.002 | 0.737 | -0.455 | 0.807 | 0.024 | 0.023 | 0.023 | 0 | 0.51 | 0.63 | 0.54 | 0.43 | 0.41 | 0.25 |
| 1 | 4685672 | rs875807 | -0.161 | 0.78 | -0.429 | 0.617 | -0.019 | 0.016 | 0.016 | 0 | 0.64 | 0.77 | 0.68 | 0.71 | 0.61 | 0.52 |
| 1 | 4686211 | rs11580863 | 0.563 | 0.953 | -0.896 | 0.515 | -0.101 | 0.021 | 0.021 | 0.835 | 0.29 | 0.05 | 0.25 | 0.19 | 0.37 | 0.37 |
| 1 | 4687814 | rs4654576 | 0.133 | 0.705 | -0.55 | 0.556 | 0.056 | -0.023 | -0.023 | 0 | 0.95 | 0.88 | 0.93 | 0.99 | 0.89 | 0.86 |
| 1 | 4693931 | rs6687814 | -1.074 | 0.365 | -0.907 | 0.521 | -0.002 | -0.001 | -0.001 | 0 | 0.34 | 0.32 | 0.3 | 0.7 | 0.27 | 0.55 |
| 1 | 4697300 | rs2101195 | -1.15 | 0.571 | -0.853 | 0.631 | 0.002 | 0.009 | 0.009 | 0.282 | 0.33 | 0.34 | 0.28 | 0.61 | 0.35 | 0.62 |
| 1 | 4698559 | rs12122754 | -0.154 | 0.696 | -1.165 | 1.041 | 0.011 | -0.01 | -0.01 | 3.025 | 0.22 | 0.09 | 0.12 | 0.22 | 0.19 | 0.34 |
| 1 | 4701258 | rs10915575 | 1.416 | 1.362 | -1.421 | 0.953 | -0.021 | -0.027 | -0.027 | 0 | 0.18 | 0.11 | 0.26 | 0.31 | 0.24 | 0.12 |
| 1 | 4704955 | rs2035453 | -0.519 | 0.405 | -1.439 | 0.804 | -0.001 | -0.008 | -0.008 | 0 | 0.36 | 0.67 | 0.32 | 0.56 | 0.37 | 0.55 |
| 1 | 4710486 | rs10915577 | 0.486 | 0.177 | -2.275 | 0.348 | -0.058 | 0.012 | 0.012 | 0.087 | 0.42 | 0.13 | 0.6 | 0.32 | 0.54 | 0.28 |
| 1 | 4717340 | rs900501 | 0.086 | 0.012 | -1.671 | 0.099 | -0.019 | -0.046 | -0.046 | 0.088 | 0.17 | 0.45 | 0.08 | 0.07 | 0.12 | 0.17 |
| 1 | 4717391 | rs1483198 | 1.016 | 0.287 | -2.183 | 0.473 | -0.068 | -0.1 | -0.1 | 0 | 0.15 | 0.02 | 0.15 | 0.32 | 0.22 | 0.3 |
| 1 | 4718551 | rs3947662 | 1.584 | 0.373 | -2.161 | -0.047 | -0.01 | -0.005 | -0.005 | 0.004 | 0.2 | 0.12 | 0.27 | 0.22 | 0.21 | 0.16 |
| 1 | 4722431 | rs2035454 | 1.238 | 0.238 | -2.237 | -0.263 | -0.003 | 0.039 | 0.039 | 0 | 0.22 | 0.14 | 0.14 | 0.01 | 0.26 | 0.11 |
| 1 | 4723046 | rs12408160 | 1.737 | 0.191 | -2.264 | -0.204 | -0.028 | -0.008 | -0.008 | 0 | 0.21 | 0.1 | 0.3 | 0.22 | 0.24 | 0.16 |
| 1 | 4723818 | rs12083131 | 0.349 | 0.204 | -2.166 | -0.104 | 0.027 | 0.068 | 0.068 | 0 | 0.19 | 0.47 | 0.13 | 0.25 | 0.21 | 0.21 |
| 1 | 4723917 | rs10915578 | 0.525 | -0.272 | -2.029 | -0.132 | 0.008 | -0.077 | -0.077 | 0 | 0.26 | 0.33 | 0.37 | 0.01 | 0.32 | 0.14 |
| 1 | 4724957 | rs17421247 | 1.242 | -0.248 | -2.071 | -0.118 | -0.046 | 0.125 | 0.125 | 0 | 0.21 | 0.03 | 0.13 | 0.01 | 0.26 | 0.11 |
| 1 | 4725585 | rs7517857 | 0.551 | -0.311 | -2.023 | 0.007 | -0.001 | -0.001 | -0.001 | 0.697 | 0.41 | 0.57 | 0.43 | 0.54 | 0.45 | 0.46 |
| 1 | 4726454 | rs12567266 | 0.361 | -0.585 | -1.636 | 0.724 | 0.042 | -0.055 | -0.055 | 0 | 0.65 | 0.79 | 0.79 | 0.48 | 0.76 | 0.52 |
| 1 | 4727243 | rs6663166 | -1.129 | -0.419 | -1.468 | 0.708 | 0.017 | 0.021 | 0.021 | 0 | 0.84 | 0.9 | 0.95 | 0.88 | 0.92 | 0.94 |
| 1 | 4730964 | rs16839422 | 0.109 | -0.028 | -1.193 | 1.281 | -0.005 | -0.114 | -0.114 | 0.231 | 0.19 | 0.02 | 0.12 | 0.34 | 0.17 | 0.34 |
| 1 | 4731689 | rs10799256 | -0.767 | -0.037 | -1.026 | 1.139 | 0.001 | -0.029 | -0.029 | 2.704 | 0.28 | 0.2 | 0.39 | 0.44 | 0.27 | 0.52 |
| 1 | 4731928 | rs11587129 | -0.679 | 0.013 | -1.277 | 1.163 | -0.01 | -0.098 | -0.098 | 4.134 | 0.27 | 0.12 | 0.39 | 0.44 | 0.27 | 0.51 |

| | | | | | | | | | | | | | | | | |
|---|---|---|---|---|---|---|---|---|---|---|---|---|---|---|---|---|
| 1 | 4735056 | rs13376219 | -1.125 | 0.077 | -0.526 | 0.625 | -0.013 | 0.106 | 0.106 | 0.049 | 0.11 | 0.25 | 0.19 | 0.3 | 0.09 | 0.21 |
| 1 | 4737693 | rs16839451 | 0.435 | 0.273 | -0.75 | 0.773 | -0.004 | -0.034 | -0.034 | 0 | 0.2 | 0.05 | 0.14 | 0.26 | 0.21 | 0.34 |
| 1 | 4738058 | rs2071990 | 0.562 | 0.24 | -0.617 | 0.818 | -0.003 | 0.089 | 0.089 | 0 | 0.05 | 0.12 | 0.15 | 0.22 | 0.05 | 0.07 |
| 1 | 4738678 | rs2071995 | 0.031 | 0.215 | -0.45 | 0.601 | 0.004 | 0.137 | 0.137 | 0 | 0.08 | 0.21 | 0.19 | 0.3 | 0.09 | 0.15 |
| 1 | 4739318 | rs10799257 | -0.625 | 0.208 | -0.514 | 0.574 | -0.005 | 0.023 | 0.023 | 0 | 0.29 | 0.32 | 0.34 | 0.56 | 0.29 | 0.5 |
| 1 | 4743704 | rs763238 | 0.855 | 0.165 | -1.02 | 0.024 | 0.031 | 0.052 | 0.052 | 0 | 0.28 | 0.16 | 0.11 | 0.01 | 0.22 | 0.14 |
| 1 | 4746636 | rs10915587 | -0.464 | 0.455 | -0.967 | 0.387 | 0.001 | 0.001 | 0.001 | 0 | 0.61 | 0.61 | 0.69 | 0.55 | 0.58 | 0.66 |
| 1 | 4748305 | rs1076944 | -0.123 | 0.342 | -1.07 | 0.076 | -0.011 | -0.012 | -0.012 | 0 | 0.38 | 0.48 | 0.53 | 0.31 | 0.35 | 0.31 |
| 1 | 4751030 | rs242052 | -0.439 | 0.817 | -1.444 | -0.079 | -0.01 | 0.062 | 0.062 | 0 | 0.21 | 0.1 | 0.12 | 0.07 | 0.19 | 0.16 |
| 1 | 4756276 | rs7518469 | -0.839 | 0.754 | -1.15 | 0.104 | 0 | -0.002 | -0.002 | 1.52 | 0.07 | 0.08 | 0.13 | 0.04 | 0.06 | 0.06 |
| 1 | 4756601 | rs3886389 | -0.624 | 0.674 | -1.003 | 0.079 | -0.003 | -0.004 | -0.004 | 0 | 0.43 | 0.48 | 0.33 | 0.39 | 0.39 | 0.51 |
| 1 | 4763189 | rs6686733 | -0.673 | 0.458 | -0.877 | 0.453 | -0.008 | 0.002 | 0.002 | 1.175 | 0.19 | 0.18 | 0.09 | 0.22 | 0.15 | 0.27 |
| 1 | 4768540 | rs6688423 | -0.544 | 0.714 | -0.836 | -0.022 | -0.001 | -0.002 | -0.002 | 0 | 0.55 | 0.56 | 0.57 | 0.55 | 0.49 | 0.74 |
| 1 | 4769825 | rs4654593 | -0.414 | 0.793 | -0.745 | -0.17 | 0.003 | -0.005 | -0.005 | 0 | 0.09 | 0.1 | 0.27 | 0.07 | 0.1 | 0.15 |
| 1 | 4772053 | rs1061968 | -0.674 | 0.497 | -0.928 | -0.07 | -0.01 | -0.018 | -0.018 | 0.083 | 0.4 | 0.52 | 0.5 | 0.66 | 0.37 | 0.58 |
| 1 | 4774438 | rs242058 | -0.584 | 0.809 | -1.45 | -0.841 | -0.008 | 0.056 | 0.056 | 0 | 0.76 | 0.9 | 0.83 | 1 | 0.77 | 0.89 |
| 1 | 4781151 | rs3820251 | -0.427 | 1.416 | -1.173 | 0.138 | -0.001 | -0.002 | -0.002 | 0.718 | 0.07 | 0.08 | 0.28 | 0.09 | 0.11 | 0.14 |
| 1 | 4783355 | rs12085854 | -1.315 | 1.946 | -1.262 | -0.175 | -0.011 | 0.006 | 0.006 | 0.629 | 0.18 | 0.18 | 0.37 | 0.13 | 0.32 | 0.35 |
| 1 | 4784556 | rs3820252 | 0.652 | 0.75 | -0.848 | -0.392 | -0.061 | 0.022 | 0.022 | 0 | 0.16 | 0.3 | 0.08 | 0.2 | 0.05 | 0.15 |
| 1 | 4786095 | rs11586740 | -0.667 | 1.425 | -1.193 | -0.672 | -0.056 | 0.05 | 0.05 | 0.163 | 0.12 | 0.02 | 0.08 | 0.03 | 0.21 | 0.09 |
| 1 | 4789179 | rs242050 | 0.264 | 1.371 | -1.038 | -0.565 | -0.001 | 0 | 0 | 0 | 0.08 | 0.05 | 0.08 | 0.03 | 0.19 | 0.08 |
| 1 | 4792184 | rs3766964 | 0.247 | 0.876 | -1.12 | -1.218 | 0.004 | -0.001 | -0.001 | 0 | 0.39 | 0.36 | 0.23 | 0.18 | 0.33 | 0.32 |
| 1 | 4795414 | rs446529 | 0.033 | 1.08 | -0.989 | -1.306 | 0 | -0.011 | -0.011 | 0.334 | 0.67 | 0.66 | 0.42 | 0.21 | 0.69 | 0.47 |
| 1 | 4799200 | rs3753702 | 0.434 | 0.701 | -1.519 | -1.433 | -0.006 | 0.053 | 0.053 | 0.019 | 0.17 | 0.04 | 0.09 | 0.06 | 0.14 | 0.32 |
| 1 | 4799270 | rs449181 | -1.018 | 0.712 | -1.301 | -0.927 | -0.003 | 0.04 | 0.04 | 0 | 0.91 | 0.91 | 0.83 | 0.28 | 0.93 | 0.85 |
| 1 | 4805434 | rs371605 | -1.248 | 0.712 | -1.284 | -0.73 | -0.001 | 0.04 | 0.04 | 0 | 0.92 | 0.93 | 0.83 | 0.3 | 0.92 | 0.85 |
| 1 | 4805674 | rs409048 | 0.475 | 0.762 | -0.873 | -1.598 | -0.003 | -0.069 | -0.069 | 0 | 0.4 | 0.39 | 0.25 | 0.07 | 0.37 | 0.38 |
| 1 | 4806207 | rs7532708 | 1.889 | 0.288 | -1.013 | -1.952 | 0.02 | 0.005 | 0.005 | 0 | 0.2 | 0.09 | 0.11 | 0.16 | 0.13 | 0.2 |
| 1 | 4806381 | rs368808 | -0.399 | 0.819 | -0.809 | -2.476 | 0.004 | 0.006 | 0.006 | 0.065 | 0.6 | 0.66 | 0.64 | 0.13 | 0.66 | 0.58 |
| 1 | 4806550 | rs11586743 | 0.003 | 0.849 | -0.804 | -2.465 | -0.01 | 0.009 | 0.009 | 0 | 0.22 | 0.22 | 0.39 | 0.08 | 0.31 | 0.36 |
| 1 | 4808454 | rs2802721 | 0.145 | 0.303 | -0.907 | -2.454 | 0 | -0.026 | -0.026 | 0 | 0.52 | 0.53 | 0.32 | 0.12 | 0.48 | 0.47 |
| 1 | 4809120 | rs2802720 | 1.466 | -0.258 | -0.918 | -2.719 | -0.006 | 0.042 | 0.042 | 2.059 | 0.29 | 0.19 | 0.22 | 0.05 | 0.3 | 0.17 |
| 1 | 4810552 | rs364642 | -1.484 | -0.085 | -1.063 | -2.603 | 0.02 | 0.027 | 0.027 | 0 | 0.72 | 0.84 | 0.7 | 0.14 | 0.8 | 0.66 |
| 1 | 4811383 | rs449154 | 1.166 | 0.07 | -1.144 | -2.536 | -0.011 | 0.023 | 0.023 | 0 | 0.1 | 0.04 | 0.09 | 0.05 | 0.14 | 0.11 |
| 1 | 4814400 | rs428001 | 0.468 | -0.021 | -0.959 | -2.417 | -0.002 | -0.063 | -0.063 | 0 | 0.4 | 0.59 | 0.26 | 0.05 | 0.35 | 0.22 |
| 1 | 4814751 | rs3766969 | 1.062 | 0.761 | -0.778 | -1.584 | -0.007 | 0.014 | 0.014 | 0.08 | 0.12 | 0.06 | 0.07 | 0.06 | 0.11 | 0.09 |
| 1 | 4818363 | rs7537345 | 0.953 | 1.223 | -0.301 | -1.592 | 0.086 | -0.069 | -0.069 | 0 | 0.12 | 0.25 | 0.33 | 0.01 | 0.25 | 0.14 |
| 1 | 4819667 | rs523141 | -0.771 | 0.686 | -0.449 | -1.319 | -0.003 | -0.003 | -0.003 | 1.028 | 0.72 | 0.69 | 0.68 | 0.31 | 0.75 | 0.7 |
| 1 | 4823492 | rs12119789 | -0.589 | 0.626 | -0.393 | -1.126 | 0.096 | 0.047 | 0.047 | 0 | 0.25 | 0.49 | 0.44 | 0.35 | 0.42 | 0.26 |
| 1 | 4824311 | rs12066439 | 1.213 | 0.709 | -0.397 | -1.273 | 0.118 | -0.074 | -0.074 | 0 | 0.12 | 0.27 | 0.34 | 0.01 | 0.27 | 0.14 |
| 1 | 4828606 | rs7518471 | -0.352 | 0.197 | -0.533 | -1.484 | 0.022 | -0.092 | -0.092 | 0.016 | 0.31 | 0.49 | 0.43 | 0.06 | 0.37 | 0.44 |
| 1 | 4830900 | rs9426483 | 1.462 | 0.011 | -0.868 | -1.794 | 0.012 | 0.088 | 0.088 | 1.224 | 0.33 | 0.15 | 0.17 | 0.05 | 0.27 | 0.16 |
| 1 | 4831076 | rs423285 | -1.275 | 0.29 | -0.689 | -1.29 | 0.001 | -0.024 | -0.024 | 0 | 0.75 | 0.82 | 0.72 | 0.49 | 0.77 | 0.72 |
| 1 | 4831320 | rs396112 | 0.175 | 0.227 | -0.597 | -1.362 | -0.006 | -0.083 | -0.083 | 0 | 0.61 | 0.55 | 0.6 | 0.09 | 0.63 | 0.46 |
| 1 | 4833498 | rs3766977 | 0.51 | 0.341 | -0.659 | -1.131 | 0.003 | 0.064 | 0.064 | 0 | 0.11 | 0.14 | 0.11 | 0.39 | 0.14 | 0.12 |
| 1 | 4833605 | rs874668 | 0.907 | 0.419 | -0.919 | -1.082 | -0.004 | 0.011 | 0.011 | 0 | 0.1 | 0.06 | 0.11 | 0.39 | 0.14 | 0.12 |
| 1 | 4835375 | rs399377 | -1.369 | 0.35 | -1.036 | -1.05 | 0 | -0.006 | -0.006 | 0.148 | 0.77 | 0.82 | 0.73 | 0.49 | 0.8 | 0.73 |
| 1 | 4835893 | rs874613 | 1.742 | 0.186 | -1.305 | -1.989 | -0.024 | 0.096 | 0.096 | 2.621 | 0.18 | 0.02 | 0.08 | 0.05 | 0.16 | 0.09 |
| 1 | 4847594 | rs11805709 | 0.805 | 0.64 | -0.388 | -1.641 | 0.157 | -0.082 | -0.082 | 0 | 0.13 | 0.39 | 0.34 | 0.01 | 0.27 | 0.13 |
| 1 | 4849384 | rs2235438 | -0.742 | 0.475 | -1.162 | -1.761 | -0.034 | 0.115 | 0.115 | 0.461 | 0.67 | 0.89 | 0.61 | 0.13 | 0.63 | 0.61 |
| 1 | 4850657 | rs875402 | -0.179 | 0.894 | -0.188 | -0.662 | 0.017 | 0.018 | 0.018 | 0 | 0.11 | 0.06 | 0.06 | 0.04 | 0.07 | 0.16 |
| 1 | 4851303 | rs2235443 | -0.467 | 0.902 | -0.185 | -0.659 | 0.024 | 0.026 | 0.026 | 0 | 0.13 | 0.06 | 0.06 | 0.05 | 0.07 | 0.32 |
| 1 | 4852907 | rs2411894 | -0.795 | 1.054 | -0.166 | -0.565 | 0.013 | -0.081 | -0.081 | 0.086 | 0.28 | 0.6 | 0.42 | 0.07 | 0.33 | 0.45 |
| 1 | 4859488 | rs17345273 | -0.358 | 0.396 | -0.086 | 0.358 | 0.006 | 0.013 | 0.013 | 0 | 0.14 | 0.09 | 0.1 | 0.05 | 0.11 | 0.09 |
| 1 | 4861160 | rs4654607 | 1.131 | -0.036 | 0.305 | -0.244 | 0.025 | 0.088 | 0.088 | 0.569 | 0.56 | 0.3 | 0.57 | 0.1 | 0.59 | 0.4 |
| 1 | 4863913 | rs760817 | -0.916 | -0.086 | 0.441 | 0.063 | -0.001 | 0.031 | 0.031 | 0 | 0.9 | 0.9 | 0.88 | 0.65 | 0.86 | 0.86 |
| 1 | 4866420 | rs12752806 | 1.471 | -0.088 | 0.4 | 0.015 | 0.033 | 0.138 | 0.138 | 0.726 | 0.55 | 0.24 | 0.57 | 0.1 | 0.59 | 0.4 |
| 1 | 4872571 | rs10799262 | 0.362 | 0.612 | 0.543 | 0.743 | 0.007 | 0.041 | 0.041 | 0 | 0.87 | 0.82 | 0.79 | 0.32 | 0.81 | 0.75 |
| 1 | 4873056 | rs10915612 | -0.203 | 0.359 | 0.32 | 0.722 | -0.001 | 0.025 | 0.025 | 0 | 0.89 | 0.9 | 0.88 | 0.65 | 0.86 | 0.86 |
| 1 | 4878871 | rs1120120 | -0.391 | 0.426 | 0.109 | 0.759 | -0.001 | -0.009 | -0.009 | 0 | 0.9 | 0.96 | 0.89 | 0.65 | 0.87 | 0.88 |
| 1 | 4886522 | rs6661090 | -1.265 | 0.704 | -0.571 | 0.382 | -0.106 | 0.034 | 0.034 | 1.778 | 0.54 | 0.93 | 0.64 | 0.56 | 0.51 | 0.66 |
| 1 | 4887910 | rs12026226 | 0.249 | 0.543 | -0.268 | 0.374 | -0.007 | -0.019 | -0.019 | 0 | 0.11 | 0.16 | 0.08 | 0.05 | 0.08 | 0.05 |
| 1 | 4888012 | rs2181699 | -0.299 | 0.54 | -0.032 | 0.392 | 0 | 0.001 | 0.001 | 0 | 0.37 | 0.37 | 0.49 | 0.35 | 0.39 | 0.48 |
| 1 | 4889912 | rs12028290 | 0.473 | 0.806 | -0.914 | 0.301 | -0.004 | -0.012 | -0.012 | 0 | 0.1 | 0.12 | 0.06 | 0.05 | 0.07 | 0.05 |
| 1 | 4892026 | rs2025377 | -0.769 | 0.81 | -0.701 | 0.522 | 0.006 | 0.028 | 0.028 | 1.625 | 0.45 | 0.81 | 0.58 | 0.51 | 0.43 | 0.62 |
| 1 | 4898833 | rs7522386 | -0.752 | 1.275 | -0.154 | 1.026 | 0.006 | 0.016 | 0.016 | 0.007 | 0.27 | 0.33 | 0.53 | 0.35 | 0.36 | 0.33 |
| 1 | 4899709 | rs11579593 | -0.045 | 0.961 | -0.242 | 1.093 | -0.005 | 0.004 | 0.004 | 0.074 | 0.08 | 0.01 | 0.05 | 0.05 | 0.07 | 0.06 |
| 1 | 4906783 | rs10915626 | -2.109 | 0.565 | 0.903 | 1.932 | -0.01 | 0.436 | 0.436 | 0.607 | 0.09 | 0.38 | 0.18 | 0.4 | 0.02 | 0.31 |
| 1 | 4914910 | rs6426440 | 0.833 | 0.949 | 0.677 | 1.887 | -0.027 | 0.005 | 0.005 | 0.255 | 0.78 | 0.74 | 0.58 | 0.78 | 0.62 | 0.79 |

| | | | | | | | | | | | | | | | | |
|---|---|---|---|---|---|---|---|---|---|---|---|---|---|---|---|---|
| 1 | 4928187 | rs10915645 | 1.327 | 0.973 | 0.713 | 1.467 | 0.064 | 0.091 | 0.091 | 0.209 | 0.73 | 0.47 | 0.45 | 0.53 | 0.62 | 0.65 |
| 1 | 4930181 | rs1886112 | 0.991 | 1.318 | -0.155 | 1.355 | 0.005 | 0.012 | 0.012 | 1.301 | 0.78 | 0.74 | 0.48 | 0.54 | 0.65 | 0.73 |
| 1 | 4941419 | rs10915652 | -0.816 | 1.163 | 0.129 | 1.404 | -0.002 | 0.006 | 0.006 | 0.073 | 0.15 | 0.19 | 0.51 | 0.32 | 0.26 | 0.43 |
| 1 | 4944214 | rs10733017 | 0.004 | 1.132 | 0.147 | 1.27 | -0.06 | -0.034 | -0.034 | 0.035 | 0.86 | 0.98 | 0.94 | 0.8 | 0.82 | 0.84 |
| 1 | 4944842 | rs7543026 | 0.194 | 0.658 | 0.042 | 0.839 | -0.006 | -0.016 | -0.016 | 4.878 | 0.2 | 0.16 | 0.17 | 0.31 | 0.2 | 0.14 |
| 1 | 4952788 | rs6664880 | -0.466 | 0.677 | 0.008 | 0.844 | 0.002 | 0.009 | 0.009 | 0 | 0.25 | 0.29 | 0.2 | 0.34 | 0.26 | 0.19 |
| 1 | 4954775 | rs4654465 | -0.097 | 0.781 | -0.237 | 0.936 | -0.004 | 0.001 | 0.001 | 0.086 | 0.16 | 0.18 | 0.51 | 0.32 | 0.26 | 0.42 |
| 1 | 4955862 | rs6426446 | 1.685 | 0.495 | -0.048 | 0.784 | 0.014 | 0.005 | 0.005 | 0 | 0.62 | 0.46 | 0.38 | 0.43 | 0.5 | 0.31 |
| 1 | 4965152 | rs17345545 | -0.818 | 0.746 | -0.221 | 0.735 | -0.044 | -0.008 | -0.008 | 5.655 | 0.2 | 0.12 | 0.1 | 0.23 | 0.24 | 0.23 |
| 1 | 4965631 | rs4654468 | -1.254 | 0.749 | -0.109 | 0.799 | 0.01 | 0.02 | 0.02 | 0.982 | 0.22 | 0.33 | 0.53 | 0.35 | 0.33 | 0.52 |
| 1 | 4968528 | rs6678459 | -0.153 | 0.742 | -0.183 | 0.82 | 0.008 | 0.025 | 0.025 | 0.174 | 0.16 | 0.21 | 0.5 | 0.32 | 0.26 | 0.46 |
| 1 | 4971507 | rs12034557 | -0.189 | 0.96 | -0.029 | 0.761 | -0.072 | -0.047 | -0.047 | 0 | 0.14 | 0.03 | 0.07 | 0.2 | 0.17 | 0.17 |
| 1 | 4973107 | rs11805753 | 0.347 | 1.043 | 0.107 | 0.753 | 0.001 | 0.019 | 0.019 | 0 | 0.14 | 0.16 | 0.15 | 0.31 | 0.15 | 0.07 |
| 1 | 4973301 | rs10915668 | -1.833 | 1.018 | 0.127 | 0.647 | -0.006 | -0.004 | -0.004 | 0 | 0.38 | 0.43 | 0.26 | 0.56 | 0.42 | 0.37 |
| 1 | 4975417 | rs10915670 | -1.242 | 1.2 | -0.408 | 0.451 | 0.003 | -0.012 | -0.012 | 0.003 | 0.1 | 0.18 | 0.05 | 0.03 | 0.11 | 0.07 |
| 1 | 4980692 | rs12035499 | -0.369 | 1.099 | -0.503 | 0.529 | -0.015 | -0.003 | -0.003 | 0 | 0.16 | 0.11 | 0.09 | 0.2 | 0.18 | 0.17 |
| 1 | 4981107 | rs11585268 | -0.482 | 1.132 | -0.486 | 0.504 | -0.002 | 0.004 | 0.004 | 0.054 | 0.07 | 0.06 | 0.02 | 0.03 | 0.05 | 0.04 |
| 1 | 4981503 | rs942245 | -1.685 | 0.646 | -0.814 | 0.323 | -0.023 | 0.001 | 0.001 | 0 | 0.55 | 0.77 | 0.37 | 0.59 | 0.48 | 0.57 |
| 1 | 4985137 | rs2093763 | 0.494 | 0.601 | -0.398 | 0.372 | -0.043 | -0.074 | -0.074 | 0.012 | 0.17 | 0.34 | 0.1 | 0.03 | 0.09 | 0.13 |
| 1 | 4985257 | rs6675013 | -0.585 | 0.657 | -0.387 | 0.408 | -0.005 | -0.013 | -0.013 | 0.916 | 0.08 | 0.18 | 0.03 | 0.03 | 0.04 | 0.04 |
| 1 | 4987424 | rs6666453 | -0.426 | 0.673 | -0.485 | 0.434 | -0.002 | 0.026 | 0.026 | 0 | 0.22 | 0.29 | 0.18 | 0.35 | 0.2 | 0.17 |
| 1 | 4988262 | rs11584727 | -0.726 | 0.754 | -0.864 | 0.459 | 0.01 | -0.008 | -0.008 | 0.07 | 0.36 | 0.63 | 0.27 | 0.38 | 0.27 | 0.28 |
| 1 | 4991218 | rs10915677 | 0.284 | 0.655 | -0.899 | 0.672 | -0.017 | -0.018 | -0.018 | 0 | 0.11 | 0.08 | 0.08 | 0.21 | 0.17 | 0.15 |
| 1 | 4992204 | rs2898853 | 1.55 | 0.61 | -1.038 | 0.415 | 0.067 | -0.031 | -0.031 | 0.017 | 0.37 | 0.12 | 0.29 | 0.38 | 0.28 | 0.38 |
| 1 | 4994693 | rs11588299 | 1.599 | 0.612 | -1.125 | 0.315 | 0.044 | -0.031 | -0.031 | 0.434 | 0.34 | 0.12 | 0.29 | 0.37 | 0.26 | 0.34 |
| 1 | 4996185 | rs10915683 | 0.919 | 0.635 | -1.101 | 0.343 | -0.002 | -0.019 | -0.019 | 0 | 0.46 | 0.23 | 0.37 | 0.56 | 0.45 | 0.47 |
| 1 | 4996226 | rs4654470 | -0.55 | 0.967 | -0.58 | 1.044 | -0.014 | -0.008 | -0.008 | 0.05 | 0.13 | 0.11 | 0.09 | 0.2 | 0.22 | 0.21 |
| 1 | 4996772 | rs12040043 | -1.602 | 0.833 | -0.383 | 1.012 | 0.004 | -0.009 | -0.009 | 0.592 | 0.23 | 0.36 | 0.11 | 0.22 | 0.24 | 0.18 |
| 1 | 4997363 | rs9426467 | -0.414 | 1.504 | -0.581 | 1.413 | -0.001 | -0.001 | -0.001 | 0 | 0.21 | 0.22 | 0.21 | 0.36 | 0.21 | 0.24 |
| 1 | 4998385 | rs9426468 | -0.732 | 0.961 | -0.682 | 1.132 | -0.011 | -0.014 | -0.014 | 0 | 0.32 | 0.31 | 0.28 | 0.55 | 0.4 | 0.38 |
| 1 | 5002053 | rs12042532 | -1.163 | 0.206 | -0.108 | 1.29 | -0.029 | 0.073 | 0.073 | 0 | 0.2 | 0.48 | 0.18 | 0.3 | 0.12 | 0.28 |
| 1 | 5003504 | rs9286971 | 0.351 | 0.466 | -0.377 | 0.969 | 0 | -0.01 | -0.01 | 0 | 0.56 | 0.47 | 0.38 | 0.33 | 0.58 | 0.51 |
| 1 | 5011402 | rs9426490 | -0.211 | 0.322 | -0.509 | 1.527 | -0.027 | -0.071 | -0.071 | 3.924 | 0.68 | 0.9 | 0.73 | 0.61 | 0.69 | 0.69 |
| 1 | 5012038 | rs7522655 | -0.124 | 0.304 | -0.341 | 1.517 | -0.006 | -0.027 | -0.027 | 0 | 0.68 | 0.79 | 0.73 | 0.61 | 0.69 | 0.69 |
| 1 | 5012301 | rs10915298 | -0.643 | 0.306 | -0.514 | 1.38 | -0.015 | 0.121 | 0.121 | 0.079 | 0.06 | 0.16 | 0.04 | 0.27 | 0.04 | 0.08 |
| 1 | 5015844 | rs6656223 | -0.098 | 0.167 | -0.831 | 1.417 | -0.014 | -0.08 | -0.08 | 1.316 | 0.66 | 0.88 | 0.72 | 0.52 | 0.68 | 0.69 |
| 1 | 5017652 | rs12029491 | -0.819 | 0.272 | -0.482 | 1.434 | -0.011 | 0.063 | 0.063 | 0.005 | 0.06 | 0.12 | 0.04 | 0.23 | 0.04 | 0.07 |
| 1 | 5019553 | rs12126932 | -0.664 | 0.391 | -0.772 | 0.906 | -0.025 | 0.032 | 0.032 | 0 | 0.08 | 0 | 0.06 | 0.01 | 0.09 | 0.08 |
| 1 | 5021200 | rs10799145 | -0.071 | 0.336 | -0.739 | 0.799 | 0 | 0.034 | 0.034 | 3.354 | 0.52 | 0.3 | 0.58 | 0.41 | 0.51 | 0.64 |
| 1 | 5023321 | rs12754637 | 0.261 | 0.144 | -0.637 | 0.976 | -0.025 | 0.114 | 0.114 | 0 | 0.22 | 0.47 | 0.28 | 0.33 | 0.22 | 0.18 |
| 1 | 5023430 | rs6702660 | -0.686 | 0.419 | -1.069 | 0.745 | -0.095 | -0.054 | -0.054 | 3.932 | 0.75 | 0.96 | 0.82 | 0.74 | 0.72 | 0.82 |
| 1 | 5025179 | rs3128678 | -0.512 | 0.437 | -1.085 | 0.579 | -0.078 | 0.016 | 0.016 | 3.917 | 0.7 | 0.9 | 0.61 | 0.75 | 0.63 | 0.77 |
| 1 | 5027479 | rs6426375 | 0.612 | -0.011 | -0.353 | 0.42 | 0.04 | -0.024 | -0.024 | 0 | 0.19 | 0.39 | 0.17 | 0.15 | 0.21 | 0.16 |
| 1 | 5031379 | rs10799149 | -0.942 | -0.424 | -1.576 | 0.479 | -0.014 | -0.084 | -0.084 | 0 | 0.51 | 0.96 | 0.39 | 0.48 | 0.43 | 0.5 |
| 1 | 5031561 | rs7518523 | -0.08 | -0.432 | -0.85 | -0.04 | -0.001 | -0.002 | -0.002 | 0 | 0.31 | 0.69 | 0.23 | 0.17 | 0.26 | 0.33 |
| 1 | 5033661 | rs7535462 | -0.119 | -0.587 | -1.034 | 0.086 | 0.013 | 0.023 | 0.023 | 5.112 | 0.28 | 0.79 | 0.21 | 0.17 | 0.2 | 0.32 |
| 1 | 5035937 | rs6665175 | -0.157 | -0.316 | -1.267 | 0.419 | 0.011 | -0.021 | -0.021 | 3.371 | 0.18 | 0.06 | 0.15 | 0.24 | 0.18 | 0.15 |
| 1 | 5036234 | rs12757902 | 0.914 | -0.237 | -0.673 | 0.328 | 0 | 0.005 | 0.005 | 2.996 | 0.18 | 0.19 | 0.14 | 0.11 | 0.15 | 0.2 |
| 1 | 5037812 | rs6666330 | -0.844 | -1.067 | -0.317 | 0.38 | -0.022 | 0.002 | 0.002 | 0 | 0.3 | 0.61 | 0.19 | 0.36 | 0.16 | 0.4 |
| 1 | 5038051 | rs12739724 | 0.635 | -0.969 | -0.672 | -0.581 | 0.01 | 0.036 | 0.036 | 6.304 | 0.22 | 0.12 | 0.15 | 0.12 | 0.17 | 0.22 |
| 1 | 5039093 | rs6657381 | 0.686 | -0.026 | -0.338 | -0.875 | -0.015 | 0.047 | 0.047 | 4.285 | 0.26 | 0.19 | 0.37 | 0.13 | 0.29 | 0.17 |
| 1 | 5039380 | rs1841241 | 0.307 | -0.051 | -0.183 | -0.795 | -0.019 | 0.045 | 0.045 | 2.496 | 0.32 | 0.27 | 0.44 | 0.15 | 0.37 | 0.2 |
| 1 | 5043734 | rs1563791 | 0.488 | 0.234 | 0.749 | -0.873 | 0.062 | -0.057 | -0.057 | 0 | 0.85 | 0.57 | 0.87 | 0.95 | 0.83 | 0.74 |
| 1 | 5047038 | rs6679312 | -0.745 | 0.284 | 0.87 | -0.846 | 0.023 | -0.012 | -0.012 | 0.016 | 0.92 | 0.89 | 0.9 | 0.95 | 0.86 | 0.85 |
| 1 | 5047163 | rs7365286 | 0.751 | 0.205 | 0.884 | -0.977 | -0.019 | 0.042 | 0.042 | 4.03 | 0.3 | 0.24 | 0.44 | 0.12 | 0.36 | 0.2 |
| 1 | 5048624 | rs7541612 | -0.805 | 0.27 | 0.613 | -0.922 | 0.019 | -0.01 | -0.01 | 0 | 0.92 | 0.9 | 0.9 | 0.95 | 0.87 | 0.85 |
| 1 | 5051232 | rs12725971 | 1.235 | -0.076 | 0.272 | -0.562 | -0.001 | -0.111 | -0.111 | 0 | 0.32 | 0.08 | 0.23 | 0.38 | 0.26 | 0.39 |
| 1 | 5051710 | rs6683624 | 0.61 | -0.077 | 0.274 | -0.486 | -0.006 | -0.121 | -0.121 | 0 | 0.24 | 0.03 | 0.21 | 0.45 | 0.21 | 0.17 |
| 1 | 5060684 | rs10158116 | -0.145 | 0.436 | 0.547 | -0.539 | 0.01 | -0.053 | -0.053 | 0.03 | 0.6 | 0.5 | 0.47 | 0.84 | 0.49 | 0.66 |
| 1 | 5061519 | rs10158288 | -0.168 | 0.375 | 0.544 | -0.53 | 0.009 | -0.058 | -0.058 | 0 | 0.6 | 0.53 | 0.48 | 0.84 | 0.49 | 0.66 |
| 1 | 5066205 | rs9662885 | 0.596 | 1.03 | 0.516 | -0.032 | 0.037 | -0.018 | -0.018 | 0 | 0.09 | 0.14 | 0.09 | 0.05 | 0.13 | 0.14 |
| 1 | 5068210 | rs10915310 | 0.401 | 0.546 | -0.158 | 0.119 | 0.01 | -0.016 | -0.016 | 0 | 0.36 | 0.29 | 0.26 | 0.43 | 0.28 | 0.5 |
| 1 | 5072295 | rs1870516 | 0.642 | 0.89 | -0.904 | -0.07 | 0.027 | -0.069 | -0.069 | 0.008 | 0.4 | 0.01 | 0.28 | 0.42 | 0.31 | 0.24 |
| 1 | 5073092 | rs9286975 | 0.127 | 1.027 | -0.475 | 0.205 | 0.112 | 0.053 | 0.053 | 0 | 0.75 | 0.58 | 0.47 | 0.69 | 0.56 | 0.67 |
| 1 | 5073658 | rs10799152 | 0.965 | 1.068 | -1.184 | -0.156 | 0.013 | 0.005 | 0.005 | 0.832 | 0.29 | 0.18 | 0.16 | 0.26 | 0.25 | 0.33 |
| 1 | 5078939 | rs10915316 | 0.248 | 1.879 | -0.91 | 0.325 | -0.001 | -0.001 | -0.001 | 0.038 | 0.52 | 0.54 | 0.55 | 0.37 | 0.59 | 0.57 |
| 1 | 5088062 | rs1459765 | 0.209 | 1.734 | -1.359 | 0.245 | 0.092 | 0.003 | 0.003 | 0 | 0.11 | 0.24 | 0.33 | 0.1 | 0.22 | 0.11 |
| 1 | 5088617 | rs2169089 | 0.069 | 0.182 | -1.99 | 0.356 | 0.026 | -0.114 | -0.114 | 0 | 0.68 | 0.98 | 0.76 | 0.57 | 0.78 | 0.78 |
| 1 | 5094435 | rs6693990 | -0.952 | 1.138 | -0.81 | -0.094 | 0.033 | -0.027 | -0.027 | 0.055 | 0.25 | 0.39 | 0.25 | 0.16 | 0.3 | 0.28 |

| | | | | | | | | | | | | | | | | |
|---|---|---|---|---|---|---|---|---|---|---|---|---|---|---|---|---|
| 1 | 5095862 | rs1459762 | -0.677 | 1.136 | -0.559 | -0.158 | 0.003 | -0.026 | -0.026 | 0.1 | 0.11 | 0.23 | 0.13 | 0.03 | 0.07 | 0.09 |
| 1 | 5103621 | rs10915324 | -0.385 | 1.562 | -0.6 | -0.169 | 0.005 | -0.001 | -0.001 | 0 | 0.14 | 0.17 | 0.12 | 0.13 | 0.24 | 0.22 |
| 1 | 5104614 | rs10799155 | 0.604 | 1.375 | -1.12 | -0.841 | 0.02 | 0.096 | 0.096 | 0.844 | 0.53 | 0.65 | 0.43 | 0.17 | 0.42 | 0.39 |
| 1 | 5113911 | rs10799157 | 1.094 | 0.951 | -1.404 | -1.472 | 0 | -0.022 | -0.022 | 0 | 0.23 | 0.22 | 0.12 | 0.02 | 0.18 | 0.18 |
| 1 | 5114102 | rs10915329 | 0.745 | 0.936 | -1.146 | -1.145 | 0.008 | 0.023 | 0.023 | 0 | 0.45 | 0.39 | 0.4 | 0.17 | 0.4 | 0.37 |
| 1 | 5124859 | rs1563792 | -0.628 | 0.901 | -1.517 | -0.424 | -0.02 | -0.04 | -0.04 | 0 | 0.74 | 0.91 | 0.65 | 0.33 | 0.72 | 0.69 |
| 1 | 5127543 | rs6603832 | -0.136 | 0.857 | -1.431 | -0.457 | -0.015 | 0.02 | 0.02 | 0.045 | 0.67 | 0.83 | 0.54 | 0.3 | 0.66 | 0.6 |
| 1 | 5136644 | rs922123 | -0.448 | 0.931 | -1.095 | 0.013 | 0 | 0.001 | 0.001 | 0.049 | 0.42 | 0.46 | 0.41 | 0.45 | 0.45 | 0.58 |
| 1 | 5140226 | rs11590026 | -0.364 | 0.75 | -1.01 | -1.027 | 0.002 | -0.043 | -0.043 | 0.27 | 0.2 | 0.21 | 0.2 | 0.01 | 0.26 | 0.14 |
| 1 | 5141293 | rs3128687 | 0.791 | 0.547 | -0.851 | -0.436 | 0.007 | 0.012 | 0.012 | 2.733 | 0.64 | 0.54 | 0.61 | 0.48 | 0.58 | 0.54 |
| 1 | 5142552 | rs10915341 | -0.342 | -0.282 | -1.26 | -1.02 | 0.04 | 0.04 | 0.04 | 0.26 | 0.41 | 0.27 | 0.3 | 0.69 | 0.31 | 0.47 |
| 1 | 5142784 | rs12047111 | -0.465 | -0.156 | -1.048 | -0.895 | 0.041 | 0.044 | 0.044 | 0 | 0.46 | 0.34 | 0.34 | 0.66 | 0.33 | 0.49 |
| 1 | 5142818 | rs2334263 | -0.412 | -0.1 | -1.027 | -0.965 | 0.032 | 0.039 | 0.039 | 0 | 0.45 | 0.34 | 0.34 | 0.67 | 0.35 | 0.49 |
| 1 | 5145066 | rs6680801 | -0.084 | 0.065 | -0.276 | -1.379 | 0.034 | -0.083 | -0.083 | 1.575 | 0.13 | 0.37 | 0.22 | 0 | 0.14 | 0.09 |
| 1 | 5145463 | rs12120144 | 1.45 | -0.205 | -1.093 | -1.722 | -0.037 | 0.018 | 0.018 | 0 | 0.25 | 0.09 | 0.21 | 0.23 | 0.29 | 0.19 |
| 1 | 5150534 | rs3101214 | 0.066 | -0.707 | -0.928 | -2.103 | -0.02 | -0.113 | -0.113 | 2.498 | 0.26 | 0.55 | 0.2 | 0.11 | 0.2 | 0.16 |
| 1 | 5152159 | rs12121092 | 0.863 | -0.727 | -1.253 | -2.255 | -0.015 | -0.048 | -0.048 | 0 | 0.18 | 0.06 | 0.3 | 0.6 | 0.18 | 0.4 |
| 1 | 5154882 | rs13374190 | 0.974 | -0.491 | -0.704 | -2.541 | -0.01 | -0.026 | -0.026 | 0 | 0.06 | 0.09 | 0.1 | 0 | 0.05 | 0.06 |
| 1 | 5156245 | rs3101225 | 1.023 | -0.444 | -1.013 | -2.513 | 0.023 | 0.037 | 0.037 | 3.678 | 0.17 | 0.08 | 0.08 | 0.11 | 0.12 | 0.13 |
| 1 | 5158857 | rs9803863 | -0.894 | -0.099 | -1.184 | -2.307 | -0.013 | 0.04 | 0.04 | 0 | 0.77 | 0.84 | 0.82 | 0.98 | 0.73 | 0.84 |
| 1 | 5160662 | rs2242157 | 0.218 | 0.136 | -1.138 | -1.724 | -0.035 | 0.117 | 0.117 | 0.232 | 0.43 | 0.21 | 0.51 | 0.82 | 0.47 | 0.6 |
| 1 | 5168663 | rs10733013 | 0.469 | -0.037 | -0.857 | -1.499 | 0.013 | 0.1 | 0.1 | 0.336 | 0.44 | 0.27 | 0.51 | 0.84 | 0.42 | 0.63 |
| 1 | 5170712 | rs7513590 | -0.168 | -0.032 | -0.165 | -1.431 | 0.005 | -0.008 | -0.008 | 0.778 | 0.1 | 0.27 | 0.17 | 0.07 | 0.09 | 0.09 |
| 1 | 5174078 | rs2611424 | -0.689 | -0.027 | -0.069 | -1.673 | 0.008 | -0.049 | -0.049 | 0 | 0.79 | 0.76 | 0.82 | 0.98 | 0.73 | 0.83 |
| 1 | 5179574 | rs10915352 | 0.7 | -0.235 | -0.16 | -1.678 | 0.01 | 0.074 | 0.074 | 0.864 | 0.49 | 0.37 | 0.54 | 0.82 | 0.47 | 0.62 |
| 1 | 5181615 | rs10737396 | -0.305 | -0.4 | -0.51 | -1.836 | -0.023 | 0.071 | 0.071 | 0.09 | 0.74 | 0.9 | 0.77 | 0.91 | 0.7 | 0.74 |
| 1 | 5183873 | rs4313443 | 1.113 | -0.632 | -0.19 | -1.786 | -0.003 | 0.103 | 0.103 | 0.188 | 0.45 | 0.24 | 0.51 | 0.79 | 0.46 | 0.62 |
| 1 | 5186247 | rs6674378 | 0.394 | -0.612 | 0.383 | -1.714 | 0 | -0.074 | -0.074 | 0 | 0.2 | 0.42 | 0.13 | 0.12 | 0.16 | 0.09 |
| 1 | 5189642 | rs17374719 | 0.045 | -0.597 | 0.094 | -1.699 | 0.004 | 0.012 | 0.012 | 3.8 | 0.2 | 0.11 | 0.15 | 0.18 | 0.14 | 0.13 |
| 1 | 5190166 | rs12083573 | 0.018 | -0.617 | 0.149 | -1.709 | 0.001 | 0.004 | 0.004 | 3.873 | 0.2 | 0.16 | 0.15 | 0.19 | 0.14 | 0.13 |
| 1 | 5192703 | rs6671889 | 1.006 | -0.457 | -0.096 | -1.807 | -0.001 | 0.141 | 0.141 | 0.111 | 0.47 | 0.23 | 0.51 | 0.79 | 0.47 | 0.64 |
| 1 | 5193026 | rs10492923 | 0.962 | -0.494 | -0.359 | -0.578 | 0.004 | 0.011 | 0.011 | 0 | 0.08 | 0.06 | 0.23 | 0.34 | 0.07 | 0.19 |
| 1 | 5195371 | rs10492925 | 0.073 | -0.736 | -0.451 | -0.88 | 0.003 | -0.026 | -0.026 | 1.892 | 0.28 | 0.16 | 0.34 | 0.45 | 0.24 | 0.27 |
| 1 | 5196739 | rs9439506 | 1.33 | -1.005 | -0.526 | -0.999 | -0.023 | -0.066 | -0.066 | 0.059 | 0.35 | 0.18 | 0.27 | 0.43 | 0.37 | 0.38 |
| 1 | 5197977 | rs10799173 | -0.861 | -1.074 | -0.319 | -1.142 | 0 | 0.039 | 0.039 | 0 | 0.73 | 0.78 | 0.8 | 0.97 | 0.71 | 0.74 |
| 1 | 5198789 | rs16840051 | 1.274 | -0.885 | -0.214 | -1.04 | -0.031 | -0.059 | -0.059 | 0.104 | 0.36 | 0.16 | 0.28 | 0.44 | 0.39 | 0.37 |
| 1 | 5200875 | rs12401618 | 0.837 | -0.723 | -0.677 | -1 | -0.005 | -0.001 | -0.001 | 0 | 0.05 | 0.06 | 0.11 | 0 | 0.08 | 0.03 |
| 1 | 5210124 | rs12121369 | 0.872 | -0.827 | -0.616 | -0.798 | 0 | -0.011 | -0.011 | 0 | 0.39 | 0.34 | 0.3 | 0.43 | 0.38 | 0.37 |
| 1 | 5211134 | rs10753353 | -0.974 | -0.724 | -1.652 | -0.765 | -0.059 | 0.033 | 0.033 | 0.061 | 0.78 | 0.94 | 0.77 | 0.88 | 0.72 | 0.68 |
| 1 | 5211338 | rs12027352 | -0.252 | -0.7 | -1.431 | -0.845 | -0.001 | -0.01 | -0.01 | 0 | 0.35 | 0.27 | 0.46 | 0.46 | 0.34 | 0.31 |
| 1 | 5215285 | rs12411059 | 0.557 | -0.791 | -1.372 | -0.13 | 0.02 | -0.037 | -0.037 | 0 | 0.09 | 0 | 0.05 | 0.08 | 0.06 | 0.16 |
| 1 | 5218601 | rs9439511 | 0.231 | -0.342 | -1.128 | -0.202 | 0.015 | -0.006 | -0.006 | 0 | 0.45 | 0.64 | 0.36 | 0.49 | 0.36 | 0.43 |
| 1 | 5222596 | rs10915365 | 0.181 | -0.203 | -1.408 | -0.112 | 0.031 | 0.009 | 0.009 | 0 | 0.51 | 0.72 | 0.4 | 0.5 | 0.43 | 0.49 |
| 1 | 5223182 | rs12026730 | -0.164 | 0.018 | -0.65 | 0.657 | -0.014 | 0.107 | 0.107 | 0 | 0.11 | 0.16 | 0.23 | 0.33 | 0.07 | 0.19 |
| 1 | 5225152 | rs4512698 | -0.424 | 0.227 | -0.952 | 0.295 | -0.009 | 0.004 | 0.004 | 0.049 | 0.06 | 0.03 | 0.12 | 0 | 0.11 | 0.03 |
| 1 | 5229311 | rs16840071 | -0.599 | -0.183 | -0.94 | 0.747 | 0.002 | 0.039 | 0.039 | 0 | 0.09 | 0.07 | 0.22 | 0.34 | 0.05 | 0.16 |
| 1 | 5231892 | rs4394703 | -0.441 | 0.073 | -1.457 | 0.158 | -0.038 | 0.067 | 0.067 | 0.188 | 0.83 | 0.93 | 0.78 | 0.97 | 0.76 | 0.77 |
| 1 | 5233284 | rs9286977 | 0.989 | 0.199 | -2.047 | 0.554 | -0.025 | -0.059 | -0.059 | 0 | 0.18 | 0.05 | 0.12 | 0.32 | 0.16 | 0.18 |
| 1 | 5236110 | rs6698702 | 0.25 | 0.198 | -1.925 | 0.408 | 0 | -0.011 | -0.011 | 1.241 | 0.07 | 0.06 | 0.06 | 0 | 0.07 | 0.05 |
| 1 | 5239931 | rs6603818 | 0.412 | 0.182 | -1.682 | 0.58 | -0.029 | 0.039 | 0.039 | 0.237 | 0.58 | 0.77 | 0.6 | 0.65 | 0.51 | 0.54 |
| 1 | 5240211 | rs7547229 | 1.216 | -0.049 | -0.446 | 0.441 | 0.004 | 0.038 | 0.038 | 0.658 | 0.45 | 0.33 | 0.33 | 0.26 | 0.45 | 0.31 |
| 1 | 5243848 | rs4641339 | 0.315 | -0.062 | 0.104 | 0.728 | -0.002 | 0.151 | 0.151 | 0 | 0.09 | 0.19 | 0.22 | 0.29 | 0.07 | 0.16 |
| 1 | 5246626 | rs17366838 | 1.354 | -0.599 | -0.016 | 0.141 | 0.01 | 0.013 | 0.013 | 2.121 | 0.18 | 0.15 | 0.15 | 0.11 | 0.16 | 0.17 |
| 1 | 5246841 | rs6669501 | 1.951 | -0.657 | 0.15 | 0.079 | -0.007 | 0.012 | 0.012 | 0 | 0.26 | 0.17 | 0.18 | 0.19 | 0.29 | 0.14 |
| 1 | 5247082 | rs10492930 | 1.457 | -0.672 | 0.493 | 0.072 | 0.001 | -0.007 | -0.007 | 0 | 0.3 | 0.31 | 0.23 | 0.26 | 0.29 | 0.21 |
| 1 | 5247355 | rs10492931 | 0.32 | -0.662 | 0.429 | 0.061 | 0.011 | 0.028 | 0.028 | 0.658 | 0.49 | 0.61 | 0.38 | 0.36 | 0.45 | 0.39 |
| 1 | 5248448 | rs12043691 | 0.702 | -0.604 | 0.238 | 0.154 | 0.002 | 0.153 | 0.153 | 0 | 0.09 | 0.19 | 0.22 | 0.28 | 0.07 | 0.16 |
| 1 | 5251569 | rs17375504 | 1.259 | -0.56 | -0.204 | 0.079 | 0.01 | 0.013 | 0.013 | 1.988 | 0.18 | 0.15 | 0.14 | 0.07 | 0.16 | 0.17 |
| 1 | 5253532 | rs12083008 | 0.945 | -0.597 | -0.276 | 0.059 | 0.008 | 0.009 | 0.009 | 2.668 | 0.2 | 0.21 | 0.14 | 0.07 | 0.16 | 0.17 |
| 1 | 5253664 | rs7536712 | -0.819 | -0.501 | -0.826 | 0.076 | -0.111 | -0.013 | -0.013 | 0 | 0.64 | 0.89 | 0.63 | 0.65 | 0.55 | 0.58 |
| 1 | 5256356 | rs1320178 | 0.152 | -0.4 | -0.71 | 0.475 | 0.001 | 0.015 | 0.015 | 0 | 0.11 | 0.09 | 0.23 | 0.29 | 0.08 | 0.23 |
| 1 | 5257229 | rs2722781 | 1.504 | -0.247 | -0.664 | -0.052 | -0.009 | 0.02 | 0.02 | 0 | 0.28 | 0.15 | 0.19 | 0.23 | 0.31 | 0.19 |
| 1 | 5259095 | rs7528871 | 0.903 | -0.089 | -0.901 | 0.213 | -0.073 | 0.086 | 0.086 | 0 | 0.18 | 0.02 | 0.23 | 0.03 | 0.26 | 0.26 |
| 1 | 5262767 | rs16840171 | 0.442 | -0.18 | -0.606 | 0.403 | -0.002 | -0.001 | -0.001 | 0 | 0.1 | 0.11 | 0.19 | 0.09 | 0.07 | 0.17 |
| 1 | 5262901 | rs2679218 | 1.173 | -0.111 | -0.664 | 0.237 | -0.037 | 0.107 | 0.107 | 3.643 | 0.37 | 0.1 | 0.25 | 0.23 | 0.39 | 0.18 |
| 1 | 5267850 | rs6603821 | -0.973 | -0.732 | -0.206 | 0.82 | 0.032 | 0.02 | 0.02 | 0.07 | 0.48 | 0.84 | 0.63 | 0.5 | 0.45 | 0.66 |
| 1 | 5270277 | rs12140099 | 0.478 | -0.826 | 0.352 | 1.249 | -0.002 | 0.003 | 0.003 | 0 | 0.27 | 0.27 | 0.28 | 0.2 | 0.32 | 0.41 |
| 1 | 5277437 | rs10915385 | 0.487 | -0.836 | 0.344 | 1.254 | -0.003 | 0.006 | 0.006 | 0 | 0.26 | 0.26 | 0.28 | 0.2 | 0.32 | 0.41 |
| 1 | 5277439 | rs2883104 | 0.78 | -0.83 | 0.039 | 1.511 | -0.003 | 0.012 | 0.012 | 0 | 0.1 | 0.11 | 0.05 | 0.25 | 0.08 | 0.12 |

| | | | | | | | | | | | | | | | | |
|---|---|---|---|---|---|---|---|---|---|---|---|---|---|---|---|---|
| 1 | 5279346 | rs12047612 | 0.751 | -0.833 | 0.039 | 1.523 | -0.004 | 0.011 | 0.011 | 0 | 0.11 | 0.11 | 0.05 | 0.25 | 0.08 | 0.13 |
| 1 | 5280502 | rs16840183 | 0.41 | -0.899 | 0.056 | 1.72 | 0 | 0.032 | 0.032 | 0 | 0.07 | 0.05 | 0.26 | 0.23 | 0.05 | 0.21 |
| 1 | 5281117 | rs7549461 | 0.6 | -0.92 | 0.126 | 1.669 | -0.001 | 0 | 0 | 0 | 0.27 | 0.32 | 0.29 | 0.2 | 0.32 | 0.41 |
| 1 | 5281421 | rs7511995 | -1.964 | -1.061 | 0.031 | 1.558 | -0.116 | 0.122 | 0.122 | 1.279 | 0.59 | 0.94 | 0.68 | 0.77 | 0.54 | 0.8 |
| 1 | 5282726 | rs16840190 | 1.293 | -1.029 | 0.097 | 1.557 | 0.002 | -0.003 | -0.003 | 2.09 | 0.09 | 0.05 | 0.04 | 0.25 | 0.07 | 0.1 |
| 1 | 5282817 | rs7546667 | 0.662 | -1.033 | 0.108 | 1.578 | -0.001 | 0 | 0 | 0.02 | 0.27 | 0.32 | 0.29 | 0.2 | 0.32 | 0.41 |
| 1 | 5283942 | rs938962 | -0.538 | -1.05 | 0.42 | 1.572 | -0.002 | 0 | 0 | 0 | 0.41 | 0.6 | 0.36 | 0.46 | 0.4 | 0.53 |
| 1 | 5284023 | rs12024316 | 1.16 | -1.028 | 0.199 | 1.656 | 0.002 | -0.003 | -0.003 | 0 | 0.09 | 0.05 | 0.04 | 0.25 | 0.07 | 0.11 |
| 1 | 5287108 | rs4259664 | -1.836 | -1.087 | 0.185 | 1.471 | 0.003 | -0.005 | -0.005 | 0 | 0.32 | 0.63 | 0.39 | 0.56 | 0.22 | 0.39 |
| 1 | 5295176 | rs677735 | -0.333 | -0.617 | -0.085 | 1.653 | 0.002 | 0.051 | 0.051 | 0 | 0.9 | 0.84 | 0.7 | 0.8 | 0.89 | 0.79 |
| 1 | 5296844 | rs2353169 | 1.267 | -0.577 | -0.279 | 1.571 | -0.116 | 0.12 | 0.12 | 1.182 | 0.42 | 0.05 | 0.32 | 0.24 | 0.47 | 0.21 |
| 1 | 5297887 | rs586165 | 0.663 | -0.375 | 0.022 | 1.475 | -0.005 | 0.062 | 0.062 | 0 | 0.8 | 0.78 | 0.66 | 0.52 | 0.82 | 0.69 |
| 1 | 5298490 | rs12026938 | 1.054 | -0.36 | -0.093 | 1.242 | 0 | 0.012 | 0.012 | 0 | 0.08 | 0.05 | 0.04 | 0.27 | 0.07 | 0.1 |
| 1 | 5302315 | rs12125575 | 1.247 | -0.456 | -0.628 | 1.073 | -0.149 | 0.17 | 0.17 | 0.462 | 0.37 | 0.05 | 0.3 | 0.13 | 0.45 | 0.19 |
| 1 | 5302405 | rs599481 | 0.873 | -0.561 | -0.173 | 1.114 | -0.031 | 0.111 | 0.111 | 0.079 | 0.74 | 0.56 | 0.63 | 0.52 | 0.79 | 0.66 |
| 1 | 5302528 | rs12066405 | 0.751 | -0.68 | -0.385 | 1.114 | -0.031 | 0.112 | 0.112 | 0.452 | 0.75 | 0.52 | 0.64 | 0.52 | 0.81 | 0.67 |
| 1 | 5304873 | rs675738 | 0.855 | -0.636 | -0.447 | 1.139 | -0.041 | 0.075 | 0.075 | 0.873 | 0.72 | 0.52 | 0.63 | 0.52 | 0.8 | 0.65 |
| 1 | 5306840 | rs12125074 | 0.223 | -0.619 | -0.611 | 1.048 | -0.016 | 0.112 | 0.112 | 0.01 | 0.84 | 0.61 | 0.68 | 0.67 | 0.89 | 0.75 |
| 1 | 5307076 | rs521430 | -0.367 | -0.456 | -0.717 | 0.612 | 0.012 | -0.002 | -0.002 | 0 | 0.47 | 0.59 | 0.34 | 0.43 | 0.4 | 0.54 |
| 1 | 5309255 | rs7534404 | 0.556 | -0.414 | -0.976 | 0.528 | -0.005 | -0.006 | -0.006 | 0.133 | 0.32 | 0.38 | 0.27 | 0.2 | 0.29 | 0.41 |
| 1 | 5309667 | rs12135834 | 0.7 | 0.17 | -1.677 | 0.162 | -0.134 | -0.059 | -0.059 | 0 | 0.37 | 0.05 | 0.34 | 0.39 | 0.5 | 0.24 |
| 1 | 5310258 | rs4465222 | 0.891 | 0.36 | -0.478 | 0.049 | -0.017 | 0 | 0 | 0 | 0.85 | 0.73 | 0.68 | 0.83 | 0.9 | 0.78 |
| 1 | 5312052 | rs580309 | -0.584 | 0.926 | -1.025 | -0.397 | 0.001 | 0.001 | 0.001 | 0 | 0.45 | 0.61 | 0.44 | 0.22 | 0.41 | 0.32 |
| 1 | 5312308 | rs670971 | 0.482 | 1.119 | -1.407 | -0.462 | -0.017 | 0.037 | 0.037 | 0 | 0.28 | 0.02 | 0.25 | 0.14 | 0.24 | 0.12 |
| 1 | 5312366 | rs494046 | -0.865 | 1.067 | -1.133 | -0.358 | 0.003 | -0.055 | -0.055 | 0.604 | 0.63 | 0.84 | 0.52 | 0.39 | 0.63 | 0.48 |
| 1 | 5313445 | rs6703948 | -1.385 | 1.123 | 0.036 | -0.253 | 0.024 | -0.035 | -0.035 | 0.524 | 0.18 | 0.53 | 0.19 | 0.11 | 0.17 | 0.2 |
| 1 | 5315481 | rs16840289 | -1.026 | 0.418 | 0.028 | 0.089 | -0.024 | 0.043 | 0.043 | 4.535 | 0.11 | 0.59 | 0.12 | 0.11 | 0.04 | 0.08 |
| 1 | 5318552 | rs12089315 | -0.738 | 1.204 | -0.411 | -0.562 | 0.01 | -0.051 | -0.051 | 0 | 0.28 | 0.3 | 0.21 | 0.05 | 0.35 | 0.21 |
| 1 | 5320534 | rs632798 | -0.727 | 0.965 | -1.004 | -0.204 | -0.025 | -0.051 | -0.051 | 4.951 | 0.76 | 0.85 | 0.75 | 0.61 | 0.77 | 0.64 |
| 1 | 5321139 | rs630153 | 0.716 | 0.023 | -1.029 | -0.158 | -0.014 | -0.01 | -0.01 | 0 | 0.36 | 0.39 | 0.28 | 0.3 | 0.34 | 0.2 |
| 1 | 5324231 | rs499416 | 0.074 | -0.102 | -0.811 | -0.105 | -0.003 | -0.003 | -0.003 | 0 | 0.61 | 0.51 | 0.53 | 0.34 | 0.7 | 0.41 |
| 1 | 5325724 | rs12139398 | 0.351 | 0.121 | -1.209 | -0.627 | -0.016 | 0.105 | 0.105 | 0.103 | 0.29 | 0.09 | 0.25 | 0 | 0.32 | 0.14 |
| 1 | 5326050 | rs536995 | -0.91 | 0.203 | -0.653 | -0.635 | 0.018 | 0.029 | 0.029 | 0 | 0.71 | 0.61 | 0.75 | 0.59 | 0.71 | 0.62 |
| 1 | 5329865 | rs579467 | 0.017 | 0.093 | -0.642 | -0.624 | 0.01 | 0.022 | 0.022 | 0 | 0.56 | 0.3 | 0.48 | 0.34 | 0.64 | 0.4 |
| 1 | 5330415 | rs495765 | 1.133 | -0.191 | -0.819 | -0.653 | 0.005 | -0.014 | -0.014 | 0.566 | 0.23 | 0.15 | 0.25 | 0.3 | 0.23 | 0.18 |
| 1 | 5333340 | rs11808981 | 1.445 | -0.213 | -0.825 | -0.653 | 0.015 | -0.018 | -0.018 | 0 | 0.12 | 0.09 | 0.26 | 0.25 | 0.07 | 0.22 |
| 1 | 5334079 | rs584938 | 1.128 | -0.169 | -0.789 | -0.64 | 0.004 | -0.01 | -0.01 | 0.039 | 0.23 | 0.16 | 0.25 | 0.3 | 0.23 | 0.18 |
| 1 | 5335599 | rs4847576 | 0.656 | -0.041 | -0.558 | -0.583 | -0.046 | 0.065 | 0.065 | 0.019 | 0.15 | 0.31 | 0.27 | 0.25 | 0.07 | 0.22 |
| 1 | 5341464 | rs17423547 | 0.256 | 0.365 | -0.719 | -0.949 | -0.002 | -0.098 | -0.098 | 0 | 0.31 | 0.32 | 0.22 | 0 | 0.32 | 0.14 |
| 1 | 5342295 | rs12141954 | -0.197 | 0.366 | -0.681 | -0.946 | 0.014 | -0.119 | -0.119 | 0 | 0.35 | 0.39 | 0.26 | 0.04 | 0.41 | 0.21 |
| 1 | 5343032 | rs604664 | -0.47 | 0.492 | -1.462 | -0.68 | -0.076 | -0.06 | -0.06 | 0 | 0.71 | 0.85 | 0.74 | 0.55 | 0.63 | 0.57 |
| 1 | 5344838 | rs623058 | -0.952 | 0.259 | -1.69 | -0.427 | -0.101 | -0.108 | -0.108 | 0 | 0.75 | 0.93 | 0.78 | 0.62 | 0.72 | 0.64 |
| 1 | 5346474 | rs557477 | -0.169 | -0.095 | -1.625 | -0.549 | 0.055 | 0.035 | 0.035 | 2.235 | 0.46 | 0.83 | 0.59 | 0.62 | 0.39 | 0.5 |
| 1 | 5347457 | rs665471 | -0.665 | -0.26 | -2.05 | -1.137 | -0.003 | 0.242 | 0.242 | 0 | 0.5 | 0.83 | 0.48 | 0.22 | 0.54 | 0.43 |
| 1 | 5347627 | rs478357 | -0.613 | 0.055 | -1.199 | -1.087 | -0.001 | 0.001 | 0.001 | 0 | 0.48 | 0.52 | 0.61 | 0.47 | 0.51 | 0.57 |
| 1 | 5347979 | rs678476 | -0.158 | 0.135 | -0.899 | -1.053 | -0.008 | -0.008 | -0.008 | 0 | 0.35 | 0.28 | 0.53 | 0.43 | 0.43 | 0.44 |
| 1 | 5349616 | rs10915406 | 0.303 | 0.551 | -0.626 | -1.5 | -0.006 | -0.019 | -0.019 | 0 | 0.21 | 0.16 | 0.17 | 0 | 0.23 | 0.19 |
| 1 | 5350428 | rs2353174 | -0.117 | 0.451 | -0.505 | -0.81 | 0 | 0 | 0 | 0 | 0.41 | 0.42 | 0.56 | 0.34 | 0.45 | 0.41 |
| 1 | 5354559 | rs12131122 | -0.071 | 0.504 | -0.579 | -0.785 | -0.002 | 0.002 | 0.002 | 0.043 | 0.4 | 0.35 | 0.55 | 0.34 | 0.45 | 0.41 |
| 1 | 5354844 | rs912988 | 0.019 | 0.515 | -0.904 | -0.737 | -0.009 | 0.07 | 0.07 | 0 | 0.34 | 0.18 | 0.33 | 0.13 | 0.37 | 0.22 |
| 1 | 5356607 | rs10799191 | -0.289 | 0.715 | -0.404 | -0.621 | 0.018 | -0.009 | -0.009 | 0.115 | 0.28 | 0.53 | 0.4 | 0.23 | 0.3 | 0.36 |
| 1 | 5357171 | rs9439550 | -0.618 | 0.259 | -0.561 | -0.672 | 0.015 | 0.01 | 0.01 | 1.695 | 0.38 | 0.71 | 0.49 | 0.3 | 0.34 | 0.38 |
| 1 | 5358384 | rs912991 | 0.869 | 0.64 | 0.404 | -0.844 | 0.017 | 0.024 | 0.024 | 0.089 | 0.46 | 0.35 | 0.41 | 0.65 | 0.43 | 0.52 |
| 1 | 5359498 | rs11579529 | -1.393 | 1.097 | 0.906 | -0.857 | 0.165 | -0.032 | -0.032 | 0 | 0.08 | 0.4 | 0.11 | 0 | 0.15 | 0.05 |
| 1 | 5369798 | rs9439527 | 0.028 | 0.838 | 0.675 | -0.99 | -0.057 | 0.013 | 0.013 | 0.021 | 0.62 | 0.87 | 0.55 | 0.69 | 0.59 | 0.6 |
| 1 | 5371159 | rs12040942 | 0.841 | 0.865 | 0.88 | -1.067 | 0 | 0.004 | 0.004 | 0.935 | 0.48 | 0.48 | 0.48 | 0.63 | 0.54 | 0.57 |
| 1 | 5374940 | rs5026665 | 1.211 | 0.214 | 0.715 | -1.028 | -0.004 | -0.002 | -0.002 | 0 | 0.44 | 0.46 | 0.39 | 0.63 | 0.38 | 0.48 |
| 1 | 5378416 | rs2148210 | 0.487 | 0.211 | 0.409 | -1.053 | -0.009 | -0.026 | -0.026 | 0 | 0.55 | 0.53 | 0.52 | 0.8 | 0.41 | 0.53 |
| 1 | 5379624 | rs4847569 | 0.533 | 0.182 | 0.405 | -1.129 | -0.008 | -0.009 | -0.009 | 0 | 0.54 | 0.5 | 0.43 | 0.7 | 0.41 | 0.52 |
| 1 | 5384158 | rs1570964 | -0.378 | 0.443 | 0.326 | -1.239 | -0.103 | 0.026 | 0.026 | 0.246 | 0.66 | 0.92 | 0.62 | 0.75 | 0.54 | 0.65 |
| 1 | 5391497 | rs9439531 | 0.855 | 0.293 | 0.424 | -1.205 | 0.07 | 0.069 | 0.069 | 0 | 0.51 | 0.25 | 0.41 | 0.7 | 0.4 | 0.52 |
| 1 | 5395996 | rs912993 | -0.078 | 0.359 | -0.301 | -1.139 | -0.029 | 0.015 | 0.015 | 0 | 0.6 | 0.85 | 0.53 | 0.7 | 0.5 | 0.55 |
| 1 | 5401364 | rs2148211 | 0.411 | 0.39 | -0.396 | -1.113 | 0 | 0.001 | 0.001 | 0 | 0.56 | 0.6 | 0.5 | 0.7 | 0.49 | 0.54 |
| 1 | 5403031 | rs10915416 | 1.328 | 0.495 | -1.439 | -1.034 | 0.024 | 0.002 | 0.002 | 0.213 | 0.4 | 0.16 | 0.37 | 0.63 | 0.37 | 0.48 |
| 1 | 5403585 | rs1326005 | -0.379 | 0.837 | -1.572 | -1.53 | -0.122 | 0.143 | 0.143 | 0 | 0.71 | 0.91 | 0.65 | 0.95 | 0.6 | 0.82 |
| 1 | 5410542 | rs4129058 | -1.023 | 0.41 | -1.204 | -0.928 | 0.003 | 0.009 | 0.009 | 0 | 0.58 | 0.75 | 0.62 | 0.3 | 0.62 | 0.48 |
| 1 | 5415884 | rs12738923 | -0.9 | -0.142 | -1.676 | 0.309 | 0.041 | -0.073 | -0.073 | 0 | 0.21 | 0.02 | 0.13 | 0.23 | 0.12 | 0.29 |
| 1 | 5421533 | rs7364349 | -0.904 | -0.138 | -1.674 | 0.311 | 0.041 | -0.073 | -0.073 | 1.961 | 0.21 | 0.02 | 0.13 | 0.23 | 0.12 | 0.3 |
| 1 | 5463937 | rs11801490 | -0.582 | 0.603 | -0.627 | -0.333 | 0.013 | 0.078 | 0.078 | 1.578 | 0.36 | 0.79 | 0.23 | 0.09 | 0.31 | 0.19 |

| | | | | | | | | | | | | | | | | |
|---|---|---|---|---|---|---|---|---|---|---|---|---|---|---|---|---|
| 1 | 5464296 | rs11578764 | -0.891 | 0.26 | 0.174 | -0.319 | -0.015 | -0.014 | -0.014 | 0 | 0.1 | 0.28 | 0.03 | 0.09 | 0.05 | 0.09 |
| 1 | 5464332 | rs7368252 | -1.274 | 0.283 | -0.276 | 0.19 | 0.022 | -0.02 | -0.02 | 0 | 0.65 | 0.8 | 0.66 | 0.36 | 0.72 | 0.58 |
| 1 | 5465673 | rs12029480 | -0.296 | 0.827 | -0.082 | 0.473 | -0.059 | -0.026 | -0.026 | 0.48 | 0.2 | 0.03 | 0.25 | 0.15 | 0.19 | 0.29 |
| 1 | 5471517 | rs7418357 | -0.971 | 0.766 | 0.617 | 0.42 | 0.018 | -0.03 | -0.03 | 0 | 0.27 | 0.51 | 0.3 | 0.16 | 0.25 | 0.31 |
| 1 | 5473808 | rs7366321 | -0.551 | 0.72 | 0.15 | 0.738 | -0.01 | -0.011 | -0.011 | 0 | 0.22 | 0.17 | 0.28 | 0.23 | 0.19 | 0.3 |
| 1 | 5475581 | rs11576458 | -0.662 | 0.803 | 0.268 | 1.037 | -0.005 | -0.006 | -0.006 | 0 | 0.1 | 0.08 | 0.28 | 0.21 | 0.16 | 0.13 |
| 1 | 5475663 | rs6603835 | -1.254 | 0.446 | 0.768 | 0.694 | -0.003 | 0.003 | 0.003 | 0 | 0.59 | 0.63 | 0.65 | 0.43 | 0.61 | 0.53 |
| 1 | 5478587 | rs11260646 | -0.346 | 0.266 | 1.23 | 0.619 | -0.014 | 0.072 | 0.072 | 0 | 0.46 | 0.34 | 0.35 | 0.21 | 0.45 | 0.41 |
| 1 | 5479861 | rs3924260 | -1.4 | 0.275 | 1.414 | 0.629 | -0.001 | -0.005 | -0.005 | 0.004 | 0.6 | 0.69 | 0.65 | 0.43 | 0.59 | 0.54 |
| 1 | 5482386 | rs11260657 | -0.044 | 0.27 | 1.013 | 0.723 | -0.001 | -0.01 | -0.01 | 0 | 0.09 | 0.06 | 0.27 | 0.21 | 0.13 | 0.13 |
| 1 | 5485177 | rs3922826 | 0.097 | 0.164 | 1.175 | 0.283 | 0 | -0.003 | -0.003 | 0 | 0.2 | 0.28 | 0.15 | 0.14 | 0.16 | 0.18 |
| 1 | 5489867 | rs12735700 | 0.865 | 0.259 | 0.997 | -0.003 | 0.001 | -0.051 | -0.051 | 0.288 | 0.21 | 0.25 | 0.17 | 0 | 0.25 | 0.06 |
| 1 | 5492043 | rs10907211 | -0.779 | 0.28 | 0.843 | 0.223 | -0.024 | -0.016 | -0.016 | 0 | 0.29 | 0.43 | 0.15 | 0.25 | 0.18 | 0.35 |
| 1 | 5504553 | rs10907214 | 1.455 | -0.149 | 0.117 | 1.059 | -0.005 | 0.038 | 0.038 | 0 | 0.31 | 0.09 | 0.29 | 0.22 | 0.32 | 0.33 |
| 1 | 5505010 | rs12562483 | 0.154 | -0.036 | 0.32 | 0.904 | 0 | 0.001 | 0.001 | 0.803 | 0.09 | 0.09 | 0.1 | 0.19 | 0.08 | 0.14 |
| 1 | 5506194 | rs12065301 | -0.146 | -0.107 | 0.017 | 0.852 | 0.003 | -0.003 | -0.003 | 0 | 0.08 | 0.04 | 0.01 | 0.15 | 0.03 | 0.17 |
| 1 | 5514194 | rs11260694 | 0.106 | 0.041 | 0.004 | 0.5 | -0.028 | 0.094 | 0.094 | 0 | 0.38 | 0.22 | 0.28 | 0.11 | 0.4 | 0.21 |
| 1 | 5521691 | rs12568815 | -0.843 | -0.629 | 0.091 | 1.014 | -0.014 | 0.015 | 0.015 | 0 | 0.19 | 0.21 | 0.1 | 0.34 | 0.08 | 0.31 |
| 1 | 5528517 | rs7536506 | 0.236 | -0.32 | -0.106 | 0.42 | -0.025 | 0.035 | 0.035 | 0 | 0.36 | 0.25 | 0.31 | 0.11 | 0.45 | 0.22 |
| 1 | 5533601 | rs7414322 | 1.348 | -1.19 | 0.232 | 1.146 | -0.05 | 0.061 | 0.061 | 0 | 0.81 | 0.63 | 0.85 | 0.64 | 0.94 | 0.7 |
| 1 | 5534326 | rs7414864 | 1.068 | -1.042 | -0.199 | 1.216 | -0.034 | 0.048 | 0.048 | 0 | 0.83 | 0.71 | 0.85 | 0.64 | 0.94 | 0.7 |
| 1 | 5536722 | rs12023817 | -1.017 | -1.143 | -0.869 | 1.318 | 0.017 | -0.023 | -0.023 | 0 | 0.14 | 0.08 | 0.11 | 0.36 | 0.02 | 0.29 |
| 1 | 5541205 | rs2012852 | 0.164 | -1.665 | -1.763 | 0.768 | 0.133 | -0.038 | -0.038 | 0.104 | 0.52 | 0.15 | 0.57 | 0.52 | 0.64 | 0.57 |
| 1 | 5542439 | rs9729575 | -0.582 | -2.462 | -1.244 | 0.241 | 0.018 | -0.003 | -0.003 | 0 | 0.61 | 0.72 | 0.43 | 0.58 | 0.38 | 0.51 |
| 1 | 5545452 | rs9729637 | 0.126 | -3.278 | -0.654 | 0.265 | -0.038 | 0.093 | 0.093 | 0 | 0.17 | 0.23 | 0.1 | 0.45 | 0.07 | 0.13 |
| 1 | 5547562 | rs12045091 | 0.457 | -3.922 | -0.275 | -0.239 | 0.006 | 0.006 | 0.006 | 0.069 | 0.27 | 0.24 | 0.19 | 0.15 | 0.18 | 0.3 |
| 1 | 5549118 | rs12030717 | 0.473 | -3.954 | -0.163 | -0.251 | -0.01 | -0.011 | -0.011 | 0.878 | 0.27 | 0.23 | 0.19 | 0.14 | 0.18 | 0.3 |
| 1 | 5549525 | rs11260698 | -1.677 | -3.94 | 0.005 | -0.351 | 0.025 | -0.028 | -0.028 | 0 | 0.62 | 0.85 | 0.44 | 0.6 | 0.39 | 0.5 |
| 1 | 5551233 | rs12120052 | 0.346 | -3.29 | 0.062 | -0.115 | 0.017 | 0.022 | 0.022 | 0.496 | 0.27 | 0.17 | 0.21 | 0.12 | 0.2 | 0.32 |
| 1 | 5553470 | rs10907219 | -0.583 | -2.98 | -0.068 | -0.111 | 0.026 | 0.061 | 0.061 | 0.574 | 0.73 | 0.89 | 0.8 | 0.87 | 0.79 | 0.68 |
| 1 | 5554049 | rs9662668 | -0.81 | -2.852 | 0.391 | -0.246 | -0.045 | 0.129 | 0.129 | 0 | 0.22 | 0.45 | 0.12 | 0.48 | 0.11 | 0.25 |
| 1 | 5555141 | rs2071953 | 1.146 | -2.802 | 0.363 | -0.26 | 0.006 | 0.004 | 0.004 | 0.076 | 0.5 | 0.44 | 0.67 | 0.41 | 0.67 | 0.42 |
| 1 | 5555419 | rs2071956 | 0.756 | -2.777 | 0.519 | -0.136 | -0.007 | -0.002 | -0.002 | 0 | 0.54 | 0.44 | 0.68 | 0.42 | 0.7 | 0.52 |
| 1 | 5555529 | rs2071957 | -0.661 | -2.751 | 0.452 | -0.138 | 0.021 | 0.07 | 0.07 | 0.602 | 0.74 | 0.94 | 0.8 | 0.88 | 0.79 | 0.67 |
| 1 | 5556067 | rs2071959 | -0.205 | -2.744 | 0.462 | -0.132 | 0.029 | 0.046 | 0.046 | 0.328 | 0.71 | 0.82 | 0.79 | 0.87 | 0.78 | 0.67 |
| 1 | 5558726 | rs17028104 | -0.642 | -2.836 | 0.652 | -0.229 | 0.004 | -0.062 | -0.062 | 0 | 0.1 | 0.27 | 0.11 | 0 | 0.1 | 0.03 |
| 1 | 5559249 | rs10796408 | 0.339 | -2.835 | 0.578 | -0.227 | 0.019 | 0.037 | 0.037 | 0.053 | 0.63 | 0.74 | 0.7 | 0.87 | 0.69 | 0.64 |
| 1 | 5561412 | rs9661310 | 0.542 | -2.845 | 0.706 | -0.203 | 0.008 | -0.008 | -0.008 | 0 | 0.14 | 0.13 | 0.07 | 0.44 | 0.09 | 0.15 |
| 1 | 5561478 | rs4845816 | 0.466 | -2.729 | 0.63 | -0.207 | 0.045 | 0.005 | 0.005 | 0.241 | 0.57 | 0.67 | 0.71 | 0.42 | 0.7 | 0.52 |
| 1 | 5563399 | rs9659669 | 1.308 | -2.748 | 0.359 | -0.199 | 0.017 | 0.009 | 0.009 | 0 | 0.46 | 0.36 | 0.61 | 0.42 | 0.61 | 0.49 |
| 1 | 5565117 | rs916374 | -1.155 | -1.812 | 0.929 | 0.466 | -0.007 | -0.058 | -0.058 | 0 | 0.11 | 0.27 | 0.1 | 0.01 | 0.1 | 0.03 |
| 1 | 5572304 | rs9729649 | -1.091 | -1.604 | 0.168 | 0.656 | -0.037 | 0.115 | 0.115 | 0 | 0.26 | 0.51 | 0.17 | 0.47 | 0.17 | 0.18 |
| 1 | 5575158 | rs878141 | 0.583 | -1.481 | -0.376 | 0.141 | 0.01 | 0.011 | 0.011 | 0 | 0.71 | 0.78 | 0.8 | 0.86 | 0.77 | 0.79 |
| 1 | 5580192 | rs10796400 | 0.539 | -1.629 | -0.239 | -0.126 | 0.008 | 0.011 | 0.011 | 0.05 | 0.71 | 0.76 | 0.83 | 0.9 | 0.76 | 0.82 |
| 1 | 5581798 | rs9660967 | 0.464 | -1.609 | -0.231 | -0.108 | 0.006 | 0.008 | 0.008 | 0.103 | 0.72 | 0.76 | 0.83 | 0.9 | 0.76 | 0.82 |
| 1 | 5594737 | rs2153900 | 0.029 | -1.564 | 0.234 | -0.155 | 0.004 | 0.007 | 0.007 | 0.638 | 0.77 | 0.83 | 0.86 | 0.9 | 0.79 | 0.83 |
| 1 | 5603850 | rs7515958 | 0.473 | -0.756 | 0.136 | 0.365 | -0.009 | -0.011 | -0.011 | 1.384 | 0.15 | 0.12 | 0.32 | 0.34 | 0.18 | 0.22 |
| 1 | 5607988 | rs12119802 | 0.89 | -0.893 | -0.583 | 0.166 | -0.005 | 0.038 | 0.038 | 1.81 | 0.19 | 0.01 | 0.1 | 0.08 | 0.12 | 0.16 |
| 1 | 5612956 | rs1418632 | 1.206 | -1.003 | -0.089 | 0.108 | -0.035 | -0.029 | -0.029 | 0.005 | 0.35 | 0.25 | 0.39 | 0.47 | 0.46 | 0.43 |
| 1 | 5613219 | rs1935766 | -0.171 | -1.036 | -0.1 | 0.095 | -0.03 | -0.052 | -0.052 | 1.504 | 0.72 | 0.66 | 0.82 | 0.9 | 0.77 | 0.78 |
| 1 | 5616830 | rs1935762 | -0.176 | -0.689 | 0.104 | 0.038 | -0.002 | -0.053 | -0.053 | 0 | 0.1 | 0.29 | 0.11 | 0 | 0.09 | 0.04 |
| 1 | 5623965 | rs9439613 | 0.792 | -0.522 | -0.808 | 0.149 | -0.026 | -0.021 | -0.021 | 0 | 0.34 | 0.26 | 0.4 | 0.48 | 0.46 | 0.42 |
| 1 | 5624529 | rs6687987 | 0.482 | -0.186 | -1.35 | -0.306 | -0.002 | 0.029 | 0.029 | 2.243 | 0.19 | 0.01 | 0.09 | 0.09 | 0.12 | 0.15 |
| 1 | 5626094 | rs9439614 | 0.355 | 0.664 | -0.57 | 0.183 | -0.074 | -0.001 | -0.001 | 0.179 | 0.28 | 0.12 | 0.32 | 0.23 | 0.39 | 0.24 |
| 1 | 5629592 | rs2891080 | -1.174 | 0.243 | 0.114 | 0.28 | 0.001 | -0.049 | -0.049 | 0 | 0.09 | 0.24 | 0.11 | 0 | 0.1 | 0.04 |
| 1 | 5635316 | rs7522344 | -0.674 | 0.357 | -0.343 | 0.631 | -0.005 | 0.01 | 0.01 | 0 | 0.38 | 0.48 | 0.36 | 0.45 | 0.41 | 0.36 |
| 1 | 5635724 | rs904525 | -0.555 | 0.354 | -0.216 | 0.615 | -0.012 | 0.026 | 0.026 | 0 | 0.36 | 0.41 | 0.35 | 0.44 | 0.39 | 0.34 |
| 1 | 5641064 | rs6603844 | -0.523 | 0.364 | -0.337 | 0.678 | 0.012 | -0.006 | -0.006 | 0 | 0.36 | 0.3 | 0.37 | 0.67 | 0.32 | 0.31 |
| 1 | 5641582 | rs1775421 | -0.359 | -0.365 | -0.607 | -0.077 | 0.021 | 0.027 | 0.027 | 0.679 | 0.17 | 0.1 | 0.08 | 0.01 | 0.1 | 0.08 |
| 1 | 5641919 | rs1775420 | -0.386 | -0.381 | -0.503 | -0.09 | 0.015 | 0.014 | 0.014 | 0.76 | 0.17 | 0.12 | 0.08 | 0.01 | 0.1 | 0.08 |
| 1 | 5642511 | rs1818038 | -0.065 | -0.122 | -0.434 | -0.198 | 0.007 | -0.002 | -0.002 | 0.553 | 0.37 | 0.33 | 0.45 | 0.67 | 0.37 | 0.34 |
| 1 | 5643758 | rs1695645 | -1.27 | 0.596 | -0.085 | -0.462 | 0.001 | -0.001 | -0.001 | 0 | 0.67 | 0.67 | 0.71 | 0.75 | 0.66 | 0.72 |
| 1 | 5646002 | rs10907200 | -0.508 | 0.139 | 0.246 | -0.234 | 0 | 0.001 | 0.001 | 0 | 0.44 | 0.56 | 0.47 | 0.34 | 0.44 | 0.51 |
| 1 | 5646264 | rs1775416 | -0.664 | -0.074 | 0.296 | -0.373 | 0 | 0 | 0 | 0 | 0.51 | 0.49 | 0.45 | 0.34 | 0.49 | 0.54 |
| 1 | 5646357 | rs12407564 | 0.924 | -0.025 | -0.289 | -0.482 | 0.006 | 0.035 | 0.035 | 0 | 0.24 | 0.04 | 0.17 | 0.13 | 0.25 | 0.18 |
| 1 | 5656069 | rs13376100 | 0.35 | 0.082 | 0.128 | -0.427 | 0 | -0.003 | -0.003 | 0 | 0.15 | 0.16 | 0.11 | 0.1 | 0.17 | 0.28 |
| 1 | 5658104 | rs1775414 | -0.845 | -0.055 | 0.557 | -0.404 | -0.011 | -0.048 | -0.048 | 0 | 0.14 | 0.25 | 0.08 | 0.02 | 0.09 | 0.1 |
| 1 | 5658267 | rs10907201 | -0.983 | -0.16 | -0.107 | -0.683 | 0 | 0.025 | 0.025 | 0 | 0.41 | 0.72 | 0.36 | 0.27 | 0.35 | 0.36 |
| 1 | 5658419 | rs12139270 | -0.281 | 0.078 | -0.687 | -0.524 | -0.004 | -0.031 | -0.031 | 0 | 0.12 | 0.18 | 0.07 | 0.02 | 0.08 | 0.1 |

| | | | | | | | | | | | | | | | | |
|---|---|---|---|---|---|---|---|---|---|---|---|---|---|---|---|---|
| 1 | 5659587 | rs950601 | 0.071 | 0.421 | -0.763 | -0.173 | -0.018 | -0.012 | -0.012 | 0 | 0.25 | 0.15 | 0.22 | 0.23 | 0.29 | 0.25 |
| 1 | 5669910 | rs1081382 | 0.045 | 0.608 | -0.455 | 0.276 | 0.013 | 0.014 | 0.014 | 0 | 0.4 | 0.5 | 0.58 | 0.49 | 0.48 | 0.31 |
| 1 | 5670530 | rs709186 | 0.164 | 0.732 | -0.128 | 0.218 | -0.015 | -0.025 | -0.025 | 0 | 0.37 | 0.26 | 0.49 | 0.48 | 0.42 | 0.3 |
| 1 | 5671786 | rs707582 | -0.075 | 0.724 | -0.119 | 0.22 | -0.006 | -0.01 | -0.01 | 0.028 | 0.4 | 0.34 | 0.51 | 0.49 | 0.45 | 0.3 |
| 1 | 5674434 | rs770718 | -1.056 | 0.578 | -0.516 | 0.343 | 0.048 | 0.026 | 0.026 | 0 | 0.64 | 0.82 | 0.78 | 0.68 | 0.73 | 0.49 |
| 1 | 5675044 | rs11260662 | 0.657 | 1.234 | -0.426 | 0.291 | -0.027 | -0.021 | -0.021 | 0 | 0.12 | 0.04 | 0.15 | 0.11 | 0.18 | 0.1 |
| 1 | 5675474 | rs1316481 | -0.553 | 0.791 | 0.045 | 0.175 | 0.009 | -0.011 | -0.011 | 0.14 | 0.09 | 0.26 | 0.05 | 0.08 | 0.06 | 0.07 |
| 1 | 5677775 | rs1775395 | -0.735 | 0.391 | -0.635 | -0.094 | 0.005 | 0.033 | 0.033 | 0 | 0.78 | 0.85 | 0.86 | 0.93 | 0.81 | 0.8 |
| 1 | 5679118 | rs9439602 | -0.273 | 0.5 | -0.172 | -0.064 | -0.003 | 0.031 | 0.031 | 0.013 | 0.45 | 0.35 | 0.41 | 0.16 | 0.47 | 0.27 |
| 1 | 5679222 | rs10907205 | 0.057 | 0.662 | -0.217 | -0.205 | 0 | -0.002 | -0.002 | 0 | 0.13 | 0.11 | 0.18 | 0.06 | 0.14 | 0.09 |
| 1 | 5679441 | rs9439603 | -0.582 | 0.72 | -0.123 | -0.243 | -0.005 | 0.059 | 0.059 | 0.028 | 0.44 | 0.31 | 0.37 | 0.16 | 0.43 | 0.26 |
| 1 | 5679808 | rs9439587 | 0.532 | 0.761 | -0.095 | -0.249 | 0.009 | 0.044 | 0.044 | 0 | 0.29 | 0.21 | 0.17 | 0.11 | 0.25 | 0.17 |
| 1 | 5680372 | rs9329417 | -0.672 | 0.689 | 0.642 | -0.15 | -0.001 | 0.001 | 0.001 | 0 | 0.45 | 0.59 | 0.45 | 0.22 | 0.5 | 0.23 |
| 1 | 5680572 | rs10157819 | 0.019 | 0.736 | 0.623 | -0.085 | 0.003 | -0.001 | -0.001 | 0 | 0.13 | 0.13 | 0.23 | 0.12 | 0.21 | 0.06 |
| 1 | 5680808 | rs9439605 | -1.771 | 0.528 | 0.033 | 0.381 | -0.053 | -0.052 | -0.052 | 1.175 | 0.79 | 0.98 | 0.82 | 0.8 | 0.75 | 0.75 |
| 1 | 5682832 | rs1695623 | 1.246 | 0.388 | 0.398 | 0.96 | -0.032 | 0.033 | 0.033 | 0.016 | 0.14 | 0.25 | 0.21 | 0.27 | 0.05 | 0.26 |
| 1 | 5684301 | rs945322 | 1.23 | 0.389 | 0.402 | 0.96 | -0.033 | 0.033 | 0.033 | 0.048 | 0.14 | 0.27 | 0.21 | 0.27 | 0.05 | 0.27 |
| 1 | 5684788 | rs12145625 | 0.723 | 0.428 | -0.193 | 1.202 | -0.032 | -0.038 | -0.038 | 0 | 0.18 | 0.07 | 0.23 | 0.26 | 0.22 | 0.21 |
| 1 | 5687879 | rs17460221 | 0.187 | 0.367 | 0.044 | 0.861 | 0.009 | -0.006 | -0.006 | 0 | 0.13 | 0.2 | 0.14 | 0.09 | 0.17 | 0.05 |
| 1 | 5688356 | rs770679 | 0.637 | 0.262 | 0.157 | 1.072 | -0.051 | 0.057 | 0.057 | 0.037 | 0.16 | 0.36 | 0.23 | 0.27 | 0.05 | 0.27 |
| 1 | 5689408 | rs11578045 | 0.024 | 1.225 | -0.5 | 1.432 | -0.036 | -0.067 | -0.067 | 0 | 0.18 | 0.06 | 0.22 | 0.33 | 0.22 | 0.2 |
| 1 | 5689702 | rs770683 | -1.047 | 1.272 | -0.504 | 0.832 | 0 | 0.05 | 0.05 | 0 | 0.81 | 0.93 | 0.8 | 0.92 | 0.79 | 0.78 |
| 1 | 5690136 | rs6663290 | -0.441 | 1.53 | -0.063 | 1.918 | -0.056 | -0.097 | -0.097 | 0.021 | 0.18 | 0.06 | 0.2 | 0.46 | 0.24 | 0.26 |
| 1 | 5691787 | rs17028422 | -0.971 | 1.546 | 1.135 | 1.442 | 0.112 | 0.09 | 0.09 | 0.041 | 0.05 | 0.37 | 0.05 | 0.05 | 0.04 | 0.06 |
| 1 | 5692728 | rs1855786 | -1.619 | 1.576 | 1.498 | 1.393 | 0.02 | 0.036 | 0.036 | 0.012 | 0.19 | 0.6 | 0.2 | 0.26 | 0.2 | 0.21 |
| 1 | 5699475 | rs10492947 | -0.508 | 1.763 | 1.27 | 1.358 | 0.017 | 0.036 | 0.036 | 0.231 | 0.12 | 0.19 | 0.15 | 0.21 | 0.16 | 0.16 |
| 1 | 5700043 | rs10465910 | -0.517 | 1.792 | 1.172 | 1.368 | 0.018 | 0.042 | 0.042 | 0 | 0.13 | 0.25 | 0.15 | 0.21 | 0.15 | 0.15 |
| 1 | 5700410 | rs10465911 | -2.297 | 1.685 | 1.071 | 1.494 | -0.058 | 0.002 | 0.002 | 0 | 0.43 | 0.88 | 0.5 | 0.62 | 0.49 | 0.47 |
| 1 | 5701323 | rs6690868 | -0.585 | 1.663 | 1.218 | 1.584 | 0.022 | 0.051 | 0.051 | 0.047 | 0.13 | 0.29 | 0.16 | 0.21 | 0.16 | 0.15 |
| 1 | 5703552 | rs17028491 | -0.209 | 1.784 | 0.418 | 1.576 | -0.024 | -0.041 | -0.041 | 0 | 0.19 | 0.09 | 0.21 | 0.37 | 0.25 | 0.24 |
| 1 | 5704836 | rs17028511 | -1.464 | 1.554 | 0.394 | 1.064 | 0.1 | 0.028 | 0.028 | 0.165 | 0.06 | 0.26 | 0.1 | 0.05 | 0.07 | 0.08 |
| 1 | 5707091 | rs971816 | -1.278 | 1.457 | -0.358 | 1 | -0.029 | 0.015 | 0.015 | 0.764 | 0.41 | 0.78 | 0.53 | 0.63 | 0.49 | 0.47 |
| 1 | 5707162 | rs971814 | -1.187 | 1.374 | -0.312 | 1.052 | -0.025 | 0.004 | 0.004 | 0.542 | 0.38 | 0.71 | 0.46 | 0.63 | 0.45 | 0.46 |
| 1 | 5707938 | rs1499395 | -1.146 | 1.16 | -0.503 | 1.154 | -0.025 | -0.037 | -0.037 | 0 | 0.37 | 0.8 | 0.49 | 0.56 | 0.41 | 0.45 |
| 1 | 5709833 | rs709184 | -0.253 | 0.812 | -0.538 | 1.382 | -0.045 | -0.057 | -0.057 | 0.208 | 0.1 | 0 | 0.11 | 0.3 | 0.13 | 0.16 |
| 1 | 5711647 | rs707589 | 0.012 | 0.278 | -0.061 | 0.912 | 0.001 | -0.001 | -0.001 | 0.249 | 0.44 | 0.38 | 0.43 | 0.5 | 0.36 | 0.29 |
| 1 | 5712831 | rs707590 | -0.868 | 0.014 | -0.007 | 0.725 | -0.015 | 0.078 | 0.078 | 1.891 | 0.55 | 0.75 | 0.57 | 0.88 | 0.5 | 0.5 |
| 1 | 5717388 | rs11260671 | -0.134 | -0.059 | 0.337 | 0.946 | 0.005 | 0.003 | 0.003 | 0.118 | 0.46 | 0.57 | 0.45 | 0.54 | 0.38 | 0.31 |
| 1 | 5719342 | rs771073 | -1.261 | -0.391 | 0.534 | 1.358 | 0 | 0.069 | 0.069 | 0 | 0.13 | 0.24 | 0.14 | 0.38 | 0.11 | 0.18 |
| 1 | 5721189 | rs12091928 | -2.493 | -1.008 | 0.785 | 1.154 | -0.065 | 0.027 | 0.027 | 5.076 | 0.12 | 0.72 | 0.06 | 0.12 | 0.01 | 0.06 |
| 1 | 5722405 | rs770721 | 0.131 | -1.203 | 0.335 | 1.071 | 0.016 | -0.122 | -0.122 | 0 | 0.32 | 0.09 | 0.28 | 0.52 | 0.24 | 0.19 |
| 1 | 5723643 | rs12038771 | -1.319 | -1.197 | 0.877 | 1.091 | 0.005 | 0.155 | 0.155 | 0.037 | 0.07 | 0.39 | 0.03 | 0.12 | 0.01 | 0.06 |
| 1 | 5724940 | rs707580 | -0.793 | -0.998 | 0.222 | 0.968 | 0.049 | 0.024 | 0.024 | 1.17 | 0.42 | 0.73 | 0.34 | 0.64 | 0.26 | 0.24 |
| 1 | 5732250 | rs7544257 | -1.343 | -0.666 | 0.938 | 0.67 | 0.037 | 0.165 | 0.165 | 2.276 | 0.06 | 0.42 | 0.04 | 0.1 | 0.01 | 0.03 |
| 1 | 5732473 | rs771071 | 1.067 | -0.592 | 0.318 | 1.161 | -0.004 | 0.015 | 0.015 | 0 | 0.1 | 0.06 | 0.12 | 0.43 | 0.09 | 0.18 |
| 1 | 5736682 | rs707593 | 0.664 | -1.049 | 0.005 | 0.311 | 0.061 | 0.19 | 0.19 | 1.162 | 0.48 | 0.22 | 0.53 | 0.07 | 0.64 | 0.52 |
| 1 | 5743509 | rs770701 | 0.139 | -0.188 | -0.298 | 0.324 | -0.046 | 0.008 | 0.008 | 0.658 | 0.08 | 0.01 | 0.07 | 0.03 | 0.14 | 0.08 |
| 1 | 5744921 | rs770702 | -0.025 | -1.326 | 0.281 | 0.264 | 0.005 | 0.008 | 0.008 | 1.658 | 0.41 | 0.38 | 0.37 | 0.73 | 0.33 | 0.28 |
| 1 | 5744963 | rs10492949 | -0.474 | -1.321 | 0.654 | 0.367 | -0.031 | 0.001 | 0.001 | 0 | 0.13 | 0.21 | 0.09 | 0.11 | 0.06 | 0.19 |
| 1 | 5745339 | rs770703 | 1.335 | -1.216 | 0.623 | 0.616 | 0.001 | 0.088 | 0.088 | 0 | 0.11 | 0.13 | 0.13 | 0.43 | 0.09 | 0.19 |
| 1 | 5747184 | rs770705 | -0.774 | -1.227 | 0.364 | 0.131 | 0.006 | 0.01 | 0.01 | 0.512 | 0.49 | 0.6 | 0.48 | 0.82 | 0.38 | 0.31 |
| 1 | 5747488 | rs770707 | -0.026 | -1.083 | 0.126 | 0.204 | -0.003 | -0.004 | -0.004 | 0.232 | 0.39 | 0.4 | 0.37 | 0.72 | 0.32 | 0.28 |
| 1 | 5748611 | rs1278832 | -1.058 | -0.828 | -0.681 | -0.064 | -0.02 | 0.12 | 0.12 | 0.535 | 0.58 | 0.8 | 0.51 | 0.86 | 0.41 | 0.51 |
| 1 | 5750013 | rs2649069 | -0.015 | -0.917 | -0.064 | 0.309 | -0.009 | -0.009 | -0.009 | 0 | 0.62 | 0.49 | 0.67 | 0.65 | 0.66 | 0.71 |
| 1 | 5758772 | rs7521197 | -0.537 | -1.108 | -0.021 | -0.033 | -0.02 | -0.008 | -0.008 | 0 | 0.18 | 0.25 | 0.22 | 0.13 | 0.08 | 0.2 |
| 1 | 5765353 | rs770693 | 1.018 | -1.276 | -0.265 | -0.277 | -0.06 | 0.117 | 0.117 | 0 | 0.39 | 0.12 | 0.43 | 0.2 | 0.48 | 0.45 |
| 1 | 5767805 | rs2649078 | 0.372 | -1.368 | -0.084 | 0.024 | 0.011 | -0.011 | -0.011 | 5.025 | 0.57 | 0.65 | 0.56 | 0.54 | 0.62 | 0.62 |
| 1 | 5774876 | rs1414380 | -0.547 | -1.02 | -0.474 | -0.52 | -0.052 | 0.037 | 0.037 | 0 | 0.66 | 0.81 | 0.65 | 0.33 | 0.59 | 0.7 |
| 1 | 5777457 | rs904534 | -0.608 | -1.032 | -0.402 | -0.524 | -0.057 | 0.032 | 0.032 | 0 | 0.66 | 0.84 | 0.71 | 0.33 | 0.59 | 0.7 |
| 1 | 5777920 | rs904532 | 0.421 | -0.538 | 0.214 | -0.744 | 0.001 | -0.019 | -0.019 | 0 | 0.42 | 0.48 | 0.48 | 0.24 | 0.49 | 0.47 |
| 1 | 5780697 | rs2244632 | 0.613 | -0.532 | 0.063 | -0.71 | -0.013 | 0.038 | 0.038 | 0.088 | 0.41 | 0.34 | 0.46 | 0.25 | 0.49 | 0.47 |
| 1 | 5782721 | rs10907212 | 1.153 | -0.744 | -0.49 | -0.519 | -0.034 | -0.085 | -0.085 | 0.002 | 0.15 | 0.05 | 0.17 | 0.51 | 0.21 | 0.28 |
| 1 | 5782769 | rs2649086 | 0.018 | -1.073 | 0.173 | -0.628 | -0.003 | -0.001 | -0.001 | 0.008 | 0.61 | 0.67 | 0.72 | 0.6 | 0.72 | 0.69 |
| 1 | 5790498 | rs2842258 | 0.325 | -1.169 | 0.618 | -1.71 | -0.002 | -0.009 | -0.009 | 0 | 0.31 | 0.26 | 0.34 | 0.04 | 0.35 | 0.28 |
| 1 | 5790655 | rs2842257 | -0.404 | -1.246 | 0.917 | -1.764 | 0.005 | -0.05 | -0.05 | 2.799 | 0.4 | 0.52 | 0.41 | 0.19 | 0.48 | 0.58 |
| 1 | 5793755 | rs12738264 | 0.469 | -1.118 | 0.914 | -1.717 | -0.003 | -0.006 | -0.006 | 0 | 0.29 | 0.24 | 0.33 | 0.04 | 0.33 | 0.27 |
| 1 | 5799631 | rs2649063 | -0.594 | -1.291 | 0.899 | -1.743 | -0.002 | -0.002 | -0.002 | 0 | 0.4 | 0.58 | 0.41 | 0.18 | 0.46 | 0.57 |
| 1 | 5800904 | rs2133496 | 0.489 | -1.253 | 0.274 | -1.796 | -0.024 | 0.073 | 0.073 | 0 | 0.28 | 0.13 | 0.31 | 0.04 | 0.33 | 0.27 |
| 1 | 5805605 | rs11260682 | -1.195 | -2.269 | 1.26 | -1.339 | -0.1 | -0.149 | -0.149 | 0 | 0.24 | 0.4 | 0.06 | 0.02 | 0.05 | 0.08 |

| | | | | | | | | | | | | | | | | |
|---|---|---|---|---|---|---|---|---|---|---|---|---|---|---|---|---|
| 1 | 5808093 | rs12059728 | -1.96 | -2.334 | 1.342 | -1.46 | 0.019 | -0.002 | -0.002 | 0 | 0.33 | 0.77 | 0.13 | 0.06 | 0.18 | 0.17 |
| 1 | 5810354 | rs4428894 | -0.616 | -2.467 | 1.572 | -1.221 | -0.015 | -0.03 | -0.03 | 0.065 | 0.16 | 0.2 | 0.02 | 0.02 | 0.03 | 0.06 |
| 1 | 5811124 | rs4433399 | -2.118 | -2.496 | 1.345 | -1.099 | -0.074 | -0.072 | -0.072 | 0 | 0.29 | 0.51 | 0.15 | 0.16 | 0.16 | 0.36 |
| 1 | 5814135 | rs17429690 | -0.028 | -2.112 | 1.058 | -0.862 | -0.026 | 0.031 | 0.031 | 0 | 0.06 | 0.01 | 0.06 | 0.01 | 0.11 | 0.08 |
| 1 | 5814154 | rs12060804 | -1.664 | -2.094 | 1.214 | -0.834 | -0.026 | 0.038 | 0.038 | 0.668 | 0.06 | 0.16 | 0.06 | 0.12 | 0.01 | 0.2 |
| 1 | 5829039 | rs12076190 | -1.893 | -2.127 | 0.356 | -0.889 | 0.049 | 0.043 | 0.043 | 2.022 | 0.3 | 0.78 | 0.17 | 0.19 | 0.18 | 0.36 |
| 1 | 5830248 | rs4072521 | -1.766 | -2.098 | 0.328 | -0.779 | 0.037 | 0.01 | 0.01 | 2.071 | 0.3 | 0.75 | 0.23 | 0.29 | 0.2 | 0.37 |
| 1 | 5831601 | rs4129514 | -1.724 | -1.908 | 0.203 | -0.684 | -0.016 | -0.002 | -0.002 | 0 | 0.28 | 0.64 | 0.22 | 0.29 | 0.2 | 0.37 |
| 1 | 5834437 | rs17028791 | -0.409 | -1.819 | 0.337 | -0.667 | -0.048 | -0.06 | -0.06 | 0 | 0.13 | 0.21 | 0.02 | 0.02 | 0.03 | 0.07 |
| 1 | 5836317 | rs11260693 | -1.018 | -1.594 | -0.797 | -0.507 | -0.012 | -0.001 | -0.001 | 0 | 0.3 | 0.64 | 0.22 | 0.32 | 0.19 | 0.38 |
| 1 | 5861105 | rs4845818 | 0.084 | -0.99 | -1.511 | -0.929 | 0.041 | 0.067 | 0.067 | 0 | 0.63 | 0.78 | 0.74 | 0.91 | 0.72 | 0.63 |
| 1 | 5870534 | rs6701947 | 0.091 | -0.985 | -1.553 | -0.95 | 0.042 | 0.074 | 0.074 | 0 | 0.63 | 0.79 | 0.75 | 0.89 | 0.71 | 0.63 |
| 1 | 5880466 | rs9308476 | -0.345 | -0.792 | -1.828 | -0.968 | 0.006 | 0.095 | 0.095 | 0 | 0.72 | 0.87 | 0.79 | 0.98 | 0.71 | 0.79 |
| 1 | 5882279 | rs9661839 | -0.673 | -0.113 | -1.81 | -0.251 | -0.001 | 0.008 | 0.008 | 0 | 0.1 | 0.11 | 0.1 | 0.23 | 0.1 | 0.25 |
| 1 | 5887140 | rs1890191 | 0.197 | -0.884 | -2.445 | -0.987 | 0.062 | 0.252 | 0.252 | 0 | 0.45 | 0.13 | 0.68 | 0.85 | 0.65 | 0.46 |
| 1 | 5897007 | rs4623725 | -0.123 | -0.381 | -2.508 | -1.059 | -0.068 | 0.115 | 0.115 | 0 | 0.58 | 0.88 | 0.64 | 0.84 | 0.49 | 0.64 |
| 1 | 5900795 | rs1698976 | 0.388 | -0.61 | -1.874 | -1.15 | -0.006 | 0.02 | 0.02 | 0 | 0.37 | 0.29 | 0.53 | 0.68 | 0.4 | 0.35 |
| 1 | 5901929 | rs1765723 | -1.029 | -0.905 | -2.466 | -1.618 | -0.028 | 0.075 | 0.075 | 0 | 0.79 | 0.97 | 0.83 | 0.91 | 0.79 | 0.75 |
| 1 | 5902023 | rs11590577 | -0.33 | -0.404 | -2.468 | -1.348 | -0.044 | -0.022 | -0.022 | 0 | 0.13 | 0.03 | 0.17 | 0.17 | 0.25 | 0.24 |
| 1 | 5902086 | rs11579676 | -0.987 | -1.131 | -2.116 | -1.686 | 0 | 0.006 | 0.006 | 0 | 0.23 | 0.23 | 0.19 | 0.18 | 0.24 | 0.18 |
| 1 | 5902367 | rs9729210 | -0.253 | -1.635 | -2.494 | -1.353 | 0.09 | 0.102 | 0.102 | 0 | 0.54 | 0.83 | 0.75 | 0.72 | 0.69 | 0.51 |
| 1 | 5904718 | rs1624858 | -0.718 | -1.574 | -2.451 | -1.576 | -0.011 | 0.11 | 0.11 | 0 | 0.67 | 0.91 | 0.76 | 0.86 | 0.68 | 0.62 |
| 1 | 5905929 | rs1765719 | -0.364 | -1.643 | -1.919 | -1.515 | 0.016 | 0.073 | 0.073 | 0 | 0.62 | 0.77 | 0.74 | 0.82 | 0.65 | 0.59 |
| 1 | 5906864 | rs2062783 | 0.21 | -1.357 | -1.937 | -1.391 | -0.054 | 0.024 | 0.024 | 0 | 0.15 | 0.03 | 0.17 | 0.11 | 0.24 | 0.12 |
| 1 | 5908733 | rs1287616 | -0.167 | -0.935 | -1.185 | -1.464 | 0.006 | 0.021 | 0.021 | 0 | 0.54 | 0.6 | 0.7 | 0.81 | 0.6 | 0.54 |
| 1 | 5913621 | rs548726 | -0.694 | -0.38 | -1.108 | -1.365 | 0.013 | 0.048 | 0.048 | 0.851 | 0.66 | 0.79 | 0.78 | 0.9 | 0.73 | 0.68 |
| 1 | 5915662 | rs10157281 | 0.852 | 0.435 | -1.134 | -1.474 | -0.007 | 0.002 | 0.002 | 1.463 | 0.15 | 0.1 | 0.07 | 0.02 | 0.14 | 0.12 |
| 1 | 5921042 | rs11121600 | 0.631 | 0.872 | -1.065 | -1.55 | -0.012 | 0.022 | 0.022 | 0.441 | 0.17 | 0.12 | 0.14 | 0.09 | 0.23 | 0.15 |
| 1 | 5923788 | rs11121648 | -1.848 | 0.186 | -0.935 | -1.807 | -0.001 | -0.002 | -0.002 | 0 | 0.57 | 0.59 | 0.49 | 0.3 | 0.55 | 0.56 |
| 1 | 5924746 | rs1287635 | -0.411 | -0.041 | -1.095 | -1.798 | 0.002 | 0.058 | 0.058 | 0 | 0.39 | 0.2 | 0.38 | 0.29 | 0.41 | 0.43 |
| 1 | 5925371 | rs1287634 | -0.289 | -0.038 | -1.148 | -1.774 | -0.006 | 0.067 | 0.067 | 0 | 0.38 | 0.16 | 0.38 | 0.28 | 0.41 | 0.43 |
| 1 | 5926507 | rs555164 | -0.299 | -0.047 | -1.152 | -1.779 | -0.005 | 0.059 | 0.059 | 0 | 0.39 | 0.16 | 0.38 | 0.28 | 0.41 | 0.43 |
| 1 | 5930209 | rs12405167 | 2.081 | -0.317 | -1.321 | -1.784 | 0.001 | -0.036 | -0.036 | 7.091 | 0.1 | 0 | 0.14 | 0.16 | 0.09 | 0.13 |
| 1 | 5932517 | rs472651 | 0.703 | -0.303 | -1.321 | -1.806 | 0.009 | -0.006 | -0.006 | 0.008 | 0.2 | 0.03 | 0.24 | 0.19 | 0.18 | 0.26 |
| 1 | 5934490 | rs868163 | -2.481 | -0.568 | -1.232 | -2.319 | -0.005 | 0.011 | 0.011 | 2.098 | 0.64 | 0.71 | 0.53 | 0.31 | 0.63 | 0.62 |
| 1 | 5934500 | rs905467 | -0.198 | -1.142 | -1.523 | -1.925 | 0.025 | -0.072 | -0.072 | 1.861 | 0.29 | 0.07 | 0.29 | 0.38 | 0.23 | 0.47 |
| 1 | 5934837 | rs1287638 | -0.596 | -1.096 | -1.295 | -1.939 | -0.003 | 0.03 | 0.03 | 3.529 | 0.48 | 0.6 | 0.34 | 0.1 | 0.49 | 0.42 |
| 1 | 5937091 | rs1622955 | -2.469 | -1.079 | -1.326 | -1.761 | -0.002 | -0.025 | -0.025 | 0.027 | 0.89 | 0.99 | 0.85 | 0.87 | 0.9 | 0.87 |
| 1 | 5937168 | rs3747992 | 1.899 | -1.266 | -1.307 | -1.871 | -0.012 | 0.019 | 0.019 | 0 | 0.36 | 0.29 | 0.46 | 0.73 | 0.42 | 0.42 |
| 1 | 5937327 | rs3747990 | 2.385 | -0.826 | -1.057 | -1.604 | -0.033 | 0.022 | 0.022 | 0 | 0.13 | 0.01 | 0.12 | 0.08 | 0.16 | 0.12 |
| 1 | 5946443 | rs12735231 | 2.115 | -0.92 | -0.679 | -1.348 | 0.006 | -0.037 | -0.037 | 0 | 0.14 | 0.29 | 0.13 | 0.09 | 0.16 | 0.11 |
| 1 | 5948946 | rs11122097 | 2.657 | -0.885 | -0.94 | -1.345 | -0.037 | 0.019 | 0.019 | 0 | 0.12 | 0.01 | 0.11 | 0.08 | 0.16 | 0.1 |
| 1 | 5956909 | rs12058375 | 1.578 | -1.097 | -0.631 | -1.308 | -0.012 | 0.04 | 0.04 | 0.143 | 0.11 | 0.2 | 0.05 | 0.18 | 0.1 | 0.19 |
| 1 | 5957601 | rs12401883 | 2.765 | -0.74 | -0.383 | -1.505 | -0.072 | 0.088 | 0.088 | 1.682 | 0.15 | 0.01 | 0.1 | 0.01 | 0.19 | 0.12 |
| 1 | 5958409 | rs11586334 | 2.416 | -0.734 | -0.239 | -1.443 | -0.008 | 0.024 | 0.024 | 0 | 0.13 | 0.05 | 0.11 | 0.09 | 0.16 | 0.11 |
| 1 | 5961870 | rs2134551 | -0.611 | -0.912 | -0.354 | -1.581 | 0.018 | 0.018 | 0.018 | 0 | 0.8 | 0.95 | 0.75 | 0.82 | 0.82 | 0.73 |
| 1 | 5965381 | rs12120967 | 2.814 | -0.832 | -0.412 | -1.538 | -0.038 | 0.018 | 0.018 | 0 | 0.12 | 0.01 | 0.11 | 0.08 | 0.16 | 0.1 |
| 1 | 5978510 | rs962662 | 0.391 | -0.875 | 0.251 | -1.581 | -0.02 | -0.044 | -0.044 | 0 | 0.63 | 0.53 | 0.61 | 0.74 | 0.66 | 0.63 |
| 1 | 5982044 | rs7549324 | 1.097 | -0.855 | 0.18 | -1.612 | -0.006 | 0.064 | 0.064 | 0 | 0.43 | 0.3 | 0.5 | 0.74 | 0.45 | 0.47 |
| 1 | 5983114 | rs7535806 | 1.161 | -0.902 | 0.362 | -1.058 | -0.005 | -0.051 | -0.051 | 0.483 | 0.16 | 0.31 | 0.13 | 0.08 | 0.16 | 0.1 |
| 1 | 6007446 | rs551207 | 1.218 | -0.758 | 0.107 | -0.072 | -0.001 | 0.002 | 0.002 | 0 | 0.84 | 0.84 | 0.78 | 0.73 | 0.85 | 0.81 |
| 1 | 6031853 | rs2047811 | -0.033 | -1.573 | -0.158 | -0.685 | -0.002 | 0.012 | 0.012 | 0 | 0.82 | 0.97 | 0.75 | 0.83 | 0.82 | 0.74 |
| 1 | 6031944 | rs11588889 | 0.714 | -1.638 | 0.031 | -0.762 | -0.019 | -0.045 | -0.045 | 0.768 | 0.63 | 0.52 | 0.62 | 0.75 | 0.66 | 0.64 |
| 1 | 6033973 | rs4908458 | 1.312 | -1.62 | -0.295 | -0.748 | -0.005 | 0.013 | 0.013 | 0 | 0.08 | 0.02 | 0.09 | 0.03 | 0.08 | 0.12 |
| 1 | 6037603 | rs11120934 | 0.408 | -1.603 | -0.167 | -0.35 | 0.011 | 0.027 | 0.027 | 0 | 0.65 | 0.74 | 0.63 | 0.75 | 0.66 | 0.64 |
| 1 | 6039254 | rs10779685 | -0.147 | -1.591 | -0.595 | -0.343 | -0.002 | 0.012 | 0.012 | 0 | 0.82 | 0.97 | 0.76 | 0.83 | 0.82 | 0.74 |
| 1 | 6043809 | rs11120961 | 1.159 | -1.622 | -0.321 | -0.135 | 0.004 | 0.018 | 0.018 | 0 | 0.16 | 0.1 | 0.12 | 0.09 | 0.16 | 0.1 |
| 1 | 6053630 | rs806111 | -0.918 | -1.576 | -0.46 | -0.094 | -0.001 | -0.005 | -0.005 | 0 | 0.38 | 0.59 | 0.34 | 0.26 | 0.35 | 0.39 |
| 1 | 6059229 | rs1295100 | -0.922 | -1.581 | -0.467 | 0.094 | -0.009 | -0.001 | -0.001 | 0 | 0.38 | 0.56 | 0.46 | 0.41 | 0.35 | 0.39 |
| 1 | 6059437 | rs1295099 | -0.193 | -1.512 | -0.399 | 0.434 | 0.007 | -0.069 | -0.069 | 0 | 0.16 | 0.04 | 0.33 | 0.32 | 0.16 | 0.27 |
| 1 | 6070870 | rs546526 | 0.746 | -1.251 | 0.003 | 0.125 | -0.007 | 0.003 | 0.003 | 0.095 | 0.4 | 0.33 | 0.39 | 0.59 | 0.45 | 0.46 |
| 1 | 6079392 | rs1295107 | -0.519 | -1.431 | -0.013 | 0.133 | -0.004 | -0.009 | -0.009 | 0.916 | 0.43 | 0.51 | 0.47 | 0.36 | 0.35 | 0.42 |
| 1 | 6085392 | rs1295105 | -0.418 | -1.149 | -0.202 | 0.507 | -0.004 | 0.023 | 0.023 | 0.01 | 0.95 | 0.98 | 0.97 | 1 | 0.95 | 0.96 |
| 1 | 6088364 | rs11121065 | 0.424 | -1.152 | 0.042 | 0.568 | -0.033 | 0.036 | 0.036 | 0.183 | 0.14 | 0.24 | 0.17 | 0.29 | 0.06 | 0.15 |
| 1 | 6088671 | rs1295103 | -0.468 | -0.948 | -0.385 | 0.394 | 0.011 | -0.033 | -0.033 | 0 | 0.06 | 0.11 | 0.16 | 0 | 0.08 | 0.02 |
| 1 | 6095199 | rs1295094 | -0.359 | -0.769 | -0.378 | 0.619 | -0.009 | 0.03 | 0.03 | 0 | 0.18 | 0.26 | 0.34 | 0.32 | 0.17 | 0.29 |
| 1 | 6098266 | rs3761923 | 0.525 | -0.774 | -0.591 | 0.673 | 0.009 | -0.009 | -0.009 | 0.692 | 0.13 | 0.14 | 0.17 | 0.32 | 0.05 | 0.13 |
| 1 | 6100313 | rs1295087 | -0.51 | -0.215 | -0.662 | 0.294 | -0.048 | 0.069 | 0.069 | 0 | 0.83 | 0.97 | 0.86 | 0.95 | 0.77 | 0.87 |
| 1 | 6104002 | rs3789563 | -0.224 | -0.163 | -0.125 | 0.849 | -0.002 | 0.066 | 0.066 | 0 | 0.53 | 0.67 | 0.43 | 0.75 | 0.47 | 0.59 |

| | | | | | | | | | | | | | | | | |
|---|---|---|---|---|---|---|---|---|---|---|---|---|---|---|---|---|
| 1 | 6108165 | rs1295085 | -0.614 | 0.508 | 0.33 | 1.331 | -0.042 | -0.024 | -0.024 | 0 | 0.73 | 0.84 | 0.7 | 0.66 | 0.65 | 0.75 |
| 1 | 6112653 | rs12047044 | -0.826 | 0.21 | 1.015 | 1.534 | 0.001 | -0.002 | -0.002 | 0 | 0.3 | 0.61 | 0.33 | 0.4 | 0.23 | 0.24 |
| 1 | 6117171 | rs3789553 | -0.937 | 0.279 | 0.937 | 1.433 | 0.002 | -0.006 | -0.006 | 0 | 0.32 | 0.62 | 0.34 | 0.39 | 0.24 | 0.25 |
| 1 | 6125084 | rs4908752 | -0.826 | 0.103 | 0.711 | 1.075 | -0.009 | 0.016 | 0.016 | 0.842 | 0.3 | 0.47 | 0.32 | 0.34 | 0.23 | 0.26 |
| 1 | 6129864 | rs1883771 | -1.185 | 0.179 | 0.374 | 0.877 | 0.076 | 0.066 | 0.066 | 1.976 | 0.05 | 0.4 | 0.04 | 0 | 0.01 | 0 |
| 1 | 6141325 | rs3789541 | -1.079 | 0.407 | -0.795 | 1.585 | -0.002 | -0.04 | -0.04 | 0 | 0.08 | 0.01 | 0.04 | 0.29 | 0.09 | 0.21 |
| 1 | 6141454 | rs12741018 | 0.254 | 0.7 | -0.972 | 0.615 | -0.003 | 0.007 | 0.007 | 0 | 0.75 | 0.75 | 0.6 | 0.88 | 0.76 | 0.82 |
| 1 | 6144914 | rs3789531 | 0.579 | 0.957 | -0.915 | 0.59 | -0.003 | 0.004 | 0.004 | 0 | 0.61 | 0.39 | 0.51 | 0.66 | 0.59 | 0.57 |
| 1 | 6149122 | rs2273042 | 0.038 | 1.015 | -1.281 | 1.135 | -0.029 | -0.035 | -0.035 | 0 | 0.07 | 0.01 | 0.06 | 0.21 | 0.14 | 0.07 |
| 1 | 6162407 | rs3827728 | -0.517 | 0.789 | -1.387 | 0.213 | 0.002 | 0.012 | 0.012 | 0.893 | 0.27 | 0.21 | 0.27 | 0.78 | 0.27 | 0.5 |
| 1 | 6162583 | rs3810993 | -1.005 | 0.435 | -1.683 | 0.113 | 0.003 | 0.016 | 0.016 | 0 | 0.61 | 0.78 | 0.64 | 0.64 | 0.65 | 0.63 |
| 1 | 6162645 | rs3810990 | 1.311 | 0.236 | -1.44 | 0.413 | -0.039 | -0.035 | -0.035 | 3.435 | 0.08 | 0.01 | 0.08 | 0.35 | 0.1 | 0.13 |
| 1 | 6169411 | rs12046716 | 0.683 | -0.022 | -1.264 | 0.769 | 0.018 | -0.017 | -0.017 | 0.735 | 0.14 | 0.09 | 0.04 | 0.23 | 0.05 | 0.18 |
| 1 | 6170353 | rs2281303 | -0.452 | -0.205 | -1.009 | 0.747 | 0.007 | 0.09 | 0.09 | 0.627 | 0.08 | 0.18 | 0.16 | 0.2 | 0.1 | 0.22 |
| 1 | 6175489 | rs1883767 | 0.273 | 0.028 | -1.072 | 0.717 | -0.028 | -0.024 | -0.024 | 0 | 0.13 | 0.07 | 0.15 | 0.56 | 0.26 | 0.2 |
| 1 | 6177898 | rs2785581 | -0.896 | -0.293 | -0.807 | 0.226 | 0.001 | -0.071 | -0.071 | 0 | 0.64 | 0.61 | 0.59 | 1 | 0.58 | 0.83 |
| 1 | 6182321 | rs2843495 | 0.418 | -1.899 | -0.824 | 0.371 | 0.066 | 0.082 | 0.082 | 0.158 | 0.44 | 0.09 | 0.37 | 0.21 | 0.29 | 0.48 |
| 1 | 6183926 | rs2843494 | -0.347 | -1.907 | -0.797 | 0.453 | 0.004 | 0.004 | 0.004 | 0 | 0.09 | 0.05 | 0.07 | 0.27 | 0.09 | 0.21 |
| 1 | 6184192 | rs2273041 | -0.938 | -1.856 | -0.678 | 0.423 | -0.001 | -0.011 | -0.011 | 0 | 0.63 | 0.61 | 0.52 | 0.83 | 0.45 | 0.77 |
| 1 | 6187086 | rs6703057 | 1.29 | -1.837 | -0.63 | 0.922 | -0.043 | -0.039 | -0.039 | 0.133 | 0.08 | 0.01 | 0.08 | 0.35 | 0.12 | 0.13 |
| 1 | 6190215 | rs9434662 | -0.407 | -1.878 | -0.317 | 0.555 | 0.007 | 0.064 | 0.064 | 0 | 0.15 | 0.17 | 0.14 | 0.63 | 0.17 | 0.3 |
| 1 | 6194018 | rs11121299 | -1.11 | -1.929 | -0.308 | 0.514 | 0.001 | 0.146 | 0.146 | 0.817 | 0.19 | 0.45 | 0.17 | 0.64 | 0.17 | 0.32 |
| 1 | 6194580 | rs3765451 | -0.462 | -2.441 | -0.593 | 0.736 | -0.013 | -0.04 | -0.04 | 0 | 0.14 | 0.06 | 0.13 | 0.64 | 0.16 | 0.3 |
| 1 | 6196869 | rs2250358 | -0.358 | -2.656 | -0.612 | -0.064 | 0.013 | 0.077 | 0.077 | 0 | 0.56 | 0.34 | 0.47 | 0.8 | 0.39 | 0.63 |
| 1 | 6199900 | rs7523555 | 1.148 | -2.6 | -0.645 | -0.209 | -0.014 | -0.08 | -0.08 | 0.239 | 0.31 | 0.3 | 0.24 | 0 | 0.15 | 0.11 |
| 1 | 6202245 | rs2273032 | 1.376 | -2.31 | -0.795 | 0.498 | -0.015 | 0.014 | 0.014 | 0 | 0.07 | 0.04 | 0.06 | 0.33 | 0.11 | 0.13 |
| 1 | 6204161 | rs17436816 | 1.463 | -2.694 | -0.985 | -0.041 | 0.082 | 0.17 | 0.17 | 4.756 | 0.28 | 0.02 | 0.21 | 0.03 | 0.12 | 0.1 |
| 1 | 6204266 | rs9435102 | -1.551 | -2.554 | -0.501 | 0.078 | -0.05 | 0.036 | 0.036 | 0.306 | 0.72 | 0.82 | 0.55 | 0.84 | 0.49 | 0.77 |
| 1 | 6204842 | rs11121328 | -2.049 | -1.819 | 0.862 | 1.027 | 0.009 | 0.018 | 0.018 | 1.368 | 0.39 | 0.65 | 0.3 | 0.78 | 0.34 | 0.62 |
| 1 | 6205349 | rs12067480 | -0.402 | -1.922 | 0.927 | 0.908 | 0.01 | 0.15 | 0.15 | 0.467 | 0.14 | 0.31 | 0.08 | 0.4 | 0.14 | 0.17 |
| 1 | 6207217 | rs1883606 | -2.231 | -1.779 | 0.119 | 0.76 | 0.053 | 0.041 | 0.041 | 1.993 | 0.45 | 0.74 | 0.24 | 0.66 | 0.34 | 0.55 |
| 1 | 6212843 | rs2311388 | -1.983 | -1.728 | 0.36 | 1.074 | 0.05 | 0.009 | 0.009 | 2.01 | 0.4 | 0.75 | 0.21 | 0.65 | 0.3 | 0.53 |
| 1 | 6212870 | rs1883604 | -1.156 | -1.366 | 1.076 | 0.662 | 0.097 | 0.004 | 0.004 | 0 | 0.07 | 0.35 | 0.06 | 0.01 | 0.08 | 0.13 |
| 1 | 6217859 | rs9434741 | -1.059 | -1.495 | 0.509 | 0.826 | -0.051 | 0.016 | 0.016 | 0.005 | 0.25 | 0.51 | 0.13 | 0.27 | 0.15 | 0.35 |
| 1 | 6221975 | rs747393 | -1.895 | -1.518 | 0.691 | 0.776 | 0.068 | 0.016 | 0.016 | 0 | 0.38 | 0.74 | 0.26 | 0.68 | 0.18 | 0.38 |
| 1 | 6222501 | rs731975 | 1.405 | -1.53 | 1.427 | 1 | -0.021 | 0.068 | 0.068 | 0 | 0.74 | 0.68 | 0.78 | 0.53 | 0.82 | 0.72 |
| 1 | 6222883 | rs4908529 | -1.415 | -1.334 | 1.67 | 1.164 | -0.025 | 0.126 | 0.126 | 0.431 | 0.18 | 0.27 | 0.07 | 0.46 | 0.12 | 0.23 |
| 1 | 6226910 | rs4908850 | -2.144 | -0.917 | 1.713 | 0.18 | -0.04 | -0.022 | -0.022 | 0 | 0.2 | 0.53 | 0.21 | 0.2 | 0.11 | 0.16 |
| 1 | 6243715 | rs1534997 | -2.43 | -0.951 | 1.567 | 0.129 | 0.011 | 0.01 | 0.01 | 0.916 | 0.25 | 0.71 | 0.22 | 0.19 | 0.12 | 0.17 |
| 1 | 6247066 | rs4908863 | -1.554 | -0.912 | 1.651 | 0.192 | -0.035 | 0.031 | 0.031 | 0 | 0.12 | 0.32 | 0.15 | 0.17 | 0.05 | 0.11 |
| 1 | 6252443 | rs11121449 | -1.528 | -0.584 | 1.562 | 0.141 | -0.007 | -0.086 | -0.086 | 0.009 | 0.13 | 0.38 | 0.07 | 0.01 | 0.06 | 0.06 |
| 1 | 6258794 | rs3789524 | -1.259 | 0.271 | 1.274 | 0.437 | 0.007 | 0.012 | 0.012 | 1.332 | 0.27 | 0.48 | 0.32 | 0.28 | 0.26 | 0.3 |
| 1 | 6263354 | rs4908864 | -1.109 | 0.357 | 1.324 | 0.442 | 0.011 | 0.011 | 0.011 | 1.349 | 0.25 | 0.39 | 0.32 | 0.28 | 0.27 | 0.28 |
| 1 | 6272813 | rs2294711 | -1.55 | -0.139 | 2.227 | 0.462 | 0.007 | 0.039 | 0.039 | 0 | 0.08 | 0.34 | 0.07 | 0.05 | 0.04 | 0.07 |
| 1 | 6276167 | rs12071731 | -2.66 | -0.258 | 2.41 | 0.4 | -0.046 | -0.016 | -0.016 | 0 | 0.17 | 0.51 | 0.1 | 0.09 | 0.08 | 0.12 |
| 1 | 6278414 | rs709209 | -2.615 | -0.134 | 2.262 | 0.044 | 0.031 | 0.058 | 0.058 | 1.466 | 0.45 | 0.74 | 0.29 | 0.26 | 0.35 | 0.4 |
| 1 | 6283062 | rs12070839 | -1.747 | -0.228 | 2.605 | 0.206 | -0.011 | 0.029 | 0.029 | 0.008 | 0.1 | 0.36 | 0.08 | 0.05 | 0.05 | 0.08 |
| 1 | 6284680 | rs1802353 | -2.59 | -0.172 | 2.626 | 0.155 | -0.028 | -0.032 | -0.032 | 5.892 | 0.16 | 0.4 | 0.1 | 0.07 | 0.09 | 0.11 |
| 1 | 6291810 | rs4908541 | -0.702 | -0.065 | 2.186 | 0.234 | 0 | -0.001 | -0.001 | 0 | 0.05 | 0.05 | 0.02 | 0.03 | 0.05 | 0.04 |
| 1 | 6293890 | rs12073932 | -1.207 | -0.031 | 2.246 | 0.271 | -0.02 | -0.009 | -0.009 | 0 | 0.08 | 0.18 | 0.05 | 0.04 | 0.04 | 0.07 |
| 1 | 6301282 | rs10779736 | 0.525 | 0.295 | 2.064 | 0.79 | 0.002 | 0.02 | 0.02 | 8.645 | 0.79 | 0.71 | 0.81 | 0.62 | 0.79 | 0.76 |
| 1 | 6306912 | rs12240078 | -0.471 | 0.243 | 1.918 | 0.723 | 0.002 | -0.004 | -0.004 | 0 | 0.13 | 0.1 | 0.04 | 0.19 | 0.08 | 0.03 |
| 1 | 6307255 | rs6577565 | 0.661 | 0.114 | 1.953 | 0.766 | -0.004 | -0.009 | -0.009 | 0 | 0.78 | 0.84 | 0.81 | 0.77 | 0.79 | 0.69 |
| 1 | 6312016 | rs4908542 | 1.392 | 0.29 | 2.189 | 0.554 | 0.01 | -0.009 | -0.009 | 0 | 0.61 | 0.27 | 0.76 | 0.59 | 0.72 | 0.69 |
| 1 | 6314479 | rs12740426 | 1.361 | 0.456 | 2.205 | 0.571 | 0.004 | -0.009 | -0.009 | 0 | 0.61 | 0.27 | 0.75 | 0.59 | 0.7 | 0.67 |
| 1 | 6314822 | rs11590458 | 0.592 | 0.446 | 1.904 | 0.584 | -0.014 | -0.035 | -0.035 | 4.412 | 0.69 | 0.81 | 0.8 | 0.62 | 0.71 | 0.71 |
| 1 | 6317121 | rs11811690 | -1.653 | 0.493 | 1.975 | 0.622 | 0.026 | 0.043 | 0.043 | 1.981 | 0.05 | 0.36 | 0.04 | 0.03 | 0.01 | 0.04 |
| 1 | 6317983 | rs4908543 | 1.029 | 0.605 | 1.448 | 0.661 | -0.002 | 0.002 | 0.002 | 2.221 | 0.62 | 0.35 | 0.76 | 0.59 | 0.7 | 0.67 |
| 1 | 6322936 | rs4908874 | 0.834 | 0.452 | 0.482 | 0.563 | 0.002 | 0 | 0 | 3.191 | 0.6 | 0.36 | 0.76 | 0.63 | 0.69 | 0.72 |
| 1 | 6330015 | rs3789512 | -1.112 | 0.274 | 0.439 | 1.024 | 0.002 | 0.137 | 0.137 | 3.881 | 0.2 | 0.38 | 0.15 | 0.46 | 0.2 | 0.16 |
| 1 | 6336557 | rs3789508 | -1.345 | -0.989 | -0.432 | 0.238 | 0.073 | 0.09 | 0.09 | 5.322 | 0.35 | 0.82 | 0.22 | 0.16 | 0.18 | 0.2 |
| 1 | 6342226 | rs3827720 | -1.078 | -1.154 | 0.267 | 0.083 | -0.024 | -0.012 | -0.012 | 0 | 0.26 | 0.66 | 0.16 | 0.15 | 0.1 | 0.16 |
| 1 | 6345134 | rs3789504 | -1.669 | -1.246 | 1.16 | -0.009 | -0.037 | -0.037 | -0.037 | 0 | 0.27 | 0.58 | 0.17 | 0.16 | 0.15 | 0.17 |
| 1 | 6345818 | rs12404358 | -0.175 | -1.211 | 0.983 | -0.016 | -0.038 | 0.016 | 0.016 | 0 | 0.07 | 0.01 | 0.06 | 0.01 | 0.1 | 0.06 |
| 1 | 6352084 | rs3789500 | -0.553 | -1.371 | 1.579 | -0.038 | -0.024 | -0.004 | -0.004 | 0 | 0.18 | 0.21 | 0.06 | 0.14 | 0.04 | 0.09 |
| 1 | 6359034 | rs17361663 | -0.272 | -1.303 | 1.541 | -0.035 | -0.016 | 0.021 | 0.021 | 0 | 0.07 | 0.03 | 0.05 | 0.01 | 0.09 | 0.06 |
| 1 | 6361437 | rs17029372 | -2.613 | -1.632 | 1.511 | -0.194 | 0.096 | 0.107 | 0.107 | 5.636 | 0.37 | 0.83 | 0.2 | 0.15 | 0.19 | 0.23 |
| 1 | 6362035 | rs7547093 | -2.698 | -1.728 | 1.511 | -0.241 | 0.112 | 0.125 | 0.125 | 7.004 | 0.38 | 0.87 | 0.2 | 0.15 | 0.19 | 0.23 |
| 1 | 6366471 | rs7543514 | -1.172 | -1.768 | 2.128 | -0.194 | -0.121 | -0.051 | -0.051 | 0.215 | 0.22 | 0.47 | 0.08 | 0.14 | 0.05 | 0.1 |

| | | | | | | | | | | | | | | | | |
|---|---|---|---|---|---|---|---|---|---|---|---|---|---|---|---|---|
| 1 | 6367219 | rs11121611 | 1.043 | -2.119 | 1.104 | -0.135 | -0.053 | -0.137 | -0.137 | 0 | 0.18 | 0.01 | 0.17 | 0.41 | 0.21 | 0.45 |
| 1 | 6379607 | rs6696861 | -2.024 | -2.13 | 2.035 | 0.086 | 0.017 | 0.046 | 0.046 | 0 | 0.07 | 0.34 | 0.04 | 0.02 | 0.02 | 0.06 |
| 1 | 6387276 | rs6577570 | -2.132 | -2.134 | 2.034 | 0.094 | 0.013 | 0.048 | 0.048 | 0.055 | 0.07 | 0.34 | 0.04 | 0.02 | 0.02 | 0.06 |
| 1 | 6399780 | rs6680950 | -1.677 | -2.138 | 2.176 | 0.06 | -0.007 | -0.021 | -0.021 | 0 | 0.11 | 0.24 | 0.08 | 0.04 | 0.1 | 0.1 |
| 1 | 6408060 | rs742505 | -2.576 | -2.191 | 2.304 | 0.042 | -0.032 | -0.002 | -0.002 | 0.12 | 0.08 | 0.29 | 0.04 | 0.02 | 0.02 | 0.05 |
| 1 | 6411024 | rs3827718 | -0.169 | -2.276 | 2.341 | 0.025 | -0.029 | -0.01 | -0.01 | 0 | 0.15 | 0.2 | 0.04 | 0.12 | 0.03 | 0.06 |
| 1 | 6414084 | rs11121699 | -2.58 | -2.214 | 2.484 | 0.012 | 0.028 | 0.027 | 0.027 | 1.935 | 0.32 | 0.75 | 0.15 | 0.15 | 0.15 | 0.18 |
| 1 | 6418218 | rs7515832 | -0.505 | -2.154 | 2.337 | 0.049 | -0.05 | -0.023 | -0.023 | 0 | 0.17 | 0.29 | 0.05 | 0.11 | 0.03 | 0.06 |
| 1 | 6447574 | rs3789463 | -1.288 | -1.995 | 2.424 | 0.083 | 0.008 | -0.013 | -0.013 | 0 | 0.1 | 0.27 | 0.09 | 0.03 | 0.1 | 0.11 |
| 1 | 6456472 | rs7524775 | -0.296 | -1.981 | 2.259 | 0.091 | -0.003 | 0.002 | 0.002 | 0.006 | 0.07 | 0.06 | 0.05 | 0.01 | 0.09 | 0.06 |
| 1 | 6461530 | rs4582826 | -1.702 | -2.267 | 2.376 | 0.1 | -0.081 | -0.061 | -0.061 | 0 | 0.26 | 0.5 | 0.11 | 0.13 | 0.09 | 0.12 |
| 1 | 6465054 | rs17026985 | -0.08 | -2.308 | 2.361 | 0.089 | -0.028 | 0.021 | 0.021 | 0.033 | 0.09 | 0.2 | 0.03 | 0.13 | 0.02 | 0.03 |
| 1 | 6474008 | rs11485227 | -2.534 | -2.403 | 2.453 | 0.058 | -0.011 | -0.003 | -0.003 | 0 | 0.25 | 0.7 | 0.14 | 0.16 | 0.13 | 0.14 |
| 1 | 6474433 | rs3791184 | -0.036 | -2.404 | 2.476 | 0.062 | -0.028 | 0.022 | 0.022 | 0.01 | 0.09 | 0.19 | 0.04 | 0.13 | 0.02 | 0.03 |
| 1 | 6475686 | rs11364 | -2.398 | -2.382 | 2.299 | 0.067 | -0.011 | -0.004 | -0.004 | 0 | 0.25 | 0.7 | 0.14 | 0.16 | 0.13 | 0.14 |
| 1 | 6481842 | rs4908892 | -3.139 | -2.639 | 1.719 | 0.283 | 0.142 | -0.023 | -0.023 | 2.123 | 0.45 | 0.88 | 0.25 | 0.57 | 0.23 | 0.32 |
| 1 | 6481916 | rs12078257 | -0.346 | -2.255 | 1.849 | -0.338 | -0.017 | 0.008 | 0.008 | 0 | 0.11 | 0.14 | 0.05 | 0.15 | 0.03 | 0.05 |
| 1 | 6483583 | rs932402 | -0.312 | -2.218 | 1.856 | -0.409 | -0.016 | 0.007 | 0.007 | 0 | 0.11 | 0.13 | 0.05 | 0.14 | 0.03 | 0.06 |
| 1 | 6487031 | rs34064241 | -1.116 | -2.227 | 2.006 | -0.403 | -0.052 | 0.025 | 0.025 | 0.495 | 0.13 | 0.32 | 0.06 | 0.14 | 0.04 | 0.06 |
| 1 | 6494399 | rs2103673 | -2.129 | -2.227 | 1.882 | -0.46 | -0.035 | -0.131 | -0.131 | 0 | 0.22 | 0.5 | 0.13 | 0.01 | 0.14 | 0.18 |
| 1 | 6494743 | rs12074379 | -0.264 | -2.064 | 1.278 | -0.103 | -0.01 | 0.005 | 0.005 | 0 | 0.1 | 0.12 | 0.05 | 0.14 | 0.04 | 0.07 |
| 1 | 6520024 | rs3138159 | -1.214 | -1.643 | 1.418 | -0.134 | 0.01 | 0.019 | 0.019 | 6.086 | 0.11 | 0.07 | 0.06 | 0.02 | 0.08 | 0.09 |
| 1 | 6523430 | rs3138156 | -1.96 | -1.662 | 1.82 | -0.183 | -0.001 | -0.041 | -0.041 | 0 | 0.09 | 0.21 | 0.05 | 0.01 | 0.07 | 0.11 |
| 1 | 6524501 | rs11800462 | -2.125 | -1.739 | 1.82 | -0.198 | -0.007 | -0.037 | -0.037 | 0 | 0.09 | 0.21 | 0.05 | 0.01 | 0.07 | 0.11 |
| 1 | 6527333 | rs14708 | -3.088 | -2.008 | 2.201 | -0.327 | -0.019 | -0.07 | -0.07 | 0 | 0.28 | 0.68 | 0.15 | 0.01 | 0.15 | 0.18 |
| 1 | 6529443 | rs730600 | -3.146 | -2.05 | 2.224 | -0.339 | -0.024 | -0.078 | -0.078 | 0.397 | 0.28 | 0.68 | 0.14 | 0.01 | 0.15 | 0.21 |
| 1 | 6530435 | rs2986753 | -2.66 | -2.004 | 2.513 | -0.317 | -0.022 | -0.075 | -0.075 | 1.114 | 0.23 | 0.36 | 0.12 | 0.01 | 0.12 | 0.18 |
| 1 | 6534781 | rs2986751 | -1.972 | -1.819 | 2.714 | -0.159 | -0.068 | -0.115 | -0.115 | 0 | 0.16 | 0.39 | 0.08 | 0 | 0.07 | 0.07 |
| 1 | 6542330 | rs1002966 | -2.149 | -1.617 | 2.592 | 0.593 | -0.065 | 0.087 | 0.087 | 2.292 | 0.16 | 0.35 | 0.08 | 0.28 | 0.06 | 0.19 |
| 1 | 6548056 | rs2986739 | -2.659 | -1.459 | 2.45 | 0.563 | -0.065 | 0.063 | 0.063 | 2.449 | 0.22 | 0.52 | 0.15 | 0.25 | 0.12 | 0.24 |
| 1 | 6550376 | rs2986742 | -2.934 | -1.513 | 1.702 | 0.522 | 0.036 | -0.01 | -0.01 | 0.378 | 0.26 | 0.79 | 0.17 | 0.25 | 0.13 | 0.23 |
| 1 | 6550505 | rs11806429 | -1.932 | -1.495 | 1.846 | 0.54 | -0.031 | 0.04 | 0.04 | 2.313 | 0.18 | 0.3 | 0.11 | 0.23 | 0.11 | 0.17 |
| 1 | 6552499 | rs9434795 | -2.318 | -1.414 | 0.823 | 0.612 | 0.027 | -0.009 | -0.009 | 3.803 | 0.26 | 0.78 | 0.18 | 0.28 | 0.13 | 0.24 |
| 1 | 6560363 | rs4908898 | -1.802 | -1.068 | 1.021 | 0.807 | -0.077 | 0.085 | 0.085 | 5.125 | 0.18 | 0.49 | 0.11 | 0.26 | 0.07 | 0.18 |
| 1 | 6568387 | rs4908554 | 0.809 | 0.027 | 0.267 | 0.759 | -0.015 | 0.008 | 0.008 | 1.634 | 0.6 | 0.29 | 0.71 | 0.37 | 0.58 | 0.57 |
| 1 | 6572175 | rs3935435 | -1.407 | 0.326 | 1.045 | 0.826 | 0.023 | 0.126 | 0.126 | 0.144 | 0.19 | 0.34 | 0.18 | 0.6 | 0.25 | 0.22 |
| 1 | 6577765 | rs4243829 | 0.805 | -0.106 | -0.211 | -0.425 | -0.061 | 0.138 | 0.138 | 1.431 | 0.24 | 0.04 | 0.38 | 0 | 0.35 | 0.12 |
| 1 | 6577871 | rs4243830 | 0.615 | -0.15 | 1.147 | 0.648 | -0.01 | 0.064 | 0.064 | 0 | 0.88 | 0.67 | 0.88 | 0.81 | 0.89 | 0.87 |
| 1 | 6579607 | rs1556035 | 1.11 | -0.206 | 1.883 | 0.604 | -0.014 | 0.079 | 0.079 | 0.087 | 0.86 | 0.45 | 0.87 | 0.81 | 0.88 | 0.86 |
| 1 | 6579843 | rs1010584 | 1.775 | -0.347 | 1.798 | 0.603 | -0.066 | 0.013 | 0.013 | 0.871 | 0.79 | 0.45 | 0.85 | 0.79 | 0.86 | 0.85 |
| 1 | 6580397 | rs12046229 | -0.445 | -0.291 | 1.749 | 0.626 | -0.003 | 0.006 | 0.006 | 0 | 0.11 | 0.14 | 0.11 | 0.16 | 0.11 | 0.14 |
| 1 | 6584420 | rs11809957 | -0.974 | -0.325 | 1.743 | 0.622 | -0.01 | 0.059 | 0.059 | 0 | 0.13 | 0.38 | 0.12 | 0.16 | 0.12 | 0.18 |
| 1 | 6585539 | rs1064835 | 1.819 | -0.456 | 1.765 | 0.569 | -0.076 | 0.009 | 0.009 | 0.47 | 0.78 | 0.48 | 0.85 | 0.79 | 0.86 | 0.81 |
| 1 | 6585804 | rs13513 | 1.621 | -0.433 | 1.749 | 0.581 | -0.068 | 0.017 | 0.017 | 0.186 | 0.79 | 0.5 | 0.86 | 0.79 | 0.86 | 0.81 |
| 1 | 6599041 | rs4908909 | 0.765 | -0.407 | 1.736 | 0.658 | -0.014 | 0.046 | 0.046 | 0 | 0.87 | 0.62 | 0.87 | 0.84 | 0.88 | 0.83 |
| 1 | 6614230 | rs10864625 | -1.594 | -0.496 | 1.639 | 0.614 | -0.022 | -0.002 | -0.002 | 0 | 0.2 | 0.28 | 0.13 | 0.18 | 0.13 | 0.17 |
| 1 | 6614535 | rs4908923 | 1.777 | -0.499 | 1.751 | 0.618 | -0.069 | 0 | 0 | 0.542 | 0.77 | 0.43 | 0.85 | 0.79 | 0.86 | 0.82 |
| 1 | 6620545 | rs12565181 | -1.697 | -0.459 | 1.142 | 0.65 | -0.031 | 0 | 0 | 0.016 | 0.2 | 0.34 | 0.13 | 0.18 | 0.13 | 0.17 |
| 1 | 6623499 | rs4908563 | 1.334 | -0.216 | 0.488 | 0.484 | -0.033 | 0.006 | 0.006 | 4.06 | 0.52 | 0.1 | 0.65 | 0.44 | 0.55 | 0.55 |
| 1 | 6631319 | rs11122096 | 0.253 | -0.463 | 1.23 | 0.412 | -0.024 | 0.017 | 0.017 | 1.708 | 0.92 | 0.67 | 0.96 | 0.94 | 0.98 | 0.98 |
| 1 | 6631431 | rs11587438 | 1.693 | -0.656 | 1.252 | 0.514 | -0.064 | -0.005 | -0.005 | 0 | 0.78 | 0.51 | 0.84 | 0.81 | 0.86 | 0.8 |
| 1 | 6632951 | rs12041318 | -1.014 | -0.207 | 0.75 | 0.572 | 0.004 | -0.002 | -0.002 | 0.033 | 0.13 | 0.1 | 0.11 | 0.13 | 0.12 | 0.18 |
| 1 | 6638064 | rs11122100 | -1.226 | -0.243 | 0.951 | 0.55 | -0.02 | 0.002 | 0.002 | 0 | 0.15 | 0.31 | 0.13 | 0.13 | 0.13 | 0.2 |
| 1 | 6639847 | rs4908932 | -0.171 | -0.085 | 0.954 | -0.126 | -0.006 | 0.009 | 0.009 | 0 | 0.13 | 0.09 | 0.27 | 0 | 0.18 | 0.07 |
| 1 | 6644723 | rs731024 | 1.178 | -0.003 | 0.313 | -0.146 | 0.021 | -0.047 | -0.047 | 0.373 | 0.43 | 0.05 | 0.31 | 0.38 | 0.32 | 0.46 |
| 1 | 6651093 | rs201171 | -0.097 | 1.211 | 0.235 | 0.181 | -0.03 | 0.031 | 0.031 | 0 | 0.05 | 0.01 | 0.07 | 0.41 | 0.14 | 0.12 |
| 1 | 6651728 | rs1556036 | 0.13 | 1.022 | 0.77 | -0.113 | -0.011 | 0.019 | 0.019 | 0 | 0.29 | 0.21 | 0.25 | 0.16 | 0.34 | 0.31 |
| 1 | 6657240 | rs7521134 | 0.126 | 1.001 | 0.885 | -0.099 | -0.023 | 0.04 | 0.04 | 0 | 0.31 | 0.17 | 0.25 | 0.16 | 0.35 | 0.32 |
| 1 | 6659872 | rs6678681 | 1.252 | 0.974 | 1.635 | -0.115 | 0.048 | 0.07 | 0.07 | 0 | 0.79 | 0.67 | 0.61 | 0.58 | 0.69 | 0.81 |
| 1 | 6661732 | rs12028517 | 1.826 | 0.93 | 1.626 | -0.127 | 0.056 | -0.007 | -0.007 | 0.386 | 0.43 | 0.13 | 0.31 | 0.38 | 0.32 | 0.49 |
| 1 | 6662913 | rs6674407 | 1.82 | 0.908 | 1.616 | -0.137 | 0.057 | -0.007 | -0.007 | 6.11 | 0.43 | 0.13 | 0.3 | 0.38 | 0.32 | 0.49 |
| 1 | 6666848 | rs481771 | 0.543 | 1.168 | 1.86 | -0.099 | -0.002 | 0.069 | 0.069 | 0 | 0.94 | 0.96 | 0.96 | 0.59 | 0.92 | 0.88 |
| 1 | 6672729 | rs11583755 | -0.183 | 1.124 | 1.93 | -0.136 | -0.006 | 0.011 | 0.011 | 0 | 0.33 | 0.28 | 0.27 | 0.2 | 0.37 | 0.32 |
| 1 | 6677064 | rs10779795 | 1.728 | 1.084 | 1.944 | -0.128 | 0.06 | 0.018 | 0.018 | 0.679 | 0.45 | 0.31 | 0.33 | 0.39 | 0.33 | 0.49 |
| 1 | 6679848 | rs2072948 | 0.214 | 1.075 | 1.976 | -0.14 | -0.057 | 0.037 | 0.037 | 0 | 0.3 | 0.09 | 0.24 | 0.15 | 0.35 | 0.3 |
| 1 | 6681275 | rs12034227 | 1.707 | 1.035 | 1.907 | -0.127 | 0.06 | 0.018 | 0.018 | 0.631 | 0.45 | 0.31 | 0.33 | 0.38 | 0.33 | 0.49 |
| 1 | 6681744 | rs3747994 | 1.2 | 0.971 | 2.08 | -0.112 | 0.057 | 0.077 | 0.077 | 0 | 0.81 | 0.74 | 0.62 | 0.58 | 0.7 | 0.81 |
| 1 | 6683240 | rs14057 | 1.676 | 1.204 | 2.011 | -0.1 | 0.057 | 0.026 | 0.026 | 0.758 | 0.46 | 0.3 | 0.34 | 0.38 | 0.34 | 0.51 |
| 1 | 6685851 | rs12127219 | 1.289 | 1.164 | 2.023 | -0.171 | 0.058 | 0.074 | 0.074 | 0 | 0.8 | 0.71 | 0.61 | 0.58 | 0.69 | 0.8 |

| | | | | | | | | | | | | | | | | |
|---|---|---|---|---|---|---|---|---|---|---|---|---|---|---|---|---|
| 1 | 6688358 | rs11122109 | -0.216 | 1.126 | 1.84 | -0.164 | -0.001 | -0.001 | -0.001 | 2.957 | 0.33 | 0.32 | 0.26 | 0.19 | 0.36 | 0.3 |
| 1 | 6689824 | rs12079872 | 1.675 | 1.147 | 1.656 | -0.161 | 0.057 | 0.003 | 0.003 | 0.469 | 0.45 | 0.19 | 0.33 | 0.39 | 0.33 | 0.49 |
| 1 | 6694207 | rs12727844 | 1.66 | 1.15 | 1.869 | -0.159 | 0.054 | 0.013 | 0.013 | 0.405 | 0.45 | 0.31 | 0.34 | 0.39 | 0.33 | 0.49 |
| 1 | 6695123 | rs1043681 | 1.177 | 1.269 | 1.735 | -0.096 | 0.059 | 0.074 | 0.074 | 0 | 0.81 | 0.73 | 0.62 | 0.58 | 0.7 | 0.79 |
| 1 | 6695244 | rs10523 | 1.294 | 1.27 | 1.731 | -0.029 | 0.059 | 0.066 | 0.066 | 0 | 0.8 | 0.71 | 0.61 | 0.63 | 0.7 | 0.8 |
| 1 | 6701978 | rs200448 | 0.763 | 1.071 | 1.803 | -0.118 | -0.004 | 0 | 0 | 0.35 | 0.58 | 0.4 | 0.64 | 0.39 | 0.56 | 0.56 |
| 1 | 6702151 | rs200447 | -0.473 | 1.058 | 1.822 | -0.17 | 0.013 | -0.041 | -0.041 | 0.238 | 0.13 | 0.13 | 0.3 | 0 | 0.22 | 0.07 |
| 1 | 6704635 | rs200458 | 0.852 | 1.005 | 1.731 | -0.137 | 0.009 | 0.155 | 0.155 | 0 | 0.92 | 0.87 | 0.96 | 0.57 | 0.92 | 0.87 |
| 1 | 6713114 | rs2235564 | -0.082 | 1.262 | 1.423 | -0.474 | 0.006 | -0.015 | -0.015 | 0 | 0.33 | 0.41 | 0.32 | 0.18 | 0.37 | 0.31 |
| 1 | 6723806 | rs6659873 | -0.123 | 1.286 | 1.368 | -0.477 | 0.011 | -0.025 | -0.025 | 0 | 0.33 | 0.44 | 0.33 | 0.17 | 0.39 | 0.31 |
| 1 | 6725427 | rs11122115 | -0.211 | 1.268 | 0.985 | -0.63 | 0.023 | -0.053 | -0.053 | 0.615 | 0.13 | 0.17 | 0.31 | 0 | 0.22 | 0.07 |
| 1 | 6732424 | rs201151 | 0.523 | 1.302 | 0.946 | -0.539 | -0.008 | 0 | 0 | 0.361 | 0.58 | 0.39 | 0.63 | 0.4 | 0.54 | 0.56 |
| 1 | 6768856 | rs11122119 | 1.496 | 1.127 | 0.75 | -0.624 | 0.068 | -0.046 | -0.046 | 0.011 | 0.44 | 0.15 | 0.33 | 0.43 | 0.3 | 0.49 |
| 1 | 6771607 | rs277672 | 0.373 | 1.145 | 0.903 | -0.345 | 0.021 | 0.018 | 0.018 | 0 | 0.62 | 0.47 | 0.65 | 0.43 | 0.53 | 0.56 |
| 1 | 6783548 | rs277679 | 0.459 | 1.101 | 0.568 | -0.328 | -0.004 | -0.004 | -0.004 | 0 | 0.6 | 0.39 | 0.65 | 0.43 | 0.54 | 0.56 |
| 1 | 6790946 | rs11122122 | 0.134 | 1.062 | 0.022 | -0.269 | -0.006 | -0.011 | -0.011 | 0 | 0.65 | 0.71 | 0.6 | 0.54 | 0.57 | 0.6 |
| 1 | 6795230 | rs447558 | 0.364 | 1.208 | -0.474 | -0.644 | -0.039 | 0.074 | 0.074 | 0 | 0.65 | 0.76 | 0.39 | 0.89 | 0.55 | 0.72 |
| 1 | 6797684 | rs449250 | 0.829 | 1.304 | -0.401 | -0.689 | -0.022 | 0.077 | 0.077 | 0 | 0.58 | 0.66 | 0.35 | 0.87 | 0.48 | 0.64 |
| 1 | 6798102 | rs2134318 | 0.871 | 1.168 | -0.087 | -0.669 | 0.009 | -0.049 | -0.049 | 0 | 0.57 | 0.46 | 0.35 | 0.87 | 0.53 | 0.67 |
| 1 | 6812518 | rs11122135 | -0.633 | 1.435 | -0.135 | -0.241 | 0 | -0.009 | -0.009 | 0 | 0.08 | 0.07 | 0.04 | 0.27 | 0.11 | 0.15 |
| 1 | 6814758 | rs277665 | 1.169 | 1.349 | -0.126 | -0.31 | -0.001 | -0.001 | -0.001 | 0 | 0.53 | 0.46 | 0.32 | 0.6 | 0.44 | 0.54 |
| 1 | 6822235 | rs656205 | 0.452 | 1.046 | -0.287 | -0.48 | 0.003 | 0.002 | 0.002 | 0.083 | 0.54 | 0.58 | 0.33 | 0.89 | 0.42 | 0.7 |
| 1 | 6826467 | rs378930 | 1.132 | 0.928 | -0.131 | -0.313 | 0.011 | 0.006 | 0.006 | 0.065 | 0.45 | 0.39 | 0.27 | 0.63 | 0.31 | 0.55 |
| 1 | 6828290 | rs11122141 | 0.098 | 0.965 | -0.107 | -0.321 | 0.001 | 0.03 | 0.03 | 0 | 0.07 | 0.09 | 0.05 | 0.27 | 0.12 | 0.14 |
| 1 | 6828608 | rs430962 | 0.397 | 1.023 | -0.097 | -0.366 | 0.004 | 0.002 | 0.002 | 0.062 | 0.54 | 0.58 | 0.33 | 0.89 | 0.42 | 0.69 |
| 1 | 6832052 | rs391674 | 0.373 | 1.013 | -0.042 | -0.37 | 0.007 | 0.019 | 0.019 | 0.08 | 0.54 | 0.59 | 0.33 | 0.89 | 0.42 | 0.7 |
| 1 | 6835761 | rs11122144 | -1.811 | 1.026 | 0.074 | -0.248 | 0.014 | 0.061 | 0.061 | 0 | 0.12 | 0.22 | 0.09 | 0.29 | 0.14 | 0.19 |
| 1 | 6846763 | rs10864639 | 0.172 | 0.953 | 0.291 | -0.156 | 0 | 0.037 | 0.037 | 0 | 0.07 | 0.08 | 0.05 | 0.27 | 0.11 | 0.13 |
| 1 | 6863342 | rs17029864 | -1.447 | 1.087 | 0.265 | -0.262 | 0.013 | 0.047 | 0.047 | 0 | 0.17 | 0.32 | 0.12 | 0.3 | 0.19 | 0.2 |
| 1 | 6870942 | rs589178 | -1.736 | 0.999 | 0.346 | -0.334 | 0.038 | 0.001 | 0.001 | 0 | 0.57 | 0.69 | 0.76 | 0.38 | 0.72 | 0.49 |
| 1 | 6873932 | rs659459 | 0.788 | 1.774 | -0.025 | -0.577 | -0.065 | 0.094 | 0.094 | 0 | 0.24 | 0.05 | 0.3 | 0.06 | 0.3 | 0.24 |
| 1 | 6877562 | rs629196 | -2.054 | 1.625 | 0.33 | -0.535 | -0.011 | 0 | 0 | 0 | 0.57 | 0.55 | 0.76 | 0.42 | 0.72 | 0.49 |
| 1 | 6882986 | rs623974 | 0.768 | 2.098 | -0.021 | -0.582 | -0.039 | 0 | 0 | 0 | 0.07 | 0 | 0.09 | 0.04 | 0.14 | 0.11 |
| 1 | 6903599 | rs12058103 | -1.636 | 2.196 | 0.129 | -0.536 | -0.006 | -0.01 | -0.01 | 0 | 0.25 | 0.23 | 0.38 | 0.35 | 0.3 | 0.2 |
| 1 | 6916210 | rs11582310 | 2.256 | 2.407 | 0.149 | -0.584 | 0 | -0.005 | -0.005 | 0 | 0.05 | 0.03 | 0.07 | 0 | 0.11 | 0.03 |
| 1 | 6929259 | rs17029968 | 0.818 | 1.947 | 0.221 | -0.489 | 0 | -0.009 | -0.009 | 0.067 | 0.08 | 0.1 | 0.06 | 0.03 | 0.06 | 0.04 |
| 1 | 6929760 | rs6688732 | -1.786 | 1.981 | 0.279 | -0.488 | -0.007 | -0.009 | -0.009 | 0 | 0.26 | 0.25 | 0.39 | 0.35 | 0.32 | 0.23 |
| 1 | 6936272 | rs7546903 | -2.723 | 1.218 | -0.101 | -0.976 | -0.014 | 0.012 | 0.012 | 1.655 | 0.67 | 0.61 | 0.78 | 0.41 | 0.74 | 0.57 |
| 1 | 6936375 | rs2063313 | 0.938 | 1.36 | -0.464 | -0.811 | -0.04 | -0.01 | -0.01 | 1.125 | 0.09 | 0.02 | 0.28 | 0.05 | 0.13 | 0.03 |
| 1 | 6944804 | rs950493 | -0.783 | 1.259 | -0.288 | -0.807 | 0.005 | 0.046 | 0.046 | 0 | 0.16 | 0.28 | 0.11 | 0.29 | 0.17 | 0.16 |
| 1 | 6968412 | rs6577393 | -1.18 | 1.254 | -0.719 | -0.963 | 0.036 | 0.08 | 0.08 | 1.168 | 0.49 | 0.22 | 0.45 | 0.33 | 0.57 | 0.49 |
| 1 | 6968465 | rs12133292 | 1.114 | 1.349 | -0.742 | -0.75 | -0.025 | -0.014 | -0.014 | 0.274 | 0.06 | 0 | 0.04 | 0.26 | 0.1 | 0.12 |
| 1 | 6972393 | rs11586867 | -1.078 | 1.159 | -0.72 | -0.91 | 0.031 | 0.047 | 0.047 | 0 | 0.46 | 0.26 | 0.47 | 0.36 | 0.6 | 0.44 |
| 1 | 6976226 | rs4908427 | -0.032 | 1.252 | -0.604 | -0.816 | 0.006 | -0.001 | -0.001 | 0 | 0.07 | 0.12 | 0.28 | 0.05 | 0.11 | 0.03 |
| 1 | 6976368 | rs4908428 | 0.574 | 1.253 | -0.694 | -0.82 | -0.023 | -0.017 | -0.017 | 0.452 | 0.05 | 0.01 | 0.2 | 0.04 | 0.07 | 0.03 |
| 1 | 6977200 | rs1892261 | -1.043 | 0.994 | -0.479 | -0.744 | 0.027 | 0.021 | 0.021 | 0 | 0.46 | 0.29 | 0.46 | 0.38 | 0.63 | 0.46 |
| 1 | 6978623 | rs12122806 | 0.163 | 1.13 | -0.507 | -0.771 | -0.001 | 0.019 | 0.019 | 0 | 0.07 | 0.08 | 0.05 | 0.23 | 0.13 | 0.13 |
| 1 | 6985069 | rs867389 | -0.408 | 1.123 | -0.625 | -0.655 | -0.059 | 0.105 | 0.105 | 1.982 | 0.39 | 0.13 | 0.36 | 0.16 | 0.45 | 0.34 |
| 1 | 6986109 | rs4908574 | -1.982 | 0.906 | -0.332 | -0.693 | 0.028 | -0.014 | -0.014 | 0 | 0.57 | 0.62 | 0.74 | 0.48 | 0.71 | 0.5 |
| 1 | 6991925 | rs6577395 | -1.012 | 1.047 | -0.485 | -0.735 | 0.02 | 0.103 | 0.103 | 3.355 | 0.47 | 0.25 | 0.44 | 0.2 | 0.53 | 0.42 |
| 1 | 6995218 | rs12043429 | 0.603 | 1.623 | -0.284 | -0.375 | -0.021 | -0.017 | -0.017 | 0 | 0.09 | 0.03 | 0.1 | 0.13 | 0.17 | 0.1 |
| 1 | 7007311 | rs6688108 | -1.648 | 1.25 | -0.105 | -0.272 | 0.009 | 0.002 | 0.002 | 0 | 0.49 | 0.44 | 0.48 | 0.42 | 0.64 | 0.47 |
| 1 | 7011805 | rs10864251 | -1.454 | 1.282 | -0.196 | -0.257 | 0.028 | 0.004 | 0.004 | 0 | 0.47 | 0.35 | 0.47 | 0.42 | 0.65 | 0.47 |
| 1 | 7014330 | rs11120796 | -0.094 | 1.505 | -0.416 | -0.134 | -0.048 | 0.024 | 0.024 | 0.014 | 0.29 | 0.14 | 0.34 | 0.19 | 0.42 | 0.25 |
| 1 | 7015633 | rs12726405 | 0.107 | 1.351 | -0.245 | 0.086 | 0.01 | 0.041 | 0.041 | 0 | 0.92 | 0.87 | 0.95 | 0.83 | 0.91 | 0.88 |
| 1 | 7017985 | rs4908581 | -0.415 | 1.359 | -0.16 | 0.049 | -0.025 | 0.022 | 0.022 | 0 | 0.33 | 0.23 | 0.48 | 0.21 | 0.47 | 0.3 |
| 1 | 7024222 | rs11120799 | 1.316 | 1.161 | -0.213 | -0.069 | 0 | -0.017 | -0.017 | 0 | 0.07 | 0.07 | 0.05 | 0 | 0.06 | 0.06 |
| 1 | 7029262 | rs17030082 | 0.733 | 1.209 | -0.232 | -0.035 | -0.001 | -0.001 | -0.001 | 0 | 0.58 | 0.63 | 0.47 | 0.62 | 0.42 | 0.58 |
| 1 | 7032168 | rs6673655 | 1.777 | 1.004 | -0.302 | -0.081 | 0.038 | -0.019 | -0.019 | 0 | 0.42 | 0.33 | 0.36 | 0.57 | 0.3 | 0.49 |
| 1 | 7035830 | rs6679134 | 0.914 | 1.437 | -0.119 | 0.219 | 0 | 0.001 | 0.001 | 0 | 0.58 | 0.56 | 0.47 | 0.59 | 0.43 | 0.58 |
| 1 | 7036802 | rs4518923 | -1.507 | 1.851 | -0.302 | 0.584 | -0.076 | -0.082 | -0.082 | 0 | 0.19 | 0.06 | 0.17 | 0.31 | 0.33 | 0.27 |
| 1 | 7041175 | rs6698901 | -1.883 | 1.444 | -0.199 | 0.602 | -0.008 | -0.006 | -0.006 | 0 | 0.41 | 0.61 | 0.45 | 0.42 | 0.48 | 0.49 |
| 1 | 7042757 | rs6703004 | -1.746 | 1.503 | -0.125 | 0.655 | 0.023 | 0.067 | 0.067 | 0 | 0.25 | 0.57 | 0.35 | 0.4 | 0.32 | 0.39 |
| 1 | 7043270 | rs11120807 | 0.082 | 1.708 | -0.009 | 0.664 | -0.007 | 0 | 0 | 0.38 | 0.09 | 0.07 | 0.09 | 0.1 | 0.17 | 0.16 |
| 1 | 7048779 | rs1317017 | -1.073 | 1.674 | 0.233 | 0.769 | 0.019 | 0.032 | 0.032 | 0 | 0.17 | 0.22 | 0.28 | 0.28 | 0.28 | 0.29 |
| 1 | 7052566 | rs6681529 | -3.184 | 1.353 | 0.099 | 0.551 | 0 | -0.046 | -0.046 | 0.591 | 0.53 | 0.9 | 0.52 | 0.44 | 0.57 | 0.53 |
| 1 | 7054607 | rs12128446 | 0.686 | 1.478 | 0.166 | 0.648 | -0.019 | -0.01 | -0.01 | 0.001 | 0.07 | 0.03 | 0.19 | 0.19 | 0.11 | 0.13 |
| 1 | 7055024 | rs12043302 | 0.531 | 1.489 | 0.527 | 0.596 | 0.082 | 0.001 | 0.001 | 0.004 | 0.09 | 0.27 | 0.11 | 0.09 | 0.16 | 0.06 |
| 1 | 7055326 | rs10128022 | -1.147 | 1.426 | 0.409 | 0.589 | 0.092 | -0.022 | -0.022 | 0.028 | 0.21 | 0.53 | 0.22 | 0.19 | 0.34 | 0.24 |

| | | | | | | | | | | | | | | | | |
|---|---|---|---|---|---|---|---|---|---|---|---|---|---|---|---|---|
| 1 | 7058450 | rs17030127 | -0.377 | 1.156 | 0.126 | 0.626 | 0.012 | -0.034 | -0.034 | 0.427 | 0.18 | 0.08 | 0.22 | 0.22 | 0.14 | 0.25 |
| 1 | 7068672 | rs2275909 | -1.643 | 1.031 | 0.239 | 0.511 | -0.001 | -0.005 | -0.005 | 0 | 0.34 | 0.68 | 0.36 | 0.33 | 0.33 | 0.34 |
| 1 | 7078364 | rs7543753 | -1.56 | 0.903 | 0.442 | 0.34 | -0.052 | -0.09 | -0.09 | 0.36 | 0.27 | 0.41 | 0.14 | 0.06 | 0.15 | 0.18 |
| 1 | 7078434 | rs6690584 | -1.27 | 0.973 | 0.605 | 0.445 | 0.067 | 0.091 | 0.091 | 1.617 | 0.23 | 0.49 | 0.34 | 0.33 | 0.3 | 0.26 |
| 1 | 7083719 | rs17030152 | -1.596 | 1.047 | 0.446 | 0.359 | 0.004 | 0.006 | 0.006 | 0 | 0.31 | 0.28 | 0.19 | 0.12 | 0.26 | 0.28 |
| 1 | 7087739 | rs11120820 | 0.189 | 1.089 | 0.38 | 0.381 | -0.016 | 0 | 0 | 0 | 0.09 | 0.07 | 0.1 | 0.1 | 0.17 | 0.16 |
| 1 | 7088261 | rs11588846 | -2.101 | 1.081 | 0.48 | 0.369 | -0.006 | -0.033 | -0.033 | 0 | 0.37 | 0.44 | 0.23 | 0.16 | 0.31 | 0.34 |
| 1 | 7088855 | rs4243823 | -1.861 | 1.123 | 0.344 | 0.444 | -0.02 | -0.009 | -0.009 | 0 | 0.4 | 0.67 | 0.43 | 0.39 | 0.46 | 0.45 |
| 1 | 7091586 | rs12048981 | -1.658 | 1.161 | 0.335 | 0.518 | 0.023 | 0.034 | 0.034 | 0.694 | 0.28 | 0.42 | 0.36 | 0.33 | 0.33 | 0.22 |
| 1 | 7093213 | rs12738895 | -2.373 | 1.462 | 0.174 | 0.673 | 0.047 | -0.024 | -0.024 | 0.106 | 0.48 | 0.87 | 0.52 | 0.45 | 0.57 | 0.51 |
| 1 | 7096700 | rs7553537 | -0.7 | 1.954 | 0.554 | 1.323 | 0.098 | 0.234 | 0.234 | 0 | 0.05 | 0.29 | 0.05 | 0.22 | 0.06 | 0.06 |
| 1 | 7098807 | rs12728534 | -0.021 | 2.16 | 0.456 | 1.197 | -0.048 | 0.001 | 0.001 | 0 | 0.11 | 0.03 | 0.12 | 0.1 | 0.22 | 0.17 |
| 1 | 7099660 | rs6686587 | -1.706 | 2.114 | 0.556 | 1.193 | 0.011 | 0.007 | 0.007 | 1.063 | 0.39 | 0.48 | 0.44 | 0.39 | 0.52 | 0.46 |
| 1 | 7105079 | rs4074964 | -0.401 | 1.895 | 0.467 | 1.202 | 0.01 | 0.026 | 0.026 | 1.406 | 0.25 | 0.36 | 0.31 | 0.29 | 0.28 | 0.3 |
| 1 | 7106963 | rs6577409 | -2.754 | 1.869 | -0.1 | 1.126 | 0.093 | 0.075 | 0.075 | 2.015 | 0.54 | 0.88 | 0.56 | 0.64 | 0.66 | 0.56 |
| 1 | 7110334 | rs4908590 | -1.873 | 2.102 | 0.071 | 1.191 | 0.029 | -0.01 | -0.01 | 0 | 0.47 | 0.67 | 0.52 | 0.45 | 0.57 | 0.48 |
| 1 | 7110510 | rs4908591 | -0.965 | 2.18 | 0.262 | 0.974 | -0.003 | 0.002 | 0.002 | 0 | 0.17 | 0.12 | 0.14 | 0.1 | 0.24 | 0.2 |
| 1 | 7112390 | rs6577410 | -2.279 | 1.816 | 0.727 | 0.938 | 0.051 | 0.041 | 0.041 | 0.283 | 0.52 | 0.67 | 0.56 | 0.62 | 0.67 | 0.55 |
| 1 | 7114033 | rs6700922 | -0.433 | 2.042 | 0.669 | 1.33 | -0.002 | 0 | 0 | 0.426 | 0.21 | 0.2 | 0.26 | 0.52 | 0.21 | 0.28 |
| 1 | 7114641 | rs12133505 | 0.534 | 2.012 | 0.279 | 1.144 | -0.022 | 0.005 | 0.005 | 0.343 | 0.15 | 0.02 | 0.09 | 0.1 | 0.13 | 0.21 |
| 1 | 7119202 | rs4908595 | -2.357 | 1.769 | 0.684 | 0.956 | 0.032 | 0.03 | 0.03 | 0.435 | 0.53 | 0.64 | 0.56 | 0.66 | 0.67 | 0.56 |
| 1 | 7119489 | rs10779671 | -2.511 | 1.758 | 0.638 | 0.951 | 0.099 | 0.085 | 0.085 | 0.789 | 0.54 | 0.89 | 0.59 | 0.66 | 0.67 | 0.56 |
| 1 | 7120915 | rs4436414 | -1.345 | 2.08 | 0.754 | 0.957 | 0 | 0.001 | 0.001 | 1.534 | 0.34 | 0.64 | 0.4 | 0.51 | 0.38 | 0.36 |
| 1 | 7126671 | rs7522545 | -2.02 | 2.116 | 0.487 | 0.799 | 0.009 | 0.009 | 0.009 | 1.513 | 0.47 | 0.62 | 0.51 | 0.59 | 0.61 | 0.54 |
| 1 | 7128056 | rs7536178 | -0.606 | 2.089 | -0.06 | 1.004 | -0.007 | -0.009 | -0.009 | 1.343 | 0.26 | 0.23 | 0.32 | 0.33 | 0.29 | 0.3 |
| 1 | 7130084 | rs4908597 | 0.067 | 1.889 | -0.79 | 0.486 | -0.023 | -0.009 | -0.009 | 0 | 0.26 | 0.15 | 0.29 | 0.26 | 0.33 | 0.37 |
| 1 | 7131192 | rs11120831 | -0.934 | 1.738 | -0.434 | 0.486 | 0.004 | 0.041 | 0.041 | 0 | 0.31 | 0.54 | 0.33 | 0.39 | 0.34 | 0.3 |
| 1 | 7135268 | rs10864271 | -0.541 | 1.369 | -1.021 | 0.282 | -0.044 | 0.025 | 0.025 | 1.815 | 0.47 | 0.17 | 0.49 | 0.35 | 0.5 | 0.57 |
| 1 | 7137637 | rs9434824 | -1.619 | 1.266 | -0.879 | 0.229 | -0.002 | -0.002 | -0.002 | 0.355 | 0.67 | 0.69 | 0.67 | 0.69 | 0.74 | 0.7 |
| 1 | 7141122 | rs9434826 | 2.804 | 0.929 | -1.175 | -0.079 | 0.003 | -0.002 | -0.002 | 0 | 0.2 | 0.14 | 0.13 | 0.26 | 0.14 | 0.13 |
| 1 | 7141327 | rs17345754 | -0.466 | 1.11 | -1.215 | -0.193 | -0.037 | -0.001 | -0.001 | 0.201 | 0.12 | 0.04 | 0.24 | 0.12 | 0.2 | 0.27 |
| 1 | 7143344 | rs2187901 | 1.043 | 0.444 | -0.931 | 0.027 | -0.001 | 0.005 | 0.005 | 0 | 0.58 | 0.32 | 0.6 | 0.42 | 0.58 | 0.65 |
| 1 | 7143807 | rs1009113 | -1.264 | 0.523 | -0.912 | 0.115 | -0.03 | -0.056 | -0.056 | 0 | 0.91 | 0.99 | 0.82 | 0.77 | 0.88 | 0.83 |
| 1 | 7144591 | rs9434459 | 2.004 | 0.509 | -0.888 | 0.106 | -0.002 | 0.001 | 0.001 | 4.658 | 0.24 | 0.22 | 0.29 | 0.31 | 0.21 | 0.26 |
| 1 | 7148455 | rs7544611 | 1.147 | 0.538 | -0.851 | 0.13 | 0.035 | 0.068 | 0.068 | 0 | 0.18 | 0.29 | 0.16 | 0.33 | 0.23 | 0.13 |
| 1 | 7150527 | rs2071917 | -0.14 | 0.516 | -0.889 | 0.08 | -0.027 | 0.035 | 0.035 | 1.956 | 0.53 | 0.81 | 0.51 | 0.65 | 0.5 | 0.43 |
| 1 | 7150577 | rs2071918 | 2.6 | 0.334 | -0.907 | -0.024 | -0.006 | 0.002 | 0.002 | 4.764 | 0.2 | 0.21 | 0.13 | 0.26 | 0.14 | 0.13 |
| 1 | 7152038 | rs10864275 | 0.795 | 0.427 | -1.255 | -0.045 | -0.006 | -0.019 | -0.019 | 0 | 0.1 | 0.13 | 0.04 | 0.02 | 0.05 | 0.05 |
| 1 | 7152069 | rs10864276 | 1.143 | 0.501 | -1.279 | -0.008 | 0 | -0.005 | -0.005 | 0 | 0.09 | 0.09 | 0.04 | 0.02 | 0.05 | 0.05 |
| 1 | 7156082 | rs11120840 | 0.214 | 0.946 | -0.936 | 0.443 | 0.022 | 0.034 | 0.034 | 0 | 0.71 | 0.59 | 0.65 | 0.44 | 0.63 | 0.7 |
| 1 | 7158561 | rs4908440 | -0.065 | 0.857 | -1.739 | 0.007 | 0.004 | 0.123 | 0.123 | 1.363 | 0.43 | 0.21 | 0.34 | 0.15 | 0.4 | 0.42 |
| 1 | 7167031 | rs6694886 | 0.187 | 0.788 | -1.26 | 0.367 | 0.012 | 0.01 | 0.01 | 1.157 | 0.44 | 0.74 | 0.32 | 0.62 | 0.39 | 0.26 |
| 1 | 7169779 | rs17030322 | -1.439 | 0.83 | -0.628 | 0.281 | 0.041 | -0.015 | -0.015 | 0 | 0.08 | 0.3 | 0.11 | 0.04 | 0.1 | 0.2 |
| 1 | 7170719 | rs17349921 | -0.295 | -0.532 | -1.454 | 0.027 | 0.071 | 0.055 | 0.055 | 0 | 0.25 | 0.11 | 0.14 | 0.22 | 0.09 | 0.3 |
| 1 | 7173895 | rs7527268 | -0.566 | 0.077 | -1.546 | 0.209 | -0.004 | 0.065 | 0.065 | 0 | 0.4 | 0.26 | 0.34 | 0.25 | 0.39 | 0.45 |
| 1 | 7177744 | rs9434469 | -0.113 | 0.045 | -1.67 | 0.193 | -0.036 | 0.066 | 0.066 | 0 | 0.32 | 0.15 | 0.3 | 0.19 | 0.35 | 0.36 |
| 1 | 7177813 | rs9434845 | -0.113 | 0.002 | -1.67 | 0.193 | -0.036 | 0.066 | 0.066 | 0 | 0.32 | 0.15 | 0.3 | 0.19 | 0.35 | 0.36 |
| 1 | 7179479 | rs7532024 | 0.92 | 0.452 | -1.181 | 0.901 | -0.007 | -0.004 | -0.004 | 0 | 0.43 | 0.5 | 0.44 | 0.42 | 0.34 | 0.38 |
| 1 | 7179721 | rs4908443 | -0.469 | 0.782 | -1.059 | 1.346 | -0.047 | -0.061 | -0.061 | 0 | 0.85 | 0.93 | 0.78 | 0.66 | 0.73 | 0.82 |
| 1 | 7186325 | rs4460663 | 0.493 | 0.784 | -0.516 | 1.544 | -0.008 | -0.009 | -0.009 | 0 | 0.65 | 0.73 | 0.53 | 0.46 | 0.57 | 0.68 |
| 1 | 7196659 | rs4908600 | 0.893 | 1.141 | -0.739 | 1.268 | -0.025 | -0.017 | -0.017 | 0 | 0.15 | 0.07 | 0.21 | 0.2 | 0.24 | 0.24 |
| 1 | 7202190 | rs970973 | 0.611 | 0.981 | -0.8 | 1.319 | -0.012 | -0.005 | -0.005 | 0 | 0.61 | 0.73 | 0.49 | 0.43 | 0.52 | 0.57 |
| 1 | 7208217 | rs12751444 | -0.231 | 1.612 | -0.946 | 1.918 | -0.065 | -0.102 | -0.102 | 0 | 0.76 | 0.93 | 0.61 | 0.64 | 0.66 | 0.76 |
| 1 | 7211045 | rs11587739 | 1.043 | 1.487 | -0.629 | 1.8 | -0.001 | -0.003 | -0.003 | 0 | 0.56 | 0.54 | 0.38 | 0.43 | 0.4 | 0.49 |
| 1 | 7211340 | rs9434482 | -0.49 | 1.685 | -1.009 | 1.762 | -0.073 | -0.064 | -0.064 | 0 | 0.78 | 0.99 | 0.67 | 0.81 | 0.72 | 0.79 |
| 1 | 7217727 | rs9434483 | -0.259 | 1.268 | -0.623 | 1.727 | -0.002 | -0.001 | -0.001 | 0 | 0.2 | 0.22 | 0.24 | 0.2 | 0.26 | 0.24 |
| 1 | 7220970 | rs4908602 | -1.066 | 1.14 | -0.596 | 1.602 | -0.001 | 0 | 0 | 0 | 0.47 | 0.52 | 0.62 | 0.56 | 0.57 | 0.51 |
| 1 | 7236810 | rs6680365 | -1.263 | 1.193 | -0.706 | 1.435 | 0 | 0 | 0 | 0.038 | 0.52 | 0.55 | 0.52 | 0.54 | 0.58 | 0.48 |
| 1 | 7241523 | rs4908445 | -0.593 | 1.863 | -1.018 | 1.638 | 0.039 | 0.04 | 0.04 | 0 | 0.23 | 0.37 | 0.3 | 0.34 | 0.34 | 0.28 |
| 1 | 7245162 | rs12735487 | 0.121 | 1.748 | -1.245 | 1.456 | -0.069 | -0.05 | -0.05 | 0.4 | 0.17 | 0.04 | 0.25 | 0.2 | 0.27 | 0.23 |
| 1 | 7245434 | rs4908605 | -0.423 | 1.254 | -0.855 | 1.104 | 0 | 0.001 | 0.001 | 0.482 | 0.48 | 0.52 | 0.52 | 0.52 | 0.58 | 0.48 |
| 1 | 7253886 | rs11581497 | -0.196 | 1.094 | -1.478 | 0.702 | -0.025 | 0.073 | 0.073 | 0 | 0.13 | 0.01 | 0.1 | 0 | 0.15 | 0.03 |
| 1 | 7256943 | rs17030562 | -0.435 | 0.027 | 0.365 | 0.258 | -0.014 | -0.017 | -0.017 | 0 | 0.07 | 0.31 | 0.01 | 0 | 0 | 0 |
| 1 | 7261994 | rs1116254 | -0.162 | -0.559 | -0.417 | 0.163 | 0.042 | 0.082 | 0.082 | 0.551 | 0.49 | 0.87 | 0.6 | 0.67 | 0.57 | 0.64 |
| 1 | 7269283 | rs4908448 | 0.716 | -0.589 | -0.185 | 0.226 | -0.024 | -0.028 | -0.028 | 0 | 0.37 | 0.23 | 0.5 | 0.57 | 0.55 | 0.6 |
| 1 | 7272250 | rs916393 | -0.229 | -0.541 | -0.263 | 0.267 | 0.026 | 0.041 | 0.041 | 0.185 | 0.48 | 0.85 | 0.55 | 0.63 | 0.56 | 0.61 |
| 1 | 7310351 | rs2067996 | 0.917 | -0.489 | 0.29 | 0.211 | 0.003 | 0 | 0 | 1.764 | 0.05 | 0.24 | 0.01 | 0 | 0 | 0 |
| 1 | 7311260 | rs4908610 | 0.148 | -0.856 | 0.255 | 0.202 | -0.005 | -0.015 | -0.015 | 0 | 0.45 | 0.53 | 0.55 | 0.71 | 0.59 | 0.73 |
| 1 | 7312172 | rs1476047 | 0.444 | -0.94 | -0.066 | 0.226 | -0.003 | 0 | 0 | 0.144 | 0.43 | 0.48 | 0.52 | 0.57 | 0.58 | 0.6 |

| | | | | | | | | | | | | | | | | |
|---|---|---|---|---|---|---|---|---|---|---|---|---|---|---|---|---|
| 1 | 7312258 | rs6701870 | -0.294 | -0.698 | -0.081 | 0.316 | -0.01 | 0.09 | 0.09 | 0 | 0.06 | 0.26 | 0.03 | 0.14 | 0 | 0.14 |
| 1 | 7314529 | rs7544004 | 0.257 | -0.731 | -0.253 | 0.229 | 0 | -0.001 | -0.001 | 1.187 | 0.46 | 0.47 | 0.53 | 0.57 | 0.6 | 0.61 |
| 1 | 7316114 | rs7549870 | -0.343 | -0.372 | -0.663 | 0.253 | 0.011 | -0.036 | -0.036 | 0 | 0.66 | 0.87 | 0.66 | 0.61 | 0.73 | 0.69 |
| 1 | 7317541 | rs6698548 | 0.117 | -0.195 | -0.645 | -0.053 | 0.002 | 0.027 | 0.027 | 0 | 0.57 | 0.65 | 0.47 | 0.26 | 0.61 | 0.37 |
| 1 | 7318448 | rs10864289 | -0.758 | -0.283 | -0.715 | 0.087 | 0.008 | 0.031 | 0.031 | 1.399 | 0.7 | 0.92 | 0.72 | 0.79 | 0.76 | 0.84 |
| 1 | 7324286 | rs11581774 | -0.296 | -0.17 | -0.515 | 0.456 | -0.001 | 0.038 | 0.038 | 0.037 | 0.3 | 0.4 | 0.37 | 0.51 | 0.31 | 0.57 |
| 1 | 7324323 | rs11120889 | -0.367 | -0.288 | -0.492 | 0.399 | -0.007 | 0.038 | 0.038 | 0.069 | 0.32 | 0.43 | 0.37 | 0.52 | 0.31 | 0.58 |
| 1 | 7324468 | rs6577430 | 0.465 | -0.285 | -0.817 | 0.128 | -0.015 | 0.013 | 0.013 | 0 | 0.35 | 0.29 | 0.32 | 0.27 | 0.45 | 0.3 |
| 1 | 7324503 | rs11120890 | -0.375 | -0.717 | -1.108 | -0.042 | 0.001 | -0.001 | -0.001 | 0 | 0.45 | 0.45 | 0.44 | 0.5 | 0.39 | 0.6 |
| 1 | 7324959 | rs1034453 | -0.224 | -1.085 | -1.272 | -0.404 | 0.03 | 0.028 | 0.028 | 1.333 | 0.27 | 0.15 | 0.13 | 0.15 | 0.12 | 0.13 |
| 1 | 7326201 | rs7517429 | -0.54 | -0.606 | -1.477 | -0.158 | -0.011 | -0.029 | -0.029 | 0 | 0.87 | 1 | 0.93 | 0.84 | 0.91 | 0.88 |
| 1 | 7326245 | rs10864290 | 0.322 | -0.418 | -1.099 | -0.168 | -0.006 | -0.001 | -0.001 | 0 | 0.59 | 0.53 | 0.63 | 0.38 | 0.65 | 0.44 |
| 1 | 7331060 | rs2412148 | -0.358 | -0.541 | -0.85 | -0.296 | -0.003 | -0.007 | -0.007 | 0.275 | 0.78 | 0.72 | 0.89 | 0.84 | 0.84 | 0.78 |
| 1 | 7331347 | rs2412147 | 0.889 | 0.461 | -0.581 | -0.419 | 0.043 | 0.041 | 0.041 | 0 | 0.17 | 0.31 | 0.37 | 0.23 | 0.24 | 0.16 |
| 1 | 7337805 | rs12745074 | 1.425 | 0.114 | -0.975 | -0.626 | 0.001 | -0.002 | -0.002 | 0 | 0.12 | 0.1 | 0.06 | 0.13 | 0.09 | 0.15 |
| 1 | 7338813 | rs7539551 | 0.932 | 0.162 | -0.936 | -0.557 | 0 | 0.053 | 0.053 | 0 | 0.17 | 0.19 | 0.27 | 0.45 | 0.19 | 0.33 |
| 1 | 7342106 | rs12409413 | -0.143 | 0.568 | -0.946 | -0.715 | -0.006 | -0.003 | -0.003 | 0.026 | 0.36 | 0.32 | 0.59 | 0.44 | 0.44 | 0.33 |
| 1 | 7348943 | rs2001142 | -1.3 | 0.343 | -1.614 | -0.851 | -0.002 | 0.113 | 0.113 | 0.301 | 0.66 | 0.95 | 0.83 | 0.84 | 0.71 | 0.78 |
| 1 | 7351434 | rs1894654 | -0.967 | 0.531 | -1.459 | -0.893 | 0.019 | 0.104 | 0.104 | 0.135 | 0.6 | 0.84 | 0.78 | 0.83 | 0.64 | 0.75 |
| 1 | 7352974 | rs2071924 | 1.029 | 1.067 | -2.032 | -0.59 | -0.036 | -0.03 | -0.03 | 0 | 0.17 | 0.08 | 0.38 | 0.27 | 0.29 | 0.18 |
| 1 | 7354117 | rs17350193 | 2.615 | 0.719 | -2.005 | -0.6 | -0.002 | 0.041 | 0.041 | 0.16 | 0.07 | 0.02 | 0.03 | 0 | 0.06 | 0.02 |
| 1 | 7356673 | rs6662782 | 0.355 | 1.122 | -1.702 | -0.346 | -0.064 | -0.031 | -0.031 | 0.062 | 0.23 | 0.08 | 0.43 | 0.27 | 0.35 | 0.19 |
| 1 | 7361338 | rs11120896 | 0.329 | 0.884 | -1.429 | -0.143 | -0.001 | 0.004 | 0.004 | 0.67 | 0.27 | 0.34 | 0.22 | 0.42 | 0.26 | 0.44 |
| 1 | 7362478 | rs9659928 | 0.228 | 1.837 | -1.634 | 0.053 | -0.061 | -0.063 | -0.063 | 2.904 | 0.14 | 0.04 | 0.3 | 0.26 | 0.23 | 0.12 |
| 1 | 7362781 | rs9660884 | -1.254 | 1.745 | -1.682 | -0.121 | -0.049 | -0.035 | -0.035 | 0 | 0.33 | 0.19 | 0.57 | 0.41 | 0.42 | 0.29 |
| 1 | 7362992 | rs12410983 | 0.75 | 1.636 | -1.703 | 0.078 | -0.022 | -0.024 | -0.024 | 0 | 0.11 | 0.01 | 0.11 | 0.13 | 0.1 | 0.13 |
| 1 | 7363310 | rs10489142 | 0.018 | 1.896 | -1.526 | 0.236 | -0.061 | -0.064 | -0.064 | 0 | 0.14 | 0.04 | 0.3 | 0.26 | 0.23 | 0.12 |
| 1 | 7368829 | rs12026592 | -0.322 | 2.003 | -1.286 | -0.041 | -0.021 | -0.026 | -0.026 | 0 | 0.21 | 0.15 | 0.26 | 0.66 | 0.29 | 0.44 |
| 1 | 7372804 | rs12404510 | 0.124 | 2.259 | -1.617 | -0.378 | -0.075 | -0.013 | -0.013 | 0 | 0.13 | 0.01 | 0.1 | 0.13 | 0.22 | 0.16 |
| 1 | 7375258 | rs12128469 | 1.902 | 1.773 | -2.285 | -0.802 | 0.009 | 0.075 | 0.075 | 0.216 | 0.37 | 0.09 | 0.41 | 0.21 | 0.3 | 0.19 |
| 1 | 7375395 | rs12402763 | 0.444 | 1.922 | -2.159 | -0.736 | -0.069 | -0.011 | -0.011 | 0.021 | 0.14 | 0.02 | 0.1 | 0.13 | 0.22 | 0.16 |
| 1 | 7376671 | rs17030707 | 0.559 | 1.561 | -0.78 | -0.924 | -0.029 | -0.031 | -0.031 | 0 | 0.16 | 0.21 | 0.13 | 0.05 | 0.09 | 0.11 |
| 1 | 7378180 | rs11120900 | -0.196 | 1.866 | -1.056 | -0.88 | -0.118 | 0.02 | 0.02 | 1.459 | 0.25 | 0.03 | 0.45 | 0.85 | 0.39 | 0.51 |
| 1 | 7379198 | rs7543711 | 0.011 | 1.067 | -0.823 | -0.869 | -0.095 | 0.054 | 0.054 | 2.73 | 0.76 | 0.96 | 0.81 | 0.84 | 0.65 | 0.73 |
| 1 | 7380062 | rs2001143 | 0.39 | 0.7 | -0.323 | -0.654 | -0.024 | -0.026 | -0.026 | 0 | 0.71 | 0.79 | 0.67 | 0.39 | 0.61 | 0.46 |
| 1 | 7380747 | rs12563394 | 0.286 | 0.742 | -0.273 | -0.562 | -0.01 | -0.011 | -0.011 | 2.935 | 0.18 | 0.16 | 0.24 | 0.58 | 0.26 | 0.43 |
| 1 | 7389944 | rs2301460 | -0.104 | 0.634 | -0.075 | -0.574 | 0.007 | 0.03 | 0.03 | 0.145 | 0.22 | 0.3 | 0.25 | 0.58 | 0.27 | 0.45 |
| 1 | 7394728 | rs12041403 | 1.4 | 0.416 | -0.719 | -0.645 | -0.029 | 0.089 | 0.089 | 0.465 | 0.51 | 0.14 | 0.51 | 0.35 | 0.51 | 0.34 |
| 1 | 7396032 | rs1029322 | 0.032 | 0.118 | -0.053 | -0.639 | -0.019 | -0.025 | -0.025 | 1.492 | 0.15 | 0.17 | 0.12 | 0.05 | 0.09 | 0.09 |
| 1 | 7405293 | rs845246 | -0.399 | 0.135 | -0.209 | -0.638 | -0.037 | 0.044 | 0.044 | 0.33 | 0.33 | 0.24 | 0.32 | 0.24 | 0.45 | 0.29 |
| 1 | 7412051 | rs845237 | -0.978 | -0.109 | -0.07 | -0.73 | 0.003 | -0.013 | -0.013 | 0 | 0.87 | 0.87 | 0.94 | 1 | 0.91 | 0.88 |
| 1 | 7412088 | rs4908624 | 1.587 | -0.356 | 0.245 | -0.779 | 0.026 | 0.046 | 0.046 | 1.443 | 0.37 | 0.21 | 0.23 | 0.17 | 0.28 | 0.16 |
| 1 | 7412961 | rs4908625 | 0.751 | -0.17 | 0.578 | -0.243 | -0.004 | 0.02 | 0.02 | 0.171 | 0.12 | 0.08 | 0.32 | 0.55 | 0.15 | 0.42 |
| 1 | 7419680 | rs845195 | 0.811 | -0.281 | 0.851 | -0.356 | -0.004 | 0.008 | 0.008 | 0.271 | 0.7 | 0.37 | 0.6 | 0.43 | 0.75 | 0.47 |
| 1 | 7419774 | rs705675 | -0.55 | -0.356 | 0.747 | -0.398 | 0.016 | 0.048 | 0.048 | 0 | 0.84 | 0.93 | 0.94 | 1 | 0.9 | 0.88 |
| 1 | 7422216 | rs4908626 | 0.351 | -0.149 | 1.272 | -0.292 | 0.021 | 0.143 | 0.143 | 0 | 0.13 | 0.22 | 0.32 | 0.56 | 0.14 | 0.42 |
| 1 | 7426875 | rs845270 | 0.877 | -0.123 | 1.171 | -0.278 | -0.005 | 0.023 | 0.023 | 0.047 | 0.71 | 0.41 | 0.61 | 0.43 | 0.75 | 0.47 |
| 1 | 7432615 | rs11579620 | 0.928 | 0.24 | 0.827 | 0.097 | 0 | 0.05 | 0.05 | 0.011 | 0.1 | 0.1 | 0.3 | 0.54 | 0.14 | 0.41 |
| 1 | 7448154 | rs2040433 | -0.262 | 0.206 | 0.981 | 0.03 | 0.017 | 0.126 | 0.126 | 0.531 | 0.15 | 0.22 | 0.32 | 0.55 | 0.16 | 0.44 |
| 1 | 7448256 | rs845232 | 0.396 | 0.166 | 1.008 | -0.064 | 0.011 | 0.056 | 0.056 | 0 | 0.74 | 0.62 | 0.62 | 0.44 | 0.75 | 0.46 |
| 1 | 7464375 | rs845202 | 0.86 | 0.052 | 0.598 | -0.111 | -0.004 | 0.02 | 0.02 | 0.671 | 0.7 | 0.41 | 0.57 | 0.42 | 0.73 | 0.43 |
| 1 | 7468614 | rs763161 | 0.309 | 0.039 | 0.669 | 0.015 | -0.001 | 0.051 | 0.051 | 0.003 | 0.75 | 0.64 | 0.6 | 0.47 | 0.76 | 0.47 |
| 1 | 7469780 | rs4908635 | 1.377 | -0.156 | 0.004 | -0.513 | -0.022 | 0.079 | 0.079 | 4.492 | 0.35 | 0.03 | 0.24 | 0.18 | 0.29 | 0.15 |
| 1 | 7478076 | rs11120917 | 0.941 | 0.24 | 0.349 | -0.537 | 0.059 | 0.006 | 0.006 | 1.695 | 0.15 | 0.26 | 0.26 | 0.22 | 0.32 | 0.11 |
| 1 | 7478924 | rs845259 | -0.334 | 0.119 | 0.242 | -0.367 | -0.014 | -0.092 | -0.092 | 0 | 0.83 | 0.54 | 0.91 | 0.97 | 0.89 | 0.89 |
| 1 | 7480054 | rs6677745 | 1.17 | 0.301 | -0.035 | -0.084 | -0.03 | -0.033 | -0.033 | 0 | 0.09 | 0.01 | 0.33 | 0.49 | 0.13 | 0.42 |
| 1 | 7482963 | rs845252 | 0.545 | 0.14 | -0.299 | -0.393 | -0.002 | -0.003 | -0.003 | 0 | 0.16 | 0.16 | 0.26 | 0.25 | 0.34 | 0.14 |
| 1 | 7483046 | rs845251 | -0.831 | -0.804 | -0.369 | -0.474 | -0.002 | -0.031 | -0.031 | 0 | 0.86 | 0.9 | 0.91 | 0.99 | 0.9 | 0.88 |
| 1 | 7485306 | rs11120921 | -0.715 | -0.679 | -0.135 | 0.075 | -0.016 | -0.07 | -0.07 | 0 | 0.23 | 0.31 | 0.09 | 0.04 | 0.15 | 0.21 |
| 1 | 7485370 | rs11120922 | -1.154 | -0.551 | -0.115 | 0.215 | -0.009 | 0.002 | 0.002 | 0 | 0.36 | 0.53 | 0.43 | 0.59 | 0.3 | 0.64 |
| 1 | 7486317 | rs12088178 | -1.409 | -0.802 | -0.208 | 0.161 | 0.004 | 0.005 | 0.005 | 0 | 0.47 | 0.54 | 0.5 | 0.6 | 0.39 | 0.65 |
| 1 | 7487277 | rs10864302 | -0.706 | -0.748 | -0.294 | 0.908 | -0.006 | 0.076 | 0.076 | 0 | 0.19 | 0.32 | 0.39 | 0.56 | 0.16 | 0.5 |
| 1 | 7487330 | rs11120923 | -0.88 | -0.591 | -0.669 | 0.889 | 0.01 | -0.012 | -0.012 | 0 | 0.28 | 0.29 | 0.44 | 0.58 | 0.23 | 0.53 |
| 1 | 7492120 | rs1193219 | -0.387 | -0.333 | -0.404 | 0.984 | 0.003 | 0.002 | 0.002 | 0 | 0.45 | 0.58 | 0.42 | 0.66 | 0.4 | 0.67 |
| 1 | 7492265 | rs1193220 | -0.329 | -0.464 | -0.362 | 0.937 | 0.004 | 0.002 | 0.002 | 0 | 0.42 | 0.65 | 0.42 | 0.7 | 0.38 | 0.67 |
| 1 | 7493231 | rs9988557 | -0.873 | -0.645 | -0.595 | 1.014 | 0.008 | 0.08 | 0.08 | 0 | 0.53 | 0.8 | 0.48 | 0.75 | 0.44 | 0.71 |
| 1 | 7497407 | rs1193224 | -0.223 | -0.514 | -0.797 | 1.333 | -0.031 | 0.013 | 0.013 | 0 | 0.34 | 0.09 | 0.3 | 0.29 | 0.34 | 0.42 |
| 1 | 7501037 | rs1193227 | -0.157 | -0.217 | -1.502 | 1.335 | -0.043 | -0.005 | -0.005 | 0 | 0.33 | 0.05 | 0.25 | 0.3 | 0.32 | 0.42 |
| 1 | 7501060 | rs12565108 | -0.898 | 0.031 | -0.347 | 1.504 | -0.006 | 0.107 | 0.107 | 0 | 0.07 | 0.13 | 0.14 | 0.26 | 0.04 | 0.22 |

| | | | | | | | | | | | | | | | | |
|---|---|---|---|---|---|---|---|---|---|---|---|---|---|---|---|---|
| 1 | 7513025 | rs6577445 | 0.4 | 0.064 | -0.824 | 0.642 | 0.007 | 0.016 | 0.016 | 0 | 0.7 | 0.79 | 0.71 | 0.79 | 0.7 | 0.66 |
| 1 | 7513252 | rs1193247 | -0.101 | 0.406 | -0.587 | 0.87 | 0.003 | 0.044 | 0.044 | 0.037 | 0.19 | 0.26 | 0.3 | 0.57 | 0.2 | 0.2 |
| 1 | 7519264 | rs1750838 | -0.295 | 0.313 | -1.805 | 0.055 | -0.062 | 0.047 | 0.047 | 0 | 0.71 | 0.93 | 0.7 | 0.81 | 0.65 | 0.65 |
| 1 | 7520129 | rs1725235 | -0.279 | 0.059 | -1.319 | -0.393 | 0.015 | 0.037 | 0.037 | 0 | 0.33 | 0.5 | 0.4 | 0.56 | 0.39 | 0.33 |
| 1 | 7520601 | rs1750837 | -0.32 | -0.306 | -1.594 | -0.744 | -0.001 | -0.004 | -0.004 | 0 | 0.45 | 0.59 | 0.47 | 0.56 | 0.49 | 0.4 |
| 1 | 7520927 | rs1750836 | -0.025 | -0.403 | -1.617 | -0.872 | 0.002 | -0.002 | -0.002 | 0 | 0.46 | 0.56 | 0.51 | 0.55 | 0.44 | 0.37 |
| 1 | 7521479 | rs11805932 | 0.254 | -0.446 | -1.353 | -0.915 | -0.01 | 0.091 | 0.091 | 0 | 0.12 | 0.3 | 0.14 | 0.26 | 0.06 | 0.23 |
| 1 | 7524342 | rs2693953 | -0.031 | -0.27 | -1.207 | -0.706 | 0.004 | 0.014 | 0.014 | 0 | 0.44 | 0.55 | 0.31 | 0.09 | 0.4 | 0.19 |
| 1 | 7524476 | rs6702132 | -1.284 | -0.346 | -1.215 | -0.42 | -0.04 | 0.006 | 0.006 | 0 | 0.66 | 0.77 | 0.41 | 0.36 | 0.59 | 0.48 |
| 1 | 7527308 | rs6663735 | 0.574 | -0.345 | -1.175 | -0.423 | -0.004 | -0.014 | -0.014 | 0.102 | 0.17 | 0.19 | 0.07 | 0.09 | 0.16 | 0.22 |
| 1 | 7528669 | rs1149331 | -0.276 | -0.376 | -1.171 | -0.475 | -0.012 | 0.044 | 0.044 | 0.091 | 0.55 | 0.59 | 0.36 | 0.18 | 0.53 | 0.26 |
| 1 | 7528846 | rs1149332 | 1.383 | -0.638 | -1.176 | -0.689 | -0.035 | 0.038 | 0.038 | 0.145 | 0.2 | 0.07 | 0.17 | 0.08 | 0.25 | 0.09 |
| 1 | 7530341 | rs1149335 | -1.465 | -0.571 | -1.182 | -0.614 | -0.051 | 0.006 | 0.006 | 0 | 0.66 | 0.85 | 0.43 | 0.35 | 0.58 | 0.48 |
| 1 | 7531338 | rs6664987 | 0.932 | -0.596 | -0.7 | -0.754 | -0.047 | -0.123 | -0.123 | 0 | 0.22 | 0.34 | 0.13 | 0 | 0.15 | 0.11 |
| 1 | 7533164 | rs1011124 | -1.312 | 0.058 | -1.099 | 0.11 | -0.094 | -0.133 | -0.133 | 0 | 0.69 | 0.93 | 0.62 | 0.49 | 0.6 | 0.53 |
| 1 | 7534464 | rs11805543 | 0.141 | -0.108 | -0.798 | -0.185 | -0.006 | -0.012 | -0.012 | 0 | 0.18 | 0.24 | 0.07 | 0.12 | 0.15 | 0.21 |
| 1 | 7536187 | rs1011127 | -0.194 | 0.227 | -0.565 | 0.157 | -0.009 | 0.001 | 0.001 | 0 | 0.6 | 0.66 | 0.59 | 0.56 | 0.61 | 0.39 |
| 1 | 7536259 | rs1149338 | -0.854 | 0.249 | -0.632 | 0.151 | -0.044 | 0.073 | 0.073 | 0 | 0.68 | 0.84 | 0.67 | 0.81 | 0.63 | 0.6 |
| 1 | 7537500 | rs1193182 | -0.971 | 0.365 | -0.933 | -0.022 | -0.076 | 0.106 | 0.106 | 0 | 0.66 | 0.89 | 0.67 | 0.81 | 0.6 | 0.67 |
| 1 | 7538437 | rs1193185 | -0.12 | 0.479 | -0.853 | 0.754 | -0.014 | 0.11 | 0.11 | 0 | 0.56 | 0.78 | 0.48 | 0.24 | 0.49 | 0.42 |
| 1 | 7540770 | rs6678279 | 0.433 | 0.327 | -0.405 | 0.783 | 0.002 | 0.066 | 0.066 | 0.151 | 0.07 | 0.17 | 0.09 | 0.17 | 0.05 | 0.13 |
| 1 | 7541345 | rs2796483 | 0.241 | 0.285 | -0.256 | 0.745 | 0.009 | 0.024 | 0.024 | 0 | 0.52 | 0.62 | 0.55 | 0.57 | 0.51 | 0.36 |
| 1 | 7543417 | rs11120943 | 0.186 | 0.291 | -0.282 | 0.75 | -0.013 | 0.011 | 0.011 | 0.059 | 0.06 | 0.14 | 0.02 | 0.08 | 0.02 | 0.15 |
| 1 | 7544964 | rs1193165 | -0.124 | 0.293 | -0.287 | 0.765 | -0.01 | 0.026 | 0.026 | 0 | 0.54 | 0.72 | 0.57 | 0.56 | 0.52 | 0.38 |
| 1 | 7548616 | rs947675 | 0.539 | 0.257 | 0.246 | 0.675 | 0.007 | 0.008 | 0.008 | 0 | 0.47 | 0.37 | 0.4 | 0.16 | 0.43 | 0.25 |
| 1 | 7549488 | rs17030993 | 1.139 | 1.056 | 0.101 | 0.593 | 0.006 | 0.029 | 0.029 | 0.105 | 0.09 | 0.18 | 0.13 | 0.16 | 0.11 | 0.18 |
| 1 | 7550126 | rs1193233 | -0.171 | 0.463 | -0.041 | 0.539 | -0.018 | 0.061 | 0.061 | 0 | 0.58 | 0.71 | 0.63 | 0.74 | 0.54 | 0.49 |
| 1 | 7551631 | rs1193230 | -0.731 | 0.495 | -0.274 | 0.466 | -0.079 | 0.129 | 0.129 | 0.099 | 0.63 | 0.87 | 0.67 | 0.81 | 0.58 | 0.64 |
| 1 | 7555322 | rs1725268 | -0.821 | 0.512 | -0.257 | 0.481 | -0.077 | 0.123 | 0.123 | 0.113 | 0.63 | 0.87 | 0.66 | 0.81 | 0.58 | 0.64 |
| 1 | 7557152 | rs10864304 | -1.159 | 0.466 | 0.574 | 0.793 | 0.058 | 0.374 | 0.374 | 0.584 | 0.08 | 0.33 | 0.17 | 0.35 | 0.09 | 0.12 |
| 1 | 7558193 | rs1725265 | -0.152 | 0.421 | 0.406 | 0.686 | 0 | 0.009 | 0.009 | 0 | 0.54 | 0.65 | 0.53 | 0.5 | 0.51 | 0.37 |
| 1 | 7561152 | rs1750851 | -0.157 | 0.269 | 0.398 | 0.568 | 0.004 | 0.034 | 0.034 | 0 | 0.56 | 0.61 | 0.43 | 0.29 | 0.48 | 0.42 |
| 1 | 7561298 | rs4908463 | -1.82 | 0.776 | 0.875 | 1.325 | 0.07 | 0.046 | 0.046 | 1.74 | 0.11 | 0.44 | 0.12 | 0.15 | 0.14 | 0.17 |
| 1 | 7563901 | rs2462925 | 0.807 | 1.009 | 0.119 | 1.524 | 0.044 | 0.188 | 0.188 | 0 | 0.92 | 0.82 | 0.84 | 0.64 | 0.86 | 0.85 |
| 1 | 7566066 | rs11120954 | 1.563 | 1.031 | -0.046 | 1.15 | 0.047 | 0.117 | 0.117 | 2.359 | 0.84 | 0.68 | 0.76 | 0.48 | 0.73 | 0.72 |
| 1 | 7567768 | rs12565334 | -0.624 | 0.894 | -0.418 | 0.694 | -0.005 | -0.006 | -0.006 | 0 | 0.12 | 0.09 | 0.16 | 0.19 | 0.17 | 0.19 |
| 1 | 7569602 | rs1193179 | 0.583 | 0.383 | -0.597 | 0.598 | -0.001 | 0.015 | 0.015 | 0.179 | 0.71 | 0.38 | 0.54 | 0.52 | 0.74 | 0.64 |
| 1 | 7572262 | rs749958 | -0.102 | -0.229 | -1.058 | 0.402 | -0.011 | 0.007 | 0.007 | 0.001 | 0.45 | 0.27 | 0.34 | 0.43 | 0.51 | 0.35 |
| 1 | 7575666 | rs7529399 | 0.241 | -0.544 | -1.074 | 0.412 | -0.02 | 0.035 | 0.035 | 0 | 0.27 | 0.11 | 0.19 | 0.15 | 0.32 | 0.2 |
| 1 | 7578333 | rs6693468 | 0.403 | -0.764 | -0.75 | 0.587 | 0.018 | 0.076 | 0.076 | 0 | 0.54 | 0.3 | 0.47 | 0.26 | 0.63 | 0.38 |
| 1 | 7578713 | rs6670316 | 0.318 | -0.875 | -0.319 | 0.69 | -0.05 | 0.122 | 0.122 | 0 | 0.22 | 0.35 | 0.27 | 0.43 | 0.11 | 0.43 |
| 1 | 7578776 | rs1750817 | -1.264 | -0.827 | -0.462 | 0.701 | -0.035 | -0.03 | -0.03 | 0.158 | 0.81 | 0.9 | 0.76 | 0.7 | 0.75 | 0.83 |
| 1 | 7580769 | rs1618346 | 0.352 | -0.67 | -0.293 | 0.376 | 0.016 | 0.058 | 0.058 | 0 | 0.54 | 0.32 | 0.47 | 0.26 | 0.63 | 0.37 |
| 1 | 7584814 | rs2478267 | -0.005 | -0.625 | -0.409 | 0.445 | -0.002 | -0.01 | -0.01 | 1.446 | 0.59 | 0.57 | 0.5 | 0.26 | 0.65 | 0.4 |
| 1 | 7590448 | rs1193167 | 0.116 | -0.441 | -1.071 | 0.362 | 0.016 | 0.06 | 0.06 | 0 | 0.55 | 0.36 | 0.41 | 0.2 | 0.63 | 0.39 |
| 1 | 7591573 | rs12723779 | 0.737 | -0.11 | -1.605 | 0.637 | -0.013 | -0.041 | -0.041 | 0.356 | 0.13 | 0 | 0.06 | 0.14 | 0.13 | 0.12 |
| 1 | 7595233 | rs2478266 | -0.695 | 0.177 | -1.069 | 0.91 | -0.004 | -0.004 | -0.004 | 0.085 | 0.82 | 0.85 | 0.77 | 0.68 | 0.76 | 0.84 |
| 1 | 7598094 | rs6577449 | -0.232 | -0.916 | -0.569 | 1.14 | -0.066 | 0.086 | 0.086 | 0 | 0.14 | 0.47 | 0.06 | 0.25 | 0.04 | 0.24 |
| 1 | 7600905 | rs12407666 | -0.198 | -0.075 | -0.996 | 0.83 | 0.005 | 0.003 | 0.003 | 0 | 0.41 | 0.3 | 0.37 | 0.34 | 0.4 | 0.39 |
| 1 | 7607505 | rs4908650 | -0.653 | -0.53 | -0.744 | 0.994 | -0.068 | 0.041 | 0.041 | 0.467 | 0.27 | 0.45 | 0.26 | 0.3 | 0.16 | 0.27 |
| 1 | 7611606 | rs4908654 | -0.313 | 0.872 | -1.722 | 0.093 | -0.064 | 0.093 | 0.093 | 0 | 0.31 | 0.14 | 0.28 | 0.17 | 0.4 | 0.32 |
| 1 | 7613745 | rs957856 | 0.28 | 0.681 | -1.073 | 0.78 | -0.013 | 0.041 | 0.041 | 0 | 0.67 | 0.47 | 0.57 | 0.55 | 0.68 | 0.76 |
| 1 | 7615909 | rs11120972 | 0.299 | 0.783 | -1.131 | 0.884 | 0.005 | 0.044 | 0.044 | 0 | 0.72 | 0.38 | 0.64 | 0.55 | 0.71 | 0.77 |
| 1 | 7616430 | rs11120973 | 0.157 | 1.328 | -1.628 | 1.122 | -0.103 | -0.101 | -0.101 | 0 | 0.82 | 0.98 | 0.76 | 0.75 | 0.75 | 0.81 |
| 1 | 7616593 | rs4908656 | -1.078 | 1.871 | -1.18 | 1.706 | -0.036 | -0.034 | -0.034 | 0.348 | 0.06 | 0.01 | 0.21 | 0.24 | 0.19 | 0.26 |
| 1 | 7616740 | rs12755955 | -0.956 | 0.852 | -1.279 | 1.118 | -0.046 | 0.013 | 0.013 | 0 | 0.3 | 0.82 | 0.3 | 0.22 | 0.35 | 0.41 |
| 1 | 7617484 | rs17031113 | -1.503 | -0.297 | 0.053 | 1.716 | -0.001 | 0.015 | 0.015 | 0 | 0.21 | 0.21 | 0.15 | 0.45 | 0.19 | 0.18 |
| 1 | 7621589 | rs6696544 | -1.9 | -0.305 | 1.217 | 1.772 | 0.023 | 0.224 | 0.224 | 0 | 0.19 | 0.49 | 0.17 | 0.52 | 0.19 | 0.2 |
| 1 | 7624855 | rs17031140 | -1.493 | -0.439 | 1.379 | 1.844 | -0.003 | 0.03 | 0.03 | 0 | 0.2 | 0.22 | 0.15 | 0.4 | 0.19 | 0.2 |
| 1 | 7634687 | rs7526052 | -1.981 | -0.48 | 1.723 | 1.584 | 0.06 | 0.117 | 0.117 | 2.382 | 0.06 | 0.6 | 0.05 | 0.03 | 0 | 0.07 |
| 1 | 7637979 | rs10492961 | -2.146 | -0.751 | 0.975 | 1.464 | 0.029 | -0.035 | -0.035 | 0.014 | 0.33 | 0.84 | 0.29 | 0.54 | 0.26 | 0.33 |
| 1 | 7644802 | rs7527087 | -1.503 | -0.682 | 1.169 | 1.353 | 0.004 | -0.01 | -0.01 | 0 | 0.26 | 0.22 | 0.28 | 0.41 | 0.21 | 0.28 |
| 1 | 7649715 | rs7519687 | -1.631 | -0.629 | 2.445 | 1.241 | -0.027 | 0.003 | 0.003 | 0.339 | 0.17 | 0.47 | 0.13 | 0.12 | 0.12 | 0.19 |
| 1 | 7667122 | rs11590447 | -1.451 | -0.659 | 2.536 | 1.166 | -0.019 | -0.036 | -0.036 | 0 | 0.16 | 0.48 | 0.13 | 0.09 | 0.13 | 0.1 |
| 1 | 7668346 | rs10864308 | -2.313 | -0.988 | 2.466 | 1.208 | 0.041 | -0.009 | -0.009 | 0.074 | 0.38 | 0.82 | 0.53 | 0.57 | 0.31 | 0.42 |
| 1 | 7669922 | rs2056675 | -1.386 | -0.663 | 2.476 | 0.818 | -0.015 | -0.03 | -0.03 | 0 | 0.15 | 0.4 | 0.12 | 0.09 | 0.13 | 0.1 |
| 1 | 7670574 | rs17031163 | -1.358 | -0.643 | 2.331 | 0.828 | -0.012 | -0.022 | -0.022 | 0.088 | 0.14 | 0.34 | 0.11 | 0.09 | 0.13 | 0.1 |
| 1 | 7681743 | rs7555394 | -1.027 | -0.583 | 2.073 | 0.906 | -0.028 | 0.017 | 0.017 | 0 | 0.17 | 0.42 | 0.13 | 0.14 | 0.13 | 0.19 |
| 1 | 7682237 | rs12139380 | -1.139 | -0.521 | 1.63 | 0.909 | -0.029 | 0.005 | 0.005 | 0 | 0.18 | 0.52 | 0.14 | 0.14 | 0.14 | 0.19 |

| | | | | | | | | | | | | | | | | |
|---|---|---|---|---|---|---|---|---|---|---|---|---|---|---|---|---|
| 1 | 7682489 | rs12139470 | -1.183 | -0.516 | 1.237 | 0.913 | -0.03 | 0.001 | 0.001 | 0 | 0.18 | 0.59 | 0.14 | 0.15 | 0.14 | 0.19 |
| 1 | 7708733 | rs12403557 | 0.437 | 0.232 | -1.129 | 0.718 | -0.052 | -0.006 | -0.006 | 0 | 0.54 | 0.14 | 0.48 | 0.48 | 0.52 | 0.49 |
| 1 | 7710308 | rs17027053 | -0.405 | -0.956 | -0.78 | 0.022 | -0.009 | -0.003 | -0.003 | 0 | 0.29 | 0.38 | 0.32 | 0.15 | 0.15 | 0.19 |
| 1 | 7712132 | rs4908663 | -0.279 | -0.219 | -1.026 | -0.718 | 0.005 | 0.08 | 0.08 | 0 | 0.65 | 0.82 | 0.81 | 0.92 | 0.69 | 0.74 |
| 1 | 7713021 | rs9434879 | -0.104 | -0.598 | -0.162 | -0.331 | -0.01 | -0.024 | -0.024 | 0.144 | 0.62 | 0.57 | 0.61 | 0.72 | 0.69 | 0.73 |
| 1 | 7715776 | rs4908665 | -0.101 | -0.573 | 0.561 | -0.496 | -0.007 | -0.078 | -0.078 | 0.395 | 0.62 | 0.57 | 0.79 | 0.92 | 0.69 | 0.69 |
| 1 | 7717080 | rs17031253 | 0.438 | -0.502 | 0.404 | -0.607 | 0.011 | 0.027 | 0.027 | 0 | 0.27 | 0.19 | 0.14 | 0.07 | 0.2 | 0.28 |
| 1 | 7718355 | rs11585947 | 0.739 | -0.04 | 0.293 | -0.276 | -0.003 | -0.003 | -0.003 | 0 | 0.06 | 0.04 | 0.08 | 0.06 | 0.09 | 0.02 |
| 1 | 7722501 | rs6662747 | -0.147 | 1.359 | 0.318 | -0.302 | 0.019 | -0.01 | -0.01 | 0 | 0.37 | 0.54 | 0.47 | 0.33 | 0.44 | 0.42 |
| 1 | 7722858 | rs870283 | 0.47 | 1.539 | 0.251 | -0.322 | -0.08 | 0.106 | 0.106 | 0.087 | 0.29 | 0.14 | 0.2 | 0.09 | 0.38 | 0.37 |
| 1 | 7727854 | rs1891215 | -0.496 | 1.822 | 0.593 | -0.37 | 0.021 | -0.013 | -0.013 | 0 | 0.39 | 0.5 | 0.47 | 0.34 | 0.43 | 0.42 |
| 1 | 7731730 | rs4908473 | -0.542 | 1.729 | 0.428 | -0.248 | 0 | 0.096 | 0.096 | 0.012 | 0.1 | 0.2 | 0.26 | 0.25 | 0.08 | 0.06 |
| 1 | 7735380 | rs6658296 | 0.692 | 1.142 | -0.211 | -0.753 | 0.018 | 0.025 | 0.025 | 0 | 0.21 | 0.1 | 0.11 | 0.14 | 0.16 | 0.2 |
| 1 | 7736097 | rs6673977 | 0.752 | 1.481 | -0.327 | -0.749 | -0.055 | 0.087 | 0.087 | 0.628 | 0.29 | 0.16 | 0.2 | 0.09 | 0.35 | 0.36 |
| 1 | 7748525 | rs3124796 | -0.279 | 1.556 | -0.143 | -0.864 | 0.014 | 0.075 | 0.075 | 1.655 | 0.53 | 0.64 | 0.72 | 0.9 | 0.61 | 0.61 |
| 1 | 7758760 | rs12756299 | 0.177 | 1.656 | -0.114 | -0.234 | -0.027 | 0.022 | 0.022 | 0.366 | 0.44 | 0.22 | 0.35 | 0.36 | 0.49 | 0.39 |
| 1 | 7760733 | rs3011931 | -0.956 | 1.525 | -0.019 | -0.192 | 0.009 | -0.033 | -0.033 | 0 | 0.94 | 0.87 | 0.96 | 1 | 0.94 | 0.97 |
| 1 | 7774071 | rs3121224 | -0.841 | 1.383 | 0.388 | -0.172 | -0.007 | -0.004 | -0.004 | 0 | 0.94 | 0.78 | 0.98 | 1 | 1 | 1 |
| 1 | 7775407 | rs3011937 | 0.074 | 1.536 | 0.349 | -0.184 | -0.002 | -0.004 | -0.004 | 0 | 0.47 | 0.46 | 0.6 | 0.63 | 0.47 | 0.57 |
| 1 | 7780128 | rs4908686 | -0.111 | 1.208 | -0.05 | -0.291 | 0.008 | 0.009 | 0.009 | 0 | 0.34 | 0.44 | 0.29 | 0.36 | 0.35 | 0.36 |
| 1 | 7784902 | rs12028339 | 0.343 | 1.103 | -0.452 | -0.378 | 0.04 | 0.097 | 0.097 | 0 | 0.39 | 0.2 | 0.34 | 0.2 | 0.3 | 0.47 |
| 1 | 7792411 | rs2301488 | 0.147 | 1.092 | -0.566 | -0.319 | 0.005 | 0.002 | 0.002 | 0 | 0.41 | 0.43 | 0.49 | 0.44 | 0.45 | 0.48 |
| 1 | 7796641 | rs2071985 | -0.48 | 1.13 | -0.482 | -0.179 | 0 | 0 | 0 | 0 | 0.48 | 0.47 | 0.67 | 0.54 | 0.48 | 0.53 |
| 1 | 7800385 | rs4908690 | -1.012 | 1.712 | -0.543 | -0.227 | 0.013 | 0.084 | 0.084 | 0 | 0.73 | 0.51 | 0.79 | 0.49 | 0.72 | 0.54 |
| 1 | 7801288 | rs2032563 | 1.999 | 1.639 | -0.845 | -0.343 | -0.018 | 0.047 | 0.047 | 0 | 0.22 | 0.02 | 0.1 | 0.13 | 0.18 | 0.28 |
| 1 | 7805208 | rs2071986 | -0.673 | 1.981 | -0.086 | 0.214 | 0.014 | 0.066 | 0.066 | 0 | 0.72 | 0.58 | 0.78 | 0.47 | 0.72 | 0.53 |
| 1 | 7811961 | rs9919220 | -0.657 | 2.104 | -0.097 | 0.163 | 0.123 | 0.2 | 0.2 | 0 | 0.09 | 0.28 | 0.32 | 0.3 | 0.2 | 0.08 |
| 1 | 7812398 | rs9919223 | -0.581 | 1.844 | -0.138 | 0.245 | 0.006 | 0.039 | 0.039 | 0 | 0.73 | 0.64 | 0.79 | 0.49 | 0.72 | 0.55 |
| 1 | 7817856 | rs2051646 | -0.441 | 1.801 | -0.268 | 0.242 | 0.062 | 0.039 | 0.039 | 0 | 0.72 | 0.57 | 0.7 | 0.42 | 0.56 | 0.52 |
| 1 | 7823085 | rs11587479 | 0.829 | 0.97 | -0.99 | -0.026 | 0.049 | -0.048 | -0.048 | 1.249 | 0.21 | 0.09 | 0.26 | 0.28 | 0.1 | 0.08 |
| 1 | 7825311 | rs707455 | -0.334 | 1.258 | -0.724 | 0.076 | -0.002 | 0.068 | 0.068 | 0 | 0.46 | 0.7 | 0.53 | 0.75 | 0.46 | 0.44 |
| 1 | 7826347 | rs697672 | -1.084 | 0.977 | -1.095 | -0.385 | 0.005 | 0.03 | 0.03 | 0 | 0.83 | 0.92 | 0.91 | 0.98 | 0.85 | 0.88 |
| 1 | 7829107 | rs1044245 | 2.933 | 0.991 | -1.396 | -0.661 | -0.04 | 0.091 | 0.091 | 1.101 | 0.15 | 0 | 0.09 | 0 | 0.11 | 0.03 |
| 1 | 7830262 | rs2153733 | -0.48 | 0.756 | -0.725 | -0.501 | 0.008 | 0.051 | 0.051 | 0 | 0.14 | 0.25 | 0.44 | 0.28 | 0.19 | 0.17 |
| 1 | 7832212 | rs707458 | 0.927 | 1.094 | -0.237 | -0.323 | 0.022 | -0.02 | -0.02 | 0 | 0.35 | 0.47 | 0.46 | 0.29 | 0.52 | 0.45 |
| 1 | 7834026 | rs2071987 | 1.471 | 1.035 | -0.204 | -0.328 | -0.013 | 0.069 | 0.069 | 0 | 0.22 | 0.38 | 0.16 | 0.39 | 0.2 | 0.32 |
| 1 | 7837842 | rs697673 | -0.602 | 0.947 | -0.3 | -0.431 | 0.026 | 0.04 | 0.04 | 0 | 0.68 | 0.79 | 0.84 | 0.96 | 0.76 | 0.84 |
| 1 | 7838691 | rs707460 | 0.246 | 0.96 | -0.359 | -0.309 | 0 | 0.002 | 0.002 | 0 | 0.42 | 0.49 | 0.67 | 0.55 | 0.54 | 0.48 |
| 1 | 7845695 | rs228729 | -1.346 | 0.591 | -0.554 | -0.441 | 0.019 | 0.086 | 0.086 | 0.973 | 0.58 | 0.78 | 0.76 | 0.76 | 0.66 | 0.66 |
| 1 | 7848557 | rs11121023 | 2.408 | 0.526 | -0.565 | -0.471 | 0 | 0.01 | 0.01 | 0 | 0.24 | 0.12 | 0.14 | 0.21 | 0.2 | 0.28 |
| 1 | 7850081 | rs10864315 | 1.433 | 0.559 | -0.536 | -0.451 | 0.045 | 0.084 | 0.084 | 0.737 | 0.4 | 0.22 | 0.21 | 0.22 | 0.29 | 0.33 |
| 1 | 7850218 | rs697686 | 0.592 | 0.609 | -0.405 | -0.376 | 0.027 | 0.046 | 0.046 | 0 | 0.29 | 0.57 | 0.26 | 0.45 | 0.37 | 0.38 |
| 1 | 7861684 | rs707467 | 1.442 | 0.597 | -0.383 | -0.51 | -0.011 | 0.054 | 0.054 | 0 | 0.23 | 0.31 | 0.16 | 0.49 | 0.2 | 0.31 |
| 1 | 7863293 | rs228642 | 0.024 | 0.594 | -0.391 | -0.55 | -0.011 | 0.024 | 0.024 | 0 | 0.61 | 0.33 | 0.49 | 0.2 | 0.56 | 0.56 |
| 1 | 7868725 | rs228666 | -0.946 | 0.609 | -0.428 | -0.463 | 0.015 | -0.008 | -0.008 | 0 | 0.32 | 0.2 | 0.5 | 0.33 | 0.31 | 0.3 |
| 1 | 7879322 | rs228690 | 1.804 | 0.668 | -0.438 | -0.459 | -0.014 | 0.001 | 0.001 | 0 | 0.07 | 0 | 0.06 | 0.04 | 0.08 | 0.07 |
| 1 | 7880683 | rs10462020 | 0.25 | 0.818 | -0.32 | -0.346 | -0.012 | 0.009 | 0.009 | 0.042 | 0.12 | 0.07 | 0.21 | 0.01 | 0.2 | 0.16 |
| 1 | 7881780 | rs10127838 | 1.962 | 0.749 | -0.336 | -0.367 | 0.001 | 0.025 | 0.025 | 0.002 | 0.24 | 0.15 | 0.15 | 0.14 | 0.19 | 0.29 |
| 1 | 7889941 | rs2640908 | 1.052 | 1.069 | -0.253 | -0.113 | -0.011 | 0.049 | 0.049 | 0 | 0.23 | 0.32 | 0.16 | 0.49 | 0.2 | 0.31 |
| 1 | 7897133 | rs10462021 | 0.102 | 1.226 | -0.404 | -0.172 | -0.012 | 0.009 | 0.009 | 0.042 | 0.12 | 0.07 | 0.21 | 0.01 | 0.2 | 0.16 |
| 1 | 7906008 | rs707472 | 0.871 | 0.584 | -0.453 | 0.16 | -0.013 | 0.056 | 0.056 | 0 | 0.23 | 0.33 | 0.16 | 0.48 | 0.2 | 0.31 |
| 1 | 7906914 | rs10864317 | 1.312 | 0.551 | -0.949 | 0.178 | 0.001 | 0.022 | 0.022 | 0.013 | 0.24 | 0.16 | 0.15 | 0.14 | 0.19 | 0.28 |
| 1 | 7908888 | rs228652 | -0.658 | 0.31 | -1.28 | 0.28 | 0.023 | -0.014 | -0.014 | 0 | 0.3 | 0.19 | 0.35 | 0.32 | 0.25 | 0.29 |
| 1 | 7910391 | rs170629 | 0.557 | 0.773 | -1.713 | 0.203 | -0.061 | 0.001 | 0.001 | 0.048 | 0.39 | 0.03 | 0.23 | 0.29 | 0.33 | 0.45 |
| 1 | 7911099 | rs228651 | 0.666 | 0.686 | -1.649 | -0.013 | -0.051 | 0.106 | 0.106 | 0.122 | 0.38 | 0.08 | 0.23 | 0.18 | 0.34 | 0.43 |
| 1 | 7916387 | rs11582983 | 0.409 | 0.237 | -1.501 | -0.294 | 0.018 | 0.022 | 0.022 | 0 | 0.24 | 0.16 | 0.12 | 0.09 | 0.16 | 0.15 |
| 1 | 7918106 | rs707476 | -0.209 | 0.549 | -1.348 | -0.335 | -0.01 | 0.079 | 0.079 | 0 | 0.69 | 0.84 | 0.69 | 0.91 | 0.68 | 0.69 |
| 1 | 7920423 | rs2859389 | 1.684 | 0.544 | -1.06 | -0.254 | 0.004 | 0.063 | 0.063 | 0 | 0.33 | 0.15 | 0.2 | 0.2 | 0.27 | 0.39 |
| 1 | 7921952 | rs531485 | 0.428 | 0.734 | -1.113 | -0.391 | -0.073 | 0.097 | 0.097 | 2.431 | 0.16 | 0.01 | 0.24 | 0.02 | 0.23 | 0.24 |
| 1 | 7924094 | rs12566535 | 0.733 | 0.596 | -0.899 | 0.226 | -0.002 | -0.001 | -0.001 | 0 | 0.23 | 0.2 | 0.13 | 0.27 | 0.15 | 0.16 |
| 1 | 7926542 | rs504560 | -0.388 | 1.121 | -0.099 | 0.077 | 0.106 | 0.073 | 0.073 | 2.222 | 0.81 | 0.51 | 0.81 | 0.73 | 0.71 | 0.86 |
| 1 | 7927981 | rs579992 | -1.307 | 0.912 | -0.68 | -0.864 | -0.002 | -0.018 | -0.018 | 0.02 | 0.86 | 0.85 | 0.92 | 0.97 | 0.9 | 0.92 |
| 1 | 7929819 | rs228725 | -1.333 | 1.228 | -0.307 | -0.557 | 0.024 | 0.114 | 0.114 | 0.164 | 0.55 | 0.81 | 0.73 | 0.74 | 0.64 | 0.58 |
| 1 | 7936272 | rs228716 | -1.282 | 1.218 | -0.144 | -0.557 | 0.022 | 0.06 | 0.06 | 0.151 | 0.54 | 0.66 | 0.72 | 0.75 | 0.64 | 0.58 |
| 1 | 7944264 | rs228703 | -0.775 | 1.446 | -0.61 | -0.935 | 0.012 | 0.117 | 0.117 | 0.28 | 0.43 | 0.22 | 0.39 | 0.04 | 0.43 | 0.47 |
| 1 | 7952427 | rs1040397 | -0.744 | 2.128 | -0.346 | -0.441 | 0.001 | 0.005 | 0.005 | 0.383 | 0.12 | 0.12 | 0.26 | 0.06 | 0.33 | 0.27 |
| 1 | 7955894 | rs170551 | -0.835 | 1.319 | -0.214 | -0.167 | -0.002 | 0.025 | 0.025 | 0.793 | 0.17 | 0.23 | 0.42 | 0.69 | 0.15 | 0.29 |
| 1 | 7961206 | rs227163 | -0.964 | 0.76 | -0.534 | -0.423 | 0.023 | 0.106 | 0.106 | 0.833 | 0.48 | 0.77 | 0.7 | 0.73 | 0.59 | 0.55 |
| 1 | 7969166 | rs6656926 | 0.076 | 0.303 | 0.396 | -0.359 | 0.038 | 0.034 | 0.034 | 0.508 | 0.06 | 0.54 | 0.04 | 0 | 0 | 0 |

| | | | | | | | | | | | | | | | | |
|---|---|---|---|---|---|---|---|---|---|---|---|---|---|---|---|---|
| 1 | 7971078 | rs679563 | 0.096 | 0.948 | -0.087 | -0.874 | -0.025 | 0.039 | 0.039 | 1.116 | 0.69 | 0.74 | 0.71 | 0.93 | 0.47 | 0.78 |
| 1 | 7973170 | rs585672 | 0.078 | 1.658 | -0.038 | -0.987 | -0.023 | 0.021 | 0.021 | 0.148 | 0.7 | 0.77 | 0.8 | 1 | 0.64 | 0.82 |
| 1 | 7982034 | rs161827 | -0.878 | 1.027 | 0.81 | -0.127 | -0.042 | 0.265 | 0.265 | 0 | 0.1 | 0.28 | 0.26 | 0.55 | 0.03 | 0.05 |
| 1 | 7989566 | rs2453021 | 1.608 | 0.78 | 0.197 | -0.694 | 0.05 | 0.131 | 0.131 | 1.226 | 0.45 | 0.07 | 0.35 | 0.3 | 0.33 | 0.43 |
| 1 | 7992615 | rs863171 | 0.873 | 0.843 | 0.691 | -0.542 | -0.011 | 0.026 | 0.026 | 0.222 | 0.59 | 0.24 | 0.43 | 0.39 | 0.45 | 0.65 |
| 1 | 7999602 | rs161811 | -0.1 | 0.86 | 0.825 | -0.392 | -0.021 | 0.233 | 0.233 | 0.017 | 0.06 | 0.15 | 0.26 | 0.55 | 0.03 | 0.05 |
| 1 | 8000797 | rs161810 | -0.643 | 0.839 | 1.525 | -0.361 | -0.038 | 0.319 | 0.319 | 0 | 0.1 | 0.35 | 0.27 | 0.54 | 0.03 | 0.05 |
| 1 | 8009763 | rs226474 | 0.233 | 0.666 | 1.034 | -0.796 | -0.04 | 0.1 | 0.1 | 1.124 | 0.65 | 0.8 | 0.7 | 0.95 | 0.45 | 0.7 |
| 1 | 8022197 | rs3766606 | -1.797 | 0.67 | 1.399 | -0.548 | 0.127 | 0.041 | 0.041 | 0.432 | 0.08 | 0.36 | 0.12 | 0.06 | 0.16 | 0.11 |
| 1 | 8024592 | rs4908488 | 0.33 | -0.273 | 0.612 | -1.078 | 0.029 | 0.062 | 0.062 | 0 | 0.22 | 0.09 | 0.21 | 0.07 | 0.15 | 0.29 |
| 1 | 8042736 | rs161801 | 0.128 | 0.324 | 1.053 | -0.775 | -0.023 | 0.004 | 0.004 | 0.819 | 0.35 | 0.29 | 0.25 | 0 | 0.43 | 0.2 |
| 1 | 8044361 | rs225119 | -1.024 | 0.235 | 1.004 | -0.76 | 0.027 | 0.106 | 0.106 | 0 | 0.48 | 0.91 | 0.63 | 0.65 | 0.59 | 0.35 |
| 1 | 8066914 | rs225100 | -1.006 | 0.292 | 0.977 | -0.723 | 0.035 | 0.105 | 0.105 | 0 | 0.48 | 0.89 | 0.63 | 0.65 | 0.59 | 0.35 |
| 1 | 8078309 | rs400736 | -0.722 | 0.284 | 0.972 | -0.701 | 0.005 | 0.078 | 0.078 | 0 | 0.45 | 0.89 | 0.62 | 0.65 | 0.58 | 0.32 |
| 1 | 8098462 | rs225133 | -0.011 | 0.311 | 1.197 | -0.608 | -0.009 | -0.001 | -0.001 | 0 | 0.38 | 0.36 | 0.53 | 0.6 | 0.45 | 0.23 |
| 1 | 8100173 | rs6577459 | -0.052 | 0.378 | 1.03 | -0.412 | 0.002 | -0.001 | -0.001 | 0 | 0.08 | 0.09 | 0.09 | 0.06 | 0.16 | 0.12 |
| 1 | 8151224 | rs11121086 | 0.501 | 0.062 | 0.687 | -0.749 | 0.118 | 0.149 | 0.149 | 0 | 0.5 | 0.22 | 0.34 | 0.28 | 0.36 | 0.64 |
| 1 | 8154547 | rs10864327 | 0.275 | 0.302 | 0.607 | -0.727 | -0.13 | 0.214 | 0.214 | 3.058 | 0.34 | 0.05 | 0.24 | 0 | 0.47 | 0.22 |
| 1 | 8168564 | rs10864330 | 0.521 | 0.261 | 0.778 | -0.711 | 0.043 | 0.048 | 0.048 | 0.093 | 0.53 | 0.36 | 0.34 | 0.29 | 0.36 | 0.56 |
| 1 | 8177028 | rs12124756 | 0.317 | 0.323 | 0.361 | -0.731 | -0.078 | 0.143 | 0.143 | 2.648 | 0.35 | 0.1 | 0.25 | 0 | 0.47 | 0.22 |
| 1 | 8177564 | rs7537362 | -0.743 | 0.397 | 0.204 | -0.655 | 0.005 | 0.016 | 0.016 | 0.83 | 0.1 | 0.11 | 0.38 | 0.7 | 0.17 | 0.22 |
| 1 | 8182535 | rs6577466 | -0.235 | 0.308 | 0.088 | -0.974 | 0.011 | -0.116 | -0.116 | 0 | 0.63 | 0.48 | 0.72 | 1 | 0.53 | 0.78 |
| 1 | 8187211 | rs4908736 | -0.642 | -0.265 | -0.007 | 0.115 | -0.042 | 0.041 | 0.041 | 0 | 0.22 | 0.32 | 0.45 | 0.7 | 0.11 | 0.26 |
| 1 | 8189854 | rs7527904 | -0.354 | 0.507 | -0.753 | -0.523 | -0.048 | 0.147 | 0.147 | 0.036 | 0.7 | 0.95 | 0.83 | 1 | 0.69 | 0.81 |
| 1 | 8191792 | rs12136685 | 0.414 | 0.153 | -0.411 | 0.218 | 0.032 | -0.014 | -0.014 | 0 | 0.58 | 0.27 | 0.42 | 0.52 | 0.65 | 0.66 |
| 1 | 8191889 | rs12144409 | 0.414 | 0.153 | -0.422 | 0.218 | 0.033 | -0.015 | -0.015 | 0.462 | 0.58 | 0.27 | 0.42 | 0.52 | 0.65 | 0.66 |
| 1 | 8192898 | rs7545284 | 1.251 | 0.398 | -0.817 | 0.108 | -0.044 | -0.014 | -0.014 | 0.234 | 0.41 | 0.18 | 0.29 | 0.4 | 0.45 | 0.36 |
| 1 | 8193294 | rs1542387 | -0.005 | 0.023 | -0.147 | -0.3 | 0.015 | 0.142 | 0.142 | 0 | 0.65 | 0.89 | 0.74 | 0.98 | 0.69 | 0.65 |
| 1 | 8195226 | rs7529511 | 0.047 | 0.118 | 0.268 | 0.056 | -0.011 | -0.012 | -0.012 | 0.006 | 0.64 | 0.53 | 0.45 | 0.64 | 0.67 | 0.71 |
| 1 | 8195724 | rs10746477 | 0.169 | 0.323 | -0.192 | -0.814 | -0.006 | -0.094 | -0.094 | 0 | 0.66 | 0.39 | 0.78 | 1 | 0.72 | 0.83 |
| 1 | 8200328 | rs9434617 | -0.419 | 1.098 | -1.051 | -0.089 | -0.019 | 0.035 | 0.035 | 0 | 0.36 | 0.74 | 0.28 | 0.23 | 0.36 | 0.25 |
| 1 | 8200578 | rs9434618 | 0.329 | 0.491 | -1.376 | -0.442 | 0.037 | -0.005 | -0.005 | 0.386 | 0.85 | 1 | 0.96 | 0.88 | 0.95 | 0.74 |
| 1 | 8200690 | rs6701300 | 0.407 | 1.086 | -1.601 | -0.817 | -0.038 | 0.239 | 0.239 | 0.261 | 0.52 | 0.89 | 0.72 | 0.8 | 0.53 | 0.47 |
| 1 | 8212425 | rs12564077 | 1.113 | 1.949 | 0.038 | 0.054 | 0.088 | -0.013 | -0.013 | 0.797 | 0.75 | 0.55 | 0.76 | 0.74 | 0.66 | 0.65 |
| 1 | 8214803 | rs2100516 | 1.099 | 2.028 | 0.216 | 0.115 | 0.085 | -0.005 | -0.005 | 0 | 0.77 | 0.64 | 0.77 | 0.74 | 0.67 | 0.63 |
| 1 | 8217087 | rs12048097 | 1 | 2.222 | -0.202 | -0.002 | -0.046 | 0.016 | 0.016 | 2.566 | 0.78 | 0.86 | 0.78 | 0.79 | 0.69 | 0.69 |
| 1 | 8217303 | rs11121117 | 0.856 | 2.307 | -0.173 | 0.028 | -0.025 | 0.005 | 0.005 | 0.781 | 0.81 | 0.86 | 0.82 | 0.79 | 0.75 | 0.71 |
| 1 | 8220106 | rs10864334 | 0.55 | 2.341 | -0.249 | 0.044 | -0.046 | 0.001 | 0.001 | 1.13 | 0.82 | 0.91 | 0.83 | 0.79 | 0.74 | 0.71 |
| 1 | 8222156 | rs17032281 | 0.907 | 1.419 | -0.321 | 0.054 | 0.054 | 0.048 | 0.048 | 0 | 0.45 | 0.25 | 0.51 | 0.33 | 0.37 | 0.34 |
| 1 | 8223832 | rs7554600 | 0.571 | 1.055 | 0.294 | 0.313 | -0.058 | -0.001 | -0.001 | 0.336 | 0.82 | 0.91 | 0.83 | 0.79 | 0.74 | 0.72 |
| 1 | 8238978 | rs11803446 | -0.441 | 0.789 | 1.001 | 0.307 | 0.062 | 0.027 | 0.027 | 0.263 | 0.07 | 0.49 | 0.04 | 0.01 | 0.02 | 0.06 |
| 1 | 8244145 | rs6663882 | 1.642 | 0.498 | 0.69 | 0.064 | 0.001 | -0.014 | -0.014 | 0.276 | 0.65 | 0.4 | 0.72 | 0.8 | 0.64 | 0.59 |
| 1 | 8256313 | rs11121127 | -0.639 | 0.876 | 0.655 | 0.597 | 0.077 | 0.028 | 0.028 | 0.168 | 0.09 | 0.17 | 0.17 | 0.26 | 0.23 | 0.21 |
| 1 | 8260834 | rs11587140 | -0.486 | 0.883 | 0.465 | 0.599 | 0.01 | -0.003 | -0.003 | 0 | 0.09 | 0.1 | 0.17 | 0.25 | 0.23 | 0.2 |
| 1 | 8264457 | rs12745477 | -0.432 | 0.886 | 0.54 | 0.592 | -0.012 | -0.014 | -0.014 | 0.574 | 0.09 | 0.07 | 0.17 | 0.25 | 0.23 | 0.21 |
| 1 | 8267513 | rs7548511 | -1.549 | 0.339 | 0.807 | 0.814 | 0.012 | -0.001 | -0.001 | 0 | 0.2 | 0.23 | 0.43 | 0.29 | 0.26 | 0.29 |
| 1 | 8268095 | rs11121129 | -1.419 | 0.358 | 0.575 | 0.823 | -0.015 | -0.001 | -0.001 | 0.058 | 0.19 | 0.14 | 0.42 | 0.29 | 0.25 | 0.3 |
| 1 | 8268715 | rs6701941 | -1.435 | 0.359 | 0.576 | 0.811 | -0.015 | 0.001 | 0.001 | 1.124 | 0.19 | 0.14 | 0.42 | 0.29 | 0.25 | 0.3 |
| 1 | 8272526 | rs425371 | -1.709 | 0.137 | 0.683 | 1.216 | 0.013 | 0.082 | 0.082 | 0.001 | 0.23 | 0.53 | 0.44 | 0.48 | 0.25 | 0.35 |
| 1 | 8273392 | rs12041676 | 0.215 | 0.622 | 0.14 | 1.486 | -0.043 | -0.013 | -0.013 | 0 | 0.07 | 0.01 | 0.12 | 0.41 | 0.18 | 0.23 |
| 1 | 8273940 | rs395847 | -0.126 | -0.009 | -0.238 | 1.227 | -0.013 | -0.025 | -0.025 | 0.294 | 0.85 | 0.99 | 0.91 | 0.87 | 0.87 | 0.88 |
| 1 | 8274018 | rs390468 | -0.981 | -0.02 | 0.466 | 1.228 | 0.018 | 0.029 | 0.029 | 0 | 0.05 | 0.2 | 0.05 | 0.04 | 0.06 | 0.06 |
| 1 | 8274807 | rs411654 | -1.04 | -0.361 | 0.799 | 1.094 | -0.034 | -0.005 | -0.005 | 0.105 | 0.18 | 0.46 | 0.11 | 0.1 | 0.11 | 0.19 |
| 1 | 8278899 | rs428180 | -1.253 | -0.623 | 0.691 | 1.09 | 0.041 | 0.107 | 0.107 | 0.822 | 0.21 | 0.44 | 0.21 | 0.43 | 0.26 | 0.44 |
| 1 | 8280517 | rs10864340 | -0.853 | -0.575 | 0.464 | 1 | 0.009 | -0.022 | -0.022 | 0.406 | 0.06 | 0.25 | 0.04 | 0.01 | 0.06 | 0.1 |
| 1 | 8281564 | rs506403 | -0.108 | -1.002 | -0.388 | 0.911 | 0.02 | -0.06 | -0.06 | 0.02 | 0.71 | 0.91 | 0.73 | 0.65 | 0.8 | 0.73 |
| 1 | 8284497 | rs11589993 | -0.217 | -0.424 | 0.078 | 1.109 | -0.026 | -0.034 | -0.034 | 0 | 0.14 | 0.07 | 0.16 | 0.43 | 0.2 | 0.34 |
| 1 | 8289660 | rs382192 | -1.244 | -0.718 | 0.334 | 0.47 | 0.008 | 0.002 | 0.002 | 2.899 | 0.33 | 0.49 | 0.29 | 0.67 | 0.34 | 0.55 |
| 1 | 8292761 | rs299499 | -0.807 | -0.845 | 0.244 | -0.074 | -0.005 | 0.025 | 0.025 | 3.472 | 0.55 | 0.55 | 0.49 | 0.81 | 0.49 | 0.69 |
| 1 | 8294762 | rs11121142 | 0.776 | -0.85 | -0.324 | -0.248 | 0.012 | 0.026 | 0.026 | 0 | 0.2 | 0.13 | 0.11 | 0.06 | 0.14 | 0.14 |
| 1 | 8295130 | rs7528376 | -0.396 | -0.671 | -0.577 | 0.079 | -0.015 | -0.015 | -0.015 | 0 | 0.33 | 0.18 | 0.35 | 0.67 | 0.34 | 0.54 |
| 1 | 8304607 | rs4908744 | 0.806 | -1.124 | -0.663 | -0.378 | -0.009 | 0.024 | 0.024 | 6.242 | 0.33 | 0.27 | 0.43 | 0.08 | 0.42 | 0.17 |
| 1 | 8313894 | rs4301655 | 0.738 | -1.952 | -0.557 | 0.194 | 0.041 | -0.014 | -0.014 | 0 | 0.63 | 0.76 | 0.68 | 0.56 | 0.79 | 0.54 |
| 1 | 8325211 | rs2661865 | 0.47 | -1.747 | -0.805 | 0.188 | 0.064 | -0.07 | -0.07 | 1.352 | 0.69 | 0.9 | 0.71 | 0.59 | 0.81 | 0.61 |
| 1 | 8338355 | rs438895 | 0.642 | -1.246 | 0.005 | -0.537 | -0.046 | -0.062 | -0.062 | 0 | 0.86 | 0.6 | 0.81 | 0.98 | 0.93 | 0.86 |
| 1 | 8352398 | rs299689 | -0.063 | -1.046 | -0.685 | -0.602 | 0.038 | 0.044 | 0.044 | 0.034 | 0.56 | 0.35 | 0.68 | 0.76 | 0.7 | 0.67 |
| 1 | 8368980 | rs10864346 | 0.016 | -1.608 | -0.27 | -1.196 | -0.056 | -0.095 | -0.095 | 0 | 0.75 | 0.61 | 0.82 | 0.97 | 0.85 | 0.79 |
| 1 | 8370088 | rs11121157 | 0.449 | -0.806 | -1.341 | -1.557 | -0.04 | 0.105 | 0.105 | 0 | 0.31 | 0.03 | 0.17 | 0.12 | 0.32 | 0.19 |
| 1 | 8373208 | rs966134 | -0.055 | -0.933 | -0.889 | -1.734 | 0.036 | 0.095 | 0.095 | 0 | 0.45 | 0.27 | 0.63 | 0.75 | 0.6 | 0.48 |

| | | | | | | | | | | | | | | | | |
|---|---|---|---|---|---|---|---|---|---|---|---|---|---|---|---|---|
| 1 | 8375913 | rs1466654 | -0.245 | -0.567 | 0.026 | -1.821 | 0.002 | 0.018 | 0.018 | 0 | 0.45 | 0.62 | 0.62 | 0.74 | 0.54 | 0.46 |
| 1 | 8382607 | rs12132135 | -0.989 | -0.844 | -0.107 | -2.008 | -0.016 | -0.038 | -0.038 | 0.51 | 0.7 | 0.5 | 0.76 | 0.81 | 0.76 | 0.79 |
| 1 | 8382681 | rs7517675 | -0.049 | -0.584 | -0.27 | -1.867 | 0.002 | 0.054 | 0.054 | 0 | 0.43 | 0.36 | 0.59 | 0.76 | 0.54 | 0.46 |
| 1 | 8385059 | rs2292242 | -0.134 | -0.724 | -0.526 | -1.841 | 0.022 | 0.084 | 0.084 | 0 | 0.44 | 0.29 | 0.6 | 0.76 | 0.58 | 0.47 |
| 1 | 8390888 | rs2289731 | 0.154 | -0.33 | -0.489 | -2.082 | 0.011 | 0.019 | 0.019 | 0.039 | 0.46 | 0.54 | 0.29 | 0.17 | 0.33 | 0.3 |
| 1 | 8395441 | rs2305016 | 0.068 | -0.112 | -0.454 | -2.289 | -0.001 | 0.044 | 0.044 | 0.179 | 0.55 | 0.28 | 0.4 | 0.29 | 0.58 | 0.39 |
| 1 | 8395560 | rs7535752 | 0.729 | -0.551 | -0.596 | -2.637 | 0.055 | 0.09 | 0.09 | 0.514 | 0.27 | 0.1 | 0.19 | 0.02 | 0.18 | 0.32 |
| 1 | 8412935 | rs8627 | 0.224 | -0.343 | -0.308 | -2.905 | -0.007 | 0.061 | 0.061 | 0.327 | 0.43 | 0.3 | 0.56 | 0.84 | 0.47 | 0.41 |
| 1 | 8422676 | rs2252865 | -0.222 | -0.504 | -0.12 | -3.119 | -0.019 | 0.035 | 0.035 | 0 | 0.78 | 0.84 | 0.78 | 0.92 | 0.68 | 0.75 |
| 1 | 8425900 | rs3753275 | 0.905 | -0.66 | 0.225 | -3.186 | -0.037 | -0.089 | -0.089 | 0 | 0.31 | 0.41 | 0.19 | 0.06 | 0.2 | 0.31 |
| 1 | 8445360 | rs3765971 | -0.149 | -0.693 | 0.77 | -3.248 | 0.028 | -0.09 | -0.09 | 0 | 0.77 | 0.65 | 0.78 | 0.92 | 0.69 | 0.78 |
| 1 | 8455712 | rs11121179 | 1.063 | -0.706 | 0.119 | -3.241 | 0.067 | 0.107 | 0.107 | 0 | 0.29 | 0.12 | 0.17 | 0.06 | 0.19 | 0.31 |
| 1 | 8488565 | rs301798 | -0.128 | -0.398 | 0.125 | -3.092 | 0.009 | -0.035 | -0.035 | 1.178 | 0.22 | 0.29 | 0.22 | 0.08 | 0.31 | 0.22 |
| 1 | 8497558 | rs2784739 | 0.675 | -0.278 | -0.62 | -3 | -0.027 | 0.103 | 0.103 | 0.023 | 0.33 | 0.03 | 0.19 | 0.11 | 0.27 | 0.22 |
| 1 | 8522553 | rs11121182 | 0.847 | -0.298 | -0.522 | -3.162 | 0.004 | 0.051 | 0.051 | 0 | 0.24 | 0.05 | 0.22 | 0.1 | 0.22 | 0.18 |
| 1 | 8554471 | rs4908501 | 0.64 | -0.099 | -0.757 | -3.463 | -0.057 | 0.083 | 0.083 | 0 | 0.2 | 0.06 | 0.19 | 0.06 | 0.32 | 0.19 |
| 1 | 8563663 | rs4908762 | -1.814 | -0.148 | -0.581 | -3.611 | 0.066 | 0.079 | 0.079 | 0 | 0.63 | 0.8 | 0.83 | 0.94 | 0.8 | 0.69 |
| 1 | 8580960 | rs7545384 | 1.366 | -0.068 | -0.615 | -3.633 | -0.068 | -0.059 | -0.059 | 0 | 0.11 | 0.02 | 0.36 | 0.77 | 0.24 | 0.25 |
| 1 | 8586563 | rs2297867 | -1.875 | -0.19 | -0.6 | -3.643 | 0.06 | 0.072 | 0.072 | 0 | 0.64 | 0.8 | 0.83 | 0.94 | 0.8 | 0.69 |
| 1 | 8598005 | rs10492966 | 1.114 | 0.104 | -0.55 | -3.468 | 0.006 | 0.043 | 0.043 | 0 | 0.24 | 0.09 | 0.22 | 0.11 | 0.22 | 0.18 |
| 1 | 8598127 | rs10492965 | 1.155 | 0.058 | -0.638 | -3.465 | 0.001 | 0.049 | 0.049 | 0 | 0.24 | 0.04 | 0.22 | 0.11 | 0.21 | 0.18 |
| 1 | 8629380 | rs10492964 | 0.492 | 0.622 | -0.603 | -3.293 | -0.095 | -0.064 | -0.064 | 0.018 | 0.14 | 0.02 | 0.38 | 0.77 | 0.3 | 0.24 |
| 1 | 8648240 | rs11121203 | 1.143 | -0.221 | -0.687 | -3.439 | -0.005 | 0.059 | 0.059 | 0 | 0.26 | 0.03 | 0.22 | 0.11 | 0.22 | 0.2 |
| 1 | 8699682 | rs10779705 | -1.806 | -0.485 | -0.683 | -3.502 | 0.093 | 0.176 | 0.176 | 0 | 0.67 | 0.91 | 0.85 | 0.94 | 0.81 | 0.65 |
| 1 | 8750591 | rs7533442 | -1.676 | -0.414 | -0.731 | -3.489 | 0.086 | 0.138 | 0.138 | 0 | 0.65 | 0.86 | 0.85 | 0.94 | 0.8 | 0.65 |
| 1 | 8751268 | rs6658881 | -1.715 | -0.41 | -0.729 | -3.487 | 0.083 | 0.135 | 0.135 | 0 | 0.66 | 0.86 | 0.85 | 0.94 | 0.8 | 0.66 |
| 1 | 8767311 | rs10864363 | 1.076 | -0.215 | -0.701 | -3.474 | -0.078 | -0.035 | -0.035 | 0.205 | 0.2 | 0.03 | 0.38 | 0.78 | 0.26 | 0.23 |
| 1 | 8767780 | rs10864364 | -1.937 | -0.265 | -0.674 | -3.487 | 0.051 | 0.077 | 0.077 | 0 | 0.73 | 0.87 | 0.86 | 0.93 | 0.81 | 0.66 |
| 1 | 8770538 | rs12125525 | 0.347 | -0.143 | -0.685 | -3.51 | -0.023 | 0.045 | 0.045 | 0 | 0.21 | 0.13 | 0.19 | 0.06 | 0.3 | 0.21 |
| 1 | 8770883 | rs7553298 | 1.008 | -0.124 | -0.755 | -3.538 | 0.003 | 0.061 | 0.061 | 0 | 0.27 | 0.1 | 0.23 | 0.1 | 0.24 | 0.19 |
| 1 | 8774451 | rs953043 | 0.524 | -0.142 | -0.751 | -3.547 | -0.047 | 0.077 | 0.077 | 0 | 0.2 | 0.05 | 0.19 | 0.06 | 0.29 | 0.21 |
| 1 | 8776719 | rs953535 | 0.843 | -0.166 | -0.665 | -3.339 | 0.004 | 0.06 | 0.06 | 0 | 0.27 | 0.1 | 0.23 | 0.1 | 0.24 | 0.21 |
| 1 | 8804237 | rs4908777 | 0.437 | 0.215 | -1.025 | -3.23 | -0.048 | 0.069 | 0.069 | 0.036 | 0.29 | 0.06 | 0.25 | 0.12 | 0.31 | 0.29 |
| 1 | 8808185 | rs12024032 | -0.344 | -0.01 | -0.798 | -3.522 | 0 | -0.006 | -0.006 | 1.13 | 0.4 | 0.4 | 0.6 | 0.82 | 0.55 | 0.43 |
| 1 | 8816438 | rs11121225 | 1.717 | -0.031 | -0.95 | -3.506 | -0.049 | -0.004 | -0.004 | 4.148 | 0.2 | 0.01 | 0.2 | 0.11 | 0.21 | 0.15 |
| 1 | 8831283 | rs11121229 | 0.462 | -0.058 | -0.851 | -3.501 | -0.005 | 0.058 | 0.058 | 0 | 0.28 | 0.13 | 0.23 | 0.11 | 0.27 | 0.23 |
| 1 | 8832499 | rs10864366 | -1.939 | -0.259 | -0.722 | -3.558 | -0.042 | -0.082 | -0.082 | 0.484 | 0.7 | 0.59 | 0.83 | 0.93 | 0.82 | 0.66 |
| 1 | 8834706 | rs11121230 | 0.7 | -0.049 | -0.763 | -3.567 | -0.053 | 0.065 | 0.065 | 0 | 0.28 | 0.05 | 0.24 | 0.11 | 0.28 | 0.24 |
| 1 | 8839300 | rs7513880 | -0.173 | -0.083 | -0.73 | -3.57 | 0.001 | -0.014 | -0.014 | 0 | 0.4 | 0.41 | 0.59 | 0.82 | 0.54 | 0.43 |
| 1 | 8845199 | rs11586056 | 1.822 | -0.092 | -0.759 | -3.578 | -0.067 | 0 | 0 | 0.034 | 0.2 | 0.01 | 0.19 | 0.08 | 0.22 | 0.14 |
| 1 | 8847380 | rs10864367 | -0.226 | -0.116 | -0.672 | -3.51 | 0.003 | -0.024 | -0.024 | 0 | 0.4 | 0.43 | 0.59 | 0.82 | 0.54 | 0.42 |
| 1 | 8856603 | rs7524437 | 1.543 | 0.118 | -1.099 | -3.553 | -0.099 | -0.066 | -0.066 | 1.951 | 0.18 | 0.02 | 0.37 | 0.77 | 0.26 | 0.22 |
| 1 | 8868406 | rs12137865 | 0.338 | 0.053 | -0.888 | -3.401 | -0.004 | 0.054 | 0.054 | 0 | 0.3 | 0.16 | 0.24 | 0.11 | 0.28 | 0.24 |
| 1 | 8871690 | rs7520572 | 0.581 | 0.114 | -0.979 | -3.373 | -0.015 | 0.063 | 0.063 | 0 | 0.28 | 0.13 | 0.24 | 0.11 | 0.28 | 0.24 |
| 1 | 8871965 | rs7546225 | 0.28 | 0.2 | -0.986 | -3.307 | 0.023 | -0.075 | -0.075 | 0.578 | 0.21 | 0.3 | 0.21 | 0.05 | 0.31 | 0.2 |
| 1 | 8878885 | rs12410893 | 0.486 | 0.104 | -1.593 | -2.954 | -0.044 | 0.06 | 0.06 | 0.007 | 0.27 | 0.05 | 0.24 | 0.11 | 0.28 | 0.24 |
| 1 | 8906301 | rs11121242 | -0.025 | 0.14 | -1.346 | -2.83 | -0.01 | 0.229 | 0.229 | 0 | 0.44 | 0.13 | 0.61 | 0.88 | 0.53 | 0.43 |
| 1 | 8910347 | rs12135416 | -0.104 | -0.082 | -1.101 | -3.037 | -0.011 | -0.005 | -0.005 | 0 | 0.18 | 0.15 | 0.19 | 0.04 | 0.28 | 0.18 |
| 1 | 8911834 | rs6677249 | -1.472 | -0.527 | -0.479 | -3.233 | -0.064 | -0.092 | -0.092 | 0 | 0.71 | 0.55 | 0.83 | 0.92 | 0.82 | 0.65 |
| 1 | 8916517 | rs2038903 | 0.741 | -0.355 | -0.736 | -3.319 | -0.096 | 0.196 | 0.196 | 0 | 0.4 | 0.1 | 0.55 | 0.87 | 0.47 | 0.33 |
| 1 | 8918795 | rs11121247 | -1.523 | -0.659 | -0.141 | -3.165 | -0.02 | -0.041 | -0.041 | 0.586 | 0.72 | 0.69 | 0.85 | 0.92 | 0.81 | 0.66 |
| 1 | 8920712 | rs2071414 | 0.84 | -0.438 | -0.49 | -2.973 | -0.029 | -0.008 | -0.008 | 0 | 0.23 | 0.12 | 0.42 | 0.74 | 0.25 | 0.28 |
| 1 | 8926329 | rs2781066 | 1.041 | -0.268 | -0.634 | -3.068 | -0.037 | 0.038 | 0.038 | 0 | 0.08 | 0.02 | 0.05 | 0 | 0.16 | 0.13 |
| 1 | 8926868 | rs2781064 | 0.062 | -0.579 | -0.215 | -3.015 | -0.038 | 0.142 | 0.142 | 0.474 | 0.42 | 0.18 | 0.39 | 0.17 | 0.56 | 0.36 |
| 1 | 8931149 | rs2274971 | 0.66 | -0.389 | -0.316 | -2.845 | -0.027 | -0.007 | -0.007 | 0 | 0.23 | 0.13 | 0.42 | 0.74 | 0.25 | 0.28 |
| 1 | 8934237 | rs3820037 | -1.196 | -0.491 | -0.024 | -2.52 | -0.05 | -0.083 | -0.083 | 0.003 | 0.71 | 0.59 | 0.84 | 0.91 | 0.81 | 0.68 |
| 1 | 8949480 | rs7539600 | -1.117 | -0.526 | -0.363 | -2.531 | 0.045 | 0.074 | 0.074 | 0 | 0.7 | 0.84 | 0.82 | 0.9 | 0.77 | 0.67 |
| 1 | 8960153 | rs12034719 | 0.399 | -0.01 | -1.397 | -1.906 | -0.014 | -0.013 | -0.013 | 0 | 0.29 | 0.13 | 0.42 | 0.69 | 0.25 | 0.29 |
| 1 | 8960387 | rs12032404 | 0.399 | -0.01 | -1.397 | -1.922 | -0.014 | -0.011 | -0.011 | 0 | 0.29 | 0.13 | 0.42 | 0.69 | 0.25 | 0.29 |
| 1 | 8969000 | rs7526401 | 0.488 | 0.271 | -1.237 | -1.768 | -0.048 | 0.048 | 0.048 | 0.547 | 0.26 | 0.11 | 0.28 | 0.15 | 0.38 | 0.2 |
| 1 | 8969219 | rs7534552 | -1.119 | 0.103 | -1.352 | -1.935 | 0.027 | 0.135 | 0.135 | 0 | 0.53 | 0.82 | 0.54 | 0.24 | 0.62 | 0.53 |
| 1 | 8972122 | rs6682376 | -1.138 | 0.082 | -0.903 | -1.706 | 0.004 | 0.03 | 0.03 | 0 | 0.33 | 0.7 | 0.31 | 0.11 | 0.32 | 0.39 |
| 1 | 8978514 | rs6665631 | -0.592 | 0.265 | -0.768 | -1.457 | 0 | -0.059 | -0.059 | 0.144 | 0.26 | 0.56 | 0.25 | 0.13 | 0.25 | 0.33 |
| 1 | 8982906 | rs4908796 | -0.133 | 0.118 | -1.153 | -0.899 | -0.002 | 0.019 | 0.019 | 0 | 0.54 | 0.58 | 0.44 | 0.26 | 0.56 | 0.58 |
| 1 | 8983768 | rs11805233 | 0.171 | 0.178 | -1.411 | -0.786 | 0.001 | 0.111 | 0.111 | 0 | 0.5 | 0.71 | 0.4 | 0.25 | 0.53 | 0.43 |
| 1 | 8985036 | rs10864370 | -0.287 | 0.399 | -1.138 | 0.062 | 0 | 0.001 | 0.001 | 0 | 0.29 | 0.24 | 0.18 | 0.26 | 0.24 | 0.37 |
| 1 | 8987259 | rs2765512 | -0.339 | 0.539 | -1.372 | 0.451 | -0.036 | 0.009 | 0.009 | 0.197 | 0.61 | 0.73 | 0.67 | 0.71 | 0.51 | 0.64 |
| 1 | 8989235 | rs6577539 | -0.673 | 0.065 | -1.365 | 0.338 | -0.034 | 0.011 | 0.011 | 0 | 0.9 | 0.99 | 0.95 | 0.95 | 0.89 | 0.84 |
| 1 | 8991141 | rs4609454 | -0.211 | -0.254 | -0.706 | -0.025 | 0.004 | -0.006 | -0.006 | 0 | 0.49 | 0.51 | 0.48 | 0.83 | 0.31 | 0.58 |

| | | | | | | | | | | | | | | | | |
|---|---|---|---|---|---|---|---|---|---|---|---|---|---|---|---|---|
| 1 | 8991873 | rs9434652 | 0.689 | -0.201 | -0.586 | -0.098 | -0.024 | 0.088 | 0.088 | 0 | 0.38 | 0.24 | 0.44 | 0.12 | 0.57 | 0.25 |
| 1 | 8994112 | rs4418632 | -0.132 | -0.359 | -0.423 | -0.103 | 0.075 | 0.121 | 0.121 | 0 | 0.5 | 0.7 | 0.5 | 0.83 | 0.32 | 0.58 |
| 1 | 8998059 | rs914997 | 0.393 | -0.728 | -0.446 | 0.026 | 0.002 | 0.006 | 0.006 | 0 | 0.79 | 0.79 | 0.89 | 0.87 | 0.84 | 0.74 |
| 1 | 8999506 | rs6688840 | 0.636 | -1.092 | -0.438 | 1.044 | -0.022 | -0.021 | -0.021 | 0 | 0.06 | 0.01 | 0.09 | 0.24 | 0.07 | 0.17 |
| 1 | 9000220 | rs12091891 | 0.392 | -1.72 | -0.476 | 1.004 | -0.007 | -0.001 | -0.001 | 0 | 0.14 | 0.14 | 0.05 | 0.08 | 0.06 | 0.09 |
| 1 | 9000628 | rs1475734 | 1.04 | -1.878 | -0.165 | 1.337 | 0.001 | -0.001 | -0.001 | 0 | 0.18 | 0.14 | 0.21 | 0.24 | 0.13 | 0.22 |
| 1 | 9000882 | rs6692694 | -0.957 | -1.842 | -0.076 | 1.013 | -0.034 | 0.026 | 0.026 | 0 | 0.25 | 0.38 | 0.19 | 0.35 | 0.1 | 0.22 |
| 1 | 9001562 | rs17032875 | -0.78 | -1.03 | -0.144 | 0.545 | -0.016 | -0.013 | -0.013 | 0 | 0.36 | 0.41 | 0.46 | 0.75 | 0.23 | 0.52 |
| 1 | 9003178 | rs6684149 | -0.452 | -0.807 | -0.298 | 0.425 | -0.013 | -0.025 | -0.025 | 0.02 | 0.07 | 0.05 | 0.02 | 0.01 | 0.02 | 0.03 |
| 1 | 9005320 | rs1409147 | -0.466 | -0.144 | -0.714 | 0.433 | -0.05 | 0.004 | 0.004 | 0.063 | 0.78 | 0.88 | 0.66 | 0.8 | 0.72 | 0.72 |
| 1 | 9009406 | rs2274327 | 1.191 | -0.477 | -1.032 | 0.413 | -0.041 | 0.078 | 0.078 | 0 | 0.37 | 0.08 | 0.38 | 0.24 | 0.42 | 0.33 |
| 1 | 9009444 | rs2274328 | 0.666 | -0.569 | 0.089 | 0.751 | 0.001 | 0.004 | 0.004 | 0 | 0.52 | 0.56 | 0.47 | 0.41 | 0.54 | 0.5 |
| 1 | 9010405 | rs17032907 | -0.3 | -0.64 | -0.165 | 0.835 | 0.012 | -0.027 | -0.027 | 0 | 0.17 | 0.18 | 0.15 | 0.4 | 0.15 | 0.19 |
| 1 | 9011586 | rs3765968 | 0.464 | -0.399 | 0.64 | 0.178 | -0.002 | 0.001 | 0.001 | 0 | 0.62 | 0.71 | 0.55 | 0.7 | 0.64 | 0.6 |
| 1 | 9011722 | rs3765966 | -0.067 | -0.103 | 0.453 | 0.198 | -0.026 | 0.024 | 0.024 | 0 | 0.69 | 0.83 | 0.57 | 0.8 | 0.66 | 0.68 |
| 1 | 9015436 | rs12021597 | 0.12 | -0.054 | 0.01 | 0.149 | -0.003 | 0.006 | 0.006 | 0.109 | 0.64 | 0.35 | 0.55 | 0.7 | 0.7 | 0.62 |
| 1 | 9016302 | rs17389460 | -0.848 | -0.085 | -0.146 | -0.502 | 0 | -0.001 | -0.001 | 0 | 0.2 | 0.17 | 0.19 | 0.17 | 0.23 | 0.22 |
| 1 | 9017091 | rs2274332 | -0.348 | 0.026 | -0.09 | -0.449 | 0.001 | -0.003 | -0.003 | 0.039 | 0.16 | 0.14 | 0.17 | 0.15 | 0.21 | 0.15 |
| 1 | 9017204 | rs2274333 | 0.222 | -0.091 | -0.556 | -0.506 | 0.004 | -0.04 | -0.04 | 0.022 | 0.39 | 0.11 | 0.39 | 0.56 | 0.29 | 0.46 |
| 1 | 9020905 | rs1832262 | -1.602 | -0.112 | -0.9 | -0.52 | -0.094 | 0.065 | 0.065 | 0.172 | 0.68 | 0.93 | 0.62 | 0.8 | 0.53 | 0.66 |
| 1 | 9021053 | rs6691526 | -0.526 | -0.095 | -0.457 | -0.258 | 0.007 | -0.015 | -0.015 | 0 | 0.17 | 0.2 | 0.18 | 0.15 | 0.21 | 0.15 |
| 1 | 9027391 | rs4466678 | -0.322 | -0.103 | -0.22 | -0.022 | -0.02 | 0.014 | 0.014 | 0 | 0.08 | 0.19 | 0.05 | 0.09 | 0.03 | 0.05 |
| 1 | 9034421 | rs3765964 | -0.268 | -0.083 | -0.639 | -0.292 | 0.008 | 0.037 | 0.037 | 0 | 0.44 | 0.32 | 0.43 | 0.68 | 0.39 | 0.52 |
| 1 | 9037273 | rs6697763 | -0.484 | -0.129 | -1.371 | -0.336 | 0.049 | 0.19 | 0.19 | 2.148 | 0.52 | 0.92 | 0.43 | 0.75 | 0.33 | 0.5 |
| 1 | 9039022 | rs6577546 | 0.663 | 0.593 | -0.661 | -0.162 | 0.001 | -0.003 | -0.003 | 0 | 0.77 | 0.82 | 0.81 | 0.84 | 0.76 | 0.83 |
| 1 | 9044562 | rs17032942 | -1.005 | 0.736 | -0.024 | 0.114 | 0.046 | 0.055 | 0.055 | 0 | 0.07 | 0.24 | 0.07 | 0.2 | 0.1 | 0.09 |
| 1 | 9045405 | rs4908802 | -0.646 | 0.78 | -0.807 | -0.414 | -0.05 | 0.055 | 0.055 | 0 | 0.71 | 0.83 | 0.58 | 0.86 | 0.54 | 0.76 |
| 1 | 9049032 | rs875148 | 0.436 | 0.242 | -0.703 | -0.627 | -0.003 | 0.034 | 0.034 | 0 | 0.41 | 0.24 | 0.4 | 0.72 | 0.34 | 0.54 |
| 1 | 9058546 | rs2895997 | -1.092 | 0.158 | -0.602 | -0.697 | -0.073 | 0.065 | 0.065 | 0 | 0.7 | 0.8 | 0.57 | 0.86 | 0.5 | 0.76 |
| 1 | 9060170 | rs4908803 | 0.649 | -0.048 | -0.644 | -0.675 | -0.002 | 0 | 0 | 0 | 0.34 | 0.28 | 0.4 | 0.65 | 0.31 | 0.48 |
| 1 | 9060789 | rs17033008 | 0.042 | -0.237 | -0.416 | -0.602 | 0.008 | -0.003 | -0.003 | 0 | 0.13 | 0.09 | 0.09 | 0.16 | 0.07 | 0.1 |
| 1 | 9061161 | rs7516862 | 0.1 | -0.197 | -0.207 | -0.522 | -0.007 | -0.046 | -0.046 | 0 | 0.16 | 0.18 | 0.07 | 0.06 | 0.11 | 0.09 |
| 1 | 9061382 | rs12047015 | 1.308 | 0.4 | -0.958 | -0.598 | -0.043 | -0.016 | -0.016 | 0 | 0.14 | 0.09 | 0.22 | 0.14 | 0.25 | 0.22 |
| 1 | 9062952 | rs17389948 | 1.306 | 0.39 | -0.96 | -0.603 | -0.042 | -0.009 | -0.009 | 0 | 0.14 | 0.09 | 0.22 | 0.13 | 0.25 | 0.22 |
| 1 | 9063238 | rs11810507 | -0.181 | 0.147 | -0.786 | -0.643 | -0.051 | -0.026 | -0.026 | 0.72 | 0.24 | 0.4 | 0.15 | 0.22 | 0.15 | 0.19 |
| 1 | 9063265 | rs12402611 | 0.257 | 0.85 | -1.266 | 0.505 | -0.029 | -0.032 | -0.032 | 0.38 | 0.06 | 0 | 0.08 | 0.14 | 0.09 | 0.03 |
| 1 | 9066695 | rs10864377 | 0.458 | 0.535 | -1.104 | 0.321 | -0.002 | -0.02 | -0.02 | 0 | 0.37 | 0.28 | 0.42 | 0.59 | 0.34 | 0.46 |
| 1 | 9067142 | rs6672506 | 0.014 | 0.71 | -0.748 | 0.637 | -0.069 | -0.008 | -0.008 | 0 | 0.16 | 0.04 | 0.22 | 0.13 | 0.27 | 0.24 |
| 1 | 9068020 | rs2039631 | -1.524 | -0.049 | 0.315 | 0.899 | -0.001 | 0.085 | 0.085 | 0 | 0.09 | 0.31 | 0.09 | 0.13 | 0.05 | 0.09 |
| 1 | 9068386 | rs7530465 | -1.146 | -0.141 | -0.625 | 0.815 | 0.018 | 0.005 | 0.005 | 0 | 0.21 | 0.76 | 0.13 | 0.15 | 0.1 | 0.15 |
| 1 | 9073801 | rs7537425 | -0.533 | 0.989 | -0.189 | 1.716 | 0.011 | 0.025 | 0.025 | 0 | 0.34 | 0.39 | 0.52 | 0.56 | 0.36 | 0.33 |
| 1 | 9080966 | rs769902 | -0.566 | 1.22 | 0.277 | 1.75 | 0.025 | -0.008 | -0.008 | 0 | 0.29 | 0.4 | 0.21 | 0.25 | 0.36 | 0.33 |
| 1 | 9098230 | rs875996 | 0.487 | 0.595 | -0.575 | 1.61 | -0.006 | -0.021 | -0.021 | 0 | 0.25 | 0.01 | 0.25 | 0.16 | 0.17 | 0.2 |
| 1 | 9100479 | rs1060998 | 0.129 | 1.244 | -0.47 | 1.398 | 0.001 | -0.001 | -0.001 | 0 | 0.16 | 0.14 | 0.17 | 0.18 | 0.28 | 0.19 |
| 1 | 9116743 | rs12080175 | 0.175 | 0.6 | -0.519 | 1.428 | 0.015 | -0.114 | -0.114 | 0.155 | 0.32 | 0.11 | 0.45 | 0.58 | 0.26 | 0.37 |
| 1 | 9118596 | rs12145292 | -0.034 | 0.424 | -0.277 | 1.345 | 0.02 | -0.018 | -0.018 | 0 | 0.35 | 0.27 | 0.46 | 0.62 | 0.25 | 0.43 |
| 1 | 9121782 | rs765617 | -0.344 | 0.926 | -0.09 | 1.275 | 0.134 | 0.104 | 0.104 | 0.212 | 0.2 | 0.48 | 0.21 | 0.29 | 0.31 | 0.25 |
| 1 | 9122198 | rs17033196 | 0.762 | 0.649 | -0.506 | 1.009 | 0.017 | 0.009 | 0.009 | 0 | 0.07 | 0.02 | 0.03 | 0.04 | 0.04 | 0.07 |
| 1 | 9129867 | rs3820034 | 1.178 | 0.677 | -0.624 | 1.119 | -0.021 | -0.004 | -0.004 | 0 | 0.26 | 0.01 | 0.25 | 0.16 | 0.2 | 0.16 |
| 1 | 9129945 | rs770041 | -1.519 | 0.792 | -0.511 | 1.047 | -0.003 | 0.154 | 0.154 | 0 | 0.64 | 0.88 | 0.78 | 0.95 | 0.65 | 0.75 |
| 1 | 9130650 | rs12736085 | 0.109 | 0.796 | -0.421 | 1.142 | 0.015 | 0.011 | 0.011 | 0.397 | 0.37 | 0.27 | 0.36 | 0.27 | 0.27 | 0.3 |
| 1 | 9132520 | rs12117043 | -0.194 | 1.31 | -0.767 | 1.309 | -0.05 | -0.042 | -0.042 | 0.987 | 0.18 | 0.11 | 0.19 | 0.3 | 0.32 | 0.26 |
| 1 | 9140078 | rs12726360 | -0.943 | 0.963 | -0.997 | 1.184 | 0.004 | -0.037 | -0.037 | 1.653 | 0.22 | 0.19 | 0.31 | 0.5 | 0.2 | 0.35 |
| 1 | 9140327 | rs12127491 | 1.209 | 0.543 | -1.523 | 0.818 | -0.035 | -0.023 | -0.023 | 0 | 0.26 | 0.01 | 0.27 | 0.18 | 0.22 | 0.14 |
| 1 | 9143332 | rs770036 | -1.294 | 0.399 | -1.294 | 0.452 | -0.035 | 0.182 | 0.182 | 0 | 0.68 | 0.92 | 0.78 | 0.99 | 0.66 | 0.74 |
| 1 | 9145013 | rs12057692 | -1.042 | 0.337 | -0.24 | 0.994 | -0.035 | 0.102 | 0.102 | 0 | 0.22 | 0.39 | 0.26 | 0.52 | 0.15 | 0.32 |
| 1 | 9147856 | rs12751466 | -0.731 | 0.438 | -0.453 | 1.087 | -0.011 | 0.049 | 0.049 | 0 | 0.19 | 0.23 | 0.19 | 0.46 | 0.15 | 0.25 |
| 1 | 9152080 | rs12738912 | -0.362 | 0.466 | -0.686 | 1.063 | 0 | -0.026 | -0.026 | 0 | 0.16 | 0.07 | 0.19 | 0.23 | 0.15 | 0.2 |
| 1 | 9155711 | rs12239637 | -0.452 | 0.402 | -0.654 | 1.067 | 0.004 | -0.003 | -0.003 | 0.062 | 0.17 | 0.11 | 0.18 | 0.22 | 0.15 | 0.2 |
| 1 | 9161619 | rs6694515 | -0.113 | 0.403 | -1.026 | 0.263 | -0.012 | 0.115 | 0.115 | 0 | 0.58 | 0.79 | 0.69 | 0.94 | 0.54 | 0.66 |
| 1 | 9164133 | rs3737669 | 0.317 | 0.193 | -0.058 | 0.579 | 0.006 | 0.012 | 0.012 | 0 | 0.52 | 0.42 | 0.55 | 0.48 | 0.54 | 0.52 |
| 1 | 9165818 | rs12753196 | 0.202 | 0.145 | -0.316 | 0.048 | -0.002 | 0 | 0 | 0.008 | 0.13 | 0.08 | 0.08 | 0.04 | 0.13 | 0.11 |
| 1 | 9170878 | rs770028 | -0.245 | 0.135 | -0.057 | 0.043 | 0.002 | -0.069 | -0.069 | 0.211 | 0.16 | 0.3 | 0.11 | 0.04 | 0.14 | 0.11 |
| 1 | 9173043 | rs6577548 | -0.888 | 0.005 | -0.437 | -0.073 | -0.027 | 0.115 | 0.115 | 0 | 0.72 | 0.89 | 0.78 | 0.98 | 0.67 | 0.77 |
| 1 | 9177317 | rs1705299 | -1.054 | -0.009 | -0.451 | -0.081 | -0.046 | 0.137 | 0.137 | 0 | 0.74 | 0.94 | 0.8 | 0.98 | 0.68 | 0.77 |
| 1 | 9182914 | rs10864390 | -0.791 | -0.002 | -0.388 | -0.052 | -0.008 | 0.055 | 0.055 | 0 | 0.72 | 0.83 | 0.78 | 0.98 | 0.67 | 0.77 |
| 1 | 9185660 | rs4908812 | 0.203 | -0.011 | -0.504 | 0.196 | 0.061 | 0.081 | 0.081 | 1.241 | 0.19 | 0.32 | 0.26 | 0.28 | 0.29 | 0.33 |
| 1 | 9190254 | rs12726928 | 0.63 | 0.062 | -1.26 | -0.356 | -0.015 | 0.049 | 0.049 | 0.045 | 0.12 | 0.04 | 0.08 | 0.03 | 0.13 | 0.11 |
| 1 | 9194614 | rs1001567 | -0.077 | 0.139 | -1.526 | -0.404 | -0.011 | 0.041 | 0.041 | 0 | 0.59 | 0.72 | 0.71 | 0.87 | 0.54 | 0.67 |

| | | | | | | | | | | | | | | | | |
|---|---|---|---|---|---|---|---|---|---|---|---|---|---|---|---|---|
| 1 | 9210262 | rs17393144 | 1.057 | 0.227 | -2.339 | -0.328 | -0.108 | 0.145 | 0.145 | 0 | 0.24 | 0.01 | 0.18 | 0.02 | 0.33 | 0.19 |
| 1 | 9211303 | rs12128240 | 1.329 | -0.081 | -2.553 | -0.259 | -0.035 | 0.036 | 0.036 | 0.181 | 0.27 | 0.02 | 0.29 | 0.18 | 0.23 | 0.16 |
| 1 | 9218721 | rs12119878 | 0.979 | 0.091 | -2.666 | -0.4 | -0.014 | 0.094 | 0.094 | 0.031 | 0.35 | 0.06 | 0.33 | 0.17 | 0.31 | 0.2 |
| 1 | 9219383 | rs6577556 | 0.119 | 0.768 | -2.183 | 0.128 | -0.077 | 0.046 | 0.046 | 0 | 0.23 | 0.04 | 0.24 | 0.17 | 0.32 | 0.27 |
| 1 | 9223160 | rs797252 | -0.918 | 0.192 | -1.695 | 0.833 | 0.014 | -0.015 | -0.015 | 0 | 0.93 | 0.95 | 0.9 | 0.81 | 0.97 | 0.96 |
| 1 | 9234143 | rs4908527 | -0.288 | 0.864 | -1.591 | 0.854 | -0.043 | -0.033 | -0.033 | 0 | 0.31 | 0.24 | 0.4 | 0.46 | 0.44 | 0.46 |
| 1 | 9238553 | rs797263 | 0.25 | 0.14 | -1.16 | 0.188 | 0 | 0 | 0 | 0.445 | 0.35 | 0.3 | 0.2 | 0.19 | 0.23 | 0.25 |
| 1 | 9245572 | rs4908817 | 1.146 | 0.223 | -1.242 | -0.033 | 0.014 | 0.029 | 0.029 | 0 | 0.22 | 0.08 | 0.17 | 0.11 | 0.16 | 0.24 |
| 1 | 9250991 | rs10864392 | -0.366 | 0.4 | -0.736 | -0.007 | -0.014 | 0.068 | 0.068 | 0 | 0.43 | 0.24 | 0.34 | 0.24 | 0.48 | 0.34 |
| 1 | 9256200 | rs4908823 | 0.607 | 0.244 | -0.274 | -0.268 | 0 | -0.009 | -0.009 | 0.551 | 0.42 | 0.45 | 0.36 | 0.23 | 0.36 | 0.44 |
| 1 | 9269638 | rs2032540 | -0.502 | -0.017 | 0.593 | 0.291 | -0.005 | 0.128 | 0.128 | 0 | 0.82 | 0.64 | 0.68 | 0.5 | 0.83 | 0.74 |
| 1 | 9274446 | rs6577561 | -0.43 | -0.003 | 0.462 | 0.277 | -0.002 | 0.196 | 0.196 | 0 | 0.81 | 0.56 | 0.67 | 0.5 | 0.83 | 0.74 |
| 1 | 9275194 | rs1009941 | 0.215 | 0.036 | -0.295 | 0.096 | -0.004 | -0.035 | -0.035 | 0.055 | 0.66 | 0.26 | 0.57 | 0.48 | 0.65 | 0.61 |
| 1 | 9278142 | rs1009940 | 0.611 | 0.118 | -0.693 | 0.09 | 0.022 | -0.041 | -0.041 | 1.42 | 0.55 | 0.2 | 0.53 | 0.51 | 0.61 | 0.48 |
| 1 | 9285721 | rs10489436 | 0.029 | -0.187 | 0.028 | 0.582 | -0.034 | 0.138 | 0.138 | 0 | 0.18 | 0.31 | 0.28 | 0.44 | 0.12 | 0.25 |
| 1 | 9292082 | rs3753271 | -0.095 | 0.368 | -0.471 | 0.338 | -0.017 | -0.034 | -0.034 | 0.261 | 0.22 | 0.14 | 0.27 | 0.41 | 0.25 | 0.13 |
| 1 | 9292282 | rs9434723 | 0.712 | 0.271 | -0.477 | 0.086 | -0.006 | -0.015 | -0.015 | 0.094 | 0.19 | 0.23 | 0.25 | 0.07 | 0.15 | 0.17 |
| 1 | 9300608 | rs2268175 | -1.099 | 0.08 | -0.165 | -0.019 | -0.008 | -0.022 | -0.022 | 0 | 0.45 | 0.75 | 0.54 | 0.48 | 0.4 | 0.31 |
| 1 | 9304731 | rs2239560 | 0.736 | 0.151 | -0.207 | 0.025 | -0.019 | -0.04 | -0.04 | 0.178 | 0.2 | 0.31 | 0.25 | 0.07 | 0.15 | 0.17 |
| 1 | 9307033 | rs7524046 | -0.019 | 0.347 | -0.709 | 0.239 | -0.04 | -0.072 | -0.072 | 0 | 0.21 | 0.11 | 0.26 | 0.4 | 0.24 | 0.14 |
| 1 | 9315847 | rs2268170 | 1.053 | 0.304 | -1.379 | -0.289 | -0.041 | 0.116 | 0.116 | 4.245 | 0.27 | 0.01 | 0.21 | 0.12 | 0.28 | 0.3 |
| 1 | 9317873 | rs3753165 | -0.358 | -0.018 | -0.735 | -0.216 | -0.002 | -0.002 | -0.002 | 0 | 0.57 | 0.55 | 0.52 | 0.53 | 0.54 | 0.42 |
| 1 | 9318401 | rs3753163 | 0.053 | 0.087 | -1.163 | -0.204 | -0.031 | -0.033 | -0.033 | 0.682 | 0.49 | 0.17 | 0.48 | 0.53 | 0.51 | 0.41 |
| 1 | 9321241 | rs2268169 | -0.062 | 0.14 | -1.29 | -0.073 | -0.063 | -0.041 | -0.041 | 0 | 0.52 | 0.11 | 0.51 | 0.54 | 0.51 | 0.42 |
| 1 | 9323910 | rs6688832 | -0.206 | -0.184 | -0.525 | -0.132 | 0.008 | 0.127 | 0.127 | 0 | 0.24 | 0.49 | 0.37 | 0.48 | 0.25 | 0.31 |
| 1 | 9324213 | rs17368528 | 1.882 | -0.282 | -1.035 | -1.23 | 0.003 | 0.033 | 0.033 | 0 | 0.16 | 0.01 | 0.11 | 0.07 | 0.13 | 0.2 |
| 1 | 9324571 | rs9434742 | 0.171 | 0.119 | -0.627 | -1.5 | 0 | -0.027 | -0.027 | 1.315 | 0.58 | 0.5 | 0.42 | 0.17 | 0.41 | 0.5 |
| 1 | 9325676 | rs1294014 | 0.541 | 0.049 | -0.53 | -1.494 | 0 | -0.014 | -0.014 | 0 | 0.25 | 0.31 | 0.23 | 0.16 | 0.27 | 0.18 |
| 1 | 9327387 | rs1294015 | -1.289 | 0.244 | -0.83 | -1.333 | -0.026 | -0.021 | -0.021 | 0 | 0.82 | 0.83 | 0.64 | 0.34 | 0.66 | 0.65 |
| 1 | 9327413 | rs11121354 | 0.676 | -0.608 | -0.563 | -1.665 | 0.012 | 0.012 | 0.012 | 0 | 0.3 | 0.21 | 0.23 | 0.02 | 0.2 | 0.22 |
| 1 | 9332703 | rs7517538 | 0.086 | -0.197 | -0.441 | -1.465 | 0.005 | 0.061 | 0.061 | 0 | 0.58 | 0.64 | 0.46 | 0.16 | 0.4 | 0.47 |
| 1 | 9334714 | rs12093524 | 0.264 | -0.073 | -0.676 | -1.359 | 0.006 | -0.004 | -0.004 | 0.128 | 0.52 | 0.41 | 0.39 | 0.16 | 0.36 | 0.46 |
| 1 | 9341038 | rs916380 | -0.03 | -0.051 | -0.975 | -1.353 | 0.029 | 0.061 | 0.061 | 0 | 0.5 | 0.4 | 0.6 | 0.72 | 0.58 | 0.63 |
| 1 | 9347278 | rs4465231 | 0.531 | 0.224 | -0.789 | -1.62 | 0.032 | 0.147 | 0.147 | 0 | 0.37 | 0.13 | 0.25 | 0.09 | 0.28 | 0.48 |
| 1 | 9349938 | rs9726385 | 0.483 | 0.965 | 0.251 | -1.031 | 0.056 | -0.029 | -0.029 | 0.121 | 0.06 | 0.18 | 0.04 | 0 | 0.06 | 0.01 |
| 1 | 9358243 | rs621846 | 0.171 | 0.71 | -0.198 | -0.947 | -0.007 | 0.007 | 0.007 | 0 | 0.66 | 0.71 | 0.59 | 0.2 | 0.57 | 0.68 |
| 1 | 9362341 | rs9662290 | 0.181 | 0.523 | -0.525 | 0.113 | 0.013 | 0.005 | 0.005 | 0 | 0.4 | 0.63 | 0.44 | 0.4 | 0.33 | 0.41 |
| 1 | 9364634 | rs1294028 | 0.384 | 0.863 | -0.427 | -0.261 | 0.004 | -0.035 | -0.035 | 0 | 0.66 | 0.56 | 0.74 | 0.84 | 0.65 | 0.67 |
| 1 | 9366852 | rs581587 | 0.182 | 1.006 | -0.631 | -0.023 | -0.007 | -0.004 | -0.004 | 0 | 0.8 | 0.83 | 0.77 | 0.8 | 0.73 | 0.75 |
| 1 | 9367908 | rs9442562 | -0.349 | 0.414 | -0.471 | -0.442 | 0.008 | 0.02 | 0.02 | 0.576 | 0.47 | 0.39 | 0.48 | 0.26 | 0.6 | 0.47 |
| 1 | 9377864 | rs926249 | -0.505 | 0.036 | 0.11 | -0.535 | -0.013 | -0.043 | -0.043 | 0 | 0.09 | 0.3 | 0.05 | 0 | 0.03 | 0.07 |
| 1 | 9379693 | rs1294046 | -0.332 | -0.467 | -0.116 | -0.921 | 0.009 | -0.006 | -0.006 | 0 | 0.67 | 0.49 | 0.54 | 0.27 | 0.66 | 0.59 |
| 1 | 9381504 | rs17377604 | -0.392 | -0.208 | -0.278 | -1.108 | -0.02 | 0.064 | 0.064 | 0.005 | 0.21 | 0.07 | 0.17 | 0.04 | 0.23 | 0.16 |
| 1 | 9382285 | rs564709 | -1.197 | -0.256 | -0.007 | -1.084 | 0.006 | -0.016 | -0.016 | 0 | 0.08 | 0.48 | 0.07 | 0 | 0.03 | 0.04 |
| 1 | 9382469 | rs9660159 | 0.401 | -0.046 | -0.653 | -0.991 | -0.063 | 0.047 | 0.047 | 0.018 | 0.15 | 0.01 | 0.14 | 0.04 | 0.2 | 0.13 |
| 1 | 9387311 | rs4908842 | 0.63 | -0.422 | -0.294 | -0.804 | -0.004 | 0.029 | 0.029 | 0 | 0.6 | 0.28 | 0.45 | 0.24 | 0.59 | 0.55 |
| 1 | 9387359 | rs4908843 | 0.427 | -0.436 | -0.396 | -0.817 | 0.005 | -0.014 | -0.014 | 0 | 0.63 | 0.52 | 0.48 | 0.24 | 0.61 | 0.55 |
| 1 | 9387444 | rs1569422 | 0.041 | -0.427 | -1.251 | -0.147 | 0 | 0.004 | 0.004 | 0.002 | 0.89 | 0.91 | 0.79 | 0.73 | 0.9 | 0.87 |
| 1 | 9389218 | rs732591 | 1.171 | -0.418 | -1.756 | -0.361 | -0.096 | 0.062 | 0.062 | 0.102 | 0.4 | 0.07 | 0.36 | 0.21 | 0.49 | 0.39 |
| 1 | 9392728 | rs11121371 | -1.322 | 0.643 | -0.845 | 0.705 | -0.021 | 0.024 | 0.024 | 0.033 | 0.18 | 0.62 | 0.27 | 0.22 | 0.13 | 0.13 |
| 1 | 9394008 | rs926247 | -0.271 | 0.614 | -0.88 | 0.727 | -0.082 | -0.149 | -0.149 | 0.188 | 0.77 | 0.95 | 0.72 | 0.47 | 0.7 | 0.66 |
| 1 | 9396660 | rs11121374 | 0.466 | 0.601 | -0.796 | 0.672 | -0.022 | 0.095 | 0.095 | 0.053 | 0.86 | 0.39 | 0.75 | 0.74 | 0.91 | 0.91 |
| 1 | 9396935 | rs12728282 | 0.5 | 0.79 | -0.753 | 0.139 | -0.004 | 0.013 | 0.013 | 0 | 0.59 | 0.31 | 0.45 | 0.24 | 0.58 | 0.53 |
| 1 | 9406979 | rs2075977 | -0.633 | 1.094 | -0.231 | 0.594 | 0.02 | 0.031 | 0.031 | 0 | 0.23 | 0.45 | 0.3 | 0.31 | 0.23 | 0.3 |
| 1 | 9412408 | rs1299374 | 0.624 | 1.521 | -0.313 | 0.135 | 0.017 | -0.045 | -0.045 | 0 | 0.86 | 0.76 | 0.84 | 0.96 | 0.83 | 0.85 |
| 1 | 9413646 | rs11121384 | 0.021 | 1.359 | -1.466 | 0.343 | -0.056 | -0.076 | -0.076 | 0.153 | 0.26 | 0.05 | 0.16 | 0.28 | 0.31 | 0.18 |
| 1 | 9415727 | rs2310919 | 0.95 | 0.187 | -1.543 | 0.127 | 0.059 | 0.023 | 0.023 | 0 | 0.34 | 0.24 | 0.34 | 0.27 | 0.2 | 0.28 |
| 1 | 9416852 | rs12760778 | 0.07 | 0.948 | -0.939 | -0.288 | 0.005 | 0.023 | 0.023 | 3.874 | 0.22 | 0.25 | 0.26 | 0.39 | 0.25 | 0.36 |
| 1 | 9416962 | rs4908852 | 0.949 | 1.503 | -1.159 | -0.4 | -0.02 | 0.012 | 0.012 | 0 | 0.56 | 0.3 | 0.52 | 0.58 | 0.52 | 0.46 |
| 1 | 9420502 | rs4908853 | -0.453 | 1.484 | -0.726 | -0.419 | 0.032 | -0.081 | -0.081 | 0.011 | 0.8 | 0.67 | 0.8 | 0.96 | 0.77 | 0.81 |
| 1 | 9422435 | rs11121389 | 0.678 | 1.802 | -1.037 | -0.215 | -0.066 | -0.073 | -0.073 | 0 | 0.11 | 0.01 | 0.09 | 0.26 | 0.16 | 0.16 |
| 1 | 9422875 | rs552230 | 0.968 | 0.971 | -0.557 | -0.169 | 0.05 | 0.018 | 0.018 | 0 | 0.39 | 0.37 | 0.37 | 0.31 | 0.21 | 0.3 |
| 1 | 9425540 | rs10864411 | -0.373 | 1.285 | -0.091 | -0.085 | 0.015 | -0.002 | -0.002 | 1.525 | 0.36 | 0.55 | 0.32 | 0.33 | 0.39 | 0.25 |
| 1 | 9427869 | rs2275457 | 0.063 | 0.994 | -0.228 | 0.06 | 0.005 | 0.092 | 0.092 | 0.319 | 0.83 | 0.37 | 0.74 | 0.68 | 0.82 | 0.75 |
| 1 | 9430583 | rs9442600 | 1.412 | -0.064 | -1.008 | -0.473 | 0.085 | 0.104 | 0.104 | 0 | 0.26 | 0.1 | 0.1 | 0.07 | 0.13 | 0.25 |
| 1 | 9431223 | rs10492967 | 0.214 | 0.596 | -0.867 | -0.134 | -0.01 | -0.035 | -0.035 | 0 | 0.24 | 0.11 | 0.2 | 0.27 | 0.27 | 0.18 |
| 1 | 9432390 | rs9308447 | -0.035 | 1.054 | -0.506 | 0.098 | 0.036 | 0.03 | 0.03 | 0.008 | 0.18 | 0.43 | 0.29 | 0.19 | 0.21 | 0.13 |
| 1 | 9432911 | rs17393334 | 0.455 | 0.924 | -1.059 | 0.33 | -0.128 | -0.127 | -0.127 | 0.007 | 0.34 | 0.02 | 0.35 | 0.4 | 0.37 | 0.29 |
| 1 | 9434137 | rs11802522 | -0.041 | 0.753 | -0.541 | 0.198 | 0.008 | 0.003 | 0.003 | 0.808 | 0.19 | 0.52 | 0.31 | 0.17 | 0.2 | 0.13 |

| | | | | | | | | | | | | | | | | |
|---|---|---|---|---|---|---|---|---|---|---|---|---|---|---|---|---|
| 1 | 9441040 | rs9729149 | 0.729 | 0.488 | -0.708 | 0.374 | 0.002 | 0.01 | 0.01 | 0.124 | 0.36 | 0.36 | 0.34 | 0.48 | 0.39 | 0.5 |
| 1 | 9446761 | rs12134602 | -0.484 | 0.449 | -0.577 | 0.484 | -0.025 | 0.001 | 0.001 | 0 | 0.44 | 0.21 | 0.36 | 0.37 | 0.42 | 0.37 |
| 1 | 9447371 | rs2095904 | -0.565 | 0.441 | -0.763 | 0.359 | -0.002 | -0.009 | -0.009 | 0 | 0.45 | 0.45 | 0.36 | 0.33 | 0.4 | 0.33 |
| 1 | 9458793 | rs6688312 | -0.731 | 0.539 | -1.153 | 0.008 | 0.008 | -0.094 | -0.094 | 0 | 0.16 | 0.4 | 0.11 | 0 | 0.16 | 0.1 |
| 1 | 9466292 | rs11121406 | -0.436 | 0.42 | -2.321 | 0.73 | -0.067 | -0.032 | -0.032 | 0 | 0.49 | 0.09 | 0.4 | 0.43 | 0.48 | 0.47 |
| 1 | 9466803 | rs12044809 | -0.134 | 0.438 | -2.284 | 0.723 | -0.078 | -0.038 | -0.038 | 0 | 0.41 | 0.04 | 0.32 | 0.36 | 0.42 | 0.39 |
| 1 | 9476781 | rs6670392 | 0.178 | -0.116 | -1.286 | 1.124 | -0.023 | 0.069 | 0.069 | 0 | 0.93 | 0.86 | 0.92 | 0.8 | 0.97 | 0.94 |
| 1 | 9483388 | rs12145857 | -0.017 | -1.588 | -1.832 | -0.077 | 0.056 | 0.043 | 0.043 | 0 | 0.26 | 0.15 | 0.12 | 0.01 | 0.11 | 0.2 |
| 1 | 9489972 | rs10864418 | 0.674 | -1.204 | -2.485 | 0.083 | -0.073 | -0.074 | -0.074 | 0 | 0.32 | 0.1 | 0.48 | 0.4 | 0.55 | 0.25 |
| 1 | 9490339 | rs11121412 | -0.406 | -1.673 | -2.508 | -0.083 | 0.151 | 0.131 | 0.131 | 0 | 0.49 | 0.8 | 0.29 | 0.32 | 0.28 | 0.43 |
| 1 | 9495571 | rs9726404 | 0.264 | -1.606 | -1.51 | -0.067 | -0.015 | 0.062 | 0.062 | 0 | 0.13 | 0.37 | 0.12 | 0.19 | 0.1 | 0.13 |
| 1 | 9497525 | rs9659326 | -0.066 | -1.781 | -1.662 | -0.163 | -0.032 | -0.025 | -0.025 | 0 | 0.41 | 0.57 | 0.26 | 0.19 | 0.21 | 0.34 |
| 1 | 9499246 | rs9787285 | 0.719 | -1.442 | -2.135 | 0.021 | -0.035 | -0.048 | -0.048 | 8.488 | 0.15 | 0.03 | 0.19 | 0.21 | 0.16 | 0.3 |
| 1 | 9501492 | rs9662546 | 0.501 | -1.887 | -2.12 | -0.448 | 0.052 | 0.033 | 0.033 | 0 | 0.28 | 0.22 | 0.15 | 0 | 0.09 | 0.21 |
| 1 | 9503008 | rs4926481 | -0.584 | -0.283 | -2.243 | 0.392 | -0.099 | -0.076 | -0.076 | 0 | 0.69 | 0.89 | 0.52 | 0.71 | 0.45 | 0.73 |
| 1 | 9507004 | rs10735516 | -1.282 | 0.71 | -2.14 | 0.498 | -0.031 | -0.064 | -0.064 | 0 | 0.82 | 0.9 | 0.77 | 0.76 | 0.77 | 0.84 |
| 1 | 9508955 | rs12034064 | 0.351 | 0.212 | -2.342 | 0.535 | -0.027 | -0.1 | -0.1 | 0 | 0.17 | 0.01 | 0.18 | 0.36 | 0.16 | 0.33 |
| 1 | 9509782 | rs10779725 | -0.828 | -0.311 | -1.502 | 0.558 | -0.047 | -0.005 | -0.005 | 0 | 0.64 | 0.56 | 0.74 | 0.68 | 0.8 | 0.7 |
| 1 | 9510088 | rs10864424 | -0.411 | -0.573 | -0.843 | 0.006 | 0.014 | 0.018 | 0.018 | 0 | 0.54 | 0.58 | 0.75 | 0.82 | 0.7 | 0.64 |
| 1 | 9510150 | rs12565445 | -0.537 | -0.157 | -0.36 | -0.114 | -0.004 | -0.004 | -0.004 | 0.058 | 0.44 | 0.52 | 0.37 | 0.71 | 0.38 | 0.53 |
| 1 | 9510526 | rs11121423 | -1.093 | -0.298 | -0.577 | -0.441 | 0.037 | 0.062 | 0.062 | 0 | 0.71 | 0.85 | 0.83 | 0.95 | 0.84 | 0.77 |
| 1 | 9511569 | rs11121424 | -0.699 | -0.218 | -0.962 | -0.792 | 0.016 | 0.065 | 0.065 | 0 | 0.71 | 0.85 | 0.81 | 0.95 | 0.78 | 0.67 |
| 1 | 9514846 | rs4926491 | -0.098 | 0.14 | -0.655 | -0.458 | 0.008 | -0.031 | -0.031 | 0.123 | 0.38 | 0.44 | 0.43 | 0.3 | 0.44 | 0.29 |
| 1 | 9525697 | rs10746490 | -1.016 | -0.24 | -1.122 | -0.492 | 0.004 | 0.028 | 0.028 | 0 | 0.74 | 0.8 | 0.78 | 0.88 | 0.8 | 0.72 |
| 1 | 9525764 | rs12042017 | 0.933 | -0.44 | -0.888 | -0.666 | -0.03 | 0.014 | 0.014 | 0.434 | 0.31 | 0.43 | 0.33 | 0.63 | 0.21 | 0.38 |
| 1 | 9526956 | rs6695925 | -1.06 | -0.777 | -0.726 | -0.347 | -0.003 | -0.02 | -0.02 | 0 | 0.74 | 0.7 | 0.76 | 0.88 | 0.8 | 0.73 |
| 1 | 9526979 | rs10864425 | 0.634 | -0.626 | -0.858 | -0.429 | 0 | -0.013 | -0.013 | 0.529 | 0.47 | 0.47 | 0.58 | 0.7 | 0.55 | 0.5 |
| 1 | 9533238 | rs6679380 | -0.198 | -0.95 | -0.707 | -0.256 | -0.003 | -0.025 | -0.025 | 0 | 0.7 | 0.65 | 0.73 | 0.88 | 0.77 | 0.72 |
| 1 | 9535303 | rs7418365 | -0.313 | -0.378 | -0.62 | 0.911 | -0.009 | -0.027 | -0.027 | 0 | 0.37 | 0.29 | 0.44 | 0.52 | 0.34 | 0.52 |
| 1 | 9539693 | rs7367253 | 0.069 | -0.2 | -1.061 | 0.218 | -0.025 | -0.022 | -0.022 | 0.572 | 0.92 | 0.98 | 0.96 | 0.88 | 0.91 | 0.92 |
| 1 | 9542612 | rs10157778 | -0.068 | -0.18 | -0.745 | 0.184 | -0.002 | 0.004 | 0.004 | 0 | 0.36 | 0.35 | 0.34 | 0.22 | 0.36 | 0.45 |
| 1 | 9549328 | rs12735814 | 1.404 | -0.102 | -1.545 | 0.148 | -0.116 | 0.209 | 0.209 | 1.372 | 0.33 | 0.03 | 0.26 | 0.06 | 0.44 | 0.13 |
| 1 | 9556061 | rs7413464 | -0.302 | -0.476 | -1.079 | 0.681 | -0.018 | 0.059 | 0.059 | 0 | 0.4 | 0.16 | 0.38 | 0.28 | 0.37 | 0.51 |
| 1 | 9562830 | rs7550613 | -0.684 | -0.442 | 0.368 | 1.803 | 0.014 | 0.275 | 0.275 | 0.268 | 0.09 | 0.29 | 0.16 | 0.45 | 0.05 | 0.13 |
| 1 | 9567789 | rs12755956 | -0.809 | -0.405 | -0.072 | 1.275 | 0.001 | 0.025 | 0.025 | 0 | 0.13 | 0.13 | 0.16 | 0.51 | 0.09 | 0.18 |
| 1 | 9576675 | rs7525571 | -1.706 | -0.58 | 1.023 | 1.082 | 0.038 | 0.208 | 0.208 | 0.157 | 0.09 | 0.51 | 0.15 | 0.15 | 0.05 | 0.23 |
| 1 | 9579964 | rs7412149 | -1.17 | -0.974 | 0.832 | 0.415 | -0.004 | 0.013 | 0.013 | 1.409 | 0.52 | 0.58 | 0.53 | 0.42 | 0.49 | 0.64 |
| 1 | 9591499 | rs12040542 | -1.772 | -1.447 | 1.622 | 0.658 | -0.031 | 0.049 | 0.049 | 1.229 | 0.2 | 0.55 | 0.16 | 0.31 | 0.14 | 0.17 |
| 1 | 9602962 | rs12023288 | -1.007 | -1.386 | 1.785 | 0.664 | 0 | 0.002 | 0.002 | 0 | 0.16 | 0.13 | 0.09 | 0.29 | 0.17 | 0.17 |
| 1 | 9607041 | rs7411273 | -1.68 | -1.393 | 2.034 | 0.662 | -0.001 | 0.08 | 0.08 | 0.176 | 0.19 | 0.46 | 0.17 | 0.31 | 0.18 | 0.18 |
| 1 | 9613871 | rs6664362 | -1.673 | -1.626 | 2.106 | 0.543 | 0.006 | 0.006 | 0.006 | 0 | 0.12 | 0.08 | 0 | 0 | 0 | 0.01 |
| 1 | 9621407 | rs4423053 | -1.062 | -0.646 | 2.575 | 0.996 | -0.012 | -0.035 | -0.035 | 0.696 | 0.16 | 0.05 | 0.08 | 0.29 | 0.17 | 0.17 |
| 1 | 9647137 | rs9430174 | 1.566 | -0.607 | 2.587 | 1.013 | -0.011 | 0.04 | 0.04 | 0.818 | 0.8 | 0.45 | 0.77 | 0.69 | 0.83 | 0.82 |
| 1 | 9656571 | rs4074744 | 1.7 | -0.646 | 2.224 | 1.086 | -0.016 | 0.025 | 0.025 | 0.609 | 0.78 | 0.43 | 0.79 | 0.69 | 0.81 | 0.79 |
| 1 | 9658707 | rs9430182 | 1.396 | -0.622 | 2.066 | 1.097 | -0.007 | 0.038 | 0.038 | 0.01 | 0.8 | 0.68 | 0.78 | 0.7 | 0.81 | 0.79 |
| 1 | 9666364 | rs7364899 | 1.322 | -0.516 | 1.482 | 1.091 | -0.011 | 0.096 | 0.096 | 0.236 | 0.81 | 0.61 | 0.8 | 0.67 | 0.84 | 0.79 |
| 1 | 9677002 | rs6540967 | 0.712 | -0.509 | 1.183 | 1.114 | 0 | -0.004 | -0.004 | 0 | 0.84 | 0.86 | 0.93 | 0.69 | 0.84 | 0.8 |
| 1 | 9677853 | rs10779727 | 1.181 | -0.46 | 0.854 | 1.079 | -0.015 | 0.106 | 0.106 | 0.65 | 0.8 | 0.51 | 0.85 | 0.67 | 0.84 | 0.79 |
| 1 | 9688512 | rs4076986 | 1.012 | -0.318 | 0.402 | 0.918 | -0.001 | 0.016 | 0.016 | 0 | 0.79 | 0.74 | 0.84 | 0.71 | 0.81 | 0.67 |
| 1 | 9693988 | rs9430506 | -1.66 | -1.138 | 0.499 | 0.52 | 0.014 | 0.018 | 0.018 | 0.385 | 0.37 | 0.67 | 0.37 | 0.83 | 0.26 | 0.52 |
| 1 | 9696690 | rs11121465 | -0.858 | -1.065 | 1.21 | 0.852 | -0.045 | 0.149 | 0.149 | 0.105 | 0.17 | 0.47 | 0.2 | 0.6 | 0.1 | 0.37 |
| 1 | 9704925 | rs4129341 | -0.844 | -2.202 | 0.729 | 0.068 | -0.06 | 0.009 | 0.009 | 0.042 | 0.27 | 0.44 | 0.34 | 0.65 | 0.16 | 0.49 |
| 1 | 9708688 | rs6540985 | -1.719 | -2.727 | 0.361 | -0.351 | 0.122 | 0.198 | 0.198 | 0.245 | 0.49 | 0.8 | 0.51 | 0.85 | 0.33 | 0.66 |
| 1 | 9711487 | rs7518793 | -0.615 | -2.602 | 0.59 | -0.301 | 0.003 | 0.004 | 0.004 | 0 | 0.28 | 0.28 | 0.11 | 0.25 | 0.24 | 0.19 |
| 1 | 9713386 | rs4240895 | -1.57 | -2.241 | 0.329 | -0.282 | 0.027 | 0.095 | 0.095 | 0.308 | 0.57 | 0.76 | 0.52 | 0.9 | 0.41 | 0.69 |
| 1 | 9713500 | rs4240896 | -1.606 | -2.228 | 0.186 | -0.298 | 0.027 | 0.107 | 0.107 | 0.44 | 0.58 | 0.77 | 0.53 | 0.92 | 0.41 | 0.69 |
| 1 | 9729599 | rs6671868 | -1.186 | -2.501 | 0.59 | 0.409 | -0.005 | 0.069 | 0.069 | 0 | 0.13 | 0.18 | 0.16 | 0.55 | 0.09 | 0.33 |
| 1 | 9733221 | rs11121472 | -2.025 | -3.028 | 0.63 | 0.061 | -0.052 | 0.01 | 0.01 | 0 | 0.28 | 0.42 | 0.37 | 0.65 | 0.14 | 0.54 |
| 1 | 9739435 | rs9430636 | -1.586 | -2.432 | 0.236 | -0.506 | -0.014 | -0.023 | -0.023 | 0 | 0.59 | 0.72 | 0.56 | 0.49 | 0.49 | 0.63 |
| 1 | 9741792 | rs4333853 | -1.743 | -2.517 | 0.189 | -0.537 | -0.015 | -0.022 | -0.022 | 0.059 | 0.61 | 0.73 | 0.55 | 0.48 | 0.49 | 0.63 |
| 1 | 9746248 | rs6677649 | -1.078 | -2.683 | 0.562 | -0.561 | -0.025 | -0.007 | -0.007 | 0.22 | 0.19 | 0.26 | 0.25 | 0.14 | 0.1 | 0.27 |
| 1 | 9748445 | rs6669509 | -1.768 | -2.514 | 0.719 | -0.614 | -0.034 | -0.046 | -0.046 | 0 | 0.63 | 0.77 | 0.56 | 0.48 | 0.49 | 0.64 |
| 1 | 9755273 | rs11801864 | -1.391 | -3.311 | 1.258 | -0.868 | -0.057 | -0.02 | -0.02 | 0.542 | 0.15 | 0.28 | 0.17 | 0.08 | 0.05 | 0.21 |
| 1 | 9760651 | rs12566637 | -1.73 | -3.737 | 1.236 | -1.067 | -0.027 | -0.02 | -0.02 | 0.06 | 0.23 | 0.27 | 0.2 | 0.13 | 0.09 | 0.46 |
| 1 | 9767817 | rs12075554 | -1.781 | -3.707 | 1.293 | -1.052 | -0.052 | -0.036 | -0.036 | 0.704 | 0.23 | 0.41 | 0.21 | 0.14 | 0.09 | 0.44 |
| 1 | 9771564 | rs6541017 | -0.455 | -3.972 | 0.845 | -1.151 | 0.009 | 0 | 0 | 0 | 0.22 | 0.22 | 0.12 | 0.24 | 0.15 | 0.15 |
| 1 | 9776339 | rs3934934 | -2.037 | -4.111 | 1.138 | -1.168 | -0.017 | -0.005 | -0.005 | 1.292 | 0.29 | 0.66 | 0.24 | 0.18 | 0.11 | 0.45 |
| 1 | 9779871 | rs9430220 | -1.897 | -3.935 | 0.856 | -1.103 | 0.05 | 0.008 | 0.008 | 0 | 0.38 | 0.79 | 0.33 | 0.36 | 0.22 | 0.53 |
| 1 | 9782556 | rs11589267 | 2.771 | -3.647 | 0.371 | -1.508 | -0.181 | 0.186 | 0.186 | 0 | 0.28 | 0.04 | 0.28 | 0 | 0.45 | 0.13 |

| | | | | | | | | | | | | | | | | |
|---|---|---|---|---|---|---|---|---|---|---|---|---|---|---|---|---|
| 1 | 9782766 | rs10864435 | -1.297 | -3.634 | 1.354 | -0.738 | -0.047 | -0.028 | -0.028 | 0 | 0.19 | 0.35 | 0.19 | 0.12 | 0.07 | 0.39 |
| 1 | 9787508 | rs1135426 | -1.127 | -3.528 | 1.173 | -0.604 | -0.015 | -0.015 | -0.015 | 1.283 | 0.92 | 0.99 | 0.82 | 0.96 | 0.9 | 0.86 |
| 1 | 9787840 | rs1135427 | 2.119 | -3.575 | 1.166 | -0.55 | -0.018 | -0.063 | -0.063 | 0.076 | 0.42 | 0.14 | 0.44 | 0.52 | 0.56 | 0.25 |
| 1 | 9789891 | rs7526958 | -1.158 | -3.5 | 1.414 | -0.585 | -0.066 | 0.012 | 0.012 | 0.618 | 0.18 | 0.63 | 0.2 | 0.19 | 0.06 | 0.33 |
| 1 | 9801341 | rs10864436 | -0.475 | -3.452 | 1.506 | -0.51 | -0.015 | 0.026 | 0.026 | 0 | 0.1 | 0.21 | 0.18 | 0.15 | 0.07 | 0.33 |
| 1 | 9804693 | rs7550295 | -0.038 | -3.423 | 1.27 | -0.495 | -0.003 | 0.069 | 0.069 | 0 | 0.09 | 0.27 | 0.16 | 0.15 | 0.05 | 0.32 |
| 1 | 9811435 | rs10779731 | -0.274 | -3.306 | 1.057 | -0.453 | -0.062 | 0.008 | 0.008 | 0.007 | 0.18 | 0.61 | 0.22 | 0.19 | 0.08 | 0.33 |
| 1 | 9842576 | rs10746492 | 0.721 | -3.216 | 0.724 | -0.488 | -0.007 | 0.033 | 0.033 | 0 | 0.63 | 0.86 | 0.66 | 0.72 | 0.65 | 0.59 |
| 1 | 9854238 | rs2119508 | -0.488 | -3.159 | 1.897 | -0.462 | -0.012 | -0.02 | -0.02 | 0.004 | 0.92 | 0.98 | 0.8 | 0.96 | 0.9 | 0.85 |
| 1 | 9862372 | rs767762 | 0.877 | -2.827 | 1.417 | -0.503 | -0.012 | -0.006 | -0.006 | 0 | 0.79 | 0.29 | 0.88 | 0.91 | 0.92 | 0.8 |
| 1 | 9885290 | rs12078414 | -1.022 | -2.785 | 0.784 | -0.374 | -0.077 | -0.059 | -0.059 | 0.24 | 0.2 | 0.58 | 0.12 | 0.09 | 0.08 | 0.19 |
| 1 | 9890158 | rs6666296 | -1.237 | -2.548 | -0.075 | -0.246 | -0.032 | -0.008 | -0.008 | 0 | 0.33 | 0.62 | 0.23 | 0.26 | 0.15 | 0.4 |
| 1 | 9897001 | rs12032209 | -0.841 | -2.115 | 0.254 | -0.172 | -0.101 | -0.041 | -0.041 | 0 | 0.21 | 0.45 | 0.16 | 0.16 | 0.08 | 0.24 |
| 1 | 9902128 | rs1933734 | 0.181 | -1.579 | 0.091 | -0.923 | 0 | -0.018 | -0.018 | 0.074 | 0.44 | 0.51 | 0.44 | 0.13 | 0.49 | 0.25 |
| 1 | 9902900 | rs10864439 | -0.325 | -2.001 | 0.193 | -1.074 | -0.017 | -0.005 | -0.005 | 0 | 0.3 | 0.33 | 0.38 | 0.77 | 0.25 | 0.47 |
| 1 | 9911291 | rs2379107 | -0.258 | -2.116 | 0.719 | -0.203 | -0.008 | 0.013 | 0.013 | 0.428 | 0.22 | 0.27 | 0.28 | 0.75 | 0.22 | 0.38 |
| 1 | 9922245 | rs1220405 | -0.434 | -1.312 | 1.325 | -0.559 | 0.002 | 0.014 | 0.014 | 0 | 0.21 | 0.66 | 0.17 | 0.28 | 0.18 | 0.39 |
| 1 | 9937041 | rs12128766 | 1.394 | -1.132 | -1.204 | -0.669 | -0.178 | 0.257 | 0.257 | 0 | 0.34 | 0.02 | 0.31 | 0.01 | 0.48 | 0.16 |
| 1 | 9942118 | rs935075 | -0.037 | -0.234 | 0.063 | 0.656 | 0.014 | 0.031 | 0.031 | 0 | 0.15 | 0.15 | 0.11 | 0.28 | 0.18 | 0.39 |
| 1 | 9987007 | rs11828 | 0.929 | -1.595 | -0.275 | 0.341 | 0.111 | -0.046 | -0.046 | 2.213 | 0.63 | 0.1 | 0.67 | 0.61 | 0.75 | 0.5 |
| 1 | 9987322 | rs17034440 | -0.867 | -1.645 | 0.686 | 0.288 | 0.033 | -0.001 | -0.001 | 0 | 0.06 | 0.16 | 0.17 | 0.04 | 0.07 | 0.13 |
| 1 | 9996707 | rs12065908 | -0.693 | -1.645 | 0.77 | 0.289 | 0.011 | -0.009 | -0.009 | 0 | 0.05 | 0.12 | 0.17 | 0.04 | 0.06 | 0.13 |
| 1 | 10008191 | rs12064822 | -0.693 | -1.649 | 0.975 | 0.551 | 0.011 | 0.116 | 0.116 | 0 | 0.05 | 0.11 | 0.17 | 0.23 | 0.06 | 0.18 |
| 1 | 10008221 | rs1220421 | 1.298 | -1.979 | 1.159 | 0.407 | 0.008 | -0.01 | -0.01 | 0.184 | 0.75 | 0.23 | 0.7 | 0.61 | 0.83 | 0.54 |
| 1 | 10013014 | rs7516611 | -1.042 | -1.91 | 1.42 | 0.433 | 0.04 | 0.167 | 0.167 | 0 | 0.06 | 0.19 | 0.19 | 0.23 | 0.09 | 0.18 |
| 1 | 10029913 | rs6683452 | 0.682 | -1.94 | 1.379 | 0.343 | -0.073 | 0.011 | 0.011 | 0.204 | 0.82 | 0.4 | 0.88 | 0.84 | 0.91 | 0.73 |
| 1 | 10046460 | rs12136213 | 1.021 | -0.838 | -0.091 | 0.243 | -0.065 | 0.026 | 0.026 | 0.076 | 0.2 | 0.07 | 0.25 | 0.08 | 0.28 | 0.14 |
| 1 | 10093457 | rs7528979 | -1.558 | -0.774 | 0.291 | 0.662 | 0.034 | -0.045 | -0.045 | 0 | 0.25 | 0.87 | 0.2 | 0.37 | 0.18 | 0.4 |
| 1 | 10134226 | rs17034493 | 0.6 | -0.904 | 1.187 | 0.529 | -0.015 | -0.008 | -0.008 | 2.322 | 0.07 | 0.11 | 0.05 | 0.05 | 0.03 | 0.02 |
| 1 | 10155283 | rs9430244 | 0.378 | -0.95 | 1.9 | 0.571 | -0.002 | 0.004 | 0.004 | 0 | 0.89 | 0.82 | 0.9 | 0.91 | 0.89 | 0.74 |
| 1 | 10165301 | rs12076922 | -1.244 | -0.987 | 1.935 | 0.591 | -0.007 | 0.118 | 0.118 | 0 | 0.12 | 0.4 | 0.11 | 0.21 | 0.07 | 0.25 |
| 1 | 10193026 | rs17034501 | -0.831 | -0.534 | 1.819 | 0.466 | 0.002 | -0.043 | -0.043 | 0 | 0.11 | 0.25 | 0.1 | 0.05 | 0.09 | 0.06 |
| 1 | 10193821 | rs6673887 | -2.176 | -0.522 | 2.158 | 0.586 | 0.044 | 0.111 | 0.111 | 1.348 | 0.1 | 0.63 | 0.1 | 0.15 | 0.09 | 0.11 |
| 1 | 10195940 | rs6671289 | -2.605 | -0.636 | 2.015 | 0.528 | 0.037 | 0.015 | 0.015 | 0.401 | 0.25 | 0.87 | 0.19 | 0.19 | 0.13 | 0.23 |
| 1 | 10204186 | rs12144133 | 1.977 | -0.635 | 1.535 | 0.676 | -0.109 | -0.12 | -0.12 | 0 | 0.46 | 0.02 | 0.41 | 0.54 | 0.48 | 0.45 |
| 1 | 10209557 | rs6696978 | 0.714 | -0.309 | 1.852 | 0.489 | -0.023 | 0.012 | 0.012 | 0 | 0.2 | 0.1 | 0.36 | 0.1 | 0.27 | 0.16 |
| 1 | 10218377 | rs2273298 | -2.298 | -0.512 | 2.069 | 0.588 | 0.011 | -0.007 | -0.007 | 0 | 0.22 | 0.8 | 0.19 | 0.3 | 0.14 | 0.21 |
| 1 | 10218439 | rs2273299 | -2.298 | -0.512 | 2.066 | 0.588 | 0.013 | -0.008 | -0.008 | 0 | 0.22 | 0.8 | 0.19 | 0.3 | 0.14 | 0.21 |
| 1 | 10230506 | rs17034563 | -1.333 | -0.43 | 2.022 | 0.58 | 0.107 | 0.033 | 0.033 | 0.47 | 0.06 | 0.39 | 0.07 | 0 | 0.05 | 0.01 |
| 1 | 10231463 | rs12092513 | -2.166 | -0.426 | 2.059 | 0.63 | 0.039 | 0.17 | 0.17 | 1.297 | 0.08 | 0.62 | 0.09 | 0.24 | 0.08 | 0.09 |
| 1 | 10232200 | rs17034569 | 0.024 | -0.427 | 1.726 | 0.399 | -0.024 | -0.022 | -0.022 | 0 | 0.09 | 0.18 | 0.06 | 0.05 | 0.03 | 0.01 |
| 1 | 10233756 | rs9783053 | -1.871 | -0.44 | 1.639 | 0.365 | -0.04 | -0.125 | -0.125 | 0 | 0.2 | 0.55 | 0.17 | 0.05 | 0.13 | 0.13 |
| 1 | 10235946 | rs2180184 | 0.021 | 0.27 | 1.464 | 0.385 | 0.066 | 0.036 | 0.036 | 0.174 | 0.26 | 0.36 | 0.37 | 0.37 | 0.38 | 0.26 |
| 1 | 10236039 | rs6541080 | -1.747 | -0.177 | 1.28 | 0.156 | -0.051 | 0.004 | 0.004 | 1.867 | 0.35 | 0.85 | 0.41 | 0.34 | 0.37 | 0.29 |
| 1 | 10236402 | rs7418410 | -2.123 | -0.142 | 1.14 | 0.201 | -0.084 | -0.014 | -0.014 | 0.075 | 0.43 | 0.93 | 0.49 | 0.41 | 0.41 | 0.36 |
| 1 | 10237960 | rs12733312 | 0.362 | -0.065 | 1.275 | 0.316 | -0.04 | -0.038 | -0.038 | 0 | 0.22 | 0.16 | 0.37 | 0.35 | 0.29 | 0.23 |
| 1 | 10238770 | rs7546642 | 0.333 | -0.117 | 1.628 | 0.27 | -0.019 | -0.044 | -0.044 | 0.375 | 0.08 | 0.21 | 0.06 | 0 | 0.03 | 0.1 |
| 1 | 10239197 | rs6682295 | -0.015 | -0.113 | 1.7 | 0.27 | -0.008 | -0.047 | -0.047 | 0.19 | 0.09 | 0.29 | 0.06 | 0 | 0.03 | 0.1 |
| 1 | 10241587 | rs4846195 | -0.247 | -0.118 | 1.612 | 0.279 | 0.014 | 0.006 | 0.006 | 0 | 0.1 | 0.33 | 0.1 | 0.06 | 0.07 | 0.03 |
| 1 | 10242654 | rs1556915 | -1.729 | -0.13 | 1.159 | 0.279 | -0.063 | -0.019 | -0.019 | 0 | 0.41 | 0.85 | 0.48 | 0.42 | 0.41 | 0.35 |
| 1 | 10257167 | rs6691645 | -0.69 | -0.126 | 1.543 | 0.087 | -0.017 | -0.068 | -0.068 | 0 | 0.11 | 0.51 | 0.1 | 0 | 0.04 | 0.1 |
| 1 | 10260592 | rs3903151 | -2.103 | -0.398 | 1.082 | 0.015 | -0.126 | 0.007 | 0.007 | 1.572 | 0.47 | 0.97 | 0.52 | 0.36 | 0.43 | 0.39 |
| 1 | 10265606 | rs4348727 | 0.132 | -0.348 | 1.372 | 0.046 | -0.018 | -0.011 | -0.011 | 0.037 | 0.23 | 0.22 | 0.33 | 0.3 | 0.31 | 0.27 |
| 1 | 10286358 | rs10492969 | -1.829 | -0.67 | 0.923 | -0.527 | -0.009 | 0.059 | 0.059 | 0.02 | 0.52 | 0.78 | 0.52 | 0.36 | 0.48 | 0.47 |
| 1 | 10290268 | rs17396382 | 0.839 | -0.594 | 0.817 | -0.434 | -0.082 | 0.05 | 0.05 | 0.096 | 0.2 | 0.05 | 0.26 | 0.09 | 0.28 | 0.17 |
| 1 | 10311877 | rs17034615 | 0.595 | -0.145 | 1.006 | -0.603 | -0.025 | -0.016 | -0.016 | 0 | 0.1 | 0.13 | 0.06 | 0.06 | 0.05 | 0.02 |
| 1 | 10322054 | rs4846209 | -0.843 | -0.176 | 1.072 | -0.604 | 0.002 | 0 | 0 | 0.035 | 0.33 | 0.34 | 0.38 | 0.35 | 0.36 | 0.28 |
| 1 | 10338598 | rs10492970 | 0.912 | -0.182 | 1.031 | -0.595 | 0.012 | 0.008 | 0.008 | 0.01 | 0.09 | 0.05 | 0.04 | 0.06 | 0.03 | 0.01 |
| 1 | 10342522 | rs17034660 | 0.831 | -0.18 | 1.031 | -0.594 | 0.011 | 0.007 | 0.007 | 0 | 0.09 | 0.06 | 0.04 | 0.06 | 0.03 | 0.01 |
| 1 | 10342629 | rs3748576 | -2.451 | -0.21 | 0.731 | -0.139 | -0.089 | -0.174 | -0.174 | 1.419 | 0.65 | 0.98 | 0.73 | 0.5 | 0.62 | 0.64 |
| 1 | 10345772 | rs12120191 | -0.808 | 0.095 | 1.144 | -0.353 | 0 | 0 | 0 | 0 | 0.33 | 0.3 | 0.38 | 0.35 | 0.36 | 0.28 |
| 1 | 10348042 | rs912963 | -1.102 | 0.091 | 1.313 | -0.355 | 0.015 | 0 | 0 | 0 | 0.34 | 0.48 | 0.4 | 0.35 | 0.36 | 0.28 |
| 1 | 10370312 | rs4846212 | -0.835 | 0.092 | 1.257 | -0.355 | -0.014 | -0.003 | -0.003 | 0 | 0.33 | 0.27 | 0.38 | 0.35 | 0.36 | 0.29 |
| 1 | 10371854 | rs17410217 | -0.781 | 0.11 | 1.254 | -0.355 | -0.011 | -0.005 | -0.005 | 0 | 0.33 | 0.26 | 0.37 | 0.35 | 0.33 | 0.28 |
| 1 | 10373503 | rs17034716 | 0.606 | 0.085 | 1.012 | -0.351 | 0.01 | 0.007 | 0.007 | 0.014 | 0.09 | 0.06 | 0.04 | 0.06 | 0.03 | 0.01 |
| 1 | 10385664 | rs2182326 | -0.857 | 0.083 | 1.069 | -0.354 | 0.005 | 0 | 0 | 0.702 | 0.33 | 0.34 | 0.38 | 0.35 | 0.36 | 0.29 |
| 1 | 10397567 | rs2297881 | 0.645 | 0.054 | 1.03 | -0.344 | 0.014 | 0.009 | 0.009 | 0 | 0.09 | 0.05 | 0.04 | 0.06 | 0.03 | 0.01 |
| 1 | 10420918 | rs3748578 | 0.119 | -0.01 | 1.239 | -0.272 | 0.023 | 0.007 | 0.007 | 0.115 | 0.24 | 0.28 | 0.33 | 0.28 | 0.31 | 0.27 |
| 1 | 10421878 | rs12125492 | 0.799 | -0.009 | 0.917 | -0.367 | -0.02 | 0.093 | 0.093 | 0.246 | 0.15 | 0 | 0.09 | 0.01 | 0.12 | 0.2 |

| | | | | | | | | | | | | | | | | |
|---|---|---|---|---|---|---|---|---|---|---|---|---|---|---|---|---|
| 1 | 10423988 | rs12403443 | 0.472 | 0.043 | 0.942 | -0.357 | -0.078 | -0.043 | -0.043 | 0 | 0.22 | 0.08 | 0.33 | 0.29 | 0.33 | 0.27 |
| 1 | 10438687 | rs1536262 | -1.934 | -0.144 | 0.801 | -0.373 | -0.017 | 0.062 | 0.062 | 0.011 | 0.53 | 0.8 | 0.52 | 0.36 | 0.49 | 0.48 |
| 1 | 10438892 | rs1002076 | -0.633 | -0.144 | 0.842 | -0.373 | 0.004 | 0.001 | 0.001 | 0.836 | 0.33 | 0.34 | 0.37 | 0.35 | 0.34 | 0.29 |
| 1 | 10441664 | rs3748581 | -1.788 | -0.145 | 0.813 | -0.372 | -0.006 | 0.063 | 0.063 | 0.012 | 0.51 | 0.72 | 0.51 | 0.36 | 0.49 | 0.49 |
| 1 | 10442867 | rs11121556 | -0.169 | 0.004 | 0.597 | -0.466 | -0.1 | 0.168 | 0.168 | 0.458 | 0.28 | 0.07 | 0.23 | 0.01 | 0.34 | 0.26 |
| 1 | 10451799 | rs4846217 | 0.463 | -0.04 | 0.729 | -0.29 | -0.005 | -0.045 | -0.045 | 3.833 | 0.17 | 0.21 | 0.11 | 0.01 | 0.12 | 0.19 |
| 1 | 10457540 | rs12409754 | -0.551 | 0.084 | 0.74 | -0.22 | 0.005 | 0.001 | 0.001 | 0.845 | 0.33 | 0.35 | 0.37 | 0.35 | 0.34 | 0.29 |
| 1 | 10463888 | rs4846219 | -1.621 | -0.014 | 0.701 | -0.307 | -0.012 | 0.091 | 0.091 | 0.215 | 0.51 | 0.8 | 0.52 | 0.35 | 0.49 | 0.46 |
| 1 | 10470846 | rs11579365 | -0.639 | 0.149 | 0.341 | -0.225 | 0.007 | -0.003 | -0.003 | 2.41 | 0.33 | 0.37 | 0.38 | 0.34 | 0.36 | 0.29 |
| 1 | 10472026 | rs4846221 | -1.534 | 0.23 | -0.156 | 0.001 | 0 | 0.031 | 0.031 | 0 | 0.49 | 0.59 | 0.5 | 0.35 | 0.48 | 0.46 |
| 1 | 10479678 | rs1889309 | -0.109 | 0.598 | -0.379 | 0.261 | -0.029 | -0.012 | -0.012 | 0.502 | 0.22 | 0.13 | 0.32 | 0.28 | 0.31 | 0.27 |
| 1 | 10484416 | rs663364 | -0.327 | 0.646 | -1.255 | 0.122 | -0.004 | 0.111 | 0.111 | 0 | 0.5 | 0.16 | 0.43 | 0.29 | 0.52 | 0.46 |
| 1 | 10485019 | rs2483679 | -1.068 | -0.735 | -0.531 | 0.293 | -0.002 | -0.004 | -0.004 | 0 | 0.56 | 0.4 | 0.6 | 0.47 | 0.53 | 0.56 |
| 1 | 10485445 | rs491880 | -1.188 | -0.886 | 0.049 | 0.262 | -0.019 | -0.021 | -0.021 | 0 | 0.58 | 0.68 | 0.61 | 0.47 | 0.49 | 0.56 |
| 1 | 10488314 | rs505141 | -1.219 | -0.826 | 0.118 | 0.301 | -0.02 | -0.02 | -0.02 | 4.275 | 0.57 | 0.69 | 0.61 | 0.47 | 0.49 | 0.59 |
| 1 | 10488555 | rs528713 | -1.413 | -0.889 | 0.199 | 0.27 | -0.046 | -0.046 | -0.046 | 0 | 0.59 | 0.8 | 0.61 | 0.47 | 0.49 | 0.59 |
| 1 | 10490881 | rs481571 | -1.205 | -0.904 | 0.191 | 0.244 | -0.018 | -0.021 | -0.021 | 0 | 0.57 | 0.68 | 0.61 | 0.46 | 0.49 | 0.59 |
| 1 | 10493415 | rs597441 | -0.471 | -0.97 | -0.001 | 0.28 | 0 | 0.001 | 0.001 | 0 | 0.46 | 0.44 | 0.55 | 0.37 | 0.46 | 0.57 |
| 1 | 10502710 | rs2847337 | 1.809 | -0.898 | 0.337 | 0.618 | 0.056 | 0.238 | 0.238 | 0.386 | 0.94 | 0.71 | 0.81 | 0.86 | 0.93 | 0.82 |
| 1 | 10510682 | rs1308520 | 1.437 | -0.885 | 0.338 | 0.621 | 0.056 | 0.228 | 0.228 | 0.241 | 0.95 | 0.75 | 0.81 | 0.86 | 0.93 | 0.82 |
| 1 | 10511544 | rs628462 | 1.645 | -0.939 | -0.061 | 0.549 | -0.13 | -0.182 | -0.182 | 0.462 | 0.34 | 0.02 | 0.31 | 0.49 | 0.39 | 0.32 |
| 1 | 10524467 | rs6667629 | 0.585 | -0.748 | 0.511 | 0.594 | -0.003 | -0.021 | -0.021 | 0 | 0.23 | 0.25 | 0.25 | 0.07 | 0.22 | 0.12 |
| 1 | 10527461 | rs2274986 | 2.14 | -0.951 | 0.694 | 0.677 | -0.025 | 0.005 | 0.005 | 1.727 | 0.07 | 0.11 | 0.02 | 0.07 | 0.02 | 0.02 |
| 1 | 10539543 | rs607941 | -0.808 | -0.904 | 0.512 | 0.703 | -0.025 | 0.017 | 0.017 | 2.527 | 0.76 | 0.88 | 0.58 | 0.81 | 0.71 | 0.74 |
| 1 | 10543836 | rs9970001 | 0.71 | -1.067 | 0.355 | 0.713 | 0.019 | -0.009 | -0.009 | 0.07 | 0.23 | 0.16 | 0.14 | 0.25 | 0.19 | 0.31 |
| 1 | 10549380 | rs670666 | 1.493 | -1.106 | 0.686 | 0.702 | -0.063 | -0.07 | -0.07 | 0 | 0.39 | 0.13 | 0.37 | 0.48 | 0.5 | 0.41 |
| 1 | 10550352 | rs673254 | 1.532 | -1.11 | 0.683 | 1.16 | -0.064 | 0.002 | 0.002 | 0.105 | 0.39 | 0.12 | 0.32 | 0.35 | 0.5 | 0.41 |
| 1 | 10554794 | rs620405 | -0.473 | -0.925 | 1.239 | 0.993 | 0.016 | -0.029 | -0.029 | 0 | 0.73 | 0.6 | 0.54 | 0.81 | 0.71 | 0.74 |
| 1 | 10555257 | rs622623 | -0.094 | -0.921 | 1.151 | 0.994 | 0.007 | -0.047 | -0.047 | 0 | 0.7 | 0.47 | 0.53 | 0.81 | 0.71 | 0.74 |
| 1 | 10556097 | rs636291 | 0.878 | -0.907 | 0.752 | 1.006 | 0.026 | 0.054 | 0.054 | 6.158 | 0.61 | 0.19 | 0.49 | 0.74 | 0.68 | 0.71 |
| 1 | 10563492 | rs2506901 | 0.299 | -0.76 | 0.884 | 1.098 | -0.014 | -0.015 | -0.015 | 0.262 | 0.26 | 0.15 | 0.48 | 0.32 | 0.29 | 0.24 |
| 1 | 10563609 | rs10864459 | 0.513 | -0.832 | 1.07 | 1.088 | -0.003 | 0.001 | 0.001 | 3.918 | 0.64 | 0.41 | 0.47 | 0.6 | 0.69 | 0.74 |
| 1 | 10563717 | rs2781230 | 0.348 | -0.684 | 0.586 | 1.084 | -0.026 | -0.004 | -0.004 | 0 | 0.25 | 0.09 | 0.36 | 0.28 | 0.25 | 0.23 |
| 1 | 10566095 | rs2506878 | 0.514 | -1.091 | 0.95 | 1.122 | -0.016 | 0 | 0 | 0 | 0.63 | 0.34 | 0.8 | 0.65 | 0.79 | 0.67 |
| 1 | 10566215 | rs616488 | 0.55 | -0.859 | 0.591 | 0.93 | -0.054 | -0.031 | -0.031 | 0 | 0.25 | 0.09 | 0.5 | 0.32 | 0.33 | 0.24 |
| 1 | 10567122 | rs604814 | 0.615 | -0.92 | 1.007 | 1.167 | 0.005 | 0.005 | 0.005 | 0 | 0.49 | 0.57 | 0.36 | 0.49 | 0.53 | 0.47 |
| 1 | 10567658 | rs2480786 | -0.926 | -0.936 | 1.048 | 1.047 | 0.009 | 0.009 | 0.009 | 0.461 | 0.76 | 0.77 | 0.86 | 0.81 | 0.83 | 0.71 |
| 1 | 10571841 | rs6540935 | 1.409 | -0.927 | 1.216 | 1.123 | -0.032 | 0.031 | 0.031 | 2.528 | 0.08 | 0.27 | 0.04 | 0.08 | 0.02 | 0.02 |
| 1 | 10573867 | rs2274985 | 1.135 | -0.545 | 1.073 | 1.327 | -0.071 | -0.044 | -0.044 | 0 | 0.39 | 0.12 | 0.37 | 0.53 | 0.51 | 0.44 |
| 1 | 10574677 | rs11587326 | 1.297 | -0.51 | 0.984 | 1.222 | -0.032 | 0.032 | 0.032 | 0.605 | 0.06 | 0.01 | 0.05 | 0 | 0.12 | 0.07 |
| 1 | 10578791 | rs11584327 | 1.079 | -0.572 | 0.996 | 1.258 | -0.071 | -0.044 | -0.044 | 0 | 0.39 | 0.13 | 0.37 | 0.53 | 0.51 | 0.44 |
| 1 | 10581658 | rs2056417 | 0.576 | -0.549 | 0.926 | 1.079 | -0.05 | 0.022 | 0.022 | 3.085 | 0.25 | 0.07 | 0.43 | 0.2 | 0.3 | 0.24 |
| 1 | 10582047 | rs2483678 | -0.799 | -0.627 | 1.014 | 1.086 | 0.023 | 0.019 | 0.019 | 0.642 | 0.7 | 0.74 | 0.86 | 0.77 | 0.83 | 0.7 |
| 1 | 10585527 | rs17035156 | 0.867 | -0.631 | 0.934 | 1.054 | 0.053 | 0.046 | 0.046 | 1.005 | 0.28 | 0.09 | 0.13 | 0.18 | 0.17 | 0.29 |
| 1 | 10597024 | rs17035189 | 0.822 | -0.491 | 0.858 | 0.862 | -0.025 | -0.02 | -0.02 | 0 | 0.42 | 0.19 | 0.37 | 0.48 | 0.49 | 0.43 |
| 1 | 10597551 | rs2480777 | 0.618 | -0.465 | 0.779 | 0.837 | -0.062 | 0.02 | 0.02 | 2.888 | 0.24 | 0.07 | 0.43 | 0.2 | 0.31 | 0.25 |
| 1 | 10607394 | rs710130 | 0.197 | -0.471 | 0.853 | 0.853 | 0.013 | 0.003 | 0.003 | 0.339 | 0.6 | 0.31 | 0.75 | 0.68 | 0.69 | 0.61 |
| 1 | 10608398 | rs11121583 | 0.68 | -0.464 | 0.901 | 0.82 | 0.024 | -0.041 | -0.041 | 7.348 | 0.08 | 0.11 | 0.06 | 0 | 0.12 | 0.07 |
| 1 | 10609505 | rs12120962 | 1.433 | -0.463 | 0.813 | 0.892 | -0.054 | -0.015 | -0.015 | 0 | 0.34 | 0.12 | 0.26 | 0.35 | 0.37 | 0.36 |
| 1 | 10613344 | rs10157111 | 0.655 | -0.482 | 0.969 | 0.882 | -0.008 | 0.002 | 0.002 | 0.019 | 0.3 | 0.36 | 0.17 | 0.32 | 0.19 | 0.33 |
| 1 | 10613660 | rs12127400 | 1.571 | -0.485 | 0.76 | 0.902 | -0.06 | -0.081 | -0.081 | 0 | 0.33 | 0.12 | 0.31 | 0.48 | 0.37 | 0.36 |
| 1 | 10616505 | rs17035305 | 0.637 | -0.355 | 0.812 | 0.963 | -0.075 | 0.009 | 0.009 | 3.01 | 0.24 | 0.04 | 0.43 | 0.2 | 0.31 | 0.24 |
| 1 | 10617924 | rs12402967 | 0.954 | -0.289 | 0.726 | 0.728 | -0.068 | 0.147 | 0.147 | 4.272 | 0.22 | 0.02 | 0.24 | 0.07 | 0.24 | 0.09 |
| 1 | 10629562 | rs12565494 | 0.217 | -0.193 | 1.05 | 0.882 | 0 | 0 | 0 | 0.048 | 0.61 | 0.36 | 0.75 | 0.68 | 0.68 | 0.6 |
| 1 | 10631237 | rs7512495 | 1.425 | -0.153 | 0.378 | 0.959 | -0.047 | -0.009 | -0.009 | 0 | 0.35 | 0.12 | 0.26 | 0.35 | 0.37 | 0.36 |
| 1 | 10638604 | rs12741973 | -1.052 | 0.825 | 1.104 | 1.005 | 0.013 | -0.001 | -0.001 | 0 | 0.32 | 0.3 | 0.54 | 0.36 | 0.43 | 0.31 |
| 1 | 10642506 | rs12083820 | -1.319 | 0.518 | 1.468 | 0.988 | 0.079 | -0.048 | -0.048 | 0 | 0.1 | 0.36 | 0.07 | 0 | 0.12 | 0.07 |
| 1 | 10643829 | rs12045923 | 1.176 | 0.246 | 0.958 | 1.375 | -0.004 | 0.012 | 0.012 | 0 | 0.66 | 0.42 | 0.45 | 0.39 | 0.67 | 0.65 |
| 1 | 10651460 | rs6540940 | 1.053 | -0.403 | 1.35 | 1.226 | -0.019 | 0.051 | 0.051 | 0.666 | 0.86 | 0.68 | 0.79 | 0.79 | 0.9 | 0.84 |
| 1 | 10653628 | rs284272 | -2.109 | -0.732 | 1.508 | 1.017 | -0.071 | -0.027 | -0.027 | 0 | 0.2 | 0.54 | 0.13 | 0.12 | 0.09 | 0.12 |
| 1 | 10666517 | rs284281 | 1.457 | -0.508 | 1.52 | 1.344 | -0.006 | 0.071 | 0.071 | 0 | 0.9 | 0.84 | 0.91 | 0.7 | 0.92 | 0.8 |
| 1 | 10672935 | rs481453 | -1.193 | -0.664 | 1.611 | 1.123 | -0.041 | -0.029 | -0.029 | 0.128 | 0.2 | 0.47 | 0.13 | 0.12 | 0.14 | 0.19 |
| 1 | 10673094 | rs3790630 | 1.747 | -0.677 | 1.429 | 1.066 | -0.025 | 0.036 | 0.036 | 0.546 | 0.7 | 0.36 | 0.78 | 0.57 | 0.78 | 0.61 |
| 1 | 10678610 | rs284237 | -1.237 | -0.581 | 1.588 | 0.492 | -0.032 | -0.075 | -0.075 | 0.034 | 0.19 | 0.36 | 0.14 | 0.06 | 0.15 | 0.19 |
| 1 | 10678843 | rs284235 | -1.286 | -0.604 | 1.577 | 0.486 | -0.04 | -0.085 | -0.085 | 0 | 0.19 | 0.4 | 0.14 | 0.06 | 0.15 | 0.19 |
| 1 | 10683683 | rs190775 | -1.183 | -0.505 | 1.617 | 0.471 | -0.029 | -0.078 | -0.078 | 0.69 | 0.19 | 0.38 | 0.14 | 0.06 | 0.15 | 0.19 |
| 1 | 10694125 | rs284301 | -1.171 | -0.503 | 1.534 | 0.444 | -0.024 | -0.066 | -0.066 | 0 | 0.19 | 0.31 | 0.13 | 0.05 | 0.15 | 0.19 |
| 1 | 10700448 | rs2128416 | 0.365 | -0.009 | 1.164 | 0.612 | -0.005 | 0.031 | 0.031 | 0 | 0.1 | 0.06 | 0.39 | 0.53 | 0.14 | 0.15 |
| 1 | 10703628 | rs729104 | -1.456 | -0.319 | 1.738 | -0.107 | 0.014 | 0.004 | 0.004 | 0 | 0.07 | 0.24 | 0.07 | 0.11 | 0.06 | 0.14 |

| | | | | | | | | | | | | | | | | |
|---|---|---|---|---|---|---|---|---|---|---|---|---|---|---|---|---|
| 1 | 10704688 | rs11587145 | -0.556 | -0.335 | 1.87 | -0.187 | 0.001 | 0 | 0 | 0 | 0.16 | 0.13 | 0.08 | 0.12 | 0.14 | 0.2 |
| 1 | 10708142 | rs778228 | -1.956 | -0.378 | 2.077 | -0.216 | -0.025 | -0.02 | -0.02 | 0 | 0.25 | 0.54 | 0.19 | 0.24 | 0.21 | 0.35 |
| 1 | 10709441 | rs284299 | -1.918 | -0.373 | 2.08 | -0.214 | -0.028 | -0.022 | -0.022 | 0.02 | 0.24 | 0.54 | 0.14 | 0.23 | 0.18 | 0.34 |
| 1 | 10714911 | rs2242287 | -1.212 | -0.423 | 1.732 | -0.335 | -0.004 | -0.002 | -0.002 | 0 | 0.19 | 0.24 | 0.15 | 0.14 | 0.18 | 0.21 |
| 1 | 10725498 | rs284322 | -0.576 | -0.279 | 1.728 | -0.261 | -0.004 | 0.002 | 0.002 | 0.149 | 0.16 | 0.22 | 0.14 | 0.15 | 0.17 | 0.19 |
| 1 | 10727773 | rs12567769 | -1.508 | -0.254 | 1.702 | 0.012 | 0.001 | -0.024 | -0.024 | 0.602 | 0.08 | 0.23 | 0.06 | 0.02 | 0.05 | 0.11 |
| 1 | 10739255 | rs284307 | -0.001 | -0.191 | 0.887 | -0.735 | 0.001 | 0.013 | 0.013 | 0 | 0.49 | 0.34 | 0.31 | 0.01 | 0.46 | 0.37 |
| 1 | 10743363 | rs284262 | -0.535 | -0.11 | 0.412 | -0.429 | -0.004 | 0.021 | 0.021 | 0 | 0.59 | 0.67 | 0.45 | 0.24 | 0.56 | 0.51 |
| 1 | 10750432 | rs516243 | -0.092 | 0.214 | -0.677 | -0.478 | -0.005 | 0.036 | 0.036 | 0 | 0.47 | 0.42 | 0.39 | 0.15 | 0.49 | 0.39 |
| 1 | 10757199 | rs4845948 | -1.41 | 0.355 | -0.806 | 0.458 | -0.011 | -0.023 | -0.023 | 0 | 0.07 | 0.01 | 0.12 | 0.08 | 0.1 | 0.04 |
| 1 | 10762432 | rs7514751 | -0.404 | 0.04 | -0.817 | 0.108 | 0.003 | 0.014 | 0.014 | 0 | 0.3 | 0.34 | 0.22 | 0.14 | 0.26 | 0.3 |
| 1 | 10765383 | rs12567505 | -1.146 | -0.485 | -0.523 | 1.08 | -0.007 | 0.01 | 0.01 | 0 | 0.14 | 0.16 | 0.15 | 0.33 | 0.08 | 0.27 |
| 1 | 10766466 | rs11804707 | -1.168 | -0.34 | -0.336 | 1.217 | -0.012 | 0.048 | 0.048 | 0.381 | 0.11 | 0.18 | 0.15 | 0.33 | 0.06 | 0.21 |
| 1 | 10767785 | rs11805515 | -1.315 | -0.59 | -0.367 | 1.159 | -0.014 | 0.02 | 0.02 | 2.95 | 0.11 | 0.17 | 0.13 | 0.31 | 0.04 | 0.21 |
| 1 | 10767902 | rs488834 | 0.455 | -0.56 | -0.198 | 1.186 | 0 | 0.065 | 0.065 | 2.628 | 0.75 | 0.44 | 0.58 | 0.53 | 0.75 | 0.74 |
| 1 | 10777260 | rs284233 | -1.618 | -1.155 | 0.922 | 0.953 | -0.089 | 0 | 0 | 0.193 | 0.26 | 0.5 | 0.2 | 0.32 | 0.1 | 0.33 |
| 1 | 10780727 | rs706007 | -1.472 | -0.882 | 1.361 | 1.068 | -0.077 | 0.006 | 0.006 | 0 | 0.2 | 0.42 | 0.19 | 0.25 | 0.08 | 0.32 |
| 1 | 10786806 | rs284255 | -1.14 | -0.567 | 0.566 | 0.755 | -0.004 | -0.013 | -0.013 | 0.14 | 0.18 | 0.18 | 0.17 | 0.06 | 0.12 | 0.14 |
| 1 | 10790797 | rs284277 | 1.173 | -0.328 | 0.329 | 0.553 | -0.002 | 0.059 | 0.059 | 0.021 | 0.6 | 0.31 | 0.44 | 0.16 | 0.63 | 0.55 |
| 1 | 10796866 | rs880315 | -1.111 | -0.271 | -0.239 | 0.978 | -0.013 | -0.004 | -0.004 | 0 | 0.35 | 0.16 | 0.51 | 0.64 | 0.36 | 0.4 |
| 1 | 10801585 | rs4845843 | -0.623 | -0.209 | -0.237 | 0.575 | 0 | 0 | 0 | 1.703 | 0.22 | 0.17 | 0.2 | 0.15 | 0.16 | 0.25 |
| 1 | 10801989 | rs4845953 | 0.653 | -0.02 | -0.292 | 0.119 | -0.001 | 0.042 | 0.042 | 0 | 0.51 | 0.62 | 0.43 | 0.15 | 0.53 | 0.45 |
| 1 | 10808587 | rs2076492 | -0.41 | -0.224 | -0.442 | 0.58 | -0.013 | -0.037 | -0.037 | 0 | 0.27 | 0.42 | 0.33 | 0.16 | 0.26 | 0.21 |
| 1 | 10811672 | rs205478 | -0.74 | -0.234 | -1.656 | 0.886 | 0.029 | -0.005 | -0.005 | 0 | 0.39 | 0.85 | 0.49 | 0.36 | 0.31 | 0.29 |
| 1 | 10813675 | rs205482 | -0.772 | -0.107 | -1.694 | 0.921 | 0.018 | 0.015 | 0.015 | 0 | 0.43 | 0.83 | 0.42 | 0.35 | 0.39 | 0.3 |
| 1 | 10814938 | rs205483 | 0.837 | 0.041 | -1.129 | 2.305 | -0.006 | -0.002 | -0.002 | 0.181 | 0.91 | 0.95 | 0.77 | 0.58 | 0.91 | 0.79 |
| 1 | 10815421 | rs205484 | -1.322 | 0.181 | -0.977 | 2.322 | -0.003 | 0.022 | 0.022 | 0 | 0.05 | 0.06 | 0.13 | 0.23 | 0.07 | 0.04 |
| 1 | 10815702 | rs205485 | -1.516 | -0.502 | -0.2 | 2.047 | -0.068 | 0.05 | 0.05 | 0 | 0.24 | 0.5 | 0.24 | 0.28 | 0.12 | 0.28 |
| 1 | 10827135 | rs205488 | 1.129 | -0.221 | -0.033 | 1.67 | -0.007 | 0.005 | 0.005 | 0 | 0.75 | 0.73 | 0.56 | 0.35 | 0.82 | 0.61 |
| 1 | 10832861 | rs205474 | 0.962 | -0.507 | 0.287 | 1.256 | -0.023 | 0.015 | 0.015 | 0 | 0.67 | 0.43 | 0.59 | 0.4 | 0.75 | 0.59 |
| 1 | 10840905 | rs17035713 | -1.489 | -0.297 | -0.156 | 1.545 | 0 | 0.124 | 0.124 | 0 | 0.08 | 0.11 | 0.2 | 0.42 | 0.08 | 0.14 |
| 1 | 10842021 | rs11121617 | 0.499 | -0.048 | -1.271 | -0.144 | -0.097 | 0.204 | 0.204 | 0 | 0.29 | 0.01 | 0.16 | 0.01 | 0.25 | 0.24 |
| 1 | 10847340 | rs6658023 | 0.684 | -0.719 | -0.47 | 0.384 | 0.022 | -0.004 | -0.004 | 0 | 0.64 | 0.32 | 0.63 | 0.61 | 0.75 | 0.47 |
| 1 | 10848699 | rs800761 | -0.086 | -0.624 | -0.228 | 0.284 | 0.006 | -0.033 | -0.033 | 0 | 0.91 | 0.79 | 0.84 | 0.96 | 0.91 | 0.85 |
| 1 | 10849645 | rs10864471 | -0.633 | -0.444 | -0.65 | 0.444 | -0.001 | 0.12 | 0.12 | 0 | 0.06 | 0.1 | 0.21 | 0.57 | 0.09 | 0.08 |
| 1 | 10850798 | rs1737617 | 0.641 | -0.564 | -0.259 | 0.599 | -0.015 | 0.008 | 0.008 | 0 | 0.67 | 0.38 | 0.62 | 0.57 | 0.76 | 0.49 |
| 1 | 10869052 | rs1750861 | -0.893 | -0.206 | -0.162 | 0.15 | -0.021 | -0.056 | -0.056 | 0 | 0.24 | 0.45 | 0.29 | 0.13 | 0.2 | 0.14 |
| 1 | 10873382 | rs1769669 | -1.03 | 0.052 | 0.083 | 0.962 | -0.026 | 0.139 | 0.139 | 0 | 0.22 | 0.53 | 0.25 | 0.56 | 0.17 | 0.36 |
| 1 | 10874522 | rs1776220 | -0.455 | 0.312 | -1.104 | 0.279 | 0.002 | 0.121 | 0.121 | 2.31 | 0.46 | 0.74 | 0.6 | 0.76 | 0.44 | 0.54 |
| 1 | 10879725 | rs6540951 | -1.103 | 0.496 | -0.764 | 0.997 | 0 | 0.016 | 0.016 | 0 | 0.13 | 0.13 | 0.21 | 0.31 | 0.12 | 0.16 |
| 1 | 10887923 | rs539790 | -0.455 | 0.515 | -0.931 | 0.857 | -0.025 | -0.016 | -0.016 | 0 | 0.08 | 0.01 | 0.03 | 0.07 | 0.08 | 0.13 |
| 1 | 10888791 | rs634478 | 0.311 | 1.058 | -0.673 | 0.571 | 0.008 | 0.004 | 0.004 | 0 | 0.86 | 0.74 | 0.65 | 0.85 | 0.81 | 0.87 |
| 1 | 10890026 | rs7555879 | -1.074 | 1.031 | -0.479 | 0.51 | 0.008 | 0.097 | 0.097 | 0 | 0.05 | 0.24 | 0.16 | 0.14 | 0.05 | 0.03 |
| 1 | 10890155 | rs661352 | -0.408 | 0.671 | -0.362 | 0.017 | 0 | -0.009 | -0.009 | 0 | 0.63 | 0.62 | 0.61 | 0.76 | 0.63 | 0.66 |
| 1 | 10890771 | rs497483 | 0.104 | -0.116 | -0.526 | -0.399 | 0.034 | 0.045 | 0.045 | 0 | 0.5 | 0.32 | 0.42 | 0.63 | 0.6 | 0.59 |
| 1 | 10891500 | rs2840299 | 0.828 | 0.2 | -0.777 | -0.355 | 0.006 | -0.032 | -0.032 | 0.054 | 0.32 | 0.18 | 0.27 | 0.45 | 0.31 | 0.42 |
| 1 | 10891777 | rs527676 | 0.231 | 0.163 | -1.028 | -0.084 | -0.009 | -0.004 | -0.004 | 0.105 | 0.13 | 0.04 | 0.04 | 0.1 | 0.1 | 0.14 |
| 1 | 10893081 | rs1292657 | 0.993 | -0.081 | -0.642 | -0.098 | 0.025 | 0.04 | 0.04 | 0 | 0.33 | 0.16 | 0.2 | 0.24 | 0.26 | 0.3 |
| 1 | 10901907 | rs555027 | 0.128 | 0.359 | -0.206 | 0.22 | 0.012 | -0.015 | -0.015 | 1.295 | 0.1 | 0.04 | 0.03 | 0.13 | 0.06 | 0.13 |
| 1 | 10907842 | rs672270 | -0.866 | 0.813 | 0.163 | 0.201 | 0.085 | -0.012 | -0.012 | 0 | 0.88 | 0.66 | 0.84 | 0.92 | 0.77 | 0.95 |
| 1 | 10913147 | rs12085982 | -0.608 | 0.564 | -0.324 | 0.265 | -0.009 | -0.003 | -0.003 | 0 | 0.21 | 0.32 | 0.35 | 0.15 | 0.2 | 0.15 |
| 1 | 10920576 | rs550887 | -0.784 | 0.999 | -1.04 | 0.376 | -0.032 | 0.01 | 0.01 | 0.047 | 0.88 | 0.92 | 0.85 | 0.94 | 0.74 | 0.92 |
| 1 | 10921228 | rs1280993 | 0.676 | 0.565 | -1.561 | 0.523 | 0.019 | -0.079 | -0.079 | 0 | 0.32 | 0.1 | 0.34 | 0.45 | 0.24 | 0.33 |
| 1 | 10922467 | rs7543046 | -0.243 | 0.669 | -1.03 | 0.284 | 0.04 | -0.095 | -0.095 | 0 | 0.24 | 0.5 | 0.38 | 0.05 | 0.33 | 0.14 |
| 1 | 10924273 | rs1280989 | 0.655 | 0.396 | -1.289 | 0.748 | 0.001 | -0.002 | -0.002 | 0 | 0.29 | 0.06 | 0.24 | 0.25 | 0.24 | 0.27 |
| 1 | 10924710 | rs1280988 | -0.161 | -0.082 | -0.724 | 0.419 | 0.001 | 0.024 | 0.024 | 0 | 0.51 | 0.41 | 0.53 | 0.58 | 0.56 | 0.46 |
| 1 | 10929928 | rs1281000 | -0.266 | 0.005 | -0.862 | 0.375 | -0.007 | 0.008 | 0.008 | 0 | 0.44 | 0.62 | 0.54 | 0.57 | 0.52 | 0.37 |
| 1 | 10935232 | rs1280996 | 0.151 | -0.423 | -1.498 | 0.274 | 0.047 | -0.08 | -0.08 | 0 | 0.61 | 0.14 | 0.57 | 0.55 | 0.73 | 0.67 |
| 1 | 10935423 | rs1280995 | -0.169 | 0.098 | -1.05 | 0.617 | -0.105 | -0.07 | -0.07 | 0 | 0.36 | 0.1 | 0.46 | 0.51 | 0.48 | 0.34 |
| 1 | 10936544 | rs7537363 | -1.785 | -0.016 | 0.399 | 0.845 | -0.053 | 0.195 | 0.195 | 0.81 | 0.13 | 0.66 | 0.2 | 0.35 | 0.06 | 0.23 |
| 1 | 10940895 | rs11586137 | -1.281 | -0.006 | 0.72 | 0.75 | -0.058 | 0.262 | 0.262 | 0.776 | 0.11 | 0.66 | 0.18 | 0.31 | 0.03 | 0.14 |
| 1 | 10947553 | rs7543908 | -0.71 | 0.003 | 0.759 | 0.646 | -0.012 | 0.018 | 0.018 | 0.274 | 0.08 | 0.46 | 0.05 | 0.02 | 0.01 | 0.06 |
| 1 | 10957719 | rs6670956 | -1.088 | 0.197 | 0.041 | 0.418 | -0.011 | -0.031 | -0.031 | 0.144 | 0.41 | 0.76 | 0.45 | 0.42 | 0.44 | 0.34 |
| 1 | 10957984 | rs12410376 | -0.513 | 0.942 | -0.226 | -0.596 | -0.061 | 0.104 | 0.104 | 0.025 | 0.29 | 0.12 | 0.26 | 0.05 | 0.4 | 0.2 |
| 1 | 10968730 | rs10864479 | 0.19 | 0.366 | -0.27 | 0.551 | 0 | -0.001 | -0.001 | 0 | 0.6 | 0.42 | 0.63 | 0.59 | 0.63 | 0.65 |
| 1 | 10969950 | rs4845962 | 0.801 | 0.555 | -0.949 | 0.067 | -0.078 | -0.106 | -0.106 | 0.036 | 0.36 | 0.08 | 0.36 | 0.47 | 0.42 | 0.38 |
| 1 | 10974329 | rs10157927 | 0.437 | 0.568 | -1.005 | -0.015 | -0.062 | 0.12 | 0.12 | 1.164 | 0.21 | 0.01 | 0.2 | 0.01 | 0.22 | 0.24 |
| 1 | 10977102 | rs6540954 | 0.788 | 0.673 | -0.868 | -0.034 | -0.053 | -0.083 | -0.083 | 0 | 0.38 | 0.14 | 0.37 | 0.48 | 0.41 | 0.41 |
| 1 | 10978316 | rs1280971 | 0.069 | 0.611 | -0.476 | 0.144 | -0.014 | 0.013 | 0.013 | 0.488 | 0.35 | 0.39 | 0.33 | 0.44 | 0.28 | 0.25 |

| | | | | | | | | | | | | | | | | |
|---|---|---|---|---|---|---|---|---|---|---|---|---|---|---|---|---|
| 1 | 10980998 | rs2387422 | 0.032 | 0.633 | -1.085 | 0.148 | 0.002 | -0.007 | -0.007 | 0 | 0.61 | 0.37 | 0.59 | 0.53 | 0.64 | 0.67 |
| 1 | 10984562 | rs1280984 | 0.451 | 0.389 | -1.975 | 0.076 | 0.013 | 0.112 | 0.112 | 2.145 | 0.26 | 0.08 | 0.18 | 0.03 | 0.21 | 0.17 |
| 1 | 10986308 | rs12748162 | -0.666 | 0.803 | -1.421 | 0.99 | -0.033 | -0.097 | -0.097 | 0 | 0.25 | 0.05 | 0.3 | 0.41 | 0.26 | 0.35 |
| 1 | 10996382 | rs9803711 | -0.095 | 0.83 | -1.111 | 1.107 | -0.018 | -0.026 | -0.026 | 0 | 0.31 | 0.26 | 0.46 | 0.46 | 0.43 | 0.33 |
| 1 | 10999600 | rs1925666 | 0.48 | -0.027 | -1.652 | 0.491 | 0.069 | 0.038 | 0.038 | 0 | 0.34 | 0.05 | 0.26 | 0.21 | 0.18 | 0.48 |
| 1 | 11002488 | rs6656310 | -0.322 | 0.037 | -0.755 | 0.386 | 0.053 | -0.084 | -0.084 | 0 | 0.79 | 0.58 | 0.84 | 0.91 | 0.74 | 0.9 |
| 1 | 11004245 | rs17036014 | 0.879 | -0.233 | -1.12 | 0.352 | 0.077 | 0.017 | 0.017 | 0 | 0.31 | 0.05 | 0.25 | 0.22 | 0.15 | 0.43 |
| 1 | 11005495 | rs6666764 | 0.526 | -0.149 | -0.878 | 0.766 | -0.066 | -0.044 | -0.044 | 0 | 0.38 | 0.18 | 0.46 | 0.43 | 0.52 | 0.4 |
| 1 | 11007159 | rs6696682 | 0.693 | -0.411 | -0.833 | 0.777 | -0.068 | -0.062 | -0.062 | 0.23 | 0.4 | 0.17 | 0.47 | 0.5 | 0.54 | 0.42 |
| 1 | 11008341 | rs10864483 | -0.346 | -1.436 | 0.142 | 1.034 | -0.003 | 0.182 | 0.182 | 0 | 0.09 | 0.37 | 0.13 | 0.21 | 0.06 | 0.05 |
| 1 | 11008695 | rs17036032 | 0.48 | -1.499 | 0.023 | 1.046 | 0.095 | 0.001 | 0.001 | 0 | 0.31 | 0.05 | 0.26 | 0.23 | 0.14 | 0.45 |
| 1 | 11009679 | rs1281013 | -0.121 | -1.511 | 0.503 | 0.979 | 0.031 | -0.016 | -0.016 | 0.468 | 0.06 | 0.2 | 0.01 | 0 | 0.04 | 0.02 |
| 1 | 11009703 | rs1281012 | -0.121 | -1.511 | 0.503 | 0.979 | 0.031 | -0.016 | -0.016 | 0 | 0.06 | 0.2 | 0.01 | 0 | 0.04 | 0.02 |
| 1 | 11011002 | rs1281008 | -0.077 | -1.458 | 0.464 | 1.051 | -0.034 | -0.022 | -0.022 | 0.046 | 0.38 | 0.42 | 0.29 | 0.24 | 0.19 | 0.48 |
| 1 | 11013286 | rs11121657 | -0.696 | -1.396 | -0.115 | 1.143 | 0.154 | -0.014 | -0.014 | 0 | 0.48 | 0.84 | 0.43 | 0.46 | 0.27 | 0.52 |
| 1 | 11017551 | rs12030940 | -0.363 | -1.094 | 0.2 | 1.146 | -0.019 | 0.091 | 0.091 | 0 | 0.07 | 0.16 | 0.1 | 0.23 | 0.06 | 0.06 |
| 1 | 11022020 | rs6685896 | -0.426 | -0.964 | 0.036 | 0.963 | 0.007 | -0.002 | -0.002 | 0.007 | 0.43 | 0.59 | 0.45 | 0.43 | 0.29 | 0.52 |
| 1 | 11024381 | rs6659326 | -0.539 | -0.971 | -0.527 | 0.959 | 0.125 | -0.033 | -0.033 | 0 | 0.46 | 0.87 | 0.47 | 0.44 | 0.29 | 0.54 |
| 1 | 11024395 | rs11121663 | -0.334 | -0.86 | -0.616 | 0.99 | 0.042 | -0.007 | -0.007 | 0 | 0.42 | 0.31 | 0.45 | 0.42 | 0.3 | 0.55 |
| 1 | 11029478 | rs12085319 | -1.085 | -1.029 | -1.297 | 1.232 | 0.039 | -0.149 | -0.149 | 0 | 0.32 | 0.95 | 0.46 | 0.46 | 0.19 | 0.38 |
| 1 | 11032827 | rs2003046 | -0.957 | -1.042 | -0.566 | 0.888 | -0.009 | -0.004 | -0.004 | 0 | 0.28 | 0.36 | 0.26 | 0.19 | 0.23 | 0.18 |
| 1 | 11038476 | rs11121667 | -1.226 | -1.138 | -0.084 | 1.009 | 0.018 | -0.002 | -0.002 | 0 | 0.32 | 0.75 | 0.28 | 0.29 | 0.24 | 0.21 |
| 1 | 11046855 | rs9430161 | -0.825 | -0.946 | 0.001 | 0.338 | -0.019 | -0.085 | -0.085 | 0.113 | 0.24 | 0.53 | 0.17 | 0.08 | 0.2 | 0.09 |
| 1 | 11048188 | rs9430163 | -0.846 | -0.945 | 0.003 | 0.339 | -0.02 | -0.085 | -0.085 | 0.069 | 0.24 | 0.53 | 0.17 | 0.08 | 0.2 | 0.09 |
| 1 | 11060075 | rs11121675 | 0.594 | -0.676 | -1.124 | 0.275 | 0.022 | 0.002 | 0.002 | 3.533 | 0.52 | 0.12 | 0.68 | 0.57 | 0.65 | 0.61 |
| 1 | 11060799 | rs7517230 | 0.967 | -0.244 | -1.608 | 0.009 | -0.112 | -0.169 | -0.169 | 4.856 | 0.25 | 0.02 | 0.39 | 0.52 | 0.33 | 0.43 |
| 1 | 11063043 | rs11121676 | 0.061 | -0.929 | -0.982 | 0.135 | 0.009 | -0.018 | -0.018 | 0.54 | 0.73 | 0.17 | 0.77 | 0.65 | 0.8 | 0.73 |
| 1 | 11072117 | rs9430335 | 0.14 | -1.431 | 0.342 | 0.403 | 0.002 | 0.002 | 0.002 | 0.338 | 0.79 | 0.83 | 0.84 | 0.87 | 0.84 | 0.81 |
| 1 | 11074250 | rs3765895 | 0.114 | -1.439 | 0.348 | 0.399 | 0.009 | 0.009 | 0.009 | 0.723 | 0.79 | 0.86 | 0.84 | 0.87 | 0.84 | 0.81 |
| 1 | 11079077 | rs2273348 | 1.018 | -1.619 | 0.613 | 0.461 | 0.073 | -0.025 | -0.025 | 0 | 0.7 | 0.18 | 0.76 | 0.67 | 0.82 | 0.68 |
| 1 | 11086332 | rs2273347 | 0.997 | -1.62 | 0.661 | 0.461 | 0.068 | -0.023 | -0.023 | 0 | 0.7 | 0.18 | 0.76 | 0.67 | 0.82 | 0.68 |
| 1 | 11086717 | rs1033638 | 0.652 | -1.629 | 1.383 | 0.451 | -0.018 | -0.018 | -0.018 | 0 | 0.74 | 0.38 | 0.8 | 0.87 | 0.84 | 0.8 |
| 1 | 11087524 | rs1782455 | 0.716 | -1.633 | 1.395 | 0.449 | 0.018 | 0.018 | 0.018 | 1.185 | 0.74 | 0.3 | 0.8 | 0.87 | 0.84 | 0.8 |
| 1 | 11090897 | rs2273346 | -0.889 | -1.626 | 1.831 | 0.459 | -0.021 | 0.103 | 0.103 | 0 | 0.05 | 0.17 | 0.05 | 0.2 | 0.02 | 0.12 |
| 1 | 11090916 | rs12711521 | 1.026 | -1.631 | 1.65 | 0.428 | 0.079 | -0.028 | -0.028 | 0 | 0.69 | 0.17 | 0.75 | 0.67 | 0.82 | 0.68 |
| 1 | 11104676 | rs3765901 | 1.132 | -1.506 | 1.353 | -0.035 | 0.076 | 0.072 | 0.072 | 1.313 | 0.65 | 0.17 | 0.72 | 0.77 | 0.77 | 0.63 |
| 1 | 11105122 | rs2273344 | 0.105 | -1.508 | 1.525 | -0.051 | 0.017 | 0.041 | 0.041 | 0.344 | 0.76 | 0.9 | 0.84 | 0.88 | 0.8 | 0.78 |
| 1 | 11107439 | rs7548659 | 1.099 | -1.736 | 1.251 | -0.007 | 0.091 | 0.052 | 0.052 | 1.513 | 0.65 | 0.17 | 0.72 | 0.74 | 0.78 | 0.63 |
| 1 | 11111629 | rs12121344 | -1.543 | -1.599 | 1.543 | 0.042 | -0.011 | 0.109 | 0.109 | 0.489 | 0.06 | 0.25 | 0.12 | 0.18 | 0.02 | 0.23 |
| 1 | 11119135 | rs743577 | 1.072 | -1.511 | 1.296 | -0.042 | 0.041 | 0.071 | 0.071 | 1.739 | 0.65 | 0.07 | 0.7 | 0.76 | 0.74 | 0.63 |
| 1 | 11125050 | rs2486920 | 0.859 | -1.466 | 1.415 | 0.021 | 0.029 | 0.04 | 0.04 | 0.484 | 0.67 | 0.2 | 0.76 | 0.78 | 0.74 | 0.72 |
| 1 | 11125866 | rs2039840 | 0.859 | -1.466 | 1.415 | 0.021 | 0.029 | 0.04 | 0.04 | 0.484 | 0.67 | 0.2 | 0.76 | 0.78 | 0.74 | 0.72 |
| 1 | 11127764 | rs2261217 | 0.87 | -1.466 | 1.415 | 0.021 | 0.03 | 0.041 | 0.041 | 0.531 | 0.67 | 0.2 | 0.76 | 0.78 | 0.74 | 0.72 |
| 1 | 11128654 | rs2791655 | 1.104 | -1.531 | 1.34 | -0.061 | 0.039 | 0.062 | 0.062 | 0.103 | 0.63 | 0.2 | 0.74 | 0.78 | 0.72 | 0.67 |
| 1 | 11129848 | rs2791653 | 0.458 | -1.344 | 1.687 | 0.181 | -0.011 | -0.062 | -0.062 | 0 | 0.72 | 0.37 | 0.83 | 0.94 | 0.77 | 0.87 |
| 1 | 11130643 | rs910660 | 0.965 | -1.342 | 1.684 | 0.182 | 0.029 | 0.04 | 0.04 | 0.661 | 0.67 | 0.2 | 0.76 | 0.78 | 0.74 | 0.72 |
| 1 | 11136838 | rs17036297 | -0.923 | -1.217 | 1.557 | 0.315 | -0.016 | 0.072 | 0.072 | 0 | 0.05 | 0.17 | 0.08 | 0.16 | 0.03 | 0.15 |
| 1 | 11159823 | rs12133922 | -0.874 | -1.201 | 1.241 | 0.323 | -0.018 | 0.06 | 0.06 | 0 | 0.05 | 0.14 | 0.07 | 0.16 | 0.03 | 0.15 |
| 1 | 11166713 | rs2536 | -0.854 | -1.195 | 1.219 | 0.016 | -0.02 | 0.023 | 0.023 | 0 | 0.05 | 0.13 | 0.07 | 0.09 | 0.02 | 0.15 |
| 1 | 11169676 | rs2275525 | -1.605 | -1.218 | 1.359 | 0.019 | -0.021 | 0.056 | 0.056 | 0.057 | 0.07 | 0.24 | 0.13 | 0.13 | 0.02 | 0.24 |
| 1 | 11198502 | rs6701524 | -0.568 | -1.313 | 1.514 | -0.107 | -0.012 | -0.099 | -0.099 | 0 | 0.27 | 0.53 | 0.16 | 0.08 | 0.24 | 0.13 |
| 1 | 11205058 | rs1057079 | 1.13 | -1.321 | 0.945 | -0.111 | 0.026 | 0.097 | 0.097 | 1.508 | 0.66 | 0.06 | 0.7 | 0.82 | 0.74 | 0.62 |
| 1 | 11248020 | rs12122605 | -1.498 | -1.313 | 0.582 | 0.222 | -0.022 | 0.152 | 0.152 | 0.037 | 0.07 | 0.22 | 0.12 | 0.24 | 0.02 | 0.25 |
| 1 | 11255008 | rs2076658 | -0.843 | -1.302 | 0.205 | -0.477 | -0.02 | 0.022 | 0.022 | 0 | 0.05 | 0.13 | 0.07 | 0.09 | 0.02 | 0.16 |
| 1 | 11255431 | rs12124598 | -0.996 | -0.78 | 0.597 | -0.112 | -0.02 | 0.023 | 0.023 | 0 | 0.05 | 0.13 | 0.07 | 0.09 | 0.02 | 0.15 |
| 1 | 11289161 | rs2076655 | 1.297 | -0.624 | 0.573 | -0.142 | 0.012 | 0.101 | 0.101 | 1.187 | 0.65 | 0.09 | 0.69 | 0.79 | 0.7 | 0.6 |
| 1 | 11318236 | rs7525957 | 0.91 | -0.457 | 0.622 | -0.217 | 0.007 | 0.048 | 0.048 | 1.043 | 0.7 | 0.22 | 0.76 | 0.88 | 0.73 | 0.78 |
| 1 | 11326788 | rs1074078 | -0.821 | -0.302 | 0.077 | -0.298 | -0.018 | 0.109 | 0.109 | 0 | 0.31 | 0.08 | 0.18 | 0.09 | 0.34 | 0.16 |
| 1 | 11340472 | rs6685950 | -1.134 | 0.085 | 0.725 | 0.351 | 0.013 | 0.073 | 0.073 | 1.753 | 0.09 | 0.45 | 0.06 | 0.11 | 0.04 | 0.15 |
| 1 | 11347092 | rs3765906 | -1.196 | 0.078 | 0.722 | 0.648 | 0.01 | 0.279 | 0.279 | 1.327 | 0.09 | 0.45 | 0.11 | 0.24 | 0.04 | 0.16 |
| 1 | 11361427 | rs2744816 | -0.921 | 0.155 | 0.458 | -0.151 | 0.026 | 0.093 | 0.093 | 1.076 | 0.08 | 0.45 | 0.06 | 0.12 | 0.05 | 0.17 |
| 1 | 11372638 | rs3010223 | -0.967 | 0.457 | -0.34 | -0.429 | -0.003 | -0.048 | -0.048 | 0 | 0.24 | 0.31 | 0.14 | 0.11 | 0.25 | 0.17 |
| 1 | 11377114 | rs2982373 | -0.777 | 0.809 | -1.591 | -0.311 | -0.063 | 0.094 | 0.094 | 0 | 0.24 | 0.03 | 0.14 | 0.11 | 0.31 | 0.17 |
| 1 | 11398365 | rs909836 | -0.495 | 0.413 | -1.634 | -0.126 | 0.057 | 0.14 | 0.14 | 0 | 0.5 | 0.85 | 0.36 | 0.24 | 0.57 | 0.37 |
| 1 | 11400409 | rs3120819 | -0.301 | -0.02 | -0.469 | -0.341 | 0.026 | 0.035 | 0.035 | 0 | 0.5 | 0.56 | 0.37 | 0.26 | 0.62 | 0.39 |
| 1 | 11401745 | rs2982384 | -0.054 | -0.235 | -0.249 | -0.58 | -0.004 | 0.064 | 0.064 | 0 | 0.4 | 0.18 | 0.31 | 0.22 | 0.57 | 0.27 |
| 1 | 11409469 | rs10864498 | 0.596 | -0.423 | -0.319 | -0.732 | 0.031 | 0.112 | 0.112 | 0 | 0.48 | 0.08 | 0.61 | 0.73 | 0.39 | 0.63 |
| 1 | 11411432 | rs2788558 | -0.345 | -0.424 | -0.437 | -0.795 | -0.09 | 0.06 | 0.06 | 0 | 0.64 | 0.95 | 0.71 | 0.81 | 0.44 | 0.76 |
| 1 | 11422737 | rs2744858 | 0.361 | -0.405 | 0.418 | -0.754 | -0.004 | -0.012 | -0.012 | 0 | 0.56 | 0.55 | 0.66 | 0.77 | 0.42 | 0.69 |

| | | | | | | | | | | | | | | | | |
|---|---|---|---|---|---|---|---|---|---|---|---|---|---|---|---|---|
| 1 | 11432744 | rs2982364 | -0.847 | -0.31 | -0.151 | -0.728 | 0.003 | 0.109 | 0.109 | 0 | 0.49 | 0.95 | 0.39 | 0.28 | 0.6 | 0.33 |
| 1 | 11435942 | rs2335404 | -0.797 | -0.161 | 0.186 | -0.617 | 0.04 | 0.129 | 0.129 | 0 | 0.49 | 0.89 | 0.38 | 0.32 | 0.59 | 0.31 |
| 1 | 11439588 | rs2788557 | -0.714 | -0.268 | 1.136 | 0.3 | 0.024 | 0.001 | 0.001 | 0 | 0.49 | 0.6 | 0.5 | 0.51 | 0.62 | 0.4 |
| 1 | 11444975 | rs3010208 | -0.582 | -0.315 | 1.403 | 0.667 | 0.002 | -0.001 | -0.001 | 0 | 0.47 | 0.53 | 0.49 | 0.47 | 0.61 | 0.36 |
| 1 | 11447805 | rs6660541 | -0.464 | -0.281 | 1.123 | 0.579 | -0.133 | 0.012 | 0.012 | 0 | 0.64 | 0.97 | 0.7 | 0.76 | 0.45 | 0.76 |
| 1 | 11456027 | rs12058244 | -0.831 | -0.319 | 1.861 | 0.514 | 0.04 | 0.02 | 0.02 | 0.184 | 0.08 | 0.36 | 0.05 | 0.05 | 0.04 | 0.11 |
| 1 | 11461601 | rs6671983 | 0.936 | -0.473 | 1.597 | 0.851 | 0.069 | -0.008 | -0.008 | 0 | 0.44 | 0.18 | 0.51 | 0.44 | 0.35 | 0.52 |
| 1 | 11468442 | rs11121728 | 0.497 | -0.397 | 1.962 | 0.975 | 0.025 | 0.011 | 0.011 | 0 | 0.52 | 0.33 | 0.54 | 0.42 | 0.41 | 0.6 |
| 1 | 11476203 | rs10864504 | 0.619 | -0.057 | 1.884 | 0.961 | 0.024 | 0.009 | 0.009 | 0 | 0.51 | 0.17 | 0.55 | 0.43 | 0.45 | 0.56 |
| 1 | 11477294 | rs10864505 | -0.098 | -0.05 | 2.091 | 0.928 | -0.015 | 0.027 | 0.027 | 0 | 0.6 | 0.65 | 0.69 | 0.74 | 0.48 | 0.75 |
| 1 | 11488203 | rs2077480 | 0.916 | 0.036 | 1.946 | 1.022 | 0.024 | -0.015 | -0.015 | 0.435 | 0.45 | 0.29 | 0.54 | 0.47 | 0.41 | 0.51 |
| 1 | 11489678 | rs877309 | -0.153 | 0.149 | 1.841 | 0.919 | -0.012 | 0.019 | 0.019 | 0 | 0.41 | 0.38 | 0.31 | 0.3 | 0.51 | 0.28 |
| 1 | 11490744 | rs10864508 | 0.546 | -0.015 | 1.255 | 0.641 | -0.009 | -0.001 | -0.001 | 0.023 | 0.57 | 0.31 | 0.63 | 0.49 | 0.5 | 0.55 |
| 1 | 11493832 | rs2922240 | 0.007 | -0.448 | 0.771 | 0.611 | -0.056 | -0.001 | -0.001 | 0 | 0.37 | 0.19 | 0.52 | 0.38 | 0.48 | 0.39 |
| 1 | 11493868 | rs11121734 | 0.873 | -0.601 | 0.794 | 0.701 | 0.019 | -0.037 | -0.037 | 0 | 0.5 | 0.1 | 0.37 | 0.45 | 0.46 | 0.45 |
| 1 | 11494573 | rs2379151 | -0.915 | -0.81 | 0.876 | 0.607 | 0.014 | -0.021 | -0.021 | 0 | 0.47 | 0.78 | 0.62 | 0.55 | 0.54 | 0.54 |
| 1 | 11496715 | rs3010217 | 0.559 | -0.727 | 1.023 | 0.644 | -0.003 | -0.038 | -0.038 | 0.224 | 0.93 | 0.82 | 0.94 | 0.99 | 0.94 | 0.97 |
| 1 | 11499286 | rs11806225 | -2.125 | -0.741 | 0.876 | 0.665 | -0.019 | 0.041 | 0.041 | 0.674 | 0.12 | 0.62 | 0.1 | 0.18 | 0.06 | 0.16 |
| 1 | 11499416 | rs3099628 | 1.758 | -0.739 | 0.514 | 0.666 | -0.027 | 0.015 | 0.015 | 1.069 | 0.88 | 0.3 | 0.89 | 0.82 | 0.94 | 0.84 |
| 1 | 11500472 | rs3124625 | 0.221 | -0.629 | -0.416 | 0.538 | -0.176 | -0.078 | -0.078 | 0 | 0.34 | 0.03 | 0.5 | 0.35 | 0.48 | 0.22 |
| 1 | 11501431 | rs11585441 | -0.983 | -0.705 | 0.05 | 0.366 | -0.007 | -0.028 | -0.028 | 0.618 | 0.07 | 0.16 | 0.05 | 0.02 | 0.04 | 0.06 |
| 1 | 11504514 | rs2379159 | -0.58 | -1.221 | -0.129 | 0.402 | 0.034 | 0.063 | 0.063 | 0.06 | 0.51 | 0.7 | 0.63 | 0.37 | 0.57 | 0.46 |
| 1 | 11508016 | rs3124627 | 0.272 | -1.215 | -0.94 | 0.351 | -0.087 | 0.008 | 0.008 | 0.656 | 0.37 | 0.1 | 0.55 | 0.32 | 0.55 | 0.26 |
| 1 | 11514206 | rs11121736 | 0.71 | -2.029 | -1.056 | -0.06 | 0.08 | 0.07 | 0.07 | 0.463 | 0.67 | 0.2 | 0.81 | 0.8 | 0.83 | 0.56 |
| 1 | 11518397 | rs12026602 | 1.313 | -2.389 | -1.483 | -0.094 | -0.04 | 0.015 | 0.015 | 0 | 0.41 | 0.11 | 0.56 | 0.32 | 0.59 | 0.27 |
| 1 | 11520653 | rs11121742 | -0.304 | -2.475 | -0.691 | -0.136 | -0.064 | -0.058 | -0.058 | 0 | 0.31 | 0.45 | 0.17 | 0.19 | 0.18 | 0.4 |
| 1 | 11525533 | rs12046038 | 0.762 | -2.236 | -1.108 | -0.05 | -0.02 | -0.103 | -0.103 | 0 | 0.25 | 0.09 | 0.25 | 0.48 | 0.23 | 0.29 |
| 1 | 11531059 | rs4846010 | 2.273 | -2.261 | -1.433 | -0.241 | -0.05 | -0.005 | -0.005 | 3.723 | 0.19 | 0.02 | 0.31 | 0.12 | 0.22 | 0.18 |
| 1 | 11531918 | rs17036930 | 0.223 | -3.341 | -0.67 | 0.429 | 0.006 | -0.101 | -0.101 | 0 | 0.24 | 0.08 | 0.22 | 0.46 | 0.19 | 0.3 |
| 1 | 11534053 | rs10159458 | -1.754 | -3.239 | -0.584 | 0.488 | 0.01 | 0.086 | 0.086 | 0 | 0.58 | 0.84 | 0.42 | 0.77 | 0.37 | 0.72 |
| 1 | 11537977 | rs2007215 | -1.758 | -3.418 | -0.159 | 0.642 | 0.069 | 0.092 | 0.092 | 0.036 | 0.55 | 0.73 | 0.4 | 0.77 | 0.34 | 0.66 |
| 1 | 11542957 | rs7555562 | 0.49 | -3.332 | -0.239 | 0.609 | -0.001 | 0 | 0 | 0 | 0.18 | 0.16 | 0.11 | 0.26 | 0.15 | 0.19 |
| 1 | 11554687 | rs7534208 | -1.66 | -3.181 | -0.112 | 0.603 | 0.081 | 0.066 | 0.066 | 0.085 | 0.54 | 0.76 | 0.4 | 0.72 | 0.34 | 0.66 |
| 1 | 11561593 | rs2817580 | 1.373 | -3.16 | -0.24 | 0.612 | 0.077 | 0.064 | 0.064 | 0.048 | 0.45 | 0.14 | 0.63 | 0.27 | 0.66 | 0.3 |
| 1 | 11574942 | rs2745265 | 1.273 | -3.184 | 0.016 | 0.594 | 0.043 | 0.038 | 0.038 | 0 | 0.47 | 0.32 | 0.64 | 0.28 | 0.65 | 0.31 |
| 1 | 11575260 | rs17036947 | -0.433 | -2.893 | 0.396 | 1.225 | 0.008 | -0.055 | -0.055 | 0 | 0.19 | 0.08 | 0.17 | 0.44 | 0.13 | 0.25 |
| 1 | 11577125 | rs2072997 | -0.491 | -3.11 | 0.27 | 1.239 | 0.012 | -0.032 | -0.032 | 0 | 0.17 | 0.09 | 0.1 | 0.44 | 0.11 | 0.21 |
| 1 | 11578973 | rs2072996 | -0.489 | -3.026 | 0.229 | 1.278 | 0.011 | -0.032 | -0.032 | 0 | 0.17 | 0.09 | 0.1 | 0.44 | 0.11 | 0.22 |
| 1 | 11584164 | rs2076469 | -1.642 | -3.273 | 0.591 | 1.204 | -0.062 | 0.083 | 0.083 | 0 | 0.29 | 0.41 | 0.14 | 0.45 | 0.16 | 0.28 |
| 1 | 11589911 | rs2076468 | -0.641 | -2.827 | -0.039 | 1.432 | -0.003 | -0.022 | -0.022 | 0 | 0.13 | 0.07 | 0.1 | 0.44 | 0.11 | 0.2 |
| 1 | 11595331 | rs2235660 | 0.587 | -3.067 | -0.055 | 1.362 | 0.014 | -0.015 | -0.015 | 0 | 0.16 | 0.13 | 0.11 | 0.25 | 0.16 | 0.2 |
| 1 | 11596455 | rs2594310 | -0.965 | -3.316 | 0.257 | 1.199 | -0.085 | -0.08 | -0.08 | 0.067 | 0.2 | 0.61 | 0.09 | 0.06 | 0.06 | 0.23 |
| 1 | 11601119 | rs2817621 | -2.381 | -3.402 | 0.123 | 1.263 | 0.117 | -0.124 | -0.124 | 0 | 0.36 | 0.91 | 0.2 | 0.51 | 0.17 | 0.45 |
| 1 | 11603063 | rs2817619 | -0.885 | -3.259 | 0.128 | 1.329 | -0.013 | -0.044 | -0.044 | 0 | 0.15 | 0.07 | 0.11 | 0.43 | 0.13 | 0.21 |
| 1 | 11605183 | rs6683331 | -0.968 | -3.245 | 0.146 | 1.194 | -0.114 | -0.088 | -0.088 | 0.534 | 0.2 | 0.51 | 0.08 | 0.08 | 0.06 | 0.23 |
| 1 | 11607932 | rs2745291 | -1.976 | -3.043 | -0.704 | 1.374 | 0.042 | -0.043 | -0.043 | 0 | 0.35 | 0.77 | 0.18 | 0.51 | 0.19 | 0.45 |
| 1 | 11619154 | rs11808316 | -0.296 | -3.147 | -1.007 | 1.206 | 0.056 | 0.007 | 0.007 | 0 | 0.25 | 0.1 | 0.11 | 0.28 | 0.18 | 0.36 |
| 1 | 11622979 | rs2817581 | -0.226 | -2.959 | -1.163 | 1.896 | -0.029 | -0.031 | -0.031 | 0 | 0.08 | 0.01 | 0.05 | 0.33 | 0.08 | 0.12 |
| 1 | 11626858 | rs2745285 | -2.054 | -3.157 | -1.222 | 1.294 | 0.125 | 0.142 | 0.142 | 0.217 | 0.52 | 0.94 | 0.3 | 0.78 | 0.31 | 0.63 |
| 1 | 11633982 | rs10864520 | -0.299 | -2.616 | -0.631 | 1.644 | 0.032 | -0.008 | -0.008 | 0.332 | 0.19 | 0.11 | 0.12 | 0.26 | 0.14 | 0.36 |
| 1 | 11641283 | rs2745276 | -1.598 | -2.425 | 0.122 | 2.143 | -0.009 | 0.123 | 0.123 | 0 | 0.13 | 0.17 | 0.07 | 0.49 | 0.08 | 0.2 |
| 1 | 11645598 | rs2745275 | -1.325 | -2.428 | 0.077 | 1.986 | -0.006 | 0.103 | 0.103 | 0 | 0.11 | 0.13 | 0.05 | 0.46 | 0.07 | 0.21 |
| 1 | 11652205 | rs2335767 | 1.365 | -2.806 | 0.33 | 1.192 | -0.001 | -0.001 | -0.001 | 0.036 | 0.59 | 0.44 | 0.73 | 0.39 | 0.71 | 0.52 |
| 1 | 11654491 | rs7530509 | 1.124 | -2.594 | 0.701 | 1.424 | -0.003 | 0.142 | 0.142 | 0 | 0.84 | 0.76 | 0.88 | 0.58 | 0.88 | 0.73 |
| 1 | 11656684 | rs761087 | 1.607 | -2.718 | 0.595 | 1.149 | 0.027 | 0.002 | 0.002 | 0 | 0.6 | 0.33 | 0.68 | 0.34 | 0.71 | 0.38 |
| 1 | 11660710 | rs761084 | 1.799 | -2.776 | 1.021 | 0.846 | -0.034 | 0.004 | 0.004 | 0 | 0.66 | 0.49 | 0.66 | 0.33 | 0.79 | 0.41 |
| 1 | 11665198 | rs11121778 | 0.754 | -2.805 | 0.982 | 0.9 | -0.018 | -0.048 | -0.048 | 0 | 0.86 | 0.92 | 0.82 | 0.55 | 0.86 | 0.79 |
| 1 | 11665575 | rs4846019 | 1.474 | -2.831 | 1.222 | 0.889 | 0.003 | 0.122 | 0.122 | 0 | 0.8 | 0.63 | 0.73 | 0.52 | 0.84 | 0.65 |
| 1 | 11670959 | rs4542213 | 1.55 | -3.016 | 1.286 | 0.848 | -0.009 | 0.002 | 0.002 | 0 | 0.67 | 0.61 | 0.68 | 0.36 | 0.8 | 0.41 |
| 1 | 11675059 | rs734737 | 1.719 | -2.985 | 1.303 | 0.731 | -0.001 | -0.001 | -0.001 | 0 | 0.66 | 0.32 | 0.65 | 0.31 | 0.79 | 0.39 |
| 1 | 11680283 | rs7550768 | 0.535 | -2.791 | 1.439 | 0.803 | 0.014 | 0.241 | 0.241 | 0 | 0.88 | 0.71 | 0.84 | 0.58 | 0.88 | 0.83 |
| 1 | 11689663 | rs17037122 | -1.061 | -2.782 | 1.034 | 0.828 | -0.03 | -0.022 | -0.022 | 1.635 | 0.21 | 0.31 | 0.18 | 0.21 | 0.1 | 0.35 |
| 1 | 11690766 | rs10779763 | 1.276 | -3.463 | 0.775 | 0.462 | -0.035 | 0.006 | 0.006 | 0.04 | 0.64 | 0.47 | 0.72 | 0.42 | 0.83 | 0.54 |
| 1 | 11691161 | rs2335768 | 1.506 | -3.006 | 0.702 | 0.645 | 0.076 | 0.047 | 0.047 | 0.561 | 0.51 | 0.36 | 0.64 | 0.3 | 0.77 | 0.37 |
| 1 | 11694773 | rs17037129 | -0.341 | -2.888 | 0.322 | 0.673 | -0.015 | -0.025 | -0.025 | 0.232 | 0.09 | 0.18 | 0.1 | 0.06 | 0.05 | 0.17 |
| 1 | 11698902 | rs6671295 | 1.072 | -2.443 | -0.154 | 0.901 | 0.156 | 0.082 | 0.082 | 0.846 | 0.5 | 0.24 | 0.61 | 0.31 | 0.78 | 0.38 |
| 1 | 11702155 | rs7526268 | 1.107 | -2.255 | -0.269 | 0.818 | 0.097 | 0.059 | 0.059 | 0.042 | 0.47 | 0.22 | 0.56 | 0.31 | 0.73 | 0.36 |
| 1 | 11703843 | rs6540987 | 0.926 | -1.5 | -0.328 | 1.069 | 0.001 | 0.149 | 0.149 | 0 | 0.85 | 0.76 | 0.81 | 0.58 | 0.87 | 0.84 |
| 1 | 11708736 | rs1055547 | 0.641 | -1.942 | -0.804 | 0.736 | 0.067 | 0.014 | 0.014 | 0.056 | 0.52 | 0.35 | 0.62 | 0.41 | 0.78 | 0.46 |

| | | | | | | | | | | | | | | | | |
|---|---|---|---|---|---|---|---|---|---|---|---|---|---|---|---|---|
| 1 | 11710561 | rs9614 | -1.309 | -1.57 | -1.032 | 1.1 | 0.001 | 0 | 0 | 0 | 0.18 | 0.14 | 0.26 | 0.45 | 0.18 | 0.23 |
| 1 | 11711485 | rs12045158 | 0.079 | -2.022 | -1.417 | 1.818 | 0.037 | -0.009 | -0.009 | 0 | 0.08 | 0 | 0.1 | 0.46 | 0.03 | 0.14 |
| 1 | 11711589 | rs3117061 | -0.832 | -1.438 | -2.003 | 0.278 | 0.021 | 0.119 | 0.119 | 0.018 | 0.55 | 0.83 | 0.66 | 0.73 | 0.45 | 0.66 |
| 1 | 11713166 | rs1883575 | -0.349 | -1.361 | -2.108 | -0.226 | 0.012 | 0.024 | 0.024 | 0 | 0.45 | 0.72 | 0.62 | 0.57 | 0.56 | 0.4 |
| 1 | 11713563 | rs909934 | -1.099 | -1.337 | -2.424 | -0.215 | 0.04 | 0.005 | 0.005 | 0 | 0.74 | 0.94 | 0.88 | 0.76 | 0.8 | 0.75 |
| 1 | 11717300 | rs12139487 | -0.039 | -0.675 | -2.188 | -0.076 | 0.005 | 0.033 | 0.033 | 0.078 | 0.19 | 0.07 | 0.3 | 0.09 | 0.18 | 0.11 |
| 1 | 11721854 | rs2294641 | -0.932 | -0.837 | -1.851 | -0.216 | 0.024 | 0.016 | 0.016 | 0 | 0.48 | 0.71 | 0.54 | 0.45 | 0.43 | 0.52 |
| 1 | 11723520 | rs3820340 | 0.241 | -0.421 | -1.669 | 0.143 | -0.001 | 0.001 | 0.001 | 0 | 0.07 | 0.05 | 0.04 | 0.18 | 0.06 | 0.15 |
| 1 | 11732932 | rs747863 | 0.204 | -0.874 | -1.729 | -0.224 | 0.022 | 0.015 | 0.015 | 0 | 0.18 | 0.1 | 0.19 | 0.15 | 0.15 | 0.21 |
| 1 | 11735652 | rs2233025 | 0.29 | -0.164 | -1.857 | 0.035 | -0.041 | -0.037 | -0.037 | 0 | 0.14 | 0.03 | 0.28 | 0.2 | 0.18 | 0.17 |
| 1 | 11736600 | rs2336030 | 0.788 | 0.188 | -2.077 | -0.04 | -0.134 | -0.121 | -0.121 | 2.591 | 0.21 | 0.03 | 0.28 | 0.52 | 0.37 | 0.34 |
| 1 | 11740358 | rs7417320 | -0.471 | -0.138 | -1.446 | -0.264 | -0.007 | -0.007 | -0.007 | 0.175 | 0.44 | 0.4 | 0.51 | 0.43 | 0.4 | 0.47 |
| 1 | 11742074 | rs2233004 | -0.823 | -0.292 | -1.649 | -0.442 | 0.029 | 0.111 | 0.111 | 0 | 0.71 | 0.9 | 0.85 | 0.93 | 0.79 | 0.82 |
| 1 | 11746500 | rs2143296 | -0.273 | 0.29 | -0.985 | 0.24 | -0.003 | -0.002 | -0.002 | 0 | 0.14 | 0.1 | 0.29 | 0.22 | 0.18 | 0.15 |
| 1 | 11751135 | rs12088541 | -0.402 | -0.337 | -0.149 | -0.056 | -0.051 | -0.012 | -0.012 | 0.182 | 0.18 | 0.56 | 0.09 | 0.15 | 0.09 | 0.09 |
| 1 | 11754168 | rs7539455 | 0.501 | 1.066 | 0.416 | 0.792 | 0.107 | 0.164 | 0.164 | 0 | 0.87 | 0.71 | 0.77 | 0.63 | 0.77 | 0.76 |
| 1 | 11761252 | rs1534991 | -0.325 | 1.158 | 0.338 | 0.222 | 0 | -0.08 | -0.08 | 0 | 0.71 | 0.58 | 0.85 | 0.9 | 0.73 | 0.73 |
| 1 | 11763677 | rs9430624 | 0.964 | 1.063 | -0.418 | 0.242 | 0.061 | 0 | 0 | 0 | 0.44 | 0.26 | 0.49 | 0.4 | 0.37 | 0.38 |
| 1 | 11764281 | rs12045323 | -0.795 | 1.389 | -0.228 | 0.495 | 0.076 | 0.15 | 0.15 | 0 | 0.19 | 0.37 | 0.28 | 0.42 | 0.26 | 0.27 |
| 1 | 11766424 | rs11121804 | 0.487 | 1.439 | -0.482 | 0.737 | 0.014 | 0.048 | 0.048 | 0 | 0.08 | 0.08 | 0.15 | 0.28 | 0.16 | 0.11 |
| 1 | 11768869 | rs6687381 | -0.393 | 0.908 | -0.694 | 0.009 | -0.004 | -0.026 | -0.026 | 0 | 0.58 | 0.56 | 0.71 | 0.83 | 0.62 | 0.58 |
| 1 | 11772340 | rs2336034 | 0 | 0.633 | -0.978 | 0.339 | -0.003 | 0.005 | 0.005 | 0 | 0.11 | 0.09 | 0.08 | 0.07 | 0.13 | 0.11 |
| 1 | 11774956 | rs9430631 | -0.147 | 0.042 | -0.92 | 0.623 | -0.003 | 0.002 | 0.002 | 0.603 | 0.41 | 0.46 | 0.48 | 0.52 | 0.35 | 0.39 |
| 1 | 11779499 | rs4846028 | 0.089 | -0.345 | -1.651 | 0.54 | 0.106 | 0.014 | 0.014 | 0 | 0.56 | 0.17 | 0.69 | 0.59 | 0.71 | 0.66 |
| 1 | 11783484 | rs4846029 | -0.32 | 0.085 | -1.369 | 0.037 | -0.002 | -0.004 | -0.004 | 0.086 | 0.46 | 0.48 | 0.56 | 0.63 | 0.4 | 0.6 |
| 1 | 11784118 | rs7529001 | 0.105 | 0.486 | -1.791 | 0.242 | -0.016 | -0.056 | -0.056 | 0 | 0.33 | 0.14 | 0.5 | 0.5 | 0.37 | 0.39 |
| 1 | 11794553 | rs12121577 | 0.076 | 0.718 | -2.056 | 0.059 | -0.046 | -0.041 | -0.041 | 0 | 0.28 | 0.05 | 0.23 | 0.32 | 0.28 | 0.18 |
| 1 | 11794676 | rs17875979 | -0.063 | 1.137 | -1.917 | -0.128 | -0.023 | -0.023 | -0.023 | 0 | 0.08 | 0.03 | 0.05 | 0.1 | 0.1 | 0.17 |
| 1 | 11797024 | rs12116971 | -0.011 | 0.739 | -1.574 | -0.938 | 0 | 0 | 0 | 0 | 0.53 | 0.45 | 0.55 | 0.4 | 0.45 | 0.52 |
| 1 | 11797186 | rs12095517 | 0.061 | 0.502 | -1.497 | -0.941 | -0.013 | 0.012 | 0.012 | 0 | 0.63 | 0.74 | 0.67 | 0.66 | 0.6 | 0.68 |
| 1 | 11802434 | rs4073574 | 0.117 | 0.548 | -1.633 | -1.157 | 0.014 | 0.053 | 0.053 | 0 | 0.58 | 0.29 | 0.74 | 0.89 | 0.63 | 0.57 |
| 1 | 11807426 | rs12752700 | 0.202 | 0.59 | -2.197 | -1.116 | 0.025 | 0.167 | 0.167 | 0 | 0.57 | 0.18 | 0.68 | 0.88 | 0.63 | 0.57 |
| 1 | 11810505 | rs8617 | 0.763 | 0.428 | -2.254 | -1.179 | 0.001 | 0.113 | 0.113 | 0 | 0.25 | 0.07 | 0.16 | 0.03 | 0.21 | 0.27 |
| 1 | 11810674 | rs1128059 | 0.537 | 0.819 | -2.137 | -0.534 | -0.057 | -0.07 | -0.07 | 1.867 | 0.26 | 0.05 | 0.29 | 0.33 | 0.31 | 0.24 |
| 1 | 11810691 | rs1128065 | 0.549 | 0.818 | -2.137 | -0.532 | -0.058 | -0.068 | -0.068 | 0 | 0.26 | 0.05 | 0.29 | 0.33 | 0.31 | 0.24 |
| 1 | 11810844 | rs11121821 | 0.427 | 0.818 | -2.077 | -0.544 | -0.028 | -0.041 | -0.041 | 0.453 | 0.27 | 0.11 | 0.29 | 0.34 | 0.31 | 0.24 |
| 1 | 11810973 | rs4075034 | 0.456 | 0.834 | -2.06 | -0.67 | -0.006 | -0.024 | -0.024 | 0.467 | 0.31 | 0.18 | 0.4 | 0.55 | 0.32 | 0.32 |
| 1 | 11811234 | rs4075033 | 0.463 | 0.936 | -2.121 | -0.603 | -0.065 | -0.096 | -0.096 | 0 | 0.27 | 0.04 | 0.34 | 0.38 | 0.32 | 0.26 |
| 1 | 11812127 | rs6661776 | -0.345 | 0.815 | -1.143 | 0.308 | -0.002 | -0.015 | -0.015 | 0 | 0.31 | 0.23 | 0.37 | 0.52 | 0.31 | 0.31 |
| 1 | 11824133 | rs4845875 | 0.278 | 0.307 | -0.909 | -0.486 | -0.001 | -0.019 | -0.019 | 0 | 0.56 | 0.45 | 0.78 | 0.88 | 0.65 | 0.6 |
| 1 | 11824303 | rs4845877 | 0.353 | -0.174 | -0.787 | -0.555 | 0.014 | 0.027 | 0.027 | 0 | 0.54 | 0.4 | 0.79 | 0.79 | 0.66 | 0.44 |
| 1 | 11825544 | rs12564559 | 0.057 | -0.091 | -1.1 | -0.569 | -0.02 | -0.019 | -0.019 | 0 | 0.13 | 0.04 | 0.09 | 0.15 | 0.09 | 0.22 |
| 1 | 11828319 | rs4845881 | 0.843 | -0.24 | -0.679 | -0.728 | -0.012 | -0.023 | -0.023 | 0.934 | 0.55 | 0.48 | 0.77 | 0.73 | 0.69 | 0.47 |
| 1 | 11836450 | rs12132479 | 0.731 | -0.495 | -1.047 | -1.008 | -0.036 | 0 | 0 | 0 | 0.2 | 0.01 | 0.08 | 0.1 | 0.18 | 0.26 |
| 1 | 11836628 | rs6699881 | 0.905 | -0.642 | 0.09 | -1.143 | -0.022 | -0.063 | -0.063 | 0 | 0.58 | 0.52 | 0.82 | 0.88 | 0.73 | 0.65 |
| 1 | 11838011 | rs10779765 | 0.896 | -0.696 | 0.652 | -1.149 | -0.022 | -0.063 | -0.063 | 0 | 0.58 | 0.52 | 0.82 | 0.88 | 0.73 | 0.65 |
| 1 | 11846252 | rs4846048 | 0.85 | -0.517 | 0.671 | -1.159 | -0.017 | -0.056 | -0.056 | 0 | 0.59 | 0.53 | 0.82 | 0.89 | 0.72 | 0.66 |
| 1 | 11847436 | rs2184226 | 0.256 | -0.542 | 0.792 | -1.167 | 0.013 | 0.02 | 0.02 | 0.014 | 0.14 | 0.08 | 0.06 | 0 | 0.07 | 0.08 |
| 1 | 11852110 | rs13306556 | -0.73 | -0.448 | 1.136 | -1.163 | 0.012 | 0.005 | 0.005 | 0 | 0.06 | 0.1 | 0.06 | 0.12 | 0.1 | 0.22 |
| 1 | 11852300 | rs1476413 | -0.109 | -0.453 | 1.169 | -1.145 | -0.019 | 0.009 | 0.009 | 0.006 | 0.27 | 0.15 | 0.14 | 0.24 | 0.28 | 0.46 |
| 1 | 11854476 | rs1801131 | -0.779 | -0.902 | 0.87 | -1.491 | -0.006 | 0.055 | 0.055 | 0 | 0.34 | 0.15 | 0.15 | 0.22 | 0.31 | 0.42 |
| 1 | 11855867 | rs6541003 | 1.398 | -1.01 | 0.776 | -1.539 | 0.005 | 0.022 | 0.022 | 0 | 0.49 | 0.43 | 0.75 | 0.78 | 0.61 | 0.49 |
| 1 | 11856378 | rs1801133 | 1.656 | -0.398 | -0.196 | -1.049 | -0.029 | -0.05 | -0.05 | 0.009 | 0.19 | 0.09 | 0.47 | 0.3 | 0.36 | 0.12 |
| 1 | 11857711 | rs1572151 | 0.846 | -0.381 | -0.078 | -0.968 | 0.012 | 0.049 | 0.049 | 0 | 0.14 | 0.06 | 0.05 | 0 | 0.07 | 0.09 |
| 1 | 11857951 | rs4846052 | 1.124 | -0.555 | -0.187 | -1.087 | -0.011 | 0.127 | 0.127 | 0.21 | 0.45 | 0.15 | 0.73 | 0.77 | 0.6 | 0.47 |
| 1 | 11860843 | rs17037390 | -1.121 | -0.447 | -0.235 | -0.991 | 0.004 | -0.019 | -0.019 | 0.246 | 0.15 | 0.22 | 0.09 | 0.13 | 0.14 | 0.19 |
| 1 | 11862047 | rs17037396 | 0.202 | -0.437 | -0.24 | -0.937 | 0.015 | 0.009 | 0.009 | 0 | 0.06 | 0.09 | 0.06 | 0.13 | 0.1 | 0.16 |
| 1 | 11862214 | rs9651118 | 2.152 | -0.676 | -0.569 | -1.169 | -0.069 | -0.095 | -0.095 | 0 | 0.18 | 0 | 0.23 | 0.41 | 0.19 | 0.3 |
| 1 | 11862778 | rs17367504 | -1.194 | -0.752 | 0.057 | -1.059 | -0.005 | 0.009 | 0.009 | 0 | 0.14 | 0.11 | 0.08 | 0.12 | 0.14 | 0.19 |
| 1 | 11863057 | rs2066470 | -0.061 | -0.736 | 0.064 | -1.048 | 0.003 | 0 | 0 | 0 | 0.06 | 0.07 | 0.05 | 0.12 | 0.1 | 0.16 |
| 1 | 11864149 | rs7553194 | -0.061 | -0.709 | 0.082 | -1.048 | 0.018 | 0.01 | 0.01 | 0 | 0.06 | 0.09 | 0.06 | 0.12 | 0.1 | 0.16 |
| 1 | 11864586 | rs3753584 | -1.177 | -0.722 | 0.075 | -1.058 | -0.003 | 0.003 | 0.003 | 0 | 0.14 | 0.13 | 0.08 | 0.13 | 0.14 | 0.19 |
| 1 | 11865804 | rs13306561 | -1.566 | -0.732 | 0.097 | -1.103 | 0.003 | -0.02 | -0.02 | 0 | 0.16 | 0.24 | 0.09 | 0.13 | 0.14 | 0.19 |
| 1 | 11867044 | rs3737964 | -0.093 | -0.784 | 0.308 | -1.169 | 0.011 | 0.042 | 0.042 | 0 | 0.66 | 0.72 | 0.84 | 0.9 | 0.74 | 0.68 |
| 1 | 11869230 | rs4846054 | 0.866 | -0.786 | 0.337 | -1.165 | -0.002 | -0.021 | -0.021 | 1.959 | 0.54 | 0.46 | 0.76 | 0.77 | 0.62 | 0.5 |
| 1 | 11870383 | rs17037425 | -1.258 | -0.768 | 0.328 | -1.136 | 0.014 | -0.006 | -0.006 | 0.516 | 0.11 | 0.21 | 0.08 | 0.13 | 0.12 | 0.18 |
| 1 | 11879685 | rs198393 | -1.607 | -0.878 | 0.367 | -1.154 | 0.002 | 0.013 | 0.013 | 0 | 0.52 | 0.62 | 0.26 | 0.23 | 0.4 | 0.52 |
| 1 | 11879699 | rs2050265 | -1.432 | -0.772 | 0.204 | -1.046 | 0.003 | -0.02 | -0.02 | 1.024 | 0.16 | 0.25 | 0.09 | 0.13 | 0.14 | 0.19 |
| 1 | 11883731 | rs12567136 | -1.055 | -0.772 | 0.192 | -1.047 | -0.007 | 0.011 | 0.011 | 0 | 0.14 | 0.11 | 0.08 | 0.13 | 0.14 | 0.19 |

| | | | | | | | | | | | | | | | | |
|---|---|---|---|---|---|---|---|---|---|---|---|---|---|---|---|---|
| 1 | 11886813 | rs2076004 | -0.088 | -0.81 | 0.197 | -1.043 | 0.016 | 0.007 | 0.007 | 0 | 0.06 | 0.09 | 0.06 | 0.12 | 0.1 | 0.16 |
| 1 | 11887303 | rs7537765 | -1.409 | -0.811 | 0.199 | -1.043 | 0.002 | -0.019 | -0.019 | 0 | 0.16 | 0.22 | 0.09 | 0.13 | 0.14 | 0.19 |
| 1 | 11892650 | rs2236797 | -0.341 | -0.769 | 0.14 | -1.016 | 0.014 | 0.006 | 0.006 | 0 | 0.06 | 0.09 | 0.06 | 0.12 | 0.11 | 0.16 |
| 1 | 11897271 | rs198405 | -0.951 | -0.887 | 0.141 | -1.102 | -0.035 | -0.121 | -0.121 | 0 | 0.25 | 0.37 | 0.11 | 0.01 | 0.11 | 0.13 |
| 1 | 11897592 | rs198406 | -1.433 | -0.914 | 0.099 | -1.13 | -0.001 | -0.008 | -0.008 | 0 | 0.51 | 0.53 | 0.24 | 0.23 | 0.4 | 0.51 |
| 1 | 11900385 | rs198411 | -0.917 | -0.914 | 0.076 | -1.13 | -0.035 | -0.121 | -0.121 | 0 | 0.25 | 0.37 | 0.11 | 0.01 | 0.11 | 0.13 |
| 1 | 11900437 | rs198412 | 0.399 | -0.929 | 0.095 | -1.139 | -0.019 | -0.079 | -0.079 | 0.264 | 0.16 | 0.28 | 0.08 | 0.01 | 0.07 | 0.09 |
| 1 | 11900825 | rs198414 | -0.717 | -0.845 | 0.098 | -1.089 | -0.035 | -0.121 | -0.121 | 0 | 0.25 | 0.37 | 0.11 | 0.01 | 0.11 | 0.13 |
| 1 | 11900838 | rs17350396 | 1.299 | -0.845 | 0.089 | -1.09 | -0.037 | -0.01 | -0.01 | 0 | 0.19 | 0.01 | 0.07 | 0.09 | 0.18 | 0.23 |
| 1 | 11901592 | rs198415 | -0.932 | -0.737 | 0.415 | -0.939 | -0.035 | -0.121 | -0.121 | 0 | 0.25 | 0.37 | 0.11 | 0.01 | 0.11 | 0.13 |
| 1 | 11904076 | rs198358 | -1.729 | -0.715 | 0.328 | -0.891 | -0.016 | -0.075 | -0.075 | 0.023 | 0.32 | 0.55 | 0.17 | 0.15 | 0.21 | 0.29 |
| 1 | 11906068 | rs5065 | -1.001 | -0.938 | 0.282 | -0.891 | -0.039 | -0.146 | -0.146 | 0.014 | 0.26 | 0.42 | 0.11 | 0.01 | 0.12 | 0.13 |
| 1 | 11909514 | rs198372 | -0.465 | -0.887 | -0.135 | -0.862 | 0.02 | 0.016 | 0.016 | 0 | 0.21 | 0.18 | 0.08 | 0.01 | 0.11 | 0.15 |
| 1 | 11913757 | rs198375 | -0.835 | -0.902 | 0.175 | -0.658 | -0.001 | -0.019 | -0.019 | 0 | 0.42 | 0.62 | 0.26 | 0.2 | 0.39 | 0.46 |
| 1 | 11914829 | rs6668352 | 0.381 | -0.672 | 0.074 | -0.843 | -0.012 | 0.035 | 0.035 | 0 | 0.28 | 0.18 | 0.15 | 0.16 | 0.28 | 0.33 |
| 1 | 11917340 | rs198388 | -1.261 | -0.687 | 0.133 | -0.773 | -0.006 | 0.053 | 0.053 | 0 | 0.55 | 0.64 | 0.27 | 0.2 | 0.43 | 0.53 |
| 1 | 11919271 | rs198389 | -0.941 | -0.661 | 0.113 | -0.777 | 0.017 | 0.074 | 0.074 | 0 | 0.5 | 0.4 | 0.24 | 0.19 | 0.41 | 0.49 |
| 1 | 11922298 | rs6668659 | -0.788 | -0.593 | 0.579 | -0.374 | 0.006 | 0.017 | 0.017 | 0 | 0.4 | 0.39 | 0.19 | 0.18 | 0.34 | 0.42 |
| 1 | 11925300 | rs6676300 | -1.338 | -0.563 | 0.508 | -0.39 | 0.026 | 0.07 | 0.07 | 0 | 0.48 | 0.63 | 0.21 | 0.21 | 0.37 | 0.46 |
| 1 | 11927056 | rs12562952 | -1.419 | -0.646 | 0.644 | -0.157 | 0.001 | 0.002 | 0.002 | 0 | 0.13 | 0.15 | 0.07 | 0.1 | 0.08 | 0.17 |
| 1 | 11928819 | rs11801879 | -1.927 | -0.545 | 0.612 | -0.121 | -0.039 | -0.102 | -0.102 | 0 | 0.17 | 0.37 | 0.05 | 0.04 | 0.07 | 0.11 |
| 1 | 11928895 | rs11803049 | -1.649 | -0.449 | 0.703 | -0.006 | 0.013 | 0.013 | 0.013 | 0.032 | 0.12 | 0.38 | 0.08 | 0.13 | 0.05 | 0.16 |
| 1 | 11936324 | rs9661078 | 0.782 | -0.159 | 0.071 | 0.207 | 0.003 | 0.008 | 0.008 | 0 | 0.39 | 0.44 | 0.34 | 0.62 | 0.42 | 0.48 |
| 1 | 11937404 | rs7554327 | -0.526 | -0.362 | -0.049 | 0.081 | -0.034 | -0.052 | -0.052 | 0 | 0.61 | 0.82 | 0.43 | 0.66 | 0.52 | 0.71 |
| 1 | 11941484 | rs2336395 | -1.114 | -0.183 | 0.103 | 0.232 | 0.002 | 0.026 | 0.026 | 0 | 0.14 | 0.12 | 0.09 | 0.07 | 0.16 | 0.15 |
| 1 | 11941936 | rs11588551 | -0.087 | -0.128 | 0.337 | 0.448 | -0.006 | -0.003 | -0.003 | 0 | 0.54 | 0.64 | 0.39 | 0.63 | 0.5 | 0.59 |
| 1 | 11947957 | rs6667637 | 1.044 | 0.071 | 0.455 | 0.554 | -0.001 | 0.002 | 0.002 | 0 | 0.38 | 0.57 | 0.4 | 0.49 | 0.34 | 0.41 |
| 1 | 11952406 | rs17037520 | 0.6 | 0.11 | 0.472 | 0.572 | 0 | -0.001 | -0.001 | 0.284 | 0.43 | 0.57 | 0.42 | 0.52 | 0.38 | 0.44 |
| 1 | 11957639 | rs12724129 | 0.136 | 0.097 | 0.468 | 0.574 | 0.007 | 0 | 0 | 0 | 0.49 | 0.62 | 0.44 | 0.56 | 0.4 | 0.51 |
| 1 | 11959216 | rs1999594 | -0.208 | 0.189 | 0.507 | 0.391 | -0.004 | -0.007 | -0.007 | 0 | 0.55 | 0.63 | 0.46 | 0.61 | 0.46 | 0.65 |
| 1 | 11969748 | rs926269 | 1.4 | 0.211 | 0.563 | 0.517 | 0 | 0.001 | 0.001 | 0 | 0.36 | 0.38 | 0.38 | 0.46 | 0.34 | 0.43 |
| 1 | 11971749 | rs11800086 | 0.956 | 0.232 | -0.022 | 0.527 | 0 | 0 | 0 | 0 | 0.38 | 0.57 | 0.4 | 0.46 | 0.35 | 0.43 |
| 1 | 11973888 | rs7554547 | 0.198 | 0.273 | -0.048 | 0.523 | -0.007 | -0.004 | -0.004 | 0 | 0.48 | 0.66 | 0.49 | 0.56 | 0.5 | 0.52 |
| 1 | 11979231 | rs6691499 | 0.288 | 0.58 | -0.514 | 0.387 | 0.001 | 0 | 0 | 0 | 0.05 | 0.06 | 0.06 | 0.07 | 0.11 | 0.07 |
| 1 | 11979556 | rs2038028 | -0.129 | 0.534 | -0.535 | 0.358 | -0.007 | -0.029 | -0.029 | 0 | 0.57 | 0.83 | 0.53 | 0.68 | 0.55 | 0.68 |
| 1 | 11980449 | rs17037529 | -1.336 | 1.007 | -0.056 | 0.862 | 0.002 | -0.005 | -0.005 | 0.017 | 0.1 | 0.14 | 0.09 | 0.1 | 0.15 | 0.09 |
| 1 | 11981270 | rs9659237 | -0.363 | 1.221 | -0.256 | 1.022 | 0.019 | 0.004 | 0.004 | 0 | 0.05 | 0.1 | 0.06 | 0.07 | 0.11 | 0.07 |
| 1 | 11982544 | rs2639453 | -0.432 | 1.075 | -0.681 | 0.988 | -0.028 | -0.061 | -0.061 | 0 | 0.83 | 0.94 | 0.62 | 0.76 | 0.78 | 0.91 |
| 1 | 12000670 | rs11802668 | -0.169 | 0.933 | -0.167 | 0.869 | 0.01 | 0.009 | 0.009 | 0.08 | 0.07 | 0.14 | 0.04 | 0.14 | 0.06 | 0.14 |
| 1 | 12000840 | rs6674770 | -1.723 | 0.657 | 0.826 | 0.927 | 0.008 | 0.043 | 0.043 | 0.304 | 0.16 | 0.58 | 0.13 | 0.26 | 0.15 | 0.24 |
| 1 | 12004999 | rs4846080 | -0.448 | 0.499 | 0.325 | 0.618 | -0.008 | 0.032 | 0.032 | 0 | 0.53 | 0.72 | 0.48 | 0.72 | 0.45 | 0.68 |
| 1 | 12008863 | rs2336379 | -1.366 | 0.327 | 0.44 | 0.652 | 0 | 0.048 | 0.048 | 0.474 | 0.17 | 0.62 | 0.13 | 0.31 | 0.15 | 0.25 |
| 1 | 12009955 | rs7529452 | 0.967 | 0.548 | 0.082 | 0.691 | -0.067 | -0.054 | -0.054 | 1.916 | 0.34 | 0.02 | 0.34 | 0.36 | 0.31 | 0.42 |
| 1 | 12010469 | rs2273285 | -0.372 | 0.639 | 1.188 | 0.785 | 0.058 | 0.138 | 0.138 | 0.016 | 0.06 | 0.3 | 0.07 | 0.13 | 0.07 | 0.08 |
| 1 | 12018290 | rs2273289 | -1.462 | 0.616 | 0.991 | 0.791 | 0.019 | 0.12 | 0.12 | 0.46 | 0.16 | 0.52 | 0.12 | 0.29 | 0.15 | 0.24 |
| 1 | 12024923 | rs7529312 | -1.389 | 0.664 | 1.354 | 0.797 | 0.049 | 0.145 | 0.145 | 0 | 0.07 | 0.39 | 0.09 | 0.15 | 0.09 | 0.12 |
| 1 | 12025353 | rs2273291 | -0.201 | 0.433 | 0.771 | 0.616 | -0.031 | 0.019 | 0.019 | 0 | 0.52 | 0.86 | 0.48 | 0.68 | 0.46 | 0.68 |
| 1 | 12027308 | rs3818157 | -0.017 | 0.436 | 1.086 | 0.642 | 0.002 | 0.001 | 0.001 | 0 | 0.49 | 0.59 | 0.45 | 0.63 | 0.46 | 0.65 |
| 1 | 12038591 | rs3753579 | 0.262 | 0.543 | 1.073 | 0.563 | 0.007 | 0.004 | 0.004 | 0 | 0.45 | 0.58 | 0.39 | 0.52 | 0.36 | 0.55 |
| 1 | 12038990 | rs3820189 | -0.12 | 0.445 | 0.954 | 0.529 | 0.012 | 0.029 | 0.029 | 0 | 0.49 | 0.72 | 0.46 | 0.67 | 0.44 | 0.64 |
| 1 | 12042755 | rs4240897 | 0.222 | 0.372 | 1.095 | 0.443 | 0 | 0 | 0 | 0 | 0.43 | 0.44 | 0.39 | 0.54 | 0.41 | 0.52 |
| 1 | 12059412 | rs2295281 | 1.563 | 1.005 | 1.05 | 0.364 | -0.011 | 0.03 | 0.03 | 0 | 0.55 | 0.29 | 0.46 | 0.39 | 0.53 | 0.44 |
| 1 | 12062205 | rs2236057 | 0.586 | 1.065 | 1.064 | 0.476 | 0.001 | 0.006 | 0.006 | 0 | 0.69 | 0.63 | 0.56 | 0.68 | 0.67 | 0.72 |
| 1 | 12070292 | rs730123 | 1.111 | 1.073 | 0.867 | 0.434 | 0 | 0.023 | 0.023 | 0 | 0.63 | 0.47 | 0.5 | 0.55 | 0.6 | 0.6 |
| 1 | 12082461 | rs11588712 | 0.135 | 1.259 | 0.425 | 0.358 | -0.027 | -0.014 | -0.014 | 0 | 0.05 | 0.02 | 0.09 | 0.07 | 0.16 | 0.1 |
| 1 | 12082881 | rs11588779 | -0.368 | 1.062 | 0.505 | 0.181 | -0.004 | 0.005 | 0.005 | 0 | 0.17 | 0.18 | 0.14 | 0.07 | 0.25 | 0.16 |
| 1 | 12082926 | rs2295283 | 0.601 | 1.039 | 0.549 | 0.233 | -0.012 | -0.034 | -0.034 | 0.379 | 0.73 | 0.8 | 0.6 | 0.55 | 0.73 | 0.61 |
| 1 | 12087749 | rs1208988 | 0.324 | 1.136 | 0.396 | 0.365 | -0.035 | -0.081 | -0.081 | 0.073 | 0.76 | 0.91 | 0.62 | 0.55 | 0.73 | 0.62 |
| 1 | 12090001 | rs4514248 | -0.6 | 1.105 | 0.582 | 0.183 | 0 | -0.008 | -0.008 | 0.026 | 0.19 | 0.26 | 0.15 | 0.1 | 0.25 | 0.16 |
| 1 | 12095868 | rs1208993 | 0.744 | 1.041 | -0.052 | 0.616 | -0.019 | -0.046 | -0.046 | 0 | 0.76 | 0.87 | 0.61 | 0.56 | 0.73 | 0.63 |
| 1 | 12097568 | rs11121850 | -0.577 | 0.989 | -0.242 | 0.709 | -0.018 | -0.061 | -0.061 | 1.09 | 0.2 | 0.08 | 0.34 | 0.46 | 0.21 | 0.36 |
| 1 | 12105048 | rs12042105 | -0.957 | 0.997 | -0.198 | 0.687 | 0.003 | 0.024 | 0.024 | 0 | 0.26 | 0.29 | 0.37 | 0.47 | 0.23 | 0.4 |
| 1 | 12117901 | rs6665486 | -0.709 | -0.014 | -0.662 | 0.455 | 0.025 | -0.012 | -0.012 | 0.498 | 0.42 | 0.66 | 0.56 | 0.44 | 0.32 | 0.44 |
| 1 | 12128498 | rs12029016 | -0.112 | 0.249 | -1.303 | 0.646 | 0.001 | 0.039 | 0.039 | 0.315 | 0.19 | 0.19 | 0.45 | 0.4 | 0.19 | 0.34 |
| 1 | 12130051 | rs4491070 | 0.001 | 0.531 | -1.521 | 0.531 | -0.012 | -0.034 | -0.034 | 0 | 0.34 | 0.21 | 0.53 | 0.4 | 0.32 | 0.44 |
| 1 | 12135526 | rs6690493 | -0.169 | 1.075 | -1.958 | 0.392 | -0.026 | 0.173 | 0.173 | 0 | 0.55 | 0.86 | 0.62 | 0.22 | 0.42 | 0.53 |
| 1 | 12153188 | rs1201110 | 0.141 | 1.407 | -1.43 | 1.174 | -0.015 | -0.051 | -0.051 | 0.07 | 0.77 | 0.85 | 0.81 | 0.46 | 0.72 | 0.71 |
| 1 | 12164071 | rs562844 | -0.321 | 0.712 | -1.545 | 0.742 | 0.054 | -0.001 | -0.001 | 0 | 0.45 | 0.82 | 0.53 | 0.41 | 0.3 | 0.47 |
| 1 | 12164078 | rs4845894 | -0.189 | 0.533 | -0.722 | 0.656 | 0.017 | -0.006 | -0.006 | 0 | 0.4 | 0.33 | 0.48 | 0.4 | 0.3 | 0.47 |

| | | | | | | | | | | | | | | | | |
|---|---|---|---|---|---|---|---|---|---|---|---|---|---|---|---|---|
| 1 | 12164090 | rs6541014 | 0.197 | 1.486 | -1.456 | 0.421 | -0.055 | 0.042 | 0.042 | 0 | 0.18 | 0.06 | 0.27 | 0.06 | 0.35 | 0.12 |
| 1 | 12166855 | rs12092682 | 0.05 | 0.894 | -0.478 | 0.73 | -0.001 | -0.001 | -0.001 | 4.245 | 0.33 | 0.34 | 0.41 | 0.33 | 0.23 | 0.37 |
| 1 | 12168523 | rs11569864 | -0.319 | 1.468 | -0.471 | 0.807 | -0.029 | -0.023 | -0.023 | 0 | 0.05 | 0.01 | 0.09 | 0.07 | 0.07 | 0.1 |
| 1 | 12183245 | rs2486316 | -1.531 | 1.832 | 0.227 | 0.682 | 0.119 | 0.009 | 0.009 | 0 | 0.06 | 0.24 | 0.1 | 0.02 | 0.14 | 0.07 |
| 1 | 12184375 | rs8179353 | -0.34 | 1.593 | -0.502 | 0.574 | -0.003 | 0.007 | 0.007 | 0 | 0.37 | 0.69 | 0.64 | 0.68 | 0.43 | 0.34 |
| 1 | 12184423 | rs17422381 | 0.014 | 1.479 | 0.186 | 1.41 | -0.013 | -0.006 | -0.006 | 0.407 | 0.32 | 0.25 | 0.56 | 0.35 | 0.4 | 0.27 |
| 1 | 12186797 | rs1148476 | -0.843 | 0.948 | 0.512 | 1.134 | 0.001 | 0.008 | 0.008 | 0 | 0.69 | 0.77 | 0.83 | 0.81 | 0.75 | 0.7 |
| 1 | 12187130 | rs6668157 | -1.128 | 0.868 | 1.049 | 1.339 | 0.065 | 0.243 | 0.243 | 0.051 | 0.08 | 0.4 | 0.09 | 0.22 | 0.1 | 0.04 |
| 1 | 12188482 | rs1201122 | -0.517 | 0.641 | 0.454 | 1.667 | 0.002 | 0.001 | 0.001 | 0 | 0.64 | 0.71 | 0.77 | 0.63 | 0.68 | 0.61 |
| 1 | 12189561 | rs535068 | -0.173 | 0.658 | 0.11 | 1.663 | -0.011 | -0.01 | -0.01 | 2.395 | 0.61 | 0.51 | 0.75 | 0.34 | 0.68 | 0.57 |
| 1 | 12194451 | rs1148472 | -0.781 | 0.698 | -0.978 | 1.338 | -0.058 | 0.041 | 0.041 | 0 | 0.64 | 0.95 | 0.83 | 0.79 | 0.67 | 0.64 |
| 1 | 12194612 | rs1148471 | 0.157 | 0.638 | -0.886 | 0.687 | 0.002 | -0.006 | -0.006 | 0 | 0.41 | 0.55 | 0.65 | 0.33 | 0.53 | 0.33 |
| 1 | 12194745 | rs3766735 | 0.152 | 0.634 | -1.48 | 0.674 | 0.016 | -0.001 | -0.001 | 0 | 0.14 | 0.06 | 0.08 | 0.11 | 0.08 | 0.23 |
| 1 | 12198179 | rs671106 | -0.159 | 0.877 | -1.676 | 0.266 | -0.055 | 0.172 | 0.172 | 0 | 0.54 | 0.9 | 0.49 | 0.76 | 0.53 | 0.48 |
| 1 | 12206755 | rs631272 | 0.313 | 1.392 | -1.85 | 0.233 | -0.014 | 0.161 | 0.161 | 1.267 | 0.51 | 0.17 | 0.29 | 0.26 | 0.53 | 0.44 |
| 1 | 12209208 | rs967403 | 0.056 | 1.197 | -0.971 | 0.199 | -0.001 | -0.008 | -0.008 | 0 | 0.53 | 0.51 | 0.33 | 0.28 | 0.5 | 0.44 |
| 1 | 12216488 | rs1148460 | 0.199 | 1.183 | -0.643 | 0.175 | -0.012 | 0.08 | 0.08 | 0.33 | 0.52 | 0.34 | 0.33 | 0.28 | 0.52 | 0.44 |
| 1 | 12224239 | rs522807 | -1.495 | 1.133 | -0.321 | 0.149 | 0.113 | 0.06 | 0.06 | 3.41 | 0.06 | 0.46 | 0.06 | 0.01 | 0.04 | 0.03 |
| 1 | 12232806 | rs496888 | -0.369 | 1.551 | -0.751 | 0.504 | 0.061 | 0.027 | 0.027 | 0 | 0.18 | 0.32 | 0.46 | 0.21 | 0.28 | 0.16 |
| 1 | 12233754 | rs976881 | -0.042 | 0.875 | -0.914 | 0.145 | 0.036 | 0.06 | 0.06 | 0.363 | 0.63 | 0.75 | 0.76 | 0.81 | 0.66 | 0.7 |
| 1 | 12240676 | rs3766730 | 1.071 | 0.562 | -1.32 | 0.507 | -0.019 | -0.033 | -0.033 | 0 | 0.18 | 0.01 | 0.07 | 0.18 | 0.14 | 0.21 |
| 1 | 12241991 | rs816050 | 0.04 | 0.492 | -1.176 | 0.537 | -0.002 | -0.007 | -0.007 | 0 | 0.21 | 0.25 | 0.14 | 0.12 | 0.2 | 0.17 |
| 1 | 12248942 | rs945439 | 0.324 | 0.576 | -1.376 | 0.571 | 0.012 | 0.023 | 0.023 | 0.347 | 0.26 | 0.2 | 0.13 | 0.15 | 0.21 | 0.27 |
| 1 | 12249031 | rs1201157 | -0.816 | 0.737 | -1.297 | 0.788 | 0.011 | -0.006 | -0.006 | 0 | 0.67 | 0.65 | 0.44 | 0.7 | 0.58 | 0.62 |
| 1 | 12249829 | rs683240 | -0.163 | 0.713 | -1.597 | 0.556 | 0.009 | 0.023 | 0.023 | 0.717 | 0.26 | 0.16 | 0.13 | 0.15 | 0.21 | 0.28 |
| 1 | 12251318 | rs666078 | -0.095 | 0.755 | -1.625 | 0.545 | 0.014 | 0.029 | 0.029 | 0.698 | 0.24 | 0.06 | 0.11 | 0.13 | 0.19 | 0.24 |
| 1 | 12251808 | rs653667 | -1.107 | 0.724 | -1.684 | 0.596 | 0.04 | 0.014 | 0.014 | 0.514 | 0.42 | 0.22 | 0.21 | 0.36 | 0.35 | 0.37 |
| 1 | 12252955 | rs1061622 | -0.667 | 1.145 | -0.967 | 0.485 | 0.012 | 0.021 | 0.021 | 0.338 | 0.26 | 0.19 | 0.13 | 0.15 | 0.22 | 0.28 |
| 1 | 12253351 | rs636964 | -0.614 | 1.195 | -1.145 | 0.491 | 0.025 | 0.011 | 0.011 | 0.358 | 0.25 | 0.13 | 0.09 | 0.15 | 0.18 | 0.24 |
| 1 | 12262298 | rs5746053 | -0.51 | 1.085 | -1.404 | 0.433 | 0.016 | 0.006 | 0.006 | 0.473 | 0.23 | 0.15 | 0.11 | 0.16 | 0.17 | 0.23 |
| 1 | 12262792 | rs5746059 | -0.439 | 1.239 | -1.857 | 0.387 | -0.048 | 0.005 | 0.005 | 0.845 | 0.23 | 0.03 | 0.11 | 0.14 | 0.22 | 0.21 |
| 1 | 12267265 | rs1061624 | -0.255 | 1.031 | -1.614 | 0.381 | 0.001 | 0.021 | 0.021 | 0 | 0.51 | 0.3 | 0.56 | 0.51 | 0.53 | 0.46 |
| 1 | 12267464 | rs5746065 | -0.853 | 0.914 | -1.148 | 0.466 | -0.004 | 0.023 | 0.023 | 0 | 0.12 | 0.16 | 0.17 | 0.2 | 0.11 | 0.15 |
| 1 | 12267999 | rs1061628 | -0.736 | 0.837 | -1.413 | -0.127 | 0.005 | 0.01 | 0.01 | 0 | 0.49 | 0.46 | 0.25 | 0.3 | 0.44 | 0.29 |
| 1 | 12268499 | rs1061631 | -0.316 | 1.017 | -1.998 | -0.212 | -0.037 | 0.055 | 0.055 | 0 | 0.25 | 0.03 | 0.09 | 0.09 | 0.22 | 0.16 |
| 1 | 12277967 | rs7552664 | -0.314 | 1.831 | -1.631 | 0.287 | -0.028 | -0.003 | -0.003 | 0 | 0.17 | 0.06 | 0.16 | 0.19 | 0.27 | 0.19 |
| 1 | 12278932 | rs7550488 | -0.256 | 1.031 | -1.936 | 0.462 | 0.017 | -0.03 | -0.03 | 0 | 0.7 | 0.82 | 0.84 | 0.6 | 0.75 | 0.78 |
| 1 | 12295188 | rs11121891 | -0.851 | 1.911 | -1.27 | -0.163 | -0.021 | 0.015 | 0.015 | 0 | 0.1 | 0.06 | 0.08 | 0 | 0.2 | 0.03 |
| 1 | 12296232 | rs235256 | -0.535 | 1.043 | -0.533 | -0.047 | -0.039 | 0.029 | 0.029 | 1.301 | 0.14 | 0.62 | 0.26 | 0.18 | 0.08 | 0.15 |
| 1 | 12305286 | rs374450 | -1.248 | 0.693 | -0.828 | -0.23 | 0 | -0.001 | -0.001 | 0 | 0.23 | 0.73 | 0.3 | 0.26 | 0.14 | 0.26 |
| 1 | 12329295 | rs3765339 | 1.972 | 0.27 | -0.418 | -0.611 | -0.025 | 0.003 | 0.003 | 0 | 0.26 | 0.01 | 0.21 | 0.2 | 0.23 | 0.2 |
| 1 | 12342599 | rs3765337 | 0.049 | 0.247 | 0.096 | -0.733 | -0.054 | 0.024 | 0.024 | 0.48 | 0.15 | 0.62 | 0.26 | 0.18 | 0.07 | 0.16 |
| 1 | 12363811 | rs7523900 | 0.609 | 0.28 | 0.433 | -0.717 | -0.015 | 0.056 | 0.056 | 0 | 0.08 | 0.25 | 0.11 | 0.16 | 0.04 | 0.09 |
| 1 | 12401868 | rs28551666 | 1.384 | 0.579 | 0.464 | -0.919 | 0.001 | -0.014 | -0.014 | 0 | 0.81 | 0.74 | 0.76 | 0.9 | 0.79 | 0.8 |
| 1 | 12418700 | rs10864552 | -2.854 | 0.67 | 0.372 | -0.893 | -0.032 | 0.041 | 0.041 | 0 | 0.93 | 1 | 0.96 | 1 | 0.92 | 0.93 |
| 1 | 12420300 | rs6660347 | -0.57 | 0.096 | 0.861 | -0.622 | -0.018 | 0.085 | 0.085 | 0.177 | 0.1 | 0.39 | 0.12 | 0.16 | 0.05 | 0.1 |
| 1 | 12429611 | rs2295338 | 0.631 | 0.126 | 0.719 | -0.622 | -0.012 | -0.027 | -0.027 | 0.035 | 0.07 | 0.11 | 0.06 | 0.02 | 0.04 | 0.07 |
| 1 | 12436431 | rs6541019 | -1.992 | 0.149 | 0.53 | -0.564 | -0.055 | 0.044 | 0.044 | 0 | 0.77 | 0.95 | 0.76 | 0.86 | 0.7 | 0.68 |
| 1 | 12436898 | rs17457661 | 0.2 | 0.271 | 0.786 | -0.578 | -0.012 | 0.003 | 0.003 | 0 | 0.11 | 0.08 | 0.09 | 0 | 0.2 | 0.05 |
| 1 | 12450829 | rs7526212 | -1.36 | -0.089 | 0.895 | -0.338 | -0.054 | 0.022 | 0.022 | 0 | 0.21 | 0.59 | 0.27 | 0.26 | 0.11 | 0.2 |
| 1 | 12463073 | rs7547740 | -1.499 | -0.139 | 0.553 | -0.316 | -0.045 | 0.059 | 0.059 | 0 | 0.72 | 0.89 | 0.74 | 0.85 | 0.67 | 0.59 |
| 1 | 12467620 | rs11801678 | -0.07 | -0.954 | 0.839 | 0.135 | -0.016 | 0.03 | 0.03 | 0 | 0.08 | 0.17 | 0.1 | 0.16 | 0.05 | 0.1 |
| 1 | 12480504 | rs6660365 | -1.26 | -0.96 | 0.584 | 0.133 | -0.057 | 0.042 | 0.042 | 0 | 0.15 | 0.53 | 0.24 | 0.18 | 0.07 | 0.17 |
| 1 | 12496021 | rs6541023 | -1.018 | -0.931 | 0.526 | 0.151 | -0.06 | 0.034 | 0.034 | 0 | 0.15 | 0.56 | 0.25 | 0.18 | 0.08 | 0.16 |
| 1 | 12546694 | rs12125606 | 1.404 | -0.771 | -0.553 | 0.277 | -0.033 | 0.055 | 0.055 | 0 | 0.26 | 0.01 | 0.21 | 0.11 | 0.24 | 0.21 |
| 1 | 12573823 | rs12129750 | -0.173 | -0.883 | 0.264 | 1.835 | -0.006 | -0.113 | -0.113 | 1.736 | 0.14 | 0 | 0.1 | 0.46 | 0.11 | 0.11 |
| 1 | 12575090 | rs11121917 | -1.773 | -0.648 | 0.945 | 1.201 | -0.035 | 0.051 | 0.051 | 0 | 0.12 | 0.35 | 0.22 | 0.18 | 0.07 | 0.15 |
| 1 | 12598434 | rs760943 | -1.161 | -0.22 | 0.362 | 1.244 | -0.011 | 0.015 | 0.015 | 0 | 0.11 | 0.15 | 0.09 | 0.18 | 0.07 | 0.12 |
| 1 | 12599422 | rs6541025 | -1.692 | -0.51 | 0.425 | 1.473 | -0.031 | 0.023 | 0.023 | 0.516 | 0.32 | 0.55 | 0.34 | 0.58 | 0.21 | 0.34 |
| 1 | 12599575 | rs7517886 | -1.551 | -0.073 | -0.242 | 1.264 | -0.002 | 0.004 | 0.004 | 1.309 | 0.34 | 0.33 | 0.26 | 0.47 | 0.3 | 0.29 |
| 1 | 12600396 | rs7521256 | -1.097 | 0.346 | -0.249 | 0.742 | 0 | -0.025 | -0.025 | 0 | 0.17 | 0.19 | 0.13 | 0.08 | 0.18 | 0.1 |
| 1 | 12602979 | rs2038118 | -1.09 | 0.539 | -0.561 | 1.388 | -0.014 | -0.029 | -0.029 | 0 | 0.15 | 0.08 | 0.25 | 0.29 | 0.19 | 0.19 |
| 1 | 12605011 | rs6541027 | 0.849 | 0.039 | -0.051 | 1.458 | -0.009 | 0.03 | 0.03 | 0 | 0.69 | 0.47 | 0.63 | 0.63 | 0.69 | 0.73 |
| 1 | 12606394 | rs4845902 | 0.338 | -0.468 | 0.568 | 1.701 | -0.04 | 0.047 | 0.047 | 0 | 0.88 | 0.76 | 0.74 | 0.76 | 0.94 | 0.89 |
| 1 | 12606725 | rs10779770 | 1.506 | -0.933 | 0.567 | 1.848 | -0.016 | 0.023 | 0.023 | 0.058 | 0.76 | 0.32 | 0.65 | 0.48 | 0.84 | 0.74 |
| 1 | 12611852 | rs12062307 | -1.521 | -0.543 | 0.324 | 1.237 | 0.002 | -0.006 | -0.006 | 0.506 | 0.28 | 0.24 | 0.16 | 0.33 | 0.33 | 0.27 |
| 1 | 12616535 | rs3923904 | -1.593 | -0.567 | 0.451 | 1.211 | 0.002 | -0.003 | -0.003 | 2.211 | 0.29 | 0.27 | 0.17 | 0.33 | 0.32 | 0.27 |
| 1 | 12619096 | rs4845904 | -1.257 | -0.566 | 0.43 | 1.235 | 0.002 | 0.006 | 0.006 | 0 | 0.25 | 0.23 | 0.15 | 0.19 | 0.27 | 0.22 |
| 1 | 12619267 | rs4846113 | -1.212 | -0.577 | 0.414 | 1.229 | 0.003 | 0.008 | 0.008 | 0 | 0.24 | 0.21 | 0.14 | 0.19 | 0.27 | 0.22 |

| | | | | | | | | | | | | | | | | |
|---|---|---|---|---|---|---|---|---|---|---|---|---|---|---|---|---|
| 1 | 12620545 | rs6541035 | -1.025 | -0.545 | 0.481 | 1.296 | 0 | 0.001 | 0.001 | 0 | 0.21 | 0.21 | 0.12 | 0.19 | 0.24 | 0.21 |
| 1 | 12625101 | rs10864570 | -1.482 | -0.486 | 0.596 | 1.272 | -0.001 | -0.001 | -0.001 | 0 | 0.25 | 0.26 | 0.14 | 0.31 | 0.29 | 0.26 |
| 1 | 12626210 | rs6541038 | -1.746 | -0.487 | 0.571 | 1.404 | -0.001 | -0.001 | -0.001 | 0.123 | 0.28 | 0.29 | 0.17 | 0.34 | 0.34 | 0.32 |
| 1 | 12626261 | rs6541039 | -1.735 | -0.482 | 0.574 | 1.346 | -0.001 | -0.001 | -0.001 | 0.035 | 0.28 | 0.29 | 0.17 | 0.33 | 0.34 | 0.32 |
| 1 | 12626549 | rs4846118 | -0.245 | -0.795 | 0.462 | 1.379 | 0.011 | -0.014 | -0.014 | 0 | 0.78 | 0.82 | 0.7 | 0.69 | 0.85 | 0.87 |
| 1 | 12627680 | rs4997711 | 1.819 | -0.856 | 0.814 | 1.339 | 0.03 | 0.071 | 0.071 | 0 | 0.49 | 0.28 | 0.5 | 0.36 | 0.52 | 0.56 |
| 1 | 12627763 | rs4385721 | 1.894 | -0.911 | 0.816 | 1.312 | 0.025 | 0.075 | 0.075 | 0 | 0.48 | 0.22 | 0.5 | 0.36 | 0.52 | 0.55 |
| 1 | 12628188 | rs4580 | -1.508 | -0.681 | 0.629 | 1.436 | -0.001 | -0.003 | -0.003 | 0 | 0.28 | 0.27 | 0.2 | 0.36 | 0.34 | 0.33 |
| 1 | 12628471 | rs4072868 | -0.524 | -0.694 | 0.454 | 1.349 | -0.06 | 0.073 | 0.073 | 0 | 0.16 | 0.03 | 0.1 | 0.02 | 0.25 | 0.19 |
| 1 | 12637160 | rs4845907 | 1.17 | -0.656 | 0.701 | 1.402 | -0.022 | 0.002 | 0.002 | 0 | 0.65 | 0.47 | 0.6 | 0.36 | 0.72 | 0.71 |
| 1 | 12639168 | rs3128461 | -0.047 | -0.641 | 0.535 | 1.442 | 0.003 | -0.007 | -0.007 | 0 | 0.78 | 0.79 | 0.71 | 0.71 | 0.8 | 0.84 |
| 1 | 12640127 | rs3128458 | 0.052 | -0.635 | 0.507 | 1.445 | -0.009 | 0.02 | 0.02 | 0.133 | 0.77 | 0.7 | 0.7 | 0.71 | 0.8 | 0.84 |
| 1 | 12640650 | rs11540058 | 0.198 | -0.614 | 0.26 | 1.394 | -0.031 | 0.043 | 0.043 | 0 | 0.12 | 0.04 | 0.08 | 0 | 0.19 | 0.09 |
| 1 | 12641488 | rs3128451 | 1.414 | -0.674 | 0.38 | 1.553 | 0.014 | 0.039 | 0.039 | 0 | 0.52 | 0.32 | 0.51 | 0.36 | 0.52 | 0.61 |
| 1 | 12641758 | rs3128449 | 0.45 | -0.337 | -0.259 | 1.429 | 0 | -0.001 | -0.001 | 0.299 | 0.68 | 0.7 | 0.65 | 0.72 | 0.66 | 0.8 |
| 1 | 12650076 | rs7535209 | 0.1 | 0.022 | -0.255 | 0.505 | -0.013 | -0.063 | -0.063 | 0.011 | 0.32 | 0.43 | 0.25 | 0.19 | 0.25 | 0.3 |
| 1 | 12654035 | rs12045736 | 0.635 | 0.124 | -0.238 | 0.768 | 0.007 | -0.01 | -0.01 | 3.718 | 0.51 | 0.19 | 0.43 | 0.48 | 0.51 | 0.58 |
| 1 | 12661179 | rs13376289 | -0.904 | 0.299 | 0.572 | 0.561 | 0.013 | -0.037 | -0.037 | 0 | 0.14 | 0.35 | 0.21 | 0.07 | 0.17 | 0.2 |
| 1 | 12663774 | rs4846127 | 1.02 | 0.279 | -0.039 | 0.754 | 0.012 | 0.014 | 0.014 | 0 | 0.5 | 0.22 | 0.51 | 0.51 | 0.5 | 0.54 |
| 1 | 12670396 | rs4846128 | -1.406 | 0.537 | -0.163 | 0.648 | 0.003 | 0.019 | 0.019 | 0 | 0.56 | 0.83 | 0.61 | 0.59 | 0.62 | 0.53 |
| 1 | 12671229 | rs4394668 | -0.854 | 0.619 | 0.051 | 0.855 | 0 | 0.007 | 0.007 | 0 | 0.16 | 0.14 | 0.21 | 0.39 | 0.2 | 0.21 |
| 1 | 12671703 | rs5022242 | -1.58 | 0.736 | 0.102 | 0.719 | 0.012 | 0.028 | 0.028 | 0 | 0.52 | 0.7 | 0.57 | 0.58 | 0.6 | 0.44 |
| 1 | 12673497 | rs6675673 | -1.268 | 0.83 | 0.532 | 0.612 | 0 | -0.024 | -0.024 | 1.553 | 0.19 | 0.16 | 0.21 | 0.02 | 0.21 | 0.16 |
| 1 | 12674271 | rs6679088 | -1.268 | 0.834 | 0.534 | 0.614 | 0 | -0.023 | -0.023 | 0 | 0.19 | 0.16 | 0.21 | 0.02 | 0.2 | 0.16 |
| 1 | 12676205 | rs11121951 | 0.109 | 0.625 | 0.292 | 0.939 | 0 | 0.086 | 0.086 | 0 | 0.06 | 0.04 | 0.15 | 0.35 | 0.11 | 0.18 |
| 1 | 12678741 | rs3010928 | 1.286 | 0.595 | 0.666 | 0.378 | 0 | -0.084 | -0.084 | 0.36 | 0.77 | 0.7 | 0.76 | 0.97 | 0.77 | 0.84 |
| 1 | 12680180 | rs6689733 | 1.194 | 0.296 | 0.691 | 0.709 | -0.01 | -0.056 | -0.056 | 0 | 0.85 | 0.8 | 0.83 | 0.98 | 0.84 | 0.93 |
| 1 | 12682574 | rs3000896 | 1.37 | 0.221 | 0.724 | 1.343 | -0.007 | 0.143 | 0.143 | 0 | 0.85 | 0.65 | 0.67 | 0.63 | 0.84 | 0.79 |
| 1 | 12682855 | rs3000895 | 0.84 | 0.168 | 0.732 | 0.879 | -0.004 | -0.024 | -0.024 | 0 | 0.9 | 0.68 | 0.81 | 0.96 | 0.94 | 0.95 |
| 1 | 12683190 | rs6699993 | 1.421 | 0.246 | -0.048 | 0.781 | 0.009 | 0.021 | 0.021 | 0 | 0.56 | 0.23 | 0.31 | 0.36 | 0.56 | 0.48 |
| 1 | 12683421 | rs3000893 | -2.172 | 0.076 | 0.301 | 0.504 | -0.021 | -0.001 | -0.001 | 0 | 0.13 | 0.32 | 0.16 | 0.06 | 0.07 | 0.16 |
| 1 | 12684324 | rs3010931 | -1.358 | 0.442 | 0.63 | 0.361 | 0.055 | 0.012 | 0.012 | 0 | 0.06 | 0.3 | 0.04 | 0 | 0.04 | 0.1 |
| 1 | 12689692 | rs12070206 | -1.198 | 0.334 | 0.116 | 0.292 | -0.022 | -0.08 | -0.08 | 0.001 | 0.13 | 0.46 | 0.07 | 0.01 | 0.07 | 0.18 |
| 1 | 12699337 | rs3000891 | 0.339 | 1.302 | -0.982 | 1.434 | -0.03 | 0.005 | 0.005 | 0.319 | 0.93 | 0.99 | 0.9 | 0.59 | 0.85 | 0.7 |
| 1 | 12709554 | rs17458785 | -0.506 | 1.072 | -1.232 | 0.824 | -0.043 | -0.04 | -0.04 | 0.018 | 0.18 | 0.06 | 0.35 | 0.34 | 0.23 | 0.35 |
| 1 | 12710882 | rs2493875 | -0.81 | 0.082 | -0.906 | 0.439 | 0.002 | -0.002 | -0.002 | 0 | 0.06 | 0.04 | 0.03 | 0.01 | 0.03 | 0.04 |
| 1 | 12718815 | rs2257017 | -1.615 | -0.237 | 0.208 | 0.317 | -0.008 | -0.007 | -0.007 | 0 | 0.11 | 0.5 | 0.09 | 0.04 | 0.05 | 0.07 |
| 1 | 12724931 | rs2027589 | -1.518 | -1.296 | -0.124 | -0.082 | -0.065 | -0.001 | -0.001 | 0.58 | 0.16 | 0.51 | 0.11 | 0.11 | 0.07 | 0.11 |
| 1 | 12731572 | rs2265709 | -1.543 | -1.312 | -0.126 | -0.065 | -0.067 | 0.014 | 0.014 | 0.164 | 0.16 | 0.52 | 0.11 | 0.12 | 0.07 | 0.11 |
| 1 | 12734774 | rs2265711 | -1.369 | -1.304 | 0.059 | -0.061 | -0.041 | 0.004 | 0.004 | 0.045 | 0.15 | 0.3 | 0.08 | 0.12 | 0.07 | 0.1 |
| 1 | 12740907 | rs1419299 | -1.313 | -1.133 | 0.138 | 0.531 | -0.017 | 0.035 | 0.035 | 0 | 0.14 | 0.18 | 0.1 | 0.28 | 0.08 | 0.1 |
| 1 | 12741240 | rs2489260 | -1.533 | -1.491 | 0.39 | 0.363 | -0.089 | 0.092 | 0.092 | 0 | 0.2 | 0.53 | 0.13 | 0.29 | 0.08 | 0.16 |
| 1 | 12743160 | rs2489262 | -1.531 | -1.525 | 0.365 | 0.369 | -0.09 | 0.088 | 0.088 | 0 | 0.2 | 0.54 | 0.14 | 0.3 | 0.08 | 0.16 |
| 1 | 12754569 | rs3010948 | 0.796 | -0.952 | 0.283 | 0.59 | 0 | 0.032 | 0.032 | 0 | 0.91 | 0.89 | 0.92 | 0.73 | 0.91 | 0.91 |
| 1 | 13859570 | rs3856278 | -1.373 | -0.249 | 0.928 | 0.855 | 0.018 | 0.032 | 0.032 | 0 | 0.13 | 0.3 | 0.12 | 0.17 | 0.15 | 0.12 |
| 1 | 13866127 | rs12034380 | -0.494 | -0.246 | 0.135 | 0.674 | 0.001 | -0.001 | -0.001 | 0 | 0.06 | 0.09 | 0.07 | 0.07 | 0.11 | 0.06 |
| 1 | 13867459 | rs183052 | -0.019 | -0.345 | 0.161 | 0.709 | -0.002 | 0.009 | 0.009 | 0 | 0.06 | 0.07 | 0.03 | 0.1 | 0.04 | 0.05 |
| 1 | 13870966 | rs444516 | -1.259 | -0.406 | 0.007 | 0.841 | 0.007 | -0.004 | -0.004 | 0 | 0.3 | 0.28 | 0.44 | 0.59 | 0.25 | 0.39 |
| 1 | 13873974 | rs452477 | -0.829 | -0.405 | 0.046 | 0.919 | -0.003 | 0.019 | 0.019 | 0 | 0.25 | 0.29 | 0.32 | 0.49 | 0.21 | 0.33 |
| 1 | 13874853 | rs1610562 | -1.389 | -0.442 | 0.186 | 0.887 | -0.013 | 0.005 | 0.005 | 0 | 0.31 | 0.39 | 0.46 | 0.59 | 0.25 | 0.4 |
| 1 | 13875610 | rs17391912 | -0.002 | -0.7 | -0.48 | 0.627 | 0.007 | -0.008 | -0.008 | 0 | 0.05 | 0.02 | 0.02 | 0.1 | 0.03 | 0.05 |
| 1 | 13882346 | rs35979056 | -0.865 | -1.07 | -0.514 | 0.719 | -0.002 | -0.015 | -0.015 | 0.752 | 0.3 | 0.28 | 0.33 | 0.42 | 0.3 | 0.31 |
| 1 | 13885203 | rs10927872 | 0.554 | -0.692 | -0.489 | 0.416 | -0.064 | 0.092 | 0.092 | 0 | 0.4 | 0.15 | 0.33 | 0.24 | 0.53 | 0.33 |
| 1 | 13905218 | rs12735797 | -0.33 | -2.248 | -0.417 | -1.128 | -0.02 | -0.124 | -0.124 | 0 | 0.41 | 0.47 | 0.33 | 0.05 | 0.33 | 0.25 |
| 1 | 13906554 | rs2361030 | 0.385 | -2.346 | -0.27 | -1.145 | -0.051 | -0.144 | -0.144 | 0.095 | 0.31 | 0.43 | 0.25 | 0.02 | 0.18 | 0.21 |
| 1 | 13907484 | rs355020 | 0.392 | -2.246 | -0.295 | -0.048 | 0 | 0.082 | 0.082 | 0 | 0.08 | 0.06 | 0.12 | 0.45 | 0.07 | 0.26 |
| 1 | 13908690 | rs355022 | -0.847 | -2.759 | -0.438 | -0.253 | 0.004 | 0.002 | 0.002 | 0 | 0.6 | 0.65 | 0.48 | 0.66 | 0.33 | 0.68 |
| 1 | 13909211 | rs12757529 | 1.057 | -2.779 | -0.549 | -0.372 | -0.024 | -0.095 | -0.095 | 0 | 0.34 | 0.38 | 0.26 | 0.07 | 0.18 | 0.22 |
| 1 | 13910152 | rs12031781 | 0.303 | -2.694 | -0.662 | -0.356 | -0.027 | 0.047 | 0.047 | 0 | 0.12 | 0.01 | 0.07 | 0.06 | 0.13 | 0.06 |
| 1 | 13912887 | rs10927903 | 1.07 | -2.667 | -0.516 | -0.324 | -0.024 | -0.096 | -0.096 | 0 | 0.33 | 0.38 | 0.25 | 0.07 | 0.18 | 0.23 |
| 1 | 13912977 | rs355027 | -0.572 | -2.691 | -0.728 | -0.077 | 0.019 | -0.035 | -0.035 | 0.18 | 0.24 | 0.22 | 0.22 | 0.52 | 0.15 | 0.4 |
| 1 | 13913180 | rs355028 | -0.411 | -2.554 | -0.612 | 0.037 | -0.007 | 0.082 | 0.082 | 0 | 0.15 | 0.24 | 0.14 | 0.5 | 0.12 | 0.31 |
| 1 | 13914230 | rs1148455 | 0.26 | -2.496 | -0.54 | 0.203 | 0.013 | -0.002 | -0.002 | 0 | 0.42 | 0.61 | 0.37 | 0.6 | 0.26 | 0.47 |
| 1 | 13914909 | rs10927904 | 0.003 | -2.434 | -0.489 | -0.066 | 0.034 | 0.057 | 0.057 | 0 | 0.46 | 0.62 | 0.37 | 0.21 | 0.31 | 0.32 |
| 1 | 13916002 | rs3845612 | 0.37 | -1.752 | -0.529 | 0.215 | 0.002 | -0.021 | -0.021 | 0.028 | 0.1 | 0.1 | 0.07 | 0.02 | 0.12 | 0.03 |
| 1 | 13920193 | rs1261025 | 0.015 | -2.04 | -0.666 | 0.225 | 0.073 | 0.037 | 0.037 | 0.448 | 0.51 | 0.71 | 0.41 | 0.65 | 0.3 | 0.5 |
| 1 | 13922117 | rs12566684 | 0.47 | -1.058 | -0.651 | 1.272 | -0.002 | -0.03 | -0.03 | 0 | 0.08 | 0.02 | 0.05 | 0.23 | 0.06 | 0.1 |
| 1 | 13922799 | rs7514217 | -0.313 | -0.654 | -0.053 | 0.724 | 0.021 | -0.006 | -0.006 | 0 | 0.65 | 0.51 | 0.52 | 0.67 | 0.43 | 0.56 |
| 1 | 13923574 | rs10927907 | -1.135 | -0.144 | 0.067 | 0.865 | 0.01 | 0.028 | 0.028 | 0.109 | 0.14 | 0.36 | 0.14 | 0.18 | 0.12 | 0.18 |

| | | | | | | | | | | | | | | | | |
|---|---|---|---|---|---|---|---|---|---|---|---|---|---|---|---|---|
| 1 | 13935667 | rs372349 | -0.039 | 0.083 | -0.486 | 0.898 | 0 | -0.003 | -0.003 | 0 | 0.36 | 0.37 | 0.34 | 0.42 | 0.34 | 0.57 |
| 1 | 13937089 | rs2487643 | -0.365 | 0.681 | -0.471 | 0.716 | 0.013 | 0.001 | 0.001 | 0.959 | 0.56 | 0.63 | 0.8 | 0.55 | 0.67 | 0.71 |
| 1 | 13937281 | rs4391657 | -0.909 | 0.275 | -0.564 | 0.726 | 0.026 | 0.032 | 0.032 | 0 | 0.71 | 0.85 | 0.89 | 0.75 | 0.77 | 0.87 |
| 1 | 13938511 | rs1412839 | -0.085 | 0.229 | -0.327 | 0.778 | 0 | -0.001 | -0.001 | 0.94 | 0.59 | 0.61 | 0.81 | 0.58 | 0.68 | 0.76 |
| 1 | 13938788 | rs3820304 | 0.758 | 0.276 | -0.734 | 0.751 | -0.101 | 0.004 | 0.004 | 0.981 | 0.19 | 0.04 | 0.45 | 0.13 | 0.33 | 0.16 |
| 1 | 13942731 | rs11880 | 0.76 | 0.302 | -0.713 | 0.764 | -0.096 | 0.007 | 0.007 | 0.94 | 0.19 | 0.05 | 0.45 | 0.13 | 0.33 | 0.15 |
| 1 | 13943528 | rs1539114 | 0.843 | 0.27 | -0.677 | 0.807 | 0.006 | -0.011 | -0.011 | 3.025 | 0.1 | 0.09 | 0.06 | 0.15 | 0.09 | 0.1 |
| 1 | 13943724 | rs1061615 | 0.767 | 0.307 | -0.686 | 0.804 | -0.101 | 0.005 | 0.005 | 0.975 | 0.19 | 0.05 | 0.45 | 0.13 | 0.33 | 0.16 |
| 1 | 13944568 | rs876931 | 0.181 | 0.291 | -0.575 | 0.803 | 0.004 | 0.005 | 0.005 | 0 | 0.48 | 0.49 | 0.42 | 0.57 | 0.43 | 0.65 |
| 1 | 13946586 | rs6703700 | 1.007 | 0.244 | -0.628 | 0.683 | 0 | 0.009 | 0.009 | 0 | 0.33 | 0.4 | 0.34 | 0.41 | 0.32 | 0.48 |
| 1 | 13946868 | rs1767676 | 0.171 | 0.739 | -0.976 | 0.585 | -0.003 | 0.001 | 0.001 | 0.12 | 0.22 | 0.17 | 0.46 | 0.16 | 0.34 | 0.22 |
| 1 | 13947905 | rs11588651 | 0.049 | 0.299 | -1.163 | 0.866 | 0.006 | -0.005 | -0.005 | 0 | 0.1 | 0.08 | 0.06 | 0.08 | 0.08 | 0.08 |
| 1 | 13948225 | rs1772635 | -0.042 | 0.649 | -1.099 | 0.868 | 0 | -0.004 | -0.004 | 0 | 0.73 | 0.85 | 0.7 | 0.69 | 0.73 | 0.89 |
| 1 | 13952864 | rs7515525 | -0.305 | 0.51 | -1.108 | 0.895 | -0.014 | 0.027 | 0.027 | 0 | 0.43 | 0.25 | 0.55 | 0.34 | 0.48 | 0.4 |
| 1 | 13954726 | rs12408369 | 0.409 | 0.286 | -1.382 | 1.162 | -0.002 | -0.025 | -0.025 | 0 | 0.08 | 0.02 | 0.06 | 0.07 | 0.09 | 0.13 |
| 1 | 13954853 | rs10803420 | -0.527 | -0.073 | -0.511 | 0.986 | 0.006 | -0.008 | -0.008 | 0 | 0.57 | 0.62 | 0.65 | 0.41 | 0.59 | 0.58 |
| 1 | 13957405 | rs12143609 | 0.308 | 0.231 | -0.827 | 0.892 | -0.033 | 0.033 | 0.033 | 0 | 0.35 | 0.21 | 0.32 | 0.21 | 0.44 | 0.24 |
| 1 | 13966218 | rs10927926 | -1.599 | 0.065 | -0.436 | 1.216 | 0.079 | 0.053 | 0.053 | 0 | 0.08 | 0.42 | 0.28 | 0.06 | 0.07 | 0.06 |
| 1 | 13971360 | rs16852640 | -0.872 | 0.083 | -0.532 | 1.225 | 0.039 | 0.023 | 0.023 | 0 | 0.06 | 0.19 | 0.26 | 0.06 | 0.07 | 0.06 |
| 1 | 13989227 | rs2486182 | -1.913 | -0.108 | -0.524 | 1.691 | 0.061 | 0.03 | 0.03 | 0 | 0.11 | 0.45 | 0.28 | 0.08 | 0.1 | 0.18 |
| 1 | 13990291 | rs2495030 | -0.953 | -0.576 | -1.112 | 1.626 | -0.047 | -0.004 | -0.004 | 0 | 0.79 | 0.99 | 0.79 | 0.81 | 0.78 | 0.82 |
| 1 | 13992846 | rs2495037 | -1.738 | -0.646 | -0.949 | 1.849 | 0.014 | 0.191 | 0.191 | 0 | 0.13 | 0.47 | 0.28 | 0.26 | 0.09 | 0.28 |
| 1 | 14001018 | rs4662010 | 0.037 | -0.615 | -1.732 | 1.29 | 0.006 | 0.013 | 0.013 | 0 | 0.71 | 0.77 | 0.7 | 0.75 | 0.73 | 0.62 |
| 1 | 14014472 | rs9660610 | 1.463 | -1.18 | -2.095 | 0.916 | 0.023 | 0.022 | 0.022 | 0.196 | 0.13 | 0.04 | 0.02 | 0.01 | 0.04 | 0.04 |
| 1 | 14020405 | rs4077841 | 0.939 | -1.106 | -2.458 | 0.784 | -0.071 | 0.038 | 0.038 | 0 | 0.23 | 0.01 | 0.11 | 0.11 | 0.27 | 0.35 |
| 1 | 14031043 | rs3806425 | 0.66 | -1 | -2.325 | 1.098 | 0.013 | 0.041 | 0.041 | 0 | 0.05 | 0.06 | 0.27 | 0.15 | 0.08 | 0.04 |
| 1 | 14033128 | rs2487622 | -1.61 | -1.141 | -2.404 | 1.068 | -0.051 | -0.047 | -0.047 | 0.05 | 0.8 | 0.99 | 0.75 | 0.79 | 0.79 | 0.79 |
| 1 | 14053474 | rs16852866 | 0.943 | -1.124 | -2.437 | 1.032 | -0.055 | -0.062 | -0.062 | 0 | 0.23 | 0.01 | 0.26 | 0.25 | 0.23 | 0.36 |
| 1 | 14080748 | rs7522689 | 0.688 | -0.992 | -2.3 | 1.098 | 0.013 | 0.039 | 0.039 | 0 | 0.05 | 0.06 | 0.27 | 0.14 | 0.08 | 0.04 |
| 1 | 14096420 | rs2359756 | 0.31 | -1.08 | -2.405 | 0.925 | 0.018 | -0.016 | -0.016 | 0 | 0.47 | 0.72 | 0.35 | 0.51 | 0.41 | 0.26 |
| 1 | 14104818 | rs11579363 | 1.47 | -0.895 | -2.252 | 1.053 | 0.019 | 0.017 | 0.017 | 0.269 | 0.14 | 0.05 | 0.03 | 0.01 | 0.04 | 0.04 |
| 1 | 14105298 | rs1203678 | -0.383 | 0.215 | -2.011 | 1.659 | -0.007 | -0.023 | -0.023 | 1.038 | 0.29 | 0.2 | 0.54 | 0.37 | 0.31 | 0.26 |
| 1 | 14109114 | rs1203651 | -0.622 | 0.228 | -2.007 | 1.651 | -0.007 | -0.024 | -0.024 | 0 | 0.31 | 0.23 | 0.54 | 0.4 | 0.33 | 0.42 |
| 1 | 14121892 | rs1203638 | 0.782 | 0.717 | -2.408 | 1.326 | -0.06 | -0.082 | -0.082 | 0 | 0.17 | 0.01 | 0.17 | 0.23 | 0.21 | 0.2 |
| 1 | 14124098 | rs2281170 | 0.886 | 0.486 | -2.307 | 0.888 | 0.001 | 0.017 | 0.017 | 0 | 0.42 | 0.62 | 0.34 | 0.28 | 0.41 | 0.17 |
| 1 | 14124110 | rs2281169 | -0.334 | 0.492 | -2.637 | 1.029 | -0.002 | 0 | 0 | 0 | 0.33 | 0.29 | 0.44 | 0.59 | 0.33 | 0.45 |
| 1 | 14126453 | rs2014788 | 0.638 | 0.623 | -3.344 | 1.049 | -0.111 | -0.099 | -0.099 | 0 | 0.54 | 0.94 | 0.61 | 0.56 | 0.51 | 0.3 |
| 1 | 14126651 | rs760914 | 0.272 | 0.844 | -2.465 | 0.54 | -0.054 | 0.054 | 0.054 | 0 | 0.75 | 0.98 | 0.79 | 0.87 | 0.75 | 0.68 |
| 1 | 14130628 | rs2281164 | -0.446 | 1.717 | -2.104 | 0.926 | -0.059 | -0.03 | -0.03 | 0 | 0.17 | 0.05 | 0.38 | 0.24 | 0.3 | 0.1 |
| 1 | 14134437 | rs2281161 | 0.313 | 1.229 | -2.215 | 1.15 | -0.05 | -0.058 | -0.058 | 0 | 0.87 | 0.99 | 0.85 | 0.85 | 0.84 | 0.77 |
| 1 | 14137801 | rs2744690 | 0.34 | 0.775 | -2.094 | 1.228 | -0.012 | -0.054 | -0.054 | 0 | 0.87 | 0.96 | 0.86 | 0.82 | 0.86 | 0.78 |
| 1 | 14138813 | rs2744687 | 0.565 | 0.554 | -2.101 | 1.233 | 0.01 | -0.02 | -0.02 | 0 | 0.86 | 0.94 | 0.86 | 0.83 | 0.87 | 0.77 |
| 1 | 14139196 | rs2245197 | 1.293 | 0.338 | -1.096 | 1.364 | -0.013 | 0.053 | 0.053 | 0.871 | 0.74 | 0.36 | 0.73 | 0.55 | 0.79 | 0.54 |
| 1 | 14139826 | rs2245218 | 0.201 | 0.331 | -1.073 | 1.444 | 0.006 | -0.057 | -0.057 | 0 | 0.85 | 0.94 | 0.85 | 0.7 | 0.85 | 0.72 |
| 1 | 14141988 | rs11588426 | 0.653 | 0.219 | -1.151 | 1.267 | -0.001 | -0.001 | -0.001 | 0 | 0.14 | 0.12 | 0.09 | 0.14 | 0.17 | 0.09 |
| 1 | 14143003 | rs1046331 | -0.259 | 0.146 | -1.05 | 1.216 | 0.01 | -0.042 | -0.042 | 0 | 0.85 | 0.93 | 0.86 | 0.71 | 0.86 | 0.73 |
| 1 | 14144694 | rs6663564 | -0.445 | 0.057 | -0.537 | 1.092 | 0.022 | 0.005 | 0.005 | 0 | 0.06 | 0.13 | 0.08 | 0.09 | 0.06 | 0.05 |
| 1 | 14145914 | rs1810474 | 0.884 | 0.009 | 0.222 | 1.07 | -0.002 | 0.072 | 0.072 | 0.355 | 0.77 | 0.54 | 0.75 | 0.63 | 0.8 | 0.68 |
| 1 | 14146935 | rs2235514 | -0.083 | 0.003 | 0.537 | 1.07 | -0.002 | 0.004 | 0.004 | 0 | 0.84 | 0.82 | 0.84 | 0.71 | 0.85 | 0.71 |
| 1 | 14148441 | rs2697971 | 0.634 | 0.045 | 0.389 | 1.039 | -0.002 | 0.016 | 0.016 | 0.192 | 0.79 | 0.73 | 0.78 | 0.62 | 0.8 | 0.67 |
| 1 | 14148790 | rs2744685 | 0.628 | 0.045 | 0.395 | 1.04 | -0.002 | 0.016 | 0.016 | 0.192 | 0.79 | 0.73 | 0.78 | 0.62 | 0.8 | 0.68 |
| 1 | 14150032 | rs2697967 | 1.352 | -0.14 | 0.139 | 0.782 | 0.003 | -0.007 | -0.007 | 1.582 | 0.69 | 0.73 | 0.75 | 0.58 | 0.74 | 0.51 |
| 1 | 14150138 | rs3820012 | -0.349 | -0.595 | 0.024 | 0.598 | -0.004 | 0.018 | 0.018 | 0 | 0.09 | 0 | 0.02 | 0.05 | 0.05 | 0.16 |
| 1 | 14150709 | rs2697964 | -0.345 | -0.6 | 0.041 | 0.647 | 0.004 | -0.002 | -0.002 | 0 | 0.84 | 0.86 | 0.86 | 0.81 | 0.85 | 0.72 |
| 1 | 14150740 | rs2697963 | -0.345 | -0.6 | 0.041 | 0.647 | 0.004 | -0.002 | -0.002 | 2.832 | 0.84 | 0.86 | 0.86 | 0.81 | 0.85 | 0.72 |
| 1 | 14150992 | rs2697962 | -0.345 | -0.6 | 0.041 | 0.647 | 0.004 | -0.002 | -0.002 | 0 | 0.84 | 0.86 | 0.86 | 0.81 | 0.85 | 0.72 |
| 1 | 14151670 | rs2744679 | 0.695 | -0.573 | -0.068 | 0.645 | 0.016 | 0.007 | 0.007 | 0 | 0.76 | 0.87 | 0.83 | 0.76 | 0.81 | 0.56 |
| 1 | 14152449 | rs12044064 | 0.844 | -0.988 | -0.414 | 0.471 | -0.028 | 0.013 | 0.013 | 0 | 0.2 | 0.07 | 0.38 | 0.17 | 0.28 | 0.14 |
| 1 | 14154723 | rs2744677 | -0.053 | -0.912 | -0.256 | 0.588 | -0.001 | -0.001 | -0.001 | 0 | 0.8 | 0.73 | 0.77 | 0.64 | 0.79 | 0.7 |
| 1 | 14155055 | rs2495090 | -0.53 | -1.48 | -0.271 | 0.364 | -0.024 | 0.008 | 0.008 | 0 | 0.33 | 0.6 | 0.33 | 0.43 | 0.24 | 0.28 |
| 1 | 14158755 | rs1881561 | 1.187 | -0.948 | -0.719 | 0.418 | -0.056 | -0.043 | -0.043 | 0.251 | 0.31 | 0.12 | 0.27 | 0.3 | 0.37 | 0.31 |
| 1 | 14158776 | rs7530376 | -0.273 | -0.308 | 0.152 | 0.818 | -0.007 | 0.011 | 0.011 | 0 | 0.21 | 0.36 | 0.29 | 0.31 | 0.21 | 0.11 |
| 1 | 14160212 | rs10927938 | -0.349 | -0.218 | -0.448 | 0.668 | -0.01 | -0.007 | -0.007 | 0 | 0.22 | 0.31 | 0.3 | 0.26 | 0.21 | 0.11 |
| 1 | 14161251 | rs2495052 | 0.461 | 0.032 | -0.924 | 0.69 | -0.001 | -0.015 | -0.015 | 0 | 0.16 | 0.21 | 0.22 | 0.11 | 0.18 | 0.05 |
| 1 | 14162023 | rs1203690 | 0.099 | 0.121 | -1.447 | 0.659 | -0.055 | -0.014 | -0.014 | 0 | 0.85 | 0.97 | 0.87 | 0.88 | 0.81 | 0.76 |
| 1 | 14164286 | rs1203683 | 0.289 | 0.23 | -0.941 | 0.752 | -0.002 | 0.013 | 0.013 | 0 | 0.65 | 0.58 | 0.6 | 0.46 | 0.63 | 0.61 |
| 1 | 14166750 | rs6702813 | -0.32 | 0.516 | -1.359 | 0.45 | -0.034 | 0.026 | 0.026 | 0 | 0.2 | 0.04 | 0.25 | 0.12 | 0.2 | 0.24 |
| 1 | 14170060 | rs1973904 | -0.375 | 0.312 | -0.356 | 0.694 | -0.012 | 0.009 | 0.009 | 0 | 0.19 | 0.39 | 0.25 | 0.25 | 0.18 | 0.11 |
| 1 | 14177608 | rs1203626 | -0.031 | 0.778 | -1.242 | 0.815 | -0.09 | -0.086 | -0.086 | 0.174 | 0.4 | 0.09 | 0.55 | 0.56 | 0.47 | 0.67 |

| | | | | | | | | | | | | | | | | |
|---|---|---|---|---|---|---|---|---|---|---|---|---|---|---|---|---|
| 1 | 14190669 | rs720035 | -0.861 | 0.767 | 0.191 | 0.633 | 0.027 | 0.032 | 0.032 | 0 | 0.55 | 0.36 | 0.69 | 0.74 | 0.65 | 0.71 |
| 1 | 14192526 | rs1203660 | 0.026 | 0.747 | 0.339 | 0.595 | -0.019 | -0.015 | -0.015 | 0 | 0.43 | 0.27 | 0.57 | 0.56 | 0.48 | 0.65 |
| 1 | 14195122 | rs2744671 | -0.485 | 1.313 | 0.073 | -0.255 | -0.002 | -0.008 | -0.008 | 0.916 | 0.45 | 0.53 | 0.64 | 0.86 | 0.47 | 0.49 |
| 1 | 14196212 | rs16853069 | -0.128 | 1.422 | -0.054 | -0.108 | 0.029 | 0.006 | 0.006 | 0 | 0.11 | 0.16 | 0.11 | 0.14 | 0.16 | 0.06 |
| 1 | 14197715 | rs12407665 | 0.527 | 1.48 | -0.401 | -0.081 | 0.002 | 0.012 | 0.012 | 0 | 0.33 | 0.22 | 0.52 | 0.72 | 0.31 | 0.43 |
| 1 | 14198107 | rs11806865 | -0.69 | 1.918 | -0.396 | 0.387 | 0.021 | -0.056 | -0.056 | 0 | 0.14 | 0.32 | 0.12 | 0.05 | 0.19 | 0.26 |
| 1 | 14198401 | rs17350949 | -0.647 | 1.835 | -1.004 | 0.819 | -0.061 | -0.038 | -0.038 | 0 | 0.1 | 0.01 | 0.1 | 0.13 | 0.16 | 0.06 |
| 1 | 14205300 | rs7521299 | -1.463 | 2.013 | -0.903 | 0.77 | -0.031 | -0.017 | -0.017 | 0 | 0.05 | 0 | 0.05 | 0.08 | 0.13 | 0.03 |
| 1 | 14205571 | rs2744656 | -0.684 | 1.935 | -0.845 | 0.628 | 0.002 | -0.003 | -0.003 | 0.41 | 0.41 | 0.41 | 0.63 | 0.66 | 0.46 | 0.51 |
| 1 | 14208571 | rs9287293 | -0.174 | 1.891 | -1.06 | 1.041 | -0.006 | 0.005 | 0.005 | 0 | 0.08 | 0.05 | 0.09 | 0.37 | 0.09 | 0.1 |
| 1 | 14210730 | rs10737914 | 0.715 | 1.797 | -1.482 | 0.504 | -0.011 | 0.107 | 0.107 | 0 | 0.51 | 0.25 | 0.29 | 0.7 | 0.48 | 0.59 |
| 1 | 14212217 | rs12090035 | 0.705 | 1.414 | -0.653 | 0.37 | -0.028 | 0.069 | 0.069 | 1.977 | 0.09 | 0.37 | 0.06 | 0.1 | 0.01 | 0.04 |
| 1 | 14214348 | rs1528787 | 1.218 | 1.702 | -0.773 | 0.44 | 0.002 | 0.049 | 0.049 | 0 | 0.44 | 0.22 | 0.21 | 0.34 | 0.41 | 0.5 |
| 1 | 14214412 | rs1528786 | 1.14 | 1.668 | -0.719 | 0.447 | 0.016 | 0.048 | 0.048 | 0 | 0.45 | 0.27 | 0.21 | 0.34 | 0.39 | 0.51 |
| 1 | 14216203 | rs2744652 | 0.443 | 1.628 | -0.77 | 0.302 | 0.01 | -0.039 | -0.039 | 0 | 0.61 | 0.56 | 0.69 | 0.79 | 0.59 | 0.65 |
| 1 | 14217674 | rs2744649 | 0.779 | 1.639 | -0.715 | 0.424 | -0.001 | -0.002 | -0.002 | 0 | 0.53 | 0.53 | 0.6 | 0.44 | 0.5 | 0.55 |
| 1 | 14218438 | rs760911 | 0.122 | 1.552 | -0.862 | 0.171 | 0.011 | 0.039 | 0.039 | 0 | 0.09 | 0.27 | 0.39 | 0.1 | 0.12 | 0.05 |
| 1 | 14220942 | rs2744647 | -0.787 | 1.757 | -1.122 | 0.07 | -0.068 | -0.004 | -0.004 | 0 | 0.88 | 0.99 | 0.93 | 0.9 | 0.81 | 0.89 |
| 1 | 14224999 | rs2038095 | 1.184 | 1.497 | -0.644 | 0.172 | 0.022 | 0.005 | 0.005 | 0 | 0.45 | 0.35 | 0.22 | 0.43 | 0.36 | 0.52 |
| 1 | 14226409 | rs6677556 | -0.172 | 1.345 | -0.591 | 0.274 | 0.023 | 0.053 | 0.053 | 0 | 0.08 | 0.27 | 0.4 | 0.11 | 0.12 | 0.04 |
| 1 | 14228077 | rs6429800 | -0.228 | 2.145 | -1.416 | 0.006 | -0.06 | 0.049 | 0.049 | 0 | 0.81 | 0.95 | 0.73 | 0.91 | 0.7 | 0.79 |
| 1 | 14228490 | rs1203709 | -0.089 | 1.008 | -1.328 | 0.228 | 0.002 | -0.003 | -0.003 | 0 | 0.33 | 0.32 | 0.46 | 0.38 | 0.33 | 0.28 |
| 1 | 14232929 | rs7543809 | 0.496 | 0.71 | -1.732 | 0.269 | 0.011 | -0.027 | -0.027 | 0 | 0.26 | 0.1 | 0.15 | 0.29 | 0.2 | 0.37 |
| 1 | 14240110 | rs12745720 | -0.371 | 0.875 | -1.788 | 0.132 | -0.02 | -0.007 | -0.007 | 0 | 0.11 | 0.04 | 0.05 | 0.12 | 0.13 | 0.16 |
| 1 | 14240350 | rs12122426 | -0.295 | 0.762 | -1.885 | -0.182 | -0.017 | 0.071 | 0.071 | 0 | 0.24 | 0.1 | 0.18 | 0.03 | 0.28 | 0.18 |
| 1 | 14243517 | rs10927953 | -0.66 | -0.444 | -1.041 | 0.416 | -0.02 | -0.009 | -0.009 | 2.379 | 0.4 | 0.56 | 0.24 | 0.64 | 0.34 | 0.33 |
| 1 | 14245737 | rs10803436 | -0.699 | -0.507 | -1.055 | 0.48 | -0.02 | -0.012 | -0.012 | 0 | 0.4 | 0.56 | 0.31 | 0.65 | 0.34 | 0.32 |
| 1 | 14249565 | rs10927957 | -0.614 | -0.7 | -1.093 | 0.917 | -0.016 | -0.003 | -0.003 | 0.158 | 0.36 | 0.54 | 0.21 | 0.35 | 0.33 | 0.27 |
| 1 | 14250674 | rs6429804 | -1.236 | -0.878 | -1.135 | 0.81 | 0 | 0 | 0 | 0 | 0.5 | 0.55 | 0.32 | 0.4 | 0.45 | 0.38 |
| 1 | 14252088 | rs12127990 | 0.268 | -1.1 | -1.056 | 0.565 | -0.021 | -0.048 | -0.048 | 0 | 0.12 | 0.16 | 0.06 | 0 | 0.07 | 0.1 |
| 1 | 14252372 | rs7542884 | -1.9 | -1.157 | -1.131 | 0.992 | -0.001 | -0.002 | -0.002 | 3.335 | 0.53 | 0.59 | 0.39 | 0.7 | 0.46 | 0.49 |
| 1 | 14252557 | rs7535860 | -0.679 | -1.259 | -1.214 | 0.901 | -0.001 | -0.006 | -0.006 | 0 | 0.29 | 0.3 | 0.22 | 0.39 | 0.31 | 0.26 |
| 1 | 14252577 | rs7543063 | -0.315 | -1.281 | -1.172 | 1.055 | -0.011 | 0.006 | 0.006 | 0 | 0.24 | 0.16 | 0.2 | 0.18 | 0.29 | 0.16 |
| 1 | 14253566 | rs10489150 | 1.362 | -1.249 | -1.164 | 1.07 | -0.013 | -0.01 | -0.01 | 1.824 | 0.12 | 0.09 | 0.14 | 0.15 | 0.18 | 0.05 |
| 1 | 14253821 | rs10927959 | 1.806 | -1.311 | -1.181 | 1.061 | -0.003 | -0.002 | -0.002 | 0 | 0.51 | 0.44 | 0.69 | 0.6 | 0.58 | 0.63 |
| 1 | 14255847 | rs7535502 | -1.308 | -1.329 | -1.044 | 1.05 | -0.007 | -0.001 | -0.001 | 0.186 | 0.29 | 0.3 | 0.13 | 0.25 | 0.21 | 0.32 |
| 1 | 14255863 | rs12123612 | -0.383 | -1.36 | -1.011 | 1.067 | 0.006 | -0.004 | -0.004 | 0 | 0.16 | 0.14 | 0.07 | 0.2 | 0.1 | 0.21 |
| 1 | 14255978 | rs7538374 | 0.508 | -1.293 | -1.115 | 1.102 | -0.008 | -0.014 | -0.014 | 0.302 | 0.13 | 0.17 | 0.06 | 0.05 | 0.11 | 0.11 |
| 1 | 14256246 | rs1005302 | 0.349 | -1.207 | -1.157 | 1.181 | 0.01 | -0.061 | -0.061 | 0 | 0.12 | 0.01 | 0.04 | 0.2 | 0.08 | 0.15 |
| 1 | 14261909 | rs2206585 | -0.241 | -1.148 | -1.018 | 1.038 | -0.001 | 0 | 0 | 0 | 0.79 | 0.74 | 0.81 | 0.85 | 0.79 | 0.95 |
| 1 | 14269713 | rs6660353 | 0.029 | -1.178 | -0.917 | 0.886 | 0.013 | 0.014 | 0.014 | 0.045 | 0.21 | 0.18 | 0.08 | 0.03 | 0.14 | 0.12 |
| 1 | 14271131 | rs12133920 | -0.38 | -1.386 | -0.923 | 0.954 | -0.001 | -0.002 | -0.002 | 0.04 | 0.16 | 0.16 | 0.07 | 0.2 | 0.09 | 0.21 |
| 1 | 14272338 | rs4661453 | 1.057 | -0.937 | -0.838 | 1.151 | 0 | -0.002 | -0.002 | 0.41 | 0.64 | 0.59 | 0.68 | 0.49 | 0.68 | 0.72 |
| 1 | 14277713 | rs10927967 | 0.149 | -1.059 | -1.034 | 0.84 | 0.031 | 0.065 | 0.065 | 0.3 | 0.2 | 0.09 | 0.08 | 0.03 | 0.14 | 0.12 |
| 1 | 14279513 | rs10927969 | 0.123 | -1.068 | -1.04 | 0.836 | 0.028 | 0.067 | 0.067 | 0.447 | 0.19 | 0.08 | 0.08 | 0.03 | 0.14 | 0.12 |
| 1 | 14280562 | rs2359896 | -0.946 | -0.886 | -0.89 | 0.983 | 0.003 | 0.002 | 0.002 | 0 | 0.87 | 0.85 | 0.84 | 0.84 | 0.81 | 0.94 |
| 1 | 14281451 | rs6678469 | -1.361 | -1.234 | -0.444 | 1.006 | 0.017 | -0.018 | -0.018 | 0.371 | 0.42 | 0.67 | 0.37 | 0.5 | 0.35 | 0.28 |
| 1 | 14284460 | rs12043211 | 0.216 | -0.063 | -1.061 | 1.081 | -0.082 | -0.106 | -0.106 | 0 | 0.21 | 0.05 | 0.39 | 0.38 | 0.24 | 0.37 |
| 1 | 14284642 | rs16853271 | 1.33 | -0.362 | -0.446 | 1.333 | 0.001 | 0.02 | 0.02 | 0 | 0.1 | 0.15 | 0.1 | 0.16 | 0.13 | 0.05 |
| 1 | 14285838 | rs6676708 | 0.291 | -0.51 | -0.605 | 1.203 | -0.016 | 0.053 | 0.053 | 1.447 | 0.19 | 0.01 | 0.14 | 0.06 | 0.15 | 0.12 |
| 1 | 14287919 | rs10803439 | -1.056 | -0.708 | -0.349 | 1.251 | -0.011 | -0.003 | -0.003 | 0 | 0.43 | 0.54 | 0.26 | 0.42 | 0.3 | 0.39 |
| 1 | 14288119 | rs12044299 | 0.236 | -0.497 | -0.366 | 1.255 | -0.003 | 0.018 | 0.018 | 0 | 0.07 | 0.05 | 0.05 | 0.28 | 0.07 | 0.1 |
| 1 | 14290847 | rs12118400 | -0.664 | -0.597 | 0.107 | 1.132 | -0.001 | 0 | 0 | 0 | 0.28 | 0.3 | 0.15 | 0.38 | 0.22 | 0.3 |
| 1 | 14291845 | rs1883645 | 0.906 | -0.277 | 0.011 | 1.195 | 0 | 0.01 | 0.01 | 0 | 0.14 | 0.18 | 0.16 | 0.28 | 0.21 | 0.07 |
| 1 | 14297110 | rs10927972 | -1.266 | -0.371 | 0.133 | 1.141 | 0 | 0 | 0 | 2.756 | 0.43 | 0.48 | 0.31 | 0.66 | 0.43 | 0.37 |
| 1 | 14297220 | rs12024045 | 0.607 | -0.262 | 0.244 | 0.978 | 0.012 | 0.078 | 0.078 | 0.012 | 0.15 | 0.32 | 0.15 | 0.28 | 0.2 | 0.07 |
| 1 | 14298014 | rs7543486 | -1.129 | -0.152 | 0.067 | 0.53 | 0.019 | -0.046 | -0.046 | 0.472 | 0.44 | 0.17 | 0.23 | 0.55 | 0.35 | 0.45 |
| 1 | 14301802 | rs12123608 | 0.4 | -0.245 | 0.139 | 0.129 | 0.007 | 0.006 | 0.006 | 1.342 | 0.19 | 0.17 | 0.09 | 0.16 | 0.15 | 0.23 |
| 1 | 14301942 | rs16853319 | 0.591 | -0.241 | -0.119 | 0.139 | -0.014 | -0.032 | -0.032 | 0.886 | 0.17 | 0.01 | 0.06 | 0.17 | 0.14 | 0.22 |
| 1 | 14302077 | rs12024854 | 0.536 | -0.137 | -0.024 | 0.191 | 0.018 | 0.014 | 0.014 | 1.812 | 0.18 | 0.08 | 0.08 | 0.15 | 0.14 | 0.22 |
| 1 | 14302142 | rs10927976 | 0.536 | -0.137 | -0.024 | 0.191 | 0.018 | 0.014 | 0.014 | 1.812 | 0.18 | 0.08 | 0.08 | 0.15 | 0.14 | 0.22 |
| 1 | 14303160 | rs2359899 | 1.806 | -0.008 | 0.181 | 0.231 | -0.02 | 0.007 | 0.007 | 0.356 | 0.38 | 0.29 | 0.61 | 0.32 | 0.42 | 0.4 |
| 1 | 14304168 | rs725397 | 1.089 | 0.262 | 0.162 | 0.413 | 0.003 | -0.034 | -0.034 | 0.264 | 0.12 | 0.17 | 0.12 | 0 | 0.19 | 0.03 |
| 1 | 14308446 | rs17351137 | 1.095 | 0.305 | 0.151 | 0.417 | 0 | 0.002 | 0.002 | 1.601 | 0.58 | 0.55 | 0.68 | 0.49 | 0.54 | 0.63 |
| 1 | 14311056 | rs4661462 | -0.156 | 0.277 | -0.342 | 0.473 | -0.022 | -0.012 | -0.012 | 0.381 | 0.92 | 0.98 | 0.92 | 0.65 | 0.89 | 0.88 |
| 1 | 14315141 | rs12072002 | 0.542 | 0.145 | -0.013 | -0.001 | 0.015 | -0.057 | -0.057 | 0 | 0.13 | 0.24 | 0.14 | 0 | 0.2 | 0.04 |
| 1 | 14320502 | rs6681501 | -0.086 | -0.104 | -0.47 | 0.267 | 0.019 | 0.007 | 0.007 | 2.458 | 0.19 | 0.12 | 0.08 | 0.17 | 0.14 | 0.23 |
| 1 | 14331940 | rs12129723 | 1.131 | 0.239 | -0.375 | 0.032 | -0.036 | -0.008 | -0.008 | 0 | 0.47 | 0.17 | 0.63 | 0.47 | 0.49 | 0.61 |
| 1 | 14334758 | rs10803448 | 0.355 | 0.4 | 0.078 | -0.134 | 0.02 | -0.06 | -0.06 | 0 | 0.15 | 0.26 | 0.17 | 0 | 0.23 | 0.04 |

| | | | | | | | | | | | | | | | | |
|---|---|---|---|---|---|---|---|---|---|---|---|---|---|---|---|---|
| 1 | 14337014 | rs10465640 | 0.333 | 0.308 | 0.538 | 0.033 | 0.019 | 0.082 | 0.082 | 0.039 | 0.79 | 0.61 | 0.81 | 0.62 | 0.74 | 0.85 |
| 1 | 14337926 | rs6689299 | -1.356 | 0.181 | 0.836 | -0.017 | -0.001 | -0.019 | -0.019 | 0 | 0.46 | 0.51 | 0.3 | 0.2 | 0.46 | 0.29 |
| 1 | 14337994 | rs6689383 | -0.886 | 0.097 | 0.753 | -0.11 | 0.009 | -0.121 | -0.121 | 0 | 0.29 | 0.5 | 0.24 | 0.04 | 0.31 | 0.08 |
| 1 | 14338328 | rs6429820 | 0.554 | 0.107 | 0.661 | 0.075 | 0.002 | -0.004 | -0.004 | 1.785 | 0.76 | 0.77 | 0.87 | 0.47 | 0.81 | 0.68 |
| 1 | 14338807 | rs12566758 | -0.202 | 0.106 | 0.677 | -0.053 | -0.003 | 0 | 0 | 0 | 0.19 | 0.2 | 0.09 | 0.2 | 0.14 | 0.22 |
| 1 | 14340185 | rs17353372 | 0.295 | 0.147 | 0.308 | -0.102 | -0.006 | -0.02 | -0.02 | 0 | 0.16 | 0.01 | 0.07 | 0.15 | 0.14 | 0.21 |
| 1 | 14340816 | rs1883647 | 0.465 | 0.141 | 0.278 | -0.104 | -0.011 | -0.013 | -0.013 | 0 | 0.15 | 0.01 | 0.06 | 0.12 | 0.12 | 0.16 |
| 1 | 14341876 | rs16853407 | 0.616 | 0.321 | 0.567 | -0.101 | 0.008 | -0.042 | -0.042 | 0 | 0.13 | 0.19 | 0.13 | 0 | 0.21 | 0.04 |
| 1 | 14346629 | rs7520101 | 0.11 | 0.289 | 0.558 | 0.026 | -0.004 | -0.049 | -0.049 | 0.292 | 0.82 | 0.9 | 0.87 | 0.66 | 0.85 | 0.87 |
| 1 | 14348533 | rs932407 | 1.117 | 0.286 | 0.926 | 0.013 | 0.001 | 0.002 | 0.002 | 9.526 | 0.66 | 0.36 | 0.71 | 0.62 | 0.64 | 0.82 |
| 1 | 14349616 | rs970723 | -0.783 | 0.29 | 0.821 | 0.015 | -0.003 | 0.051 | 0.051 | 0 | 0.95 | 0.97 | 0.96 | 0.66 | 0.95 | 0.9 |
| 1 | 14351366 | rs973857 | 1.13 | 0.259 | 1.037 | -0.039 | -0.004 | -0.009 | -0.009 | 0 | 0.65 | 0.27 | 0.7 | 0.62 | 0.64 | 0.82 |
| 1 | 14352963 | rs10927998 | 0.252 | 0.337 | 0.995 | 0.021 | 0.03 | -0.068 | -0.068 | 0 | 0.14 | 0.27 | 0.14 | 0.01 | 0.21 | 0.04 |
| 1 | 14353071 | rs3929673 | 1.011 | 0.284 | 0.554 | 0.075 | 0.001 | -0.027 | -0.027 | 1.728 | 0.61 | 0.25 | 0.72 | 0.47 | 0.6 | 0.64 |
| 1 | 14354311 | rs4579751 | -0.845 | -0.158 | 0.518 | -0.459 | -0.054 | 0.014 | 0.014 | 0 | 0.25 | 0.49 | 0.12 | 0.22 | 0.16 | 0.26 |
| 1 | 14358424 | rs7527934 | 0.464 | 0.021 | 0.508 | -0.344 | -0.003 | -0.003 | -0.003 | 0 | 0.7 | 0.71 | 0.86 | 0.69 | 0.74 | 0.65 |
| 1 | 14359224 | rs2206586 | 0.495 | 0.016 | 0.558 | -0.347 | -0.011 | -0.012 | -0.012 | 0 | 0.7 | 0.64 | 0.86 | 0.69 | 0.74 | 0.63 |
| 1 | 14359780 | rs7414562 | 1.662 | 0.419 | 0 | -0.661 | -0.038 | 0.011 | 0.011 | 1.011 | 0.42 | 0.14 | 0.64 | 0.6 | 0.43 | 0.45 |
| 1 | 14364575 | rs2359908 | 0.224 | 0.375 | 0.115 | -0.418 | 0.048 | 0.093 | 0.093 | 0 | 0.94 | 0.74 | 0.95 | 0.83 | 0.94 | 0.87 |
| 1 | 14365793 | rs2294811 | -0.096 | 0.321 | -0.022 | -0.488 | -0.017 | 0.037 | 0.037 | 0 | 0.24 | 0.01 | 0.09 | 0.13 | 0.2 | 0.24 |
| 1 | 14368289 | rs4662089 | 1.102 | 0.403 | 0.003 | -0.578 | -0.038 | 0.136 | 0.136 | 2.191 | 0.54 | 0.83 | 0.75 | 0.7 | 0.51 | 0.54 |
| 1 | 14368784 | rs7514368 | 0.056 | 0.396 | -0.001 | -0.582 | 0.001 | 0.011 | 0.011 | 0 | 0.72 | 0.92 | 0.88 | 0.7 | 0.74 | 0.64 |
| 1 | 14372058 | rs12061818 | -0.36 | 0.394 | 0.033 | -0.586 | 0.018 | -0.109 | -0.109 | 0 | 0.2 | 0.33 | 0.15 | 0 | 0.24 | 0.09 |
| 1 | 14372163 | rs10754907 | 1.297 | 0.4 | -0.015 | -0.574 | -0.001 | 0.026 | 0.026 | 1.067 | 0.52 | 0.59 | 0.73 | 0.7 | 0.51 | 0.54 |
| 1 | 14376799 | rs12748863 | 0.063 | 0.28 | -0.292 | -0.779 | -0.019 | 0.038 | 0.038 | 1.865 | 0.17 | 0.09 | 0.13 | 0 | 0.22 | 0.09 |
| 1 | 14379546 | rs761161 | -0.184 | 0.417 | -0.211 | -0.673 | 0.011 | -0.042 | -0.042 | 0 | 0.2 | 0.42 | 0.17 | 0.07 | 0.23 | 0.15 |
| 1 | 14385135 | rs10737918 | 0.703 | 0.483 | -0.398 | -0.278 | 0.004 | 0.02 | 0.02 | 0.624 | 0.51 | 0.43 | 0.71 | 0.66 | 0.51 | 0.5 |
| 1 | 14391685 | rs6701426 | -0.693 | -0.037 | -0.508 | -0.327 | -0.021 | -0.026 | -0.026 | 0 | 0.28 | 0.36 | 0.13 | 0.15 | 0.19 | 0.31 |
| 1 | 14394225 | rs715340 | 0.512 | 0.272 | -1.21 | -0.083 | -0.03 | -0.034 | -0.034 | 0 | 0.11 | 0.01 | 0.1 | 0.11 | 0.11 | 0.04 |
| 1 | 14394291 | rs7538925 | -0.375 | 0.214 | -1.808 | -0.836 | -0.016 | 0.055 | 0.055 | 0.241 | 0.26 | 0.13 | 0.16 | 0.11 | 0.24 | 0.27 |
| 1 | 14394717 | rs715339 | 0.401 | 0.309 | -0.702 | -1.084 | 0.011 | 0.025 | 0.025 | 0 | 0.12 | 0.29 | 0.14 | 0.13 | 0.12 | 0.04 |
| 1 | 14397803 | rs742545 | 0.255 | -0.446 | -1.006 | -0.57 | -0.006 | 0.043 | 0.043 | 2.896 | 0.15 | 0.16 | 0.17 | 0.51 | 0.1 | 0.17 |
| 1 | 14399627 | rs12746681 | 0.365 | -0.285 | -0.599 | -0.601 | 0.01 | 0.034 | 0.034 | 0.158 | 0.14 | 0.32 | 0.14 | 0.15 | 0.12 | 0.09 |
| 1 | 14401350 | rs2246175 | -0.69 | -0.359 | -0.846 | -0.799 | 0.014 | -0.019 | -0.019 | 0 | 0.41 | 0.31 | 0.32 | 0.52 | 0.34 | 0.31 |
| 1 | 14407802 | rs4661470 | 1.452 | 0.343 | -1.427 | -0.915 | -0.052 | -0.001 | -0.001 | 0.82 | 0.12 | 0.02 | 0.31 | 0.07 | 0.21 | 0.25 |
| 1 | 14413237 | rs12139399 | -0.372 | -0.417 | -0.821 | -1.001 | 0.003 | -0.025 | -0.025 | 0 | 0.34 | 0.34 | 0.29 | 0.13 | 0.36 | 0.14 |
| 1 | 14418186 | rs10928003 | -0.747 | -0.607 | -0.86 | -0.847 | 0 | 0.001 | 0.001 | 0 | 0.42 | 0.41 | 0.4 | 0.67 | 0.36 | 0.34 |
| 1 | 14418448 | rs10928004 | -0.696 | -0.591 | -0.805 | -0.84 | -0.001 | -0.002 | -0.002 | 0 | 0.41 | 0.4 | 0.4 | 0.67 | 0.36 | 0.34 |
| 1 | 14433775 | rs10928009 | -0.356 | -0.556 | -0.322 | -0.773 | -0.003 | 0.009 | 0.009 | 0 | 0.33 | 0.33 | 0.29 | 0.55 | 0.28 | 0.31 |
| 1 | 14439042 | rs7515917 | -1.096 | -0.896 | -1.208 | -1.252 | -0.012 | 0.002 | 0.002 | 0.079 | 0.64 | 0.67 | 0.49 | 0.67 | 0.51 | 0.44 |
| 1 | 14440141 | rs6664213 | -0.627 | -0.359 | -0.722 | -0.653 | -0.011 | 0.029 | 0.029 | 0 | 0.34 | 0.52 | 0.31 | 0.54 | 0.28 | 0.31 |
| 1 | 14440876 | rs6664316 | -0.813 | 0.117 | -1.001 | -0.168 | -0.016 | 0.026 | 0.026 | 0 | 0.37 | 0.46 | 0.28 | 0.54 | 0.28 | 0.33 |
| 1 | 14441384 | rs7519796 | -0.201 | 1.139 | -1.523 | 0.211 | -0.036 | -0.063 | -0.063 | 0.409 | 0.84 | 0.95 | 0.77 | 0.41 | 0.81 | 0.82 |
| 1 | 14441738 | rs10928016 | -0.771 | 0.614 | -1.637 | -0.092 | -0.105 | 0.018 | 0.018 | 0 | 0.35 | 0.11 | 0.41 | 0.69 | 0.42 | 0.33 |
| 1 | 14444425 | rs2157565 | -0.686 | -0.584 | -0.457 | -0.518 | -0.042 | -0.075 | -0.075 | 5.973 | 0.23 | 0.37 | 0.17 | 0.09 | 0.15 | 0.08 |
| 1 | 14444796 | rs16853600 | -0.431 | -1.026 | -0.442 | -1.025 | -0.011 | -0.04 | -0.04 | 3.486 | 0.2 | 0.2 | 0.09 | 0.02 | 0.12 | 0.15 |
| 1 | 14448184 | rs4662105 | 0.226 | -1.011 | -0.823 | -1 | 0.037 | 0.096 | 0.096 | 0 | 0.52 | 0.72 | 0.68 | 0.74 | 0.46 | 0.51 |
| 1 | 14448571 | rs4662110 | 1.014 | -1.046 | -0.85 | -0.996 | -0.025 | -0.016 | -0.016 | 4.739 | 0.37 | 0.49 | 0.55 | 0.68 | 0.33 | 0.48 |
| 1 | 14449136 | rs2359942 | 0.45 | -0.6 | -0.941 | -0.805 | 0.035 | 0.145 | 0.145 | 0 | 0.53 | 0.83 | 0.69 | 0.74 | 0.46 | 0.51 |
| 1 | 14457230 | rs717538 | 0.235 | -0.361 | -1.033 | -0.725 | 0.014 | 0.09 | 0.09 | 0 | 0.57 | 0.77 | 0.69 | 0.76 | 0.47 | 0.55 |
| 1 | 14462146 | rs4661474 | 0.248 | -0.359 | -1.053 | -0.605 | 0.015 | 0.069 | 0.069 | 0 | 0.56 | 0.78 | 0.69 | 0.7 | 0.47 | 0.55 |
| 1 | 14463067 | rs6671072 | -0.908 | -0.194 | -1.436 | -0.616 | -0.174 | -0.04 | -0.04 | 0.576 | 0.74 | 0.97 | 0.78 | 0.72 | 0.58 | 0.67 |
| 1 | 14464167 | rs10803459 | 0.109 | -0.404 | -1.118 | -0.596 | 0.03 | 0.104 | 0.104 | 0 | 0.53 | 0.74 | 0.69 | 0.71 | 0.44 | 0.54 |
| 1 | 14467720 | rs4661477 | 0.14 | -0.383 | -1.139 | -0.658 | 0.031 | 0.113 | 0.113 | 0 | 0.53 | 0.73 | 0.68 | 0.72 | 0.45 | 0.54 |
| 1 | 14468557 | rs1934656 | -0.304 | -0.134 | -0.804 | -0.544 | -0.028 | 0.037 | 0.037 | 0 | 0.63 | 0.78 | 0.73 | 0.73 | 0.53 | 0.51 |
| 1 | 14473926 | rs12758112 | 0.732 | -0.971 | -0.292 | -0.202 | -0.034 | 0.081 | 0.081 | 0 | 0.2 | 0.31 | 0.32 | 0.59 | 0.13 | 0.16 |
| 1 | 14476490 | rs17353931 | -0.315 | -0.854 | -0.373 | -0.554 | 0.005 | 0.002 | 0.002 | 0 | 0.48 | 0.49 | 0.42 | 0.1 | 0.4 | 0.38 |
| 1 | 14477944 | rs929125 | -0.211 | -0.88 | -0.301 | -0.566 | 0.016 | 0.092 | 0.092 | 0 | 0.46 | 0.34 | 0.41 | 0.1 | 0.4 | 0.38 |
| 1 | 14487284 | rs10928043 | 0.229 | -0.683 | -0.039 | -0.598 | -0.001 | -0.002 | -0.002 | 0 | 0.25 | 0.19 | 0.19 | 0.02 | 0.26 | 0.37 |
| 1 | 14489962 | rs7546077 | 0.467 | -0.797 | 0.047 | -0.014 | 0.021 | 0.146 | 0.146 | 0 | 0.12 | 0.34 | 0.2 | 0.32 | 0.12 | 0.03 |
| 1 | 14490497 | rs9803674 | -0.813 | -0.798 | 0.027 | -0.023 | 0.004 | -0.005 | -0.005 | 0 | 0.38 | 0.53 | 0.39 | 0.36 | 0.4 | 0.43 |
| 1 | 14492478 | rs10803470 | 0.754 | -0.775 | -0.02 | -0.002 | 0.011 | 0.092 | 0.092 | 0 | 0.11 | 0.24 | 0.2 | 0.32 | 0.11 | 0.03 |
| 1 | 14493845 | rs7541978 | -1.427 | -0.55 | -0.316 | -0.032 | -0.114 | 0.018 | 0.018 | 0 | 0.62 | 0.91 | 0.64 | 0.69 | 0.54 | 0.59 |
| 1 | 14495003 | rs10489144 | 0.651 | -0.5 | 0.41 | 0.903 | -0.004 | 0.008 | 0.008 | 0 | 0.1 | 0.1 | 0.18 | 0.32 | 0.11 | 0.03 |
| 1 | 14495162 | rs6692372 | 0.195 | -0.501 | 0.399 | 0.903 | 0.008 | 0.081 | 0.081 | 0 | 0.11 | 0.21 | 0.19 | 0.32 | 0.11 | 0.03 |
| 1 | 14495224 | rs12092197 | -0.299 | -0.394 | 0.034 | 0.92 | -0.053 | 0.056 | 0.056 | 0 | 0.26 | 0.49 | 0.26 | 0.35 | 0.17 | 0.19 |
| 1 | 14496787 | rs12568548 | -0.174 | -0.297 | -0.651 | 1.017 | -0.022 | 0.021 | 0.021 | 0 | 0.24 | 0.3 | 0.23 | 0.33 | 0.15 | 0.15 |
| 1 | 14500883 | rs10928048 | 0.024 | 0.766 | -1.024 | 1.196 | -0.003 | 0.011 | 0.011 | 0 | 0.1 | 0.1 | 0.19 | 0.32 | 0.11 | 0.03 |
| 1 | 14501296 | rs16853705 | -0.187 | 0.782 | -1.018 | 1.202 | 0.001 | 0.031 | 0.031 | 0 | 0.11 | 0.13 | 0.19 | 0.32 | 0.11 | 0.03 |

| | | | | | | | | | | | | | | | | |
|---|---|---|---|---|---|---|---|---|---|---|---|---|---|---|---|---|
| 1 | 14507057 | rs10928050 | -0.813 | 1.008 | -1.36 | 1.192 | -0.043 | -0.069 | -0.069 | 0 | 0.15 | 0.05 | 0.2 | 0.36 | 0.19 | 0.06 |
| 1 | 14507529 | rs526188 | 0.314 | 0.752 | -1.711 | 0.753 | 0.014 | 0.06 | 0.06 | 0 | 0.52 | 0.7 | 0.51 | 0.27 | 0.61 | 0.46 |
| 1 | 14507542 | rs675997 | -0.297 | 0.528 | -1.777 | 0.936 | 0.005 | 0.005 | 0.005 | 0 | 0.67 | 0.73 | 0.8 | 0.7 | 0.74 | 0.59 |
| 1 | 14507739 | rs10754914 | -0.878 | 0.495 | -1.896 | 0.423 | -0.013 | 0.04 | 0.04 | 0 | 0.52 | 0.27 | 0.44 | 0.62 | 0.46 | 0.47 |
| 1 | 14507803 | rs10803474 | -0.389 | 0.107 | -1.748 | 0.507 | 0.027 | 0.024 | 0.024 | 0 | 0.45 | 0.26 | 0.36 | 0.62 | 0.34 | 0.45 |
| 1 | 14509276 | rs577987 | -0.141 | -0.682 | -0.759 | 0.394 | -0.035 | -0.019 | -0.019 | 0 | 0.14 | 0.3 | 0.17 | 0.09 | 0.07 | 0.11 |
| 1 | 14509334 | rs12736419 | 0.237 | -0.965 | -0.881 | 0.358 | 0.003 | -0.045 | -0.045 | 0.064 | 0.39 | 0.19 | 0.35 | 0.48 | 0.31 | 0.43 |
| 1 | 14509688 | rs574360 | 0.311 | -1.024 | -0.488 | 0.328 | -0.044 | -0.023 | -0.023 | 0 | 0.15 | 0.36 | 0.18 | 0.09 | 0.07 | 0.11 |
| 1 | 14510164 | rs12736835 | -0.002 | -1.186 | -0.162 | 0.25 | 0.007 | -0.013 | -0.013 | 3.987 | 0.41 | 0.31 | 0.36 | 0.48 | 0.32 | 0.43 |
| 1 | 14511098 | rs12563497 | 0.965 | -1.626 | -0.424 | 0.405 | 0.012 | -0.015 | -0.015 | 0.127 | 0.14 | 0.1 | 0.07 | 0.29 | 0.05 | 0.18 |
| 1 | 14511349 | rs517269 | -1.433 | -1.592 | -0.375 | 0.329 | -0.003 | 0 | 0 | 1.079 | 0.57 | 0.6 | 0.54 | 0.57 | 0.45 | 0.56 |
| 1 | 14512986 | rs565765 | -1.635 | -1.619 | -0.311 | 0.316 | -0.019 | -0.006 | -0.006 | 0 | 0.59 | 0.73 | 0.56 | 0.57 | 0.45 | 0.56 |
| 1 | 14513609 | rs657359 | 0.188 | -1.595 | -0.392 | 0.256 | -0.028 | -0.029 | -0.029 | 0 | 0.18 | 0.43 | 0.19 | 0.1 | 0.13 | 0.13 |
| 1 | 14515925 | rs6679368 | 1.032 | -1.553 | -0.574 | 0.273 | -0.033 | 0.037 | 0.037 | 0 | 0.26 | 0.12 | 0.28 | 0.15 | 0.27 | 0.23 |
| 1 | 14518979 | rs641343 | 0.029 | -1.542 | -0.469 | 0.472 | 0.016 | -0.023 | -0.023 | 0.029 | 0.56 | 0.8 | 0.52 | 0.49 | 0.65 | 0.55 |
| 1 | 14518991 | rs1339367 | 0.638 | -1.446 | -0.034 | 0.293 | -0.013 | -0.032 | -0.032 | 0 | 0.11 | 0.22 | 0.14 | 0.05 | 0.1 | 0.07 |
| 1 | 14525934 | rs653504 | 0.273 | -1.62 | -0.182 | 0.231 | -0.067 | -0.104 | -0.104 | 0 | 0.79 | 0.54 | 0.8 | 0.96 | 0.91 | 0.86 |
| 1 | 14536470 | rs614719 | 1.819 | -1.202 | -0.611 | 0.285 | -0.076 | 0.022 | 0.022 | 0 | 0.31 | 0.08 | 0.21 | 0.25 | 0.44 | 0.41 |
| 1 | 14539152 | rs647287 | 0.955 | -1.126 | -0.342 | 0.494 | -0.011 | 0.012 | 0.012 | 0 | 0.43 | 0.28 | 0.32 | 0.39 | 0.57 | 0.47 |
| 1 | 14547727 | rs2027315 | 0.317 | -1.009 | -0.731 | 0.547 | 0 | -0.023 | -0.023 | 0.944 | 0.32 | 0.26 | 0.44 | 0.48 | 0.29 | 0.36 |
| 1 | 14558018 | rs12742276 | 0.914 | -0.991 | -1.275 | 0.65 | -0.072 | -0.007 | -0.007 | 0.849 | 0.26 | 0.02 | 0.36 | 0.16 | 0.26 | 0.22 |
| 1 | 14567691 | rs12024171 | 0.347 | -0.598 | -0.775 | 0.512 | -0.008 | -0.055 | -0.055 | 0 | 0.29 | 0.16 | 0.42 | 0.48 | 0.28 | 0.36 |
| 1 | 14575084 | rs10157045 | -0.206 | -0.909 | -0.48 | 0.474 | 0.04 | -0.035 | -0.035 | 0 | 0.42 | 0.12 | 0.4 | 0.39 | 0.32 | 0.4 |
| 1 | 14575256 | rs10158007 | -1.098 | -1.025 | -0.148 | 0.455 | 0.002 | 0.012 | 0.012 | 0 | 0.58 | 0.75 | 0.63 | 0.6 | 0.38 | 0.5 |
| 1 | 14575842 | rs6662041 | 0.288 | -0.61 | 0.497 | 0.754 | -0.025 | 0.036 | 0.036 | 0 | 0.1 | 0.35 | 0.17 | 0.09 | 0.03 | 0.07 |
| 1 | 14575992 | rs12403204 | 0.258 | -0.612 | 0.497 | 0.753 | -0.026 | 0.035 | 0.035 | 0 | 0.1 | 0.35 | 0.17 | 0.09 | 0.03 | 0.07 |
| 1 | 14577474 | rs4631686 | -0.595 | -0.589 | 0.45 | 0.745 | -0.023 | 0.06 | 0.06 | 0.018 | 0.15 | 0.45 | 0.2 | 0.2 | 0.06 | 0.11 |
| 1 | 14579742 | rs6668555 | -1.072 | -0.296 | -0.025 | 0.818 | -0.018 | 0.15 | 0.15 | 0.236 | 0.23 | 0.44 | 0.28 | 0.46 | 0.2 | 0.27 |
| 1 | 14583472 | rs12403871 | 0.687 | -0.28 | -0.657 | 0.403 | 0.011 | 0.092 | 0.092 | 0.684 | 0.28 | 0.1 | 0.15 | 0.03 | 0.23 | 0.17 |
| 1 | 14585648 | rs10928056 | -0.056 | -0.054 | -0.733 | 0.932 | 0.004 | 0.035 | 0.035 | 0 | 0.07 | 0.03 | 0.22 | 0.39 | 0.05 | 0.17 |
| 1 | 14586879 | rs4662139 | -0.361 | -0.021 | -0.472 | 0.666 | 0.05 | 0.037 | 0.037 | 0 | 0.5 | 0.7 | 0.59 | 0.62 | 0.35 | 0.5 |
| 1 | 14594027 | rs10754916 | 0.354 | 0.054 | 0.227 | 1.137 | -0.006 | 0.057 | 0.057 | 0 | 0.84 | 0.79 | 0.61 | 0.48 | 0.87 | 0.69 |
| 1 | 14595692 | rs10928060 | 0.31 | 0.406 | 0.146 | 0.842 | -0.033 | 0.006 | 0.006 | 1.405 | 0.34 | 0.25 | 0.34 | 0.36 | 0.51 | 0.37 |
| 1 | 14603198 | rs7555815 | 0.567 | 0.393 | 0.248 | 0.743 | 0 | -0.019 | -0.019 | 0.053 | 0.48 | 0.47 | 0.26 | 0.12 | 0.33 | 0.3 |
| 1 | 14620369 | rs16853880 | -0.218 | 0.483 | 0.308 | 0.873 | 0.023 | 0.005 | 0.005 | 0.209 | 0.1 | 0.39 | 0.1 | 0.04 | 0.05 | 0.06 |
| 1 | 14621842 | rs1416628 | 0.106 | 0.883 | -0.065 | 1.047 | -0.029 | 0.001 | 0.001 | 1.148 | 0.35 | 0.24 | 0.33 | 0.41 | 0.5 | 0.39 |
| 1 | 14622770 | rs1538534 | -0.121 | 0.992 | 0.316 | 0.623 | 0.007 | -0.003 | -0.003 | 0 | 0.08 | 0.16 | 0.07 | 0.04 | 0.05 | 0.06 |
| 1 | 14623470 | rs6668724 | -0.236 | 1.113 | -0.174 | 0.901 | 0.003 | 0.022 | 0.022 | 0 | 0.09 | 0.16 | 0.17 | 0.15 | 0.08 | 0.08 |
| 1 | 14625432 | rs7527299 | 0.08 | 1.35 | -0.504 | 1.036 | -0.004 | -0.004 | -0.004 | 0.014 | 0.7 | 0.73 | 0.55 | 0.39 | 0.58 | 0.48 |
| 1 | 14629281 | rs10927407 | -0.829 | 0.942 | 0.218 | 1.022 | 0.04 | 0.094 | 0.094 | 0 | 0.07 | 0.29 | 0.06 | 0.09 | 0.04 | 0.03 |
| 1 | 14629413 | rs1416626 | 0.22 | 1.274 | 0.061 | 1.138 | -0.002 | -0.002 | -0.002 | 0 | 0.65 | 0.68 | 0.7 | 0.51 | 0.57 | 0.6 |
| 1 | 14629449 | rs1416625 | 0.312 | 0.87 | -0.603 | 0.747 | 0 | -0.002 | -0.002 | 0.074 | 0.26 | 0.29 | 0.49 | 0.24 | 0.27 | 0.31 |
| 1 | 14629604 | rs1416624 | -0.01 | 0.773 | -0.744 | 0.761 | 0 | 0.002 | 0.002 | 0 | 0.55 | 0.51 | 0.59 | 0.33 | 0.45 | 0.5 |
| 1 | 14631625 | rs12143153 | 1.147 | 0.497 | -0.812 | 0.561 | 0.022 | 0.18 | 0.18 | 0 | 0.27 | 0.04 | 0.09 | 0.01 | 0.18 | 0.19 |
| 1 | 14633150 | rs2027316 | 0.506 | 0.57 | -0.494 | 0.391 | 0.015 | 0.021 | 0.021 | 0 | 0.34 | 0.24 | 0.16 | 0.01 | 0.26 | 0.22 |
| 1 | 14635852 | rs946481 | -0.471 | 0.622 | -0.261 | 0.566 | 0.023 | 0.054 | 0.054 | 0 | 0.51 | 0.71 | 0.31 | 0.35 | 0.4 | 0.33 |
| 1 | 14638612 | rs3737643 | -0.009 | 0.745 | -0.009 | 0.84 | 0.046 | 0.103 | 0.103 | 0 | 0.08 | 0.33 | 0.06 | 0.09 | 0.04 | 0.05 |
| 1 | 14649876 | rs1416623 | 0.039 | 0.833 | -0.503 | 0.861 | 0.002 | 0.026 | 0.026 | 0.014 | 0.07 | 0.12 | 0.09 | 0.25 | 0.1 | 0.06 |
| 1 | 14652782 | rs2149507 | 0.416 | 1.036 | -0.978 | 0.711 | -0.022 | -0.033 | -0.033 | 0 | 0.34 | 0.27 | 0.59 | 0.52 | 0.48 | 0.39 |
| 1 | 14654660 | rs820629 | -0.858 | 0.928 | -0.63 | 0.81 | -0.008 | -0.063 | -0.063 | 0 | 0.88 | 0.99 | 0.93 | 0.77 | 0.91 | 0.78 |
| 1 | 14657100 | rs820626 | -1.148 | 0.721 | -0.719 | 0.609 | 0.016 | -0.011 | -0.011 | 0 | 0.28 | 0.36 | 0.23 | 0.24 | 0.3 | 0.36 |
| 1 | 14661294 | rs820623 | -1.013 | 0.682 | -0.737 | 0.677 | 0.009 | -0.008 | -0.008 | 0 | 0.27 | 0.46 | 0.2 | 0.24 | 0.26 | 0.35 |
| 1 | 14663626 | rs703801 | -1.713 | 0.684 | -0.886 | 0.694 | 0 | -0.002 | -0.002 | 0 | 0.34 | 0.63 | 0.27 | 0.38 | 0.29 | 0.39 |
| 1 | 14666142 | rs703797 | -0.122 | 0.603 | -1.146 | 0.896 | 0.022 | -0.047 | -0.047 | 0 | 0.17 | 0.09 | 0.1 | 0.35 | 0.11 | 0.29 |
| 1 | 14668801 | rs2487751 | -0.96 | 0.758 | -1.123 | 1.006 | -0.006 | -0.025 | -0.025 | 0 | 0.26 | 0.15 | 0.33 | 0.33 | 0.25 | 0.37 |
| 1 | 14670296 | rs2789740 | 1.138 | 0.721 | -1.183 | 0.897 | -0.009 | -0.003 | -0.003 | 0 | 0.06 | 0.02 | 0.03 | 0.05 | 0.06 | 0.07 |
| 1 | 14670764 | rs2789742 | 0.577 | 0.708 | -1.165 | 0.891 | -0.002 | 0.002 | 0.002 | 0 | 0.08 | 0.06 | 0.03 | 0.05 | 0.06 | 0.07 |
| 1 | 14672479 | rs768725 | -0.085 | 1.356 | -1.377 | 0.923 | -0.007 | -0.035 | -0.035 | 0.763 | 0.23 | 0.17 | 0.36 | 0.47 | 0.31 | 0.48 |
| 1 | 14673025 | rs2789746 | -0.712 | 1.355 | -1.374 | 0.906 | -0.012 | -0.043 | -0.043 | 0.012 | 0.31 | 0.2 | 0.38 | 0.52 | 0.37 | 0.56 |
| 1 | 14676909 | rs7355173 | 0.348 | 1.409 | -1.52 | 0.251 | -0.028 | 0.018 | 0.018 | 0 | 0.23 | 0.15 | 0.21 | 0.17 | 0.31 | 0.16 |
| 1 | 14677729 | rs12134023 | 1.171 | 1.379 | -1.536 | 0.257 | -0.004 | 0.001 | 0.001 | 0 | 0.17 | 0.13 | 0.13 | 0.18 | 0.19 | 0.12 |
| 1 | 14678726 | rs3897492 | -0.335 | 1.081 | -1.339 | 0.439 | 0.029 | 0.072 | 0.072 | 0 | 0.46 | 0.66 | 0.55 | 0.22 | 0.65 | 0.42 |
| 1 | 14679601 | rs12239349 | 0.634 | 1.008 | -1.251 | 0.497 | 0.13 | -0.057 | -0.057 | 0 | 0.11 | 0.32 | 0.13 | 0 | 0.17 | 0.08 |
| 1 | 14680899 | rs4113453 | -0.679 | 0.311 | -1.869 | 0.762 | 0.077 | -0.019 | -0.019 | 0.158 | 0.58 | 0.87 | 0.76 | 0.46 | 0.71 | 0.54 |
| 1 | 14688707 | rs12129808 | 1.215 | 0.508 | -1.802 | 0.284 | 0.011 | 0.204 | 0.204 | 0 | 0.3 | 0.04 | 0.12 | 0.01 | 0.22 | 0.18 |
| 1 | 14692271 | rs820671 | 0.114 | 0.335 | -1.526 | 1.104 | -0.009 | 0.138 | 0.138 | 0 | 0.91 | 0.87 | 0.72 | 0.49 | 0.94 | 0.91 |
| 1 | 14692598 | rs10927414 | 1.141 | 0.604 | -1.927 | 0.624 | -0.071 | 0.173 | 0.173 | 0 | 0.32 | 0.06 | 0.19 | 0.07 | 0.38 | 0.23 |
| 1 | 14692672 | rs7538154 | 0.624 | 0.912 | -1.485 | 1.095 | -0.002 | 0.017 | 0.017 | 0 | 0.83 | 0.83 | 0.68 | 0.41 | 0.89 | 0.83 |
| 1 | 14697489 | rs12407887 | 0.915 | 1.124 | -1.216 | 0.647 | 0.008 | 0.022 | 0.022 | 0 | 0.07 | 0.17 | 0.07 | 0.16 | 0.08 | 0.11 |

| | | | | | | | | | | | | | | | | |
|---|---|---|---|---|---|---|---|---|---|---|---|---|---|---|---|---|
| 1 | 14698609 | rs6429654 | 0.031 | 1.079 | -1.147 | 0.901 | -0.006 | 0.115 | 0.115 | 0 | 0.93 | 0.87 | 0.92 | 0.68 | 0.96 | 0.92 |
| 1 | 14700829 | rs10803271 | -0.383 | 1.282 | -1.001 | 0.182 | 0.029 | -0.003 | -0.003 | 0 | 0.18 | 0.48 | 0.15 | 0.19 | 0.17 | 0.16 |
| 1 | 14702567 | rs7519457 | 0.543 | 1.31 | -1.188 | -0.257 | 0.03 | -0.063 | -0.063 | 0 | 0.1 | 0.28 | 0.08 | 0 | 0.09 | 0.06 |
| 1 | 14703298 | rs1556083 | 0.238 | 1.392 | -2.299 | 0.246 | 0 | 0.034 | 0.034 | 0 | 0.9 | 0.87 | 0.91 | 0.7 | 0.92 | 0.88 |
| 1 | 14703854 | rs11590846 | -0.015 | 1.178 | -2.717 | -0.172 | -0.058 | 0.153 | 0.153 | 0 | 0.28 | 0.05 | 0.3 | 0.03 | 0.32 | 0.21 |
| 1 | 14704441 | rs6689141 | 0.272 | 1.091 | -1.953 | 0.767 | 0 | 0.044 | 0.044 | 0 | 0.9 | 0.86 | 0.91 | 0.7 | 0.92 | 0.88 |
| 1 | 14704676 | rs735832 | 0.367 | 1.086 | -1.96 | 0.765 | 0.001 | 0.072 | 0.072 | 0 | 0.89 | 0.8 | 0.9 | 0.7 | 0.92 | 0.88 |
| 1 | 14715919 | rs6686602 | -0.344 | 1.358 | -2.888 | -0.239 | -0.092 | 0.149 | 0.149 | 0 | 0.69 | 0.94 | 0.67 | 0.92 | 0.6 | 0.71 |
| 1 | 14718957 | rs7527767 | -0.283 | 0.893 | -2.457 | 0.346 | 0.041 | 0.028 | 0.028 | 0 | 0.4 | 0.17 | 0.3 | 0.75 | 0.3 | 0.53 |
| 1 | 14721450 | rs3932686 | -0.692 | 2.212 | -2.3 | 0.591 | -0.067 | -0.004 | -0.004 | 0.052 | 0.19 | 0.08 | 0.23 | 0.19 | 0.34 | 0.12 |
| 1 | 14726049 | rs943918 | 0.164 | 1.888 | -2.574 | 0.825 | -0.016 | -0.026 | -0.026 | 0 | 0.16 | 0.08 | 0.21 | 0.25 | 0.19 | 0.14 |
| 1 | 14726396 | rs12122566 | 0.397 | 1.115 | -3.141 | 0.244 | 0.013 | 0.08 | 0.08 | 0 | 0.13 | 0 | 0.06 | 0 | 0.11 | 0.11 |
| 1 | 14726485 | rs12141361 | 0.194 | 1.162 | -3.19 | 0.92 | 0.003 | -0.034 | -0.034 | 0 | 0.14 | 0 | 0.11 | 0.12 | 0.15 | 0.17 |
| 1 | 14732019 | rs12405445 | 0.313 | 1.294 | -2.926 | 0.868 | -0.008 | -0.019 | -0.019 | 0.413 | 0.16 | 0.1 | 0.21 | 0.25 | 0.18 | 0.15 |
| 1 | 14733533 | rs12410249 | -0.55 | 1.927 | -3.059 | 0.603 | -0.042 | -0.001 | -0.001 | 0 | 0.22 | 0.12 | 0.36 | 0.23 | 0.38 | 0.13 |
| 1 | 14740155 | rs3928343 | -0.445 | 2.469 | -3.018 | 0.873 | -0.073 | -0.022 | -0.022 | 0.556 | 0.19 | 0.07 | 0.23 | 0.24 | 0.37 | 0.1 |
| 1 | 14742723 | rs10754855 | -0.406 | 1.943 | -3.367 | 0.507 | -0.001 | 0.028 | 0.028 | 0 | 0.88 | 0.94 | 0.88 | 0.94 | 0.91 | 0.85 |
| 1 | 14743972 | rs4573512 | -0.443 | 0.737 | -2.189 | 0.271 | 0.024 | -0.015 | -0.015 | 0.159 | 0.36 | 0.49 | 0.31 | 0.34 | 0.44 | 0.19 |
| 1 | 14747769 | rs6665391 | -0.732 | 0.064 | -2.13 | 0.04 | 0.012 | 0.05 | 0.05 | 0 | 0.48 | 0.62 | 0.38 | 0.38 | 0.54 | 0.31 |
| 1 | 14755827 | rs10754856 | -0.881 | -0.248 | -1.972 | 0.576 | 0.004 | -0.019 | -0.019 | 0 | 0.56 | 0.68 | 0.51 | 0.53 | 0.59 | 0.37 |
| 1 | 14757240 | rs12130873 | 1.993 | -0.252 | -2.064 | 0.418 | -0.011 | 0.032 | 0.032 | 0 | 0.12 | 0 | 0.06 | 0.03 | 0.12 | 0.11 |
| 1 | 14760634 | rs2183102 | -0.28 | -0.326 | -1.952 | 0.401 | 0.003 | -0.004 | -0.004 | 0 | 0.38 | 0.55 | 0.33 | 0.35 | 0.44 | 0.19 |
| 1 | 14761482 | rs4661493 | 1.893 | -0.718 | -1.326 | 0.308 | -0.011 | 0.033 | 0.033 | 0 | 0.12 | 0 | 0.06 | 0.03 | 0.11 | 0.11 |
| 1 | 14771398 | rs12401776 | 2.653 | -1.064 | -1.573 | 0.686 | -0.003 | -0.05 | -0.05 | 0.548 | 0.08 | 0 | 0.08 | 0.22 | 0.07 | 0.13 |
| 1 | 14772774 | rs10803278 | -0.592 | -0.539 | -1.555 | 0.499 | -0.008 | 0.007 | 0.007 | 0 | 0.51 | 0.67 | 0.49 | 0.47 | 0.51 | 0.33 |
| 1 | 14775805 | rs6658316 | -0.595 | -0.571 | -0.964 | 0.407 | -0.004 | 0.008 | 0.008 | 0 | 0.4 | 0.35 | 0.4 | 0.4 | 0.41 | 0.25 |
| 1 | 14776392 | rs6658744 | 0.636 | -0.551 | -0.968 | 0.468 | -0.056 | 0.058 | 0.058 | 0 | 0.3 | 0.07 | 0.24 | 0.23 | 0.36 | 0.16 |
| 1 | 14780811 | rs6429661 | 0.44 | -0.669 | -1.006 | 0.374 | -0.015 | -0.003 | -0.003 | 1.586 | 0.24 | 0.33 | 0.22 | 0.2 | 0.17 | 0.19 |
| 1 | 14782380 | rs6682103 | -1.002 | -0.485 | -1.567 | 0.482 | -0.038 | -0.036 | -0.036 | 0 | 0.58 | 0.77 | 0.51 | 0.47 | 0.55 | 0.36 |
| 1 | 14786269 | rs12122476 | 2.362 | -0.365 | -1.548 | 0.426 | 0.005 | -0.007 | -0.007 | 4.738 | 0.1 | 0.01 | 0.04 | 0.1 | 0.06 | 0.07 |
| 1 | 14787410 | rs16850285 | 2.461 | -0.265 | -1.397 | 0.761 | 0.015 | -0.022 | -0.022 | 0 | 0.08 | 0.03 | 0.1 | 0.27 | 0.07 | 0.11 |
| 1 | 14788274 | rs17401833 | -0.005 | -0.005 | -1.076 | -0.177 | 0 | -0.016 | -0.016 | 0 | 0.24 | 0.26 | 0.19 | 0.07 | 0.29 | 0.12 |
| 1 | 14789535 | rs12564154 | 1.339 | -0.049 | -1.034 | 0.017 | 0.016 | 0.003 | 0.003 | 0 | 0.18 | 0.06 | 0.07 | 0.16 | 0.12 | 0.11 |
| 1 | 14790075 | rs12116420 | 1.339 | -0.049 | -1.034 | 0.017 | 0.016 | 0.003 | 0.003 | 0 | 0.18 | 0.06 | 0.07 | 0.16 | 0.13 | 0.11 |
| 1 | 14790631 | rs4661496 | -0.699 | 0.202 | -1.109 | 0.138 | -0.026 | -0.023 | -0.023 | 0 | 0.55 | 0.77 | 0.46 | 0.43 | 0.49 | 0.39 |
| 1 | 14790842 | rs10927440 | -0.213 | 1.227 | -0.383 | 0.268 | 0.025 | -0.032 | -0.032 | 0.075 | 0.18 | 0.29 | 0.18 | 0.08 | 0.24 | 0.11 |
| 1 | 14792535 | rs2480054 | 0.472 | 1.123 | -0.511 | 0.31 | 0.002 | -0.011 | -0.011 | 0.035 | 0.59 | 0.5 | 0.53 | 0.7 | 0.57 | 0.58 |
| 1 | 14796432 | rs2480057 | -0.495 | 1.282 | -0.478 | 0.525 | 0.002 | 0.008 | 0.008 | 0 | 0.92 | 1 | 0.88 | 0.97 | 0.96 | 0.88 |
| 1 | 14800247 | rs2486774 | -0.781 | 2.479 | 0.068 | 1.101 | 0.004 | 0.008 | 0.008 | 0 | 0.28 | 0.25 | 0.34 | 0.42 | 0.36 | 0.2 |
| 1 | 14800369 | rs2486772 | -2.263 | 2.669 | 0.328 | 0.9 | 0.024 | 0.069 | 0.069 | 0 | 0.51 | 0.63 | 0.6 | 0.89 | 0.62 | 0.42 |
| 1 | 14806750 | rs2486764 | -1.285 | 2.398 | 0.325 | 1.242 | 0.027 | 0.011 | 0.011 | 0 | 0.32 | 0.45 | 0.53 | 0.6 | 0.42 | 0.26 |
| 1 | 14807686 | rs2486762 | -0.951 | 2.417 | 0.245 | 1.291 | 0.008 | 0.018 | 0.018 | 0 | 0.29 | 0.28 | 0.38 | 0.47 | 0.36 | 0.21 |
| 1 | 14808756 | rs2480061 | -1.117 | 2.354 | 0.236 | 1.26 | 0.014 | 0.008 | 0.008 | 0 | 0.31 | 0.35 | 0.51 | 0.59 | 0.4 | 0.26 |
| 1 | 14819757 | rs12129734 | -0.764 | 2.416 | 0.113 | 1.172 | -0.032 | -0.017 | -0.017 | 0 | 0.12 | 0.06 | 0.1 | 0.17 | 0.24 | 0.13 |
| 1 | 14821610 | rs12142754 | -0.625 | 2.418 | 0.112 | 1.162 | -0.032 | -0.018 | -0.018 | 0 | 0.11 | 0.05 | 0.1 | 0.16 | 0.24 | 0.13 |
| 1 | 14826029 | rs12733612 | -0.019 | 2.206 | 0.126 | 1.134 | -0.029 | 0.079 | 0.079 | 0.054 | 0.78 | 0.6 | 0.57 | 0.4 | 0.81 | 0.79 |
| 1 | 14830866 | rs10803284 | -0.876 | 2.412 | 0.139 | 0.984 | -0.029 | -0.016 | -0.016 | 0.789 | 0.11 | 0.07 | 0.1 | 0.17 | 0.23 | 0.13 |
| 1 | 14830984 | rs10803285 | 1.333 | 2.347 | 0.102 | 0.851 | 0.003 | -0.028 | -0.028 | 0 | 0.56 | 0.47 | 0.41 | 0.21 | 0.51 | 0.66 |
| 1 | 14831066 | rs10733135 | -0.051 | 1.069 | -0.449 | 0.982 | 0.001 | -0.002 | -0.002 | 0 | 0.72 | 0.75 | 0.68 | 0.54 | 0.79 | 0.85 |
| 1 | 14837953 | rs10927452 | -0.715 | 1.555 | -0.925 | 0.82 | -0.02 | -0.023 | -0.023 | 10.757 | 0.09 | 0.06 | 0.08 | 0.18 | 0.17 | 0.1 |
| 1 | 14838731 | rs10927453 | 0.963 | 1.46 | -0.874 | 0.713 | 0.013 | 0.042 | 0.042 | 0 | 0.6 | 0.28 | 0.45 | 0.22 | 0.59 | 0.66 |
| 1 | 14841189 | rs11807837 | -0.598 | 1.384 | -0.844 | 0.823 | 0.017 | -0.065 | -0.065 | 0 | 0.77 | 0.87 | 0.73 | 0.56 | 0.82 | 0.85 |
| 1 | 14842486 | rs12733185 | 0.581 | 1.607 | -1.036 | -0.007 | 0.003 | 0.015 | 0.015 | 0 | 0.58 | 0.35 | 0.43 | 0.2 | 0.56 | 0.65 |
| 1 | 14845459 | rs12138326 | -0.502 | 1.67 | -1.433 | 0.199 | -0.066 | -0.023 | -0.023 | 0 | 0.18 | 0.07 | 0.17 | 0.2 | 0.31 | 0.12 |
| 1 | 14849088 | rs3892594 | 0.535 | 1.137 | -1.285 | -0.146 | -0.007 | 0.116 | 0.116 | 1.132 | 0.51 | 0.27 | 0.35 | 0.2 | 0.48 | 0.64 |
| 1 | 14849996 | rs6673698 | -0.594 | 2.026 | -1.529 | -0.062 | -0.085 | -0.019 | -0.019 | 0 | 0.22 | 0.07 | 0.18 | 0.24 | 0.33 | 0.12 |
| 1 | 14850351 | rs7526119 | 0.116 | 2.394 | -0.829 | 0.028 | 0.061 | 0.001 | 0.001 | 0.413 | 0.89 | 0.78 | 0.79 | 0.83 | 0.77 | 0.85 |
| 1 | 14853297 | rs12044865 | 0.275 | 1.41 | -0.982 | 0.024 | -0.009 | 0.05 | 0.05 | 0 | 0.46 | 0.27 | 0.38 | 0.36 | 0.5 | 0.52 |
| 1 | 14857643 | rs2105239 | 0.054 | 1.296 | -0.992 | 0.007 | 0.008 | 0.059 | 0.059 | 0 | 0.51 | 0.3 | 0.4 | 0.38 | 0.5 | 0.53 |
| 1 | 14861570 | rs10927459 | 0.781 | 1.635 | -1.004 | 0.108 | 0.029 | 0.065 | 0.065 | 0 | 0.34 | 0.21 | 0.23 | 0.16 | 0.26 | 0.39 |
| 1 | 14863561 | rs1890784 | 0.845 | 1.81 | -1.124 | 0.15 | -0.068 | -0.053 | -0.053 | 0 | 0.1 | 0.01 | 0.13 | 0.2 | 0.22 | 0.11 |
| 1 | 14870568 | rs10927460 | -0.196 | 0.931 | -0.886 | 0.108 | 0.007 | -0.021 | -0.021 | 0 | 0.2 | 0.27 | 0.18 | 0.18 | 0.2 | 0.24 |
| 1 | 14876032 | rs4385705 | 0.092 | 1.073 | -1.433 | 0.331 | -0.008 | 0.018 | 0.018 | 0 | 0.47 | 0.33 | 0.38 | 0.43 | 0.49 | 0.5 |
| 1 | 14883844 | rs17403250 | -0.583 | 1.438 | -1.245 | 0.089 | -0.001 | -0.003 | -0.003 | 0 | 0.21 | 0.18 | 0.25 | 0.07 | 0.27 | 0.19 |
| 1 | 14890071 | rs10927466 | -0.472 | 1.34 | -1.763 | 0.739 | -0.035 | -0.08 | -0.08 | 0.007 | 0.24 | 0.08 | 0.4 | 0.58 | 0.28 | 0.35 |
| 1 | 14901191 | rs10927468 | -0.39 | 1.149 | -1.169 | 0.688 | 0.001 | 0.009 | 0.009 | 0.179 | 0.35 | 0.44 | 0.33 | 0.37 | 0.34 | 0.33 |
| 1 | 14902252 | rs7527439 | -0.061 | 1.489 | -1.067 | 0.562 | 0.009 | -0.001 | -0.001 | 1.495 | 0.06 | 0.06 | 0.08 | 0.04 | 0.12 | 0.07 |
| 1 | 14908073 | rs2103694 | -0.128 | 0.553 | -0.258 | -0.043 | 0 | -0.015 | -0.015 | 0 | 0.15 | 0.13 | 0.1 | 0.02 | 0.14 | 0.12 |
| 1 | 14910920 | rs6685406 | 0.131 | 1.027 | -0.602 | 0.203 | -0.039 | -0.053 | -0.053 | 0.361 | 0.22 | 0.14 | 0.49 | 0.42 | 0.36 | 0.34 |

| | | | | | | | | | | | | | | | | |
|---|---|---|---|---|---|---|---|---|---|---|---|---|---|---|---|---|
| 1 | 14915147 | rs4661292 | -0.839 | 0.21 | -0.266 | 0.232 | -0.012 | 0.006 | 0.006 | 0.11 | 0.69 | 0.49 | 0.73 | 0.65 | 0.76 | 0.72 |
| 1 | 14934220 | rs10803289 | -0.904 | 0.14 | -0.13 | 0.184 | -0.017 | 0.004 | 0.004 | 0.037 | 0.69 | 0.6 | 0.73 | 0.68 | 0.76 | 0.71 |
| 1 | 14938803 | rs693191 | -0.956 | 0.122 | 0.202 | 0.105 | 0.017 | 0.076 | 0.076 | 1.631 | 0.08 | 0.42 | 0.05 | 0.07 | 0.02 | 0.11 |
| 1 | 14947920 | rs11579756 | -0.125 | 0.272 | -0.166 | 0.047 | 0.013 | 0.012 | 0.012 | 0 | 0.37 | 0.64 | 0.2 | 0.34 | 0.26 | 0.34 |
| 1 | 14961083 | rs16850635 | -0.225 | 0.613 | -0.093 | -0.212 | 0.012 | -0.052 | -0.052 | 0.033 | 0.32 | 0.45 | 0.21 | 0.18 | 0.32 | 0.34 |
| 1 | 14965581 | rs2294887 | 0.038 | 0.311 | -0.118 | -0.172 | -0.003 | -0.007 | -0.007 | 0 | 0.28 | 0.25 | 0.18 | 0.21 | 0.22 | 0.31 |
| 1 | 14967039 | rs10927475 | 0.469 | 0.294 | -0.504 | -0.261 | 0.005 | 0.014 | 0.014 | 0 | 0.23 | 0.13 | 0.15 | 0.21 | 0.18 | 0.31 |
| 1 | 14972976 | rs34645579 | 0.587 | 0.296 | -0.503 | -0.27 | 0.004 | 0.012 | 0.012 | 0 | 0.22 | 0.12 | 0.15 | 0.21 | 0.18 | 0.27 |
| 1 | 14978291 | rs6699411 | -0.416 | 0.208 | -0.278 | -0.337 | 0.002 | 0.003 | 0.003 | 0 | 0.33 | 0.3 | 0.19 | 0.25 | 0.24 | 0.38 |
| 1 | 14981920 | rs761191 | -0.295 | -0.007 | -0.701 | -0.412 | -0.002 | -0.001 | -0.001 | 0 | 0.39 | 0.59 | 0.6 | 0.57 | 0.51 | 0.44 |
| 1 | 14994487 | rs1028389 | 0.445 | 0.041 | -0.746 | -0.342 | 0.008 | 0.014 | 0.014 | 0 | 0.25 | 0.21 | 0.15 | 0.21 | 0.19 | 0.28 |
| 1 | 14997933 | rs3820066 | 0.871 | -0.242 | -0.031 | -0.417 | 0.017 | 0.065 | 0.065 | 2.515 | 0.36 | 0.26 | 0.27 | 0.19 | 0.29 | 0.18 |
| 1 | 14998872 | rs6701296 | -0.298 | 0.026 | 0.267 | 0.151 | 0.001 | -0.001 | -0.001 | 0.004 | 0.47 | 0.43 | 0.36 | 0.55 | 0.47 | 0.61 |
| 1 | 15009937 | rs2871581 | -0.16 | 0.06 | 0.633 | 0.351 | 0 | 0 | 0 | 0 | 0.47 | 0.4 | 0.37 | 0.53 | 0.5 | 0.65 |
| 1 | 15018660 | rs3737686 | -0.174 | -0.025 | 0.569 | 0.396 | -0.014 | -0.046 | -0.046 | 0 | 0.45 | 0.18 | 0.36 | 0.49 | 0.51 | 0.6 |
| 1 | 15032340 | rs7534010 | -1.153 | -0.024 | 0.343 | 0.226 | -0.013 | 0.128 | 0.128 | 1.189 | 0.59 | 0.82 | 0.7 | 0.79 | 0.58 | 0.77 |
| 1 | 15033508 | rs4661514 | 0.5 | 0.08 | 0.214 | 0.308 | -0.072 | -0.071 | -0.071 | 0 | 0.36 | 0.13 | 0.28 | 0.48 | 0.43 | 0.54 |
| 1 | 15036694 | rs10803304 | 0.307 | 0.099 | 0.23 | 0.309 | -0.069 | -0.071 | -0.071 | 0 | 0.38 | 0.16 | 0.3 | 0.49 | 0.44 | 0.54 |
| 1 | 15039411 | rs7516095 | -0.721 | 0.103 | 0.283 | 0.488 | -0.028 | 0.051 | 0.051 | 0.047 | 0.2 | 0.39 | 0.39 | 0.26 | 0.16 | 0.21 |
| 1 | 15048326 | rs12758392 | -0.969 | 0.88 | -0.451 | 0.756 | 0.005 | 0.003 | 0.003 | 0 | 0.12 | 0.1 | 0.12 | 0.18 | 0.26 | 0.1 |
| 1 | 15048871 | rs12045403 | 1.129 | 0.329 | -0.669 | 0.395 | 0.034 | 0.09 | 0.09 | 1.338 | 0.38 | 0.17 | 0.29 | 0.22 | 0.33 | 0.19 |
| 1 | 15050524 | rs4661517 | -0.973 | 0.465 | -0.7 | 0.401 | 0.033 | 0.136 | 0.136 | 2.995 | 0.57 | 0.84 | 0.68 | 0.78 | 0.62 | 0.79 |
| 1 | 15056175 | rs2050126 | 0.503 | 0.946 | -0.505 | 0.561 | -0.058 | 0.013 | 0.013 | 0 | 0.29 | 0.1 | 0.19 | 0.28 | 0.31 | 0.41 |
| 1 | 15058013 | rs7532344 | -0.067 | 0.433 | 0.341 | 0.425 | 0.018 | 0.153 | 0.153 | 0 | 0.89 | 0.63 | 0.67 | 0.75 | 0.88 | 0.86 |
| 1 | 15067407 | rs1997762 | -0.172 | 0.539 | 0.313 | -0.426 | 0.003 | 0.006 | 0.006 | 0 | 0.51 | 0.55 | 0.35 | 0.23 | 0.43 | 0.36 |
| 1 | 15068997 | rs6660333 | 0.902 | 0.328 | 0.148 | -0.505 | -0.049 | 0.056 | 0.056 | 0.453 | 0.36 | 0.09 | 0.24 | 0.19 | 0.35 | 0.22 |
| 1 | 15072614 | rs6429674 | 0.285 | 0.199 | 0.322 | -0.45 | -0.001 | -0.005 | -0.005 | 0 | 0.43 | 0.51 | 0.34 | 0.25 | 0.4 | 0.3 |
| 1 | 15076305 | rs10927530 | -0.583 | 0.223 | 0.169 | -0.368 | 0 | 0.007 | 0.007 | 0 | 0.12 | 0.08 | 0.06 | 0.04 | 0.03 | 0.14 |
| 1 | 15077567 | rs6661767 | -0.353 | 0.692 | 0.113 | -0.501 | 0.003 | 0.038 | 0.038 | 0.04 | 0.51 | 0.5 | 0.38 | 0.24 | 0.46 | 0.33 |
| 1 | 15080560 | rs4357518 | -0.304 | 1.004 | -0.422 | -0.077 | -0.001 | 0.004 | 0.004 | 1.385 | 0.44 | 0.38 | 0.34 | 0.22 | 0.53 | 0.27 |
| 1 | 15089767 | rs4543785 | -0.083 | 1.032 | -0.412 | 0.146 | 0.006 | -0.014 | -0.014 | 0.05 | 0.38 | 0.39 | 0.31 | 0.21 | 0.49 | 0.25 |
| 1 | 15093562 | rs732469 | 0.674 | 0.758 | -0.065 | 0.84 | 0.003 | 0.087 | 0.087 | 0.244 | 0.92 | 0.74 | 0.68 | 0.82 | 0.94 | 0.88 |
| 1 | 15101472 | rs4661303 | -0.956 | 1.101 | -0.342 | 0.55 | -0.004 | 0.004 | 0.004 | 0 | 0.11 | 0.13 | 0.08 | 0.06 | 0.12 | 0.12 |
| 1 | 15102393 | rs4661529 | -1.46 | 1.337 | -0.129 | 0.347 | 0.008 | -0.015 | -0.015 | 0.087 | 0.15 | 0.22 | 0.11 | 0.06 | 0.19 | 0.15 |
| 1 | 15111927 | rs804125 | -0.078 | 1.225 | 0.232 | 0.211 | -0.008 | -0.002 | -0.002 | 0 | 0.73 | 0.61 | 0.67 | 0.68 | 0.75 | 0.77 |
| 1 | 15117060 | rs1721829 | -0.525 | 1.078 | 0.461 | 0.115 | 0 | -0.003 | -0.003 | 0 | 0.79 | 0.78 | 0.79 | 0.77 | 0.78 | 0.84 |
| 1 | 15118081 | rs1721828 | 0.624 | 0.927 | 0.346 | -0.149 | -0.001 | -0.008 | -0.008 | 0 | 0.66 | 0.63 | 0.54 | 0.7 | 0.7 | 0.69 |
| 1 | 15120526 | rs1737352 | 0.941 | -0.069 | 0.086 | -0.807 | -0.009 | 0.058 | 0.058 | 0.023 | 0.43 | 0.3 | 0.53 | 0.69 | 0.57 | 0.59 |
| 1 | 15123100 | rs7525851 | 1.609 | 0.913 | -0.043 | -0.691 | -0.008 | 0.072 | 0.072 | 0 | 0.31 | 0.15 | 0.26 | 0.1 | 0.35 | 0.31 |
| 1 | 15123189 | rs1737350 | -1.121 | 0.956 | 0.383 | -0.693 | 0.035 | 0.005 | 0.005 | 0.014 | 0.26 | 0.45 | 0.41 | 0.67 | 0.3 | 0.37 |
| 1 | 15124318 | rs1737349 | 0.259 | 0.384 | 0.486 | -0.615 | 0.014 | 0.033 | 0.033 | 0 | 0.52 | 0.62 | 0.57 | 0.63 | 0.63 | 0.6 |
| 1 | 15125699 | rs804130 | -0.153 | 0.372 | 0.435 | -0.611 | -0.001 | 0.011 | 0.011 | 0 | 0.64 | 0.68 | 0.61 | 0.71 | 0.67 | 0.64 |
| 1 | 15127627 | rs12568933 | -0.488 | 0.128 | 0.55 | 0.598 | 0.003 | 0.049 | 0.049 | 0 | 0.22 | 0.35 | 0.26 | 0.42 | 0.24 | 0.32 |
| 1 | 15127743 | rs12563293 | -0.4 | 0.168 | 0.57 | 0.922 | 0.004 | 0.034 | 0.034 | 0 | 0.21 | 0.34 | 0.24 | 0.36 | 0.23 | 0.32 |
| 1 | 15133668 | rs804136 | -0.697 | 0.665 | -0.082 | 0.463 | -0.031 | 0.024 | 0.024 | 0.662 | 0.7 | 0.79 | 0.71 | 0.71 | 0.65 | 0.72 |
| 1 | 15138738 | rs3845590 | -1.189 | 0.617 | 0.444 | 0.771 | -0.008 | -0.023 | -0.023 | 0 | 0.31 | 0.69 | 0.45 | 0.51 | 0.35 | 0.3 |
| 1 | 15142300 | rs6658818 | -0.31 | 1.032 | 0.502 | 0.291 | 0.043 | -0.056 | -0.056 | 0 | 0.17 | 0.37 | 0.21 | 0.04 | 0.24 | 0.16 |
| 1 | 15144305 | rs10489354 | -0.188 | 1.146 | -0.413 | 0.245 | 0.061 | -0.076 | -0.076 | 0 | 0.17 | 0.36 | 0.21 | 0.03 | 0.24 | 0.14 |
| 1 | 15150914 | rs804134 | 0.139 | 0.431 | -1.525 | 0.644 | 0.023 | 0.03 | 0.03 | 0 | 0.36 | 0.1 | 0.26 | 0.31 | 0.3 | 0.44 |
| 1 | 15152755 | rs12127407 | -0.41 | 0.747 | -0.828 | 1.066 | -0.007 | -0.003 | -0.003 | 0.041 | 0.06 | 0.03 | 0.12 | 0.07 | 0.07 | 0.08 |
| 1 | 15154630 | rs12134149 | -0.396 | -0.257 | -0.942 | 0.676 | 0.044 | 0.028 | 0.028 | 4.974 | 0.37 | 0.3 | 0.16 | 0.24 | 0.24 | 0.24 |
| 1 | 15154825 | rs12123938 | -0.247 | -0.491 | -0.657 | 0.584 | 0.025 | 0.004 | 0.004 | 7.028 | 0.36 | 0.31 | 0.16 | 0.3 | 0.24 | 0.23 |
| 1 | 15156736 | rs2103117 | -1.683 | 0.814 | 0.48 | 0.408 | 0.044 | -0.046 | -0.046 | 0 | 0.08 | 0.17 | 0.16 | 0.01 | 0.1 | 0.04 |
| 1 | 15165021 | rs7546946 | -1.761 | 0.495 | 0.429 | 0.297 | -0.02 | -0.079 | -0.079 | 0 | 0.13 | 0.38 | 0.07 | 0 | 0.07 | 0.07 |
| 1 | 15167003 | rs11585327 | -1.752 | 0.573 | 0.418 | 0.297 | 0.002 | -0.108 | -0.108 | 0.032 | 0.16 | 0.4 | 0.2 | 0.01 | 0.12 | 0.09 |
| 1 | 15182153 | rs3856273 | -1.272 | 0.898 | -0.093 | 0.891 | 0.014 | 0.015 | 0.015 | 0.018 | 0.14 | 0.2 | 0.13 | 0.21 | 0.17 | 0.13 |
| 1 | 15183309 | rs804129 | 0.04 | 0.796 | -0.276 | 0.55 | -0.001 | 0 | 0 | 0.399 | 0.63 | 0.71 | 0.57 | 0.64 | 0.65 | 0.62 |
| 1 | 15185229 | rs2103116 | -0.274 | 0.427 | -0.554 | -0.76 | 0.017 | 0.083 | 0.083 | 0 | 0.45 | 0.73 | 0.3 | 0.1 | 0.38 | 0.29 |
| 1 | 15188019 | rs10803320 | -0.249 | 0.384 | -0.574 | -0.788 | 0.016 | 0.076 | 0.076 | 0 | 0.45 | 0.72 | 0.29 | 0.08 | 0.38 | 0.29 |
| 1 | 15189088 | rs4661533 | -0.659 | 0.335 | -0.851 | -0.824 | -0.004 | 0.094 | 0.094 | 0 | 0.54 | 0.75 | 0.39 | 0.18 | 0.45 | 0.33 |
| 1 | 15189231 | rs4661534 | -0.51 | 0.41 | -0.758 | -1.094 | 0.009 | 0.04 | 0.04 | 0 | 0.21 | 0.11 | 0.16 | 0.11 | 0.18 | 0.15 |
| 1 | 15189299 | rs3845596 | -0.863 | 0.435 | -0.787 | -1.074 | -0.003 | 0.099 | 0.099 | 0 | 0.54 | 0.74 | 0.36 | 0.14 | 0.45 | 0.31 |
| 1 | 15189693 | rs3845598 | 0.656 | 0.462 | -0.166 | -1.1 | -0.004 | -0.097 | -0.097 | 1.563 | 0.31 | 0.38 | 0.18 | 0.03 | 0.27 | 0.16 |
| 1 | 15190264 | rs3765380 | -0.846 | 0.33 | 0.275 | -0.958 | -0.012 | -0.073 | -0.073 | 0 | 0.21 | 0.31 | 0.17 | 0.06 | 0.17 | 0.12 |
| 1 | 15191252 | rs12734818 | 0.554 | 0.527 | -0.005 | -0.751 | -0.003 | 0.002 | 0.002 | 0.176 | 0.06 | 0.04 | 0.04 | 0.06 | 0.07 | 0.14 |
| 1 | 15192285 | rs2462190 | 0.036 | 1.173 | 0.269 | -0.176 | -0.001 | -0.003 | -0.003 | 3.013 | 0.29 | 0.26 | 0.49 | 0.58 | 0.25 | 0.37 |
| 1 | 15192559 | rs2803399 | -0.304 | 1.332 | 0.047 | -0.517 | -0.009 | -0.07 | -0.07 | 0 | 0.62 | 0.41 | 0.8 | 0.95 | 0.72 | 0.83 |
| 1 | 15192740 | rs2807568 | -0.072 | 1.373 | 0.032 | -0.513 | 0.001 | -0.047 | -0.047 | 0.008 | 0.59 | 0.4 | 0.75 | 0.93 | 0.68 | 0.81 |
| 1 | 15192916 | rs4661539 | 0.721 | 1.233 | -0.014 | -0.403 | -0.007 | -0.003 | -0.003 | 0.229 | 0.35 | 0.32 | 0.33 | 0.4 | 0.47 | 0.46 |

| | | | | | | | | | | | | | | | | |
|---|---|---|---|---|---|---|---|---|---|---|---|---|---|---|---|---|
| 1 | 15193585 | rs2807545 | -0.555 | 1.178 | 0.098 | -0.474 | -0.007 | -0.066 | -0.066 | 0 | 0.63 | 0.62 | 0.81 | 0.95 | 0.71 | 0.83 |
| 1 | 15194562 | rs7416912 | 1.135 | 1.741 | -0.396 | -0.268 | -0.023 | 0.01 | 0.01 | 0 | 0.31 | 0.08 | 0.2 | 0.25 | 0.31 | 0.32 |
| 1 | 15194652 | rs2803397 | 0.891 | 0.516 | -0.542 | 0.193 | -0.068 | -0.114 | -0.114 | 0.021 | 0.22 | 0.02 | 0.42 | 0.34 | 0.23 | 0.29 |
| 1 | 15197814 | rs12564579 | 1.094 | 0.626 | -0.425 | -0.499 | 0 | 0 | 0 | 3.963 | 0.33 | 0.33 | 0.31 | 0.34 | 0.42 | 0.45 |
| 1 | 15202204 | rs4661540 | 0.239 | 0.143 | -0.571 | -0.242 | 0.006 | -0.043 | -0.043 | 0 | 0.26 | 0.16 | 0.46 | 0.55 | 0.2 | 0.34 |
| 1 | 15207952 | rs2690022 | -0.294 | -0.297 | -0.226 | -0.208 | -0.015 | -0.08 | -0.08 | 0 | 0.67 | 0.66 | 0.85 | 0.95 | 0.76 | 0.82 |
| 1 | 15208212 | rs2801178 | 0.171 | -0.062 | -0.114 | 0.114 | -0.008 | -0.026 | -0.026 | 0.307 | 0.21 | 0.16 | 0.44 | 0.59 | 0.2 | 0.31 |
| 1 | 15211824 | rs2690024 | 0.137 | -0.911 | 0.152 | -0.392 | -0.024 | -0.095 | -0.095 | 0.424 | 0.73 | 0.65 | 0.86 | 0.95 | 0.8 | 0.85 |
| 1 | 15216378 | rs10927567 | -0.746 | -0.784 | 0.33 | 0.112 | -0.025 | -0.032 | -0.032 | 0 | 0.17 | 0.29 | 0.1 | 0.1 | 0.13 | 0.07 |
| 1 | 15216489 | rs12029721 | -0.357 | -0.954 | 0.389 | 0.659 | -0.015 | 0.039 | 0.039 | 0 | 0.1 | 0.14 | 0.21 | 0.18 | 0.05 | 0.11 |
| 1 | 15226055 | rs2807555 | 1.115 | -1.115 | 1.316 | 0.653 | 0.064 | 0.142 | 0.142 | 0.008 | 0.93 | 0.66 | 0.94 | 0.87 | 0.95 | 0.91 |
| 1 | 15229101 | rs4661543 | 0.262 | -1.214 | 0.776 | 0.483 | -0.003 | -0.001 | -0.001 | 9.781 | 0.85 | 0.87 | 0.9 | 0.87 | 0.87 | 0.91 |
| 1 | 15234029 | rs2807562 | -0.39 | -1.595 | 0.756 | 0.039 | -0.027 | -0.044 | -0.044 | 0.917 | 0.28 | 0.47 | 0.28 | 0.18 | 0.22 | 0.23 |
| 1 | 15244761 | rs2489772 | 0.616 | -1.633 | 0.583 | 0.616 | -0.017 | 0.035 | 0.035 | 0 | 0.66 | 0.61 | 0.77 | 0.56 | 0.74 | 0.56 |
| 1 | 15249739 | rs1404061 | -0.12 | -1.738 | 0.302 | 0.012 | -0.018 | -0.032 | -0.032 | 0 | 0.62 | 0.58 | 0.7 | 0.85 | 0.78 | 0.83 |
| 1 | 15257211 | rs12141413 | -0.66 | -1.387 | 0.347 | 0.807 | -0.013 | 0.154 | 0.154 | 0 | 0.17 | 0.55 | 0.18 | 0.4 | 0.1 | 0.35 |
| 1 | 15257426 | rs12568821 | 0.165 | -1.341 | 0.099 | 0.897 | -0.01 | -0.045 | -0.045 | 0 | 0.12 | 0.06 | 0.13 | 0.26 | 0.09 | 0.27 |
| 1 | 15258425 | rs10927583 | -1.101 | -1.335 | -0.077 | 0.719 | -0.001 | 0.002 | 0.002 | 0 | 0.34 | 0.65 | 0.29 | 0.47 | 0.26 | 0.47 |
| 1 | 15264270 | rs12410007 | 0.257 | -1.877 | -0.326 | 0.165 | -0.033 | -0.039 | -0.039 | 0 | 0.26 | 0.29 | 0.26 | 0.12 | 0.14 | 0.13 |
| 1 | 15265404 | rs6693394 | -1.138 | -1.778 | -1.285 | 0.402 | -0.054 | -0.156 | -0.156 | 0 | 0.58 | 0.95 | 0.47 | 0.49 | 0.38 | 0.51 |
| 1 | 15267837 | rs12741828 | 0.006 | -1.301 | -0.711 | 0.485 | -0.014 | 0.006 | 0.006 | 0 | 0.13 | 0.08 | 0.1 | 0.09 | 0.15 | 0.1 |
| 1 | 15273097 | rs7546786 | -1.018 | -1.378 | 0.062 | 0.653 | 0.003 | 0.031 | 0.031 | 0 | 0.27 | 0.49 | 0.22 | 0.28 | 0.24 | 0.32 |
| 1 | 15273115 | rs6429695 | -1.409 | -1.946 | -0.573 | 0.446 | -0.068 | -0.135 | -0.135 | 0.993 | 0.58 | 0.94 | 0.48 | 0.53 | 0.39 | 0.51 |
| 1 | 15273181 | rs6694029 | 0.104 | -2.06 | 0.226 | 0.375 | -0.075 | -0.015 | -0.015 | 0 | 0.29 | 0.44 | 0.26 | 0.26 | 0.14 | 0.15 |
| 1 | 15276526 | rs16851049 | -0.339 | -2.105 | 0.409 | 0.096 | 0.006 | 0.008 | 0.008 | 0.081 | 0.16 | 0.12 | 0.22 | 0.13 | 0.13 | 0.22 |
| 1 | 15277710 | rs4661548 | -1.604 | -2.535 | 0.606 | -0.176 | -0.009 | -0.004 | -0.004 | 0 | 0.66 | 0.74 | 0.49 | 0.43 | 0.46 | 0.47 |
| 1 | 15278427 | rs10927593 | 0.348 | -2.802 | 0.631 | -0.646 | -0.069 | -0.08 | -0.08 | 0 | 0.28 | 0.63 | 0.12 | 0.11 | 0.09 | 0.11 |
| 1 | 15280823 | rs7522410 | 0 | -2.646 | 0.55 | -0.468 | -0.033 | -0.024 | -0.024 | 0 | 0.23 | 0.09 | 0.14 | 0.21 | 0.23 | 0.13 |
| 1 | 15283217 | rs7514440 | -0.834 | -2.726 | 0.463 | -0.488 | 0.005 | 0.012 | 0.012 | 1.085 | 0.3 | 0.77 | 0.21 | 0.21 | 0.24 | 0.18 |
| 1 | 15286096 | rs12068752 | -1.342 | -2.455 | 0.67 | -0.372 | -0.014 | -0.016 | -0.016 | 0.037 | 0.06 | 0.68 | 0.07 | 0 | 0 | 0.01 |
| 1 | 15286805 | rs6429696 | -1.619 | -2.383 | 0.382 | -0.374 | -0.196 | -0.121 | -0.121 | 0 | 0.68 | 0.98 | 0.52 | 0.39 | 0.48 | 0.51 |
| 1 | 15294031 | rs10754875 | -0.456 | -1.93 | -0.088 | -0.478 | -0.026 | -0.024 | -0.024 | 0 | 0.44 | 0.55 | 0.35 | 0.3 | 0.26 | 0.37 |
| 1 | 15303919 | rs10927611 | -0.551 | -0.913 | -0.115 | -0.487 | 0.006 | -0.054 | -0.054 | 0.262 | 0.21 | 0.29 | 0.13 | 0.07 | 0.18 | 0.1 |
| 1 | 15303970 | rs12084008 | -0.539 | -0.907 | -0.112 | -0.484 | 0.007 | -0.054 | -0.054 | 0.517 | 0.21 | 0.29 | 0.13 | 0.07 | 0.18 | 0.1 |
| 1 | 15304859 | rs4661553 | 0.076 | -1.029 | -0.333 | -0.561 | 0.001 | 0.037 | 0.037 | 0 | 0.13 | 0.08 | 0.2 | 0.05 | 0.12 | 0.24 |
| 1 | 15307478 | rs7522712 | -1.157 | -1.001 | -0.31 | -0.456 | 0 | -0.028 | -0.028 | 0 | 0.34 | 0.33 | 0.33 | 0.12 | 0.31 | 0.35 |
| 1 | 15315176 | rs7531035 | -0.123 | -0.804 | -0.642 | -0.247 | -0.004 | -0.012 | -0.012 | 0 | 0.16 | 0.15 | 0.1 | 0 | 0.16 | 0.06 |
| 1 | 15317488 | rs12409399 | -0.344 | -0.803 | -1.236 | -0.01 | 0.007 | 0.005 | 0.005 | 0 | 0.39 | 0.65 | 0.46 | 0.28 | 0.28 | 0.45 |
| 1 | 15320285 | rs7525238 | -1.338 | -0.851 | -1.933 | 0.226 | -0.166 | -0.129 | -0.129 | 3.87 | 0.63 | 0.97 | 0.5 | 0.46 | 0.49 | 0.6 |
| 1 | 15320449 | rs4661310 | -0.141 | -0.826 | -0.964 | 0.137 | 0 | 0.056 | 0.056 | 0 | 0.31 | 0.13 | 0.29 | 0.18 | 0.27 | 0.32 |
| 1 | 15323416 | rs1316257 | -1.007 | -1.23 | -0.51 | 0.554 | 0.012 | 0.009 | 0.009 | 0 | 0.43 | 0.58 | 0.22 | 0.28 | 0.24 | 0.41 |
| 1 | 15328151 | rs7529561 | -0.468 | -1.399 | -0.622 | 0.788 | 0.025 | -0.025 | -0.025 | 0 | 0.24 | 0.12 | 0.24 | 0.32 | 0.18 | 0.36 |
| 1 | 15328697 | rs16851112 | 0.282 | -1.587 | -0.768 | 0.811 | 0.023 | -0.016 | -0.016 | 1.73 | 0.16 | 0.07 | 0.19 | 0.16 | 0.11 | 0.2 |
| 1 | 15329987 | rs10754876 | -1.74 | -1.732 | -1.042 | 0.595 | 0.021 | -0.055 | -0.055 | 0 | 0.56 | 0.86 | 0.45 | 0.44 | 0.37 | 0.53 |
| 1 | 15330300 | rs10803338 | -0.239 | -1.833 | -0.209 | 0.499 | -0.012 | -0.066 | -0.066 | 0.145 | 0.28 | 0.35 | 0.16 | 0.03 | 0.18 | 0.16 |
| 1 | 15331802 | rs10927627 | -1.66 | -1.244 | -0.654 | 0.786 | -0.027 | -0.058 | -0.058 | 0 | 0.58 | 0.85 | 0.48 | 0.47 | 0.42 | 0.54 |
| 1 | 15332823 | rs2311975 | -1.704 | -1.283 | -0.952 | 0.757 | -0.109 | -0.16 | -0.16 | 2.256 | 0.58 | 0.96 | 0.49 | 0.46 | 0.42 | 0.54 |
| 1 | 15334456 | rs6665532 | -0.327 | -1.354 | -0.2 | 0.436 | -0.006 | -0.017 | -0.017 | 0 | 0.25 | 0.28 | 0.14 | 0.11 | 0.18 | 0.12 |
| 1 | 15334782 | rs6681053 | -0.966 | -0.933 | 0.445 | 0.982 | 0.029 | 0.077 | 0.077 | 0 | 0.13 | 0.31 | 0.14 | 0.23 | 0.13 | 0.17 |
| 1 | 15335548 | rs9783012 | 0.221 | -1.27 | -0.09 | 1.085 | 0.025 | -0.056 | -0.056 | 0 | 0.12 | 0.04 | 0.15 | 0.24 | 0.1 | 0.25 |
| 1 | 15336720 | rs6672536 | -0.815 | -1.299 | 0.642 | 0.478 | -0.008 | 0.001 | 0.001 | 0 | 0.13 | 0.33 | 0.11 | 0.13 | 0.1 | 0.08 |
| 1 | 15337966 | rs7554543 | -0.294 | -1.434 | 0.591 | 0.031 | -0.001 | -0.105 | -0.105 | 0 | 0.21 | 0.31 | 0.14 | 0 | 0.17 | 0.08 |
| 1 | 15338903 | rs11578349 | -0.917 | -1.359 | 0.839 | 0.329 | 0.021 | 0.022 | 0.022 | 0.074 | 0.08 | 0.31 | 0.09 | 0.09 | 0.06 | 0.06 |
| 1 | 15339960 | rs6429703 | 0.675 | -1.296 | 0.512 | 0.526 | -0.005 | 0 | 0 | 0 | 0.83 | 0.8 | 0.81 | 0.83 | 0.87 | 0.84 |
| 1 | 15341055 | rs4661561 | -0.562 | -1.29 | 0.431 | 0.48 | 0.006 | 0.01 | 0.01 | 0 | 0.92 | 0.95 | 0.95 | 1 | 0.93 | 0.94 |
| 1 | 15341413 | rs4661563 | 1.783 | -1.934 | -0.438 | 0.199 | -0.127 | 0.143 | 0.143 | 0 | 0.39 | 0.08 | 0.49 | 0.23 | 0.49 | 0.51 |
| 1 | 15343094 | rs10157110 | 0.656 | -1.341 | -1.084 | -0.381 | -0.007 | 0.006 | 0.006 | 0 | 0.23 | 0.14 | 0.35 | 0.24 | 0.21 | 0.26 |
| 1 | 15344403 | rs869712 | -0.769 | -1.539 | -0.394 | -0.169 | 0.001 | -0.023 | -0.023 | 0 | 0.59 | 0.51 | 0.58 | 0.69 | 0.54 | 0.56 |
| 1 | 15344936 | rs938252 | -0.52 | -1.372 | 0.327 | -0.114 | -0.001 | 0.043 | 0.043 | 0 | 0.89 | 0.58 | 0.88 | 0.88 | 0.93 | 0.92 |
| 1 | 15346330 | rs10927631 | 0.537 | -1.282 | -0.404 | -0.486 | 0.005 | -0.031 | -0.031 | 0 | 0.12 | 0.14 | 0.12 | 0 | 0.18 | 0.07 |
| 1 | 15347640 | rs6667220 | 0.459 | -1.647 | -0.456 | -0.19 | 0.036 | -0.024 | -0.024 | 0.015 | 0.64 | 0.76 | 0.84 | 0.54 | 0.73 | 0.7 |
| 1 | 15347711 | rs6658629 | -0.06 | -1.617 | -0.509 | 0.036 | 0.016 | -0.023 | -0.023 | 0.032 | 0.24 | 0.18 | 0.07 | 0.4 | 0.12 | 0.21 |
| 1 | 15351283 | rs17413451 | 0.655 | -0.774 | -0.44 | -0.389 | 0.01 | -0.036 | -0.036 | 5.17 | 0.09 | 0.12 | 0.1 | 0 | 0.13 | 0.06 |
| 1 | 15352820 | rs6684496 | -0.781 | -1.207 | -0.556 | -0.504 | -0.003 | 0.015 | 0.015 | 0 | 0.49 | 0.64 | 0.55 | 0.46 | 0.47 | 0.42 |
| 1 | 15354399 | rs11803332 | -0.276 | -0.897 | -0.562 | -0.566 | 0.005 | 0.009 | 0.009 | 0 | 0.3 | 0.5 | 0.33 | 0.3 | 0.3 | 0.22 |
| 1 | 15356223 | rs6679504 | -0.878 | -0.863 | -0.892 | -0.549 | 0 | 0.018 | 0.018 | 0 | 0.5 | 0.65 | 0.55 | 0.46 | 0.47 | 0.42 |
| 1 | 15357505 | rs2175771 | 1.441 | -0.559 | -1.767 | -0.678 | -0.095 | 0.064 | 0.064 | 0 | 0.28 | 0.04 | 0.31 | 0.16 | 0.37 | 0.33 |
| 1 | 15359967 | rs4520412 | -1.215 | -0.666 | -1.549 | -0.527 | 0 | 0.003 | 0.003 | 0.057 | 0.77 | 0.75 | 0.87 | 0.63 | 0.84 | 0.74 |
| 1 | 15360808 | rs12129618 | 0.834 | -0.431 | -2.109 | -0.63 | -0.078 | 0.103 | 0.103 | 0 | 0.39 | 0.12 | 0.5 | 0.24 | 0.47 | 0.47 |

| | | | | | | | | | | | | | | | | |
|---|---|---|---|---|---|---|---|---|---|---|---|---|---|---|---|---|
| 1 | 15363223 | rs6674129 | -0.991 | 0.137 | -1.883 | -0.267 | -0.024 | -0.028 | -0.028 | 0.063 | 0.52 | 0.9 | 0.56 | 0.46 | 0.54 | 0.45 |
| 1 | 15365648 | rs12139386 | -0.21 | 1.204 | -1.436 | -0.82 | 0.003 | -0.011 | -0.011 | 0.007 | 0.18 | 0.23 | 0.17 | 0.07 | 0.25 | 0.12 |
| 1 | 15368553 | rs10927653 | 0.498 | 1.272 | -1.056 | -0.472 | 0.004 | -0.026 | -0.026 | 0 | 0.16 | 0.19 | 0.2 | 0.09 | 0.16 | 0.18 |
| 1 | 15371332 | rs6704367 | -0.249 | 1.287 | -0.895 | 0.041 | 0.014 | 0.021 | 0.021 | 0 | 0.06 | 0.15 | 0.06 | 0.08 | 0.07 | 0.06 |
| 1 | 15371724 | rs578771 | -0.498 | 1.51 | -1.606 | -0.189 | -0.07 | 0.024 | 0.024 | 0 | 0.78 | 0.95 | 0.71 | 0.85 | 0.68 | 0.7 |
| 1 | 15371782 | rs12757909 | -0.723 | 0.934 | -0.875 | -0.063 | 0.006 | -0.04 | -0.04 | 0 | 0.08 | 0.19 | 0.05 | 0 | 0.06 | 0.04 |
| 1 | 15373038 | rs10754879 | -0.494 | 0.824 | -1.395 | 0.101 | -0.004 | 0.015 | 0.015 | 0 | 0.43 | 0.26 | 0.47 | 0.36 | 0.57 | 0.41 |
| 1 | 15373378 | rs10927657 | 0.018 | -0.129 | -0.591 | -0.702 | -0.016 | -0.073 | -0.073 | 0.017 | 0.44 | 0.47 | 0.23 | 0.06 | 0.34 | 0.22 |
| 1 | 15376091 | rs10803341 | -1.264 | 0.548 | -0.466 | 0.759 | 0.017 | -0.07 | -0.07 | 0 | 0.2 | 0.08 | 0.19 | 0.71 | 0.18 | 0.29 |
| 1 | 15377687 | rs761288 | -2.021 | 0.277 | 0.138 | 0.479 | -0.039 | -0.009 | -0.009 | 0 | 0.35 | 0.51 | 0.28 | 0.75 | 0.25 | 0.42 |
| 1 | 15386628 | rs2073091 | -1.545 | 0.333 | 0.164 | 0.743 | 0.004 | -0.055 | -0.055 | 0.101 | 0.21 | 0.12 | 0.19 | 0.62 | 0.17 | 0.29 |
| 1 | 15388673 | rs3795757 | -0.484 | 1.24 | -0.002 | 0.619 | -0.12 | 0.008 | 0.008 | 0 | 0.83 | 0.98 | 0.55 | 0.94 | 0.65 | 0.79 |
| 1 | 15389862 | rs2076566 | -2.275 | 0.023 | 1.026 | 1.22 | -0.018 | 0.014 | 0.014 | 0 | 0.29 | 0.63 | 0.25 | 0.72 | 0.22 | 0.34 |
| 1 | 15392450 | rs12057431 | -1.576 | 0.077 | 1.08 | 0.953 | 0.046 | 0.116 | 0.116 | 0.087 | 0.06 | 0.35 | 0.04 | 0.07 | 0.01 | 0.08 |
| 1 | 15395881 | rs9943244 | -0.289 | 0.249 | 0.748 | 1.236 | -0.02 | -0.023 | -0.023 | 0 | 0.1 | 0.04 | 0.13 | 0.53 | 0.11 | 0.18 |
| 1 | 15401158 | rs16851241 | -0.9 | 0.173 | 0.706 | 1.178 | -0.002 | 0.006 | 0.006 | 0 | 0.14 | 0.16 | 0.07 | 0.25 | 0.1 | 0.16 |
| 1 | 15402156 | rs4661572 | -2.438 | 0.208 | 0.781 | 1.139 | 0.005 | 0.007 | 0.007 | 0 | 0.36 | 0.67 | 0.3 | 0.77 | 0.28 | 0.42 |
| 1 | 15403275 | rs742662 | -1.138 | 0.169 | 0.596 | 1.172 | -0.036 | 0.055 | 0.055 | 0 | 0.15 | 0.5 | 0.09 | 0.18 | 0.1 | 0.09 |
| 1 | 15405489 | rs1000313 | -1.963 | 0.331 | 0.082 | 1.024 | -0.019 | -0.075 | -0.075 | 0.065 | 0.29 | 0.5 | 0.23 | 0.9 | 0.23 | 0.4 |
| 1 | 15406696 | rs2175774 | -0.95 | 0.354 | 0.103 | 1.197 | 0.015 | 0.215 | 0.215 | 0 | 0.16 | 0.44 | 0.17 | 0.64 | 0.13 | 0.26 |
| 1 | 15411661 | rs6669698 | -1.709 | 0.28 | 0.087 | 1.204 | -0.027 | -0.019 | -0.019 | 0 | 0.27 | 0.48 | 0.23 | 0.82 | 0.22 | 0.33 |
| 1 | 15412415 | rs10927670 | -1.234 | 0.387 | 0.187 | 1.351 | -0.003 | 0.097 | 0.097 | 0 | 0.19 | 0.44 | 0.18 | 0.73 | 0.13 | 0.31 |
| 1 | 15414413 | rs4661575 | -1.288 | 0.667 | -0.252 | 0.832 | -0.007 | -0.044 | -0.044 | 0 | 0.38 | 0.57 | 0.29 | 0.9 | 0.31 | 0.47 |
| 1 | 15416326 | rs10803352 | -0.792 | 0.796 | -0.711 | 1.067 | -0.027 | 0.069 | 0.069 | 0 | 0.28 | 0.42 | 0.15 | 0.4 | 0.23 | 0.3 |
| 1 | 15424731 | rs3752934 | 0.597 | 1.784 | -1.343 | 0.669 | -0.044 | 0.003 | 0.003 | 0 | 0.87 | 0.92 | 0.71 | 0.89 | 0.81 | 0.8 |
| 1 | 15427117 | rs532075 | -0.476 | 1.983 | -1.923 | 0.879 | -0.041 | -0.065 | -0.065 | 0.011 | 0.29 | 0.13 | 0.38 | 0.55 | 0.35 | 0.27 |
| 1 | 15427982 | rs502918 | -0.71 | 1.89 | -1.581 | -0.117 | -0.006 | -0.028 | -0.028 | 0 | 0.59 | 0.42 | 0.61 | 0.84 | 0.59 | 0.59 |
| 1 | 15428508 | rs476859 | -0.03 | 1.685 | -1.865 | -1.014 | 0.02 | 0.069 | 0.069 | 0 | 0.43 | 0.26 | 0.49 | 0.8 | 0.37 | 0.44 |
| 1 | 15430895 | rs6695209 | 0.724 | 1.876 | -1.384 | -0.989 | 0.027 | -0.012 | -0.012 | 0.072 | 0.1 | 0.15 | 0.1 | 0.06 | 0.16 | 0.16 |
| 1 | 15431657 | rs494983 | -0.196 | 1.484 | -1.157 | -1.703 | 0.004 | -0.019 | -0.019 | 3.876 | 0.6 | 0.45 | 0.63 | 0.79 | 0.57 | 0.48 |
| 1 | 15435257 | rs496567 | 0.828 | 1.298 | -1.385 | -1.61 | 0.016 | -0.04 | -0.04 | 0 | 0.27 | 0.2 | 0.27 | 0.52 | 0.24 | 0.18 |
| 1 | 15437520 | rs10927678 | 0.989 | 1.463 | -0.906 | -1.796 | 0.007 | 0.026 | 0.026 | 0 | 0.43 | 0.33 | 0.46 | 0.66 | 0.4 | 0.51 |
| 1 | 15441275 | rs1055412 | 0.016 | 1.336 | -0.703 | -1.81 | -0.001 | -0.001 | -0.001 | 0.012 | 0.63 | 0.43 | 0.61 | 0.7 | 0.64 | 0.7 |
| 1 | 15441926 | rs910105 | -0.004 | 1.335 | -0.687 | -1.81 | 0.001 | -0.009 | -0.009 | 0.118 | 0.63 | 0.44 | 0.61 | 0.7 | 0.64 | 0.7 |
| 1 | 15442058 | rs2235789 | 1.06 | 1.241 | -0.435 | -1.868 | 0.015 | 0.055 | 0.055 | 0 | 0.46 | 0.28 | 0.45 | 0.65 | 0.43 | 0.49 |
| 1 | 15442550 | rs11807820 | -0.684 | 1.391 | -0.226 | -1.885 | 0.038 | -0.031 | -0.031 | 0 | 0.1 | 0.46 | 0.12 | 0 | 0.08 | 0.04 |
| 1 | 15442795 | rs2235788 | 0.765 | 1.401 | -0.371 | -1.872 | 0.004 | 0.01 | 0.01 | 0 | 0.51 | 0.47 | 0.49 | 0.64 | 0.44 | 0.5 |
| 1 | 15444288 | rs2235784 | 0.208 | 1.484 | -0.464 | -1.846 | 0.001 | -0.035 | -0.035 | 0 | 0.6 | 0.46 | 0.58 | 0.71 | 0.62 | 0.7 |
| 1 | 15444487 | rs2235783 | 0.199 | 1.485 | -0.464 | -1.846 | 0.001 | -0.035 | -0.035 | 0 | 0.6 | 0.46 | 0.58 | 0.71 | 0.62 | 0.7 |
| 1 | 15446168 | rs13375820 | 0.309 | 1.413 | -0.407 | -1.795 | 0.022 | -0.032 | -0.032 | 0.039 | 0.14 | 0.24 | 0.14 | 0.06 | 0.19 | 0.2 |
| 1 | 15455192 | rs10803355 | 0.073 | 1.228 | -0.489 | -1.551 | -0.001 | -0.001 | -0.001 | 0 | 0.26 | 0.27 | 0.35 | 0.34 | 0.31 | 0.22 |
| 1 | 15456919 | rs12046866 | 1.98 | 0.997 | -0.755 | -1.572 | 0.015 | -0.064 | -0.064 | 0 | 0.39 | 0.07 | 0.4 | 0.57 | 0.34 | 0.46 |
| 1 | 15466832 | rs2206761 | 0.611 | 1.086 | -0.068 | -1.466 | 0.013 | 0.006 | 0.006 | 0 | 0.67 | 0.38 | 0.59 | 0.59 | 0.57 | 0.58 |
| 1 | 15467263 | rs6429711 | -0.369 | 1.057 | -0.144 | -1.344 | 0.025 | 0.036 | 0.036 | 0.024 | 0.8 | 0.67 | 0.69 | 0.65 | 0.75 | 0.79 |
| 1 | 15471992 | rs761301 | -1.071 | 0.762 | -0.279 | -1.471 | -0.009 | -0.01 | -0.01 | 0 | 0.38 | 0.53 | 0.38 | 0.37 | 0.34 | 0.33 |
| 1 | 15477865 | rs1883761 | -0.613 | 1.118 | 0.03 | -1.535 | 0.048 | -0.033 | -0.033 | 0 | 0.85 | 0.75 | 0.86 | 0.94 | 0.75 | 0.78 |
| 1 | 15478066 | rs1883759 | -0.533 | 1.108 | -0.056 | -1.539 | 0.093 | -0.043 | -0.043 | 0 | 0.84 | 0.62 | 0.85 | 0.94 | 0.75 | 0.78 |
| 1 | 15479283 | rs1076726 | -0.554 | 1.116 | -0.073 | -1.542 | 0.107 | -0.042 | -0.042 | 0 | 0.85 | 0.62 | 0.85 | 0.94 | 0.75 | 0.78 |
| 1 | 15482932 | rs4661582 | -0.537 | 1.036 | -0.23 | -1.264 | -0.018 | -0.012 | -0.012 | 0 | 0.31 | 0.27 | 0.34 | 0.39 | 0.39 | 0.45 |
| 1 | 15489884 | rs10489452 | 0.478 | 0.908 | -0.151 | -1.256 | 0.012 | 0.049 | 0.049 | 0.01 | 0.19 | 0.34 | 0.25 | 0.35 | 0.18 | 0.25 |
| 1 | 15492032 | rs7540957 | 1.066 | 0.835 | -0.176 | -1.555 | 0.002 | -0.067 | -0.067 | 0 | 0.65 | 0.47 | 0.73 | 0.92 | 0.6 | 0.67 |
| 1 | 15495620 | rs16851354 | 0.702 | 0.827 | -0.149 | -1.41 | 0.015 | 0.057 | 0.057 | 0 | 0.19 | 0.36 | 0.26 | 0.34 | 0.18 | 0.25 |
| 1 | 15498452 | rs4436387 | 0.119 | 0.698 | -0.16 | -1.476 | -0.022 | -0.105 | -0.105 | 0.046 | 0.76 | 0.41 | 0.81 | 0.97 | 0.81 | 0.87 |
| 1 | 15500332 | rs16851358 | 0.096 | 0.466 | -0.309 | -1.404 | -0.024 | -0.084 | -0.084 | 0 | 0.2 | 0.31 | 0.14 | 0.02 | 0.15 | 0.11 |
| 1 | 15504403 | rs10927698 | 1.646 | 0.507 | -0.729 | -1.194 | -0.025 | 0.009 | 0.009 | 0 | 0.44 | 0.1 | 0.47 | 0.58 | 0.42 | 0.41 |
| 1 | 15507197 | rs12145963 | 0.096 | 0.454 | -0.191 | -1.645 | -0.009 | 0.032 | 0.032 | 0 | 0.34 | 0.25 | 0.23 | 0.08 | 0.31 | 0.3 |
| 1 | 15507952 | rs4661594 | 0.945 | 0.488 | 0.033 | -1.677 | 0.021 | 0.054 | 0.054 | 0 | 0.46 | 0.37 | 0.6 | 0.88 | 0.42 | 0.49 |
| 1 | 15510203 | rs1079314 | 1.347 | 0.476 | 0.027 | -1.365 | 0.006 | 0.004 | 0.004 | 0 | 0.38 | 0.29 | 0.54 | 0.6 | 0.38 | 0.36 |
| 1 | 15512222 | rs11588628 | 0.516 | 0.429 | 0.157 | -1.482 | 0.006 | 0.027 | 0.027 | 0 | 0.53 | 0.4 | 0.66 | 0.63 | 0.59 | 0.58 |
| 1 | 15516299 | rs10927709 | -0.036 | 0.476 | 0.295 | -1.608 | 0.008 | -0.124 | -0.124 | 0 | 0.62 | 0.51 | 0.71 | 0.94 | 0.64 | 0.7 |
| 1 | 15518792 | rs16851387 | 0.247 | 0.489 | 0.349 | -1.62 | 0.001 | -0.037 | -0.037 | 0.344 | 0.18 | 0.24 | 0.13 | 0.03 | 0.22 | 0.17 |
| 1 | 15519417 | rs742671 | -0.605 | 0.61 | 0.188 | -1.658 | 0.015 | -0.077 | -0.077 | 0.71 | 0.8 | 0.73 | 0.82 | 0.97 | 0.77 | 0.89 |
| 1 | 15520583 | rs6692220 | -1.201 | 0.592 | -0.025 | -1.652 | -0.003 | -0.022 | -0.022 | 0 | 0.91 | 0.89 | 0.93 | 0.97 | 0.9 | 0.91 |
| 1 | 15520987 | rs10927713 | 1.823 | 0.622 | -0.402 | -1.605 | -0.054 | 0.083 | 0.083 | 0 | 0.14 | 0 | 0.09 | 0.03 | 0.13 | 0.19 |
| 1 | 15528487 | rs2050233 | -0.504 | 0.613 | -0.239 | -1.618 | 0.031 | -0.117 | -0.117 | 1.023 | 0.79 | 0.58 | 0.8 | 0.97 | 0.77 | 0.89 |
| 1 | 15529536 | rs4661599 | 1.007 | 0.555 | -0.187 | -1.615 | 0.025 | 0.109 | 0.109 | 0 | 0.46 | 0.32 | 0.59 | 0.91 | 0.42 | 0.53 |
| 1 | 15530260 | rs7536482 | -0.109 | 0.562 | -0.242 | -1.581 | 0.02 | -0.134 | -0.134 | 0.059 | 0.65 | 0.56 | 0.71 | 0.94 | 0.64 | 0.71 |
| 1 | 15530336 | rs12028875 | 1.454 | 0.748 | -0.534 | -0.854 | 0.029 | -0.03 | -0.03 | 0 | 0.08 | 0.02 | 0.05 | 0.31 | 0.05 | 0.12 |
| 1 | 15532130 | rs10927720 | -0.529 | 1.043 | -0.386 | -1.043 | 0.001 | -0.027 | -0.027 | 0 | 0.21 | 0.25 | 0.15 | 0.07 | 0.24 | 0.22 |

| | | | | | | | | | | | | | | | | |
|---|---|---|---|---|---|---|---|---|---|---|---|---|---|---|---|---|
| 1 | 15537122 | rs12070469 | -0.655 | 1.198 | -0.676 | -0.91 | -0.008 | 0.008 | 0.008 | 0 | 0.36 | 0.22 | 0.26 | 0.36 | 0.38 | 0.31 |
| 1 | 15539483 | rs1316533 | 0.824 | 1.271 | -0.509 | -1.019 | 0 | -0.001 | -0.001 | 0 | 0.56 | 0.57 | 0.63 | 0.61 | 0.51 | 0.62 |
| 1 | 15539582 | rs12117872 | 1.684 | 1.171 | -0.751 | -1.143 | -0.026 | 0.074 | 0.074 | 0 | 0.14 | 0.02 | 0.08 | 0.04 | 0.12 | 0.2 |
| 1 | 15539953 | rs1316277 | 0.652 | 1.184 | -0.711 | -1.137 | 0.011 | -0.027 | -0.027 | 0 | 0.21 | 0.13 | 0.16 | 0.33 | 0.16 | 0.1 |
| 1 | 15539998 | rs1316276 | -1.287 | 1.322 | -0.645 | -1.154 | 0.046 | -0.028 | -0.028 | 0 | 0.92 | 0.75 | 0.91 | 0.98 | 0.89 | 0.89 |
| 1 | 15540685 | rs6429728 | 0.118 | 1.311 | -0.693 | -1.17 | 0.023 | -0.071 | -0.071 | 0.016 | 0.77 | 0.68 | 0.79 | 0.94 | 0.66 | 0.71 |
| 1 | 15540734 | rs6429729 | 0.841 | 1.319 | -0.694 | -1.163 | -0.006 | 0.009 | 0.009 | 0 | 0.57 | 0.61 | 0.64 | 0.61 | 0.5 | 0.61 |
| 1 | 15541412 | rs3753315 | 0.682 | 1.221 | -0.676 | -1.102 | 0.004 | -0.061 | -0.061 | 0 | 0.2 | 0.07 | 0.16 | 0.33 | 0.16 | 0.1 |
| 1 | 15541537 | rs3753314 | -0.929 | 1.265 | -0.63 | -1.111 | 0.061 | -0.028 | -0.028 | 0.324 | 0.91 | 0.69 | 0.9 | 0.98 | 0.89 | 0.9 |
| 1 | 15542968 | rs10803365 | 0.645 | 1.181 | -0.671 | -1.092 | 0.009 | -0.043 | -0.043 | 0.007 | 0.2 | 0.1 | 0.16 | 0.33 | 0.16 | 0.1 |
| 1 | 15543354 | rs17405463 | 1.828 | 1.315 | -0.78 | -1.134 | -0.031 | 0.031 | 0.031 | 0 | 0.08 | 0.01 | 0.07 | 0.03 | 0.12 | 0.07 |
| 1 | 15544366 | rs12057512 | 0.472 | 0.957 | -0.644 | -1.021 | 0.004 | -0.01 | -0.01 | 0 | 0.21 | 0.16 | 0.17 | 0.32 | 0.17 | 0.1 |
| 1 | 15546077 | rs3753313 | 0.252 | 0.955 | -0.73 | -1.134 | -0.058 | 0.142 | 0.142 | 0 | 0.29 | 0.07 | 0.2 | 0.08 | 0.35 | 0.37 |
| 1 | 15546761 | rs7681 | 1.162 | 0.805 | -0.694 | -1.151 | -0.007 | -0.008 | -0.008 | 0 | 0.41 | 0.52 | 0.55 | 0.58 | 0.36 | 0.43 |
| 1 | 15546825 | rs549 | 0.235 | 0.878 | -0.858 | -1.122 | -0.015 | 0.016 | 0.016 | 0 | 0.62 | 0.69 | 0.71 | 0.9 | 0.54 | 0.53 |
| 1 | 15548397 | rs12124285 | 1.59 | 0.942 | -1.513 | -0.942 | -0.032 | 0.072 | 0.072 | 0 | 0.14 | 0.01 | 0.08 | 0.04 | 0.12 | 0.19 |
| 1 | 15550525 | rs10803369 | -0.916 | 0.923 | -1.296 | -0.88 | 0.001 | 0.006 | 0.006 | 0 | 0.54 | 0.45 | 0.4 | 0.42 | 0.54 | 0.52 |
| 1 | 15558938 | rs6429732 | -0.242 | 0.854 | -2.073 | -0.666 | 0.011 | -0.007 | -0.007 | 0 | 0.27 | 0.33 | 0.31 | 0.18 | 0.28 | 0.21 |
| 1 | 15560900 | rs767255 | -0.258 | 0.89 | -2.028 | -0.536 | -0.016 | 0.016 | 0.016 | 0 | 0.42 | 0.34 | 0.4 | 0.31 | 0.43 | 0.4 |
| 1 | 15561181 | rs12034146 | 0.121 | 1.116 | -2.17 | -0.21 | -0.006 | -0.005 | -0.005 | 0.269 | 0.3 | 0.25 | 0.31 | 0.31 | 0.31 | 0.38 |
| 1 | 15562593 | rs1780593 | 0.288 | 1.316 | -2.461 | -0.275 | -0.062 | 0.07 | 0.07 | 0 | 0.14 | 0.01 | 0.1 | 0.03 | 0.21 | 0.18 |
| 1 | 15562759 | rs1985917 | 0.691 | 1.211 | -2.366 | -0.168 | 0.036 | -0.049 | -0.049 | 0 | 0.24 | 0.09 | 0.31 | 0.33 | 0.19 | 0.22 |
| 1 | 15566803 | rs4644485 | -0.008 | 2.093 | -1.689 | -0.324 | -0.05 | 0.045 | 0.045 | 0 | 0.86 | 0.96 | 0.88 | 0.94 | 0.78 | 0.89 |
| 1 | 15569521 | rs16851463 | -0.731 | 1.438 | -1.151 | -0.007 | 0.008 | 0.001 | 0.001 | 0 | 0.13 | 0.22 | 0.21 | 0.1 | 0.14 | 0.14 |
| 1 | 15569822 | rs11585832 | -0.91 | 0.805 | -1.247 | -0.276 | 0 | -0.021 | -0.021 | 0 | 0.11 | 0.09 | 0.09 | 0 | 0.12 | 0.14 |
| 1 | 15571315 | rs12038868 | -0.755 | 0.139 | -1.989 | -0.328 | 0.008 | 0.018 | 0.018 | 0 | 0.23 | 0.01 | 0.12 | 0.14 | 0.2 | 0.37 |
| 1 | 15572133 | rs1752020 | 0.336 | -0.325 | -1.441 | -1.039 | -0.009 | -0.041 | -0.041 | 0 | 0.73 | 0.73 | 0.76 | 0.95 | 0.8 | 0.79 |
| 1 | 15577022 | rs1752005 | 0.65 | 0.194 | -1.958 | -1.787 | -0.067 | 0.129 | 0.129 | 2.847 | 0.32 | 0.1 | 0.22 | 0.04 | 0.37 | 0.3 |
| 1 | 15583819 | rs16851480 | -0.508 | -0.313 | -1.333 | -1.08 | 0.008 | -0.018 | -0.018 | 0 | 0.63 | 0.51 | 0.7 | 0.71 | 0.61 | 0.58 |
| 1 | 15584079 | rs943558 | -0.508 | -0.313 | -1.333 | -1.08 | 0.008 | -0.018 | -0.018 | 0 | 0.63 | 0.51 | 0.7 | 0.71 | 0.61 | 0.58 |
| 1 | 15585613 | rs7520597 | 0.653 | -0.498 | -1.376 | -1.234 | -0.003 | 0.084 | 0.084 | 0 | 0.2 | 0.07 | 0.16 | 0.03 | 0.18 | 0.11 |
| 1 | 15588759 | rs6676098 | 0.257 | -0.498 | -1.34 | -1.206 | 0.055 | 0.358 | 0.358 | 0.347 | 0.06 | 0.39 | 0.08 | 0.25 | 0.01 | 0.13 |
| 1 | 15591641 | rs12144738 | 0.676 | -0.241 | -1.477 | -1.212 | -0.038 | 0.046 | 0.046 | 0 | 0.2 | 0.05 | 0.19 | 0.12 | 0.21 | 0.13 |
| 1 | 15591929 | rs10927753 | -0.682 | -0.38 | -1.464 | -0.646 | -0.01 | 0.091 | 0.091 | 3.29 | 0.57 | 0.76 | 0.69 | 0.82 | 0.59 | 0.49 |
| 1 | 15593666 | rs4661617 | 0.857 | -0.428 | -1.236 | -0.639 | -0.027 | -0.068 | -0.068 | 0 | 0.3 | 0.13 | 0.42 | 0.55 | 0.37 | 0.22 |
| 1 | 15599160 | rs6703580 | -0.246 | -0.02 | -1.035 | -0.405 | -0.013 | 0.086 | 0.086 | 3.222 | 0.56 | 0.67 | 0.67 | 0.94 | 0.57 | 0.49 |
| 1 | 15600049 | rs10927757 | -0.258 | 0.044 | -0.922 | -0.339 | -0.005 | 0.089 | 0.089 | 3.185 | 0.55 | 0.64 | 0.67 | 0.89 | 0.58 | 0.44 |
| 1 | 15600085 | rs10927758 | -0.239 | 0.01 | -0.885 | -0.332 | -0.006 | 0.08 | 0.08 | 3.106 | 0.55 | 0.63 | 0.65 | 0.81 | 0.57 | 0.44 |
| 1 | 15602330 | rs752626 | 0.164 | -0.226 | -0.367 | -0.537 | -0.034 | 0.005 | 0.005 | 2.466 | 0.35 | 0.48 | 0.29 | 0.36 | 0.26 | 0.26 |
| 1 | 15606453 | rs10927764 | -0.07 | 0.695 | -0.605 | -0.627 | -0.023 | 0.091 | 0.091 | 1.905 | 0.51 | 0.2 | 0.62 | 0.7 | 0.57 | 0.45 |
| 1 | 15606723 | rs7553794 | -0.558 | 0.709 | -0.166 | -0.723 | 0.028 | 0.157 | 0.157 | 0 | 0.14 | 0.49 | 0.16 | 0.26 | 0.1 | 0.27 |
| 1 | 15609117 | rs572953 | 0.879 | 0.8 | -0.22 | -1.057 | -0.004 | 0.019 | 0.019 | 0.178 | 0.34 | 0.25 | 0.22 | 0.04 | 0.33 | 0.28 |
| 1 | 15610004 | rs10927765 | 0.463 | 0.837 | -0.388 | -0.904 | 0.022 | 0.094 | 0.094 | 0 | 0.22 | 0.46 | 0.4 | 0.61 | 0.28 | 0.19 |
| 1 | 15610609 | rs1737344 | -0.752 | 0.902 | -0.67 | -0.877 | -0.042 | 0.085 | 0.085 | 2.232 | 0.65 | 0.92 | 0.71 | 0.82 | 0.66 | 0.73 |
| 1 | 15613478 | rs536364 | -0.349 | 0.824 | -0.204 | -0.62 | 0.008 | 0.029 | 0.029 | 0.025 | 0.54 | 0.58 | 0.59 | 0.65 | 0.59 | 0.46 |
| 1 | 15616139 | rs486557 | 0.093 | 0.89 | -0.571 | -0.864 | -0.008 | 0.057 | 0.057 | 0.153 | 0.33 | 0.21 | 0.21 | 0.04 | 0.33 | 0.26 |
| 1 | 15626718 | rs10803378 | -1.348 | 0.952 | 1.132 | -0.05 | 0.135 | 0.142 | 0.142 | 1.473 | 0.07 | 0.52 | 0.09 | 0.07 | 0.06 | 0.09 |
| 1 | 15643218 | rs477590 | 0.293 | 0.999 | 0.468 | 0.062 | -0.002 | 0.013 | 0.013 | 0 | 0.65 | 0.28 | 0.75 | 0.75 | 0.68 | 0.75 |
| 1 | 15644729 | rs7554324 | -0.217 | 1.128 | -0.072 | -0.132 | -0.002 | 0.004 | 0.004 | 0.027 | 0.37 | 0.64 | 0.49 | 0.31 | 0.38 | 0.56 |
| 1 | 15647828 | rs7518884 | 0.343 | 0.902 | -0.591 | -0.187 | -0.027 | 0.039 | 0.039 | 0 | 0.31 | 0.14 | 0.4 | 0.25 | 0.32 | 0.53 |
| 1 | 15649031 | rs504775 | 0.699 | 0.676 | -0.626 | -0.18 | -0.006 | -0.041 | -0.041 | 0 | 0.31 | 0.12 | 0.33 | 0.39 | 0.33 | 0.23 |
| 1 | 15651062 | rs3765356 | 0.347 | 0.694 | -0.46 | -0.807 | 0.001 | -0.046 | -0.046 | 0 | 0.31 | 0.38 | 0.36 | 0.07 | 0.35 | 0.4 |
| 1 | 15653563 | rs3765357 | 0.874 | 0.747 | -1.029 | -0.812 | -0.029 | 0.068 | 0.068 | 0 | 0.26 | 0.13 | 0.32 | 0.05 | 0.33 | 0.34 |
| 1 | 15655304 | rs567419 | 0.781 | 0.5 | -1.267 | -0.477 | 0.063 | -0.13 | -0.13 | 0.012 | 0.28 | 0.02 | 0.17 | 0.42 | 0.18 | 0.21 |
| 1 | 15668276 | rs763821 | -0.216 | 1.017 | -0.591 | -1.836 | 0.001 | -0.098 | -0.098 | 0.415 | 0.41 | 0.48 | 0.41 | 0.04 | 0.4 | 0.37 |
| 1 | 15668319 | rs12126178 | -0.225 | 0.041 | -0.662 | -1.584 | 0.004 | -0.014 | -0.014 | 0.065 | 0.13 | 0.1 | 0.04 | 0 | 0.05 | 0.03 |
| 1 | 15671483 | rs486089 | -0.161 | 0.218 | -0.541 | -1.574 | 0.05 | 0.094 | 0.094 | 0 | 0.48 | 0.28 | 0.43 | 0.89 | 0.35 | 0.43 |
| 1 | 15680451 | rs2473358 | -0.318 | 0.997 | -0.712 | -0.74 | -0.002 | 0.009 | 0.009 | 0 | 0.46 | 0.62 | 0.47 | 0.23 | 0.46 | 0.46 |
| 1 | 15686006 | rs1751998 | -0.063 | 1.163 | -1.623 | -0.56 | -0.102 | 0.146 | 0.146 | 0 | 0.59 | 0.9 | 0.59 | 0.79 | 0.54 | 0.56 |
| 1 | 15687059 | rs4661330 | -1.226 | 1.477 | -0.667 | 0.226 | 0.019 | 0.013 | 0.013 | 0 | 0.19 | 0.26 | 0.21 | 0.29 | 0.29 | 0.2 |
| 1 | 15687814 | rs542008 | -1.159 | 0.905 | -0.654 | 0.019 | -0.003 | 0.002 | 0.002 | 0 | 0.28 | 0.68 | 0.25 | 0.25 | 0.29 | 0.34 |
| 1 | 15691540 | rs10803382 | -1.142 | 0.853 | -0.575 | -0.187 | 0.007 | -0.019 | -0.019 | 0 | 0.28 | 0.59 | 0.23 | 0.19 | 0.29 | 0.33 |
| 1 | 15697183 | rs876295 | -1.282 | 0.59 | -0.791 | -0.119 | -0.001 | 0.013 | 0.013 | 0 | 0.39 | 0.6 | 0.29 | 0.28 | 0.33 | 0.37 |
| 1 | 15702130 | rs17368978 | -0.298 | -0.033 | -0.439 | -0.26 | -0.001 | -0.016 | -0.016 | 0 | 0.07 | 0.08 | 0.04 | 0 | 0.05 | 0.07 |
| 1 | 15702434 | rs17368999 | -0.298 | -0.033 | -0.439 | -0.26 | -0.001 | -0.016 | -0.016 | 0 | 0.07 | 0.08 | 0.04 | 0 | 0.04 | 0.07 |
| 1 | 15705072 | rs736185 | -0.483 | -0.008 | -0.555 | -0.342 | -0.038 | -0.039 | -0.039 | 0 | 0.12 | 0.01 | 0.09 | 0.15 | 0.15 | 0.25 |
| 1 | 15708540 | rs1010294 | -1.781 | -0.456 | -0.214 | -0.568 | 0.052 | 0.056 | 0.056 | 0 | 0.39 | 0.77 | 0.31 | 0.26 | 0.26 | 0.37 |
| 1 | 15723843 | rs2294490 | 1.66 | -0.43 | -1.028 | -0.319 | -0.097 | -0.116 | -0.116 | 0.257 | 0.36 | 0.1 | 0.29 | 0.45 | 0.45 | 0.32 |
| 1 | 15724033 | rs2253372 | -0.781 | -0.439 | -1.213 | -0.356 | -0.017 | -0.019 | -0.019 | 0 | 0.76 | 0.91 | 0.58 | 0.71 | 0.74 | 0.69 |

| | | | | | | | | | | | | | | | | |
|---|---|---|---|---|---|---|---|---|---|---|---|---|---|---|---|---|
| 1 | 15726566 | rs742361 | -0.621 | -0.51 | -0.848 | -0.393 | 0.012 | 0.022 | 0.022 | 0 | 0.32 | 0.18 | 0.19 | 0.26 | 0.26 | 0.36 |
| 1 | 15726791 | rs742362 | 0.409 | -0.477 | -0.826 | -0.37 | 0.005 | 0.004 | 0.004 | 0 | 0.21 | 0.17 | 0.11 | 0.11 | 0.11 | 0.11 |
| 1 | 15728289 | rs2473344 | -0.537 | -0.344 | -1.165 | -0.213 | -0.012 | -0.014 | -0.014 | 0.023 | 0.76 | 0.89 | 0.58 | 0.71 | 0.74 | 0.69 |
| 1 | 15732793 | rs733887 | 1.498 | -0.245 | -1.344 | -0.228 | -0.161 | -0.117 | -0.117 | 0 | 0.36 | 0.06 | 0.3 | 0.55 | 0.47 | 0.33 |
| 1 | 15733394 | rs2092324 | -0.051 | -0.231 | -0.956 | -0.209 | -0.01 | -0.027 | -0.027 | 0 | 0.7 | 0.56 | 0.52 | 0.78 | 0.7 | 0.68 |
| 1 | 15739016 | rs2496325 | -1.169 | -0.556 | -1.765 | 0.034 | 0.069 | 0.003 | 0.003 | 0.054 | 0.5 | 0.94 | 0.37 | 0.38 | 0.35 | 0.38 |
| 1 | 15744271 | rs2496320 | -0.534 | -0.776 | -0.559 | -0.139 | -0.061 | -0.074 | -0.074 | 2.634 | 0.24 | 0.44 | 0.14 | 0.11 | 0.12 | 0.08 |
| 1 | 15745380 | rs7525279 | -1.149 | -1.015 | -0.879 | -0.49 | 0.021 | 0.017 | 0.017 | 0 | 0.49 | 0.53 | 0.36 | 0.39 | 0.39 | 0.39 |
| 1 | 15750217 | rs2473355 | -0.286 | -0.848 | -2.153 | -0.62 | -0.049 | 0.052 | 0.052 | 1.631 | 0.56 | 0.91 | 0.57 | 0.31 | 0.41 | 0.48 |
| 1 | 15752716 | rs3765369 | 0.309 | -1.015 | -1.868 | -0.847 | -0.005 | -0.001 | -0.001 | 0 | 0.19 | 0.16 | 0.22 | 0.05 | 0.23 | 0.11 |
| 1 | 15753398 | rs3765370 | 1.686 | -0.515 | -2.101 | -0.079 | -0.032 | 0.02 | 0.02 | 0.017 | 0.07 | 0 | 0.08 | 0.51 | 0.11 | 0.07 |
| 1 | 15757044 | rs2473357 | -1.097 | -1.195 | -2.01 | -1.171 | 0.015 | 0.15 | 0.15 | 0 | 0.42 | 0.87 | 0.4 | 0.14 | 0.34 | 0.22 |
| 1 | 15763340 | rs6693417 | -0.435 | -1.066 | -1.129 | -1.108 | 0.005 | 0.01 | 0.01 | 0 | 0.52 | 0.57 | 0.42 | 0.32 | 0.39 | 0.42 |
| 1 | 15768304 | rs10436957 | 0.658 | -0.816 | -0.909 | -1.013 | 0.017 | -0.051 | -0.051 | 0.124 | 0.19 | 0.23 | 0.21 | 0.06 | 0.21 | 0.11 |
| 1 | 15772933 | rs555015 | -1.003 | -0.392 | -1.018 | -0.615 | -0.032 | 0.007 | 0.007 | 0 | 0.64 | 0.81 | 0.52 | 0.34 | 0.48 | 0.48 |
| 1 | 15786195 | rs4661627 | 0.587 | -0.31 | -0.53 | -0.651 | 0.036 | 0.07 | 0.07 | 0.105 | 0.51 | 0.35 | 0.61 | 0.73 | 0.64 | 0.66 |
| 1 | 15788789 | rs7547591 | -0.322 | -0.496 | -0.196 | -0.658 | 0 | -0.002 | -0.002 | 0 | 0.38 | 0.37 | 0.25 | 0.24 | 0.26 | 0.29 |
| 1 | 15790639 | rs11579489 | 1.145 | -0.117 | -0.216 | -0.496 | 0.01 | -0.032 | -0.032 | 0 | 0.07 | 0.16 | 0.06 | 0 | 0.06 | 0.05 |
| 1 | 15796714 | rs7556116 | 0.233 | -0.844 | -0.55 | -0.585 | -0.003 | -0.001 | -0.001 | 0 | 0.72 | 0.72 | 0.78 | 0.77 | 0.81 | 0.74 |
| 1 | 15796808 | rs10927791 | 1.156 | -0.772 | -1.345 | -0.877 | -0.076 | -0.018 | -0.018 | 0 | 0.42 | 0.03 | 0.56 | 0.69 | 0.62 | 0.56 |
| 1 | 15808767 | rs3820071 | 0.395 | 0.032 | -0.765 | -0.534 | 0.046 | 0.171 | 0.171 | 0.061 | 0.13 | 0.28 | 0.35 | 0.57 | 0.24 | 0.22 |
| 1 | 15808872 | rs3766160 | 0.365 | 0.018 | -0.775 | -0.539 | 0.043 | 0.168 | 0.168 | 0.017 | 0.14 | 0.28 | 0.29 | 0.56 | 0.24 | 0.21 |
| 1 | 15810624 | rs12130370 | -0.813 | -0.199 | -0.432 | -0.821 | -0.013 | -0.011 | -0.011 | 0.266 | 0.41 | 0.74 | 0.53 | 0.61 | 0.43 | 0.4 |
| 1 | 15814186 | rs6667862 | -0.693 | -0.249 | -0.402 | -0.965 | -0.008 | 0.067 | 0.067 | 0.042 | 0.5 | 0.78 | 0.56 | 0.67 | 0.52 | 0.44 |
| 1 | 15821673 | rs4646092 | 0.047 | -0.328 | 0.052 | -1.047 | -0.005 | -0.083 | -0.083 | 0 | 0.28 | 0.44 | 0.19 | 0.06 | 0.22 | 0.2 |
| 1 | 15823061 | rs4661636 | 1.905 | -0.41 | -0.138 | -1.133 | -0.119 | 0.112 | 0.112 | 0.36 | 0.31 | 0.01 | 0.31 | 0.1 | 0.36 | 0.36 |
| 1 | 15825195 | rs6685648 | 0.876 | -0.295 | 0.163 | -0.959 | 0.004 | 0.049 | 0.049 | 0.064 | 0.22 | 0.29 | 0.31 | 0.6 | 0.3 | 0.24 |
| 1 | 15829187 | rs4233533 | 0.919 | -0.298 | 0.238 | -0.867 | -0.002 | -0.005 | -0.005 | 0 | 0.22 | 0.22 | 0.3 | 0.59 | 0.3 | 0.24 |
| 1 | 15832543 | rs1052576 | -0.819 | -0.356 | 0.209 | -1.068 | -0.005 | 0.048 | 0.048 | 0.011 | 0.5 | 0.74 | 0.5 | 0.65 | 0.51 | 0.44 |
| 1 | 15834360 | rs2020902 | 1.346 | -0.562 | 0.329 | -1.244 | -0.022 | 0.052 | 0.052 | 0 | 0.16 | 0.09 | 0.11 | 0.03 | 0.14 | 0.1 |
| 1 | 15844057 | rs1862710 | -1.368 | -0.437 | 0.322 | -1.166 | -0.05 | 0.025 | 0.025 | 0.972 | 0.54 | 0.89 | 0.53 | 0.65 | 0.53 | 0.45 |
| 1 | 15850603 | rs4645983 | -0.17 | -0.423 | 0.386 | -1.146 | -0.005 | -0.081 | -0.081 | 0 | 0.28 | 0.44 | 0.19 | 0.06 | 0.22 | 0.19 |
| 1 | 15850613 | rs1052571 | -0.816 | -0.421 | 0.377 | -1.148 | -0.005 | 0.048 | 0.048 | 0.011 | 0.5 | 0.74 | 0.5 | 0.65 | 0.51 | 0.44 |
| 1 | 15852704 | rs4263990 | -0.808 | -0.424 | 0.375 | -1.15 | -0.004 | 0.049 | 0.049 | 0.016 | 0.5 | 0.74 | 0.5 | 0.65 | 0.51 | 0.44 |
| 1 | 15857134 | rs10927796 | -0.82 | -0.421 | 0.377 | -1.133 | -0.005 | 0.052 | 0.052 | 0.024 | 0.5 | 0.74 | 0.5 | 0.66 | 0.51 | 0.44 |
| 1 | 15860803 | rs11583306 | -0.163 | -0.423 | 0.351 | -1.112 | -0.005 | -0.091 | -0.091 | 0 | 0.28 | 0.43 | 0.19 | 0.05 | 0.22 | 0.19 |
| 1 | 15865604 | rs10927797 | -0.81 | -0.408 | 0.283 | -1.271 | -0.01 | 0.061 | 0.061 | 0.033 | 0.51 | 0.78 | 0.5 | 0.66 | 0.51 | 0.43 |
| 1 | 15894959 | rs1467513 | 0.808 | -0.271 | 0.223 | -1.1 | 0.006 | 0.067 | 0.067 | 0.058 | 0.22 | 0.31 | 0.31 | 0.61 | 0.3 | 0.24 |
| 1 | 15909744 | rs11580170 | -0.444 | -0.379 | 0.321 | -1.301 | -0.013 | -0.104 | -0.104 | 0 | 0.33 | 0.55 | 0.21 | 0.06 | 0.23 | 0.2 |
| 1 | 15910003 | rs6429759 | -0.961 | -0.367 | 0.043 | -1.157 | -0.03 | 0.02 | 0.02 | 1.899 | 0.55 | 0.83 | 0.52 | 0.64 | 0.53 | 0.45 |
| 1 | 15918974 | rs1547032 | 0.387 | -0.394 | 0.451 | -1.189 | -0.008 | -0.027 | -0.027 | 0 | 0.07 | 0.14 | 0.06 | 0 | 0.06 | 0.1 |
| 1 | 15920454 | rs556258 | -2.032 | -0.477 | 0.085 | -1.386 | -0.115 | 0.086 | 0.086 | 0.166 | 0.71 | 0.99 | 0.7 | 0.91 | 0.68 | 0.66 |
| 1 | 15923530 | rs6675252 | -1.437 | -0.189 | 0.41 | -1.313 | -0.045 | -0.023 | -0.023 | 0 | 0.58 | 0.91 | 0.56 | 0.66 | 0.59 | 0.48 |
| 1 | 15930869 | rs16851855 | -0.652 | -0.285 | 0.854 | -1.558 | -0.004 | -0.101 | -0.101 | 0 | 0.33 | 0.44 | 0.22 | 0.03 | 0.27 | 0.21 |
| 1 | 15951743 | rs6697813 | 0.182 | -0.335 | 0.868 | -1.554 | -0.002 | -0.006 | -0.006 | 0.742 | 0.26 | 0.25 | 0.15 | 0.03 | 0.21 | 0.13 |
| 1 | 15975237 | rs1980607 | -0.801 | -0.305 | 1.148 | -1.5 | -0.006 | -0.06 | -0.06 | 5.829 | 0.35 | 0.61 | 0.23 | 0.06 | 0.27 | 0.21 |
| 1 | 15986547 | rs3766163 | -0.608 | -0.236 | 1.118 | -1.496 | -0.004 | -0.098 | -0.098 | 0 | 0.33 | 0.45 | 0.22 | 0.03 | 0.27 | 0.21 |
| 1 | 15990552 | rs927207 | -0.608 | -0.267 | 1.118 | -1.56 | -0.004 | -0.098 | -0.098 | 0.451 | 0.33 | 0.45 | 0.22 | 0.03 | 0.27 | 0.21 |
| 1 | 15993338 | rs10492987 | -1.391 | -0.23 | 0.872 | -1.546 | 0.001 | 0.016 | 0.016 | 0.49 | 0.58 | 0.75 | 0.54 | 0.68 | 0.61 | 0.48 |
| 1 | 16001588 | rs12746773 | -1.04 | -0.628 | 1.513 | -2.244 | -0.016 | -0.119 | -0.119 | 0 | 0.34 | 0.46 | 0.2 | 0.03 | 0.24 | 0.21 |
| 1 | 16015784 | rs7546138 | -0.833 | -0.611 | 1.126 | -2.257 | -0.013 | -0.05 | -0.05 | 0 | 0.11 | 0.43 | 0.07 | 0 | 0.06 | 0.08 |
| 1 | 16041606 | rs12745403 | -0.41 | -0.273 | 0.361 | -2.04 | -0.008 | -0.095 | -0.095 | 0 | 0.26 | 0.37 | 0.15 | 0.03 | 0.19 | 0.12 |
| 1 | 16043157 | rs2280115 | -0.41 | -0.273 | 0.361 | -2.04 | -0.008 | -0.095 | -0.095 | 0.055 | 0.26 | 0.37 | 0.15 | 0.03 | 0.19 | 0.12 |
| 1 | 16044401 | rs16851970 | -0.415 | -0.272 | 0.362 | -2.04 | -0.008 | -0.095 | -0.095 | 0 | 0.26 | 0.37 | 0.15 | 0.03 | 0.19 | 0.12 |
| 1 | 16055039 | rs12724755 | -0.051 | -0.605 | 0.181 | -2.036 | 0.007 | 0.004 | 0.004 | 0 | 0.26 | 0.21 | 0.19 | 0.04 | 0.19 | 0.16 |
| 1 | 16055931 | rs7539884 | -0.982 | -0.799 | 0.469 | -2.083 | 0.009 | 0.005 | 0.005 | 0.508 | 0.35 | 0.71 | 0.26 | 0.07 | 0.21 | 0.17 |
| 1 | 16056364 | rs3817269 | -0.953 | -0.79 | 0.473 | -2.085 | 0.008 | -0.015 | -0.015 | 0 | 0.34 | 0.71 | 0.26 | 0.04 | 0.21 | 0.17 |
| 1 | 16063560 | rs11577707 | 1.633 | -0.61 | 0.179 | -2.34 | -0.059 | 0.089 | 0.089 | 0.006 | 0.14 | 0.01 | 0.12 | 0.05 | 0.17 | 0.07 |
| 1 | 16063698 | rs6701500 | -1.555 | -1.059 | 0.8 | -1.908 | 0.023 | 0.036 | 0.036 | 0.52 | 0.4 | 0.72 | 0.29 | 0.08 | 0.27 | 0.24 |
| 1 | 16068364 | rs35365738 | -0.707 | -0.88 | 0.385 | -2.122 | 0.008 | 0.064 | 0.064 | 0 | 0.36 | 0.21 | 0.27 | 0.04 | 0.33 | 0.22 |
| 1 | 16069644 | rs11584945 | -0.485 | -0.72 | 0.161 | -2.062 | 0.006 | 0.013 | 0.013 | 0.47 | 0.32 | 0.23 | 0.17 | 0.03 | 0.29 | 0.18 |
| 1 | 16075906 | rs10927848 | -1.132 | -0.572 | -0.122 | -1.927 | 0.011 | 0.01 | 0.01 | 0.17 | 0.42 | 0.67 | 0.28 | 0.07 | 0.35 | 0.24 |
| 1 | 16076705 | rs10927849 | -1.071 | -0.561 | -0.123 | -1.921 | 0.006 | -0.008 | -0.008 | 0 | 0.42 | 0.63 | 0.28 | 0.07 | 0.35 | 0.24 |
| 1 | 16079818 | rs6681468 | -0.763 | -0.695 | -0.091 | -1.633 | -0.032 | -0.05 | -0.05 | 0 | 0.12 | 0.32 | 0.07 | 0.01 | 0.08 | 0.11 |
| 1 | 16080171 | rs12725198 | -0.348 | -0.611 | -0.258 | -1.659 | 0.001 | -0.011 | -0.011 | 0 | 0.26 | 0.21 | 0.16 | 0.07 | 0.22 | 0.21 |
| 1 | 16082127 | rs34108989 | -1.091 | -0.704 | -0.408 | -1.387 | -0.007 | -0.019 | -0.019 | 0.156 | 0.36 | 0.42 | 0.25 | 0.14 | 0.28 | 0.3 |
| 1 | 16082976 | rs35609485 | 0.4 | -0.527 | -0.75 | -1.288 | -0.014 | 0.022 | 0.022 | 0 | 0.12 | 0 | 0.08 | 0.05 | 0.1 | 0.07 |
| 1 | 16087164 | rs867321 | -0.125 | -0.642 | -0.709 | -1.044 | 0.005 | -0.002 | -0.002 | 0 | 0.18 | 0.05 | 0.12 | 0.14 | 0.16 | 0.12 |
| 1 | 16087260 | rs938295 | -1.15 | -0.913 | -0.038 | -1.19 | -0.038 | -0.04 | -0.04 | 0 | 0.26 | 0.49 | 0.21 | 0.14 | 0.22 | 0.18 |

| | | | | | | | | | | | | | | | | |
|---|---|---|---|---|---|---|---|---|---|---|---|---|---|---|---|---|
| 1 | 16089356 | rs12083988 | -1.692 | -1.097 | -0.347 | -1.557 | -0.018 | -0.041 | -0.041 | 0 | 0.4 | 0.52 | 0.3 | 0.15 | 0.35 | 0.22 |
| 1 | 16095316 | rs2271545 | 1.22 | -1.058 | -0.27 | -1.566 | -0.007 | -0.014 | -0.014 | 0 | 0.62 | 0.38 | 0.69 | 0.83 | 0.65 | 0.76 |
| 1 | 16097237 | rs10803398 | 0.097 | -1.016 | -0.08 | -1.566 | -0.013 | -0.102 | -0.102 | 0 | 0.81 | 0.6 | 0.83 | 1 | 0.81 | 0.91 |
| 1 | 16098866 | rs10927852 | 0.38 | -0.878 | -0.298 | -1.4 | -0.011 | -0.027 | -0.027 | 0 | 0.63 | 0.57 | 0.71 | 0.83 | 0.65 | 0.76 |
| 1 | 16102124 | rs12730950 | 0.334 | -0.85 | -0.503 | -1.279 | -0.017 | -0.027 | -0.027 | 0 | 0.63 | 0.55 | 0.71 | 0.8 | 0.65 | 0.76 |
| 1 | 16110815 | rs11576404 | 0.678 | -0.111 | -1.564 | -1.466 | -0.001 | 0.019 | 0.019 | 0 | 0.41 | 0.29 | 0.46 | 0.74 | 0.42 | 0.41 |
| 1 | 16112795 | rs7663 | 0.492 | -0.042 | -1.564 | -1.651 | 0.008 | 0.022 | 0.022 | 0 | 0.43 | 0.36 | 0.47 | 0.74 | 0.42 | 0.41 |
| 1 | 16122018 | rs12026094 | 0.474 | 0.328 | -1.649 | -1.475 | -0.052 | 0.066 | 0.066 | 0 | 0.2 | 0.07 | 0.23 | 0.02 | 0.23 | 0.34 |
| 1 | 16123199 | rs12026701 | 0.473 | 0.05 | -1.636 | -1.174 | -0.015 | -0.034 | -0.034 | 0 | 0.39 | 0.14 | 0.43 | 0.68 | 0.39 | 0.33 |
| 1 | 16124438 | rs12569079 | 0.724 | 0.032 | -2.056 | -1.185 | -0.105 | -0.125 | -0.125 | 1.239 | 0.35 | 0.03 | 0.42 | 0.69 | 0.39 | 0.32 |
| 1 | 16126207 | rs11589957 | 0.286 | 0.139 | -1.972 | -2.089 | -0.094 | 0.181 | 0.181 | 0 | 0.26 | 0.01 | 0.26 | 0 | 0.28 | 0.32 |
| 1 | 16129320 | rs2013019 | -0.933 | -0.091 | -0.955 | -2.427 | -0.001 | -0.052 | -0.052 | 0 | 0.51 | 0.49 | 0.42 | 0.1 | 0.44 | 0.5 |
| 1 | 16142960 | rs2862162 | -0.048 | -0.168 | -0.96 | -2.544 | -0.066 | 0.198 | 0.198 | 0 | 0.34 | 0.08 | 0.32 | 0.09 | 0.35 | 0.42 |
| 1 | 16149301 | rs6677321 | -0.061 | 0.009 | -0.893 | -2.477 | -0.067 | 0.203 | 0.203 | 0 | 0.35 | 0.08 | 0.33 | 0.09 | 0.35 | 0.43 |
| 1 | 16161836 | rs4661661 | -0.334 | -0.112 | -0.74 | -2.517 | 0.049 | -0.057 | -0.057 | 0 | 0.11 | 0.37 | 0.1 | 0.01 | 0.09 | 0.07 |
| 1 | 16188681 | rs6701290 | 0.078 | -0.223 | -0.375 | -2.206 | -0.008 | 0.011 | 0.011 | 0 | 0.85 | 0.8 | 0.74 | 0.77 | 0.89 | 0.9 |
| 1 | 16200163 | rs904910 | 0.307 | 0.059 | -0.437 | -2.304 | -0.005 | 0.189 | 0.189 | 0 | 0.33 | 0.08 | 0.31 | 0.09 | 0.31 | 0.42 |
| 1 | 16201502 | rs16852052 | 0.001 | -0.023 | 0.283 | -2.367 | 0.09 | -0.018 | -0.018 | 0.63 | 0.06 | 0.2 | 0.06 | 0 | 0.08 | 0.03 |
| 1 | 16207862 | rs16852063 | 1.105 | 0 | 0.253 | -2.365 | 0.004 | 0.01 | 0.01 | 0 | 0.08 | 0.06 | 0.06 | 0.08 | 0.07 | 0.18 |
| 1 | 16212219 | rs4661670 | 0.862 | 0.042 | 0.371 | -2.329 | -0.003 | 0 | 0 | 0 | 0.06 | 0.06 | 0.04 | 0.01 | 0.06 | 0.05 |
| 1 | 16242620 | rs12063905 | 0.443 | 0.018 | 0.352 | -2.329 | -0.022 | -0.048 | -0.048 | 0 | 0.08 | 0.02 | 0.24 | 0.22 | 0.1 | 0.09 |
| 1 | 16251571 | rs4661345 | -0.879 | 0.019 | 0.367 | -2.329 | 0.106 | -0.078 | -0.078 | 0 | 0.13 | 0.38 | 0.12 | 0.01 | 0.14 | 0.08 |
| 1 | 16256007 | rs848209 | -2.072 | 0.033 | -0.109 | -2.273 | -0.12 | -0.021 | -0.021 | 0 | 0.64 | 0.98 | 0.71 | 0.32 | 0.56 | 0.59 |
| 1 | 16260916 | rs848211 | 0.364 | 0.089 | 0.001 | -2.157 | -0.022 | -0.048 | -0.048 | 0.584 | 0.08 | 0.02 | 0.24 | 0.22 | 0.1 | 0.09 |
| 1 | 16267147 | rs848215 | 0.364 | 0.089 | 0.001 | -2.149 | -0.022 | -0.048 | -0.048 | 0.502 | 0.08 | 0.02 | 0.24 | 0.23 | 0.1 | 0.09 |
| 1 | 16275247 | rs2268341 | -0.211 | 0.048 | 0.034 | -2.239 | 0.011 | 0.132 | 0.132 | 0 | 0.4 | 0.21 | 0.36 | 0.09 | 0.37 | 0.46 |
| 1 | 16277647 | rs848189 | -0.995 | 0.082 | -0.027 | -2.255 | 0.012 | 0.027 | 0.027 | 0 | 0.2 | 0.37 | 0.32 | 0.23 | 0.18 | 0.13 |
| 1 | 16285342 | rs848194 | -1.892 | -0.054 | -0.37 | -2.126 | -0.08 | 0.011 | 0.011 | 0 | 0.64 | 0.93 | 0.71 | 0.31 | 0.56 | 0.59 |
| 1 | 16299312 | rs10927875 | 0.059 | 0.108 | -0.268 | -2.096 | 0.005 | 0.056 | 0.056 | 0 | 0.36 | 0.22 | 0.31 | 0.01 | 0.32 | 0.28 |
| 1 | 16302982 | rs698893 | 0.37 | 0.225 | -0.237 | -2.092 | -0.003 | 0.078 | 0.078 | 0 | 0.54 | 0.72 | 0.44 | 0.77 | 0.56 | 0.64 |
| 1 | 16312138 | rs1763632 | -0.372 | 0.143 | -0.21 | -1.855 | -0.022 | -0.048 | -0.048 | 0 | 0.92 | 0.98 | 0.76 | 0.78 | 0.9 | 0.92 |
| 1 | 16314726 | rs17383551 | 0.384 | 0.189 | 0.013 | -1.879 | 0.009 | -0.028 | -0.028 | 1.202 | 0.07 | 0.1 | 0.05 | 0 | 0.06 | 0.05 |
| 1 | 16319232 | rs848307 | -0.364 | 0.138 | -0.053 | -1.834 | -0.022 | -0.047 | -0.047 | 0.111 | 0.92 | 0.98 | 0.85 | 0.77 | 0.9 | 0.92 |
| 1 | 16324233 | rs1739822 | -0.533 | 0.31 | 0.079 | -1.787 | 0.007 | 0.009 | 0.009 | 0.957 | 0.15 | 0.13 | 0.08 | 0.08 | 0.1 | 0.19 |
| 1 | 16327422 | rs1739828 | -1.082 | 0.32 | 0.127 | -1.786 | 0.004 | -0.009 | -0.009 | 0.12 | 0.19 | 0.17 | 0.22 | 0.29 | 0.18 | 0.23 |
| 1 | 16331909 | rs1763613 | -0.567 | 0.334 | 0.107 | -1.774 | 0 | -0.007 | -0.007 | 0 | 0.13 | 0.14 | 0.21 | 0.29 | 0.16 | 0.21 |
| 1 | 16332571 | rs1763612 | 0.811 | 0.33 | 0.07 | -1.75 | 0.004 | 0.013 | 0.013 | 0 | 0.08 | 0.05 | 0.06 | 0.08 | 0.07 | 0.16 |
| 1 | 16336654 | rs11578845 | 1.34 | 0.32 | 0.379 | -1.956 | 0 | 0.006 | 0.006 | 0 | 0.39 | 0.35 | 0.31 | 0.67 | 0.42 | 0.41 |
| 1 | 16340761 | rs1048334 | -0.247 | 0.39 | 0.402 | -1.979 | -0.009 | 0.025 | 0.025 | 0.007 | 0.62 | 0.72 | 0.46 | 0.76 | 0.59 | 0.64 |
| 1 | 16341354 | rs1048245 | -0.256 | 0.389 | 0.419 | -1.98 | -0.01 | 0.026 | 0.026 | 0.01 | 0.62 | 0.73 | 0.46 | 0.76 | 0.59 | 0.64 |
| 1 | 16341525 | rs1739839 | 0.892 | 0.397 | 0.414 | -1.946 | 0.004 | 0.008 | 0.008 | 0 | 0.08 | 0.07 | 0.06 | 0.08 | 0.07 | 0.16 |
| 1 | 16344858 | rs945419 | 0.241 | 0.408 | 0.349 | -1.966 | -0.006 | 0.055 | 0.055 | 0 | 0.56 | 0.71 | 0.45 | 0.76 | 0.57 | 0.63 |
| 1 | 16345735 | rs1544131 | 0.164 | 0.404 | 0.298 | -1.896 | -0.006 | -0.016 | -0.016 | 0.015 | 0.1 | 0.29 | 0.09 | 0.09 | 0.07 | 0.16 |
| 1 | 16346507 | rs3754322 | -0.533 | 0.547 | -0.283 | -1.848 | -0.022 | -0.042 | -0.042 | 0 | 0.17 | 0.09 | 0.19 | 0.23 | 0.19 | 0.11 |
| 1 | 16348412 | rs945425 | -0.441 | 0.542 | -0.251 | -1.888 | -0.004 | 0.013 | 0.013 | 0 | 0.66 | 0.73 | 0.6 | 0.98 | 0.68 | 0.69 |
| 1 | 16348729 | rs1889785 | 0.407 | 0.38 | 0.032 | -1.591 | 0.008 | 0.047 | 0.047 | 0 | 0.48 | 0.58 | 0.39 | 0.76 | 0.49 | 0.58 |
| 1 | 16352937 | rs1010069 | 0.338 | 0.435 | -0.395 | -1.538 | 0.017 | 0.109 | 0.109 | 0 | 0.49 | 0.67 | 0.4 | 0.76 | 0.49 | 0.58 |
| 1 | 16367339 | rs9442193 | 0.127 | 0.558 | -0.177 | -1.447 | 0.005 | 0.039 | 0.039 | 0 | 0.41 | 0.22 | 0.49 | 0.31 | 0.44 | 0.51 |
| 1 | 16368324 | rs10803410 | 0.297 | 0.382 | 0.011 | -1.71 | 0.002 | -0.016 | -0.016 | 0 | 0.43 | 0.42 | 0.42 | 0.68 | 0.49 | 0.46 |
| 1 | 16368977 | rs2007471 | 1.173 | 0.635 | 0.248 | -1.624 | -0.005 | 0 | 0 | 0 | 0.31 | 0.19 | 0.35 | 0.67 | 0.36 | 0.4 |
| 1 | 16371932 | rs6604909 | 0.346 | 0.414 | 0.988 | -1.854 | -0.035 | -0.118 | -0.118 | 0 | 0.85 | 0.46 | 0.9 | 1 | 0.94 | 0.94 |
| 1 | 16373124 | rs5257 | 1.476 | 0.417 | 0.714 | -1.889 | -0.003 | -0.117 | -0.117 | 0 | 0.77 | 0.31 | 0.85 | 0.99 | 0.83 | 0.89 |
| 1 | 16376831 | rs10803412 | 1.472 | 0.208 | 0.709 | -1.88 | -0.014 | -0.115 | -0.115 | 0 | 0.77 | 0.32 | 0.85 | 0.99 | 0.84 | 0.89 |
| 1 | 16378215 | rs5252 | -1.082 | 0.628 | 0.196 | -1.498 | 0.018 | -0.043 | -0.043 | 0.171 | 0.09 | 0.21 | 0.07 | 0.01 | 0.1 | 0.05 |
| 1 | 16393357 | rs9442235 | -0.479 | 0.457 | -0.064 | -2.187 | -0.001 | -0.09 | -0.09 | 0 | 0.54 | 0.58 | 0.53 | 0.02 | 0.5 | 0.4 |
| 1 | 16395147 | rs10927902 | -0.978 | 0.4 | 0.045 | -2.536 | 0.011 | -0.132 | -0.132 | 0.95 | 0.22 | 0.45 | 0.15 | 0 | 0.19 | 0.09 |
| 1 | 16398375 | rs9442200 | -0.593 | 0.05 | -0.225 | -2.488 | 0.034 | 0.112 | 0.112 | 0 | 0.5 | 0.66 | 0.51 | 0.08 | 0.39 | 0.45 |
| 1 | 16413291 | rs2064595 | -1.582 | 0.015 | 0.101 | -2.167 | 0.001 | -0.049 | -0.049 | 0 | 0.29 | 0.52 | 0.24 | 0.16 | 0.27 | 0.29 |
| 1 | 16414088 | rs4661351 | 0.106 | 0.303 | -0.219 | -2.272 | 0.012 | 0.035 | 0.035 | 0.468 | 0.53 | 0.61 | 0.5 | 0.82 | 0.61 | 0.46 |
| 1 | 16422010 | rs28757713 | -0.112 | 0.468 | -0.502 | -2.013 | -0.004 | -0.01 | -0.01 | 1.033 | 0.13 | 0.11 | 0.1 | 0.01 | 0.17 | 0.07 |
| 1 | 16442908 | rs4661702 | -0.233 | -0.406 | -0.877 | -2.087 | 0.017 | 0.173 | 0.173 | 0 | 0.63 | 0.89 | 0.68 | 0.86 | 0.71 | 0.55 |
| 1 | 16446349 | rs924201 | 0.505 | -0.634 | -0.73 | -1.797 | 0.048 | 0.15 | 0.15 | 0.053 | 0.49 | 0.16 | 0.53 | 0.75 | 0.6 | 0.53 |
| 1 | 16446980 | rs17416878 | 0.107 | -0.684 | -0.975 | -2.215 | -0.007 | 0.223 | 0.223 | 0.045 | 0.38 | 0.08 | 0.32 | 0.03 | 0.34 | 0.32 |
| 1 | 16449746 | rs11260711 | 0.203 | -0.82 | -0.191 | -1.808 | 0.012 | 0.103 | 0.103 | 0 | 0.36 | 0.13 | 0.32 | 0.15 | 0.31 | 0.34 |
| 1 | 16451767 | rs3754334 | 0.215 | -0.842 | -0.377 | -1.794 | 0.005 | 0.121 | 0.121 | 0 | 0.36 | 0.11 | 0.31 | 0.15 | 0.32 | 0.35 |
| 1 | 16452998 | rs3820610 | 0.343 | -0.898 | -0.476 | -1.815 | 0.057 | 0.126 | 0.126 | 0 | 0.54 | 0.16 | 0.54 | 0.72 | 0.64 | 0.55 |
| 1 | 16459745 | rs10907223 | -1.61 | -0.875 | 0.757 | -1.578 | -0.029 | 0.041 | 0.041 | 6.283 | 0.12 | 0.48 | 0.08 | 0.11 | 0.05 | 0.12 |
| 1 | 16460541 | rs6603855 | -1.985 | -0.904 | 0.507 | -1.618 | -0.017 | -0.004 | -0.004 | 0 | 0.16 | 0.79 | 0.14 | 0.12 | 0.07 | 0.13 |
| 1 | 16464673 | rs2230597 | 0.171 | -0.988 | 0.383 | -2.088 | 0.004 | 0.089 | 0.089 | 0 | 0.46 | 0.34 | 0.43 | 0.15 | 0.42 | 0.44 |

| | | | | | | | | | | | | | | | | |
|---|---|---|---|---|---|---|---|---|---|---|---|---|---|---|---|---|
| 1 | 16466003 | rs11260745 | -1.208 | -1.219 | 0.915 | -1.778 | -0.022 | 0.062 | 0.062 | 0 | 0.11 | 0.4 | 0.08 | 0.11 | 0.06 | 0.16 |
| 1 | 16467924 | rs3768293 | -0.124 | -1.129 | 0.946 | -1.934 | 0.003 | -0.019 | -0.019 | 4.899 | 0.49 | 0.48 | 0.52 | 0.84 | 0.58 | 0.59 |
| 1 | 16474701 | rs6603867 | -0.038 | -1.079 | 1.234 | -2.085 | -0.001 | -0.065 | -0.065 | 0.976 | 0.51 | 0.47 | 0.45 | 0.07 | 0.4 | 0.34 |
| 1 | 16478654 | rs1472408 | 0.026 | -1.152 | 1.274 | -2.111 | 0.017 | 0.092 | 0.092 | 0 | 0.46 | 0.38 | 0.52 | 0.89 | 0.59 | 0.6 |
| 1 | 16482976 | rs6603883 | -0.251 | -1.234 | 1.509 | -2.149 | 0.004 | -0.032 | -0.032 | 0.382 | 0.5 | 0.57 | 0.54 | 0.89 | 0.6 | 0.6 |
| 1 | 16486793 | rs904106 | 1.729 | -1.044 | 1.53 | -1.626 | 0.059 | 0.101 | 0.101 | 0 | 0.93 | 0.63 | 0.9 | 0.89 | 0.96 | 0.89 |
| 1 | 16495846 | rs4661715 | 1.434 | -1.011 | 0.848 | -1.562 | -0.02 | 0.022 | 0.022 | 0 | 0.87 | 0.7 | 0.91 | 0.85 | 0.9 | 0.82 |
| 1 | 16495998 | rs924206 | 1.434 | -1.011 | 0.85 | -1.562 | -0.02 | 0.023 | 0.023 | 0 | 0.87 | 0.7 | 0.91 | 0.85 | 0.9 | 0.83 |
| 1 | 16504549 | rs7538966 | -0.051 | -1.047 | 0.726 | -1.854 | -0.001 | -0.024 | -0.024 | 0 | 0.47 | 0.66 | 0.49 | 0.87 | 0.61 | 0.49 |
| 1 | 16514369 | rs4661721 | -0.531 | -1.038 | 0.356 | -2.114 | -0.025 | -0.063 | -0.063 | 3.212 | 0.18 | 0.34 | 0.17 | 0.08 | 0.11 | 0.07 |
| 1 | 16530364 | rs221047 | 0.831 | -0.756 | 0.688 | -1.906 | 0.008 | -0.027 | -0.027 | 0 | 0.84 | 0.68 | 0.86 | 0.86 | 0.84 | 0.88 |
| 1 | 16591072 | rs6698317 | -0.723 | 0.726 | 0.974 | -1.963 | 0.061 | 0.017 | 0.017 | 0 | 0.09 | 0.1 | 0.13 | 0.17 | 0.23 | 0.1 |
| 1 | 17462261 | rs877648 | 1.482 | -2.198 | 0.212 | -0.831 | 0.116 | 0.089 | 0.089 | 0.637 | 0.5 | 0.22 | 0.48 | 0.32 | 0.63 | 0.57 |
| 1 | 17473895 | rs6586503 | 1.698 | -0.96 | 1.208 | -0.12 | 0.016 | -0.015 | -0.015 | 0.074 | 0.7 | 0.33 | 0.74 | 0.37 | 0.77 | 0.61 |
| 1 | 17476831 | rs12085482 | -0.294 | -1.048 | 1.174 | -0.126 | -0.041 | -0.01 | -0.01 | 0 | 0.1 | 0.16 | 0.03 | 0.06 | 0.01 | 0.11 |
| 1 | 17486722 | rs4620585 | 1.695 | -0.905 | 0.763 | -0.113 | 0.018 | -0.014 | -0.014 | 0.117 | 0.67 | 0.35 | 0.68 | 0.37 | 0.71 | 0.56 |
| 1 | 17487951 | rs2454163 | 1.05 | -0.2 | 0.74 | 0.267 | 0 | 0.009 | 0.009 | 0 | 0.83 | 0.84 | 0.8 | 0.43 | 0.78 | 0.71 |
| 1 | 17490249 | rs2454170 | 0.725 | 0.212 | -0.904 | 0.437 | -0.004 | -0.025 | -0.025 | 0 | 0.84 | 0.92 | 0.81 | 0.43 | 0.78 | 0.72 |
| 1 | 17501898 | rs2977258 | -0.873 | 0.178 | -0.845 | -1.225 | -0.001 | -0.037 | -0.037 | 1.442 | 0.3 | 0.28 | 0.21 | 0.01 | 0.34 | 0.15 |
| 1 | 17503210 | rs2977265 | -0.853 | 0.087 | -0.796 | -1.266 | -0.001 | -0.055 | -0.055 | 0 | 0.31 | 0.34 | 0.22 | 0.01 | 0.34 | 0.18 |
| 1 | 17510703 | rs2935923 | -0.898 | -0.009 | -0.118 | -1.31 | -0.001 | -0.102 | -0.102 | 0 | 0.31 | 0.4 | 0.22 | 0.01 | 0.33 | 0.18 |
| 1 | 17518160 | rs12744275 | 0.661 | -0.292 | -1.369 | -1.585 | 0.03 | 0.056 | 0.056 | 0 | 0.29 | 0.03 | 0.12 | 0.15 | 0.21 | 0.16 |
| 1 | 17521274 | rs3003469 | -0.258 | -0.012 | -0.706 | -1.322 | -0.044 | 0.064 | 0.064 | 0 | 0.2 | 0.08 | 0.18 | 0.13 | 0.28 | 0.12 |
| 1 | 17522821 | rs754171 | -1.064 | 0.151 | -0.48 | -1.525 | 0.003 | -0.043 | -0.043 | 0 | 0.26 | 0.31 | 0.21 | 0.04 | 0.33 | 0.15 |
| 1 | 17523194 | rs2935941 | -0.299 | 0.325 | -0.829 | -1.405 | -0.015 | 0.027 | 0.027 | 0 | 0.19 | 0.16 | 0.16 | 0.04 | 0.27 | 0.12 |
| 1 | 17523806 | rs873319 | -0.275 | 0.335 | -0.865 | -1.4 | -0.017 | 0.029 | 0.029 | 0 | 0.19 | 0.15 | 0.16 | 0.04 | 0.27 | 0.12 |
| 1 | 17525965 | rs3003473 | -0.873 | 0.257 | -0.916 | -1.43 | -0.018 | 0.041 | 0.041 | 0 | 0.27 | 0.21 | 0.2 | 0.07 | 0.34 | 0.19 |
| 1 | 17531582 | rs2293910 | 0.176 | 0.331 | -0.482 | -1.216 | 0.002 | -0.017 | -0.017 | 0 | 0.13 | 0.12 | 0.1 | 0.04 | 0.17 | 0.1 |
| 1 | 17534310 | rs4384275 | -1.047 | 0.077 | -0.127 | -1.392 | -0.003 | -0.052 | -0.052 | 0 | 0.3 | 0.38 | 0.19 | 0.12 | 0.32 | 0.26 |
| 1 | 17536952 | rs2977272 | 0.705 | -0.105 | -0.799 | -1.695 | -0.045 | 0.137 | 0.137 | 0 | 0.32 | 0.05 | 0.22 | 0.09 | 0.32 | 0.15 |
| 1 | 17538952 | rs2977278 | 0.383 | -0.382 | 0.264 | -1.151 | -0.004 | -0.041 | -0.041 | 0.018 | 0.76 | 0.75 | 0.84 | 0.95 | 0.76 | 0.81 |
| 1 | 17539073 | rs4920358 | -0.876 | -0.404 | 0.52 | -1.16 | -0.019 | -0.053 | -0.053 | 0.286 | 0.27 | 0.57 | 0.2 | 0.1 | 0.25 | 0.19 |
| 1 | 17540695 | rs6667138 | -1.704 | -0.087 | 1.208 | -0.851 | 0.055 | -0.033 | -0.033 | 0 | 0.11 | 0.47 | 0.11 | 0.02 | 0.09 | 0.07 |
| 1 | 17541949 | rs2977283 | -2.008 | -0.338 | 0.958 | -0.68 | -0.009 | -0.018 | -0.018 | 0 | 0.29 | 0.64 | 0.44 | 0.2 | 0.23 | 0.22 |
| 1 | 17543077 | rs3115788 | -0.958 | -0.387 | 0.729 | -0.464 | -0.006 | 0.03 | 0.03 | 0.036 | 0.09 | 0.29 | 0.09 | 0.13 | 0.06 | 0.11 |
| 1 | 17545283 | rs12401313 | -1.371 | -0.574 | 0.43 | -0.575 | -0.007 | -0.003 | -0.003 | 0 | 0.21 | 0.21 | 0.36 | 0.17 | 0.14 | 0.22 |
| 1 | 17545548 | rs3003479 | -1.589 | -0.526 | 0.298 | -0.588 | -0.062 | -0.062 | -0.062 | 0 | 0.26 | 0.5 | 0.38 | 0.17 | 0.15 | 0.22 |
| 1 | 17550121 | rs2293912 | -1.181 | 0.361 | -0.421 | 0.136 | -0.021 | -0.026 | -0.026 | 0.006 | 0.11 | 0.05 | 0.31 | 0.16 | 0.14 | 0.16 |
| 1 | 17550314 | rs2293914 | -1.019 | 0.186 | -0.249 | 0.268 | 0 | 0.006 | 0.006 | 0 | 0.1 | 0.11 | 0.31 | 0.2 | 0.11 | 0.13 |
| 1 | 17550601 | rs3003482 | 0.034 | 0.678 | 0.065 | 1.071 | 0.055 | 0.235 | 0.235 | 0 | 0.86 | 0.61 | 0.85 | 0.62 | 0.81 | 0.71 |
| 1 | 17551943 | rs16823950 | 0.492 | 0.331 | -0.771 | 0.654 | -0.005 | -0.025 | -0.025 | 0.5 | 0.22 | 0.07 | 0.18 | 0.25 | 0.21 | 0.14 |
| 1 | 17552875 | rs2977290 | -0.398 | 0.691 | -0.32 | 0.651 | 0.039 | 0.238 | 0.238 | 0 | 0.87 | 0.64 | 0.87 | 0.59 | 0.83 | 0.71 |
| 1 | 17553091 | rs12024174 | -0.466 | 0.278 | -0.746 | 0.394 | 0.022 | -0.013 | -0.013 | 0 | 0.4 | 0.36 | 0.52 | 0.44 | 0.31 | 0.28 |
| 1 | 17554853 | rs10127662 | -0.274 | 1.181 | -0.826 | -0.228 | -0.008 | -0.004 | -0.004 | 0.122 | 0.13 | 0.08 | 0.09 | 0.01 | 0.17 | 0.13 |
| 1 | 17555931 | rs12091488 | -0.499 | 0.545 | -0.607 | -0.576 | -0.021 | -0.067 | -0.067 | 0 | 0.29 | 0.57 | 0.21 | 0.11 | 0.22 | 0.29 |
| 1 | 17557039 | rs10888010 | -1.198 | 0.929 | -0.434 | 0.122 | 0.002 | 0.008 | 0.008 | 0 | 0.35 | 0.59 | 0.35 | 0.45 | 0.43 | 0.51 |
| 1 | 17557133 | rs3003406 | -1.099 | 0.847 | 0.303 | 0.214 | 0.015 | 0.027 | 0.027 | 0 | 0.33 | 0.4 | 0.32 | 0.37 | 0.43 | 0.5 |
| 1 | 17557178 | rs2977235 | -0.962 | 0.941 | 0.276 | 0.252 | 0.007 | 0.046 | 0.046 | 2.445 | 0.28 | 0.41 | 0.28 | 0.3 | 0.33 | 0.43 |
| 1 | 17559196 | rs4268393 | -1.162 | 0.888 | 0.63 | 0.345 | 0.002 | -0.012 | -0.012 | 0 | 0.25 | 0.46 | 0.22 | 0.18 | 0.27 | 0.37 |
| 1 | 17559676 | rs3750297 | -1.397 | 0.624 | 0.346 | 0.045 | 0.001 | -0.014 | -0.014 | 0 | 0.27 | 0.32 | 0.24 | 0.14 | 0.32 | 0.31 |
| 1 | 17560646 | rs12026071 | -0.068 | 0.673 | 0.283 | 0.057 | -0.023 | 0.008 | 0.008 | 0.223 | 0.15 | 0.06 | 0.15 | 0.11 | 0.23 | 0.21 |
| 1 | 17560972 | rs11203339 | -2.211 | 0.098 | 0.263 | -0.237 | -0.007 | -0.013 | -0.013 | 0.018 | 0.4 | 0.57 | 0.33 | 0.21 | 0.37 | 0.51 |
| 1 | 17562823 | rs12028888 | -0.174 | 0.214 | -0.274 | -0.705 | -0.038 | 0.046 | 0.046 | 0.736 | 0.22 | 0.06 | 0.25 | 0.11 | 0.3 | 0.26 |
| 1 | 17563967 | rs2977306 | -2.016 | 0.538 | 0.714 | -0.23 | 0.005 | 0.022 | 0.022 | 0.053 | 0.37 | 0.53 | 0.4 | 0.44 | 0.42 | 0.59 |
| 1 | 17564990 | rs2977310 | -2.31 | 0.344 | 0.603 | -0.38 | 0.001 | -0.005 | -0.005 | 0 | 0.46 | 0.61 | 0.44 | 0.45 | 0.48 | 0.64 |
| 1 | 17569949 | rs2977313 | -0.92 | 0.573 | 0.875 | -0.162 | 0.025 | 0.062 | 0.062 | 0 | 0.12 | 0.28 | 0.12 | 0.19 | 0.15 | 0.14 |
| 1 | 17573439 | rs2977225 | -1.322 | 0.579 | 0.871 | -0.22 | -0.001 | -0.001 | -0.001 | 1.651 | 0.25 | 0.22 | 0.26 | 0.34 | 0.3 | 0.36 |
| 1 | 17575155 | rs12404500 | 3.113 | 0.644 | 0.512 | -0.556 | -0.096 | -0.097 | -0.097 | 0.084 | 0.46 | 0.03 | 0.39 | 0.51 | 0.43 | 0.32 |
| 1 | 17576328 | rs2977227 | -2.477 | 0.743 | 0.978 | -0.445 | 0 | 0.002 | 0.002 | 0.249 | 0.39 | 0.59 | 0.43 | 0.44 | 0.45 | 0.6 |
| 1 | 17576499 | rs2977228 | -0.614 | 0.78 | 0.991 | -0.385 | 0.018 | 0.016 | 0.016 | 0 | 0.15 | 0.19 | 0.13 | 0.19 | 0.22 | 0.13 |
| 1 | 17578750 | rs6673643 | -2.676 | 0.916 | 0.948 | -0.407 | -0.004 | -0.009 | -0.009 | 0 | 0.42 | 0.64 | 0.43 | 0.44 | 0.46 | 0.64 |
| 1 | 17579783 | rs12025184 | -0.503 | 0.863 | 1.123 | -0.368 | 0.008 | 0.003 | 0.003 | 0.183 | 0.12 | 0.29 | 0.15 | 0.1 | 0.15 | 0.24 |
| 1 | 17580568 | rs4920581 | 0.46 | 0.869 | 1.019 | -0.336 | -0.001 | -0.03 | -0.03 | 0 | 0.11 | 0.03 | 0.13 | 0.15 | 0.09 | 0.23 |
| 1 | 17586134 | rs3750300 | 0.318 | 0.867 | 1.023 | -0.331 | 0.003 | -0.008 | -0.008 | 0.627 | 0.11 | 0.08 | 0.14 | 0.16 | 0.09 | 0.23 |
| 1 | 17588162 | rs3003426 | -1.801 | 0.852 | 1.118 | -0.356 | -0.016 | 0.046 | 0.046 | 2.197 | 0.3 | 0.58 | 0.34 | 0.38 | 0.28 | 0.56 |
| 1 | 17589048 | rs3003427 | -0.291 | 1.031 | 1.324 | -0.357 | 0.02 | -0.022 | -0.022 | 0 | 0.92 | 0.82 | 0.91 | 0.95 | 0.89 | 0.96 |
| 1 | 17591077 | rs2977269 | -1.377 | 1.163 | 1.224 | -0.104 | 0.004 | 0.015 | 0.015 | 0.154 | 0.23 | 0.26 | 0.26 | 0.29 | 0.25 | 0.32 |
| 1 | 17592324 | rs2272627 | -0.445 | 1.049 | 0.884 | -0.107 | 0.009 | 0.005 | 0.005 | 0.449 | 0.11 | 0.23 | 0.15 | 0.1 | 0.15 | 0.24 |
| 1 | 17593316 | rs2272629 | 0.328 | 1.132 | 0.419 | -0.18 | 0.002 | -0.026 | -0.026 | 0 | 0.1 | 0.03 | 0.12 | 0.15 | 0.09 | 0.24 |

| | | | | | | | | | | | | | | | | |
|---|---|---|---|---|---|---|---|---|---|---|---|---|---|---|---|---|
| 1 | 17595811 | rs10888012 | -1.406 | 1.253 | 0.612 | -0.211 | 0.03 | 0.027 | 0.027 | 0.673 | 0.23 | 0.4 | 0.26 | 0.23 | 0.32 | 0.33 |
| 1 | 17597721 | rs4595409 | -1.311 | 1.286 | 0.598 | -0.683 | 0.003 | -0.038 | -0.038 | 0 | 0.26 | 0.43 | 0.27 | 0.15 | 0.27 | 0.32 |
| 1 | 17599444 | rs2977293 | -0.781 | 1.713 | 0.439 | -0.448 | -0.012 | -0.001 | -0.001 | 0 | 0.11 | 0.05 | 0.14 | 0.11 | 0.23 | 0.11 |
| 1 | 17600437 | rs2977295 | -1.053 | 1.433 | 0.859 | -0.714 | -0.001 | -0.012 | -0.012 | 0 | 0.19 | 0.32 | 0.2 | 0.18 | 0.17 | 0.15 |
| 1 | 17601083 | rs2293917 | -2.219 | 1.578 | 1.212 | -0.623 | 0.025 | -0.028 | -0.028 | 0 | 0.3 | 0.58 | 0.34 | 0.25 | 0.39 | 0.2 |
| 1 | 17601165 | rs11585357 | 0.172 | 1.589 | 1.175 | -0.619 | -0.046 | 0.04 | 0.04 | 0 | 0.07 | 0.01 | 0.1 | 0 | 0.18 | 0.04 |
| 1 | 17602191 | rs3003434 | -1.433 | 1.575 | 1.225 | -0.618 | 0.007 | -0.006 | -0.006 | 0 | 0.22 | 0.5 | 0.23 | 0.25 | 0.21 | 0.16 |
| 1 | 17605774 | rs11585118 | 0.115 | 1.633 | 1 | -0.567 | -0.046 | 0.04 | 0.04 | 0 | 0.07 | 0.01 | 0.1 | 0 | 0.17 | 0.04 |
| 1 | 17610899 | rs3750304 | 0.002 | 1.657 | 0.846 | -0.957 | 0.001 | -0.012 | -0.012 | 0.186 | 0.23 | 0.3 | 0.26 | 0.2 | 0.22 | 0.35 |
| 1 | 17622654 | rs7548900 | -0.718 | 1.607 | 0.855 | -1.01 | -0.009 | 0.011 | 0.011 | 0 | 0.32 | 0.29 | 0.38 | 0.19 | 0.41 | 0.38 |
| 1 | 17622897 | rs7542629 | 0.451 | 1.579 | 0.695 | -1.001 | -0.009 | 0.025 | 0.025 | 0 | 0.21 | 0.06 | 0.25 | 0.12 | 0.19 | 0.33 |
| 1 | 17629004 | rs2477142 | -2.002 | 1.458 | 0.776 | -1.033 | 0.02 | 0.07 | 0.07 | 0 | 0.53 | 0.81 | 0.61 | 0.38 | 0.58 | 0.52 |
| 1 | 17635396 | rs1886302 | -1.286 | 0.915 | 0.917 | -0.848 | 0 | 0.003 | 0.003 | 0 | 0.41 | 0.32 | 0.45 | 0.31 | 0.36 | 0.49 |
| 1 | 17636321 | rs2477131 | 0.164 | 1.01 | 0.587 | -0.718 | -0.011 | -0.039 | -0.039 | 0.187 | 0.75 | 0.59 | 0.73 | 0.87 | 0.81 | 0.66 |
| 1 | 17647475 | rs10788664 | -0.561 | 1.023 | 0.124 | -0.614 | -0.001 | -0.001 | -0.001 | 0 | 0.39 | 0.34 | 0.39 | 0.38 | 0.31 | 0.43 |
| 1 | 17649796 | rs10888024 | 0.899 | 1.361 | 0.259 | -0.634 | 0 | 0.009 | 0.009 | 0.753 | 0.53 | 0.48 | 0.54 | 0.56 | 0.57 | 0.52 |
| 1 | 17655407 | rs1748041 | 0.274 | 1.36 | 0.298 | -0.637 | -0.007 | -0.01 | -0.01 | 0 | 0.6 | 0.61 | 0.61 | 0.62 | 0.69 | 0.58 |
| 1 | 17657534 | rs11203366 | 0.929 | 1.332 | 0.333 | -0.64 | 0.001 | 0.016 | 0.016 | 0.96 | 0.52 | 0.45 | 0.54 | 0.56 | 0.58 | 0.52 |
| 1 | 17657616 | rs11203367 | 0.872 | 1.335 | 0.336 | -0.634 | 0 | 0.009 | 0.009 | 0.753 | 0.53 | 0.48 | 0.55 | 0.56 | 0.57 | 0.52 |
| 1 | 17662662 | rs1748033 | 0.46 | 1.376 | 0.316 | -0.629 | -0.007 | -0.017 | -0.017 | 0.016 | 0.57 | 0.54 | 0.59 | 0.62 | 0.66 | 0.57 |
| 1 | 17666326 | rs1635576 | -0.659 | 0.758 | 0.186 | -0.283 | 0.02 | -0.002 | -0.002 | 0 | 0.05 | 0.14 | 0.05 | 0.05 | 0.1 | 0.05 |
| 1 | 17674108 | rs2240339 | 0.842 | 0.521 | -0.441 | -0.52 | -0.001 | 0 | 0 | 0 | 0.38 | 0.34 | 0.41 | 0.56 | 0.35 | 0.45 |
| 1 | 17674185 | rs2240338 | 0.842 | 0.521 | -0.441 | -0.52 | -0.001 | 0 | 0 | 0 | 0.38 | 0.34 | 0.41 | 0.56 | 0.35 | 0.45 |
| 1 | 17674222 | rs2240337 | 0.466 | 0.471 | -0.403 | -0.641 | -0.005 | -0.009 | -0.009 | 0 | 0.12 | 0.13 | 0.13 | 0.07 | 0.08 | 0.02 |
| 1 | 17674537 | rs2240335 | 0.499 | 0.529 | -0.593 | -0.298 | 0 | 0 | 0 | 0.203 | 0.39 | 0.35 | 0.41 | 0.56 | 0.35 | 0.45 |
| 1 | 17674821 | rs12746451 | -1.446 | 1.097 | -0.818 | 0.015 | -0.004 | -0.011 | -0.011 | 0.037 | 0.08 | 0.19 | 0.06 | 0.05 | 0.07 | 0.02 |
| 1 | 17681277 | rs6683201 | 0.266 | 1.641 | -1.501 | 0.283 | -0.088 | -0.047 | -0.047 | 0 | 0.81 | 0.92 | 0.7 | 0.77 | 0.7 | 0.71 |
| 1 | 17683526 | rs1635564 | 0.486 | 1.408 | -1.847 | 0.052 | -0.112 | -0.054 | -0.054 | 0.722 | 0.75 | 0.98 | 0.69 | 0.79 | 0.7 | 0.66 |
| 1 | 17687027 | rs1635561 | -0.321 | -0.085 | -1.507 | 0.24 | 0.007 | -0.003 | -0.003 | 0 | 0.3 | 0.23 | 0.14 | 0.3 | 0.21 | 0.23 |
| 1 | 17693711 | rs1204892 | -0.504 | -0.19 | -1.431 | 0.324 | -0.001 | -0.001 | -0.001 | 0 | 0.24 | 0.22 | 0.14 | 0.31 | 0.19 | 0.19 |
| 1 | 17694701 | rs1204890 | -0.746 | -0.376 | -0.731 | 0.266 | -0.042 | 0.027 | 0.027 | 1.201 | 0.27 | 0.48 | 0.17 | 0.3 | 0.19 | 0.2 |
| 1 | 17695153 | rs12732819 | -0.343 | -0.313 | -0.956 | 0.308 | -0.008 | 0.003 | 0.003 | 1.656 | 0.23 | 0.21 | 0.13 | 0.27 | 0.16 | 0.19 |
| 1 | 17707739 | rs1204898 | 0.199 | 0.148 | -0.473 | -0.121 | 0.005 | -0.045 | -0.045 | 0 | 0.76 | 0.61 | 0.73 | 0.82 | 0.7 | 0.79 |
| 1 | 17707798 | rs10788668 | 0.207 | 0.024 | -0.046 | 0.182 | 0.013 | 0.061 | 0.061 | 0 | 0.77 | 0.6 | 0.73 | 0.59 | 0.7 | 0.73 |
| 1 | 17708095 | rs2181867 | -0.013 | -0.113 | -0.276 | 0.087 | -0.001 | 0.002 | 0.002 | 0 | 0.23 | 0.2 | 0.2 | 0.31 | 0.23 | 0.23 |
| 1 | 17714864 | rs6696295 | -0.701 | -0.318 | -0.58 | -0.119 | 0.042 | -0.035 | -0.035 | 0.041 | 0.28 | 0.15 | 0.25 | 0.32 | 0.2 | 0.27 |
| 1 | 17715081 | rs2526842 | -1.015 | -0.372 | -0.077 | -0.171 | -0.014 | 0.004 | 0.004 | 0 | 0.32 | 0.4 | 0.29 | 0.36 | 0.24 | 0.28 |
| 1 | 17719061 | rs10458538 | 0.955 | -0.369 | -0.343 | -0.088 | 0.011 | -0.072 | -0.072 | 1.944 | 0.11 | 0.01 | 0.08 | 0.27 | 0.09 | 0.15 |
| 1 | 17719843 | rs6695214 | -0.579 | -0.218 | -0.09 | -0.273 | -0.001 | 0.011 | 0.011 | 0 | 0.64 | 0.49 | 0.63 | 0.53 | 0.61 | 0.65 |
| 1 | 17720615 | rs10458537 | -0.594 | -0.226 | -0.094 | -0.277 | -0.001 | 0.012 | 0.012 | 0 | 0.64 | 0.49 | 0.63 | 0.53 | 0.61 | 0.65 |
| 1 | 17722363 | rs7538876 | 1.094 | -0.139 | -0.077 | -0.289 | 0.003 | -0.006 | -0.006 | 0 | 0.34 | 0.34 | 0.37 | 0.17 | 0.37 | 0.37 |
| 1 | 17723126 | rs2526833 | -0.645 | -0.116 | 0.087 | -0.298 | -0.003 | -0.008 | -0.008 | 0.582 | 0.66 | 0.72 | 0.66 | 0.52 | 0.61 | 0.65 |
| 1 | 17728119 | rs2526830 | -0.745 | -0.155 | 0.072 | -0.241 | -0.002 | -0.005 | -0.005 | 1.182 | 0.66 | 0.69 | 0.65 | 0.54 | 0.61 | 0.66 |
| 1 | 17739313 | rs942458 | -0.613 | -0.193 | 0.155 | -0.244 | 0 | 0.003 | 0.003 | 0 | 0.65 | 0.63 | 0.64 | 0.55 | 0.61 | 0.68 |
| 1 | 17739586 | rs942457 | 0.967 | -0.15 | 0.081 | -0.218 | 0.007 | -0.016 | -0.016 | 0 | 0.35 | 0.37 | 0.37 | 0.16 | 0.37 | 0.39 |
| 1 | 17753639 | rs6688886 | 0.978 | -0.091 | -0.049 | -0.262 | 0.015 | -0.03 | -0.03 | 0 | 0.34 | 0.43 | 0.37 | 0.17 | 0.35 | 0.39 |
| 1 | 17759245 | rs2489611 | -0.747 | -0.116 | -0.024 | -0.305 | -0.004 | -0.011 | -0.011 | 0 | 0.66 | 0.72 | 0.66 | 0.55 | 0.61 | 0.69 |
| 1 | 17763312 | rs2800686 | -0.747 | -0.116 | -0.024 | -0.305 | -0.004 | -0.011 | -0.011 | 0.143 | 0.66 | 0.72 | 0.65 | 0.55 | 0.61 | 0.69 |
| 1 | 17773159 | rs2883272 | -0.364 | -0.439 | 0.166 | -0.552 | 0.019 | 0.027 | 0.027 | 0 | 0.27 | 0.18 | 0.22 | 0.05 | 0.2 | 0.21 |
| 1 | 17778571 | rs2526822 | -0.214 | 0.401 | 0.371 | 0.229 | -0.002 | 0.004 | 0.004 | 0 | 0.38 | 0.54 | 0.27 | 0.34 | 0.36 | 0.41 |
| 1 | 17791882 | rs12078935 | -0.001 | 0.338 | 0.403 | 0.239 | -0.001 | 0 | 0 | 0.121 | 0.59 | 0.46 | 0.71 | 0.66 | 0.59 | 0.58 |
| 1 | 17796711 | rs4284283 | 0.235 | 0.399 | 0.279 | 0.319 | 0 | -0.014 | -0.014 | 3.564 | 0.36 | 0.37 | 0.23 | 0.13 | 0.37 | 0.36 |
| 1 | 17815433 | rs4387213 | 0.222 | 0.306 | 0.323 | -0.238 | 0 | -0.027 | -0.027 | 0.582 | 0.34 | 0.32 | 0.21 | 0.09 | 0.36 | 0.35 |
| 1 | 17818579 | rs7534298 | 0.122 | 0.193 | 0.437 | -0.166 | 0.005 | -0.051 | -0.051 | 0 | 0.36 | 0.44 | 0.24 | 0.13 | 0.37 | 0.38 |
| 1 | 17820742 | rs6689677 | -0.129 | 0.165 | 0.44 | -0.038 | 0 | -0.021 | -0.021 | 0.354 | 0.62 | 0.51 | 0.75 | 0.75 | 0.62 | 0.57 |
| 1 | 17829956 | rs11203414 | 0.305 | 0.143 | 0.54 | -0.161 | 0.003 | -0.049 | -0.049 | 0 | 0.3 | 0.32 | 0.19 | 0.07 | 0.31 | 0.31 |
| 1 | 17843099 | rs4083558 | 0.128 | 0.122 | 0.574 | -0.179 | 0 | -0.033 | -0.033 | 0.121 | 0.33 | 0.32 | 0.21 | 0.08 | 0.34 | 0.31 |
| 1 | 17846922 | rs2244444 | -0.039 | -0.2 | 0.754 | 0.024 | -0.001 | 0 | 0 | 0 | 0.59 | 0.46 | 0.68 | 0.75 | 0.56 | 0.57 |
| 1 | 17857601 | rs2244300 | 0.625 | -0.524 | -0.159 | -0.109 | -0.008 | 0.036 | 0.036 | 0 | 0.44 | 0.34 | 0.51 | 0.71 | 0.46 | 0.46 |
| 1 | 17858638 | rs7549022 | -0.103 | -0.194 | -0.017 | 0.038 | -0.011 | 0.027 | 0.027 | 0 | 0.59 | 0.41 | 0.59 | 0.76 | 0.54 | 0.58 |
| 1 | 17864514 | rs2245148 | -0.375 | -0.145 | 0.018 | 0.002 | 0.006 | -0.055 | -0.055 | 0.357 | 0.62 | 0.54 | 0.65 | 0.89 | 0.57 | 0.65 |
| 1 | 17869179 | rs2477735 | 0.18 | -0.033 | -0.257 | 0.133 | 0.009 | -0.027 | -0.027 | 0 | 0.05 | 0.12 | 0.05 | 0.01 | 0.07 | 0.07 |
| 1 | 17872028 | rs4920374 | 0.048 | -0.142 | -0.27 | 0.141 | -0.009 | 0.025 | 0.025 | 0 | 0.56 | 0.46 | 0.57 | 0.76 | 0.49 | 0.51 |
| 1 | 17872110 | rs2248697 | -0.24 | 0.458 | -0.718 | 0.383 | 0.007 | -0.027 | -0.027 | 4.892 | 0.05 | 0.1 | 0.05 | 0.01 | 0.07 | 0.07 |
| 1 | 17874843 | rs4920375 | 0.257 | 0.33 | -1.724 | 1.011 | -0.137 | -0.179 | -0.179 | 0.998 | 0.34 | 0.03 | 0.42 | 0.51 | 0.39 | 0.31 |
| 1 | 17876389 | rs2249161 | -1.17 | -0.075 | -0.303 | 1.638 | 0.001 | 0.164 | 0.164 | 0 | 0.14 | 0.59 | 0.16 | 0.35 | 0.15 | 0.1 |
| 1 | 17878619 | rs1408953 | -1.03 | -0.491 | -0.636 | 1.377 | 0.011 | -0.02 | -0.02 | 0.002 | 0.37 | 0.69 | 0.28 | 0.5 | 0.32 | 0.24 |
| 1 | 17885856 | rs12039118 | 0.38 | -0.674 | -0.525 | 1.646 | -0.02 | 0.05 | 0.05 | 0.155 | 0.1 | 0.17 | 0.07 | 0.21 | 0.05 | 0.04 |
| 1 | 17887963 | rs2256114 | -1.589 | -0.018 | -0.888 | 1.307 | -0.062 | -0.064 | -0.064 | 0 | 0.41 | 0.85 | 0.37 | 0.5 | 0.44 | 0.28 |

| | | | | | | | | | | | | | | | | |
|---|---|---|---|---|---|---|---|---|---|---|---|---|---|---|---|---|
| 1 | 17892396 | rs12725975 | 1.773 | -0.762 | -1.359 | 1.124 | -0.114 | 0.034 | 0.034 | 0 | 0.35 | 0.05 | 0.5 | 0.25 | 0.43 | 0.24 |
| 1 | 17892816 | rs2256787 | -0.986 | -1.818 | -0.715 | 1.075 | 0.011 | 0.014 | 0.014 | 0 | 0.88 | 0.93 | 0.92 | 0.93 | 0.94 | 0.78 |
| 1 | 17895744 | rs2257142 | -0.418 | -1.189 | 0.729 | 1.631 | 0.008 | -0.054 | -0.054 | 0 | 0.21 | 0.4 | 0.15 | 0.11 | 0.22 | 0.17 |
| 1 | 17899151 | rs12049589 | -0.211 | -1.162 | 0.718 | 1.7 | -0.015 | 0.108 | 0.108 | 0.037 | 0.12 | 0.17 | 0.1 | 0.37 | 0.08 | 0.06 |
| 1 | 17899719 | rs2246282 | 0.117 | -1.152 | 0.575 | 1.335 | -0.002 | -0.001 | -0.001 | 0.287 | 0.07 | 0.05 | 0.05 | 0.07 | 0.09 | 0.04 |
| 1 | 17903298 | rs6702679 | -0.996 | -1.071 | 0.539 | 1.428 | -0.008 | 0.032 | 0.032 | 0 | 0.27 | 0.58 | 0.19 | 0.37 | 0.22 | 0.2 |
| 1 | 17905190 | rs12066034 | 0.002 | -1.091 | 0.189 | 1.005 | -0.023 | 0.071 | 0.071 | 0.099 | 0.1 | 0.29 | 0.06 | 0.17 | 0.05 | 0.04 |
| 1 | 17907505 | rs11203423 | 0.137 | -0.885 | -0.591 | 0.872 | -0.001 | 0 | 0 | 0 | 0.66 | 0.36 | 0.78 | 0.65 | 0.71 | 0.76 |
| 1 | 17909412 | rs12756041 | -1.028 | -0.818 | -0.426 | 0.856 | -0.019 | -0.004 | -0.004 | 0 | 0.2 | 0.18 | 0.09 | 0.14 | 0.11 | 0.26 |
| 1 | 17915833 | rs12404333 | -0.675 | 0.034 | -0.488 | 1.036 | 0.001 | 0.018 | 0.018 | 0 | 0.22 | 0.28 | 0.47 | 0.34 | 0.26 | 0.35 |
| 1 | 17916382 | rs4920379 | 0.44 | -0.227 | -0.792 | 0.579 | 0.011 | -0.016 | -0.016 | 0.202 | 0.46 | 0.26 | 0.55 | 0.55 | 0.4 | 0.36 |
| 1 | 17916962 | rs10788679 | -0.633 | -0.079 | -0.751 | 0.529 | 0.015 | -0.071 | -0.071 | 0.622 | 0.64 | 0.59 | 0.75 | 0.94 | 0.57 | 0.66 |
| 1 | 17918600 | rs4920380 | -0.017 | 0.219 | -1.377 | 0.977 | -0.011 | -0.049 | -0.049 | 0 | 0.42 | 0.21 | 0.55 | 0.52 | 0.4 | 0.35 |
| 1 | 17920155 | rs1193339 | 0.761 | -0.031 | -0.628 | 1.454 | -0.026 | 0.174 | 0.174 | 0 | 0.87 | 0.66 | 0.93 | 0.66 | 0.93 | 0.86 |
| 1 | 17928259 | rs12724758 | -0.112 | -0.017 | -1.835 | 0.176 | -0.025 | 0.111 | 0.111 | 0.505 | 0.26 | 0.01 | 0.16 | 0.06 | 0.21 | 0.38 |
| 1 | 17954411 | rs12060961 | -1.046 | 0.774 | -0.486 | 0.328 | 0.035 | -0.009 | -0.009 | 0.065 | 0.14 | 0.38 | 0.12 | 0.1 | 0.14 | 0.14 |
| 1 | 17954486 | rs12059707 | -0.765 | 0.749 | -0.496 | 0.314 | 0.015 | 0.001 | 0.001 | 0 | 0.09 | 0.17 | 0.09 | 0.08 | 0.13 | 0.11 |
| 1 | 17955297 | rs12117763 | 0.281 | 0.136 | -1.282 | 0.647 | 0.016 | -0.075 | -0.075 | 5.186 | 0.15 | 0.01 | 0.19 | 0.19 | 0.13 | 0.15 |
| 1 | 17958240 | rs12562076 | -1.701 | 0.278 | -0.572 | 0.504 | 0.02 | -0.037 | -0.037 | 2.506 | 0.17 | 0.48 | 0.15 | 0.1 | 0.16 | 0.13 |
| 1 | 17964617 | rs7522176 | -0.931 | 0.275 | -0.528 | 0.5 | -0.032 | 0.027 | 0.027 | 0 | 0.11 | 0.06 | 0.1 | 0.07 | 0.16 | 0.11 |
| 1 | 17965194 | rs2297916 | -1.646 | 1.924 | -0.608 | 0.459 | 0.092 | 0.006 | 0.006 | 0.781 | 0.06 | 0.21 | 0.07 | 0.01 | 0.13 | 0.08 |
| 1 | 17968988 | rs6665682 | -0.923 | 1.632 | -0.673 | 0.443 | 0.004 | -0.004 | -0.004 | 0 | 0.11 | 0.16 | 0.09 | 0.07 | 0.14 | 0.12 |
| 1 | 17972321 | rs3766307 | 0.854 | 1.498 | -1.04 | 0.999 | -0.009 | -0.043 | -0.043 | 0 | 0.07 | 0 | 0.03 | 0.26 | 0.06 | 0.03 |
| 1 | 17976560 | rs6703771 | -1.915 | 2.466 | -0.086 | 0.837 | 0.116 | 0.021 | 0.021 | 0.073 | 0.17 | 0.42 | 0.22 | 0.17 | 0.29 | 0.19 |
| 1 | 17977378 | rs3766309 | -1.328 | 2.53 | -0.038 | 0.839 | 0.044 | 0.007 | 0.007 | 0 | 0.14 | 0.27 | 0.19 | 0.12 | 0.18 | 0.16 |
| 1 | 17979118 | rs10888046 | -1.995 | 2.757 | 0.029 | 0.728 | 0.123 | 0.008 | 0.008 | 0 | 0.2 | 0.49 | 0.49 | 0.18 | 0.32 | 0.23 |
| 1 | 18006107 | rs12737453 | -0.289 | 2.384 | 0.419 | 0.82 | 0.003 | 0 | 0 | 0 | 0.09 | 0.12 | 0.11 | 0.09 | 0.18 | 0.09 |
| 1 | 18006550 | rs12742948 | -1.346 | 2.447 | 0.421 | 0.823 | 0.069 | 0.049 | 0.049 | 0 | 0.13 | 0.4 | 0.18 | 0.17 | 0.18 | 0.11 |
| 1 | 18013602 | rs12753636 | -0.659 | 2.375 | 0.39 | 0.777 | 0.067 | -0.014 | -0.014 | 0 | 0.11 | 0.29 | 0.11 | 0.09 | 0.17 | 0.09 |
| 1 | 18017522 | rs871087 | 0.092 | 2.371 | 0.413 | 0.786 | 0.008 | -0.003 | -0.003 | 0 | 0.91 | 0.87 | 0.89 | 0.91 | 0.83 | 0.91 |
| 1 | 18017704 | rs871089 | 0.092 | 2.371 | 0.413 | 0.786 | 0.008 | -0.003 | -0.003 | 0 | 0.91 | 0.87 | 0.89 | 0.91 | 0.83 | 0.91 |
| 1 | 18019524 | rs10788681 | 0.522 | 2.425 | 0.412 | 0.758 | 0.057 | -0.024 | -0.024 | 0.047 | 0.88 | 0.65 | 0.88 | 0.91 | 0.82 | 0.91 |
| 1 | 18019957 | rs2134482 | 0.835 | 2.212 | 0.063 | 0.322 | -0.001 | 0.039 | 0.039 | 0 | 0.2 | 0.11 | 0.11 | 0.08 | 0.2 | 0.11 |
| 1 | 18023365 | rs2270978 | 0.57 | 2.342 | 0.843 | 0.866 | 0.083 | 0.026 | 0.026 | 0 | 0.88 | 0.59 | 0.87 | 0.84 | 0.82 | 0.91 |
| 1 | 18023690 | rs2270976 | 0.342 | 2.294 | 0.798 | 0.862 | 0.061 | -0.02 | -0.02 | 0 | 0.88 | 0.67 | 0.87 | 0.9 | 0.82 | 0.92 |
| 1 | 18024794 | rs1316637 | 1.344 | 1.991 | 0.187 | 1.028 | -0.007 | -0.051 | -0.051 | 0.149 | 0.47 | 0.13 | 0.49 | 0.54 | 0.44 | 0.35 |
| 1 | 18026435 | rs12139706 | 1.285 | 2.253 | 1.01 | 1.746 | 0.05 | 0.029 | 0.029 | 0 | 0.85 | 0.53 | 0.81 | 0.82 | 0.82 | 0.87 |
| 1 | 18031533 | rs12063085 | -1.217 | 2.348 | 1.314 | 1.87 | 0.006 | -0.009 | -0.009 | 0 | 0.18 | 0.16 | 0.13 | 0.19 | 0.13 | 0.15 |
| 1 | 18033499 | rs7522419 | -1.794 | 2.583 | 1.674 | 2.022 | 0.033 | 0.12 | 0.12 | 0 | 0.21 | 0.41 | 0.18 | 0.32 | 0.24 | 0.22 |
| 1 | 18034588 | rs7528432 | 1.883 | 2.362 | 1.828 | 2.094 | 0.033 | -0.003 | -0.003 | 0.625 | 0.8 | 0.64 | 0.75 | 0.75 | 0.67 | 0.78 |
| 1 | 18034640 | rs16830549 | -1.239 | 2.355 | 1.849 | 2.115 | -0.033 | 0.169 | 0.169 | 0 | 0.13 | 0.31 | 0.11 | 0.24 | 0.08 | 0.11 |
| 1 | 18038560 | rs4920624 | 1.884 | 2.472 | 1.548 | 2.133 | 0.038 | 0.015 | 0.015 | 0 | 0.81 | 0.63 | 0.63 | 0.69 | 0.67 | 0.78 |
| 1 | 18039182 | rs6664873 | 0.668 | 2.457 | 1.542 | 1.906 | 0.029 | 0.003 | 0.003 | 0 | 0.88 | 0.72 | 0.82 | 0.82 | 0.81 | 0.88 |
| 1 | 18040004 | rs2185324 | -1.357 | 1.387 | 1.374 | 1.746 | 0.002 | 0 | 0 | 0 | 0.08 | 0.09 | 0.2 | 0.08 | 0.15 | 0.1 |
| 1 | 18042035 | rs1553400 | -1.825 | 1.364 | 1.083 | 1.785 | 0.003 | 0.016 | 0.016 | 0 | 0.19 | 0.25 | 0.3 | 0.25 | 0.22 | 0.2 |
| 1 | 18044374 | rs6687077 | 1.744 | 1.58 | 1.337 | 1.305 | 0.049 | -0.083 | -0.083 | 0.006 | 0.83 | 0.52 | 0.66 | 0.94 | 0.81 | 0.77 |
| 1 | 18046287 | rs4920626 | 1.861 | 1.573 | 1.07 | 1.95 | 0.013 | 0.029 | 0.029 | 0 | 0.72 | 0.45 | 0.55 | 0.65 | 0.72 | 0.66 |
| 1 | 18048668 | rs16830671 | -1.395 | 1.336 | 0.534 | 2.278 | -0.002 | -0.058 | -0.058 | 0 | 0.12 | 0.03 | 0.11 | 0.29 | 0.11 | 0.15 |
| 1 | 18049872 | rs11581535 | -0.437 | 1.583 | 0.651 | 2.131 | -0.034 | 0.002 | 0.002 | 0 | 0.08 | 0.01 | 0.03 | 0.07 | 0.09 | 0.07 |
| 1 | 18052645 | rs260514 | 2.422 | 1.362 | 1.621 | 2.805 | 0.047 | 0.413 | 0.413 | 0.435 | 0.94 | 0.63 | 0.93 | 0.68 | 0.96 | 0.86 |
| 1 | 18059492 | rs1890747 | -2.464 | 1.628 | 1.609 | 2.707 | 0.014 | 0.097 | 0.097 | 0.621 | 0.17 | 0.63 | 0.39 | 0.34 | 0.19 | 0.21 |
| 1 | 18060264 | rs11584149 | -1.701 | 1.654 | 1.451 | 2.78 | -0.051 | -0.007 | -0.007 | 0.023 | 0.11 | 0.02 | 0.31 | 0.11 | 0.18 | 0.18 |
| 1 | 18060436 | rs11584168 | -2.183 | 1.636 | 1.798 | 2.743 | 0.003 | 0.017 | 0.017 | 0 | 0.13 | 0.16 | 0.35 | 0.29 | 0.19 | 0.2 |
| 1 | 18065643 | rs2077398 | -2.593 | 1.793 | 1.912 | 2.755 | 0.076 | 0.15 | 0.15 | 0.567 | 0.16 | 0.54 | 0.39 | 0.34 | 0.23 | 0.22 |
| 1 | 18074702 | rs11203452 | -1.934 | 1.798 | 1.846 | 2.747 | -0.012 | -0.013 | -0.013 | 0 | 0.13 | 0.1 | 0.35 | 0.27 | 0.22 | 0.23 |
| 1 | 18076398 | rs6658566 | -2.669 | 1.765 | 1.884 | 2.734 | 0.043 | 0.1 | 0.1 | 0.251 | 0.17 | 0.55 | 0.39 | 0.33 | 0.23 | 0.24 |
| 1 | 18084290 | rs1610410 | -2.428 | 1.631 | 1.165 | 3.319 | -0.018 | -0.032 | -0.032 | 0.93 | 0.17 | 0.14 | 0.36 | 0.58 | 0.25 | 0.4 |
| 1 | 18085290 | rs1604625 | -1.642 | 1.693 | 1.04 | 3.202 | -0.036 | -0.042 | -0.042 | 0 | 0.15 | 0.08 | 0.37 | 0.28 | 0.26 | 0.18 |
| 1 | 18090975 | rs16830722 | -1.011 | 1.478 | 1.097 | 3.303 | -0.002 | 0.053 | 0.053 | 0 | 0.05 | 0.03 | 0.03 | 0.28 | 0.03 | 0.19 |
| 1 | 18094337 | rs11203456 | -1.841 | 2.171 | 1.033 | 3.339 | -0.028 | -0.028 | -0.028 | 0 | 0.17 | 0.13 | 0.41 | 0.34 | 0.35 | 0.22 |
| 1 | 18098553 | rs1977251 | -1.456 | 2.125 | 1.503 | 3.161 | 0.018 | 0.295 | 0.295 | 0.174 | 0.07 | 0.31 | 0.06 | 0.28 | 0.03 | 0.19 |
| 1 | 18099283 | rs10888060 | -1.885 | 2.417 | 1.058 | 2.543 | 0.086 | 0.071 | 0.071 | 0 | 0.17 | 0.3 | 0.41 | 0.29 | 0.35 | 0.22 |
| 1 | 18110701 | rs260519 | -1.293 | 1.466 | 0.613 | 1.726 | 0 | -0.027 | -0.027 | 0 | 0.37 | 0.45 | 0.49 | 0.78 | 0.41 | 0.39 |
| 1 | 18122009 | rs4141983 | -1.999 | 2.517 | 1.405 | 3.599 | 0.082 | 0.102 | 0.102 | 0 | 0.23 | 0.41 | 0.54 | 0.61 | 0.33 | 0.31 |
| 1 | 18123332 | rs4920385 | -1.881 | 2.525 | 1.411 | 3.964 | 0.079 | 0.231 | 0.231 | 0 | 0.1 | 0.29 | 0.42 | 0.35 | 0.15 | 0.21 |
| 1 | 18123957 | rs1847235 | -1.905 | 2.524 | 1.373 | 3.964 | 0.096 | 0.268 | 0.268 | 0 | 0.1 | 0.32 | 0.42 | 0.34 | 0.15 | 0.21 |
| 1 | 18124527 | rs16830757 | -1.102 | 2.507 | 1.289 | 3.908 | -0.03 | -0.031 | -0.031 | 0 | 0.13 | 0.04 | 0.11 | 0.26 | 0.18 | 0.09 |
| 1 | 18125179 | rs7552919 | -1.98 | 2.174 | 1.394 | 3.887 | 0.102 | 0.286 | 0.286 | 0 | 0.1 | 0.32 | 0.42 | 0.35 | 0.15 | 0.21 |
| 1 | 18128532 | rs4920642 | -1.79 | 2.063 | 1.667 | 3.789 | 0.194 | 0.271 | 0.271 | 0 | 0.06 | 0.35 | 0.19 | 0.12 | 0.11 | 0.03 |

| | | | | | | | | | | | | | | | | |
|---|---|---|---|---|---|---|---|---|---|---|---|---|---|---|---|---|
| 1 | 18132601 | rs560514 | -2.074 | 1.766 | 1.563 | 2.802 | -0.001 | -0.001 | -0.001 | 0 | 0.43 | 0.48 | 0.62 | 0.73 | 0.54 | 0.46 |
| 1 | 18135882 | rs561698 | -2.242 | 1.572 | 1.708 | 2.701 | -0.009 | 0.038 | 0.038 | 0 | 0.44 | 0.67 | 0.65 | 0.75 | 0.53 | 0.45 |
| 1 | 18136684 | rs533374 | -2.01 | 1.584 | 1.718 | 2.674 | 0.016 | -0.009 | -0.009 | 0 | 0.39 | 0.49 | 0.6 | 0.69 | 0.48 | 0.44 |
| 1 | 18149510 | rs694214 | -1.524 | 1.594 | 1.633 | 3.067 | -0.072 | -0.025 | -0.025 | 0.451 | 0.33 | 0.14 | 0.32 | 0.43 | 0.44 | 0.25 |
| 1 | 18149566 | rs2296035 | -1.882 | 1.58 | 1.594 | 2.51 | -0.002 | 0 | 0 | 0 | 0.37 | 0.32 | 0.58 | 0.69 | 0.47 | 0.43 |
| 1 | 18150330 | rs619993 | -2.261 | 1.569 | 1.278 | 2.371 | -0.009 | 0.032 | 0.032 | 0 | 0.43 | 0.65 | 0.65 | 0.75 | 0.52 | 0.45 |
| 1 | 18152582 | rs478698 | -0.671 | 1.298 | 0.708 | 2.096 | -0.055 | -0.022 | -0.022 | 1.207 | 0.14 | 0.02 | 0.1 | 0.18 | 0.19 | 0.16 |
| 1 | 18155405 | rs648305 | -1.471 | 1.343 | 0.843 | 2.096 | -0.04 | 0.012 | 0.012 | 0 | 0.25 | 0.16 | 0.25 | 0.24 | 0.34 | 0.2 |
| 1 | 18158156 | rs7545559 | -1.296 | 1.204 | 1.198 | 2.247 | -0.001 | 0.171 | 0.171 | 0 | 0.08 | 0.16 | 0.19 | 0.38 | 0.07 | 0.08 |
| 1 | 18160092 | rs11203472 | -1.388 | 1.536 | 0.757 | 2.029 | -0.011 | -0.008 | -0.008 | 0.028 | 0.1 | 0.07 | 0.15 | 0.14 | 0.14 | 0.07 |
| 1 | 18161596 | rs4920645 | 1.3 | 0.426 | 0.993 | 1.907 | -0.018 | 0.002 | 0.002 | 0 | 0.66 | 0.68 | 0.56 | 0.33 | 0.58 | 0.68 |
| 1 | 18163922 | rs495347 | 1.149 | 0.382 | 0.902 | 1.852 | 0.007 | -0.002 | -0.002 | 0.163 | 0.66 | 0.37 | 0.53 | 0.32 | 0.62 | 0.69 |
| 1 | 18164285 | rs491778 | 0.445 | 0.538 | 0.249 | 1.374 | 0.017 | -0.007 | -0.007 | 0 | 0.74 | 0.71 | 0.72 | 0.77 | 0.64 | 0.78 |
| 1 | 18167395 | rs510504 | 0.236 | 0.035 | -0.443 | 0.908 | -0.023 | 0.089 | 0.089 | 0 | 0.56 | 0.77 | 0.73 | 0.75 | 0.51 | 0.61 |
| 1 | 18170886 | rs647325 | -0.797 | -0.631 | -0.022 | 1.148 | -0.022 | 0.028 | 0.028 | 0 | 0.3 | 0.63 | 0.54 | 0.5 | 0.21 | 0.17 |
| 1 | 18175932 | rs539889 | -0.223 | -0.158 | -0.238 | 0.293 | -0.009 | -0.021 | -0.021 | 0.111 | 0.19 | 0.24 | 0.22 | 0.1 | 0.18 | 0.07 |
| 1 | 18179816 | rs697760 | -0.399 | 0.065 | -0.21 | 0.356 | -0.008 | -0.034 | -0.034 | 0.533 | 0.21 | 0.34 | 0.23 | 0.1 | 0.2 | 0.08 |
| 1 | 18190225 | rs615917 | -0.027 | 0.145 | -1.231 | 1.122 | -0.054 | 0.065 | 0.065 | 0 | 0.35 | 0.14 | 0.28 | 0.19 | 0.47 | 0.3 |
| 1 | 18195801 | rs542963 | -0.652 | 0.45 | -1.391 | 0.54 | 0.039 | 0.14 | 0.14 | 0 | 0.61 | 0.79 | 0.76 | 0.87 | 0.68 | 0.79 |
| 1 | 18196800 | rs11203223 | -0.138 | 0.87 | -0.588 | 1.224 | 0.008 | 0.015 | 0.015 | 0 | 0.17 | 0.18 | 0.21 | 0.37 | 0.2 | 0.21 |
| 1 | 18202181 | rs7538231 | -0.062 | 0.89 | -0.742 | 1.224 | -0.011 | -0.011 | -0.011 | 0 | 0.17 | 0.14 | 0.2 | 0.37 | 0.2 | 0.21 |
| 1 | 18210028 | rs638425 | 0.486 | 0.434 | -1.117 | 0.739 | -0.015 | 0.013 | 0.013 | 0 | 0.16 | 0.01 | 0.12 | 0.11 | 0.17 | 0.07 |
| 1 | 18212230 | rs597865 | -0.76 | 1.413 | 0.085 | 1.537 | 0.132 | 0.237 | 0.237 | 0 | 0.07 | 0.24 | 0.39 | 0.35 | 0.14 | 0.17 |
| 1 | 18212707 | rs579985 | 0.014 | 1.623 | -0.037 | 0.824 | -0.036 | 0.006 | 0.006 | 0 | 0.76 | 0.83 | 0.8 | 0.79 | 0.57 | 0.64 |
| 1 | 18219827 | rs477558 | 0.798 | 1.199 | 0.397 | 0.804 | 0.006 | -0.008 | -0.008 | 0.192 | 0.65 | 0.45 | 0.72 | 0.76 | 0.52 | 0.48 |
| 1 | 18225521 | rs12037123 | 0.248 | 1.601 | 0.365 | 1.252 | 0.056 | 0.181 | 0.181 | 0 | 0.05 | 0.08 | 0.21 | 0.44 | 0.14 | 0.1 |
| 1 | 18228861 | rs679002 | -0.22 | 1.542 | 0.588 | 1.031 | -0.002 | -0.002 | -0.002 | 1.723 | 0.74 | 0.75 | 0.81 | 0.91 | 0.61 | 0.79 |
| 1 | 18231349 | rs620547 | 0.312 | 1.533 | 0.589 | 1.05 | 0.01 | 0.005 | 0.005 | 0 | 0.68 | 0.66 | 0.59 | 0.46 | 0.47 | 0.6 |
| 1 | 18232418 | rs1537772 | -0.276 | 1.484 | 0.755 | 0.972 | -0.003 | 0 | 0 | 0.557 | 0.12 | 0.11 | 0.05 | 0.12 | 0.07 | 0.24 |
| 1 | 18234374 | rs12071205 | -0.005 | 1.449 | 0.71 | 0.598 | -0.002 | -0.002 | -0.002 | 0 | 0.26 | 0.25 | 0.19 | 0.09 | 0.39 | 0.21 |
| 1 | 18236459 | rs1339934 | -1.387 | 1.429 | 0.797 | 0.585 | -0.003 | -0.012 | -0.012 | 0.04 | 0.45 | 0.5 | 0.47 | 0.67 | 0.6 | 0.67 |
| 1 | 18240000 | rs11203232 | -1.137 | 1.4 | 0.769 | 0.665 | 0.026 | 0.076 | 0.076 | 0.052 | 0.19 | 0.29 | 0.28 | 0.56 | 0.21 | 0.46 |
| 1 | 18242493 | rs12089138 | 0.042 | 1.738 | 0.718 | 0.608 | -0.002 | 0.008 | 0.008 | 0 | 0.25 | 0.22 | 0.15 | 0.1 | 0.34 | 0.19 |
| 1 | 18244707 | rs6686929 | -1.505 | 1.749 | 0.792 | 0.601 | 0 | 0.005 | 0.005 | 0 | 0.43 | 0.4 | 0.46 | 0.65 | 0.59 | 0.55 |
| 1 | 18245365 | rs12083022 | 0.045 | 1.747 | 0.593 | 0.605 | -0.002 | 0.008 | 0.008 | 0 | 0.25 | 0.22 | 0.15 | 0.1 | 0.34 | 0.19 |
| 1 | 18245540 | rs7541126 | -0.555 | 1.844 | 0.229 | 0.762 | -0.018 | -0.022 | -0.022 | 0 | 0.15 | 0.08 | 0.26 | 0.55 | 0.19 | 0.34 |
| 1 | 18247287 | rs12033416 | -0.972 | 1.796 | 0.238 | 0.77 | -0.035 | -0.043 | -0.043 | 0.771 | 0.18 | 0.08 | 0.3 | 0.55 | 0.26 | 0.36 |
| 1 | 18261837 | rs11203242 | -0.555 | 1.607 | 0.082 | 0.607 | 0.013 | -0.059 | -0.059 | 0 | 0.29 | 0.5 | 0.4 | 0.12 | 0.39 | 0.32 |
| 1 | 18268281 | rs651743 | 0.146 | 1.476 | -0.839 | 0.496 | 0.004 | -0.012 | -0.012 | 0.042 | 0.34 | 0.7 | 0.3 | 0.04 | 0.29 | 0.28 |
| 1 | 18268968 | rs2095518 | 0.556 | 1.057 | -1.44 | 0.271 | 0.013 | 0.006 | 0.006 | 0 | 0.26 | 0.19 | 0.2 | 0.2 | 0.15 | 0.23 |
| 1 | 18271363 | rs1572345 | 0.193 | 0.987 | -1.6 | 0.418 | -0.085 | 0.015 | 0.015 | 0 | 0.36 | 0.04 | 0.24 | 0.31 | 0.44 | 0.33 |
| 1 | 18272751 | rs661977 | -0.008 | 0.923 | -1.423 | 0.387 | -0.07 | 0.089 | 0.089 | 0 | 0.59 | 0.88 | 0.5 | 0.24 | 0.46 | 0.49 |
| 1 | 18273842 | rs7551015 | 0.087 | 1.201 | -0.478 | 0.547 | 0.003 | 0 | 0 | 0 | 0.69 | 0.66 | 0.62 | 0.35 | 0.71 | 0.61 |
| 1 | 18274898 | rs1192586 | 0.281 | 1.122 | -0.113 | 0.36 | 0 | -0.018 | -0.018 | 0 | 0.68 | 0.57 | 0.52 | 0.3 | 0.72 | 0.58 |
| 1 | 18277480 | rs659810 | -0.374 | 1.25 | -0.498 | 1.127 | -0.025 | 0.063 | 0.063 | 0.177 | 0.95 | 0.99 | 0.81 | 0.57 | 0.86 | 0.82 |
| 1 | 18278257 | rs673567 | 0.824 | 0.853 | -0.341 | 0.262 | 0.007 | -0.008 | -0.008 | 0 | 0.71 | 0.7 | 0.56 | 0.29 | 0.65 | 0.51 |
| 1 | 18278422 | rs12408792 | -0.504 | 0.344 | -1.083 | -0.015 | 0.001 | 0.102 | 0.102 | 0 | 0.3 | 0.03 | 0.2 | 0.17 | 0.27 | 0.2 |
| 1 | 18283857 | rs1192579 | 0.742 | 0.644 | -0.008 | 0.326 | 0.027 | 0.092 | 0.092 | 0 | 0.81 | 0.53 | 0.57 | 0.68 | 0.79 | 0.83 |
| 1 | 18285258 | rs11203247 | -0.16 | 0.715 | -0.058 | 0.277 | 0.006 | -0.008 | -0.008 | 0 | 0.14 | 0.16 | 0.22 | 0.14 | 0.19 | 0.08 |
| 1 | 18297407 | rs2095842 | 0.374 | 0.42 | -0.188 | 0.258 | 0.002 | 0.045 | 0.045 | 0 | 0.23 | 0.12 | 0.17 | 0.16 | 0.22 | 0.09 |
| 1 | 18297688 | rs871644 | 0.397 | 0.384 | -0.186 | 0.256 | 0.003 | 0.05 | 0.05 | 0 | 0.22 | 0.12 | 0.17 | 0.13 | 0.22 | 0.09 |
| 1 | 18305292 | rs1192630 | 0.538 | 0.249 | -0.018 | 0.369 | -0.023 | 0.09 | 0.09 | 0.039 | 0.94 | 0.82 | 0.81 | 0.83 | 0.99 | 0.93 |
| 1 | 18307682 | rs7414500 | 0.171 | 0.349 | 0.067 | 0.183 | -0.015 | 0.024 | 0.024 | 0 | 0.79 | 0.64 | 0.69 | 0.78 | 0.83 | 0.76 |
| 1 | 18310576 | rs1192607 | 0.342 | 0.857 | 0.148 | 0.317 | 0.068 | 0.235 | 0.235 | 0 | 0.08 | 0.18 | 0.1 | 0.44 | 0.14 | 0.15 |
| 1 | 18311140 | rs6685120 | 1.353 | 0.764 | -0.227 | -0.738 | 0.019 | 0.162 | 0.162 | 0 | 0.45 | 0.18 | 0.32 | 0.19 | 0.44 | 0.45 |
| 1 | 18312084 | rs7340046 | 0.083 | 0.756 | -0.171 | -0.742 | -0.009 | -0.065 | -0.065 | 0 | 0.2 | 0.33 | 0.15 | 0.06 | 0.17 | 0.17 |
| 1 | 18313386 | rs7533732 | 1.006 | 0.44 | -1.004 | -0.777 | 0.016 | 0.163 | 0.163 | 0 | 0.45 | 0.18 | 0.32 | 0.17 | 0.44 | 0.45 |
| 1 | 18314621 | rs11588830 | 0.282 | 0.886 | -1.402 | -0.6 | -0.003 | 0.025 | 0.025 | 0 | 0.24 | 0.17 | 0.25 | 0.16 | 0.25 | 0.1 |
| 1 | 18317841 | rs12726822 | 0.868 | 0.683 | -1.839 | -0.686 | -0.074 | 0.174 | 0.174 | 0.014 | 0.34 | 0.03 | 0.27 | 0.11 | 0.35 | 0.29 |
| 1 | 18322825 | rs1192613 | -0.4 | 0.605 | -0.842 | -0.22 | -0.002 | -0.02 | -0.02 | 0.069 | 0.28 | 0.34 | 0.29 | 0.19 | 0.28 | 0.18 |
| 1 | 18323910 | rs11203252 | 0.162 | 0.426 | -0.406 | 0.237 | -0.001 | -0.003 | -0.003 | 5.558 | 0.46 | 0.49 | 0.32 | 0.34 | 0.41 | 0.39 |
| 1 | 18325257 | rs12022529 | 0.516 | 0.331 | -1.036 | 0.599 | -0.052 | -0.087 | -0.087 | 0.243 | 0.23 | 0.02 | 0.23 | 0.3 | 0.2 | 0.24 |
| 1 | 18325601 | rs1192614 | -0.87 | 0.278 | -0.903 | 0.745 | 0.001 | -0.003 | -0.003 | 0 | 0.89 | 0.88 | 0.92 | 0.85 | 0.92 | 0.88 |
| 1 | 18325885 | rs11203253 | 1.646 | 0.382 | -0.674 | 0.73 | 0.008 | 0.015 | 0.015 | 0 | 0.32 | 0.26 | 0.23 | 0.21 | 0.28 | 0.13 |
| 1 | 18326099 | rs12743154 | 0.704 | 0.634 | -0.709 | 0.711 | -0.028 | -0.011 | -0.011 | 0 | 0.16 | 0.09 | 0.34 | 0.21 | 0.28 | 0.28 |
| 1 | 18326623 | rs11203254 | 1.173 | 0.571 | -0.427 | 0.611 | -0.007 | -0.003 | -0.003 | 0 | 0.13 | 0.1 | 0.07 | 0.12 | 0.14 | 0.23 |
| 1 | 18327209 | rs1192616 | 0.042 | 0.496 | -0.18 | 0.603 | -0.003 | -0.003 | -0.003 | 0 | 0.72 | 0.7 | 0.57 | 0.64 | 0.63 | 0.62 |
| 1 | 18327378 | rs1192617 | 0.562 | 0.443 | -0.136 | 0.556 | 0.004 | 0.005 | 0.005 | 0 | 0.58 | 0.53 | 0.49 | 0.52 | 0.49 | 0.39 |
| 1 | 18331242 | rs1192610 | -1.046 | 0.06 | -0.13 | 0.737 | 0.019 | 0.151 | 0.151 | 0 | 0.5 | 0.81 | 0.45 | 0.73 | 0.42 | 0.55 |

| | | | | | | | | | | | | | | | | |
|---|---|---|---|---|---|---|---|---|---|---|---|---|---|---|---|---|
| 1 | 18331332 | rs1890789 | -0.267 | -0.322 | 0.67 | 1.085 | -0.015 | 0.003 | 0.003 | 0 | 0.31 | 0.36 | 0.3 | 0.36 | 0.22 | 0.28 |
| 1 | 18331592 | rs4433436 | -0.319 | -0.327 | 0.665 | 1.337 | -0.014 | 0.019 | 0.019 | 0 | 0.31 | 0.36 | 0.3 | 0.47 | 0.22 | 0.3 |
| 1 | 18335916 | rs6678740 | 0.303 | -0.038 | 0.976 | 1.386 | -0.01 | 0.008 | 0.008 | 0 | 0.86 | 0.79 | 0.92 | 0.82 | 0.9 | 0.89 |
| 1 | 18336903 | rs1192618 | 0.864 | -0.237 | 1.431 | 1.416 | -0.029 | -0.015 | -0.015 | 0 | 0.84 | 0.73 | 0.89 | 0.88 | 0.91 | 0.84 |
| 1 | 18338598 | rs6686302 | -0.112 | -0.34 | 1.611 | 1.395 | 0.003 | 0.016 | 0.016 | 0 | 0.25 | 0.32 | 0.27 | 0.37 | 0.24 | 0.25 |
| 1 | 18339748 | rs6586487 | 0.128 | -0.347 | 1.617 | 1.396 | -0.004 | -0.004 | -0.004 | 0 | 0.24 | 0.18 | 0.25 | 0.3 | 0.23 | 0.22 |
| 1 | 18339791 | rs1192625 | 0.302 | -0.364 | 1.663 | 1.424 | 0.018 | -0.035 | -0.035 | 1.481 | 0.87 | 0.92 | 0.9 | 0.78 | 0.91 | 0.82 |
| 1 | 18340025 | rs17472583 | 0.126 | -0.342 | 1.668 | 1.406 | -0.004 | -0.004 | -0.004 | 0.716 | 0.24 | 0.18 | 0.25 | 0.3 | 0.23 | 0.22 |
| 1 | 18340225 | rs11203256 | -0.706 | -0.343 | 1.746 | 1.381 | 0.002 | 0.038 | 0.038 | 0.014 | 0.31 | 0.41 | 0.34 | 0.48 | 0.28 | 0.35 |
| 1 | 18342964 | rs794237 | -0.23 | -0.459 | 1.626 | 1.367 | 0.012 | 0.001 | 0.001 | 0 | 0.89 | 0.92 | 0.96 | 1 | 0.94 | 0.94 |
| 1 | 18343025 | rs12058798 | 1.146 | -0.189 | 1.697 | 1.457 | -0.02 | -0.015 | -0.015 | 0 | 0.13 | 0.09 | 0.15 | 0.15 | 0.16 | 0.22 |
| 1 | 18343551 | rs875294 | -1.334 | -0.113 | 1.976 | 1.285 | -0.009 | 0.062 | 0.062 | 0 | 0.46 | 0.68 | 0.5 | 0.75 | 0.45 | 0.61 |
| 1 | 18349764 | rs2355459 | 1.495 | -0.199 | 1.847 | 1.656 | -0.004 | 0.028 | 0.028 | 0.021 | 0.65 | 0.43 | 0.69 | 0.56 | 0.7 | 0.49 |
| 1 | 18349821 | rs4532860 | 0.889 | -0.22 | 1.943 | 1.683 | -0.026 | 0.009 | 0.009 | 0 | 0.84 | 0.73 | 0.92 | 0.83 | 0.9 | 0.76 |
| 1 | 18351999 | rs4539169 | 0.747 | -0.223 | 1.864 | 1.664 | 0.006 | 0.003 | 0.003 | 0.006 | 0.14 | 0.18 | 0.17 | 0.16 | 0.16 | 0.17 |
| 1 | 18352458 | rs2355462 | -0.06 | -0.22 | 1.778 | 1.674 | 0.017 | 0.03 | 0.03 | 0 | 0.18 | 0.25 | 0.23 | 0.28 | 0.19 | 0.25 |
| 1 | 18353743 | rs7550969 | 1.961 | -0.109 | 1.778 | 1.602 | 0.011 | 0.072 | 0.072 | 0 | 0.44 | 0.26 | 0.45 | 0.25 | 0.48 | 0.34 |
| 1 | 18355150 | rs12132348 | 2.165 | -0.16 | 1.544 | 1.577 | -0.089 | 0.064 | 0.064 | 0.1 | 0.42 | 0.03 | 0.43 | 0.25 | 0.47 | 0.34 |
| 1 | 18355412 | rs6669252 | 0.302 | -0.029 | 1.847 | 1.645 | 0.005 | -0.01 | -0.01 | 0 | 0.87 | 0.85 | 0.95 | 1 | 0.92 | 0.91 |
| 1 | 18355580 | rs6586490 | 0.762 | 0.047 | 1.881 | 1.716 | 0.007 | 0.004 | 0.004 | 0.003 | 0.14 | 0.16 | 0.17 | 0.17 | 0.16 | 0.15 |
| 1 | 18359580 | rs12067141 | -1.216 | 0.027 | 1.993 | 1.745 | 0.029 | 0.14 | 0.14 | 0 | 0.25 | 0.47 | 0.27 | 0.44 | 0.24 | 0.43 |
| 1 | 18359901 | rs7553424 | 0.497 | 0.001 | 2.01 | 1.716 | 0 | -0.004 | -0.004 | 0 | 0.76 | 0.73 | 0.79 | 0.83 | 0.79 | 0.79 |
| 1 | 18360161 | rs16861481 | -2.107 | 0.011 | 2.029 | 1.751 | 0.008 | 0.129 | 0.129 | 0.615 | 0.09 | 0.26 | 0.09 | 0.21 | 0.07 | 0.19 |
| 1 | 18364893 | rs870042 | 0.536 | 0.031 | 1.941 | 1.738 | 0.012 | 0.032 | 0.032 | 0 | 0.15 | 0.2 | 0.18 | 0.26 | 0.17 | 0.17 |
| 1 | 18365279 | rs946374 | 0.162 | -0.005 | 1.914 | 1.655 | -0.038 | -0.071 | -0.071 | 0.004 | 0.88 | 0.73 | 0.95 | 1 | 0.94 | 0.9 |
| 1 | 18366795 | rs10887986 | -1.072 | 0.048 | 1.817 | 1.725 | 0.026 | 0.124 | 0.124 | 0 | 0.24 | 0.42 | 0.27 | 0.44 | 0.24 | 0.43 |
| 1 | 18370015 | rs7523606 | 1.192 | 0.063 | 1.658 | 1.647 | -0.009 | 0 | 0 | 0 | 0.68 | 0.45 | 0.74 | 0.7 | 0.73 | 0.61 |
| 1 | 18370771 | rs12032253 | 0.554 | 0.174 | 1.363 | 1.569 | -0.012 | -0.014 | -0.014 | 0 | 0.19 | 0.15 | 0.24 | 0.33 | 0.23 | 0.18 |
| 1 | 18373009 | rs12034254 | 1.48 | 0.203 | 1.299 | 1.391 | -0.004 | 0.012 | 0.012 | 0 | 0.65 | 0.43 | 0.69 | 0.58 | 0.69 | 0.52 |
| 1 | 18374940 | rs1551477 | 0.445 | 0.446 | 0.953 | 1.278 | 0.006 | 0 | 0 | 0 | 0.15 | 0.18 | 0.17 | 0.15 | 0.17 | 0.17 |
| 1 | 18377790 | rs1031567 | 1.611 | 0.259 | 0.908 | 1.227 | 0.012 | 0.086 | 0.086 | 0 | 0.47 | 0.22 | 0.46 | 0.26 | 0.48 | 0.35 |
| 1 | 18378569 | rs10887988 | 0.233 | 0.318 | 1.115 | 1.172 | 0.008 | -0.006 | -0.006 | 0 | 0.16 | 0.23 | 0.14 | 0.16 | 0.17 | 0.25 |
| 1 | 18379379 | rs4920419 | 0.247 | 0.327 | 1.12 | 1.143 | 0.008 | -0.008 | -0.008 | 0 | 0.16 | 0.23 | 0.14 | 0.16 | 0.17 | 0.24 |
| 1 | 18379766 | rs7534337 | 0.437 | 0.338 | 1.11 | 1.109 | 0.014 | -0.016 | -0.016 | 0 | 0.15 | 0.22 | 0.14 | 0.11 | 0.16 | 0.19 |
| 1 | 18391974 | rs9286088 | -1.925 | 0.331 | 0.969 | 1.149 | 0.022 | 0.089 | 0.089 | 0 | 0.52 | 0.79 | 0.55 | 0.73 | 0.53 | 0.64 |
| 1 | 18392608 | rs4920421 | 0.313 | 0.447 | 0.774 | 1.141 | 0.007 | -0.01 | -0.01 | 0 | 0.15 | 0.18 | 0.13 | 0.11 | 0.16 | 0.19 |
| 1 | 18405073 | rs17486629 | 0.457 | -0.005 | 0.552 | 1.47 | -0.085 | -0.085 | -0.085 | 0 | 0.17 | 0.03 | 0.24 | 0.32 | 0.22 | 0.19 |
| 1 | 18405541 | rs11260973 | -1.407 | -0.079 | 0.78 | 1.505 | -0.002 | -0.007 | -0.007 | 0 | 0.31 | 0.33 | 0.36 | 0.49 | 0.32 | 0.31 |
| 1 | 18408767 | rs9660584 | 1.574 | -0.005 | 0.279 | 1.467 | 0 | -0.003 | -0.003 | 0 | 0.63 | 0.32 | 0.57 | 0.38 | 0.66 | 0.61 |
| 1 | 18412381 | rs897496 | -0.557 | 0.507 | -0.678 | 0.945 | -0.064 | -0.077 | -0.077 | 1.35 | 0.84 | 0.97 | 0.77 | 0.67 | 0.81 | 0.83 |
| 1 | 18414524 | rs16861538 | -0.907 | 0.178 | -0.106 | 0.496 | -0.011 | -0.04 | -0.04 | 0 | 0.1 | 0.18 | 0.07 | 0 | 0.07 | 0.05 |
| 1 | 18425011 | rs9804128 | -1.106 | 0.054 | -0.115 | 0.35 | -0.02 | -0.022 | -0.022 | 0 | 0.28 | 0.58 | 0.22 | 0.2 | 0.26 | 0.28 |
| 1 | 18425601 | rs4920428 | 1.199 | 0.296 | -1 | 0.556 | -0.062 | -0.031 | -0.031 | 0 | 0.52 | 0.08 | 0.52 | 0.49 | 0.51 | 0.53 |
| 1 | 18427031 | rs12740505 | 1.145 | 0.211 | -0.729 | 0.438 | -0.075 | -0.043 | -0.043 | 0.961 | 0.53 | 0.07 | 0.52 | 0.5 | 0.51 | 0.55 |
| 1 | 18427615 | rs9659352 | 0.008 | 0.107 | -0.134 | -0.149 | -0.005 | -0.034 | -0.034 | 0 | 0.83 | 0.79 | 0.92 | 0.94 | 0.83 | 0.82 |
| 1 | 18429575 | rs930851 | 0.747 | 0.138 | -0.543 | -0.516 | -0.009 | -0.003 | -0.003 | 1.675 | 0.39 | 0.42 | 0.41 | 0.63 | 0.26 | 0.57 |
| 1 | 18433355 | rs12133062 | 0.583 | 0.892 | -1.188 | -0.695 | -0.063 | 0.128 | 0.128 | 0.792 | 0.2 | 0.02 | 0.15 | 0 | 0.24 | 0.11 |
| 1 | 18438239 | rs4920295 | 0.212 | 0.698 | -0.639 | -0.207 | 0.003 | 0.003 | 0.003 | 0 | 0.77 | 0.72 | 0.69 | 0.71 | 0.66 | 0.83 |
| 1 | 18438351 | rs1342569 | 0.281 | 0.746 | -0.467 | -0.54 | 0.024 | -0.003 | -0.003 | 0 | 0.9 | 0.76 | 0.9 | 0.91 | 0.85 | 0.93 |
| 1 | 18438930 | rs10157701 | -0.468 | 0.762 | -0.943 | -0.528 | -0.03 | -0.003 | -0.003 | 0 | 0.09 | 0.03 | 0.08 | 0.08 | 0.15 | 0.07 |
| 1 | 18447551 | rs1858589 | 0.376 | 0.767 | -0.431 | -0.81 | -0.007 | -0.014 | -0.014 | 1.891 | 0.22 | 0.15 | 0.17 | 0.01 | 0.28 | 0.12 |
| 1 | 18449524 | rs2355729 | 1.247 | 0.324 | -1.055 | -0.761 | -0.084 | -0.135 | -0.135 | 0 | 0.33 | 0.03 | 0.39 | 0.58 | 0.29 | 0.56 |
| 1 | 18450321 | rs12735521 | 0.464 | 0.247 | -0.239 | -0.833 | 0.014 | -0.055 | -0.055 | 2.274 | 0.25 | 0.31 | 0.19 | 0.1 | 0.31 | 0.15 |
| 1 | 18450595 | rs12072266 | -0.623 | 0.023 | 0.081 | -1.014 | -0.004 | -0.048 | -0.048 | 0 | 0.44 | 0.5 | 0.29 | 0.11 | 0.42 | 0.29 |
| 1 | 18453918 | rs16861613 | 1.032 | -0.111 | 0.119 | -1.007 | 0.037 | 0.042 | 0.042 | 0 | 0.16 | 0.07 | 0.07 | 0 | 0.1 | 0.1 |
| 1 | 18454964 | rs1858587 | -0.26 | -0.04 | 0.52 | -0.967 | -0.001 | -0.002 | -0.002 | 0 | 0.32 | 0.37 | 0.33 | 0.25 | 0.32 | 0.25 |
| 1 | 18455424 | rs11261067 | 0.541 | 0.061 | 0.214 | -0.898 | 0.009 | -0.038 | -0.038 | 0.04 | 0.26 | 0.28 | 0.21 | 0.11 | 0.32 | 0.16 |
| 1 | 18456382 | rs4920435 | 1.108 | -0.117 | -0.412 | -0.805 | -0.068 | -0.042 | -0.042 | 0 | 0.36 | 0.07 | 0.42 | 0.64 | 0.33 | 0.57 |
| 1 | 18457568 | rs12033458 | 1.17 | -0.107 | -0.5 | -0.8 | -0.089 | -0.061 | -0.061 | 0 | 0.36 | 0.05 | 0.42 | 0.64 | 0.33 | 0.57 |
| 1 | 18458372 | rs11577112 | -0.014 | -0.134 | 0.001 | -0.776 | 0.003 | -0.009 | -0.009 | 0.012 | 0.2 | 0.2 | 0.1 | 0.06 | 0.15 | 0.17 |
| 1 | 18460469 | rs12131391 | 0.607 | 0.314 | 0.269 | -0.824 | 0.013 | -0.075 | -0.075 | 0.253 | 0.2 | 0.22 | 0.18 | 0 | 0.25 | 0.1 |
| 1 | 18460526 | rs9725656 | 0.275 | -0.026 | 0.607 | -0.77 | 0.011 | -0.031 | -0.031 | 0 | 0.61 | 0.55 | 0.62 | 0.8 | 0.56 | 0.75 |
| 1 | 18461490 | rs12036744 | 0.93 | -0.07 | 0.231 | -0.475 | 0.001 | -0.01 | -0.01 | 0 | 0.22 | 0.17 | 0.37 | 0.49 | 0.23 | 0.47 |
| 1 | 18467153 | rs2005010 | 0.185 | 0.139 | 0.205 | -0.541 | -0.02 | -0.025 | -0.025 | 1.732 | 0.41 | 0.27 | 0.52 | 0.52 | 0.51 | 0.57 |
| 1 | 18467265 | rs2005334 | -0.337 | 0.02 | 0.208 | -0.577 | -0.004 | 0.066 | 0.066 | 0 | 0.77 | 0.45 | 0.69 | 0.65 | 0.79 | 0.8 |
| 1 | 18468093 | rs876032 | -1.125 | 0.029 | 0.344 | -0.551 | 0 | 0.023 | 0.023 | 0 | 0.81 | 0.65 | 0.76 | 0.74 | 0.83 | 0.86 |
| 1 | 18470009 | rs753305 | 0.647 | -0.045 | 0.307 | -0.548 | 0.009 | 0.029 | 0.029 | 3.469 | 0.37 | 0.29 | 0.18 | 0.13 | 0.28 | 0.22 |
| 1 | 18471278 | rs10907331 | -0.405 | 0.044 | 0.316 | -0.487 | 0.009 | -0.023 | -0.023 | 0 | 0.63 | 0.57 | 0.6 | 0.78 | 0.56 | 0.77 |
| 1 | 18471650 | rs9730142 | 1.136 | 0.061 | 0.217 | -0.632 | 0.004 | -0.007 | -0.007 | 0 | 0.22 | 0.22 | 0.15 | 0.15 | 0.21 | 0.17 |

| | | | | | | | | | | | | | | | | |
|---|---|---|---|---|---|---|---|---|---|---|---|---|---|---|---|---|
| 1 | 18473343 | rs9329469 | -1.095 | 0.046 | 0.294 | -0.562 | 0.021 | 0.126 | 0.126 | 0 | 0.06 | 0.26 | 0.08 | 0.13 | 0.05 | 0.08 |
| 1 | 18475260 | rs12742611 | 1.33 | -0.044 | 0.086 | -0.634 | 0.04 | 0.042 | 0.042 | 0 | 0.17 | 0.04 | 0.06 | 0.06 | 0.12 | 0.12 |
| 1 | 18475498 | rs4920440 | -0.584 | -0.025 | 0.148 | -0.529 | -0.009 | 0.002 | 0.002 | 0.255 | 0.63 | 0.68 | 0.68 | 0.62 | 0.67 | 0.77 |
| 1 | 18475523 | rs2027530 | -0.215 | -1.467 | -0.272 | -0.449 | 0.002 | 0.013 | 0.013 | 0.21 | 0.4 | 0.33 | 0.59 | 0.73 | 0.4 | 0.6 |
| 1 | 18476187 | rs9727014 | -0.612 | -0.762 | -0.02 | 0.165 | -0.001 | -0.001 | -0.001 | 0 | 0.41 | 0.43 | 0.45 | 0.54 | 0.35 | 0.58 |
| 1 | 18479417 | rs12564645 | 0.138 | -0.731 | -0.858 | 0.107 | -0.01 | 0.082 | 0.082 | 0.019 | 0.24 | 0.39 | 0.39 | 0.51 | 0.24 | 0.46 |
| 1 | 18480012 | rs9662795 | 0.511 | -1.026 | -2.233 | 0.056 | 0.028 | 0.053 | 0.053 | 0 | 0.18 | 0.04 | 0.07 | 0.06 | 0.15 | 0.16 |
| 1 | 18480091 | rs9662299 | 0.464 | -1.028 | -2.235 | 0.055 | 0.029 | 0.055 | 0.055 | 0 | 0.19 | 0.04 | 0.07 | 0.06 | 0.15 | 0.16 |
| 1 | 18481003 | rs10907335 | -0.59 | -0.894 | -2.29 | 0.029 | 0.049 | -0.008 | -0.008 | 0 | 0.43 | 0.29 | 0.42 | 0.55 | 0.36 | 0.64 |
| 1 | 18483129 | rs16861648 | 0.096 | -0.457 | -2.521 | 0.405 | 0.02 | -0.056 | -0.056 | 0.661 | 0.23 | 0.12 | 0.33 | 0.46 | 0.2 | 0.46 |
| 1 | 18483918 | rs9659675 | -0.179 | -0.1 | -2.281 | -0.147 | 0.018 | 0.06 | 0.06 | 0.084 | 0.31 | 0.02 | 0.17 | 0.19 | 0.19 | 0.15 |
| 1 | 18489319 | rs12076097 | -0.824 | 0.715 | -1.228 | 0.024 | 0.012 | -0.035 | -0.035 | 0.32 | 0.07 | 0.15 | 0.05 | 0 | 0.07 | 0.08 |
| 1 | 18489949 | rs4920299 | -0.598 | 1.189 | -2.424 | 0.326 | -0.051 | -0.074 | -0.074 | 0 | 0.31 | 0.1 | 0.49 | 0.64 | 0.36 | 0.44 |
| 1 | 18490269 | rs11260939 | -0.409 | 1.581 | -2.67 | 0.34 | -0.093 | -0.164 | -0.164 | 0 | 0.26 | 0.05 | 0.44 | 0.54 | 0.32 | 0.31 |
| 1 | 18490551 | rs11260940 | -0.205 | 0.932 | -2.558 | -0.603 | -0.003 | 0.026 | 0.026 | 0 | 0.44 | 0.37 | 0.32 | 0.27 | 0.45 | 0.39 |
| 1 | 18493076 | rs11802233 | 0.507 | 0.567 | -2.546 | -0.662 | 0.021 | 0.09 | 0.09 | 0 | 0.29 | 0.08 | 0.14 | 0.13 | 0.19 | 0.26 |
| 1 | 18496159 | rs17435169 | 1.143 | 0.76 | -2.872 | -0.993 | -0.04 | 0.153 | 0.153 | 0 | 0.25 | 0.03 | 0.15 | 0.01 | 0.27 | 0.15 |
| 1 | 18497278 | rs16861658 | 0.535 | 0.505 | -2.038 | -0.906 | 0.021 | 0.14 | 0.14 | 0.002 | 0.05 | 0.26 | 0.1 | 0.17 | 0.05 | 0.06 |
| 1 | 18499322 | rs10907288 | -0.523 | 0.361 | -1.656 | -1.114 | -0.034 | 0.03 | 0.03 | 0.808 | 0.33 | 0.56 | 0.28 | 0.33 | 0.24 | 0.28 |
| 1 | 18501662 | rs9726406 | -0.691 | 0.613 | -1.536 | -1.088 | 0.008 | 0.013 | 0.013 | 0 | 0.62 | 0.61 | 0.45 | 0.41 | 0.59 | 0.58 |
| 1 | 18503402 | rs749917 | 0.848 | 2.238 | -0.932 | -0.351 | -0.072 | -0.066 | -0.066 | 0 | 0.1 | 0.02 | 0.43 | 0.49 | 0.26 | 0.24 |
| 1 | 18503419 | rs749918 | 0.262 | 2.264 | -0.246 | 0.04 | 0.019 | 0.015 | 0.015 | 0 | 0.07 | 0.19 | 0.07 | 0.06 | 0.08 | 0.17 |
| 1 | 18504396 | rs755967 | -1.065 | 2.208 | -0.19 | 0.105 | -0.001 | -0.001 | -0.001 | 2.716 | 0.46 | 0.61 | 0.4 | 0.44 | 0.5 | 0.51 |
| 1 | 18505196 | rs11260950 | -0.504 | 2.125 | 0.057 | -0.53 | -0.03 | 0.046 | 0.046 | 0 | 0.34 | 0.22 | 0.25 | 0.09 | 0.44 | 0.3 |
| 1 | 18505429 | rs16861677 | -1.155 | 1.517 | 0.429 | -0.287 | 0.07 | 0.03 | 0.03 | 0.01 | 0.1 | 0.4 | 0.1 | 0.06 | 0.09 | 0.09 |
| 1 | 18510033 | rs9728569 | 0.564 | 2.015 | -0.593 | 0.104 | -0.02 | 0.027 | 0.027 | 0 | 0.65 | 0.3 | 0.71 | 0.83 | 0.57 | 0.72 |
| 1 | 18512328 | rs10907289 | 0.741 | 1.856 | 0.301 | 0.995 | 0.068 | 0.12 | 0.12 | 0.015 | 0.9 | 0.46 | 0.89 | 0.84 | 0.89 | 0.85 |
| 1 | 18516703 | rs12567796 | 0.188 | 1.387 | -1.046 | 0.517 | -0.112 | 0.034 | 0.034 | 0 | 0.3 | 0.04 | 0.16 | 0.24 | 0.33 | 0.37 |
| 1 | 18516989 | rs1539051 | -0.178 | 0.734 | -1.211 | 0.837 | 0.054 | -0.039 | -0.039 | 0 | 0.43 | 0.77 | 0.56 | 0.47 | 0.32 | 0.34 |
| 1 | 18517072 | rs10752432 | 0.36 | 0.264 | -0.608 | 1.027 | -0.001 | 0.012 | 0.012 | 0 | 0.14 | 0.18 | 0.36 | 0.15 | 0.13 | 0.1 |
| 1 | 18521689 | rs767654 | 1.145 | 0.173 | 0.18 | 1.163 | 0.004 | -0.004 | -0.004 | 0 | 0.32 | 0.32 | 0.19 | 0.32 | 0.33 | 0.38 |
| 1 | 18521755 | rs12023073 | 0.404 | 0.027 | 0.222 | 1.349 | -0.014 | 0.005 | 0.005 | 0 | 0.3 | 0.14 | 0.29 | 0.25 | 0.38 | 0.38 |
| 1 | 18523331 | rs4920453 | 1.111 | 0.147 | 0.195 | 1.187 | -0.037 | 0.014 | 0.014 | 0 | 0.31 | 0.15 | 0.17 | 0.33 | 0.34 | 0.38 |
| 1 | 18527213 | rs4920459 | 0.644 | -0.95 | 0.602 | 1.197 | 0.006 | 0.051 | 0.051 | 0 | 0.46 | 0.34 | 0.31 | 0.29 | 0.51 | 0.38 |
| 1 | 18527267 | rs4920460 | -0.08 | -1.007 | 0.768 | 1.668 | 0.014 | 0.125 | 0.125 | 0 | 0.87 | 0.74 | 0.59 | 0.57 | 0.83 | 0.64 |
| 1 | 18528026 | rs9725311 | 0.542 | -1.265 | 0.424 | 1.601 | -0.016 | -0.047 | -0.047 | 1.921 | 0.75 | 0.82 | 0.51 | 0.41 | 0.72 | 0.54 |
| 1 | 18528848 | rs1033190 | 0.357 | -1.338 | 0.734 | 1.474 | 0.029 | 0.039 | 0.039 | 0 | 0.26 | 0.16 | 0.17 | 0.12 | 0.21 | 0.17 |
| 1 | 18533281 | rs4920461 | -0.353 | -1.329 | 0.721 | 1.484 | -0.008 | -0.006 | -0.006 | 0.007 | 0.87 | 0.76 | 0.88 | 0.84 | 0.89 | 0.8 |
| 1 | 18543784 | rs11260958 | -0.723 | -1.399 | 0.856 | 1.692 | 0.023 | -0.008 | -0.008 | 0 | 0.39 | 0.24 | 0.53 | 0.51 | 0.34 | 0.38 |
| 1 | 18544840 | rs4310409 | -0.732 | -1.399 | 0.856 | 1.698 | 0.006 | 0.002 | 0.002 | 0 | 0.39 | 0.3 | 0.53 | 0.54 | 0.34 | 0.38 |
| 1 | 18545932 | rs11578597 | 1.234 | -1.501 | 0.817 | 1.689 | 0.009 | 0.024 | 0.024 | 0 | 0.47 | 0.43 | 0.34 | 0.28 | 0.55 | 0.44 |
| 1 | 18546610 | rs12753104 | 0.163 | -1.442 | 0.619 | 1.705 | -0.043 | -0.088 | -0.088 | 3.723 | 0.12 | 0.01 | 0.35 | 0.4 | 0.14 | 0.21 |
| 1 | 18550683 | rs12033929 | 0.502 | -1.37 | 0.645 | 1.521 | -0.014 | -0.003 | -0.003 | 0 | 0.11 | 0.05 | 0.11 | 0.15 | 0.13 | 0.1 |
| 1 | 18550837 | rs1913592 | 0.473 | -1.347 | 0.741 | 1.542 | -0.025 | 0.002 | 0.002 | 0 | 0.6 | 0.53 | 0.44 | 0.5 | 0.66 | 0.61 |
| 1 | 18554356 | rs3738087 | 0.262 | -1.394 | 0.308 | 1.444 | -0.015 | 0.049 | 0.049 | 0.043 | 0.26 | 0.05 | 0.24 | 0.18 | 0.29 | 0.22 |
| 1 | 18554711 | rs3738089 | -0.383 | -1.515 | 0.184 | 1.821 | -0.039 | -0.031 | -0.031 | 0 | 0.17 | 0.03 | 0.18 | 0.22 | 0.19 | 0.2 |
| 1 | 18554886 | rs12032745 | 0.391 | -1.432 | 0.2 | 1.837 | -0.01 | 0.043 | 0.043 | 0.332 | 0.26 | 0.05 | 0.24 | 0.18 | 0.28 | 0.22 |
| 1 | 18556959 | rs17436172 | 1.356 | -1.724 | 0.561 | 1.979 | 0 | 0.002 | 0.002 | 0 | 0.44 | 0.29 | 0.5 | 0.45 | 0.5 | 0.44 |
| 1 | 18558575 | rs17436219 | 0.252 | -1.602 | 0.445 | 2.008 | -0.003 | 0.001 | 0.001 | 0.35 | 0.26 | 0.14 | 0.26 | 0.27 | 0.33 | 0.22 |
| 1 | 18559122 | rs6664613 | -0.21 | -1.911 | 0.645 | 2.15 | 0 | 0 | 0 | 0.475 | 0.3 | 0.28 | 0.23 | 0.44 | 0.27 | 0.36 |
| 1 | 18561590 | rs12041734 | 0.377 | -1.888 | 0.061 | 2.116 | 0.001 | -0.014 | -0.014 | 0.311 | 0.25 | 0.07 | 0.21 | 0.28 | 0.27 | 0.22 |
| 1 | 18563449 | rs11260963 | 0.701 | -1.79 | 0.024 | 2.189 | -0.011 | -0.01 | -0.01 | 0 | 0.57 | 0.44 | 0.61 | 0.33 | 0.62 | 0.46 |
| 1 | 18564669 | rs1341306 | -1.431 | -1.369 | 0.272 | 2.362 | -0.013 | 0.489 | 0.489 | 0.335 | 0.09 | 0.45 | 0.29 | 0.33 | 0.03 | 0.15 |
| 1 | 18566065 | rs1341309 | -0.932 | -0.768 | -1.053 | 1.443 | -0.005 | -0.045 | -0.045 | 0.044 | 0.09 | 0.02 | 0.13 | 0.23 | 0.1 | 0.16 |
| 1 | 18566851 | rs2883821 | 0.632 | -0.988 | -0.828 | 1.36 | -0.038 | 0.01 | 0.01 | 0.325 | 0.77 | 0.61 | 0.87 | 0.7 | 0.89 | 0.87 |
| 1 | 18569510 | rs12566407 | -0.839 | -0.183 | -1.54 | 1.401 | 0 | -0.029 | -0.029 | 0 | 0.24 | 0.17 | 0.22 | 0.35 | 0.23 | 0.2 |
| 1 | 18572901 | rs109490 | 0.182 | -0.478 | -1.894 | 1.008 | 0.089 | -0.032 | -0.032 | 1.218 | 0.56 | 0.22 | 0.75 | 0.57 | 0.74 | 0.68 |
| 1 | 18578075 | rs1003741 | 0.098 | 1.002 | -2.347 | 0.978 | -0.114 | -0.117 | -0.117 | 0.327 | 0.26 | 0.04 | 0.51 | 0.51 | 0.36 | 0.47 |
| 1 | 18581074 | rs223179 | -0.497 | 0.364 | -2.588 | 0.344 | 0.008 | -0.027 | -0.027 | 0 | 0.39 | 0.16 | 0.47 | 0.63 | 0.29 | 0.6 |
| 1 | 18585845 | rs223188 | 0.052 | -0.309 | -1.45 | -0.551 | -0.01 | -0.077 | -0.077 | 0.084 | 0.23 | 0.3 | 0.11 | 0.06 | 0.2 | 0.11 |
| 1 | 18586420 | rs371505 | 0.409 | -0.305 | -1.405 | -0.551 | 0.006 | 0.046 | 0.046 | 0 | 0.21 | 0.13 | 0.09 | 0.05 | 0.2 | 0.1 |
| 1 | 18589620 | rs419971 | -0.706 | -0.38 | -0.729 | -1 | -0.003 | -0.037 | -0.037 | 0.323 | 0.44 | 0.5 | 0.29 | 0.13 | 0.37 | 0.24 |
| 1 | 18597809 | rs4920468 | 0.807 | -0.116 | -0.968 | -0.427 | 0.007 | -0.042 | -0.042 | 0 | 0.07 | 0 | 0.23 | 0.18 | 0.04 | 0.15 |
| 1 | 18600824 | rs11801569 | -0.112 | 0.108 | -0.371 | -1.206 | 0.005 | -0.08 | -0.08 | 1.175 | 0.21 | 0.38 | 0.22 | 0.05 | 0.2 | 0.08 |
| 1 | 18605945 | rs11260970 | -0.533 | 0.162 | -0.142 | -1.111 | -0.003 | -0.063 | -0.063 | 0 | 0.44 | 0.48 | 0.33 | 0.13 | 0.41 | 0.26 |
| 1 | 18611016 | rs223192 | -1.779 | 0.51 | 1.372 | -0.44 | 0.035 | -0.058 | -0.058 | 0 | 0.14 | 0.53 | 0.15 | 0.04 | 0.14 | 0.12 |
| 1 | 18612669 | rs6670195 | 0.809 | 0.552 | 1.071 | -0.42 | 0.045 | -0.04 | -0.04 | 0 | 0.89 | 0.71 | 0.87 | 0.97 | 0.86 | 0.91 |
| 1 | 18615392 | rs6603893 | 1.478 | 0.538 | 0.198 | -0.517 | 0 | -0.037 | -0.037 | 0 | 0.79 | 0.24 | 0.81 | 0.96 | 0.82 | 0.88 |
| 1 | 18630106 | rs1533345 | -1.282 | 0.735 | 0.411 | -0.202 | 0.069 | -0.002 | -0.002 | 0.094 | 0.12 | 0.32 | 0.14 | 0.06 | 0.14 | 0.12 |

| | | | | | | | | | | | | | | | | |
|---|---|---|---|---|---|---|---|---|---|---|---|---|---|---|---|---|
| 1 | 18631175 | rs12134352 | 1.248 | 0.58 | -0.751 | -0.683 | -0.102 | 0.145 | 0.145 | 0 | 0.58 | 0.08 | 0.61 | 0.76 | 0.61 | 0.6 |
| 1 | 18633683 | rs761584 | 1.247 | 0.574 | -0.649 | -0.686 | -0.082 | 0.146 | 0.146 | 0 | 0.58 | 0.09 | 0.61 | 0.76 | 0.61 | 0.61 |
| 1 | 18643247 | rs4920474 | -0.854 | 0.499 | -0.047 | -0.867 | 0.006 | -0.03 | -0.03 | 0.922 | 0.24 | 0.26 | 0.19 | 0.07 | 0.25 | 0.17 |
| 1 | 18643577 | rs6663855 | 1.247 | 0.655 | 0.433 | -0.919 | 0.018 | -0.063 | -0.063 | 0.02 | 0.67 | 0.49 | 0.68 | 0.91 | 0.65 | 0.68 |
| 1 | 18643632 | rs4920477 | 0.314 | 0.308 | 0.248 | -0.561 | 0.069 | -0.024 | -0.024 | 1.103 | 0.94 | 0.85 | 0.88 | 0.99 | 0.9 | 0.87 |
| 1 | 18646231 | rs1040623 | 0.283 | -0.483 | -0.171 | 0.027 | -0.021 | 0.041 | 0.041 | 0 | 0.72 | 0.53 | 0.74 | 0.45 | 0.82 | 0.66 |
| 1 | 18655339 | rs4920478 | 0.816 | -0.144 | -1.112 | -0.113 | -0.103 | 0.013 | 0.013 | 0.271 | 0.31 | 0.05 | 0.34 | 0.25 | 0.43 | 0.26 |
| 1 | 18656738 | rs573262 | -0.11 | -0.24 | -0.734 | 0.219 | -0.063 | -0.021 | -0.021 | 0 | 0.19 | 0.56 | 0.12 | 0.13 | 0.08 | 0.12 |
| 1 | 18659641 | rs552749 | -0.173 | 0.433 | -1.405 | 0.626 | -0.018 | -0.021 | -0.021 | 0.107 | 0.6 | 0.78 | 0.59 | 0.57 | 0.58 | 0.49 |
| 1 | 18661583 | rs536540 | -0.634 | -0.607 | -1.28 | 0.605 | 0.029 | 0 | 0 | 0.956 | 0.34 | 0.74 | 0.27 | 0.35 | 0.15 | 0.24 |
| 1 | 18664626 | rs517004 | 0.533 | -0.249 | -0.414 | 0.909 | 0.035 | 0.074 | 0.074 | 0 | 0.94 | 0.83 | 0.94 | 0.89 | 0.95 | 0.93 |
| 1 | 18668501 | rs11260984 | 0.565 | 0.035 | -1.171 | 1.145 | -0.017 | 0.041 | 0.041 | 0 | 0.52 | 0.21 | 0.43 | 0.39 | 0.58 | 0.43 |
| 1 | 18672774 | rs542634 | 0.044 | -0.252 | -0.278 | 1.411 | -0.007 | -0.004 | -0.004 | 0 | 0.3 | 0.32 | 0.34 | 0.34 | 0.3 | 0.39 |
| 1 | 18676961 | rs555115 | 0.834 | -0.102 | 0.07 | 1.348 | 0.009 | 0.01 | 0.01 | 0 | 0.2 | 0.26 | 0.16 | 0.3 | 0.2 | 0.29 |
| 1 | 18678170 | rs16861827 | 0.32 | -0.091 | 0.444 | 1.132 | 0 | 0 | 0 | 0 | 0.11 | 0.11 | 0.24 | 0.13 | 0.13 | 0.11 |
| 1 | 18680330 | rs539132 | -0.019 | -0.159 | 0.893 | 1.243 | -0.019 | 0.012 | 0.012 | 0 | 0.85 | 0.8 | 0.86 | 0.83 | 0.9 | 0.83 |
| 1 | 18685202 | rs10489573 | 0.633 | -0.069 | 0.235 | 0.598 | -0.017 | 0.097 | 0.097 | 0 | 0.26 | 0.06 | 0.31 | 0.08 | 0.28 | 0.24 |
| 1 | 18685434 | rs525448 | 0.518 | -0.244 | 0.463 | 1.038 | 0.012 | 0.001 | 0.001 | 0 | 0.3 | 0.44 | 0.24 | 0.3 | 0.27 | 0.35 |
| 1 | 18691614 | rs12079178 | 0.114 | -0.382 | 0.643 | 1.175 | -0.012 | 0.021 | 0.021 | 0 | 0.14 | 0.19 | 0.15 | 0.21 | 0.1 | 0.17 |
| 1 | 18693245 | rs7543948 | -0.753 | -0.202 | 0.844 | 1.101 | 0 | -0.002 | -0.002 | 0 | 0.7 | 0.62 | 0.76 | 0.72 | 0.75 | 0.65 |
| 1 | 18693707 | rs11260991 | 0.865 | -0.002 | 0.685 | 0.884 | -0.012 | 0.055 | 0.055 | 0 | 0.27 | 0.14 | 0.33 | 0.04 | 0.32 | 0.24 |
| 1 | 18694036 | rs12123650 | 1.263 | -0.022 | 0.832 | 1.019 | 0 | 0.003 | 0.003 | 0 | 0.29 | 0.29 | 0.24 | 0.36 | 0.33 | 0.23 |
| 1 | 18696215 | rs882916 | -0.933 | 0.071 | 0.577 | 1.04 | 0 | 0.002 | 0.002 | 0 | 0.71 | 0.77 | 0.79 | 0.73 | 0.76 | 0.65 |
| 1 | 18697149 | rs12035573 | 0.991 | 0.323 | -0.187 | 1.671 | -0.041 | -0.054 | -0.054 | 0 | 0.27 | 0.12 | 0.26 | 0.37 | 0.35 | 0.23 |
| 1 | 18697680 | rs10907305 | 0.991 | 0.323 | -0.228 | 1.671 | -0.048 | -0.063 | -0.063 | 0 | 0.27 | 0.11 | 0.26 | 0.37 | 0.35 | 0.23 |
| 1 | 18700256 | rs884129 | 0.98 | 0.326 | -0.323 | 1.672 | -0.055 | -0.071 | -0.071 | 0 | 0.27 | 0.1 | 0.26 | 0.37 | 0.35 | 0.23 |
| 1 | 18701899 | rs10465918 | -0.873 | 0.07 | 0.362 | 1.849 | -0.008 | 0.023 | 0.023 | 0 | 0.15 | 0.17 | 0.19 | 0.31 | 0.1 | 0.17 |
| 1 | 18703672 | rs2271861 | -0.343 | 0.475 | -0.267 | 1.853 | -0.001 | 0.029 | 0.029 | 0 | 0.07 | 0.09 | 0.11 | 0.2 | 0.06 | 0.21 |
| 1 | 18704862 | rs1043459 | -0.967 | 0.617 | -0.424 | 1.951 | 0.001 | 0.002 | 0.002 | 0.397 | 0.42 | 0.77 | 0.56 | 0.61 | 0.44 | 0.58 |
| 1 | 18705694 | rs12566510 | -0.522 | 0.705 | 0.182 | 2.529 | 0 | 0.248 | 0.248 | 0 | 0.09 | 0.2 | 0.12 | 0.42 | 0.06 | 0.31 |
| 1 | 18707467 | rs4920489 | 0.31 | 0.585 | -0.042 | 2.385 | -0.058 | 0.074 | 0.074 | 0 | 0.23 | 0.06 | 0.31 | 0.11 | 0.32 | 0.28 |
| 1 | 18708402 | rs1577608 | -0.543 | 0.534 | 0.247 | 2.453 | 0.008 | 0.311 | 0.311 | 0 | 0.09 | 0.26 | 0.12 | 0.42 | 0.06 | 0.3 |
| 1 | 18709741 | rs7538502 | -1.053 | 0.475 | 0.015 | 1.613 | -0.011 | 0.075 | 0.075 | 0 | 0.46 | 0.7 | 0.61 | 0.76 | 0.47 | 0.71 |
| 1 | 18714272 | rs12040552 | 0.059 | 0.374 | 0.026 | 1.92 | -0.004 | 0.047 | 0.047 | 0 | 0.07 | 0.08 | 0.05 | 0.24 | 0.05 | 0.26 |
| 1 | 18716278 | rs7538442 | 0.015 | 0.864 | -0.208 | 1.511 | -0.028 | -0.013 | -0.013 | 0.351 | 0.23 | 0.14 | 0.35 | 0.25 | 0.32 | 0.19 |
| 1 | 18718305 | rs11260998 | -1.161 | 0.737 | 0.085 | 1.536 | -0.001 | 0.016 | 0.016 | 0.302 | 0.38 | 0.48 | 0.4 | 0.38 | 0.35 | 0.45 |
| 1 | 18718530 | rs12041781 | -0.206 | 0.504 | -0.106 | 1.921 | 0.01 | -0.009 | -0.009 | 0 | 0.14 | 0.09 | 0.06 | 0.34 | 0.09 | 0.3 |
| 1 | 18719844 | rs4920492 | 0.927 | 0.888 | -0.318 | 1.497 | 0.001 | 0.008 | 0.008 | 0 | 0.26 | 0.12 | 0.18 | 0.23 | 0.25 | 0.33 |
| 1 | 18722491 | rs1336129 | -1.964 | 0.692 | -0.197 | 1.352 | 0.002 | -0.003 | -0.003 | 0 | 0.73 | 0.76 | 0.79 | 0.72 | 0.79 | 0.69 |
| 1 | 18722692 | rs11810587 | 0.942 | 0.708 | -0.23 | 1.408 | 0.002 | 0.001 | 0.001 | 0 | 0.1 | 0.12 | 0.17 | 0.3 | 0.09 | 0.1 |
| 1 | 18722713 | rs1336130 | -0.661 | 0.716 | 0.055 | 1.37 | -0.002 | -0.004 | -0.004 | 0.543 | 0.4 | 0.39 | 0.53 | 0.43 | 0.44 | 0.29 |
| 1 | 18722846 | rs1533622 | 1.286 | 0.717 | 0.054 | 1.402 | 0.001 | 0.026 | 0.026 | 0 | 0.26 | 0.13 | 0.18 | 0.18 | 0.25 | 0.34 |
| 1 | 18723444 | rs7539612 | -0.662 | 0.773 | -0.014 | 1.353 | -0.003 | -0.005 | -0.005 | 0 | 0.39 | 0.38 | 0.53 | 0.43 | 0.44 | 0.29 |
| 1 | 18724101 | rs12138463 | 0.27 | 0.822 | -0.169 | 1.373 | -0.004 | 0.005 | 0.005 | 0 | 0.3 | 0.27 | 0.35 | 0.14 | 0.35 | 0.18 |
| 1 | 18728581 | rs7512723 | -0.086 | 0.771 | -0.064 | 1.087 | -0.016 | -0.021 | -0.021 | 1.091 | 0.38 | 0.25 | 0.51 | 0.43 | 0.44 | 0.28 |
| 1 | 18730305 | rs7523920 | -0.864 | 0.77 | -0.003 | 1.017 | 0.001 | -0.001 | -0.001 | 0.02 | 0.67 | 0.58 | 0.72 | 0.69 | 0.69 | 0.65 |
| 1 | 18731662 | rs7522460 | -1.508 | 0.46 | -0.276 | 1.042 | 0.011 | -0.006 | -0.006 | 0 | 0.75 | 0.79 | 0.86 | 0.73 | 0.83 | 0.7 |
| 1 | 18732980 | rs867538 | 0.119 | 0.649 | -0.439 | 0.683 | -0.007 | 0.001 | 0.001 | 0 | 0.29 | 0.22 | 0.22 | 0.25 | 0.28 | 0.39 |
| 1 | 18733717 | rs6695244 | -0.52 | 0.84 | -0.276 | 0.724 | -0.001 | -0.001 | -0.001 | 0 | 0.47 | 0.52 | 0.5 | 0.39 | 0.58 | 0.5 |
| 1 | 18733958 | rs1336125 | -0.316 | 0.732 | -0.295 | 0.88 | 0.022 | 0.004 | 0.004 | 0 | 0.51 | 0.35 | 0.49 | 0.48 | 0.4 | 0.6 |
| 1 | 18736393 | rs4543765 | 0.72 | 0.688 | -0.26 | 0.871 | -0.001 | 0.009 | 0.009 | 0 | 0.24 | 0.28 | 0.34 | 0.33 | 0.22 | 0.27 |
| 1 | 18736536 | rs9662941 | 1.16 | 0.763 | -0.409 | 0.853 | 0.005 | 0.006 | 0.006 | 0 | 0.22 | 0.12 | 0.15 | 0.15 | 0.19 | 0.36 |
| 1 | 18747095 | rs11261002 | -0.553 | 1.104 | -0.147 | 0.566 | 0.002 | -0.004 | -0.004 | 0 | 0.58 | 0.46 | 0.55 | 0.57 | 0.51 | 0.65 |
| 1 | 18750431 | rs12128025 | -0.634 | 1.138 | -0.257 | 0.583 | 0 | -0.001 | -0.001 | 0 | 0.58 | 0.55 | 0.56 | 0.57 | 0.51 | 0.64 |
| 1 | 18751864 | rs869080 | 0.8 | 1.157 | -0.299 | 0.607 | -0.003 | -0.001 | -0.001 | 1.132 | 0.17 | 0.16 | 0.28 | 0.15 | 0.31 | 0.12 |
| 1 | 18759377 | rs2222257 | -0.576 | 1.104 | -0.324 | 0.578 | -0.002 | 0 | 0 | 0 | 0.57 | 0.55 | 0.55 | 0.57 | 0.49 | 0.63 |
| 1 | 18761893 | rs1850750 | 1.088 | 1.335 | -0.413 | 0.641 | 0.011 | 0.003 | 0.003 | 0 | 0.06 | 0.08 | 0.07 | 0.1 | 0.09 | 0.02 |
| 1 | 18765153 | rs17437569 | 1.338 | 0.95 | -0.547 | 0.451 | -0.008 | 0.018 | 0.018 | 0 | 0.15 | 0.09 | 0.19 | 0.04 | 0.17 | 0.13 |
| 1 | 18765308 | rs1336128 | -1.286 | 0.751 | -0.366 | 0.608 | -0.005 | -0.001 | -0.001 | 0.32 | 0.74 | 0.72 | 0.82 | 0.72 | 0.79 | 0.74 |
| 1 | 18766761 | rs1568339 | 0.02 | 0.984 | -0.525 | 0.52 | 0.003 | 0.008 | 0.008 | 0 | 0.5 | 0.42 | 0.48 | 0.54 | 0.44 | 0.6 |
| 1 | 18766850 | rs1568340 | -0.854 | 0.54 | -0.252 | 0.486 | -0.038 | -0.023 | -0.023 | 0.013 | 0.71 | 0.43 | 0.77 | 0.71 | 0.78 | 0.76 |
| 1 | 18767081 | rs12136062 | -0.349 | 0.399 | -0.613 | 0.483 | 0.018 | 0 | 0 | 0 | 0.52 | 0.36 | 0.5 | 0.47 | 0.59 | 0.58 |
| 1 | 18767738 | rs6677903 | 0.701 | -0.186 | -0.476 | 0.04 | 0 | 0 | 0 | 0 | 0.15 | 0.15 | 0.13 | 0.08 | 0.13 | 0.32 |
| 1 | 18767777 | rs16861969 | 0.62 | -0.041 | -0.616 | 0.211 | -0.014 | -0.022 | -0.022 | 0 | 0.06 | 0.02 | 0.17 | 0.21 | 0.08 | 0.05 |
| 1 | 18767791 | rs2946523 | -1.793 | -0.246 | -0.391 | -0.078 | 0.002 | 0.002 | 0.002 | 1.035 | 0.44 | 0.49 | 0.53 | 0.45 | 0.45 | 0.66 |
| 1 | 18768011 | rs3007722 | -0.341 | -0.244 | -0.47 | -0.085 | 0.004 | -0.018 | -0.018 | 0.228 | 0.16 | 0.14 | 0.2 | 0.28 | 0.14 | 0.23 |
| 1 | 18768277 | rs2946524 | -0.965 | -0.27 | -0.505 | -0.108 | 0.002 | -0.002 | -0.002 | 0.044 | 0.31 | 0.29 | 0.33 | 0.32 | 0.27 | 0.48 |
| 1 | 18769575 | rs12408368 | 1.357 | -0.224 | -0.673 | -0.099 | -0.005 | 0.028 | 0.028 | 0 | 0.14 | 0.05 | 0.13 | 0.04 | 0.13 | 0.25 |
| 1 | 18770671 | rs2992749 | -1.171 | -0.223 | -0.438 | -0.094 | -0.008 | 0 | 0 | 0 | 0.33 | 0.42 | 0.36 | 0.32 | 0.29 | 0.52 |

| | | | | | | | | | | | | | | | | |
|---|---|---|---|---|---|---|---|---|---|---|---|---|---|---|---|---|
| 1 | 18772971 | rs2946532 | -1.271 | -0.071 | -0.674 | 0.015 | 0.016 | 0.029 | 0.029 | 0.307 | 0.3 | 0.44 | 0.42 | 0.42 | 0.33 | 0.42 |
| 1 | 18774783 | rs2946534 | -0.869 | -0.551 | -1.334 | -0.366 | 0.023 | 0.024 | 0.024 | 0 | 0.53 | 0.79 | 0.52 | 0.39 | 0.41 | 0.68 |
| 1 | 18776939 | rs2992746 | 1.134 | -0.403 | -1.086 | -0.146 | -0.007 | 0.036 | 0.036 | 0.02 | 0.16 | 0.1 | 0.15 | 0.03 | 0.16 | 0.12 |
| 1 | 18781288 | rs2184708 | 0.909 | -0.131 | -1.106 | -0.013 | -0.035 | -0.037 | -0.037 | 0.061 | 0.07 | 0 | 0.17 | 0.18 | 0.09 | 0.06 |
| 1 | 18781486 | rs4920497 | 0.404 | -0.646 | -0.843 | -0.838 | -0.001 | -0.008 | -0.008 | 0 | 0.59 | 0.53 | 0.54 | 0.26 | 0.61 | 0.56 |
| 1 | 18785466 | rs10907312 | 1.111 | -0.667 | -0.98 | -1.425 | -0.04 | 0.092 | 0.092 | 0 | 0.38 | 0.19 | 0.35 | 0.16 | 0.46 | 0.18 |
| 1 | 18791334 | rs12747775 | 1.279 | -0.518 | -0.9 | -1.686 | -0.026 | 0.075 | 0.075 | 0.678 | 0.32 | 0.16 | 0.27 | 0.13 | 0.32 | 0.14 |
| 1 | 18792300 | rs12144656 | 1.555 | -0.733 | -1.064 | -1.779 | -0.007 | 0.006 | 0.006 | 0 | 0.15 | 0.06 | 0.14 | 0.08 | 0.13 | 0.1 |
| 1 | 18793520 | rs3007727 | -1.644 | -1.17 | -0.806 | -2.002 | 0.005 | 0.005 | 0.005 | 0.499 | 0.78 | 0.83 | 0.86 | 0.93 | 0.81 | 0.62 |
| 1 | 18793636 | rs11261014 | 1.849 | -1.189 | -0.538 | -2.016 | -0.01 | 0.033 | 0.033 | 0.507 | 0.34 | 0.24 | 0.27 | 0.13 | 0.32 | 0.14 |
| 1 | 18795255 | rs3007729 | -0.008 | -1.08 | -0.173 | -1.784 | 0.012 | 0.026 | 0.026 | 0 | 0.29 | 0.44 | 0.38 | 0.66 | 0.36 | 0.38 |
| 1 | 18797100 | rs3007730 | -1.727 | -1.027 | -0.301 | -1.843 | -0.023 | -0.049 | -0.049 | 0 | 0.87 | 0.81 | 0.93 | 0.99 | 0.91 | 0.74 |
| 1 | 18797703 | rs2992736 | -0.399 | -0.784 | -0.341 | -1.692 | -0.004 | -0.009 | -0.009 | 0 | 0.33 | 0.43 | 0.46 | 0.71 | 0.3 | 0.34 |
| 1 | 18800547 | rs2992734 | -0.46 | -1.037 | -0.674 | -1.902 | 0.002 | -0.018 | -0.018 | 0.001 | 0.7 | 0.71 | 0.81 | 0.94 | 0.71 | 0.62 |
| 1 | 18800911 | rs3007733 | -0.344 | -0.989 | -0.709 | -1.919 | 0.012 | 0.034 | 0.034 | 0.013 | 0.61 | 0.68 | 0.73 | 0.88 | 0.61 | 0.49 |
| 1 | 18805328 | rs11261017 | 0.885 | -1.133 | -0.588 | -2.111 | -0.013 | 0.077 | 0.077 | 0.927 | 0.44 | 0.3 | 0.4 | 0.19 | 0.46 | 0.25 |
| 1 | 18805564 | rs11261020 | 0.975 | -1.157 | -0.566 | -2.122 | -0.027 | 0.117 | 0.117 | 0.756 | 0.43 | 0.24 | 0.39 | 0.19 | 0.46 | 0.25 |
| 1 | 18805698 | rs1360920 | 0.471 | -1.134 | -0.381 | -2.087 | -0.001 | 0.053 | 0.053 | 0 | 0.17 | 0.32 | 0.32 | 0.7 | 0.15 | 0.24 |
| 1 | 18805988 | rs1360918 | 0.88 | -1.208 | -0.387 | -2.115 | 0 | 0.007 | 0.007 | 0 | 0.19 | 0.19 | 0.34 | 0.62 | 0.28 | 0.22 |
| 1 | 18806585 | rs10796456 | 0.767 | -1.32 | -0.275 | -2.094 | 0.001 | 0.021 | 0.021 | 0.427 | 0.45 | 0.39 | 0.46 | 0.27 | 0.48 | 0.3 |
| 1 | 18807117 | rs7528206 | 2.35 | -1.272 | -0.558 | -2.238 | -0.001 | -0.001 | -0.001 | 0 | 0.12 | 0.12 | 0.12 | 0.04 | 0.12 | 0.06 |
| 1 | 18808069 | rs7528572 | 2.35 | -1.272 | -0.558 | -2.238 | -0.001 | -0.001 | -0.001 | 0.318 | 0.12 | 0.12 | 0.12 | 0.04 | 0.12 | 0.06 |
| 1 | 18808292 | rs2992753 | -0.645 | -1.202 | -0.517 | -2.203 | 0.01 | 0.024 | 0.024 | 0.054 | 0.62 | 0.66 | 0.73 | 0.9 | 0.61 | 0.48 |
| 1 | 18809916 | rs12061708 | 1.896 | -0.964 | -0.662 | -2.062 | -0.03 | 0.038 | 0.038 | 0.276 | 0.27 | 0.17 | 0.25 | 0.15 | 0.33 | 0.13 |
| 1 | 18812176 | rs872355 | 1.378 | -0.951 | -0.695 | -2.105 | -0.002 | 0.003 | 0.003 | 0 | 0.36 | 0.32 | 0.34 | 0.26 | 0.36 | 0.23 |
| 1 | 18813026 | rs3007720 | -0.1 | -0.941 | -0.452 | -2.023 | -0.005 | -0.03 | -0.03 | 0.811 | 0.29 | 0.43 | 0.46 | 0.74 | 0.27 | 0.35 |
| 1 | 18813445 | rs4542191 | 0.13 | -0.778 | -0.614 | -1.673 | -0.006 | 0.068 | 0.068 | 0 | 0.14 | 0.24 | 0.31 | 0.67 | 0.13 | 0.21 |
| 1 | 18814565 | rs10907313 | -0.922 | -0.987 | -1.08 | -2.01 | -0.003 | 0.052 | 0.052 | 0 | 0.75 | 0.86 | 0.8 | 0.94 | 0.7 | 0.67 |
| 1 | 18814596 | rs11577512 | 0.76 | -0.688 | -1.392 | -1.918 | -0.066 | 0.111 | 0.111 | 0 | 0.34 | 0.14 | 0.36 | 0.14 | 0.4 | 0.25 |
| 1 | 18815334 | rs4920502 | -0.518 | -1.001 | -0.81 | -0.707 | 0.003 | 0.003 | 0.003 | 0 | 0.81 | 0.85 | 0.77 | 0.86 | 0.85 | 0.81 |
| 1 | 18816616 | rs4531264 | 1.267 | -1.109 | -0.654 | -0.565 | 0.01 | -0.015 | -0.015 | 0.06 | 0.21 | 0.17 | 0.14 | 0.53 | 0.13 | 0.28 |
| 1 | 18817181 | rs10907314 | 1.135 | -1.034 | -0.686 | -0.735 | -0.045 | 0.06 | 0.06 | 0.017 | 0.34 | 0.19 | 0.35 | 0.1 | 0.49 | 0.19 |
| 1 | 18821416 | rs6686713 | 0.817 | -0.203 | 0.155 | -0.015 | -0.007 | -0.024 | -0.024 | 0.059 | 0.67 | 0.5 | 0.71 | 0.73 | 0.7 | 0.6 |
| 1 | 18823903 | rs17351864 | 1.175 | -0.436 | -0.323 | -0.033 | -0.014 | -0.016 | -0.016 | 0 | 0.12 | 0.01 | 0.2 | 0.09 | 0.1 | 0.14 |
| 1 | 18825203 | rs909948 | -1.156 | -0.317 | 0.182 | -0.078 | 0.006 | -0.08 | -0.08 | 0 | 0.12 | 0.41 | 0.1 | 0 | 0.08 | 0.21 |
| 1 | 18829672 | rs4920327 | 0.876 | -0.092 | -0.15 | 0.048 | 0.007 | -0.027 | -0.027 | 0 | 0.66 | 0.43 | 0.7 | 0.75 | 0.65 | 0.58 |
| 1 | 18834756 | rs11261027 | -0.109 | -0.23 | -0.167 | 0.41 | -0.002 | -0.029 | -0.029 | 0 | 0.18 | 0.17 | 0.09 | 0.07 | 0.15 | 0.14 |
| 1 | 18835437 | rs2883890 | -0.519 | -0.927 | 0.214 | 1.583 | 0.031 | -0.03 | -0.03 | 0.364 | 0.22 | 0.1 | 0.2 | 0.28 | 0.12 | 0.26 |
| 1 | 18835504 | rs2745304 | -0.903 | -0.87 | 0.333 | 1.504 | -0.011 | 0.019 | 0.019 | 0 | 0.19 | 0.15 | 0.19 | 0.05 | 0.25 | 0.25 |
| 1 | 18836431 | rs2745303 | 0.617 | -1.278 | 0.304 | 1.545 | 0.017 | 0.008 | 0.008 | 0 | 0.1 | 0.07 | 0.09 | 0.05 | 0.07 | 0.04 |
| 1 | 18839126 | rs2745302 | -2.293 | -1.387 | 0.718 | 1.631 | 0.067 | 0.033 | 0.033 | 0 | 0.49 | 0.71 | 0.43 | 0.37 | 0.34 | 0.48 |
| 1 | 18839394 | rs2294656 | -2.115 | -1.224 | 0.812 | 1.496 | -0.003 | 0.006 | 0.006 | 0 | 0.21 | 0.47 | 0.14 | 0.17 | 0.16 | 0.26 |
| 1 | 18850054 | rs9439709 | -0.635 | -1.328 | -0.011 | 2.086 | 0.022 | -0.108 | -0.108 | 0 | 0.23 | 0.04 | 0.27 | 0.43 | 0.16 | 0.22 |
| 1 | 18851113 | rs2816025 | -1.567 | -1.433 | 0.203 | 2.055 | 0.002 | -0.003 | -0.003 | 0 | 0.41 | 0.44 | 0.39 | 0.46 | 0.32 | 0.36 |
| 1 | 18855616 | rs2076778 | -0.743 | -1.738 | 0.545 | 2.105 | -0.056 | 0.012 | 0.012 | 0 | 0.15 | 0.32 | 0.06 | 0.11 | 0.08 | 0.17 |
| 1 | 18857598 | rs2816030 | -1.257 | -1.586 | -0.015 | 2.163 | -0.086 | 0.147 | 0.147 | 0 | 0.24 | 0.52 | 0.17 | 0.44 | 0.14 | 0.23 |
| 1 | 18863502 | rs12401403 | -0.769 | -0.977 | -0.81 | 1.643 | 0.034 | -0.023 | -0.023 | 1.707 | 0.36 | 0.25 | 0.3 | 0.48 | 0.24 | 0.31 |
| 1 | 18876408 | rs2745327 | 0.595 | 0.54 | -0.749 | 1.901 | -0.015 | -0.027 | -0.027 | 0 | 0.86 | 0.9 | 0.8 | 0.7 | 0.82 | 0.77 |
| 1 | 18880442 | rs2745325 | 1.393 | 0.675 | -1.239 | 1.755 | -0.057 | 0.017 | 0.017 | 0 | 0.55 | 0.15 | 0.54 | 0.37 | 0.54 | 0.63 |
| 1 | 18882231 | rs725387 | -0.536 | 0.598 | -0.971 | 1.934 | 0 | 0 | 0 | 0.078 | 0.26 | 0.66 | 0.25 | 0.29 | 0.28 | 0.15 |
| 1 | 18882489 | rs10907320 | -0.32 | 0.477 | -0.879 | 1.689 | 0.01 | -0.037 | -0.037 | 0 | 0.22 | 0.6 | 0.19 | 0.08 | 0.23 | 0.11 |
| 1 | 18885736 | rs2745322 | -0.106 | 0.522 | -0.966 | 1.813 | -0.023 | -0.043 | -0.043 | 0.243 | 0.86 | 0.92 | 0.8 | 0.58 | 0.82 | 0.78 |
| 1 | 18891233 | rs11261034 | -0.226 | 0.521 | 0.216 | 1.505 | 0.034 | -0.057 | -0.057 | 0.047 | 0.2 | 0.41 | 0.18 | 0.08 | 0.23 | 0.11 |
| 1 | 18893763 | rs2816048 | 1.225 | 0.442 | 0.208 | 1.408 | -0.047 | 0.022 | 0.022 | 0 | 0.55 | 0.16 | 0.54 | 0.38 | 0.54 | 0.64 |
| 1 | 18895806 | rs6698315 | 1.144 | 0.482 | 0.196 | 1.427 | -0.025 | 0.02 | 0.02 | 0 | 0.56 | 0.27 | 0.55 | 0.38 | 0.54 | 0.64 |
| 1 | 18897505 | rs12131648 | -0.35 | 0.861 | 0.912 | 1.071 | 0.066 | -0.023 | -0.023 | 0 | 0.06 | 0.25 | 0.07 | 0 | 0.11 | 0.01 |
| 1 | 18904707 | rs7549273 | -1.106 | 0.799 | 0.685 | 1.551 | 0.018 | 0.186 | 0.186 | 0.007 | 0.14 | 0.36 | 0.17 | 0.35 | 0.15 | 0.23 |
| 1 | 18908865 | rs12566790 | -1.058 | 1.125 | 0.13 | 0.859 | -0.016 | -0.018 | -0.018 | 0 | 0.08 | 0.03 | 0.07 | 0.14 | 0.1 | 0.16 |
| 1 | 18913558 | rs4920334 | -0.29 | 0.949 | -0.343 | -0.332 | -0.016 | 0.161 | 0.161 | 0.036 | 0.49 | 0.24 | 0.35 | 0.18 | 0.5 | 0.39 |
| 1 | 18914423 | rs2816065 | 0.192 | 0.81 | 0.441 | -0.361 | 0.001 | 0.016 | 0.016 | 0 | 0.47 | 0.4 | 0.61 | 0.79 | 0.47 | 0.53 |
| 1 | 18929842 | rs4920520 | -0.307 | 0.33 | 0.88 | -0.652 | 0 | -0.039 | -0.039 | 0 | 0.53 | 0.49 | 0.45 | 0.15 | 0.5 | 0.34 |
| 1 | 18940380 | rs4920522 | -0.573 | 0.365 | 0.895 | -0.556 | -0.029 | 0.15 | 0.15 | 0 | 0.57 | 0.8 | 0.49 | 0.2 | 0.52 | 0.42 |
| 1 | 18943584 | rs11799871 | -0.573 | 0.365 | 0.902 | -0.587 | -0.031 | 0.165 | 0.165 | 0 | 0.57 | 0.8 | 0.49 | 0.18 | 0.52 | 0.42 |
| 1 | 18945536 | rs6664982 | -1.708 | 0.446 | 1.328 | -0.636 | 0.039 | 0.025 | 0.025 | 0.198 | 0.12 | 0.43 | 0.17 | 0.14 | 0.11 | 0.16 |
| 1 | 18952112 | rs17352100 | 0.015 | 0.262 | 1.191 | -0.722 | 0.013 | 0.092 | 0.092 | 0 | 0.48 | 0.33 | 0.33 | 0.07 | 0.43 | 0.33 |
| 1 | 18956404 | rs766324 | -0.812 | 0.054 | 0.696 | -0.753 | -0.048 | 0.134 | 0.134 | 0 | 0.62 | 0.84 | 0.5 | 0.19 | 0.55 | 0.42 |
| 1 | 18972776 | rs9439713 | 0.532 | 0.193 | -0.137 | -1.056 | -0.038 | 0.117 | 0.117 | 0 | 0.37 | 0.16 | 0.28 | 0.03 | 0.4 | 0.24 |
| 1 | 18978482 | rs1339062 | -0.508 | 0.386 | -0.55 | -0.532 | 0.002 | 0.037 | 0.037 | 0 | 0.55 | 0.65 | 0.48 | 0.18 | 0.55 | 0.32 |
| 1 | 18979320 | rs6695765 | -0.622 | 0.498 | -1.257 | -0.075 | -0.012 | 0.065 | 0.065 | 0 | 0.58 | 0.83 | 0.56 | 0.33 | 0.56 | 0.41 |

| | | | | | | | | | | | | | | | | |
|---|---|---|---|---|---|---|---|---|---|---|---|---|---|---|---|---|
| 1 | 18990793 | rs4920338 | -0.118 | 0.487 | -1.189 | -0.303 | -0.005 | 0.193 | 0.193 | 0 | 0.48 | 0.78 | 0.43 | 0.2 | 0.44 | 0.33 |
| 1 | 18994150 | rs2743218 | 0.086 | 0.404 | -1.177 | -0.202 | 0 | 0.073 | 0.073 | 0 | 0.38 | 0.73 | 0.39 | 0.18 | 0.38 | 0.24 |
| 1 | 18999794 | rs16862125 | -0.716 | 0.425 | -1.527 | -0.04 | -0.017 | -0.056 | -0.056 | 0 | 0.11 | 0.05 | 0.18 | 0.3 | 0.12 | 0.19 |
| 1 | 18999831 | rs2236832 | -0.538 | 0.214 | -1.453 | 0.049 | -0.004 | 0.139 | 0.139 | 0.388 | 0.5 | 0.85 | 0.53 | 0.32 | 0.48 | 0.35 |
| 1 | 19003593 | rs11576839 | -0.604 | 0.096 | -1.046 | 0.471 | -0.007 | 0.001 | 0.001 | 0 | 0.07 | 0.02 | 0.03 | 0.1 | 0.07 | 0.1 |
| 1 | 19006152 | rs618941 | -0.048 | -0.101 | -1.086 | 1.896 | 0.001 | 0.019 | 0.019 | 0 | 0.91 | 0.9 | 0.89 | 0.68 | 0.92 | 0.85 |
| 1 | 19015628 | rs574590 | 1.273 | -0.669 | -0.653 | 1.355 | -0.01 | 0.003 | 0.003 | 0 | 0.69 | 0.61 | 0.73 | 0.58 | 0.75 | 0.63 |
| 1 | 19017855 | rs2743208 | 0.275 | -0.612 | -0.656 | 1.513 | -0.008 | 0.031 | 0.031 | 0 | 0.87 | 0.85 | 0.88 | 0.67 | 0.89 | 0.83 |
| 1 | 19018131 | rs2076021 | -0.584 | -0.606 | -0.668 | 1.508 | 0.004 | -0.039 | -0.039 | 0 | 0.92 | 0.99 | 0.91 | 0.73 | 0.93 | 0.86 |
| 1 | 19018432 | rs2076019 | 0.187 | -0.632 | -0.629 | 1.528 | 0.007 | -0.009 | -0.009 | 0 | 0.09 | 0.05 | 0.09 | 0.26 | 0.06 | 0.19 |
| 1 | 19019351 | rs12741080 | -1.572 | -0.75 | -0.564 | 1.41 | 0.009 | -0.004 | -0.004 | 0 | 0.25 | 0.19 | 0.22 | 0.36 | 0.18 | 0.36 |
| 1 | 19021373 | rs585075 | 0.836 | -0.973 | -0.773 | 1.134 | 0.006 | -0.031 | -0.031 | 2.326 | 0.32 | 0.21 | 0.24 | 0.37 | 0.3 | 0.28 |
| 1 | 19021544 | rs12760805 | -0.789 | -1.476 | -0.453 | 1.218 | -0.005 | 0.033 | 0.033 | 0 | 0.15 | 0.05 | 0.12 | 0.09 | 0.13 | 0.17 |
| 1 | 19022911 | rs12563055 | 0.336 | -1.321 | -0.379 | 1.324 | 0.013 | -0.03 | -0.03 | 0 | 0.09 | 0.01 | 0.08 | 0.26 | 0.06 | 0.2 |
| 1 | 19025822 | rs2282701 | -0.363 | -1.45 | -0.222 | 1.333 | -0.017 | 0.178 | 0.178 | 0.596 | 0.11 | 0.35 | 0.11 | 0.25 | 0.06 | 0.1 |
| 1 | 19027239 | rs2743201 | -1.731 | -1.517 | -0.386 | 1.11 | -0.018 | 0.008 | 0.008 | 0 | 0.37 | 0.64 | 0.39 | 0.6 | 0.26 | 0.47 |
| 1 | 19030396 | rs764289 | -0.703 | -1.204 | -1.065 | 0.799 | 0.024 | -0.036 | -0.036 | 0.01 | 0.13 | 0.07 | 0.12 | 0.34 | 0.08 | 0.24 |
| 1 | 19033575 | rs589463 | 0.253 | -1.363 | -0.882 | 0.065 | -0.031 | -0.052 | -0.052 | 0 | 0.63 | 0.47 | 0.77 | 0.81 | 0.74 | 0.65 |
| 1 | 19035393 | rs501080 | 0.063 | -1.259 | -1.253 | 0.193 | -0.001 | 0 | 0 | 0 | 0.76 | 0.72 | 0.73 | 0.75 | 0.78 | 0.7 |
| 1 | 19036394 | rs17352309 | 0.36 | -1.195 | -1.606 | -0.254 | 0.007 | 0.007 | 0.007 | 0.017 | 0.07 | 0.03 | 0.04 | 0.03 | 0.03 | 0.07 |
| 1 | 19041598 | rs6670751 | 0.447 | -0.893 | -1.723 | -0.009 | -0.017 | 0.066 | 0.066 | 0.254 | 0.38 | 0.15 | 0.38 | 0.22 | 0.55 | 0.4 |
| 1 | 19044877 | rs624761 | 0.248 | -0.119 | -1.262 | 0.172 | 0 | 0.007 | 0.007 | 0 | 0.8 | 0.79 | 0.78 | 0.81 | 0.8 | 0.81 |
| 1 | 19049615 | rs2236817 | -0.471 | -0.25 | -1.573 | 0.389 | 0.017 | -0.026 | -0.026 | 2.989 | 0.16 | 0.07 | 0.06 | 0.3 | 0.11 | 0.26 |
| 1 | 19049946 | rs2236815 | 0.316 | 0.651 | -0.891 | 0.546 | 0.028 | 0.047 | 0.047 | 0 | 0.71 | 0.64 | 0.41 | 0.55 | 0.6 | 0.7 |
| 1 | 19056382 | rs2743189 | 0.006 | -0.117 | -0.65 | -0.648 | -0.025 | -0.106 | -0.106 | 0 | 0.37 | 0.41 | 0.17 | 0.08 | 0.27 | 0.3 |
| 1 | 19057480 | rs84353 | -0.491 | 0.145 | -0.847 | -0.756 | 0.031 | 0.072 | 0.072 | 0 | 0.49 | 0.61 | 0.3 | 0.13 | 0.43 | 0.35 |
| 1 | 19057553 | rs17352351 | 0.273 | -0.073 | -1.004 | -0.776 | 0.036 | 0.097 | 0.097 | 0 | 0.36 | 0.16 | 0.15 | 0.08 | 0.3 | 0.3 |
| 1 | 19057730 | rs2236805 | -0.121 | 0.125 | -1.223 | -0.527 | 0.033 | 0.15 | 0.15 | 0 | 0.46 | 0.16 | 0.27 | 0.23 | 0.41 | 0.35 |
| 1 | 19057941 | rs2236804 | -0.857 | -0.201 | -0.643 | 0.453 | -0.011 | 0.083 | 0.083 | 0.031 | 0.11 | 0.22 | 0.1 | 0.25 | 0.07 | 0.03 |
| 1 | 19058918 | rs2236801 | 0.383 | 0.074 | -1.434 | 0.141 | -0.019 | -0.015 | -0.015 | 0 | 0.06 | 0 | 0.06 | 0.08 | 0.06 | 0.04 |
| 1 | 19058929 | rs2236800 | 0.237 | -0.004 | -1.721 | 0.066 | 0.017 | 0.028 | 0.028 | 0 | 0.47 | 0.36 | 0.48 | 0.4 | 0.46 | 0.41 |
| 1 | 19059471 | rs9439731 | -0.38 | -0.293 | -1.74 | -0.18 | 0.036 | 0.075 | 0.075 | 0 | 0.43 | 0.18 | 0.48 | 0.24 | 0.35 | 0.26 |
| 1 | 19060653 | rs12144695 | -0.744 | 0.227 | -1.157 | 0.432 | 0.014 | 0.013 | 0.013 | 0 | 0.13 | 0.05 | 0.11 | 0.1 | 0.1 | 0.1 |
| 1 | 19072926 | rs485874 | 0.098 | 0.769 | -0.915 | 0.748 | -0.014 | -0.014 | -0.014 | 0 | 0.65 | 0.72 | 0.52 | 0.45 | 0.53 | 0.64 |
| 1 | 19074980 | rs12121503 | 0.4 | 0.678 | -1.089 | 1.006 | -0.012 | -0.028 | -0.028 | 0 | 0.05 | 0 | 0.09 | 0.1 | 0.06 | 0.04 |
| 1 | 19077825 | rs626600 | 1.202 | 0.163 | -1.566 | 0.238 | 0.005 | 0.131 | 0.131 | 0.009 | 0.3 | 0.06 | 0.12 | 0 | 0.24 | 0.14 |
| 1 | 19080055 | rs479064 | 0.018 | -0.053 | -0.367 | 0.4 | -0.011 | 0.003 | 0.003 | 1.014 | 0.57 | 0.36 | 0.39 | 0.33 | 0.47 | 0.59 |
| 1 | 19080506 | rs1160680 | 0.539 | -0.163 | -0.492 | 0.033 | 0.005 | 0.005 | 0.005 | 0.372 | 0.5 | 0.41 | 0.4 | 0.42 | 0.47 | 0.52 |
| 1 | 19087996 | rs17354622 | -0.067 | -0.146 | -0.383 | 0.031 | 0.001 | 0.086 | 0.086 | 0.013 | 0.19 | 0.05 | 0.1 | 0.03 | 0.15 | 0.17 |
| 1 | 19088899 | rs669138 | -1.056 | -0.191 | 0.066 | 0.032 | -0.02 | -0.017 | -0.017 | 0 | 0.71 | 0.81 | 0.58 | 0.52 | 0.61 | 0.68 |
| 1 | 19094738 | rs7414734 | 1.237 | -0.224 | 0.266 | 0.008 | 0.005 | -0.003 | -0.003 | 0 | 0.38 | 0.33 | 0.28 | 0.3 | 0.31 | 0.41 |
| 1 | 19096297 | rs16862248 | 1.219 | 0.504 | 0.013 | 0.239 | -0.001 | 0.002 | 0.002 | 0 | 0.18 | 0.17 | 0.4 | 0.42 | 0.25 | 0.18 |
| 1 | 19100057 | rs6603908 | -1.006 | 0.498 | 0.008 | 0.21 | -0.002 | -0.003 | -0.003 | 0 | 0.95 | 0.96 | 0.94 | 0.92 | 0.92 | 0.96 |
| 1 | 19100178 | rs2789337 | -0.025 | 0.61 | 0.016 | 0.126 | 0.006 | -0.053 | -0.053 | 0 | 0.11 | 0.35 | 0.13 | 0.02 | 0.1 | 0.09 |
| 1 | 19100436 | rs2789334 | -0.658 | 0.626 | -0.002 | 0.184 | 0.024 | -0.02 | -0.02 | 0 | 0.21 | 0.38 | 0.16 | 0.19 | 0.22 | 0.26 |
| 1 | 19102509 | rs2841087 | 0.975 | 0.198 | 0.023 | 0.123 | 0.011 | 0.018 | 0.018 | 0 | 0.59 | 0.34 | 0.67 | 0.74 | 0.68 | 0.63 |
| 1 | 19104381 | rs4920344 | 0.251 | 0.424 | -0.107 | 0.628 | 0.004 | -0.005 | -0.005 | 0 | 0.89 | 0.95 | 0.94 | 0.89 | 0.91 | 0.85 |
| 1 | 19105580 | rs6697715 | 0.869 | 0.507 | 0.112 | 1.084 | 0.002 | 0.003 | 0.003 | 0 | 0.69 | 0.66 | 0.73 | 0.61 | 0.7 | 0.64 |
| 1 | 19106254 | rs11261065 | 0.337 | 0.756 | -0.221 | 1.703 | 0.004 | -0.019 | -0.019 | 0.21 | 0.18 | 0.12 | 0.38 | 0.34 | 0.21 | 0.11 |
| 1 | 19107476 | rs6667983 | -0.138 | 0.823 | -0.282 | 1.728 | 0.001 | 0.008 | 0.008 | 0 | 0.94 | 0.94 | 0.94 | 0.91 | 0.92 | 0.95 |
| 1 | 19108355 | rs12565650 | 0.459 | 0.858 | -0.305 | 1.66 | 0.001 | -0.029 | -0.029 | 0 | 0.17 | 0.11 | 0.37 | 0.34 | 0.2 | 0.11 |
| 1 | 19119606 | rs7549129 | 0.195 | 0.955 | -0.092 | 1.364 | 0.056 | 0.131 | 0.131 | 0 | 0.88 | 0.72 | 0.82 | 0.72 | 0.84 | 0.91 |
| 1 | 19119851 | rs7533384 | -0.597 | 0.826 | 0.035 | 1.108 | 0 | 0.118 | 0.118 | 0.07 | 0.2 | 0.42 | 0.15 | 0.4 | 0.17 | 0.35 |
| 1 | 19119910 | rs11261070 | 1.145 | 0.192 | -1.545 | 0.588 | -0.024 | 0.152 | 0.152 | 0 | 0.38 | 0.14 | 0.27 | 0.11 | 0.39 | 0.42 |
| 1 | 19122339 | rs11261071 | -0.026 | -0.26 | -1.345 | 1.233 | 0.027 | 0.048 | 0.048 | 0.018 | 0.3 | 0.17 | 0.39 | 0.14 | 0.28 | 0.13 |
| 1 | 19123033 | rs12566799 | -0.05 | -0.213 | -1.336 | 1.322 | 0.001 | 0.039 | 0.039 | 0.073 | 0.08 | 0.09 | 0.09 | 0.4 | 0.09 | 0.3 |
| 1 | 19124368 | rs11261073 | -0.301 | -0.09 | -1.326 | 0.492 | 0.026 | 0.049 | 0.049 | 0.052 | 0.3 | 0.16 | 0.38 | 0.14 | 0.28 | 0.12 |
| 1 | 19130827 | rs4074733 | -0.318 | -0.235 | -1.195 | 0.863 | 0.02 | -0.006 | -0.006 | 0 | 0.28 | 0.2 | 0.24 | 0.24 | 0.16 | 0.1 |
| 1 | 19131620 | rs4075768 | 0.35 | -0.156 | -0.94 | 0.227 | -0.004 | -0.009 | -0.009 | 0 | 0.64 | 0.6 | 0.62 | 0.72 | 0.7 | 0.79 |
| 1 | 19133990 | rs6603912 | 0.982 | 0.282 | -0.66 | -0.56 | 0.006 | -0.05 | -0.05 | 0 | 0.41 | 0.31 | 0.21 | 0.01 | 0.31 | 0.42 |
| 1 | 19134260 | rs7411058 | 0.257 | 0.96 | -0.724 | 0.466 | -0.014 | -0.022 | -0.022 | 0 | 0.11 | 0.08 | 0.24 | 0.37 | 0.21 | 0.29 |
| 1 | 19136610 | rs11261075 | -0.21 | 0.587 | -0.341 | 0.649 | 0.003 | -0.011 | -0.011 | 0 | 0.29 | 0.23 | 0.26 | 0.4 | 0.21 | 0.15 |
| 1 | 19138295 | rs7521204 | 0.319 | 0.691 | -0.038 | 0.581 | -0.002 | -0.008 | -0.008 | 0 | 0.64 | 0.59 | 0.58 | 0.28 | 0.67 | 0.71 |
| 1 | 19145099 | rs12026106 | 0.451 | 0.558 | -0.582 | 0.264 | 0.007 | -0.01 | -0.01 | 0 | 0.48 | 0.25 | 0.49 | 0.51 | 0.57 | 0.69 |
| 1 | 19145241 | rs12748059 | 1.119 | 0.549 | -0.728 | 0.113 | 0.038 | -0.015 | -0.015 | 0 | 0.22 | 0.03 | 0.21 | 0.23 | 0.14 | 0.11 |
| 1 | 19147761 | rs12125880 | -0.514 | 0.425 | -0.888 | -0.394 | -0.008 | -0.01 | -0.01 | 0 | 0.39 | 0.28 | 0.5 | 0.55 | 0.47 | 0.53 |
| 1 | 19147846 | rs6686865 | -0.49 | 0.426 | -0.884 | -0.389 | -0.008 | -0.013 | -0.013 | 0 | 0.39 | 0.29 | 0.5 | 0.55 | 0.47 | 0.48 |
| 1 | 19149527 | rs11261081 | -0.518 | 0.374 | -0.54 | -0.453 | -0.001 | 0.001 | 0.001 | 3.111 | 0.66 | 0.63 | 0.66 | 0.62 | 0.71 | 0.74 |
| 1 | 19153969 | rs6684577 | 0.207 | 0.414 | -0.514 | -0.506 | 0.008 | -0.005 | -0.005 | 0.402 | 0.49 | 0.35 | 0.5 | 0.51 | 0.56 | 0.62 |

| | | | | | | | | | | | | | | | | |
|---|---|---|---|---|---|---|---|---|---|---|---|---|---|---|---|---|
| 1 | 19154065 | rs6674964 | -0.227 | 0.49 | -0.653 | -0.83 | -0.026 | -0.062 | -0.062 | 0 | 0.23 | 0.15 | 0.35 | 0.5 | 0.31 | 0.41 |
| 1 | 19155969 | rs6691420 | -0.165 | 0.695 | -0.491 | -0.697 | 0.02 | 0.044 | 0.044 | 0 | 0.2 | 0.24 | 0.34 | 0.44 | 0.33 | 0.4 |
| 1 | 19157933 | rs6603918 | -1.308 | 0.869 | -0.303 | -0.709 | -0.007 | -0.014 | -0.014 | 0.171 | 0.85 | 0.82 | 0.89 | 0.91 | 0.88 | 0.82 |
| 1 | 19159657 | rs11261084 | -0.9 | 1.498 | -0.527 | -0.476 | 0.001 | -0.002 | -0.002 | 0 | 0.59 | 0.58 | 0.64 | 0.54 | 0.69 | 0.64 |
| 1 | 19160236 | rs4920550 | 1.402 | 1.577 | -0.537 | -0.332 | 0.007 | 0.031 | 0.031 | 0 | 0.16 | 0.04 | 0.14 | 0.1 | 0.15 | 0.11 |
| 1 | 19160523 | rs12038239 | -0.652 | 1.629 | -0.556 | -0.281 | 0.028 | 0.059 | 0.059 | 0 | 0.2 | 0.26 | 0.35 | 0.44 | 0.33 | 0.4 |
| 1 | 19163075 | rs12035855 | 0.156 | 1.381 | -0.579 | -0.95 | 0.018 | 0.05 | 0.05 | 0 | 0.22 | 0.36 | 0.28 | 0.57 | 0.27 | 0.31 |
| 1 | 19164527 | rs12034674 | 0.156 | 1.381 | -0.584 | -0.95 | 0.02 | 0.054 | 0.054 | 0 | 0.22 | 0.36 | 0.28 | 0.57 | 0.27 | 0.31 |
| 1 | 19166893 | rs6662276 | 0.894 | 0.962 | -0.378 | -0.8 | 0.001 | -0.002 | -0.002 | 0 | 0.07 | 0.06 | 0.03 | 0.11 | 0.06 | 0.19 |
| 1 | 19167371 | rs3935570 | 0.581 | 0.988 | -0.457 | -0.957 | -0.001 | -0.014 | -0.014 | 0 | 0.3 | 0.34 | 0.17 | 0.12 | 0.26 | 0.31 |
| 1 | 19170078 | rs7534618 | 0.905 | 0.933 | -0.796 | -0.926 | -0.014 | -0.029 | -0.029 | 0 | 0.3 | 0.23 | 0.25 | 0.46 | 0.34 | 0.28 |
| 1 | 19170946 | rs12408808 | -0.372 | 0.888 | -0.738 | -0.944 | -0.008 | -0.002 | -0.002 | 0 | 0.58 | 0.56 | 0.65 | 0.57 | 0.7 | 0.51 |
| 1 | 19175593 | rs4920561 | 0.273 | 1.206 | -1.409 | -0.694 | -0.012 | -0.003 | -0.003 | 0.003 | 0.38 | 0.28 | 0.27 | 0.57 | 0.4 | 0.48 |
| 1 | 19176908 | rs4076838 | 0.062 | 0.985 | -1.435 | -1.248 | 0.021 | 0.098 | 0.098 | 1.418 | 0.52 | 0.27 | 0.52 | 0.2 | 0.58 | 0.46 |
| 1 | 19179104 | rs4920564 | -0.025 | 1.186 | -1.506 | -1.114 | -0.034 | 0.119 | 0.119 | 0 | 0.69 | 0.86 | 0.75 | 0.96 | 0.64 | 0.83 |
| 1 | 19188233 | rs35217608 | -0.081 | 1.092 | -1.095 | -0.884 | -0.011 | -0.003 | -0.003 | 0.065 | 0.19 | 0.12 | 0.16 | 0.01 | 0.25 | 0.07 |
| 1 | 19188456 | rs28410948 | -0.659 | 0.722 | -0.924 | -0.544 | -0.015 | 0.012 | 0.012 | 0.348 | 0.64 | 0.45 | 0.74 | 0.48 | 0.71 | 0.72 |
| 1 | 19191326 | rs9662598 | -0.592 | 0.392 | -0.716 | -1.023 | 0 | -0.011 | -0.011 | 0 | 0.31 | 0.28 | 0.28 | 0.04 | 0.33 | 0.21 |
| 1 | 19194995 | rs28652778 | 0.082 | 0.697 | -0.709 | -0.899 | -0.028 | 0.052 | 0.052 | 0 | 0.19 | 0.09 | 0.15 | 0.01 | 0.24 | 0.07 |
| 1 | 19197849 | rs3935824 | 0.726 | 0.091 | -0.381 | -0.642 | -0.004 | 0 | 0 | 0 | 0.26 | 0.28 | 0.12 | 0.25 | 0.13 | 0.19 |
| 1 | 19198064 | rs1138333 | -0.253 | 0.141 | -0.143 | -0.908 | -0.001 | -0.036 | -0.036 | 1.022 | 0.26 | 0.22 | 0.22 | 0.04 | 0.28 | 0.14 |
| 1 | 19198895 | rs9117 | 0.483 | 0.158 | -0.237 | -0.907 | -0.032 | 0.057 | 0.057 | 0 | 0.19 | 0.06 | 0.18 | 0 | 0.25 | 0.09 |
| 1 | 19199221 | rs11740 | 0.573 | 0.504 | -0.066 | -0.636 | -0.013 | 0.045 | 0.045 | 0.989 | 0.43 | 0.2 | 0.55 | 0.66 | 0.45 | 0.61 |
| 1 | 19200180 | rs7555171 | -0.411 | 0.274 | 0.369 | -0.087 | -0.003 | -0.001 | -0.001 | 0 | 0.69 | 0.61 | 0.79 | 0.69 | 0.72 | 0.75 |
| 1 | 19201919 | rs6695033 | -0.131 | 0.212 | 0.472 | -0.06 | 0.008 | -0.02 | -0.02 | 0 | 0.06 | 0.2 | 0.04 | 0 | 0.03 | 0.02 |
| 1 | 19201956 | rs2230708 | -0.536 | 0.102 | 0.485 | -0.119 | 0 | -0.001 | -0.001 | 0.04 | 0.73 | 0.71 | 0.81 | 0.71 | 0.74 | 0.76 |
| 1 | 19202507 | rs6426802 | 0.05 | 0.187 | 0.149 | -0.062 | -0.015 | -0.011 | -0.011 | 0 | 0.65 | 0.46 | 0.77 | 0.71 | 0.71 | 0.74 |
| 1 | 19205672 | rs28497538 | -0.385 | 0.686 | 0.066 | 0.717 | 0.028 | 0.046 | 0.046 | 0.055 | 0.31 | 0.5 | 0.47 | 0.36 | 0.39 | 0.45 |
| 1 | 19210816 | rs7365362 | 0.629 | 0.427 | -0.086 | 0.805 | -0.001 | 0.045 | 0.045 | 0.054 | 0.78 | 0.33 | 0.61 | 0.72 | 0.78 | 0.73 |
| 1 | 19215418 | rs4912075 | -1.179 | 0.52 | -1.169 | 0.557 | -0.024 | 0.044 | 0.044 | 0 | 0.45 | 0.91 | 0.47 | 0.56 | 0.42 | 0.5 |
| 1 | 19215440 | rs4911985 | 0.158 | 0.403 | -0.851 | 0.239 | 0 | -0.013 | -0.013 | 0 | 0.22 | 0.25 | 0.25 | 0.08 | 0.22 | 0.16 |
| 1 | 19220748 | rs16862341 | 0.116 | 0.315 | -0.933 | 0.298 | -0.001 | 0.01 | 0.01 | 0 | 0.63 | 0.51 | 0.68 | 0.6 | 0.61 | 0.52 |
| 1 | 19221842 | rs16862344 | -0.59 | 0.357 | -0.925 | -0.044 | -0.006 | 0.02 | 0.02 | 0 | 0.07 | 0.19 | 0.07 | 0.05 | 0.04 | 0.07 |
| 1 | 19222020 | rs1009971 | -0.59 | 0.357 | -0.925 | -0.044 | -0.006 | 0.02 | 0.02 | 0 | 0.07 | 0.19 | 0.07 | 0.05 | 0.05 | 0.07 |
| 1 | 19224486 | rs7554974 | -1.077 | 0.353 | -0.908 | -0.06 | 0 | 0.058 | 0.058 | 0 | 0.08 | 0.31 | 0.08 | 0.11 | 0.04 | 0.08 |
| 1 | 19230263 | rs4912044 | -0.81 | 0.388 | -1.08 | -0.082 | -0.057 | 0.065 | 0.065 | 0 | 0.81 | 0.99 | 0.85 | 0.9 | 0.76 | 0.81 |
| 1 | 19231017 | rs2295227 | -0.374 | 0.309 | -0.203 | 0.59 | -0.005 | -0.01 | -0.01 | 0 | 0.05 | 0.09 | 0.05 | 0.02 | 0.04 | 0.01 |
| 1 | 19235517 | rs6673711 | -1.19 | 0.343 | -0.378 | 0.513 | -0.005 | -0.031 | -0.031 | 1.227 | 0.25 | 0.3 | 0.26 | 0.11 | 0.25 | 0.17 |
| 1 | 19237107 | rs2024724 | -1.162 | 0.584 | -0.578 | 0.881 | 0.001 | -0.014 | -0.014 | 2.86 | 0.07 | 0.19 | 0.15 | 0.05 | 0.07 | 0.04 |
| 1 | 19238942 | rs16862363 | -0.499 | 0.522 | -1.342 | 1.062 | 0.009 | -0.033 | -0.033 | 0.236 | 0.36 | 0.16 | 0.33 | 0.43 | 0.33 | 0.2 |
| 1 | 19241904 | rs16862370 | -1.036 | 0.898 | -0.614 | 1.386 | 0.007 | 0.032 | 0.032 | 0.284 | 0.13 | 0.22 | 0.35 | 0.23 | 0.12 | 0.08 |
| 1 | 19243589 | rs3748586 | 0.893 | 1.101 | -0.493 | 1.251 | -0.004 | 0.017 | 0.017 | 0.003 | 0.59 | 0.38 | 0.37 | 0.27 | 0.6 | 0.56 |
| 1 | 19244678 | rs4912056 | -0.758 | 1.191 | -0.501 | 1.197 | -0.012 | 0.004 | 0.004 | 0.054 | 0.19 | 0.1 | 0.44 | 0.19 | 0.17 | 0.15 |
| 1 | 19255671 | rs6698642 | -1.028 | 1.537 | -0.001 | 1.693 | 0.077 | 0.106 | 0.106 | 2.255 | 0.13 | 0.59 | 0.14 | 0.21 | 0.15 | 0.13 |
| 1 | 19255754 | rs6674337 | -1.59 | 1.607 | -0.075 | 1.827 | 0.035 | 0.07 | 0.07 | 1.181 | 0.18 | 0.67 | 0.19 | 0.34 | 0.2 | 0.19 |
| 1 | 19258375 | rs17387963 | 2.053 | 1.413 | -0.778 | 1.483 | -0.021 | 0.083 | 0.083 | 0 | 0.43 | 0.04 | 0.26 | 0.22 | 0.44 | 0.53 |
| 1 | 19258723 | rs16862387 | -1.746 | 1.447 | -0.708 | 1.525 | -0.068 | -0.035 | -0.035 | 0 | 0.38 | 0.87 | 0.62 | 0.47 | 0.38 | 0.31 |
| 1 | 19261241 | rs992660 | -1.294 | 1.471 | -0.651 | 1.492 | -0.028 | -0.001 | -0.001 | 1.171 | 0.33 | 0.76 | 0.57 | 0.34 | 0.33 | 0.27 |
| 1 | 19263783 | rs4912087 | -0.239 | 1.56 | -0.621 | 1.438 | -0.002 | -0.035 | -0.035 | 0.872 | 0.83 | 0.92 | 0.88 | 0.71 | 0.82 | 0.84 |
| 1 | 19274525 | rs2024727 | 0.593 | 1.451 | 0.372 | 1.043 | 0.029 | -0.062 | -0.062 | 0 | 0.82 | 0.6 | 0.9 | 0.94 | 0.83 | 0.95 |
| 1 | 19284564 | rs4912013 | -1.299 | 1.353 | 0.385 | 1.124 | 0.016 | -0.015 | -0.015 | 0 | 0.37 | 0.59 | 0.64 | 0.74 | 0.4 | 0.43 |
| 1 | 19291757 | rs17388255 | -0.823 | 1.375 | 0.423 | 0.825 | 0.005 | -0.059 | -0.059 | 0.019 | 0.19 | 0.29 | 0.09 | 0.04 | 0.18 | 0.04 |
| 1 | 19293249 | rs6680748 | -0.911 | 1.397 | 0.509 | 0.827 | 0.006 | -0.073 | -0.073 | 0 | 0.2 | 0.36 | 0.1 | 0.04 | 0.18 | 0.04 |
| 1 | 19293290 | rs16862410 | -0.323 | 1.378 | 0.107 | 0.99 | -0.07 | -0.101 | -0.101 | 0 | 0.26 | 0.08 | 0.53 | 0.66 | 0.3 | 0.29 |
| 1 | 19297205 | rs17384241 | -1.977 | 1.123 | 0.454 | 1.016 | 0.044 | 0.112 | 0.112 | 0 | 0.56 | 0.81 | 0.75 | 0.8 | 0.59 | 0.49 |
| 1 | 19319245 | rs6426610 | -2.02 | 1.047 | 0.423 | 1.017 | 0.03 | 0.1 | 0.1 | 0 | 0.54 | 0.75 | 0.74 | 0.8 | 0.57 | 0.48 |
| 1 | 19320209 | rs6697292 | -0.682 | 1.041 | 0.787 | 1.255 | 0.006 | 0.067 | 0.067 | 0 | 0.12 | 0.29 | 0.16 | 0.27 | 0.11 | 0.16 |
| 1 | 19321462 | rs17384449 | -1.152 | 1.048 | 0.882 | 1.254 | 0.006 | 0.006 | 0.006 | 0 | 0.42 | 0.41 | 0.58 | 0.52 | 0.46 | 0.32 |
| 1 | 19326813 | rs6699706 | -0.624 | 1.06 | 0.882 | 1.238 | 0.007 | 0.071 | 0.071 | 0 | 0.12 | 0.29 | 0.16 | 0.27 | 0.11 | 0.16 |
| 1 | 19327310 | rs2147949 | -2.124 | 1.011 | 0.789 | 1.188 | 0.023 | 0.07 | 0.07 | 0 | 0.54 | 0.71 | 0.74 | 0.8 | 0.57 | 0.48 |
| 1 | 19335792 | rs7531434 | -1.248 | 1.004 | 0.439 | 0.947 | -0.003 | -0.001 | -0.001 | 0.431 | 0.42 | 0.53 | 0.57 | 0.56 | 0.45 | 0.32 |
| 1 | 19339070 | rs6426630 | -0.242 | 0.966 | 0.43 | 1.042 | -0.006 | 0.006 | 0.006 | 0 | 0.09 | 0.16 | 0.11 | 0.13 | 0.05 | 0.07 |
| 1 | 19340317 | rs16862453 | -0.394 | 1.098 | 0.163 | 0.942 | -0.044 | -0.081 | -0.081 | 0 | 0.24 | 0.09 | 0.49 | 0.56 | 0.3 | 0.26 |
| 1 | 19341553 | rs6681972 | -1.216 | 0.969 | 0.517 | 0.72 | -0.017 | -0.13 | -0.13 | 0 | 0.23 | 0.54 | 0.14 | 0.01 | 0.16 | 0.06 |
| 1 | 19344501 | rs7531834 | -1.306 | 1.147 | 0.511 | 0.739 | -0.001 | -0.103 | -0.103 | 0 | 0.24 | 0.59 | 0.22 | 0.01 | 0.19 | 0.07 |
| 1 | 19344586 | rs7545211 | -2.061 | 0.821 | -0.898 | -0.228 | -0.001 | 0.019 | 0.019 | 0 | 0.75 | 0.99 | 0.86 | 0.84 | 0.78 | 0.69 |
| 1 | 19348605 | rs6687825 | -0.484 | 1.401 | 0.23 | 0.198 | 0 | -0.097 | -0.097 | 0 | 0.17 | 0.27 | 0.15 | 0 | 0.14 | 0.02 |
| 1 | 19358015 | rs912845 | -0.552 | 1.405 | 0.256 | 0.2 | 0.001 | -0.11 | -0.11 | 0.471 | 0.18 | 0.28 | 0.15 | 0 | 0.14 | 0.02 |
| 1 | 19365552 | rs10492996 | -0.229 | 1.572 | -0.275 | 0.371 | -0.082 | -0.101 | -0.101 | 0 | 0.23 | 0.08 | 0.41 | 0.56 | 0.33 | 0.27 |

| | | | | | | | | | | | | | | | | |
|---|---|---|---|---|---|---|---|---|---|---|---|---|---|---|---|---|
| 1 | 19375841 | rs6683868 | -0.021 | 1.383 | 0.063 | 0.122 | 0.032 | -0.045 | -0.045 | 0.496 | 0.08 | 0.17 | 0.05 | 0 | 0.08 | 0.01 |
| 1 | 19376017 | rs4912036 | -1.72 | 1.283 | -0.494 | -0.012 | 0.062 | 0.173 | 0.173 | 0 | 0.51 | 0.83 | 0.72 | 0.78 | 0.57 | 0.45 |
| 1 | 19380992 | rs7549904 | -1.378 | 1.263 | -0.242 | 0.582 | 0.016 | -0.022 | -0.022 | 0.079 | 0.36 | 0.71 | 0.49 | 0.47 | 0.26 | 0.18 |
| 1 | 19382691 | rs10799662 | -1.72 | 1.537 | -0.307 | 0.172 | 0.099 | 0.122 | 0.122 | 0 | 0.49 | 0.76 | 0.71 | 0.77 | 0.62 | 0.55 |
| 1 | 19386358 | rs942186 | -0.266 | 1.863 | 0.085 | 0.278 | 0.014 | -0.023 | -0.023 | 0 | 0.05 | 0.12 | 0.04 | 0 | 0.09 | 0.04 |
| 1 | 19391177 | rs10916867 | -0.101 | 2.384 | 0.525 | -0.004 | 0.009 | -0.067 | -0.067 | 0 | 0.16 | 0.21 | 0.16 | 0 | 0.17 | 0.03 |
| 1 | 19393033 | rs10753508 | -1.055 | 2.526 | 1.054 | -0.067 | 0.07 | -0.153 | -0.153 | 0 | 0.22 | 0.49 | 0.22 | 0 | 0.27 | 0.05 |
| 1 | 19401405 | rs2281748 | -1.235 | 2.618 | 1.302 | 0.299 | -0.037 | -0.033 | -0.033 | 0 | 0.24 | 0.18 | 0.46 | 0.61 | 0.41 | 0.36 |
| 1 | 19405237 | rs2146866 | -2.305 | 2.544 | 1.533 | 0.386 | -0.002 | -0.004 | -0.004 | 0 | 0.43 | 0.6 | 0.7 | 0.75 | 0.6 | 0.45 |
| 1 | 19408380 | rs12164540 | -1.362 | 2.856 | 1.593 | 0.846 | -0.006 | -0.004 | -0.004 | 6.815 | 0.3 | 0.33 | 0.43 | 0.36 | 0.41 | 0.37 |
| 1 | 19415304 | rs1043899 | -2.169 | 2.707 | 1.66 | 0.366 | -0.01 | -0.026 | -0.026 | 0.21 | 0.42 | 0.51 | 0.68 | 0.76 | 0.58 | 0.48 |
| 1 | 19437860 | rs2313509 | -0.52 | 2.737 | 1.71 | 0.396 | -0.041 | -0.05 | -0.05 | 0.056 | 0.19 | 0.12 | 0.41 | 0.63 | 0.3 | 0.33 |
| 1 | 19437923 | rs12740975 | -0.503 | 2.738 | 1.701 | 0.396 | -0.051 | -0.062 | -0.062 | 0 | 0.19 | 0.1 | 0.41 | 0.63 | 0.3 | 0.33 |
| 1 | 19446587 | rs6693149 | 0 | 2.64 | 1.667 | 0.358 | 0.018 | -0.047 | -0.047 | 0.027 | 0.17 | 0.29 | 0.2 | 0.07 | 0.17 | 0.09 |
| 1 | 19457257 | rs7547104 | -2.425 | 2.523 | 1.742 | 0.352 | 0.002 | -0.003 | -0.003 | 0.264 | 0.44 | 0.58 | 0.7 | 0.8 | 0.6 | 0.54 |
| 1 | 19476581 | rs942182 | 0.362 | 2.702 | 1.471 | 0.394 | -0.012 | 0.006 | 0.006 | 0 | 0.15 | 0.1 | 0.15 | 0 | 0.17 | 0.02 |
| 1 | 19500551 | rs1009806 | -0.506 | 2.826 | 1.473 | 0.433 | -0.041 | -0.05 | -0.05 | 0 | 0.19 | 0.12 | 0.41 | 0.63 | 0.3 | 0.33 |
| 1 | 19528847 | rs12121004 | -0.525 | 2.784 | 1.571 | 0.378 | 0.022 | -0.019 | -0.019 | 6.749 | 0.06 | 0.09 | 0.06 | 0 | 0.12 | 0.03 |
| 1 | 19531534 | rs12758524 | 0.3 | 2.751 | 1.552 | 0.355 | -0.013 | 0.007 | 0.007 | 0 | 0.15 | 0.11 | 0.15 | 0 | 0.17 | 0.02 |
| 1 | 19537447 | rs10158374 | -1.618 | 2.737 | 1.772 | 0.656 | 0.094 | -0.022 | -0.022 | 0.012 | 0.08 | 0.34 | 0.07 | 0 | 0.11 | 0.05 |
| 1 | 19537755 | rs10158092 | -2.224 | 2.741 | 1.771 | 0.698 | 0.08 | 0.07 | 0.07 | 0.874 | 0.09 | 0.36 | 0.1 | 0.1 | 0.12 | 0.14 |
| 1 | 19545053 | rs12130051 | -0.961 | 2.979 | 1.433 | 0.645 | 0.053 | -0.032 | -0.032 | 0 | 0.06 | 0.2 | 0.06 | 0 | 0.11 | 0.05 |
| 1 | 19549864 | rs2073105 | -0.541 | 2.936 | 1.339 | 0.764 | -0.027 | -0.028 | -0.028 | 0 | 0.18 | 0.13 | 0.29 | 0.68 | 0.26 | 0.34 |
| 1 | 19555060 | rs12565770 | -0.141 | 2.615 | 1.158 | 0.631 | 0.001 | 0.002 | 0.002 | 0 | 0.06 | 0.1 | 0.08 | 0.09 | 0.1 | 0.1 |
| 1 | 19556408 | rs3762395 | -2.287 | 2.287 | 1.591 | 0.518 | 0.065 | 0.069 | 0.069 | 1.099 | 0.1 | 0.38 | 0.1 | 0.13 | 0.12 | 0.16 |
| 1 | 19563900 | rs16862632 | -1.269 | 2.338 | 1.436 | 0.535 | 0.02 | 0.013 | 0.013 | 0.172 | 0.07 | 0.16 | 0.08 | 0.08 | 0.1 | 0.07 |
| 1 | 19565338 | rs709682 | -1.89 | 2.316 | 1.406 | 0.538 | 0.063 | 0.066 | 0.066 | 0.404 | 0.1 | 0.32 | 0.1 | 0.12 | 0.12 | 0.13 |
| 1 | 19578192 | rs2294942 | 0.409 | 1.584 | 0.437 | -0.326 | -0.001 | 0.002 | 0.002 | 0 | 0.57 | 0.42 | 0.7 | 0.83 | 0.63 | 0.74 |
| 1 | 19584004 | rs1042380 | -0.812 | 1.624 | 0.243 | -0.505 | 0.086 | -0.022 | -0.022 | 0 | 0.18 | 0.49 | 0.18 | 0.1 | 0.23 | 0.15 |
| 1 | 19585207 | rs2294939 | -0.817 | 1.72 | 0.35 | -0.526 | 0.008 | 0.004 | 0.004 | 0.224 | 0.07 | 0.09 | 0.09 | 0.1 | 0.13 | 0.14 |
| 1 | 19586294 | rs1063984 | -0.978 | 1.72 | 0.357 | -0.51 | 0.086 | -0.018 | -0.018 | 0 | 0.18 | 0.49 | 0.19 | 0.11 | 0.23 | 0.15 |
| 1 | 19586360 | rs1063985 | 0.681 | 1.182 | 0.42 | -0.519 | 0.051 | -0.075 | -0.075 | 0 | 0.12 | 0.41 | 0.09 | 0 | 0.1 | 0.01 |
| 1 | 19586571 | rs1063986 | 0.898 | 1.181 | 0.518 | -0.519 | 0.032 | -0.07 | -0.07 | 0 | 0.11 | 0.31 | 0.09 | 0 | 0.1 | 0.01 |
| 1 | 19586684 | rs7551095 | 0.647 | 1.175 | 0.356 | -0.522 | 0.05 | -0.076 | -0.076 | 0.018 | 0.12 | 0.41 | 0.09 | 0 | 0.1 | 0.01 |
| 1 | 19590043 | rs2179660 | -1.064 | 0.999 | -0.35 | -0.3 | 0.039 | 0.052 | 0.052 | 0.317 | 0.75 | 0.93 | 0.85 | 0.87 | 0.85 | 0.85 |
| 1 | 19602979 | rs4912064 | -0.71 | 0.921 | -0.254 | 0.052 | 0.02 | -0.017 | -0.017 | 0.726 | 0.86 | 0.93 | 0.89 | 0.85 | 0.91 | 0.92 |
| 1 | 19608769 | rs1738026 | -1.191 | 0.903 | -0.059 | 0.026 | 0.015 | -0.012 | -0.012 | 0 | 0.87 | 0.98 | 0.92 | 0.9 | 0.92 | 0.94 |
| 1 | 19611241 | rs1738023 | 0.159 | 1.22 | 0.522 | -0.036 | -0.013 | 0.013 | 0.013 | 0 | 0.81 | 0.64 | 0.86 | 0.85 | 0.86 | 0.9 |
| 1 | 19613412 | rs12405546 | 0.773 | 2.084 | 0.656 | 0.6 | 0.049 | 0.129 | 0.129 | 0 | 0.1 | 0.16 | 0.29 | 0.38 | 0.2 | 0.11 |
| 1 | 19618601 | rs9729157 | -0.66 | 1.289 | 0.108 | -0.251 | -0.009 | -0.013 | -0.013 | 0 | 0.62 | 0.57 | 0.75 | 0.69 | 0.66 | 0.62 |
| 1 | 19627876 | rs12407355 | 0.829 | 1.381 | -0.272 | -0.525 | -0.011 | -0.011 | -0.011 | 0 | 0.21 | 0.18 | 0.51 | 0.6 | 0.38 | 0.33 |
| 1 | 19627931 | rs12132200 | 1.841 | 1.2 | -0.372 | -0.763 | 0.019 | 0.024 | 0.024 | 0 | 0.29 | 0.23 | 0.16 | 0.08 | 0.18 | 0.27 |
| 1 | 19632332 | rs3828102 | -0.441 | 1.413 | -0.293 | -0.594 | 0.001 | 0 | 0 | 0 | 0.34 | 0.35 | 0.59 | 0.62 | 0.48 | 0.35 |
| 1 | 19635011 | rs1043657 | 1.567 | 1.171 | -0.892 | -0.624 | -0.013 | 0.042 | 0.042 | 0 | 0.1 | 0.03 | 0.06 | 0 | 0.1 | 0.01 |
| 1 | 19636041 | rs11808495 | 0.838 | 0.975 | -0.723 | -0.631 | 0.019 | 0.021 | 0.021 | 0 | 0.29 | 0.24 | 0.16 | 0.08 | 0.18 | 0.28 |
| 1 | 19638309 | rs2227295 | -1.204 | 1.276 | -0.962 | -0.181 | -0.005 | -0.04 | -0.04 | 0 | 0.72 | 0.84 | 0.61 | 0.57 | 0.72 | 0.73 |
| 1 | 19638883 | rs2473808 | -0.984 | 1.309 | -0.632 | -0.037 | -0.02 | -0.027 | -0.027 | 0 | 0.66 | 0.83 | 0.56 | 0.58 | 0.64 | 0.73 |
| 1 | 19640676 | rs859218 | -0.944 | 1.353 | -0.617 | -0.016 | -0.02 | -0.027 | -0.027 | 0.108 | 0.65 | 0.84 | 0.56 | 0.58 | 0.64 | 0.73 |
| 1 | 19642326 | rs2073098 | 0.293 | 1.498 | -0.752 | -0.133 | -0.011 | 0.004 | 0.004 | 0 | 0.26 | 0.21 | 0.22 | 0.24 | 0.31 | 0.36 |
| 1 | 19645673 | rs4912069 | 0.333 | 0.747 | -0.476 | -0.198 | 0.032 | 0.033 | 0.033 | 0 | 0.34 | 0.25 | 0.29 | 0.2 | 0.29 | 0.32 |
| 1 | 19647578 | rs12410435 | 0.061 | 0.833 | -0.506 | 0.031 | -0.039 | 0.074 | 0.074 | 0 | 0.46 | 0.14 | 0.38 | 0.27 | 0.5 | 0.55 |
| 1 | 19650220 | rs3790775 | 0.891 | 0.667 | -0.554 | 0.043 | -0.007 | 0.07 | 0.07 | 0.01 | 0.29 | 0.21 | 0.19 | 0.08 | 0.29 | 0.33 |
| 1 | 19658506 | rs3748756 | 0.771 | 0.677 | -0.378 | 0.24 | -0.009 | -0.035 | -0.035 | 0 | 0.18 | 0.06 | 0.24 | 0.19 | 0.22 | 0.23 |
| 1 | 19666808 | rs2268813 | 0.208 | 0.676 | 0.062 | 0.323 | -0.002 | -0.01 | -0.01 | 0.062 | 0.22 | 0.24 | 0.28 | 0.33 | 0.26 | 0.27 |
| 1 | 19674070 | rs214327 | -0.488 | 0.633 | -0.314 | 0.24 | 0 | 0.002 | 0.002 | 0 | 0.53 | 0.65 | 0.48 | 0.41 | 0.55 | 0.59 |
| 1 | 19678473 | rs214336 | -1.282 | 0.59 | -0.588 | 0.415 | 0.012 | 0.007 | 0.007 | 0 | 0.67 | 0.84 | 0.71 | 0.65 | 0.7 | 0.73 |
| 1 | 19679556 | rs2268805 | 0.338 | 0.795 | -0.18 | 0.526 | -0.001 | -0.002 | -0.002 | 0.568 | 0.22 | 0.21 | 0.26 | 0.19 | 0.25 | 0.23 |
| 1 | 19679873 | rs214338 | 0.11 | 0.512 | -0.493 | 0.344 | -0.004 | 0.04 | 0.04 | 1.104 | 0.47 | 0.75 | 0.58 | 0.63 | 0.43 | 0.43 |
| 1 | 19680532 | rs12751502 | 0.384 | 1.158 | -0.014 | 0.758 | -0.006 | -0.006 | -0.006 | 0 | 0.19 | 0.11 | 0.23 | 0.18 | 0.23 | 0.22 |
| 1 | 19680544 | rs17477654 | 0.74 | 0.886 | -0.115 | 0.608 | -0.048 | 0.077 | 0.077 | 0 | 0.23 | 0.06 | 0.16 | 0.09 | 0.26 | 0.33 |
| 1 | 19681647 | rs214344 | -1.087 | 0.886 | 0.547 | 0.981 | 0.037 | 0.111 | 0.111 | 0.3 | 0.21 | 0.45 | 0.32 | 0.38 | 0.23 | 0.24 |
| 1 | 19682429 | rs214346 | -1.05 | 0.439 | 0.588 | 0.857 | 0.003 | 0.003 | 0.003 | 0 | 0.39 | 0.51 | 0.53 | 0.56 | 0.41 | 0.4 |
| 1 | 19683301 | rs169957 | -1.048 | 0.473 | 0.603 | 0.874 | 0.003 | 0.004 | 0.004 | 0 | 0.39 | 0.51 | 0.53 | 0.56 | 0.41 | 0.4 |
| 1 | 19684540 | rs214348 | -0.717 | 0.486 | 0.624 | 0.875 | -0.023 | -0.044 | -0.044 | 0 | 0.35 | 0.13 | 0.48 | 0.56 | 0.38 | 0.4 |
| 1 | 19698026 | rs10753559 | 1.436 | 0.418 | 0.787 | 0.816 | 0.019 | 0.013 | 0.013 | 0 | 0.37 | 0.19 | 0.31 | 0.36 | 0.33 | 0.28 |
| 1 | 19706751 | rs6663076 | -1.725 | 0.45 | 0.949 | 1 | 0.061 | 0.174 | 0.174 | 0.644 | 0.06 | 0.36 | 0.07 | 0.13 | 0.03 | 0.06 |
| 1 | 19713004 | rs1472566 | 0.178 | 0.478 | 0.474 | 0.87 | -0.022 | 0.009 | 0.009 | 0 | 0.64 | 0.72 | 0.63 | 0.67 | 0.61 | 0.54 |
| 1 | 19713910 | rs6672756 | 1.06 | 0.521 | 0.189 | 0.827 | -0.058 | 0.067 | 0.067 | 0 | 0.21 | 0.06 | 0.14 | 0.09 | 0.26 | 0.32 |
| 1 | 19724079 | rs10917443 | 0.613 | 0.404 | 0.341 | 0.764 | -0.015 | -0.054 | -0.054 | 0 | 0.21 | 0.09 | 0.27 | 0.34 | 0.25 | 0.25 |

| | | | | | | | | | | | | | | | | |
|---|---|---|---|---|---|---|---|---|---|---|---|---|---|---|---|---|
| 1 | 19725033 | rs7513800 | -0.15 | 0.371 | 0.392 | 0.742 | -0.066 | 0.016 | 0.016 | 0 | 0.66 | 0.79 | 0.63 | 0.71 | 0.58 | 0.56 |
| 1 | 19727896 | rs1566531 | 1.455 | 0.344 | 0.561 | 0.642 | 0.02 | 0.01 | 0.01 | 0.01 | 0.37 | 0.18 | 0.31 | 0.36 | 0.33 | 0.26 |
| 1 | 19730181 | rs4912086 | 0.755 | 0.38 | 0.556 | 0.682 | -0.009 | -0.043 | -0.043 | 1.42 | 0.2 | 0.12 | 0.28 | 0.43 | 0.24 | 0.29 |
| 1 | 19730346 | rs7515937 | -0.132 | 0.362 | 0.603 | 0.671 | -0.059 | 0.113 | 0.113 | 0 | 0.65 | 0.78 | 0.78 | 0.94 | 0.58 | 0.61 |
| 1 | 19733320 | rs881247 | 0.683 | 0.463 | 0.322 | 0.731 | -0.009 | -0.045 | -0.045 | 0 | 0.2 | 0.12 | 0.28 | 0.43 | 0.24 | 0.31 |
| 1 | 19738518 | rs12026907 | 0.941 | 0.408 | 0.516 | 0.574 | 0.01 | 0.009 | 0.009 | 0 | 0.39 | 0.28 | 0.32 | 0.36 | 0.33 | 0.27 |
| 1 | 19751333 | rs12036866 | 1.152 | 0.392 | 0.463 | 0.574 | 0.013 | 0.008 | 0.008 | 0 | 0.37 | 0.24 | 0.32 | 0.36 | 0.34 | 0.29 |
| 1 | 19755204 | rs6661421 | -1.007 | 0.536 | 0.121 | 0.626 | -0.025 | -0.014 | -0.014 | 0 | 0.76 | 0.82 | 0.82 | 0.75 | 0.72 | 0.61 |
| 1 | 19757646 | rs2088828 | 0.689 | 0.269 | 0.123 | 0.653 | -0.018 | -0.017 | -0.017 | 0 | 0.21 | 0.1 | 0.26 | 0.23 | 0.27 | 0.22 |
| 1 | 19762466 | rs4912094 | 0.707 | 0.269 | 0.101 | 0.654 | -0.031 | -0.029 | -0.029 | 0.006 | 0.21 | 0.07 | 0.26 | 0.24 | 0.27 | 0.22 |
| 1 | 19766326 | rs12754186 | -0.343 | 0.221 | 0.556 | 0.604 | 0.033 | 0.005 | 0.005 | 0 | 0.14 | 0.48 | 0.23 | 0.14 | 0.14 | 0.08 |
| 1 | 19769243 | rs10458381 | 1.411 | 0.175 | 0.259 | 0.613 | -0.025 | -0.029 | -0.029 | 0 | 0.36 | 0.04 | 0.31 | 0.36 | 0.34 | 0.25 |
| 1 | 19770968 | rs10917452 | -0.475 | 0.527 | 0.538 | 0.813 | 0.011 | 0.008 | 0.008 | 0.139 | 0.32 | 0.57 | 0.49 | 0.38 | 0.39 | 0.25 |
| 1 | 19771925 | rs12091047 | 1.154 | 0.496 | 0.26 | 0.778 | 0.04 | 0.036 | 0.036 | 0.045 | 0.42 | 0.18 | 0.32 | 0.36 | 0.35 | 0.33 |
| 1 | 19772847 | rs10492998 | 0.983 | 0.749 | 0.143 | 0.731 | -0.034 | 0.001 | 0.001 | 0 | 0.19 | 0.06 | 0.14 | 0.13 | 0.23 | 0.33 |
| 1 | 19775113 | rs4912096 | 0.493 | 0.718 | 0.165 | 0.741 | -0.005 | -0.004 | -0.004 | 0 | 0.21 | 0.2 | 0.28 | 0.24 | 0.26 | 0.22 |
| 1 | 19785334 | rs2088824 | -0.071 | 0.75 | 0.278 | 0.781 | 0.014 | -0.002 | -0.002 | 0 | 0.69 | 0.39 | 0.64 | 0.7 | 0.65 | 0.65 |
| 1 | 19791086 | rs4481878 | -0.388 | 0.613 | 0.247 | 1.26 | 0.004 | 0.03 | 0.03 | 0 | 0.8 | 0.71 | 0.72 | 0.63 | 0.77 | 0.72 |
| 1 | 19792562 | rs12755497 | 0.895 | 0.417 | -0.34 | 0.815 | -0.009 | 0.039 | 0.039 | 0.895 | 0.39 | 0.08 | 0.32 | 0.32 | 0.37 | 0.34 |
| 1 | 19793984 | rs999974 | 0.104 | 0.405 | 0.039 | 0.843 | 0.001 | 0.06 | 0.06 | 0 | 0.73 | 0.53 | 0.69 | 0.6 | 0.74 | 0.68 |
| 1 | 19797248 | rs1002365 | -0.009 | 0.254 | 0.167 | 1.023 | 0.011 | 0.011 | 0.011 | 0.293 | 0.48 | 0.32 | 0.43 | 0.39 | 0.53 | 0.52 |
| 1 | 19797740 | rs4912099 | 0.21 | 0.232 | 0.249 | 1.05 | 0.003 | 0.004 | 0.004 | 0.447 | 0.46 | 0.19 | 0.41 | 0.39 | 0.53 | 0.52 |
| 1 | 19798007 | rs4912100 | -0.358 | 0.29 | 0.342 | 0.943 | 0.007 | 0.02 | 0.02 | 0.533 | 0.53 | 0.67 | 0.48 | 0.53 | 0.54 | 0.63 |
| 1 | 19800954 | rs4912105 | 0.046 | -0.021 | -0.115 | 0.59 | 0.027 | -0.005 | -0.005 | 0 | 0.92 | 0.57 | 0.96 | 1 | 0.98 | 0.99 |
| 1 | 19806259 | rs924519 | 0.048 | -0.008 | -0.619 | 0.761 | 0.027 | 0.01 | 0.01 | 0.117 | 0.51 | 0.36 | 0.59 | 0.51 | 0.61 | 0.57 |
| 1 | 19806692 | rs10917459 | 0.872 | -0.028 | 0.021 | 0.962 | -0.032 | 0.096 | 0.096 | 1.648 | 0.89 | 0.4 | 0.92 | 0.82 | 0.98 | 0.88 |
| 1 | 19809171 | rs6664461 | -0.079 | 0.325 | -1.067 | 0.428 | -0.008 | 0.043 | 0.043 | 0 | 0.39 | 0.28 | 0.26 | 0.24 | 0.41 | 0.44 |
| 1 | 19809758 | rs10917461 | -0.055 | -0.019 | -1.438 | 0.735 | 0.014 | -0.007 | -0.007 | 3.067 | 0.28 | 0.17 | 0.28 | 0.31 | 0.22 | 0.22 |
| 1 | 19822813 | rs16822413 | 0.27 | 0.083 | -1.854 | 1.015 | -0.019 | -0.109 | -0.109 | 0 | 0.24 | 0.02 | 0.14 | 0.41 | 0.26 | 0.24 |
| 1 | 19824008 | rs2314146 | -0.86 | 0.061 | -1.492 | 0.593 | 0.013 | -0.016 | -0.016 | 0 | 0.59 | 0.53 | 0.6 | 0.71 | 0.52 | 0.6 |
| 1 | 19831939 | rs11807599 | 0.017 | -0.222 | -1.237 | 0.455 | -0.003 | -0.005 | -0.005 | 0 | 0.23 | 0.3 | 0.26 | 0.17 | 0.18 | 0.15 |
| 1 | 19837932 | rs12087005 | 0.946 | -0.173 | -1.078 | 0.444 | -0.015 | 0.007 | 0.007 | 0 | 0.12 | 0.17 | 0.19 | 0.14 | 0.08 | 0.22 |
| 1 | 19841174 | rs10799824 | 0.16 | -0.249 | -1.053 | 0.389 | -0.001 | -0.009 | -0.009 | 0 | 0.18 | 0.26 | 0.19 | 0.13 | 0.14 | 0.11 |
| 1 | 19849656 | rs9783019 | -0.779 | -0.204 | -1.182 | 0.438 | -0.018 | -0.008 | -0.008 | 2.523 | 0.32 | 0.5 | 0.41 | 0.29 | 0.23 | 0.34 |
| 1 | 19851031 | rs12726742 | 1.336 | 0.125 | -1.826 | -0.142 | -0.139 | 0.02 | 0.02 | 0.005 | 0.24 | 0.03 | 0.27 | 0.17 | 0.39 | 0.26 |
| 1 | 19858807 | rs4350219 | -1.222 | -0.141 | -1.382 | 0.042 | 0.006 | -0.051 | -0.051 | 0 | 0.74 | 0.83 | 0.75 | 0.46 | 0.73 | 0.73 |
| 1 | 19861027 | rs4912110 | -1.422 | -0.14 | -1.512 | 0.052 | -0.009 | -0.131 | -0.131 | 1.04 | 0.75 | 0.93 | 0.68 | 0.47 | 0.74 | 0.75 |
| 1 | 19876025 | rs4508039 | 0.543 | 0.307 | -1.841 | -0.141 | 0.019 | 0.017 | 0.017 | 0.164 | 0.52 | 0.09 | 0.61 | 0.64 | 0.63 | 0.55 |
| 1 | 19876438 | rs4912122 | 0.588 | 0.302 | -1.846 | -0.143 | 0.017 | 0.014 | 0.014 | 0.228 | 0.52 | 0.07 | 0.61 | 0.64 | 0.63 | 0.55 |
| 1 | 19882170 | rs10917493 | -0.259 | 0.027 | -1.223 | -0.201 | 0.018 | 0.021 | 0.021 | 0.362 | 0.49 | 0.59 | 0.49 | 0.35 | 0.57 | 0.56 |
| 1 | 19889959 | rs1572823 | -1.182 | 0.105 | -1.026 | -0.301 | -0.034 | -0.029 | -0.029 | 0 | 0.77 | 0.68 | 0.87 | 0.89 | 0.86 | 0.82 |
| 1 | 19897768 | rs12132648 | 0.727 | 0.602 | -0.805 | -0.005 | 0.009 | -0.013 | -0.013 | 0.132 | 0.06 | 0.07 | 0.06 | 0.02 | 0.1 | 0.08 |
| 1 | 19914083 | rs12142530 | 0.439 | 0.503 | -0.424 | -0.03 | 0.005 | -0.015 | -0.015 | 0 | 0.06 | 0.06 | 0.05 | 0.01 | 0.08 | 0.07 |
| 1 | 19924662 | rs2792043 | -1.192 | 0.587 | -0.511 | -0.084 | -0.054 | 0.114 | 0.114 | 0 | 0.55 | 0.92 | 0.63 | 0.74 | 0.54 | 0.55 |
| 1 | 19927553 | rs860847 | -0.4 | 0.625 | -0.416 | -0.053 | 0.099 | -0.023 | -0.023 | 1.154 | 0.09 | 0.52 | 0.1 | 0.02 | 0.1 | 0.08 |
| 1 | 19945189 | rs1770489 | -0.963 | 0.597 | -0.441 | -0.026 | 0.001 | -0.004 | -0.004 | 0 | 0.23 | 0.71 | 0.2 | 0.14 | 0.24 | 0.29 |
| 1 | 19947256 | rs10489858 | 0.579 | 0.59 | -0.466 | -0.014 | 0.001 | 0.002 | 0.002 | 0.051 | 0.14 | 0.18 | 0.1 | 0.12 | 0.14 | 0.2 |
| 1 | 19947973 | rs1320978 | -1.299 | 0.566 | -0.555 | -0.035 | -0.055 | 0.112 | 0.112 | 0 | 0.55 | 0.92 | 0.63 | 0.74 | 0.54 | 0.56 |
| 1 | 19953165 | rs2281237 | 0.535 | 0.462 | -0.437 | 0.126 | 0.001 | 0.002 | 0.002 | 0.04 | 0.14 | 0.18 | 0.1 | 0.12 | 0.14 | 0.2 |
| 1 | 19954014 | rs880554 | 0.836 | 0.407 | -0.529 | 0.119 | 0.02 | -0.126 | -0.126 | 0.257 | 0.27 | 0.07 | 0.32 | 0.52 | 0.23 | 0.23 |
| 1 | 19962874 | rs7523079 | -0.955 | 0.662 | -0.752 | 0.157 | -0.013 | -0.004 | -0.004 | 0 | 0.23 | 0.78 | 0.27 | 0.21 | 0.25 | 0.28 |
| 1 | 19963047 | rs7523192 | -0.449 | 0.663 | -0.962 | 0.203 | 0.028 | 0.021 | 0.021 | 1.304 | 0.2 | 0.36 | 0.2 | 0.19 | 0.21 | 0.26 |
| 1 | 19968058 | rs12408663 | 0.683 | 0.598 | -1.737 | 0.306 | -0.013 | -0.19 | -0.19 | 0 | 0.27 | 0.01 | 0.3 | 0.52 | 0.23 | 0.23 |
| 1 | 19969350 | rs6702295 | 0.301 | 0.559 | -1.072 | 0.364 | 0.012 | 0.007 | 0.007 | 0 | 0.06 | 0.1 | 0.1 | 0.07 | 0.08 | 0.05 |
| 1 | 19973358 | rs7522389 | -0.822 | 0.497 | -1.409 | 0.272 | -0.026 | 0.016 | 0.016 | 2.629 | 0.23 | 0.84 | 0.26 | 0.16 | 0.25 | 0.28 |
| 1 | 19973564 | rs7535952 | -0.862 | 0.46 | -1.328 | 0.157 | -0.015 | 0.008 | 0.008 | 2.164 | 0.27 | 0.76 | 0.3 | 0.19 | 0.29 | 0.33 |
| 1 | 19974669 | rs2314331 | -0.128 | -0.073 | -1.431 | 0.327 | -0.013 | -0.182 | -0.182 | 2.467 | 0.75 | 0.99 | 0.69 | 0.51 | 0.79 | 0.75 |
| 1 | 19976782 | rs1115163 | 0.834 | 0.076 | -0.387 | -0.35 | -0.002 | -0.002 | -0.002 | 0.074 | 0.45 | 0.58 | 0.46 | 0.15 | 0.5 | 0.32 |
| 1 | 19978895 | rs753612 | 0.468 | 0.054 | -0.397 | -0.085 | -0.004 | -0.008 | -0.008 | 0 | 0.63 | 0.82 | 0.59 | 0.38 | 0.63 | 0.57 |
| 1 | 19981791 | rs2294630 | 0.915 | 0.033 | -0.284 | -0.15 | -0.009 | 0.078 | 0.078 | 0 | 0.47 | 0.7 | 0.48 | 0.17 | 0.51 | 0.35 |
| 1 | 19984461 | rs3087566 | 0.392 | -0.174 | 0.09 | 0.052 | -0.004 | 0.017 | 0.017 | 0 | 0.67 | 0.65 | 0.64 | 0.45 | 0.7 | 0.7 |
| 1 | 19988894 | rs1977101 | 1.176 | -0.759 | 0.299 | 0.377 | 0.005 | -0.039 | -0.039 | 0 | 0.15 | 0.05 | 0.11 | 0.21 | 0.12 | 0.19 |
| 1 | 19992513 | rs1805054 | 1.056 | -0.763 | 0.346 | 0.374 | -0.002 | 0.004 | 0.004 | 0 | 0.16 | 0.2 | 0.12 | 0.21 | 0.12 | 0.19 |
| 1 | 19999645 | rs6699866 | 0.093 | -0.749 | 0.296 | 0.356 | 0.01 | -0.018 | -0.018 | 0 | 0.74 | 0.79 | 0.65 | 0.45 | 0.79 | 0.74 |
| 1 | 20002518 | rs9659997 | 1.1 | -0.799 | 0.428 | 0.14 | 0.012 | 0.013 | 0.013 | 0 | 0.52 | 0.44 | 0.49 | 0.2 | 0.65 | 0.51 |
| 1 | 20006471 | rs10917512 | 1.005 | -0.62 | 0.658 | 0.279 | -0.01 | 0.04 | 0.04 | 0 | 0.17 | 0.27 | 0.14 | 0.29 | 0.12 | 0.21 |
| 1 | 20008600 | rs10799831 | 0.056 | -0.707 | 0.341 | 0.183 | 0.023 | -0.052 | -0.052 | 0 | 0.74 | 0.9 | 0.7 | 0.46 | 0.79 | 0.72 |
| 1 | 20008880 | rs9064 | 0.612 | -0.7 | 0.771 | 0.155 | -0.027 | 0.028 | 0.028 | 0 | 0.69 | 0.5 | 0.65 | 0.42 | 0.77 | 0.68 |
| 1 | 20014739 | rs7533981 | 1.132 | -0.688 | 0.664 | 0.134 | 0.057 | 0.122 | 0.122 | 0 | 0.53 | 0.28 | 0.53 | 0.21 | 0.66 | 0.53 |

| | | | | | | | | | | | | | | | | |
|---|---|---|---|---|---|---|---|---|---|---|---|---|---|---|---|---|
| 1 | 20015142 | rs2143292 | 1.317 | -0.509 | 0.562 | 0.159 | 0.007 | -0.034 | -0.034 | 0 | 0.15 | 0.05 | 0.11 | 0.21 | 0.12 | 0.16 |
| 1 | 20015655 | rs3813987 | -0.842 | -0.575 | 0.431 | -0.058 | 0.002 | -0.017 | -0.017 | 0 | 0.28 | 0.19 | 0.32 | 0.67 | 0.28 | 0.31 |
| 1 | 20015955 | rs10159394 | 0.96 | -0.326 | 1.256 | 0.541 | 0.004 | 0.004 | 0.004 | 0 | 0.58 | 0.65 | 0.58 | 0.28 | 0.67 | 0.56 |
| 1 | 20018077 | rs11582254 | -0.199 | -0.176 | 0.695 | 0.649 | 0.007 | -0.029 | -0.029 | 0 | 0.78 | 0.87 | 0.71 | 0.58 | 0.8 | 0.76 |
| 1 | 20018195 | rs3748748 | 0.546 | -0.192 | 0.627 | 0.64 | -0.004 | 0.011 | 0.011 | 0 | 0.16 | 0.22 | 0.13 | 0.25 | 0.12 | 0.15 |
| 1 | 20020044 | rs6689496 | -0.182 | -0.029 | 0.038 | 0.643 | 0.004 | -0.014 | -0.014 | 0.703 | 0.79 | 0.88 | 0.71 | 0.58 | 0.8 | 0.76 |
| 1 | 20020466 | rs1883567 | 0.013 | 0.116 | -0.627 | 0.726 | -0.032 | 0.104 | 0.104 | 0 | 0.75 | 0.53 | 0.68 | 0.58 | 0.81 | 0.76 |
| 1 | 20020994 | rs4515815 | 0.165 | 0.616 | -1.678 | -0.165 | 0.054 | 0.076 | 0.076 | 0 | 0.57 | 0.19 | 0.54 | 0.28 | 0.67 | 0.59 |
| 1 | 20026572 | rs3934191 | 0.386 | 1.063 | -1.721 | -0.19 | -0.016 | 0.083 | 0.083 | 0 | 0.45 | 0.12 | 0.5 | 0.32 | 0.57 | 0.5 |
| 1 | 20027404 | rs10799834 | -0.093 | 1.365 | -0.678 | -0.028 | 0.009 | 0.032 | 0.032 | 0 | 0.27 | 0.45 | 0.31 | 0.63 | 0.28 | 0.33 |
| 1 | 20030565 | rs6657561 | 0.504 | 1.632 | -1.062 | -0.733 | 0.024 | 0.14 | 0.14 | 0 | 0.48 | 0.26 | 0.41 | 0.14 | 0.41 | 0.43 |
| 1 | 20036890 | rs4912019 | 0.522 | 1.619 | -0.571 | -0.641 | -0.001 | -0.003 | -0.003 | 0.02 | 0.52 | 0.42 | 0.45 | 0.18 | 0.46 | 0.46 |
| 1 | 20039820 | rs6704156 | 0.598 | 1.424 | -0.235 | -0.281 | 0.006 | 0.046 | 0.046 | 0 | 0.74 | 0.82 | 0.84 | 0.84 | 0.73 | 0.77 |
| 1 | 20041973 | rs11589338 | 0.007 | 1.674 | -0.348 | -0.463 | -0.003 | -0.012 | -0.012 | 0 | 0.1 | 0.1 | 0.05 | 0 | 0.1 | 0.07 |
| 1 | 20047186 | rs3934190 | 1.215 | 1.708 | 0.236 | 0.128 | 0.008 | -0.006 | -0.006 | 0.868 | 0.62 | 0.45 | 0.69 | 0.36 | 0.61 | 0.61 |
| 1 | 20049085 | rs7539563 | 0.892 | 1.933 | 0.371 | -0.035 | 0.01 | -0.016 | -0.016 | 0.462 | 0.81 | 0.67 | 0.83 | 0.81 | 0.77 | 0.8 |
| 1 | 20049351 | rs7539841 | 0.897 | 2.187 | -0.11 | -0.006 | -0.033 | 0.038 | 0.038 | 0 | 0.82 | 0.92 | 0.8 | 0.9 | 0.76 | 0.76 |
| 1 | 20052571 | rs3929667 | -0.568 | 1.944 | -0.009 | 0.045 | -0.003 | -0.001 | -0.001 | 0 | 0.13 | 0.13 | 0.12 | 0.18 | 0.18 | 0.06 |
| 1 | 20056631 | rs4486471 | -0.531 | 1.082 | -0.07 | -0.06 | -0.001 | -0.003 | -0.003 | 2.495 | 0.4 | 0.39 | 0.5 | 0.74 | 0.41 | 0.48 |
| 1 | 20056759 | rs4326647 | -1.126 | 1.324 | -0.541 | -0.13 | 0.019 | 0.061 | 0.061 | 0 | 0.51 | 0.67 | 0.57 | 0.74 | 0.52 | 0.55 |
| 1 | 20057205 | rs10917527 | -0.754 | 1.313 | -0.267 | 0.126 | 0.02 | -0.047 | -0.047 | 0 | 0.1 | 0.16 | 0.05 | 0 | 0.11 | 0.07 |
| 1 | 20061456 | rs3850533 | -0.971 | 0.509 | -0.376 | 0.798 | -0.005 | 0.002 | 0.002 | 0 | 0.35 | 0.6 | 0.2 | 0.38 | 0.25 | 0.33 |
| 1 | 20061862 | rs1002103 | -0.898 | 0.579 | -0.293 | 1.352 | -0.001 | 0.012 | 0.012 | 0 | 0.1 | 0.14 | 0.13 | 0.33 | 0.07 | 0.06 |
| 1 | 20064575 | rs1782469 | -1.028 | 0.873 | 0.381 | 1.249 | 0.058 | -0.041 | -0.041 | 0.074 | 0.21 | 0.4 | 0.21 | 0.13 | 0.27 | 0.15 |
| 1 | 20066490 | rs6426825 | -0.7 | 0.866 | 0.822 | 1.138 | 0.098 | 0.013 | 0.013 | 0.011 | 0.09 | 0.32 | 0.14 | 0.05 | 0.13 | 0.07 |
| 1 | 20080310 | rs1473689 | -1.043 | 0.577 | 0.174 | 1.163 | -0.006 | -0.007 | -0.007 | 2.312 | 0.18 | 0.19 | 0.32 | 0.45 | 0.25 | 0.18 |
| 1 | 20081064 | rs1983602 | 0.272 | -0.11 | -0.339 | 1.201 | 0.022 | -0.038 | -0.038 | 0.07 | 0.16 | 0.1 | 0.03 | 0.25 | 0.08 | 0.27 |
| 1 | 20082849 | rs10917532 | -0.359 | 0.077 | -0.358 | 0.931 | -0.017 | -0.022 | -0.022 | 0 | 0.15 | 0.11 | 0.21 | 0.41 | 0.21 | 0.15 |
| 1 | 20083197 | rs12133589 | -1.353 | -0.134 | -0.471 | 0.765 | -0.046 | -0.002 | -0.002 | 0.214 | 0.29 | 0.17 | 0.29 | 0.41 | 0.34 | 0.2 |
| 1 | 20087406 | rs1473688 | -1.386 | -0.165 | -0.443 | 0.734 | -0.022 | 0.009 | 0.009 | 0 | 0.31 | 0.23 | 0.27 | 0.4 | 0.35 | 0.27 |
| 1 | 20090156 | rs2294634 | -0.24 | -0.116 | -0.503 | 0.729 | -0.029 | -0.027 | -0.027 | 0 | 0.16 | 0.1 | 0.21 | 0.36 | 0.21 | 0.14 |
| 1 | 20091859 | rs16822850 | -0.235 | -0.041 | -0.55 | 0.618 | -0.003 | -0.005 | -0.005 | 0 | 0.15 | 0.11 | 0.06 | 0 | 0.14 | 0.11 |
| 1 | 20096176 | rs1266438 | -1.86 | -0.11 | -0.272 | 1.025 | -0.014 | 0.021 | 0.021 | 0 | 0.57 | 0.76 | 0.47 | 0.69 | 0.51 | 0.6 |
| 1 | 20097941 | rs10917536 | -1.254 | 0.249 | 0.025 | 0.634 | -0.001 | 0 | 0 | 0.027 | 0.33 | 0.39 | 0.38 | 0.43 | 0.37 | 0.31 |
| 1 | 20101034 | rs6426826 | -1.393 | 0.35 | -0.529 | 0.707 | -0.001 | 0.005 | 0.005 | 0.187 | 0.41 | 0.65 | 0.44 | 0.44 | 0.43 | 0.33 |
| 1 | 20109839 | rs6669059 | -0.343 | 0.528 | -1.038 | 0.637 | -0.008 | -0.006 | -0.006 | 0 | 0.1 | 0.06 | 0.07 | 0 | 0.15 | 0.05 |
| 1 | 20131771 | rs2314757 | 0.718 | -0.129 | -1.946 | 0.326 | -0.13 | 0.163 | 0.163 | 0.149 | 0.38 | 0.02 | 0.37 | 0.11 | 0.49 | 0.43 |
| 1 | 20138436 | rs7513369 | 0.688 | -0.18 | -1.28 | 0.849 | 0.008 | 0.019 | 0.019 | 0 | 0.76 | 0.81 | 0.69 | 0.76 | 0.77 | 0.86 |
| 1 | 20140036 | rs1317209 | -0.97 | -0.692 | -0.874 | 1.3 | -0.018 | 0.02 | 0.02 | 0 | 0.25 | 0.3 | 0.27 | 0.37 | 0.18 | 0.27 |
| 1 | 20140340 | rs3767216 | -0.917 | -0.072 | -0.863 | 0.96 | -0.004 | -0.006 | -0.006 | 2.213 | 0.4 | 0.57 | 0.35 | 0.63 | 0.33 | 0.36 |
| 1 | 20141528 | rs1541185 | -0.423 | 0.535 | -1.145 | 1.296 | -0.003 | -0.002 | -0.002 | 0.132 | 0.21 | 0.25 | 0.38 | 0.33 | 0.24 | 0.28 |
| 1 | 20141920 | rs12064796 | -0.713 | -0.101 | -1.195 | 0.832 | -0.002 | 0.008 | 0.008 | 0 | 0.52 | 0.46 | 0.51 | 0.63 | 0.48 | 0.48 |
| 1 | 20142434 | rs3820329 | 0.14 | 0.803 | -1.061 | 1.222 | 0.01 | 0.054 | 0.054 | 0 | 0.16 | 0.27 | 0.17 | 0.22 | 0.21 | 0.25 |
| 1 | 20142681 | rs3806310 | -0.414 | 0.825 | -1.245 | 0.992 | -0.012 | -0.019 | -0.019 | 0 | 0.3 | 0.29 | 0.44 | 0.5 | 0.37 | 0.36 |
| 1 | 20144143 | rs12405608 | 0.444 | 0.644 | -1.605 | 1.297 | -0.001 | -0.039 | -0.039 | 0 | 0.07 | 0.01 | 0.24 | 0.19 | 0.05 | 0.18 |
| 1 | 20144332 | rs17396847 | 0.126 | 1.075 | -1.329 | 0.874 | -0.052 | 0.013 | 0.013 | 0 | 0.26 | 0.16 | 0.21 | 0.31 | 0.38 | 0.22 |
| 1 | 20146417 | rs9651026 | -0.354 | 0.955 | -0.736 | 1.035 | 0.011 | 0.145 | 0.145 | 0 | 0.12 | 0.27 | 0.13 | 0.28 | 0.12 | 0.18 |
| 1 | 20147302 | rs10917549 | 0.107 | 1.014 | -1.512 | -0.119 | -0.034 | 0.078 | 0.078 | 0 | 0.24 | 0.06 | 0.23 | 0.07 | 0.26 | 0.17 |
| 1 | 20148249 | rs10799839 | 0.366 | 0.7 | -0.939 | -0.121 | -0.002 | -0.007 | -0.007 | 0 | 0.83 | 0.81 | 0.83 | 0.89 | 0.84 | 0.67 |
| 1 | 20149058 | rs1832574 | 0.114 | 0.809 | -1.617 | 0.052 | -0.028 | -0.08 | -0.08 | 0.359 | 0.25 | 0.07 | 0.18 | 0.58 | 0.23 | 0.37 |
| 1 | 20150998 | rs4655232 | -0.269 | 0.691 | -1.06 | -0.002 | -0.001 | 0.004 | 0.004 | 0 | 0.29 | 0.27 | 0.2 | 0.58 | 0.24 | 0.45 |
| 1 | 20158241 | rs4509550 | -0.412 | 0.783 | -0.71 | -0.178 | 0 | 0 | 0 | 0 | 0.4 | 0.35 | 0.41 | 0.73 | 0.38 | 0.63 |
| 1 | 20165383 | rs12137940 | -0.116 | 0.824 | -0.527 | -0.092 | 0.001 | -0.003 | -0.003 | 0 | 0.06 | 0.09 | 0.13 | 0.06 | 0.06 | 0.09 |
| 1 | 20171860 | rs6426833 | 0.524 | 1.143 | -0.238 | -0.407 | 0.028 | -0.011 | -0.011 | 0 | 0.65 | 0.46 | 0.49 | 0.76 | 0.55 | 0.69 |
| 1 | 20171884 | rs7532822 | 0.647 | 1.441 | -0.25 | -0.501 | 0.032 | 0.032 | 0.032 | 0 | 0.8 | 0.51 | 0.62 | 0.82 | 0.8 | 0.75 |
| 1 | 20172956 | rs10799844 | 1.141 | 0.897 | -0.368 | -0.712 | 0.021 | -0.022 | -0.022 | 0 | 0.42 | 0.25 | 0.39 | 0.55 | 0.39 | 0.58 |
| 1 | 20173263 | rs10917557 | 1.397 | 0.847 | -0.45 | -0.805 | 0.01 | -0.032 | -0.032 | 0 | 0.36 | 0.25 | 0.37 | 0.55 | 0.34 | 0.53 |
| 1 | 20173858 | rs10737482 | 0.181 | 0.877 | -0.249 | -0.907 | 0.014 | -0.023 | -0.023 | 0.626 | 0.72 | 0.68 | 0.63 | 0.9 | 0.62 | 0.83 |
| 1 | 20175112 | rs12145220 | 0.4 | 0.731 | -0.43 | -0.815 | 0 | -0.004 | -0.004 | 0 | 0.57 | 0.47 | 0.64 | 0.76 | 0.66 | 0.74 |
| 1 | 20183342 | rs6690967 | 0.614 | 0.785 | -0.146 | -0.652 | -0.001 | 0.036 | 0.036 | 0 | 0.48 | 0.31 | 0.57 | 0.6 | 0.48 | 0.64 |
| 1 | 20186581 | rs12121573 | -0.521 | 0.699 | 0.32 | -0.911 | -0.003 | 0.003 | 0.003 | 0 | 0.29 | 0.3 | 0.28 | 0.19 | 0.34 | 0.24 |
| 1 | 20187321 | rs7531128 | 0.58 | 0.672 | 0.39 | -0.945 | -0.007 | -0.027 | -0.027 | 0 | 0.07 | 0.11 | 0.03 | 0 | 0.04 | 0.05 |
| 1 | 20188458 | rs16823060 | -0.674 | 0.67 | 0.621 | -0.93 | 0.006 | -0.012 | -0.012 | 0 | 0.29 | 0.38 | 0.3 | 0.19 | 0.34 | 0.24 |
| 1 | 20190672 | rs16823061 | 0.03 | 0.505 | 0.925 | -1.031 | 0.01 | -0.016 | -0.016 | 0 | 0.15 | 0.27 | 0.22 | 0.07 | 0.14 | 0.1 |
| 1 | 20191436 | rs4654897 | 1.108 | 0.583 | 0.863 | -1.046 | -0.012 | 0 | 0 | 0 | 0.16 | 0.11 | 0.13 | 0.13 | 0.24 | 0.11 |
| 1 | 20193622 | rs1416819 | 0.736 | 0.62 | 0.767 | -1.001 | 0.042 | 0.027 | 0.027 | 0.251 | 0.06 | 0.11 | 0.18 | 0.07 | 0.12 | 0.07 |
| 1 | 20200990 | rs4654903 | -1.538 | 0.566 | 0.708 | -1.021 | -0.007 | 0.019 | 0.019 | 0 | 0.44 | 0.65 | 0.52 | 0.25 | 0.47 | 0.33 |
| 1 | 20233086 | rs10916668 | 1.315 | 0.577 | 0.55 | -0.949 | -0.008 | 0.007 | 0.007 | 0 | 0.05 | 0.01 | 0.12 | 0.02 | 0.06 | 0.08 |
| 1 | 20236731 | rs17401735 | -0.942 | 0.565 | 0.677 | -1.016 | 0.016 | -0.007 | -0.007 | 6.193 | 0.15 | 0.21 | 0.35 | 0.09 | 0.2 | 0.17 |

| | | | | | | | | | | | | | | | | |
|---|---|---|---|---|---|---|---|---|---|---|---|---|---|---|---|---|
| 1 | 20237760 | rs10492999 | 1.096 | 0.571 | 0.487 | -0.998 | -0.015 | 0.002 | 0.002 | 0 | 0.05 | 0 | 0.12 | 0.02 | 0.06 | 0.08 |
| 1 | 20238343 | rs1046548 | -1.604 | 0.753 | 0.545 | -1.031 | -0.001 | 0.014 | 0.014 | 0 | 0.45 | 0.62 | 0.54 | 0.24 | 0.52 | 0.4 |
| 1 | 20238466 | rs17401847 | 0.286 | 0.665 | 0.726 | -0.473 | 0.024 | 0.008 | 0.008 | 0.282 | 0.06 | 0.06 | 0.19 | 0.07 | 0.14 | 0.12 |
| 1 | 20240590 | rs1111872 | 0.188 | 0.466 | 0.7 | -0.447 | -0.004 | -0.024 | -0.024 | 0 | 0.07 | 0.1 | 0.04 | 0 | 0.06 | 0.05 |
| 1 | 20244408 | rs2066130 | -1.414 | 0.652 | 0.568 | -0.485 | -0.006 | -0.013 | -0.013 | 0 | 0.34 | 0.54 | 0.44 | 0.25 | 0.33 | 0.31 |
| 1 | 20245708 | rs7531863 | -0.576 | 0.667 | 0.444 | -0.487 | -0.003 | -0.045 | -0.045 | 0 | 0.26 | 0.41 | 0.39 | 0.09 | 0.24 | 0.23 |
| 1 | 20247674 | rs12028511 | 0.553 | 0.934 | -0.21 | 0.12 | -0.031 | -0.037 | -0.037 | 0 | 0.1 | 0.01 | 0.05 | 0.11 | 0.11 | 0.08 |
| 1 | 20248709 | rs1108974 | -0.658 | 0.859 | 0.307 | 0.1 | -0.003 | 0.021 | 0.021 | 0.139 | 0.1 | 0.13 | 0.16 | 0.18 | 0.12 | 0.13 |
| 1 | 20254017 | rs12066827 | -1.039 | 0.612 | 1.053 | -0.005 | 0.1 | 0.178 | 0.178 | 0.478 | 0.07 | 0.49 | 0.1 | 0.18 | 0.05 | 0.06 |
| 1 | 20258473 | rs1093733 | -0.173 | 0.247 | 0.661 | -0.338 | 0.017 | 0.043 | 0.043 | 0 | 0.82 | 0.9 | 0.83 | 0.97 | 0.88 | 0.9 |
| 1 | 20260536 | rs1796915 | 1.412 | 0.149 | 0.01 | -0.855 | 0.031 | 0.067 | 0.067 | 0 | 0.57 | 0.28 | 0.63 | 0.7 | 0.67 | 0.72 |
| 1 | 20262931 | rs1796918 | 0.636 | 0.153 | -0.002 | -0.852 | -0.017 | -0.022 | -0.022 | 0 | 0.73 | 0.38 | 0.73 | 0.8 | 0.83 | 0.85 |
| 1 | 20263901 | rs17410800 | 0.412 | 0.324 | -0.542 | -0.844 | -0.008 | 0.037 | 0.037 | 0.21 | 0.16 | 0.04 | 0.1 | 0.09 | 0.16 | 0.13 |
| 1 | 20272338 | rs12138723 | 0.142 | 0.264 | -0.678 | -0.678 | -0.005 | 0.082 | 0.082 | 0 | 0.18 | 0 | 0.15 | 0.02 | 0.12 | 0.1 |
| 1 | 20278924 | rs12132830 | 0.372 | 0.249 | -0.654 | -0.65 | 0.001 | 0.069 | 0.069 | 0 | 0.16 | 0 | 0.14 | 0.02 | 0.09 | 0.09 |
| 1 | 20279405 | rs818968 | 0.745 | 0.536 | -0.682 | -0.611 | -0.007 | 0.083 | 0.083 | 0 | 0.57 | 0.26 | 0.6 | 0.71 | 0.57 | 0.64 |
| 1 | 20281969 | rs1796924 | 0.682 | 0.564 | -0.605 | -0.54 | -0.006 | 0.063 | 0.063 | 0 | 0.57 | 0.3 | 0.6 | 0.68 | 0.57 | 0.62 |
| 1 | 20283943 | rs818963 | -0.616 | 0.351 | -0.545 | 0.058 | 0 | 0.005 | 0.005 | 0 | 0.37 | 0.38 | 0.32 | 0.25 | 0.29 | 0.33 |
| 1 | 20285487 | rs12408941 | 0.821 | 0.393 | -1.052 | 0.107 | -0.012 | -0.072 | -0.072 | 0.064 | 0.17 | 0 | 0.17 | 0.23 | 0.11 | 0.2 |
| 1 | 20286563 | rs10916679 | 1.357 | 0.329 | -1.729 | 0.429 | -0.013 | -0.044 | -0.044 | 0 | 0.2 | 0.05 | 0.13 | 0.21 | 0.25 | 0.12 |
| 1 | 20287919 | rs4655010 | -0.198 | 0.346 | -0.902 | 0.764 | 0 | 0 | 0 | 0.077 | 0.34 | 0.36 | 0.3 | 0.28 | 0.26 | 0.3 |
| 1 | 20287981 | rs2795480 | -1.945 | 0.421 | -0.827 | 0.755 | 0.004 | -0.006 | -0.006 | 0 | 0.78 | 0.86 | 0.72 | 0.67 | 0.83 | 0.71 |
| 1 | 20292006 | rs12023742 | -0.008 | 0.676 | -0.696 | 0.596 | -0.014 | -0.002 | -0.002 | 0 | 0.35 | 0.21 | 0.42 | 0.32 | 0.39 | 0.45 |
| 1 | 20292378 | rs13375942 | 1.044 | 0.694 | -0.703 | 0.585 | 0.005 | -0.004 | -0.004 | 0 | 0.18 | 0.2 | 0.25 | 0.14 | 0.28 | 0.25 |
| 1 | 20293055 | rs17411112 | 1.052 | 0.752 | -0.933 | 0.45 | -0.022 | 0.044 | 0.044 | 0 | 0.15 | 0.09 | 0.11 | 0.03 | 0.15 | 0.09 |
| 1 | 20293791 | rs6672057 | -1.112 | 0.697 | -0.9 | 0.517 | 0.002 | 0.001 | 0.001 | 0 | 0.52 | 0.57 | 0.55 | 0.42 | 0.55 | 0.55 |
| 1 | 20295022 | rs4655020 | -0.517 | 0.792 | -1.044 | 0.502 | 0.025 | -0.029 | -0.029 | 0 | 0.35 | 0.5 | 0.38 | 0.24 | 0.44 | 0.34 |
| 1 | 20296319 | rs17402685 | -0.521 | 0.797 | -1.041 | 0.505 | 0.024 | -0.029 | -0.029 | 0.114 | 0.35 | 0.5 | 0.38 | 0.24 | 0.44 | 0.35 |
| 1 | 20300544 | rs10916683 | 0.559 | 0.819 | -1.692 | 0.517 | -0.016 | 0.036 | 0.036 | 0 | 0.16 | 0.1 | 0.23 | 0.04 | 0.16 | 0.11 |
| 1 | 20301964 | rs11677 | 0.712 | 0.559 | -1.526 | 0.993 | -0.011 | -0.06 | -0.06 | 0 | 0.17 | 0 | 0.16 | 0.18 | 0.11 | 0.2 |
| 1 | 20304857 | rs2307246 | 0.692 | 0.827 | -1.439 | 0.537 | -0.008 | 0.015 | 0.015 | 0 | 0.16 | 0.1 | 0.11 | 0.03 | 0.21 | 0.1 |
| 1 | 20304926 | rs4744 | 0.692 | 0.827 | -1.439 | 0.537 | -0.008 | 0.015 | 0.015 | 0 | 0.16 | 0.1 | 0.11 | 0.03 | 0.21 | 0.1 |
| 1 | 20305887 | rs3753827 | 0.276 | 0.675 | -1.531 | 0.879 | -0.006 | 0.03 | 0.03 | 0.112 | 0.45 | 0.31 | 0.39 | 0.42 | 0.48 | 0.5 |
| 1 | 20311676 | rs12568139 | 0.777 | 0.309 | -1.717 | 1.478 | 0.004 | -0.088 | -0.088 | 1.046 | 0.14 | 0 | 0.04 | 0.26 | 0.05 | 0.13 |
| 1 | 20316055 | rs12021819 | 0.981 | 1.031 | -1.367 | 1.671 | 0.002 | 0.014 | 0.014 | 0 | 0.1 | 0.08 | 0.09 | 0.21 | 0.18 | 0.16 |
| 1 | 20342503 | rs1093661 | 0.425 | 0.511 | -1.494 | 1.067 | -0.007 | 0 | 0 | 0 | 0.28 | 0.24 | 0.29 | 0.27 | 0.35 | 0.31 |
| 1 | 20373391 | rs12410635 | -0.176 | 0.361 | -1.005 | 0.659 | 0.005 | 0.006 | 0.006 | 0 | 0.25 | 0.26 | 0.37 | 0.36 | 0.28 | 0.36 |
| 1 | 20380145 | rs617363 | -1.006 | -0.275 | -0.864 | 0.427 | 0.023 | 0.026 | 0.026 | 0.066 | 0.48 | 0.7 | 0.58 | 0.61 | 0.39 | 0.53 |
| 1 | 20387608 | rs12405285 | 1.169 | -0.282 | -0.808 | 0.431 | -0.002 | -0.003 | -0.003 | 0 | 0.13 | 0.1 | 0.12 | 0.29 | 0.16 | 0.25 |
| 1 | 20390322 | rs17354650 | 1.956 | -0.548 | -1.188 | 0.202 | -0.054 | 0.036 | 0.036 | 0 | 0.22 | 0.02 | 0.15 | 0.09 | 0.24 | 0.14 |
| 1 | 20392941 | rs1891318 | 1.606 | 0.344 | -0.667 | 0.417 | -0.001 | 0.044 | 0.044 | 0.022 | 0.42 | 0.11 | 0.31 | 0.31 | 0.42 | 0.34 |
| 1 | 20394713 | rs606980 | 0.895 | 0.324 | -0.333 | 0.419 | -0.035 | 0.031 | 0.031 | 4.5 | 0.22 | 0.37 | 0.21 | 0.25 | 0.11 | 0.17 |
| 1 | 20395401 | rs11573185 | 0.808 | 0.317 | -0.206 | 0.392 | -0.04 | 0.038 | 0.038 | 1.235 | 0.53 | 0.19 | 0.54 | 0.38 | 0.54 | 0.45 |
| 1 | 20395497 | rs10916703 | 1.494 | 0.204 | 0.216 | 0.343 | 0.013 | -0.038 | -0.038 | 0 | 0.08 | 0.26 | 0.06 | 0 | 0.06 | 0.05 |
| 1 | 20396415 | rs11573191 | 1.241 | 0.243 | 0.189 | 0.376 | 0.005 | 0.021 | 0.021 | 0 | 0.14 | 0.16 | 0.13 | 0.29 | 0.17 | 0.25 |
| 1 | 20396648 | rs3806311 | 2.459 | 0.172 | 0.051 | 0.362 | 0.002 | -0.078 | -0.078 | 0.06 | 0.12 | 0 | 0.14 | 0.24 | 0.05 | 0.1 |
| 1 | 20396674 | rs668630 | 0.447 | 0.177 | 0.062 | 0.331 | -0.01 | -0.068 | -0.068 | 0 | 0.14 | 0.36 | 0.16 | 0.03 | 0.1 | 0.08 |
| 1 | 20396733 | rs3806313 | 2.363 | 0.229 | -0.012 | 0.374 | 0.01 | -0.07 | -0.07 | 0.746 | 0.12 | 0.02 | 0.14 | 0.24 | 0.05 | 0.12 |
| 1 | 20397240 | rs656110 | 0.39 | 0.319 | 0.203 | 0.375 | 0.05 | 0.023 | 0.023 | 0 | 0.68 | 0.49 | 0.61 | 0.58 | 0.57 | 0.53 |
| 1 | 20402767 | rs11573219 | 1.605 | 0.269 | -0.183 | 0.075 | 0.023 | 0.048 | 0.048 | 0 | 0.42 | 0.13 | 0.3 | 0.31 | 0.41 | 0.33 |
| 1 | 20403098 | rs11573222 | 1.618 | 0.267 | -0.219 | 0.073 | 0.013 | 0.043 | 0.043 | 0 | 0.42 | 0.12 | 0.3 | 0.31 | 0.41 | 0.33 |
| 1 | 20404055 | rs3767227 | 1.12 | 0.406 | 0.006 | 0.381 | 0.005 | 0.028 | 0.028 | 0 | 0.14 | 0.13 | 0.13 | 0.33 | 0.18 | 0.26 |
| 1 | 20405962 | rs11573236 | 1.636 | 0.412 | -0.032 | 0.34 | -0.003 | 0.007 | 0.007 | 0 | 0.08 | 0.07 | 0.14 | 0.04 | 0.09 | 0.09 |
| 1 | 20406586 | rs11573238 | -0.042 | 0.328 | -0.057 | 0.364 | 0.007 | 0.021 | 0.021 | 0 | 0.22 | 0.25 | 0.28 | 0.36 | 0.27 | 0.35 |
| 1 | 20409374 | rs525380 | 0.315 | 0.239 | -0.218 | 0.341 | 0.047 | 0.043 | 0.043 | 0 | 0.68 | 0.53 | 0.61 | 0.54 | 0.55 | 0.52 |
| 1 | 20410795 | rs660414 | 0.332 | 0.209 | -0.487 | 0.244 | -0.022 | -0.012 | -0.012 | 0.252 | 0.26 | 0.39 | 0.32 | 0.23 | 0.16 | 0.2 |
| 1 | 20411332 | rs2020887 | 1.793 | 0.175 | -1.484 | 0.054 | 0.037 | 0.002 | 0.002 | 0 | 0.36 | 0.1 | 0.26 | 0.31 | 0.34 | 0.31 |
| 1 | 20414239 | rs11573272 | 1.544 | 0.31 | -1.367 | 0.197 | 0.02 | 0.04 | 0.04 | 0.004 | 0.4 | 0.12 | 0.29 | 0.3 | 0.4 | 0.31 |
| 1 | 20416794 | rs11573288 | 1.806 | 0.402 | -1.17 | 0.273 | 0.002 | -0.033 | -0.033 | 0 | 0.13 | 0.01 | 0.14 | 0.13 | 0.06 | 0.13 |
| 1 | 20418207 | rs11573298 | 1.047 | 0.654 | -1.114 | 0.356 | -0.032 | -0.09 | -0.09 | 0 | 0.15 | 0.04 | 0.12 | 0.39 | 0.18 | 0.23 |
| 1 | 20423099 | rs6699362 | 0.961 | 0.469 | -0.943 | -0.072 | 0.045 | 0.133 | 0.133 | 0 | 0.48 | 0.21 | 0.35 | 0.25 | 0.44 | 0.4 |
| 1 | 20424167 | rs656755 | -1.364 | 0.997 | -0.655 | 0.205 | 0.01 | 0 | 0 | 0 | 0.9 | 0.88 | 0.87 | 0.89 | 0.81 | 0.86 |
| 1 | 20427541 | rs4655155 | 1.24 | 0.609 | -1.478 | -0.162 | -0.019 | -0.006 | -0.006 | 0 | 0.36 | 0.17 | 0.4 | 0.38 | 0.38 | 0.31 |
| 1 | 20429861 | rs492738 | 0.129 | 0.525 | -1.226 | -0.117 | -0.024 | -0.037 | -0.037 | 0.56 | 0.63 | 0.85 | 0.77 | 0.62 | 0.62 | 0.65 |
| 1 | 20434402 | rs599367 | -0.681 | -0.905 | -0.986 | 0.302 | 0.023 | 0.002 | 0.002 | 0 | 0.44 | 0.65 | 0.25 | 0.39 | 0.29 | 0.39 |
| 1 | 20437404 | rs12059492 | -0.018 | -0.962 | -0.825 | -0.638 | 0.004 | 0.009 | 0.009 | 0.256 | 0.27 | 0.23 | 0.24 | 0.14 | 0.26 | 0.3 |
| 1 | 20437540 | rs11576937 | 0.791 | -1.175 | -0.688 | -0.269 | -0.002 | 0.052 | 0.052 | 0 | 0.13 | 0.15 | 0.2 | 0.43 | 0.07 | 0.08 |
| 1 | 20437563 | rs7518058 | -1.059 | -1.266 | -0.869 | -0.753 | -0.075 | 0.003 | 0.003 | 0 | 0.69 | 0.9 | 0.59 | 0.72 | 0.56 | 0.66 |
| 1 | 20437764 | rs7551550 | -0.256 | -1.125 | -0.443 | -0.893 | 0.01 | 0.061 | 0.061 | 0.032 | 0.49 | 0.66 | 0.37 | 0.22 | 0.48 | 0.45 |

| | | | | | | | | | | | | | | | | |
|---|---|---|---|---|---|---|---|---|---|---|---|---|---|---|---|---|
| 1 | 20441633 | rs10916711 | 0.425 | -0.96 | -0.638 | -0.971 | -0.001 | 0.031 | 0.031 | 0.068 | 0.33 | 0.25 | 0.29 | 0.13 | 0.33 | 0.3 |
| 1 | 20442054 | rs584367 | -1.325 | -0.555 | -0.716 | -0.613 | -0.131 | -0.041 | -0.041 | 0 | 0.77 | 0.97 | 0.7 | 0.76 | 0.59 | 0.75 |
| 1 | 20442311 | rs12045689 | 0.149 | -0.857 | 0.378 | 0.388 | 0.005 | 0 | 0 | 0 | 0.15 | 0.13 | 0.2 | 0.45 | 0.08 | 0.13 |
| 1 | 20442553 | rs679667 | -0.023 | -0.929 | 0.882 | -0.547 | -0.012 | 0.002 | 0.002 | 0 | 0.13 | 0.4 | 0.08 | 0.1 | 0.07 | 0.12 |
| 1 | 20442691 | rs3736979 | -0.53 | -0.747 | 0.158 | -0.441 | -0.017 | -0.032 | -0.032 | 0 | 0.35 | 0.52 | 0.38 | 0.74 | 0.27 | 0.29 |
| 1 | 20455220 | rs12722987 | -0.856 | -0.35 | -0.603 | -0.235 | -0.004 | -0.012 | -0.012 | 0 | 0.19 | 0.19 | 0.16 | 0.09 | 0.17 | 0.15 |
| 1 | 20457340 | rs12725043 | -0.717 | -0.237 | -1.09 | -0.192 | 0.011 | 0.023 | 0.023 | 0 | 0.18 | 0.1 | 0.16 | 0.08 | 0.17 | 0.15 |
| 1 | 20464891 | rs631134 | -0.515 | 0.003 | -1.322 | 0.269 | 0.007 | -0.035 | -0.035 | 0 | 0.23 | 0.12 | 0.33 | 0.62 | 0.18 | 0.3 |
| 1 | 20466830 | rs12065685 | -0.895 | -0.042 | -0.903 | -0.261 | -0.003 | -0.016 | -0.016 | 0 | 0.19 | 0.18 | 0.15 | 0.07 | 0.12 | 0.17 |
| 1 | 20467993 | rs35231462 | -0.718 | -0.099 | -0.934 | -0.236 | 0.012 | 0.02 | 0.02 | 0 | 0.16 | 0.12 | 0.14 | 0.07 | 0.1 | 0.13 |
| 1 | 20470148 | rs6657574 | -0.664 | 0.549 | -1.104 | -0.552 | -0.016 | 0.019 | 0.019 | 0 | 0.11 | 0.07 | 0.1 | 0.02 | 0.15 | 0.08 |
| 1 | 20470767 | rs6684376 | -0.84 | 0.457 | -0.975 | -0.598 | -0.004 | -0.006 | -0.006 | 0 | 0.12 | 0.11 | 0.11 | 0.02 | 0.16 | 0.08 |
| 1 | 20472808 | rs7522524 | -0.764 | 0.342 | -1.241 | -0.577 | 0.023 | -0.007 | -0.007 | 1.097 | 0.15 | 0.06 | 0.08 | 0.15 | 0.11 | 0.26 |
| 1 | 20473179 | rs11582551 | -2.319 | 0.103 | -0.522 | -0.704 | -0.002 | -0.006 | -0.006 | 1.991 | 0.29 | 0.31 | 0.19 | 0.17 | 0.26 | 0.35 |
| 1 | 20474907 | rs3738126 | -0.741 | 0.066 | -0.494 | -0.613 | -0.005 | -0.001 | -0.001 | 0 | 0.16 | 0.12 | 0.09 | 0.16 | 0.11 | 0.26 |
| 1 | 20476391 | rs818571 | -1.452 | -0.148 | 0.016 | -0.138 | -0.014 | 0.06 | 0.06 | 0 | 0.29 | 0.47 | 0.37 | 0.62 | 0.27 | 0.31 |
| 1 | 20479395 | rs10753481 | -1.238 | -0.826 | 0.267 | -0.37 | 0 | -0.029 | -0.029 | 2.144 | 0.16 | 0.22 | 0.13 | 0.02 | 0.18 | 0.09 |
| 1 | 20485770 | rs7546956 | -0.648 | -0.837 | 0.381 | -0.386 | -0.011 | 0.015 | 0.015 | 0 | 0.08 | 0.02 | 0.07 | 0.02 | 0.13 | 0.07 |
| 1 | 20487743 | rs876171 | -2.176 | -1.215 | -0.289 | -0.5 | -0.011 | 0.023 | 0.023 | 0 | 0.6 | 0.74 | 0.49 | 0.72 | 0.58 | 0.67 |
| 1 | 20490518 | rs6426616 | -2.186 | -1.076 | -0.29 | -0.505 | -0.01 | 0.025 | 0.025 | 0 | 0.6 | 0.75 | 0.49 | 0.73 | 0.58 | 0.67 |
| 1 | 20491842 | rs7515752 | -0.633 | -0.96 | -0.299 | -0.528 | 0.002 | -0.037 | -0.037 | 2.265 | 0.16 | 0.23 | 0.15 | 0.02 | 0.22 | 0.09 |
| 1 | 20498914 | rs2301472 | 0.411 | -1.004 | -0.499 | -0.387 | -0.013 | -0.014 | -0.014 | 0 | 0.09 | 0.03 | 0.15 | 0.49 | 0.14 | 0.15 |
| 1 | 20499033 | rs2301474 | -0.856 | -1.204 | -0.164 | -0.602 | 0.004 | -0.002 | -0.002 | 0 | 0.36 | 0.31 | 0.27 | 0.66 | 0.32 | 0.53 |
| 1 | 20499168 | rs2301475 | -0.856 | -1.204 | -0.164 | -0.602 | 0.004 | -0.002 | -0.002 | 0 | 0.36 | 0.31 | 0.27 | 0.66 | 0.32 | 0.53 |
| 1 | 20501442 | rs10916718 | -1.65 | -1.227 | -0.039 | -0.612 | 0.001 | 0.008 | 0.008 | 0 | 0.33 | 0.4 | 0.36 | 0.55 | 0.4 | 0.29 |
| 1 | 20501582 | rs12139100 | -0.309 | -1.088 | 0.095 | 0.056 | -0.002 | -0.008 | -0.008 | 0 | 0.26 | 0.28 | 0.12 | 0.18 | 0.18 | 0.39 |
| 1 | 20501974 | rs2301478 | -0.405 | -0.738 | -0.221 | 0.351 | -0.021 | -0.068 | -0.068 | 0.345 | 0.13 | 0.03 | 0.18 | 0.49 | 0.19 | 0.15 |
| 1 | 20503183 | rs11588698 | -0.349 | -0.564 | 0.225 | -0.009 | 0.002 | -0.041 | -0.041 | 0 | 0.13 | 0.21 | 0.1 | 0.01 | 0.14 | 0.11 |
| 1 | 20503710 | rs10753483 | -1.019 | -0.529 | 0.152 | 0.788 | 0.001 | -0.001 | -0.001 | 0 | 0.66 | 0.41 | 0.6 | 0.68 | 0.58 | 0.76 |
| 1 | 20518226 | rs3179690 | 0.447 | -0.822 | -0.023 | 2.051 | -0.041 | -0.048 | -0.048 | 0 | 0.08 | 0.01 | 0.11 | 0.28 | 0.14 | 0.1 |
| 1 | 20519808 | rs1061588 | -0.198 | -1.242 | 0.692 | 1.435 | -0.059 | 0.037 | 0.037 | 0 | 0.31 | 0.43 | 0.28 | 0.41 | 0.19 | 0.4 |
| 1 | 20520732 | rs7542020 | -0.763 | -1.232 | 0.689 | 1.409 | 0.013 | -0.005 | -0.005 | 0.168 | 0.37 | 0.61 | 0.33 | 0.41 | 0.22 | 0.47 |
| 1 | 20524056 | rs10799613 | -0.941 | -1.015 | 0.725 | 1.081 | 0.026 | -0.018 | -0.018 | 0.276 | 0.39 | 0.62 | 0.33 | 0.47 | 0.22 | 0.5 |
| 1 | 20530208 | rs3820320 | -0.747 | -1.285 | 0.462 | 1.137 | -0.032 | 0.009 | 0.009 | 0.081 | 0.36 | 0.54 | 0.3 | 0.46 | 0.19 | 0.43 |
| 1 | 20533678 | rs719683 | -0.801 | -0.691 | 0.239 | 1.434 | -0.004 | 0 | 0 | 0 | 0.4 | 0.56 | 0.34 | 0.46 | 0.29 | 0.45 |
| 1 | 20541370 | rs10799615 | 0.592 | -0.485 | 0.197 | 1.055 | 0.002 | -0.002 | -0.002 | 0 | 0.61 | 0.44 | 0.67 | 0.44 | 0.76 | 0.59 |
| 1 | 20544835 | rs2213801 | -0.098 | -0.055 | -0.463 | 0.686 | 0.002 | -0.004 | -0.004 | 0 | 0.7 | 0.72 | 0.7 | 0.55 | 0.78 | 0.61 |
| 1 | 20551745 | rs2097532 | 0.13 | 0.255 | -0.996 | 0.453 | -0.013 | -0.012 | -0.012 | 5.857 | 0.62 | 0.53 | 0.73 | 0.63 | 0.67 | 0.56 |
| 1 | 20556029 | rs12135822 | -0.847 | 0.312 | -1.454 | 0.302 | -0.012 | 0.014 | 0.014 | 0 | 0.14 | 0.01 | 0.08 | 0.09 | 0.13 | 0.11 |
| 1 | 20557542 | rs11585587 | -0.57 | 0.46 | -1.791 | 0.257 | -0.019 | -0.003 | -0.003 | 0 | 0.26 | 0.14 | 0.3 | 0.29 | 0.29 | 0.27 |
| 1 | 20557582 | rs10916727 | 0.077 | -0.344 | -2.373 | 0.295 | 0.02 | -0.098 | -0.098 | 0 | 0.19 | 0.07 | 0.22 | 0.49 | 0.14 | 0.3 |
| 1 | 20561434 | rs7514659 | 0.213 | -0.224 | -0.637 | -0.358 | 0 | -0.029 | -0.029 | 0 | 0.11 | 0.12 | 0.08 | 0 | 0.14 | 0.1 |
| 1 | 20562389 | rs4655165 | -1.394 | -0.372 | -0.351 | -0.423 | -0.006 | 0.028 | 0.028 | 0.253 | 0.59 | 0.63 | 0.63 | 0.78 | 0.56 | 0.67 |
| 1 | 20570287 | rs10916732 | 0.885 | -0.373 | -0.591 | -0.115 | 0.022 | -0.105 | -0.105 | 0 | 0.14 | 0.01 | 0.18 | 0.41 | 0.08 | 0.19 |
| 1 | 20571807 | rs4655166 | 0.089 | -0.281 | -0.37 | -0.277 | -0.002 | -0.008 | -0.008 | 0.763 | 0.12 | 0.12 | 0.09 | 0 | 0.14 | 0.07 |
| 1 | 20572454 | rs6426623 | 0.295 | -0.464 | -0.351 | -0.317 | 0.002 | -0.002 | -0.002 | 0 | 0.26 | 0.26 | 0.3 | 0.28 | 0.27 | 0.25 |
| 1 | 20572621 | rs2157508 | 0.167 | -0.466 | -0.245 | -0.318 | 0.005 | -0.005 | -0.005 | 0 | 0.27 | 0.35 | 0.3 | 0.28 | 0.27 | 0.25 |
| 1 | 20575549 | rs10799617 | 0.078 | -0.495 | -0.083 | -0.294 | 0.001 | 0 | 0 | 0 | 0.27 | 0.35 | 0.31 | 0.32 | 0.27 | 0.24 |
| 1 | 20575902 | rs10489156 | -0.773 | -0.587 | -0.144 | -0.377 | 0.003 | -0.015 | -0.015 | 0 | 0.38 | 0.45 | 0.39 | 0.35 | 0.4 | 0.32 |
| 1 | 20577954 | rs10916736 | -0.116 | -0.513 | -0.356 | -0.557 | -0.001 | -0.002 | -0.002 | 0 | 0.31 | 0.28 | 0.22 | 0.35 | 0.29 | 0.25 |
| 1 | 20580487 | rs7541616 | -1.11 | -0.481 | -0.308 | -0.593 | -0.002 | 0.005 | 0.005 | 0 | 0.47 | 0.57 | 0.33 | 0.37 | 0.48 | 0.45 |
| 1 | 20582685 | rs736665 | 0.17 | -0.464 | -0.678 | -0.408 | 0 | -0.003 | -0.003 | 0 | 0.27 | 0.23 | 0.2 | 0.35 | 0.26 | 0.24 |
| 1 | 20583238 | rs743982 | 0.757 | -0.316 | -0.714 | 0.07 | 0.021 | -0.04 | -0.04 | 0 | 0.14 | 0.03 | 0.27 | 0.4 | 0.08 | 0.2 |
| 1 | 20584645 | rs11800772 | -0.731 | 0.011 | -0.667 | -0.383 | -0.005 | -0.003 | -0.003 | 0.09 | 0.15 | 0.16 | 0.1 | 0.02 | 0.19 | 0.19 |
| 1 | 20585364 | rs12067030 | 0.045 | -0.055 | -0.974 | -0.706 | -0.001 | -0.004 | -0.004 | 1.071 | 0.15 | 0.11 | 0.11 | 0 | 0.16 | 0.06 |
| 1 | 20585773 | rs17354986 | -0.95 | 0.285 | -1.109 | -0.716 | -0.041 | 0.068 | 0.068 | 0 | 0.17 | 0.07 | 0.12 | 0.04 | 0.23 | 0.18 |
| 1 | 20588698 | rs10916742 | -1.309 | -0.065 | -1.826 | -0.975 | -0.106 | -0.015 | -0.015 | 0 | 0.73 | 0.93 | 0.65 | 0.7 | 0.64 | 0.8 |
| 1 | 20591780 | rs1894662 | -1.485 | -0.116 | -1.807 | -1.03 | -0.095 | 0.052 | 0.052 | 0 | 0.75 | 0.94 | 0.68 | 0.86 | 0.66 | 0.84 |
| 1 | 20592887 | rs763202 | -0.19 | -0.721 | -0.976 | -0.524 | 0.011 | 0.019 | 0.019 | 0 | 0.49 | 0.61 | 0.43 | 0.42 | 0.49 | 0.46 |
| 1 | 20594695 | rs6678323 | -0.835 | -0.62 | -1.323 | -0.378 | -0.084 | 0.072 | 0.072 | 0 | 0.66 | 0.82 | 0.63 | 0.8 | 0.56 | 0.7 |
| 1 | 20601012 | rs4654842 | 1.031 | -0.764 | -1.194 | -0.138 | 0.021 | -0.043 | -0.043 | 0 | 0.14 | 0.07 | 0.18 | 0.41 | 0.07 | 0.25 |
| 1 | 20604959 | rs10916749 | -0.141 | -0.501 | -0.757 | -0.333 | 0 | 0 | 0 | 2.772 | 0.41 | 0.39 | 0.35 | 0.38 | 0.37 | 0.42 |
| 1 | 20605863 | rs11582590 | -1.495 | -0.246 | -0.145 | -0.504 | -0.012 | -0.078 | -0.078 | 0 | 0.18 | 0.4 | 0.16 | 0.05 | 0.12 | 0.13 |
| 1 | 20606320 | rs6683970 | -1.067 | 1.125 | -0.589 | 0.057 | 0.027 | 0.01 | 0.01 | 0 | 0.33 | 0.48 | 0.46 | 0.33 | 0.4 | 0.39 |
| 1 | 20607376 | rs6426628 | -1.163 | 0.988 | -1.504 | -0.805 | -0.02 | 0.205 | 0.205 | 0.635 | 0.61 | 0.89 | 0.79 | 0.92 | 0.63 | 0.77 |
| 1 | 20608200 | rs2142629 | -0.705 | 0.855 | -0.648 | -0.983 | 0 | -0.038 | -0.038 | 0 | 0.5 | 0.46 | 0.63 | 0.85 | 0.53 | 0.61 |
| 1 | 20613063 | rs4422977 | 0.899 | 1.134 | -0.996 | -0.33 | -0.042 | -0.097 | -0.097 | 0.053 | 0.14 | 0.01 | 0.17 | 0.49 | 0.18 | 0.2 |
| 1 | 20613458 | rs1076624 | -0.596 | 0.897 | -0.312 | -0.906 | -0.02 | -0.166 | -0.166 | 0 | 0.23 | 0.47 | 0.14 | 0 | 0.17 | 0.14 |
| 1 | 20617419 | rs12738826 | -0.034 | 0.85 | -0.185 | -0.383 | -0.011 | -0.039 | -0.039 | 0 | 0.18 | 0.09 | 0.29 | 0.37 | 0.23 | 0.25 |

| | | | | | | | | | | | | | | | | |
|---|---|---|---|---|---|---|---|---|---|---|---|---|---|---|---|---|
| 1 | 20620175 | rs12044035 | 0.907 | 0.978 | -0.128 | -0.426 | 0.002 | -0.007 | -0.007 | 0.055 | 0.14 | 0.08 | 0.2 | 0.56 | 0.12 | 0.2 |
| 1 | 20620323 | rs2223225 | -1.247 | 0.79 | -0.234 | -0.982 | 0.002 | 0.004 | 0.004 | 0 | 0.36 | 0.55 | 0.44 | 0.36 | 0.42 | 0.39 |
| 1 | 20623631 | rs12080654 | 0.582 | 0.782 | 0.468 | -1.005 | 0.034 | -0.056 | -0.056 | 0 | 0.1 | 0.35 | 0.07 | 0 | 0.1 | 0.08 |
| 1 | 20624063 | rs10916754 | -0.464 | 0.599 | 0.371 | -1.149 | 0.007 | -0.022 | -0.022 | 0.014 | 0.25 | 0.19 | 0.14 | 0.57 | 0.19 | 0.24 |
| 1 | 20625054 | rs10753494 | -2.353 | 0.615 | 0.339 | -1.144 | -0.013 | 0.111 | 0.111 | 0 | 0.58 | 0.75 | 0.57 | 0.93 | 0.58 | 0.62 |
| 1 | 20630834 | rs10916755 | -1.707 | 0.745 | 0.174 | -1.074 | 0.002 | -0.009 | -0.009 | 0 | 0.43 | 0.65 | 0.37 | 0.56 | 0.42 | 0.39 |
| 1 | 20632407 | rs4654849 | -0.153 | 0.739 | 0.109 | -1.064 | -0.012 | -0.094 | -0.094 | 0 | 0.23 | 0.07 | 0.21 | 0.56 | 0.23 | 0.25 |
| 1 | 20633829 | rs6676380 | -1.727 | 0.81 | 0.292 | -1.163 | -0.003 | -0.029 | -0.029 | 0 | 0.47 | 0.4 | 0.58 | 0.93 | 0.53 | 0.55 |
| 1 | 20635722 | rs11809335 | -0.379 | 0.67 | 0.3 | -1.18 | -0.001 | -0.118 | -0.118 | 0 | 0.19 | 0.6 | 0.17 | 0 | 0.2 | 0.14 |
| 1 | 20636038 | rs12749098 | -0.568 | 0.753 | 0.246 | -1.155 | -0.001 | -0.013 | -0.013 | 0 | 0.24 | 0.2 | 0.32 | 0.38 | 0.26 | 0.27 |
| 1 | 20636232 | rs12095205 | 0.614 | 0.785 | 0.327 | -1.165 | 0.029 | -0.077 | -0.077 | 0 | 0.12 | 0.37 | 0.09 | 0 | 0.12 | 0.09 |
| 1 | 20637374 | rs10916761 | -2.504 | 0.796 | 0.104 | -1.109 | -0.059 | 0.266 | 0.266 | 0 | 0.61 | 0.93 | 0.69 | 0.93 | 0.65 | 0.64 |
| 1 | 20644752 | rs10916762 | -0.269 | 0.833 | 0.548 | -0.496 | 0.01 | 0.043 | 0.043 | 0.502 | 0.17 | 0.21 | 0.3 | 0.37 | 0.25 | 0.26 |
| 1 | 20648985 | rs2064073 | -1.97 | 0.751 | 0.615 | -0.511 | -0.002 | -0.009 | -0.009 | 0.391 | 0.41 | 0.63 | 0.55 | 0.91 | 0.43 | 0.48 |
| 1 | 20650550 | rs17353088 | -0.6 | 0.702 | 0.624 | -0.498 | -0.003 | -0.005 | -0.005 | 0 | 0.08 | 0.07 | 0.13 | 0.01 | 0.14 | 0.04 |
| 1 | 20652798 | rs4590651 | -0.197 | 0.695 | 0.762 | -0.442 | 0.024 | 0.166 | 0.166 | 0.056 | 0.17 | 0.39 | 0.22 | 0.36 | 0.17 | 0.23 |
| 1 | 20656824 | rs2072752 | 0.549 | 0.717 | 0.486 | -0.45 | -0.053 | -0.091 | -0.091 | 0 | 0.12 | 0.01 | 0.19 | 0.36 | 0.17 | 0.23 |
| 1 | 20657352 | rs2072753 | -1.992 | 0.725 | 0.579 | -0.485 | -0.001 | -0.025 | -0.025 | 0.907 | 0.41 | 0.62 | 0.56 | 0.91 | 0.43 | 0.48 |
| 1 | 20662938 | rs10916769 | 0.052 | 0.779 | 0.31 | -0.366 | 0 | 0.024 | 0.024 | 0 | 0.15 | 0.16 | 0.2 | 0.36 | 0.17 | 0.23 |
| 1 | 20671961 | rs11582960 | -1.076 | 0.84 | 0.264 | -0.362 | 0.002 | 0.009 | 0.009 | 0 | 0.23 | 0.24 | 0.35 | 0.35 | 0.32 | 0.28 |
| 1 | 20680718 | rs2072751 | -0.846 | 0.759 | 0.221 | -0.143 | 0 | 0.006 | 0.006 | 0 | 0.2 | 0.21 | 0.18 | 0.29 | 0.22 | 0.24 |
| 1 | 20681571 | rs12034781 | 0.396 | 0.539 | 0.126 | -0.126 | -0.001 | -0.015 | -0.015 | 0 | 0.14 | 0.1 | 0.26 | 0.59 | 0.12 | 0.21 |
| 1 | 20683946 | rs230177 | -1.353 | 0.543 | 0.269 | -0.482 | 0.002 | -0.136 | -0.136 | 1.08 | 0.25 | 0.58 | 0.24 | 0.01 | 0.28 | 0.16 |
| 1 | 20684705 | rs230175 | -2.005 | 0.501 | 0.226 | 0.125 | 0.004 | 0 | 0 | 0 | 0.39 | 0.75 | 0.44 | 0.43 | 0.39 | 0.35 |
| 1 | 20687285 | rs12045792 | 0.517 | 0.561 | 0.227 | 0.188 | -0.002 | -0.045 | -0.045 | 0.551 | 0.14 | 0.06 | 0.18 | 0.42 | 0.12 | 0.19 |
| 1 | 20695831 | rs1886740 | 0.274 | 0.589 | 0.36 | 0.175 | 0.026 | -0.122 | -0.122 | 0 | 0.82 | 0.44 | 0.8 | 0.99 | 0.79 | 0.87 |
| 1 | 20701277 | rs10159448 | -0.161 | 0.437 | -0.028 | 0.124 | 0.003 | 0.034 | 0.034 | 0 | 0.2 | 0.26 | 0.25 | 0.35 | 0.19 | 0.22 |
| 1 | 20703026 | rs10916777 | -0.202 | 0.468 | -0.088 | 0.129 | 0.002 | 0.022 | 0.022 | 0 | 0.2 | 0.27 | 0.23 | 0.34 | 0.19 | 0.21 |
| 1 | 20705117 | rs10916779 | -0.344 | 0.101 | -0.221 | 0.135 | -0.029 | -0.106 | -0.106 | 0.515 | 0.18 | 0.02 | 0.22 | 0.42 | 0.18 | 0.22 |
| 1 | 20707153 | rs10916780 | -0.502 | 0.116 | -0.192 | 0.097 | -0.007 | -0.064 | -0.064 | 0 | 0.19 | 0.06 | 0.22 | 0.42 | 0.19 | 0.22 |
| 1 | 20717539 | rs7538084 | 0.241 | -0.155 | 0.171 | -0.352 | -0.008 | -0.092 | -0.092 | 0.114 | 0.17 | 0.3 | 0.1 | 0 | 0.14 | 0.1 |
| 1 | 20717657 | rs12045307 | -0.091 | -0.036 | 0.041 | -0.01 | -0.001 | 0 | 0 | 0 | 0.13 | 0.11 | 0.18 | 0.29 | 0.15 | 0.21 |
| 1 | 20729212 | rs879484 | 0.315 | 0.093 | -0.023 | 0.005 | -0.028 | -0.039 | -0.039 | 0 | 0.1 | 0.03 | 0.16 | 0.26 | 0.15 | 0.21 |
| 1 | 20732082 | rs10916786 | -1.079 | 0.022 | -0.035 | -0.329 | 0.001 | -0.006 | -0.006 | 0 | 0.39 | 0.44 | 0.52 | 0.75 | 0.4 | 0.44 |
| 1 | 20732420 | rs7523815 | -1.395 | 0.061 | -0.451 | -0.183 | 0.004 | 0.068 | 0.068 | 0.55 | 0.57 | 0.77 | 0.62 | 0.75 | 0.55 | 0.53 |
| 1 | 20733376 | rs10753498 | -1.256 | 0.084 | -0.417 | -0.161 | 0.003 | 0.015 | 0.015 | 0.553 | 0.56 | 0.63 | 0.61 | 0.76 | 0.56 | 0.53 |
| 1 | 20735361 | rs7533344 | 1.48 | 0.035 | -0.632 | -0.168 | 0.017 | 0.074 | 0.074 | 0.045 | 0.4 | 0.2 | 0.3 | 0.23 | 0.37 | 0.41 |
| 1 | 20735452 | rs6426635 | 0.667 | 0.361 | -0.257 | 0.186 | 0.012 | -0.018 | -0.018 | 0 | 0.66 | 0.49 | 0.73 | 0.73 | 0.63 | 0.65 |
| 1 | 20736532 | rs10916790 | 1.009 | 1.141 | 0.178 | 0.78 | 0.023 | 0.029 | 0.029 | 0 | 0.83 | 0.69 | 0.73 | 0.73 | 0.74 | 0.69 |
| 1 | 20740016 | rs10799633 | -0.711 | 1.065 | -0.009 | 0.654 | -0.002 | -0.014 | -0.014 | 0.019 | 0.29 | 0.22 | 0.41 | 0.52 | 0.26 | 0.29 |
| 1 | 20746568 | rs764151 | -0.202 | 1.34 | -0.093 | 0.63 | 0.023 | -0.012 | -0.012 | 0 | 0.94 | 0.85 | 0.8 | 0.94 | 0.87 | 0.88 |
| 1 | 20750561 | rs7537381 | 0.88 | 1.241 | -0.169 | 0.755 | -0.001 | 0.001 | 0.001 | 0 | 0.55 | 0.46 | 0.31 | 0.23 | 0.49 | 0.4 |
| 1 | 20758192 | rs10753501 | -0.618 | 1.273 | -0.096 | 1.307 | 0.065 | 0.219 | 0.219 | 0 | 0.18 | 0.44 | 0.31 | 0.45 | 0.22 | 0.2 |
| 1 | 20762133 | rs7513147 | 0.32 | 0.733 | -0.712 | 0.637 | -0.001 | -0.003 | -0.003 | 0.331 | 0.52 | 0.51 | 0.36 | 0.26 | 0.45 | 0.38 |
| 1 | 20766279 | rs12044712 | -1.31 | 0.428 | -0.829 | 1.008 | 0.006 | -0.008 | -0.008 | 0.261 | 0.14 | 0.12 | 0.25 | 0.25 | 0.07 | 0.33 |
| 1 | 20767524 | rs10916799 | -1.518 | 0.379 | -0.262 | 1.397 | -0.007 | 0.019 | 0.019 | 0.255 | 0.33 | 0.58 | 0.42 | 0.57 | 0.27 | 0.51 |
| 1 | 20769184 | rs750070 | -1.822 | 0.165 | 0.684 | 1.447 | -0.008 | 0.01 | 0.01 | 0.979 | 0.35 | 0.55 | 0.41 | 0.61 | 0.28 | 0.48 |
| 1 | 20770061 | rs11577932 | -1.523 | 0.151 | 0.396 | 1.315 | -0.001 | 0 | 0 | 0.083 | 0.41 | 0.63 | 0.3 | 0.39 | 0.4 | 0.28 |
| 1 | 20770887 | rs10799636 | -1.921 | 0.457 | 0.39 | 1.342 | 0 | 0.004 | 0.004 | 0 | 0.45 | 0.66 | 0.51 | 0.6 | 0.43 | 0.5 |
| 1 | 20772487 | rs721360 | -1.446 | 0.365 | 0.492 | 1.403 | -0.008 | 0.005 | 0.005 | 4.064 | 0.14 | 0.12 | 0.23 | 0.2 | 0.08 | 0.27 |
| 1 | 20772817 | rs7414922 | -1.941 | -0.047 | 0.35 | 1.022 | 0.007 | 0.028 | 0.028 | 0 | 0.5 | 0.67 | 0.53 | 0.61 | 0.47 | 0.53 |
| 1 | 20773115 | rs10916800 | -0.445 | 0.245 | 0.27 | 1.039 | -0.017 | -0.02 | -0.02 | 0 | 0.22 | 0.27 | 0.21 | 0.42 | 0.34 | 0.22 |
| 1 | 20773962 | rs4654861 | -0.387 | 0.085 | 0.185 | 0.79 | 0.005 | 0.01 | 0.01 | 0.231 | 0.11 | 0.07 | 0.05 | 0.04 | 0.08 | 0.07 |
| 1 | 20775350 | rs11799849 | -1.805 | 0.01 | 0.268 | 1.057 | 0 | -0.001 | -0.001 | 0 | 0.45 | 0.56 | 0.51 | 0.54 | 0.45 | 0.5 |
| 1 | 20779003 | rs11812029 | 0.261 | 0.667 | -0.17 | 1.099 | 0.017 | -0.023 | -0.023 | 1.459 | 0.12 | 0.1 | 0.28 | 0.21 | 0.07 | 0.22 |
| 1 | 20785310 | rs7541314 | -0.25 | 0.714 | -0.062 | 0.984 | 0.001 | -0.002 | -0.002 | 0 | 0.15 | 0.21 | 0.09 | 0.24 | 0.1 | 0.18 |
| 1 | 20792099 | rs618041 | 0.284 | 0.744 | 0.573 | 1.086 | 0.023 | 0.145 | 0.145 | 0.076 | 0.06 | 0.22 | 0.06 | 0.21 | 0.06 | 0.13 |
| 1 | 20793015 | rs6663450 | -1.192 | 0.683 | 0.885 | 0.697 | 0.002 | 0 | 0 | 0 | 0.36 | 0.32 | 0.47 | 0.59 | 0.4 | 0.47 |
| 1 | 20798595 | rs12741740 | -0.949 | 0.756 | 0.722 | 0.776 | -0.003 | 0.008 | 0.008 | 0 | 0.32 | 0.36 | 0.48 | 0.54 | 0.31 | 0.4 |
| 1 | 20801589 | rs576436 | -1.954 | 0.254 | 0.193 | 0.631 | 0.012 | 0.148 | 0.148 | 0 | 0.49 | 0.84 | 0.62 | 0.69 | 0.45 | 0.59 |
| 1 | 20803771 | rs433816 | 1.252 | 0.046 | 0.05 | 0.067 | 0.004 | 0.009 | 0.009 | 0 | 0.43 | 0.37 | 0.28 | 0.31 | 0.33 | 0.4 |
| 1 | 20804065 | rs97922 | -0.209 | 0.257 | 0.192 | 0.434 | 0.008 | -0.016 | -0.016 | 0 | 0.75 | 0.69 | 0.8 | 0.87 | 0.68 | 0.81 |
| 1 | 20813488 | rs1253918 | 0.48 | 0.576 | 0.031 | 0.766 | 0.007 | 0.034 | 0.034 | 2.395 | 0.79 | 0.7 | 0.55 | 0.43 | 0.76 | 0.58 |
| 1 | 20814522 | rs1152984 | -0.151 | 0.59 | 0.03 | 0.692 | -0.005 | -0.001 | -0.001 | 0 | 0.83 | 0.81 | 0.6 | 0.54 | 0.8 | 0.65 |
| 1 | 20818935 | rs1152986 | 1.032 | 0.383 | -0.338 | 0.466 | -0.002 | 0.001 | 0.001 | 0.689 | 0.64 | 0.64 | 0.48 | 0.4 | 0.63 | 0.45 |
| 1 | 20821965 | rs12076947 | -0.301 | 0.316 | -0.503 | 0.311 | 0.001 | 0.002 | 0.002 | 0 | 0.2 | 0.17 | 0.12 | 0.14 | 0.18 | 0.19 |
| 1 | 20826910 | rs529974 | -0.458 | 0.371 | -0.72 | -0.058 | -0.008 | 0.069 | 0.069 | 0 | 0.35 | 0.24 | 0.2 | 0.09 | 0.36 | 0.2 |
| 1 | 20827031 | rs531651 | 0.388 | 0.364 | -0.743 | -0.088 | -0.021 | 0.029 | 0.029 | 0 | 0.24 | 0.19 | 0.17 | 0.06 | 0.32 | 0.12 |
| 1 | 20827216 | rs595980 | 1.656 | 0.416 | -1.086 | -0.085 | -0.052 | 0.093 | 0.093 | 0.059 | 0.16 | 0.03 | 0.1 | 0 | 0.2 | 0.05 |

| | | | | | | | | | | | | | | | | |
|---|---|---|---|---|---|---|---|---|---|---|---|---|---|---|---|---|
| 1 | 20830127 | rs17412284 | 1.124 | 0.382 | -1.009 | 0.012 | -0.035 | -0.023 | -0.023 | 0.051 | 0.06 | 0 | 0.05 | 0.05 | 0.12 | 0.07 |
| 1 | 20832674 | rs12566937 | -0.292 | 0.332 | -0.942 | 0.02 | 0.007 | 0.013 | 0.013 | 0 | 0.2 | 0.15 | 0.12 | 0.14 | 0.17 | 0.19 |
| 1 | 20851834 | rs7544348 | -0.482 | 0.293 | -0.995 | 0.475 | 0.012 | -0.055 | -0.055 | 0 | 0.22 | 0.13 | 0.3 | 0.36 | 0.19 | 0.31 |
| 1 | 20853195 | rs476622 | 1.581 | 0.306 | -0.971 | 0.356 | -0.03 | 0.061 | 0.061 | 0.035 | 0.17 | 0.07 | 0.11 | 0 | 0.2 | 0.05 |
| 1 | 20855056 | rs10916806 | 0.495 | 0.302 | -0.904 | 0.525 | 0.018 | -0.021 | -0.021 | 0 | 0.15 | 0.13 | 0.25 | 0.31 | 0.07 | 0.24 |
| 1 | 20857630 | rs2000366 | 1.476 | 0.272 | -0.92 | -0.135 | -0.01 | 0.028 | 0.028 | 0 | 0.24 | 0.13 | 0.16 | 0.06 | 0.27 | 0.17 |
| 1 | 20859701 | rs12074436 | 1.59 | 0.59 | -0.649 | 0.306 | 0.027 | 0.025 | 0.025 | 0 | 0.11 | 0.05 | 0.03 | 0 | 0.05 | 0.06 |
| 1 | 20862106 | rs616145 | 1.319 | 0.735 | -0.303 | 0.494 | 0.006 | -0.002 | -0.002 | 0 | 0.48 | 0.38 | 0.48 | 0.55 | 0.44 | 0.54 |
| 1 | 20865879 | rs10493000 | 0.134 | 0.728 | -0.562 | 0.766 | 0.006 | -0.008 | -0.008 | 0 | 0.17 | 0.14 | 0.26 | 0.3 | 0.1 | 0.26 |
| 1 | 20866735 | rs645496 | 1.135 | 0.704 | -0.733 | -0.107 | -0.012 | 0.019 | 0.019 | 0 | 0.18 | 0.14 | 0.11 | 0 | 0.19 | 0.05 |
| 1 | 20872694 | rs667357 | -0.882 | 0.443 | -0.7 | 0.039 | 0.004 | 0.021 | 0.021 | 0 | 0.39 | 0.37 | 0.29 | 0.24 | 0.36 | 0.36 |
| 1 | 20872901 | rs666501 | -0.501 | 0.461 | -0.794 | 0.01 | -0.006 | 0.07 | 0.07 | 0.003 | 0.34 | 0.23 | 0.21 | 0.05 | 0.34 | 0.18 |
| 1 | 20877812 | rs12072199 | -0.728 | 0.401 | -0.453 | 0.42 | -0.011 | 0.008 | 0.008 | 0 | 0.22 | 0.28 | 0.18 | 0.24 | 0.19 | 0.31 |
| 1 | 20882523 | rs6689263 | -0.759 | 1.04 | -0.059 | 0.891 | 0.029 | 0.11 | 0.11 | 0 | 0.1 | 0.23 | 0.14 | 0.24 | 0.12 | 0.23 |
| 1 | 20883203 | rs12565883 | -0.838 | 1.252 | -0.287 | 0.796 | 0.03 | 0.068 | 0.068 | 0 | 0.23 | 0.35 | 0.24 | 0.36 | 0.29 | 0.34 |
| 1 | 20883270 | rs12046455 | 0.482 | 0.816 | -0.794 | 0.945 | -0.005 | -0.109 | -0.109 | 0.92 | 0.14 | 0.01 | 0.28 | 0.45 | 0.16 | 0.27 |
| 1 | 20883422 | rs675397 | -0.223 | 1.073 | -0.443 | 0.546 | -0.002 | 0.002 | 0.002 | 0 | 0.39 | 0.53 | 0.28 | 0.41 | 0.38 | 0.34 |
| 1 | 20883816 | rs677214 | 1.017 | 1.194 | 0.47 | 0.337 | 0.028 | -0.024 | -0.024 | 0 | 0.84 | 0.64 | 0.87 | 0.93 | 0.82 | 0.8 |
| 1 | 20884833 | rs529553 | -0.929 | 1.257 | 0.619 | 0.367 | 0.033 | -0.007 | -0.007 | 0 | 0.09 | 0.18 | 0.09 | 0.04 | 0.14 | 0.11 |
| 1 | 20893965 | rs472055 | -0.862 | 1.259 | 1.069 | -0.156 | 0.001 | -0.002 | -0.002 | 0 | 0.54 | 0.55 | 0.57 | 0.79 | 0.56 | 0.67 |
| 1 | 20895874 | rs10916815 | 1.24 | 1.368 | 0.91 | -0.31 | 0.004 | -0.008 | -0.008 | 0 | 0.09 | 0.21 | 0.1 | 0.08 | 0.08 | 0.08 |
| 1 | 20899016 | rs1890005 | 0.167 | 1.417 | 1.021 | 0.166 | 0 | -0.001 | -0.001 | 0 | 0.38 | 0.42 | 0.52 | 0.53 | 0.35 | 0.4 |
| 1 | 20900123 | rs639400 | -0.221 | 1.453 | 0.944 | 0.234 | 0.046 | -0.006 | -0.006 | 0.056 | 0.09 | 0.2 | 0.09 | 0.04 | 0.14 | 0.1 |
| 1 | 20904743 | rs1152991 | -0.008 | 1.461 | 1.157 | 0.31 | 0.005 | 0.006 | 0.006 | 0 | 0.6 | 0.49 | 0.5 | 0.51 | 0.54 | 0.59 |
| 1 | 20904858 | rs1253795 | 1.523 | 1.558 | 1.224 | -0.044 | 0.057 | 0.004 | 0.004 | 0 | 0.86 | 0.6 | 0.9 | 0.93 | 0.82 | 0.86 |
| 1 | 20905098 | rs622179 | -0.491 | 1.126 | 0.882 | -0.151 | 0.001 | -0.02 | -0.02 | 0 | 0.14 | 0.37 | 0.14 | 0.09 | 0.11 | 0.13 |
| 1 | 20907051 | rs640880 | -0.282 | 1.214 | 0.493 | -0.045 | 0.033 | -0.011 | -0.011 | 0 | 0.89 | 0.72 | 0.93 | 0.97 | 0.89 | 0.94 |
| 1 | 20907307 | rs652536 | -0.282 | 1.214 | 0.493 | -0.045 | 0.033 | -0.011 | -0.011 | 0 | 0.89 | 0.72 | 0.93 | 0.97 | 0.89 | 0.94 |
| 1 | 20907830 | rs485716 | -0.226 | 1.211 | 0.492 | -0.046 | 0.031 | -0.012 | -0.012 | 0.021 | 0.89 | 0.72 | 0.93 | 0.97 | 0.89 | 0.94 |
| 1 | 20910358 | rs485833 | -0.281 | 1.218 | 0.495 | -0.043 | 0.033 | -0.011 | -0.011 | 0 | 0.89 | 0.72 | 0.93 | 0.97 | 0.89 | 0.94 |
| 1 | 20911404 | rs678980 | -0.281 | 1.218 | 0.495 | -0.043 | 0.033 | -0.011 | -0.011 | 0.02 | 0.89 | 0.72 | 0.93 | 0.97 | 0.89 | 0.94 |
| 1 | 20911830 | rs650264 | -0.281 | 1.218 | 0.495 | -0.043 | 0.033 | -0.011 | -0.011 | 0.129 | 0.89 | 0.72 | 0.93 | 0.97 | 0.89 | 0.94 |
| 1 | 20914451 | rs6690069 | 0 | 1.254 | 0.041 | -0.069 | 0.002 | -0.007 | -0.007 | 0.014 | 0.11 | 0.12 | 0.06 | 0.03 | 0.12 | 0.08 |
| 1 | 20915172 | rs532545 | 1.363 | 1.164 | -0.579 | -0.326 | -0.014 | 0.136 | 0.136 | 0 | 0.34 | 0.06 | 0.31 | 0.12 | 0.3 | 0.22 |
| 1 | 20915441 | rs12726436 | 0.379 | 1.334 | -0.656 | -0.275 | -0.027 | -0.011 | -0.011 | 0 | 0.07 | 0 | 0.04 | 0.05 | 0.11 | 0.08 |
| 1 | 20915701 | rs2072671 | 1.399 | 1.47 | -0.717 | -0.359 | -0.054 | 0.144 | 0.144 | 0.1 | 0.34 | 0.07 | 0.31 | 0.12 | 0.31 | 0.23 |
| 1 | 20916791 | rs818202 | -0.396 | 1.269 | -0.264 | -0.283 | 0.001 | -0.01 | -0.01 | 0 | 0.62 | 0.41 | 0.5 | 0.27 | 0.61 | 0.46 |
| 1 | 20917206 | rs471760 | 1.063 | 1.102 | -1.098 | -0.256 | -0.01 | 0.163 | 0.163 | 0.248 | 0.35 | 0.08 | 0.3 | 0.12 | 0.3 | 0.2 |
| 1 | 20918912 | rs10916824 | 1.342 | 1.157 | -0.46 | 0.352 | -0.02 | -0.01 | -0.01 | 0 | 0.07 | 0 | 0.07 | 0.07 | 0.07 | 0.09 |
| 1 | 20924483 | rs818197 | 0.801 | 1.134 | 0.188 | 0.707 | -0.003 | 0.027 | 0.027 | 0 | 0.83 | 0.66 | 0.88 | 0.77 | 0.82 | 0.68 |
| 1 | 20928154 | rs577042 | 0.851 | 1.119 | 0.186 | 0.7 | -0.004 | 0.026 | 0.026 | 0 | 0.82 | 0.66 | 0.88 | 0.77 | 0.82 | 0.68 |
| 1 | 20928284 | rs4655226 | 0.583 | 1.419 | 0.284 | 0.785 | 0.026 | 0.032 | 0.032 | 0 | 0.06 | 0.14 | 0.18 | 0.16 | 0.11 | 0.12 |
| 1 | 20928916 | rs12022958 | 1.019 | 1.088 | -0.224 | 0.273 | -0.036 | 0.078 | 0.078 | 0 | 0.22 | 0.01 | 0.18 | 0.12 | 0.21 | 0.16 |
| 1 | 20930035 | rs517483 | 0.479 | 1.106 | -0.119 | 0.335 | -0.003 | 0.028 | 0.028 | 0 | 0.83 | 0.66 | 0.88 | 0.77 | 0.82 | 0.68 |
| 1 | 20930381 | rs10799647 | -0.258 | 1.015 | -0.04 | 0.173 | 0.008 | 0.042 | 0.042 | 0 | 0.34 | 0.21 | 0.22 | 0.15 | 0.31 | 0.21 |
| 1 | 20933278 | rs12036909 | -0.579 | 0.993 | 0.197 | 0.342 | 0.005 | 0.016 | 0.016 | 0 | 0.39 | 0.28 | 0.31 | 0.36 | 0.39 | 0.48 |
| 1 | 20933341 | rs10916827 | -0.579 | 0.993 | 0.197 | 0.342 | 0.005 | 0.016 | 0.016 | 0 | 0.39 | 0.28 | 0.31 | 0.36 | 0.39 | 0.48 |
| 1 | 20933596 | rs12036993 | -0.056 | 0.965 | 0.212 | 0.208 | 0.003 | 0.018 | 0.018 | 0 | 0.32 | 0.28 | 0.27 | 0.18 | 0.34 | 0.25 |
| 1 | 20933893 | rs580032 | 0.845 | 0.982 | 0.457 | 1.051 | -0.004 | 0.103 | 0.103 | 0.133 | 0.91 | 0.81 | 0.95 | 0.69 | 0.94 | 0.76 |
| 1 | 20934213 | rs11579252 | -0.057 | 1.129 | 0.464 | 0.766 | 0.006 | -0.017 | -0.017 | 0 | 0.12 | 0.22 | 0.12 | 0.05 | 0.15 | 0.1 |
| 1 | 20934283 | rs527912 | 1.565 | 1.231 | 0.62 | 0.998 | 0.009 | 0.026 | 0.026 | 0.02 | 0.4 | 0.27 | 0.43 | 0.29 | 0.38 | 0.29 |
| 1 | 20934796 | rs1689924 | 0.828 | 1.135 | 0.609 | 1.041 | -0.001 | -0.002 | -0.002 | 0 | 0.52 | 0.5 | 0.54 | 0.34 | 0.53 | 0.42 |
| 1 | 20935358 | rs558946 | 0.399 | 1.119 | 0.972 | 1.071 | -0.001 | -0.003 | -0.003 | 0 | 0.61 | 0.61 | 0.58 | 0.36 | 0.63 | 0.44 |
| 1 | 20936248 | rs477155 | 0.756 | 1.208 | 0.975 | 1.16 | 0.014 | 0.094 | 0.094 | 0.26 | 0.86 | 0.78 | 0.76 | 0.49 | 0.84 | 0.59 |
| 1 | 20940463 | rs3738130 | -0.489 | 1.162 | 0.979 | 1.047 | 0.014 | -0.022 | -0.022 | 0 | 0.2 | 0.32 | 0.14 | 0.07 | 0.22 | 0.15 |
| 1 | 20943139 | rs12072318 | -0.571 | 1.158 | 1.036 | 1.045 | 0.005 | -0.01 | -0.01 | 0 | 0.2 | 0.23 | 0.12 | 0.07 | 0.22 | 0.15 |
| 1 | 20943195 | rs12404655 | 0.982 | 1.1 | 1.042 | 0.969 | -0.003 | 0.089 | 0.089 | 0 | 0.24 | 0.07 | 0.18 | 0.13 | 0.22 | 0.16 |
| 1 | 20945644 | rs10916832 | 0.909 | 1.145 | 1.134 | 1.166 | -0.013 | 0.127 | 0.127 | 1.248 | 0.88 | 0.74 | 0.91 | 0.66 | 0.93 | 0.71 |
| 1 | 20945686 | rs10916833 | 0.909 | 1.145 | 1.134 | 1.166 | -0.013 | 0.127 | 0.127 | 1.248 | 0.88 | 0.74 | 0.91 | 0.66 | 0.93 | 0.71 |
| 1 | 20946756 | rs1614627 | 0.775 | 1.146 | 1.135 | 1.166 | -0.011 | 0.134 | 0.134 | 1.107 | 0.89 | 0.75 | 0.91 | 0.66 | 0.93 | 0.71 |
| 1 | 20951359 | rs6687370 | -0.551 | 1.19 | 1.027 | 1.064 | 0.005 | -0.01 | -0.01 | 0 | 0.2 | 0.24 | 0.12 | 0.07 | 0.22 | 0.15 |
| 1 | 20957431 | rs10799655 | 0.926 | 1.195 | 0.985 | 1.118 | 0.029 | 0.17 | 0.17 | 0.031 | 0.83 | 0.68 | 0.73 | 0.5 | 0.83 | 0.58 |
| 1 | 20959421 | rs10916840 | 1.624 | 1.206 | 0.968 | 1.088 | 0.001 | 0.006 | 0.006 | 0 | 0.41 | 0.36 | 0.44 | 0.29 | 0.41 | 0.29 |
| 1 | 20965442 | rs3738133 | -1.168 | 1.26 | 0.977 | 1.228 | -0.011 | -0.024 | -0.024 | 0 | 0.26 | 0.21 | 0.19 | 0.36 | 0.29 | 0.39 |
| 1 | 20968011 | rs3102071 | -0.149 | 1.256 | 1.015 | 1.238 | 0.049 | 0.077 | 0.077 | 0 | 0.9 | 0.74 | 0.8 | 0.81 | 0.9 | 0.87 |
| 1 | 20972111 | rs3738136 | 0.035 | 1.264 | 0.963 | 1.254 | -0.003 | -0.006 | -0.006 | 0 | 0.06 | 0.01 | 0.06 | 0.27 | 0.05 | 0.24 |
| 1 | 20977000 | rs1043424 | -1.185 | 1.283 | 1.005 | 1.243 | -0.008 | -0.019 | -0.019 | 0 | 0.27 | 0.25 | 0.19 | 0.36 | 0.29 | 0.4 |
| 1 | 20978058 | rs650616 | 1.596 | 1.209 | 1.056 | 1.1 | 0.011 | 0.001 | 0.001 | 0.454 | 0.64 | 0.42 | 0.6 | 0.42 | 0.6 | 0.43 |
| 1 | 20982151 | rs3738140 | -0.357 | 1.2 | 0.962 | 1.092 | -0.004 | 0.002 | 0.002 | 0 | 0.18 | 0.18 | 0.12 | 0.08 | 0.22 | 0.16 |

| | | | | | | | | | | | | | | | | |
|---|---|---|---|---|---|---|---|---|---|---|---|---|---|---|---|---|
| 1 | 20986078 | rs623817 | 1.529 | 1.155 | 0.908 | 1.034 | 0.005 | 0.019 | 0.019 | 0 | 0.41 | 0.32 | 0.44 | 0.29 | 0.41 | 0.28 |
| 1 | 20986557 | rs10916846 | 0.526 | 1.249 | 0.643 | 0.921 | 0 | 0.004 | 0.004 | 0 | 0.09 | 0.07 | 0.08 | 0.04 | 0.07 | 0.08 |
| 1 | 20988911 | rs3753834 | 0.554 | 1.279 | 0.648 | 0.929 | 0.001 | -0.001 | -0.001 | 0 | 0.1 | 0.09 | 0.04 | 0.04 | 0.15 | 0.08 |
| 1 | 20990905 | rs10799659 | -0.735 | 1.253 | 0.604 | 0.925 | -0.002 | 0.001 | 0.001 | 0.001 | 0.19 | 0.2 | 0.12 | 0.08 | 0.22 | 0.16 |
| 1 | 20993151 | rs617367 | 0.447 | 1.351 | 0.636 | 1.093 | 0.02 | 0.09 | 0.09 | 0 | 0.84 | 0.76 | 0.75 | 0.56 | 0.82 | 0.64 |
| 1 | 20993162 | rs3818392 | -0.815 | 1.328 | 0.499 | 0.864 | 0.001 | 0.019 | 0.019 | 0 | 0.42 | 0.39 | 0.31 | 0.25 | 0.41 | 0.36 |
| 1 | 20994952 | rs2170336 | 0.014 | 1.559 | 0.37 | 1.005 | 0.005 | -0.003 | -0.003 | 0.238 | 0.11 | 0.13 | 0.05 | 0.03 | 0.15 | 0.08 |
| 1 | 20996555 | rs1125090 | 0.658 | 1.381 | 0.457 | 1.397 | 0.003 | 0.239 | 0.239 | 0.207 | 0.95 | 0.72 | 0.96 | 0.79 | 0.99 | 0.88 |
| 1 | 20996757 | rs1570663 | -0.936 | 1.877 | -0.162 | 0.817 | -0.004 | -0.004 | -0.004 | 0 | 0.17 | 0.14 | 0.27 | 0.24 | 0.2 | 0.24 |
| 1 | 20997340 | rs10157351 | -0.925 | 1.584 | -0.648 | 0.353 | 0.011 | 0.003 | 0.003 | 0 | 0.51 | 0.44 | 0.67 | 0.54 | 0.57 | 0.48 |
| 1 | 20999804 | rs12118760 | 0.739 | 1.763 | -1.318 | -0.489 | -0.076 | 0.087 | 0.087 | 0 | 0.22 | 0.01 | 0.18 | 0.06 | 0.25 | 0.13 |
| 1 | 21002613 | rs10916848 | -1.021 | 1.648 | -1.212 | -0.418 | 0.034 | 0.058 | 0.058 | 0 | 0.54 | 0.25 | 0.51 | 0.42 | 0.6 | 0.49 |
| 1 | 21002726 | rs12025916 | 0.767 | 1.334 | -1.111 | -0.189 | 0.004 | 0.07 | 0.07 | 0 | 0.16 | 0.04 | 0.11 | 0.08 | 0.15 | 0.09 |
| 1 | 21008104 | rs4654866 | -0.279 | 1.281 | -0.985 | 0.038 | -0.003 | 0.001 | 0.001 | 0 | 0.3 | 0.23 | 0.29 | 0.34 | 0.3 | 0.33 |
| 1 | 21009586 | rs669410 | -0.49 | 1.009 | -1.092 | 0.115 | -0.002 | -0.002 | -0.002 | 0.02 | 0.54 | 0.44 | 0.51 | 0.42 | 0.61 | 0.47 |
| 1 | 21009800 | rs4655235 | -0.166 | 0.927 | -1.765 | -0.191 | -0.077 | 0.151 | 0.151 | 0 | 0.3 | 0.01 | 0.19 | 0.08 | 0.29 | 0.2 |
| 1 | 21010824 | rs581635 | -0.942 | 0.611 | -0.485 | 0.819 | -0.014 | 0.071 | 0.071 | 0 | 0.23 | 0.49 | 0.23 | 0.35 | 0.19 | 0.22 |
| 1 | 21011182 | rs2296223 | -1.379 | 0.539 | -0.192 | 1.246 | -0.004 | 0.154 | 0.154 | 0 | 0.07 | 0.15 | 0.11 | 0.33 | 0.04 | 0.09 |
| 1 | 21013546 | rs4143111 | -1.098 | 0.471 | -0.199 | 0.884 | -0.021 | 0.039 | 0.039 | 0 | 0.12 | 0.02 | 0.08 | 0.03 | 0.13 | 0.13 |
| 1 | 21015685 | rs10799660 | -0.678 | 1.296 | -0.204 | 1.444 | -0.005 | 0.015 | 0.015 | 0 | 0.12 | 0.1 | 0.27 | 0.37 | 0.18 | 0.28 |
| 1 | 21018485 | rs628265 | -2.059 | 0.687 | -0.087 | 0.568 | -0.009 | 0.108 | 0.108 | 0 | 0.47 | 0.73 | 0.46 | 0.78 | 0.42 | 0.58 |
| 1 | 21020128 | rs610719 | -0.764 | 0.698 | 0.097 | 0.699 | -0.001 | -0.008 | -0.008 | 0 | 0.21 | 0.26 | 0.23 | 0.09 | 0.18 | 0.15 |
| 1 | 21020868 | rs12728802 | -1.363 | 0.839 | 0.245 | 0.74 | 0.006 | -0.017 | -0.017 | 0.491 | 0.06 | 0.09 | 0.05 | 0 | 0.1 | 0.07 |
| 1 | 21022402 | rs16824489 | -1.77 | 0.744 | 0.257 | 0.751 | -0.011 | -0.019 | -0.019 | 0 | 0.37 | 0.53 | 0.26 | 0.7 | 0.27 | 0.48 |
| 1 | 21022653 | rs683936 | -0.082 | 0.754 | 0.247 | 0.757 | 0.013 | 0.017 | 0.017 | 0 | 0.05 | 0.17 | 0.15 | 0.08 | 0.06 | 0.03 |
| 1 | 21024460 | rs16824496 | -1.519 | 0.592 | 0.283 | 1.231 | -0.022 | 0.013 | 0.013 | 0 | 0.32 | 0.53 | 0.23 | 0.33 | 0.22 | 0.4 |
| 1 | 21024743 | rs521424 | -1.958 | 0.591 | 0.14 | 1.173 | 0.002 | -0.009 | -0.009 | 0 | 0.38 | 0.69 | 0.4 | 0.41 | 0.33 | 0.46 |
| 1 | 21031042 | rs2296225 | -1.179 | 0.749 | 0.036 | 1.486 | -0.002 | 0.018 | 0.018 | 0 | 0.09 | 0.1 | 0.06 | 0.29 | 0.09 | 0.26 |
| 1 | 21032041 | rs752555 | 0.682 | 0.269 | 0.021 | 0.847 | 0.01 | 0.014 | 0.014 | 0 | 0.64 | 0.39 | 0.55 | 0.78 | 0.74 | 0.67 |
| 1 | 21035149 | rs6426645 | 2.2 | 0.534 | -0.289 | 0.622 | -0.12 | 0.262 | 0.262 | 0 | 0.4 | 0.04 | 0.33 | 0.05 | 0.48 | 0.29 |
| 1 | 21035578 | rs12042868 | -2.04 | 0.464 | 0.335 | 0.964 | 0.066 | 0.297 | 0.297 | 0 | 0.14 | 0.42 | 0.25 | 0.5 | 0.17 | 0.2 |
| 1 | 21040151 | rs10916851 | -1.552 | 0.369 | 0.639 | 0.934 | 0.002 | 0.167 | 0.167 | 0.001 | 0.16 | 0.39 | 0.18 | 0.46 | 0.15 | 0.15 |
| 1 | 21045508 | rs6668854 | -0.825 | 0.57 | 0.501 | 0.956 | -0.011 | -0.01 | -0.01 | 0 | 0.09 | 0.06 | 0.15 | 0.71 | 0.13 | 0.3 |
| 1 | 21047404 | rs3738144 | -2.15 | 0.458 | 1.124 | 0.869 | 0.059 | 0.039 | 0.039 | 2.932 | 0.09 | 0.35 | 0.11 | 0.1 | 0.07 | 0.12 |
| 1 | 21047616 | rs637990 | 2.22 | 0.519 | 0.645 | 0.286 | -0.059 | 0.2 | 0.2 | 0 | 0.38 | 0.1 | 0.31 | 0.08 | 0.42 | 0.29 |
| 1 | 21047725 | rs659046 | 0.189 | 0.534 | 0.775 | 0.275 | 0.014 | 0.027 | 0.027 | 0 | 0.62 | 0.7 | 0.63 | 0.89 | 0.68 | 0.75 |
| 1 | 21050958 | rs592060 | -0.989 | 0.335 | 0.533 | 0.117 | -0.058 | 0.139 | 0.139 | 0 | 0.78 | 0.98 | 0.76 | 0.98 | 0.77 | 0.83 |
| 1 | 21052693 | rs10916852 | -1.645 | 0.421 | 0.781 | 0.277 | 0.066 | 0.039 | 0.039 | 2.071 | 0.09 | 0.34 | 0.11 | 0.1 | 0.07 | 0.1 |
| 1 | 21060356 | rs951805 | 2.195 | 0.482 | 0.749 | 0.194 | 0.002 | 0.093 | 0.093 | 0 | 0.37 | 0.22 | 0.32 | 0.04 | 0.39 | 0.25 |
| 1 | 21062001 | rs710311 | 1.515 | 0.524 | 0.758 | 0.236 | 0 | -0.013 | -0.013 | 0 | 0.44 | 0.56 | 0.4 | 0.17 | 0.42 | 0.35 |
| 1 | 21067001 | rs12123092 | 1.453 | 0.553 | 0.86 | 0.223 | 0 | -0.012 | -0.012 | 0 | 0.46 | 0.47 | 0.42 | 0.17 | 0.46 | 0.36 |
| 1 | 21071967 | rs2274119 | 0.579 | 0.598 | 0.718 | 0.517 | -0.032 | 0.001 | 0.001 | 0.87 | 0.06 | 0.01 | 0.05 | 0.29 | 0.09 | 0.26 |
| 1 | 21087927 | rs7552264 | -0.738 | 0.557 | 0.819 | 0.332 | 0.001 | 0.004 | 0.004 | 0 | 0.11 | 0.12 | 0.18 | 0.72 | 0.17 | 0.35 |
| 1 | 21091861 | rs1985278 | -0.498 | 0.496 | 0.846 | 0.29 | 0.007 | 0.088 | 0.088 | 0 | 0.4 | 0.22 | 0.39 | 0.11 | 0.36 | 0.29 |
| 1 | 21105994 | rs495522 | 0.318 | 0.499 | 0.852 | 0.286 | 0.007 | 0.037 | 0.037 | 0 | 0.59 | 0.69 | 0.6 | 0.88 | 0.64 | 0.71 |
| 1 | 21114833 | rs3856173 | 1.546 | 0.333 | 0.951 | 0.294 | -0.002 | -0.022 | -0.022 | 0.61 | 0.42 | 0.44 | 0.39 | 0.17 | 0.42 | 0.35 |
| 1 | 21140385 | rs7520481 | -1.658 | 0.372 | 1.025 | 0.321 | -0.004 | -0.039 | -0.039 | 0.076 | 0.58 | 0.52 | 0.6 | 0.83 | 0.59 | 0.64 |
| 1 | 21145660 | rs935918 | 1.921 | 0.314 | 1.114 | 0.306 | 0.005 | 0.104 | 0.104 | 0 | 0.38 | 0.23 | 0.33 | 0.04 | 0.38 | 0.26 |
| 1 | 21152640 | rs10799665 | -1.504 | 0.205 | 0.777 | 0.035 | -0.006 | -0.034 | -0.034 | 0 | 0.56 | 0.52 | 0.64 | 0.83 | 0.64 | 0.64 |
| 1 | 21163308 | rs3767239 | -0.032 | 0.343 | 0.952 | 0.523 | -0.007 | 0.026 | 0.026 | 0 | 0.08 | 0.08 | 0.07 | 0.3 | 0.13 | 0.28 |
| 1 | 21180181 | rs2305463 | -1.071 | 0.311 | 0.98 | 0.308 | 0.008 | 0.051 | 0.051 | 0 | 0.52 | 0.37 | 0.57 | 0.83 | 0.55 | 0.61 |
| 1 | 21203898 | rs4654726 | -1.471 | 0.245 | 0.972 | 0.28 | -0.004 | -0.041 | -0.041 | 3.172 | 0.58 | 0.52 | 0.6 | 0.83 | 0.59 | 0.63 |
| 1 | 21224253 | rs4654729 | -1.045 | 0.198 | 1.106 | 0.25 | 0.002 | 0.002 | 0.002 | 0 | 0.52 | 0.4 | 0.57 | 0.83 | 0.55 | 0.61 |
| 1 | 21227083 | rs2053981 | 1.096 | 0.203 | 0.98 | 0.252 | -0.008 | 0.124 | 0.124 | 0 | 0.44 | 0.28 | 0.39 | 0.17 | 0.45 | 0.38 |
| 1 | 21234923 | rs4654887 | 1.296 | 0.208 | 0.979 | 0.25 | 0.003 | 0.013 | 0.013 | 0 | 0.42 | 0.36 | 0.39 | 0.17 | 0.41 | 0.37 |
| 1 | 21276854 | rs3767248 | 1.612 | 0.07 | 0.925 | 0.461 | -0.015 | 0.064 | 0.064 | 0 | 0.38 | 0.22 | 0.38 | 0.21 | 0.43 | 0.35 |
| 1 | 21306161 | rs12021529 | -0.463 | 0.058 | 1.006 | 0.499 | -0.001 | -0.026 | -0.026 | 0 | 0.44 | 0.22 | 0.5 | 0.53 | 0.42 | 0.35 |
| 1 | 21317524 | rs10916919 | 1.952 | 0.067 | 0.957 | 0.483 | -0.018 | 0.118 | 0.118 | 0 | 0.36 | 0.19 | 0.33 | 0.07 | 0.4 | 0.26 |
| 1 | 21324491 | rs2874367 | -1.041 | 0.098 | 0.982 | 0.512 | 0.003 | 0.055 | 0.055 | 0 | 0.52 | 0.35 | 0.56 | 0.77 | 0.53 | 0.62 |
| 1 | 21324933 | rs1110240 | 0.041 | 0.085 | 0.942 | 0.522 | -0.008 | -0.007 | -0.007 | 0.374 | 0.08 | 0.06 | 0.06 | 0.24 | 0.11 | 0.27 |
| 1 | 21332812 | rs12130664 | 1.806 | 0.079 | 0.903 | 0.51 | -0.01 | 0.102 | 0.102 | 0 | 0.37 | 0.22 | 0.33 | 0.09 | 0.4 | 0.28 |
| 1 | 21337313 | rs4654732 | 0.748 | -0.012 | 0.913 | 0.456 | -0.077 | -0.114 | -0.114 | 0.082 | 0.27 | 0.04 | 0.38 | 0.43 | 0.33 | 0.26 |
| 1 | 21339644 | rs6681064 | 0.831 | 0.364 | 1.26 | 0.661 | 0.002 | 0.016 | 0.016 | 0 | 0.43 | 0.39 | 0.39 | 0.23 | 0.43 | 0.37 |
| 1 | 21395133 | rs12567861 | -0.191 | 0.418 | 1.071 | 0.63 | 0.004 | -0.017 | -0.017 | 0 | 0.45 | 0.25 | 0.5 | 0.53 | 0.42 | 0.35 |
| 1 | 21525228 | rs1609558 | 0.423 | 0.374 | 0.739 | 0.318 | -0.001 | 0.02 | 0.02 | 0 | 0.46 | 0.43 | 0.41 | 0.22 | 0.47 | 0.38 |
| 1 | 21534055 | rs12128470 | 0.827 | 0.446 | 0.395 | -0.121 | -0.024 | 0.131 | 0.131 | 0 | 0.4 | 0.23 | 0.35 | 0.08 | 0.44 | 0.28 |
| 1 | 21534237 | rs12131134 | 0.827 | 0.446 | 0.395 | -0.121 | -0.024 | 0.131 | 0.131 | 0 | 0.4 | 0.23 | 0.35 | 0.08 | 0.44 | 0.28 |
| 1 | 21541735 | rs7539288 | -1.454 | 0.251 | 1.019 | 0.257 | -0.025 | -0.079 | -0.079 | 0 | 0.12 | 0.5 | 0.06 | 0 | 0.04 | 0.03 |
| 1 | 21544601 | rs3026913 | -2.262 | 0.301 | 1.1 | 0.56 | -0.018 | 0.037 | 0.037 | 2.404 | 0.19 | 0.6 | 0.14 | 0.22 | 0.16 | 0.28 |

| | | | | | | | | | | | | | | | | |
|---|---|---|---|---|---|---|---|---|---|---|---|---|---|---|---|---|
| 1 | 21544735 | rs3026912 | -0.307 | 0.346 | 0.919 | 0.585 | 0.005 | 0.046 | 0.046 | 1.557 | 0.07 | 0.08 | 0.07 | 0.23 | 0.11 | 0.26 |
| 1 | 21551457 | rs2038089 | -0.89 | 0.391 | 0.373 | 0.085 | -0.007 | 0.02 | 0.02 | 0 | 0.41 | 0.72 | 0.38 | 0.31 | 0.45 | 0.48 |
| 1 | 21551550 | rs2038090 | 1.202 | 0.605 | 0.114 | -0.192 | -0.029 | 0.045 | 0.045 | 0 | 0.21 | 0.14 | 0.24 | 0.08 | 0.29 | 0.2 |
| 1 | 21570426 | rs212508 | -0.847 | 0.31 | 0.674 | 0.176 | 0.024 | 0.021 | 0.021 | 6.132 | 0.06 | 0.3 | 0.02 | 0.01 | 0 | 0.01 |
| 1 | 21571601 | rs12562197 | 0.943 | 0.415 | -0.304 | -0.025 | -0.057 | 0.027 | 0.027 | 0 | 0.22 | 0.03 | 0.28 | 0.13 | 0.27 | 0.18 |
| 1 | 21575045 | rs212515 | -0.723 | 0.634 | -0.528 | 0.183 | -0.001 | 0.014 | 0.014 | 0 | 0.43 | 0.36 | 0.42 | 0.32 | 0.44 | 0.37 |
| 1 | 21575456 | rs212516 | -0.837 | 0.651 | -0.534 | 0.19 | 0.008 | 0.038 | 0.038 | 2.15 | 0.14 | 0.33 | 0.12 | 0.19 | 0.13 | 0.18 |
| 1 | 21576796 | rs9426750 | 0.732 | 0.748 | -1.097 | 0.239 | -0.025 | -0.03 | -0.03 | 0 | 0.08 | 0.03 | 0.09 | 0.19 | 0.13 | 0.16 |
| 1 | 21581021 | rs3026886 | 1.41 | 0.708 | -1.153 | 0.215 | -0.034 | 0.014 | 0.014 | 0 | 0.21 | 0.02 | 0.27 | 0.13 | 0.24 | 0.15 |
| 1 | 21581979 | rs3026883 | 0.491 | 0.744 | -1.159 | 0.205 | -0.056 | 0.068 | 0.068 | 0.631 | 0.29 | 0.02 | 0.3 | 0.13 | 0.32 | 0.19 |
| 1 | 21582425 | rs212522 | 0.657 | 0.843 | -1.012 | 0.102 | -0.026 | -0.03 | -0.03 | 0 | 0.08 | 0.03 | 0.1 | 0.18 | 0.13 | 0.16 |
| 1 | 21583311 | rs212524 | -1.45 | 0.505 | -1.238 | 0.214 | -0.037 | 0.141 | 0.141 | 0 | 0.59 | 0.86 | 0.67 | 0.77 | 0.61 | 0.68 |
| 1 | 21584941 | rs212526 | -1.471 | 0.519 | -1.27 | -0.106 | -0.042 | 0.216 | 0.216 | 0 | 0.59 | 0.87 | 0.68 | 0.9 | 0.61 | 0.69 |
| 1 | 21585014 | rs212527 | 0.751 | 0.675 | -1.071 | 0.143 | 0.004 | 0.013 | 0.013 | 1.597 | 0.09 | 0.09 | 0.1 | 0.18 | 0.12 | 0.17 |
| 1 | 21587849 | rs2282714 | 1.154 | 0.646 | -1.005 | 0.283 | -0.012 | -0.053 | -0.053 | 0 | 0.26 | 0.16 | 0.43 | 0.42 | 0.31 | 0.36 |
| 1 | 21592392 | rs212539 | -1.234 | 0.709 | -0.989 | -0.751 | -0.032 | 0.159 | 0.159 | 0 | 0.62 | 0.86 | 0.68 | 0.91 | 0.61 | 0.73 |
| 1 | 21593117 | rs212540 | -1.105 | 0.696 | -0.978 | -0.742 | -0.032 | 0.162 | 0.162 | 0 | 0.61 | 0.86 | 0.68 | 0.9 | 0.6 | 0.72 |
| 1 | 21593773 | rs212541 | -0.012 | 0.045 | -0.855 | -1.104 | 0.026 | 0.138 | 0.138 | 0 | 0.45 | 0.74 | 0.58 | 0.9 | 0.41 | 0.59 |
| 1 | 21598377 | rs212549 | 0.538 | -0.082 | 0.036 | -0.232 | -0.008 | 0.083 | 0.083 | 0.445 | 0.06 | 0.04 | 0.13 | 0.45 | 0.09 | 0.19 |
| 1 | 21600677 | rs4654916 | 0.078 | -0.401 | 0.425 | -0.612 | -0.024 | -0.07 | -0.07 | 0.022 | 0.13 | 0.27 | 0.09 | 0.03 | 0.1 | 0.09 |
| 1 | 21621536 | rs213051 | -2.123 | -0.462 | -0.112 | -0.626 | 0.031 | -0.028 | -0.028 | 0.001 | 0.38 | 0.76 | 0.31 | 0.47 | 0.29 | 0.37 |
| 1 | 21625145 | rs10916958 | 0.186 | -0.381 | -0.301 | -1.396 | -0.001 | -0.017 | -0.017 | 0 | 0.19 | 0.16 | 0.17 | 0.01 | 0.24 | 0.15 |
| 1 | 21627370 | rs213057 | 0.829 | -0.389 | -0.296 | -1.029 | 0.007 | 0.127 | 0.127 | 0.443 | 0.06 | 0.07 | 0.14 | 0.4 | 0.08 | 0.23 |
| 1 | 21628927 | rs213058 | -2.341 | 0.053 | -1.189 | -1.675 | -0.121 | -0.178 | -0.178 | 0 | 0.74 | 0.96 | 0.55 | 0.55 | 0.62 | 0.73 |
| 1 | 21631472 | rs2051031 | 2.053 | 0.1 | -0.983 | -1.832 | -0.012 | 0.019 | 0.019 | 0.895 | 0.12 | 0.04 | 0.12 | 0 | 0.19 | 0.06 |
| 1 | 21631663 | rs10916959 | 1.221 | -0.269 | -1.13 | -1.93 | -0.027 | 0.128 | 0.128 | 0 | 0.26 | 0.02 | 0.11 | 0.12 | 0.19 | 0.26 |
| 1 | 21632470 | rs3026845 | 1.985 | 0.101 | -0.852 | -2.102 | -0.011 | 0.063 | 0.063 | 0 | 0.11 | 0.01 | 0.03 | 0 | 0.08 | 0.05 |
| 1 | 21635529 | rs3026840 | 0.644 | 0.233 | 0.575 | -1.585 | 0 | -0.005 | -0.005 | 0.029 | 0.06 | 0.05 | 0.04 | 0.01 | 0.05 | 0.09 |
| 1 | 21636746 | rs12756690 | 0.123 | 0.212 | 0.499 | -1.653 | -0.01 | 0.082 | 0.082 | 0 | 0.13 | 0.01 | 0.06 | 0 | 0.1 | 0.08 |
| 1 | 21639167 | rs213010 | -1.569 | 0.592 | 1.191 | -1.483 | 0.015 | 0.038 | 0.038 | 0.895 | 0.29 | 0.54 | 0.34 | 0.39 | 0.34 | 0.38 |
| 1 | 21639935 | rs3026837 | -0.228 | 0.538 | 1.226 | -1.478 | -0.008 | 0 | 0 | 0.676 | 0.19 | 0.12 | 0.17 | 0.01 | 0.26 | 0.15 |
| 1 | 21640718 | rs12116832 | -0.211 | 0.541 | 1.227 | -1.477 | -0.008 | 0 | 0 | 0 | 0.19 | 0.12 | 0.16 | 0.01 | 0.25 | 0.15 |
| 1 | 21641522 | rs84853 | -1.811 | 0.534 | 1.299 | -1.44 | -0.024 | -0.007 | -0.007 | 0 | 0.43 | 0.82 | 0.43 | 0.39 | 0.44 | 0.46 |
| 1 | 21654735 | rs213032 | -1.328 | 0.689 | 0.112 | -1.345 | 0.011 | -0.027 | -0.027 | 0.049 | 0.32 | 0.74 | 0.32 | 0.39 | 0.26 | 0.42 |
| 1 | 21657162 | rs213039 | -1.482 | 0.4 | -0.097 | -1.289 | -0.031 | -0.019 | -0.019 | 0.621 | 0.44 | 0.82 | 0.45 | 0.39 | 0.47 | 0.47 |
| 1 | 21657239 | rs4654918 | -0.474 | 0.068 | 0.615 | -1.417 | -0.004 | -0.031 | -0.031 | 0 | 0.08 | 0.12 | 0.06 | 0.01 | 0.07 | 0.1 |
| 1 | 21664475 | rs3026815 | -0.547 | 0.214 | 0.457 | -1.432 | 0.029 | 0.088 | 0.088 | 0 | 0.34 | 0.2 | 0.24 | 0.1 | 0.26 | 0.37 |
| 1 | 21678600 | rs213006 | -1.027 | 0.211 | 1.108 | -1.17 | 0.005 | -0.026 | -0.026 | 0 | 0.44 | 0.41 | 0.34 | 0.13 | 0.31 | 0.38 |
| 1 | 21681636 | rs2745252 | -1.711 | 1.317 | 2.241 | -0.408 | 0.01 | -0.054 | -0.054 | 2.668 | 0.1 | 0.54 | 0.1 | 0 | 0.06 | 0.03 |
| 1 | 21682361 | rs10916967 | -0.36 | 1.294 | 2.153 | -0.449 | -0.006 | -0.03 | -0.03 | 0 | 0.1 | 0.14 | 0.04 | 0 | 0.07 | 0.05 |
| 1 | 21682434 | rs2796345 | 1.603 | 1.303 | 2.073 | -0.449 | -0.021 | -0.098 | -0.098 | 0 | 0.81 | 0.37 | 0.87 | 1 | 0.87 | 0.92 |
| 1 | 21685445 | rs16825276 | -1.889 | 1.634 | 2.01 | 0.084 | 0.029 | 0.138 | 0.138 | 0 | 0.11 | 0.24 | 0.12 | 0.31 | 0.12 | 0.22 |
| 1 | 21687154 | rs12744958 | 1.196 | 2.145 | 1.988 | -0.244 | -0.002 | 0 | 0 | 0.008 | 0.68 | 0.36 | 0.7 | 0.71 | 0.62 | 0.69 |
| 1 | 21687854 | rs2796354 | 1.505 | 2.159 | 1.956 | -0.629 | -0.011 | 0.011 | 0.011 | 1.009 | 0.64 | 0.35 | 0.72 | 0.79 | 0.56 | 0.68 |
| 1 | 21688141 | rs2682362 | 0.896 | 1.608 | 1.986 | -0.637 | 0.027 | -0.15 | -0.15 | 0.062 | 0.72 | 0.46 | 0.78 | 1 | 0.67 | 0.87 |
| 1 | 21697469 | rs785198 | 1.949 | 0.603 | 2.211 | 0.309 | 0.038 | 0.103 | 0.103 | 0 | 0.86 | 0.57 | 0.9 | 0.8 | 0.86 | 0.8 |
| 1 | 21705378 | rs4614227 | -1.478 | 0.637 | 1.584 | 0.211 | -0.004 | -0.035 | -0.035 | 0.373 | 0.07 | 0.17 | 0.05 | 0 | 0.06 | 0.02 |
| 1 | 21709489 | rs7530779 | -1.647 | 0.78 | 1.414 | 0.281 | 0.003 | 0.029 | 0.029 | 0 | 0.18 | 0.42 | 0.1 | 0.21 | 0.16 | 0.24 |
| 1 | 21711731 | rs10916972 | -2.111 | 0.781 | 1.361 | 0.201 | -0.034 | -0.131 | -0.131 | 0.072 | 0.23 | 0.44 | 0.12 | 0 | 0.16 | 0.1 |
| 1 | 21725824 | rs12124123 | -1.297 | 0.689 | 0.034 | 0.035 | 0.001 | -0.005 | -0.005 | 0 | 0.07 | 0.07 | 0.04 | 0 | 0.06 | 0.02 |
| 1 | 21736327 | rs12750103 | -1.319 | 0.826 | 0.58 | 0.012 | 0.017 | -0.044 | -0.044 | 0.184 | 0.08 | 0.29 | 0.07 | 0 | 0.08 | 0.08 |
| 1 | 21785981 | rs4654938 | 0.819 | 1.477 | 0.94 | -0.013 | 0.029 | -0.013 | -0.013 | 0 | 0.77 | 0.38 | 0.75 | 0.81 | 0.73 | 0.89 |
| 1 | 21795388 | rs1827293 | 1.278 | 1.623 | 0.356 | 0.231 | -0.078 | -0.03 | -0.03 | 0 | 0.59 | 0.14 | 0.48 | 0.58 | 0.54 | 0.48 |
| 1 | 21796616 | rs7537914 | 0.292 | 1.609 | 0.595 | 0.18 | 0.046 | -0.012 | -0.012 | 0 | 0.78 | 0.44 | 0.76 | 0.81 | 0.73 | 0.89 |
| 1 | 21805048 | rs4654947 | -0.024 | 1.711 | 0.962 | 0.132 | 0.072 | 0.026 | 0.026 | 0 | 0.82 | 0.54 | 0.79 | 0.81 | 0.77 | 0.92 |
| 1 | 21815660 | rs904927 | -1.082 | 1.467 | 1.146 | 0.108 | 0.001 | -0.027 | -0.027 | 0 | 0.22 | 0.35 | 0.14 | 0.14 | 0.2 | 0.34 |
| 1 | 21821757 | rs1780324 | 1.486 | 1.364 | 0.668 | 0.187 | -0.029 | 0.057 | 0.057 | 0 | 0.58 | 0.24 | 0.55 | 0.71 | 0.55 | 0.53 |
| 1 | 21823292 | rs1697421 | 1.597 | 1.675 | 0.383 | 0.257 | -0.068 | -0.067 | -0.067 | 0 | 0.56 | 0.07 | 0.44 | 0.52 | 0.51 | 0.46 |
| 1 | 21823629 | rs1697420 | 1.794 | 1.604 | 1.001 | 0.691 | 0.067 | 0.254 | 0.254 | 2.633 | 0.89 | 0.48 | 0.66 | 0.72 | 0.88 | 0.9 |
| 1 | 21824108 | rs1780321 | 1.047 | 1.461 | 0.678 | 0.377 | 0.026 | 0.042 | 0.042 | 0 | 0.69 | 0.5 | 0.57 | 0.55 | 0.68 | 0.46 |
| 1 | 21826566 | rs6667242 | -1.686 | 1.24 | 0.589 | 0.801 | 0.047 | 0.183 | 0.183 | 0.356 | 0.08 | 0.25 | 0.29 | 0.28 | 0.1 | 0.11 |
| 1 | 21837755 | rs16825455 | -1.552 | 1.328 | 0.662 | 0.794 | 0.069 | 0.222 | 0.222 | 0.394 | 0.08 | 0.33 | 0.3 | 0.29 | 0.1 | 0.11 |
| 1 | 21839690 | rs975000 | 0.262 | 0.461 | -0.528 | 0.218 | 0.05 | -0.059 | -0.059 | 0 | 0.35 | 0.17 | 0.37 | 0.48 | 0.28 | 0.43 |
| 1 | 21840129 | rs869179 | 0.118 | 0.798 | -0.938 | 0.201 | 0.024 | -0.104 | -0.104 | 2.935 | 0.37 | 0.11 | 0.42 | 0.47 | 0.33 | 0.45 |
| 1 | 21842892 | rs12116444 | 0.189 | 1.15 | -0.342 | -0.192 | 0.01 | -0.057 | -0.057 | 0 | 0.13 | 0.22 | 0.1 | 0 | 0.13 | 0.03 |
| 1 | 21847548 | rs12145027 | 0.255 | 0.483 | -1.101 | 0.382 | 0.02 | -0.053 | -0.053 | 0 | 0.24 | 0.08 | 0.38 | 0.37 | 0.21 | 0.44 |
| 1 | 21851312 | rs1256323 | 0.372 | 0.845 | -1.55 | -0.324 | -0.016 | 0.082 | 0.082 | 0.369 | 0.2 | 0.05 | 0.1 | 0 | 0.18 | 0.04 |
| 1 | 21856119 | rs1697407 | -1.082 | 0.57 | -0.944 | -0.508 | -0.004 | -0.025 | -0.025 | 0 | 0.95 | 0.84 | 0.96 | 1 | 0.96 | 0.99 |
| 1 | 21859507 | rs10465542 | 0.234 | 0.788 | -0.559 | -0.299 | 0.041 | 0.176 | 0.176 | 0 | 0.2 | 0.49 | 0.28 | 0.49 | 0.21 | 0.27 |

| | | | | | | | | | | | | | | | | |
|---|---|---|---|---|---|---|---|---|---|---|---|---|---|---|---|---|
| 1 | 21861656 | rs2840331 | 1.549 | 0.812 | -0.835 | -0.45 | 0.002 | -0.028 | -0.028 | 0 | 0.29 | 0.17 | 0.4 | 0.36 | 0.3 | 0.45 |
| 1 | 21863691 | rs4021228 | 0.767 | 0.677 | -0.909 | -0.538 | 0.004 | 0.076 | 0.076 | 0 | 0.5 | 0.22 | 0.5 | 0.34 | 0.48 | 0.48 |
| 1 | 21864206 | rs10917006 | 0.337 | 0.858 | -0.141 | -0.546 | 0.021 | -0.031 | -0.031 | 0 | 0.06 | 0.23 | 0.11 | 0 | 0.06 | 0.03 |
| 1 | 21866625 | rs12565845 | 1.35 | 0.829 | -0.205 | -0.489 | -0.043 | -0.075 | -0.075 | 0 | 0.12 | 0.01 | 0.15 | 0.51 | 0.15 | 0.24 |
| 1 | 21866829 | rs1256341 | 0.902 | 0.796 | 0.046 | -0.759 | 0.027 | 0.065 | 0.065 | 4.848 | 0.37 | 0.23 | 0.2 | 0.15 | 0.3 | 0.11 |
| 1 | 21869773 | rs12127343 | 0.685 | 0.79 | 0.179 | -0.786 | -0.002 | 0.002 | 0.002 | 2.21 | 0.42 | 0.38 | 0.63 | 0.84 | 0.44 | 0.69 |
| 1 | 21873002 | rs12141569 | 0.205 | 0.782 | 0.08 | -0.76 | -0.002 | 0.057 | 0.057 | 3.21 | 0.48 | 0.61 | 0.68 | 0.84 | 0.5 | 0.71 |
| 1 | 21875916 | rs885814 | 1.849 | 0.792 | -0.096 | -0.694 | -0.002 | -0.027 | -0.027 | 0.046 | 0.28 | 0.14 | 0.36 | 0.33 | 0.29 | 0.45 |
| 1 | 21876492 | rs1256342 | 1.359 | 0.872 | -0.176 | -0.731 | -0.019 | 0.012 | 0.012 | 0 | 0.09 | 0.06 | 0.08 | 0.03 | 0.16 | 0.16 |
| 1 | 21877074 | rs885813 | 0.269 | 0.787 | 0.067 | -0.711 | -0.001 | 0.112 | 0.112 | 1.753 | 0.46 | 0.27 | 0.29 | 0.16 | 0.45 | 0.27 |
| 1 | 21885121 | rs1780326 | 0.15 | 0.574 | -0.131 | -0.865 | -0.006 | 0.117 | 0.117 | 1.987 | 0.43 | 0.23 | 0.26 | 0.14 | 0.42 | 0.25 |
| 1 | 21885195 | rs3767155 | 1.543 | 0.557 | -0.036 | -0.829 | 0.003 | -0.021 | -0.021 | 0 | 0.32 | 0.19 | 0.37 | 0.35 | 0.33 | 0.47 |
| 1 | 21887290 | rs1767430 | 0.158 | 0.588 | -0.195 | -0.824 | -0.005 | 0.115 | 0.115 | 2.015 | 0.43 | 0.24 | 0.27 | 0.14 | 0.42 | 0.25 |
| 1 | 21889340 | rs1767429 | 0.821 | 0.588 | 0 | -0.903 | -0.003 | -0.002 | -0.002 | 1.209 | 0.41 | 0.53 | 0.43 | 0.37 | 0.39 | 0.51 |
| 1 | 21889506 | rs1256336 | 0.345 | 0.571 | 0.039 | -1.197 | -0.005 | -0.087 | -0.087 | 2.079 | 0.3 | 0.35 | 0.15 | 0.02 | 0.25 | 0.08 |
| 1 | 21889635 | rs1780316 | -1.259 | 0.452 | -0.14 | -1.019 | -0.003 | -0.032 | -0.032 | 0 | 0.93 | 0.85 | 0.95 | 0.98 | 0.94 | 0.96 |
| 1 | 21890386 | rs1256335 | -0.208 | 0.469 | -0.163 | -1.033 | 0.012 | -0.044 | -0.044 | 0 | 0.84 | 0.76 | 0.86 | 0.98 | 0.78 | 0.79 |
| 1 | 21894214 | rs3767150 | 0.728 | 0.4 | -0.207 | -0.473 | -0.002 | 0.143 | 0.143 | 0 | 0.14 | 0.27 | 0.27 | 0.45 | 0.12 | 0.23 |
| 1 | 21899491 | rs1780320 | 0.926 | 0.644 | -0.451 | -0.708 | -0.055 | 0.088 | 0.088 | 0.052 | 0.24 | 0.1 | 0.17 | 0 | 0.3 | 0.05 |
| 1 | 21900420 | rs2275370 | -0.151 | 0.345 | 0.219 | -0.143 | -0.004 | -0.005 | -0.005 | 0 | 0.74 | 0.7 | 0.83 | 0.79 | 0.75 | 0.74 |
| 1 | 21901457 | rs1697404 | -0.581 | 0.653 | 0.673 | 0.317 | -0.011 | 0.137 | 0.137 | 0 | 0.86 | 0.74 | 0.74 | 0.55 | 0.89 | 0.78 |
| 1 | 21902950 | rs1780329 | -0.601 | 0.728 | 0.641 | 0.333 | -0.007 | 0.119 | 0.119 | 0 | 0.2 | 0.37 | 0.32 | 0.47 | 0.18 | 0.26 |
| 1 | 21904267 | rs1697406 | -1.349 | 0.523 | 0.063 | -0.201 | 0.032 | -0.003 | -0.003 | 0 | 0.78 | 0.86 | 0.91 | 0.8 | 0.86 | 0.93 |
| 1 | 21904374 | rs1772719 | -0.688 | 0.722 | 0.114 | -0.368 | 0.002 | -0.029 | -0.029 | 1.252 | 0.18 | 0.22 | 0.13 | 0.02 | 0.21 | 0.22 |
| 1 | 21904529 | rs2242420 | 1.202 | 0.39 | -0.231 | -0.111 | 0.041 | -0.012 | -0.012 | 2.237 | 0.21 | 0.09 | 0.09 | 0.2 | 0.14 | 0.07 |
| 1 | 21907604 | rs3121206 | -0.623 | 0.496 | -0.354 | -0.322 | 0.026 | 0.017 | 0.017 | 0.81 | 0.67 | 0.73 | 0.83 | 0.78 | 0.76 | 0.87 |
| 1 | 21910890 | rs12124283 | 0.399 | 1.42 | -0.782 | 0.169 | -0.09 | -0.078 | -0.078 | 0 | 0.44 | 0.1 | 0.3 | 0.45 | 0.49 | 0.58 |
| 1 | 21914362 | rs1256279 | 0.945 | 1.467 | -0.435 | 0.213 | -0.008 | -0.032 | -0.032 | 0.296 | 0.33 | 0.21 | 0.22 | 0.43 | 0.35 | 0.37 |
| 1 | 21917234 | rs10917021 | 0.57 | 1.495 | -0.357 | 0.47 | 0.009 | 0.069 | 0.069 | 0.316 | 0.08 | 0.12 | 0.29 | 0.25 | 0.1 | 0.14 |
| 1 | 21921058 | rs829391 | -0.186 | 1.335 | -0.285 | 0.376 | 0.003 | 0.016 | 0.016 | 2.893 | 0.82 | 0.76 | 0.64 | 0.73 | 0.79 | 0.78 |
| 1 | 21921736 | rs7515386 | 1.115 | 0.654 | -0.219 | -0.195 | 0.039 | 0.014 | 0.014 | 9.661 | 0.28 | 0.19 | 0.22 | 0.24 | 0.18 | 0.16 |
| 1 | 21925603 | rs10917023 | 0.237 | 1.359 | 0.217 | -0.205 | 0.011 | -0.014 | -0.014 | 0 | 0.08 | 0.17 | 0.06 | 0.04 | 0.07 | 0.05 |
| 1 | 21927848 | rs12759525 | -0.565 | 1.335 | 0.045 | -0.261 | 0.008 | -0.053 | -0.053 | 0.705 | 0.21 | 0.29 | 0.16 | 0.05 | 0.25 | 0.26 |
| 1 | 21933193 | rs829370 | -0.953 | 1.254 | -0.342 | -0.186 | 0 | 0.025 | 0.025 | 0.02 | 0.9 | 0.93 | 0.94 | 0.98 | 0.89 | 0.93 |
| 1 | 21935467 | rs12130950 | 0.389 | 1.494 | -0.143 | -0.06 | 0.006 | -0.01 | -0.01 | 0 | 0.1 | 0.11 | 0.08 | 0.03 | 0.14 | 0.19 |
| 1 | 21938375 | rs2275364 | 0.802 | 1.571 | 0.034 | -0.146 | 0.004 | -0.017 | -0.017 | 0 | 0.31 | 0.33 | 0.25 | 0.13 | 0.36 | 0.23 |
| 1 | 21940555 | rs2275363 | 0.338 | 1.395 | -0.013 | 0.389 | -0.007 | -0.004 | -0.004 | 0 | 0.39 | 0.36 | 0.28 | 0.42 | 0.44 | 0.43 |
| 1 | 21943136 | rs829417 | -0.262 | 1.435 | 0.086 | 0.346 | 0.013 | 0.005 | 0.005 | 3.681 | 0.22 | 0.26 | 0.45 | 0.26 | 0.3 | 0.35 |
| 1 | 21943504 | rs829418 | -1.029 | 1.012 | -0.138 | 0.111 | 0.004 | 0.028 | 0.028 | 0 | 0.87 | 0.92 | 0.93 | 0.98 | 0.87 | 0.89 |
| 1 | 21946396 | rs12134357 | 0.336 | 1.142 | 0.145 | 0.335 | 0.007 | -0.002 | -0.002 | 0 | 0.38 | 0.29 | 0.34 | 0.37 | 0.31 | 0.31 |
| 1 | 21949462 | rs829406 | 1.218 | 0.666 | 0.417 | 0.507 | -0.053 | 0.024 | 0.024 | 7.784 | 0.27 | 0.36 | 0.23 | 0.33 | 0.13 | 0.11 |
| 1 | 21953287 | rs7354914 | 0.416 | 1.03 | 0.376 | 0.631 | 0.007 | 0.031 | 0.031 | 0.084 | 0.08 | 0.1 | 0.25 | 0.23 | 0.11 | 0.13 |
| 1 | 21953304 | rs1531829 | -0.693 | 1.382 | 0.464 | 0.556 | 0.005 | 0.012 | 0.012 | 0.355 | 0.27 | 0.28 | 0.43 | 0.54 | 0.33 | 0.41 |
| 1 | 21953436 | rs829412 | -1.134 | 1.422 | 0.264 | 0.291 | 0.01 | -0.056 | -0.056 | 0.055 | 0.87 | 0.75 | 0.91 | 0.98 | 0.83 | 0.91 |
| 1 | 21954294 | rs4654974 | 1.239 | 1.283 | -0.017 | 0.312 | -0.035 | 0.084 | 0.084 | 0 | 0.34 | 0.16 | 0.27 | 0.11 | 0.4 | 0.37 |
| 1 | 21958536 | rs11591167 | 1.5 | 1.225 | -0.181 | 0.459 | -0.05 | -0.069 | -0.069 | 0 | 0.12 | 0.01 | 0.05 | 0.26 | 0.14 | 0.24 |
| 1 | 21958909 | rs16825746 | 1.892 | 1.033 | -0.152 | 0.425 | 0.06 | -0.051 | -0.051 | 0 | 0.23 | 0.09 | 0.18 | 0.33 | 0.12 | 0.08 |
| 1 | 21959015 | rs1557087 | -0.778 | 1.317 | 0.079 | 0.354 | 0.022 | -0.085 | -0.085 | 0 | 0.85 | 0.61 | 0.9 | 0.98 | 0.82 | 0.91 |
| 1 | 21961581 | rs12045276 | 0.396 | 1.351 | -0.675 | 0.543 | -0.024 | -0.042 | -0.042 | 4.722 | 0.16 | 0.07 | 0.08 | 0.3 | 0.18 | 0.3 |
| 1 | 21966833 | rs12142081 | 0.666 | 1.186 | -0.661 | 0.415 | -0.052 | 0.094 | 0.094 | 0 | 0.38 | 0.18 | 0.29 | 0.15 | 0.44 | 0.43 |
| 1 | 21972480 | rs6426721 | 0.99 | 1.598 | -0.19 | 0.727 | 0.014 | 0.011 | 0.011 | 0.47 | 0.59 | 0.48 | 0.47 | 0.44 | 0.53 | 0.46 |
| 1 | 21974208 | rs2128699 | 0.554 | 1.258 | -0.082 | 0.569 | 0.008 | 0.009 | 0.009 | 0.171 | 0.72 | 0.56 | 0.56 | 0.69 | 0.71 | 0.71 |
| 1 | 21979230 | rs2625408 | -0.299 | 1.065 | -0.416 | 0.171 | -0.001 | -0.005 | -0.005 | 0 | 0.66 | 0.65 | 0.67 | 0.83 | 0.63 | 0.68 |
| 1 | 21983611 | rs829404 | -0.158 | 0.834 | -0.516 | 0.017 | -0.001 | 0.001 | 0.001 | 0.021 | 0.46 | 0.43 | 0.71 | 0.65 | 0.5 | 0.51 |
| 1 | 21988563 | rs829373 | 1.416 | 1.178 | -0.444 | 0.074 | 0.004 | -0.003 | -0.003 | 0.324 | 0.17 | 0.19 | 0.26 | 0.14 | 0.22 | 0.26 |
| 1 | 21988622 | rs829374 | -0.885 | 0.845 | -0.594 | -0.466 | -0.001 | -0.031 | -0.031 | 0 | 0.7 | 0.71 | 0.85 | 0.97 | 0.7 | 0.74 |
| 1 | 21996848 | rs829398 | 0.315 | 0.534 | -0.611 | -0.216 | -0.001 | 0 | 0 | 0 | 0.56 | 0.5 | 0.49 | 0.45 | 0.59 | 0.66 |
| 1 | 21997282 | rs3738091 | 0.346 | 0.613 | -0.675 | -0.184 | -0.003 | -0.014 | -0.014 | 0 | 0.31 | 0.24 | 0.47 | 0.49 | 0.34 | 0.3 |
| 1 | 21998761 | rs829400 | -1.474 | 0.756 | -1.111 | 0.115 | 0.005 | 0.009 | 0.009 | 0 | 0.94 | 0.88 | 0.98 | 0.94 | 0.96 | 0.99 |
| 1 | 21998838 | rs12036106 | 1.505 | 1.079 | -1.629 | 0.61 | -0.034 | -0.005 | -0.005 | 0.047 | 0.06 | 0 | 0.25 | 0.29 | 0.1 | 0.12 |
| 1 | 22000026 | rs3818942 | 1.682 | 0.597 | -1.462 | 0.225 | -0.007 | 0.022 | 0.022 | 0 | 0.11 | 0.13 | 0.07 | 0.28 | 0.07 | 0.16 |
| 1 | 22001981 | rs12135009 | 0.616 | 0.812 | -1.446 | 0.224 | 0.006 | 0.006 | 0.006 | 0 | 0.34 | 0.3 | 0.18 | 0.31 | 0.3 | 0.38 |
| 1 | 22003793 | rs829396 | 0.729 | 0.975 | -1.553 | 0.371 | -0.014 | 0.005 | 0.005 | 0 | 0.18 | 0.08 | 0.25 | 0.13 | 0.23 | 0.25 |
| 1 | 22006721 | rs3767132 | 0.02 | 0.972 | -1.599 | 0.33 | -0.054 | -0.044 | -0.044 | 0.09 | 0.14 | 0.03 | 0.37 | 0.24 | 0.19 | 0.16 |
| 1 | 22008798 | rs12404648 | 0.866 | 0.327 | -1.754 | -0.003 | -0.006 | -0.051 | -0.051 | 0.138 | 0.16 | 0.01 | 0.14 | 0.19 | 0.08 | 0.09 |
| 1 | 22013771 | rs10493008 | -0.537 | 0.419 | -1.217 | 0.005 | -0.012 | 0.005 | 0.005 | 0 | 0.4 | 0.34 | 0.67 | 0.64 | 0.41 | 0.44 |
| 1 | 22020152 | rs3767133 | 1.403 | -0.087 | -1.03 | -0.137 | -0.03 | 0.128 | 0.128 | 0 | 0.12 | 0.27 | 0.08 | 0.28 | 0.07 | 0.19 |
| 1 | 22021862 | rs16825823 | 1.758 | -0.054 | -1.274 | -0.129 | 0.005 | 0.002 | 0.002 | 0 | 0.1 | 0.06 | 0.06 | 0.29 | 0.06 | 0.19 |
| 1 | 22033705 | rs333171 | 1.338 | -0.148 | -1.769 | -0.793 | -0.047 | 0.192 | 0.192 | 0 | 0.3 | 0.03 | 0.15 | 0.02 | 0.33 | 0.26 |

| | | | | | | | | | | | | | | | | |
|---|---|---|---|---|---|---|---|---|---|---|---|---|---|---|---|---|
| 1 | 22040294 | rs333188 | -0.335 | -0.225 | -1.617 | -0.815 | -0.049 | 0.026 | 0.026 | 0 | 0.72 | 0.58 | 0.87 | 0.66 | 0.82 | 0.68 |
| 1 | 22041073 | rs1018102 | 0.144 | -0.151 | -1.649 | -0.862 | 0.013 | -0.007 | -0.007 | 0 | 0.4 | 0.52 | 0.68 | 0.63 | 0.41 | 0.41 |
| 1 | 22042643 | rs333185 | -1.12 | -0.308 | -1.727 | -0.851 | -0.044 | 0.084 | 0.084 | 0 | 0.8 | 0.65 | 0.89 | 0.66 | 0.85 | 0.71 |
| 1 | 22046558 | rs10799708 | -0.64 | -0.239 | -1.73 | -0.754 | 0.003 | 0.004 | 0.004 | 0 | 0.5 | 0.55 | 0.74 | 0.71 | 0.52 | 0.45 |
| 1 | 22048664 | rs12030578 | 2.014 | 0.027 | -1.83 | -0.516 | -0.038 | 0.012 | 0.012 | 0 | 0.06 | 0.01 | 0.39 | 0.37 | 0.14 | 0.12 |
| 1 | 22049667 | rs6426724 | 0.638 | -0.526 | -1.657 | -1.458 | 0.03 | 0.018 | 0.018 | 0 | 0.56 | 0.39 | 0.74 | 0.66 | 0.63 | 0.63 |
| 1 | 22052814 | rs6702312 | 2.516 | -0.495 | -2.087 | -1.461 | -0.066 | 0.095 | 0.095 | 0 | 0.18 | 0.01 | 0.09 | 0.03 | 0.22 | 0.24 |
| 1 | 22054427 | rs16825896 | 1.11 | -1.182 | -1.364 | -0.797 | 0.068 | 0.042 | 0.042 | 0 | 0.55 | 0.25 | 0.73 | 0.66 | 0.62 | 0.63 |
| 1 | 22076705 | rs2315931 | 0.82 | -1.395 | -1.695 | -0.459 | -0.036 | -0.008 | -0.008 | 0 | 0.62 | 0.6 | 0.82 | 0.67 | 0.71 | 0.67 |
| 1 | 22080775 | rs333204 | 1.623 | -1.422 | -1.905 | -0.48 | 0.013 | 0.045 | 0.045 | 0 | 0.11 | 0.03 | 0.06 | 0 | 0.11 | 0.02 |
| 1 | 22088379 | rs16825927 | 1.448 | -1.31 | -1.805 | -0.387 | -0.004 | 0.019 | 0.019 | 0 | 0.11 | 0.08 | 0.06 | 0.28 | 0.07 | 0.21 |
| 1 | 22104971 | rs10733030 | 0.545 | -1.197 | -1.808 | -0.386 | -0.034 | -0.024 | -0.024 | 0 | 0.62 | 0.54 | 0.82 | 0.74 | 0.71 | 0.67 |
| 1 | 22109370 | rs10917042 | 0.139 | -0.357 | -1.352 | 0.397 | -0.063 | -0.124 | -0.124 | 0 | 0.22 | 0.03 | 0.53 | 0.59 | 0.22 | 0.2 |
| 1 | 22127992 | rs7518006 | 0.461 | -0.94 | -0.528 | 0.553 | -0.049 | 0.009 | 0.009 | 0 | 0.78 | 0.59 | 0.88 | 0.77 | 0.84 | 0.7 |
| 1 | 22138198 | rs4654986 | -0.138 | -0.502 | -1.35 | 0.773 | 0.001 | -0.001 | -0.001 | 2.186 | 0.14 | 0.09 | 0.08 | 0.29 | 0.07 | 0.17 |
| 1 | 22140348 | rs13374837 | 0.065 | -0.444 | -1.54 | 0.739 | 0.022 | -0.019 | -0.019 | 0.371 | 0.17 | 0.09 | 0.16 | 0.18 | 0.12 | 0.07 |
| 1 | 22141983 | rs2010397 | -0.462 | -0.403 | -1.314 | 0.426 | 0.011 | 0.002 | 0.002 | 0.223 | 0.32 | 0.29 | 0.17 | 0.31 | 0.24 | 0.26 |
| 1 | 22150120 | rs3736360 | -0.727 | 0.55 | -0.956 | 0.722 | -0.023 | -0.005 | -0.005 | 0 | 0.15 | 0.07 | 0.15 | 0.16 | 0.22 | 0.32 |
| 1 | 22154987 | rs10917052 | -1.537 | 0.151 | -1.233 | 0.286 | -0.021 | 0.05 | 0.05 | 0.168 | 0.27 | 0.11 | 0.2 | 0.16 | 0.3 | 0.51 |
| 1 | 22158120 | rs2270701 | -0.304 | 0.205 | -1.243 | 0.305 | -0.033 | -0.012 | -0.012 | 0 | 0.14 | 0.04 | 0.15 | 0.16 | 0.2 | 0.32 |
| 1 | 22158363 | rs4654771 | 1.618 | 0.201 | -1.911 | 0.449 | -0.102 | -0.113 | -0.113 | 0 | 0.45 | 0.03 | 0.38 | 0.44 | 0.48 | 0.28 |
| 1 | 22160043 | rs2229493 | -0.478 | 0.267 | -0.853 | 0.47 | -0.014 | -0.062 | -0.062 | 0 | 0.11 | 0.17 | 0.05 | 0 | 0.07 | 0.11 |
| 1 | 22160148 | rs7547731 | 1.874 | 0.185 | -0.828 | 0.6 | -0.042 | 0.07 | 0.07 | 0.401 | 0.5 | 0.08 | 0.46 | 0.65 | 0.48 | 0.33 |
| 1 | 22165987 | rs2291827 | 0.197 | 0.276 | -0.786 | 0.634 | -0.062 | -0.036 | -0.036 | 0.262 | 0.13 | 0.01 | 0.15 | 0.16 | 0.19 | 0.3 |
| 1 | 22168216 | rs7556412 | 0.935 | 0.071 | 0.274 | 0.543 | 0.001 | -0.033 | -0.033 | 0 | 0.67 | 0.46 | 0.58 | 0.81 | 0.68 | 0.63 |
| 1 | 22168801 | rs2229491 | -0.443 | 0.061 | 0.347 | 0.527 | -0.014 | -0.066 | -0.066 | 0 | 0.11 | 0.18 | 0.05 | 0 | 0.07 | 0.11 |
| 1 | 22174426 | rs2290501 | -2.009 | 0.092 | 0.362 | 0.517 | -0.002 | -0.031 | -0.031 | 12.26 | 0.34 | 0.54 | 0.26 | 0.16 | 0.33 | 0.5 |
| 1 | 22174600 | rs3767138 | -0.883 | 0.31 | 0.437 | 0.661 | 0.027 | 0.03 | 0.03 | 0.487 | 0.85 | 0.67 | 0.68 | 0.81 | 0.82 | 0.84 |
| 1 | 22176831 | rs2290500 | -0.18 | 0.315 | 0.624 | 0.666 | 0.01 | 0.003 | 0.003 | 0 | 0.15 | 0.16 | 0.16 | 0.17 | 0.19 | 0.3 |
| 1 | 22180548 | rs12117402 | -0.091 | 0.247 | 0.506 | 0.795 | 0.015 | 0.093 | 0.093 | 1.742 | 0.8 | 0.57 | 0.63 | 0.62 | 0.8 | 0.78 |
| 1 | 22183739 | rs2305562 | 1.222 | 0.227 | 0.236 | 0.698 | -0.002 | -0.002 | -0.002 | 0 | 0.57 | 0.4 | 0.46 | 0.47 | 0.54 | 0.39 |
| 1 | 22184552 | rs4654991 | -0.705 | 0.211 | -0.009 | 0.65 | -0.015 | -0.04 | -0.04 | 0 | 0.18 | 0.06 | 0.18 | 0.35 | 0.19 | 0.35 |
| 1 | 22191454 | rs897471 | 1.209 | 0.475 | 0.819 | 0.426 | 0.009 | -0.055 | -0.055 | 0 | 0.74 | 0.59 | 0.78 | 0.89 | 0.73 | 0.53 |
| 1 | 22198190 | rs11586789 | -1.29 | -0.146 | 1.42 | 0.638 | 0.048 | 0.005 | 0.005 | 4.067 | 0.06 | 0.27 | 0.05 | 0 | 0.02 | 0.01 |
| 1 | 22199245 | rs2229482 | 1.7 | -0.013 | 1.699 | 1.296 | 0.006 | 0.002 | 0.002 | 1.624 | 0.59 | 0.21 | 0.41 | 0.41 | 0.65 | 0.52 |
| 1 | 22199821 | rs2454290 | 1.741 | -0.011 | 1.774 | 1.055 | -0.044 | -0.148 | -0.148 | 0 | 0.75 | 0.42 | 0.87 | 0.97 | 0.86 | 0.73 |
| 1 | 22204506 | rs747546 | -2.179 | 0.209 | 2.474 | 1.376 | -0.007 | -0.043 | -0.043 | 0.356 | 0.14 | 0.29 | 0.13 | 0.04 | 0.09 | 0.14 |
| 1 | 22216279 | rs3767141 | -1.303 | 0.273 | 1.668 | 2.037 | -0.033 | -0.057 | -0.057 | 0.058 | 0.26 | 0.15 | 0.45 | 0.37 | 0.3 | 0.27 |
| 1 | 22224102 | rs11810496 | -1.041 | 0.327 | 1.539 | 2.061 | -0.072 | -0.122 | -0.122 | 0.615 | 0.23 | 0.06 | 0.42 | 0.38 | 0.28 | 0.27 |
| 1 | 22230207 | rs2501255 | -1.923 | 0.24 | 1.893 | 2.263 | 0 | 0 | 0 | 5.616 | 0.36 | 0.68 | 0.57 | 0.66 | 0.32 | 0.38 |
| 1 | 22233670 | rs1545593 | -1.877 | 0.244 | 1.567 | 1.97 | 0.001 | -0.005 | -0.005 | 0.152 | 0.35 | 0.65 | 0.57 | 0.72 | 0.33 | 0.38 |
| 1 | 22271839 | rs7544500 | 1.581 | 0.659 | 1.771 | 1.944 | 0.003 | -0.013 | -0.013 | 0 | 0.65 | 0.43 | 0.45 | 0.37 | 0.65 | 0.61 |
| 1 | 22281759 | rs10917074 | 1.655 | 0.634 | 1.91 | 2.243 | 0.024 | 0.175 | 0.175 | 3.295 | 0.87 | 0.59 | 0.84 | 0.76 | 0.87 | 0.88 |
| 1 | 22282619 | rs7529220 | 2.012 | 0.647 | 1.985 | 2.245 | 0.03 | 0.164 | 0.164 | 7.994 | 0.86 | 0.52 | 0.82 | 0.76 | 0.86 | 0.88 |
| 1 | 22283991 | rs4655001 | 1.549 | 0.621 | 1.789 | 2.136 | 0.035 | -0.014 | -0.014 | 0 | 0.88 | 0.59 | 0.86 | 0.94 | 0.88 | 0.93 |
| 1 | 22284495 | rs11589754 | -0.743 | 0.524 | 1.497 | 2.415 | -0.037 | -0.086 | -0.086 | 0.828 | 0.21 | 0.09 | 0.38 | 0.4 | 0.22 | 0.23 |
| 1 | 22290156 | rs8179387 | -0.772 | 0.788 | 1.256 | 2.73 | -0.033 | -0.058 | -0.058 | 0 | 0.19 | 0.07 | 0.41 | 0.41 | 0.21 | 0.21 |
| 1 | 22296307 | rs10917079 | -0.783 | 0.788 | 1.428 | 2.788 | -0.012 | -0.019 | -0.019 | 0.012 | 0.19 | 0.11 | 0.41 | 0.39 | 0.21 | 0.21 |
| 1 | 22308178 | rs7553024 | 1.499 | 0.701 | 1.769 | 2.841 | 0.008 | 0.371 | 0.371 | 1.119 | 0.94 | 0.71 | 0.94 | 0.65 | 0.97 | 0.91 |
| 1 | 22308807 | rs10917097 | 1.899 | 0.752 | 1.943 | 2.727 | 0.006 | 0.343 | 0.343 | 2.313 | 0.89 | 0.51 | 0.9 | 0.73 | 0.93 | 0.94 |
| 1 | 22321550 | rs12405048 | -0.934 | 1.012 | 0.534 | 2.448 | -0.086 | -0.128 | -0.128 | 0 | 0.2 | 0.02 | 0.41 | 0.4 | 0.21 | 0.23 |
| 1 | 22336305 | rs9187 | -1.134 | 1.038 | 0.842 | 2.442 | 0.014 | 0.034 | 0.034 | 1.731 | 0.2 | 0.23 | 0.41 | 0.4 | 0.22 | 0.22 |
| 1 | 22345093 | rs3117048 | 1.435 | 0.991 | 1.028 | 2.147 | 0.028 | 0.055 | 0.055 | 0.521 | 0.71 | 0.62 | 0.48 | 0.48 | 0.68 | 0.72 |
| 1 | 22354110 | rs3768574 | -0.753 | 0.465 | 0.589 | 1.799 | -0.013 | 0.032 | 0.032 | 0 | 0.09 | 0.12 | 0.09 | 0.18 | 0.05 | 0.1 |
| 1 | 22354866 | rs2473295 | -0.056 | 0.127 | -0.015 | 0.842 | 0.011 | 0.035 | 0.035 | 0.706 | 0.76 | 0.84 | 0.75 | 0.89 | 0.74 | 0.88 |
| 1 | 22361845 | rs2473277 | 0.341 | 0.519 | -0.865 | 0.192 | -0.037 | 0.014 | 0.014 | 0 | 0.55 | 0.82 | 0.41 | 0.36 | 0.52 | 0.6 |
| 1 | 22373624 | rs2501276 | 0.265 | 0.283 | -0.826 | -0.257 | -0.001 | -0.004 | -0.004 | 0 | 0.08 | 0.07 | 0.05 | 0.02 | 0.09 | 0.14 |
| 1 | 22375065 | rs2501275 | -0.478 | -0.001 | -0.3 | -0.314 | -0.02 | -0.055 | -0.055 | 0.509 | 0.74 | 0.68 | 0.74 | 0.86 | 0.75 | 0.88 |
| 1 | 22390072 | rs2473322 | -1.504 | -0.169 | -0.364 | -0.457 | 0.005 | -0.005 | -0.005 | 0.051 | 0.91 | 0.95 | 0.97 | 1 | 0.94 | 0.94 |
| 1 | 22414070 | rs11800014 | 0.823 | -0.141 | -0.387 | -0.477 | -0.003 | -0.018 | -0.018 | 0 | 0.16 | 0.21 | 0.23 | 0.13 | 0.19 | 0.06 |
| 1 | 22415680 | rs10917148 | -1.502 | -0.141 | -0.41 | -0.473 | 0.005 | -0.005 | -0.005 | 1.885 | 0.91 | 0.95 | 0.97 | 1 | 0.94 | 0.94 |
| 1 | 22423543 | rs4076750 | 0.613 | -0.157 | -0.369 | -0.468 | -0.006 | -0.023 | -0.023 | 0.34 | 0.17 | 0.24 | 0.24 | 0.13 | 0.19 | 0.07 |
| 1 | 22426187 | rs1534949 | 0.828 | -0.078 | -0.394 | -0.528 | -0.001 | -0.002 | -0.002 | 3.163 | 0.57 | 0.48 | 0.41 | 0.37 | 0.58 | 0.6 |
| 1 | 22432587 | rs10489437 | 1.157 | -0.206 | -0.561 | -0.512 | 0.007 | -0.047 | -0.047 | 1.878 | 0.08 | 0 | 0.08 | 0.18 | 0.05 | 0.09 |
| 1 | 22442025 | rs1474646 | -0.334 | -0.138 | -0.564 | -0.665 | -0.003 | -0.061 | -0.061 | 0 | 0.75 | 0.77 | 0.86 | 1 | 0.76 | 0.89 |
| 1 | 22443887 | rs1046310 | 1.185 | -0.1 | -0.543 | -0.631 | 0.001 | -0.007 | -0.007 | 0 | 0.5 | 0.53 | 0.34 | 0.2 | 0.54 | 0.51 |
| 1 | 22445991 | rs3765351 | 1.193 | -0.141 | -0.72 | -0.625 | 0.002 | 0.001 | 0.001 | 0 | 0.5 | 0.5 | 0.35 | 0.2 | 0.55 | 0.52 |
| 1 | 22448217 | rs12131703 | 0.307 | -0.477 | -0.7 | -1.023 | -0.005 | -0.087 | -0.087 | 0.165 | 0.15 | 0.34 | 0.11 | 0.02 | 0.13 | 0.21 |
| 1 | 22450487 | rs2235529 | 1.244 | -0.351 | -0.889 | -0.573 | -0.059 | -0.103 | -0.103 | 0.088 | 0.13 | 0.01 | 0.28 | 0.48 | 0.14 | 0.21 |

| | | | | | | | | | | | | | | | | |
|---|---|---|---|---|---|---|---|---|---|---|---|---|---|---|---|---|
| 1 | 22451845 | rs7526484 | -0.002 | -0.172 | -0.638 | -0.81 | 0.007 | 0.015 | 0.015 | 0.145 | 0.73 | 0.79 | 0.75 | 0.87 | 0.74 | 0.85 |
| 1 | 22454001 | rs2235526 | 0.918 | -0.125 | -0.634 | -0.759 | -0.001 | -0.013 | -0.013 | 0 | 0.16 | 0.2 | 0.23 | 0.13 | 0.19 | 0.07 |
| 1 | 22455142 | rs7544210 | 1.025 | -0.133 | -0.907 | -0.894 | -0.001 | -0.016 | -0.016 | 0 | 0.51 | 0.5 | 0.4 | 0.18 | 0.56 | 0.54 |
| 1 | 22472732 | rs12042083 | -0.2 | 0.081 | -0.539 | -0.415 | -0.001 | 0 | 0 | 0 | 0.24 | 0.21 | 0.44 | 0.81 | 0.2 | 0.3 |
| 1 | 22477493 | rs7542242 | 0.237 | 0.29 | -0.952 | -0.411 | -0.001 | -0.001 | -0.001 | 0 | 0.37 | 0.38 | 0.22 | 0.27 | 0.33 | 0.37 |
| 1 | 22480312 | rs4623666 | 0.043 | 0.02 | -0.948 | -0.419 | 0 | 0.001 | 0.001 | 0 | 0.5 | 0.5 | 0.62 | 0.63 | 0.41 | 0.45 |
| 1 | 22487036 | rs2744721 | -0.599 | -0.102 | -1.525 | -0.827 | -0.029 | -0.037 | -0.037 | 0 | 0.67 | 0.74 | 0.51 | 0.56 | 0.58 | 0.61 |
| 1 | 22488940 | rs2807362 | 1.766 | -0.221 | -1.494 | -1.218 | -0.005 | 0.031 | 0.031 | 0 | 0.07 | 0.01 | 0.02 | 0 | 0.06 | 0.06 |
| 1 | 22491984 | rs2807357 | -0.254 | -0.408 | -1.206 | -1.372 | -0.007 | -0.074 | -0.074 | 0.009 | 0.44 | 0.46 | 0.23 | 0.06 | 0.37 | 0.27 |
| 1 | 22493672 | rs2807356 | 0.14 | 0.076 | -1.536 | -1.083 | -0.011 | 0.102 | 0.102 | 0 | 0.74 | 0.88 | 0.86 | 0.92 | 0.73 | 0.8 |
| 1 | 22493824 | rs12405138 | 0.27 | -0.102 | -1.464 | -1.152 | 0.002 | 0.031 | 0.031 | 1.686 | 0.17 | 0.09 | 0.29 | 0.08 | 0.17 | 0.08 |
| 1 | 22493977 | rs2807355 | -0.75 | -0.075 | -1.326 | -0.796 | 0.003 | -0.003 | -0.003 | 0 | 0.86 | 0.87 | 0.92 | 0.87 | 0.87 | 0.85 |
| 1 | 22494791 | rs12747516 | 0.357 | 0.11 | -1.47 | -0.977 | 0 | 0.044 | 0.044 | 3.472 | 0.2 | 0.09 | 0.12 | 0 | 0.19 | 0.05 |
| 1 | 22495215 | rs17360017 | 1.2 | 0.161 | -1.392 | -0.554 | -0.019 | -0.007 | -0.007 | 0 | 0.23 | 0.17 | 0.21 | 0.23 | 0.35 | 0.33 |
| 1 | 22496703 | rs909813 | -0.645 | -0.222 | -0.872 | -0.634 | -0.002 | -0.004 | -0.004 | 0 | 0.73 | 0.71 | 0.64 | 0.84 | 0.78 | 0.78 |
| 1 | 22498103 | rs4655030 | 0.462 | -0.298 | -0.524 | -0.727 | -0.008 | 0.008 | 0.008 | 0 | 0.32 | 0.29 | 0.27 | 0.21 | 0.45 | 0.34 |
| 1 | 22498765 | rs2179378 | 1.611 | -0.448 | -0.543 | -0.537 | 0.002 | -0.048 | -0.048 | 0 | 0.27 | 0.15 | 0.27 | 0.48 | 0.23 | 0.22 |
| 1 | 22499222 | rs1014986 | 0.82 | -0.434 | -0.503 | -0.691 | -0.002 | 0.006 | 0.006 | 0 | 0.16 | 0.12 | 0.09 | 0 | 0.17 | 0.1 |
| 1 | 22499530 | rs17360053 | 1.756 | -0.395 | -0.569 | -0.632 | -0.031 | 0.003 | 0.003 | 0 | 0.23 | 0.16 | 0.21 | 0.21 | 0.35 | 0.3 |
| 1 | 22500228 | rs12568557 | 1.329 | -0.331 | -0.579 | -0.444 | 0.002 | 0.026 | 0.026 | 0 | 0.06 | 0.04 | 0.04 | 0.24 | 0.05 | 0.2 |
| 1 | 22501628 | rs10917165 | 0.048 | -0.579 | -0.412 | -0.59 | 0.015 | 0.013 | 0.013 | 0 | 0.42 | 0.25 | 0.4 | 0.65 | 0.36 | 0.43 |
| 1 | 22501846 | rs742356 | 1.483 | -0.586 | -0.376 | -0.492 | -0.001 | -0.075 | -0.075 | 0 | 0.27 | 0.12 | 0.3 | 0.52 | 0.23 | 0.23 |
| 1 | 22503667 | rs2744705 | -0.633 | -0.544 | -0.063 | -0.577 | -0.009 | -0.031 | -0.031 | 0 | 0.72 | 0.58 | 0.66 | 0.83 | 0.77 | 0.78 |
| 1 | 22503776 | rs10737465 | 0.181 | -0.499 | -0.066 | -0.548 | -0.004 | -0.005 | -0.005 | 0 | 0.6 | 0.53 | 0.6 | 0.71 | 0.67 | 0.55 |
| 1 | 22506253 | rs2982285 | 0.074 | -0.3 | -0.101 | -0.626 | 0.006 | -0.02 | -0.02 | 0 | 0.4 | 0.44 | 0.32 | 0.2 | 0.5 | 0.37 |
| 1 | 22506851 | rs2744708 | 0.318 | -0.234 | -0.267 | -0.598 | -0.022 | 0.036 | 0.036 | 0 | 0.38 | 0.3 | 0.3 | 0.2 | 0.51 | 0.37 |
| 1 | 22509251 | rs2744712 | -0.242 | -0.095 | 0.02 | -0.444 | -0.009 | -0.037 | -0.037 | 0 | 0.72 | 0.57 | 0.64 | 0.83 | 0.77 | 0.79 |
| 1 | 22512847 | rs2807376 | -0.167 | 0.093 | -0.081 | -0.037 | 0.003 | -0.01 | -0.01 | 2.681 | 0.72 | 0.48 | 0.61 | 0.79 | 0.74 | 0.79 |
| 1 | 22514236 | rs10753531 | 0.355 | -0.211 | -0.373 | -0.491 | 0.01 | 0.024 | 0.024 | 0 | 0.65 | 0.67 | 0.82 | 0.87 | 0.73 | 0.57 |
| 1 | 22515992 | rs7531383 | 1.406 | -0.133 | -0.143 | -0.687 | 0.003 | -0.07 | -0.07 | 0 | 0.11 | 0.25 | 0.09 | 0 | 0.09 | 0.02 |
| 1 | 22518382 | rs2998273 | 2.251 | -0.151 | -0.49 | -0.737 | -0.017 | 0.031 | 0.031 | 0 | 0.06 | 0 | 0.03 | 0 | 0.06 | 0.06 |
| 1 | 22518578 | rs11579887 | 1.912 | -0.231 | -0.26 | -0.699 | -0.001 | -0.049 | -0.049 | 1.274 | 0.1 | 0.19 | 0.08 | 0 | 0.09 | 0.02 |
| 1 | 22522927 | rs12734979 | 1.436 | -0.08 | -0.994 | -0.443 | -0.017 | -0.044 | -0.044 | 0 | 0.24 | 0.09 | 0.3 | 0.47 | 0.25 | 0.22 |
| 1 | 22523466 | rs16826741 | 1.395 | -0.335 | -0.959 | -0.232 | 0.041 | -0.042 | -0.042 | 0 | 0.08 | 0.01 | 0.01 | 0.14 | 0.01 | 0.23 |
| 1 | 22529868 | rs2744720 | 0.084 | -0.154 | -0.709 | -0.262 | -0.004 | 0 | 0 | 0 | 0.63 | 0.43 | 0.6 | 0.67 | 0.75 | 0.56 |
| 1 | 22531553 | rs17356059 | 1.596 | -0.051 | -1.131 | -0.321 | -0.061 | -0.02 | -0.02 | 0.014 | 0.23 | 0.08 | 0.19 | 0.2 | 0.32 | 0.25 |
| 1 | 22533106 | rs2807348 | 0.427 | -0.312 | -0.824 | -0.412 | 0.043 | -0.007 | -0.007 | 0 | 0.61 | 0.28 | 0.59 | 0.67 | 0.74 | 0.56 |
| 1 | 22533608 | rs2807347 | -1.074 | -0.151 | -0.66 | -0.329 | -0.011 | 0.004 | 0.004 | 0 | 0.82 | 0.72 | 0.71 | 0.82 | 0.86 | 0.78 |
| 1 | 22534242 | rs3765340 | 0.797 | -0.151 | -1.342 | -0.22 | -0.09 | -0.126 | -0.126 | 0.284 | 0.3 | 0.02 | 0.36 | 0.42 | 0.31 | 0.26 |
| 1 | 22535413 | rs7513455 | 0.389 | -0.07 | -0.834 | -0.515 | -0.008 | -0.009 | -0.009 | 0 | 0.23 | 0.26 | 0.12 | 0.14 | 0.18 | 0.22 |
| 1 | 22535811 | rs2473247 | -0.438 | 0.069 | -1.139 | -0.738 | 0 | 0.025 | 0.025 | 0 | 0.69 | 0.72 | 0.71 | 0.83 | 0.67 | 0.71 |
| 1 | 22537071 | rs2473249 | -0.143 | 0.004 | -0.952 | -0.688 | 0.011 | 0.021 | 0.021 | 0 | 0.5 | 0.65 | 0.4 | 0.36 | 0.47 | 0.5 |
| 1 | 22537649 | rs2473250 | 0.266 | -0.03 | -0.772 | -0.685 | -0.029 | -0.036 | -0.036 | 0 | 0.26 | 0.48 | 0.13 | 0.14 | 0.19 | 0.24 |
| 1 | 22538597 | rs2473252 | -0.118 | -0.162 | -0.852 | -0.69 | 0.015 | 0.002 | 0.002 | 0.568 | 0.46 | 0.65 | 0.43 | 0.61 | 0.37 | 0.45 |
| 1 | 22538787 | rs2473253 | 0.007 | -0.175 | -0.836 | -0.693 | -0.023 | -0.023 | -0.023 | 0 | 0.28 | 0.59 | 0.14 | 0.14 | 0.17 | 0.24 |
| 1 | 22539813 | rs12080095 | 2.053 | -0.107 | -0.831 | -0.62 | -0.026 | -0.061 | -0.061 | 4.084 | 0.18 | 0.07 | 0.3 | 0.46 | 0.21 | 0.21 |
| 1 | 22542445 | rs2505722 | -0.632 | -0.065 | -0.831 | -0.653 | -0.013 | 0.002 | 0.002 | 0 | 0.55 | 0.66 | 0.42 | 0.37 | 0.52 | 0.52 |
| 1 | 22547290 | rs7552560 | 1.644 | -0.046 | -0.849 | -0.675 | -0.02 | -0.007 | -0.007 | 0.38 | 0.24 | 0.15 | 0.27 | 0.22 | 0.3 | 0.26 |
| 1 | 22547352 | rs1569583 | 0.996 | -0.053 | -0.726 | -0.665 | -0.009 | -0.014 | -0.014 | 1.615 | 0.3 | 0.19 | 0.38 | 0.46 | 0.34 | 0.27 |
| 1 | 22549729 | rs2744734 | -0.858 | -0.293 | -0.764 | -0.628 | -0.004 | -0.005 | -0.005 | 0 | 0.74 | 0.71 | 0.73 | 0.83 | 0.75 | 0.77 |
| 1 | 22551813 | rs2744738 | -0.736 | -0.114 | -0.698 | -0.643 | -0.002 | 0.001 | 0.001 | 0 | 0.55 | 0.6 | 0.39 | 0.39 | 0.53 | 0.55 |
| 1 | 22551925 | rs2505711 | -0.501 | -0.139 | -0.673 | -0.616 | 0.007 | 0.013 | 0.013 | 0 | 0.5 | 0.6 | 0.36 | 0.39 | 0.46 | 0.51 |
| 1 | 22553478 | rs12749135 | 1.608 | -0.097 | -0.854 | -0.559 | -0.043 | -0.027 | -0.027 | 0 | 0.23 | 0.07 | 0.23 | 0.24 | 0.29 | 0.25 |
| 1 | 22555993 | rs2491210 | -0.023 | -0.266 | -0.469 | -0.594 | -0.031 | -0.03 | -0.03 | 0 | 0.26 | 0.38 | 0.12 | 0.15 | 0.17 | 0.26 |
| 1 | 22564787 | rs12128603 | 0.078 | -0.159 | -0.207 | -0.343 | 0.017 | 0.143 | 0.143 | 1.975 | 0.09 | 0.33 | 0.06 | 0.15 | 0.05 | 0.2 |
| 1 | 22565371 | rs2744741 | 0.367 | -0.04 | -0.262 | -0.369 | 0.018 | 0.026 | 0.026 | 0 | 0.31 | 0.34 | 0.43 | 0.59 | 0.36 | 0.41 |
| 1 | 22568747 | rs2744744 | -0.105 | -0.155 | -0.445 | -0.503 | 0.011 | 0.022 | 0.022 | 0 | 0.48 | 0.52 | 0.65 | 0.8 | 0.6 | 0.64 |
| 1 | 22569226 | rs932371 | 2.098 | -0.025 | -0.67 | -0.437 | -0.04 | -0.06 | -0.06 | 0 | 0.18 | 0.07 | 0.33 | 0.44 | 0.24 | 0.19 |
| 1 | 22572123 | rs7553227 | -0.389 | -0.181 | -0.641 | -0.549 | 0.002 | 0.002 | 0.002 | 0 | 0.39 | 0.61 | 0.38 | 0.35 | 0.42 | 0.49 |
| 1 | 22573259 | rs10489160 | 1.916 | 0.022 | -0.719 | -0.467 | -0.03 | -0.018 | -0.018 | 0.475 | 0.17 | 0.06 | 0.21 | 0.21 | 0.24 | 0.22 |
| 1 | 22574426 | rs2744750 | 0.543 | 0.101 | -0.565 | -0.416 | 0.017 | 0.025 | 0.025 | 0 | 0.3 | 0.31 | 0.42 | 0.59 | 0.35 | 0.4 |
| 1 | 22577290 | rs12239548 | 2.041 | 0.112 | -0.975 | -0.482 | -0.033 | -0.042 | -0.042 | 0.054 | 0.18 | 0.06 | 0.21 | 0.26 | 0.24 | 0.24 |
| 1 | 22577425 | rs2807338 | -0.165 | 0.058 | -1.047 | -0.728 | 0.042 | 0.081 | 0.081 | 0 | 0.49 | 0.64 | 0.66 | 0.8 | 0.6 | 0.64 |
| 1 | 22578063 | rs2807339 | -0.819 | 0.157 | -1.272 | -0.611 | 0.031 | 0.035 | 0.035 | 0 | 0.63 | 0.74 | 0.75 | 0.8 | 0.73 | 0.71 |
| 1 | 22579798 | rs2744759 | -0.457 | 0.596 | -1.428 | -0.198 | -0.003 | -0.002 | -0.002 | 1.907 | 0.39 | 0.34 | 0.45 | 0.5 | 0.41 | 0.46 |
| 1 | 22585236 | rs3845519 | 0.484 | 1.32 | -1.914 | 0.452 | -0.105 | -0.105 | -0.105 | 0 | 0.23 | 0.04 | 0.37 | 0.43 | 0.3 | 0.21 |
| 1 | 22587878 | rs4655036 | 0.187 | 1.274 | -1.716 | 0.29 | -0.037 | 0.018 | 0.018 | 0 | 0.37 | 0.18 | 0.55 | 0.75 | 0.46 | 0.41 |
| 1 | 22588355 | rs4654790 | 0.412 | 1.35 | -1.665 | 0.687 | -0.034 | -0.042 | -0.042 | 0.023 | 0.23 | 0.11 | 0.25 | 0.32 | 0.31 | 0.28 |
| 1 | 22588993 | rs2473237 | -0.316 | 0.75 | -1.353 | 0.232 | 0.007 | 0.001 | 0.001 | 0 | 0.75 | 0.89 | 0.77 | 0.82 | 0.78 | 0.77 |

| | | | | | | | | | | | | | | | | |
|---|---|---|---|---|---|---|---|---|---|---|---|---|---|---|---|---|
| 1 | 22591456 | rs2505714 | -0.714 | 0.795 | -0.811 | -0.141 | 0.004 | -0.038 | -0.038 | 0 | 0.79 | 0.78 | 0.89 | 0.96 | 0.74 | 0.89 |
| 1 | 22593385 | rs736473 | 0.018 | 0.668 | -1.03 | 0.404 | -0.078 | 0.061 | 0.061 | 0 | 0.62 | 0.87 | 0.58 | 0.71 | 0.51 | 0.72 |
| 1 | 22594695 | rs12741884 | 1.061 | 0.704 | -1.066 | 0.435 | -0.069 | -0.022 | -0.022 | 0 | 0.17 | 0.02 | 0.3 | 0.22 | 0.21 | 0.15 |
| 1 | 22594956 | rs6682404 | 0.647 | 0.338 | -0.529 | -0.105 | 0.001 | -0.074 | -0.074 | 0 | 0.13 | 0.21 | 0.1 | 0 | 0.13 | 0.08 |
| 1 | 22595026 | rs962770 | -0.261 | 0.225 | -0.372 | -0.053 | -0.002 | -0.002 | -0.002 | 0 | 0.65 | 0.42 | 0.77 | 0.69 | 0.65 | 0.66 |
| 1 | 22596600 | rs2473241 | 0.178 | 0.843 | -0.458 | -0.07 | -0.007 | 0.039 | 0.039 | 0 | 0.2 | 0.23 | 0.41 | 0.52 | 0.12 | 0.32 |
| 1 | 22598534 | rs6681709 | 1.468 | 0.822 | -0.602 | -0.394 | -0.004 | -0.003 | -0.003 | 2.892 | 0.22 | 0.23 | 0.11 | 0.17 | 0.21 | 0.3 |
| 1 | 22599878 | rs2143100 | 0.439 | 0.682 | -0.534 | -0.437 | 0.002 | -0.015 | -0.015 | 1.031 | 0.33 | 0.36 | 0.2 | 0.17 | 0.36 | 0.33 |
| 1 | 22602254 | rs12406679 | 1.883 | 0.803 | -0.804 | -0.224 | -0.059 | -0.044 | -0.044 | 0 | 0.13 | 0.01 | 0.18 | 0.24 | 0.19 | 0.09 |
| 1 | 22603552 | rs760915 | -0.916 | 0.747 | -0.682 | -0.305 | 0.004 | 0.014 | 0.014 | 0 | 0.67 | 0.71 | 0.8 | 0.94 | 0.66 | 0.74 |
| 1 | 22607557 | rs12753092 | 1.38 | 1.104 | -0.786 | 0.449 | -0.059 | -0.05 | -0.05 | 0 | 0.13 | 0.01 | 0.18 | 0.25 | 0.19 | 0.09 |
| 1 | 22612690 | rs12737805 | 0.585 | 0.86 | -0.611 | 0.297 | -0.002 | 0.002 | 0.002 | 0 | 0.22 | 0.17 | 0.12 | 0.17 | 0.23 | 0.26 |
| 1 | 22614646 | rs17356374 | 0.875 | 0.924 | -0.595 | 0.315 | -0.001 | 0 | 0 | 0 | 0.21 | 0.17 | 0.12 | 0.17 | 0.22 | 0.26 |
| 1 | 22615740 | rs12734427 | 0.758 | 0.867 | -0.671 | 0.255 | -0.001 | 0.002 | 0.002 | 0 | 0.22 | 0.16 | 0.12 | 0.16 | 0.23 | 0.26 |
| 1 | 22616978 | rs880467 | 0.625 | 0.994 | -0.859 | 0.302 | -0.008 | 0.017 | 0.017 | 0 | 0.24 | 0.09 | 0.12 | 0.16 | 0.25 | 0.31 |
| 1 | 22622513 | rs10917196 | -0.25 | 1.037 | -0.76 | 0.346 | -0.002 | -0.001 | -0.001 | 0 | 0.55 | 0.54 | 0.65 | 0.51 | 0.57 | 0.45 |
| 1 | 22625246 | rs10799746 | 0.01 | 1.405 | -0.38 | 0.614 | 0.041 | 0.093 | 0.093 | 0 | 0.17 | 0.43 | 0.28 | 0.34 | 0.17 | 0.07 |
| 1 | 22629057 | rs6426746 | -0.749 | 1.154 | -1.234 | 0.089 | -0.002 | -0.025 | -0.025 | 0 | 0.83 | 0.86 | 0.89 | 0.94 | 0.82 | 0.85 |
| 1 | 22636808 | rs4589080 | -0.374 | 1.249 | -2.11 | 0.049 | -0.026 | 0.165 | 0.165 | 0 | 0.46 | 0.08 | 0.6 | 0.82 | 0.53 | 0.52 |
| 1 | 22638050 | rs4655040 | 0.576 | 0.886 | -1.752 | 0.027 | 0.052 | 0.077 | 0.077 | 0 | 0.42 | 0.23 | 0.26 | 0.19 | 0.34 | 0.4 |
| 1 | 22645889 | rs8179370 | -0.504 | 1.745 | -1.103 | 1.168 | -0.008 | -0.037 | -0.037 | 0 | 0.14 | 0.07 | 0.21 | 0.29 | 0.12 | 0.21 |
| 1 | 22648411 | rs6690148 | -0.966 | 1.922 | -0.723 | 1.151 | -0.001 | 0.002 | 0.002 | 0 | 0.23 | 0.22 | 0.43 | 0.48 | 0.28 | 0.39 |
| 1 | 22649994 | rs10917209 | -0.94 | 1.901 | -0.754 | 1.027 | -0.019 | -0.028 | -0.028 | 0.883 | 0.23 | 0.17 | 0.42 | 0.53 | 0.28 | 0.39 |
| 1 | 22653884 | rs12137132 | 0.363 | 0.443 | -0.696 | 0.254 | 0.02 | -0.007 | -0.007 | 0.085 | 0.38 | 0.22 | 0.28 | 0.35 | 0.32 | 0.47 |
| 1 | 22654254 | rs4655043 | -0.51 | 0.322 | -0.307 | -0.066 | 0.013 | -0.02 | -0.02 | 0.151 | 0.66 | 0.63 | 0.63 | 0.74 | 0.58 | 0.75 |
| 1 | 22656863 | rs11580249 | -1.103 | -0.01 | -0.201 | -0.297 | -0.008 | 0.015 | 0.015 | 0 | 0.8 | 0.63 | 0.74 | 0.75 | 0.83 | 0.8 |
| 1 | 22658301 | rs12128312 | 1.599 | -0.056 | -0.289 | -0.299 | -0.025 | -0.064 | -0.064 | 0 | 0.15 | 0.01 | 0.22 | 0.24 | 0.13 | 0.19 |
| 1 | 22659375 | rs4655044 | -1.031 | 0.147 | 0.598 | 0.1 | 0.004 | -0.015 | -0.015 | 0.015 | 0.67 | 0.61 | 0.64 | 0.73 | 0.63 | 0.75 |
| 1 | 22664003 | rs12407338 | -0.842 | 0.139 | 0.593 | 0.227 | -0.006 | -0.023 | -0.023 | 0 | 0.28 | 0.36 | 0.28 | 0.2 | 0.25 | 0.31 |
| 1 | 22665400 | rs10917216 | -0.081 | 0.256 | 0.565 | -0.01 | 0.039 | 0.12 | 0.12 | 0 | 0.51 | 0.26 | 0.58 | 0.73 | 0.57 | 0.63 |
| 1 | 22669656 | rs7553482 | -1.064 | 0.349 | 0.134 | 0.015 | -0.005 | 0.041 | 0.041 | 0.512 | 0.65 | 0.74 | 0.62 | 0.73 | 0.6 | 0.71 |
| 1 | 22670664 | rs12078899 | -0.864 | 0.532 | 0.436 | 0.42 | 0.015 | 0.024 | 0.024 | 0.475 | 0.52 | 0.66 | 0.35 | 0.37 | 0.42 | 0.51 |
| 1 | 22674370 | rs10917219 | -0.256 | 0.841 | -0.187 | 0.759 | 0.001 | -0.008 | -0.008 | 0.145 | 0.38 | 0.18 | 0.27 | 0.37 | 0.34 | 0.42 |
| 1 | 22682366 | rs12742784 | -0.648 | 0.591 | 0.269 | 0.726 | -0.017 | 0.009 | 0.009 | 0 | 0.31 | 0.38 | 0.25 | 0.37 | 0.19 | 0.38 |
| 1 | 22683405 | rs10917220 | -1.434 | 0.635 | 0.621 | 0.373 | 0.017 | 0.112 | 0.112 | 0 | 0.57 | 0.8 | 0.64 | 0.79 | 0.57 | 0.62 |
| 1 | 22685251 | rs10753536 | -0.788 | 0.524 | 1.12 | 0.301 | -0.001 | 0 | 0 | 0 | 0.43 | 0.55 | 0.35 | 0.39 | 0.4 | 0.43 |
| 1 | 22688484 | rs4655052 | -0.8 | 0.351 | 0.992 | 0.361 | -0.019 | -0.027 | -0.027 | 0 | 0.13 | 0.12 | 0.27 | 0.39 | 0.17 | 0.18 |
| 1 | 22690923 | rs6682619 | -1.138 | 0.313 | 1.141 | 0.299 | 0 | -0.003 | -0.003 | 0.16 | 0.38 | 0.61 | 0.49 | 0.76 | 0.31 | 0.5 |
| 1 | 22691058 | rs6680239 | -0.39 | 0.285 | 1.328 | 0.698 | 0.056 | 0.096 | 0.096 | 0 | 0.22 | 0.51 | 0.36 | 0.62 | 0.23 | 0.28 |
| 1 | 22698447 | rs7524102 | -0.528 | 0.239 | 1.386 | 0.561 | -0.019 | 0.003 | 0.003 | 0 | 0.24 | 0.34 | 0.14 | 0.24 | 0.17 | 0.21 |
| 1 | 22701924 | rs4531265 | -0.174 | 0.299 | 1.191 | 0.476 | -0.001 | -0.001 | -0.001 | 0.009 | 0.32 | 0.34 | 0.49 | 0.55 | 0.42 | 0.4 |
| 1 | 22702280 | rs12561997 | 0.588 | 0.401 | 0.996 | 0.052 | -0.001 | 0.042 | 0.042 | 0 | 0.16 | 0.26 | 0.11 | 0.23 | 0.11 | 0.15 |
| 1 | 22702858 | rs6696981 | 0.669 | 0.402 | 0.996 | 0.053 | 0.001 | 0.046 | 0.046 | 0 | 0.15 | 0.26 | 0.1 | 0.22 | 0.11 | 0.15 |
| 1 | 22706434 | rs6684375 | -0.605 | 0.4 | 0.996 | 0.051 | -0.015 | -0.001 | -0.001 | 0 | 0.24 | 0.31 | 0.14 | 0.23 | 0.16 | 0.21 |
| 1 | 22706534 | rs6684479 | 0.509 | 0.5 | 0.996 | 0.123 | -0.001 | 0.043 | 0.043 | 0 | 0.16 | 0.26 | 0.12 | 0.23 | 0.12 | 0.15 |
| 1 | 22713451 | rs6690260 | 0.234 | 0.495 | 0.729 | 0.005 | -0.001 | -0.005 | -0.005 | 0 | 0.28 | 0.3 | 0.25 | 0.14 | 0.3 | 0.29 |
| 1 | 22718261 | rs6704135 | -1.533 | 0.577 | 0.404 | 0.232 | 0.007 | -0.011 | -0.011 | 0.144 | 0.6 | 0.78 | 0.57 | 0.58 | 0.59 | 0.67 |
| 1 | 22718991 | rs6426752 | -0.429 | 0.531 | 0.52 | 0.339 | 0.034 | 0.106 | 0.106 | 0.175 | 0.23 | 0.42 | 0.28 | 0.41 | 0.23 | 0.26 |
| 1 | 22722694 | rs6667212 | -0.476 | 0.472 | 0.354 | 0.325 | 0.002 | -0.003 | -0.003 | 0 | 0.16 | 0.18 | 0.21 | 0.15 | 0.17 | 0.15 |
| 1 | 22725224 | rs10917223 | 0.147 | 0.027 | -0.042 | 0.068 | -0.007 | 0.021 | 0.021 | 0 | 0.27 | 0.24 | 0.25 | 0.17 | 0.3 | 0.35 |
| 1 | 22725825 | rs2185341 | 0.069 | 0.097 | -0.132 | 0.068 | -0.002 | 0.004 | 0.004 | 0 | 0.28 | 0.32 | 0.25 | 0.16 | 0.31 | 0.36 |
| 1 | 22731269 | rs7543680 | -0.236 | 0.468 | -0.489 | 0.199 | -0.008 | 0.009 | 0.009 | 0 | 0.27 | 0.42 | 0.16 | 0.27 | 0.22 | 0.28 |
| 1 | 22735906 | rs4655059 | 0.127 | 0.479 | -1.343 | 0.407 | -0.038 | -0.022 | -0.022 | 0.694 | 0.43 | 0.11 | 0.58 | 0.56 | 0.48 | 0.37 |
| 1 | 22742075 | rs4589081 | -0.534 | -0.177 | -0.545 | -0.344 | -0.029 | -0.106 | -0.106 | 0 | 0.23 | 0.35 | 0.22 | 0 | 0.12 | 0.16 |
| 1 | 22742985 | rs12145722 | 0.792 | -0.372 | -0.256 | -0.05 | -0.041 | -0.055 | -0.055 | 0.277 | 0.69 | 0.51 | 0.57 | 0.79 | 0.8 | 0.75 |
| 1 | 22744686 | rs4515757 | 0.51 | -0.343 | -0.016 | 0.125 | -0.003 | 0.058 | 0.058 | 0 | 0.92 | 0.81 | 0.79 | 0.76 | 0.93 | 0.9 |
| 1 | 22746686 | rs10799754 | 0.807 | -0.419 | -0.129 | 0.039 | -0.039 | -0.053 | -0.053 | 0 | 0.67 | 0.51 | 0.56 | 0.79 | 0.78 | 0.73 |
| 1 | 22753127 | rs12089317 | 0.97 | -0.629 | 0.219 | -0.052 | 0.032 | 0.034 | 0.034 | 0.165 | 0.66 | 0.25 | 0.55 | 0.76 | 0.78 | 0.73 |
| 1 | 22760325 | rs7530165 | 0.658 | -0.668 | 0.244 | -0.07 | 0.017 | 0.013 | 0.013 | 0.176 | 0.7 | 0.26 | 0.64 | 0.9 | 0.84 | 0.81 |
| 1 | 22760900 | rs11591062 | 0.669 | -0.138 | -1.039 | 0.121 | -0.108 | -0.114 | -0.114 | 0 | 0.39 | 0.03 | 0.23 | 0.49 | 0.4 | 0.43 |
| 1 | 22768784 | rs2078450 | -0.29 | -0.135 | -0.343 | 0.336 | 0.029 | -0.048 | -0.048 | 0 | 0.7 | 0.58 | 0.75 | 0.87 | 0.59 | 0.66 |
| 1 | 22774266 | rs1890775 | -0.264 | 0.032 | -0.719 | 0.448 | 0.016 | -0.023 | -0.023 | 0 | 0.31 | 0.74 | 0.58 | 0.4 | 0.24 | 0.25 |
| 1 | 22778456 | rs4655064 | 0.275 | -0.035 | -0.192 | 0.469 | -0.008 | 0.087 | 0.087 | 0 | 0.25 | 0.46 | 0.46 | 0.36 | 0.23 | 0.23 |
| 1 | 22780927 | rs12133700 | 0.606 | 0.302 | -0.534 | 0.227 | -0.001 | 0.079 | 0.079 | 0.503 | 0.23 | 0.09 | 0.13 | 0.01 | 0.25 | 0.16 |
| 1 | 22784335 | rs1934482 | 0.262 | 0.266 | -0.537 | 0.281 | -0.047 | 0.069 | 0.069 | 0 | 0.28 | 0.13 | 0.2 | 0.11 | 0.37 | 0.3 |
| 1 | 22787711 | rs10493014 | 0.403 | 0.266 | -0.544 | 0.281 | -0.082 | 0.076 | 0.076 | 0.014 | 0.27 | 0.06 | 0.2 | 0.11 | 0.37 | 0.31 |
| 1 | 22795459 | rs4369203 | 0.256 | 0.253 | -0.505 | 0.293 | -0.011 | 0.03 | 0.03 | 0 | 0.27 | 0.59 | 0.47 | 0.36 | 0.23 | 0.23 |
| 1 | 22806281 | rs1011380 | -0.563 | 0.271 | -0.685 | 0.299 | -0.082 | 0.075 | 0.075 | 0 | 0.73 | 0.94 | 0.8 | 0.89 | 0.63 | 0.69 |
| 1 | 22807171 | rs2007200 | 1.406 | 0.177 | -0.572 | 0.477 | -0.063 | -0.03 | -0.03 | 0.475 | 0.36 | 0.02 | 0.16 | 0.32 | 0.36 | 0.39 |

| | | | | | | | | | | | | | | | | |
|---|---|---|---|---|---|---|---|---|---|---|---|---|---|---|---|---|
| 1 | 22835677 | rs209729 | -0.747 | 0.195 | -0.571 | 0.146 | -0.052 | 0.043 | 0.043 | 0 | 0.78 | 0.94 | 0.84 | 0.89 | 0.72 | 0.69 |
| 1 | 22837849 | rs209727 | -0.333 | 0.182 | -0.535 | 0.169 | 0.013 | -0.027 | -0.027 | 4.695 | 0.36 | 0.76 | 0.62 | 0.4 | 0.32 | 0.27 |
| 1 | 22847552 | rs209719 | -0.301 | 0.168 | -0.653 | 0.131 | -0.027 | 0.037 | 0.037 | 0 | 0.75 | 0.91 | 0.74 | 0.85 | 0.71 | 0.66 |
| 1 | 22851434 | rs10917246 | 0.822 | 0.255 | -0.583 | 0.072 | 0.008 | -0.028 | -0.028 | 0 | 0.44 | 0.19 | 0.32 | 0.54 | 0.4 | 0.44 |
| 1 | 22855441 | rs12137696 | 0.209 | 0.535 | -0.781 | 0.096 | 0.002 | -0.121 | -0.121 | 0.67 | 0.28 | 0.38 | 0.3 | 0 | 0.29 | 0.19 |
| 1 | 22863680 | rs12756490 | 0.464 | 0.371 | -1.437 | 0.461 | -0.007 | -0.022 | -0.022 | 0 | 0.41 | 0.2 | 0.21 | 0.5 | 0.39 | 0.43 |
| 1 | 22871312 | rs209714 | -0.178 | -0.363 | -1.55 | -0.223 | 0.046 | 0.066 | 0.066 | 0 | 0.79 | 0.93 | 0.92 | 0.99 | 0.86 | 0.86 |
| 1 | 22872687 | rs766910 | -0.429 | -0.006 | -1.718 | 0.281 | -0.007 | -0.036 | -0.036 | 0 | 0.2 | 0.14 | 0.27 | 0.32 | 0.2 | 0.24 |
| 1 | 22875879 | rs209710 | 0.418 | -0.441 | -1.419 | -0.214 | -0.002 | -0.003 | -0.003 | 0 | 0.63 | 0.65 | 0.61 | 0.32 | 0.67 | 0.61 |
| 1 | 22877240 | rs11584687 | 0.689 | 0.156 | -1.124 | -0.555 | 0.021 | -0.052 | -0.052 | 0 | 0.32 | 0.38 | 0.25 | 0.11 | 0.45 | 0.39 |
| 1 | 22877813 | rs209709 | -0.404 | 0.217 | -0.685 | 0.137 | -0.025 | 0.088 | 0.088 | 0 | 0.81 | 0.67 | 0.77 | 0.36 | 0.85 | 0.81 |
| 1 | 22885766 | rs7521074 | 0.463 | 0.687 | -0.34 | 0.031 | 0.004 | 0.045 | 0.045 | 0 | 0.72 | 0.53 | 0.71 | 0.37 | 0.71 | 0.76 |
| 1 | 22891409 | rs209736 | -1.41 | 0.431 | -0.08 | -0.319 | 0.018 | 0.018 | 0.018 | 0 | 0.4 | 0.67 | 0.35 | 0.32 | 0.31 | 0.51 |
| 1 | 22899011 | rs209741 | -1.703 | 0.625 | -0.295 | -0.472 | 0.003 | 0.092 | 0.092 | 0 | 0.46 | 0.73 | 0.4 | 0.3 | 0.4 | 0.56 |
| 1 | 22906247 | rs649754 | -1.427 | 0.443 | 0.214 | -0.395 | -0.001 | -0.009 | -0.009 | 0 | 0.26 | 0.66 | 0.23 | 0.19 | 0.23 | 0.34 |
| 1 | 22916864 | rs671485 | -1.653 | 0.274 | 0.847 | -0.333 | 0.04 | 0.012 | 0.012 | 0.8 | 0.15 | 0.55 | 0.19 | 0.12 | 0.15 | 0.12 |
| 1 | 22919904 | rs209696 | -1.428 | -0.217 | 0.752 | -0.641 | -0.009 | -0.032 | -0.032 | 0.559 | 0.4 | 0.39 | 0.35 | 0.17 | 0.27 | 0.49 |
| 1 | 22921503 | rs209694 | -2.078 | -0.152 | 0.945 | -0.513 | 0.02 | 0.054 | 0.054 | 0 | 0.43 | 0.71 | 0.49 | 0.25 | 0.35 | 0.43 |
| 1 | 22922800 | rs209690 | -0.207 | -0.13 | 0.664 | -0.506 | -0.013 | -0.009 | -0.009 | 1.817 | 0.16 | 0.21 | 0.13 | 0.12 | 0.11 | 0.22 |
| 1 | 22924464 | rs12404556 | -1.082 | -0.157 | 1.124 | -0.485 | 0 | -0.006 | -0.006 | 0.151 | 0.2 | 0.19 | 0.3 | 0.1 | 0.2 | 0.16 |
| 1 | 22926021 | rs999800 | -1.727 | -0.397 | 1.245 | -0.528 | -0.041 | -0.069 | -0.069 | 1.228 | 0.32 | 0.48 | 0.35 | 0.16 | 0.24 | 0.25 |
| 1 | 22929376 | rs209697 | -0.699 | -0.255 | 1.241 | -0.364 | -0.014 | -0.014 | -0.014 | 0 | 0.18 | 0.28 | 0.15 | 0.12 | 0.13 | 0.16 |
| 1 | 22932162 | rs12128121 | -0.082 | -0.335 | 1.121 | -0.437 | 0.027 | 0.065 | 0.065 | 0 | 0.16 | 0.03 | 0.2 | 0.06 | 0.11 | 0.15 |
| 1 | 22934644 | rs209705 | -0.359 | -0.355 | 1.217 | -0.396 | 0.001 | 0 | 0 | 0.07 | 0.16 | 0.18 | 0.14 | 0.14 | 0.13 | 0.16 |
| 1 | 22936072 | rs209704 | -2.203 | -0.313 | 1.443 | -0.347 | -0.008 | -0.014 | -0.014 | 0 | 0.41 | 0.48 | 0.47 | 0.27 | 0.35 | 0.38 |
| 1 | 22937094 | rs12128546 | -1.068 | -0.393 | 0.935 | -0.274 | 0.003 | 0.021 | 0.021 | 0 | 0.23 | 0.13 | 0.31 | 0.13 | 0.2 | 0.22 |
| 1 | 22937818 | rs10917268 | -0.111 | -0.493 | 1.15 | -0.229 | 0.001 | -0.001 | -0.001 | 0 | 0.13 | 0.14 | 0.12 | 0.14 | 0.09 | 0.12 |
| 1 | 22952386 | rs186037 | -1.005 | -0.469 | 1.171 | -0.21 | -0.04 | -0.001 | -0.001 | 0.218 | 0.2 | 0.39 | 0.16 | 0.18 | 0.14 | 0.17 |
| 1 | 22952823 | rs158762 | -2.078 | -0.709 | 0.922 | -0.447 | 0.021 | 0.028 | 0.028 | 0.344 | 0.33 | 0.74 | 0.21 | 0.19 | 0.22 | 0.22 |
| 1 | 22954372 | rs158761 | -1.922 | -0.156 | 0.482 | 0.409 | 0.002 | 0.072 | 0.072 | 3.965 | 0.48 | 0.82 | 0.61 | 0.61 | 0.43 | 0.65 |
| 1 | 22958318 | rs587585 | -0.783 | -0.127 | 0.597 | 0.451 | -0.005 | 0.006 | 0.006 | 0 | 0.18 | 0.44 | 0.13 | 0.15 | 0.15 | 0.19 |
| 1 | 22965438 | rs172378 | -1.555 | -0.044 | -0.24 | 0.341 | 0.015 | 0.048 | 0.048 | 1.27 | 0.45 | 0.79 | 0.6 | 0.61 | 0.4 | 0.63 |
| 1 | 22970908 | rs12073436 | -0.285 | -0.154 | -0.4 | 0.228 | 0.019 | 0.059 | 0.059 | 1.077 | 0.14 | 0.03 | 0.04 | 0.05 | 0.1 | 0.2 |
| 1 | 22971892 | rs294185 | -1.576 | -0.266 | -0.09 | 0.956 | -0.032 | -0.041 | -0.041 | 0.099 | 0.4 | 0.89 | 0.5 | 0.62 | 0.38 | 0.43 |
| 1 | 22974928 | rs294179 | -2.053 | -0.236 | -0.123 | 0.955 | -0.05 | 0.053 | 0.053 | 4.237 | 0.5 | 0.92 | 0.53 | 0.68 | 0.47 | 0.6 |
| 1 | 22977883 | rs12734877 | -1.576 | -0.255 | 0.64 | 0.879 | 0.002 | 0 | 0 | 1.404 | 0.4 | 0.47 | 0.41 | 0.46 | 0.42 | 0.53 |
| 1 | 22978646 | rs17433222 | -0.071 | -0.256 | 0.887 | 1.12 | -0.025 | 0.013 | 0.013 | 2.541 | 0.2 | 0.1 | 0.34 | 0.14 | 0.29 | 0.18 |
| 1 | 22981876 | rs12756603 | -1.141 | -0.391 | 0.974 | 1.171 | -0.031 | 0.025 | 0.025 | 0 | 0.2 | 0.37 | 0.08 | 0.3 | 0.14 | 0.35 |
| 1 | 22982066 | rs12722729 | -1.141 | -0.391 | 0.974 | 1.171 | -0.031 | 0.025 | 0.025 | 0 | 0.2 | 0.37 | 0.08 | 0.3 | 0.14 | 0.35 |
| 1 | 22986403 | rs631090 | -1.046 | -0.413 | 1.183 | 1.117 | -0.031 | 0.132 | 0.132 | 2.991 | 0.13 | 0.53 | 0.14 | 0.26 | 0.08 | 0.14 |
| 1 | 22986631 | rs623183 | -1.314 | -0.521 | 0.901 | 1.018 | -0.047 | 0.02 | 0.02 | 0 | 0.23 | 0.38 | 0.15 | 0.28 | 0.16 | 0.31 |
| 1 | 22989745 | rs292010 | -0.121 | -0.463 | 0.583 | 1.105 | -0.018 | 0.072 | 0.072 | 0 | 0.1 | 0.17 | 0.17 | 0.22 | 0.07 | 0.13 |
| 1 | 22993335 | rs292005 | -2.35 | -0.285 | 0.237 | 1.037 | -0.086 | 0.026 | 0.026 | 2.737 | 0.51 | 0.93 | 0.6 | 0.69 | 0.48 | 0.62 |
| 1 | 22996571 | rs7549747 | -1.789 | -0.195 | 0.319 | 0.94 | -0.014 | 0.01 | 0.01 | 2.731 | 0.41 | 0.78 | 0.44 | 0.45 | 0.41 | 0.49 |
| 1 | 22997379 | rs2031749 | 0.077 | -0.094 | 0.385 | 0.708 | 0.01 | 0.014 | 0.014 | 0 | 0.12 | 0.09 | 0.04 | 0.05 | 0.08 | 0.21 |
| 1 | 23007175 | rs17486657 | 0.316 | 0.126 | -0.251 | 0.275 | 0 | -0.001 | -0.001 | 0 | 0.27 | 0.24 | 0.33 | 0.21 | 0.33 | 0.2 |
| 1 | 23016103 | rs17433871 | -0.078 | 0.64 | -0.482 | 0.156 | -0.011 | 0.001 | 0.001 | 0.932 | 0.16 | 0.13 | 0.2 | 0.15 | 0.28 | 0.15 |
| 1 | 23026293 | rs294214 | -0.945 | -0.419 | -1.292 | 0.2 | -0.039 | -0.135 | -0.135 | 0.078 | 0.72 | 0.93 | 0.62 | 0.48 | 0.71 | 0.62 |
| 1 | 23031451 | rs6676345 | -0.931 | -0.168 | -0.374 | 0.051 | 0.008 | 0.021 | 0.021 | 0 | 0.36 | 0.46 | 0.28 | 0.5 | 0.37 | 0.34 |
| 1 | 23033257 | rs294218 | -0.334 | -0.425 | -0.019 | -0.296 | 0.019 | 0.022 | 0.022 | 0 | 0.49 | 0.58 | 0.48 | 0.35 | 0.64 | 0.37 |
| 1 | 23039909 | rs2043969 | 0.056 | -0.275 | 0.005 | -0.253 | -0.006 | 0.024 | 0.024 | 0 | 0.17 | 0.19 | 0.13 | 0.37 | 0.13 | 0.19 |
| 1 | 23042127 | rs6695566 | -0.279 | -0.218 | 0.482 | -0.511 | 0 | -0.075 | -0.075 | 0 | 0.57 | 0.49 | 0.63 | 0.93 | 0.45 | 0.72 |
| 1 | 23042318 | rs12074571 | 0.598 | -0.207 | 0.404 | -0.477 | -0.027 | -0.014 | -0.014 | 0 | 0.16 | 0.19 | 0.05 | 0.1 | 0.07 | 0.1 |
| 1 | 23044828 | rs734445 | -0.687 | -0.229 | 0.169 | -0.601 | 0.004 | -0.063 | -0.063 | 0 | 0.63 | 0.64 | 0.66 | 0.93 | 0.46 | 0.75 |
| 1 | 23045943 | rs2869513 | 0.907 | -0.025 | -0.096 | -0.37 | -0.001 | -0.124 | -0.124 | 1.876 | 0.38 | 0.05 | 0.47 | 0.49 | 0.31 | 0.54 |
| 1 | 23050262 | rs882024 | 0.374 | 0.529 | 0.194 | -0.71 | -0.007 | 0.297 | 0.297 | 0.024 | 0.52 | 0.1 | 0.57 | 0.88 | 0.4 | 0.72 |
| 1 | 23050352 | rs882025 | 0.069 | 0.524 | 0.352 | -0.714 | 0.006 | 0.081 | 0.081 | 0 | 0.55 | 0.34 | 0.59 | 0.95 | 0.41 | 0.71 |
| 1 | 23055360 | rs4655093 | 0.182 | 0.519 | -0.018 | -0.6 | -0.052 | 0.258 | 0.258 | 0 | 0.54 | 0.11 | 0.6 | 0.87 | 0.45 | 0.73 |
| 1 | 23057190 | rs12732926 | 1.323 | 0.932 | 0.055 | -0.679 | -0.02 | 0.03 | 0.03 | 0 | 0.23 | 0.13 | 0.23 | 0.01 | 0.34 | 0.15 |
| 1 | 23062908 | rs10917284 | 0.419 | 0.264 | -0.148 | -0.348 | 0.049 | 0.028 | 0.028 | 0.013 | 0.51 | 0.09 | 0.38 | 0.59 | 0.35 | 0.61 |
| 1 | 23063930 | rs4263970 | -0.333 | 0.338 | -0.018 | -0.773 | 0.011 | -0.104 | -0.104 | 0 | 0.6 | 0.56 | 0.66 | 0.96 | 0.46 | 0.75 |
| 1 | 23068761 | rs6426763 | -0.593 | -0.078 | -0.311 | -0.042 | 0.019 | -0.029 | -0.029 | 0 | 0.72 | 0.78 | 0.8 | 0.48 | 0.82 | 0.57 |
| 1 | 23069649 | rs6673115 | -0.194 | 0.308 | -0.17 | -0.506 | 0.01 | 0.057 | 0.057 | 0 | 0.32 | 0.2 | 0.16 | 0.08 | 0.28 | 0.19 |
| 1 | 23072420 | rs876685 | -1.13 | 0.279 | 1.071 | 0.296 | 0.081 | 0.105 | 0.105 | 1.285 | 0.08 | 0.4 | 0.25 | 0.14 | 0.11 | 0.09 |
| 1 | 23080351 | rs4233292 | -0.002 | -0.189 | 0.428 | -0.069 | -0.054 | -0.098 | -0.098 | 0.785 | 0.83 | 0.6 | 0.92 | 0.94 | 0.93 | 0.91 |
| 1 | 23086096 | rs893964 | -0.343 | 0.599 | -0.625 | 0.645 | 0.012 | -0.044 | -0.044 | 0 | 0.26 | 0.1 | 0.18 | 0.6 | 0.18 | 0.37 |
| 1 | 23086883 | rs1561625 | -1.642 | 1.374 | 0.11 | 0.706 | 0.084 | 0.068 | 0.068 | 0 | 0.1 | 0.28 | 0.18 | 0.21 | 0.13 | 0.13 |
| 1 | 23094421 | rs4654814 | -0.822 | 1.257 | -0.517 | -0.062 | -0.024 | 0.023 | 0.023 | 0 | 0.19 | 0.12 | 0.28 | 0.02 | 0.3 | 0.1 |
| 1 | 23094454 | rs4655107 | -0.72 | 1.309 | -0.516 | -0.055 | -0.019 | 0.024 | 0.024 | 0 | 0.18 | 0.11 | 0.25 | 0.02 | 0.26 | 0.1 |

| | | | | | | | | | | | | | | | | |
|---|---|---|---|---|---|---|---|---|---|---|---|---|---|---|---|---|
| 1 | 23096818 | rs7516175 | -0.289 | 0.96 | -0.039 | -0.015 | 0.001 | -0.03 | -0.03 | 0 | 0.09 | 0.12 | 0.15 | 0.01 | 0.1 | 0.2 |
| 1 | 23098982 | rs12410046 | -0.833 | 0.809 | -0.325 | 1.077 | -0.021 | -0.025 | -0.025 | 0 | 0.08 | 0.01 | 0.2 | 0.41 | 0.08 | 0.08 |
| 1 | 23101688 | rs1561624 | 1.85 | 0.64 | 0.548 | 0.951 | -0.006 | 0.142 | 0.142 | 0.163 | 0.83 | 0.65 | 0.68 | 0.45 | 0.86 | 0.83 |
| 1 | 23102856 | rs6692683 | 1.535 | 0.097 | 0.637 | 0.61 | 0.005 | 0.004 | 0.004 | 0.053 | 0.68 | 0.32 | 0.68 | 0.25 | 0.8 | 0.71 |
| 1 | 23106213 | rs2817909 | 0.806 | -0.151 | 1.127 | 0.748 | -0.084 | 0.021 | 0.021 | 3.874 | 0.78 | 0.56 | 0.87 | 0.75 | 0.91 | 0.79 |
| 1 | 23109291 | rs2817901 | 0.673 | -0.15 | 1.312 | 0.743 | -0.023 | 0.005 | 0.005 | 3.199 | 0.8 | 0.74 | 0.88 | 0.75 | 0.91 | 0.79 |
| 1 | 23112369 | rs10799762 | -0.753 | -0.06 | 1.117 | 0.876 | -0.001 | 0.032 | 0.032 | 0.044 | 0.1 | 0.09 | 0.21 | 0.54 | 0.1 | 0.14 |
| 1 | 23115939 | rs11801544 | 0.073 | -0.23 | 1.138 | 0.803 | -0.035 | 0.017 | 0.017 | 2.242 | 0.13 | 0.14 | 0.03 | 0.2 | 0.03 | 0.13 |
| 1 | 23117460 | rs10799764 | -0.793 | 0.063 | 1.208 | 0.98 | 0.005 | -0.001 | -0.001 | 0 | 0.1 | 0.05 | 0.21 | 0.5 | 0.1 | 0.14 |
| 1 | 23118173 | rs893961 | -1.373 | 0.053 | 1.215 | 0.719 | -0.001 | 0.056 | 0.056 | 0.266 | 0.21 | 0.28 | 0.38 | 0.54 | 0.21 | 0.31 |
| 1 | 23122280 | rs4655110 | 0.773 | 0.107 | 1.075 | 0.62 | 0.001 | 0.059 | 0.059 | 0 | 0.83 | 0.65 | 0.75 | 0.69 | 0.85 | 0.67 |
| 1 | 23134218 | rs10465543 | -0.409 | 0.105 | 1.363 | 0.342 | -0.001 | -0.002 | -0.002 | 0 | 0.26 | 0.17 | 0.23 | 0.2 | 0.17 | 0.33 |
| 1 | 23137761 | rs12723359 | 1.402 | -0.04 | 1.399 | 0.542 | 0.025 | 0.027 | 0.027 | 0 | 0.6 | 0.37 | 0.55 | 0.28 | 0.72 | 0.5 |
| 1 | 23140700 | rs2869511 | 1.32 | -0.11 | 1.282 | 0.59 | 0.054 | 0.048 | 0.048 | 0 | 0.59 | 0.24 | 0.54 | 0.29 | 0.71 | 0.51 |
| 1 | 23142496 | rs7536195 | -0.477 | 0.04 | 1.185 | 0.724 | -0.001 | 0.047 | 0.047 | 0 | 0.08 | 0.05 | 0.16 | 0.51 | 0.12 | 0.16 |
| 1 | 23143338 | rs6660612 | -1.388 | 0.655 | 1.016 | 0.379 | -0.015 | 0.076 | 0.076 | 0.105 | 0.46 | 0.27 | 0.48 | 0.79 | 0.46 | 0.56 |
| 1 | 23143647 | rs751022 | -0.069 | 0.641 | 0.956 | 0.39 | -0.006 | -0.015 | -0.015 | 0 | 0.26 | 0.23 | 0.24 | 0.2 | 0.18 | 0.32 |
| 1 | 23144006 | rs893966 | -0.457 | 0.903 | 1.068 | 0.438 | 0.015 | 0.075 | 0.075 | 0 | 0.17 | 0.3 | 0.22 | 0.51 | 0.22 | 0.2 |
| 1 | 23145603 | rs4390147 | 0.129 | 0.957 | 1.106 | 0.465 | 0.028 | 0.13 | 0.13 | 0 | 0.83 | 0.64 | 0.78 | 0.5 | 0.79 | 0.8 |
| 1 | 23146497 | rs6662622 | 0.089 | 0.959 | 1.062 | 0.467 | 0.019 | 0.097 | 0.097 | 0 | 0.83 | 0.69 | 0.78 | 0.5 | 0.79 | 0.8 |
| 1 | 23152969 | rs10917312 | 0.916 | 0.938 | 0.681 | 0.021 | 0 | -0.009 | -0.009 | 0 | 0.53 | 0.52 | 0.47 | 0.68 | 0.57 | 0.45 |
| 1 | 23155231 | rs12090415 | 0.898 | 0.94 | 0.557 | 0.021 | -0.001 | 0.015 | 0.015 | 0 | 0.54 | 0.59 | 0.48 | 0.69 | 0.57 | 0.45 |
| 1 | 23156102 | rs12027585 | 0.704 | 1.205 | 0.276 | -0.101 | -0.022 | -0.006 | -0.006 | 0 | 0.12 | 0.05 | 0.21 | 0.11 | 0.17 | 0.15 |
| 1 | 23158122 | rs10753545 | -1.269 | 0.883 | 0.542 | 0.316 | -0.023 | -0.018 | -0.018 | 0 | 0.4 | 0.76 | 0.51 | 0.53 | 0.47 | 0.34 |
| 1 | 23164184 | rs12731889 | 1.967 | 0.861 | -0.037 | 0.193 | 0.013 | 0.139 | 0.139 | 0.109 | 0.34 | 0.07 | 0.17 | 0.1 | 0.32 | 0.19 |
| 1 | 23169503 | rs4655128 | 0.257 | 0.766 | 0.09 | 0.061 | -0.032 | 0.012 | 0.012 | 0 | 0.7 | 0.49 | 0.71 | 0.68 | 0.82 | 0.61 |
| 1 | 23171706 | rs11800828 | 0.697 | 0.968 | -0.153 | 0.189 | 0 | 0 | 0 | 0.051 | 0.53 | 0.49 | 0.51 | 0.54 | 0.56 | 0.4 |
| 1 | 23177189 | rs4655131 | -1.011 | 0.767 | -0.156 | 0.46 | -0.003 | 0.01 | 0.01 | 0 | 0.4 | 0.37 | 0.53 | 0.83 | 0.41 | 0.57 |
| 1 | 23183130 | rs4654824 | -1.187 | 1.509 | -0.07 | 1.394 | 0.01 | 0.027 | 0.027 | 0.688 | 0.3 | 0.34 | 0.39 | 0.47 | 0.32 | 0.44 |
| 1 | 23184206 | rs7413155 | 0.852 | 1.275 | -0.408 | 1.393 | -0.021 | -0.053 | -0.053 | 0 | 0.76 | 0.84 | 0.69 | 0.52 | 0.69 | 0.64 |
| 1 | 23207135 | rs11807221 | -1.311 | 0.116 | 0.287 | 1.515 | -0.015 | 0.035 | 0.035 | 0 | 0.15 | 0.23 | 0.21 | 0.59 | 0.09 | 0.16 |
| 1 | 23212740 | rs309476 | 0.086 | 0.262 | -0.039 | 0.075 | 0.003 | -0.023 | -0.023 | 0 | 0.42 | 0.48 | 0.53 | 0.25 | 0.5 | 0.41 |
| 1 | 23219187 | rs116119 | 0.123 | 0.05 | 0.428 | -0.033 | -0.036 | 0.07 | 0.07 | 0 | 0.39 | 0.22 | 0.5 | 0.25 | 0.5 | 0.4 |
| 1 | 23225322 | rs309543 | -0.93 | -0.006 | 0.7 | -0.165 | -0.038 | 0.166 | 0.166 | 0.461 | 0.6 | 0.83 | 0.75 | 0.88 | 0.62 | 0.62 |
| 1 | 23227166 | rs2675494 | -0.123 | -0.126 | 0.746 | -0.276 | -0.014 | 0.092 | 0.092 | 0 | 0.44 | 0.68 | 0.52 | 0.25 | 0.5 | 0.43 |
| 1 | 23233600 | rs309500 | -0.113 | 0.334 | 0.395 | 0.309 | -0.041 | 0.062 | 0.062 | 0 | 0.46 | 0.8 | 0.56 | 0.36 | 0.53 | 0.4 |
| 1 | 23238023 | rs309495 | -1.191 | 0.271 | 0.227 | -0.017 | -0.022 | 0.129 | 0.129 | 0 | 0.63 | 0.82 | 0.77 | 0.88 | 0.66 | 0.61 |
| 1 | 23247395 | rs744834 | 0.212 | 0.122 | 0.377 | -0.043 | -0.034 | 0.07 | 0.07 | 0 | 0.37 | 0.21 | 0.29 | 0.2 | 0.48 | 0.2 |
| 1 | 23267090 | rs722993 | -0.886 | 0.715 | 0.73 | 0.415 | -0.011 | -0.017 | -0.017 | 1.669 | 0.28 | 0.25 | 0.48 | 0.56 | 0.37 | 0.41 |
| 1 | 23274948 | rs309539 | -1.287 | 0.463 | 0.545 | -0.081 | 0.024 | 0.096 | 0.096 | 0 | 0.57 | 0.75 | 0.75 | 0.85 | 0.67 | 0.58 |
| 1 | 23278632 | rs620514 | 0.053 | 0.449 | 0.464 | -0.035 | 0.003 | -0.022 | -0.022 | 0 | 0.29 | 0.49 | 0.27 | 0.29 | 0.3 | 0.16 |
| 1 | 23283076 | rs4654828 | -0.536 | 0.487 | 0.138 | 0.147 | 0.059 | 0.101 | 0.101 | 0 | 0.47 | 0.21 | 0.49 | 0.78 | 0.67 | 0.38 |
| 1 | 23288580 | rs7543064 | -0.994 | 0.204 | 0.458 | 0.489 | 0.007 | 0.089 | 0.089 | 0.616 | 0.17 | 0.37 | 0.37 | 0.3 | 0.18 | 0.25 |
| 1 | 23288828 | rs6426775 | -0.903 | -0.777 | -0.636 | -0.28 | -0.063 | -0.026 | -0.026 | 0.689 | 0.74 | 0.54 | 0.8 | 0.81 | 0.86 | 0.7 |
| 1 | 23289397 | rs10799776 | 0.612 | -0.786 | -0.755 | -0.291 | 0.065 | -0.098 | -0.098 | 0 | 0.56 | 0.07 | 0.41 | 0.52 | 0.67 | 0.45 |
| 1 | 23291090 | rs925826 | -0.791 | -0.693 | -0.427 | -0.377 | -0.059 | -0.027 | -0.027 | 0 | 0.74 | 0.55 | 0.8 | 0.82 | 0.84 | 0.7 |
| 1 | 23294839 | rs648322 | 0.32 | -0.813 | -0.545 | -0.52 | -0.003 | -0.002 | -0.002 | 0 | 0.39 | 0.4 | 0.38 | 0.55 | 0.49 | 0.33 |
| 1 | 23312025 | rs4655141 | -0.61 | -1.165 | -0.285 | -0.598 | -0.045 | -0.034 | -0.034 | 0 | 0.71 | 0.57 | 0.59 | 0.83 | 0.82 | 0.52 |
| 1 | 23345051 | rs561136 | -1.736 | -1.23 | -0.197 | -0.615 | 0.032 | 0.041 | 0.041 | 0 | 0.77 | 0.86 | 0.83 | 0.88 | 0.84 | 0.7 |
| 1 | 23376541 | rs10917347 | 1.813 | -1.051 | -0.555 | -0.645 | -0.042 | -0.122 | -0.122 | 0 | 0.15 | 0.01 | 0.21 | 0.49 | 0.19 | 0.18 |
| 1 | 23381116 | rs7543975 | 0.01 | -1.634 | -0.139 | -0.691 | 0.005 | 0.007 | 0.007 | 0.049 | 0.34 | 0.29 | 0.22 | 0.25 | 0.31 | 0.24 |
| 1 | 23409538 | rs2235546 | 1.127 | -1.77 | -0.243 | -0.674 | -0.147 | -0.138 | -0.138 | 0 | 0.35 | 0.04 | 0.34 | 0.55 | 0.49 | 0.32 |
| 1 | 23413458 | rs580878 | -0.93 | -1.819 | -0.117 | -0.727 | 0.029 | 0.013 | 0.013 | 0 | 0.74 | 0.81 | 0.62 | 0.81 | 0.83 | 0.56 |
| 1 | 23415102 | rs683893 | -0.933 | -1.815 | -0.126 | -0.736 | 0.029 | 0.013 | 0.013 | 0 | 0.74 | 0.81 | 0.62 | 0.81 | 0.83 | 0.56 |
| 1 | 23418153 | rs10799790 | -0.968 | -1.696 | -0.077 | -0.677 | 0.031 | 0.012 | 0.012 | 0 | 0.74 | 0.84 | 0.62 | 0.81 | 0.83 | 0.56 |
| 1 | 23419261 | rs477717 | -0.813 | -1.684 | -0.02 | -0.671 | -0.022 | -0.011 | -0.011 | 0 | 0.73 | 0.64 | 0.59 | 0.81 | 0.83 | 0.56 |
| 1 | 23419374 | rs3765407 | -0.727 | -1.685 | -0.006 | -0.671 | -0.041 | -0.021 | -0.021 | 0.775 | 0.72 | 0.59 | 0.59 | 0.81 | 0.83 | 0.56 |
| 1 | 23419383 | rs477830 | -0.812 | -1.678 | -0.053 | -0.668 | -0.022 | -0.011 | -0.011 | 0 | 0.73 | 0.64 | 0.59 | 0.81 | 0.83 | 0.56 |
| 1 | 23419855 | rs1983965 | -0.984 | -1.66 | -0.233 | -0.659 | 0.03 | 0.012 | 0.012 | 0 | 0.74 | 0.84 | 0.62 | 0.81 | 0.83 | 0.56 |
| 1 | 23429122 | rs1208932 | 0.849 | -1.434 | 0.146 | -0.417 | 0.032 | 0.009 | 0.009 | 0 | 0.49 | 0.32 | 0.47 | 0.56 | 0.55 | 0.4 |
| 1 | 23431777 | rs765803 | -0.744 | -1.456 | 0.172 | -0.449 | 0 | -0.001 | -0.001 | 0.056 | 0.85 | 0.85 | 0.72 | 0.82 | 0.89 | 0.65 |
| 1 | 23432346 | rs1208930 | 0.466 | -1.534 | 0.336 | -0.49 | -0.014 | 0.092 | 0.092 | 0.012 | 0.73 | 0.56 | 0.57 | 0.56 | 0.73 | 0.53 |
| 1 | 23435538 | rs16827981 | 0.938 | -1.435 | 0.346 | -0.482 | 0.002 | -0.041 | -0.041 | 0.286 | 0.24 | 0.22 | 0.09 | 0 | 0.18 | 0.13 |
| 1 | 23435669 | rs2274687 | 0.735 | -1.358 | 0.344 | -0.41 | 0.02 | 0.078 | 0.078 | 0 | 0.11 | 0.19 | 0.15 | 0.25 | 0.16 | 0.11 |
| 1 | 23464974 | rs10917351 | 0.763 | -1.351 | 0.347 | -0.393 | 0.02 | 0.082 | 0.082 | 0 | 0.11 | 0.21 | 0.15 | 0.26 | 0.16 | 0.11 |
| 1 | 23472341 | rs3845521 | 0.763 | -1.351 | 0.347 | -0.393 | 0.02 | 0.082 | 0.082 | 0 | 0.11 | 0.2 | 0.15 | 0.26 | 0.16 | 0.11 |
| 1 | 23501354 | rs643492 | -0.261 | -1.41 | 0.314 | -0.429 | -0.006 | -0.017 | -0.017 | 0 | 0.37 | 0.5 | 0.26 | 0.26 | 0.35 | 0.25 |
| 1 | 23510850 | rs1738468 | -0.261 | -1.41 | 0.298 | -0.429 | -0.006 | -0.017 | -0.017 | 0 | 0.37 | 0.51 | 0.27 | 0.26 | 0.35 | 0.25 |
| 1 | 23513980 | rs641032 | -0.316 | -1.407 | 0.214 | -0.448 | -0.006 | -0.018 | -0.018 | 0 | 0.38 | 0.51 | 0.27 | 0.26 | 0.35 | 0.25 |

| | | | | | | | | | | | | | | | | |
|---|---|---|---|---|---|---|---|---|---|---|---|---|---|---|---|---|
| 1 | 23519477 | rs623988 | -0.303 | -1.398 | 0.22 | -0.429 | -0.007 | -0.019 | -0.019 | 0 | 0.38 | 0.5 | 0.26 | 0.26 | 0.35 | 0.25 |
| 1 | 23537555 | rs627304 | -1.48 | -1.798 | 0.141 | -0.406 | 0.036 | 0.004 | 0.004 | 0 | 0.51 | 0.65 | 0.51 | 0.45 | 0.42 | 0.6 |
| 1 | 23539010 | rs10917360 | -0.367 | -1.832 | 0.312 | -0.437 | -0.001 | -0.002 | -0.002 | 0 | 0.39 | 0.4 | 0.35 | 0.19 | 0.25 | 0.49 |
| 1 | 23540125 | rs16828047 | 0.421 | -1.812 | 0.269 | -0.427 | 0.008 | -0.009 | -0.009 | 0.171 | 0.13 | 0.11 | 0.26 | 0.19 | 0.09 | 0.35 |
| 1 | 23540543 | rs16828054 | 0.943 | -1.796 | 0.267 | -0.446 | 0 | -0.047 | -0.047 | 0 | 0.24 | 0.24 | 0.07 | 0 | 0.16 | 0.14 |
| 1 | 23547194 | rs10917362 | 1.156 | -1.431 | 0.809 | 0.457 | -0.013 | 0.067 | 0.067 | 0 | 0.74 | 0.64 | 0.57 | 0.55 | 0.73 | 0.54 |
| 1 | 23548871 | rs10917363 | -0.156 | -1.053 | 0.786 | 0.499 | 0.022 | 0.086 | 0.086 | 0 | 0.88 | 0.77 | 0.84 | 0.73 | 0.83 | 0.89 |
| 1 | 23560000 | rs4654837 | 0.508 | -1.165 | 0.478 | 0.466 | 0 | -0.007 | -0.007 | 0 | 0.84 | 0.89 | 0.86 | 0.7 | 0.83 | 0.83 |
| 1 | 23565599 | rs1740482 | -1.869 | -1.253 | 1.101 | 0.572 | -0.018 | 0.112 | 0.112 | 0.142 | 0.19 | 0.48 | 0.37 | 0.36 | 0.16 | 0.25 |
| 1 | 23577073 | rs1535180 | -1.763 | -1.345 | 1.156 | 0.45 | -0.022 | 0.077 | 0.077 | 0 | 0.2 | 0.38 | 0.44 | 0.3 | 0.17 | 0.26 |
| 1 | 23577666 | rs2144732 | -0.394 | -1.536 | 0.812 | 0.25 | 0.011 | -0.002 | -0.002 | 0 | 0.88 | 0.9 | 0.94 | 0.88 | 0.9 | 0.76 |
| 1 | 23587590 | rs9658850 | -1.851 | -1.19 | 1.282 | 0.456 | -0.002 | 0.009 | 0.009 | 0 | 0.19 | 0.19 | 0.4 | 0.25 | 0.16 | 0.28 |
| 1 | 23604370 | rs628465 | -2.224 | -1.51 | 1.846 | 0.353 | -0.005 | 0.002 | 0.002 | 2.253 | 0.21 | 0.62 | 0.45 | 0.22 | 0.18 | 0.22 |
| 1 | 23631625 | rs655600 | -2.225 | -1.577 | 2.083 | 0.336 | -0.005 | 0.004 | 0.004 | 2.227 | 0.21 | 0.62 | 0.45 | 0.22 | 0.17 | 0.23 |
| 1 | 23639009 | rs1535179 | -1.368 | -1.589 | 1.289 | 0.358 | 0.004 | -0.047 | -0.047 | 0 | 0.09 | 0.01 | 0.28 | 0.13 | 0.05 | 0.19 |
| 1 | 23667751 | rs2103911 | 2.986 | -1.647 | 1.204 | 0.45 | 0.031 | -0.089 | -0.089 | 0 | 0.67 | 0.13 | 0.48 | 0.59 | 0.73 | 0.51 |
| 1 | 23669770 | rs3753250 | -1.871 | -1.443 | 1.304 | 0.324 | -0.002 | -0.002 | -0.002 | 0 | 0.16 | 0.14 | 0.4 | 0.22 | 0.18 | 0.22 |
| 1 | 23686798 | rs601786 | 0.369 | -1.25 | 1.088 | 0.346 | 0.007 | 0.003 | 0.003 | 0 | 0.06 | 0.06 | 0.11 | 0.1 | 0.12 | 0.04 |
| 1 | 23687827 | rs644749 | -2.46 | -1.406 | 1.234 | 0.343 | 0.002 | 0.043 | 0.043 | 0.289 | 0.2 | 0.33 | 0.24 | 0.27 | 0.19 | 0.31 |
| 1 | 23688178 | rs646193 | 0.588 | -1.742 | 1.293 | 0.214 | -0.059 | -0.005 | -0.005 | 0 | 0.86 | 0.56 | 0.89 | 0.91 | 0.92 | 0.81 |
| 1 | 23688933 | rs2849028 | 2.947 | -1.751 | 1.158 | 0.209 | 0.041 | -0.031 | -0.031 | 0 | 0.65 | 0.19 | 0.65 | 0.63 | 0.73 | 0.5 |
| 1 | 23689083 | rs3795308 | -1.002 | -1.73 | 1.137 | 0.035 | 0.011 | -0.05 | -0.05 | 1.051 | 0.1 | 0.02 | 0.11 | 0.18 | 0.07 | 0.27 |
| 1 | 23696152 | rs665977 | 0.666 | -1.703 | 1.172 | 0.029 | -0.063 | -0.005 | -0.005 | 0 | 0.86 | 0.51 | 0.89 | 0.91 | 0.91 | 0.81 |
| 1 | 23700811 | rs9887921 | -1.912 | -1.359 | 0.9 | 0.143 | -0.03 | 0.027 | 0.027 | 0.184 | 0.14 | 0.25 | 0.13 | 0.18 | 0.06 | 0.28 |
| 1 | 23701312 | rs1740481 | 0.393 | -1.281 | 0.687 | 0.121 | 0.007 | 0 | 0 | 0 | 0.06 | 0.06 | 0.11 | 0.09 | 0.12 | 0.05 |
| 1 | 23708989 | rs10917375 | -2.091 | -1.355 | 0.746 | 0.15 | -0.039 | 0.037 | 0.037 | 0.14 | 0.14 | 0.3 | 0.12 | 0.18 | 0.07 | 0.28 |
| 1 | 23710177 | rs686546 | 0.242 | -1.205 | 0.327 | 0.053 | 0.015 | 0.005 | 0.005 | 0.786 | 0.89 | 0.91 | 0.94 | 0.91 | 0.92 | 0.8 |
| 1 | 23710475 | rs2298632 | 0.616 | -1.159 | -0.436 | -0.569 | -0.149 | -0.048 | -0.048 | 0 | 0.31 | 0.07 | 0.52 | 0.66 | 0.49 | 0.24 |
| 1 | 23711376 | rs2092933 | 0.75 | -1.366 | 0.674 | 0.207 | -0.024 | -0.008 | -0.008 | 0 | 0.87 | 0.76 | 0.93 | 0.91 | 0.91 | 0.8 |
| 1 | 23714448 | rs1198447 | 0.999 | -1.37 | 0.844 | 0.205 | -0.057 | -0.023 | -0.023 | 0 | 0.86 | 0.62 | 0.92 | 0.91 | 0.91 | 0.8 |
| 1 | 23714665 | rs1212034 | 1.508 | -1.359 | 0.816 | 0.188 | -0.063 | 0.037 | 0.037 | 0.748 | 0.79 | 0.56 | 0.83 | 0.73 | 0.86 | 0.56 |
| 1 | 23723935 | rs7516962 | -1.279 | -1.056 | 0.658 | 0.457 | -0.015 | 0.125 | 0.125 | 0.389 | 0.09 | 0.28 | 0.08 | 0.25 | 0.04 | 0.29 |
| 1 | 23724183 | rs2275355 | 1.507 | -1.328 | -0.242 | 0.145 | 0.038 | -0.038 | -0.038 | 0.193 | 0.6 | 0.09 | 0.57 | 0.55 | 0.71 | 0.49 |
| 1 | 23734350 | rs1767141 | 0.075 | -0.961 | 0.12 | 0.3 | 0.007 | -0.008 | -0.008 | 2.007 | 0.91 | 0.92 | 0.85 | 0.86 | 0.92 | 0.86 |
| 1 | 23735241 | rs1198423 | 0.027 | -0.881 | 0.76 | 0.09 | 0.013 | -0.005 | -0.005 | 0 | 0.9 | 0.76 | 0.87 | 0.88 | 0.88 | 0.86 |
| 1 | 23743296 | rs6682786 | -0.544 | -1.112 | 1.465 | 0.225 | -0.011 | 0.078 | 0.078 | 0 | 0.06 | 0.09 | 0.09 | 0.24 | 0.02 | 0.27 |
| 1 | 23745228 | rs3737646 | -1.459 | -1.36 | 1.261 | -0.021 | -0.054 | 0.042 | 0.042 | 1.068 | 0.17 | 0.39 | 0.16 | 0.24 | 0.07 | 0.31 |
| 1 | 23748440 | rs7515139 | -0.32 | -1.336 | 1.576 | -0.002 | -0.004 | 0.167 | 0.167 | 0 | 0.06 | 0.17 | 0.09 | 0.24 | 0.02 | 0.28 |
| 1 | 23755513 | rs1198430 | -0.093 | -1.355 | 1.419 | 0 | 0.001 | -0.003 | -0.003 | 0 | 0.9 | 0.84 | 0.88 | 0.88 | 0.89 | 0.88 |
| 1 | 23756695 | rs1198433 | 1.224 | -1.422 | 1.671 | -0.011 | -0.021 | 0.103 | 0.103 | 1.946 | 0.83 | 0.44 | 0.76 | 0.64 | 0.87 | 0.6 |
| 1 | 23765472 | rs1767148 | -0.304 | -1.395 | 1.534 | -0.028 | -0.001 | -0.003 | -0.003 | 0 | 0.9 | 0.84 | 0.88 | 0.88 | 0.89 | 0.88 |
| 1 | 23767669 | rs12117542 | 1.14 | -1.383 | 0.908 | -0.056 | 0.01 | 0.037 | 0.037 | 0.216 | 0.07 | 0.01 | 0.04 | 0 | 0.04 | 0.03 |
| 1 | 23778362 | rs2077066 | 1.966 | -1.552 | 0.612 | -0.119 | 0.055 | -0.034 | -0.034 | 7.795 | 0.54 | 0.09 | 0.61 | 0.55 | 0.67 | 0.48 |
| 1 | 23794914 | rs7522616 | 1.999 | -1.418 | 1.239 | 0.239 | 0.05 | -0.032 | -0.032 | 0 | 0.72 | 0.17 | 0.71 | 0.64 | 0.83 | 0.57 |
| 1 | 23805293 | rs2742958 | -0.923 | -1.017 | 1.117 | -0.3 | 0.065 | 0.061 | 0.061 | 9.399 | 0.05 | 0.41 | 0.04 | 0 | 0 | 0 |
| 1 | 23824242 | rs2811963 | 0.529 | -0.678 | 0.526 | -0.148 | -0.009 | -0.013 | -0.013 | 0 | 0.88 | 0.64 | 0.71 | 0.89 | 0.92 | 0.88 |
| 1 | 23831840 | rs12562830 | -0.472 | -0.715 | -0.325 | 0.247 | 0.013 | -0.038 | -0.038 | 0 | 0.07 | 0.01 | 0.1 | 0.26 | 0.05 | 0.32 |
| 1 | 23839769 | rs2282720 | 0.322 | -0.734 | -0.237 | -0.566 | -0.008 | -0.004 | -0.004 | 0 | 0.38 | 0.36 | 0.45 | 0.56 | 0.53 | 0.33 |
| 1 | 23845288 | rs3218177 | -0.183 | -0.864 | -0.544 | -0.649 | 0.03 | -0.051 | -0.051 | 0.527 | 0.09 | 0.01 | 0.11 | 0.22 | 0.06 | 0.36 |
| 1 | 23847464 | rs2075995 | 0.693 | -1.036 | -0.719 | -0.784 | -0.159 | -0.115 | -0.115 | 0 | 0.32 | 0.06 | 0.41 | 0.6 | 0.51 | 0.3 |
| 1 | 23848042 | rs2038027 | -0.343 | -1.039 | -0.18 | -0.831 | -0.001 | -0.03 | -0.03 | 0 | 0.46 | 0.5 | 0.56 | 0.83 | 0.57 | 0.67 |
| 1 | 23858317 | rs3218121 | 1.553 | -1.334 | -0.772 | -1.025 | 0.008 | 0.016 | 0.016 | 8.004 | 0.11 | 0.04 | 0.04 | 0 | 0.07 | 0.09 |
| 1 | 23859521 | rs6667575 | 1.05 | -0.206 | -0.962 | -0.985 | -0.043 | -0.143 | -0.143 | 0 | 0.25 | 0.03 | 0.32 | 0.6 | 0.27 | 0.24 |
| 1 | 23884369 | rs2071494 | -0.155 | 0.104 | -0.322 | -0.768 | -0.015 | 0.02 | 0.02 | 0 | 0.09 | 0.1 | 0.09 | 0.18 | 0.05 | 0.37 |
| 1 | 23884661 | rs1050096 | -0.01 | 0.404 | -0.323 | -1.61 | -0.008 | -0.009 | -0.009 | 0.039 | 0.72 | 0.72 | 0.64 | 0.91 | 0.63 | 0.76 |
| 1 | 23884780 | rs2920 | 0.291 | 1.426 | -0.089 | -0.899 | 0.023 | 0.013 | 0.013 | 1.044 | 0.14 | 0.19 | 0.16 | 0.17 | 0.2 | 0.08 |
| 1 | 23885498 | rs11574 | -0.404 | 1.467 | -0.236 | -1.11 | -0.027 | 0.029 | 0.029 | 0.978 | 0.93 | 0.99 | 0.74 | 0.97 | 0.76 | 0.92 |
| 1 | 23886935 | rs1555025 | -0.052 | 0.309 | 0.104 | -1.207 | -0.001 | 0.003 | 0.003 | 1.923 | 0.54 | 0.61 | 0.53 | 0.81 | 0.52 | 0.36 |
| 1 | 23902416 | rs2811946 | 0.093 | 0.271 | -0.37 | -1.12 | 0.003 | 0.036 | 0.036 | 0.067 | 0.51 | 0.64 | 0.5 | 0.78 | 0.55 | 0.48 |
| 1 | 23904345 | rs10917395 | 0.523 | 0.992 | -0.839 | -0.779 | 0.016 | 0.079 | 0.079 | 1.972 | 0.5 | 0.36 | 0.44 | 0.78 | 0.48 | 0.45 |
| 1 | 23907702 | rs4649099 | -0.319 | 1.157 | -1.766 | -0.335 | -0.073 | 0.102 | 0.102 | 2.486 | 0.39 | 0.11 | 0.47 | 0.21 | 0.44 | 0.45 |
| 1 | 23926256 | rs2864889 | -0.897 | 1.493 | -0.876 | 0.576 | 0.029 | 0.037 | 0.037 | 0 | 0.33 | 0.56 | 0.59 | 0.55 | 0.4 | 0.53 |
| 1 | 23927112 | rs10917397 | -0.607 | 1.778 | -0.713 | 0.794 | -0.001 | 0 | 0 | 0 | 0.19 | 0.19 | 0.47 | 0.36 | 0.25 | 0.26 |
| 1 | 23927929 | rs4649100 | -0.632 | 1.768 | -0.592 | 0.79 | 0.011 | 0.016 | 0.016 | 0 | 0.19 | 0.25 | 0.49 | 0.36 | 0.25 | 0.26 |
| 1 | 23937567 | rs2811986 | 0.048 | 0.848 | -1.135 | -0.532 | 0.034 | 0.067 | 0.067 | 2.34 | 0.59 | 0.33 | 0.69 | 0.95 | 0.69 | 0.63 |
| 1 | 23941735 | rs12040769 | 0.367 | 1.2 | -1.629 | -0.257 | -0.011 | 0.075 | 0.075 | 1.123 | 0.44 | 0.15 | 0.44 | 0.67 | 0.59 | 0.48 |
| 1 | 23942802 | rs6677615 | -0.267 | 0.45 | -0.718 | -0.753 | -0.005 | -0.012 | -0.012 | 0 | 0.38 | 0.41 | 0.42 | 0.28 | 0.31 | 0.4 |
| 1 | 23943905 | rs4649102 | 0.543 | 0.978 | -0.914 | -0.078 | -0.008 | -0.005 | -0.005 | 0.257 | 0.74 | 0.49 | 0.72 | 0.76 | 0.76 | 0.8 |
| 1 | 23945443 | rs2864884 | 0.96 | 0.803 | -0.924 | -0.333 | -0.001 | 0.003 | 0.003 | 0 | 0.6 | 0.4 | 0.43 | 0.49 | 0.61 | 0.56 |

| | | | | | | | | | | | | | | | | |
|---|---|---|---|---|---|---|---|---|---|---|---|---|---|---|---|---|
| 1 | 23950147 | rs2811981 | -0.777 | 0.483 | -0.644 | -0.509 | -0.016 | -0.017 | -0.017 | 0 | 0.89 | 0.88 | 0.92 | 0.95 | 0.93 | 0.92 |
| 1 | 23950177 | rs12746273 | 0.16 | 1.317 | -0.904 | -0.61 | -0.015 | 0.005 | 0.005 | 0 | 0.1 | 0.06 | 0.12 | 0.08 | 0.23 | 0.1 |
| 1 | 23951114 | rs2813875 | 0.124 | 0.764 | -0.44 | -0.942 | -0.033 | -0.113 | -0.113 | 0.124 | 0.64 | 0.59 | 0.78 | 0.95 | 0.78 | 0.74 |
| 1 | 23957318 | rs2811956 | 0.449 | 0.805 | -0.582 | -0.594 | 0.008 | 0.01 | 0.01 | 0 | 0.52 | 0.64 | 0.48 | 0.57 | 0.6 | 0.52 |
| 1 | 23964809 | rs12760004 | -0.09 | 1.185 | -1.098 | -0.283 | -0.082 | 0.044 | 0.044 | 0.025 | 0.13 | 0.01 | 0.13 | 0.06 | 0.25 | 0.13 |
| 1 | 23965829 | rs10917404 | -0.31 | 0.847 | -0.988 | -0.273 | -0.017 | -0.092 | -0.092 | 0 | 0.25 | 0.08 | 0.4 | 0.43 | 0.25 | 0.32 |
| 1 | 23970650 | rs4649104 | 1.054 | 0.652 | -0.856 | -0.305 | 0 | 0 | 0 | 0.059 | 0.56 | 0.61 | 0.38 | 0.53 | 0.44 | 0.51 |
| 1 | 23972740 | rs2864098 | 1.295 | 0.714 | -1.355 | -0.17 | -0.016 | -0.085 | -0.085 | 0 | 0.28 | 0.11 | 0.2 | 0.51 | 0.27 | 0.35 |
| 1 | 23973908 | rs7527966 | -0.01 | 1 | -1.544 | -0.369 | -0.071 | 0.054 | 0.054 | 0.007 | 0.15 | 0.03 | 0.13 | 0.05 | 0.27 | 0.1 |
| 1 | 23974031 | rs10753557 | 1.195 | 0.747 | -1.355 | -0.249 | -0.008 | -0.07 | -0.07 | 0.018 | 0.29 | 0.13 | 0.21 | 0.5 | 0.27 | 0.35 |
| 1 | 23985309 | rs6684096 | 0.364 | 0.614 | -1.452 | -0.129 | -0.094 | -0.081 | -0.081 | 0 | 0.39 | 0.11 | 0.35 | 0.51 | 0.49 | 0.41 |
| 1 | 23994072 | rs7549155 | 0.309 | 0.325 | -0.693 | -0.583 | 0.016 | -0.001 | -0.001 | 0 | 0.45 | 0.39 | 0.24 | 0.05 | 0.34 | 0.26 |
| 1 | 23997640 | rs6673098 | 0.691 | 0.318 | -0.706 | -0.541 | -0.104 | -0.093 | -0.093 | 0 | 0.39 | 0.08 | 0.35 | 0.51 | 0.48 | 0.37 |
| 1 | 23998136 | rs6679217 | -0.578 | 0.372 | -0.502 | -0.522 | 0.009 | -0.02 | -0.02 | 0 | 0.54 | 0.56 | 0.76 | 0.93 | 0.64 | 0.73 |
| 1 | 24004202 | rs2864704 | -0.766 | 0.736 | 0.286 | 0.151 | 0.008 | 0.128 | 0.128 | 0 | 0.13 | 0.21 | 0.38 | 0.42 | 0.16 | 0.35 |
| 1 | 24005465 | rs10917411 | -0.752 | 0.702 | -0.494 | 0.466 | -0.072 | -0.115 | -0.115 | 0 | 0.24 | 0.07 | 0.3 | 0.53 | 0.38 | 0.31 |
| 1 | 24009544 | rs10799802 | -0.768 | 0.679 | -0.512 | 0.303 | -0.075 | -0.117 | -0.117 | 0 | 0.24 | 0.06 | 0.31 | 0.53 | 0.38 | 0.29 |
| 1 | 24018224 | rs3753270 | -0.523 | 0.913 | 0.277 | 0.27 | 0.046 | 0.234 | 0.234 | 0 | 0.14 | 0.35 | 0.21 | 0.42 | 0.18 | 0.43 |
| 1 | 24022437 | rs2280437 | -0.924 | 0.899 | 0.018 | 0.452 | -0.032 | -0.053 | -0.053 | 0 | 0.26 | 0.15 | 0.33 | 0.57 | 0.38 | 0.29 |
| 1 | 24043985 | rs985086 | 0.759 | 1.171 | 0.198 | 0.087 | -0.002 | -0.002 | -0.002 | 0.147 | 0.67 | 0.71 | 0.61 | 0.52 | 0.55 | 0.64 |
| 1 | 24049595 | rs561733 | 0.554 | 1.303 | 0.152 | 0.057 | 0.016 | 0.019 | 0.019 | 0 | 0.77 | 0.75 | 0.66 | 0.54 | 0.63 | 0.75 |
| 1 | 24062987 | rs10917420 | 1.549 | 1.051 | 0.51 | -0.066 | -0.008 | -0.006 | -0.006 | 0 | 0.61 | 0.34 | 0.56 | 0.51 | 0.54 | 0.61 |
| 1 | 24072355 | rs550850 | -1.147 | 1.536 | 0.462 | 0.256 | -0.005 | 0.007 | 0.007 | 0 | 0.94 | 0.97 | 0.89 | 0.82 | 0.92 | 0.87 |
| 1 | 24077451 | rs2235541 | 1.093 | 1.366 | 0.119 | 0.235 | -0.017 | -0.018 | -0.018 | 0 | 0.12 | 0 | 0.11 | 0.09 | 0.09 | 0.15 |
| 1 | 24083494 | rs2076345 | 0.274 | 0.738 | 0.009 | -0.194 | -0.02 | 0.014 | 0.014 | 0 | 0.25 | 0.09 | 0.24 | 0.27 | 0.31 | 0.14 |
| 1 | 24083649 | rs2076346 | 0.696 | 1.048 | 0.115 | -0.285 | -0.031 | -0.049 | -0.049 | 0 | 0.18 | 0.09 | 0.21 | 0.52 | 0.26 | 0.43 |
| 1 | 24085063 | rs10917423 | 1.305 | 1.269 | 0.229 | 0.028 | -0.039 | 0.1 | 0.1 | 0 | 0.16 | 0.01 | 0.14 | 0 | 0.2 | 0.07 |
| 1 | 24088128 | rs1045017 | 1.439 | 0.996 | 0.681 | 0.173 | 0.029 | 0.059 | 0.059 | 0 | 0.34 | 0.15 | 0.31 | 0 | 0.26 | 0.19 |
| 1 | 24100223 | rs7514394 | 1.322 | 1.313 | 1.445 | 1.458 | 0.002 | 0.005 | 0.005 | 0 | 0.63 | 0.43 | 0.6 | 0.55 | 0.59 | 0.69 |
| 1 | 24102080 | rs11591202 | -0.421 | 1.359 | 1.48 | 1.477 | -0.015 | 0.017 | 0.017 | 0 | 0.26 | 0.14 | 0.26 | 0.27 | 0.33 | 0.14 |
| 1 | 24114129 | rs4237 | 0.804 | 1.363 | 1.513 | 1.485 | -0.021 | -0.029 | -0.029 | 0 | 0.68 | 0.82 | 0.63 | 0.55 | 0.59 | 0.71 |
| 1 | 24128467 | rs11714 | -0.123 | 1.368 | 1.552 | 1.498 | 0.006 | 0.057 | 0.057 | 0 | 0.93 | 0.93 | 0.88 | 0.82 | 0.92 | 0.85 |
| 1 | 24129126 | rs2076343 | -0.431 | 1.354 | 1.406 | 1.417 | -0.021 | 0.023 | 0.023 | 0 | 0.27 | 0.09 | 0.26 | 0.28 | 0.33 | 0.14 |
| 1 | 24133612 | rs1018396 | 1.361 | 1.466 | 1.495 | 1.526 | 0.002 | 0.012 | 0.012 | 0 | 0.63 | 0.43 | 0.6 | 0.51 | 0.59 | 0.7 |
| 1 | 24134721 | rs719400 | -0.433 | 1.637 | 0.397 | 1.6 | -0.005 | 0.025 | 0.025 | 0 | 0.94 | 0.97 | 0.89 | 0.78 | 0.93 | 0.84 |
| 1 | 24136545 | rs2473375 | 0.987 | 1.628 | 0.432 | 1.542 | -0.005 | -0.015 | -0.015 | 0 | 0.67 | 0.72 | 0.63 | 0.51 | 0.6 | 0.7 |
| 1 | 24148365 | rs2179395 | 0.917 | 1.525 | 0.402 | 1.553 | -0.02 | -0.041 | -0.041 | 0.001 | 0.67 | 0.8 | 0.63 | 0.51 | 0.59 | 0.7 |
| 1 | 24149315 | rs2502979 | 1.18 | 1.53 | 0.353 | 1.556 | -0.001 | -0.001 | -0.001 | 0 | 0.65 | 0.64 | 0.62 | 0.51 | 0.59 | 0.7 |
| 1 | 24170458 | rs10917430 | -0.405 | 1.782 | -0.262 | 1.563 | -0.027 | 0.007 | 0.007 | 0 | 0.25 | 0.09 | 0.25 | 0.28 | 0.33 | 0.14 |
| 1 | 24174093 | rs10917431 | -1.014 | 1.749 | -0.222 | 1.52 | -0.03 | -0.053 | -0.053 | 0 | 0.31 | 0.18 | 0.36 | 0.5 | 0.42 | 0.3 |
| 1 | 24193875 | rs3003329 | -0.315 | 1.607 | -0.072 | 1.734 | 0.006 | 0.064 | 0.064 | 0 | 0.07 | 0.12 | 0.12 | 0.22 | 0.09 | 0.16 |
| 1 | 24196401 | rs3123554 | 0.868 | 1.835 | -0.253 | 1.652 | -0.029 | -0.051 | -0.051 | 0 | 0.7 | 0.79 | 0.64 | 0.5 | 0.59 | 0.7 |
| 1 | 24199182 | rs2501424 | 0.89 | 1.796 | 0.19 | 1.606 | 0 | 0.002 | 0.002 | 0 | 0.69 | 0.65 | 0.63 | 0.5 | 0.59 | 0.7 |
| 1 | 24200800 | rs1105 | 0.857 | 1.794 | 0.003 | 1.623 | -0.029 | -0.051 | -0.051 | 0 | 0.7 | 0.79 | 0.64 | 0.5 | 0.59 | 0.7 |
| 1 | 24201162 | rs2229579 | 0.513 | 1.819 | -0.008 | 1.674 | -0.029 | 0.003 | 0.003 | 0 | 0.05 | 0 | 0.11 | 0.21 | 0.09 | 0.16 |
| 1 | 24201262 | rs2502993 | 0.858 | 1.786 | 0.004 | 1.612 | -0.029 | -0.051 | -0.051 | 1.926 | 0.7 | 0.79 | 0.64 | 0.5 | 0.59 | 0.7 |
| 1 | 24201357 | rs4649124 | 0.858 | 1.786 | 0.004 | 1.612 | -0.029 | -0.051 | -0.051 | 0 | 0.7 | 0.79 | 0.64 | 0.5 | 0.59 | 0.7 |
| 1 | 24201643 | rs2501431 | 0.865 | 1.781 | 0.002 | 1.61 | -0.029 | -0.052 | -0.052 | 5.329 | 0.7 | 0.79 | 0.64 | 0.5 | 0.59 | 0.7 |
| 1 | 24206985 | rs6689530 | 0.865 | 1.356 | 0.678 | 1.901 | -0.002 | 0.042 | 0.042 | 0.007 | 0.87 | 0.8 | 0.87 | 0.76 | 0.87 | 0.85 |
| 1 | 24209378 | rs12733278 | 1.514 | 1.624 | 1.081 | 1.742 | 0.019 | -0.008 | -0.008 | 3.044 | 0.79 | 0.69 | 0.77 | 0.22 | 0.75 | 0.6 |
| 1 | 24209971 | rs4625225 | 1.514 | 1.624 | 1.081 | 1.742 | 0.019 | -0.008 | -0.008 | 0 | 0.79 | 0.69 | 0.77 | 0.22 | 0.75 | 0.6 |
| 1 | 24215130 | rs16828926 | -0.717 | 1.23 | 0.759 | 1.82 | -0.036 | -0.078 | -0.078 | 0 | 0.12 | 0.03 | 0.12 | 0.36 | 0.14 | 0.18 |
| 1 | 24216861 | rs2502959 | 0.813 | 1.229 | 0.844 | 1.819 | 0 | 0.023 | 0.023 | 0.016 | 0.86 | 0.82 | 0.86 | 0.64 | 0.86 | 0.82 |
| 1 | 24221834 | rs9424398 | 0.828 | 1.157 | 0.806 | 1.684 | 0 | 0.005 | 0.005 | 0.272 | 0.84 | 0.82 | 0.86 | 0.63 | 0.83 | 0.8 |
| 1 | 24264244 | rs2501430 | 0.594 | 1.115 | 0.483 | 1.233 | 0.022 | 0.069 | 0.069 | 0.358 | 0.87 | 0.57 | 0.77 | 0.79 | 0.83 | 0.89 |
| 1 | 24286885 | rs9424346 | -0.769 | 1.033 | -0.52 | 1.129 | -0.078 | -0.039 | -0.039 | 0 | 0.33 | 0.08 | 0.29 | 0.55 | 0.5 | 0.06 |
| 1 | 24294587 | rs35233638 | 0.819 | 0.447 | -0.266 | 0.799 | -0.003 | 0.007 | 0.007 | 0 | 0.53 | 0.54 | 0.65 | 0.45 | 0.43 | 0.85 |
| 1 | 24365512 | rs17184588 | -0.217 | 0.979 | -1.119 | 1.211 | -0.175 | -0.142 | -0.142 | 0 | 0.28 | 0.02 | 0.32 | 0.36 | 0.48 | 0.19 |
| 1 | 24372971 | rs12401982 | -0.158 | 0.76 | -0.744 | 1.52 | -0.004 | 0.073 | 0.073 | 0 | 0.06 | 0.16 | 0.08 | 0.17 | 0.05 | 0.04 |
| 1 | 24377326 | rs11586125 | -0.208 | -0.216 | -0.632 | 0.443 | -0.03 | 0.011 | 0.011 | 0 | 0.36 | 0.45 | 0.21 | 0.59 | 0.23 | 0.49 |
| 1 | 24379601 | rs4649163 | -1.074 | -0.087 | -0.263 | 0.13 | -0.008 | -0.145 | -0.145 | 0 | 0.65 | 0.54 | 0.57 | 0.95 | 0.73 | 0.7 |
| 1 | 24382298 | rs6424147 | -1.226 | -0.212 | 0.093 | -0.051 | -0.003 | -0.024 | -0.024 | 0 | 0.94 | 0.92 | 0.94 | 0.99 | 0.94 | 0.93 |
| 1 | 24383819 | rs7540491 | -0.875 | -0.22 | 0.108 | -0.069 | -0.008 | -0.153 | -0.153 | 0.383 | 0.65 | 0.53 | 0.57 | 0.95 | 0.73 | 0.7 |
| 1 | 24385997 | rs7549149 | -0.232 | -0.425 | 0.814 | 0.326 | -0.024 | 0.004 | 0.004 | 0 | 0.36 | 0.54 | 0.17 | 0.41 | 0.23 | 0.47 |
| 1 | 24386125 | rs11249061 | -0.197 | -0.413 | 0.62 | 0.335 | -0.002 | -0.001 | -0.001 | 0.996 | 0.35 | 0.31 | 0.15 | 0.41 | 0.23 | 0.47 |
| 1 | 24388417 | rs11811060 | -0.003 | -0.535 | 0.627 | 0.199 | 0.037 | -0.025 | -0.025 | 0 | 0.32 | 0.2 | 0.13 | 0.41 | 0.2 | 0.39 |
| 1 | 24389035 | rs4648930 | 0.457 | -0.456 | 1.016 | 0.155 | 0.028 | -0.053 | -0.053 | 1.358 | 0.89 | 0.5 | 0.88 | 0.99 | 0.91 | 0.84 |
| 1 | 24389213 | rs4648931 | 0.625 | -0.516 | 0.971 | 0.387 | 0.002 | 0.087 | 0.087 | 0 | 0.87 | 0.73 | 0.9 | 0.64 | 0.9 | 0.82 |
| 1 | 24392063 | rs4456048 | -0.319 | -0.961 | 1.058 | 0.232 | -0.046 | -0.028 | -0.028 | 0.039 | 0.29 | 0.48 | 0.16 | 0.2 | 0.14 | 0.31 |

| | | | | | | | | | | | | | | | | |
|---|---|---|---|---|---|---|---|---|---|---|---|---|---|---|---|---|
| 1 | 24393563 | rs16829071 | -0.33 | -0.962 | 1.062 | 0.231 | -0.047 | -0.029 | -0.029 | 0 | 0.29 | 0.49 | 0.16 | 0.2 | 0.14 | 0.31 |
| 1 | 24395575 | rs4649171 | 1.55 | -0.681 | 0.629 | -0.005 | -0.006 | 0.087 | 0.087 | 0.283 | 0.3 | 0.16 | 0.35 | 0.06 | 0.25 | 0.29 |
| 1 | 24396500 | rs6656699 | -0.756 | -0.22 | 0.503 | 0.123 | 0.007 | 0.034 | 0.034 | 0 | 0.51 | 0.69 | 0.51 | 0.62 | 0.57 | 0.48 |
| 1 | 24406647 | rs4649175 | 0.439 | 0.011 | 0.619 | 0.164 | 0.044 | 0.048 | 0.048 | 0 | 0.23 | 0.32 | 0.34 | 0.43 | 0.44 | 0.24 |
| 1 | 24408414 | rs4313343 | 0.425 | 0.009 | 0.623 | 0.163 | 0.078 | 0.086 | 0.086 | 0 | 0.23 | 0.37 | 0.35 | 0.43 | 0.44 | 0.24 |
| 1 | 24409165 | rs4320728 | 0.253 | -0.182 | 0.567 | 0.079 | 0.037 | 0.018 | 0.018 | 0 | 0.27 | 0.19 | 0.14 | 0.19 | 0.13 | 0.25 |
| 1 | 24409191 | rs4320729 | 0.894 | -0.13 | 0.468 | 0.013 | -0.145 | -0.148 | -0.148 | 0.106 | 0.2 | 0.02 | 0.32 | 0.43 | 0.44 | 0.18 |
| 1 | 24412447 | rs4276861 | -1.18 | -0.155 | 0.599 | -0.081 | 0.009 | 0.098 | 0.098 | 0 | 0.56 | 0.75 | 0.54 | 0.93 | 0.61 | 0.64 |
| 1 | 24413159 | rs6424152 | 0.44 | -0.034 | 0.521 | 0.145 | 0.08 | 0.093 | 0.093 | 0 | 0.23 | 0.36 | 0.35 | 0.46 | 0.45 | 0.24 |
| 1 | 24415047 | rs4378147 | -1.103 | 0.068 | 0.404 | -0.544 | 0.021 | 0.041 | 0.041 | 0 | 0.56 | 0.65 | 0.7 | 0.93 | 0.65 | 0.65 |
| 1 | 24415723 | rs4649183 | -0.052 | -0.123 | 0.354 | -0.42 | 0.037 | -0.007 | -0.007 | 0 | 0.28 | 0.36 | 0.39 | 0.73 | 0.49 | 0.39 |
| 1 | 24416133 | rs4276859 | 0.033 | -0.337 | 0.107 | -0.328 | 0.035 | 0.027 | 0.027 | 0 | 0.26 | 0.19 | 0.13 | 0.18 | 0.11 | 0.24 |
| 1 | 24417415 | rs6700245 | -0.379 | -0.463 | -0.087 | -1.093 | 0.023 | 0.021 | 0.021 | 0.175 | 0.3 | 0.24 | 0.13 | 0 | 0.18 | 0.27 |
| 1 | 24417868 | rs4487973 | -0.341 | -0.138 | -0.217 | -0.603 | -0.001 | 0.004 | 0.004 | 0 | 0.53 | 0.63 | 0.5 | 0.55 | 0.58 | 0.41 |
| 1 | 24418768 | rs7556176 | -0.312 | 0.701 | -0.063 | -1.027 | 0.012 | -0.01 | -0.01 | 0 | 0.37 | 0.47 | 0.32 | 0.35 | 0.47 | 0.22 |
| 1 | 24419496 | rs4233050 | -0.382 | 0.463 | 0.041 | -0.997 | 0.011 | -0.008 | -0.008 | 0 | 0.38 | 0.45 | 0.32 | 0.35 | 0.5 | 0.23 |
| 1 | 24421474 | rs6678540 | 0.14 | 0.704 | -0.476 | -0.863 | -0.008 | 0 | 0 | 0.105 | 0.31 | 0.27 | 0.29 | 0.34 | 0.48 | 0.22 |
| 1 | 24422626 | rs12568789 | -0.358 | 0.153 | -0.177 | -1.091 | -0.016 | -0.092 | -0.092 | 4.78 | 0.19 | 0.26 | 0.09 | 0.02 | 0.12 | 0.19 |
| 1 | 24424241 | rs3885698 | 0.006 | -0.017 | -0.393 | -1.085 | 0 | 0.001 | 0.001 | 0.905 | 0.39 | 0.63 | 0.36 | 0.32 | 0.4 | 0.26 |
| 1 | 24425730 | rs7545471 | 0.001 | -0.02 | -0.385 | -1.086 | -0.001 | 0 | 0 | 0.878 | 0.39 | 0.63 | 0.36 | 0.31 | 0.4 | 0.26 |
| 1 | 24431956 | rs3934861 | -0.502 | -0.067 | -0.615 | -1.515 | -0.012 | 0.025 | 0.025 | 0 | 0.57 | 0.69 | 0.41 | 0.32 | 0.49 | 0.36 |
| 1 | 24434595 | rs11249170 | -1.195 | -0.315 | -0.383 | -1.784 | -0.009 | 0.002 | 0.002 | 0 | 0.68 | 0.78 | 0.58 | 0.7 | 0.62 | 0.78 |
| 1 | 24437063 | rs7534564 | 0.635 | -0.591 | 0.097 | -1.799 | -0.045 | -0.099 | -0.099 | 0.377 | 0.19 | 0.33 | 0.09 | 0 | 0.09 | 0.14 |
| 1 | 24442907 | rs6686007 | 0.078 | 0.062 | -0.083 | -1.752 | 0.002 | -0.021 | -0.021 | 0.139 | 0.43 | 0.5 | 0.57 | 0.74 | 0.43 | 0.65 |
| 1 | 24446128 | rs10794665 | 0.748 | 0.03 | -0.549 | -1.898 | -0.09 | -0.008 | -0.008 | 2.507 | 0.32 | 0.07 | 0.4 | 0.73 | 0.39 | 0.55 |
| 1 | 24447116 | rs3795302 | 0.547 | 0.03 | -0.223 | -1.909 | -0.01 | 0.007 | 0.007 | 0 | 0.34 | 0.28 | 0.55 | 0.73 | 0.41 | 0.55 |
| 1 | 24447214 | rs3795300 | -1.136 | 0.002 | -0.359 | -1.86 | -0.016 | -0.002 | -0.002 | 0 | 0.73 | 0.8 | 0.57 | 0.73 | 0.62 | 0.8 |
| 1 | 24448084 | rs17852648 | 1.632 | 0.035 | -0.53 | -1.799 | -0.035 | -0.021 | -0.021 | 0.257 | 0.21 | 0.08 | 0.22 | 0.26 | 0.31 | 0.19 |
| 1 | 24453078 | rs11577442 | 0.348 | -0.35 | -0.245 | -1.869 | 0.004 | -0.011 | -0.011 | 0 | 0.4 | 0.41 | 0.32 | 0.3 | 0.44 | 0.3 |
| 1 | 24454688 | rs16829204 | 0.225 | -0.823 | -0.331 | -2.292 | 0.007 | -0.033 | -0.033 | 0 | 0.29 | 0.24 | 0.15 | 0 | 0.17 | 0.18 |
| 1 | 24463563 | rs4486393 | 0.135 | -0.6 | -0.487 | -2.408 | 0.001 | -0.052 | -0.052 | 0 | 0.32 | 0.3 | 0.16 | 0 | 0.22 | 0.22 |
| 1 | 24465113 | rs10903022 | 0.668 | -0.529 | -0.565 | -2.197 | -0.025 | 0.009 | 0.009 | 0 | 0.29 | 0.26 | 0.52 | 0.76 | 0.36 | 0.53 |
| 1 | 24465309 | rs10794642 | 0.686 | -0.537 | -0.569 | -2.201 | -0.025 | 0.008 | 0.008 | 0 | 0.29 | 0.26 | 0.52 | 0.77 | 0.36 | 0.53 |
| 1 | 24466035 | rs4648935 | 0.172 | -0.529 | -0.501 | -2.367 | -0.02 | 0.02 | 0.02 | 0 | 0.36 | 0.34 | 0.58 | 0.76 | 0.47 | 0.56 |
| 1 | 24469814 | rs4292900 | 1.136 | -1.071 | -0.404 | -2.394 | 0.055 | 0.047 | 0.047 | 0 | 0.52 | 0.36 | 0.73 | 0.98 | 0.67 | 0.72 |
| 1 | 24471200 | rs4448470 | 1.149 | -1.119 | -0.427 | -2.398 | 0.084 | 0.126 | 0.126 | 0 | 0.52 | 0.31 | 0.73 | 0.98 | 0.67 | 0.72 |
| 1 | 24471256 | rs7514676 | -1.364 | -1.149 | -0.076 | -2.26 | -0.018 | -0.048 | -0.048 | 0 | 0.91 | 0.84 | 0.95 | 0.98 | 0.95 | 0.92 |
| 1 | 24472083 | rs7513249 | -0.491 | -1.147 | -0.222 | -2.285 | -0.054 | -0.152 | -0.152 | 0.017 | 0.38 | 0.49 | 0.17 | 0.02 | 0.2 | 0.21 |
| 1 | 24475177 | rs4648940 | -1.075 | -0.653 | -0.392 | -1.312 | 0.007 | 0.03 | 0.03 | 0.013 | 0.9 | 0.8 | 0.92 | 0.8 | 0.89 | 0.75 |
| 1 | 24476370 | rs4330872 | -0.779 | -1.003 | -0.397 | -1.573 | -0.035 | -0.107 | -0.107 | 0 | 0.8 | 0.71 | 0.9 | 1 | 0.87 | 0.88 |
| 1 | 24476886 | rs11579657 | 1.715 | -0.679 | -0.808 | -1.463 | -0.049 | 0.038 | 0.038 | 0 | 0.17 | 0.03 | 0.28 | 0.08 | 0.2 | 0.09 |
| 1 | 24479274 | rs4649191 | 1.382 | -0.656 | -0.227 | -0.95 | -0.002 | -0.009 | -0.009 | 0 | 0.54 | 0.53 | 0.58 | 0.29 | 0.57 | 0.45 |
| 1 | 24480905 | rs10903034 | 1.382 | -0.596 | -0.221 | -0.939 | -0.002 | -0.01 | -0.01 | 0.74 | 0.54 | 0.53 | 0.58 | 0.28 | 0.57 | 0.46 |
| 1 | 24481940 | rs10903035 | 0.529 | -0.613 | -0.263 | -0.932 | -0.011 | 0.03 | 0.03 | 0 | 0.63 | 0.54 | 0.64 | 0.46 | 0.68 | 0.65 |
| 1 | 24482004 | rs12028945 | 0.983 | -0.16 | -0.326 | -0.546 | -0.028 | -0.042 | -0.042 | 0 | 0.1 | 0.01 | 0.05 | 0.18 | 0.1 | 0.19 |
| 1 | 24482474 | rs11249006 | -0.735 | -0.186 | -0.255 | -0.637 | -0.015 | -0.02 | -0.02 | 0 | 0.26 | 0.2 | 0.33 | 0.7 | 0.3 | 0.42 |
| 1 | 24485941 | rs4649195 | 0.419 | -0.242 | -0.454 | -0.52 | -0.009 | 0.025 | 0.025 | 0 | 0.66 | 0.45 | 0.64 | 0.51 | 0.72 | 0.71 |
| 1 | 24486955 | rs1416835 | 0.735 | -0.266 | -0.289 | -0.894 | 0.001 | 0.001 | 0.001 | 0 | 0.57 | 0.44 | 0.59 | 0.36 | 0.61 | 0.51 |
| 1 | 24488450 | rs7552086 | -0.557 | 0.102 | -0.265 | -0.376 | 0.003 | 0.014 | 0.014 | 0 | 0.3 | 0.31 | 0.35 | 0.63 | 0.3 | 0.45 |
| 1 | 24495722 | rs4649197 | -0.6 | 0.145 | -0.522 | -1.054 | 0.027 | 0.059 | 0.059 | 0 | 0.54 | 0.58 | 0.57 | 0.92 | 0.63 | 0.8 |
| 1 | 24498911 | rs7520329 | -0.646 | 0.207 | -1.199 | -0.795 | 0.006 | 0.046 | 0.046 | 0 | 0.3 | 0.61 | 0.36 | 0.58 | 0.3 | 0.54 |
| 1 | 24506861 | rs3897440 | 0.099 | -0.048 | -1.335 | -1.044 | 0.033 | 0.131 | 0.131 | 0.108 | 0.52 | 0.71 | 0.48 | 0.16 | 0.4 | 0.3 |
| 1 | 24507774 | rs6683433 | 0.113 | -0.142 | -1.505 | -0.908 | 0.064 | 0.121 | 0.121 | 0.034 | 0.52 | 0.68 | 0.46 | 0.17 | 0.34 | 0.29 |
| 1 | 24508710 | rs7524076 | -0.254 | 0.305 | -1.742 | -0.743 | -0.075 | 0.069 | 0.069 | 0.07 | 0.64 | 0.9 | 0.78 | 0.8 | 0.58 | 0.74 |
| 1 | 24512804 | rs11249020 | -0.615 | -0.347 | -1.286 | -0.343 | 0.032 | -0.028 | -0.028 | 0 | 0.8 | 0.93 | 0.9 | 0.8 | 0.86 | 0.89 |
| 1 | 24512881 | rs11249021 | -0.616 | -0.346 | -1.287 | -0.205 | 0.036 | -0.064 | -0.064 | 0.462 | 0.8 | 0.93 | 0.9 | 0.65 | 0.86 | 0.88 |
| 1 | 24516880 | rs11587500 | 0.377 | -0.334 | -1.173 | -0.375 | 0.004 | 0.104 | 0.104 | 0.491 | 0.51 | 0.28 | 0.28 | 0.13 | 0.51 | 0.55 |
| 1 | 24518206 | rs10794648 | 1.052 | -0.97 | -1.009 | 0.377 | -0.042 | 0.003 | 0.003 | 0.21 | 0.63 | 0.49 | 0.79 | 0.63 | 0.75 | 0.71 |
| 1 | 24518643 | rs7552167 | 0.385 | -0.111 | -1.747 | 0.641 | 0.03 | -0.028 | -0.028 | 1.07 | 0.8 | 0.88 | 0.9 | 0.77 | 0.85 | 0.87 |
| 1 | 24531634 | rs946662 | -0.21 | 0.417 | -1.923 | 0.195 | 0 | 0.066 | 0.066 | 0 | 0.58 | 0.79 | 0.55 | 0.71 | 0.58 | 0.53 |
| 1 | 24532144 | rs16860738 | -0.12 | 0.795 | -1.527 | 0.818 | -0.004 | 0.021 | 0.021 | 0 | 0.12 | 0.15 | 0.24 | 0.57 | 0.16 | 0.19 |
| 1 | 24535178 | rs12061941 | -0.291 | 0.168 | -0.633 | -0.001 | -0.038 | -0.029 | -0.029 | 0 | 0.18 | 0.35 | 0.1 | 0.13 | 0.1 | 0.19 |
| 1 | 24540521 | rs7539076 | 1.041 | 0.165 | -0.085 | -0.129 | 0.066 | 0.045 | 0.045 | 0.826 | 0.94 | 0.61 | 0.95 | 1 | 0.98 | 0.98 |
| 1 | 24552835 | rs11249039 | 0.241 | -0.066 | -1.477 | -0.171 | 0.036 | -0.038 | -0.038 | 0 | 0.4 | 0.17 | 0.28 | 0.58 | 0.31 | 0.37 |
| 1 | 24556756 | rs10903062 | 0.278 | 0.193 | -1.668 | -0.349 | -0.026 | -0.068 | -0.068 | 0 | 0.4 | 0.09 | 0.28 | 0.58 | 0.35 | 0.35 |
| 1 | 24557760 | rs11249045 | 0.143 | 0.507 | -2.106 | -0.696 | -0.081 | -0.137 | -0.137 | 0 | 0.19 | 0.02 | 0.44 | 0.42 | 0.23 | 0.43 |
| 1 | 24567435 | rs12131684 | 0.469 | 0.357 | -2.04 | -1.035 | -0.056 | 0.165 | 0.165 | 0.054 | 0.29 | 0.05 | 0.2 | 0 | 0.38 | 0.18 |
| 1 | 24568090 | rs4648955 | 0.433 | 0.352 | -2.042 | -1.037 | -0.055 | 0.167 | 0.167 | 0.032 | 0.3 | 0.05 | 0.2 | 0 | 0.38 | 0.18 |
| 1 | 24572806 | rs7517309 | -0.63 | 0.206 | -1.988 | -0.844 | 0.031 | -0.009 | -0.009 | 0 | 0.52 | 0.13 | 0.66 | 0.43 | 0.64 | 0.64 |

| | | | | | | | | | | | | | | | | |
|---|---|---|---|---|---|---|---|---|---|---|---|---|---|---|---|---|
| 1 | 24575345 | rs7526505 | 0.085 | 0.481 | -1.7 | -0.372 | -0.062 | -0.137 | -0.137 | 0 | 0.17 | 0.03 | 0.43 | 0.45 | 0.2 | 0.45 |
| 1 | 24579435 | rs6688902 | 0.148 | 0.896 | -1.001 | -0.23 | -0.079 | -0.084 | -0.084 | 0 | 0.12 | 0.01 | 0.42 | 0.56 | 0.21 | 0.43 |
| 1 | 24581493 | rs11249057 | -1.25 | 0.014 | -0.312 | -0.842 | 0.004 | 0.004 | 0.004 | 0 | 0.31 | 0.41 | 0.55 | 0.56 | 0.35 | 0.49 |
| 1 | 24582162 | rs12088212 | -0.814 | -0.037 | -0.298 | -0.848 | -0.063 | -0.119 | -0.119 | 0 | 0.19 | 0.54 | 0.14 | 0 | 0.13 | 0.07 |
| 1 | 24582304 | rs12144166 | 1.333 | -0.27 | -0.623 | -1.214 | -0.002 | 0.054 | 0.054 | 0 | 0.56 | 0.13 | 0.35 | 0.43 | 0.56 | 0.48 |
| 1 | 24588518 | rs12139592 | 0.982 | -0.393 | -1.364 | -1.94 | -0.089 | 0.081 | 0.081 | 0.063 | 0.34 | 0.02 | 0.26 | 0.31 | 0.37 | 0.37 |
| 1 | 24590691 | rs6424169 | -0.262 | -0.43 | -0.326 | -1.813 | -0.042 | -0.103 | -0.103 | 4.303 | 0.18 | 0.3 | 0.1 | 0 | 0.14 | 0.06 |
| 1 | 24602795 | rs4233036 | -0.641 | -0.149 | 0.15 | -1.729 | 0.019 | -0.086 | -0.086 | 0 | 0.87 | 0.61 | 0.89 | 1 | 0.87 | 0.96 |
| 1 | 24604438 | rs12063430 | -0.121 | 0.151 | 0.252 | -1.801 | 0.012 | -0.006 | -0.006 | 4.11 | 0.31 | 0.4 | 0.53 | 0.59 | 0.35 | 0.48 |
| 1 | 24607419 | rs7535789 | -1.162 | 0.119 | 0.181 | -1.811 | 0.014 | -0.067 | -0.067 | 0.238 | 0.89 | 0.71 | 0.9 | 1 | 0.87 | 0.96 |
| 1 | 24609935 | rs12026525 | 1.013 | 0.055 | -0.291 | -1.708 | -0.081 | 0.076 | 0.076 | 0 | 0.53 | 0.09 | 0.32 | 0.41 | 0.48 | 0.49 |
| 1 | 24610584 | rs12121909 | 0.662 | 0.169 | -0.071 | -1.728 | -0.028 | 0.027 | 0.027 | 0 | 0.6 | 0.27 | 0.34 | 0.41 | 0.51 | 0.51 |
| 1 | 24616294 | rs1572786 | 0.621 | 0.205 | -0.085 | -1.688 | -0.005 | 0.003 | 0.003 | 0 | 0.6 | 0.36 | 0.35 | 0.44 | 0.51 | 0.51 |
| 1 | 24618605 | rs4648960 | 0.831 | 0.25 | -0.302 | -1.632 | 0.009 | 0.064 | 0.064 | 0 | 0.53 | 0.24 | 0.32 | 0.43 | 0.46 | 0.48 |
| 1 | 24620801 | rs12405071 | 0.662 | 0.32 | -0.258 | -1.629 | -0.023 | -0.017 | -0.017 | 1.34 | 0.24 | 0.18 | 0.5 | 0.56 | 0.3 | 0.45 |
| 1 | 24624870 | rs4648967 | 0.767 | 0.35 | -0.329 | -1.656 | 0.005 | 0.046 | 0.046 | 0 | 0.55 | 0.28 | 0.34 | 0.43 | 0.47 | 0.48 |
| 1 | 24625549 | rs4641259 | 0.529 | 0.571 | -0.199 | -1.526 | -0.007 | -0.005 | -0.005 | 0 | 0.25 | 0.25 | 0.51 | 0.57 | 0.3 | 0.46 |
| 1 | 24633812 | rs12080352 | -0.805 | 0.009 | -0.513 | -1.616 | 0.001 | 0.008 | 0.008 | 0 | 0.47 | 0.42 | 0.63 | 0.56 | 0.53 | 0.53 |
| 1 | 24634151 | rs4648973 | 1.197 | -0.423 | -1.144 | -1.558 | -0.018 | 0 | 0 | 1.343 | 0.21 | 0.01 | 0.11 | 0.14 | 0.17 | 0.13 |
| 1 | 24640314 | rs621535 | -0.34 | -0.299 | -1.581 | -1.408 | 0.001 | -0.049 | -0.049 | 0 | 0.26 | 0.71 | 0.18 | 0 | 0.22 | 0.1 |
| 1 | 24645356 | rs2248219 | -0.551 | 0.085 | -1.588 | -0.511 | 0.019 | 0.027 | 0.027 | 0 | 0.73 | 0.88 | 0.8 | 0.71 | 0.77 | 0.69 |
| 1 | 24653792 | rs483202 | -0.628 | 0.156 | -1.666 | -0.265 | 0.025 | 0.048 | 0.048 | 0.338 | 0.63 | 0.9 | 0.72 | 0.67 | 0.66 | 0.71 |
| 1 | 24657860 | rs2486669 | -0.486 | 1.794 | -0.19 | 0.601 | 0.207 | 0.188 | 0.188 | 0.507 | 0.08 | 0.34 | 0.15 | 0.19 | 0.16 | 0.15 |
| 1 | 24659435 | rs10903078 | 0.137 | 0.861 | -1.205 | 0.472 | -0.001 | -0.098 | -0.098 | 0.021 | 0.3 | 0.09 | 0.36 | 0.38 | 0.32 | 0.25 |
| 1 | 24662506 | rs6659209 | -0.057 | 1.079 | -0.009 | 0.154 | 0.101 | 0.023 | 0.023 | 0.936 | 0.1 | 0.45 | 0.12 | 0.06 | 0.13 | 0.07 |
| 1 | 24667300 | rs12568599 | 0.306 | 0.992 | -0.812 | 0.058 | -0.05 | 0.008 | 0.008 | 0 | 0.18 | 0.06 | 0.15 | 0.11 | 0.21 | 0.25 |
| 1 | 24670264 | rs4648975 | -0.031 | 0.523 | -0.426 | 0.437 | -0.016 | -0.058 | -0.058 | 0 | 0.25 | 0.1 | 0.37 | 0.43 | 0.3 | 0.24 |
| 1 | 24671287 | rs4648859 | -0.906 | 0.204 | -0.083 | -0.033 | 0.004 | -0.086 | -0.086 | 0 | 0.18 | 0.66 | 0.21 | 0.03 | 0.17 | 0.2 |
| 1 | 24672641 | rs6675501 | -1.255 | 0.056 | -0.002 | 0.161 | -0.004 | 0.01 | 0.01 | 0 | 0.44 | 0.73 | 0.59 | 0.42 | 0.49 | 0.42 |
| 1 | 24684116 | rs7527465 | -0.687 | 0.016 | 0.837 | -0.002 | 0.036 | -0.073 | -0.073 | 1.764 | 0.12 | 0.35 | 0.13 | 0.01 | 0.13 | 0.06 |
| 1 | 24684728 | rs11249092 | -0.035 | 0.061 | 0.615 | 0.017 | -0.008 | 0.002 | 0.002 | 0 | 0.09 | 0.06 | 0.1 | 0 | 0.12 | 0.05 |
| 1 | 24687653 | rs532147 | -1.411 | -0.162 | 0.178 | -0.331 | -0.041 | 0.074 | 0.074 | 0 | 0.71 | 0.87 | 0.8 | 0.88 | 0.68 | 0.7 |
| 1 | 24690080 | rs486575 | -0.008 | -0.344 | 0.541 | -0.382 | -0.002 | 0.125 | 0.125 | 0.06 | 0.51 | 0.3 | 0.58 | 0.83 | 0.52 | 0.53 |
| 1 | 24695114 | rs7535482 | -1.473 | 0.578 | 1.562 | 0.436 | 0.029 | 0.049 | 0.049 | 0 | 0.28 | 0.41 | 0.47 | 0.42 | 0.36 | 0.22 |
| 1 | 24695151 | rs7537875 | -1.452 | 0.58 | 1.578 | 0.437 | 0.026 | 0.044 | 0.044 | 0 | 0.28 | 0.39 | 0.46 | 0.42 | 0.36 | 0.22 |
| 1 | 24705688 | rs10489438 | 0.04 | 0.491 | 1.287 | 0.309 | -0.01 | 0.011 | 0.011 | 0 | 0.09 | 0.05 | 0.09 | 0 | 0.11 | 0.05 |
| 1 | 24706292 | rs1142057 | -1.486 | 0.456 | 1.776 | 0.398 | 0.012 | 0.042 | 0.042 | 0 | 0.29 | 0.31 | 0.48 | 0.44 | 0.35 | 0.29 |
| 1 | 24706410 | rs3765431 | -1.576 | 0.441 | 1.958 | 0.391 | 0.009 | 0.046 | 0.046 | 0.156 | 0.31 | 0.48 | 0.51 | 0.44 | 0.36 | 0.29 |
| 1 | 24712169 | rs2244762 | -0.848 | 0.52 | 1.617 | 0.58 | 0.03 | 0.006 | 0.006 | 0 | 0.29 | 0.4 | 0.32 | 0.32 | 0.35 | 0.26 |
| 1 | 24718075 | rs2294520 | -0.336 | 0.328 | 1.533 | 0.517 | -0.005 | -0.06 | -0.06 | 0 | 0.24 | 0.1 | 0.38 | 0.46 | 0.28 | 0.29 |
| 1 | 24720003 | rs501748 | -2.303 | 0.196 | 1.512 | 0.454 | -0.02 | 0.106 | 0.106 | 0 | 0.59 | 0.91 | 0.75 | 0.79 | 0.63 | 0.59 |
| 1 | 24725076 | rs577168 | -0.737 | 0.309 | 1.698 | 0.508 | 0.021 | 0.003 | 0.003 | 0 | 0.29 | 0.38 | 0.32 | 0.32 | 0.35 | 0.27 |
| 1 | 24735363 | rs10489440 | -0.016 | 0.348 | 1.666 | 0.466 | 0.006 | -0.033 | -0.033 | 0.091 | 0.09 | 0.15 | 0.09 | 0.01 | 0.11 | 0.08 |
| 1 | 24738491 | rs12123880 | -0.391 | 0.329 | 1.865 | 0.473 | 0.001 | -0.014 | -0.014 | 0 | 0.25 | 0.2 | 0.39 | 0.44 | 0.28 | 0.29 |
| 1 | 24742112 | rs10489441 | 0.202 | 0.358 | 1.66 | 0.464 | 0.003 | -0.024 | -0.024 | 0 | 0.09 | 0.13 | 0.09 | 0 | 0.11 | 0.06 |
| 1 | 24748310 | rs6686862 | 0.166 | 0.358 | 1.66 | 0.464 | 0.002 | -0.021 | -0.021 | 0.44 | 0.09 | 0.13 | 0.09 | 0.01 | 0.11 | 0.08 |
| 1 | 24751197 | rs4648983 | -2.285 | 0.331 | 1.511 | 0.503 | -0.016 | 0.092 | 0.092 | 0 | 0.59 | 0.91 | 0.76 | 0.78 | 0.63 | 0.58 |
| 1 | 24751920 | rs6670829 | -1.929 | 0.328 | 1.744 | 0.503 | 0.025 | 0.059 | 0.059 | 0 | 0.56 | 0.72 | 0.73 | 0.77 | 0.63 | 0.57 |
| 1 | 24753310 | rs2235552 | -0.522 | 0.339 | 1.544 | 0.532 | -0.002 | 0.012 | 0.012 | 0 | 0.26 | 0.25 | 0.4 | 0.44 | 0.28 | 0.29 |
| 1 | 24755281 | rs10903086 | -1.079 | 0.392 | 1.597 | 0.514 | 0.003 | -0.001 | -0.001 | 3.957 | 0.32 | 0.66 | 0.36 | 0.34 | 0.35 | 0.29 |
| 1 | 24757806 | rs6676449 | 0.064 | 0.35 | 1.568 | 0.493 | -0.002 | -0.011 | -0.011 | 0.502 | 0.09 | 0.09 | 0.09 | 0 | 0.11 | 0.06 |
| 1 | 24759685 | rs3765888 | -0.232 | 0.47 | 1.318 | 0.575 | 0.027 | 0.046 | 0.046 | 0 | 0.23 | 0.54 | 0.26 | 0.33 | 0.24 | 0.21 |
| 1 | 24764689 | rs1008485 | -0.758 | 0.874 | 1.062 | 0.805 | -0.001 | 0.016 | 0.016 | 0 | 0.27 | 0.26 | 0.42 | 0.43 | 0.29 | 0.29 |
| 1 | 24775000 | rs4648988 | -0.585 | 1.041 | 0.887 | 0.809 | 0.007 | 0.001 | 0.001 | 0.078 | 0.25 | 0.27 | 0.26 | 0.31 | 0.35 | 0.24 |
| 1 | 24779855 | rs2272934 | -0.721 | 0.998 | 0.813 | 1.128 | -0.004 | -0.026 | -0.026 | 0.39 | 0.26 | 0.16 | 0.44 | 0.46 | 0.28 | 0.3 |
| 1 | 24781247 | rs2281188 | -1.806 | 1.063 | 0.971 | 1.088 | 0 | -0.002 | -0.002 | 0 | 0.45 | 0.51 | 0.64 | 0.76 | 0.51 | 0.47 |
| 1 | 24785393 | rs2272935 | -0.917 | 1.085 | -0.338 | 1.257 | -0.006 | -0.03 | -0.03 | 0 | 0.25 | 0.16 | 0.44 | 0.45 | 0.28 | 0.29 |
| 1 | 24788950 | rs6688953 | -0.424 | 1.224 | -0.103 | 0.931 | 0.003 | -0.014 | -0.014 | 0 | 0.06 | 0.09 | 0.05 | 0.02 | 0.08 | 0.11 |
| 1 | 24796838 | rs12117331 | -1.184 | 1.37 | -0.102 | 1.274 | 0.017 | 0.04 | 0.04 | 0.25 | 0.15 | 0.22 | 0.3 | 0.25 | 0.19 | 0.2 |
| 1 | 24798383 | rs195729 | -1.013 | 1.647 | -0.883 | 1.197 | -0.064 | 0.002 | 0.002 | 1.059 | 0.35 | 0.13 | 0.6 | 0.68 | 0.52 | 0.39 |
| 1 | 24813155 | rs195704 | -0.966 | 2.061 | -0.658 | 1.794 | -0.061 | -0.064 | -0.064 | 5.778 | 0.17 | 0.06 | 0.2 | 0.36 | 0.31 | 0.27 |
| 1 | 24822500 | rs9424374 | -1.276 | 1.15 | -1.035 | 1.478 | 0.063 | -0.078 | -0.078 | 0 | 0.28 | 0.89 | 0.25 | 0.41 | 0.18 | 0.21 |
| 1 | 24828929 | rs11805096 | -0.903 | 1.168 | 0.271 | 1.678 | 0.033 | 0.244 | 0.244 | 1.618 | 0.07 | 0.31 | 0.13 | 0.26 | 0.06 | 0.07 |
| 1 | 24832734 | rs7540336 | -1 | 1.155 | 0.285 | 1.672 | 0.029 | 0.234 | 0.234 | 1.226 | 0.07 | 0.31 | 0.13 | 0.26 | 0.06 | 0.07 |
| 1 | 24838493 | rs196402 | -1.938 | 1.205 | 0.357 | 1.245 | -0.035 | 0.025 | 0.025 | 0 | 0.39 | 0.72 | 0.42 | 0.73 | 0.46 | 0.48 |
| 1 | 24857086 | rs185511 | -0.979 | 1.375 | 1.444 | 1.318 | 0 | 0 | 0 | 1.848 | 0.27 | 0.23 | 0.33 | 0.68 | 0.32 | 0.35 |
| 1 | 24859665 | rs196430 | -0.992 | 1.368 | 1.458 | 1.336 | 0 | 0.001 | 0.001 | 1.847 | 0.27 | 0.25 | 0.33 | 0.68 | 0.32 | 0.37 |
| 1 | 24861399 | rs11801547 | -1.489 | 1.515 | 1.445 | 1.252 | 0.032 | -0.004 | -0.004 | 0 | 0.06 | 0.25 | 0.03 | 0 | 0.02 | 0.03 |
| 1 | 24861704 | rs196432 | -1.79 | 1.562 | 1.559 | 1.219 | 0.033 | -0.009 | -0.009 | 5.338 | 0.34 | 0.47 | 0.4 | 0.73 | 0.46 | 0.47 |

| | | | | | | | | | | | | | | | | |
|---|---|---|---|---|---|---|---|---|---|---|---|---|---|---|---|---|
| 1 | 24862643 | rs196433 | -2.351 | 1.591 | 1.589 | 1.219 | -0.035 | 0.03 | 0.03 | 0 | 0.4 | 0.72 | 0.43 | 0.73 | 0.48 | 0.51 |
| 1 | 24863285 | rs11231 | -0.241 | 1.616 | 1.596 | 1.243 | 0.06 | -0.001 | -0.001 | 0 | 0.07 | 0.19 | 0.07 | 0.05 | 0.14 | 0.11 |
| 1 | 24863530 | rs10489444 | -1.301 | 1.433 | 1.592 | 1.298 | 0.045 | 0.009 | 0.009 | 0.668 | 0.05 | 0.25 | 0.03 | 0 | 0.02 | 0.03 |
| 1 | 24864576 | rs7540973 | -2.187 | 1.373 | 1.74 | 1.269 | 0.036 | -0.007 | -0.007 | 0 | 0.08 | 0.41 | 0.04 | 0 | 0.02 | 0.03 |
| 1 | 24871730 | rs196426 | -1.689 | 1.422 | 1.273 | 1.29 | 0.001 | -0.001 | -0.001 | 0.461 | 0.35 | 0.44 | 0.39 | 0.7 | 0.38 | 0.39 |
| 1 | 24873860 | rs12095513 | -0.041 | 1.423 | 0.887 | 1.631 | 0 | 0.042 | 0.042 | 1.71 | 0.15 | 0.16 | 0.17 | 0.38 | 0.16 | 0.18 |
| 1 | 24874674 | rs16829859 | 0.025 | 1.771 | 0.698 | 1.612 | -0.005 | -0.01 | -0.01 | 0 | 0.15 | 0.11 | 0.18 | 0.4 | 0.19 | 0.18 |
| 1 | 24875490 | rs2809964 | -1.479 | 1.512 | 0.653 | 0.78 | 0.02 | -0.019 | -0.019 | 1.199 | 0.31 | 0.45 | 0.27 | 0.34 | 0.37 | 0.37 |
| 1 | 24876383 | rs2776747 | -0.128 | 1.548 | 0.614 | 0.813 | 0.004 | 0.006 | 0.006 | 0 | 0.13 | 0.13 | 0.17 | 0.28 | 0.16 | 0.18 |
| 1 | 24877458 | rs12065278 | -0.996 | 1.215 | 0.575 | 0.422 | 0.01 | -0.061 | -0.061 | 0 | 0.19 | 0.36 | 0.11 | 0.06 | 0.22 | 0.18 |
| 1 | 24878789 | rs12069273 | -1.06 | 1.23 | 0.543 | 0.388 | 0.01 | -0.061 | -0.061 | 0 | 0.19 | 0.36 | 0.11 | 0.06 | 0.22 | 0.19 |
| 1 | 24881126 | rs6424099 | 0.863 | 0.671 | -0.178 | 0.183 | -0.013 | -0.012 | -0.012 | 0.081 | 0.44 | 0.52 | 0.47 | 0.29 | 0.31 | 0.39 |
| 1 | 24884460 | rs3893315 | 0.601 | 0.724 | -0.543 | 0.009 | 0.011 | 0.066 | 0.066 | 0.743 | 0.61 | 0.77 | 0.62 | 0.28 | 0.41 | 0.45 |
| 1 | 24886818 | rs12122463 | -0.602 | 0.603 | -0.299 | 0.167 | 0 | 0.01 | 0.01 | 0.29 | 0.21 | 0.29 | 0.12 | 0.21 | 0.21 | 0.21 |
| 1 | 24887920 | rs2481930 | 1.105 | 0.614 | -0.308 | 0.053 | -0.002 | -0.004 | -0.004 | 2.515 | 0.58 | 0.57 | 0.71 | 0.77 | 0.64 | 0.69 |
| 1 | 24888585 | rs4610948 | 1.105 | 0.614 | -0.303 | 0.053 | -0.002 | -0.004 | -0.004 | 0.218 | 0.58 | 0.57 | 0.71 | 0.77 | 0.64 | 0.69 |
| 1 | 24894997 | rs1929301 | -0.698 | 1.377 | -0.012 | 0.316 | -0.001 | -0.001 | -0.001 | 0 | 0.22 | 0.25 | 0.15 | 0.23 | 0.29 | 0.19 |
| 1 | 24898453 | rs6424106 | -0.061 | 1.363 | 0.087 | 0.34 | 0.009 | 0.047 | 0.047 | 0 | 0.08 | 0.18 | 0.06 | 0.14 | 0.08 | 0.06 |
| 1 | 24901121 | rs4648998 | 1.081 | 1.259 | 0.086 | 0.33 | -0.005 | 0.027 | 0.027 | 0.007 | 0.53 | 0.39 | 0.59 | 0.39 | 0.52 | 0.53 |
| 1 | 24904255 | rs12129478 | -0.729 | 0.375 | -0.562 | 0.075 | -0.014 | -0.008 | -0.008 | 1.1 | 0.77 | 0.71 | 0.83 | 0.77 | 0.8 | 0.79 |
| 1 | 24907415 | rs4073051 | -0.361 | 0.202 | -0.574 | -0.103 | 0.014 | 0.02 | 0.02 | 0 | 0.59 | 0.71 | 0.52 | 0.8 | 0.67 | 0.67 |
| 1 | 24908959 | rs3924436 | 0.471 | 0.303 | -0.49 | 0.036 | 0.004 | -0.005 | -0.005 | 0 | 0.39 | 0.44 | 0.34 | 0.41 | 0.38 | 0.4 |
| 1 | 24910319 | rs11249138 | 0.511 | 0.398 | -0.381 | 0.019 | 0 | 0 | 0 | 0 | 0.4 | 0.43 | 0.49 | 0.61 | 0.44 | 0.52 |
| 1 | 24917456 | rs9328948 | 0.81 | 0.846 | -0.413 | 0.416 | 0.004 | 0.026 | 0.026 | 0 | 0.06 | 0.1 | 0.05 | 0.12 | 0.07 | 0.09 |
| 1 | 24922129 | rs10903095 | -0.715 | 0.631 | -0.645 | 0.321 | -0.005 | -0.004 | -0.004 | 0 | 0.31 | 0.69 | 0.28 | 0.39 | 0.44 | 0.34 |
| 1 | 24927001 | rs4425931 | 0.067 | 0.871 | -0.384 | 0.384 | 0.062 | 0.014 | 0.014 | 0 | 0.14 | 0.22 | 0.14 | 0.17 | 0.24 | 0.17 |
| 1 | 24927561 | rs4553122 | -0.044 | 0.06 | -0.641 | 0.084 | 0 | -0.001 | -0.001 | 0 | 0.57 | 0.51 | 0.52 | 0.59 | 0.56 | 0.58 |
| 1 | 24929676 | rs4649002 | -0.222 | 0.026 | -0.419 | 0.067 | -0.007 | -0.004 | -0.004 | 0 | 0.59 | 0.7 | 0.54 | 0.59 | 0.56 | 0.58 |
| 1 | 24932733 | rs11249141 | 0.382 | -1.011 | -0.851 | -0.025 | 0.084 | 0.004 | 0.004 | 0 | 0.44 | 0.25 | 0.38 | 0.42 | 0.32 | 0.41 |
| 1 | 24947948 | rs4649005 | 1.237 | -1.349 | -1.819 | 0.065 | -0.015 | -0.137 | -0.137 | 0 | 0.24 | 0.02 | 0.32 | 0.4 | 0.21 | 0.37 |
| 1 | 24955492 | rs4648870 | -0.441 | -1.249 | -1.199 | -0.195 | -0.02 | 0.007 | 0.007 | 0 | 0.76 | 0.97 | 0.69 | 0.8 | 0.77 | 0.74 |
| 1 | 24970623 | rs4617346 | 0.573 | -1.237 | -0.298 | 0.012 | -0.039 | -0.047 | -0.047 | 0 | 0.12 | 0.24 | 0.04 | 0.02 | 0.04 | 0.01 |
| 1 | 25012493 | rs6691770 | 0.322 | -1.234 | -0.279 | 0.221 | -0.059 | -0.008 | -0.008 | 0 | 0.13 | 0.36 | 0.11 | 0.13 | 0.04 | 0.02 |
| 1 | 25015418 | rs4648873 | -0.165 | -1.624 | -0.858 | 0.201 | 0.049 | 0.019 | 0.019 | 0.306 | 0.53 | 0.4 | 0.72 | 0.59 | 0.74 | 0.64 |
| 1 | 25015514 | rs4649013 | -0.228 | -1.223 | -0.952 | 0.011 | -0.008 | -0.004 | -0.004 | 0.977 | 0.39 | 0.69 | 0.46 | 0.57 | 0.54 | 0.37 |
| 1 | 25015530 | rs4649014 | -0.509 | -1.242 | -1.076 | -0.151 | 0.052 | 0.12 | 0.12 | 0 | 0.49 | 0.77 | 0.56 | 0.76 | 0.59 | 0.43 |
| 1 | 25021037 | rs2885174 | 0.365 | 0.021 | -1.083 | 0.519 | -0.043 | -0.051 | -0.051 | 0 | 0.24 | 0.21 | 0.38 | 0.55 | 0.38 | 0.33 |
| 1 | 25022055 | rs11249163 | 0.516 | 0.092 | -1.022 | 0.025 | 0 | -0.001 | -0.001 | 0.117 | 0.12 | 0.11 | 0.09 | 0.14 | 0.16 | 0.12 |
| 1 | 25029118 | rs7543969 | 0.146 | -0.509 | -0.694 | -0.177 | 0.002 | -0.033 | -0.033 | 0 | 0.65 | 0.59 | 0.59 | 0.8 | 0.64 | 0.68 |
| 1 | 25029488 | rs11249167 | 0.786 | 0.02 | -0.51 | 0.357 | -0.113 | -0.103 | -0.103 | 0 | 0.21 | 0.05 | 0.32 | 0.32 | 0.38 | 0.23 |
| 1 | 25051153 | rs1002706 | 0.637 | 0.032 | -0.504 | 0.37 | -0.09 | -0.08 | -0.08 | 0 | 0.22 | 0.11 | 0.34 | 0.32 | 0.37 | 0.22 |
| 1 | 25069939 | rs2185151 | -0.165 | -0.006 | -0.423 | 0.405 | 0.015 | -0.013 | -0.013 | 0 | 0.29 | 0.34 | 0.38 | 0.21 | 0.33 | 0.32 |
| 1 | 25075801 | rs1890449 | -0.777 | -0.213 | -0.461 | 0.124 | 0.045 | 0.011 | 0.011 | 1.312 | 0.48 | 0.36 | 0.7 | 0.52 | 0.7 | 0.54 |
| 1 | 25080522 | rs7525139 | 0.172 | -0.211 | -0.379 | 0.127 | -0.05 | 0.045 | 0.045 | 0 | 0.26 | 0.08 | 0.34 | 0.14 | 0.3 | 0.31 |
| 1 | 25136713 | rs10903104 | -0.193 | -1.372 | -0.047 | 0.135 | 0.045 | 0.011 | 0.011 | 1.312 | 0.48 | 0.36 | 0.7 | 0.52 | 0.7 | 0.55 |
| 1 | 25140825 | rs3890756 | 0.271 | -1.391 | 0.032 | 0.126 | 0.033 | 0.009 | 0.009 | 0.338 | 0.48 | 0.38 | 0.27 | 0.48 | 0.3 | 0.46 |
| 1 | 25148686 | rs10903106 | -0.175 | -1.323 | -0.252 | 0.158 | 0.044 | 0.011 | 0.011 | 1.139 | 0.48 | 0.35 | 0.7 | 0.52 | 0.7 | 0.54 |
| 1 | 25158500 | rs6667299 | -0.18 | -1.271 | -0.515 | 0.182 | 0.114 | 0.019 | 0.019 | 2.317 | 0.48 | 0.24 | 0.68 | 0.51 | 0.7 | 0.58 |
| 1 | 25169634 | rs12439 | 0.107 | -1.21 | -0.648 | 0.213 | 0.005 | 0.002 | 0.002 | 1.03 | 0.48 | 0.46 | 0.28 | 0.49 | 0.3 | 0.42 |
| 1 | 25185786 | rs4295833 | -0.819 | -0.76 | -1.399 | 0.002 | 0.009 | 0.06 | 0.06 | 0 | 0.18 | 0.06 | 0.28 | 0.09 | 0.12 | 0.18 |
| 1 | 25186716 | rs7523543 | -0.034 | -0.981 | -1.235 | 0.228 | -0.002 | 0 | 0 | 0 | 0.34 | 0.55 | 0.21 | 0.41 | 0.29 | 0.4 |
| 1 | 25187878 | rs11249186 | 0.46 | -0.552 | -2.066 | 0.335 | -0.128 | -0.081 | -0.081 | 0 | 0.29 | 0.09 | 0.3 | 0.32 | 0.39 | 0.33 |
| 1 | 25187952 | rs11249187 | 0.449 | -0.554 | -2.066 | 0.334 | -0.128 | -0.081 | -0.081 | 0 | 0.29 | 0.09 | 0.3 | 0.32 | 0.39 | 0.33 |
| 1 | 25188064 | rs6681012 | 0.196 | -0.587 | -1.907 | 0.272 | 0.016 | 0.023 | 0.023 | 0 | 0.33 | 0.19 | 0.47 | 0.27 | 0.33 | 0.26 |
| 1 | 25188940 | rs10903108 | 1.047 | 0.108 | -1.813 | 0.566 | -0.092 | -0.063 | -0.063 | 0 | 0.2 | 0.07 | 0.27 | 0.28 | 0.33 | 0.27 |
| 1 | 25193155 | rs3131513 | -0.356 | -0.141 | -1.812 | 0.259 | -0.03 | 0.004 | 0.004 | 0.02 | 0.67 | 0.86 | 0.54 | 0.7 | 0.6 | 0.74 |
| 1 | 25193199 | rs3131514 | 0.858 | -0.359 | -0.814 | 0.571 | 0.038 | 0.033 | 0.033 | 0 | 0.24 | 0.33 | 0.28 | 0.35 | 0.28 | 0.28 |
| 1 | 25194299 | rs2361150 | 0.204 | -1.576 | -0.393 | 0.473 | -0.022 | -0.034 | -0.034 | 0 | 0.39 | 0.49 | 0.27 | 0.26 | 0.3 | 0.44 |
| 1 | 25194700 | rs2206047 | -0.158 | -1.532 | -0.402 | 0.539 | 0.036 | -0.016 | -0.016 | 2.276 | 0.62 | 0.81 | 0.55 | 0.63 | 0.64 | 0.7 |
| 1 | 25196370 | rs3131518 | -0.112 | -1.711 | -0.519 | 0.359 | 0.032 | 0.034 | 0.034 | 0 | 0.46 | 0.61 | 0.29 | 0.29 | 0.32 | 0.45 |
| 1 | 25197155 | rs4648881 | 0.307 | -2.194 | -0.152 | 0.551 | 0 | 0.002 | 0.002 | 0 | 0.44 | 0.49 | 0.38 | 0.37 | 0.5 | 0.35 |
| 1 | 25199722 | rs3131522 | 0.456 | -2.146 | -0.05 | 0.483 | -0.043 | -0.009 | -0.009 | 0.019 | 0.25 | 0.37 | 0.18 | 0.26 | 0.16 | 0.43 |
| 1 | 25207629 | rs4649033 | 1.422 | -2.013 | -0.48 | 0.529 | -0.163 | -0.104 | -0.104 | 0 | 0.31 | 0.02 | 0.31 | 0.35 | 0.47 | 0.3 |
| 1 | 25213379 | rs12027805 | 1.039 | -2.224 | -0.507 | 0.501 | -0.002 | 0.003 | 0.003 | 0 | 0.16 | 0.1 | 0.12 | 0.15 | 0.2 | 0.07 |
| 1 | 25213803 | rs7551358 | 0.329 | -2.486 | -0.294 | 0.627 | -0.009 | 0.007 | 0.007 | 0 | 0.24 | 0.22 | 0.16 | 0.35 | 0.16 | 0.44 |
| 1 | 25215750 | rs10903113 | 0.176 | -2.464 | 0.056 | 0.298 | -0.029 | -0.002 | -0.002 | 0.007 | 0.64 | 0.53 | 0.5 | 0.39 | 0.71 | 0.43 |
| 1 | 25215989 | rs11578143 | -0.088 | -2.57 | 0.017 | 0.114 | -0.018 | -0.091 | -0.091 | 0 | 0.29 | 0.31 | 0.15 | 0.02 | 0.23 | 0.12 |
| 1 | 25216494 | rs6657823 | 0.673 | -2.365 | -0.161 | 0.513 | 0.012 | -0.023 | -0.023 | 0 | 0.18 | 0.12 | 0.17 | 0.46 | 0.13 | 0.46 |
| 1 | 25217346 | rs4649036 | -0.568 | -2.362 | -0.028 | 0.488 | -0.005 | 0.002 | 0.002 | 0 | 0.83 | 0.76 | 0.66 | 0.85 | 0.84 | 0.88 |

| | | | | | | | | | | | | | | | | |
|---|---|---|---|---|---|---|---|---|---|---|---|---|---|---|---|---|
| 1 | 25220541 | rs9438856 | 0.463 | -2.199 | -0.061 | 0.533 | 0.019 | -0.052 | -0.052 | 0 | 0.18 | 0.08 | 0.17 | 0.46 | 0.14 | 0.46 |
| 1 | 25220590 | rs9438873 | 0.333 | -2.183 | -0.032 | 0.533 | 0.005 | -0.004 | -0.004 | 0 | 0.19 | 0.15 | 0.18 | 0.46 | 0.14 | 0.46 |
| 1 | 25222379 | rs6663310 | -0.468 | -2.151 | -0.124 | 0.456 | -0.004 | 0.001 | 0.001 | 0 | 0.4 | 0.53 | 0.32 | 0.49 | 0.34 | 0.55 |
| 1 | 25224010 | rs7553295 | -0.971 | -2.331 | -0.327 | 0.356 | 0.085 | 0.05 | 0.05 | 0.126 | 0.44 | 0.79 | 0.46 | 0.3 | 0.24 | 0.29 |
| 1 | 25224321 | rs9438859 | -0.463 | -2.19 | -0.089 | 0.398 | -0.005 | -0.002 | -0.002 | 0 | 0.31 | 0.35 | 0.31 | 0.31 | 0.27 | 0.31 |
| 1 | 25227677 | rs13157 | -0.91 | -1.874 | 0.39 | 0.079 | -0.074 | -0.083 | -0.083 | 0.584 | 0.16 | 0.45 | 0.07 | 0.04 | 0.06 | 0.07 |
| 1 | 25239289 | rs16830141 | 0.07 | -1.778 | 0.344 | 0.078 | -0.007 | -0.026 | -0.026 | 0 | 0.09 | 0.28 | 0.05 | 0.03 | 0.05 | 0.05 |
| 1 | 25240341 | rs2236850 | -1.304 | -1.509 | -0.51 | 0.155 | -0.036 | -0.022 | -0.022 | 0 | 0.6 | 0.94 | 0.5 | 0.35 | 0.44 | 0.38 |
| 1 | 25241056 | rs2282718 | -0.29 | -1.279 | -0.063 | 0.192 | 0.001 | 0.001 | 0.001 | 0.002 | 0.46 | 0.47 | 0.41 | 0.33 | 0.38 | 0.33 |
| 1 | 25241116 | rs9438876 | -0.746 | 0.239 | -0.042 | 0.122 | -0.002 | 0.009 | 0.009 | 0 | 0.54 | 0.59 | 0.5 | 0.27 | 0.49 | 0.37 |
| 1 | 25243554 | rs2236851 | 0.198 | 0.215 | -0.673 | 0.103 | -0.053 | -0.04 | -0.04 | 19.757 | 0.09 | 0.01 | 0.07 | 0.12 | 0.15 | 0.12 |
| 1 | 25243767 | rs2236852 | -0.007 | 0.361 | -0.128 | 0.181 | 0.001 | 0 | 0 | 4.584 | 0.48 | 0.53 | 0.51 | 0.57 | 0.45 | 0.63 |
| 1 | 25251424 | rs742230 | -0.121 | -0.042 | -0.308 | 0.274 | 0 | -0.001 | -0.001 | 0.385 | 0.49 | 0.46 | 0.58 | 0.55 | 0.58 | 0.61 |
| 1 | 25251938 | rs6662768 | -0.292 | -0.131 | -0.001 | 0.132 | -0.038 | -0.04 | -0.04 | 0.128 | 0.07 | 0.18 | 0.01 | 0 | 0.01 | 0.01 |
| 1 | 25273200 | rs7551188 | -0.105 | 0.332 | -0.397 | 0.19 | 0.002 | 0.002 | 0.002 | 0 | 0.45 | 0.41 | 0.59 | 0.52 | 0.47 | 0.53 |
| 1 | 25277982 | rs11249206 | -0.195 | 0.225 | -0.286 | -0.153 | -0.003 | 0.019 | 0.019 | 0 | 0.47 | 0.31 | 0.66 | 0.64 | 0.52 | 0.54 |
| 1 | 25279180 | rs2135756 | -0.328 | 0.194 | -0.247 | -0.178 | 0.001 | 0.018 | 0.018 | 0 | 0.49 | 0.54 | 0.67 | 0.64 | 0.52 | 0.53 |
| 1 | 25281015 | rs4478762 | 0.5 | -0.094 | -0.85 | -0.481 | -0.009 | -0.027 | -0.027 | 0 | 0.17 | 0.23 | 0.1 | 0.05 | 0.13 | 0.26 |
| 1 | 25282961 | rs1848185 | -0.601 | 0.237 | -0.536 | -0.026 | -0.088 | -0.046 | -0.046 | 0.079 | 0.69 | 0.93 | 0.81 | 0.69 | 0.65 | 0.79 |
| 1 | 25289427 | rs11249209 | -0.412 | 0.411 | -0.047 | -0.025 | 0.008 | -0.042 | -0.042 | 0.304 | 0.09 | 0.18 | 0.07 | 0.02 | 0.12 | 0.06 |
| 1 | 25294264 | rs7523412 | 0.416 | 0.443 | -0.068 | -0.096 | -0.026 | 0.076 | 0.076 | 0 | 0.54 | 0.22 | 0.71 | 0.69 | 0.54 | 0.68 |
| 1 | 25294878 | rs10903118 | 0.329 | 0.536 | -0.002 | -0.051 | -0.027 | 0.065 | 0.065 | 0 | 0.55 | 0.24 | 0.71 | 0.69 | 0.54 | 0.67 |
| 1 | 25295580 | rs7529070 | 0.373 | 0.531 | 0.074 | -0.056 | -0.007 | 0.015 | 0.015 | 0 | 0.56 | 0.42 | 0.73 | 0.69 | 0.55 | 0.68 |
| 1 | 25297184 | rs11249215 | 0.655 | 0.516 | -0.254 | 0.16 | 0.002 | 0.018 | 0.018 | 0.1 | 0.49 | 0.23 | 0.62 | 0.53 | 0.52 | 0.63 |
| 1 | 25297217 | rs11249216 | -0.567 | 1.013 | -0.549 | -0.035 | 0.038 | -0.042 | -0.042 | 0.273 | 0.1 | 0.37 | 0.08 | 0.01 | 0.12 | 0.07 |
| 1 | 25301869 | rs1609996 | 0.415 | 0.83 | -0.797 | 0.029 | -0.006 | 0.007 | 0.007 | 0 | 0.58 | 0.59 | 0.75 | 0.68 | 0.54 | 0.68 |
| 1 | 25305607 | rs6690896 | -0.078 | 0.979 | -1.718 | 0.405 | -0.092 | -0.011 | -0.011 | 0 | 0.68 | 0.93 | 0.82 | 0.71 | 0.67 | 0.74 |
| 1 | 25306562 | rs12719821 | 0.052 | 0.656 | -1.005 | 0.486 | 0.001 | 0 | 0 | 0 | 0.52 | 0.44 | 0.64 | 0.45 | 0.55 | 0.51 |
| 1 | 25308276 | rs311484 | -0.232 | 0.676 | -0.962 | 0.065 | 0.004 | -0.038 | -0.038 | 0 | 0.42 | 0.55 | 0.33 | 0.17 | 0.43 | 0.33 |
| 1 | 25314894 | rs432169 | -0.504 | -0.37 | -1.334 | 0.696 | 0.038 | -0.032 | -0.032 | 6.138 | 0.35 | 0.2 | 0.26 | 0.43 | 0.26 | 0.37 |
| 1 | 25319751 | rs311437 | -0.967 | -0.873 | -1.859 | 1.136 | 0.028 | -0.07 | -0.07 | 0.823 | 0.26 | 0.86 | 0.31 | 0.36 | 0.15 | 0.38 |
| 1 | 25321597 | rs9438878 | 0.132 | -0.821 | -1.563 | 1.102 | -0.004 | 0.003 | 0.003 | 0 | 0.11 | 0.15 | 0.09 | 0.15 | 0.09 | 0.19 |
| 1 | 25321765 | rs9438879 | 0.132 | -0.821 | -1.563 | 1.102 | -0.004 | 0.003 | 0.003 | 0 | 0.11 | 0.15 | 0.09 | 0.15 | 0.09 | 0.18 |
| 1 | 25322574 | rs9438880 | 0.072 | -0.918 | -1.783 | 0.931 | 0.015 | -0.014 | -0.014 | 4.229 | 0.76 | 0.87 | 0.84 | 0.77 | 0.79 | 0.78 |
| 1 | 25322808 | rs9438881 | -0.252 | -1.003 | -1.76 | 0.932 | -0.013 | -0.044 | -0.044 | 1.444 | 0.79 | 0.97 | 0.89 | 0.78 | 0.82 | 0.79 |
| 1 | 25323047 | rs11249227 | 0.558 | -0.929 | -0.613 | 1.157 | -0.003 | 0.077 | 0.077 | 0 | 0.73 | 0.62 | 0.69 | 0.58 | 0.77 | 0.61 |
| 1 | 25325837 | rs4649042 | 1.395 | -1.37 | 0.046 | 1.135 | 0.004 | -0.006 | -0.006 | 0 | 0.61 | 0.35 | 0.57 | 0.46 | 0.63 | 0.44 |
| 1 | 25329752 | rs11249231 | 0.231 | -1.359 | 0.368 | 1.272 | 0.004 | 0.15 | 0.15 | 0 | 0.1 | 0.21 | 0.12 | 0.37 | 0.1 | 0.28 |
| 1 | 25331411 | rs311469 | 0.247 | -1.402 | 0.454 | 1.226 | 0.003 | 0.009 | 0.009 | 0 | 0.75 | 0.81 | 0.73 | 0.83 | 0.76 | 0.72 |
| 1 | 25331468 | rs311470 | -0.456 | -1.425 | 0.434 | 1.187 | 0.018 | 0.065 | 0.065 | 0 | 0.81 | 0.93 | 0.89 | 1 | 0.81 | 0.85 |
| 1 | 25333062 | rs12403486 | -0.506 | -1.153 | 0.837 | 1.504 | -0.007 | 0.148 | 0.148 | 0.061 | 0.13 | 0.33 | 0.15 | 0.37 | 0.13 | 0.28 |
| 1 | 25333264 | rs12058910 | 0.15 | -1.144 | 0.656 | 1.539 | 0.021 | 0.078 | 0.078 | 0.351 | 0.06 | 0.13 | 0.16 | 0.17 | 0.05 | 0.14 |
| 1 | 25335222 | rs12117144 | -0.941 | -1.509 | 0.514 | 1.384 | -0.051 | 0.095 | 0.095 | 2.814 | 0.22 | 0.49 | 0.18 | 0.38 | 0.15 | 0.32 |
| 1 | 25338799 | rs10489161 | -1.308 | -1.471 | 0.368 | 1.278 | -0.02 | 0.054 | 0.054 | 5.007 | 0.27 | 0.62 | 0.34 | 0.54 | 0.19 | 0.45 |
| 1 | 25344155 | rs10903126 | -1.446 | -1.473 | -0.054 | 1.104 | 0.02 | -0.024 | -0.024 | 0 | 0.3 | 0.78 | 0.33 | 0.43 | 0.2 | 0.46 |
| 1 | 25344494 | rs11802335 | 1.033 | -1.438 | -0.754 | 0.668 | -0.002 | -0.004 | -0.004 | 1.043 | 0.23 | 0.2 | 0.13 | 0.15 | 0.2 | 0.11 |
| 1 | 25347577 | rs11249243 | 0.633 | -1.41 | -0.936 | 0.737 | -0.003 | -0.04 | -0.04 | 0.351 | 0.12 | 0.09 | 0.17 | 0.19 | 0.07 | 0.18 |
| 1 | 25347711 | rs11249244 | 0.958 | -1.878 | -1.401 | 0.296 | -0.024 | -0.07 | -0.07 | 0 | 0.41 | 0.06 | 0.53 | 0.57 | 0.6 | 0.39 |
| 1 | 25348657 | rs6700387 | 0.645 | -1.666 | -0.622 | 0.397 | -0.015 | 0.042 | 0.042 | 0.014 | 0.11 | 0.25 | 0.1 | 0.16 | 0.1 | 0.13 |
| 1 | 25351581 | rs4320727 | 0.659 | -1.468 | -0.762 | 0.307 | 0.016 | -0.004 | -0.004 | 0 | 0.43 | 0.26 | 0.56 | 0.54 | 0.62 | 0.47 |
| 1 | 25354020 | rs11249245 | 0.53 | -1.498 | -0.83 | 0.14 | -0.009 | -0.017 | -0.017 | 0 | 0.15 | 0.11 | 0.2 | 0.19 | 0.13 | 0.18 |
| 1 | 25356719 | rs311425 | -0.621 | -1.851 | -0.24 | -0.16 | -0.015 | -0.086 | -0.086 | 0.676 | 0.68 | 0.66 | 0.85 | 1 | 0.8 | 0.86 |
| 1 | 25358211 | rs311426 | 0.155 | -1.824 | -0.39 | -0.063 | 0.095 | 0.112 | 0.112 | 0.051 | 0.54 | 0.27 | 0.71 | 0.74 | 0.7 | 0.64 |
| 1 | 25359929 | rs387240 | -0.761 | -2.043 | -0.499 | -0.244 | 0.021 | 0.03 | 0.03 | 0 | 0.7 | 0.8 | 0.71 | 0.86 | 0.73 | 0.8 |
| 1 | 25363656 | rs452044 | -0.001 | -1.992 | -0.431 | -0.141 | 0.012 | 0.01 | 0.01 | 0 | 0.64 | 0.64 | 0.7 | 0.72 | 0.72 | 0.65 |
| 1 | 25364652 | rs1702312 | 0.209 | -0.573 | -0.452 | -0.236 | -0.001 | 0.028 | 0.028 | 1.522 | 0.53 | 0.67 | 0.61 | 0.72 | 0.58 | 0.62 |
| 1 | 25366182 | rs380497 | -1.543 | -0.62 | -0.574 | -0.239 | 0.017 | 0.02 | 0.02 | 0 | 0.8 | 0.89 | 0.91 | 0.91 | 0.85 | 0.82 |
| 1 | 25366987 | rs404100 | 1.185 | -0.707 | -0.496 | -0.051 | -0.016 | -0.032 | -0.032 | 0 | 0.37 | 0.21 | 0.47 | 0.49 | 0.47 | 0.46 |
| 1 | 25368768 | rs427848 | -1.471 | -0.656 | -0.96 | -0.036 | 0.037 | 0.046 | 0.046 | 0 | 0.79 | 0.93 | 0.91 | 0.91 | 0.85 | 0.82 |
| 1 | 25370521 | rs380939 | -0.883 | -0.677 | -1.078 | -0.127 | 0.056 | 0.083 | 0.083 | 0.104 | 0.73 | 0.95 | 0.9 | 0.91 | 0.84 | 0.8 |
| 1 | 25375468 | rs4649058 | -0.409 | -0.156 | -0.733 | 0.388 | 0.022 | 0.035 | 0.035 | 0.521 | 0.34 | 0.43 | 0.49 | 0.44 | 0.39 | 0.31 |
| 1 | 25375583 | rs11248971 | 0.785 | -0.172 | -0.698 | 0.341 | -0.006 | 0.037 | 0.037 | 0.768 | 0.15 | 0.43 | 0.13 | 0.19 | 0.11 | 0.19 |
| 1 | 25377908 | rs4082732 | 1.833 | -0.168 | -0.846 | 0.333 | -0.005 | -0.031 | -0.031 | 0 | 0.11 | 0.03 | 0.2 | 0.13 | 0.14 | 0.08 |
| 1 | 25378444 | rs369427 | 0.717 | -0.198 | -0.792 | 0.437 | 0.001 | 0.001 | 0.001 | 0.506 | 0.47 | 0.56 | 0.38 | 0.54 | 0.46 | 0.57 |
| 1 | 25378513 | rs11248974 | 0.633 | 0.013 | -0.665 | 0.57 | 0.034 | 0.009 | 0.009 | 0.444 | 0.21 | 0.29 | 0.32 | 0.25 | 0.27 | 0.18 |
| 1 | 25379360 | rs1702311 | 1.642 | -0.015 | -0.76 | 0.473 | -0.051 | -0.042 | -0.042 | 4.787 | 0.32 | 0.08 | 0.24 | 0.33 | 0.34 | 0.38 |
| 1 | 25380023 | rs7520145 | 0.569 | 0.044 | -0.624 | 0.388 | 0.023 | 0.006 | 0.006 | 0.285 | 0.21 | 0.28 | 0.33 | 0.26 | 0.27 | 0.18 |
| 1 | 25381063 | rs9438886 | -0.226 | -0.082 | -0.583 | 0.32 | -0.027 | -0.028 | -0.028 | 0 | 0.64 | 0.42 | 0.76 | 0.78 | 0.75 | 0.69 |
| 1 | 25382556 | rs12403541 | 1.183 | 0.282 | -0.351 | 0.548 | -0.006 | 0.004 | 0.004 | 0 | 0.14 | 0.17 | 0.23 | 0.15 | 0.16 | 0.08 |

| | | | | | | | | | | | | | | | | |
|---|---|---|---|---|---|---|---|---|---|---|---|---|---|---|---|---|
| 1 | 25382618 | rs1997927 | -0.037 | 0.243 | -0.284 | 0.481 | 0.011 | 0.008 | 0.008 | 0.509 | 0.35 | 0.41 | 0.42 | 0.44 | 0.38 | 0.38 |
| 1 | 25384533 | rs4649060 | -0.913 | 0.274 | -0.468 | 0.473 | 0.003 | 0.017 | 0.017 | 0.327 | 0.49 | 0.64 | 0.65 | 0.58 | 0.55 | 0.46 |
| 1 | 25390827 | rs311448 | 0.706 | 0.433 | -0.813 | 0.361 | -0.013 | 0.009 | 0.009 | 0 | 0.44 | 0.15 | 0.45 | 0.55 | 0.46 | 0.51 |
| 1 | 25391629 | rs12031599 | 0.516 | 0.511 | -0.789 | 0.474 | -0.009 | -0.003 | -0.003 | 0 | 0.08 | 0.06 | 0.21 | 0.24 | 0.14 | 0.15 |
| 1 | 25393816 | rs311454 | -0.321 | 0.052 | -0.702 | 0.314 | 0.005 | 0.007 | 0.007 | 0 | 0.69 | 0.74 | 0.68 | 0.85 | 0.7 | 0.81 |
| 1 | 25395879 | rs12126466 | 0.735 | -0.696 | -0.804 | 0.372 | -0.034 | -0.021 | -0.021 | 0 | 0.27 | 0.34 | 0.17 | 0.2 | 0.16 | 0.21 |
| 1 | 25396833 | rs365865 | -0.544 | -0.464 | -0.887 | 0.401 | -0.089 | 0.01 | 0.01 | 0 | 0.74 | 0.9 | 0.66 | 0.77 | 0.62 | 0.73 |
| 1 | 25397349 | rs10903024 | 0.775 | -0.47 | -0.819 | 0.339 | -0.035 | -0.022 | -0.022 | 0 | 0.27 | 0.34 | 0.17 | 0.2 | 0.16 | 0.21 |
| 1 | 25397967 | rs4568778 | 0.259 | -0.246 | -1.495 | 0.402 | -0.04 | -0.003 | -0.003 | 0 | 0.6 | 0.81 | 0.44 | 0.61 | 0.47 | 0.65 |
| 1 | 25399474 | rs11248979 | -0.582 | -0.413 | -1.639 | 0.577 | 0.099 | -0.001 | -0.001 | 0 | 0.44 | 0.79 | 0.41 | 0.41 | 0.31 | 0.37 |
| 1 | 25401437 | rs427614 | 0.441 | 0.392 | -1.439 | 0.312 | 0.015 | -0.021 | -0.021 | 0.117 | 0.64 | 0.57 | 0.71 | 0.79 | 0.53 | 0.6 |
| 1 | 25401760 | rs6693687 | 1.844 | 0.459 | -1.551 | 0.533 | -0.054 | -0.092 | -0.092 | 0 | 0.25 | 0.03 | 0.37 | 0.32 | 0.25 | 0.25 |
| 1 | 25401850 | rs380337 | 0.367 | 0.469 | -1.524 | 0.429 | 0.015 | -0.021 | -0.021 | 0 | 0.64 | 0.57 | 0.71 | 0.79 | 0.53 | 0.59 |
| 1 | 25403405 | rs401668 | -1.874 | 0.61 | -1.579 | 0.162 | -0.088 | 0.071 | 0.071 | 0 | 0.86 | 0.99 | 0.89 | 0.99 | 0.76 | 0.8 |
| 1 | 25403783 | rs443523 | 0.84 | 0.335 | -1.483 | 0.287 | -0.008 | -0.009 | -0.009 | 0 | 0.34 | 0.4 | 0.21 | 0.26 | 0.25 | 0.24 |
| 1 | 25406674 | rs6702929 | -0.212 | 0.421 | -1.725 | 0.34 | 0.001 | 0.001 | 0.001 | 0 | 0.5 | 0.53 | 0.34 | 0.39 | 0.43 | 0.33 |
| 1 | 25407854 | rs10493026 | 1.192 | 0.595 | -2.034 | 0.423 | -0.008 | -0.017 | -0.017 | 0 | 0.14 | 0.04 | 0.11 | 0.13 | 0.18 | 0.09 |
| 1 | 25408685 | rs12081177 | 0.934 | 0.736 | -1.732 | 0.603 | -0.001 | -0.025 | -0.025 | 0 | 0.06 | 0.11 | 0.04 | 0.04 | 0.04 | 0.12 |
| 1 | 25409607 | rs311477 | -0.329 | 0.799 | -1.977 | 0.641 | -0.012 | -0.006 | -0.006 | 1.194 | 0.58 | 0.83 | 0.4 | 0.44 | 0.48 | 0.5 |
| 1 | 25413466 | rs7533661 | -0.354 | 0.886 | -1.046 | 0.395 | 0.043 | 0.173 | 0.173 | 0.487 | 0.1 | 0.38 | 0.18 | 0.26 | 0.08 | 0.19 |
| 1 | 25418010 | rs311461 | -0.303 | 0.94 | -1.261 | 0.239 | 0.069 | -0.013 | -0.013 | 0 | 0.76 | 0.65 | 0.71 | 0.79 | 0.68 | 0.71 |
| 1 | 25421777 | rs11578154 | 0.555 | 0.708 | -2.643 | -0.026 | 0.007 | -0.005 | -0.005 | 0 | 0.11 | 0.06 | 0.13 | 0.14 | 0.14 | 0.18 |
| 1 | 25421990 | rs11588172 | 0.128 | 1.06 | -2.449 | -0.035 | -0.04 | -0.003 | -0.003 | 0 | 0.53 | 0.14 | 0.39 | 0.39 | 0.51 | 0.49 |
| 1 | 25424889 | rs803314 | -0.752 | 1.277 | -2.377 | -1.017 | -0.097 | 0.077 | 0.077 | 0 | 0.27 | 0.05 | 0.2 | 0.12 | 0.33 | 0.29 |
| 1 | 25429473 | rs706844 | 0.027 | 0.983 | -2.027 | -1.47 | -0.024 | 0.017 | 0.017 | 2.901 | 0.75 | 0.83 | 0.8 | 0.95 | 0.63 | 0.65 |
| 1 | 25434242 | rs803313 | 0.503 | 0.924 | -2.104 | -1.576 | -0.038 | 0.044 | 0.044 | 1.328 | 0.7 | 0.83 | 0.78 | 0.95 | 0.58 | 0.65 |
| 1 | 25447125 | rs6692950 | 0.574 | -0.068 | -1.592 | -2.077 | -0.04 | -0.052 | -0.052 | 0 | 0.32 | 0.5 | 0.52 | 0.81 | 0.28 | 0.38 |
| 1 | 25452671 | rs2753319 | 0.146 | -0.39 | -1.464 | -1.984 | -0.008 | -0.037 | -0.037 | 0 | 0.4 | 0.43 | 0.29 | 0.17 | 0.31 | 0.33 |
| 1 | 25457038 | rs803304 | 0.157 | -0.394 | -1.49 | -1.973 | -0.008 | -0.048 | -0.048 | 4.262 | 0.4 | 0.47 | 0.3 | 0.17 | 0.32 | 0.31 |
| 1 | 25460648 | rs2660367 | 0.034 | -0.445 | -1.458 | -2.007 | -0.026 | -0.023 | -0.023 | 0.121 | 0.07 | 0.21 | 0.04 | 0 | 0.01 | 0.01 |
| 1 | 25461078 | rs6673342 | 0.959 | -0.475 | -1.582 | -2.031 | -0.101 | 0.06 | 0.06 | 0 | 0.3 | 0.03 | 0.23 | 0.14 | 0.3 | 0.26 |
| 1 | 25462079 | rs11804893 | 0.084 | -0.436 | -1.513 | -2.006 | -0.006 | -0.045 | -0.045 | 0.025 | 0.39 | 0.45 | 0.29 | 0.14 | 0.31 | 0.27 |
| 1 | 25467161 | rs7530932 | -0.501 | -0.755 | -1.02 | -1.564 | -0.012 | -0.01 | -0.01 | 0 | 0.07 | 0.28 | 0.04 | 0 | 0.01 | 0.01 |
| 1 | 25469913 | rs2064251 | -0.171 | -0.37 | -1.567 | -1.452 | 0.049 | 0.057 | 0.057 | 0.75 | 0.39 | 0.67 | 0.54 | 0.82 | 0.31 | 0.38 |
| 1 | 25486203 | rs926461 | 0.107 | 0.688 | -0.905 | -0.926 | 0.032 | -0.069 | -0.069 | 0.486 | 0.83 | 0.7 | 0.84 | 0.96 | 0.72 | 0.74 |
| 1 | 25495966 | rs7527379 | -0.357 | 0.628 | -1.43 | -0.297 | 0.018 | -0.003 | -0.003 | 0 | 0.23 | 0.6 | 0.23 | 0.23 | 0.29 | 0.41 |
| 1 | 25499030 | rs742390 | -0.224 | 0.623 | -2.123 | 0.043 | -0.064 | 0.001 | 0.001 | 0.28 | 0.35 | 0.13 | 0.32 | 0.34 | 0.36 | 0.36 |
| 1 | 25519623 | rs4520361 | -0.392 | -0.241 | -1.851 | 0.24 | 0.068 | 0.038 | 0.038 | 0 | 0.46 | 0.77 | 0.35 | 0.38 | 0.34 | 0.3 |
| 1 | 25521090 | rs12736858 | -0.174 | -1.159 | -1.299 | -0.209 | -0.018 | -0.021 | -0.021 | 0.905 | 0.26 | 0.23 | 0.17 | 0.09 | 0.11 | 0.13 |
| 1 | 25525711 | rs742391 | -0.022 | -0.836 | -0.848 | 0.02 | -0.003 | -0.008 | -0.008 | 0 | 0.2 | 0.12 | 0.12 | 0.28 | 0.21 | 0.13 |
| 1 | 25525881 | rs742393 | -0.131 | -0.811 | -0.862 | 0.006 | 0.001 | -0.008 | -0.008 | 0 | 0.2 | 0.12 | 0.11 | 0.28 | 0.18 | 0.12 |
| 1 | 25527944 | rs7415034 | -0.871 | -1.285 | 0.277 | -0.486 | -0.053 | -0.106 | -0.106 | 0.003 | 0.2 | 0.52 | 0.13 | 0.07 | 0.12 | 0.04 |
| 1 | 25529038 | rs2375278 | -0.275 | -1.368 | -0.159 | -0.066 | 0.019 | -0.04 | -0.04 | 0 | 0.79 | 0.96 | 0.78 | 0.8 | 0.83 | 0.9 |
| 1 | 25555672 | rs10903036 | -1.198 | -1.513 | 2.396 | -0.238 | 0.004 | -0.004 | -0.004 | 0 | 0.07 | 0.65 | 0.07 | 0 | 0.01 | 0 |
| 1 | 25570081 | rs1043879 | 0.182 | -1.278 | 0.95 | -1.131 | -0.031 | 0.195 | 0.195 | 0 | 0.29 | 0.01 | 0.21 | 0.03 | 0.27 | 0.07 |
| 1 | 25687901 | rs1053438 | 0.014 | -1.271 | 1.061 | -1.005 | -0.02 | 0.073 | 0.073 | 0.65 | 0.46 | 0.1 | 0.49 | 0.76 | 0.44 | 0.68 |
| 1 | 25688276 | rs9689 | 0.018 | -1.294 | 1.277 | -1.016 | 0.011 | 0.047 | 0.047 | 0.014 | 0.51 | 0.64 | 0.48 | 0.24 | 0.55 | 0.32 |
| 1 | 25695835 | rs1293259 | 0.824 | -1.422 | 0.475 | -0.988 | 0.015 | -0.04 | -0.04 | 0 | 0.18 | 0.04 | 0.22 | 0.21 | 0.15 | 0.09 |
| 1 | 25753051 | rs11249248 | 0.139 | -0.765 | -0.095 | -0.507 | -0.001 | 0.09 | 0.09 | 0.507 | 0.53 | 0.81 | 0.49 | 0.23 | 0.55 | 0.32 |
| 1 | 25753638 | rs926438 | 0.139 | -0.765 | -0.095 | -0.507 | -0.001 | 0.09 | 0.09 | 0.507 | 0.53 | 0.81 | 0.49 | 0.23 | 0.55 | 0.32 |
| 1 | 25760920 | rs11802413 | 0.141 | -0.761 | -0.176 | -0.466 | 0.004 | 0.077 | 0.077 | 0.297 | 0.53 | 0.76 | 0.48 | 0.23 | 0.55 | 0.32 |
| 1 | 25768937 | rs10903129 | 0.147 | -0.755 | -0.174 | -0.463 | 0.004 | 0.078 | 0.078 | 0.333 | 0.53 | 0.76 | 0.48 | 0.23 | 0.55 | 0.32 |
| 1 | 25889422 | rs12096438 | -0.544 | 0.172 | -0.104 | 0.516 | -0.003 | -0.001 | -0.001 | 0 | 0.52 | 0.32 | 0.51 | 0.44 | 0.52 | 0.44 |
| 1 | 25889632 | rs6687605 | -0.549 | 0.144 | -0.117 | 0.502 | -0.003 | -0.001 | -0.001 | 0 | 0.52 | 0.32 | 0.51 | 0.44 | 0.52 | 0.44 |
| 1 | 25889672 | rs6688931 | -0.483 | 0.245 | -0.071 | 0.51 | -0.001 | 0 | 0 | 0 | 0.51 | 0.28 | 0.49 | 0.44 | 0.49 | 0.44 |
| 1 | 25894853 | rs11563 | 0.283 | 0.232 | -0.054 | 0.513 | -0.004 | -0.002 | -0.002 | 0 | 0.48 | 0.71 | 0.48 | 0.55 | 0.47 | 0.56 |
| 1 | 25895599 | rs3924234 | 0.23 | 0.193 | -0.177 | 0.52 | -0.003 | 0.002 | 0.002 | 0 | 0.49 | 0.75 | 0.5 | 0.56 | 0.5 | 0.54 |
| 1 | 25900215 | rs4659356 | -0.645 | -0.675 | 0.053 | 0.719 | 0.015 | 0.001 | 0.001 | 0 | 0.51 | 0.41 | 0.45 | 0.48 | 0.4 | 0.43 |
| 1 | 25900240 | rs4659357 | -0.177 | -0.501 | -0.338 | 0.658 | 0.001 | 0.002 | 0.002 | 0 | 0.17 | 0.13 | 0.28 | 0.13 | 0.18 | 0.06 |
| 1 | 25908241 | rs10902661 | -0.353 | -0.514 | 0.175 | 1.107 | -0.032 | 0.026 | 0.026 | 0 | 0.33 | 0.42 | 0.38 | 0.42 | 0.23 | 0.36 |
| 1 | 25932186 | rs4488019 | -0.163 | -0.422 | 0.42 | 0.612 | -0.071 | 0.085 | 0.085 | 0.799 | 0.18 | 0.01 | 0.13 | 0.02 | 0.29 | 0.1 |
| 1 | 25941936 | rs3754168 | -1.181 | -0.921 | 1.305 | 1.19 | -0.005 | 0.196 | 0.196 | 4.198 | 0.07 | 0.57 | 0.21 | 0.21 | 0.01 | 0.08 |
| 1 | 25942685 | rs3754166 | -1.265 | -0.929 | 1.302 | 1.186 | -0.01 | 0.188 | 0.188 | 4.214 | 0.08 | 0.57 | 0.21 | 0.21 | 0.01 | 0.08 |
| 1 | 25946094 | rs3754164 | -1.169 | -0.897 | 1.316 | 1.202 | -0.005 | 0.196 | 0.196 | 4.262 | 0.07 | 0.57 | 0.21 | 0.21 | 0.01 | 0.08 |
| 1 | 25968690 | rs3767908 | -1.046 | -0.86 | 0.285 | 1.244 | -0.006 | 0.213 | 0.213 | 5.12 | 0.07 | 0.59 | 0.21 | 0.23 | 0.01 | 0.08 |
| 1 | 25977004 | rs3767904 | -0.197 | -0.877 | -0.648 | 1.427 | 0.009 | -0.026 | -0.026 | 0 | 0.26 | 0.09 | 0.3 | 0.35 | 0.24 | 0.23 |
| 1 | 25981789 | rs3014714 | -0.187 | -0.896 | -0.634 | 1.456 | 0.011 | -0.016 | -0.016 | 0 | 0.26 | 0.18 | 0.32 | 0.34 | 0.25 | 0.24 |
| 1 | 25991356 | rs3014712 | -0.786 | -0.631 | -1.345 | 1.199 | 0.061 | -0.087 | -0.087 | 0 | 0.37 | 0.89 | 0.54 | 0.58 | 0.26 | 0.33 |
| 1 | 25999517 | rs2982313 | -0.778 | -0.736 | -1.396 | 1.204 | 0.072 | -0.099 | -0.099 | 0 | 0.37 | 0.91 | 0.54 | 0.57 | 0.26 | 0.37 |

| | | | | | | | | | | | | | | | | |
|---|---|---|---|---|---|---|---|---|---|---|---|---|---|---|---|---|
| 1 | 26003756 | rs3767891 | -1.507 | -0.335 | -0.622 | 1.482 | -0.011 | 0.087 | 0.087 | 0 | 0.05 | 0.13 | 0.18 | 0.29 | 0.04 | 0.12 |
| 1 | 26008659 | rs11247595 | 0.589 | -0.431 | -1.699 | 0.573 | -0.051 | -0.007 | -0.007 | 2.535 | 0.46 | 0.04 | 0.5 | 0.52 | 0.5 | 0.52 |
| 1 | 26010611 | rs2786882 | -1.058 | -0.742 | -0.305 | 0.48 | -0.012 | 0.026 | 0.026 | 6.745 | 0.14 | 0.63 | 0.25 | 0.2 | 0.09 | 0.1 |
| 1 | 26013252 | rs2744778 | -0.05 | -0.77 | 0.566 | 0.435 | 0.012 | -0.035 | -0.035 | 0 | 0.07 | 0.19 | 0.04 | 0 | 0.05 | 0.01 |
| 1 | 26020784 | rs2744780 | -1.744 | -0.884 | 0.394 | 0.387 | -0.025 | 0.008 | 0.008 | 4.41 | 0.16 | 0.65 | 0.26 | 0.21 | 0.1 | 0.12 |
| 1 | 26040514 | rs2744761 | -1.434 | -0.782 | 0.624 | 0.387 | 0.032 | 0.059 | 0.059 | 1.842 | 0.13 | 0.53 | 0.27 | 0.21 | 0.11 | 0.12 |
| 1 | 26057309 | rs2744768 | -1.913 | -0.876 | 0.627 | 0.373 | -0.044 | 0.003 | 0.003 | 4.763 | 0.16 | 0.65 | 0.27 | 0.21 | 0.09 | 0.14 |
| 1 | 26061266 | rs2786875 | -0.306 | -0.756 | 0.557 | 0.333 | -0.007 | -0.041 | -0.041 | 0.338 | 0.08 | 0.14 | 0.04 | 0 | 0.05 | 0.02 |
| 1 | 26069415 | rs3767880 | 0.045 | -1.347 | 0.378 | 0.025 | 0.028 | 0 | 0 | 0 | 0.23 | 0.11 | 0.1 | 0.29 | 0.16 | 0.23 |
| 1 | 26069507 | rs807267 | -0.36 | -1.233 | 0.757 | -0.024 | 0.019 | -0.048 | -0.048 | 0 | 0.09 | 0.25 | 0.06 | 0 | 0.07 | 0.02 |
| 1 | 26070909 | rs3767879 | 1.588 | -1.378 | 0.107 | -0.092 | -0.115 | -0.08 | -0.08 | 0.391 | 0.42 | 0.02 | 0.48 | 0.49 | 0.49 | 0.5 |
| 1 | 26075377 | rs2066995 | 0.483 | -1.282 | 1.07 | -0.121 | 0.004 | -0.007 | -0.007 | 0 | 0.77 | 0.65 | 0.64 | 0.75 | 0.73 | 0.74 |
| 1 | 26080217 | rs2281000 | -0.525 | -1.194 | 1.301 | -0.153 | 0.023 | 0.11 | 0.11 | 0 | 0.06 | 0.15 | 0.23 | 0.25 | 0.09 | 0.13 |
| 1 | 26081051 | rs807262 | 1.302 | -1.453 | 1.182 | -0.359 | -0.006 | -0.025 | -0.025 | 0.578 | 0.72 | 0.41 | 0.62 | 0.74 | 0.72 | 0.72 |
| 1 | 26087744 | rs4659378 | 1.554 | -1.656 | 0.786 | -0.395 | 0.01 | 0.06 | 0.06 | 1.257 | 0.64 | 0.17 | 0.55 | 0.74 | 0.66 | 0.72 |
| 1 | 26090072 | rs807270 | -0.571 | -1.652 | 0.774 | -0.41 | -0.033 | 0.08 | 0.08 | 0 | 0.85 | 0.97 | 0.89 | 0.99 | 0.82 | 0.86 |
| 1 | 26092248 | rs3820515 | 1.413 | -1.612 | 0.862 | -0.38 | 0.007 | 0.037 | 0.037 | 1.218 | 0.65 | 0.23 | 0.57 | 0.75 | 0.66 | 0.72 |
| 1 | 26097731 | rs2072748 | 1.441 | -2.204 | 0.386 | -0.431 | 0.021 | 0.018 | 0.018 | 0 | 0.48 | 0.32 | 0.5 | 0.48 | 0.51 | 0.35 |
| 1 | 26098107 | rs2072747 | 1.015 | -2.033 | 0.382 | -0.428 | 0 | -0.019 | -0.019 | 1.274 | 0.67 | 0.38 | 0.57 | 0.74 | 0.66 | 0.71 |
| 1 | 26098350 | rs807263 | -0.869 | -1.728 | 0.316 | 0.028 | 0.016 | 0.028 | 0.028 | 0 | 0.15 | 0.34 | 0.29 | 0.24 | 0.16 | 0.15 |
| 1 | 26104973 | rs1007108 | -0.028 | -1.587 | -0.389 | -0.028 | 0.036 | -0.015 | -0.015 | 0 | 0.19 | 0.05 | 0.07 | 0.26 | 0.13 | 0.35 |
| 1 | 26106769 | rs12043011 | -0.011 | -1.569 | -0.38 | -0.032 | 0.035 | -0.016 | -0.016 | 0 | 0.19 | 0.05 | 0.07 | 0.26 | 0.13 | 0.35 |
| 1 | 26116076 | rs807253 | 0.687 | -0.577 | 0.089 | -0.588 | 0.008 | -0.07 | -0.07 | 1.105 | 0.77 | 0.66 | 0.86 | 0.97 | 0.75 | 0.86 |
| 1 | 26117323 | rs807252 | 0.71 | -0.606 | 0.845 | -0.776 | 0.025 | -0.085 | -0.085 | 3.68 | 0.79 | 0.68 | 0.87 | 0.97 | 0.77 | 0.84 |
| 1 | 26118299 | rs6694897 | 0.75 | -0.373 | -0.46 | -0.733 | -0.032 | 0.074 | 0.074 | 0.096 | 0.39 | 0.11 | 0.6 | 0.3 | 0.42 | 0.53 |
| 1 | 26120869 | rs807247 | 0.467 | 0.224 | -0.736 | -0.249 | -0.015 | -0.094 | -0.094 | 3.111 | 0.29 | 0.03 | 0.17 | 0.34 | 0.27 | 0.18 |
| 1 | 26121077 | rs6696298 | 0.642 | 0.178 | -0.171 | -0.256 | 0 | 0.015 | 0.015 | 0 | 0.52 | 0.63 | 0.7 | 0.63 | 0.52 | 0.7 |
| 1 | 26121555 | rs807245 | -0.215 | 0.215 | -0.145 | -0.253 | 0.025 | -0.067 | -0.067 | 0 | 0.81 | 0.67 | 0.87 | 0.97 | 0.79 | 0.89 |
| 1 | 26125175 | rs7524154 | 0.446 | 0.235 | -0.065 | -0.224 | 0 | 0.012 | 0.012 | 0 | 0.53 | 0.63 | 0.7 | 0.64 | 0.53 | 0.74 |
| 1 | 26131021 | rs2072749 | 0.445 | 0.337 | -0.268 | -0.12 | 0.006 | -0.034 | -0.034 | 0.19 | 0.29 | 0.17 | 0.2 | 0.33 | 0.27 | 0.16 |
| 1 | 26131459 | rs11247735 | 0.433 | 0.296 | 0.097 | -0.114 | -0.001 | -0.002 | -0.002 | 0.935 | 0.48 | 0.48 | 0.7 | 0.61 | 0.48 | 0.7 |
| 1 | 26138136 | rs760598 | 0.225 | 0.151 | 0.525 | -0.369 | 0.007 | -0.099 | -0.099 | 2.12 | 0.77 | 0.41 | 0.83 | 0.97 | 0.77 | 0.9 |
| 1 | 26140573 | rs2294228 | 0.31 | 0.083 | 0.695 | -0.393 | 0 | 0.003 | 0.003 | 0.468 | 0.76 | 0.21 | 0.83 | 0.97 | 0.78 | 0.9 |
| 1 | 26144674 | rs6872 | 0.976 | 0.003 | 0.179 | -0.325 | -0.035 | -0.106 | -0.106 | 1.973 | 0.24 | 0.02 | 0.18 | 0.31 | 0.22 | 0.14 |
| 1 | 26146449 | rs2294226 | 0.823 | 0 | 0.169 | -0.336 | -0.042 | -0.104 | -0.104 | 6.441 | 0.26 | 0.02 | 0.2 | 0.31 | 0.27 | 0.14 |
| 1 | 26148011 | rs2049919 | -0.718 | -0.061 | 0.9 | -0.396 | 0.007 | -0.045 | -0.045 | 0 | 0.81 | 0.8 | 0.88 | 0.98 | 0.78 | 0.9 |
| 1 | 26155356 | rs10902694 | -0.525 | -0.061 | 1.141 | -0.396 | 0.014 | -0.077 | -0.077 | 0 | 0.8 | 0.71 | 0.86 | 0.98 | 0.77 | 0.9 |
| 1 | 26158962 | rs15045 | -0.533 | -0.065 | 1.139 | -0.398 | 0.014 | -0.077 | -0.077 | 0 | 0.8 | 0.71 | 0.86 | 0.98 | 0.77 | 0.9 |
| 1 | 26167843 | rs213632 | -0.368 | -0.055 | 1.325 | -0.404 | 0.012 | -0.102 | -0.102 | 0 | 0.23 | 0.57 | 0.16 | 0.03 | 0.23 | 0.1 |
| 1 | 26172111 | rs213627 | -0.104 | -0.067 | 1.417 | -0.405 | 0.022 | -0.132 | -0.132 | 0 | 0.22 | 0.43 | 0.14 | 0.02 | 0.23 | 0.1 |
| 1 | 26188625 | rs213637 | 0.258 | -0.017 | 1.3 | -0.344 | -0.028 | 0.026 | 0.026 | 0 | 0.56 | 0.8 | 0.68 | 0.67 | 0.51 | 0.76 |
| 1 | 26207244 | rs213621 | 0.442 | -0.106 | 1.343 | -0.39 | 0.002 | 0.033 | 0.033 | 0 | 0.56 | 0.79 | 0.7 | 0.68 | 0.53 | 0.76 |
| 1 | 26210791 | rs159530 | 0.425 | -0.109 | 1.099 | -0.391 | -0.003 | -0.008 | -0.008 | 0 | 0.55 | 0.54 | 0.69 | 0.68 | 0.53 | 0.76 |
| 1 | 26211773 | rs159528 | 1.065 | -0.13 | 1.058 | -0.403 | 0.025 | 0.068 | 0.068 | 0 | 0.5 | 0.17 | 0.65 | 0.67 | 0.53 | 0.76 |
| 1 | 26218084 | rs2095426 | 0.499 | -0.116 | 1.065 | -0.397 | -0.003 | -0.01 | -0.01 | 2.388 | 0.54 | 0.53 | 0.68 | 0.68 | 0.53 | 0.76 |
| 1 | 26224633 | rs12037513 | -0.598 | -0.064 | 0.799 | -0.035 | 0.011 | 0.05 | 0.05 | 0.304 | 0.29 | 0.5 | 0.19 | 0.38 | 0.31 | 0.3 |
| 1 | 26235705 | rs807057 | 0.902 | -0.508 | 0.554 | -0.401 | 0.051 | 0.068 | 0.068 | 0 | 0.51 | 0.19 | 0.66 | 0.68 | 0.56 | 0.77 |
| 1 | 26242882 | rs807061 | -0.024 | -0.577 | 1.255 | -0.047 | 0.003 | -0.001 | -0.001 | 0 | 0.69 | 0.29 | 0.74 | 0.71 | 0.73 | 0.85 |
| 1 | 26266697 | rs10902707 | -0.171 | -0.311 | 0.739 | -0.074 | -0.014 | -0.005 | -0.005 | 0 | 0.7 | 0.41 | 0.75 | 0.77 | 0.73 | 0.79 |
| 1 | 26274156 | rs10902709 | 0.626 | -0.671 | 0.208 | -0.258 | 0.044 | 0.045 | 0.045 | 0 | 0.49 | 0.15 | 0.67 | 0.71 | 0.61 | 0.69 |
| 1 | 26276335 | rs2782810 | -0.481 | -0.672 | 0.98 | -0.145 | -0.019 | -0.03 | -0.03 | 0.062 | 0.79 | 0.69 | 0.91 | 0.93 | 0.87 | 0.9 |
| 1 | 26284884 | rs10751731 | 0.947 | -0.898 | 0.604 | -0.632 | 0.069 | 0.153 | 0.153 | 0.274 | 0.5 | 0.15 | 0.69 | 0.78 | 0.64 | 0.62 |
| 1 | 26286840 | rs2782814 | 0.842 | -0.908 | 1.018 | -0.595 | -0.003 | 0.012 | 0.012 | 0.03 | 0.21 | 0.3 | 0.18 | 0.21 | 0.2 | 0.18 |
| 1 | 26297909 | rs1429936 | 0.941 | -1.009 | 0.962 | -0.585 | 0.074 | 0.079 | 0.079 | 0.125 | 0.51 | 0.09 | 0.69 | 0.71 | 0.66 | 0.68 |
| 1 | 26301699 | rs298442 | 1.022 | -0.969 | 1.051 | -0.58 | -0.001 | 0.001 | 0.001 | 0.04 | 0.2 | 0.19 | 0.17 | 0.21 | 0.2 | 0.14 |
| 1 | 26304999 | rs298443 | 0.82 | -0.959 | 0.704 | -0.569 | -0.003 | 0.017 | 0.017 | 0.07 | 0.22 | 0.35 | 0.19 | 0.22 | 0.2 | 0.14 |
| 1 | 26306701 | rs298447 | 0.369 | -0.93 | 0.489 | -0.558 | 0.025 | 0.045 | 0.045 | 0 | 0.22 | 0.12 | 0.08 | 0.07 | 0.14 | 0.17 |
| 1 | 26314564 | rs2245979 | -0.269 | -0.876 | 0.635 | -0.532 | -0.03 | -0.051 | -0.051 | 0 | 0.75 | 0.62 | 0.88 | 0.93 | 0.86 | 0.83 |
| 1 | 26324520 | rs11247841 | 0.596 | -0.642 | -0.491 | -0.4 | 0.071 | 0.075 | 0.075 | 0.076 | 0.51 | 0.09 | 0.68 | 0.71 | 0.66 | 0.69 |
| 1 | 26344477 | rs3008244 | -0.441 | -0.655 | -0.128 | -0.528 | 0.019 | 0.03 | 0.03 | 0 | 0.76 | 0.81 | 0.92 | 0.94 | 0.86 | 0.83 |
| 1 | 26345027 | rs12074420 | 1.184 | -0.79 | -0.207 | -0.605 | -0.017 | 0.036 | 0.036 | 0 | 0.19 | 0.08 | 0.11 | 0.06 | 0.2 | 0.07 |
| 1 | 26348049 | rs4659362 | 1.29 | -0.384 | -0.543 | -0.41 | -0.025 | 0.012 | 0.012 | 0 | 0.23 | 0.08 | 0.18 | 0.19 | 0.3 | 0.11 |
| 1 | 26351382 | rs17163470 | 0.636 | -1.15 | -0.255 | 0.136 | -0.025 | -0.055 | -0.055 | 0 | 0.1 | 0.03 | 0.3 | 0.35 | 0.14 | 0.24 |
| 1 | 26362194 | rs2802315 | 0.268 | -1.638 | 1.051 | -0.736 | -0.006 | -0.019 | -0.019 | 0.181 | 0.56 | 0.5 | 0.81 | 0.87 | 0.7 | 0.76 |
| 1 | 26365374 | rs4659404 | 0.485 | -1.645 | 1.203 | -0.737 | 0.071 | 0.078 | 0.078 | 0.742 | 0.53 | 0.2 | 0.73 | 0.75 | 0.69 | 0.75 |
| 1 | 26371788 | rs3121763 | -0.527 | -1.649 | 1.334 | -0.788 | -0.053 | -0.055 | -0.055 | 0.05 | 0.75 | 0.6 | 0.89 | 0.93 | 0.89 | 0.83 |
| 1 | 26376607 | rs3008428 | -0.844 | -1.634 | 1.277 | -0.78 | 0.025 | 0.025 | 0.025 | 0 | 0.77 | 0.83 | 0.93 | 0.93 | 0.89 | 0.83 |
| 1 | 26377880 | rs1135253 | 0.413 | -1.588 | 1.216 | -0.776 | 0 | 0 | 0 | 0 | 0.53 | 0.42 | 0.81 | 0.77 | 0.68 | 0.74 |
| 1 | 26378002 | rs1135251 | 0.647 | -1.565 | 1.094 | -0.764 | 0.075 | 0.095 | 0.095 | 0.338 | 0.52 | 0.19 | 0.77 | 0.77 | 0.67 | 0.74 |

| | | | | | | | | | | | | | | | | |
|---|---|---|---|---|---|---|---|---|---|---|---|---|---|---|---|---|
| 1 | 26380805 | rs7553840 | -0.27 | -1.548 | 1.061 | -0.705 | -0.077 | -0.026 | -0.026 | 0 | 0.73 | 0.51 | 0.91 | 0.82 | 0.89 | 0.82 |
| 1 | 26383875 | rs2275947 | 0.005 | -0.609 | 1.057 | -0.523 | -0.005 | -0.022 | -0.022 | 0 | 0.25 | 0.55 | 0.15 | 0.17 | 0.22 | 0.08 |
| 1 | 26387423 | rs2997447 | 0.12 | -0.616 | 1.093 | -0.783 | -0.01 | -0.089 | -0.089 | 1.274 | 0.23 | 0.42 | 0.13 | 0.04 | 0.21 | 0.06 |
| 1 | 26394997 | rs3008226 | 0.527 | -0.737 | 1.704 | -0.685 | -0.006 | -0.03 | -0.03 | 0 | 0.2 | 0.33 | 0.11 | 0.11 | 0.19 | 0.08 |
| 1 | 26402820 | rs3008419 | 0.34 | -0.757 | 1.681 | -0.662 | -0.013 | -0.039 | -0.039 | 0 | 0.22 | 0.45 | 0.15 | 0.12 | 0.18 | 0.08 |
| 1 | 26403098 | rs3008418 | 0.4 | -0.744 | 1.687 | -0.656 | -0.011 | -0.037 | -0.037 | 0 | 0.22 | 0.45 | 0.15 | 0.12 | 0.18 | 0.08 |
| 1 | 26411918 | rs12069719 | 0.668 | -0.742 | 1.588 | -0.731 | -0.002 | -0.026 | -0.026 | 0.287 | 0.19 | 0.35 | 0.12 | 0.11 | 0.18 | 0.07 |
| 1 | 26431337 | rs2802314 | 0.607 | -0.743 | 1.622 | -0.755 | 0.004 | -0.106 | -0.106 | 0.68 | 0.2 | 0.44 | 0.13 | 0.02 | 0.18 | 0.08 |
| 1 | 26447739 | rs1981135 | -0.759 | -1.844 | 0.413 | -1.331 | 0.074 | 0.076 | 0.076 | 0 | 0.69 | 0.8 | 0.89 | 0.93 | 0.87 | 0.67 |
| 1 | 26448447 | rs17257107 | 1.558 | -1.661 | 0.452 | -1.508 | 0.001 | -0.024 | -0.024 | 0 | 0.17 | 0.19 | 0.11 | 0.02 | 0.18 | 0.06 |
| 1 | 26450009 | rs17163588 | 1.334 | -1.628 | 0.302 | -1.491 | 0.013 | -0.078 | -0.078 | 0.336 | 0.18 | 0.33 | 0.12 | 0.02 | 0.18 | 0.06 |
| 1 | 26450101 | rs17257120 | 1.537 | -1.605 | -0.645 | -1.48 | 0.001 | -0.024 | -0.024 | 0 | 0.17 | 0.19 | 0.11 | 0.02 | 0.18 | 0.06 |
| 1 | 26471743 | rs1335759 | 0.196 | -1.461 | -1.615 | -1.469 | 0.117 | 0.21 | 0.21 | 0.333 | 0.48 | 0.19 | 0.76 | 0.87 | 0.69 | 0.59 |
| 1 | 26474439 | rs4281320 | 0.063 | 0.246 | -1.45 | -0.897 | -0.074 | 0.083 | 0.083 | 0.055 | 0.17 | 0.02 | 0.14 | 0.02 | 0.19 | 0.11 |
| 1 | 26475521 | rs868551 | 0.066 | -0.111 | -0.62 | -0.218 | -0.009 | -0.036 | -0.036 | 0.939 | 0.23 | 0.59 | 0.14 | 0.11 | 0.19 | 0.08 |
| 1 | 26481589 | rs12034664 | -0.365 | 0.007 | -1.418 | 0.763 | -0.032 | -0.107 | -0.107 | 0.187 | 0.14 | 0.01 | 0.39 | 0.56 | 0.15 | 0.44 |
| 1 | 26483381 | rs12565228 | 0.948 | 0.546 | -2.176 | -0.074 | -0.102 | -0.057 | -0.057 | 0.02 | 0.19 | 0.03 | 0.22 | 0.27 | 0.31 | 0.12 |
| 1 | 26484201 | rs4992499 | -0.017 | 0.116 | -2.129 | -0.063 | 0.01 | -0.035 | -0.035 | 0 | 0.12 | 0.09 | 0.4 | 0.57 | 0.11 | 0.44 |
| 1 | 26485013 | rs3806382 | 1.065 | -0.11 | -2.268 | -0.207 | -0.103 | -0.051 | -0.051 | 0.569 | 0.21 | 0.04 | 0.22 | 0.27 | 0.31 | 0.12 |
| 1 | 26486880 | rs10751733 | -0.342 | -0.883 | -1.862 | -1.111 | 0.004 | 0.001 | 0.001 | 0 | 0.59 | 0.42 | 0.75 | 0.9 | 0.64 | 0.66 |
| 1 | 26488947 | rs12069762 | 1.354 | -0.962 | -1.731 | -1.156 | 0.001 | 0.018 | 0.018 | 0 | 0.22 | 0.16 | 0.1 | 0.01 | 0.2 | 0.08 |
| 1 | 26491355 | rs11247860 | -0.239 | -0.936 | -1.715 | -1.135 | 0.006 | 0.009 | 0.009 | 0 | 0.58 | 0.42 | 0.75 | 0.89 | 0.64 | 0.65 |
| 1 | 26493171 | rs10794523 | 0.591 | -1.027 | -1.768 | -1.111 | 0.001 | 0.049 | 0.049 | 0 | 0.45 | 0.32 | 0.35 | 0.33 | 0.53 | 0.2 |
| 1 | 26493240 | rs10794524 | -0.138 | -0.921 | -1.666 | -1.108 | 0.015 | 0.05 | 0.05 | 0 | 0.58 | 0.35 | 0.75 | 0.89 | 0.64 | 0.65 |
| 1 | 26495550 | rs3762460 | 0.181 | -1.12 | -1.283 | -1.166 | 0.006 | 0.003 | 0.003 | 0 | 0.35 | 0.28 | 0.52 | 0.65 | 0.32 | 0.53 |
| 1 | 26496370 | rs2232646 | 0.639 | -1.089 | -1.372 | -1.467 | -0.016 | 0.105 | 0.105 | 0 | 0.31 | 0.08 | 0.17 | 0.1 | 0.3 | 0.23 |
| 1 | 26496455 | rs2232648 | 1.625 | -1.07 | -1.047 | -1.418 | -0.089 | -0.028 | -0.028 | 0 | 0.24 | 0.04 | 0.21 | 0.27 | 0.32 | 0.13 |
| 1 | 26504036 | rs17356937 | 1.626 | -1.165 | -0.96 | -1.418 | -0.005 | 0.063 | 0.063 | 0 | 0.21 | 0.12 | 0.09 | 0.01 | 0.19 | 0.07 |
| 1 | 26505265 | rs2783633 | -0.041 | -1.163 | -0.279 | -1.424 | -0.001 | -0.001 | -0.001 | 0 | 0.38 | 0.63 | 0.55 | 0.64 | 0.3 | 0.53 |
| 1 | 26508293 | rs3819033 | 0.876 | -0.994 | -0.263 | -1.276 | -0.007 | -0.089 | -0.089 | 0.7 | 0.12 | 0.01 | 0.41 | 0.59 | 0.1 | 0.45 |
| 1 | 26511139 | rs2297711 | 0.989 | -1.436 | -0.668 | -1.841 | -0.015 | 0.025 | 0.025 | 0 | 0.18 | 0.03 | 0.1 | 0.1 | 0.15 | 0.22 |
| 1 | 26511700 | rs2783680 | -0.598 | -1.783 | -0.505 | -2.033 | 0.006 | -0.069 | -0.069 | 0.949 | 0.64 | 0.47 | 0.78 | 0.97 | 0.6 | 0.84 |
| 1 | 26516845 | rs4659411 | -0.713 | -1.829 | -0.582 | -2.102 | -0.003 | 0.01 | 0.01 | 0 | 0.55 | 0.45 | 0.61 | 0.72 | 0.45 | 0.77 |
| 1 | 26517667 | rs11247864 | 0.487 | -1.732 | -0.488 | -1.891 | -0.004 | -0.029 | -0.029 | 0 | 0.37 | 0.44 | 0.28 | 0.12 | 0.31 | 0.27 |
| 1 | 26517773 | rs11247865 | 1.885 | -1.692 | -0.529 | -1.895 | 0.002 | -0.056 | -0.056 | 0 | 0.2 | 0.27 | 0.1 | 0.01 | 0.19 | 0.06 |
| 1 | 26517891 | rs11247867 | 2.014 | -1.671 | -0.585 | -1.885 | 0 | -0.032 | -0.032 | 0 | 0.19 | 0.2 | 0.09 | 0.01 | 0.19 | 0.06 |
| 1 | 26519614 | rs17163647 | 1.24 | -1.654 | -0.648 | -1.847 | 0.01 | 0.004 | 0.004 | 0 | 0.16 | 0.08 | 0.17 | 0.12 | 0.12 | 0.21 |
| 1 | 26522723 | rs12732367 | 1.82 | -1.541 | -0.748 | -1.793 | -0.061 | -0.001 | -0.001 | 0 | 0.28 | 0.1 | 0.23 | 0.28 | 0.33 | 0.14 |
| 1 | 26524706 | rs9970046 | -0.952 | -1.626 | -0.539 | -1.697 | -0.013 | 0.068 | 0.068 | 0.188 | 0.85 | 0.82 | 0.66 | 0.42 | 0.9 | 0.51 |
| 1 | 26526439 | rs17257155 | 1.943 | -1.514 | -0.89 | -1.723 | -0.001 | -0.016 | -0.016 | 0 | 0.19 | 0.18 | 0.09 | 0.01 | 0.18 | 0.17 |
| 1 | 26527163 | rs11247870 | 0.552 | -1.346 | -1.138 | -1.623 | -0.062 | 0.005 | 0.005 | 0.628 | 0.44 | 0.12 | 0.41 | 0.39 | 0.46 | 0.29 |
| 1 | 26527255 | rs12145714 | -0.71 | -1.275 | -0.571 | -1.553 | 0.016 | -0.113 | -0.113 | 1.426 | 0.22 | 0.39 | 0.15 | 0.02 | 0.25 | 0.15 |
| 1 | 26527863 | rs17163674 | 0.338 | -1.135 | -1.015 | -1.03 | -0.01 | 0.01 | 0.01 | 0 | 0.16 | 0.01 | 0.08 | 0.1 | 0.12 | 0.17 |
| 1 | 26546723 | rs17162102 | -0.637 | -1.146 | -0.562 | -1.036 | -0.003 | -0.016 | -0.016 | 3.43 | 0.15 | 0.22 | 0.07 | 0.02 | 0.13 | 0.09 |
| 1 | 26547375 | rs4659413 | -0.866 | -1.523 | -0.461 | -0.776 | -0.007 | 0.06 | 0.06 | 0 | 0.88 | 0.88 | 0.69 | 0.46 | 0.93 | 0.61 |
| 1 | 26552921 | rs10493028 | -0.301 | -1.513 | -0.489 | -0.794 | 0.001 | 0.054 | 0.054 | 0 | 0.14 | 0.03 | 0.05 | 0.02 | 0.13 | 0.09 |
| 1 | 26560745 | rs3795687 | 0.523 | -1.795 | 0.001 | -1.074 | -0.018 | 0.003 | 0.003 | 0.084 | 0.18 | 0.33 | 0.22 | 0.13 | 0.17 | 0.23 |
| 1 | 26561856 | rs10493030 | -0.387 | -1.843 | 0.024 | -1.079 | 0 | -0.003 | -0.003 | 1.173 | 0.17 | 0.2 | 0.11 | 0.02 | 0.19 | 0.11 |
| 1 | 26568165 | rs17163749 | -0.387 | -1.843 | 0.024 | -1.079 | 0 | -0.003 | -0.003 | 0 | 0.17 | 0.2 | 0.11 | 0.02 | 0.19 | 0.11 |
| 1 | 26569611 | rs6598948 | -1.441 | -1.887 | 0.038 | -1.081 | 0.02 | -0.026 | -0.026 | 0.441 | 0.89 | 0.97 | 0.7 | 0.46 | 0.93 | 0.61 |
| 1 | 26570009 | rs17257169 | -0.05 | -1.862 | 0.045 | -1.068 | -0.013 | 0.051 | 0.051 | 0 | 0.15 | 0.11 | 0.1 | 0.02 | 0.19 | 0.11 |
| 1 | 26576032 | rs10902724 | -1.108 | -1.844 | 0.053 | -1.062 | -0.005 | 0.055 | 0.055 | 0 | 0.88 | 0.88 | 0.7 | 0.46 | 0.93 | 0.61 |
| 1 | 26596080 | rs7550997 | -0.012 | -1.879 | 0.035 | -1.081 | 0.001 | -0.012 | -0.012 | 0.945 | 0.14 | 0.2 | 0.11 | 0.02 | 0.19 | 0.1 |
| 1 | 26596709 | rs12096059 | 1.42 | -1.89 | 0.049 | -1.087 | 0.022 | 0.025 | 0.025 | 0.359 | 0.55 | 0.34 | 0.41 | 0.31 | 0.63 | 0.29 |
| 1 | 26597673 | rs11247894 | -0.543 | -1.822 | -0.218 | -1.447 | 0.001 | 0.001 | 0.001 | 0 | 0.6 | 0.62 | 0.52 | 0.41 | 0.6 | 0.45 |
| 1 | 26606174 | rs10902732 | 0.208 | -1.749 | -0.257 | -1.46 | 0.001 | -0.012 | -0.012 | 0.073 | 0.14 | 0.2 | 0.11 | 0.02 | 0.19 | 0.1 |
| 1 | 26607726 | rs7544 | 0.208 | -1.749 | -0.251 | -1.46 | 0.001 | -0.014 | -0.014 | 0 | 0.14 | 0.2 | 0.11 | 0.02 | 0.19 | 0.1 |
| 1 | 26612460 | rs11247900 | 0.221 | -1.646 | -0.197 | -1.409 | 0 | -0.003 | -0.003 | 0 | 0.17 | 0.2 | 0.11 | 0.02 | 0.19 | 0.09 |
| 1 | 26615616 | rs4233458 | 1.231 | -1.623 | -0.191 | -1.396 | 0.008 | -0.049 | -0.049 | 0 | 0.28 | 0.26 | 0.17 | 0.05 | 0.33 | 0.16 |
| 1 | 26619649 | rs6598955 | -0.905 | -1.702 | -0.119 | -1.27 | -0.032 | -0.038 | -0.038 | 0 | 0.33 | 0.54 | 0.29 | 0.15 | 0.3 | 0.31 |
| 1 | 26621101 | rs6681452 | -0.079 | -1.678 | -0.002 | -1.268 | -0.002 | -0.038 | -0.038 | 2.69 | 0.17 | 0.27 | 0.11 | 0.02 | 0.19 | 0.09 |
| 1 | 26622911 | rs7541416 | 0.949 | -1.68 | 0.04 | -1.244 | -0.02 | 0.003 | 0.003 | 0 | 0.16 | 0.22 | 0.17 | 0.13 | 0.11 | 0.21 |
| 1 | 26627478 | rs2276712 | 0.849 | -1.405 | -0.208 | -1.064 | -0.014 | 0.023 | 0.023 | 0 | 0.7 | 0.59 | 0.57 | 0.44 | 0.74 | 0.49 |
| 1 | 26646726 | rs1071849 | 0.809 | -1.339 | -0.266 | -1.057 | -0.008 | 0.015 | 0.015 | 0 | 0.71 | 0.63 | 0.58 | 0.44 | 0.74 | 0.5 |
| 1 | 26646730 | rs17645 | 0.809 | -1.339 | -0.266 | -1.057 | -0.008 | 0.015 | 0.015 | 0 | 0.71 | 0.63 | 0.58 | 0.44 | 0.74 | 0.5 |
| 1 | 26648034 | rs4526602 | 0.896 | -1.381 | -0.216 | -1.077 | -0.01 | 0.014 | 0.014 | 0 | 0.7 | 0.62 | 0.58 | 0.45 | 0.74 | 0.5 |
| 1 | 26648088 | rs17163866 | -0.111 | -1.374 | -0.205 | -1.075 | 0 | -0.005 | -0.005 | 0 | 0.17 | 0.2 | 0.11 | 0.02 | 0.19 | 0.09 |
| 1 | 26648328 | rs12568163 | 0.875 | -1.368 | -0.245 | -1.067 | -0.004 | 0.057 | 0.057 | 0 | 0.12 | 0.12 | 0.31 | 0.54 | 0.07 | 0.41 |
| 1 | 26648381 | rs17163868 | -0.285 | -1.291 | -0.218 | -1.103 | -0.003 | -0.049 | -0.049 | 0 | 0.17 | 0.29 | 0.11 | 0.02 | 0.19 | 0.09 |

| | | | | | | | | | | | | | | | | |
|---|---|---|---|---|---|---|---|---|---|---|---|---|---|---|---|---|
| 1 | 26655208 | rs11247916 | -0.02 | -0.965 | -0.241 | -0.682 | -0.016 | 0.042 | 0.042 | 0 | 0.57 | 0.67 | 0.38 | 0.18 | 0.53 | 0.43 |
| 1 | 26658391 | rs4659431 | -0.202 | -0.859 | -0.258 | -0.547 | -0.001 | 0.04 | 0.04 | 0 | 0.84 | 0.83 | 0.64 | 0.45 | 0.83 | 0.57 |
| 1 | 26663362 | rs10751735 | -0.131 | -0.805 | -0.272 | -0.805 | -0.016 | 0.04 | 0.04 | 0 | 0.57 | 0.66 | 0.38 | 0.18 | 0.53 | 0.43 |
| 1 | 26664968 | rs11247919 | 0.481 | -0.954 | -0.215 | -0.959 | 0 | -0.067 | -0.067 | 0.841 | 0.41 | 0.44 | 0.22 | 0.06 | 0.43 | 0.22 |
| 1 | 26671084 | rs12562454 | -0.011 | -0.61 | -0.581 | -0.481 | -0.001 | 0 | 0 | 0 | 0.18 | 0.15 | 0.19 | 0.15 | 0.19 | 0.23 |
| 1 | 26673466 | rs7535656 | 0.041 | -0.587 | -0.586 | -0.47 | -0.001 | 0 | 0 | 0.037 | 0.17 | 0.15 | 0.19 | 0.15 | 0.19 | 0.23 |
| 1 | 26677721 | rs11582416 | 0.704 | 0.221 | -0.631 | -0.149 | -0.029 | 0.017 | 0.017 | 0.043 | 0.26 | 0.13 | 0.25 | 0.23 | 0.28 | 0.14 |
| 1 | 26679041 | rs12031946 | -0.026 | 0.016 | -0.709 | 0.026 | 0.021 | -0.052 | -0.052 | 0 | 0.14 | 0.07 | 0.33 | 0.56 | 0.1 | 0.38 |
| 1 | 26679968 | rs6691909 | -0.721 | -0.128 | -0.083 | -0.383 | 0.005 | 0.01 | 0.01 | 0 | 0.49 | 0.47 | 0.67 | 0.87 | 0.5 | 0.73 |
| 1 | 26681306 | rs4233461 | -0.679 | -0.22 | -0.021 | -0.533 | 0.007 | 0.068 | 0.068 | 0 | 0.54 | 0.38 | 0.68 | 0.86 | 0.53 | 0.73 |
| 1 | 26687561 | rs4584380 | 0.866 | -0.219 | -0.199 | -0.51 | -0.055 | 0.027 | 0.027 | 0.006 | 0.31 | 0.13 | 0.43 | 0.28 | 0.34 | 0.25 |
| 1 | 26689220 | rs12030833 | -0.433 | -0.235 | -0.148 | -0.525 | 0.002 | -0.003 | -0.003 | 0 | 0.23 | 0.25 | 0.25 | 0.58 | 0.19 | 0.47 |
| 1 | 26690481 | rs7547496 | -1.219 | -0.355 | -0.015 | -0.569 | 0.011 | 0.09 | 0.09 | 0 | 0.61 | 0.76 | 0.75 | 0.9 | 0.63 | 0.81 |
| 1 | 26691174 | rs11247933 | 0.914 | -0.325 | -0.104 | -0.541 | -0.055 | 0.026 | 0.026 | 0.008 | 0.31 | 0.12 | 0.42 | 0.28 | 0.34 | 0.25 |
| 1 | 26694245 | rs10794531 | -1.698 | -0.349 | 0.648 | 0.255 | -0.01 | 0.013 | 0.013 | 0.46 | 0.3 | 0.59 | 0.32 | 0.62 | 0.29 | 0.57 |
| 1 | 26694260 | rs10794532 | -1.1 | -0.326 | -0.344 | 0.409 | 0.002 | -0.001 | -0.001 | 0 | 0.23 | 0.24 | 0.25 | 0.58 | 0.19 | 0.47 |
| 1 | 26709040 | rs11247940 | -0.328 | -0.043 | -0.684 | 0.968 | 0.005 | 0.015 | 0.015 | 0 | 0.13 | 0.12 | 0.25 | 0.54 | 0.09 | 0.32 |
| 1 | 26709504 | rs7529342 | 0.532 | 0.385 | -1.268 | 0.54 | -0.051 | -0.037 | -0.037 | 0.103 | 0.28 | 0.1 | 0.36 | 0.32 | 0.36 | 0.31 |
| 1 | 26710499 | rs12084041 | -0.995 | 0.429 | -0.983 | 0.233 | -0.001 | -0.031 | -0.031 | 0 | 0.57 | 0.48 | 0.72 | 0.94 | 0.63 | 0.86 |
| 1 | 26712597 | rs12137896 | -0.007 | 0.5 | -1.121 | 0.669 | -0.006 | 0.009 | 0.009 | 0 | 0.08 | 0.07 | 0.11 | 0.03 | 0.14 | 0.14 |
| 1 | 26714161 | rs12043608 | -0.081 | 0.368 | -1.068 | 1.053 | 0.006 | -0.025 | -0.025 | 0 | 0.11 | 0.08 | 0.23 | 0.61 | 0.08 | 0.31 |
| 1 | 26715232 | rs4436378 | -0.416 | 0.423 | -0.776 | 0.179 | 0.003 | 0.015 | 0.015 | 0 | 0.5 | 0.47 | 0.56 | 0.88 | 0.47 | 0.71 |
| 1 | 26716136 | rs9438620 | -0.032 | 0.549 | -0.846 | 0.759 | 0 | 0.005 | 0.005 | 0 | 0.38 | 0.41 | 0.29 | 0.46 | 0.35 | 0.26 |
| 1 | 26719115 | rs11247946 | -0.556 | 0.256 | -1.512 | 0.333 | 0.024 | -0.025 | -0.025 | 0 | 0.34 | 0.19 | 0.39 | 0.64 | 0.36 | 0.42 |
| 1 | 26719264 | rs11587947 | 1.043 | 0.538 | -2.23 | -0.399 | -0.071 | 0.111 | 0.111 | 0 | 0.2 | 0.02 | 0.09 | 0.01 | 0.18 | 0.04 |
| 1 | 26721287 | rs12722898 | 0.48 | 0.467 | -1.948 | -0.739 | 0.047 | 0.201 | 0.201 | 0 | 0.51 | 0.21 | 0.49 | 0.15 | 0.55 | 0.46 |
| 1 | 26722384 | rs9438621 | -0.093 | 0.255 | -1.963 | -0.761 | -0.016 | 0.011 | 0.011 | 0.045 | 0.84 | 0.34 | 0.86 | 0.8 | 0.91 | 0.87 |
| 1 | 26727205 | rs4360511 | 0.404 | 0.323 | -2.131 | -1.009 | -0.02 | 0.173 | 0.173 | 0 | 0.45 | 0.16 | 0.42 | 0.15 | 0.49 | 0.43 |
| 1 | 26735773 | rs12122703 | 0.317 | 0.161 | -1.194 | -1.093 | 0.015 | 0.177 | 0.177 | 0 | 0.47 | 0.21 | 0.43 | 0.16 | 0.5 | 0.44 |
| 1 | 26741285 | rs11247954 | 0.562 | -0.321 | -1.5 | -0.897 | 0.009 | -0.091 | -0.091 | 0 | 0.12 | 0 | 0.19 | 0.59 | 0.1 | 0.22 |
| 1 | 26743903 | rs6683792 | 0.761 | -0.089 | -1.056 | -1.296 | 0.012 | -0.042 | -0.042 | 0 | 0.28 | 0.49 | 0.38 | 0.15 | 0.31 | 0.39 |
| 1 | 26744087 | rs11581746 | 1.595 | -0.07 | -1.316 | -1.391 | -0.045 | 0.11 | 0.11 | 0 | 0.21 | 0.06 | 0.1 | 0.01 | 0.18 | 0.04 |
| 1 | 26746199 | rs4274112 | 0.599 | 0.011 | -0.689 | -1.363 | 0.002 | -0.007 | -0.007 | 0 | 0.28 | 0.31 | 0.44 | 0.15 | 0.32 | 0.41 |
| 1 | 26746327 | rs6598964 | -0.408 | -0.163 | -0.876 | -1.39 | 0.014 | -0.006 | -0.006 | 0 | 0.63 | 0.79 | 0.65 | 0.34 | 0.6 | 0.6 |
| 1 | 26746807 | rs12728900 | 0.047 | -0.206 | -0.617 | -1.581 | -0.002 | -0.067 | -0.067 | 0 | 0.36 | 0.44 | 0.19 | 0.14 | 0.27 | 0.16 |
| 1 | 26748208 | rs11247955 | 0.473 | -0.008 | -0.64 | -1.968 | -0.015 | 0.145 | 0.145 | 0 | 0.46 | 0.23 | 0.42 | 0.11 | 0.5 | 0.29 |
| 1 | 26752129 | rs35015532 | 1.049 | 0.235 | -0.671 | -2.013 | -0.012 | 0.023 | 0.023 | 0 | 0.24 | 0.08 | 0.37 | 0.13 | 0.24 | 0.26 |
| 1 | 26756209 | rs6697410 | 0.112 | 0.564 | -0.725 | -2.126 | -0.017 | 0.014 | 0.014 | 0.002 | 0.81 | 0.85 | 0.85 | 0.86 | 0.74 | 0.85 |
| 1 | 26756856 | rs17261915 | 1.257 | 0.531 | -0.742 | -2.131 | -0.031 | 0.011 | 0.011 | 0 | 0.23 | 0.03 | 0.36 | 0.13 | 0.24 | 0.26 |
| 1 | 26758602 | rs3811462 | -0.082 | 0.598 | -0.709 | -2.081 | -0.009 | 0.009 | 0.009 | 0 | 0.82 | 0.85 | 0.86 | 0.86 | 0.78 | 0.85 |
| 1 | 26759149 | rs3811461 | 0.938 | 0.366 | -0.546 | -2.149 | -0.038 | 0.029 | 0.029 | 0 | 0.28 | 0.48 | 0.31 | 0.71 | 0.23 | 0.49 |
| 1 | 26765541 | rs9438549 | 0.761 | 0.369 | -0.548 | -2.156 | 0.003 | -0.011 | -0.011 | 0 | 0.73 | 0.71 | 0.81 | 0.85 | 0.69 | 0.78 |
| 1 | 26767367 | rs11247958 | 1.024 | 0.064 | -0.384 | -2.103 | -0.027 | 0.026 | 0.026 | 0 | 0.27 | 0.4 | 0.3 | 0.71 | 0.23 | 0.49 |
| 1 | 26770999 | rs11584995 | 2.385 | 0.031 | -0.607 | -2.131 | -0.071 | 0.103 | 0.103 | 0 | 0.2 | 0.02 | 0.13 | 0.01 | 0.27 | 0.04 |
| 1 | 26773400 | rs12042656 | 1.032 | 0.028 | -0.576 | -2.115 | -0.003 | -0.005 | -0.005 | 0 | 0.08 | 0.07 | 0.04 | 0.02 | 0.04 | 0.07 |
| 1 | 26775223 | rs9438551 | 0.023 | -0.004 | -0.55 | -2.131 | 0.012 | 0.006 | 0.006 | 0 | 0.37 | 0.64 | 0.35 | 0.73 | 0.28 | 0.56 |
| 1 | 26779270 | rs4659374 | 0.108 | 0.019 | -0.536 | -2.118 | -0.009 | -0.007 | -0.007 | 0.29 | 0.36 | 0.58 | 0.44 | 0.76 | 0.28 | 0.56 |
| 1 | 26786627 | rs3816539 | 0.981 | 0.023 | -0.541 | -2.114 | -0.034 | 0.034 | 0.034 | 0 | 0.27 | 0.48 | 0.31 | 0.71 | 0.23 | 0.49 |
| 1 | 26792358 | rs2245071 | 0.854 | 0.042 | -0.534 | -2.102 | -0.016 | -0.007 | -0.007 | 0.157 | 0.09 | 0.16 | 0.13 | 0.05 | 0.05 | 0.07 |
| 1 | 26797654 | rs1055323 | 2.12 | 0.085 | -0.651 | -2.115 | -0.049 | 0.082 | 0.082 | 0 | 0.21 | 0.07 | 0.13 | 0.01 | 0.27 | 0.04 |
| 1 | 26797905 | rs11590999 | 2.12 | 0.085 | -0.651 | -2.115 | -0.049 | 0.082 | 0.082 | 0 | 0.21 | 0.07 | 0.13 | 0.01 | 0.27 | 0.04 |
| 1 | 26802465 | rs17340420 | 2.126 | 0.081 | -0.653 | -2.116 | -0.049 | 0.081 | 0.081 | 0 | 0.21 | 0.07 | 0.13 | 0.01 | 0.27 | 0.04 |
| 1 | 26802548 | rs12741800 | 1.006 | 0.339 | -0.474 | -1.87 | -0.015 | 0.146 | 0.146 | 0 | 0.46 | 0.21 | 0.52 | 0.14 | 0.51 | 0.29 |
| 1 | 26812892 | rs2494401 | 0.889 | 0.403 | -0.501 | -1.909 | -0.003 | -0.005 | -0.005 | 1.428 | 0.08 | 0.07 | 0.04 | 0.02 | 0.04 | 0.07 |
| 1 | 26849312 | rs4970466 | 0.689 | 0.638 | -0.259 | -1.53 | -0.031 | 0.06 | 0.06 | 0 | 0.27 | 0.49 | 0.35 | 0.64 | 0.24 | 0.49 |
| 1 | 26855401 | rs11247963 | -0.457 | 0.689 | -0.664 | -1.32 | -0.007 | 0.008 | 0.008 | 0 | 0.17 | 0.15 | 0.13 | 0.14 | 0.22 | 0.14 |
| 1 | 26862939 | rs737465 | -0.457 | 0.316 | -0.881 | -1.286 | 0.005 | 0.022 | 0.022 | 0 | 0.7 | 0.92 | 0.66 | 0.79 | 0.76 | 0.7 |
| 1 | 26865971 | rs1002487 | 0.461 | 0.339 | -0.267 | -1.228 | 0.002 | 0.005 | 0.005 | 0 | 0.06 | 0.04 | 0.03 | 0.13 | 0.06 | 0.09 |
| 1 | 26872832 | rs17162190 | 1.038 | 0.17 | -0.321 | -1.5 | 0.017 | -0.046 | -0.046 | 0 | 0.23 | 0.13 | 0.3 | 0.64 | 0.22 | 0.5 |
| 1 | 26873245 | rs2278978 | -0.743 | 0.145 | -0.334 | -1.759 | 0.004 | 0.015 | 0.015 | 0 | 0.71 | 0.92 | 0.66 | 0.79 | 0.76 | 0.71 |
| 1 | 26877708 | rs2278979 | 0.158 | 0.077 | 0.088 | -1.419 | 0.009 | 0.006 | 0.006 | 0 | 0.3 | 0.19 | 0.33 | 0.77 | 0.28 | 0.58 |
| 1 | 26884864 | rs3790645 | -0.178 | 0.169 | 0.058 | -1.291 | -0.022 | -0.012 | -0.012 | 0 | 0.33 | 0.45 | 0.36 | 0.76 | 0.29 | 0.59 |
| 1 | 26887592 | rs1064196 | -0.161 | 0.199 | 0.126 | -1.276 | -0.024 | -0.013 | -0.013 | 0 | 0.33 | 0.51 | 0.36 | 0.76 | 0.29 | 0.59 |
| 1 | 26899125 | rs282177 | -0.288 | 0.192 | -0.445 | -1.366 | -0.022 | -0.02 | -0.02 | 0 | 0.35 | 0.52 | 0.41 | 0.76 | 0.29 | 0.65 |
| 1 | 26900708 | rs2229714 | 0.247 | 0.625 | -0.765 | -1.507 | -0.003 | 0.002 | 0.002 | 0 | 0.09 | 0.1 | 0.08 | 0.01 | 0.14 | 0.06 |
| 1 | 26901090 | rs11113 | 0.484 | 0.626 | -1.005 | -1.826 | -0.005 | 0.026 | 0.026 | 0 | 0.59 | 0.68 | 0.58 | 0.71 | 0.58 | 0.68 |
| 1 | 26901319 | rs7726 | 0.366 | 0.402 | -1.003 | -1.535 | -0.002 | 0 | 0 | 0 | 0.32 | 0.31 | 0.37 | 0.71 | 0.26 | 0.63 |
| 1 | 26902388 | rs6668958 | -0.421 | 0.724 | -1.873 | -1.635 | -0.039 | 0.009 | 0.009 | 0 | 0.14 | 0.04 | 0.11 | 0.14 | 0.2 | 0.15 |
| 1 | 26903570 | rs406985 | -0.329 | 0.582 | -1.837 | -1.598 | 0.01 | 0.086 | 0.086 | 0 | 0.39 | 0.16 | 0.51 | 0.77 | 0.39 | 0.66 |

| | | | | | | | | | | | | | | | | |
|---|---|---|---|---|---|---|---|---|---|---|---|---|---|---|---|---|
| 1 | 26906586 | rs4970503 | -0.802 | 1.26 | -1.343 | -1.544 | -0.005 | 0.053 | 0.053 | 0.316 | 0.34 | 0.73 | 0.43 | 0.19 | 0.29 | 0.26 |
| 1 | 26909666 | rs566421 | 1.392 | 0.901 | -1.664 | -1.269 | 0.021 | 0.036 | 0.036 | 0 | 0.43 | 0.14 | 0.28 | 0.69 | 0.41 | 0.52 |
| 1 | 26909765 | rs6598858 | -1.237 | 1.06 | -0.72 | -1.657 | 0.017 | -0.058 | -0.058 | 0.12 | 0.4 | 0.43 | 0.46 | 0.18 | 0.45 | 0.43 |
| 1 | 26919414 | rs12408288 | -0.443 | 0.886 | -0.573 | -1.622 | -0.071 | 0.104 | 0.104 | 0.584 | 0.29 | 0.12 | 0.29 | 0.09 | 0.37 | 0.27 |
| 1 | 26924308 | rs17162247 | -1.085 | 0.386 | 0.622 | -0.893 | 0.06 | 0.074 | 0.074 | 0.26 | 0.07 | 0.46 | 0.07 | 0.05 | 0.03 | 0.03 |
| 1 | 26929002 | rs17162257 | -1.673 | 0.389 | 0.497 | -0.829 | -0.031 | 0.051 | 0.051 | 1.197 | 0.1 | 0.68 | 0.27 | 0.12 | 0.03 | 0.05 |
| 1 | 26940681 | rs12137317 | 0.392 | 0.547 | 0.13 | -0.817 | -0.03 | -0.015 | -0.015 | 0 | 0.06 | 0.01 | 0.16 | 0.06 | 0.08 | 0.06 |
| 1 | 26945712 | rs4970480 | -0.927 | 0.229 | 0.442 | -1.136 | 0.052 | 0.172 | 0.172 | 1.257 | 0.06 | 0.48 | 0.24 | 0.07 | 0.01 | 0.02 |
| 1 | 26988026 | rs11809030 | 0.205 | 0.566 | -0.957 | -2.432 | -0.002 | -0.075 | -0.075 | 0 | 0.29 | 0.29 | 0.19 | 0.01 | 0.29 | 0.1 |
| 1 | 27020012 | rs4466675 | -0.156 | 0.079 | -0.995 | -3.219 | -0.001 | -0.038 | -0.038 | 0 | 0.39 | 0.41 | 0.21 | 0.06 | 0.36 | 0.33 |
| 1 | 27021684 | rs6598860 | 0.381 | 0.056 | -1.216 | -3.493 | -0.011 | -0.079 | -0.079 | 0 | 0.29 | 0.34 | 0.13 | 0.03 | 0.23 | 0.24 |
| 1 | 27068202 | rs4970481 | -1.151 | 0.486 | -1.476 | -3.757 | -0.027 | 0.039 | 0.039 | 0.3 | 0.68 | 0.82 | 0.8 | 1 | 0.61 | 0.89 |
| 1 | 27093657 | rs11247596 | 0.915 | 0.309 | -1.569 | -3.8 | -0.021 | 0.149 | 0.149 | 0.248 | 0.27 | 0.05 | 0.1 | 0.03 | 0.23 | 0.25 |
| 1 | 27161670 | rs11247599 | -0.457 | 0.231 | -1.584 | -3.847 | -0.038 | 0.067 | 0.067 | 0 | 0.64 | 0.77 | 0.73 | 0.97 | 0.5 | 0.79 |
| 1 | 27164335 | rs7527776 | -1.502 | 0.221 | -1.614 | -3.859 | -0.12 | 0.169 | 0.169 | 0 | 0.74 | 0.98 | 0.83 | 1 | 0.63 | 0.89 |
| 1 | 27169484 | rs35650645 | 0.513 | 0.252 | -1.577 | -3.814 | -0.017 | 0.013 | 0.013 | 0 | 0.09 | 0.04 | 0.08 | 0.03 | 0.13 | 0.1 |
| 1 | 27180829 | rs737524 | -0.608 | 0.073 | -1.478 | -3.934 | -0.001 | -0.036 | -0.036 | 0.516 | 0.45 | 0.44 | 0.24 | 0.06 | 0.38 | 0.34 |
| 1 | 27182261 | rs12747620 | 0.207 | 0.647 | -1.58 | -3.549 | -0.004 | 0.003 | 0.003 | 0 | 0.09 | 0.09 | 0.09 | 0.04 | 0.13 | 0.1 |
| 1 | 27182984 | rs10902646 | -1 | 0.692 | -1.618 | -3.64 | -0.034 | 0.03 | 0.03 | 0 | 0.74 | 0.83 | 0.82 | 1 | 0.63 | 0.89 |
| 1 | 27220521 | rs11810321 | 0.038 | 1.005 | -1.442 | -3.401 | 0.028 | -0.038 | -0.038 | 0 | 0.1 | 0.21 | 0.1 | 0.02 | 0.13 | 0.11 |
| 1 | 27234682 | rs12724052 | -0.088 | 0.997 | -1.373 | -3.404 | 0.027 | -0.044 | -0.044 | 0 | 0.1 | 0.22 | 0.1 | 0.02 | 0.13 | 0.11 |
| 1 | 27236212 | rs17162330 | 0.037 | 1.003 | -1.407 | -3.401 | 0.029 | -0.038 | -0.038 | 1.108 | 0.1 | 0.21 | 0.1 | 0.02 | 0.13 | 0.11 |
| 1 | 27242009 | rs11581460 | 0.862 | 0.741 | -1.577 | -3.668 | -0.018 | -0.011 | -0.011 | 0 | 0.24 | 0.2 | 0.18 | 0 | 0.36 | 0.11 |
| 1 | 27247822 | rs36109400 | -0.1 | 1.029 | -1.677 | -3.291 | 0.029 | -0.038 | -0.038 | 0 | 0.1 | 0.21 | 0.1 | 0.02 | 0.13 | 0.11 |
| 1 | 27278553 | rs3010109 | -0.017 | 0.839 | -1.888 | -3.231 | 0.032 | 0.151 | 0.151 | 0.833 | 0.27 | 0.01 | 0.09 | 0.03 | 0.17 | 0.2 |
| 1 | 27312816 | rs4970469 | 0.761 | 1.699 | -1.081 | -2.301 | -0.034 | 0.055 | 0.055 | 0.391 | 0.1 | 0 | 0.07 | 0 | 0.14 | 0.02 |
| 1 | 27347994 | rs11576827 | -0.233 | 1.673 | -0.771 | -1.675 | -0.004 | -0.017 | -0.017 | 0 | 0.26 | 0.21 | 0.18 | 0.02 | 0.35 | 0.14 |
| 1 | 27358792 | rs2993066 | -1.244 | 1.177 | -0.978 | -1.327 | -0.011 | -0.036 | -0.036 | 2.539 | 0.07 | 0.19 | 0.05 | 0.01 | 0.04 | 0.01 |
| 1 | 27367693 | rs525621 | 0.776 | 1.2 | -1.02 | -1.316 | -0.011 | -0.029 | -0.029 | 0 | 0.93 | 0.88 | 0.96 | 0.99 | 0.97 | 0.99 |
| 1 | 27372924 | rs568533 | 0.882 | 1.142 | -1.019 | -0.933 | -0.019 | 0.049 | 0.049 | 0.102 | 0.93 | 0.83 | 0.94 | 0.85 | 0.96 | 0.99 |
| 1 | 27374218 | rs537951 | 1.624 | 1.267 | -1.091 | -0.975 | -0.007 | 0.016 | 0.016 | 0 | 0.88 | 0.8 | 0.9 | 0.82 | 0.89 | 0.85 |
| 1 | 27375717 | rs2256674 | 0.298 | 1.202 | -1.184 | -0.925 | -0.007 | -0.018 | -0.018 | 0.689 | 0.94 | 0.93 | 0.96 | 1 | 0.97 | 0.99 |
| 1 | 27384315 | rs10430146 | 0.044 | 1.494 | -1.47 | -1.604 | -0.069 | 0.128 | 0.128 | 0.5 | 0.25 | 0.03 | 0.18 | 0.03 | 0.31 | 0.15 |
| 1 | 27384989 | rs11805628 | -1.823 | 1.378 | -0.566 | -0.835 | 0.002 | -0.012 | -0.012 | 0.202 | 0.11 | 0.12 | 0.11 | 0.04 | 0.13 | 0.17 |
| 1 | 27392946 | rs7537450 | -1.23 | 1.686 | -0.745 | -0.664 | 0.029 | -0.019 | -0.019 | 0 | 0.08 | 0.14 | 0.07 | 0.04 | 0.12 | 0.2 |
| 1 | 27397604 | rs12410656 | -0.789 | 1.325 | -1.158 | -0.546 | -0.016 | 0.002 | 0.002 | 0 | 0.06 | 0 | 0.11 | 0.04 | 0.08 | 0.04 |
| 1 | 27403761 | rs484677 | 0.848 | 1.711 | -0.8 | -0.897 | 0.016 | -0.068 | -0.068 | 0 | 0.68 | 0.62 | 0.78 | 0.89 | 0.61 | 0.71 |
| 1 | 27403974 | rs482812 | 0.5 | 1.833 | -1.088 | -0.917 | -0.059 | 0.071 | 0.071 | 0 | 0.81 | 0.95 | 0.88 | 0.92 | 0.71 | 0.86 |
| 1 | 27417917 | rs12125221 | 0.416 | 1.26 | -1.857 | -0.416 | -0.013 | -0.108 | -0.108 | 0.248 | 0.16 | 0.01 | 0.4 | 0.62 | 0.12 | 0.06 |
| 1 | 27426219 | rs5810 | -1.186 | 0.925 | -0.915 | -1.207 | 0.032 | -0.049 | -0.049 | 0 | 0.23 | 0.34 | 0.2 | 0.12 | 0.36 | 0.35 |
| 1 | 27426796 | rs4266911 | -0.582 | 0.678 | -1.027 | -1.346 | 0.026 | 0.039 | 0.039 | 0.008 | 0.92 | 0.99 | 0.98 | 1 | 0.96 | 1 |
| 1 | 27437217 | rs12123077 | -1.287 | 0.809 | -0.713 | -1.266 | 0.041 | -0.062 | -0.062 | 0 | 0.23 | 0.36 | 0.19 | 0.11 | 0.35 | 0.34 |
| 1 | 27467984 | rs11247611 | 0.102 | 0.701 | -0.523 | -1.356 | -0.025 | 0.007 | 0.007 | 0.012 | 0.07 | 0 | 0.05 | 0.06 | 0.09 | 0.2 |
| 1 | 27477549 | rs6696706 | -1.2 | 0.696 | -0.353 | -1.4 | 0.014 | -0.028 | -0.028 | 0 | 0.21 | 0.25 | 0.19 | 0.12 | 0.33 | 0.34 |
| 1 | 27488299 | rs12402451 | -1.103 | 0.718 | -0.415 | -1.421 | -0.015 | 0.014 | 0.014 | 0.32 | 0.21 | 0.2 | 0.19 | 0.12 | 0.34 | 0.34 |
| 1 | 27493679 | rs12751422 | -0.861 | 0.123 | -0.387 | -2.183 | -0.005 | -0.014 | -0.014 | 0.233 | 0.58 | 0.59 | 0.45 | 0.21 | 0.7 | 0.72 |
| 1 | 27530146 | rs11247623 | -1.404 | 0.368 | -0.236 | -1.984 | 0.037 | -0.043 | -0.043 | 5.072 | 0.28 | 0.45 | 0.29 | 0.17 | 0.44 | 0.39 |
| 1 | 27537101 | rs6687674 | -0.66 | 0.571 | -0.608 | -1.933 | -0.029 | 0.02 | 0.02 | 0.182 | 0.21 | 0.2 | 0.23 | 0.15 | 0.37 | 0.33 |
| 1 | 27540993 | rs17162492 | 1.428 | 0.285 | -1.045 | -1.307 | -0.011 | -0.052 | -0.052 | 2.408 | 0.2 | 0 | 0.09 | 0.22 | 0.14 | 0.2 |
| 1 | 27543523 | rs10751722 | 0.713 | 0.484 | 0.118 | -1.335 | 0.051 | 0.004 | 0.004 | 0.213 | 0.91 | 0.54 | 0.91 | 0.99 | 0.92 | 0.94 |
| 1 | 27548657 | rs11247626 | 1.115 | 0.207 | -0.124 | -1.173 | 0.026 | 0.036 | 0.036 | 0.005 | 0.35 | 0.14 | 0.5 | 0.78 | 0.26 | 0.24 |
| 1 | 27558522 | rs10902652 | 0.315 | 0.506 | 0.056 | -1.045 | 0.071 | 0.025 | 0.025 | 0.17 | 0.92 | 0.56 | 0.91 | 0.99 | 0.92 | 0.95 |
| 1 | 27586900 | rs6598879 | 0.551 | 0.589 | -0.158 | -0.988 | 0.02 | -0.031 | -0.031 | 1.978 | 0.81 | 0.51 | 0.83 | 0.87 | 0.77 | 0.74 |
| 1 | 27595862 | rs11589265 | -0.085 | 0.957 | -0.828 | -0.906 | -0.06 | -0.013 | -0.013 | 0 | 0.09 | 0.02 | 0.14 | 0.06 | 0.25 | 0.11 |
| 1 | 27647082 | rs4970518 | 0.462 | 0.432 | -1.521 | -1.045 | 0.012 | 0.112 | 0.112 | 0 | 0.31 | 0.09 | 0.19 | 0.06 | 0.28 | 0.38 |
| 1 | 27660340 | rs12758154 | -1.006 | 0.497 | -1.479 | -0.841 | -0.039 | 0.012 | 0.012 | 0.608 | 0.28 | 0.2 | 0.26 | 0.21 | 0.46 | 0.39 |
| 1 | 27661037 | rs12565684 | -0.724 | 0.556 | -1.418 | -0.722 | -0.042 | 0.005 | 0.005 | 0.451 | 0.25 | 0.15 | 0.23 | 0.21 | 0.41 | 0.35 |
| 1 | 27662666 | rs9203 | -0.788 | 0.588 | -1.459 | -0.411 | -0.039 | -0.007 | -0.007 | 0 | 0.25 | 0.15 | 0.23 | 0.29 | 0.41 | 0.4 |
| 1 | 27679692 | rs11247639 | -0.941 | 0.193 | -1.308 | -0.24 | 0.013 | 0.015 | 0.015 | 0 | 0.59 | 0.34 | 0.43 | 0.25 | 0.68 | 0.74 |
| 1 | 27679797 | rs3813795 | -0.935 | 0.205 | -1.408 | -0.235 | 0.008 | 0.009 | 0.009 | 0 | 0.59 | 0.35 | 0.43 | 0.25 | 0.68 | 0.73 |
| 1 | 27682126 | rs12742921 | -0.058 | 0.723 | -2.027 | -0.181 | -0.053 | 0.006 | 0.006 | 0.085 | 0.09 | 0.01 | 0.12 | 0.04 | 0.19 | 0.11 |
| 1 | 27686621 | rs17162551 | 0.39 | -0.035 | -1.945 | -0.313 | -0.012 | -0.022 | -0.022 | 0 | 0.07 | 0.09 | 0.02 | 0 | 0.02 | 0.01 |
| 1 | 27688633 | rs1138294 | -0.245 | 0.391 | -2.25 | 0.445 | -0.053 | -0.09 | -0.09 | 0 | 0.29 | 0.1 | 0.21 | 0.54 | 0.38 | 0.36 |
| 1 | 27689763 | rs4525067 | 0.098 | 0.452 | -1.496 | 0.222 | -0.087 | 0.057 | 0.057 | 0.051 | 0.25 | 0.02 | 0.19 | 0.1 | 0.29 | 0.14 |
| 1 | 27705565 | rs2474287 | -1.364 | -0.617 | 0.501 | 1.286 | -0.036 | 0.059 | 0.059 | 0.694 | 0.19 | 0.61 | 0.07 | 0.4 | 0.13 | 0.29 |
| 1 | 27709020 | rs2504779 | -1.447 | -0.695 | 1.62 | 1.271 | -0.037 | 0.059 | 0.059 | 0.702 | 0.19 | 0.61 | 0.08 | 0.4 | 0.14 | 0.3 |
| 1 | 27720178 | rs2504786 | -2.347 | -0.666 | 1.613 | 0.944 | 0.013 | -0.061 | -0.061 | 0 | 0.37 | 0.89 | 0.23 | 0.46 | 0.33 | 0.38 |
| 1 | 27731651 | rs11548323 | -0.604 | -0.678 | 2.49 | 0.899 | -0.013 | 0.023 | 0.023 | 0 | 0.15 | 0.11 | 0.1 | 0.05 | 0.18 | 0.08 |
| 1 | 27744513 | rs1005413 | -1.278 | -1.439 | 2.826 | 1.168 | -0.029 | 0.043 | 0.043 | 0.777 | 0.19 | 0.63 | 0.07 | 0.4 | 0.14 | 0.29 |

| | | | | | | | | | | | | | | | | |
|---|---|---|---|---|---|---|---|---|---|---|---|---|---|---|---|---|
| 1 | 27772009 | rs17371357 | -0.685 | -1.402 | 3.141 | 1.183 | 0.009 | 0.019 | 0.019 | 2.669 | 0.12 | 0.05 | 0.03 | 0 | 0.12 | 0.03 |
| 1 | 27773902 | rs2474301 | -1.867 | -1.394 | 3.128 | 1.23 | -0.029 | 0.048 | 0.048 | 0.716 | 0.19 | 0.62 | 0.07 | 0.41 | 0.14 | 0.29 |
| 1 | 27775196 | rs2480224 | -0.802 | -1.367 | 3.024 | 1.237 | -0.006 | -0.043 | -0.043 | 3.923 | 0.13 | 0.16 | 0.04 | 0 | 0.12 | 0.03 |
| 1 | 27782390 | rs7531313 | 1.009 | -1.36 | 2.992 | 1.247 | -0.029 | 0.113 | 0.113 | 1.39 | 0.84 | 0.71 | 0.95 | 0.6 | 0.86 | 0.71 |
| 1 | 27786813 | rs6668823 | 1.272 | -1.358 | 2.903 | 1.247 | -0.039 | 0.149 | 0.149 | 1.13 | 0.83 | 0.61 | 0.94 | 0.6 | 0.86 | 0.71 |
| 1 | 27836014 | rs4075401 | -2.049 | -1.369 | 2.343 | 1.326 | -0.038 | 0.078 | 0.078 | 2.265 | 0.18 | 0.66 | 0.1 | 0.38 | 0.11 | 0.3 |
| 1 | 27859036 | rs17361679 | -1.469 | -0.975 | 1.672 | 0.683 | -0.003 | 0.021 | 0.021 | 1.884 | 0.13 | 0.07 | 0.05 | 0 | 0.13 | 0.03 |
| 1 | 27862451 | rs17361686 | -0.474 | -0.938 | 1.65 | 0.742 | -0.022 | 0.047 | 0.047 | 1.267 | 0.08 | 0.01 | 0.04 | 0 | 0.12 | 0.02 |
| 1 | 27890533 | rs12146138 | -1.442 | -0.738 | 1.953 | 0.701 | 0.029 | -0.056 | -0.056 | 0 | 0.1 | 0.26 | 0.06 | 0 | 0.11 | 0.01 |
| 1 | 27930898 | rs6677501 | -2.622 | -1.044 | 2.174 | 0.795 | 0.007 | -0.002 | -0.002 | 0.126 | 0.07 | 0.58 | 0.07 | 0 | 0.02 | 0 |
| 1 | 27931698 | rs4908343 | 1.544 | -1.649 | 1.603 | 0.751 | 0.116 | -0.051 | -0.051 | 0.001 | 0.6 | 0.08 | 0.8 | 0.6 | 0.8 | 0.56 |
| 1 | 27933024 | rs11809483 | -2.713 | -1.012 | 2.363 | 0.607 | 0.005 | -0.004 | -0.004 | 0.03 | 0.07 | 0.58 | 0.07 | 0 | 0.02 | 0 |
| 1 | 27937456 | rs6598891 | -2.688 | -1.014 | 2.362 | 0.606 | 0.005 | -0.004 | -0.004 | 0 | 0.07 | 0.58 | 0.07 | 0 | 0.02 | 0 |
| 1 | 27945359 | rs17162676 | -2.219 | -0.946 | 2.092 | 0.636 | 0.056 | 0.041 | 0.041 | 0.316 | 0.06 | 0.44 | 0.05 | 0 | 0.02 | 0 |
| 1 | 27958245 | rs12726763 | -1.418 | -1.023 | 1.75 | 0.809 | -0.038 | -0.011 | -0.011 | 0.829 | 0.14 | 0.31 | 0.06 | 0.06 | 0.08 | 0.06 |
| 1 | 27971502 | rs1292081 | -1.813 | -0.838 | 2.012 | 1.021 | -0.026 | 0.045 | 0.045 | 4.175 | 0.22 | 0.59 | 0.11 | 0.36 | 0.16 | 0.26 |
| 1 | 27973748 | rs1292076 | -1.601 | -0.944 | 2.528 | 1.314 | 0.006 | 0.379 | 0.379 | 5.558 | 0.07 | 0.4 | 0.05 | 0.37 | 0 | 0.25 |
| 1 | 27975384 | rs73494 | -1.877 | -0.959 | 2.121 | 1.307 | -0.005 | 0.376 | 0.376 | 7.662 | 0.08 | 0.55 | 0.06 | 0.37 | 0 | 0.25 |
| 1 | 27975910 | rs475570 | -2.001 | -0.958 | 2.113 | 1.298 | -0.059 | 0.209 | 0.209 | 4.712 | 0.13 | 0.55 | 0.07 | 0.37 | 0.02 | 0.25 |
| 1 | 27978664 | rs12087211 | -1.581 | -0.868 | 1.651 | 1.342 | -0.04 | 0.244 | 0.244 | 1.759 | 0.11 | 0.39 | 0.06 | 0.37 | 0.02 | 0.25 |
| 1 | 27996472 | rs12732199 | -0.245 | 0.251 | 0.306 | 0.197 | -0.133 | 0.091 | 0.091 | 0.345 | 0.2 | 0.01 | 0.15 | 0.06 | 0.32 | 0.16 |
| 1 | 28011854 | rs12743229 | -0.851 | -0.035 | 0.609 | 0.419 | 0.005 | 0.04 | 0.04 | 0.031 | 0.15 | 0.05 | 0.12 | 0.02 | 0.15 | 0.07 |
| 1 | 28017518 | rs2808422 | 1.213 | -0.082 | 1.639 | 0.794 | 0.073 | 0.392 | 0.392 | 0.752 | 0.93 | 0.53 | 0.94 | 0.73 | 0.98 | 0.9 |
| 1 | 28028200 | rs4908360 | 1.384 | -0.056 | 1.839 | 0.806 | 0.072 | 0.389 | 0.389 | 3.665 | 0.93 | 0.39 | 0.92 | 0.73 | 0.98 | 0.9 |
| 1 | 28028261 | rs4908335 | 1.384 | -0.056 | 1.839 | 0.806 | 0.072 | 0.389 | 0.389 | 3.665 | 0.93 | 0.39 | 0.92 | 0.73 | 0.98 | 0.9 |
| 1 | 28051611 | rs2272795 | -1.446 | -0.194 | 1.774 | 0.776 | 0.033 | 0.315 | 0.315 | 0.13 | 0.06 | 0.42 | 0.06 | 0.27 | 0.01 | 0.1 |
| 1 | 28054814 | rs12022270 | -1.446 | -0.194 | 1.793 | 0.776 | 0.033 | 0.315 | 0.315 | 0.13 | 0.06 | 0.42 | 0.06 | 0.27 | 0.01 | 0.1 |
| 1 | 28056342 | rs1767523 | -1.23 | -0.508 | 0.886 | 0.435 | -0.137 | 0.189 | 0.189 | 0.384 | 0.6 | 0.97 | 0.66 | 0.85 | 0.52 | 0.59 |
| 1 | 28060574 | rs6670793 | -2.177 | -0.433 | 1.935 | 1.298 | 0.006 | 0.279 | 0.279 | 2.188 | 0.08 | 0.62 | 0.09 | 0.27 | 0.01 | 0.1 |
| 1 | 28071836 | rs1569432 | -1.711 | -0.432 | 1.899 | 1.298 | 0.045 | 0.368 | 0.368 | 1.427 | 0.07 | 0.58 | 0.08 | 0.27 | 0.01 | 0.1 |
| 1 | 28084571 | rs10902670 | -1.711 | -0.432 | 1.899 | 1.298 | 0.045 | 0.368 | 0.368 | 1.427 | 0.07 | 0.58 | 0.08 | 0.27 | 0.01 | 0.1 |
| 1 | 28084735 | rs10794504 | -1.711 | -0.432 | 1.899 | 1.298 | 0.045 | 0.368 | 0.368 | 1.427 | 0.07 | 0.58 | 0.08 | 0.27 | 0.01 | 0.1 |
| 1 | 28157871 | rs2072750 | -2.039 | -0.148 | 0.586 | 1.409 | 0.006 | 0.269 | 0.269 | 3.295 | 0.09 | 0.62 | 0.09 | 0.27 | 0.02 | 0.1 |
| 1 | 28209305 | rs3766398 | 0.655 | 0.226 | -1.309 | 1.201 | 0.004 | -0.054 | -0.054 | 0 | 0.66 | 0.14 | 0.51 | 0.51 | 0.63 | 0.69 |
| 1 | 28209366 | rs3766400 | 0.768 | 0.161 | -1.367 | 1.167 | -0.006 | -0.091 | -0.091 | 2.017 | 0.65 | 0.07 | 0.5 | 0.5 | 0.63 | 0.69 |
| 1 | 28211403 | rs1467465 | 0.501 | 0.225 | -0.261 | 1.325 | 0.014 | 0.011 | 0.011 | 0.126 | 0.74 | 0.47 | 0.74 | 0.63 | 0.67 | 0.69 |
| 1 | 28212806 | rs11399 | -0.194 | 0.41 | -0.912 | 1.259 | -0.018 | -0.042 | -0.042 | 0.324 | 0.1 | 0.02 | 0.15 | 0.13 | 0.12 | 0.08 |
| 1 | 28212975 | rs6564 | -0.962 | 0.421 | -0.914 | 0.869 | 0.001 | -0.002 | -0.002 | 0 | 0.42 | 0.62 | 0.39 | 0.5 | 0.38 | 0.33 |
| 1 | 28213157 | rs6565 | -0.968 | 0.42 | -0.914 | 0.87 | 0.001 | -0.002 | -0.002 | 0 | 0.42 | 0.62 | 0.39 | 0.5 | 0.38 | 0.33 |
| 1 | 28218100 | rs7356 | -0.976 | 0.415 | -0.906 | 0.908 | 0.001 | -0.003 | -0.003 | 2.633 | 0.42 | 0.63 | 0.39 | 0.49 | 0.38 | 0.31 |
| 1 | 28220484 | rs17257252 | 0.827 | 0.445 | -1.089 | 1.018 | -0.073 | 0.036 | 0.036 | 0.004 | 0.23 | 0.01 | 0.12 | 0.08 | 0.23 | 0.13 |
| 1 | 28234999 | rs2474470 | -0.788 | 0.621 | -0.852 | 0.916 | 0.005 | -0.009 | -0.009 | 0 | 0.42 | 0.69 | 0.39 | 0.49 | 0.38 | 0.3 |
| 1 | 28251700 | rs11247723 | -0.256 | 0.656 | -1.003 | 0.953 | -0.069 | -0.041 | -0.041 | 0 | 0.18 | 0.04 | 0.27 | 0.27 | 0.27 | 0.16 |
| 1 | 28257209 | rs6676158 | -0.045 | 0.599 | -0.963 | 0.961 | 0.001 | -0.003 | -0.003 | 0 | 0.64 | 0.35 | 0.63 | 0.43 | 0.71 | 0.71 |
| 1 | 28261902 | rs3737807 | 1.12 | 0.489 | -0.809 | 0.909 | -0.006 | 0.1 | 0.1 | 2.554 | 0.44 | 0.25 | 0.35 | 0.17 | 0.44 | 0.56 |
| 1 | 28263202 | rs4253940 | -0.032 | 0.483 | -0.683 | 0.971 | 0.001 | -0.002 | -0.002 | 0 | 0.64 | 0.37 | 0.64 | 0.43 | 0.71 | 0.71 |
| 1 | 28267869 | rs1010082 | 1.021 | 0.527 | -0.838 | 0.677 | -0.009 | -0.014 | -0.014 | 0 | 0.47 | 0.26 | 0.42 | 0.52 | 0.47 | 0.66 |
| 1 | 28268573 | rs10902678 | -0.034 | 0.48 | -0.635 | 0.472 | -0.018 | -0.014 | -0.014 | 0.093 | 0.67 | 0.48 | 0.71 | 0.79 | 0.75 | 0.82 |
| 1 | 28273016 | rs4908379 | 0.032 | 0.613 | -0.738 | 0.372 | -0.012 | 0 | 0 | 0 | 0.64 | 0.52 | 0.67 | 0.71 | 0.69 | 0.81 |
| 1 | 28274984 | rs10902682 | 0.09 | 0.737 | -0.601 | 0.279 | -0.001 | -0.008 | -0.008 | 0 | 0.6 | 0.57 | 0.57 | 0.78 | 0.64 | 0.79 |
| 1 | 28276389 | rs4256835 | 0.169 | 0.891 | -0.634 | 0.376 | 0.016 | 0 | 0 | 4.396 | 0.18 | 0.24 | 0.29 | 0.22 | 0.26 | 0.17 |
| 1 | 28277669 | rs5027651 | 0.695 | 0.734 | -1.002 | 0.577 | 0.014 | -0.011 | -0.011 | 0 | 0.42 | 0.3 | 0.27 | 0.56 | 0.38 | 0.61 |
| 1 | 28281924 | rs3813805 | -0.179 | 0.702 | -1.111 | 0.655 | -0.023 | -0.038 | -0.038 | 0 | 0.27 | 0.07 | 0.26 | 0.21 | 0.27 | 0.18 |
| 1 | 28282292 | rs3813803 | -0.158 | 0.728 | -1.108 | 0.671 | -0.027 | -0.039 | -0.039 | 0 | 0.26 | 0.07 | 0.26 | 0.21 | 0.27 | 0.18 |
| 1 | 28292739 | rs4243820 | 0.622 | 1.165 | 0.109 | 0.931 | -0.002 | -0.007 | -0.007 | 0 | 0.64 | 0.36 | 0.64 | 0.51 | 0.63 | 0.62 |
| 1 | 28298951 | rs6699701 | -0.738 | 1.34 | 0.675 | 1.161 | 0.005 | 0.052 | 0.052 | 0 | 0.26 | 0.36 | 0.29 | 0.53 | 0.28 | 0.34 |
| 1 | 28311224 | rs12040932 | 1.511 | 1.175 | 1.113 | 1.288 | 0.005 | 0.101 | 0.101 | 0.257 | 0.48 | 0.18 | 0.45 | 0.26 | 0.44 | 0.47 |
| 1 | 28313993 | rs11247734 | -0.648 | 1.171 | 1.127 | 1.3 | 0.007 | 0.048 | 0.048 | 0 | 0.27 | 0.37 | 0.29 | 0.53 | 0.29 | 0.34 |
| 1 | 28344980 | rs12144891 | -0.406 | 1.192 | 1.1 | 1.225 | 0.006 | -0.001 | -0.001 | 0 | 0.23 | 0.21 | 0.21 | 0.26 | 0.26 | 0.23 |
| 1 | 28352647 | rs11247739 | -0.395 | 1.189 | 1.132 | 1.223 | 0.006 | -0.001 | -0.001 | 0 | 0.23 | 0.21 | 0.21 | 0.26 | 0.26 | 0.23 |
| 1 | 28369279 | rs2476547 | -1.951 | 1.183 | 1.13 | 1.232 | 0.005 | 0.101 | 0.101 | 0.257 | 0.52 | 0.81 | 0.55 | 0.74 | 0.56 | 0.53 |
| 1 | 28377053 | rs10902684 | -0.531 | 1.265 | 1.094 | 1.267 | 0.005 | -0.001 | -0.001 | 0.344 | 0.23 | 0.21 | 0.21 | 0.26 | 0.26 | 0.23 |
| 1 | 28390596 | rs3753864 | 0.247 | 1.449 | 0.454 | 1.437 | 0.01 | 0.01 | 0.01 | 0.023 | 0.14 | 0.12 | 0.18 | 0.26 | 0.19 | 0.2 |
| 1 | 28416855 | rs2871807 | -1.454 | 1.276 | 0.403 | 1.327 | 0.02 | 0.093 | 0.093 | 1.356 | 0.11 | 0.3 | 0.13 | 0.27 | 0.1 | 0.11 |
| 1 | 28420384 | rs2486674 | -0.075 | 1.261 | -0.215 | 1.078 | -0.008 | -0.036 | -0.036 | 1.568 | 0.19 | 0.09 | 0.19 | 0.22 | 0.21 | 0.19 |
| 1 | 28426001 | rs11247752 | -1.185 | 1.952 | 0.14 | 1.361 | -0.006 | -0.012 | -0.012 | 0 | 0.25 | 0.17 | 0.28 | 0.63 | 0.32 | 0.41 |
| 1 | 28431100 | rs2481981 | -1.161 | 1.625 | 0.171 | 0.919 | -0.024 | 0.019 | 0.019 | 0.878 | 0.37 | 0.3 | 0.37 | 0.77 | 0.49 | 0.5 |
| 1 | 28432399 | rs536132 | -1.507 | 1.726 | 0.564 | 0.868 | 0.001 | 0.012 | 0.012 | 0.803 | 0.43 | 0.65 | 0.44 | 0.77 | 0.56 | 0.57 |
| 1 | 28432549 | rs6598919 | -1.042 | 1.821 | 0.449 | 1.265 | 0.035 | -0.016 | -0.016 | 0 | 0.33 | 0.44 | 0.38 | 0.77 | 0.46 | 0.48 |

| | | | | | | | | | | | | | | | | |
|---|---|---|---|---|---|---|---|---|---|---|---|---|---|---|---|---|
| 1 | 28461640 | rs10794512 | -1.196 | 0.603 | 0.32 | 1.77 | -0.023 | 0.058 | 0.058 | 0 | 0.24 | 0.58 | 0.19 | 0.33 | 0.18 | 0.17 |
| 1 | 28461661 | rs10751727 | -1.938 | 0.774 | -0.394 | 1.21 | 0.01 | 0.058 | 0.058 | 0.16 | 0.49 | 0.8 | 0.35 | 0.62 | 0.46 | 0.51 |
| 1 | 28464178 | rs12088946 | -0.535 | 0.547 | 0.376 | 1.064 | -0.013 | 0.059 | 0.059 | 0 | 0.21 | 0.39 | 0.17 | 0.26 | 0.17 | 0.17 |
| 1 | 28466221 | rs12065041 | -0.126 | 0.596 | 0.192 | 0.84 | -0.044 | 0.09 | 0.09 | 0 | 0.19 | 0.05 | 0.14 | 0 | 0.24 | 0.11 |
| 1 | 28471443 | rs313153 | -1.457 | -0.153 | 0.371 | 0.855 | -0.035 | 0.005 | 0.005 | 0.138 | 0.29 | 0.57 | 0.19 | 0.25 | 0.18 | 0.15 |
| 1 | 28509799 | rs6668505 | 0.738 | 0.218 | 0.998 | 0.442 | 0.002 | -0.016 | -0.016 | 0.204 | 0.06 | 0.08 | 0.04 | 0 | 0.08 | 0 |
| 1 | 28523238 | rs17162990 | 0.982 | 0.319 | 0.591 | 0.46 | -0.035 | -0.038 | -0.038 | 0 | 0.08 | 0 | 0.08 | 0.16 | 0.11 | 0.09 |
| 1 | 28535233 | rs510379 | -2.219 | 0.657 | 1.017 | -0.04 | 0.002 | 0.001 | 0.001 | 0 | 0.39 | 0.67 | 0.29 | 0.1 | 0.36 | 0.22 |
| 1 | 28538204 | rs490633 | -1.716 | 0.654 | 1.136 | -0.044 | -0.022 | -0.086 | -0.086 | 0 | 0.33 | 0.5 | 0.23 | 0.1 | 0.29 | 0.22 |
| 1 | 28547077 | rs489259 | -1.792 | 0.531 | 1.332 | -0.156 | -0.004 | -0.134 | -0.134 | 0 | 0.17 | 0.5 | 0.12 | 0 | 0.12 | 0.05 |
| 1 | 28550448 | rs4908393 | -2.183 | 0.562 | 0.979 | -0.142 | -0.013 | -0.052 | -0.052 | 0.171 | 0.34 | 0.59 | 0.25 | 0.1 | 0.29 | 0.22 |
| 1 | 28551450 | rs7518571 | -1.883 | 0.547 | 0.996 | -0.152 | -0.017 | -0.058 | -0.058 | 0 | 0.3 | 0.58 | 0.22 | 0.1 | 0.26 | 0.21 |
| 1 | 28555426 | rs567086 | -2.565 | 0.564 | 0.964 | -0.14 | 0.002 | 0.001 | 0.001 | 0 | 0.39 | 0.67 | 0.29 | 0.1 | 0.36 | 0.22 |
| 1 | 28563084 | rs2078348 | -0.829 | 0.523 | 1.127 | -0.154 | -0.003 | 0.026 | 0.026 | 0 | 0.22 | 0.14 | 0.17 | 0.1 | 0.25 | 0.17 |
| 1 | 28564252 | rs8559 | -2.3 | 0.684 | 0.824 | -0.898 | 0.01 | 0.064 | 0.064 | 0 | 0.4 | 0.74 | 0.29 | 0.1 | 0.37 | 0.22 |
| 1 | 28564279 | rs9508 | -1.802 | 0.641 | 0.84 | -0.863 | 0.002 | -0.006 | -0.006 | 0 | 0.34 | 0.66 | 0.25 | 0.1 | 0.29 | 0.22 |
| 1 | 28581064 | rs11805172 | 0.621 | 0.727 | 0.795 | -0.7 | 0.001 | -0.017 | -0.017 | 3.532 | 0.06 | 0.09 | 0.05 | 0 | 0.08 | 0 |
| 1 | 28589578 | rs508772 | -2.301 | 0.789 | 0.222 | -1.195 | -0.022 | 0.121 | 0.121 | 0.026 | 0.55 | 0.75 | 0.4 | 0.11 | 0.53 | 0.29 |
| 1 | 28596084 | rs543240 | -1.842 | 0.806 | 0.457 | -1.011 | 0.008 | 0.026 | 0.026 | 0 | 0.4 | 0.71 | 0.29 | 0.11 | 0.37 | 0.21 |
| 1 | 28605319 | rs11247766 | -1.532 | 0.864 | 0.407 | -0.97 | -0.007 | -0.048 | -0.048 | 0.111 | 0.36 | 0.6 | 0.28 | 0.11 | 0.35 | 0.21 |
| 1 | 28607890 | rs1053107 | -0.682 | 1.359 | 0.286 | -0.561 | -0.002 | -0.033 | -0.033 | 0 | 0.06 | 0.13 | 0.04 | 0 | 0.05 | 0.01 |
| 1 | 28636693 | rs12040403 | 1.282 | 1.038 | -0.296 | -0.778 | 0.013 | 0.238 | 0.238 | 3.552 | 0.5 | 0.14 | 0.52 | 0.85 | 0.39 | 0.58 |
| 1 | 28640773 | rs7523070 | 0.427 | 1.251 | 0.303 | -0.875 | 0 | -0.03 | -0.03 | 0 | 0.67 | 0.64 | 0.65 | 0.94 | 0.59 | 0.83 |
| 1 | 28648893 | rs17163135 | -0.965 | 1.321 | 0.66 | -0.733 | 0.113 | 0.017 | 0.017 | 0 | 0.09 | 0.28 | 0.09 | 0.09 | 0.17 | 0.22 |
| 1 | 28649402 | rs11247771 | -1.937 | 1.068 | 0.983 | -0.976 | 0.008 | 0.042 | 0.042 | 0.635 | 0.44 | 0.68 | 0.36 | 0.14 | 0.46 | 0.38 |
| 1 | 28650377 | rs6598927 | -1.042 | 0.985 | 1.016 | -0.976 | 0.003 | -0.007 | -0.007 | 0 | 0.34 | 0.37 | 0.31 | 0.06 | 0.34 | 0.17 |
| 1 | 28659696 | rs10794514 | -1.768 | 1.058 | 0.906 | -0.956 | 0.013 | 0.063 | 0.063 | 0.883 | 0.45 | 0.72 | 0.37 | 0.14 | 0.46 | 0.38 |
| 1 | 28691734 | rs4908404 | -1.749 | 1.041 | 0.918 | -1.128 | 0.01 | 0.09 | 0.09 | 0.112 | 0.46 | 0.72 | 0.42 | 0.06 | 0.5 | 0.19 |
| 1 | 28696091 | rs11247790 | -1.117 | 0.932 | 0.878 | -1.036 | -0.003 | -0.029 | -0.029 | 0.006 | 0.37 | 0.63 | 0.31 | 0.07 | 0.35 | 0.17 |
| 1 | 28740421 | rs11587757 | 2.319 | 0.825 | 0.651 | -0.383 | -0.075 | -0.062 | -0.062 | 0 | 0.41 | 0.04 | 0.41 | 0.46 | 0.32 | 0.42 |
| 1 | 28815282 | rs7518859 | -0.54 | 0.53 | 1.434 | -0.976 | 0.069 | 0.028 | 0.028 | 0 | 0.06 | 0.34 | 0.04 | 0 | 0.04 | 0.01 |
| 1 | 28824065 | rs10158229 | -0.493 | 0.53 | 1.434 | -0.976 | 0.072 | 0.03 | 0.03 | 0 | 0.06 | 0.34 | 0.04 | 0 | 0.04 | 0.01 |
| 1 | 28828893 | rs11804512 | -0.493 | 0.53 | 1.434 | -0.976 | 0.072 | 0.03 | 0.03 | 0 | 0.06 | 0.34 | 0.04 | 0 | 0.04 | 0.01 |
| 1 | 28833494 | rs2840758 | -1.472 | 0.514 | 1.371 | -0.986 | 0.006 | 0.002 | 0.002 | 0 | 0.36 | 0.65 | 0.31 | 0.07 | 0.35 | 0.17 |
| 1 | 28848588 | rs10915206 | -0.88 | 0.655 | 1.126 | -0.921 | 0.005 | 0.02 | 0.02 | 0 | 0.29 | 0.26 | 0.25 | 0.07 | 0.31 | 0.16 |
| 1 | 28861636 | rs2066726 | -0.91 | 0.698 | 1.157 | -0.913 | 0 | -0.005 | -0.005 | 0.029 | 0.29 | 0.31 | 0.25 | 0.07 | 0.31 | 0.16 |
| 1 | 28864120 | rs10799123 | -0.836 | 0.696 | 1.104 | -0.914 | 0.003 | 0.018 | 0.018 | 0 | 0.28 | 0.27 | 0.25 | 0.07 | 0.31 | 0.16 |
| 1 | 28864435 | rs2227977 | -0.405 | 0.69 | 1.121 | -0.888 | 0.034 | -0.002 | -0.002 | 0.008 | 0.06 | 0.25 | 0.04 | 0 | 0.04 | 0.01 |
| 1 | 28874969 | rs12090855 | -0.73 | 0.665 | 0.866 | -0.898 | 0.018 | -0.018 | -0.018 | 1.704 | 0.07 | 0.27 | 0.05 | 0 | 0.04 | 0.01 |
| 1 | 28882973 | rs12139243 | -1.485 | 0.64 | 0.828 | -0.909 | -0.012 | -0.089 | -0.089 | 2.134 | 0.34 | 0.51 | 0.29 | 0.08 | 0.35 | 0.17 |
| 1 | 28893572 | rs9426420 | 0.644 | 0.887 | -0.293 | -0.914 | -0.027 | 0.057 | 0.057 | 0 | 0.1 | 0 | 0.05 | 0 | 0.09 | 0.03 |
| 1 | 28901455 | rs9426442 | -1.104 | 0.913 | -0.271 | -0.902 | 0.001 | 0.034 | 0.034 | 0 | 0.27 | 0.2 | 0.24 | 0.07 | 0.3 | 0.16 |
| 1 | 28925538 | rs34749063 | 0.076 | 1.075 | -0.639 | -0.728 | -0.04 | -0.026 | -0.026 | 0 | 0.12 | 0.01 | 0.17 | 0.08 | 0.21 | 0.08 |
| 1 | 28939138 | rs6695883 | -0.963 | 0.982 | -0.623 | -0.804 | -0.023 | 0.06 | 0.06 | 0 | 0.24 | 0.09 | 0.22 | 0.08 | 0.3 | 0.12 |
| 1 | 28971281 | rs12142346 | -0.643 | 1.034 | -0.751 | -0.779 | -0.067 | 0.03 | 0.03 | 0 | 0.22 | 0.02 | 0.22 | 0.08 | 0.29 | 0.12 |
| 1 | 28990922 | rs9426298 | 0.792 | 1.031 | -0.906 | -0.852 | -0.01 | 0.067 | 0.067 | 0 | 0.69 | 0.17 | 0.75 | 0.86 | 0.68 | 0.77 |
| 1 | 29004746 | rs12746205 | 1.618 | 0.928 | -0.991 | -0.921 | -0.069 | 0.076 | 0.076 | 0.064 | 0.44 | 0.1 | 0.54 | 0.77 | 0.39 | 0.65 |
| 1 | 29013516 | rs4654361 | 1.542 | 0.91 | -0.901 | -0.93 | -0.027 | 0.093 | 0.093 | 0.016 | 0.46 | 0.17 | 0.54 | 0.77 | 0.39 | 0.65 |
| 1 | 29037793 | rs10799119 | 1.584 | 0.842 | -0.909 | -0.929 | -0.062 | 0.091 | 0.091 | 0.012 | 0.45 | 0.12 | 0.54 | 0.77 | 0.4 | 0.65 |
| 1 | 29038610 | rs6682357 | 1.559 | 0.835 | -0.913 | -0.933 | -0.061 | 0.093 | 0.093 | 0 | 0.46 | 0.12 | 0.54 | 0.77 | 0.4 | 0.65 |
| 1 | 29054752 | rs10799120 | 1.456 | 0.824 | -0.544 | -0.938 | -0.012 | 0.081 | 0.081 | 0.122 | 0.47 | 0.25 | 0.55 | 0.78 | 0.4 | 0.65 |
| 1 | 29060898 | rs3753693 | 1.581 | 0.83 | -0.598 | -1.019 | -0.03 | 0.083 | 0.083 | 0.529 | 0.44 | 0.16 | 0.54 | 0.79 | 0.39 | 0.63 |
| 1 | 29070856 | rs17340752 | -0.616 | 0.844 | 0.172 | -1.228 | -0.041 | 0.124 | 0.124 | 0 | 0.21 | 0.02 | 0.14 | 0 | 0.24 | 0.1 |
| 1 | 29074983 | rs10915194 | 1.557 | 0.903 | 0.318 | -1.046 | -0.025 | 0.081 | 0.081 | 0.392 | 0.44 | 0.16 | 0.54 | 0.79 | 0.39 | 0.63 |
| 1 | 29080166 | rs7520756 | -1.736 | 0.953 | -0.072 | -1.004 | -0.005 | 0.079 | 0.079 | 0.574 | 0.54 | 0.79 | 0.45 | 0.21 | 0.61 | 0.37 |
| 1 | 29107895 | rs528265 | -1.658 | 1.075 | -0.252 | -1.094 | -0.007 | 0.142 | 0.142 | 0.151 | 0.56 | 0.81 | 0.46 | 0.1 | 0.63 | 0.38 |
| 1 | 29123347 | rs1338064 | -0.402 | 1.097 | -0.015 | -1.162 | 0 | -0.026 | -0.026 | 0 | 0.18 | 0.19 | 0.1 | 0 | 0.14 | 0.07 |
| 1 | 29136686 | rs569356 | 0.133 | 1.346 | -0.089 | -1.143 | -0.024 | 0.022 | 0.022 | 0 | 0.07 | 0.02 | 0.08 | 0 | 0.13 | 0.06 |
| 1 | 29138975 | rs1042114 | -0.221 | 1.358 | -0.088 | -1.146 | -0.021 | 0.018 | 0.018 | 0 | 0.93 | 0.98 | 0.92 | 1 | 0.87 | 0.94 |
| 1 | 29139756 | rs2236861 | 0.059 | 1.191 | -0.129 | -1.198 | -0.012 | 0.035 | 0.035 | 0.336 | 0.19 | 0.05 | 0.2 | 0.1 | 0.26 | 0.12 |
| 1 | 29145188 | rs678849 | 1.401 | 1.146 | -0.138 | -0.766 | 0.01 | 0.098 | 0.098 | 0 | 0.5 | 0.22 | 0.35 | 0.61 | 0.51 | 0.66 |
| 1 | 29149361 | rs6669447 | -1.362 | 1.148 | 0.249 | -1.067 | -0.006 | -0.013 | -0.013 | 1.013 | 0.39 | 0.56 | 0.4 | 0.23 | 0.36 | 0.25 |
| 1 | 29151183 | rs797397 | 1.701 | 1.132 | 0.183 | -1.235 | -0.009 | 0.156 | 0.156 | 0 | 0.52 | 0.18 | 0.49 | 0.8 | 0.51 | 0.71 |
| 1 | 29151245 | rs4654322 | -1.45 | 1.074 | 0.209 | -1.165 | 0.007 | 0.017 | 0.017 | 0 | 0.39 | 0.66 | 0.42 | 0.2 | 0.36 | 0.25 |
| 1 | 29161609 | rs2236857 | -0.786 | 1.092 | 0.288 | -1.29 | -0.001 | -0.013 | -0.013 | 0 | 0.33 | 0.34 | 0.35 | 0.09 | 0.29 | 0.15 |
| 1 | 29162568 | rs760588 | -1.293 | 1.122 | 0.36 | -1.267 | -0.005 | -0.02 | -0.02 | 0 | 0.39 | 0.47 | 0.4 | 0.18 | 0.36 | 0.25 |
| 1 | 29169559 | rs3766951 | -1.237 | 1.058 | 0.289 | -1.258 | 0 | -0.003 | -0.003 | 1.483 | 0.38 | 0.41 | 0.39 | 0.19 | 0.36 | 0.25 |
| 1 | 29170005 | rs618886 | 0.036 | 1.055 | 0.141 | -1.539 | 0.048 | -0.053 | -0.053 | 0 | 0.91 | 0.77 | 0.9 | 1 | 0.87 | 0.94 |
| 1 | 29172715 | rs595725 | 1.687 | 1.212 | -0.937 | -1.489 | -0.045 | 0.171 | 0.171 | 0 | 0.45 | 0.17 | 0.47 | 0.87 | 0.45 | 0.66 |

| | | | | | | | | | | | | | | | | |
|---|---|---|---|---|---|---|---|---|---|---|---|---|---|---|---|---|
| 1 | 29174044 | rs545247 | 1.752 | 1.212 | -0.963 | -1.489 | -0.076 | 0.191 | 0.191 | 0 | 0.44 | 0.1 | 0.46 | 0.87 | 0.45 | 0.66 |
| 1 | 29174946 | rs529520 | 1.547 | 1.296 | -0.733 | -1.24 | 0.009 | 0.192 | 0.192 | 0 | 0.51 | 0.23 | 0.51 | 0.87 | 0.51 | 0.7 |
| 1 | 29175052 | rs499062 | 0.257 | 1.283 | -0.741 | -1.287 | -0.004 | -0.085 | -0.085 | 0 | 0.76 | 0.35 | 0.81 | 1 | 0.79 | 0.91 |
| 1 | 29175373 | rs581111 | 0.462 | 1.22 | -0.939 | -1.202 | 0 | 0.009 | 0.009 | 0.453 | 0.71 | 0.28 | 0.79 | 0.92 | 0.73 | 0.81 |
| 1 | 29176213 | rs12749204 | -0.472 | 0.974 | -1.113 | -0.732 | 0 | 0.074 | 0.074 | 0.305 | 0.2 | 0.04 | 0.27 | 0.05 | 0.22 | 0.11 |
| 1 | 29181525 | rs508448 | 0.469 | 1.305 | -0.773 | -0.334 | 0.001 | -0.02 | -0.02 | 0 | 0.59 | 0.48 | 0.62 | 0.77 | 0.45 | 0.64 |
| 1 | 29190138 | rs4654327 | -0.149 | 0.429 | -1.373 | -0.781 | 0.006 | 0.073 | 0.073 | 0 | 0.45 | 0.29 | 0.65 | 0.76 | 0.55 | 0.69 |
| 1 | 29194818 | rs204069 | -0.608 | -0.799 | -1.658 | -1.428 | -0.017 | 0.082 | 0.082 | 0 | 0.68 | 0.84 | 0.82 | 0.91 | 0.68 | 0.85 |
| 1 | 29194901 | rs1485471 | 1.571 | -0.703 | -2.427 | -1.68 | -0.145 | 0.109 | 0.109 | 0 | 0.28 | 0.02 | 0.31 | 0.07 | 0.38 | 0.29 |
| 1 | 29200219 | rs7511894 | 0.247 | -1.218 | -2.362 | -1.787 | 0.038 | 0.139 | 0.139 | 0 | 0.46 | 0.19 | 0.69 | 0.77 | 0.59 | 0.66 |
| 1 | 29203907 | rs157198 | 0.689 | -1.484 | -1.655 | -1.869 | -0.023 | -0.005 | -0.005 | 0 | 0.14 | 0.17 | 0.08 | 0.14 | 0.09 | 0.18 |
| 1 | 29223377 | rs10915207 | 0.4 | -1.446 | -1.679 | -1.9 | 0.028 | 0.139 | 0.139 | 0 | 0.46 | 0.18 | 0.69 | 0.77 | 0.58 | 0.68 |
| 1 | 29252322 | rs12122962 | 1.766 | -1.606 | -2 | -2.175 | -0.04 | 0.077 | 0.077 | 0 | 0.13 | 0.01 | 0.06 | 0 | 0.15 | 0.06 |
| 1 | 29268716 | rs12038347 | 1.101 | -1.542 | -1.905 | -0.78 | 0 | 0.009 | 0.009 | 0 | 0.09 | 0.1 | 0.18 | 0.35 | 0.1 | 0.24 |
| 1 | 29288008 | rs10915212 | 0.395 | -1.636 | -1.119 | -0.52 | -0.034 | -0.07 | -0.07 | 0 | 0.11 | 0.29 | 0.07 | 0 | 0.07 | 0.02 |
| 1 | 29289616 | rs2985319 | 1.536 | -1.637 | -1.048 | -0.473 | -0.018 | 0.092 | 0.092 | 0 | 0.05 | 0.11 | 0.05 | 0.23 | 0.02 | 0.15 |
| 1 | 29331189 | rs10429865 | 1.837 | -1.74 | -1.353 | -0.923 | -0.112 | 0.178 | 0.178 | 0 | 0.36 | 0.06 | 0.32 | 0.08 | 0.42 | 0.32 |
| 1 | 29335280 | rs16837840 | 1.386 | -1.653 | -1.262 | -0.801 | -0.003 | -0.007 | -0.007 | 1.592 | 0.09 | 0.07 | 0.18 | 0.36 | 0.1 | 0.24 |
| 1 | 29363985 | rs2486200 | -1.094 | -1.674 | -0.723 | -0.862 | -0.001 | 0.001 | 0.001 | 0 | 0.28 | 0.64 | 0.34 | 0.49 | 0.21 | 0.32 |
| 1 | 29429004 | rs12045777 | 1.19 | -1.609 | -0.924 | -0.906 | -0.065 | -0.026 | -0.026 | 0 | 0.45 | 0.13 | 0.5 | 0.43 | 0.52 | 0.56 |
| 1 | 29443723 | rs2249138 | -0.61 | -1.664 | -0.465 | -0.724 | 0.036 | 0.25 | 0.25 | 0 | 0.88 | 0.74 | 0.75 | 0.51 | 0.87 | 0.7 |
| 1 | 29444544 | rs569716 | 0.821 | -1.72 | -0.247 | -0.762 | -0.024 | -0.057 | -0.057 | 0 | 0.1 | 0.24 | 0.06 | 0 | 0.07 | 0.01 |
| 1 | 29445514 | rs553253 | 1.242 | -1.397 | -0.512 | -0.845 | 0.007 | -0.01 | -0.01 | 0.324 | 0.61 | 0.37 | 0.61 | 0.5 | 0.68 | 0.65 |
| 1 | 29446811 | rs2452788 | 0.4 | -1.655 | 0.048 | -0.114 | -0.004 | 0.129 | 0.129 | 0 | 0.82 | 0.68 | 0.73 | 0.56 | 0.86 | 0.7 |
| 1 | 29446847 | rs575675 | 0.497 | -1.874 | -0.025 | -0.283 | -0.025 | -0.062 | -0.062 | 0.489 | 0.1 | 0.28 | 0.07 | 0 | 0.07 | 0.01 |
| 1 | 29461133 | rs12402749 | 0.894 | -1.785 | -0.404 | 0.202 | 0 | 0.003 | 0.003 | 0 | 0.09 | 0.12 | 0.1 | 0.36 | 0.09 | 0.25 |
| 1 | 29464954 | rs12032470 | 2.064 | -1.896 | -0.668 | -0.477 | -0.165 | 0.194 | 0.194 | 0.002 | 0.36 | 0.05 | 0.32 | 0.08 | 0.45 | 0.31 |
| 1 | 29475648 | rs2230679 | -0.662 | -1.973 | -0.092 | -0.4 | 0.028 | 0.258 | 0.258 | 0 | 0.89 | 0.72 | 0.75 | 0.56 | 0.88 | 0.69 |
| 1 | 29488047 | rs10915234 | 0.997 | -2.005 | -0.306 | -0.39 | 0.003 | 0.015 | 0.015 | 0 | 0.1 | 0.12 | 0.17 | 0.41 | 0.09 | 0.25 |
| 1 | 29518293 | rs3103882 | -0.712 | -1.99 | -0.205 | -0.387 | 0.009 | 0.146 | 0.146 | 0 | 0.89 | 0.82 | 0.75 | 0.56 | 0.88 | 0.69 |
| 1 | 29519778 | rs2640464 | -0.694 | -1.987 | -0.267 | -0.386 | 0.033 | 0.26 | 0.26 | 0 | 0.89 | 0.72 | 0.73 | 0.56 | 0.88 | 0.69 |
| 1 | 29521705 | rs12738007 | 2.218 | -2.016 | -0.332 | -0.435 | -0.166 | 0.19 | 0.19 | 0 | 0.36 | 0.04 | 0.32 | 0.08 | 0.45 | 0.31 |
| 1 | 29533232 | rs12743830 | 1.683 | -1.862 | -0.161 | -0.306 | 0.003 | -0.004 | -0.004 | 0 | 0.06 | 0.04 | 0.02 | 0 | 0.02 | 0 |
| 1 | 29542637 | rs1128400 | -0.683 | -1.841 | -0.132 | -0.278 | 0.034 | 0.263 | 0.263 | 0 | 0.89 | 0.72 | 0.73 | 0.56 | 0.88 | 0.69 |
| 1 | 29549996 | rs2319014 | 0.889 | -1.768 | -0.152 | -0.231 | 0.002 | 0.015 | 0.015 | 0 | 0.1 | 0.12 | 0.17 | 0.41 | 0.09 | 0.25 |
| 1 | 29556158 | rs2640467 | -0.335 | -1.912 | -0.028 | -0.314 | 0.006 | 0.025 | 0.025 | 0 | 0.33 | 0.3 | 0.21 | 0.08 | 0.32 | 0.13 |
| 1 | 29556399 | rs2819606 | -0.015 | -2.005 | 0.12 | -0.156 | -0.011 | 0.148 | 0.148 | 0 | 0.82 | 0.69 | 0.72 | 0.56 | 0.86 | 0.69 |
| 1 | 29558468 | rs2503011 | 0.979 | -1.969 | 0.262 | -0.155 | 0.012 | -0.03 | -0.03 | 0 | 0.06 | 0.23 | 0.06 | 0 | 0.06 | 0.01 |
| 1 | 29562765 | rs12409260 | 0.83 | -1.987 | 0.128 | -0.135 | 0.004 | 0.009 | 0.009 | 0 | 0.1 | 0.12 | 0.17 | 0.41 | 0.09 | 0.25 |
| 1 | 29569212 | rs7349137 | 1.141 | -2.063 | 0.23 | -0.137 | -0.015 | 0.012 | 0.012 | 0.213 | 0.7 | 0.38 | 0.65 | 0.56 | 0.78 | 0.68 |
| 1 | 29571758 | rs12057666 | 0.592 | -1.521 | 0.264 | 0.13 | 0.006 | -0.032 | -0.032 | 0 | 0.07 | 0.25 | 0.06 | 0 | 0.06 | 0.01 |
| 1 | 29571788 | rs1113735 | -0.055 | -1.264 | -0.195 | 0.404 | 0.004 | 0.078 | 0.078 | 0 | 0.8 | 0.73 | 0.67 | 0.57 | 0.79 | 0.68 |
| 1 | 29575605 | rs1032425 | 0.709 | -1.155 | -0.398 | 0.19 | -0.008 | -0.024 | -0.024 | 0 | 0.18 | 0.21 | 0.11 | 0.08 | 0.17 | 0.1 |
| 1 | 29579826 | rs2131556 | 0.733 | -0.836 | -0.617 | 1.041 | 0 | 0.017 | 0.017 | 0 | 0.78 | 0.78 | 0.65 | 0.5 | 0.77 | 0.72 |
| 1 | 29581381 | rs12401971 | -0.401 | -0.514 | -0.788 | 1.332 | 0.004 | 0.006 | 0.006 | 0 | 0.09 | 0.1 | 0.15 | 0.34 | 0.07 | 0.18 |
| 1 | 29586703 | rs12404526 | -0.438 | -0.484 | -0.903 | 1.024 | 0.011 | -0.063 | -0.063 | 0.165 | 0.11 | 0.02 | 0.29 | 0.65 | 0.1 | 0.32 |
| 1 | 29603999 | rs10799128 | 0.335 | -0.161 | -0.412 | 0.515 | 0.045 | -0.008 | -0.008 | 0 | 0.85 | 0.68 | 0.8 | 0.85 | 0.8 | 0.88 |
| 1 | 29616318 | rs2076620 | -0.027 | -0.063 | -0.881 | 0.485 | -0.017 | 0.001 | 0.001 | 0.312 | 0.12 | 0.07 | 0.19 | 0.16 | 0.18 | 0.14 |
| 1 | 29630455 | rs2295061 | -0.639 | -0.479 | -0.694 | 0.547 | -0.023 | 0.092 | 0.092 | 0.087 | 0.18 | 0.35 | 0.33 | 0.64 | 0.16 | 0.38 |
| 1 | 29630956 | rs3737606 | -0.584 | -0.404 | -0.79 | 0.43 | 0.028 | -0.053 | -0.053 | 0 | 0.21 | 0.52 | 0.27 | 0.17 | 0.24 | 0.25 |
| 1 | 29631050 | rs7524056 | -0.97 | -0.693 | -2.172 | 0.058 | -0.047 | 0.119 | 0.119 | 0.011 | 0.54 | 0.94 | 0.63 | 0.81 | 0.45 | 0.65 |
| 1 | 29631909 | rs2235937 | -0.479 | -1.25 | -1.509 | 0.58 | 0.015 | -0.017 | -0.017 | 0 | 0.31 | 0.22 | 0.38 | 0.49 | 0.24 | 0.41 |
| 1 | 29640621 | rs11811458 | -0.3 | -1.709 | -0.922 | 0.241 | -0.038 | -0.023 | -0.023 | 0 | 0.22 | 0.3 | 0.09 | 0.17 | 0.14 | 0.23 |
| 1 | 29642318 | rs2179795 | -0.817 | -0.966 | -1.839 | 0.499 | -0.025 | -0.052 | -0.052 | 0 | 0.2 | 0.14 | 0.36 | 0.51 | 0.3 | 0.32 |
| 1 | 29646127 | rs2007350 | -1.523 | -1.218 | -2.064 | 0.095 | 0.017 | 0.035 | 0.035 | 0 | 0.5 | 0.92 | 0.5 | 0.66 | 0.45 | 0.54 |
| 1 | 29649189 | rs12749276 | 0.113 | -0.745 | -1.611 | -0.354 | 0.044 | 0.103 | 0.103 | 0 | 0.2 | 0.07 | 0.04 | 0.01 | 0.13 | 0.21 |
| 1 | 29655656 | rs12402313 | 0.367 | -0.407 | -1.624 | -0.089 | -0.059 | -0.059 | -0.059 | 0 | 0.1 | 0.01 | 0.31 | 0.54 | 0.15 | 0.28 |
| 1 | 29655746 | rs10489445 | 0.367 | -0.407 | -1.624 | -0.089 | -0.059 | -0.059 | -0.059 | 0 | 0.1 | 0.01 | 0.31 | 0.54 | 0.15 | 0.28 |
| 1 | 29658494 | rs7530986 | -0.653 | -0.458 | -1.011 | -0.036 | 0.002 | -0.002 | -0.002 | 0 | 0.4 | 0.35 | 0.42 | 0.56 | 0.38 | 0.55 |
| 1 | 29671081 | rs212315 | -0.735 | -0.339 | -0.948 | -0.023 | 0.001 | -0.001 | -0.001 | 0 | 0.42 | 0.41 | 0.45 | 0.53 | 0.4 | 0.54 |
| 1 | 29678592 | rs761429 | -0.064 | -0.074 | -1.425 | -1 | -0.054 | 0.232 | 0.232 | 0.058 | 0.58 | 0.87 | 0.67 | 0.93 | 0.49 | 0.79 |
| 1 | 29678774 | rs761430 | 0.315 | 0.115 | -0.747 | -0.98 | 0 | -0.032 | -0.032 | 0 | 0.36 | 0.38 | 0.33 | 0.06 | 0.44 | 0.16 |
| 1 | 29684158 | rs4654305 | 0.637 | 0.442 | -0.577 | -0.182 | -0.004 | -0.002 | -0.002 | 0 | 0.92 | 0.96 | 0.96 | 0.9 | 0.9 | 0.92 |
| 1 | 29685790 | rs6426356 | 0.046 | 0.24 | -0.435 | -0.397 | -0.003 | 0.004 | 0.004 | 0 | 0.6 | 0.66 | 0.37 | 0.35 | 0.5 | 0.41 |
| 1 | 29694741 | rs6694460 | 0.022 | 0.201 | -0.429 | -0.397 | -0.001 | 0.024 | 0.024 | 0 | 0.57 | 0.7 | 0.38 | 0.35 | 0.49 | 0.42 |
| 1 | 29695231 | rs1325280 | 0.197 | 0.167 | -0.928 | -0.37 | 0.025 | 0.089 | 0.089 | 0 | 0.51 | 0.23 | 0.31 | 0.35 | 0.47 | 0.37 |
| 1 | 29707412 | rs1886651 | 0.176 | 0.414 | -1.291 | -0.25 | -0.001 | 0.085 | 0.085 | 0 | 0.5 | 0.24 | 0.35 | 0.33 | 0.51 | 0.38 |
| 1 | 29718655 | rs4654337 | -0.231 | 0.565 | -1.425 | -0.374 | -0.024 | 0.05 | 0.05 | 0 | 0.42 | 0.24 | 0.53 | 0.24 | 0.51 | 0.48 |
| 1 | 29719475 | rs212306 | -0.13 | 0.42 | -1.147 | -0.14 | -0.009 | 0.006 | 0.006 | 0 | 0.6 | 0.35 | 0.56 | 0.67 | 0.6 | 0.58 |

| | | | | | | | | | | | | | | | | |
|---|---|---|---|---|---|---|---|---|---|---|---|---|---|---|---|---|
| 1 | 29722553 | rs212290 | 0.306 | 0.162 | -1.538 | -0.241 | 0.008 | -0.063 | -0.063 | 0.336 | 0.36 | 0.17 | 0.32 | 0.51 | 0.36 | 0.46 |
| 1 | 29732223 | rs6671744 | -0.317 | 0.434 | -0.505 | -0.571 | -0.001 | -0.016 | -0.016 | 0 | 0.27 | 0.3 | 0.31 | 0.16 | 0.28 | 0.15 |
| 1 | 29735095 | rs1886650 | -0.629 | 0.239 | -0.809 | -0.589 | 0.004 | 0.005 | 0.005 | 0 | 0.34 | 0.65 | 0.33 | 0.17 | 0.29 | 0.32 |
| 1 | 29742743 | rs4654339 | -0.153 | 0.392 | -0.868 | -0.747 | -0.001 | 0.009 | 0.009 | 0 | 0.78 | 0.78 | 0.86 | 0.98 | 0.78 | 0.88 |
| 1 | 29743408 | rs6657309 | 0.15 | 0.19 | -0.608 | 0.578 | -0.002 | 0.018 | 0.018 | 0 | 0.76 | 0.69 | 0.67 | 0.46 | 0.77 | 0.75 |
| 1 | 29743630 | rs1408857 | 0.017 | 0.251 | -1.021 | -0.323 | -0.006 | 0.023 | 0.023 | 0 | 0.35 | 0.15 | 0.43 | 0.28 | 0.37 | 0.49 |
| 1 | 29744397 | rs8179300 | 0.848 | 0.249 | -0.666 | -0.291 | -0.009 | -0.017 | -0.017 | 0.083 | 0.12 | 0.06 | 0.15 | 0.15 | 0.13 | 0.31 |
| 1 | 29750827 | rs12134081 | 0.186 | 0.361 | -0.622 | -0.24 | 0.001 | 0 | 0 | 0 | 0.1 | 0.09 | 0.2 | 0.14 | 0.13 | 0.13 |
| 1 | 29751247 | rs912772 | 0.889 | 0.227 | -0.268 | -0.425 | -0.006 | 0.021 | 0.021 | 0 | 0.38 | 0.24 | 0.38 | 0.69 | 0.39 | 0.37 |
| 1 | 29753125 | rs12085934 | 0.517 | 0.167 | -0.217 | -0.177 | -0.01 | 0.042 | 0.042 | 0 | 0.26 | 0.07 | 0.23 | 0.15 | 0.25 | 0.36 |
| 1 | 29754510 | rs10915271 | 0.756 | 0.54 | -0.65 | -0.224 | -0.056 | 0.089 | 0.089 | 0 | 0.36 | 0.08 | 0.2 | 0.17 | 0.4 | 0.26 |
| 1 | 29755818 | rs16838233 | -0.829 | 0.648 | -0.111 | 0.221 | 0.002 | 0 | 0 | 0 | 0.11 | 0.1 | 0.2 | 0.14 | 0.14 | 0.12 |
| 1 | 29761148 | rs16838255 | -0.448 | 0.812 | -0.673 | 0.347 | -0.004 | -0.004 | -0.004 | 0 | 0.09 | 0.05 | 0.2 | 0.14 | 0.13 | 0.12 |
| 1 | 29765644 | rs4654342 | -0.538 | 0.991 | -0.809 | 0.434 | -0.028 | -0.029 | -0.029 | 0.795 | 0.07 | 0.02 | 0.19 | 0.15 | 0.13 | 0.08 |
| 1 | 29766120 | rs6685390 | -0.205 | 0.272 | -0.534 | 0.367 | -0.019 | 0.03 | 0.03 | 0 | 0.3 | 0.61 | 0.34 | 0.58 | 0.22 | 0.25 |
| 1 | 29770078 | rs1408858 | 0.268 | 0.325 | -1.073 | -0.564 | 0.013 | 0.027 | 0.027 | 0 | 0.25 | 0.11 | 0.22 | 0.19 | 0.23 | 0.31 |
| 1 | 29773040 | rs4654344 | -0.538 | 0.77 | -0.373 | -1.128 | 0.001 | -0.076 | -0.076 | 0 | 0.62 | 0.62 | 0.79 | 0.95 | 0.64 | 0.73 |
| 1 | 29779305 | rs4654347 | 0.974 | 0.212 | -0.031 | -0.856 | 0.014 | 0.03 | 0.03 | 0 | 0.16 | 0.08 | 0.16 | 0.05 | 0.1 | 0.13 |
| 1 | 29780573 | rs7521309 | -1.091 | 0.213 | -0.669 | -1.579 | -0.007 | 0.149 | 0.149 | 0 | 0.59 | 0.86 | 0.71 | 0.82 | 0.63 | 0.54 |
| 1 | 29781221 | rs4654349 | -0.964 | 0.125 | 0.191 | -1.595 | 0.024 | 0.059 | 0.059 | 0 | 0.55 | 0.64 | 0.58 | 0.25 | 0.59 | 0.43 |
| 1 | 29781530 | rs9426359 | 1.407 | 0.282 | 0.022 | -1.436 | 0.013 | 0.024 | 0.024 | 0 | 0.16 | 0.08 | 0.16 | 0.05 | 0.1 | 0.13 |
| 1 | 29783755 | rs4654351 | 0.149 | 0.85 | -0.524 | -2.305 | 0.011 | 0.12 | 0.12 | 0.508 | 0.42 | 0.12 | 0.41 | 0.2 | 0.39 | 0.36 |
| 1 | 29784150 | rs12758197 | 0.239 | 1.215 | 0.134 | -1.973 | 0.083 | -0.033 | -0.033 | 0 | 0.11 | 0.2 | 0.14 | 0.05 | 0.22 | 0.08 |
| 1 | 29785574 | rs912773 | 0.028 | 0.451 | -0.168 | -2.15 | 0.042 | 0.087 | 0.087 | 0 | 0.51 | 0.68 | 0.57 | 0.79 | 0.62 | 0.62 |
| 1 | 29789248 | rs10915288 | 0.659 | 1.072 | -0.359 | -2.064 | 0.007 | -0.013 | -0.013 | 1.626 | 0.37 | 0.42 | 0.44 | 0.74 | 0.46 | 0.54 |
| 1 | 29790682 | rs10915293 | 0.529 | 1.056 | -0.697 | -2.067 | -0.013 | 0.024 | 0.024 | 0 | 0.38 | 0.32 | 0.43 | 0.74 | 0.46 | 0.54 |
| 1 | 29792303 | rs766385 | -0.668 | 0.764 | -0.949 | -2.141 | -0.018 | 0.028 | 0.028 | 0 | 0.77 | 0.9 | 0.77 | 0.84 | 0.73 | 0.77 |
| 1 | 29793205 | rs927727 | -0.765 | 1.052 | -0.578 | -1.531 | -0.017 | 0.022 | 0.022 | 0 | 0.63 | 0.67 | 0.56 | 0.24 | 0.53 | 0.44 |
| 1 | 29794648 | rs11804085 | -0.165 | 1.053 | -0.391 | -1.752 | 0 | -0.004 | -0.004 | 0 | 0.46 | 0.55 | 0.46 | 0.14 | 0.43 | 0.38 |
| 1 | 29794833 | rs9426362 | 0.147 | 0.663 | 0.219 | -1.766 | -0.019 | -0.062 | -0.062 | 0 | 0.12 | 0.24 | 0.08 | 0 | 0.1 | 0.03 |
| 1 | 29796218 | rs4654314 | 0.439 | 0.991 | 0.575 | -1.955 | 0.072 | -0.063 | -0.063 | 0.389 | 0.89 | 0.64 | 0.88 | 0.98 | 0.86 | 0.95 |
| 1 | 29796645 | rs9426458 | -0.444 | 0.776 | 0.559 | -2 | -0.034 | -0.087 | -0.087 | 0 | 0.15 | 0.29 | 0.07 | 0 | 0.09 | 0.02 |
| 1 | 29799510 | rs1325281 | -0.388 | 0.969 | 0.327 | -1.912 | 0.004 | -0.006 | -0.006 | 0 | 0.94 | 0.91 | 0.92 | 0.98 | 0.89 | 0.96 |
| 1 | 29800081 | rs9426464 | -0.152 | 0.698 | 0.424 | -1.883 | -0.02 | -0.065 | -0.065 | 0 | 0.14 | 0.2 | 0.07 | 0 | 0.09 | 0.02 |
| 1 | 29801368 | rs10753340 | 1.17 | 0.655 | 0.123 | -2.395 | 0.006 | 0.063 | 0.063 | 0 | 0.51 | 0.39 | 0.59 | 0.9 | 0.57 | 0.76 |
| 1 | 29801397 | rs6694109 | -0.027 | 0.64 | 0.238 | -2.42 | -0.011 | 0.045 | 0.045 | 0.29 | 0.66 | 0.72 | 0.68 | 0.9 | 0.67 | 0.79 |
| 1 | 29802337 | rs1983855 | -0.21 | 0.488 | 0.329 | -2.345 | -0.027 | -0.099 | -0.099 | 0 | 0.14 | 0.33 | 0.09 | 0 | 0.1 | 0.03 |
| 1 | 29804812 | rs9426466 | 0.886 | 0.592 | 0.199 | -2.382 | 0.019 | 0.025 | 0.025 | 0 | 0.2 | 0.09 | 0.15 | 0.08 | 0.11 | 0.14 |
| 1 | 29805775 | rs4147252 | -0.572 | 0.543 | 0.447 | -2.305 | -0.001 | -0.021 | -0.021 | 0 | 0.36 | 0.37 | 0.32 | 0.09 | 0.31 | 0.2 |
| 1 | 29806634 | rs6426365 | 0.803 | 0.492 | 0.528 | -1.984 | 0.005 | -0.077 | -0.077 | 0 | 0.87 | 0.49 | 0.89 | 0.99 | 0.9 | 0.97 |
| 1 | 29810250 | rs11808245 | -0.16 | 0.628 | -0.136 | -1.921 | 0 | -0.012 | -0.012 | 0 | 0.1 | 0.11 | 0.06 | 0.01 | 0.1 | 0.03 |
| 1 | 29812405 | rs9426275 | -0.488 | 0.573 | -0.07 | -1.895 | -0.001 | -0.006 | -0.006 | 0.295 | 0.34 | 0.31 | 0.3 | 0.09 | 0.29 | 0.21 |
| 1 | 29813268 | rs11586865 | 0.348 | 0.513 | -0.256 | -1.756 | -0.004 | 0.011 | 0.011 | 0 | 0.16 | 0.1 | 0.21 | 0.07 | 0.2 | 0.18 |
| 1 | 29815645 | rs4078312 | -1.192 | 0.344 | -0.037 | -1.496 | -0.003 | -0.021 | -0.021 | 0.322 | 0.06 | 0.24 | 0.05 | 0.01 | 0.05 | 0.01 |
| 1 | 29827798 | rs6664690 | -0.959 | 0.368 | -0.19 | -1.657 | -0.01 | -0.079 | -0.079 | 1.45 | 0.39 | 0.53 | 0.32 | 0.09 | 0.29 | 0.2 |
| 1 | 29835996 | rs9426279 | -0.1 | -0.096 | -0.219 | -1.554 | -0.036 | -0.123 | -0.123 | 0 | 0.22 | 0.34 | 0.1 | 0 | 0.14 | 0.08 |
| 1 | 29852167 | rs7519921 | 0.356 | 0.344 | -0.67 | -0.492 | -0.033 | -0.073 | -0.073 | 0 | 0.13 | 0.03 | 0.08 | 0.34 | 0.14 | 0.29 |
| 1 | 29859210 | rs7537024 | -1.01 | -0.039 | -0.307 | -0.614 | 0.02 | 0.011 | 0.011 | 0.147 | 0.4 | 0.68 | 0.22 | 0.36 | 0.29 | 0.37 |
| 1 | 29865107 | rs6426343 | -0.996 | -0.058 | -0.122 | -0.623 | -0.003 | -0.002 | -0.002 | 1.263 | 0.38 | 0.43 | 0.19 | 0.36 | 0.29 | 0.37 |
| 1 | 29867522 | rs6688164 | 0.991 | 0.179 | -0.973 | -0.117 | -0.008 | 0.035 | 0.035 | 0 | 0.46 | 0.21 | 0.55 | 0.38 | 0.48 | 0.48 |
| 1 | 29867677 | rs6677824 | -1.603 | 0.186 | -1.087 | -0.102 | 0 | 0.008 | 0.008 | 0.028 | 0.62 | 0.68 | 0.61 | 0.7 | 0.6 | 0.62 |
| 1 | 29868073 | rs6700641 | -0.579 | 0.102 | -1.126 | -0.181 | 0.019 | 0 | 0 | 0 | 0.38 | 0.46 | 0.36 | 0.63 | 0.42 | 0.46 |
| 1 | 29869915 | rs6605085 | 0.479 | 0.224 | -1.955 | -0.014 | -0.019 | -0.077 | -0.077 | 0 | 0.18 | 0.03 | 0.09 | 0.34 | 0.16 | 0.31 |
| 1 | 29870454 | rs7418865 | 1.037 | 0.19 | -1.689 | -0.348 | -0.026 | 0.015 | 0.015 | 0.444 | 0.36 | 0.13 | 0.37 | 0.3 | 0.4 | 0.38 |
| 1 | 29870952 | rs28716253 | -0.462 | 0.078 | -0.971 | -0.197 | -0.005 | -0.038 | -0.038 | 0 | 0.15 | 0.18 | 0.07 | 0 | 0.1 | 0.07 |
| 1 | 30033473 | rs28526732 | -0.657 | 0.546 | -0.941 | 0.686 | 0.003 | 0.015 | 0.015 | 0 | 0.27 | 0.31 | 0.18 | 0.39 | 0.27 | 0.34 |
| 1 | 30039872 | rs28589362 | -0.799 | 0.333 | -1.887 | 0.429 | -0.045 | 0.12 | 0.12 | 0.118 | 0.54 | 0.85 | 0.44 | 0.71 | 0.46 | 0.49 |
| 1 | 30063197 | rs1932397 | 0.693 | 0.456 | -1.614 | 0.7 | -0.036 | -0.084 | -0.084 | 0 | 0.19 | 0.03 | 0.16 | 0.34 | 0.18 | 0.13 |
| 1 | 30064009 | rs6603927 | 0.411 | -0.021 | -0.86 | 0.561 | -0.003 | 0.008 | 0.008 | 0 | 0.59 | 0.45 | 0.71 | 0.45 | 0.66 | 0.68 |
| 1 | 30067017 | rs10915035 | 0.331 | -0.199 | -0.901 | 0.552 | -0.003 | 0.003 | 0.003 | 0 | 0.16 | 0.17 | 0.15 | 0.15 | 0.15 | 0.26 |
| 1 | 30073328 | rs10914201 | 1.364 | -0.088 | -1.13 | 0.433 | -0.112 | 0.08 | 0.08 | 0 | 0.37 | 0.11 | 0.44 | 0.23 | 0.47 | 0.33 |
| 1 | 30073828 | rs9729975 | 0.188 | 0.32 | -0.518 | 0.751 | -0.001 | -0.003 | -0.003 | 0.728 | 0.09 | 0.09 | 0.07 | 0.21 | 0.08 | 0.08 |
| 1 | 30077312 | rs10914306 | -0.463 | 0.241 | -0.412 | 0.841 | -0.001 | -0.006 | -0.006 | 0.042 | 0.23 | 0.21 | 0.2 | 0.41 | 0.23 | 0.32 |
| 1 | 30086662 | rs10798895 | -1.492 | 0.245 | 0.681 | 0.872 | 0 | 0.018 | 0.018 | 0 | 0.8 | 0.77 | 0.83 | 0.66 | 0.82 | 0.87 |
| 1 | 30090260 | rs4949283 | -1.016 | 0.289 | 0.76 | 0.88 | 0.006 | 0.126 | 0.126 | 0 | 0.77 | 0.54 | 0.78 | 0.6 | 0.81 | 0.87 |
| 1 | 30093351 | rs16832695 | 0.749 | 0.514 | 0.557 | 0.826 | -0.025 | -0.033 | -0.033 | 0 | 0.08 | 0 | 0.04 | 0.09 | 0.1 | 0.12 |
| 1 | 30093568 | rs9729003 | 1.432 | 0.956 | 0.257 | 0.431 | -0.04 | -0.021 | -0.021 | 0 | 0.1 | 0.01 | 0.18 | 0.12 | 0.1 | 0.15 |
| 1 | 30097294 | rs9283399 | -1.02 | 1.223 | 0.447 | 0.389 | 0.004 | 0.012 | 0.012 | 0 | 0.36 | 0.45 | 0.44 | 0.5 | 0.38 | 0.46 |
| 1 | 30098510 | rs271321 | 1.396 | 1.256 | 0.386 | 0.344 | -0.003 | -0.045 | -0.045 | 0 | 0.36 | 0.36 | 0.35 | 0.07 | 0.35 | 0.29 |

| | | | | | | | | | | | | | | | | |
|---|---|---|---|---|---|---|---|---|---|---|---|---|---|---|---|---|
| 1 | 30098743 | rs9286944 | -0.277 | 1.389 | 0.362 | 0.453 | -0.003 | -0.006 | -0.006 | 0 | 0.23 | 0.19 | 0.22 | 0.32 | 0.26 | 0.32 |
| 1 | 30101569 | rs9286945 | -0.683 | 1.534 | 0.492 | 0.38 | 0.036 | 0.029 | 0.029 | 0 | 0.21 | 0.41 | 0.33 | 0.34 | 0.27 | 0.2 |
| 1 | 30101765 | rs10914758 | 1.417 | 1.411 | 0.387 | 0.252 | 0.013 | 0.158 | 0.158 | 0 | 0.34 | 0.13 | 0.32 | 0.07 | 0.34 | 0.3 |
| 1 | 30102340 | rs271316 | 0.078 | 1.268 | 0.529 | 0.452 | 0.001 | 0 | 0 | 1.823 | 0.57 | 0.68 | 0.68 | 0.46 | 0.61 | 0.5 |
| 1 | 30103858 | rs4949611 | 0.157 | 1.262 | 0.385 | 0.389 | 0.056 | 0.077 | 0.077 | 0 | 0.07 | 0.18 | 0.11 | 0.21 | 0.12 | 0.05 |
| 1 | 30109482 | rs502545 | 0.274 | 0.736 | 0.093 | -0.048 | 0.008 | -0.016 | -0.016 | 0 | 0.56 | 0.75 | 0.68 | 0.49 | 0.61 | 0.5 |
| 1 | 30114164 | rs9439752 | -0.41 | 0.77 | -0.189 | -0.055 | 0.006 | 0.019 | 0.019 | 0.101 | 0.26 | 0.1 | 0.13 | 0.21 | 0.23 | 0.33 |
| 1 | 30118583 | rs9286955 | 1.231 | 0.72 | -0.215 | -0.236 | -0.017 | 0.166 | 0.166 | 0 | 0.34 | 0.09 | 0.35 | 0.07 | 0.36 | 0.31 |
| 1 | 30120287 | rs271383 | -0.93 | 0.973 | 0.284 | 0.203 | -0.008 | 0.028 | 0.028 | 0 | 0.77 | 0.47 | 0.85 | 0.73 | 0.83 | 0.89 |
| 1 | 30127106 | rs446335 | -1.257 | 0.941 | 0.346 | 0.243 | -0.003 | 0.002 | 0.002 | 0 | 0.78 | 0.81 | 0.88 | 0.73 | 0.8 | 0.88 |
| 1 | 30133502 | rs199613 | -0.959 | 1.027 | 0.115 | 0.193 | 0.006 | -0.001 | -0.001 | 0 | 0.41 | 0.45 | 0.52 | 0.62 | 0.43 | 0.56 |
| 1 | 30134961 | rs11589926 | -0.258 | 1.177 | -0.389 | 0.296 | 0 | -0.001 | -0.001 | 0 | 0.22 | 0.09 | 0.15 | 0.21 | 0.2 | 0.31 |
| 1 | 30137055 | rs271363 | 0.484 | 1.172 | -0.474 | 0.322 | -0.02 | -0.024 | -0.024 | 0.531 | 0.17 | 0.13 | 0.35 | 0.41 | 0.24 | 0.25 |
| 1 | 30142718 | rs10915104 | 0.401 | 1.04 | -0.724 | 0.037 | 0.024 | -0.046 | -0.046 | 0.004 | 0.15 | 0.02 | 0.09 | 0.2 | 0.09 | 0.23 |
| 1 | 30143707 | rs12122023 | 1.15 | 1.06 | -0.613 | 0.095 | 0.011 | 0.105 | 0.105 | 0 | 0.35 | 0.21 | 0.34 | 0.08 | 0.34 | 0.32 |
| 1 | 30148652 | rs166604 | -1.336 | 1.073 | -0.589 | 0.104 | -0.011 | 0.013 | 0.013 | 0 | 0.77 | 0.76 | 0.88 | 0.73 | 0.79 | 0.88 |
| 1 | 30149249 | rs9661274 | 1.163 | 1.067 | -0.607 | 0.081 | 0.008 | 0.093 | 0.093 | 0 | 0.35 | 0.22 | 0.33 | 0.07 | 0.35 | 0.31 |
| 1 | 30155341 | rs12564818 | 0.156 | 1.16 | -0.513 | 0.134 | -0.001 | -0.002 | -0.002 | 0 | 0.22 | 0.28 | 0.15 | 0.21 | 0.19 | 0.32 |
| 1 | 30156975 | rs271344 | -0.143 | 1.347 | -0.915 | 0.231 | 0.025 | 0.061 | 0.061 | 0 | 0.2 | 0.27 | 0.39 | 0.45 | 0.25 | 0.24 |
| 1 | 30157559 | rs387232 | -0.011 | 0.777 | -0.871 | 0.512 | -0.001 | 0 | 0 | 0.055 | 0.55 | 0.49 | 0.73 | 0.53 | 0.59 | 0.56 |
| 1 | 30162007 | rs9729887 | -0.938 | 1.235 | -1.23 | 0.809 | -0.014 | -0.01 | -0.01 | 0.006 | 0.25 | 0.15 | 0.25 | 0.24 | 0.28 | 0.44 |
| 1 | 30162368 | rs271392 | -0.942 | 1.223 | -0.703 | 0.852 | 0.008 | 0.005 | 0.005 | 0 | 0.17 | 0.32 | 0.35 | 0.29 | 0.2 | 0.19 |
| 1 | 30162535 | rs9726412 | 0.587 | 0.677 | -0.746 | 1.061 | -0.007 | 0.006 | 0.006 | 0 | 0.61 | 0.36 | 0.64 | 0.5 | 0.63 | 0.41 |
| 1 | 30163578 | rs9442467 | -0.599 | 1.133 | -1.029 | 1.576 | -0.026 | -0.013 | -0.013 | 0 | 0.05 | 0.01 | 0.2 | 0.21 | 0.1 | 0.18 |
| 1 | 30165211 | rs10753234 | 0.428 | 1.225 | -0.685 | 1.647 | 0.006 | -0.002 | -0.002 | 0 | 0.68 | 0.42 | 0.77 | 0.67 | 0.65 | 0.69 |
| 1 | 30165913 | rs11584888 | 0.286 | 1.187 | -1.692 | 1.407 | -0.056 | -0.062 | -0.062 | 0.243 | 0.23 | 0.03 | 0.32 | 0.25 | 0.24 | 0.22 |
| 1 | 30166174 | rs12408008 | -0.446 | 0.993 | -1.665 | 1.128 | -0.036 | -0.141 | -0.141 | 0.262 | 0.33 | 0.1 | 0.57 | 0.51 | 0.33 | 0.46 |
| 1 | 30171995 | rs12137141 | -0.065 | 0.563 | -0.762 | 0.525 | -0.014 | -0.078 | -0.078 | 0 | 0.14 | 0.25 | 0.05 | 0 | 0.08 | 0.1 |
| 1 | 30172044 | rs722497 | 0.324 | 0.455 | -0.795 | 0.48 | 0 | 0.179 | 0.179 | 0 | 0.52 | 0.84 | 0.3 | 0.19 | 0.43 | 0.31 |
| 1 | 30172441 | rs1441837 | -0.36 | 0.379 | -0.364 | 0.558 | 0.087 | 0.302 | 0.302 | 0.019 | 0.05 | 0.43 | 0.06 | 0.14 | 0.03 | 0.08 |
| 1 | 30173101 | rs1441834 | 0.98 | 0.34 | -0.507 | 0.549 | 0.002 | 0.006 | 0.006 | 0 | 0.39 | 0.61 | 0.25 | 0.19 | 0.35 | 0.22 |
| 1 | 30174552 | rs10914187 | 1.177 | 0.375 | -0.4 | 0.563 | -0.004 | -0.019 | -0.019 | 0.278 | 0.37 | 0.56 | 0.25 | 0.19 | 0.35 | 0.21 |
| 1 | 30175592 | rs12141268 | 0.356 | 0.275 | -0.199 | 0.609 | 0.012 | -0.06 | -0.06 | 0 | 0.1 | 0.26 | 0.05 | 0 | 0.07 | 0.05 |
| 1 | 30183014 | rs894220 | 0.492 | -0.081 | -0.286 | 0.4 | 0.007 | 0.025 | 0.025 | 0 | 0.46 | 0.58 | 0.4 | 0.28 | 0.42 | 0.36 |
| 1 | 30184426 | rs4259710 | 0.133 | -0.157 | -0.282 | 0.754 | 0.029 | -0.003 | -0.003 | 0 | 0.5 | 0.82 | 0.43 | 0.44 | 0.43 | 0.39 |
| 1 | 30185088 | rs894216 | -0.104 | -0.306 | 0.101 | 0.97 | -0.012 | 0.028 | 0.028 | 0 | 0.37 | 0.48 | 0.49 | 0.53 | 0.32 | 0.43 |
| 1 | 30185467 | rs12118066 | -1.26 | -0.291 | 0.336 | 0.987 | 0.027 | 0.019 | 0.019 | 0.367 | 0.05 | 0.3 | 0.03 | 0 | 0.01 | 0.01 |
| 1 | 30185662 | rs1866967 | 0.458 | -0.388 | 0.175 | 0.993 | 0.024 | 0.019 | 0.019 | 0 | 0.51 | 0.82 | 0.37 | 0.39 | 0.43 | 0.4 |
| 1 | 30186956 | rs4949268 | 1.204 | -0.355 | 0.182 | 0.726 | -0.001 | -0.003 | -0.003 | 0.362 | 0.4 | 0.4 | 0.31 | 0.24 | 0.36 | 0.28 |
| 1 | 30191202 | rs7534377 | -0.56 | -0.085 | 0.194 | 0.842 | -0.005 | -0.005 | -0.005 | 0 | 0.89 | 0.91 | 0.88 | 0.76 | 0.76 | 0.87 |
| 1 | 30198580 | rs2376723 | 0.043 | 0.099 | -0.048 | 0.879 | -0.022 | -0.018 | -0.018 | 0 | 0.85 | 0.9 | 0.81 | 0.76 | 0.74 | 0.82 |
| 1 | 30204943 | rs12125419 | -0.169 | -0.706 | 0.678 | 1.229 | -0.007 | 0.032 | 0.032 | 0 | 0.07 | 0.06 | 0.02 | 0.24 | 0.01 | 0.22 |
| 1 | 30206771 | rs1441831 | 0.738 | -1.329 | 0.367 | 0.858 | -0.009 | 0.049 | 0.049 | 0 | 0.38 | 0.25 | 0.22 | 0.22 | 0.39 | 0.22 |
| 1 | 30211197 | rs10914266 | 0.021 | -1.334 | 0.392 | 0.969 | -0.001 | -0.009 | -0.009 | 0 | 0.53 | 0.52 | 0.34 | 0.38 | 0.56 | 0.34 |
| 1 | 30213676 | rs7411511 | 0.807 | -1.363 | 0.285 | 1.129 | 0 | -0.007 | -0.007 | 0 | 0.4 | 0.44 | 0.23 | 0.23 | 0.39 | 0.22 |
| 1 | 30213840 | rs10914271 | -0.89 | -1.427 | 0.319 | 1.154 | -0.026 | -0.003 | -0.003 | 0 | 0.62 | 0.77 | 0.38 | 0.63 | 0.59 | 0.56 |
| 1 | 30214062 | rs1441839 | 0.781 | -1.365 | 0.264 | 1.037 | -0.001 | 0.013 | 0.013 | 0 | 0.39 | 0.35 | 0.22 | 0.23 | 0.39 | 0.22 |
| 1 | 30217585 | rs4082047 | 1.191 | -1.326 | -0.075 | 1.008 | -0.109 | 0.037 | 0.037 | 0 | 0.32 | 0.06 | 0.53 | 0.23 | 0.39 | 0.39 |
| 1 | 30219493 | rs1348198 | 0.026 | -1.238 | 0.181 | 0.924 | -0.001 | -0.008 | -0.008 | 0 | 0.53 | 0.52 | 0.34 | 0.39 | 0.56 | 0.34 |
| 1 | 30221061 | rs10914307 | 0.675 | -1.23 | 0.165 | 0.908 | 0 | 0.008 | 0.008 | 0 | 0.41 | 0.4 | 0.23 | 0.24 | 0.4 | 0.22 |
| 1 | 30222525 | rs920600 | -0.917 | -1.21 | -0.011 | 1.013 | -0.007 | 0.002 | 0.002 | 0 | 0.62 | 0.67 | 0.36 | 0.64 | 0.58 | 0.57 |
| 1 | 30225125 | rs17262734 | 0.897 | -0.954 | -0.555 | 0.747 | 0.001 | -0.005 | -0.005 | 0 | 0.13 | 0.16 | 0.11 | 0.15 | 0.17 | 0.12 |
| 1 | 30225137 | rs12566608 | 0.901 | -1.03 | -0.588 | 0.66 | -0.006 | 0.093 | 0.093 | 0 | 0.35 | 0.21 | 0.2 | 0.07 | 0.34 | 0.14 |
| 1 | 30226168 | rs16832876 | 0.018 | -0.53 | -1.04 | 1.107 | 0.001 | 0.011 | 0.011 | 0 | 0.17 | 0.21 | 0.13 | 0.32 | 0.22 | 0.2 |
| 1 | 30226532 | rs17341169 | 1.186 | -0.489 | -1.541 | 0.611 | -0.083 | 0.072 | 0.072 | 0 | 0.19 | 0.01 | 0.1 | 0.06 | 0.24 | 0.11 |
| 1 | 30232687 | rs10493041 | 0.499 | -0.609 | -0.959 | 1.065 | -0.001 | -0.001 | -0.001 | 0 | 0.14 | 0.12 | 0.1 | 0.14 | 0.16 | 0.12 |
| 1 | 30235650 | rs9727539 | -0.891 | -0.922 | -0.731 | 1.227 | -0.036 | 0.071 | 0.071 | 0.026 | 0.24 | 0.39 | 0.22 | 0.4 | 0.19 | 0.33 |
| 1 | 30238984 | rs6684453 | -0.884 | -0.92 | -0.73 | 1.228 | -0.036 | 0.071 | 0.071 | 0.015 | 0.24 | 0.39 | 0.22 | 0.39 | 0.19 | 0.36 |
| 1 | 30244847 | rs12145142 | -0.016 | -0.798 | -0.905 | 1.128 | -0.037 | 0.041 | 0.041 | 0 | 0.08 | 0.18 | 0.03 | 0.13 | 0.02 | 0.23 |
| 1 | 30245413 | rs12145316 | 0.54 | -0.653 | -1.695 | 0.396 | -0.137 | 0.043 | 0.043 | 0 | 0.31 | 0.06 | 0.49 | 0.23 | 0.4 | 0.39 |
| 1 | 30246262 | rs2121954 | 0.755 | -0.762 | -1.514 | 0.574 | -0.109 | 0.087 | 0.087 | 0 | 0.37 | 0.06 | 0.41 | 0.24 | 0.44 | 0.44 |
| 1 | 30255396 | rs10753253 | -0.571 | -1.263 | -0.269 | 0.729 | -0.053 | 0.08 | 0.08 | 0.249 | 0.25 | 0.53 | 0.21 | 0.51 | 0.15 | 0.32 |
| 1 | 30257664 | rs17262748 | 0.967 | -1.166 | -0.384 | 0.143 | 0.019 | -0.056 | -0.056 | 0 | 0.2 | 0.31 | 0.13 | 0.06 | 0.2 | 0.12 |
| 1 | 30258944 | rs6425840 | 1.015 | -1.113 | -1.08 | -0.017 | -0.115 | 0.139 | 0.139 | 0 | 0.43 | 0.06 | 0.45 | 0.25 | 0.52 | 0.46 |
| 1 | 30259725 | rs1868302 | -1.252 | -0.882 | -0.736 | 0.002 | 0 | 0.014 | 0.014 | 0 | 0.89 | 0.93 | 0.89 | 0.97 | 0.87 | 0.91 |
| 1 | 30261280 | rs2376727 | -0.326 | -0.958 | -0.492 | 0.257 | -0.002 | 0.014 | 0.014 | 0 | 0.37 | 0.42 | 0.23 | 0.4 | 0.32 | 0.38 |
| 1 | 30264883 | rs12117184 | 0.672 | -0.634 | -0.121 | -0.045 | -0.002 | -0.005 | -0.005 | 0 | 0.12 | 0.13 | 0.07 | 0.14 | 0.11 | 0.08 |
| 1 | 30274404 | rs16832918 | -0.035 | -0.503 | -0.454 | -0.201 | 0.006 | -0.08 | -0.08 | 0.446 | 0.25 | 0.39 | 0.15 | 0.05 | 0.22 | 0.12 |
| 1 | 30277330 | rs4949476 | 0.892 | -0.241 | -0.995 | -0.271 | -0.043 | 0.057 | 0.057 | 0 | 0.3 | 0.12 | 0.37 | 0.2 | 0.41 | 0.27 |

| | | | | | | | | | | | | | | | | |
|---|---|---|---|---|---|---|---|---|---|---|---|---|---|---|---|---|
| 1 | 30277668 | rs4949477 | 0.284 | -0.465 | -0.701 | -0.582 | 0 | 0.036 | 0.036 | 0 | 0.43 | 0.35 | 0.39 | 0.18 | 0.55 | 0.35 |
| 1 | 30277785 | rs10914474 | -0.342 | -0.427 | -0.715 | -0.12 | -0.007 | 0.014 | 0.014 | 0.293 | 0.73 | 0.8 | 0.73 | 0.77 | 0.74 | 0.73 |
| 1 | 30278193 | rs1868305 | -0.021 | -0.379 | -0.587 | 0.362 | 0.002 | -0.012 | -0.012 | 0 | 0.67 | 0.51 | 0.73 | 0.75 | 0.71 | 0.64 |
| 1 | 30278550 | rs10914475 | 0.485 | -0.399 | -0.569 | 0.497 | -0.003 | 0.003 | 0.003 | 0 | 0.16 | 0.12 | 0.23 | 0.16 | 0.15 | 0.19 |
| 1 | 30280472 | rs13374873 | 0.305 | -0.692 | -0.477 | 0.701 | 0.005 | 0.015 | 0.015 | 0.02 | 0.6 | 0.53 | 0.61 | 0.53 | 0.61 | 0.65 |
| 1 | 30284090 | rs12030673 | 0.145 | -1.027 | -0.663 | 0.505 | -0.006 | 0.031 | 0.031 | 0 | 0.45 | 0.32 | 0.56 | 0.68 | 0.56 | 0.47 |
| 1 | 30284327 | rs7524204 | 1.34 | -1.127 | -0.79 | 0.542 | -0.084 | 0.061 | 0.061 | 0 | 0.29 | 0.09 | 0.34 | 0.2 | 0.41 | 0.27 |
| 1 | 30284674 | rs6678054 | 0.227 | -1.566 | -0.292 | 0.489 | -0.008 | 0.101 | 0.101 | 0.179 | 0.91 | 0.58 | 0.86 | 0.87 | 0.98 | 0.82 |
| 1 | 30285005 | rs4949483 | -0.072 | -1.636 | -0.28 | 0.449 | -0.016 | -0.027 | -0.027 | 0.478 | 0.94 | 0.92 | 0.97 | 1 | 0.96 | 0.94 |
| 1 | 30285530 | rs10798894 | 1.032 | -1.589 | -0.318 | 0.742 | -0.016 | -0.03 | -0.03 | 0 | 0.11 | 0.05 | 0.07 | 0.35 | 0.13 | 0.14 |
| 1 | 30285848 | rs4949485 | 1.221 | -1.572 | -0.259 | 0.569 | -0.034 | 0.059 | 0.059 | 0 | 0.32 | 0.16 | 0.35 | 0.2 | 0.43 | 0.29 |
| 1 | 30288207 | rs2166884 | 1.003 | -1.847 | -0.744 | 0.324 | -0.093 | 0.086 | 0.086 | 0 | 0.24 | 0.04 | 0.29 | 0.1 | 0.28 | 0.2 |
| 1 | 30297793 | rs993245 | 0.376 | -1.602 | -0.538 | 0.636 | -0.105 | 0.175 | 0.175 | 0.013 | 0.45 | 0.05 | 0.37 | 0.18 | 0.42 | 0.35 |
| 1 | 30297930 | rs10157715 | 0.453 | -1.419 | -0.289 | 0.835 | 0.001 | 0.079 | 0.079 | 0 | 0.12 | 0.17 | 0.11 | 0.48 | 0.13 | 0.18 |
| 1 | 30301308 | rs12748731 | 0.543 | -1.389 | -0.258 | 0.513 | -0.012 | 0.086 | 0.086 | 0 | 0.92 | 0.66 | 0.87 | 0.87 | 0.98 | 0.82 |
| 1 | 30307366 | rs17341224 | 0.129 | -1.393 | -0.423 | 0.39 | 0.007 | 0.092 | 0.092 | 0 | 0.48 | 0.34 | 0.39 | 0.16 | 0.41 | 0.39 |
| 1 | 30307903 | rs6426048 | -0.013 | -1.236 | -0.362 | 0.488 | 0.025 | 0.137 | 0.137 | 1.496 | 0.06 | 0.32 | 0.12 | 0.13 | 0.01 | 0.14 |
| 1 | 30307979 | rs1979088 | 0.4 | -0.963 | -1.078 | 0.659 | -0.015 | 0.026 | 0.026 | 0.108 | 0.38 | 0.29 | 0.44 | 0.27 | 0.48 | 0.31 |
| 1 | 30308744 | rs749663 | 1.065 | -0.644 | -1.682 | 0.66 | -0.081 | 0.103 | 0.103 | 0 | 0.29 | 0.1 | 0.35 | 0.15 | 0.41 | 0.29 |
| 1 | 30308981 | rs749665 | 1.108 | -0.484 | -1.246 | 1.176 | -0.01 | 0.069 | 0.069 | 0.18 | 0.06 | 0.02 | 0.04 | 0.45 | 0.09 | 0.12 |
| 1 | 30310092 | rs12045961 | 0.654 | -0.56 | -1.375 | 1.085 | -0.005 | 0.008 | 0.008 | 1.057 | 0.11 | 0.07 | 0.06 | 0.41 | 0.1 | 0.12 |
| 1 | 30310877 | rs895785 | 0.643 | -0.592 | -1.444 | 1.068 | -0.021 | -0.054 | -0.054 | 0.551 | 0.11 | 0.03 | 0.06 | 0.42 | 0.1 | 0.16 |
| 1 | 30311124 | rs895786 | -0.891 | -1.446 | -2.066 | 0.453 | 0.058 | -0.012 | -0.012 | 0 | 0.43 | 0.14 | 0.18 | 0.52 | 0.31 | 0.37 |
| 1 | 30311769 | rs12046763 | -0.131 | -1.527 | -2.132 | 0.4 | -0.002 | -0.074 | -0.074 | 0.005 | 0.19 | 0.1 | 0.1 | 0.43 | 0.17 | 0.16 |
| 1 | 30314005 | rs12145293 | 1.239 | -1.47 | -2.079 | 0.304 | 0.005 | 0.024 | 0.024 | 0 | 0.08 | 0.04 | 0.04 | 0.01 | 0.07 | 0.01 |
| 1 | 30315354 | rs12142061 | 1.431 | -1.283 | -2.405 | 0.06 | -0.051 | 0.016 | 0.016 | 0 | 0.16 | 0.07 | 0.25 | 0.13 | 0.2 | 0.24 |
| 1 | 30317648 | rs10798896 | -0.894 | -1.277 | -2.278 | 0.129 | 0.058 | -0.016 | -0.016 | 0 | 0.44 | 0.14 | 0.18 | 0.52 | 0.31 | 0.37 |
| 1 | 30317917 | rs12120617 | -0.086 | -1.339 | -2.311 | 0.118 | 0.06 | 0.103 | 0.103 | 0 | 0.32 | 0.08 | 0.12 | 0.1 | 0.21 | 0.22 |
| 1 | 30318753 | rs10798897 | -0.047 | -1.661 | -2.394 | -0.064 | 0.017 | 0.023 | 0.023 | 0 | 0.59 | 0.17 | 0.47 | 0.62 | 0.66 | 0.67 |
| 1 | 30319889 | rs921197 | 0.649 | -1.66 | -2.403 | -0.062 | -0.023 | 0.174 | 0.174 | 0.475 | 0.48 | 0.14 | 0.42 | 0.2 | 0.55 | 0.52 |
| 1 | 30322046 | rs7512834 | -0.571 | -1.39 | -2.107 | -0.037 | 0.007 | -0.005 | -0.005 | 0.084 | 0.68 | 0.26 | 0.52 | 0.62 | 0.72 | 0.68 |
| 1 | 30323808 | rs6663901 | 1.098 | -0.618 | -1.558 | -0.069 | -0.068 | 0.075 | 0.075 | 0.178 | 0.25 | 0.12 | 0.34 | 0.1 | 0.41 | 0.31 |
| 1 | 30324462 | rs10798898 | 0.115 | -0.575 | -1.506 | 0.44 | 0.003 | -0.014 | -0.014 | 0.699 | 0.22 | 0.21 | 0.13 | 0.34 | 0.17 | 0.2 |
| 1 | 30328390 | rs6425687 | 0.622 | -0.739 | -1.223 | -0.329 | 0.004 | -0.035 | -0.035 | 1.205 | 0.28 | 0.3 | 0.32 | 0.1 | 0.37 | 0.25 |
| 1 | 30330967 | rs6669972 | 0.658 | -1.043 | -1.506 | -0.048 | -0.045 | -0.093 | -0.093 | 0.005 | 0.23 | 0.05 | 0.17 | 0.39 | 0.25 | 0.19 |
| 1 | 30332121 | rs10914549 | 0.142 | -1.334 | -0.723 | -0.337 | -0.015 | 0.066 | 0.066 | 2.549 | 0.08 | 0.31 | 0.23 | 0.11 | 0.02 | 0.02 |
| 1 | 30344788 | rs10914556 | 0.239 | -1.631 | -1.532 | -0.382 | -0.042 | -0.084 | -0.084 | 3.066 | 0.41 | 0.09 | 0.22 | 0.55 | 0.36 | 0.34 |
| 1 | 30346741 | rs7513406 | 0.566 | -1.716 | -1.542 | -0.619 | -0.04 | 0.081 | 0.081 | 0 | 0.41 | 0.19 | 0.43 | 0.27 | 0.54 | 0.56 |
| 1 | 30348274 | rs6669732 | 0.537 | -1.716 | -1.476 | -0.62 | -0.025 | 0.067 | 0.067 | 0 | 0.41 | 0.25 | 0.44 | 0.27 | 0.54 | 0.56 |
| 1 | 30348925 | rs12134222 | 1.584 | -1.612 | -1.622 | -0.601 | -0.04 | -0.054 | -0.054 | 1.595 | 0.14 | 0.04 | 0.14 | 0.17 | 0.17 | 0.31 |
| 1 | 30349276 | rs1448337 | 1.594 | -1.617 | -1.624 | -0.602 | -0.04 | -0.054 | -0.054 | 1.292 | 0.14 | 0.04 | 0.14 | 0.17 | 0.17 | 0.31 |
| 1 | 30351032 | rs12407932 | 0.58 | -1.713 | -1.376 | -0.727 | -0.055 | -0.025 | -0.025 | 0.879 | 0.16 | 0.44 | 0.28 | 0.12 | 0.1 | 0.06 |
| 1 | 30351529 | rs4949509 | 1.696 | -1.197 | -1.556 | -0.864 | -0.062 | 0.083 | 0.083 | 0 | 0.23 | 0.06 | 0.28 | 0.11 | 0.33 | 0.22 |
| 1 | 30353687 | rs10458370 | 0.103 | -1.624 | -1.498 | -0.737 | 0.001 | -0.002 | -0.002 | 2.814 | 0.4 | 0.38 | 0.45 | 0.6 | 0.36 | 0.29 |
| 1 | 30354850 | rs12039019 | 0.044 | -1.12 | -1.494 | -0.644 | 0.031 | 0.028 | 0.028 | 0 | 0.64 | 0.31 | 0.7 | 0.83 | 0.76 | 0.77 |
| 1 | 30357612 | rs2053806 | 0.093 | -0.013 | -1.343 | -0.776 | 0.006 | -0.005 | -0.005 | 0 | 0.36 | 0.46 | 0.41 | 0.34 | 0.42 | 0.43 |
| 1 | 30361690 | rs11582334 | 1.214 | 0.309 | -1.553 | -1.217 | -0.035 | 0.046 | 0.046 | 0 | 0.08 | 0 | 0.06 | 0.01 | 0.09 | 0.09 |
| 1 | 30362080 | rs2377151 | 1.651 | 0.163 | -1.584 | -1.298 | 0.006 | 0.022 | 0.022 | 0 | 0.07 | 0.02 | 0.05 | 0 | 0.07 | 0.01 |
| 1 | 30364629 | rs12407360 | 1.734 | 0.067 | -1.833 | -1.286 | -0.003 | -0.085 | -0.085 | 0 | 0.21 | 0.08 | 0.3 | 0.56 | 0.19 | 0.22 |
| 1 | 30365461 | rs12082727 | 0.632 | 0.127 | -1.848 | -1.56 | -0.015 | 0.017 | 0.017 | 0 | 0.2 | 0.14 | 0.17 | 0.1 | 0.28 | 0.16 |
| 1 | 30365605 | rs12083826 | 0.694 | 0.122 | -1.724 | -1.562 | -0.005 | 0.005 | 0.005 | 0 | 0.2 | 0.17 | 0.18 | 0.1 | 0.28 | 0.16 |
| 1 | 30365857 | rs7540340 | -0.268 | 0.209 | -0.732 | -1.32 | 0.002 | -0.005 | -0.005 | 0 | 0.3 | 0.43 | 0.38 | 0.33 | 0.3 | 0.46 |
| 1 | 30367970 | rs11587046 | -0.285 | 0.351 | -0.365 | -1.506 | -0.004 | -0.028 | -0.028 | 0 | 0.18 | 0.29 | 0.27 | 0.13 | 0.15 | 0.17 |
| 1 | 30368883 | rs895796 | -1.414 | 0.377 | -0.337 | -1.59 | 0.002 | -0.047 | -0.047 | 0 | 0.82 | 0.77 | 0.92 | 0.99 | 0.85 | 0.87 |
| 1 | 30369220 | rs752685 | 0.313 | 0.381 | -0.404 | -1.524 | -0.003 | -0.032 | -0.032 | 0 | 0.64 | 0.46 | 0.67 | 0.83 | 0.7 | 0.67 |
| 1 | 30369659 | rs10753261 | 1.029 | 0.413 | -0.777 | -1.342 | -0.002 | 0.063 | 0.063 | 0 | 0.41 | 0.19 | 0.39 | 0.72 | 0.39 | 0.5 |
| 1 | 30371100 | rs10798913 | -0.381 | 0.558 | -0.275 | -1.141 | 0.004 | -0.075 | -0.075 | 0 | 0.77 | 0.7 | 0.89 | 0.98 | 0.8 | 0.85 |
| 1 | 30374868 | rs4949230 | -0.475 | 0.373 | 0.168 | -0.659 | 0 | -0.018 | -0.018 | 1.086 | 0.46 | 0.5 | 0.53 | 0.25 | 0.5 | 0.38 |
| 1 | 30380136 | rs873894 | -0.447 | 0.294 | -0.135 | -0.679 | -0.023 | 0.076 | 0.076 | 0 | 0.43 | 0.19 | 0.51 | 0.24 | 0.49 | 0.38 |
| 1 | 30381489 | rs10914615 | 1.008 | 0.087 | -0.057 | -0.511 | -0.02 | -0.045 | -0.045 | 0 | 0.39 | 0.22 | 0.24 | 0.51 | 0.37 | 0.31 |
| 1 | 30382007 | rs2084436 | -0.935 | 0.235 | -0.191 | -0.434 | -0.011 | 0.028 | 0.028 | 1.073 | 0.85 | 0.68 | 0.78 | 0.76 | 0.86 | 0.69 |
| 1 | 30383408 | rs1448343 | 0.712 | 0.429 | -0.457 | -0.642 | -0.028 | 0.022 | 0.022 | 0 | 0.11 | 0.04 | 0.11 | 0.04 | 0.2 | 0.06 |
| 1 | 30392970 | rs12401689 | 0.907 | 0.123 | -0.462 | -0.596 | -0.009 | 0.046 | 0.046 | 0 | 0.24 | 0.02 | 0.32 | 0.12 | 0.23 | 0.24 |
| 1 | 30393790 | rs268955 | -0.809 | 0.062 | -0.185 | -0.332 | -0.011 | 0.025 | 0.025 | 1.395 | 0.85 | 0.69 | 0.78 | 0.75 | 0.87 | 0.69 |
| 1 | 30394090 | rs12063790 | 0.132 | 0.195 | -0.314 | -0.405 | 0.016 | 0.044 | 0.044 | 0 | 0.34 | 0.15 | 0.42 | 0.2 | 0.31 | 0.33 |
| 1 | 30398797 | rs155637 | -0.699 | 0.311 | -0.175 | -0.656 | -0.012 | 0.04 | 0.04 | 0.05 | 0.84 | 0.66 | 0.75 | 0.71 | 0.86 | 0.66 |
| 1 | 30399109 | rs10798921 | 0.494 | 0.254 | -0.141 | -0.783 | 0.012 | 0.046 | 0.046 | 0 | 0.31 | 0.19 | 0.38 | 0.07 | 0.29 | 0.28 |
| 1 | 30399154 | rs198271 | 0.811 | 0.157 | 0.033 | -0.526 | -0.001 | -0.001 | -0.001 | 0 | 0.32 | 0.23 | 0.19 | 0.59 | 0.31 | 0.22 |
| 1 | 30399634 | rs155635 | -0.463 | 0.007 | 0.044 | -0.361 | -0.013 | 0.045 | 0.045 | 0.069 | 0.84 | 0.66 | 0.74 | 0.7 | 0.86 | 0.66 |

| | | | | | | | | | | | | | | | | |
|---|---|---|---|---|---|---|---|---|---|---|---|---|---|---|---|---|
| 1 | 30401563 | rs12074244 | -0.389 | 0.252 | -0.231 | -0.96 | 0.005 | 0.03 | 0.03 | 0 | 0.52 | 0.42 | 0.48 | 0.07 | 0.55 | 0.43 |
| 1 | 30403470 | rs4478859 | 0.625 | -0.125 | -0.116 | 0.001 | -0.018 | -0.007 | -0.007 | 0 | 0.7 | 0.49 | 0.7 | 0.77 | 0.73 | 0.75 |
| 1 | 30405460 | rs448357 | 0.727 | -0.313 | -0.12 | -0.772 | -0.004 | -0.038 | -0.038 | 0 | 0.6 | 0.54 | 0.74 | 0.86 | 0.63 | 0.58 |
| 1 | 30407575 | rs174838 | 0.377 | -0.759 | 0.065 | -1.103 | -0.008 | -0.028 | -0.028 | 0 | 0.73 | 0.73 | 0.82 | 0.9 | 0.79 | 0.67 |
| 1 | 30410570 | rs4949233 | 0.604 | -1.094 | 0.484 | -0.905 | -0.021 | -0.024 | -0.024 | 0 | 0.72 | 0.66 | 0.78 | 0.81 | 0.8 | 0.59 |
| 1 | 30413903 | rs10798927 | 1.717 | -1.319 | 0.294 | -0.9 | 0.044 | 0.095 | 0.095 | 0 | 0.51 | 0.16 | 0.58 | 0.7 | 0.64 | 0.43 |
| 1 | 30414182 | rs16833076 | -0.619 | -1.328 | 0.556 | -0.894 | 0.009 | -0.05 | -0.05 | 0 | 0.19 | 0.4 | 0.18 | 0.08 | 0.17 | 0.14 |
| 1 | 30417143 | rs10914662 | 1.659 | -1.334 | 0.54 | -0.897 | 0.051 | 0.093 | 0.093 | 0 | 0.52 | 0.24 | 0.59 | 0.7 | 0.64 | 0.43 |
| 1 | 30417560 | rs10798928 | 0.504 | -1.545 | 0.448 | -1.023 | -0.02 | -0.016 | -0.016 | 0.267 | 0.68 | 0.66 | 0.76 | 0.78 | 0.79 | 0.56 |
| 1 | 30419370 | rs11583644 | 0.526 | -1.387 | 0.628 | -0.928 | -0.04 | -0.029 | -0.029 | 0 | 0.7 | 0.59 | 0.76 | 0.77 | 0.8 | 0.57 |
| 1 | 30419456 | rs11583665 | 1.669 | -1.384 | 0.587 | -0.93 | 0.052 | 0.093 | 0.093 | 0 | 0.52 | 0.24 | 0.59 | 0.7 | 0.64 | 0.43 |
| 1 | 30422596 | rs4949238 | -0.837 | -1.243 | 0.777 | -0.819 | 0.008 | -0.041 | -0.041 | 0.41 | 0.18 | 0.36 | 0.17 | 0.08 | 0.16 | 0.14 |
| 1 | 30426014 | rs12407717 | -0.182 | -1.032 | 0.381 | -0.729 | 0.006 | 0.001 | 0.001 | 0.049 | 0.08 | 0.09 | 0.07 | 0.06 | 0.13 | 0.03 |
| 1 | 30430366 | rs267700 | 1.257 | -1.582 | 0.318 | -1.006 | 0.001 | 0 | 0 | 0 | 0.58 | 0.47 | 0.72 | 0.78 | 0.7 | 0.49 |
| 1 | 30431560 | rs1009080 | 1.455 | -1.588 | -0.002 | -0.973 | 0.071 | 0.101 | 0.101 | 0.097 | 0.54 | 0.11 | 0.51 | 0.7 | 0.69 | 0.43 |
| 1 | 30437268 | rs6694545 | 1.458 | -1.419 | 0.224 | -0.847 | 0.041 | 0.029 | 0.029 | 0.087 | 0.64 | 0.12 | 0.62 | 0.72 | 0.76 | 0.43 |
| 1 | 30439760 | rs12401298 | 0.171 | -1.304 | 0.238 | -0.845 | -0.027 | -0.007 | -0.007 | 0 | 0.07 | 0.03 | 0.06 | 0.06 | 0.11 | 0.03 |
| 1 | 30444979 | rs17263039 | -0.044 | -1.603 | 0.105 | -1.164 | -0.016 | 0.034 | 0.034 | 0 | 0.18 | 0.02 | 0.14 | 0.08 | 0.13 | 0.27 |
| 1 | 30445050 | rs10753284 | 1.265 | -1.201 | 0.451 | -1.042 | 0.039 | 0.072 | 0.072 | 0 | 0.58 | 0.18 | 0.54 | 0.77 | 0.7 | 0.42 |
| 1 | 30445631 | rs267708 | 1.128 | -1.187 | 0.416 | -1.036 | -0.012 | -0.033 | -0.033 | 1.277 | 0.61 | 0.51 | 0.68 | 0.82 | 0.71 | 0.47 |
| 1 | 30459412 | rs904300 | 1.116 | -1.12 | -0.216 | -1.004 | 0.029 | 0.067 | 0.067 | 0.006 | 0.59 | 0.14 | 0.54 | 0.77 | 0.7 | 0.46 |
| 1 | 30459797 | rs4949241 | 0.279 | -0.552 | -0.078 | -0.791 | -0.034 | -0.014 | -0.014 | 0 | 0.06 | 0.01 | 0.06 | 0.05 | 0.11 | 0.03 |
| 1 | 30472960 | rs409226 | 0.605 | -1.16 | -0.135 | -1.27 | -0.001 | -0.011 | -0.011 | 0 | 0.52 | 0.53 | 0.64 | 0.83 | 0.63 | 0.56 |
| 1 | 30478996 | rs452132 | 0.637 | -1.423 | -0.023 | -1.061 | 0.009 | 0.014 | 0.014 | 0 | 0.52 | 0.46 | 0.68 | 0.79 | 0.66 | 0.54 |
| 1 | 30485510 | rs373316 | 0.191 | -1.602 | -0.021 | -1.171 | -0.016 | -0.034 | -0.034 | 0 | 0.6 | 0.49 | 0.66 | 0.78 | 0.72 | 0.48 |
| 1 | 30487342 | rs453149 | 0.194 | -1.579 | 0.067 | -1.175 | -0.011 | -0.026 | -0.026 | 0 | 0.6 | 0.43 | 0.64 | 0.79 | 0.72 | 0.48 |
| 1 | 30494591 | rs1603724 | 1.257 | -0.82 | -0.743 | -1.643 | -0.162 | -0.072 | -0.072 | 0 | 0.27 | 0.03 | 0.42 | 0.75 | 0.43 | 0.3 |
| 1 | 30496615 | rs10914821 | 1.298 | -0.83 | -0.553 | -1.641 | -0.047 | 0.001 | 0.001 | 0 | 0.26 | 0.13 | 0.46 | 0.77 | 0.43 | 0.31 |
| 1 | 30496978 | rs267721 | 0.079 | -0.873 | -0.494 | -1.691 | 0.001 | -0.038 | -0.038 | 0 | 0.33 | 0.3 | 0.19 | 0.02 | 0.28 | 0.19 |
| 1 | 30497242 | rs267720 | 0.026 | -0.865 | -0.606 | -1.691 | -0.001 | -0.033 | -0.033 | 0 | 0.34 | 0.35 | 0.22 | 0.04 | 0.28 | 0.19 |
| 1 | 30506995 | rs11588815 | -0.141 | -0.785 | -0.797 | -1.4 | 0.018 | 0.024 | 0.024 | 0 | 0.63 | 0.27 | 0.61 | 0.73 | 0.74 | 0.49 |
| 1 | 30509439 | rs7511843 | 0.308 | -0.766 | -0.587 | -1.337 | 0.028 | 0.05 | 0.05 | 0.393 | 0.55 | 0.28 | 0.57 | 0.71 | 0.68 | 0.43 |
| 1 | 30510262 | rs904304 | 0.205 | -0.762 | -0.61 | -1.336 | 0.003 | 0.005 | 0.005 | 0.219 | 0.56 | 0.43 | 0.58 | 0.71 | 0.68 | 0.43 |
| 1 | 30513725 | rs12145453 | 0.735 | -0.638 | -0.822 | -1.52 | 0.031 | 0.133 | 0.133 | 0 | 0.28 | 0.06 | 0.13 | 0 | 0.24 | 0.15 |
| 1 | 30516776 | rs10753300 | 0.451 | -0.352 | -0.359 | -0.402 | -0.006 | 0.002 | 0.002 | 0.024 | 0.57 | 0.47 | 0.58 | 0.59 | 0.68 | 0.41 |
| 1 | 30519784 | rs924772 | -0.487 | -0.176 | -0.309 | -0.602 | -0.022 | -0.044 | -0.044 | 20.855 | 0.31 | 0.55 | 0.22 | 0.14 | 0.21 | 0.35 |
| 1 | 30521571 | rs370325 | -0.443 | 0.299 | -0.068 | -0.486 | 0.003 | -0.014 | -0.014 | 1.299 | 0.2 | 0.45 | 0.2 | 0.16 | 0.17 | 0.31 |
| 1 | 30522300 | rs1391489 | -0.437 | 0.296 | -0.07 | -0.395 | 0.004 | -0.011 | -0.011 | 0.012 | 0.2 | 0.45 | 0.18 | 0.16 | 0.17 | 0.31 |
| 1 | 30523923 | rs1503772 | 0.579 | 0.777 | -0.491 | 0.052 | -0.037 | -0.017 | -0.017 | 0.814 | 0.54 | 0.17 | 0.44 | 0.55 | 0.53 | 0.39 |
| 1 | 30524419 | rs3131419 | -0.675 | 0.76 | -0.35 | 0.109 | 0.029 | 0.018 | 0.018 | 0 | 0.76 | 0.63 | 0.62 | 0.72 | 0.7 | 0.71 |
| 1 | 30525355 | rs437441 | -0.707 | 0.937 | -0.643 | -0.033 | 0.009 | 0.001 | 0.001 | 0 | 0.46 | 0.72 | 0.57 | 0.43 | 0.51 | 0.68 |
| 1 | 30528143 | rs372125 | -0.213 | 1.18 | 0.687 | -0.324 | -0.003 | -0.003 | -0.003 | 0.255 | 0.73 | 0.45 | 0.7 | 0.73 | 0.74 | 0.67 |
| 1 | 30529226 | rs12125183 | 0.287 | 1.19 | 0.646 | -0.325 | 0 | -0.003 | -0.003 | 0 | 0.23 | 0.22 | 0.27 | 0.24 | 0.24 | 0.32 |
| 1 | 30529658 | rs1846217 | -0.141 | 1.179 | 0.708 | -0.277 | -0.003 | -0.017 | -0.017 | 1.53 | 0.73 | 0.44 | 0.7 | 0.78 | 0.74 | 0.66 |
| 1 | 30530417 | rs438226 | 0.586 | 0.901 | -0.038 | -0.732 | 0.016 | 0.064 | 0.064 | 0 | 0.28 | 0.2 | 0.18 | 0.1 | 0.23 | 0.17 |
| 1 | 30530885 | rs391479 | -1.683 | 0.831 | -0.324 | -1.232 | 0.023 | -0.096 | -0.096 | 0.001 | 0.76 | 0.6 | 0.82 | 0.97 | 0.7 | 0.91 |
| 1 | 30531572 | rs404169 | 0.098 | 0.83 | -0.325 | -1.227 | 0.001 | 0.024 | 0.024 | 0 | 0.52 | 0.41 | 0.45 | 0.33 | 0.47 | 0.49 |
| 1 | 30532095 | rs1966006 | -1.667 | 0.87 | -0.326 | -1.206 | 0.023 | -0.096 | -0.096 | 0 | 0.76 | 0.6 | 0.82 | 0.97 | 0.7 | 0.91 |
| 1 | 30533883 | rs16833203 | 0.371 | 0.798 | -0.296 | -0.963 | -0.001 | -0.019 | -0.019 | 0 | 0.24 | 0.2 | 0.36 | 0.62 | 0.24 | 0.38 |
| 1 | 30534426 | rs870859 | 0.897 | 0.841 | -0.322 | -0.911 | 0.001 | -0.005 | -0.005 | 0 | 0.24 | 0.23 | 0.28 | 0.25 | 0.24 | 0.37 |
| 1 | 30536711 | rs746758 | -0.381 | 0.689 | -0.337 | -1.092 | -0.003 | 0.051 | 0.051 | 0 | 0.47 | 0.34 | 0.64 | 0.87 | 0.48 | 0.76 |
| 1 | 30538254 | rs12074770 | 0.65 | 0.359 | -0.817 | -0.792 | -0.029 | -0.009 | -0.009 | 0 | 0.22 | 0.12 | 0.25 | 0.23 | 0.26 | 0.4 |
| 1 | 30538346 | rs267658 | 0.957 | 0.099 | -0.714 | -0.818 | 0.02 | 0.037 | 0.037 | 0 | 0.26 | 0.17 | 0.14 | 0.16 | 0.2 | 0.14 |
| 1 | 30540710 | rs267660 | 0.888 | -0.016 | -0.531 | -0.96 | 0.016 | 0.018 | 0.018 | 0.044 | 0.26 | 0.2 | 0.14 | 0.12 | 0.18 | 0.11 |
| 1 | 30543026 | rs1149064 | -0.301 | -0.102 | -0.544 | -1.036 | 0.067 | 0.069 | 0.069 | 0 | 0.49 | 0.75 | 0.42 | 0.38 | 0.42 | 0.54 |
| 1 | 30545515 | rs1228828 | -1.489 | 0.434 | -0.77 | -1.572 | -0.049 | 0.118 | 0.118 | 0 | 0.78 | 0.97 | 0.86 | 1 | 0.75 | 0.93 |
| 1 | 30546997 | rs267667 | -0.02 | 0.594 | -0.182 | -1.437 | -0.005 | -0.023 | -0.023 | 0 | 0.24 | 0.62 | 0.29 | 0.18 | 0.24 | 0.41 |
| 1 | 30548348 | rs267669 | -1.079 | 0.538 | 0.161 | -1.546 | 0.001 | -0.067 | -0.067 | 0 | 0.78 | 0.73 | 0.84 | 1 | 0.75 | 0.93 |
| 1 | 30550982 | rs267674 | -0.365 | 0.464 | -0.243 | -1.591 | -0.012 | -0.129 | -0.129 | 5.737 | 0.73 | 0.43 | 0.82 | 1 | 0.75 | 0.93 |
| 1 | 30554684 | rs369715 | 0.182 | 0.627 | -0.315 | -1.441 | -0.002 | 0.033 | 0.033 | 0.293 | 0.57 | 0.36 | 0.66 | 0.81 | 0.52 | 0.53 |
| 1 | 30562066 | rs6661562 | 1.096 | 0.227 | -0.542 | -1.407 | 0.013 | 0.025 | 0.025 | 0 | 0.28 | 0.2 | 0.15 | 0.13 | 0.21 | 0.13 |
| 1 | 30562263 | rs7530417 | -0.049 | 0.265 | -0.244 | -1.393 | -0.006 | -0.013 | -0.013 | 0.962 | 0.48 | 0.54 | 0.29 | 0.34 | 0.4 | 0.51 |
| 1 | 30566926 | rs16833246 | 0.885 | 0.288 | -0.454 | -1.237 | 0.004 | 0.008 | 0.008 | 1.423 | 0.18 | 0.12 | 0.12 | 0.21 | 0.2 | 0.39 |
| 1 | 30572812 | rs1746272 | 0.341 | 0.654 | -0.44 | -1.017 | -0.002 | -0.002 | -0.002 | 0.015 | 0.27 | 0.27 | 0.48 | 0.62 | 0.28 | 0.37 |
| 1 | 30574158 | rs2076977 | -0.305 | 0.597 | -0.349 | -1.126 | -0.01 | -0.056 | -0.056 | 0 | 0.58 | 0.49 | 0.73 | 0.79 | 0.57 | 0.54 |
| 1 | 30574413 | rs11807362 | 0.994 | 0.655 | -0.507 | -1.069 | 0.004 | 0.019 | 0.019 | 0 | 0.31 | 0.22 | 0.24 | 0.17 | 0.29 | 0.17 |
| 1 | 30576228 | rs6691680 | 0.534 | 0.848 | -0.995 | -0.91 | -0.017 | -0.019 | -0.019 | 0 | 0.26 | 0.18 | 0.48 | 0.59 | 0.28 | 0.37 |
| 1 | 30576573 | rs12742458 | 0.392 | 0.702 | -1.066 | -0.987 | -0.012 | -0.035 | -0.035 | 0 | 0.34 | 0.53 | 0.28 | 0.18 | 0.29 | 0.18 |
| 1 | 30578594 | rs10915027 | 0.608 | 1.107 | -1.988 | -0.551 | -0.056 | -0.005 | -0.005 | 0 | 0.2 | 0.06 | 0.32 | 0.2 | 0.22 | 0.24 |

| | | | | | | | | | | | | | | | | |
|---|---|---|---|---|---|---|---|---|---|---|---|---|---|---|---|---|
| 1 | 30578618 | rs10799077 | 0.053 | 0.904 | -2.236 | -0.545 | 0.003 | -0.022 | -0.022 | 0 | 0.64 | 0.27 | 0.62 | 0.58 | 0.65 | 0.75 |
| 1 | 30578922 | rs10915029 | 0.332 | 0.847 | -2.041 | -1.242 | 0.005 | 0.162 | 0.162 | 0.011 | 0.5 | 0.3 | 0.48 | 0.78 | 0.5 | 0.6 |
| 1 | 30579227 | rs926980 | -1.155 | 0.692 | -1.984 | -1.084 | -0.032 | 0.087 | 0.087 | 0.468 | 0.8 | 0.93 | 0.88 | 1 | 0.79 | 0.92 |
| 1 | 30581567 | rs2376907 | -0.206 | 0.561 | -1.219 | -0.162 | 0.012 | 0.043 | 0.043 | 0 | 0.35 | 0.19 | 0.33 | 0.75 | 0.35 | 0.47 |
| 1 | 30590082 | rs10799079 | -0.166 | 0.588 | -1.346 | -0.268 | 0.018 | 0.075 | 0.075 | 0 | 0.36 | 0.19 | 0.31 | 0.79 | 0.35 | 0.47 |
| 1 | 30600504 | rs7541381 | -1.09 | 0.671 | -0.572 | -0.014 | -0.013 | -0.015 | -0.015 | 0 | 0.15 | 0.7 | 0.21 | 0.09 | 0.11 | 0.06 |
| 1 | 30604682 | rs16833316 | -1.29 | 0.695 | -0.423 | -0.05 | -0.001 | -0.022 | -0.022 | 4.201 | 0.1 | 0.59 | 0.09 | 0.04 | 0.06 | 0.02 |
| 1 | 30605323 | rs10157877 | -1.29 | 0.695 | -0.423 | -0.05 | -0.001 | -0.022 | -0.022 | 0 | 0.1 | 0.59 | 0.09 | 0.04 | 0.06 | 0.02 |
| 1 | 30610536 | rs2376487 | -1.019 | 1.224 | -1.524 | 0.629 | -0.028 | -0.08 | -0.08 | 0 | 0.35 | 0.83 | 0.41 | 0.47 | 0.4 | 0.18 |
| 1 | 30617856 | rs1575072 | -0.447 | 0.906 | -2.015 | 0.364 | -0.065 | -0.03 | -0.03 | 0 | 0.35 | 0.2 | 0.39 | 0.42 | 0.45 | 0.67 |
| 1 | 30620480 | rs2146850 | -1.311 | 0.499 | -1.165 | 0.238 | 0.003 | 0.02 | 0.02 | 0 | 0.47 | 0.57 | 0.58 | 0.74 | 0.54 | 0.46 |
| 1 | 30627619 | rs1321117 | -1.326 | 0.409 | -0.917 | 0.222 | 0.003 | 0.028 | 0.028 | 0 | 0.47 | 0.57 | 0.59 | 0.75 | 0.54 | 0.46 |
| 1 | 30635302 | rs3934202 | 0.056 | 0.437 | -0.496 | 0.402 | -0.004 | 0.01 | 0.01 | 0 | 0.25 | 0.2 | 0.4 | 0.47 | 0.24 | 0.14 |
| 1 | 30636313 | rs10799083 | -1.289 | 0.628 | -0.496 | 0.267 | 0.031 | 0.067 | 0.067 | 0 | 0.44 | 0.67 | 0.63 | 0.71 | 0.41 | 0.55 |
| 1 | 30636813 | rs12142439 | -0.073 | 0.625 | -0.435 | 0.271 | 0.016 | 0.027 | 0.027 | 0.569 | 0.19 | 0.43 | 0.23 | 0.21 | 0.17 | 0.41 |
| 1 | 30637397 | rs6678179 | -0.637 | 0.929 | -0.181 | 0.557 | 0.056 | 0.002 | 0.002 | 0.536 | 0.2 | 0.43 | 0.16 | 0.21 | 0.27 | 0.24 |
| 1 | 30639318 | rs7530575 | 0.43 | 0.892 | -0.238 | 0.556 | 0.015 | -0.026 | -0.026 | 0 | 0.69 | 0.3 | 0.49 | 0.47 | 0.68 | 0.79 |
| 1 | 30641610 | rs6425865 | -1.357 | 0.884 | 0.064 | 0.49 | 0.108 | 0.03 | 0.03 | 1.175 | 0.06 | 0.52 | 0.09 | 0 | 0.06 | 0.01 |
| 1 | 30644096 | rs6701005 | -0.981 | 0.886 | 0.05 | 0.491 | 0.162 | 0.072 | 0.072 | 0.588 | 0.05 | 0.43 | 0.09 | 0 | 0.06 | 0 |
| 1 | 30645412 | rs6687452 | 1.581 | 0.918 | -0.025 | 0.506 | 0.002 | 0.017 | 0.017 | 0 | 0.53 | 0.45 | 0.42 | 0.2 | 0.46 | 0.54 |
| 1 | 30645986 | rs10493043 | -0.165 | 0.834 | -0.109 | 0.528 | -0.015 | -0.006 | -0.006 | 0 | 0.17 | 0.18 | 0.1 | 0.22 | 0.22 | 0.24 |
| 1 | 30654053 | rs16833375 | -1.644 | 1.071 | 0.279 | 1.061 | 0.036 | 0.014 | 0.014 | 0.848 | 0.06 | 0.14 | 0.07 | 0.06 | 0.07 | 0.07 |
| 1 | 30655212 | rs12049424 | 0.643 | 0.568 | 0.119 | 1.402 | 0.037 | -0.003 | -0.003 | 0 | 0.12 | 0.01 | 0.05 | 0.08 | 0.08 | 0.08 |
| 1 | 30655709 | rs4949550 | 0.443 | 0.61 | 0.252 | 1.966 | -0.008 | 0.007 | 0.007 | 0 | 0.76 | 0.81 | 0.66 | 0.72 | 0.77 | 0.86 |
| 1 | 30656664 | rs2377060 | 0.217 | 0.691 | 0.481 | 1.924 | 0.004 | 0.238 | 0.238 | 0 | 0.86 | 0.47 | 0.6 | 0.57 | 0.85 | 0.94 |
| 1 | 30657535 | rs10799084 | -0.47 | 0.627 | 0.402 | 1.58 | -0.003 | -0.008 | -0.008 | 0 | 0.9 | 0.94 | 0.75 | 0.74 | 0.85 | 0.94 |
| 1 | 30660403 | rs7523165 | -1.268 | 0.743 | 0.352 | 0.905 | -0.004 | 0.003 | 0.003 | 1.604 | 0.28 | 0.71 | 0.29 | 0.27 | 0.26 | 0.49 |
| 1 | 30660839 | rs12072928 | -0.455 | 0.721 | 0.361 | 0.83 | 0.016 | -0.003 | -0.003 | 0 | 0.2 | 0.39 | 0.24 | 0.22 | 0.21 | 0.39 |
| 1 | 30663554 | rs10915040 | 0.747 | 0.302 | 0.505 | 1.34 | 0.013 | -0.028 | -0.028 | 0 | 0.67 | 0.64 | 0.5 | 0.23 | 0.57 | 0.58 |
| 1 | 30669469 | rs10799085 | -0.541 | 0.225 | 0.513 | 1.848 | -0.025 | -0.051 | -0.051 | 0.136 | 0.24 | 0.15 | 0.23 | 0.59 | 0.32 | 0.32 |
| 1 | 30675881 | rs4381243 | -0.782 | 0.203 | 0.598 | 1.855 | 0.018 | 0.029 | 0.029 | 0 | 0.31 | 0.42 | 0.27 | 0.62 | 0.36 | 0.41 |
| 1 | 30687128 | rs16833396 | -1.399 | -0.293 | 1.091 | 2.063 | 0 | 0 | 0 | 0 | 0.21 | 0.19 | 0.38 | 0.21 | 0.12 | 0.29 |
| 1 | 30687827 | rs16833405 | -1.626 | -0.294 | 1.295 | 2.041 | -0.001 | -0.004 | -0.004 | 0 | 0.23 | 0.28 | 0.39 | 0.21 | 0.16 | 0.29 |
| 1 | 30691713 | rs13375139 | -2.12 | -0.452 | 1.717 | 2.474 | -0.014 | 0.076 | 0.076 | 0.191 | 0.29 | 0.44 | 0.42 | 0.43 | 0.18 | 0.38 |
| 1 | 30705639 | rs462649 | -0.113 | -0.523 | 1.273 | 2.158 | 0.003 | -0.026 | -0.026 | 0 | 0.07 | 0.02 | 0.04 | 0.1 | 0.06 | 0.04 |
| 1 | 30706530 | rs909623 | -2.534 | -0.545 | 1.994 | 2.572 | -0.007 | 0.071 | 0.071 | 3.916 | 0.22 | 0.62 | 0.36 | 0.4 | 0.16 | 0.3 |
| 1 | 30711813 | rs6661218 | -2.139 | -0.535 | 2.012 | 2.834 | -0.001 | 0.224 | 0.224 | 0 | 0.05 | 0.15 | 0.17 | 0.57 | 0.05 | 0.05 |
| 1 | 30716625 | rs7529543 | -2.346 | -0.537 | 2.048 | 2.833 | 0.002 | 0.263 | 0.263 | 0 | 0.05 | 0.18 | 0.17 | 0.57 | 0.05 | 0.05 |
| 1 | 30718102 | rs12401958 | -2.055 | -0.544 | 2.143 | 2.68 | 0.01 | 0.001 | 0.001 | 4.211 | 0.17 | 0.5 | 0.33 | 0.16 | 0.11 | 0.27 |
| 1 | 30720094 | rs12060888 | -2.052 | -0.563 | 2.132 | 2.671 | 0.009 | 0 | 0 | 0 | 0.17 | 0.48 | 0.33 | 0.17 | 0.11 | 0.27 |
| 1 | 30723490 | rs1998012 | -0.804 | -0.647 | 2.161 | 2.615 | -0.029 | -0.058 | -0.058 | 0 | 0.13 | 0.39 | 0.05 | 0.03 | 0.03 | 0.04 |
| 1 | 30723600 | rs1555042 | -0.915 | -0.589 | 1.896 | 2.645 | -0.007 | -0.017 | -0.017 | 0.169 | 0.09 | 0.04 | 0.26 | 0.17 | 0.06 | 0.21 |
| 1 | 30728368 | rs6425919 | -2.365 | -0.599 | 2.11 | 2.718 | 0.002 | 0.167 | 0.167 | 1.326 | 0.18 | 0.41 | 0.46 | 0.53 | 0.12 | 0.29 |
| 1 | 30729050 | rs11802750 | -2.42 | -0.588 | 1.877 | 2.71 | -0.026 | 0.083 | 0.083 | 0.755 | 0.29 | 0.53 | 0.48 | 0.56 | 0.15 | 0.32 |
| 1 | 30730173 | rs7524879 | -1.717 | -0.694 | 1.43 | 2.83 | -0.003 | 0.016 | 0.016 | 0 | 0.2 | 0.24 | 0.22 | 0.56 | 0.14 | 0.14 |
| 1 | 30731138 | rs10158763 | -0.936 | -0.672 | 1.37 | 2.68 | -0.005 | 0.016 | 0.016 | 0 | 0.16 | 0.23 | 0.2 | 0.35 | 0.09 | 0.11 |
| 1 | 30731688 | rs6673391 | -0.974 | -0.674 | 1.369 | 2.658 | -0.005 | 0.014 | 0.014 | 0 | 0.16 | 0.23 | 0.2 | 0.35 | 0.09 | 0.11 |
| 1 | 30732553 | rs6679932 | -1.676 | -0.719 | 0.907 | 2.012 | -0.011 | 0.013 | 0.013 | 0 | 0.13 | 0.2 | 0.28 | 0.2 | 0.06 | 0.23 |
| 1 | 30734036 | rs4949565 | -2.513 | -0.5 | 0.75 | 2.01 | -0.011 | -0.019 | -0.019 | 0 | 0.35 | 0.58 | 0.52 | 0.77 | 0.25 | 0.39 |
| 1 | 30734163 | rs6676025 | -1.26 | -0.929 | 0.012 | 2.01 | 0.003 | -0.027 | -0.027 | 2.116 | 0.12 | 0.05 | 0.27 | 0.22 | 0.06 | 0.22 |
| 1 | 30737443 | rs1008196 | -1.278 | -0.856 | 0.582 | 1.852 | -0.035 | 0.015 | 0.015 | 0 | 0.21 | 0.4 | 0.45 | 0.75 | 0.1 | 0.26 |
| 1 | 30739134 | rs2142716 | -1.123 | -1.05 | 0.92 | 1.36 | 0.038 | -0.025 | -0.025 | 0.16 | 0.09 | 0.34 | 0.07 | 0.02 | 0.05 | 0.14 |
| 1 | 30740493 | rs12035857 | -1.236 | -1.141 | 0.773 | 2.089 | -0.086 | 0.2 | 0.2 | 1.022 | 0.19 | 0.46 | 0.43 | 0.56 | 0.06 | 0.21 |
| 1 | 30745268 | rs11582464 | -1.548 | -0.684 | -0.065 | 1.834 | 0.005 | 0.01 | 0.01 | 0.349 | 0.05 | 0.03 | 0.07 | 0.19 | 0.02 | 0.17 |
| 1 | 30755532 | rs58245 | -0.597 | -0.567 | -0.316 | 0.597 | 0.004 | 0.039 | 0.039 | 0.672 | 0.36 | 0.69 | 0.21 | 0.11 | 0.32 | 0.23 |
| 1 | 30756008 | rs12040153 | -0.129 | -0.98 | 0.215 | 0.357 | -0.032 | -0.011 | -0.011 | 1.849 | 0.15 | 0.16 | 0.05 | 0.09 | 0.05 | 0.15 |
| 1 | 30756280 | rs396577 | -1.247 | -1.077 | -0.747 | 0.294 | -0.029 | -0.07 | -0.07 | 0 | 0.73 | 0.93 | 0.48 | 0.67 | 0.73 | 0.61 |
| 1 | 30758871 | rs457252 | -0.752 | -1.061 | 0.322 | 0.143 | -0.001 | -0.087 | -0.087 | 0.751 | 0.33 | 0.47 | 0.19 | 0.09 | 0.31 | 0.24 |
| 1 | 30758928 | rs455494 | -1.015 | -1.134 | 0.13 | 0.347 | 0.008 | 0.106 | 0.106 | 1.349 | 0.37 | 0.84 | 0.22 | 0.17 | 0.32 | 0.24 |
| 1 | 30759956 | rs463128 | 0.284 | -0.935 | 0.281 | 0.277 | 0.044 | -0.097 | -0.097 | 0.011 | 0.18 | 0.35 | 0.11 | 0.01 | 0.24 | 0.09 |
| 1 | 30761304 | rs460911 | 0.295 | -0.923 | 0.403 | 0.315 | 0.018 | -0.026 | -0.026 | 0 | 0.18 | 0.22 | 0.12 | 0.06 | 0.25 | 0.09 |
| 1 | 30762830 | rs1122816 | 0.777 | -1.221 | -0.199 | 0.318 | -0.022 | -0.058 | -0.058 | 0 | 0.26 | 0.16 | 0.22 | 0.48 | 0.35 | 0.28 |
| 1 | 30766261 | rs12135910 | 0.381 | -1.238 | -0.042 | -0.078 | -0.029 | -0.013 | -0.013 | 1.408 | 0.14 | 0.2 | 0.06 | 0.1 | 0.06 | 0.11 |
| 1 | 30787227 | rs411238 | 0.273 | -0.889 | -0.528 | -0.226 | 0.027 | -0.095 | -0.095 | 0 | 0.2 | 0.33 | 0.13 | 0 | 0.24 | 0.1 |
| 1 | 30787475 | rs6672668 | 0.504 | -0.966 | -1.538 | -0.12 | -0.008 | -0.051 | -0.051 | 0 | 0.28 | 0.18 | 0.32 | 0.56 | 0.34 | 0.32 |
| 1 | 30788183 | rs12057521 | 0.446 | -0.965 | -1.532 | 0.096 | -0.005 | -0.029 | -0.029 | 0 | 0.28 | 0.21 | 0.3 | 0.47 | 0.34 | 0.31 |
| 1 | 30788475 | rs433737 | -0.75 | -1.049 | -1.698 | 0.001 | 0.011 | -0.046 | -0.046 | 0 | 0.52 | 0.84 | 0.48 | 0.57 | 0.58 | 0.42 |
| 1 | 30788889 | rs1555045 | -0.021 | -1.071 | -1.214 | 0.691 | -0.016 | -0.084 | -0.084 | 0.261 | 0.29 | 0.1 | 0.32 | 0.56 | 0.34 | 0.32 |
| 1 | 30791516 | rs10799096 | -0.481 | -0.863 | -1.122 | 0.681 | 0.026 | -0.018 | -0.018 | 0 | 0.41 | 0.3 | 0.36 | 0.61 | 0.37 | 0.39 |

| | | | | | | | | | | | | | | | | |
|---|---|---|---|---|---|---|---|---|---|---|---|---|---|---|---|---|
| 1 | 30791525 | rs10915079 | -0.515 | -0.789 | -1.082 | 0.575 | 0.02 | -0.027 | -0.027 | 0 | 0.4 | 0.3 | 0.35 | 0.51 | 0.37 | 0.38 |
| 1 | 30798436 | rs7528451 | 0.088 | -0.704 | -1.388 | 0.129 | 0.056 | 0.039 | 0.039 | 0.03 | 0.49 | 0.28 | 0.45 | 0.52 | 0.64 | 0.35 |
| 1 | 30799299 | rs2007451 | 0.23 | -0.832 | -1.546 | -0.036 | 0.082 | 0.044 | 0.044 | 0 | 0.51 | 0.22 | 0.45 | 0.52 | 0.67 | 0.38 |
| 1 | 30803478 | rs6425951 | 0.403 | -0.365 | -0.768 | 0.026 | 0.068 | 0.103 | 0.103 | 0 | 0.23 | 0.37 | 0.34 | 0.48 | 0.43 | 0.26 |
| 1 | 30807154 | rs375526 | -0.628 | -1.229 | -0.259 | 0.134 | 0.003 | 0.021 | 0.021 | 0.02 | 0.85 | 0.91 | 0.93 | 0.89 | 0.88 | 0.8 |
| 1 | 30808141 | rs10915081 | 0.504 | -1.62 | -0.107 | -0.013 | 0.008 | -0.001 | -0.001 | 0.008 | 0.62 | 0.41 | 0.54 | 0.58 | 0.76 | 0.5 |
| 1 | 30808552 | rs4949578 | 0.699 | -1.656 | -0.096 | -0.036 | 0.049 | -0.014 | -0.014 | 0 | 0.61 | 0.25 | 0.52 | 0.51 | 0.76 | 0.5 |
| 1 | 30809179 | rs453756 | 1.178 | -1.712 | 0.399 | -0.062 | 0.006 | -0.011 | -0.011 | 1 | 0.17 | 0.11 | 0.12 | 0.26 | 0.12 | 0.25 |
| 1 | 30809347 | rs760655 | 0.516 | -1.744 | 0.631 | -0.079 | -0.011 | -0.002 | -0.002 | 0 | 0.64 | 0.46 | 0.78 | 0.63 | 0.76 | 0.55 |
| 1 | 30809473 | rs375489 | 1.193 | -1.732 | 0.519 | -0.072 | 0.007 | -0.012 | -0.012 | 0 | 0.17 | 0.11 | 0.12 | 0.26 | 0.12 | 0.25 |
| 1 | 30810150 | rs4949256 | 0.622 | -1.717 | 0.739 | -0.078 | -0.017 | 0 | 0 | 0.302 | 0.63 | 0.51 | 0.54 | 0.58 | 0.75 | 0.5 |
| 1 | 30827240 | rs762798 | 0.61 | -1.681 | 0.685 | 0.1 | 0.032 | -0.017 | -0.017 | 0 | 0.63 | 0.26 | 0.68 | 0.57 | 0.76 | 0.54 |
| 1 | 30827563 | rs10799099 | 0.61 | -1.681 | 0.685 | 0.1 | 0.032 | -0.017 | -0.017 | 0 | 0.63 | 0.26 | 0.68 | 0.57 | 0.76 | 0.54 |
| 1 | 30830505 | rs636797 | -0.789 | -1.548 | 0.65 | 0.086 | 0.007 | -0.005 | -0.005 | 0 | 0.86 | 0.91 | 0.84 | 0.84 | 0.9 | 0.86 |
| 1 | 30843915 | rs674001 | 0.356 | -1.582 | 0.783 | 0.09 | -0.04 | 0.008 | 0.008 | 1.263 | 0.8 | 0.41 | 0.78 | 0.77 | 0.9 | 0.85 |
| 1 | 30844120 | rs642819 | -0.668 | -1.466 | 0.419 | 0.099 | 0.005 | -0.011 | -0.011 | 0 | 0.86 | 0.93 | 0.83 | 0.84 | 0.9 | 0.86 |
| 1 | 30845037 | rs658840 | 0.815 | -1.184 | 0.612 | 0.327 | 0.001 | 0 | 0 | 1.825 | 0.17 | 0.14 | 0.11 | 0.17 | 0.13 | 0.3 |
| 1 | 30850174 | rs675696 | 0.607 | -1.152 | 0.754 | 0.352 | -0.011 | 0.013 | 0.013 | 0 | 0.18 | 0.26 | 0.12 | 0.19 | 0.14 | 0.32 |
| 1 | 30850797 | rs6704397 | 0.088 | -1.222 | 1.105 | 0.319 | -0.033 | 0.012 | 0.012 | 0.288 | 0.21 | 0.53 | 0.14 | 0.19 | 0.14 | 0.32 |
| 1 | 30850902 | rs6696645 | -0.57 | -1.537 | 0.706 | 0.088 | -0.01 | -0.004 | -0.004 | 0 | 0.63 | 0.6 | 0.74 | 0.71 | 0.8 | 0.83 |
| 1 | 30854069 | rs1329112 | -0.818 | -1.531 | 0.583 | 0.075 | 0.055 | 0.032 | 0.032 | 3.285 | 0.66 | 0.89 | 0.53 | 0.76 | 0.8 | 0.84 |
| 1 | 30855815 | rs750148 | 0.082 | -1.548 | 0.673 | 0.066 | 0.008 | 0.001 | 0.001 | 0.23 | 0.47 | 0.58 | 0.4 | 0.49 | 0.66 | 0.51 |
| 1 | 30855956 | rs750150 | -0.749 | -1.526 | 0.58 | 0.075 | 0.055 | 0.004 | 0.004 | 2.903 | 0.65 | 0.86 | 0.53 | 0.69 | 0.8 | 0.83 |
| 1 | 30858250 | rs1887916 | -0.809 | -1.508 | 0.628 | 0.058 | 0.031 | 0.022 | 0.022 | 2.699 | 0.65 | 0.77 | 0.52 | 0.76 | 0.8 | 0.84 |
| 1 | 30862461 | rs4949262 | 0 | -1.357 | 0.532 | 0.047 | -0.005 | 0 | 0 | 0.19 | 0.47 | 0.51 | 0.4 | 0.5 | 0.67 | 0.51 |
| 1 | 30873469 | rs3867305 | -0.158 | -1.528 | 0.305 | -0.153 | -0.014 | -0.014 | -0.014 | 0 | 0.55 | 0.52 | 0.77 | 0.71 | 0.73 | 0.58 |
| 1 | 30875110 | rs6699099 | -0.543 | -1.417 | 0.291 | -0.081 | 0.013 | 0.018 | 0.018 | 0.437 | 0.59 | 0.64 | 0.79 | 0.79 | 0.74 | 0.64 |
| 1 | 30875987 | rs6656841 | 0.021 | -1.045 | 0.034 | -0.202 | 0.022 | -0.005 | -0.005 | 0.196 | 0.37 | 0.54 | 0.64 | 0.67 | 0.5 | 0.38 |
| 1 | 30878285 | rs6656216 | 1.13 | -0.981 | -0.168 | -0.139 | -0.126 | -0.14 | -0.14 | 0 | 0.23 | 0.02 | 0.21 | 0.44 | 0.43 | 0.28 |
| 1 | 30878363 | rs642696 | -0.283 | -1.202 | 0.071 | -0.191 | -0.031 | -0.013 | -0.013 | 0 | 0.62 | 0.42 | 0.5 | 0.7 | 0.81 | 0.86 |
| 1 | 30882036 | rs12071101 | 0.54 | -1.114 | -0.139 | -0.185 | -0.014 | -0.01 | -0.01 | 0 | 0.23 | 0.46 | 0.15 | 0.21 | 0.18 | 0.36 |
| 1 | 30884114 | rs6666421 | 0.54 | -1.114 | -0.139 | -0.166 | -0.014 | 0.031 | 0.031 | 0.152 | 0.23 | 0.46 | 0.15 | 0.28 | 0.18 | 0.37 |
| 1 | 30885210 | rs650676 | -0.333 | -1.192 | -0.241 | -0.28 | 0.003 | -0.001 | -0.001 | 0 | 0.53 | 0.51 | 0.76 | 0.79 | 0.68 | 0.6 |
| 1 | 30885349 | rs12029866 | 0.9 | -1.172 | -0.26 | -0.271 | -0.002 | -0.001 | -0.001 | 0 | 0.21 | 0.22 | 0.13 | 0.21 | 0.18 | 0.36 |
| 1 | 30885525 | rs646198 | -1.336 | -1.212 | -0.374 | -0.569 | 0.015 | 0.02 | 0.02 | 3.744 | 0.75 | 0.76 | 0.9 | 0.93 | 0.86 | 0.95 |
| 1 | 30887732 | rs6676701 | -0.262 | -0.324 | -0.359 | -0.113 | 0 | 0.004 | 0.004 | 4.417 | 0.45 | 0.58 | 0.35 | 0.73 | 0.57 | 0.65 |
| 1 | 30890053 | rs12407723 | 0.198 | -0.467 | -0.519 | 0.048 | -0.017 | 0.044 | 0.044 | 2.349 | 0.1 | 0.19 | 0.39 | 0.23 | 0.06 | 0.1 |
| 1 | 30890532 | rs12407007 | 0.198 | -0.467 | -0.519 | 0.048 | -0.017 | 0.044 | 0.044 | 2.349 | 0.1 | 0.19 | 0.39 | 0.23 | 0.06 | 0.1 |
| 1 | 30890587 | rs686393 | -0.674 | 0.045 | -0.597 | 0.233 | 0.001 | 0.034 | 0.034 | 3.459 | 0.46 | 0.65 | 0.35 | 0.72 | 0.55 | 0.65 |
| 1 | 30891217 | rs591858 | -0.801 | -0.117 | -0.647 | 0.61 | 0.011 | -0.002 | -0.002 | 0 | 0.24 | 0.18 | 0.11 | 0.24 | 0.18 | 0.27 |
| 1 | 30892426 | rs10489163 | -0.916 | 0.046 | -0.281 | 0.859 | -0.008 | 0.045 | 0.045 | 0 | 0.24 | 0.29 | 0.27 | 0.52 | 0.22 | 0.33 |
| 1 | 30893256 | rs728338 | -0.445 | 0.029 | -0.456 | 0.408 | -0.047 | -0.137 | -0.137 | 0 | 0.21 | 0.02 | 0.25 | 0.39 | 0.21 | 0.35 |
| 1 | 30894160 | rs606404 | -1.821 | -0.094 | -0.045 | 0.12 | 0.031 | 0.256 | 0.256 | 0 | 0.49 | 0.76 | 0.56 | 0.91 | 0.43 | 0.65 |
| 1 | 30894530 | rs654075 | -1.819 | -0.097 | -0.037 | 0.118 | 0.03 | 0.26 | 0.26 | 0 | 0.49 | 0.76 | 0.56 | 0.91 | 0.43 | 0.65 |
| 1 | 30898855 | rs10915113 | 0.115 | -0.217 | 0.417 | 0.352 | -0.047 | -0.132 | -0.132 | 0 | 0.18 | 0.02 | 0.19 | 0.39 | 0.2 | 0.27 |
| 1 | 30908689 | rs1085906 | -1.564 | -0.187 | 0.123 | 0.13 | 0.03 | 0.239 | 0.239 | 0 | 0.49 | 0.74 | 0.56 | 0.9 | 0.43 | 0.65 |
| 1 | 30918166 | rs11577262 | -0.539 | 0.003 | -0.122 | 0.82 | 0.004 | -0.067 | -0.067 | 0 | 0.18 | 0.07 | 0.22 | 0.44 | 0.17 | 0.3 |
| 1 | 30918453 | rs744528 | 1.576 | -0.221 | 0.001 | -0.125 | -0.015 | 0.086 | 0.086 | 0 | 0.45 | 0.35 | 0.41 | 0.1 | 0.48 | 0.4 |
| 1 | 30919030 | rs657816 | 0.358 | -0.177 | 0.493 | 0.386 | -0.017 | 0.072 | 0.072 | 0 | 0.79 | 0.73 | 0.74 | 0.48 | 0.84 | 0.71 |
| 1 | 30919378 | rs12117449 | -0.309 | -0.405 | 0.387 | -0.126 | -0.008 | -0.008 | -0.008 | 0 | 0.32 | 0.26 | 0.33 | 0.38 | 0.41 | 0.42 |
| 1 | 30920818 | rs1001704 | -0.242 | -0.365 | 0.374 | -0.105 | -0.018 | -0.026 | -0.026 | 0 | 0.32 | 0.23 | 0.33 | 0.47 | 0.41 | 0.43 |
| 1 | 30920920 | rs686964 | -0.098 | -0.451 | 0.22 | -0.149 | 0.023 | -0.095 | -0.095 | 0 | 0.8 | 0.93 | 0.77 | 0.57 | 0.84 | 0.72 |
| 1 | 30923786 | rs12047343 | 1.39 | -0.437 | 0.085 | -0.308 | -0.006 | 0.103 | 0.103 | 0 | 0.44 | 0.3 | 0.39 | 0.1 | 0.43 | 0.29 |
| 1 | 30925952 | rs667153 | 0.439 | -0.427 | -0.051 | -0.199 | 0.037 | -0.092 | -0.092 | 0 | 0.77 | 0.91 | 0.78 | 0.51 | 0.84 | 0.71 |
| 1 | 30933736 | rs6673363 | 1.586 | -0.361 | -0.072 | -0.338 | -0.001 | 0.059 | 0.059 | 0 | 0.42 | 0.32 | 0.38 | 0.1 | 0.41 | 0.29 |
| 1 | 30933984 | rs262623 | 0.856 | -0.367 | -0.809 | -0.336 | 0.051 | 0.052 | 0.052 | 2.948 | 0.94 | 0.5 | 0.94 | 1 | 1 | 1 |
| 1 | 30938590 | rs262597 | -0.104 | -0.295 | -1.627 | -0.284 | -0.11 | -0.087 | -0.087 | 0 | 0.31 | 0.06 | 0.32 | 0.33 | 0.41 | 0.42 |
| 1 | 30940894 | rs10915123 | 1.553 | -0.22 | -1.543 | -0.55 | -0.016 | 0.142 | 0.142 | 0 | 0.4 | 0.21 | 0.37 | 0.09 | 0.41 | 0.29 |
| 1 | 30944362 | rs1552319 | 0.099 | -0.052 | -1.578 | -0.102 | -0.116 | -0.001 | -0.001 | 0 | 0.67 | 0.9 | 0.53 | 0.73 | 0.56 | 0.58 |
| 1 | 30945974 | rs12406008 | -0.715 | 0.426 | -1.134 | 0.434 | 0.009 | -0.022 | -0.022 | 0 | 0.19 | 0.17 | 0.23 | 0.52 | 0.15 | 0.33 |
| 1 | 30962474 | rs415314 | -0.722 | 0.698 | -1.156 | 0.348 | 0.035 | 0.021 | 0.021 | 0 | 0.29 | 0.54 | 0.49 | 0.39 | 0.37 | 0.32 |
| 1 | 30981662 | rs16833801 | -0.59 | 0.578 | -1.398 | 0.402 | -0.027 | -0.143 | -0.143 | 0.184 | 0.19 | 0.01 | 0.25 | 0.52 | 0.19 | 0.37 |
| 1 | 30985076 | rs3104429 | 0.73 | 0.568 | -1.387 | 0.393 | -0.006 | 0.01 | 0.01 | 0 | 0.77 | 0.74 | 0.73 | 0.37 | 0.8 | 0.65 |
| 1 | 30985651 | rs10915140 | -0.373 | 1.014 | -1.061 | 0.686 | -0.127 | -0.053 | -0.053 | 0 | 0.23 | 0.07 | 0.43 | 0.3 | 0.36 | 0.29 |
| 1 | 30985821 | rs262634 | 0.32 | 1.039 | -1.004 | 0.703 | 0.001 | 0 | 0 | 0.058 | 0.74 | 0.73 | 0.56 | 0.6 | 0.64 | 0.7 |
| 1 | 30988344 | rs1484849 | -0.723 | 0.428 | -0.318 | 0.795 | 0.078 | 0.073 | 0.073 | 3.301 | 0.05 | 0.49 | 0.06 | 0 | 0 | 0 |
| 1 | 30988365 | rs1484850 | -1.955 | 0.268 | -0.786 | 0.748 | -0.041 | -0.016 | -0.016 | 0 | 0.17 | 0.71 | 0.1 | 0.17 | 0.08 | 0.07 |
| 1 | 30988530 | rs379605 | -0.419 | 1.079 | -1.402 | 1.149 | -0.003 | -0.012 | -0.012 | 0 | 0.39 | 0.64 | 0.41 | 0.48 | 0.42 | 0.32 |
| 1 | 30989639 | rs12144776 | -0.878 | 1.186 | -0.573 | 1.585 | -0.035 | -0.025 | -0.025 | 0 | 0.13 | 0.08 | 0.19 | 0.17 | 0.18 | 0.2 |

| | | | | | | | | | | | | | | | | |
|---|---|---|---|---|---|---|---|---|---|---|---|---|---|---|---|---|
| 1 | 30993005 | rs396648 | -1.141 | 1.084 | -0.53 | 1.701 | 0.002 | 0.038 | 0.038 | 0 | 0.24 | 0.69 | 0.3 | 0.33 | 0.25 | 0.26 |
| 1 | 30998096 | rs6692098 | -0.996 | 1.316 | -0.487 | 1.404 | 0.014 | 0.021 | 0.021 | 0 | 0.13 | 0.2 | 0.25 | 0.16 | 0.2 | 0.07 |
| 1 | 30998579 | rs10915147 | 0.846 | 1.246 | -1.372 | 1.588 | -0.033 | -0.034 | -0.034 | 0 | 0.59 | 0.13 | 0.53 | 0.49 | 0.61 | 0.68 |
| 1 | 31000290 | rs2019878 | 1.387 | 0.713 | -0.065 | 2.712 | 0.002 | 0.288 | 0.288 | 0.305 | 0.94 | 0.76 | 0.84 | 0.65 | 0.95 | 0.94 |
| 1 | 31000635 | rs7540267 | 1.625 | -0.156 | 0.625 | 2.353 | -0.04 | 0.058 | 0.058 | 2.016 | 0.81 | 0.36 | 0.76 | 0.59 | 0.92 | 0.91 |
| 1 | 31007037 | rs12127272 | -1.679 | -0.186 | 0.332 | 2.366 | -0.014 | -0.002 | -0.002 | 0.275 | 0.33 | 0.51 | 0.32 | 0.34 | 0.23 | 0.16 |
| 1 | 31008215 | rs7517833 | -1.648 | -0.204 | -0.511 | 2.213 | 0.006 | -0.003 | -0.003 | 0 | 0.51 | 0.53 | 0.5 | 0.54 | 0.45 | 0.46 |
| 1 | 31009394 | rs10915149 | 0.222 | -0.213 | 0.419 | 2.154 | 0.001 | 0 | 0 | 0 | 0.19 | 0.25 | 0.23 | 0.25 | 0.23 | 0.33 |
| 1 | 31010302 | rs1994239 | 1.646 | -0.212 | 0.474 | 2.224 | -0.014 | 0.076 | 0.076 | 0 | 0.46 | 0.22 | 0.46 | 0.31 | 0.53 | 0.53 |
| 1 | 31011165 | rs4949270 | 0.912 | -0.223 | 0.853 | 2.182 | -0.01 | 0.014 | 0.014 | 0.66 | 0.66 | 0.43 | 0.7 | 0.56 | 0.76 | 0.86 |
| 1 | 31012257 | rs10915150 | -0.772 | -0.293 | 1.624 | 2.113 | -0.004 | 0.022 | 0.022 | 0 | 0.28 | 0.39 | 0.26 | 0.42 | 0.27 | 0.35 |
| 1 | 31013426 | rs9628644 | 0.749 | -0.355 | 1.724 | 2.017 | -0.034 | -0.043 | -0.043 | 0.012 | 0.08 | 0.13 | 0.02 | 0 | 0.02 | 0 |
| 1 | 31016564 | rs12035623 | 0.261 | -0.355 | 1.76 | 2.077 | -0.001 | 0.003 | 0.003 | 0 | 0.19 | 0.13 | 0.2 | 0.36 | 0.24 | 0.3 |
| 1 | 31022237 | rs904257 | 1.368 | -0.582 | 1.802 | 2.082 | 0.043 | -0.016 | -0.016 | 0.019 | 0.6 | 0.26 | 0.57 | 0.4 | 0.68 | 0.6 |
| 1 | 31023291 | rs3104434 | 1.49 | -0.562 | 1.853 | 2.102 | -0.012 | 0.203 | 0.203 | 0 | 0.92 | 0.74 | 0.82 | 0.6 | 0.93 | 0.89 |
| 1 | 31024967 | rs2377368 | -0.267 | -0.551 | 1.731 | 2.101 | 0.006 | -0.029 | -0.029 | 0.973 | 0.05 | 0 | 0.13 | 0.18 | 0.04 | 0.06 |
| 1 | 31027917 | rs2377367 | -1.419 | -0.555 | 1.821 | 2.06 | -0.012 | 0.219 | 0.219 | 0 | 0.08 | 0.19 | 0.16 | 0.34 | 0.06 | 0.07 |
| 1 | 31028135 | rs924267 | -1.419 | -0.555 | 1.818 | 2.06 | -0.013 | 0.214 | 0.214 | 0 | 0.08 | 0.19 | 0.16 | 0.34 | 0.06 | 0.07 |
| 1 | 31034663 | rs3104425 | 1.282 | -0.555 | 1.698 | 2.035 | -0.021 | 0.01 | 0.01 | 0 | 0.62 | 0.47 | 0.59 | 0.4 | 0.68 | 0.6 |
| 1 | 31045509 | rs3104435 | -0.155 | -0.619 | 1.381 | 1.909 | -0.001 | 0.003 | 0.003 | 0 | 0.78 | 0.84 | 0.78 | 0.75 | 0.76 | 0.66 |
| 1 | 31049746 | rs4445518 | 1.803 | -0.703 | 1.246 | 1.953 | -0.029 | 0.064 | 0.064 | 0 | 0.43 | 0.15 | 0.46 | 0.29 | 0.52 | 0.53 |
| 1 | 31058914 | rs525813 | 0.467 | -0.665 | 1.273 | 1.955 | -0.001 | 0.003 | 0.003 | 0 | 0.18 | 0.19 | 0.13 | 0.12 | 0.16 | 0.07 |
| 1 | 31060480 | rs472061 | -1.497 | -0.649 | 1.346 | 1.987 | 0 | 0 | 0 | 0 | 0.49 | 0.53 | 0.38 | 0.53 | 0.43 | 0.41 |
| 1 | 31061327 | rs610679 | -0.923 | -0.727 | 1.486 | 2.002 | -0.03 | 0.02 | 0.02 | 0.134 | 0.31 | 0.49 | 0.2 | 0.33 | 0.21 | 0.1 |
| 1 | 31062295 | rs530973 | -1.451 | -0.833 | 0.856 | 1.822 | 0.033 | -0.025 | -0.025 | 0 | 0.38 | 0.75 | 0.35 | 0.51 | 0.25 | 0.16 |
| 1 | 31063390 | rs485447 | 0.543 | -0.524 | 0.735 | 1.336 | -0.003 | 0.005 | 0.005 | 0 | 0.17 | 0.12 | 0.11 | 0.1 | 0.17 | 0.06 |
| 1 | 31064203 | rs788871 | 0.729 | -0.897 | 0.628 | 1.15 | 0.005 | -0.001 | -0.001 | 0.773 | 0.59 | 0.35 | 0.6 | 0.4 | 0.73 | 0.64 |
| 1 | 31070465 | rs1575513 | -0.179 | -0.789 | 0.983 | 0.807 | -0.039 | -0.084 | -0.084 | 0.293 | 0.13 | 0.35 | 0.07 | 0 | 0.06 | 0.02 |
| 1 | 31076399 | rs10914150 | 0.022 | -0.479 | 0.353 | 0.547 | 0.005 | -0.001 | -0.001 | 0 | 0.62 | 0.57 | 0.73 | 0.66 | 0.59 | 0.75 |
| 1 | 31076994 | rs7551357 | 0.257 | -0.371 | 0.123 | 0.563 | -0.001 | 0.003 | 0.003 | 0 | 0.26 | 0.28 | 0.31 | 0.42 | 0.26 | 0.36 |
| 1 | 31080612 | rs4949279 | -0.249 | -0.38 | 0.255 | 0.523 | 0.003 | -0.001 | -0.001 | 0 | 0.63 | 0.61 | 0.74 | 0.67 | 0.59 | 0.75 |
| 1 | 31080962 | rs548273 | 0.174 | -0.329 | 0.196 | 0.537 | 0.002 | 0.026 | 0.026 | 0 | 0.53 | 0.64 | 0.54 | 0.39 | 0.55 | 0.58 |
| 1 | 31083478 | rs544789 | -0.744 | -0.447 | 0.411 | 0.52 | 0.002 | 0.136 | 0.136 | 0 | 0.06 | 0.25 | 0.06 | 0.19 | 0.03 | 0.03 |
| 1 | 31086685 | rs666750 | -0.271 | -0.525 | 0.363 | 0.493 | -0.025 | 0.045 | 0.045 | 0 | 0.32 | 0.5 | 0.36 | 0.61 | 0.29 | 0.39 |
| 1 | 31086765 | rs666322 | 0.439 | -0.445 | 0.425 | 0.516 | -0.003 | 0.009 | 0.009 | 0 | 0.26 | 0.33 | 0.32 | 0.42 | 0.26 | 0.36 |
| 1 | 31087850 | rs10914158 | 1.269 | -0.452 | 0.622 | 0.454 | 0.002 | -0.044 | -0.044 | 0 | 0.15 | 0.17 | 0.09 | 0 | 0.2 | 0.17 |
| 1 | 31088967 | rs7518010 | 0.571 | -0.449 | 0.6 | 0.482 | 0.002 | -0.018 | -0.018 | 0.221 | 0.25 | 0.25 | 0.31 | 0.42 | 0.26 | 0.36 |
| 1 | 31091447 | rs474814 | 0.184 | -0.468 | 0.637 | 0.401 | -0.006 | 0.064 | 0.064 | 0 | 0.55 | 0.68 | 0.55 | 0.3 | 0.53 | 0.58 |
| 1 | 31099501 | rs11589302 | 1.472 | -0.467 | 0.627 | 0.381 | 0.001 | 0.018 | 0.018 | 0 | 0.37 | 0.29 | 0.42 | 0.25 | 0.33 | 0.39 |
| 1 | 31099929 | rs1188414 | 0.221 | -0.403 | 0.475 | 0.45 | -0.003 | 0.037 | 0.037 | 0 | 0.55 | 0.58 | 0.54 | 0.3 | 0.53 | 0.58 |
| 1 | 31101134 | rs1201394 | -1.99 | -0.134 | 0.024 | 0.539 | -0.035 | -0.035 | -0.035 | 0 | 0.83 | 0.97 | 0.87 | 0.86 | 0.82 | 0.95 |
| 1 | 31108443 | rs585889 | 0.015 | -0.065 | 0.152 | 0.605 | 0 | 0 | 0 | 0.06 | 0.64 | 0.65 | 0.76 | 0.72 | 0.59 | 0.77 |
| 1 | 31110186 | rs2104311 | 0.801 | 0.018 | 0.051 | 0.567 | 0.006 | -0.006 | -0.006 | 0 | 0.16 | 0.24 | 0.11 | 0.14 | 0.22 | 0.18 |
| 1 | 31113232 | rs1882027 | 0.865 | -0.008 | 0.04 | 0.372 | 0.001 | -0.041 | -0.041 | 0 | 0.15 | 0.18 | 0.1 | 0.01 | 0.2 | 0.17 |
| 1 | 31113723 | rs904255 | 0.45 | -0.311 | 0.21 | 0.859 | -0.006 | -0.014 | -0.014 | 0 | 0.44 | 0.47 | 0.46 | 0.34 | 0.38 | 0.52 |
| 1 | 31115997 | rs4508063 | -1.221 | -0.183 | -0.184 | 0.689 | -0.001 | 0.003 | 0.003 | 0 | 0.77 | 0.88 | 0.85 | 0.86 | 0.81 | 0.94 |
| 1 | 31116014 | rs10914170 | 0.878 | -0.161 | 0 | 0.585 | 0.001 | -0.013 | -0.013 | 1.135 | 0.37 | 0.38 | 0.45 | 0.25 | 0.36 | 0.4 |
| 1 | 31116813 | rs10914172 | -0.291 | 0.047 | -0.239 | 0.847 | -0.008 | -0.01 | -0.01 | 0.04 | 0.35 | 0.22 | 0.36 | 0.41 | 0.41 | 0.48 |
| 1 | 31118741 | rs522294 | -0.23 | 0.294 | 0.285 | 0.786 | 0.006 | -0.004 | -0.004 | 0 | 0.25 | 0.32 | 0.23 | 0.21 | 0.29 | 0.36 |
| 1 | 31125281 | rs1628852 | 0.161 | 0.282 | 0.185 | 0.862 | -0.019 | 0.005 | 0.005 | 0.033 | 0.23 | 0.14 | 0.22 | 0.19 | 0.29 | 0.35 |
| 1 | 31133242 | rs1188432 | 0.12 | 0.281 | 0.603 | 0.888 | 0.004 | -0.001 | -0.001 | 2.678 | 0.23 | 0.2 | 0.22 | 0.2 | 0.3 | 0.35 |
| 1 | 31139078 | rs1983822 | -0.931 | -0.123 | 0.7 | 1.22 | -0.036 | 0.035 | 0.035 | 1.585 | 0.21 | 0.66 | 0.41 | 0.32 | 0.15 | 0.14 |
| 1 | 31151452 | rs1408151 | -0.887 | -0.298 | 0.722 | 1.063 | -0.006 | -0.011 | -0.011 | 1.993 | 0.24 | 0.25 | 0.36 | 0.12 | 0.2 | 0.35 |
| 1 | 31151500 | rs1188390 | 0.36 | 0.283 | 0.158 | 1.287 | 0.005 | -0.001 | -0.001 | 0 | 0.23 | 0.2 | 0.22 | 0.2 | 0.29 | 0.33 |
| 1 | 31159619 | rs1188453 | 0.359 | 0.287 | 0.158 | 1.27 | 0.005 | -0.002 | -0.002 | 0 | 0.23 | 0.2 | 0.22 | 0.19 | 0.29 | 0.33 |
| 1 | 31163876 | rs1188445 | -1.213 | 0.154 | 0.034 | 1.268 | -0.041 | 0.057 | 0.057 | 1.851 | 0.19 | 0.61 | 0.28 | 0.34 | 0.13 | 0.29 |
| 1 | 31164624 | rs1188444 | -0.919 | 0.259 | -0.591 | 1.105 | 0 | -0.039 | -0.039 | 0 | 0.43 | 0.77 | 0.52 | 0.53 | 0.42 | 0.67 |
| 1 | 31167561 | rs716486 | 0.73 | 0.431 | -0.707 | 0.823 | -0.049 | 0.015 | 0.015 | 0.022 | 0.29 | 0.05 | 0.19 | 0.18 | 0.3 | 0.41 |
| 1 | 31167998 | rs1033867 | 0.651 | 0.561 | -0.981 | 0.458 | -0.02 | 0.06 | 0.06 | 0.056 | 0.2 | 0.1 | 0.12 | 0.08 | 0.25 | 0.1 |
| 1 | 31168162 | rs1072498 | 0.61 | 0.63 | -0.431 | 0.512 | -0.01 | 0.016 | 0.016 | 0 | 0.36 | 0.26 | 0.49 | 0.28 | 0.41 | 0.58 |
| 1 | 31168219 | rs1830705 | -0.286 | 0.588 | -0.068 | 0.665 | -0.007 | -0.003 | -0.003 | 0 | 0.57 | 0.49 | 0.64 | 0.44 | 0.66 | 0.68 |
| 1 | 31168270 | rs1188443 | -0.199 | 0.528 | 0.856 | 0.765 | -0.011 | -0.066 | -0.066 | 0 | 0.11 | 0.28 | 0.08 | 0 | 0.08 | 0.04 |
| 1 | 31176942 | rs1149023 | -1.039 | 0.598 | 0.805 | 1.11 | -0.002 | 0.21 | 0.21 | 0.702 | 0.15 | 0.5 | 0.25 | 0.33 | 0.1 | 0.1 |
| 1 | 31178663 | rs12750278 | -0.609 | 0.643 | 0.836 | 1.042 | 0.033 | 0.024 | 0.024 | 0 | 0.06 | 0.18 | 0.3 | 0.1 | 0.11 | 0.17 |
| 1 | 31183812 | rs7555715 | -0.362 | 0.439 | 0.772 | 1.004 | -0.003 | 0.001 | 0.001 | 0 | 0.26 | 0.26 | 0.41 | 0.18 | 0.37 | 0.27 |
| 1 | 31184218 | rs1188394 | -1.568 | 0.419 | 0.768 | 1.063 | -0.008 | -0.019 | -0.019 | 0 | 0.41 | 0.75 | 0.65 | 0.52 | 0.47 | 0.35 |
| 1 | 31184368 | rs1188395 | -1.585 | 0.42 | 0.769 | 1.064 | -0.008 | -0.019 | -0.019 | 0 | 0.41 | 0.74 | 0.65 | 0.52 | 0.47 | 0.37 |
| 1 | 31192304 | rs1149040 | -1.563 | 0.419 | 0.781 | 1.066 | -0.023 | -0.044 | -0.044 | 1.399 | 0.41 | 0.82 | 0.66 | 0.52 | 0.47 | 0.37 |
| 1 | 31197045 | rs1149045 | -0.441 | 0.37 | 0.917 | 1.088 | 0.001 | 0.17 | 0.17 | 0 | 0.12 | 0.27 | 0.23 | 0.42 | 0.1 | 0.1 |

| | | | | | | | | | | | | | | | | |
|---|---|---|---|---|---|---|---|---|---|---|---|---|---|---|---|---|
| 1 | 31197799 | rs3828051 | -0.714 | 0.411 | 0.683 | 1.091 | 0.016 | -0.031 | -0.031 | 0.216 | 0.29 | 0.53 | 0.46 | 0.27 | 0.37 | 0.27 |
| 1 | 31198733 | rs1149048 | -1.487 | 0.553 | -0.567 | 0.914 | -0.034 | -0.044 | -0.044 | 0 | 0.42 | 0.89 | 0.68 | 0.6 | 0.47 | 0.37 |
| 1 | 31206637 | rs3795438 | 0.467 | 1.184 | -0.445 | 1.013 | -0.085 | 0.078 | 0.078 | 0.602 | 0.17 | 0.01 | 0.1 | 0.08 | 0.27 | 0.1 |
| 1 | 31206806 | rs3795437 | -0.569 | 1.062 | -0.026 | 1.2 | 0.032 | 0.109 | 0.109 | 2.994 | 0.25 | 0.45 | 0.31 | 0.5 | 0.31 | 0.17 |
| 1 | 31210667 | rs1188355 | -0.427 | 0.901 | -0.382 | 0.454 | -0.006 | 0.02 | 0.02 | 0 | 0.17 | 0.14 | 0.3 | 0.08 | 0.2 | 0.18 |
| 1 | 31211282 | rs1188357 | -0.9 | 0.797 | -0.576 | 0.458 | 0.003 | -0.006 | -0.006 | 0 | 0.66 | 0.74 | 0.77 | 0.67 | 0.66 | 0.57 |
| 1 | 31212376 | rs1188360 | -0.667 | 0.809 | -0.974 | 0.543 | 0.002 | -0.005 | -0.005 | 0 | 0.63 | 0.42 | 0.7 | 0.46 | 0.66 | 0.55 |
| 1 | 31215364 | rs1050663 | -0.59 | 1.018 | -1.143 | 0.398 | 0 | 0.007 | 0.007 | 0.315 | 0.56 | 0.62 | 0.69 | 0.67 | 0.53 | 0.54 |
| 1 | 31216969 | rs11585511 | -0.692 | 1.457 | -1.557 | 0.008 | -0.066 | 0.114 | 0.114 | 0.532 | 0.24 | 0.03 | 0.13 | 0.08 | 0.3 | 0.12 |
| 1 | 31222316 | rs1188350 | -0.561 | 0.8 | -0.729 | 0.621 | -0.001 | -0.002 | -0.002 | 0 | 0.39 | 0.62 | 0.58 | 0.31 | 0.36 | 0.27 |
| 1 | 31226340 | rs1188347 | -0.408 | 0.717 | -0.686 | 0.723 | 0.008 | 0.036 | 0.036 | 1.401 | 0.46 | 0.68 | 0.64 | 0.69 | 0.53 | 0.55 |
| 1 | 31231333 | rs10914193 | -0.469 | 0.815 | -0.221 | 0.887 | 0.007 | -0.005 | -0.005 | 1.522 | 0.42 | 0.55 | 0.55 | 0.61 | 0.48 | 0.48 |
| 1 | 31231680 | rs3762297 | -0.464 | 1.59 | 0.206 | 1.635 | 0.028 | 0.048 | 0.048 | 0 | 0.14 | 0.34 | 0.42 | 0.26 | 0.17 | 0.13 |
| 1 | 31239262 | rs12124742 | 0.304 | 1.474 | -0.615 | 2.179 | -0.049 | -0.047 | -0.047 | 0 | 0.25 | 0.07 | 0.13 | 0.25 | 0.3 | 0.37 |
| 1 | 31241312 | rs12736883 | -0.121 | 1.467 | -0.642 | 2.176 | -0.05 | -0.064 | -0.064 | 0 | 0.11 | 0.03 | 0.38 | 0.33 | 0.18 | 0.13 |
| 1 | 31247236 | rs878467 | -0.658 | 1.227 | -0.489 | 1.987 | -0.053 | -0.035 | -0.035 | 0 | 0.37 | 0.19 | 0.52 | 0.56 | 0.47 | 0.49 |
| 1 | 31248562 | rs10798806 | -0.697 | 1.216 | -0.487 | 1.772 | -0.051 | -0.009 | -0.009 | 0 | 0.38 | 0.18 | 0.52 | 0.61 | 0.48 | 0.5 |
| 1 | 31258259 | rs11802256 | -0.239 | 0.748 | -0.674 | 1.788 | -0.012 | -0.045 | -0.045 | 0 | 0.31 | 0.2 | 0.14 | 0.34 | 0.3 | 0.33 |
| 1 | 31258827 | rs7516769 | -0.135 | 0.24 | -0.657 | 1.028 | -0.001 | 0.015 | 0.015 | 0 | 0.43 | 0.6 | 0.68 | 0.36 | 0.46 | 0.33 |
| 1 | 31264978 | rs10914206 | -0.064 | -0.244 | -0.538 | 0.569 | -0.023 | -0.01 | -0.01 | 0 | 0.42 | 0.46 | 0.21 | 0.4 | 0.31 | 0.3 |
| 1 | 31267335 | rs11578558 | 0.872 | -0.682 | -0.161 | 0.336 | -0.009 | -0.017 | -0.017 | 0 | 0.22 | 0.26 | 0.12 | 0.08 | 0.16 | 0.09 |
| 1 | 31267470 | rs12726862 | 0.648 | -0.628 | -0.228 | 0.494 | 0.006 | 0.007 | 0.007 | 0.707 | 0.17 | 0.1 | 0.09 | 0.14 | 0.14 | 0.18 |
| 1 | 31272279 | rs10798809 | 0.405 | 0.138 | -0.553 | 0.435 | -0.017 | -0.029 | -0.029 | 0.846 | 0.25 | 0.2 | 0.56 | 0.48 | 0.32 | 0.28 |
| 1 | 31272450 | rs16834077 | 1.103 | 0.216 | -0.615 | 0.19 | 0.01 | 0.013 | 0.013 | 0.75 | 0.17 | 0.1 | 0.09 | 0.14 | 0.14 | 0.18 |
| 1 | 31278481 | rs6425686 | -1.379 | 0.533 | -0.766 | 0.073 | 0 | 0.017 | 0.017 | 0.451 | 0.78 | 0.87 | 0.88 | 0.9 | 0.77 | 0.75 |
| 1 | 31282326 | rs4949181 | -1.362 | 0.53 | -0.767 | 0.071 | 0 | 0.017 | 0.017 | 0.468 | 0.78 | 0.86 | 0.88 | 0.9 | 0.77 | 0.75 |
| 1 | 31286796 | rs12038524 | -0.209 | 0.727 | -0.777 | 0.455 | 0.009 | -0.009 | -0.009 | 0 | 0.39 | 0.24 | 0.46 | 0.51 | 0.34 | 0.43 |
| 1 | 31289265 | rs1407887 | 1.705 | 0.53 | -1.159 | 0.315 | 0.022 | 0.112 | 0.112 | 2.119 | 0.25 | 0.04 | 0.07 | 0.04 | 0.17 | 0.15 |
| 1 | 31291197 | rs12401326 | 1.068 | 0.677 | -0.835 | 0.699 | 0.003 | -0.003 | -0.003 | 0.871 | 0.09 | 0.07 | 0.26 | 0.17 | 0.24 | 0.1 |
| 1 | 31292641 | rs16834081 | 1.068 | 0.471 | -1.018 | 0.653 | 0.006 | -0.034 | -0.034 | 0.103 | 0.14 | 0.06 | 0.29 | 0.39 | 0.1 | 0.14 |
| 1 | 31294767 | rs1407891 | -1.331 | 0.396 | -0.383 | 0.464 | -0.011 | -0.001 | -0.001 | 0.291 | 0.75 | 0.65 | 0.85 | 0.83 | 0.77 | 0.74 |
| 1 | 31296585 | rs4949309 | 0.828 | 0.594 | -0.762 | 0.54 | 0.055 | 0.039 | 0.039 | 0 | 0.1 | 0.17 | 0.26 | 0.17 | 0.24 | 0.1 |
| 1 | 31297519 | rs746700 | -1.392 | 0.257 | -1.233 | 0.524 | 0.002 | -0.001 | -0.001 | 0 | 0.76 | 0.76 | 0.86 | 0.83 | 0.77 | 0.74 |
| 1 | 31298769 | rs2377570 | -1.606 | 0.363 | -1.281 | 0.478 | 0.002 | 0.01 | 0.01 | 0.765 | 0.77 | 0.85 | 0.88 | 0.9 | 0.77 | 0.75 |
| 1 | 31302186 | rs12117057 | 1.247 | 0.331 | -1.195 | 0.248 | 0.02 | 0.027 | 0.027 | 0 | 0.16 | 0.08 | 0.1 | 0.11 | 0.15 | 0.15 |
| 1 | 31306486 | rs4949310 | -0.318 | 0.466 | -1.063 | 0.451 | 0.011 | -0.028 | -0.028 | 0 | 0.64 | 0.73 | 0.6 | 0.5 | 0.67 | 0.62 |
| 1 | 31306541 | rs10914217 | 0.888 | 0.868 | -1.153 | 0.507 | -0.003 | -0.005 | -0.005 | 0 | 0.08 | 0.07 | 0.07 | 0.01 | 0.08 | 0.13 |
| 1 | 31309717 | rs3923872 | -0.102 | 0.771 | -1.036 | 0.795 | 0.005 | -0.034 | -0.034 | 0 | 0.66 | 0.73 | 0.6 | 0.5 | 0.68 | 0.61 |
| 1 | 31311418 | rs4551635 | -0.01 | 0.936 | -0.949 | 1.047 | 0.008 | -0.031 | -0.031 | 0 | 0.71 | 0.73 | 0.63 | 0.52 | 0.75 | 0.68 |
| 1 | 31313417 | rs12756105 | 0.348 | 1.46 | -1.152 | 1.039 | -0.044 | -0.031 | -0.031 | 4.358 | 0.14 | 0.07 | 0.28 | 0.17 | 0.26 | 0.1 |
| 1 | 31315191 | rs10914220 | -0.761 | 0.82 | -0.746 | 1.475 | -0.002 | -0.003 | -0.003 | 0.222 | 0.83 | 0.79 | 0.9 | 0.91 | 0.85 | 0.83 |
| 1 | 31315286 | rs12029838 | 0.714 | 0.587 | -0.821 | 1.537 | 0.015 | -0.018 | -0.018 | 2.985 | 0.1 | 0.06 | 0.05 | 0.21 | 0.09 | 0.16 |
| 1 | 31315402 | rs4949315 | -0.57 | 1.133 | -0.612 | 1.45 | -0.003 | -0.004 | -0.004 | 0.04 | 0.83 | 0.79 | 0.91 | 0.91 | 0.85 | 0.83 |
| 1 | 31316029 | rs10914221 | -0.304 | 1.219 | -0.613 | 1.72 | -0.001 | 0.025 | 0.025 | 0 | 0.12 | 0.12 | 0.29 | 0.38 | 0.09 | 0.16 |
| 1 | 31317480 | rs10914222 | -0.657 | 2.06 | -0.043 | 1.623 | 0.307 | 0.359 | 0.359 | 1.315 | 0.06 | 0.5 | 0.13 | 0.25 | 0.16 | 0.05 |
| 1 | 31317530 | rs7530095 | 0.235 | 1.506 | -1.617 | 0.852 | -0.061 | 0.002 | 0.002 | 0.919 | 0.75 | 0.98 | 0.68 | 0.81 | 0.71 | 0.88 |
| 1 | 31320651 | rs2488230 | -0.523 | 1.142 | -1.492 | 0.349 | -0.006 | 0.045 | 0.045 | 0.491 | 0.36 | 0.24 | 0.45 | 0.21 | 0.38 | 0.34 |
| 1 | 31321577 | rs6703029 | 0.153 | 1.327 | -1.643 | 0.312 | -0.012 | 0.01 | 0.01 | 0 | 0.11 | 0.06 | 0.08 | 0.09 | 0.13 | 0.16 |
| 1 | 31323909 | rs12405481 | 0.54 | 0.725 | -1.692 | 0.116 | 0.008 | -0.004 | -0.004 | 0 | 0.11 | 0.09 | 0.26 | 0.13 | 0.07 | 0.15 |
| 1 | 31324881 | rs2506967 | -1.105 | 0.151 | -1.409 | 0.173 | -0.002 | -0.001 | -0.001 | 0 | 0.36 | 0.36 | 0.47 | 0.36 | 0.33 | 0.35 |
| 1 | 31325090 | rs947645 | 0.305 | 0.21 | -1.302 | 0.243 | 0.009 | -0.005 | -0.005 | 0.458 | 0.18 | 0.21 | 0.2 | 0.13 | 0.25 | 0.14 |
| 1 | 31326201 | rs2796210 | -0.603 | 0.259 | -1.29 | 0.281 | -0.001 | -0.002 | -0.002 | 0 | 0.3 | 0.32 | 0.46 | 0.27 | 0.32 | 0.29 |
| 1 | 31326966 | rs6680621 | 0.325 | 0.48 | -1.155 | 0.476 | 0.114 | 0.201 | 0.201 | 0.256 | 0.2 | 0.43 | 0.36 | 0.5 | 0.3 | 0.34 |
| 1 | 31327011 | rs2491144 | -0.403 | 0.493 | -1.219 | 0.37 | -0.001 | -0.002 | -0.002 | 0 | 0.3 | 0.33 | 0.45 | 0.27 | 0.32 | 0.29 |
| 1 | 31330181 | rs2488233 | -1.609 | -0.005 | -1.581 | -0.343 | -0.001 | -0.001 | -0.001 | 0 | 0.84 | 0.86 | 0.93 | 0.87 | 0.85 | 0.96 |
| 1 | 31330844 | rs3935266 | -0.03 | -0.032 | -1.716 | -0.426 | 0.002 | 0.01 | 0.01 | 0.678 | 0.31 | 0.25 | 0.44 | 0.24 | 0.33 | 0.18 |
| 1 | 31335379 | rs2796208 | 0.762 | -0.144 | -2.052 | -0.567 | 0.002 | 0.155 | 0.155 | 0 | 0.4 | 0.12 | 0.24 | 0.15 | 0.38 | 0.39 |
| 1 | 31336439 | rs2506969 | 0.932 | -0.044 | -1.989 | -0.755 | -0.022 | 0.14 | 0.14 | 0 | 0.35 | 0.12 | 0.22 | 0.09 | 0.37 | 0.33 |
| 1 | 31337417 | rs12408228 | 0.087 | 0.181 | -1.473 | -0.337 | -0.02 | -0.012 | -0.012 | 0 | 0.22 | 0.27 | 0.34 | 0.17 | 0.17 | 0.12 |
| 1 | 31339570 | rs4949320 | -0.173 | 0.974 | -1.158 | 0.228 | 0.06 | 0.116 | 0.116 | 0 | 0.22 | 0.36 | 0.34 | 0.55 | 0.3 | 0.46 |
| 1 | 31344952 | rs2488238 | 0.234 | 0.698 | -2.047 | -0.162 | -0.033 | 0.118 | 0.118 | 0 | 0.51 | 0.17 | 0.45 | 0.25 | 0.53 | 0.44 |
| 1 | 31350566 | rs2491133 | -0.576 | 0.704 | -1.699 | 0.334 | 0.005 | 0.024 | 0.024 | 0.915 | 0.66 | 0.87 | 0.7 | 0.77 | 0.69 | 0.83 |
| 1 | 31351513 | rs1891419 | 0.606 | 1.261 | -2.071 | 0.159 | -0.097 | -0.022 | -0.022 | 0 | 0.19 | 0.03 | 0.28 | 0.11 | 0.3 | 0.25 |
| 1 | 31352419 | rs2807013 | -0.394 | 1.041 | -1.17 | 0.58 | 0.015 | 0.025 | 0.025 | 0.44 | 0.62 | 0.69 | 0.52 | 0.26 | 0.68 | 0.68 |
| 1 | 31353409 | rs12406720 | 0.204 | 0.289 | -0.709 | 0.391 | 0.002 | 0.347 | 0.347 | 0.176 | 0.14 | 0.44 | 0.24 | 0.57 | 0.11 | 0.23 |
| 1 | 31362224 | rs10753239 | -0.286 | 0.722 | -1.575 | 0.296 | -0.068 | -0.006 | -0.006 | 0 | 0.57 | 0.86 | 0.64 | 0.69 | 0.49 | 0.79 |
| 1 | 31372509 | rs7368024 | 0.273 | -0.013 | -0.572 | -0.334 | 0.019 | 0.062 | 0.062 | 0 | 0.5 | 0.33 | 0.51 | 0.66 | 0.51 | 0.67 |
| 1 | 31372686 | rs7529390 | -0.38 | 0.254 | 0.265 | -0.007 | 0.004 | 0.002 | 0.002 | 0 | 0.08 | 0.13 | 0.06 | 0.1 | 0.1 | 0.17 |
| 1 | 31372734 | rs12085929 | 0.36 | -0.029 | 0.372 | -0.143 | -0.012 | 0.055 | 0.055 | 0 | 0.1 | 0.38 | 0.11 | 0.13 | 0.04 | 0.05 |

| | | | | | | | | | | | | | | | | |
|---|---|---|---|---|---|---|---|---|---|---|---|---|---|---|---|---|
| 1 | 31377743 | rs4949324 | -0.664 | -0.116 | -0.23 | -0.35 | 0.032 | 0.053 | 0.053 | 0 | 0.62 | 0.78 | 0.61 | 0.75 | 0.64 | 0.86 |
| 1 | 31378286 | rs10158813 | 1.002 | 0.383 | 0.17 | -0.61 | 0.006 | -0.051 | -0.051 | 0 | 0.12 | 0.13 | 0.06 | 0 | 0.11 | 0.18 |
| 1 | 31380174 | rs4303117 | -1.006 | 0.048 | 0.538 | -0.798 | -0.024 | -0.067 | -0.067 | 0 | 0.6 | 0.51 | 0.62 | 0.87 | 0.66 | 0.87 |
| 1 | 31387330 | rs6680835 | -0.107 | 0.585 | 0.032 | -1.201 | -0.021 | 0.069 | 0.069 | 0.117 | 0.37 | 0.18 | 0.33 | 0.01 | 0.36 | 0.34 |
| 1 | 31408056 | rs7514514 | -1.878 | 0.529 | -0.039 | -1.075 | 0.001 | 0.119 | 0.119 | 0.858 | 0.66 | 0.95 | 0.67 | 0.87 | 0.67 | 0.91 |
| 1 | 31413972 | rs3795433 | 0.11 | 0.919 | 0.677 | 0.371 | -0.02 | -0.014 | -0.014 | 0.557 | 0.06 | 0 | 0.13 | 0.27 | 0.08 | 0.18 |
| 1 | 31417458 | rs12406355 | -0.95 | 0.954 | 0.976 | 0.294 | -0.003 | 0.006 | 0.006 | 1.642 | 0.38 | 0.3 | 0.47 | 0.35 | 0.38 | 0.37 |
| 1 | 31424889 | rs6672383 | -1.311 | 0.916 | 1.331 | 0.268 | -0.033 | -0.009 | -0.009 | 0 | 0.45 | 0.8 | 0.49 | 0.55 | 0.48 | 0.58 |
| 1 | 31426815 | rs2275741 | -0.884 | 0.994 | 1.195 | 0.325 | -0.003 | 0.005 | 0.005 | 1.895 | 0.37 | 0.29 | 0.47 | 0.35 | 0.38 | 0.36 |
| 1 | 31429785 | rs6696300 | -0.2 | 0.75 | 1.234 | -0.477 | -0.005 | 0.014 | 0.014 | 0.588 | 0.29 | 0.18 | 0.33 | 0 | 0.28 | 0.22 |
| 1 | 31464327 | rs1999811 | -1.36 | 0.024 | 1.097 | -0.09 | 0.02 | 0.204 | 0.204 | 0.247 | 0.53 | 0.83 | 0.62 | 0.84 | 0.55 | 0.73 |
| 1 | 31593640 | rs4245633 | -0.904 | -0.423 | 2.632 | 0.806 | -0.037 | -0.014 | -0.014 | 0 | 0.21 | 0.38 | 0.26 | 0.15 | 0.12 | 0.08 |
| 1 | 31606783 | rs2185419 | -1.169 | -0.309 | 1.501 | 0.86 | -0.029 | -0.015 | -0.015 | 0 | 0.23 | 0.65 | 0.29 | 0.15 | 0.12 | 0.08 |
| 1 | 31671059 | rs6677966 | 1.042 | 0.984 | 1.126 | 2.136 | 0.075 | 0.239 | 0.239 | 2.797 | 0.84 | 0.53 | 0.74 | 0.58 | 0.78 | 0.65 |
| 1 | 31687659 | rs6680246 | 1.125 | 0.392 | 0.588 | 1.23 | 0.004 | 0.049 | 0.049 | 2.515 | 0.55 | 0.37 | 0.49 | 0.18 | 0.56 | 0.32 |
| 1 | 31687958 | rs6680565 | 1.33 | 0.199 | 0.616 | 1.174 | 0.009 | 0.077 | 0.077 | 1.267 | 0.54 | 0.31 | 0.5 | 0.2 | 0.57 | 0.33 |
| 1 | 31688111 | rs6672056 | -2.038 | -0.069 | 0.484 | 1.054 | -0.018 | 0.202 | 0.202 | 0.157 | 0.53 | 0.82 | 0.7 | 0.89 | 0.47 | 0.72 |
| 1 | 31692035 | rs12755429 | -0.399 | -0.236 | 0.423 | 1.672 | 0.013 | -0.069 | -0.069 | 1.561 | 0.25 | 0.01 | 0.23 | 0.26 | 0.18 | 0.26 |
| 1 | 31699002 | rs10914332 | -1.571 | -0.178 | 1.479 | 1.898 | 0.023 | 0.048 | 0.048 | 0.11 | 0.13 | 0.36 | 0.12 | 0.22 | 0.16 | 0.24 |
| 1 | 31700412 | rs12087046 | -2.408 | -0.02 | 1.549 | 2.042 | 0.016 | 0.055 | 0.055 | 0.724 | 0.2 | 0.63 | 0.17 | 0.41 | 0.23 | 0.38 |
| 1 | 31704333 | rs6699372 | -2.248 | -0.236 | 1.416 | 2.018 | 0.01 | 0.048 | 0.048 | 0 | 0.2 | 0.47 | 0.33 | 0.27 | 0.21 | 0.32 |
| 1 | 31706636 | rs2377728 | -0.88 | -0.536 | 1.074 | 1.984 | 0.014 | -0.108 | -0.108 | 0 | 0.3 | 0.1 | 0.35 | 0.46 | 0.26 | 0.4 |
| 1 | 31707857 | rs12026718 | -0.469 | -0.535 | 1.033 | 1.993 | 0.005 | -0.164 | -0.164 | 0 | 0.27 | 0.02 | 0.33 | 0.41 | 0.22 | 0.27 |
| 1 | 31707971 | rs6701667 | 1.777 | -0.568 | 1.581 | 2.023 | 0.013 | 0.033 | 0.033 | 0.071 | 0.54 | 0.35 | 0.49 | 0.34 | 0.57 | 0.32 |
| 1 | 31719450 | rs10798838 | 1.48 | -0.585 | 1.693 | 1.975 | -0.013 | -0.021 | -0.021 | 0 | 0.57 | 0.45 | 0.52 | 0.34 | 0.62 | 0.33 |
| 1 | 31722059 | rs6658964 | -0.799 | -0.598 | 1.704 | 2 | -0.01 | 0.037 | 0.037 | 0 | 0.06 | 0.08 | 0.06 | 0.22 | 0.01 | 0.21 |
| 1 | 31728206 | rs10914337 | -0.799 | -0.598 | 1.704 | 1.976 | -0.01 | 0.031 | 0.031 | 0 | 0.06 | 0.08 | 0.06 | 0.21 | 0.01 | 0.21 |
| 1 | 31731208 | rs10753245 | 1.542 | -0.617 | 1.746 | 1.973 | -0.013 | -0.022 | -0.022 | 1.926 | 0.57 | 0.44 | 0.51 | 0.34 | 0.62 | 0.33 |
| 1 | 31732892 | rs2292165 | -2.386 | -0.625 | 1.808 | 1.997 | 0.01 | 0.037 | 0.037 | 0 | 0.14 | 0.29 | 0.14 | 0.21 | 0.17 | 0.26 |
| 1 | 31741577 | rs4949373 | 1.571 | -0.625 | 1.808 | 1.997 | -0.013 | -0.022 | -0.022 | 0.716 | 0.57 | 0.44 | 0.51 | 0.34 | 0.62 | 0.33 |
| 1 | 31741777 | rs10493047 | -1.048 | -0.627 | 1.835 | 1.996 | 0.03 | -0.036 | -0.036 | 0.952 | 0.07 | 0.14 | 0.08 | 0 | 0.15 | 0.03 |
| 1 | 31747393 | rs6425730 | 0.257 | -0.628 | 1.867 | 1.996 | 0.03 | -0.071 | -0.071 | 0.003 | 0.72 | 0.85 | 0.67 | 0.55 | 0.79 | 0.59 |
| 1 | 31753884 | rs7555364 | -1.337 | -0.626 | 1.874 | 2.031 | 0.034 | -0.042 | -0.042 | 0.842 | 0.08 | 0.19 | 0.08 | 0 | 0.15 | 0.06 |
| 1 | 31758467 | rs4949377 | -2.435 | -0.626 | 1.87 | 2.032 | 0.01 | 0.037 | 0.037 | 0 | 0.14 | 0.29 | 0.14 | 0.21 | 0.17 | 0.26 |
| 1 | 31769025 | rs10914344 | 1.565 | -0.605 | 1.842 | 2 | -0.011 | -0.023 | -0.023 | 0.033 | 0.57 | 0.44 | 0.51 | 0.34 | 0.61 | 0.33 |
| 1 | 31770944 | rs6679393 | 1.604 | -0.6 | 1.844 | 1.972 | -0.011 | -0.04 | -0.04 | 0 | 0.57 | 0.44 | 0.51 | 0.19 | 0.61 | 0.32 |
| 1 | 31773496 | rs10914347 | -2.449 | -0.6 | 1.852 | 1.996 | 0.016 | 0.055 | 0.055 | 0.004 | 0.14 | 0.33 | 0.15 | 0.22 | 0.17 | 0.26 |
| 1 | 31788313 | rs7523438 | 1.49 | -0.601 | 1.854 | 1.996 | -0.003 | -0.006 | -0.006 | 0 | 0.58 | 0.53 | 0.52 | 0.35 | 0.61 | 0.33 |
| 1 | 31793256 | rs4949385 | 1.61 | -0.601 | 1.852 | 1.997 | -0.011 | -0.04 | -0.04 | 0 | 0.57 | 0.44 | 0.51 | 0.18 | 0.61 | 0.32 |
| 1 | 31795460 | rs11576621 | 1.601 | -0.602 | 1.851 | 1.996 | -0.011 | -0.04 | -0.04 | 0 | 0.57 | 0.44 | 0.51 | 0.18 | 0.61 | 0.32 |
| 1 | 31805314 | rs12564155 | -0.399 | -0.598 | 1.856 | 1.998 | 0.023 | -0.112 | -0.112 | 0.622 | 0.28 | 0.07 | 0.33 | 0.43 | 0.22 | 0.41 |
| 1 | 31810681 | rs11582342 | 1.602 | -0.598 | 1.853 | 1.997 | -0.011 | -0.04 | -0.04 | 0 | 0.57 | 0.44 | 0.51 | 0.18 | 0.61 | 0.32 |
| 1 | 31819128 | rs10798844 | 1.094 | -0.551 | 1.857 | 1.996 | -0.001 | 0 | 0 | 0 | 0.63 | 0.57 | 0.57 | 0.4 | 0.63 | 0.54 |
| 1 | 31822636 | rs10798845 | 1.601 | -0.585 | 1.853 | 1.998 | -0.011 | -0.041 | -0.041 | 0.146 | 0.57 | 0.44 | 0.51 | 0.18 | 0.61 | 0.32 |
| 1 | 31829873 | rs3795432 | -2.703 | -0.573 | 1.853 | 2.004 | 0.017 | 0.122 | 0.122 | 0.035 | 0.15 | 0.43 | 0.16 | 0.38 | 0.17 | 0.27 |
| 1 | 31837089 | rs12139997 | -1.061 | -0.569 | 1.836 | 2 | 0.03 | -0.036 | -0.036 | 0 | 0.07 | 0.14 | 0.08 | 0 | 0.15 | 0.03 |
| 1 | 31841619 | rs7532845 | -0.82 | -0.577 | 1.802 | 2.006 | -0.023 | 0.076 | 0.076 | 0 | 0.06 | 0.12 | 0.07 | 0.22 | 0.01 | 0.21 |
| 1 | 31846206 | rs10914367 | 0.259 | -0.588 | 1.756 | 1.972 | 0.027 | -0.077 | -0.077 | 0 | 0.72 | 0.87 | 0.67 | 0.57 | 0.78 | 0.59 |
| 1 | 31847419 | rs12401792 | -0.769 | -0.582 | 1.751 | 1.965 | 0.031 | -0.114 | -0.114 | 0.035 | 0.31 | 0.08 | 0.35 | 0.43 | 0.26 | 0.43 |
| 1 | 31850657 | rs6425744 | 0.252 | -0.633 | 1.746 | 1.933 | 0.029 | -0.064 | -0.064 | 0 | 0.72 | 0.87 | 0.67 | 0.6 | 0.78 | 0.59 |
| 1 | 31851555 | rs6694857 | -1.221 | -0.628 | 1.811 | 1.93 | 0.016 | -0.025 | -0.025 | 0 | 0.07 | 0.11 | 0.06 | 0 | 0.15 | 0.03 |
| 1 | 31851875 | rs4949393 | 1.601 | -0.58 | 1.718 | 1.944 | -0.01 | -0.034 | -0.034 | 0 | 0.57 | 0.44 | 0.5 | 0.2 | 0.61 | 0.32 |
| 1 | 31854882 | rs12562824 | -2.833 | -0.549 | 1.698 | 1.95 | 0.019 | 0.144 | 0.144 | 0.027 | 0.15 | 0.42 | 0.16 | 0.4 | 0.17 | 0.26 |
| 1 | 31858784 | rs12122184 | -2.751 | -0.544 | 1.689 | 1.886 | 0.024 | 0.091 | 0.091 | 0.051 | 0.14 | 0.43 | 0.16 | 0.24 | 0.17 | 0.26 |
| 1 | 31864450 | rs6683793 | -1.641 | -0.512 | 1.573 | 1.823 | 0.041 | -0.044 | -0.044 | 0.918 | 0.08 | 0.24 | 0.09 | 0 | 0.15 | 0.05 |
| 1 | 31869669 | rs6659255 | -1.602 | -0.521 | 1.51 | 1.903 | 0.046 | 0.134 | 0.134 | 0 | 0.1 | 0.36 | 0.14 | 0.24 | 0.13 | 0.25 |
| 1 | 31871407 | rs2889753 | -0.709 | -0.619 | 1.313 | 1.931 | 0.029 | -0.097 | -0.097 | 0.015 | 0.31 | 0.07 | 0.35 | 0.4 | 0.25 | 0.42 |
| 1 | 31872581 | rs10914374 | -1.911 | -0.534 | 1.604 | 1.89 | 0.029 | 0.103 | 0.103 | 0.166 | 0.15 | 0.45 | 0.16 | 0.24 | 0.17 | 0.26 |
| 1 | 31873525 | rs1566960 | -1.864 | -0.533 | 1.605 | 1.891 | 0.031 | 0.106 | 0.106 | 0.192 | 0.14 | 0.44 | 0.16 | 0.24 | 0.17 | 0.26 |
| 1 | 31883925 | rs4478858 | -1.538 | -0.454 | 1.421 | 1.881 | -0.003 | -0.017 | -0.017 | 0.86 | 0.48 | 0.53 | 0.52 | 0.8 | 0.44 | 0.68 |
| 1 | 31885859 | rs10914375 | -1.556 | -0.454 | 1.412 | 1.881 | -0.002 | -0.014 | -0.014 | 0 | 0.48 | 0.53 | 0.52 | 0.8 | 0.44 | 0.68 |
| 1 | 31898132 | rs4949400 | 1.339 | -0.451 | 1.008 | 1.865 | 0.033 | 0.188 | 0.188 | 0.045 | 0.5 | 0.15 | 0.45 | 0.2 | 0.56 | 0.33 |
| 1 | 31898234 | rs4949402 | 1.546 | -0.441 | 1.013 | 1.834 | 0.033 | 0.108 | 0.108 | 0.026 | 0.5 | 0.15 | 0.45 | 0.36 | 0.56 | 0.33 |
| 1 | 31902418 | rs10047085 | -1.031 | -0.319 | 1.071 | 1.715 | -0.003 | 0.004 | 0.004 | 0 | 0.39 | 0.47 | 0.43 | 0.41 | 0.38 | 0.46 |
| 1 | 31909124 | rs10914387 | 0.987 | -0.694 | 1.321 | 1.449 | 0.125 | 0.108 | 0.108 | 0.074 | 0.55 | 0.15 | 0.53 | 0.21 | 0.73 | 0.4 |
| 1 | 31910482 | rs6702129 | 0.691 | -0.774 | 1.693 | 1.407 | 0.019 | 0.016 | 0.016 | 0 | 0.61 | 0.63 | 0.58 | 0.21 | 0.74 | 0.42 |
| 1 | 31915790 | rs7548805 | 1.087 | -0.676 | 1.49 | 1.429 | 0.123 | 0.116 | 0.116 | 0.035 | 0.55 | 0.14 | 0.53 | 0.21 | 0.72 | 0.4 |
| 1 | 31921177 | rs7546536 | 1.097 | -0.692 | 1.503 | 1.753 | 0.126 | 0.119 | 0.119 | 0.054 | 0.55 | 0.12 | 0.53 | 0.21 | 0.72 | 0.41 |
| 1 | 31928387 | rs2105217 | 0.884 | -0.693 | 1.356 | 1.721 | -0.024 | -0.006 | -0.006 | 0 | 0.58 | 0.48 | 0.57 | 0.38 | 0.73 | 0.46 |

| | | | | | | | | | | | | | | | | |
|---|---|---|---|---|---|---|---|---|---|---|---|---|---|---|---|---|
| 1 | 31934343 | rs10798854 | 0.756 | -0.704 | 1.323 | 1.717 | 0.008 | 0.002 | 0.002 | 0 | 0.6 | 0.61 | 0.59 | 0.38 | 0.73 | 0.46 |
| 1 | 31949720 | rs10914410 | -0.526 | -0.572 | 0.838 | 2.196 | -0.03 | -0.107 | -0.107 | 1.148 | 0.32 | 0.07 | 0.36 | 0.47 | 0.32 | 0.37 |
| 1 | 31953898 | rs4949416 | -1.152 | -0.581 | 1.266 | 2.496 | -0.001 | 0.219 | 0.219 | 0.212 | 0.05 | 0.19 | 0.05 | 0.27 | 0.03 | 0.18 |
| 1 | 31955533 | rs10798859 | -0.499 | -0.612 | 1.262 | 2.418 | -0.03 | -0.107 | -0.107 | 0 | 0.32 | 0.07 | 0.35 | 0.47 | 0.32 | 0.38 |
| 1 | 31972756 | rs1316509 | 1.295 | -0.594 | 1.742 | 2.389 | 0.008 | 0.029 | 0.029 | 0.056 | 0.54 | 0.4 | 0.39 | 0.21 | 0.58 | 0.29 |
| 1 | 31973440 | rs1574104 | 0.817 | -0.589 | 1.762 | 2.475 | 0 | -0.002 | -0.002 | 0 | 0.59 | 0.59 | 0.45 | 0.49 | 0.6 | 0.5 |
| 1 | 31977350 | rs10914420 | 1.281 | -0.5 | 1.569 | 2.091 | -0.006 | 0.024 | 0.024 | 0.799 | 0.45 | 0.39 | 0.34 | 0.21 | 0.5 | 0.18 |
| 1 | 31982479 | rs6684549 | -1.466 | -0.508 | 1.619 | 2.132 | 0.001 | -0.024 | -0.024 | 0 | 0.53 | 0.5 | 0.64 | 0.79 | 0.5 | 0.8 |
| 1 | 31987662 | rs10914430 | -0.69 | -0.554 | 1.485 | 2.152 | 0.023 | 0.019 | 0.019 | 0 | 0.42 | 0.28 | 0.42 | 0.64 | 0.38 | 0.59 |
| 1 | 31990783 | rs6657902 | -1.478 | -0.548 | 1.53 | 2.137 | -0.009 | 0.096 | 0.096 | 0.397 | 0.53 | 0.33 | 0.63 | 0.79 | 0.5 | 0.8 |
| 1 | 31992493 | rs6667261 | 1.463 | -0.501 | 1.872 | 2.146 | -0.033 | 0.134 | 0.134 | 1.673 | 0.43 | 0.26 | 0.32 | 0.1 | 0.5 | 0.17 |
| 1 | 31993870 | rs10493048 | -0.825 | -0.488 | 1.738 | 2.172 | -0.013 | -0.022 | -0.022 | 1.907 | 0.1 | 0.06 | 0.21 | 0.15 | 0.12 | 0.21 |
| 1 | 31995973 | rs6662915 | 0.631 | -0.493 | 1.686 | 2.18 | 0.025 | 0.041 | 0.041 | 0.34 | 0.56 | 0.67 | 0.56 | 0.25 | 0.62 | 0.39 |
| 1 | 32001500 | rs6425763 | 0.66 | -0.484 | 1.639 | 2.131 | 0.027 | 0.024 | 0.024 | 0.252 | 0.55 | 0.68 | 0.57 | 0.37 | 0.62 | 0.39 |
| 1 | 32002287 | rs4245634 | 0.151 | -0.516 | 1.275 | 2.259 | -0.002 | -0.027 | -0.027 | 0 | 0.61 | 0.88 | 0.62 | 0.64 | 0.65 | 0.61 |
| 1 | 32002794 | rs10914436 | -1.231 | -0.498 | 1.452 | 2.263 | 0.067 | 0.409 | 0.409 | 0.327 | 0.06 | 0.41 | 0.07 | 0.28 | 0.03 | 0.22 |
| 1 | 32004898 | rs4341399 | 0.171 | -0.506 | 1.462 | 2.278 | 0.005 | -0.016 | -0.016 | 0.07 | 0.61 | 0.87 | 0.62 | 0.65 | 0.64 | 0.61 |
| 1 | 32007672 | rs6425766 | 1.367 | -0.438 | 1.459 | 1.992 | -0.043 | 0.151 | 0.151 | 1.346 | 0.43 | 0.23 | 0.32 | 0.1 | 0.49 | 0.17 |
| 1 | 32009650 | rs10798873 | -1.556 | -0.341 | 1.407 | 2.185 | 0.067 | 0.409 | 0.409 | 0.327 | 0.06 | 0.41 | 0.07 | 0.28 | 0.03 | 0.22 |
| 1 | 32012957 | rs4245636 | -0.715 | -0.389 | 0.507 | 1.948 | 0.001 | -0.001 | -0.001 | 0.19 | 0.39 | 0.2 | 0.42 | 0.6 | 0.39 | 0.57 |
| 1 | 32015974 | rs4949441 | 0.564 | -0.366 | 0.561 | 1.946 | 0.008 | 0.009 | 0.009 | 0.017 | 0.53 | 0.55 | 0.54 | 0.36 | 0.63 | 0.45 |
| 1 | 32017993 | rs4949213 | 0.883 | -0.289 | 0.864 | 2.2 | 0.002 | 0.023 | 0.023 | 0.722 | 0.47 | 0.27 | 0.4 | 0.45 | 0.54 | 0.38 |
| 1 | 32019153 | rs6670560 | 1 | -0.293 | 0.752 | 2.098 | -0.024 | 0.131 | 0.131 | 1 | 0.46 | 0.14 | 0.37 | 0.25 | 0.54 | 0.37 |
| 1 | 32022031 | rs4949444 | 0.479 | -0.422 | 0.719 | 2.195 | 0.001 | 0 | 0 | 0 | 0.53 | 0.52 | 0.56 | 0.59 | 0.63 | 0.46 |
| 1 | 32031672 | rs2992028 | 0.373 | -0.45 | 0.561 | 2.023 | -0.001 | 0.001 | 0.001 | 0 | 0.42 | 0.53 | 0.51 | 0.57 | 0.56 | 0.39 |
| 1 | 32031843 | rs2377894 | -0.65 | -0.259 | 0.654 | 1.671 | 0.078 | 0.097 | 0.097 | 0.174 | 0.06 | 0.29 | 0.17 | 0.12 | 0.09 | 0.07 |
| 1 | 32036927 | rs9425884 | 0.94 | -0.288 | -0.671 | 1.453 | -0.111 | 0.043 | 0.043 | 0.045 | 0.35 | 0.11 | 0.31 | 0.24 | 0.48 | 0.3 |
| 1 | 32038716 | rs10914443 | -0.229 | -1.158 | -0.426 | 1.454 | -0.011 | 0.009 | 0.009 | 0 | 0.38 | 0.48 | 0.44 | 0.34 | 0.34 | 0.42 |
| 1 | 32038787 | rs3008406 | -1.057 | -0.573 | -0.696 | 1.533 | -0.018 | 0.037 | 0.037 | 0 | 0.63 | 0.7 | 0.58 | 0.83 | 0.5 | 0.7 |
| 1 | 32042268 | rs3738008 | -0.972 | -0.123 | -0.888 | 1.545 | -0.018 | -0.033 | -0.033 | 0 | 0.06 | 0 | 0.04 | 0.18 | 0.08 | 0.04 |
| 1 | 32047632 | rs945213 | 0.269 | -0.202 | -1.057 | 1.344 | -0.005 | -0.008 | -0.008 | 0 | 0.66 | 0.75 | 0.72 | 0.56 | 0.65 | 0.66 |
| 1 | 32048052 | rs1107550 | 0.371 | -0.333 | -0.396 | 1.025 | 0.007 | 0.024 | 0.024 | 0 | 0.72 | 0.62 | 0.76 | 0.53 | 0.73 | 0.69 |
| 1 | 32054094 | rs10914446 | 0.717 | -0.909 | -0.145 | 0.781 | -0.01 | 0.014 | 0.014 | 0.519 | 0.68 | 0.38 | 0.56 | 0.36 | 0.71 | 0.62 |
| 1 | 32062066 | rs4949448 | 1.557 | -1.182 | -1.133 | -0.125 | -0.151 | 0.167 | 0.167 | 0.187 | 0.4 | 0.05 | 0.28 | 0.12 | 0.48 | 0.23 |
| 1 | 32070234 | rs6681414 | 1.07 | -1.115 | -1.201 | -0.314 | -0.024 | 0.136 | 0.136 | 0.345 | 0.47 | 0.08 | 0.47 | 0.25 | 0.56 | 0.55 |
| 1 | 32075036 | rs10914453 | 0.411 | -1.318 | -0.53 | -0.433 | -0.009 | -0.012 | -0.012 | 0 | 0.6 | 0.46 | 0.53 | 0.29 | 0.65 | 0.63 |
| 1 | 32081785 | rs3753614 | 1.123 | -1.12 | -0.802 | -0.479 | 0.002 | 0.143 | 0.143 | 0.09 | 0.49 | 0.09 | 0.47 | 0.24 | 0.59 | 0.56 |
| 1 | 32084904 | rs1056526 | 1.234 | -1.154 | -0.775 | -0.488 | -0.019 | 0.147 | 0.147 | 0.086 | 0.47 | 0.08 | 0.47 | 0.24 | 0.56 | 0.55 |
| 1 | 32085637 | rs10914455 | 0.549 | -1.18 | -0.534 | -0.508 | -0.008 | -0.061 | -0.061 | 3.762 | 0.11 | 0.2 | 0.04 | 0.01 | 0.07 | 0.07 |
| 1 | 32086089 | rs10753257 | 0.53 | -1.184 | -0.533 | -0.504 | 0.023 | 0.05 | 0.05 | 0 | 0.58 | 0.28 | 0.51 | 0.24 | 0.63 | 0.62 |
| 1 | 32092525 | rs2271933 | 0.522 | -1.164 | -0.52 | -0.502 | 0.023 | 0.046 | 0.046 | 0 | 0.58 | 0.28 | 0.51 | 0.24 | 0.63 | 0.62 |
| 1 | 32092735 | rs2176807 | 1.166 | -1.085 | -0.585 | -0.47 | -0.015 | 0.141 | 0.141 | 0.055 | 0.47 | 0.08 | 0.47 | 0.24 | 0.57 | 0.54 |
| 1 | 32093561 | rs6703093 | 1.172 | -1.09 | -0.546 | -0.473 | -0.017 | 0.143 | 0.143 | 0.004 | 0.47 | 0.09 | 0.47 | 0.24 | 0.56 | 0.54 |
| 1 | 32094261 | rs10914457 | 1.985 | -1.034 | -0.67 | -0.562 | -0.124 | 0.14 | 0.14 | 0 | 0.37 | 0.06 | 0.24 | 0.12 | 0.44 | 0.22 |
| 1 | 32105192 | rs7529064 | 1.294 | -1.407 | -0.356 | -0.291 | -0.017 | 0.138 | 0.138 | 0.042 | 0.47 | 0.08 | 0.47 | 0.24 | 0.56 | 0.54 |
| 1 | 32108579 | rs7524405 | 0.459 | -1.459 | 0.162 | -0.322 | -0.011 | -0.016 | -0.016 | 0 | 0.6 | 0.51 | 0.53 | 0.29 | 0.64 | 0.62 |
| 1 | 32109849 | rs7549590 | 1.166 | -1.386 | -0.065 | -0.313 | 0.009 | 0.195 | 0.195 | 0.022 | 0.48 | 0.09 | 0.47 | 0.22 | 0.57 | 0.54 |
| 1 | 32110813 | rs7542583 | 1.071 | -1.389 | 0.053 | -0.314 | 0.041 | 0.129 | 0.129 | 0 | 0.49 | 0.23 | 0.48 | 0.22 | 0.57 | 0.54 |
| 1 | 32117779 | rs2139539 | 1.174 | -1.377 | -0.362 | -0.311 | 0.013 | 0.194 | 0.194 | 0.048 | 0.48 | 0.09 | 0.47 | 0.22 | 0.57 | 0.54 |
| 1 | 32124597 | rs16834652 | -0.359 | -1.249 | 0.108 | -0.231 | 0.011 | -0.035 | -0.035 | 0 | 0.12 | 0.41 | 0.06 | 0.05 | 0.07 | 0.07 |
| 1 | 32125943 | rs10914462 | -0.971 | -1.387 | -0.008 | -0.112 | -0.001 | -0.003 | -0.003 | 0 | 0.4 | 0.52 | 0.48 | 0.69 | 0.42 | 0.34 |
| 1 | 32127953 | rs2271928 | -0.831 | -1.383 | 0.001 | -0.121 | -0.001 | 0 | 0 | 0 | 0.39 | 0.52 | 0.47 | 0.64 | 0.4 | 0.31 |
| 1 | 32135495 | rs7537646 | 1.338 | -1.351 | -0.464 | -0.121 | 0.066 | 0.198 | 0.198 | 0 | 0.5 | 0.11 | 0.46 | 0.23 | 0.6 | 0.6 |
| 1 | 32135773 | rs2292989 | -0.084 | -1.372 | -0.164 | -0.036 | -0.001 | -0.016 | -0.016 | 0 | 0.31 | 0.24 | 0.31 | 0.6 | 0.34 | 0.28 |
| 1 | 32139635 | rs909002 | 0.928 | -0.982 | -0.093 | -0.412 | 0.067 | 0.144 | 0.144 | 0 | 0.51 | 0.2 | 0.6 | 0.25 | 0.58 | 0.6 |
| 1 | 32142354 | rs4949459 | 0.834 | -0.736 | -0.576 | -0.654 | 0.017 | 0.129 | 0.129 | 0.075 | 0.46 | 0.18 | 0.57 | 0.24 | 0.55 | 0.59 |
| 1 | 32146764 | rs2297684 | 0.326 | -0.538 | -0.114 | -0.518 | -0.008 | -0.043 | -0.043 | 0.293 | 0.6 | 0.49 | 0.65 | 0.23 | 0.62 | 0.61 |
| 1 | 32155911 | rs6425780 | 0.292 | -0.299 | -0.203 | -0.965 | 0.02 | 0.122 | 0.122 | 0 | 0.5 | 0.69 | 0.34 | 0.15 | 0.56 | 0.32 |
| 1 | 32156940 | rs2297680 | -0.224 | -0.206 | -0.285 | -0.544 | -0.001 | -0.004 | -0.004 | 0 | 0.13 | 0.08 | 0.35 | 0.15 | 0.11 | 0.4 |
| 1 | 32157531 | rs2297676 | 0.939 | -0.406 | 0.231 | -0.848 | -0.015 | 0.086 | 0.086 | 0 | 0.43 | 0.3 | 0.29 | 0.12 | 0.53 | 0.27 |
| 1 | 32158374 | rs3818788 | -0.271 | -0.448 | 0.391 | -0.423 | 0.03 | 0.027 | 0.027 | 0 | 0.59 | 0.77 | 0.68 | 0.36 | 0.65 | 0.67 |
| 1 | 32160918 | rs6656079 | -0.22 | -0.408 | 0.283 | -0.549 | 0.041 | 0.081 | 0.081 | 0 | 0.54 | 0.81 | 0.67 | 0.36 | 0.65 | 0.39 |
| 1 | 32165059 | rs6657383 | 0.891 | -0.349 | 1.047 | -0.804 | -0.017 | 0.049 | 0.049 | 0 | 0.4 | 0.3 | 0.29 | 0.16 | 0.53 | 0.26 |
| 1 | 32165495 | rs2228552 | -0.186 | -0.419 | 1.17 | -0.82 | 0.052 | 0.123 | 0.123 | 0 | 0.52 | 0.26 | 0.46 | 0.27 | 0.65 | 0.57 |
| 1 | 32168879 | rs10493049 | -1.313 | -0.28 | 1.84 | -0.659 | -0.007 | -0.032 | -0.032 | 0.032 | 0.15 | 0.37 | 0.21 | 0.13 | 0.12 | 0.33 |
| 1 | 32169415 | rs4949463 | -1.117 | -0.273 | 1.936 | -0.655 | 0 | -0.024 | -0.024 | 0 | 0.14 | 0.23 | 0.19 | 0.11 | 0.12 | 0.32 |
| 1 | 32179261 | rs7524140 | -1.324 | -0.215 | 1.533 | -0.6 | -0.001 | -0.006 | -0.006 | 0.356 | 0.14 | 0.27 | 0.19 | 0.16 | 0.12 | 0.32 |
| 1 | 32181220 | rs10914472 | 0.966 | -0.418 | 0.848 | -0.846 | -0.057 | 0.112 | 0.112 | 0 | 0.37 | 0.18 | 0.28 | 0.16 | 0.53 | 0.24 |
| 1 | 32195819 | rs1889571 | -1.234 | -0.167 | 1.335 | -0.281 | -0.002 | 0.017 | 0.017 | 0.915 | 0.13 | 0.04 | 0.17 | 0.12 | 0.11 | 0.32 |

| | | | | | | | | | | | | | | | | |
|---|---|---|---|---|---|---|---|---|---|---|---|---|---|---|---|---|
| 1 | 32203218 | rs2271932 | 0.039 | -0.118 | 1.326 | -0.249 | -0.009 | -0.003 | -0.003 | 0 | 0.06 | 0.02 | 0.15 | 0.08 | 0.07 | 0.08 |
| 1 | 32204683 | rs2050256 | -2.118 | -0.068 | 1.801 | -0.29 | -0.005 | -0.007 | -0.007 | 0 | 0.19 | 0.5 | 0.23 | 0.21 | 0.16 | 0.34 |
| 1 | 32212514 | rs12035217 | -2.33 | 0.061 | 1.832 | -0.192 | 0.092 | 0.146 | 0.146 | 3.154 | 0.06 | 0.53 | 0.07 | 0.12 | 0.05 | 0.03 |
| 1 | 32213655 | rs12037019 | -1.814 | 0.068 | 1.877 | -0.188 | 0.1 | 0.153 | 0.153 | 1.832 | 0.05 | 0.42 | 0.06 | 0.12 | 0.05 | 0.03 |
| 1 | 32215178 | rs12037400 | -2.043 | 0.065 | 1.653 | -0.189 | 0.087 | 0.14 | 0.14 | 3.956 | 0.07 | 0.54 | 0.07 | 0.12 | 0.05 | 0.04 |
| 1 | 32242206 | rs877674 | -0.603 | 0.165 | 0.279 | -0.071 | -0.025 | -0.025 | -0.025 | 0.201 | 0.11 | 0.01 | 0.32 | 0.16 | 0.1 | 0.34 |
| 1 | 32252615 | rs6656287 | -1.217 | 1.411 | 0.867 | 0.005 | 0.195 | 0.022 | 0.022 | 0.288 | 0.09 | 0.5 | 0.13 | 0.04 | 0.18 | 0.03 |
| 1 | 32256086 | rs13426 | -0.31 | 0.972 | -0.507 | -0.174 | -0.001 | -0.016 | -0.016 | 0 | 0.72 | 0.87 | 0.74 | 0.43 | 0.74 | 0.76 |
| 1 | 32266026 | rs552124 | -0.339 | 0.99 | -0.463 | 0.496 | 0.007 | -0.053 | -0.053 | 0 | 0.71 | 0.83 | 0.74 | 0.56 | 0.73 | 0.79 |
| 1 | 32277199 | rs2184202 | -0.563 | 1.403 | 0.272 | 0.466 | 0.024 | 0.034 | 0.034 | 0 | 0.49 | 0.61 | 0.61 | 0.56 | 0.56 | 0.41 |
| 1 | 32278308 | rs751137 | -1.234 | 1.035 | -0.034 | 0.249 | -0.002 | -0.003 | -0.003 | 0.158 | 0.68 | 0.64 | 0.67 | 0.72 | 0.67 | 0.75 |
| 1 | 32279629 | rs6669563 | 0.628 | 1.361 | 0.239 | 0.253 | -0.055 | -0.059 | -0.059 | 0 | 0.37 | 0.19 | 0.5 | 0.42 | 0.44 | 0.28 |
| 1 | 32280610 | rs6664445 | -1.387 | 1.349 | 0.176 | 0.218 | -0.002 | -0.003 | -0.003 | 0.012 | 0.69 | 0.64 | 0.67 | 0.72 | 0.67 | 0.75 |
| 1 | 32285400 | rs11806757 | 0.275 | 1.422 | 0.075 | 0.252 | -0.003 | -0.003 | -0.003 | 0.026 | 0.11 | 0.11 | 0.09 | 0.14 | 0.12 | 0.13 |
| 1 | 32290078 | rs10798893 | 0.54 | 1.444 | 0.046 | 0.458 | -0.1 | -0.076 | -0.076 | 0 | 0.37 | 0.13 | 0.49 | 0.37 | 0.44 | 0.25 |
| 1 | 32296525 | rs4949473 | 0.585 | 1.416 | 0.309 | 0.538 | -0.009 | -0.01 | -0.01 | 1.994 | 0.34 | 0.29 | 0.47 | 0.38 | 0.4 | 0.25 |
| 1 | 32300130 | rs11589992 | -0.235 | 1.392 | 0.309 | 0.56 | -0.005 | 0.017 | 0.017 | 0 | 0.53 | 0.31 | 0.53 | 0.58 | 0.51 | 0.6 |
| 1 | 32302108 | rs11578323 | -0.277 | 1.378 | 0.437 | 0.552 | -0.003 | 0.006 | 0.006 | 0.117 | 0.53 | 0.37 | 0.29 | 0.49 | 0.51 | 0.59 |
| 1 | 32302791 | rs10914486 | -0.11 | 1.317 | 0.553 | 0.565 | 0.039 | 0.035 | 0.035 | 0 | 0.12 | 0.34 | 0.11 | 0.12 | 0.12 | 0.13 |
| 1 | 32307399 | rs12733547 | 0.514 | 1.134 | 0.183 | 0.541 | 0.04 | 0.014 | 0.014 | 0 | 0.19 | 0.07 | 0.06 | 0.21 | 0.11 | 0.34 |
| 1 | 32307489 | rs2025600 | -0.118 | 1.54 | 0.164 | 0.573 | 0.006 | 0.012 | 0.012 | 0 | 0.52 | 0.34 | 0.26 | 0.48 | 0.45 | 0.58 |
| 1 | 32311335 | rs12565351 | -0.221 | 1.742 | -0.05 | 0.612 | 0.004 | 0 | 0 | 0.014 | 0.12 | 0.12 | 0.12 | 0.12 | 0.19 | 0.14 |
| 1 | 32315147 | rs7518553 | 0.568 | 1.239 | -0.119 | 0.581 | -0.001 | 0 | 0 | 0 | 0.3 | 0.32 | 0.2 | 0.28 | 0.29 | 0.23 |
| 1 | 32322733 | rs12565681 | -0.029 | 0.614 | -0.493 | 0.499 | 0.012 | -0.038 | -0.038 | 0 | 0.36 | 0.2 | 0.24 | 0.46 | 0.32 | 0.37 |
| 1 | 32336098 | rs671101 | -1.154 | 0.083 | 0.284 | 0.053 | -0.034 | 0.012 | 0.012 | 0 | 0.32 | 0.4 | 0.32 | 0.4 | 0.21 | 0.46 |
| 1 | 32336784 | rs11805047 | -0.31 | 0.09 | 0.191 | 0.043 | -0.005 | 0.007 | 0.007 | 0 | 0.12 | 0.17 | 0.09 | 0.15 | 0.11 | 0.1 |
| 1 | 32342268 | rs687234 | -1.474 | -0.006 | 0.539 | 0.008 | -0.015 | 0.003 | 0.003 | 0 | 0.34 | 0.63 | 0.34 | 0.4 | 0.21 | 0.47 |
| 1 | 32342334 | rs687605 | -1.245 | -0.002 | 0.544 | 0.01 | -0.04 | 0.017 | 0.017 | 0 | 0.32 | 0.43 | 0.33 | 0.4 | 0.21 | 0.46 |
| 1 | 32343368 | rs594540 | 0.211 | 0.038 | 0.116 | -0.01 | 0.04 | -0.04 | -0.04 | 0 | 0.18 | 0.05 | 0.21 | 0.24 | 0.1 | 0.34 |
| 1 | 32347171 | rs597987 | -1.366 | 0.061 | 0.33 | -0.258 | -0.021 | 0.004 | 0.004 | 0 | 0.34 | 0.64 | 0.34 | 0.4 | 0.21 | 0.43 |
| 1 | 32371442 | rs16834765 | 0.631 | 0.067 | 0.516 | -0.21 | -0.02 | -0.064 | -0.064 | 0.371 | 0.12 | 0.24 | 0.06 | 0 | 0.06 | 0.01 |
| 1 | 32371523 | rs682654 | -1.38 | -0.164 | 0.205 | -0.253 | -0.052 | 0.041 | 0.041 | 1.649 | 0.26 | 0.63 | 0.31 | 0.39 | 0.14 | 0.34 |
| 1 | 32372026 | rs646689 | -0.176 | -0.142 | 0.2 | -0.256 | 0.001 | 0.028 | 0.028 | 0.75 | 0.1 | 0.12 | 0.21 | 0.39 | 0.08 | 0.31 |
| 1 | 32372881 | rs639233 | -1.316 | -0.146 | -0.018 | -0.242 | -0.049 | 0.038 | 0.038 | 1.663 | 0.26 | 0.64 | 0.31 | 0.4 | 0.14 | 0.34 |
| 1 | 32378540 | rs10493050 | 0.931 | 0.201 | -0.639 | -0.558 | 0.016 | 0.04 | 0.04 | 0.052 | 0.11 | 0.03 | 0.04 | 0 | 0.06 | 0.01 |
| 1 | 32390324 | rs563978 | -1.51 | 0.94 | -0.873 | -0.766 | -0.059 | 0.102 | 0.102 | 0 | 0.77 | 0.95 | 0.84 | 0.92 | 0.68 | 0.83 |
| 1 | 32410482 | rs648718 | -1.003 | 0.467 | -0.783 | -0.686 | -0.149 | 0.192 | 0.192 | 0.682 | 0.67 | 0.94 | 0.78 | 0.92 | 0.52 | 0.81 |
| 1 | 32419044 | rs9792944 | 0.573 | 0.187 | 0.092 | 0.063 | -0.008 | 0.038 | 0.038 | 0.774 | 0.11 | 0.03 | 0.06 | 0 | 0.09 | 0.01 |
| 1 | 32424756 | rs3886437 | -0.747 | 0.122 | 0.726 | -0.011 | -0.004 | -0.102 | -0.102 | 0.097 | 0.16 | 0.39 | 0.12 | 0 | 0.11 | 0.04 |
| 1 | 32427388 | rs4949486 | -0.874 | 0.106 | 0.831 | -0.018 | -0.002 | -0.13 | -0.13 | 0 | 0.17 | 0.48 | 0.12 | 0 | 0.11 | 0.04 |
| 1 | 32435958 | rs687860 | -1.229 | 0.011 | 0.348 | 0.712 | 0.006 | -0.009 | -0.009 | 0 | 0.16 | 0.15 | 0.25 | 0.38 | 0.12 | 0.39 |
| 1 | 32437268 | rs624017 | -1.635 | -0.001 | 0.47 | 0.701 | -0.003 | 0.008 | 0.008 | 0 | 0.2 | 0.23 | 0.28 | 0.38 | 0.14 | 0.41 |
| 1 | 32438234 | rs680328 | -1.843 | -0.292 | 0.844 | 0.708 | -0.032 | 0.043 | 0.043 | 0 | 0.22 | 0.36 | 0.31 | 0.42 | 0.13 | 0.41 |
| 1 | 32456015 | rs7541800 | -1.038 | -0.264 | 1.496 | 0.534 | -0.004 | 0.011 | 0.011 | 0 | 0.07 | 0.23 | 0.03 | 0.04 | 0.03 | 0.03 |
| 1 | 32466331 | rs16834799 | -0.603 | -0.131 | 1.419 | 0.687 | -0.017 | -0.009 | -0.009 | 1.071 | 0.11 | 0.03 | 0.06 | 0.13 | 0.13 | 0.1 |
| 1 | 32501215 | rs12094507 | -2.422 | -0.474 | 2.301 | 0.768 | 0.002 | -0.003 | -0.003 | 0 | 0.3 | 0.69 | 0.36 | 0.4 | 0.23 | 0.43 |
| 1 | 32547587 | rs16834858 | -0.704 | -0.393 | 2.54 | 0.884 | 0.056 | 0.259 | 0.259 | 0 | 0.06 | 0.26 | 0.22 | 0.24 | 0.06 | 0.29 |
| 1 | 32557486 | rs744455 | -1.029 | -0.437 | 2.595 | 0.879 | 0.052 | 0.262 | 0.262 | 0.004 | 0.06 | 0.29 | 0.23 | 0.24 | 0.06 | 0.28 |
| 1 | 32567617 | rs2282517 | -1.029 | -0.438 | 2.6 | 0.91 | 0.049 | 0.26 | 0.26 | 0.004 | 0.06 | 0.29 | 0.23 | 0.24 | 0.06 | 0.28 |
| 1 | 32661059 | rs16834922 | -1.734 | -0.411 | 3.328 | 0.887 | 0.059 | 0.271 | 0.271 | 0.48 | 0.07 | 0.39 | 0.25 | 0.23 | 0.06 | 0.29 |
| 1 | 32664866 | rs16834926 | -0.881 | -0.396 | 3.154 | 0.894 | 0.028 | 0.191 | 0.191 | 0 | 0.06 | 0.27 | 0.23 | 0.23 | 0.06 | 0.29 |
| 1 | 32667609 | rs1407134 | -0.177 | -0.454 | 3.075 | 0.866 | -0.004 | -0.001 | -0.001 | 0.232 | 0.05 | 0.05 | 0.02 | 0.04 | 0.04 | 0.04 |
| 1 | 32671030 | rs2295116 | -0.346 | -0.335 | 2.693 | 0.919 | 0.006 | 0.122 | 0.122 | 0 | 0.05 | 0.13 | 0.21 | 0.23 | 0.06 | 0.28 |
| 1 | 32672908 | rs3903683 | -0.868 | -0.344 | 2.544 | 0.915 | 0.042 | 0.225 | 0.225 | 0 | 0.06 | 0.31 | 0.24 | 0.23 | 0.06 | 0.29 |
| 1 | 32686594 | rs3738001 | -1.604 | 0.162 | 1.621 | 1.516 | 0.022 | 0.186 | 0.186 | 1.361 | 0.1 | 0.53 | 0.27 | 0.23 | 0.07 | 0.29 |
| 1 | 32689857 | rs589034 | -1.875 | -0.136 | 0.012 | 1.244 | 0.053 | -0.008 | -0.008 | 0 | 0.41 | 0.89 | 0.5 | 0.69 | 0.31 | 0.52 |
| 1 | 32711360 | rs669538 | -2.098 | -0.229 | -0.203 | 1.016 | 0.065 | -0.011 | -0.011 | 0 | 0.44 | 0.94 | 0.5 | 0.69 | 0.31 | 0.51 |
| 1 | 32723355 | rs35021967 | 0.283 | 0.577 | 0.072 | 1.486 | -0.037 | -0.008 | -0.008 | 0 | 0.06 | 0.01 | 0.13 | 0.33 | 0.13 | 0.09 |
| 1 | 32729702 | rs1004420 | -1.122 | 0.174 | 0.628 | 1.097 | -0.036 | 0.083 | 0.083 | 0.759 | 0.23 | 0.47 | 0.32 | 0.35 | 0.17 | 0.38 |
| 1 | 32743866 | rs695161 | -1.054 | 0.415 | 0.011 | -0.278 | -0.007 | 0.015 | 0.015 | 0 | 0.56 | 0.7 | 0.64 | 0.98 | 0.53 | 0.63 |
| 1 | 32756439 | rs1741981 | -1.049 | 0.349 | 0.095 | 0.454 | 0.017 | -0.057 | -0.057 | 0.029 | 0.37 | 0.86 | 0.26 | 0.49 | 0.34 | 0.27 |
| 1 | 32831543 | rs476153 | -1.22 | 0.282 | 0.447 | -0.249 | -0.033 | 0.069 | 0.069 | 1.57 | 0.87 | 0.98 | 0.92 | 1 | 0.86 | 0.96 |
| 1 | 32862797 | rs9970917 | -0.819 | 0.236 | 0.308 | 0.008 | -0.004 | -0.023 | -0.023 | 0.007 | 0.11 | 0.56 | 0.08 | 0.04 | 0.06 | 0.07 |
| 1 | 32866717 | rs587149 | 0.606 | 0.56 | -0.765 | -0.051 | -0.044 | 0.171 | 0.171 | 0.845 | 0.26 | 0.02 | 0.12 | 0 | 0.22 | 0.05 |
| 1 | 32875280 | rs16834981 | -0.351 | 0.318 | 0.148 | 1.526 | -0.027 | -0.017 | -0.017 | 0 | 0.07 | 0 | 0.04 | 0.33 | 0.08 | 0.09 |
| 1 | 32879586 | rs10914555 | -0.839 | 0.181 | 0.55 | 0.448 | 0.022 | -0.082 | -0.082 | 0 | 0.13 | 0.31 | 0.1 | 0 | 0.14 | 0.04 |
| 1 | 32904611 | rs11588809 | -0.271 | -0.026 | 0.008 | -0.099 | 0.019 | -0.028 | -0.028 | 13.321 | 0.12 | 0.22 | 0.1 | 0.05 | 0.14 | 0.07 |
| 1 | 32913807 | rs12144474 | -0.145 | -0.005 | -0.115 | -0.089 | 0.014 | -0.023 | -0.023 | 0 | 0.11 | 0.21 | 0.1 | 0.05 | 0.14 | 0.05 |
| 1 | 32942662 | rs4970524 | -0.489 | -0.21 | -0.236 | 0.124 | -0.005 | 0.009 | 0.009 | 0 | 0.07 | 0.17 | 0.05 | 0.05 | 0.06 | 0.02 |

| | | | | | | | | | | | | | | | | |
|---|---|---|---|---|---|---|---|---|---|---|---|---|---|---|---|---|
| 1 | 32960336 | rs12140988 | -0.494 | -0.267 | -0.284 | 0.073 | -0.012 | -0.005 | -0.005 | 0 | 0.07 | 0.2 | 0.05 | 0.05 | 0.06 | 0.02 |
| 1 | 32973271 | rs12141725 | -0.511 | -0.198 | -0.356 | 0.055 | -0.003 | -0.005 | -0.005 | 0.24 | 0.08 | 0.13 | 0.04 | 0.05 | 0.08 | 0.02 |
| 1 | 32975903 | rs2453244 | -1.012 | -0.522 | -0.522 | -0.079 | 0 | 0.001 | 0.001 | 0 | 0.51 | 0.51 | 0.51 | 0.55 | 0.43 | 0.5 |
| 1 | 32993522 | rs16835005 | 0.451 | -0.244 | -0.677 | 0.044 | 0.007 | 0.055 | 0.055 | 0 | 0.2 | 0.19 | 0.36 | 0.5 | 0.19 | 0.4 |
| 1 | 33012259 | rs7516064 | -0.395 | -0.294 | -0.74 | -0.025 | -0.01 | -0.021 | -0.021 | 0 | 0.13 | 0.23 | 0.07 | 0.05 | 0.11 | 0.05 |
| 1 | 33030461 | rs11588497 | 0.521 | -0.292 | -0.742 | -0.014 | 0.012 | 0.076 | 0.076 | 0 | 0.19 | 0.2 | 0.36 | 0.5 | 0.19 | 0.4 |
| 1 | 33032412 | rs9425886 | -0.761 | -0.306 | -1.198 | -0.061 | -0.082 | 0.025 | 0.025 | 0 | 0.66 | 0.86 | 0.6 | 0.7 | 0.53 | 0.63 |
| 1 | 33034876 | rs10914577 | -1.57 | -0.484 | 0.043 | 0.145 | -0.007 | -0.015 | -0.015 | 0.513 | 0.13 | 0.2 | 0.06 | 0.05 | 0.11 | 0.06 |
| 1 | 33051960 | rs12025475 | -1.598 | -0.47 | -0.031 | 0.158 | -0.005 | -0.004 | -0.004 | 0 | 0.13 | 0.19 | 0.06 | 0.11 | 0.1 | 0.05 |
| 1 | 33055668 | rs12124774 | -1.491 | -0.444 | -0.265 | 0.139 | -0.005 | -0.036 | -0.036 | 0 | 0.13 | 0.19 | 0.06 | 0.03 | 0.1 | 0.03 |
| 1 | 33062554 | rs6678903 | -1.554 | -0.443 | -0.268 | 0.169 | -0.01 | -0.008 | -0.008 | 0 | 0.14 | 0.23 | 0.07 | 0.11 | 0.11 | 0.05 |
| 1 | 33066299 | rs2294815 | -1.401 | -0.794 | -0.065 | -0.032 | -0.039 | -0.034 | -0.034 | 2.867 | 0.17 | 0.3 | 0.24 | 0.12 | 0.09 | 0.11 |
| 1 | 33068754 | rs422215 | -1.14 | -0.825 | -1.395 | -0.268 | -0.099 | 0.01 | 0.01 | 0.431 | 0.53 | 0.97 | 0.5 | 0.38 | 0.47 | 0.45 |
| 1 | 33070778 | rs12136921 | -1.077 | -1.396 | -0.431 | -0.446 | 0.012 | -0.019 | -0.019 | 0 | 0.07 | 0.28 | 0.04 | 0.04 | 0.04 | 0.04 |
| 1 | 33100815 | rs3954228 | -1.852 | -1.56 | -0.239 | -0.335 | 0.045 | 0.132 | 0.132 | 0.121 | 0.07 | 0.4 | 0.21 | 0.12 | 0.03 | 0.06 |
| 1 | 33116544 | rs389409 | 2.232 | -1.74 | 0.131 | -0.422 | -0.029 | 0.053 | 0.053 | 0.683 | 0.88 | 0.49 | 0.74 | 0.87 | 0.96 | 0.92 |
| 1 | 33119593 | rs2934766 | 1.602 | -1.737 | 0.148 | -0.42 | 0.048 | 0.118 | 0.118 | 0.204 | 0.93 | 0.59 | 0.78 | 0.89 | 0.97 | 0.94 |
| 1 | 33134774 | rs359955 | 1.349 | -2.043 | 0.785 | -0.652 | 0.101 | 0.092 | 0.092 | 0.276 | 0.57 | 0.19 | 0.64 | 0.77 | 0.75 | 0.77 |
| 1 | 33134989 | rs359956 | 2.162 | -2.049 | 0.801 | -0.576 | -0.032 | 0.05 | 0.05 | 0.603 | 0.88 | 0.49 | 0.74 | 0.87 | 0.96 | 0.92 |
| 1 | 33146501 | rs360511 | 1.442 | -2.021 | 0.799 | -0.565 | 0.049 | 0.12 | 0.12 | 0.204 | 0.93 | 0.59 | 0.78 | 0.89 | 0.97 | 0.94 |
| 1 | 33148935 | rs16835131 | -0.092 | -2.285 | 0.41 | -0.711 | 0.018 | 0.004 | 0.004 | 0 | 0.13 | 0.07 | 0.04 | 0.1 | 0.1 | 0.09 |
| 1 | 33160878 | rs360042 | 1.314 | -2.146 | 0.36 | -0.642 | -0.022 | 0.017 | 0.017 | 0 | 0.92 | 0.78 | 0.77 | 0.87 | 0.97 | 0.92 |
| 1 | 33170413 | rs9425961 | 1.463 | -2.36 | 0.297 | -0.633 | 0.121 | 0.085 | 0.085 | 0.36 | 0.56 | 0.19 | 0.64 | 0.76 | 0.76 | 0.77 |
| 1 | 33179156 | rs419145 | 1.49 | -1.479 | 0.638 | -0.091 | -0.032 | 0.006 | 0.006 | 0 | 0.82 | 0.66 | 0.73 | 0.82 | 0.86 | 0.84 |
| 1 | 33187046 | rs359952 | 1.671 | -1.164 | 0.824 | 0.121 | 0.004 | 0.134 | 0.134 | 0.112 | 0.92 | 0.61 | 0.77 | 0.84 | 0.97 | 0.94 |
| 1 | 33203461 | rs2184931 | -0.922 | -1.098 | -0.187 | 0.196 | 0.008 | -0.009 | -0.009 | 0 | 0.07 | 0.03 | 0.06 | 0.17 | 0.04 | 0.06 |
| 1 | 33215380 | rs644879 | -0.32 | -1.181 | -0.689 | 0.051 | -0.036 | 0.02 | 0.02 | 4.936 | 0.38 | 0.22 | 0.37 | 0.31 | 0.43 | 0.41 |
| 1 | 33221282 | rs360490 | -0.372 | -0.443 | -0.792 | -0.313 | 0.031 | 0.036 | 0.036 | 0.069 | 0.37 | 0.2 | 0.18 | 0.23 | 0.29 | 0.34 |
| 1 | 33225668 | rs12561762 | 0.126 | -0.606 | -0.99 | -0.446 | 0.003 | 0.019 | 0.019 | 0.166 | 0.28 | 0.03 | 0.11 | 0.19 | 0.2 | 0.22 |
| 1 | 33232525 | rs12022461 | 0.508 | -0.573 | -1.009 | -0.285 | -0.02 | -0.041 | -0.041 | 0.208 | 0.24 | 0.01 | 0.1 | 0.23 | 0.18 | 0.21 |
| 1 | 33235651 | rs12730560 | -0.523 | -0.503 | -0.467 | -0.995 | 0.012 | -0.081 | -0.081 | 0.309 | 0.14 | 0.27 | 0.23 | 0 | 0.13 | 0.08 |
| 1 | 33241289 | rs16866009 | 0.404 | -0.666 | 0.328 | -1.087 | -0.004 | -0.019 | -0.019 | 0 | 0.08 | 0.07 | 0.03 | 0 | 0.03 | 0 |
| 1 | 33256884 | rs1765223 | -0.559 | -0.78 | 0.342 | -1.1 | 0.008 | 0.036 | 0.036 | 0.017 | 0.39 | 0.3 | 0.32 | 0.23 | 0.32 | 0.29 |
| 1 | 33257919 | rs10914614 | 0.91 | -0.837 | 0.852 | -1.146 | 0.005 | 0.012 | 0.012 | 0.095 | 0.49 | 0.46 | 0.6 | 0.74 | 0.58 | 0.58 |
| 1 | 33275981 | rs10798918 | 1.028 | -0.808 | 0.747 | -1.126 | -0.014 | -0.04 | -0.04 | 0 | 0.25 | 0.01 | 0.1 | 0.23 | 0.18 | 0.21 |
| 1 | 33277080 | rs10914621 | -1.016 | -0.787 | 0.816 | -1.119 | 0.04 | -0.06 | -0.06 | 0 | 0.1 | 0.29 | 0.21 | 0 | 0.12 | 0.07 |
| 1 | 33282392 | rs16835176 | -1.388 | -0.795 | 0.746 | -1.126 | 0.012 | -0.077 | -0.077 | 0 | 0.14 | 0.25 | 0.22 | 0 | 0.13 | 0.07 |
| 1 | 33289388 | rs4951788 | 0.745 | -0.837 | 0.764 | -1.141 | 0.005 | 0.013 | 0.013 | 0.88 | 0.53 | 0.44 | 0.61 | 0.73 | 0.6 | 0.58 |
| 1 | 33298822 | rs10914625 | 1.805 | 0.169 | -0.351 | -0.644 | -0.12 | -0.148 | -0.148 | 0 | 0.24 | 0.03 | 0.44 | 0.46 | 0.36 | 0.39 |
| 1 | 33314700 | rs4951774 | -0.671 | 0.406 | 0.004 | -0.566 | -0.008 | -0.036 | -0.036 | 0.149 | 0.64 | 0.52 | 0.67 | 0.77 | 0.7 | 0.71 |
| 1 | 33320601 | rs6673337 | 1.011 | 0.56 | -0.027 | -0.545 | 0.001 | 0.001 | 0.001 | 0 | 0.11 | 0.09 | 0.06 | 0.04 | 0.1 | 0.14 |
| 1 | 33324136 | rs9500 | -0.711 | 0.297 | -0.157 | -0.674 | -0.006 | -0.023 | -0.023 | 0 | 0.64 | 0.56 | 0.67 | 0.77 | 0.69 | 0.71 |
| 1 | 33326172 | rs1300114 | 1.996 | 0.279 | -0.208 | -0.684 | -0.013 | -0.039 | -0.039 | 1.506 | 0.25 | 0.01 | 0.11 | 0.23 | 0.19 | 0.21 |
| 1 | 33328165 | rs3480 | 0.175 | 0.281 | -0.194 | -0.682 | 0 | -0.001 | -0.001 | 1.756 | 0.53 | 0.46 | 0.61 | 0.73 | 0.59 | 0.57 |
| 1 | 33335039 | rs1746661 | 1.909 | 0.355 | -0.186 | -0.57 | -0.008 | -0.038 | -0.038 | 0.092 | 0.25 | 0.01 | 0.11 | 0.23 | 0.19 | 0.21 |
| 1 | 33348442 | rs35452140 | 0.346 | 1.11 | -0.328 | -0.711 | -0.001 | -0.001 | -0.001 | 0 | 0.38 | 0.35 | 0.5 | 0.59 | 0.42 | 0.56 |
| 1 | 33350183 | rs11811702 | -0.042 | 0.907 | -0.161 | -0.192 | -0.019 | -0.014 | -0.014 | 0 | 0.56 | 0.52 | 0.54 | 0.71 | 0.71 | 0.72 |
| 1 | 33355923 | rs1284373 | 1.112 | 1.241 | -0.551 | 0.036 | 0.009 | -0.015 | -0.015 | 0 | 0.3 | 0.01 | 0.21 | 0.23 | 0.21 | 0.22 |
| 1 | 33364095 | rs942242 | -0.687 | 1.379 | -0.475 | 0.031 | 0.044 | 0.014 | 0.014 | 0 | 0.64 | 0.88 | 0.6 | 0.7 | 0.72 | 0.72 |
| 1 | 33401773 | rs194637 | 1.04 | 1.556 | -0.538 | 0.092 | -0.018 | -0.059 | -0.059 | 1.761 | 0.3 | 0.03 | 0.24 | 0.33 | 0.25 | 0.22 |
| 1 | 33403481 | rs194640 | 1.04 | 1.544 | -0.538 | 0.092 | -0.022 | -0.059 | -0.059 | 1.627 | 0.3 | 0.03 | 0.24 | 0.33 | 0.25 | 0.22 |
| 1 | 33404367 | rs171837 | 1.008 | 1.547 | -0.538 | 0.106 | -0.021 | -0.018 | -0.018 | 0 | 0.3 | 0.03 | 0.24 | 0.27 | 0.25 | 0.22 |
| 1 | 33405972 | rs194641 | 1.008 | 1.547 | -0.534 | 0.106 | -0.017 | -0.015 | -0.015 | 0.177 | 0.3 | 0.03 | 0.24 | 0.28 | 0.25 | 0.22 |
| 1 | 33408138 | rs1149128 | -0.728 | 1.655 | -0.344 | 0.069 | 0.095 | 0.017 | 0.017 | 0.013 | 0.05 | 0.29 | 0.19 | 0 | 0.08 | 0.07 |
| 1 | 33409551 | rs194644 | 0.056 | 1.688 | -0.37 | 0.064 | -0.005 | -0.007 | -0.007 | 0.047 | 0.38 | 0.43 | 0.45 | 0.33 | 0.34 | 0.3 |
| 1 | 33417785 | rs194648 | 1.03 | 1.671 | -0.392 | 0.065 | -0.021 | -0.018 | -0.018 | 0 | 0.3 | 0.03 | 0.24 | 0.27 | 0.25 | 0.22 |
| 1 | 33419206 | rs194652 | 0.25 | 1.71 | -0.301 | 0.116 | -0.002 | -0.003 | -0.003 | 0 | 0.38 | 0.39 | 0.45 | 0.33 | 0.35 | 0.3 |
| 1 | 33437042 | rs194663 | 0.187 | 1.694 | -0.354 | 0.131 | -0.002 | -0.006 | -0.006 | 0 | 0.39 | 0.43 | 0.46 | 0.32 | 0.37 | 0.31 |
| 1 | 33448220 | rs4412566 | 0.28 | 1.65 | -0.375 | 0.298 | 0 | 0 | 0 | 0 | 0.34 | 0.34 | 0.42 | 0.33 | 0.32 | 0.25 |
| 1 | 33448509 | rs17511157 | 0.106 | 2.147 | -0.625 | 0.544 | -0.013 | -0.012 | -0.012 | 0 | 0.06 | 0.03 | 0.2 | 0.1 | 0.14 | 0.09 |
| 1 | 33460871 | rs10737365 | -1.116 | 1.298 | -1.121 | 0.47 | -0.136 | -0.117 | -0.117 | 0 | 0.65 | 0.95 | 0.59 | 0.44 | 0.51 | 0.56 |
| 1 | 33472924 | rs1534892 | 1.06 | 1.356 | -0.431 | 0.72 | -0.061 | -0.048 | -0.048 | 0 | 0.27 | 0.02 | 0.38 | 0.25 | 0.28 | 0.28 |
| 1 | 33472942 | rs1534893 | 0.299 | 1.314 | 0.206 | 0.695 | -0.01 | -0.002 | -0.002 | 0 | 0.33 | 0.47 | 0.45 | 0.32 | 0.3 | 0.3 |
| 1 | 33475426 | rs998664 | 1.093 | 1.336 | 0.041 | 0.678 | -0.061 | -0.048 | -0.048 | 0.024 | 0.27 | 0.02 | 0.38 | 0.26 | 0.28 | 0.29 |
| 1 | 33496639 | rs10753271 | 1.103 | 1.289 | -0.22 | 0.681 | -0.062 | -0.048 | -0.048 | 0.317 | 0.27 | 0.02 | 0.38 | 0.26 | 0.28 | 0.29 |
| 1 | 33540443 | rs3863724 | 1.126 | 0.868 | -0.349 | 0.538 | -0.041 | 0.012 | 0.012 | 0 | 0.28 | 0.02 | 0.31 | 0.21 | 0.27 | 0.29 |
| 1 | 33543052 | rs10429857 | 0.351 | 0.935 | -0.146 | 0.599 | -0.011 | -0.03 | -0.03 | 0 | 0.34 | 0.4 | 0.36 | 0.23 | 0.3 | 0.3 |
| 1 | 33565513 | rs804429 | -0.124 | 0.77 | -0.383 | 0.23 | -0.015 | 0.008 | 0.008 | 0 | 0.19 | 0.08 | 0.07 | 0.16 | 0.18 | 0.22 |
| 1 | 33585070 | rs638130 | 0.61 | 1.004 | -0.437 | 0.3 | -0.002 | -0.002 | -0.002 | 0 | 0.32 | 0.31 | 0.35 | 0.27 | 0.28 | 0.29 |

| | | | | | | | | | | | | | | | | |
|---|---|---|---|---|---|---|---|---|---|---|---|---|---|---|---|---|
| 1 | 33592887 | rs664182 | 0.502 | 1.027 | -0.687 | 0.316 | -0.027 | -0.026 | -0.026 | 0 | 0.33 | 0.45 | 0.37 | 0.27 | 0.28 | 0.29 |
| 1 | 33595212 | rs84314 | -1.029 | 1.276 | -1.687 | 0.464 | -0.117 | -0.073 | -0.073 | 0 | 0.6 | 0.94 | 0.5 | 0.42 | 0.48 | 0.52 |
| 1 | 33600115 | rs17542433 | -0.029 | 1.487 | -1.07 | 0.555 | -0.044 | 0.01 | 0.01 | 0.023 | 0.12 | 0.01 | 0.06 | 0.06 | 0.14 | 0.12 |
| 1 | 33611890 | rs689187 | -0.324 | 0.583 | -0.158 | 0.46 | -0.016 | -0.001 | -0.001 | 0 | 0.71 | 0.57 | 0.76 | 0.68 | 0.74 | 0.68 |
| 1 | 33612508 | rs3766813 | 0.699 | 0.548 | -0.309 | 0.465 | -0.032 | 0.017 | 0.017 | 0 | 0.25 | 0.03 | 0.18 | 0.2 | 0.25 | 0.29 |
| 1 | 33614162 | rs3912145 | 0.648 | 0.547 | -0.193 | 0.472 | 0.007 | 0.006 | 0.006 | 0 | 0.26 | 0.14 | 0.19 | 0.26 | 0.25 | 0.29 |
| 1 | 33622706 | rs4481817 | 0.685 | 0.635 | -0.516 | 0.423 | -0.04 | 0.011 | 0.011 | 0 | 0.25 | 0.02 | 0.18 | 0.2 | 0.25 | 0.29 |
| 1 | 33631069 | rs2306257 | 0.779 | 0.624 | -0.511 | 0.39 | -0.01 | 0.032 | 0.032 | 0 | 0.25 | 0.05 | 0.18 | 0.2 | 0.25 | 0.29 |
| 1 | 33641256 | rs2927970 | 0.898 | 0.022 | -0.635 | 0.357 | 0.022 | -0.008 | -0.008 | 0 | 0.46 | 0.07 | 0.63 | 0.59 | 0.61 | 0.47 |
| 1 | 33642312 | rs603186 | -0.62 | 0.333 | -0.038 | 0.673 | 0.007 | 0.03 | 0.03 | 0 | 0.75 | 0.9 | 0.81 | 0.8 | 0.76 | 0.72 |
| 1 | 33644352 | rs680960 | -0.378 | 0.321 | 0.218 | 0.667 | -0.002 | -0.003 | -0.003 | 0 | 0.73 | 0.71 | 0.79 | 0.8 | 0.75 | 0.72 |
| 1 | 33646950 | rs622407 | -0.387 | 0.334 | 0.224 | 0.673 | -0.002 | -0.003 | -0.003 | 0 | 0.73 | 0.71 | 0.8 | 0.79 | 0.75 | 0.72 |
| 1 | 33684101 | rs1498319 | 0.208 | 0.159 | 0.397 | 0.628 | -0.032 | -0.011 | -0.011 | 0.062 | 0.67 | 0.54 | 0.74 | 0.69 | 0.75 | 0.69 |
| 1 | 33684733 | rs6686546 | 1.579 | 0.334 | 0.593 | 0.742 | 0.017 | 0.056 | 0.056 | 0.144 | 0.93 | 0.62 | 0.91 | 0.96 | 0.97 | 0.97 |
| 1 | 33690698 | rs12045968 | 0.454 | 0.334 | 0.306 | 0.678 | 0.002 | -0.014 | -0.014 | 0 | 0.23 | 0.07 | 0.17 | 0.27 | 0.22 | 0.27 |
| 1 | 33694810 | rs7540951 | 0.149 | 0.361 | 0.527 | 0.675 | -0.025 | -0.007 | -0.007 | 1.07 | 0.7 | 0.52 | 0.77 | 0.7 | 0.73 | 0.69 |
| 1 | 33710429 | rs7534724 | -0.073 | 0.525 | 0.019 | 0.66 | 0.009 | 0.006 | 0.006 | 0 | 0.27 | 0.21 | 0.22 | 0.27 | 0.26 | 0.28 |
| 1 | 33719642 | rs1566230 | 0.197 | 0.643 | -0.167 | 0.596 | -0.03 | 0.009 | 0.009 | 0 | 0.24 | 0.05 | 0.2 | 0.2 | 0.26 | 0.28 |
| 1 | 33723652 | rs10914671 | -0.332 | 0.736 | -0.003 | 0.629 | 0.001 | 0.01 | 0.01 | 0 | 0.76 | 0.79 | 0.79 | 0.84 | 0.74 | 0.71 |
| 1 | 33732192 | rs6425813 | 0.047 | 0.765 | 0.298 | 0.593 | -0.001 | -0.006 | -0.006 | 3.039 | 0.73 | 0.71 | 0.78 | 0.83 | 0.7 | 0.69 |
| 1 | 33735361 | rs4077439 | -0.359 | 0.903 | 0.187 | 0.778 | -0.001 | 0 | 0 | 0.097 | 0.76 | 0.79 | 0.79 | 0.78 | 0.75 | 0.7 |
| 1 | 33749027 | rs10914677 | -0.01 | 0.95 | 0.493 | 0.754 | 0.008 | -0.004 | -0.004 | 0 | 0.73 | 0.64 | 0.77 | 0.76 | 0.7 | 0.69 |
| 1 | 33755543 | rs12748225 | 0.24 | 0.964 | 0.135 | 0.777 | -0.008 | -0.007 | -0.007 | 0 | 0.22 | 0.13 | 0.18 | 0.23 | 0.26 | 0.29 |
| 1 | 33760197 | rs12723104 | 0.612 | 1.018 | -0.057 | 0.875 | -0.067 | -0.078 | -0.078 | 0 | 0.19 | 0.01 | 0.17 | 0.26 | 0.24 | 0.29 |
| 1 | 33760743 | rs4641276 | -0.872 | 1.162 | -0.141 | 0.845 | -0.058 | -0.064 | -0.064 | 0.477 | 0.8 | 0.97 | 0.84 | 0.76 | 0.76 | 0.72 |
| 1 | 33761201 | rs7527638 | 0.627 | 1.067 | 0.077 | 1.011 | 0.006 | 0.017 | 0.017 | 0 | 0.58 | 0.29 | 0.71 | 0.62 | 0.62 | 0.58 |
| 1 | 33765272 | rs1132790 | -0.94 | 1.005 | 0.036 | 0.957 | 0.007 | 0.037 | 0.037 | 0 | 0.42 | 0.74 | 0.29 | 0.38 | 0.4 | 0.45 |
| 1 | 33766706 | rs7523119 | -0.83 | 0.971 | 0.068 | 0.949 | 0.005 | 0.027 | 0.027 | 0 | 0.41 | 0.67 | 0.28 | 0.37 | 0.39 | 0.45 |
| 1 | 33770526 | rs4653002 | -0.983 | 0.929 | 0.186 | 0.94 | 0.009 | 0.049 | 0.049 | 0.098 | 0.42 | 0.79 | 0.3 | 0.37 | 0.39 | 0.45 |
| 1 | 33776431 | rs4653015 | -0.112 | 1.232 | 0.361 | 0.886 | -0.026 | -0.094 | -0.094 | 0 | 0.34 | 0.53 | 0.2 | 0.09 | 0.23 | 0.32 |
| 1 | 33779043 | rs16835351 | -0.728 | 1.253 | -0.049 | 1.491 | 0.056 | 0.051 | 0.051 | 0.063 | 0.39 | 0.85 | 0.24 | 0.25 | 0.28 | 0.35 |
| 1 | 33785978 | rs6686500 | -0.395 | 1.254 | -0.046 | 1.513 | 0.028 | 0.024 | 0.024 | 0 | 0.37 | 0.71 | 0.23 | 0.26 | 0.28 | 0.35 |
| 1 | 33788884 | rs7542777 | 1.308 | 1.319 | -0.146 | 1.576 | -0.01 | -0.035 | -0.035 | 0 | 0.2 | 0.02 | 0.14 | 0.21 | 0.19 | 0.27 |
| 1 | 33789382 | rs7264 | -0.309 | 1.147 | -0.079 | 1.586 | 0.023 | 0.021 | 0.021 | 0 | 0.37 | 0.69 | 0.21 | 0.25 | 0.28 | 0.35 |
| 1 | 33789968 | rs1130800 | 1.482 | 1.21 | -0.11 | 1.57 | -0.036 | -0.039 | -0.039 | 0 | 0.2 | 0.02 | 0.14 | 0.21 | 0.19 | 0.27 |
| 1 | 33790496 | rs11554674 | 1.28 | 1.215 | -0.092 | 1.589 | -0.057 | -0.033 | -0.033 | 0 | 0.21 | 0.02 | 0.17 | 0.21 | 0.24 | 0.27 |
| 1 | 33795572 | rs10914684 | 1.86 | 1.172 | -0.079 | 1.646 | -0.09 | 0.013 | 0.013 | 0.028 | 0.35 | 0.03 | 0.41 | 0.21 | 0.37 | 0.25 |
| 1 | 33796554 | rs4361958 | -1.187 | 1.202 | -0.081 | 1.599 | 0.006 | -0.044 | -0.044 | 0 | 0.42 | 0.81 | 0.3 | 0.52 | 0.4 | 0.46 |
| 1 | 33819562 | rs4233199 | -1.576 | 1.216 | -0.097 | 1.524 | -0.053 | -0.053 | -0.053 | 0 | 0.6 | 0.85 | 0.4 | 0.48 | 0.51 | 0.63 |
| 1 | 33819814 | rs4653080 | -0.546 | 1.205 | -0.151 | 1.518 | 0.026 | 0.025 | 0.025 | 0 | 0.38 | 0.7 | 0.21 | 0.25 | 0.28 | 0.35 |
| 1 | 33820134 | rs12026290 | 1.259 | 1.19 | -0.27 | 1.543 | -0.05 | -0.03 | -0.03 | 0 | 0.21 | 0.02 | 0.17 | 0.21 | 0.23 | 0.27 |
| 1 | 33832865 | rs6425816 | -2.002 | 1.183 | -0.263 | 1.537 | -0.029 | 0.033 | 0.033 | 0 | 0.64 | 0.88 | 0.49 | 0.77 | 0.63 | 0.74 |
| 1 | 33850890 | rs12127097 | 0.474 | 0.987 | 0.001 | 1.388 | 0 | -0.028 | -0.028 | 0 | 0.27 | 0.28 | 0.14 | 0.04 | 0.19 | 0.25 |
| 1 | 33861853 | rs1535917 | -1.446 | 1.007 | 0.015 | 1.439 | 0.042 | 0.223 | 0.223 | 0.599 | 0.16 | 0.57 | 0.18 | 0.5 | 0.2 | 0.21 |
| 1 | 33865068 | rs6685849 | 0.314 | 1.001 | 0.252 | 1.412 | 0.016 | -0.011 | -0.011 | 0 | 0.08 | 0.32 | 0.07 | 0.05 | 0.06 | 0.08 |
| 1 | 33867447 | rs12137943 | 0.51 | 1.011 | -0.792 | 1.353 | 0.006 | -0.003 | -0.003 | 0 | 0.26 | 0.23 | 0.13 | 0.04 | 0.19 | 0.25 |
| 1 | 33872593 | rs6661037 | -0.348 | 1.029 | -0.762 | 1.389 | 0.023 | 0.116 | 0.116 | 0 | 0.47 | 0.25 | 0.3 | 0.25 | 0.43 | 0.52 |
| 1 | 33875477 | rs7525843 | -0.023 | 0.977 | -0.747 | 1.368 | 0.003 | 0.152 | 0.152 | 0.77 | 0.11 | 0.54 | 0.1 | 0.21 | 0.09 | 0.11 |
| 1 | 33877808 | rs6663355 | -0.752 | 0.981 | -0.757 | 1.37 | 0.006 | -0.014 | -0.014 | 0.784 | 0.51 | 0.26 | 0.37 | 0.56 | 0.54 | 0.63 |
| 1 | 33886888 | rs10914706 | -0.094 | 0.907 | -0.661 | 1.353 | 0.003 | 0.154 | 0.154 | 0.77 | 0.11 | 0.53 | 0.1 | 0.21 | 0.09 | 0.11 |
| 1 | 33893830 | rs4653135 | 0.537 | 0.938 | -0.976 | 1.19 | 0.012 | 0.038 | 0.038 | 0 | 0.24 | 0.14 | 0.11 | 0.04 | 0.19 | 0.25 |
| 1 | 33901751 | rs7544329 | 0.843 | 0.87 | -0.958 | 1.269 | 0.016 | 0.054 | 0.054 | 0 | 0.21 | 0.11 | 0.1 | 0.04 | 0.16 | 0.24 |
| 1 | 33905230 | rs10914716 | 1.148 | 1.095 | -1.11 | 1.46 | -0.051 | -0.03 | -0.03 | 0 | 0.21 | 0.02 | 0.17 | 0.21 | 0.24 | 0.27 |
| 1 | 33910453 | rs1123295 | -0.648 | 1.149 | -0.927 | 1.282 | 0 | -0.001 | -0.001 | 0 | 0.37 | 0.62 | 0.35 | 0.7 | 0.44 | 0.5 |
| 1 | 33911647 | rs12564600 | 0.372 | 1.442 | -1.456 | 1.408 | -0.032 | -0.008 | -0.008 | 0 | 0.05 | 0.01 | 0.08 | 0.3 | 0.12 | 0.11 |
| 1 | 33915751 | rs2336240 | -0.892 | 0.937 | -1.471 | 0.867 | -0.042 | 0.097 | 0.097 | 0 | 0.6 | 0.89 | 0.55 | 0.81 | 0.6 | 0.78 |
| 1 | 33920466 | rs6669405 | -0.002 | 0.242 | -0.978 | 0.864 | 0 | -0.004 | -0.004 | 0 | 0.3 | 0.29 | 0.21 | 0.42 | 0.28 | 0.33 |
| 1 | 33920586 | rs12032996 | 0.297 | -0.043 | -1.096 | 0.772 | 0.001 | -0.014 | -0.014 | 4.965 | 0.19 | 0.02 | 0.2 | 0.2 | 0.14 | 0.3 |
| 1 | 33921878 | rs10737370 | -0.291 | 0.385 | -0.414 | 0.68 | 0 | 0 | 0 | 0 | 0.44 | 0.52 | 0.36 | 0.58 | 0.54 | 0.6 |
| 1 | 33928758 | rs12565140 | -0.117 | 1.26 | -0.474 | 0.962 | -0.04 | -0.062 | -0.062 | 0 | 0.11 | 0.02 | 0.19 | 0.36 | 0.15 | 0.29 |
| 1 | 33930301 | rs9662700 | -0.315 | 1.007 | -0.032 | 0.741 | 0.005 | 0.015 | 0.015 | 0 | 0.58 | 0.64 | 0.53 | 0.74 | 0.61 | 0.66 |
| 1 | 33934824 | rs10914731 | 0.074 | 1.014 | -0.061 | 0.754 | -0.044 | -0.065 | -0.065 | 0.154 | 0.12 | 0.02 | 0.2 | 0.37 | 0.16 | 0.29 |
| 1 | 33936347 | rs6425820 | 0.73 | 1.006 | 0.342 | 0.683 | 0 | -0.008 | -0.008 | 0.023 | 0.43 | 0.44 | 0.3 | 0.28 | 0.43 | 0.33 |
| 1 | 33937135 | rs10798953 | -1.102 | 0.56 | 0.351 | 0.436 | 0.007 | 0.015 | 0.015 | 0 | 0.71 | 0.89 | 0.62 | 0.75 | 0.73 | 0.74 |
| 1 | 33945591 | rs10798956 | -1.076 | 0.546 | 0.433 | 0.415 | 0.007 | 0.01 | 0.01 | 0 | 0.69 | 0.76 | 0.61 | 0.75 | 0.72 | 0.74 |
| 1 | 33945685 | rs10493052 | -0.67 | 0.554 | 0.339 | 0.509 | -0.002 | -0.028 | -0.028 | 0 | 0.12 | 0.15 | 0.08 | 0.01 | 0.13 | 0.09 |
| 1 | 33947275 | rs16835511 | -0.293 | 0.685 | 0.401 | 0.538 | -0.008 | 0.021 | 0.021 | 0 | 0.1 | 0.07 | 0.05 | 0.01 | 0.12 | 0.11 |
| 1 | 33947467 | rs7531293 | 0.915 | 0.754 | 0.507 | 0.543 | 0 | -0.002 | -0.002 | 0.367 | 0.45 | 0.45 | 0.32 | 0.28 | 0.45 | 0.29 |
| 1 | 33957130 | rs3795419 | 0.142 | 0.89 | 0.143 | 0.485 | -0.029 | -0.077 | -0.077 | 0.852 | 0.13 | 0.02 | 0.21 | 0.37 | 0.16 | 0.28 |

| | | | | | | | | | | | | | | | | |
|---|---|---|---|---|---|---|---|---|---|---|---|---|---|---|---|---|
| 1 | 33957773 | rs9662260 | -0.949 | 0.879 | 0.146 | 0.478 | 0.01 | 0.01 | 0.01 | 0 | 0.71 | 0.85 | 0.62 | 0.74 | 0.74 | 0.74 |
| 1 | 33960062 | rs9943259 | -0.99 | 0.878 | 0.146 | 0.478 | 0.009 | 0.009 | 0.009 | 0 | 0.71 | 0.85 | 0.62 | 0.74 | 0.74 | 0.74 |
| 1 | 33964984 | rs12046307 | 0.122 | 0.943 | 0.129 | 0.514 | -0.029 | -0.077 | -0.077 | 0 | 0.13 | 0.02 | 0.21 | 0.37 | 0.16 | 0.28 |
| 1 | 33978546 | rs12402882 | 0.134 | 0.933 | 0.332 | 0.465 | 0.002 | 0.005 | 0.005 | 0 | 0.56 | 0.62 | 0.4 | 0.36 | 0.57 | 0.56 |
| 1 | 33980084 | rs3816922 | 0.223 | 0.951 | 0.148 | 0.479 | -0.03 | -0.075 | -0.075 | 6.796 | 0.13 | 0.02 | 0.22 | 0.39 | 0.16 | 0.26 |
| 1 | 33981351 | rs1690549 | -1.043 | 0.945 | 0.173 | 0.462 | 0.007 | 0.009 | 0.009 | 0 | 0.72 | 0.85 | 0.61 | 0.74 | 0.73 | 0.73 |
| 1 | 33981769 | rs3889480 | 0.25 | 0.992 | 0.094 | 0.503 | -0.03 | -0.073 | -0.073 | 0 | 0.12 | 0.02 | 0.22 | 0.39 | 0.16 | 0.25 |
| 1 | 33983019 | rs10798959 | -0.976 | 0.985 | 0.207 | 0.513 | -0.002 | -0.005 | -0.005 | 0.34 | 0.23 | 0.21 | 0.28 | 0.39 | 0.28 | 0.37 |
| 1 | 33984970 | rs2131776 | 1.094 | 1.021 | 0.148 | 0.506 | 0.009 | 0.014 | 0.014 | 0.028 | 0.43 | 0.37 | 0.24 | 0.35 | 0.37 | 0.31 |
| 1 | 33987778 | rs923880 | -0.446 | 1.181 | 0.081 | 0.531 | 0.016 | 0.16 | 0.16 | 0.359 | 0.18 | 0.4 | 0.28 | 0.39 | 0.2 | 0.33 |
| 1 | 33987836 | rs7532550 | 0.035 | 1.488 | -1.1 | 0.581 | -0.001 | -0.006 | -0.006 | 0.06 | 0.09 | 0.08 | 0.04 | 0.01 | 0.09 | 0.1 |
| 1 | 33992370 | rs2794593 | -0.247 | 1.468 | -1.076 | 0.629 | 0.002 | 0.027 | 0.027 | 0 | 0.63 | 0.75 | 0.58 | 0.73 | 0.65 | 0.67 |
| 1 | 33993669 | rs10798962 | -0.104 | 1.318 | -1.051 | 0.661 | -0.007 | 0.014 | 0.014 | 0 | 0.61 | 0.72 | 0.55 | 0.72 | 0.59 | 0.59 |
| 1 | 33999657 | rs1532307 | -0.095 | 1.476 | -1.111 | 0.891 | -0.017 | -0.04 | -0.04 | 0 | 0.14 | 0.05 | 0.24 | 0.3 | 0.2 | 0.28 |
| 1 | 33999809 | rs923879 | -0.181 | 1.456 | -1.141 | 0.808 | -0.012 | 0.028 | 0.028 | 0 | 0.6 | 0.76 | 0.53 | 0.64 | 0.57 | 0.59 |
| 1 | 34001296 | rs10429856 | -0.245 | 1.293 | -1.149 | 1.118 | -0.041 | -0.088 | -0.088 | 0 | 0.14 | 0.02 | 0.24 | 0.34 | 0.19 | 0.28 |
| 1 | 34004912 | rs12729472 | -0.311 | 1.293 | -1.036 | 1.13 | -0.001 | 0.012 | 0.012 | 0.097 | 0.15 | 0.13 | 0.24 | 0.33 | 0.19 | 0.31 |
| 1 | 34008297 | rs1000418 | -0.31 | 1.303 | -1.081 | 1.114 | -0.002 | 0.007 | 0.007 | 0.145 | 0.15 | 0.14 | 0.25 | 0.34 | 0.2 | 0.31 |
| 1 | 34010708 | rs12136254 | -0.162 | 1.355 | -1.415 | 1.131 | -0.015 | -0.043 | -0.043 | 0 | 0.15 | 0.09 | 0.24 | 0.34 | 0.19 | 0.31 |
| 1 | 34013242 | rs2794591 | -0.17 | 1.13 | -1.151 | 0.556 | 0.005 | 0.005 | 0.005 | 0 | 0.71 | 0.85 | 0.62 | 0.7 | 0.74 | 0.76 |
| 1 | 34019467 | rs2794606 | -0.154 | 1.089 | -1.248 | 0.545 | -0.016 | 0.019 | 0.019 | 0 | 0.32 | 0.12 | 0.4 | 0.31 | 0.33 | 0.32 |
| 1 | 34023749 | rs915108 | 0.458 | 0.929 | -0.347 | 0.399 | -0.001 | -0.006 | -0.006 | 0 | 0.86 | 0.71 | 0.84 | 0.84 | 0.82 | 0.92 |
| 1 | 34023779 | rs1318411 | 1.171 | 1.082 | -0.794 | 0.542 | -0.04 | -0.019 | -0.019 | 0 | 0.57 | 0.18 | 0.63 | 0.42 | 0.51 | 0.51 |
| 1 | 34025030 | rs16835593 | 0.464 | 1.628 | -0.929 | 0.952 | -0.04 | -0.032 | -0.032 | 0 | 0.06 | 0.01 | 0.09 | 0.22 | 0.15 | 0.16 |
| 1 | 34026479 | rs933044 | -0.149 | 1.496 | -0.276 | 0.826 | 0.001 | -0.046 | -0.046 | 0 | 0.11 | 0.21 | 0.07 | 0.01 | 0.09 | 0.08 |
| 1 | 34028430 | rs1493995 | -0.096 | 1.407 | -0.249 | 0.76 | -0.004 | 0.007 | 0.007 | 0.083 | 0.1 | 0.06 | 0.06 | 0 | 0.08 | 0.08 |
| 1 | 34029070 | rs923882 | 0.759 | 1.293 | -0.078 | 0.755 | -0.023 | 0.038 | 0.038 | 1.064 | 0.75 | 0.51 | 0.8 | 0.63 | 0.75 | 0.68 |
| 1 | 34029349 | rs923881 | 0.646 | 1.33 | -0.11 | 0.793 | -0.039 | -0.03 | -0.03 | 0 | 0.06 | 0.01 | 0.09 | 0.23 | 0.15 | 0.1 |
| 1 | 34029462 | rs7367222 | 2.276 | 1.276 | -0.068 | 0.759 | -0.04 | -0.013 | -0.013 | 0 | 0.55 | 0.18 | 0.62 | 0.4 | 0.51 | 0.49 |
| 1 | 34030514 | rs1493994 | 1.733 | 1.356 | -0.252 | 0.803 | 0.007 | 0.003 | 0.003 | 3.792 | 0.59 | 0.43 | 0.66 | 0.41 | 0.51 | 0.5 |
| 1 | 34034649 | rs5007951 | -0.876 | 1.232 | -0.128 | 0.634 | -0.011 | 0.004 | 0.004 | 0.742 | 0.18 | 0.26 | 0.15 | 0.2 | 0.19 | 0.27 |
| 1 | 34039844 | rs10914746 | -1.251 | 1.471 | -0.183 | 0.571 | -0.004 | -0.004 | -0.004 | 0.005 | 0.18 | 0.16 | 0.16 | 0.24 | 0.26 | 0.19 |
| 1 | 34041191 | rs6701268 | -1.126 | 1.448 | -0.249 | 0.583 | -0.003 | -0.05 | -0.05 | 0 | 0.2 | 0.25 | 0.1 | 0.03 | 0.16 | 0.29 |
| 1 | 34042245 | rs16835617 | 0.109 | 1.8 | -0.425 | 0.937 | -0.039 | -0.032 | -0.032 | 0.022 | 0.07 | 0.01 | 0.09 | 0.24 | 0.16 | 0.13 |
| 1 | 34044616 | rs6425830 | 0.139 | 0.516 | -0.66 | 0.282 | -0.012 | -0.006 | -0.006 | 0 | 0.3 | 0.36 | 0.17 | 0.23 | 0.23 | 0.22 |
| 1 | 34046066 | rs292809 | -1.416 | 1.349 | -0.867 | 0.252 | -0.005 | 0.007 | 0.007 | 0 | 0.35 | 0.73 | 0.26 | 0.3 | 0.35 | 0.35 |
| 1 | 34048281 | rs389724 | -0.6 | 0.817 | -0.623 | -0.155 | -0.057 | -0.116 | -0.116 | 0 | 0.18 | 0.55 | 0.1 | 0.02 | 0.08 | 0.19 |
| 1 | 34050753 | rs16835640 | -0.159 | 1.636 | -0.756 | 0.408 | 0.01 | 0.013 | 0.013 | 0 | 0.09 | 0.1 | 0.04 | 0.01 | 0.05 | 0.03 |
| 1 | 34051242 | rs11586497 | -0.359 | 2.298 | -0.829 | 0.586 | -0.035 | -0.036 | -0.036 | 0.286 | 0.1 | 0.03 | 0.13 | 0.21 | 0.28 | 0.16 |
| 1 | 34052305 | rs12727575 | -0.666 | 2.155 | -0.824 | 0.578 | -0.065 | -0.019 | -0.019 | 0.749 | 0.19 | 0.12 | 0.17 | 0.21 | 0.33 | 0.19 |
| 1 | 34054862 | rs2781790 | -0.569 | 1.482 | -0.385 | 1.107 | 0.027 | 0.106 | 0.106 | 0 | 0.17 | 0.26 | 0.41 | 0.53 | 0.22 | 0.36 |
| 1 | 34063516 | rs10914749 | -0.049 | 1.123 | -0.786 | 0.348 | 0 | 0.02 | 0.02 | 0 | 0.6 | 0.66 | 0.74 | 0.62 | 0.65 | 0.71 |
| 1 | 34065436 | rs10914750 | -0.884 | 0.367 | -0.2 | -0.182 | -0.022 | -0.038 | -0.038 | 0 | 0.24 | 0.49 | 0.13 | 0.19 | 0.15 | 0.25 |
| 1 | 34068364 | rs1493988 | 0.518 | 0.744 | -0.059 | -0.342 | 0.002 | 0.01 | 0.01 | 0.061 | 0.51 | 0.4 | 0.7 | 0.55 | 0.58 | 0.63 |
| 1 | 34068723 | rs2131774 | -0.496 | 0.693 | -0.089 | -0.301 | 0.008 | 0.015 | 0.015 | 0 | 0.65 | 0.68 | 0.79 | 0.73 | 0.72 | 0.74 |
| 1 | 34071525 | rs1874045 | 0.589 | 0.512 | -0.284 | -0.453 | -0.009 | 0.022 | 0.022 | 0 | 0.44 | 0.32 | 0.68 | 0.55 | 0.58 | 0.58 |
| 1 | 34076550 | rs6672528 | 0.433 | 0.408 | 0.139 | -0.073 | -0.014 | 0.003 | 0.003 | 0 | 0.12 | 0.06 | 0.05 | 0.14 | 0.13 | 0.11 |
| 1 | 34078820 | rs17357341 | 0.39 | 0.407 | 0.158 | -0.044 | -0.012 | 0.002 | 0.002 | 0 | 0.12 | 0.07 | 0.06 | 0.14 | 0.14 | 0.11 |
| 1 | 34082279 | rs10914755 | -1.303 | 0.246 | 0.405 | -0.03 | -0.022 | -0.014 | -0.014 | 0 | 0.23 | 0.47 | 0.12 | 0.25 | 0.15 | 0.24 |
| 1 | 34089394 | rs4652956 | 0.795 | -0.349 | -0.03 | -0.295 | -0.015 | 0.051 | 0.051 | 2.434 | 0.14 | 0.01 | 0.23 | 0.05 | 0.19 | 0.11 |
| 1 | 34090679 | rs16835683 | 0.725 | -0.439 | 0.003 | -0.225 | 0.003 | 0.003 | 0.003 | 0 | 0.11 | 0.07 | 0.04 | 0.15 | 0.08 | 0.11 |
| 1 | 34091591 | rs12023541 | -1.232 | -0.595 | 0.538 | -0.254 | -0.049 | 0.007 | 0.007 | 2.397 | 0.25 | 0.65 | 0.12 | 0.35 | 0.11 | 0.24 |
| 1 | 34098756 | rs603893 | 0.074 | -0.515 | 0.516 | -0.359 | -0.04 | -0.001 | -0.001 | 0 | 0.1 | 0.26 | 0.09 | 0.14 | 0.02 | 0.12 |
| 1 | 34101727 | rs6701479 | 0.266 | -0.052 | 0.531 | -0.301 | -0.001 | -0.004 | -0.004 | 0.123 | 0.16 | 0.13 | 0.27 | 0.05 | 0.21 | 0.16 |
| 1 | 34102537 | rs292813 | 0.376 | -0.132 | 0.478 | 0.14 | 0 | -0.001 | -0.001 | 0 | 0.11 | 0.12 | 0.17 | 0.15 | 0.09 | 0.08 |
| 1 | 34109200 | rs1690553 | 0.716 | -0.119 | 0.342 | 0.227 | -0.021 | -0.042 | -0.042 | 0 | 0.09 | 0.01 | 0.03 | 0.14 | 0.08 | 0.08 |
| 1 | 34114077 | rs2439650 | 0.082 | -0.035 | 0.133 | 0.144 | -0.001 | 0 | 0 | 0 | 0.12 | 0.12 | 0.04 | 0.14 | 0.08 | 0.09 |
| 1 | 34129477 | rs508546 | -0.479 | -0.015 | 0.021 | 0.123 | 0 | -0.001 | -0.001 | 0 | 0.89 | 0.89 | 0.96 | 0.87 | 0.92 | 0.91 |
| 1 | 34136452 | rs4653334 | 0.492 | -0.156 | -0.209 | -0.113 | 0.018 | 0.033 | 0.033 | 0.363 | 0.15 | 0.06 | 0.26 | 0.07 | 0.17 | 0.11 |
| 1 | 34137334 | rs10798977 | 1.252 | -0.758 | 0.03 | -0.563 | 0.006 | 0.005 | 0.005 | 0 | 0.49 | 0.49 | 0.54 | 0.77 | 0.61 | 0.58 |
| 1 | 34137448 | rs529714 | -1.34 | -0.663 | -0.512 | -0.482 | -0.027 | -0.031 | -0.031 | 0.42 | 0.91 | 0.99 | 0.98 | 0.91 | 0.9 | 0.94 |
| 1 | 34149885 | rs625757 | -1.274 | -0.665 | -0.513 | -0.385 | -0.026 | -0.04 | -0.04 | 0 | 0.91 | 0.99 | 0.98 | 0.91 | 0.9 | 0.93 |
| 1 | 34150551 | rs649214 | 1.441 | -0.567 | -0.168 | -0.364 | 0.046 | 0.07 | 0.07 | 0 | 0.5 | 0.72 | 0.54 | 0.73 | 0.54 | 0.54 |
| 1 | 34151260 | rs12023377 | -0.216 | -0.508 | -0.38 | -0.284 | 0.033 | 0.043 | 0.043 | 0 | 0.21 | 0.08 | 0.15 | 0.11 | 0.13 | 0.2 |
| 1 | 34155501 | rs732889 | 1.067 | -0.128 | -0.333 | -0.129 | 0.014 | 0.023 | 0.023 | 0 | 0.52 | 0.65 | 0.52 | 0.65 | 0.53 | 0.51 |
| 1 | 34156533 | rs668310 | -0.751 | 0.092 | 0.077 | 0.435 | -0.034 | -0.05 | -0.05 | 0 | 0.91 | 0.99 | 0.97 | 0.84 | 0.89 | 0.9 |
| 1 | 34158482 | rs12757320 | 0.828 | 0.433 | -0.161 | 0.92 | 0.027 | 0.038 | 0.038 | 0 | 0.53 | 0.27 | 0.42 | 0.33 | 0.38 | 0.38 |
| 1 | 34164286 | rs1887918 | 0.736 | 0.456 | -0.152 | 0.989 | 0.018 | 0.024 | 0.024 | 0 | 0.54 | 0.28 | 0.43 | 0.38 | 0.41 | 0.39 |
| 1 | 34173973 | rs612279 | -0.283 | 0.453 | -0.694 | 1.005 | -0.039 | -0.037 | -0.037 | 0 | 0.94 | 0.99 | 0.95 | 0.89 | 0.84 | 0.91 |

| | | | | | | | | | | | | | | | | |
|---|---|---|---|---|---|---|---|---|---|---|---|---|---|---|---|---|
| 1 | 34175351 | rs16835734 | 0.347 | 0.13 | -0.324 | 1.052 | -0.004 | -0.003 | -0.003 | 3.742 | 0.08 | 0.03 | 0.05 | 0.06 | 0.08 | 0.04 |
| 1 | 34175928 | rs10914766 | 0.251 | 0.136 | -0.341 | 1.02 | 0.009 | 0.011 | 0.011 | 0 | 0.28 | 0.35 | 0.45 | 0.45 | 0.38 | 0.4 |
| 1 | 34178064 | rs562189 | -1.062 | 0.342 | -1.533 | 0.747 | -0.072 | -0.011 | -0.011 | 0 | 0.44 | 0.85 | 0.55 | 0.62 | 0.47 | 0.58 |
| 1 | 34178508 | rs565953 | -0.897 | 0.615 | -1.397 | 0.921 | -0.03 | 0.005 | 0.005 | 0 | 0.42 | 0.72 | 0.53 | 0.62 | 0.46 | 0.55 |
| 1 | 34180842 | rs16835742 | -0.867 | 0.375 | -0.885 | 0.735 | -0.035 | -0.01 | -0.01 | 0 | 0.11 | 0.01 | 0.07 | 0.07 | 0.12 | 0.15 |
| 1 | 34186193 | rs528059 | -1.061 | 0.147 | 0.11 | 0.973 | -0.02 | 0.077 | 0.077 | 0 | 0.1 | 0.31 | 0.19 | 0.18 | 0.04 | 0.13 |
| 1 | 34187258 | rs16835754 | -0.686 | 0.299 | 0.299 | 1.261 | 0.027 | 0.103 | 0.103 | 0 | 0.21 | 0.44 | 0.22 | 0.36 | 0.22 | 0.46 |
| 1 | 34202162 | rs652391 | -1.139 | 0.231 | 0.235 | 1.387 | -0.001 | -0.005 | -0.005 | 0.703 | 0.22 | 0.76 | 0.26 | 0.35 | 0.24 | 0.33 |
| 1 | 34205134 | rs12117137 | 1.673 | 0.917 | 0.139 | 1.303 | -0.015 | -0.015 | -0.015 | 0 | 0.7 | 0.23 | 0.56 | 0.61 | 0.67 | 0.58 |
| 1 | 34205859 | rs481776 | -1.438 | 0.139 | 1.032 | 0.968 | -0.011 | 0.006 | 0.006 | 0 | 0.24 | 0.51 | 0.36 | 0.26 | 0.21 | 0.34 |
| 1 | 34207470 | rs475622 | -0.884 | 0.308 | 0.643 | 0.929 | 0 | -0.006 | -0.006 | 0.789 | 0.2 | 0.69 | 0.23 | 0.21 | 0.19 | 0.26 |
| 1 | 34208399 | rs580682 | -1.356 | -0.19 | -0.194 | 0.285 | -0.162 | -0.151 | -0.151 | 0 | 0.62 | 0.97 | 0.65 | 0.48 | 0.49 | 0.8 |
| 1 | 34210532 | rs4578181 | -0.941 | 0.317 | 0.727 | 0.286 | 0 | -0.007 | -0.007 | 7.533 | 0.21 | 0.68 | 0.24 | 0.21 | 0.2 | 0.27 |
| 1 | 34210880 | rs10914770 | -0.6 | 0.347 | 0.835 | 0.305 | 0.035 | 0.017 | 0.017 | 0.008 | 0.18 | 0.46 | 0.21 | 0.21 | 0.19 | 0.25 |
| 1 | 34213109 | rs6682875 | -1.195 | 0.476 | 0.655 | 0.339 | 0.013 | 0.002 | 0.002 | 0 | 0.22 | 0.66 | 0.39 | 0.24 | 0.25 | 0.36 |
| 1 | 34215907 | rs9943211 | -0.623 | 0.5 | 0.794 | 0.345 | 0.041 | 0.023 | 0.023 | 0 | 0.18 | 0.46 | 0.21 | 0.21 | 0.2 | 0.27 |
| 1 | 34217931 | rs989661 | -0.677 | 0.51 | 0.441 | 0.335 | 0.018 | 0.002 | 0.002 | 0 | 0.18 | 0.59 | 0.22 | 0.2 | 0.19 | 0.26 |
| 1 | 34221810 | rs490109 | -0.7 | -0.115 | 0.097 | 0.066 | -0.028 | -0.025 | -0.025 | 0 | 0.27 | 0.43 | 0.16 | 0.19 | 0.2 | 0.35 |
| 1 | 34227380 | rs527315 | -1.369 | -0.707 | -0.01 | -0.158 | -0.03 | -0.019 | -0.019 | 0 | 0.39 | 0.53 | 0.32 | 0.27 | 0.24 | 0.48 |
| 1 | 34232280 | rs479056 | -0.607 | -0.398 | 0.266 | -0.02 | -0.026 | -0.003 | -0.003 | 0.337 | 0.27 | 0.41 | 0.28 | 0.25 | 0.2 | 0.46 |
| 1 | 34233391 | rs1372279 | 0.271 | -0.53 | 0.058 | -0.118 | 0.011 | 0.035 | 0.035 | 0.868 | 0.12 | 0.06 | 0.17 | 0.02 | 0.08 | 0.17 |
| 1 | 34238283 | rs474474 | -1.268 | -0.466 | 0.092 | -0.055 | -0.032 | -0.014 | -0.014 | 0 | 0.33 | 0.53 | 0.32 | 0.27 | 0.24 | 0.48 |
| 1 | 34238852 | rs479938 | 0.026 | -0.415 | 0.055 | -0.052 | -0.015 | 0.051 | 0.051 | 0 | 0.14 | 0.28 | 0.1 | 0.23 | 0.1 | 0.27 |
| 1 | 34241397 | rs10798986 | -0.601 | -0.27 | -0.808 | -0.299 | -0.015 | -0.026 | -0.026 | 0 | 0.2 | 0.27 | 0.23 | 0.08 | 0.14 | 0.22 |
| 1 | 34243635 | rs524787 | -0.107 | 0.512 | -1.579 | -0.054 | -0.081 | 0.024 | 0.024 | 0 | 0.7 | 0.93 | 0.79 | 0.81 | 0.62 | 0.81 |
| 1 | 34248064 | rs476463 | -0.756 | -0.031 | -0.506 | -0.238 | -0.031 | -0.067 | -0.067 | 0.05 | 0.19 | 0.38 | 0.12 | 0.06 | 0.13 | 0.25 |
| 1 | 34248836 | rs4652974 | 1.178 | -0.204 | -0.594 | -0.159 | 0.031 | 0.023 | 0.023 | 0 | 0.58 | 0.31 | 0.55 | 0.64 | 0.6 | 0.55 |
| 1 | 34249018 | rs507641 | -1.305 | -0.48 | -0.524 | -0.457 | -0.026 | -0.104 | -0.104 | 0 | 0.32 | 0.47 | 0.22 | 0.11 | 0.29 | 0.34 |
| 1 | 34251312 | rs472363 | -1.387 | -0.442 | -0.021 | -0.477 | -0.018 | -0.085 | -0.085 | 2.46 | 0.33 | 0.44 | 0.21 | 0.12 | 0.3 | 0.35 |
| 1 | 34251481 | rs12133088 | -0.123 | -0.178 | -0.138 | -0.25 | -0.004 | 0.018 | 0.018 | 0.092 | 0.13 | 0.07 | 0.09 | 0.04 | 0.17 | 0.1 |
| 1 | 34251754 | rs12128339 | -0.049 | -0.169 | -0.304 | -0.245 | -0.036 | 0.057 | 0.057 | 0.071 | 0.12 | 0.01 | 0.09 | 0.04 | 0.17 | 0.1 |
| 1 | 34254416 | rs6425838 | -0.884 | -0.228 | -0.277 | -0.629 | -0.013 | -0.019 | -0.019 | 1.309 | 0.3 | 0.4 | 0.26 | 0.25 | 0.29 | 0.15 |
| 1 | 34254701 | rs1441521 | -0.193 | 0.113 | -0.651 | -0.345 | -0.002 | 0 | 0 | 0 | 0.45 | 0.34 | 0.44 | 0.42 | 0.43 | 0.34 |
| 1 | 34257471 | rs10737373 | -0.578 | 0.125 | -1.122 | -0.371 | -0.031 | -0.051 | -0.051 | 0 | 0.68 | 0.85 | 0.6 | 0.61 | 0.64 | 0.61 |
| 1 | 34260948 | rs6690935 | -0.414 | -0.475 | -0.134 | -0.597 | -0.004 | -0.071 | -0.071 | 0 | 0.12 | 0.28 | 0.09 | 0 | 0.08 | 0.1 |
| 1 | 34265384 | rs12737807 | 0.484 | -0.279 | -0.794 | -0.599 | -0.033 | 0.038 | 0.038 | 0 | 0.43 | 0.16 | 0.59 | 0.66 | 0.52 | 0.59 |
| 1 | 34270187 | rs12408037 | 0.41 | -0.316 | 0.084 | -0.635 | 0.007 | 0.028 | 0.028 | 0 | 0.52 | 0.58 | 0.64 | 0.7 | 0.53 | 0.61 |
| 1 | 34270498 | rs10737374 | -0.767 | -0.472 | -0.099 | -0.968 | -0.053 | 0.071 | 0.071 | 0 | 0.84 | 0.96 | 0.86 | 0.96 | 0.69 | 0.92 |
| 1 | 34271055 | rs10914783 | 0.802 | -0.703 | -0.144 | -1.11 | -0.053 | -0.092 | -0.092 | 0.241 | 0.17 | 0.33 | 0.07 | 0 | 0.04 | 0.08 |
| 1 | 34278533 | rs1325242 | 0.355 | -0.761 | -0.007 | -0.828 | 0.008 | 0.005 | 0.005 | 0 | 0.64 | 0.33 | 0.77 | 0.96 | 0.64 | 0.83 |
| 1 | 34282691 | rs3845473 | 0.489 | -0.624 | -0.286 | -0.654 | -0.092 | -0.068 | -0.068 | 0.031 | 0.72 | 0.93 | 0.72 | 0.66 | 0.58 | 0.7 |
| 1 | 34283024 | rs4653352 | -0.299 | -0.667 | -0.316 | -0.66 | -0.017 | -0.072 | -0.072 | 0.026 | 0.14 | 0.02 | 0.15 | 0.3 | 0.12 | 0.23 |
| 1 | 34287512 | rs2281595 | 1.31 | -0.861 | -0.25 | -0.622 | 0.004 | 0.029 | 0.029 | 0 | 0.47 | 0.19 | 0.61 | 0.63 | 0.52 | 0.59 |
| 1 | 34288935 | rs12096149 | 0.835 | -0.856 | 0.209 | -0.58 | -0.053 | -0.101 | -0.101 | 0 | 0.18 | 0.32 | 0.08 | 0 | 0.05 | 0.1 |
| 1 | 34292015 | rs10914786 | -0.333 | -0.945 | -0.068 | -0.564 | 0.005 | -0.069 | -0.069 | 0 | 0.19 | 0.05 | 0.17 | 0.3 | 0.13 | 0.38 |
| 1 | 34293848 | rs1015266 | 1.014 | -0.934 | 0.061 | -0.646 | 0.002 | 0.002 | 0.002 | 0 | 0.6 | 0.56 | 0.66 | 0.62 | 0.52 | 0.53 |
| 1 | 34294874 | rs10914789 | 0.138 | -1.021 | 0.316 | -0.691 | -0.01 | -0.126 | -0.126 | 0 | 0.66 | 0.51 | 0.8 | 0.95 | 0.64 | 0.83 |
| 1 | 34306705 | rs1321623 | -0.788 | -1.017 | 0.242 | -0.678 | -0.01 | -0.001 | -0.001 | 0 | 0.37 | 0.4 | 0.24 | 0.3 | 0.19 | 0.46 |
| 1 | 34307751 | rs987954 | 0.386 | -1.019 | 0.299 | -0.651 | -0.011 | -0.072 | -0.072 | 0 | 0.63 | 0.53 | 0.8 | 0.92 | 0.64 | 0.82 |
| 1 | 34313983 | rs2180745 | 1.633 | -0.633 | -0.025 | -0.327 | -0.013 | -0.017 | -0.017 | 0 | 0.41 | 0.3 | 0.6 | 0.55 | 0.5 | 0.46 |
| 1 | 34322237 | rs753893 | -1.138 | -0.024 | -0.511 | -0.575 | -0.072 | -0.021 | -0.021 | 0.679 | 0.87 | 0.99 | 0.91 | 0.9 | 0.75 | 0.93 |
| 1 | 34325440 | rs6425853 | 0.195 | -0.188 | -0.396 | -0.643 | 0.016 | -0.003 | -0.003 | 0.75 | 0.21 | 0.14 | 0.05 | 0.01 | 0.09 | 0.13 |
| 1 | 34330067 | rs10379 | -0.912 | 0.385 | -0.648 | -0.169 | -0.072 | -0.023 | -0.023 | 0 | 0.87 | 0.99 | 0.91 | 0.9 | 0.75 | 0.93 |
| 1 | 34332933 | rs1108929 | -0.912 | 0.385 | -0.648 | -0.169 | -0.072 | -0.023 | -0.023 | 0 | 0.87 | 0.99 | 0.91 | 0.9 | 0.75 | 0.93 |
| 1 | 34340467 | rs10799000 | 0.032 | -0.014 | -0.813 | 0.016 | -0.014 | -0.069 | -0.069 | 0 | 0.16 | 0.02 | 0.18 | 0.26 | 0.13 | 0.23 |
| 1 | 34342399 | rs6425854 | 0.043 | -0.646 | -0.254 | -0.177 | -0.051 | -0.051 | -0.051 | 0 | 0.22 | 0.34 | 0.17 | 0.08 | 0.09 | 0.12 |
| 1 | 34345249 | rs10914802 | 1.747 | -0.537 | 0.153 | -0.308 | 0 | -0.002 | -0.002 | 0.079 | 0.45 | 0.42 | 0.53 | 0.52 | 0.53 | 0.51 |
| 1 | 34347846 | rs12058145 | 0.296 | -0.494 | -0.165 | -0.389 | -0.035 | -0.082 | -0.082 | 0 | 0.2 | 0.29 | 0.06 | 0.01 | 0.08 | 0.1 |
| 1 | 34349815 | rs4653358 | -1.036 | -0.213 | -0.752 | -0.117 | -0.07 | -0.007 | -0.007 | 0 | 0.87 | 0.99 | 0.91 | 0.9 | 0.75 | 0.91 |
| 1 | 34350918 | rs16836066 | 0.343 | -0.742 | -0.573 | -0.214 | -0.003 | -0.005 | -0.005 | 0 | 0.17 | 0.12 | 0.06 | 0.05 | 0.04 | 0.11 |
| 1 | 34353184 | rs12568620 | 0.34 | -0.737 | -0.571 | -0.201 | -0.006 | -0.008 | -0.008 | 0 | 0.17 | 0.15 | 0.06 | 0.05 | 0.04 | 0.11 |
| 1 | 34353648 | rs10914803 | 1.466 | 0.094 | -0.558 | -0.139 | -0.001 | 0 | 0 | 0 | 0.45 | 0.44 | 0.48 | 0.51 | 0.43 | 0.47 |
| 1 | 34358476 | rs10799004 | 0.286 | 0.347 | -0.949 | -0.447 | -0.003 | 0.039 | 0.039 | 0 | 0.68 | 0.8 | 0.77 | 0.85 | 0.65 | 0.81 |
| 1 | 34362665 | rs10799006 | -1.038 | 0.762 | -1.62 | -0.263 | -0.078 | -0.001 | -0.001 | 0.212 | 0.87 | 0.99 | 0.84 | 0.91 | 0.72 | 0.94 |
| 1 | 34363830 | rs12029918 | 1.067 | 0.671 | -1.552 | -0.067 | -0.098 | -0.092 | -0.092 | 0 | 0.42 | 0.06 | 0.49 | 0.56 | 0.41 | 0.49 |
| 1 | 34366411 | rs7528354 | 0.941 | 0.59 | -1.492 | -0.058 | -0.067 | -0.066 | -0.066 | 0 | 0.43 | 0.09 | 0.49 | 0.56 | 0.41 | 0.5 |
| 1 | 34369245 | rs1570089 | -0.7 | 0.709 | -0.603 | 0.281 | 0 | 0.005 | 0.005 | 0 | 0.24 | 0.29 | 0.24 | 0.28 | 0.22 | 0.32 |
| 1 | 34375199 | rs911216 | -0.533 | 1.138 | -0.304 | -0.104 | 0.009 | -0.027 | -0.027 | 0 | 0.21 | 0.27 | 0.14 | 0.09 | 0.23 | 0.28 |
| 1 | 34377214 | rs11580955 | -0.224 | 1.683 | -0.72 | -0.194 | -0.032 | 0.012 | 0.012 | 0 | 0.05 | 0 | 0.12 | 0.02 | 0.14 | 0.04 |

| | | | | | | | | | | | | | | | | |
|---|---|---|---|---|---|---|---|---|---|---|---|---|---|---|---|---|
| 1 | 34382423 | rs7533958 | -1.355 | 1.336 | -0.696 | -0.337 | 0.018 | 0.031 | 0.031 | 0 | 0.56 | 0.71 | 0.42 | 0.28 | 0.63 | 0.47 |
| 1 | 34383554 | rs12124122 | -0.1 | 1.082 | -0.849 | -0.352 | -0.037 | -0.05 | -0.05 | 1.357 | 0.17 | 0.03 | 0.08 | 0.15 | 0.15 | 0.12 |
| 1 | 34386710 | rs12031839 | -0.881 | 0.91 | -0.112 | -0.243 | 0.015 | -0.071 | -0.071 | 0 | 0.18 | 0.37 | 0.21 | 0.05 | 0.25 | 0.11 |
| 1 | 34387864 | rs6697504 | -1.378 | 0.7 | -0.241 | 0.177 | 0.018 | 0.005 | 0.005 | 0 | 0.47 | 0.7 | 0.35 | 0.37 | 0.47 | 0.44 |
| 1 | 34392843 | rs10914816 | -0.205 | 1.144 | 0.2 | 0.637 | -0.038 | -0.002 | -0.002 | 0 | 0.07 | 0.01 | 0.13 | 0.04 | 0.15 | 0.06 |
| 1 | 34394259 | rs10914817 | -0.403 | 1.107 | 0.228 | 0.632 | -0.026 | 0.008 | 0.008 | 0 | 0.08 | 0.03 | 0.13 | 0.04 | 0.16 | 0.06 |
| 1 | 34394682 | rs7555410 | -0.447 | 0.852 | 0.525 | 0.443 | -0.008 | -0.033 | -0.033 | 0 | 0.2 | 0.25 | 0.09 | 0.07 | 0.14 | 0.17 |
| 1 | 34396256 | rs10799010 | -0.959 | 0.739 | 0.93 | 0.571 | 0 | -0.062 | -0.062 | 0.065 | 0.29 | 0.47 | 0.25 | 0.15 | 0.3 | 0.25 |
| 1 | 34398046 | rs6665339 | -0.936 | 0.832 | 0.989 | 0.609 | 0 | -0.067 | -0.067 | 0 | 0.29 | 0.43 | 0.22 | 0.12 | 0.29 | 0.24 |
| 1 | 34398141 | rs6425863 | -1.312 | 0.86 | 0.889 | 0.855 | -0.001 | 0.001 | 0.001 | 0.051 | 0.37 | 0.63 | 0.3 | 0.35 | 0.36 | 0.32 |
| 1 | 34418942 | rs504471 | -0.533 | 0.992 | 0.588 | 0.792 | 0 | -0.003 | -0.003 | 0 | 0.38 | 0.64 | 0.27 | 0.24 | 0.37 | 0.29 |
| 1 | 34424393 | rs549048 | -1.005 | 0.969 | -0.28 | 0.708 | 0.023 | 0.011 | 0.011 | 0 | 0.51 | 0.75 | 0.42 | 0.47 | 0.57 | 0.46 |
| 1 | 34424462 | rs549944 | -0.482 | 0.907 | -0.679 | 0.019 | -0.016 | 0.053 | 0.053 | 0 | 0.32 | 0.22 | 0.21 | 0.09 | 0.34 | 0.24 |
| 1 | 34428365 | rs483083 | -1.03 | -0.574 | 0.036 | 1.238 | -0.041 | 0.052 | 0.052 | 0 | 0.23 | 0.61 | 0.25 | 0.4 | 0.12 | 0.34 |
| 1 | 34430533 | rs545337 | -0.722 | -0.424 | -0.824 | 0.19 | -0.059 | 0.207 | 0.207 | 0 | 0.57 | 0.84 | 0.55 | 0.85 | 0.47 | 0.75 |
| 1 | 34432484 | rs12128266 | -0.133 | -0.742 | -0.889 | 0.72 | -0.003 | 0.012 | 0.012 | 0.834 | 0.08 | 0.1 | 0.04 | 0.18 | 0.03 | 0.1 |
| 1 | 34437596 | rs6663957 | 0.305 | -0.853 | -0.444 | 1.401 | 0.001 | 0.041 | 0.041 | 0 | 0.08 | 0.06 | 0.21 | 0.36 | 0.03 | 0.23 |
| 1 | 34438881 | rs502199 | -0.721 | -0.863 | -0.06 | 1.335 | -0.004 | 0.07 | 0.07 | 0.357 | 0.48 | 0.69 | 0.5 | 0.67 | 0.43 | 0.66 |
| 1 | 34441126 | rs1536110 | -0.974 | -0.982 | 0.751 | 1.313 | 0.001 | 0.019 | 0.019 | 0 | 0.47 | 0.55 | 0.49 | 0.72 | 0.41 | 0.61 |
| 1 | 34443688 | rs749733 | -0.124 | -1.038 | 0.8 | 1.321 | -0.04 | -0.034 | -0.034 | 0 | 0.36 | 0.17 | 0.41 | 0.55 | 0.38 | 0.51 |
| 1 | 34444230 | rs12127584 | 0.177 | -1.026 | 0.824 | 1.327 | -0.013 | 0.045 | 0.045 | 0.852 | 0.08 | 0.14 | 0.04 | 0.18 | 0.03 | 0.1 |
| 1 | 34444494 | rs560563 | -1.023 | -0.996 | 1.103 | 1.262 | -0.011 | 0.041 | 0.041 | 0.004 | 0.1 | 0.3 | 0.06 | 0.13 | 0.05 | 0.14 |
| 1 | 34445919 | rs12072000 | 0.812 | -0.978 | 0.977 | 1.324 | -0.095 | 0.13 | 0.13 | 0 | 0.27 | 0.03 | 0.16 | 0.06 | 0.34 | 0.17 |
| 1 | 34446139 | rs4408120 | 0.449 | -0.971 | 1.051 | 1.327 | 0.002 | -0.031 | -0.031 | 1.07 | 0.29 | 0.27 | 0.18 | 0.06 | 0.34 | 0.17 |
| 1 | 34451605 | rs10914827 | -0.956 | -0.978 | 1.006 | 1.454 | -0.001 | 0.024 | 0.024 | 0 | 0.14 | 0.14 | 0.27 | 0.55 | 0.06 | 0.34 |
| 1 | 34453282 | rs10449236 | -0.003 | -0.982 | 0.982 | 1.455 | -0.014 | -0.019 | -0.019 | 0 | 0.33 | 0.25 | 0.25 | 0.36 | 0.37 | 0.36 |
| 1 | 34456109 | rs520226 | -0.404 | -0.95 | 1.263 | 1.474 | -0.015 | 0.138 | 0.138 | 0 | 0.11 | 0.23 | 0.22 | 0.4 | 0.04 | 0.28 |
| 1 | 34460961 | rs2358755 | -1.081 | -1.469 | 1.043 | 0.928 | 0 | 0.005 | 0.005 | 0.402 | 0.5 | 0.45 | 0.29 | 0.28 | 0.46 | 0.37 |
| 1 | 34463451 | rs533861 | -0.677 | -1.449 | 1.266 | 1.2 | -0.012 | 0.327 | 0.327 | 0 | 0.13 | 0.47 | 0.25 | 0.39 | 0.04 | 0.3 |
| 1 | 34465455 | rs9425977 | -0.367 | -1.471 | 1.169 | 1.145 | 0.004 | -0.012 | -0.012 | 0 | 0.13 | 0.1 | 0.04 | 0.01 | 0.07 | 0.03 |
| 1 | 34470308 | rs487472 | -1.923 | -1.426 | 1.183 | 1.206 | -0.027 | 0.034 | 0.034 | 0.155 | 0.62 | 0.78 | 0.53 | 0.66 | 0.5 | 0.65 |
| 1 | 34472159 | rs7515286 | -0.049 | -1.354 | 1.125 | 1.168 | 0 | 0.041 | 0.041 | 0.001 | 0.1 | 0.1 | 0.21 | 0.38 | 0.04 | 0.29 |
| 1 | 34472274 | rs7529498 | -0.285 | -1.357 | 1.161 | 1.166 | -0.01 | 0.209 | 0.209 | 0 | 0.11 | 0.26 | 0.23 | 0.38 | 0.04 | 0.29 |
| 1 | 34473767 | rs12085816 | 0.91 | -1.391 | 0.819 | 1.073 | -0.073 | 0.112 | 0.112 | 0.002 | 0.27 | 0.09 | 0.16 | 0.06 | 0.33 | 0.14 |
| 1 | 34473786 | rs9286948 | -1.094 | -1.549 | 0.917 | 1.169 | -0.001 | -0.001 | -0.001 | 0 | 0.2 | 0.21 | 0.08 | 0.11 | 0.12 | 0.11 |
| 1 | 34474234 | rs10914838 | -0.199 | -1.175 | 0.875 | 1.365 | 0.001 | 0.039 | 0.039 | 0 | 0.1 | 0.08 | 0.21 | 0.38 | 0.04 | 0.29 |
| 1 | 34475200 | rs561207 | 0.456 | -0.869 | 0.961 | 0.798 | 0.001 | -0.027 | -0.027 | 0.371 | 0.28 | 0.3 | 0.17 | 0.07 | 0.29 | 0.14 |
| 1 | 34475772 | rs7541778 | -1.042 | -0.806 | 0.957 | 1.237 | -0.011 | 0.09 | 0.09 | 0 | 0.16 | 0.24 | 0.27 | 0.48 | 0.08 | 0.35 |
| 1 | 34477581 | rs1325258 | -0.957 | -0.731 | 0.915 | 1.278 | -0.005 | 0.052 | 0.052 | 0 | 0.16 | 0.2 | 0.27 | 0.47 | 0.08 | 0.36 |
| 1 | 34477893 | rs10914839 | -0.834 | -0.773 | 0.901 | 1.096 | 0.002 | 0.023 | 0.023 | 0 | 0.51 | 0.42 | 0.46 | 0.6 | 0.41 | 0.56 |
| 1 | 34479922 | rs2358756 | -1.504 | -0.916 | 0.943 | 1.048 | 0.007 | -0.03 | -0.03 | 1.724 | 0.24 | 0.16 | 0.31 | 0.59 | 0.16 | 0.47 |
| 1 | 34486225 | rs12563211 | -0.709 | -0.906 | 1.036 | 1.122 | 0 | -0.005 | -0.005 | 0 | 0.13 | 0.13 | 0.08 | 0.19 | 0.1 | 0.13 |
| 1 | 34488115 | rs10914845 | 1.968 | -0.895 | 0.926 | 0.994 | -0.142 | 0.123 | 0.123 | 0 | 0.36 | 0.05 | 0.38 | 0.19 | 0.54 | 0.27 |
| 1 | 34488578 | rs1465782 | -1.695 | -0.76 | 1.235 | 1.177 | -0.007 | 0.041 | 0.041 | 0 | 0.57 | 0.69 | 0.58 | 0.75 | 0.41 | 0.66 |
| 1 | 34490495 | rs10159305 | 0.857 | -0.71 | 1.057 | 1.151 | -0.004 | -0.072 | -0.072 | 0 | 0.27 | 0.3 | 0.14 | 0 | 0.22 | 0.09 |
| 1 | 34490934 | rs10914846 | 0.357 | -0.603 | 0.817 | 1.298 | 0 | 0.048 | 0.048 | 1.158 | 0.09 | 0.09 | 0.22 | 0.34 | 0.05 | 0.27 |
| 1 | 34491558 | rs2164581 | -1.819 | -0.601 | 0.82 | 1.281 | 0.001 | -0.005 | -0.005 | 0 | 0.24 | 0.25 | 0.32 | 0.59 | 0.16 | 0.47 |
| 1 | 34495028 | rs1550164 | -0.729 | -0.567 | 0.74 | 1.258 | -0.005 | -0.02 | -0.02 | 0.543 | 0.12 | 0.07 | 0.07 | 0.19 | 0.1 | 0.13 |
| 1 | 34496094 | rs10914850 | 0.035 | -0.544 | 0.714 | 1.274 | -0.002 | 0.079 | 0.079 | 0.638 | 0.1 | 0.14 | 0.23 | 0.34 | 0.05 | 0.27 |
| 1 | 34496322 | rs10914852 | 0.182 | -0.542 | 0.671 | 1.275 | 0.001 | 0.022 | 0.022 | 0.768 | 0.09 | 0.08 | 0.22 | 0.34 | 0.05 | 0.27 |
| 1 | 34499892 | rs6701264 | 0.74 | -0.636 | 0.766 | 1.038 | -0.009 | -0.098 | -0.098 | 0.098 | 0.27 | 0.34 | 0.15 | 0 | 0.23 | 0.09 |
| 1 | 34499921 | rs12035559 | -0.018 | -0.549 | 0.142 | 1.183 | 0.001 | 0.041 | 0.041 | 0.918 | 0.11 | 0.09 | 0.23 | 0.4 | 0.05 | 0.34 |
| 1 | 34502161 | rs12062460 | 0.872 | -0.495 | 0.121 | 0.969 | 0.006 | 0.077 | 0.077 | 0.261 | 0.25 | 0.13 | 0.13 | 0 | 0.23 | 0.09 |
| 1 | 34505115 | rs2625684 | -0.257 | -0.407 | 0.229 | 1.081 | -0.002 | 0.062 | 0.062 | 0 | 0.64 | 0.75 | 0.79 | 0.8 | 0.66 | 0.78 |
| 1 | 34514486 | rs3131529 | -0.167 | -0.63 | 0.297 | 1.166 | 0.005 | 0.01 | 0.01 | 0.179 | 0.62 | 0.61 | 0.78 | 0.74 | 0.7 | 0.79 |
| 1 | 34515330 | rs899378 | 1.056 | -0.692 | 0.37 | 1.274 | 0 | 0.016 | 0.016 | 0 | 0.43 | 0.34 | 0.43 | 0.26 | 0.62 | 0.41 |
| 1 | 34516663 | rs12127677 | -0.591 | 0.236 | 0.445 | 1.211 | -0.002 | 0.001 | 0.001 | 0.468 | 0.06 | 0.05 | 0.25 | 0.15 | 0.11 | 0.18 |
| 1 | 34523335 | rs9425897 | -1.082 | -0.445 | 0.761 | 1.293 | 0.037 | 0.051 | 0.051 | 0 | 0.5 | 0.68 | 0.55 | 0.61 | 0.35 | 0.51 |
| 1 | 34523743 | rs12043450 | -0.1 | -0.418 | 0.658 | 1.277 | 0.001 | 0.02 | 0.02 | 0.19 | 0.12 | 0.1 | 0.3 | 0.34 | 0.05 | 0.27 |
| 1 | 34524849 | rs6425872 | -0.443 | -0.208 | 0.59 | 0.501 | -0.014 | -0.056 | -0.056 | 0 | 0.34 | 0.44 | 0.2 | 0.13 | 0.28 | 0.14 |
| 1 | 34525105 | rs6700561 | 0.445 | -0.207 | 0.402 | 0.481 | 0.01 | 0.042 | 0.042 | 1.981 | 0.26 | 0.16 | 0.14 | 0 | 0.24 | 0.08 |
| 1 | 34533702 | rs1865892 | -0.198 | 0.102 | -0.289 | 0.109 | -0.023 | -0.075 | -0.075 | 0.001 | 0.34 | 0.51 | 0.21 | 0.13 | 0.28 | 0.14 |
| 1 | 34541724 | rs9425986 | -0.44 | 0.473 | -0.879 | 0.41 | -0.022 | 0.004 | 0.004 | 0 | 0.24 | 0.15 | 0.4 | 0.2 | 0.32 | 0.22 |
| 1 | 34546694 | rs7551631 | -1.327 | -0.301 | -0.426 | 0.733 | 0.003 | 0.002 | 0.002 | 0 | 0.46 | 0.64 | 0.63 | 0.57 | 0.44 | 0.38 |
| 1 | 34556046 | rs10914862 | 0.424 | -0.299 | -0.553 | 0.655 | -0.025 | -0.008 | -0.008 | 0.06 | 0.2 | 0.12 | 0.38 | 0.2 | 0.29 | 0.18 |
| 1 | 34564086 | rs9425991 | -1.644 | -0.485 | -0.56 | 0.748 | 0.012 | 0.018 | 0.018 | 0 | 0.48 | 0.69 | 0.66 | 0.61 | 0.43 | 0.38 |
| 1 | 34564936 | rs6665353 | -0.567 | -0.545 | -0.223 | 0.717 | 0.01 | -0.017 | -0.017 | 0.05 | 0.3 | 0.57 | 0.49 | 0.25 | 0.33 | 0.24 |
| 1 | 34565359 | rs12048816 | -0.429 | -0.826 | -0.288 | 0.984 | 0.015 | -0.026 | -0.026 | 0 | 0.18 | 0.12 | 0.19 | 0.37 | 0.11 | 0.15 |
| 1 | 34569267 | rs2196541 | 0.308 | -1.122 | 0.102 | 0.844 | -0.045 | 0.058 | 0.058 | 0 | 0.73 | 0.58 | 0.84 | 0.61 | 0.84 | 0.55 |

| | | | | | | | | | | | | | | | | |
|---|---|---|---|---|---|---|---|---|---|---|---|---|---|---|---|---|
| 1 | 34571206 | rs720212 | -0.694 | -1.062 | 0.289 | 0.409 | 0.003 | 0.014 | 0.014 | 0 | 0.4 | 0.36 | 0.64 | 0.68 | 0.39 | 0.49 |
| 1 | 34574691 | rs1382600 | -0.418 | -1.139 | 0.35 | 0.349 | 0.004 | 0.017 | 0.017 | 0 | 0.37 | 0.24 | 0.58 | 0.64 | 0.36 | 0.43 |
| 1 | 34583840 | rs1065966 | 0.081 | -1.254 | 0.881 | 0.284 | -0.001 | -0.024 | -0.024 | 1.2 | 0.94 | 0.8 | 0.97 | 1 | 0.98 | 0.96 |
| 1 | 34586161 | rs11581276 | -0.057 | -1.356 | 1.082 | 0.069 | 0.016 | -0.005 | -0.005 | 0 | 0.25 | 0.21 | 0.11 | 0.31 | 0.16 | 0.39 |
| 1 | 34590285 | rs7551988 | -0.293 | -1.194 | 1.089 | 0.219 | 0.002 | 0.031 | 0.031 | 0 | 0.36 | 0.2 | 0.61 | 0.69 | 0.36 | 0.38 |
| 1 | 34591069 | rs6425876 | -0.383 | -1.069 | 1.114 | 0.19 | 0.045 | 0.017 | 0.017 | 1 | 0.95 | 0.73 | 0.96 | 1 | 0.98 | 0.96 |
| 1 | 34602870 | rs9426003 | 0.53 | -1.166 | 0.406 | 0.506 | -0.053 | -0.065 | -0.065 | 0.257 | 0.23 | 0.06 | 0.4 | 0.24 | 0.27 | 0.21 |
| 1 | 34604567 | rs9662615 | -0.811 | -1.188 | 0.494 | 0.363 | -0.013 | 0.068 | 0.068 | 0 | 0.44 | 0.17 | 0.64 | 0.68 | 0.45 | 0.55 |
| 1 | 34606984 | rs771150 | 0.78 | -1.876 | 0.798 | 0.36 | -0.04 | 0.017 | 0.017 | 0.187 | 0.72 | 0.39 | 0.85 | 0.6 | 0.87 | 0.63 |
| 1 | 34609394 | rs705210 | 0.909 | -1.891 | 0.909 | 0.331 | -0.055 | -0.021 | -0.021 | 0 | 0.86 | 0.48 | 0.93 | 0.96 | 0.95 | 0.85 |
| 1 | 34610222 | rs771145 | -0.068 | -1.819 | 1.082 | 0.266 | -0.051 | 0.041 | 0.041 | 0 | 0.78 | 0.66 | 0.91 | 0.61 | 0.9 | 0.69 |
| 1 | 34613920 | rs796405 | 0.218 | -1.813 | 0.92 | 0.27 | -0.086 | 0.062 | 0.062 | 0 | 0.76 | 0.48 | 0.89 | 0.61 | 0.9 | 0.7 |
| 1 | 34613986 | rs771138 | -0.401 | -1.613 | 0.498 | 0.227 | -0.041 | -0.022 | -0.022 | 0 | 0.38 | 0.12 | 0.61 | 0.59 | 0.39 | 0.49 |
| 1 | 34615576 | rs4653395 | 0.65 | -1.538 | 0.565 | 0.239 | -0.007 | 0.002 | 0.002 | 0.007 | 0.69 | 0.37 | 0.83 | 0.6 | 0.81 | 0.63 |
| 1 | 34616045 | rs771132 | 1.143 | -1.372 | 0.732 | -0.226 | 0.014 | -0.132 | -0.132 | 0 | 0.4 | 0.48 | 0.3 | 0.03 | 0.51 | 0.2 |
| 1 | 34616871 | rs11590546 | -0.284 | -1.588 | 0.282 | 0.18 | -0.064 | 0.083 | 0.083 | 0 | 0.26 | 0.46 | 0.23 | 0.45 | 0.14 | 0.27 |
| 1 | 34617256 | rs12755908 | 1.25 | -1.451 | -0.499 | 0.155 | -0.01 | 0.035 | 0.035 | 0 | 0.4 | 0.25 | 0.32 | 0.25 | 0.54 | 0.29 |
| 1 | 34620770 | rs10914880 | 0.836 | -1.416 | -0.582 | 0.16 | 0.032 | 0.019 | 0.019 | 0 | 0.6 | 0.31 | 0.62 | 0.27 | 0.76 | 0.51 |
| 1 | 34620986 | rs12145664 | 0.172 | -1.241 | -1.138 | 0.372 | -0.036 | -0.038 | -0.038 | 0.087 | 0.27 | 0.07 | 0.39 | 0.26 | 0.29 | 0.37 |
| 1 | 34622349 | rs4652992 | 0.636 | -0.926 | -0.605 | 0.595 | 0 | 0.024 | 0.024 | 0 | 0.08 | 0.07 | 0.09 | 0.24 | 0.07 | 0.15 |
| 1 | 34625898 | rs7552457 | 0.544 | -0.462 | -0.951 | 0.097 | -0.04 | -0.047 | -0.047 | 0 | 0.19 | 0.08 | 0.39 | 0.28 | 0.25 | 0.26 |
| 1 | 34626340 | rs10493055 | 0.822 | -0.307 | -0.882 | -0.519 | -0.061 | 0.112 | 0.112 | 0.075 | 0.36 | 0.16 | 0.21 | 0 | 0.46 | 0.21 |
| 1 | 34627298 | rs771124 | -0.098 | -0.388 | -0.237 | 1.05 | -0.002 | -0.017 | -0.017 | 0 | 0.89 | 0.92 | 0.89 | 0.61 | 0.91 | 0.85 |
| 1 | 34627949 | rs771122 | 0.334 | -0.219 | -0.188 | 1.136 | -0.018 | 0.026 | 0.026 | 0 | 0.79 | 0.76 | 0.84 | 0.61 | 0.88 | 0.85 |
| 1 | 34643837 | rs750572 | -1.336 | 0.907 | 0.211 | 0.478 | 0.037 | -0.007 | -0.007 | 0 | 0.09 | 0.27 | 0.17 | 0.1 | 0.12 | 0.13 |
| 1 | 34653822 | rs2792928 | -0.987 | 0.481 | 0.471 | -0.075 | -0.015 | -0.077 | -0.077 | 0 | 0.22 | 0.38 | 0.19 | 0.08 | 0.2 | 0.07 |
| 1 | 34656128 | rs1771374 | -1.426 | -0.096 | 1.242 | -0.351 | -0.018 | -0.013 | -0.013 | 0 | 0.12 | 0.28 | 0.2 | 0.1 | 0.07 | 0.19 |
| 1 | 34659156 | rs1771365 | -1.404 | -0.051 | 1.236 | -0.358 | 0 | -0.011 | -0.011 | 0 | 0.1 | 0.28 | 0.21 | 0.08 | 0.07 | 0.16 |
| 1 | 34667863 | rs12746310 | -0.863 | -0.326 | 1.586 | -0.512 | -0.011 | 0.087 | 0.087 | 1.31 | 0.22 | 0.06 | 0.22 | 0.07 | 0.2 | 0.18 |
| 1 | 34671355 | rs297804 | -1.791 | -0.315 | 2.025 | -0.506 | -0.009 | -0.034 | -0.034 | 0 | 0.31 | 0.63 | 0.35 | 0.14 | 0.23 | 0.27 |
| 1 | 34674814 | rs297802 | -1.636 | -0.261 | 1.585 | -0.513 | -0.019 | -0.069 | -0.069 | 0 | 0.29 | 0.36 | 0.27 | 0.08 | 0.22 | 0.26 |
| 1 | 34679066 | rs4652997 | 0.178 | -0.194 | 0.483 | -0.231 | -0.01 | -0.007 | -0.007 | 0 | 0.46 | 0.28 | 0.63 | 0.51 | 0.54 | 0.59 |
| 1 | 34680319 | rs719769 | -0.752 | -1.19 | 0.241 | -0.686 | 0.004 | 0.046 | 0.046 | 0 | 0.71 | 0.88 | 0.81 | 0.83 | 0.77 | 0.9 |
| 1 | 34682856 | rs1023325 | -0.238 | -1.396 | 0.46 | -0.714 | 0.018 | 0.041 | 0.041 | 0 | 0.65 | 0.78 | 0.77 | 0.79 | 0.72 | 0.76 |
| 1 | 34683153 | rs1023324 | 0.027 | -1.498 | 1.252 | -0.76 | -0.008 | -0.015 | -0.015 | 0.744 | 0.62 | 0.55 | 0.75 | 0.78 | 0.71 | 0.74 |
| 1 | 34684463 | rs705207 | -0.424 | -1.469 | 1.363 | -0.734 | 0.009 | 0.018 | 0.018 | 0 | 0.67 | 0.75 | 0.79 | 0.79 | 0.75 | 0.79 |
| 1 | 34684621 | rs1061074 | -0.786 | -1.503 | 1.332 | -0.736 | 0.005 | 0.035 | 0.035 | 0 | 0.71 | 0.86 | 0.8 | 0.83 | 0.76 | 0.9 |
| 1 | 34686130 | rs12043879 | 0.202 | -1.033 | 1.778 | -0.391 | 0.002 | 0.004 | 0.004 | 0 | 0.19 | 0.19 | 0.13 | 0.27 | 0.2 | 0.17 |
| 1 | 34697488 | rs1414472 | -0.13 | -1.461 | 1.369 | -0.686 | 0.003 | -0.007 | -0.007 | 1.326 | 0.77 | 0.59 | 0.76 | 0.77 | 0.79 | 0.8 |
| 1 | 34706158 | rs10914890 | 0.417 | -1.975 | 1.178 | -0.884 | -0.016 | -0.026 | -0.026 | 1.297 | 0.74 | 0.59 | 0.81 | 0.78 | 0.79 | 0.72 |
| 1 | 34714693 | rs2795634 | -0.479 | -1.676 | 1.25 | -0.726 | 0.014 | 0.007 | 0.007 | 0 | 0.83 | 0.71 | 0.84 | 0.79 | 0.81 | 0.92 |
| 1 | 34723163 | rs771410 | -0.647 | -1.645 | 1.089 | -0.711 | 0.005 | 0.002 | 0.002 | 0 | 0.84 | 0.81 | 0.85 | 0.79 | 0.81 | 0.92 |
| 1 | 34723928 | rs12128858 | -0.055 | -0.826 | 0.849 | 0.206 | -0.01 | -0.005 | -0.005 | 0 | 0.08 | 0.04 | 0.23 | 0.26 | 0.1 | 0.17 |
| 1 | 34726959 | rs7512138 | -0.325 | -0.864 | 0.815 | 0.259 | 0.022 | 0.083 | 0.083 | 0 | 0.25 | 0.36 | 0.35 | 0.41 | 0.28 | 0.2 |
| 1 | 34727536 | rs10799032 | 0.697 | -0.712 | 0.782 | -0.196 | 0 | -0.01 | -0.01 | 0.005 | 0.15 | 0.14 | 0.1 | 0.05 | 0.14 | 0.06 |
| 1 | 34727579 | rs771401 | -0.213 | -0.554 | 0.52 | -0.565 | 0.01 | 0.041 | 0.041 | 0 | 0.69 | 0.78 | 0.81 | 0.88 | 0.72 | 0.82 |
| 1 | 34727902 | rs771400 | -0.126 | -0.7 | 0.547 | -0.512 | 0.035 | 0.047 | 0.047 | 0 | 0.71 | 0.81 | 0.85 | 0.88 | 0.79 | 0.85 |
| 1 | 34727990 | rs6425885 | 0.557 | -0.707 | 0.886 | -0.561 | -0.023 | -0.051 | -0.051 | 0 | 0.62 | 0.51 | 0.79 | 0.85 | 0.72 | 0.61 |
| 1 | 34734066 | rs10914894 | 1.223 | -0.809 | 0.243 | -0.557 | -0.046 | 0.04 | 0.04 | 0 | 0.43 | 0.2 | 0.53 | 0.62 | 0.5 | 0.5 |
| 1 | 34735938 | rs10914896 | 1.256 | -0.7 | 0.171 | -0.528 | 0.018 | 0.006 | 0.006 | 0.116 | 0.18 | 0.21 | 0.23 | 0.22 | 0.22 | 0.11 |
| 1 | 34736287 | rs1414470 | 1.126 | -0.754 | 0.089 | -0.532 | -0.034 | 0.05 | 0.05 | 0 | 0.44 | 0.22 | 0.54 | 0.63 | 0.5 | 0.5 |
| 1 | 34741605 | rs880017 | 0.382 | -0.79 | -0.099 | -0.512 | -0.023 | -0.061 | -0.061 | 0 | 0.64 | 0.53 | 0.8 | 0.87 | 0.73 | 0.65 |
| 1 | 34748645 | rs6677202 | 0.46 | -0.784 | -0.177 | -0.51 | -0.034 | -0.067 | -0.067 | 0 | 0.63 | 0.5 | 0.8 | 0.85 | 0.73 | 0.62 |
| 1 | 34750936 | rs771390 | -0.539 | -0.668 | -0.544 | -0.439 | 0.036 | 0.066 | 0.066 | 0.18 | 0.73 | 0.85 | 0.85 | 0.88 | 0.79 | 0.85 |
| 1 | 34754967 | rs4653013 | 0.044 | -0.537 | -0.059 | -0.124 | 0.008 | -0.002 | -0.002 | 0 | 0.23 | 0.53 | 0.29 | 0.25 | 0.23 | 0.14 |
| 1 | 34756509 | rs6704508 | -0.734 | -0.527 | -0.336 | -0.138 | 0 | 0.003 | 0.003 | 2.155 | 0.31 | 0.67 | 0.32 | 0.27 | 0.29 | 0.35 |
| 1 | 34764974 | rs1404350 | 0.86 | -0.294 | -0.292 | -0.035 | 0.035 | 0.093 | 0.093 | 0 | 0.55 | 0.27 | 0.6 | 0.74 | 0.63 | 0.48 |
| 1 | 34783684 | rs1541221 | -0.789 | -0.317 | -0.805 | -0.051 | 0.033 | 0.037 | 0.037 | 0 | 0.78 | 0.87 | 0.93 | 0.98 | 0.86 | 0.78 |
| 1 | 34790866 | rs12130968 | 0.719 | -0.149 | -0.982 | 0.063 | 0.033 | 0.131 | 0.131 | 0 | 0.54 | 0.14 | 0.61 | 0.73 | 0.62 | 0.61 |
| 1 | 34793830 | rs10914912 | 0.82 | -0.15 | -0.935 | -0.758 | -0.036 | 0.095 | 0.095 | 0.423 | 0.48 | 0.12 | 0.34 | 0.34 | 0.55 | 0.37 |
| 1 | 34796092 | rs10914915 | 0.787 | 0.191 | -0.753 | -0.476 | -0.002 | 0.016 | 0.016 | 0 | 0.47 | 0.46 | 0.42 | 0.32 | 0.52 | 0.36 |
| 1 | 34797348 | rs1361675 | -0.633 | 1.256 | -0.683 | 0.547 | 0.007 | 0 | 0 | 0.41 | 0.3 | 0.27 | 0.49 | 0.68 | 0.33 | 0.4 |
| 1 | 34804499 | rs1342160 | -0.38 | 1.021 | -0.825 | 0.316 | 0.002 | 0.022 | 0.022 | 0 | 0.47 | 0.63 | 0.48 | 0.64 | 0.44 | 0.5 |
| 1 | 34806304 | rs4652836 | -0.166 | 0.952 | -0.564 | 1.064 | -0.002 | 0.028 | 0.028 | 0 | 0.29 | 0.44 | 0.32 | 0.32 | 0.27 | 0.38 |
| 1 | 34809587 | rs10799047 | 0.325 | 0.418 | -0.368 | 0.725 | -0.011 | 0.048 | 0.048 | 0 | 0.78 | 0.61 | 0.81 | 0.68 | 0.83 | 0.8 |
| 1 | 34809903 | rs7416554 | 1.18 | 0.414 | -0.433 | 0.713 | -0.016 | 0.088 | 0.088 | 0 | 0.48 | 0.16 | 0.48 | 0.35 | 0.54 | 0.42 |
| 1 | 34812189 | rs17500033 | 0.792 | 1.536 | -0.793 | 0.364 | -0.027 | 0.025 | 0.025 | 0 | 0.06 | 0.02 | 0.06 | 0 | 0.13 | 0.04 |
| 1 | 34812948 | rs10914926 | 0.42 | 1.348 | -0.743 | 0.356 | -0.024 | -0.066 | -0.066 | 1.112 | 0.15 | 0.03 | 0.14 | 0.32 | 0.16 | 0.12 |
| 1 | 34815453 | rs6660749 | 0.426 | 1.397 | 0.181 | 0.531 | 0 | -0.003 | -0.003 | 0 | 0.27 | 0.21 | 0.29 | 0.28 | 0.24 | 0.37 |

| | | | | | | | | | | | | | | | | |
|---|---|---|---|---|---|---|---|---|---|---|---|---|---|---|---|---|
| 1 | 34815894 | rs2359229 | -0.59 | 1.362 | 0.498 | 0.471 | -0.003 | 0.006 | 0.006 | 1.522 | 0.37 | 0.38 | 0.36 | 0.27 | 0.39 | 0.4 |
| 1 | 34823560 | rs2049420 | 1.35 | 1.313 | 0.501 | 0.577 | 0.012 | 0.032 | 0.032 | 0 | 0.44 | 0.33 | 0.46 | 0.36 | 0.43 | 0.35 |
| 1 | 34823622 | rs2140682 | 1.041 | 1.178 | 0.661 | 0.938 | 0.011 | 0.009 | 0.009 | 0 | 0.51 | 0.49 | 0.55 | 0.65 | 0.44 | 0.5 |
| 1 | 34828319 | rs12117975 | 0.808 | 1.457 | 0.432 | 0.8 | 0 | 0.002 | 0.002 | 0.098 | 0.61 | 0.39 | 0.57 | 0.38 | 0.65 | 0.46 |
| 1 | 34830640 | rs7537783 | -0.545 | 1.479 | 0.501 | 0.811 | -0.007 | -0.004 | -0.004 | 0 | 0.33 | 0.29 | 0.39 | 0.61 | 0.35 | 0.46 |
| 1 | 34835613 | rs12565280 | 1.373 | 0.359 | -0.406 | 0.192 | 0.042 | -0.093 | -0.093 | 0 | 0.18 | 0.07 | 0.21 | 0.53 | 0.12 | 0.22 |
| 1 | 34839501 | rs705203 | -0.552 | 0.363 | -0.456 | 0.078 | -0.02 | -0.004 | -0.004 | 0 | 0.7 | 0.77 | 0.68 | 0.73 | 0.64 | 0.59 |
| 1 | 34844208 | rs17380703 | -0.041 | 0.355 | -0.355 | 0.079 | 0 | 0.011 | 0.011 | 1.008 | 0.19 | 0.12 | 0.11 | 0.01 | 0.19 | 0.08 |
| 1 | 34844836 | rs705197 | 0.596 | 0.36 | -0.119 | 0.416 | -0.04 | 0.071 | 0.071 | 0 | 0.21 | 0.28 | 0.23 | 0.55 | 0.12 | 0.22 |
| 1 | 34866656 | rs2294193 | 0.219 | 0.63 | -0.17 | 0.191 | 0.003 | 0.062 | 0.062 | 0.008 | 0.49 | 0.59 | 0.45 | 0.2 | 0.51 | 0.37 |
| 1 | 34871620 | rs732699 | 0.652 | 0.478 | -0.195 | -0.183 | -0.015 | 0.05 | 0.05 | 3.209 | 0.39 | 0.24 | 0.38 | 0.18 | 0.46 | 0.2 |
| 1 | 34872737 | rs2092028 | -0.419 | 0.419 | -0.297 | -0.006 | 0.005 | -0.001 | -0.001 | 0 | 0.7 | 0.69 | 0.68 | 0.75 | 0.67 | 0.59 |
| 1 | 34879050 | rs7530700 | 0.614 | 0.518 | -0.343 | 0.019 | 0.005 | -0.034 | -0.034 | 0.041 | 0.4 | 0.42 | 0.39 | 0.19 | 0.47 | 0.2 |
| 1 | 34883744 | rs6693232 | 0.117 | 0.167 | -0.854 | 0.487 | 0.025 | -0.05 | -0.05 | 0 | 0.25 | 0.17 | 0.26 | 0.57 | 0.2 | 0.29 |
| 1 | 34886654 | rs2294190 | -0.198 | 0.475 | -0.781 | 0.288 | 0.002 | 0.012 | 0.012 | 0 | 0.05 | 0.08 | 0.16 | 0.09 | 0.06 | 0.04 |
| 1 | 34889492 | rs4653029 | -0.03 | -0.086 | -1.625 | 0.318 | 0.032 | -0.043 | -0.043 | 0 | 0.24 | 0.09 | 0.3 | 0.27 | 0.18 | 0.29 |
| 1 | 34895009 | rs17385253 | 1.323 | 0.233 | -1.804 | 0.247 | -0.011 | -0.042 | -0.042 | 0 | 0.24 | 0.05 | 0.15 | 0.31 | 0.27 | 0.15 |
| 1 | 34896603 | rs7533470 | 1.116 | 0.66 | -1.253 | 0.103 | 0.006 | 0.008 | 0.008 | 0 | 0.26 | 0.34 | 0.22 | 0.34 | 0.35 | 0.16 |
| 1 | 34900172 | rs2142534 | -0.968 | 0.641 | -1.082 | -0.242 | 0 | -0.001 | -0.001 | 0 | 0.67 | 0.7 | 0.53 | 0.6 | 0.64 | 0.51 |
| 1 | 34902910 | rs4653032 | 0.552 | 0.037 | -1.038 | -0.272 | 0.016 | 0.011 | 0.011 | 0 | 0.28 | 0.13 | 0.28 | 0.26 | 0.24 | 0.3 |
| 1 | 34903822 | rs4653033 | 0.357 | 0.018 | -0.788 | -0.358 | -0.001 | -0.001 | -0.001 | 0 | 0.31 | 0.34 | 0.3 | 0.3 | 0.24 | 0.3 |
| 1 | 34907148 | rs10753308 | -0.947 | 0.717 | -0.603 | -0.027 | 0.005 | 0.025 | 0.025 | 0 | 0.73 | 0.7 | 0.56 | 0.6 | 0.69 | 0.5 |
| 1 | 34915051 | rs6681948 | -0.648 | 0.437 | -0.951 | -0.297 | 0.007 | 0.009 | 0.009 | 0 | 0.45 | 0.59 | 0.35 | 0.26 | 0.29 | 0.32 |
| 1 | 34915394 | rs12134720 | 0.828 | 0.64 | -0.764 | -0.126 | 0.01 | -0.004 | -0.004 | 1.169 | 0.06 | 0.1 | 0.08 | 0.03 | 0.12 | 0.05 |
| 1 | 34916121 | rs4653036 | -0.034 | 0.553 | -0.651 | -0.058 | -0.005 | -0.006 | -0.006 | 0.601 | 0.67 | 0.35 | 0.46 | 0.55 | 0.55 | 0.5 |
| 1 | 34919266 | rs10914953 | 1.531 | 0.485 | -0.58 | -0.109 | 0.015 | 0.021 | 0.021 | 0 | 0.3 | 0.22 | 0.16 | 0.28 | 0.28 | 0.21 |
| 1 | 34919755 | rs12132086 | 0.681 | 0.52 | -0.552 | -0.112 | 0.019 | -0.009 | -0.009 | 0 | 0.06 | 0.13 | 0.08 | 0.03 | 0.12 | 0.05 |
| 1 | 34921075 | rs2075933 | 0.213 | 0.488 | -0.522 | -0.115 | 0.023 | 0.026 | 0.026 | 0 | 0.36 | 0.2 | 0.3 | 0.27 | 0.27 | 0.29 |
| 1 | 34921613 | rs12035126 | 1.738 | 0.485 | -0.533 | -0.106 | 0.011 | 0.023 | 0.023 | 0.36 | 0.29 | 0.16 | 0.15 | 0.26 | 0.27 | 0.15 |
| 1 | 34922823 | rs4653037 | 1.095 | 0.274 | -0.794 | -0.239 | 0.005 | -0.024 | -0.024 | 0.206 | 0.25 | 0.07 | 0.24 | 0.27 | 0.19 | 0.2 |
| 1 | 34924642 | rs1028282 | 1.935 | 0.324 | -0.74 | -0.21 | 0.004 | 0.024 | 0.024 | 0.433 | 0.28 | 0.07 | 0.14 | 0.26 | 0.27 | 0.15 |
| 1 | 34924784 | rs1016090 | 0.904 | 0.366 | -0.675 | -0.215 | -0.002 | 0.007 | 0.007 | 0 | 0.37 | 0.38 | 0.25 | 0.32 | 0.42 | 0.25 |
| 1 | 34925069 | rs1033371 | -0.134 | 0.294 | -0.788 | -0.209 | -0.001 | 0 | 0 | 0.231 | 0.33 | 0.36 | 0.34 | 0.32 | 0.31 | 0.31 |
| 1 | 34925601 | rs7514397 | 1.422 | 0.472 | -1.667 | -0.11 | 0.016 | 0.051 | 0.051 | 0 | 0.11 | 0.04 | 0.05 | 0 | 0.09 | 0.08 |
| 1 | 34928441 | rs7544781 | 0.367 | 0.498 | -1.683 | -0.07 | -0.005 | -0.01 | -0.01 | 0 | 0.54 | 0.24 | 0.39 | 0.53 | 0.46 | 0.4 |
| 1 | 34930461 | rs4653042 | -0.419 | 0.611 | -1.879 | -0.019 | 0.029 | 0.025 | 0.025 | 0 | 0.39 | 0.2 | 0.31 | 0.31 | 0.29 | 0.37 |
| 1 | 34931680 | rs10799060 | 0.283 | 0.598 | -1.8 | 0.057 | -0.008 | -0.014 | -0.014 | 0 | 0.53 | 0.17 | 0.39 | 0.59 | 0.45 | 0.36 |
| 1 | 34935035 | rs7549758 | 0.923 | 0.706 | -1.788 | 0.024 | -0.047 | 0.077 | 0.077 | 0 | 0.17 | 0.02 | 0.12 | 0.03 | 0.21 | 0.14 |
| 1 | 34936568 | rs798076 | 1.535 | 0.647 | -1.722 | 0.129 | 0.005 | 0.02 | 0.02 | 0.019 | 0.28 | 0.09 | 0.14 | 0.27 | 0.27 | 0.14 |
| 1 | 34936992 | rs798075 | 0.855 | 0.595 | -1.145 | 0.694 | 0.002 | 0.068 | 0.068 | 0 | 0.47 | 0.25 | 0.28 | 0.35 | 0.5 | 0.28 |
| 1 | 34937352 | rs798074 | 0.424 | 1.033 | -0.094 | 1.165 | 0.009 | 0.027 | 0.027 | 0 | 0.73 | 0.3 | 0.53 | 0.64 | 0.69 | 0.56 |
| 1 | 34944591 | rs41416648 | -1.287 | 0.964 | 0.348 | 1.093 | 0.042 | 0.022 | 0.022 | 0 | 0.13 | 0.29 | 0.19 | 0.14 | 0.15 | 0.04 |
| 1 | 34958515 | rs3845481 | -1.061 | 1.007 | 0.984 | 1.758 | 0.095 | 0.236 | 0.236 | 0 | 0.1 | 0.28 | 0.17 | 0.29 | 0.14 | 0.04 |
| 1 | 34962675 | rs2359106 | -0.784 | 0.724 | 0.686 | 2.08 | 0.002 | -0.003 | -0.003 | 0 | 0.1 | 0.12 | 0.05 | 0.21 | 0.06 | 0.11 |
| 1 | 34962846 | rs3845483 | -0.234 | 0.578 | 0.559 | 1.044 | 0.001 | 0.001 | 0.001 | 0 | 0.58 | 0.6 | 0.76 | 0.79 | 0.67 | 0.67 |
| 1 | 34967241 | rs4652841 | 0.768 | 0.442 | 0.226 | 0.921 | -0.051 | -0.104 | -0.104 | 0 | 0.41 | 0.07 | 0.51 | 0.48 | 0.46 | 0.57 |
| 1 | 34978409 | rs12027125 | 0.224 | 0.411 | 0.326 | 0.987 | -0.002 | 0.024 | 0.024 | 0 | 0.08 | 0.13 | 0.05 | 0.2 | 0.06 | 0.09 |
| 1 | 34982041 | rs6694043 | -0.973 | 0.354 | 0.488 | 0.934 | 0.02 | 0.09 | 0.09 | 0.102 | 0.18 | 0.61 | 0.24 | 0.29 | 0.18 | 0.12 |
| 1 | 34987339 | rs1539374 | 0.644 | 0.333 | -0.102 | 0.911 | -0.06 | -0.097 | -0.097 | 0 | 0.41 | 0.08 | 0.52 | 0.48 | 0.49 | 0.58 |
| 1 | 34988954 | rs10493058 | 0.106 | 0.232 | 0.075 | 0.97 | 0 | -0.001 | -0.001 | 0 | 0.42 | 0.47 | 0.33 | 0.59 | 0.47 | 0.42 |
| 1 | 34992037 | rs4653052 | -1.328 | 0.355 | -0.472 | 0.549 | -0.081 | 0.202 | 0.202 | 0 | 0.7 | 0.96 | 0.81 | 0.98 | 0.7 | 0.81 |
| 1 | 34995974 | rs4595337 | -0.55 | 0.581 | 0.23 | 0.987 | 0.031 | 0.014 | 0.014 | 0 | 0.13 | 0.26 | 0.21 | 0.17 | 0.13 | 0.15 |
| 1 | 34996413 | rs10737381 | -0.438 | 0.274 | -0.101 | 0.79 | -0.042 | 0.093 | 0.093 | 0.372 | 0.57 | 0.86 | 0.68 | 0.72 | 0.59 | 0.51 |
| 1 | 34997944 | rs1418483 | 0.53 | 0.247 | 0.289 | 0.886 | 0.005 | 0.005 | 0.005 | 0.303 | 0.4 | 0.27 | 0.46 | 0.6 | 0.45 | 0.34 |
| 1 | 35013586 | rs12136079 | 0.409 | 0.302 | 0.19 | 0.949 | -0.009 | -0.005 | -0.005 | 0.018 | 0.42 | 0.49 | 0.48 | 0.63 | 0.45 | 0.33 |
| 1 | 35029632 | rs6425900 | -0.65 | 0.402 | 0.16 | 0.903 | -0.018 | 0.039 | 0.039 | 0.021 | 0.6 | 0.8 | 0.69 | 0.76 | 0.6 | 0.53 |
| 1 | 35038958 | rs947671 | -0.595 | 0.455 | 0.2 | 0.856 | 0.001 | -0.002 | -0.002 | 0 | 0.15 | 0.13 | 0.18 | 0.12 | 0.15 | 0.19 |
| 1 | 35040371 | rs6661959 | -0.239 | 0.107 | 0.213 | 0.816 | -0.004 | 0.035 | 0.035 | 0 | 0.56 | 0.65 | 0.64 | 0.86 | 0.56 | 0.64 |
| 1 | 35040991 | rs6665029 | -0.217 | 0.091 | 0.155 | 1.362 | 0.002 | -0.002 | -0.002 | 0 | 0.12 | 0.08 | 0.15 | 0.23 | 0.1 | 0.35 |
| 1 | 35042287 | rs6677231 | -0.5 | 0.275 | 0.488 | 1.306 | 0.016 | 0.021 | 0.021 | 0.024 | 0.07 | 0.11 | 0.2 | 0.11 | 0.07 | 0.13 |
| 1 | 35042445 | rs6668930 | -0.906 | 0.289 | 0.395 | 1.437 | 0 | 0.001 | 0.001 | 0 | 0.11 | 0.09 | 0.21 | 0.18 | 0.12 | 0.2 |
| 1 | 35042498 | rs6671294 | -0.668 | 0.224 | 0.846 | 1.201 | -0.015 | 0.039 | 0.039 | 0 | 0.18 | 0.32 | 0.21 | 0.25 | 0.18 | 0.14 |
| 1 | 35044934 | rs7530276 | -0.774 | 0.248 | 1.05 | 1.237 | 0.035 | 0.094 | 0.094 | 2.368 | 0.08 | 0.3 | 0.15 | 0.13 | 0.05 | 0.11 |
| 1 | 35049252 | rs10799071 | 0.053 | 0.24 | 0.507 | 0.933 | -0.004 | -0.011 | -0.011 | 0 | 0.43 | 0.5 | 0.45 | 0.66 | 0.35 | 0.37 |
| 1 | 35049332 | rs10914983 | 0.656 | 0.208 | 0.254 | 1.039 | -0.002 | -0.066 | -0.066 | 0 | 0.35 | 0.2 | 0.27 | 0.5 | 0.31 | 0.26 |
| 1 | 35050334 | rs12074036 | 0.191 | 0.377 | 0.082 | 0.685 | 0.004 | 0.01 | 0.01 | 0 | 0.44 | 0.61 | 0.45 | 0.65 | 0.36 | 0.37 |
| 1 | 35055281 | rs10914986 | -1.017 | 0.23 | -0.387 | 0.615 | -0.133 | 0.137 | 0.137 | 0 | 0.68 | 0.97 | 0.73 | 0.93 | 0.59 | 0.8 |
| 1 | 35058101 | rs728448 | -0.625 | 0.11 | -0.135 | 0.656 | -0.034 | 0.005 | 0.005 | 0 | 0.23 | 0.4 | 0.27 | 0.26 | 0.18 | 0.39 |
| 1 | 35058148 | rs728447 | -0.067 | 0.504 | 0.128 | 0.927 | -0.001 | -0.006 | -0.006 | 5.639 | 0.44 | 0.55 | 0.44 | 0.66 | 0.39 | 0.41 |

| | | | | | | | | | | | | | | | | |
|---|---|---|---|---|---|---|---|---|---|---|---|---|---|---|---|---|
| 1 | 35065969 | rs7528705 | -1.17 | 0.59 | 0.509 | 1.247 | 0.053 | 0.259 | 0.259 | 0 | 0.05 | 0.21 | 0.03 | 0.27 | 0.04 | 0.21 |
| 1 | 35069285 | rs1886340 | 0.261 | 0.789 | 0.109 | 0.325 | 0.001 | 0.007 | 0.007 | 0 | 0.47 | 0.55 | 0.55 | 0.67 | 0.47 | 0.5 |
| 1 | 35072298 | rs4653061 | -0.01 | 0.786 | -0.274 | 0.081 | -0.059 | 0.24 | 0.24 | 0 | 0.44 | 0.14 | 0.38 | 0.05 | 0.46 | 0.27 |
| 1 | 35084074 | rs7544557 | -0.553 | 0.768 | 0.109 | 0.315 | 0.008 | 0.025 | 0.025 | 0 | 0.52 | 0.41 | 0.45 | 0.35 | 0.53 | 0.49 |
| 1 | 35085870 | rs6695510 | -0.621 | 0.784 | 0.168 | 0.315 | -0.001 | -0.004 | -0.004 | 0.344 | 0.53 | 0.52 | 0.46 | 0.35 | 0.53 | 0.49 |
| 1 | 35086365 | rs716289 | -1.384 | 0.72 | 0.105 | 0.231 | 0.02 | 0.138 | 0.138 | 0 | 0.09 | 0.27 | 0.07 | 0.3 | 0.08 | 0.22 |
| 1 | 35088484 | rs4653067 | 0.028 | 0.742 | 0.007 | 0.218 | -0.041 | 0.175 | 0.175 | 0 | 0.43 | 0.17 | 0.39 | 0.12 | 0.45 | 0.3 |
| 1 | 35090340 | rs6702527 | -1.775 | 0.514 | 0.48 | 0.687 | 0.049 | 0.175 | 0.175 | 0.934 | 0.09 | 0.34 | 0.06 | 0.22 | 0.07 | 0.2 |
| 1 | 35092406 | rs12059222 | -0.755 | 0.856 | -0.093 | 0.301 | -0.003 | -0.006 | -0.006 | 0 | 0.37 | 0.41 | 0.3 | 0.35 | 0.37 | 0.36 |
| 1 | 35093593 | rs2093880 | -0.4 | 0.594 | -0.151 | 0.255 | -0.005 | 0.056 | 0.056 | 0 | 0.07 | 0.17 | 0.05 | 0.19 | 0.05 | 0.19 |
| 1 | 35094871 | rs6425904 | -0.803 | 0.613 | -0.283 | -0.16 | 0 | -0.007 | -0.007 | 0.02 | 0.2 | 0.22 | 0.2 | 0.07 | 0.18 | 0.22 |
| 1 | 35095368 | rs7548924 | -1.53 | 0.701 | -0.314 | -0.158 | -0.005 | -0.013 | -0.013 | 0 | 0.27 | 0.36 | 0.27 | 0.26 | 0.25 | 0.41 |
| 1 | 35095609 | rs10914996 | -1.734 | 0.851 | -0.379 | -0.069 | 0 | 0 | 0 | 8.314 | 0.43 | 0.44 | 0.42 | 0.32 | 0.42 | 0.47 |
| 1 | 35095652 | rs12132123 | 1.029 | 0.927 | -0.541 | -0.033 | -0.003 | 0.009 | 0.009 | 0.044 | 0.16 | 0.06 | 0.15 | 0.06 | 0.16 | 0.06 |
| 1 | 35113184 | rs612353 | -1.564 | 0.761 | -0.439 | -0.208 | 0 | -0.001 | -0.001 | 0 | 0.46 | 0.49 | 0.42 | 0.33 | 0.44 | 0.5 |
| 1 | 35114289 | rs4652852 | -0.972 | 0.773 | -0.612 | -0.265 | 0.002 | 0.017 | 0.017 | 0.125 | 0.34 | 0.26 | 0.25 | 0.25 | 0.32 | 0.35 |
| 1 | 35114411 | rs491000 | -0.296 | 0.871 | -0.653 | -0.394 | -0.001 | -0.009 | -0.009 | 0 | 0.11 | 0.15 | 0.16 | 0.08 | 0.12 | 0.15 |
| 1 | 35114647 | rs12140922 | -0.375 | 0.839 | -0.68 | -0.181 | -0.003 | 0.024 | 0.024 | 0.233 | 0.25 | 0.17 | 0.19 | 0.06 | 0.25 | 0.14 |
| 1 | 35115542 | rs17561354 | 0.287 | 0.734 | -0.756 | -0.153 | 0 | 0.022 | 0.022 | 0 | 0.1 | 0.04 | 0.03 | 0 | 0.07 | 0.08 |
| 1 | 35118140 | rs6425908 | -0.571 | 1.046 | -0.756 | -0.168 | -0.02 | 0.023 | 0.023 | 0 | 0.27 | 0.13 | 0.22 | 0.16 | 0.31 | 0.22 |
| 1 | 35118282 | rs9787098 | -0.743 | 0.889 | -0.542 | 0.129 | -0.008 | 0.012 | 0.012 | 0 | 0.37 | 0.54 | 0.38 | 0.59 | 0.31 | 0.64 |
| 1 | 35118335 | rs509788 | 0.131 | 0.783 | -0.689 | 0.139 | -0.009 | 0.012 | 0.012 | 0.631 | 0.42 | 0.33 | 0.37 | 0.36 | 0.43 | 0.35 |
| 1 | 35122523 | rs10915010 | -0.401 | 0.67 | -0.591 | 0.526 | 0.001 | 0.006 | 0.006 | 0 | 0.28 | 0.33 | 0.23 | 0.33 | 0.31 | 0.31 |
| 1 | 35127074 | rs912537 | -0.582 | 0.627 | -0.466 | 0.542 | 0.003 | 0.015 | 0.015 | 0 | 0.29 | 0.35 | 0.23 | 0.42 | 0.31 | 0.37 |
| 1 | 35133818 | rs12024912 | -0.212 | 0.739 | -0.62 | 0.581 | -0.045 | -0.079 | -0.079 | 0 | 0.25 | 0.08 | 0.2 | 0.42 | 0.31 | 0.36 |
| 1 | 35142875 | rs12142734 | -0.34 | 0.711 | -0.574 | 0.576 | -0.003 | -0.007 | -0.007 | 0 | 0.27 | 0.21 | 0.21 | 0.42 | 0.31 | 0.37 |
| 1 | 35152490 | rs1177663 | 0.519 | 0.222 | -0.424 | 0.193 | -0.044 | -0.047 | -0.047 | 0.095 | 0.25 | 0.35 | 0.38 | 0.12 | 0.2 | 0.19 |
| 1 | 35159799 | rs12136236 | 1.089 | 0.346 | -0.841 | 0.274 | -0.054 | -0.009 | -0.009 | 0.44 | 0.17 | 0.02 | 0.16 | 0.12 | 0.21 | 0.09 |
| 1 | 35160136 | rs16836901 | 0.599 | 0.537 | -0.717 | 0.141 | 0 | -0.02 | -0.02 | 0.472 | 0.08 | 0.09 | 0.15 | 0 | 0.09 | 0.05 |
| 1 | 35160674 | rs10915021 | -0.434 | 0.514 | -0.47 | 0.2 | 0.01 | -0.011 | -0.011 | 0 | 0.12 | 0.32 | 0.18 | 0.09 | 0.11 | 0.12 |
| 1 | 35161802 | rs4320749 | -0.506 | 0.422 | -0.394 | 0.157 | 0.004 | 0.008 | 0.008 | 0.325 | 0.59 | 0.68 | 0.48 | 0.76 | 0.59 | 0.72 |
| 1 | 35162901 | rs1336625 | 0.284 | 0.501 | -0.694 | 0.175 | 0 | 0.028 | 0.028 | 0 | 0.47 | 0.37 | 0.3 | 0.67 | 0.48 | 0.6 |
| 1 | 35164392 | rs10493063 | 0.135 | 0.721 | -0.849 | 0.331 | 0.001 | -0.034 | -0.034 | 0 | 0.08 | 0.08 | 0.14 | 0 | 0.09 | 0.05 |
| 1 | 35170315 | rs17473524 | -0.676 | 0.663 | -0.896 | 0.32 | -0.002 | -0.014 | -0.014 | 0 | 0.13 | 0.16 | 0.16 | 0.09 | 0.11 | 0.12 |
| 1 | 35182232 | rs1741961 | -0.196 | 0.992 | -1.564 | 0.506 | -0.025 | 0.013 | 0.013 | 0 | 0.75 | 0.88 | 0.7 | 0.79 | 0.69 | 0.85 |
| 1 | 35183638 | rs684524 | -0.15 | 0.984 | -1.553 | 0.503 | -0.018 | 0.011 | 0.011 | 0 | 0.75 | 0.84 | 0.69 | 0.79 | 0.7 | 0.85 |
| 1 | 35185845 | rs10489658 | 0.621 | 0.727 | -1.878 | 0.465 | 0 | 0.003 | 0.003 | 0 | 0.41 | 0.59 | 0.48 | 0.37 | 0.41 | 0.42 |
| 1 | 35190138 | rs1741953 | -0.128 | 0.619 | -2.27 | 0.712 | 0.001 | -0.033 | -0.033 | 1.819 | 0.3 | 0.12 | 0.2 | 0.33 | 0.28 | 0.36 |
| 1 | 35198358 | rs666919 | 0.06 | 0.516 | -2.382 | 0.637 | -0.018 | -0.059 | -0.059 | 0 | 0.77 | 0.99 | 0.85 | 0.75 | 0.79 | 0.87 |
| 1 | 35208698 | rs9426007 | -0.246 | 1.014 | -2.63 | 0.267 | -0.051 | 0.034 | 0.034 | 0.321 | 0.29 | 0.08 | 0.13 | 0.21 | 0.34 | 0.29 |
| 1 | 35215989 | rs2064899 | -0.485 | 0.445 | -2.281 | -0.128 | -0.001 | -0.015 | -0.015 | 0 | 0.61 | 0.41 | 0.6 | 0.83 | 0.59 | 0.81 |
| 1 | 35216973 | rs9331222 | 0.261 | -0.114 | -1.998 | -0.231 | 0.005 | 0.019 | 0.019 | 0.009 | 0.46 | 0.38 | 0.45 | 0.68 | 0.55 | 0.5 |
| 1 | 35217376 | rs585268 | -1.12 | -0.796 | -1.716 | -0.107 | 0.018 | -0.03 | -0.03 | 0 | 0.42 | 0.77 | 0.39 | 0.53 | 0.38 | 0.43 |
| 1 | 35218077 | rs692923 | -0.279 | -0.659 | -1.19 | -0.236 | -0.001 | -0.029 | -0.029 | 0 | 0.27 | 0.55 | 0.21 | 0.2 | 0.27 | 0.29 |
| 1 | 35223345 | rs3738345 | -0.114 | -0.587 | -0.753 | -0.298 | -0.013 | -0.082 | -0.082 | 1.279 | 0.28 | 0.48 | 0.24 | 0.11 | 0.25 | 0.16 |
| 1 | 35224312 | rs2093185 | -1.007 | -0.529 | -0.969 | 0.017 | 0.053 | -0.017 | -0.017 | 0 | 0.47 | 0.83 | 0.42 | 0.52 | 0.39 | 0.43 |
| 1 | 35227673 | rs755931 | -0.912 | -0.176 | -0.442 | -0.384 | -0.005 | 0.108 | 0.108 | 0 | 0.07 | 0.3 | 0.13 | 0.13 | 0.04 | 0.06 |
| 1 | 35231378 | rs10489656 | -0.862 | 0.074 | -1.526 | -0.922 | -0.001 | -0.007 | -0.007 | 1.458 | 0.37 | 0.41 | 0.25 | 0.14 | 0.36 | 0.37 |
| 1 | 35239992 | rs639126 | 0.021 | 0.436 | -1.331 | -0.63 | -0.003 | -0.006 | -0.006 | 0.055 | 0.2 | 0.69 | 0.24 | 0.16 | 0.2 | 0.07 |
| 1 | 35246297 | rs1764405 | -1.51 | 0.477 | -2.03 | -0.773 | -0.113 | -0.146 | -0.146 | 2.429 | 0.66 | 0.99 | 0.53 | 0.49 | 0.67 | 0.49 |
| 1 | 35251669 | rs13774 | 0.752 | 0.599 | -1.898 | -0.916 | -0.019 | 0.053 | 0.053 | 0.867 | 0.14 | 0.03 | 0.11 | 0 | 0.14 | 0.05 |
| 1 | 35253376 | rs483721 | 0.368 | 0.846 | -1.268 | -0.733 | 0.023 | -0.035 | -0.035 | 0 | 0.21 | 0.48 | 0.16 | 0.14 | 0.22 | 0.08 |
| 1 | 35256125 | rs1741969 | -1.424 | 0.692 | -1.72 | -0.517 | -0.098 | -0.134 | -0.134 | 0 | 0.67 | 0.96 | 0.53 | 0.49 | 0.66 | 0.5 |
| 1 | 35257416 | rs4653091 | -0.314 | 0.409 | -1.402 | -0.476 | 0.021 | 0.021 | 0.021 | 0 | 0.36 | 0.27 | 0.24 | 0.32 | 0.32 | 0.3 |
| 1 | 35265770 | rs564226 | -0.483 | 0.898 | -0.695 | -0.698 | 0.007 | -0.031 | -0.031 | 0 | 0.31 | 0.58 | 0.27 | 0.17 | 0.32 | 0.2 |
| 1 | 35268368 | rs1764392 | -0.901 | 0.812 | -0.171 | -0.636 | -0.001 | 0 | 0 | 0 | 0.54 | 0.56 | 0.4 | 0.49 | 0.52 | 0.47 |
| 1 | 35269895 | rs512161 | -0.573 | 0.702 | 0.006 | -0.821 | 0.024 | 0.008 | 0.008 | 0 | 0.48 | 0.63 | 0.36 | 0.48 | 0.44 | 0.36 |
| 1 | 35272080 | rs6689695 | 0.552 | 0.513 | 0.082 | -1.163 | -0.011 | 0.013 | 0.013 | 0 | 0.07 | 0.09 | 0.06 | 0.1 | 0.06 | 0.07 |
| 1 | 35282297 | rs1407135 | 0.496 | 0.89 | 0.272 | -1.307 | -0.001 | -0.003 | -0.003 | 0 | 0.55 | 0.51 | 0.73 | 0.57 | 0.53 | 0.64 |
| 1 | 35289873 | rs1741972 | -0.963 | 0.98 | -0.134 | -1.203 | 0 | 0.008 | 0.008 | 0 | 0.93 | 0.94 | 0.96 | 0.9 | 0.93 | 0.92 |
| 1 | 35299572 | rs12064230 | 0.554 | 1.007 | -0.122 | -1.263 | 0.002 | -0.016 | -0.016 | 0 | 0.05 | 0.1 | 0.04 | 0 | 0.07 | 0.1 |
| 1 | 35301579 | rs6689081 | -1.023 | 1.34 | -0.024 | -1.852 | 0.001 | 0.004 | 0.004 | 0 | 0.51 | 0.61 | 0.43 | 0.15 | 0.54 | 0.37 |
| 1 | 35302617 | rs11264125 | -0.472 | 1.349 | -0.038 | -1.889 | -0.002 | -0.017 | -0.017 | 0.026 | 0.19 | 0.2 | 0.23 | 0.02 | 0.24 | 0.2 |
| 1 | 35303369 | rs11264139 | -2.119 | 1.32 | -0.167 | -2.13 | -0.007 | -0.057 | -0.057 | 0 | 0.66 | 0.95 | 0.53 | 0.46 | 0.71 | 0.56 |
| 1 | 35304381 | rs12239410 | -0.13 | 1.083 | -0.055 | -2.163 | -0.005 | -0.021 | -0.021 | 0 | 0.31 | 0.43 | 0.19 | 0.13 | 0.3 | 0.17 |
| 1 | 35307121 | rs4653104 | -0.428 | 0.686 | -0.381 | -1.663 | 0 | -0.005 | -0.005 | 0 | 0.2 | 0.19 | 0.23 | 0.1 | 0.22 | 0.18 |
| 1 | 35312631 | rs11802507 | -0.245 | 0.511 | -0.357 | -1.377 | -0.013 | 0.015 | 0.015 | 0 | 0.11 | 0.13 | 0.05 | 0.3 | 0.07 | 0.14 |
| 1 | 35314841 | rs12128624 | -0.597 | 0.886 | -0.774 | -1.977 | -0.004 | 0.094 | 0.094 | 0 | 0.65 | 0.85 | 0.5 | 0.23 | 0.67 | 0.53 |
| 1 | 35316497 | rs12076918 | -0.036 | 1.073 | -0.812 | -2.288 | -0.001 | -0.058 | -0.058 | 0 | 0.51 | 0.47 | 0.39 | 0.18 | 0.49 | 0.44 |

| | | | | | | | | | | | | | | | | |
|---|---|---|---|---|---|---|---|---|---|---|---|---|---|---|---|---|
| 1 | 35317472 | rs2025005 | -0.049 | 0.468 | -0.889 | -1.9 | 0.019 | 0.129 | 0.129 | 0 | 0.48 | 0.29 | 0.39 | 0.16 | 0.57 | 0.42 |
| 1 | 35321242 | rs14103 | -0.38 | 0.383 | -0.761 | -1.262 | 0.001 | -0.002 | -0.002 | 3.192 | 0.09 | 0.09 | 0.07 | 0.15 | 0.07 | 0.1 |
| 1 | 35322503 | rs4653107 | 0.058 | 0.248 | -0.549 | -1.69 | -0.004 | -0.069 | -0.069 | 5.978 | 0.85 | 0.79 | 0.92 | 1 | 0.87 | 0.96 |
| 1 | 35328032 | rs4653108 | 0.825 | 0.149 | -1.133 | -1.807 | -0.011 | 0.02 | 0.02 | 0 | 0.23 | 0.02 | 0.42 | 0.79 | 0.2 | 0.38 |
| 1 | 35341982 | rs7541937 | 0.904 | 0.39 | -0.646 | -2.111 | -0.002 | -0.049 | -0.049 | 0 | 0.44 | 0.45 | 0.38 | 0.04 | 0.47 | 0.37 |
| 1 | 35342095 | rs11264126 | -0.878 | 0.803 | -0.878 | -1.465 | -0.005 | 0.056 | 0.056 | 0 | 0.45 | 0.32 | 0.43 | 0.76 | 0.47 | 0.52 |
| 1 | 35364457 | rs4259608 | -0.145 | 0.718 | -1.781 | -0.873 | -0.007 | 0.101 | 0.101 | 0 | 0.28 | 0.08 | 0.47 | 0.92 | 0.25 | 0.46 |
| 1 | 35367290 | rs4652867 | -0.349 | 0.773 | -1.861 | -0.719 | -0.014 | 0.039 | 0.039 | 0 | 0.32 | 0.07 | 0.34 | 0.79 | 0.3 | 0.41 |
| 1 | 35368431 | rs11264172 | 0.049 | 0.702 | -1.947 | -0.635 | 0.013 | 0.02 | 0.02 | 0 | 0.46 | 0.32 | 0.39 | 0.37 | 0.38 | 0.61 |
| 1 | 35368760 | rs6686484 | 0.344 | -0.258 | -1.483 | -1.396 | -0.002 | -0.103 | -0.103 | 0 | 0.61 | 0.4 | 0.76 | 1 | 0.64 | 0.81 |
| 1 | 35368771 | rs11264173 | -0.164 | -0.36 | -0.553 | -0.899 | -0.001 | -0.051 | -0.051 | 0 | 0.35 | 0.32 | 0.33 | 0.03 | 0.38 | 0.23 |
| 1 | 35369014 | rs10493064 | 0.012 | -0.929 | -0.333 | 0.608 | 0 | 0.023 | 0.023 | 0 | 0.15 | 0.13 | 0.31 | 0.45 | 0.15 | 0.28 |
| 1 | 35373374 | rs3587 | 0.774 | -0.89 | -0.474 | 0.619 | -0.09 | -0.101 | -0.101 | 0.441 | 0.38 | 0.09 | 0.32 | 0.48 | 0.46 | 0.51 |
| 1 | 35373878 | rs7555884 | 0.046 | -1.013 | 0.298 | 0.813 | 0.01 | -0.006 | -0.006 | 0.451 | 0.55 | 0.36 | 0.4 | 0.48 | 0.59 | 0.64 |
| 1 | 35378934 | rs4652869 | 0.489 | -0.809 | -0.054 | 1.179 | -0.06 | -0.055 | -0.055 | 0 | 0.38 | 0.1 | 0.31 | 0.45 | 0.44 | 0.51 |
| 1 | 35379119 | rs6694435 | -0.31 | -1.26 | 1.129 | 1.179 | 0.009 | 0.011 | 0.011 | 0 | 0.23 | 0.18 | 0.23 | 0.03 | 0.19 | 0.11 |
| 1 | 35379493 | rs10908403 | -0.238 | -1.258 | 1.189 | 1.21 | 0.005 | -0.009 | -0.009 | 0.424 | 0.22 | 0.21 | 0.23 | 0.03 | 0.18 | 0.11 |
| 1 | 35381295 | rs6686482 | 1.675 | -0.738 | 2.239 | 2.649 | -0.028 | 0.134 | 0.134 | 0.682 | 0.79 | 0.55 | 0.78 | 0.5 | 0.88 | 0.82 |
| 1 | 35403677 | rs10437318 | -1.352 | -1.015 | 2.065 | 2.07 | -0.008 | -0.04 | -0.04 | 0 | 0.08 | 0.2 | 0.02 | 0 | 0.03 | 0.03 |
| 1 | 35428428 | rs11263859 | -2.38 | -1.165 | 2.46 | 2.519 | -0.046 | 0.079 | 0.079 | 0.039 | 0.16 | 0.41 | 0.07 | 0.25 | 0.05 | 0.14 |
| 1 | 35441614 | rs12562704 | -1.51 | -1.09 | 2.54 | 2.627 | -0.005 | 0.102 | 0.102 | 0 | 0.06 | 0.07 | 0.06 | 0.38 | 0.03 | 0.09 |
| 1 | 35444037 | rs4072108 | -3.408 | -1.231 | 3.309 | 2.574 | -0.037 | 0.087 | 0.087 | 0.05 | 0.13 | 0.73 | 0.12 | 0.38 | 0.03 | 0.12 |
| 1 | 35446830 | rs11263872 | -1.46 | -1.233 | 3.277 | 2.578 | -0.006 | 0.114 | 0.114 | 0 | 0.06 | 0.09 | 0.06 | 0.38 | 0.03 | 0.09 |
| 1 | 35473850 | rs12086023 | -1.46 | -1.234 | 3.481 | 2.577 | -0.006 | 0.114 | 0.114 | 0 | 0.06 | 0.09 | 0.06 | 0.38 | 0.03 | 0.09 |
| 1 | 35484861 | rs11263938 | -2.941 | -1.236 | 3.92 | 2.576 | -0.036 | 0.283 | 0.283 | 0 | 0.11 | 0.57 | 0.1 | 0.38 | 0.03 | 0.12 |
| 1 | 35509416 | rs11263952 | -3.492 | -1.192 | 3.995 | 2.577 | -0.034 | 0.087 | 0.087 | 1.224 | 0.13 | 0.73 | 0.12 | 0.38 | 0.04 | 0.13 |
| 1 | 35544502 | rs11264015 | -1.668 | -1.176 | 4.054 | 2.627 | -0.007 | 0.131 | 0.131 | 0 | 0.06 | 0.13 | 0.07 | 0.39 | 0.03 | 0.09 |
| 1 | 35546991 | rs12048974 | 1.212 | -1.178 | 3.524 | 3.021 | 0 | 0 | 0 | 13.738 | 0.87 | 0.85 | 0.75 | 0.74 | 0.92 | 0.93 |
| 1 | 35562965 | rs2971408 | 1.381 | -1.179 | 3.428 | 3.027 | -0.009 | 0.011 | 0.011 | 0 | 0.86 | 0.8 | 0.74 | 0.74 | 0.92 | 0.93 |
| 1 | 35586432 | rs10908374 | 2.438 | -1.183 | 3.301 | 2.827 | -0.045 | 0.053 | 0.053 | 0.098 | 0.77 | 0.36 | 0.64 | 0.35 | 0.89 | 0.79 |
| 1 | 35628789 | rs6425939 | 2.017 | -1.128 | 2.398 | 2.902 | -0.058 | 0.097 | 0.097 | 0 | 0.79 | 0.56 | 0.66 | 0.37 | 0.91 | 0.84 |
| 1 | 35633937 | rs2971400 | -1.832 | -0.86 | 1.639 | 2.87 | 0.004 | -0.005 | -0.005 | 0 | 0.14 | 0.15 | 0.14 | 0.26 | 0.07 | 0.08 |
| 1 | 35776603 | rs12075713 | -2.204 | -0.682 | 2.946 | 3.31 | 0.001 | 0.162 | 0.162 | 0 | 0.06 | 0.18 | 0.32 | 0.67 | 0.03 | 0.12 |
| 1 | 35950705 | rs1203138 | -0.054 | -0.763 | 3.829 | 2.409 | -0.016 | 0.048 | 0.048 | 0.41 | 0.89 | 0.98 | 0.94 | 1 | 0.88 | 0.99 |
| 1 | 35961737 | rs2486297 | -1.935 | -0.949 | 4.14 | 2.404 | 0 | -0.015 | -0.015 | 0 | 0.09 | 0.11 | 0.06 | 0 | 0.06 | 0.04 |
| 1 | 36038775 | rs6703909 | -2.075 | -0.945 | 4.285 | 2.45 | 0.002 | -0.013 | -0.013 | 0 | 0.1 | 0.11 | 0.06 | 0 | 0.07 | 0.05 |
| 1 | 36162133 | rs960629 | 2.618 | -1.092 | 3.008 | 1.642 | -0.04 | -0.033 | -0.033 | 0.015 | 0.76 | 0.48 | 0.59 | 0.27 | 0.87 | 0.76 |
| 1 | 36167456 | rs12563583 | -2.646 | -1.367 | 3.039 | 1.633 | -0.025 | 0.02 | 0.02 | 0.935 | 0.19 | 0.3 | 0.38 | 0.67 | 0.12 | 0.24 |
| 1 | 36178041 | rs13375070 | -1.27 | -1.25 | 3.168 | 1.528 | -0.001 | -0.015 | -0.015 | 0 | 0.09 | 0.22 | 0.05 | 0.07 | 0.04 | 0.06 |
| 1 | 36180972 | rs3795501 | -2.27 | -1.52 | 3.397 | 1.501 | -0.018 | 0.066 | 0.066 | 0 | 0.07 | 0.1 | 0.31 | 0.67 | 0.02 | 0.09 |
| 1 | 36195372 | rs11264194 | -2.847 | -1.564 | 3.891 | 1.472 | -0.03 | 0.047 | 0.047 | 0.016 | 0.16 | 0.32 | 0.37 | 0.67 | 0.09 | 0.14 |
| 1 | 36200821 | rs6660890 | -1.29 | -1.571 | 4.003 | 1.479 | 0.001 | -0.011 | -0.011 | 0 | 0.09 | 0.26 | 0.05 | 0.07 | 0.04 | 0.06 |
| 1 | 36218675 | rs12089175 | -1.322 | -1.571 | 4.227 | 1.479 | 0.013 | -0.003 | -0.003 | 0 | 0.09 | 0.31 | 0.05 | 0.07 | 0.04 | 0.06 |
| 1 | 36225948 | rs7537203 | -3.449 | -1.596 | 4.344 | 1.47 | -0.009 | -0.007 | -0.007 | 0.094 | 0.25 | 0.67 | 0.43 | 0.74 | 0.14 | 0.2 |
| 1 | 36253394 | rs7513938 | -2.484 | -1.6 | 4.259 | 1.5 | -0.013 | 0.165 | 0.165 | 0.005 | 0.07 | 0.23 | 0.33 | 0.67 | 0.02 | 0.07 |
| 1 | 36256339 | rs12076840 | -1.301 | -1.601 | 4.244 | 1.486 | 0.004 | -0.009 | -0.009 | 4.29 | 0.09 | 0.28 | 0.06 | 0.07 | 0.04 | 0.06 |
| 1 | 36276399 | rs12044203 | -2.54 | -1.585 | 4.001 | 1.518 | -0.014 | 0.167 | 0.167 | 0 | 0.07 | 0.23 | 0.33 | 0.67 | 0.02 | 0.07 |
| 1 | 36281336 | rs727005 | -2.932 | -1.527 | 3.928 | 1.507 | -0.031 | 0.055 | 0.055 | 0.003 | 0.16 | 0.34 | 0.37 | 0.67 | 0.1 | 0.12 |
| 1 | 36326196 | rs6682769 | -1.719 | -1.511 | 3.71 | 1.478 | -0.007 | 0.011 | 0.011 | 0.558 | 0.11 | 0.36 | 0.07 | 0.11 | 0.05 | 0.07 |
| 1 | 36444957 | rs4351606 | -0.985 | -0.987 | 3.567 | 1.297 | -0.006 | -0.009 | -0.009 | 0 | 0.09 | 0.12 | 0.04 | 0.1 | 0.04 | 0.19 |
| 1 | 36488199 | rs274732 | 1.11 | -0.867 | 3.462 | 1.211 | -0.002 | 0 | 0 | 0.077 | 0.9 | 0.93 | 0.94 | 1 | 0.91 | 0.94 |
| 1 | 36549157 | rs3754076 | 2.323 | -0.973 | 3.871 | 3.188 | -0.016 | 0.253 | 0.253 | 0.438 | 0.84 | 0.46 | 0.62 | 0.52 | 0.9 | 0.88 |
| 1 | 36553673 | rs399393 | 0.913 | -0.966 | 3.876 | 3.161 | -0.001 | -0.012 | -0.012 | 0.443 | 0.9 | 0.91 | 0.94 | 1 | 0.91 | 0.94 |
| 1 | 36555758 | rs3767703 | -0.657 | -0.849 | 3.786 | 3.164 | -0.022 | -0.057 | -0.057 | 0 | 0.09 | 0.01 | 0.06 | 0.23 | 0.07 | 0.25 |
| 1 | 36565617 | rs274754 | 2.245 | -0.814 | 3.32 | 2.683 | 0.006 | 0.007 | 0.007 | 0.084 | 0.68 | 0.33 | 0.52 | 0.26 | 0.78 | 0.59 |
| 1 | 36571920 | rs96067 | 1.839 | -0.838 | 3.095 | 2.687 | -0.01 | 0.04 | 0.04 | 0 | 0.72 | 0.61 | 0.63 | 0.5 | 0.8 | 0.62 |
| 1 | 36572161 | rs274759 | 0.398 | -0.818 | 2.775 | 2.698 | 0.002 | 0.019 | 0.019 | 0 | 0.9 | 0.87 | 0.93 | 0.79 | 0.92 | 0.76 |
| 1 | 36581792 | rs274133 | 0.865 | -0.83 | 2.61 | 2.544 | -0.003 | 0.017 | 0.017 | 0 | 0.81 | 0.77 | 0.68 | 0.75 | 0.86 | 0.87 |
| 1 | 36581978 | rs274132 | 0.71 | -0.822 | 2.477 | 2.547 | 0 | 0.001 | 0.001 | 0 | 0.82 | 0.82 | 0.69 | 0.75 | 0.86 | 0.87 |
| 1 | 36585077 | rs12074848 | 1.735 | -0.415 | 1.95 | 2.011 | 0.02 | 0.06 | 0.06 | 0 | 0.6 | 0.25 | 0.46 | 0.27 | 0.69 | 0.59 |
| 1 | 36591612 | rs12123086 | -1.165 | -0.478 | 1.791 | 1.871 | 0 | 0.003 | 0.003 | 0 | 0.12 | 0.11 | 0.07 | 0.08 | 0.11 | 0.28 |
| 1 | 36599547 | rs4653160 | 1.364 | -0.573 | 2.029 | 2.227 | 0.004 | -0.008 | -0.008 | 0 | 0.66 | 0.33 | 0.52 | 0.44 | 0.73 | 0.53 |
| 1 | 36622730 | rs272834 | 0.03 | -0.696 | 1.689 | 1.815 | -0.023 | 0.06 | 0.06 | 0 | 0.83 | 0.96 | 0.9 | 0.93 | 0.82 | 0.67 |
| 1 | 36637708 | rs795047 | 1.116 | -0.159 | 0.978 | 1.253 | 0 | -0.001 | -0.001 | 0 | 0.58 | 0.47 | 0.51 | 0.51 | 0.66 | 0.46 |
| 1 | 36653540 | rs6680302 | 1.176 | 0.211 | -0.291 | 0.451 | 0.009 | -0.011 | -0.011 | 0.233 | 0.53 | 0.29 | 0.44 | 0.5 | 0.56 | 0.45 |
| 1 | 36653677 | rs522468 | 0.299 | 0.814 | -0.741 | -0.31 | -0.068 | 0.077 | 0.077 | 0 | 0.8 | 0.95 | 0.86 | 0.93 | 0.73 | 0.66 |
| 1 | 36690870 | rs272814 | -0.171 | -0.452 | -0.225 | -0.633 | -0.063 | -0.092 | -0.092 | 0 | 0.27 | 0.48 | 0.14 | 0.1 | 0.12 | 0.13 |
| 1 | 36752433 | rs6425977 | 0.214 | -0.633 | 0.416 | -0.249 | -0.063 | -0.11 | -0.11 | 0 | 0.73 | 0.52 | 0.86 | 0.95 | 0.88 | 0.87 |
| 1 | 36755635 | rs4653173 | 0.371 | -0.748 | 0.844 | -0.304 | -0.074 | -0.047 | -0.047 | 0 | 0.71 | 0.44 | 0.8 | 0.83 | 0.88 | 0.87 |

| | | | | | | | | | | | | | | | | |
|---|---|---|---|---|---|---|---|---|---|---|---|---|---|---|---|---|
| 1 | 36756491 | rs4653175 | 0.72 | -0.398 | -0.042 | -0.919 | -0.035 | -0.029 | -0.029 | 0 | 0.19 | 0.05 | 0.17 | 0.22 | 0.21 | 0.19 |
| 1 | 36764770 | rs10908292 | -0.084 | -0.397 | -0.103 | -0.912 | 0.068 | 0.119 | 0.119 | 3.965 | 0.95 | 0.52 | 0.95 | 0.95 | 1 | 0.99 |
| 1 | 36765912 | rs9659735 | 1.047 | -0.208 | -0.573 | -1.412 | 0.004 | 0.043 | 0.043 | 0 | 0.24 | 0.34 | 0.47 | 0.65 | 0.25 | 0.22 |
| 1 | 36773405 | rs35343437 | 0.404 | -0.316 | -0.137 | -1.594 | -0.067 | 0.045 | 0.045 | 0 | 0.38 | 0.12 | 0.56 | 0.72 | 0.44 | 0.39 |
| 1 | 36787294 | rs10796881 | -0.996 | -0.262 | -0.229 | -1.627 | -0.072 | -0.114 | -0.114 | 0 | 0.72 | 0.51 | 0.86 | 0.95 | 0.88 | 0.87 |
| 1 | 36789546 | rs12116935 | 0.632 | -0.171 | -0.301 | -1.575 | -0.097 | 0.147 | 0.147 | 0 | 0.29 | 0.06 | 0.22 | 0.08 | 0.42 | 0.45 |
| 1 | 36795430 | rs4653178 | 0.899 | 0.011 | -0.159 | -1.547 | -0.018 | -0.006 | -0.006 | 0.757 | 0.19 | 0.11 | 0.17 | 0.21 | 0.21 | 0.19 |
| 1 | 36807481 | rs3795498 | 1.161 | 0.157 | -0.192 | -1.537 | -0.001 | 0.005 | 0.005 | 0 | 0.22 | 0.17 | 0.14 | 0.14 | 0.23 | 0.21 |
| 1 | 36818791 | rs6701311 | 0.573 | -0.148 | -0.63 | -1.966 | -0.002 | -0.028 | -0.028 | 0.984 | 0.28 | 0.33 | 0.12 | 0.04 | 0.12 | 0.12 |
| 1 | 36820927 | rs11263875 | 1.363 | -0.178 | -1.253 | -1.985 | 0.046 | 0.109 | 0.109 | 0 | 0.22 | 0.05 | 0.1 | 0.04 | 0.12 | 0.13 |
| 1 | 36822312 | rs11583322 | 0.57 | 0.49 | -1.276 | -1.769 | -0.101 | 0.05 | 0.05 | 0 | 0.26 | 0.07 | 0.25 | 0.15 | 0.42 | 0.27 |
| 1 | 36833080 | rs6680494 | 0.346 | -0.013 | -0.555 | -1.463 | -0.055 | -0.107 | -0.107 | 0.01 | 0.29 | 0.44 | 0.13 | 0.04 | 0.13 | 0.13 |
| 1 | 36834478 | rs11588213 | 0.284 | 0.052 | -0.597 | -1.488 | -0.014 | 0.034 | 0.034 | 0 | 0.22 | 0.12 | 0.13 | 0.08 | 0.23 | 0.17 |
| 1 | 36835804 | rs6690378 | 0.394 | -0.188 | -0.58 | -1.479 | -0.051 | -0.104 | -0.104 | 2.797 | 0.3 | 0.45 | 0.13 | 0.04 | 0.14 | 0.13 |
| 1 | 36847766 | rs4653184 | -0.573 | 1.391 | -0.725 | -0.271 | -0.026 | 0.033 | 0.033 | 0 | 0.22 | 0.13 | 0.15 | 0.09 | 0.26 | 0.18 |
| 1 | 36858680 | rs10796884 | 0.26 | 1.1 | -0.134 | -0.645 | -0.001 | 0.002 | 0.002 | 0 | 0.58 | 0.41 | 0.36 | 0.22 | 0.61 | 0.6 |
| 1 | 36860826 | rs16822801 | -1.082 | 1.458 | 0.238 | -0.371 | 0.002 | -0.007 | -0.007 | 0.085 | 0.28 | 0.35 | 0.22 | 0.27 | 0.3 | 0.32 |
| 1 | 36861854 | rs4653185 | -1.082 | 1.458 | 0.237 | -0.371 | 0.002 | -0.007 | -0.007 | 0.237 | 0.28 | 0.35 | 0.22 | 0.27 | 0.3 | 0.32 |
| 1 | 36861993 | rs884925 | 1.414 | 1.277 | 0.243 | -0.559 | 0.001 | 0.215 | 0.215 | 0.65 | 0.4 | 0.11 | 0.21 | 0.12 | 0.36 | 0.39 |
| 1 | 36863456 | rs1126984 | 1.418 | 1.267 | 0.239 | -0.563 | 0 | 0.214 | 0.214 | 0.604 | 0.4 | 0.11 | 0.21 | 0.12 | 0.36 | 0.39 |
| 1 | 36867363 | rs6696875 | -1.146 | 1.357 | 0.397 | 0.833 | 0.002 | 0.022 | 0.022 | 0.57 | 0.29 | 0.37 | 0.29 | 0.54 | 0.3 | 0.37 |
| 1 | 36872665 | rs12070076 | -0.276 | 1.288 | 0.729 | 0.79 | 0.023 | 0.036 | 0.036 | 0.017 | 0.17 | 0.33 | 0.16 | 0.26 | 0.21 | 0.27 |
| 1 | 36873176 | rs6696510 | -0.323 | 1.258 | 0.716 | 0.776 | 0.021 | 0.034 | 0.034 | 0.039 | 0.18 | 0.33 | 0.16 | 0.26 | 0.21 | 0.27 |
| 1 | 36874242 | rs16822823 | -0.272 | 1.274 | 0.723 | 0.784 | 0.023 | 0.036 | 0.036 | 0.017 | 0.17 | 0.33 | 0.16 | 0.26 | 0.21 | 0.27 |
| 1 | 36875406 | rs16822826 | 0.181 | 1.287 | 1.038 | 0.852 | -0.005 | -0.031 | -0.031 | 0 | 0.14 | 0.35 | 0.16 | 0.1 | 0.15 | 0.17 |
| 1 | 36878024 | rs883805 | -0.317 | 1.343 | 1.177 | 0.792 | 0.017 | 0.04 | 0.04 | 0.082 | 0.16 | 0.31 | 0.16 | 0.26 | 0.2 | 0.27 |
| 1 | 36886117 | rs2275477 | -0.533 | 1.211 | 1.929 | 0.728 | 0.008 | 0.028 | 0.028 | 0.129 | 0.17 | 0.32 | 0.15 | 0.26 | 0.21 | 0.27 |
| 1 | 36887163 | rs2282353 | 0.097 | 1.133 | 1.797 | 0.609 | -0.042 | 0.015 | 0.015 | 0 | 0.59 | 0.76 | 0.47 | 0.69 | 0.54 | 0.72 |
| 1 | 36888462 | rs1416840 | 0.426 | 1.209 | 1.787 | 0.624 | 0.008 | -0.031 | -0.031 | 0 | 0.12 | 0.07 | 0.13 | 0.31 | 0.09 | 0.09 |
| 1 | 36890010 | rs10157686 | -0.775 | 1.208 | 1.791 | 0.62 | -0.045 | -0.031 | -0.031 | 0.149 | 0.71 | 0.84 | 0.53 | 0.69 | 0.63 | 0.76 |
| 1 | 36897803 | rs6696986 | 1.763 | 1.203 | 1.81 | 0.605 | -0.001 | 0.193 | 0.193 | 0 | 0.38 | 0.11 | 0.2 | 0.12 | 0.33 | 0.4 |
| 1 | 36910277 | rs953517 | -0.774 | 1.18 | 1.816 | 0.594 | -0.029 | -0.021 | -0.021 | 3.039 | 0.71 | 0.79 | 0.52 | 0.69 | 0.63 | 0.76 |
| 1 | 36914747 | rs10493076 | 0.367 | 1.109 | 1.786 | 0.591 | 0.01 | -0.035 | -0.035 | 0 | 0.13 | 0.07 | 0.13 | 0.31 | 0.09 | 0.09 |
| 1 | 36921371 | rs1538634 | 0.919 | 1.574 | 1.76 | 0.696 | -0.047 | -0.052 | -0.052 | 0 | 0.24 | 0.07 | 0.45 | 0.25 | 0.3 | 0.22 |
| 1 | 36921533 | rs2275480 | 0.434 | 1.571 | 1.719 | 0.668 | 0.009 | -0.031 | -0.031 | 0 | 0.13 | 0.08 | 0.13 | 0.31 | 0.09 | 0.09 |
| 1 | 36923009 | rs12023003 | 1.925 | 1.541 | 1.703 | 0.649 | -0.004 | 0.194 | 0.194 | 0 | 0.38 | 0.11 | 0.2 | 0.12 | 0.33 | 0.4 |
| 1 | 36923032 | rs16823042 | -0.079 | 1.449 | 1.907 | 0.712 | -0.007 | 0.048 | 0.048 | 0.108 | 0.17 | 0.34 | 0.15 | 0.26 | 0.18 | 0.27 |
| 1 | 36928903 | rs10796885 | 0.376 | 1.589 | 1.502 | 0.82 | -0.007 | -0.017 | -0.017 | 0 | 0.28 | 0.23 | 0.47 | 0.32 | 0.33 | 0.23 |
| 1 | 36929988 | rs3806289 | 0.098 | 1.346 | 1.306 | 0.876 | 0.005 | -0.002 | -0.002 | 0.315 | 0.13 | 0.1 | 0.13 | 0.31 | 0.09 | 0.08 |
| 1 | 36937059 | rs3917981 | 0.158 | 1.535 | 1.519 | 0.706 | -0.023 | 0.014 | 0.014 | 0 | 0.6 | 0.74 | 0.4 | 0.45 | 0.57 | 0.71 |
| 1 | 36937065 | rs3917980 | 1.639 | 1.453 | 1.382 | 0.58 | 0 | 0.149 | 0.149 | 0 | 0.37 | 0.17 | 0.2 | 0.13 | 0.34 | 0.41 |
| 1 | 36937392 | rs3917979 | -0.618 | 1.197 | 1.68 | 0.71 | -0.011 | 0.015 | 0.015 | 0 | 0.19 | 0.33 | 0.16 | 0.25 | 0.2 | 0.3 |
| 1 | 36941751 | rs3917950 | -0.577 | 0.922 | 1.057 | 0.756 | 0.011 | -0.047 | -0.047 | 0 | 0.19 | 0.08 | 0.13 | 0.33 | 0.23 | 0.22 |
| 1 | 36945559 | rs3917925 | 1.172 | 1.257 | 1.064 | 0.969 | -0.012 | 0.001 | 0.001 | 0 | 0.39 | 0.15 | 0.47 | 0.39 | 0.4 | 0.51 |
| 1 | 36945653 | rs3917924 | 1.237 | 1.263 | 1.042 | 0.405 | -0.017 | 0.178 | 0.178 | 0 | 0.39 | 0.14 | 0.44 | 0.13 | 0.39 | 0.46 |
| 1 | 36957831 | rs6667450 | 1.224 | 1.215 | 1.249 | 1.203 | 0 | 0.036 | 0.036 | 1.129 | 0.73 | 0.39 | 0.76 | 0.43 | 0.73 | 0.65 |
| 1 | 36960626 | rs11263901 | -1.548 | 0.808 | 0.953 | 1.495 | 0.039 | 0.452 | 0.452 | 0.135 | 0.1 | 0.49 | 0.1 | 0.48 | 0.07 | 0.08 |
| 1 | 36975893 | rs4462104 | -0.976 | 0.695 | 0.077 | 0.623 | -0.006 | -0.053 | -0.053 | 0.084 | 0.24 | 0.27 | 0.17 | 0.07 | 0.21 | 0.31 |
| 1 | 36976253 | rs4653193 | 0.94 | 0.838 | -1.026 | 0.667 | -0.092 | -0.101 | -0.101 | 0 | 0.22 | 0.02 | 0.25 | 0.28 | 0.27 | 0.19 |
| 1 | 36977565 | rs4233207 | -1.508 | 0.733 | -1.187 | 0.608 | -0.047 | 0.105 | 0.105 | 0.673 | 0.62 | 0.88 | 0.56 | 0.74 | 0.61 | 0.58 |
| 1 | 36980804 | rs4078260 | 0.377 | 0.711 | -0.921 | 0.693 | -0.036 | 0.051 | 0.051 | 0 | 0.12 | 0.01 | 0.1 | 0.06 | 0.16 | 0.21 |
| 1 | 36984734 | rs11804163 | -1.127 | 0.689 | -0.375 | 1.25 | -0.026 | 0.051 | 0.051 | 3.254 | 0.15 | 0.62 | 0.18 | 0.2 | 0.1 | 0.03 |
| 1 | 36986143 | rs7515998 | 0.437 | 0.703 | -0.615 | 1.242 | 0.005 | 0.04 | 0.04 | 0 | 0.07 | 0.01 | 0.04 | 0 | 0.08 | 0.01 |
| 1 | 36986518 | rs7530283 | -1.783 | 0.661 | -0.684 | 1.46 | 0.005 | -0.084 | -0.084 | 0.3 | 0.35 | 0.83 | 0.32 | 0.5 | 0.33 | 0.35 |
| 1 | 36988601 | rs6425991 | -0.625 | 0.629 | -0.259 | 1.472 | 0.018 | 0.164 | 0.164 | 0.923 | 0.12 | 0.45 | 0.15 | 0.2 | 0.1 | 0.03 |
| 1 | 36990036 | rs4345779 | 0.802 | 0.711 | -1.53 | 1.134 | -0.065 | -0.077 | -0.077 | 2.656 | 0.24 | 0.01 | 0.2 | 0.24 | 0.25 | 0.16 |
| 1 | 36990158 | rs4576633 | -0.957 | 0.747 | -1.266 | 1.099 | -0.039 | -0.016 | -0.016 | 0 | 0.59 | 0.83 | 0.47 | 0.57 | 0.57 | 0.5 |
| 1 | 37000090 | rs7526895 | 0.513 | 0.823 | -1.802 | 0.988 | -0.088 | -0.051 | -0.051 | 0 | 0.26 | 0.02 | 0.17 | 0.22 | 0.27 | 0.13 |
| 1 | 37000226 | rs4503315 | -0.719 | 0.045 | -1.54 | 0.29 | 0.018 | 0.084 | 0.084 | 0 | 0.49 | 0.76 | 0.52 | 0.56 | 0.45 | 0.3 |
| 1 | 37000433 | rs4074399 | -0.784 | -0.015 | -1.554 | 0.373 | -0.023 | -0.015 | -0.015 | 0.086 | 0.17 | 0.72 | 0.13 | 0.07 | 0.08 | 0.08 |
| 1 | 37001923 | rs10752589 | -1.555 | -0.072 | -2.125 | 0.091 | -0.054 | -0.007 | -0.007 | 0 | 0.7 | 0.93 | 0.66 | 0.71 | 0.7 | 0.6 |
| 1 | 37003273 | rs17498436 | 1.041 | 0.086 | -2.157 | 0.106 | -0.11 | -0.138 | -0.138 | 0.411 | 0.31 | 0.04 | 0.39 | 0.49 | 0.37 | 0.22 |
| 1 | 37005756 | rs3924538 | -1.215 | 0.009 | -1.482 | -0.131 | 0.008 | 0.025 | 0.025 | 0.885 | 0.37 | 0.67 | 0.24 | 0.19 | 0.33 | 0.38 |
| 1 | 37007987 | rs12726247 | -0.42 | 0.428 | -1.34 | 0.071 | 0.001 | -0.049 | -0.049 | 0 | 0.24 | 0.31 | 0.14 | 0.15 | 0.25 | 0.3 |
| 1 | 37013031 | rs12072868 | -0.48 | 0.435 | -1.505 | 0.074 | 0 | -0.03 | -0.03 | 0 | 0.25 | 0.27 | 0.14 | 0.15 | 0.25 | 0.3 |
| 1 | 37019556 | rs11263915 | -0.073 | 0.338 | -1.657 | 0.787 | 0 | -0.015 | -0.015 | 0 | 0.37 | 0.39 | 0.44 | 0.41 | 0.38 | 0.23 |
| 1 | 37020422 | rs12031548 | -0.642 | 0.387 | -1.864 | 0.846 | -0.004 | -0.033 | -0.033 | 0.121 | 0.08 | 0.01 | 0.07 | 0.14 | 0.04 | 0.09 |
| 1 | 37021076 | rs6425992 | -0.064 | 0.737 | -2.582 | 0.721 | -0.185 | -0.103 | -0.103 | 0 | 0.65 | 0.98 | 0.67 | 0.63 | 0.54 | 0.7 |
| 1 | 37022310 | rs4026414 | -0.114 | -0.089 | -1.582 | 0.345 | 0.002 | 0.04 | 0.04 | 0.057 | 0.51 | 0.65 | 0.56 | 0.71 | 0.51 | 0.65 |

| | | | | | | | | | | | | | | | | |
|---|---|---|---|---|---|---|---|---|---|---|---|---|---|---|---|---|
| 1 | 37033128 | rs4026409 | -0.887 | -0.935 | -0.574 | 1.104 | -0.016 | 0.041 | 0.041 | 1.274 | 0.25 | 0.34 | 0.31 | 0.38 | 0.19 | 0.13 |
| 1 | 37035588 | rs6698957 | -0.159 | -0.887 | -1.057 | 1 | 0.01 | 0.066 | 0.066 | 3.967 | 0.19 | 0.04 | 0.15 | 0.05 | 0.16 | 0.04 |
| 1 | 37040374 | rs7540413 | -1.091 | -0.899 | -0.891 | 1.025 | 0 | -0.002 | -0.002 | 0 | 0.33 | 0.68 | 0.37 | 0.38 | 0.22 | 0.18 |
| 1 | 37043894 | rs7538665 | -1.017 | -0.679 | -0.433 | 1.508 | -0.06 | 0.145 | 0.145 | 0.504 | 0.2 | 0.47 | 0.12 | 0.35 | 0.1 | 0.17 |
| 1 | 37055734 | rs6682952 | -0.293 | -0.399 | -0.935 | 2.108 | 0.002 | 0 | 0 | 0 | 0.1 | 0.05 | 0.09 | 0.09 | 0.1 | 0.04 |
| 1 | 37058354 | rs4653204 | -0.729 | -0.129 | -1.046 | 1.624 | 0.005 | 0.03 | 0.03 | 0 | 0.48 | 0.41 | 0.71 | 0.64 | 0.59 | 0.56 |
| 1 | 37063921 | rs4653207 | -0.893 | -0.284 | -0.817 | 1.569 | 0 | 0.004 | 0.004 | 0 | 0.52 | 0.5 | 0.71 | 0.64 | 0.55 | 0.55 |
| 1 | 37070248 | rs12063741 | -1.212 | -0.037 | -0.874 | 1.86 | -0.007 | 0.063 | 0.063 | 0 | 0.19 | 0.32 | 0.31 | 0.33 | 0.17 | 0.32 |
| 1 | 37073451 | rs1022629 | -1.115 | -0.07 | -0.931 | 1.143 | 0.004 | 0.013 | 0.013 | 0 | 0.53 | 0.46 | 0.74 | 0.72 | 0.6 | 0.58 |
| 1 | 37074351 | rs4653210 | -0.87 | -0.475 | -0.561 | 1.241 | -0.001 | -0.002 | -0.002 | 0 | 0.14 | 0.18 | 0.09 | 0.08 | 0.14 | 0.17 |
| 1 | 37084522 | rs7533162 | -0.894 | -1.246 | -1.087 | 1.75 | 0.019 | 0.002 | 0.002 | 0 | 0.55 | 0.67 | 0.41 | 0.4 | 0.39 | 0.3 |
| 1 | 37089450 | rs12131598 | -0.938 | -1.323 | -0.995 | 2.335 | 0.011 | -0.018 | -0.018 | 1.721 | 0.57 | 0.83 | 0.42 | 0.51 | 0.4 | 0.39 |
| 1 | 37089923 | rs7532617 | -0.613 | -1.54 | -1.029 | 2.163 | 0.03 | 0.064 | 0.064 | 0 | 0.45 | 0.68 | 0.27 | 0.27 | 0.35 | 0.28 |
| 1 | 37093026 | rs761719 | -0.824 | -1.766 | -0.785 | 2.408 | 0.068 | 0.02 | 0.02 | 0.52 | 0.49 | 0.82 | 0.48 | 0.61 | 0.35 | 0.53 |
| 1 | 37093821 | rs743015 | -1.448 | -1.752 | -0.793 | 2.34 | -0.024 | 0.22 | 0.22 | 0 | 0.59 | 0.9 | 0.65 | 0.91 | 0.41 | 0.66 |
| 1 | 37096184 | rs1474850 | -0.826 | -1.823 | -0.142 | 2.443 | 0.029 | 0.025 | 0.025 | 0 | 0.49 | 0.67 | 0.57 | 0.68 | 0.35 | 0.52 |
| 1 | 37096594 | rs6702202 | -0.548 | -1.853 | -0.078 | 2.502 | -0.004 | -0.007 | -0.007 | 0.374 | 0.44 | 0.56 | 0.36 | 0.31 | 0.35 | 0.27 |
| 1 | 37115263 | rs7355025 | 0.376 | -1.811 | -0.149 | 2.52 | -0.008 | 0.05 | 0.05 | 0 | 0.09 | 0.07 | 0.05 | 0.24 | 0.05 | 0.12 |
| 1 | 37122035 | rs2983117 | -1.476 | -1.83 | -0.29 | 1.956 | -0.022 | 0.241 | 0.241 | 0 | 0.6 | 0.94 | 0.64 | 0.91 | 0.38 | 0.66 |
| 1 | 37126512 | rs219002 | -1.108 | -1.72 | -0.202 | 2.014 | 0.029 | 0.039 | 0.039 | 0 | 0.56 | 0.79 | 0.43 | 0.56 | 0.37 | 0.41 |
| 1 | 37136423 | rs1076773 | -1.381 | -0.931 | 0.039 | 1.893 | 0.016 | -0.051 | -0.051 | 0 | 0.19 | 0.11 | 0.41 | 0.53 | 0.13 | 0.36 |
| 1 | 37136762 | rs218990 | -1.755 | -1.364 | 0.178 | 1.58 | 0.011 | 0.012 | 0.012 | 0 | 0.45 | 0.55 | 0.57 | 0.88 | 0.35 | 0.54 |
| 1 | 37143732 | rs1395457 | 0.005 | -1.266 | 0.258 | 1.596 | -0.005 | 0.023 | 0.023 | 0 | 0.25 | 0.38 | 0.16 | 0.34 | 0.22 | 0.14 |
| 1 | 37152471 | rs10430444 | 0.642 | -1.192 | -0.545 | 1.789 | 0.011 | -0.017 | -0.017 | 0 | 0.08 | 0.01 | 0.16 | 0.3 | 0.05 | 0.07 |
| 1 | 37154993 | rs535175 | -1.079 | -1.118 | -0.581 | 1.69 | 0.022 | 0.054 | 0.054 | 1.009 | 0.38 | 0.8 | 0.39 | 0.68 | 0.28 | 0.22 |
| 1 | 37163878 | rs2983118 | -1.831 | -0.441 | -0.07 | 1.744 | 0.014 | -0.05 | -0.05 | 4.808 | 0.41 | 0.86 | 0.5 | 0.58 | 0.32 | 0.49 |
| 1 | 37165467 | rs12755038 | -1.881 | -0.113 | 0.297 | 1.778 | -0.001 | 0 | 0 | 0 | 0.12 | 0.16 | 0.28 | 0.22 | 0.1 | 0.31 |
| 1 | 37170745 | rs218984 | -0.728 | -0.082 | 0.251 | 1.698 | -0.001 | 0.002 | 0.002 | 0.712 | 0.28 | 0.67 | 0.18 | 0.39 | 0.22 | 0.15 |
| 1 | 37174773 | rs4652921 | 0.299 | -0.047 | 0.336 | 1.703 | -0.004 | 0.034 | 0.034 | 0.085 | 0.73 | 0.49 | 0.83 | 0.61 | 0.78 | 0.84 |
| 1 | 37178411 | rs12063010 | -1.739 | -0.055 | 0.343 | 1.669 | -0.004 | 0.009 | 0.009 | 0 | 0.12 | 0.21 | 0.18 | 0.18 | 0.1 | 0.32 |
| 1 | 37178768 | rs7548051 | -1.954 | -0.055 | 0.345 | 1.669 | -0.001 | 0.019 | 0.019 | 0 | 0.12 | 0.29 | 0.19 | 0.18 | 0.1 | 0.32 |
| 1 | 37182786 | rs1116900 | -1.749 | -0.119 | 0.337 | 1.642 | -0.011 | 0.006 | 0.006 | 0 | 0.12 | 0.19 | 0.18 | 0.19 | 0.08 | 0.31 |
| 1 | 37185190 | rs1395455 | 1.525 | -0.125 | 0.33 | 1.661 | 0.004 | -0.073 | -0.073 | 0 | 0.61 | 0.2 | 0.61 | 0.42 | 0.68 | 0.52 |
| 1 | 37185336 | rs1395453 | 0.361 | -0.338 | 0.343 | 1.662 | 0 | 0.013 | 0.013 | 0.11 | 0.73 | 0.41 | 0.82 | 0.61 | 0.77 | 0.84 |
| 1 | 37189396 | rs906227 | 0.331 | -0.337 | 0.323 | 1.66 | 0.001 | 0.036 | 0.036 | 0.127 | 0.73 | 0.48 | 0.82 | 0.61 | 0.77 | 0.84 |
| 1 | 37192741 | rs1002656 | 1.451 | -0.194 | 0.011 | 1.115 | 0.012 | -0.003 | -0.003 | 0 | 0.61 | 0.24 | 0.61 | 0.35 | 0.69 | 0.52 |
| 1 | 37193645 | rs16823426 | -1.622 | -0.704 | 0.145 | 0.953 | -0.009 | 0.003 | 0.003 | 0 | 0.11 | 0.13 | 0.17 | 0.19 | 0.08 | 0.31 |
| 1 | 37195557 | rs16823431 | -1.128 | -0.578 | -0.338 | 0.921 | -0.001 | 0.002 | 0.002 | 0 | 0.09 | 0.09 | 0.2 | 0.19 | 0.1 | 0.3 |
| 1 | 37198352 | rs7549591 | -0.974 | -0.159 | -0.329 | 0.164 | -0.001 | 0 | 0 | 0 | 0.37 | 0.55 | 0.45 | 0.64 | 0.32 | 0.41 |
| 1 | 37200680 | rs11576073 | 1.331 | -0.364 | -0.389 | 0.161 | 0.006 | 0.027 | 0.027 | 0 | 0.55 | 0.3 | 0.42 | 0.35 | 0.59 | 0.52 |
| 1 | 37201769 | rs10796891 | 0.328 | -0.16 | -0.731 | -0.362 | 0.002 | -0.021 | -0.021 | 0 | 0.93 | 0.88 | 0.87 | 0.99 | 0.9 | 0.93 |
| 1 | 37210112 | rs494022 | 1.474 | -0.307 | -0.85 | -0.358 | -0.021 | 0.017 | 0.017 | 0.222 | 0.53 | 0.07 | 0.41 | 0.38 | 0.59 | 0.48 |
| 1 | 37220721 | rs1472823 | -0.945 | -0.525 | -0.095 | 0.323 | 0.02 | -0.016 | -0.016 | 0 | 0.32 | 0.71 | 0.42 | 0.61 | 0.22 | 0.45 |
| 1 | 37240886 | rs525798 | -1.243 | -0.536 | -0.298 | 0.187 | 0.053 | -0.028 | -0.028 | 0 | 0.35 | 0.83 | 0.43 | 0.62 | 0.22 | 0.44 |
| 1 | 37250088 | rs11801494 | -1.322 | -0.001 | 0.607 | -0.318 | -0.003 | 0.006 | 0.006 | 0 | 0.06 | 0.05 | 0.08 | 0.01 | 0.09 | 0.06 |
| 1 | 37270283 | rs493106 | -2.125 | 0.054 | 0.935 | -0.222 | 0.001 | -0.026 | -0.026 | 0 | 0.13 | 0.36 | 0.13 | 0.15 | 0.13 | 0.15 |
| 1 | 37280785 | rs528137 | -2.365 | 0.085 | 1.016 | -0.29 | 0.004 | -0.045 | -0.045 | 1.108 | 0.13 | 0.42 | 0.14 | 0.1 | 0.13 | 0.14 |
| 1 | 37280827 | rs2135596 | -1.27 | 0.097 | 0.98 | -0.334 | -0.004 | 0.008 | 0.008 | 0 | 0.09 | 0.08 | 0.11 | 0.06 | 0.13 | 0.12 |
| 1 | 37285123 | rs476227 | -2.146 | 0.102 | 0.975 | -0.332 | 0.008 | -0.05 | -0.05 | 0 | 0.13 | 0.37 | 0.13 | 0.06 | 0.13 | 0.12 |
| 1 | 37294722 | rs544932 | -1.825 | 0.024 | 0.844 | -0.324 | 0.009 | -0.041 | -0.041 | 0.159 | 0.12 | 0.3 | 0.12 | 0.06 | 0.13 | 0.12 |
| 1 | 37297307 | rs942973 | -1.971 | 0.05 | 0.761 | -0.312 | 0.003 | -0.038 | -0.038 | 0 | 0.12 | 0.31 | 0.13 | 0.08 | 0.13 | 0.15 |
| 1 | 37302038 | rs572688 | -2.053 | -0.092 | 0.723 | -0.336 | 0 | -0.038 | -0.038 | 2.496 | 0.13 | 0.34 | 0.11 | 0.07 | 0.14 | 0.13 |
| 1 | 37303737 | rs484017 | -1.763 | -0.203 | 0.682 | -0.349 | -0.02 | -0.052 | -0.052 | 0 | 0.17 | 0.36 | 0.12 | 0.08 | 0.15 | 0.16 |
| 1 | 37308444 | rs522480 | -1.428 | -0.035 | -0.034 | -0.268 | -0.005 | -0.017 | -0.017 | 0 | 0.16 | 0.22 | 0.1 | 0.08 | 0.15 | 0.16 |
| 1 | 37324935 | rs12067006 | -0.98 | 0.083 | 0.237 | -0.674 | -0.01 | -0.165 | -0.165 | 0.492 | 0.27 | 0.48 | 0.25 | 0.01 | 0.29 | 0.18 |
| 1 | 37326083 | rs4652931 | -0.944 | -0.266 | 0.09 | 0.17 | 0.012 | -0.015 | -0.015 | 0 | 0.29 | 0.21 | 0.3 | 0.28 | 0.26 | 0.26 |
| 1 | 37327225 | rs1392744 | 1.036 | -0.645 | 1.145 | -0.029 | -0.063 | 0.016 | 0.016 | 0 | 0.86 | 0.5 | 0.91 | 0.82 | 0.95 | 0.85 |
| 1 | 37335085 | rs574214 | 0.843 | -0.626 | 1.401 | -0.213 | -0.023 | -0.059 | -0.059 | 0 | 0.89 | 0.8 | 0.94 | 0.99 | 0.93 | 0.95 |
| 1 | 37349733 | rs3820271 | 1.739 | -0.814 | 2.313 | -0.019 | -0.055 | -0.014 | -0.014 | 0 | 0.84 | 0.44 | 0.89 | 0.85 | 0.92 | 0.85 |
| 1 | 37376745 | rs3767065 | 2.036 | -0.969 | 2.734 | -0.158 | -0.05 | -0.031 | -0.031 | 0 | 0.82 | 0.39 | 0.86 | 0.85 | 0.89 | 0.84 |
| 1 | 37384630 | rs471710 | 1.126 | -0.954 | 2.379 | -0.162 | -0.011 | -0.044 | -0.044 | 0 | 0.87 | 0.86 | 0.92 | 0.99 | 0.89 | 0.93 |
| 1 | 37391420 | rs1027599 | 1.758 | -0.955 | 2.458 | -0.161 | -0.039 | -0.023 | -0.023 | 0.657 | 0.84 | 0.69 | 0.9 | 0.86 | 0.9 | 0.85 |
| 1 | 37395244 | rs1334804 | 1.753 | -1.171 | 2.16 | -0.556 | -0.032 | -0.029 | -0.029 | 1.194 | 0.79 | 0.65 | 0.88 | 0.84 | 0.88 | 0.81 |
| 1 | 37396339 | rs1334802 | 1.3 | -1.027 | 1.795 | -0.606 | 0.006 | -0.018 | -0.018 | 0 | 0.82 | 0.87 | 0.92 | 0.99 | 0.88 | 0.9 |
| 1 | 37398436 | rs4653233 | 1.499 | -0.429 | 1.707 | -0.676 | -0.001 | -0.047 | -0.047 | 0.845 | 0.81 | 0.82 | 0.88 | 0.99 | 0.84 | 0.89 |
| 1 | 37402001 | rs7517274 | 0.249 | -0.306 | -0.146 | -1.292 | -0.14 | 0.073 | 0.073 | 0 | 0.33 | 0.01 | 0.23 | 0.15 | 0.36 | 0.26 |
| 1 | 37403014 | rs481047 | 0.789 | -0.205 | 0.659 | -1.473 | -0.018 | -0.115 | -0.115 | 0 | 0.78 | 0.69 | 0.9 | 0.99 | 0.84 | 0.89 |
| 1 | 37403567 | rs1160751 | 1.017 | -0.201 | -0.067 | -1.85 | 0.016 | 0.071 | 0.071 | 0 | 0.56 | 0.32 | 0.72 | 0.82 | 0.66 | 0.69 |
| 1 | 37408081 | rs16823687 | -0.327 | -0.226 | -0.544 | -2.216 | 0.01 | -0.003 | -0.003 | 0.011 | 0.26 | 0.23 | 0.11 | 0.03 | 0.17 | 0.29 |

| | | | | | | | | | | | | | | | | |
|---|---|---|---|---|---|---|---|---|---|---|---|---|---|---|---|---|
| 1 | 37409665 | rs3767080 | 0.787 | 0.402 | -1.368 | -1.304 | -0.079 | -0.15 | -0.15 | 1.122 | 0.3 | 0.02 | 0.27 | 0.6 | 0.28 | 0.27 |
| 1 | 37411578 | rs550115 | -0.663 | 0.745 | 0.116 | -0.385 | -0.063 | -0.088 | -0.088 | 0.044 | 0.84 | 0.96 | 0.59 | 0.69 | 0.68 | 0.75 |
| 1 | 37411893 | rs1762067 | -1.479 | 0.187 | 0.84 | -0.809 | -0.013 | -0.012 | -0.012 | 0.746 | 0.1 | 0.36 | 0.07 | 0.05 | 0.05 | 0.24 |
| 1 | 37416269 | rs3767087 | -1.057 | 0.38 | -0.449 | -0.717 | -0.003 | -0.003 | -0.003 | 0.013 | 0.08 | 0.07 | 0.05 | 0.05 | 0.05 | 0.16 |
| 1 | 37419841 | rs535620 | 0.671 | 1.016 | -0.269 | -0.678 | 0.013 | -0.09 | -0.09 | 0.008 | 0.72 | 0.54 | 0.82 | 0.93 | 0.7 | 0.81 |
| 1 | 37430266 | rs6426003 | -1.014 | 1.173 | -0.103 | -0.592 | 0.01 | -0.04 | -0.04 | 0 | 0.1 | 0.18 | 0.06 | 0.01 | 0.08 | 0.08 |
| 1 | 37430810 | rs488959 | -1.956 | 1.812 | 0.705 | -1.069 | 0.026 | -0.081 | -0.081 | 0 | 0.25 | 0.62 | 0.28 | 0.08 | 0.28 | 0.29 |
| 1 | 37431392 | rs6693285 | -1.06 | 1.576 | 0.945 | -1.179 | 0.043 | -0.058 | -0.058 | 0 | 0.12 | 0.39 | 0.08 | 0.01 | 0.08 | 0.08 |
| 1 | 37435475 | rs3753771 | -1.957 | 1.556 | 0.829 | -1.189 | -0.04 | 0.098 | 0.098 | 0.26 | 0.27 | 0.85 | 0.29 | 0.08 | 0.28 | 0.29 |
| 1 | 37437776 | rs575605 | -0.894 | 0.885 | -0.539 | -2.098 | -0.005 | 0.177 | 0.177 | 0 | 0.65 | 0.99 | 0.81 | 0.95 | 0.69 | 0.75 |
| 1 | 37456220 | rs16823744 | -0.795 | 0.511 | 0.523 | -2.222 | 0 | -0.061 | -0.061 | 0 | 0.22 | 0.73 | 0.19 | 0.01 | 0.21 | 0.26 |
| 1 | 37459950 | rs9628638 | -1.491 | 0.586 | 0.214 | -2.562 | -0.022 | 0.029 | 0.029 | 1.193 | 0.38 | 0.73 | 0.31 | 0.03 | 0.38 | 0.37 |
| 1 | 37478063 | rs12138633 | -0.736 | 0.776 | 0.52 | -2.439 | 0.033 | -0.082 | -0.082 | 2.501 | 0.2 | 0.22 | 0.2 | 0.01 | 0.25 | 0.12 |
| 1 | 37480144 | rs3753776 | -0.838 | 0.707 | 0.247 | -1.306 | -0.005 | -0.007 | -0.007 | 0 | 0.29 | 0.2 | 0.27 | 0.58 | 0.34 | 0.31 |
| 1 | 37483391 | rs563293 | 0.204 | 0.623 | 0.887 | -1.376 | -0.007 | -0.097 | -0.097 | 0.537 | 0.16 | 0.35 | 0.09 | 0.01 | 0.12 | 0.23 |
| 1 | 37488337 | rs522481 | -1.559 | 0.615 | 0.891 | -1.383 | -0.001 | -0.014 | -0.014 | 0 | 0.46 | 0.65 | 0.38 | 0.59 | 0.47 | 0.54 |
| 1 | 37489380 | rs6686296 | -0.776 | 0.618 | 0.952 | -1.382 | -0.01 | -0.015 | -0.015 | 0.184 | 0.29 | 0.19 | 0.27 | 0.58 | 0.34 | 0.31 |
| 1 | 37494602 | rs485719 | 0.051 | 0.612 | 0.953 | -1.359 | -0.004 | -0.111 | -0.111 | 0 | 0.17 | 0.42 | 0.11 | 0.01 | 0.14 | 0.23 |
| 1 | 37506171 | rs864584 | 0.306 | 0.585 | 0.824 | -1.362 | -0.003 | -0.043 | -0.043 | 0 | 0.15 | 0.2 | 0.07 | 0.01 | 0.12 | 0.23 |
| 1 | 37509352 | rs2859741 | -1.601 | 0.716 | 0.575 | -1.332 | 0.009 | -0.013 | -0.013 | 0 | 0.47 | 0.69 | 0.39 | 0.58 | 0.48 | 0.54 |
| 1 | 37517643 | rs640665 | 0.125 | 0.514 | 0.366 | -0.706 | 0.023 | -0.045 | -0.045 | 0.015 | 0.3 | 0.15 | 0.32 | 0.32 | 0.23 | 0.44 |
| 1 | 37525639 | rs550273 | -1.016 | 0.139 | 0.655 | -0.813 | -0.002 | -0.054 | -0.054 | 0 | 0.6 | 0.47 | 0.66 | 0.85 | 0.58 | 0.8 |
| 1 | 37526481 | rs521696 | -1.4 | 0.15 | -0.059 | -0.818 | -0.039 | 0.152 | 0.152 | 0 | 0.64 | 0.88 | 0.72 | 0.91 | 0.59 | 0.81 |
| 1 | 37530824 | rs1334792 | -0.498 | 0.284 | -0.395 | -0.056 | -0.005 | -0.087 | -0.087 | 0 | 0.14 | 0.02 | 0.29 | 0.42 | 0.14 | 0.28 |
| 1 | 37530961 | rs1334794 | -0.498 | 0.284 | -0.395 | -0.056 | -0.005 | -0.087 | -0.087 | 0.313 | 0.14 | 0.02 | 0.29 | 0.42 | 0.14 | 0.28 |
| 1 | 37532771 | rs12135742 | -0.012 | 0.241 | -0.37 | -0.031 | -0.007 | -0.071 | -0.071 | 0 | 0.12 | 0.02 | 0.2 | 0.42 | 0.12 | 0.25 |
| 1 | 37534052 | rs849943 | -0.274 | 0.248 | -0.147 | -0.059 | -0.008 | 0.129 | 0.129 | 0 | 0.13 | 0.28 | 0.23 | 0.48 | 0.13 | 0.25 |
| 1 | 37534255 | rs6702349 | -0.376 | 0.3 | -0.973 | -0.008 | -0.062 | -0.052 | -0.052 | 0 | 0.43 | 0.11 | 0.35 | 0.47 | 0.44 | 0.53 |
| 1 | 37537709 | rs218422 | 0.558 | 1.154 | -0.629 | 0.038 | 0.012 | 0.025 | 0.025 | 0 | 0.72 | 0.64 | 0.75 | 0.39 | 0.7 | 0.64 |
| 1 | 37538771 | rs218419 | -0.303 | 0.68 | -0.901 | 0.416 | 0 | -0.001 | -0.001 | 0 | 0.29 | 0.3 | 0.36 | 0.25 | 0.3 | 0.42 |
| 1 | 37544223 | rs218428 | 0.52 | 0.672 | -0.696 | 1.474 | -0.009 | -0.024 | -0.024 | 0 | 0.87 | 0.93 | 0.85 | 0.42 | 0.82 | 0.75 |
| 1 | 37548473 | rs218396 | -1.187 | 0.138 | 0.225 | 1.229 | 0.029 | 0.268 | 0.268 | 6.363 | 0.06 | 0.59 | 0.23 | 0.17 | 0.01 | 0.04 |
| 1 | 37551870 | rs11263972 | 0.064 | -0.004 | -0.92 | -0.489 | -0.017 | 0.225 | 0.225 | 0.034 | 0.44 | 0.12 | 0.33 | 0.04 | 0.53 | 0.36 |
| 1 | 37561638 | rs849952 | 0.094 | -0.25 | -0.588 | -0.12 | 0.016 | 0.179 | 0.179 | 0.05 | 0.47 | 0.16 | 0.34 | 0.03 | 0.55 | 0.35 |
| 1 | 37567439 | rs3856135 | -0.006 | -0.491 | 0.005 | -0.225 | -0.002 | -0.027 | -0.027 | 0 | 0.16 | 0.25 | 0.07 | 0.05 | 0.13 | 0.1 |
| 1 | 37581882 | rs10908321 | 0.237 | -0.215 | -0.097 | -0.528 | -0.003 | -0.042 | -0.042 | 0 | 0.55 | 0.44 | 0.36 | 0.08 | 0.6 | 0.43 |
| 1 | 37585973 | rs215770 | -0.187 | -0.537 | -0.918 | -0.458 | -0.018 | 0.099 | 0.099 | 0 | 0.72 | 0.95 | 0.67 | 0.86 | 0.74 | 0.72 |
| 1 | 37596760 | rs4653247 | 0.442 | -1.321 | -1.367 | -0.366 | 0.043 | -0.077 | -0.077 | 0 | 0.19 | 0.01 | 0.23 | 0.24 | 0.12 | 0.22 |
| 1 | 37606291 | rs215791 | 0.412 | -0.575 | -0.485 | -0.955 | 0.01 | -0.051 | -0.051 | 1.319 | 0.32 | 0.55 | 0.25 | 0.06 | 0.39 | 0.22 |
| 1 | 37611513 | rs12131641 | 0.287 | -0.006 | -1.061 | -1.132 | -0.067 | 0.065 | 0.065 | 0 | 0.14 | 0.01 | 0.1 | 0.03 | 0.2 | 0.15 |
| 1 | 37611583 | rs215789 | -0.69 | -0.185 | -1.883 | -1.551 | -0.044 | 0.056 | 0.056 | 0.227 | 0.72 | 0.91 | 0.67 | 0.82 | 0.69 | 0.69 |
| 1 | 37614806 | rs11263989 | 0.615 | -0.073 | -1.257 | -1.484 | -0.003 | -0.009 | -0.009 | 0 | 0.15 | 0.11 | 0.11 | 0.03 | 0.2 | 0.15 |
| 1 | 37617809 | rs2884796 | -0.533 | -0.198 | -1.215 | -1.627 | -0.007 | 0.067 | 0.067 | 0 | 0.68 | 0.82 | 0.66 | 0.82 | 0.68 | 0.67 |
| 1 | 37622070 | rs4653252 | 0.58 | -0.215 | -0.625 | -1.594 | -0.002 | -0.022 | -0.022 | 0 | 0.43 | 0.47 | 0.49 | 0.76 | 0.41 | 0.46 |
| 1 | 37631514 | rs12141503 | 0.601 | -0.037 | -0.847 | -1.449 | -0.005 | -0.003 | -0.003 | 0.001 | 0.15 | 0.11 | 0.11 | 0.03 | 0.2 | 0.15 |
| 1 | 37636117 | rs998176 | 0.66 | -0.228 | -0.36 | -1.388 | 0 | -0.009 | -0.009 | 0 | 0.41 | 0.38 | 0.49 | 0.73 | 0.41 | 0.46 |
| 1 | 37637422 | rs7518123 | 0.537 | -0.31 | -0.276 | -1.641 | -0.001 | -0.025 | -0.025 | 0 | 0.43 | 0.42 | 0.59 | 0.85 | 0.42 | 0.5 |
| 1 | 37641047 | rs218388 | 1.033 | -0.434 | -0.025 | -1.65 | -0.016 | -0.024 | -0.024 | 0 | 0.1 | 0.17 | 0.05 | 0.03 | 0.06 | 0.06 |
| 1 | 37649899 | rs215833 | 0.196 | -0.441 | -0.166 | -1.772 | -0.009 | -0.068 | -0.068 | 0 | 0.58 | 0.5 | 0.69 | 0.88 | 0.63 | 0.66 |
| 1 | 37655620 | rs12123283 | 0.542 | 0.719 | -0.335 | -1.34 | -0.06 | 0.025 | 0.025 | 0 | 0.1 | 0.01 | 0.09 | 0.04 | 0.17 | 0.13 |
| 1 | 37657963 | rs150571 | 1.248 | 0.497 | -0.164 | -1.317 | 0.006 | 0.01 | 0.01 | 0.031 | 0.08 | 0.04 | 0.05 | 0.03 | 0.06 | 0.07 |
| 1 | 37663143 | rs215820 | -1.307 | 0.558 | 0.271 | -1.445 | 0.006 | -0.061 | -0.061 | 0.009 | 0.25 | 0.39 | 0.16 | 0.07 | 0.24 | 0.22 |
| 1 | 37663172 | rs12743241 | -0.925 | 0.558 | 0.215 | -1.459 | -0.032 | 0.068 | 0.068 | 0 | 0.22 | 0.1 | 0.13 | 0.07 | 0.23 | 0.22 |
| 1 | 37683919 | rs215862 | -0.565 | 0.431 | 0.502 | -1.523 | -0.025 | -0.05 | -0.05 | 2.795 | 0.15 | 0.42 | 0.09 | 0.06 | 0.1 | 0.25 |
| 1 | 37692018 | rs17462022 | 1.544 | 0.446 | 0.502 | -1.512 | 0.001 | -0.001 | -0.001 | 0 | 0.08 | 0 | 0.04 | 0.06 | 0.06 | 0.07 |
| 1 | 37703332 | rs11264003 | 0.539 | 0.351 | 1.107 | -1.562 | -0.036 | -0.067 | -0.067 | 0 | 0.84 | 0.42 | 0.9 | 0.94 | 0.9 | 0.76 |
| 1 | 37704048 | rs12140004 | -0.297 | 0.746 | 0.761 | -1.673 | -0.08 | 0.07 | 0.07 | 0 | 0.13 | 0.01 | 0.1 | 0.02 | 0.2 | 0.15 |
| 1 | 37707456 | rs6679108 | 0.355 | 0.626 | 0.977 | -1.633 | -0.035 | -0.065 | -0.065 | 0.106 | 0.84 | 0.49 | 0.9 | 0.94 | 0.9 | 0.76 |
| 1 | 37708339 | rs1566599 | 0.485 | 0.529 | 0.945 | -1.614 | -0.043 | -0.052 | -0.052 | 0 | 0.83 | 0.48 | 0.89 | 0.92 | 0.9 | 0.79 |
| 1 | 37708832 | rs1540721 | -1.04 | 0.551 | 0.954 | -1.595 | -0.014 | 0.005 | 0.005 | 0 | 0.9 | 0.71 | 0.92 | 0.93 | 0.94 | 0.89 |
| 1 | 37710590 | rs16823912 | 0.076 | -0.198 | 0.512 | -2.394 | -0.041 | -0.07 | -0.07 | 0.01 | 0.31 | 0.4 | 0.17 | 0.07 | 0.2 | 0.22 |
| 1 | 37716037 | rs872157 | 1.094 | -0.183 | 0.414 | -2.466 | 0.013 | 0.091 | 0.091 | 0 | 0.5 | 0.32 | 0.69 | 0.9 | 0.52 | 0.54 |
| 1 | 37716718 | rs6691915 | 0.457 | -0.148 | 0.387 | -2.463 | 0.032 | -0.056 | -0.056 | 0 | 0.83 | 0.71 | 0.87 | 0.97 | 0.75 | 0.79 |
| 1 | 37717219 | rs2047706 | 1.546 | -0.03 | 0.184 | -1.396 | -0.005 | 0.013 | 0.013 | 0.164 | 0.06 | 0 | 0.04 | 0.34 | 0.05 | 0.09 |
| 1 | 37717520 | rs2047707 | -1.058 | 0.004 | 0.221 | -1.552 | -0.021 | 0.018 | 0.018 | 0 | 0.9 | 0.93 | 0.91 | 0.97 | 0.8 | 0.88 |
| 1 | 37718209 | rs4384192 | 0.264 | -0.129 | 0.168 | -1.597 | -0.001 | -0.052 | -0.052 | 0 | 0.62 | 0.61 | 0.77 | 0.97 | 0.6 | 0.65 |
| 1 | 37724275 | rs10908330 | 1.768 | -0.05 | 0.247 | -1.918 | -0.02 | -0.051 | -0.051 | 0 | 0.29 | 0.16 | 0.55 | 0.58 | 0.35 | 0.35 |
| 1 | 37725041 | rs1542885 | 0.134 | 0 | 0.072 | -1.989 | 0 | -0.055 | -0.055 | 0 | 0.62 | 0.6 | 0.77 | 0.97 | 0.6 | 0.65 |
| 1 | 37730550 | rs12037190 | 0.809 | 0.11 | 0.145 | -1.86 | 0.015 | 0.067 | 0.067 | 0 | 0.51 | 0.37 | 0.7 | 0.9 | 0.55 | 0.59 |

| | | | | | | | | | | | | | | | | |
|---|---|---|---|---|---|---|---|---|---|---|---|---|---|---|---|---|
| 1 | 37748214 | rs823926 | -0.538 | 0.344 | -0.015 | -1.808 | 0.017 | -0.04 | -0.04 | 0 | 0.15 | 0.26 | 0.08 | 0.03 | 0.18 | 0.13 |
| 1 | 37749227 | rs624650 | -0.732 | 0.308 | -0.275 | -1.88 | 0.055 | 0.119 | 0.119 | 0 | 0.67 | 0.87 | 0.84 | 0.93 | 0.73 | 0.67 |
| 1 | 37752850 | rs301992 | -0.566 | 0.585 | 0 | -1.672 | 0.039 | -0.067 | -0.067 | 0 | 0.15 | 0.36 | 0.09 | 0.03 | 0.18 | 0.13 |
| 1 | 37757131 | rs16823951 | 1.642 | 0.648 | -0.31 | -1.558 | -0.01 | -0.021 | -0.021 | 0.108 | 0.07 | 0 | 0.05 | 0.06 | 0.07 | 0.06 |
| 1 | 37764586 | rs1888698 | -0.502 | 0.275 | -0.328 | -1.352 | -0.001 | -0.009 | -0.009 | 6.58 | 0.66 | 0.65 | 0.85 | 0.81 | 0.79 | 0.73 |
| 1 | 37765920 | rs10908337 | 0.84 | 0.23 | -0.284 | -1.965 | 0.013 | 0.065 | 0.065 | 0 | 0.47 | 0.33 | 0.71 | 0.72 | 0.55 | 0.55 |
| 1 | 37769074 | rs1888695 | -0.38 | 0.112 | -0.316 | -1.922 | -0.039 | -0.055 | -0.055 | 0 | 0.65 | 0.58 | 0.84 | 0.81 | 0.79 | 0.73 |
| 1 | 37773050 | rs7516603 | 0.256 | 0.549 | -0.596 | -1.853 | 0.008 | -0.052 | -0.052 | 0 | 0.27 | 0.29 | 0.17 | 0.03 | 0.3 | 0.25 |
| 1 | 37778661 | rs1331335 | 1.062 | 0.103 | -0.958 | -1.227 | -0.009 | -0.01 | -0.01 | 0.474 | 0.07 | 0 | 0.05 | 0.05 | 0.08 | 0.05 |
| 1 | 37782841 | rs963295 | -1.208 | -0.205 | -1.023 | -1.49 | 0.057 | 0.069 | 0.069 | 0.093 | 0.77 | 0.94 | 0.91 | 1 | 0.84 | 0.75 |
| 1 | 37782874 | rs1411931 | -0.08 | 0.216 | -0.765 | -1.597 | -0.017 | -0.115 | -0.115 | 0 | 0.59 | 0.5 | 0.79 | 0.97 | 0.66 | 0.62 |
| 1 | 37782902 | rs949586 | 0.373 | 0.258 | -0.94 | -1.235 | 0.014 | 0.022 | 0.022 | 0 | 0.48 | 0.41 | 0.58 | 0.69 | 0.58 | 0.52 |
| 1 | 37785527 | rs10908340 | -0.106 | 0.725 | -1.136 | -1.429 | -0.011 | 0.044 | 0.044 | 0.163 | 0.39 | 0.35 | 0.69 | 0.92 | 0.51 | 0.52 |
| 1 | 37787733 | rs761158 | -1.398 | -0.02 | -1.764 | -1.647 | 0.02 | 0.148 | 0.148 | 0.388 | 0.78 | 0.99 | 0.92 | 0.99 | 0.83 | 0.78 |
| 1 | 37796436 | rs4653277 | -0.112 | 0.713 | -1.406 | -1.185 | 0.001 | -0.004 | -0.004 | 0 | 0.45 | 0.5 | 0.62 | 0.46 | 0.51 | 0.45 |
| 1 | 37796608 | rs11582151 | -0.134 | 1.451 | -0.804 | -0.42 | -0.029 | -0.042 | -0.042 | 0.187 | 0.09 | 0.04 | 0.32 | 0.39 | 0.14 | 0.17 |
| 1 | 37809549 | rs727670 | -0.199 | 1 | -0.395 | -0.22 | 0.008 | 0.015 | 0.015 | 0 | 0.32 | 0.42 | 0.3 | 0.53 | 0.33 | 0.34 |
| 1 | 37824340 | rs11809018 | -0.898 | 0.622 | -0.314 | -0.171 | -0.003 | -0.011 | -0.011 | 0 | 0.15 | 0.26 | 0.12 | 0.1 | 0.15 | 0.12 |
| 1 | 37824508 | rs1004766 | 1.129 | 0.521 | -0.664 | -0.342 | -0.016 | 0.112 | 0.112 | 0.041 | 0.53 | 0.11 | 0.55 | 0.37 | 0.53 | 0.56 |
| 1 | 37826908 | rs765807 | 0.057 | 0.649 | -0.564 | -0.254 | 0.006 | -0.014 | -0.014 | 0.027 | 0.1 | 0.15 | 0.09 | 0.04 | 0.14 | 0.11 |
| 1 | 37853334 | rs1569661 | -0.073 | 0.532 | 0.064 | -0.158 | 0.003 | 0.018 | 0.018 | 0 | 0.72 | 0.64 | 0.74 | 0.51 | 0.69 | 0.7 |
| 1 | 37855834 | rs16824043 | -1.106 | 0.613 | 0.045 | -0.466 | -0.002 | -0.007 | -0.007 | 0.298 | 0.12 | 0.19 | 0.05 | 0.05 | 0.07 | 0.06 |
| 1 | 37859876 | rs2745487 | 0.052 | 0.713 | 0.224 | -0.442 | 0.011 | -0.019 | -0.019 | 0 | 0.1 | 0.16 | 0.09 | 0.04 | 0.14 | 0.12 |
| 1 | 37861046 | rs2745486 | -0.534 | 0.649 | 0.252 | -0.29 | 0.018 | 0.074 | 0.074 | 0 | 0.79 | 0.64 | 0.78 | 0.46 | 0.75 | 0.76 |
| 1 | 37861476 | rs2745485 | 0.353 | 0.61 | 0.204 | -0.361 | -0.005 | 0.013 | 0.013 | 0 | 0.72 | 0.48 | 0.74 | 0.39 | 0.73 | 0.74 |
| 1 | 37864911 | rs876465 | 1.034 | 0.663 | -0.296 | -0.412 | -0.005 | 0.044 | 0.044 | 0 | 0.61 | 0.22 | 0.65 | 0.35 | 0.59 | 0.61 |
| 1 | 37865131 | rs876466 | 0.456 | 0.635 | -0.088 | -0.27 | 0.007 | 0.006 | 0.006 | 0 | 0.68 | 0.42 | 0.69 | 0.44 | 0.61 | 0.64 |
| 1 | 37865954 | rs875361 | 1.234 | -0.149 | -1.276 | -0.992 | -0.077 | 0.018 | 0.018 | 3.355 | 0.37 | 0.1 | 0.57 | 0.38 | 0.46 | 0.29 |
| 1 | 37866343 | rs2745481 | -0.439 | -0.247 | -0.981 | -0.861 | -0.018 | -0.097 | -0.097 | 0 | 0.75 | 0.44 | 0.84 | 0.94 | 0.78 | 0.77 |
| 1 | 37866643 | rs2745480 | -0.444 | -0.284 | -1.044 | -0.691 | -0.001 | -0.004 | -0.004 | 0 | 0.17 | 0.2 | 0.13 | 0.08 | 0.16 | 0.14 |
| 1 | 37867275 | rs16824060 | 0.49 | -0.236 | -1.103 | -0.534 | -0.027 | 0.042 | 0.042 | 0 | 0.06 | 0.01 | 0.09 | 0.47 | 0.07 | 0.07 |
| 1 | 37878610 | rs10908348 | -0.015 | -0.534 | -1.098 | -0.815 | -0.046 | -0.073 | -0.073 | 0 | 0.15 | 0.69 | 0.09 | 0.01 | 0.06 | 0.15 |
| 1 | 37882114 | rs2811630 | -0.55 | 0.11 | -1.167 | -0.38 | -0.017 | -0.031 | -0.031 | 0 | 0.2 | 0.19 | 0.23 | 0.51 | 0.24 | 0.19 |
| 1 | 37887745 | rs10493079 | 1.305 | -0.257 | -1.367 | -0.634 | 0.036 | 0.065 | 0.065 | 0 | 0.17 | 0.04 | 0.05 | 0.02 | 0.08 | 0.25 |
| 1 | 37888656 | rs16824119 | 0.471 | -0.209 | -1.389 | -0.569 | -0.023 | -0.022 | -0.022 | 0 | 0.1 | 0.04 | 0.12 | 0.44 | 0.09 | 0.07 |
| 1 | 37889172 | rs4428850 | 0.478 | -0.178 | -1.39 | -0.659 | -0.005 | -0.031 | -0.031 | 0 | 0.62 | 0.78 | 0.71 | 0.47 | 0.68 | 0.55 |
| 1 | 37891045 | rs12139812 | 1.325 | -0.154 | -1.877 | -1.258 | -0.02 | -0.039 | -0.039 | 0 | 0.13 | 0 | 0.15 | 0.15 | 0.14 | 0.14 |
| 1 | 37892154 | rs12734239 | -0.938 | -0.154 | -1.603 | -1.514 | 0.001 | 0.178 | 0.178 | 0.38 | 0.53 | 0.77 | 0.32 | 0.13 | 0.45 | 0.62 |
| 1 | 37892550 | rs4653291 | 0.565 | 0.247 | -1.347 | -1.22 | -0.003 | -0.002 | -0.002 | 0.329 | 0.11 | 0.13 | 0.12 | 0.03 | 0.15 | 0.13 |
| 1 | 37892624 | rs11264045 | 0.101 | 0.051 | -0.899 | -1.023 | -0.02 | -0.109 | -0.109 | 0.011 | 0.24 | 0.48 | 0.14 | 0.06 | 0.22 | 0.24 |
| 1 | 37897737 | rs10752596 | -0.128 | 0.811 | -0.772 | -0.538 | 0.002 | -0.004 | -0.004 | 0 | 0.68 | 0.48 | 0.67 | 0.59 | 0.69 | 0.52 |
| 1 | 37900173 | rs10796907 | -0.068 | 1.436 | -1.109 | -0.61 | -0.048 | -0.022 | -0.022 | 0.474 | 0.78 | 0.85 | 0.64 | 0.58 | 0.64 | 0.67 |
| 1 | 37902331 | rs10908350 | 0.426 | 1.119 | -1.236 | -0.836 | 0.002 | 0.022 | 0.022 | 0 | 0.49 | 0.42 | 0.48 | 0.22 | 0.55 | 0.35 |
| 1 | 37907433 | rs6685425 | -0.784 | 0.996 | -0.909 | -0.825 | 0.004 | -0.041 | -0.041 | 0.689 | 0.78 | 0.69 | 0.83 | 0.92 | 0.79 | 0.65 |
| 1 | 37909488 | rs10493078 | 1.173 | 0.918 | -1.245 | -1.181 | -0.005 | -0.09 | -0.09 | 0 | 0.3 | 0.33 | 0.17 | 0.03 | 0.24 | 0.25 |
| 1 | 37912389 | rs11264055 | 0.179 | 1.355 | -1.583 | -0.748 | -0.095 | -0.085 | -0.085 | 0 | 0.29 | 0.09 | 0.49 | 0.51 | 0.43 | 0.31 |
| 1 | 37913283 | rs2484556 | -0.336 | 1.36 | -1.226 | -0.691 | -0.003 | -0.003 | -0.003 | 0.314 | 0.77 | 0.8 | 0.63 | 0.59 | 0.64 | 0.68 |
| 1 | 37914557 | rs3105708 | 1.55 | 1.086 | -1.146 | -0.922 | 0.001 | -0.055 | -0.055 | 0 | 0.3 | 0.29 | 0.16 | 0.03 | 0.23 | 0.24 |
| 1 | 37918320 | rs1395720 | 0.484 | 1.493 | -1.592 | -0.638 | -0.028 | -0.029 | -0.029 | 1.267 | 0.62 | 0.78 | 0.42 | 0.43 | 0.46 | 0.61 |
| 1 | 37922859 | rs2660569 | 1.287 | 1.325 | -1.384 | -0.882 | -0.01 | -0.106 | -0.106 | 0 | 0.44 | 0.54 | 0.22 | 0.06 | 0.33 | 0.51 |
| 1 | 37931786 | rs2245872 | 1.262 | 1.305 | -1.426 | -0.879 | 0.002 | -0.064 | -0.064 | 2.168 | 0.45 | 0.57 | 0.23 | 0.06 | 0.33 | 0.52 |
| 1 | 37939094 | rs6681785 | -0.698 | 1.949 | -1.207 | -0.109 | -0.013 | 0.022 | 0.022 | 0.311 | 0.31 | 0.16 | 0.43 | 0.72 | 0.33 | 0.35 |
| 1 | 37939173 | rs2492301 | 1.154 | 2.148 | -1.136 | -0.105 | 0.002 | -0.009 | -0.009 | 0.091 | 0.61 | 0.58 | 0.45 | 0.22 | 0.54 | 0.58 |
| 1 | 37941953 | rs1395718 | 0.258 | 2.476 | -1.2 | 0.223 | -0.008 | -0.013 | -0.013 | 0 | 0.17 | 0.12 | 0.26 | 0.38 | 0.21 | 0.27 |
| 1 | 37951754 | rs215206 | 0.41 | 2.325 | -1.164 | 0.383 | -0.021 | -0.003 | -0.003 | 0.005 | 0.71 | 0.74 | 0.56 | 0.66 | 0.61 | 0.71 |
| 1 | 37957376 | rs6700586 | -1.374 | 2.18 | -1.296 | 0.343 | 0.013 | -0.014 | -0.014 | 0 | 0.43 | 0.33 | 0.64 | 0.55 | 0.59 | 0.54 |
| 1 | 37961613 | rs2294873 | 0.299 | 2.353 | -1.313 | 0.355 | -0.048 | -0.044 | -0.044 | 0 | 0.13 | 0.02 | 0.18 | 0.18 | 0.17 | 0.16 |
| 1 | 37967774 | rs215864 | -1.05 | 1.606 | -1.408 | 0.367 | -0.054 | -0.062 | -0.062 | 0 | 0.72 | 0.53 | 0.84 | 0.9 | 0.84 | 0.65 |
| 1 | 37973832 | rs12125547 | 1.739 | 1.711 | -1.698 | 0.192 | 0.005 | 0.013 | 0.013 | 0 | 0.08 | 0.01 | 0.04 | 0.04 | 0.06 | 0.13 |
| 1 | 37975279 | rs215214 | -1.506 | 1.678 | -1.715 | 0.176 | 0.006 | 0.015 | 0.015 | 0 | 0.91 | 0.99 | 0.95 | 0.96 | 0.94 | 0.87 |
| 1 | 37991458 | rs11264067 | -0.829 | 1.833 | -1.692 | 0.303 | -0.004 | -0.009 | -0.009 | 0.267 | 0.41 | 0.27 | 0.61 | 0.61 | 0.53 | 0.45 |
| 1 | 37996750 | rs7545493 | -1.055 | 1.498 | -1.59 | 0.212 | -0.045 | -0.081 | -0.081 | 0 | 0.72 | 0.47 | 0.84 | 0.97 | 0.84 | 0.68 |
| 1 | 37999227 | rs12119795 | 0.459 | 1.777 | -1.968 | 0.304 | -0.063 | -0.056 | -0.056 | 0.017 | 0.13 | 0.02 | 0.18 | 0.17 | 0.17 | 0.16 |
| 1 | 38000185 | rs7515306 | -1.355 | 1.403 | -1.508 | -0.206 | -0.018 | -0.041 | -0.041 | 1.395 | 0.76 | 0.73 | 0.87 | 0.97 | 0.84 | 0.67 |
| 1 | 38018536 | rs11264072 | 1.332 | 2.096 | -2.784 | -0.003 | -0.047 | 0.012 | 0.012 | 0 | 0.1 | 0.03 | 0.19 | 0.06 | 0.17 | 0.04 |
| 1 | 38019747 | rs2273013 | 1.248 | 1.899 | -2.803 | 0.084 | -0.018 | -0.086 | -0.086 | 0 | 0.21 | 0.06 | 0.29 | 0.39 | 0.22 | 0.25 |
| 1 | 38024054 | rs4653300 | -0.927 | 1.813 | -2.722 | 0.016 | 0.03 | 0.005 | 0.005 | 0 | 0.44 | 0.35 | 0.64 | 0.62 | 0.59 | 0.49 |
| 1 | 38028374 | rs4653301 | -0.132 | 1.9 | -2.601 | 0.149 | -0.079 | -0.07 | -0.07 | 0.24 | 0.34 | 0.2 | 0.56 | 0.57 | 0.47 | 0.33 |
| 1 | 38030696 | rs6619 | -0.145 | 1.908 | -2.596 | 0.153 | -0.071 | -0.063 | -0.063 | 0 | 0.34 | 0.22 | 0.56 | 0.57 | 0.47 | 0.33 |

| | | | | | | | | | | | | | | | | |
|---|---|---|---|---|---|---|---|---|---|---|---|---|---|---|---|---|
| 1 | 38031972 | rs3856137 | 1.623 | 1.667 | -2.615 | 0.157 | 0.007 | 0.012 | 0.012 | 0 | 0.08 | 0.03 | 0.05 | 0.04 | 0.06 | 0.12 |
| 1 | 38035286 | rs10908355 | -0.423 | 1.809 | -2.353 | 0.214 | 0.028 | 0.004 | 0.004 | 0 | 0.44 | 0.33 | 0.63 | 0.62 | 0.58 | 0.49 |
| 1 | 38051585 | rs7535432 | 1.152 | 1.633 | -2.897 | 0.283 | 0.016 | -0.048 | -0.048 | 0 | 0.28 | 0.08 | 0.2 | 0.35 | 0.25 | 0.18 |
| 1 | 38078300 | rs4652964 | -0.277 | 1.64 | -2.917 | 0.243 | -0.036 | -0.004 | -0.004 | 0.001 | 0.42 | 0.21 | 0.59 | 0.68 | 0.53 | 0.44 |
| 1 | 38086578 | rs12023270 | 1.099 | 1.104 | -2.86 | 0.299 | 0.008 | -0.027 | -0.027 | 0 | 0.28 | 0.07 | 0.21 | 0.28 | 0.25 | 0.2 |
| 1 | 38089114 | rs12144790 | -0.059 | 1.489 | -2.059 | 0.718 | -0.016 | -0.106 | -0.106 | 0 | 0.29 | 0.07 | 0.33 | 0.44 | 0.28 | 0.37 |
| 1 | 38092723 | rs4074961 | -0.619 | 1.317 | -1.975 | 0.711 | -0.045 | -0.054 | -0.054 | 0.001 | 0.41 | 0.21 | 0.52 | 0.49 | 0.46 | 0.41 |
| 1 | 38098036 | rs4233215 | -0.011 | 0.961 | -1.977 | 0.848 | -0.009 | -0.12 | -0.12 | 0.009 | 0.27 | 0.05 | 0.32 | 0.45 | 0.23 | 0.38 |
| 1 | 38104340 | rs4653309 | 0.04 | 1.361 | -1.075 | 0.187 | 0.063 | -0.01 | -0.01 | 2.195 | 0.85 | 0.42 | 0.81 | 0.83 | 0.77 | 0.92 |
| 1 | 38105212 | rs11264079 | 0.119 | 1.232 | -1.269 | -0.184 | -0.039 | -0.026 | -0.026 | 6.643 | 0.4 | 0.24 | 0.56 | 0.52 | 0.47 | 0.47 |
| 1 | 38109157 | rs4653311 | 0.09 | 1.077 | -1.179 | 0.09 | -0.032 | -0.042 | -0.042 | 0.693 | 0.15 | 0.08 | 0.2 | 0.28 | 0.24 | 0.13 |
| 1 | 38109660 | rs7521055 | -0.867 | 0.394 | -0.874 | -0.428 | -0.026 | -0.016 | -0.016 | 0.297 | 0.79 | 0.67 | 0.9 | 0.83 | 0.88 | 0.66 |
| 1 | 38109752 | rs7552071 | 0.953 | 0.413 | -0.695 | -0.5 | 0.001 | -0.01 | -0.01 | 1.022 | 0.6 | 0.33 | 0.66 | 0.52 | 0.61 | 0.51 |
| 1 | 38111706 | rs4653312 | -0.848 | 0.533 | -0.94 | -0.449 | -0.026 | -0.015 | -0.015 | 0 | 0.79 | 0.69 | 0.9 | 0.83 | 0.88 | 0.66 |
| 1 | 38116724 | rs12742985 | -0.193 | 0.68 | -1.271 | -0.375 | -0.095 | -0.087 | -0.087 | 0 | 0.22 | 0.12 | 0.26 | 0.36 | 0.35 | 0.24 |
| 1 | 38117142 | rs11264081 | -1.175 | 0.558 | -1.407 | -0.556 | 0.057 | -0.003 | -0.003 | 0 | 0.47 | 0.75 | 0.59 | 0.5 | 0.66 | 0.47 |
| 1 | 38121934 | rs10908366 | -0.696 | 1.279 | -1.556 | -0.316 | 0.042 | 0.026 | 0.026 | 0 | 0.3 | 0.46 | 0.44 | 0.42 | 0.5 | 0.27 |
| 1 | 38125758 | rs11578786 | 0.474 | 1.117 | -1.781 | -0.178 | -0.008 | 0.014 | 0.014 | 0 | 0.52 | 0.3 | 0.6 | 0.52 | 0.54 | 0.54 |
| 1 | 38127154 | rs6687329 | 0.344 | 0.457 | -1.705 | -0.76 | 0.012 | 0.053 | 0.053 | 0 | 0.57 | 0.36 | 0.75 | 0.8 | 0.65 | 0.63 |
| 1 | 38134606 | rs10908367 | -0.774 | 0.932 | -0.963 | -0.127 | 0.05 | 0.135 | 0.135 | 0 | 0.15 | 0.44 | 0.21 | 0.33 | 0.18 | 0.18 |
| 1 | 38139551 | rs11585866 | 0.001 | 0.633 | -2.031 | -0.646 | -0.006 | 0.064 | 0.064 | 0.424 | 0.17 | 0.06 | 0.11 | 0.03 | 0.18 | 0.04 |
| 1 | 38139870 | rs12135570 | 0.213 | 0.162 | -2.221 | -0.4 | 0.008 | -0.112 | -0.112 | 0.597 | 0.35 | 0.06 | 0.32 | 0.41 | 0.29 | 0.45 |
| 1 | 38144268 | rs10908368 | 0.046 | 0.591 | -2.129 | -0.376 | 0.023 | -0.04 | -0.04 | 0 | 0.48 | 0.08 | 0.4 | 0.41 | 0.39 | 0.45 |
| 1 | 38147166 | rs510208 | -0.441 | 0.4 | -1.507 | -1.071 | -0.003 | -0.027 | -0.027 | 0.029 | 0.23 | 0.21 | 0.14 | 0.03 | 0.27 | 0.14 |
| 1 | 38149193 | rs686083 | -0.578 | 0.33 | -0.788 | -1.103 | 0.016 | -0.121 | -0.121 | 0 | 0.24 | 0.39 | 0.16 | 0.03 | 0.27 | 0.15 |
| 1 | 38151705 | rs525379 | -0.602 | 0.317 | -0.794 | -1.109 | 0.015 | -0.121 | -0.121 | 0.007 | 0.24 | 0.39 | 0.16 | 0.03 | 0.27 | 0.15 |
| 1 | 38156902 | rs3762352 | 1.013 | -0.334 | -0.589 | -1.195 | 0.001 | 0 | 0 | 0 | 0.28 | 0.28 | 0.36 | 0.67 | 0.17 | 0.35 |
| 1 | 38157717 | rs652109 | -0.683 | -0.234 | -0.555 | -1.228 | 0.003 | 0.003 | 0.003 | 0.095 | 0.51 | 0.53 | 0.51 | 0.7 | 0.44 | 0.5 |
| 1 | 38157844 | rs651626 | 0.774 | -0.292 | -0.646 | -1.047 | 0 | -0.004 | -0.004 | 0 | 0.1 | 0.1 | 0.07 | 0.18 | 0.11 | 0.05 |
| 1 | 38163387 | rs11264087 | 1.693 | -0.366 | -0.82 | -1.026 | 0.01 | -0.169 | -0.169 | 0.02 | 0.23 | 0.01 | 0.26 | 0.44 | 0.16 | 0.33 |
| 1 | 38169578 | rs3790623 | 1.711 | 0.176 | -0.643 | -1.274 | -0.026 | 0.021 | 0.021 | 0 | 0.26 | 0.16 | 0.26 | 0.2 | 0.29 | 0.26 |
| 1 | 38174185 | rs557897 | -0.382 | 0.19 | -0.914 | -1.202 | 0.021 | 0.065 | 0.065 | 0.069 | 0.51 | 0.66 | 0.52 | 0.7 | 0.44 | 0.5 |
| 1 | 38174987 | rs7822 | -1.161 | 0.56 | -1.645 | -1.48 | -0.072 | 0.119 | 0.119 | 0 | 0.81 | 0.98 | 0.82 | 0.95 | 0.78 | 0.78 |
| 1 | 38184881 | rs7533921 | 1.101 | -0.224 | -0.928 | -0.876 | 0.032 | -0.057 | -0.057 | 0 | 0.26 | 0.16 | 0.28 | 0.44 | 0.17 | 0.33 |
| 1 | 38185723 | rs6670599 | 0.059 | 0.686 | -0.579 | -1.199 | -0.011 | 0.029 | 0.029 | 0 | 0.08 | 0.02 | 0.04 | 0.03 | 0.11 | 0.03 |
| 1 | 38185814 | rs12124242 | -0.474 | 0.688 | -0.532 | -1.227 | -0.004 | -0.009 | -0.009 | 0 | 0.14 | 0.11 | 0.08 | 0 | 0.16 | 0.11 |
| 1 | 38186189 | rs6671088 | -0.57 | 0.691 | -0.483 | -1.225 | 0.004 | -0.054 | -0.054 | 0 | 0.14 | 0.18 | 0.09 | 0 | 0.16 | 0.11 |
| 1 | 38186294 | rs1008879 | -0.57 | 0.691 | -0.483 | -1.225 | 0.004 | -0.054 | -0.054 | 0.75 | 0.14 | 0.18 | 0.09 | 0 | 0.16 | 0.11 |
| 1 | 38188195 | rs6426030 | -1.463 | 0.709 | -0.533 | -1.222 | -0.002 | -0.026 | -0.026 | 0 | 0.22 | 0.23 | 0.14 | 0.03 | 0.26 | 0.14 |
| 1 | 38188740 | rs12405650 | 0.02 | 0.812 | -0.572 | -1.172 | -0.011 | 0.029 | 0.029 | 0 | 0.08 | 0.02 | 0.05 | 0.03 | 0.11 | 0.03 |
| 1 | 38188768 | rs17464871 | 0.02 | 0.812 | -0.572 | -1.172 | -0.011 | 0.029 | 0.029 | 0 | 0.08 | 0.02 | 0.04 | 0.03 | 0.11 | 0.03 |
| 1 | 38188787 | rs17511304 | 0.02 | 0.812 | -0.572 | -1.172 | -0.011 | 0.029 | 0.029 | 0 | 0.08 | 0.02 | 0.04 | 0.03 | 0.11 | 0.03 |
| 1 | 38188841 | rs12121777 | -0.507 | 0.777 | -0.259 | -0.83 | 0.004 | 0.01 | 0.01 | 0 | 0.21 | 0.17 | 0.11 | 0 | 0.2 | 0.12 |
| 1 | 38189142 | rs1010805 | -0.697 | 0.786 | -0.233 | -0.819 | 0.003 | -0.021 | -0.021 | 0 | 0.22 | 0.23 | 0.13 | 0 | 0.2 | 0.13 |
| 1 | 38192296 | rs6686248 | -0.609 | 0.946 | -0.462 | -0.809 | -0.001 | -0.011 | -0.011 | 0 | 0.22 | 0.2 | 0.13 | 0 | 0.22 | 0.12 |
| 1 | 38196026 | rs10908376 | -0.453 | 0.839 | -1.255 | -0.7 | 0.001 | 0.071 | 0.071 | 0 | 0.21 | 0.1 | 0.11 | 0 | 0.2 | 0.12 |
| 1 | 38197629 | rs12127877 | -0.27 | 1.057 | -0.938 | -0.597 | 0 | 0 | 0 | 0 | 0.19 | 0.16 | 0.12 | 0 | 0.19 | 0.14 |
| 1 | 38199542 | rs7549842 | -0.266 | 1.597 | -0.94 | -0.416 | -0.014 | 0.021 | 0.021 | 0 | 0.17 | 0.13 | 0.18 | 0.09 | 0.19 | 0.24 |
| 1 | 38199716 | rs659573 | 0.901 | 1.454 | -1.013 | -0.007 | 0.04 | -0.044 | -0.044 | 0 | 0.36 | 0.17 | 0.38 | 0.42 | 0.34 | 0.41 |
| 1 | 38201371 | rs631432 | -0.226 | 1.425 | -0.395 | -0.104 | 0.003 | -0.004 | -0.004 | 0 | 0.58 | 0.46 | 0.6 | 0.53 | 0.54 | 0.65 |
| 1 | 38206797 | rs663428 | -0.584 | 2.127 | 0.289 | -0.142 | -0.006 | -0.008 | -0.008 | 0 | 0.71 | 0.65 | 0.86 | 0.71 | 0.8 | 0.83 |
| 1 | 38207825 | rs483201 | 1.428 | 2.093 | 0.372 | -0.144 | 0.028 | -0.023 | -0.023 | 0 | 0.37 | 0.24 | 0.39 | 0.44 | 0.34 | 0.42 |
| 1 | 38222805 | rs508992 | 1.314 | 2.109 | 0.371 | -0.161 | 0.006 | -0.006 | -0.006 | 8.347 | 0.39 | 0.33 | 0.4 | 0.42 | 0.34 | 0.41 |
| 1 | 38224635 | rs565645 | 0.564 | 2.209 | 0.288 | 0.247 | 0 | 0 | 0 | 0 | 0.48 | 0.49 | 0.62 | 0.42 | 0.54 | 0.57 |
| 1 | 38231069 | rs10157024 | -0.161 | 1.747 | 0.263 | 0.254 | 0.035 | 0.274 | 0.274 | 6.201 | 0.05 | 0.35 | 0.04 | 0.28 | 0.01 | 0.03 |
| 1 | 38243785 | rs489408 | -0.401 | 2.035 | -0.024 | 0.094 | -0.102 | 0.114 | 0.114 | 0 | 0.38 | 0.12 | 0.38 | 0.2 | 0.47 | 0.32 |
| 1 | 38247673 | rs12118489 | -0.062 | 2.074 | -0.045 | 0.065 | -0.087 | -0.034 | -0.034 | 0 | 0.16 | 0.04 | 0.29 | 0.2 | 0.31 | 0.19 |
| 1 | 38247738 | rs4308943 | -0.965 | 1.869 | 0.024 | 0.063 | -0.025 | -0.067 | -0.067 | 0 | 0.38 | 0.18 | 0.51 | 0.49 | 0.56 | 0.47 |
| 1 | 38254611 | rs664939 | 0.352 | 2.059 | 0.124 | -0.248 | 0.003 | 0.018 | 0.018 | 0 | 0.26 | 0.16 | 0.13 | 0 | 0.23 | 0.16 |
| 1 | 38258644 | rs12563037 | 0.508 | 2.06 | -0.023 | -0.039 | -0.071 | -0.079 | -0.079 | 0.001 | 0.21 | 0.04 | 0.21 | 0.28 | 0.28 | 0.28 |
| 1 | 38260503 | rs2306627 | -0.184 | 2.122 | 0.05 | -0.039 | 0.013 | 0.001 | 0.001 | 0.207 | 0.17 | 0.23 | 0.3 | 0.21 | 0.28 | 0.19 |
| 1 | 38268397 | rs3845498 | -1.138 | 2.108 | 0.072 | -0.016 | -0.012 | -0.04 | -0.04 | 1.11 | 0.38 | 0.26 | 0.51 | 0.48 | 0.56 | 0.47 |
| 1 | 38275802 | rs473279 | -2.007 | 2.172 | 0.289 | -0.017 | -0.013 | -0.007 | -0.007 | 0 | 0.65 | 0.68 | 0.62 | 0.75 | 0.71 | 0.63 |
| 1 | 38279987 | rs3748682 | 0.707 | 2.128 | 0.139 | -0.013 | -0.03 | -0.055 | -0.055 | 1.924 | 0.19 | 0.06 | 0.15 | 0.26 | 0.24 | 0.28 |
| 1 | 38289383 | rs12751325 | 0.669 | 1.664 | 0.194 | 0.031 | -0.051 | -0.049 | -0.049 | 0 | 0.21 | 0.06 | 0.21 | 0.26 | 0.28 | 0.28 |
| 1 | 38309322 | rs12745294 | -1.607 | 1.61 | 0.189 | 0.02 | 0.048 | -0.029 | -0.029 | 1.165 | 0.61 | 0.3 | 0.6 | 0.48 | 0.72 | 0.62 |
| 1 | 38318929 | rs12743834 | -1.061 | 1.636 | 0.263 | 0.02 | -0.006 | -0.027 | -0.027 | 0 | 0.38 | 0.28 | 0.51 | 0.48 | 0.57 | 0.47 |
| 1 | 38338795 | rs11488569 | -1.584 | 1.572 | 0.472 | 0.012 | 0.044 | -0.027 | -0.027 | 0 | 0.61 | 0.3 | 0.6 | 0.48 | 0.73 | 0.62 |
| 1 | 38345896 | rs2306427 | -1.592 | 1.574 | 0.486 | 0.013 | 0.043 | -0.028 | -0.028 | 0 | 0.61 | 0.3 | 0.6 | 0.48 | 0.72 | 0.62 |

| | | | | | | | | | | | | | | | | |
|---|---|---|---|---|---|---|---|---|---|---|---|---|---|---|---|---|
| 1 | 38347417 | rs12742756 | -0.626 | 1.626 | 0.348 | 0.002 | -0.062 | 0.068 | 0.068 | 0.555 | 0.44 | 0.08 | 0.3 | 0.26 | 0.44 | 0.43 |
| 1 | 38349400 | rs12728438 | 0.65 | 1.597 | 0.371 | 0.025 | -0.051 | -0.054 | -0.054 | 0 | 0.21 | 0.06 | 0.21 | 0.26 | 0.28 | 0.29 |
| 1 | 38349606 | rs9329570 | -1.981 | 1.366 | 0.54 | -0.127 | -0.031 | -0.013 | -0.013 | 1.751 | 0.65 | 0.6 | 0.63 | 0.75 | 0.74 | 0.64 |
| 1 | 38366907 | rs28739509 | 0.726 | 1.136 | -0.018 | -0.002 | -0.049 | -0.067 | -0.067 | 0.039 | 0.2 | 0.07 | 0.2 | 0.28 | 0.25 | 0.29 |
| 1 | 38380285 | rs12733578 | -1.36 | -0.376 | 0.253 | 0.007 | 0.045 | -0.027 | -0.027 | 0.253 | 0.6 | 0.3 | 0.59 | 0.5 | 0.72 | 0.61 |
| 1 | 38388708 | rs12723117 | -1.33 | -0.415 | 0.161 | 0.016 | 0.06 | -0.042 | -0.042 | 0 | 0.6 | 0.25 | 0.59 | 0.48 | 0.72 | 0.61 |
| 1 | 38398588 | rs17465420 | -0.443 | -0.189 | -0.026 | 0.077 | -0.002 | 0.031 | 0.031 | 0 | 0.46 | 0.09 | 0.44 | 0.32 | 0.58 | 0.47 |
| 1 | 38401933 | rs12739026 | 0.114 | -0.173 | -0.009 | 0.084 | -0.019 | -0.027 | -0.027 | 2.374 | 0.1 | 0.06 | 0.13 | 0.18 | 0.13 | 0.14 |
| 1 | 38407014 | rs28435150 | -0.462 | -0.235 | -0.214 | 0.084 | 0.001 | 0.034 | 0.034 | 0 | 0.46 | 0.09 | 0.44 | 0.32 | 0.58 | 0.47 |
| 1 | 38411350 | rs11485595 | 0.716 | -0.133 | -0.202 | 0.104 | -0.124 | -0.075 | -0.075 | 0.054 | 0.24 | 0.07 | 0.35 | 0.32 | 0.43 | 0.33 |
| 1 | 38415315 | rs16824518 | -1.724 | -0.181 | 0.034 | 0.051 | -0.024 | -0.008 | -0.008 | 0.146 | 0.63 | 0.63 | 0.62 | 0.76 | 0.73 | 0.63 |
| 1 | 38421508 | rs1383243 | -1.005 | 0.063 | -0.348 | 0.16 | 0.01 | 0.158 | 0.158 | 0 | 0.77 | 0.98 | 0.91 | 1 | 0.85 | 0.86 |
| 1 | 38423588 | rs1050257 | -1.005 | 0.063 | -0.348 | 0.16 | 0.01 | 0.158 | 0.158 | 0 | 0.77 | 0.98 | 0.91 | 1 | 0.85 | 0.86 |
| 1 | 38442547 | rs10890216 | -0.999 | 0.07 | -0.345 | 0.163 | 0.011 | 0.159 | 0.159 | 0 | 0.77 | 0.97 | 0.91 | 1 | 0.85 | 0.85 |
| 1 | 38449910 | rs11210870 | 1.416 | 0.135 | -0.159 | 0.249 | 0.018 | 0.006 | 0.006 | 0 | 0.36 | 0.25 | 0.37 | 0.24 | 0.28 | 0.37 |
| 1 | 38452581 | rs12138115 | 0.447 | 0.378 | -0.207 | -0.343 | 0.016 | 0.021 | 0.021 | 0 | 0.49 | 0.4 | 0.51 | 0.68 | 0.42 | 0.52 |
| 1 | 38453041 | rs12044531 | 0.478 | 0.381 | -0.375 | -0.335 | -0.116 | -0.067 | -0.067 | 0 | 0.24 | 0.08 | 0.35 | 0.31 | 0.42 | 0.33 |
| 1 | 38453576 | rs10890285 | 0.54 | 0.223 | 0.06 | 0.297 | 0.007 | -0.007 | -0.007 | 0 | 0.39 | 0.31 | 0.38 | 0.5 | 0.29 | 0.39 |
| 1 | 38456106 | rs4072980 | 0.115 | 0.444 | -0.742 | 0.026 | 0.029 | 0.037 | 0.037 | 0 | 0.49 | 0.57 | 0.51 | 0.68 | 0.42 | 0.52 |
| 1 | 38463504 | rs7366048 | 0.207 | 0.557 | -0.979 | 0.078 | 0.047 | 0.058 | 0.058 | 0 | 0.5 | 0.69 | 0.52 | 0.68 | 0.42 | 0.52 |
| 1 | 38465315 | rs4634868 | -0.62 | 0.057 | -0.612 | -1.301 | -0.031 | -0.084 | -0.084 | 1.903 | 0.33 | 0.44 | 0.24 | 0.12 | 0.26 | 0.26 |
| 1 | 38469500 | rs6600436 | -0.54 | 0.464 | -0.886 | -1.543 | 0.002 | 0.058 | 0.058 | 0 | 0.48 | 0.64 | 0.64 | 0.81 | 0.49 | 0.55 |
| 1 | 38482356 | rs11211383 | -0.77 | 0.882 | -1.253 | -1.865 | -0.049 | 0.139 | 0.139 | 0 | 0.27 | 0.08 | 0.2 | 0.06 | 0.28 | 0.29 |
| 1 | 38488170 | rs35867479 | -0.515 | 0.312 | -1.044 | -1.62 | 0.021 | 0.079 | 0.079 | 0.03 | 0.2 | 0.07 | 0.07 | 0.06 | 0.12 | 0.15 |
| 1 | 38492904 | rs12745452 | 0.066 | 0.296 | -1.063 | -1.655 | 0.021 | 0.041 | 0.041 | 0.439 | 0.15 | 0.07 | 0.05 | 0.07 | 0.08 | 0.13 |
| 1 | 38499643 | rs12743979 | -0.715 | 0.207 | -0.958 | -1.667 | 0.022 | 0.056 | 0.056 | 0.037 | 0.2 | 0.09 | 0.07 | 0.06 | 0.12 | 0.15 |
| 1 | 38501187 | rs1037976 | -1.457 | 0.101 | -0.5 | -1.821 | -0.004 | -0.015 | -0.015 | 0.796 | 0.29 | 0.36 | 0.15 | 0.15 | 0.21 | 0.25 |
| 1 | 38507495 | rs569791 | -1.713 | -0.15 | -0.89 | -2.208 | -0.11 | 0.287 | 0.287 | 0.117 | 0.56 | 0.97 | 0.63 | 0.95 | 0.47 | 0.57 |
| 1 | 38517981 | rs11205722 | -1.123 | 0.02 | -0.76 | -2.123 | 0.004 | 0.006 | 0.006 | 0.022 | 0.28 | 0.31 | 0.15 | 0.15 | 0.21 | 0.25 |
| 1 | 38527331 | rs12723955 | -0.8 | -0.088 | -0.587 | -2.037 | -0.017 | -0.023 | -0.023 | 0 | 0.25 | 0.36 | 0.13 | 0.16 | 0.16 | 0.23 |
| 1 | 38529500 | rs11205858 | 0.137 | -0.198 | -0.047 | -1.95 | 0.018 | -0.023 | -0.023 | 0.425 | 0.26 | 0.37 | 0.48 | 0.79 | 0.3 | 0.35 |
| 1 | 38531745 | rs12750006 | -0.61 | -0.469 | 1.034 | -1.184 | -0.013 | -0.033 | -0.033 | 0.053 | 0.19 | 0.28 | 0.1 | 0.08 | 0.13 | 0.16 |
| 1 | 38533001 | rs12733163 | -1.305 | -0.467 | 1.034 | -1.185 | -0.028 | -0.041 | -0.041 | 2.002 | 0.25 | 0.41 | 0.14 | 0.15 | 0.17 | 0.23 |
| 1 | 38539961 | rs560654 | -1.515 | -0.31 | 0.647 | -1.06 | -0.056 | -0.081 | -0.081 | 0.078 | 0.3 | 0.51 | 0.16 | 0.15 | 0.17 | 0.24 |
| 1 | 38541256 | rs12566302 | -0.213 | -0.323 | 0.718 | -1.068 | 0.007 | -0.001 | -0.001 | 0.977 | 0.2 | 0.21 | 0.07 | 0.07 | 0.09 | 0.14 |
| 1 | 38541428 | rs12074522 | -0.579 | -0.28 | 0.341 | -1.044 | -0.002 | 0.013 | 0.013 | 0.137 | 0.07 | 0.19 | 0.04 | 0.07 | 0.04 | 0.08 |
| 1 | 38550105 | rs571881 | -0.747 | -0.059 | -0.604 | -0.909 | -0.019 | -0.029 | -0.029 | 0.041 | 0.31 | 0.64 | 0.16 | 0.15 | 0.17 | 0.24 |
| 1 | 38559049 | rs486563 | 0.64 | 1.108 | -0.598 | -0.536 | 0.003 | -0.015 | -0.015 | 0 | 0.88 | 0.87 | 0.89 | 0.95 | 0.85 | 0.86 |
| 1 | 38559352 | rs1108182 | -1.012 | 1.353 | -0.463 | -0.761 | -0.001 | -0.031 | -0.031 | 5.785 | 0.37 | 0.52 | 0.49 | 0.83 | 0.4 | 0.56 |
| 1 | 38559533 | rs544700 | 1.156 | 1.854 | -0.794 | 0.46 | -0.046 | -0.055 | -0.055 | 0 | 0.78 | 0.86 | 0.63 | 0.3 | 0.72 | 0.77 |
| 1 | 38560968 | rs523864 | 0.493 | 0.975 | -0.413 | 0.393 | 0.003 | 0.01 | 0.01 | 0 | 0.79 | 0.85 | 0.86 | 0.93 | 0.8 | 0.9 |
| 1 | 38563923 | rs578107 | 1.246 | 1.455 | 0.523 | 1.348 | 0.012 | 0.016 | 0.016 | 0 | 0.7 | 0.5 | 0.61 | 0.39 | 0.67 | 0.66 |
| 1 | 38568328 | rs11206585 | -1.581 | 1.242 | 0.995 | 1.584 | 0.003 | -0.003 | -0.003 | 0.598 | 0.16 | 0.31 | 0.18 | 0.15 | 0.17 | 0.13 |
| 1 | 38570853 | rs2615811 | 1.729 | 1.242 | 0.633 | 1.522 | 0.006 | 0.056 | 0.056 | 0 | 0.55 | 0.25 | 0.52 | 0.3 | 0.57 | 0.42 |
| 1 | 38591115 | rs12404747 | -1.386 | 0.994 | 0.569 | 2.374 | -0.001 | 0.071 | 0.071 | 0.185 | 0.09 | 0.1 | 0.25 | 0.52 | 0.12 | 0.29 |
| 1 | 38595864 | rs6662282 | -1.983 | 1.186 | 0.713 | 1.418 | 0.002 | 0.084 | 0.084 | 0 | 0.42 | 0.74 | 0.47 | 0.82 | 0.42 | 0.58 |
| 1 | 38596210 | rs6665139 | -0.897 | 0.681 | 0.863 | 1.05 | 0 | -0.002 | -0.002 | 0 | 0.27 | 0.25 | 0.16 | 0.12 | 0.23 | 0.17 |
| 1 | 38601480 | rs6600309 | -1.423 | 0.788 | 1.121 | 1.268 | 0.018 | 0.094 | 0.094 | 0 | 0.05 | 0.25 | 0.02 | 0.09 | 0.02 | 0.09 |
| 1 | 38609325 | rs4593783 | -0.988 | 1.116 | 0.591 | 0.452 | -0.017 | 0.034 | 0.034 | 0 | 0.32 | 0.21 | 0.26 | 0.06 | 0.39 | 0.38 |
| 1 | 38616871 | rs883220 | -0.632 | 1.401 | 0.823 | 0.378 | 0.001 | 0.001 | 0.001 | 0 | 0.23 | 0.16 | 0.13 | 0.01 | 0.22 | 0.22 |
| 1 | 38623285 | rs12083795 | -0.213 | 1.068 | 1.375 | -0.077 | -0.008 | -0.045 | -0.045 | 0 | 0.1 | 0.2 | 0.07 | 0 | 0.06 | 0.02 |
| 1 | 38630862 | rs6693777 | -0.26 | 1.02 | 1.504 | -0.102 | -0.01 | -0.054 | -0.054 | 0 | 0.1 | 0.21 | 0.07 | 0 | 0.06 | 0.02 |
| 1 | 38636296 | rs10889314 | 0.722 | 0.455 | 2.164 | 1.005 | -0.017 | -0.077 | -0.077 | 0 | 0.84 | 0.58 | 0.88 | 0.97 | 0.88 | 0.95 |
| 1 | 38636712 | rs12138920 | 2.241 | 0.587 | 1.647 | 1.044 | -0.042 | 0.137 | 0.137 | 0 | 0.43 | 0.1 | 0.28 | 0.2 | 0.41 | 0.33 |
| 1 | 38639208 | rs7544568 | 0.098 | 0.65 | 1.807 | 1.175 | -0.003 | 0.032 | 0.032 | 0 | 0.22 | 0.11 | 0.13 | 0.01 | 0.22 | 0.22 |
| 1 | 38644359 | rs6600333 | -0.43 | 0.644 | 2.094 | 1.52 | 0.019 | 0.195 | 0.195 | 0 | 0.93 | 0.82 | 0.93 | 0.77 | 0.93 | 0.93 |
| 1 | 38645708 | rs898982 | -0.16 | 0.641 | 2.132 | 1.518 | 0.021 | 0.209 | 0.209 | 0 | 0.93 | 0.76 | 0.93 | 0.77 | 0.93 | 0.93 |
| 1 | 38650917 | rs17534243 | 0.055 | 0.748 | 1.954 | 1.076 | -0.008 | 0.01 | 0.01 | 0 | 0.19 | 0.12 | 0.11 | 0.01 | 0.21 | 0.21 |
| 1 | 38651444 | rs6666216 | 0.279 | 0.683 | 2.058 | 1.101 | -0.004 | -0.053 | -0.053 | 0 | 0.09 | 0.2 | 0.06 | 0 | 0.06 | 0.01 |
| 1 | 38656716 | rs11208403 | 0.159 | 0.662 | 2.048 | 1.091 | -0.004 | -0.057 | -0.057 | 0 | 0.1 | 0.26 | 0.07 | 0 | 0.06 | 0.01 |
| 1 | 38657551 | rs748229 | -1.421 | 0.739 | 2.169 | 1.343 | -0.015 | 0.039 | 0.039 | 0 | 0.41 | 0.71 | 0.55 | 0.75 | 0.41 | 0.58 |
| 1 | 38659743 | rs7543360 | -1.032 | 0.704 | 2.233 | 1.369 | 0.005 | 0.011 | 0.011 | 0 | 0.36 | 0.75 | 0.42 | 0.71 | 0.28 | 0.49 |
| 1 | 38660593 | rs1993804 | -0.854 | 0.715 | 2.444 | 1.843 | -0.002 | 0 | 0 | 0 | 0.33 | 0.6 | 0.38 | 0.35 | 0.27 | 0.45 |
| 1 | 38666914 | rs898979 | 0.45 | 0.789 | 1.891 | 1.938 | 0.034 | -0.026 | -0.026 | 1.272 | 0.92 | 0.85 | 0.85 | 0.96 | 0.86 | 0.9 |
| 1 | 38667704 | rs898978 | 1.803 | 0.682 | 1.668 | 1.779 | -0.026 | 0.177 | 0.177 | 0.139 | 0.5 | 0.13 | 0.39 | 0.22 | 0.49 | 0.39 |
| 1 | 38668093 | rs898977 | 0.923 | 0.688 | 1.968 | 1.706 | 0.022 | -0.041 | -0.041 | 0.999 | 0.84 | 0.74 | 0.79 | 0.93 | 0.76 | 0.86 |
| 1 | 38672209 | rs12075354 | 0.686 | 0.635 | 1.408 | 1.456 | -0.001 | 0 | 0 | 0 | 0.66 | 0.35 | 0.6 | 0.29 | 0.72 | 0.53 |
| 1 | 38675803 | rs745526 | -0.79 | 0.429 | 0.54 | 0.896 | -0.037 | 0.002 | 0.002 | 0 | 0.1 | 0.01 | 0.06 | 0.04 | 0.12 | 0.12 |
| 1 | 38681026 | rs6683126 | -1.11 | 0.332 | 1.242 | 0.847 | 0.005 | -0.087 | -0.087 | 0 | 0.3 | 0.41 | 0.24 | 0.06 | 0.26 | 0.29 |

| | | | | | | | | | | | | | | | | |
|---|---|---|---|---|---|---|---|---|---|---|---|---|---|---|---|---|
| 1 | 38682207 | rs7537377 | -0.832 | 0.24 | 1.015 | 1.197 | -0.011 | -0.001 | -0.001 | 0 | 0.16 | 0.15 | 0.08 | 0.12 | 0.13 | 0.07 |
| 1 | 38685317 | rs6600353 | 0.115 | 0.289 | 0.988 | 1.027 | 0 | -0.004 | -0.004 | 0 | 0.81 | 0.83 | 0.9 | 0.91 | 0.82 | 0.83 |
| 1 | 38686908 | rs6675079 | -0.853 | 0.203 | 1.34 | 1.857 | -0.024 | 0.056 | 0.056 | 0 | 0.15 | 0.28 | 0.26 | 0.34 | 0.13 | 0.18 |
| 1 | 38687060 | rs6685558 | -1.447 | 0.094 | 1.446 | 1.848 | -0.036 | 0.061 | 0.061 | 0 | 0.17 | 0.34 | 0.26 | 0.35 | 0.13 | 0.18 |
| 1 | 38694395 | rs1807832 | -0.911 | 0.182 | 1.594 | 2.166 | -0.029 | 0.158 | 0.158 | 0 | 0.15 | 0.32 | 0.28 | 0.47 | 0.12 | 0.29 |
| 1 | 38695702 | rs16824807 | -1.435 | 0.159 | 1.634 | 2.16 | -0.035 | 0.181 | 0.181 | 0 | 0.18 | 0.44 | 0.29 | 0.47 | 0.14 | 0.29 |
| 1 | 38697702 | rs12409212 | -1.427 | 0.161 | 1.635 | 2.161 | -0.035 | 0.182 | 0.182 | 0 | 0.18 | 0.44 | 0.29 | 0.47 | 0.14 | 0.28 |
| 1 | 38705915 | rs872595 | -1.478 | 0.566 | 1.597 | 1.917 | 0.042 | 0.148 | 0.148 | 0 | 0.06 | 0.29 | 0.13 | 0.17 | 0.05 | 0.25 |
| 1 | 38715252 | rs6702480 | -1.203 | 1.199 | 1.097 | 1.985 | 0.037 | 0.06 | 0.06 | 0 | 0.29 | 0.42 | 0.4 | 0.59 | 0.38 | 0.36 |
| 1 | 38715353 | rs6600367 | -1.145 | 1.193 | 1.121 | 1.982 | 0.02 | 0.032 | 0.032 | 0 | 0.28 | 0.34 | 0.39 | 0.59 | 0.38 | 0.36 |
| 1 | 38721479 | rs12732447 | -1.163 | 1.174 | 1.09 | 1.98 | 0.021 | 0.034 | 0.034 | 0 | 0.28 | 0.34 | 0.39 | 0.59 | 0.38 | 0.36 |
| 1 | 38721663 | rs11802770 | -2.154 | 0.808 | 0.138 | 1.763 | 0.001 | 0.086 | 0.086 | 0.004 | 0.49 | 0.67 | 0.5 | 0.7 | 0.52 | 0.58 |
| 1 | 38723544 | rs10889902 | -0.788 | 1.052 | -0.382 | 2.294 | -0.04 | -0.034 | -0.034 | 0 | 0.09 | 0.01 | 0.25 | 0.43 | 0.16 | 0.24 |
| 1 | 38726435 | rs12022969 | -1.05 | 0.582 | -0.083 | 1.434 | 0.006 | 0.006 | 0.006 | 0.016 | 0.2 | 0.19 | 0.09 | 0.11 | 0.14 | 0.22 |
| 1 | 38729507 | rs12031152 | -0.95 | 0.635 | -0.322 | 1.441 | 0.01 | 0.016 | 0.016 | 0 | 0.2 | 0.17 | 0.09 | 0.11 | 0.15 | 0.22 |
| 1 | 38731622 | rs12028799 | -1.824 | 1.254 | 0.08 | 2.207 | 0.014 | 0.117 | 0.117 | 0 | 0.1 | 0.19 | 0.13 | 0.4 | 0.12 | 0.28 |
| 1 | 38733788 | rs2480697 | -0.897 | 1.222 | -0.605 | 1.258 | 0.036 | 0.02 | 0.02 | 0 | 0.49 | 0.76 | 0.53 | 0.56 | 0.55 | 0.45 |
| 1 | 38734768 | rs10789332 | -0.729 | 1.437 | -0.03 | 1.017 | 0.006 | 0.003 | 0.003 | 0.041 | 0.41 | 0.54 | 0.49 | 0.52 | 0.48 | 0.45 |
| 1 | 38736874 | rs6698091 | -1.096 | 1.652 | 0.031 | 0.987 | 0.02 | 0.007 | 0.007 | 0.158 | 0.37 | 0.4 | 0.58 | 0.38 | 0.51 | 0.32 |
| 1 | 38738214 | rs12075234 | -0.613 | 1.6 | 0.404 | 0.919 | -0.011 | 0.002 | 0.002 | 0.547 | 0.36 | 0.31 | 0.54 | 0.67 | 0.42 | 0.58 |
| 1 | 38747960 | rs1543632 | -1.769 | 1.337 | 0.126 | 0.568 | -0.09 | 0.272 | 0.272 | 0.303 | 0.62 | 0.95 | 0.76 | 0.98 | 0.57 | 0.76 |
| 1 | 38749134 | rs16824833 | -0.535 | 1.594 | 1.368 | 1.54 | 0.041 | 0.07 | 0.07 | 0.32 | 0.08 | 0.23 | 0.18 | 0.1 | 0.13 | 0.05 |
| 1 | 38750042 | rs12565767 | -0.46 | 1.731 | 1.513 | 1.727 | 0.054 | 0.122 | 0.122 | 0 | 0.1 | 0.18 | 0.29 | 0.29 | 0.17 | 0.19 |
| 1 | 38750228 | rs6600376 | -1.458 | 1.698 | 1.292 | 1.645 | 0.108 | 0.23 | 0.23 | 0.038 | 0.18 | 0.4 | 0.47 | 0.39 | 0.29 | 0.24 |
| 1 | 38752476 | rs1329967 | -1.466 | 1.692 | 0.757 | 1.584 | 0.033 | 0.051 | 0.051 | 0 | 0.18 | 0.26 | 0.46 | 0.6 | 0.29 | 0.32 |
| 1 | 38752491 | rs11586720 | 0.118 | 1.687 | 0.321 | 0.872 | 0.005 | 0.011 | 0.011 | 0 | 0.06 | 0.08 | 0.16 | 0.1 | 0.13 | 0.05 |
| 1 | 38763942 | rs12083721 | -1.356 | 1.479 | 0.461 | 0.665 | 0 | 0.003 | 0.003 | 0 | 0.23 | 0.29 | 0.39 | 0.28 | 0.22 | 0.25 |
| 1 | 38764346 | rs12091117 | -0.949 | 1.394 | 0.431 | 0.687 | 0.001 | 0.002 | 0.002 | 0 | 0.17 | 0.14 | 0.12 | 0.14 | 0.17 | 0.11 |
| 1 | 38768561 | rs1034091 | -0.852 | 1.158 | 1.229 | 0.544 | -0.046 | -0.102 | -0.102 | 1.788 | 0.14 | 0.38 | 0.05 | 0.01 | 0.04 | 0.07 |
| 1 | 38769077 | rs1778040 | 1.273 | 1.06 | 0.82 | 0.084 | 0.05 | 0.083 | 0.083 | 0 | 0.51 | 0.28 | 0.56 | 0.27 | 0.61 | 0.45 |
| 1 | 38769144 | rs12748758 | 0.433 | 1.151 | 0.166 | 0.22 | -0.017 | -0.032 | -0.032 | 0 | 0.07 | 0.03 | 0.08 | 0.14 | 0.11 | 0.05 |
| 1 | 38769523 | rs2149935 | 0.956 | 1.634 | -0.094 | -0.172 | -0.05 | 0.048 | 0.048 | 0 | 0.1 | 0.02 | 0.09 | 0 | 0.21 | 0.06 |
| 1 | 38772343 | rs11210514 | 0.153 | 0.885 | 0.284 | 0.208 | 0.012 | 0.001 | 0.001 | 0 | 0.12 | 0.1 | 0.03 | 0.01 | 0.04 | 0.07 |
| 1 | 38774882 | rs1778037 | -0.246 | 1.032 | 0.464 | 0.378 | 0.003 | -0.017 | -0.017 | 0 | 0.73 | 0.77 | 0.69 | 0.42 | 0.76 | 0.58 |
| 1 | 38782783 | rs1778048 | -0.38 | 1.047 | 0.283 | 0.385 | 0.002 | -0.066 | -0.066 | 0 | 0.74 | 0.85 | 0.7 | 0.41 | 0.76 | 0.58 |
| 1 | 38790638 | rs1778041 | 0.104 | 1.089 | 0.26 | 0.151 | -0.002 | -0.038 | -0.038 | 0 | 0.27 | 0.18 | 0.33 | 0.6 | 0.27 | 0.42 |
| 1 | 38791901 | rs869720 | 0.37 | 0.783 | 0.903 | -0.011 | -0.017 | -0.002 | -0.002 | 3.241 | 0.63 | 0.57 | 0.66 | 0.34 | 0.72 | 0.51 |
| 1 | 38791931 | rs913864 | 1.317 | 0.931 | 1.182 | -0.14 | 0.015 | 0.166 | 0.166 | 0 | 0.49 | 0.25 | 0.47 | 0.08 | 0.54 | 0.41 |
| 1 | 38791968 | rs913863 | 0.847 | 0.944 | 1.229 | -0.035 | -0.002 | -0.005 | -0.005 | 0.176 | 0.56 | 0.42 | 0.57 | 0.25 | 0.65 | 0.47 |
| 1 | 38794068 | rs17477313 | -0.162 | 0.91 | 0.757 | 0.005 | -0.006 | -0.007 | -0.007 | 0 | 0.06 | 0.04 | 0.07 | 0.09 | 0.06 | 0.03 |
| 1 | 38796920 | rs2637848 | -0.122 | 1.05 | 1.048 | -0.08 | 0.111 | 0.066 | 0.066 | 0.321 | 0.91 | 0.57 | 0.78 | 0.93 | 0.9 | 0.87 |
| 1 | 38799191 | rs2637846 | -0.295 | 1.045 | 0.772 | -0.161 | -0.016 | 0.051 | 0.051 | 0 | 0.33 | 0.17 | 0.4 | 0.77 | 0.37 | 0.51 |
| 1 | 38801278 | rs12073718 | 0.635 | 1.043 | 0.777 | -0.151 | -0.015 | -0.005 | -0.005 | 7.493 | 0.09 | 0.08 | 0.01 | 0.06 | 0.01 | 0.07 |
| 1 | 38806630 | rs6679637 | 0.816 | 1.369 | 0.848 | 0.065 | -0.001 | -0.001 | -0.001 | 0.667 | 0.06 | 0.07 | 0.05 | 0.08 | 0.11 | 0.05 |
| 1 | 38809520 | rs2798795 | 0.418 | 1.163 | 0.687 | 0.103 | -0.014 | -0.081 | -0.081 | 4.813 | 0.25 | 0.1 | 0.32 | 0.61 | 0.26 | 0.44 |
| 1 | 38814355 | rs2637876 | 0.496 | 1.124 | 1.104 | 0.227 | 0.074 | 0.076 | 0.076 | 0.476 | 0.88 | 0.5 | 0.71 | 0.84 | 0.87 | 0.85 |
| 1 | 38829224 | rs2040073 | -1.546 | 1.443 | 0.531 | 0.085 | -0.02 | -0.008 | -0.008 | 0 | 0.27 | 0.73 | 0.41 | 0.31 | 0.39 | 0.26 |
| 1 | 38835477 | rs12726098 | 0.487 | 1.525 | 0.314 | 0.169 | -0.001 | -0.001 | -0.001 | 0 | 0.06 | 0.06 | 0.07 | 0.08 | 0.14 | 0.05 |
| 1 | 38836186 | rs2798798 | 0.199 | 1.247 | -0.109 | 0.033 | 0.016 | -0.1 | -0.1 | 0 | 0.26 | 0.09 | 0.36 | 0.6 | 0.23 | 0.43 |
| 1 | 38843016 | rs4278342 | -0.627 | 1.79 | 0.613 | 0.252 | 0.05 | 0.022 | 0.022 | 0 | 0.89 | 0.79 | 0.81 | 0.86 | 0.81 | 0.82 |
| 1 | 38845391 | rs17537940 | 0.014 | 1.747 | 0.237 | 0.294 | -0.007 | -0.006 | -0.006 | 1.019 | 0.06 | 0.05 | 0.09 | 0.1 | 0.07 | 0.02 |
| 1 | 38848996 | rs11210654 | 1.577 | 1.411 | 0.115 | 0.092 | -0.02 | -0.009 | -0.009 | 0 | 0.6 | 0.25 | 0.52 | 0.57 | 0.48 | 0.61 |
| 1 | 38850304 | rs4284229 | 0.03 | 0.771 | -0.096 | -0.3 | 0 | -0.004 | -0.004 | 0.689 | 0.24 | 0.19 | 0.31 | 0.56 | 0.15 | 0.38 |
| 1 | 38850339 | rs4266846 | -0.296 | 0.836 | 0.019 | -0.304 | 0.116 | 0.042 | 0.042 | 0 | 0.82 | 0.64 | 0.73 | 0.78 | 0.68 | 0.77 |
| 1 | 38853024 | rs7521522 | 0.899 | 0.876 | -0.44 | -0.28 | 0.002 | 0.053 | 0.053 | 0 | 0.14 | 0.08 | 0.08 | 0.03 | 0.12 | 0.12 |
| 1 | 38858428 | rs12061862 | 0.101 | 0.903 | -0.447 | -0.209 | -0.002 | -0.017 | -0.017 | 0.014 | 0.21 | 0.33 | 0.17 | 0.13 | 0.19 | 0.14 |
| 1 | 38858540 | rs12740005 | 1.743 | 0.491 | -1.256 | -0.007 | -0.018 | 0.154 | 0.154 | 0 | 0.37 | 0.12 | 0.25 | 0.09 | 0.34 | 0.24 |
| 1 | 38865601 | rs6690257 | 0.801 | 1.125 | -0.789 | 0.253 | -0.023 | 0.024 | 0.024 | 0.287 | 0.6 | 0.23 | 0.55 | 0.65 | 0.48 | 0.62 |
| 1 | 38876635 | rs12132779 | 0.919 | 1.217 | -0.758 | 0.826 | 0.004 | 0.047 | 0.047 | 0 | 0.57 | 0.27 | 0.35 | 0.31 | 0.6 | 0.37 |
| 1 | 38888749 | rs7540859 | -0.358 | 0.857 | -0.898 | 1.325 | 0.015 | -0.073 | -0.073 | 0.347 | 0.8 | 0.93 | 0.81 | 0.65 | 0.84 | 0.65 |
| 1 | 38890659 | rs10890223 | -0.241 | 0.841 | -0.797 | 1.317 | 0.02 | -0.056 | -0.056 | 0 | 0.79 | 0.89 | 0.81 | 0.65 | 0.84 | 0.65 |
| 1 | 38900680 | rs4970591 | -1.018 | 1.451 | 0.015 | 0.998 | 0.101 | 0.04 | 0.04 | 0 | 0.26 | 0.49 | 0.51 | 0.33 | 0.42 | 0.34 |
| 1 | 38904931 | rs11586142 | 1.652 | 1.011 | -0.341 | 0.649 | 0.02 | 0.097 | 0.097 | 0.025 | 0.43 | 0.27 | 0.28 | 0.12 | 0.38 | 0.25 |
| 1 | 38906637 | rs12567295 | -0.041 | 1.297 | -0.389 | 0.756 | -0.012 | -0.007 | -0.007 | 3.605 | 0.18 | 0.16 | 0.42 | 0.25 | 0.31 | 0.29 |
| 1 | 38908461 | rs2256357 | 0.223 | 0.38 | 0.092 | 1.208 | 0.044 | -0.06 | -0.06 | 0 | 0.7 | 0.88 | 0.79 | 0.45 | 0.81 | 0.58 |
| 1 | 38910848 | rs1932362 | -1.259 | 1.133 | 0.688 | 1.073 | 0.066 | 0.044 | 0.044 | 0.01 | 0.07 | 0.32 | 0.08 | 0.07 | 0.1 | 0.05 |
| 1 | 38915251 | rs1006812 | -1.11 | 1.165 | 0.39 | 1.396 | 0.027 | 0.055 | 0.055 | 0 | 0.1 | 0.2 | 0.13 | 0.22 | 0.15 | 0.27 |
| 1 | 38916976 | rs11210795 | -2.014 | 0.54 | 0.879 | 1.627 | -0.014 | 0.089 | 0.089 | 0 | 0.29 | 0.54 | 0.27 | 0.52 | 0.23 | 0.34 |
| 1 | 38917810 | rs882000 | -1.153 | 0.647 | 0.48 | 1.121 | -0.007 | 0.016 | 0.016 | 0 | 0.54 | 0.61 | 0.44 | 0.57 | 0.53 | 0.48 |

| | | | | | | | | | | | | | | | | |
|---|---|---|---|---|---|---|---|---|---|---|---|---|---|---|---|---|
| 1 | 38919423 | rs2252153 | -0.063 | 0.249 | 0.345 | 1.029 | 0.018 | 0.017 | 0.017 | 0 | 0.35 | 0.19 | 0.24 | 0.23 | 0.32 | 0.17 |
| 1 | 38920889 | rs16825062 | -0.175 | 0.274 | 0.456 | 1.046 | -0.001 | -0.001 | -0.001 | 0 | 0.08 | 0.1 | 0.06 | 0.09 | 0.07 | 0.05 |
| 1 | 38922053 | rs2251803 | -0.461 | 0.399 | 0.623 | 1.179 | 0.014 | 0.007 | 0.007 | 0 | 0.44 | 0.34 | 0.3 | 0.33 | 0.39 | 0.24 |
| 1 | 38928492 | rs2248323 | -0.458 | 0.377 | 0.556 | 0.921 | -0.012 | -0.059 | -0.059 | 0.274 | 0.12 | 0.17 | 0.05 | 0 | 0.07 | 0.17 |
| 1 | 38929864 | rs2494441 | -0.872 | 0.506 | 0.448 | 0.867 | 0.002 | 0.036 | 0.036 | 0 | 0.56 | 0.58 | 0.34 | 0.21 | 0.43 | 0.36 |
| 1 | 38931261 | rs2247973 | -0.99 | -0.083 | 0.631 | 0.983 | 0.035 | 0.03 | 0.03 | 0 | 0.53 | 0.6 | 0.33 | 0.3 | 0.33 | 0.36 |
| 1 | 38931492 | rs2247965 | -1.139 | 0.373 | 0.998 | 1.114 | -0.013 | -0.07 | -0.07 | 0 | 0.13 | 0.26 | 0.05 | 0 | 0.07 | 0.13 |
| 1 | 38932405 | rs12564999 | -0.944 | 0.615 | 0.706 | 1.736 | 0.031 | 0.173 | 0.173 | 0 | 0.06 | 0.19 | 0.07 | 0.25 | 0.07 | 0.22 |
| 1 | 38936977 | rs1329817 | -0.708 | 0.425 | 0.359 | 1.548 | -0.014 | 0.023 | 0.023 | 0 | 0.36 | 0.47 | 0.25 | 0.4 | 0.24 | 0.43 |
| 1 | 38940464 | rs2244275 | -0.844 | 0.35 | -0.443 | 1.263 | 0.008 | -0.01 | -0.01 | 0.285 | 0.38 | 0.62 | 0.29 | 0.41 | 0.25 | 0.44 |
| 1 | 38954125 | rs2992434 | 0.807 | 0.447 | -0.513 | 1.415 | -0.007 | -0.017 | -0.017 | 0 | 0.73 | 0.82 | 0.73 | 0.68 | 0.68 | 0.68 |
| 1 | 38957463 | rs3011205 | 0.708 | 0.435 | 0.206 | 1.377 | -0.023 | -0.046 | -0.046 | 0 | 0.74 | 0.89 | 0.74 | 0.69 | 0.69 | 0.69 |
| 1 | 38962135 | rs1932364 | 1.012 | 0.253 | 1.02 | 1.327 | 0.003 | 0.014 | 0.014 | 0 | 0.71 | 0.64 | 0.72 | 0.68 | 0.68 | 0.69 |
| 1 | 38984339 | rs2184611 | -1.545 | -0.486 | 1.1 | 0.926 | 0.004 | 0.006 | 0.006 | 0 | 0.31 | 0.72 | 0.26 | 0.2 | 0.25 | 0.24 |
| 1 | 38989193 | rs12062702 | -1.668 | -0.621 | 1.384 | 0.879 | -0.014 | 0.02 | 0.02 | 0.11 | 0.17 | 0.5 | 0.19 | 0.17 | 0.13 | 0.17 |
| 1 | 38994115 | rs11590421 | -0.519 | -0.367 | 1.383 | 0.842 | -0.025 | 0.072 | 0.072 | 0 | 0.17 | 0.07 | 0.09 | 0.02 | 0.19 | 0.12 |
| 1 | 38998839 | rs4970551 | -1.67 | -0.401 | 1.535 | 0.857 | -0.006 | 0.082 | 0.082 | 0 | 0.13 | 0.46 | 0.15 | 0.17 | 0.08 | 0.13 |
| 1 | 39000106 | rs16825164 | -0.229 | -0.367 | 1.305 | 0.855 | -0.02 | -0.004 | -0.004 | 0 | 0.06 | 0.01 | 0.1 | 0.05 | 0.07 | 0.08 |
| 1 | 39010828 | rs2992439 | -2.082 | -0.473 | 1.024 | 0.95 | 0.009 | 0.019 | 0.019 | 0 | 0.34 | 0.73 | 0.26 | 0.24 | 0.27 | 0.25 |
| 1 | 39029775 | rs10890297 | 1.688 | -0.537 | 1.492 | 1.096 | -0.015 | -0.041 | -0.041 | 0 | 0.85 | 0.53 | 0.86 | 0.92 | 0.92 | 0.85 |
| 1 | 39033922 | rs4469689 | 1.858 | -0.52 | 0.871 | 1.199 | -0.013 | 0 | 0 | 0 | 0.83 | 0.26 | 0.82 | 0.8 | 0.91 | 0.89 |
| 1 | 39036468 | rs12567715 | -0.92 | -0.579 | 0.367 | 2.08 | -0.002 | 0.106 | 0.106 | 0 | 0.07 | 0.05 | 0.16 | 0.52 | 0.06 | 0.18 |
| 1 | 39039169 | rs16825217 | -1.524 | -0.495 | 0.522 | 1.42 | 0.031 | 0.181 | 0.181 | 0.482 | 0.06 | 0.35 | 0.05 | 0.17 | 0.01 | 0.01 |
| 1 | 39041234 | rs10789446 | 0.252 | -0.636 | -1.4 | 0.894 | -0.043 | -0.066 | -0.066 | 0 | 0.45 | 0.06 | 0.54 | 0.31 | 0.56 | 0.48 |
| 1 | 39041667 | rs4970603 | 0.12 | -0.348 | -0.99 | 0.461 | -0.002 | -0.013 | -0.013 | 0 | 0.55 | 0.56 | 0.52 | 0.11 | 0.53 | 0.43 |
| 1 | 39042285 | rs11210993 | 0.448 | -0.02 | -2.06 | 0.645 | -0.155 | -0.134 | -0.134 | 9.315 | 0.32 | 0.05 | 0.44 | 0.34 | 0.42 | 0.44 |
| 1 | 39046200 | rs12048178 | 0.127 | 0.086 | -1.978 | -0.205 | -0.138 | 0.092 | 0.092 | 0.446 | 0.38 | 0.07 | 0.5 | 0.77 | 0.44 | 0.54 |
| 1 | 39047119 | rs4970605 | 0.051 | -0.402 | -0.806 | -0.266 | -0.033 | -0.094 | -0.094 | 0 | 0.67 | 0.58 | 0.83 | 0.88 | 0.79 | 0.76 |
| 1 | 39050786 | rs1001450 | -1.031 | 0.103 | -0.757 | 0.476 | -0.003 | 0.005 | 0.005 | 0 | 0.29 | 0.35 | 0.15 | 0.41 | 0.24 | 0.19 |
| 1 | 39054557 | rs2985755 | -0.853 | 0.534 | -0.367 | 0.273 | 0.01 | 0.035 | 0.035 | 0.278 | 0.22 | 0.38 | 0.14 | 0.3 | 0.22 | 0.3 |
| 1 | 39060495 | rs12026014 | 0.97 | 0.725 | -0.908 | -0.403 | 0.012 | 0.02 | 0.02 | 0 | 0.45 | 0.3 | 0.54 | 0.58 | 0.44 | 0.4 |
| 1 | 39060900 | rs3011179 | -0.414 | 1.184 | -1.224 | -1.231 | -0.034 | 0.008 | 0.008 | 0 | 0.9 | 1 | 0.85 | 0.9 | 0.91 | 0.91 |
| 1 | 39065974 | rs2985763 | 0.58 | 1.101 | -0.757 | -1.49 | -0.01 | -0.008 | -0.008 | 4.051 | 0.65 | 0.68 | 0.66 | 0.95 | 0.59 | 0.76 |
| 1 | 39071272 | rs2985774 | -0.934 | 0.527 | -1.12 | -1.523 | -0.023 | 0.174 | 0.174 | 0.049 | 0.48 | 0.81 | 0.42 | 0.21 | 0.47 | 0.35 |
| 1 | 39076855 | rs1925379 | -0.1 | 0.078 | -0.741 | -1.335 | -0.004 | 0.005 | 0.005 | 0 | 0.09 | 0.15 | 0.06 | 0.16 | 0.08 | 0.12 |
| 1 | 39077100 | rs1925382 | -1.29 | 0.151 | -0.691 | -1.483 | 0 | -0.001 | -0.001 | 0 | 0.15 | 0.19 | 0.08 | 0.16 | 0.14 | 0.16 |
| 1 | 39078789 | rs3011196 | -1.192 | -0.016 | 0.347 | -1.57 | 0.033 | -0.045 | -0.045 | 0 | 0.09 | 0.28 | 0.04 | 0 | 0.06 | 0.04 |
| 1 | 39080431 | rs17482558 | 0.601 | 0.189 | -0.103 | -1.809 | -0.035 | 0.047 | 0.047 | 0 | 0.1 | 0.02 | 0.09 | 0.02 | 0.13 | 0.07 |
| 1 | 39081829 | rs4970606 | -1.268 | 0.177 | -0.115 | -1.81 | -0.02 | 0.007 | 0.007 | 0 | 0.24 | 0.12 | 0.17 | 0.18 | 0.27 | 0.24 |
| 1 | 39087604 | rs12037910 | -0.336 | 0.124 | 0.441 | -1.858 | 0.002 | -0.037 | -0.037 | 1.983 | 0.13 | 0.18 | 0.1 | 0.02 | 0.14 | 0.07 |
| 1 | 39090577 | rs658866 | -1.301 | 0.174 | 1.021 | -1.83 | -0.008 | -0.007 | -0.007 | 0 | 0.21 | 0.28 | 0.16 | 0.17 | 0.21 | 0.39 |
| 1 | 39092056 | rs7548027 | -1.395 | 0.14 | 1.122 | -1.846 | -0.001 | -0.001 | -0.001 | 0 | 0.22 | 0.24 | 0.17 | 0.17 | 0.21 | 0.39 |
| 1 | 39092449 | rs4970609 | -1.954 | 0.026 | 0.904 | -2.215 | -0.005 | -0.021 | -0.021 | 0 | 0.36 | 0.6 | 0.26 | 0.2 | 0.32 | 0.47 |
| 1 | 39103961 | rs17482903 | 0.659 | 0.206 | 0.929 | -1.931 | -0.008 | 0.03 | 0.03 | 0.179 | 0.06 | 0.01 | 0.07 | 0 | 0.07 | 0.19 |
| 1 | 39119083 | rs10082063 | -0.695 | 0.44 | 0.493 | -2.157 | -0.002 | 0.005 | 0.005 | 0 | 0.82 | 0.87 | 0.86 | 0.98 | 0.83 | 0.95 |
| 1 | 39123603 | rs724321 | -1.173 | 0.31 | 0.808 | -2.018 | -0.004 | -0.007 | -0.007 | 0.254 | 0.35 | 0.45 | 0.27 | 0.29 | 0.34 | 0.46 |
| 1 | 39125390 | rs4970612 | -0.468 | 0.504 | 0.349 | -2.04 | -0.001 | -0.047 | -0.047 | 0 | 0.75 | 0.75 | 0.8 | 0.98 | 0.76 | 0.92 |
| 1 | 39125973 | rs4079876 | 0.299 | 0.415 | 0.386 | -2.069 | -0.012 | 0.01 | 0.01 | 0.092 | 0.17 | 0.25 | 0.15 | 0.26 | 0.15 | 0.32 |
| 1 | 39126938 | rs730219 | 1.86 | 0.381 | 0.119 | -2.008 | -0.046 | 0.011 | 0.011 | 0.434 | 0.38 | 0.14 | 0.53 | 0.67 | 0.41 | 0.44 |
| 1 | 39127471 | rs1971467 | -0.561 | 0.335 | 0.182 | -2.118 | 0 | -0.042 | -0.042 | 0 | 0.76 | 0.74 | 0.81 | 0.98 | 0.77 | 0.91 |
| 1 | 39129295 | rs6687438 | -0.158 | 0.533 | 0.069 | -1.987 | -0.007 | -0.02 | -0.02 | 0 | 0.22 | 0.16 | 0.18 | 0.27 | 0.27 | 0.38 |
| 1 | 39130160 | rs4474214 | -1.177 | 0.511 | 0.123 | -2.008 | -0.004 | -0.002 | -0.002 | 0 | 0.32 | 0.65 | 0.27 | 0.3 | 0.27 | 0.42 |
| 1 | 39133299 | rs6665540 | -0.11 | 0.519 | 0.13 | -1.994 | -0.006 | -0.014 | -0.014 | 0.109 | 0.22 | 0.16 | 0.18 | 0.26 | 0.27 | 0.38 |
| 1 | 39134880 | rs6687034 | -1.521 | 0.662 | -0.044 | -2.013 | -0.002 | 0.006 | 0.006 | 0 | 0.38 | 0.66 | 0.32 | 0.31 | 0.39 | 0.47 |
| 1 | 39135748 | rs4970614 | 0.134 | 0.479 | -0.137 | -1.976 | 0.008 | -0.076 | -0.076 | 0.214 | 0.15 | 0.43 | 0.11 | 0.04 | 0.1 | 0.1 |
| 1 | 39139260 | rs10789461 | -0.993 | 0.739 | -1.03 | -1.763 | 0.009 | 0.02 | 0.02 | 0.076 | 0.39 | 0.74 | 0.3 | 0.32 | 0.37 | 0.47 |
| 1 | 39143056 | rs11585248 | 0.697 | 0.637 | -0.8 | -0.921 | -0.055 | 0.071 | 0.071 | 0.223 | 0.46 | 0.12 | 0.55 | 0.66 | 0.5 | 0.51 |
| 1 | 39159100 | rs17484306 | -1.903 | 0.733 | 0.09 | -0.319 | -0.009 | 0 | 0 | 0 | 0.1 | 0.18 | 0.08 | 0.08 | 0.06 | 0.23 |
| 1 | 39160381 | rs4970619 | 2.083 | 0.935 | 0.888 | -0.657 | 0.013 | -0.068 | -0.068 | 0.633 | 0.74 | 0.39 | 0.73 | 0.89 | 0.75 | 0.71 |
| 1 | 39163072 | rs17556209 | -0.603 | 1.009 | 0.844 | -0.582 | -0.008 | -0.014 | -0.014 | 0.109 | 0.05 | 0.01 | 0.02 | 0.05 | 0.04 | 0.13 |
| 1 | 39166802 | rs10789469 | -0.128 | 1.031 | 0.972 | -0.603 | 0.104 | -0.056 | -0.056 | 2.02 | 0.9 | 0.67 | 0.83 | 0.99 | 0.83 | 0.97 |
| 1 | 39169671 | rs12409323 | -2.198 | 0.808 | 0.294 | -0.432 | -0.02 | 0.009 | 0.009 | 0 | 0.08 | 0.18 | 0.07 | 0.08 | 0.03 | 0.23 |
| 1 | 39172288 | rs6665159 | -0.362 | 0.901 | 0.169 | -0.393 | -0.008 | -0.007 | -0.007 | 0 | 0.05 | 0.01 | 0.02 | 0.04 | 0.04 | 0.13 |
| 1 | 39182130 | rs6663716 | -1.382 | 0.86 | 0.315 | -0.408 | -0.004 | 0.025 | 0.025 | 0 | 0.08 | 0.22 | 0.04 | 0.07 | 0.05 | 0.14 |
| 1 | 39192498 | rs6680382 | -2.085 | 0.69 | 0.294 | 0.136 | -0.047 | 0.122 | 0.122 | 0 | 0.19 | 0.41 | 0.14 | 0.37 | 0.1 | 0.34 |
| 1 | 39194716 | rs597563 | -2.121 | 0.614 | -0.592 | -0.792 | 0.046 | 0.084 | 0.084 | 0 | 0.23 | 0.06 | 0.13 | 0.06 | 0.12 | 0.32 |
| 1 | 39196638 | rs4595336 | -2.241 | 0.582 | -0.53 | -0.79 | 0.006 | -0.002 | -0.002 | 0.022 | 0.23 | 0.2 | 0.15 | 0.06 | 0.12 | 0.32 |
| 1 | 39202183 | rs3118213 | 1.194 | 1.216 | -0.33 | -0.603 | 0 | -0.018 | -0.018 | 0.42 | 0.76 | 0.73 | 0.76 | 0.93 | 0.76 | 0.72 |
| 1 | 39204184 | rs679143 | 1.199 | 1.598 | -0.524 | -0.533 | -0.022 | 0.023 | 0.023 | 0 | 0.75 | 0.79 | 0.75 | 0.94 | 0.7 | 0.71 |

| | | | | | | | | | | | | | | | | |
|---|---|---|---|---|---|---|---|---|---|---|---|---|---|---|---|---|
| 1 | 39209490 | rs7547720 | -1.289 | 1.421 | -0.366 | -0.539 | 0.005 | -0.024 | -0.024 | 0.286 | 0.18 | 0.24 | 0.13 | 0.09 | 0.2 | 0.23 |
| 1 | 39211558 | rs2182146 | -0.46 | 1.67 | -0.504 | -0.307 | -0.036 | -0.026 | -0.026 | 2.87 | 0.06 | 0.01 | 0.03 | 0.1 | 0.09 | 0.07 |
| 1 | 39213604 | rs2377375 | 0.153 | 1.296 | -0.317 | -0.343 | -0.023 | 0.035 | 0.035 | 0 | 0.32 | 0.46 | 0.51 | 0.58 | 0.26 | 0.35 |
| 1 | 39213862 | rs11211260 | -0.081 | 2.296 | -0.633 | -0.65 | -0.088 | 0.002 | 0.002 | 0 | 0.15 | 0.04 | 0.13 | 0.14 | 0.28 | 0.15 |
| 1 | 39214152 | rs942759 | -0.223 | 2.187 | -0.556 | -0.716 | 0.005 | -0.002 | -0.002 | 0 | 0.42 | 0.51 | 0.61 | 0.59 | 0.45 | 0.43 |
| 1 | 39215163 | rs1412912 | -0.202 | 2.184 | -0.517 | -0.717 | 0.003 | -0.001 | -0.001 | 0 | 0.42 | 0.49 | 0.61 | 0.59 | 0.45 | 0.43 |
| 1 | 39217732 | rs675766 | -1.4 | 1.562 | -1.095 | -1.427 | 0.034 | 0.113 | 0.113 | 0.019 | 0.71 | 0.88 | 0.87 | 1 | 0.77 | 0.83 |
| 1 | 39219090 | rs594797 | 0.276 | 1.383 | -0.181 | -1.269 | -0.02 | 0.096 | 0.096 | 0 | 0.14 | 0.28 | 0.14 | 0.27 | 0.11 | 0.19 |
| 1 | 39221925 | rs4970625 | 0.406 | 1.63 | -0.232 | -1.241 | -0.008 | -0.004 | -0.004 | 0 | 0.4 | 0.39 | 0.56 | 0.58 | 0.44 | 0.39 |
| 1 | 39224652 | rs2483119 | -0.788 | 1.583 | -0.233 | -1.588 | -0.001 | 0.046 | 0.046 | 0 | 0.58 | 0.7 | 0.78 | 1 | 0.59 | 0.72 |
| 1 | 39225303 | rs4147130 | 0.959 | 1.693 | -0.07 | -1.248 | 0.004 | -0.018 | -0.018 | 0 | 0.29 | 0.25 | 0.38 | 0.61 | 0.3 | 0.3 |
| 1 | 39228906 | rs12567070 | 1.141 | 1.636 | -0.139 | -1.279 | -0.027 | -0.111 | -0.111 | 0 | 0.27 | 0.06 | 0.36 | 0.61 | 0.29 | 0.29 |
| 1 | 39229313 | rs12043383 | -0.992 | 1.435 | 0.087 | -1.137 | 0.021 | 0.057 | 0.057 | 0.088 | 0.29 | 0.47 | 0.42 | 0.39 | 0.3 | 0.43 |
| 1 | 39238597 | rs4970627 | 0.197 | 0.39 | -0.557 | -1.373 | -0.005 | 0.003 | 0.003 | 0 | 0.32 | 0.36 | 0.36 | 0.68 | 0.29 | 0.38 |
| 1 | 39242855 | rs12046969 | 1.201 | 0.462 | -0.826 | -1.287 | -0.026 | -0.081 | -0.081 | 0 | 0.2 | 0.08 | 0.32 | 0.64 | 0.25 | 0.3 |
| 1 | 39247489 | rs12729493 | 1.452 | 0.939 | -1.122 | -1.214 | -0.088 | -0.089 | -0.089 | 16.664 | 0.14 | 0.01 | 0.3 | 0.24 | 0.21 | 0.2 |
| 1 | 39248701 | rs2151266 | -1.36 | 0.522 | -1.211 | -1.653 | 0.025 | 0.111 | 0.111 | 0 | 0.69 | 0.85 | 0.87 | 1 | 0.74 | 0.78 |
| 1 | 39248886 | rs2209477 | 0.051 | 0.647 | -0.768 | -1.546 | 0 | -0.025 | -0.025 | 0.203 | 0.19 | 0.22 | 0.16 | 0.09 | 0.21 | 0.19 |
| 1 | 39250964 | rs1925421 | -1.377 | 0.69 | -1.096 | -1.467 | 0.013 | 0.082 | 0.082 | 0 | 0.73 | 0.85 | 0.87 | 1 | 0.74 | 0.79 |
| 1 | 39255293 | rs10789490 | -0.903 | 0.665 | -0.808 | -0.756 | 0.04 | 0.112 | 0.112 | 0 | 0.48 | 0.71 | 0.53 | 0.79 | 0.41 | 0.56 |
| 1 | 39258674 | rs6675193 | -0.862 | 0.164 | -0.73 | -0.781 | -0.029 | -0.011 | -0.011 | 0 | 0.27 | 0.54 | 0.25 | 0.24 | 0.19 | 0.27 |
| 1 | 39266840 | rs12026925 | -0.232 | 0.302 | -1.223 | -0.348 | 0.008 | 0.086 | 0.086 | 0 | 0.09 | 0.24 | 0.14 | 0.19 | 0.08 | 0.15 |
| 1 | 39274125 | rs9326146 | 0.3 | 0.366 | -2.157 | -1.082 | -0.035 | 0.251 | 0.251 | 0 | 0.36 | 0.05 | 0.16 | 0.01 | 0.37 | 0.25 |
| 1 | 39278261 | rs4970630 | -0.621 | 0.152 | -1.631 | 0.098 | -0.002 | 0.016 | 0.016 | 0 | 0.79 | 0.76 | 0.75 | 0.58 | 0.83 | 0.74 |
| 1 | 39278828 | rs9439072 | 0.364 | 0.048 | -1.998 | 0.489 | -0.02 | -0.102 | -0.102 | 0.012 | 0.26 | 0.08 | 0.26 | 0.42 | 0.29 | 0.38 |
| 1 | 39279037 | rs16825524 | 0.466 | -0.247 | -2.105 | 0.327 | 0.026 | 0.003 | 0.003 | 4.518 | 0.16 | 0.06 | 0.04 | 0.14 | 0.1 | 0.08 |
| 1 | 39279324 | rs1184068 | -0.465 | 0.282 | -1.16 | 0.471 | 0 | -0.004 | -0.004 | 0 | 0.34 | 0.31 | 0.33 | 0.42 | 0.37 | 0.41 |
| 1 | 39291708 | rs6690335 | -0.591 | 0.219 | -0.851 | 0.88 | -0.004 | 0.037 | 0.037 | 0 | 0.23 | 0.33 | 0.3 | 0.41 | 0.21 | 0.27 |
| 1 | 39295972 | rs9438970 | -0.154 | 0.418 | -1.616 | 0.508 | -0.045 | -0.107 | -0.107 | 0.003 | 0.81 | 0.95 | 0.8 | 0.58 | 0.76 | 0.71 |
| 1 | 39299306 | rs16825572 | 1.325 | -0.167 | -1.679 | -0.056 | 0.007 | -0.007 | -0.007 | 0.725 | 0.14 | 0.08 | 0.06 | 0 | 0.1 | 0.06 |
| 1 | 39302993 | rs7527128 | 1.391 | -0.489 | -1.537 | 0.171 | -0.073 | 0.046 | 0.046 | 0 | 0.38 | 0.13 | 0.49 | 0.27 | 0.45 | 0.33 |
| 1 | 39305403 | rs17499125 | 0.81 | -0.463 | -1.539 | 0.468 | -0.076 | -0.04 | -0.04 | 0 | 0.15 | 0.01 | 0.11 | 0.13 | 0.21 | 0.08 |
| 1 | 39309107 | rs11211460 | 1.309 | -0.374 | -1.436 | 0.401 | -0.039 | 0.067 | 0.067 | 0 | 0.41 | 0.18 | 0.51 | 0.27 | 0.52 | 0.39 |
| 1 | 39328203 | rs34393348 | 0.516 | -0.462 | -1.615 | 0.478 | 0.08 | -0.01 | -0.01 | 0 | 0.53 | 0.15 | 0.61 | 0.4 | 0.68 | 0.41 |
| 1 | 39339449 | rs12127091 | 1.321 | -0.355 | -1.509 | 0.554 | -0.07 | 0.052 | 0.052 | 0 | 0.39 | 0.15 | 0.48 | 0.27 | 0.45 | 0.31 |
| 1 | 39340282 | rs880303 | 1.321 | -0.355 | -1.509 | 0.554 | -0.07 | 0.052 | 0.052 | 0 | 0.39 | 0.15 | 0.48 | 0.27 | 0.45 | 0.31 |
| 1 | 39340607 | rs16825625 | 1.594 | -0.558 | -1.579 | 0.784 | 0.027 | -0.01 | -0.01 | 0 | 0.11 | 0.02 | 0.05 | 0.08 | 0.07 | 0.06 |
| 1 | 39340862 | rs874243 | 0.986 | -0.248 | -1.438 | 0.67 | -0.078 | -0.048 | -0.048 | 0.23 | 0.13 | 0.01 | 0.12 | 0.13 | 0.22 | 0.08 |
| 1 | 39350742 | rs4970633 | 1.357 | -0.238 | -1.192 | 0.645 | -0.066 | -0.05 | -0.05 | 0 | 0.11 | 0.01 | 0.08 | 0.13 | 0.18 | 0.06 |
| 1 | 39352083 | rs4491033 | 0.83 | -0.375 | -1.119 | 0.666 | -0.009 | 0.032 | 0.032 | 0 | 0.42 | 0.38 | 0.5 | 0.27 | 0.46 | 0.35 |
| 1 | 39352271 | rs2147914 | 1.048 | -0.232 | -1.178 | 0.741 | -0.07 | 0.053 | 0.053 | 0 | 0.39 | 0.15 | 0.48 | 0.27 | 0.45 | 0.31 |
| 1 | 39354517 | rs4414011 | 0.203 | -0.232 | -1.178 | 0.741 | 0.068 | 0.01 | 0.01 | 0 | 0.53 | 0.32 | 0.61 | 0.4 | 0.67 | 0.39 |
| 1 | 39354602 | rs10890491 | 0.558 | -0.43 | -1.073 | 0.658 | -0.07 | 0.053 | 0.053 | 0 | 0.39 | 0.15 | 0.48 | 0.27 | 0.45 | 0.31 |
| 1 | 39372843 | rs9438989 | 0.908 | -0.074 | -0.682 | 0.829 | -0.065 | 0.007 | 0.007 | 0 | 0.17 | 0.03 | 0.24 | 0.14 | 0.22 | 0.19 |
| 1 | 39381576 | rs6675870 | -0.247 | -1.156 | 0.24 | 1.264 | -0.059 | 0.036 | 0.036 | 0 | 0.21 | 0.41 | 0.19 | 0.29 | 0.12 | 0.22 |
| 1 | 39385086 | rs7522863 | 0.397 | -0.883 | 0.498 | 1.681 | 0.005 | 0.171 | 0.171 | 0 | 0.92 | 0.8 | 0.9 | 0.73 | 0.95 | 0.86 |
| 1 | 39387082 | rs1925676 | -0.772 | -1.062 | 0.25 | 1.262 | -0.022 | 0.025 | 0.025 | 0.049 | 0.31 | 0.56 | 0.32 | 0.66 | 0.25 | 0.46 |
| 1 | 39388838 | rs9438994 | -0.54 | -2.269 | 0.12 | 1.082 | -0.055 | 0.005 | 0.005 | 0 | 0.3 | 0.6 | 0.17 | 0.29 | 0.14 | 0.28 |
| 1 | 39396070 | rs6656771 | 1.346 | -2.212 | -0.562 | 1.026 | -0.001 | 0.165 | 0.165 | 0 | 0.43 | 0.05 | 0.45 | 0.18 | 0.58 | 0.31 |
| 1 | 39396424 | rs12042694 | -0.686 | -2.361 | -0.249 | 1.088 | -0.022 | 0.007 | 0.007 | 0 | 0.32 | 0.54 | 0.4 | 0.67 | 0.27 | 0.5 |
| 1 | 39398869 | rs10749676 | 0.729 | -2.373 | -0.223 | 1.062 | -0.006 | -0.005 | -0.005 | 0 | 0.64 | 0.43 | 0.55 | 0.26 | 0.68 | 0.42 |
| 1 | 39406152 | rs12047330 | -0.341 | -2.432 | 0.258 | 0.203 | 0.015 | -0.002 | -0.002 | 0.059 | 0.27 | 0.2 | 0.24 | 0.26 | 0.24 | 0.27 |
| 1 | 39406644 | rs6422258 | -0.875 | -2.41 | 0.077 | 0.216 | 0.032 | 0.013 | 0.013 | 0 | 0.83 | 0.9 | 0.93 | 0.92 | 0.91 | 0.88 |
| 1 | 39417403 | rs11810357 | 1.714 | -2.288 | 0.186 | 0.239 | -0.013 | -0.029 | -0.029 | 0 | 0.09 | 0.09 | 0.04 | 0 | 0.04 | 0.02 |
| 1 | 39423848 | rs4508001 | -0.345 | -2.217 | 0.291 | 0.284 | 0.01 | 0.003 | 0.003 | 0 | 0.06 | 0.18 | 0.06 | 0.07 | 0.05 | 0.07 |
| 1 | 39424634 | rs4512638 | 1.205 | -2.177 | 0.076 | 0.324 | 0.051 | 0.012 | 0.012 | 0 | 0.5 | 0.36 | 0.64 | 0.59 | 0.62 | 0.55 |
| 1 | 39425489 | rs4475740 | -0.205 | -1.62 | 0.282 | 0.534 | -0.053 | 0.068 | 0.068 | 0 | 0.82 | 0.73 | 0.85 | 0.7 | 0.92 | 0.75 |
| 1 | 39430262 | rs7548458 | -0.404 | -1.415 | 0.258 | 0.606 | 0.019 | 0.259 | 0.259 | 2.463 | 0.05 | 0.3 | 0.12 | 0.22 | 0.01 | 0.14 |
| 1 | 39431045 | rs6665494 | 0.335 | -1.119 | -0.443 | -0.245 | 0.021 | 0.032 | 0.032 | 0.051 | 0.67 | 0.8 | 0.83 | 0.87 | 0.72 | 0.76 |
| 1 | 39438889 | rs4660446 | 0.011 | -0.399 | -0.713 | -0.466 | 0.068 | 0.239 | 0.239 | 1.454 | 0.47 | 0.18 | 0.34 | 0.11 | 0.64 | 0.29 |
| 1 | 39439538 | rs7537437 | 0.702 | 0.165 | -0.82 | -0.686 | -0.012 | 0.037 | 0.037 | 0 | 0.22 | 0.16 | 0.11 | 0.03 | 0.23 | 0.17 |
| 1 | 39447322 | rs3934859 | -0.926 | -0.181 | -0.812 | -0.693 | -0.015 | 0.013 | 0.013 | 0 | 0.9 | 0.95 | 0.85 | 0.99 | 0.85 | 0.87 |
| 1 | 39451857 | rs6696013 | -0.897 | -0.259 | -0.812 | -0.692 | -0.017 | 0.015 | 0.015 | 0 | 0.9 | 0.95 | 0.84 | 0.99 | 0.84 | 0.87 |
| 1 | 39455470 | rs12756255 | 1.561 | -0.493 | -0.922 | -0.674 | 0.027 | 0.027 | 0.027 | 0 | 0.05 | 0 | 0.01 | 0 | 0.02 | 0.02 |
| 1 | 39457859 | rs4660652 | 0.453 | 0.031 | 0.155 | -0.02 | -0.011 | -0.021 | -0.021 | 0 | 0.57 | 0.32 | 0.53 | 0.47 | 0.53 | 0.58 |
| 1 | 39458110 | rs7553434 | 1.28 | -0.188 | 0.1 | -0.304 | 0.024 | -0.022 | -0.022 | 0 | 0.32 | 0.23 | 0.33 | 0.47 | 0.14 | 0.46 |
| 1 | 39472585 | rs10888621 | 0.304 | 0.262 | 0.185 | -0.375 | -0.004 | -0.002 | -0.002 | 0 | 0.27 | 0.26 | 0.24 | 0.11 | 0.39 | 0.11 |
| 1 | 39477635 | rs7538228 | -1.21 | 0.16 | 0.274 | -0.271 | -0.065 | 0.004 | 0.004 | 0.729 | 0.63 | 0.6 | 0.64 | 0.4 | 0.84 | 0.46 |
| 1 | 39492462 | rs3768324 | 0.652 | 0.294 | -0.003 | -0.283 | -0.043 | -0.025 | -0.025 | 0 | 0.25 | 0.16 | 0.23 | 0.29 | 0.3 | 0.25 |

| | | | | | | | | | | | | | | | | |
|---|---|---|---|---|---|---|---|---|---|---|---|---|---|---|---|---|
| 1 | 39498491 | rs6663155 | -1.318 | 0.282 | -0.013 | -0.285 | 0.012 | -0.005 | -0.005 | 0.105 | 0.64 | 0.7 | 0.65 | 0.4 | 0.84 | 0.47 |
| 1 | 39507161 | rs10888650 | 0.623 | 0.346 | -0.401 | -0.252 | -0.032 | -0.018 | -0.018 | 0.393 | 0.26 | 0.18 | 0.24 | 0.29 | 0.3 | 0.25 |
| 1 | 39511058 | rs3901310 | 0.375 | -0.436 | -0.553 | 0.085 | -0.136 | 0.061 | 0.061 | 0.098 | 0.24 | 0.01 | 0.21 | 0.11 | 0.38 | 0.11 |
| 1 | 39518006 | rs10465799 | -0.258 | -1.167 | 0.012 | 0.562 | -0.047 | 0.008 | 0.008 | 2.299 | 0.61 | 0.44 | 0.62 | 0.46 | 0.83 | 0.52 |
| 1 | 39525201 | rs6692557 | -0.377 | -1.066 | -0.625 | 0.643 | -0.044 | 0.009 | 0.009 | 1.05 | 0.63 | 0.59 | 0.64 | 0.46 | 0.84 | 0.52 |
| 1 | 39528071 | rs9786967 | -0.403 | -0.957 | -0.886 | 0.206 | -0.024 | 0 | 0 | 0 | 0.63 | 0.65 | 0.65 | 0.4 | 0.84 | 0.47 |
| 1 | 39542129 | rs12142033 | -0.201 | 1.061 | -1.028 | 0.411 | -0.029 | 0.031 | 0.031 | 0 | 0.22 | 0.16 | 0.17 | 0.1 | 0.34 | 0.09 |
| 1 | 39545269 | rs9659165 | -0.55 | 0.621 | -0.591 | 1.215 | 0.004 | 0.16 | 0.16 | 0.161 | 0.06 | 0.11 | 0.21 | 0.33 | 0.04 | 0.12 |
| 1 | 39548798 | rs260974 | -0.614 | 0.353 | -0.639 | 0.211 | -0.017 | -0.047 | -0.047 | 0.332 | 0.17 | 0.24 | 0.08 | 0.06 | 0.11 | 0.12 |
| 1 | 39550621 | rs7546735 | -0.465 | 1.01 | -1.594 | 0.137 | -0.076 | -0.002 | -0.002 | 0 | 0.2 | 0.01 | 0.24 | 0.12 | 0.27 | 0.08 |
| 1 | 39551621 | rs260969 | -0.171 | 0.943 | -2.322 | 0.41 | -0.008 | -0.063 | -0.063 | 0 | 0.32 | 0.11 | 0.4 | 0.5 | 0.3 | 0.65 |
| 1 | 39555643 | rs11205632 | 0.444 | 1.652 | -2.386 | 0.014 | -0.009 | 0.003 | 0.003 | 0 | 0.06 | 0.04 | 0.04 | 0 | 0.13 | 0.05 |
| 1 | 39557688 | rs4660401 | -0.498 | 1.878 | -2.484 | 0.247 | -0.036 | 0.012 | 0.012 | 0.72 | 0.42 | 0.21 | 0.52 | 0.63 | 0.48 | 0.72 |
| 1 | 39557772 | rs260964 | -1.282 | 1.212 | -2.049 | -0.64 | 0.005 | -0.147 | -0.147 | 0.704 | 0.75 | 0.48 | 0.8 | 0.99 | 0.74 | 0.87 |
| 1 | 39568765 | rs2282232 | -0.79 | 1.337 | -1.308 | -0.743 | -0.005 | -0.001 | -0.001 | 0.235 | 0.63 | 0.76 | 0.5 | 0.33 | 0.65 | 0.33 |
| 1 | 39569571 | rs2282231 | 0.69 | 1.07 | -1.45 | -0.716 | 0.014 | 0.015 | 0.015 | 2.655 | 0.25 | 0.15 | 0.23 | 0.17 | 0.2 | 0.08 |
| 1 | 39571211 | rs13374459 | -0.174 | 0.972 | -0.573 | -0.823 | 0.004 | -0.04 | -0.04 | 0 | 0.37 | 0.57 | 0.36 | 0.17 | 0.41 | 0.19 |
| 1 | 39571830 | rs905380 | 0.448 | 0.958 | -0.461 | -0.804 | 0 | 0 | 0 | 0 | 0.27 | 0.24 | 0.15 | 0.16 | 0.25 | 0.14 |
| 1 | 39581931 | rs7550056 | 0.756 | 0.872 | -0.011 | -0.797 | -0.007 | 0.002 | 0.002 | 1.802 | 0.41 | 0.56 | 0.56 | 0.67 | 0.41 | 0.68 |
| 1 | 39583686 | rs11580989 | 0.061 | 0.813 | 0.044 | -0.804 | -0.013 | 0.058 | 0.058 | 0 | 0.31 | 0.14 | 0.18 | 0.16 | 0.36 | 0.19 |
| 1 | 39590824 | rs11205665 | 1.205 | 0.907 | -0.166 | -0.22 | -0.003 | -0.019 | -0.019 | 0.436 | 0.09 | 0.02 | 0.04 | 0.15 | 0.08 | 0.07 |
| 1 | 39626381 | rs656619 | 1.16 | 0.266 | 0.154 | -0.39 | -0.059 | -0.07 | -0.07 | 0.197 | 0.12 | 0.27 | 0.03 | 0 | 0.01 | 0 |
| 1 | 39644067 | rs560372 | 2.018 | 0.265 | 0.161 | -0.39 | -0.038 | -0.043 | -0.043 | 0.013 | 0.1 | 0.14 | 0.02 | 0 | 0 | 0 |
| 1 | 39647038 | rs1537818 | 0.313 | 0.369 | -0.107 | -0.463 | -0.017 | 0.001 | 0.001 | 0.921 | 0.63 | 0.48 | 0.68 | 0.67 | 0.71 | 0.74 |
| 1 | 39662654 | rs6700698 | 1.08 | 0.452 | -0.446 | -0.054 | -0.076 | -0.091 | -0.091 | 0 | 0.15 | 0.02 | 0.34 | 0.39 | 0.21 | 0.41 |
| 1 | 39692941 | rs495879 | 0.492 | 0.166 | -0.243 | -0.052 | -0.051 | 0.014 | 0.014 | 2.364 | 0.18 | 0.38 | 0.08 | 0.16 | 0.07 | 0.05 |
| 1 | 39695155 | rs10788933 | -0.061 | 0.141 | -0.363 | -0.012 | 0.018 | -0.006 | -0.006 | 2.059 | 0.64 | 0.27 | 0.67 | 0.67 | 0.73 | 0.8 |
| 1 | 39703847 | rs16837533 | 2.497 | 0.367 | -0.539 | -0.03 | -0.036 | -0.053 | -0.053 | 0 | 0.14 | 0.01 | 0.17 | 0.17 | 0.16 | 0.07 |
| 1 | 39705458 | rs950805 | 0.215 | 0.307 | -0.431 | 0.042 | 0.018 | -0.022 | -0.022 | 0 | 0.64 | 0.27 | 0.67 | 0.54 | 0.72 | 0.79 |
| 1 | 39705874 | rs12120111 | 1.951 | 0.32 | -0.676 | -0.401 | -0.013 | 0.102 | 0.102 | 0 | 0.24 | 0.07 | 0.15 | 0 | 0.29 | 0.11 |
| 1 | 39731550 | rs4660214 | 2.296 | 0.319 | -0.232 | -0.12 | -0.053 | -0.064 | -0.064 | 0 | 0.16 | 0.01 | 0.21 | 0.18 | 0.19 | 0.08 |
| 1 | 39744607 | rs2252410 | -1.236 | 0.173 | 0.199 | 0.36 | -0.059 | 0.041 | 0.041 | 2.732 | 0.84 | 0.62 | 0.93 | 0.82 | 0.95 | 0.96 |
| 1 | 39748921 | rs2275188 | 0.148 | 0.178 | 0.216 | 0.318 | 0.017 | -0.012 | -0.012 | 0 | 0.64 | 0.27 | 0.67 | 0.65 | 0.72 | 0.8 |
| 1 | 39749884 | rs2039597 | 2.04 | 0.404 | 0.063 | 0.375 | -0.043 | -0.059 | -0.059 | 0.668 | 0.11 | 0.02 | 0.25 | 0.32 | 0.14 | 0.38 |
| 1 | 39758955 | rs2484749 | -0.607 | -0.137 | 0.242 | 0.485 | -0.01 | 0.009 | 0.009 | 2.789 | 0.67 | 0.58 | 0.72 | 0.67 | 0.76 | 0.87 |
| 1 | 39763242 | rs2275187 | -0.064 | -0.12 | 0.244 | 0.474 | 0.019 | -0.016 | -0.016 | 0.049 | 0.64 | 0.27 | 0.67 | 0.6 | 0.72 | 0.79 |
| 1 | 39772180 | rs3120282 | -1.68 | -0.152 | 0.218 | 0.47 | -0.06 | -0.059 | -0.059 | 0.435 | 0.87 | 0.62 | 0.95 | 0.94 | 0.97 | 0.99 |
| 1 | 39785641 | rs3116389 | -1.049 | -0.185 | 0.383 | 0.582 | -0.078 | 0.111 | 0.111 | 2.894 | 0.83 | 0.52 | 0.92 | 0.76 | 0.95 | 0.95 |
| 1 | 39797055 | rs16826069 | 2.283 | -0.264 | 0.194 | 0.607 | -0.055 | -0.087 | -0.087 | 0 | 0.16 | 0.01 | 0.25 | 0.26 | 0.2 | 0.15 |
| 1 | 39812822 | rs2761764 | -2.948 | -0.159 | 0.15 | 0.488 | -0.032 | -0.021 | -0.021 | 0 | 0.91 | 0.89 | 0.98 | 0.94 | 1 | 0.99 |
| 1 | 39835817 | rs2296172 | 2.388 | -0.142 | 0.099 | 0.499 | -0.054 | -0.041 | -0.041 | 0 | 0.16 | 0.01 | 0.2 | 0.16 | 0.19 | 0.08 |
| 1 | 39867508 | rs2451672 | 2.682 | -0.158 | 0.114 | 0.503 | -0.04 | -0.028 | -0.028 | 0 | 0.1 | 0.14 | 0.02 | 0.06 | 0.01 | 0 |
| 1 | 39871926 | rs616630 | 1.843 | -0.164 | 0.105 | 0.473 | -0.026 | 0.016 | 0.016 | 1.353 | 0.13 | 0.14 | 0.04 | 0.16 | 0.03 | 0.04 |
| 1 | 39879517 | rs587523 | 2.68 | -0.159 | 0.134 | 0.452 | -0.04 | -0.046 | -0.046 | 0.041 | 0.1 | 0.14 | 0.02 | 0 | 0 | 0 |
| 1 | 39894839 | rs645061 | -1.931 | 0.048 | -0.045 | 0.462 | -0.07 | -0.078 | -0.078 | 0 | 0.86 | 0.98 | 0.66 | 0.62 | 0.79 | 0.59 |
| 1 | 39897271 | rs599823 | 0.241 | -0.545 | 0.708 | 0.387 | -0.062 | 0.048 | 0.048 | 3.028 | 0.18 | 0.5 | 0.11 | 0.17 | 0.09 | 0.12 |
| 1 | 39908506 | rs587404 | -0.38 | -0.525 | 0.424 | 0.374 | -0.014 | 0.017 | 0.017 | 4.169 | 0.34 | 0.51 | 0.32 | 0.35 | 0.28 | 0.2 |
| 1 | 39913351 | rs2296173 | 2.009 | -0.518 | 0.408 | 0.377 | -0.054 | -0.043 | -0.043 | 0 | 0.16 | 0.01 | 0.21 | 0.16 | 0.2 | 0.08 |
| 1 | 39919234 | rs687848 | -0.933 | -0.576 | 0.452 | 0.279 | 0.011 | 0.003 | 0.003 | 2.297 | 0.59 | 0.6 | 0.46 | 0.35 | 0.58 | 0.31 |
| 1 | 39926824 | rs4660762 | 0.183 | -0.574 | 0.398 | 0.097 | -0.014 | -0.022 | -0.022 | 0.379 | 0.57 | 0.43 | 0.75 | 0.81 | 0.62 | 0.77 |
| 1 | 39929240 | rs7554206 | 0.204 | -0.192 | 0.863 | 0.816 | -0.012 | 0.01 | 0.01 | 4.294 | 0.66 | 0.52 | 0.69 | 0.69 | 0.72 | 0.8 |
| 1 | 39945297 | rs2275767 | 0.495 | -0.135 | 0.398 | 0.497 | 0.018 | -0.027 | -0.027 | 0 | 0.62 | 0.28 | 0.59 | 0.58 | 0.66 | 0.79 |
| 1 | 39952745 | rs710915 | 1.003 | -0.255 | 0.742 | 0.675 | -0.047 | 0.111 | 0.111 | 0.935 | 0.13 | 0.21 | 0.1 | 0.3 | 0.03 | 0.04 |
| 1 | 39963873 | rs2004330 | 1.171 | -0.366 | 0.361 | 0.306 | -0.044 | -0.025 | -0.025 | 0 | 0.43 | 0.17 | 0.47 | 0.56 | 0.42 | 0.74 |
| 1 | 39973646 | rs782869 | 2.245 | -0.514 | 1.01 | 0.216 | -0.051 | -0.053 | -0.053 | 0.138 | 0.1 | 0.16 | 0.02 | 0 | 0 | 0 |
| 1 | 39991588 | rs3738676 | 0.559 | -0.361 | 0.713 | 0.197 | 0.026 | 0.071 | 0.071 | 0 | 0.26 | 0.36 | 0.36 | 0.44 | 0.37 | 0.16 |
| 1 | 39992808 | rs1180341 | 0.805 | -0.537 | 0.524 | 0.002 | 0.006 | -0.024 | -0.024 | 0 | 0.56 | 0.27 | 0.55 | 0.55 | 0.53 | 0.78 |
| 1 | 39993432 | rs1180343 | 1.61 | -0.548 | 0.443 | -0.051 | -0.048 | -0.048 | -0.048 | 0.065 | 0.44 | 0.17 | 0.47 | 0.55 | 0.43 | 0.76 |
| 1 | 40035686 | rs17513135 | 1.703 | 0.463 | 0.301 | 0.496 | -0.068 | 0 | 0 | 0 | 0.16 | 0.01 | 0.11 | 0.1 | 0.24 | 0.05 |
| 1 | 40037902 | rs2242500 | 0.023 | 0.456 | 0.417 | 0.496 | -0.002 | 0.001 | 0.001 | 0 | 0.24 | 0.19 | 0.25 | 0.42 | 0.34 | 0.14 |
| 1 | 40049184 | rs11206378 | 0.193 | 0.485 | 0.542 | 0.547 | 0.005 | 0.022 | 0.022 | 0.871 | 0.69 | 0.36 | 0.68 | 0.57 | 0.62 | 0.86 |
| 1 | 40050568 | rs7539279 | -0.157 | 0.488 | 0.55 | 0.552 | 0.024 | -0.067 | -0.067 | 0.792 | 0.72 | 0.46 | 0.77 | 0.88 | 0.65 | 0.89 |
| 1 | 40051616 | rs2243761 | 0.647 | 0.489 | 0.166 | 0.505 | -0.015 | -0.039 | -0.039 | 0 | 0.65 | 0.27 | 0.63 | 0.57 | 0.54 | 0.8 |
| 1 | 40082002 | rs1180320 | -1.341 | 0.757 | 0.236 | 0.837 | -0.04 | -0.095 | -0.095 | 0 | 0.36 | 0.86 | 0.37 | 0.45 | 0.45 | 0.2 |
| 1 | 40083666 | rs12567371 | -0.325 | 0.753 | 0.172 | 0.906 | 0.001 | 0.086 | 0.086 | 0 | 0.07 | 0.09 | 0.13 | 0.31 | 0.11 | 0.09 |
| 1 | 40089595 | rs12030495 | -0.972 | 0.647 | 0.104 | 0.665 | -0.011 | -0.027 | -0.027 | 0.113 | 0.35 | 0.75 | 0.35 | 0.42 | 0.45 | 0.19 |
| 1 | 40091119 | rs11206478 | -0.202 | -0.067 | 0.511 | 0.402 | 0.027 | -0.037 | -0.037 | 0 | 0.08 | 0.28 | 0.07 | 0.02 | 0.1 | 0.02 |
| 1 | 40095493 | rs6694831 | -1.324 | -0.283 | 0.805 | 0.848 | 0.007 | 0.12 | 0.12 | 1.196 | 0.09 | 0.31 | 0.1 | 0.17 | 0.1 | 0.05 |
| 1 | 40096053 | rs1148945 | -0.04 | -0.681 | 0.369 | 0.492 | -0.002 | 0 | 0 | 0 | 0.41 | 0.4 | 0.32 | 0.4 | 0.33 | 0.49 |

| | | | | | | | | | | | | | | | | |
|---|---|---|---|---|---|---|---|---|---|---|---|---|---|---|---|---|
| 1 | 40104584 | rs4660892 | 1.294 | -0.352 | 0.113 | -0.039 | 0.02 | -0.068 | -0.068 | 0 | 0.22 | 0.31 | 0.25 | 0.06 | 0.3 | 0.1 |
| 1 | 40104814 | rs784622 | -0.67 | -0.507 | -0.201 | 0.321 | 0.04 | 0.006 | 0.006 | 0 | 0.73 | 0.86 | 0.77 | 0.75 | 0.86 | 0.68 |
| 1 | 40106095 | rs1180280 | 0.755 | -0.471 | -0.348 | 0.405 | 0.066 | 0.109 | 0.109 | 0 | 0.36 | 0.11 | 0.12 | 0.2 | 0.23 | 0.42 |
| 1 | 40112610 | rs784619 | -1.466 | -0.325 | -0.238 | 0.552 | 0.013 | -0.035 | -0.035 | 0 | 0.88 | 0.93 | 0.8 | 0.76 | 0.93 | 0.75 |
| 1 | 40120085 | rs1180275 | 1.592 | -0.702 | -0.169 | -0.259 | -0.013 | -0.017 | -0.017 | 0 | 0.16 | 0.17 | 0.09 | 0.09 | 0.11 | 0.09 |
| 1 | 40125704 | rs3851911 | 1.478 | -0.572 | -0.592 | -0.23 | -0.003 | 0.026 | 0.026 | 0.183 | 0.19 | 0.11 | 0.09 | 0.09 | 0.17 | 0.3 |
| 1 | 40132795 | rs873917 | 0.04 | -0.068 | -0.423 | 0.085 | 0.06 | 0.096 | 0.096 | 0 | 0.77 | 0.6 | 0.56 | 0.46 | 0.69 | 0.66 |
| 1 | 40138123 | rs1533479 | -0.199 | -0.228 | -0.626 | -0.019 | 0.037 | -0.01 | -0.01 | 0 | 0.32 | 0.25 | 0.24 | 0.36 | 0.21 | 0.22 |
| 1 | 40139553 | rs784600 | -0.76 | 0.209 | -0.671 | -0.723 | 0.004 | -0.08 | -0.08 | 0.293 | 0.58 | 0.53 | 0.64 | 0.93 | 0.48 | 0.78 |
| 1 | 40143327 | rs6663003 | 1.308 | 0.387 | -0.626 | -0.629 | 0 | 0.01 | 0.01 | 0 | 0.19 | 0.17 | 0.1 | 0.07 | 0.18 | 0.23 |
| 1 | 40144097 | rs11206608 | 0.444 | 0.531 | -0.766 | -0.316 | 0.003 | 0.002 | 0.002 | 0 | 0.5 | 0.44 | 0.59 | 0.55 | 0.47 | 0.5 |
| 1 | 40145297 | rs6703320 | 0.247 | 0.536 | -0.772 | -0.313 | 0.003 | 0.002 | 0.002 | 0 | 0.5 | 0.44 | 0.59 | 0.55 | 0.47 | 0.51 |
| 1 | 40145616 | rs12039225 | 1.619 | 0.57 | -0.819 | -0.26 | -0.015 | -0.08 | -0.08 | 0 | 0.11 | 0.01 | 0.3 | 0.47 | 0.09 | 0.32 |
| 1 | 40146253 | rs784629 | -0.298 | 0.891 | -0.439 | 0.074 | 0.015 | 0.014 | 0.014 | 0 | 0.91 | 0.92 | 0.83 | 0.82 | 0.83 | 0.87 |
| 1 | 40149642 | rs784634 | -0.298 | 0.891 | -0.439 | 0.074 | 0.015 | 0.014 | 0.014 | 0 | 0.91 | 0.92 | 0.83 | 0.82 | 0.83 | 0.87 |
| 1 | 40150156 | rs9662128 | -0.312 | 0.855 | -0.588 | -0.953 | -0.004 | -0.009 | -0.009 | 2.188 | 0.41 | 0.48 | 0.24 | 0.27 | 0.35 | 0.36 |
| 1 | 40152117 | rs2248221 | 0.465 | 1.876 | -0.732 | -0.661 | 0 | -0.015 | -0.015 | 0 | 0.49 | 0.5 | 0.59 | 0.85 | 0.4 | 0.59 |
| 1 | 40152676 | rs785108 | 0.433 | 2.523 | -0.507 | -0.082 | 0.009 | -0.039 | -0.039 | 0.751 | 0.66 | 0.66 | 0.68 | 0.92 | 0.53 | 0.81 |
| 1 | 40153141 | rs477913 | -0.154 | 2.271 | -0.805 | 0.075 | 0.021 | 0.027 | 0.027 | 0 | 0.05 | 0.11 | 0.07 | 0.11 | 0.12 | 0.07 |
| 1 | 40155623 | rs564249 | -0.304 | 1.561 | -0.781 | -0.392 | 0.002 | -0.06 | -0.06 | 0.007 | 0.28 | 0.38 | 0.27 | 0.08 | 0.32 | 0.15 |
| 1 | 40161013 | rs704771 | 0.294 | 1.815 | -1.187 | 0.236 | -0.015 | 0.01 | 0.01 | 0 | 0.93 | 0.96 | 0.93 | 1 | 0.86 | 0.96 |
| 1 | 40164552 | rs7549005 | -0.566 | 1.144 | -1.262 | 0.535 | 0.005 | -0.005 | -0.005 | 0 | 0.38 | 0.28 | 0.37 | 0.42 | 0.34 | 0.47 |
| 1 | 40168587 | rs12565115 | -0.114 | 1.035 | -1.986 | 0.814 | -0.028 | -0.097 | -0.097 | 0 | 0.17 | 0.01 | 0.18 | 0.27 | 0.14 | 0.15 |
| 1 | 40168865 | rs17561616 | -0.26 | 1.597 | -1.829 | 0.485 | 0.014 | -0.02 | -0.02 | 0 | 0.07 | 0.12 | 0.08 | 0.01 | 0.11 | 0.18 |
| 1 | 40171226 | rs17561720 | -0.034 | 1.842 | -2.421 | 0.035 | -0.05 | 0.048 | 0.048 | 0.021 | 0.13 | 0.02 | 0.1 | 0.04 | 0.19 | 0.08 |
| 1 | 40183922 | rs481423 | 0.745 | 1.078 | -2.009 | -0.684 | -0.041 | 0.041 | 0.041 | 1.579 | 0.31 | 0.37 | 0.22 | 0.51 | 0.19 | 0.27 |
| 1 | 40184733 | rs704782 | 0.078 | 1.022 | -2.003 | -0.433 | -0.024 | -0.016 | -0.016 | 0 | 0.43 | 0.41 | 0.31 | 0.66 | 0.24 | 0.37 |
| 1 | 40188113 | rs704784 | -0.808 | 1.252 | -2.127 | -0.851 | -0.001 | -0.008 | -0.008 | 0 | 0.63 | 0.62 | 0.55 | 0.79 | 0.54 | 0.68 |
| 1 | 40190102 | rs12406805 | 1.825 | 1.245 | -2.054 | -0.749 | -0.023 | -0.093 | -0.093 | 0 | 0.15 | 0.01 | 0.18 | 0.27 | 0.12 | 0.14 |
| 1 | 40193772 | rs11579412 | 1.585 | 0.665 | -1.725 | -1.03 | 0.031 | 0.032 | 0.032 | 1.672 | 0.18 | 0.1 | 0.07 | 0.04 | 0.09 | 0.04 |
| 1 | 40194903 | rs539853 | 0.887 | 0.651 | -1.749 | -1 | -0.061 | 0.095 | 0.095 | 0.305 | 0.23 | 0.38 | 0.12 | 0.39 | 0.11 | 0.21 |
| 1 | 40195563 | rs7536524 | 0.134 | 0.99 | -1.54 | -1.341 | -0.017 | 0.01 | 0.01 | 0 | 0.17 | 0.13 | 0.13 | 0.01 | 0.23 | 0.24 |
| 1 | 40195574 | rs704785 | -1.098 | 0.705 | -1.427 | -1.041 | 0.021 | -0.03 | -0.03 | 0 | 0.82 | 0.65 | 0.66 | 0.83 | 0.8 | 0.85 |
| 1 | 40199854 | rs704787 | -0.288 | 0.499 | -1.465 | -1.098 | -0.001 | -0.019 | -0.019 | 0 | 0.09 | 0.1 | 0.05 | 0.02 | 0.1 | 0.1 |
| 1 | 40203722 | rs12074147 | -0.226 | 0.323 | -1.22 | -0.726 | -0.001 | 0 | 0 | 0 | 0.29 | 0.23 | 0.29 | 0.25 | 0.36 | 0.41 |
| 1 | 40207169 | rs497333 | -1.13 | 0.35 | -0.871 | -0.462 | 0.054 | 0.12 | 0.12 | 1.94 | 0.77 | 0.54 | 0.54 | 0.46 | 0.68 | 0.64 |
| 1 | 40216993 | rs501523 | 1.222 | -0.015 | -1.407 | -0.895 | -0.01 | 0.007 | 0.007 | 0 | 0.12 | 0.05 | 0.14 | 0 | 0.16 | 0.09 |
| 1 | 40219065 | rs1046988 | 0.611 | -0.386 | -1.386 | -0.766 | -0.008 | 0.004 | 0.004 | 0.318 | 0.28 | 0.17 | 0.28 | 0.23 | 0.36 | 0.41 |
| 1 | 40219280 | rs14528 | 0.061 | -0.243 | -1.218 | -0.842 | 0.004 | 0.011 | 0.011 | 0.333 | 0.55 | 0.37 | 0.35 | 0.37 | 0.45 | 0.5 |
| 1 | 40223140 | rs1053846 | 0.774 | 0.863 | -1.025 | -0.338 | -0.067 | 0.023 | 0.023 | 3.043 | 0.15 | 0.02 | 0.12 | 0.09 | 0.23 | 0.33 |
| 1 | 40242315 | rs914962 | 0.254 | 0.692 | -1.124 | -0.057 | -0.009 | -0.002 | -0.002 | 0 | 0.35 | 0.6 | 0.27 | 0.32 | 0.27 | 0.22 |
| 1 | 40255872 | rs230280 | -0.315 | 1.876 | -0.438 | 0.9 | 0.028 | 0.057 | 0.057 | 1.952 | 0.79 | 0.73 | 0.55 | 0.46 | 0.66 | 0.63 |
| 1 | 40258993 | rs230284 | 0.831 | 1.229 | -0.265 | 0.911 | -0.001 | -0.001 | -0.001 | 0 | 0.54 | 0.54 | 0.57 | 0.4 | 0.56 | 0.58 |
| 1 | 40275071 | rs7531010 | 0.614 | 0.381 | -0.574 | 0.735 | -0.017 | -0.056 | -0.056 | 0 | 0.17 | 0.23 | 0.07 | 0 | 0.1 | 0.04 |
| 1 | 40275549 | rs2676697 | 0.175 | 1.124 | -0.913 | 1.736 | -0.004 | -0.031 | -0.031 | 0 | 0.87 | 0.93 | 0.8 | 0.67 | 0.87 | 0.83 |
| 1 | 40286005 | rs10889022 | -0.599 | 1.007 | -0.499 | 1.272 | 0.001 | 0.069 | 0.069 | 0 | 0.18 | 0.27 | 0.24 | 0.58 | 0.22 | 0.22 |
| 1 | 40288051 | rs1999574 | 0.384 | 1.107 | -1.236 | 0.376 | 0.002 | 0.028 | 0.028 | 0 | 0.37 | 0.25 | 0.47 | 0.31 | 0.35 | 0.42 |
| 1 | 40289812 | rs1999573 | 0.023 | 0.925 | -1.387 | 0.562 | -0.023 | 0.059 | 0.059 | 0.042 | 0.48 | 0.11 | 0.55 | 0.36 | 0.55 | 0.54 |
| 1 | 40290277 | rs2087796 | -0.067 | 0.663 | -1.281 | 0.256 | 0.018 | -0.028 | -0.028 | 0 | 0.68 | 0.29 | 0.66 | 0.46 | 0.76 | 0.75 |
| 1 | 40290597 | rs230271 | -0.712 | 0.769 | -1.369 | 0.209 | 0.005 | 0.078 | 0.078 | 0.133 | 0.51 | 0.87 | 0.46 | 0.65 | 0.45 | 0.46 |
| 1 | 40294186 | rs6600301 | -0.659 | 1.006 | -1.479 | 0.379 | -0.018 | 0.213 | 0.213 | 0.053 | 0.5 | 0.92 | 0.47 | 0.81 | 0.44 | 0.48 |
| 1 | 40302463 | rs6693041 | -0.331 | 0.439 | -0.824 | 0.832 | -0.007 | 0.018 | 0.018 | 0 | 0.18 | 0.22 | 0.25 | 0.29 | 0.16 | 0.22 |
| 1 | 40303103 | rs6693076 | -0.936 | 0.315 | 0.178 | 1.059 | 0 | 0 | 0 | 1.403 | 0.39 | 0.62 | 0.41 | 0.59 | 0.39 | 0.46 |
| 1 | 40306865 | rs1043843 | 1.426 | 0.305 | 0.402 | 1.108 | -0.021 | 0.027 | 0.027 | 0 | 0.16 | 0.1 | 0.12 | 0.01 | 0.19 | 0.21 |
| 1 | 40306898 | rs7315 | -1.062 | 0.295 | 0.383 | 1.104 | 0 | 0 | 0 | 1.227 | 0.38 | 0.62 | 0.41 | 0.59 | 0.39 | 0.46 |
| 1 | 40307719 | rs230310 | -1.638 | 0.017 | -0.494 | 0.407 | -0.061 | -0.087 | -0.087 | 0.497 | 0.83 | 0.98 | 0.64 | 0.68 | 0.77 | 0.75 |
| 1 | 40324097 | rs6680378 | -1.228 | 0.027 | 0.419 | 0.344 | -0.013 | 0.027 | 0.027 | 0 | 0.52 | 0.9 | 0.47 | 0.59 | 0.44 | 0.48 |
| 1 | 40324210 | rs17570439 | 1.478 | 0.032 | 0.42 | 0.328 | -0.002 | -0.024 | -0.024 | 0 | 0.16 | 0.19 | 0.13 | 0.01 | 0.19 | 0.21 |
| 1 | 40343432 | rs10789049 | -1.114 | 0.058 | 0.206 | 0.313 | 0.005 | 0.033 | 0.033 | 0.048 | 0.51 | 0.9 | 0.46 | 0.59 | 0.43 | 0.48 |
| 1 | 40357268 | rs3131691 | -0.515 | 0.041 | 0.402 | -0.013 | -0.048 | -0.008 | -0.008 | 0 | 0.31 | 0.45 | 0.3 | 0.29 | 0.21 | 0.24 |
| 1 | 40362066 | rs3134615 | 1.238 | 0.218 | -0.008 | -0.401 | -0.057 | 0.06 | 0.06 | 0 | 0.17 | 0.06 | 0.12 | 0.04 | 0.22 | 0.23 |
| 1 | 40364803 | rs3134613 | -1.307 | 0.01 | -0.114 | -0.482 | -0.06 | -0.066 | -0.066 | 0 | 0.54 | 0.9 | 0.47 | 0.5 | 0.47 | 0.51 |
| 1 | 40368905 | rs7548524 | -0.6 | -0.271 | 0.586 | -0.111 | 0.043 | 0.56 | 0.56 | 0.58 | 0.06 | 0.44 | 0.16 | 0.45 | 0.01 | 0.08 |
| 1 | 40373232 | rs6657848 | 1.363 | -0.072 | 0.291 | -0.587 | -0.038 | -0.078 | -0.078 | 0 | 0.13 | 0.25 | 0.05 | 0 | 0.05 | 0.01 |
| 1 | 40384968 | rs12127002 | 0.207 | 0.081 | -0.298 | -0.472 | -0.012 | 0.052 | 0.052 | 0.406 | 0.12 | 0.02 | 0.11 | 0 | 0.1 | 0.14 |
| 1 | 40389420 | rs3131644 | -0.996 | 0.053 | -0.205 | -0.399 | 0.009 | -0.043 | -0.043 | 2.911 | 0.64 | 0.77 | 0.59 | 0.5 | 0.66 | 0.56 |
| 1 | 40391752 | rs7542907 | 0.635 | 0.197 | 0.016 | -0.731 | 0.02 | -0.155 | -0.155 | 0 | 0.22 | 0.49 | 0.15 | 0.01 | 0.23 | 0.23 |
| 1 | 40393195 | rs7514618 | 0.178 | 0.125 | -0.401 | -0.922 | -0.029 | 0.173 | 0.173 | 0.595 | 0.33 | 0.08 | 0.24 | 0 | 0.33 | 0.17 |
| 1 | 40395169 | rs3103780 | 0.342 | 0.027 | -0.422 | -0.628 | 0.001 | -0.017 | -0.017 | 0 | 0.27 | 0.75 | 0.29 | 0.41 | 0.25 | 0.31 |

| | | | | | | | | | | | | | | | | |
|---|---|---|---|---|---|---|---|---|---|---|---|---|---|---|---|---|
| 1 | 40399734 | rs12566893 | 0.714 | -0.052 | -0.313 | -0.382 | -0.023 | -0.09 | -0.09 | 0 | 0.36 | 0.07 | 0.39 | 0.57 | 0.31 | 0.47 |
| 1 | 40403414 | rs11207167 | 0.225 | 0.218 | 0.192 | -0.233 | -0.041 | 0.159 | 0.159 | 0.349 | 0.34 | 0.1 | 0.26 | 0 | 0.38 | 0.17 |
| 1 | 40413098 | rs3131706 | -0.172 | -0.092 | 0.638 | -0.102 | 0 | -0.095 | -0.095 | 0 | 0.61 | 0.54 | 0.66 | 0.99 | 0.57 | 0.78 |
| 1 | 40418589 | rs3131704 | -0.886 | -0.044 | -0.19 | -0.047 | -0.074 | 0.157 | 0.157 | 0 | 0.69 | 0.93 | 0.71 | 0.98 | 0.59 | 0.79 |
| 1 | 40433771 | rs3103778 | -0.143 | 0.125 | 0.065 | 0.383 | -0.005 | 0.015 | 0.015 | 0.136 | 0.53 | 0.29 | 0.53 | 0.57 | 0.51 | 0.72 |
| 1 | 40437831 | rs11207226 | -0.507 | 0.19 | 1.028 | 1.387 | -0.004 | -0.011 | -0.011 | 0 | 0.05 | 0.13 | 0.01 | 0 | 0.01 | 0 |
| 1 | 40437923 | rs11207227 | -0.654 | 0.22 | 0.363 | 1.693 | -0.022 | 0.028 | 0.028 | 0 | 0.29 | 0.54 | 0.39 | 0.42 | 0.2 | 0.37 |
| 1 | 40458920 | rs4660371 | -1.56 | 0.175 | 0.17 | 1.365 | -0.02 | 0.005 | 0.005 | 0 | 0.14 | 0.24 | 0.09 | 0.17 | 0.06 | 0.28 |
| 1 | 40462662 | rs4233484 | 0.761 | 0.342 | 0.926 | 1.434 | 0.035 | 0.019 | 0.019 | 0 | 0.87 | 0.68 | 0.76 | 0.87 | 0.87 | 0.92 |
| 1 | 40499302 | rs11207370 | -1.339 | 0.364 | 1.213 | 1.404 | 0.003 | -0.002 | -0.002 | 0 | 0.14 | 0.17 | 0.24 | 0.12 | 0.14 | 0.12 |
| 1 | 40504550 | rs3806207 | -0.524 | 0.182 | 1.46 | 1.789 | -0.009 | 0.026 | 0.026 | 0 | 0.13 | 0.26 | 0.09 | 0.25 | 0.08 | 0.2 |
| 1 | 40513710 | rs3131683 | 2.336 | 0.108 | 1.57 | 2.003 | 0.027 | -0.041 | -0.041 | 0 | 0.58 | 0.25 | 0.54 | 0.46 | 0.67 | 0.45 |
| 1 | 40523328 | rs3122416 | 1.664 | 0.118 | 1.735 | 2.025 | -0.006 | 0.027 | 0.027 | 0.466 | 0.71 | 0.45 | 0.64 | 0.6 | 0.77 | 0.68 |
| 1 | 40539076 | rs1126973 | 1.373 | 0.122 | 1.749 | 2.018 | 0.001 | 0.016 | 0.016 | 0.404 | 0.73 | 0.6 | 0.67 | 0.63 | 0.77 | 0.69 |
| 1 | 40543019 | rs6681804 | -1.09 | 0.12 | 1.521 | 1.967 | -0.014 | 0.009 | 0.009 | 0 | 0.12 | 0.17 | 0.1 | 0.14 | 0.1 | 0.21 |
| 1 | 40547950 | rs3131660 | 2.191 | 0.017 | 1.345 | 1.998 | 0 | -0.04 | -0.04 | 0 | 0.65 | 0.24 | 0.55 | 0.56 | 0.68 | 0.55 |
| 1 | 40551745 | rs6694800 | 0.505 | 0.483 | 1.263 | 2.053 | -0.01 | -0.014 | -0.014 | 7.473 | 0.06 | 0.02 | 0.03 | 0.11 | 0.06 | 0.07 |
| 1 | 40553333 | rs7513553 | -1.852 | 0.459 | 1.551 | 2.015 | 0.022 | 0.087 | 0.087 | 0 | 0.2 | 0.36 | 0.19 | 0.39 | 0.21 | 0.37 |
| 1 | 40553444 | rs12118846 | -0.862 | 0.462 | 1.573 | 2.027 | 0.024 | 0.085 | 0.085 | 0.379 | 0.11 | 0.23 | 0.08 | 0.22 | 0.13 | 0.23 |
| 1 | 40555321 | rs3131651 | 2.649 | 0.446 | 1.638 | 2.013 | -0.006 | -0.039 | -0.039 | 0 | 0.6 | 0.24 | 0.52 | 0.45 | 0.63 | 0.48 |
| 1 | 40560399 | rs4660390 | 2.054 | 0.489 | 1.678 | 1.941 | -0.004 | 0.005 | 0.005 | 0.659 | 0.7 | 0.45 | 0.6 | 0.66 | 0.75 | 0.7 |
| 1 | 40560831 | rs11207493 | -0.09 | 0.499 | 1.625 | 1.943 | 0.004 | 0.01 | 0.01 | 0 | 0.09 | 0.13 | 0.12 | 0.18 | 0.09 | 0.14 |
| 1 | 40561471 | rs10889147 | 0.845 | 0.509 | 1.686 | 1.915 | 0.006 | 0.017 | 0.017 | 0.132 | 0.87 | 0.72 | 0.78 | 0.88 | 0.89 | 0.9 |
| 1 | 40561911 | rs6600312 | 2.41 | 0.498 | 1.69 | 1.941 | -0.002 | -0.006 | -0.006 | 2.867 | 0.67 | 0.34 | 0.56 | 0.64 | 0.68 | 0.57 |
| 1 | 40563700 | rs4660159 | 0.176 | 0.718 | 1.902 | 2.172 | 0.04 | 0.069 | 0.069 | 0.035 | 0.92 | 0.78 | 0.8 | 0.88 | 0.91 | 0.94 |
| 1 | 40564994 | rs6600317 | 1.277 | 0.716 | 1.915 | 2.174 | 0.038 | 0.079 | 0.079 | 0.698 | 0.83 | 0.61 | 0.68 | 0.7 | 0.83 | 0.8 |
| 1 | 40565508 | rs7524958 | 0.937 | 0.72 | 1.849 | 2.153 | 0.094 | 0.159 | 0.159 | 0 | 0.88 | 0.58 | 0.75 | 0.84 | 0.84 | 0.81 |
| 1 | 40573134 | rs12074195 | -0.696 | 0.765 | 1.703 | 2.074 | 0.008 | 0.038 | 0.038 | 0 | 0.1 | 0.23 | 0.15 | 0.22 | 0.09 | 0.14 |
| 1 | 40584798 | rs10493089 | 0.124 | 0.906 | 1.61 | 2.125 | -0.001 | -0.002 | -0.002 | 0 | 0.08 | 0.07 | 0.13 | 0.17 | 0.08 | 0.16 |
| 1 | 40586438 | rs12035579 | 2.584 | 0.851 | 1.618 | 2.126 | -0.048 | -0.07 | -0.07 | 0.008 | 0.63 | 0.13 | 0.59 | 0.5 | 0.63 | 0.5 |
| 1 | 40598775 | rs4492609 | -2.02 | 0.847 | 1.709 | 2.094 | 0.038 | -0.025 | -0.025 | 0 | 0.19 | 0.56 | 0.22 | 0.12 | 0.23 | 0.19 |
| 1 | 40601636 | rs6600318 | -2.534 | 0.836 | 1.704 | 2.092 | -0.003 | 0 | 0 | 1.112 | 0.27 | 0.7 | 0.35 | 0.29 | 0.31 | 0.36 |
| 1 | 40624273 | rs9725035 | -1.274 | 0.829 | 1.718 | 2.079 | 0.018 | -0.048 | -0.048 | 0.422 | 0.14 | 0.29 | 0.15 | 0.06 | 0.14 | 0.04 |
| 1 | 40626918 | rs10493090 | 0.598 | 0.727 | 1.65 | 2.086 | -0.001 | 0 | 0 | 0 | 0.07 | 0.07 | 0.13 | 0.17 | 0.08 | 0.16 |
| 1 | 40631460 | rs11584362 | -1.04 | 0.634 | 1.619 | 2.094 | -0.001 | 0 | 0 | 0 | 0.15 | 0.14 | 0.18 | 0.17 | 0.14 | 0.2 |
| 1 | 40636387 | rs4600036 | -0.017 | 0.762 | 1.648 | 2.091 | 0.009 | -0.022 | -0.022 | 0 | 0.14 | 0.19 | 0.16 | 0.06 | 0.18 | 0.05 |
| 1 | 40646447 | rs11207626 | 2.072 | 0.75 | 1.65 | 2.128 | -0.033 | -0.038 | -0.038 | 0 | 0.64 | 0.22 | 0.59 | 0.51 | 0.61 | 0.49 |
| 1 | 40676406 | rs9438936 | 0.18 | 0.777 | 1.648 | 2.134 | 0 | 0.01 | 0.01 | 0.911 | 0.08 | 0.08 | 0.04 | 0.21 | 0.06 | 0.15 |
| 1 | 40679709 | rs6600321 | -2.649 | 0.77 | 1.642 | 2.115 | -0.055 | -0.062 | -0.062 | 0.375 | 0.38 | 0.89 | 0.43 | 0.49 | 0.39 | 0.51 |
| 1 | 40708853 | rs10889209 | -1.969 | 0.807 | 1.555 | 2.1 | -0.005 | 0.001 | 0.001 | 0 | 0.29 | 0.7 | 0.37 | 0.29 | 0.33 | 0.39 |
| 1 | 40714010 | rs12126903 | 0.114 | 0.885 | 0.884 | 2.067 | 0 | -0.006 | -0.006 | 0 | 0.08 | 0.07 | 0.04 | 0.15 | 0.06 | 0.19 |
| 1 | 40722795 | rs926830 | -0.183 | 0.974 | 0.799 | 2.247 | 0.008 | 0.004 | 0.004 | 0 | 0.19 | 0.25 | 0.19 | 0.22 | 0.23 | 0.18 |
| 1 | 40723618 | rs3765483 | -0.097 | 1.034 | -0.289 | 2.116 | 0.002 | 0.014 | 0.014 | 0 | 0.08 | 0.12 | 0.12 | 0.17 | 0.08 | 0.17 |
| 1 | 40735817 | rs16827109 | 0.603 | 1.035 | -0.292 | 2.101 | -0.002 | -0.002 | -0.002 | 0 | 0.06 | 0.07 | 0.12 | 0.17 | 0.07 | 0.16 |
| 1 | 40736487 | rs6682701 | -1.621 | 1.016 | -0.335 | 2.228 | -0.012 | -0.002 | -0.002 | 0 | 0.38 | 0.75 | 0.49 | 0.63 | 0.37 | 0.56 |
| 1 | 40740206 | rs2284446 | -0.472 | 0.843 | 0.087 | 2.536 | 0 | -0.001 | -0.001 | 0 | 0.08 | 0.07 | 0.14 | 0.21 | 0.06 | 0.2 |
| 1 | 40741899 | rs6676644 | -0.788 | 0.793 | -0.281 | 2.413 | -0.013 | -0.03 | -0.03 | 0 | 0.87 | 0.98 | 0.93 | 0.85 | 0.86 | 0.93 |
| 1 | 40742366 | rs12757549 | -0.412 | 0.883 | 0.288 | 2.132 | 0.005 | 0.017 | 0.017 | 0 | 0.07 | 0.12 | 0.13 | 0.17 | 0.08 | 0.17 |
| 1 | 40753985 | rs11207778 | -1.223 | 0.749 | -0.028 | 1.746 | 0.004 | 0.004 | 0.004 | 0.019 | 0.07 | 0.16 | 0.06 | 0.08 | 0.08 | 0.13 |
| 1 | 40759064 | rs10489431 | -1.16 | 0.669 | -0.417 | 1.667 | 0 | -0.004 | -0.004 | 1.005 | 0.08 | 0.06 | 0.14 | 0.2 | 0.06 | 0.19 |
| 1 | 40759231 | rs10489432 | -1.16 | 0.669 | -0.417 | 1.667 | 0 | -0.004 | -0.004 | 0 | 0.08 | 0.06 | 0.14 | 0.2 | 0.06 | 0.19 |
| 1 | 40773289 | rs2075560 | -1.028 | 0.143 | 0.21 | 1.741 | -0.002 | 0.007 | 0.007 | 0 | 0.06 | 0.07 | 0.05 | 0.18 | 0.03 | 0.12 |
| 1 | 40775229 | rs209914 | -0.507 | 0.012 | -0.912 | 0.539 | -0.01 | 0.19 | 0.19 | 2.317 | 0.63 | 0.92 | 0.69 | 0.95 | 0.64 | 0.81 |
| 1 | 40779841 | rs2273195 | -1.546 | -0.079 | -0.411 | 0.695 | -0.037 | 0.029 | 0.029 | 0 | 0.11 | 0.22 | 0.07 | 0.2 | 0.04 | 0.17 |
| 1 | 40788961 | rs578039 | -1.442 | 0.116 | -0.019 | 0.551 | -0.007 | 0.04 | 0.04 | 0 | 0.14 | 0.42 | 0.11 | 0.17 | 0.09 | 0.16 |
| 1 | 40789927 | rs11207870 | 0.178 | 0.983 | -0.738 | 1.311 | -0.043 | -0.001 | -0.001 | 0 | 0.07 | 0.01 | 0.23 | 0.44 | 0.18 | 0.1 |
| 1 | 40800486 | rs10889285 | -1.014 | 0.327 | -0.532 | 0.82 | -0.01 | 0.043 | 0.043 | 0 | 0.12 | 0.23 | 0.18 | 0.25 | 0.1 | 0.2 |
| 1 | 40809669 | rs432803 | 0.657 | 0.108 | -0.714 | -0.294 | 0.052 | 0.078 | 0.078 | 0.117 | 0.5 | 0.58 | 0.25 | 0.14 | 0.35 | 0.42 |
| 1 | 40814917 | rs734922 | -0.908 | 0.454 | -0.452 | 0.013 | 0.002 | 0.01 | 0.01 | 0.036 | 0.14 | 0.13 | 0.11 | 0.04 | 0.12 | 0.13 |
| 1 | 40823379 | rs388538 | -1.209 | 0.37 | -0.005 | -0.027 | -0.014 | -0.07 | -0.07 | 0 | 0.16 | 0.4 | 0.14 | 0.04 | 0.12 | 0.11 |
| 1 | 40827598 | rs3013462 | 0.005 | 0.831 | -0.663 | 0.359 | -0.034 | -0.014 | -0.014 | 0 | 0.58 | 0.8 | 0.56 | 0.6 | 0.46 | 0.54 |
| 1 | 40838976 | rs176627 | -0.267 | 1.424 | -0.688 | 0.303 | -0.002 | -0.007 | -0.007 | 0 | 0.39 | 0.28 | 0.36 | 0.39 | 0.42 | 0.42 |
| 1 | 40845981 | rs209580 | 0.268 | 1.278 | -0.867 | 0.457 | -0.013 | 0.026 | 0.026 | 0 | 0.31 | 0.2 | 0.23 | 0.19 | 0.37 | 0.27 |
| 1 | 40847491 | rs209583 | -0.429 | 1.287 | -0.866 | 0.446 | -0.006 | -0.012 | -0.012 | 0 | 0.38 | 0.23 | 0.36 | 0.39 | 0.42 | 0.42 |
| 1 | 40848821 | rs6656226 | -0.138 | 1.63 | -0.806 | 1.121 | -0.004 | -0.001 | -0.001 | 0.073 | 0.07 | 0.09 | 0.06 | 0.19 | 0.09 | 0.11 |
| 1 | 40865633 | rs2982506 | -0.124 | 1.598 | 0.163 | 0.931 | 0.055 | -0.017 | -0.017 | 0.013 | 0.1 | 0.33 | 0.11 | 0.04 | 0.13 | 0.06 |
| 1 | 40866288 | rs2982508 | -1.116 | 1.393 | 0.38 | 1.161 | 0.036 | 0.039 | 0.039 | 0 | 0.14 | 0.47 | 0.15 | 0.11 | 0.15 | 0.1 |
| 1 | 40869272 | rs3013456 | 0.256 | 1.401 | 0.399 | 1.153 | -0.004 | 0.003 | 0.003 | 0 | 0.08 | 0.09 | 0.09 | 0.04 | 0.13 | 0.06 |
| 1 | 40870545 | rs16827275 | -0.318 | 1.313 | 0.272 | 1.241 | 0.016 | -0.032 | -0.032 | 0 | 0.09 | 0.05 | 0.13 | 0.2 | 0.07 | 0.15 |

| | | | | | | | | | | | | | | | | |
|---|---|---|---|---|---|---|---|---|---|---|---|---|---|---|---|---|
| 1 | 40871497 | rs209599 | -0.539 | 1.431 | 0.314 | 1.155 | -0.012 | 0.008 | 0.008 | 0 | 0.94 | 0.97 | 0.92 | 0.98 | 0.88 | 0.94 |
| 1 | 40872623 | rs2982510 | -1.077 | 0.679 | 0.382 | 1.221 | 0.049 | 0.073 | 0.073 | 0 | 0.2 | 0.53 | 0.2 | 0.29 | 0.24 | 0.22 |
| 1 | 40885832 | rs11208057 | 1.352 | 0.359 | -0.198 | 1.083 | 0.054 | 0.022 | 0.022 | 1.83 | 0.34 | 0.16 | 0.35 | 0.3 | 0.2 | 0.3 |
| 1 | 40897492 | rs209618 | 0.053 | 0.274 | 0.134 | 1.182 | 0.023 | 0.003 | 0.003 | 0.404 | 0.5 | 0.4 | 0.47 | 0.5 | 0.4 | 0.45 |
| 1 | 40916218 | rs209607 | 0.468 | 0.171 | 0.338 | 1.508 | 0.008 | -0.003 | -0.003 | 0.03 | 0.44 | 0.56 | 0.54 | 0.52 | 0.29 | 0.51 |
| 1 | 40921145 | rs2982500 | 0.245 | 0.208 | 0.044 | 1.41 | 0.017 | 0.017 | 0.017 | 0.113 | 0.47 | 0.43 | 0.55 | 0.68 | 0.3 | 0.53 |
| 1 | 40921238 | rs12568089 | -1.18 | 0.527 | 0.115 | 1.616 | -0.004 | 0.023 | 0.023 | 0 | 0.08 | 0.14 | 0.14 | 0.21 | 0.04 | 0.18 |
| 1 | 40922062 | rs10489434 | 0.142 | 0.661 | -0.001 | 1.616 | -0.004 | 0.002 | 0.002 | 0.397 | 0.07 | 0.06 | 0.04 | 0.04 | 0.13 | 0.08 |
| 1 | 40922377 | rs10489435 | -0.435 | 0.61 | -0.005 | 1.683 | -0.003 | 0.032 | 0.032 | 0 | 0.05 | 0.07 | 0.12 | 0.21 | 0.02 | 0.17 |
| 1 | 40924019 | rs7535790 | -1.28 | 0.523 | 0.084 | 1.429 | -0.025 | 0.101 | 0.101 | 0 | 0.24 | 0.52 | 0.24 | 0.51 | 0.17 | 0.28 |
| 1 | 40925551 | rs2036198 | -0.265 | 0.759 | -0.702 | 0.956 | -0.012 | 0.015 | 0.015 | 0.38 | 0.1 | 0.06 | 0.05 | 0.03 | 0.14 | 0.07 |
| 1 | 40930624 | rs2982502 | -1.561 | 0.605 | -0.68 | 0.991 | 0.008 | -0.021 | -0.021 | 0 | 0.35 | 0.73 | 0.26 | 0.47 | 0.33 | 0.27 |
| 1 | 40930890 | rs3969615 | 0.207 | 0.834 | -0.761 | 0.77 | 0 | 0 | 0 | 0 | 0.56 | 0.43 | 0.65 | 0.57 | 0.53 | 0.61 |
| 1 | 40931003 | rs736861 | 0.472 | 0.621 | -1.248 | 0.71 | 0 | -0.018 | -0.018 | 1.449 | 0.44 | 0.16 | 0.57 | 0.47 | 0.39 | 0.53 |
| 1 | 40945022 | rs4076058 | -0.48 | 0.182 | -0.223 | 0.243 | -0.019 | -0.039 | -0.039 | 0 | 0.08 | 0.12 | 0.03 | 0 | 0.05 | 0.01 |
| 1 | 40948669 | rs7534068 | -0.364 | -0.237 | -0.335 | 0.165 | 0.029 | 0.001 | 0.001 | 0 | 0.51 | 0.65 | 0.53 | 0.5 | 0.37 | 0.55 |
| 1 | 40961449 | rs3795348 | 0.387 | -0.156 | -0.315 | 0.064 | 0 | -0.003 | -0.003 | 0 | 0.05 | 0.06 | 0.03 | 0.02 | 0.03 | 0.06 |
| 1 | 40966167 | rs7511983 | 0.672 | -0.178 | -0.31 | 0.051 | -0.018 | -0.039 | -0.039 | 0 | 0.08 | 0.12 | 0.03 | 0 | 0.04 | 0.01 |
| 1 | 40966183 | rs7512061 | -1.313 | -0.148 | -0.718 | 0.018 | 0.01 | 0.026 | 0.026 | 0 | 0.89 | 0.73 | 0.8 | 0.85 | 0.88 | 0.86 |
| 1 | 40969063 | rs6666654 | -0.103 | -0.293 | -0.163 | 0.001 | 0.005 | -0.004 | -0.004 | 0 | 0.07 | 0.17 | 0.04 | 0.03 | 0.03 | 0.06 |
| 1 | 40974156 | rs3795347 | 0.64 | -0.717 | -0.384 | -0.245 | 0.015 | 0.026 | 0.026 | 0 | 0.31 | 0.21 | 0.15 | 0.09 | 0.22 | 0.19 |
| 1 | 40974799 | rs12068587 | -0.919 | -0.723 | -0.427 | -0.23 | 0.036 | 0.016 | 0.016 | 0 | 0.51 | 0.71 | 0.54 | 0.57 | 0.38 | 0.56 |
| 1 | 40980731 | rs11208299 | -0.894 | -0.734 | -0.432 | -0.235 | 0.036 | 0.006 | 0.006 | 0 | 0.51 | 0.7 | 0.54 | 0.5 | 0.38 | 0.55 |
| 1 | 40981230 | rs3795344 | 1.088 | -0.733 | -0.432 | -0.235 | 0 | -0.003 | -0.003 | 0.053 | 0.05 | 0.06 | 0.03 | 0.02 | 0.03 | 0.06 |
| 1 | 40997033 | rs2362438 | 1.692 | 0.01 | -0.838 | -0.445 | -0.04 | -0.008 | -0.008 | 0 | 0.07 | 0 | 0.05 | 0.04 | 0.17 | 0.06 |
| 1 | 40998406 | rs1000370 | -0.733 | -0.136 | -0.767 | -0.392 | 0.035 | 0.016 | 0.016 | 0 | 0.51 | 0.71 | 0.54 | 0.57 | 0.38 | 0.56 |
| 1 | 40998811 | rs6659361 | 1.929 | -0.175 | -0.78 | -0.412 | 0.002 | 0.073 | 0.073 | 0 | 0.22 | 0.06 | 0.11 | 0.03 | 0.18 | 0.16 |
| 1 | 40998902 | rs1001400 | -0.046 | -0.164 | -0.769 | -0.399 | 0.002 | 0.192 | 0.192 | 0 | 0.2 | 0.5 | 0.39 | 0.47 | 0.15 | 0.37 |
| 1 | 41006154 | rs16865682 | 1.632 | -0.182 | -0.701 | -0.408 | -0.012 | -0.029 | -0.029 | 0 | 0.08 | 0.09 | 0.03 | 0 | 0.04 | 0.01 |
| 1 | 41008259 | rs11208360 | -0.656 | -0.187 | -0.382 | -0.413 | 0.03 | 0.016 | 0.016 | 0 | 0.5 | 0.66 | 0.54 | 0.57 | 0.38 | 0.56 |
| 1 | 41010378 | rs7524182 | 1.808 | -0.163 | -0.397 | -0.421 | 0.005 | 0.074 | 0.074 | 0 | 0.22 | 0.07 | 0.11 | 0.03 | 0.18 | 0.16 |
| 1 | 41010454 | rs11208363 | -0.147 | -0.107 | -0.296 | -0.383 | -0.002 | 0.167 | 0.167 | 0 | 0.2 | 0.44 | 0.39 | 0.54 | 0.15 | 0.38 |
| 1 | 41013015 | rs2297755 | 1.189 | -0.078 | -0.25 | -0.355 | 0 | 0.006 | 0.006 | 0 | 0.13 | 0.11 | 0.34 | 0.45 | 0.11 | 0.31 |
| 1 | 41013260 | rs2297756 | -0.614 | -0.034 | -0.282 | -0.331 | 0.038 | 0.016 | 0.016 | 0 | 0.51 | 0.71 | 0.54 | 0.57 | 0.37 | 0.56 |
| 1 | 41021976 | rs4660429 | -0.518 | -0.081 | -0.281 | -0.338 | 0.048 | 0.008 | 0.008 | 0 | 0.48 | 0.69 | 0.52 | 0.56 | 0.35 | 0.53 |
| 1 | 41032059 | rs7537003 | -0.428 | -0.091 | -0.34 | -0.21 | -0.004 | 0.004 | 0.004 | 0 | 0.4 | 0.56 | 0.48 | 0.56 | 0.31 | 0.56 |
| 1 | 41042068 | rs4660436 | 0.174 | 0.034 | -0.625 | -0.305 | -0.029 | -0.069 | -0.069 | 0 | 0.23 | 0.12 | 0.39 | 0.49 | 0.3 | 0.42 |
| 1 | 41050952 | rs4660438 | -0.771 | -0.309 | -0.312 | -0.652 | 0.003 | 0.001 | 0.001 | 0 | 0.53 | 0.62 | 0.58 | 0.55 | 0.53 | 0.66 |
| 1 | 41051233 | rs16865687 | -0.395 | -0.288 | -0.241 | -0.795 | -0.004 | -0.014 | -0.014 | 0.341 | 0.09 | 0.12 | 0.05 | 0.02 | 0.05 | 0.08 |
| 1 | 41053260 | rs6677621 | -0.812 | -0.383 | -0.165 | -0.787 | -0.01 | 0.005 | 0.005 | 0 | 0.09 | 0.2 | 0.05 | 0.06 | 0.05 | 0.08 |
| 1 | 41057570 | rs16827403 | 1.447 | -0.352 | -0.394 | -0.904 | -0.001 | 0.032 | 0.032 | 0 | 0.16 | 0.07 | 0.09 | 0.02 | 0.15 | 0.16 |
| 1 | 41061519 | rs7526314 | -0.532 | -0.53 | 0.093 | -0.987 | -0.025 | -0.077 | -0.077 | 0 | 0.31 | 0.53 | 0.17 | 0.08 | 0.22 | 0.26 |
| 1 | 41073802 | rs618144 | 0.481 | -0.484 | 0.085 | -0.654 | -0.006 | -0.002 | -0.002 | 0 | 0.46 | 0.43 | 0.47 | 0.46 | 0.5 | 0.34 |
| 1 | 41075209 | rs500586 | -0.15 | -0.349 | -0.113 | -0.667 | 0.011 | -0.021 | -0.021 | 0 | 0.63 | 0.77 | 0.6 | 0.52 | 0.68 | 0.52 |
| 1 | 41083947 | rs12067173 | 0.784 | -0.331 | -0.186 | -0.76 | -0.003 | 0.024 | 0.024 | 0 | 0.19 | 0.12 | 0.11 | 0.06 | 0.19 | 0.19 |
| 1 | 41088577 | rs17412556 | 0.405 | -0.046 | -0.226 | -0.696 | -0.003 | 0 | 0 | 0 | 0.12 | 0.11 | 0.07 | 0.04 | 0.14 | 0.11 |
| 1 | 41088676 | rs12117629 | 0.079 | -0.246 | -0.402 | -0.851 | 0.003 | 0.008 | 0.008 | 0 | 0.08 | 0.02 | 0.04 | 0.02 | 0.06 | 0.06 |
| 1 | 41090250 | rs7520333 | -0.859 | -0.085 | -0.276 | -1 | -0.006 | 0.043 | 0.043 | 0 | 0.42 | 0.71 | 0.4 | 0.18 | 0.45 | 0.34 |
| 1 | 41091127 | rs570671 | -1.19 | 0.16 | -0.718 | -0.543 | 0.021 | -0.075 | -0.075 | 0 | 0.74 | 0.9 | 0.68 | 0.54 | 0.79 | 0.53 |
| 1 | 41093830 | rs661221 | 0.587 | 0.444 | -0.4 | -0.475 | 0.007 | -0.04 | -0.04 | 0.16 | 0.13 | 0.21 | 0.09 | 0.04 | 0.14 | 0.11 |
| 1 | 41104475 | rs3767955 | 1.548 | 0.462 | -0.468 | -0.427 | -0.048 | -0.036 | -0.036 | 0 | 0.29 | 0.09 | 0.27 | 0.32 | 0.33 | 0.18 |
| 1 | 41105238 | rs12562229 | 0.727 | 0.399 | 0.029 | -0.464 | 0.002 | -0.012 | -0.012 | 1.245 | 0.24 | 0.3 | 0.26 | 0.16 | 0.26 | 0.21 |
| 1 | 41106583 | rs4660439 | 0.638 | 0.29 | 0.01 | -0.52 | -0.001 | -0.002 | -0.002 | 0 | 0.23 | 0.23 | 0.27 | 0.09 | 0.27 | 0.23 |
| 1 | 41106774 | rs7524868 | 1.627 | 0.372 | -0.269 | -0.481 | -0.04 | -0.028 | -0.028 | 0 | 0.29 | 0.11 | 0.27 | 0.32 | 0.33 | 0.19 |
| 1 | 41108094 | rs664350 | 2.077 | 0.338 | -0.231 | -0.515 | 0.054 | -0.033 | -0.033 | 7.545 | 0.08 | 0.21 | 0.06 | 0.01 | 0.12 | 0.07 |
| 1 | 41108149 | rs11208590 | -0.476 | 0.301 | -0.219 | -0.524 | -0.003 | -0.009 | -0.009 | 0 | 0.34 | 0.61 | 0.34 | 0.18 | 0.33 | 0.26 |
| 1 | 41109562 | rs7542025 | -0.559 | 0.361 | -0.197 | -0.486 | 0.003 | -0.018 | -0.018 | 0 | 0.64 | 0.71 | 0.61 | 0.5 | 0.66 | 0.45 |
| 1 | 41115608 | rs2780944 | -0.177 | 0.52 | -0.035 | -0.567 | -0.001 | -0.006 | -0.006 | 0 | 0.31 | 0.65 | 0.33 | 0.18 | 0.31 | 0.24 |
| 1 | 41127976 | rs627178 | 1.839 | 0.489 | -0.102 | -0.442 | 0.039 | -0.029 | -0.029 | 0 | 0.08 | 0.14 | 0.06 | 0.01 | 0.12 | 0.07 |
| 1 | 41139341 | rs12044426 | 1.269 | 0.197 | -0.451 | -0.264 | -0.042 | -0.051 | -0.051 | 0.139 | 0.29 | 0.14 | 0.25 | 0.4 | 0.39 | 0.25 |
| 1 | 41140985 | rs545218 | -1.067 | -0.028 | -0.506 | -0.274 | 0.009 | -0.077 | -0.077 | 0 | 0.73 | 0.92 | 0.65 | 0.55 | 0.78 | 0.56 |
| 1 | 41154407 | rs965949 | 0.323 | 0.087 | -0.15 | -0.242 | 0.013 | 0 | 0 | 0.859 | 0.51 | 0.41 | 0.52 | 0.47 | 0.62 | 0.45 |
| 1 | 41170670 | rs11208727 | 1.967 | 0.077 | 0.136 | -0.158 | -0.007 | 0.019 | 0.019 | 0.563 | 0.06 | 0.2 | 0.04 | 0.05 | 0.03 | 0.01 |
| 1 | 41204569 | rs1057925 | 0.503 | 0.147 | 0.114 | -0.268 | 0.009 | 0 | 0 | 0 | 0.51 | 0.42 | 0.51 | 0.47 | 0.59 | 0.44 |
| 1 | 41225520 | rs10158384 | 2.029 | 0.102 | 0.35 | -0.177 | 0.01 | 0.047 | 0.047 | 0.341 | 0.06 | 0.23 | 0.04 | 0.05 | 0.03 | 0.01 |
| 1 | 41226602 | rs11208830 | 0.822 | 0.365 | -0.206 | -0.22 | 0.002 | -0.011 | -0.011 | 0 | 0.22 | 0.24 | 0.26 | 0.08 | 0.23 | 0.2 |
| 1 | 41233031 | rs3767951 | 0.834 | 0.119 | -0.394 | -0.242 | 0.013 | -0.064 | -0.064 | 0 | 0.27 | 0.1 | 0.35 | 0.45 | 0.22 | 0.45 |
| 1 | 41237388 | rs753753 | 0.389 | 0.052 | -0.185 | -0.061 | 0.002 | -0.011 | -0.011 | 0.171 | 0.37 | 0.26 | 0.43 | 0.48 | 0.33 | 0.53 |
| 1 | 41240519 | rs823673 | -1.136 | 0.282 | -0.296 | -0.147 | 0.001 | -0.015 | -0.015 | 2.307 | 0.8 | 0.77 | 0.77 | 0.95 | 0.8 | 0.81 |

| | | | | | | | | | | | | | | | | |
|---|---|---|---|---|---|---|---|---|---|---|---|---|---|---|---|---|
| 1 | 41245354 | rs2769255 | 0.026 | 0.445 | -0.501 | 0.151 | 0 | -0.002 | -0.002 | 0 | 0.41 | 0.36 | 0.48 | 0.5 | 0.36 | 0.53 |
| 1 | 41245859 | rs2251413 | -0.106 | 0.097 | -0.579 | -0.057 | -0.007 | -0.027 | -0.027 | 0.359 | 0.67 | 0.59 | 0.7 | 0.89 | 0.72 | 0.76 |
| 1 | 41248488 | rs2769256 | -0.1 | 0.098 | -0.59 | -0.09 | -0.008 | -0.033 | -0.033 | 0.019 | 0.67 | 0.59 | 0.71 | 0.91 | 0.72 | 0.78 |
| 1 | 41253185 | rs709688 | 0.698 | 0.349 | -0.817 | 1.019 | 0.001 | 0.042 | 0.042 | 0 | 0.92 | 0.96 | 0.74 | 0.53 | 0.93 | 0.78 |
| 1 | 41255387 | rs12723678 | 0.246 | 0.21 | -0.788 | 0.378 | -0.002 | -0.003 | -0.003 | 0 | 0.11 | 0.15 | 0.09 | 0.04 | 0.09 | 0.09 |
| 1 | 41260168 | rs2769258 | -0.278 | 0.295 | -0.897 | 0.038 | 0.005 | 0.065 | 0.065 | 0 | 0.49 | 0.31 | 0.61 | 0.72 | 0.47 | 0.49 |
| 1 | 41267258 | rs4660464 | -0.19 | 0.179 | -0.889 | 0.004 | 0.003 | 0.058 | 0.058 | 0 | 0.51 | 0.34 | 0.64 | 0.73 | 0.55 | 0.54 |
| 1 | 41272533 | rs727145 | 0.506 | 0.396 | -0.617 | -0.047 | -0.005 | 0.01 | 0.01 | 0 | 0.35 | 0.31 | 0.3 | 0.19 | 0.39 | 0.33 |
| 1 | 41276073 | rs11208958 | -0.294 | 0.659 | -0.551 | 0.976 | 0.001 | 0.067 | 0.067 | 0 | 0.16 | 0.19 | 0.31 | 0.49 | 0.19 | 0.24 |
| 1 | 41276188 | rs878042 | 0.5 | 0.649 | -0.544 | 0.931 | 0.006 | 0.084 | 0.084 | 0 | 0.81 | 0.71 | 0.64 | 0.46 | 0.77 | 0.74 |
| 1 | 41285087 | rs4660468 | 0.59 | 0.397 | -0.711 | 0.696 | -0.015 | 0.015 | 0.015 | 0 | 0.66 | 0.53 | 0.56 | 0.47 | 0.7 | 0.56 |
| 1 | 41286353 | rs4660469 | 0.201 | 0.527 | -1.084 | 0.128 | 0 | 0.003 | 0.003 | 0 | 0.6 | 0.56 | 0.52 | 0.5 | 0.57 | 0.58 |
| 1 | 41286883 | rs3754173 | 0.508 | 0.76 | -0.799 | 0.053 | -0.004 | -0.023 | -0.023 | 0 | 0.59 | 0.51 | 0.48 | 0.21 | 0.6 | 0.42 |
| 1 | 41290235 | rs3767941 | -0.468 | 0.637 | -0.461 | -0.068 | -0.015 | -0.027 | -0.027 | 0 | 0.28 | 0.56 | 0.4 | 0.2 | 0.18 | 0.21 |
| 1 | 41291531 | rs3767942 | -1.17 | 0.631 | -0.544 | 0.268 | -0.001 | -0.001 | -0.001 | 0 | 0.39 | 0.6 | 0.49 | 0.47 | 0.29 | 0.46 |
| 1 | 41293001 | rs6697721 | 1.425 | 0.892 | -1.049 | 0.038 | 0.002 | 0.029 | 0.029 | 0 | 0.35 | 0.16 | 0.29 | 0.26 | 0.38 | 0.25 |
| 1 | 41293419 | rs3767943 | 0.697 | 0.836 | -0.627 | -0.002 | -0.001 | 0 | 0 | 0 | 0.2 | 0.24 | 0.36 | 0.21 | 0.15 | 0.22 |
| 1 | 41293649 | rs4660176 | 1.241 | 0.712 | -0.795 | 0.03 | 0.005 | 0.03 | 0.03 | 0 | 0.36 | 0.17 | 0.29 | 0.26 | 0.38 | 0.25 |
| 1 | 41293664 | rs3820524 | 1.242 | 0.714 | -0.794 | 0.03 | 0.005 | 0.028 | 0.028 | 0 | 0.36 | 0.17 | 0.29 | 0.26 | 0.38 | 0.25 |
| 1 | 41295542 | rs1041239 | 0.189 | 0.466 | -0.892 | -0.196 | 0.006 | -0.023 | -0.023 | 0.507 | 0.11 | 0.06 | 0.08 | 0.25 | 0.1 | 0.27 |
| 1 | 41297411 | rs963377 | -0.676 | 0.539 | -0.654 | -1.273 | 0.006 | 0.006 | 0.006 | 0 | 0.5 | 0.59 | 0.64 | 0.4 | 0.55 | 0.49 |
| 1 | 41298510 | rs874847 | 0.047 | 0.521 | -0.712 | -1.392 | 0.001 | -0.032 | -0.032 | 0 | 0.24 | 0.36 | 0.43 | 0.13 | 0.2 | 0.22 |
| 1 | 41298652 | rs913378 | -0.016 | 0.972 | -0.965 | -1.345 | 0.005 | -0.022 | -0.022 | 0 | 0.7 | 0.68 | 0.77 | 0.84 | 0.65 | 0.77 |
| 1 | 41300952 | rs729590 | 0.47 | 0.805 | -1.25 | -1.056 | -0.024 | 0.023 | 0.023 | 0 | 0.12 | 0.08 | 0.28 | 0.06 | 0.17 | 0.1 |
| 1 | 41306895 | rs10889654 | -0.925 | 0.344 | -1.365 | -1.09 | 0.024 | 0.031 | 0.031 | 0 | 0.79 | 0.91 | 0.84 | 0.93 | 0.85 | 0.74 |
| 1 | 41308352 | rs4660471 | -0.015 | 0.629 | -1.552 | -0.965 | 0.066 | 0.107 | 0.107 | 0.136 | 0.52 | 0.19 | 0.69 | 0.78 | 0.66 | 0.62 |
| 1 | 41315679 | rs11209015 | 0.371 | 0.281 | -0.653 | -0.182 | -0.039 | 0.084 | 0.084 | 0 | 0.24 | 0.39 | 0.25 | 0.58 | 0.21 | 0.31 |
| 1 | 41315867 | rs12403717 | -0.603 | 0.578 | -1.096 | -0.334 | 0.002 | -0.013 | -0.013 | 0.832 | 0.42 | 0.54 | 0.59 | 0.78 | 0.48 | 0.44 |
| 1 | 41317020 | rs11209020 | -0.583 | -0.42 | -1.416 | 0.258 | 0.005 | -0.004 | -0.004 | 0.153 | 0.32 | 0.31 | 0.28 | 0.6 | 0.26 | 0.36 |
| 1 | 41318440 | rs4660473 | -0.207 | -0.114 | -1.86 | 0.422 | 0.05 | -0.057 | -0.057 | 0 | 0.3 | 0.12 | 0.27 | 0.62 | 0.25 | 0.37 |
| 1 | 41319146 | rs12564627 | -1.138 | 0.968 | -1.86 | 0.453 | -0.06 | -0.052 | -0.052 | 0.844 | 0.11 | 0.01 | 0.15 | 0.15 | 0.15 | 0.06 |
| 1 | 41320774 | rs17357593 | -0.03 | 0.97 | -1.942 | 0.013 | -0.021 | 0.033 | 0.033 | 0 | 0.07 | 0.03 | 0.03 | 0 | 0.09 | 0.12 |
| 1 | 41323882 | rs913382 | -0.187 | 0.795 | -2.393 | -0.424 | -0.008 | 0.089 | 0.089 | 0 | 0.58 | 0.81 | 0.7 | 0.78 | 0.45 | 0.63 |
| 1 | 41325801 | rs11209041 | 0.737 | 0.382 | -2.014 | 0.616 | -0.006 | 0.135 | 0.135 | 0.09 | 0.09 | 0.14 | 0.37 | 0.49 | 0.07 | 0.13 |
| 1 | 41338665 | rs1359893 | 0.849 | -0.387 | -1.484 | -0.148 | -0.011 | -0.005 | -0.005 | 0 | 0.3 | 0.31 | 0.16 | 0.23 | 0.16 | 0.3 |
| 1 | 41339538 | rs11209068 | 0.847 | -0.107 | -1.155 | -0.33 | -0.002 | -0.013 | -0.013 | 0 | 0.08 | 0.06 | 0.05 | 0 | 0.11 | 0.12 |
| 1 | 41343061 | rs11209075 | 0.219 | -0.164 | -0.856 | -0.462 | 0.002 | -0.037 | -0.037 | 0 | 0.13 | 0.16 | 0.1 | 0 | 0.15 | 0.13 |
| 1 | 41343608 | rs913389 | 0.164 | -0.627 | -1.047 | -0.58 | 0.002 | 0.003 | 0.003 | 0.474 | 0.42 | 0.42 | 0.54 | 0.73 | 0.25 | 0.44 |
| 1 | 41346428 | rs35809276 | 0.135 | -0.235 | -0.625 | -0.613 | 0.01 | -0.044 | -0.044 | 0 | 0.09 | 0.14 | 0.05 | 0 | 0.1 | 0.13 |
| 1 | 41347228 | rs728859 | -0.275 | 0.431 | -0.672 | -0.682 | -0.009 | -0.008 | -0.008 | 2.961 | 0.87 | 0.87 | 0.89 | 1 | 0.82 | 0.85 |
| 1 | 41348072 | rs6600354 | 0.817 | 0.634 | -1.189 | -1.439 | 0.001 | 0.087 | 0.087 | 0 | 0.26 | 0.1 | 0.14 | 0.01 | 0.26 | 0.18 |
| 1 | 41351613 | rs10158668 | 0.375 | 0.581 | -0.708 | -1.079 | -0.003 | -0.001 | -0.001 | 0 | 0.42 | 0.51 | 0.5 | 0.43 | 0.4 | 0.46 |
| 1 | 41354681 | rs4660481 | 1.1 | 0.03 | 0.274 | -1.045 | -0.04 | -0.005 | -0.005 | 0.02 | 0.25 | 0.38 | 0.33 | 0.26 | 0.11 | 0.27 |
| 1 | 41356226 | rs6696977 | -0.521 | 0.386 | -0.259 | -1.256 | -0.069 | 0.16 | 0.16 | 0 | 0.6 | 0.88 | 0.72 | 0.83 | 0.44 | 0.64 |
| 1 | 41365445 | rs12120974 | 0.573 | 0.413 | 0.152 | -1.18 | -0.003 | 0.002 | 0.002 | 0 | 0.24 | 0.23 | 0.19 | 0.13 | 0.31 | 0.29 |
| 1 | 41366059 | rs12026763 | 1.64 | 0.29 | 0.375 | -1.176 | 0.019 | -0.002 | -0.002 | 0 | 0.27 | 0.2 | 0.34 | 0.33 | 0.2 | 0.26 |
| 1 | 41367667 | rs913392 | -0.405 | 0.344 | 0.522 | -1.259 | -0.031 | 0.08 | 0.08 | 0 | 0.26 | 0.46 | 0.31 | 0.53 | 0.21 | 0.26 |
| 1 | 41368215 | rs7528422 | 0.506 | 0.201 | 0.655 | -1.213 | -0.023 | -0.034 | -0.034 | 6.303 | 0.44 | 0.5 | 0.58 | 0.82 | 0.29 | 0.46 |
| 1 | 41369334 | rs4660484 | -2.08 | 0.288 | 0.249 | -1.523 | -0.035 | 0.104 | 0.104 | 0 | 0.83 | 0.98 | 0.92 | 0.99 | 0.84 | 0.91 |
| 1 | 41370066 | rs12133860 | 1.39 | 0.382 | 0.357 | -1.434 | -0.021 | 0.023 | 0.023 | 0.005 | 0.2 | 0.14 | 0.17 | 0.08 | 0.31 | 0.25 |
| 1 | 41374443 | rs3856250 | 1.517 | 0.069 | 0.304 | -1.283 | 0.015 | -0.008 | -0.008 | 0 | 0.11 | 0.07 | 0.22 | 0.53 | 0.08 | 0.16 |
| 1 | 41378076 | rs6674863 | 1.37 | 0.114 | 0.385 | -1.357 | -0.002 | -0.001 | -0.001 | 0 | 0.29 | 0.3 | 0.34 | 0.34 | 0.2 | 0.28 |
| 1 | 41382006 | rs17357745 | 1.191 | 0.318 | -0.121 | -1.309 | -0.034 | 0.039 | 0.039 | 0 | 0.12 | 0.05 | 0.11 | 0 | 0.17 | 0.13 |
| 1 | 41383838 | rs6698251 | -0.475 | -0.071 | -0.102 | -1.161 | 0.022 | -0.055 | -0.055 | 0.036 | 0.7 | 0.83 | 0.49 | 0.51 | 0.76 | 0.68 |
| 1 | 41384630 | rs11209170 | -0.215 | -0.27 | 0.06 | -1.064 | 0.01 | -0.034 | -0.034 | 0 | 0.35 | 0.6 | 0.27 | 0.16 | 0.41 | 0.34 |
| 1 | 41385086 | rs11805475 | 1.591 | -0.186 | 0.146 | -1.13 | -0.021 | 0.023 | 0.023 | 0 | 0.18 | 0.11 | 0.14 | 0.12 | 0.26 | 0.19 |
| 1 | 41386871 | rs7515340 | 0.015 | -0.147 | -0.191 | -1.051 | 0.01 | -0.056 | -0.056 | 0 | 0.34 | 0.59 | 0.26 | 0.13 | 0.4 | 0.32 |
| 1 | 41387003 | rs7554973 | -1.286 | -0.017 | -0.561 | -0.959 | -0.019 | 0.045 | 0.045 | 0 | 0.59 | 0.79 | 0.75 | 0.65 | 0.56 | 0.65 |
| 1 | 41390175 | rs10789250 | -0.587 | 0.182 | -0.429 | -1.135 | 0.002 | 0.062 | 0.062 | 0 | 0.56 | 0.63 | 0.41 | 0.12 | 0.53 | 0.42 |
| 1 | 41395072 | rs1887542 | 0.505 | 0.394 | -0.259 | -1.03 | 0.001 | -0.088 | -0.088 | 0.118 | 0.25 | 0.42 | 0.23 | 0.11 | 0.27 | 0.26 |
| 1 | 41395698 | rs12083349 | 0.988 | 0.913 | -0.518 | -0.771 | -0.002 | -0.018 | -0.018 | 0 | 0.12 | 0.12 | 0.1 | 0 | 0.14 | 0.1 |
| 1 | 41399738 | rs2361654 | 0.049 | 0.868 | -0.359 | -0.307 | -0.01 | 0.074 | 0.074 | 0 | 0.28 | 0.43 | 0.49 | 0.44 | 0.23 | 0.33 |
| 1 | 41402143 | rs4660492 | -1.13 | 0.912 | -0.438 | -0.345 | -0.013 | -0.013 | -0.013 | 0 | 0.7 | 0.79 | 0.8 | 0.61 | 0.64 | 0.66 |
| 1 | 41402399 | rs11209209 | 1.179 | 1.238 | -1.156 | -0.613 | 0.004 | -0.01 | -0.01 | 0 | 0.17 | 0.19 | 0.21 | 0.11 | 0.25 | 0.21 |
| 1 | 41404596 | rs12043667 | -0.494 | 0.882 | -1.405 | 0.006 | -0.007 | -0.029 | -0.029 | 0.223 | 0.28 | 0.22 | 0.47 | 0.34 | 0.28 | 0.31 |
| 1 | 41406460 | rs11209213 | -0.903 | 1.063 | -1.414 | 0.231 | -0.003 | 0 | 0 | 0.403 | 0.52 | 0.41 | 0.57 | 0.4 | 0.44 | 0.44 |
| 1 | 41412547 | rs11582127 | -0.666 | 1.504 | -1.444 | 1.471 | -0.025 | -0.083 | -0.083 | 0 | 0.21 | 0.07 | 0.46 | 0.35 | 0.21 | 0.22 |
| 1 | 41420985 | rs6671627 | 1.125 | 1.88 | -0.865 | 1.193 | 0.039 | 0.038 | 0.038 | 0 | 0.93 | 0.89 | 0.84 | 0.84 | 0.85 | 0.88 |
| 1 | 41425864 | rs12067645 | 0.108 | 0.699 | -0.858 | 0.339 | -0.002 | -0.005 | -0.005 | 0 | 0.36 | 0.35 | 0.26 | 0.16 | 0.23 | 0.33 |

| | | | | | | | | | | | | | | | | |
|---|---|---|---|---|---|---|---|---|---|---|---|---|---|---|---|---|
| 1 | 41425927 | rs12723637 | 0.212 | 1.797 | -1.392 | 0.836 | -0.047 | -0.069 | -0.069 | 1.411 | 0.15 | 0.04 | 0.29 | 0.29 | 0.17 | 0.13 |
| 1 | 41433516 | rs7533665 | -0.191 | 2.226 | -1.242 | 0.431 | 0.002 | 0.021 | 0.021 | 0 | 0.71 | 0.71 | 0.73 | 0.56 | 0.69 | 0.6 |
| 1 | 41437417 | rs10789262 | -0.175 | 2.309 | -1.15 | 0.426 | 0.004 | 0.028 | 0.028 | 0 | 0.71 | 0.68 | 0.72 | 0.56 | 0.69 | 0.6 |
| 1 | 41442786 | rs4440845 | 0.623 | 1.529 | -1.26 | 0.042 | 0.013 | 0.026 | 0.026 | 0.752 | 0.44 | 0.27 | 0.42 | 0.39 | 0.35 | 0.34 |
| 1 | 41447366 | rs16827810 | 1.679 | 1.454 | -1.267 | -0.068 | 0.005 | -0.002 | -0.002 | 0 | 0.28 | 0.23 | 0.13 | 0.1 | 0.18 | 0.18 |
| 1 | 41449629 | rs4132440 | -0.794 | 1.473 | -1.16 | 0.097 | 0.003 | 0.132 | 0.132 | 0 | 0.85 | 0.73 | 0.8 | 0.67 | 0.84 | 0.78 |
| 1 | 41449859 | rs7523029 | 0.815 | 1.412 | -1.013 | 0.031 | 0.017 | -0.028 | -0.028 | 0 | 0.18 | 0.33 | 0.18 | 0.12 | 0.29 | 0.14 |
| 1 | 41452880 | rs12044786 | -0.218 | 1.247 | -1.147 | 0.195 | 0.025 | 0.109 | 0.109 | 0 | 0.17 | 0.25 | 0.23 | 0.41 | 0.25 | 0.25 |
| 1 | 41469666 | rs11209342 | -0.536 | 1.669 | -1.888 | 0.05 | -0.097 | -0.107 | -0.107 | 0 | 0.29 | 0.09 | 0.54 | 0.44 | 0.41 | 0.32 |
| 1 | 41485077 | rs11209363 | 1.059 | 2.336 | -2.009 | 0.079 | -0.03 | -0.06 | -0.06 | 4.198 | 0.1 | 0.01 | 0.25 | 0.28 | 0.11 | 0.1 |
| 1 | 41486245 | rs1138293 | 0.518 | 2.95 | -2.097 | 0.016 | -0.065 | 0.005 | 0.005 | 0.014 | 0.11 | 0.01 | 0.13 | 0.05 | 0.2 | 0.09 |
| 1 | 41486408 | rs3738369 | 1.108 | 2.479 | -2.067 | 0.054 | -0.026 | -0.059 | -0.059 | 0 | 0.1 | 0.01 | 0.25 | 0.28 | 0.11 | 0.11 |
| 1 | 41487337 | rs17357954 | 0.907 | 2.479 | -2.065 | 0.054 | -0.059 | -0.002 | -0.002 | 0 | 0.1 | 0.01 | 0.14 | 0.05 | 0.2 | 0.1 |
| 1 | 41488722 | rs11209368 | 0.794 | 2.461 | -2.059 | 0.061 | -0.017 | -0.023 | -0.023 | 0 | 0.09 | 0 | 0.15 | 0.12 | 0.1 | 0.12 |
| 1 | 41493111 | rs3157 | -1.244 | 2.256 | -2.15 | 0.156 | -0.094 | -0.138 | -0.138 | 0.266 | 0.33 | 0.11 | 0.55 | 0.54 | 0.44 | 0.37 |
| 1 | 41508575 | rs11209391 | -0.036 | 2.091 | -1.639 | -0.015 | 0.01 | 0.075 | 0.075 | 0 | 0.23 | 0.58 | 0.27 | 0.4 | 0.25 | 0.31 |
| 1 | 41509674 | rs2749418 | 0.479 | 2.13 | -1.711 | -0.011 | -0.099 | -0.108 | -0.108 | 8.477 | 0.71 | 0.9 | 0.46 | 0.55 | 0.58 | 0.72 |
| 1 | 41510708 | rs2268680 | 1.135 | 2.008 | -1.839 | -0.151 | -0.028 | -0.025 | -0.025 | 0.934 | 0.08 | 0 | 0.04 | 0.1 | 0.08 | 0.05 |
| 1 | 41524538 | rs4660184 | 0.526 | 2.011 | -1.639 | -0.124 | -0.052 | -0.056 | -0.056 | 0 | 0.68 | 0.82 | 0.45 | 0.55 | 0.57 | 0.64 |
| 1 | 41552300 | rs6685340 | 3.146 | 1.695 | -1.758 | -0.329 | 0.008 | 0.169 | 0.169 | 0.5 | 0.27 | 0.03 | 0.11 | 0.03 | 0.19 | 0.16 |
| 1 | 41554088 | rs12022978 | 1.399 | 1.984 | -1.672 | -0.147 | -0.029 | -0.064 | -0.064 | 0.267 | 0.11 | 0.01 | 0.25 | 0.28 | 0.11 | 0.1 |
| 1 | 41562243 | rs2024860 | -1.3 | 2.036 | -1.661 | -0.151 | -0.059 | -0.003 | -0.003 | 0 | 0.9 | 0.99 | 0.86 | 0.95 | 0.8 | 0.91 |
| 1 | 41563086 | rs3819842 | 0.514 | 2.028 | -1.597 | -0.045 | 0.052 | 0.133 | 0.133 | 0 | 0.19 | 0.48 | 0.26 | 0.29 | 0.23 | 0.27 |
| 1 | 41582006 | rs213765 | 0.439 | 1.929 | -2.073 | 0.414 | -0.065 | -0.001 | -0.001 | 0 | 0.81 | 0.93 | 0.7 | 0.83 | 0.7 | 0.78 |
| 1 | 41615358 | rs213769 | 1.613 | 1.634 | -2.065 | 0.4 | -0.009 | -0.013 | -0.013 | 0 | 0.51 | 0.83 | 0.4 | 0.43 | 0.45 | 0.48 |
| 1 | 41634956 | rs11802778 | -0.153 | 1.619 | -1.954 | 0.428 | 0.011 | 0.023 | 0.023 | 0 | 0.19 | 0.07 | 0.06 | 0.13 | 0.13 | 0.2 |
| 1 | 41650222 | rs12410899 | 0.255 | 1.869 | -1.525 | 0.765 | -0.017 | -0.02 | -0.02 | 3.573 | 0.09 | 0 | 0.15 | 0.12 | 0.1 | 0.12 |
| 1 | 41702942 | rs10493094 | 0.225 | 1.869 | -1.525 | 0.765 | -0.017 | -0.02 | -0.02 | 0 | 0.1 | 0 | 0.15 | 0.12 | 0.1 | 0.12 |
| 1 | 41744807 | rs12143111 | -1.027 | 1.958 | -1.527 | 0.76 | -0.111 | -0.114 | -0.114 | 0 | 0.28 | 0.1 | 0.54 | 0.44 | 0.42 | 0.3 |
| 1 | 41748424 | rs11209792 | -1.15 | 1.963 | -0.929 | 1.432 | 0.018 | 0.168 | 0.168 | 0 | 0.08 | 0.16 | 0.15 | 0.47 | 0.12 | 0.12 |
| 1 | 41755556 | rs11209799 | 1.728 | 1.612 | -0.698 | 1.378 | 0 | 0.004 | 0.004 | 4.13 | 0.51 | 0.39 | 0.37 | 0.32 | 0.47 | 0.44 |
| 1 | 41761220 | rs12030183 | -1.048 | 1.913 | -0.585 | 1.339 | -0.006 | -0.006 | -0.006 | 0 | 0.21 | 0.21 | 0.27 | 0.49 | 0.31 | 0.26 |
| 1 | 41763018 | rs6660666 | 1.361 | 1.837 | -0.446 | 1.287 | 0.017 | 0.061 | 0.061 | 1.557 | 0.75 | 0.58 | 0.72 | 0.51 | 0.68 | 0.71 |
| 1 | 41764175 | rs12037657 | -0.093 | 1.868 | -0.648 | 1.464 | -0.025 | -0.026 | -0.026 | 0 | 0.11 | 0.09 | 0.11 | 0.35 | 0.21 | 0.1 |
| 1 | 41785792 | rs2154317 | -1.556 | 1.582 | 0.122 | 1.458 | 0.025 | 0.068 | 0.068 | 1.68 | 0.23 | 0.36 | 0.48 | 0.5 | 0.36 | 0.26 |
| 1 | 41790077 | rs16828092 | 0.176 | 1.623 | 0.028 | 1.507 | -0.024 | -0.048 | -0.048 | 0 | 0.09 | 0.01 | 0.35 | 0.15 | 0.14 | 0.16 |
| 1 | 41794403 | rs17362424 | 0.242 | 1.663 | 0.043 | 1.806 | -0.066 | -0.047 | -0.047 | 0 | 0.1 | 0.01 | 0.11 | 0.19 | 0.22 | 0.07 |
| 1 | 41799160 | rs4660187 | -1.173 | 1.534 | 0.149 | 1.532 | 0.018 | 0.049 | 0.049 | 1.655 | 0.24 | 0.36 | 0.47 | 0.5 | 0.36 | 0.26 |
| 1 | 41817502 | rs4660190 | 0.972 | 2.092 | -0.67 | 1.523 | 0.018 | 0.048 | 0.048 | 0.945 | 0.73 | 0.4 | 0.5 | 0.48 | 0.62 | 0.71 |
| 1 | 41821570 | rs4660526 | 0.175 | 1.964 | -0.773 | 1.384 | -0.026 | 0.02 | 0.02 | 0 | 0.06 | 0.02 | 0.12 | 0.29 | 0.07 | 0.27 |
| 1 | 41821779 | rs4660527 | 1.314 | 1.74 | -0.779 | 1.149 | 0.013 | 0.007 | 0.007 | 1.009 | 0.52 | 0.29 | 0.35 | 0.38 | 0.45 | 0.53 |
| 1 | 41830257 | rs11585393 | -0.876 | 1.723 | -1.141 | 0.708 | -0.04 | 0.029 | 0.029 | 2.769 | 0.24 | 0.15 | 0.47 | 0.15 | 0.27 | 0.29 |
| 1 | 41831047 | rs7539614 | 1.357 | 1.891 | -1.881 | 0.349 | -0.064 | 0.109 | 0.109 | 0.591 | 0.57 | 0.08 | 0.4 | 0.76 | 0.57 | 0.53 |
| 1 | 41839083 | rs1317558 | -0.052 | 1.704 | -1.375 | 0.231 | -0.002 | -0.017 | -0.017 | 0 | 0.18 | 0.18 | 0.08 | 0.05 | 0.11 | 0.32 |
| 1 | 41839822 | rs4660531 | 1.35 | 1.677 | -2.409 | 0.697 | -0.038 | 0.032 | 0.032 | 1.075 | 0.31 | 0.09 | 0.2 | 0.17 | 0.33 | 0.28 |
| 1 | 41841986 | rs6690527 | -0.279 | 1.306 | -1.753 | 0.556 | 0.019 | 0.031 | 0.031 | 0 | 0.44 | 0.63 | 0.24 | 0.18 | 0.29 | 0.4 |
| 1 | 41842350 | rs4660192 | 0.706 | 1.634 | -1.944 | 0.743 | -0.021 | -0.002 | -0.002 | 0.048 | 0.42 | 0.22 | 0.6 | 0.35 | 0.51 | 0.38 |
| 1 | 41844293 | rs11209971 | 0.457 | 1.572 | -1.739 | 0.727 | 0.003 | -0.013 | -0.013 | 2.526 | 0.66 | 0.29 | 0.74 | 0.44 | 0.69 | 0.45 |
| 1 | 41849735 | rs10889959 | 0.185 | 1.284 | -1.472 | 0.08 | -0.02 | 0.076 | 0.076 | 0 | 0.45 | 0.22 | 0.67 | 0.63 | 0.54 | 0.51 |
| 1 | 41850182 | rs10749805 | 0.193 | 1.288 | -1.47 | 0.082 | -0.02 | 0.075 | 0.075 | 0 | 0.45 | 0.22 | 0.67 | 0.63 | 0.53 | 0.51 |
| 1 | 41854357 | rs2364546 | 0.777 | 0.769 | -1.216 | -0.462 | 0.013 | 0.044 | 0.044 | 0.232 | 0.31 | 0.23 | 0.22 | 0.04 | 0.3 | 0.19 |
| 1 | 41861383 | rs2996016 | -0.126 | 0.871 | -0.985 | 0.1 | 0.003 | 0.044 | 0.044 | 0 | 0.93 | 0.89 | 0.64 | 0.81 | 0.92 | 0.88 |
| 1 | 41862125 | rs10889978 | -0.152 | 0.568 | -1.397 | -0.366 | 0.018 | 0.008 | 0.008 | 0.062 | 0.24 | 0.09 | 0.43 | 0.19 | 0.18 | 0.38 |
| 1 | 41864364 | rs3845572 | 1.312 | 0.714 | -0.375 | -0.369 | -0.035 | -0.05 | -0.05 | 0 | 0.28 | 0.45 | 0.15 | 0.13 | 0.18 | 0.09 |
| 1 | 41866315 | rs7536490 | 1.579 | 0.733 | -0.454 | -0.36 | 0.021 | 0.033 | 0.033 | 0 | 0.26 | 0.1 | 0.13 | 0.13 | 0.17 | 0.08 |
| 1 | 41867980 | rs2971987 | -0.294 | 0.906 | -0.435 | -0.34 | 0.014 | 0.021 | 0.021 | 0 | 0.53 | 0.42 | 0.73 | 0.83 | 0.69 | 0.66 |
| 1 | 41872078 | rs2997338 | -0.013 | 1.118 | -0.447 | -0.162 | 0.043 | 0.057 | 0.057 | 0 | 0.2 | 0.28 | 0.21 | 0.64 | 0.34 | 0.36 |
| 1 | 41875951 | rs2997341 | 0.038 | 1.23 | -0.391 | -0.005 | 0.034 | 0.06 | 0.06 | 0.047 | 0.19 | 0.28 | 0.21 | 0.59 | 0.32 | 0.34 |
| 1 | 41878988 | rs2801962 | 0.037 | 0.845 | -0.669 | 0.132 | 0.025 | 0.094 | 0.094 | 0 | 0.51 | 0.66 | 0.71 | 0.82 | 0.62 | 0.58 |
| 1 | 41879081 | rs12402711 | -0.199 | 1.105 | 0.043 | 0.728 | 0.03 | 0.179 | 0.179 | 0 | 0.06 | 0.19 | 0.37 | 0.22 | 0.08 | 0.12 |
| 1 | 41880343 | rs2810583 | 0.072 | 1.018 | -0.701 | 0.279 | 0.005 | 0.011 | 0.011 | 0 | 0.77 | 0.81 | 0.86 | 0.93 | 0.8 | 0.65 |
| 1 | 41883722 | rs3856254 | 0.964 | 0.915 | -0.986 | 0.348 | 0.007 | 0.013 | 0.013 | 1.342 | 0.48 | 0.19 | 0.36 | 0.37 | 0.45 | 0.23 |
| 1 | 41889513 | rs12133441 | -0.752 | 1.194 | -0.799 | 0.414 | -0.003 | 0.002 | 0.002 | 0 | 0.36 | 0.38 | 0.27 | 0.31 | 0.45 | 0.36 |
| 1 | 41895497 | rs2810569 | -0.355 | 1.226 | -1.137 | 0.842 | -0.012 | -0.008 | -0.008 | 0 | 0.07 | 0.04 | 0.29 | 0.24 | 0.09 | 0.17 |
| 1 | 41903038 | rs12086029 | -0.835 | 0.996 | -1.073 | 0.454 | -0.021 | -0.018 | -0.018 | 0 | 0.3 | 0.26 | 0.41 | 0.45 | 0.4 | 0.37 |
| 1 | 41904021 | rs7517186 | -0.985 | 1.1 | -1.186 | 0.392 | -0.031 | -0.019 | -0.019 | 0 | 0.31 | 0.26 | 0.42 | 0.45 | 0.43 | 0.36 |
| 1 | 41904551 | rs12065902 | 0.5 | 0.8 | -1.148 | 0.257 | 0.008 | 0.012 | 0.012 | 0 | 0.14 | 0.11 | 0.09 | 0.06 | 0.1 | 0.17 |
| 1 | 41909856 | rs7534137 | 0.58 | 0.785 | -1.204 | 0.241 | 0.012 | 0.017 | 0.017 | 0 | 0.15 | 0.11 | 0.1 | 0.06 | 0.12 | 0.2 |
| 1 | 41910825 | rs11210142 | -1 | 0.973 | -1.091 | 0.525 | -0.026 | -0.021 | -0.021 | 0 | 0.3 | 0.26 | 0.42 | 0.45 | 0.4 | 0.37 |

| | | | | | | | | | | | | | | | | |
|---|---|---|---|---|---|---|---|---|---|---|---|---|---|---|---|---|
| 1 | 41922577 | rs4660540 | -0.353 | -0.184 | -0.553 | 0.287 | -0.004 | 0.001 | 0.001 | 0 | 0.2 | 0.17 | 0.3 | 0.18 | 0.11 | 0.32 |
| 1 | 41925050 | rs2483692 | -1.272 | 0.288 | -0.714 | 0.404 | -0.018 | -0.009 | -0.009 | 0 | 0.35 | 0.26 | 0.47 | 0.49 | 0.46 | 0.56 |
| 1 | 41927706 | rs9645433 | 0.579 | -0.402 | -0.794 | -0.064 | 0.01 | 0.015 | 0.015 | 1.677 | 0.11 | 0.09 | 0.04 | 0.02 | 0.07 | 0.02 |
| 1 | 41929950 | rs12138576 | 0.352 | -0.358 | -0.725 | -0.03 | 0.002 | -0.003 | -0.003 | 0 | 0.11 | 0.15 | 0.04 | 0.02 | 0.07 | 0.02 |
| 1 | 41935450 | rs1537522 | 0.229 | -0.181 | -0.411 | 0.009 | 0.085 | 0.049 | 0.049 | 0.873 | 0.83 | 0.43 | 0.8 | 0.68 | 0.69 | 0.81 |
| 1 | 41940687 | rs12069358 | -0.485 | -1.039 | -0.565 | 0.054 | -0.005 | -0.019 | -0.019 | 0.873 | 0.18 | 0.14 | 0.13 | 0.43 | 0.25 | 0.26 |
| 1 | 41941115 | rs11210250 | 0.901 | -0.997 | -0.658 | 0.1 | -0.045 | -0.057 | -0.057 | 0 | 0.1 | 0.01 | 0.1 | 0.35 | 0.2 | 0.18 |
| 1 | 41941599 | rs10890041 | 0.149 | -1.234 | -0.842 | -0.145 | 0.005 | 0.068 | 0.068 | 0 | 0.14 | 0.01 | 0.05 | 0.02 | 0.09 | 0.08 |
| 1 | 41941726 | rs2492076 | 1.347 | -1.394 | -0.532 | -0.158 | 0.086 | 0.01 | 0.01 | 0.131 | 0.48 | 0.17 | 0.7 | 0.47 | 0.6 | 0.45 |
| 1 | 41944690 | rs5800 | -0.539 | -1.37 | -0.713 | -0.265 | 0.027 | 0.029 | 0.029 | 0 | 0.28 | 0.22 | 0.12 | 0.1 | 0.15 | 0.27 |
| 1 | 41950905 | rs11572340 | 0.552 | -0.763 | -1.145 | -0.171 | -0.048 | 0.077 | 0.077 | 0.385 | 0.25 | 0.02 | 0.14 | 0.06 | 0.22 | 0.07 |
| 1 | 41952597 | rs3754287 | -0.137 | -0.587 | -0.679 | 0.3 | -0.004 | 0.005 | 0.005 | 0 | 0.16 | 0.21 | 0.13 | 0.16 | 0.14 | 0.14 |
| 1 | 41953745 | rs12718439 | -1.415 | -0.526 | -1.139 | 0.17 | -0.018 | 0.001 | 0.001 | 1.878 | 0.66 | 0.8 | 0.44 | 0.68 | 0.61 | 0.45 |
| 1 | 41954444 | rs4660541 | 0.318 | -0.734 | -1.246 | 0.144 | 0.005 | 0.07 | 0.07 | 0.156 | 0.33 | 0.13 | 0.18 | 0.13 | 0.26 | 0.16 |
| 1 | 41957439 | rs6671507 | -0.816 | -0.275 | -0.879 | 0.106 | -0.006 | 0.051 | 0.051 | 0 | 0.32 | 0.56 | 0.25 | 0.54 | 0.34 | 0.29 |
| 1 | 41962681 | rs747459 | -0.359 | -0.243 | -1.62 | -0.515 | -0.008 | 0.089 | 0.089 | 0 | 0.71 | 0.88 | 0.84 | 0.97 | 0.74 | 0.88 |
| 1 | 41969327 | rs2991345 | 0.575 | -0.45 | -1.632 | -0.38 | -0.005 | 0.069 | 0.069 | 1.622 | 0.52 | 0.76 | 0.71 | 0.61 | 0.48 | 0.72 |
| 1 | 41971212 | rs11210339 | -1.394 | -0.977 | -1.486 | 0.037 | 0.024 | -0.027 | -0.027 | 0 | 0.36 | 0.82 | 0.27 | 0.5 | 0.32 | 0.29 |
| 1 | 41971227 | rs12043801 | 0 | -0.629 | -1.605 | 0.225 | -0.004 | -0.005 | -0.005 | 0 | 0.16 | 0.14 | 0.14 | 0.43 | 0.22 | 0.18 |
| 1 | 41972251 | rs7542858 | 0.25 | -0.523 | -1.67 | 0.439 | -0.001 | 0.006 | 0.006 | 0.059 | 0.06 | 0.07 | 0.04 | 0.14 | 0.06 | 0.12 |
| 1 | 41973908 | rs1969749 | 0.265 | -0.579 | -1.985 | 0.039 | 0.031 | -0.033 | -0.033 | 0 | 0.5 | 0.08 | 0.65 | 0.48 | 0.58 | 0.69 |
| 1 | 41976529 | rs11809423 | 0.005 | -1.24 | -0.769 | -0.408 | -0.048 | -0.057 | -0.057 | 0 | 0.16 | 0.24 | 0.04 | 0.01 | 0.02 | 0.13 |
| 1 | 41984658 | rs600401 | -0.625 | -1.099 | -0.521 | -1.536 | 0.01 | -0.157 | -0.157 | 0.227 | 0.61 | 0.5 | 0.67 | 0.99 | 0.55 | 0.71 |
| 1 | 41990867 | rs941970 | 0.056 | -1.226 | -0.497 | -1.646 | -0.006 | 0.225 | 0.225 | 0.792 | 0.49 | 0.21 | 0.41 | 0.02 | 0.48 | 0.39 |
| 1 | 41998561 | rs12117690 | 0.934 | -0.977 | -0.805 | -1.605 | -0.071 | 0.021 | 0.021 | 0.139 | 0.16 | 0.04 | 0.14 | 0.08 | 0.26 | 0.21 |
| 1 | 42001301 | rs10890105 | 0.78 | -1.416 | 0.358 | -0.678 | -0.002 | 0.018 | 0.018 | 0 | 0.2 | 0.23 | 0.16 | 0.39 | 0.19 | 0.19 |
| 1 | 42001444 | rs638859 | 1.448 | -1.521 | -0.043 | -0.894 | 0.001 | -0.072 | -0.072 | 0 | 0.34 | 0.31 | 0.28 | 0.01 | 0.41 | 0.29 |
| 1 | 42001831 | rs640688 | 0.781 | -1.55 | 0.154 | -0.755 | -0.001 | -0.001 | -0.001 | 0 | 0.48 | 0.45 | 0.62 | 0.48 | 0.47 | 0.44 |
| 1 | 42006005 | rs2475839 | 0.685 | -1.555 | 0.223 | -0.706 | 0.007 | -0.005 | -0.005 | 0.653 | 0.57 | 0.57 | 0.47 | 0.48 | 0.65 | 0.58 |
| 1 | 42008521 | rs10493102 | 1.305 | -1.513 | 0.118 | -0.713 | -0.083 | 0.042 | 0.042 | 0.148 | 0.15 | 0.02 | 0.14 | 0.05 | 0.26 | 0.21 |
| 1 | 42021616 | rs2759253 | -0.109 | -1.69 | 0.313 | -0.718 | -0.061 | 0.055 | 0.055 | 0.714 | 0.74 | 0.61 | 0.6 | 0.53 | 0.9 | 0.79 |
| 1 | 42027561 | rs1001925 | 0.815 | -1.59 | 0.28 | -0.741 | -0.012 | 0.01 | 0.01 | 0 | 0.18 | 0.16 | 0.14 | 0.05 | 0.25 | 0.2 |
| 1 | 42029134 | rs2181276 | 1.614 | -1.628 | 0.317 | -0.741 | 0.022 | -0.019 | -0.019 | 1.13 | 0.14 | 0.07 | 0.34 | 0.46 | 0.05 | 0.14 |
| 1 | 42031065 | rs1570359 | -0.757 | -1.638 | 0.523 | -0.756 | 0.01 | 0.037 | 0.037 | 0.462 | 0.3 | 0.38 | 0.42 | 0.52 | 0.32 | 0.32 |
| 1 | 42031693 | rs7524240 | -0.721 | -1.638 | 0.523 | -0.756 | 0.01 | 0.037 | 0.037 | 0.478 | 0.29 | 0.37 | 0.42 | 0.52 | 0.32 | 0.33 |
| 1 | 42033410 | rs7527409 | -0.028 | -1.572 | 0.568 | -0.692 | -0.03 | 0.024 | 0.024 | 0.939 | 0.25 | 0.29 | 0.39 | 0.47 | 0.1 | 0.22 |
| 1 | 42036859 | rs1885653 | 0.363 | -1.534 | 0.53 | -0.782 | -0.007 | 0.082 | 0.082 | 0 | 0.24 | 0.35 | 0.19 | 0.46 | 0.23 | 0.28 |
| 1 | 42041479 | rs1815257 | 1.796 | -1.533 | 0.413 | -0.811 | -0.067 | 0.133 | 0.133 | 0 | 0.32 | 0.09 | 0.26 | 0.02 | 0.41 | 0.3 |
| 1 | 42041806 | rs10749832 | -1.932 | -1.534 | 0.42 | -0.811 | -0.053 | 0.108 | 0.108 | 0 | 0.68 | 0.87 | 0.73 | 0.98 | 0.59 | 0.7 |
| 1 | 42042317 | rs1570355 | -0.631 | -1.4 | 0.524 | -0.69 | 0.001 | 0.001 | 0.001 | 0.969 | 0.42 | 0.55 | 0.34 | 0.51 | 0.48 | 0.48 |
| 1 | 42050366 | rs2146315 | 0.12 | -1.714 | 0.545 | -0.427 | -0.007 | 0.028 | 0.028 | 0 | 0.26 | 0.32 | 0.19 | 0.43 | 0.23 | 0.32 |
| 1 | 42062667 | rs6600382 | -0.504 | -1.963 | 0.915 | -0.19 | -0.087 | -0.091 | -0.091 | 4.155 | 0.27 | 0.43 | 0.19 | 0.11 | 0.12 | 0.2 |
| 1 | 42071438 | rs6700352 | -1.017 | -2.008 | 0.908 | -0.203 | -0.052 | 0.028 | 0.028 | 0.732 | 0.32 | 0.57 | 0.45 | 0.57 | 0.16 | 0.26 |
| 1 | 42075267 | rs12740722 | 0.423 | -2.033 | 0.921 | -0.236 | 0.023 | 0.005 | 0.005 | 2.687 | 0.2 | 0.11 | 0.14 | 0.01 | 0.07 | 0.12 |
| 1 | 42076607 | rs17363834 | 0.223 | -1.863 | 0.685 | -0.588 | -0.023 | -0.009 | -0.009 | 0.116 | 0.41 | 0.29 | 0.31 | 0.42 | 0.49 | 0.49 |
| 1 | 42076915 | rs7515066 | -1.938 | -1.829 | 0.602 | -0.739 | -0.075 | 0.138 | 0.138 | 0 | 0.73 | 0.95 | 0.76 | 1 | 0.65 | 0.74 |
| 1 | 42079046 | rs7515948 | -0.865 | -1.629 | 0.746 | -0.367 | -0.002 | -0.004 | -0.004 | 2.339 | 0.6 | 0.6 | 0.46 | 0.5 | 0.56 | 0.6 |
| 1 | 42083369 | rs7519983 | 0.011 | -1.485 | 0.715 | -0.285 | 0.014 | 0.016 | 0.016 | 0.002 | 0.4 | 0.5 | 0.32 | 0.48 | 0.49 | 0.48 |
| 1 | 42091634 | rs11210501 | 0.077 | -1.574 | 0.542 | -0.116 | -0.023 | -0.002 | -0.002 | 0 | 0.38 | 0.3 | 0.3 | 0.43 | 0.48 | 0.47 |
| 1 | 42091708 | rs1109255 | -0.646 | -1.596 | 0.026 | -0.13 | -0.022 | 0.024 | 0.024 | 0 | 0.55 | 0.78 | 0.45 | 0.43 | 0.52 | 0.58 |
| 1 | 42099583 | rs4083593 | -0.259 | -1.652 | 0.044 | -0.091 | -0.004 | 0.023 | 0.023 | 0.403 | 0.45 | 0.76 | 0.36 | 0.42 | 0.52 | 0.48 |
| 1 | 42102877 | rs11807949 | -0.386 | -1.741 | 0.093 | 0.023 | -0.013 | 0 | 0 | 0.16 | 0.26 | 0.24 | 0.37 | 0.55 | 0.12 | 0.25 |
| 1 | 42105182 | rs10789406 | -0.547 | -1.804 | 0.18 | -0.031 | -0.001 | -0.001 | -0.001 | 0 | 0.56 | 0.43 | 0.44 | 0.58 | 0.59 | 0.6 |
| 1 | 42106935 | rs11210508 | -0.331 | -1.742 | 0.168 | 0.055 | 0.001 | -0.004 | -0.004 | 0 | 0.43 | 0.39 | 0.36 | 0.51 | 0.56 | 0.49 |
| 1 | 42110888 | rs6696511 | -1.243 | -1.667 | 0.172 | 0.005 | -0.025 | 0.079 | 0.079 | 0 | 0.64 | 0.8 | 0.68 | 0.9 | 0.58 | 0.56 |
| 1 | 42113215 | rs7541348 | -0.573 | -1.591 | 0.243 | 0.079 | 0.005 | 0.006 | 0.006 | 0 | 0.55 | 0.62 | 0.45 | 0.58 | 0.59 | 0.57 |
| 1 | 42117737 | rs12141084 | 0.097 | -1.149 | 0.169 | -0.141 | -0.002 | -0.001 | -0.001 | 0 | 0.36 | 0.36 | 0.3 | 0.41 | 0.45 | 0.39 |
| 1 | 42120011 | rs10890160 | -0.729 | -1.178 | -0.099 | -0.581 | 0.009 | -0.164 | -0.164 | 0.027 | 0.62 | 0.48 | 0.65 | 0.98 | 0.57 | 0.65 |
| 1 | 42122117 | rs943376 | -1.029 | -1.011 | -0.452 | -0.496 | 0.004 | -0.09 | -0.09 | 0.634 | 0.66 | 0.63 | 0.68 | 1 | 0.6 | 0.69 |
| 1 | 42133853 | rs943372 | -0.168 | -1.589 | -0.273 | 0.4 | -0.073 | 0.029 | 0.029 | 0.024 | 0.29 | 0.36 | 0.33 | 0.45 | 0.1 | 0.22 |
| 1 | 42139393 | rs17372813 | -0.073 | 0.21 | -0.447 | 0.555 | -0.017 | 0.025 | 0.025 | 0 | 0.14 | 0.04 | 0.12 | 0.07 | 0.19 | 0.1 |
| 1 | 42141252 | rs17364300 | 1.017 | 0.543 | -0.48 | 0.451 | -0.018 | 0.021 | 0.021 | 0 | 0.07 | 0.01 | 0.05 | 0 | 0.09 | 0.02 |
| 1 | 42143899 | rs11801716 | -1.011 | 0.197 | 0.37 | 0.549 | 0.042 | -0.01 | -0.01 | 0 | 0.07 | 0.33 | 0.05 | 0.01 | 0.06 | 0.01 |
| 1 | 42146295 | rs731862 | -0.882 | 0.375 | -0.307 | 1.041 | -0.04 | -0.017 | -0.017 | 0 | 0.23 | 0.05 | 0.18 | 0.21 | 0.31 | 0.33 |
| 1 | 42165629 | rs12134005 | 0.043 | 0.402 | -0.266 | 1.237 | -0.023 | -0.037 | -0.037 | 0.049 | 0.09 | 0.02 | 0.08 | 0.15 | 0.12 | 0.05 |
| 1 | 42180082 | rs2077354 | -1.207 | 0.109 | 1.511 | 1.816 | 0.046 | 0.367 | 0.367 | 2.34 | 0.05 | 0.63 | 0.08 | 0.24 | 0.01 | 0.04 |
| 1 | 42187478 | rs710243 | -0.69 | -0.569 | 0.936 | 1.341 | 0.086 | 0.102 | 0.102 | 0 | 0.47 | 0.86 | 0.54 | 0.66 | 0.35 | 0.51 |
| 1 | 42189456 | rs6600388 | -1.354 | -0.602 | 1.059 | 1.333 | 0.034 | 0.32 | 0.32 | 2.258 | 0.06 | 0.61 | 0.1 | 0.23 | 0.02 | 0.04 |
| 1 | 42191999 | rs7537131 | -1.504 | -0.604 | 1.017 | 1.331 | 0.008 | 0.255 | 0.255 | 2.24 | 0.07 | 0.68 | 0.11 | 0.23 | 0.02 | 0.04 |

| | | | | | | | | | | | | | | | | |
|---|---|---|---|---|---|---|---|---|---|---|---|---|---|---|---|---|
| 1 | 42198576 | rs710239 | -0.232 | -0.699 | 0.924 | 1.262 | 0.037 | -0.042 | -0.042 | 0 | 0.39 | 0.18 | 0.43 | 0.43 | 0.32 | 0.47 |
| 1 | 42201943 | rs6667709 | -0.238 | -0.528 | 0.885 | 1.317 | -0.01 | -0.098 | -0.098 | 0 | 0.39 | 0.1 | 0.43 | 0.48 | 0.36 | 0.45 |
| 1 | 42202714 | rs710235 | -2.346 | -0.556 | 0.95 | 1.332 | -0.032 | 0.105 | 0.105 | 3.699 | 0.08 | 0.77 | 0.11 | 0.22 | 0.02 | 0.04 |
| 1 | 42202857 | rs710234 | -1.524 | -0.548 | 1.249 | 1.338 | 0.052 | 0.39 | 0.39 | 4.073 | 0.06 | 0.5 | 0.09 | 0.22 | 0.01 | 0.03 |
| 1 | 42204406 | rs710231 | -0.734 | -0.533 | 1.177 | 1.331 | 0.056 | 0.107 | 0.107 | 0 | 0.46 | 0.86 | 0.53 | 0.69 | 0.36 | 0.49 |
| 1 | 42206473 | rs2668939 | -1.181 | -0.487 | 1.293 | 1.369 | 0.06 | 0.393 | 0.393 | 2.589 | 0.05 | 0.4 | 0.08 | 0.22 | 0.01 | 0.03 |
| 1 | 42218817 | rs4660203 | 1.34 | -0.49 | 1.305 | 1.367 | 0.059 | 0.405 | 0.405 | 2.05 | 0.94 | 0.52 | 0.91 | 0.78 | 1 | 0.97 |
| 1 | 42219601 | rs2099757 | 2.339 | -0.568 | 1.337 | 1.327 | -0.01 | 0.261 | 0.261 | 6.216 | 0.92 | 0.43 | 0.9 | 0.78 | 1 | 0.97 |
| 1 | 42223133 | rs12746339 | -0.2 | -0.656 | 0.899 | 1.276 | 0.013 | -0.021 | -0.021 | 4.279 | 0.39 | 0.28 | 0.43 | 0.47 | 0.34 | 0.45 |
| 1 | 42225580 | rs2055953 | 0.517 | -0.647 | 0.902 | 1.281 | 0.058 | 0.106 | 0.106 | 0 | 0.53 | 0.17 | 0.46 | 0.31 | 0.64 | 0.52 |
| 1 | 42228019 | rs2165303 | 1.468 | -0.601 | 0.967 | 1.311 | 0.036 | 0.335 | 0.335 | 1.277 | 0.93 | 0.49 | 0.9 | 0.78 | 0.98 | 0.96 |
| 1 | 42232085 | rs6693668 | 0.469 | -0.618 | 0.662 | 1.177 | 0.056 | 0.102 | 0.102 | 0 | 0.54 | 0.18 | 0.46 | 0.31 | 0.64 | 0.52 |
| 1 | 42234315 | rs10789409 | 1.439 | -0.609 | 0.94 | 1.361 | 0.005 | 0.248 | 0.248 | 0.987 | 0.93 | 0.35 | 0.92 | 0.77 | 0.97 | 0.96 |
| 1 | 42236004 | rs6680824 | 0.784 | -0.84 | 0.131 | 0.45 | -0.033 | 0.052 | 0.052 | 0 | 0.39 | 0.17 | 0.41 | 0.28 | 0.55 | 0.46 |
| 1 | 42238233 | rs1533184 | -0.092 | -0.061 | -0.014 | 0.105 | -0.002 | 0.074 | 0.074 | 0 | 0.48 | 0.31 | 0.4 | 0.31 | 0.54 | 0.5 |
| 1 | 42239717 | rs765040 | 0.167 | -0.115 | -0.169 | 0.292 | -0.051 | 0.152 | 0.152 | 0 | 0.45 | 0.14 | 0.38 | 0.24 | 0.53 | 0.49 |
| 1 | 42242159 | rs349430 | -0.659 | 0.071 | -0.304 | 0.83 | -0.038 | -0.041 | -0.041 | 0 | 0.87 | 0.94 | 0.69 | 0.65 | 0.79 | 0.89 |
| 1 | 42247715 | rs402379 | 0.124 | 0.102 | -0.186 | 0.284 | -0.054 | 0.183 | 0.183 | 0 | 0.46 | 0.16 | 0.38 | 0.23 | 0.51 | 0.44 |
| 1 | 42250012 | rs7548246 | -1.488 | 0.041 | 0.415 | 0.357 | -0.02 | 0.166 | 0.166 | 1.417 | 0.08 | 0.63 | 0.1 | 0.23 | 0.04 | 0.05 |
| 1 | 42253788 | rs2492604 | -1.757 | 0.034 | 0.395 | 0.353 | -0.034 | 0.107 | 0.107 | 2.551 | 0.09 | 0.71 | 0.11 | 0.23 | 0.04 | 0.05 |
| 1 | 42255767 | rs400187 | -0.647 | 0.016 | 0.289 | 0.265 | -0.026 | 0.012 | 0.012 | 0 | 0.54 | 0.83 | 0.49 | 0.46 | 0.55 | 0.49 |
| 1 | 42256262 | rs11210527 | 0.159 | 0.73 | 0.011 | -0.063 | 0.018 | 0.235 | 0.235 | 0.27 | 0.16 | 0.48 | 0.17 | 0.43 | 0.17 | 0.28 |
| 1 | 42259133 | rs17373758 | 1.393 | 0.769 | -0.181 | -0.104 | 0.004 | 0.083 | 0.083 | 0 | 0.15 | 0.01 | 0.05 | 0.02 | 0.09 | 0.08 |
| 1 | 42260474 | rs11210529 | 0.859 | 0.859 | -0.078 | -0.091 | -0.01 | -0.009 | -0.009 | 0 | 0.17 | 0.1 | 0.15 | 0.27 | 0.23 | 0.25 |
| 1 | 42261077 | rs1073926 | 1.503 | 0.798 | -0.164 | -0.463 | -0.068 | 0.108 | 0.108 | 0 | 0.2 | 0.04 | 0.25 | 0 | 0.29 | 0.13 |
| 1 | 42261889 | rs10493098 | 0.367 | 0.86 | 0.058 | -0.253 | -0.002 | 0.002 | 0.002 | 0 | 0.39 | 0.32 | 0.32 | 0.03 | 0.4 | 0.23 |
| 1 | 42271163 | rs2786480 | -0.359 | 0.855 | -0.667 | 0.246 | 0.006 | 0.019 | 0.019 | 0 | 0.6 | 0.69 | 0.53 | 0.37 | 0.62 | 0.49 |
| 1 | 42273653 | rs12563071 | 0.611 | 0.93 | -1.251 | -0.031 | -0.057 | 0.005 | 0.005 | 0 | 0.3 | 0.02 | 0.19 | 0.18 | 0.31 | 0.26 |
| 1 | 42274690 | rs647368 | 0.298 | 0.996 | -1.409 | -0.454 | -0.068 | 0.108 | 0.108 | 0 | 0.41 | 0.17 | 0.43 | 0.82 | 0.42 | 0.6 |
| 1 | 42275134 | rs606850 | 0.359 | 0.976 | -1.418 | -0.464 | -0.1 | 0.1 | 0.1 | 0.014 | 0.4 | 0.12 | 0.42 | 0.81 | 0.42 | 0.6 |
| 1 | 42277396 | rs349443 | 0.687 | 0.871 | -0.557 | -0.564 | -0.008 | -0.041 | -0.041 | 0 | 0.36 | 0.47 | 0.3 | 0.18 | 0.36 | 0.27 |
| 1 | 42281302 | rs401869 | -0.811 | 0.788 | -0.445 | -0.576 | -0.007 | -0.134 | -0.134 | 0.653 | 0.77 | 0.66 | 0.73 | 1 | 0.76 | 0.86 |
| 1 | 42284761 | rs694236 | 0.373 | 0.853 | -0.668 | -0.557 | -0.068 | 0.108 | 0.108 | 0 | 0.41 | 0.17 | 0.42 | 0.81 | 0.42 | 0.6 |
| 1 | 42285999 | rs2492603 | -1.046 | 0.8 | -0.35 | -0.558 | -0.004 | -0.094 | -0.094 | 0 | 0.79 | 0.74 | 0.75 | 1 | 0.78 | 0.87 |
| 1 | 42296145 | rs168382 | 0.053 | 0.762 | -0.438 | -0.494 | -0.029 | 0.091 | 0.091 | 0.055 | 0.43 | 0.28 | 0.45 | 0.81 | 0.45 | 0.6 |
| 1 | 42297236 | rs12562508 | 0.856 | 0.703 | -0.209 | -0.564 | -0.002 | -0.017 | -0.017 | 0 | 0.35 | 0.4 | 0.3 | 0.19 | 0.35 | 0.27 |
| 1 | 42298324 | rs611628 | 0.237 | 0.746 | -0.233 | -0.503 | -0.09 | 0.119 | 0.119 | 0.306 | 0.42 | 0.13 | 0.44 | 0.81 | 0.45 | 0.6 |
| 1 | 42299909 | rs2786484 | 0.005 | 0.483 | -0.14 | -0.524 | 0.015 | -0.061 | -0.061 | 0 | 0.89 | 0.58 | 0.84 | 1 | 0.95 | 0.9 |
| 1 | 42301595 | rs16828739 | 0.619 | 0.819 | -0.825 | 0.363 | -0.022 | -0.076 | -0.076 | 0 | 0.16 | 0.03 | 0.08 | 0.31 | 0.16 | 0.24 |
| 1 | 42317819 | rs633520 | -0.072 | 1.209 | -0.888 | -0.042 | -0.042 | -0.014 | -0.014 | 9.19 | 0.79 | 0.87 | 0.66 | 0.24 | 0.73 | 0.63 |
| 1 | 42325122 | rs10493096 | 1.093 | 1.137 | -0.235 | 0.079 | -0.021 | -0.109 | -0.109 | 0 | 0.21 | 0.31 | 0.25 | 0 | 0.18 | 0.13 |
| 1 | 42327814 | rs582123 | 0.994 | 1.202 | -0.544 | 0.032 | -0.006 | -0.052 | -0.052 | 0 | 0.35 | 0.49 | 0.31 | 0.16 | 0.35 | 0.26 |
| 1 | 42330585 | rs12734651 | -0.003 | 0.856 | -0.611 | 0.159 | -0.009 | -0.081 | -0.081 | 0 | 0.38 | 0.15 | 0.2 | 0.56 | 0.33 | 0.52 |
| 1 | 42335347 | rs783432 | -1.264 | 0.78 | -0.597 | 0.152 | -0.027 | -0.131 | -0.131 | 0 | 0.79 | 0.64 | 0.75 | 1 | 0.81 | 0.87 |
| 1 | 42350837 | rs2993120 | 0.108 | 1.306 | -0.571 | 0.904 | 0.002 | 0.002 | 0.002 | 0 | 0.39 | 0.39 | 0.53 | 0.47 | 0.47 | 0.34 |
| 1 | 42354233 | rs12759780 | 0.862 | 1.39 | -0.45 | 2.05 | 0.006 | 0.09 | 0.09 | 0.327 | 0.91 | 0.9 | 0.74 | 0.53 | 0.85 | 0.8 |
| 1 | 42355482 | rs7523641 | -1.044 | 1.374 | 0.471 | 1.603 | 0.046 | 0.305 | 0.305 | 0 | 0.12 | 0.35 | 0.1 | 0.42 | 0.13 | 0.26 |
| 1 | 42356812 | rs12059980 | -1.285 | 0.871 | 0.845 | 1.379 | 0.065 | 0.033 | 0.033 | 0.117 | 0.06 | 0.4 | 0.04 | 0 | 0.03 | 0.01 |
| 1 | 42360713 | rs811461 | 0.254 | -0.304 | -0.105 | 0.591 | 0.033 | 0.016 | 0.016 | 0 | 0.31 | 0.13 | 0.31 | 0.27 | 0.19 | 0.27 |
| 1 | 42361102 | rs783628 | -0.883 | -0.154 | -0.513 | 0.2 | -0.041 | -0.102 | -0.102 | 0 | 0.7 | 0.9 | 0.59 | 0.56 | 0.63 | 0.59 |
| 1 | 42366129 | rs12407004 | 1.767 | -0.39 | 0.013 | 0.047 | 0 | -0.008 | -0.008 | 0 | 0.09 | 0.06 | 0.05 | 0.04 | 0.06 | 0.1 |
| 1 | 42366988 | rs783622 | 0.06 | -0.388 | 0.178 | 0.057 | -0.002 | 0.001 | 0.001 | 0 | 0.55 | 0.48 | 0.51 | 0.52 | 0.54 | 0.48 |
| 1 | 42368704 | rs2764839 | -0.617 | -0.322 | 0.108 | 0.147 | 0 | 0.006 | 0.006 | 0 | 0.66 | 0.63 | 0.57 | 0.56 | 0.63 | 0.59 |
| 1 | 42378784 | rs2786485 | 0.595 | -0.47 | 0.084 | -0.245 | 0.019 | -0.005 | -0.005 | 0.089 | 0.23 | 0.16 | 0.28 | 0.23 | 0.13 | 0.18 |
| 1 | 42386080 | rs4660582 | 0.073 | -0.509 | 0.4 | -0.439 | 0.002 | -0.012 | -0.012 | 0 | 0.44 | 0.47 | 0.31 | 0.7 | 0.52 | 0.55 |
| 1 | 42390317 | rs4660584 | 0.747 | -0.505 | 0.205 | -0.433 | 0.017 | -0.006 | -0.006 | 0 | 0.23 | 0.15 | 0.28 | 0.24 | 0.12 | 0.18 |
| 1 | 42397435 | rs6687206 | 0.229 | -0.193 | 0.268 | -0.11 | -0.087 | 0.045 | 0.045 | 0 | 0.4 | 0.11 | 0.29 | 0.7 | 0.52 | 0.55 |
| 1 | 42408070 | rs7515635 | 0.053 | -0.398 | 0.842 | -0.082 | -0.008 | 0.042 | 0.042 | 0 | 0.43 | 0.3 | 0.31 | 0.7 | 0.53 | 0.55 |
| 1 | 42417182 | rs10890172 | 0.125 | -0.432 | 0.847 | -0.084 | -0.017 | 0.051 | 0.051 | 0 | 0.42 | 0.23 | 0.3 | 0.7 | 0.53 | 0.52 |
| 1 | 42418570 | rs12404436 | 0.352 | -0.611 | 0.95 | -0.252 | 0.052 | 0.019 | 0.019 | 0 | 0.31 | 0.15 | 0.31 | 0.27 | 0.17 | 0.31 |
| 1 | 42424594 | rs653944 | -0.068 | -0.593 | 1.257 | -0.257 | -0.002 | -0.012 | -0.012 | 0 | 0.45 | 0.54 | 0.33 | 0.69 | 0.55 | 0.54 |
| 1 | 42424713 | rs6677053 | 0.429 | -0.445 | 1.008 | 0.12 | -0.005 | -0.011 | -0.011 | 2.485 | 0.09 | 0.07 | 0.05 | 0.16 | 0.12 | 0.23 |
| 1 | 42441288 | rs4660590 | 0.227 | -0.497 | 1.02 | -0.04 | -0.009 | 0.039 | 0.039 | 0 | 0.42 | 0.28 | 0.31 | 0.69 | 0.53 | 0.52 |
| 1 | 42444975 | rs654961 | -0.037 | -0.488 | 0.778 | -0.016 | 0.007 | 0.036 | 0.036 | 0 | 0.46 | 0.68 | 0.33 | 0.69 | 0.55 | 0.54 |
| 1 | 42445113 | rs604145 | -1.573 | -0.109 | 0.795 | -0.007 | -0.005 | 0.033 | 0.033 | 0 | 0.78 | 0.88 | 0.65 | 0.97 | 0.72 | 0.85 |
| 1 | 42466251 | rs348142 | -0.184 | -0.323 | 0.86 | -0.013 | 0.031 | 0.085 | 0.085 | 0 | 0.51 | 0.74 | 0.36 | 0.69 | 0.56 | 0.59 |
| 1 | 42470901 | rs348139 | -2.012 | -0.398 | 0.96 | -0.131 | -0.023 | 0.067 | 0.067 | 0 | 0.8 | 0.93 | 0.66 | 0.97 | 0.73 | 0.86 |
| 1 | 42472711 | rs3856257 | 0.158 | -0.429 | 0.991 | -0.122 | 0.006 | 0.1 | 0.1 | 0 | 0.47 | 0.17 | 0.32 | 0.7 | 0.54 | 0.58 |
| 1 | 42476676 | rs4660210 | -1.672 | -0.437 | 1.171 | -0.129 | 0.003 | -0.066 | -0.066 | 0 | 0.78 | 0.65 | 0.64 | 0.97 | 0.73 | 0.86 |

| | | | | | | | | | | | | | | | | |
|---|---|---|---|---|---|---|---|---|---|---|---|---|---|---|---|---|
| 1 | 42484367 | rs4660596 | 1.092 | -0.525 | 0.605 | 0.115 | -0.137 | -0.129 | -0.129 | 4.172 | 0.34 | 0.02 | 0.24 | 0.41 | 0.41 | 0.29 |
| 1 | 42486843 | rs6683904 | 0.429 | -0.56 | 0.821 | -0.256 | 0.031 | 0.007 | 0.007 | 0 | 0.28 | 0.18 | 0.3 | 0.27 | 0.17 | 0.26 |
| 1 | 42487523 | rs6689536 | -1.31 | -0.52 | 0.601 | -0.234 | 0.01 | -0.125 | -0.125 | 0 | 0.76 | 0.54 | 0.63 | 0.97 | 0.71 | 0.84 |
| 1 | 42489031 | rs1024278 | 0.044 | -0.289 | -0.26 | 0.011 | -0.043 | 0.086 | 0.086 | 0 | 0.45 | 0.1 | 0.3 | 0.69 | 0.53 | 0.58 |
| 1 | 42513929 | rs4660600 | -1.339 | -0.093 | 0.699 | -0.138 | 0.008 | -0.072 | -0.072 | 0 | 0.77 | 0.63 | 0.63 | 0.97 | 0.7 | 0.83 |
| 1 | 42514452 | rs11210568 | 0.487 | -0.214 | 0.264 | 0.239 | -0.076 | -0.102 | -0.102 | 0 | 0.37 | 0.1 | 0.26 | 0.48 | 0.4 | 0.31 |
| 1 | 42534375 | rs6691697 | 0.272 | -0.262 | 0.831 | 0.101 | 0.006 | 0.011 | 0.011 | 0 | 0.4 | 0.47 | 0.29 | 0.49 | 0.4 | 0.31 |
| 1 | 42547061 | rs10157817 | 0.156 | -0.267 | 0.771 | 0.107 | -0.001 | -0.002 | -0.002 | 0 | 0.42 | 0.52 | 0.29 | 0.49 | 0.4 | 0.31 |
| 1 | 42551252 | rs9943167 | -0.318 | -0.369 | 0.669 | 0.082 | 0.007 | 0.079 | 0.079 | 0 | 0.51 | 0.63 | 0.34 | 0.69 | 0.51 | 0.59 |
| 1 | 42570803 | rs1016867 | 0.836 | -0.574 | 0.945 | 0.285 | 0.035 | -0.022 | -0.022 | 0 | 0.59 | 0.24 | 0.63 | 0.4 | 0.7 | 0.44 |
| 1 | 42571439 | rs1024909 | 0.672 | -0.72 | 0.747 | 0.261 | -0.003 | 0.103 | 0.103 | 0.31 | 0.75 | 0.49 | 0.67 | 0.43 | 0.75 | 0.51 |
| 1 | 42577444 | rs6672120 | 0.549 | -0.581 | 0.493 | 0.055 | 0.014 | 0.015 | 0.015 | 0 | 0.66 | 0.3 | 0.83 | 0.77 | 0.74 | 0.63 |
| 1 | 42580407 | rs11210587 | -0.274 | -0.294 | 0.204 | -0.026 | 0 | 0 | 0 | 0 | 0.2 | 0.25 | 0.3 | 0.37 | 0.18 | 0.23 |
| 1 | 42591695 | rs4660608 | -0.257 | -0.282 | 0.153 | -0.025 | -0.001 | -0.006 | -0.006 | 3.079 | 0.2 | 0.22 | 0.29 | 0.36 | 0.18 | 0.23 |
| 1 | 42598964 | rs7542228 | -1.06 | -0.297 | 0.097 | -0.037 | 0.01 | 0.003 | 0.003 | 0.183 | 0.38 | 0.66 | 0.37 | 0.41 | 0.24 | 0.3 |
| 1 | 42609867 | rs883062 | 1.397 | -0.247 | -0.581 | 0.06 | -0.062 | -0.044 | -0.044 | 0 | 0.47 | 0.07 | 0.54 | 0.36 | 0.56 | 0.39 |
| 1 | 42616910 | rs4660610 | 0.228 | -0.315 | -0.201 | 0.136 | -0.003 | -0.026 | -0.026 | 0 | 0.15 | 0.12 | 0.26 | 0.4 | 0.16 | 0.2 |
| 1 | 42618740 | rs12069432 | 0.458 | -0.293 | 0.005 | 0.126 | -0.002 | 0 | 0 | 0 | 0.14 | 0.14 | 0.09 | 0.16 | 0.19 | 0.3 |
| 1 | 42620154 | rs885846 | 1.314 | -0.312 | -0.223 | 0.145 | -0.013 | -0.069 | -0.069 | 0.894 | 0.52 | 0.08 | 0.57 | 0.41 | 0.6 | 0.42 |
| 1 | 42621645 | rs3818727 | -1.034 | -0.177 | 0.414 | 0.421 | 0.063 | 0.169 | 0.169 | 0.056 | 0.07 | 0.25 | 0.23 | 0.35 | 0.11 | 0.13 |
| 1 | 42627280 | rs1877477 | -0.349 | -0.279 | 0.444 | 0.463 | -0.013 | 0.05 | 0.05 | 0.019 | 0.16 | 0.25 | 0.25 | 0.41 | 0.13 | 0.2 |
| 1 | 42635107 | rs7546673 | -0.506 | 0.052 | -0.054 | 0.127 | -0.01 | 0.001 | 0.001 | 0.127 | 0.26 | 0.21 | 0.15 | 0.26 | 0.26 | 0.36 |
| 1 | 42638163 | rs12045477 | 0.971 | -0.128 | -0.912 | -0.185 | -0.044 | -0.138 | -0.138 | 0 | 0.27 | 0.02 | 0.17 | 0.35 | 0.28 | 0.24 |
| 1 | 42641141 | rs1877478 | 1.115 | -0.333 | -0.655 | -0.331 | 0.033 | 0.065 | 0.065 | 0 | 0.13 | 0.01 | 0.02 | 0.01 | 0.05 | 0.11 |
| 1 | 42641625 | rs651674 | -0.282 | -0.358 | -0.351 | -0.342 | -0.003 | 0.002 | 0.002 | 0.178 | 0.27 | 0.26 | 0.16 | 0.24 | 0.26 | 0.36 |
| 1 | 42644770 | rs686247 | -0.479 | -0.303 | 0.217 | -0.381 | 0.061 | 0.084 | 0.084 | 0 | 0.92 | 0.77 | 0.72 | 0.78 | 0.88 | 0.84 |
| 1 | 42657196 | rs1139978 | 2.139 | -0.35 | -0.241 | -0.627 | -0.018 | 0.049 | 0.049 | 0 | 0.11 | 0.02 | 0.07 | 0 | 0.13 | 0.05 |
| 1 | 42664238 | rs585320 | 1.041 | -0.532 | 0.564 | -0.329 | 0.02 | -0.001 | -0.001 | 0.007 | 0.14 | 0.19 | 0.11 | 0.15 | 0.2 | 0.28 |
| 1 | 42680327 | rs343360 | 0.585 | -0.609 | 0.616 | -0.361 | 0.007 | -0.025 | -0.025 | 0 | 0.71 | 0.51 | 0.83 | 0.76 | 0.74 | 0.64 |
| 1 | 42693597 | rs343376 | -1.196 | -0.61 | 0.614 | -0.365 | 0.036 | -0.003 | -0.003 | 0.602 | 0.85 | 0.76 | 0.89 | 0.85 | 0.79 | 0.72 |
| 1 | 42702344 | rs608277 | 0.078 | -0.521 | 0.309 | -0.318 | 0.007 | -0.009 | -0.009 | 0 | 0.73 | 0.64 | 0.84 | 0.76 | 0.74 | 0.64 |
| 1 | 42727950 | rs510157 | 0.403 | -0.527 | 0.335 | -0.321 | -0.001 | -0.011 | -0.011 | 0 | 0.7 | 0.39 | 0.82 | 0.76 | 0.74 | 0.64 |
| 1 | 42737932 | rs10789416 | 0.076 | -0.507 | -0.195 | -0.312 | 0.007 | -0.009 | -0.009 | 0 | 0.73 | 0.64 | 0.84 | 0.76 | 0.74 | 0.64 |
| 1 | 42738366 | rs12402117 | 1.419 | -0.518 | -0.221 | -0.316 | 0.023 | -0.015 | -0.015 | 0 | 0.11 | 0 | 0.04 | 0.09 | 0.06 | 0.08 |
| 1 | 42748731 | rs3754280 | 1.109 | -0.423 | -0.019 | -0.137 | -0.048 | -0.011 | -0.011 | 0.103 | 0.49 | 0.12 | 0.52 | 0.5 | 0.57 | 0.37 |
| 1 | 42768969 | rs10736423 | -0.161 | -0.402 | -0.001 | -0.257 | 0.006 | -0.025 | -0.025 | 0 | 0.71 | 0.51 | 0.83 | 0.76 | 0.74 | 0.64 |
| 1 | 42797765 | rs1566584 | 0.284 | 0.082 | 0.443 | -0.091 | -0.002 | -0.012 | -0.012 | 0 | 0.15 | 0.14 | 0.03 | 0.01 | 0.05 | 0.1 |
| 1 | 42798204 | rs2494366 | -0.85 | 0.986 | 0.54 | 0.498 | 0.111 | 0.145 | 0.145 | 0.339 | 0.09 | 0.37 | 0.29 | 0.22 | 0.12 | 0.16 |
| 1 | 42806191 | rs913426 | -0.129 | 0.446 | -0.131 | 0.158 | -0.001 | -0.001 | -0.001 | 0 | 0.43 | 0.59 | 0.51 | 0.44 | 0.5 | 0.34 |
| 1 | 42823768 | rs1536651 | 0.56 | 0.187 | -0.258 | 0.207 | -0.125 | -0.094 | -0.094 | 0 | 0.32 | 0.1 | 0.46 | 0.33 | 0.47 | 0.25 |
| 1 | 42825442 | rs17369290 | 0.733 | -0.147 | -0.632 | -0.222 | 0.03 | 0.014 | 0.014 | 0 | 0.16 | 0.01 | 0.06 | 0.11 | 0.13 | 0.13 |
| 1 | 42835396 | rs6679533 | -0.064 | 0.231 | 0.32 | 0.095 | 0.001 | -0.001 | -0.001 | 0 | 0.47 | 0.4 | 0.43 | 0.57 | 0.4 | 0.62 |
| 1 | 42850403 | rs11210628 | 0.504 | 0.558 | 0.634 | -0.268 | -0.003 | -0.043 | -0.043 | 0.968 | 0.52 | 0.49 | 0.39 | 0.12 | 0.47 | 0.3 |
| 1 | 42857663 | rs17378108 | 0.302 | 0.933 | 0.71 | 0.146 | 0.025 | 0.081 | 0.081 | 0 | 0.09 | 0.14 | 0.14 | 0.23 | 0.17 | 0.09 |
| 1 | 42864868 | rs4660630 | 0.124 | 0.938 | 0.954 | 0.118 | 0.065 | 0.152 | 0.152 | 0 | 0.1 | 0.22 | 0.14 | 0.22 | 0.18 | 0.1 |
| 1 | 42876423 | rs7413966 | 0.226 | 0.759 | 0.786 | 0.191 | 0.004 | 0.002 | 0.002 | 0 | 0.61 | 0.52 | 0.52 | 0.34 | 0.64 | 0.42 |
| 1 | 42876696 | rs7418303 | -0.064 | 0.763 | 0.347 | 0.193 | -0.019 | -0.012 | -0.012 | 0.002 | 0.63 | 0.7 | 0.54 | 0.34 | 0.65 | 0.42 |
| 1 | 42880516 | rs1055055 | 0.157 | 0.863 | 0.016 | 0.241 | 0.005 | 0.002 | 0.002 | 0 | 0.61 | 0.52 | 0.52 | 0.34 | 0.64 | 0.42 |
| 1 | 42882296 | rs4660632 | -0.295 | 0.978 | -0.218 | 0.69 | -0.008 | -0.005 | -0.005 | 0 | 0.08 | 0.06 | 0.1 | 0.18 | 0.13 | 0.06 |
| 1 | 42901082 | rs3748845 | -0.823 | 0.896 | 0.274 | 0.685 | 0.027 | 0.092 | 0.092 | 0 | 0.11 | 0.24 | 0.13 | 0.21 | 0.15 | 0.11 |
| 1 | 42904304 | rs7529199 | -0.359 | 0.751 | -0.152 | 0.533 | 0.011 | -0.026 | -0.026 | 0.211 | 0.28 | 0.23 | 0.16 | 0.43 | 0.19 | 0.47 |
| 1 | 42908848 | rs4660226 | -0.411 | 0.528 | -0.323 | 0.548 | -0.015 | -0.034 | -0.034 | 0 | 0.14 | 0.07 | 0.13 | 0.16 | 0.17 | 0.12 |
| 1 | 42913695 | rs1002701 | -1.611 | 0.38 | -0.438 | -0.228 | -0.007 | 0.009 | 0.009 | 0 | 0.63 | 0.57 | 0.59 | 0.61 | 0.66 | 0.72 |
| 1 | 42918754 | rs12039696 | -0.988 | 0.366 | -0.323 | -0.307 | 0.001 | 0 | 0 | 0 | 0.42 | 0.45 | 0.3 | 0.6 | 0.35 | 0.57 |
| 1 | 42933906 | rs12121532 | 1.423 | 0.321 | -0.066 | -0.347 | -0.084 | 0.13 | 0.13 | 0.039 | 0.2 | 0.02 | 0.28 | 0.01 | 0.31 | 0.15 |
| 1 | 42999325 | rs4660229 | 0.079 | 0.234 | -0.021 | -0.309 | -0.008 | -0.024 | -0.024 | 0 | 0.14 | 0.08 | 0.13 | 0.16 | 0.16 | 0.12 |
| 1 | 43009616 | rs12032119 | 0.372 | 0.18 | -0.024 | -0.312 | 0.006 | -0.016 | -0.016 | 0 | 0.27 | 0.24 | 0.16 | 0.43 | 0.19 | 0.45 |
| 1 | 43024213 | rs11210669 | 1.496 | 0.237 | -0.157 | -0.365 | -0.078 | 0.125 | 0.125 | 0 | 0.19 | 0.02 | 0.27 | 0.01 | 0.3 | 0.15 |
| 1 | 43029788 | rs4660646 | 0.145 | 0.279 | -0.123 | -0.322 | -0.016 | -0.038 | -0.038 | 0 | 0.14 | 0.06 | 0.13 | 0.14 | 0.16 | 0.12 |
| 1 | 43034105 | rs12033025 | 0.909 | -0.059 | -0.241 | -0.007 | 0.013 | 0.005 | 0.005 | 0 | 0.15 | 0.13 | 0.06 | 0.17 | 0.05 | 0.22 |
| 1 | 43036700 | rs2762699 | 1.703 | -0.035 | 0.503 | -0.006 | -0.002 | -0.011 | -0.011 | 0.007 | 0.36 | 0.23 | 0.4 | 0.39 | 0.35 | 0.28 |
| 1 | 43042125 | rs2762697 | -0.275 | 0.015 | 0.426 | 0.006 | -0.017 | 0.033 | 0.033 | 0 | 0.65 | 0.75 | 0.59 | 0.85 | 0.54 | 0.73 |
| 1 | 43042703 | rs12049046 | 0.046 | -0.019 | 0.275 | -0.006 | 0.011 | -0.041 | -0.041 | 0.953 | 0.27 | 0.18 | 0.16 | 0.46 | 0.19 | 0.45 |
| 1 | 43045757 | rs1016544 | -0.137 | -0.001 | 0.378 | -0.083 | 0.011 | -0.039 | -0.039 | 0.427 | 0.64 | 0.53 | 0.58 | 0.85 | 0.54 | 0.73 |
| 1 | 43050396 | rs2809658 | -0.225 | 0.005 | 0.344 | -0.08 | -0.006 | 0.013 | 0.013 | 0 | 0.64 | 0.69 | 0.59 | 0.85 | 0.54 | 0.73 |
| 1 | 43073504 | rs12025456 | -0.389 | -0.165 | 0.24 | 0.143 | -0.004 | 0.004 | 0.004 | 0 | 0.28 | 0.35 | 0.16 | 0.44 | 0.19 | 0.49 |
| 1 | 43081293 | rs12081092 | 0.407 | 0.216 | 0.226 | -0.082 | 0.001 | -0.004 | -0.004 | 0 | 0.32 | 0.24 | 0.32 | 0.16 | 0.32 | 0.29 |
| 1 | 43101562 | rs6600418 | -1.441 | 0.35 | 0.101 | 0.183 | -0.011 | -0.012 | -0.012 | 0 | 0.61 | 0.74 | 0.56 | 0.6 | 0.6 | 0.7 |
| 1 | 43102844 | rs16829822 | 0.109 | 0.32 | -0.16 | 0.298 | -0.025 | -0.054 | -0.054 | 0.143 | 0.17 | 0.09 | 0.12 | 0.29 | 0.15 | 0.29 |

| | | | | | | | | | | | | | | | | |
|---|---|---|---|---|---|---|---|---|---|---|---|---|---|---|---|---|
| 1 | 43104952 | rs1951918 | -0.155 | 0.297 | 0.023 | 0.287 | -0.003 | -0.015 | -0.015 | 1.529 | 0.18 | 0.19 | 0.13 | 0.29 | 0.15 | 0.34 |
| 1 | 43108188 | rs12036818 | 0.972 | 0.41 | -0.154 | 0.448 | -0.021 | -0.072 | -0.072 | 1.042 | 0.12 | 0.02 | 0.09 | 0.31 | 0.11 | 0.21 |
| 1 | 43110276 | rs7523056 | -1.648 | 0.279 | 0.006 | 0.247 | -0.021 | -0.005 | -0.005 | 0 | 0.62 | 0.78 | 0.57 | 0.64 | 0.6 | 0.7 |
| 1 | 43118882 | rs11210691 | -0.484 | 0.244 | 0.285 | 0.209 | 0.015 | 0.014 | 0.014 | 0 | 0.46 | 0.28 | 0.41 | 0.47 | 0.44 | 0.57 |
| 1 | 43123855 | rs12145700 | 0.755 | 0.249 | 0.5 | 0.037 | 0.002 | 0.07 | 0.07 | 0.143 | 0.31 | 0.13 | 0.31 | 0.16 | 0.32 | 0.31 |
| 1 | 43144021 | rs912074 | -0.846 | 0.21 | 0.705 | 0.033 | -0.002 | -0.009 | -0.009 | 0 | 0.46 | 0.52 | 0.46 | 0.32 | 0.47 | 0.43 |
| 1 | 43155062 | rs11210696 | 0.193 | 0.161 | 0.768 | -0.004 | -0.001 | 0.037 | 0.037 | 0 | 0.34 | 0.23 | 0.34 | 0.15 | 0.34 | 0.38 |
| 1 | 43158510 | rs10493113 | 0.971 | 0.158 | 0.804 | 0.014 | -0.001 | -0.002 | -0.002 | 0.084 | 0.1 | 0.07 | 0.1 | 0.17 | 0.13 | 0.06 |
| 1 | 43169007 | rs11210698 | -0.475 | 0.074 | 0.746 | -0.032 | 0.001 | 0.009 | 0.009 | 0 | 0.4 | 0.33 | 0.41 | 0.32 | 0.43 | 0.36 |
| 1 | 43200526 | rs912075 | -0.645 | 0.321 | 0.626 | -0.343 | 0 | -0.004 | -0.004 | 0 | 0.6 | 0.52 | 0.62 | 0.67 | 0.6 | 0.65 |
| 1 | 43202095 | rs7548008 | -0.194 | 0.342 | 0.635 | -0.26 | -0.003 | 0.014 | 0.014 | 0 | 0.52 | 0.36 | 0.45 | 0.45 | 0.48 | 0.59 |
| 1 | 43202636 | rs10890212 | -1.487 | 0.321 | 0.677 | -0.274 | 0.003 | 0.004 | 0.004 | 0 | 0.68 | 0.62 | 0.66 | 0.67 | 0.64 | 0.71 |
| 1 | 43208561 | rs11590362 | 1.599 | 0.569 | 0.365 | -0.406 | -0.012 | 0.005 | 0.005 | 0.018 | 0.05 | 0.01 | 0.05 | 0.04 | 0.08 | 0.12 |
| 1 | 43212111 | rs6882 | -0.81 | 0.438 | 0.516 | -0.477 | -0.005 | -0.006 | -0.006 | 1.595 | 0.65 | 0.52 | 0.66 | 0.71 | 0.68 | 0.78 |
| 1 | 43213610 | rs7552138 | -2.137 | 0.467 | 0.558 | -0.461 | -0.001 | -0.001 | -0.001 | 0 | 0.75 | 0.75 | 0.71 | 0.71 | 0.73 | 0.85 |
| 1 | 43213797 | rs3738500 | 1.299 | 0.646 | 0.489 | -0.402 | 0.002 | 0.03 | 0.03 | 0 | 0.07 | 0.07 | 0.16 | 0.22 | 0.12 | 0.06 |
| 1 | 43215930 | rs11581921 | 1.502 | 0.702 | 0.449 | -0.432 | -0.012 | 0.005 | 0.005 | 0.018 | 0.05 | 0.01 | 0.05 | 0.04 | 0.08 | 0.12 |
| 1 | 43223489 | rs6700677 | 0.423 | 0.536 | 0.502 | -0.37 | 0.001 | -0.013 | -0.013 | 0 | 0.08 | 0.11 | 0.04 | 0 | 0.04 | 0.06 |
| 1 | 43230559 | rs2816594 | 0.234 | 0.8 | 0.182 | -0.392 | -0.001 | -0.001 | -0.001 | 0 | 0.33 | 0.29 | 0.32 | 0.66 | 0.29 | 0.44 |
| 1 | 43232798 | rs11210712 | 1.218 | 0.837 | 0.052 | -0.449 | -0.038 | 0.085 | 0.085 | 0 | 0.19 | 0.06 | 0.26 | 0.01 | 0.25 | 0.17 |
| 1 | 43238031 | rs17385526 | 1.245 | 0.848 | 0.043 | -0.444 | -0.047 | 0.097 | 0.097 | 0 | 0.19 | 0.05 | 0.26 | 0.01 | 0.25 | 0.17 |
| 1 | 43254108 | rs719676 | 0.699 | 0.698 | 0.022 | -0.424 | 0.014 | -0.062 | -0.062 | 0.031 | 0.31 | 0.07 | 0.16 | 0.44 | 0.22 | 0.36 |
| 1 | 43261781 | rs7547717 | -0.209 | 0.84 | 0.059 | -0.414 | 0.005 | 0.004 | 0.004 | 0 | 0.4 | 0.28 | 0.34 | 0.65 | 0.34 | 0.42 |
| 1 | 43270928 | rs323719 | -0.207 | 1.098 | -0.236 | -0.488 | -0.078 | 0.175 | 0.175 | 0 | 0.27 | 0.07 | 0.27 | 0.01 | 0.3 | 0.23 |
| 1 | 43271033 | rs323720 | -0.867 | 1.091 | -0.047 | -0.502 | 0.005 | -0.095 | -0.095 | 0 | 0.32 | 0.42 | 0.31 | 0.01 | 0.32 | 0.28 |
| 1 | 43272246 | rs11210721 | 0.559 | 1.287 | -0.183 | -0.283 | -0.066 | 0.117 | 0.117 | 0 | 0.19 | 0.04 | 0.25 | 0.01 | 0.25 | 0.17 |
| 1 | 43278242 | rs1108734 | 0.559 | 1.28 | -0.183 | -0.269 | -0.063 | 0.117 | 0.117 | 0 | 0.19 | 0.04 | 0.25 | 0.01 | 0.25 | 0.17 |
| 1 | 43285544 | rs6658712 | 1.185 | 1.332 | -0.542 | -0.144 | 0 | -0.001 | -0.001 | 0 | 0.35 | 0.31 | 0.37 | 0.33 | 0.39 | 0.32 |
| 1 | 43296173 | rs33950227 | 0.093 | 1.626 | -1.142 | -0.054 | -0.031 | 0.077 | 0.077 | 0 | 0.19 | 0.09 | 0.26 | 0.01 | 0.25 | 0.17 |
| 1 | 43296195 | rs33953680 | -0.255 | 1.594 | -1.169 | 0.019 | -0.092 | 0.174 | 0.174 | 0 | 0.26 | 0.02 | 0.27 | 0.01 | 0.3 | 0.19 |
| 1 | 43298871 | rs4660671 | -0.124 | 1.197 | -0.228 | 0.996 | -0.001 | 0.019 | 0.019 | 0.273 | 0.09 | 0.06 | 0.17 | 0.25 | 0.11 | 0.22 |
| 1 | 43305753 | rs12034000 | 0.638 | 1.206 | 0.166 | 0.831 | -0.002 | 0 | 0 | 0.897 | 0.26 | 0.26 | 0.16 | 0.34 | 0.2 | 0.31 |
| 1 | 43306128 | rs11210729 | -0.141 | 1.498 | 0.152 | 1.34 | 0.013 | 0.042 | 0.042 | 0.953 | 0.09 | 0.14 | 0.3 | 0.22 | 0.14 | 0.18 |
| 1 | 43310639 | rs12406279 | -0.084 | 1.207 | -0.328 | 0.428 | -0.01 | 0.002 | 0.002 | 0 | 0.13 | 0.08 | 0.22 | 0 | 0.17 | 0.15 |
| 1 | 43317793 | rs9960 | -0.43 | 1.185 | -0.269 | 0.415 | 0.002 | -0.041 | -0.041 | 0 | 0.14 | 0.16 | 0.21 | 0 | 0.16 | 0.14 |
| 1 | 43319272 | rs4660234 | 0.467 | 1.142 | -0.339 | 0.486 | -0.026 | 0.036 | 0.036 | 0 | 0.1 | 0.04 | 0.17 | 0 | 0.13 | 0.12 |
| 1 | 43321476 | rs6685920 | 0.479 | 1.141 | -0.287 | 0.485 | -0.017 | 0.02 | 0.02 | 0 | 0.11 | 0.07 | 0.16 | 0 | 0.13 | 0.12 |
| 1 | 43326098 | rs12136910 | 0.141 | 1.335 | -0.311 | 0.99 | 0 | 0.028 | 0.028 | 0.593 | 0.09 | 0.11 | 0.28 | 0.23 | 0.13 | 0.16 |
| 1 | 43326208 | rs12022524 | 0.678 | 0.88 | -0.443 | 0.284 | 0.005 | 0.003 | 0.003 | 0 | 0.28 | 0.17 | 0.21 | 0.34 | 0.25 | 0.38 |
| 1 | 43341662 | rs1601240 | -0.234 | 0.47 | -0.501 | -0.119 | -0.015 | 0.084 | 0.084 | 0 | 0.44 | 0.26 | 0.27 | 0.2 | 0.46 | 0.33 |
| 1 | 43353134 | rs6674019 | -0.309 | 0.598 | -0.687 | -0.054 | -0.001 | 0.032 | 0.032 | 0.248 | 0.47 | 0.43 | 0.29 | 0.19 | 0.46 | 0.33 |
| 1 | 43357803 | rs2792600 | -0.101 | 0.814 | -0.944 | 0.556 | 0.005 | 0.079 | 0.079 | 0.615 | 0.49 | 0.33 | 0.28 | 0.19 | 0.45 | 0.34 |
| 1 | 43366891 | rs841860 | 0.926 | 0.848 | -0.108 | 1.826 | 0.017 | 0.142 | 0.142 | 0 | 0.83 | 0.64 | 0.75 | 0.39 | 0.86 | 0.75 |
| 1 | 43367726 | rs12564800 | -0.527 | 0.775 | -0.63 | 1.279 | -0.011 | 0.023 | 0.023 | 0 | 0.37 | 0.26 | 0.47 | 0.3 | 0.39 | 0.39 |
| 1 | 43367875 | rs841859 | -0.69 | 0.729 | -0.934 | 1.331 | -0.01 | -0.006 | -0.006 | 0.061 | 0.15 | 0.08 | 0.17 | 0.14 | 0.14 | 0.26 |
| 1 | 43368150 | rs11210751 | 0.305 | 0.695 | -0.835 | 1.309 | -0.023 | 0.051 | 0.051 | 0.133 | 0.22 | 0.08 | 0.31 | 0.11 | 0.21 | 0.26 |
| 1 | 43374498 | rs6699129 | 1.167 | 0.403 | -0.847 | 1.321 | -0.111 | 0.034 | 0.034 | 0 | 0.42 | 0.03 | 0.29 | 0.24 | 0.5 | 0.31 |
| 1 | 43380982 | rs841837 | -0.129 | 0.551 | -0.654 | 1.027 | 0.001 | 0.009 | 0.009 | 0 | 0.08 | 0.08 | 0.14 | 0.14 | 0.08 | 0.18 |
| 1 | 43381026 | rs1098782 | -0.701 | 0.434 | -0.755 | 0.922 | -0.015 | -0.018 | -0.018 | 0 | 0.52 | 0.85 | 0.66 | 0.48 | 0.47 | 0.62 |
| 1 | 43398085 | rs3820546 | 0.589 | 0.465 | -1.679 | 0.785 | -0.111 | -0.027 | -0.027 | 0.06 | 0.44 | 0.03 | 0.36 | 0.32 | 0.55 | 0.37 |
| 1 | 43398396 | rs3820548 | 0.216 | 0.012 | -0.753 | 1.021 | -0.002 | 0.008 | 0.008 | 0 | 0.66 | 0.62 | 0.66 | 0.51 | 0.71 | 0.66 |
| 1 | 43399167 | rs841858 | 0.473 | 0.181 | -0.936 | 1.137 | -0.019 | -0.011 | -0.011 | 0 | 0.16 | 0.11 | 0.2 | 0.2 | 0.17 | 0.16 |
| 1 | 43399686 | rs3754219 | 1.394 | 0.373 | -1.687 | 0.835 | -0.098 | -0.066 | -0.066 | 0.121 | 0.42 | 0.04 | 0.33 | 0.36 | 0.47 | 0.4 |
| 1 | 43400352 | rs3729548 | 1.442 | 0.67 | -1.706 | 0.844 | -0.076 | -0.065 | -0.065 | 0 | 0.41 | 0.04 | 0.28 | 0.35 | 0.45 | 0.37 |
| 1 | 43401438 | rs841853 | 0.451 | 0.436 | -0.456 | 0.882 | -0.006 | -0.016 | -0.016 | 0 | 0.68 | 0.51 | 0.62 | 0.78 | 0.7 | 0.77 |
| 1 | 43401499 | rs841852 | 0.084 | 0.331 | -0.021 | 0.974 | 0.012 | -0.007 | -0.007 | 1.075 | 0.21 | 0.31 | 0.23 | 0.24 | 0.2 | 0.2 |
| 1 | 43402545 | rs841848 | 0.056 | 0.324 | 0.031 | 0.971 | 0.007 | -0.006 | -0.006 | 0 | 0.22 | 0.29 | 0.23 | 0.24 | 0.2 | 0.2 |
| 1 | 43408966 | rs1385129 | -0.089 | 0.274 | -0.149 | 0.796 | -0.003 | -0.016 | -0.016 | 0 | 0.24 | 0.15 | 0.32 | 0.32 | 0.22 | 0.24 |
| 1 | 43410859 | rs751210 | -1.477 | 0.223 | -0.154 | 0.661 | -0.001 | -0.06 | -0.06 | 0 | 0.43 | 0.89 | 0.55 | 0.47 | 0.33 | 0.43 |
| 1 | 43411831 | rs1770807 | -1.486 | 0.221 | -0.155 | 0.66 | 0 | -0.059 | -0.059 | 0 | 0.43 | 0.89 | 0.55 | 0.47 | 0.33 | 0.43 |
| 1 | 43412259 | rs2086856 | 1.176 | 0.221 | -0.155 | 0.66 | -0.001 | -0.06 | -0.06 | 0.455 | 0.57 | 0.11 | 0.45 | 0.53 | 0.67 | 0.57 |
| 1 | 43414200 | rs710221 | -1.547 | 0.117 | 0.314 | 0.558 | 0.037 | -0.012 | -0.012 | 0.11 | 0.46 | 0.79 | 0.53 | 0.54 | 0.4 | 0.47 |
| 1 | 43415288 | rs2297977 | 0.698 | 0.099 | 0.629 | 0.557 | 0.003 | 0.001 | 0.001 | 0 | 0.2 | 0.26 | 0.27 | 0.23 | 0.21 | 0.18 |
| 1 | 43415346 | rs710220 | -0.548 | 0.062 | 1.103 | 0.555 | -0.015 | 0.005 | 0.005 | 0 | 0.3 | 0.44 | 0.35 | 0.35 | 0.23 | 0.28 |
| 1 | 43416133 | rs17387775 | 0.047 | 0.09 | 1.045 | 0.565 | -0.001 | -0.004 | -0.004 | 0 | 0.13 | 0.06 | 0.15 | 0.17 | 0.16 | 0.2 |
| 1 | 43418026 | rs2297972 | 0.618 | -0.017 | 1.119 | 0.642 | 0.009 | 0 | 0 | 0 | 0.67 | 0.27 | 0.62 | 0.64 | 0.77 | 0.72 |
| 1 | 43419737 | rs12406072 | -1.445 | -0.051 | 1.188 | 0.605 | -0.061 | 0.02 | 0.02 | 5.935 | 0.13 | 0.42 | 0.1 | 0.12 | 0.02 | 0.1 |
| 1 | 43421524 | rs3768045 | -0.899 | -0.031 | 0.82 | 0.615 | -0.043 | 0.005 | 0.005 | 2.668 | 0.11 | 0.19 | 0.08 | 0.12 | 0.02 | 0.1 |
| 1 | 43436051 | rs841572 | -1.448 | 0.27 | 0.872 | 0.507 | -0.002 | -0.012 | -0.012 | 0 | 0.42 | 0.72 | 0.56 | 0.52 | 0.4 | 0.52 |

| | | | | | | | | | | | | | | | | |
|---|---|---|---|---|---|---|---|---|---|---|---|---|---|---|---|---|
| 1 | 43437675 | rs2367161 | -0.003 | 0.33 | 0.571 | 0.545 | -0.036 | -0.034 | -0.034 | 0 | 0.12 | 0.02 | 0.16 | 0.11 | 0.11 | 0.16 |
| 1 | 43440201 | rs3738516 | -1.595 | 0.375 | 0.663 | 0.581 | -0.015 | 0.015 | 0.015 | 0 | 0.24 | 0.51 | 0.3 | 0.28 | 0.21 | 0.35 |
| 1 | 43447091 | rs7413414 | 1.133 | 0.47 | 0.168 | 0.412 | 0 | -0.012 | -0.012 | 0.968 | 0.54 | 0.23 | 0.45 | 0.48 | 0.58 | 0.47 |
| 1 | 43451901 | rs7515271 | 1.113 | 0.476 | 0.149 | 0.391 | 0 | -0.006 | -0.006 | 0 | 0.54 | 0.26 | 0.46 | 0.49 | 0.58 | 0.47 |
| 1 | 43458250 | rs4660697 | -0.081 | 0.506 | 0.134 | 0.305 | -0.002 | 0.009 | 0.009 | 0 | 0.7 | 0.61 | 0.66 | 0.6 | 0.71 | 0.66 |
| 1 | 43458458 | rs12404679 | -0.828 | 0.747 | -0.114 | 0.277 | 0.005 | -0.016 | -0.016 | 0 | 0.15 | 0.25 | 0.19 | 0.09 | 0.13 | 0.19 |
| 1 | 43460561 | rs841833 | 0.598 | 0.374 | -0.599 | 0.318 | 0 | 0 | 0 | 0 | 0.78 | 0.82 | 0.79 | 0.81 | 0.8 | 0.79 |
| 1 | 43467068 | rs2792599 | 1.529 | 0.293 | -0.763 | 0.579 | 0.021 | -0.048 | -0.048 | 0 | 0.71 | 0.18 | 0.62 | 0.53 | 0.81 | 0.64 |
| 1 | 43468555 | rs841568 | -0.583 | 0.356 | -0.497 | 0.769 | 0.005 | 0.054 | 0.054 | 0 | 0.13 | 0.17 | 0.3 | 0.36 | 0.13 | 0.27 |
| 1 | 43471182 | rs9326132 | -1.008 | 0.233 | -1.253 | 0.42 | 0 | 0 | 0 | 0.905 | 0.5 | 0.8 | 0.49 | 0.46 | 0.42 | 0.48 |
| 1 | 43471613 | rs13375629 | -0.473 | 0.022 | -0.771 | 0.333 | -0.024 | 0.003 | 0.003 | 0.255 | 0.08 | 0.13 | 0.02 | 0.09 | 0.01 | 0.07 |
| 1 | 43523549 | rs17388983 | -0.183 | 0.92 | -1.355 | -0.309 | -0.072 | 0.105 | 0.105 | 0 | 0.83 | 0.99 | 0.9 | 1 | 0.8 | 0.96 |
| 1 | 43529875 | rs11210780 | 0.937 | 0.245 | -2.051 | 0.057 | -0.027 | -0.057 | -0.057 | 5.226 | 0.53 | 0.04 | 0.44 | 0.37 | 0.62 | 0.56 |
| 1 | 43530154 | rs588662 | -1.108 | -0.006 | -0.706 | -0.051 | -0.039 | 0.144 | 0.144 | 2.165 | 0.19 | 0.56 | 0.38 | 0.59 | 0.1 | 0.25 |
| 1 | 43535013 | rs515781 | -0.436 | 0.183 | -0.271 | 0.191 | -0.001 | 0.101 | 0.101 | 0.396 | 0.07 | 0.07 | 0.22 | 0.53 | 0.07 | 0.17 |
| 1 | 43536477 | rs607464 | -0.209 | 0.15 | -0.167 | 0.093 | -0.035 | 0.105 | 0.105 | 0 | 0.76 | 0.9 | 0.84 | 1 | 0.73 | 0.83 |
| 1 | 43548419 | rs583682 | -1.011 | 0.303 | 0.387 | 0.527 | 0.004 | 0.137 | 0.137 | 0.399 | 0.08 | 0.1 | 0.26 | 0.54 | 0.09 | 0.18 |
| 1 | 43557168 | rs1754144 | -0.897 | 0.278 | 0.484 | 0.518 | 0.003 | 0.147 | 0.147 | 0.586 | 0.08 | 0.09 | 0.22 | 0.53 | 0.09 | 0.18 |
| 1 | 43565492 | rs578703 | -1.658 | 0.12 | 1.31 | -0.233 | 0.065 | 0.097 | 0.097 | 9.738 | 0.06 | 0.54 | 0.16 | 0.05 | 0.02 | 0.09 |
| 1 | 43568133 | rs7512851 | -1.465 | 0.119 | 1.598 | -0.235 | 0.102 | 0.111 | 0.111 | 8.072 | 0.05 | 0.48 | 0.12 | 0.04 | 0.02 | 0.09 |
| 1 | 43587725 | rs578598 | -2.691 | 0.262 | 2.092 | -0.316 | 0.033 | -0.004 | -0.004 | 0.01 | 0.08 | 0.65 | 0.14 | 0.01 | 0.05 | 0.02 |
| 1 | 43595231 | rs16830355 | -1.804 | 0.263 | 2.325 | -0.312 | 0.07 | 0.045 | 0.045 | 2.843 | 0.06 | 0.35 | 0.11 | 0.01 | 0.04 | 0.01 |
| 1 | 43597014 | rs585655 | -3.608 | 0.22 | 2.431 | -0.373 | -0.005 | -0.041 | -0.041 | 0 | 0.1 | 0.75 | 0.15 | 0.01 | 0.06 | 0.1 |
| 1 | 43598129 | rs3903911 | -1.606 | 0.237 | 2.546 | -0.331 | 0.053 | 0.04 | 0.04 | 2.778 | 0.05 | 0.28 | 0.11 | 0.01 | 0.04 | 0.01 |
| 1 | 43600931 | rs490876 | -2.926 | 0.054 | 2.434 | 0.064 | -0.003 | 0.189 | 0.189 | 4.159 | 0.13 | 0.52 | 0.31 | 0.32 | 0.09 | 0.25 |
| 1 | 43602321 | rs558404 | -1.442 | 0.06 | 2.319 | 0.072 | 0.002 | 0.2 | 0.2 | 0 | 0.08 | 0.23 | 0.2 | 0.31 | 0.05 | 0.23 |
| 1 | 43606731 | rs1012120 | -0.205 | 0.069 | 2.152 | 0.08 | 0.009 | 0.007 | 0.007 | 0 | 0.05 | 0.01 | 0.18 | 0.32 | 0.04 | 0.15 |
| 1 | 43608078 | rs528664 | -0.615 | 0.144 | 1.855 | -0.478 | 0.005 | -0.11 | -0.11 | 2.615 | 0.31 | 0.45 | 0.27 | 0.01 | 0.34 | 0.16 |
| 1 | 43609205 | rs559834 | 0.038 | 0.149 | 1.718 | -0.478 | -0.047 | 0.135 | 0.135 | 0 | 0.25 | 0.06 | 0.15 | 0 | 0.28 | 0.14 |
| 1 | 43610910 | rs626842 | -1.687 | -0.061 | 1.618 | -0.661 | 0.015 | 0.101 | 0.101 | 0.168 | 0.47 | 0.86 | 0.51 | 0.33 | 0.42 | 0.4 |
| 1 | 43611005 | rs627263 | 0.189 | -0.053 | 1.615 | -0.658 | -0.048 | 0.134 | 0.134 | 0 | 0.25 | 0.06 | 0.15 | 0 | 0.28 | 0.14 |
| 1 | 43611230 | rs641350 | -0.93 | -0.063 | 1.77 | -0.658 | -0.003 | -0.015 | -0.015 | 0 | 0.36 | 0.56 | 0.43 | 0.32 | 0.37 | 0.3 |
| 1 | 43612919 | rs12030306 | 0.055 | -0.024 | 1.763 | -0.544 | 0.004 | 0.038 | 0.038 | 0 | 0.05 | 0.04 | 0.19 | 0.32 | 0.04 | 0.15 |
| 1 | 43615173 | rs10890239 | -0.576 | -0.034 | 1.806 | -0.549 | -0.006 | 0.193 | 0.193 | 0 | 0.06 | 0.21 | 0.2 | 0.32 | 0.05 | 0.15 |
| 1 | 43615668 | rs584672 | -0.819 | -0.021 | 1.705 | -0.602 | -0.009 | -0.049 | -0.049 | 2.044 | 0.11 | 0.3 | 0.08 | 0.01 | 0.05 | 0.1 |
| 1 | 43621759 | rs589819 | -1.512 | -0.024 | 1.672 | -0.63 | -0.04 | 0.086 | 0.086 | 0 | 0.16 | 0.39 | 0.24 | 0.32 | 0.08 | 0.24 |
| 1 | 43632536 | rs7163 | 0.183 | 0.154 | 1.461 | -0.751 | -0.033 | 0.091 | 0.091 | 0 | 0.26 | 0.13 | 0.15 | 0 | 0.29 | 0.14 |
| 1 | 43636692 | rs1010621 | 0.157 | 0.145 | 1.468 | -0.755 | -0.073 | 0.156 | 0.156 | 0 | 0.25 | 0.04 | 0.14 | 0 | 0.29 | 0.14 |
| 1 | 43640544 | rs11210802 | 0.18 | 0.147 | 1.6 | -0.76 | -0.033 | 0.091 | 0.091 | 0 | 0.26 | 0.13 | 0.15 | 0 | 0.29 | 0.15 |
| 1 | 43645411 | rs621559 | -1.564 | 0.009 | 1.823 | -0.696 | 0.008 | 0.207 | 0.207 | 0.909 | 0.1 | 0.35 | 0.26 | 0.32 | 0.07 | 0.24 |
| 1 | 43649508 | rs663824 | -1.741 | -0.338 | 1.375 | -0.941 | 0.029 | 0.153 | 0.153 | 0 | 0.5 | 0.84 | 0.51 | 0.32 | 0.41 | 0.41 |
| 1 | 43663376 | rs513009 | 1.255 | -0.284 | 1.636 | -0.871 | 0.005 | 0.225 | 0.225 | 0.865 | 0.89 | 0.55 | 0.74 | 0.68 | 0.93 | 0.76 |
| 1 | 43664218 | rs603123 | -0.11 | -0.231 | 1.542 | -0.943 | -0.006 | -0.026 | -0.026 | 0 | 0.29 | 0.28 | 0.21 | 0 | 0.32 | 0.17 |
| 1 | 43666454 | rs680343 | -0.212 | -0.245 | 1.009 | -1.096 | 0.01 | -0.002 | -0.002 | 0 | 0.14 | 0.08 | 0.04 | 0 | 0.05 | 0.01 |
| 1 | 43666683 | rs12026626 | 0.286 | -0.216 | 0.978 | -1.02 | 0.01 | -0.003 | -0.003 | 0 | 0.05 | 0.01 | 0.19 | 0.32 | 0.05 | 0.14 |
| 1 | 43669850 | rs4660248 | -0.705 | -0.213 | 1.053 | -1.082 | -0.011 | 0.022 | 0.022 | 0 | 0.75 | 0.79 | 0.85 | 1 | 0.73 | 0.86 |
| 1 | 43671182 | rs620418 | -2.037 | -0.233 | 1.151 | -0.993 | 0.006 | 0.13 | 0.13 | 2.195 | 0.14 | 0.51 | 0.32 | 0.32 | 0.11 | 0.27 |
| 1 | 43674974 | rs2453414 | -0.265 | -0.24 | 1.283 | -1.033 | -0.005 | -0.036 | -0.036 | 0 | 0.15 | 0.15 | 0.04 | 0 | 0.05 | 0.01 |
| 1 | 43681872 | rs611261 | 1.693 | -0.04 | 1.378 | -0.752 | 0.016 | 0.15 | 0.15 | 2.129 | 0.86 | 0.54 | 0.68 | 0.68 | 0.89 | 0.73 |
| 1 | 43687464 | rs2781236 | 1.393 | 0.145 | 1.189 | -0.861 | 0.107 | 0.03 | 0.03 | 0.446 | 0.94 | 0.69 | 0.88 | 1 | 0.94 | 0.89 |
| 1 | 43688077 | rs11210815 | 0.104 | 0.15 | 0.367 | -0.9 | -0.028 | 0.065 | 0.065 | 0 | 0.25 | 0.17 | 0.15 | 0 | 0.28 | 0.13 |
| 1 | 43693532 | rs594389 | -0.105 | 0.074 | 0.302 | -0.769 | -0.005 | 0.062 | 0.062 | 0 | 0.68 | 0.8 | 0.64 | 0.86 | 0.7 | 0.63 |
| 1 | 43695653 | rs622762 | -0.702 | -0.595 | 0.407 | -0.601 | -0.024 | -0.003 | -0.003 | 3.568 | 0.2 | 0.3 | 0.09 | 0.22 | 0.09 | 0.13 |
| 1 | 43696080 | rs522909 | -0.702 | -0.595 | 0.407 | -0.601 | -0.024 | -0.003 | -0.003 | 0.006 | 0.2 | 0.3 | 0.1 | 0.22 | 0.09 | 0.13 |
| 1 | 43697968 | rs3120045 | 0.777 | -0.686 | 0.091 | -0.598 | 0.103 | 0.066 | 0.066 | 0.179 | 0.49 | 0.18 | 0.5 | 0.59 | 0.66 | 0.51 |
| 1 | 43699813 | rs3820588 | -0.559 | -0.527 | 0.566 | -0.608 | -0.008 | -0.031 | -0.031 | 0.164 | 0.73 | 0.73 | 0.68 | 0.89 | 0.77 | 0.84 |
| 1 | 43707220 | rs3898794 | 1.105 | -0.192 | 0.304 | -0.584 | -0.064 | -0.088 | -0.088 | 1.64 | 0.32 | 0.09 | 0.33 | 0.56 | 0.41 | 0.48 |
| 1 | 43707283 | rs10493115 | -0.336 | -0.259 | 0.487 | -0.76 | -0.035 | 0.045 | 0.045 | 0 | 0.16 | 0.38 | 0.16 | 0.25 | 0.12 | 0.28 |
| 1 | 43707556 | rs3862226 | 1.274 | -0.211 | 0.394 | -0.73 | -0.064 | -0.09 | -0.09 | 0 | 0.32 | 0.09 | 0.33 | 0.56 | 0.41 | 0.48 |
| 1 | 43708658 | rs1760668 | -0.469 | -0.408 | 0.554 | -0.743 | -0.022 | -0.001 | -0.001 | 0 | 0.19 | 0.22 | 0.08 | 0.21 | 0.06 | 0.12 |
| 1 | 43709967 | rs1760671 | 0.538 | -0.405 | 0.542 | -0.744 | 0.019 | 0.008 | 0.008 | 1.772 | 0.13 | 0.1 | 0.02 | 0 | 0.02 | 0.01 |
| 1 | 43723986 | rs529113 | 0.538 | -0.373 | 0.484 | -0.747 | 0 | -0.018 | -0.018 | 1.37 | 0.13 | 0.13 | 0.02 | 0 | 0.02 | 0.01 |
| 1 | 43735196 | rs11210824 | 0.557 | -0.27 | 0.521 | -0.702 | -0.006 | -0.027 | -0.027 | 0.41 | 0.11 | 0.14 | 0.12 | 0.04 | 0.09 | 0.18 |
| 1 | 43737296 | rs7536340 | 0.633 | -0.188 | 0.446 | -0.643 | -0.021 | 0.084 | 0.084 | 0 | 0.06 | 0.14 | 0.02 | 0.2 | 0.03 | 0.1 |
| 1 | 43741713 | rs2991996 | 0.318 | -0.189 | 0.349 | -0.734 | -0.014 | -0.021 | -0.021 | 0 | 0.38 | 0.24 | 0.43 | 0.4 | 0.43 | 0.3 |
| 1 | 43746448 | rs4660727 | -0.76 | -0.216 | 0.455 | -0.744 | 0.01 | -0.022 | -0.022 | 0.489 | 0.65 | 0.57 | 0.56 | 0.81 | 0.59 | 0.77 |
| 1 | 43748051 | rs1763696 | 0.102 | 0.552 | 0.46 | -0.298 | 0.019 | 0.025 | 0.025 | 0.049 | 0.5 | 0.26 | 0.45 | 0.4 | 0.46 | 0.31 |
| 1 | 43749755 | rs3120123 | 0.372 | 0.624 | -0.104 | 0.286 | 0.008 | 0.029 | 0.029 | 0.39 | 0.46 | 0.31 | 0.46 | 0.26 | 0.44 | 0.25 |
| 1 | 43751701 | rs6697017 | -0.27 | 0.851 | -0.137 | 0.687 | 0.039 | 0.033 | 0.033 | 0 | 0.13 | 0.35 | 0.19 | 0.09 | 0.12 | 0.2 |

| | | | | | | | | | | | | | | | | |
|---|---|---|---|---|---|---|---|---|---|---|---|---|---|---|---|---|
| 1 | 43751977 | rs2784466 | 0.167 | 0.953 | -0.775 | 0.576 | -0.001 | 0 | 0 | 0.275 | 0.59 | 0.46 | 0.51 | 0.41 | 0.51 | 0.48 |
| 1 | 43757762 | rs34783763 | -0.64 | 0.936 | -1.171 | 0.775 | -0.001 | -0.005 | -0.005 | 0.162 | 0.14 | 0.1 | 0.16 | 0.12 | 0.14 | 0.22 |
| 1 | 43757955 | rs6690113 | 0.231 | 1.113 | -1.716 | 0.933 | -0.109 | -0.113 | -0.113 | 0 | 0.28 | 0.05 | 0.28 | 0.47 | 0.37 | 0.37 |
| 1 | 43763086 | rs11210831 | -0.491 | 1.104 | -1.943 | 1.192 | -0.041 | -0.061 | -0.061 | 0 | 0.3 | 0.13 | 0.32 | 0.46 | 0.37 | 0.45 |
| 1 | 43766426 | rs3768046 | -0.674 | 0.116 | -2.272 | 0.24 | -0.025 | 0.089 | 0.089 | 0 | 0.66 | 0.81 | 0.64 | 0.87 | 0.63 | 0.9 |
| 1 | 43767138 | rs7527092 | -0.214 | -0.591 | -1.417 | 0.434 | 0.001 | -0.006 | -0.006 | 0.94 | 0.44 | 0.49 | 0.5 | 0.71 | 0.49 | 0.75 |
| 1 | 43774437 | rs1467809 | -1.062 | -0.582 | -1.148 | 0.53 | -0.025 | 0.08 | 0.08 | 0 | 0.66 | 0.81 | 0.65 | 0.84 | 0.63 | 0.89 |
| 1 | 43777852 | rs1999595 | -1.063 | -0.91 | -1.089 | 0.399 | -0.014 | -0.003 | -0.003 | 0 | 0.77 | 0.69 | 0.85 | 0.84 | 0.86 | 0.86 |
| 1 | 43779564 | rs3120276 | -0.858 | -0.662 | -0.934 | 0.533 | -0.078 | 0.158 | 0.158 | 0 | 0.66 | 0.89 | 0.64 | 0.88 | 0.63 | 0.9 |
| 1 | 43779869 | rs2991990 | -0.748 | -0.669 | -0.93 | 0.536 | -0.068 | 0.14 | 0.14 | 0 | 0.65 | 0.86 | 0.64 | 0.86 | 0.63 | 0.9 |
| 1 | 43784956 | rs1199039 | 0.558 | -0.7 | -0.904 | 0.52 | -0.039 | 0.11 | 0.11 | 0 | 0.36 | 0.18 | 0.36 | 0.15 | 0.38 | 0.11 |
| 1 | 43787437 | rs11210834 | 1.184 | -0.393 | -0.566 | 1.13 | -0.005 | -0.009 | -0.009 | 0 | 0.2 | 0.16 | 0.24 | 0.4 | 0.24 | 0.32 |
| 1 | 43800001 | rs1199038 | 0.785 | -0.475 | -0.646 | 1.006 | -0.052 | 0.169 | 0.169 | 0 | 0.35 | 0.15 | 0.36 | 0.08 | 0.37 | 0.08 |
| 1 | 43803120 | rs839993 | 0.814 | -0.506 | -0.562 | 0.993 | -0.02 | 0.099 | 0.099 | 0 | 0.36 | 0.24 | 0.37 | 0.11 | 0.37 | 0.09 |
| 1 | 43812075 | rs1760670 | 0.83 | -0.564 | -0.253 | 0.938 | -0.034 | 0.147 | 0.147 | 0 | 0.36 | 0.19 | 0.36 | 0.08 | 0.37 | 0.08 |
| 1 | 43826099 | rs2069385 | 1.369 | -0.672 | -0.536 | 0.887 | -0.029 | 0.037 | 0.037 | 0 | 0.07 | 0.02 | 0.04 | 0 | 0.08 | 0.05 |
| 1 | 43826312 | rs710251 | -0.669 | -0.569 | -0.355 | 1.075 | 0.011 | -0.022 | -0.022 | 1.298 | 0.61 | 0.69 | 0.64 | 0.52 | 0.64 | 0.47 |
| 1 | 43831545 | rs3768048 | 1.151 | 0.211 | -0.853 | 0.874 | -0.028 | -0.045 | -0.045 | 0 | 0.08 | 0.01 | 0.15 | 0.13 | 0.09 | 0.11 |
| 1 | 43841744 | rs11587504 | 0.471 | 0.265 | -0.617 | -0.513 | -0.097 | 0.177 | 0.177 | 0 | 0.33 | 0.08 | 0.36 | 0.08 | 0.37 | 0.08 |
| 1 | 43845658 | rs11210839 | 0.128 | 0.293 | -0.568 | -0.477 | -0.023 | 0.12 | 0.12 | 0 | 0.36 | 0.23 | 0.37 | 0.11 | 0.37 | 0.09 |
| 1 | 43855546 | rs839752 | 0.185 | 0.412 | -0.49 | -0.405 | -0.021 | 0.106 | 0.106 | 0 | 0.36 | 0.24 | 0.37 | 0.11 | 0.37 | 0.09 |
| 1 | 43872067 | rs839769 | 0.139 | 0.267 | -0.451 | -0.346 | -0.084 | 0.193 | 0.193 | 0 | 0.37 | 0.15 | 0.39 | 0.08 | 0.42 | 0.09 |
| 1 | 43886321 | rs2782642 | 0.022 | 0.266 | -0.444 | -0.345 | -0.044 | 0.126 | 0.126 | 0 | 0.38 | 0.24 | 0.39 | 0.11 | 0.42 | 0.1 |
| 1 | 43886494 | rs2782643 | 0.195 | 0.599 | -0.456 | -0.359 | -0.102 | 0.203 | 0.203 | 0 | 0.36 | 0.1 | 0.38 | 0.08 | 0.42 | 0.09 |
| 1 | 43887146 | rs2782644 | 0.195 | 0.599 | -0.456 | -0.359 | -0.102 | 0.203 | 0.203 | 0 | 0.36 | 0.1 | 0.38 | 0.08 | 0.42 | 0.09 |
| 1 | 43888602 | rs2782645 | 0.007 | 0.611 | -0.459 | -0.353 | -0.044 | 0.126 | 0.126 | 0 | 0.38 | 0.23 | 0.39 | 0.11 | 0.42 | 0.1 |
| 1 | 43901916 | rs2842180 | 0.037 | 0.613 | -0.591 | -0.38 | -0.069 | 0.179 | 0.179 | 0 | 0.37 | 0.19 | 0.39 | 0.08 | 0.42 | 0.09 |
| 1 | 43917409 | rs2251804 | 0.071 | 0.611 | -0.603 | -0.405 | -0.086 | 0.205 | 0.205 | 0 | 0.37 | 0.15 | 0.35 | 0.08 | 0.43 | 0.1 |
| 1 | 43917637 | rs2251802 | 0.104 | 0.611 | -0.586 | -0.386 | -0.071 | 0.177 | 0.177 | 0 | 0.37 | 0.19 | 0.39 | 0.08 | 0.42 | 0.1 |
| 1 | 43926305 | rs2782652 | -0.388 | 0.705 | -0.563 | -0.313 | 0.002 | -0.03 | -0.03 | 0 | 0.42 | 0.53 | 0.42 | 0.12 | 0.45 | 0.16 |
| 1 | 43938801 | rs7413861 | -0.486 | 1.047 | -1.186 | -0.238 | -0.084 | 0.135 | 0.135 | 0 | 0.68 | 0.85 | 0.63 | 0.9 | 0.61 | 0.9 |
| 1 | 43957515 | rs6687571 | 0.265 | 0.947 | -0.699 | 0.134 | 0.008 | 0.003 | 0.003 | 1.654 | 0.3 | 0.33 | 0.34 | 0.59 | 0.34 | 0.46 |
| 1 | 43960479 | rs2039529 | 0.541 | 0.965 | -0.655 | 0.114 | -0.011 | -0.007 | -0.007 | 0.229 | 0.28 | 0.24 | 0.33 | 0.59 | 0.33 | 0.45 |
| 1 | 43964954 | rs867605 | 0.981 | 0.415 | -0.795 | -0.056 | 0.042 | 0.062 | 0.062 | 0.236 | 0.35 | 0.15 | 0.41 | 0.27 | 0.31 | 0.42 |
| 1 | 43974764 | rs882464 | -0.848 | 0.365 | -0.323 | 0.273 | 0.025 | 0.002 | 0.002 | 0.476 | 0.63 | 0.71 | 0.54 | 0.61 | 0.69 | 0.58 |
| 1 | 43976157 | rs2819341 | 0.944 | 0.306 | -0.406 | 0.236 | 0.055 | 0.021 | 0.021 | 1.101 | 0.36 | 0.18 | 0.46 | 0.33 | 0.31 | 0.42 |
| 1 | 43978837 | rs11210858 | 0.802 | 0.452 | -0.379 | 0.177 | 0.015 | 0.023 | 0.023 | 2 | 0.3 | 0.36 | 0.29 | 0.52 | 0.33 | 0.45 |
| 1 | 43981265 | rs1413938 | 0.767 | 0.441 | -0.254 | -0.007 | 0.025 | 0.016 | 0.016 | 2.16 | 0.31 | 0.42 | 0.3 | 0.59 | 0.33 | 0.45 |
| 1 | 43982527 | rs11210860 | 0.51 | 0.483 | -0.278 | -0.294 | -0.045 | 0.132 | 0.132 | 0.218 | 0.31 | 0.14 | 0.24 | 0.03 | 0.36 | 0.12 |
| 1 | 44005928 | rs1889587 | 1.727 | 0.583 | -1.157 | -0.284 | -0.017 | 0.068 | 0.068 | 0 | 0.12 | 0.03 | 0.05 | 0 | 0.09 | 0.06 |
| 1 | 44007648 | rs2842189 | -0.654 | 1.037 | -0.997 | 0.162 | -0.046 | 0.056 | 0.056 | 0.236 | 0.71 | 0.92 | 0.77 | 0.83 | 0.64 | 0.87 |
| 1 | 44009033 | rs2782640 | -0.104 | 0.858 | -0.625 | 0.355 | -0.001 | 0.01 | 0.01 | 0.055 | 0.66 | 0.66 | 0.75 | 0.73 | 0.64 | 0.86 |
| 1 | 44010571 | rs6695915 | -1.461 | 1.143 | -0.29 | 0.347 | 0.057 | 0.097 | 0.097 | 0 | 0.07 | 0.21 | 0.06 | 0.21 | 0.09 | 0.07 |
| 1 | 44013355 | rs2782641 | -0.512 | 0.795 | -0.516 | 0.414 | -0.008 | 0.016 | 0.016 | 0 | 0.68 | 0.7 | 0.72 | 0.76 | 0.63 | 0.87 |
| 1 | 44024484 | rs11580074 | -1.295 | 0.413 | -0.75 | 0.198 | 0.032 | 0.033 | 0.033 | 0.043 | 0.64 | 0.7 | 0.57 | 0.75 | 0.74 | 0.58 |
| 1 | 44035352 | rs1065771 | 1.537 | 1.042 | -1.085 | 1.082 | -0.107 | -0.091 | -0.091 | 0 | 0.18 | 0.04 | 0.24 | 0.29 | 0.31 | 0.3 |
| 1 | 44037124 | rs16830922 | 1.532 | 0.79 | -0.425 | 1.267 | -0.009 | 0.021 | 0.021 | 0 | 0.12 | 0.2 | 0.07 | 0.15 | 0.07 | 0.14 |
| 1 | 44044468 | rs11210874 | 1.279 | 0.923 | -0.475 | 1.62 | -0.055 | -0.068 | -0.068 | 0.184 | 0.2 | 0.09 | 0.24 | 0.42 | 0.32 | 0.34 |
| 1 | 44045294 | rs752963 | 1.565 | 0.827 | -0.389 | 1.414 | -0.009 | 0.041 | 0.041 | 0 | 0.12 | 0.21 | 0.06 | 0.18 | 0.07 | 0.12 |
| 1 | 44045465 | rs2004899 | -1.125 | 0.537 | -0.366 | 0.838 | -0.018 | -0.024 | -0.024 | 0 | 0.63 | 0.49 | 0.55 | 0.77 | 0.75 | 0.58 |
| 1 | 44045919 | rs2077652 | 1.899 | 0.552 | -0.425 | 0.848 | -0.007 | 0 | 0 | 0 | 0.12 | 0.2 | 0.06 | 0.08 | 0.07 | 0.11 |
| 1 | 44050747 | rs10789437 | 1.656 | 0.569 | -0.578 | 0.893 | -0.077 | -0.108 | -0.108 | 0 | 0.19 | 0.07 | 0.24 | 0.48 | 0.32 | 0.35 |
| 1 | 44050856 | rs3828150 | -1.429 | 0.502 | -0.374 | 0.853 | 0.042 | 0.054 | 0.054 | 0.019 | 0.65 | 0.77 | 0.56 | 0.77 | 0.75 | 0.58 |
| 1 | 44051834 | rs501299 | 0.713 | 0.522 | -0.61 | 0.858 | -0.07 | 0.018 | 0.018 | 0.18 | 0.3 | 0.06 | 0.23 | 0.21 | 0.35 | 0.12 |
| 1 | 44058034 | rs2117316 | 1.628 | 0.587 | -0.566 | 0.889 | -0.075 | -0.108 | -0.108 | 0 | 0.19 | 0.07 | 0.24 | 0.48 | 0.32 | 0.35 |
| 1 | 44058265 | rs3828151 | -0.077 | 0.538 | -0.202 | 0.816 | 0.028 | 0.02 | 0.02 | 0 | 0.33 | 0.46 | 0.32 | 0.56 | 0.4 | 0.46 |
| 1 | 44066424 | rs596522 | 0.662 | 0.568 | -0.539 | 0.761 | -0.073 | 0.014 | 0.014 | 0.187 | 0.3 | 0.06 | 0.23 | 0.21 | 0.35 | 0.12 |
| 1 | 44071221 | rs631248 | -0.929 | 0.35 | -0.304 | 0.781 | 0.004 | 0.004 | 0.004 | 0.272 | 0.61 | 0.43 | 0.54 | 0.76 | 0.74 | 0.58 |
| 1 | 44071546 | rs11210886 | 1.794 | 0.374 | -0.329 | 0.844 | -0.01 | -0.008 | -0.008 | 0.115 | 0.12 | 0.24 | 0.06 | 0.07 | 0.07 | 0.11 |
| 1 | 44071796 | rs618369 | -0.825 | 0.347 | -0.792 | 0.53 | 0 | -0.001 | -0.001 | 0 | 0.54 | 0.6 | 0.51 | 0.69 | 0.67 | 0.51 |
| 1 | 44072018 | rs1065772 | 0.969 | 0.354 | -1.398 | 0.493 | -0.128 | -0.15 | -0.15 | 0 | 0.22 | 0.04 | 0.27 | 0.47 | 0.36 | 0.33 |
| 1 | 44072066 | rs10890266 | 0.882 | 0.254 | -1.398 | 0.421 | -0.133 | -0.153 | -0.153 | 1.556 | 0.23 | 0.03 | 0.28 | 0.47 | 0.36 | 0.34 |
| 1 | 44072420 | rs539096 | -1.167 | -1.024 | -1.073 | 0.3 | 0.05 | 0.058 | 0.058 | 0.162 | 0.66 | 0.86 | 0.58 | 0.77 | 0.74 | 0.57 |
| 1 | 44076019 | rs11210887 | -0.062 | -1.246 | -1.439 | 0.343 | 0.083 | 0.083 | 0.083 | 0.992 | 0.49 | 0.09 | 0.48 | 0.69 | 0.66 | 0.46 |
| 1 | 44087496 | rs673485 | -1.03 | -1.192 | -1.388 | 0.329 | 0.022 | 0.04 | 0.04 | 0.175 | 0.66 | 0.95 | 0.59 | 0.77 | 0.73 | 0.56 |
| 1 | 44123191 | rs10789439 | -0.247 | -1.057 | -1.244 | 0.432 | -0.032 | -0.049 | -0.049 | 1.8 | 0.36 | 0.77 | 0.34 | 0.53 | 0.39 | 0.45 |
| 1 | 44129209 | rs6658300 | -0.788 | -0.982 | -1.427 | 0.069 | 0.004 | 0.082 | 0.082 | 0 | 0.63 | 0.76 | 0.56 | 0.81 | 0.67 | 0.56 |
| 1 | 44131794 | rs489319 | 0.49 | -1.666 | -0.608 | 0.473 | 0.026 | -0.02 | -0.02 | 0 | 0.27 | 0.5 | 0.25 | 0.22 | 0.31 | 0.12 |
| 1 | 44133016 | rs607062 | -0.937 | -1.635 | -0.8 | 0.441 | 0.055 | 0.142 | 0.142 | 1.661 | 0.59 | 0.95 | 0.56 | 0.79 | 0.7 | 0.57 |

| | | | | | | | | | | | | | | | | |
|---|---|---|---|---|---|---|---|---|---|---|---|---|---|---|---|---|
| 1 | 44133028 | rs2296757 | 1.119 | -1.662 | -0.838 | 0.486 | -0.116 | -0.159 | -0.159 | 0.51 | 0.19 | 0.02 | 0.23 | 0.49 | 0.31 | 0.35 |
| 1 | 44137257 | rs586339 | -0.85 | -1.885 | -0.711 | 0.501 | 0.055 | 0.142 | 0.142 | 1.661 | 0.59 | 0.95 | 0.56 | 0.79 | 0.7 | 0.57 |
| 1 | 44151015 | rs6697354 | -0.614 | -1.836 | -0.345 | 0.42 | 0.024 | 0.021 | 0.021 | 0.664 | 0.54 | 0.63 | 0.51 | 0.71 | 0.68 | 0.53 |
| 1 | 44174081 | rs7528454 | -0.085 | -1.839 | -0.213 | 0.417 | 0.044 | 0.051 | 0.051 | 0.201 | 0.31 | 0.46 | 0.31 | 0.49 | 0.39 | 0.42 |
| 1 | 44200656 | rs12119146 | 1.288 | -1.957 | -0.3 | 0.204 | -0.125 | -0.156 | -0.156 | 0.052 | 0.18 | 0.02 | 0.25 | 0.52 | 0.32 | 0.35 |
| 1 | 44202733 | rs3791040 | 0.482 | -1.963 | -0.206 | 0.209 | 0.007 | -0.012 | -0.012 | 0.101 | 0.27 | 0.6 | 0.24 | 0.2 | 0.3 | 0.12 |
| 1 | 44202991 | rs3791041 | -0.721 | -1.975 | -0.211 | 0.201 | 0.025 | 0.025 | 0.025 | 0.576 | 0.55 | 0.65 | 0.52 | 0.72 | 0.69 | 0.54 |
| 1 | 44208122 | rs10890276 | 0.492 | -1.959 | -0.51 | 0.209 | -0.143 | -0.158 | -0.158 | 0 | 0.27 | 0.06 | 0.28 | 0.52 | 0.39 | 0.42 |
| 1 | 44209075 | rs1472661 | 0.542 | -1.734 | -0.308 | 0.318 | 0.017 | -0.01 | -0.01 | 0 | 0.26 | 0.52 | 0.23 | 0.2 | 0.3 | 0.12 |
| 1 | 44226657 | rs7520053 | 0.659 | -1.721 | -0.366 | 0.324 | 0.022 | -0.003 | -0.003 | 0.144 | 0.25 | 0.42 | 0.23 | 0.2 | 0.3 | 0.12 |
| 1 | 44245670 | rs7546818 | 0.632 | -1.714 | -0.366 | 0.373 | 0.063 | 0.119 | 0.119 | 0.255 | 0.23 | 0.41 | 0.27 | 0.52 | 0.33 | 0.36 |
| 1 | 44248272 | rs12354267 | 0.555 | -2.152 | -0.062 | 0.692 | -0.004 | 0 | 0 | 0 | 0.24 | 0.23 | 0.21 | 0.19 | 0.3 | 0.12 |
| 1 | 44249344 | rs12760274 | 1.109 | -2.04 | -0.047 | 0.628 | -0.047 | 0.169 | 0.169 | 0 | 0.27 | 0.03 | 0.2 | 0 | 0.32 | 0.08 |
| 1 | 44249770 | rs6692652 | 0.958 | -2.126 | 0.008 | 0.847 | -0.075 | -0.08 | -0.08 | 0.464 | 0.29 | 0.16 | 0.29 | 0.52 | 0.39 | 0.45 |
| 1 | 44250381 | rs4660750 | -1.785 | -2.42 | -0.033 | 0.287 | 0.054 | 0.099 | 0.099 | 0 | 0.82 | 0.98 | 0.82 | 0.91 | 0.89 | 0.79 |
| 1 | 44252908 | rs3791051 | 1.048 | -2.414 | 0.09 | 0.314 | -0.047 | -0.054 | -0.054 | 0 | 0.29 | 0.19 | 0.29 | 0.54 | 0.39 | 0.44 |
| 1 | 44253015 | rs3828139 | -0.314 | -2.434 | 0.195 | 0.307 | 0 | 0.002 | 0.002 | 0 | 0.49 | 0.47 | 0.5 | 0.38 | 0.49 | 0.34 |
| 1 | 44261195 | rs11210924 | 1.063 | -2.237 | 0.238 | -0.071 | -0.087 | -0.068 | -0.068 | 0 | 0.3 | 0.14 | 0.29 | 0.58 | 0.4 | 0.45 |
| 1 | 44263415 | rs12401813 | 0.881 | -2.279 | 0.233 | -0.098 | -0.026 | 0.071 | 0.071 | 0 | 0.29 | 0.11 | 0.22 | 0.14 | 0.35 | 0.12 |
| 1 | 44264435 | rs975761 | 1.01 | -2.176 | 0.299 | -0.043 | -0.02 | -0.015 | -0.015 | 0.008 | 0.31 | 0.28 | 0.3 | 0.58 | 0.4 | 0.45 |
| 1 | 44266310 | rs976312 | 1.069 | -2.174 | 0.283 | -0.042 | -0.044 | -0.034 | -0.034 | 0 | 0.3 | 0.19 | 0.29 | 0.58 | 0.4 | 0.45 |
| 1 | 44272761 | rs246778 | -1.137 | -2.206 | 0.305 | -0.106 | -0.06 | -0.093 | -0.093 | 0 | 0.78 | 0.68 | 0.8 | 0.91 | 0.89 | 0.78 |
| 1 | 44274699 | rs246776 | -0.252 | -2.137 | 0.297 | -0.061 | -0.001 | 0.005 | 0.005 | 0 | 0.47 | 0.46 | 0.5 | 0.33 | 0.48 | 0.33 |
| 1 | 44278722 | rs37469 | -0.252 | -2.137 | 0.297 | -0.061 | -0.001 | 0.005 | 0.005 | 0 | 0.47 | 0.46 | 0.5 | 0.33 | 0.48 | 0.32 |
| 1 | 44290530 | rs37458 | 0.566 | -1.971 | 0.282 | -0.022 | -0.001 | -0.027 | -0.027 | 0 | 0.31 | 0.31 | 0.24 | 0.14 | 0.35 | 0.11 |
| 1 | 44312109 | rs3011221 | -1.004 | -2.067 | 0.305 | 0 | -0.039 | -0.062 | -0.062 | 0.107 | 0.78 | 0.77 | 0.81 | 0.91 | 0.89 | 0.77 |
| 1 | 44319373 | rs3011225 | 0.558 | -2.155 | 0.206 | 0.044 | 0.116 | 0.093 | 0.093 | 0.368 | 0.57 | 0.18 | 0.51 | 0.72 | 0.74 | 0.55 |
| 1 | 44345292 | rs2527776 | 1.066 | -2.293 | 0.167 | 0.016 | 0.138 | 0.02 | 0.02 | 0.981 | 0.51 | 0.1 | 0.58 | 0.59 | 0.69 | 0.5 |
| 1 | 44365457 | rs783304 | 0.149 | -2.472 | 0.425 | -0.096 | -0.073 | -0.044 | -0.044 | 0 | 0.27 | 0.45 | 0.23 | 0.23 | 0.15 | 0.28 |
| 1 | 44366250 | rs3791101 | 1.546 | -2.409 | 0.313 | 0.008 | -0.077 | 0.074 | 0.074 | 0.018 | 0.24 | 0.02 | 0.19 | 0.11 | 0.3 | 0.09 |
| 1 | 44373439 | rs2428963 | 0.431 | -2.812 | 1.534 | 0.739 | 0.068 | 0.099 | 0.099 | 0 | 0.19 | 0.02 | 0.17 | 0.1 | 0.11 | 0.23 |
| 1 | 44379957 | rs6674176 | 1.214 | -2.831 | 1.468 | 0.747 | 0.138 | 0.09 | 0.09 | 0.711 | 0.6 | 0.18 | 0.66 | 0.74 | 0.8 | 0.64 |
| 1 | 44381246 | rs2486011 | 0.205 | -2.798 | 1.546 | 0.764 | -0.045 | 0.032 | 0.032 | 0 | 0.83 | 0.58 | 0.86 | 0.83 | 0.91 | 0.87 |
| 1 | 44386615 | rs3120803 | 0.13 | -2.798 | 1.544 | 0.765 | -0.036 | 0.021 | 0.021 | 0 | 0.83 | 0.64 | 0.86 | 0.84 | 0.91 | 0.87 |
| 1 | 44393533 | rs4031031 | 1.576 | -2.803 | 1.573 | 0.77 | 0.008 | 0.003 | 0.003 | 0.412 | 0.56 | 0.36 | 0.63 | 0.6 | 0.74 | 0.6 |
| 1 | 44395786 | rs2108202 | 1.582 | -2.826 | 1.545 | 0.76 | 0.153 | 0.01 | 0.01 | 0.934 | 0.56 | 0.14 | 0.63 | 0.61 | 0.76 | 0.61 |
| 1 | 44396215 | rs1859726 | 0.355 | -2.796 | 1.563 | 0.653 | 0.071 | 0.094 | 0.094 | 0 | 0.19 | 0.03 | 0.17 | 0.09 | 0.11 | 0.23 |
| 1 | 44399608 | rs1548634 | -0.14 | -2.788 | 1.411 | 0.657 | -0.083 | -0.114 | -0.114 | 0 | 0.22 | 0.32 | 0.19 | 0.09 | 0.11 | 0.23 |
| 1 | 44401384 | rs2242637 | -0.247 | -2.702 | 1.203 | 0.73 | -0.022 | 0.027 | 0.027 | 0 | 0.85 | 0.74 | 0.87 | 0.83 | 0.91 | 0.88 |
| 1 | 44411589 | rs4448553 | 1.333 | -2.633 | 1.058 | 0.755 | 0.093 | 0.017 | 0.017 | 0.986 | 0.58 | 0.27 | 0.63 | 0.61 | 0.76 | 0.61 |
| 1 | 44416090 | rs7556565 | 1.372 | -2.655 | 0.946 | 0.72 | 0.14 | -0.023 | -0.023 | 0 | 0.58 | 0.09 | 0.62 | 0.62 | 0.76 | 0.61 |
| 1 | 44420248 | rs2486013 | -0.181 | -2.933 | 1.322 | 0.943 | -0.029 | 0.016 | 0.016 | 0 | 0.84 | 0.71 | 0.87 | 0.83 | 0.92 | 0.91 |
| 1 | 44426823 | rs2301992 | -1.029 | -2.884 | 1.434 | 0.941 | 0.052 | 0.122 | 0.122 | 0 | 0.06 | 0.16 | 0.04 | 0.12 | 0.04 | 0.11 |
| 1 | 44438396 | rs783308 | -0.418 | -2.938 | 1.406 | 0.913 | -0.028 | -0.02 | -0.02 | 0 | 0.24 | 0.21 | 0.22 | 0.21 | 0.15 | 0.3 |
| 1 | 44438961 | rs4221 | -0.664 | -2.937 | 1.273 | 0.913 | -0.035 | -0.026 | -0.026 | 0 | 0.26 | 0.57 | 0.25 | 0.21 | 0.15 | 0.3 |
| 1 | 44438974 | rs7161 | 1.668 | -2.981 | 1.186 | 0.917 | 0.144 | -0.025 | -0.025 | 0 | 0.54 | 0.06 | 0.6 | 0.61 | 0.74 | 0.6 |
| 1 | 44440146 | rs4660761 | 1.272 | -2.986 | 1.201 | 0.87 | 0.17 | 0.076 | 0.076 | 1.651 | 0.6 | 0.09 | 0.65 | 0.73 | 0.8 | 0.68 |
| 1 | 44441533 | rs2286243 | -1.07 | -2.96 | 1.227 | 1.012 | -0.001 | 0.041 | 0.041 | 0 | 0.06 | 0.08 | 0.05 | 0.18 | 0.05 | 0.09 |
| 1 | 44442521 | rs12410334 | 1.247 | -2.963 | 1.196 | 1.007 | 0.17 | 0.076 | 0.076 | 1.679 | 0.6 | 0.09 | 0.65 | 0.74 | 0.8 | 0.69 |
| 1 | 44442930 | rs2428954 | -0.138 | -2.799 | 1.346 | 0.944 | -0.092 | -0.138 | -0.138 | 0.102 | 0.22 | 0.41 | 0.2 | 0.09 | 0.12 | 0.22 |
| 1 | 44445667 | rs3762423 | -1.259 | -2.755 | 1.33 | 0.975 | 0.005 | 0.098 | 0.098 | 0 | 0.07 | 0.19 | 0.06 | 0.18 | 0.05 | 0.09 |
| 1 | 44446651 | rs2486005 | -0.687 | -2.738 | 0.971 | 0.975 | -0.019 | -0.006 | -0.006 | 0.126 | 0.29 | 0.6 | 0.26 | 0.27 | 0.17 | 0.31 |
| 1 | 44468235 | rs2248829 | -0.254 | -2.22 | 0.675 | 0.795 | -0.04 | 0.024 | 0.024 | 0.396 | 0.17 | 0.32 | 0.14 | 0.17 | 0.08 | 0.1 |
| 1 | 44469031 | rs1362153 | -1.105 | -2.233 | 0.406 | 0.787 | -0.033 | -0.001 | -0.001 | 0 | 0.22 | 0.6 | 0.17 | 0.17 | 0.11 | 0.14 |
| 1 | 44475408 | rs2486002 | -0.353 | -1.918 | -0.558 | 1.02 | -0.04 | 0.04 | 0.04 | 0.268 | 0.17 | 0.37 | 0.15 | 0.17 | 0.08 | 0.1 |
| 1 | 44475586 | rs1766967 | -0.287 | -1.906 | -1.02 | 1.025 | -0.024 | 0.016 | 0.016 | 0.357 | 0.16 | 0.24 | 0.14 | 0.17 | 0.08 | 0.1 |
| 1 | 44478007 | rs2486000 | 0.031 | -1.306 | -1.631 | 0.989 | 0.042 | 0.024 | 0.024 | 0 | 0.79 | 0.96 | 0.76 | 0.78 | 0.84 | 0.79 |
| 1 | 44486239 | rs9429225 | 0.386 | -1.237 | -1.45 | 0.861 | 0.034 | -0.011 | -0.011 | 0 | 0.76 | 0.83 | 0.75 | 0.62 | 0.85 | 0.77 |
| 1 | 44487280 | rs6429644 | -1.064 | -1.652 | -1.739 | 0.256 | 0.063 | 0.066 | 0.066 | 5.335 | 0.19 | 0.86 | 0.13 | 0.1 | 0.07 | 0.06 |
| 1 | 44487950 | rs11585120 | -0.678 | -1.559 | -1.331 | 0.298 | -0.018 | -0.02 | -0.02 | 0 | 0.15 | 0.74 | 0.12 | 0.08 | 0.06 | 0.06 |
| 1 | 44493795 | rs7523468 | -0.007 | -0.689 | -2.21 | 0.714 | -0.011 | -0.057 | -0.057 | 0 | 0.63 | 0.82 | 0.52 | 0.4 | 0.61 | 0.49 |
| 1 | 44499242 | rs1978195 | 0.477 | -0.796 | -2.555 | 0.498 | -0.028 | -0.027 | -0.027 | 0.296 | 0.33 | 0.21 | 0.32 | 0.33 | 0.41 | 0.28 |
| 1 | 44502163 | rs16831558 | -0.376 | -1.062 | -1.704 | 0.67 | -0.008 | 0.045 | 0.045 | 0 | 0.16 | 0.25 | 0.27 | 0.27 | 0.15 | 0.24 |
| 1 | 44503040 | rs2485993 | -1.144 | -1.479 | -1.836 | 0.318 | -0.022 | 0.036 | 0.036 | 0 | 0.75 | 0.93 | 0.83 | 0.83 | 0.73 | 0.76 |
| 1 | 44508195 | rs1476670 | -2.088 | -1.531 | -1.948 | 0.154 | -0.064 | 0.107 | 0.107 | 0 | 0.82 | 0.98 | 0.88 | 1 | 0.79 | 0.94 |
| 1 | 44509551 | rs751442 | -0.689 | -1.686 | -1.605 | 0.241 | -0.002 | 0.02 | 0.02 | 0 | 0.72 | 0.86 | 0.83 | 0.8 | 0.73 | 0.76 |
| 1 | 44511409 | rs6693689 | 0.261 | -0.651 | -1.804 | 0.444 | 0.037 | 0.047 | 0.047 | 0 | 0.13 | 0.19 | 0.27 | 0.33 | 0.19 | 0.36 |
| 1 | 44512756 | rs12240184 | 0.788 | -1.002 | -1.822 | 0.207 | 0.036 | 0.047 | 0.047 | 0.133 | 0.14 | 0.06 | 0.04 | 0 | 0.05 | 0.02 |
| 1 | 44515696 | rs2057968 | 0.122 | -0.756 | -1.697 | 0.454 | 0.016 | 0.093 | 0.093 | 3.477 | 0.46 | 0.24 | 0.57 | 0.73 | 0.6 | 0.67 |

| | | | | | | | | | | | | | | | | |
|---|---|---|---|---|---|---|---|---|---|---|---|---|---|---|---|---|
| 1 | 44517189 | rs2158821 | 0.754 | -0.594 | -1.648 | 0.5 | -0.047 | -0.045 | -0.045 | 0 | 0.35 | 0.21 | 0.49 | 0.5 | 0.5 | 0.45 |
| 1 | 44517409 | rs1007795 | 0.754 | -0.594 | -1.648 | 0.5 | -0.047 | -0.045 | -0.045 | 1.49 | 0.35 | 0.21 | 0.49 | 0.49 | 0.5 | 0.46 |
| 1 | 44522274 | rs942756 | -0.451 | -0.571 | -1.192 | 0.251 | 0.012 | 0.028 | 0.028 | 0 | 0.59 | 0.67 | 0.78 | 0.73 | 0.66 | 0.73 |
| 1 | 44524676 | rs12031261 | 1.463 | 0.242 | -1.018 | 0.553 | 0.007 | -0.011 | -0.011 | 0.052 | 0.18 | 0.13 | 0.19 | 0.15 | 0.18 | 0.07 |
| 1 | 44529157 | rs12759480 | -0.043 | 0.492 | -0.648 | 0.518 | -0.008 | -0.017 | -0.017 | 0 | 0.33 | 0.31 | 0.47 | 0.56 | 0.4 | 0.51 |
| 1 | 44532725 | rs7519215 | -0.087 | 0.81 | -0.374 | 0.446 | 0.046 | -0.012 | -0.012 | 0 | 0.07 | 0.23 | 0.07 | 0.07 | 0.1 | 0.08 |
| 1 | 44541586 | rs2057969 | -0.011 | 0.702 | -0.336 | 0.445 | 0.029 | -0.009 | -0.009 | 0 | 0.07 | 0.23 | 0.04 | 0.11 | 0.07 | 0.08 |
| 1 | 44541961 | rs11210951 | 0.307 | 0.473 | -0.838 | 0.209 | -0.002 | 0.018 | 0.018 | 0 | 0.13 | 0.09 | 0.05 | 0.06 | 0.11 | 0.08 |
| 1 | 44541996 | rs4660770 | -0.352 | 0.372 | -1.046 | 0.455 | -0.021 | 0.011 | 0.011 | 0 | 0.46 | 0.22 | 0.52 | 0.55 | 0.47 | 0.59 |
| 1 | 44554457 | rs6698333 | -0.513 | 0.805 | -0.454 | 0.721 | 0.03 | 0.026 | 0.026 | 0 | 0.35 | 0.5 | 0.59 | 0.49 | 0.43 | 0.46 |
| 1 | 44559401 | rs2025615 | -0.696 | 0.051 | -0.857 | 0.748 | 0.033 | 0.008 | 0.008 | 0 | 0.4 | 0.22 | 0.4 | 0.32 | 0.33 | 0.37 |
| 1 | 44568322 | rs509261 | -0.715 | 0.059 | -0.223 | 0.73 | 0.032 | 0.311 | 0.311 | 0 | 0.93 | 0.82 | 0.92 | 0.66 | 0.92 | 0.86 |
| 1 | 44571434 | rs1291050 | -0.587 | 0.051 | -0.097 | 0.726 | 0.04 | 0.341 | 0.341 | 0 | 0.93 | 0.78 | 0.92 | 0.66 | 0.92 | 0.86 |
| 1 | 44571545 | rs12048838 | -0.288 | -0.029 | -0.36 | 0.591 | 0.021 | -0.025 | -0.025 | 0 | 0.35 | 0.17 | 0.36 | 0.32 | 0.3 | 0.33 |
| 1 | 44578559 | rs663818 | 1.24 | 0.223 | -0.003 | 0.026 | 0.003 | 0.066 | 0.066 | 0 | 0.49 | 0.38 | 0.48 | 0.22 | 0.51 | 0.42 |
| 1 | 44587132 | rs6657162 | -0.177 | 0.293 | -0.224 | 0.246 | -0.001 | -0.001 | -0.001 | 0 | 0.31 | 0.31 | 0.41 | 0.66 | 0.34 | 0.36 |
| 1 | 44587512 | rs6690437 | -1.015 | 0.124 | -0.232 | 0.228 | 0.003 | 0.006 | 0.006 | 0 | 0.41 | 0.39 | 0.35 | 0.59 | 0.38 | 0.42 |
| 1 | 44587895 | rs2485646 | 1.807 | 0.032 | -1.022 | -0.612 | -0.079 | 0.065 | 0.065 | 0 | 0.28 | 0.08 | 0.41 | 0.17 | 0.34 | 0.33 |
| 1 | 44595047 | rs11210969 | 0.541 | -0.441 | -0.96 | -0.196 | -0.045 | -0.005 | -0.005 | 0.018 | 0.25 | 0.12 | 0.26 | 0.2 | 0.3 | 0.25 |
| 1 | 44595410 | rs2485652 | 2.046 | -0.413 | -0.422 | -0.353 | -0.04 | -0.063 | -0.063 | 0.255 | 0.32 | 0.09 | 0.26 | 0.42 | 0.4 | 0.26 |
| 1 | 44597200 | rs2995018 | -0.108 | -0.476 | -0.055 | -0.417 | 0.001 | 0.004 | 0.004 | 0 | 0.63 | 0.68 | 0.66 | 0.71 | 0.72 | 0.54 |
| 1 | 44598760 | rs2254746 | -0.147 | -0.476 | -0.055 | -0.417 | 0.001 | 0.004 | 0.004 | 0 | 0.37 | 0.32 | 0.34 | 0.29 | 0.28 | 0.46 |
| 1 | 44602159 | rs6662888 | 0.696 | -0.446 | -0.148 | -0.375 | -0.036 | 0 | 0 | 0 | 0.25 | 0.14 | 0.26 | 0.2 | 0.29 | 0.26 |
| 1 | 44602422 | rs11581177 | 2.396 | -0.38 | -0.257 | -0.658 | -0.062 | -0.014 | -0.014 | 0 | 0.14 | 0.01 | 0.13 | 0.1 | 0.2 | 0.06 |
| 1 | 44603506 | rs2429060 | -0.552 | -0.389 | -0.103 | -0.75 | 0.023 | 0.053 | 0.053 | 0 | 0.41 | 0.71 | 0.38 | 0.3 | 0.3 | 0.47 |
| 1 | 44610760 | rs6675129 | 0.142 | -0.477 | 0.092 | -0.685 | 0.001 | -0.004 | -0.004 | 2.341 | 0.28 | 0.3 | 0.28 | 0.25 | 0.3 | 0.28 |
| 1 | 44617687 | rs6429532 | -0.975 | -0.666 | 0.399 | -0.824 | 0.003 | 0.004 | 0.004 | 0 | 0.38 | 0.66 | 0.4 | 0.27 | 0.34 | 0.34 |
| 1 | 44623775 | rs803379 | -2.182 | -0.555 | 0.146 | -0.784 | -0.078 | -0.146 | -0.146 | 0 | 0.69 | 0.96 | 0.75 | 0.51 | 0.61 | 0.76 |
| 1 | 44656429 | rs11576516 | 0.315 | -0.467 | 0.2 | -0.789 | -0.001 | -0.002 | -0.002 | 0.005 | 0.27 | 0.25 | 0.24 | 0.2 | 0.27 | 0.23 |
| 1 | 44666345 | rs803371 | -2.07 | -0.49 | 0.157 | -0.792 | -0.041 | -0.098 | -0.098 | 0 | 0.67 | 0.92 | 0.74 | 0.51 | 0.6 | 0.74 |
| 1 | 44681113 | rs1291169 | -0.106 | -0.485 | 0.377 | -0.84 | -0.01 | -0.021 | -0.021 | 0 | 0.3 | 0.45 | 0.27 | 0.21 | 0.28 | 0.24 |
| 1 | 44699848 | rs17414912 | 0.908 | -0.463 | 0.275 | -0.905 | -0.02 | -0.016 | -0.016 | 0.002 | 0.14 | 0.07 | 0.28 | 0.24 | 0.18 | 0.44 |
| 1 | 44709218 | rs11589379 | 2.003 | -0.496 | 0.244 | -0.914 | -0.049 | -0.029 | -0.029 | 0 | 0.16 | 0.01 | 0.12 | 0.13 | 0.19 | 0.06 |
| 1 | 44728001 | rs10489168 | 0.658 | -0.279 | 0.084 | -0.788 | -0.001 | -0.009 | -0.009 | 0 | 0.06 | 0.07 | 0.02 | 0 | 0.05 | 0.05 |
| 1 | 44745624 | rs459358 | 0.047 | -0.322 | 0.187 | -0.82 | -0.01 | -0.018 | -0.018 | 0 | 0.31 | 0.58 | 0.28 | 0.21 | 0.28 | 0.24 |
| 1 | 44760103 | rs6699067 | 1.043 | -0.331 | 0.088 | -0.823 | -0.022 | -0.024 | -0.024 | 0 | 0.06 | 0.01 | 0.09 | 0.08 | 0.08 | 0.12 |
| 1 | 44764677 | rs325154 | -0.785 | -0.506 | 0.329 | -0.812 | -0.003 | -0.006 | -0.006 | 0 | 0.72 | 0.71 | 0.74 | 0.79 | 0.72 | 0.76 |
| 1 | 44765646 | rs12058010 | 1.617 | -0.522 | 0.167 | -0.879 | 0.022 | 0.082 | 0.082 | 0.481 | 0.16 | 0.03 | 0.05 | 0 | 0.07 | 0 |
| 1 | 44772626 | rs325145 | 0.269 | -0.45 | -0.325 | -0.831 | 0.007 | 0.009 | 0.009 | 0 | 0.62 | 0.31 | 0.6 | 0.73 | 0.66 | 0.69 |
| 1 | 44776360 | rs4958 | -0.028 | -0.367 | -0.806 | -0.79 | -0.025 | 0.036 | 0.036 | 0 | 0.29 | 0.1 | 0.33 | 0.23 | 0.26 | 0.43 |
| 1 | 44793760 | rs11210982 | 1.174 | -0.331 | -0.888 | -0.769 | -0.028 | -0.066 | -0.066 | 0 | 0.38 | 0.15 | 0.28 | 0.49 | 0.45 | 0.3 |
| 1 | 44800498 | rs12123509 | 2.164 | 0.158 | -0.925 | -0.701 | -0.078 | 0.16 | 0.16 | 0.026 | 0.24 | 0.01 | 0.15 | 0 | 0.3 | 0.09 |
| 1 | 44832896 | rs226072 | 0.088 | 0.496 | -0.221 | -0.001 | -0.046 | 0.033 | 0.033 | 0 | 0.29 | 0.06 | 0.32 | 0.23 | 0.25 | 0.42 |
| 1 | 44840084 | rs226075 | 1.102 | 0.801 | 0.894 | 0.631 | 0.001 | -0.004 | -0.004 | 0 | 0.68 | 0.31 | 0.59 | 0.73 | 0.66 | 0.72 |
| 1 | 44844958 | rs4660275 | -0.142 | 0.892 | 0.44 | 0.637 | 0.019 | 0.002 | 0.002 | 0 | 0.21 | 0.25 | 0.25 | 0.2 | 0.27 | 0.2 |
| 1 | 44855878 | rs3862229 | -0.098 | 0.951 | 0.443 | 0.679 | 0.019 | 0.005 | 0.005 | 0 | 0.19 | 0.21 | 0.26 | 0.2 | 0.28 | 0.22 |
| 1 | 44855891 | rs3862230 | -1.409 | 0.602 | 0.606 | 0.581 | 0 | -0.002 | -0.002 | 0 | 0.32 | 0.7 | 0.32 | 0.25 | 0.33 | 0.28 |
| 1 | 44856052 | rs3888585 | -0.394 | 0.835 | 0.527 | 0.57 | 0.049 | 0.026 | 0.026 | 0 | 0.23 | 0.39 | 0.38 | 0.22 | 0.33 | 0.26 |
| 1 | 44858169 | rs3915854 | -0.86 | 0.011 | -0.088 | 0.591 | -0.068 | 0.179 | 0.179 | 1.293 | 0.2 | 0.55 | 0.13 | 0.43 | 0.08 | 0.16 |
| 1 | 44858313 | rs619199 | 1.127 | 0.009 | -0.364 | 0.322 | -0.039 | 0.158 | 0.158 | 0.002 | 0.48 | 0.1 | 0.41 | 0.27 | 0.54 | 0.54 |
| 1 | 44859885 | rs7516912 | -1.261 | -0.43 | 0.205 | 0.99 | -0.047 | 0.119 | 0.119 | 0.651 | 0.19 | 0.45 | 0.12 | 0.43 | 0.07 | 0.15 |
| 1 | 44860224 | rs6429539 | -1.129 | 0.088 | -0.328 | 1.477 | -0.025 | 0.318 | 0.318 | 0.051 | 0.09 | 0.31 | 0.07 | 0.43 | 0.03 | 0.16 |
| 1 | 44878549 | rs11210992 | 0.106 | 0.398 | -1.629 | 0.164 | -0.016 | 0.182 | 0.182 | 0 | 0.45 | 0.09 | 0.5 | 0.23 | 0.57 | 0.45 |
| 1 | 44879221 | rs4660794 | -0.319 | 0.308 | -0.956 | 0.238 | 0 | -0.005 | -0.005 | 0 | 0.37 | 0.56 | 0.37 | 0.57 | 0.39 | 0.43 |
| 1 | 44885123 | rs3866643 | 0.346 | 0.5 | -1.401 | -0.236 | -0.016 | 0.078 | 0.078 | 0.126 | 0.36 | 0.16 | 0.3 | 0.19 | 0.4 | 0.22 |
| 1 | 44887362 | rs11210996 | 0.069 | 0.311 | -0.741 | 0.716 | 0.004 | 0.161 | 0.161 | 0.086 | 0.09 | 0.19 | 0.08 | 0.35 | 0.1 | 0.24 |
| 1 | 44889509 | rs1340647 | -0.764 | -0.197 | -0.925 | 0.02 | -0.012 | -0.045 | -0.045 | 0.229 | 0.83 | 0.84 | 0.93 | 1 | 0.91 | 0.9 |
| 1 | 44891488 | rs12022308 | 0.477 | 0.223 | -1.162 | 0.356 | -0.073 | -0.067 | -0.067 | 0 | 0.15 | 0.03 | 0.23 | 0.19 | 0.22 | 0.07 |
| 1 | 44902285 | rs6429540 | 0.385 | -0.124 | -0.65 | 0.033 | 0.001 | 0.004 | 0.004 | 0.048 | 0.69 | 0.68 | 0.81 | 0.8 | 0.73 | 0.59 |
| 1 | 44911812 | rs7553056 | 1.105 | -0.2 | 0.288 | 0.617 | 0.014 | 0.004 | 0.004 | 0.107 | 0.57 | 0.27 | 0.7 | 0.57 | 0.63 | 0.52 |
| 1 | 44912073 | rs7518522 | 1.083 | -0.08 | 0.343 | 0.843 | 0.006 | -0.025 | -0.025 | 0 | 0.57 | 0.28 | 0.6 | 0.52 | 0.62 | 0.52 |
| 1 | 44916216 | rs6668996 | -0.726 | -0.267 | 0.704 | 0.484 | 0.009 | -0.002 | -0.002 | 0 | 0.32 | 0.44 | 0.25 | 0.37 | 0.33 | 0.29 |
| 1 | 44920992 | rs1417371 | 0.366 | -0.244 | 0.205 | 0.025 | -0.018 | 0.009 | 0.009 | 0 | 0.92 | 0.83 | 0.95 | 0.91 | 0.96 | 0.86 |
| 1 | 44922683 | rs12746640 | 0.029 | -0.098 | 0.145 | 0.056 | -0.016 | 0.056 | 0.056 | 0.063 | 0.24 | 0.11 | 0.15 | 0.09 | 0.24 | 0.26 |
| 1 | 44923473 | rs11211003 | 0.337 | -0.065 | -0.146 | -0.162 | 0.01 | 0.003 | 0.003 | 0.196 | 0.3 | 0.26 | 0.13 | 0.26 | 0.22 | 0.28 |
| 1 | 44924989 | rs12730333 | 0.818 | 0.582 | -0.145 | -0.225 | -0.043 | 0.07 | 0.07 | 0.266 | 0.19 | 0.05 | 0.14 | 0 | 0.23 | 0.07 |
| 1 | 44925227 | rs4660800 | -0.324 | 0.602 | -0.052 | -0.174 | 0 | -0.036 | -0.036 | 0 | 0.27 | 0.31 | 0.19 | 0.09 | 0.26 | 0.31 |
| 1 | 44925508 | rs6664827 | 0.004 | 0.602 | -0.393 | -0.142 | -0.009 | 0.031 | 0.031 | 0 | 0.24 | 0.21 | 0.17 | 0.08 | 0.25 | 0.26 |
| 1 | 44929865 | rs1890948 | -0.557 | 0.592 | -0.242 | 0.059 | -0.004 | 0.036 | 0.036 | 0 | 0.34 | 0.28 | 0.23 | 0.3 | 0.33 | 0.34 |

| | | | | | | | | | | | | | | | | |
|---|---|---|---|---|---|---|---|---|---|---|---|---|---|---|---|---|
| 1 | 44934995 | rs2185500 | -0.587 | 0.599 | -0.278 | 0.062 | -0.001 | 0.028 | 0.028 | 0 | 0.34 | 0.3 | 0.24 | 0.3 | 0.33 | 0.34 |
| 1 | 44937974 | rs11211006 | -0.635 | 0.879 | 0.27 | 0.726 | 0.02 | 0.1 | 0.1 | 0 | 0.17 | 0.22 | 0.35 | 0.47 | 0.21 | 0.36 |
| 1 | 44939696 | rs4660806 | -1.096 | 0.491 | 0.36 | 0.245 | -0.033 | -0.079 | -0.079 | 0.407 | 0.13 | 0.32 | 0.04 | 0 | 0.06 | 0.09 |
| 1 | 44951085 | rs781783 | 0.716 | 0.792 | 0.026 | 0.529 | -0.001 | 0.022 | 0.022 | 0 | 0.83 | 0.82 | 0.67 | 0.55 | 0.82 | 0.63 |
| 1 | 44951245 | rs9804150 | 0.132 | 0.172 | -0.821 | -0.031 | -0.037 | 0.011 | 0.011 | 0 | 0.37 | 0.13 | 0.23 | 0.33 | 0.35 | 0.21 |
| 1 | 44956714 | rs12077687 | -0.233 | -0.071 | 0.047 | 0.254 | -0.003 | -0.025 | -0.025 | 0 | 0.09 | 0.14 | 0.02 | 0 | 0.06 | 0.05 |
| 1 | 44957986 | rs7354900 | 1.046 | -0.197 | -0.099 | 0.276 | -0.049 | -0.039 | -0.039 | 0 | 0.83 | 0.41 | 0.92 | 0.9 | 0.93 | 0.76 |
| 1 | 44965723 | rs6429541 | -0.592 | 0.571 | -0.771 | 0.423 | 0.001 | -0.003 | -0.003 | 0 | 0.37 | 0.3 | 0.42 | 0.41 | 0.33 | 0.2 |
| 1 | 44972309 | rs781775 | -0.256 | 0.543 | -0.617 | 0.176 | -0.005 | -0.003 | -0.003 | 0 | 0.24 | 0.26 | 0.23 | 0.2 | 0.22 | 0.13 |
| 1 | 44972725 | rs7550459 | -0.214 | 1.062 | -1.072 | -0.553 | -0.039 | 0.041 | 0.041 | 0.06 | 0.24 | 0.17 | 0.26 | 0.09 | 0.36 | 0.22 |
| 1 | 44974119 | rs781770 | 0.456 | 1.02 | -0.405 | -0.514 | -0.007 | -0.005 | -0.005 | 0.006 | 0.25 | 0.28 | 0.24 | 0.2 | 0.22 | 0.13 |
| 1 | 44974796 | rs12138940 | 0.528 | 1.687 | -0.741 | -0.557 | -0.055 | -0.016 | -0.016 | 0 | 0.1 | 0.03 | 0.16 | 0.1 | 0.22 | 0.14 |
| 1 | 44976806 | rs781790 | 0.252 | 1.377 | -0.474 | -0.741 | -0.014 | -0.023 | -0.023 | 0.056 | 0.29 | 0.38 | 0.25 | 0.2 | 0.25 | 0.13 |
| 1 | 44979498 | rs16832018 | 0.635 | 1.067 | -2.147 | -0.613 | 0.009 | -0.123 | -0.123 | 0.071 | 0.22 | 0.09 | 0.35 | 0.47 | 0.18 | 0.32 |
| 1 | 44979966 | rs6704115 | -0.185 | 1.257 | -2.155 | -0.743 | -0.054 | -0.058 | -0.058 | 0 | 0.31 | 0.12 | 0.47 | 0.61 | 0.36 | 0.47 |
| 1 | 44980081 | rs2120823 | 0.305 | 1.264 | -1.768 | -0.564 | -0.005 | 0.004 | 0.004 | 0 | 0.13 | 0.1 | 0.06 | 0 | 0.12 | 0.06 |
| 1 | 44984932 | rs12749754 | 0.146 | 1.212 | -1.772 | -0.515 | -0.003 | 0.009 | 0.009 | 0 | 0.14 | 0.12 | 0.06 | 0.13 | 0.12 | 0.14 |
| 1 | 44988433 | rs272531 | 1.58 | 0.783 | -2.249 | -0.851 | 0.021 | 0.154 | 0.154 | 0.002 | 0.28 | 0.01 | 0.13 | 0.07 | 0.23 | 0.2 |
| 1 | 44989160 | rs3842990 | 0.226 | 1.357 | -1.964 | -0.454 | -0.046 | -0.116 | -0.116 | 2.005 | 0.3 | 0.1 | 0.45 | 0.55 | 0.33 | 0.34 |
| 1 | 44991278 | rs270724 | 0.198 | 1.201 | -1.463 | -0.251 | -0.048 | -0.018 | -0.018 | 0 | 0.09 | 0.03 | 0.2 | 0.07 | 0.18 | 0.15 |
| 1 | 44991991 | rs12749503 | -0.276 | 0.512 | -0.812 | -0.476 | 0 | -0.039 | -0.039 | 0.113 | 0.16 | 0.17 | 0.08 | 0 | 0.16 | 0.13 |
| 1 | 44994835 | rs782135 | -0.189 | 0.693 | -0.981 | -0.103 | 0.003 | -0.016 | -0.016 | 0.062 | 0.42 | 0.22 | 0.65 | 0.57 | 0.58 | 0.47 |
| 1 | 44997314 | rs6672462 | 1.031 | 1.117 | -1.097 | -0.504 | -0.013 | 0.025 | 0.025 | 0 | 0.2 | 0.17 | 0.2 | 0.1 | 0.29 | 0.09 |
| 1 | 44998464 | rs10890308 | 0.117 | 0.522 | -0.503 | -0.549 | -0.005 | -0.041 | -0.041 | 0 | 0.54 | 0.47 | 0.23 | 0.13 | 0.4 | 0.44 |
| 1 | 44999614 | rs746733 | 1.171 | 0.328 | -1.034 | -0.446 | 0.049 | 0.171 | 0.171 | 0.142 | 0.3 | 0.04 | 0.11 | 0.05 | 0.23 | 0.23 |
| 1 | 45003255 | rs156653 | -0.066 | 0.897 | -0.711 | 0.676 | -0.051 | -0.074 | -0.074 | 0.045 | 0.81 | 0.92 | 0.58 | 0.55 | 0.73 | 0.65 |
| 1 | 45007595 | rs12065045 | 0.972 | 0.471 | 0.23 | 0.119 | 0.002 | -0.049 | -0.049 | 0 | 0.36 | 0.28 | 0.18 | 0.03 | 0.28 | 0.24 |
| 1 | 45009801 | rs3896232 | 1.047 | 0.387 | 0.523 | 0.26 | -0.018 | -0.046 | -0.046 | 0 | 0.34 | 0.38 | 0.17 | 0.14 | 0.25 | 0.24 |
| 1 | 45012273 | rs272564 | -0.58 | 0.877 | 0.897 | 0.661 | 0.017 | 0.031 | 0.031 | 0 | 0.22 | 0.28 | 0.44 | 0.6 | 0.29 | 0.39 |
| 1 | 45013415 | rs272565 | -0.792 | 0.737 | 0.882 | 0.718 | 0.001 | 0.004 | 0.004 | 0 | 0.31 | 0.32 | 0.46 | 0.5 | 0.35 | 0.44 |
| 1 | 45019240 | rs175222 | -0.634 | 0.283 | 0.762 | 0.688 | 0.006 | -0.004 | -0.004 | 0.132 | 0.57 | 0.43 | 0.67 | 0.53 | 0.67 | 0.56 |
| 1 | 45023415 | rs270765 | 0.699 | 0.351 | 0.259 | 0.726 | -0.016 | -0.069 | -0.069 | 2.065 | 0.13 | 0.02 | 0.07 | 0.23 | 0.1 | 0.16 |
| 1 | 45024342 | rs270764 | 0.394 | 0.241 | 0.636 | 0.592 | 0.016 | 0.083 | 0.083 | 0 | 0.45 | 0.2 | 0.22 | 0.23 | 0.39 | 0.39 |
| 1 | 45028820 | rs1889759 | -0.555 | 0.316 | 1.194 | 0.647 | 0.043 | 0.101 | 0.101 | 0 | 0.11 | 0.47 | 0.12 | 0.16 | 0.09 | 0.03 |
| 1 | 45033371 | rs755597 | 0.836 | 0.365 | 0.905 | 0.448 | 0.016 | -0.031 | -0.031 | 0 | 0.07 | 0.13 | 0.06 | 0.02 | 0.08 | 0.02 |
| 1 | 45035621 | rs10158231 | 0.955 | 0.232 | 1.328 | 0.503 | -0.014 | -0.044 | -0.044 | 0 | 0.36 | 0.5 | 0.28 | 0.19 | 0.3 | 0.39 |
| 1 | 45039205 | rs69141 | -0.09 | 0.542 | 1.006 | 0.776 | -0.013 | -0.004 | -0.004 | 0 | 0.58 | 0.65 | 0.42 | 0.44 | 0.49 | 0.56 |
| 1 | 45050047 | rs270709 | 0.189 | 0.574 | 0.381 | 0.768 | -0.012 | 0.005 | 0.005 | 0 | 0.55 | 0.28 | 0.38 | 0.4 | 0.5 | 0.53 |
| 1 | 45050958 | rs272529 | 0.413 | 0.952 | -0.433 | 1.186 | -0.063 | -0.099 | -0.099 | 0 | 0.15 | 0.02 | 0.38 | 0.38 | 0.19 | 0.33 |
| 1 | 45053760 | rs928685 | 0.647 | 0.861 | -0.251 | 0.421 | -0.003 | 0.029 | 0.029 | 0 | 0.3 | 0.25 | 0.3 | 0.16 | 0.33 | 0.25 |
| 1 | 45054866 | rs270716 | 0.341 | 1.064 | -0.156 | 1.318 | 0.017 | 0.132 | 0.132 | 0 | 0.82 | 0.63 | 0.59 | 0.41 | 0.81 | 0.65 |
| 1 | 45056293 | rs156650 | -0.214 | 0.807 | -1.305 | 1.12 | -0.021 | -0.025 | -0.025 | 0 | 0.2 | 0.05 | 0.14 | 0.22 | 0.21 | 0.16 |
| 1 | 45060687 | rs270703 | 0.362 | 0.793 | -0.805 | 1.763 | 0.001 | -0.016 | -0.016 | 0 | 0.85 | 0.9 | 0.64 | 0.39 | 0.85 | 0.63 |
| 1 | 45067768 | rs10890311 | -1.155 | 0.798 | -0.8 | 1.727 | 0.002 | -0.011 | -0.011 | 0.681 | 0.19 | 0.21 | 0.28 | 0.57 | 0.16 | 0.48 |
| 1 | 45072074 | rs12129788 | 0.253 | 1.187 | -1.372 | 0.635 | -0.054 | 0.111 | 0.111 | 0 | 0.21 | 0.05 | 0.19 | 0 | 0.3 | 0.13 |
| 1 | 45076929 | rs11582764 | -0.087 | 1.004 | -1.135 | 1.65 | 0.019 | -0.018 | -0.018 | 0.069 | 0.08 | 0.01 | 0.15 | 0.44 | 0.06 | 0.25 |
| 1 | 45078152 | rs272547 | -0.564 | 1.78 | -1.227 | 0.569 | -0.037 | -0.025 | -0.025 | 0 | 0.14 | 0.08 | 0.17 | 0.19 | 0.19 | 0.11 |
| 1 | 45081298 | rs459088 | 0.811 | 1.373 | -0.549 | 0.575 | 0.026 | 0.068 | 0.068 | 0.446 | 0.95 | 0.76 | 0.97 | 0.85 | 0.96 | 0.79 |
| 1 | 45082340 | rs11585508 | 1.243 | 1.096 | -0.438 | -0.344 | 0.001 | 0.035 | 0.035 | 0.005 | 0.56 | 0.34 | 0.37 | 0.23 | 0.59 | 0.38 |
| 1 | 45083937 | rs270728 | 0.948 | 0.222 | 0.076 | -0.033 | -0.029 | 0.088 | 0.088 | 0.029 | 0.77 | 0.56 | 0.64 | 0.37 | 0.79 | 0.57 |
| 1 | 45088053 | rs270729 | -0.227 | 0.415 | -0.482 | -0.497 | -0.102 | 0.037 | 0.037 | 0 | 0.28 | 0.09 | 0.31 | 0.18 | 0.44 | 0.19 |
| 1 | 45098461 | rs995089 | 1.515 | 0.105 | -0.365 | -0.585 | 0.058 | 0.189 | 0.189 | 0.135 | 0.4 | 0.14 | 0.25 | 0.04 | 0.33 | 0.34 |
| 1 | 45101478 | rs2236219 | 0.897 | 0.137 | -0.016 | -0.517 | 0.01 | 0.008 | 0.008 | 0 | 0.49 | 0.49 | 0.39 | 0.2 | 0.38 | 0.39 |
| 1 | 45116985 | rs1051664 | 1.141 | 0.066 | -0.103 | -0.54 | 0.041 | 0.087 | 0.087 | 0 | 0.46 | 0.36 | 0.27 | 0.16 | 0.36 | 0.33 |
| 1 | 45123135 | rs11211021 | 0.027 | 0.252 | -0.043 | -0.427 | -0.019 | 0.028 | 0.028 | 0.02 | 0.64 | 0.79 | 0.65 | 0.81 | 0.51 | 0.8 |
| 1 | 45125356 | rs3850853 | 0.642 | 0.266 | -0.008 | -0.654 | 0 | -0.002 | -0.002 | 0 | 0.55 | 0.61 | 0.45 | 0.24 | 0.47 | 0.41 |
| 1 | 45131401 | rs6667593 | -0.749 | 0.559 | 0.303 | -0.288 | 0.011 | 0.025 | 0.025 | 0.108 | 0.15 | 0.6 | 0.21 | 0.17 | 0.14 | 0.09 |
| 1 | 45133561 | rs6683133 | 0.658 | 0.702 | 0.234 | -0.25 | 0.009 | -0.019 | -0.019 | 0 | 0.62 | 0.63 | 0.51 | 0.23 | 0.54 | 0.42 |
| 1 | 45139252 | rs4660818 | 1.361 | 0.612 | 0.305 | -0.144 | 0.004 | 0.002 | 0.002 | 0 | 0.52 | 0.58 | 0.43 | 0.23 | 0.41 | 0.36 |
| 1 | 45161845 | rs12022162 | 0.235 | 1.215 | -0.089 | 0.42 | 0.002 | -0.001 | -0.001 | 0 | 0.73 | 0.8 | 0.69 | 0.78 | 0.61 | 0.84 |
| 1 | 45166383 | rs11585861 | -0.225 | 0.768 | 0.076 | 0.631 | -0.003 | 0.039 | 0.039 | 0 | 0.08 | 0.21 | 0.14 | 0.15 | 0.06 | 0.04 |
| 1 | 45170012 | rs12088087 | -0.591 | 0.614 | -0.072 | 0.149 | -0.003 | -0.022 | -0.022 | 0 | 0.41 | 0.46 | 0.53 | 0.76 | 0.41 | 0.48 |
| 1 | 45175925 | rs10890313 | -1.545 | 0.584 | 0.675 | 1.011 | -0.005 | 0.096 | 0.096 | 0 | 0.2 | 0.5 | 0.29 | 0.33 | 0.14 | 0.21 |
| 1 | 45184306 | rs11211024 | -1.293 | 0.563 | 0.981 | 1.089 | 0.041 | 0.098 | 0.098 | 0 | 0.14 | 0.41 | 0.21 | 0.16 | 0.12 | 0.12 |
| 1 | 45186056 | rs2983715 | -1.847 | 1.091 | 1.052 | 1.569 | 0.037 | 0.143 | 0.143 | 0.027 | 0.18 | 0.3 | 0.35 | 0.45 | 0.25 | 0.3 |
| 1 | 45196677 | rs11803622 | -1.191 | 0.933 | 1.051 | 1.464 | -0.005 | 0.038 | 0.038 | 0 | 0.17 | 0.21 | 0.26 | 0.29 | 0.13 | 0.18 |
| 1 | 45213915 | rs6704436 | -1.091 | 0.932 | 1.088 | 1.464 | 0 | -0.001 | -0.001 | 0.014 | 0.17 | 0.14 | 0.25 | 0.28 | 0.14 | 0.18 |
| 1 | 45224998 | rs4342887 | -1.137 | 0.869 | 1.312 | 1.485 | -0.003 | 0.028 | 0.028 | 0 | 0.17 | 0.18 | 0.25 | 0.29 | 0.14 | 0.18 |
| 1 | 45226084 | rs1140279 | -1.098 | 0.866 | 1.144 | 1.484 | 0 | -0.001 | -0.001 | 0.061 | 0.17 | 0.14 | 0.25 | 0.29 | 0.14 | 0.18 |

| | | | | | | | | | | | | | | | | |
|---|---|---|---|---|---|---|---|---|---|---|---|---|---|---|---|---|
| 1 | 45228580 | rs12746270 | -1.646 | 0.791 | 1.586 | 1.446 | -0.012 | 0.069 | 0.069 | 0 | 0.19 | 0.48 | 0.28 | 0.29 | 0.14 | 0.18 |
| 1 | 45237119 | rs12136048 | -2.725 | 0.536 | 1.424 | 1.511 | -0.001 | -0.003 | -0.003 | 0 | 0.35 | 0.6 | 0.46 | 0.49 | 0.36 | 0.35 |
| 1 | 45241113 | rs11211031 | -2.745 | 0.534 | 1.423 | 1.543 | -0.001 | -0.004 | -0.004 | 0 | 0.35 | 0.6 | 0.46 | 0.49 | 0.36 | 0.35 |
| 1 | 45244692 | rs10465818 | -2.86 | 0.554 | 1.432 | 1.547 | 0 | -0.001 | -0.001 | 0 | 0.35 | 0.6 | 0.46 | 0.49 | 0.36 | 0.33 |
| 1 | 45258141 | rs12057238 | -1.397 | 0.569 | 1.607 | 1.515 | 0.04 | 0.032 | 0.032 | 0 | 0.13 | 0.38 | 0.17 | 0.16 | 0.18 | 0.14 |
| 1 | 45259295 | rs12059964 | -2.352 | 0.596 | 1.609 | 1.573 | 0.001 | 0.026 | 0.026 | 0 | 0.3 | 0.55 | 0.42 | 0.45 | 0.31 | 0.31 |
| 1 | 45264545 | rs6676749 | -0.959 | 0.583 | 1.547 | 1.557 | -0.001 | 0.009 | 0.009 | 0 | 0.17 | 0.15 | 0.25 | 0.29 | 0.14 | 0.17 |
| 1 | 45275551 | rs11211037 | -2.845 | 0.494 | 1.963 | 2.157 | 0 | -0.001 | -0.001 | 0.043 | 0.34 | 0.65 | 0.45 | 0.61 | 0.32 | 0.33 |
| 1 | 45279813 | rs11211038 | -1.01 | 0.57 | 1.653 | 1.523 | -0.001 | 0.013 | 0.013 | 0 | 0.17 | 0.16 | 0.25 | 0.29 | 0.14 | 0.18 |
| 1 | 45291908 | rs11211040 | -1.011 | 0.572 | 1.699 | 1.524 | -0.001 | 0.013 | 0.013 | 0 | 0.17 | 0.16 | 0.25 | 0.29 | 0.14 | 0.17 |
| 1 | 45292866 | rs2295996 | -0.912 | 0.6 | 1.712 | 1.438 | -0.017 | -0.028 | -0.028 | 0.842 | 0.12 | 0.08 | 0.17 | 0.2 | 0.17 | 0.15 |
| 1 | 45293518 | rs7525308 | -2.593 | 0.567 | 1.799 | 1.422 | 0 | 0.005 | 0.005 | 0 | 0.33 | 0.6 | 0.44 | 0.49 | 0.32 | 0.32 |
| 1 | 45293752 | rs2295997 | -1.096 | 0.537 | 1.71 | 0.943 | -0.004 | 0.043 | 0.043 | 0 | 0.18 | 0.22 | 0.25 | 0.29 | 0.14 | 0.17 |
| 1 | 45301735 | rs11573562 | -1.245 | 0.541 | 1.621 | 0.949 | -0.006 | 0.048 | 0.048 | 0 | 0.18 | 0.25 | 0.25 | 0.3 | 0.15 | 0.17 |
| 1 | 45311749 | rs6429550 | -1.031 | 0.513 | 1.559 | 0.935 | -0.001 | 0.007 | 0.007 | 0 | 0.18 | 0.14 | 0.35 | 0.29 | 0.15 | 0.18 |
| 1 | 45350028 | rs12139143 | -1.013 | 0.492 | 1.537 | 0.886 | 0 | 0.003 | 0.003 | 0 | 0.18 | 0.14 | 0.35 | 0.29 | 0.14 | 0.17 |
| 1 | 45358527 | rs263975 | -2.604 | 0.503 | 1.468 | 0.849 | -0.001 | 0.003 | 0.003 | 0 | 0.34 | 0.61 | 0.54 | 0.45 | 0.32 | 0.31 |
| 1 | 45363887 | rs608649 | -1.642 | 0.616 | 1.224 | 0.753 | 0.018 | -0.003 | -0.003 | 2.866 | 0.16 | 0.41 | 0.18 | 0.16 | 0.17 | 0.14 |
| 1 | 45369086 | rs1572379 | -1.664 | 0.617 | 1.213 | 0.761 | 0.027 | 0.009 | 0.009 | 0 | 0.16 | 0.45 | 0.18 | 0.17 | 0.18 | 0.14 |
| 1 | 45400653 | rs17393012 | -0.281 | 0.964 | 0.737 | 0.501 | -0.011 | 0.005 | 0.005 | 0 | 0.1 | 0.07 | 0.06 | 0.01 | 0.15 | 0.09 |
| 1 | 45400827 | rs17390824 | -0.753 | 0.804 | 0.831 | 0.728 | -0.006 | -0.003 | -0.003 | 2.788 | 0.1 | 0.11 | 0.15 | 0.16 | 0.17 | 0.15 |
| 1 | 45416900 | rs12407565 | -0.926 | 0.819 | 0.899 | 0.729 | 0.024 | 0.014 | 0.014 | 0 | 0.11 | 0.2 | 0.16 | 0.16 | 0.18 | 0.15 |
| 1 | 45417184 | rs452989 | -1.789 | 0.714 | 0.678 | 0.439 | 0.002 | -0.008 | -0.008 | 0.295 | 0.4 | 0.63 | 0.59 | 0.44 | 0.38 | 0.34 |
| 1 | 45432392 | rs17390983 | -0.725 | 0.73 | 0.644 | 0.443 | 0.003 | 0.001 | 0.001 | 0 | 0.11 | 0.15 | 0.15 | 0.16 | 0.17 | 0.15 |
| 1 | 45444038 | rs11556200 | -0.989 | 0.525 | 0.766 | 0.301 | -0.024 | 0.008 | 0.008 | 0 | 0.29 | 0.4 | 0.43 | 0.28 | 0.2 | 0.18 |
| 1 | 45444996 | rs264025 | -1.085 | 0.527 | 0.712 | 0.301 | -0.023 | 0.009 | 0.009 | 0 | 0.3 | 0.46 | 0.43 | 0.28 | 0.2 | 0.18 |
| 1 | 45451477 | rs364482 | -0.49 | 0.515 | 0.713 | 0.29 | -0.053 | 0.058 | 0.058 | 0 | 0.23 | 0.46 | 0.39 | 0.28 | 0.13 | 0.16 |
| 1 | 45457278 | rs4399199 | -1.006 | 0.514 | 0.711 | 0.298 | -0.017 | 0.004 | 0.004 | 0 | 0.29 | 0.5 | 0.43 | 0.27 | 0.2 | 0.18 |
| 1 | 45466927 | rs6666806 | 1.013 | 0.85 | 0.67 | 0.348 | 0 | 0.003 | 0.003 | 0 | 0.68 | 0.42 | 0.5 | 0.59 | 0.71 | 0.71 |
| 1 | 45468606 | rs8681 | 0.64 | 0.854 | 0.704 | 0.345 | -0.011 | 0.005 | 0.005 | 0 | 0.72 | 0.54 | 0.57 | 0.71 | 0.79 | 0.76 |
| 1 | 45474008 | rs7541207 | -0.544 | 0.837 | 0.88 | 0.337 | -0.001 | -0.005 | -0.005 | 0 | 0.24 | 0.16 | 0.41 | 0.29 | 0.21 | 0.24 |
| 1 | 45476489 | rs3806405 | -0.905 | 0.835 | 0.888 | 0.337 | -0.01 | 0.006 | 0.006 | 0 | 0.28 | 0.43 | 0.43 | 0.29 | 0.21 | 0.24 |
| 1 | 45479772 | rs11211066 | -0.574 | 0.946 | 0.109 | 0.377 | -0.001 | -0.005 | -0.005 | 0 | 0.24 | 0.16 | 0.41 | 0.29 | 0.21 | 0.24 |
| 1 | 45480829 | rs13948 | -0.574 | 0.927 | 0.109 | 0.377 | -0.001 | -0.005 | -0.005 | 6.425 | 0.24 | 0.16 | 0.41 | 0.29 | 0.21 | 0.24 |
| 1 | 45506964 | rs2202152 | -0.574 | 1.194 | -0.51 | -0.36 | -0.007 | 0.003 | 0.003 | 0 | 0.88 | 0.9 | 0.86 | 0.86 | 0.84 | 0.96 |
| 1 | 45511301 | rs11211073 | 1.347 | 1.383 | -1.395 | -1.069 | -0.013 | 0.211 | 0.211 | 0 | 0.37 | 0.1 | 0.22 | 0.03 | 0.34 | 0.34 |
| 1 | 45513219 | rs4660834 | -0.632 | 1.45 | -0.926 | -0.832 | -0.006 | 0.005 | 0.005 | 0 | 0.87 | 0.86 | 0.86 | 0.87 | 0.84 | 0.94 |
| 1 | 45513552 | rs6688318 | -0.547 | 1.433 | -0.584 | -0.841 | 0.009 | -0.01 | -0.01 | 0 | 0.87 | 0.77 | 0.85 | 0.87 | 0.84 | 0.94 |
| 1 | 45523377 | rs7552211 | 0.258 | 1.603 | -0.048 | -1.015 | 0.101 | -0.032 | -0.032 | 0.687 | 0.09 | 0.32 | 0.06 | 0 | 0.09 | 0.06 |
| 1 | 45523893 | rs3850855 | 0.766 | 1.573 | -0.021 | -1.031 | 0.026 | -0.054 | -0.054 | 0 | 0.78 | 0.41 | 0.78 | 0.87 | 0.74 | 0.87 |
| 1 | 45529693 | rs16832411 | 0.299 | 1.553 | 0.079 | -0.983 | 0.074 | -0.041 | -0.041 | 0 | 0.09 | 0.27 | 0.06 | 0 | 0.09 | 0.06 |
| 1 | 45576594 | rs2997399 | -0.411 | 1.077 | -0.553 | -1.045 | -0.028 | 0.057 | 0.057 | 0 | 0.89 | 0.99 | 0.91 | 1 | 0.84 | 0.89 |
| 1 | 45585207 | rs346733 | -1.003 | 1.058 | -0.339 | -1.032 | 0.038 | -0.067 | -0.067 | 0 | 0.23 | 0.56 | 0.21 | 0.13 | 0.26 | 0.13 |
| 1 | 45585348 | rs346732 | 0.601 | 1.062 | -0.381 | -1.002 | -0.011 | 0.006 | 0.006 | 0 | 0.12 | 0.11 | 0.14 | 0.13 | 0.17 | 0.06 |
| 1 | 45610193 | rs346717 | 0.605 | 1.079 | -0.262 | -0.515 | 0.001 | -0.002 | -0.002 | 0.044 | 0.12 | 0.14 | 0.14 | 0.13 | 0.17 | 0.07 |
| 1 | 45612038 | rs4660838 | 1.132 | 1.04 | -0.286 | -0.539 | -0.028 | 0.038 | 0.038 | 0 | 0.1 | 0 | 0.08 | 0.02 | 0.11 | 0.03 |
| 1 | 45627630 | rs346724 | 0.443 | 1.119 | -0.22 | -0.501 | 0.004 | -0.004 | -0.004 | 0 | 0.12 | 0.17 | 0.15 | 0.13 | 0.17 | 0.06 |
| 1 | 45645069 | rs3009969 | -1.233 | 1.137 | -0.402 | -0.543 | 0.029 | -0.068 | -0.068 | 0.259 | 0.23 | 0.56 | 0.21 | 0.14 | 0.26 | 0.13 |
| 1 | 45663038 | rs4660839 | 0.579 | 1.105 | -0.85 | -0.479 | -0.002 | 0 | 0 | 0 | 0.12 | 0.12 | 0.14 | 0.13 | 0.17 | 0.06 |
| 1 | 45699200 | rs17192059 | 1.405 | 1.101 | -0.972 | -0.475 | -0.005 | 0.01 | 0.01 | 0 | 0.09 | 0.04 | 0.08 | 0.02 | 0.11 | 0.03 |
| 1 | 45716285 | rs3121730 | 0.275 | 1.007 | -1.134 | -0.61 | 0.012 | -0.012 | -0.012 | 3.148 | 0.7 | 0.84 | 0.53 | 0.41 | 0.72 | 0.57 |
| 1 | 45740469 | rs1938297 | -0.423 | 1.259 | -0.925 | -0.703 | -0.003 | 0.007 | 0.007 | 0 | 0.9 | 0.93 | 0.92 | 0.99 | 0.88 | 0.91 |
| 1 | 45757124 | rs11211091 | -0.038 | 1.217 | -0.454 | -0.285 | 0.094 | -0.031 | -0.031 | 0 | 0.09 | 0.44 | 0.07 | 0 | 0.08 | 0.06 |
| 1 | 45763650 | rs4233496 | -1.108 | 1.207 | -0.647 | -0.235 | 0.018 | -0.037 | -0.037 | 0 | 0.23 | 0.61 | 0.23 | 0.17 | 0.27 | 0.14 |
| 1 | 45764803 | rs6429564 | -0.653 | 0.613 | -0.592 | -0.312 | 0.024 | -0.107 | -0.107 | 1.53 | 0.19 | 0.5 | 0.17 | 0.03 | 0.19 | 0.1 |
| 1 | 45765523 | rs4484960 | 0.074 | 0.619 | -1.072 | 0.563 | -0.027 | -0.04 | -0.04 | 0 | 0.16 | 0.03 | 0.31 | 0.18 | 0.14 | 0.13 |
| 1 | 45766451 | rs12094184 | -0.352 | 0.498 | -0.573 | 0.792 | -0.002 | 0.039 | 0.039 | 0 | 0.12 | 0.2 | 0.16 | 0.31 | 0.12 | 0.17 |
| 1 | 45791154 | rs10890324 | 0.595 | 0.268 | -0.796 | 0.144 | 0.006 | 0.01 | 0.01 | 0 | 0.7 | 0.25 | 0.78 | 0.81 | 0.75 | 0.76 |
| 1 | 45796354 | rs3219492 | -0.007 | 0.072 | -0.688 | 0.281 | 0.004 | -0.003 | -0.003 | 0.959 | 0.15 | 0.11 | 0.36 | 0.16 | 0.1 | 0.1 |
| 1 | 45798555 | rs3219487 | -0.564 | -0.027 | -0.791 | 0.217 | -0.004 | -0.009 | -0.009 | 0 | 0.91 | 0.98 | 0.94 | 0.91 | 0.92 | 0.93 |
| 1 | 45804050 | rs3219472 | -1.048 | -0.035 | -0.568 | 0.19 | -0.002 | 0.003 | 0.003 | 1.077 | 0.28 | 0.25 | 0.48 | 0.35 | 0.24 | 0.26 |
| 1 | 45810091 | rs9429072 | -1.342 | -0.175 | 0.236 | 0.18 | -0.009 | 0.013 | 0.013 | 1.689 | 0.31 | 0.61 | 0.51 | 0.39 | 0.25 | 0.26 |
| 1 | 45810923 | rs3893383 | 0.292 | -0.119 | 0.264 | 0.194 | 0.009 | 0.076 | 0.076 | 0 | 0.11 | 0.29 | 0.11 | 0.19 | 0.1 | 0.12 |
| 1 | 45819489 | rs9429076 | -1.164 | -0.147 | 0.203 | 0.131 | -0.009 | 0.013 | 0.013 | 1.689 | 0.31 | 0.61 | 0.5 | 0.38 | 0.25 | 0.26 |
| 1 | 45872662 | rs781226 | 0.832 | -0.213 | 0.613 | 0.074 | -0.011 | 0.012 | 0.012 | 1.666 | 0.69 | 0.37 | 0.64 | 0.62 | 0.76 | 0.72 |
| 1 | 45879088 | rs6657284 | -0.926 | -0.216 | 0.641 | 0.068 | -0.005 | 0.004 | 0.004 | 0.479 | 0.29 | 0.29 | 0.35 | 0.35 | 0.24 | 0.27 |
| 1 | 45890155 | rs12139364 | 1.95 | -0.292 | 0.269 | -0.222 | -0.004 | 0.01 | 0.01 | 0 | 0.47 | 0.24 | 0.49 | 0.58 | 0.53 | 0.54 |
| 1 | 45896779 | rs1771549 | -0.502 | -0.253 | 0.094 | -0.244 | -0.008 | 0.044 | 0.044 | 0 | 0.22 | 0.17 | 0.15 | 0.04 | 0.23 | 0.19 |
| 1 | 45903488 | rs10493120 | -0.471 | -0.297 | 0.071 | -0.255 | -0.013 | 0.067 | 0.067 | 0 | 0.22 | 0.13 | 0.14 | 0.04 | 0.22 | 0.19 |

| | | | | | | | | | | | | | | | | |
|---|---|---|---|---|---|---|---|---|---|---|---|---|---|---|---|---|
| 1 | 45909994 | rs1006216 | -0.615 | -0.322 | 0.248 | -0.226 | 0.007 | -0.01 | -0.01 | 0 | 0.28 | 0.23 | 0.33 | 0.35 | 0.23 | 0.27 |
| 1 | 45927848 | rs10789463 | -1.97 | -0.294 | 0.286 | -0.201 | 0.004 | 0.011 | 0.011 | 0 | 0.51 | 0.58 | 0.48 | 0.38 | 0.46 | 0.46 |
| 1 | 45955868 | rs4660863 | -0.628 | -0.313 | 0.209 | -0.186 | 0.011 | -0.009 | -0.009 | 0 | 0.28 | 0.22 | 0.33 | 0.32 | 0.22 | 0.29 |
| 1 | 45957290 | rs1494813 | 0.293 | -0.097 | 0.054 | -0.404 | -0.027 | 0.064 | 0.064 | 0.02 | 0.75 | 0.86 | 0.84 | 0.94 | 0.7 | 0.78 |
| 1 | 45962409 | rs11211124 | -0.469 | -0.098 | 0.053 | -0.403 | 0.009 | -0.008 | -0.008 | 0 | 0.28 | 0.22 | 0.32 | 0.32 | 0.22 | 0.29 |
| 1 | 45964766 | rs12042481 | 0.947 | -0.076 | 0.021 | -0.446 | -0.014 | 0.025 | 0.025 | 0 | 0.12 | 0.04 | 0.07 | 0.05 | 0.15 | 0.1 |
| 1 | 45970446 | rs12029322 | -0.495 | -0.065 | 0.07 | -0.392 | 0.009 | -0.008 | -0.008 | 0 | 0.28 | 0.22 | 0.32 | 0.32 | 0.22 | 0.29 |
| 1 | 45971759 | rs10789465 | 1.934 | -0.067 | 0.243 | -0.379 | -0.004 | 0.001 | 0.001 | 0 | 0.45 | 0.37 | 0.5 | 0.6 | 0.48 | 0.48 |
| 1 | 45973928 | rs2275276 | 1.934 | -0.067 | 0.24 | -0.379 | -0.004 | 0.001 | 0.001 | 0 | 0.45 | 0.37 | 0.5 | 0.6 | 0.48 | 0.48 |
| 1 | 45976147 | rs882803 | 0.621 | -0.02 | 0.175 | -0.395 | 0.012 | -0.056 | -0.056 | 0.051 | 0.73 | 0.59 | 0.82 | 0.91 | 0.69 | 0.77 |
| 1 | 45977685 | rs2065708 | 0.398 | 0.002 | 0.114 | -0.4 | -0.02 | 0.058 | 0.058 | 0 | 0.18 | 0.07 | 0.11 | 0.05 | 0.22 | 0.15 |
| 1 | 45984353 | rs10736426 | 0.726 | -0.015 | 0.169 | -0.387 | 0.012 | -0.055 | -0.055 | 0 | 0.73 | 0.59 | 0.82 | 0.91 | 0.69 | 0.77 |
| 1 | 45987574 | rs17522918 | 0.753 | -0.116 | 0.108 | -0.382 | 0.02 | 0.003 | 0.003 | 0.653 | 0.15 | 0.1 | 0.21 | 0.13 | 0.08 | 0.09 |
| 1 | 46009818 | rs868351 | -0.306 | -0.037 | 0.127 | -0.394 | 0.013 | -0.009 | -0.009 | 0 | 0.28 | 0.21 | 0.32 | 0.3 | 0.21 | 0.29 |
| 1 | 46009946 | rs512026 | 1.663 | 0.032 | 0.079 | -0.333 | -0.005 | 0.001 | 0.001 | 0 | 0.45 | 0.36 | 0.5 | 0.61 | 0.48 | 0.48 |
| 1 | 46020795 | rs666720 | 1.627 | 0.006 | -0.833 | -0.442 | -0.006 | 0.022 | 0.022 | 0 | 0.45 | 0.63 | 0.51 | 0.66 | 0.46 | 0.5 |
| 1 | 46021556 | rs12049503 | -0.27 | -0.073 | -0.861 | -0.419 | 0.007 | -0.008 | -0.008 | 1.231 | 0.27 | 0.24 | 0.31 | 0.3 | 0.21 | 0.29 |
| 1 | 46024701 | rs534534 | 0.107 | 0.032 | -1.134 | -0.786 | -0.057 | 0.17 | 0.17 | 0 | 0.67 | 0.88 | 0.8 | 0.99 | 0.62 | 0.78 |
| 1 | 46032974 | rs2088102 | 1.47 | 0.056 | -0.943 | -0.689 | -0.001 | 0.003 | 0.003 | 0 | 0.45 | 0.52 | 0.52 | 0.65 | 0.48 | 0.48 |
| 1 | 46037357 | rs518216 | -0.415 | 0.046 | -0.989 | -0.72 | -0.011 | 0.023 | 0.023 | 0.004 | 0.77 | 0.72 | 0.73 | 0.66 | 0.86 | 0.7 |
| 1 | 46039612 | rs2934857 | 1.188 | 0.153 | -0.886 | -0.568 | -0.001 | 0.002 | 0.002 | 0 | 0.45 | 0.52 | 0.52 | 0.65 | 0.48 | 0.48 |
| 1 | 46050561 | rs11579176 | 0.439 | 0.273 | -0.926 | -0.554 | -0.02 | 0.082 | 0.082 | 0 | 0.2 | 0.06 | 0.12 | 0.01 | 0.24 | 0.13 |
| 1 | 46078854 | rs2230658 | -0.202 | 0.234 | -0.899 | -0.498 | -0.002 | -0.016 | -0.016 | 0 | 0.67 | 0.73 | 0.65 | 0.66 | 0.72 | 0.61 |
| 1 | 46081282 | rs2991986 | -0.197 | 0.238 | -0.897 | -0.496 | -0.002 | -0.016 | -0.016 | 0 | 0.67 | 0.73 | 0.65 | 0.66 | 0.72 | 0.61 |
| 1 | 46084383 | rs1053941 | 0.519 | 0.229 | -0.951 | -0.573 | -0.027 | 0.093 | 0.093 | 0 | 0.19 | 0.05 | 0.12 | 0.01 | 0.23 | 0.13 |
| 1 | 46086329 | rs11590549 | 0.102 | 0.211 | -0.722 | -0.571 | 0 | -0.046 | -0.046 | 0 | 0.22 | 0.31 | 0.15 | 0.04 | 0.25 | 0.13 |
| 1 | 46094027 | rs1250 | -0.128 | 0.15 | -0.692 | -0.432 | -0.004 | -0.02 | -0.02 | 0 | 0.33 | 0.26 | 0.35 | 0.35 | 0.28 | 0.4 |
| 1 | 46100322 | rs10890333 | -0.173 | 0.223 | -0.788 | -0.443 | -0.003 | -0.016 | -0.016 | 0 | 0.34 | 0.27 | 0.35 | 0.34 | 0.29 | 0.42 |
| 1 | 46101323 | rs4431884 | -0.285 | 0.177 | -0.769 | -0.443 | 0 | -0.001 | -0.001 | 0 | 0.35 | 0.38 | 0.36 | 0.34 | 0.29 | 0.42 |
| 1 | 46125880 | rs3811436 | -0.165 | 0.211 | -0.905 | -0.446 | -0.001 | -0.005 | -0.005 | 0 | 0.33 | 0.27 | 0.35 | 0.32 | 0.28 | 0.41 |
| 1 | 46153203 | rs3013595 | -1.394 | 0.206 | -0.832 | -0.475 | 0 | -0.004 | -0.004 | 0.207 | 0.56 | 0.58 | 0.49 | 0.35 | 0.53 | 0.54 |
| 1 | 46153548 | rs11211163 | -0.088 | 0.173 | -0.851 | -0.574 | -0.002 | -0.016 | -0.016 | 0.1 | 0.34 | 0.27 | 0.35 | 0.34 | 0.29 | 0.42 |
| 1 | 46158331 | rs10789470 | -0.088 | 0.173 | -0.827 | -0.574 | -0.002 | -0.015 | -0.015 | 0 | 0.34 | 0.27 | 0.35 | 0.34 | 0.29 | 0.42 |
| 1 | 46166709 | rs11211169 | -0.043 | 0.171 | -0.831 | -0.582 | -0.002 | -0.003 | -0.003 | 0 | 0.33 | 0.27 | 0.35 | 0.31 | 0.28 | 0.41 |
| 1 | 46188914 | rs28550303 | -0.043 | 0.171 | -0.874 | -0.572 | -0.003 | -0.003 | -0.003 | 0 | 0.33 | 0.25 | 0.35 | 0.31 | 0.28 | 0.41 |
| 1 | 46195375 | rs28375469 | -0.043 | 0.171 | -0.875 | -0.575 | -0.002 | -0.004 | -0.004 | 0.646 | 0.33 | 0.25 | 0.35 | 0.31 | 0.28 | 0.41 |
| 1 | 46196813 | rs28433203 | 0.505 | 0.157 | -0.864 | -0.603 | -0.021 | 0.088 | 0.088 | 0 | 0.2 | 0.06 | 0.12 | 0.01 | 0.24 | 0.12 |
| 1 | 46202283 | rs28719889 | -0.028 | 0.16 | -0.854 | -0.579 | -0.002 | 0.001 | 0.001 | 0 | 0.33 | 0.25 | 0.35 | 0.3 | 0.28 | 0.41 |
| 1 | 46206399 | rs28647556 | -1.522 | 0.157 | -0.857 | -0.589 | 0 | -0.006 | -0.006 | 0 | 0.56 | 0.56 | 0.5 | 0.35 | 0.53 | 0.54 |
| 1 | 46216575 | rs6429576 | -1.522 | 0.157 | -0.85 | -0.589 | 0 | -0.005 | -0.005 | 0.831 | 0.56 | 0.54 | 0.49 | 0.35 | 0.53 | 0.54 |
| 1 | 46221440 | rs10430123 | -0.1 | 0.233 | -0.913 | -0.538 | -0.003 | -0.016 | -0.016 | 0.652 | 0.33 | 0.25 | 0.35 | 0.34 | 0.29 | 0.42 |
| 1 | 46245423 | rs11582058 | 0.223 | 0.109 | -1.107 | -0.689 | -0.015 | 0.1 | 0.1 | 0 | 0.23 | 0.07 | 0.14 | 0.02 | 0.27 | 0.14 |
| 1 | 46248645 | rs4626927 | 1.38 | -0.127 | -0.997 | -0.636 | 0.007 | -0.005 | -0.005 | 0.772 | 0.41 | 0.42 | 0.48 | 0.65 | 0.46 | 0.42 |
| 1 | 46267923 | rs7525281 | -0.569 | -0.03 | -1.062 | -0.654 | -0.022 | 0.089 | 0.089 | 0 | 0.77 | 0.92 | 0.86 | 0.98 | 0.72 | 0.86 |
| 1 | 46306583 | rs10789477 | -0.569 | -0.03 | -1.064 | -0.654 | -0.024 | 0.093 | 0.093 | 0 | 0.77 | 0.92 | 0.87 | 0.98 | 0.72 | 0.86 |
| 1 | 46321843 | rs6429582 | 1.378 | 0.102 | -1.044 | -0.676 | 0.003 | -0.005 | -0.005 | 0.014 | 0.41 | 0.43 | 0.48 | 0.65 | 0.45 | 0.42 |
| 1 | 46340554 | rs11211211 | -0.615 | 0.103 | -1.044 | -0.676 | -0.024 | 0.093 | 0.093 | 0 | 0.77 | 0.92 | 0.87 | 0.98 | 0.72 | 0.86 |
| 1 | 46342337 | rs6682683 | -0.049 | 0.079 | -1.051 | -0.709 | -0.005 | -0.004 | -0.004 | 0 | 0.36 | 0.47 | 0.38 | 0.3 | 0.27 | 0.44 |
| 1 | 46406767 | rs11211230 | 1.381 | 0.08 | -1.032 | -0.689 | 0.004 | -0.006 | -0.006 | 1.132 | 0.41 | 0.43 | 0.48 | 0.66 | 0.45 | 0.42 |
| 1 | 46411146 | rs11211232 | 0.098 | 0.06 | -1 | -0.698 | 0.005 | 0.005 | 0.005 | 0 | 0.34 | 0.23 | 0.36 | 0.29 | 0.27 | 0.44 |
| 1 | 46425500 | rs4420029 | -0.292 | 0.083 | -0.983 | -0.698 | 0 | -0.035 | -0.035 | 0 | 0.75 | 0.68 | 0.84 | 0.95 | 0.71 | 0.86 |
| 1 | 46440252 | rs10749861 | -0.593 | 0.091 | -1.038 | -0.709 | -0.024 | 0.094 | 0.094 | 0 | 0.77 | 0.92 | 0.87 | 0.98 | 0.72 | 0.86 |
| 1 | 46461587 | rs4660905 | -0.013 | 0.075 | -1.024 | -0.708 | -0.006 | -0.006 | -0.006 | 0.046 | 0.36 | 0.47 | 0.38 | 0.29 | 0.27 | 0.43 |
| 1 | 46465662 | rs6661500 | 0.082 | 0.077 | -1.005 | -0.707 | -0.002 | -0.002 | -0.002 | 0.024 | 0.35 | 0.44 | 0.38 | 0.29 | 0.27 | 0.43 |
| 1 | 46476587 | rs11211247 | -0.573 | 0.115 | -1.053 | -0.711 | -0.024 | 0.094 | 0.094 | 0 | 0.77 | 0.92 | 0.87 | 0.98 | 0.72 | 0.86 |
| 1 | 46493460 | rs1707336 | 1.306 | 0.095 | -1.009 | -0.674 | 0.003 | -0.007 | -0.007 | 0 | 0.41 | 0.43 | 0.48 | 0.66 | 0.45 | 0.43 |
| 1 | 46495024 | rs3795318 | 0.445 | 0.143 | -1.121 | -0.718 | -0.039 | 0.116 | 0.116 | 0 | 0.22 | 0.04 | 0.14 | 0.02 | 0.28 | 0.14 |
| 1 | 46498375 | rs1622208 | -0.347 | 0.09 | -1.077 | -0.703 | 0.007 | 0.007 | 0.007 | 0 | 0.66 | 0.75 | 0.64 | 0.71 | 0.73 | 0.56 |
| 1 | 46500993 | rs1052610 | 2.491 | 0.47 | -1.7 | -1.238 | -0.033 | 0.054 | 0.054 | 0 | 0.09 | 0.01 | 0.07 | 0 | 0.1 | 0.02 |
| 1 | 46502836 | rs1768809 | 0.697 | 0.368 | -1.405 | -0.97 | -0.001 | -0.001 | -0.001 | 0.01 | 0.48 | 0.49 | 0.53 | 0.68 | 0.57 | 0.45 |
| 1 | 46503217 | rs1768808 | 0.222 | 0.178 | -0.996 | -0.404 | 0.001 | -0.005 | -0.005 | 0 | 0.61 | 0.71 | 0.47 | 0.62 | 0.65 | 0.53 |
| 1 | 46521517 | rs785468 | -0.044 | 0.49 | -1.352 | -0.48 | -0.004 | -0.001 | -0.001 | 0 | 0.65 | 0.59 | 0.63 | 0.69 | 0.73 | 0.56 |
| 1 | 46544768 | rs1085243 | 0.981 | 0.498 | -1.352 | -0.512 | 0.003 | -0.003 | -0.003 | 0 | 0.42 | 0.48 | 0.49 | 0.64 | 0.45 | 0.43 |
| 1 | 46558927 | rs9429187 | 0.409 | 0.507 | -1.386 | -0.55 | -0.01 | 0.067 | 0.067 | 0 | 0.24 | 0.11 | 0.14 | 0.01 | 0.28 | 0.14 |
| 1 | 46568836 | rs785520 | 0.903 | 0.662 | -1.328 | -0.46 | 0.003 | -0.003 | -0.003 | 0.504 | 0.42 | 0.48 | 0.49 | 0.64 | 0.45 | 0.42 |
| 1 | 46598273 | rs1707303 | 0.229 | 0.256 | -1.159 | -0.106 | -0.003 | 0 | 0 | 0 | 0.66 | 0.59 | 0.63 | 0.69 | 0.73 | 0.57 |
| 1 | 46600917 | rs1707302 | 0.139 | 0.568 | -0.84 | 0.176 | 0 | -0.001 | -0.001 | 1.82 | 0.64 | 0.61 | 0.6 | 0.71 | 0.68 | 0.52 |
| 1 | 46609736 | rs785475 | 0.396 | 0.52 | -0.657 | 0.843 | -0.013 | 0.077 | 0.077 | 0 | 0.82 | 0.74 | 0.72 | 0.5 | 0.87 | 0.63 |
| 1 | 46621740 | rs7512966 | -0.77 | 0.419 | -0.439 | 0.749 | -0.035 | 0.093 | 0.093 | 0.033 | 0.21 | 0.45 | 0.31 | 0.38 | 0.14 | 0.46 |

| | | | | | | | | | | | | | | | | |
|---|---|---|---|---|---|---|---|---|---|---|---|---|---|---|---|---|
| 1 | 46633874 | rs3855959 | -1.044 | 0.882 | -0.613 | 0.34 | 0 | 0.008 | 0.008 | 0 | 0.6 | 0.6 | 0.56 | 0.43 | 0.59 | 0.7 |
| 1 | 46635209 | rs3845301 | -0.8 | 0.835 | -0.498 | 0.416 | 0.001 | 0.006 | 0.006 | 0 | 0.38 | 0.53 | 0.44 | 0.39 | 0.39 | 0.55 |
| 1 | 46645524 | rs3820082 | -0.557 | 0.184 | -0.308 | 0.621 | -0.002 | 0.014 | 0.014 | 0 | 0.18 | 0.21 | 0.28 | 0.33 | 0.15 | 0.45 |
| 1 | 46649046 | rs10890384 | -0.08 | 0.304 | -0.372 | 0.178 | -0.01 | 0.002 | 0.002 | 0 | 0.14 | 0.11 | 0.25 | 0.12 | 0.15 | 0.21 |
| 1 | 46656964 | rs11584024 | 1.126 | 0.088 | -0.401 | -0.154 | 0.002 | 0.126 | 0.126 | 0 | 0.2 | 0.02 | 0.1 | 0 | 0.17 | 0.15 |
| 1 | 46658343 | rs2292485 | -0.752 | -0.078 | -0.225 | -0.158 | 0.013 | 0.052 | 0.052 | 0 | 0.4 | 0.33 | 0.4 | 0.17 | 0.35 | 0.43 |
| 1 | 46659104 | rs7527668 | -0.687 | -0.096 | -0.166 | -0.167 | 0.002 | -0.001 | -0.001 | 0 | 0.4 | 0.46 | 0.4 | 0.17 | 0.35 | 0.43 |
| 1 | 46664249 | rs28364741 | 1.212 | 0.074 | -0.503 | 0.228 | 0.009 | -0.102 | -0.102 | 0 | 0.22 | 0.01 | 0.28 | 0.38 | 0.18 | 0.12 |
| 1 | 46677918 | rs12141928 | 0.094 | 0.279 | -0.181 | -0.43 | 0.002 | -0.001 | -0.001 | 0 | 0.16 | 0.18 | 0.31 | 0.16 | 0.19 | 0.22 |
| 1 | 46691621 | rs953070 | 0.655 | 0.191 | -0.358 | -0.556 | 0.018 | -0.042 | -0.042 | 0 | 0.78 | 0.46 | 0.66 | 0.84 | 0.77 | 0.72 |
| 1 | 46705106 | rs12141269 | 1.031 | 0.227 | -1.248 | -0.517 | -0.028 | -0.023 | -0.023 | 0 | 0.09 | 0.04 | 0.28 | 0.15 | 0.15 | 0.13 |
| 1 | 46718905 | rs10789488 | 0.272 | -0.07 | -1.111 | -0.522 | 0.017 | -0.028 | -0.028 | 6.285 | 0.77 | 0.46 | 0.65 | 0.83 | 0.75 | 0.7 |
| 1 | 46721155 | rs11211262 | 0.386 | -0.3 | -1.315 | -0.526 | 0.033 | -0.005 | -0.005 | 0.796 | 0.51 | 0.3 | 0.52 | 0.52 | 0.61 | 0.29 |
| 1 | 46722389 | rs9793263 | 0.386 | -0.34 | -1.442 | -0.524 | 0.012 | 0.036 | 0.036 | 0 | 0.57 | 0.33 | 0.54 | 0.73 | 0.6 | 0.68 |
| 1 | 46722939 | rs17102086 | 1.529 | -0.031 | -1.487 | -0.622 | -0.032 | -0.067 | -0.067 | 0 | 0.13 | 0.04 | 0.27 | 0.41 | 0.17 | 0.19 |
| 1 | 46726057 | rs2295465 | 1.047 | 0.257 | -0.985 | -0.52 | 0.03 | -0.092 | -0.092 | 0 | 0.24 | 0.02 | 0.12 | 0.28 | 0.15 | 0.45 |
| 1 | 46731694 | rs4660918 | -0.179 | 0.232 | -0.908 | -0.511 | 0.026 | 0 | 0 | 0 | 0.8 | 0.63 | 0.66 | 0.81 | 0.78 | 0.76 |
| 1 | 46734792 | rs12125573 | 0.602 | 0.232 | -0.921 | -0.512 | 0.004 | -0.012 | -0.012 | 0 | 0.75 | 0.34 | 0.64 | 0.79 | 0.75 | 0.69 |
| 1 | 46743900 | rs1048771 | 1.091 | 0.201 | -0.96 | -0.493 | 0.04 | -0.089 | -0.089 | 0 | 0.24 | 0.03 | 0.12 | 0.28 | 0.14 | 0.41 |
| 1 | 46746164 | rs11542623 | 1.091 | 0.154 | -0.956 | -0.493 | 0.042 | -0.084 | -0.084 | 0.089 | 0.24 | 0.03 | 0.12 | 0.28 | 0.14 | 0.41 |
| 1 | 46747301 | rs12142240 | 1.305 | 0.923 | -0.343 | -0.626 | -0.027 | 0.016 | 0.016 | 0 | 0.18 | 0.12 | 0.14 | 0.01 | 0.29 | 0.08 |
| 1 | 46768408 | rs1547180 | 0.334 | 0.416 | -0.076 | 0.073 | 0.023 | 0.003 | 0.003 | 0 | 0.81 | 0.64 | 0.66 | 0.81 | 0.79 | 0.76 |
| 1 | 46781602 | rs4660921 | 0.699 | 0.444 | -0.213 | 0.081 | 0.011 | -0.014 | -0.014 | 0 | 0.77 | 0.45 | 0.65 | 0.79 | 0.76 | 0.7 |
| 1 | 46805222 | rs4660344 | 0.325 | 0.199 | -1.505 | 0.166 | 0 | -0.08 | -0.08 | 0 | 0.35 | 0.14 | 0.36 | 0.52 | 0.34 | 0.23 |
| 1 | 46806480 | rs17361749 | 1.314 | 0.627 | -1.638 | -0.402 | -0.025 | 0.019 | 0.019 | 0 | 0.17 | 0.12 | 0.15 | 0.02 | 0.28 | 0.09 |
| 1 | 46807597 | rs10489770 | 0.769 | 0.464 | -1.828 | -0.474 | -0.019 | -0.012 | -0.012 | 0.097 | 0.1 | 0.04 | 0.26 | 0.15 | 0.12 | 0.14 |
| 1 | 46809419 | rs7515284 | 1.321 | 0.223 | -1.308 | -0.516 | -0.015 | 0.005 | 0.005 | 0 | 0.18 | 0.13 | 0.15 | 0.02 | 0.27 | 0.09 |
| 1 | 46810530 | rs3737744 | 0.94 | 0.18 | -1.339 | -0.463 | 0.004 | -0.087 | -0.087 | 0.809 | 0.35 | 0.14 | 0.36 | 0.52 | 0.34 | 0.23 |
| 1 | 46810670 | rs17361763 | 1.284 | 0.193 | -1.319 | -0.471 | -0.015 | 0.005 | 0.005 | 0 | 0.18 | 0.13 | 0.15 | 0.02 | 0.27 | 0.09 |
| 1 | 46815154 | rs5013330 | 1.284 | 0.193 | -1.319 | -0.471 | -0.015 | 0.005 | 0.005 | 0 | 0.18 | 0.13 | 0.15 | 0.02 | 0.27 | 0.09 |
| 1 | 46827456 | rs9865 | 0.868 | -0.142 | -1.513 | -0.668 | 0.057 | -0.077 | -0.077 | 0 | 0.25 | 0.06 | 0.13 | 0.28 | 0.15 | 0.44 |
| 1 | 46828700 | rs17361833 | 1.412 | -0.109 | -1.505 | -0.652 | -0.015 | 0.005 | 0.005 | 0 | 0.18 | 0.13 | 0.15 | 0.02 | 0.27 | 0.09 |
| 1 | 46830430 | rs10252 | -0.178 | -0.275 | -1.109 | -0.741 | -0.001 | -0.004 | -0.004 | 0 | 0.44 | 0.51 | 0.28 | 0.3 | 0.41 | 0.53 |
| 1 | 46830447 | rs12062 | 1.455 | -0.234 | -1.161 | -0.726 | -0.015 | 0.005 | 0.005 | 0 | 0.18 | 0.13 | 0.15 | 0.02 | 0.27 | 0.09 |
| 1 | 46834775 | rs17361887 | 1.653 | -0.235 | -1.182 | -0.73 | -0.057 | 0.057 | 0.057 | 0 | 0.17 | 0.07 | 0.14 | 0.01 | 0.27 | 0.08 |
| 1 | 46843216 | rs3863641 | -0.004 | -0.221 | -1.078 | -0.645 | 0.001 | 0.02 | 0.02 | 0 | 0.45 | 0.39 | 0.29 | 0.3 | 0.42 | 0.54 |
| 1 | 46865671 | rs3766246 | 0.364 | -0.217 | -1.171 | -0.571 | 0.052 | -0.013 | -0.013 | 1.436 | 0.54 | 0.24 | 0.51 | 0.53 | 0.63 | 0.34 |
| 1 | 46870761 | rs324420 | 0.179 | -0.094 | -0.925 | -0.48 | 0.053 | 0.003 | 0.003 | 0 | 0.16 | 0.37 | 0.35 | 0.18 | 0.21 | 0.2 |
| 1 | 46875067 | rs11576941 | 0.888 | -0.042 | -1.22 | -0.558 | 0.007 | -0.094 | -0.094 | 0 | 0.33 | 0.13 | 0.35 | 0.51 | 0.31 | 0.21 |
| 1 | 46885940 | rs4660928 | 1.129 | 0.593 | -0.516 | -0.631 | 0.006 | -0.028 | -0.028 | 0 | 0.75 | 0.44 | 0.61 | 0.83 | 0.72 | 0.74 |
| 1 | 46895641 | rs1571138 | 0.278 | 0.74 | -0.561 | -0.368 | 0.048 | 0.005 | 0.005 | 0 | 0.83 | 0.63 | 0.65 | 0.82 | 0.79 | 0.73 |
| 1 | 46897698 | rs1535737 | 0.091 | 0.038 | -1.038 | -0.518 | 0.007 | 0.022 | 0.022 | 0 | 0.54 | 0.46 | 0.44 | 0.76 | 0.43 | 0.55 |
| 1 | 46902461 | rs661902 | -0.575 | -0.277 | -1.222 | -0.331 | 0.013 | 0.045 | 0.045 | 0 | 0.41 | 0.63 | 0.53 | 0.78 | 0.37 | 0.57 |
| 1 | 46905802 | rs11211276 | 0.066 | -0.415 | -1.472 | 0.313 | -0.005 | -0.042 | -0.042 | 2.19 | 0.16 | 0.1 | 0.25 | 0.25 | 0.2 | 0.31 |
| 1 | 46912162 | rs606884 | -0.495 | -0.945 | -1.382 | 0.291 | -0.001 | 0 | 0 | 0 | 0.29 | 0.61 | 0.31 | 0.53 | 0.19 | 0.32 |
| 1 | 46913300 | rs4660347 | 0.123 | -0.837 | -1.4 | 0.342 | -0.007 | -0.046 | -0.046 | 0 | 0.17 | 0.1 | 0.25 | 0.26 | 0.2 | 0.32 |
| 1 | 46920331 | rs7532468 | 0.969 | -0.966 | -0.308 | 0.522 | 0.067 | 0.126 | 0.126 | 0.665 | 0.07 | 0.32 | 0.07 | 0.09 | 0.04 | 0.08 |
| 1 | 46923102 | rs2031248 | -0.619 | -0.991 | -0.282 | 0.513 | -0.001 | 0.051 | 0.051 | 0 | 0.22 | 0.33 | 0.28 | 0.31 | 0.23 | 0.36 |
| 1 | 46925594 | rs10890398 | -0.595 | -1.376 | -0.953 | 0.318 | 0.021 | -0.026 | -0.026 | 0 | 0.29 | 0.24 | 0.3 | 0.48 | 0.19 | 0.34 |
| 1 | 46927103 | rs942256 | -0.916 | -1.294 | -0.843 | 0.197 | 0.02 | 0.012 | 0.012 | 0 | 0.39 | 0.28 | 0.48 | 0.7 | 0.36 | 0.56 |
| 1 | 46928349 | rs11211282 | -1.473 | -1.09 | -1.289 | 0.135 | 0.021 | 0.132 | 0.132 | 0 | 0.53 | 0.81 | 0.6 | 0.8 | 0.43 | 0.7 |
| 1 | 46938837 | rs2031247 | -0.376 | -0.003 | -1.041 | 0.046 | -0.015 | 0.001 | 0.001 | 0 | 0.25 | 0.15 | 0.22 | 0.22 | 0.26 | 0.32 |
| 1 | 46954587 | rs1998545 | 0.435 | -0.205 | -1.241 | 0.176 | 0.002 | -0.02 | -0.02 | 0.443 | 0.69 | 0.79 | 0.61 | 0.66 | 0.67 | 0.71 |
| 1 | 46967890 | rs2025384 | -1.183 | -0.79 | -0.038 | 0.352 | -0.059 | -0.017 | -0.017 | 0.337 | 0.28 | 0.45 | 0.18 | 0.24 | 0.17 | 0.28 |
| 1 | 46975773 | rs6683116 | -1.255 | -0.638 | 0.394 | 0.33 | -0.008 | 0 | 0 | 0.168 | 0.31 | 0.63 | 0.24 | 0.31 | 0.19 | 0.31 |
| 1 | 46978497 | rs2209172 | 0.796 | -0.826 | 0.467 | 0.252 | -0.009 | -0.014 | -0.014 | 0 | 0.19 | 0.17 | 0.12 | 0.24 | 0.22 | 0.27 |
| 1 | 46978885 | rs2209171 | 0.645 | -0.78 | 0.898 | 0.291 | -0.034 | -0.024 | -0.024 | 0 | 0.07 | 0.23 | 0.02 | 0.02 | 0.01 | 0.07 |
| 1 | 46981536 | rs6429601 | 0.179 | -0.609 | 0.457 | 0.416 | 0.009 | 0.119 | 0.119 | 0 | 0.82 | 0.74 | 0.68 | 0.52 | 0.81 | 0.76 |
| 1 | 46982455 | rs942249 | 1.076 | -0.577 | 0.612 | 0.214 | -0.006 | 0.045 | 0.045 | 0 | 0.73 | 0.49 | 0.64 | 0.48 | 0.76 | 0.69 |
| 1 | 46986641 | rs7519040 | -0.81 | -0.755 | 0.933 | 0.196 | 0.008 | 0.02 | 0.02 | 0.331 | 0.22 | 0.43 | 0.39 | 0.26 | 0.23 | 0.2 |
| 1 | 46987718 | rs1267308 | -0.217 | -0.756 | 0.998 | 0.16 | -0.002 | 0 | 0 | 0.513 | 0.82 | 0.85 | 0.71 | 0.83 | 0.77 | 0.85 |
| 1 | 46989742 | rs1267305 | -2.731 | -0.949 | 0.071 | -0.288 | -0.031 | 0.085 | 0.085 | 0 | 0.86 | 0.99 | 0.8 | 1 | 0.87 | 0.96 |
| 1 | 46989792 | rs942255 | -0.862 | -1.435 | 0.92 | 0.278 | -0.002 | -0.001 | -0.001 | 0 | 0.21 | 0.41 | 0.35 | 0.22 | 0.22 | 0.16 |
| 1 | 46990044 | rs12408243 | 2.409 | -1.615 | 0.525 | -0.065 | -0.09 | 0.234 | 0.234 | 0.534 | 0.4 | 0.12 | 0.4 | 0.01 | 0.46 | 0.27 |
| 1 | 46990457 | rs1267303 | 1.36 | -1.297 | 0.766 | 0.552 | -0.01 | 0.054 | 0.054 | 0 | 0.72 | 0.47 | 0.57 | 0.54 | 0.73 | 0.74 |
| 1 | 46990681 | rs7542172 | 0.153 | -1.162 | 0.563 | 0.631 | -0.016 | -0.048 | -0.048 | 0 | 0.2 | 0.1 | 0.11 | 0.26 | 0.2 | 0.28 |
| 1 | 46998009 | rs7542296 | -0.78 | -1.32 | 0.821 | 0.694 | 0.002 | 0.003 | 0.003 | 0.435 | 0.21 | 0.34 | 0.31 | 0.21 | 0.21 | 0.15 |
| 1 | 47000566 | rs3737741 | -1.913 | -1.335 | 0.531 | 0.686 | -0.004 | -0.007 | -0.007 | 0 | 0.44 | 0.66 | 0.52 | 0.49 | 0.47 | 0.51 |
| 1 | 47002889 | rs952947 | 0.222 | -1.173 | 0.568 | 0.415 | -0.018 | -0.075 | -0.075 | 0.013 | 0.75 | 0.82 | 0.84 | 0.47 | 0.72 | 0.53 |

| | | | | | | | | | | | | | | | | |
|---|---|---|---|---|---|---|---|---|---|---|---|---|---|---|---|---|
| 1 | 47007844 | rs2476163 | -1.759 | -1.228 | 0.681 | 0.087 | -0.003 | 0.013 | 0.013 | 0 | 0.46 | 0.59 | 0.55 | 0.68 | 0.5 | 0.53 |
| 1 | 47010027 | rs1258033 | -0.587 | -1.074 | 0.324 | 0.271 | 0.001 | 0.02 | 0.02 | 0 | 0.27 | 0.37 | 0.33 | 0.4 | 0.29 | 0.4 |
| 1 | 47011762 | rs885125 | -1.864 | -1.307 | 0.275 | 0.124 | 0.001 | 0.034 | 0.034 | 0 | 0.47 | 0.64 | 0.51 | 0.67 | 0.48 | 0.52 |
| 1 | 47012144 | rs885127 | 0.935 | -1.488 | -0.003 | -0.495 | -0.004 | -0.004 | -0.004 | 0.821 | 0.65 | 0.4 | 0.69 | 0.3 | 0.68 | 0.48 |
| 1 | 47013180 | rs10890404 | 0.87 | -1.493 | 0.033 | -0.46 | -0.011 | -0.004 | -0.004 | 0 | 0.67 | 0.4 | 0.72 | 0.3 | 0.73 | 0.49 |
| 1 | 47023531 | rs8602 | -0.833 | -1.325 | -0.608 | -0.242 | -0.015 | -0.035 | -0.035 | 0.005 | 0.24 | 0.1 | 0.34 | 0.28 | 0.28 | 0.21 |
| 1 | 47028115 | rs6673787 | 1.571 | -1.307 | -0.537 | -0.203 | -0.007 | -0.022 | -0.022 | 0 | 0.05 | 0.14 | 0.02 | 0 | 0.03 | 0.01 |
| 1 | 47031340 | rs7543083 | 1.761 | -1.578 | -1.002 | -0.33 | 0.055 | 0.017 | 0.017 | 0.776 | 0.55 | 0.09 | 0.52 | 0.67 | 0.66 | 0.68 |
| 1 | 47035643 | rs1054743 | 0.442 | -1.523 | -0.961 | -0.3 | -0.02 | -0.078 | -0.078 | 1.273 | 0.13 | 0.01 | 0.19 | 0.22 | 0.11 | 0.11 |
| 1 | 47042612 | rs2297850 | 2.353 | -1.408 | -0.387 | -0.378 | -0.198 | 0.111 | 0.111 | 0.653 | 0.34 | 0.04 | 0.39 | 0.12 | 0.48 | 0.24 |
| 1 | 47047441 | rs3766236 | -1.117 | -1.453 | 0.171 | -0.739 | 0.008 | 0.007 | 0.007 | 0.054 | 0.37 | 0.34 | 0.38 | 0.27 | 0.3 | 0.3 |
| 1 | 47049747 | rs3766235 | 0.417 | -1.447 | 0.211 | -0.749 | -0.002 | -0.003 | -0.003 | 0.416 | 0.05 | 0.09 | 0.06 | 0.03 | 0.03 | 0.08 |
| 1 | 47056347 | rs3766231 | -0.31 | -1.309 | 0.47 | -0.513 | 0.015 | 0.011 | 0.011 | 0.06 | 0.31 | 0.21 | 0.32 | 0.24 | 0.27 | 0.22 |
| 1 | 47062900 | rs6429606 | -0.583 | -1.284 | 0.671 | -0.515 | -0.008 | -0.007 | -0.007 | 0 | 0.33 | 0.35 | 0.33 | 0.24 | 0.27 | 0.22 |
| 1 | 47063085 | rs7556481 | -0.351 | -1.313 | 0.673 | -0.484 | 0.009 | 0.007 | 0.007 | 0.028 | 0.31 | 0.24 | 0.32 | 0.24 | 0.27 | 0.22 |
| 1 | 47066383 | rs11211319 | -1.622 | -1.271 | 0.753 | -0.668 | -0.006 | 0.045 | 0.045 | 0.054 | 0.54 | 0.35 | 0.5 | 0.93 | 0.5 | 0.71 |
| 1 | 47068584 | rs12022855 | -1.662 | -1.266 | 0.776 | -0.666 | -0.002 | -0.02 | -0.02 | 0 | 0.55 | 0.43 | 0.51 | 0.93 | 0.5 | 0.71 |
| 1 | 47076498 | rs10890408 | -1.595 | -1.197 | 0.291 | -0.633 | -0.001 | -0.05 | -0.05 | 0 | 0.56 | 0.47 | 0.52 | 0.93 | 0.5 | 0.71 |
| 1 | 47080665 | rs11211328 | -0.307 | -1.555 | 0.203 | -0.545 | 0.008 | 0.005 | 0.005 | 0.07 | 0.31 | 0.24 | 0.33 | 0.25 | 0.27 | 0.22 |
| 1 | 47080679 | rs6671527 | -1.602 | -1.353 | 0.227 | -1.152 | 0.002 | -0.075 | -0.075 | 0 | 0.57 | 0.56 | 0.52 | 0.93 | 0.48 | 0.71 |
| 1 | 47087968 | rs6669160 | 0.698 | -1.107 | 0.337 | -0.441 | 0.049 | 0.088 | 0.088 | 0.949 | 0.6 | 0.27 | 0.55 | 0.13 | 0.7 | 0.62 |
| 1 | 47089617 | rs6675720 | 0.698 | -1.107 | 0.337 | -0.441 | 0.049 | 0.088 | 0.088 | 0.949 | 0.6 | 0.27 | 0.55 | 0.13 | 0.7 | 0.62 |
| 1 | 47099346 | rs723333 | 1.425 | -1.019 | 1.908 | 0.554 | 0.044 | 0.574 | 0.574 | 5.456 | 0.93 | 0.53 | 0.93 | 0.57 | 0.98 | 0.85 |
| 1 | 47101015 | rs724309 | 1.398 | -1.016 | 1.909 | 0.576 | 0.047 | 0.591 | 0.591 | 5.192 | 0.93 | 0.54 | 0.93 | 0.53 | 0.98 | 0.85 |
| 1 | 47105000 | rs1258061 | -0.306 | -1.044 | 1.891 | 0.263 | -0.006 | 0.013 | 0.013 | 0 | 0.18 | 0.15 | 0.12 | 0.07 | 0.22 | 0.29 |
| 1 | 47109089 | rs1745377 | 0.669 | -1.072 | 1.89 | 0.139 | 0.033 | 0.032 | 0.032 | 0.591 | 0.59 | 0.35 | 0.53 | 0.2 | 0.69 | 0.59 |
| 1 | 47121502 | rs631368 | -0.76 | -1.066 | 2.032 | 0.145 | 0.058 | 0.574 | 0.574 | 2.323 | 0.06 | 0.4 | 0.06 | 0.42 | 0.01 | 0.15 |
| 1 | 47138819 | rs614486 | -0.866 | -1.072 | 2.054 | 0.134 | 0.024 | 0.024 | 0.024 | 0.152 | 0.4 | 0.65 | 0.46 | 0.8 | 0.31 | 0.41 |
| 1 | 47139103 | rs1025806 | -0.26 | -1.078 | 2.07 | 0.144 | 0.02 | 0.016 | 0.016 | 0.018 | 0.34 | 0.18 | 0.35 | 0.34 | 0.3 | 0.22 |
| 1 | 47141718 | rs12794 | -0.221 | -1.073 | 2.028 | 0.14 | -0.006 | 0.013 | 0.013 | 0 | 0.83 | 0.85 | 0.88 | 0.93 | 0.78 | 0.71 |
| 1 | 47142538 | rs1048380 | -0.22 | -1.08 | 2.114 | 0.16 | 0.022 | 0.018 | 0.018 | 0.056 | 0.34 | 0.14 | 0.35 | 0.33 | 0.3 | 0.22 |
| 1 | 47153122 | rs3766222 | -0.852 | -1.074 | 1.997 | 0.152 | 0.055 | 0.573 | 0.573 | 2.572 | 0.06 | 0.4 | 0.06 | 0.42 | 0.01 | 0.15 |
| 1 | 47154272 | rs2304745 | -0.224 | -1.073 | 1.451 | 0.135 | 0.022 | 0.015 | 0.015 | 0.056 | 0.34 | 0.13 | 0.35 | 0.34 | 0.3 | 0.22 |
| 1 | 47169210 | rs2083486 | -0.785 | -0.962 | 1.368 | 0.185 | 0.002 | 0.002 | 0.002 | 0.408 | 0.41 | 0.59 | 0.46 | 0.8 | 0.31 | 0.41 |
| 1 | 47174149 | rs10890412 | 1.726 | -0.961 | 1.288 | 0.166 | -0.083 | 0.135 | 0.135 | 0 | 0.41 | 0.13 | 0.41 | 0.14 | 0.48 | 0.3 |
| 1 | 47182206 | rs3737731 | -0.248 | -1.147 | 1.346 | 0.146 | 0.024 | 0.02 | 0.02 | 0.432 | 0.36 | 0.21 | 0.36 | 0.34 | 0.3 | 0.22 |
| 1 | 47184196 | rs3766211 | -0.779 | -1.121 | 0.997 | 0.072 | 0.005 | 0.005 | 0.005 | 0.408 | 0.41 | 0.59 | 0.46 | 0.8 | 0.31 | 0.41 |
| 1 | 47188150 | rs4660957 | 1.425 | -0.768 | 0.412 | -0.186 | -0.048 | 0.237 | 0.237 | 0 | 0.44 | 0.13 | 0.48 | 0.12 | 0.51 | 0.31 |
| 1 | 47201798 | rs723690 | -1.193 | -1.305 | 0.575 | 0.269 | 0.031 | 0.052 | 0.052 | 0 | 0.41 | 0.64 | 0.41 | 0.82 | 0.3 | 0.42 |
| 1 | 47208081 | rs1971518 | -0.751 | -0.965 | -0.276 | -0.067 | 0.014 | -0.014 | -0.014 | 0.578 | 0.35 | 0.24 | 0.28 | 0.37 | 0.29 | 0.22 |
| 1 | 47213386 | rs594387 | 0.796 | -1.33 | -0.252 | -1.145 | 0.021 | 0.017 | 0.017 | 0 | 0.59 | 0.35 | 0.61 | 0.11 | 0.78 | 0.43 |
| 1 | 47214785 | rs686053 | -0.179 | -1.374 | -0.014 | -1.128 | -0.038 | -0.027 | -0.027 | 0 | 0.78 | 0.69 | 0.71 | 0.15 | 0.86 | 0.62 |
| 1 | 47219823 | rs720413 | -1.628 | -0.037 | 0.032 | -0.778 | -0.003 | 0.018 | 0.018 | 1.332 | 0.41 | 0.67 | 0.28 | 0.15 | 0.4 | 0.38 |
| 1 | 47220059 | rs6662321 | -1.622 | -0.624 | 0.516 | -0.299 | -0.028 | -0.066 | -0.066 | 0 | 0.12 | 0.3 | 0.06 | 0.01 | 0.07 | 0.16 |
| 1 | 47234335 | rs11583588 | -1.537 | -0.359 | -0.694 | -0.245 | 0.012 | 0.101 | 0.101 | 0.272 | 0.16 | 0.02 | 0.06 | 0.01 | 0.09 | 0.16 |
| 1 | 47259138 | rs863915 | 1.569 | -0.421 | 1.01 | -0.213 | 0.012 | -0.043 | -0.043 | 0.393 | 0.9 | 0.64 | 0.92 | 0.98 | 0.95 | 0.9 |
| 1 | 47263223 | rs6677350 | -1.504 | -0.688 | 1.328 | -0.455 | -0.013 | -0.03 | -0.03 | 0 | 0.07 | 0.07 | 0.01 | 0 | 0.02 | 0.06 |
| 1 | 47263434 | rs837403 | 1.51 | -0.706 | 1.336 | -0.465 | 0.01 | -0.045 | -0.045 | 2.49 | 0.9 | 0.62 | 0.92 | 0.97 | 0.95 | 0.9 |
| 1 | 47264133 | rs632645 | 2.494 | -0.896 | 1.169 | 0.045 | -0.026 | -0.019 | -0.019 | 0 | 0.69 | 0.45 | 0.8 | 0.69 | 0.8 | 0.65 |
| 1 | 47264495 | rs632133 | 1.297 | -0.899 | 1.166 | 0.044 | 0.006 | -0.049 | -0.049 | 0 | 0.9 | 0.61 | 0.92 | 0.97 | 0.95 | 0.9 |
| 1 | 47264724 | rs2297813 | -0.149 | -0.804 | 1.055 | 0.762 | -0.003 | 0.005 | 0.005 | 0 | 0.06 | 0.03 | 0.15 | 0.17 | 0.08 | 0.05 |
| 1 | 47265273 | rs3766209 | -0.071 | -0.81 | 1.05 | 0.759 | -0.023 | -0.026 | -0.026 | 0 | 0.05 | 0.01 | 0.15 | 0.17 | 0.08 | 0.05 |
| 1 | 47265665 | rs837402 | 1.104 | -0.815 | 1.061 | 0.887 | 0.006 | -0.049 | -0.049 | 0.01 | 0.9 | 0.61 | 0.92 | 0.98 | 0.95 | 0.9 |
| 1 | 47265677 | rs837401 | 1.104 | -0.815 | 1.061 | 0.887 | 0.006 | -0.049 | -0.049 | 1.297 | 0.9 | 0.61 | 0.92 | 0.97 | 0.95 | 0.9 |
| 1 | 47265776 | rs837400 | 3.032 | -0.549 | 1.236 | 1.305 | 0.011 | -0.011 | -0.011 | 0 | 0.6 | 0.3 | 0.61 | 0.44 | 0.65 | 0.6 |
| 1 | 47266422 | rs837398 | 2.258 | -0.486 | 1.227 | 1.33 | -0.023 | -0.017 | -0.017 | 0 | 0.69 | 0.45 | 0.8 | 0.69 | 0.79 | 0.65 |
| 1 | 47266590 | rs12142787 | -0.184 | -0.451 | 1.211 | 1.36 | -0.014 | 0.012 | 0.012 | 0 | 0.06 | 0.01 | 0.15 | 0.26 | 0.08 | 0.06 |
| 1 | 47268041 | rs681840 | 1.531 | -0.503 | 1.797 | 1.289 | -0.003 | -0.011 | -0.011 | 0 | 0.79 | 0.65 | 0.81 | 0.75 | 0.81 | 0.77 |
| 1 | 47268526 | rs3766208 | -1.179 | -0.51 | 1.79 | 1.264 | -0.015 | -0.023 | -0.023 | 0 | 0.1 | 0.04 | 0.18 | 0.26 | 0.13 | 0.11 |
| 1 | 47269292 | rs751027 | 1.469 | -0.523 | 1.801 | 1.298 | -0.003 | -0.022 | -0.022 | 1.259 | 0.79 | 0.63 | 0.81 | 0.76 | 0.81 | 0.77 |
| 1 | 47271538 | rs4646481 | -1.177 | -0.487 | 1.886 | 1.302 | -0.016 | -0.023 | -0.023 | 0 | 0.1 | 0.04 | 0.18 | 0.26 | 0.13 | 0.11 |
| 1 | 47272185 | rs2405335 | -2.79 | -0.497 | 1.907 | 1.282 | -0.018 | -0.052 | -0.052 | 0.016 | 0.28 | 0.44 | 0.19 | 0.24 | 0.22 | 0.28 |
| 1 | 47272785 | rs1890250 | -1.014 | -0.485 | 1.862 | 1.289 | -0.025 | 0.001 | 0.001 | 0 | 0.16 | 0.07 | 0.14 | 0.21 | 0.19 | 0.18 |
| 1 | 47275241 | rs2065996 | -1.393 | -0.525 | 2.177 | 1.269 | 0.001 | -0.001 | -0.001 | 0.378 | 0.15 | 0.21 | 0.15 | 0.21 | 0.18 | 0.18 |
| 1 | 47275770 | rs10890422 | -4.047 | -0.644 | 2.499 | 1.156 | 0.008 | -0.029 | -0.029 | 0 | 0.44 | 0.85 | 0.43 | 0.56 | 0.38 | 0.43 |
| 1 | 47277217 | rs4646483 | -1.461 | -0.541 | 2.55 | 1.245 | -0.007 | -0.008 | -0.008 | 0 | 0.11 | 0.08 | 0.19 | 0.26 | 0.13 | 0.11 |
| 1 | 47278662 | rs4646484 | -2.389 | -0.548 | 2.59 | 1.241 | 0.004 | 0.152 | 0.152 | 0 | 0.16 | 0.45 | 0.23 | 0.29 | 0.13 | 0.16 |
| 1 | 47278997 | rs4646486 | -0.285 | -0.531 | 2.508 | 1.233 | 0.004 | 0.005 | 0.005 | 0 | 0.1 | 0.13 | 0.11 | 0.21 | 0.12 | 0.17 |
| 1 | 47279175 | rs4646487 | -1.622 | -0.542 | 2.524 | 1.225 | -0.002 | -0.007 | -0.007 | 0 | 0.17 | 0.2 | 0.11 | 0.21 | 0.14 | 0.23 |

| | | | | | | | | | | | | | | | | |
|---|---|---|---|---|---|---|---|---|---|---|---|---|---|---|---|---|
| 1 | 47280550 | rs11576162 | -0.758 | -0.538 | 2.468 | 1.239 | -0.012 | -0.017 | -0.017 | 0 | 0.12 | 0.05 | 0.1 | 0.22 | 0.12 | 0.19 |
| 1 | 47280859 | rs2297810 | -2.488 | -0.602 | 2.677 | 1.224 | -0.001 | 0.142 | 0.142 | 0 | 0.17 | 0.46 | 0.23 | 0.28 | 0.13 | 0.17 |
| 1 | 47280884 | rs4646491 | -1.303 | -0.603 | 2.68 | 1.224 | -0.014 | -0.03 | -0.03 | 0.001 | 0.11 | 0.04 | 0.18 | 0.26 | 0.13 | 0.11 |
| 1 | 47281183 | rs4646492 | -0.735 | -0.603 | 2.68 | 1.22 | -0.012 | -0.017 | -0.017 | 0 | 0.12 | 0.05 | 0.1 | 0.22 | 0.12 | 0.19 |
| 1 | 47281302 | rs4646493 | -2.066 | -0.607 | 2.713 | 1.226 | -0.018 | -0.024 | -0.024 | 0 | 0.21 | 0.35 | 0.13 | 0.22 | 0.14 | 0.24 |
| 1 | 47281780 | rs3766198 | -3.614 | -0.605 | 2.691 | 1.221 | 0.006 | -0.013 | -0.013 | 0 | 0.36 | 0.67 | 0.35 | 0.5 | 0.27 | 0.42 |
| 1 | 47282772 | rs2297809 | -1.438 | -0.598 | 2.673 | 1.229 | 0.004 | 0.047 | 0.047 | 0 | 0.11 | 0.13 | 0.19 | 0.25 | 0.13 | 0.11 |
| 1 | 47284526 | rs3766196 | -0.26 | -0.599 | 2.663 | 1.223 | -0.016 | -0.019 | -0.019 | 0.988 | 0.1 | 0.05 | 0.1 | 0.22 | 0.12 | 0.19 |
| 1 | 47284923 | rs12059860 | -1.265 | -0.605 | 2.674 | 1.219 | -0.02 | -0.038 | -0.038 | 0.071 | 0.08 | 0.12 | 0.01 | 0 | 0.02 | 0.06 |
| 1 | 47285150 | rs3766194 | -1.37 | -0.603 | 2.647 | 1.225 | -0.014 | -0.032 | -0.032 | 1.595 | 0.11 | 0.04 | 0.18 | 0.26 | 0.13 | 0.11 |
| 1 | 47286528 | rs11583378 | -0.258 | -0.603 | 2.685 | 1.224 | -0.013 | -0.015 | -0.015 | 0.02 | 0.1 | 0.05 | 0.1 | 0.22 | 0.12 | 0.19 |
| 1 | 47288951 | rs7535312 | -1.377 | -0.59 | 2.662 | 1.229 | -0.014 | -0.032 | -0.032 | 0 | 0.11 | 0.04 | 0.18 | 0.26 | 0.13 | 0.11 |
| 1 | 47289636 | rs6429613 | -3.354 | -0.604 | 2.66 | 1.196 | 0.002 | -0.01 | -0.01 | 0 | 0.34 | 0.67 | 0.34 | 0.5 | 0.27 | 0.39 |
| 1 | 47289913 | rs13373822 | -1.362 | -0.617 | 2.712 | 1.158 | 0.003 | -0.045 | -0.045 | 0 | 0.09 | 0.17 | 0.07 | 0 | 0.1 | 0.03 |
| 1 | 47309265 | rs10749865 | -3.278 | -0.633 | 2.546 | 1.282 | -0.008 | 0.032 | 0.032 | 0 | 0.33 | 0.54 | 0.33 | 0.55 | 0.28 | 0.39 |
| 1 | 47326226 | rs6679068 | -3.519 | -0.567 | 2.539 | 1.247 | 0.004 | -0.007 | -0.007 | 0 | 0.35 | 0.68 | 0.34 | 0.47 | 0.28 | 0.38 |
| 1 | 47329647 | rs7528956 | -2.823 | -0.673 | 2.416 | 1.326 | -0.007 | 0.025 | 0.025 | 0 | 0.31 | 0.4 | 0.31 | 0.41 | 0.27 | 0.37 |
| 1 | 47339113 | rs2211012 | -2.707 | -0.655 | 2.465 | 1.481 | -0.014 | 0.065 | 0.065 | 0 | 0.28 | 0.39 | 0.28 | 0.5 | 0.22 | 0.34 |
| 1 | 47374039 | rs6687264 | -1.935 | -0.629 | 2.634 | 1.618 | -0.008 | -0.001 | -0.001 | 0.061 | 0.19 | 0.21 | 0.12 | 0.26 | 0.13 | 0.26 |
| 1 | 47392054 | rs11211402 | 2.323 | -0.644 | 2.676 | 1.612 | -0.045 | 0.012 | 0.012 | 1.156 | 0.76 | 0.42 | 0.84 | 0.69 | 0.86 | 0.74 |
| 1 | 47392596 | rs11587251 | -0.905 | -0.633 | 2.779 | 1.622 | 0.001 | 0.001 | 0.001 | 0 | 0.12 | 0.11 | 0.06 | 0.09 | 0.08 | 0.16 |
| 1 | 47394096 | rs4491016 | -2.029 | -0.619 | 2.741 | 1.615 | -0.008 | -0.001 | -0.001 | 0 | 0.19 | 0.21 | 0.12 | 0.26 | 0.13 | 0.26 |
| 1 | 47396903 | rs9333025 | -0.727 | -0.628 | 2.683 | 1.64 | 0.001 | 0.03 | 0.03 | 0 | 0.07 | 0.08 | 0.15 | 0.19 | 0.09 | 0.07 |
| 1 | 47397089 | rs4660979 | -2.021 | -0.625 | 2.699 | 1.642 | -0.008 | -0.001 | -0.001 | 0 | 0.19 | 0.21 | 0.12 | 0.26 | 0.13 | 0.26 |
| 1 | 47399034 | rs9333016 | -0.665 | -0.651 | 2.644 | 1.633 | 0 | 0.031 | 0.031 | 0 | 0.07 | 0.08 | 0.15 | 0.19 | 0.08 | 0.07 |
| 1 | 47399497 | rs4660980 | -2.139 | -0.649 | 2.63 | 1.631 | -0.005 | -0.002 | -0.002 | 0.513 | 0.2 | 0.21 | 0.13 | 0.26 | 0.14 | 0.26 |
| 1 | 47404186 | rs9332998 | -2.055 | -0.597 | 2.267 | 1.756 | -0.005 | 0 | 0 | 0 | 0.19 | 0.21 | 0.13 | 0.26 | 0.14 | 0.26 |
| 1 | 47406178 | rs9332982 | -0.58 | -0.522 | 2.011 | 1.751 | 0 | 0.024 | 0.024 | 0 | 0.07 | 0.07 | 0.15 | 0.19 | 0.09 | 0.07 |
| 1 | 47411821 | rs12022394 | -0.408 | -0.518 | 1.846 | 1.753 | -0.025 | -0.029 | -0.029 | 0 | 0.06 | 0.01 | 0.15 | 0.19 | 0.09 | 0.07 |
| 1 | 47413786 | rs11211406 | 2.523 | -0.688 | 2.082 | 1.716 | -0.011 | 0.018 | 0.018 | 0.359 | 0.68 | 0.35 | 0.68 | 0.49 | 0.77 | 0.67 |
| 1 | 47416441 | rs10789496 | 1.526 | -0.554 | 2.176 | 1.824 | 0.007 | 0.232 | 0.232 | 0.974 | 0.89 | 0.55 | 0.81 | 0.75 | 0.91 | 0.92 |
| 1 | 47441156 | rs6692958 | -1.781 | -0.47 | 1.497 | 1.587 | -0.006 | -0.033 | -0.033 | 0 | 0.25 | 0.16 | 0.31 | 0.51 | 0.27 | 0.3 |
| 1 | 47449065 | rs11211414 | -1.866 | -0.479 | 1.458 | 1.676 | 0.001 | -0.028 | -0.028 | 1.761 | 0.25 | 0.16 | 0.26 | 0.39 | 0.25 | 0.28 |
| 1 | 47449486 | rs6680773 | -1.662 | -0.446 | 1.509 | 1.647 | -0.005 | -0.016 | -0.016 | 0.179 | 0.23 | 0.18 | 0.31 | 0.52 | 0.27 | 0.33 |
| 1 | 47467796 | rs10890438 | -2.335 | -0.586 | 1.629 | 1.576 | 0.005 | -0.005 | -0.005 | 0 | 0.34 | 0.68 | 0.35 | 0.57 | 0.27 | 0.37 |
| 1 | 47481616 | rs9727862 | 0.393 | -0.673 | 1.127 | 1.368 | -0.009 | 0.05 | 0.05 | 0 | 0.82 | 0.7 | 0.86 | 0.69 | 0.89 | 0.64 |
| 1 | 47481640 | rs9726786 | -2.261 | -0.489 | 1.377 | 1.666 | 0.012 | 0.008 | 0.008 | 0.281 | 0.36 | 0.68 | 0.37 | 0.67 | 0.29 | 0.4 |
| 1 | 47487081 | rs12403250 | -0.847 | -0.43 | 1.4 | 1.491 | -0.001 | 0.01 | 0.01 | 0 | 0.05 | 0.05 | 0.08 | 0.16 | 0.04 | 0.04 |
| 1 | 47489786 | rs2036462 | -1.686 | 0.002 | 1.636 | 1.316 | 0.005 | -0.006 | -0.006 | 0 | 0.3 | 0.39 | 0.37 | 0.69 | 0.3 | 0.54 |
| 1 | 47490974 | rs1553391 | -2.381 | -0.019 | 1.672 | 1.278 | 0.001 | -0.063 | -0.063 | 2.316 | 0.38 | 0.39 | 0.46 | 0.93 | 0.37 | 0.64 |
| 1 | 47496738 | rs9793989 | -1.927 | 0.078 | 1.72 | 1.685 | 0 | 0.019 | 0.019 | 0 | 0.09 | 0.11 | 0.11 | 0.24 | 0.07 | 0.1 |
| 1 | 47504717 | rs2292059 | -1.474 | 0.083 | 1.543 | 1.69 | -0.016 | -0.042 | -0.042 | 0 | 0.08 | 0.03 | 0.1 | 0.24 | 0.07 | 0.1 |
| 1 | 47509513 | rs9699826 | -1.948 | 0.077 | 1.65 | 1.674 | -0.002 | 0.025 | 0.025 | 0 | 0.09 | 0.14 | 0.11 | 0.23 | 0.07 | 0.12 |
| 1 | 47548974 | rs12026359 | -1.31 | 0.068 | 1.748 | 1.653 | -0.013 | -0.019 | -0.019 | 0 | 0.2 | 0.12 | 0.32 | 0.69 | 0.24 | 0.5 |
| 1 | 47551273 | rs11211436 | -1.322 | 0.082 | 1.772 | 1.655 | -0.007 | -0.013 | -0.013 | 0 | 0.21 | 0.15 | 0.32 | 0.69 | 0.24 | 0.5 |
| 1 | 47565477 | rs11582907 | -1.829 | 0.083 | 1.798 | 1.675 | -0.001 | 0.023 | 0.023 | 0 | 0.09 | 0.11 | 0.11 | 0.24 | 0.07 | 0.12 |
| 1 | 47572502 | rs1847225 | -1.42 | 0.145 | 1.363 | 1.726 | -0.009 | -0.013 | -0.013 | 0 | 0.21 | 0.16 | 0.33 | 0.65 | 0.25 | 0.5 |
| 1 | 47587253 | rs12040022 | -1.367 | 0.401 | 0.704 | 1.849 | -0.011 | -0.015 | -0.015 | 0.113 | 0.21 | 0.15 | 0.33 | 0.65 | 0.25 | 0.5 |
| 1 | 47594935 | rs12564103 | -1.753 | -0.062 | 0.708 | 2.124 | -0.008 | -0.052 | -0.052 | 0 | 0.09 | 0.01 | 0.1 | 0.29 | 0.07 | 0.12 |
| 1 | 47608197 | rs2056898 | -2.391 | 0.028 | 0.407 | 2.553 | 0.001 | -0.007 | -0.007 | 0 | 0.27 | 0.21 | 0.24 | 0.55 | 0.2 | 0.53 |
| 1 | 47620126 | rs1832992 | -1.625 | -0.035 | 0.284 | 2.628 | -0.015 | -0.062 | -0.062 | 0 | 0.1 | 0.01 | 0.1 | 0.31 | 0.08 | 0.26 |
| 1 | 47627246 | rs3737772 | -1.089 | 0.06 | 0.545 | 2.441 | -0.002 | 0.039 | 0.039 | 0 | 0.15 | 0.17 | 0.18 | 0.31 | 0.12 | 0.28 |
| 1 | 47637999 | rs12034515 | -0.83 | 0.017 | 0.499 | 2.39 | -0.001 | 0.004 | 0.004 | 0 | 0.14 | 0.12 | 0.13 | 0.25 | 0.11 | 0.27 |
| 1 | 47639520 | rs6695624 | -1.807 | 0 | 0.544 | 2.398 | -0.01 | 0.076 | 0.076 | 0 | 0.21 | 0.3 | 0.22 | 0.42 | 0.16 | 0.45 |
| 1 | 47641041 | rs2798353 | -1.241 | 0.272 | -0.049 | 1.779 | -0.011 | 0.051 | 0.051 | 0 | 0.19 | 0.22 | 0.18 | 0.34 | 0.13 | 0.32 |
| 1 | 47645995 | rs2245122 | 0.312 | 0.686 | -0.732 | 1.208 | -0.01 | -0.004 | -0.004 | 0 | 0.58 | 0.64 | 0.58 | 0.63 | 0.59 | 0.51 |
| 1 | 47654724 | rs3737770 | -0.282 | 0.622 | -1.079 | 2.071 | -0.024 | -0.051 | -0.051 | 1.595 | 0.08 | 0 | 0.03 | 0.27 | 0.07 | 0.17 |
| 1 | 47655816 | rs2248907 | 0.234 | 1.116 | -1.668 | 2.142 | -0.059 | -0.063 | -0.063 | 0 | 0.87 | 0.98 | 0.76 | 0.8 | 0.82 | 0.9 |
| 1 | 47657141 | rs12184246 | 1.774 | 0.193 | -1.583 | 1.719 | 0.017 | 0.108 | 0.108 | 0.34 | 0.6 | 0.19 | 0.55 | 0.21 | 0.69 | 0.49 |
| 1 | 47661564 | rs4926718 | 0.766 | 0.278 | -1.488 | 2.079 | -0.017 | -0.059 | -0.059 | 0.002 | 0.82 | 0.91 | 0.65 | 0.59 | 0.78 | 0.73 |
| 1 | 47661620 | rs7531972 | -1.141 | 0.167 | -0.279 | 2.063 | -0.027 | 0.111 | 0.111 | 0 | 0.2 | 0.46 | 0.1 | 0.35 | 0.11 | 0.25 |
| 1 | 47666146 | rs17103015 | -0.276 | 0.122 | 0.253 | 1.771 | -0.015 | -0.052 | -0.052 | 0 | 0.09 | 0.22 | 0.04 | 0 | 0.04 | 0.07 |
| 1 | 47669325 | rs12760654 | -0.525 | 0.487 | -0.362 | 1.588 | -0.015 | -0.006 | -0.006 | 0 | 0.17 | 0.12 | 0.18 | 0.21 | 0.2 | 0.13 |
| 1 | 47673296 | rs13373881 | -0.539 | 0.529 | -0.399 | 1.377 | -0.001 | -0.014 | -0.014 | 0 | 0.07 | 0.07 | 0.05 | 0 | 0.07 | 0.03 |
| 1 | 47673802 | rs1008455 | 0.537 | 0.545 | -0.365 | 1.934 | 0.008 | 0.085 | 0.085 | 0 | 0.95 | 0.93 | 0.69 | 0.76 | 0.89 | 0.93 |
| 1 | 47683464 | rs2070930 | 0.642 | 0.533 | -0.401 | 1.948 | -0.026 | -0.049 | -0.049 | 0 | 0.08 | 0 | 0.04 | 0.33 | 0.08 | 0.2 |
| 1 | 47686533 | rs2249636 | -1.526 | 0.411 | -0.049 | 1.374 | 0.001 | -0.004 | -0.004 | 0 | 0.33 | 0.65 | 0.27 | 0.7 | 0.29 | 0.48 |
| 1 | 47710154 | rs2758738 | 0.714 | -0.795 | 0.115 | 0.597 | -0.074 | -0.125 | -0.125 | 0.691 | 0.19 | 0.54 | 0.09 | 0 | 0.1 | 0.09 |
| 1 | 47726087 | rs13376679 | 0.074 | -0.199 | 0.301 | -0.192 | -0.018 | 0.068 | 0.068 | 0.432 | 0.32 | 0.18 | 0.22 | 0.03 | 0.29 | 0.18 |

| | | | | | | | | | | | | | | | | |
|---|---|---|---|---|---|---|---|---|---|---|---|---|---|---|---|---|
| 1 | 47745506 | rs7532148 | 0.212 | 0.087 | 0.058 | 0.674 | -0.01 | 0.007 | 0.007 | 0.023 | 0.12 | 0.12 | 0.32 | 0.54 | 0.16 | 0.26 |
| 1 | 47747445 | rs3122619 | -1.127 | -0.1 | 0.052 | 0.543 | -0.035 | 0.014 | 0.014 | 0 | 0.49 | 0.88 | 0.59 | 0.57 | 0.46 | 0.44 |
| 1 | 47766943 | rs6697248 | -0.473 | -0.121 | 0.245 | 0.56 | 0.015 | 0.106 | 0.106 | 0.626 | 0.15 | 0.3 | 0.34 | 0.54 | 0.16 | 0.26 |
| 1 | 47767914 | rs3125630 | -1.112 | -0.08 | 0.23 | 0.549 | -0.03 | 0.014 | 0.014 | 0 | 0.49 | 0.88 | 0.59 | 0.57 | 0.46 | 0.44 |
| 1 | 47806087 | rs12123269 | 0.23 | -0.363 | 0.36 | 0.577 | -0.005 | -0.008 | -0.008 | 0 | 0.32 | 0.25 | 0.22 | 0.03 | 0.29 | 0.19 |
| 1 | 47842777 | rs1044457 | 0.862 | -0.272 | -0.811 | 0.873 | -0.117 | 0.235 | 0.235 | 0.208 | 0.48 | 0.03 | 0.4 | 0.22 | 0.54 | 0.45 |
| 1 | 47844017 | rs3088062 | 0.29 | -0.269 | -0.79 | 0.943 | -0.008 | 0.013 | 0.013 | 0.113 | 0.11 | 0.12 | 0.31 | 0.55 | 0.15 | 0.25 |
| 1 | 47844464 | rs6795 | 0.736 | -0.306 | -0.944 | 0.908 | -0.112 | 0.232 | 0.232 | 0.233 | 0.48 | 0.04 | 0.4 | 0.22 | 0.54 | 0.45 |
| 1 | 47868005 | rs11579058 | -0.664 | -0.189 | 0.51 | 1.27 | -0.007 | 0.015 | 0.015 | 0.369 | 0.14 | 0.19 | 0.06 | 0.26 | 0.08 | 0.24 |
| 1 | 47869316 | rs7555040 | 1.518 | -0.252 | 0.017 | 0.959 | -0.025 | -0.015 | -0.015 | 1.314 | 0.11 | 0.01 | 0.14 | 0.11 | 0.15 | 0.06 |
| 1 | 47870271 | rs7528075 | 0.136 | 0.243 | 0.545 | 0.66 | -0.015 | 0.042 | 0.042 | 0 | 0.29 | 0.17 | 0.19 | 0.03 | 0.29 | 0.13 |
| 1 | 47873684 | rs2405922 | 1.477 | 0.465 | 0.597 | 0.99 | -0.05 | 0.04 | 0.04 | 0.555 | 0.5 | 0.07 | 0.49 | 0.45 | 0.55 | 0.56 |
| 1 | 47891361 | rs7525184 | -0.502 | 0.41 | 0.932 | 1.295 | -0.011 | 0.041 | 0.041 | 0 | 0.38 | 0.23 | 0.24 | 0.21 | 0.36 | 0.36 |
| 1 | 47893929 | rs2820974 | 1.053 | 0.402 | 1.509 | 1.267 | -0.009 | 0.023 | 0.023 | 0 | 0.54 | 0.37 | 0.52 | 0.4 | 0.56 | 0.58 |
| 1 | 47902817 | rs3738316 | 0.428 | 0.545 | 1.766 | 1.623 | -0.008 | 0.019 | 0.019 | 0 | 0.1 | 0.13 | 0.06 | 0.18 | 0.08 | 0.24 |
| 1 | 47924275 | rs497606 | -1.089 | 0.579 | 1.853 | 1.232 | -0.001 | -0.001 | -0.001 | 0 | 0.43 | 0.52 | 0.26 | 0.32 | 0.36 | 0.41 |
| 1 | 47927545 | rs6695969 | -0.058 | 0.542 | 1.409 | 1.115 | -0.014 | 0.109 | 0.109 | 0.474 | 0.12 | 0.47 | 0.09 | 0.19 | 0.08 | 0.24 |
| 1 | 47932550 | rs543640 | -0.764 | 0.546 | 1.41 | 1.163 | 0.012 | 0.022 | 0.022 | 0 | 0.37 | 0.43 | 0.39 | 0.46 | 0.35 | 0.23 |
| 1 | 47933238 | rs2098 | 0.248 | 0.567 | 0.506 | 0.88 | -0.015 | 0.061 | 0.061 | 0 | 0.3 | 0.16 | 0.17 | 0.03 | 0.28 | 0.17 |
| 1 | 47936696 | rs505399 | -1.534 | 0.546 | 0.383 | 0.891 | -0.059 | 0.102 | 0.102 | 0.179 | 0.5 | 0.91 | 0.48 | 0.65 | 0.43 | 0.47 |
| 1 | 47938129 | rs10789509 | -0.038 | 0.938 | 0.563 | 0.563 | -0.014 | 0.051 | 0.051 | 0 | 0.3 | 0.17 | 0.17 | 0.03 | 0.28 | 0.17 |
| 1 | 47943032 | rs6588397 | 0.805 | -0.299 | 1.024 | 0.932 | -0.036 | -0.011 | -0.011 | 0 | 0.42 | 0.51 | 0.31 | 0.36 | 0.21 | 0.31 |
| 1 | 47948764 | rs2065945 | 0.627 | 0.506 | 0.334 | 0.612 | -0.118 | 0.149 | 0.149 | 0 | 0.29 | 0.06 | 0.27 | 0.04 | 0.43 | 0.22 |
| 1 | 47948798 | rs2065944 | -0.229 | 0.595 | 0.848 | 0.747 | 0.057 | 0.129 | 0.129 | 0 | 0.88 | 0.78 | 0.66 | 0.41 | 0.81 | 0.7 |
| 1 | 47953757 | rs11578572 | -0.49 | 0.558 | 0.673 | 0.636 | 0.017 | 0.179 | 0.179 | 1.429 | 0.45 | 0.18 | 0.39 | 0.07 | 0.59 | 0.36 |
| 1 | 47955790 | rs515146 | 0.653 | 1.292 | 1.556 | 1.899 | -0.006 | -0.006 | -0.006 | 0.012 | 0.82 | 0.87 | 0.67 | 0.62 | 0.78 | 0.75 |
| 1 | 47960253 | rs12134032 | -0.182 | 1.276 | 1.523 | 2.063 | -0.002 | -0.005 | -0.005 | 0 | 0.09 | 0.05 | 0.08 | 0.24 | 0.1 | 0.2 |
| 1 | 47960631 | rs12123884 | -0.306 | 1.304 | 1.581 | 2.085 | 0 | 0.029 | 0.029 | 0 | 0.09 | 0.07 | 0.12 | 0.26 | 0.1 | 0.21 |
| 1 | 47981663 | rs822886 | -0.583 | 0.38 | 1.346 | 1.114 | 0 | -0.022 | -0.022 | 0 | 0.17 | 0.2 | 0.16 | 0.06 | 0.17 | 0.16 |
| 1 | 47986452 | rs2506985 | -0.207 | 0.399 | 1.288 | 1.275 | 0.008 | 0.136 | 0.136 | 0 | 0.09 | 0.21 | 0.1 | 0.32 | 0.1 | 0.23 |
| 1 | 47990749 | rs1024108 | -0.69 | 0.404 | 0.498 | 1.267 | -0.003 | -0.08 | -0.08 | 2.576 | 0.2 | 0.43 | 0.18 | 0.06 | 0.17 | 0.16 |
| 1 | 47993332 | rs606852 | -1.154 | 0.429 | -0.56 | 1.186 | 0.002 | -0.015 | -0.015 | 0 | 0.3 | 0.75 | 0.29 | 0.37 | 0.27 | 0.38 |
| 1 | 48003121 | rs518063 | -0.489 | 0.076 | -1.038 | 0.812 | 0.035 | 0.068 | 0.068 | 0 | 0.47 | 0.75 | 0.61 | 0.63 | 0.44 | 0.54 |
| 1 | 48004776 | rs2485439 | -1.576 | -0.454 | 0.204 | 1.278 | 0.001 | 0.169 | 0.169 | 1.804 | 0.07 | 0.41 | 0.11 | 0.14 | 0.01 | 0.14 |
| 1 | 48005984 | rs12024835 | -0.062 | -0.574 | 0.045 | 1.153 | -0.003 | -0.004 | -0.004 | 0.12 | 0.09 | 0.11 | 0.08 | 0.06 | 0.12 | 0.05 |
| 1 | 48006898 | rs10158159 | -0.05 | -0.574 | 0.039 | 1.152 | -0.003 | -0.003 | -0.003 | 0.211 | 0.09 | 0.1 | 0.08 | 0.06 | 0.12 | 0.05 |
| 1 | 48009686 | rs518057 | -1.829 | -1.019 | 0.248 | 1.539 | -0.001 | 0 | 0 | 0.23 | 0.23 | 0.74 | 0.28 | 0.31 | 0.2 | 0.28 |
| 1 | 48011533 | rs10789511 | -1.647 | -1.382 | 0.665 | 2.078 | -0.032 | 0.137 | 0.137 | 0.107 | 0.2 | 0.49 | 0.15 | 0.42 | 0.16 | 0.25 |
| 1 | 48013721 | rs11584269 | 0.578 | -0.788 | -0.277 | 1.455 | -0.05 | 0.006 | 0.006 | 0 | 0.17 | 0.03 | 0.12 | 0.12 | 0.22 | 0.23 |
| 1 | 48015258 | rs694308 | -1.838 | -1.055 | -0.319 | 1.048 | -0.004 | -0.004 | -0.004 | 0.202 | 0.72 | 0.79 | 0.78 | 0.95 | 0.66 | 0.83 |
| 1 | 48015566 | rs12028069 | -0.498 | -1.261 | -0.294 | 1.074 | -0.027 | 0.078 | 0.078 | 0 | 0.54 | 0.21 | 0.37 | 0.3 | 0.49 | 0.56 |
| 1 | 48015862 | rs12124180 | 0.836 | -1.16 | -0.389 | 0.994 | -0.058 | -0.012 | -0.012 | 0 | 0.21 | 0.09 | 0.21 | 0.19 | 0.3 | 0.25 |
| 1 | 48016511 | rs12743540 | 0.878 | -1.158 | -0.388 | 0.96 | -0.058 | 0.012 | 0.012 | 0 | 0.21 | 0.09 | 0.21 | 0.15 | 0.3 | 0.25 |
| 1 | 48017067 | rs502392 | -1.721 | -1.153 | -0.2 | 1.117 | 0.002 | -0.002 | -0.002 | 0.049 | 0.68 | 0.71 | 0.55 | 0.74 | 0.61 | 0.7 |
| 1 | 48018359 | rs12121534 | 1.755 | -1.093 | -0.562 | 1.105 | -0.017 | -0.018 | -0.018 | 0 | 0.06 | 0.03 | 0.06 | 0.13 | 0.09 | 0.04 |
| 1 | 48019222 | rs17420782 | 1.019 | -1.205 | -0.557 | 0.991 | 0.046 | 0.089 | 0.089 | 0 | 0.25 | 0.08 | 0.14 | 0 | 0.16 | 0.16 |
| 1 | 48020922 | rs978246 | -0.467 | -0.593 | 0.008 | 1.824 | -0.012 | -0.023 | -0.023 | 0.017 | 0.27 | 0.21 | 0.24 | 0.42 | 0.31 | 0.4 |
| 1 | 48022068 | rs6660504 | 0.59 | -0.374 | -0.102 | 1.851 | -0.04 | -0.036 | -0.036 | 1.2 | 0.19 | 0.11 | 0.22 | 0.32 | 0.28 | 0.22 |
| 1 | 48025824 | rs4926901 | -0.015 | -0.386 | -0.101 | 1.895 | -0.061 | -0.086 | -0.086 | 0.216 | 0.23 | 0.13 | 0.5 | 0.46 | 0.34 | 0.4 |
| 1 | 48029727 | rs6656330 | 0.718 | -0.774 | 0.188 | 1.827 | 0 | 0.043 | 0.043 | 1.734 | 0.09 | 0.12 | 0.26 | 0.26 | 0.07 | 0.16 |
| 1 | 48030249 | rs7550183 | 1.568 | -1.162 | 0.312 | 1.423 | -0.005 | 0.136 | 0.136 | 0 | 0.47 | 0.24 | 0.35 | 0.04 | 0.51 | 0.32 |
| 1 | 48033804 | rs12131166 | 0.537 | -1.237 | 0.446 | 1.689 | 0 | -0.007 | -0.007 | 0 | 0.1 | 0.1 | 0.06 | 0.21 | 0.11 | 0.08 |
| 1 | 48034735 | rs1876599 | 1.155 | -1.293 | 0.416 | 1.762 | -0.029 | 0.058 | 0.058 | 1.232 | 0.77 | 0.67 | 0.59 | 0.44 | 0.86 | 0.62 |
| 1 | 48034778 | rs2457066 | 0.215 | -1.23 | 0.283 | 1.648 | -0.033 | 0.035 | 0.035 | 0 | 0.91 | 0.85 | 0.95 | 0.74 | 0.96 | 0.82 |
| 1 | 48035813 | rs10890490 | 1.068 | -0.998 | 0.146 | 1.531 | -0.003 | -0.003 | -0.003 | 0.25 | 0.18 | 0.14 | 0.24 | 0.25 | 0.23 | 0.22 |
| 1 | 48040577 | rs11211563 | -0.582 | -1.019 | 0.089 | 1.925 | -0.001 | 0.003 | 0.003 | 1.286 | 0.15 | 0.14 | 0.29 | 0.29 | 0.1 | 0.28 |
| 1 | 48041606 | rs2457050 | 0.786 | -0.92 | -0.06 | 1.501 | -0.021 | 0.038 | 0.038 | 0.743 | 0.73 | 0.55 | 0.62 | 0.35 | 0.79 | 0.59 |
| 1 | 48046320 | rs2507007 | -0.977 | -1.028 | -0.711 | 1.39 | 0.019 | -0.031 | -0.031 | 0 | 0.29 | 0.76 | 0.41 | 0.53 | 0.19 | 0.34 |
| 1 | 48048060 | rs1605672 | -0.405 | -1.065 | -0.638 | 0.419 | 0.001 | 0 | 0 | 0.009 | 0.4 | 0.62 | 0.32 | 0.39 | 0.45 | 0.38 |
| 1 | 48050570 | rs12094838 | 0.733 | -0.622 | -1.215 | 0.234 | -0.029 | 0.078 | 0.078 | 0.605 | 0.28 | 0.12 | 0.26 | 0.05 | 0.36 | 0.26 |
| 1 | 48052723 | rs12134806 | 0.581 | -0.762 | -1.437 | 0.091 | -0.005 | -0.032 | -0.032 | 0 | 0.39 | 0.21 | 0.58 | 0.43 | 0.38 | 0.39 |
| 1 | 48053426 | rs11576372 | 0.838 | -0.053 | -1.162 | -0.015 | -0.078 | 0.076 | 0.076 | 0 | 0.23 | 0.08 | 0.22 | 0 | 0.37 | 0.18 |
| 1 | 48060089 | rs1108866 | 0.999 | -1.056 | -0.461 | 0.77 | -0.066 | 0.125 | 0.125 | 0.221 | 0.83 | 0.32 | 0.92 | 0.59 | 0.93 | 0.75 |
| 1 | 48063350 | rs10890492 | -0.035 | -0.459 | -0.57 | 0.369 | 0.002 | 0.003 | 0.003 | 0 | 0.6 | 0.39 | 0.62 | 0.69 | 0.63 | 0.58 |
| 1 | 48065019 | rs12742768 | -1.21 | -0.606 | -0.598 | 0.42 | -0.019 | 0.071 | 0.071 | 0 | 0.14 | 0.23 | 0.14 | 0.29 | 0.1 | 0.2 |
| 1 | 48066171 | rs12058525 | -0.851 | -0.595 | -0.804 | 0.385 | -0.006 | 0.004 | 0.004 | 0 | 0.12 | 0.13 | 0.1 | 0.13 | 0.09 | 0.12 |
| 1 | 48070955 | rs12047730 | -0.512 | -0.53 | -0.776 | 0.417 | 0.003 | -0.004 | -0.004 | 0 | 0.11 | 0.11 | 0.1 | 0.13 | 0.09 | 0.12 |
| 1 | 48071156 | rs4554761 | -0.316 | -0.494 | -1.348 | -0.125 | 0.018 | 0.081 | 0.081 | 0 | 0.38 | 0.18 | 0.26 | 0.16 | 0.34 | 0.3 |
| 1 | 48081679 | rs12734920 | 0.262 | 0.029 | -1.235 | 0.182 | 0.004 | -0.006 | -0.006 | 0 | 0.09 | 0.08 | 0.09 | 0.13 | 0.07 | 0.07 |

| | | | | | | | | | | | | | | | | |
|---|---|---|---|---|---|---|---|---|---|---|---|---|---|---|---|---|
| 1 | 48081838 | rs11577726 | 1.02 | -0.015 | -1.024 | -0.178 | -0.002 | -0.069 | -0.069 | 0.175 | 0.3 | 0.33 | 0.18 | 0 | 0.27 | 0.18 |
| 1 | 48082491 | rs6660568 | -1.117 | 0.94 | -1.106 | 1.136 | -0.016 | 0.033 | 0.033 | 0 | 0.12 | 0.18 | 0.1 | 0.2 | 0.09 | 0.12 |
| 1 | 48085268 | rs4926916 | -0.502 | 1.513 | -1.85 | 0.983 | -0.064 | -0.071 | -0.071 | 2.54 | 0.24 | 0.1 | 0.49 | 0.42 | 0.36 | 0.31 |
| 1 | 48090146 | rs12403651 | -0.288 | 1.587 | -1.719 | 1.144 | -0.043 | -0.05 | -0.05 | 0 | 0.23 | 0.12 | 0.51 | 0.44 | 0.35 | 0.29 |
| 1 | 48098297 | rs17455819 | 0.255 | 0.944 | -2.143 | 0.721 | 0.003 | -0.033 | -0.033 | 0 | 0.48 | 0.1 | 0.44 | 0.4 | 0.45 | 0.35 |
| 1 | 48098406 | rs2506991 | 0.094 | 1.054 | -1.501 | 0.602 | -0.018 | -0.013 | -0.013 | 0 | 0.57 | 0.32 | 0.56 | 0.44 | 0.51 | 0.46 |
| 1 | 48100829 | rs2506987 | 0.426 | 1.12 | -1.536 | 0.618 | -0.006 | -0.026 | -0.026 | 1.138 | 0.54 | 0.21 | 0.55 | 0.49 | 0.52 | 0.43 |
| 1 | 48102121 | rs931937 | -0.252 | 1.317 | -1.401 | 0.283 | -0.05 | 0.002 | 0.002 | 0 | 0.63 | 0.79 | 0.49 | 0.66 | 0.6 | 0.71 |
| 1 | 48102145 | rs1876598 | -0.102 | 1.069 | -0.442 | -0.262 | -0.007 | -0.076 | -0.076 | 0.573 | 0.17 | 0.43 | 0.12 | 0.12 | 0.1 | 0.14 |
| 1 | 48102692 | rs2457058 | 1.079 | 1.01 | 0.132 | -0.225 | -0.017 | -0.059 | -0.059 | 0.283 | 0.33 | 0.46 | 0.22 | 0.11 | 0.34 | 0.24 |
| 1 | 48102984 | rs2457057 | 0.079 | 0.948 | 0.209 | -0.264 | 0.053 | 0.14 | 0.14 | 0 | 0.47 | 0.65 | 0.29 | 0.22 | 0.4 | 0.31 |
| 1 | 48103092 | rs10157025 | 1.071 | 1.012 | 0.339 | -0.266 | -0.021 | -0.072 | -0.072 | 0 | 0.34 | 0.44 | 0.21 | 0.1 | 0.34 | 0.24 |
| 1 | 48103170 | rs2457054 | -0.221 | 1.074 | 0.321 | 0.106 | -0.014 | 0.034 | 0.034 | 0 | 0.59 | 0.63 | 0.48 | 0.7 | 0.58 | 0.65 |
| 1 | 48105228 | rs12410488 | -0.856 | 1.137 | 0.689 | 0.569 | -0.017 | 0.031 | 0.031 | 0 | 0.08 | 0.02 | 0.05 | 0 | 0.06 | 0.05 |
| 1 | 48106092 | rs2506986 | -0.995 | 1.086 | 1.184 | 0.68 | -0.004 | -0.014 | -0.014 | 0.314 | 0.15 | 0.25 | 0.1 | 0.12 | 0.09 | 0.2 |
| 1 | 48108539 | rs6699040 | -0.648 | 1.196 | 1.051 | 0.906 | 0.008 | 0.134 | 0.134 | 0 | 0.1 | 0.11 | 0.21 | 0.48 | 0.09 | 0.26 |
| 1 | 48109225 | rs4926923 | -1.099 | 1.108 | 0.934 | 0.747 | -0.002 | -0.003 | -0.003 | 0.033 | 0.15 | 0.2 | 0.09 | 0.12 | 0.09 | 0.19 |
| 1 | 48115746 | rs12402179 | -0.23 | 1.264 | 0.683 | 0.728 | -0.005 | -0.006 | -0.006 | 0 | 0.06 | 0.05 | 0.2 | 0.18 | 0.06 | 0.06 |
| 1 | 48118188 | rs12407853 | -0.987 | 1.16 | 0.93 | 0.547 | 0 | 0 | 0 | 0 | 0.15 | 0.18 | 0.09 | 0.12 | 0.09 | 0.21 |
| 1 | 48118483 | rs4633322 | -0.746 | 1.143 | 0.905 | 0.553 | 0.008 | 0.01 | 0.01 | 0 | 0.13 | 0.1 | 0.07 | 0.11 | 0.08 | 0.18 |
| 1 | 48119989 | rs893762 | -0.295 | 1.133 | 0.865 | 0.613 | 0.003 | -0.001 | -0.001 | 0 | 0.11 | 0.07 | 0.06 | 0.12 | 0.07 | 0.11 |
| 1 | 48120087 | rs893763 | -0.593 | 1.122 | 1.153 | 0.608 | -0.005 | -0.002 | -0.002 | 0 | 0.12 | 0.16 | 0.07 | 0.13 | 0.07 | 0.14 |
| 1 | 48131369 | rs12029435 | -0.737 | 1.423 | 1.242 | 0.67 | 0.007 | 0.071 | 0.071 | 0 | 0.11 | 0.1 | 0.23 | 0.48 | 0.11 | 0.25 |
| 1 | 48133693 | rs11211587 | -0.726 | 1.149 | 1.337 | 0.584 | -0.011 | -0.005 | -0.005 | 0 | 0.13 | 0.26 | 0.08 | 0.13 | 0.07 | 0.18 |
| 1 | 48161625 | rs12031428 | -1.023 | 1.187 | 1.119 | 0.239 | 0 | 0 | 0 | 0 | 0.3 | 0.3 | 0.39 | 0.32 | 0.29 | 0.22 |
| 1 | 48168375 | rs11211590 | -0.756 | 1.446 | 1.302 | 0.489 | 0.054 | 0.239 | 0.239 | 0.088 | 0.12 | 0.23 | 0.18 | 0.49 | 0.11 | 0.29 |
| 1 | 48174867 | rs1439916 | 1.764 | 1.294 | 1.429 | 0.424 | -0.005 | -0.001 | -0.001 | 0.041 | 0.71 | 0.45 | 0.57 | 0.35 | 0.81 | 0.59 |
| 1 | 48182595 | rs12038526 | -1.255 | 0.926 | 1.113 | 0.717 | 0.018 | 0.124 | 0.124 | 0 | 0.19 | 0.25 | 0.21 | 0.49 | 0.15 | 0.32 |
| 1 | 48189253 | rs7530242 | -1.357 | 0.853 | 1.068 | 0.637 | 0.013 | 0.091 | 0.091 | 0 | 0.2 | 0.27 | 0.37 | 0.53 | 0.16 | 0.36 |
| 1 | 48192754 | rs1017475 | -0.554 | 1.103 | 0.35 | 0.027 | -0.025 | -0.02 | -0.02 | 0.165 | 0.21 | 0.16 | 0.25 | 0.31 | 0.25 | 0.18 |
| 1 | 48193205 | rs4926601 | -1.845 | 1.311 | 0.844 | 0.544 | -0.011 | 0 | 0 | 0 | 0.29 | 0.55 | 0.42 | 0.65 | 0.18 | 0.41 |
| 1 | 48196590 | rs12022200 | -1.487 | 1.958 | 0.414 | 0.862 | 0.007 | 0.006 | 0.006 | 0 | 0.26 | 0.31 | 0.26 | 0.35 | 0.24 | 0.21 |
| 1 | 48209873 | rs1439920 | -1.165 | 1.612 | 0.158 | 1.066 | 0.006 | 0.029 | 0.029 | 0 | 0.24 | 0.28 | 0.25 | 0.57 | 0.21 | 0.26 |
| 1 | 48213641 | rs2165193 | -1.316 | 1.631 | 0.319 | 1.254 | 0.043 | 0.097 | 0.097 | 0 | 0.13 | 0.25 | 0.3 | 0.34 | 0.16 | 0.14 |
| 1 | 48215379 | rs11579242 | -0.766 | 1.607 | 0.1 | 0.832 | -0.031 | 0.042 | 0.042 | 1 | 0.16 | 0.06 | 0.16 | 0.09 | 0.18 | 0.09 |
| 1 | 48216629 | rs12082556 | 1.182 | 1.621 | -0.08 | -0.43 | 0 | -0.071 | -0.071 | 0 | 0.43 | 0.44 | 0.27 | 0.03 | 0.46 | 0.39 |
| 1 | 48216886 | rs1561572 | -0.175 | 1.868 | 0.187 | 0.246 | 0.014 | 0.166 | 0.166 | 0 | 0.86 | 0.71 | 0.85 | 0.44 | 0.84 | 0.72 |
| 1 | 48216976 | rs2165194 | -1.087 | 1.74 | -0.19 | 0.347 | -0.01 | 0.016 | 0.016 | 0 | 0.4 | 0.22 | 0.5 | 0.37 | 0.38 | 0.26 |
| 1 | 48220454 | rs11584807 | 0.233 | 1.786 | -0.246 | 0.357 | -0.021 | 0.039 | 0.039 | 3.202 | 0.15 | 0.05 | 0.14 | 0.09 | 0.17 | 0.07 |
| 1 | 48222050 | rs10430126 | 0.014 | 1.773 | -0.136 | 0.422 | -0.001 | -0.021 | -0.021 | 0 | 0.22 | 0.12 | 0.33 | 0.28 | 0.2 | 0.16 |
| 1 | 48224494 | rs6588490 | -1.066 | 2.097 | 0.605 | 1.266 | 0.001 | 0.011 | 0.011 | 0 | 0.42 | 0.44 | 0.24 | 0.29 | 0.36 | 0.2 |
| 1 | 48227135 | rs7543124 | -1.85 | 2.448 | 1.217 | 1.019 | 0.012 | -0.019 | -0.019 | 0 | 0.57 | 0.61 | 0.74 | 0.97 | 0.6 | 0.59 |
| 1 | 48227943 | rs1561573 | -1.217 | 2.464 | 1.418 | 1.662 | -0.001 | 0.001 | 0.001 | 0 | 0.47 | 0.54 | 0.52 | 0.44 | 0.45 | 0.27 |
| 1 | 48233438 | rs12409657 | -0.077 | 2.238 | 1.696 | 2.182 | -0.002 | -0.001 | -0.001 | 0 | 0.21 | 0.15 | 0.36 | 0.4 | 0.24 | 0.23 |
| 1 | 48233732 | rs12038078 | -2.426 | 1.932 | 2.032 | 2.036 | 0.032 | 0.076 | 0.076 | 0.727 | 0.06 | 0.21 | 0.26 | 0.13 | 0.06 | 0.07 |
| 1 | 48234949 | rs6701938 | -1.321 | 1.76 | 2.001 | 1.69 | 0.012 | 0.102 | 0.102 | 0 | 0.48 | 0.34 | 0.51 | 0.92 | 0.51 | 0.47 |
| 1 | 48236963 | rs6657202 | -1.753 | 1.767 | 2.229 | 1.611 | 0.015 | 0.083 | 0.083 | 0 | 0.54 | 0.72 | 0.74 | 0.99 | 0.54 | 0.46 |
| 1 | 48240005 | rs2282361 | -1.119 | 1.706 | 1.911 | 1.932 | -0.018 | 0.048 | 0.048 | 0 | 0.46 | 0.32 | 0.34 | 0.61 | 0.47 | 0.35 |
| 1 | 48254782 | rs6588505 | -0.818 | 1.776 | 1.969 | 2.21 | -0.008 | 0.042 | 0.042 | 0 | 0.28 | 0.42 | 0.33 | 0.48 | 0.24 | 0.23 |
| 1 | 48255008 | rs6588506 | -1.502 | 1.978 | 2.101 | 2.169 | 0.014 | 0.041 | 0.041 | 0 | 0.56 | 0.66 | 0.63 | 0.79 | 0.58 | 0.43 |
| 1 | 48257485 | rs6675571 | -1.702 | 1.945 | 2.125 | 2.012 | 0.024 | 0.085 | 0.085 | 0 | 0.58 | 0.79 | 0.76 | 0.98 | 0.59 | 0.47 |
| 1 | 48262619 | rs4927069 | -0.439 | 2.102 | 2.232 | 2.259 | 0.027 | 0.014 | 0.014 | 0 | 0.3 | 0.4 | 0.39 | 0.32 | 0.34 | 0.22 |
| 1 | 48282376 | rs12024706 | -2.271 | 1.878 | 2.44 | 2.526 | 0.046 | 0.399 | 0.399 | 0.049 | 0.05 | 0.33 | 0.21 | 0.37 | 0.02 | 0.12 |
| 1 | 48285768 | rs6671927 | -0.116 | 1.658 | 2.213 | 2.289 | -0.011 | 0.01 | 0.01 | 0 | 0.31 | 0.15 | 0.22 | 0.26 | 0.31 | 0.15 |
| 1 | 48286563 | rs12134443 | 1.796 | 1.679 | 2.238 | 2.197 | 0.024 | 0.137 | 0.137 | 0 | 0.34 | 0.13 | 0.18 | 0.02 | 0.31 | 0.5 |
| 1 | 48288016 | rs11211611 | 0.296 | 1.703 | 2.315 | 2.246 | -0.002 | -0.002 | -0.002 | 0.52 | 0.27 | 0.22 | 0.3 | 0.3 | 0.33 | 0.2 |
| 1 | 48290860 | rs11211613 | -0.059 | 1.714 | 2.416 | 2.242 | -0.004 | -0.007 | -0.007 | 0 | 0.68 | 0.59 | 0.77 | 0.74 | 0.69 | 0.85 |
| 1 | 48295710 | rs7525331 | 2.496 | 1.704 | 2.335 | 2.291 | 0.062 | 0.447 | 0.447 | 0.22 | 0.93 | 0.58 | 0.71 | 0.59 | 0.96 | 0.85 |
| 1 | 48309587 | rs10890507 | -0.055 | 1.68 | 2.048 | 2.066 | 0.018 | 0.015 | 0.015 | 0 | 0.29 | 0.31 | 0.38 | 0.35 | 0.35 | 0.24 |
| 1 | 48331522 | rs7536189 | -1.505 | 1.591 | 1.922 | 1.959 | 0.006 | 0.007 | 0.007 | 0 | 0.62 | 0.67 | 0.68 | 0.83 | 0.68 | 0.47 |
| 1 | 48335067 | rs4926664 | -0.578 | 1.613 | 1.844 | 2.062 | -0.001 | 0.001 | 0.001 | 0 | 0.35 | 0.39 | 0.39 | 0.5 | 0.34 | 0.27 |
| 1 | 48341005 | rs10493130 | 0.417 | 1.63 | 1.86 | 2.014 | -0.035 | -0.016 | -0.016 | 0.001 | 0.25 | 0.12 | 0.15 | 0.24 | 0.29 | 0.16 |
| 1 | 48343224 | rs12048888 | -0.348 | 1.639 | 1.823 | 1.957 | 0.006 | -0.029 | -0.029 | 0 | 0.33 | 0.29 | 0.3 | 0.48 | 0.33 | 0.23 |
| 1 | 48347013 | rs17103887 | 0.025 | 1.72 | 1.914 | 1.803 | -0.004 | -0.004 | -0.004 | 0 | 0.27 | 0.2 | 0.29 | 0.32 | 0.33 | 0.21 |
| 1 | 48359634 | rs11211624 | -1.792 | 1.461 | 2.55 | 2.521 | 0.041 | 0.173 | 0.173 | 0.057 | 0.06 | 0.12 | 0.13 | 0.34 | 0.1 | 0.14 |
| 1 | 48362713 | rs2088361 | -1.558 | 1.428 | 2.745 | 2.531 | -0.001 | 0 | 0 | 0 | 0.37 | 0.66 | 0.51 | 0.58 | 0.37 | 0.34 |
| 1 | 48370525 | rs1556981 | 2.691 | 1.149 | 3.035 | 2.541 | 0.111 | 0.374 | 0.374 | 0.799 | 0.88 | 0.56 | 0.75 | 0.62 | 0.87 | 0.8 |
| 1 | 48380390 | rs6684538 | 1.793 | 1.137 | 2.966 | 2.516 | -0.008 | 0.049 | 0.049 | 0.067 | 0.58 | 0.18 | 0.38 | 0.37 | 0.61 | 0.6 |
| 1 | 48386888 | rs1497095 | -0.828 | 1.117 | 3.064 | 2.497 | 0.001 | 0 | 0 | 0 | 0.3 | 0.27 | 0.42 | 0.26 | 0.28 | 0.18 |

| | | | | | | | | | | | | | | | | |
|---|---|---|---|---|---|---|---|---|---|---|---|---|---|---|---|---|
| 1 | 48387427 | rs10493131 | -0.332 | 1.118 | 2.99 | 2.494 | -0.001 | -0.007 | -0.007 | 0 | 0.26 | 0.08 | 0.31 | 0.23 | 0.26 | 0.14 |
| 1 | 48391668 | rs1361443 | -1.558 | 1.127 | 3.02 | 2.509 | 0.005 | -0.007 | -0.007 | 0.041 | 0.37 | 0.43 | 0.56 | 0.6 | 0.38 | 0.33 |
| 1 | 48392033 | rs12061891 | -1.604 | 1.151 | 2.983 | 2.523 | 0.044 | 0.169 | 0.169 | 0.045 | 0.06 | 0.13 | 0.13 | 0.34 | 0.11 | 0.15 |
| 1 | 48401697 | rs6681669 | -0.564 | 1.151 | 2.812 | 2.46 | -0.003 | -0.002 | -0.002 | 0 | 0.27 | 0.28 | 0.33 | 0.23 | 0.26 | 0.14 |
| 1 | 48406460 | rs11211632 | 1.822 | 1.146 | 2.534 | 2.461 | -0.004 | 0.05 | 0.05 | 0.076 | 0.58 | 0.21 | 0.39 | 0.37 | 0.61 | 0.6 |
| 1 | 48409812 | rs17103939 | -1.742 | 1.244 | 2.338 | 2.458 | 0.059 | 0.202 | 0.202 | 0.015 | 0.07 | 0.16 | 0.13 | 0.33 | 0.1 | 0.15 |
| 1 | 48411729 | rs1603530 | -0.785 | 0.311 | 1.964 | 1.671 | 0.003 | 0.002 | 0.002 | 0 | 0.28 | 0.27 | 0.34 | 0.23 | 0.26 | 0.12 |
| 1 | 48442182 | rs11205430 | -1.722 | 0.388 | 1.659 | 1.818 | -0.004 | 0.011 | 0.011 | 0 | 0.35 | 0.59 | 0.42 | 0.43 | 0.35 | 0.35 |
| 1 | 48454212 | rs10888594 | -1.547 | 0.499 | 1.412 | 1.818 | -0.013 | 0.034 | 0.034 | 0 | 0.32 | 0.47 | 0.3 | 0.36 | 0.33 | 0.21 |
| 1 | 48457543 | rs6676465 | -1.569 | 0.421 | 1.229 | 1.828 | -0.014 | 0.027 | 0.027 | 0 | 0.32 | 0.51 | 0.3 | 0.36 | 0.33 | 0.21 |
| 1 | 48459907 | rs3814006 | -0.712 | 0.571 | 0.3 | 1.386 | -0.051 | -0.071 | -0.071 | 0 | 0.23 | 0.01 | 0.22 | 0.25 | 0.28 | 0.12 |
| 1 | 48480638 | rs12137714 | -0.799 | 0.585 | 0.818 | 1.389 | -0.003 | 0.001 | 0.001 | 0 | 0.26 | 0.28 | 0.26 | 0.25 | 0.28 | 0.13 |
| 1 | 48481192 | rs17104042 | -1.12 | 0.58 | 0.718 | 1.351 | -0.005 | 0.016 | 0.016 | 0 | 0.28 | 0.39 | 0.34 | 0.32 | 0.3 | 0.18 |
| 1 | 48483197 | rs1693258 | 0.406 | 0.448 | -0.041 | 1.021 | 0.016 | -0.011 | -0.011 | 0 | 0.55 | 0.42 | 0.37 | 0.53 | 0.42 | 0.43 |
| 1 | 48485165 | rs1390972 | -1.684 | 0.314 | 1.055 | 1.462 | 0.033 | 0.21 | 0.21 | 0.158 | 0.06 | 0.33 | 0.03 | 0.21 | 0.01 | 0.1 |
| 1 | 48486172 | rs1693246 | -0.002 | 0.865 | 0.491 | 1.326 | -0.064 | -0.017 | -0.017 | 0 | 0.69 | 0.86 | 0.63 | 0.67 | 0.6 | 0.66 |
| 1 | 48492057 | rs303928 | 0.164 | 0.783 | 0.555 | 1.352 | -0.055 | -0.061 | -0.061 | 11.169 | 0.67 | 0.84 | 0.62 | 0.47 | 0.6 | 0.58 |
| 1 | 48496056 | rs6588639 | -1.286 | 0.536 | 1.517 | 1.487 | -0.003 | 0.006 | 0.006 | 2.035 | 0.06 | 0.15 | 0.08 | 0.09 | 0.05 | 0.16 |
| 1 | 48500746 | rs722656 | 1.162 | 0.701 | 1.527 | 1.276 | 0.078 | 0.049 | 0.049 | 0.255 | 0.52 | 0.23 | 0.41 | 0.34 | 0.39 | 0.35 |
| 1 | 48509053 | rs748142 | -0.074 | 0.991 | 1.719 | 1.299 | 0.003 | 0.001 | 0.001 | 0 | 0.67 | 0.73 | 0.63 | 0.63 | 0.61 | 0.63 |
| 1 | 48512119 | rs7549671 | 1.128 | 0.862 | 1.81 | 1.302 | 0.078 | 0.047 | 0.047 | 0.332 | 0.52 | 0.21 | 0.41 | 0.35 | 0.4 | 0.35 |
| 1 | 48512650 | rs6690390 | 0.044 | 0.996 | 1.807 | 1.33 | 0.044 | 0.069 | 0.069 | 0 | 0.07 | 0.2 | 0.11 | 0.2 | 0.14 | 0.09 |
| 1 | 48512766 | rs6588648 | -1.405 | 1.003 | 1.831 | 1.222 | 0.049 | 0.06 | 0.06 | 1.017 | 0.11 | 0.35 | 0.18 | 0.25 | 0.18 | 0.21 |
| 1 | 48515250 | rs1039131 | -1.063 | 1.011 | 1.511 | 1.311 | -0.002 | 0.011 | 0.011 | 1.522 | 0.05 | 0.13 | 0.07 | 0.09 | 0.05 | 0.15 |
| 1 | 48516401 | rs513108 | 0.1 | 1.044 | 1.449 | 1.334 | 0.001 | 0 | 0 | 1.53 | 0.65 | 0.72 | 0.61 | 0.64 | 0.58 | 0.6 |
| 1 | 48517657 | rs11582141 | -0.528 | 0.919 | 0.799 | 1.122 | 0.055 | 0.032 | 0.032 | 0 | 0.11 | 0.28 | 0.12 | 0.17 | 0.16 | 0.09 |
| 1 | 48519912 | rs302691 | 0.194 | 0.913 | -0.097 | 1.007 | -0.048 | -0.03 | -0.03 | 0 | 0.64 | 0.79 | 0.57 | 0.55 | 0.58 | 0.49 |
| 1 | 48524180 | rs6679531 | 0.077 | 0.889 | -0.811 | 0.647 | 0.043 | 0.127 | 0.127 | 0.364 | 0.47 | 0.18 | 0.31 | 0.27 | 0.35 | 0.41 |
| 1 | 48526339 | rs11205439 | 1.158 | 1.124 | -0.635 | 0.51 | -0.012 | 0.039 | 0.039 | 0 | 0.28 | 0.17 | 0.32 | 0.1 | 0.35 | 0.23 |
| 1 | 48527879 | rs12080303 | 0.632 | 1.121 | -1.137 | 0.492 | 0.035 | 0.155 | 0.155 | 0.895 | 0.44 | 0.18 | 0.18 | 0.17 | 0.32 | 0.37 |
| 1 | 48528242 | rs302681 | 0.675 | 1.044 | -0.513 | 1.221 | -0.002 | -0.018 | -0.018 | 0.668 | 0.34 | 0.5 | 0.35 | 0.23 | 0.37 | 0.34 |
| 1 | 48532672 | rs502819 | -0.423 | 1.134 | -0.435 | 1.615 | -0.004 | -0.001 | -0.001 | 0.002 | 0.18 | 0.16 | 0.32 | 0.4 | 0.28 | 0.22 |
| 1 | 48535182 | rs602024 | -0.84 | 1.18 | -0.282 | 1.33 | 0.005 | -0.017 | -0.017 | 0 | 0.66 | 0.58 | 0.63 | 0.76 | 0.6 | 0.71 |
| 1 | 48537376 | rs679282 | -0.828 | 1.481 | -0.39 | 1.912 | -0.003 | 0.006 | 0.006 | 0 | 0.18 | 0.17 | 0.32 | 0.48 | 0.28 | 0.25 |
| 1 | 48539143 | rs526274 | -0.527 | 1.351 | -0.954 | 0.976 | 0.003 | 0.003 | 0.003 | 0 | 0.33 | 0.35 | 0.49 | 0.64 | 0.38 | 0.36 |
| 1 | 48546195 | rs638965 | -0.547 | 1.015 | -1.635 | 0.291 | 0.026 | 0.082 | 0.082 | 0 | 0.57 | 0.22 | 0.73 | 0.77 | 0.62 | 0.66 |
| 1 | 48549313 | rs12036718 | 0.192 | 1.605 | -1.589 | 0.686 | -0.051 | -0.039 | -0.039 | 0 | 0.08 | 0.01 | 0.14 | 0.16 | 0.17 | 0.17 |
| 1 | 48549673 | rs560004 | 0.087 | 1.07 | -1.522 | 0.568 | -0.004 | 0.017 | 0.017 | 0 | 0.41 | 0.18 | 0.55 | 0.62 | 0.38 | 0.4 |
| 1 | 48556330 | rs4466696 | -1.009 | 1.079 | -0.736 | 1.181 | 0.008 | 0.057 | 0.057 | 0 | 0.19 | 0.26 | 0.21 | 0.42 | 0.24 | 0.31 |
| 1 | 48556821 | rs472771 | -0.585 | 0.894 | -0.927 | 0.98 | 0.004 | 0 | 0 | 0 | 0.62 | 0.49 | 0.4 | 0.63 | 0.59 | 0.63 |
| 1 | 48559463 | rs1007657 | 0.267 | 0.693 | -0.9 | 1.028 | -0.005 | 0.084 | 0.084 | 0 | 0.5 | 0.27 | 0.34 | 0.33 | 0.5 | 0.38 |
| 1 | 48564267 | rs17468833 | 0.118 | 0.67 | -0.873 | 1.033 | 0.019 | 0.07 | 0.07 | 0 | 0.52 | 0.3 | 0.36 | 0.32 | 0.55 | 0.42 |
| 1 | 48567117 | rs17428691 | 1.872 | 0.558 | -1.123 | 0.734 | 0.007 | 0.11 | 0.11 | 0.034 | 0.17 | 0.02 | 0.08 | 0.01 | 0.11 | 0.12 |
| 1 | 48568400 | rs17468972 | 1.515 | 0.413 | -1.131 | 0.787 | -0.045 | 0.1 | 0.1 | 0 | 0.35 | 0.05 | 0.18 | 0.17 | 0.33 | 0.2 |
| 1 | 48568672 | rs10888596 | 0.367 | 0.39 | -1.053 | 0.724 | 0.014 | 0.038 | 0.038 | 0 | 0.53 | 0.31 | 0.37 | 0.36 | 0.55 | 0.49 |
| 1 | 48570133 | rs2073293 | 0.976 | 0.515 | -0.662 | 1.109 | -0.01 | -0.006 | -0.006 | 0.021 | 0.18 | 0.14 | 0.27 | 0.2 | 0.26 | 0.2 |
| 1 | 48570466 | rs12047002 | 1.243 | 0.433 | -0.198 | 1.152 | -0.003 | 0.006 | 0.006 | 0 | 0.21 | 0.18 | 0.32 | 0.43 | 0.16 | 0.23 |
| 1 | 48575218 | rs12126234 | 0.828 | 0.547 | -0.555 | 0.615 | -0.042 | -0.025 | -0.025 | 0 | 0.11 | 0 | 0.13 | 0.13 | 0.15 | 0.18 |
| 1 | 48588004 | rs214203 | -0.1 | 0.531 | -0.29 | 0.741 | -0.003 | -0.011 | -0.011 | 1.237 | 0.42 | 0.67 | 0.63 | 0.56 | 0.44 | 0.42 |
| 1 | 48594282 | rs11205442 | 1.359 | 0.4 | -0.552 | 0.655 | -0.033 | 0.007 | 0.007 | 0.113 | 0.36 | 0.08 | 0.18 | 0.26 | 0.32 | 0.19 |
| 1 | 48606878 | rs12741362 | -0.125 | 0.392 | -0.187 | 0.764 | -0.007 | 0 | 0 | 0.07 | 0.45 | 0.64 | 0.66 | 0.41 | 0.5 | 0.5 |
| 1 | 48609038 | rs2134477 | 0.247 | 0.397 | -0.479 | 0.784 | -0.015 | 0.05 | 0.05 | 0.054 | 0.49 | 0.17 | 0.32 | 0.61 | 0.48 | 0.5 |
| 1 | 48622095 | rs12044066 | 2.3 | 0.373 | -0.865 | 0.587 | -0.066 | -0.088 | -0.088 | 0 | 0.21 | 0.02 | 0.09 | 0.26 | 0.19 | 0.1 |
| 1 | 48630438 | rs10788882 | -0.259 | 0.354 | -0.557 | 0.429 | -0.002 | -0.002 | -0.002 | 0 | 0.74 | 0.72 | 0.5 | 0.6 | 0.67 | 0.64 |
| 1 | 48646553 | rs214193 | -0.602 | 0.368 | -0.717 | 0.443 | -0.116 | -0.124 | -0.124 | 0 | 0.76 | 0.95 | 0.51 | 0.63 | 0.67 | 0.71 |
| 1 | 48647124 | rs11205449 | 1.226 | 0.344 | -0.743 | 0.381 | -0.098 | -0.095 | -0.095 | 0 | 0.33 | 0.04 | 0.22 | 0.38 | 0.35 | 0.33 |
| 1 | 48665598 | rs823849 | -0.307 | 0.419 | -0.446 | 0.539 | -0.01 | -0.014 | -0.014 | 0.244 | 0.75 | 0.74 | 0.5 | 0.6 | 0.67 | 0.64 |
| 1 | 48666398 | rs639297 | -0.324 | 0.425 | -0.553 | 0.542 | -0.054 | -0.078 | -0.078 | 0 | 0.75 | 0.87 | 0.5 | 0.6 | 0.67 | 0.64 |
| 1 | 48671147 | rs12136440 | 0.88 | 0.474 | -0.632 | 0.56 | -0.101 | -0.118 | -0.118 | 0.183 | 0.19 | 0.05 | 0.48 | 0.38 | 0.31 | 0.24 |
| 1 | 48672401 | rs1722710 | 0.684 | 0.288 | -0.1 | 0.41 | -0.002 | 0.002 | 0.002 | 0.385 | 0.56 | 0.56 | 0.36 | 0.4 | 0.51 | 0.52 |
| 1 | 48673385 | rs1524736 | -0.211 | 0.31 | -0.357 | 0.428 | -0.048 | -0.053 | -0.053 | 0 | 0.35 | 0.14 | 0.58 | 0.58 | 0.44 | 0.35 |
| 1 | 48682034 | rs10788883 | 0.352 | 0.29 | -0.277 | 0.353 | -0.004 | 0.034 | 0.034 | 0 | 0.51 | 0.3 | 0.34 | 0.39 | 0.5 | 0.42 |
| 1 | 48685435 | rs926979 | 0.342 | 0.194 | -0.384 | 0.307 | 0.096 | 0.171 | 0.171 | 0.007 | 0.54 | 0.17 | 0.72 | 0.79 | 0.67 | 0.63 |
| 1 | 48686096 | rs1524732 | 0.588 | 0.187 | -0.378 | 0.333 | 0.002 | -0.001 | -0.001 | 0 | 0.17 | 0.18 | 0.13 | 0.2 | 0.15 | 0.12 |
| 1 | 48687848 | rs6695408 | 0.194 | 0.215 | -0.386 | 0.337 | -0.02 | 0.042 | 0.042 | 0 | 0.52 | 0.23 | 0.36 | 0.61 | 0.51 | 0.52 |
| 1 | 48691564 | rs12139099 | 0.954 | 0.298 | -0.317 | 0.393 | -0.033 | -0.026 | -0.026 | 0 | 0.37 | 0.14 | 0.26 | 0.61 | 0.37 | 0.43 |
| 1 | 48694259 | rs764656 | 2.503 | 0.317 | -0.42 | 0.534 | -0.048 | -0.089 | -0.089 | 0 | 0.21 | 0.03 | 0.18 | 0.28 | 0.19 | 0.11 |
| 1 | 48695081 | rs212989 | 1.396 | 0.176 | 0.21 | 0.493 | -0.029 | -0.12 | -0.12 | 0 | 0.21 | 0.31 | 0.12 | 0.01 | 0.16 | 0.18 |
| 1 | 48700325 | rs10489600 | 0.333 | 0.195 | -0.079 | 0.542 | -0.019 | -0.021 | -0.021 | 0 | 0.53 | 0.25 | 0.43 | 0.44 | 0.5 | 0.48 |

| | | | | | | | | | | | | | | | | |
|---|---|---|---|---|---|---|---|---|---|---|---|---|---|---|---|---|
| 1 | 48705391 | rs11205459 | 1.34 | 0.23 | -0.281 | 0.484 | -0.104 | -0.103 | -0.103 | 0.399 | 0.35 | 0.04 | 0.31 | 0.4 | 0.36 | 0.33 |
| 1 | 48708228 | rs850763 | 1.364 | 0.455 | 0.193 | 0.655 | 0.017 | 0.02 | 0.02 | 1.31 | 0.2 | 0.11 | 0.1 | 0.01 | 0.16 | 0.18 |
| 1 | 48709019 | rs6670286 | 0.698 | 0.549 | 0.117 | 0.751 | -0.03 | -0.031 | -0.031 | 2.762 | 0.19 | 0.12 | 0.4 | 0.35 | 0.31 | 0.24 |
| 1 | 48713394 | rs986027 | -0.547 | 0.163 | 0.264 | 0.563 | -0.031 | -0.006 | -0.006 | 0.128 | 0.73 | 0.44 | 0.83 | 0.75 | 0.81 | 0.66 |
| 1 | 48721158 | rs212964 | 0.195 | 0.47 | 0.229 | 0.647 | 0.026 | -0.001 | -0.001 | 0.001 | 0.58 | 0.32 | 0.53 | 0.59 | 0.65 | 0.54 |
| 1 | 48762661 | rs17104339 | 1.391 | 0.699 | 0.174 | 0.41 | 0.004 | -0.005 | -0.005 | 0 | 0.12 | 0.09 | 0.05 | 0 | 0.09 | 0.08 |
| 1 | 48764419 | rs1056042 | 1.394 | 0.53 | 0.004 | 0.344 | -0.144 | -0.058 | -0.058 | 0 | 0.4 | 0.03 | 0.33 | 0.35 | 0.43 | 0.34 |
| 1 | 48765945 | rs6666372 | -0.095 | 0.49 | 0.147 | 0.362 | -0.003 | -0.028 | -0.028 | 4.119 | 0.37 | 0.57 | 0.44 | 0.18 | 0.34 | 0.34 |
| 1 | 48774515 | rs1608695 | 1.359 | 0.52 | -0.019 | 0.282 | -0.118 | -0.036 | -0.036 | 0.427 | 0.4 | 0.04 | 0.34 | 0.35 | 0.43 | 0.33 |
| 1 | 48785665 | rs1418105 | 1.297 | 0.506 | 0.609 | 0.105 | -0.02 | -0.077 | -0.077 | 0 | 0.18 | 0.21 | 0.1 | 0.01 | 0.13 | 0.18 |
| 1 | 48802740 | rs11205475 | 1.201 | 0.458 | 0.273 | 0.251 | -0.02 | -0.077 | -0.077 | 0 | 0.18 | 0.21 | 0.1 | 0.01 | 0.13 | 0.18 |
| 1 | 48820294 | rs11205476 | -0.509 | 0.624 | -0.157 | 0.403 | 0.009 | 0.036 | 0.036 | 0 | 0.38 | 0.66 | 0.46 | 0.21 | 0.34 | 0.43 |
| 1 | 48828015 | rs1877719 | 0.299 | 0.787 | -0.192 | 0.795 | 0.019 | 0.178 | 0.178 | 0 | 0.9 | 0.78 | 0.84 | 0.56 | 0.85 | 0.85 |
| 1 | 48839644 | rs12145298 | 1.213 | 0.883 | -0.556 | 0.544 | -0.052 | 0.05 | 0.05 | 0 | 0.16 | 0.01 | 0.14 | 0.08 | 0.2 | 0.22 |
| 1 | 48857313 | rs3767616 | 0.705 | 0.71 | -0.307 | 0.587 | -0.017 | 0.009 | 0.009 | 0 | 0.16 | 0.1 | 0.32 | 0.17 | 0.19 | 0.14 |
| 1 | 48859681 | rs10158605 | 0 | 0.637 | -0.052 | 0.557 | 0.01 | -0.021 | -0.021 | 0 | 0.19 | 0.31 | 0.34 | 0.17 | 0.21 | 0.2 |
| 1 | 48891957 | rs3767621 | 0.881 | 0.617 | -0.433 | 0.529 | -0.032 | 0.053 | 0.053 | 0.714 | 0.52 | 0.12 | 0.39 | 0.35 | 0.51 | 0.41 |
| 1 | 48946572 | rs2787875 | 0.634 | 0.507 | 0.004 | 0.493 | -0.015 | 0.089 | 0.089 | 0.958 | 0.89 | 0.64 | 0.74 | 0.81 | 0.93 | 0.81 |
| 1 | 48954476 | rs2253233 | 0.286 | 0.687 | -0.092 | 0.608 | 0.045 | 0.034 | 0.034 | 0 | 0.79 | 0.61 | 0.76 | 0.7 | 0.7 | 0.73 |
| 1 | 48957540 | rs1338319 | -0.453 | 0.656 | -0.082 | 0.579 | -0.007 | -0.021 | -0.021 | 0 | 0.94 | 0.79 | 0.96 | 1 | 0.97 | 0.96 |
| 1 | 48959912 | rs2798107 | 0.277 | 0.743 | -0.118 | 0.64 | 0.022 | 0.02 | 0.02 | 0 | 0.16 | 0.18 | 0.2 | 0.3 | 0.27 | 0.25 |
| 1 | 48961903 | rs2798105 | 1.295 | 0.468 | -0.188 | 0.469 | -0.021 | -0.061 | -0.061 | 0 | 0.18 | 0.24 | 0.06 | 0.02 | 0.12 | 0.18 |
| 1 | 48965209 | rs12045951 | 0.457 | 0.608 | -0.21 | 0.503 | -0.001 | -0.002 | -0.002 | 0 | 0.62 | 0.41 | 0.7 | 0.68 | 0.6 | 0.57 |
| 1 | 48965583 | rs12032082 | 2.317 | 0.668 | -0.996 | 0.5 | -0.092 | -0.058 | -0.058 | 0 | 0.24 | 0.02 | 0.21 | 0.22 | 0.24 | 0.1 |
| 1 | 48966113 | rs2803252 | 0.262 | 1.058 | -0.333 | 0.614 | -0.018 | -0.106 | -0.106 | 0 | 0.65 | 0.48 | 0.8 | 0.95 | 0.7 | 0.68 |
| 1 | 48967002 | rs12098127 | 0.05 | 1.112 | -0.329 | 0.674 | 0.002 | -0.026 | -0.026 | 0.399 | 0.17 | 0.27 | 0.13 | 0.03 | 0.18 | 0.14 |
| 1 | 48972810 | rs1343124 | -0.111 | 1.145 | -0.521 | 0.691 | 0.004 | -0.004 | -0.004 | 0 | 0.72 | 0.29 | 0.79 | 0.68 | 0.74 | 0.61 |
| 1 | 48973321 | rs11583202 | 0.586 | 1.281 | -0.682 | 0.696 | -0.071 | 0.038 | 0.038 | 0 | 0.17 | 0.01 | 0.17 | 0.08 | 0.24 | 0.25 |
| 1 | 48974933 | rs1539530 | 0.544 | 0.979 | 0.04 | 1.173 | 0.002 | -0.029 | -0.029 | 0.174 | 0.59 | 0.21 | 0.66 | 0.47 | 0.59 | 0.56 |
| 1 | 48977584 | rs11205499 | -1.163 | 1.24 | -0.354 | 1.406 | 0.025 | 0.07 | 0.07 | 0 | 0.09 | 0.2 | 0.15 | 0.28 | 0.15 | 0.12 |
| 1 | 48986545 | rs2354463 | 0.424 | 0.546 | -0.955 | 0.78 | 0.032 | 0.034 | 0.034 | 0.175 | 0.57 | 0.25 | 0.73 | 0.7 | 0.69 | 0.53 |
| 1 | 48991061 | rs12127061 | -0.176 | 0.197 | -0.605 | 0.68 | 0.009 | 0.017 | 0.017 | 0 | 0.3 | 0.2 | 0.17 | 0.22 | 0.23 | 0.23 |
| 1 | 48992947 | rs12029855 | 1.716 | 0.345 | -1.645 | 0.583 | -0.082 | -0.018 | -0.018 | 0 | 0.27 | 0.02 | 0.15 | 0.18 | 0.28 | 0.13 |
| 1 | 48993335 | rs2354467 | 0.345 | -0.122 | -0.933 | 0.581 | 0.018 | 0.019 | 0.019 | 0.008 | 0.55 | 0.33 | 0.43 | 0.65 | 0.64 | 0.63 |
| 1 | 48998361 | rs2246105 | -0.849 | -0.075 | -0.755 | 0.698 | -0.016 | 0.004 | 0.004 | 0 | 0.87 | 0.8 | 0.65 | 0.86 | 0.92 | 0.87 |
| 1 | 48998748 | rs2803261 | -0.338 | -0.042 | -0.528 | 0.704 | 0.02 | 0.062 | 0.062 | 0 | 0.29 | 0.5 | 0.33 | 0.6 | 0.39 | 0.39 |
| 1 | 48999555 | rs2798111 | 0.289 | 0.029 | -0.818 | 0.502 | 0.007 | 0.012 | 0.012 | 0 | 0.24 | 0.28 | 0.26 | 0.39 | 0.34 | 0.33 |
| 1 | 49001369 | rs1539525 | -0.471 | -0.028 | -0.888 | 0.517 | 0.01 | 0.012 | 0.012 | 0 | 0.5 | 0.45 | 0.39 | 0.47 | 0.57 | 0.6 |
| 1 | 49001780 | rs2246557 | 0.951 | -0.281 | -1.724 | 0.532 | -0.075 | -0.056 | -0.056 | 0 | 0.46 | 0.07 | 0.46 | 0.61 | 0.57 | 0.42 |
| 1 | 49003131 | rs4926743 | 0.807 | -0.28 | -1.303 | 0.677 | 0.012 | -0.012 | -0.012 | 0 | 0.07 | 0.05 | 0.32 | 0.14 | 0.04 | 0.11 |
| 1 | 49006567 | rs2354468 | 0.386 | -0.286 | -0.926 | 0.872 | -0.026 | 0.012 | 0.012 | 0 | 0.66 | 0.58 | 0.76 | 0.57 | 0.76 | 0.55 |
| 1 | 49006786 | rs7515225 | 1.934 | 0.197 | -0.97 | 0.564 | -0.001 | 0.002 | 0.002 | 0 | 0.32 | 0.29 | 0.19 | 0.16 | 0.3 | 0.22 |
| 1 | 49007034 | rs12079703 | 0.192 | 0.416 | -1.247 | 0.41 | -0.002 | -0.01 | -0.01 | 0 | 0.09 | 0.09 | 0.03 | 0 | 0.1 | 0.09 |
| 1 | 49016360 | rs2803264 | 0.29 | 0.292 | -1.197 | 0.733 | 0.001 | 0.03 | 0.03 | 0 | 0.87 | 0.77 | 0.61 | 0.72 | 0.85 | 0.83 |
| 1 | 49019450 | rs2803266 | -0.477 | 0.379 | -1.179 | 0.717 | 0.044 | 0.217 | 0.217 | 0 | 0.9 | 0.72 | 0.69 | 0.59 | 0.88 | 0.83 |
| 1 | 49024709 | rs2803268 | 2.805 | 0.416 | -1.78 | 0.44 | -0.032 | -0.054 | -0.054 | 1.819 | 0.28 | 0.03 | 0.16 | 0.21 | 0.24 | 0.13 |
| 1 | 49025436 | rs11205506 | 0.464 | 0.418 | -1.646 | 0.538 | -0.078 | -0.08 | -0.08 | 0.576 | 0.14 | 0.02 | 0.2 | 0.26 | 0.27 | 0.21 |
| 1 | 49031238 | rs2803270 | 0.432 | 0.387 | -1.478 | 0.456 | 0.027 | 0.038 | 0.038 | 0 | 0.75 | 0.69 | 0.48 | 0.34 | 0.61 | 0.62 |
| 1 | 49033734 | rs11802042 | 0.103 | -0.322 | -1.39 | 0.037 | 0.02 | 0.012 | 0.012 | 0 | 0.22 | 0.14 | 0.09 | 0.01 | 0.13 | 0.17 |
| 1 | 49036512 | rs2798095 | 1.23 | -0.289 | -1.512 | 0.173 | 0.003 | 0.028 | 0.028 | 0.338 | 0.54 | 0.33 | 0.53 | 0.55 | 0.61 | 0.42 |
| 1 | 49040384 | rs2798125 | 0.395 | -0.038 | -1.264 | 0.029 | 0.004 | 0.006 | 0.006 | 0 | 0.63 | 0.59 | 0.54 | 0.54 | 0.66 | 0.64 |
| 1 | 49040423 | rs17104522 | -0.024 | -0.035 | -1.32 | 0.024 | 0.003 | 0.082 | 0.082 | 0 | 0.09 | 0.2 | 0.08 | 0.25 | 0.12 | 0.15 |
| 1 | 49042349 | rs2803273 | 3.008 | -0.254 | -1.612 | -0.185 | -0.089 | 0.007 | 0.007 | 0 | 0.32 | 0.05 | 0.26 | 0.19 | 0.37 | 0.15 |
| 1 | 49042952 | rs10493134 | 1.468 | -0.248 | -1.405 | -0.389 | -0.023 | -0.009 | -0.009 | 0 | 0.05 | 0.01 | 0.06 | 0.06 | 0.1 | 0.09 |
| 1 | 49043429 | rs2803274 | 1.33 | -0.239 | -1.286 | -0.503 | -0.006 | 0.045 | 0.045 | 0 | 0.53 | 0.25 | 0.46 | 0.29 | 0.53 | 0.48 |
| 1 | 49043877 | rs6665839 | -1.171 | -0.297 | -1.087 | -0.381 | -0.023 | 0.018 | 0.018 | 0 | 0.21 | 0.4 | 0.11 | 0.25 | 0.18 | 0.24 |
| 1 | 49047578 | rs2803275 | -0.365 | -0.337 | -1.071 | -0.429 | 0.012 | 0.161 | 0.161 | 0 | 0.81 | 0.59 | 0.73 | 0.56 | 0.84 | 0.81 |
| 1 | 49049758 | rs11205510 | 0.283 | -0.243 | -1.214 | -0.426 | -0.003 | -0.018 | -0.018 | 0 | 0.07 | 0.07 | 0.02 | 0 | 0.05 | 0.08 |
| 1 | 49051568 | rs2798120 | 1.047 | -0.226 | -1.327 | -0.482 | 0 | 0.002 | 0.002 | 0 | 0.57 | 0.41 | 0.63 | 0.56 | 0.65 | 0.57 |
| 1 | 49052263 | rs2627754 | -0.607 | -0.472 | -0.593 | -1.142 | 0.001 | -0.032 | -0.032 | 0.061 | 0.31 | 0.27 | 0.15 | 0.01 | 0.26 | 0.31 |
| 1 | 49052423 | rs11579440 | -0.624 | -0.566 | -0.648 | -0.965 | -0.013 | 0.043 | 0.043 | 0 | 0.14 | 0.27 | 0.11 | 0.31 | 0.16 | 0.1 |
| 1 | 49054581 | rs12074072 | -1.027 | -0.703 | -1.039 | -1.731 | 0 | -0.014 | -0.014 | 0 | 0.13 | 0.14 | 0.03 | 0.01 | 0.06 | 0.18 |
| 1 | 49056695 | rs7543402 | -1.916 | -0.751 | -1.07 | -1.789 | 0.004 | -0.036 | -0.036 | 0.94 | 0.46 | 0.43 | 0.31 | 0.05 | 0.35 | 0.53 |
| 1 | 49059615 | rs319960 | -1.419 | -0.84 | -1.151 | -1.789 | 0.009 | 0.003 | 0.003 | 0 | 0.38 | 0.33 | 0.29 | 0.04 | 0.3 | 0.34 |
| 1 | 49064482 | rs6696631 | -0.136 | -0.346 | -0.943 | -1.373 | -0.008 | -0.021 | -0.021 | 0 | 0.08 | 0.11 | 0.03 | 0.01 | 0.04 | 0.08 |
| 1 | 49067014 | rs12127450 | -0.237 | -0.04 | -1.27 | -1.299 | -0.013 | 0.027 | 0.027 | 0 | 0.16 | 0.06 | 0.11 | 0 | 0.17 | 0.16 |
| 1 | 49067379 | rs319978 | -0.697 | -0.184 | -1.059 | -0.892 | -0.001 | -0.02 | -0.02 | 0 | 0.21 | 0.2 | 0.18 | 0.04 | 0.16 | 0.25 |
| 1 | 49070646 | rs6701767 | -0.177 | 0.027 | -1.255 | -0.826 | -0.006 | 0.072 | 0.072 | 0 | 0.2 | 0.09 | 0.12 | 0.07 | 0.15 | 0.24 |
| 1 | 49073954 | rs10888607 | -0.542 | -0.076 | -0.96 | -0.188 | -0.003 | 0.067 | 0.067 | 0 | 0.89 | 0.87 | 0.7 | 0.6 | 0.9 | 0.86 |

| | | | | | | | | | | | | | | | | |
|---|---|---|---|---|---|---|---|---|---|---|---|---|---|---|---|---|
| 1 | 49074468 | rs11811953 | 0.153 | -0.223 | -0.53 | -0.702 | -0.012 | -0.067 | -0.067 | 0 | 0.16 | 0.22 | 0.08 | 0 | 0.14 | 0.19 |
| 1 | 49075971 | rs320018 | 0.986 | 0.292 | -0.52 | -0.863 | -0.005 | 0.003 | 0.003 | 0 | 0.2 | 0.19 | 0.21 | 0.22 | 0.29 | 0.33 |
| 1 | 49076126 | rs320019 | 0.673 | 0.382 | -0.425 | -0.91 | 0.009 | -0.006 | -0.006 | 0.331 | 0.22 | 0.26 | 0.21 | 0.22 | 0.31 | 0.34 |
| 1 | 49076656 | rs320021 | 0.791 | 0.228 | 0.111 | -0.378 | -0.014 | 0.024 | 0.024 | 0 | 0.69 | 0.6 | 0.6 | 0.6 | 0.73 | 0.63 |
| 1 | 49077356 | rs4926745 | 1.717 | 0.178 | -0.176 | -0.485 | -0.001 | -0.008 | -0.008 | 0 | 0.26 | 0.22 | 0.27 | 0.32 | 0.3 | 0.08 |
| 1 | 49080074 | rs320025 | -1.153 | 0.353 | 0.681 | 0.33 | -0.015 | -0.117 | -0.117 | 0.46 | 0.2 | 0.35 | 0.1 | 0.01 | 0.17 | 0.25 |
| 1 | 49083071 | rs10888608 | 0.956 | 0.184 | 0.038 | 0.712 | 0.019 | -0.024 | -0.024 | 0 | 0.7 | 0.31 | 0.6 | 0.61 | 0.77 | 0.68 |
| 1 | 49086889 | rs320031 | 0.794 | 0.941 | 0.097 | 0.762 | -0.025 | 0.009 | 0.009 | 0 | 0.76 | 0.88 | 0.62 | 0.32 | 0.7 | 0.7 |
| 1 | 49092039 | rs11205517 | -1.463 | 0.82 | 1.433 | 0.562 | 0.043 | -0.01 | -0.01 | 0 | 0.11 | 0.44 | 0.1 | 0.04 | 0.08 | 0.05 |
| 1 | 49102047 | rs319986 | 0.708 | 0.862 | 1.252 | 0.787 | -0.026 | -0.007 | -0.007 | 0 | 0.75 | 0.88 | 0.61 | 0.35 | 0.68 | 0.67 |
| 1 | 49106416 | rs1999709 | 0.835 | 0.867 | 1.467 | 0.898 | -0.007 | -0.005 | -0.005 | 0.579 | 0.74 | 0.81 | 0.61 | 0.37 | 0.67 | 0.67 |
| 1 | 49107826 | rs12058997 | -1.382 | 0.865 | 1.451 | 0.764 | 0.019 | -0.025 | -0.025 | 0.013 | 0.08 | 0.21 | 0.08 | 0.03 | 0.07 | 0.2 |
| 1 | 49113247 | rs17373849 | -0.987 | 0.835 | 1.627 | 0.77 | 0.003 | -0.006 | -0.006 | 0 | 0.07 | 0.12 | 0.06 | 0.04 | 0.05 | 0.03 |
| 1 | 49113622 | rs319952 | 1.824 | 0.87 | 1.911 | 0.819 | -0.01 | 0.032 | 0.032 | 0 | 0.63 | 0.27 | 0.49 | 0.33 | 0.6 | 0.54 |
| 1 | 49121231 | rs319993 | 1.125 | 0.835 | 1.948 | 0.813 | 0.001 | 0 | 0 | 0 | 0.71 | 0.68 | 0.57 | 0.37 | 0.65 | 0.62 |
| 1 | 49123567 | rs7556311 | -1.275 | 0.832 | 2.061 | 0.807 | 0.065 | 0.034 | 0.034 | 0 | 0.08 | 0.36 | 0.07 | 0.04 | 0.05 | 0.03 |
| 1 | 49126778 | rs320003 | 1.884 | 0.815 | 2.225 | 0.802 | -0.009 | 0.043 | 0.043 | 0 | 0.62 | 0.25 | 0.49 | 0.33 | 0.6 | 0.54 |
| 1 | 49127186 | rs320004 | 1.705 | 0.815 | 2.166 | 0.803 | -0.006 | 0.019 | 0.019 | 0 | 0.63 | 0.31 | 0.51 | 0.33 | 0.6 | 0.59 |
| 1 | 49129465 | rs320010 | 0.277 | 0.973 | 1.746 | 1.331 | 0.032 | -0.055 | -0.055 | 4.251 | 0.81 | 0.92 | 0.79 | 0.6 | 0.88 | 0.76 |
| 1 | 49133886 | rs397260 | 1.667 | 0.964 | 1.8 | 1.147 | 0.006 | -0.016 | -0.016 | 0 | 0.64 | 0.57 | 0.52 | 0.33 | 0.6 | 0.54 |
| 1 | 49139531 | rs387244 | 1.072 | 0.903 | 1.435 | 1.326 | -0.031 | -0.027 | -0.027 | 0 | 0.73 | 0.88 | 0.61 | 0.37 | 0.66 | 0.68 |
| 1 | 49145514 | rs319959 | -1.901 | 1.177 | 1.416 | 1.232 | -0.015 | 0.059 | 0.059 | 0.158 | 0.42 | 0.73 | 0.64 | 0.66 | 0.47 | 0.53 |
| 1 | 49148942 | rs11205521 | -0.358 | 0.963 | 1.613 | 1.512 | -0.035 | -0.031 | -0.031 | 0.354 | 0.08 | 0.04 | 0.19 | 0.2 | 0.21 | 0.08 |
| 1 | 49151886 | rs319981 | -1.344 | 0.775 | 1.719 | 0.87 | -0.007 | 0.007 | 0.007 | 0.111 | 0.37 | 0.73 | 0.45 | 0.6 | 0.41 | 0.45 |
| 1 | 49164735 | rs12096159 | -1.825 | 0.545 | 2.25 | 0.853 | 0.045 | 0.03 | 0.03 | 0 | 0.07 | 0.3 | 0.04 | 0.01 | 0.03 | 0.08 |
| 1 | 49174405 | rs320050 | 2.019 | 0.505 | 1.504 | 0.891 | -0.005 | 0.012 | 0.012 | 0.575 | 0.91 | 0.53 | 0.94 | 0.97 | 0.97 | 0.86 |
| 1 | 49179498 | rs17104710 | -1.413 | 0.664 | 0.422 | 0.904 | -0.018 | -0.033 | -0.033 | 0.287 | 0.06 | 0.14 | 0.02 | 0 | 0.03 | 0.12 |
| 1 | 49184327 | rs7535594 | 0.988 | 1.254 | -0.797 | 1.337 | -0.042 | -0.056 | -0.056 | 0 | 0.78 | 0.92 | 0.68 | 0.63 | 0.76 | 0.58 |
| 1 | 49189775 | rs4926528 | -0.791 | 1.29 | -0.683 | 1.19 | -0.039 | -0.042 | -0.042 | 0.112 | 0.07 | 0.01 | 0.18 | 0.24 | 0.13 | 0.13 |
| 1 | 49193129 | rs2291335 | -0.967 | 1.265 | -0.643 | 1.182 | -0.019 | -0.022 | -0.022 | 0.057 | 0.08 | 0.03 | 0.19 | 0.24 | 0.13 | 0.14 |
| 1 | 49196334 | rs10888618 | -1.046 | 0.714 | -0.498 | 1.116 | 0.005 | -0.019 | -0.019 | 0.205 | 0.26 | 0.17 | 0.43 | 0.65 | 0.27 | 0.43 |
| 1 | 49206305 | rs6588054 | -1.075 | 0.481 | -0.22 | 0.799 | -0.017 | 0.004 | 0.004 | 0 | 0.24 | 0.37 | 0.44 | 0.74 | 0.22 | 0.43 |
| 1 | 49216371 | rs11586980 | 0.159 | 0.143 | -1.614 | -0.316 | 0.033 | 0.163 | 0.163 | 0.051 | 0.33 | 0.05 | 0.16 | 0.11 | 0.23 | 0.26 |
| 1 | 49219384 | rs1484412 | 0.584 | 0.713 | -1.598 | -0.928 | -0.035 | 0.218 | 0.218 | 0.122 | 0.47 | 0.18 | 0.41 | 0.03 | 0.5 | 0.39 |
| 1 | 49221356 | rs12096331 | -0.933 | 0.151 | -0.511 | 0.233 | -0.052 | 0.144 | 0.144 | 0 | 0.19 | 0.31 | 0.21 | 0.47 | 0.11 | 0.28 |
| 1 | 49222904 | rs1934409 | 0.766 | 0.114 | -0.191 | 0.215 | -0.07 | 0.211 | 0.211 | 0 | 0.81 | 0.6 | 0.78 | 0.54 | 0.89 | 0.72 |
| 1 | 49232075 | rs1316396 | -0.074 | 0.106 | -0.374 | 0.64 | 0.009 | -0.026 | -0.026 | 0 | 0.08 | 0.02 | 0.05 | 0.3 | 0.07 | 0.19 |
| 1 | 49234327 | rs12739065 | -0.135 | 0.102 | -0.353 | 0.638 | 0.015 | -0.024 | -0.024 | 0.013 | 0.09 | 0.02 | 0.05 | 0.3 | 0.07 | 0.2 |
| 1 | 49238750 | rs6664435 | 0.735 | 0.298 | -0.274 | 0.433 | 0.001 | 0.007 | 0.007 | 0 | 0.71 | 0.76 | 0.64 | 0.29 | 0.71 | 0.54 |
| 1 | 49241421 | rs1934379 | -0.386 | 0.289 | 0.2 | 0.36 | -0.013 | 0.012 | 0.012 | 0 | 0.1 | 0.17 | 0.15 | 0.16 | 0.06 | 0.08 |
| 1 | 49242966 | rs4926754 | 0.656 | 0.309 | 0.157 | 0.365 | 0.004 | 0.021 | 0.021 | 0 | 0.7 | 0.78 | 0.63 | 0.27 | 0.71 | 0.52 |
| 1 | 49249604 | rs4926757 | 0.883 | 0.231 | 0.665 | 0.387 | -0.007 | 0.034 | 0.034 | 0 | 0.68 | 0.5 | 0.52 | 0.41 | 0.72 | 0.67 |
| 1 | 49252456 | rs4926759 | 1.299 | 0.267 | 0.743 | 0.24 | -0.009 | -0.062 | -0.062 | 0.157 | 0.59 | 0.47 | 0.48 | 0.11 | 0.65 | 0.44 |
| 1 | 49261163 | rs7523675 | -0.07 | 0.28 | 0.788 | 0.306 | -0.012 | 0.012 | 0.012 | 0 | 0.08 | 0.11 | 0.1 | 0.14 | 0.03 | 0.06 |
| 1 | 49262542 | rs11205542 | 1.462 | 0.223 | 0.965 | 0.228 | 0.01 | 0.017 | 0.017 | 0 | 0.58 | 0.35 | 0.46 | 0.11 | 0.65 | 0.45 |
| 1 | 49279412 | rs2355686 | 0.928 | 0.159 | 1.141 | 0.3 | -0.031 | 0.061 | 0.061 | 0.328 | 0.89 | 0.45 | 0.82 | 0.84 | 0.96 | 0.93 |
| 1 | 49296228 | rs1934401 | 1.315 | 0.093 | 1.567 | 0.223 | -0.014 | -0.073 | -0.073 | 0 | 0.61 | 0.51 | 0.48 | 0.12 | 0.66 | 0.46 |
| 1 | 49315866 | rs4926765 | 0.594 | 0.054 | 1.595 | 0.233 | -0.019 | 0.055 | 0.055 | 0.032 | 0.91 | 0.62 | 0.83 | 0.85 | 0.97 | 0.95 |
| 1 | 49317239 | rs10788896 | -0.157 | 0.04 | 1.653 | 0.227 | -0.014 | 0.043 | 0.043 | 0 | 0.93 | 0.8 | 0.85 | 0.85 | 0.97 | 0.95 |
| 1 | 49317658 | rs7530277 | -1.145 | 0.048 | 1.581 | 0.144 | 0.002 | 0.021 | 0.021 | 0 | 0.33 | 0.19 | 0.38 | 0.72 | 0.34 | 0.49 |
| 1 | 49325290 | rs2153326 | 0.846 | 0.041 | 1.593 | 0.172 | -0.026 | 0.072 | 0.072 | 0.329 | 0.89 | 0.44 | 0.82 | 0.84 | 0.96 | 0.94 |
| 1 | 49335330 | rs6667780 | -1.338 | 0.055 | 1.46 | 0.176 | 0.001 | 0.002 | 0.002 | 0.364 | 0.36 | 0.6 | 0.4 | 0.72 | 0.31 | 0.48 |
| 1 | 49336456 | rs5017079 | -0.237 | 0.099 | 1.198 | 0.333 | -0.007 | -0.089 | -0.089 | 0.208 | 0.21 | 0.03 | 0.25 | 0.44 | 0.18 | 0.29 |
| 1 | 49346828 | rs4457622 | -0.117 | 0.083 | 1.127 | 0.487 | -0.016 | -0.004 | -0.004 | 0 | 0.05 | 0.01 | 0.16 | 0.38 | 0.06 | 0.18 |
| 1 | 49361990 | rs3121525 | 0.436 | 0.118 | 1.282 | 0.389 | 0.007 | 0.119 | 0.119 | 0 | 0.91 | 0.64 | 0.83 | 0.84 | 0.96 | 0.93 |
| 1 | 49365708 | rs3127565 | -1.562 | 0.195 | 1.263 | 0.378 | -0.007 | -0.007 | -0.007 | 0 | 0.33 | 0.4 | 0.37 | 0.72 | 0.29 | 0.47 |
| 1 | 49369206 | rs3121532 | -1.654 | 0.179 | 1.254 | 0.373 | -0.013 | -0.016 | -0.016 | 0.818 | 0.34 | 0.47 | 0.39 | 0.73 | 0.29 | 0.48 |
| 1 | 49374736 | rs3121537 | 0.366 | 0.29 | 1.207 | 0.532 | 0.018 | 0.185 | 0.185 | 0 | 0.91 | 0.79 | 0.91 | 0.71 | 0.88 | 0.84 |
| 1 | 49376820 | rs1343424 | -0.602 | 0.603 | 0.858 | 0.486 | -0.015 | -0.078 | -0.078 | 0 | 0.23 | 0.04 | 0.3 | 0.43 | 0.21 | 0.32 |
| 1 | 49378281 | rs17105150 | -0.428 | 0.839 | 1.207 | 0.283 | 0.017 | -0.007 | -0.007 | 0.69 | 0.11 | 0.17 | 0.05 | 0.08 | 0.14 | 0.06 |
| 1 | 49385466 | rs1108427 | -0.376 | 0.849 | 1.16 | 0.289 | 0.013 | -0.006 | -0.006 | 0 | 0.11 | 0.15 | 0.05 | 0.08 | 0.14 | 0.06 |
| 1 | 49390336 | rs11205572 | -0.202 | 0.778 | 1.184 | 0.281 | -0.012 | 0.018 | 0.018 | 0 | 0.07 | 0.15 | 0.12 | 0.15 | 0.04 | 0.06 |
| 1 | 49395184 | rs2040070 | -0.346 | 0.825 | 1.14 | 0.265 | 0.007 | -0.004 | -0.004 | 0 | 0.11 | 0.13 | 0.05 | 0.08 | 0.14 | 0.06 |
| 1 | 49403365 | rs17105276 | -0.834 | 0.803 | 1.255 | 0.252 | 0.055 | -0.009 | -0.009 | 0 | 0.12 | 0.31 | 0.06 | 0.08 | 0.15 | 0.07 |
| 1 | 49407295 | rs10493140 | -1.719 | 0.58 | 1.017 | 0.188 | 0.008 | 0.014 | 0.014 | 0 | 0.22 | 0.44 | 0.18 | 0.23 | 0.19 | 0.15 |
| 1 | 49416462 | rs12091032 | -1.011 | 0.61 | 0.99 | 0.268 | -0.004 | 0.077 | 0.077 | 0.492 | 0.11 | 0.51 | 0.14 | 0.15 | 0.04 | 0.08 |
| 1 | 49419688 | rs10888636 | -1.753 | 0.611 | 0.929 | 0.234 | 0.003 | 0.005 | 0.005 | 0.524 | 0.21 | 0.59 | 0.18 | 0.23 | 0.18 | 0.14 |
| 1 | 49428303 | rs12097797 | -0.889 | 0.346 | 0.89 | 0.315 | 0.023 | 0.119 | 0.119 | 0.239 | 0.1 | 0.42 | 0.13 | 0.15 | 0.04 | 0.08 |
| 1 | 49436107 | rs17105335 | -0.299 | 0.547 | 0.428 | 0.295 | 0 | 0 | 0 | 0 | 0.08 | 0.07 | 0.02 | 0.08 | 0.08 | 0.05 |

| | | | | | | | | | | | | | | | | |
|---|---|---|---|---|---|---|---|---|---|---|---|---|---|---|---|---|
| 1 | 49454215 | rs1543562 | 1.232 | 0.473 | 0.4 | -0.394 | -0.005 | -0.054 | -0.054 | 1.234 | 0.59 | 0.52 | 0.58 | 0.17 | 0.61 | 0.48 |
| 1 | 49461406 | rs3118220 | 0.509 | 0.446 | 0.599 | -0.386 | 0.002 | 0.009 | 0.009 | 0 | 0.69 | 0.74 | 0.62 | 0.27 | 0.69 | 0.53 |
| 1 | 49478333 | rs3118226 | 1.435 | 0.395 | 0.775 | -0.416 | 0.005 | 0.019 | 0.019 | 0.077 | 0.58 | 0.4 | 0.56 | 0.19 | 0.61 | 0.47 |
| 1 | 49511424 | rs3934240 | -0.4 | 0.331 | 0.607 | 0.27 | -0.013 | -0.075 | -0.075 | 0 | 0.2 | 0.04 | 0.26 | 0.42 | 0.19 | 0.28 |
| 1 | 49527451 | rs10888647 | 1.036 | 0.282 | 0.645 | 0.594 | -0.001 | 0 | 0 | 0 | 0.7 | 0.75 | 0.63 | 0.3 | 0.69 | 0.53 |
| 1 | 49545134 | rs632493 | 1.732 | 0.26 | 0.71 | 0.564 | -0.001 | 0.059 | 0.059 | 0 | 0.62 | 0.73 | 0.61 | 0.18 | 0.62 | 0.47 |
| 1 | 49554476 | rs589436 | -0.089 | 0.327 | 0.644 | 0.682 | -0.036 | -0.032 | -0.032 | 0 | 0.08 | 0 | 0.1 | 0.27 | 0.12 | 0.15 |
| 1 | 49580287 | rs6687090 | 0.334 | 0.207 | 0.843 | 0.493 | 0.012 | 0.084 | 0.084 | 0.07 | 0.07 | 0.25 | 0.1 | 0.15 | 0.04 | 0.06 |
| 1 | 49581006 | rs590503 | -0.056 | 0.261 | 0.834 | 0.591 | -0.017 | 0.001 | 0.001 | 0 | 0.92 | 0.97 | 0.9 | 0.68 | 0.89 | 0.87 |
| 1 | 49589847 | rs657452 | 1.549 | -0.371 | 0.531 | 0.02 | 0.006 | 0.013 | 0.013 | 0 | 0.59 | 0.42 | 0.54 | 0.26 | 0.63 | 0.46 |
| 1 | 49627502 | rs513360 | 0.844 | -0.227 | 0.801 | 0.178 | 0 | -0.011 | -0.011 | 0 | 0.69 | 0.65 | 0.62 | 0.26 | 0.69 | 0.51 |
| 1 | 49664304 | rs11591141 | 1.504 | -0.35 | 0.804 | 0.157 | -0.001 | -0.011 | -0.011 | 1.793 | 0.62 | 0.44 | 0.58 | 0.2 | 0.63 | 0.45 |
| 1 | 49709123 | rs4244635 | -0.259 | -0.326 | 0.893 | 0.169 | 0.001 | -0.011 | -0.011 | 0 | 0.93 | 0.84 | 0.96 | 0.94 | 0.93 | 0.96 |
| 1 | 49762418 | rs1431638 | 1.678 | -0.359 | 1.141 | 0.146 | 0.008 | 0.044 | 0.044 | 0.059 | 0.61 | 0.32 | 0.56 | 0.2 | 0.62 | 0.46 |
| 1 | 49765438 | rs241501 | -0.117 | -0.345 | 1.029 | 0.29 | -0.024 | -0.092 | -0.092 | 0 | 0.8 | 0.97 | 0.74 | 0.57 | 0.81 | 0.7 |
| 1 | 49782325 | rs375899 | 0.616 | -0.339 | 1.04 | 0.207 | 0 | -0.003 | -0.003 | 0 | 0.06 | 0.08 | 0.03 | 0.06 | 0.07 | 0.05 |
| 1 | 49799987 | rs241467 | 0.846 | -0.368 | 1.067 | 0.208 | 0 | 0.012 | 0.012 | 0 | 0.71 | 0.84 | 0.64 | 0.25 | 0.7 | 0.52 |
| 1 | 49800692 | rs241468 | -0.688 | -0.379 | 0.905 | 0.297 | -0.028 | 0.017 | 0.017 | 0 | 0.85 | 0.99 | 0.9 | 0.93 | 0.86 | 0.85 |
| 1 | 49828663 | rs7531656 | 0.951 | -0.216 | 0.96 | 0.212 | -0.01 | -0.025 | -0.025 | 0.702 | 0.68 | 0.52 | 0.6 | 0.25 | 0.7 | 0.52 |
| 1 | 49943664 | rs6679263 | 0.497 | -0.163 | 1.037 | 0.041 | 0.005 | 0.038 | 0.038 | 0 | 0.08 | 0.19 | 0.11 | 0.13 | 0.05 | 0.06 |
| 1 | 49949006 | rs1933271 | 1.606 | -0.134 | 1.033 | 0.03 | 0.01 | 0.044 | 0.044 | 0.079 | 0.61 | 0.32 | 0.56 | 0.19 | 0.63 | 0.46 |
| 1 | 49958961 | rs1185222 | 0.615 | -0.108 | 1.01 | 0.045 | 0.001 | 0.012 | 0.012 | 0 | 0.7 | 0.8 | 0.63 | 0.25 | 0.7 | 0.52 |
| 1 | 49990335 | rs12240044 | 0.418 | -0.093 | 1.048 | 0.073 | 0.022 | 0.007 | 0.007 | 0 | 0.07 | 0.19 | 0.04 | 0.06 | 0.06 | 0.04 |
| 1 | 50046741 | rs4926538 | -0.338 | -0.2 | 1.044 | 0.023 | -0.006 | 0.045 | 0.045 | 0 | 0.11 | 0.29 | 0.12 | 0.16 | 0.07 | 0.07 |
| 1 | 50049490 | rs1999875 | -1.518 | -0.192 | 1.061 | 0.018 | -0.003 | 0.034 | 0.034 | 0 | 0.18 | 0.48 | 0.15 | 0.21 | 0.14 | 0.12 |
| 1 | 50053885 | rs11587574 | 0.885 | -0.217 | 1.039 | 0.126 | -0.025 | -0.026 | -0.026 | 0 | 0.08 | 0.01 | 0.09 | 0.31 | 0.11 | 0.12 |
| 1 | 50057166 | rs2883976 | 0.114 | -0.22 | 1.039 | 0.113 | 0.008 | 0.048 | 0.048 | 0 | 0.09 | 0.28 | 0.12 | 0.13 | 0.05 | 0.07 |
| 1 | 50081573 | rs10888663 | 0.498 | -0.235 | 1.042 | 0.106 | 0.006 | 0.039 | 0.039 | 0.019 | 0.08 | 0.19 | 0.11 | 0.13 | 0.05 | 0.07 |
| 1 | 50081599 | rs11205633 | 0.628 | -0.223 | 0.999 | 0.076 | 0.003 | 0.013 | 0.013 | 0 | 0.7 | 0.81 | 0.64 | 0.25 | 0.7 | 0.52 |
| 1 | 50189321 | rs6660206 | 0.301 | -0.052 | 1.488 | 0.509 | 0.026 | 0 | 0 | 0.068 | 0.07 | 0.2 | 0.04 | 0.06 | 0.07 | 0.04 |
| 1 | 50192150 | rs1474910 | 0.155 | -0.045 | 1.485 | 0.568 | -0.018 | -0.088 | -0.088 | 0 | 0.2 | 0.04 | 0.27 | 0.45 | 0.19 | 0.3 |
| 1 | 50243854 | rs3899856 | 1.639 | 0.046 | 1.536 | 0.583 | 0.013 | 0.067 | 0.067 | 0.037 | 0.58 | 0.32 | 0.56 | 0.17 | 0.61 | 0.46 |
| 1 | 50312200 | rs4582848 | 0.763 | 0.071 | 1.508 | 0.66 | -0.01 | -0.027 | -0.027 | 0 | 0.68 | 0.52 | 0.6 | 0.25 | 0.69 | 0.51 |
| 1 | 50335528 | rs10736388 | 0.485 | 0.084 | 1.515 | 0.665 | 0.003 | 0.011 | 0.011 | 0 | 0.7 | 0.8 | 0.64 | 0.25 | 0.7 | 0.52 |
| 1 | 50359987 | rs10888672 | 1.517 | 0.075 | 1.263 | 0.665 | 0.013 | 0.067 | 0.067 | 0.037 | 0.58 | 0.32 | 0.56 | 0.16 | 0.61 | 0.46 |
| 1 | 50379476 | rs4926545 | 0.533 | 0.26 | 1.303 | 0.795 | 0.002 | 0.011 | 0.011 | 0 | 0.7 | 0.8 | 0.63 | 0.25 | 0.7 | 0.52 |
| 1 | 50391140 | rs11803306 | 0.494 | 0.278 | 1.315 | 0.823 | 0 | -0.003 | -0.003 | 0 | 0.06 | 0.08 | 0.03 | 0.06 | 0.07 | 0.04 |
| 1 | 50426542 | rs7518987 | -0.06 | 0.482 | 1.311 | 1.223 | 0.012 | 0.048 | 0.048 | 0 | 0.08 | 0.19 | 0.11 | 0.13 | 0.05 | 0.07 |
| 1 | 50456837 | rs11576945 | 0.071 | 0.812 | 1.205 | 1.695 | -0.025 | -0.026 | -0.026 | 0 | 0.08 | 0 | 0.09 | 0.32 | 0.11 | 0.12 |
| 1 | 50521604 | rs4388744 | -1.232 | 0.834 | 1.235 | 1.483 | 0.001 | 0.011 | 0.011 | 0 | 0.3 | 0.21 | 0.37 | 0.76 | 0.3 | 0.48 |
| 1 | 50546810 | rs4926547 | 1.17 | 0.785 | 1.198 | 1.434 | -0.013 | -0.033 | -0.033 | 0 | 0.67 | 0.52 | 0.58 | 0.24 | 0.69 | 0.52 |
| 1 | 50559162 | rs11579246 | -0.039 | 1.377 | 0.7 | 1.819 | -0.007 | -0.004 | -0.004 | 0.14 | 0.09 | 0.03 | 0.1 | 0.32 | 0.11 | 0.12 |
| 1 | 50566266 | rs10888679 | -1.319 | 1.341 | 0.753 | 1.379 | 0 | 0.009 | 0.009 | 0 | 0.29 | 0.2 | 0.38 | 0.76 | 0.29 | 0.48 |
| 1 | 50567676 | rs11205682 | -0.36 | 1.427 | 0.682 | 1.383 | 0.019 | 0.049 | 0.049 | 0 | 0.07 | 0.2 | 0.1 | 0.12 | 0.05 | 0.04 |
| 1 | 50577654 | rs2480677 | -0.661 | 1.332 | 0.728 | 1.618 | -0.001 | 0.045 | 0.045 | 1.114 | 0.13 | 0.11 | 0.12 | 0.43 | 0.16 | 0.12 |
| 1 | 50580823 | rs4078484 | -0.518 | 1.382 | 0.859 | 1.619 | 0.013 | 0.078 | 0.078 | 0 | 0.19 | 0.43 | 0.3 | 0.35 | 0.19 | 0.27 |
| 1 | 50582172 | rs967582 | -1.611 | 1.385 | 0.74 | 1.607 | 0.002 | -0.033 | -0.033 | 0.142 | 0.32 | 0.54 | 0.42 | 0.78 | 0.35 | 0.38 |
| 1 | 50589798 | rs9436444 | -0.894 | 1.28 | 0.848 | 1.651 | 0.008 | 0.058 | 0.058 | 0 | 0.19 | 0.36 | 0.3 | 0.35 | 0.19 | 0.28 |
| 1 | 50598368 | rs4307603 | -0.92 | 1.192 | 0.64 | 1.254 | 0.001 | 0.04 | 0.04 | 0 | 0.25 | 0.36 | 0.35 | 0.36 | 0.26 | 0.36 |
| 1 | 50602495 | rs6588376 | -0.501 | 1.453 | 0.684 | 1.518 | -0.005 | 0.064 | 0.064 | 0 | 0.2 | 0.25 | 0.3 | 0.36 | 0.19 | 0.35 |
| 1 | 50605337 | rs3902720 | -1.555 | 1.45 | 0.761 | 1.505 | 0.01 | -0.031 | -0.031 | 0.32 | 0.28 | 0.41 | 0.41 | 0.78 | 0.33 | 0.47 |
| 1 | 50613676 | rs4412638 | -1.133 | 1.463 | 0.488 | 1.519 | -0.001 | 0.034 | 0.034 | 0 | 0.26 | 0.35 | 0.35 | 0.36 | 0.26 | 0.36 |
| 1 | 50620945 | rs17105974 | -0.782 | 1.345 | 0.148 | 0.739 | -0.001 | -0.012 | -0.012 | 0.009 | 0.06 | 0.12 | 0.06 | 0 | 0.07 | 0.01 |
| 1 | 50652318 | rs2494875 | 1.563 | 1.118 | 1.584 | 0.635 | 0.093 | 0.06 | 0.06 | 0 | 0.95 | 0.58 | 0.96 | 1 | 0.97 | 0.98 |
| 1 | 50661411 | rs2480683 | 2.432 | 1.091 | 1.979 | 0.469 | -0.01 | -0.039 | -0.039 | 0.274 | 0.91 | 0.37 | 0.94 | 1 | 0.97 | 0.98 |
| 1 | 50666515 | rs2494876 | 2.54 | 1.16 | 1.861 | 1.036 | 0.005 | 0.224 | 0.224 | 1.904 | 0.89 | 0.34 | 0.88 | 0.65 | 0.9 | 0.88 |
| 1 | 50676278 | rs2494878 | 2.205 | 1.122 | 2.067 | 0.013 | 0.012 | -0.019 | -0.019 | 0 | 0.92 | 0.51 | 0.95 | 1 | 0.97 | 0.98 |
| 1 | 50679059 | rs2480687 | 1.902 | 1.122 | 2.092 | 0.013 | 0.052 | 0.018 | 0.018 | 1.06 | 0.93 | 0.54 | 0.95 | 1 | 0.97 | 0.99 |
| 1 | 50684153 | rs2494881 | 2.523 | 1.12 | 1.884 | 0.012 | -0.03 | -0.055 | -0.055 | 0 | 0.9 | 0.36 | 0.94 | 1 | 0.97 | 0.98 |
| 1 | 50716267 | rs4437920 | 2.123 | 1.03 | 1.819 | 0.085 | 0.092 | 0.174 | 0.174 | 0.493 | 0.93 | 0.58 | 0.94 | 0.82 | 0.93 | 0.86 |
| 1 | 50722989 | rs3001644 | -2.319 | 1.131 | 1.084 | 0.132 | 0.022 | 0.055 | 0.055 | 1.321 | 0.1 | 0.6 | 0.09 | 0.17 | 0.09 | 0.15 |
| 1 | 50735395 | rs12030672 | -0.222 | 1.093 | 0.277 | -0.163 | -0.063 | -0.104 | -0.104 | 0 | 0.2 | 0.01 | 0.29 | 0.65 | 0.23 | 0.35 |
| 1 | 50755278 | rs2938099 | 1.245 | 0.406 | 0.993 | -0.75 | -0.028 | -0.019 | -0.019 | 0.008 | 0.86 | 0.35 | 0.91 | 0.9 | 0.92 | 0.84 |
| 1 | 50765982 | rs1875635 | 1.118 | 0.7 | 1.064 | -0.877 | -0.003 | -0.022 | -0.022 | 0 | 0.8 | 0.48 | 0.85 | 0.84 | 0.76 | 0.73 |
| 1 | 50779946 | rs2356868 | 0.834 | 0.76 | 0.85 | -0.768 | 0.008 | 0 | 0 | 1.223 | 0.83 | 0.54 | 0.87 | 0.84 | 0.8 | 0.74 |
| 1 | 50800516 | rs6689749 | -0.131 | 0.873 | 0.208 | -1.36 | -0.001 | 0.043 | 0.043 | 0 | 0.55 | 0.59 | 0.33 | 0.13 | 0.49 | 0.44 |
| 1 | 50802855 | rs4491095 | 0.045 | 0.846 | 0.428 | -1.449 | 0.01 | -0.006 | -0.006 | 0 | 0.49 | 0.55 | 0.28 | 0.09 | 0.39 | 0.36 |
| 1 | 50814015 | rs1393632 | -0.15 | 0.689 | 0.474 | -1.492 | 0.009 | 0.08 | 0.08 | 0 | 0.44 | 0.35 | 0.67 | 0.87 | 0.52 | 0.55 |
| 1 | 50822880 | rs2653891 | -0.115 | 0.673 | 0.406 | -1.515 | 0.007 | 0.105 | 0.105 | 0 | 0.44 | 0.31 | 0.67 | 0.87 | 0.52 | 0.55 |

| | | | | | | | | | | | | | | | | |
|---|---|---|---|---|---|---|---|---|---|---|---|---|---|---|---|---|
| 1 | 50838990 | rs6660621 | 0.036 | 1.236 | 0.26 | -1.502 | -0.038 | 0.087 | 0.087 | 0 | 0.41 | 0.26 | 0.63 | 0.84 | 0.49 | 0.58 |
| 1 | 50844173 | rs2455636 | 0.466 | 1.191 | 1.106 | -1.632 | 0.068 | 0.114 | 0.114 | 0 | 0.89 | 0.68 | 0.88 | 0.84 | 0.83 | 0.83 |
| 1 | 50871847 | rs1502856 | -0.09 | 1.144 | 1.074 | -1.764 | 0 | 0.057 | 0.057 | 0 | 0.44 | 0.34 | 0.66 | 0.84 | 0.53 | 0.62 |
| 1 | 50874830 | rs2784143 | 0.746 | 1.017 | 1.674 | -1.83 | 0.045 | 0.096 | 0.096 | 0.486 | 0.88 | 0.69 | 0.86 | 0.84 | 0.83 | 0.85 |
| 1 | 50882159 | rs1278537 | -0.248 | 0.997 | 1.431 | -1.833 | 0.004 | 0.064 | 0.064 | 0 | 0.55 | 0.66 | 0.34 | 0.16 | 0.46 | 0.37 |
| 1 | 50900021 | rs1278516 | -0.241 | 0.97 | 1.689 | -1.635 | 0.017 | 0.032 | 0.032 | 0.326 | 0.06 | 0.08 | 0.07 | 0.08 | 0.1 | 0.06 |
| 1 | 50935031 | rs17106219 | -1.107 | 0.872 | 2.088 | -1.688 | 0.103 | 0.139 | 0.139 | 0.279 | 0.07 | 0.37 | 0.1 | 0.08 | 0.1 | 0.07 |
| 1 | 50945526 | rs3789576 | -0.227 | 0.888 | 1.773 | -1.68 | 0.017 | 0.032 | 0.032 | 0.326 | 0.06 | 0.08 | 0.07 | 0.08 | 0.1 | 0.06 |
| 1 | 50970475 | rs10493149 | -0.833 | 1.174 | 1.724 | -1.863 | 0.071 | 0.101 | 0.101 | 0.231 | 0.07 | 0.25 | 0.09 | 0.08 | 0.1 | 0.07 |
| 1 | 50972564 | rs1149795 | -0.728 | 0.765 | 1.075 | -1.865 | 0.04 | 0.009 | 0.009 | 0 | 0.83 | 0.98 | 0.94 | 0.91 | 0.9 | 0.68 |
| 1 | 51039300 | rs17383290 | 0.111 | 0.613 | 0.933 | -2.863 | 0.013 | 0.033 | 0.033 | 0 | 0.55 | 0.61 | 0.34 | 0.15 | 0.43 | 0.37 |
| 1 | 51092829 | rs12120719 | -1.103 | 0.812 | 0.715 | -2.64 | 0.017 | 0.01 | 0.01 | 0 | 0.07 | 0.21 | 0.07 | 0.07 | 0.09 | 0.06 |
| 1 | 51118253 | rs10788930 | -0.743 | 0.826 | 0.673 | -2.606 | 0.088 | 0.139 | 0.139 | 0 | 0.09 | 0.34 | 0.14 | 0.15 | 0.15 | 0.13 |
| 1 | 51121198 | rs11205753 | 0.808 | 0.117 | -0.073 | -2.813 | 0.022 | 0.085 | 0.085 | 0 | 0.14 | 0.02 | 0.07 | 0 | 0.11 | 0.28 |
| 1 | 51174330 | rs11205760 | 1.111 | 1.397 | 0.286 | -2.455 | -0.114 | -0.065 | -0.065 | 2.369 | 0.17 | 0.03 | 0.53 | 0.79 | 0.36 | 0.29 |
| 1 | 51199193 | rs10127531 | 0.735 | 0.253 | 0.723 | -2.415 | 0.042 | 0.135 | 0.135 | 0 | 0.55 | 0.76 | 0.33 | 0.15 | 0.41 | 0.36 |
| 1 | 51220127 | rs11205768 | 0.735 | 0.253 | 0.807 | -2.415 | 0.04 | 0.124 | 0.124 | 0 | 0.55 | 0.76 | 0.33 | 0.15 | 0.41 | 0.36 |
| 1 | 51242141 | rs12084054 | 0.183 | 0.297 | 1.042 | -2.368 | 0.009 | 0.017 | 0.017 | 0.048 | 0.06 | 0.1 | 0.07 | 0.08 | 0.1 | 0.06 |
| 1 | 51257608 | rs7522611 | 0.765 | 0.276 | 0.956 | -2.417 | 0.022 | 0.125 | 0.125 | 0 | 0.56 | 0.77 | 0.35 | 0.15 | 0.43 | 0.36 |
| 1 | 51280132 | rs6692340 | -0.124 | 0.134 | 1.096 | -2.452 | 0.028 | 0.028 | 0.028 | 0 | 0.82 | 0.91 | 0.94 | 0.93 | 0.9 | 0.64 |
| 1 | 51312619 | rs3747997 | 0.247 | 0.235 | 1.052 | -2.396 | 0.047 | 0.049 | 0.049 | 0 | 0.16 | 0.02 | 0.06 | 0.07 | 0.1 | 0.34 |
| 1 | 51347213 | rs12088617 | -0.323 | 0.331 | 1.668 | -2.176 | 0.066 | 0.081 | 0.081 | 0.022 | 0.07 | 0.24 | 0.09 | 0.08 | 0.1 | 0.07 |
| 1 | 51362441 | rs12076849 | -0.138 | 0.336 | 1.676 | -2.173 | 0.057 | 0.071 | 0.071 | 0.052 | 0.06 | 0.23 | 0.09 | 0.08 | 0.1 | 0.07 |
| 1 | 51403595 | rs4926873 | -0.761 | 0.334 | 1.402 | -2.218 | 0.019 | 0.087 | 0.087 | 0 | 0.44 | 0.27 | 0.65 | 0.85 | 0.57 | 0.64 |
| 1 | 51440093 | rs12855 | -0.253 | 0.463 | 1.281 | -2.084 | 0.115 | 0.134 | 0.134 | 0.118 | 0.07 | 0.36 | 0.09 | 0.08 | 0.1 | 0.06 |
| 1 | 51462439 | rs2487824 | 0.449 | 0.439 | 0.747 | -2.16 | 0.023 | 0.094 | 0.094 | 0 | 0.55 | 0.68 | 0.34 | 0.15 | 0.43 | 0.36 |
| 1 | 51470242 | rs11588271 | 0.832 | 0.362 | 0.67 | -2.231 | 0.041 | 0.055 | 0.055 | 0 | 0.5 | 0.41 | 0.26 | 0.14 | 0.33 | 0.34 |
| 1 | 51475488 | rs3813634 | 0.25 | 0.375 | 0.572 | -2.194 | 0.01 | 0.055 | 0.055 | 0 | 0.57 | 0.67 | 0.34 | 0.22 | 0.43 | 0.42 |
| 1 | 51483564 | rs11205800 | -1.284 | 0.68 | 1.411 | -1.395 | 0.019 | 0.073 | 0.073 | 0 | 0.09 | 0.3 | 0.07 | 0.13 | 0.06 | 0.11 |
| 1 | 51484626 | rs11205801 | -1.196 | 0.681 | 1.308 | -1.395 | 0.011 | 0.057 | 0.057 | 0 | 0.08 | 0.25 | 0.06 | 0.13 | 0.06 | 0.11 |
| 1 | 51500769 | rs17106389 | -2.049 | 0.677 | 1.323 | -1.414 | 0.012 | 0.059 | 0.059 | 0.346 | 0.15 | 0.55 | 0.15 | 0.21 | 0.16 | 0.17 |
| 1 | 51528697 | rs4131141 | -0.895 | 0.733 | 1.257 | -1.371 | 0.117 | 0.129 | 0.129 | 0.32 | 0.07 | 0.31 | 0.09 | 0.07 | 0.11 | 0.07 |
| 1 | 51544465 | rs7541663 | -2.1 | 0.736 | 1.043 | -1.296 | 0.013 | 0.081 | 0.081 | 0.61 | 0.13 | 0.59 | 0.11 | 0.23 | 0.12 | 0.18 |
| 1 | 51558856 | rs7530673 | -1.525 | 0.805 | 1.822 | -1.301 | 0.162 | 0.163 | 0.163 | 0.465 | 0.06 | 0.4 | 0.06 | 0.09 | 0.08 | 0.07 |
| 1 | 51570046 | rs4926879 | 0.431 | 0.244 | 1.514 | -1.624 | 0.05 | 0.04 | 0.04 | 0 | 0.8 | 0.89 | 0.9 | 0.91 | 0.87 | 0.61 |
| 1 | 51589655 | rs6588399 | 1.222 | 0.671 | 1.305 | -1.412 | 0 | 0 | 0 | 0 | 0.75 | 0.32 | 0.83 | 0.77 | 0.82 | 0.75 |
| 1 | 51616379 | rs12074413 | -2.561 | 0.83 | 1.285 | -1.278 | 0.055 | 0.079 | 0.079 | 0.861 | 0.12 | 0.5 | 0.13 | 0.18 | 0.14 | 0.12 |
| 1 | 51631884 | rs11205821 | -1.305 | 0.82 | 1.217 | -1.246 | 0.129 | 0.157 | 0.157 | 0.891 | 0.06 | 0.4 | 0.05 | 0.09 | 0.05 | 0.07 |
| 1 | 51632964 | rs12744230 | 0.412 | 1.669 | 0.298 | -1.347 | -0.133 | -0.083 | -0.083 | 0 | 0.2 | 0.02 | 0.52 | 0.74 | 0.33 | 0.32 |
| 1 | 51684980 | rs12740598 | -0.97 | 1.207 | 1.102 | -1.041 | 0.07 | 0.003 | 0.003 | 0.27 | 0.05 | 0.24 | 0.05 | 0 | 0.06 | 0.04 |
| 1 | 51699726 | rs4926884 | -0.681 | 1.283 | 0.73 | -1.071 | -0.001 | 0 | 0 | 0 | 0.08 | 0.08 | 0.07 | 0.08 | 0.11 | 0.07 |
| 1 | 51753722 | rs3748615 | -0.73 | 1.115 | 0.594 | -1.097 | -0.004 | 0 | 0 | 3.573 | 0.09 | 0.1 | 0.05 | 0.09 | 0.08 | 0.08 |
| 1 | 51753812 | rs635590 | 0.027 | 0.873 | 0.332 | -1.181 | 0.017 | -0.01 | -0.01 | 0 | 0.14 | 0.01 | 0.09 | 0.09 | 0.13 | 0.32 |
| 1 | 51754726 | rs12734773 | -0.896 | 1.351 | 0.739 | -1.001 | -0.004 | -0.002 | -0.002 | 0 | 0.09 | 0.1 | 0.05 | 0.06 | 0.08 | 0.06 |
| 1 | 51762087 | rs12074459 | -2.145 | 1.398 | 0.769 | -1.028 | 0.007 | -0.009 | -0.009 | 0 | 0.2 | 0.5 | 0.16 | 0.17 | 0.19 | 0.15 |
| 1 | 51781211 | rs12134299 | -2.111 | 1.72 | 0.7 | -1.018 | 0.02 | -0.002 | -0.002 | 0 | 0.2 | 0.59 | 0.2 | 0.16 | 0.27 | 0.14 |
| 1 | 51804731 | rs11205836 | -1.927 | 1.49 | 0.794 | -1.034 | 0.01 | 0.002 | 0.002 | 0.013 | 0.18 | 0.6 | 0.14 | 0.12 | 0.19 | 0.12 |
| 1 | 51814274 | rs6659736 | -2.331 | 1.45 | 1.116 | -1.089 | 0.025 | -0.111 | -0.111 | 0.185 | 0.19 | 0.55 | 0.16 | 0.01 | 0.2 | 0.09 |
| 1 | 51826921 | rs17567 | -2.348 | 1.439 | 1.163 | -1.099 | 0.022 | -0.106 | -0.106 | 0.011 | 0.19 | 0.57 | 0.16 | 0.01 | 0.2 | 0.09 |
| 1 | 51837737 | rs1105681 | -2.4 | 1.436 | 1.228 | -1.101 | 0.007 | -0.079 | -0.079 | 0.153 | 0.2 | 0.62 | 0.16 | 0.01 | 0.2 | 0.09 |
| 1 | 51869248 | rs12756284 | -0.11 | 1.436 | 1.446 | -1.11 | 0.001 | -0.011 | -0.011 | 0 | 0.06 | 0.08 | 0.05 | 0 | 0.08 | 0.01 |
| 1 | 51884383 | rs11205843 | -0.602 | 1.418 | 1.773 | -1.114 | -0.004 | -0.005 | -0.005 | 7.512 | 0.06 | 0.17 | 0.03 | 0.01 | 0.03 | 0.04 |
| 1 | 51903837 | rs10888721 | -2.841 | 1.417 | 1.727 | -1.117 | 0.007 | -0.079 | -0.079 | 0.316 | 0.2 | 0.62 | 0.16 | 0.01 | 0.2 | 0.09 |
| 1 | 51908788 | rs12565491 | -0.132 | 1.417 | 1.692 | -1.107 | -0.005 | -0.005 | -0.005 | 0.592 | 0.05 | 0.09 | 0.03 | 0.01 | 0.03 | 0.04 |
| 1 | 51919013 | rs1149789 | 0.377 | 1.838 | 1.094 | -0.936 | -0.12 | -0.128 | -0.128 | 0 | 0.19 | 0.01 | 0.46 | 0.69 | 0.31 | 0.29 |
| 1 | 51923614 | rs10788934 | -2.548 | 1.357 | 1.342 | -1.132 | 0.007 | -0.079 | -0.079 | 0.482 | 0.2 | 0.62 | 0.16 | 0.01 | 0.2 | 0.09 |
| 1 | 51939197 | rs10493155 | -2.579 | 1.076 | 0.995 | -1.119 | 0.019 | -0.107 | -0.107 | 0 | 0.2 | 0.57 | 0.16 | 0.01 | 0.2 | 0.09 |
| 1 | 51957970 | rs6668872 | -1.446 | 1.072 | 0.956 | -1.106 | 0.005 | -0.031 | -0.031 | 0 | 0.1 | 0.24 | 0.07 | 0.01 | 0.1 | 0.06 |
| 1 | 51992336 | rs12139552 | 0.123 | 0.69 | 0.263 | -1.31 | 0.028 | -0.081 | -0.081 | 0 | 0.16 | 0.01 | 0.07 | 0.2 | 0.11 | 0.38 |
| 1 | 51995495 | rs7543230 | -0.376 | 0.713 | 0.638 | -1.247 | 0.003 | 0.002 | 0.002 | 0.115 | 0.05 | 0.06 | 0.03 | 0.13 | 0.03 | 0.07 |
| 1 | 52003541 | rs12061410 | -2.138 | 0.745 | 0.639 | -1.261 | 0.002 | 0 | 0 | 0 | 0.22 | 0.52 | 0.17 | 0.14 | 0.2 | 0.16 |
| 1 | 52010203 | rs10399889 | -0.229 | 0.993 | 0.288 | -0.975 | 0.021 | -0.031 | -0.031 | 0 | 0.06 | 0.12 | 0.05 | 0 | 0.08 | 0.01 |
| 1 | 52015111 | rs10493157 | -0.229 | 0.993 | 0.288 | -0.975 | 0.021 | -0.031 | -0.031 | 0.895 | 0.06 | 0.12 | 0.05 | 0 | 0.07 | 0.01 |
| 1 | 52015192 | rs10493158 | -0.229 | 1.01 | 0.055 | -0.975 | 0.006 | -0.018 | -0.018 | 0.45 | 0.06 | 0.1 | 0.05 | 0 | 0.08 | 0.01 |
| 1 | 52016713 | rs6588412 | 0.002 | 0.918 | -0.115 | -1.054 | -0.032 | 0.062 | 0.062 | 0 | 0.61 | 0.72 | 0.4 | 0.12 | 0.52 | 0.31 |
| 1 | 52024718 | rs12071920 | -0.596 | 0.385 | 0.693 | -0.472 | 0.021 | 0.013 | 0.013 | 0 | 0.06 | 0.27 | 0.03 | 0 | 0.01 | 0 |
| 1 | 52026384 | rs7534689 | -0.079 | 0.403 | 0.343 | -1.047 | 0.003 | 0.014 | 0.014 | 0 | 0.58 | 0.66 | 0.35 | 0.14 | 0.46 | 0.32 |
| 1 | 52033122 | rs12038297 | -1.567 | 0.015 | 0.537 | -1.059 | -0.058 | 0.007 | 0.007 | 0 | 0.18 | 0.51 | 0.12 | 0.12 | 0.09 | 0.13 |
| 1 | 52035067 | rs10888725 | -1.416 | 0.017 | 0.592 | -0.96 | -0.045 | 0.064 | 0.064 | 0 | 0.13 | 0.4 | 0.09 | 0.12 | 0.05 | 0.11 |

| | | | | | | | | | | | | | | | | |
|---|---|---|---|---|---|---|---|---|---|---|---|---|---|---|---|---|
| 1 | 52088992 | rs12058362 | 0.187 | 0.124 | 0.407 | -0.946 | -0.013 | -0.02 | -0.02 | 0 | 0.05 | 0.07 | 0.01 | 0 | 0.01 | 0 |
| 1 | 52097580 | rs6695853 | -0.231 | -0.816 | 0.558 | -1.561 | -0.056 | -0.053 | -0.053 | 0 | 0.36 | 0.5 | 0.18 | 0.12 | 0.19 | 0.11 |
| 1 | 52112526 | rs11205875 | 0.352 | -0.826 | 0.748 | -1.287 | -0.024 | -0.034 | -0.034 | 0.3 | 0.06 | 0.14 | 0.02 | 0 | 0.01 | 0.02 |
| 1 | 52135979 | rs11205877 | -0.283 | -0.934 | 1.202 | -1.341 | -0.04 | -0.038 | -0.038 | 0 | 0.36 | 0.53 | 0.18 | 0.12 | 0.19 | 0.11 |
| 1 | 52144929 | rs11205878 | 0.527 | -0.828 | 1.193 | -1.293 | -0.026 | -0.036 | -0.036 | 0.044 | 0.07 | 0.14 | 0.02 | 0 | 0.01 | 0.02 |
| 1 | 52163537 | rs12074299 | 0.075 | -0.387 | 1.044 | -1.17 | -0.006 | -0.051 | -0.051 | 0 | 0.3 | 0.32 | 0.14 | 0.01 | 0.19 | 0.09 |
| 1 | 52165225 | rs12130751 | 1.461 | -0.132 | 0.282 | -0.775 | -0.094 | -0.163 | -0.163 | 0 | 0.2 | 0.02 | 0.5 | 0.61 | 0.29 | 0.26 |
| 1 | 52221714 | rs10489546 | -0.036 | -0.648 | 0.472 | -1.504 | -0.055 | -0.055 | -0.055 | 0 | 0.34 | 0.5 | 0.18 | 0.12 | 0.19 | 0.12 |
| 1 | 52252006 | rs856614 | -0.596 | -0.444 | 0.066 | -1.667 | -0.002 | 0.065 | 0.065 | 0 | 0.69 | 0.8 | 0.83 | 0.99 | 0.71 | 0.52 |
| 1 | 52253793 | rs12097772 | -0.205 | -0.964 | 0.385 | -0.964 | -0.051 | -0.043 | -0.043 | 0 | 0.33 | 0.49 | 0.18 | 0.13 | 0.19 | 0.12 |
| 1 | 52261130 | rs856613 | 0.437 | -0.928 | 0.355 | -0.884 | -0.061 | -0.128 | -0.128 | 0 | 0.2 | 0.46 | 0.12 | 0.01 | 0.1 | 0.07 |
| 1 | 52263082 | rs1417364 | -0.275 | -1.089 | 0.097 | -0.783 | -0.062 | -0.016 | -0.016 | 0 | 0.28 | 0.61 | 0.21 | 0.14 | 0.12 | 0.09 |
| 1 | 52264064 | rs1770791 | -0.998 | -1.005 | -0.287 | -1.043 | 0.04 | 0.042 | 0.042 | 0.007 | 0.44 | 0.67 | 0.37 | 0.15 | 0.24 | 0.27 |
| 1 | 52266350 | rs2273198 | 0.553 | -1.278 | -0.272 | -1.077 | 0.001 | -0.003 | -0.003 | 0 | 0.13 | 0.06 | 0.08 | 0.09 | 0.02 | 0.06 |
| 1 | 52293570 | rs2077725 | 0.14 | -0.917 | 0.298 | -1.769 | 0.002 | -0.048 | -0.048 | 0 | 0.67 | 0.4 | 0.77 | 0.93 | 0.64 | 0.5 |
| 1 | 52316785 | rs6676789 | 0.779 | -1.147 | 0.241 | -1.979 | 0.014 | 0.102 | 0.102 | 0.056 | 0.51 | 0.31 | 0.69 | 0.93 | 0.55 | 0.45 |
| 1 | 52322129 | rs11585468 | 0.977 | -1.123 | -0.021 | -2.094 | -0.063 | 0.125 | 0.125 | 0 | 0.2 | 0.02 | 0.14 | 0.02 | 0.26 | 0.26 |
| 1 | 52324228 | rs2842576 | -1.943 | -1.109 | 0.02 | -2.041 | 0.03 | 0.048 | 0.048 | 0.86 | 0.84 | 0.91 | 0.92 | 1 | 0.91 | 0.95 |
| 1 | 52343297 | rs6663305 | -0.448 | -1.066 | 0.367 | -2.054 | 0.015 | -0.095 | -0.095 | 0.247 | 0.31 | 0.43 | 0.21 | 0.07 | 0.37 | 0.5 |
| 1 | 52345760 | rs2842586 | -0.432 | -1.07 | 0.06 | -2.083 | -0.026 | 0.234 | 0.234 | 0 | 0.64 | 0.91 | 0.78 | 0.98 | 0.64 | 0.69 |
| 1 | 52347038 | rs10888738 | 1.531 | -1.096 | 0.381 | -1.836 | 0.033 | 0.134 | 0.134 | 0 | 0.53 | 0.28 | 0.68 | 0.85 | 0.6 | 0.48 |
| 1 | 52348349 | rs11205901 | -0.964 | -0.918 | 0.319 | -1.679 | -0.04 | 0.077 | 0.077 | 0 | 0.81 | 0.92 | 0.85 | 0.99 | 0.73 | 0.77 |
| 1 | 52353449 | rs3001778 | -0.645 | -0.856 | 0.151 | -1.748 | -0.021 | 0.048 | 0.048 | 0 | 0.79 | 0.89 | 0.85 | 0.94 | 0.72 | 0.66 |
| 1 | 52373863 | rs1045633 | -0.187 | -0.908 | 0.707 | -1.588 | -0.043 | -0.105 | -0.105 | 3.079 | 0.16 | 0.28 | 0.06 | 0 | 0.07 | 0.03 |
| 1 | 52375677 | rs2244156 | -1.406 | -0.988 | 0.89 | -1.684 | -0.029 | -0.101 | -0.101 | 0.052 | 0.24 | 0.56 | 0.1 | 0 | 0.12 | 0.13 |
| 1 | 52380808 | rs2842573 | -0.039 | -0.926 | 1.003 | -1.647 | 0.039 | -0.018 | -0.018 | 0 | 0.07 | 0.28 | 0.04 | 0 | 0.05 | 0.09 |
| 1 | 52396943 | rs12566236 | 0.422 | -0.893 | 0.907 | -1.795 | 0.03 | -0.078 | -0.078 | 1.203 | 0.71 | 0.57 | 0.71 | 0.89 | 0.63 | 0.65 |
| 1 | 52400097 | rs10888740 | 1.647 | -0.965 | 0.997 | -1.811 | 0.023 | 0.11 | 0.11 | 0.146 | 0.56 | 0.34 | 0.69 | 0.85 | 0.6 | 0.52 |
| 1 | 52402956 | rs10888742 | -0.086 | -0.582 | 0.798 | -1.606 | 0.06 | -0.004 | -0.004 | 0 | 0.93 | 0.68 | 0.95 | 1 | 0.95 | 0.91 |
| 1 | 52407430 | rs4503295 | 0.536 | -0.579 | 0.798 | -1.609 | 0.073 | -0.052 | -0.052 | 0 | 0.9 | 0.65 | 0.92 | 1 | 0.9 | 0.9 |
| 1 | 52412005 | rs7514201 | 0.257 | -0.603 | 0.831 | -1.579 | 0.027 | -0.024 | -0.024 | 0 | 0.07 | 0.21 | 0.03 | 0 | 0.05 | 0.09 |
| 1 | 52416078 | rs2809942 | -0.103 | -0.599 | 0.55 | -1.577 | 0.06 | -0.004 | -0.004 | 0 | 0.93 | 0.68 | 0.95 | 1 | 0.95 | 0.91 |
| 1 | 52417538 | rs2809944 | 0.575 | -0.491 | -0.4 | -1.466 | -0.03 | 0.021 | 0.021 | 0.228 | 0.19 | 0.09 | 0.21 | 0.1 | 0.27 | 0.24 |
| 1 | 52423167 | rs10888743 | 1.179 | -0.221 | -0.557 | -1.111 | -0.087 | 0.033 | 0.033 | 0 | 0.36 | 0.08 | 0.5 | 0.78 | 0.38 | 0.35 |
| 1 | 52431393 | rs1167169 | 0.107 | -0.429 | -0.136 | -1.033 | 0.006 | 0.002 | 0.002 | 0 | 0.09 | 0.04 | 0.04 | 0 | 0.05 | 0.04 |
| 1 | 52431812 | rs10888744 | 0.926 | -0.257 | 0.1 | -0.188 | 0.015 | -0.039 | -0.039 | 0 | 0.35 | 0.23 | 0.39 | 0.52 | 0.31 | 0.35 |
| 1 | 52466087 | rs1167174 | -1.892 | -0.236 | 1.763 | -0.119 | 0 | 0.038 | 0.038 | 1.71 | 0.15 | 0.5 | 0.14 | 0.18 | 0.09 | 0.21 |
| 1 | 52486104 | rs7512076 | 0.314 | -0.194 | 1.338 | -0.091 | 0.001 | -0.021 | -0.021 | 0.355 | 0.05 | 0.09 | 0.03 | 0 | 0.05 | 0.08 |
| 1 | 52498370 | rs2809917 | -2.282 | -0.243 | 1.562 | -0.105 | -0.016 | 0.029 | 0.029 | 1.167 | 0.16 | 0.41 | 0.14 | 0.18 | 0.09 | 0.21 |
| 1 | 52499939 | rs17106988 | -0.246 | 0.102 | 1.091 | -0.01 | 0.004 | -0.031 | -0.031 | 0 | 0.06 | 0.15 | 0.04 | 0 | 0.05 | 0.08 |
| 1 | 52630296 | rs7512531 | 1.471 | -0.66 | 2.199 | -0.255 | -0.025 | -0.006 | -0.006 | 0 | 0.81 | 0.34 | 0.84 | 0.83 | 0.92 | 0.81 |
| 1 | 52658156 | rs11205940 | 1.78 | -0.702 | 2.072 | -0.275 | -0.003 | 0 | 0 | 0 | 0.8 | 0.25 | 0.84 | 0.83 | 0.92 | 0.81 |
| 1 | 52696159 | rs6657731 | 0.282 | -0.649 | 2.33 | -0.31 | -0.009 | -0.04 | -0.04 | 4.991 | 0.09 | 0.15 | 0.04 | 0 | 0.05 | 0.09 |
| 1 | 52700196 | rs4272565 | -1.175 | -0.692 | 2.646 | -0.276 | -0.029 | 0.016 | 0.016 | 0 | 0.14 | 0.31 | 0.11 | 0.16 | 0.06 | 0.19 |
| 1 | 52708067 | rs1502689 | -2.061 | -0.963 | 2.768 | -0.167 | -0.035 | -0.011 | -0.011 | 0 | 0.2 | 0.7 | 0.15 | 0.16 | 0.08 | 0.19 |
| 1 | 52718881 | rs10493164 | 0.571 | -0.746 | 2.509 | -0.588 | -0.001 | -0.018 | -0.018 | 0 | 0.09 | 0.11 | 0.04 | 0 | 0.05 | 0.09 |
| 1 | 52744754 | rs11205949 | -1.655 | -0.757 | 2.417 | -0.592 | -0.035 | -0.01 | -0.01 | 0 | 0.18 | 0.7 | 0.14 | 0.16 | 0.06 | 0.19 |
| 1 | 52754871 | rs1288706 | 1.381 | 0.048 | 1.934 | -0.68 | 0.006 | 0 | 0 | 0 | 0.07 | 0.05 | 0.11 | 0.04 | 0.18 | 0.07 |
| 1 | 52754974 | rs7541658 | 0.994 | -0.232 | 1.951 | -0.675 | 0.003 | 0 | 0 | 0 | 0.08 | 0.06 | 0.03 | 0 | 0.05 | 0.09 |
| 1 | 52796250 | rs2762821 | 0.695 | -0.923 | 1.379 | -1.231 | 0.003 | 0.053 | 0.053 | 0 | 0.19 | 0.09 | 0.14 | 0.04 | 0.19 | 0.14 |
| 1 | 52811581 | rs7517338 | -1.566 | -0.879 | 1.769 | -1.178 | -0.029 | -0.008 | -0.008 | 0.712 | 0.18 | 0.68 | 0.14 | 0.16 | 0.06 | 0.2 |
| 1 | 52814170 | rs7546223 | 0.591 | -0.838 | 1.812 | -1.141 | -0.008 | -0.032 | -0.032 | 0 | 0.09 | 0.15 | 0.04 | 0 | 0.05 | 0.1 |
| 1 | 52816959 | rs17364363 | 0.922 | -0.624 | 1.748 | -1.105 | -0.001 | -0.017 | -0.017 | 0 | 0.08 | 0.09 | 0.03 | 0 | 0.05 | 0.1 |
| 1 | 52818188 | rs7513934 | -1.447 | -0.57 | 1.079 | -1.484 | -0.061 | 0.178 | 0.178 | 0 | 0.59 | 0.91 | 0.39 | 0.17 | 0.48 | 0.52 |
| 1 | 52836139 | rs9633423 | 1.913 | -0.531 | 0.375 | -0.785 | -0.143 | -0.141 | -0.141 | 0 | 0.31 | 0.02 | 0.5 | 0.6 | 0.38 | 0.26 |
| 1 | 52836736 | rs10157619 | -1.798 | -0.177 | 1.169 | -0.5 | -0.004 | 0 | 0 | 2.453 | 0.19 | 0.77 | 0.15 | 0.16 | 0.06 | 0.2 |
| 1 | 52843465 | rs7537734 | -0.46 | -0.101 | 1.227 | -0.477 | -0.035 | 0.004 | 0.004 | 0 | 0.13 | 0.3 | 0.11 | 0.13 | 0.06 | 0.14 |
| 1 | 52888119 | rs17107212 | 1.913 | 0.359 | 0.122 | -0.428 | -0.038 | -0.056 | -0.056 | 0 | 0.09 | 0.01 | 0.08 | 0.2 | 0.11 | 0.12 |
| 1 | 52889891 | rs17107217 | -0.756 | 0.147 | 0.644 | -0.593 | -0.034 | -0.089 | -0.089 | 0 | 0.13 | 0.55 | 0.07 | 0 | 0.05 | 0.11 |
| 1 | 52897395 | rs10493166 | 1.952 | 0.163 | 0.557 | -0.585 | -0.038 | -0.056 | -0.056 | 0.977 | 0.09 | 0.01 | 0.08 | 0.2 | 0.11 | 0.12 |
| 1 | 52901752 | rs17373092 | 0.894 | 0.148 | 0.684 | -0.593 | -0.001 | -0.018 | -0.018 | 0 | 0.08 | 0.08 | 0.04 | 0 | 0.05 | 0.11 |
| 1 | 52908280 | rs970917 | -1.849 | 0.02 | 0.811 | -0.662 | 0.018 | 0.014 | 0.014 | 0.896 | 0.29 | 0.78 | 0.19 | 0.24 | 0.19 | 0.31 |
| 1 | 52953766 | rs6682550 | -1.495 | 0.216 | 0.893 | -0.671 | 0.016 | 0.007 | 0.007 | 1.11 | 0.27 | 0.78 | 0.18 | 0.24 | 0.17 | 0.29 |
| 1 | 52955458 | rs10493168 | 1.982 | 0.18 | 0.835 | -0.654 | -0.038 | -0.056 | -0.056 | 0 | 0.09 | 0.01 | 0.08 | 0.2 | 0.11 | 0.12 |
| 1 | 52961888 | rs11205964 | 1.238 | 0.174 | 0.854 | -0.657 | 0.005 | 0.005 | 0.005 | 0 | 0.07 | 0.04 | 0.03 | 0 | 0.05 | 0.11 |
| 1 | 52996289 | rs7512677 | 0.21 | 0.215 | 0.843 | -0.768 | 0.005 | 0.029 | 0.029 | 0 | 0.16 | 0.25 | 0.12 | 0.23 | 0.14 | 0.2 |
| 1 | 53008039 | rs11800551 | 0.967 | 0.208 | 0.667 | -0.775 | -0.002 | -0.018 | -0.018 | 0.165 | 0.08 | 0.08 | 0.04 | 0 | 0.05 | 0.11 |
| 1 | 53017445 | rs1493365 | -1.573 | -0.051 | 0.569 | -0.632 | 0.022 | 0.014 | 0.014 | 1.466 | 0.28 | 0.78 | 0.18 | 0.21 | 0.16 | 0.3 |
| 1 | 53039603 | rs17373814 | 0.415 | 0.207 | 0.508 | -0.566 | -0.011 | -0.045 | -0.045 | 0 | 0.08 | 0.13 | 0.04 | 0 | 0.05 | 0.11 |

| | | | | | | | | | | | | | | | | |
|---|---|---|---|---|---|---|---|---|---|---|---|---|---|---|---|---|
| 1 | 53045440 | rs17365357 | -0.384 | 0.229 | 0.428 | -0.558 | -0.005 | -0.045 | -0.045 | 0 | 0.11 | 0.24 | 0.06 | 0.03 | 0.09 | 0.17 |
| 1 | 53050461 | rs6679249 | 0.056 | 0.414 | 0.378 | -0.621 | 0.074 | -0.05 | -0.05 | 1.131 | 0.1 | 0.33 | 0.12 | 0 | 0.14 | 0.07 |
| 1 | 53051993 | rs12737517 | 1.226 | 0.314 | -0.702 | -0.639 | -0.014 | -0.052 | -0.052 | 0 | 0.43 | 0.16 | 0.46 | 0.47 | 0.48 | 0.4 |
| 1 | 53056187 | rs12138419 | 0.731 | 0.391 | -0.543 | -1.019 | -0.003 | -0.007 | -0.007 | 0.953 | 0.07 | 0.06 | 0.03 | 0 | 0.06 | 0.02 |
| 1 | 53059106 | rs835337 | 0.29 | 0.303 | -0.494 | -1.028 | -0.01 | -0.006 | -0.006 | 0 | 0.19 | 0.12 | 0.2 | 0.2 | 0.25 | 0.19 |
| 1 | 53061766 | rs835338 | 0.23 | 0.297 | -0.792 | -1.031 | -0.023 | -0.014 | -0.014 | 0.129 | 0.2 | 0.11 | 0.2 | 0.2 | 0.25 | 0.19 |
| 1 | 53064034 | rs835342 | 1.48 | 0.166 | -0.515 | -1.2 | 0.005 | -0.017 | -0.017 | 0.007 | 0.59 | 0.43 | 0.68 | 0.77 | 0.58 | 0.62 |
| 1 | 53067083 | rs1416094 | -1.614 | 0.398 | -0.592 | -1.108 | -0.042 | -0.057 | -0.057 | 0 | 0.87 | 0.97 | 0.88 | 0.8 | 0.84 | 0.87 |
| 1 | 53073327 | rs946154 | 0.725 | 0.345 | -0.318 | -1.16 | 0.018 | -0.022 | -0.022 | 0 | 0.7 | 0.56 | 0.74 | 0.8 | 0.68 | 0.79 |
| 1 | 53074412 | rs1047619 | 0.142 | 0.101 | -0.486 | -1.28 | -0.049 | -0.091 | -0.091 | 0 | 0.14 | 0.38 | 0.06 | 0 | 0.06 | 0.08 |
| 1 | 53074532 | rs1047635 | 1.692 | 0.135 | -0.663 | -1.272 | -0.005 | 0.037 | 0.037 | 0 | 0.56 | 0.32 | 0.67 | 0.77 | 0.58 | 0.64 |
| 1 | 53076663 | rs7529595 | -0.126 | 0.209 | -0.984 | -0.856 | -0.057 | -0.031 | -0.031 | 0 | 0.78 | 0.93 | 0.78 | 0.77 | 0.7 | 0.74 |
| 1 | 53081321 | rs6588432 | -1.396 | 0.128 | -0.899 | -0.79 | -0.043 | -0.056 | -0.056 | 0 | 0.87 | 0.97 | 0.88 | 0.8 | 0.84 | 0.87 |
| 1 | 53081585 | rs6671552 | 0.316 | -0.222 | -0.529 | -1.041 | -0.031 | -0.074 | -0.074 | 0 | 0.12 | 0.29 | 0.06 | 0 | 0.06 | 0.08 |
| 1 | 53085337 | rs6672470 | 0.978 | -0.255 | -0.655 | -1.059 | -0.017 | -0.049 | -0.049 | 0.065 | 0.11 | 0.14 | 0.04 | 0 | 0.06 | 0.08 |
| 1 | 53094986 | rs2050695 | -0.862 | -0.201 | -1.195 | -1.366 | -0.104 | 0.06 | 0.06 | 0 | 0.63 | 0.9 | 0.67 | 0.75 | 0.48 | 0.67 |
| 1 | 53095582 | rs11585624 | 0.875 | -0.389 | -0.981 | -1.416 | -0.003 | -0.057 | -0.057 | 0.786 | 0.1 | 0.18 | 0.09 | 0 | 0.09 | 0.04 |
| 1 | 53102133 | rs11205983 | 0.501 | -0.888 | -1.143 | -1.156 | -0.082 | 0.177 | 0.177 | 0 | 0.16 | 0.39 | 0.35 | 0.34 | 0.04 | 0.22 |
| 1 | 53103490 | rs523451 | -0.121 | -0.815 | -1.271 | -1.143 | -0.071 | 0.113 | 0.113 | 0.062 | 0.2 | 0.57 | 0.39 | 0.36 | 0.08 | 0.26 |
| 1 | 53103888 | rs11205985 | 0.192 | -0.816 | -1.312 | -1.222 | -0.017 | -0.09 | -0.09 | 0.145 | 0.21 | 0.26 | 0.12 | 0.01 | 0.16 | 0.12 |
| 1 | 53106057 | rs554870 | -0.892 | -0.8 | -1.421 | -1.184 | 0.006 | -0.007 | -0.007 | 0 | 0.29 | 0.71 | 0.47 | 0.36 | 0.17 | 0.3 |
| 1 | 53110002 | rs17374258 | 0.245 | -0.643 | -1.416 | -1.16 | 0.001 | -0.035 | -0.035 | 0 | 0.2 | 0.16 | 0.12 | 0 | 0.16 | 0.11 |
| 1 | 53111724 | rs6682749 | 1.418 | -0.302 | -1.428 | -1.225 | -0.011 | 0.023 | 0.023 | 0 | 0.27 | 0.17 | 0.2 | 0 | 0.34 | 0.21 |
| 1 | 53111782 | rs936632 | -1.721 | 0.178 | -1.306 | -0.802 | -0.019 | -0.026 | -0.026 | 0.07 | 0.88 | 0.92 | 0.88 | 0.71 | 0.83 | 0.87 |
| 1 | 53114318 | rs6692566 | -0.594 | -0.266 | -0.831 | -0.224 | 0 | 0.021 | 0.021 | 0 | 0.42 | 0.24 | 0.28 | 0.36 | 0.4 | 0.39 |
| 1 | 53114736 | rs706497 | -0.673 | 0.44 | -0.818 | -0.628 | 0.001 | -0.019 | -0.019 | 0 | 0.39 | 0.49 | 0.59 | 0.69 | 0.37 | 0.52 |
| 1 | 53115200 | rs625829 | -0.921 | 0.175 | -0.967 | -0.656 | -0.002 | -0.037 | -0.037 | 2.287 | 0.14 | 0.15 | 0.09 | 0 | 0.12 | 0.04 |
| 1 | 53125600 | rs681865 | 0.073 | 0.339 | -1.444 | -0.794 | -0.028 | 0.146 | 0.146 | 0 | 0.43 | 0.78 | 0.33 | 0.14 | 0.46 | 0.33 |
| 1 | 53126155 | rs269324 | 0.138 | 0.16 | -1.666 | -0.489 | 0.008 | 0.001 | 0.001 | 0 | 0.37 | 0.13 | 0.54 | 0.65 | 0.32 | 0.47 |
| 1 | 53135933 | rs269316 | 1.119 | -0.038 | -0.928 | -0.884 | -0.019 | 0.04 | 0.04 | 0 | 0.31 | 0.15 | 0.22 | 0.01 | 0.36 | 0.31 |
| 1 | 53143271 | rs269303 | 0.443 | -0.061 | 0.072 | -0.887 | 0.003 | -0.071 | -0.071 | 0 | 0.37 | 0.48 | 0.25 | 0.15 | 0.39 | 0.34 |
| 1 | 53153044 | rs407856 | 0.202 | -0.166 | 0.153 | -0.937 | 0.002 | 0.094 | 0.094 | 0 | 0.4 | 0.79 | 0.3 | 0.15 | 0.4 | 0.34 |
| 1 | 53153432 | rs443751 | 0.972 | 0.098 | 0.077 | -0.929 | -0.034 | 0.082 | 0.082 | 0 | 0.29 | 0.16 | 0.22 | 0.01 | 0.37 | 0.31 |
| 1 | 53155867 | rs428366 | 0.968 | 0.1 | 0.099 | -0.929 | -0.029 | 0.069 | 0.069 | 0 | 0.3 | 0.18 | 0.22 | 0.01 | 0.37 | 0.31 |
| 1 | 53159824 | rs451821 | 0.479 | -0.493 | 0.62 | -0.209 | 0.02 | 0.014 | 0.014 | 0 | 0.71 | 0.54 | 0.46 | 0.33 | 0.68 | 0.59 |
| 1 | 53161620 | rs448788 | 1.588 | -1.033 | 0.344 | -0.592 | -0.098 | 0.249 | 0.249 | 0.326 | 0.4 | 0.09 | 0.28 | 0.02 | 0.49 | 0.37 |
| 1 | 53164220 | rs375945 | 0.775 | -1.346 | 1.007 | -0.616 | 0.001 | -0.031 | -0.031 | 0.136 | 0.58 | 0.57 | 0.34 | 0.16 | 0.53 | 0.41 |
| 1 | 53173052 | rs440871 | 1.479 | -1.207 | 1.176 | -0.678 | -0.012 | 0.008 | 0.008 | 0 | 0.42 | 0.31 | 0.28 | 0.02 | 0.5 | 0.38 |
| 1 | 53180565 | rs6703127 | 1.692 | -1.155 | 0.952 | -0.652 | -0.087 | 0.244 | 0.244 | 0.37 | 0.4 | 0.09 | 0.28 | 0.02 | 0.5 | 0.42 |
| 1 | 53180716 | rs6703281 | 1.063 | -1.258 | 0.936 | -0.709 | 0.026 | 0.104 | 0.104 | 0.067 | 0.52 | 0.33 | 0.34 | 0.02 | 0.57 | 0.44 |
| 1 | 53185631 | rs374849 | 1.511 | -1.099 | 0.833 | -0.567 | -0.087 | 0.244 | 0.244 | 0.37 | 0.4 | 0.09 | 0.27 | 0.02 | 0.5 | 0.42 |
| 1 | 53203586 | rs11205998 | -0.778 | -1.149 | 1.699 | 0.572 | 0.065 | 0.479 | 0.479 | 1.152 | 0.06 | 0.37 | 0.09 | 0.38 | 0.01 | 0.08 |
| 1 | 53203664 | rs6661196 | -1.332 | -1.101 | 1.582 | 0.409 | 0 | -0.002 | -0.002 | 0 | 0.52 | 0.51 | 0.62 | 0.61 | 0.48 | 0.49 |
| 1 | 53217335 | rs4926930 | -1.822 | -1.217 | 1.386 | 0.613 | -0.021 | 0.319 | 0.319 | 0.466 | 0.54 | 0.91 | 0.67 | 0.98 | 0.43 | 0.51 |
| 1 | 53218380 | rs11206006 | -0.401 | -1.432 | 1.273 | 0.98 | -0.086 | -0.144 | -0.144 | 0.051 | 0.24 | 0.03 | 0.48 | 0.41 | 0.34 | 0.34 |
| 1 | 53234874 | rs6664794 | -0.498 | -1.494 | 1.449 | 0.832 | 0.04 | 0.394 | 0.394 | 0.052 | 0.05 | 0.31 | 0.08 | 0.37 | 0.01 | 0.07 |
| 1 | 53245849 | rs7526275 | -2.011 | -1.49 | 1.45 | 0.825 | -0.012 | 0.321 | 0.321 | 0.461 | 0.54 | 0.91 | 0.67 | 0.98 | 0.43 | 0.51 |
| 1 | 53250915 | rs7523761 | -1.6 | -1.487 | 1.448 | 0.827 | 0.021 | 0.03 | 0.03 | 0.013 | 0.49 | 0.6 | 0.59 | 0.61 | 0.42 | 0.43 |
| 1 | 53259950 | rs11206019 | 0.993 | -1.367 | 1.402 | 1.069 | -0.059 | -0.041 | -0.041 | 0.072 | 0.09 | 0.01 | 0.12 | 0.11 | 0.22 | 0.11 |
| 1 | 53288239 | rs1048058 | -0.546 | -1.421 | 1.639 | 1.078 | 0.047 | 0.409 | 0.409 | 0.053 | 0.05 | 0.31 | 0.09 | 0.37 | 0.01 | 0.07 |
| 1 | 53318851 | rs555741 | 1.449 | -1.644 | 1.599 | 0.827 | 0.026 | 0.041 | 0.041 | 0.185 | 0.49 | 0.39 | 0.34 | 0.02 | 0.59 | 0.49 |
| 1 | 53320274 | rs480299 | 0.515 | -1.693 | 1.585 | 0.881 | -0.046 | -0.02 | -0.02 | 0 | 0.73 | 0.9 | 0.44 | 0.23 | 0.65 | 0.59 |
| 1 | 53323492 | rs10788945 | -0.94 | -1.464 | 1.415 | 0.893 | -0.007 | -0.042 | -0.042 | 1.405 | 0.87 | 0.99 | 0.9 | 0.84 | 0.88 | 0.89 |
| 1 | 53333116 | rs557847 | 0.757 | -1.225 | 1.755 | 0.946 | 0.026 | 0.041 | 0.041 | 0.185 | 0.49 | 0.39 | 0.34 | 0.02 | 0.59 | 0.49 |
| 1 | 53340824 | rs1672909 | 0.778 | -1.219 | 1.603 | 0.949 | 0.032 | 0.086 | 0.086 | 0.257 | 0.49 | 0.33 | 0.33 | 0.02 | 0.59 | 0.49 |
| 1 | 53343880 | rs512723 | 1.066 | -1.244 | 1.252 | 0.948 | -0.014 | 0.308 | 0.308 | 0.266 | 0.45 | 0.1 | 0.33 | 0.02 | 0.58 | 0.49 |
| 1 | 53374666 | rs562641 | -0.416 | -1.239 | 0.887 | 1.132 | -0.035 | 0.002 | 0.002 | 0 | 0.68 | 0.89 | 0.43 | 0.26 | 0.63 | 0.59 |
| 1 | 53396842 | rs1242330 | 1.12 | -1.2 | -0.228 | 1.115 | 0 | 0.27 | 0.27 | 0.221 | 0.46 | 0.12 | 0.33 | 0.02 | 0.57 | 0.49 |
| 1 | 53402552 | rs11206043 | -0.277 | -0.878 | -0.248 | 1.371 | -0.045 | 0 | 0 | 0 | 0.68 | 0.89 | 0.43 | 0.26 | 0.63 | 0.59 |
| 1 | 53438130 | rs11585570 | 0.178 | -0.711 | -0.351 | 1.355 | -0.092 | -0.123 | -0.123 | 0 | 0.29 | 0.04 | 0.49 | 0.37 | 0.36 | 0.34 |
| 1 | 53448372 | rs7544995 | -1.245 | -0.869 | 0.281 | 1.407 | 0.042 | -0.041 | -0.041 | 0 | 0.26 | 0.79 | 0.39 | 0.38 | 0.1 | 0.19 |
| 1 | 53460091 | rs10493170 | -0.324 | -0.067 | 0.258 | 1.579 | 0.001 | -0.029 | -0.029 | 0 | 0.07 | 0 | 0.02 | 0.21 | 0.04 | 0.04 |
| 1 | 53489668 | rs17107624 | -0.881 | 0.025 | 0.624 | 1.943 | 0.004 | 0.19 | 0.19 | 0 | 0.08 | 0.23 | 0.29 | 0.38 | 0.06 | 0.11 |
| 1 | 53495130 | rs10437066 | -1.654 | -1.074 | 0.561 | 1.496 | 0.06 | -0.052 | -0.052 | 0.566 | 0.27 | 0.81 | 0.39 | 0.38 | 0.1 | 0.2 |
| 1 | 53497421 | rs12075895 | -0.886 | -1.05 | 1.13 | 1.495 | -0.014 | 0.118 | 0.118 | 0 | 0.09 | 0.21 | 0.03 | 0.22 | 0.04 | 0.04 |
| 1 | 53502973 | rs17107710 | -1.523 | -1.091 | 1.591 | 1.47 | 0.026 | 0.309 | 0.309 | 0 | 0.11 | 0.45 | 0.32 | 0.34 | 0.07 | 0.11 |
| 1 | 53513437 | rs1288362 | 1.157 | -1.106 | -0.14 | 0.916 | -0.105 | 0.054 | 0.054 | 0.102 | 0.37 | 0.04 | 0.3 | 0.26 | 0.46 | 0.27 |
| 1 | 53514728 | rs3766763 | -1.499 | -1.181 | 0.226 | 1.029 | 0.066 | -0.053 | -0.053 | 0.054 | 0.27 | 0.81 | 0.39 | 0.38 | 0.1 | 0.21 |
| 1 | 53516742 | rs13496 | -0.077 | -1.2 | 0.274 | 1.027 | 0.002 | 0.161 | 0.161 | 0 | 0.08 | 0.23 | 0.29 | 0.34 | 0.06 | 0.11 |

| | | | | | | | | | | | | | | | | |
|---|---|---|---|---|---|---|---|---|---|---|---|---|---|---|---|---|
| 1 | 53516999 | rs8871 | 0.035 | -1.186 | 0.28 | 1.033 | 0.004 | 0.168 | 0.168 | 0 | 0.08 | 0.23 | 0.29 | 0.34 | 0.06 | 0.11 |
| 1 | 53520126 | rs7550236 | -1.543 | -1.211 | 0.015 | 1.013 | 0.069 | -0.053 | -0.053 | 0 | 0.28 | 0.81 | 0.39 | 0.38 | 0.1 | 0.21 |
| 1 | 53523904 | rs10493172 | 0.715 | -1.146 | 0.029 | 1.077 | -0.003 | -0.028 | -0.028 | 0 | 0.06 | 0 | 0.02 | 0.21 | 0.04 | 0.04 |
| 1 | 53525162 | rs12059934 | -1.402 | -1.154 | 0.032 | 1.015 | 0.068 | -0.053 | -0.053 | 0 | 0.28 | 0.82 | 0.39 | 0.38 | 0.1 | 0.21 |
| 1 | 53532117 | rs10158910 | -0.553 | -0.914 | 0.495 | 1.355 | 0.024 | 0.314 | 0.314 | 0 | 0.1 | 0.38 | 0.31 | 0.38 | 0.06 | 0.1 |
| 1 | 53532609 | rs11580093 | 0.414 | -1.067 | 0.402 | 1.207 | 0.023 | 0.022 | 0.022 | 0 | 0.13 | 0.09 | 0.05 | 0.04 | 0.03 | 0.09 |
| 1 | 53534248 | rs17107814 | 0.255 | -1.009 | 0.63 | 1.323 | 0.002 | 0.009 | 0.009 | 0 | 0.07 | 0.04 | 0.02 | 0.21 | 0.04 | 0.04 |
| 1 | 53535175 | rs1531699 | 0.365 | -0.957 | 0.649 | 1.32 | 0.004 | -0.005 | -0.005 | 0 | 0.07 | 0.05 | 0.02 | 0.2 | 0.04 | 0.04 |
| 1 | 53535478 | rs1288386 | 0.982 | -1.148 | 0.436 | 0.908 | -0.055 | -0.066 | -0.066 | 0.055 | 0.37 | 0.12 | 0.34 | 0.55 | 0.43 | 0.41 |
| 1 | 53535499 | rs17107831 | 0.694 | -1.037 | 0.464 | 0.903 | 0.022 | 0.022 | 0.022 | 0 | 0.13 | 0.09 | 0.05 | 0.04 | 0.05 | 0.09 |
| 1 | 53535918 | rs2275454 | 0.597 | -0.848 | 0.536 | 1.081 | 0.006 | -0.007 | -0.007 | 0.263 | 0.07 | 0.05 | 0.02 | 0.2 | 0.04 | 0.04 |
| 1 | 53537266 | rs1769316 | 0.502 | -0.616 | 1.039 | 0.916 | 0 | -0.01 | -0.01 | 0 | 0.42 | 0.5 | 0.65 | 0.74 | 0.47 | 0.48 |
| 1 | 53542077 | rs7524881 | -0.1 | -0.487 | 0.758 | 1.164 | -0.001 | -0.001 | -0.001 | 0 | 0.31 | 0.27 | 0.19 | 0.29 | 0.18 | 0.36 |
| 1 | 53544439 | rs1288388 | 1.083 | -0.365 | 0.997 | 1.463 | 0.082 | 0.097 | 0.097 | 0 | 0.57 | 0.15 | 0.48 | 0.29 | 0.73 | 0.5 |
| 1 | 53544807 | rs17107864 | 1.12 | -0.3 | 1.004 | 1.551 | -0.001 | -0.029 | -0.029 | 0 | 0.06 | 0 | 0.02 | 0.17 | 0.04 | 0.04 |
| 1 | 53551065 | rs2282340 | 1.141 | -0.309 | 1.211 | 1.696 | -0.002 | -0.027 | -0.027 | 0 | 0.06 | 0 | 0.02 | 0.17 | 0.04 | 0.04 |
| 1 | 53551293 | rs1288397 | 1.699 | -0.33 | 0.919 | 1.551 | -0.065 | 0.169 | 0.169 | 0 | 0.34 | 0.04 | 0.25 | 0.11 | 0.39 | 0.22 |
| 1 | 53553522 | rs11581058 | 0.387 | -0.679 | 1.362 | 1.644 | 0.03 | 0.049 | 0.049 | 0 | 0.27 | 0.19 | 0.08 | 0.03 | 0.17 | 0.16 |
| 1 | 53553754 | rs1288401 | 1.073 | -0.698 | 1.391 | 1.681 | 0.08 | 0.096 | 0.096 | 0 | 0.57 | 0.15 | 0.48 | 0.3 | 0.73 | 0.5 |
| 1 | 53554305 | rs2306459 | -0.52 | -0.703 | 1.497 | 1.691 | -0.008 | 0.375 | 0.375 | 0 | 0.12 | 0.39 | 0.14 | 0.41 | 0.05 | 0.21 |
| 1 | 53554406 | rs11583585 | 0.359 | -0.632 | 1.358 | 1.575 | 0.028 | 0.045 | 0.045 | 0 | 0.27 | 0.19 | 0.08 | 0.04 | 0.17 | 0.16 |
| 1 | 53557484 | rs1288404 | -0.24 | -0.635 | 1.299 | 1.6 | 0.009 | -0.001 | -0.001 | 0 | 0.71 | 0.74 | 0.65 | 0.71 | 0.79 | 0.7 |
| 1 | 53567546 | rs3766775 | 0.747 | -0.685 | 1.409 | 1.627 | 0.045 | 0.054 | 0.054 | 0 | 0.59 | 0.29 | 0.5 | 0.3 | 0.73 | 0.49 |
| 1 | 53567721 | rs11206093 | -0.199 | -0.733 | 1.372 | 1.642 | 0.019 | 0.011 | 0.011 | 0 | 0.31 | 0.23 | 0.11 | 0.19 | 0.21 | 0.21 |
| 1 | 53568162 | rs11206099 | -0.188 | -0.732 | 1.358 | 1.643 | 0.023 | 0.013 | 0.013 | 0 | 0.31 | 0.21 | 0.11 | 0.19 | 0.21 | 0.21 |
| 1 | 53568980 | rs2242274 | -0.192 | -0.749 | 1.341 | 1.635 | 0.021 | 0.012 | 0.012 | 0 | 0.31 | 0.21 | 0.11 | 0.19 | 0.21 | 0.21 |
| 1 | 53575575 | rs1288369 | 1.19 | 0.036 | 0.547 | 1.351 | -0.05 | 0.101 | 0.101 | 0 | 0.39 | 0.16 | 0.26 | 0.31 | 0.47 | 0.22 |
| 1 | 53576592 | rs1288371 | -0.222 | 0.16 | 1.22 | 1.435 | 0.013 | -0.005 | -0.005 | 0.972 | 0.74 | 0.74 | 0.53 | 0.75 | 0.69 | 0.63 |
| 1 | 53577284 | rs1288372 | 0.401 | 0.093 | 0.991 | 1.762 | 0.001 | 0.004 | 0.004 | 0 | 0.67 | 0.38 | 0.49 | 0.61 | 0.68 | 0.59 |
| 1 | 53579109 | rs17108022 | -0.663 | 0.268 | 0.642 | 2.108 | -0.014 | 0.297 | 0.297 | 0.168 | 0.05 | 0.16 | 0.38 | 0.42 | 0.02 | 0.23 |
| 1 | 53579182 | rs3766781 | 0.008 | 0.897 | 0.202 | 1.211 | -0.001 | -0.001 | -0.001 | 0 | 0.22 | 0.26 | 0.2 | 0.26 | 0.29 | 0.36 |
| 1 | 53586380 | rs936642 | 0.188 | 0.44 | -0.374 | 0.644 | 0.006 | 0.064 | 0.064 | 0 | 0.21 | 0.07 | 0.11 | 0.07 | 0.16 | 0.17 |
| 1 | 53587179 | rs6694834 | 0.791 | 0.397 | -0.848 | 0.262 | -0.019 | 0.003 | 0.003 | 0 | 0.39 | 0.29 | 0.57 | 0.51 | 0.42 | 0.21 |
| 1 | 53588494 | rs1679977 | -0.089 | 0.557 | -1.048 | -0.094 | -0.011 | 0.133 | 0.133 | 0 | 0.37 | 0.15 | 0.23 | 0.12 | 0.36 | 0.25 |
| 1 | 53590423 | rs3766792 | 0.416 | 0.573 | -0.36 | 0.322 | -0.021 | -0.006 | -0.006 | 1.357 | 0.16 | 0.34 | 0.15 | 0.21 | 0.14 | 0.47 |
| 1 | 53590630 | rs3766793 | -0.484 | 0.589 | -0.311 | 0.311 | -0.034 | 0.011 | 0.011 | 0.096 | 0.23 | 0.38 | 0.17 | 0.36 | 0.17 | 0.53 |
| 1 | 53591463 | rs3766794 | 0.562 | 1.406 | -0.715 | 0.728 | -0.046 | -0.06 | -0.06 | 0 | 0.1 | 0.02 | 0.34 | 0.23 | 0.15 | 0.09 |
| 1 | 53598126 | rs10888767 | 0.965 | 1.134 | -0.564 | 0.04 | 0.013 | 0.018 | 0.018 | 0 | 0.47 | 0.57 | 0.35 | 0.31 | 0.37 | 0.25 |
| 1 | 53600078 | rs3737989 | 0.42 | 1.126 | -0.307 | 0.249 | 0.011 | 0.059 | 0.059 | 0 | 0.13 | 0.13 | 0.37 | 0.29 | 0.16 | 0.06 |
| 1 | 53600350 | rs1288354 | -1.745 | 1.032 | -0.243 | 0.07 | -0.007 | -0.039 | -0.039 | 0.649 | 0.88 | 0.85 | 0.96 | 1 | 0.94 | 0.9 |
| 1 | 53605884 | rs1288367 | -0.978 | 1.02 | 0.034 | 0.36 | -0.008 | 0.038 | 0.038 | 0 | 0.82 | 0.63 | 0.86 | 0.65 | 0.82 | 0.46 |
| 1 | 53606080 | rs11206109 | 0.545 | 1.035 | -0.075 | 0.334 | -0.001 | -0.004 | -0.004 | 0.064 | 0.27 | 0.21 | 0.41 | 0.28 | 0.26 | 0.16 |
| 1 | 53608425 | rs34418279 | 1.313 | 1.054 | -0.547 | 0.249 | 0.034 | 0.026 | 0.026 | 0.096 | 0.41 | 0.22 | 0.26 | 0.28 | 0.3 | 0.18 |
| 1 | 53610372 | rs1320594 | -0.689 | 1.122 | -0.515 | 0.186 | 0.003 | -0.001 | -0.001 | 0.377 | 0.43 | 0.51 | 0.57 | 0.36 | 0.42 | 0.26 |
| 1 | 53611699 | rs1679933 | -0.209 | 1.176 | -0.517 | 0.219 | 0.004 | 0.002 | 0.002 | 2.074 | 0.26 | 0.28 | 0.24 | 0.43 | 0.38 | 0.63 |
| 1 | 53615541 | rs10888768 | 0.129 | 1.161 | -0.524 | 0.228 | 0.005 | 0.002 | 0.002 | 2.302 | 0.72 | 0.7 | 0.74 | 0.58 | 0.61 | 0.37 |
| 1 | 53619482 | rs7556421 | -0.682 | 1.204 | -0.63 | 0.183 | -0.003 | 0.031 | 0.031 | 0 | 0.5 | 0.67 | 0.62 | 0.63 | 0.5 | 0.72 |
| 1 | 53621345 | rs1288344 | -0.64 | 1.217 | -0.612 | 0.188 | 0.045 | -0.01 | -0.01 | 0 | 0.9 | 0.78 | 0.88 | 0.91 | 0.8 | 0.91 |
| 1 | 53622093 | rs1288347 | -1.269 | 1.175 | -0.62 | 0.172 | 0 | 0 | 0 | 0 | 0.59 | 0.59 | 0.69 | 0.7 | 0.6 | 0.79 |
| 1 | 53626467 | rs12119155 | -0.475 | 1.182 | -0.621 | 0.139 | -0.003 | 0.005 | 0.005 | 0 | 0.47 | 0.45 | 0.6 | 0.61 | 0.5 | 0.72 |
| 1 | 53628562 | rs1288339 | 0.687 | 1.185 | -0.629 | 0.143 | 0 | -0.01 | -0.01 | 0 | 0.13 | 0.13 | 0.09 | 0.07 | 0.11 | 0.07 |
| 1 | 53629585 | rs6695567 | 1.083 | 1.253 | -0.627 | 0.196 | -0.027 | 0.019 | 0.019 | 2.669 | 0.57 | 0.76 | 0.6 | 0.5 | 0.42 | 0.27 |
| 1 | 53631029 | rs12140961 | 0.763 | 1.32 | -0.58 | 0.201 | 0.034 | -0.054 | -0.054 | 0.012 | 0.73 | 0.59 | 0.71 | 0.84 | 0.62 | 0.81 |
| 1 | 53631049 | rs1288331 | -1.064 | 0.989 | -0.598 | 0.385 | 0.001 | 0.008 | 0.008 | 0 | 0.35 | 0.29 | 0.31 | 0.41 | 0.38 | 0.65 |
| 1 | 53631805 | rs874348 | -1.468 | 1.079 | 0.066 | 0.463 | 0.003 | 0 | 0 | 0 | 0.42 | 0.52 | 0.54 | 0.35 | 0.4 | 0.25 |
| 1 | 53631902 | rs874349 | -0.288 | 1.083 | -0.009 | 0.898 | 0.008 | 0.028 | 0.028 | 0 | 0.25 | 0.36 | 0.38 | 0.27 | 0.22 | 0.15 |
| 1 | 53633248 | rs1298238 | -0.816 | 1.017 | -0.465 | 0.856 | -0.001 | -0.002 | -0.002 | 0 | 0.37 | 0.61 | 0.47 | 0.35 | 0.32 | 0.21 |
| 1 | 53641911 | rs7531337 | -0.963 | 0.574 | -0.17 | 1.104 | 0.006 | 0.025 | 0.025 | 0.022 | 0.24 | 0.57 | 0.53 | 0.27 | 0.24 | 0.13 |
| 1 | 53643272 | rs11206121 | -0.61 | 0.465 | 0.069 | 1.046 | 0.039 | 0.072 | 0.072 | 1.315 | 0.07 | 0.5 | 0.06 | 0.05 | 0.01 | 0 |
| 1 | 53644967 | rs6588471 | -0.107 | 0.335 | -1.077 | 0.732 | -0.106 | 0.094 | 0.094 | 0 | 0.32 | 0.09 | 0.21 | 0.18 | 0.44 | 0.31 |
| 1 | 53646557 | rs3102044 | -1.409 | -0.14 | -0.76 | 1.123 | 0.044 | -0.039 | -0.039 | 0 | 0.62 | 0.82 | 0.77 | 0.52 | 0.7 | 0.49 |
| 1 | 53660993 | rs7554022 | -0.694 | -0.109 | -0.477 | 1.158 | -0.026 | 0.026 | 0.026 | 0 | 0.37 | 0.26 | 0.38 | 0.75 | 0.52 | 0.22 |
| 1 | 53666965 | rs3766759 | -0.959 | -0.139 | -0.394 | 1.226 | -0.015 | 0.046 | 0.046 | 0.052 | 0.39 | 0.28 | 0.39 | 0.81 | 0.53 | 0.26 |
| 1 | 53674542 | rs11578832 | 0.443 | 0.156 | -0.541 | 1.502 | -0.053 | 0.013 | 0.013 | 0 | 0.16 | 0.02 | 0.15 | 0.08 | 0.21 | 0.07 |
| 1 | 53675049 | rs370493 | -2.35 | -0.322 | -0.994 | 1.08 | -0.06 | 0.17 | 0.17 | 0.512 | 0.64 | 0.98 | 0.78 | 0.93 | 0.68 | 0.53 |
| 1 | 53675532 | rs12737375 | 0.751 | -0.303 | -0.907 | 1.83 | -0.083 | -0.097 | -0.097 | 2.529 | 0.2 | 0.06 | 0.18 | 0.49 | 0.31 | 0.14 |
| 1 | 53676448 | rs1799821 | -0.801 | -0.314 | -0.847 | 1.888 | -0.025 | 0.021 | 0.021 | 0 | 0.38 | 0.26 | 0.38 | 0.72 | 0.52 | 0.22 |
| 1 | 53679229 | rs1799822 | 0.684 | -0.29 | -0.887 | 1.89 | -0.053 | 0.013 | 0.013 | 0 | 0.16 | 0.02 | 0.15 | 0.09 | 0.21 | 0.07 |
| 1 | 53680090 | rs1056425 | 0.579 | -0.204 | -0.808 | 2.01 | -0.083 | -0.09 | -0.09 | 0 | 0.2 | 0.06 | 0.18 | 0.47 | 0.31 | 0.14 |

| | | | | | | | | | | | | | | | | |
|---|---|---|---|---|---|---|---|---|---|---|---|---|---|---|---|---|
| 1 | 53682678 | rs17108140 | 0.236 | -0.334 | -0.036 | 1.87 | -0.021 | -0.045 | -0.045 | 0 | 0.19 | 0.29 | 0.11 | 0.06 | 0.13 | 0.21 |
| 1 | 53684324 | rs3736118 | 0.607 | -0.295 | -0.247 | 1.88 | -0.083 | -0.095 | -0.095 | 1.06 | 0.2 | 0.06 | 0.18 | 0.5 | 0.31 | 0.14 |
| 1 | 53687589 | rs6663121 | -1.374 | -0.326 | 0.032 | 1.861 | -0.008 | 0.061 | 0.061 | 0 | 0.44 | 0.72 | 0.37 | 0.82 | 0.45 | 0.4 |
| 1 | 53694388 | rs2275087 | 0.512 | -0.196 | 0.118 | 1.973 | -0.033 | -0.036 | -0.036 | 0 | 0.2 | 0.12 | 0.18 | 0.48 | 0.32 | 0.14 |
| 1 | 53696051 | rs10788949 | 0.552 | -0.181 | 0.134 | 1.98 | -0.054 | -0.059 | -0.059 | 0.181 | 0.2 | 0.07 | 0.18 | 0.48 | 0.32 | 0.14 |
| 1 | 53697139 | rs6673692 | -0.913 | -0.22 | 0.286 | 1.954 | -0.008 | -0.011 | -0.011 | 0.437 | 0.4 | 0.31 | 0.36 | 0.58 | 0.53 | 0.22 |
| 1 | 53701581 | rs2297656 | 0.046 | 0.067 | -0.251 | 2.179 | -0.022 | -0.03 | -0.03 | 0 | 0.22 | 0.17 | 0.19 | 0.56 | 0.32 | 0.18 |
| 1 | 53707953 | rs17785382 | 0.19 | -0.21 | -0.303 | 2.371 | -0.065 | -0.053 | -0.053 | 0 | 0.23 | 0.09 | 0.24 | 0.47 | 0.4 | 0.17 |
| 1 | 53708850 | rs3737984 | -0.017 | -0.243 | -0.455 | 2.422 | -0.003 | -0.003 | -0.003 | 0 | 0.25 | 0.31 | 0.3 | 0.63 | 0.41 | 0.18 |
| 1 | 53712727 | rs5174 | 0.457 | -0.251 | -0.963 | 2.445 | -0.143 | 0.122 | 0.122 | 0.075 | 0.21 | 0.02 | 0.18 | 0.03 | 0.4 | 0.15 |
| 1 | 53713336 | rs10788950 | -1.853 | -0.329 | -1.001 | 2.296 | -0.024 | 0.147 | 0.147 | 0.179 | 0.62 | 0.94 | 0.54 | 0.87 | 0.67 | 0.49 |
| 1 | 53713549 | rs11206127 | -0.343 | -0.271 | -1.059 | 2.409 | -0.013 | -0.004 | -0.004 | 0.539 | 0.27 | 0.25 | 0.29 | 0.7 | 0.41 | 0.22 |
| 1 | 53714139 | rs12031155 | 0.163 | -0.267 | -1.062 | 2.427 | -0.094 | -0.057 | -0.057 | 8.951 | 0.23 | 0.07 | 0.22 | 0.42 | 0.4 | 0.17 |
| 1 | 53716416 | rs3737983 | -0.285 | -0.284 | -1.065 | 2.143 | -0.017 | -0.005 | -0.005 | 0.276 | 0.27 | 0.25 | 0.29 | 0.7 | 0.41 | 0.22 |
| 1 | 53722752 | rs2297663 | 0.182 | -0.287 | -1.08 | 2.159 | -0.096 | -0.099 | -0.099 | 0.212 | 0.23 | 0.07 | 0.24 | 0.53 | 0.39 | 0.18 |
| 1 | 53732315 | rs2297660 | -0.1 | -0.254 | -1.135 | 2.057 | -0.031 | -0.028 | -0.028 | 0 | 0.25 | 0.2 | 0.28 | 0.63 | 0.4 | 0.18 |
| 1 | 53736613 | rs7542607 | -0.114 | -0.254 | -1.135 | 2.057 | -0.032 | -0.028 | -0.028 | 0 | 0.25 | 0.2 | 0.28 | 0.63 | 0.4 | 0.18 |
| 1 | 53746134 | rs1288518 | -0.149 | -0.228 | -1.153 | 1.904 | 0.007 | 0.003 | 0.003 | 0 | 0.25 | 0.28 | 0.27 | 0.69 | 0.37 | 0.24 |
| 1 | 53748286 | rs1288516 | -0.736 | -0.745 | -1.815 | 1.238 | 0.079 | -0.077 | -0.077 | 0 | 0.4 | 0.89 | 0.33 | 0.35 | 0.3 | 0.31 |
| 1 | 53751610 | rs4623641 | -0.014 | -0.003 | -1.043 | 1.708 | -0.093 | -0.122 | -0.122 | 0.329 | 0.21 | 0.05 | 0.22 | 0.58 | 0.35 | 0.2 |
| 1 | 53752134 | rs12035926 | -0.331 | -0.102 | -1.096 | 1.659 | -0.103 | -0.136 | -0.136 | 0.718 | 0.24 | 0.05 | 0.23 | 0.58 | 0.37 | 0.21 |
| 1 | 53753718 | rs1002358 | 0.505 | -1.218 | -0.192 | 1.373 | -0.087 | 0.089 | 0.089 | 0 | 0.76 | 0.42 | 0.8 | 0.4 | 0.9 | 0.78 |
| 1 | 53754891 | rs17108202 | -0.307 | -0.901 | -0.577 | 0.625 | -0.058 | -0.044 | -0.044 | 0 | 0.32 | 0.18 | 0.29 | 0.6 | 0.42 | 0.3 |
| 1 | 53755679 | rs12039021 | -0.265 | -1.003 | -0.604 | 0.612 | -0.098 | -0.09 | -0.09 | 2.182 | 0.31 | 0.11 | 0.29 | 0.6 | 0.41 | 0.35 |
| 1 | 53756870 | rs1288503 | -1.541 | -1.408 | -1.02 | 0.065 | -0.058 | 0.171 | 0.171 | 0 | 0.62 | 0.86 | 0.77 | 0.92 | 0.61 | 0.48 |
| 1 | 53757154 | rs1288502 | -0.851 | -1.491 | -0.478 | 0.065 | 0 | 0.13 | 0.13 | 0 | 0.48 | 0.69 | 0.66 | 0.87 | 0.49 | 0.41 |
| 1 | 53762189 | rs1288497 | -0.52 | -1.526 | 0.156 | -0.042 | -0.049 | -0.079 | -0.079 | 0 | 0.18 | 0.44 | 0.13 | 0.03 | 0.13 | 0.13 |
| 1 | 53762226 | rs4926968 | 0.113 | -1.596 | 0.242 | 0.052 | -0.055 | 0.05 | 0.05 | 0 | 0.19 | 0.35 | 0.36 | 0.29 | 0.09 | 0.17 |
| 1 | 53762781 | rs11206139 | -0.74 | -1.462 | 0.223 | -0.374 | -0.003 | 0.027 | 0.027 | 0 | 0.47 | 0.44 | 0.63 | 0.91 | 0.49 | 0.41 |
| 1 | 53763374 | rs1288495 | -1.62 | -2.087 | 0.369 | 0.225 | 0.058 | -0.013 | -0.013 | 0.017 | 0.35 | 0.77 | 0.49 | 0.33 | 0.23 | 0.31 |
| 1 | 53763851 | rs2297824 | 0.361 | -2.041 | 0.827 | 0.206 | -0.042 | 0.102 | 0.102 | 0 | 0.12 | 0.34 | 0.13 | 0.27 | 0.09 | 0.15 |
| 1 | 53764635 | rs2782500 | -0.86 | -2.078 | 0.637 | 0.249 | -0.005 | -0.004 | -0.004 | 0 | 0.2 | 0.22 | 0.34 | 0.08 | 0.14 | 0.16 |
| 1 | 53765434 | rs11206140 | 0.133 | -1.852 | 0.371 | 0.027 | -0.008 | -0.035 | -0.035 | 0 | 0.22 | 0.16 | 0.18 | 0.58 | 0.23 | 0.2 |
| 1 | 53766972 | rs867884 | -1.448 | -1.854 | 0.229 | -0.398 | -0.003 | -0.008 | -0.008 | 0 | 0.45 | 0.52 | 0.54 | 0.66 | 0.38 | 0.37 |
| 1 | 53771202 | rs10493173 | -0.173 | -1.478 | -0.321 | -0.691 | -0.007 | -0.02 | -0.02 | 0.186 | 0.15 | 0.16 | 0.1 | 0.02 | 0.12 | 0.07 |
| 1 | 53774534 | rs1288487 | -1.013 | -1.11 | -0.685 | -0.468 | -0.048 | 0.307 | 0.307 | 0 | 0.56 | 0.95 | 0.67 | 0.93 | 0.46 | 0.49 |
| 1 | 53777543 | rs2788033 | -1.092 | -1.245 | 0.654 | 0.518 | -0.035 | 0.003 | 0.003 | 0 | 0.23 | 0.59 | 0.22 | 0.2 | 0.2 | 0.08 |
| 1 | 53777631 | rs2788032 | -0.239 | -1.106 | 0.897 | 0.907 | -0.044 | -0.043 | -0.043 | 0 | 0.14 | 0.35 | 0.09 | 0.07 | 0.1 | 0.22 |
| 1 | 53777739 | rs2788031 | -0.548 | -1.272 | 0.326 | 0.896 | -0.07 | 0.04 | 0.04 | 0.011 | 0.17 | 0.6 | 0.14 | 0.24 | 0.08 | 0.23 |
| 1 | 53778904 | rs1288479 | -0.924 | -0.77 | -0.383 | 0.826 | 0.025 | -0.01 | -0.01 | 0 | 0.38 | 0.66 | 0.3 | 0.59 | 0.3 | 0.41 |
| 1 | 53787360 | rs872315 | 0.048 | -1.266 | -0.637 | 0.066 | 0.011 | 0.071 | 0.071 | 0.693 | 0.12 | 0 | 0.04 | 0 | 0.09 | 0.02 |
| 1 | 53787633 | rs1288520 | -1.023 | -0.85 | -0.511 | 0.371 | 0.002 | 0 | 0 | 0.328 | 0.4 | 0.67 | 0.24 | 0.42 | 0.33 | 0.4 |
| 1 | 53787673 | rs1288521 | -0.664 | -0.928 | 0.356 | 0.309 | -0.001 | 0 | 0 | 0.035 | 0.34 | 0.38 | 0.2 | 0.39 | 0.28 | 0.35 |
| 1 | 53792651 | rs3820198 | -1.421 | -0.949 | 0.069 | 0.216 | 0.02 | -0.033 | -0.033 | 1.73 | 0.5 | 0.93 | 0.3 | 0.48 | 0.39 | 0.45 |
| 1 | 53797580 | rs2782491 | -1.424 | -1.014 | 0.035 | 0.195 | 0.019 | -0.03 | -0.03 | 0 | 0.5 | 0.93 | 0.3 | 0.48 | 0.39 | 0.44 |
| 1 | 53803639 | rs12083302 | -0.459 | -1 | 0.638 | 0.039 | 0.001 | -0.001 | -0.001 | 0 | 0.06 | 0.19 | 0.04 | 0.03 | 0.05 | 0.05 |
| 1 | 53807803 | rs2782493 | -1.488 | -1.042 | 0.669 | 0.167 | 0.058 | 0.061 | 0.061 | 0.543 | 0.5 | 0.83 | 0.29 | 0.4 | 0.39 | 0.46 |
| 1 | 53808658 | rs1416096 | -0.432 | -1.107 | 0.808 | 0.152 | -0.002 | -0.004 | -0.004 | 0.099 | 0.38 | 0.45 | 0.23 | 0.35 | 0.33 | 0.37 |
| 1 | 53808693 | rs1416095 | -1.564 | -1.084 | 0.572 | 0.134 | 0.016 | 0.014 | 0.014 | 0.691 | 0.51 | 0.93 | 0.3 | 0.43 | 0.39 | 0.46 |
| 1 | 53808916 | rs17108355 | -0.773 | -1.086 | 0.71 | 0.103 | -0.013 | -0.006 | -0.006 | 0 | 0.05 | 0.09 | 0.02 | 0.02 | 0.01 | 0.03 |
| 1 | 53810030 | rs12075186 | -1.193 | -1.084 | 0.8 | 0.104 | -0.026 | -0.013 | -0.013 | 0 | 0.07 | 0.17 | 0.03 | 0.02 | 0.01 | 0.03 |
| 1 | 53810614 | rs12075385 | -1.218 | -1.08 | 0.803 | 0.106 | -0.025 | -0.012 | -0.012 | 0 | 0.06 | 0.17 | 0.03 | 0.02 | 0.01 | 0.03 |
| 1 | 53813684 | rs10888780 | -0.051 | -0.917 | 1.055 | 0.306 | -0.008 | 0.006 | 0.006 | 0 | 0.64 | 0.68 | 0.78 | 0.68 | 0.65 | 0.62 |
| 1 | 53813961 | rs10888781 | 1.388 | -0.762 | 0.391 | 0.053 | -0.032 | -0.07 | -0.07 | 0.022 | 0.39 | 0.13 | 0.38 | 0.54 | 0.41 | 0.34 |
| 1 | 53813973 | rs10888782 | 1.364 | -0.777 | 0.384 | 0.046 | -0.031 | -0.069 | -0.069 | 0 | 0.39 | 0.13 | 0.38 | 0.54 | 0.41 | 0.34 |
| 1 | 53814038 | rs7535922 | 0.683 | -0.821 | 0.542 | 0.031 | 0.001 | 0.001 | 0.001 | 0 | 0.47 | 0.51 | 0.43 | 0.59 | 0.46 | 0.39 |
| 1 | 53815537 | rs4926976 | 0.678 | -0.827 | 0.515 | -0.062 | 0.002 | 0.003 | 0.003 | 0 | 0.47 | 0.54 | 0.42 | 0.6 | 0.45 | 0.39 |
| 1 | 53817135 | rs6658926 | 1.361 | -0.813 | 0.428 | 0.04 | -0.024 | -0.065 | -0.065 | 0 | 0.39 | 0.14 | 0.37 | 0.53 | 0.4 | 0.36 |
| 1 | 53817220 | rs6588474 | -0.576 | -0.737 | 0.447 | 0.031 | 0.001 | -0.001 | -0.001 | 0 | 0.66 | 0.64 | 0.76 | 0.67 | 0.64 | 0.62 |
| 1 | 53821821 | rs11206148 | -0.004 | -0.598 | 0.504 | 0.253 | -0.013 | -0.029 | -0.029 | 0 | 0.22 | 0.32 | 0.36 | 0.1 | 0.19 | 0.23 |
| 1 | 53824168 | rs10888783 | 1.503 | -0.89 | 0.201 | -0.024 | -0.037 | -0.08 | -0.08 | 0 | 0.33 | 0.12 | 0.36 | 0.5 | 0.38 | 0.28 |
| 1 | 53824428 | rs12135186 | 0.211 | -0.749 | 0.135 | 0.021 | -0.014 | -0.052 | -0.052 | 0 | 0.21 | 0.33 | 0.35 | 0.06 | 0.19 | 0.22 |
| 1 | 53827481 | rs12143675 | 0.79 | -0.674 | -0.128 | 0.057 | -0.004 | 0.042 | 0.042 | 0 | 0.18 | 0.08 | 0.33 | 0.06 | 0.19 | 0.22 |
| 1 | 53829729 | rs1999897 | 0.181 | -0.436 | 0.255 | 0.374 | -0.002 | 0.002 | 0.002 | 0 | 0.59 | 0.62 | 0.75 | 0.63 | 0.62 | 0.55 |
| 1 | 53831095 | rs11206152 | 0.373 | -0.284 | -0.404 | 0.447 | 0 | 0.005 | 0.005 | 0 | 0.57 | 0.42 | 0.74 | 0.62 | 0.62 | 0.55 |
| 1 | 53832824 | rs3207053 | 0.377 | -0.263 | -0.393 | 0.445 | 0 | 0.006 | 0.006 | 0 | 0.57 | 0.42 | 0.74 | 0.64 | 0.62 | 0.55 |
| 1 | 53837187 | rs6588477 | 0.32 | -0.351 | -0.428 | 1.253 | -0.005 | 0.069 | 0.069 | 0 | 0.88 | 0.85 | 0.93 | 0.55 | 0.9 | 0.76 |
| 1 | 53842099 | rs1316436 | 1.081 | -0.172 | -1.524 | 0.879 | -0.177 | -0.118 | -0.118 | 2.589 | 0.28 | 0.03 | 0.39 | 0.3 | 0.46 | 0.29 |
| 1 | 53844161 | rs12746317 | 1.201 | -0.074 | -1.149 | 0.823 | -0.162 | -0.134 | -0.134 | 0 | 0.3 | 0.03 | 0.37 | 0.37 | 0.45 | 0.29 |

| | | | | | | | | | | | | | | | | |
|---|---|---|---|---|---|---|---|---|---|---|---|---|---|---|---|---|
| 1 | 53845440 | rs2185077 | 0.033 | -0.178 | -0.35 | 0.702 | 0.005 | 0.199 | 0.199 | 0 | 0.87 | 0.74 | 0.88 | 0.5 | 0.88 | 0.67 |
| 1 | 53845985 | rs17108398 | -0.006 | -0.702 | -0.201 | 0.256 | 0.005 | -0.045 | -0.045 | 0.151 | 0.37 | 0.42 | 0.2 | 0.04 | 0.28 | 0.12 |
| 1 | 53846598 | rs11587217 | -0.006 | -0.702 | -0.198 | 0.256 | 0.004 | -0.048 | -0.048 | 0 | 0.37 | 0.42 | 0.2 | 0.04 | 0.28 | 0.12 |
| 1 | 53852469 | rs11206153 | 0.848 | -0.655 | -0.581 | 0.482 | -0.046 | -0.069 | -0.069 | 0 | 0.45 | 0.1 | 0.65 | 0.41 | 0.55 | 0.46 |
| 1 | 53864019 | rs17371030 | 1.305 | -0.576 | -0.409 | 0.458 | 0.014 | 0.028 | 0.028 | 0 | 0.15 | 0.07 | 0.28 | 0.04 | 0.1 | 0.17 |
| 1 | 53868891 | rs1288590 | -0.884 | -0.723 | 0.121 | 0.51 | -0.001 | 0 | 0 | 0.256 | 0.49 | 0.5 | 0.32 | 0.51 | 0.43 | 0.45 |
| 1 | 53870466 | rs4926992 | -0.549 | -0.064 | 0.697 | 0.314 | -0.01 | -0.08 | -0.08 | 0 | 0.41 | 0.48 | 0.49 | 0.09 | 0.39 | 0.34 |
| 1 | 53870831 | rs1288588 | -2.067 | -0.045 | 0.282 | 0.641 | -0.131 | -0.151 | -0.151 | 0 | 0.63 | 0.97 | 0.66 | 0.62 | 0.57 | 0.65 |
| 1 | 53874063 | rs12039843 | -0.514 | 0.125 | 0.642 | 0.898 | -0.022 | -0.039 | -0.039 | 0 | 0.15 | 0.1 | 0.11 | 0.49 | 0.16 | 0.27 |
| 1 | 53876149 | rs1342970 | -0.04 | -0.09 | 0.882 | 0.737 | -0.05 | -0.047 | -0.047 | 0 | 0.29 | 0.45 | 0.38 | 0.12 | 0.19 | 0.31 |
| 1 | 53877199 | rs11206154 | -0.568 | 0.005 | 1.053 | 0.768 | 0.024 | 0.085 | 0.085 | 0 | 0.12 | 0.18 | 0.12 | 0.49 | 0.15 | 0.27 |
| 1 | 53878651 | rs6656566 | -0.162 | -0.064 | 0.863 | 0.738 | -0.051 | -0.079 | -0.079 | 0 | 0.32 | 0.45 | 0.39 | 0.07 | 0.2 | 0.24 |
| 1 | 53880369 | rs6685296 | -0.487 | 0.201 | 0.433 | 1.027 | -0.013 | -0.005 | -0.005 | 0 | 0.12 | 0.1 | 0.1 | 0.49 | 0.14 | 0.27 |
| 1 | 53881081 | rs6686055 | -1.567 | 0.198 | 0.513 | 1.004 | -0.049 | -0.041 | -0.041 | 0 | 0.6 | 0.82 | 0.64 | 0.62 | 0.55 | 0.65 |
| 1 | 53883126 | rs2000239 | -2.138 | 0.3 | 1.897 | 1.758 | 0.001 | 0.036 | 0.036 | 0.205 | 0.3 | 0.56 | 0.25 | 0.57 | 0.26 | 0.34 |
| 1 | 53896599 | rs12023145 | -1.582 | -0.459 | 2.039 | 1.508 | 0.004 | 0.036 | 0.036 | 0 | 0.12 | 0.22 | 0.09 | 0.33 | 0.11 | 0.17 |
| 1 | 53901837 | rs1766643 | -2.321 | -0.512 | 1.936 | 1.5 | 0.031 | 0.018 | 0.018 | 4.137 | 0.27 | 0.78 | 0.17 | 0.26 | 0.17 | 0.28 |
| 1 | 53913080 | rs9436924 | -2.737 | -0.504 | 1.876 | 1.519 | 0.048 | -0.08 | -0.08 | 0 | 0.33 | 0.88 | 0.2 | 0.56 | 0.2 | 0.39 |
| 1 | 53924150 | rs1288627 | -2.833 | -0.525 | 2.272 | 1.37 | 0.017 | -0.025 | -0.025 | 0 | 0.35 | 0.74 | 0.22 | 0.52 | 0.27 | 0.4 |
| 1 | 53924164 | rs12035140 | -0.381 | -0.529 | 2.423 | 1.386 | -0.007 | 0.077 | 0.077 | 0 | 0.06 | 0.13 | 0.02 | 0.31 | 0.03 | 0.12 |
| 1 | 53926037 | rs1288628 | -2.562 | -0.564 | 2.523 | 1.378 | -0.011 | -0.001 | -0.001 | 0 | 0.26 | 0.68 | 0.13 | 0.37 | 0.15 | 0.25 |
| 1 | 53927649 | rs943521 | -3.036 | -0.51 | 2.537 | 1.365 | 0.008 | -0.017 | -0.017 | 0 | 0.36 | 0.72 | 0.22 | 0.52 | 0.28 | 0.41 |
| 1 | 53930551 | rs943514 | -0.623 | -0.521 | 2.88 | 1.353 | -0.062 | -0.064 | -0.064 | 0 | 0.12 | 0.26 | 0.05 | 0.03 | 0.03 | 0.07 |
| 1 | 53934091 | rs1288637 | -3.072 | -0.517 | 2.813 | 1.372 | 0.01 | -0.029 | -0.029 | 0 | 0.36 | 0.74 | 0.22 | 0.53 | 0.28 | 0.41 |
| 1 | 53936383 | rs11579521 | -2.016 | -0.496 | 2.868 | 1.381 | -0.001 | 0.004 | 0.004 | 0 | 0.2 | 0.21 | 0.16 | 0.47 | 0.23 | 0.32 |
| 1 | 53936718 | rs11579575 | -0.14 | -0.45 | 2.408 | 1.377 | -0.034 | -0.033 | -0.033 | 0 | 0.08 | 0.02 | 0.09 | 0.16 | 0.13 | 0.16 |
| 1 | 53942583 | rs11581093 | -0.575 | -0.452 | 2.484 | 1.367 | 0.033 | 0.037 | 0.037 | 0 | 0.1 | 0.15 | 0.1 | 0.17 | 0.13 | 0.16 |
| 1 | 53946485 | rs1288622 | -2.075 | -1.131 | 2.834 | 0.982 | -0.045 | -0.124 | -0.124 | 0 | 0.19 | 0.5 | 0.1 | 0.03 | 0.11 | 0.11 |
| 1 | 53946718 | rs12027731 | -0.987 | -1.14 | 2.772 | 0.995 | 0.001 | 0.184 | 0.184 | 0.162 | 0.08 | 0.32 | 0.04 | 0.31 | 0.03 | 0.12 |
| 1 | 53948733 | rs1298638 | -2.298 | -1.13 | 2.794 | 0.919 | -0.034 | -0.088 | -0.088 | 0.81 | 0.21 | 0.59 | 0.11 | 0.05 | 0.12 | 0.14 |
| 1 | 53952141 | rs1288617 | -0.901 | -1.095 | 2.164 | 0.921 | -0.054 | -0.056 | -0.056 | 1.461 | 0.12 | 0.23 | 0.04 | 0.03 | 0.03 | 0.08 |
| 1 | 53958815 | rs11206165 | -0.756 | -1.07 | 1.932 | 0.928 | 0.061 | -0.024 | -0.024 | 0 | 0.07 | 0.24 | 0.05 | 0 | 0.08 | 0.05 |
| 1 | 53963481 | rs10749690 | 1.995 | -1.058 | 0.854 | 1.013 | 0.076 | 0 | 0 | 0.931 | 0.7 | 0.11 | 0.85 | 0.62 | 0.84 | 0.74 |
| 1 | 53965176 | rs943520 | 1.095 | -0.938 | 1.187 | 0.611 | -0.062 | -0.069 | -0.069 | 0.037 | 0.87 | 0.75 | 0.95 | 0.97 | 0.97 | 0.92 |
| 1 | 53965546 | rs943519 | 1.095 | -0.938 | 1.117 | 0.612 | -0.055 | -0.057 | -0.057 | 1.026 | 0.87 | 0.77 | 0.95 | 0.97 | 0.97 | 0.92 |
| 1 | 53967454 | rs12727041 | 1.109 | -0.913 | 0.988 | 0.635 | -0.035 | -0.039 | -0.039 | 0 | 0.87 | 0.8 | 0.95 | 0.96 | 0.96 | 0.92 |
| 1 | 53972049 | rs2296761 | -1.075 | -0.384 | 0.772 | 0.831 | -0.02 | -0.035 | -0.035 | 0 | 0.08 | 0.24 | 0.04 | 0.04 | 0.02 | 0.07 |
| 1 | 53978119 | rs6588480 | 1.074 | -0.238 | 0.811 | 1.44 | -0.002 | 0.217 | 0.217 | 0 | 0.86 | 0.65 | 0.71 | 0.56 | 0.86 | 0.79 |
| 1 | 53996718 | rs542238 | 1.668 | -0.387 | 1.395 | 1.43 | 0.021 | 0.359 | 0.359 | 0.963 | 0.86 | 0.53 | 0.72 | 0.52 | 0.85 | 0.79 |
| 1 | 53999509 | rs3887651 | -1.784 | -0.392 | 1.73 | 1.465 | 0.063 | 0.622 | 0.622 | 1.912 | 0.05 | 0.35 | 0.21 | 0.48 | 0.02 | 0.15 |
| 1 | 54000726 | rs7520584 | 2.042 | -0.483 | 1.681 | 1.501 | 0.001 | 0.171 | 0.171 | 0.586 | 0.87 | 0.72 | 0.74 | 0.52 | 0.86 | 0.79 |
| 1 | 54001983 | rs545514 | 1.743 | -0.478 | 1.579 | 1.601 | 0.02 | 0.324 | 0.324 | 0.027 | 0.9 | 0.73 | 0.77 | 0.49 | 0.88 | 0.83 |
| 1 | 54003748 | rs12739933 | 0.534 | -0.433 | 0.462 | 0.484 | 0.004 | 0.001 | 0.001 | 0.001 | 0.37 | 0.42 | 0.43 | 0.4 | 0.41 | 0.34 |
| 1 | 54005060 | rs11206169 | 0.659 | -0.431 | 0.304 | 0.486 | -0.006 | -0.004 | -0.004 | 0.025 | 0.35 | 0.35 | 0.43 | 0.38 | 0.42 | 0.29 |
| 1 | 54007863 | rs12403311 | 0.26 | -0.864 | -0.171 | -0.017 | 0.029 | 0.086 | 0.086 | 1.037 | 0.3 | 0.12 | 0.13 | 0.01 | 0.22 | 0.38 |
| 1 | 54009376 | rs9700109 | 0.837 | -1.071 | -0.774 | -0.117 | -0.048 | 0.152 | 0.152 | 5.121 | 0.23 | 0.01 | 0.1 | 0 | 0.21 | 0.25 |
| 1 | 54010926 | rs11206170 | 0.247 | -0.736 | -0.509 | 0.139 | -0.001 | -0.001 | -0.001 | 0 | 0.59 | 0.45 | 0.63 | 0.48 | 0.65 | 0.36 |
| 1 | 54014492 | rs563403 | -0.023 | -0.224 | -0.7 | 0.04 | -0.001 | -0.004 | -0.004 | 0.209 | 0.95 | 0.97 | 0.96 | 1 | 0.91 | 0.96 |
| 1 | 54016353 | rs17108782 | -0.289 | -0.248 | -0.708 | 0.017 | -0.003 | 0.031 | 0.031 | 0 | 0.23 | 0.19 | 0.2 | 0.09 | 0.25 | 0.08 |
| 1 | 54018483 | rs11206172 | -0.444 | -0.739 | -0.817 | 0.878 | 0.005 | -0.033 | -0.033 | 0 | 0.19 | 0.16 | 0.31 | 0.56 | 0.15 | 0.28 |
| 1 | 54020542 | rs12141264 | -0.471 | -1.235 | -0.916 | 0.491 | -0.022 | 0.122 | 0.122 | 0.086 | 0.19 | 0.01 | 0.09 | 0 | 0.16 | 0.17 |
| 1 | 54024808 | rs4575030 | -1.486 | -1.72 | -0.248 | 0.222 | 0.004 | 0.006 | 0.006 | 0 | 0.52 | 0.54 | 0.51 | 0.78 | 0.39 | 0.56 |
| 1 | 54024840 | rs12035088 | -0.132 | -1.869 | 0.082 | 0.599 | -0.018 | 0.173 | 0.173 | 0 | 0.1 | 0.17 | 0.25 | 0.57 | 0.06 | 0.24 |
| 1 | 54026880 | rs529674 | -0.687 | -2.196 | 0.796 | 0.782 | -0.042 | 0.049 | 0.049 | 0.129 | 0.31 | 0.49 | 0.16 | 0.46 | 0.23 | 0.3 |
| 1 | 54027035 | rs4548383 | -0.412 | -2.275 | 0.805 | 0.751 | -0.046 | -0.028 | -0.028 | 0.145 | 0.25 | 0.46 | 0.18 | 0.19 | 0.16 | 0.26 |
| 1 | 54029642 | rs4927007 | 1.307 | -2.046 | 0.246 | 0.891 | -0.002 | -0.017 | -0.017 | 0.106 | 0.44 | 0.07 | 0.66 | 0.35 | 0.61 | 0.44 |
| 1 | 54035689 | rs10788957 | 0.421 | -1.97 | 0.904 | 0.843 | 0.007 | 0.007 | 0.007 | 1.868 | 0.78 | 0.24 | 0.84 | 0.86 | 0.92 | 0.86 |
| 1 | 54038536 | rs10493174 | -0.741 | -1.845 | 0.868 | 1.049 | -0.041 | 0.043 | 0.043 | 0.61 | 0.19 | 0.02 | 0.09 | 0.12 | 0.19 | 0.15 |
| 1 | 54042714 | rs2950250 | 1.725 | -1.931 | 1.143 | 0.906 | 0.052 | 0.099 | 0.099 | 0 | 0.55 | 0.23 | 0.74 | 0.76 | 0.69 | 0.62 |
| 1 | 54045180 | rs12089978 | -0.41 | -1.962 | 1.719 | 0.819 | -0.046 | -0.079 | -0.079 | 0 | 0.22 | 0.44 | 0.16 | 0.07 | 0.11 | 0.22 |
| 1 | 54046556 | rs941129 | -0.623 | -1.884 | 1.495 | 0.82 | -0.044 | 0.058 | 0.058 | 1.92 | 0.21 | 0.03 | 0.09 | 0.12 | 0.2 | 0.16 |
| 1 | 54051898 | rs2948045 | -1.762 | -1.91 | 1.841 | 0.989 | 0.018 | 0.008 | 0.008 | 0 | 0.43 | 0.36 | 0.27 | 0.63 | 0.32 | 0.41 |
| 1 | 54061978 | rs3013754 | -1.866 | -1.975 | 1.632 | 0.911 | 0.042 | 0.02 | 0.02 | 0 | 0.47 | 0.6 | 0.27 | 0.62 | 0.32 | 0.41 |
| 1 | 54064240 | rs10159298 | -1.109 | -1.949 | 1.677 | 0.846 | -0.01 | -0.048 | -0.048 | 0.384 | 0.24 | 0.32 | 0.1 | 0.13 | 0.2 | 0.15 |
| 1 | 54068016 | rs4927012 | 0.546 | -2.042 | 1.596 | 0.943 | -0.008 | 0.01 | 0.01 | 1.302 | 0.75 | 0.31 | 0.8 | 0.52 | 0.88 | 0.73 |
| 1 | 54071428 | rs17382457 | -1.475 | -2.074 | 1.474 | 0.869 | 0.03 | 0.12 | 0.12 | 0 | 0.4 | 0.21 | 0.17 | 0.19 | 0.31 | 0.34 |
| 1 | 54072471 | rs4927015 | 1.912 | -1.736 | 1.674 | 0.588 | 0.018 | 0.018 | 0.018 | 0 | 0.48 | 0.31 | 0.67 | 0.39 | 0.59 | 0.55 |
| 1 | 54072816 | rs2948040 | 0.817 | -2.04 | 1.845 | 0.851 | -0.012 | 0.045 | 0.045 | 0.092 | 0.75 | 0.6 | 0.77 | 0.51 | 0.79 | 0.79 |
| 1 | 54074729 | rs2948048 | 0.82 | -2.032 | 1.798 | 0.785 | -0.003 | 0.011 | 0.011 | 0.105 | 0.74 | 0.66 | 0.77 | 0.57 | 0.79 | 0.79 |

| | | | | | | | | | | | | | | | | |
|---|---|---|---|---|---|---|---|---|---|---|---|---|---|---|---|---|
| 1 | 54095360 | rs1880861 | 0.404 | -2.094 | 1.683 | 0.63 | 0.004 | 0.077 | 0.077 | 0.877 | 0.17 | 0.02 | 0.08 | 0.04 | 0.1 | 0.21 |
| 1 | 54102943 | rs17382914 | 0.022 | -2.161 | 2.039 | 0.592 | -0.016 | -0.05 | -0.05 | 0 | 0.19 | 0.24 | 0.1 | 0.05 | 0.11 | 0.31 |
| 1 | 54108777 | rs10888801 | 2.025 | -2.232 | 2.049 | 0.63 | 0.072 | 0.012 | 0.012 | 0 | 0.5 | 0.3 | 0.63 | 0.5 | 0.67 | 0.46 |
| 1 | 54108906 | rs10888802 | 1.159 | -2.229 | 1.773 | 0.632 | -0.033 | 0.079 | 0.079 | 0.615 | 0.7 | 0.53 | 0.73 | 0.54 | 0.77 | 0.76 |
| 1 | 54111733 | rs4926611 | 1.912 | -2.236 | 1.728 | 0.627 | 0.071 | -0.05 | -0.05 | 0 | 0.49 | 0.08 | 0.6 | 0.5 | 0.66 | 0.47 |
| 1 | 54114033 | rs7529819 | 1.178 | -2.22 | 1.756 | 0.639 | -0.03 | 0.071 | 0.071 | 0.627 | 0.7 | 0.54 | 0.73 | 0.54 | 0.78 | 0.76 |
| 1 | 54120873 | rs2141079 | 1.897 | -1.818 | 0.739 | 0.53 | -0.031 | -0.077 | -0.077 | 0 | 0.45 | 0.07 | 0.55 | 0.5 | 0.59 | 0.46 |
| 1 | 54121818 | rs12031794 | -0.909 | -1.387 | 1.139 | 0.96 | -0.015 | -0.054 | -0.054 | 0 | 0.19 | 0.15 | 0.25 | 0.4 | 0.24 | 0.1 |
| 1 | 54129591 | rs2950241 | 1.77 | -1.882 | 1.578 | 0.919 | -0.011 | 0.012 | 0.012 | 0.727 | 0.67 | 0.38 | 0.71 | 0.6 | 0.76 | 0.76 |
| 1 | 54131753 | rs4927025 | -0.834 | -1.822 | 1.821 | 0.884 | -0.001 | 0.029 | 0.029 | 0.013 | 0.19 | 0.19 | 0.22 | 0.34 | 0.22 | 0.09 |
| 1 | 54152703 | rs2948042 | 1.83 | -1.986 | 1.769 | 0.845 | 0.044 | -0.045 | -0.045 | 0.175 | 0.67 | 0.23 | 0.72 | 0.65 | 0.76 | 0.75 |
| 1 | 54153635 | rs2950245 | 0.76 | -1.993 | 1.967 | 0.843 | -0.054 | -0.053 | -0.053 | 1.468 | 0.87 | 0.62 | 0.94 | 0.96 | 0.97 | 0.86 |
| 1 | 54154726 | rs882879 | 1.741 | -1.982 | 1.771 | 0.849 | 0.016 | -0.017 | -0.017 | 0 | 0.68 | 0.33 | 0.72 | 0.65 | 0.75 | 0.75 |
| 1 | 54163854 | rs1780394 | 0.504 | -1.99 | 1.827 | 0.659 | -0.028 | -0.034 | -0.034 | 0 | 0.88 | 0.85 | 0.96 | 0.97 | 0.97 | 0.86 |
| 1 | 54174607 | rs11206202 | -1.337 | -1.53 | 1.783 | 1.077 | -0.012 | 0.11 | 0.11 | 0.056 | 0.21 | 0.41 | 0.23 | 0.29 | 0.22 | 0.11 |
| 1 | 54187449 | rs11579257 | -0.445 | -1.775 | 1.342 | 1.086 | -0.004 | 0 | 0 | 0 | 0.16 | 0.2 | 0.18 | 0.27 | 0.11 | 0.2 |
| 1 | 54189668 | rs12043385 | -0.331 | -1.769 | 1.337 | 1.089 | 0.013 | -0.008 | -0.008 | 0.204 | 0.16 | 0.09 | 0.17 | 0.25 | 0.11 | 0.19 |
| 1 | 54190045 | rs7547962 | -0.445 | -1.77 | 1.359 | 1.088 | -0.007 | 0.001 | 0.001 | 0 | 0.16 | 0.23 | 0.19 | 0.27 | 0.11 | 0.2 |
| 1 | 54190695 | rs797906 | -2.228 | -1.882 | 1.207 | 1.027 | 0 | -0.002 | -0.002 | 0 | 0.39 | 0.59 | 0.42 | 0.52 | 0.33 | 0.32 |
| 1 | 54191958 | rs797908 | -0.329 | -1.799 | 1.071 | 0.964 | 0.013 | -0.008 | -0.008 | 6.296 | 0.16 | 0.09 | 0.17 | 0.25 | 0.11 | 0.19 |
| 1 | 54194992 | rs702491 | 0.797 | -1.619 | 0.954 | 1.134 | -0.006 | 0.012 | 0.012 | 0 | 0.8 | 0.73 | 0.82 | 0.83 | 0.8 | 0.88 |
| 1 | 54196929 | rs808860 | 1.471 | -1.738 | 0.462 | 0.794 | 0.028 | 0.006 | 0.006 | 0 | 0.54 | 0.36 | 0.56 | 0.43 | 0.67 | 0.66 |
| 1 | 54197688 | rs702489 | 0.618 | -1.295 | 0.458 | 0.634 | 0 | -0.001 | -0.001 | 0.824 | 0.82 | 0.84 | 0.77 | 0.82 | 0.88 | 0.79 |
| 1 | 54198562 | rs797913 | -1.253 | -0.469 | 1.049 | 0.395 | 0.062 | -0.016 | -0.016 | 0 | 0.09 | 0.32 | 0.03 | 0.04 | 0.09 | 0.06 |
| 1 | 54200662 | rs4927033 | -0.661 | -0.304 | 0.301 | 0.323 | 0.002 | -0.002 | -0.002 | 0 | 0.11 | 0.11 | 0.06 | 0.01 | 0.12 | 0.11 |
| 1 | 54211771 | rs1725982 | -0.892 | -0.108 | 0.275 | 0.576 | 0.001 | -0.022 | -0.022 | 0 | 0.1 | 0.23 | 0.11 | 0.08 | 0.09 | 0.12 |
| 1 | 54222571 | rs1183394 | 0.906 | -0.081 | -0.227 | 1.044 | -0.018 | -0.059 | -0.059 | 0 | 0.89 | 0.98 | 0.9 | 0.75 | 0.9 | 0.85 |
| 1 | 54234215 | rs3753420 | -1.301 | -0.615 | 0.41 | -0.073 | -0.022 | -0.059 | -0.059 | 0.687 | 0.23 | 0.42 | 0.14 | 0.1 | 0.19 | 0.08 |
| 1 | 54234580 | rs17536223 | -0.814 | -0.622 | 0.326 | -0.1 | 0.018 | 0.039 | 0.039 | 0 | 0.17 | 0.06 | 0.08 | 0.1 | 0.14 | 0.06 |
| 1 | 54258880 | rs2273139 | -0.858 | -0.627 | 0.323 | -0.102 | 0.019 | 0.039 | 0.039 | 0 | 0.17 | 0.06 | 0.08 | 0.1 | 0.14 | 0.06 |
| 1 | 54264996 | rs1181180 | -1.944 | -0.653 | 0.476 | -0.117 | -0.031 | -0.076 | -0.076 | 5.557 | 0.24 | 0.49 | 0.15 | 0.1 | 0.19 | 0.07 |
| 1 | 54269087 | rs1181147 | -1.932 | -0.653 | 0.486 | -0.117 | -0.03 | -0.074 | -0.074 | 0.356 | 0.24 | 0.49 | 0.15 | 0.1 | 0.19 | 0.07 |
| 1 | 54272330 | rs1181145 | -1.187 | -0.31 | 0.572 | -0.124 | 0.053 | 0.003 | 0.003 | 1.971 | 0.07 | 0.39 | 0.07 | 0 | 0.05 | 0.01 |
| 1 | 54289020 | rs10489461 | 0.113 | -0.355 | 0.534 | -0.15 | -0.011 | 0.041 | 0.041 | 0 | 0.1 | 0.01 | 0.05 | 0 | 0.12 | 0.02 |
| 1 | 54292026 | rs1181186 | -1.348 | -0.263 | 0.737 | 0.109 | 0.047 | -0.002 | -0.002 | 0 | 0.07 | 0.4 | 0.07 | 0 | 0.05 | 0.01 |
| 1 | 54314043 | rs4354491 | 0.041 | 0.107 | -0.833 | 0.428 | -0.024 | -0.006 | -0.006 | 0 | 0.21 | 0.05 | 0.2 | 0.18 | 0.2 | 0.33 |
| 1 | 54320383 | rs1181094 | 0.462 | 0.475 | -0.308 | -0.025 | 0.034 | 0.068 | 0.068 | 0 | 0.28 | 0.45 | 0.27 | 0.56 | 0.38 | 0.43 |
| 1 | 54322418 | rs17109454 | 0.711 | 0.615 | -0.447 | -0.14 | 0.007 | -0.005 | -0.005 | 0 | 0.19 | 0.05 | 0.18 | 0.17 | 0.15 | 0.27 |
| 1 | 54328025 | rs6588492 | -0.593 | 0.559 | -0.25 | -0.183 | 0.008 | 0.178 | 0.178 | 0 | 0.51 | 0.76 | 0.5 | 0.85 | 0.57 | 0.71 |
| 1 | 54333594 | rs3766468 | 0.42 | 0.563 | -0.104 | -0.152 | 0.01 | 0.022 | 0.022 | 0 | 0.29 | 0.58 | 0.27 | 0.57 | 0.38 | 0.43 |
| 1 | 54341947 | rs10788964 | 0.417 | 0.568 | -0.101 | -0.15 | 0.009 | 0.021 | 0.021 | 0 | 0.29 | 0.58 | 0.27 | 0.57 | 0.38 | 0.43 |
| 1 | 54343342 | rs7548793 | 0.529 | 0.57 | -0.086 | -0.149 | 0.03 | 0.059 | 0.059 | 0.007 | 0.28 | 0.45 | 0.26 | 0.57 | 0.38 | 0.43 |
| 1 | 54354165 | rs926456 | -0.694 | 0.654 | -0.143 | -0.168 | -0.006 | 0.146 | 0.146 | 0 | 0.49 | 0.71 | 0.49 | 0.84 | 0.53 | 0.69 |
| 1 | 54354658 | rs2294508 | 0.231 | 0.689 | -0.438 | -0.098 | -0.002 | 0.003 | 0.003 | 0 | 0.22 | 0.28 | 0.24 | 0.27 | 0.19 | 0.28 |
| 1 | 54355041 | rs10888814 | 0.238 | 0.692 | -0.437 | -0.097 | -0.002 | 0.004 | 0.004 | 0 | 0.22 | 0.28 | 0.24 | 0.27 | 0.19 | 0.28 |
| 1 | 54361288 | rs2268181 | 0.651 | 0.789 | -0.849 | -0.126 | -0.008 | 0.013 | 0.013 | 0 | 0.19 | 0.07 | 0.19 | 0.17 | 0.18 | 0.27 |
| 1 | 54368253 | rs731828 | 0.01 | 0.923 | -0.144 | 0.123 | 0.013 | 0.045 | 0.045 | 0 | 0.29 | 0.47 | 0.28 | 0.58 | 0.36 | 0.42 |
| 1 | 54369730 | rs4926616 | -0.15 | 0.404 | -0.407 | -0.546 | -0.005 | -0.046 | -0.046 | 0 | 0.61 | 0.57 | 0.53 | 0.83 | 0.66 | 0.71 |
| 1 | 54375348 | rs17109582 | -0.005 | 0.093 | -0.203 | -0.675 | -0.009 | -0.06 | -0.06 | 0 | 0.14 | 0.18 | 0.08 | 0 | 0.11 | 0.03 |
| 1 | 54375570 | rs2235544 | 0.2 | 0.075 | -0.92 | -0.43 | -0.06 | -0.063 | -0.063 | 0.329 | 0.39 | 0.16 | 0.64 | 0.46 | 0.52 | 0.67 |
| 1 | 54375701 | rs11206244 | 0.573 | -0.022 | -1.072 | -0.727 | -0.033 | 0.075 | 0.075 | 0 | 0.31 | 0.16 | 0.52 | 0.15 | 0.37 | 0.49 |
| 1 | 54376730 | rs12095080 | 0.451 | -0.221 | -0.507 | -0.502 | -0.01 | -0.063 | -0.063 | 0 | 0.14 | 0.21 | 0.08 | 0 | 0.11 | 0.03 |
| 1 | 54381406 | rs12063777 | -0.803 | -0.714 | -0.336 | -0.628 | -0.018 | -0.035 | -0.035 | 2.41 | 0.24 | 0.58 | 0.11 | 0.06 | 0.13 | 0.06 |
| 1 | 54382383 | rs12065313 | -0.753 | -0.722 | -0.035 | -0.633 | -0.05 | -0.082 | -0.082 | 0 | 0.24 | 0.5 | 0.1 | 0.06 | 0.13 | 0.06 |
| 1 | 54386271 | rs6696913 | -1.073 | -0.83 | -0.076 | -0.682 | 0.036 | 0.042 | 0.042 | 0 | 0.27 | 0.77 | 0.13 | 0.06 | 0.13 | 0.06 |
| 1 | 54388307 | rs10888823 | -0.751 | -0.83 | -0.017 | -0.682 | -0.034 | -0.058 | -0.058 | 0 | 0.24 | 0.55 | 0.11 | 0.06 | 0.13 | 0.06 |
| 1 | 54389200 | rs10047036 | -1.074 | -0.83 | -0.027 | -0.682 | 0.036 | 0.042 | 0.042 | 0 | 0.27 | 0.77 | 0.13 | 0.06 | 0.13 | 0.06 |
| 1 | 54389471 | rs12067895 | -0.769 | -0.765 | -0.062 | -0.649 | -0.043 | -0.071 | -0.071 | 0 | 0.24 | 0.53 | 0.11 | 0.06 | 0.13 | 0.06 |
| 1 | 54413422 | rs7526168 | -0.901 | -0.532 | -0.055 | -0.614 | -0.032 | -0.071 | -0.071 | 0 | 0.24 | 0.54 | 0.11 | 0.06 | 0.14 | 0.06 |
| 1 | 54415698 | rs12079858 | 0.677 | -0.284 | -0.37 | -0.541 | -0.001 | -0.005 | -0.005 | 0 | 0.13 | 0.09 | 0.07 | 0 | 0.12 | 0.03 |
| 1 | 54443867 | rs11206266 | -0.543 | -0.369 | 0.148 | -0.285 | -0.041 | -0.002 | -0.002 | 0 | 0.11 | 0.35 | 0.02 | 0.05 | 0.02 | 0.03 |
| 1 | 54450490 | rs638749 | -0.154 | -0.533 | -0.479 | -0.14 | 0.005 | 0.013 | 0.013 | 0 | 0.42 | 0.34 | 0.45 | 0.4 | 0.58 | 0.68 |
| 1 | 54468166 | rs620907 | -0.906 | -0.57 | -0.562 | -0.116 | 0.042 | 0.016 | 0.016 | 0.255 | 0.52 | 0.71 | 0.49 | 0.47 | 0.61 | 0.72 |
| 1 | 54474472 | rs6672989 | 0.686 | -0.593 | -0.865 | -0.11 | 0.024 | 0.005 | 0.005 | 0.108 | 0.51 | 0.13 | 0.51 | 0.53 | 0.41 | 0.3 |
| 1 | 54482523 | rs1537323 | 0.014 | -0.499 | -0.423 | -0.059 | 0 | -0.001 | -0.001 | 0 | 0.59 | 0.58 | 0.54 | 0.6 | 0.43 | 0.32 |
| 1 | 54482915 | rs11804837 | 0.023 | -0.499 | -0.423 | -0.058 | 0 | -0.001 | -0.001 | 0 | 0.59 | 0.57 | 0.54 | 0.6 | 0.43 | 0.32 |
| 1 | 54484513 | rs7528535 | 0.622 | 0.534 | -1.032 | 0.108 | -0.054 | -0.053 | -0.053 | 0 | 0.08 | 0.01 | 0.14 | 0.26 | 0.22 | 0.15 |
| 1 | 54487417 | rs12047169 | 0.877 | -0.622 | -1.369 | 0.205 | -0.105 | -0.069 | -0.069 | 0 | 0.38 | 0.05 | 0.45 | 0.34 | 0.38 | 0.24 |
| 1 | 54494702 | rs11206274 | -1.064 | -0.898 | -0.202 | 0.34 | -0.063 | 0.065 | 0.065 | 0.855 | 0.17 | 0.63 | 0.12 | 0.27 | 0.05 | 0.1 |

| | | | | | | | | | | | | | | | | |
|---|---|---|---|---|---|---|---|---|---|---|---|---|---|---|---|---|
| 1 | 54497040 | rs1547467 | -0.649 | -1.112 | -0.459 | 0.296 | -0.102 | -0.047 | -0.047 | 0 | 0.66 | 0.9 | 0.58 | 0.61 | 0.44 | 0.34 |
| 1 | 54505437 | rs3766466 | -0.358 | -1.191 | 0.301 | 0.35 | 0.005 | 0.028 | 0.028 | 1.683 | 0.09 | 0.43 | 0.06 | 0.05 | 0.03 | 0.05 |
| 1 | 54522307 | rs3820112 | -0.938 | -1.169 | 0.646 | 0.416 | -0.044 | 0.188 | 0.188 | 0 | 0.13 | 0.48 | 0.09 | 0.27 | 0.04 | 0.1 |
| 1 | 54546449 | rs7556423 | -0.185 | -1.085 | 0.593 | 0.404 | 0.059 | 0.083 | 0.083 | 0 | 0.06 | 0.33 | 0.04 | 0.05 | 0.02 | 0.05 |
| 1 | 54557290 | rs12046885 | 1.371 | -1.114 | 0.564 | 0.252 | 0.019 | 0.023 | 0.023 | 1.275 | 0.41 | 0.23 | 0.45 | 0.35 | 0.34 | 0.22 |
| 1 | 54562444 | rs7524477 | -0.678 | -1.082 | 0.5 | 0.321 | 0.057 | 0.39 | 0.39 | 0.278 | 0.07 | 0.41 | 0.07 | 0.25 | 0.03 | 0.08 |
| 1 | 54563156 | rs6588502 | -0.15 | -0.966 | 0.183 | 0.254 | -0.047 | -0.016 | -0.016 | 0.011 | 0.6 | 0.9 | 0.57 | 0.6 | 0.4 | 0.33 |
| 1 | 54563522 | rs7538004 | 0.017 | -0.889 | 0.268 | 0.279 | 0.06 | 0.085 | 0.085 | 0 | 0.05 | 0.3 | 0.03 | 0.05 | 0.02 | 0.05 |
| 1 | 54572243 | rs10788972 | -0.696 | -0.833 | -0.16 | 0.267 | -0.11 | -0.046 | -0.046 | 0 | 0.66 | 0.9 | 0.58 | 0.6 | 0.44 | 0.35 |
| 1 | 54575746 | rs4927055 | 1.085 | -0.736 | 0.038 | 0.351 | 0.016 | 0.021 | 0.021 | 1.354 | 0.41 | 0.17 | 0.45 | 0.35 | 0.34 | 0.22 |
| 1 | 54576912 | rs12093973 | -0.714 | -0.57 | 0.557 | 0.414 | 0.077 | 0.097 | 0.097 | 0.167 | 0.06 | 0.37 | 0.04 | 0.05 | 0.02 | 0.05 |
| 1 | 54584278 | rs11579628 | -0.143 | -0.264 | 0.005 | 0.377 | -0.069 | -0.003 | -0.003 | 0.848 | 0.62 | 0.87 | 0.6 | 0.6 | 0.51 | 0.35 |
| 1 | 54585415 | rs12063422 | 0.65 | -0.867 | 0.443 | 0.341 | 0.016 | 0.025 | 0.025 | 0.146 | 0.51 | 0.41 | 0.52 | 0.38 | 0.43 | 0.35 |
| 1 | 54586538 | rs12065056 | 0.086 | -0.88 | 0.515 | 0.39 | -0.002 | 0.021 | 0.021 | 0.046 | 0.57 | 0.72 | 0.56 | 0.61 | 0.45 | 0.41 |
| 1 | 54588110 | rs7516442 | 1.035 | -0.855 | 0.318 | 0.376 | 0.022 | 0.041 | 0.041 | 1.22 | 0.44 | 0.27 | 0.51 | 0.33 | 0.39 | 0.24 |
| 1 | 54588354 | rs7530816 | -0.32 | -0.858 | 0.057 | 0.378 | -0.065 | -0.004 | -0.004 | 0 | 0.63 | 0.84 | 0.61 | 0.61 | 0.46 | 0.43 |
| 1 | 54588398 | rs7516735 | -0.281 | -0.86 | 0.219 | 0.377 | -0.035 | 0.005 | 0.005 | 0.019 | 0.62 | 0.77 | 0.6 | 0.61 | 0.46 | 0.42 |
| 1 | 54588636 | rs12095916 | -0.091 | -0.769 | 0.364 | 0.364 | -0.034 | 0.146 | 0.146 | 0 | 0.1 | 0.26 | 0.06 | 0.24 | 0.01 | 0.07 |
| 1 | 54590188 | rs6683426 | 0.861 | -1.226 | -0.231 | 0.011 | 0.01 | 0.006 | 0.006 | 0.875 | 0.34 | 0.26 | 0.2 | 0.26 | 0.27 | 0.18 |
| 1 | 54591322 | rs612368 | -0.358 | -0.689 | -0.015 | 0.16 | 0.011 | -0.006 | -0.006 | 0 | 0.46 | 0.3 | 0.44 | 0.49 | 0.58 | 0.69 |
| 1 | 54592897 | rs608395 | -0.415 | -0.654 | -0.104 | 0.177 | 0.007 | -0.002 | -0.002 | 0 | 0.47 | 0.37 | 0.45 | 0.49 | 0.58 | 0.7 |
| 1 | 54598275 | rs630538 | -0.558 | -0.4 | -0.33 | 0.182 | -0.005 | -0.005 | -0.005 | 0.187 | 0.55 | 0.5 | 0.42 | 0.41 | 0.63 | 0.66 |
| 1 | 54601232 | rs2026110 | -0.331 | -0.594 | -0.024 | -0.136 | -0.025 | -0.102 | -0.102 | 0 | 0.78 | 0.65 | 0.89 | 0.98 | 0.86 | 0.95 |
| 1 | 54601281 | rs2026111 | 0.953 | -0.846 | -0.154 | 0.227 | -0.031 | 0.014 | 0.014 | 0 | 0.65 | 0.56 | 0.72 | 0.59 | 0.72 | 0.65 |
| 1 | 54602648 | rs913811 | -0.116 | -0.492 | -0.05 | 0.615 | 0.001 | 0.041 | 0.041 | 0 | 0.09 | 0.08 | 0.16 | 0.38 | 0.12 | 0.13 |
| 1 | 54602977 | rs12078845 | -1.439 | -0.511 | 0.057 | 0.442 | -0.009 | 0.024 | 0.024 | 0 | 0.26 | 0.32 | 0.26 | 0.62 | 0.25 | 0.46 |
| 1 | 54606522 | rs12078646 | -0.455 | -0.825 | 0.475 | 0.491 | -0.032 | 0.256 | 0.256 | 0.006 | 0.09 | 0.26 | 0.13 | 0.43 | 0.03 | 0.27 |
| 1 | 54606804 | rs3766465 | 0.733 | -0.526 | 0.377 | -0.024 | -0.005 | -0.007 | -0.007 | 0.605 | 0.85 | 0.77 | 0.79 | 0.87 | 0.85 | 0.84 |
| 1 | 54607157 | rs671539 | 0.19 | -0.493 | 0.361 | -0.063 | -0.001 | -0.018 | -0.018 | 0 | 0.88 | 0.84 | 0.81 | 0.97 | 0.85 | 0.9 |
| 1 | 54608352 | rs1571541 | 0.355 | -0.868 | 0.289 | -0.36 | -0.003 | -0.022 | -0.022 | 0 | 0.22 | 0.19 | 0.1 | 0.04 | 0.17 | 0.17 |
| 1 | 54609652 | rs599597 | 0.329 | -0.801 | 0.283 | -0.225 | -0.015 | -0.006 | -0.006 | 0 | 0.86 | 0.77 | 0.91 | 0.88 | 0.88 | 0.9 |
| 1 | 54611295 | rs617067 | -0.082 | -0.556 | 0.292 | -0.297 | -0.011 | -0.047 | -0.047 | 0 | 0.24 | 0.27 | 0.22 | 0.07 | 0.2 | 0.21 |
| 1 | 54613369 | rs17109915 | 1.099 | -0.551 | 0.27 | -0.31 | -0.002 | -0.034 | -0.034 | 0 | 0.16 | 0.12 | 0.07 | 0 | 0.15 | 0.08 |
| 1 | 54615232 | rs4927065 | 0.156 | -0.57 | 0.231 | -0.471 | -0.011 | -0.061 | -0.061 | 0.045 | 0.69 | 0.68 | 0.74 | 0.93 | 0.77 | 0.75 |
| 1 | 54615337 | rs585715 | 0.536 | -0.551 | 0.065 | -0.461 | 0.001 | 0.001 | 0.001 | 0 | 0.08 | 0.05 | 0.03 | 0.04 | 0.03 | 0.11 |
| 1 | 54615558 | rs6699826 | 0.761 | -0.54 | 0.073 | -0.464 | 0.001 | -0.037 | -0.037 | 1.946 | 0.21 | 0.18 | 0.09 | 0 | 0.17 | 0.11 |
| 1 | 54616753 | rs1999175 | -0.178 | -0.294 | 0.005 | -0.266 | 0 | -0.003 | -0.003 | 0 | 0.1 | 0.1 | 0.17 | 0.07 | 0.08 | 0.15 |
| 1 | 54619106 | rs12096978 | -0.502 | -0.621 | -0.112 | -0.128 | -0.002 | 0 | 0 | 0 | 0.22 | 0.23 | 0.17 | 0.26 | 0.2 | 0.35 |
| 1 | 54619337 | rs657783 | -0.205 | -0.646 | 0.028 | 0.24 | -0.012 | 0.034 | 0.034 | 0 | 0.9 | 0.88 | 0.9 | 0.72 | 0.97 | 0.89 |
| 1 | 54619404 | rs7519887 | 0.714 | 0.407 | -0.441 | -0.086 | -0.068 | 0.031 | 0.031 | 0 | 0.11 | 0.02 | 0.13 | 0.07 | 0.23 | 0.14 |
| 1 | 54622694 | rs17109929 | 1.109 | -0.525 | -0.157 | -0.116 | 0.007 | -0.012 | -0.012 | 0 | 0.21 | 0.13 | 0.09 | 0 | 0.17 | 0.12 |
| 1 | 54625292 | rs660514 | -0.015 | -0.52 | -0.064 | -0.017 | -0.026 | -0.008 | -0.008 | 2.414 | 0.67 | 0.59 | 0.8 | 0.72 | 0.8 | 0.78 |
| 1 | 54627830 | rs618465 | -0.441 | -0.451 | 0 | 0.138 | -0.025 | 0.057 | 0.057 | 0.022 | 0.88 | 0.8 | 0.89 | 0.74 | 0.89 | 0.87 |
| 1 | 54633847 | rs17109983 | 0.608 | -0.422 | -0.051 | 0.187 | -0.027 | 0.135 | 0.135 | 1.064 | 0.06 | 0.15 | 0.05 | 0.28 | 0.01 | 0.08 |
| 1 | 54634941 | rs686106 | 1.339 | -0.18 | -0.174 | -0.069 | -0.002 | -0.004 | -0.004 | 1.776 | 0.43 | 0.34 | 0.49 | 0.44 | 0.52 | 0.41 |
| 1 | 54636689 | rs7552121 | -0.043 | -0.494 | -0.049 | 0.076 | 0.008 | -0.029 | -0.029 | 1.95 | 0.12 | 0.08 | 0.14 | 0.27 | 0.12 | 0.11 |
| 1 | 54636977 | rs10493175 | 0.835 | -0.477 | 0.002 | -0.073 | 0.003 | 0 | 0 | 0.277 | 0.23 | 0.13 | 0.1 | 0 | 0.21 | 0.13 |
| 1 | 54637167 | rs625643 | -0.263 | -0.351 | -0.295 | 0.009 | -0.022 | -0.012 | -0.012 | 0 | 0.68 | 0.6 | 0.82 | 0.72 | 0.75 | 0.62 |
| 1 | 54655563 | rs11206297 | 0.513 | 0.001 | -0.712 | 0.078 | 0.032 | -0.001 | -0.001 | 0 | 0.52 | 0.38 | 0.56 | 0.49 | 0.61 | 0.53 |
| 1 | 54655895 | rs10888837 | 0.53 | 0.011 | -0.745 | 0.101 | 0.037 | 0 | 0 | 0.671 | 0.51 | 0.39 | 0.56 | 0.5 | 0.61 | 0.53 |
| 1 | 54660515 | rs12120284 | 1.631 | 0.727 | -0.893 | 0.544 | 0.03 | 0.048 | 0.048 | 0 | 0.48 | 0.24 | 0.34 | 0.35 | 0.42 | 0.31 |
| 1 | 54661150 | rs946448 | -1.063 | 1.732 | -0.515 | 0.8 | -0.023 | 0.008 | 0.008 | 0 | 0.24 | 0.21 | 0.32 | 0.26 | 0.35 | 0.48 |
| 1 | 54662346 | rs7525757 | -1.1 | 1.736 | -0.513 | 0.802 | -0.022 | 0.008 | 0.008 | 0 | 0.24 | 0.22 | 0.32 | 0.26 | 0.35 | 0.48 |
| 1 | 54664921 | rs4926623 | -1.408 | 1.163 | 0.085 | 1.372 | 0.011 | 0.097 | 0.097 | 0 | 0.21 | 0.37 | 0.47 | 0.49 | 0.2 | 0.33 |
| 1 | 54665124 | rs946447 | 1.311 | 0.651 | 0.337 | 1.118 | -0.029 | -0.005 | -0.005 | 0 | 0.87 | 0.61 | 0.83 | 0.92 | 0.91 | 0.65 |
| 1 | 54673097 | rs10888838 | -0.482 | 1.358 | -0.008 | 1.898 | -0.009 | 0.009 | 0.009 | 0.246 | 0.12 | 0.09 | 0.21 | 0.47 | 0.13 | 0.25 |
| 1 | 54679912 | rs4927071 | 1.373 | 1.248 | 0.625 | 1.861 | 0.003 | 0.091 | 0.091 | 0 | 0.8 | 0.68 | 0.64 | 0.49 | 0.8 | 0.48 |
| 1 | 54680058 | rs4927072 | -0.254 | 1.59 | 0.599 | 1.877 | 0 | 0 | 0 | 0.008 | 0.1 | 0.09 | 0.18 | 0.08 | 0.18 | 0.06 |
| 1 | 54689461 | rs10888843 | -1.291 | 1.823 | 0.515 | 1.955 | 0.006 | 0.039 | 0.039 | 0.004 | 0.3 | 0.58 | 0.27 | 0.54 | 0.29 | 0.38 |
| 1 | 54690037 | rs2282332 | -0.44 | 1.516 | 0.711 | 1.999 | -0.006 | -0.009 | -0.009 | 0 | 0.13 | 0.1 | 0.11 | 0.37 | 0.12 | 0.24 |
| 1 | 54690954 | rs3088379 | -1.287 | 1.445 | 0.537 | 1.634 | -0.004 | 0.008 | 0.008 | 0.504 | 0.42 | 0.59 | 0.4 | 0.31 | 0.45 | 0.51 |
| 1 | 54700707 | rs686930 | -1.621 | 1.237 | 0.015 | 1.483 | 0.012 | -0.01 | -0.01 | 0.004 | 0.5 | 0.83 | 0.67 | 0.46 | 0.55 | 0.78 |
| 1 | 54702869 | rs3766453 | -1.002 | 0.903 | 0.298 | 0.905 | -0.001 | 0.001 | 0.001 | 0.579 | 0.2 | 0.23 | 0.22 | 0.1 | 0.2 | 0.51 |
| 1 | 54718770 | rs687050 | -0.656 | 0.222 | 0.183 | 0.927 | 0.001 | 0.01 | 0.01 | 0 | 0.57 | 0.64 | 0.54 | 0.33 | 0.54 | 0.77 |
| 1 | 54730036 | rs3766431 | 0.096 | 0.187 | -0.172 | 0.728 | 0.006 | -0.016 | -0.016 | 0 | 0.38 | 0.4 | 0.4 | 0.23 | 0.41 | 0.48 |
| 1 | 54736800 | rs9662034 | -0.067 | 0.532 | -0.322 | 1.198 | -0.001 | 0.007 | 0.007 | 0 | 0.58 | 0.63 | 0.64 | 0.37 | 0.52 | 0.76 |
| 1 | 54741767 | rs603901 | -0.065 | 0.716 | -0.814 | 0.276 | -0.004 | 0.023 | 0.023 | 0 | 0.58 | 0.75 | 0.72 | 0.67 | 0.52 | 0.74 |
| 1 | 54743408 | rs11206315 | -1.05 | 0.828 | -0.392 | 0.803 | 0.004 | 0.062 | 0.062 | 0 | 0.15 | 0.21 | 0.12 | 0.29 | 0.17 | 0.15 |
| 1 | 54745567 | rs3753406 | -0.978 | 0.117 | -0.711 | -0.568 | 0.009 | 0.007 | 0.007 | 0 | 0.32 | 0.28 | 0.4 | 0.18 | 0.22 | 0.5 |

| | | | | | | | | | | | | | | | | |
|---|---|---|---|---|---|---|---|---|---|---|---|---|---|---|---|---|
| 1 | 54750625 | rs9326028 | -0.567 | -0.08 | -0.586 | -0.673 | -0.004 | -0.028 | -0.028 | 0 | 0.18 | 0.15 | 0.09 | 0.02 | 0.13 | 0.15 |
| 1 | 54751900 | rs10788977 | -1.427 | 0.691 | -0.709 | -0.728 | 0.005 | 0.009 | 0.009 | 0 | 0.33 | 0.28 | 0.43 | 0.16 | 0.28 | 0.45 |
| 1 | 54753512 | rs1416558 | -0.384 | 1.914 | -0.195 | -0.832 | 0.077 | -0.029 | -0.029 | 0 | 0.07 | 0.17 | 0.06 | 0.01 | 0.13 | 0.13 |
| 1 | 54753840 | rs608547 | -0.33 | 1.92 | -0.279 | -1.667 | 0.038 | -0.094 | -0.094 | 0 | 0.24 | 0.34 | 0.2 | 0.05 | 0.34 | 0.22 |
| 1 | 54755091 | rs6697414 | -1.108 | 0.963 | -0.463 | -1.895 | -0.01 | -0.037 | -0.037 | 0 | 0.34 | 0.59 | 0.43 | 0.17 | 0.26 | 0.59 |
| 1 | 54756342 | rs4927081 | -0.009 | 0.416 | -0.766 | -1.953 | -0.001 | -0.007 | -0.007 | 2.28 | 0.61 | 0.58 | 0.53 | 0.74 | 0.66 | 0.55 |
| 1 | 54758639 | rs12040312 | -0.258 | 0.222 | -0.896 | -1.522 | 0.016 | -0.005 | -0.005 | 0 | 0.06 | 0.01 | 0.23 | 0.09 | 0.02 | 0.18 |
| 1 | 54760066 | rs4927084 | -0.02 | 0.156 | -0.818 | -2.137 | -0.007 | -0.024 | -0.024 | 0.602 | 0.58 | 0.57 | 0.55 | 0.83 | 0.69 | 0.62 |
| 1 | 54762317 | rs11206321 | 0.099 | 0.357 | -0.892 | -2.267 | -0.001 | -0.041 | -0.041 | 0.038 | 0.35 | 0.37 | 0.32 | 0.11 | 0.36 | 0.36 |
| 1 | 54762350 | rs6678619 | 0.595 | 0.447 | -1.133 | -1.123 | -0.024 | -0.072 | -0.072 | 2.727 | 0.22 | 0.13 | 0.21 | 0.52 | 0.23 | 0.26 |
| 1 | 54765289 | rs11206326 | 0.313 | -0.254 | -1.154 | -1.051 | 0.024 | -0.016 | -0.016 | 0 | 0.06 | 0 | 0.22 | 0.08 | 0.02 | 0.18 |
| 1 | 54765727 | rs10888844 | -1.192 | -0.394 | -1.414 | -1.425 | -0.002 | -0.006 | -0.006 | 0.047 | 0.71 | 0.67 | 0.76 | 0.9 | 0.63 | 0.88 |
| 1 | 54766101 | rs4927086 | -0.201 | -1.257 | -1.693 | -0.856 | 0.024 | 0.178 | 0.178 | 0.624 | 0.37 | 0.08 | 0.25 | 0.13 | 0.3 | 0.41 |
| 1 | 54766816 | rs7523947 | -0.887 | -1.268 | -1.662 | -0.836 | 0.037 | 0.058 | 0.058 | 0.006 | 0.46 | 0.27 | 0.49 | 0.22 | 0.32 | 0.59 |
| 1 | 54767767 | rs12730301 | -0.439 | -0.642 | -2.476 | -0.732 | 0.024 | 0.178 | 0.178 | 0.624 | 0.37 | 0.08 | 0.25 | 0.13 | 0.3 | 0.4 |
| 1 | 54769520 | rs6694281 | -0.627 | -0.602 | -2.466 | -0.37 | 0.038 | 0.054 | 0.054 | 0.053 | 0.39 | 0.12 | 0.33 | 0.33 | 0.31 | 0.44 |
| 1 | 54769756 | rs17392923 | -0.028 | -0.1 | -2.146 | -0.441 | 0 | 0.101 | 0.101 | 0 | 0.23 | 0.04 | 0.17 | 0.11 | 0.2 | 0.34 |
| 1 | 54770909 | rs6695534 | -0.172 | 0.112 | -1.912 | 0.204 | -0.01 | 0.029 | 0.029 | 0 | 0.61 | 0.81 | 0.58 | 0.71 | 0.6 | 0.76 |
| 1 | 54775203 | rs4927088 | -1.081 | -0.251 | -1.138 | 1.006 | -0.008 | 0.035 | 0.035 | 0.012 | 0.2 | 0.26 | 0.22 | 0.46 | 0.15 | 0.22 |
| 1 | 54784716 | rs3927580 | -1.753 | -0.387 | -0.226 | 1.391 | -0.035 | 0.033 | 0.033 | 0 | 0.3 | 0.44 | 0.29 | 0.42 | 0.22 | 0.33 |
| 1 | 54785109 | rs17101268 | 0.042 | -0.388 | -0.184 | 1.378 | 0.005 | -0.008 | -0.008 | 0.021 | 0.1 | 0.05 | 0.06 | 0.13 | 0.08 | 0.12 |
| 1 | 54787049 | rs12021758 | -2.252 | -0.446 | 0.381 | 1.443 | -0.004 | 0.004 | 0.004 | 0.136 | 0.34 | 0.64 | 0.32 | 0.54 | 0.26 | 0.34 |
| 1 | 54794819 | rs4927092 | -2.082 | -0.541 | -0.003 | 1.598 | -0.02 | 0.023 | 0.023 | 0.733 | 0.27 | 0.65 | 0.25 | 0.54 | 0.15 | 0.27 |
| 1 | 54794930 | rs12564882 | -2.082 | -0.541 | -0.003 | 1.598 | -0.02 | 0.023 | 0.023 | 0.733 | 0.27 | 0.65 | 0.25 | 0.54 | 0.15 | 0.27 |
| 1 | 54794985 | rs12728736 | -0.485 | -0.542 | -0.039 | 1.585 | 0.001 | -0.03 | -0.03 | 0 | 0.1 | 0 | 0.05 | 0.12 | 0.07 | 0.12 |
| 1 | 54796767 | rs1556528 | -1.264 | -0.535 | 0.012 | 1.603 | -0.042 | 0.126 | 0.126 | 0.542 | 0.17 | 0.6 | 0.2 | 0.43 | 0.08 | 0.15 |
| 1 | 54798752 | rs10788979 | -1.692 | -0.298 | -0.024 | 1.199 | -0.04 | 0.052 | 0.052 | 0.185 | 0.25 | 0.54 | 0.15 | 0.37 | 0.15 | 0.24 |
| 1 | 54801454 | rs12065118 | -0.112 | 0.065 | -0.209 | 1.082 | -0.002 | -0.001 | -0.001 | 0 | 0.05 | 0.07 | 0.2 | 0.03 | 0.04 | 0.09 |
| 1 | 54810970 | rs12092071 | -1.3 | 0.068 | 0.027 | 1.173 | 0 | 0.042 | 0.042 | 0 | 0.2 | 0.38 | 0.33 | 0.24 | 0.19 | 0.18 |
| 1 | 54818099 | rs2076571 | -1.671 | 0.635 | -0.961 | 0.99 | -0.063 | -0.05 | -0.05 | 0 | 0.56 | 0.97 | 0.66 | 0.62 | 0.57 | 0.43 |
| 1 | 54821429 | rs7552342 | -0.781 | 1.331 | -0.195 | 1.074 | 0.013 | 0.065 | 0.065 | 0 | 0.29 | 0.55 | 0.26 | 0.44 | 0.29 | 0.23 |
| 1 | 54826187 | rs10489565 | 1.005 | 1.604 | -0.92 | 1.158 | -0.006 | -0.026 | -0.026 | 0.208 | 0.13 | 0.08 | 0.12 | 0.27 | 0.15 | 0.08 |
| 1 | 54827554 | rs2073109 | -1.833 | 1.613 | -1.029 | 1.106 | 0.029 | 0.006 | 0.006 | 0 | 0.53 | 0.88 | 0.64 | 0.58 | 0.56 | 0.41 |
| 1 | 54830569 | rs2317387 | 0.035 | 1.779 | -0.612 | 1.082 | 0.091 | 0.052 | 0.052 | 0 | 0.1 | 0.25 | 0.11 | 0.16 | 0.16 | 0.08 |
| 1 | 54831801 | rs3855975 | -1.588 | 1.23 | -1.528 | 0.988 | -0.021 | 0.005 | 0.005 | 0 | 0.53 | 0.88 | 0.63 | 0.58 | 0.53 | 0.41 |
| 1 | 54836906 | rs12024740 | 0.537 | 0.869 | -1.106 | 1.283 | -0.021 | -0.075 | -0.075 | 0 | 0.12 | 0.03 | 0.11 | 0.27 | 0.11 | 0.15 |
| 1 | 54852994 | rs11206344 | -1.628 | 0.967 | -1.011 | 1.013 | -0.004 | 0.117 | 0.117 | 0 | 0.5 | 0.78 | 0.66 | 0.68 | 0.5 | 0.46 |
| 1 | 54854079 | rs6690450 | -1.699 | 0.99 | -1.028 | 0.954 | -0.013 | 0.184 | 0.184 | 0 | 0.51 | 0.86 | 0.67 | 0.75 | 0.51 | 0.47 |
| 1 | 54869816 | rs213496 | -1.459 | 0.854 | -0.574 | 1.001 | 0.02 | 0.01 | 0.01 | 0 | 0.48 | 0.64 | 0.3 | 0.54 | 0.39 | 0.36 |
| 1 | 54883003 | rs4927101 | 1.634 | 1.136 | -0.571 | 1.082 | -0.045 | 0.018 | 0.018 | 0 | 0.46 | 0.14 | 0.63 | 0.4 | 0.5 | 0.55 |
| 1 | 54890956 | rs1883783 | 0.688 | 0.988 | -0.049 | 1.23 | -0.023 | -0.042 | -0.042 | 0.16 | 0.66 | 0.82 | 0.77 | 0.6 | 0.62 | 0.71 |
| 1 | 54902861 | rs4927113 | 0.24 | 1.096 | -0.078 | 1.026 | -0.008 | -0.004 | -0.004 | 0 | 0.74 | 0.81 | 0.84 | 0.73 | 0.7 | 0.78 |
| 1 | 54905582 | rs11206355 | -1.247 | 0.938 | 0.335 | 1.228 | -0.022 | 0.084 | 0.084 | 0 | 0.21 | 0.5 | 0.48 | 0.35 | 0.18 | 0.21 |
| 1 | 54907935 | rs1543369 | -1.5 | 0.966 | 0.272 | 1.234 | -0.006 | 0.053 | 0.053 | 0 | 0.23 | 0.6 | 0.51 | 0.41 | 0.22 | 0.22 |
| 1 | 54908153 | rs12758577 | 2.125 | 0.907 | -0.069 | 0.991 | -0.046 | 0.021 | 0.021 | 0 | 0.39 | 0.08 | 0.26 | 0.33 | 0.39 | 0.49 |
| 1 | 54918666 | rs12746454 | 2.113 | 0.942 | -0.054 | 1.007 | -0.045 | 0.022 | 0.022 | 0 | 0.39 | 0.08 | 0.26 | 0.33 | 0.39 | 0.49 |
| 1 | 54922003 | rs4927123 | 0.685 | 0.813 | 0.149 | 0.559 | -0.005 | 0.014 | 0.014 | 0 | 0.61 | 0.63 | 0.76 | 0.73 | 0.61 | 0.71 |
| 1 | 54928493 | rs7550485 | -1.486 | 0.932 | 1.078 | 1.306 | -0.056 | -0.012 | -0.012 | 0 | 0.15 | 0.32 | 0.08 | 0.14 | 0.08 | 0.12 |
| 1 | 54930970 | rs213473 | 1.468 | 0.996 | 0.398 | 1.28 | -0.015 | 0.012 | 0.012 | 0 | 0.84 | 0.76 | 0.89 | 0.8 | 0.88 | 0.87 |
| 1 | 54938567 | rs1999205 | 0.586 | 1.285 | -0.001 | 1.188 | -0.009 | 0.013 | 0.013 | 0 | 0.86 | 0.88 | 0.88 | 0.95 | 0.81 | 0.78 |
| 1 | 54943026 | rs382327 | 1.804 | 1.34 | 0.269 | 1.549 | 0.033 | 0.08 | 0.08 | 0 | 0.74 | 0.43 | 0.45 | 0.53 | 0.66 | 0.65 |
| 1 | 54943752 | rs6656290 | -1.234 | 0.752 | 0.454 | 1.018 | 0.003 | 0.073 | 0.073 | 1.154 | 0.1 | 0.5 | 0.06 | 0.1 | 0.05 | 0.05 |
| 1 | 54946847 | rs190963 | -1.061 | 0.849 | -0.015 | 0.664 | -0.019 | -0.007 | -0.007 | 0 | 0.23 | 0.32 | 0.13 | 0.21 | 0.16 | 0.14 |
| 1 | 54953523 | rs4927133 | 1.098 | 1.19 | -0.067 | 1.249 | -0.001 | 0.006 | 0.006 | 0 | 0.79 | 0.73 | 0.84 | 0.73 | 0.78 | 0.76 |
| 1 | 54960989 | rs904218 | -0.652 | 1.294 | 0.013 | 0.983 | 0.017 | 0.002 | 0.002 | 0 | 0.07 | 0.16 | 0.11 | 0.05 | 0.09 | 0.14 |
| 1 | 54961163 | rs6697657 | -1.43 | 0.668 | 0.613 | 0.825 | -0.06 | -0.064 | -0.064 | 0.114 | 0.18 | 0.43 | 0.08 | 0.06 | 0.1 | 0.11 |
| 1 | 54961630 | rs299894 | 0.617 | 1.084 | 0.112 | 1.504 | 0.005 | 0.072 | 0.072 | 0 | 0.88 | 0.85 | 0.71 | 0.46 | 0.81 | 0.9 |
| 1 | 54967890 | rs1888182 | 1.848 | 1.589 | -0.573 | 0.801 | -0.001 | 0.157 | 0.157 | 0.063 | 0.48 | 0.11 | 0.39 | 0.31 | 0.4 | 0.52 |
| 1 | 54968494 | rs299881 | 1.447 | 1.344 | -0.236 | 1.065 | -0.002 | 0.002 | 0.002 | 0 | 0.71 | 0.36 | 0.55 | 0.35 | 0.67 | 0.73 |
| 1 | 54983931 | rs4927135 | -0.6 | 1.265 | -0.418 | 1.84 | 0.005 | 0.004 | 0.004 | 0 | 0.08 | 0.05 | 0.29 | 0.25 | 0.08 | 0.04 |
| 1 | 54984107 | rs4927134 | -1.441 | 1.646 | -0.099 | 1.723 | 0.005 | 0.083 | 0.083 | 0 | 0.14 | 0.24 | 0.34 | 0.3 | 0.15 | 0.08 |
| 1 | 54986334 | rs1105690 | 1.706 | 1.641 | -1.221 | 1.091 | -0.038 | 0.079 | 0.079 | 0.551 | 0.45 | 0.02 | 0.26 | 0.32 | 0.37 | 0.46 |
| 1 | 54992939 | rs12752888 | -0.641 | 1.482 | -0.453 | 0.772 | 0.001 | -0.036 | -0.036 | 0.001 | 0.24 | 0.26 | 0.2 | 0.04 | 0.26 | 0.22 |
| 1 | 55006557 | rs7548057 | -1.044 | 0.878 | -0.494 | 1.146 | -0.001 | -0.001 | -0.001 | 0 | 0.33 | 0.61 | 0.2 | 0.22 | 0.23 | 0.26 |
| 1 | 55009913 | rs380841 | 0.153 | 1.559 | -0.947 | 1.372 | -0.06 | -0.042 | -0.042 | 0 | 0.9 | 0.98 | 0.84 | 0.88 | 0.81 | 0.81 |
| 1 | 55012359 | rs10888857 | -0.374 | 0.632 | -0.901 | 1.714 | 0.03 | -0.033 | -0.033 | 0.028 | 0.09 | 0.03 | 0.18 | 0.23 | 0.05 | 0.04 |
| 1 | 55012825 | rs7515747 | -0.892 | 0.968 | -0.843 | 1.518 | -0.002 | -0.006 | -0.006 | 0 | 0.09 | 0.04 | 0.18 | 0.1 | 0.1 | 0.14 |
| 1 | 55014160 | rs299872 | 0.537 | 1.384 | -0.696 | 1.587 | -0.015 | -0.011 | -0.011 | 0.008 | 0.75 | 0.22 | 0.8 | 0.76 | 0.73 | 0.84 |
| 1 | 55014492 | rs3753013 | -1.277 | 0.304 | -0.137 | 1.381 | -0.047 | 0.011 | 0.011 | 0 | 0.27 | 0.41 | 0.4 | 0.32 | 0.17 | 0.18 |

| | | | | | | | | | | | | | | | | |
|---|---|---|---|---|---|---|---|---|---|---|---|---|---|---|---|---|
| 1 | 55022557 | rs2133003 | -1.268 | 2.026 | 0.349 | 1.162 | 0.139 | 0.008 | 0.008 | 0 | 0.13 | 0.29 | 0.16 | 0.17 | 0.27 | 0.08 |
| 1 | 55033582 | rs405731 | -0.645 | -0.136 | 0.171 | 1.187 | -0.027 | 0.009 | 0.009 | 0 | 0.31 | 0.62 | 0.31 | 0.4 | 0.15 | 0.36 |
| 1 | 55033869 | rs300280 | -1.391 | -0.114 | -0.573 | 0.89 | -0.035 | -0.05 | -0.05 | 0 | 0.66 | 0.88 | 0.62 | 0.61 | 0.63 | 0.7 |
| 1 | 55035348 | rs300278 | 0.071 | -0.157 | -0.277 | 0.892 | -0.075 | 0.059 | 0.059 | 0 | 0.35 | 0.17 | 0.33 | 0.26 | 0.51 | 0.31 |
| 1 | 55035395 | rs300277 | -0.468 | -0.325 | 0.254 | 1.063 | -0.011 | 0.007 | 0.007 | 0 | 0.42 | 0.32 | 0.43 | 0.44 | 0.55 | 0.58 |
| 1 | 55044940 | rs1755588 | -2.042 | -0.302 | 1.035 | 1.221 | 0.114 | 0.253 | 0.253 | 4.164 | 0.05 | 0.42 | 0.06 | 0.16 | 0.02 | 0.04 |
| 1 | 55045983 | rs300272 | -0.219 | -0.437 | 0.505 | 0.967 | -0.041 | 0.099 | 0.099 | 0.219 | 0.42 | 0.14 | 0.44 | 0.27 | 0.61 | 0.54 |
| 1 | 55046039 | rs300271 | -1.619 | -0.612 | 0.503 | 1.17 | -0.028 | -0.04 | -0.04 | 1.397 | 0.7 | 0.86 | 0.72 | 0.63 | 0.71 | 0.74 |
| 1 | 55046392 | rs300269 | 0.207 | -0.686 | 0.432 | 1.028 | -0.097 | 0.131 | 0.131 | 0.008 | 0.37 | 0.13 | 0.36 | 0.2 | 0.53 | 0.28 |
| 1 | 55046535 | rs300267 | -1.025 | -0.727 | 0.747 | 1.012 | 0.003 | -0.001 | -0.001 | 0 | 0.63 | 0.56 | 0.53 | 0.41 | 0.62 | 0.6 |
| 1 | 55048190 | rs17110749 | 1.062 | -0.978 | 0.784 | 1.045 | -0.064 | -0.063 | -0.063 | 0.186 | 0.19 | 0.24 | 0.08 | 0.03 | 0.05 | 0.06 |
| 1 | 55049806 | rs17110757 | 0.344 | -0.99 | 0.816 | 1.138 | -0.094 | 0.019 | 0.019 | 0 | 0.23 | 0.44 | 0.22 | 0.2 | 0.08 | 0.12 |
| 1 | 55059295 | rs1172779 | -1.311 | -0.402 | 1.157 | 1.475 | -0.003 | 0.118 | 0.118 | 0 | 0.07 | 0.32 | 0.11 | 0.18 | 0.04 | 0.26 |
| 1 | 55060362 | rs1702003 | -1.962 | -0.415 | 1.166 | 1.469 | -0.014 | 0.124 | 0.124 | 0.126 | 0.1 | 0.54 | 0.12 | 0.18 | 0.04 | 0.26 |
| 1 | 55060548 | rs1702002 | -1.817 | -0.391 | 1.095 | 1.48 | 0.003 | 0.161 | 0.161 | 0.065 | 0.09 | 0.48 | 0.12 | 0.18 | 0.04 | 0.26 |
| 1 | 55069986 | rs746503 | -0.386 | -0.18 | 0.476 | 1.688 | 0 | 0.002 | 0.002 | 0 | 0.61 | 0.49 | 0.56 | 0.43 | 0.64 | 0.45 |
| 1 | 55073396 | rs12027962 | 1.218 | -0.151 | 0.239 | 1.677 | -0.089 | -0.069 | -0.069 | 0 | 0.3 | 0.06 | 0.3 | 0.35 | 0.31 | 0.27 |
| 1 | 55076430 | rs733035 | -0.365 | -0.163 | 0.4 | 1.508 | 0 | -0.001 | -0.001 | 0 | 0.61 | 0.48 | 0.55 | 0.35 | 0.63 | 0.41 |
| 1 | 55078046 | rs1655528 | 1.623 | -0.277 | 0.448 | 1.909 | -0.023 | 0.331 | 0.331 | 0.609 | 0.87 | 0.53 | 0.71 | 0.6 | 0.92 | 0.69 |
| 1 | 55085141 | rs17395160 | 0.278 | 0.052 | -0.853 | 1.129 | -0.063 | -0.029 | -0.029 | 0 | 0.22 | 0.02 | 0.18 | 0.2 | 0.26 | 0.19 |
| 1 | 55085499 | rs11206396 | -1.081 | -0.197 | -0.3 | 1.313 | 0.017 | -0.003 | -0.003 | 0 | 0.51 | 0.7 | 0.53 | 0.49 | 0.5 | 0.67 |
| 1 | 55089317 | rs2060386 | -0.332 | -0.432 | -0.582 | 1.239 | 0.037 | 0.006 | 0.006 | 0 | 0.39 | 0.11 | 0.28 | 0.34 | 0.37 | 0.27 |
| 1 | 55089708 | rs2117263 | -0.34 | -0.396 | -0.578 | 1.244 | 0.042 | 0.006 | 0.006 | 0 | 0.4 | 0.1 | 0.28 | 0.34 | 0.37 | 0.27 |
| 1 | 55092069 | rs12039865 | -0.972 | -0.258 | -0.248 | 1.292 | 0.008 | 0.105 | 0.105 | 0 | 0.07 | 0.17 | 0.25 | 0.24 | 0.09 | 0.24 |
| 1 | 55103945 | rs10736390 | -0.964 | -0.336 | -0.075 | 1.048 | 0.036 | 0.006 | 0.006 | 0 | 0.51 | 0.68 | 0.52 | 0.55 | 0.45 | 0.45 |
| 1 | 55106807 | rs1655524 | -0.15 | -0.312 | -0.243 | 0.603 | -0.001 | -0.003 | -0.003 | 3.397 | 0.77 | 0.75 | 0.77 | 0.86 | 0.73 | 0.79 |
| 1 | 55107019 | rs1368882 | 1.044 | -0.131 | -0.639 | 0.743 | -0.025 | -0.029 | -0.029 | 1.415 | 0.33 | 0.23 | 0.46 | 0.48 | 0.35 | 0.53 |
| 1 | 55107055 | rs957132 | 0.927 | -0.347 | -0.856 | -0.101 | 0.033 | 0.082 | 0.082 | 0.007 | 0.14 | 0 | 0.05 | 0.03 | 0.07 | 0.08 |
| 1 | 55115501 | rs1655523 | 0.975 | 0.059 | -0.876 | -0.228 | -0.019 | 0.052 | 0.052 | 0 | 0.31 | 0.18 | 0.3 | 0.21 | 0.29 | 0.49 |
| 1 | 55115608 | rs1701999 | -0.139 | -0.123 | -0.186 | -0.401 | -0.008 | -0.075 | -0.075 | 0 | 0.76 | 0.51 | 0.76 | 0.89 | 0.76 | 0.84 |
| 1 | 55117000 | rs752468 | 1.125 | -0.103 | 0.008 | -0.357 | -0.002 | 0 | 0 | 0 | 0.33 | 0.27 | 0.34 | 0.21 | 0.3 | 0.54 |
| 1 | 55117520 | rs10888859 | 1.081 | -0.199 | 0.098 | -0.427 | 0.007 | 0.004 | 0.004 | 0 | 0.16 | 0.14 | 0.2 | 0.06 | 0.09 | 0.08 |
| 1 | 55117879 | rs1655522 | -0.612 | -0.211 | 0.442 | -0.489 | -0.024 | -0.067 | -0.067 | 0.673 | 0.81 | 0.66 | 0.75 | 0.94 | 0.87 | 0.91 |
| 1 | 55118162 | rs7533275 | 0.752 | -0.383 | 0.276 | -0.594 | -0.016 | -0.018 | -0.018 | 0 | 0.24 | 0.15 | 0.23 | 0.63 | 0.33 | 0.26 |
| 1 | 55119515 | rs1655519 | 1.256 | -0.301 | 0.355 | -0.256 | 0.013 | 0.01 | 0.01 | 0 | 0.36 | 0.43 | 0.38 | 0.42 | 0.33 | 0.57 |
| 1 | 55119534 | rs17399810 | 0.295 | -0.304 | 0.366 | -0.326 | -0.014 | 0.089 | 0.089 | 0 | 0.22 | 0.09 | 0.14 | 0 | 0.22 | 0.08 |
| 1 | 55120469 | rs2570770 | 1.25 | -0.299 | 0.294 | -0.312 | 0.013 | 0.017 | 0.017 | 0 | 0.36 | 0.43 | 0.51 | 0.46 | 0.33 | 0.58 |
| 1 | 55124388 | rs473223 | -0.402 | -0.014 | 0.031 | -0.235 | -0.044 | 0.14 | 0.14 | 0 | 0.78 | 0.98 | 0.86 | 1 | 0.78 | 0.92 |
| 1 | 55129742 | rs689258 | 0.455 | -0.113 | 0.443 | -0.277 | 0.014 | 0.024 | 0.024 | 0 | 0.61 | 0.69 | 0.74 | 0.88 | 0.68 | 0.84 |
| 1 | 55130931 | rs2078273 | 0.152 | 0.556 | 0.614 | 0.013 | 0.03 | -0.009 | -0.009 | 0 | 0.44 | 0.19 | 0.31 | 0.53 | 0.43 | 0.33 |
| 1 | 55131844 | rs501260 | -0.685 | 0.774 | 0.444 | -0.022 | -0.024 | 0.023 | 0.023 | 0 | 0.84 | 0.9 | 0.85 | 1 | 0.76 | 0.94 |
| 1 | 55132925 | rs643893 | 1.016 | 0.564 | 0.74 | 0.192 | 0 | 0.002 | 0.002 | 0 | 0.31 | 0.29 | 0.48 | 0.38 | 0.28 | 0.55 |
| 1 | 55134487 | rs478513 | 0.954 | 0.705 | 0.425 | 0.263 | 0.001 | 0.006 | 0.006 | 0 | 0.31 | 0.31 | 0.48 | 0.38 | 0.28 | 0.55 |
| 1 | 55136529 | rs480963 | 0.982 | 0.763 | 0.266 | 0.276 | -0.003 | -0.006 | -0.006 | 3.469 | 0.31 | 0.27 | 0.47 | 0.37 | 0.29 | 0.57 |
| 1 | 55138294 | rs598130 | 1.143 | 0.699 | 0.184 | 0.27 | -0.001 | 0.001 | 0.001 | 0 | 0.29 | 0.27 | 0.28 | 0.3 | 0.24 | 0.48 |
| 1 | 55139618 | rs571199 | 0.282 | 1.178 | -0.322 | 0.475 | 0.054 | 0.063 | 0.063 | 0 | 0.4 | 0.24 | 0.2 | 0.23 | 0.33 | 0.22 |
| 1 | 55139741 | rs570218 | -0.161 | 1.942 | -0.192 | 0.776 | -0.02 | 0.024 | 0.024 | 0 | 0.88 | 0.93 | 0.86 | 1 | 0.8 | 0.94 |
| 1 | 55143328 | rs12137621 | -1.442 | 1.378 | 0.192 | 1.394 | 0.076 | 0.153 | 0.153 | 0 | 0.12 | 0.26 | 0.36 | 0.32 | 0.21 | 0.16 |
| 1 | 55147547 | rs1560854 | -0.577 | 0.352 | 0.107 | 0.467 | -0.037 | -0.068 | -0.068 | 0 | 0.14 | 0.28 | 0.08 | 0.04 | 0.08 | 0.1 |
| 1 | 55149254 | rs744748 | -0.193 | 0.64 | -0.548 | 0.685 | -0.018 | -0.035 | -0.035 | 0.009 | 0.35 | 0.28 | 0.43 | 0.58 | 0.44 | 0.43 |
| 1 | 55149582 | rs6659406 | 0.036 | 0.728 | -0.707 | 0.709 | -0.024 | 0.062 | 0.062 | 0.283 | 0.24 | 0.12 | 0.17 | 0.08 | 0.28 | 0.2 |
| 1 | 55149681 | rs879062 | 0.335 | 0.656 | -0.635 | 1.31 | 0 | 0.03 | 0.03 | 0 | 0.1 | 0.15 | 0.26 | 0.31 | 0.16 | 0.18 |
| 1 | 55151126 | rs605970 | 0.616 | 0.928 | -1.199 | 1.321 | -0.063 | -0.055 | -0.055 | 0 | 0.15 | 0.03 | 0.2 | 0.18 | 0.24 | 0.21 |
| 1 | 55152223 | rs17110938 | 0.477 | 0.447 | -0.743 | 1.34 | 0.001 | 0.046 | 0.046 | 0 | 0.1 | 0.15 | 0.26 | 0.31 | 0.16 | 0.18 |
| 1 | 55156881 | rs6588528 | -0.039 | 0.173 | -0.56 | 1.168 | -0.013 | 0.054 | 0.054 | 0 | 0.17 | 0.28 | 0.28 | 0.3 | 0.18 | 0.16 |
| 1 | 55161844 | rs6671173 | 0.354 | 0.346 | -1.016 | 0.739 | -0.029 | -0.016 | -0.016 | 0 | 0.18 | 0.07 | 0.14 | 0.26 | 0.21 | 0.3 |
| 1 | 55168084 | rs644955 | -0.96 | -0.014 | -0.547 | 0.822 | 0.004 | 0.074 | 0.074 | 0 | 0.52 | 0.71 | 0.54 | 0.76 | 0.49 | 0.65 |
| 1 | 55171439 | rs1981583 | -0.339 | -0.212 | -0.462 | -0.176 | 0.001 | 0.02 | 0.02 | 0.441 | 0.36 | 0.3 | 0.23 | 0.17 | 0.33 | 0.4 |
| 1 | 55182300 | rs1147990 | -0.399 | -0.396 | -0.343 | -0.973 | 0 | -0.084 | -0.084 | 0.395 | 0.55 | 0.5 | 0.61 | 0.97 | 0.55 | 0.75 |
| 1 | 55183132 | rs3766415 | -0.111 | -0.549 | -0.091 | -0.84 | 0.004 | 0.02 | 0.02 | 0.342 | 0.09 | 0.17 | 0.03 | 0.13 | 0.06 | 0.22 |
| 1 | 55185204 | rs1911230 | -0.111 | -0.549 | -0.091 | -0.84 | 0.004 | 0.02 | 0.02 | 0.342 | 0.09 | 0.17 | 0.03 | 0.13 | 0.06 | 0.22 |
| 1 | 55189185 | rs10489662 | -0.111 | -0.549 | -0.091 | -0.84 | 0.004 | 0.02 | 0.02 | 0.342 | 0.09 | 0.17 | 0.03 | 0.13 | 0.06 | 0.22 |
| 1 | 55191644 | rs1180937 | -1.166 | -0.84 | -0.087 | -1.128 | -0.021 | -0.049 | -0.049 | 0 | 0.79 | 0.63 | 0.81 | 0.92 | 0.86 | 0.89 |
| 1 | 55196099 | rs4926653 | -0.639 | -0.978 | 0.12 | -0.958 | -0.008 | 0.003 | 0.003 | 0 | 0.56 | 0.6 | 0.61 | 0.97 | 0.55 | 0.75 |
| 1 | 55197149 | rs3815226 | 0.775 | -1.158 | -0.279 | -0.538 | -0.005 | -0.122 | -0.122 | 0.132 | 0.19 | 0.03 | 0.36 | 0.53 | 0.2 | 0.25 |
| 1 | 55203135 | rs3737829 | -0.12 | -0.947 | 0.062 | -0.853 | 0.017 | 0.042 | 0.042 | 1.139 | 0.1 | 0.28 | 0.03 | 0.13 | 0.06 | 0.22 |
| 1 | 55211397 | rs4927155 | -0.424 | -1.061 | -0.184 | -0.742 | 0.002 | -0.024 | -0.024 | 0 | 0.49 | 0.51 | 0.59 | 0.89 | 0.49 | 0.56 |
| 1 | 55212984 | rs6695931 | -0.589 | -0.918 | 0.068 | -0.122 | 0.011 | 0.039 | 0.039 | 1.781 | 0.1 | 0.3 | 0.03 | 0.13 | 0.06 | 0.21 |
| 1 | 55217907 | rs2873551 | -0.36 | -0.932 | -0.431 | -0.043 | 0.003 | -0.001 | -0.001 | 0.705 | 0.36 | 0.28 | 0.48 | 0.58 | 0.38 | 0.25 |

| | | | | | | | | | | | | | | | | |
|---|---|---|---|---|---|---|---|---|---|---|---|---|---|---|---|---|
| 1 | 55218155 | rs2108035 | 0.123 | -0.916 | -0.512 | 0.204 | 0.006 | -0.027 | -0.027 | 0 | 0.24 | 0.17 | 0.38 | 0.59 | 0.27 | 0.3 |
| 1 | 55218254 | rs1180964 | 0.423 | -1.362 | -0.546 | -0.103 | -0.008 | -0.014 | -0.014 | 0.225 | 0.23 | 0.2 | 0.12 | 0.1 | 0.16 | 0.13 |
| 1 | 55219540 | rs12046989 | 0.645 | -1.255 | -0.618 | -0.023 | 0.001 | 0 | 0 | 1.603 | 0.19 | 0.17 | 0.35 | 0.48 | 0.18 | 0.14 |
| 1 | 55220911 | rs2527848 | 0.096 | -1.387 | -0.759 | -0.295 | 0.009 | 0.007 | 0.007 | 0 | 0.31 | 0.21 | 0.17 | 0.23 | 0.26 | 0.3 |
| 1 | 55222080 | rs2764822 | -1.467 | -1.169 | -0.807 | -1.054 | 0.02 | -0.084 | -0.084 | 0 | 0.69 | 0.58 | 0.65 | 0.96 | 0.58 | 0.74 |
| 1 | 55224751 | rs11577368 | 1.044 | -0.943 | -1.083 | -0.73 | -0.002 | -0.001 | -0.001 | 0.393 | 0.19 | 0.06 | 0.13 | 0.18 | 0.15 | 0.17 |
| 1 | 55226288 | rs1180954 | -0.05 | -0.691 | -0.633 | -0.863 | -0.012 | -0.029 | -0.029 | 0 | 0.28 | 0.25 | 0.15 | 0.1 | 0.25 | 0.15 |
| 1 | 55240823 | rs1180959 | -0.399 | -0.091 | -0.62 | -0.764 | 0.003 | -0.059 | -0.059 | 0.095 | 0.66 | 0.6 | 0.64 | 0.96 | 0.57 | 0.73 |
| 1 | 55241940 | rs1180957 | -0.357 | -0.211 | -0.666 | -0.737 | -0.002 | -0.037 | -0.037 | 0 | 0.64 | 0.58 | 0.63 | 0.95 | 0.56 | 0.72 |
| 1 | 55243319 | rs10888878 | -0.274 | -0.702 | -0.202 | 0.167 | -0.018 | 0.083 | 0.083 | 0 | 0.08 | 0.18 | 0.09 | 0.22 | 0.04 | 0.11 |
| 1 | 55243590 | rs10888879 | 0.44 | -0.63 | -0.302 | 0.124 | -0.009 | 0.025 | 0.025 | 0 | 0.06 | 0.09 | 0.07 | 0.12 | 0.04 | 0.06 |
| 1 | 55249154 | rs628667 | -1.141 | -0.704 | -0.5 | 0.506 | -0.012 | 0.026 | 0.026 | 0 | 0.34 | 0.39 | 0.29 | 0.51 | 0.23 | 0.31 |
| 1 | 55250684 | rs2286202 | -0.66 | -0.88 | -0.591 | 0.453 | -0.063 | 0.138 | 0.138 | 0 | 0.21 | 0.36 | 0.21 | 0.49 | 0.1 | 0.23 |
| 1 | 55252514 | rs11581256 | 0.213 | -1.057 | -0.77 | 0.39 | -0.056 | 0.078 | 0.078 | 0 | 0.13 | 0.02 | 0.08 | 0.02 | 0.13 | 0.09 |
| 1 | 55259667 | rs1147988 | 0.059 | -1.993 | -0.969 | 0.219 | 0.003 | -0.005 | -0.005 | 0 | 0.71 | 0.72 | 0.75 | 0.57 | 0.76 | 0.83 |
| 1 | 55261752 | rs886738 | 0.714 | -1.957 | -0.758 | 0.336 | 0.022 | -0.003 | -0.003 | 1.566 | 0.55 | 0.65 | 0.59 | 0.47 | 0.63 | 0.72 |
| 1 | 55262166 | rs880755 | 0.152 | -1.833 | -0.279 | 0.735 | 0.003 | 0.001 | 0.001 | 0.129 | 0.26 | 0.25 | 0.32 | 0.24 | 0.22 | 0.4 |
| 1 | 55265268 | rs12123513 | -0.572 | -1.787 | -0.045 | 1.355 | -0.023 | 0.029 | 0.029 | 0 | 0.15 | 0.04 | 0.16 | 0.13 | 0.12 | 0.15 |
| 1 | 55266797 | rs671108 | 0.301 | -1.908 | -0.257 | 1.707 | 0.003 | -0.099 | -0.099 | 0 | 0.22 | 0.03 | 0.23 | 0.26 | 0.24 | 0.07 |
| 1 | 55267509 | rs689322 | -0.021 | -1.975 | 0.313 | 1.764 | 0.005 | -0.01 | -0.01 | 1.211 | 0.25 | 0.24 | 0.24 | 0.48 | 0.24 | 0.16 |
| 1 | 55270030 | rs11206431 | -0.867 | -1.896 | 0.67 | 1.74 | 0 | -0.007 | -0.007 | 0 | 0.18 | 0.25 | 0.17 | 0.2 | 0.12 | 0.17 |
| 1 | 55273580 | rs9782980 | 1.282 | -1.863 | 0.786 | 1.811 | 0 | -0.001 | -0.001 | 1.127 | 0.53 | 0.5 | 0.46 | 0.27 | 0.59 | 0.66 |
| 1 | 55278841 | rs616315 | -0.374 | -2.013 | 1.06 | 1.924 | -0.014 | 0.015 | 0.015 | 0 | 0.28 | 0.37 | 0.37 | 0.31 | 0.28 | 0.09 |
| 1 | 55281847 | rs631666 | 0.12 | -2.076 | 1.065 | 1.99 | 0.011 | -0.024 | -0.024 | 0 | 0.24 | 0.17 | 0.27 | 0.45 | 0.23 | 0.11 |
| 1 | 55281991 | rs630895 | -1.375 | -2.148 | 1.438 | 1.949 | -0.003 | -0.001 | -0.001 | 0 | 0.43 | 0.49 | 0.43 | 0.55 | 0.35 | 0.25 |
| 1 | 55294722 | rs4926659 | 1.142 | -2.167 | 1.411 | 2.051 | 0.01 | 0.005 | 0.005 | 0.002 | 0.57 | 0.44 | 0.56 | 0.38 | 0.65 | 0.74 |
| 1 | 55299139 | rs141014 | -0.938 | -2.179 | 1.418 | 2.055 | 0.011 | 0.02 | 0.02 | 0 | 0.17 | 0.39 | 0.2 | 0.22 | 0.12 | 0.19 |
| 1 | 55301220 | rs682615 | 0.183 | -2.226 | 1.54 | 2.041 | 0.016 | -0.014 | -0.014 | 0.16 | 0.7 | 0.76 | 0.73 | 0.66 | 0.74 | 0.91 |
| 1 | 55301309 | rs1180855 | -1.175 | -2.24 | 1.57 | 2.053 | 0.004 | 0.082 | 0.082 | 0 | 0.18 | 0.45 | 0.21 | 0.31 | 0.12 | 0.19 |
| 1 | 55302719 | rs1544908 | 1.274 | -2.221 | 1.519 | 1.961 | -0.008 | 0.035 | 0.035 | 0 | 0.78 | 0.54 | 0.76 | 0.69 | 0.85 | 0.8 |
| 1 | 55304970 | rs7549251 | 1.537 | -2.189 | 1.445 | 1.443 | 0.025 | 0.024 | 0.024 | 0.057 | 0.52 | 0.39 | 0.52 | 0.38 | 0.62 | 0.72 |
| 1 | 55306618 | rs4927164 | 1.363 | -1.982 | 1.789 | 1.634 | 0.012 | 0.027 | 0.027 | 0 | 0.83 | 0.61 | 0.81 | 0.81 | 0.87 | 0.88 |
| 1 | 55307693 | rs17111100 | 0.08 | -2.132 | 1.783 | 1.685 | 0.021 | -0.043 | -0.043 | 0 | 0.24 | 0.15 | 0.28 | 0.44 | 0.22 | 0.08 |
| 1 | 55308221 | rs12026851 | -0.335 | -2.15 | 1.843 | 1.675 | -0.015 | 0.02 | 0.02 | 0 | 0.28 | 0.32 | 0.29 | 0.44 | 0.24 | 0.12 |
| 1 | 55309354 | rs642998 | -1.433 | -2.139 | 1.893 | 1.664 | 0.007 | -0.002 | -0.002 | 0 | 0.38 | 0.62 | 0.43 | 0.59 | 0.32 | 0.21 |
| 1 | 55311452 | rs11206451 | 0.074 | -2.111 | 1.825 | 1.677 | 0.013 | -0.025 | -0.025 | 0 | 0.25 | 0.17 | 0.28 | 0.44 | 0.22 | 0.08 |
| 1 | 55312440 | rs687912 | -1.023 | -2.135 | 1.829 | 1.641 | -0.026 | 0.015 | 0.015 | 0 | 0.36 | 0.46 | 0.42 | 0.58 | 0.31 | 0.17 |
| 1 | 55316322 | rs7374 | -1.37 | -2.137 | 1.786 | 1.639 | 0.002 | -0.002 | -0.002 | 2.403 | 0.38 | 0.6 | 0.43 | 0.58 | 0.32 | 0.18 |
| 1 | 55317685 | rs1138030 | 0.195 | -2.114 | 1.682 | 1.678 | 0.019 | -0.042 | -0.042 | 0.189 | 0.24 | 0.15 | 0.27 | 0.44 | 0.22 | 0.08 |
| 1 | 55318108 | rs667222 | -0.895 | -2.026 | 1.77 | 1.655 | -0.007 | 0.003 | 0.003 | 0 | 0.36 | 0.37 | 0.42 | 0.58 | 0.32 | 0.18 |
| 1 | 55319902 | rs718265 | 0.622 | -2.026 | 1.77 | 1.655 | -0.007 | 0.003 | 0.003 | 0 | 0.64 | 0.63 | 0.59 | 0.42 | 0.68 | 0.82 |
| 1 | 55326345 | rs4927168 | 0.797 | -1.689 | 1.551 | 1.364 | -0.037 | 0.123 | 0.123 | 0.02 | 0.88 | 0.63 | 0.83 | 0.71 | 0.96 | 0.87 |
| 1 | 55326933 | rs6676123 | -0.149 | -1.891 | 1.583 | 1.274 | -0.025 | 0.011 | 0.011 | 0 | 0.32 | 0.34 | 0.32 | 0.55 | 0.29 | 0.13 |
| 1 | 55333453 | rs642737 | 0.585 | -1.88 | 1.585 | 1.278 | -0.036 | -0.015 | -0.015 | 0 | 0.59 | 0.55 | 0.52 | 0.37 | 0.67 | 0.83 |
| 1 | 55333525 | rs600491 | 1.299 | -1.865 | 1.259 | 1.011 | 0.094 | 0.198 | 0.198 | 0.032 | 0.49 | 0.26 | 0.38 | 0.15 | 0.59 | 0.66 |
| 1 | 55338679 | rs7551288 | 1.297 | -1.879 | 0.831 | 1.002 | 0.108 | 0.228 | 0.228 | 0.205 | 0.5 | 0.19 | 0.38 | 0.16 | 0.59 | 0.69 |
| 1 | 55339086 | rs6686859 | 0.217 | -1.904 | 0.953 | 1.094 | -0.018 | 0.021 | 0.021 | 0.067 | 0.28 | 0.29 | 0.31 | 0.39 | 0.26 | 0.1 |
| 1 | 55341915 | rs638944 | 1.163 | -1.878 | 0.407 | 1.068 | 0.06 | 0.015 | 0.015 | 1.065 | 0.53 | 0.34 | 0.39 | 0.39 | 0.61 | 0.72 |
| 1 | 55350023 | rs601902 | -0.272 | -1.625 | -0.13 | 0.643 | 0.01 | -0.025 | -0.025 | 0 | 0.92 | 0.9 | 0.87 | 0.99 | 0.91 | 0.89 |
| 1 | 55354335 | rs4927176 | 1.172 | -1.516 | -0.512 | 0.67 | 0.078 | 0.191 | 0.191 | 0.07 | 0.48 | 0.23 | 0.38 | 0.14 | 0.59 | 0.65 |
| 1 | 55369645 | rs12037220 | -0.383 | -0.968 | -0.668 | 0.883 | 0.002 | -0.004 | -0.004 | 0 | 0.28 | 0.23 | 0.31 | 0.39 | 0.25 | 0.14 |
| 1 | 55379939 | rs619895 | -0.763 | -0.143 | -0.68 | 0.516 | 0.001 | 0 | 0 | 0.16 | 0.07 | 0.05 | 0.15 | 0.07 | 0.04 | 0.05 |
| 1 | 55387637 | rs678358 | -0.207 | 0.342 | -0.799 | -0.275 | -0.014 | 0.015 | 0.015 | 0 | 0.39 | 0.32 | 0.33 | 0.08 | 0.5 | 0.29 |
| 1 | 55391268 | rs35526112 | 0.26 | -0.022 | -0.984 | 0.09 | -0.006 | 0.001 | 0.001 | 0 | 0.41 | 0.4 | 0.37 | 0.49 | 0.29 | 0.53 |
| 1 | 55394648 | rs2114580 | -0.166 | -0.167 | -0.683 | 0.307 | 0.002 | 0.125 | 0.125 | 0 | 0.75 | 0.51 | 0.67 | 0.52 | 0.77 | 0.77 |
| 1 | 55397406 | rs604778 | 0.003 | -0.215 | -0.873 | 0.306 | 0 | 0.048 | 0.048 | 0 | 0.72 | 0.48 | 0.65 | 0.43 | 0.72 | 0.67 |
| 1 | 55403411 | rs626353 | 0.351 | -0.071 | -1.416 | 0.024 | -0.11 | 0.162 | 0.162 | 0 | 0.36 | 0.09 | 0.31 | 0.09 | 0.5 | 0.29 |
| 1 | 55404022 | rs1889209 | 0.115 | -0.139 | -0.597 | 0.646 | -0.002 | 0.028 | 0.028 | 0 | 0.75 | 0.39 | 0.84 | 0.61 | 0.76 | 0.74 |
| 1 | 55405850 | rs11206476 | 0.692 | -0.079 | -1.408 | -0.367 | -0.099 | 0.216 | 0.216 | 0 | 0.33 | 0.02 | 0.22 | 0.04 | 0.3 | 0.28 |
| 1 | 55413361 | rs7549203 | -0.286 | 0.201 | -0.837 | 0.193 | -0.015 | -0.013 | -0.013 | 0 | 0.66 | 0.72 | 0.71 | 0.63 | 0.58 | 0.7 |
| 1 | 55422254 | rs2114574 | -0.037 | 0.215 | -0.467 | 0.142 | 0.038 | 0.083 | 0.083 | 0 | 0.72 | 0.5 | 0.77 | 0.56 | 0.68 | 0.59 |
| 1 | 55424178 | rs6588539 | -0.516 | 0.39 | -0.526 | 0.226 | -0.017 | -0.026 | -0.026 | 0.063 | 0.76 | 0.77 | 0.78 | 0.56 | 0.68 | 0.59 |
| 1 | 55432798 | rs7531509 | 0.349 | 0.245 | -0.433 | 0.361 | -0.004 | 0.022 | 0.022 | 0 | 0.26 | 0.29 | 0.47 | 0.51 | 0.24 | 0.28 |
| 1 | 55434648 | rs2317951 | 0.1 | 0.099 | -0.658 | -0.147 | 0.007 | 0.038 | 0.038 | 0 | 0.39 | 0.28 | 0.25 | 0.13 | 0.33 | 0.32 |
| 1 | 55445123 | rs2059688 | 0.662 | -0.424 | -0.389 | 0.899 | 0.019 | -0.013 | -0.013 | 0 | 0.65 | 0.33 | 0.73 | 0.48 | 0.77 | 0.68 |
| 1 | 55445958 | rs1863669 | 0.625 | -0.434 | -0.326 | 0.895 | 0.012 | -0.009 | -0.009 | 0 | 0.66 | 0.34 | 0.74 | 0.48 | 0.77 | 0.69 |
| 1 | 55446671 | rs2253466 | 0.453 | -0.478 | -0.039 | 0.933 | -0.016 | 0.018 | 0.018 | 0 | 0.69 | 0.41 | 0.75 | 0.49 | 0.77 | 0.73 |
| 1 | 55457815 | rs2495513 | 0.12 | -0.509 | 0.478 | 1.453 | -0.001 | 0.005 | 0.005 | 0 | 0.81 | 0.82 | 0.89 | 0.64 | 0.8 | 0.82 |
| 1 | 55460736 | rs1003323 | 1.301 | -0.816 | 1.167 | 1.916 | 0.033 | 0.003 | 0.003 | 0.019 | 0.69 | 0.22 | 0.73 | 0.31 | 0.77 | 0.73 |

| | | | | | | | | | | | | | | | | |
|---|---|---|---|---|---|---|---|---|---|---|---|---|---|---|---|---|
| 1 | 55462380 | rs9887913 | -2.125 | -0.769 | 1.233 | 1.919 | -0.064 | 0.187 | 0.187 | 2.711 | 0.13 | 0.6 | 0.15 | 0.33 | 0.03 | 0.2 |
| 1 | 55463086 | rs11206502 | -1.116 | -0.864 | 1.669 | 1.694 | -0.045 | 0.052 | 0.052 | 3.626 | 0.1 | 0.33 | 0.14 | 0.12 | 0.02 | 0.07 |
| 1 | 55463860 | rs1003769 | 1.037 | -0.707 | 1.469 | 1.321 | -0.008 | -0.002 | -0.002 | 0 | 0.68 | 0.33 | 0.69 | 0.31 | 0.76 | 0.75 |
| 1 | 55464320 | rs1003767 | 1.037 | -0.707 | 1.469 | 1.321 | -0.008 | -0.002 | -0.002 | 1.311 | 0.68 | 0.33 | 0.69 | 0.31 | 0.76 | 0.75 |
| 1 | 55464743 | rs2500340 | 0.869 | -0.716 | 1.7 | 1.396 | -0.035 | 0.004 | 0.004 | 0 | 0.69 | 0.5 | 0.7 | 0.32 | 0.76 | 0.75 |
| 1 | 55465594 | rs759914 | 0.999 | -0.69 | 1.238 | 1.377 | -0.007 | -0.002 | -0.002 | 0 | 0.68 | 0.34 | 0.73 | 0.31 | 0.76 | 0.75 |
| 1 | 55472365 | rs4927186 | -1.311 | 0.132 | 0.393 | 0.632 | -0.013 | 0.056 | 0.056 | 0 | 0.5 | 0.81 | 0.31 | 0.38 | 0.51 | 0.49 |
| 1 | 55473043 | rs6687801 | -1.021 | 0.31 | 0.6 | 0.12 | -0.007 | 0.037 | 0.037 | 0 | 0.42 | 0.65 | 0.26 | 0.22 | 0.44 | 0.5 |
| 1 | 55474325 | rs6682884 | -0.827 | 0.436 | 0.68 | 0.19 | 0.01 | -0.009 | -0.009 | 3.192 | 0.36 | 0.52 | 0.23 | 0.33 | 0.4 | 0.23 |
| 1 | 55475966 | rs6588543 | -0.236 | 0.435 | 0.776 | 0.15 | -0.013 | 0.026 | 0.026 | 0 | 0.24 | 0.38 | 0.14 | 0.34 | 0.21 | 0.18 |
| 1 | 55477327 | rs2495510 | -1.167 | 0.749 | 0.888 | 0.294 | 0.034 | -0.096 | -0.096 | 0 | 0.18 | 0.6 | 0.25 | 0.02 | 0.22 | 0.15 |
| 1 | 55477635 | rs10888894 | -0.817 | 0.514 | 0.126 | 0.277 | -0.002 | 0.045 | 0.045 | 0 | 0.27 | 0.42 | 0.53 | 0.61 | 0.29 | 0.43 |
| 1 | 55477821 | rs12093385 | -0.352 | 0.372 | -0.23 | -0.253 | 0.007 | 0.022 | 0.022 | 0 | 0.15 | 0.09 | 0.07 | 0.05 | 0.12 | 0.14 |
| 1 | 55479198 | rs12075602 | -0.759 | 0.539 | 0.136 | -0.187 | 0.005 | -0.079 | -0.079 | 0.104 | 0.13 | 0.42 | 0.09 | 0.01 | 0.11 | 0.22 |
| 1 | 55480211 | rs2182832 | -0.75 | 0.569 | -0.75 | -0.099 | -0.001 | 0.054 | 0.054 | 0 | 0.47 | 0.77 | 0.59 | 0.58 | 0.46 | 0.43 |
| 1 | 55480504 | rs2149038 | -0.603 | 0.43 | -0.589 | -0.57 | 0.001 | 0.01 | 0.01 | 0 | 0.5 | 0.52 | 0.66 | 0.67 | 0.53 | 0.53 |
| 1 | 55481116 | rs2149037 | -0.09 | 0.856 | -1.015 | -0.726 | -0.011 | 0.004 | 0.004 | 0 | 0.32 | 0.28 | 0.31 | 0.29 | 0.45 | 0.29 |
| 1 | 55482091 | rs2479396 | 0.16 | 0.121 | -1.315 | -0.678 | 0.011 | -0.019 | -0.019 | 0.488 | 0.66 | 0.78 | 0.57 | 0.44 | 0.67 | 0.65 |
| 1 | 55486064 | rs2479394 | 0.297 | -0.442 | -1.238 | -1.322 | -0.008 | 0.003 | 0.003 | 0 | 0.65 | 0.6 | 0.59 | 0.44 | 0.72 | 0.57 |
| 1 | 55486361 | rs11206508 | 0.463 | -0.229 | -1.405 | -1.314 | 0.001 | -0.013 | -0.013 | 0.022 | 0.18 | 0.14 | 0.08 | 0 | 0.13 | 0.12 |
| 1 | 55494682 | rs2495499 | 0.591 | -0.094 | -1.533 | -1.362 | -0.06 | 0.021 | 0.021 | 0 | 0.34 | 0.22 | 0.36 | 0.3 | 0.48 | 0.37 |
| 1 | 55495744 | rs2479417 | -0.123 | -0.303 | -1.446 | -1.361 | 0.032 | 0.046 | 0.046 | 0 | 0.52 | 0.37 | 0.48 | 0.35 | 0.65 | 0.39 |
| 1 | 55496039 | rs11206510 | -0.071 | -0.097 | -1.182 | -0.898 | 0.006 | 0.013 | 0.013 | 0.073 | 0.18 | 0.13 | 0.12 | 0.04 | 0.17 | 0.03 |
| 1 | 55496556 | rs2495495 | -0.345 | -0.069 | -1.271 | -0.123 | -0.001 | -0.098 | -0.098 | 0.036 | 0.85 | 0.97 | 0.7 | 0.5 | 0.87 | 0.84 |
| 1 | 55496648 | rs12739979 | -0.817 | 0.023 | -0.745 | -0.701 | 0 | -0.022 | -0.022 | 0 | 0.18 | 0.28 | 0.22 | 0.13 | 0.23 | 0.2 |
| 1 | 55500206 | rs2495488 | -0.116 | 0.427 | -0.192 | -0.59 | 0.066 | 0.048 | 0.048 | 0 | 0.94 | 0.76 | 0.86 | 0.92 | 0.93 | 0.97 |
| 1 | 55500429 | rs2182833 | 0.134 | 0.095 | -0.022 | -0.579 | -0.018 | 0.029 | 0.029 | 0 | 0.23 | 0.3 | 0.31 | 0.67 | 0.18 | 0.23 |
| 1 | 55504650 | rs2479409 | -0.011 | 0.354 | -0.311 | -0.793 | 0.002 | 0.013 | 0.013 | 0 | 0.65 | 0.75 | 0.43 | 0.33 | 0.68 | 0.72 |
| 1 | 55505668 | rs11583680 | -0.087 | 0.598 | -0.02 | -1.109 | -0.037 | -0.012 | -0.012 | 0 | 0.1 | 0.01 | 0.07 | 0.1 | 0.13 | 0.16 |
| 1 | 55509872 | rs4927193 | -0.714 | 0.607 | 0.468 | -1.149 | 0.003 | -0.01 | -0.01 | 0.001 | 0.13 | 0.22 | 0.1 | 0.1 | 0.13 | 0.15 |
| 1 | 55509900 | rs2479411 | -0.09 | 0.552 | 0.719 | -1.169 | 0.027 | 0.033 | 0.033 | 0 | 0.06 | 0.2 | 0.12 | 0.06 | 0.05 | 0.02 |
| 1 | 55512549 | rs499718 | 1.302 | 0.497 | 0.938 | -1.184 | 0.009 | -0.007 | -0.007 | 1.497 | 0.81 | 0.48 | 0.77 | 0.83 | 0.82 | 0.82 |
| 1 | 55512995 | rs2495478 | -0.166 | 0.539 | 0.858 | -0.976 | 0.065 | 0.092 | 0.092 | 0 | 0.06 | 0.32 | 0.12 | 0.07 | 0.05 | 0.02 |
| 1 | 55516004 | rs11206514 | 0.496 | 1.05 | -0.95 | -1.399 | -0.01 | 0.096 | 0.096 | 0 | 0.61 | 0.21 | 0.64 | 0.84 | 0.6 | 0.68 |
| 1 | 55517344 | rs572512 | 0.447 | 0.86 | -1.223 | -1.656 | -0.011 | 0.07 | 0.07 | 0.5 | 0.38 | 0.19 | 0.56 | 0.8 | 0.35 | 0.46 |
| 1 | 55518316 | rs2483205 | -0.628 | 0.873 | -0.338 | -1.675 | 0 | -0.023 | -0.023 | 0 | 0.46 | 0.51 | 0.32 | 0.2 | 0.45 | 0.44 |
| 1 | 55518467 | rs2495477 | -1.209 | -0.014 | -0.437 | -1.737 | 0.068 | 0.148 | 0.148 | 0 | 0.5 | 0.75 | 0.3 | 0.23 | 0.4 | 0.51 |
| 1 | 55518682 | rs2479413 | -1.241 | 0.483 | 0.24 | -1.227 | -0.006 | -0.044 | -0.044 | 0.647 | 0.34 | 0.49 | 0.23 | 0.2 | 0.34 | 0.45 |
| 1 | 55520864 | rs557435 | 0.738 | 0.304 | 0.689 | -1.221 | -0.005 | -0.062 | -0.062 | 0.017 | 0.81 | 0.7 | 0.89 | 0.96 | 0.8 | 0.89 |
| 1 | 55521109 | rs693668 | 1.472 | 0.087 | 0.654 | -1.378 | 0.044 | 0.078 | 0.078 | 0 | 0.55 | 0.31 | 0.73 | 0.78 | 0.63 | 0.58 |
| 1 | 55524197 | rs540796 | 0.399 | 0.364 | 1.298 | -1.189 | 0.019 | -0.058 | -0.058 | 0 | 0.88 | 0.77 | 0.89 | 0.99 | 0.82 | 0.91 |
| 1 | 55524237 | rs562556 | 0.399 | 0.364 | 1.298 | -1.189 | 0.019 | -0.058 | -0.058 | 0 | 0.88 | 0.77 | 0.89 | 0.99 | 0.82 | 0.9 |
| 1 | 55524842 | rs568052 | 1.392 | 0.284 | 1.578 | -1.269 | -0.002 | -0.014 | -0.014 | 0 | 0.66 | 0.37 | 0.79 | 0.87 | 0.71 | 0.76 |
| 1 | 55526428 | rs11206517 | -1.123 | 0.412 | 1.799 | -1.121 | 0.011 | 0.048 | 0.048 | 0.077 | 0.06 | 0.21 | 0.03 | 0.04 | 0.03 | 0.03 |
| 1 | 55529187 | rs505151 | 1.346 | 0.35 | 1.864 | -1.123 | 0.016 | 0.061 | 0.061 | 0.579 | 0.94 | 0.71 | 0.96 | 0.96 | 0.97 | 0.97 |
| 1 | 55529828 | rs662145 | 2.103 | 0.351 | 1.954 | -1.184 | 0.015 | -0.043 | -0.043 | 0 | 0.79 | 0.39 | 0.86 | 0.91 | 0.77 | 0.84 |
| 1 | 55538462 | rs17410294 | 0.116 | 0.471 | 1.907 | -1.125 | -0.032 | 0.03 | 0.03 | 0 | 0.08 | 0.03 | 0.08 | 0 | 0.16 | 0.07 |
| 1 | 55538552 | rs10493176 | 0.042 | 0.37 | 1.773 | -1.237 | -0.003 | 0.033 | 0.033 | 0 | 0.1 | 0 | 0.04 | 0.06 | 0.11 | 0.16 |
| 1 | 55541174 | rs487230 | 2.182 | 0.457 | 2.059 | -1.176 | 0.002 | -0.01 | -0.01 | 0 | 0.8 | 0.26 | 0.85 | 0.92 | 0.77 | 0.85 |
| 1 | 55555285 | rs548852 | 2.198 | 0.456 | 2.122 | -1.179 | 0.001 | -0.009 | -0.009 | 0 | 0.8 | 0.26 | 0.85 | 0.92 | 0.77 | 0.84 |
| 1 | 55557067 | rs1165284 | 1.155 | 0.47 | 2.158 | -1.175 | 0.106 | -0.062 | -0.062 | 0 | 0.86 | 0.5 | 0.88 | 0.96 | 0.81 | 0.87 |
| 1 | 55567410 | rs615652 | -0.048 | 0.474 | 2.057 | -1.171 | 0.021 | -0.036 | -0.036 | 0 | 0.91 | 0.86 | 0.92 | 1 | 0.84 | 0.93 |
| 1 | 55590465 | rs17111652 | -1.771 | 0.441 | 2.328 | -1.166 | 0.009 | 0.042 | 0.042 | 0.095 | 0.07 | 0.28 | 0.04 | 0.05 | 0.04 | 0.04 |
| 1 | 55590828 | rs502793 | 1.36 | 0.493 | 2.106 | -1.179 | 0.087 | -0.07 | -0.07 | 0 | 0.85 | 0.46 | 0.87 | 0.96 | 0.81 | 0.87 |
| 1 | 55608005 | rs1165226 | 1.993 | 0.322 | 2.057 | -1.251 | 0.041 | 0.152 | 0.152 | 0 | 0.55 | 0.18 | 0.7 | 0.79 | 0.6 | 0.62 |
| 1 | 55630204 | rs1024305 | 1.433 | 0.672 | 2.494 | -0.844 | 0.104 | -0.067 | -0.067 | 0.017 | 0.86 | 0.5 | 0.87 | 0.96 | 0.81 | 0.87 |
| 1 | 55638075 | rs1165222 | 0.196 | 0.682 | 1.921 | -0.838 | 0.014 | -0.025 | -0.025 | 0 | 0.91 | 0.87 | 0.92 | 1 | 0.84 | 0.93 |
| 1 | 55638546 | rs1475701 | -1.162 | 0.594 | 1.697 | -0.817 | -0.009 | 0.014 | 0.014 | 0 | 0.05 | 0.13 | 0.03 | 0.04 | 0.03 | 0.03 |
| 1 | 55673420 | rs6703366 | -1.254 | 0.798 | 1.417 | -0.756 | 0.023 | -0.015 | -0.015 | 0 | 0.12 | 0.19 | 0.1 | 0.04 | 0.2 | 0.1 |
| 1 | 55688016 | rs11206524 | 1.698 | 0.837 | 1.535 | -0.742 | 0.073 | -0.041 | -0.041 | 0.295 | 0.83 | 0.54 | 0.87 | 0.92 | 0.77 | 0.87 |
| 1 | 55696884 | rs7550296 | -0.018 | 0.86 | 1.163 | -0.714 | 0.032 | -0.036 | -0.036 | 0.384 | 0.92 | 0.85 | 0.93 | 1 | 0.84 | 0.95 |
| 1 | 55699023 | rs10788998 | 1.436 | 0.811 | 1.191 | -0.717 | 0.034 | -0.034 | -0.034 | 0 | 0.83 | 0.36 | 0.86 | 0.92 | 0.77 | 0.88 |
| 1 | 55711465 | rs4927205 | -0.365 | 0.279 | 0.608 | -0.874 | 0.016 | 0.042 | 0.042 | 0.016 | 0.1 | 0.02 | 0.05 | 0.05 | 0.1 | 0.13 |
| 1 | 55712335 | rs11206536 | 0.423 | 0.135 | 0.465 | -0.792 | 0.005 | 0.003 | 0.003 | 0 | 0.15 | 0.12 | 0.11 | 0.1 | 0.07 | 0.08 |
| 1 | 55713628 | rs4927207 | -0.577 | 0.2 | 0.509 | -0.747 | 0.034 | 0.031 | 0.031 | 0 | 0.24 | 0.13 | 0.16 | 0.16 | 0.17 | 0.21 |
| 1 | 55716497 | rs2647286 | -0.924 | 0.194 | 0.638 | -0.694 | -0.01 | -0.007 | -0.007 | 0.1 | 0.32 | 0.61 | 0.21 | 0.19 | 0.19 | 0.25 |
| 1 | 55719127 | rs2802865 | 0.037 | 0.78 | -0.288 | -0.774 | -0.022 | 0.014 | 0.014 | 0 | 0.9 | 0.94 | 0.92 | 1 | 0.81 | 0.94 |
| 1 | 55719166 | rs7551981 | 0.841 | 0.662 | -0.392 | -0.633 | 0.009 | 0.063 | 0.063 | 0 | 0.58 | 0.32 | 0.71 | 0.81 | 0.62 | 0.68 |
| 1 | 55725176 | rs12409228 | 0.121 | 0.136 | -1.53 | -0.112 | 0.042 | -0.003 | -0.003 | 0 | 0.44 | 0.07 | 0.42 | 0.68 | 0.38 | 0.31 |

| | | | | | | | | | | | | | | | | |
|---|---|---|---|---|---|---|---|---|---|---|---|---|---|---|---|---|
| 1 | 55727195 | rs4926674 | 0.032 | 0.049 | -0.786 | -0.11 | 0.063 | 0.055 | 0.055 | 0.428 | 0.45 | 0.13 | 0.41 | 0.69 | 0.37 | 0.29 |
| 1 | 55730327 | rs6662286 | -0.007 | 0.043 | -0.525 | -0.379 | 0.003 | 0.008 | 0.008 | 0 | 0.9 | 0.96 | 0.97 | 0.99 | 0.91 | 0.87 |
| 1 | 55735734 | rs12048828 | 1.252 | -0.394 | -0.74 | 0.003 | -0.006 | -0.002 | -0.002 | 0 | 0.14 | 0.09 | 0.09 | 0.18 | 0.04 | 0.04 |
| 1 | 55737455 | rs6588553 | 0.516 | 0.505 | -0.372 | -0.061 | -0.032 | 0.029 | 0.029 | 0.196 | 0.35 | 0.2 | 0.44 | 0.23 | 0.44 | 0.51 |
| 1 | 55738663 | rs12086676 | -0.579 | 0.466 | -0.335 | -0.077 | 0.13 | 0.018 | 0.018 | 0 | 0.12 | 0.45 | 0.13 | 0.08 | 0.17 | 0.09 |
| 1 | 55738797 | rs6680729 | 0.38 | 0.511 | -0.713 | 0.077 | -0.011 | 0.01 | 0.01 | 0.067 | 0.35 | 0.34 | 0.44 | 0.24 | 0.44 | 0.51 |
| 1 | 55743297 | rs1874774 | 0.387 | 0.679 | -0.801 | -0.076 | 0 | 0.007 | 0.007 | 0.32 | 0.76 | 0.73 | 0.85 | 0.91 | 0.78 | 0.73 |
| 1 | 55743352 | rs9787299 | 0.796 | 0.268 | -1.131 | -0.512 | -0.044 | 0.046 | 0.046 | 0 | 0.32 | 0.13 | 0.44 | 0.23 | 0.4 | 0.39 |
| 1 | 55743519 | rs1874776 | 0.049 | 0.07 | -0.728 | -0.446 | -0.001 | -0.011 | -0.011 | 0 | 0.75 | 0.65 | 0.84 | 0.91 | 0.78 | 0.73 |
| 1 | 55749478 | rs17111847 | 0.784 | -0.494 | -0.118 | -0.353 | -0.05 | -0.137 | -0.137 | 0 | 0.2 | 0.43 | 0.09 | 0.02 | 0.13 | 0.06 |
| 1 | 55751821 | rs4927219 | 0.659 | -0.485 | -0.325 | -0.242 | 0.039 | 0.083 | 0.083 | 0 | 0.55 | 0.27 | 0.67 | 0.78 | 0.66 | 0.66 |
| 1 | 55767547 | rs4927222 | -0.032 | -0.599 | -0.049 | -0.236 | -0.029 | -0.001 | -0.001 | 0 | 0.75 | 0.49 | 0.75 | 0.71 | 0.79 | 0.67 |
| 1 | 55770565 | rs1948635 | -0.054 | -0.612 | 0.034 | -0.235 | -0.022 | 0.002 | 0.002 | 0.902 | 0.76 | 0.57 | 0.75 | 0.72 | 0.79 | 0.67 |
| 1 | 55771636 | rs6680609 | 0.884 | -0.699 | -0.102 | -0.351 | 0.075 | 0.136 | 0.136 | 0 | 0.54 | 0.13 | 0.66 | 0.7 | 0.66 | 0.61 |
| 1 | 55773756 | rs13353041 | 1.029 | -0.66 | -0.016 | -0.335 | -0.025 | -0.093 | -0.093 | 1.691 | 0.2 | 0.28 | 0.08 | 0.02 | 0.14 | 0.07 |
| 1 | 55775534 | rs1566208 | 1.283 | -0.058 | -0.476 | -0.162 | -0.141 | 0.003 | 0.003 | 0 | 0.28 | 0.02 | 0.4 | 0.22 | 0.38 | 0.38 |
| 1 | 55775665 | rs6669253 | 1.192 | 0.46 | 0.134 | 0.308 | -0.044 | -0.155 | -0.155 | 1.74 | 0.21 | 0.02 | 0.2 | 0.4 | 0.23 | 0.19 |
| 1 | 55776971 | rs12071251 | -1.187 | 0.637 | 0.763 | 0.158 | 0.091 | 0.151 | 0.151 | 0 | 0.08 | 0.33 | 0.19 | 0.27 | 0.14 | 0.13 |
| 1 | 55781281 | rs207154 | -0.017 | 0.67 | 0.171 | 0.092 | 0.015 | 0.009 | 0.009 | 0 | 0.06 | 0.09 | 0.14 | 0.1 | 0.1 | 0.05 |
| 1 | 55786637 | rs10888914 | 1.502 | 0.571 | 0.144 | 0.102 | 0.049 | 0.062 | 0.062 | 0 | 0.84 | 0.39 | 0.63 | 0.72 | 0.8 | 0.85 |
| 1 | 55798691 | rs2802884 | -0.383 | -0.696 | -0.539 | -0.28 | -0.019 | -0.012 | -0.012 | 0.874 | 0.29 | 0.32 | 0.15 | 0.14 | 0.15 | 0.19 |
| 1 | 55799770 | rs207164 | 0.962 | -0.279 | -0.383 | 0.083 | 0.006 | 0.036 | 0.036 | 0.724 | 0.23 | 0.28 | 0.38 | 0.36 | 0.26 | 0.19 |
| 1 | 55804902 | rs207155 | -0.913 | -0.015 | -0.556 | -0.119 | -0.017 | 0.019 | 0.019 | 0 | 0.93 | 0.98 | 0.95 | 1 | 0.91 | 0.97 |
| 1 | 55806063 | rs6694538 | 1.43 | -0.082 | -0.449 | -0.142 | -0.026 | -0.065 | -0.065 | 0 | 0.2 | 0.07 | 0.35 | 0.36 | 0.25 | 0.19 |
| 1 | 55806861 | rs12146081 | 0.79 | 0.197 | -0.382 | -0.065 | -0.009 | 0.013 | 0.013 | 0.408 | 0.4 | 0.35 | 0.38 | 0.4 | 0.48 | 0.54 |
| 1 | 55807259 | rs207149 | -0.242 | 0.157 | -0.372 | -0.064 | 0.022 | 0.014 | 0.014 | 0.409 | 0.51 | 0.61 | 0.53 | 0.5 | 0.4 | 0.38 |
| 1 | 55811215 | rs207138 | -0.32 | 0.146 | -0.351 | -0.029 | 0.023 | 0.01 | 0.01 | 0.482 | 0.52 | 0.61 | 0.54 | 0.5 | 0.4 | 0.38 |
| 1 | 55812880 | rs12117114 | 0.869 | 0.188 | -0.375 | -0.004 | -0.006 | 0.009 | 0.009 | 0.189 | 0.4 | 0.36 | 0.38 | 0.4 | 0.48 | 0.55 |
| 1 | 55814164 | rs207136 | -0.674 | -0.228 | -0.308 | -0.221 | -0.009 | -0.034 | -0.034 | 0 | 0.8 | 0.68 | 0.87 | 0.9 | 0.85 | 0.9 |
| 1 | 55820099 | rs880385 | 0.226 | -0.091 | -0.23 | -0.091 | -0.009 | -0.001 | -0.001 | 0 | 0.54 | 0.62 | 0.5 | 0.52 | 0.51 | 0.62 |
| 1 | 55820264 | rs880386 | 0.176 | 0.21 | -0.324 | -0.171 | -0.029 | 0.004 | 0.004 | 0 | 0.21 | 0.6 | 0.29 | 0.23 | 0.14 | 0.17 |
| 1 | 55820637 | rs2746689 | 0.427 | 0.238 | -0.487 | -0.227 | -0.002 | -0.003 | -0.003 | 0 | 0.7 | 0.41 | 0.63 | 0.72 | 0.73 | 0.79 |
| 1 | 55820713 | rs2179853 | -0.577 | 0.07 | -0.385 | -0.281 | -0.013 | -0.029 | -0.029 | 0 | 0.77 | 0.71 | 0.87 | 0.91 | 0.86 | 0.9 |
| 1 | 55828641 | rs7555489 | -0.481 | 0.268 | -0.444 | -0.169 | 0.005 | -0.02 | -0.02 | 0.18 | 0.62 | 0.72 | 0.57 | 0.52 | 0.63 | 0.65 |
| 1 | 55832031 | rs12141667 | 1.346 | 0.37 | -0.184 | -0.265 | -0.041 | 0.048 | 0.048 | 0.158 | 0.31 | 0.11 | 0.19 | 0.26 | 0.36 | 0.42 |
| 1 | 55833351 | rs910270 | -1.244 | 0.023 | -0.232 | -0.493 | -0.017 | -0.024 | -0.024 | 0 | 0.7 | 0.84 | 0.65 | 0.68 | 0.68 | 0.72 |
| 1 | 55834309 | rs7520205 | 0.258 | 0.183 | -0.209 | -1.017 | -0.015 | -0.085 | -0.085 | 0 | 0.26 | 0.45 | 0.3 | 0.14 | 0.23 | 0.16 |
| 1 | 55835253 | rs7523391 | -0.339 | 0.072 | -0.45 | -0.833 | 0.004 | -0.008 | -0.008 | 0 | 0.6 | 0.63 | 0.56 | 0.52 | 0.63 | 0.64 |
| 1 | 55839117 | rs12758665 | -0.161 | -0.435 | 0.107 | -0.63 | -0.005 | -0.036 | -0.036 | 1.203 | 0.26 | 0.32 | 0.31 | 0.14 | 0.22 | 0.17 |
| 1 | 55842956 | rs17112041 | -0.33 | 0.099 | 0.24 | 0.07 | 0.004 | 0.025 | 0.025 | 0 | 0.09 | 0.16 | 0.06 | 0.14 | 0.09 | 0.13 |
| 1 | 55852987 | rs4926684 | 0.961 | 0.027 | -0.21 | -0.041 | 0.017 | -0.033 | -0.033 | 0 | 0.22 | 0.15 | 0.16 | 0.32 | 0.22 | 0.37 |
| 1 | 55860561 | rs11206577 | 1.002 | -0.01 | -0.136 | -0.232 | -0.025 | 0.001 | 0.001 | 0.075 | 0.43 | 0.22 | 0.58 | 0.39 | 0.47 | 0.32 |
| 1 | 55862783 | rs761484 | 0.55 | 0.062 | -0.202 | -0.229 | 0.002 | -0.013 | -0.013 | 0 | 0.64 | 0.4 | 0.63 | 0.75 | 0.65 | 0.71 |
| 1 | 55872780 | rs2015252 | 0.861 | 0.255 | -0.22 | -0.354 | -0.008 | -0.014 | -0.014 | 0 | 0.42 | 0.25 | 0.48 | 0.43 | 0.44 | 0.33 |
| 1 | 55874284 | rs761483 | -1.032 | 0.204 | -0.784 | -0.389 | -0.006 | -0.005 | -0.005 | 0.444 | 0.82 | 0.87 | 0.89 | 0.98 | 0.81 | 0.88 |
| 1 | 55874426 | rs2092651 | 1.106 | 0.208 | -0.821 | -0.361 | -0.044 | -0.06 | -0.06 | 0.518 | 0.38 | 0.08 | 0.43 | 0.4 | 0.41 | 0.29 |
| 1 | 55884720 | rs6588558 | 0.44 | 0.185 | -0.681 | -0.319 | 0.004 | 0.006 | 0.006 | 0.337 | 0.52 | 0.54 | 0.69 | 0.56 | 0.53 | 0.4 |
| 1 | 55894893 | rs6673648 | -0.243 | 0.755 | -0.712 | -0.455 | -0.002 | -0.029 | -0.029 | 0 | 0.81 | 0.84 | 0.89 | 0.98 | 0.81 | 0.86 |
| 1 | 55903443 | rs7516169 | 0.388 | 0.629 | -1.473 | -0.082 | -0.04 | -0.071 | -0.071 | 0 | 0.37 | 0.09 | 0.42 | 0.38 | 0.4 | 0.27 |
| 1 | 55903668 | rs10789001 | -0.223 | -0.672 | -1.113 | -0.472 | 0.026 | 0.04 | 0.04 | 0 | 0.64 | 0.7 | 0.88 | 0.83 | 0.8 | 0.84 |
| 1 | 55907494 | rs4927235 | 0.426 | -0.514 | -0.792 | -0.39 | 0.024 | 0.021 | 0.021 | 0 | 0.36 | 0.49 | 0.54 | 0.46 | 0.54 | 0.38 |
| 1 | 55909238 | rs12128309 | 0.155 | -0.811 | -1.034 | -0.455 | 0 | -0.005 | -0.005 | 0 | 0.12 | 0.04 | 0.07 | 0.1 | 0.1 | 0.17 |
| 1 | 55909305 | rs10888920 | 0.083 | -1.282 | -0.923 | -0.311 | 0.011 | 0.01 | 0.01 | 0 | 0.53 | 0.55 | 0.74 | 0.59 | 0.68 | 0.67 |
| 1 | 55930462 | rs10888923 | -0.149 | -1.221 | -0.403 | -0.239 | -0.008 | 0.042 | 0.042 | 0 | 0.09 | 0.17 | 0.06 | 0.14 | 0.07 | 0.15 |
| 1 | 55931362 | rs987029 | 1.271 | -1.413 | 0.079 | -0.353 | -0.038 | -0.035 | -0.035 | 0 | 0.32 | 0.21 | 0.5 | 0.42 | 0.52 | 0.34 |
| 1 | 55932300 | rs769891 | 0.409 | -1.456 | 0.177 | -0.383 | 0.01 | 0.016 | 0.016 | 0 | 0.39 | 0.37 | 0.21 | 0.25 | 0.25 | 0.18 |
| 1 | 55934389 | rs12568687 | -0.07 | -1.113 | 0.331 | -0.163 | -0.006 | 0.032 | 0.032 | 0 | 0.08 | 0.13 | 0.05 | 0.14 | 0.07 | 0.17 |
| 1 | 55935340 | rs12075626 | 0.157 | -1.12 | 0.495 | -0.166 | 0.002 | -0.007 | -0.007 | 0 | 0.07 | 0.06 | 0.05 | 0.14 | 0.07 | 0.17 |
| 1 | 55935941 | rs4142664 | -0.024 | -1.165 | 0.702 | -0.156 | 0 | -0.004 | -0.004 | 0 | 0.42 | 0.53 | 0.57 | 0.57 | 0.59 | 0.51 |
| 1 | 55938887 | rs7532502 | 0.456 | -1.139 | 0.428 | -0.164 | 0.005 | -0.009 | -0.009 | 1.19 | 0.07 | 0.05 | 0.04 | 0.09 | 0.05 | 0.14 |
| 1 | 55939994 | rs1499678 | 0.802 | -1.013 | 0.522 | -0.144 | -0.013 | -0.012 | -0.012 | 0 | 0.33 | 0.29 | 0.51 | 0.43 | 0.52 | 0.34 |
| 1 | 55948307 | rs7546894 | 0.836 | -1.48 | 0.362 | -0.197 | -0.067 | -0.075 | -0.075 | 0 | 0.35 | 0.12 | 0.48 | 0.42 | 0.53 | 0.34 |
| 1 | 55948478 | rs1780526 | 0.598 | -2.121 | 0.622 | -0.599 | 0 | 0.003 | 0.003 | 0 | 0.23 | 0.21 | 0.11 | 0.12 | 0.17 | 0.17 |
| 1 | 55949987 | rs630446 | 0.524 | -2.429 | 0.857 | -0.73 | -0.029 | -0.004 | -0.004 | 1.555 | 0.24 | 0.33 | 0.16 | 0.23 | 0.14 | 0.18 |
| 1 | 55950886 | rs1321119 | -1.656 | -2.388 | 0.676 | -0.691 | -0.003 | -0.02 | -0.02 | 0 | 0.86 | 0.95 | 0.79 | 0.83 | 0.87 | 0.73 |
| 1 | 55952835 | rs411611 | -0.523 | -2.475 | 1.215 | -0.758 | -0.003 | -0.005 | -0.005 | 0 | 0.44 | 0.44 | 0.24 | 0.3 | 0.27 | 0.33 |
| 1 | 55953290 | rs2649629 | 1.601 | -2.446 | 0.749 | -0.883 | -0.038 | 0.035 | 0.035 | 0 | 0.16 | 0.01 | 0.06 | 0.11 | 0.13 | 0.12 |
| 1 | 55954627 | rs379213 | 0.898 | -2.478 | 1.09 | -0.812 | -0.027 | -0.011 | -0.011 | 0.183 | 0.21 | 0.26 | 0.14 | 0.12 | 0.11 | 0.1 |
| 1 | 55963519 | rs1780542 | 1.098 | -2.435 | 1.32 | -0.83 | -0.004 | 0.016 | 0.016 | 0 | 0.2 | 0.15 | 0.1 | 0.12 | 0.17 | 0.14 |

| | | | | | | | | | | | | | | | | |
|---|---|---|---|---|---|---|---|---|---|---|---|---|---|---|---|---|
| 1 | 55963649 | rs12563278 | -0.929 | -2.45 | 1.47 | -0.798 | 0.019 | 0.128 | 0.128 | 0 | 0.05 | 0.19 | 0.05 | 0.2 | 0.06 | 0.15 |
| 1 | 55966764 | rs1165390 | -0.753 | -2.44 | 1.327 | -0.806 | -0.002 | -0.025 | -0.025 | 0 | 0.77 | 0.68 | 0.88 | 0.78 | 0.82 | 0.77 |
| 1 | 55967144 | rs11206611 | 1.265 | -2.478 | 1.229 | -0.762 | 0.048 | 0.038 | 0.038 | 0 | 0.18 | 0.08 | 0.11 | 0.07 | 0.09 | 0.06 |
| 1 | 55968289 | rs1780522 | 0.269 | -2.491 | 1.175 | -0.753 | -0.013 | -0.002 | -0.002 | 11.609 | 0.62 | 0.57 | 0.62 | 0.66 | 0.72 | 0.61 |
| 1 | 55968450 | rs1695938 | 0.269 | -2.491 | 1.175 | -0.684 | -0.013 | -0.006 | -0.006 | 0.681 | 0.62 | 0.56 | 0.64 | 0.71 | 0.72 | 0.61 |
| 1 | 55968954 | rs356116 | -1.606 | -2.384 | 0.908 | -0.676 | -0.002 | -0.03 | -0.03 | 0.421 | 0.85 | 0.94 | 0.78 | 0.79 | 0.85 | 0.69 |
| 1 | 55972500 | rs356119 | 0.558 | -2.443 | 1.097 | -0.718 | -0.032 | -0.017 | -0.017 | 0 | 0.23 | 0.33 | 0.15 | 0.12 | 0.13 | 0.08 |
| 1 | 55972596 | rs189593 | -1.59 | -2.428 | 0.955 | -0.484 | 0.005 | -0.016 | -0.016 | 0.566 | 0.85 | 0.93 | 0.72 | 0.81 | 0.86 | 0.72 |
| 1 | 55973486 | rs1695941 | 0.179 | -2.449 | 1.291 | -0.536 | -0.013 | -0.004 | -0.004 | 0.772 | 0.6 | 0.54 | 0.65 | 0.66 | 0.73 | 0.64 |
| 1 | 55976706 | rs1740122 | 0.398 | -2.382 | 1.369 | -0.661 | 0.002 | -0.03 | -0.03 | 0 | 0.23 | 0.34 | 0.12 | 0.16 | 0.19 | 0.17 |
| 1 | 55983525 | rs1695960 | 0.738 | -2.397 | 1.391 | -0.671 | 0 | 0.009 | 0.009 | 0 | 0.21 | 0.17 | 0.1 | 0.12 | 0.18 | 0.14 |
| 1 | 55984309 | rs2133555 | -0.293 | -2.525 | 1.442 | -0.624 | -0.024 | -0.028 | -0.028 | 0.047 | 0.29 | 0.53 | 0.14 | 0.29 | 0.21 | 0.35 |
| 1 | 55987794 | rs1780541 | 0.113 | -2.565 | 1.234 | -0.636 | 0.003 | 0.003 | 0.003 | 0 | 0.6 | 0.6 | 0.71 | 0.69 | 0.73 | 0.64 |
| 1 | 55989646 | rs356081 | -1.442 | -2.552 | 1.141 | -0.643 | 0 | 0.002 | 0.002 | 0 | 0.85 | 0.83 | 0.85 | 0.78 | 0.83 | 0.75 |
| 1 | 55989704 | rs1078876 | 1.203 | -2.615 | 1.178 | -0.676 | -0.018 | -0.025 | -0.025 | 0.934 | 0.38 | 0.24 | 0.6 | 0.48 | 0.55 | 0.41 |
| 1 | 55989750 | rs1078877 | 0.831 | -2.586 | 1.19 | -0.664 | -0.001 | -0.033 | -0.033 | 0 | 0.21 | 0.25 | 0.11 | 0.11 | 0.18 | 0.13 |
| 1 | 55991345 | rs92447 | -0.058 | -2.509 | 0.918 | -0.597 | -0.021 | 0.053 | 0.053 | 0 | 0.06 | 0.21 | 0.03 | 0.13 | 0.01 | 0.15 |
| 1 | 55992755 | rs356085 | -0.048 | -2.544 | 0.953 | -0.613 | -0.002 | -0.026 | -0.026 | 0 | 0.63 | 0.53 | 0.74 | 0.67 | 0.66 | 0.63 |
| 1 | 55993095 | rs6588561 | 1.289 | -2.515 | 0.776 | -0.603 | -0.035 | -0.05 | -0.05 | 1.447 | 0.37 | 0.17 | 0.59 | 0.43 | 0.53 | 0.34 |
| 1 | 55994302 | rs356086 | -0.049 | -2.474 | 0.601 | -0.539 | -0.02 | 0.062 | 0.062 | 0 | 0.06 | 0.21 | 0.03 | 0.13 | 0.01 | 0.14 |
| 1 | 55995298 | rs1995229 | 1.293 | -2.304 | -0.077 | -0.511 | -0.064 | -0.08 | -0.08 | 0 | 0.36 | 0.12 | 0.58 | 0.44 | 0.54 | 0.34 |
| 1 | 55995469 | rs7529841 | 0.747 | -2.475 | -0.035 | -0.553 | 0.041 | 0.034 | 0.034 | 0 | 0.19 | 0.09 | 0.12 | 0.07 | 0.09 | 0.07 |
| 1 | 55998435 | rs168549 | 0.019 | -2.473 | 0.161 | -0.497 | 0.005 | 0.003 | 0.003 | 0 | 0.64 | 0.65 | 0.73 | 0.72 | 0.73 | 0.66 |
| 1 | 56005177 | rs1740113 | 1.113 | -2.47 | -0.057 | -0.527 | 0.022 | 0.026 | 0.026 | 0 | 0.15 | 0.09 | 0.04 | 0.07 | 0.08 | 0.24 |
| 1 | 56013803 | rs356103 | 0.8 | -1.481 | -0.139 | -0.288 | -0.014 | -0.018 | -0.018 | 0.119 | 0.25 | 0.17 | 0.5 | 0.54 | 0.34 | 0.35 |
| 1 | 56015915 | rs1392127 | 0.396 | -1.541 | -0.123 | -0.351 | -0.001 | -0.003 | -0.003 | 0.315 | 0.16 | 0.16 | 0.15 | 0.05 | 0.14 | 0.09 |
| 1 | 56021061 | rs9436973 | -2.109 | -1.493 | -0.389 | -0.065 | -0.023 | -0.025 | -0.025 | 0.407 | 0.82 | 0.89 | 0.53 | 0.57 | 0.69 | 0.79 |
| 1 | 56023333 | rs9787362 | 1.652 | -1.531 | -0.286 | -0.045 | 0.004 | 0.014 | 0.014 | 0.427 | 0.19 | 0.2 | 0.47 | 0.43 | 0.31 | 0.21 |
| 1 | 56026844 | rs17112378 | 2.965 | -1.599 | -0.402 | -0.041 | 0.003 | -0.005 | -0.005 | 0 | 0.06 | 0.03 | 0.03 | 0.11 | 0.03 | 0.05 |
| 1 | 56027175 | rs6588562 | 0.864 | -1.523 | -0.369 | -0.136 | -0.027 | 0.049 | 0.049 | 0 | 0.33 | 0.15 | 0.24 | 0.27 | 0.38 | 0.2 |
| 1 | 56033892 | rs10888934 | 2.003 | -1.328 | -0.322 | 0.01 | -0.043 | -0.053 | -0.053 | 0 | 0.17 | 0.08 | 0.46 | 0.43 | 0.31 | 0.2 |
| 1 | 56037629 | rs1499672 | -0.633 | -1.651 | -0.045 | -0.125 | 0.004 | -0.002 | -0.002 | 0 | 0.58 | 0.51 | 0.43 | 0.44 | 0.53 | 0.45 |
| 1 | 56038407 | rs1566399 | -1.513 | -1.655 | 0.107 | -0.107 | -0.045 | -0.064 | -0.064 | 0 | 0.77 | 0.66 | 0.86 | 0.9 | 0.87 | 0.71 |
| 1 | 56041961 | rs4927255 | -0.04 | -1.639 | -0.202 | -0.068 | -0.028 | 0.01 | 0.01 | 0 | 0.41 | 0.27 | 0.59 | 0.59 | 0.48 | 0.47 |
| 1 | 56042915 | rs904604 | -0.323 | -1.569 | -0.198 | -0.275 | -0.023 | 0.008 | 0.008 | 0.277 | 0.56 | 0.32 | 0.41 | 0.44 | 0.53 | 0.45 |
| 1 | 56043876 | rs12127175 | -0.107 | -1.589 | -0.395 | -0.277 | 0.011 | 0.043 | 0.043 | 0.112 | 0.49 | 0.4 | 0.33 | 0.39 | 0.49 | 0.39 |
| 1 | 56046320 | rs4927257 | 0.069 | -1.646 | -0.507 | -0.239 | 0.003 | 0.072 | 0.072 | 0.005 | 0.47 | 0.26 | 0.32 | 0.39 | 0.49 | 0.39 |
| 1 | 56047603 | rs7535298 | -0.508 | -1.7 | -0.359 | -0.431 | -0.03 | -0.036 | -0.036 | 0.005 | 0.68 | 0.44 | 0.75 | 0.85 | 0.82 | 0.66 |
| 1 | 56050016 | rs1695945 | 0.862 | -1.518 | -0.995 | -0.443 | -0.012 | 0.074 | 0.074 | 0 | 0.29 | 0.02 | 0.17 | 0.18 | 0.24 | 0.31 |
| 1 | 56054865 | rs10493194 | 1.652 | -1.614 | -0.817 | -0.533 | -0.006 | -0.01 | -0.01 | 0.136 | 0.19 | 0.22 | 0.11 | 0.11 | 0.13 | 0.17 |
| 1 | 56056400 | rs4926690 | 0.827 | -1.552 | -0.968 | -0.158 | -0.024 | -0.06 | -0.06 | 0 | 0.48 | 0.07 | 0.6 | 0.57 | 0.57 | 0.51 |
| 1 | 56057691 | rs6680818 | 1.795 | -1.452 | -0.823 | -0.168 | -0.008 | -0.013 | -0.013 | 0 | 0.19 | 0.22 | 0.11 | 0.11 | 0.13 | 0.17 |
| 1 | 56057849 | rs1499662 | -0.394 | -1.484 | -0.293 | -0.16 | 0.051 | 0.037 | 0.037 | 0.231 | 0.85 | 0.58 | 0.82 | 0.87 | 0.83 | 0.89 |
| 1 | 56059223 | rs7526426 | -0.211 | -1.443 | -0.425 | -0.148 | 0.006 | 0.015 | 0.015 | 0.158 | 0.64 | 0.33 | 0.69 | 0.76 | 0.69 | 0.66 |
| 1 | 56060308 | rs11577328 | 0.798 | -1.043 | -0.262 | 0.043 | 0.057 | 0.027 | 0.027 | 0.032 | 0.05 | 0.15 | 0.09 | 0.08 | 0.13 | 0.05 |
| 1 | 56067651 | rs4927258 | 0.855 | -1.208 | -0.476 | 0.043 | -0.005 | 0.003 | 0.003 | 0 | 0.16 | 0.17 | 0.09 | 0.19 | 0.12 | 0.15 |
| 1 | 56068394 | rs11807964 | 1.993 | -0.671 | -0.673 | 0.334 | -0.117 | -0.124 | -0.124 | 0 | 0.17 | 0.02 | 0.43 | 0.38 | 0.32 | 0.21 |
| 1 | 56069516 | rs1499665 | 1.57 | -0.899 | -0.629 | -0.604 | 0.001 | 0.001 | 0.001 | 0 | 0.2 | 0.19 | 0.16 | 0.18 | 0.14 | 0.35 |
| 1 | 56070382 | rs6704127 | 0.298 | -0.663 | -0.475 | -0.418 | 0.016 | 0.05 | 0.05 | 0.601 | 0.55 | 0.3 | 0.66 | 0.71 | 0.59 | 0.69 |
| 1 | 56075918 | rs7555929 | 1.317 | 0.758 | 0.01 | 0.574 | -0.117 | -0.135 | -0.135 | 0 | 0.17 | 0.02 | 0.43 | 0.51 | 0.32 | 0.21 |
| 1 | 56078416 | rs10749702 | 0.715 | 0.706 | 0.064 | 0.474 | 0.009 | 0.007 | 0.007 | 0.019 | 0.2 | 0.19 | 0.47 | 0.63 | 0.34 | 0.4 |
| 1 | 56087509 | rs709396 | -0.506 | 0.273 | 0.05 | 0.585 | 0.064 | 0.115 | 0.115 | 0.031 | 0.15 | 0.48 | 0.2 | 0.21 | 0.16 | 0.22 |
| 1 | 56090323 | rs709397 | -0.625 | 0.089 | -0.819 | 0.389 | -0.041 | -0.036 | -0.036 | 0 | 0.65 | 0.77 | 0.49 | 0.47 | 0.55 | 0.73 |
| 1 | 56099371 | rs6700588 | -0.943 | -0.779 | -0.282 | -0.047 | -0.011 | -0.026 | -0.026 | 0 | 0.68 | 0.67 | 0.78 | 0.89 | 0.75 | 0.65 |
| 1 | 56101822 | rs1392140 | -0.939 | -0.948 | 0.285 | 0.162 | -0.012 | -0.028 | -0.028 | 0 | 0.68 | 0.67 | 0.78 | 0.88 | 0.75 | 0.65 |
| 1 | 56102040 | rs1499676 | 1.171 | -1.102 | 0.041 | 0.032 | -0.01 | 0.017 | 0.017 | 0 | 0.11 | 0.03 | 0.05 | 0.12 | 0.11 | 0.06 |
| 1 | 56106506 | rs12405865 | 1.26 | -1.178 | -0.487 | -0.314 | -0.161 | -0.167 | -0.167 | 0 | 0.25 | 0.02 | 0.45 | 0.53 | 0.37 | 0.26 |
| 1 | 56106677 | rs6669312 | 0.937 | -1.2 | -0.152 | -0.366 | 0.027 | 0.016 | 0.016 | 0.281 | 0.05 | 0.12 | 0.06 | 0.08 | 0.1 | 0.08 |
| 1 | 56108604 | rs1165472 | 1.25 | -1.321 | 0.018 | -0.414 | -0.005 | -0.018 | -0.018 | 0.031 | 0.32 | 0.31 | 0.18 | 0.11 | 0.25 | 0.34 |
| 1 | 56112774 | rs10443215 | -0.443 | -1.221 | 0.035 | -0.353 | 0.017 | 0.043 | 0.043 | 0 | 0.44 | 0.37 | 0.72 | 0.83 | 0.58 | 0.42 |
| 1 | 56118105 | rs17112591 | 0.699 | -0.899 | 0.37 | -0.136 | 0.044 | 0.023 | 0.023 | 1.748 | 0.05 | 0.14 | 0.07 | 0.08 | 0.11 | 0.08 |
| 1 | 56119747 | rs7540786 | 0.664 | -1.414 | 0.289 | -0.41 | 0.008 | 0.002 | 0.002 | 0 | 0.35 | 0.71 | 0.38 | 0.3 | 0.35 | 0.42 |
| 1 | 56120519 | rs9436515 | 0.762 | -1.404 | 0.24 | -0.427 | -0.004 | -0.037 | -0.037 | 0.134 | 0.33 | 0.51 | 0.23 | 0.19 | 0.34 | 0.39 |
| 1 | 56123166 | rs9436516 | -1.359 | -1.683 | 0.132 | -0.389 | -0.04 | 0.033 | 0.033 | 3.493 | 0.14 | 0.68 | 0.22 | 0.15 | 0.06 | 0.14 |
| 1 | 56133634 | rs1955318 | 1.294 | -1.501 | 0.097 | 0.306 | 0.01 | -0.003 | -0.003 | 0 | 0.26 | 0.27 | 0.47 | 0.21 | 0.38 | 0.24 |
| 1 | 56143209 | rs7518530 | 0.034 | -1.553 | 0.035 | 0.372 | 0.006 | 0.044 | 0.044 | 0 | 0.62 | 0.71 | 0.78 | 0.7 | 0.62 | 0.63 |
| 1 | 56152725 | rs2767503 | 0.024 | -1.679 | 0.159 | 0.52 | 0.005 | 0.021 | 0.021 | 0 | 0.5 | 0.45 | 0.56 | 0.25 | 0.54 | 0.37 |
| 1 | 56157154 | rs12565309 | 1.942 | -1.739 | 0.076 | 0.713 | 0.004 | -0.105 | -0.105 | 0.813 | 0.24 | 0.01 | 0.11 | 0.33 | 0.18 | 0.3 |
| 1 | 56157953 | rs2767511 | -1.004 | -1.744 | 0.125 | 0.731 | 0.021 | 0.058 | 0.058 | 6.894 | 0.94 | 0.89 | 0.93 | 0.76 | 0.88 | 0.86 |

| | | | | | | | | | | | | | | | | |
|---|---|---|---|---|---|---|---|---|---|---|---|---|---|---|---|---|
| 1 | 56158236 | rs2767512 | -0.443 | -1.722 | 0.204 | 0.755 | -0.003 | -0.003 | -0.003 | 0 | 0.79 | 0.78 | 0.85 | 0.7 | 0.76 | 0.7 |
| 1 | 56160151 | rs2819483 | -0.458 | -1.721 | 0.191 | 0.755 | -0.006 | -0.005 | -0.005 | 0.037 | 0.79 | 0.79 | 0.85 | 0.7 | 0.76 | 0.7 |
| 1 | 56161430 | rs1849680 | 1.784 | -1.695 | 0.23 | 0.772 | 0.017 | -0.023 | -0.023 | 0.605 | 0.26 | 0.13 | 0.14 | 0.33 | 0.21 | 0.3 |
| 1 | 56161500 | rs4927261 | 1.318 | -1.696 | 0.267 | 0.788 | -0.016 | 0.031 | 0.031 | 0 | 0.28 | 0.32 | 0.29 | 0.43 | 0.21 | 0.33 |
| 1 | 56163458 | rs9326037 | 1.678 | -1.627 | -0.061 | 0.623 | 0.017 | -0.019 | -0.019 | 1.069 | 0.26 | 0.16 | 0.14 | 0.33 | 0.21 | 0.3 |
| 1 | 56179614 | rs2671245 | 0.551 | -1.652 | -0.038 | 0.549 | -0.01 | 0.038 | 0.038 | 0.019 | 0.42 | 0.32 | 0.53 | 0.23 | 0.52 | 0.3 |
| 1 | 56183163 | rs1183736 | 0.12 | -1.493 | -0.068 | 0.681 | 0.008 | 0.035 | 0.035 | 0 | 0.49 | 0.4 | 0.55 | 0.27 | 0.54 | 0.37 |
| 1 | 56185027 | rs3862237 | 1.666 | -1.417 | -0.366 | 0.787 | 0.018 | -0.033 | -0.033 | 0 | 0.25 | 0.09 | 0.14 | 0.32 | 0.21 | 0.29 |
| 1 | 56186258 | rs1165486 | -0.254 | -1.174 | -0.256 | 0.602 | -0.001 | -0.003 | -0.003 | 0 | 0.5 | 0.52 | 0.51 | 0.27 | 0.54 | 0.38 |
| 1 | 56195659 | rs4111070 | 0.011 | -1.458 | -0.38 | 0.687 | 0 | 0 | 0 | 0.013 | 0.26 | 0.29 | 0.28 | 0.3 | 0.3 | 0.19 |
| 1 | 56201024 | rs2767499 | -0.248 | -1.749 | -0.23 | 0.812 | 0.04 | 0.003 | 0.003 | 0.629 | 0.48 | 0.4 | 0.72 | 0.54 | 0.68 | 0.48 |
| 1 | 56204848 | rs1184810 | -0.193 | -2.27 | -0.34 | 0.557 | -0.032 | -0.093 | -0.093 | 0.068 | 0.75 | 0.56 | 0.84 | 0.92 | 0.84 | 0.77 |
| 1 | 56207532 | rs17112735 | 1.536 | -2.278 | -0.39 | 0.584 | 0.076 | -0.093 | -0.093 | 0 | 0.26 | 0.06 | 0.12 | 0.37 | 0.16 | 0.29 |
| 1 | 56219293 | rs1165500 | 0.213 | -1.686 | -0.038 | 1.079 | -0.026 | 0.017 | 0.017 | 1.358 | 0.39 | 0.19 | 0.66 | 0.28 | 0.54 | 0.39 |
| 1 | 56222118 | rs1165495 | -0.23 | -1.694 | 0.006 | 1.136 | -0.002 | 0 | 0 | 0 | 0.44 | 0.48 | 0.72 | 0.45 | 0.65 | 0.41 |
| 1 | 56240015 | rs749330 | 0.3 | -1.697 | -0.014 | 1.135 | 0.006 | 0.001 | 0.001 | 0.526 | 0.52 | 0.49 | 0.28 | 0.46 | 0.32 | 0.53 |
| 1 | 56240702 | rs12092608 | -0.259 | -1.562 | -0.246 | 1.048 | -0.017 | 0.017 | 0.017 | 1.167 | 0.61 | 0.74 | 0.34 | 0.72 | 0.46 | 0.61 |
| 1 | 56247216 | rs3122551 | -0.54 | -2.538 | 0.114 | 0.636 | -0.024 | -0.033 | -0.033 | 0 | 0.78 | 0.71 | 0.77 | 0.93 | 0.85 | 0.82 |
| 1 | 56251667 | rs3122554 | -0.613 | -2.546 | 0.115 | 0.632 | -0.019 | -0.026 | -0.026 | 0.058 | 0.79 | 0.72 | 0.77 | 0.93 | 0.85 | 0.82 |
| 1 | 56268037 | rs4474302 | 0.061 | -3.13 | 0.041 | 1.025 | 0.003 | 0.004 | 0.004 | 1.788 | 0.42 | 0.38 | 0.5 | 0.26 | 0.52 | 0.41 |
| 1 | 56269821 | rs4367814 | 1.011 | -3.286 | -0.325 | 1.474 | 0.02 | 0.009 | 0.009 | 0.423 | 0.3 | 0.21 | 0.17 | 0.24 | 0.21 | 0.33 |
| 1 | 56287506 | rs4512692 | -0.694 | -3.09 | -0.575 | 0.828 | 0.024 | 0.041 | 0.041 | 0 | 0.75 | 0.87 | 0.89 | 0.99 | 0.82 | 0.88 |
| 1 | 56287632 | rs10493191 | 0.435 | -3.285 | -0.173 | 0.584 | -0.019 | -0.056 | -0.056 | 0 | 0.1 | 0.21 | 0.04 | 0 | 0.04 | 0.09 |
| 1 | 56287724 | rs6588574 | 1.365 | -3.536 | -0.182 | 0.774 | 0.066 | -0.006 | -0.006 | 0.48 | 0.32 | 0.16 | 0.16 | 0.29 | 0.23 | 0.37 |
| 1 | 56288116 | rs12077235 | 1.022 | -3.471 | -0.133 | 0.809 | 0.044 | -0.027 | -0.027 | 0 | 0.35 | 0.26 | 0.18 | 0.45 | 0.29 | 0.39 |
| 1 | 56288744 | rs4433442 | 0.314 | -3.467 | -0.222 | 0.897 | -0.014 | -0.007 | -0.007 | 0 | 0.6 | 0.61 | 0.65 | 0.69 | 0.74 | 0.68 |
| 1 | 56289523 | rs4129669 | 1.615 | -3.078 | -0.339 | 1.104 | -0.161 | -0.024 | -0.024 | 0 | 0.2 | 0.03 | 0.37 | 0.17 | 0.42 | 0.19 |
| 1 | 56299527 | rs10493190 | 0.524 | -3.287 | -0.03 | 0.507 | -0.016 | -0.046 | -0.046 | 0 | 0.09 | 0.15 | 0.03 | 0 | 0.04 | 0.09 |
| 1 | 56320873 | rs7349129 | -1.056 | -3.417 | -0.14 | 0.424 | 0.023 | 0.008 | 0.008 | 0 | 0.78 | 0.85 | 0.94 | 0.99 | 0.93 | 0.92 |
| 1 | 56321074 | rs10493186 | 1.121 | -3.688 | -0.217 | 0.516 | 0.046 | -0.041 | -0.041 | 0 | 0.33 | 0.22 | 0.14 | 0.45 | 0.27 | 0.37 |
| 1 | 56325957 | rs4540693 | 0.878 | -2.808 | 0.255 | 1.023 | 0.053 | 0.006 | 0.006 | 0.135 | 0.25 | 0.35 | 0.47 | 0.24 | 0.46 | 0.27 |
| 1 | 56328238 | rs6657176 | -0.849 | -2.908 | 0.887 | 1.266 | -0.002 | 0.001 | 0.001 | 0 | 0.19 | 0.2 | 0.21 | 0.25 | 0.2 | 0.2 |
| 1 | 56333935 | rs10888941 | -0.435 | -2.896 | 0.884 | 1.212 | -0.002 | -0.038 | -0.038 | 0 | 0.16 | 0.18 | 0.13 | 0.01 | 0.18 | 0.15 |
| 1 | 56340495 | rs17113009 | -1.37 | -2.934 | 1.225 | 1.325 | 0.085 | 0.27 | 0.27 | 0.098 | 0.06 | 0.38 | 0.15 | 0.24 | 0.07 | 0.12 |
| 1 | 56342433 | rs17113019 | -0.921 | -2.931 | 1.228 | 1.316 | 0.109 | 0.214 | 0.214 | 0.107 | 0.05 | 0.37 | 0.04 | 0.19 | 0.06 | 0.04 |
| 1 | 56343290 | rs7516605 | -0.069 | -2.974 | 1.012 | 1.191 | -0.002 | -0.035 | -0.035 | 0.003 | 0.14 | 0.16 | 0.11 | 0 | 0.16 | 0.15 |
| 1 | 56345824 | rs4130692 | -0.257 | -2.978 | 0.984 | 1.202 | -0.005 | -0.011 | -0.011 | 0 | 0.16 | 0.2 | 0.22 | 0.05 | 0.16 | 0.26 |
| 1 | 56359597 | rs6681658 | -1.603 | -3.068 | 0.829 | 1.385 | 0.007 | -0.011 | -0.011 | 0 | 0.28 | 0.69 | 0.36 | 0.51 | 0.26 | 0.34 |
| 1 | 56387285 | rs12144400 | 1.01 | -3.382 | 0.016 | 0.886 | 0.106 | -0.036 | -0.036 | 0 | 0.27 | 0.01 | 0.09 | 0.18 | 0.16 | 0.26 |
| 1 | 56394673 | rs11206686 | -1.151 | -1.567 | 1.246 | 1.29 | -0.002 | -0.066 | -0.066 | 0 | 0.2 | 0.43 | 0.15 | 0.08 | 0.24 | 0.21 |
| 1 | 56398037 | rs4573554 | 1.208 | -1.592 | 1.331 | 1.293 | -0.024 | -0.042 | -0.042 | 0 | 0.77 | 0.46 | 0.72 | 0.87 | 0.75 | 0.71 |
| 1 | 56401121 | rs11206687 | 1.524 | -1.528 | -0.163 | 0.903 | 0.03 | -0.046 | -0.046 | 1.741 | 0.71 | 0.23 | 0.63 | 0.57 | 0.71 | 0.65 |
| 1 | 56407227 | rs11206690 | -0.631 | -1.219 | -0.357 | 0.276 | 0.002 | -0.024 | -0.024 | 0.464 | 0.16 | 0.15 | 0.11 | 0 | 0.18 | 0.18 |
| 1 | 56412081 | rs12081053 | 1.023 | -1.287 | -0.364 | 0.753 | -0.012 | 0.024 | 0.024 | 0.389 | 0.71 | 0.38 | 0.64 | 0.48 | 0.7 | 0.64 |
| 1 | 56413260 | rs6662998 | -0.65 | -1.884 | -0.573 | -0.411 | -0.022 | -0.117 | -0.117 | 0.672 | 0.35 | 0.42 | 0.18 | 0.01 | 0.25 | 0.28 |
| 1 | 56413519 | rs4614300 | 0.029 | -2.291 | -0.824 | 0.535 | -0.01 | 0.006 | 0.006 | 0.063 | 0.22 | 0.24 | 0.14 | 0.39 | 0.13 | 0.2 |
| 1 | 56420527 | rs7540718 | -0.533 | -2.404 | -1.29 | 0.213 | 0.027 | -0.003 | -0.003 | 0 | 0.62 | 0.83 | 0.31 | 0.6 | 0.37 | 0.52 |
| 1 | 56429031 | rs2192102 | 0.261 | -3.105 | -0.451 | 0.237 | -0.04 | 0.063 | 0.063 | 0 | 0.09 | 0.23 | 0.05 | 0.14 | 0.03 | 0.16 |
| 1 | 56432184 | rs12409063 | 0.285 | -3.111 | -0.431 | 0.235 | -0.038 | 0.064 | 0.064 | 0 | 0.09 | 0.21 | 0.04 | 0.14 | 0.03 | 0.16 |
| 1 | 56434711 | rs10789012 | -0.766 | -4.309 | -0.823 | -0.117 | 0.127 | 0.001 | 0.001 | 0 | 0.47 | 0.76 | 0.27 | 0.52 | 0.26 | 0.45 |
| 1 | 56439836 | rs10789014 | -0.745 | -4.254 | -0.809 | -0.14 | 0.057 | -0.037 | -0.037 | 0.05 | 0.58 | 0.85 | 0.31 | 0.46 | 0.35 | 0.42 |
| 1 | 56439909 | rs10489777 | 0.029 | -5.025 | -1.105 | -0.51 | 0.021 | -0.005 | -0.005 | 0.005 | 0.35 | 0.24 | 0.22 | 0.39 | 0.28 | 0.31 |
| 1 | 56444449 | rs11206704 | 0.661 | -4.917 | -0.138 | -0.743 | -0.057 | -0.047 | -0.047 | 0.725 | 0.22 | 0.34 | 0.06 | 0.08 | 0.08 | 0.11 |
| 1 | 56446105 | rs12029607 | -0.459 | -5.379 | -0.463 | -0.714 | -0.046 | -0.019 | -0.019 | 0 | 0.34 | 0.6 | 0.14 | 0.22 | 0.15 | 0.2 |
| 1 | 56448410 | rs11206709 | -0.13 | -5.415 | -0.447 | -0.738 | -0.05 | -0.045 | -0.045 | 5.871 | 0.3 | 0.63 | 0.12 | 0.08 | 0.1 | 0.2 |
| 1 | 56455182 | rs12074486 | -0.139 | -5.424 | -0.432 | -0.732 | -0.052 | -0.045 | -0.045 | 0 | 0.31 | 0.63 | 0.12 | 0.08 | 0.1 | 0.2 |
| 1 | 56493106 | rs1416127 | -0.454 | -5.039 | -0.693 | -0.665 | 0.037 | 0.06 | 0.06 | 0 | 0.46 | 0.35 | 0.28 | 0.28 | 0.41 | 0.38 |
| 1 | 56502227 | rs11206718 | 0.346 | -5.106 | -0.804 | -0.711 | -0.055 | 0.124 | 0.124 | 0.648 | 0.35 | 0.03 | 0.24 | 0.14 | 0.37 | 0.24 |
| 1 | 56505301 | rs2404393 | -1.819 | -5.131 | -0.598 | -0.676 | -0.055 | -0.075 | -0.075 | 0 | 0.73 | 0.84 | 0.53 | 0.5 | 0.52 | 0.66 |
| 1 | 56506564 | rs11206722 | -0.217 | -5.155 | -0.68 | -0.686 | 0.028 | 0.066 | 0.066 | 0 | 0.44 | 0.31 | 0.42 | 0.28 | 0.43 | 0.32 |
| 1 | 56510794 | rs12031810 | 2.141 | -5.153 | -0.793 | -0.684 | 0.024 | -0.034 | -0.034 | 0.314 | 0.07 | 0 | 0.01 | 0.13 | 0.03 | 0.07 |
| 1 | 56519023 | rs4278395 | -0.023 | -5.121 | -0.771 | -0.672 | 0.01 | 0.099 | 0.099 | 0.062 | 0.42 | 0.12 | 0.4 | 0.28 | 0.43 | 0.32 |
| 1 | 56525934 | rs7525207 | -1.004 | -4.768 | -0.505 | -0.552 | 0.021 | 0.033 | 0.033 | 0 | 0.71 | 0.66 | 0.53 | 0.49 | 0.53 | 0.68 |
| 1 | 56545484 | rs6702275 | 0.359 | -5.047 | -0.818 | -0.624 | -0.063 | 0.056 | 0.056 | 0 | 0.31 | 0.03 | 0.34 | 0.19 | 0.33 | 0.26 |
| 1 | 56562921 | rs1412215 | -1.073 | -4.885 | -0.286 | -0.466 | 0.022 | 0.011 | 0.011 | 0 | 0.71 | 0.64 | 0.52 | 0.41 | 0.53 | 0.55 |
| 1 | 56564716 | rs12025982 | 1.611 | -4.805 | -0.594 | -0.497 | -0.109 | -0.119 | -0.119 | 1.256 | 0.23 | 0.04 | 0.45 | 0.45 | 0.46 | 0.32 |
| 1 | 56569364 | rs12117303 | 1.731 | -4.762 | -0.476 | -0.532 | 0.026 | 0.053 | 0.053 | 0 | 0.21 | 0.02 | 0.11 | 0.08 | 0.16 | 0.14 |
| 1 | 56574395 | rs7512228 | 1.215 | -4.808 | -0.235 | -0.555 | -0.007 | -0.013 | -0.013 | 1.327 | 0.24 | 0.26 | 0.13 | 0.08 | 0.17 | 0.14 |
| 1 | 56577257 | rs7543532 | 1.251 | -4.781 | -0.177 | -0.568 | 0.005 | 0.001 | 0.001 | 0.07 | 0.24 | 0.19 | 0.12 | 0.08 | 0.17 | 0.14 |

| | | | | | | | | | | | | | | | | |
|---|---|---|---|---|---|---|---|---|---|---|---|---|---|---|---|---|
| 1 | 56586286 | rs6588607 | 1.051 | -4.266 | -0.263 | -0.516 | 0.014 | -0.002 | -0.002 | 0 | 0.27 | 0.2 | 0.14 | 0.24 | 0.19 | 0.3 |
| 1 | 56591630 | rs10888957 | 0.066 | -4.432 | 0.166 | -0.513 | -0.052 | 0.004 | 0.004 | 0.349 | 0.39 | 0.47 | 0.17 | 0.37 | 0.22 | 0.37 |
| 1 | 56594863 | rs4926709 | 0.403 | -4.438 | 0.174 | -0.515 | 0.048 | -0.014 | -0.014 | 0 | 0.36 | 0.21 | 0.15 | 0.37 | 0.22 | 0.37 |
| 1 | 56602531 | rs1229948 | 0.873 | -4.186 | 0.19 | -0.684 | -0.026 | -0.027 | -0.027 | 0 | 0.05 | 0.12 | 0.02 | 0 | 0.01 | 0.07 |
| 1 | 56606206 | rs778124 | -0.764 | -4.015 | -0.086 | -0.824 | 0.027 | 0.041 | 0.041 | 0 | 0.48 | 0.39 | 0.42 | 0.32 | 0.38 | 0.42 |
| 1 | 56615809 | rs1757915 | -0.302 | -3.864 | -0.068 | -0.802 | -0.005 | -0.034 | -0.034 | 0 | 0.4 | 0.4 | 0.4 | 0.19 | 0.36 | 0.34 |
| 1 | 56635806 | rs778382 | -1.654 | -3.25 | -0.663 | -0.37 | -0.065 | -0.056 | -0.056 | 0 | 0.78 | 0.89 | 0.58 | 0.58 | 0.57 | 0.75 |
| 1 | 56642082 | rs778129 | -0.16 | -3.778 | -0.847 | -0.651 | 0.175 | 0.003 | 0.003 | 0 | 0.46 | 0.79 | 0.2 | 0.39 | 0.21 | 0.5 |
| 1 | 56642998 | rs1331862 | 1.77 | -3.761 | -0.814 | -0.611 | 0.035 | -0.027 | -0.027 | 0 | 0.24 | 0.11 | 0.13 | 0.25 | 0.18 | 0.34 |
| 1 | 56647459 | rs778419 | -0.592 | -3.818 | -1.048 | -1.041 | 0.006 | 0.048 | 0.048 | 0 | 0.64 | 0.72 | 0.47 | 0.22 | 0.43 | 0.51 |
| 1 | 56647914 | rs778420 | 0.426 | -3.892 | -0.408 | -1.084 | 0.028 | 0.025 | 0.025 | 0.302 | 0.05 | 0.28 | 0.02 | 0 | 0.01 | 0.05 |
| 1 | 56651430 | rs1149646 | -0.673 | -3.913 | -0.451 | -1.1 | 0.004 | 0.065 | 0.065 | 0 | 0.64 | 0.78 | 0.47 | 0.23 | 0.43 | 0.51 |
| 1 | 56657399 | rs1149648 | -0.674 | -3.913 | -0.45 | -1.167 | 0.004 | 0.078 | 0.078 | 0 | 0.64 | 0.78 | 0.47 | 0.2 | 0.43 | 0.51 |
| 1 | 56660979 | rs11206753 | 2.739 | -4.028 | -0.492 | -1.301 | 0.036 | 0.06 | 0.06 | 0 | 0.21 | 0.07 | 0.11 | 0.05 | 0.16 | 0.15 |
| 1 | 56666118 | rs17113488 | 0.861 | -4.122 | -0.708 | -1.361 | -0.026 | -0.027 | -0.027 | 0 | 0.05 | 0.13 | 0.02 | 0 | 0.01 | 0.06 |
| 1 | 56666809 | rs778432 | 1.779 | -4.105 | -0.674 | -1.35 | -0.082 | -0.143 | -0.143 | 0.576 | 0.27 | 0.43 | 0.13 | 0.05 | 0.16 | 0.19 |
| 1 | 56668472 | rs7554047 | -0.12 | -4.083 | -0.645 | -1.337 | 0 | -0.059 | -0.059 | 1.391 | 0.3 | 0.31 | 0.33 | 0.01 | 0.24 | 0.24 |
| 1 | 56670460 | rs697582 | 0.283 | -3.655 | -0.485 | -1.097 | 0.056 | 0.053 | 0.053 | 0.305 | 0.05 | 0.33 | 0.02 | 0 | 0.01 | 0.05 |
| 1 | 56681024 | rs706375 | -1.922 | -3.64 | -0.764 | -0.956 | 0.023 | 0.019 | 0.019 | 0 | 0.79 | 0.79 | 0.57 | 0.57 | 0.58 | 0.78 |
| 1 | 56686298 | rs928389 | 1.325 | -3.675 | -0.89 | -0.894 | -0.038 | -0.034 | -0.034 | 0 | 0.11 | 0.01 | 0.09 | 0.18 | 0.12 | 0.08 |
| 1 | 56687749 | rs7547680 | -0.856 | -3.949 | -0.909 | -1.044 | 0.039 | 0.007 | 0.007 | 0 | 0.39 | 0.29 | 0.35 | 0.32 | 0.28 | 0.51 |
| 1 | 56689034 | rs706371 | 1.913 | -3.982 | -0.853 | -1.071 | -0.084 | -0.139 | -0.139 | 0 | 0.28 | 0.44 | 0.13 | 0.05 | 0.17 | 0.19 |
| 1 | 56694460 | rs706356 | -0.851 | -3.937 | -0.91 | -1.042 | -0.007 | -0.011 | -0.011 | 0 | 0.67 | 0.75 | 0.48 | 0.37 | 0.45 | 0.7 |
| 1 | 56703505 | rs778406 | -0.964 | -4.037 | -0.951 | -1.092 | -0.065 | 0.037 | 0.037 | 0.004 | 0.23 | 0.58 | 0.07 | 0.32 | 0.06 | 0.38 |
| 1 | 56703833 | rs12121243 | 0.798 | -4.02 | -0.956 | -1.097 | -0.008 | 0.079 | 0.079 | 0 | 0.22 | 0.1 | 0.3 | 0.01 | 0.23 | 0.18 |
| 1 | 56707637 | rs778413 | -0.419 | -4.065 | -1.082 | -1.17 | 0.023 | 0.001 | 0.001 | 0.012 | 0.6 | 0.67 | 0.4 | 0.37 | 0.34 | 0.58 |
| 1 | 56715332 | rs2793694 | 0.565 | -3.845 | -0.797 | -0.896 | 0.024 | 0.025 | 0.025 | 0 | 0.63 | 0.32 | 0.64 | 0.81 | 0.74 | 0.52 |
| 1 | 56715711 | rs6687944 | 0.256 | -3.748 | -0.812 | -0.881 | 0.006 | -0.029 | -0.029 | 0 | 0.28 | 0.27 | 0.32 | 0.01 | 0.23 | 0.24 |
| 1 | 56724055 | rs2793660 | 1.553 | -3.725 | -0.64 | -0.819 | 0.06 | -0.019 | -0.019 | 0 | 0.27 | 0.1 | 0.13 | 0.22 | 0.19 | 0.34 |
| 1 | 56725516 | rs2793662 | 0.592 | -3.704 | -0.542 | -0.853 | 0.04 | -0.013 | -0.013 | 0.104 | 0.39 | 0.19 | 0.22 | 0.41 | 0.31 | 0.42 |
| 1 | 56727938 | rs947007 | 0.586 | -3.688 | -0.493 | -0.85 | 0.034 | -0.008 | -0.008 | 0.929 | 0.39 | 0.26 | 0.22 | 0.4 | 0.31 | 0.41 |
| 1 | 56731680 | rs4927309 | 0.64 | -3.69 | -0.772 | -0.871 | -0.061 | -0.038 | -0.038 | 0.057 | 0.31 | 0.21 | 0.45 | 0.59 | 0.46 | 0.34 |
| 1 | 56736013 | rs2792792 | -0.128 | -3.438 | -0.578 | -0.74 | -0.024 | -0.057 | -0.057 | 1.507 | 0.67 | 0.47 | 0.66 | 0.86 | 0.74 | 0.73 |
| 1 | 56744527 | rs2096086 | -1.774 | -3.358 | -0.511 | -0.666 | -0.082 | -0.141 | -0.141 | 2.513 | 0.74 | 0.58 | 0.87 | 0.95 | 0.83 | 0.86 |
| 1 | 56745004 | rs1581995 | -0.028 | -2.606 | -0.55 | -0.219 | -0.006 | -0.029 | -0.029 | 0 | 0.16 | 0.09 | 0.11 | 0.34 | 0.15 | 0.28 |
| 1 | 56751380 | rs2211292 | 0.491 | -2.31 | -0.382 | -0.053 | 0.043 | 0.04 | 0.04 | 0 | 0.27 | 0.33 | 0.35 | 0.46 | 0.43 | 0.29 |
| 1 | 56751839 | rs17113634 | 1.4 | -2.502 | -0.518 | -0.107 | 0.018 | -0.012 | -0.012 | 4.902 | 0.08 | 0.03 | 0.09 | 0.12 | 0.04 | 0.08 |
| 1 | 56754223 | rs12755760 | 1.624 | -0.582 | -0.715 | -0.05 | -0.111 | 0.017 | 0.017 | 0.053 | 0.15 | 0.02 | 0.25 | 0.08 | 0.34 | 0.14 |
| 1 | 56756491 | rs2000141 | 0.518 | -0.867 | -0.141 | -0.129 | -0.005 | -0.006 | -0.006 | 2.905 | 0.15 | 0.18 | 0.11 | 0.18 | 0.11 | 0.09 |
| 1 | 56756907 | rs2793673 | 0.675 | -1.208 | -0.238 | -0.328 | -0.001 | -0.007 | -0.007 | 0 | 0.55 | 0.56 | 0.5 | 0.43 | 0.59 | 0.58 |
| 1 | 56758973 | rs2994802 | 0.564 | -1.202 | -0.243 | -0.338 | 0.004 | 0.08 | 0.08 | 0 | 0.11 | 0.18 | 0.09 | 0.38 | 0.09 | 0.15 |
| 1 | 56759031 | rs2793677 | 1.877 | -1.23 | -0.179 | -0.324 | -0.001 | -0.001 | -0.001 | 0 | 0.29 | 0.23 | 0.15 | 0.24 | 0.2 | 0.37 |
| 1 | 56760177 | rs12563597 | -0.086 | -1.237 | -0.168 | -0.341 | 0.043 | -0.037 | -0.037 | 0 | 0.33 | 0.4 | 0.38 | 0.27 | 0.45 | 0.24 |
| 1 | 56760199 | rs2000142 | 0.92 | -1.246 | -0.313 | -0.376 | -0.022 | -0.018 | -0.018 | 0 | 0.14 | 0.01 | 0.11 | 0.18 | 0.11 | 0.11 |
| 1 | 56760429 | rs2793679 | 1.037 | -1.215 | -0.268 | -0.358 | -0.002 | 0.013 | 0.013 | 0 | 0.47 | 0.38 | 0.4 | 0.32 | 0.55 | 0.55 |
| 1 | 56760616 | rs2000144 | -0.375 | -1.326 | -1.134 | -0.336 | -0.013 | -0.027 | -0.027 | 0 | 0.71 | 0.74 | 0.62 | 0.62 | 0.7 | 0.69 |
| 1 | 56762391 | rs12070360 | 1.657 | -1.343 | -1.208 | -0.83 | -0.026 | 0.034 | 0.034 | 1.494 | 0.08 | 0.18 | 0.1 | 0.11 | 0.05 | 0.08 |
| 1 | 56768672 | rs947252 | 1.925 | -1.234 | -1.551 | -1.01 | 0.062 | 0.118 | 0.118 | 0 | 0.25 | 0.05 | 0.12 | 0.07 | 0.18 | 0.16 |
| 1 | 56770269 | rs1341353 | -1.091 | -0.981 | -1.591 | -0.98 | 0.051 | 0.283 | 0.283 | 0 | 0.54 | 0.8 | 0.58 | 0.92 | 0.6 | 0.64 |
| 1 | 56771709 | rs3005884 | -0.642 | -0.835 | -1.25 | -0.148 | -0.048 | -0.074 | -0.074 | 0.797 | 0.74 | 0.87 | 0.55 | 0.6 | 0.7 | 0.65 |
| 1 | 56775821 | rs12066561 | 0.046 | -0.567 | -0.769 | 0.097 | -0.02 | -0.02 | -0.02 | 0 | 0.16 | 0.31 | 0.13 | 0.16 | 0.1 | 0.07 |
| 1 | 56777781 | rs3005888 | -0.144 | -0.018 | -0.568 | 0.053 | -0.007 | 0.01 | 0.01 | 0.019 | 0.26 | 0.24 | 0.24 | 0.18 | 0.32 | 0.26 |
| 1 | 56783425 | rs2994555 | 0.412 | 0.08 | -0.653 | 0.048 | 0.008 | -0.014 | -0.014 | 0.959 | 0.19 | 0.23 | 0.21 | 0.11 | 0.27 | 0.17 |
| 1 | 56783945 | rs10493204 | 1.394 | 0.058 | -0.666 | 0.033 | 0.004 | -0.012 | -0.012 | 0.011 | 0.08 | 0 | 0.03 | 0.07 | 0.04 | 0.09 |
| 1 | 56785889 | rs2994553 | -0.795 | 0.051 | -0.628 | 0.049 | -0.003 | -0.005 | -0.005 | 0 | 0.31 | 0.3 | 0.26 | 0.37 | 0.34 | 0.48 |
| 1 | 56786595 | rs3005897 | 0.031 | 0.092 | -1.122 | 0.07 | -0.001 | 0 | 0 | 1.161 | 0.22 | 0.22 | 0.23 | 0.3 | 0.3 | 0.37 |
| 1 | 56790383 | rs3005906 | -0.611 | 0.094 | -1.16 | 0.045 | 0.012 | 0.02 | 0.02 | 0 | 0.67 | 0.44 | 0.43 | 0.45 | 0.64 | 0.66 |
| 1 | 56790773 | rs12138574 | 1.344 | 0.143 | -1.434 | 0.058 | 0.066 | 0.143 | 0.143 | 0 | 0.37 | 0.15 | 0.17 | 0.08 | 0.3 | 0.17 |
| 1 | 56791479 | rs2994547 | 0.263 | 0.268 | -1.309 | 0.316 | -0.013 | 0.002 | 0.002 | 0 | 0.58 | 0.3 | 0.4 | 0.38 | 0.6 | 0.55 |
| 1 | 56794965 | rs3005911 | -0.292 | 0.323 | -1.29 | 0.309 | -0.005 | -0.01 | -0.01 | 4.25 | 0.66 | 0.35 | 0.43 | 0.46 | 0.65 | 0.66 |
| 1 | 56797332 | rs17113759 | -0.101 | 0.643 | -0.325 | 1.004 | 0.019 | 0.006 | 0.006 | 0.052 | 0.09 | 0.03 | 0.03 | 0.07 | 0.04 | 0.12 |
| 1 | 56797558 | rs17113760 | -1.097 | 0.635 | -0.305 | 1.095 | 0.01 | -0.015 | -0.015 | 0 | 0.14 | 0.13 | 0.05 | 0.27 | 0.07 | 0.31 |
| 1 | 56797754 | rs3005914 | -0.352 | 0.834 | -0.447 | 1.051 | -0.016 | 0 | 0 | 0 | 0.9 | 1 | 0.97 | 0.93 | 0.93 | 0.88 |
| 1 | 56798308 | rs2994565 | -0.137 | 1.029 | -0.368 | 1.011 | -0.002 | -0.001 | -0.001 | 0 | 0.18 | 0.19 | 0.25 | 0.19 | 0.27 | 0.2 |
| 1 | 56799912 | rs7556596 | 0.818 | 1.059 | -0.455 | 0.977 | 0.004 | 0.01 | 0.01 | 0 | 0.66 | 0.62 | 0.69 | 0.56 | 0.63 | 0.47 |
| 1 | 56800736 | rs10159099 | 0.632 | 0.497 | -0.33 | 0.984 | -0.024 | 0.109 | 0.109 | 0 | 0.82 | 0.65 | 0.92 | 0.76 | 0.9 | 0.67 |
| 1 | 56801686 | rs2994570 | -0.151 | 1.338 | -1.692 | 0.08 | -0.148 | 0.072 | 0.072 | 0 | 0.29 | 0.03 | 0.34 | 0.13 | 0.41 | 0.22 |
| 1 | 56814694 | rs4912413 | -0.275 | 1.078 | -1.037 | 0.365 | -0.003 | 0.013 | 0.013 | 0 | 0.64 | 0.44 | 0.59 | 0.48 | 0.68 | 0.52 |
| 1 | 56826219 | rs7515200 | -0.328 | 0.911 | -0.137 | 0.285 | -0.001 | -0.002 | -0.002 | 0 | 0.66 | 0.71 | 0.62 | 0.48 | 0.68 | 0.52 |

| | | | | | | | | | | | | | | | | |
|---|---|---|---|---|---|---|---|---|---|---|---|---|---|---|---|---|
| 1 | 56829396 | rs17113792 | 0.327 | 0.864 | 0.053 | -0.043 | 0.003 | -0.03 | -0.03 | 0 | 0.09 | 0.12 | 0.06 | 0 | 0.1 | 0.07 |
| 1 | 56836116 | rs12123862 | -0.376 | 0.839 | 0.106 | -0.046 | -0.003 | 0.024 | 0.024 | 0 | 0.65 | 0.58 | 0.6 | 0.48 | 0.68 | 0.52 |
| 1 | 56844621 | rs4600084 | 0.344 | 0.748 | 0.136 | -0.005 | 0.008 | 0.02 | 0.02 | 0.975 | 0.54 | 0.31 | 0.52 | 0.43 | 0.58 | 0.44 |
| 1 | 56844652 | rs17113799 | 0.256 | 0.712 | 0.172 | -0.004 | 0.002 | -0.022 | -0.022 | 4.779 | 0.09 | 0.11 | 0.06 | 0.03 | 0.1 | 0.06 |
| 1 | 56846350 | rs7553068 | -0.029 | 0.71 | 0.163 | -0.005 | 0.012 | -0.048 | -0.048 | 0 | 0.1 | 0.24 | 0.08 | 0.03 | 0.1 | 0.06 |
| 1 | 56846395 | rs12564524 | -0.068 | 1.421 | -0.263 | -0.1 | 0 | -0.026 | -0.026 | 4.244 | 0.55 | 0.5 | 0.48 | 0.24 | 0.51 | 0.3 |
| 1 | 56848481 | rs9436540 | 0.025 | 1.712 | -0.72 | -0.096 | -0.06 | -0.056 | -0.056 | 0 | 0.11 | 0.01 | 0.12 | 0.16 | 0.18 | 0.21 |
| 1 | 56848954 | rs7538333 | -0.256 | 1.115 | -0.187 | -0.731 | -0.038 | 0.085 | 0.085 | 0.619 | 0.57 | 0.82 | 0.51 | 0.3 | 0.53 | 0.29 |
| 1 | 56849539 | rs6588624 | -0.116 | 1.195 | -0.211 | -0.693 | -0.026 | 0.083 | 0.083 | 0.3 | 0.56 | 0.75 | 0.5 | 0.31 | 0.53 | 0.3 |
| 1 | 56850686 | rs11206801 | -0.597 | 1.04 | 0.155 | -0.15 | 0.04 | 0.002 | 0.002 | 0 | 0.11 | 0.35 | 0.09 | 0.09 | 0.12 | 0.07 |
| 1 | 56852267 | rs4431886 | -0.884 | 0.981 | 0.15 | -0.178 | 0.035 | -0.008 | -0.008 | 0 | 0.13 | 0.45 | 0.09 | 0.09 | 0.13 | 0.07 |
| 1 | 56856989 | rs9437001 | -0.253 | 0.675 | -0.956 | -0.351 | -0.004 | 0.004 | 0.004 | 0 | 0.46 | 0.16 | 0.54 | 0.48 | 0.6 | 0.48 |
| 1 | 56860086 | rs12408513 | 0.654 | 0.419 | -0.518 | -0.998 | -0.01 | 0.05 | 0.05 | 0 | 0.24 | 0.15 | 0.22 | 0.04 | 0.3 | 0.1 |
| 1 | 56860216 | rs1931074 | -0.15 | 0.627 | -0.342 | -0.691 | -0.006 | -0.005 | -0.005 | 0.21 | 0.1 | 0.12 | 0.27 | 0.24 | 0.12 | 0.1 |
| 1 | 56860456 | rs1931073 | -0.136 | 0.631 | -0.34 | -0.734 | -0.005 | -0.005 | -0.005 | 0 | 0.1 | 0.12 | 0.27 | 0.24 | 0.12 | 0.1 |
| 1 | 56867523 | rs3005916 | 0.401 | 0.449 | 0.139 | -0.848 | -0.004 | -0.002 | -0.002 | 1.046 | 0.18 | 0.09 | 0.14 | 0.18 | 0.16 | 0.24 |
| 1 | 56867989 | rs7549365 | 0.966 | 0.585 | 0.077 | -0.876 | -0.022 | 0.109 | 0.109 | 0 | 0.27 | 0.12 | 0.19 | 0 | 0.34 | 0.12 |
| 1 | 56870834 | rs1931078 | -0.185 | 0.618 | 0.26 | -0.716 | -0.018 | 0.066 | 0.066 | 0 | 0.42 | 0.34 | 0.49 | 0.28 | 0.48 | 0.26 |
| 1 | 56872357 | rs2991265 | -0.287 | 0.571 | 0.216 | -0.77 | 0.002 | 0.094 | 0.094 | 0.349 | 0.5 | 0.35 | 0.62 | 0.23 | 0.5 | 0.4 |
| 1 | 56872794 | rs17469007 | 0.926 | 0.637 | -0.389 | -0.755 | 0.001 | 0.015 | 0.015 | 0 | 0.29 | 0.23 | 0.36 | 0 | 0.34 | 0.13 |
| 1 | 56873274 | rs12035812 | 1.489 | 0.594 | -0.415 | -0.766 | -0.003 | 0.013 | 0.013 | 0 | 0.25 | 0.29 | 0.15 | 0.44 | 0.19 | 0.36 |
| 1 | 56873524 | rs3005922 | 0.813 | 0.547 | -0.412 | -0.742 | -0.01 | -0.007 | -0.007 | 0.323 | 0.19 | 0.07 | 0.14 | 0.18 | 0.16 | 0.27 |
| 1 | 56875043 | rs17428956 | 0.727 | 0.63 | -0.471 | -0.665 | -0.013 | -0.013 | -0.013 | 0.36 | 0.09 | 0.09 | 0.12 | 0.23 | 0.12 | 0.08 |
| 1 | 56877509 | rs11206803 | 0.959 | 0.514 | -0.516 | -0.701 | 0.004 | 0 | 0 | 0 | 0.52 | 0.37 | 0.5 | 0.44 | 0.53 | 0.49 |
| 1 | 56881763 | rs884277 | -0.448 | 0.577 | -0.711 | -0.449 | -0.017 | -0.014 | -0.014 | 0.063 | 0.29 | 0.24 | 0.27 | 0.41 | 0.29 | 0.35 |
| 1 | 56885111 | rs12093027 | 0.791 | 0.538 | -0.908 | -0.981 | -0.002 | 0.002 | 0.002 | 0 | 0.34 | 0.36 | 0.23 | 0.66 | 0.31 | 0.45 |
| 1 | 56888166 | rs7367993 | -0.508 | 0.555 | -0.225 | 0.193 | -0.003 | 0.013 | 0.013 | 0 | 0.22 | 0.33 | 0.16 | 0.28 | 0.17 | 0.29 |
| 1 | 56895392 | rs2780859 | 0.494 | 0.437 | -0.234 | -0.125 | -0.008 | -0.068 | -0.068 | 0.216 | 0.81 | 0.74 | 0.73 | 0.94 | 0.81 | 0.86 |
| 1 | 56898998 | rs4363444 | -0.215 | 0.484 | -0.563 | 0.25 | -0.001 | 0.001 | 0.001 | 0 | 0.45 | 0.5 | 0.48 | 0.54 | 0.43 | 0.35 |
| 1 | 56900270 | rs875153 | -0.326 | 0.081 | -0.893 | -0.092 | 0.005 | 0.159 | 0.159 | 0 | 0.5 | 0.81 | 0.63 | 0.27 | 0.45 | 0.31 |
| 1 | 56901453 | rs1953116 | 0.327 | 0.054 | -0.311 | -0.291 | -0.004 | -0.057 | -0.057 | 0.021 | 0.38 | 0.37 | 0.26 | 0.15 | 0.34 | 0.16 |
| 1 | 56903392 | rs7550882 | 0.232 | -0.2 | 0.061 | 0.15 | 0.005 | 0.005 | 0.005 | 0 | 0.51 | 0.4 | 0.39 | 0.51 | 0.43 | 0.4 |
| 1 | 56903435 | rs914836 | -1.55 | -0.176 | 0.541 | -0.123 | 0.039 | -0.014 | -0.014 | 0.474 | 0.08 | 0.33 | 0.22 | 0.02 | 0.08 | 0.07 |
| 1 | 56904162 | rs914838 | -0.622 | 0.186 | -0.258 | -0.435 | -0.021 | 0.058 | 0.058 | 0.071 | 0.88 | 0.98 | 0.94 | 0.97 | 0.89 | 0.91 |
| 1 | 56904480 | rs6701827 | 0.495 | 0.152 | -0.076 | -0.453 | 0.002 | -0.041 | -0.041 | 0 | 0.83 | 0.78 | 0.74 | 0.96 | 0.81 | 0.88 |
| 1 | 56904885 | rs6666317 | 0.93 | 0.144 | -0.073 | -0.451 | 0.007 | -0.033 | -0.033 | 0.446 | 0.8 | 0.75 | 0.69 | 0.9 | 0.76 | 0.8 |
| 1 | 56905072 | rs2780293 | -0.498 | 0.027 | -0.217 | -0.422 | -0.003 | -0.016 | -0.016 | 0 | 0.9 | 0.89 | 0.95 | 0.97 | 0.94 | 0.94 |
| 1 | 56906274 | rs7537229 | 0.001 | 0.053 | -0.371 | -0.393 | -0.006 | 0.038 | 0.038 | 0.07 | 0.88 | 0.96 | 0.94 | 0.97 | 0.89 | 0.92 |
| 1 | 56908494 | rs3892374 | -0.03 | 0.068 | -0.384 | -0.353 | -0.008 | 0.036 | 0.036 | 0.085 | 0.88 | 0.96 | 0.93 | 0.96 | 0.88 | 0.91 |
| 1 | 56909550 | rs6694114 | 0.874 | -0.381 | -0.806 | -0.62 | 0.053 | 0.063 | 0.063 | 0 | 0.43 | 0.72 | 0.33 | 0.34 | 0.31 | 0.22 |
| 1 | 56914341 | rs12736058 | 1.367 | -0.095 | 0.322 | 0.006 | 0.001 | 0.006 | 0.006 | 0 | 0.52 | 0.42 | 0.32 | 0.23 | 0.45 | 0.28 |
| 1 | 56914815 | rs12137498 | -0.321 | -0.057 | 0.285 | -0.071 | 0 | -0.005 | -0.005 | 0 | 0.06 | 0.06 | 0.2 | 0 | 0.09 | 0.04 |
| 1 | 56915820 | rs1889145 | 1.162 | 0.288 | 0.294 | -0.072 | -0.003 | -0.008 | -0.008 | 0 | 0.8 | 0.63 | 0.68 | 0.77 | 0.78 | 0.85 |
| 1 | 56917952 | rs11206813 | 1.453 | 0.37 | 0.274 | -0.144 | -0.001 | -0.001 | -0.001 | 0 | 0.54 | 0.5 | 0.36 | 0.42 | 0.46 | 0.31 |
| 1 | 56922122 | rs11206815 | 0.974 | 0.378 | 0.328 | -0.122 | 0 | -0.007 | -0.007 | 0 | 0.82 | 0.81 | 0.72 | 0.96 | 0.78 | 0.88 |
| 1 | 56929525 | rs7368386 | 1.105 | 0.455 | 0.348 | -0.061 | 0.001 | -0.058 | -0.058 | 0 | 0.81 | 0.7 | 0.71 | 0.96 | 0.78 | 0.88 |
| 1 | 56932035 | rs11206816 | -0.207 | 0.419 | 0.303 | -0.069 | -0.006 | -0.018 | -0.018 | 0 | 0.27 | 0.23 | 0.34 | 0.47 | 0.29 | 0.46 |
| 1 | 56933544 | rs7418187 | 1.21 | 0.412 | 0.293 | -0.062 | -0.019 | 0.007 | 0.007 | 0 | 0.57 | 0.66 | 0.39 | 0.49 | 0.5 | 0.42 |
| 1 | 56938218 | rs6588634 | -0.115 | 0.416 | 0.273 | -0.067 | 0 | 0.002 | 0.002 | 0 | 0.88 | 0.88 | 0.92 | 0.96 | 0.87 | 0.93 |
| 1 | 56941387 | rs4912291 | -0.618 | 0.469 | 0.285 | 0.077 | -0.026 | -0.046 | -0.046 | 0.001 | 0.3 | 0.2 | 0.52 | 0.47 | 0.37 | 0.51 |
| 1 | 56955386 | rs4638151 | 0.333 | 0.43 | 0.355 | 0.039 | 0.008 | 0.013 | 0.013 | 0.066 | 0.7 | 0.6 | 0.58 | 0.45 | 0.67 | 0.45 |
| 1 | 56960123 | rs11206830 | -0.136 | 0.193 | 0.288 | 0.443 | -0.025 | 0.25 | 0.25 | 0 | 0.06 | 0.17 | 0.17 | 0.43 | 0.03 | 0.22 |
| 1 | 56961354 | rs10888976 | 0.162 | 0.191 | 0.293 | 0.445 | -0.027 | 0.28 | 0.28 | 0 | 0.93 | 0.81 | 0.83 | 0.57 | 0.97 | 0.77 |
| 1 | 56961756 | rs4628 | 1.649 | 0.383 | 0.292 | 0.337 | 0.001 | -0.005 | -0.005 | 6.598 | 0.55 | 0.45 | 0.35 | 0.42 | 0.48 | 0.33 |
| 1 | 56965001 | rs11206833 | 1.877 | 0.523 | 0.091 | 0.199 | -0.058 | 0.154 | 0.154 | 0.029 | 0.53 | 0.12 | 0.47 | 0.7 | 0.5 | 0.5 |
| 1 | 56966350 | rs17114046 | -0.054 | 0.684 | 0.011 | 0.253 | -0.001 | -0.041 | -0.041 | 0 | 0.1 | 0.2 | 0.08 | 0.02 | 0.12 | 0.06 |
| 1 | 56968477 | rs4912314 | -2.278 | 0.736 | 0.38 | 0.229 | -0.013 | -0.022 | -0.022 | 0.012 | 0.26 | 0.65 | 0.35 | 0.24 | 0.25 | 0.27 |
| 1 | 56968755 | rs7528118 | 0.051 | 0.999 | -0.03 | 0.271 | -0.101 | 0.068 | 0.068 | 0.008 | 0.19 | 0.01 | 0.16 | 0.06 | 0.25 | 0.23 |
| 1 | 56974091 | rs1319274 | -0.58 | 1.611 | 0.018 | 0.198 | -0.007 | 0.007 | 0.007 | 0 | 0.19 | 0.13 | 0.13 | 0.03 | 0.26 | 0.12 |
| 1 | 56974816 | rs11576602 | -1.542 | 1.654 | 0.029 | 0.186 | -0.025 | 0.026 | 0.026 | 0 | 0.28 | 0.22 | 0.28 | 0.06 | 0.38 | 0.29 |
| 1 | 56977819 | rs3738570 | 0.12 | 1.651 | -0.02 | 0.221 | -0.045 | 0.006 | 0.006 | 1.408 | 0.07 | 0.01 | 0.13 | 0.03 | 0.13 | 0.18 |
| 1 | 56979076 | rs2404716 | -1.766 | 1.634 | 0.075 | 0.208 | 0.012 | -0.02 | -0.02 | 0.441 | 0.19 | 0.21 | 0.22 | 0.08 | 0.27 | 0.35 |
| 1 | 56981625 | rs12081589 | -1.331 | 1.656 | 0.183 | 0.254 | -0.017 | 0.015 | 0.015 | 0.05 | 0.13 | 0.37 | 0.2 | 0.18 | 0.09 | 0.12 |
| 1 | 56991040 | rs11206835 | -1.095 | 1.759 | 0.091 | -0.09 | -0.03 | 0.029 | 0.029 | 0.368 | 0.12 | 0.07 | 0.3 | 0.03 | 0.19 | 0.25 |
| 1 | 56991294 | rs6674131 | 0.85 | 1.506 | 0.025 | -0.068 | -0.044 | 0.065 | 0.065 | 0 | 0.11 | 0 | 0.07 | 0 | 0.14 | 0.06 |
| 1 | 57007539 | rs1930760 | -2.322 | 1.554 | 0.14 | 0.409 | 0.011 | 0.021 | 0.021 | 0.752 | 0.32 | 0.57 | 0.55 | 0.54 | 0.36 | 0.55 |
| 1 | 57012791 | rs11206838 | -1.192 | 1.583 | 0.118 | 0.375 | 0.01 | -0.026 | -0.026 | 0 | 0.21 | 0.23 | 0.36 | 0.06 | 0.31 | 0.31 |
| 1 | 57013198 | rs1041602 | 0.901 | 1.595 | 0.075 | 0.451 | -0.044 | 0.066 | 0.066 | 0 | 0.11 | 0 | 0.07 | 0 | 0.14 | 0.06 |
| 1 | 57016862 | rs11206841 | -0.698 | 1.579 | 0.147 | 0.536 | -0.026 | 0.29 | 0.29 | 0.32 | 0.11 | 0.34 | 0.19 | 0.48 | 0.05 | 0.24 |

| | | | | | | | | | | | | | | | | |
|---|---|---|---|---|---|---|---|---|---|---|---|---|---|---|---|---|
| 1 | 57016912 | rs10493211 | -1.561 | 1.688 | -0.115 | 0.435 | 0.001 | -0.012 | -0.012 | 0 | 0.23 | 0.25 | 0.38 | 0.09 | 0.34 | 0.43 |
| 1 | 57017739 | rs1261411 | -1.163 | 1.153 | -0.319 | 0.702 | -0.001 | 0 | 0 | 2.356 | 0.41 | 0.61 | 0.49 | 0.54 | 0.45 | 0.64 |
| 1 | 57020903 | rs17416299 | -1.19 | 0.784 | -0.275 | 0.634 | -0.001 | -0.004 | -0.004 | 0 | 0.11 | 0.09 | 0.2 | 0.03 | 0.14 | 0.13 |
| 1 | 57022282 | rs7553307 | 0.516 | 0.492 | -0.007 | 1.066 | -0.003 | 0.01 | 0.01 | 0 | 0.88 | 0.8 | 0.89 | 0.78 | 0.89 | 0.61 |
| 1 | 57024560 | rs1777283 | 1.419 | 0.258 | 0.591 | 1.705 | -0.013 | 0.105 | 0.105 | 0 | 0.83 | 0.64 | 0.85 | 0.62 | 0.87 | 0.5 |
| 1 | 57036017 | rs2065553 | 0.053 | 0.199 | 0.598 | 1.671 | -0.003 | -0.006 | -0.006 | 0.249 | 0.91 | 0.84 | 0.9 | 0.88 | 0.92 | 0.73 |
| 1 | 57042335 | rs933001 | 1.131 | 0.094 | 0.101 | 1.603 | -0.025 | 0.097 | 0.097 | 0 | 0.5 | 0.15 | 0.6 | 0.3 | 0.45 | 0.3 |
| 1 | 57042954 | rs7539051 | -0.872 | 0.129 | 0.297 | 1.292 | -0.021 | 0.122 | 0.122 | 0 | 0.07 | 0.16 | 0.05 | 0.25 | 0.03 | 0.21 |
| 1 | 57049977 | rs4475779 | 1.16 | -0.238 | 0.7 | 1.178 | -0.05 | 0.081 | 0.081 | 0 | 0.8 | 0.56 | 0.84 | 0.6 | 0.88 | 0.48 |
| 1 | 57057316 | rs17408450 | 0.669 | -0.124 | 0.543 | 1.195 | -0.044 | 0.032 | 0.032 | 0 | 0.24 | 0.11 | 0.2 | 0.15 | 0.38 | 0.15 |
| 1 | 57066988 | rs7521034 | -1.068 | -0.094 | 0.56 | 1.425 | 0.009 | -0.006 | -0.006 | 0.071 | 0.1 | 0.11 | 0.05 | 0.26 | 0.03 | 0.24 |
| 1 | 57068107 | rs6663578 | -0.521 | -0.235 | 0.612 | 1.059 | 0.004 | 0.015 | 0.015 | 0 | 0.43 | 0.38 | 0.3 | 0.32 | 0.46 | 0.38 |
| 1 | 57081328 | rs12137489 | -0.444 | -0.689 | 1.152 | 1.216 | -0.042 | 0.003 | 0.003 | 1.173 | 0.09 | 0.23 | 0.04 | 0.07 | 0.02 | 0.13 |
| 1 | 57088605 | rs2404987 | 0.313 | -0.627 | 0.777 | 1.03 | -0.037 | 0.032 | 0.032 | 0 | 0.39 | 0.21 | 0.29 | 0.26 | 0.48 | 0.28 |
| 1 | 57093183 | rs6679902 | -1.25 | -0.566 | 0.954 | 1.054 | -0.055 | -0.014 | -0.014 | 0.103 | 0.13 | 0.38 | 0.06 | 0.09 | 0.05 | 0.16 |
| 1 | 57095904 | rs2404990 | 1.225 | -0.659 | 1.075 | 1.239 | -0.057 | 0.034 | 0.034 | 0.882 | 0.78 | 0.37 | 0.83 | 0.65 | 0.87 | 0.55 |
| 1 | 57100380 | rs10889000 | -0.529 | -0.613 | 0.97 | 1.258 | -0.01 | 0.08 | 0.08 | 0 | 0.9 | 0.76 | 0.89 | 0.74 | 0.92 | 0.71 |
| 1 | 57103219 | rs11206883 | 0.861 | -0.628 | 1.152 | 1.25 | -0.036 | 0.02 | 0.02 | 0.239 | 0.81 | 0.73 | 0.86 | 0.68 | 0.87 | 0.56 |
| 1 | 57103362 | rs11206884 | 0.572 | -0.576 | 1.122 | 1.17 | 0.005 | -0.015 | -0.015 | 0 | 0.36 | 0.41 | 0.28 | 0.16 | 0.46 | 0.18 |
| 1 | 57103965 | rs2404993 | 0.567 | -0.612 | 1.123 | 1.171 | 0.005 | -0.015 | -0.015 | 0 | 0.36 | 0.41 | 0.29 | 0.16 | 0.46 | 0.18 |
| 1 | 57108423 | rs6588640 | -0.659 | -0.392 | 1.296 | 1.372 | -0.006 | 0.026 | 0.026 | 0 | 0.91 | 0.88 | 0.9 | 0.77 | 0.92 | 0.71 |
| 1 | 57109660 | rs4584419 | 0.182 | -0.604 | 1.134 | 1.177 | -0.005 | 0.021 | 0.021 | 0 | 0.44 | 0.42 | 0.31 | 0.16 | 0.53 | 0.22 |
| 1 | 57114671 | rs10749709 | 0.975 | -0.543 | 0.927 | 1.24 | -0.006 | -0.012 | -0.012 | 0.779 | 0.35 | 0.29 | 0.54 | 0.42 | 0.34 | 0.24 |
| 1 | 57116909 | rs11206887 | 0.469 | -0.557 | 0.962 | 1.242 | -0.023 | 0.045 | 0.045 | 0.072 | 0.79 | 0.73 | 0.83 | 0.58 | 0.84 | 0.45 |
| 1 | 57117248 | rs11206889 | 0.238 | -0.559 | 0.968 | 1.237 | -0.004 | 0.026 | 0.026 | 0 | 0.44 | 0.41 | 0.3 | 0.16 | 0.5 | 0.21 |
| 1 | 57121883 | rs4912408 | 0.929 | -0.448 | 0.783 | 1.261 | -0.006 | -0.012 | -0.012 | 0.578 | 0.35 | 0.29 | 0.54 | 0.42 | 0.34 | 0.24 |
| 1 | 57126919 | rs2746347 | 0.013 | -0.445 | 0.798 | 1.282 | 0.014 | 0.077 | 0.077 | 0 | 0.11 | 0.27 | 0.13 | 0.26 | 0.11 | 0.3 |
| 1 | 57139370 | rs2746339 | 0.225 | -0.445 | 0.687 | 1.22 | 0 | -0.001 | -0.001 | 0 | 0.45 | 0.44 | 0.65 | 0.65 | 0.46 | 0.54 |
| 1 | 57159136 | rs1418442 | 0.527 | -0.428 | 0.339 | 1.071 | -0.019 | 0.065 | 0.065 | 0 | 0.39 | 0.23 | 0.27 | 0.16 | 0.46 | 0.21 |
| 1 | 57170229 | rs932447 | 0.519 | -0.363 | 0.053 | 1.097 | -0.008 | 0.036 | 0.036 | 0 | 0.41 | 0.33 | 0.28 | 0.16 | 0.46 | 0.21 |
| 1 | 57175158 | rs857148 | -0.236 | 0.37 | -0.695 | 1.292 | -0.014 | -0.001 | -0.001 | 0.001 | 0.43 | 0.29 | 0.63 | 0.61 | 0.47 | 0.53 |
| 1 | 57177244 | rs4912411 | 0.504 | 0.406 | -0.807 | 1.335 | -0.024 | -0.027 | -0.027 | 0.03 | 0.33 | 0.16 | 0.52 | 0.38 | 0.35 | 0.23 |
| 1 | 57177789 | rs857147 | -0.081 | 0.253 | -0.82 | 1.022 | -0.004 | -0.001 | -0.001 | 0 | 0.43 | 0.37 | 0.64 | 0.61 | 0.46 | 0.52 |
| 1 | 57178926 | rs3738567 | -0.666 | -0.152 | -0.862 | 1.139 | 0.041 | -0.015 | -0.015 | 0 | 0.42 | 0.75 | 0.35 | 0.54 | 0.27 | 0.64 |
| 1 | 57180867 | rs706466 | -0.215 | 0.49 | -0.554 | 1.17 | 0.001 | -0.001 | -0.001 | 0 | 0.49 | 0.33 | 0.63 | 0.48 | 0.48 | 0.48 |
| 1 | 57181382 | rs706468 | -1.117 | 0.532 | -1.432 | 1.047 | -0.066 | 0.007 | 0.007 | 0.007 | 0.74 | 0.98 | 0.77 | 0.78 | 0.67 | 0.77 |
| 1 | 57191659 | rs11206908 | 0.49 | 0.675 | -1.365 | 1.566 | -0.002 | 0.029 | 0.029 | 0.414 | 0.34 | 0.07 | 0.52 | 0.23 | 0.31 | 0.3 |
| 1 | 57191924 | rs857138 | -0.377 | 0.815 | -0.573 | 2.106 | -0.006 | -0.017 | -0.017 | 0 | 0.14 | 0.09 | 0.09 | 0.24 | 0.15 | 0.18 |
| 1 | 57192848 | rs10489618 | -0.669 | 0.612 | -0.087 | 1.81 | 0.035 | 0.258 | 0.258 | 0 | 0.09 | 0.38 | 0.06 | 0.3 | 0.05 | 0.19 |
| 1 | 57201871 | rs1832899 | -0.068 | 0.3 | -1.4 | 1.168 | 0.004 | -0.035 | -0.035 | 1.486 | 0.17 | 0.01 | 0.11 | 0.17 | 0.13 | 0.22 |
| 1 | 57203011 | rs6658790 | 0.227 | 0.226 | -0.835 | 0.17 | -0.003 | -0.161 | -0.161 | 0 | 0.22 | 0.41 | 0.12 | 0 | 0.18 | 0.06 |
| 1 | 57203501 | rs1935562 | 0.2 | 0.589 | -0.793 | 0.162 | 0.02 | -0.157 | -0.157 | 0 | 0.22 | 0.43 | 0.13 | 0 | 0.2 | 0.06 |
| 1 | 57205595 | rs857124 | -1.895 | 0.985 | -1.758 | -0.141 | -0.094 | 0.034 | 0.034 | 0.007 | 0.75 | 0.98 | 0.75 | 0.88 | 0.66 | 0.78 |
| 1 | 57207074 | rs1935555 | -0.268 | 0.885 | -1.668 | 0.124 | 0 | -0.025 | -0.025 | 0 | 0.34 | 0.52 | 0.33 | 0.24 | 0.32 | 0.15 |
| 1 | 57207221 | rs10889014 | 1.242 | 0.931 | -1.675 | 0.294 | -0.009 | -0.016 | -0.016 | 0 | 0.12 | 0.08 | 0.21 | 0.23 | 0.14 | 0.09 |
| 1 | 57209695 | rs1156845 | 1.464 | 0.995 | -1.577 | 0.881 | -0.005 | -0.032 | -0.032 | 0 | 0.11 | 0.05 | 0.24 | 0.2 | 0.12 | 0.14 |
| 1 | 57211399 | rs857120 | -0.301 | 0.894 | -1.159 | 1.049 | 0.007 | 0.058 | 0.058 | 0.435 | 0.13 | 0.35 | 0.07 | 0.23 | 0.09 | 0.27 |
| 1 | 57211976 | rs857119 | -0.792 | 0.836 | -1.13 | 0.973 | -0.006 | -0.002 | -0.002 | 0 | 0.46 | 0.49 | 0.45 | 0.36 | 0.35 | 0.46 |
| 1 | 57215026 | rs6666850 | 0.544 | 0.801 | -0.742 | 0.769 | 0.046 | -0.03 | -0.03 | 0 | 0.24 | 0.43 | 0.28 | 0.23 | 0.27 | 0.2 |
| 1 | 57216376 | rs6696458 | 0.206 | 0.64 | -0.907 | 0.947 | -0.005 | 0.092 | 0.092 | 0.106 | 0.18 | 0.3 | 0.28 | 0.41 | 0.16 | 0.23 |
| 1 | 57217116 | rs857111 | -0.978 | 0.572 | -0.992 | 0.848 | -0.01 | 0.011 | 0.011 | 0 | 0.58 | 0.66 | 0.48 | 0.41 | 0.5 | 0.53 |
| 1 | 57221878 | rs2298127 | 0.552 | 0.354 | -0.766 | 0.943 | -0.008 | -0.021 | -0.021 | 0 | 0.16 | 0.12 | 0.41 | 0.42 | 0.18 | 0.22 |
| 1 | 57222599 | rs857107 | -0.857 | 0.538 | -0.786 | 0.86 | 0.001 | -0.001 | -0.001 | 0.779 | 0.39 | 0.42 | 0.23 | 0.41 | 0.29 | 0.43 |
| 1 | 57231991 | rs12410615 | 0.295 | 0.627 | -0.515 | 1.304 | -0.009 | -0.032 | -0.032 | 0 | 0.16 | 0.09 | 0.41 | 0.42 | 0.18 | 0.28 |
| 1 | 57234065 | rs1935561 | -0.616 | 0.534 | -0.17 | 1.346 | 0.037 | 0.222 | 0.222 | 0.438 | 0.05 | 0.23 | 0.04 | 0.23 | 0.04 | 0.06 |
| 1 | 57238019 | rs857163 | 0.136 | 0.16 | -0.773 | 0.736 | 0.022 | 0.017 | 0.017 | 0.245 | 0.17 | 0.15 | 0.06 | 0.09 | 0.07 | 0.04 |
| 1 | 57241234 | rs1557145 | -0.777 | 0.671 | 0.034 | 1.383 | 0.069 | 0.369 | 0.369 | 1.166 | 0.1 | 0.32 | 0.21 | 0.47 | 0.09 | 0.2 |
| 1 | 57241867 | rs857160 | -1.846 | 0.948 | -0.969 | 0.756 | -0.053 | 0.123 | 0.123 | 0.095 | 0.77 | 0.99 | 0.79 | 0.96 | 0.71 | 0.79 |
| 1 | 57246662 | rs1774822 | -0.069 | 0.898 | -0.687 | 0.834 | -0.011 | -0.014 | -0.014 | 0 | 0.57 | 0.38 | 0.49 | 0.44 | 0.53 | 0.41 |
| 1 | 57246968 | rs1569746 | 0.583 | 0.721 | -1.112 | 0.799 | -0.04 | 0.104 | 0.104 | 0.461 | 0.37 | 0.09 | 0.36 | 0.16 | 0.37 | 0.2 |
| 1 | 57249558 | rs1774825 | 0.162 | 0.831 | -0.555 | 1.457 | 0.001 | -0.005 | -0.005 | 0 | 0.19 | 0.23 | 0.13 | 0.26 | 0.17 | 0.22 |
| 1 | 57252494 | rs6658389 | -1.079 | 0.937 | -0.303 | 1.777 | 0.003 | 0.138 | 0.138 | 0 | 0.16 | 0.24 | 0.23 | 0.49 | 0.12 | 0.21 |
| 1 | 57257817 | rs2298130 | -0.603 | 0.819 | -0.878 | 1.419 | 0 | 0.105 | 0.105 | 0 | 0.17 | 0.36 | 0.16 | 0.35 | 0.14 | 0.17 |
| 1 | 57267946 | rs1361730 | 1.003 | 0.713 | -2.358 | 0.185 | -0.046 | 0.069 | 0.069 | 0 | 0.23 | 0.01 | 0.16 | 0.09 | 0.26 | 0.26 |
| 1 | 57269521 | rs6662617 | 0.041 | 1.398 | -2.079 | 0.149 | -0.06 | 0.13 | 0.13 | 0 | 0.39 | 0.14 | 0.28 | 0.15 | 0.45 | 0.36 |
| 1 | 57270093 | rs17114382 | -0.319 | 1.654 | -1.508 | 1.287 | -0.019 | -0.065 | -0.065 | 0 | 0.16 | 0.05 | 0.23 | 0.31 | 0.15 | 0.15 |
| 1 | 57273017 | rs12408283 | 0.038 | 1.816 | -1.538 | 1.118 | -0.035 | 0.011 | 0.011 | 0 | 0.08 | 0.02 | 0.06 | 0.05 | 0.12 | 0.07 |
| 1 | 57273172 | rs947639 | 0.618 | 1.435 | -0.94 | 1.304 | 0 | 0.001 | 0.001 | 0 | 0.77 | 0.8 | 0.72 | 0.68 | 0.78 | 0.81 |
| 1 | 57275487 | rs10889018 | -1.042 | 1.133 | -0.599 | 1.818 | 0.018 | 0.243 | 0.243 | 0 | 0.08 | 0.22 | 0.2 | 0.35 | 0.07 | 0.15 |

| | | | | | | | | | | | | | | | | |
|---|---|---|---|---|---|---|---|---|---|---|---|---|---|---|---|---|
| 1 | 57275564 | rs10749710 | 0.451 | 1.244 | -1.363 | 1.437 | -0.025 | -0.11 | -0.11 | 0 | 0.79 | 0.93 | 0.84 | 0.56 | 0.84 | 0.69 |
| 1 | 57278215 | rs1935559 | -0.517 | 0.984 | -1.317 | 0.351 | 0.031 | 0.068 | 0.068 | 5.779 | 0.47 | 0.34 | 0.64 | 0.75 | 0.56 | 0.74 |
| 1 | 57282318 | rs6673223 | 0.01 | 0.488 | -1.485 | 0.069 | 0.015 | 0.027 | 0.027 | 0 | 0.31 | 0.19 | 0.25 | 0.18 | 0.27 | 0.18 |
| 1 | 57283181 | rs11206923 | -0.199 | 0.863 | -0.864 | -0.039 | 0.085 | 0.047 | 0.047 | 1.27 | 0.08 | 0.31 | 0.1 | 0.06 | 0.12 | 0.02 |
| 1 | 57286856 | rs6672416 | 0.829 | 1.028 | -1.597 | -0.345 | -0.034 | 0.049 | 0.049 | 0.009 | 0.11 | 0.01 | 0.09 | 0.03 | 0.16 | 0.05 |
| 1 | 57290659 | rs1418440 | 0.496 | 0.925 | -1.329 | -0.001 | -0.006 | 0.004 | 0.004 | 0 | 0.43 | 0.26 | 0.46 | 0.56 | 0.5 | 0.58 |
| 1 | 57291520 | rs12048846 | 0.922 | 0.915 | -1.97 | 0.039 | -0.07 | -0.098 | -0.098 | 0 | 0.37 | 0.07 | 0.37 | 0.5 | 0.38 | 0.56 |
| 1 | 57291548 | rs947636 | -0.133 | 0.729 | -1.378 | -0.031 | 0.021 | 0.046 | 0.046 | 0.011 | 0.31 | 0.16 | 0.25 | 0.18 | 0.27 | 0.19 |
| 1 | 57297388 | rs7544695 | -0.955 | 0.916 | -1.313 | 0.116 | -0.021 | 0.015 | 0.015 | 0.048 | 0.09 | 0.01 | 0.05 | 0.03 | 0.1 | 0.1 |
| 1 | 57301685 | rs12032625 | -0.28 | 0.816 | -0.988 | 0.386 | -0.001 | -0.001 | -0.001 | 0.056 | 0.23 | 0.23 | 0.2 | 0.16 | 0.22 | 0.23 |
| 1 | 57304135 | rs761964 | 0.483 | 1.532 | -0.89 | 0.962 | 0.004 | 0.043 | 0.043 | 0 | 0.89 | 0.9 | 0.6 | 0.31 | 0.85 | 0.71 |
| 1 | 57309137 | rs737189 | -0.333 | 1.272 | -0.892 | -0.235 | -0.001 | -0.034 | -0.034 | 0 | 0.22 | 0.14 | 0.14 | 0 | 0.21 | 0.05 |
| 1 | 57309227 | rs706469 | -1.358 | 1.19 | -0.99 | 0.431 | -0.009 | -0.02 | -0.02 | 0 | 0.4 | 0.79 | 0.52 | 0.4 | 0.39 | 0.38 |
| 1 | 57314696 | rs1033637 | 0.606 | 1.343 | -0.998 | -0.121 | -0.014 | 0.163 | 0.163 | 0 | 0.53 | 0.2 | 0.56 | 0.88 | 0.49 | 0.81 |
| 1 | 57316268 | rs1033729 | 0.863 | 1.342 | -0.98 | 0.05 | 0.008 | 0.171 | 0.171 | 0.23 | 0.5 | 0.23 | 0.54 | 0.84 | 0.48 | 0.66 |
| 1 | 57317559 | rs10489623 | 0.116 | 1.376 | -1.05 | 0.522 | -0.013 | 0.049 | 0.049 | 0 | 0.06 | 0.03 | 0.29 | 0.38 | 0.14 | 0.18 |
| 1 | 57318178 | rs647962 | 0.492 | 1.626 | -0.435 | 0.261 | -0.005 | 0.069 | 0.069 | 0.126 | 0.54 | 0.31 | 0.56 | 0.92 | 0.54 | 0.82 |
| 1 | 57324512 | rs662377 | 0.073 | 1.468 | 0.364 | 0.176 | -0.021 | 0.057 | 0.057 | 0.195 | 0.61 | 0.8 | 0.62 | 0.92 | 0.56 | 0.83 |
| 1 | 57333759 | rs601662 | 0.518 | 1.43 | 0.373 | 0.157 | -0.007 | 0.045 | 0.045 | 0.396 | 0.56 | 0.38 | 0.59 | 0.92 | 0.56 | 0.84 |
| 1 | 57340381 | rs1885002 | -0.63 | 1.473 | 0.273 | 0.338 | 0 | 0.054 | 0.054 | 0.019 | 0.08 | 0.12 | 0.31 | 0.34 | 0.15 | 0.19 |
| 1 | 57340727 | rs652785 | 1.456 | 1.488 | 0.143 | 0.275 | 0.044 | -0.008 | -0.008 | 0 | 0.44 | 0.18 | 0.25 | 0.5 | 0.34 | 0.48 |
| 1 | 57347939 | rs706476 | -0.024 | 1.435 | 0.292 | 0.247 | -0.07 | 0.17 | 0.17 | 0.232 | 0.61 | 0.9 | 0.63 | 0.92 | 0.56 | 0.83 |
| 1 | 57349614 | rs10489624 | 1.359 | 1.348 | 0.238 | 0.3 | 0.041 | -0.008 | -0.008 | 0 | 0.44 | 0.18 | 0.25 | 0.5 | 0.34 | 0.48 |
| 1 | 57359928 | rs6696924 | 1.282 | 0.962 | 0.309 | -0.093 | 0.005 | -0.004 | -0.004 | 0 | 0.35 | 0.3 | 0.29 | 0.48 | 0.24 | 0.48 |
| 1 | 57360450 | rs6699859 | -0.627 | 1.62 | -0.016 | -0.414 | -0.022 | 0.172 | 0.172 | 0.91 | 0.67 | 0.91 | 0.66 | 1 | 0.65 | 0.9 |
| 1 | 57362787 | rs3768215 | 0.105 | 1.722 | 0.262 | -0.302 | 0.015 | 0.147 | 0.147 | 2.33 | 0.07 | 0.12 | 0.31 | 0.35 | 0.14 | 0.16 |
| 1 | 57364041 | rs1360151 | 0.6 | 1.793 | -0.361 | -0.271 | 0.018 | 0 | 0 | 0 | 0.08 | 0.06 | 0.05 | 0.08 | 0.14 | 0.08 |
| 1 | 57364812 | rs679350 | 1.608 | 1.425 | -0.641 | -0.305 | 0.021 | -0.066 | -0.066 | 0 | 0.33 | 0.08 | 0.18 | 0.48 | 0.23 | 0.48 |
| 1 | 57371133 | rs12116668 | 1.492 | 1.417 | -0.67 | -0.323 | 0.005 | 0.007 | 0.007 | 0 | 0.12 | 0.07 | 0.07 | 0.08 | 0.12 | 0.02 |
| 1 | 57372591 | rs619545 | -0.454 | 1.47 | -0.881 | -0.451 | -0.01 | 0.209 | 0.209 | 0.79 | 0.68 | 0.96 | 0.65 | 1 | 0.71 | 0.91 |
| 1 | 57372917 | rs1411009 | 0.616 | 1.453 | -0.57 | 0.168 | -0.024 | -0.072 | -0.072 | 2.423 | 0.15 | 0.05 | 0.34 | 0.39 | 0.22 | 0.21 |
| 1 | 57374534 | rs624298 | -0.409 | 1.842 | 0.016 | -0.264 | 0.027 | -0.017 | -0.017 | 0 | 0.82 | 0.6 | 0.85 | 0.87 | 0.76 | 0.93 |
| 1 | 57377172 | rs638919 | 1.459 | 1.721 | -0.132 | -0.218 | 0.012 | -0.037 | -0.037 | 0 | 0.35 | 0.2 | 0.34 | 0.47 | 0.27 | 0.47 |
| 1 | 57379401 | rs11206934 | 1.143 | 1.819 | -0.267 | -0.172 | -0.06 | -0.099 | -0.099 | 0.053 | 0.15 | 0.01 | 0.26 | 0.35 | 0.21 | 0.18 |
| 1 | 57379639 | rs1754538 | -0.368 | 1.65 | 0.016 | -0.298 | -0.023 | -0.017 | -0.017 | 0 | 0.1 | 0.26 | 0.13 | 0.04 | 0.05 | 0.16 |
| 1 | 57383358 | rs17300936 | 1.523 | 1.845 | 0.011 | -0.322 | -0.014 | -0.013 | -0.013 | 0 | 0.05 | 0.02 | 0.04 | 0.06 | 0.12 | 0.07 |
| 1 | 57383683 | rs1774889 | -0.394 | 1.049 | 0.069 | -0.365 | -0.023 | -0.017 | -0.017 | 0 | 0.1 | 0.26 | 0.13 | 0.04 | 0.05 | 0.16 |
| 1 | 57389872 | rs2149735 | -1.105 | 1.166 | 0.302 | -0.493 | -0.002 | -0.002 | -0.002 | 0 | 0.48 | 0.49 | 0.47 | 0.41 | 0.57 | 0.47 |
| 1 | 57395251 | rs605648 | -1.65 | 1.118 | 0.332 | -0.553 | -0.003 | -0.029 | -0.029 | 0.838 | 0.87 | 0.78 | 0.89 | 0.93 | 0.86 | 0.97 |
| 1 | 57396014 | rs591730 | 1.585 | 1.12 | -0.255 | -0.546 | 0.021 | -0.055 | -0.055 | 0 | 0.37 | 0.19 | 0.41 | 0.52 | 0.29 | 0.5 |
| 1 | 57399622 | rs652553 | -1.769 | 1.125 | -0.221 | -0.543 | 0.002 | 0.013 | 0.013 | 0.017 | 0.87 | 0.92 | 0.9 | 0.93 | 0.86 | 0.97 |
| 1 | 57400984 | rs17301153 | 0.615 | 1.325 | -0.54 | -0.554 | 0.008 | 0.035 | 0.035 | 0 | 0.2 | 0.1 | 0.11 | 0 | 0.2 | 0.09 |
| 1 | 57405721 | rs725330 | -0.274 | 0.767 | -0.367 | -0.301 | -0.016 | -0.003 | -0.003 | 1.248 | 0.42 | 0.27 | 0.42 | 0.4 | 0.54 | 0.35 |
| 1 | 57407484 | rs626457 | 0.637 | 0.946 | -0.713 | 0.077 | 0.02 | -0.056 | -0.056 | 0.01 | 0.37 | 0.19 | 0.41 | 0.52 | 0.29 | 0.5 |
| 1 | 57413768 | rs582317 | -0.653 | 0.968 | -0.7 | -0.038 | 0.016 | -0.03 | -0.03 | 0 | 0.64 | 0.75 | 0.53 | 0.6 | 0.67 | 0.77 |
| 1 | 57416335 | rs658285 | 0.331 | 1.594 | -0.894 | 0.113 | 0.002 | -0.019 | -0.019 | 0 | 0.33 | 0.19 | 0.3 | 0.39 | 0.3 | 0.55 |
| 1 | 57417592 | rs1960384 | -0.202 | 1.681 | -0.85 | 0.095 | 0.045 | -0.01 | -0.01 | 0 | 0.14 | 0.17 | 0.14 | 0.12 | 0.27 | 0.1 |
| 1 | 57421166 | rs620092 | -0.373 | 1.096 | -0.902 | 0.216 | 0 | 0 | 0 | 0 | 0.48 | 0.48 | 0.46 | 0.51 | 0.57 | 0.69 |
| 1 | 57428381 | rs637253 | 0.041 | 0.714 | -1.201 | 0.478 | 0 | -0.011 | -0.011 | 0 | 0.32 | 0.25 | 0.31 | 0.39 | 0.3 | 0.52 |
| 1 | 57428866 | rs600876 | -0.213 | 0.295 | -1.412 | 0.126 | 0.027 | -0.014 | -0.014 | 0.77 | 0.39 | 0.19 | 0.32 | 0.4 | 0.33 | 0.61 |
| 1 | 57430805 | rs2236217 | 0.07 | -0.041 | -1.092 | 0.403 | 0.004 | -0.004 | -0.004 | 0.005 | 0.18 | 0.18 | 0.31 | 0.37 | 0.1 | 0.22 |
| 1 | 57433095 | rs1984266 | -0.845 | 0.392 | -1.66 | -0.245 | 0.004 | 0.036 | 0.036 | 0 | 0.52 | 0.81 | 0.58 | 0.59 | 0.51 | 0.43 |
| 1 | 57438919 | rs1754511 | -0.729 | 0.354 | -0.492 | 0.284 | -0.002 | 0.109 | 0.109 | 0 | 0.23 | 0.41 | 0.36 | 0.48 | 0.17 | 0.29 |
| 1 | 57440242 | rs6668180 | -0.352 | 0.265 | -0.311 | 0.394 | -0.022 | 0.069 | 0.069 | 0 | 0.18 | 0.32 | 0.32 | 0.32 | 0.11 | 0.21 |
| 1 | 57441363 | rs1623113 | -1.444 | 0.136 | 0.265 | 0.305 | -0.014 | 0.104 | 0.104 | 0.446 | 0.27 | 0.56 | 0.38 | 0.48 | 0.18 | 0.32 |
| 1 | 57444818 | rs947655 | 0.508 | 0.302 | 0.113 | 0.291 | 0.015 | 0.03 | 0.03 | 0 | 0.05 | 0.09 | 0.04 | 0.16 | 0.07 | 0.08 |
| 1 | 57463326 | rs595056 | -1.382 | 0.179 | 0.754 | 0.305 | -0.007 | 0.098 | 0.098 | 0.089 | 0.26 | 0.59 | 0.38 | 0.49 | 0.18 | 0.32 |
| 1 | 57467935 | rs11206955 | 0.072 | 0.146 | 0.791 | 0.321 | -0.02 | 0.098 | 0.098 | 0 | 0.13 | 0.24 | 0.29 | 0.32 | 0.07 | 0.17 |
| 1 | 57469961 | rs515493 | 0.317 | 0.216 | 0.631 | 0.294 | 0.033 | 0.059 | 0.059 | 0 | 0.06 | 0.13 | 0.04 | 0.16 | 0.07 | 0.08 |
| 1 | 57473273 | rs661586 | -0.013 | 0.214 | 0.633 | 0.293 | 0.049 | 0.082 | 0.082 | 0 | 0.07 | 0.19 | 0.05 | 0.16 | 0.07 | 0.08 |
| 1 | 57481828 | rs6668269 | -1.385 | 0.338 | 0.513 | -0.13 | 0.005 | -0.001 | -0.001 | 0 | 0.33 | 0.71 | 0.44 | 0.32 | 0.26 | 0.29 |
| 1 | 57481883 | rs17114816 | -0.738 | 0.325 | 0.793 | -0.097 | -0.027 | 0.087 | 0.087 | 0 | 0.19 | 0.36 | 0.3 | 0.32 | 0.11 | 0.24 |
| 1 | 57483505 | rs3820580 | 0.811 | 0.205 | -0.327 | -0.486 | -0.02 | -0.044 | -0.044 | 0 | 0.31 | 0.09 | 0.34 | 0.41 | 0.34 | 0.32 |
| 1 | 57490074 | rs706360 | 0.823 | 0.264 | -0.408 | -0.204 | -0.001 | 0.022 | 0.022 | 0 | 0.12 | 0.13 | 0.27 | 0.32 | 0.07 | 0.17 |
| 1 | 57493551 | rs706363 | -0.78 | 0.568 | -0.425 | -0.152 | 0.06 | 0.126 | 0.126 | 0 | 0.93 | 0.79 | 0.91 | 0.75 | 0.86 | 0.83 |
| 1 | 57501277 | rs17451521 | 1.909 | 0.211 | -1.126 | -0.538 | -0.011 | 0.035 | 0.035 | 0 | 0.1 | 0.04 | 0.09 | 0 | 0.13 | 0.04 |
| 1 | 57506698 | rs12033358 | 1.125 | 0.037 | -1.173 | -0.475 | -0.06 | 0.196 | 0.196 | 1.282 | 0.3 | 0.03 | 0.13 | 0.02 | 0.27 | 0.24 |
| 1 | 57507014 | rs3768203 | -0.102 | 0.181 | -0.724 | -0.425 | 0.019 | 0.031 | 0.031 | 2.272 | 0.5 | 0.4 | 0.44 | 0.35 | 0.38 | 0.5 |
| 1 | 57517439 | rs3820578 | 0.151 | 0.296 | -1.365 | -0.176 | 0.015 | 0.012 | 0.012 | 1.06 | 0.43 | 0.16 | 0.41 | 0.35 | 0.35 | 0.42 |

| | | | | | | | | | | | | | | | | |
|---|---|---|---|---|---|---|---|---|---|---|---|---|---|---|---|---|
| 1 | 57535630 | rs476744 | -0.036 | 0.383 | -0.869 | -0.771 | 0 | -0.02 | -0.02 | 0 | 0.65 | 0.67 | 0.79 | 0.86 | 0.71 | 0.63 |
| 1 | 57536582 | rs17418461 | 0.605 | 1.122 | -0.752 | -0.894 | 0.012 | -0.021 | -0.021 | 0 | 0.05 | 0.1 | 0.05 | 0 | 0.08 | 0.04 |
| 1 | 57537652 | rs6657073 | 0.875 | 0.782 | -1.537 | -1.635 | -0.039 | 0.174 | 0.174 | 0 | 0.26 | 0.03 | 0.18 | 0 | 0.29 | 0.14 |
| 1 | 57540909 | rs3768190 | -0.215 | 0.641 | -1.35 | -1.184 | 0 | -0.009 | -0.009 | 0 | 0.4 | 0.39 | 0.51 | 0.47 | 0.46 | 0.43 |
| 1 | 57541619 | rs3729546 | 0.644 | 1.006 | -0.82 | -0.642 | 0.017 | 0.154 | 0.154 | 0 | 0.11 | 0.22 | 0.32 | 0.46 | 0.17 | 0.28 |
| 1 | 57542783 | rs486706 | -1.685 | 1.243 | -1.163 | -0.877 | -0.037 | -0.02 | -0.02 | 0.145 | 0.94 | 0.99 | 0.97 | 0.94 | 0.92 | 0.9 |
| 1 | 57551569 | rs514412 | -0.812 | 1.132 | -0.691 | -1.186 | 0.008 | -0.054 | -0.054 | 0 | 0.78 | 0.67 | 0.83 | 0.94 | 0.8 | 0.84 |
| 1 | 57555391 | rs11206971 | -0.172 | 1.106 | -0.64 | -1.274 | 0.019 | 0.087 | 0.087 | 0.008 | 0.21 | 0.33 | 0.38 | 0.5 | 0.24 | 0.34 |
| 1 | 57556458 | rs474194 | -1.369 | 1.131 | -0.451 | -1.326 | 0.015 | 0 | 0 | 0 | 0.92 | 0.87 | 0.95 | 0.94 | 0.92 | 0.92 |
| 1 | 57557620 | rs1359735 | 1.148 | 0.765 | -0.888 | -1.613 | -0.015 | 0.164 | 0.164 | 0 | 0.34 | 0.08 | 0.19 | 0.06 | 0.34 | 0.33 |
| 1 | 57560649 | rs517412 | -0.245 | 0.769 | -0.743 | -1.556 | -0.002 | -0.009 | -0.009 | 0 | 0.57 | 0.51 | 0.67 | 0.59 | 0.59 | 0.69 |
| 1 | 57560931 | rs573307 | -0.981 | 0.611 | -0.618 | -1.49 | 0.013 | -0.091 | -0.091 | 0 | 0.75 | 0.51 | 0.71 | 0.93 | 0.79 | 0.83 |
| 1 | 57562190 | rs482373 | -1.321 | 0.515 | -0.923 | -1.446 | -0.002 | 0.011 | 0.011 | 0 | 0.68 | 0.72 | 0.75 | 0.89 | 0.68 | 0.7 |
| 1 | 57562656 | rs543144 | 0.683 | 0.91 | -1.506 | -1.293 | -0.023 | -0.028 | -0.028 | 0 | 0.25 | 0.09 | 0.19 | 0.37 | 0.29 | 0.2 |
| 1 | 57563074 | rs554795 | 0.35 | 1.13 | -0.867 | -0.615 | 0.002 | 0.074 | 0.074 | 0.062 | 0.24 | 0.08 | 0.1 | 0 | 0.21 | 0.18 |
| 1 | 57565437 | rs490765 | 0.963 | 1.478 | -0.137 | 0.327 | -0.011 | 0.023 | 0.023 | 0 | 0.05 | 0.01 | 0.05 | 0 | 0.09 | 0.02 |
| 1 | 57568923 | rs11206973 | -0.318 | 1.392 | 0.354 | 0.774 | 0.02 | 0.158 | 0.158 | 0 | 0.1 | 0.16 | 0.31 | 0.53 | 0.15 | 0.29 |
| 1 | 57571980 | rs561503 | 0.024 | 1.565 | 0.439 | 0.448 | 0.002 | -0.026 | -0.026 | 1.489 | 0.82 | 0.81 | 0.88 | 0.99 | 0.8 | 0.89 |
| 1 | 57572121 | rs473562 | 1.507 | 1.582 | 0.814 | 0.343 | 0.011 | 0 | 0 | 0 | 0.64 | 0.43 | 0.46 | 0.43 | 0.6 | 0.54 |
| 1 | 57575069 | rs489049 | 0.26 | 0.807 | 0.991 | -0.172 | 0.007 | -0.05 | -0.05 | 0 | 0.81 | 0.74 | 0.88 | 0.99 | 0.8 | 0.89 |
| 1 | 57575777 | rs563670 | 1.435 | 0.803 | 0.992 | 0.314 | 0.004 | -0.011 | -0.011 | 0 | 0.68 | 0.46 | 0.69 | 0.77 | 0.71 | 0.65 |
| 1 | 57580829 | rs522532 | 0.152 | 0.854 | 0.783 | 0.271 | -0.001 | -0.017 | -0.017 | 0 | 0.82 | 0.8 | 0.89 | 0.99 | 0.8 | 0.89 |
| 1 | 57584721 | rs534080 | 0.75 | 0.799 | 0.674 | 0.396 | 0.004 | -0.019 | -0.019 | 0 | 0.68 | 0.47 | 0.7 | 0.79 | 0.71 | 0.66 |
| 1 | 57587533 | rs12029757 | -0.723 | 0.446 | 0.664 | 0.804 | -0.011 | -0.04 | -0.04 | 0 | 0.24 | 0.35 | 0.15 | 0.13 | 0.21 | 0.19 |
| 1 | 57595310 | rs553508 | -0.812 | 0.14 | 1.321 | 0.754 | -0.017 | -0.064 | -0.064 | 0 | 0.22 | 0.55 | 0.25 | 0.09 | 0.17 | 0.21 |
| 1 | 57599834 | rs519401 | -0.539 | 0.113 | 1.128 | 0.737 | -0.001 | -0.008 | -0.008 | 0 | 0.19 | 0.15 | 0.21 | 0.08 | 0.17 | 0.21 |
| 1 | 57602383 | rs563921 | -1.395 | -0.057 | 1.022 | 1.012 | 0.023 | -0.011 | -0.011 | 0 | 0.25 | 0.83 | 0.29 | 0.23 | 0.19 | 0.25 |
| 1 | 57602868 | rs11206975 | -1.146 | 0.019 | 1.134 | 0.562 | 0.083 | 0.135 | 0.135 | 1.667 | 0.05 | 0.55 | 0.14 | 0.05 | 0.01 | 0.02 |
| 1 | 57607081 | rs484064 | -0.493 | 0.177 | 0.284 | 0.435 | 0.003 | 0 | 0 | 0 | 0.18 | 0.13 | 0.12 | 0.03 | 0.16 | 0.2 |
| 1 | 57614330 | rs546583 | 0.229 | 0.316 | -0.778 | 0.429 | -0.067 | -0.005 | -0.005 | 0.041 | 0.37 | 0.05 | 0.22 | 0.23 | 0.42 | 0.14 |
| 1 | 57615514 | rs17115220 | -0.281 | 0.504 | -0.286 | 0.141 | 0.002 | -0.032 | -0.032 | 0 | 0.1 | 0.15 | 0.06 | 0.03 | 0.09 | 0.17 |
| 1 | 57623047 | rs10789045 | 0.296 | -0.024 | -0.83 | -0.028 | -0.021 | 0.126 | 0.126 | 2.423 | 0.37 | 0.09 | 0.19 | 0.12 | 0.37 | 0.13 |
| 1 | 57625649 | rs10889028 | -0.916 | -0.023 | -0.457 | 0.488 | -0.011 | -0.006 | -0.006 | 0 | 0.6 | 0.73 | 0.38 | 0.52 | 0.61 | 0.55 |
| 1 | 57627203 | rs1831870 | -0.501 | -0.192 | 0.346 | 0.33 | -0.019 | 0.009 | 0.009 | 0.298 | 0.52 | 0.79 | 0.33 | 0.52 | 0.49 | 0.51 |
| 1 | 57629549 | rs6670988 | 1.269 | -0.174 | 0.075 | 0.173 | -0.089 | 0.056 | 0.056 | 1.309 | 0.27 | 0.02 | 0.17 | 0.11 | 0.34 | 0.1 |
| 1 | 57633496 | rs7533282 | -1.062 | -0.104 | 0.612 | 0.22 | 0.053 | -0.058 | -0.058 | 0 | 0.11 | 0.46 | 0.12 | 0 | 0.12 | 0.06 |
| 1 | 57641475 | rs7530049 | -1.478 | -0.092 | 0.712 | 0.404 | -0.007 | -0.013 | -0.013 | 1.392 | 0.19 | 0.68 | 0.19 | 0.12 | 0.15 | 0.24 |
| 1 | 57642394 | rs149766 | 1.14 | -0.184 | 0.372 | 0.169 | -0.005 | 0.057 | 0.057 | 0.337 | 0.1 | 0.01 | 0.06 | 0 | 0.07 | 0.04 |
| 1 | 57643701 | rs10489464 | 0.525 | -0.131 | 0.791 | 0.396 | 0 | 0.016 | 0.016 | 0 | 0.08 | 0.29 | 0.04 | 0.06 | 0.03 | 0.17 |
| 1 | 57645164 | rs7542527 | 0.737 | -0.125 | 0.704 | 0.4 | 0.003 | 0.018 | 0.018 | 0 | 0.08 | 0.23 | 0.04 | 0.06 | 0.02 | 0.15 |
| 1 | 57645301 | rs1504589 | 0.53 | -0.247 | 0.092 | 0.366 | -0.081 | -0.081 | -0.081 | 0.005 | 0.4 | 0.07 | 0.19 | 0.39 | 0.45 | 0.32 |
| 1 | 57646661 | rs9970802 | -0.751 | -0.284 | 0.166 | 0.389 | -0.005 | -0.023 | -0.023 | 0 | 0.58 | 0.76 | 0.39 | 0.51 | 0.61 | 0.55 |
| 1 | 57649626 | rs155312 | 0.118 | -0.045 | -0.064 | 0.389 | -0.016 | -0.019 | -0.019 | 0 | 0.46 | 0.15 | 0.45 | 0.46 | 0.52 | 0.49 |
| 1 | 57651497 | rs197655 | -1.496 | -0.057 | 0.407 | 0.396 | 0.035 | 0.234 | 0.234 | 1.35 | 0.11 | 0.48 | 0.35 | 0.23 | 0.1 | 0.11 |
| 1 | 57651809 | rs17482980 | 0.19 | -0.334 | -0.068 | -0.012 | -0.001 | -0.021 | -0.021 | 1.247 | 0.1 | 0.11 | 0.05 | 0 | 0.09 | 0.09 |
| 1 | 57652069 | rs155309 | -0.055 | -0.281 | -0.173 | 0.052 | -0.034 | 0.129 | 0.129 | 6.194 | 0.4 | 0.21 | 0.25 | 0.12 | 0.43 | 0.13 |
| 1 | 57656501 | rs155298 | 0.737 | -0.276 | -0.142 | 0.219 | -0.054 | 0.04 | 0.04 | 0.597 | 0.29 | 0.15 | 0.17 | 0.17 | 0.35 | 0.1 |
| 1 | 57657301 | rs155297 | -1.101 | -0.346 | 0.663 | 0.552 | 0.011 | 0.187 | 0.187 | 0.679 | 0.13 | 0.49 | 0.33 | 0.23 | 0.09 | 0.13 |
| 1 | 57658036 | rs155294 | -0.992 | -0.38 | 0.685 | 0.566 | 0.004 | 0.199 | 0.199 | 1.212 | 0.13 | 0.49 | 0.33 | 0.23 | 0.07 | 0.12 |
| 1 | 57659614 | rs688382 | -1.015 | -0.385 | 0.685 | 0.565 | -0.002 | 0.197 | 0.197 | 1.32 | 0.13 | 0.49 | 0.33 | 0.23 | 0.07 | 0.11 |
| 1 | 57660822 | rs495875 | -0.48 | -0.441 | 0.865 | 0.491 | -0.003 | 0.079 | 0.079 | 0.59 | 0.16 | 0.36 | 0.37 | 0.23 | 0.12 | 0.1 |
| 1 | 57662143 | rs483412 | 0.227 | -0.528 | 0.559 | 0.369 | -0.001 | 0.001 | 0.001 | 0.083 | 0.33 | 0.3 | 0.19 | 0.17 | 0.38 | 0.14 |
| 1 | 57662361 | rs480836 | -0.131 | -0.534 | 0.611 | 0.436 | 0.002 | -0.007 | -0.007 | 0.015 | 0.37 | 0.41 | 0.26 | 0.25 | 0.4 | 0.17 |
| 1 | 57663346 | rs561703 | 0.222 | -0.518 | 0.583 | 0.423 | -0.001 | 0.002 | 0.002 | 0.083 | 0.33 | 0.3 | 0.19 | 0.17 | 0.38 | 0.14 |
| 1 | 57665337 | rs932508 | -1.088 | -0.564 | 0.675 | 0.463 | -0.012 | 0.015 | 0.015 | 0.559 | 0.22 | 0.36 | 0.38 | 0.23 | 0.15 | 0.14 |
| 1 | 57665590 | rs197614 | -0.882 | -0.568 | 0.684 | 0.448 | 0 | 0 | 0 | 0 | 0.49 | 0.63 | 0.56 | 0.4 | 0.5 | 0.26 |
| 1 | 57666761 | rs264040 | 0.885 | -0.262 | 0.411 | 0.566 | -0.066 | 0.022 | 0.022 | 0 | 0.26 | 0.1 | 0.16 | 0.17 | 0.35 | 0.1 |
| 1 | 57666901 | rs1847537 | -0.267 | -0.256 | 0.76 | 0.804 | -0.006 | 0.043 | 0.043 | 0.721 | 0.13 | 0.28 | 0.31 | 0.15 | 0.1 | 0.07 |
| 1 | 57672097 | rs10489465 | 0.018 | -0.377 | 0.785 | 0.704 | -0.012 | 0.008 | 0.008 | 0 | 0.11 | 0.22 | 0.3 | 0.1 | 0.09 | 0.09 |
| 1 | 57672148 | rs17115362 | -1.267 | -0.193 | 0.753 | 0.559 | -0.001 | -0.006 | -0.006 | 0 | 0.08 | 0.09 | 0.06 | 0.01 | 0.09 | 0.16 |
| 1 | 57674906 | rs155284 | -0.649 | -0.058 | 0.571 | 0.43 | 0 | 0.013 | 0.013 | 1.41 | 0.48 | 0.43 | 0.54 | 0.37 | 0.52 | 0.35 |
| 1 | 57675484 | rs197103 | -0.489 | -0.066 | 0.578 | 0.436 | -0.005 | 0.005 | 0.005 | 2.145 | 0.46 | 0.44 | 0.52 | 0.37 | 0.5 | 0.33 |
| 1 | 57677319 | rs197105 | -0.858 | -0.163 | 0.887 | 0.388 | -0.005 | 0.024 | 0.024 | 2.151 | 0.52 | 0.79 | 0.57 | 0.36 | 0.52 | 0.36 |
| 1 | 57677559 | rs502350 | -0.071 | -0.159 | 0.916 | 0.403 | -0.023 | 0.034 | 0.034 | 2.714 | 0.41 | 0.24 | 0.45 | 0.28 | 0.46 | 0.28 |
| 1 | 57679080 | rs197606 | -2.305 | -0.145 | 0.958 | 0.367 | 0.02 | 0.045 | 0.045 | 1.662 | 0.1 | 0.55 | 0.11 | 0.08 | 0.06 | 0.1 |
| 1 | 57679608 | rs197603 | -0.107 | -0.135 | 0.846 | 0.333 | -0.002 | 0.078 | 0.078 | 2.046 | 0.41 | 0.27 | 0.24 | 0.17 | 0.43 | 0.17 |
| 1 | 57683417 | rs197597 | -0.538 | -0.161 | 0.874 | 0.354 | 0.003 | 0.05 | 0.05 | 2.188 | 0.47 | 0.67 | 0.31 | 0.23 | 0.46 | 0.23 |
| 1 | 57685971 | rs197617 | -0.432 | -0.361 | 0.634 | 0.393 | 0 | -0.001 | -0.001 | 0 | 0.6 | 0.44 | 0.64 | 0.57 | 0.59 | 0.5 |
| 1 | 57697059 | rs657471 | -1.197 | 0.032 | 0.754 | 0.623 | -0.001 | -0.018 | -0.018 | 0 | 0.7 | 0.73 | 0.74 | 0.81 | 0.68 | 0.64 |

| | | | | | | | | | | | | | | | | |
|---|---|---|---|---|---|---|---|---|---|---|---|---|---|---|---|---|
| 1 | 57704356 | rs169100 | -0.628 | 0.056 | 0.725 | 0.589 | 0.011 | -0.013 | -0.013 | 0.148 | 0.05 | 0.22 | 0.03 | 0 | 0.03 | 0.06 |
| 1 | 57705206 | rs584383 | 0.111 | 0.042 | 0.656 | 0.618 | -0.018 | -0.009 | -0.009 | 0.981 | 0.57 | 0.16 | 0.6 | 0.57 | 0.56 | 0.44 |
| 1 | 57705650 | rs17115430 | -0.998 | 0.059 | 1.156 | 1.043 | -0.001 | 0.004 | 0.004 | 0 | 0.07 | 0.09 | 0.05 | 0.18 | 0.08 | 0.14 |
| 1 | 57706112 | rs197628 | -1.094 | -0.045 | 1.211 | 0.689 | 0.036 | 0.009 | 0.009 | 0.142 | 0.06 | 0.28 | 0.04 | 0 | 0.03 | 0.06 |
| 1 | 57708548 | rs197627 | -0.717 | -0.11 | 0.68 | 0.838 | 0.001 | 0 | 0 | 0.373 | 0.58 | 0.34 | 0.71 | 0.53 | 0.6 | 0.48 |
| 1 | 57713829 | rs659069 | -0.515 | -0.105 | 0.489 | 0.796 | -0.006 | -0.016 | -0.016 | 0.225 | 0.56 | 0.62 | 0.68 | 0.47 | 0.56 | 0.43 |
| 1 | 57718030 | rs548897 | 0.315 | -0.138 | 0.193 | 0.828 | -0.137 | -0.07 | -0.07 | 0 | 0.42 | 0.03 | 0.26 | 0.36 | 0.45 | 0.26 |
| 1 | 57721731 | rs12132898 | -0.255 | 0.028 | 0.338 | 0.615 | -0.001 | 0.025 | 0.025 | 0 | 0.05 | 0.04 | 0.04 | 0.29 | 0.06 | 0.13 |
| 1 | 57722095 | rs197644 | -0.408 | 0.015 | 0.426 | 0.594 | -0.002 | -0.007 | -0.007 | 0.362 | 0.56 | 0.57 | 0.68 | 0.47 | 0.56 | 0.44 |
| 1 | 57726537 | rs197636 | -1.072 | 0.062 | 0.392 | 0.605 | -0.001 | 0.007 | 0.007 | 0.635 | 0.63 | 0.63 | 0.76 | 0.82 | 0.63 | 0.58 |
| 1 | 57727250 | rs7529737 | 0.672 | 0.082 | 0.233 | 0.616 | -0.005 | -0.008 | -0.008 | 0 | 0.39 | 0.3 | 0.26 | 0.47 | 0.4 | 0.51 |
| 1 | 57734544 | rs2551484 | -1.205 | 0.166 | 0.008 | 0.643 | -0.109 | 0.003 | 0.003 | 1.903 | 0.72 | 0.98 | 0.86 | 0.82 | 0.66 | 0.9 |
| 1 | 57737411 | rs584147 | 0.288 | 0.06 | 1.098 | 0.639 | 0.005 | -0.002 | -0.002 | 1.518 | 0.48 | 0.41 | 0.6 | 0.47 | 0.4 | 0.37 |
| 1 | 57741163 | rs2764664 | -0.476 | 0.057 | 1.144 | 0.637 | -0.023 | -0.013 | -0.013 | 0.53 | 0.57 | 0.81 | 0.71 | 0.53 | 0.46 | 0.44 |
| 1 | 57742783 | rs2764665 | 0.25 | 0.177 | 1.062 | 0.705 | 0.005 | -0.001 | -0.001 | 0 | 0.48 | 0.41 | 0.6 | 0.47 | 0.4 | 0.39 |
| 1 | 57744259 | rs596784 | -0.631 | 0.203 | 0.925 | 0.708 | -0.017 | 0.064 | 0.064 | 2.545 | 0.58 | 0.71 | 0.71 | 0.82 | 0.5 | 0.55 |
| 1 | 57749426 | rs492781 | -0.505 | -0.116 | 1.045 | 0.392 | -0.002 | 0 | 0 | 3.19 | 0.54 | 0.52 | 0.63 | 0.47 | 0.43 | 0.41 |
| 1 | 57751761 | rs643382 | -0.161 | -0.19 | 0.748 | 0.392 | 0.006 | 0.003 | 0.003 | 4.043 | 0.51 | 0.43 | 0.64 | 0.52 | 0.41 | 0.41 |
| 1 | 57756980 | rs595513 | -0.078 | -0.149 | 0.02 | 0.359 | 0.035 | 0.01 | 0.01 | 3.67 | 0.48 | 0.3 | 0.63 | 0.52 | 0.41 | 0.41 |
| 1 | 57771854 | rs12564258 | -1.305 | 0.497 | -0.11 | 0.309 | 0.034 | 0.03 | 0.03 | 0.594 | 0.1 | 0.43 | 0.1 | 0.06 | 0.09 | 0.08 |
| 1 | 57779221 | rs12402993 | 0.055 | 0.647 | -1.631 | 0.196 | -0.013 | 0.084 | 0.084 | 0.903 | 0.45 | 0.13 | 0.26 | 0.74 | 0.44 | 0.5 |
| 1 | 57780247 | rs17422457 | 0.228 | 0.595 | -0.793 | 0.041 | 0.01 | -0.028 | -0.028 | 0 | 0.05 | 0.08 | 0.04 | 0 | 0.04 | 0.04 |
| 1 | 57780302 | rs267643 | -1.102 | 0.062 | -0.405 | -0.248 | -0.023 | -0.051 | -0.051 | 0 | 0.16 | 0.28 | 0.08 | 0.04 | 0.11 | 0.24 |
| 1 | 57780985 | rs267642 | -0.225 | 0.032 | -0.471 | -0.188 | -0.015 | -0.011 | -0.011 | 0 | 0.1 | 0.12 | 0.04 | 0.04 | 0.07 | 0.18 |
| 1 | 57782780 | rs1149629 | -0.381 | 0.098 | -0.532 | 0.975 | 0.005 | 0.014 | 0.014 | 0 | 0.1 | 0.06 | 0.11 | 0.31 | 0.08 | 0.26 |
| 1 | 57784144 | rs746937 | -0.442 | 0.335 | -0.575 | -0.188 | -0.001 | 0 | 0 | 0 | 0.42 | 0.36 | 0.54 | 0.9 | 0.46 | 0.65 |
| 1 | 57799785 | rs6693181 | 0.8 | -0.081 | -0.877 | -0.762 | 0.015 | 0.016 | 0.016 | 0 | 0.16 | 0.11 | 0.1 | 0.01 | 0.1 | 0.1 |
| 1 | 57800885 | rs4912163 | 0.829 | 0.075 | -0.399 | -1.047 | -0.003 | 0.078 | 0.078 | 0.656 | 0.35 | 0.25 | 0.19 | 0.09 | 0.33 | 0.12 |
| 1 | 57801287 | rs1424471 | 1.023 | 0.066 | -0.055 | -1.028 | 0.066 | -0.02 | -0.02 | 0 | 0.06 | 0.18 | 0.14 | 0.01 | 0.09 | 0.07 |
| 1 | 57807897 | rs10889035 | -0.389 | 0.122 | -0.027 | -1.019 | 0 | -0.025 | -0.025 | 0.347 | 0.58 | 0.57 | 0.46 | 0.11 | 0.55 | 0.36 |
| 1 | 57812260 | rs12139429 | 0.428 | 0.057 | -0.002 | -1.034 | 0.001 | -0.039 | -0.039 | 0 | 0.41 | 0.46 | 0.34 | 0.09 | 0.42 | 0.19 |
| 1 | 57814771 | rs11800574 | 1.307 | -0.521 | -0.236 | -1.327 | -0.002 | -0.04 | -0.04 | 0 | 0.13 | 0.14 | 0.15 | 0.01 | 0.12 | 0.07 |
| 1 | 57820539 | rs1424465 | -0.673 | -0.565 | -0.17 | -1.375 | 0 | -0.036 | -0.036 | 0 | 0.58 | 0.59 | 0.44 | 0.1 | 0.52 | 0.35 |
| 1 | 57823951 | rs12565636 | -0.52 | -0.658 | -0.165 | -1.39 | 0.003 | -0.06 | -0.06 | 0.429 | 0.56 | 0.55 | 0.41 | 0.1 | 0.5 | 0.28 |
| 1 | 57831854 | rs7535763 | -0.14 | -0.701 | 0.053 | -1.386 | 0 | 0.02 | 0.02 | 0 | 0.44 | 0.44 | 0.28 | 0.1 | 0.41 | 0.25 |
| 1 | 57837089 | rs2216808 | 0.102 | -0.716 | 0.051 | -1.394 | 0 | 0.051 | 0.051 | 0 | 0.42 | 0.4 | 0.25 | 0.09 | 0.39 | 0.2 |
| 1 | 57841662 | rs10047071 | -0.184 | -0.712 | 0.05 | -1.391 | 0.001 | 0.022 | 0.022 | 0 | 0.45 | 0.44 | 0.28 | 0.1 | 0.41 | 0.26 |
| 1 | 57845605 | rs17424216 | 0.865 | -0.572 | -0.009 | -1.536 | -0.01 | 0.044 | 0.044 | 0 | 0.26 | 0.17 | 0.18 | 0.09 | 0.29 | 0.12 |
| 1 | 57846394 | rs17424278 | 1.341 | -0.599 | -0.009 | -1.552 | 0.002 | 0.064 | 0.064 | 0 | 0.15 | 0.05 | 0.07 | 0.01 | 0.11 | 0.09 |
| 1 | 57846985 | rs3909556 | 0.571 | -0.592 | 0.015 | -1.54 | 0.001 | -0.039 | -0.039 | 0 | 0.28 | 0.42 | 0.2 | 0.09 | 0.29 | 0.12 |
| 1 | 57847075 | rs10493218 | 1.439 | -0.581 | -0.037 | -1.514 | -0.004 | -0.04 | -0.04 | 0 | 0.13 | 0.14 | 0.14 | 0.01 | 0.11 | 0.07 |
| 1 | 57850717 | rs11805025 | -0.086 | -0.597 | -0.04 | -1.534 | -0.015 | -0.112 | -0.112 | 0.031 | 0.3 | 0.43 | 0.23 | 0.01 | 0.22 | 0.16 |
| 1 | 57850914 | rs12405761 | -0.23 | -0.586 | -0.228 | -1.525 | 0 | -0.01 | -0.01 | 0 | 0.45 | 0.47 | 0.3 | 0.1 | 0.42 | 0.26 |
| 1 | 57858443 | rs11588115 | 1.718 | -0.023 | -0.946 | -1.422 | -0.037 | 0.013 | 0.013 | 0.063 | 0.08 | 0.01 | 0.08 | 0.03 | 0.11 | 0.06 |
| 1 | 57858457 | rs10493220 | 1.764 | 0.061 | -1.088 | -0.772 | -0.024 | -0.023 | -0.023 | 0 | 0.06 | 0 | 0.23 | 0.27 | 0.08 | 0.21 |
| 1 | 57869430 | rs17115750 | 0.093 | 0.608 | -0.954 | -1.266 | -0.024 | 0.094 | 0.094 | 0 | 0.42 | 0.31 | 0.31 | 0.1 | 0.45 | 0.24 |
| 1 | 57871326 | rs269045 | -0.003 | 0.956 | -0.938 | -0.561 | -0.016 | 0.061 | 0.061 | 0.061 | 0.59 | 0.73 | 0.41 | 0.11 | 0.51 | 0.31 |
| 1 | 57872825 | rs144823 | -0.739 | 1.223 | -1.323 | 0.076 | -0.101 | -0.109 | -0.109 | 0 | 0.8 | 0.98 | 0.78 | 0.63 | 0.69 | 0.81 |
| 1 | 57880555 | rs269062 | -0.104 | 0.489 | -0.55 | -0.325 | 0.008 | 0.017 | 0.017 | 0 | 0.47 | 0.52 | 0.37 | 0.2 | 0.37 | 0.26 |
| 1 | 57883320 | rs574726 | -0.265 | 0.489 | -0.464 | -0.417 | 0.036 | 0.129 | 0.129 | 0 | 0.49 | 0.71 | 0.33 | 0.06 | 0.38 | 0.27 |
| 1 | 57884310 | rs2991515 | -0.385 | 0.567 | -0.43 | -0.314 | 0.013 | 0.036 | 0.036 | 0 | 0.5 | 0.57 | 0.33 | 0.24 | 0.41 | 0.25 |
| 1 | 57904853 | rs11207036 | -0.852 | 0.548 | 0.073 | 0.265 | 0.021 | 0.102 | 0.102 | 0 | 0.25 | 0.53 | 0.27 | 0.43 | 0.28 | 0.43 |
| 1 | 57904964 | rs11207037 | -0.411 | 0.601 | 0.112 | 0.3 | 0.035 | 0.178 | 0.178 | 0 | 0.21 | 0.49 | 0.23 | 0.42 | 0.24 | 0.37 |
| 1 | 57909581 | rs2781321 | 0.547 | 0.401 | -0.654 | -0.022 | -0.012 | -0.003 | -0.003 | 0.023 | 0.47 | 0.21 | 0.62 | 0.56 | 0.54 | 0.55 |
| 1 | 57913227 | rs12033655 | 0.549 | 0.81 | -0.678 | -0.293 | 0.007 | -0.036 | -0.036 | 0 | 0.36 | 0.26 | 0.52 | 0.56 | 0.34 | 0.52 |
| 1 | 57913272 | rs2860532 | -0.326 | 0.543 | -0.362 | -0.623 | 0 | 0.003 | 0.003 | 0 | 0.72 | 0.68 | 0.78 | 0.6 | 0.71 | 0.69 |
| 1 | 57914164 | rs12044871 | -0.643 | 0.265 | -0.213 | -0.441 | 0.01 | -0.008 | -0.008 | 0 | 0.67 | 0.64 | 0.75 | 0.8 | 0.56 | 0.76 |
| 1 | 57917398 | rs4279875 | -0.806 | 0.302 | -0.294 | -0.448 | -0.008 | -0.012 | -0.012 | 2.328 | 0.69 | 0.69 | 0.78 | 0.97 | 0.59 | 0.82 |
| 1 | 57917478 | rs1334240 | 0.728 | 0.271 | -0.358 | -0.463 | -0.142 | 0.156 | 0.156 | 0.003 | 0.29 | 0.04 | 0.2 | 0.03 | 0.41 | 0.18 |
| 1 | 57923439 | rs11207039 | 0.824 | -0.226 | 0.01 | -0.61 | 0.02 | -0.02 | -0.02 | 2.134 | 0.39 | 0.28 | 0.53 | 0.58 | 0.36 | 0.46 |
| 1 | 57928401 | rs9436130 | 0.935 | -0.119 | 0.126 | -0.641 | -0.091 | 0.126 | 0.126 | 0 | 0.29 | 0.12 | 0.21 | 0.03 | 0.4 | 0.17 |
| 1 | 57933076 | rs2406067 | -0.29 | -0.382 | 0.291 | -0.762 | 0.016 | 0.034 | 0.034 | 0 | 0.5 | 0.42 | 0.34 | 0.24 | 0.57 | 0.33 |
| 1 | 57933955 | rs12407896 | 0.338 | -0.501 | 0.289 | -0.829 | -0.055 | 0.219 | 0.219 | 0.295 | 0.41 | 0.15 | 0.28 | 0.06 | 0.51 | 0.23 |
| 1 | 57934034 | rs10493223 | 0.786 | -0.55 | 0.311 | -0.793 | -0.012 | -0.048 | -0.048 | 0 | 0.1 | 0.02 | 0.1 | 0.19 | 0.09 | 0.22 |
| 1 | 57936193 | rs6686072 | -0.439 | -0.437 | 0.361 | -0.912 | 0.016 | 0.034 | 0.034 | 0 | 0.5 | 0.42 | 0.34 | 0.24 | 0.57 | 0.33 |
| 1 | 57937562 | rs12132685 | 0.921 | -0.34 | -0.029 | -1.026 | -0.105 | 0.143 | 0.143 | 0 | 0.3 | 0.12 | 0.2 | 0.03 | 0.42 | 0.18 |
| 1 | 57940990 | rs2805873 | 0.465 | -0.401 | 0.524 | -0.568 | 0.028 | 0.013 | 0.013 | 1.999 | 0.93 | 0.51 | 0.94 | 0.99 | 0.98 | 0.96 |
| 1 | 57945943 | rs9436131 | -0.258 | -0.377 | 0.335 | -0.418 | -0.023 | -0.024 | -0.024 | 0 | 0.22 | 0.53 | 0.16 | 0.18 | 0.15 | 0.13 |
| 1 | 57956702 | rs2691465 | -1.152 | -0.316 | 0.376 | -0.026 | 0.006 | 0.144 | 0.144 | 0.849 | 0.08 | 0.53 | 0.09 | 0.17 | 0.03 | 0.07 |

| | | | | | | | | | | | | | | | | |
|---|---|---|---|---|---|---|---|---|---|---|---|---|---|---|---|---|
| 1 | 57957657 | rs2691466 | -0.678 | -0.405 | -0.171 | 0.285 | 0.019 | -0.014 | -0.014 | 0 | 0.61 | 0.69 | 0.47 | 0.51 | 0.64 | 0.49 |
| 1 | 57962373 | rs1341318 | -0.081 | 0.245 | 0.634 | 0.517 | -0.014 | -0.003 | -0.003 | 0 | 0.69 | 0.21 | 0.71 | 0.74 | 0.67 | 0.69 |
| 1 | 57966528 | rs1341320 | 0.614 | 0.32 | 0.893 | 0.561 | 0.033 | 0.075 | 0.075 | 1.054 | 0.89 | 0.45 | 0.86 | 0.89 | 0.92 | 0.83 |
| 1 | 57975111 | rs2691437 | 0.703 | 0.26 | 0.283 | 0.544 | 0.027 | -0.003 | -0.003 | 0 | 0.22 | 0.49 | 0.17 | 0.18 | 0.25 | 0.17 |
| 1 | 57975197 | rs17427990 | 1.421 | 0.308 | -0.65 | 0.549 | -0.045 | -0.007 | -0.007 | 0 | 0.18 | 0.07 | 0.13 | 0.15 | 0.25 | 0.14 |
| 1 | 57979583 | rs10443216 | -0.229 | 0.444 | -0.779 | 0.56 | -0.02 | 0.002 | 0.002 | 0 | 0.67 | 0.25 | 0.55 | 0.74 | 0.63 | 0.66 |
| 1 | 57987253 | rs10493228 | -0.183 | -0.093 | -0.993 | 0.782 | 0.02 | -0.021 | -0.021 | 0.359 | 0.38 | 0.13 | 0.3 | 0.4 | 0.34 | 0.34 |
| 1 | 57992118 | rs1357785 | 0.488 | 0.418 | -0.15 | 0.972 | 0.004 | 0.003 | 0.003 | 0 | 0.78 | 0.7 | 0.6 | 0.75 | 0.74 | 0.8 |
| 1 | 57992413 | rs1357787 | 0.478 | 0.415 | 0.113 | 0.97 | 0.015 | 0.013 | 0.013 | 0 | 0.78 | 0.63 | 0.6 | 0.75 | 0.74 | 0.8 |
| 1 | 57996397 | rs2793626 | -0.16 | -0.154 | 0.362 | 0.904 | -0.007 | -0.002 | -0.002 | 0 | 0.3 | 0.31 | 0.17 | 0.26 | 0.22 | 0.42 |
| 1 | 58002491 | rs2691441 | -0.446 | -0.214 | 0.366 | 0.834 | -0.026 | -0.013 | -0.013 | 0.238 | 0.33 | 0.51 | 0.35 | 0.26 | 0.23 | 0.43 |
| 1 | 58003131 | rs1524713 | 0.263 | -0.277 | 0.534 | 0.804 | 0.021 | 0.013 | 0.013 | 0 | 0.64 | 0.29 | 0.63 | 0.74 | 0.77 | 0.57 |
| 1 | 58019733 | rs12733015 | -0.075 | 0.006 | 0.424 | 0.983 | -0.008 | -0.011 | -0.011 | 0 | 0.18 | 0.13 | 0.22 | 0.26 | 0.23 | 0.18 |
| 1 | 58021206 | rs6687355 | 1.397 | -0.009 | -0.188 | 0.979 | -0.066 | -0.019 | -0.019 | 0 | 0.47 | 0.07 | 0.4 | 0.48 | 0.53 | 0.38 |
| 1 | 58037504 | rs7530027 | -0.534 | 0.394 | 0.371 | 1.198 | -0.033 | -0.048 | -0.048 | 0 | 0.17 | 0.07 | 0.22 | 0.26 | 0.23 | 0.18 |
| 1 | 58050139 | rs10493230 | -0.674 | 0.484 | 0.048 | 1.143 | -0.067 | -0.082 | -0.082 | 0 | 0.19 | 0.05 | 0.21 | 0.27 | 0.27 | 0.21 |
| 1 | 58055485 | rs11583494 | -0.436 | 0.46 | -0.101 | 1.092 | -0.079 | -0.124 | -0.124 | 0.437 | 0.18 | 0.03 | 0.21 | 0.36 | 0.26 | 0.2 |
| 1 | 58058589 | rs11207070 | -0.981 | 0.419 | 0.37 | 0.96 | 0.019 | 0.047 | 0.047 | 0 | 0.23 | 0.33 | 0.26 | 0.37 | 0.3 | 0.28 |
| 1 | 58068119 | rs7528953 | 0.451 | 0.343 | 0.083 | 0.593 | -0.004 | -0.003 | -0.003 | 0.338 | 0.78 | 0.28 | 0.79 | 0.79 | 0.82 | 0.71 |
| 1 | 58077023 | rs1534047 | -0.198 | 0.473 | 0.345 | 0.679 | 0.014 | 0.022 | 0.022 | 0.805 | 0.17 | 0.34 | 0.18 | 0.2 | 0.16 | 0.3 |
| 1 | 58079143 | rs1341322 | -0.296 | 0.66 | -0.623 | 0.873 | -0.022 | -0.042 | -0.042 | 0.638 | 0.19 | 0.11 | 0.23 | 0.38 | 0.25 | 0.24 |
| 1 | 58084162 | rs10789050 | 0.911 | 0.449 | -0.404 | 0.598 | -0.01 | 0.001 | 0.001 | 0.285 | 0.6 | 0.3 | 0.56 | 0.43 | 0.56 | 0.43 |
| 1 | 58086486 | rs6687594 | -0.119 | -0.032 | -0.906 | -0.442 | 0.013 | 0.059 | 0.059 | 0 | 0.37 | 0.2 | 0.4 | 0.13 | 0.31 | 0.42 |
| 1 | 58095791 | rs10889059 | 0.103 | -0.257 | -0.869 | -0.613 | 0.006 | 0.115 | 0.115 | 0 | 0.32 | 0.07 | 0.35 | 0.1 | 0.26 | 0.32 |
| 1 | 58096379 | rs1983454 | 0.587 | -0.678 | -0.311 | -0.689 | 0.001 | 0.038 | 0.038 | 0 | 0.45 | 0.29 | 0.44 | 0.69 | 0.57 | 0.43 |
| 1 | 58099707 | rs1539053 | 0.609 | -0.702 | -0.161 | -0.701 | 0.005 | 0.035 | 0.035 | 0 | 0.46 | 0.34 | 0.45 | 0.69 | 0.57 | 0.43 |
| 1 | 58104110 | rs7531119 | -0.223 | -0.67 | 0.23 | -0.726 | -0.008 | 0.005 | 0.005 | 0 | 0.81 | 0.7 | 0.82 | 0.79 | 0.83 | 0.76 |
| 1 | 58106887 | rs7548624 | 0.627 | -0.703 | 0.271 | -0.756 | 0.004 | 0.022 | 0.022 | 0 | 0.47 | 0.39 | 0.45 | 0.69 | 0.57 | 0.42 |
| 1 | 58111045 | rs4484949 | 0.187 | -0.674 | 0.156 | -0.738 | -0.016 | -0.062 | -0.062 | 0.152 | 0.35 | 0.44 | 0.38 | 0.1 | 0.26 | 0.33 |
| 1 | 58120489 | rs11588804 | 0.889 | -0.44 | -0.248 | -0.923 | -0.054 | -0.106 | -0.106 | 2.493 | 0.14 | 0.01 | 0.16 | 0.59 | 0.18 | 0.29 |
| 1 | 58121100 | rs10889061 | 0.299 | -0.454 | -0.093 | -1.139 | -0.004 | -0.023 | -0.023 | 0 | 0.35 | 0.38 | 0.38 | 0.1 | 0.26 | 0.33 |
| 1 | 58124563 | rs6672391 | 0.245 | -0.553 | 0.044 | -1.196 | -0.004 | -0.038 | -0.038 | 0.185 | 0.64 | 0.57 | 0.64 | 0.18 | 0.65 | 0.47 |
| 1 | 58129636 | rs4244011 | 1.221 | -0.549 | -0.158 | -1.183 | 0.006 | -0.057 | -0.057 | 0.028 | 0.3 | 0.35 | 0.28 | 0.08 | 0.39 | 0.14 |
| 1 | 58131026 | rs4912274 | 1.486 | -0.519 | -0.394 | -1.168 | -0.054 | 0.101 | 0.101 | 0 | 0.28 | 0.12 | 0.27 | 0.08 | 0.39 | 0.14 |
| 1 | 58131280 | rs4522021 | 1.425 | -0.745 | -0.663 | -1.418 | 0.019 | 0.087 | 0.087 | 0.018 | 0.19 | 0.03 | 0.11 | 0.04 | 0.13 | 0.07 |
| 1 | 58136781 | rs7526173 | 0.996 | -0.418 | -0.32 | -1.002 | -0.008 | -0.012 | -0.012 | 0.15 | 0.3 | 0.29 | 0.28 | 0.07 | 0.39 | 0.14 |
| 1 | 58138896 | rs12566928 | 0.548 | -0.347 | -0.66 | -0.835 | -0.011 | 0.007 | 0.007 | 0 | 0.18 | 0.04 | 0.29 | 0.12 | 0.17 | 0.22 |
| 1 | 58139294 | rs4568858 | -0.027 | -0.258 | -0.317 | -0.768 | 0.003 | 0.034 | 0.034 | 0 | 0.51 | 0.4 | 0.55 | 0.69 | 0.47 | 0.51 |
| 1 | 58141486 | rs3126031 | -1.567 | -0.383 | -0.425 | -0.848 | -0.027 | 0.039 | 0.039 | 0 | 0.73 | 0.8 | 0.74 | 0.94 | 0.63 | 0.86 |
| 1 | 58143696 | rs3131754 | -0.03 | -0.35 | -0.41 | -0.81 | -0.022 | 0.011 | 0.011 | 0 | 0.21 | 0.37 | 0.19 | 0.24 | 0.16 | 0.34 |
| 1 | 58160124 | rs3131772 | -0.946 | 0.279 | -1.33 | -0.743 | -0.044 | 0.089 | 0.089 | 0 | 0.72 | 0.85 | 0.75 | 0.94 | 0.63 | 0.85 |
| 1 | 58162855 | rs3126019 | -0.699 | 0.004 | -1.341 | -0.023 | 0.051 | -0.01 | -0.01 | 0 | 0.36 | 0.84 | 0.5 | 0.28 | 0.23 | 0.36 |
| 1 | 58164124 | rs10889065 | -0.717 | 1.028 | -0.34 | 0.719 | 0.033 | 0.084 | 0.084 | 1.306 | 0.25 | 0.37 | 0.31 | 0.48 | 0.33 | 0.26 |
| 1 | 58170218 | rs3131780 | -0.975 | 0.855 | 0.472 | 0.72 | 0.031 | 0.084 | 0.084 | 1.736 | 0.27 | 0.57 | 0.33 | 0.46 | 0.33 | 0.33 |
| 1 | 58178715 | rs716417 | -0.389 | 0.909 | 0.196 | 0.72 | -0.004 | -0.002 | -0.002 | 0 | 0.23 | 0.2 | 0.29 | 0.46 | 0.32 | 0.26 |
| 1 | 58181165 | rs12136964 | 1.181 | 0.814 | -1.142 | 0.381 | 0.056 | 0.038 | 0.038 | 1.09 | 0.42 | 0.14 | 0.44 | 0.39 | 0.31 | 0.51 |
| 1 | 58182151 | rs1575512 | -1.337 | 1.016 | -1.084 | 0.374 | -0.111 | 0.035 | 0.035 | 0 | 0.78 | 0.99 | 0.85 | 0.91 | 0.67 | 0.89 |
| 1 | 58185042 | rs12030998 | 0.452 | 1.243 | -0.831 | 0.795 | -0.035 | -0.03 | -0.03 | 2.041 | 0.06 | 0.01 | 0.06 | 0.11 | 0.12 | 0.11 |
| 1 | 58186694 | rs3118024 | -0.871 | 0.715 | -0.527 | 0.765 | 0 | -0.037 | -0.037 | 0 | 0.24 | 0.81 | 0.29 | 0.41 | 0.22 | 0.29 |
| 1 | 58187880 | rs3131723 | -0.381 | 0.809 | -0.444 | 0.833 | 0.015 | 0.13 | 0.13 | 0.378 | 0.21 | 0.53 | 0.26 | 0.41 | 0.21 | 0.22 |
| 1 | 58192123 | rs3131726 | -1.117 | 1.123 | -1.198 | 0.314 | -0.021 | 0.028 | 0.028 | 0 | 0.81 | 0.9 | 0.85 | 0.96 | 0.71 | 0.86 |
| 1 | 58192889 | rs3118027 | -0.339 | 0.788 | -1.171 | 0.861 | -0.059 | -0.031 | -0.031 | 0 | 0.65 | 0.86 | 0.64 | 0.62 | 0.54 | 0.67 |
| 1 | 58194530 | rs7354949 | -1.197 | 0.097 | -0.013 | 0.863 | -0.037 | 0.023 | 0.023 | 0 | 0.15 | 0.56 | 0.13 | 0.14 | 0.07 | 0.1 |
| 1 | 58202297 | rs1886139 | -1.26 | -0.416 | -0.261 | 0.954 | -0.002 | -0.006 | -0.006 | 2.267 | 0.46 | 0.55 | 0.58 | 0.55 | 0.49 | 0.5 |
| 1 | 58203986 | rs3118036 | -1.369 | -0.244 | -0.322 | 0.933 | 0.003 | 0.031 | 0.031 | 0.036 | 0.3 | 0.54 | 0.29 | 0.5 | 0.27 | 0.33 |
| 1 | 58208267 | rs4912173 | -0.781 | -0.295 | -0.249 | 1.048 | 0.001 | 0.057 | 0.057 | 0 | 0.09 | 0.07 | 0.14 | 0.42 | 0.12 | 0.13 |
| 1 | 58209883 | rs3118045 | -1.414 | -0.32 | 0.015 | 0.985 | -0.024 | 0.065 | 0.065 | 0 | 0.27 | 0.5 | 0.24 | 0.49 | 0.2 | 0.29 |
| 1 | 58214656 | rs10889070 | -0.885 | -0.694 | -0.202 | 0.64 | 0.004 | -0.026 | -0.026 | 0 | 0.33 | 0.36 | 0.42 | 0.13 | 0.38 | 0.28 |
| 1 | 58215415 | rs2224823 | -2.135 | -0.81 | 0.274 | 0.688 | -0.001 | 0.001 | 0.001 | 0 | 0.57 | 0.57 | 0.63 | 0.59 | 0.57 | 0.47 |
| 1 | 58217055 | rs7540400 | 0.552 | -0.73 | 0.314 | 0.505 | -0.015 | -0.009 | -0.009 | 0 | 0.13 | 0.17 | 0.06 | 0.07 | 0.08 | 0.05 |
| 1 | 58217784 | rs1323816 | -1.854 | -0.64 | 0.472 | 0.857 | 0.012 | -0.012 | -0.012 | 0 | 0.41 | 0.67 | 0.33 | 0.6 | 0.35 | 0.29 |
| 1 | 58225872 | rs1323821 | -1.758 | -0.72 | 0.312 | 0.789 | 0.007 | -0.001 | -0.001 | 0 | 0.43 | 0.67 | 0.52 | 0.6 | 0.42 | 0.36 |
| 1 | 58226592 | rs1323822 | -1.162 | -0.738 | 0.7 | 0.802 | 0.014 | 0.295 | 0.295 | 0 | 0.13 | 0.41 | 0.17 | 0.49 | 0.12 | 0.14 |
| 1 | 58226786 | rs1323824 | -1.48 | -0.738 | 0.372 | 0.79 | 0 | 0 | 0 | 0 | 0.27 | 0.67 | 0.23 | 0.56 | 0.21 | 0.18 |
| 1 | 58227195 | rs3131741 | -2.321 | -0.621 | 0.427 | 0.792 | -0.01 | 0.017 | 0.017 | 0 | 0.55 | 0.69 | 0.57 | 0.64 | 0.49 | 0.4 |
| 1 | 58228412 | rs17472345 | 0.521 | -0.57 | 0.232 | 0.809 | -0.043 | 0.078 | 0.078 | 0 | 0.17 | 0.06 | 0.29 | 0.02 | 0.22 | 0.17 |
| 1 | 58231158 | rs1886140 | -2.248 | -0.667 | 0.257 | 0.787 | -0.004 | 0.01 | 0.01 | 0 | 0.55 | 0.61 | 0.56 | 0.64 | 0.49 | 0.39 |
| 1 | 58232509 | rs1323826 | -0.54 | -0.701 | 0.213 | 1.027 | 0.062 | -0.006 | -0.006 | 0.012 | 0.27 | 0.09 | 0.1 | 0.29 | 0.15 | 0.14 |
| 1 | 58234014 | rs3118058 | -2.205 | -0.607 | 0.156 | 0.713 | 0 | 0.004 | 0.004 | 0 | 0.56 | 0.6 | 0.61 | 0.65 | 0.56 | 0.46 |

| | | | | | | | | | | | | | | | | |
|---|---|---|---|---|---|---|---|---|---|---|---|---|---|---|---|---|
| 1 | 58234133 | rs3131743 | -1.877 | -1.378 | 0.249 | 1.11 | -0.01 | 0 | 0 | 0 | 0.4 | 0.48 | 0.31 | 0.63 | 0.34 | 0.29 |
| 1 | 58239661 | rs3118061 | -0.48 | -1.165 | -0.692 | 0.606 | 0.041 | 0.07 | 0.07 | 0 | 0.22 | 0.07 | 0.09 | 0.1 | 0.14 | 0.09 |
| 1 | 58247772 | rs1924267 | -0.46 | -0.467 | -0.151 | 1.299 | -0.025 | -0.031 | -0.031 | 0 | 0.16 | 0.05 | 0.14 | 0.26 | 0.17 | 0.13 |
| 1 | 58251060 | rs1581762 | 1.414 | -0.583 | 0.263 | 1.537 | -0.001 | -0.001 | -0.001 | 0.473 | 0.65 | 0.67 | 0.64 | 0.36 | 0.61 | 0.7 |
| 1 | 58256909 | rs1323836 | 0.607 | -0.804 | 0.313 | 1.582 | -0.003 | 0.04 | 0.04 | 0 | 0.82 | 0.8 | 0.8 | 0.63 | 0.8 | 0.86 |
| 1 | 58259463 | rs1570595 | 1.324 | -0.966 | 0.014 | 1.301 | 0.006 | -0.035 | -0.035 | 0.935 | 0.66 | 0.82 | 0.48 | 0.61 | 0.63 | 0.69 |
| 1 | 58260994 | rs7552143 | -1.292 | -1.014 | 0.729 | 0.718 | -0.004 | -0.032 | -0.032 | 0 | 0.36 | 0.53 | 0.49 | 0.27 | 0.34 | 0.3 |
| 1 | 58269042 | rs2406284 | 0.129 | -0.868 | 0.232 | 0.882 | -0.006 | -0.005 | -0.005 | 0 | 0.85 | 0.95 | 0.81 | 0.84 | 0.8 | 0.92 |
| 1 | 58272916 | rs12140411 | 0.232 | -0.8 | 0.192 | 0.682 | -0.027 | 0.072 | 0.072 | 0 | 0.16 | 0.02 | 0.31 | 0.02 | 0.16 | 0.17 |
| 1 | 58279107 | rs11207112 | 2.147 | -0.999 | 0.539 | 0.919 | -0.016 | 0.006 | 0.006 | 0 | 0.46 | 0.31 | 0.31 | 0.43 | 0.46 | 0.53 |
| 1 | 58280509 | rs852749 | -1.077 | -1.001 | 0.538 | 0.995 | -0.01 | 0.073 | 0.073 | 0 | 0.32 | 0.08 | 0.5 | 0.15 | 0.36 | 0.25 |
| 1 | 58282044 | rs2897366 | -0.203 | -1.005 | 0.757 | 1.03 | 0.032 | 0.069 | 0.069 | 0 | 0.17 | 0.43 | 0.17 | 0.28 | 0.17 | 0.12 |
| 1 | 58283618 | rs852757 | -2.565 | -1.15 | 0.27 | 0.978 | -0.158 | -0.097 | -0.097 | 0 | 0.57 | 0.96 | 0.71 | 0.58 | 0.54 | 0.55 |
| 1 | 58286508 | rs852766 | -1.524 | -1.011 | 0.474 | 1.059 | 0.014 | 0.012 | 0.012 | 0 | 0.36 | 0.21 | 0.53 | 0.29 | 0.36 | 0.34 |
| 1 | 58288317 | rs852771 | -0.836 | -0.986 | 0.034 | 0.882 | -0.021 | 0.041 | 0.041 | 0 | 0.22 | 0.06 | 0.29 | 0.12 | 0.28 | 0.15 |
| 1 | 58292621 | rs706411 | -2.349 | -0.938 | 0.063 | 0.903 | -0.02 | 0.003 | 0.003 | 0 | 0.55 | 0.69 | 0.7 | 0.57 | 0.54 | 0.46 |
| 1 | 58302102 | rs852787 | -0.68 | -0.417 | -0.115 | 0.85 | -0.019 | 0.024 | 0.024 | 0 | 0.16 | 0.03 | 0.26 | 0.09 | 0.2 | 0.11 |
| 1 | 58303993 | rs2057225 | 0.014 | -0.266 | 0.1 | 1.008 | 0.006 | 0.002 | 0.002 | 0 | 0.07 | 0.07 | 0.08 | 0.1 | 0.12 | 0.11 |
| 1 | 58305551 | rs852788 | -0.438 | -0.448 | -0.056 | 0.859 | -0.075 | 0.008 | 0.008 | 0 | 0.18 | 0.03 | 0.26 | 0.1 | 0.26 | 0.11 |
| 1 | 58314187 | rs723714 | 0.491 | -0.542 | 0.206 | 0.862 | 0.011 | -0.002 | -0.002 | 0 | 0.09 | 0.02 | 0.21 | 0.04 | 0.07 | 0.13 |
| 1 | 58317833 | rs706412 | -2.432 | -0.754 | 0.531 | 0.979 | -0.001 | 0.002 | 0.002 | 0 | 0.53 | 0.58 | 0.67 | 0.58 | 0.55 | 0.44 |
| 1 | 58321909 | rs852801 | -1.54 | -0.746 | 0.506 | 1.002 | -0.027 | 0.002 | 0.002 | 0 | 0.38 | 0.27 | 0.4 | 0.39 | 0.46 | 0.24 |
| 1 | 58323943 | rs852802 | -1.488 | -0.773 | 0.402 | 0.975 | -0.055 | -0.004 | -0.004 | 2.303 | 0.38 | 0.18 | 0.4 | 0.38 | 0.46 | 0.24 |
| 1 | 58324779 | rs852806 | -2.353 | -0.731 | 0.385 | 1.003 | -0.001 | 0.002 | 0.002 | 0 | 0.53 | 0.58 | 0.67 | 0.58 | 0.55 | 0.44 |
| 1 | 58343983 | rs2207986 | -0.378 | -0.636 | -0.894 | 0.886 | -0.04 | 0.047 | 0.047 | 0 | 0.21 | 0.08 | 0.27 | 0.1 | 0.28 | 0.12 |
| 1 | 58345238 | rs10489669 | 0.466 | -0.452 | -0.945 | 0.918 | -0.007 | 0.011 | 0.011 | 0 | 0.11 | 0.01 | 0.23 | 0.06 | 0.1 | 0.09 |
| 1 | 58346207 | rs1202822 | 0.049 | -0.317 | -0.83 | 0.982 | -0.035 | -0.049 | -0.049 | 0.043 | 0.13 | 0.06 | 0.11 | 0.28 | 0.15 | 0.12 |
| 1 | 58349005 | rs1202824 | -1.809 | -0.497 | -0.553 | 0.871 | -0.003 | 0.005 | 0.005 | 0 | 0.56 | 0.66 | 0.7 | 0.6 | 0.56 | 0.44 |
| 1 | 58352715 | rs6670853 | -0.952 | -0.325 | -0.659 | 0.336 | -0.045 | 0.064 | 0.064 | 0 | 0.23 | 0.09 | 0.27 | 0.09 | 0.3 | 0.13 |
| 1 | 58355969 | rs1202829 | -0.055 | -0.469 | -0.758 | 0.276 | -0.034 | 0.052 | 0.052 | 0 | 0.17 | 0.05 | 0.28 | 0.08 | 0.19 | 0.16 |
| 1 | 58359310 | rs1202832 | -2.294 | -0.565 | -0.471 | 0.424 | -0.006 | 0.068 | 0.068 | 0 | 0.51 | 0.75 | 0.65 | 0.34 | 0.53 | 0.39 |
| 1 | 58363656 | rs12402927 | -0.996 | -0.608 | -0.485 | 0.411 | -0.017 | 0.053 | 0.053 | 0 | 0.24 | 0.14 | 0.28 | 0.09 | 0.3 | 0.13 |
| 1 | 58368541 | rs1202844 | -2.25 | -0.505 | -0.55 | 0.432 | -0.002 | 0.039 | 0.039 | 0 | 0.49 | 0.63 | 0.63 | 0.32 | 0.53 | 0.39 |
| 1 | 58369027 | rs1202847 | -0.082 | -0.229 | -0.901 | 0.367 | -0.025 | 0.019 | 0.019 | 0 | 0.15 | 0.07 | 0.28 | 0.1 | 0.19 | 0.16 |
| 1 | 58369662 | rs17579464 | -0.756 | -0.107 | -0.783 | 0.437 | -0.012 | 0.021 | 0.021 | 0 | 0.2 | 0.12 | 0.26 | 0.06 | 0.28 | 0.12 |
| 1 | 58371042 | rs12116562 | -0.114 | 0.171 | -1.238 | 0.531 | -0.012 | 0.027 | 0.027 | 0 | 0.11 | 0.04 | 0.2 | 0.04 | 0.12 | 0.08 |
| 1 | 58371727 | rs12406340 | -0.116 | 0.195 | -1.179 | 0.76 | -0.011 | 0.043 | 0.043 | 0 | 0.07 | 0.12 | 0.05 | 0.22 | 0.04 | 0.08 |
| 1 | 58372040 | rs1202794 | -0.231 | 0.524 | -1.591 | 0.355 | -0.043 | 0.015 | 0.015 | 0.007 | 0.14 | 0.06 | 0.23 | 0.1 | 0.19 | 0.16 |
| 1 | 58375966 | rs1884487 | -0.484 | 0.199 | -1.786 | 0.189 | 0.021 | -0.02 | -0.02 | 0 | 0.41 | 0.66 | 0.42 | 0.42 | 0.37 | 0.27 |
| 1 | 58376247 | rs1202802 | -0.684 | 0.469 | -1.572 | 0.418 | -0.013 | -0.007 | -0.007 | 0 | 0.56 | 0.74 | 0.67 | 0.5 | 0.58 | 0.43 |
| 1 | 58381521 | rs12125562 | -0.818 | 0.84 | -1.301 | 0.229 | -0.04 | 0.044 | 0.044 | 0 | 0.11 | 0.02 | 0.32 | 0.02 | 0.16 | 0.14 |
| 1 | 58387807 | rs1936413 | -0.103 | 0.731 | -1.072 | 0.025 | -0.004 | -0.025 | -0.025 | 0 | 0.57 | 0.51 | 0.62 | 0.2 | 0.64 | 0.35 |
| 1 | 58392765 | rs1202762 | -0.881 | 0.964 | -0.031 | -0.112 | 0.065 | 0.133 | 0.133 | 0 | 0.09 | 0.36 | 0.06 | 0.13 | 0.1 | 0.08 |
| 1 | 58394752 | rs1202780 | -1.468 | 0.778 | -0.158 | 0.298 | 0.024 | 0.143 | 0.143 | 0 | 0.12 | 0.36 | 0.08 | 0.19 | 0.11 | 0.15 |
| 1 | 58396783 | rs12135814 | 0.379 | 0.539 | -0.521 | 1.437 | -0.016 | -0.038 | -0.038 | 0 | 0.08 | 0.01 | 0.1 | 0.39 | 0.06 | 0.08 |
| 1 | 58398753 | rs1202769 | -0.425 | -0.334 | -0.322 | 0.517 | 0.001 | 0.038 | 0.038 | 0 | 0.47 | 0.7 | 0.58 | 0.26 | 0.56 | 0.39 |
| 1 | 58403623 | rs1857380 | 0.427 | -0.405 | -0.282 | 0.341 | -0.035 | 0.07 | 0.07 | 0 | 0.36 | 0.23 | 0.51 | 0.13 | 0.46 | 0.27 |
| 1 | 58413055 | rs12028282 | 0.56 | -0.483 | -0.486 | 0.052 | -0.027 | 0.037 | 0.037 | 0 | 0.3 | 0.22 | 0.48 | 0.13 | 0.42 | 0.26 |
| 1 | 58414715 | rs1202788 | -0.693 | -0.69 | -0.457 | 0.079 | -0.028 | 0.07 | 0.07 | 0.308 | 0.43 | 0.74 | 0.56 | 0.21 | 0.54 | 0.39 |
| 1 | 58420835 | rs1857373 | 0.272 | -0.8 | -0.258 | 0.005 | 0.042 | 0.068 | 0.068 | 0 | 0.89 | 0.6 | 0.94 | 0.88 | 0.89 | 0.9 |
| 1 | 58431135 | rs1343635 | -0.026 | -0.6 | -0.265 | -0.006 | 0.016 | -0.046 | -0.046 | 0.332 | 0.64 | 0.45 | 0.46 | 0.89 | 0.48 | 0.72 |
| 1 | 58432755 | rs7532719 | 0.11 | -0.604 | -0.271 | -0.013 | -0.05 | 0.098 | 0.098 | 0 | 0.34 | 0.2 | 0.52 | 0.03 | 0.47 | 0.21 |
| 1 | 58434572 | rs1557252 | -0.488 | -0.233 | 0.137 | 0.161 | 0.06 | 0.053 | 0.053 | 0 | 0.93 | 0.74 | 0.95 | 0.92 | 0.91 | 0.93 |
| 1 | 58441860 | rs1187983 | 1.365 | -0.295 | -0.509 | 0.133 | -0.047 | -0.073 | -0.073 | 0.061 | 0.14 | 0.01 | 0.07 | 0.28 | 0.12 | 0.3 |
| 1 | 58448771 | rs2764669 | -0.461 | -0.366 | -0.308 | 0.074 | 0.108 | 0.097 | 0.097 | 0 | 0.93 | 0.67 | 0.94 | 0.92 | 0.91 | 0.93 |
| 1 | 58455538 | rs1187997 | 0.326 | -0.466 | 0.012 | 0.078 | 0.029 | 0.011 | 0.011 | 0.017 | 0.9 | 0.65 | 0.8 | 0.93 | 0.89 | 0.94 |
| 1 | 58462929 | rs2406418 | 0.29 | -0.712 | -0.171 | 0.58 | -0.002 | 0 | 0 | 0 | 0.58 | 0.38 | 0.46 | 0.62 | 0.43 | 0.65 |
| 1 | 58463540 | rs1749777 | 1.477 | -0.598 | -0.406 | 0.766 | -0.079 | -0.075 | -0.075 | 0.521 | 0.15 | 0.01 | 0.09 | 0.29 | 0.16 | 0.35 |
| 1 | 58463763 | rs1749778 | 0.188 | -0.484 | -0.076 | 0.58 | 0.002 | -0.007 | -0.007 | 0 | 0.6 | 0.56 | 0.45 | 0.72 | 0.42 | 0.76 |
| 1 | 58466016 | rs1187988 | -0.304 | -0.426 | -0.063 | 0.61 | 0.018 | 0.003 | 0.003 | 0 | 0.67 | 0.59 | 0.48 | 0.66 | 0.46 | 0.7 |
| 1 | 58467889 | rs1343633 | -1.452 | -0.434 | 0.057 | 0.676 | 0.022 | 0.211 | 0.211 | 0.015 | 0.12 | 0.3 | 0.2 | 0.34 | 0.13 | 0.17 |
| 1 | 58471091 | rs1325449 | -0.437 | -0.507 | -0.036 | 0.57 | -0.035 | 0.073 | 0.073 | 0 | 0.66 | 0.79 | 0.63 | 0.96 | 0.52 | 0.81 |
| 1 | 58472546 | rs1797115 | 1.017 | -0.561 | 0.076 | 0.57 | 0.009 | 0.014 | 0.014 | 0 | 0.51 | 0.37 | 0.42 | 0.62 | 0.39 | 0.64 |
| 1 | 58482029 | rs11207153 | -1.416 | -0.58 | 0.136 | 0.591 | 0.024 | 0.148 | 0.148 | 0 | 0.12 | 0.33 | 0.19 | 0.27 | 0.13 | 0.1 |
| 1 | 58482341 | rs1952046 | 0.462 | -0.553 | -0.073 | 0.608 | -0.004 | 0.001 | 0.001 | 0.312 | 0.57 | 0.37 | 0.45 | 0.63 | 0.43 | 0.65 |
| 1 | 58483220 | rs7539068 | -0.644 | -0.017 | -0.544 | 0.407 | -0.021 | 0.032 | 0.032 | 0 | 0.4 | 0.29 | 0.37 | 0.29 | 0.5 | 0.28 |
| 1 | 58484116 | rs1188002 | 0.199 | -0.15 | -0.229 | 0.377 | 0.005 | -0.072 | -0.072 | 2.415 | 0.6 | 0.51 | 0.59 | 0.93 | 0.52 | 0.78 |
| 1 | 58484277 | rs1925419 | 0.303 | -0.08 | -0.302 | 0.429 | 0.044 | 0.124 | 0.124 | 0 | 0.05 | 0.16 | 0.04 | 0.22 | 0.1 | 0.06 |
| 1 | 58484869 | rs1188003 | 0.306 | -0.155 | -0.495 | 0.502 | -0.021 | 0.003 | 0.003 | 0 | 0.58 | 0.3 | 0.58 | 0.63 | 0.45 | 0.66 |

| | | | | | | | | | | | | | | | | |
|---|---|---|---|---|---|---|---|---|---|---|---|---|---|---|---|---|
| 1 | 58488751 | rs1325454 | 0.513 | 0.146 | -0.356 | 0.607 | 0.012 | 0.044 | 0.044 | 0 | 0.51 | 0.28 | 0.54 | 0.67 | 0.41 | 0.65 |
| 1 | 58489443 | rs1998739 | -0.038 | 0.409 | 0.008 | 1.373 | 0.026 | 0.085 | 0.085 | 0 | 0.14 | 0.22 | 0.13 | 0.43 | 0.19 | 0.29 |
| 1 | 58500409 | rs12728627 | -0.581 | 0.534 | -0.077 | 0.614 | -0.081 | 0.063 | 0.063 | 0 | 0.28 | 0.09 | 0.41 | 0.14 | 0.43 | 0.25 |
| 1 | 58502280 | rs6701501 | -0.349 | 0.588 | -0.101 | 0.62 | -0.131 | 0.058 | 0.058 | 0 | 0.25 | 0.03 | 0.4 | 0.1 | 0.42 | 0.22 |
| 1 | 58512049 | rs4110981 | -0.893 | 0.481 | 0.118 | 0.541 | -0.01 | 0.009 | 0.009 | 0 | 0.32 | 0.3 | 0.48 | 0.18 | 0.48 | 0.24 |
| 1 | 58520106 | rs4912296 | -0.913 | -0.876 | 0.918 | 1.228 | -0.007 | 0.055 | 0.055 | 0 | 0.09 | 0.17 | 0.05 | 0.15 | 0.05 | 0.05 |
| 1 | 58520500 | rs11591081 | 0.336 | -0.967 | 0.682 | 1.134 | 0.005 | -0.001 | -0.001 | 0 | 0.16 | 0.12 | 0.29 | 0.13 | 0.2 | 0.18 |
| 1 | 58520774 | rs11207160 | -1.034 | -0.896 | 0.386 | 1.052 | -0.031 | 0.026 | 0.026 | 0 | 0.28 | 0.51 | 0.36 | 0.28 | 0.25 | 0.29 |
| 1 | 58521203 | rs11207161 | 0.294 | -1.063 | 0.409 | 0.646 | -0.027 | -0.008 | -0.008 | 0 | 0.8 | 0.69 | 0.87 | 0.84 | 0.88 | 0.84 |
| 1 | 58523346 | rs2764676 | 0.967 | -0.769 | 0.309 | 0.725 | 0.033 | -0.003 | -0.003 | 0 | 0.51 | 0.32 | 0.5 | 0.57 | 0.57 | 0.56 |
| 1 | 58523985 | rs2806403 | 1.128 | -1.667 | 0.513 | 0.823 | -0.013 | 0.005 | 0.005 | 0 | 0.72 | 0.62 | 0.75 | 0.69 | 0.84 | 0.72 |
| 1 | 58526016 | rs2808822 | 1.116 | -1.714 | 0.716 | 0.836 | -0.058 | 0.031 | 0.031 | 0 | 0.73 | 0.38 | 0.76 | 0.71 | 0.86 | 0.72 |
| 1 | 58526235 | rs10443242 | -0.899 | -1.782 | 0.612 | 0.876 | -0.018 | 0.018 | 0.018 | 0 | 0.19 | 0.3 | 0.17 | 0.2 | 0.14 | 0.28 |
| 1 | 58526758 | rs11207162 | 0.069 | -1.792 | -0.036 | 0.871 | 0.007 | -0.011 | -0.011 | 0 | 0.13 | 0.1 | 0.11 | 0.18 | 0.11 | 0.13 |
| 1 | 58526854 | rs2025617 | 1.207 | -1.761 | -0.085 | 0.86 | 0.084 | -0.032 | -0.032 | 0 | 0.48 | 0.17 | 0.52 | 0.56 | 0.62 | 0.52 |
| 1 | 58528820 | rs6657936 | -0.935 | -1.453 | -0.409 | 1.277 | 0.022 | -0.025 | -0.025 | 0 | 0.19 | 0.19 | 0.2 | 0.29 | 0.11 | 0.23 |
| 1 | 58531627 | rs6698482 | -0.15 | -0.987 | -0.286 | 0.916 | 0 | 0 | 0 | 0 | 0.25 | 0.18 | 0.23 | 0.15 | 0.24 | 0.2 |
| 1 | 58532755 | rs12569191 | -0.376 | -0.499 | -0.28 | 1.304 | 0.006 | -0.01 | -0.01 | 0 | 0.06 | 0.08 | 0.08 | 0.11 | 0 | 0.03 |
| 1 | 58536417 | rs2764683 | 0.634 | -0.367 | 0.224 | 1.466 | -0.031 | 0.077 | 0.077 | 0 | 0.84 | 0.69 | 0.84 | 0.69 | 0.9 | 0.93 |
| 1 | 58537361 | rs2806405 | 0.084 | 0.034 | 0.297 | 1.411 | 0.006 | 0.014 | 0.014 | 0 | 0.63 | 0.47 | 0.64 | 0.59 | 0.6 | 0.52 |
| 1 | 58544805 | rs2764687 | 0.197 | -0.119 | 0.4 | 1.702 | 0.002 | 0.004 | 0.004 | 0 | 0.63 | 0.43 | 0.63 | 0.51 | 0.59 | 0.49 |
| 1 | 58556598 | rs873989 | -0.082 | -0.142 | 0.911 | 1.527 | -0.015 | 0.236 | 0.236 | 0.291 | 0.18 | 0.43 | 0.2 | 0.41 | 0.19 | 0.34 |
| 1 | 58561954 | rs1320555 | -1.256 | -0.746 | 1.475 | 1.55 | -0.005 | 0.005 | 0.005 | 1.357 | 0.27 | 0.68 | 0.29 | 0.49 | 0.24 | 0.48 |
| 1 | 58574668 | rs2788712 | -1.152 | -0.624 | 1.487 | 1.483 | -0.013 | 0.184 | 0.184 | 0 | 0.17 | 0.38 | 0.35 | 0.48 | 0.14 | 0.2 |
| 1 | 58574925 | rs1416347 | -0.792 | -0.615 | 1.452 | 1.568 | -0.001 | 0.019 | 0.019 | 0 | 0.13 | 0.16 | 0.32 | 0.39 | 0.11 | 0.12 |
| 1 | 58575317 | rs1416346 | -0.631 | -0.851 | 1.505 | 1.821 | -0.03 | 0.075 | 0.075 | 0.487 | 0.22 | 0.51 | 0.24 | 0.3 | 0.22 | 0.35 |
| 1 | 58578612 | rs1342876 | -1.354 | -0.907 | 1.305 | 1.832 | -0.016 | 0.053 | 0.053 | 0 | 0.17 | 0.23 | 0.33 | 0.49 | 0.11 | 0.21 |
| 1 | 58582979 | rs1767574 | -1.701 | -0.943 | 1.486 | 1.569 | -0.013 | -0.031 | -0.031 | 0 | 0.36 | 0.55 | 0.49 | 0.8 | 0.34 | 0.53 |
| 1 | 58586505 | rs12040062 | -1.112 | -0.686 | 1.771 | 1.709 | -0.011 | 0.16 | 0.16 | 0 | 0.2 | 0.36 | 0.25 | 0.44 | 0.22 | 0.45 |
| 1 | 58591848 | rs17116983 | -0.066 | -0.658 | 1.692 | 1.721 | 0.011 | 0.108 | 0.108 | 0 | 0.13 | 0.22 | 0.15 | 0.37 | 0.19 | 0.29 |
| 1 | 58602197 | rs17116988 | -0.023 | -0.66 | 1.599 | 1.724 | 0.01 | 0.102 | 0.102 | 0 | 0.12 | 0.2 | 0.15 | 0.37 | 0.19 | 0.29 |
| 1 | 58602918 | rs6688878 | -0.253 | -0.554 | 1.522 | 1.594 | -0.005 | -0.021 | -0.021 | 0 | 0.07 | 0.08 | 0.09 | 0 | 0.05 | 0.08 |
| 1 | 58610928 | rs6667829 | -1.257 | -0.562 | 1.636 | 1.662 | -0.006 | 0.131 | 0.131 | 0 | 0.21 | 0.36 | 0.26 | 0.45 | 0.24 | 0.47 |
| 1 | 58617269 | rs2806399 | -0.972 | -0.443 | 1.253 | 1.58 | -0.002 | 0.059 | 0.059 | 0.033 | 0.18 | 0.31 | 0.34 | 0.41 | 0.15 | 0.12 |
| 1 | 58630042 | rs11802821 | 0.196 | -0.441 | 1.124 | 1.534 | 0.002 | 0.056 | 0.056 | 0 | 0.12 | 0.15 | 0.15 | 0.38 | 0.19 | 0.29 |
| 1 | 58666076 | rs7524008 | 1.314 | -0.516 | 1.189 | 1.416 | -0.018 | 0.12 | 0.12 | 0.36 | 0.8 | 0.45 | 0.65 | 0.56 | 0.85 | 0.79 |
| 1 | 58674799 | rs572888 | 0.526 | -0.359 | 0.319 | 1.079 | -0.01 | 0.035 | 0.035 | 0 | 0.84 | 0.71 | 0.75 | 0.69 | 0.87 | 0.89 |
| 1 | 58677864 | rs7511972 | -0.686 | -0.062 | -0.386 | 0.845 | 0 | -0.011 | -0.011 | 0.18 | 0.07 | 0.06 | 0.09 | 0 | 0.05 | 0.08 |
| 1 | 58687680 | rs12146090 | -0.096 | 0.284 | -1.219 | 0.297 | -0.174 | 0.148 | 0.148 | 0 | 0.33 | 0.05 | 0.28 | 0.09 | 0.49 | 0.21 |
| 1 | 58700416 | rs553133 | -0.326 | -0.521 | -0.495 | 0.736 | 0.005 | 0.006 | 0.006 | 0 | 0.51 | 0.48 | 0.47 | 0.29 | 0.63 | 0.48 |
| 1 | 58704644 | rs4912184 | -0.383 | -0.54 | -0.171 | 0.69 | 0.048 | 0.065 | 0.065 | 0 | 0.52 | 0.68 | 0.48 | 0.29 | 0.63 | 0.48 |
| 1 | 58707111 | rs494214 | -0.559 | -0.765 | 0.101 | 0.958 | 0.012 | -0.011 | -0.011 | 0 | 0.58 | 0.63 | 0.52 | 0.48 | 0.68 | 0.75 |
| 1 | 58714427 | rs614373 | -0.512 | -0.079 | -0.435 | 0.496 | -0.043 | 0.058 | 0.058 | 0 | 0.7 | 0.85 | 0.75 | 0.8 | 0.6 | 0.78 |
| 1 | 58722463 | rs12132541 | 0.814 | -0.113 | -0.568 | 0.39 | -0.005 | 0.013 | 0.013 | 0 | 0.13 | 0.07 | 0.27 | 0.08 | 0.13 | 0.19 |
| 1 | 58725716 | rs639241 | 0.093 | -0.239 | 0.778 | 0.329 | -0.019 | 0.116 | 0.116 | 0 | 0.57 | 0.76 | 0.48 | 0.72 | 0.47 | 0.58 |
| 1 | 58726579 | rs555213 | 0.749 | -0.161 | 0.69 | 0.299 | -0.009 | -0.015 | -0.015 | 0 | 0.46 | 0.43 | 0.34 | 0.65 | 0.34 | 0.49 |
| 1 | 58730127 | rs574802 | -0.099 | -0.168 | 0.885 | 0.342 | 0.01 | -0.073 | -0.073 | 0.038 | 0.2 | 0.36 | 0.36 | 0.08 | 0.18 | 0.24 |
| 1 | 58733050 | rs552352 | -0.727 | -0.162 | 0.639 | 0.496 | 0 | 0 | 0 | 0 | 0.94 | 0.93 | 0.94 | 0.94 | 0.91 | 0.96 |
| 1 | 58742150 | rs689231 | 0.635 | -0.174 | 0.662 | 0.474 | 0.013 | -0.069 | -0.069 | 0 | 0.73 | 0.45 | 0.57 | 0.86 | 0.73 | 0.72 |
| 1 | 58742470 | rs496009 | 0.666 | -0.175 | 0.139 | 0.474 | 0 | -0.013 | -0.013 | 0.015 | 0.73 | 0.34 | 0.56 | 0.86 | 0.73 | 0.72 |
| 1 | 58743154 | rs17117233 | 1.168 | -0.406 | -0.653 | 0.36 | 0.03 | -0.086 | -0.086 | 0 | 0.36 | 0.12 | 0.27 | 0.44 | 0.29 | 0.22 |
| 1 | 58745403 | rs545696 | -0.127 | 0.168 | -0.638 | 0.135 | 0.002 | 0.002 | 0.002 | 0 | 0.94 | 0.89 | 0.94 | 0.94 | 0.91 | 0.96 |
| 1 | 58747747 | rs635459 | -0.098 | 0.168 | -0.737 | 0.135 | 0 | 0 | 0 | 2.871 | 0.94 | 0.91 | 0.94 | 0.94 | 0.91 | 0.96 |
| 1 | 58759203 | rs6665082 | 0.632 | 0.248 | -0.814 | 0.144 | -0.007 | 0.012 | 0.012 | 0.034 | 0.13 | 0.07 | 0.27 | 0.08 | 0.14 | 0.2 |
| 1 | 58767977 | rs12125333 | 0.476 | 0.301 | -1.156 | 0.111 | -0.174 | 0.034 | 0.034 | 0 | 0.28 | 0.04 | 0.23 | 0.2 | 0.43 | 0.22 |
| 1 | 58768044 | rs1924563 | 0.098 | 0.032 | -1.272 | -0.079 | 0.014 | 0.039 | 0.039 | 3.27 | 0.2 | 0.07 | 0.3 | 0.1 | 0.15 | 0.32 |
| 1 | 58768957 | rs4912187 | 0.015 | 0.236 | -0.255 | 0.53 | 0.05 | -0.005 | -0.005 | 0.119 | 0.33 | 0.49 | 0.36 | 0.33 | 0.46 | 0.27 |
| 1 | 58773286 | rs12076949 | 0.005 | 0.354 | -0.739 | -0.123 | -0.003 | 0.069 | 0.069 | 0.162 | 0.52 | 0.38 | 0.55 | 0.16 | 0.49 | 0.55 |
| 1 | 58783971 | rs10493241 | 0.899 | 0.154 | -0.857 | 0.442 | 0.01 | 0.021 | 0.021 | 2.655 | 0.17 | 0.09 | 0.3 | 0.1 | 0.12 | 0.33 |
| 1 | 58784150 | rs10493240 | 1.037 | 0.324 | -0.394 | 0.692 | 0.015 | 0.157 | 0.157 | 0 | 0.19 | 0.35 | 0.21 | 0.41 | 0.2 | 0.2 |
| 1 | 58789870 | rs10127537 | 0.356 | 0.428 | -0.896 | 0.183 | -0.002 | 0.006 | 0.006 | 0 | 0.23 | 0.31 | 0.23 | 0.64 | 0.21 | 0.27 |
| 1 | 58790501 | rs11207196 | 0.957 | 0.746 | -1.084 | -0.242 | 0.006 | -0.002 | -0.002 | 8.116 | 0.28 | 0.23 | 0.13 | 0.04 | 0.22 | 0.16 |
| 1 | 58792030 | rs12726314 | -0.454 | 1.626 | -0.959 | 0.544 | 0.103 | -0.003 | -0.003 | 0 | 0.09 | 0.32 | 0.12 | 0.03 | 0.15 | 0.07 |
| 1 | 58794017 | rs1213640 | -1.216 | 1.786 | -2.2 | 0.598 | -0.096 | -0.025 | -0.025 | 0 | 0.78 | 0.97 | 0.79 | 0.81 | 0.7 | 0.84 |
| 1 | 58796131 | rs12730076 | -0.025 | 1.812 | -1.416 | 0.882 | -0.037 | 0.014 | 0.014 | 0 | 0.07 | 0.01 | 0.11 | 0.03 | 0.15 | 0.07 |
| 1 | 58800475 | rs12567514 | 0.58 | 1.428 | -1.397 | 1.425 | -0.058 | -0.107 | -0.107 | 0.272 | 0.16 | 0.01 | 0.17 | 0.37 | 0.2 | 0.15 |
| 1 | 58801506 | rs1924569 | -1.54 | 1.391 | -0.432 | 1.568 | 0.052 | 0.221 | 0.221 | 0.292 | 0.13 | 0.46 | 0.13 | 0.27 | 0.16 | 0.1 |
| 1 | 58802060 | rs1213646 | -0.873 | 1.643 | -1.781 | 0.759 | -0.097 | 0.039 | 0.039 | 0 | 0.65 | 0.97 | 0.52 | 0.78 | 0.61 | 0.68 |
| 1 | 58806965 | rs1886306 | -1.108 | 2.007 | -0.602 | 0.762 | 0.008 | -0.039 | -0.039 | 0 | 0.09 | 0.13 | 0.1 | 0 | 0.14 | 0.12 |

| | | | | | | | | | | | | | | | | |
|---|---|---|---|---|---|---|---|---|---|---|---|---|---|---|---|---|
| 1 | 58807475 | rs2794651 | -1.718 | 2.248 | -0.38 | 0.922 | 0.046 | 0.02 | 0.02 | 0 | 0.17 | 0.28 | 0.23 | 0.19 | 0.25 | 0.17 |
| 1 | 58810057 | rs12748486 | -1.337 | 2.406 | -0.456 | 0.952 | 0.022 | -0.03 | -0.03 | 0 | 0.08 | 0.17 | 0.14 | 0 | 0.19 | 0.12 |
| 1 | 58810472 | rs947407 | -1.065 | 2.091 | -0.233 | 1.259 | 0.079 | 0.077 | 0.077 | 0 | 0.11 | 0.3 | 0.18 | 0.2 | 0.18 | 0.04 |
| 1 | 58810918 | rs1213656 | -0.754 | 1.565 | -0.656 | 1.361 | 0 | 0.027 | 0.027 | 0 | 0.11 | 0.17 | 0.08 | 0.3 | 0.12 | 0.16 |
| 1 | 58811029 | rs12404503 | 0.024 | 1.083 | -1.239 | 0.374 | 0.01 | 0.081 | 0.081 | 0 | 0.22 | 0.07 | 0.11 | 0.06 | 0.16 | 0.27 |
| 1 | 58811791 | rs6686674 | 1.18 | 1.057 | -0.794 | 0.526 | -0.014 | 0.01 | 0.01 | 0 | 0.36 | 0.4 | 0.55 | 0.51 | 0.32 | 0.34 |
| 1 | 58817583 | rs12145683 | 0.726 | 1.014 | -0.87 | 0.309 | 0.001 | 0.01 | 0.01 | 0.67 | 0.11 | 0.11 | 0.05 | 0 | 0.07 | 0.03 |
| 1 | 58819774 | rs4244014 | 1.105 | 0.935 | -0.972 | 0.285 | 0.007 | 0.009 | 0.009 | 0.024 | 0.53 | 0.37 | 0.65 | 0.41 | 0.47 | 0.4 |
| 1 | 58821318 | rs1318371 | 1.005 | 0.906 | -0.797 | 0.275 | -0.001 | 0.018 | 0.018 | 0.509 | 0.55 | 0.3 | 0.65 | 0.58 | 0.5 | 0.48 |
| 1 | 58824110 | rs2018765 | 0.205 | 1.239 | -0.695 | 0.377 | -0.023 | -0.004 | -0.004 | 0 | 0.18 | 0.1 | 0.13 | 0.19 | 0.24 | 0.3 |
| 1 | 58825601 | rs17496442 | 1.022 | 1.213 | -0.734 | 0.264 | 0.003 | 0.055 | 0.055 | 1.321 | 0.17 | 0.04 | 0.08 | 0.06 | 0.1 | 0.11 |
| 1 | 58826547 | rs11580730 | -0.716 | 1.233 | -0.427 | 0.244 | -0.002 | 0.004 | 0.004 | 0 | 0.3 | 0.31 | 0.19 | 0.2 | 0.31 | 0.33 |
| 1 | 58827079 | rs6695605 | -0.18 | 1.242 | -0.412 | 0.287 | 0.011 | 0.074 | 0.074 | 0.183 | 0.3 | 0.18 | 0.16 | 0.07 | 0.2 | 0.14 |
| 1 | 58829981 | rs12404972 | 0.145 | 1.265 | -0.399 | 0.314 | 0.004 | -0.001 | -0.001 | 0 | 0.19 | 0.21 | 0.14 | 0.2 | 0.24 | 0.3 |
| 1 | 58833588 | rs1213664 | 0.63 | 1.308 | -0.312 | 0.578 | 0.048 | 0.193 | 0.193 | 0 | 0.09 | 0.34 | 0.11 | 0.23 | 0.07 | 0.08 |
| 1 | 58833758 | rs4912192 | -0.197 | 1.412 | -0.483 | 0.577 | 0.008 | 0.095 | 0.095 | 0 | 0.4 | 0.28 | 0.23 | 0.19 | 0.32 | 0.23 |
| 1 | 58833911 | rs12028180 | 1.002 | 1.534 | -0.719 | 0.44 | 0 | 0.027 | 0.027 | 0 | 0.07 | 0.12 | 0.1 | 0.23 | 0.07 | 0.08 |
| 1 | 58834026 | rs11580092 | 1.828 | 1.582 | -0.84 | 0.382 | -0.031 | -0.079 | -0.079 | 0 | 0.33 | 0.03 | 0.52 | 0.34 | 0.34 | 0.28 |
| 1 | 58838246 | rs11580545 | 1.126 | 1.612 | -0.708 | 0.17 | -0.021 | -0.009 | -0.009 | 0 | 0.07 | 0.02 | 0.06 | 0.08 | 0.12 | 0.14 |
| 1 | 58851991 | rs338941 | -1.653 | 1.084 | -1.135 | 0.163 | -0.034 | -0.023 | -0.023 | 0 | 0.68 | 0.98 | 0.48 | 0.75 | 0.66 | 0.74 |
| 1 | 58853893 | rs338935 | -0.68 | 1.343 | -0.539 | 0.484 | 0.057 | 0.058 | 0.058 | 0 | 0.13 | 0.32 | 0.11 | 0.22 | 0.19 | 0.2 |
| 1 | 58857598 | rs849492 | -0.68 | 1.024 | -0.97 | 0.236 | -0.02 | 0.009 | 0.009 | 0 | 0.28 | 0.59 | 0.16 | 0.34 | 0.18 | 0.28 |
| 1 | 58857666 | rs338934 | -0.303 | 1.241 | -2.099 | 0.421 | -0.003 | -0.007 | -0.007 | 0 | 0.27 | 0.29 | 0.17 | 0.33 | 0.26 | 0.35 |
| 1 | 58862010 | rs338926 | -0.875 | 0.768 | -2.54 | 0.491 | -0.035 | -0.014 | -0.014 | 1.731 | 0.65 | 0.98 | 0.47 | 0.76 | 0.66 | 0.72 |
| 1 | 58864093 | rs12038113 | -0.673 | 0.849 | -1.773 | 0.663 | 0.002 | -0.051 | -0.051 | 0 | 0.25 | 0.18 | 0.14 | 0.34 | 0.2 | 0.28 |
| 1 | 58865399 | rs338919 | -0.516 | 1.274 | -1.084 | -0.182 | 0.041 | -0.069 | -0.069 | 0.175 | 0.33 | 0.49 | 0.29 | 0.19 | 0.43 | 0.3 |
| 1 | 58872128 | rs849483 | -0.748 | 1.09 | -0.719 | 0.513 | 0.005 | 0.099 | 0.099 | 0 | 0.08 | 0.18 | 0.06 | 0.26 | 0.1 | 0.12 |
| 1 | 58872522 | rs10465839 | -0.302 | 1.963 | -0.222 | -0.125 | 0.02 | -0.041 | -0.041 | 0 | 0.29 | 0.3 | 0.23 | 0.17 | 0.34 | 0.24 |
| 1 | 58873906 | rs338911 | -0.995 | 0.898 | -0.16 | -0.107 | 0.011 | 0.022 | 0.022 | 0.036 | 0.32 | 0.62 | 0.28 | 0.46 | 0.36 | 0.26 |
| 1 | 58874059 | rs849484 | 0.219 | 0.808 | -0.153 | -0.038 | 0.007 | 0.087 | 0.087 | 0 | 0.07 | 0.13 | 0.06 | 0.29 | 0.1 | 0.11 |
| 1 | 58875165 | rs849485 | -0.949 | 0.614 | 0.294 | -0.081 | -0.032 | 0.121 | 0.121 | 0 | 0.13 | 0.37 | 0.08 | 0.29 | 0.07 | 0.2 |
| 1 | 58875644 | rs338905 | 0.365 | 0.753 | -0.023 | -0.176 | -0.089 | -0.029 | -0.029 | 0 | 0.15 | 0.01 | 0.17 | 0.16 | 0.24 | 0.04 |
| 1 | 58877966 | rs11207224 | 0.62 | 0.218 | -0.196 | -0.884 | -0.006 | -0.064 | -0.064 | 0 | 0.33 | 0.29 | 0.16 | 0.03 | 0.26 | 0.22 |
| 1 | 58878270 | rs338901 | -1.122 | 0.105 | -0.652 | -0.677 | -0.114 | -0.093 | -0.093 | 0.344 | 0.65 | 0.92 | 0.42 | 0.45 | 0.59 | 0.49 |
| 1 | 58884192 | rs706416 | -0.89 | 0.223 | -0.459 | -0.584 | -0.052 | -0.029 | -0.029 | 0.207 | 0.61 | 0.84 | 0.4 | 0.46 | 0.57 | 0.47 |
| 1 | 58885233 | rs706417 | -0.33 | 0.218 | -0.092 | -0.34 | -0.018 | 0.075 | 0.075 | 0 | 0.11 | 0.21 | 0.07 | 0.33 | 0.07 | 0.15 |
| 1 | 58894050 | rs849469 | -0.668 | 0.278 | 0.257 | -0.384 | 0.003 | 0.15 | 0.15 | 0 | 0.11 | 0.29 | 0.07 | 0.32 | 0.08 | 0.19 |
| 1 | 58902488 | rs706428 | -0.741 | 0.3 | -0.186 | -0.535 | -0.02 | 0.011 | 0.011 | 0 | 0.59 | 0.75 | 0.39 | 0.44 | 0.57 | 0.46 |
| 1 | 58908819 | rs706439 | -0.007 | 0.798 | -0.045 | 0.124 | 0.013 | -0.017 | -0.017 | 2.091 | 0.6 | 0.47 | 0.46 | 0.68 | 0.49 | 0.71 |
| 1 | 58909250 | rs706440 | -0.19 | 0.372 | 0.121 | 0.633 | -0.005 | 0.05 | 0.05 | 0 | 0.12 | 0.17 | 0.17 | 0.32 | 0.09 | 0.14 |
| 1 | 58910442 | rs706444 | -0.502 | 0.369 | -0.072 | 0.592 | -0.006 | 0.071 | 0.071 | 0 | 0.16 | 0.28 | 0.21 | 0.33 | 0.12 | 0.24 |
| 1 | 58911040 | rs706445 | -0.081 | 0.478 | -0.683 | 0.262 | -0.002 | 0 | 0 | 0.032 | 0.77 | 0.75 | 0.62 | 0.8 | 0.74 | 0.81 |
| 1 | 58915024 | rs10889092 | 0.622 | 0.044 | -0.928 | 0.323 | 0.031 | 0.045 | 0.045 | 1.375 | 0.58 | 0.28 | 0.37 | 0.28 | 0.63 | 0.4 |
| 1 | 58915624 | rs706450 | 0.64 | -0.011 | -0.306 | 0.208 | 0.017 | 0.039 | 0.039 | 0 | 0.08 | 0.28 | 0.06 | 0.1 | 0.06 | 0.07 |
| 1 | 58915956 | rs849477 | -0.679 | -0.158 | -0.521 | 1.134 | -0.003 | -0.079 | -0.079 | 0 | 0.87 | 0.97 | 0.62 | 0.56 | 0.9 | 0.85 |
| 1 | 58916818 | rs822165 | 1.045 | -0.002 | -1.019 | 0.248 | -0.009 | 0.071 | 0.071 | 0 | 0.46 | 0.34 | 0.29 | 0.19 | 0.49 | 0.32 |
| 1 | 58917256 | rs12126555 | -0.165 | 0.002 | -0.867 | 0.053 | 0.001 | -0.019 | -0.019 | 0 | 0.26 | 0.24 | 0.19 | 0.07 | 0.28 | 0.12 |
| 1 | 58918699 | rs706453 | -0.18 | -0.076 | -0.838 | 0.447 | -0.002 | -0.084 | -0.084 | 0 | 0.82 | 0.94 | 0.58 | 0.34 | 0.83 | 0.6 |
| 1 | 58919090 | rs706454 | -0.164 | -0.089 | -0.58 | 0.432 | -0.002 | -0.084 | -0.084 | 0 | 0.82 | 0.94 | 0.58 | 0.34 | 0.83 | 0.59 |
| 1 | 58920920 | rs706457 | -0.516 | -0.291 | -0.524 | -0.199 | -0.001 | -0.045 | -0.045 | 0 | 0.35 | 0.39 | 0.23 | 0.06 | 0.34 | 0.15 |
| 1 | 58923511 | rs697587 | -0.606 | -0.325 | -0.266 | 0.516 | -0.004 | -0.036 | -0.036 | 0 | 0.88 | 0.95 | 0.81 | 0.51 | 0.88 | 0.67 |
| 1 | 58925978 | rs6666833 | -0.544 | -0.627 | -0.292 | 0.222 | -0.004 | -0.029 | -0.029 | 0.873 | 0.38 | 0.39 | 0.25 | 0.14 | 0.35 | 0.22 |
| 1 | 58926395 | rs12040053 | 2.189 | -0.519 | -0.071 | -0.059 | -0.024 | 0.036 | 0.036 | 0.176 | 0.32 | 0.22 | 0.23 | 0.18 | 0.36 | 0.33 |
| 1 | 58931305 | rs338248 | -0.227 | -0.482 | 0.134 | 0.057 | -0.005 | -0.093 | -0.093 | 0 | 0.32 | 0.53 | 0.22 | 0.02 | 0.29 | 0.16 |
| 1 | 58933274 | rs697590 | 1.02 | -0.468 | -0.217 | 0.139 | 0.011 | 0.044 | 0.044 | 0 | 0.49 | 0.36 | 0.55 | 0.36 | 0.53 | 0.45 |
| 1 | 58933458 | rs11207233 | 0.258 | -0.077 | -0.403 | 0.081 | -0.006 | -0.056 | -0.056 | 0 | 0.2 | 0.25 | 0.12 | 0.01 | 0.16 | 0.13 |
| 1 | 58934155 | rs338241 | -1.428 | 0.267 | -0.331 | 0.944 | -0.006 | -0.011 | -0.011 | 0 | 0.94 | 1 | 0.86 | 0.71 | 0.95 | 0.91 |
| 1 | 58934242 | rs7544701 | 0.23 | 0.382 | -0.14 | 0.338 | -0.004 | 0.02 | 0.02 | 0 | 0.13 | 0.08 | 0.09 | 0.01 | 0.13 | 0.03 |
| 1 | 58936384 | rs697592 | 0.157 | 0.367 | -0.015 | 0.653 | -0.001 | 0.004 | 0.004 | 0 | 0.1 | 0.12 | 0.3 | 0.18 | 0.09 | 0.1 |
| 1 | 58939502 | rs12118488 | -0.207 | 0.423 | 0.063 | 0.289 | -0.001 | -0.018 | -0.018 | 0 | 0.2 | 0.25 | 0.12 | 0.05 | 0.18 | 0.15 |
| 1 | 58939634 | rs12407003 | 0.74 | 0.513 | -0.299 | 0.885 | -0.038 | -0.029 | -0.029 | 0.5 | 0.08 | 0.02 | 0.18 | 0.41 | 0.13 | 0.29 |
| 1 | 58940596 | rs338230 | -1.65 | 0.737 | -0.438 | 0.649 | -0.042 | 0.055 | 0.055 | 0.37 | 0.52 | 0.87 | 0.59 | 0.36 | 0.49 | 0.39 |
| 1 | 58943495 | rs10889097 | 2.092 | 0.738 | -0.447 | 0.835 | -0.013 | 0.054 | 0.054 | 0 | 0.4 | 0.12 | 0.23 | 0.23 | 0.38 | 0.32 |
| 1 | 58949510 | rs10489815 | 0.394 | 0.552 | -0.129 | 0.606 | -0.001 | -0.032 | -0.032 | 0 | 0.18 | 0.23 | 0.11 | 0.05 | 0.18 | 0.15 |
| 1 | 58949987 | rs10489816 | 1.005 | 0.597 | -0.074 | 0.53 | -0.01 | 0.03 | 0.03 | 0 | 0.12 | 0.08 | 0.07 | 0 | 0.11 | 0.04 |
| 1 | 58971831 | rs3087585 | -0.855 | 0.623 | 0.301 | 0.553 | -0.006 | -0.108 | -0.108 | 0 | 0.33 | 0.45 | 0.21 | 0.05 | 0.29 | 0.19 |
| 1 | 58974163 | rs11581844 | 1.561 | 0.638 | 0.2 | 0.585 | -0.031 | 0.056 | 0.056 | 0.527 | 0.48 | 0.14 | 0.41 | 0.64 | 0.51 | 0.61 |
| 1 | 58978397 | rs6656025 | 1.103 | 0.638 | 0.38 | 0.603 | 0.009 | 0.053 | 0.053 | 0 | 0.6 | 0.27 | 0.75 | 0.81 | 0.66 | 0.7 |
| 1 | 58996203 | rs2271004 | 0.832 | 0.637 | 0.407 | 0.75 | 0 | 0.001 | 0.001 | 0 | 0.12 | 0.12 | 0.34 | 0.17 | 0.15 | 0.09 |

| | | | | | | | | | | | | | | | | |
|---|---|---|---|---|---|---|---|---|---|---|---|---|---|---|---|---|
| 1 | 58999651 | rs17117678 | 0 | 0.616 | 0.465 | 0.588 | -0.003 | -0.062 | -0.062 | 0 | 0.15 | 0.22 | 0.09 | 0 | 0.11 | 0.04 |
| 1 | 59002221 | rs7537470 | 0.005 | 0.615 | 0.464 | 0.599 | -0.003 | -0.004 | -0.004 | 0 | 0.15 | 0.23 | 0.1 | 0.14 | 0.12 | 0.1 |
| 1 | 59024435 | rs659061 | -1.007 | 0.526 | 0.167 | 0.266 | 0.004 | 0.011 | 0.011 | 0.652 | 0.35 | 0.68 | 0.23 | 0.19 | 0.3 | 0.24 |
| 1 | 59040719 | rs232840 | -0.916 | 0.366 | 0.169 | 0.224 | -0.014 | -0.044 | -0.044 | 0.106 | 0.22 | 0.52 | 0.14 | 0.14 | 0.16 | 0.15 |
| 1 | 59041092 | rs6683669 | 0.555 | 0.341 | 0.351 | 0.21 | 0.038 | 0.118 | 0.118 | 0 | 0.07 | 0.32 | 0.05 | 0.14 | 0.05 | 0.11 |
| 1 | 59041436 | rs9583 | 0.027 | 0.343 | 0.202 | 0.207 | -0.005 | -0.063 | -0.063 | 3.485 | 0.15 | 0.23 | 0.09 | 0 | 0.11 | 0.04 |
| 1 | 59042181 | rs14008 | 0.137 | 0.333 | 0.036 | 0.375 | 0.052 | 0.14 | 0.14 | 0.004 | 0.07 | 0.34 | 0.05 | 0.14 | 0.05 | 0.11 |
| 1 | 59047175 | rs232828 | -2.02 | 0.693 | -1.57 | 0.313 | -0.062 | -0.032 | -0.032 | 0 | 0.65 | 0.99 | 0.81 | 0.77 | 0.69 | 0.69 |
| 1 | 59056198 | rs12756352 | 0.158 | 1.139 | -1.082 | 0.603 | -0.087 | -0.115 | -0.115 | 0 | 0.18 | 0.02 | 0.27 | 0.43 | 0.27 | 0.31 |
| 1 | 59056326 | rs6664314 | 0.304 | 1.07 | -0.878 | 0.425 | -0.003 | 0.002 | 0.002 | 0 | 0.16 | 0.12 | 0.11 | 0.05 | 0.2 | 0.15 |
| 1 | 59057933 | rs6587830 | -1.265 | 0.885 | -0.469 | 0.41 | 0.008 | -0.001 | -0.001 | 0.002 | 0.42 | 0.46 | 0.44 | 0.48 | 0.52 | 0.52 |
| 1 | 59062268 | rs11207270 | -0.891 | 0.689 | -1.029 | 0.526 | -0.066 | -0.075 | -0.075 | 0 | 0.31 | 0.12 | 0.37 | 0.46 | 0.37 | 0.42 |
| 1 | 59073430 | rs11207272 | -0.658 | 0.31 | -0.194 | 0.665 | -0.036 | 0.036 | 0.036 | 0.575 | 0.18 | 0.48 | 0.11 | 0.21 | 0.11 | 0.18 |
| 1 | 59075102 | rs12745094 | 1.232 | 0.707 | -0.879 | 0.901 | -0.036 | -0.047 | -0.047 | 0 | 0.11 | 0.03 | 0.14 | 0.22 | 0.11 | 0.08 |
| 1 | 59079248 | rs232812 | 0.821 | 0.784 | -0.875 | 0.932 | -0.016 | -0.019 | -0.019 | 0 | 0.09 | 0.03 | 0.18 | 0.18 | 0.17 | 0.26 |
| 1 | 59079359 | rs232813 | -1.239 | 0.159 | -0.66 | 0.558 | -0.002 | 0.003 | 0.003 | 0 | 0.38 | 0.58 | 0.35 | 0.52 | 0.34 | 0.5 |
| 1 | 59081533 | rs170827 | -2.349 | 0.178 | -1.572 | 0.266 | -0.106 | -0.033 | -0.033 | 0.222 | 0.7 | 0.98 | 0.75 | 0.81 | 0.68 | 0.72 |
| 1 | 59084139 | rs232816 | -1.417 | 0.13 | -1.188 | 0.143 | -0.013 | -0.005 | -0.005 | 0 | 0.56 | 0.73 | 0.66 | 0.61 | 0.58 | 0.56 |
| 1 | 59086221 | rs2073094 | 0.437 | 0.228 | -1.642 | 0.195 | -0.085 | -0.096 | -0.096 | 0.688 | 0.3 | 0.07 | 0.38 | 0.41 | 0.36 | 0.22 |
| 1 | 59098228 | rs232915 | -1.237 | 0.265 | -1.394 | 0.254 | -0.041 | -0.039 | -0.039 | 0 | 0.57 | 0.83 | 0.67 | 0.6 | 0.58 | 0.57 |
| 1 | 59110167 | rs7527235 | 1.478 | -0.285 | -1.149 | -0.004 | 0.039 | 0.004 | 0.004 | 0 | 0.37 | 0.11 | 0.24 | 0.31 | 0.29 | 0.27 |
| 1 | 59114438 | rs11207282 | 0.404 | -0.066 | -0.471 | 0.003 | -0.001 | -0.012 | -0.012 | 0 | 0.69 | 0.38 | 0.64 | 0.72 | 0.67 | 0.62 |
| 1 | 59120650 | rs232857 | -0.756 | -0.203 | -0.598 | -0.053 | -0.011 | 0.017 | 0.017 | 0 | 0.88 | 0.92 | 0.91 | 0.94 | 0.87 | 0.93 |
| 1 | 59122871 | rs232851 | -0.774 | -0.305 | -0.401 | -0.096 | 0.03 | 0.035 | 0.035 | 0 | 0.49 | 0.72 | 0.67 | 0.63 | 0.59 | 0.62 |
| 1 | 59125683 | rs232777 | -0.064 | -0.322 | -0.275 | -0.098 | -0.003 | 0.011 | 0.011 | 0 | 0.19 | 0.46 | 0.27 | 0.21 | 0.2 | 0.31 |
| 1 | 59127086 | rs7523134 | 1.19 | -0.345 | -0.457 | -0.099 | 0.047 | 0.015 | 0.015 | 0 | 0.39 | 0.13 | 0.24 | 0.31 | 0.28 | 0.3 |
| 1 | 59131311 | rs232790 | -0.614 | -0.209 | -0.347 | -0.058 | 0 | -0.004 | -0.004 | 0.004 | 0.88 | 0.85 | 0.91 | 0.94 | 0.87 | 0.93 |
| 1 | 59147926 | rs12139511 | -0.772 | -0.192 | 0.215 | 0.01 | 0.017 | 0.019 | 0.019 | 0 | 0.49 | 0.66 | 0.66 | 0.63 | 0.59 | 0.62 |
| 1 | 59151958 | rs17118106 | 1.25 | -0.148 | -0.023 | 0.052 | -0.044 | -0.057 | -0.057 | 4.343 | 0.11 | 0.01 | 0.15 | 0.21 | 0.19 | 0.27 |
| 1 | 59173358 | rs4912375 | 0.58 | 0.008 | 0.194 | 0.079 | -0.053 | -0.058 | -0.058 | 0 | 0.29 | 0.12 | 0.43 | 0.39 | 0.38 | 0.24 |
| 1 | 59174026 | rs4912202 | 0.848 | -0.024 | 0.383 | 0.064 | 0.002 | 0.007 | 0.007 | 0 | 0.16 | 0.23 | 0.19 | 0.21 | 0.2 | 0.31 |
| 1 | 59175142 | rs12073211 | 0.376 | -0.042 | 0.268 | 0.059 | -0.008 | -0.008 | -0.008 | 0 | 0.3 | 0.25 | 0.45 | 0.39 | 0.38 | 0.24 |
| 1 | 59175979 | rs10158502 | 0.547 | -0.06 | 0.111 | 0.021 | 0.019 | 0.034 | 0.034 | 0 | 0.52 | 0.28 | 0.34 | 0.39 | 0.42 | 0.43 |
| 1 | 59180394 | rs17264915 | 0.078 | -0.242 | -0.31 | 0.092 | 0.007 | 0.053 | 0.053 | 3.864 | 0.18 | 0.1 | 0.09 | 0.06 | 0.13 | 0.1 |
| 1 | 59181597 | rs999157 | -1.513 | 0.026 | -0.433 | 0.287 | -0.024 | 0.004 | 0.004 | 0 | 0.84 | 0.9 | 0.66 | 0.88 | 0.75 | 0.83 |
| 1 | 59186777 | rs11583201 | 0.556 | -0.333 | 0.182 | 0.253 | -0.007 | 0.009 | 0.009 | 0 | 0.19 | 0.33 | 0.22 | 0.24 | 0.22 | 0.31 |
| 1 | 59203121 | rs2760480 | -0.191 | -0.261 | 0.164 | 0.373 | 0.027 | 0.015 | 0.015 | 0 | 0.66 | 0.57 | 0.52 | 0.62 | 0.6 | 0.69 |
| 1 | 59205102 | rs1969772 | 1.552 | -0.171 | -0.293 | 0.428 | 0.002 | -0.011 | -0.011 | 0 | 0.3 | 0.19 | 0.24 | 0.35 | 0.28 | 0.34 |
| 1 | 59209416 | rs12137541 | -0.554 | -0.11 | -0.374 | 0.46 | 0.019 | 0.032 | 0.032 | 3.396 | 0.34 | 0.25 | 0.27 | 0.28 | 0.32 | 0.37 |
| 1 | 59215791 | rs2811876 | -0.484 | 0.188 | 0.614 | 0.411 | 0.006 | 0.014 | 0.014 | 0 | 0.64 | 0.63 | 0.51 | 0.55 | 0.62 | 0.69 |
| 1 | 59221647 | rs10889118 | 1.481 | 0.212 | -0.247 | 0.427 | 0.006 | -0.015 | -0.015 | 0 | 0.3 | 0.12 | 0.25 | 0.34 | 0.27 | 0.29 |
| 1 | 59227624 | rs11207303 | 1.22 | 0.259 | -0.297 | 0.501 | -0.041 | -0.005 | -0.005 | 0 | 0.36 | 0.08 | 0.29 | 0.33 | 0.38 | 0.32 |
| 1 | 59228062 | rs2811881 | -0.461 | 0.113 | 0.05 | 0.426 | -0.03 | -0.053 | -0.053 | 1.091 | 0.67 | 0.8 | 0.52 | 0.54 | 0.62 | 0.68 |
| 1 | 59230033 | rs6677589 | 0.207 | 0.172 | 0.685 | 0.541 | -0.001 | 0.011 | 0.011 | 0 | 0.64 | 0.43 | 0.67 | 0.58 | 0.65 | 0.6 |
| 1 | 59230228 | rs2811883 | 0.438 | -0.327 | 0.557 | 0.37 | -0.028 | -0.004 | -0.004 | 0.138 | 0.16 | 0.32 | 0.14 | 0.16 | 0.11 | 0.26 |
| 1 | 59231160 | rs1007271 | 0.416 | -0.012 | 0.23 | 0.325 | 0.012 | -0.034 | -0.034 | 0 | 0.63 | 0.17 | 0.66 | 0.6 | 0.71 | 0.65 |
| 1 | 59231953 | rs2760488 | -0.961 | 0.437 | 0.171 | 0.354 | -0.002 | -0.003 | -0.003 | 0 | 0.81 | 0.81 | 0.83 | 0.74 | 0.81 | 0.9 |
| 1 | 59240954 | rs12738233 | 0.785 | 0.149 | 0.155 | 0.335 | -0.072 | -0.047 | -0.047 | 0 | 0.15 | 0.03 | 0.37 | 0.2 | 0.25 | 0.22 |
| 1 | 59242028 | rs2760496 | 0.106 | -0.755 | 0.627 | 0.533 | 0.028 | -0.078 | -0.078 | 0 | 0.37 | 0.13 | 0.25 | 0.51 | 0.31 | 0.55 |
| 1 | 59251097 | rs2760501 | -0.887 | -1.095 | 0.534 | 0.582 | 0.048 | 0.013 | 0.013 | 6.654 | 0.19 | 0.86 | 0.22 | 0.17 | 0.09 | 0.25 |
| 1 | 59253695 | rs2984915 | 0.564 | -1.101 | 0.599 | 0.544 | -0.01 | 0.001 | 0.001 | 5.823 | 0.81 | 0.27 | 0.85 | 0.84 | 0.92 | 0.75 |
| 1 | 59258879 | rs11207306 | 1.107 | -0.934 | 0.396 | 0.599 | -0.104 | -0.149 | -0.149 | 0.323 | 0.26 | 0.01 | 0.2 | 0.39 | 0.29 | 0.33 |
| 1 | 59271489 | rs3015315 | -0.797 | -1.249 | 1.62 | 0.456 | 0 | -0.002 | -0.002 | 0.382 | 0.34 | 0.61 | 0.42 | 0.22 | 0.31 | 0.27 |
| 1 | 59277926 | rs2999134 | -0.702 | -1.14 | 0.98 | 0.484 | 0.001 | 0.008 | 0.008 | 0 | 0.33 | 0.66 | 0.43 | 0.21 | 0.3 | 0.27 |
| 1 | 59283810 | rs2984904 | -0.007 | -0.968 | -0.064 | 0.567 | -0.023 | 0.014 | 0.014 | 0 | 0.26 | 0.12 | 0.35 | 0.21 | 0.3 | 0.26 |
| 1 | 59286691 | rs11207307 | 0.239 | -0.978 | -0.15 | 0.514 | 0.021 | 0.003 | 0.003 | 0 | 0.11 | 0.04 | 0.06 | 0.14 | 0.07 | 0.25 |
| 1 | 59301218 | rs1815274 | 1.073 | -0.941 | -0.239 | 0.528 | 0.04 | 0.036 | 0.036 | 0.053 | 0.55 | 0.16 | 0.5 | 0.63 | 0.63 | 0.48 |
| 1 | 59304719 | rs2984908 | -0.262 | -0.246 | -0.45 | 0.759 | -0.013 | 0.003 | 0.003 | 0 | 0.27 | 0.16 | 0.35 | 0.23 | 0.3 | 0.26 |
| 1 | 59304910 | rs942124 | 0.559 | -0.489 | -0.02 | 0.802 | -0.028 | -0.008 | -0.008 | 0 | 0.87 | 0.85 | 0.93 | 0.86 | 0.93 | 0.75 |
| 1 | 59321745 | rs9436174 | -0.084 | -0.289 | -0.139 | 0.74 | 0.003 | 0.045 | 0.045 | 0 | 0.37 | 0.22 | 0.33 | 0.24 | 0.31 | 0.38 |
| 1 | 59328135 | rs6684750 | -0.481 | -0.473 | 0.335 | 1.088 | 0 | 0.007 | 0.007 | 0 | 0.69 | 0.76 | 0.68 | 0.74 | 0.73 | 0.64 |
| 1 | 59330763 | rs12133457 | 0.492 | -0.234 | -0.269 | 1.088 | -0.054 | -0.024 | -0.024 | 0 | 0.24 | 0.12 | 0.29 | 0.2 | 0.33 | 0.2 |
| 1 | 59330863 | rs6679571 | 0.316 | -0.408 | -0.237 | 0.348 | 0.003 | 0.008 | 0.008 | 0 | 0.26 | 0.14 | 0.16 | 0.21 | 0.21 | 0.32 |
| 1 | 59332599 | rs4326648 | -0.821 | -0.137 | -0.101 | 0.546 | 0.004 | -0.019 | -0.019 | 0.428 | 0.64 | 0.64 | 0.65 | 0.76 | 0.66 | 0.61 |
| 1 | 59342968 | rs2716129 | -0.297 | 0.573 | -0.338 | 1.021 | 0.03 | 0.06 | 0.06 | 0 | 0.3 | 0.42 | 0.44 | 0.53 | 0.4 | 0.29 |
| 1 | 59347149 | rs11207319 | 0.164 | 0.415 | -0.71 | 0.778 | 0.032 | 0.008 | 0.008 | 0 | 0.52 | 0.66 | 0.5 | 0.5 | 0.36 | 0.58 |
| 1 | 59347543 | rs1536159 | 0.211 | 0.326 | -0.228 | 0.681 | -0.002 | 0 | 0 | 0 | 0.49 | 0.54 | 0.48 | 0.48 | 0.33 | 0.6 |
| 1 | 59348528 | rs2764908 | -0.516 | 1.293 | -0.107 | 0.63 | 0.071 | 0.011 | 0.011 | 0 | 0.09 | 0.27 | 0.22 | 0.06 | 0.13 | 0.12 |
| 1 | 59349570 | rs12239208 | 0.234 | 0.417 | 0.024 | 0.556 | -0.025 | -0.004 | -0.004 | 0 | 0.1 | 0.14 | 0.04 | 0.06 | 0.03 | 0.13 |

| | | | | | | | | | | | | | | | | |
|---|---|---|---|---|---|---|---|---|---|---|---|---|---|---|---|---|
| 1 | 59351446 | rs2492321 | -0.812 | 0.396 | 0.047 | 0.443 | 0.063 | -0.011 | -0.011 | 0 | 0.12 | 0.27 | 0.32 | 0.07 | 0.19 | 0.23 |
| 1 | 59352323 | rs11207322 | 0.399 | -0.07 | 0.618 | 0.297 | 0.048 | -0.013 | -0.013 | 0 | 0.42 | 0.3 | 0.34 | 0.49 | 0.25 | 0.47 |
| 1 | 59353441 | rs2716107 | -0.808 | -0.045 | 0.606 | 0.309 | -0.066 | 0.015 | 0.015 | 0 | 0.59 | 0.87 | 0.69 | 0.58 | 0.47 | 0.71 |
| 1 | 59355900 | rs12117564 | 1.038 | -0.063 | 0.614 | 0.305 | 0.026 | -0.054 | -0.054 | 0 | 0.35 | 0.18 | 0.4 | 0.43 | 0.28 | 0.46 |
| 1 | 59357252 | rs11207324 | 0.4 | -0.056 | 0.605 | 0.305 | -0.018 | -0.002 | -0.002 | 0 | 0.1 | 0.13 | 0.04 | 0.06 | 0.03 | 0.05 |
| 1 | 59360403 | rs2011623 | -0.366 | -0.053 | 0.862 | 0.297 | 0.079 | 0.023 | 0.023 | 0 | 0.1 | 0.36 | 0.22 | 0.07 | 0.14 | 0.19 |
| 1 | 59364918 | rs6658104 | 0.421 | -0.159 | 0.945 | 0.249 | 0.017 | -0.006 | -0.006 | 0 | 0.42 | 0.34 | 0.42 | 0.5 | 0.31 | 0.51 |
| 1 | 59371517 | rs11207327 | -1.96 | -0.243 | 0.96 | 0.16 | 0.001 | -0.006 | -0.006 | 0 | 0.19 | 0.32 | 0.26 | 0.13 | 0.19 | 0.25 |
| 1 | 59379043 | rs17096897 | -0.092 | -0.003 | 0.742 | 0.047 | -0.042 | 0.013 | 0.013 | 0 | 0.08 | 0.01 | 0.19 | 0.04 | 0.14 | 0.23 |
| 1 | 59389845 | rs10159430 | 0.545 | -0.01 | 0.822 | 0.195 | -0.012 | 0.046 | 0.046 | 0 | 0.06 | 0.17 | 0.05 | 0.08 | 0.02 | 0.03 |
| 1 | 59394773 | rs2716116 | -1.015 | 1.094 | 0.083 | 0.039 | -0.065 | -0.063 | -0.063 | 0 | 0.86 | 0.99 | 0.88 | 0.73 | 0.78 | 0.85 |
| 1 | 59395523 | rs2764914 | 0.018 | 0.936 | 0.054 | 0.118 | -0.019 | 0.014 | 0.014 | 0 | 0.57 | 0.82 | 0.7 | 0.56 | 0.46 | 0.73 |
| 1 | 59396262 | rs6664131 | 1.533 | 0.721 | 0.057 | -0.006 | -0.006 | 0.005 | 0.005 | 0 | 0.38 | 0.57 | 0.44 | 0.45 | 0.29 | 0.46 |
| 1 | 59397257 | rs6667591 | 1.474 | 0.744 | 0.053 | -0.065 | -0.009 | 0.014 | 0.014 | 0 | 0.05 | 0.07 | 0.03 | 0.08 | 0.02 | 0.03 |
| 1 | 59397745 | rs6681035 | -1.38 | 0.742 | 0.044 | -0.069 | -0.004 | -0.007 | -0.007 | 0 | 0.19 | 0.26 | 0.26 | 0.12 | 0.17 | 0.27 |
| 1 | 59402566 | rs4912205 | 1.256 | 0.628 | -0.078 | -0.077 | -0.089 | 0.034 | 0.034 | 0 | 0.26 | 0.02 | 0.16 | 0.15 | 0.3 | 0.12 |
| 1 | 59406024 | rs6677752 | 0.734 | 0.214 | 0.349 | -0.254 | 0.024 | -0.009 | -0.009 | 0 | 0.38 | 0.2 | 0.43 | 0.56 | 0.3 | 0.5 |
| 1 | 59413234 | rs11803507 | 0.427 | 0.219 | 0.021 | -0.152 | -0.012 | 0.038 | 0.038 | 0 | 0.05 | 0.11 | 0.03 | 0.08 | 0.02 | 0.03 |
| 1 | 59414592 | rs11207344 | 0.007 | 0.199 | 0.043 | -0.162 | 0.014 | 0.098 | 0.098 | 0 | 0.06 | 0.27 | 0.03 | 0.08 | 0.02 | 0.03 |
| 1 | 59415757 | rs7413192 | -0.246 | 0.702 | -1.379 | -0.476 | -0.09 | 0.012 | 0.012 | 0 | 0.42 | 0.09 | 0.38 | 0.39 | 0.48 | 0.39 |
| 1 | 59422789 | rs2764928 | 0.081 | 0.171 | -1.149 | -1.289 | 0.006 | 0.036 | 0.036 | 0.902 | 0.28 | 0.74 | 0.19 | 0.14 | 0.23 | 0.18 |
| 1 | 59423103 | rs3949912 | 0.183 | 0.17 | -1.662 | -1.566 | -0.018 | 0.159 | 0.159 | 0.704 | 0.55 | 0.19 | 0.63 | 0.81 | 0.62 | 0.66 |
| 1 | 59435688 | rs7543260 | 0.401 | -0.171 | -0.344 | -1.427 | 0 | -0.006 | -0.006 | 1.558 | 0.23 | 0.25 | 0.12 | 0.14 | 0.21 | 0.17 |
| 1 | 59435778 | rs2764933 | -0.911 | -0.335 | -0.263 | -1.769 | 0.011 | 0.116 | 0.116 | 0 | 0.49 | 0.67 | 0.42 | 0.19 | 0.44 | 0.28 |
| 1 | 59438203 | rs2716128 | 0.079 | -0.338 | -0.375 | -1.769 | 0.006 | 0.029 | 0.029 | 0.285 | 0.23 | 0.16 | 0.27 | 0.05 | 0.22 | 0.11 |
| 1 | 59443050 | rs7537319 | -1.695 | -0.221 | -0.665 | -1.57 | -0.134 | -0.054 | -0.054 | 0.071 | 0.65 | 0.94 | 0.5 | 0.33 | 0.54 | 0.42 |
| 1 | 59443353 | rs6661726 | 0.589 | -0.039 | -0.151 | -1.324 | 0 | -0.004 | -0.004 | 0.497 | 0.13 | 0.01 | 0.06 | 0.11 | 0.1 | 0.13 |
| 1 | 59456803 | rs11207361 | 0.171 | 0.493 | 0.986 | -1.146 | 0.01 | -0.021 | -0.021 | 0 | 0.22 | 0.33 | 0.12 | 0.15 | 0.21 | 0.16 |
| 1 | 59464697 | rs2985884 | -1.092 | 0.472 | 0.758 | -1.2 | -0.007 | 0.109 | 0.109 | 0 | 0.45 | 0.76 | 0.39 | 0.21 | 0.42 | 0.28 |
| 1 | 59466192 | rs2764900 | -1.259 | 0.465 | 0.748 | -1.204 | -0.003 | 0.057 | 0.057 | 0 | 0.45 | 0.69 | 0.38 | 0.21 | 0.42 | 0.28 |
| 1 | 59470439 | rs11207367 | 0.498 | 0.441 | 0.886 | -1.242 | -0.004 | 0.004 | 0.004 | 0 | 0.21 | 0.19 | 0.11 | 0.14 | 0.22 | 0.17 |
| 1 | 59470494 | rs10158505 | -1.107 | 0.372 | 0.885 | -1.232 | 0.002 | -0.033 | -0.033 | 0 | 0.43 | 0.47 | 0.36 | 0.21 | 0.42 | 0.28 |
| 1 | 59472397 | rs2716140 | -1.193 | 0.316 | 0.571 | -1.234 | -0.001 | 0.016 | 0.016 | 0 | 0.44 | 0.58 | 0.37 | 0.21 | 0.42 | 0.28 |
| 1 | 59474027 | rs1812402 | -1.145 | 0.321 | 0.059 | -1.231 | -0.001 | 0.094 | 0.094 | 0.004 | 0.45 | 0.74 | 0.39 | 0.21 | 0.42 | 0.28 |
| 1 | 59482626 | rs3010336 | -0.195 | 0.686 | -0.832 | -1.212 | 0 | -0.02 | -0.02 | 0 | 0.21 | 0.23 | 0.25 | 0.07 | 0.22 | 0.11 |
| 1 | 59486551 | rs12748496 | -0.467 | 0.716 | -1.043 | -0.549 | -0.006 | 0.015 | 0.015 | 0 | 0.24 | 0.18 | 0.13 | 0.13 | 0.23 | 0.18 |
| 1 | 59489803 | rs9436591 | -0.632 | -0.058 | -1.451 | -0.938 | 0.058 | 0.16 | 0.16 | 0 | 0.42 | 0.73 | 0.39 | 0.12 | 0.34 | 0.24 |
| 1 | 59498925 | rs17530657 | -0.585 | 0.087 | -1.536 | -0.943 | -0.04 | 0.033 | 0.033 | 0 | 0.07 | 0.01 | 0.19 | 0.01 | 0.12 | 0.06 |
| 1 | 59523179 | rs17118649 | -0.567 | -0.552 | -0.851 | -1.033 | -0.009 | -0.025 | -0.025 | 0.058 | 0.3 | 0.38 | 0.18 | 0.11 | 0.22 | 0.25 |
| 1 | 59526466 | rs6687397 | 0.787 | -0.614 | -0.249 | -0.343 | 0.004 | 0.001 | 0.001 | 0 | 0.57 | 0.42 | 0.61 | 0.56 | 0.63 | 0.47 |
| 1 | 59526860 | rs10789073 | -0.431 | -0.604 | 0.138 | -0.199 | -0.003 | 0.017 | 0.017 | 0.302 | 0.84 | 0.83 | 0.85 | 0.75 | 0.85 | 0.72 |
| 1 | 59531321 | rs12733019 | -0.406 | -0.606 | 0.138 | -0.2 | -0.003 | 0.017 | 0.017 | 0.282 | 0.84 | 0.83 | 0.85 | 0.75 | 0.85 | 0.72 |
| 1 | 59533996 | rs4128414 | 0.078 | -0.702 | 0.392 | -0.307 | 0.003 | 0.004 | 0.004 | 0 | 0.27 | 0.3 | 0.24 | 0.18 | 0.21 | 0.25 |
| 1 | 59537214 | rs7536474 | -0.925 | -0.856 | 0.33 | -0.458 | 0 | 0.006 | 0.006 | 0 | 0.16 | 0.17 | 0.22 | 0.11 | 0.16 | 0.09 |
| 1 | 59544435 | rs4424536 | 1.732 | -0.798 | 0.269 | -0.39 | -0.03 | -0.021 | -0.021 | 0 | 0.39 | 0.24 | 0.37 | 0.46 | 0.47 | 0.33 |
| 1 | 59551895 | rs1331748 | 0.225 | -0.736 | 0.228 | -0.349 | -0.001 | 0.021 | 0.021 | 0.351 | 0.16 | 0.22 | 0.15 | 0.25 | 0.16 | 0.32 |
| 1 | 59552704 | rs9970281 | 0.336 | -0.769 | 0.202 | -0.372 | 0.02 | 0.028 | 0.028 | 0 | 0.27 | 0.22 | 0.23 | 0.18 | 0.21 | 0.26 |
| 1 | 59556763 | rs3010318 | 0.148 | -0.615 | -0.046 | -0.617 | -0.014 | 0.013 | 0.013 | 0 | 0.87 | 0.94 | 0.79 | 0.89 | 0.83 | 0.91 |
| 1 | 59559189 | rs11207402 | 0.106 | -0.573 | 0.053 | -0.52 | -0.001 | 0.001 | 0.001 | 0.419 | 0.16 | 0.13 | 0.15 | 0.24 | 0.16 | 0.32 |
| 1 | 59567263 | rs3010323 | -0.219 | -0.609 | -0.039 | -0.749 | 0.007 | -0.008 | -0.008 | 0 | 0.75 | 0.78 | 0.85 | 1 | 0.77 | 0.87 |
| 1 | 59569829 | rs3010325 | 1.086 | -0.434 | 0.049 | -0.226 | -0.001 | -0.003 | -0.003 | 0.047 | 0.48 | 0.49 | 0.46 | 0.51 | 0.53 | 0.47 |
| 1 | 59573341 | rs2068743 | -0.317 | -0.43 | -0.171 | -0.589 | 0.004 | -0.025 | -0.025 | 0 | 0.25 | 0.25 | 0.15 | 0 | 0.23 | 0.13 |
| 1 | 59577136 | rs914462 | 0.029 | -0.774 | -0.231 | -0.647 | 0.005 | 0.001 | 0.001 | 0 | 0.7 | 0.76 | 0.86 | 0.91 | 0.79 | 0.91 |
| 1 | 59578582 | rs10493260 | -0.05 | -0.958 | -0.667 | -0.716 | 0.001 | 0.017 | 0.017 | 1.483 | 0.15 | 0.09 | 0.1 | 0.13 | 0.12 | 0.14 |
| 1 | 59578785 | rs1331746 | -0.175 | -1.103 | -0.585 | -0.813 | 0.006 | 0.001 | 0.001 | 0 | 0.69 | 0.76 | 0.86 | 0.91 | 0.79 | 0.87 |
| 1 | 59580485 | rs17118788 | 1.19 | -0.536 | -0.116 | -0.306 | -0.018 | -0.052 | -0.052 | 0 | 0.24 | 0.13 | 0.32 | 0.41 | 0.26 | 0.22 |
| 1 | 59588128 | rs332230 | 0.75 | -0.588 | -0.117 | -0.347 | 0.013 | 0.017 | 0.017 | 0 | 0.3 | 0.21 | 0.14 | 0.09 | 0.21 | 0.14 |
| 1 | 59588984 | rs2985500 | -0.109 | -0.604 | 0.277 | -0.35 | 0.001 | 0.001 | 0.001 | 0 | 0.45 | 0.5 | 0.52 | 0.5 | 0.53 | 0.65 |
| 1 | 59589399 | rs17118797 | 1.237 | -0.6 | 0.255 | -0.347 | -0.027 | -0.072 | -0.072 | 0 | 0.24 | 0.11 | 0.32 | 0.41 | 0.26 | 0.22 |
| 1 | 59592256 | rs632986 | -0.048 | -0.67 | 0.691 | -0.376 | -0.006 | -0.004 | -0.004 | 0 | 0.54 | 0.35 | 0.46 | 0.5 | 0.47 | 0.35 |
| 1 | 59593315 | rs617922 | 0.026 | -0.66 | 0.597 | -0.372 | 0.004 | 0.003 | 0.003 | 0 | 0.56 | 0.47 | 0.47 | 0.5 | 0.47 | 0.35 |
| 1 | 59597553 | rs2989870 | 0.534 | -0.619 | 0.252 | -0.355 | -0.004 | -0.008 | -0.008 | 0 | 0.15 | 0.22 | 0.11 | 0.13 | 0.11 | 0.14 |
| 1 | 59601907 | rs182816 | -1.439 | -0.527 | -0.025 | -0.299 | -0.003 | -0.008 | -0.008 | 0 | 0.76 | 0.78 | 0.67 | 0.59 | 0.74 | 0.78 |
| 1 | 59603757 | rs4912389 | 1.037 | -0.609 | -0.341 | -0.299 | -0.127 | -0.15 | -0.15 | 0 | 0.26 | 0.03 | 0.37 | 0.43 | 0.38 | 0.49 |
| 1 | 59606640 | rs11207409 | 0.345 | -0.357 | 0.46 | -0.043 | 0.002 | -0.004 | -0.004 | 0 | 0.55 | 0.35 | 0.53 | 0.45 | 0.58 | 0.55 |
| 1 | 59608025 | rs1475629 | -1.223 | 0.023 | 0.486 | 0.342 | 0.007 | 0.001 | 0.001 | 0 | 0.81 | 0.8 | 0.7 | 0.77 | 0.78 | 0.84 |
| 1 | 59610174 | rs6666629 | -0.238 | -0.353 | 1.045 | 0.147 | -0.001 | -0.045 | -0.045 | 0.073 | 0.1 | 0.39 | 0.07 | 0 | 0.05 | 0.03 |
| 1 | 59614192 | rs17118875 | 0.681 | -0.298 | 1.012 | 0.177 | -0.012 | -0.035 | -0.035 | 0 | 0.07 | 0.12 | 0.05 | 0 | 0.04 | 0.03 |
| 1 | 59618816 | rs6676724 | -0.559 | 0.092 | 1.091 | 0.307 | 0.007 | -0.028 | -0.028 | 0 | 0.15 | 0.24 | 0.12 | 0.07 | 0.16 | 0.17 |

| | | | | | | | | | | | | | | | | |
|---|---|---|---|---|---|---|---|---|---|---|---|---|---|---|---|---|
| 1 | 59619571 | rs474103 | 1.506 | 0.58 | 0.933 | 0.621 | 0.015 | -0.001 | -0.001 | 0.59 | 0.67 | 0.39 | 0.62 | 0.66 | 0.55 | 0.59 |
| 1 | 59623181 | rs615743 | 0.584 | -0.117 | 0.954 | 1.011 | -0.027 | 0.021 | 0.021 | 0.909 | 0.11 | 0.14 | 0.03 | 0.19 | 0.03 | 0.08 |
| 1 | 59630555 | rs12139438 | 1.937 | -0.019 | 0.8 | 0.945 | -0.027 | -0.029 | -0.029 | 1.207 | 0.56 | 0.25 | 0.59 | 0.46 | 0.52 | 0.52 |
| 1 | 59633846 | rs3908770 | -1.005 | 0.298 | 0.853 | 0.743 | 0.022 | -0.015 | -0.015 | 0 | 0.27 | 0.35 | 0.27 | 0.2 | 0.4 | 0.31 |
| 1 | 59635870 | rs3850542 | -0.023 | 0.301 | 0.866 | 0.733 | 0.012 | -0.037 | -0.037 | 0 | 0.15 | 0.31 | 0.13 | 0.07 | 0.16 | 0.17 |
| 1 | 59642102 | rs11207416 | 0.576 | 0.353 | -0.259 | 0.873 | -0.073 | -0.021 | -0.021 | 0 | 0.12 | 0.03 | 0.14 | 0.13 | 0.24 | 0.15 |
| 1 | 59646978 | rs12733512 | 0.895 | -0.494 | -0.247 | 0.895 | -0.059 | -0.02 | -0.02 | 0 | 0.11 | 0.04 | 0.13 | 0.13 | 0.23 | 0.16 |
| 1 | 59647365 | rs11207422 | 1.353 | -0.906 | -0.572 | 0.923 | -0.024 | 0.041 | 0.041 | 0 | 0.47 | 0.23 | 0.58 | 0.38 | 0.49 | 0.5 |
| 1 | 59649116 | rs9660067 | -0.547 | -0.117 | -0.003 | 1.011 | 0.035 | 0.024 | 0.024 | 0 | 0.17 | 0.26 | 0.23 | 0.26 | 0.27 | 0.25 |
| 1 | 59653630 | rs3850543 | -1.044 | -0.624 | -0.221 | 0.532 | -0.002 | 0 | 0 | 0.069 | 0.33 | 0.34 | 0.22 | 0.32 | 0.34 | 0.21 |
| 1 | 59654019 | rs12753566 | 0.635 | -0.479 | -0.251 | 0.6 | -0.042 | -0.016 | -0.016 | 0 | 0.13 | 0.07 | 0.14 | 0.16 | 0.24 | 0.14 |
| 1 | 59655643 | rs17119044 | 0.876 | -0.645 | -0.187 | 0.524 | -0.005 | -0.024 | -0.024 | 0 | 0.1 | 0.13 | 0.04 | 0 | 0.06 | 0.02 |
| 1 | 59662719 | rs12047394 | 0.68 | -0.704 | -0.138 | 0.88 | -0.01 | 0.017 | 0.017 | 3.931 | 0.09 | 0.18 | 0.03 | 0.16 | 0.04 | 0.04 |
| 1 | 59665542 | rs331635 | 1.343 | -0.686 | 0.384 | 0.732 | 0.003 | 0.007 | 0.007 | 0.249 | 0.65 | 0.41 | 0.73 | 0.51 | 0.63 | 0.69 |
| 1 | 59665770 | rs10493258 | 1.045 | -1 | 0.583 | 0.215 | -0.022 | -0.053 | -0.053 | 0.805 | 0.09 | 0.18 | 0.04 | 0 | 0.06 | 0.01 |
| 1 | 59669708 | rs12758288 | -0.415 | -0.988 | 0.568 | 0.226 | 0.027 | -0.018 | -0.018 | 0.178 | 0.22 | 0.32 | 0.18 | 0.17 | 0.3 | 0.15 |
| 1 | 59680579 | rs331633 | 1.04 | -1.736 | -0.773 | -0.158 | 0.042 | 0.103 | 0.103 | 0 | 0.18 | 0.02 | 0.06 | 0 | 0.11 | 0.11 |
| 1 | 59685265 | rs7542194 | -1.086 | -1.188 | -0.305 | 0.373 | 0.032 | -0.007 | -0.007 | 0 | 0.31 | 0.42 | 0.21 | 0.33 | 0.34 | 0.21 |
| 1 | 59688948 | rs10493256 | 0.602 | -1.098 | -0.503 | 0.41 | -0.036 | -0.015 | -0.015 | 0.764 | 0.13 | 0.07 | 0.14 | 0.17 | 0.24 | 0.15 |
| 1 | 59693470 | rs7540474 | -0.462 | -2.884 | -0.056 | 0.3 | -0.045 | -0.021 | -0.021 | 2.297 | 0.19 | 0.3 | 0.06 | 0.16 | 0.1 | 0.06 |
| 1 | 59694611 | rs7531917 | 0.527 | -3.028 | -0.65 | 0.272 | 0.014 | 0.01 | 0.01 | 0 | 0.46 | 0.6 | 0.55 | 0.65 | 0.63 | 0.58 |
| 1 | 59698062 | rs1037038 | 1.217 | -2.812 | -1.046 | 0.031 | -0.029 | -0.044 | -0.044 | 0 | 0.13 | 0.03 | 0.25 | 0.13 | 0.15 | 0.2 |
| 1 | 59702797 | rs6700125 | 0.753 | -2.753 | -0.75 | 0.277 | 0.077 | -0.01 | -0.01 | 2.679 | 0.52 | 0.36 | 0.67 | 0.5 | 0.68 | 0.58 |
| 1 | 59703738 | rs6703534 | -0.148 | -2.775 | -0.669 | 0.263 | -0.057 | -0.084 | -0.084 | 0.053 | 0.64 | 0.6 | 0.82 | 0.85 | 0.76 | 0.81 |
| 1 | 59703982 | rs6690993 | 0.544 | -2.765 | -0.646 | 0.382 | 0.014 | 0.013 | 0.013 | 0 | 0.54 | 0.5 | 0.72 | 0.68 | 0.68 | 0.7 |
| 1 | 59704822 | rs7547161 | 0.093 | -2.791 | -0.676 | 0.366 | 0.009 | -0.028 | -0.028 | 0 | 0.28 | 0.26 | 0.38 | 0.5 | 0.26 | 0.41 |
| 1 | 59705447 | rs6587852 | -0.229 | -2.789 | -0.671 | 0.359 | -0.02 | -0.035 | -0.035 | 0 | 0.64 | 0.66 | 0.82 | 0.85 | 0.76 | 0.81 |
| 1 | 59707689 | rs12034496 | 0.665 | -2.711 | -1.061 | 0.409 | 0.012 | -0.036 | -0.036 | 0.163 | 0.23 | 0.18 | 0.31 | 0.33 | 0.2 | 0.36 |
| 1 | 59708325 | rs6683082 | 1.581 | -2.708 | -1.052 | 0.412 | 0.006 | 0 | 0 | 0 | 0.3 | 0.33 | 0.41 | 0.36 | 0.48 | 0.34 |
| 1 | 59714992 | rs1470407 | -0.629 | -2.807 | -1.187 | 0.352 | -0.067 | -0.041 | -0.041 | 0 | 0.73 | 0.62 | 0.84 | 0.83 | 0.84 | 0.86 |
| 1 | 59715273 | rs11207432 | -1.023 | -2.889 | -1.016 | -0.256 | 0.031 | -0.022 | -0.022 | 0 | 0.43 | 0.3 | 0.43 | 0.48 | 0.37 | 0.53 |
| 1 | 59716008 | rs12743706 | -0.557 | -2.634 | -1.61 | -1.378 | 0.012 | 0.029 | 0.029 | 0 | 0.2 | 0.11 | 0.12 | 0.15 | 0.17 | 0.16 |
| 1 | 59722872 | rs17478612 | 0.241 | -2.349 | -1.542 | -1.008 | -0.001 | 0 | 0 | 0.142 | 0.14 | 0.05 | 0.08 | 0.15 | 0.14 | 0.05 |
| 1 | 59729342 | rs751101 | 0.031 | -2.202 | -1.081 | -1.064 | -0.015 | -0.012 | -0.012 | 0 | 0.22 | 0.04 | 0.27 | 0.14 | 0.21 | 0.27 |
| 1 | 59731843 | rs333683 | 0.328 | -1.606 | 0.14 | -0.761 | -0.083 | -0.133 | -0.133 | 0.324 | 0.78 | 0.66 | 0.91 | 1 | 0.9 | 0.91 |
| 1 | 59743914 | rs333662 | 0.439 | -1.641 | -0.093 | -0.838 | -0.047 | -0.102 | -0.102 | 0.636 | 0.71 | 0.65 | 0.86 | 1 | 0.84 | 0.88 |
| 1 | 59744841 | rs333661 | -0.667 | -1.905 | 0.213 | -0.581 | -0.03 | 0.006 | 0.006 | 0 | 0.06 | 0.12 | 0.06 | 0.04 | 0.02 | 0.08 |
| 1 | 59745790 | rs7532046 | -0.324 | -2.122 | 0.179 | -0.751 | 0.004 | 0.006 | 0.006 | 0 | 0.14 | 0.13 | 0.05 | 0.07 | 0.11 | 0.09 |
| 1 | 59754855 | rs863123 | 0.192 | -1.85 | -0.018 | -0.045 | 0 | 0.014 | 0.014 | 0 | 0.21 | 0.21 | 0.26 | 0.39 | 0.22 | 0.36 |
| 1 | 59757049 | rs835365 | -1.218 | -2.115 | 0.567 | -0.124 | -0.008 | 0.005 | 0.005 | 0 | 0.37 | 0.54 | 0.41 | 0.52 | 0.35 | 0.51 |
| 1 | 59782017 | rs835401 | -1.236 | -2.011 | 0.538 | 0.103 | -0.005 | 0.008 | 0.008 | 0 | 0.37 | 0.55 | 0.41 | 0.48 | 0.35 | 0.51 |
| 1 | 59791970 | rs17480463 | 0.272 | -1.337 | 1.069 | 0.063 | 0.002 | 0.007 | 0.007 | 0 | 0.14 | 0.12 | 0.05 | 0.07 | 0.11 | 0.1 |
| 1 | 59794792 | rs11207437 | 0.682 | -0.559 | 1.062 | 0.039 | 0.011 | 0 | 0 | 0 | 0.12 | 0.16 | 0.18 | 0.15 | 0.13 | 0.12 |
| 1 | 59796808 | rs9787234 | 0.852 | -0.548 | 1.084 | 0.074 | -0.038 | -0.111 | -0.111 | 0 | 0.19 | 0.03 | 0.27 | 0.34 | 0.23 | 0.36 |
| 1 | 59796838 | rs835378 | -1.457 | -0.559 | 1.101 | 0.069 | -0.012 | 0.09 | 0.09 | 0 | 0.5 | 0.79 | 0.59 | 0.61 | 0.49 | 0.63 |
| 1 | 59800996 | rs835385 | -1.248 | -0.563 | 1.431 | 0.294 | -0.005 | 0.014 | 0.014 | 0 | 0.35 | 0.55 | 0.41 | 0.4 | 0.35 | 0.51 |
| 1 | 59812293 | rs835356 | -0.681 | -0.565 | 1.914 | 0.29 | -0.004 | 0.025 | 0.025 | 0 | 0.16 | 0.41 | 0.13 | 0.12 | 0.13 | 0.15 |
| 1 | 59814228 | rs835361 | -1.206 | -0.549 | 1.83 | 0.277 | 0.001 | -0.003 | -0.003 | 0 | 0.36 | 0.63 | 0.42 | 0.47 | 0.36 | 0.51 |
| 1 | 59832762 | rs7539697 | -0.572 | -0.313 | 1.802 | 0.257 | -0.009 | 0.076 | 0.076 | 0 | 0.27 | 0.44 | 0.36 | 0.39 | 0.27 | 0.41 |
| 1 | 59841785 | rs949827 | -0.608 | -0.334 | 1.685 | 0.262 | -0.015 | -0.034 | -0.034 | 0 | 0.27 | 0.13 | 0.3 | 0.32 | 0.31 | 0.44 |
| 1 | 59844471 | rs835435 | -1.518 | 0 | 1.97 | 0.295 | -0.007 | -0.037 | -0.037 | 0 | 0.37 | 0.68 | 0.41 | 0.43 | 0.4 | 0.47 |
| 1 | 59850907 | rs2173033 | 0.209 | -0.194 | 2.007 | 0.311 | -0.002 | -0.022 | -0.022 | 0 | 0.18 | 0.1 | 0.25 | 0.34 | 0.2 | 0.36 |
| 1 | 59853319 | rs7534016 | 0.227 | -0.194 | 1.667 | 0.311 | -0.046 | -0.123 | -0.123 | 0.495 | 0.18 | 0.01 | 0.23 | 0.34 | 0.2 | 0.36 |
| 1 | 59860445 | rs835426 | -0.954 | -0.236 | 1.844 | 0.282 | -0.014 | 0.017 | 0.017 | 0 | 0.28 | 0.53 | 0.31 | 0.34 | 0.23 | 0.46 |
| 1 | 59869604 | rs628177 | -1.349 | -0.207 | 1.186 | 0.334 | 0.015 | -0.037 | -0.037 | 0 | 0.35 | 0.7 | 0.4 | 0.41 | 0.33 | 0.5 |
| 1 | 59875451 | rs11576871 | 0.034 | -0.143 | 0.984 | 0.351 | 0 | 0.006 | 0.006 | 0 | 0.2 | 0.19 | 0.25 | 0.34 | 0.19 | 0.37 |
| 1 | 59877775 | rs17119433 | -0.619 | -0.155 | 0.469 | 0.356 | -0.008 | -0.008 | -0.008 | 0 | 0.14 | 0.36 | 0.14 | 0.07 | 0.13 | 0.12 |
| 1 | 59881472 | rs657746 | -0.776 | 0.029 | -0.733 | 0.3 | 0.013 | -0.031 | -0.031 | 0 | 0.35 | 0.7 | 0.4 | 0.41 | 0.33 | 0.49 |
| 1 | 59891290 | rs10493264 | -0.106 | 0.472 | -1.584 | 0.328 | -0.029 | -0.057 | -0.057 | 0 | 0.39 | 0.17 | 0.42 | 0.4 | 0.41 | 0.35 |
| 1 | 59891821 | rs7520470 | 0.062 | 0.892 | -1.695 | 0.575 | -0.033 | -0.058 | -0.058 | 0 | 0.36 | 0.13 | 0.4 | 0.38 | 0.39 | 0.36 |
| 1 | 59892655 | rs946574 | -0.115 | 1.156 | -1.129 | 0.208 | 0.013 | -0.033 | -0.033 | 0.2 | 0.18 | 0.26 | 0.17 | 0.08 | 0.19 | 0.16 |
| 1 | 59894323 | rs1416788 | -0.62 | 1.175 | -0.389 | 0.029 | 0.017 | -0.021 | -0.021 | 0 | 0.61 | 0.69 | 0.66 | 0.48 | 0.64 | 0.57 |
| 1 | 59900041 | rs1879689 | 0.406 | 1.349 | 0.228 | 0.061 | 0.003 | 0 | 0 | 0.062 | 0.43 | 0.49 | 0.48 | 0.39 | 0.44 | 0.4 |
| 1 | 59901944 | rs4912396 | 0.062 | 1.187 | 0.511 | 0.069 | 0.004 | 0.005 | 0.005 | 1.047 | 0.49 | 0.53 | 0.54 | 0.35 | 0.57 | 0.46 |
| 1 | 59904180 | rs11207448 | 0.444 | 1.33 | 0.476 | 0.067 | 0.038 | -0.036 | -0.036 | 0 | 0.09 | 0.12 | 0.1 | 0.02 | 0.14 | 0.08 |
| 1 | 59909614 | rs10889136 | 0.768 | 0.975 | 0.572 | 0.277 | -0.078 | -0.016 | -0.016 | 0 | 0.38 | 0.12 | 0.41 | 0.34 | 0.42 | 0.38 |
| 1 | 59912718 | rs12145590 | 0.959 | 1.046 | 0.411 | 0.406 | -0.127 | 0.042 | 0.042 | 0.959 | 0.36 | 0.03 | 0.31 | 0.21 | 0.37 | 0.34 |
| 1 | 59913542 | rs2177072 | 0.737 | 0.877 | 0.564 | 0.374 | -0.005 | 0.01 | 0.01 | 0 | 0.06 | 0.07 | 0.06 | 0.1 | 0.04 | 0.03 |
| 1 | 59924256 | rs7529551 | 0.624 | 1.045 | 0.907 | 0.454 | -0.065 | 0.046 | 0.046 | 0 | 0.4 | 0.16 | 0.43 | 0.26 | 0.47 | 0.38 |

| | | | | | | | | | | | | | | | | |
|---|---|---|---|---|---|---|---|---|---|---|---|---|---|---|---|---|
| 1 | 59925666 | rs12046441 | 1.042 | 0.279 | 1.034 | 0.329 | 0.001 | 0.006 | 0.006 | 0.255 | 0.19 | 0.2 | 0.25 | 0.49 | 0.17 | 0.21 |
| 1 | 59936406 | rs2407515 | 0.814 | 0.437 | 0.799 | 0.174 | -0.001 | 0.001 | 0.001 | 0 | 0.41 | 0.37 | 0.45 | 0.27 | 0.46 | 0.38 |
| 1 | 59940664 | rs6663897 | 0.608 | 0.382 | 0.873 | 0.168 | 0.002 | -0.01 | -0.01 | 0 | 0.43 | 0.52 | 0.46 | 0.27 | 0.45 | 0.38 |
| 1 | 59946306 | rs12563198 | 0.48 | 0.305 | 0.66 | 0.18 | -0.011 | 0.003 | 0.003 | 0 | 0.08 | 0.11 | 0.06 | 0.1 | 0.04 | 0.06 |
| 1 | 59946350 | rs626733 | 0.679 | 0.296 | 0.694 | 0.146 | -0.001 | 0.001 | 0.001 | 0 | 0.42 | 0.53 | 0.48 | 0.26 | 0.44 | 0.38 |
| 1 | 59951705 | rs1188236 | 0.178 | 0.296 | 0.521 | 0.169 | -0.003 | 0.029 | 0.029 | 0 | 0.51 | 0.67 | 0.54 | 0.37 | 0.48 | 0.44 |
| 1 | 59956092 | rs4258244 | 0.184 | 0.305 | 0.489 | 0.149 | -0.002 | 0.03 | 0.03 | 0 | 0.51 | 0.66 | 0.53 | 0.37 | 0.48 | 0.44 |
| 1 | 59964314 | rs11808336 | 0.411 | 0.355 | 0.906 | 0.363 | -0.002 | -0.001 | -0.001 | 0.017 | 0.07 | 0.07 | 0.05 | 0.1 | 0.04 | 0.06 |
| 1 | 59969263 | rs592484 | -0.173 | 0.252 | -0.18 | 0.026 | -0.001 | 0.026 | 0.026 | 0 | 0.37 | 0.66 | 0.46 | 0.22 | 0.39 | 0.23 |
| 1 | 59971351 | rs10789077 | 0.859 | 0.303 | -0.226 | 0.02 | -0.02 | 0.041 | 0.041 | 0 | 0.27 | 0.19 | 0.36 | 0.13 | 0.34 | 0.17 |
| 1 | 59973874 | rs4912402 | 1.101 | 0.281 | -0.483 | 0.061 | 0.004 | -0.121 | -0.121 | 0 | 0.16 | 0.03 | 0.22 | 0.55 | 0.14 | 0.19 |
| 1 | 59977019 | rs621517 | -0.351 | 0.597 | 0.121 | -0.391 | 0.001 | 0.017 | 0.017 | 0 | 0.36 | 0.63 | 0.45 | 0.22 | 0.37 | 0.23 |
| 1 | 59985298 | rs10889144 | 0.452 | 0.829 | -0.659 | -0.861 | -0.002 | 0 | 0 | 0 | 0.23 | 0.18 | 0.13 | 0 | 0.24 | 0.11 |
| 1 | 59990791 | rs12732719 | -0.606 | 0.197 | 0.7 | -0.91 | 0.003 | -0.025 | -0.025 | 0 | 0.11 | 0.42 | 0.07 | 0.05 | 0.05 | 0.12 |
| 1 | 59992986 | rs6692151 | -0.512 | 0.097 | 0.02 | -1.791 | -0.014 | 0.04 | 0.04 | 0 | 0.58 | 0.69 | 0.35 | 0.17 | 0.48 | 0.3 |
| 1 | 60006785 | rs7527689 | -0.442 | 0.474 | 0.097 | -1.195 | 0.005 | 0.014 | 0.014 | 0 | 0.15 | 0.28 | 0.13 | 0.16 | 0.12 | 0.35 |
| 1 | 60007275 | rs2798652 | -0.044 | 1.022 | -0.385 | -1.384 | 0.04 | -0.029 | -0.029 | 0 | 0.2 | 0.3 | 0.34 | 0.14 | 0.27 | 0.28 |
| 1 | 60007658 | rs11811141 | -0.255 | 0.934 | -0.216 | -1.289 | 0.024 | 0.045 | 0.045 | 0 | 0.16 | 0.43 | 0.14 | 0.16 | 0.12 | 0.35 |
| 1 | 60007922 | rs12069567 | -0.255 | 0.934 | -0.216 | -1.289 | 0.024 | 0.045 | 0.045 | 0 | 0.16 | 0.43 | 0.14 | 0.16 | 0.12 | 0.35 |
| 1 | 60015280 | rs12562305 | 1.245 | 0.65 | -0.477 | -1.564 | 0.001 | -0.003 | -0.003 | 0.953 | 0.11 | 0.06 | 0.09 | 0.09 | 0.09 | 0.05 |
| 1 | 60019782 | rs2272052 | 1.055 | 0.44 | -0.212 | -1.591 | 0.001 | -0.009 | -0.009 | 0 | 0.05 | 0.2 | 0.02 | 0.02 | 0.02 | 0.05 |
| 1 | 60023941 | rs1440575 | -0.312 | 0.087 | -0.382 | -1.791 | 0.004 | 0.09 | 0.09 | 0 | 0.51 | 0.74 | 0.57 | 0.36 | 0.47 | 0.65 |
| 1 | 60030451 | rs2685541 | 0.746 | 0.528 | 0.563 | -0.983 | 0.035 | 0.016 | 0.016 | 0 | 0.12 | 0.26 | 0.31 | 0.14 | 0.14 | 0.22 |
| 1 | 60045220 | rs4497239 | -0.333 | 0.492 | 0.345 | -0.963 | 0.025 | 0.032 | 0.032 | 0.009 | 0.13 | 0.26 | 0.12 | 0.14 | 0.11 | 0.32 |
| 1 | 60059135 | rs6674107 | -2.145 | 0.356 | -0.126 | -1.159 | -0.04 | -0.046 | -0.046 | 0 | 0.56 | 0.96 | 0.63 | 0.38 | 0.53 | 0.7 |
| 1 | 60061007 | rs9436596 | 0.324 | 0.386 | 0.688 | -0.676 | -0.012 | 0 | 0 | 0 | 0.13 | 0.08 | 0.29 | 0.14 | 0.16 | 0.25 |
| 1 | 60065474 | rs12740006 | -1.447 | 0.239 | 0.962 | -0.738 | -0.01 | -0.007 | -0.007 | 0 | 0.19 | 0.31 | 0.15 | 0.16 | 0.13 | 0.37 |
| 1 | 60065520 | rs12757929 | -0.164 | 0.29 | 0.907 | -0.707 | -0.002 | -0.004 | -0.004 | 0 | 0.11 | 0.11 | 0.11 | 0.14 | 0.09 | 0.32 |
| 1 | 60066022 | rs12724194 | 0.043 | 0.295 | 0.897 | -0.705 | -0.004 | -0.007 | -0.007 | 0 | 0.11 | 0.09 | 0.11 | 0.14 | 0.09 | 0.32 |
| 1 | 60069296 | rs11207488 | -0.102 | 0.289 | 1.173 | -0.708 | 0.02 | 0.036 | 0.036 | 0.557 | 0.11 | 0.22 | 0.12 | 0.14 | 0.09 | 0.32 |
| 1 | 60071746 | rs4571987 | -0.597 | 0.294 | 1.759 | -0.741 | 0.014 | -0.003 | -0.003 | 0 | 0.86 | 0.81 | 0.7 | 0.86 | 0.84 | 0.75 |
| 1 | 60099282 | rs6690357 | 1.566 | 0.203 | 1.61 | -0.79 | 0.008 | 0.021 | 0.021 | 0 | 0.65 | 0.23 | 0.52 | 0.7 | 0.7 | 0.37 |
| 1 | 60100662 | rs7524379 | 0.197 | 0.436 | 1.954 | -0.502 | 0.014 | 0.006 | 0.006 | 0 | 0.05 | 0.26 | 0.04 | 0 | 0.02 | 0.03 |
| 1 | 60128805 | rs7540838 | -0.18 | 0.482 | 1.305 | -0.453 | 0.004 | 0 | 0 | 0.072 | 0.86 | 0.85 | 0.7 | 0.85 | 0.83 | 0.76 |
| 1 | 60140439 | rs4578243 | 1.2 | 0.398 | 1.4 | -0.538 | -0.023 | -0.047 | -0.047 | 0 | 0.75 | 0.51 | 0.65 | 0.84 | 0.79 | 0.67 |
| 1 | 60146753 | rs4912219 | 0.169 | 1.017 | 1.033 | -0.268 | 0.007 | -0.002 | -0.002 | 0 | 0.78 | 0.78 | 0.61 | 0.79 | 0.73 | 0.7 |
| 1 | 60157178 | rs12732614 | 1.295 | 0.669 | 0.642 | 0.008 | -0.005 | -0.03 | -0.03 | 0.325 | 0.62 | 0.29 | 0.44 | 0.57 | 0.65 | 0.48 |
| 1 | 60161256 | rs4357555 | -0.99 | 0.768 | 0.534 | -0.076 | -0.04 | -0.026 | -0.026 | 0.467 | 0.13 | 0.09 | 0.35 | 0.17 | 0.19 | 0.22 |
| 1 | 60164209 | rs4244022 | 1.581 | 1.374 | 0.628 | 0.221 | -0.054 | -0.051 | -0.051 | 0 | 0.6 | 0.21 | 0.39 | 0.52 | 0.53 | 0.46 |
| 1 | 60168554 | rs6661666 | -0.725 | 1.794 | 0.79 | 0.484 | 0.043 | -0.002 | -0.002 | 0 | 0.16 | 0.24 | 0.39 | 0.14 | 0.29 | 0.26 |
| 1 | 60173734 | rs6685309 | 0.31 | 0.916 | 0.778 | 0.767 | 0.001 | 0.088 | 0.088 | 0 | 0.81 | 0.67 | 0.81 | 0.67 | 0.83 | 0.7 |
| 1 | 60183673 | rs1933213 | 0.62 | 0.89 | 0.121 | 0.733 | 0.009 | 0.066 | 0.066 | 0 | 0.84 | 0.69 | 0.68 | 0.76 | 0.85 | 0.71 |
| 1 | 60184557 | rs2152938 | 0.397 | 0.998 | -0.85 | 0.126 | 0.012 | 0.055 | 0.055 | 0 | 0.5 | 0.3 | 0.47 | 0.65 | 0.55 | 0.5 |
| 1 | 60196337 | rs17557490 | -0.821 | 1.469 | -1.308 | 0.524 | -0.054 | -0.048 | -0.048 | 0.533 | 0.1 | 0.01 | 0.35 | 0.14 | 0.16 | 0.24 |
| 1 | 60198430 | rs1890212 | -0.499 | 1.026 | -1.371 | -0.14 | -0.009 | 0.02 | 0.02 | 0 | 0.41 | 0.69 | 0.45 | 0.81 | 0.49 | 0.43 |
| 1 | 60199591 | rs9436185 | -1.182 | 0.951 | -1.54 | 0.178 | -0.001 | -0.001 | -0.001 | 0 | 0.12 | 0.14 | 0.06 | 0.13 | 0.11 | 0.22 |
| 1 | 60201195 | rs2897708 | 0.105 | 0.839 | -2.188 | 0.269 | -0.032 | -0.002 | -0.002 | 0 | 0.39 | 0.1 | 0.46 | 0.36 | 0.41 | 0.27 |
| 1 | 60205784 | rs4915905 | -0.694 | 0.964 | -2.01 | 0.526 | -0.006 | 0.018 | 0.018 | 0.679 | 0.1 | 0.17 | 0.04 | 0.19 | 0.08 | 0.21 |
| 1 | 60210649 | rs1004772 | -0.027 | 1.072 | -2.129 | 0.404 | -0.023 | -0.013 | -0.013 | 0 | 0.86 | 0.98 | 0.91 | 0.85 | 0.81 | 0.78 |
| 1 | 60213519 | rs2179662 | 0.061 | 1.018 | -2.089 | 0.362 | -0.001 | 0 | 0 | 0 | 0.84 | 0.83 | 0.88 | 0.88 | 0.8 | 0.79 |
| 1 | 60215711 | rs9726956 | 0.015 | 0.84 | -2.672 | 0.634 | -0.066 | 0.015 | 0.015 | 0.364 | 0.45 | 0.12 | 0.37 | 0.39 | 0.44 | 0.51 |
| 1 | 60222380 | rs4915686 | 0.11 | 0.766 | -2.235 | 0.814 | -0.049 | -0.033 | -0.033 | 0 | 0.52 | 0.13 | 0.38 | 0.52 | 0.49 | 0.54 |
| 1 | 60223460 | rs590704 | 0.078 | 0.898 | -1.265 | 0.664 | -0.014 | -0.014 | -0.014 | 0.241 | 0.38 | 0.47 | 0.41 | 0.28 | 0.31 | 0.21 |
| 1 | 60224206 | rs1933216 | 0.019 | 1.751 | -1.272 | 0.684 | 0.007 | 0 | 0 | 0 | 0.12 | 0.14 | 0.32 | 0.11 | 0.27 | 0.22 |
| 1 | 60226496 | rs654165 | -0.024 | 0.542 | -0.946 | 0.663 | -0.017 | -0.011 | -0.011 | 0 | 0.43 | 0.47 | 0.35 | 0.34 | 0.35 | 0.35 |
| 1 | 60233747 | rs1007266 | -0.12 | 0.599 | -1.081 | 1.003 | 0.001 | -0.006 | -0.006 | 0.128 | 0.15 | 0.13 | 0.08 | 0.29 | 0.14 | 0.08 |
| 1 | 60238762 | rs7554648 | 0.311 | -0.919 | -2.034 | 0.997 | 0.073 | -0.064 | -0.064 | 0 | 0.18 | 0.01 | 0.07 | 0.21 | 0.07 | 0.27 |
| 1 | 60240601 | rs674949 | -1.001 | -0.126 | -2.548 | 0.598 | -0.046 | -0.005 | -0.005 | 0 | 0.79 | 0.95 | 0.71 | 0.83 | 0.76 | 0.83 |
| 1 | 60243182 | rs9436605 | 0.177 | 0.621 | -2.081 | 0.197 | -0.045 | 0.076 | 0.076 | 0.037 | 0.23 | 0.13 | 0.28 | 0.04 | 0.33 | 0.19 |
| 1 | 60257931 | rs9436188 | 0.063 | 1.189 | -1.642 | 0.446 | 0.006 | -0.001 | -0.001 | 0.09 | 0.15 | 0.18 | 0.28 | 0.1 | 0.28 | 0.17 |
| 1 | 60259626 | rs6672451 | -0.656 | 1 | -0.606 | 0.465 | 0.088 | -0.008 | -0.008 | 0 | 0.2 | 0.45 | 0.31 | 0.17 | 0.32 | 0.17 |
| 1 | 60275095 | rs11207520 | -0.13 | 1.002 | -0.149 | 0.475 | 0.096 | 0.001 | 0.001 | 0 | 0.17 | 0.34 | 0.28 | 0.14 | 0.3 | 0.17 |
| 1 | 60279540 | rs1933218 | 0.397 | 0.953 | -0.302 | 0.425 | -0.017 | 0.017 | 0.017 | 0 | 0.14 | 0.11 | 0.19 | 0.04 | 0.23 | 0.16 |
| 1 | 60302414 | rs11207523 | -0.897 | 0.784 | -0.017 | 0.555 | 0.009 | 0.03 | 0.03 | 0 | 0.15 | 0.46 | 0.16 | 0.15 | 0.16 | 0.08 |
| 1 | 60327117 | rs11207526 | -0.991 | 0.773 | 0.006 | 0.572 | 0.002 | 0.026 | 0.026 | 0 | 0.16 | 0.47 | 0.16 | 0.16 | 0.16 | 0.08 |
| 1 | 60327434 | rs10493269 | 0.438 | 0.756 | 0.184 | 0.563 | -0.016 | -0.005 | -0.005 | 0 | 0.09 | 0.2 | 0.05 | 0.05 | 0.06 | 0.08 |
| 1 | 60328613 | rs2235798 | -0.109 | 0.753 | 0.186 | 0.564 | -0.013 | 0.044 | 0.044 | 0 | 0.11 | 0.37 | 0.05 | 0.09 | 0.07 | 0.08 |
| 1 | 60352377 | rs12064386 | 0.021 | 0.76 | 0.19 | 0.567 | -0.009 | 0.046 | 0.046 | 0 | 0.11 | 0.35 | 0.05 | 0.09 | 0.08 | 0.08 |
| 1 | 60357969 | rs10789082 | 1.194 | 0.562 | -0.23 | 0.694 | -0.061 | -0.078 | -0.078 | 0 | 0.2 | 0.02 | 0.22 | 0.24 | 0.24 | 0.34 |

| | | | | | | | | | | | | | | | | |
|---|---|---|---|---|---|---|---|---|---|---|---|---|---|---|---|---|
| 1 | 60358982 | rs2280273 | 0.083 | 0.554 | -0.092 | 0.646 | -0.012 | 0.042 | 0.042 | 0 | 0.11 | 0.35 | 0.05 | 0.09 | 0.08 | 0.08 |
| 1 | 60359156 | rs4388726 | 0.647 | 0.557 | -0.1 | 0.645 | -0.014 | -0.005 | -0.005 | 0 | 0.09 | 0.17 | 0.04 | 0.05 | 0.06 | 0.08 |
| 1 | 60360027 | rs1408479 | -0.257 | 0.569 | -0.093 | 0.644 | 0.039 | 0.034 | 0.034 | 0 | 0.12 | 0.4 | 0.13 | 0.11 | 0.12 | 0.08 |
| 1 | 60360195 | rs11207535 | 0.332 | 0.571 | -0.095 | 0.645 | 0.034 | 0.03 | 0.03 | 0 | 0.1 | 0.3 | 0.12 | 0.11 | 0.12 | 0.08 |
| 1 | 60361109 | rs2147397 | -0.87 | 0.56 | -0.11 | 0.642 | 0.011 | 0.032 | 0.032 | 0 | 0.15 | 0.44 | 0.16 | 0.16 | 0.16 | 0.09 |
| 1 | 60363358 | rs9659919 | 0.92 | 0.538 | -0.113 | 0.615 | -0.007 | -0.008 | -0.008 | 0 | 0.08 | 0.12 | 0.04 | 0.04 | 0.06 | 0.07 |
| 1 | 60366337 | rs11572307 | 1.165 | 0.56 | -0.112 | 0.623 | 0.001 | 0.002 | 0.002 | 0 | 0.08 | 0.09 | 0.03 | 0.04 | 0.06 | 0.07 |
| 1 | 60366386 | rs11572305 | 1.165 | 0.56 | -0.112 | 0.623 | 0.001 | 0.002 | 0.002 | 0 | 0.08 | 0.09 | 0.03 | 0.04 | 0.06 | 0.07 |
| 1 | 60367240 | rs2280275 | -0.875 | 0.55 | -0.077 | 0.628 | 0.013 | 0.029 | 0.029 | 0 | 0.15 | 0.45 | 0.16 | 0.15 | 0.17 | 0.09 |
| 1 | 60372721 | rs11572285 | 0.308 | 0.554 | -0.081 | 0.626 | 0.014 | 0.006 | 0.006 | 0 | 0.1 | 0.24 | 0.13 | 0.09 | 0.11 | 0.08 |
| 1 | 60372958 | rs3754205 | 0.455 | 0.549 | -0.073 | 0.627 | 0.026 | 0.057 | 0.057 | 0 | 0.05 | 0.21 | 0.03 | 0.07 | 0.06 | 0.01 |
| 1 | 60373666 | rs2271800 | -0.954 | 0.497 | 0.025 | 0.6 | 0.003 | 0.018 | 0.018 | 0.114 | 0.16 | 0.51 | 0.16 | 0.15 | 0.16 | 0.09 |
| 1 | 60373775 | rs1155002 | -1.163 | 0.529 | -0.154 | 0.474 | -0.012 | 0.06 | 0.06 | 0 | 0.59 | 0.78 | 0.69 | 0.68 | 0.61 | 0.84 |
| 1 | 60378836 | rs11572255 | 1.438 | 0.708 | -0.037 | 0.389 | -0.008 | 0.006 | 0.006 | 0 | 0.08 | 0.06 | 0.17 | 0.04 | 0.15 | 0.03 |
| 1 | 60378853 | rs11572254 | -0.219 | 0.66 | -0.018 | 0.379 | 0.004 | -0.019 | -0.019 | 1.003 | 0.17 | 0.25 | 0.22 | 0.09 | 0.21 | 0.11 |
| 1 | 60379433 | rs11572250 | 1.725 | 0.644 | 0.083 | 0.387 | -0.042 | 0.026 | 0.026 | 0 | 0.08 | 0.02 | 0.17 | 0.04 | 0.15 | 0.03 |
| 1 | 60382682 | rs7515289 | -1.078 | 0.617 | 0.117 | 0.377 | -0.002 | -0.041 | -0.041 | 0.313 | 0.23 | 0.52 | 0.25 | 0.15 | 0.26 | 0.11 |
| 1 | 60383109 | rs11572237 | -0.975 | 0.633 | 0.115 | 0.385 | 0.003 | -0.041 | -0.041 | 0 | 0.22 | 0.46 | 0.25 | 0.15 | 0.26 | 0.11 |
| 1 | 60383417 | rs11572235 | 1.284 | 0.645 | 0.129 | 0.398 | -0.006 | -0.008 | -0.008 | 0 | 0.08 | 0.11 | 0.04 | 0.03 | 0.06 | 0.07 |
| 1 | 60385461 | rs11572227 | 1.284 | 0.64 | 0.112 | 0.394 | -0.006 | -0.006 | -0.006 | 0 | 0.08 | 0.11 | 0.04 | 0.03 | 0.06 | 0.07 |
| 1 | 60388674 | rs10493270 | 1.434 | 0.606 | -0.109 | 0.517 | -0.019 | -0.05 | -0.05 | 0 | 0.1 | 0.01 | 0.17 | 0.14 | 0.1 | 0.13 |
| 1 | 60390033 | rs11572201 | 1.439 | 0.582 | 0.098 | 0.322 | -0.027 | 0.011 | 0.011 | 0 | 0.08 | 0.03 | 0.17 | 0.05 | 0.15 | 0.03 |
| 1 | 60391946 | rs3754206 | -0.607 | 0.574 | 0.137 | 0.319 | 0.02 | -0.029 | -0.029 | 0 | 0.19 | 0.42 | 0.22 | 0.12 | 0.22 | 0.11 |
| 1 | 60394522 | rs12729119 | 1.616 | 0.547 | -0.059 | 0.33 | -0.042 | 0.027 | 0.027 | 0 | 0.08 | 0.02 | 0.17 | 0.04 | 0.15 | 0.03 |
| 1 | 60402488 | rs12743907 | 0.573 | 0.517 | -0.072 | 0.346 | -0.01 | 0.036 | 0.036 | 0 | 0.1 | 0.28 | 0.04 | 0.08 | 0.08 | 0.08 |
| 1 | 60407502 | rs11207545 | -0.478 | 0.594 | -0.195 | 0.336 | 0.02 | -0.047 | -0.047 | 0 | 0.19 | 0.41 | 0.22 | 0.11 | 0.22 | 0.1 |
| 1 | 60408975 | rs12731852 | -0.791 | 0.576 | -0.251 | 0.34 | 0.008 | -0.037 | -0.037 | 0 | 0.22 | 0.44 | 0.24 | 0.15 | 0.26 | 0.11 |
| 1 | 60413925 | rs11207546 | 0.55 | 0.282 | -0.748 | 0.566 | -0.072 | -0.156 | -0.156 | 0.002 | 0.3 | 0.02 | 0.4 | 0.38 | 0.32 | 0.47 |
| 1 | 60421762 | rs7537739 | -0.131 | 0.135 | 0.098 | 0.396 | 0.006 | 0.003 | 0.003 | 0 | 0.11 | 0.1 | 0.06 | 0.08 | 0.1 | 0.08 |
| 1 | 60427947 | rs12072111 | -0.942 | -0.036 | 0.27 | 0.402 | -0.041 | 0.016 | 0.016 | 0 | 0.13 | 0.36 | 0.05 | 0.11 | 0.09 | 0.08 |
| 1 | 60430348 | rs10493273 | 0.517 | 0.252 | -0.4 | 0.43 | -0.001 | 0 | 0 | 0 | 0.08 | 0.1 | 0.04 | 0.05 | 0.06 | 0.08 |
| 1 | 60437812 | rs890291 | 0.079 | 0.171 | -0.176 | 0.392 | -0.004 | -0.017 | -0.017 | 0.453 | 0.12 | 0.18 | 0.14 | 0.07 | 0.11 | 0.18 |
| 1 | 60440185 | rs2554297 | -0.876 | -0.131 | -0.237 | 0.429 | 0.011 | 0.017 | 0.017 | 0.048 | 0.67 | 0.75 | 0.8 | 0.72 | 0.74 | 0.85 |
| 1 | 60446474 | rs11800976 | 0.085 | -0.038 | -0.14 | 0.585 | 0 | 0 | 0 | 0 | 0.18 | 0.16 | 0.18 | 0.22 | 0.17 | 0.13 |
| 1 | 60446789 | rs7411728 | 0.764 | -0.012 | -0.273 | 0.614 | 0.017 | -0.039 | -0.039 | 0 | 0.13 | 0.01 | 0.08 | 0.17 | 0.08 | 0.1 |
| 1 | 60447656 | rs2807616 | -1.347 | -0.08 | -0.263 | 0.538 | 0.008 | -0.006 | -0.006 | 0.412 | 0.7 | 0.82 | 0.81 | 0.65 | 0.74 | 0.82 |
| 1 | 60448209 | rs2807619 | 0.552 | -0.137 | -0.261 | 0.514 | -0.014 | -0.03 | -0.03 | 0 | 0.13 | 0.19 | 0.15 | 0.05 | 0.1 | 0.2 |
| 1 | 60454069 | rs4915842 | 0.683 | 0.155 | -0.808 | 0.87 | -0.032 | -0.014 | -0.014 | 0 | 0.33 | 0.23 | 0.45 | 0.35 | 0.45 | 0.48 |
| 1 | 60455797 | rs10493275 | -0.175 | -0.109 | -0.734 | 0.991 | 0.003 | -0.004 | -0.004 | 0 | 0.14 | 0.12 | 0.09 | 0.2 | 0.07 | 0.11 |
| 1 | 60460443 | rs10889167 | -0.054 | -0.138 | -0.715 | 0.641 | 0.02 | 0.015 | 0.015 | 0 | 0.4 | 0.33 | 0.29 | 0.38 | 0.26 | 0.22 |
| 1 | 60460733 | rs642307 | -0.126 | 0.896 | -0.732 | 0.65 | 0.004 | 0.016 | 0.016 | 0 | 0.5 | 0.64 | 0.45 | 0.5 | 0.54 | 0.68 |
| 1 | 60461958 | rs4915847 | -0.109 | 1.033 | -0.68 | 0.525 | 0.014 | 0.059 | 0.059 | 0 | 0.2 | 0.29 | 0.35 | 0.48 | 0.2 | 0.19 |
| 1 | 60463036 | rs919055 | 0.872 | 1.032 | -0.598 | 0.631 | -0.01 | 0.073 | 0.073 | 0 | 0.1 | 0.16 | 0.16 | 0.34 | 0.06 | 0.11 |
| 1 | 60464168 | rs649859 | 0.874 | 0.501 | -0.166 | 0.286 | -0.03 | -0.026 | -0.026 | 0 | 0.13 | 0.2 | 0.08 | 0.05 | 0.11 | 0.19 |
| 1 | 60466814 | rs626251 | -0.879 | 0.363 | -0.815 | 0.044 | 0.005 | 0.04 | 0.04 | 0 | 0.52 | 0.61 | 0.62 | 0.63 | 0.45 | 0.45 |
| 1 | 60472097 | rs11207565 | 1.611 | 0.751 | -1.204 | -0.375 | -0.037 | 0.051 | 0.051 | 0.06 | 0.16 | 0.06 | 0.19 | 0 | 0.27 | 0.16 |
| 1 | 60474983 | rs681724 | -1.082 | 0.605 | -0.777 | 0.376 | -0.002 | 0.04 | 0.04 | 0 | 0.85 | 0.83 | 0.82 | 0.57 | 0.89 | 0.85 |
| 1 | 60476423 | rs630572 | -0.173 | 0.644 | -0.712 | 0.359 | 0.003 | 0.015 | 0.015 | 0 | 0.74 | 0.72 | 0.61 | 0.43 | 0.74 | 0.78 |
| 1 | 60477322 | rs17119990 | 1.873 | 0.749 | -0.635 | 0.311 | -0.009 | -0.007 | -0.007 | 0 | 0.1 | 0.11 | 0.2 | 0.14 | 0.14 | 0.07 |
| 1 | 60478610 | rs681128 | -0.297 | 0.235 | -0.458 | -0.043 | 0.001 | 0.005 | 0.005 | 0 | 0.5 | 0.52 | 0.47 | 0.33 | 0.54 | 0.64 |
| 1 | 60479803 | rs675493 | 0.225 | 0.251 | -0.509 | 0.037 | 0.001 | 0.033 | 0.033 | 0 | 0.42 | 0.32 | 0.35 | 0.26 | 0.48 | 0.59 |
| 1 | 60484684 | rs657373 | 0.769 | 0.151 | -0.561 | -0.017 | -0.001 | 0.002 | 0.002 | 0 | 0.31 | 0.29 | 0.29 | 0.24 | 0.39 | 0.56 |
| 1 | 60490458 | rs713567 | 1.971 | 0.144 | -0.455 | -0.022 | 0 | 0.041 | 0.041 | 0 | 0.21 | 0.12 | 0.13 | 0.02 | 0.19 | 0.09 |
| 1 | 60494956 | rs642294 | 0.633 | 0.269 | -0.5 | -0.118 | 0.008 | 0.017 | 0.017 | 0 | 0.37 | 0.32 | 0.27 | 0.26 | 0.4 | 0.56 |
| 1 | 60496313 | rs634094 | 0.648 | 0.25 | -0.429 | -0.126 | 0.005 | 0.01 | 0.01 | 0 | 0.37 | 0.37 | 0.27 | 0.26 | 0.4 | 0.55 |
| 1 | 60502624 | rs17120030 | 1.4 | 0.384 | -0.368 | 0.051 | -0.001 | -0.006 | -0.006 | 0 | 0.24 | 0.24 | 0.17 | 0.09 | 0.21 | 0.13 |
| 1 | 60503762 | rs17560663 | 1.98 | 0.397 | -0.82 | -0.103 | -0.048 | 0.115 | 0.115 | 0 | 0.2 | 0.01 | 0.13 | 0.02 | 0.19 | 0.09 |
| 1 | 60504967 | rs17120032 | 1.575 | 0.188 | -0.613 | 0.263 | 0 | 0.002 | 0.002 | 0 | 0.11 | 0.09 | 0.11 | 0.07 | 0.11 | 0.11 |
| 1 | 60505480 | rs12081664 | 1.874 | 0.188 | -0.561 | 0.216 | -0.008 | -0.033 | -0.033 | 0 | 0.19 | 0.22 | 0.12 | 0.02 | 0.12 | 0.09 |
| 1 | 60505578 | rs680390 | -0.703 | 0.603 | -0.547 | 0.986 | 0 | 0.003 | 0.003 | 0 | 0.88 | 0.92 | 0.8 | 0.7 | 0.9 | 0.9 |
| 1 | 60506744 | rs10493277 | 2.127 | 0.881 | -0.709 | 0.938 | -0.034 | -0.035 | -0.035 | 0 | 0.08 | 0.03 | 0.2 | 0.22 | 0.13 | 0.07 |
| 1 | 60529363 | rs640639 | 0.321 | 1.012 | -0.619 | 0.696 | 0.004 | 0.021 | 0.021 | 0 | 0.43 | 0.31 | 0.29 | 0.35 | 0.45 | 0.48 |
| 1 | 60531017 | rs7543186 | 1.769 | 0.861 | -0.839 | 0.596 | 0.004 | -0.021 | -0.021 | 1.672 | 0.14 | 0.09 | 0.1 | 0.26 | 0.16 | 0.31 |
| 1 | 60543334 | rs17120097 | 2.838 | 1.028 | -0.779 | 0.696 | -0.042 | -0.042 | -0.042 | 0 | 0.08 | 0.03 | 0.21 | 0.28 | 0.14 | 0.08 |
| 1 | 60543950 | rs675310 | 0.586 | 0.86 | -0.86 | 0.557 | 0.003 | -0.006 | -0.006 | 0.094 | 0.23 | 0.22 | 0.16 | 0.3 | 0.24 | 0.43 |
| 1 | 60547426 | rs685952 | -0.203 | 0.897 | -0.226 | 0.525 | 0.033 | 0.102 | 0.102 | 0.103 | 0.83 | 0.6 | 0.74 | 0.69 | 0.83 | 0.87 |
| 1 | 60563524 | rs12071650 | 1.117 | 0.798 | -0.195 | 0.548 | 0.009 | 0.017 | 0.017 | 0 | 0.31 | 0.22 | 0.26 | 0.11 | 0.25 | 0.33 |
| 1 | 60575925 | rs677574 | -0.321 | 0.926 | -0.249 | 0.664 | 0.014 | 0.015 | 0.015 | 0 | 0.84 | 0.67 | 0.77 | 0.77 | 0.85 | 0.9 |
| 1 | 60577184 | rs608556 | 0.066 | 1.015 | -0.452 | 0.71 | 0.034 | 0.066 | 0.066 | 0 | 0.75 | 0.59 | 0.56 | 0.49 | 0.7 | 0.81 |

| | | | | | | | | | | | | | | | | |
|---|---|---|---|---|---|---|---|---|---|---|---|---|---|---|---|---|
| 1 | 60581154 | rs6677345 | 1.407 | 0.811 | -0.769 | 0.475 | 0.024 | 0.05 | 0.05 | 0 | 0.3 | 0.15 | 0.23 | 0.1 | 0.24 | 0.3 |
| 1 | 60585118 | rs12028437 | 1.428 | 0.835 | -0.656 | 0.872 | -0.004 | -0.03 | -0.03 | 0 | 0.14 | 0.08 | 0.11 | 0.26 | 0.19 | 0.31 |
| 1 | 60585522 | rs665062 | 0.974 | 1.096 | 0.05 | 1.28 | 0.048 | 0.097 | 0.097 | 0 | 0.73 | 0.52 | 0.53 | 0.45 | 0.69 | 0.78 |
| 1 | 60589775 | rs634161 | 0.836 | 0.881 | 0.173 | 0.639 | 0.024 | 0.059 | 0.059 | 0 | 0.81 | 0.54 | 0.74 | 0.73 | 0.83 | 0.86 |
| 1 | 60594980 | rs87061 | 0.574 | 0.926 | -0.32 | 0.559 | 0.011 | -0.005 | -0.005 | 0 | 0.54 | 0.57 | 0.48 | 0.49 | 0.44 | 0.74 |
| 1 | 60597914 | rs645024 | 0.457 | 1.215 | -0.278 | 0.563 | 0.01 | 0.023 | 0.023 | 0.035 | 0.76 | 0.66 | 0.65 | 0.54 | 0.68 | 0.79 |
| 1 | 60599251 | rs680811 | 0.598 | 1.185 | -0.267 | 0.272 | 0.026 | 0.044 | 0.044 | 0.023 | 0.75 | 0.59 | 0.64 | 0.58 | 0.68 | 0.79 |
| 1 | 60601856 | rs656464 | 1.569 | 0.827 | -0.198 | 0.109 | -0.005 | 0.001 | 0.001 | 0 | 0.39 | 0.36 | 0.34 | 0.45 | 0.28 | 0.52 |
| 1 | 60607178 | rs1462935 | 0.877 | 1.049 | -0.283 | 0.204 | -0.029 | -0.037 | -0.037 | 0 | 0.12 | 0.06 | 0.15 | 0.24 | 0.17 | 0.08 |
| 1 | 60607633 | rs583614 | 0.273 | 1.017 | -0.243 | 0.193 | 0.018 | 0.027 | 0.027 | 0 | 0.87 | 0.68 | 0.79 | 0.81 | 0.85 | 0.87 |
| 1 | 60609043 | rs609462 | 0.749 | 1.014 | -0.224 | 0.197 | 0.013 | 0.054 | 0.054 | 0.025 | 0.85 | 0.65 | 0.76 | 0.76 | 0.84 | 0.85 |
| 1 | 60611291 | rs656442 | 1.682 | 0.973 | -0.326 | 0.141 | -0.001 | -0.001 | -0.001 | 0.831 | 0.39 | 0.35 | 0.34 | 0.45 | 0.28 | 0.54 |
| 1 | 60613082 | rs17120264 | 0.408 | 1.022 | -0.367 | 0.221 | -0.034 | -0.026 | -0.026 | 6.56 | 0.08 | 0.01 | 0.19 | 0.15 | 0.14 | 0.1 |
| 1 | 60614595 | rs657909 | 0.883 | 0.749 | -0.33 | 0.219 | 0.007 | 0.027 | 0.027 | 0.012 | 0.75 | 0.62 | 0.65 | 0.57 | 0.73 | 0.79 |
| 1 | 60628446 | rs12130292 | 0.397 | 0.769 | -0.38 | 0.163 | -0.019 | 0.006 | 0.006 | 0 | 0.1 | 0.05 | 0.12 | 0.07 | 0.14 | 0.2 |
| 1 | 60631000 | rs922356 | 0.036 | 0.773 | -0.32 | 0.191 | 0.007 | 0.043 | 0.043 | 0 | 0.09 | 0.21 | 0.19 | 0.18 | 0.1 | 0.12 |
| 1 | 60637233 | rs12079264 | -1.285 | 0.734 | -0.397 | 0.135 | 0.011 | 0.004 | 0.004 | 0 | 0.49 | 0.48 | 0.52 | 0.48 | 0.53 | 0.25 |
| 1 | 60648441 | rs2035174 | -1.828 | 0.762 | -0.318 | 0.174 | -0.018 | 0.007 | 0.007 | 0 | 0.59 | 0.55 | 0.64 | 0.54 | 0.67 | 0.45 |
| 1 | 60650062 | rs11584678 | 1.356 | 0.815 | -0.36 | 0.173 | -0.059 | -0.056 | -0.056 | 0 | 0.1 | 0.01 | 0.12 | 0.19 | 0.16 | 0.06 |
| 1 | 60659543 | rs6587877 | -1.938 | 0.791 | -0.259 | 0.166 | 0 | -0.001 | -0.001 | 0 | 0.61 | 0.63 | 0.65 | 0.55 | 0.67 | 0.45 |
| 1 | 60659719 | rs10443220 | -0.174 | 0.809 | -0.692 | 0.176 | -0.002 | -0.002 | -0.002 | 0 | 0.17 | 0.18 | 0.17 | 0.13 | 0.17 | 0.22 |
| 1 | 60663224 | rs12030859 | 1.751 | 0.799 | -1.09 | 0.14 | 0 | -0.01 | -0.01 | 0.02 | 0.06 | 0 | 0.02 | 0.06 | 0.04 | 0.09 |
| 1 | 60666641 | rs6661520 | -0.775 | 0.976 | -0.988 | 0.161 | -0.004 | 0.003 | 0.003 | 0 | 0.42 | 0.39 | 0.48 | 0.42 | 0.51 | 0.24 |
| 1 | 60667702 | rs12125049 | 0.217 | 1.024 | -1.451 | -0.008 | -0.018 | 0.027 | 0.027 | 0 | 0.14 | 0.05 | 0.13 | 0.07 | 0.16 | 0.2 |
| 1 | 60667904 | rs1381888 | 1.142 | 1.122 | -1.233 | -0.026 | -0.079 | 0.099 | 0.099 | 0 | 0.21 | 0.01 | 0.16 | 0.05 | 0.24 | 0.05 |
| 1 | 60691588 | rs17120400 | -0.837 | 1.039 | -0.674 | 0.435 | 0.008 | 0.024 | 0.024 | 0.018 | 0.11 | 0.23 | 0.2 | 0.19 | 0.13 | 0.13 |
| 1 | 60701724 | rs10515373 | 0.143 | 1.146 | -0.95 | 0.287 | -0.02 | -0.038 | -0.038 | 0.002 | 0.15 | 0.09 | 0.15 | 0.4 | 0.18 | 0.15 |
| 1 | 60702323 | rs3906402 | -1.489 | 0.961 | -0.949 | -0.092 | -0.01 | 0.004 | 0.004 | 0 | 0.6 | 0.55 | 0.59 | 0.57 | 0.64 | 0.52 |
| 1 | 60704278 | rs7535541 | -0.109 | 1.044 | -1.134 | -0.014 | -0.051 | -0.068 | -0.068 | 0 | 0.17 | 0.07 | 0.25 | 0.37 | 0.24 | 0.17 |
| 1 | 60730938 | rs6662682 | -0.461 | 0.972 | -1.301 | -0.134 | -0.005 | 0.015 | 0.015 | 0 | 0.2 | 0.16 | 0.25 | 0.14 | 0.22 | 0.32 |
| 1 | 60739301 | rs10493281 | 0.4 | 0.977 | -1.419 | -0.107 | -0.035 | 0.028 | 0.028 | 0 | 0.15 | 0.01 | 0.12 | 0.07 | 0.15 | 0.26 |
| 1 | 60752854 | rs12741643 | -0.469 | 1.023 | -1.205 | 0.013 | -0.012 | -0.008 | -0.008 | 2.397 | 0.38 | 0.29 | 0.32 | 0.43 | 0.42 | 0.2 |
| 1 | 60756493 | rs6587879 | -0.48 | 0.793 | -0.615 | 0.004 | 0.05 | 0.096 | 0.096 | 0.283 | 0.94 | 0.69 | 0.96 | 0.94 | 0.97 | 0.91 |
| 1 | 60758915 | rs4259695 | -0.148 | 0.994 | -0.88 | 0.081 | 0.001 | 0.013 | 0.013 | 0 | 0.23 | 0.23 | 0.16 | 0.41 | 0.23 | 0.18 |
| 1 | 60762564 | rs4147294 | -0.383 | 0.559 | -0.772 | 0.224 | -0.016 | 0.006 | 0.006 | 0.066 | 0.63 | 0.59 | 0.59 | 0.58 | 0.66 | 0.49 |
| 1 | 60782286 | rs298174 | -0.35 | 1.463 | -0.799 | -0.282 | -0.01 | -0.001 | -0.001 | 0.003 | 0.24 | 0.23 | 0.32 | 0.06 | 0.4 | 0.25 |
| 1 | 60787556 | rs298169 | 0.728 | 0.181 | -0.807 | -0.045 | 0.029 | -0.011 | -0.011 | 0 | 0.38 | 0.25 | 0.39 | 0.45 | 0.27 | 0.51 |
| 1 | 60794707 | rs454776 | 0.651 | 0.201 | -0.365 | -0.498 | -0.026 | 0.035 | 0.035 | 0 | 0.16 | 0.07 | 0.14 | 0 | 0.23 | 0.07 |
| 1 | 60804975 | rs7415271 | 0.915 | 0.163 | -0.249 | -0.376 | 0.007 | -0.005 | -0.005 | 0 | 0.2 | 0.22 | 0.29 | 0.11 | 0.34 | 0.24 |
| 1 | 60805150 | rs298167 | 0.699 | -0.076 | -0.145 | -0.382 | 0.008 | 0.042 | 0.042 | 0 | 0.54 | 0.37 | 0.51 | 0.78 | 0.41 | 0.63 |
| 1 | 60806703 | rs298166 | 0.569 | -0.069 | -0.14 | -0.389 | -0.002 | -0.023 | -0.023 | 1.605 | 0.56 | 0.51 | 0.53 | 0.8 | 0.41 | 0.64 |
| 1 | 60815316 | rs12137879 | 0.922 | -0.078 | -0.153 | -0.327 | 0.006 | -0.012 | -0.012 | 0 | 0.16 | 0.11 | 0.13 | 0.37 | 0.14 | 0.13 |
| 1 | 60822957 | rs7536342 | 1.017 | -0.118 | -0.138 | -0.366 | 0.007 | -0.004 | -0.004 | 0.241 | 0.2 | 0.22 | 0.29 | 0.12 | 0.33 | 0.24 |
| 1 | 60824982 | rs412924 | 0.772 | -0.026 | -0.186 | -0.429 | -0.006 | 0.029 | 0.029 | 0 | 0.15 | 0.07 | 0.11 | 0 | 0.18 | 0.06 |
| 1 | 60828648 | rs1501246 | 1.027 | 0.016 | -0.204 | -0.401 | 0.007 | -0.004 | -0.004 | 0 | 0.2 | 0.22 | 0.29 | 0.12 | 0.33 | 0.24 |
| 1 | 60833726 | rs298142 | 0.716 | -0.036 | -0.158 | -0.129 | 0.001 | 0.008 | 0.008 | 0 | 0.16 | 0.13 | 0.12 | 0.09 | 0.19 | 0.09 |
| 1 | 60834294 | rs1354085 | -0.615 | -0.422 | -0.112 | 0.124 | -0.007 | 0.104 | 0.104 | 0 | 0.94 | 0.8 | 0.95 | 0.89 | 0.98 | 0.94 |
| 1 | 60838146 | rs298147 | 0.255 | -0.478 | -0.215 | 0.022 | -0.002 | 0.032 | 0.032 | 0 | 0.54 | 0.36 | 0.52 | 0.68 | 0.45 | 0.61 |
| 1 | 60840443 | rs407173 | 0.123 | -0.334 | -0.243 | -0.38 | -0.007 | 0.007 | 0.007 | 0 | 0.37 | 0.39 | 0.41 | 0.19 | 0.52 | 0.33 |
| 1 | 60848538 | rs298128 | 0.181 | 0.124 | -0.365 | -0.442 | 0.006 | 0.01 | 0.01 | 0 | 0.38 | 0.25 | 0.4 | 0.4 | 0.32 | 0.51 |
| 1 | 60862487 | rs11207599 | 0.478 | 0.046 | -1.837 | -0.214 | -0.025 | -0.08 | -0.08 | 1.037 | 0.32 | 0.02 | 0.36 | 0.38 | 0.25 | 0.4 |
| 1 | 60869362 | rs403304 | 0.365 | 0.58 | -0.98 | -1.086 | 0.012 | -0.176 | -0.176 | 0 | 0.67 | 0.48 | 0.76 | 1 | 0.64 | 0.89 |
| 1 | 60872496 | rs10493285 | -0.266 | 0.757 | -1.464 | -1.022 | -0.049 | 0.065 | 0.065 | 0 | 0.12 | 0.02 | 0.13 | 0.01 | 0.14 | 0.12 |
| 1 | 60874772 | rs298155 | -0.904 | -0.465 | -1.413 | -1.195 | 0.007 | 0.01 | 0.01 | 0 | 0.35 | 0.31 | 0.34 | 0.2 | 0.24 | 0.37 |
| 1 | 60876146 | rs7529292 | 1.023 | -1.151 | -1.665 | -0.769 | 0.001 | -0.003 | -0.003 | 0 | 0.26 | 0.22 | 0.28 | 0.52 | 0.18 | 0.22 |
| 1 | 60876186 | rs657300 | 1.064 | -1.181 | -1.954 | -0.786 | 0.036 | 0.019 | 0.019 | 0 | 0.2 | 0.1 | 0.19 | 0.15 | 0.1 | 0.23 |
| 1 | 60879175 | rs298160 | -0.035 | -0.956 | -1.709 | -0.641 | 0.022 | 0.068 | 0.068 | 0 | 0.33 | 0.17 | 0.32 | 0.16 | 0.24 | 0.34 |
| 1 | 60882355 | rs17120672 | 1.138 | -1.342 | -0.962 | 0.196 | 0.003 | 0 | 0 | 0.878 | 0.17 | 0.15 | 0.15 | 0.14 | 0.11 | 0.04 |
| 1 | 60887538 | rs11207605 | -0.195 | -0.632 | -1.202 | 0.127 | 0.001 | 0.025 | 0.025 | 0 | 0.73 | 0.35 | 0.73 | 0.53 | 0.71 | 0.56 |
| 1 | 60888482 | rs10493287 | 0.372 | -0.639 | -1.56 | 0.043 | 0.037 | 0.017 | 0.017 | 1.655 | 0.55 | 0.19 | 0.57 | 0.39 | 0.6 | 0.52 |
| 1 | 60888999 | rs7551363 | -0.115 | -1.418 | -0.859 | 0.233 | -0.058 | 0.039 | 0.039 | 1.11 | 0.27 | 0.37 | 0.3 | 0.39 | 0.13 | 0.35 |
| 1 | 60889219 | rs2062629 | 0.785 | -1.209 | -1.226 | 0.084 | 0.044 | 0.007 | 0.007 | 0.338 | 0.21 | 0.03 | 0.14 | 0.14 | 0.12 | 0.09 |
| 1 | 60891642 | rs11207606 | 0.107 | -1.445 | -1.038 | 0.157 | 0.078 | -0.021 | -0.021 | 0 | 0.41 | 0.23 | 0.37 | 0.56 | 0.28 | 0.4 |
| 1 | 60897332 | rs167229 | 1.614 | -0.496 | -1.184 | -0.41 | -0.086 | 0.087 | 0.087 | 0 | 0.21 | 0.07 | 0.2 | 0 | 0.38 | 0.06 |
| 1 | 60907460 | rs1501228 | 1.193 | -0.568 | -1.275 | -0.053 | -0.029 | -0.024 | -0.024 | 0 | 0.16 | 0.07 | 0.21 | 0.18 | 0.22 | 0.18 |
| 1 | 60925043 | rs4595390 | 0.796 | -0.873 | -0.11 | -0.133 | 0.096 | -0.053 | -0.053 | 0 | 0.41 | 0.14 | 0.37 | 0.56 | 0.29 | 0.39 |
| 1 | 60927512 | rs4915693 | 1.468 | -0.699 | -0.044 | -0.128 | -0.064 | 0.034 | 0.034 | 0 | 0.23 | 0.13 | 0.22 | 0.14 | 0.37 | 0.09 |
| 1 | 60939827 | rs4644526 | 0.202 | -0.796 | 0.32 | -0.076 | -0.002 | 0 | 0 | 0.135 | 0.5 | 0.49 | 0.48 | 0.54 | 0.33 | 0.49 |
| 1 | 60944959 | rs12404114 | 0.132 | -0.893 | 0.21 | -0.017 | -0.006 | 0.005 | 0.005 | 0.293 | 0.39 | 0.21 | 0.44 | 0.34 | 0.6 | 0.27 |

| | | | | | | | | | | | | | | | | |
|---|---|---|---|---|---|---|---|---|---|---|---|---|---|---|---|---|
| 1 | 60952106 | rs4075690 | 1.291 | -0.564 | 0.23 | 0.088 | -0.053 | -0.051 | -0.051 | 0 | 0.1 | 0.01 | 0.17 | 0.16 | 0.18 | 0.14 |
| 1 | 60962168 | rs4915695 | 1.905 | -0.908 | 0.049 | 0.112 | 0.043 | -0.018 | -0.018 | 0.128 | 0.21 | 0.03 | 0.22 | 0.2 | 0.12 | 0.11 |
| 1 | 60966964 | rs7355119 | 0.722 | -0.016 | 0.892 | 0.248 | -0.001 | -0.001 | -0.001 | 0 | 0.26 | 0.29 | 0.28 | 0.29 | 0.38 | 0.21 |
| 1 | 60975097 | rs11806181 | -0.211 | -0.201 | 1.038 | 0.091 | 0.084 | -0.012 | -0.012 | 3.813 | 0.09 | 0.3 | 0.12 | 0.02 | 0.07 | 0.12 |
| 1 | 60983620 | rs7522454 | 0.648 | -0.623 | 0.536 | 0.169 | 0.119 | -0.031 | -0.031 | 0 | 0.49 | 0.14 | 0.4 | 0.53 | 0.34 | 0.48 |
| 1 | 60995104 | rs11207631 | -0.211 | -0.502 | 1.108 | -0.286 | -0.02 | -0.06 | -0.06 | 4.113 | 0.13 | 0.37 | 0.15 | 0.05 | 0.04 | 0.15 |
| 1 | 60998195 | rs10889182 | 0.937 | -0.47 | 0.899 | -0.164 | 0.111 | 0.021 | 0.021 | 2.371 | 0.5 | 0.19 | 0.45 | 0.56 | 0.34 | 0.57 |
| 1 | 60999977 | rs10889183 | -0.072 | -0.45 | 1.097 | -0.207 | -0.016 | -0.065 | -0.065 | 0.153 | 0.87 | 0.58 | 0.85 | 0.95 | 0.95 | 0.85 |
| 1 | 61007182 | rs11207633 | 0.887 | -0.442 | 0.942 | -0.129 | 0.11 | 0.011 | 0.011 | 2.034 | 0.51 | 0.17 | 0.45 | 0.56 | 0.34 | 0.57 |
| 1 | 61009286 | rs6587887 | 0.642 | -0.266 | 0.933 | -0.15 | -0.114 | -0.059 | -0.059 | 0.647 | 0.3 | 0.13 | 0.3 | 0.31 | 0.52 | 0.16 |
| 1 | 61014021 | rs7522761 | 0.917 | -0.282 | 0.921 | -0.117 | 0.106 | 0.012 | 0.012 | 2.034 | 0.51 | 0.17 | 0.45 | 0.56 | 0.34 | 0.57 |
| 1 | 61023058 | rs6680587 | -0.701 | 0.007 | 0.822 | -0.256 | 0.056 | 0.033 | 0.033 | 0 | 0.06 | 0.22 | 0.07 | 0.09 | 0.07 | 0.12 |
| 1 | 61037015 | rs1461356 | 0.35 | -0.063 | 0.874 | -0.248 | -0.006 | -0.009 | -0.009 | 0 | 0.6 | 0.44 | 0.6 | 0.63 | 0.38 | 0.68 |
| 1 | 61037311 | rs12563618 | -0.009 | -0.007 | 0.671 | -0.221 | -0.024 | -0.006 | -0.006 | 0 | 0.34 | 0.31 | 0.35 | 0.37 | 0.55 | 0.26 |
| 1 | 61041875 | rs12565755 | 0.914 | 0.203 | 0.298 | -0.217 | -0.042 | -0.05 | -0.05 | 0 | 0.08 | 0.02 | 0.09 | 0.28 | 0.15 | 0.1 |
| 1 | 61061996 | rs3896178 | 0.937 | -0.617 | 0.19 | -0.432 | 0.097 | -0.055 | -0.055 | 0.076 | 0.37 | 0.07 | 0.32 | 0.57 | 0.25 | 0.48 |
| 1 | 61062474 | rs484743 | 0.132 | 0.676 | 1.059 | 0.306 | -0.01 | 0.006 | 0.006 | 0.102 | 0.66 | 0.68 | 0.65 | 0.72 | 0.42 | 0.79 |
| 1 | 61066603 | rs898729 | 0.559 | 0.723 | 0.791 | 0.263 | -0.001 | -0.003 | -0.003 | 0 | 0.62 | 0.37 | 0.51 | 0.63 | 0.39 | 0.69 |
| 1 | 61084790 | rs537110 | -1.033 | 1.161 | -0.275 | -0.167 | -0.05 | 0.041 | 0.041 | 0 | 0.79 | 0.88 | 0.79 | 1 | 0.63 | 0.94 |
| 1 | 61092456 | rs814197 | 0.662 | 0.761 | 0.341 | 0.259 | 0.002 | -0.01 | -0.01 | 0.05 | 0.61 | 0.43 | 0.56 | 0.66 | 0.43 | 0.59 |
| 1 | 61099717 | rs496946 | 0.353 | 0.575 | 0.522 | 0.313 | 0.014 | -0.004 | -0.004 | 0 | 0.64 | 0.55 | 0.6 | 0.66 | 0.48 | 0.61 |
| 1 | 61111774 | rs497005 | 0.082 | 0.732 | -0.081 | 0.435 | -0.038 | -0.006 | -0.006 | 0 | 0.68 | 0.73 | 0.62 | 0.66 | 0.5 | 0.7 |
| 1 | 61121928 | rs473368 | -0.082 | 0.717 | -0.136 | 0.432 | -0.063 | -0.015 | -0.015 | 0 | 0.69 | 0.8 | 0.62 | 0.66 | 0.5 | 0.7 |
| 1 | 61122317 | rs7555033 | 1.403 | 0.319 | -0.291 | 0.763 | 0.013 | -0.014 | -0.014 | 0 | 0.09 | 0.06 | 0.09 | 0.19 | 0.04 | 0.14 |
| 1 | 61123028 | rs550049 | 0.026 | 0.849 | -0.15 | 0.534 | -0.031 | -0.007 | -0.007 | 0.485 | 0.69 | 0.72 | 0.62 | 0.66 | 0.5 | 0.69 |
| 1 | 61129046 | rs4916077 | -0.836 | 0.994 | -0.006 | -0.114 | 0.072 | -0.138 | -0.138 | 0.186 | 0.78 | 0.6 | 0.76 | 1 | 0.65 | 0.94 |
| 1 | 61129957 | rs1020863 | -0.059 | 0.306 | -0.886 | 0.567 | 0.009 | -0.088 | -0.088 | 6.136 | 0.14 | 0.03 | 0.06 | 0.48 | 0.12 | 0.19 |
| 1 | 61132661 | rs17120892 | 0.309 | 0.069 | -0.686 | 0.724 | -0.003 | 0.002 | 0.002 | 0 | 0.05 | 0.06 | 0.07 | 0.11 | 0.01 | 0.13 |
| 1 | 61133462 | rs4915699 | 0.128 | 0.133 | -0.753 | 0.298 | 0.019 | 0.054 | 0.054 | 0 | 0.49 | 0.31 | 0.56 | 0.39 | 0.42 | 0.5 |
| 1 | 61137283 | rs10493291 | -0.05 | -0.323 | -0.745 | 0.187 | 0.002 | -0.011 | -0.011 | 0 | 0.35 | 0.27 | 0.43 | 0.48 | 0.36 | 0.45 |
| 1 | 61140736 | rs12404699 | 0.332 | -0.344 | -0.258 | 0.082 | -0.003 | 0.01 | 0.01 | 0 | 0.05 | 0.03 | 0.04 | 0 | 0.04 | 0.05 |
| 1 | 61141016 | rs10493034 | -1.61 | -0.54 | 0.255 | 0.046 | -0.02 | 0.025 | 0.025 | 0.305 | 0.13 | 0.53 | 0.13 | 0.1 | 0.07 | 0.14 |
| 1 | 61143095 | rs11207648 | -0.987 | -0.474 | 0.275 | 0.062 | 0.018 | -0.003 | -0.003 | 0 | 0.09 | 0.34 | 0.09 | 0.03 | 0.06 | 0.07 |
| 1 | 61144642 | rs1381166 | -0.652 | -0.498 | -0.08 | 0.018 | 0.008 | -0.008 | -0.008 | 0 | 0.44 | 0.62 | 0.52 | 0.52 | 0.42 | 0.51 |
| 1 | 61149086 | rs2219615 | -0.393 | -0.228 | -0.77 | -0.231 | 0.004 | -0.044 | -0.044 | 0 | 0.27 | 0.23 | 0.21 | 0.46 | 0.21 | 0.37 |
| 1 | 61157162 | rs1461353 | -0.897 | -0.177 | -1.248 | -0.45 | -0.019 | -0.051 | -0.051 | 0 | 0.35 | 0.23 | 0.27 | 0.52 | 0.36 | 0.44 |
| 1 | 61162222 | rs1381170 | 0.812 | -0.295 | -0.948 | -0.464 | -0.01 | 0.015 | 0.015 | 0 | 0.27 | 0.28 | 0.37 | 0.42 | 0.21 | 0.37 |
| 1 | 61162380 | rs17120999 | 0.37 | -0.066 | -1.227 | -0.452 | 0.004 | 0.015 | 0.015 | 1.945 | 0.2 | 0.16 | 0.08 | 0.01 | 0.14 | 0.12 |
| 1 | 61170302 | rs12038037 | -0.336 | 0.144 | -0.716 | 0.277 | -0.008 | 0.062 | 0.062 | 0 | 0.06 | 0.04 | 0.08 | 0.4 | 0.06 | 0.23 |
| 1 | 61182251 | rs11207656 | -0.689 | -0.308 | -0.524 | 0.18 | -0.004 | 0.006 | 0.006 | 0 | 0.3 | 0.31 | 0.45 | 0.66 | 0.3 | 0.47 |
| 1 | 61184923 | rs12039857 | 1.017 | -0.169 | -0.301 | 0.485 | -0.021 | 0.06 | 0.06 | 0 | 0.06 | 0.09 | 0.1 | 0.29 | 0.03 | 0.18 |
| 1 | 61185400 | rs11207657 | -0.382 | -0.07 | -0.289 | 0.491 | -0.001 | 0.009 | 0.009 | 0 | 0.16 | 0.17 | 0.24 | 0.4 | 0.22 | 0.26 |
| 1 | 61192181 | rs12045994 | 0.651 | 0.118 | -0.435 | -0.157 | 0.003 | 0.053 | 0.053 | 0.979 | 0.47 | 0.32 | 0.4 | 0.41 | 0.49 | 0.4 |
| 1 | 61193225 | rs13376444 | 0.279 | -0.048 | 0.262 | -0.033 | 0.011 | 0.39 | 0.39 | 0.175 | 0.07 | 0.29 | 0.07 | 0.4 | 0.03 | 0.15 |
| 1 | 61196715 | rs1022265 | 1.449 | -0.04 | 0.389 | -0.125 | 0.014 | 0.039 | 0.039 | 0.605 | 0.38 | 0.28 | 0.27 | 0.3 | 0.3 | 0.32 |
| 1 | 61198872 | rs17121048 | 1.302 | -0.062 | 0.406 | -0.116 | -0.026 | 0.078 | 0.078 | 0 | 0.07 | 0.12 | 0.11 | 0.29 | 0.03 | 0.18 |
| 1 | 61202901 | rs1563954 | -1.443 | 0.596 | 0.672 | -0.029 | 0.001 | 0.011 | 0.011 | 0.202 | 0.42 | 0.63 | 0.49 | 0.66 | 0.41 | 0.45 |
| 1 | 61206485 | rs10889197 | -0.752 | 0.614 | 0.6 | -0.032 | -0.003 | -0.009 | -0.009 | 0 | 0.79 | 0.8 | 0.77 | 0.95 | 0.73 | 0.78 |
| 1 | 61212213 | rs1010910 | -0.061 | 0.607 | 0.551 | -0.012 | -0.049 | 0.053 | 0.053 | 0 | 0.71 | 0.85 | 0.66 | 0.84 | 0.56 | 0.69 |
| 1 | 61212569 | rs6661743 | 0.396 | 0.627 | 0.751 | 0.146 | 0.014 | 0.011 | 0.011 | 0 | 0.64 | 0.44 | 0.58 | 0.44 | 0.52 | 0.56 |
| 1 | 61214988 | rs12749586 | 1.945 | 0.73 | 0.693 | 0.142 | 0.001 | 0.074 | 0.074 | 0.542 | 0.36 | 0.08 | 0.25 | 0.3 | 0.3 | 0.32 |
| 1 | 61216080 | rs867914 | 0.218 | 0.674 | 0.635 | 0.089 | -0.001 | -0.013 | -0.013 | 0 | 0.18 | 0.16 | 0.14 | 0 | 0.14 | 0.16 |
| 1 | 61216954 | rs12133031 | 0.811 | 0.778 | 0.54 | 0.089 | -0.001 | -0.015 | -0.015 | 0.186 | 0.19 | 0.08 | 0.3 | 0.19 | 0.17 | 0.26 |
| 1 | 61222399 | rs11207664 | 0.529 | 0.856 | 0.458 | 0.234 | 0.001 | 0.002 | 0.002 | 0 | 0.1 | 0.11 | 0.1 | 0.13 | 0.15 | 0.09 |
| 1 | 61223973 | rs4915707 | 0.705 | 0.893 | 0.609 | 0.298 | 0.017 | 0.103 | 0.103 | 0 | 0.16 | 0.33 | 0.17 | 0.38 | 0.22 | 0.16 |
| 1 | 61225064 | rs11583359 | -0.65 | 0.948 | 0.723 | 0.117 | 0.001 | -0.014 | -0.014 | 0 | 0.28 | 0.32 | 0.25 | 0.13 | 0.29 | 0.26 |
| 1 | 61225613 | rs1355911 | -1.015 | 0.889 | 0.519 | 0.176 | -0.006 | 0.017 | 0.017 | 0 | 0.44 | 0.32 | 0.53 | 0.3 | 0.44 | 0.5 |
| 1 | 61227108 | rs310162 | -1.963 | 0.879 | 0.209 | 0.388 | 0.011 | 0.048 | 0.048 | 0.157 | 0.62 | 0.8 | 0.73 | 0.69 | 0.67 | 0.65 |
| 1 | 61229194 | rs7553940 | 0.568 | 0.972 | 0.254 | 0.358 | 0 | -0.001 | -0.001 | 0 | 0.11 | 0.09 | 0.1 | 0.13 | 0.16 | 0.1 |
| 1 | 61229363 | rs310163 | -0.467 | 1.033 | 0.456 | 0.369 | -0.001 | 0.003 | 0.003 | 0 | 0.35 | 0.57 | 0.46 | 0.56 | 0.37 | 0.4 |
| 1 | 61229730 | rs310164 | 0.583 | 1.063 | 0.45 | 0.508 | 0.005 | 0.042 | 0.042 | 0 | 0.17 | 0.26 | 0.15 | 0.35 | 0.22 | 0.13 |
| 1 | 61229925 | rs10889198 | -0.302 | 1.031 | 0.399 | 0.329 | 0.003 | -0.029 | -0.029 | 0.189 | 0.32 | 0.39 | 0.45 | 0.18 | 0.29 | 0.35 |
| 1 | 61231047 | rs310166 | -0.316 | 1.071 | -0.135 | 0.357 | 0.003 | 0.018 | 0.018 | 0 | 0.25 | 0.29 | 0.23 | 0.48 | 0.36 | 0.22 |
| 1 | 61233723 | rs310170 | 0.858 | 1.066 | -0.675 | 0.373 | 0 | 0.017 | 0.017 | 0 | 0.15 | 0.2 | 0.15 | 0.36 | 0.22 | 0.12 |
| 1 | 61234270 | rs6676070 | 0.578 | 0.867 | -1.259 | 0.064 | 0 | -0.002 | -0.002 | 0 | 0.19 | 0.16 | 0.3 | 0.2 | 0.16 | 0.24 |
| 1 | 61235684 | rs425181 | -0.779 | 1.1 | -1.403 | -0.237 | 0.03 | 0.067 | 0.067 | 0.501 | 0.61 | 0.88 | 0.76 | 0.67 | 0.7 | 0.65 |
| 1 | 61255619 | rs12403020 | -0.51 | 1.605 | -0.58 | 0.256 | -0.017 | 0.016 | 0.016 | 0.05 | 0.12 | 0.04 | 0.15 | 0 | 0.14 | 0.13 |
| 1 | 61260443 | rs310177 | 1.527 | 1.779 | -0.279 | 0.544 | 0.03 | -0.022 | -0.022 | 0 | 0.94 | 0.9 | 0.87 | 1 | 0.87 | 0.94 |
| 1 | 61264038 | rs12404270 | -2.19 | 2.263 | 0.164 | 1.708 | 0.008 | 0.099 | 0.099 | 0 | 0.08 | 0.11 | 0.27 | 0.45 | 0.14 | 0.23 |

| | | | | | | | | | | | | | | | | |
|---|---|---|---|---|---|---|---|---|---|---|---|---|---|---|---|---|
| 1 | 61264342 | rs6650097 | -0.168 | 1.63 | -0.428 | 1.263 | 0.02 | 0.011 | 0.011 | 0 | 0.65 | 0.79 | 0.67 | 0.35 | 0.74 | 0.52 |
| 1 | 61265635 | rs2211560 | 0.799 | 1.296 | 0.179 | 0.857 | -0.003 | 0.106 | 0.106 | 0 | 0.49 | 0.31 | 0.43 | 0.19 | 0.53 | 0.32 |
| 1 | 61274442 | rs647554 | -0.178 | 1.215 | 0.025 | 0.978 | -0.005 | 0.065 | 0.065 | 0 | 0.63 | 0.81 | 0.63 | 0.27 | 0.68 | 0.44 |
| 1 | 61296240 | rs648787 | -2.012 | 1.107 | 0.995 | 0.937 | 0.053 | 0.011 | 0.011 | 1.502 | 0.15 | 0.56 | 0.3 | 0.16 | 0.19 | 0.21 |
| 1 | 61298261 | rs7516519 | -0.391 | 0.983 | 0.685 | 1.285 | 0 | 0.029 | 0.029 | 0 | 0.06 | 0.06 | 0.12 | 0.18 | 0.06 | 0.14 |
| 1 | 61298399 | rs476077 | -0.079 | 1.012 | 0.608 | 1.224 | 0.03 | 0.023 | 0.023 | 0 | 0.63 | 0.82 | 0.66 | 0.35 | 0.73 | 0.51 |
| 1 | 61303553 | rs11808039 | -1.577 | 1.062 | 0.738 | 1.179 | 0.053 | 0.099 | 0.099 | 0.415 | 0.12 | 0.35 | 0.21 | 0.26 | 0.16 | 0.26 |
| 1 | 61304565 | rs17303103 | 0.924 | 1.018 | 0.425 | 1.282 | -0.01 | 0.079 | 0.079 | 0 | 0.27 | 0.01 | 0.12 | 0.11 | 0.17 | 0.24 |
| 1 | 61313338 | rs788040 | 0.356 | 0.834 | 0.363 | 1.214 | 0.036 | 0.091 | 0.091 | 0 | 0.58 | 0.8 | 0.61 | 0.28 | 0.69 | 0.43 |
| 1 | 61318179 | rs805499 | -0.202 | 0.824 | 0.307 | 1.163 | 0.032 | -0.007 | -0.007 | 0 | 0.65 | 0.86 | 0.74 | 0.7 | 0.76 | 0.58 |
| 1 | 61320892 | rs310159 | -0.405 | 0.788 | 0.218 | 0.981 | 0.044 | -0.03 | -0.03 | 1.547 | 0.62 | 0.9 | 0.73 | 0.66 | 0.74 | 0.55 |
| 1 | 61321009 | rs6587905 | -1.269 | 1.026 | 0.712 | 1.145 | 0.054 | 0.1 | 0.1 | 1.804 | 0.09 | 0.37 | 0.17 | 0.21 | 0.1 | 0.17 |
| 1 | 61329593 | rs581503 | -0.495 | 1.246 | 0.407 | 1.348 | -0.013 | -0.002 | -0.002 | 0.693 | 0.08 | 0.03 | 0.15 | 0.08 | 0.11 | 0.11 |
| 1 | 61332973 | rs6657429 | 0.841 | 1.113 | 0.484 | 1.067 | 0 | -0.003 | -0.003 | 0 | 0.46 | 0.49 | 0.42 | 0.38 | 0.55 | 0.2 |
| 1 | 61334177 | rs17375290 | 0.777 | 1.073 | 0.432 | 1.102 | 0.005 | 0.142 | 0.142 | 0.16 | 0.32 | 0.02 | 0.13 | 0.09 | 0.2 | 0.31 |
| 1 | 61335153 | rs7532404 | 1.462 | 1.216 | 0.541 | 1.135 | -0.005 | -0.012 | -0.012 | 0.05 | 0.4 | 0.31 | 0.29 | 0.03 | 0.45 | 0.12 |
| 1 | 61336529 | rs10493298 | 1.046 | 1.249 | 0.556 | 1.189 | -0.005 | -0.034 | -0.034 | 0 | 0.43 | 0.38 | 0.33 | 0.03 | 0.51 | 0.14 |
| 1 | 61337677 | rs310174 | -0.363 | 0.588 | 0.269 | 1.177 | 0.029 | 0.013 | 0.013 | 0.116 | 0.43 | 0.29 | 0.32 | 0.34 | 0.32 | 0.55 |
| 1 | 61341375 | rs421735 | 0.026 | 0.438 | -0.094 | 1.226 | 0.055 | -0.019 | -0.019 | 1.159 | 0.39 | 0.11 | 0.27 | 0.33 | 0.27 | 0.46 |
| 1 | 61345009 | rs17121177 | -0.02 | 0.427 | -0.053 | 1.254 | 0.004 | -0.001 | -0.001 | 0.18 | 0.08 | 0.06 | 0.15 | 0.28 | 0.06 | 0.21 |
| 1 | 61353698 | rs599989 | -0.332 | 0.593 | -0.272 | 0.645 | 0.026 | 0.047 | 0.047 | 0.047 | 0.4 | 0.28 | 0.2 | 0.09 | 0.27 | 0.43 |
| 1 | 61355010 | rs1447184 | -1.427 | 1.225 | 0.207 | 1.343 | 0.014 | 0.068 | 0.068 | 0 | 0.08 | 0.11 | 0.22 | 0.24 | 0.12 | 0.14 |
| 1 | 61355371 | rs600411 | -1.526 | 0.8 | 0.136 | 0.653 | 0.035 | -0.011 | -0.011 | 0 | 0.05 | 0.2 | 0.05 | 0 | 0.05 | 0.09 |
| 1 | 61359395 | rs10493299 | -0.337 | 0.98 | -0.427 | 0.688 | -0.004 | 0.004 | 0.004 | 0.172 | 0.38 | 0.36 | 0.59 | 0.68 | 0.43 | 0.52 |
| 1 | 61359835 | rs11207680 | -1.579 | 0.723 | -0.268 | 1.874 | -0.002 | 0.087 | 0.087 | 0 | 0.13 | 0.17 | 0.31 | 0.53 | 0.14 | 0.32 |
| 1 | 61363707 | rs2044548 | -0.216 | -0.037 | 0.349 | 1.631 | 0.009 | -0.041 | -0.041 | 0 | 0.1 | 0.29 | 0.06 | 0.05 | 0.08 | 0.05 |
| 1 | 61364921 | rs12079568 | 0.574 | 0.086 | -0.322 | 1.09 | 0.001 | 0.008 | 0.008 | 0 | 0.34 | 0.3 | 0.28 | 0.08 | 0.34 | 0.3 |
| 1 | 61365837 | rs12565296 | -0.533 | 0.281 | -0.69 | 1.469 | -0.007 | 0.277 | 0.277 | 1.937 | 0.08 | 0.34 | 0.21 | 0.38 | 0.05 | 0.21 |
| 1 | 61368955 | rs591979 | -0.119 | 0.317 | -0.902 | 0.865 | 0.004 | 0.018 | 0.018 | 0 | 0.49 | 0.6 | 0.54 | 0.37 | 0.49 | 0.43 |
| 1 | 61376144 | rs573733 | -0.098 | 0.595 | -1.588 | 0.92 | -0.153 | -0.134 | -0.134 | 0 | 0.58 | 0.96 | 0.66 | 0.6 | 0.57 | 0.67 |
| 1 | 61376228 | rs2121598 | -0.708 | 1.064 | -1.074 | 1.245 | -0.029 | -0.003 | -0.003 | 1.096 | 0.05 | 0 | 0.11 | 0.23 | 0.07 | 0.09 |
| 1 | 61376522 | rs655460 | -0.148 | 1.279 | -1.226 | 0.713 | -0.063 | 0.022 | 0.022 | 0.011 | 0.79 | 0.95 | 0.77 | 0.92 | 0.74 | 0.86 |
| 1 | 61377927 | rs11207682 | -0.031 | 1.305 | 0.376 | 0.804 | 0.035 | -0.018 | -0.018 | 0 | 0.06 | 0.14 | 0.13 | 0.03 | 0.1 | 0.04 |
| 1 | 61380205 | rs17121250 | -0.581 | 1.074 | 0.839 | 1.055 | -0.018 | 0.091 | 0.091 | 0.037 | 0.08 | 0.23 | 0.09 | 0.22 | 0.05 | 0.23 |
| 1 | 61382409 | rs10889201 | -0.261 | 1.056 | 0.838 | 0.996 | 0.033 | -0.027 | -0.027 | 0.031 | 0.08 | 0.21 | 0.13 | 0.03 | 0.1 | 0.05 |
| 1 | 61383717 | rs4915717 | -0.516 | 1.559 | 0.182 | 1.033 | 0.064 | 0.067 | 0.067 | 0 | 0.83 | 0.64 | 0.8 | 0.74 | 0.74 | 0.9 |
| 1 | 61383945 | rs4915719 | 0.221 | 1.33 | -0.14 | 0.87 | 0.01 | 0.011 | 0.011 | 0.001 | 0.5 | 0.6 | 0.54 | 0.64 | 0.49 | 0.71 |
| 1 | 61384107 | rs12062121 | 0.778 | 0.848 | -1.03 | 0.941 | 0.017 | -0.013 | -0.013 | 0 | 0.42 | 0.27 | 0.39 | 0.61 | 0.39 | 0.67 |
| 1 | 61384237 | rs12062201 | -0.391 | 0.164 | -1.042 | 0.845 | -0.006 | 0.023 | 0.023 | 0.014 | 0.07 | 0.07 | 0.07 | 0.22 | 0.05 | 0.25 |
| 1 | 61386502 | rs6673313 | -0.591 | 0.533 | -1.117 | 0.246 | -0.004 | 0.08 | 0.08 | 0.002 | 0.33 | 0.05 | 0.26 | 0.1 | 0.25 | 0.19 |
| 1 | 61389213 | rs6587909 | -0.554 | 0.143 | -0.865 | 0.471 | 0.036 | 0.057 | 0.057 | 0 | 0.3 | 0.09 | 0.24 | 0.1 | 0.22 | 0.18 |
| 1 | 61396954 | rs11207685 | -1.668 | 0.769 | 0.504 | 1.264 | 0.03 | 0.145 | 0.145 | 0 | 0.1 | 0.39 | 0.14 | 0.1 | 0.07 | 0.07 |
| 1 | 61398759 | rs607777 | -2.177 | 0.394 | 1.169 | 1.496 | -0.031 | 0.002 | 0.002 | 0.564 | 0.16 | 0.6 | 0.17 | 0.18 | 0.1 | 0.24 |
| 1 | 61403867 | rs632853 | -1.717 | 0.81 | -0.093 | 1.522 | -0.118 | -0.115 | -0.115 | 0 | 0.45 | 0.97 | 0.52 | 0.54 | 0.46 | 0.48 |
| 1 | 61408445 | rs11207686 | -1.013 | 1.273 | 0.777 | 1.554 | 0.122 | 0.09 | 0.09 | 0 | 0.12 | 0.38 | 0.22 | 0.16 | 0.18 | 0.16 |
| 1 | 61409064 | rs1414277 | -0.989 | 1.361 | 0.838 | 1.435 | 0.131 | 0.094 | 0.094 | 0 | 0.12 | 0.38 | 0.22 | 0.15 | 0.18 | 0.14 |
| 1 | 61409566 | rs11207687 | -1.545 | 1.386 | 0.941 | 1.519 | 0.134 | 0.087 | 0.087 | 0.01 | 0.13 | 0.4 | 0.24 | 0.19 | 0.18 | 0.19 |
| 1 | 61412131 | rs17121331 | -0.344 | 0.959 | 0.317 | 1.581 | -0.005 | -0.04 | -0.04 | 7.455 | 0.18 | 0.08 | 0.16 | 0.18 | 0.19 | 0.08 |
| 1 | 61413691 | rs529823 | -2.059 | 1.083 | 1.106 | 1.91 | 0.002 | 0.003 | 0.003 | 0.02 | 0.35 | 0.59 | 0.48 | 0.6 | 0.39 | 0.51 |
| 1 | 61414438 | rs537075 | -1.869 | 1.034 | 1.689 | 2.344 | 0.054 | 0.19 | 0.19 | 2.092 | 0.06 | 0.33 | 0.1 | 0.2 | 0.04 | 0.09 |
| 1 | 61417729 | rs1572312 | -0.685 | 1.048 | 1.463 | 2.265 | 0.001 | 0.008 | 0.008 | 0.867 | 0.1 | 0.09 | 0.07 | 0.03 | 0.11 | 0.11 |
| 1 | 61423517 | rs2361817 | 1.712 | 1.096 | 1.937 | 2.295 | 0.097 | 0.253 | 0.253 | 3.289 | 0.93 | 0.52 | 0.89 | 0.8 | 0.95 | 0.91 |
| 1 | 61425824 | rs12030799 | -0.995 | 1.027 | 1.344 | 2.598 | 0.003 | 0.072 | 0.072 | 0.375 | 0.07 | 0.07 | 0.1 | 0.3 | 0.09 | 0.32 |
| 1 | 61435075 | rs12038942 | -1.748 | 0.973 | 1.399 | 2.575 | -0.001 | 0.017 | 0.017 | 1.783 | 0.18 | 0.21 | 0.19 | 0.34 | 0.2 | 0.43 |
| 1 | 61447172 | rs1032622 | -1.245 | 0.425 | 1.758 | 3.043 | -0.009 | 0.014 | 0.014 | 0 | 0.21 | 0.3 | 0.2 | 0.18 | 0.19 | 0.11 |
| 1 | 61449494 | rs12140465 | -2.154 | -0.134 | 1.703 | 2.687 | 0.06 | 0.002 | 0.002 | 0.055 | 0.39 | 0.76 | 0.31 | 0.66 | 0.25 | 0.43 |
| 1 | 61456362 | rs10082014 | -1.771 | 0.07 | 1.701 | 2.653 | -0.04 | 0.265 | 0.265 | 0.673 | 0.1 | 0.35 | 0.11 | 0.33 | 0.03 | 0.07 |
| 1 | 61463426 | rs12028122 | -1.144 | 0.674 | 1.175 | 2.391 | 0.002 | 0.048 | 0.048 | 0 | 0.08 | 0.16 | 0.07 | 0.23 | 0.09 | 0.08 |
| 1 | 61465289 | rs1283256 | 1.874 | 0.551 | 1.042 | 2.436 | 0.013 | 0.035 | 0.035 | 0 | 0.55 | 0.28 | 0.58 | 0.65 | 0.59 | 0.52 |
| 1 | 61465314 | rs8179472 | -1.931 | 0.163 | 1.061 | 2.731 | -0.012 | 0.182 | 0.182 | 0 | 0.13 | 0.26 | 0.28 | 0.52 | 0.1 | 0.31 |
| 1 | 61467602 | rs17121462 | -1.868 | 0.171 | 1.065 | 2.415 | -0.01 | 0.098 | 0.098 | 0 | 0.13 | 0.26 | 0.26 | 0.29 | 0.1 | 0.28 |
| 1 | 61470318 | rs870752 | -1.238 | 0.061 | 0.77 | 2.296 | 0.013 | 0.023 | 0.023 | 0 | 0.21 | 0.1 | 0.09 | 0.11 | 0.19 | 0.19 |
| 1 | 61472174 | rs12072258 | -1.781 | 0.062 | 1.394 | 2.744 | -0.017 | 0.18 | 0.18 | 0.87 | 0.13 | 0.41 | 0.21 | 0.32 | 0.07 | 0.12 |
| 1 | 61476762 | rs1498109 | 0.87 | -0.012 | 0.439 | 1.828 | 0.009 | 0.131 | 0.131 | 0 | 0.37 | 0.13 | 0.2 | 0.05 | 0.34 | 0.24 |
| 1 | 61477603 | rs1419336 | 0.727 | 0.237 | 0.605 | 2.238 | 0.007 | 0.016 | 0.016 | 0 | 0.5 | 0.34 | 0.48 | 0.43 | 0.48 | 0.44 |
| 1 | 61483042 | rs1779865 | -0.05 | 0.194 | 0.761 | 2.436 | 0.007 | 0.008 | 0.008 | 0 | 0.59 | 0.54 | 0.55 | 0.5 | 0.55 | 0.49 |
| 1 | 61484997 | rs2173346 | -1.281 | 0.19 | 0.728 | 2.469 | -0.017 | 0.151 | 0.151 | 0.096 | 0.12 | 0.36 | 0.25 | 0.25 | 0.07 | 0.12 |
| 1 | 61485568 | rs2173345 | -0.443 | 0.121 | 0.574 | 2.421 | -0.01 | 0.061 | 0.061 | 0 | 0.09 | 0.23 | 0.21 | 0.18 | 0.06 | 0.07 |
| 1 | 61493285 | rs1777533 | 0.132 | -0.024 | 0.838 | 2.522 | 0.001 | 0.004 | 0.004 | 0 | 0.6 | 0.6 | 0.57 | 0.5 | 0.56 | 0.49 |

| | | | | | | | | | | | | | | | | |
|---|---|---|---|---|---|---|---|---|---|---|---|---|---|---|---|---|
| 1 | 61522880 | rs12565220 | -2.327 | -0.093 | 1.238 | 3.25 | 0.049 | 0.224 | 0.224 | 0.013 | 0.06 | 0.25 | 0.21 | 0.28 | 0.08 | 0.07 |
| 1 | 61528110 | rs6587910 | -1.599 | -0.295 | 1.17 | 2.652 | -0.032 | 0.018 | 0.018 | 0.566 | 0.07 | 0.24 | 0.05 | 0.04 | 0.02 | 0.02 |
| 1 | 61567939 | rs964091 | -2.481 | -0.174 | 1.268 | 3.616 | 0.053 | 0.372 | 0.372 | 0 | 0.06 | 0.25 | 0.12 | 0.44 | 0.07 | 0.14 |
| 1 | 61571392 | rs10493302 | -3.432 | -0.278 | 1.346 | 3.686 | -0.004 | 0.228 | 0.228 | 0 | 0.15 | 0.38 | 0.19 | 0.51 | 0.13 | 0.28 |
| 1 | 61574202 | rs4915728 | 1.592 | -0.326 | 1.341 | 3.67 | -0.008 | -0.001 | -0.001 | 0 | 0.92 | 0.88 | 0.94 | 0.95 | 0.94 | 0.9 |
| 1 | 61582546 | rs6693597 | -0.671 | -0.527 | 1.172 | 3.66 | 0.009 | 0.022 | 0.022 | 0 | 0.2 | 0.11 | 0.08 | 0.1 | 0.13 | 0.21 |
| 1 | 61607034 | rs2008968 | -0.524 | -0.496 | 1.607 | 3.536 | 0.015 | 0.021 | 0.021 | 0 | 0.09 | 0.2 | 0.12 | 0.06 | 0.12 | 0.2 |
| 1 | 61615536 | rs334707 | 2.76 | -0.77 | 1.512 | 3.881 | -0.049 | 0.22 | 0.22 | 0 | 0.86 | 0.64 | 0.82 | 0.57 | 0.93 | 0.75 |
| 1 | 61619056 | rs334701 | 2.046 | -0.754 | 1.401 | 3.73 | -0.018 | 0.014 | 0.014 | 0 | 0.89 | 0.83 | 0.9 | 0.86 | 0.93 | 0.89 |
| 1 | 61622287 | rs334713 | 1.499 | -0.748 | 1.349 | 3.691 | -0.002 | 0.021 | 0.021 | 0 | 0.94 | 0.81 | 0.95 | 0.95 | 0.96 | 0.9 |
| 1 | 61622755 | rs334712 | 2.163 | -0.756 | 1.336 | 3.747 | -0.019 | 0.055 | 0.055 | 0 | 0.9 | 0.81 | 0.92 | 0.81 | 0.94 | 0.89 |
| 1 | 61625310 | rs334711 | 3.398 | -0.747 | 1.515 | 3.703 | -0.048 | 0.265 | 0.265 | 0.063 | 0.84 | 0.53 | 0.82 | 0.55 | 0.91 | 0.75 |
| 1 | 61625872 | rs334710 | 1.996 | -0.879 | 1.435 | 3.815 | 0.014 | 0.187 | 0.187 | 0 | 0.93 | 0.73 | 0.92 | 0.74 | 0.96 | 0.84 |
| 1 | 61629453 | rs168022 | 2.545 | -0.88 | 1.439 | 3.77 | -0.028 | 0.061 | 0.061 | 0 | 0.89 | 0.73 | 0.9 | 0.77 | 0.94 | 0.81 |
| 1 | 61632029 | rs334723 | 1.25 | -0.865 | 1.419 | 3.733 | 0 | 0.007 | 0.007 | 0 | 0.94 | 0.84 | 0.95 | 0.96 | 0.96 | 0.9 |
| 1 | 61639382 | rs334717 | 2.385 | -0.835 | 1.083 | 3.01 | -0.039 | 0.082 | 0.082 | 0 | 0.86 | 0.69 | 0.85 | 0.68 | 0.93 | 0.75 |
| 1 | 61646425 | rs914735 | -1.233 | -0.739 | 0.936 | 2.978 | 0.015 | 0.171 | 0.171 | 0.044 | 0.07 | 0.23 | 0.22 | 0.28 | 0.05 | 0.16 |
| 1 | 61647103 | rs12091215 | -1.43 | -0.764 | 1.043 | 2.949 | 0.023 | 0.216 | 0.216 | 0 | 0.09 | 0.36 | 0.23 | 0.28 | 0.05 | 0.2 |
| 1 | 61648634 | rs10789093 | -2.309 | -0.857 | 0.696 | 2.848 | -0.035 | 0.102 | 0.102 | 0 | 0.24 | 0.42 | 0.33 | 0.47 | 0.17 | 0.39 |
| 1 | 61652197 | rs12142891 | -0.589 | -0.716 | 0.139 | 2.304 | -0.022 | 0.001 | 0.001 | 0.051 | 0.1 | 0.01 | 0.06 | 0.05 | 0.09 | 0.16 |
| 1 | 61659786 | rs12126115 | -1.388 | -0.674 | -0.296 | 1.868 | -0.027 | 0.021 | 0.021 | 0 | 0.31 | 0.51 | 0.36 | 0.59 | 0.21 | 0.42 |
| 1 | 61668784 | rs10889215 | -1.376 | -0.91 | -0.56 | 1.92 | 0.004 | -0.007 | -0.007 | 0 | 0.21 | 0.73 | 0.31 | 0.37 | 0.1 | 0.16 |
| 1 | 61670344 | rs441374 | -1.056 | -1.329 | -0.821 | 1.657 | 0.009 | 0.002 | 0.002 | 0 | 0.23 | 0.13 | 0.22 | 0.26 | 0.17 | 0.11 |
| 1 | 61671468 | rs12118221 | 0.776 | -1.244 | -0.725 | 1.315 | 0.028 | -0.012 | -0.012 | 0 | 0.33 | 0.41 | 0.38 | 0.25 | 0.5 | 0.34 |
| 1 | 61674052 | rs332823 | -0.696 | -1.192 | -0.798 | 1.09 | -0.032 | -0.005 | -0.005 | 0.08 | 0.57 | 0.78 | 0.48 | 0.63 | 0.47 | 0.44 |
| 1 | 61674963 | rs334668 | -0.265 | -1.348 | -0.49 | 1.15 | 0.043 | -0.012 | -0.012 | 0 | 0.44 | 0.31 | 0.35 | 0.4 | 0.33 | 0.46 |
| 1 | 61678557 | rs332819 | -2.003 | -1.259 | -0.634 | 1.225 | -0.14 | -0.064 | -0.064 | 1.051 | 0.74 | 0.97 | 0.69 | 0.75 | 0.58 | 0.67 |
| 1 | 61678783 | rs333151 | 0.058 | -1.676 | -0.198 | 0.91 | 0.024 | 0.043 | 0.043 | 0 | 0.25 | 0.52 | 0.19 | 0.28 | 0.25 | 0.23 |
| 1 | 61679283 | rs332818 | -1.547 | -1.8 | -0.895 | 0.925 | -0.109 | -0.152 | -0.152 | 0.047 | 0.64 | 0.93 | 0.6 | 0.5 | 0.44 | 0.57 |
| 1 | 61680057 | rs333149 | -0.986 | -1.551 | -0.03 | 0.881 | 0.025 | 0.067 | 0.067 | 0.041 | 0.3 | 0.56 | 0.3 | 0.52 | 0.32 | 0.3 |
| 1 | 61681474 | rs11207714 | 1.378 | -1.521 | 0.023 | 0.819 | -0.001 | 0.004 | 0.004 | 0 | 0.1 | 0.1 | 0.08 | 0.04 | 0.11 | 0.11 |
| 1 | 61681500 | rs333146 | 0.14 | -1.358 | 0.571 | 1.01 | 0.021 | 0.029 | 0.029 | 0 | 0.13 | 0.21 | 0.12 | 0.23 | 0.15 | 0.12 |
| 1 | 61684005 | rs333142 | 1.424 | -1.317 | 0.391 | 1.029 | -0.011 | -0.021 | -0.021 | 0 | 0.08 | 0.03 | 0.04 | 0.23 | 0.07 | 0.1 |
| 1 | 61684518 | rs12734797 | 0.002 | -2.075 | 0.305 | 0.557 | 0.006 | 0.002 | 0.002 | 0 | 0.21 | 0.13 | 0.22 | 0.13 | 0.11 | 0.26 |
| 1 | 61694210 | rs6678065 | 0.901 | -1.514 | 0.221 | 0.568 | -0.019 | -0.022 | -0.022 | 0 | 0.05 | 0 | 0.08 | 0.26 | 0.05 | 0.07 |
| 1 | 61704202 | rs17377253 | 0.956 | -1.614 | 0.246 | 0.267 | 0.002 | -0.008 | -0.008 | 0.364 | 0.1 | 0.05 | 0.04 | 0 | 0.07 | 0.04 |
| 1 | 61704221 | rs6669982 | -0.71 | -1.473 | 0.632 | 0.638 | -0.025 | 0.161 | 0.161 | 0.21 | 0.14 | 0.45 | 0.13 | 0.31 | 0.05 | 0.06 |
| 1 | 61706073 | rs7540527 | 0.969 | -0.851 | -0.315 | 0.213 | -0.002 | 0.021 | 0.021 | 0 | 0.09 | 0.03 | 0.08 | 0.03 | 0.08 | 0.17 |
| 1 | 61712185 | rs17121858 | 0.469 | -0.743 | -0.198 | 0.427 | -0.011 | -0.011 | -0.011 | 0 | 0.11 | 0.09 | 0.11 | 0.22 | 0.15 | 0.12 |
| 1 | 61725652 | rs332837 | 0.134 | -1.276 | -0.325 | -0.012 | 0.026 | 0.049 | 0.049 | 0.157 | 0.37 | 0.24 | 0.19 | 0.12 | 0.28 | 0.22 |
| 1 | 61728597 | rs2065841 | -0.761 | -0.944 | -0.024 | 0.379 | 0.017 | 0.076 | 0.076 | 0.008 | 0.2 | 0.57 | 0.17 | 0.39 | 0.18 | 0.22 |
| 1 | 61731440 | rs6684051 | 0.825 | -0.942 | 0.139 | 0.395 | 0 | 0.036 | 0.036 | 0 | 0.11 | 0.18 | 0.08 | 0.21 | 0.07 | 0.11 |
| 1 | 61731652 | rs17121875 | -0.422 | -0.927 | -0.218 | 0.404 | 0.054 | 0.144 | 0.144 | 0 | 0.09 | 0.32 | 0.06 | 0.24 | 0.09 | 0.16 |
| 1 | 61733856 | rs7544358 | 0.073 | -0.837 | -0.181 | 0.395 | 0.079 | 0.141 | 0.141 | 0 | 0.08 | 0.32 | 0.06 | 0.18 | 0.08 | 0.1 |
| 1 | 61738270 | rs437021 | 1.239 | -0.971 | -0.869 | 0.187 | -0.124 | -0.162 | -0.162 | 0 | 0.34 | 0.05 | 0.56 | 0.46 | 0.45 | 0.4 |
| 1 | 61739394 | rs332830 | 0.282 | -1.267 | -0.269 | -0.762 | 0.034 | 0.074 | 0.074 | 0.02 | 0.54 | 0.65 | 0.73 | 0.85 | 0.63 | 0.62 |
| 1 | 61751410 | rs2806425 | 0.53 | 0.243 | -0.932 | -0.222 | -0.02 | 0.104 | 0.104 | 0 | 0.57 | 0.24 | 0.71 | 0.75 | 0.53 | 0.7 |
| 1 | 61755978 | rs2806426 | 1.026 | -0.293 | 0.112 | 0.045 | -0.022 | 0.037 | 0.037 | 0 | 0.88 | 0.69 | 0.84 | 0.88 | 0.94 | 0.81 |
| 1 | 61757024 | rs2243874 | 1.174 | -0.097 | 0.668 | -0.537 | 0.034 | -0.07 | -0.07 | 2.006 | 0.82 | 0.49 | 0.84 | 0.9 | 0.78 | 0.83 |
| 1 | 61759942 | rs11207721 | -0.198 | -1.05 | 0.098 | -0.828 | -0.046 | -0.019 | -0.019 | 0.009 | 0.29 | 0.38 | 0.14 | 0.29 | 0.19 | 0.18 |
| 1 | 61762362 | rs1414265 | 1.222 | -0.895 | -0.29 | -0.791 | 0.039 | 0.101 | 0.101 | 0 | 0.49 | 0.11 | 0.69 | 0.62 | 0.56 | 0.7 |
| 1 | 61774969 | rs2806436 | 0.094 | -0.477 | -0.002 | -0.761 | -0.01 | 0.006 | 0.006 | 0.007 | 0.89 | 0.92 | 0.92 | 0.93 | 0.83 | 0.9 |
| 1 | 61776742 | rs2363876 | -0.861 | -0.848 | 0.498 | -0.923 | -0.002 | -0.038 | -0.038 | 0 | 0.28 | 0.27 | 0.17 | 0.04 | 0.28 | 0.09 |
| 1 | 61781139 | rs2782541 | -0.404 | -0.75 | 0.099 | -0.859 | -0.003 | 0.001 | 0.001 | 0 | 0.91 | 0.91 | 0.91 | 0.95 | 0.84 | 0.93 |
| 1 | 61784017 | rs2067774 | 0.553 | -0.853 | 0.115 | -0.909 | 0.028 | 0.056 | 0.056 | 0 | 0.16 | 0.06 | 0.06 | 0 | 0.12 | 0.03 |
| 1 | 61786398 | rs2806440 | -0.115 | -0.668 | 0.269 | -0.841 | 0.007 | -0.003 | -0.003 | 0 | 0.9 | 0.88 | 0.89 | 0.95 | 0.83 | 0.93 |
| 1 | 61791863 | rs6691768 | 1.475 | -1.114 | 0.322 | -0.907 | 0.048 | 0.093 | 0.093 | 0.058 | 0.54 | 0.16 | 0.69 | 0.69 | 0.59 | 0.74 |
| 1 | 61792008 | rs6691941 | 0.552 | -1.225 | 0.071 | -1.097 | 0.002 | 0.09 | 0.09 | 0 | 0.71 | 0.85 | 0.83 | 0.94 | 0.71 | 0.91 |
| 1 | 61792661 | rs6694984 | -0.515 | -1.131 | 0.219 | -0.924 | 0.04 | 0.036 | 0.036 | 0 | 0.27 | 0.19 | 0.13 | 0.2 | 0.18 | 0.06 |
| 1 | 61803889 | rs7521242 | 1.087 | -0.134 | 0.475 | -0.916 | -0.002 | 0.046 | 0.046 | 0.21 | 0.49 | 0.41 | 0.57 | 0.67 | 0.46 | 0.62 |
| 1 | 61810515 | rs766488 | -0.529 | -0.006 | 0.827 | -0.998 | -0.004 | -0.108 | -0.108 | 0 | 0.34 | 0.51 | 0.29 | 0.08 | 0.35 | 0.16 |
| 1 | 61812773 | rs9436636 | -0.817 | 0.051 | 0.639 | -0.974 | -0.006 | -0.035 | -0.035 | 0 | 0.41 | 0.52 | 0.33 | 0.26 | 0.4 | 0.17 |
| 1 | 61825664 | rs6587917 | -0.298 | 0.256 | 0.135 | -0.904 | 0.006 | -0.013 | -0.013 | 0 | 0.9 | 0.9 | 0.9 | 0.95 | 0.86 | 0.81 |
| 1 | 61826395 | rs2180200 | 0.162 | 0.224 | 0.121 | -0.913 | 0.005 | -0.011 | -0.011 | 0 | 0.87 | 0.88 | 0.8 | 0.95 | 0.8 | 0.78 |
| 1 | 61831890 | rs1125777 | 0.666 | -0.017 | -0.179 | -0.674 | -0.008 | 0.108 | 0.108 | 0.359 | 0.48 | 0.25 | 0.55 | 0.71 | 0.45 | 0.62 |
| 1 | 61831938 | rs12568010 | -0.278 | -0.009 | -0.048 | 0.19 | -0.013 | 0.021 | 0.021 | 0 | 0.09 | 0.19 | 0.07 | 0.17 | 0.06 | 0.03 |
| 1 | 61835924 | rs12568891 | -0.832 | 0.251 | -0.702 | 0.743 | -0.006 | -0.075 | -0.075 | 0 | 0.15 | 0.06 | 0.22 | 0.34 | 0.16 | 0.29 |
| 1 | 61837588 | rs7533561 | -0.338 | 0.151 | -0.656 | 0.504 | 0.013 | 0.031 | 0.031 | 0 | 0.16 | 0.08 | 0.05 | 0.05 | 0.11 | 0.06 |
| 1 | 61838509 | rs10489911 | -0.475 | 0.615 | -0.397 | 1.09 | -0.01 | -0.047 | -0.047 | 0 | 0.09 | 0.05 | 0.19 | 0.29 | 0.09 | 0.12 |

| | | | | | | | | | | | | | | | | |
|---|---|---|---|---|---|---|---|---|---|---|---|---|---|---|---|---|
| 1 | 61844272 | rs11807521 | -1.319 | 0.983 | 0.132 | 0.737 | 0.13 | 0.095 | 0.095 | 0.016 | 0.08 | 0.36 | 0.13 | 0.09 | 0.11 | 0.05 |
| 1 | 61851210 | rs6686758 | -0.966 | 1.256 | -0.794 | 1.03 | -0.04 | -0.063 | -0.063 | 0 | 0.13 | 0.09 | 0.29 | 0.32 | 0.2 | 0.14 |
| 1 | 61859802 | rs2474388 | -0.15 | 1.341 | -1.519 | -0.58 | -0.069 | 0.191 | 0.191 | 0 | 0.6 | 0.87 | 0.71 | 0.95 | 0.51 | 0.63 |
| 1 | 61861024 | rs11581697 | 0.864 | 0.808 | -1.357 | -0.26 | -0.006 | -0.002 | -0.002 | 0 | 0.28 | 0.26 | 0.19 | 0.27 | 0.19 | 0.32 |
| 1 | 61864231 | rs1474872 | -0.668 | 0.902 | -1.687 | -0.298 | -0.158 | 0.283 | 0.283 | 0 | 0.6 | 0.96 | 0.72 | 0.95 | 0.52 | 0.63 |
| 1 | 61864763 | rs10489908 | 1.108 | 1.122 | -2.14 | -0.74 | -0.103 | 0.175 | 0.175 | 0 | 0.25 | 0.02 | 0.13 | 0 | 0.3 | 0.08 |
| 1 | 61865995 | rs17356825 | 1.183 | 0.994 | -1.836 | -0.49 | -0.023 | 0.005 | 0.005 | 0 | 0.06 | 0 | 0.04 | 0.04 | 0.07 | 0.14 |
| 1 | 61869118 | rs2474382 | 0.209 | 0.802 | -1.079 | -0.057 | 0.001 | -0.006 | -0.006 | 2.266 | 0.29 | 0.31 | 0.38 | 0.56 | 0.29 | 0.28 |
| 1 | 61871356 | rs7535730 | 0.709 | 0.775 | -1.064 | 0.095 | 0.009 | -0.003 | -0.003 | 0.691 | 0.26 | 0.16 | 0.18 | 0.27 | 0.18 | 0.34 |
| 1 | 61872541 | rs10889221 | 0.479 | 0.913 | -1.173 | 0.021 | -0.008 | 0.006 | 0.006 | 0 | 0.28 | 0.12 | 0.26 | 0.22 | 0.29 | 0.32 |
| 1 | 61873246 | rs2474379 | -0.399 | 0.883 | -1.145 | -0.446 | -0.082 | 0.1 | 0.1 | 0 | 0.83 | 0.95 | 0.85 | 0.99 | 0.73 | 0.89 |
| 1 | 61877445 | rs2092867 | -0.071 | 0.644 | -1.111 | -0.484 | -0.098 | 0.202 | 0.202 | 0.07 | 0.7 | 0.95 | 0.8 | 0.99 | 0.66 | 0.85 |
| 1 | 61880686 | rs2499533 | -0.106 | 0.519 | -1.11 | -0.485 | -0.092 | 0.201 | 0.201 | 0.131 | 0.7 | 0.95 | 0.81 | 0.99 | 0.66 | 0.85 |
| 1 | 61880714 | rs2474373 | 0.153 | -0.145 | -0.157 | 0.273 | -0.039 | 0.134 | 0.134 | 0 | 0.2 | 0.36 | 0.39 | 0.5 | 0.15 | 0.22 |
| 1 | 61890465 | rs11207738 | 0.436 | -0.119 | -0.537 | 0.404 | 0.013 | -0.082 | -0.082 | 0.668 | 0.18 | 0.07 | 0.29 | 0.43 | 0.16 | 0.23 |
| 1 | 61891184 | rs17265796 | 0.971 | -0.077 | -0.617 | 0.414 | -0.025 | -0.007 | -0.007 | 1.775 | 0.07 | 0 | 0.04 | 0.06 | 0.09 | 0.1 |
| 1 | 61894709 | rs2207792 | -0.056 | -0.226 | 0.053 | 0.379 | -0.015 | -0.048 | -0.048 | 0 | 0.63 | 0.48 | 0.69 | 0.88 | 0.68 | 0.85 |
| 1 | 61895257 | rs6587924 | 0.252 | 0.216 | 0.259 | 0.838 | -0.005 | 0.019 | 0.019 | 0 | 0.44 | 0.23 | 0.38 | 0.44 | 0.52 | 0.62 |
| 1 | 61920769 | rs2474358 | 0.373 | -0.378 | -1.242 | 0.289 | 0.05 | 0.088 | 0.088 | 0 | 0.76 | 0.95 | 0.88 | 0.9 | 0.83 | 0.77 |
| 1 | 61929717 | rs2474363 | -1.067 | 0.105 | -0.043 | 1.549 | 0.022 | 0.299 | 0.299 | 0.308 | 0.16 | 0.44 | 0.31 | 0.46 | 0.15 | 0.22 |
| 1 | 61930062 | rs2474359 | 0.447 | -0.263 | -0.213 | 1.472 | -0.004 | 0.003 | 0.003 | 0 | 0.85 | 0.84 | 0.82 | 0.82 | 0.88 | 0.79 |
| 1 | 61931141 | rs2474366 | -0.454 | -0.255 | 0.272 | 2.154 | 0.003 | 0.051 | 0.051 | 0 | 0.09 | 0.09 | 0.21 | 0.3 | 0.11 | 0.18 |
| 1 | 61931351 | rs9436212 | 0.387 | -0.324 | 0.576 | 2.05 | -0.006 | 0.018 | 0.018 | 0 | 0.86 | 0.76 | 0.8 | 0.8 | 0.86 | 0.76 |
| 1 | 61949463 | rs4281350 | -0.993 | -0.212 | 1.285 | 2.228 | -0.003 | 0.08 | 0.08 | 0 | 0.12 | 0.3 | 0.24 | 0.22 | 0.09 | 0.17 |
| 1 | 61952574 | rs4915746 | 1.408 | -0.227 | 1.149 | 2.183 | 0.001 | 0.131 | 0.131 | 1.535 | 0.77 | 0.46 | 0.53 | 0.49 | 0.79 | 0.66 |
| 1 | 61954169 | rs4915748 | 0.391 | -0.1 | 0.316 | 1.046 | -0.007 | 0.015 | 0.015 | 0 | 0.38 | 0.32 | 0.27 | 0.26 | 0.42 | 0.35 |
| 1 | 61961953 | rs3890764 | 0.638 | -0.069 | 0.624 | 1.751 | 0.017 | 0.254 | 0.254 | 0 | 0.91 | 0.7 | 0.83 | 0.72 | 0.91 | 0.87 |
| 1 | 61966582 | rs6682077 | 0.213 | -0.137 | -0.3 | 1.354 | -0.05 | -0.005 | -0.005 | 0 | 0.6 | 0.2 | 0.51 | 0.69 | 0.57 | 0.74 |
| 1 | 61977016 | rs2499515 | -0.012 | -0.149 | -0.14 | 1.338 | -0.01 | 0.005 | 0.005 | 0 | 0.62 | 0.4 | 0.57 | 0.71 | 0.57 | 0.74 |
| 1 | 61977281 | rs4915591 | 1.271 | -0.066 | -1.415 | 1.226 | -0.093 | -0.101 | -0.101 | 0 | 0.22 | 0.02 | 0.19 | 0.34 | 0.31 | 0.28 |
| 1 | 61978694 | rs7512200 | 0.024 | 0.088 | -1.055 | 0.998 | -0.057 | -0.002 | -0.002 | 0 | 0.61 | 0.18 | 0.51 | 0.7 | 0.57 | 0.74 |
| 1 | 61987970 | rs2246514 | -0.901 | 0.212 | -0.078 | 1.227 | -0.002 | -0.003 | -0.003 | 0 | 0.47 | 0.55 | 0.52 | 0.63 | 0.39 | 0.6 |
| 1 | 61998177 | rs1890265 | -0.004 | 0.189 | -0.462 | 1.444 | 0.006 | -0.045 | -0.045 | 0 | 0.14 | 0.07 | 0.17 | 0.31 | 0.11 | 0.17 |
| 1 | 62000281 | rs1933299 | -0.2 | 0.127 | 0.233 | 1.482 | -0.013 | 0.079 | 0.079 | 0 | 0.15 | 0.3 | 0.23 | 0.34 | 0.11 | 0.17 |
| 1 | 62002888 | rs1933301 | -0.322 | 0.286 | 0.191 | 1.428 | -0.004 | 0.007 | 0.007 | 0 | 0.33 | 0.28 | 0.34 | 0.26 | 0.34 | 0.44 |
| 1 | 62003946 | rs7541526 | -1.229 | 0.276 | 0.218 | 1.306 | -0.004 | 0.004 | 0.004 | 0.202 | 0.46 | 0.37 | 0.51 | 0.57 | 0.45 | 0.62 |
| 1 | 62022914 | rs1933314 | 0.035 | 0.256 | -0.505 | 1.352 | 0.012 | -0.049 | -0.049 | 0 | 0.16 | 0.08 | 0.18 | 0.36 | 0.12 | 0.18 |
| 1 | 62023059 | rs1933315 | -1.013 | 0.32 | -0.248 | 1.069 | 0 | 0.004 | 0.004 | 0.478 | 0.49 | 0.55 | 0.57 | 0.6 | 0.46 | 0.61 |
| 1 | 62027313 | rs1933316 | -0.276 | 0.146 | -0.467 | 1.239 | 0.002 | -0.028 | -0.028 | 0 | 0.16 | 0.1 | 0.15 | 0.36 | 0.14 | 0.32 |
| 1 | 62027586 | rs6661015 | 0.033 | 0.03 | -0.379 | 0.663 | 0 | -0.001 | -0.001 | 0 | 0.3 | 0.35 | 0.32 | 0.17 | 0.23 | 0.3 |
| 1 | 62032669 | rs11207774 | -0.031 | 0.068 | -0.139 | 0.67 | 0.001 | 0.004 | 0.004 | 0 | 0.3 | 0.3 | 0.17 | 0.14 | 0.24 | 0.31 |
| 1 | 62034999 | rs2365259 | 1.121 | -0.176 | -0.052 | 0.496 | 0.002 | -0.006 | -0.006 | 0 | 0.45 | 0.36 | 0.49 | 0.47 | 0.59 | 0.33 |
| 1 | 62048979 | rs2185040 | -0.45 | -0.133 | -0.038 | 0.529 | 0.008 | -0.001 | -0.001 | 0.002 | 0.92 | 0.92 | 0.97 | 1 | 0.96 | 0.96 |
| 1 | 62050490 | rs1933323 | -0.283 | -0.248 | 0.332 | 0.607 | -0.006 | 0.036 | 0.036 | 0 | 0.2 | 0.34 | 0.19 | 0.34 | 0.18 | 0.35 |
| 1 | 62053966 | rs2365263 | 0.936 | -0.386 | 0.137 | 0.368 | 0.055 | -0.014 | -0.014 | 0.078 | 0.5 | 0.18 | 0.56 | 0.47 | 0.61 | 0.34 |
| 1 | 62059647 | rs12143476 | 0.816 | 0.182 | 0.042 | 0.268 | 0 | -0.001 | -0.001 | 0.024 | 0.58 | 0.39 | 0.57 | 0.66 | 0.57 | 0.5 |
| 1 | 62059660 | rs10493308 | -0.331 | 0.177 | 0.393 | 0.28 | -0.013 | -0.004 | -0.004 | 0 | 0.76 | 0.82 | 0.82 | 0.81 | 0.7 | 0.66 |
| 1 | 62067741 | rs6660502 | 0.329 | 0.127 | 0.604 | 0.296 | 0.01 | 0.019 | 0.019 | 0 | 0.1 | 0.29 | 0.07 | 0.1 | 0.09 | 0.13 |
| 1 | 62068302 | rs2029857 | 0.516 | 0.017 | 0.662 | 0.45 | -0.002 | -0.005 | -0.005 | 0 | 0.58 | 0.37 | 0.57 | 0.48 | 0.6 | 0.47 |
| 1 | 62068667 | rs17122481 | 1.521 | 0.151 | 0.761 | 0.503 | 0.003 | -0.009 | -0.009 | 0.037 | 0.16 | 0.13 | 0.07 | 0.27 | 0.1 | 0.2 |
| 1 | 62068999 | rs7519218 | 1.661 | 0.126 | 0.68 | 0.491 | -0.039 | 0.02 | 0.02 | 0 | 0.4 | 0.12 | 0.49 | 0.38 | 0.46 | 0.29 |
| 1 | 62070183 | rs4915593 | -0.222 | -0.001 | 0.812 | 0.39 | -0.025 | -0.011 | -0.011 | 0 | 0.76 | 0.88 | 0.82 | 0.81 | 0.7 | 0.67 |
| 1 | 62071167 | rs11207785 | 1.024 | -0.002 | 0.755 | 0.394 | 0.006 | 0.007 | 0.007 | 0 | 0.5 | 0.44 | 0.56 | 0.48 | 0.56 | 0.43 |
| 1 | 62071180 | rs4915763 | 1.393 | -0.018 | 0.588 | 0.269 | 0.006 | -0.04 | -0.04 | 0 | 0.15 | 0.06 | 0.06 | 0.27 | 0.1 | 0.2 |
| 1 | 62071712 | rs17122491 | 0.787 | -0.017 | 0.624 | 0.271 | -0.005 | 0.007 | 0.007 | 0 | 0.17 | 0.18 | 0.07 | 0.27 | 0.1 | 0.2 |
| 1 | 62075577 | rs12029654 | -0.038 | 0.066 | 0.175 | 0.24 | -0.007 | 0.001 | 0.001 | 0 | 0.24 | 0.26 | 0.1 | 0.27 | 0.14 | 0.24 |
| 1 | 62078347 | rs7545391 | -0.338 | 0.276 | -0.073 | -0.402 | -0.046 | -0.066 | -0.066 | 0 | 0.27 | 0.5 | 0.3 | 0.18 | 0.18 | 0.24 |
| 1 | 62080228 | rs12068634 | -0.256 | 0.618 | -0.213 | -0.377 | -0.016 | -0.037 | -0.037 | 0 | 0.26 | 0.49 | 0.17 | 0.16 | 0.22 | 0.33 |
| 1 | 62080405 | rs11207789 | -0.863 | 0.684 | -0.15 | -0.488 | 0.002 | 0.014 | 0.014 | 0 | 0.37 | 0.66 | 0.26 | 0.16 | 0.35 | 0.37 |
| 1 | 62080668 | rs17377908 | 1.524 | 0.689 | -0.237 | -0.445 | -0.014 | -0.013 | -0.013 | 0 | 0.15 | 0.02 | 0.08 | 0.15 | 0.14 | 0.21 |
| 1 | 62082162 | rs6687792 | 0.336 | 0.758 | -0.325 | -0.581 | -0.003 | -0.012 | -0.012 | 0.24 | 0.16 | 0.13 | 0.11 | 0 | 0.17 | 0.08 |
| 1 | 62082189 | rs6699079 | 1.129 | 0.651 | -0.098 | -0.355 | 0.001 | -0.03 | -0.03 | 0 | 0.11 | 0.12 | 0.08 | 0 | 0.12 | 0.04 |
| 1 | 62082312 | rs6675169 | 1.211 | 0.593 | -0.123 | -0.307 | -0.017 | 0.045 | 0.045 | 0 | 0.46 | 0.3 | 0.66 | 0.68 | 0.51 | 0.42 |
| 1 | 62082800 | rs7530003 | -0.855 | 0.579 | -0.141 | -0.285 | 0 | -0.002 | -0.002 | 0 | 0.37 | 0.62 | 0.39 | 0.22 | 0.35 | 0.37 |
| 1 | 62083960 | rs9629017 | -1.186 | 0.677 | -0.028 | -0.132 | -0.008 | 0.011 | 0.011 | 0 | 0.53 | 0.64 | 0.48 | 0.39 | 0.5 | 0.59 |
| 1 | 62084503 | rs6694989 | 0.438 | 0.54 | 0.182 | -0.147 | -0.01 | -0.015 | -0.015 | 0.018 | 0.6 | 0.52 | 0.78 | 0.69 | 0.66 | 0.4 |
| 1 | 62089261 | rs1933293 | 0.101 | 0.525 | 0.438 | -0.639 | -0.01 | -0.063 | -0.063 | 0 | 0.7 | 0.49 | 0.76 | 0.83 | 0.72 | 0.62 |
| 1 | 62096235 | rs870675 | -0.443 | 0.524 | 0.407 | -0.654 | -0.005 | -0.055 | -0.055 | 0 | 0.29 | 0.4 | 0.2 | 0.08 | 0.27 | 0.35 |
| 1 | 62102307 | rs12139976 | 0.675 | 0.52 | -0.092 | -0.693 | -0.045 | -0.006 | -0.006 | 0.144 | 0.41 | 0.14 | 0.4 | 0.6 | 0.49 | 0.35 |

| | | | | | | | | | | | | | | | | |
|---|---|---|---|---|---|---|---|---|---|---|---|---|---|---|---|---|
| 1 | 62103029 | rs10889245 | 0.566 | 0.021 | -0.207 | -0.341 | 0.009 | 0.016 | 0.016 | 0 | 0.64 | 0.3 | 0.61 | 0.78 | 0.73 | 0.61 |
| 1 | 62103414 | rs6686620 | -0.874 | 0.109 | -0.706 | -0.301 | -0.024 | -0.089 | -0.089 | 0 | 0.34 | 0.51 | 0.36 | 0.12 | 0.27 | 0.36 |
| 1 | 62107558 | rs4915769 | -0.902 | 0.813 | -1.836 | -0.095 | -0.019 | 0.053 | 0.053 | 0 | 0.86 | 0.98 | 0.91 | 0.97 | 0.86 | 0.82 |
| 1 | 62108681 | rs11207795 | 0.216 | 0.823 | -1.858 | 0.242 | -0.063 | -0.139 | -0.139 | 0 | 0.25 | 0.01 | 0.39 | 0.61 | 0.3 | 0.32 |
| 1 | 62109502 | rs10736398 | 0.323 | 0.612 | -0.713 | -0.314 | -0.004 | -0.038 | -0.038 | 0.144 | 0.72 | 0.61 | 0.82 | 0.93 | 0.74 | 0.69 |
| 1 | 62119809 | rs10889249 | -0.389 | 0.498 | 0.181 | -0.091 | -0.001 | -0.008 | -0.008 | 0 | 0.45 | 0.54 | 0.45 | 0.69 | 0.42 | 0.65 |
| 1 | 62120764 | rs12140608 | 0.307 | 0.929 | -0.825 | -0.584 | -0.078 | 0.07 | 0.07 | 4.772 | 0.24 | 0.05 | 0.2 | 0.07 | 0.32 | 0.25 |
| 1 | 62121303 | rs6700526 | 0.907 | 0.885 | -0.779 | 0.092 | -0.017 | -0.031 | -0.031 | 0 | 0.17 | 0.09 | 0.2 | 0.4 | 0.23 | 0.26 |
| 1 | 62124388 | rs4915770 | 0.585 | 0.53 | -0.143 | -0.131 | -0.025 | -0.005 | -0.005 | 0 | 0.21 | 0.4 | 0.28 | 0.19 | 0.15 | 0.07 |
| 1 | 62126262 | rs11207801 | 0.217 | 0.217 | -0.705 | -0.385 | 0 | 0.002 | 0.002 | 0.003 | 0.37 | 0.21 | 0.34 | 0.35 | 0.37 | 0.47 |
| 1 | 62126948 | rs6669826 | -0.364 | 0.029 | -0.139 | -0.786 | 0.002 | -0.061 | -0.061 | 1.234 | 0.79 | 0.72 | 0.81 | 0.98 | 0.77 | 0.81 |
| 1 | 62129132 | rs6587943 | -0.357 | -0.173 | -0.345 | -0.479 | -0.017 | -0.024 | -0.024 | 0 | 0.4 | 0.29 | 0.36 | 0.5 | 0.45 | 0.6 |
| 1 | 62130167 | rs6694023 | -0.671 | -0.613 | -0.381 | -0.342 | -0.028 | -0.012 | -0.012 | 0.05 | 0.26 | 0.26 | 0.15 | 0.19 | 0.17 | 0.34 |
| 1 | 62131807 | rs10493310 | 1.381 | -0.482 | -1.045 | -0.363 | -0.151 | -0.182 | -0.182 | 0.116 | 0.32 | 0.03 | 0.45 | 0.49 | 0.38 | 0.32 |
| 1 | 62133112 | rs2365465 | 0.299 | -0.474 | -0.229 | -0.201 | 0 | -0.024 | -0.024 | 0 | 0.14 | 0.15 | 0.1 | 0 | 0.12 | 0.03 |
| 1 | 62134264 | rs11207804 | 1.045 | -0.151 | -0.029 | 0.028 | 0.023 | 0.05 | 0.05 | 0 | 0.16 | 0.22 | 0.2 | 0.3 | 0.2 | 0.13 |
| 1 | 62140388 | rs1390450 | 0.237 | -0.611 | 0.079 | 0.043 | -0.028 | 0.004 | 0.004 | 0.319 | 0.13 | 0.25 | 0.11 | 0.15 | 0.07 | 0.02 |
| 1 | 62146438 | rs11207808 | -0.548 | -0.536 | 0.175 | -0.643 | -0.008 | -0.069 | -0.069 | 0.648 | 0.27 | 0.62 | 0.21 | 0.02 | 0.25 | 0.21 |
| 1 | 62165212 | rs11207811 | 0.328 | -0.592 | -0.437 | -0.436 | -0.004 | 0.007 | 0.007 | 0.038 | 0.26 | 0.28 | 0.3 | 0.36 | 0.24 | 0.45 |
| 1 | 62177027 | rs1779267 | -0.62 | 0.131 | -0.139 | -0.817 | -0.006 | -0.067 | -0.067 | 0 | 0.27 | 0.62 | 0.21 | 0.02 | 0.25 | 0.21 |
| 1 | 62179196 | rs2799626 | -0.439 | 0.26 | -0.437 | -0.755 | 0.001 | -0.106 | -0.106 | 0 | 0.24 | 0.46 | 0.19 | 0.02 | 0.25 | 0.21 |
| 1 | 62189541 | rs17312971 | 0.754 | 0.278 | -1.149 | -0.321 | -0.053 | 0.025 | 0.025 | 0 | 0.16 | 0.03 | 0.12 | 0.08 | 0.24 | 0.13 |
| 1 | 62207254 | rs7551399 | -0.989 | -0.034 | -0.774 | -0.169 | -0.004 | 0.002 | 0.002 | 0 | 0.89 | 0.9 | 0.96 | 0.87 | 0.95 | 0.84 |
| 1 | 62210481 | rs6685551 | 0.554 | 0.537 | -1.208 | -0.069 | -0.016 | -0.023 | -0.023 | 0 | 0.24 | 0.17 | 0.29 | 0.34 | 0.31 | 0.16 |
| 1 | 62212822 | rs12408292 | 0.228 | 0.175 | -1.207 | -0.358 | -0.005 | -0.006 | -0.006 | 0 | 0.36 | 0.25 | 0.44 | 0.44 | 0.4 | 0.5 |
| 1 | 62212909 | rs3790580 | 0.222 | 0.163 | -1.298 | 0.053 | -0.011 | -0.041 | -0.041 | 3.42 | 0.1 | 0.01 | 0.16 | 0.16 | 0.09 | 0.12 |
| 1 | 62224365 | rs7549691 | 0.945 | 0.443 | -1.174 | -0.187 | 0.006 | -0.067 | -0.067 | 0 | 0.35 | 0.08 | 0.38 | 0.43 | 0.28 | 0.48 |
| 1 | 62225073 | rs11807037 | 0.572 | 0.961 | -0.796 | -0.268 | 0.004 | 0.024 | 0.024 | 0 | 0.21 | 0.22 | 0.17 | 0.37 | 0.25 | 0.15 |
| 1 | 62232130 | rs11207827 | 0.645 | 0.986 | -0.639 | -0.257 | 0.018 | 0.069 | 0.069 | 0 | 0.19 | 0.31 | 0.17 | 0.37 | 0.23 | 0.15 |
| 1 | 62238413 | rs1892022 | 0.703 | 0.71 | -0.074 | -1.027 | 0.001 | -0.039 | -0.039 | 0 | 0.58 | 0.54 | 0.59 | 0.86 | 0.54 | 0.66 |
| 1 | 62242938 | rs7528744 | 0.43 | 1.315 | -0.095 | -0.335 | 0.005 | -0.052 | -0.052 | 1.407 | 0.91 | 0.69 | 0.9 | 1 | 0.91 | 0.96 |
| 1 | 62244279 | rs11207830 | 0.348 | 1.31 | -0.458 | -0.281 | 0.009 | -0.022 | -0.022 | 0 | 0.73 | 0.61 | 0.68 | 0.87 | 0.69 | 0.75 |
| 1 | 62248195 | rs1954436 | 0.062 | 1.6 | -0.6 | 0.854 | -0.007 | 0.014 | 0.014 | 0 | 0.13 | 0.11 | 0.11 | 0.44 | 0.2 | 0.16 |
| 1 | 62249367 | rs12038135 | 0.957 | 1.516 | -0.557 | 0.349 | -0.02 | 0.033 | 0.033 | 0 | 0.58 | 0.22 | 0.47 | 0.35 | 0.5 | 0.6 |
| 1 | 62253106 | rs11207837 | 1.377 | 1.347 | -0.669 | 0.198 | -0.023 | 0.215 | 0.215 | 0 | 0.51 | 0.09 | 0.42 | 0.22 | 0.47 | 0.55 |
| 1 | 62253483 | rs3762321 | 1.234 | 1.194 | -1.163 | -0.239 | -0.032 | -0.021 | -0.021 | 0 | 0.12 | 0.01 | 0.13 | 0.16 | 0.1 | 0.23 |
| 1 | 62257036 | rs1286823 | 0.581 | 1.33 | -0.847 | -0.051 | -0.028 | 0.02 | 0.02 | 0.022 | 0.08 | 0.01 | 0.06 | 0.02 | 0.12 | 0.03 |
| 1 | 62257223 | rs3790577 | 0.739 | 1.112 | -0.753 | 0.004 | -0.022 | -0.007 | -0.007 | 0.363 | 0.58 | 0.22 | 0.57 | 0.45 | 0.5 | 0.61 |
| 1 | 62261168 | rs7516332 | 0.747 | 1.124 | -0.782 | 0.008 | -0.033 | -0.011 | -0.011 | 0 | 0.58 | 0.18 | 0.56 | 0.45 | 0.49 | 0.61 |
| 1 | 62263416 | rs4915778 | 1.272 | 1.145 | -0.987 | -0.004 | -0.021 | 0.107 | 0.107 | 0 | 0.51 | 0.09 | 0.53 | 0.32 | 0.47 | 0.56 |
| 1 | 62270355 | rs10889256 | 0.901 | 1.592 | -0.293 | 0.103 | 0.019 | 0.013 | 0.013 | 0 | 0.61 | 0.48 | 0.61 | 0.48 | 0.5 | 0.63 |
| 1 | 62270494 | rs1286830 | -0.011 | 1.111 | -0.286 | -0.312 | 0.003 | -0.037 | -0.037 | 0.176 | 0.84 | 0.77 | 0.8 | 0.93 | 0.83 | 0.81 |
| 1 | 62271962 | rs1286831 | -0.011 | 1.111 | -0.401 | -0.312 | 0.002 | -0.029 | -0.029 | 0 | 0.84 | 0.78 | 0.81 | 0.93 | 0.83 | 0.81 |
| 1 | 62284504 | rs1286809 | -0.003 | 1.016 | -0.775 | -0.448 | -0.001 | 0.003 | 0.003 | 0 | 0.55 | 0.4 | 0.54 | 0.25 | 0.52 | 0.42 |
| 1 | 62289710 | rs1286816 | 0.5 | 0.915 | -0.331 | -0.694 | 0.006 | -0.048 | -0.048 | 0.054 | 0.78 | 0.62 | 0.78 | 0.88 | 0.79 | 0.75 |
| 1 | 62305682 | rs17122796 | 0.703 | 1.397 | -0.84 | -0.259 | -0.032 | -0.037 | -0.037 | 0.018 | 0.06 | 0.01 | 0.13 | 0.22 | 0.11 | 0.05 |
| 1 | 62306819 | rs1572670 | 0.521 | 0.577 | -0.972 | -1.28 | 0.013 | 0.042 | 0.042 | 1.119 | 0.5 | 0.41 | 0.65 | 0.83 | 0.61 | 0.42 |
| 1 | 62309562 | rs17371694 | 0.834 | 0.172 | -1.06 | -1.431 | 0.01 | 0.082 | 0.082 | 0 | 0.21 | 0.08 | 0.1 | 0.05 | 0.12 | 0.25 |
| 1 | 62309796 | rs12045057 | 0.945 | 0.16 | -1.045 | -1.374 | -0.019 | -0.033 | -0.033 | 0.549 | 0.1 | 0 | 0.05 | 0.11 | 0.06 | 0.09 |
| 1 | 62312654 | rs2365498 | 1.5 | 0.409 | -1.478 | -1.682 | -0.034 | -0.105 | -0.105 | 0 | 0.23 | 0.08 | 0.27 | 0.54 | 0.24 | 0.24 |
| 1 | 62313142 | rs2788616 | 0.991 | 0.254 | -0.76 | -1.773 | -0.011 | -0.061 | -0.061 | 0 | 0.76 | 0.5 | 0.78 | 0.88 | 0.81 | 0.66 |
| 1 | 62324451 | rs7515512 | -0.344 | 0.378 | -0.649 | -1.626 | 0.034 | -0.01 | -0.01 | 0 | 0.16 | 0.36 | 0.19 | 0.11 | 0.18 | 0.24 |
| 1 | 62324742 | rs11207848 | -0.626 | 0.253 | -0.768 | -1.901 | -0.006 | -0.038 | -0.038 | 0 | 0.38 | 0.48 | 0.27 | 0.17 | 0.32 | 0.52 |
| 1 | 62337807 | rs6698337 | 0.192 | -0.046 | -0.487 | -1.374 | 0.001 | -0.012 | -0.012 | 2.753 | 0.39 | 0.34 | 0.45 | 0.16 | 0.58 | 0.37 |
| 1 | 62340726 | rs12096192 | 0.795 | -0.231 | -0.329 | -1.711 | 0.002 | -0.034 | -0.034 | 0 | 0.63 | 0.63 | 0.82 | 0.87 | 0.65 | 0.53 |
| 1 | 62341157 | rs4598521 | 0.475 | -0.259 | -0.338 | -1.756 | -0.004 | -0.036 | -0.036 | 0 | 0.68 | 0.67 | 0.83 | 0.87 | 0.71 | 0.58 |
| 1 | 62342230 | rs6685516 | 0.441 | -0.283 | -0.484 | -1.671 | 0.002 | 0.007 | 0.007 | 0 | 0.75 | 0.78 | 0.88 | 0.87 | 0.79 | 0.66 |
| 1 | 62344107 | rs10889262 | -0.383 | -0.441 | -0.539 | -1.608 | 0.004 | -0.013 | -0.013 | 0 | 0.59 | 0.57 | 0.56 | 0.73 | 0.46 | 0.59 |
| 1 | 62345961 | rs4313427 | 0.196 | -0.386 | -0.362 | -1.077 | 0.01 | 0.025 | 0.025 | 0 | 0.85 | 0.92 | 0.92 | 0.98 | 0.88 | 0.82 |
| 1 | 62347976 | rs4540685 | -0.136 | -0.269 | -0.308 | -1.003 | 0.011 | 0.003 | 0.003 | 0.005 | 0.86 | 0.9 | 0.94 | 0.87 | 0.91 | 0.73 |
| 1 | 62356590 | rs9326052 | 1.228 | -0.431 | -0.207 | -0.734 | -0.011 | -0.003 | -0.003 | 0.031 | 0.59 | 0.5 | 0.68 | 0.57 | 0.67 | 0.33 |
| 1 | 62357771 | rs10889266 | -1.277 | 0.075 | 0.157 | -0.741 | 0.012 | -0.019 | -0.019 | 0 | 0.32 | 0.53 | 0.28 | 0.3 | 0.3 | 0.44 |
| 1 | 62363350 | rs2498997 | -0.869 | -0.409 | 0.074 | -0.29 | -0.006 | 0.027 | 0.027 | 0 | 0.21 | 0.42 | 0.22 | 0.3 | 0.16 | 0.31 |
| 1 | 62365217 | rs1412622 | 0.284 | -0.636 | -0.121 | -0.408 | -0.033 | -0.004 | -0.004 | 0 | 0.61 | 0.47 | 0.65 | 0.41 | 0.69 | 0.63 |
| 1 | 62366878 | rs2499002 | 0.454 | -0.155 | 0.063 | -0.975 | -0.007 | -0.081 | -0.081 | 0 | 0.76 | 0.7 | 0.73 | 1 | 0.76 | 0.85 |
| 1 | 62375977 | rs1332636 | 0.599 | -0.138 | -0.134 | -0.965 | -0.016 | -0.157 | -0.157 | 0 | 0.73 | 0.46 | 0.69 | 1 | 0.71 | 0.85 |
| 1 | 62379774 | rs2499006 | 0.464 | -0.278 | -0.533 | -0.981 | 0.004 | -0.004 | -0.004 | 0.998 | 0.64 | 0.66 | 0.68 | 0.86 | 0.65 | 0.59 |
| 1 | 62380124 | rs4915789 | -0.56 | 0.238 | -0.646 | -1.093 | 0.025 | -0.023 | -0.023 | 0 | 0.7 | 0.6 | 0.67 | 0.8 | 0.61 | 0.76 |
| 1 | 62380298 | rs1056513 | 0.251 | -1.135 | -1.97 | -1.171 | 0.055 | -0.032 | -0.032 | 0.014 | 0.54 | 0.09 | 0.52 | 0.57 | 0.63 | 0.56 |

| | | | | | | | | | | | | | | | | |
|---|---|---|---|---|---|---|---|---|---|---|---|---|---|---|---|---|
| 1 | 62380658 | rs11207864 | 0.272 | -1.284 | -2.032 | -1.669 | -0.04 | 0.057 | 0.057 | 0 | 0.27 | 0.12 | 0.4 | 0.77 | 0.3 | 0.3 |
| 1 | 62387652 | rs11207865 | 0.91 | -1.178 | -2.082 | -0.954 | -0.054 | -0.083 | -0.083 | 0.717 | 0.21 | 0.06 | 0.21 | 0.5 | 0.28 | 0.25 |
| 1 | 62392494 | rs4915790 | 0.978 | -1.384 | -2.411 | -2.057 | -0.013 | 0.154 | 0.154 | 0 | 0.23 | 0.02 | 0.23 | 0 | 0.2 | 0.18 |
| 1 | 62393394 | rs2251182 | -1.172 | -1.524 | -1.719 | -2.06 | -0.002 | -0.002 | -0.002 | 0 | 0.87 | 0.87 | 0.92 | 0.81 | 0.85 | 0.79 |
| 1 | 62400487 | rs10889271 | 0.492 | -0.966 | -1.677 | -2.229 | -0.054 | 0.162 | 0.162 | 0 | 0.4 | 0.19 | 0.48 | 0.78 | 0.42 | 0.36 |
| 1 | 62408549 | rs7530734 | 0.43 | -0.75 | -0.744 | -2.172 | -0.035 | 0.163 | 0.163 | 0 | 0.42 | 0.23 | 0.49 | 0.79 | 0.42 | 0.42 |
| 1 | 62422442 | rs11585194 | 0.302 | -0.687 | -0.365 | -2.165 | -0.004 | 0.019 | 0.019 | 0 | 0.4 | 0.25 | 0.38 | 0.02 | 0.41 | 0.35 |
| 1 | 62433100 | rs2498967 | 0.065 | -0.608 | -0.684 | -2.13 | 0.008 | 0.101 | 0.101 | 0 | 0.43 | 0.32 | 0.4 | 0.02 | 0.42 | 0.38 |
| 1 | 62435141 | rs2498968 | -0.969 | -0.243 | -0.052 | -1.383 | -0.001 | 0 | 0 | 0 | 0.87 | 0.87 | 0.92 | 0.8 | 0.85 | 0.8 |
| 1 | 62448245 | rs7545272 | -0.42 | -0.37 | 0.1 | -0.51 | 0.006 | 0.033 | 0.033 | 0 | 0.26 | 0.42 | 0.23 | 0.45 | 0.26 | 0.41 |
| 1 | 62456014 | rs1134767 | -1.144 | -0.016 | 0.081 | -0.778 | 0.003 | -0.004 | -0.004 | 0 | 0.43 | 0.55 | 0.32 | 0.47 | 0.41 | 0.56 |
| 1 | 62457787 | rs12728194 | -0.242 | -0.08 | 0.552 | -0.293 | -0.006 | -0.015 | -0.015 | 0.733 | 0.23 | 0.24 | 0.2 | 0.44 | 0.24 | 0.38 |
| 1 | 62458463 | rs10493316 | 0.872 | -0.022 | 0.47 | -0.162 | -0.015 | 0.144 | 0.144 | 0 | 0.06 | 0.11 | 0.05 | 0.38 | 0.02 | 0.19 |
| 1 | 62468855 | rs7526836 | 0.517 | 0.177 | 0.309 | -0.42 | -0.008 | 0.007 | 0.007 | 0 | 0.17 | 0.15 | 0.16 | 0.06 | 0.21 | 0.22 |
| 1 | 62479403 | rs2183995 | -0.952 | 0.445 | 0.968 | -0.019 | 0.106 | 0.291 | 0.291 | 0.126 | 0.13 | 0.44 | 0.15 | 0.44 | 0.15 | 0.34 |
| 1 | 62486381 | rs6697273 | 0.568 | -0.453 | -0.031 | -0.99 | 0.042 | 0.05 | 0.05 | 0 | 0.24 | 0.12 | 0.23 | 0.02 | 0.15 | 0.21 |
| 1 | 62488071 | rs11590158 | 0.712 | -0.498 | -0.102 | -1.022 | 0.049 | 0.065 | 0.065 | 0 | 0.22 | 0.1 | 0.23 | 0.02 | 0.14 | 0.18 |
| 1 | 62492215 | rs1332630 | -0.082 | -0.256 | -0.003 | -1.232 | -0.013 | 0.017 | 0.017 | 0 | 0.41 | 0.67 | 0.3 | 0.35 | 0.4 | 0.53 |
| 1 | 62498825 | rs7537323 | -0.514 | -0.236 | 0.281 | -1.286 | 0.013 | 0.009 | 0.009 | 0 | 0.42 | 0.61 | 0.42 | 0.36 | 0.32 | 0.57 |
| 1 | 62499673 | rs11585888 | 1.446 | -0.22 | 0.248 | -1.45 | -0.015 | 0.077 | 0.077 | 0 | 0.25 | 0.14 | 0.14 | 0.01 | 0.24 | 0.12 |
| 1 | 62500555 | rs7520924 | -0.709 | 0.377 | 0.389 | -0.937 | -0.023 | 0.008 | 0.008 | 0 | 0.67 | 0.8 | 0.57 | 0.34 | 0.56 | 0.69 |
| 1 | 62506473 | rs4325166 | 0.28 | 0.373 | 0.414 | -1.051 | -0.003 | -0.112 | -0.112 | 0.177 | 0.54 | 0.51 | 0.4 | 0.03 | 0.41 | 0.34 |
| 1 | 62511933 | rs13374203 | -0.324 | -0.08 | 0.085 | -0.68 | -0.001 | 0 | 0 | 0 | 0.66 | 0.68 | 0.55 | 0.28 | 0.54 | 0.64 |
| 1 | 62517305 | rs12117302 | 1.156 | 0.387 | -0.661 | -0.423 | -0.05 | -0.037 | -0.037 | 0.396 | 0.13 | 0.03 | 0.1 | 0.13 | 0.24 | 0.11 |
| 1 | 62523223 | rs12072096 | 0.045 | -0.744 | -0.57 | -0.831 | 0.048 | 0.049 | 0.049 | 0 | 0.29 | 0.12 | 0.26 | 0.15 | 0.17 | 0.35 |
| 1 | 62534938 | rs6674622 | 0.207 | -0.567 | 0.224 | -1.194 | -0.001 | 0.006 | 0.006 | 0 | 0.43 | 0.59 | 0.33 | 0.21 | 0.42 | 0.48 |
| 1 | 62538372 | rs6699078 | -0.29 | -0.387 | 0.524 | -1.181 | 0.004 | -0.018 | -0.018 | 0 | 0.66 | 0.59 | 0.56 | 0.19 | 0.6 | 0.66 |
| 1 | 62555219 | rs17123057 | -0.408 | -0.846 | 1.003 | -1.334 | 0.041 | 0.073 | 0.073 | 0 | 0.05 | 0.24 | 0.04 | 0.09 | 0.04 | 0.01 |
| 1 | 62555619 | rs4255414 | -0.377 | -1.247 | 0.761 | -1.649 | -0.022 | -0.048 | -0.048 | 0 | 0.34 | 0.38 | 0.31 | 0.13 | 0.2 | 0.35 |
| 1 | 62560928 | rs10127438 | 0.188 | -1.253 | 0.937 | -1.663 | -0.002 | -0.012 | -0.012 | 0.077 | 0.29 | 0.25 | 0.28 | 0.13 | 0.18 | 0.35 |
| 1 | 62568680 | rs7535670 | -0.241 | -1.376 | 0.886 | -1.71 | -0.019 | -0.043 | -0.043 | 0.636 | 0.33 | 0.38 | 0.3 | 0.13 | 0.19 | 0.43 |
| 1 | 62575427 | rs2476188 | -0.686 | -1.165 | 0.817 | -1.838 | 0.001 | -0.037 | -0.037 | 0 | 0.67 | 0.61 | 0.56 | 0.17 | 0.62 | 0.66 |
| 1 | 62575911 | rs6688017 | -0.207 | -1.177 | 0.793 | -1.764 | -0.034 | -0.068 | -0.068 | 0 | 0.33 | 0.43 | 0.3 | 0.13 | 0.19 | 0.43 |
| 1 | 62578944 | rs2476190 | 0.19 | -0.843 | 0.853 | -1.64 | -0.002 | -0.086 | -0.086 | 0 | 0.61 | 0.46 | 0.52 | 0.05 | 0.58 | 0.49 |
| 1 | 62579672 | rs1475564 | -0.785 | -0.695 | 0.264 | -1.334 | -0.002 | 0.007 | 0.007 | 0.063 | 0.76 | 0.81 | 0.79 | 0.9 | 0.69 | 0.84 |
| 1 | 62603544 | rs6680526 | -0.262 | -1.269 | 0.223 | -1.013 | -0.012 | -0.003 | -0.003 | 0 | 0.36 | 0.59 | 0.5 | 0.72 | 0.21 | 0.5 |
| 1 | 62606693 | rs2476197 | -0.794 | -1.109 | 0.003 | -1.166 | -0.033 | 0.016 | 0.016 | 0 | 0.73 | 0.92 | 0.79 | 0.83 | 0.67 | 0.8 |
| 1 | 62607786 | rs11207909 | 0.452 | -1.315 | 0.23 | -1.151 | -0.058 | 0.036 | 0.036 | 0 | 0.28 | 0.44 | 0.45 | 0.65 | 0.17 | 0.34 |
| 1 | 62614453 | rs2666494 | 0.263 | -1.27 | -0.129 | -1.093 | 0.007 | 0.015 | 0.015 | 0 | 0.63 | 0.67 | 0.71 | 0.76 | 0.62 | 0.56 |
| 1 | 62615684 | rs2481672 | -0.759 | -1.141 | -0.334 | -1.069 | -0.014 | 0.019 | 0.019 | 0 | 0.73 | 0.88 | 0.78 | 0.83 | 0.67 | 0.79 |
| 1 | 62632571 | rs2481677 | 1.447 | -1.046 | -0.571 | -1.282 | -0.108 | 0.154 | 0.154 | 0 | 0.32 | 0.09 | 0.24 | 0.02 | 0.42 | 0.21 |
| 1 | 62633581 | rs2457831 | 0.778 | -1.084 | -0.457 | -1.223 | -0.067 | 0.183 | 0.183 | 0 | 0.39 | 0.16 | 0.27 | 0.03 | 0.47 | 0.27 |
| 1 | 62634834 | rs12095754 | 0.526 | -1.361 | -0.149 | -1.259 | -0.021 | 0.009 | 0.009 | 0.058 | 0.08 | 0.11 | 0.05 | 0.09 | 0.03 | 0.24 |
| 1 | 62635017 | rs6697095 | 1.984 | -1.671 | -0.463 | -1.469 | 0.019 | 0.098 | 0.098 | 0 | 0.16 | 0.01 | 0.21 | 0 | 0.09 | 0.14 |
| 1 | 62635058 | rs6684324 | 0.461 | -1.526 | 0.176 | -0.959 | 0.009 | 0.338 | 0.338 | 0.291 | 0.09 | 0.31 | 0.24 | 0.6 | 0.06 | 0.17 |
| 1 | 62635081 | rs2666504 | -0.806 | -1.407 | -0.069 | -0.87 | -0.001 | -0.001 | -0.001 | 2.591 | 0.53 | 0.52 | 0.5 | 0.38 | 0.48 | 0.6 |
| 1 | 62635116 | rs2457832 | 0.116 | -0.809 | -0.377 | -0.793 | -0.001 | 0.001 | 0.001 | 0 | 0.26 | 0.25 | 0.21 | 0.19 | 0.32 | 0.23 |
| 1 | 62644118 | rs2481703 | -1.07 | 0.39 | -0.841 | -1.211 | -0.025 | 0.079 | 0.079 | 0.121 | 0.63 | 0.77 | 0.59 | 0.89 | 0.54 | 0.77 |
| 1 | 62646182 | rs12759978 | -0.651 | 0.54 | -1.121 | -0.89 | -0.014 | 0.002 | 0.002 | 0 | 0.2 | 0.13 | 0.14 | 0.17 | 0.25 | 0.41 |
| 1 | 62657530 | rs12116810 | -0.361 | 0.889 | -1.279 | -0.554 | -0.001 | -0.001 | -0.001 | 0 | 0.27 | 0.27 | 0.53 | 0.58 | 0.36 | 0.32 |
| 1 | 62663868 | rs10889296 | -0.274 | 0.896 | -1.269 | -0.477 | 0.008 | 0.139 | 0.139 | 0.463 | 0.12 | 0.2 | 0.24 | 0.58 | 0.11 | 0.22 |
| 1 | 62666518 | rs12137088 | -0.418 | 0.504 | -1.245 | -0.807 | -0.003 | 0.001 | 0.001 | 0 | 0.66 | 0.7 | 0.79 | 0.69 | 0.71 | 0.59 |
| 1 | 62672546 | rs2457829 | -0.984 | 0.766 | -1.258 | -1.28 | -0.049 | -0.053 | -0.053 | 0 | 0.91 | 0.99 | 0.94 | 0.85 | 0.84 | 0.77 |
| 1 | 62673037 | rs7542665 | -0.592 | 0.989 | -0.826 | -1.093 | 0.004 | 0.005 | 0.005 | 0 | 0.62 | 0.67 | 0.65 | 0.32 | 0.68 | 0.55 |
| 1 | 62673225 | rs7533274 | -0.436 | 1.134 | -0.864 | -0.961 | -0.003 | -0.008 | -0.008 | 2.743 | 0.58 | 0.6 | 0.62 | 0.32 | 0.65 | 0.51 |
| 1 | 62673287 | rs2457828 | 1.95 | 1.397 | -1.236 | -0.911 | -0.016 | 0.016 | 0.016 | 0 | 0.13 | 0.06 | 0.13 | 0 | 0.17 | 0.04 |
| 1 | 62674507 | rs12729017 | 1.903 | 1.089 | -1.476 | -0.88 | -0.033 | 0.149 | 0.149 | 0 | 0.25 | 0.03 | 0.16 | 0.04 | 0.26 | 0.19 |
| 1 | 62674547 | rs11207932 | -0.383 | 0.931 | -0.67 | -0.246 | 0.023 | 0.083 | 0.083 | 0.001 | 0.1 | 0.25 | 0.11 | 0.21 | 0.08 | 0.2 |
| 1 | 62675709 | rs7551777 | -0.007 | 1.024 | -0.91 | -0.2 | 0.018 | 0.067 | 0.067 | 0.2 | 0.08 | 0.18 | 0.1 | 0.21 | 0.08 | 0.2 |
| 1 | 62676091 | rs11207933 | -0.754 | 1.012 | -0.945 | -0.224 | 0.003 | 0.031 | 0.031 | 0 | 0.13 | 0.19 | 0.12 | 0.28 | 0.12 | 0.22 |
| 1 | 62677050 | rs12137401 | 1.061 | 1.246 | -1.04 | -0.321 | -0.05 | 0.049 | 0.049 | 0 | 0.14 | 0.08 | 0.3 | 0.07 | 0.25 | 0.08 |
| 1 | 62677118 | rs12130050 | 0.346 | 1.195 | -1.022 | -0.287 | -0.064 | 0.08 | 0.08 | 0 | 0.19 | 0.08 | 0.32 | 0.07 | 0.29 | 0.1 |
| 1 | 62677723 | rs11207936 | 0.088 | 0.058 | -0.546 | 0.178 | 0.035 | 0.095 | 0.095 | 1.038 | 0.07 | 0.18 | 0.1 | 0.21 | 0.07 | 0.18 |
| 1 | 62680981 | rs6704042 | -0.001 | -0.775 | -0.822 | 0.068 | -0.024 | 0.006 | 0.006 | 0 | 0.62 | 0.62 | 0.73 | 0.51 | 0.79 | 0.63 |
| 1 | 62687104 | rs4019803 | 0.171 | 0.385 | -0.859 | -0.186 | 0.005 | -0.001 | -0.001 | 1.274 | 0.26 | 0.32 | 0.31 | 0.24 | 0.37 | 0.24 |
| 1 | 62692092 | rs6683592 | -0.532 | -0.039 | -1.105 | -0.509 | 0 | 0 | 0 | 0 | 0.34 | 0.51 | 0.44 | 0.67 | 0.3 | 0.45 |
| 1 | 62693352 | rs11577415 | 1.079 | 0.014 | -1.276 | 0.197 | -0.002 | 0.061 | 0.061 | 0 | 0.05 | 0.02 | 0.13 | 0.38 | 0.04 | 0.04 |
| 1 | 62701741 | rs12144282 | 1.294 | -0.011 | -1.397 | -0.999 | -0.058 | -0.011 | -0.011 | 0.752 | 0.19 | 0.07 | 0.27 | 0.14 | 0.32 | 0.19 |
| 1 | 62702720 | rs6688986 | 0.414 | -0.154 | -1.433 | -1.104 | -0.06 | 0.015 | 0.015 | 0 | 0.24 | 0.08 | 0.29 | 0.14 | 0.35 | 0.21 |

| | | | | | | | | | | | | | | | | |
|---|---|---|---|---|---|---|---|---|---|---|---|---|---|---|---|---|
| 1 | 62703236 | rs1050334 | 1.352 | -0.127 | -1.421 | -1.085 | -0.058 | -0.011 | -0.011 | 0 | 0.19 | 0.07 | 0.27 | 0.14 | 0.32 | 0.19 |
| 1 | 62707613 | rs11207944 | -0.153 | -0.518 | -1.433 | -1.345 | -0.036 | 0.072 | 0.072 | 0 | 0.34 | 0.18 | 0.35 | 0.14 | 0.42 | 0.26 |
| 1 | 62708339 | rs2067994 | -0.358 | -0.702 | -0.825 | -1.441 | 0 | -0.008 | -0.008 | 0 | 0.4 | 0.44 | 0.38 | 0.14 | 0.45 | 0.35 |
| 1 | 62711941 | rs6664672 | 0.883 | -0.625 | -0.572 | -1.621 | -0.003 | -0.017 | -0.017 | 0.068 | 0.38 | 0.37 | 0.37 | 0.84 | 0.34 | 0.4 |
| 1 | 62713246 | rs10889315 | -0.064 | -0.608 | -0.651 | -1.707 | 0.004 | 0.055 | 0.055 | 0 | 0.41 | 0.3 | 0.34 | 0.14 | 0.42 | 0.4 |
| 1 | 62714918 | rs12122645 | -1.211 | -0.664 | -0.679 | -1.606 | -0.006 | -0.031 | -0.031 | 0 | 0.63 | 0.62 | 0.74 | 0.84 | 0.67 | 0.77 |
| 1 | 62716323 | rs2457809 | -0.664 | -0.654 | -0.345 | -1.654 | 0.023 | 0.016 | 0.016 | 0 | 0.26 | 0.35 | 0.44 | 0.7 | 0.32 | 0.4 |
| 1 | 62716353 | rs2457808 | -1.004 | -0.467 | -0.237 | -1.656 | -0.005 | -0.027 | -0.027 | 0 | 0.63 | 0.63 | 0.74 | 0.84 | 0.67 | 0.77 |
| 1 | 62717759 | rs4915605 | 0.378 | 0.151 | -0.299 | -1.245 | 0.036 | -0.024 | -0.024 | 0 | 0.06 | 0.11 | 0.21 | 0 | 0.15 | 0.13 |
| 1 | 62719839 | rs6692004 | 1.389 | -0.201 | -0.444 | -1.257 | 0.006 | -0.006 | -0.006 | 0 | 0.1 | 0.1 | 0.06 | 0 | 0.07 | 0.03 |
| 1 | 62723207 | rs2057634 | 0.209 | 0.136 | -0.232 | -1.18 | 0.056 | -0.03 | -0.03 | 0 | 0.06 | 0.16 | 0.22 | 0 | 0.14 | 0.13 |
| 1 | 62723566 | rs958802 | 0.389 | -0.205 | -0.777 | -1.498 | -0.077 | 0.091 | 0.091 | 2.397 | 0.35 | 0.05 | 0.25 | 0.12 | 0.39 | 0.32 |
| 1 | 62724753 | rs12062559 | 0.322 | -0.262 | -0.4 | -1.527 | 0.006 | 0.028 | 0.028 | 0 | 0.15 | 0.21 | 0.21 | 0.72 | 0.16 | 0.26 |
| 1 | 62726134 | rs6685907 | 0.862 | -0.276 | -0.367 | -1.571 | 0.02 | -0.016 | -0.016 | 0 | 0.06 | 0.07 | 0.21 | 0 | 0.14 | 0.13 |
| 1 | 62727596 | rs2765241 | -1.621 | -0.892 | -0.09 | -1.677 | 0.007 | 0.022 | 0.022 | 0 | 0.71 | 0.78 | 0.81 | 0.84 | 0.74 | 0.79 |
| 1 | 62728784 | rs2666472 | -0.901 | -0.83 | -0.027 | -1.636 | -0.035 | -0.075 | -0.075 | 0 | 0.62 | 0.55 | 0.79 | 0.84 | 0.74 | 0.74 |
| 1 | 62728861 | rs2260581 | -0.901 | -0.83 | -0.019 | -1.636 | -0.036 | -0.077 | -0.077 | 0 | 0.62 | 0.55 | 0.79 | 0.84 | 0.74 | 0.74 |
| 1 | 62728918 | rs2262110 | -0.892 | -0.829 | -0.035 | -1.635 | -0.035 | -0.075 | -0.075 | 0 | 0.62 | 0.55 | 0.79 | 0.84 | 0.74 | 0.74 |
| 1 | 62732421 | rs11207949 | 0.041 | 0.343 | 0.505 | -0.669 | 0.083 | 0.134 | 0.134 | 0.056 | 0.12 | 0.21 | 0.4 | 0.59 | 0.27 | 0.12 |
| 1 | 62732945 | rs11579934 | -0.692 | -0.252 | 0.398 | -0.984 | -0.003 | -0.005 | -0.005 | 0 | 0.48 | 0.49 | 0.37 | 0.25 | 0.39 | 0.65 |
| 1 | 62734089 | rs17123306 | -0.308 | -0.275 | 0.431 | -0.855 | -0.031 | 0.035 | 0.035 | 0 | 0.09 | 0.21 | 0.04 | 0.14 | 0.04 | 0.17 |
| 1 | 62737766 | rs17123307 | -0.632 | -0.296 | 0.425 | -0.871 | -0.045 | 0.02 | 0.02 | 0.093 | 0.12 | 0.24 | 0.12 | 0.14 | 0.04 | 0.12 |
| 1 | 62740065 | rs12127930 | -0.944 | -0.179 | 0.116 | -0.93 | 0.012 | 0.038 | 0.038 | 0 | 0.53 | 0.64 | 0.39 | 0.25 | 0.44 | 0.61 |
| 1 | 62740930 | rs2666508 | -1.745 | 0.003 | -0.06 | -0.933 | 0.029 | 0.129 | 0.129 | 0 | 0.65 | 0.88 | 0.79 | 0.85 | 0.7 | 0.76 |
| 1 | 62741049 | rs11207955 | -0.196 | -0.092 | 0.087 | -0.988 | -0.006 | -0.044 | -0.044 | 0 | 0.41 | 0.46 | 0.28 | 0.12 | 0.38 | 0.4 |
| 1 | 62743398 | rs12138544 | -0.232 | -0.089 | 0.049 | -0.986 | -0.003 | -0.024 | -0.024 | 0.192 | 0.41 | 0.53 | 0.29 | 0.12 | 0.38 | 0.4 |
| 1 | 62745206 | rs2765251 | -1.762 | -0.095 | -0.393 | -0.982 | 0.029 | 0.129 | 0.129 | 0 | 0.65 | 0.88 | 0.79 | 0.85 | 0.7 | 0.76 |
| 1 | 62750051 | rs4915810 | -0.257 | 0.105 | -0.348 | -1.071 | -0.059 | 0.056 | 0.056 | 0 | 0.39 | 0.21 | 0.52 | 0.24 | 0.5 | 0.54 |
| 1 | 62753383 | rs2765255 | -0.26 | -0.163 | -0.407 | -1.053 | -0.019 | 0.066 | 0.066 | 0 | 0.41 | 0.17 | 0.32 | 0.24 | 0.4 | 0.61 |
| 1 | 62757670 | rs2765261 | 1.137 | -0.271 | 0.273 | -1.058 | 0.001 | -0.018 | -0.018 | 0.175 | 0.41 | 0.58 | 0.26 | 0.1 | 0.29 | 0.23 |
| 1 | 62758183 | rs4915811 | 0.268 | -0.083 | 0.038 | -1.019 | 0.073 | 0.09 | 0.09 | 0 | 0.18 | 0.28 | 0.42 | 0.65 | 0.29 | 0.2 |
| 1 | 62759369 | rs2765263 | 1.208 | -0.163 | 0.054 | -1.058 | 0.025 | 0.034 | 0.034 | 0 | 0.39 | 0.33 | 0.24 | 0.1 | 0.29 | 0.23 |
| 1 | 62760352 | rs1886830 | 0.238 | -0.15 | 0.059 | -1.046 | 0.002 | 0.013 | 0.013 | 0 | 0.57 | 0.59 | 0.65 | 0.76 | 0.57 | 0.43 |
| 1 | 62760625 | rs12082238 | 1.151 | -0.024 | -0.533 | -1.098 | -0.014 | -0.046 | -0.046 | 0 | 0.37 | 0.39 | 0.23 | 0.1 | 0.28 | 0.22 |
| 1 | 62760802 | rs10465842 | -0.567 | -0.04 | -1.432 | -1.027 | -0.103 | 0.122 | 0.122 | 0.062 | 0.66 | 0.96 | 0.77 | 0.89 | 0.61 | 0.66 |
| 1 | 62765049 | rs1886834 | -0.813 | -0.046 | -1.437 | -1.053 | -0.092 | 0.113 | 0.113 | 0.486 | 0.69 | 0.96 | 0.79 | 0.9 | 0.64 | 0.65 |
| 1 | 62766406 | rs10493320 | 1.198 | -0.007 | -0.88 | -0.75 | 0.025 | 0.053 | 0.053 | 0 | 0.32 | 0.18 | 0.21 | 0.1 | 0.25 | 0.2 |
| 1 | 62773268 | rs10493321 | 0.112 | 0.677 | -1.148 | -0.577 | -0.002 | 0.004 | 0.004 | 0 | 0.19 | 0.13 | 0.22 | 0.14 | 0.2 | 0.17 |
| 1 | 62773907 | rs11207957 | 1.379 | 0.896 | -1.551 | -0.646 | 0.034 | 0.09 | 0.09 | 0 | 0.29 | 0.11 | 0.19 | 0.1 | 0.24 | 0.15 |
| 1 | 62774469 | rs7512821 | 0.418 | 1.059 | -1.175 | -0.131 | 0.011 | 0.057 | 0.057 | 0 | 0.53 | 0.29 | 0.61 | 0.76 | 0.53 | 0.43 |
| 1 | 62776978 | rs2366208 | -0.62 | 1.212 | -1.26 | 0.223 | -0.004 | -0.003 | -0.003 | 0 | 0.89 | 0.9 | 0.8 | 0.86 | 0.83 | 0.84 |
| 1 | 62778298 | rs4915816 | -0.467 | 1.018 | -1.627 | -0.061 | -0.003 | 0.074 | 0.074 | 0 | 0.57 | 0.8 | 0.59 | 0.74 | 0.58 | 0.64 |
| 1 | 62778382 | rs12758910 | 0.215 | 0.774 | -1.66 | -0.083 | 0.033 | 0.03 | 0.03 | 0 | 0.29 | 0.12 | 0.13 | 0.18 | 0.24 | 0.32 |
| 1 | 62778757 | rs1555831 | 0.907 | 0.75 | -1.651 | -0.298 | 0.037 | 0.123 | 0.123 | 0 | 0.33 | 0.12 | 0.18 | 0.05 | 0.24 | 0.31 |
| 1 | 62788511 | rs4915818 | 0.799 | 1.334 | -0.743 | 1.078 | -0.001 | 0 | 0 | 0 | 0.63 | 0.36 | 0.67 | 0.4 | 0.6 | 0.68 |
| 1 | 62791652 | rs4323745 | -0.347 | 1.59 | -0.57 | 1.302 | 0.029 | 0.033 | 0.033 | 0 | 0.18 | 0.27 | 0.22 | 0.36 | 0.3 | 0.21 |
| 1 | 62791745 | rs2366210 | 1.127 | 1.547 | -0.851 | 1 | -0.013 | 0.029 | 0.029 | 0 | 0.58 | 0.24 | 0.63 | 0.29 | 0.52 | 0.61 |
| 1 | 62797677 | rs4915824 | 0.491 | 1.692 | -1.416 | 1.285 | -0.021 | -0.058 | -0.058 | 0.251 | 0.1 | 0.01 | 0.19 | 0.23 | 0.13 | 0.09 |
| 1 | 62799451 | rs10749732 | -0.303 | 1.761 | -0.578 | 1.805 | -0.006 | 0.018 | 0.018 | 0 | 0.85 | 0.83 | 0.9 | 0.74 | 0.86 | 0.9 |
| 1 | 62803060 | rs6675251 | -1.111 | 2.448 | -0.906 | 1.993 | 0.046 | 0.047 | 0.047 | 0.005 | 0.21 | 0.29 | 0.46 | 0.5 | 0.39 | 0.38 |
| 1 | 62803211 | rs6690755 | 0.338 | 0.902 | -1.579 | 1.374 | -0.001 | 0.002 | 0.002 | 0 | 0.83 | 0.84 | 0.88 | 0.74 | 0.83 | 0.87 |
| 1 | 62803277 | rs10889320 | -0.17 | 1.211 | -1.716 | 1.578 | -0.057 | -0.019 | -0.019 | 0.121 | 0.11 | 0.01 | 0.16 | 0.11 | 0.2 | 0.13 |
| 1 | 62803803 | rs12028582 | -0.051 | 0.364 | -1.166 | 1.738 | 0.001 | -0.016 | -0.016 | 0 | 0.87 | 0.92 | 0.89 | 0.74 | 0.85 | 0.93 |
| 1 | 62804719 | rs4915827 | -1.061 | 0.42 | -0.909 | 1.585 | 0.001 | -0.025 | -0.025 | 0.02 | 0.16 | 0.13 | 0.14 | 0.36 | 0.15 | 0.24 |
| 1 | 62808765 | rs12403723 | -0.793 | 0.007 | -0.904 | 0.541 | 0.023 | 0.053 | 0.053 | 2.035 | 0.18 | 0.1 | 0.15 | 0 | 0.11 | 0.07 |
| 1 | 62811748 | rs12404736 | -1.404 | -0.004 | -0.181 | 0.563 | -0.01 | -0.038 | -0.038 | 0.268 | 0.28 | 0.33 | 0.23 | 0.09 | 0.21 | 0.22 |
| 1 | 62818217 | rs6675459 | -1.493 | -0.564 | 0.3 | 1.157 | 0.013 | 0.002 | 0.002 | 0 | 0.36 | 0.62 | 0.39 | 0.31 | 0.23 | 0.37 |
| 1 | 62819067 | rs7541112 | -0.142 | -0.66 | 0.543 | 1.162 | 0.011 | -0.006 | -0.006 | 0 | 0.15 | 0.1 | 0.09 | 0.19 | 0.05 | 0.11 |
| 1 | 62819617 | rs11207960 | -0.477 | 0.399 | 0.15 | 1.473 | -0.073 | -0.082 | -0.082 | 0 | 0.87 | 0.97 | 0.84 | 0.66 | 0.76 | 0.93 |
| 1 | 62820001 | rs17123525 | -1.207 | -0.284 | 0.905 | 0.99 | -0.037 | -0.007 | -0.007 | 0 | 0.34 | 0.46 | 0.37 | 0.29 | 0.23 | 0.41 |
| 1 | 62824790 | rs3861941 | -1.17 | -0.284 | 0.099 | 1.001 | -0.008 | -0.001 | -0.001 | 0 | 0.35 | 0.61 | 0.4 | 0.31 | 0.23 | 0.41 |
| 1 | 62833738 | rs12042483 | -0.722 | 0.281 | 0.16 | 0.785 | -0.009 | -0.023 | -0.023 | 0 | 0.23 | 0.39 | 0.14 | 0.2 | 0.2 | 0.23 |
| 1 | 62836976 | rs6587971 | -0.53 | 0.568 | -0.7 | 0.763 | -0.001 | 0.042 | 0.042 | 0 | 0.56 | 0.75 | 0.47 | 0.69 | 0.56 | 0.67 |
| 1 | 62842677 | rs997532 | -0.185 | 0.92 | -0.366 | 1.051 | -0.002 | -0.011 | -0.011 | 0 | 0.27 | 0.21 | 0.31 | 0.48 | 0.26 | 0.31 |
| 1 | 62843575 | rs7417430 | 0.482 | 0.866 | 0.055 | 0.629 | -0.004 | 0.023 | 0.023 | 0 | 0.64 | 0.48 | 0.61 | 0.51 | 0.68 | 0.69 |
| 1 | 62848090 | rs3949904 | 0.405 | 0.868 | 0.125 | 0.63 | -0.002 | 0.022 | 0.022 | 0 | 0.64 | 0.53 | 0.62 | 0.56 | 0.68 | 0.69 |
| 1 | 62848113 | rs1473752 | -1.068 | 0.849 | -0.064 | 0.532 | -0.016 | 0.037 | 0.037 | 0 | 0.92 | 0.99 | 0.93 | 1 | 0.92 | 0.99 |
| 1 | 62851933 | rs6587975 | 0.407 | 0.804 | 0.718 | 0.464 | 0.011 | -0.098 | -0.098 | 0.836 | 0.86 | 0.64 | 0.79 | 1 | 0.89 | 0.97 |
| 1 | 62854808 | rs2104448 | -1.242 | 1.014 | 0.657 | 0.642 | 0.084 | 0.102 | 0.102 | 0 | 0.08 | 0.35 | 0.16 | 0.07 | 0.07 | 0.03 |

| | | | | | | | | | | | | | | | | |
|---|---|---|---|---|---|---|---|---|---|---|---|---|---|---|---|---|
| 1 | 62856695 | rs10889325 | 0.509 | 1.001 | 0.825 | 0.629 | 0.012 | -0.027 | -0.027 | 2.067 | 0.85 | 0.73 | 0.78 | 0.93 | 0.85 | 0.96 |
| 1 | 62866854 | rs4439380 | 0.034 | 1.106 | 0.653 | 0.826 | 0 | -0.024 | -0.024 | 2.886 | 0.71 | 0.81 | 0.82 | 0.62 | 0.73 | 0.78 |
| 1 | 62867790 | rs10889328 | 0.764 | 1.211 | 0.635 | 0.769 | 0.032 | -0.023 | -0.023 | 0.025 | 0.83 | 0.5 | 0.71 | 0.91 | 0.84 | 0.92 |
| 1 | 62875313 | rs1334806 | -0.658 | 1.038 | 0.244 | 0.9 | -0.013 | -0.061 | -0.061 | 0.159 | 0.76 | 0.91 | 0.83 | 0.59 | 0.75 | 0.82 |
| 1 | 62892082 | rs13375306 | 0.484 | 1.057 | -0.128 | 0.303 | -0.042 | 0.046 | 0.046 | 0 | 0.43 | 0.18 | 0.42 | 0.35 | 0.44 | 0.59 |
| 1 | 62896419 | rs635897 | -0.177 | 1.125 | 0.008 | 0.363 | 0.021 | 0.006 | 0.006 | 0 | 0.52 | 0.67 | 0.64 | 0.44 | 0.54 | 0.66 |
| 1 | 62898142 | rs10493324 | 0.186 | 1.066 | -0.42 | 0.152 | -0.015 | 0.063 | 0.063 | 0 | 0.46 | 0.23 | 0.43 | 0.35 | 0.45 | 0.6 |
| 1 | 62905893 | rs10493322 | -0.917 | 1.153 | -0.227 | -0.249 | 0.013 | -0.03 | -0.03 | 0 | 0.3 | 0.36 | 0.35 | 0.18 | 0.3 | 0.43 |
| 1 | 62912919 | rs10158897 | -0.94 | 1.008 | 0.147 | -0.308 | 0.013 | -0.03 | -0.03 | 0.03 | 0.3 | 0.36 | 0.35 | 0.18 | 0.3 | 0.43 |
| 1 | 62916796 | rs583609 | -1.085 | 0.803 | 0.435 | -0.183 | -0.005 | 0.006 | 0.006 | 1.131 | 0.31 | 0.69 | 0.39 | 0.23 | 0.3 | 0.47 |
| 1 | 62921586 | rs3913007 | -0.893 | 0.967 | 0.433 | -0.161 | 0.013 | -0.03 | -0.03 | 5.596 | 0.3 | 0.36 | 0.35 | 0.18 | 0.3 | 0.43 |
| 1 | 62953373 | rs10493326 | 0.977 | 1.021 | -0.232 | -0.517 | -0.042 | 0.016 | 0.016 | 0 | 0.19 | 0.05 | 0.23 | 0.15 | 0.26 | 0.21 |
| 1 | 62980607 | rs10889337 | -0.792 | 0.885 | -0.121 | -0.579 | 0.002 | -0.012 | -0.012 | 0.001 | 0.33 | 0.58 | 0.38 | 0.23 | 0.31 | 0.48 |
| 1 | 62997573 | rs7550306 | 1.839 | 1.603 | -0.164 | -0.684 | -0.04 | 0.033 | 0.033 | 0 | 0.06 | 0.01 | 0.17 | 0 | 0.18 | 0.05 |
| 1 | 63020816 | rs10159255 | -0.428 | 1.372 | -0.178 | -0.656 | 0.015 | -0.045 | -0.045 | 0.45 | 0.31 | 0.45 | 0.35 | 0.18 | 0.3 | 0.43 |
| 1 | 63044464 | rs11207993 | 1.043 | 1.497 | -0.202 | -0.747 | -0.041 | 0.047 | 0.047 | 0.529 | 0.1 | 0.04 | 0.17 | 0 | 0.17 | 0.09 |
| 1 | 63055280 | rs12048208 | 0.708 | 1.38 | 0.034 | -0.762 | 0.002 | -0.002 | -0.002 | 0.121 | 0.16 | 0.14 | 0.09 | 0.17 | 0.11 | 0.27 |
| 1 | 63056112 | rs1748197 | -1.101 | 1.326 | 0.033 | -0.759 | -0.003 | 0.016 | 0.016 | 1.323 | 0.33 | 0.69 | 0.37 | 0.2 | 0.3 | 0.47 |
| 1 | 63072265 | rs10789117 | -1.123 | 1.377 | 0.026 | -0.756 | -0.004 | 0.01 | 0.01 | 1.417 | 0.34 | 0.69 | 0.39 | 0.23 | 0.31 | 0.47 |
| 1 | 63083289 | rs2031373 | -1.756 | 1.287 | -0.032 | -0.757 | 0.031 | 0.026 | 0.026 | 1.575 | 0.53 | 0.76 | 0.62 | 0.38 | 0.58 | 0.67 |
| 1 | 63090858 | rs6587980 | -1.181 | 1.243 | 0.456 | -0.65 | 0.002 | -0.011 | -0.011 | 0 | 0.33 | 0.6 | 0.38 | 0.23 | 0.31 | 0.46 |
| 1 | 63103217 | rs1570694 | -0.942 | 1.269 | 0.457 | -0.873 | 0.015 | -0.045 | -0.045 | 0.472 | 0.31 | 0.46 | 0.35 | 0.18 | 0.3 | 0.43 |
| 1 | 63145439 | rs11208004 | -0.819 | 1.293 | 0.489 | -0.861 | 0.015 | -0.034 | -0.034 | 0 | 0.3 | 0.38 | 0.34 | 0.18 | 0.3 | 0.42 |
| 1 | 63156098 | rs1168113 | 0.937 | 1.24 | 0.752 | -0.793 | -0.004 | 0.011 | 0.011 | 1.398 | 0.67 | 0.32 | 0.63 | 0.78 | 0.69 | 0.49 |
| 1 | 63163073 | rs12136083 | -0.932 | 1.231 | 0.821 | -0.812 | 0.012 | -0.027 | -0.027 | 0 | 0.3 | 0.32 | 0.34 | 0.17 | 0.3 | 0.43 |
| 1 | 63191577 | rs4915853 | 0.893 | 1.333 | 0.683 | -0.922 | -0.013 | 0.037 | 0.037 | 1.831 | 0.62 | 0.29 | 0.61 | 0.78 | 0.61 | 0.48 |
| 1 | 63191777 | rs12130333 | 0.149 | 1.366 | 0.688 | -0.947 | -0.033 | 0.047 | 0.047 | 0.043 | 0.14 | 0.09 | 0.23 | 0.01 | 0.18 | 0.16 |
| 1 | 63199272 | rs4630152 | 0.057 | 1.122 | 0.8 | -0.799 | -0.005 | -0.005 | -0.005 | 0.013 | 0.75 | 0.36 | 0.84 | 0.8 | 0.79 | 0.63 |
| 1 | 63212679 | rs12141183 | -0.23 | 0.892 | 0.785 | -0.979 | 0.02 | -0.07 | -0.07 | 0 | 0.16 | 0.37 | 0.26 | 0.02 | 0.14 | 0.19 |
| 1 | 63214147 | rs12143139 | 0.166 | 0.911 | 0.771 | -0.986 | -0.018 | 0.039 | 0.039 | 0.191 | 0.14 | 0.07 | 0.22 | 0.02 | 0.14 | 0.15 |
| 1 | 63221191 | rs6663281 | -0.483 | 0.886 | 0.48 | -1.034 | -0.042 | 0 | 0 | 0 | 0.66 | 0.82 | 0.76 | 0.68 | 0.59 | 0.58 |
| 1 | 63231401 | rs11807368 | 0.646 | 0.883 | 0.565 | -1.025 | 0.044 | 0.056 | 0.056 | 0 | 0.49 | 0.27 | 0.49 | 0.67 | 0.44 | 0.39 |
| 1 | 63246546 | rs17123817 | 0.145 | 0.845 | 0.704 | -0.928 | 0.027 | -0.073 | -0.073 | 2.848 | 0.16 | 0.42 | 0.27 | 0.02 | 0.14 | 0.19 |
| 1 | 63248080 | rs17123830 | 0.065 | 0.844 | 0.716 | -0.929 | 0.032 | -0.081 | -0.081 | 0 | 0.16 | 0.47 | 0.27 | 0.02 | 0.14 | 0.19 |
| 1 | 63248396 | rs17388170 | 0.641 | 0.842 | 0.865 | -0.93 | 0.005 | -0.024 | -0.024 | 0 | 0.13 | 0.22 | 0.25 | 0.02 | 0.13 | 0.15 |
| 1 | 63250266 | rs12067646 | 0.113 | 0.857 | 0.813 | -0.923 | 0.031 | -0.08 | -0.08 | 0 | 0.16 | 0.45 | 0.27 | 0.02 | 0.14 | 0.19 |
| 1 | 63252766 | rs6587988 | 0.098 | 0.87 | 0.718 | -0.917 | 0.026 | -0.084 | -0.084 | 0.351 | 0.16 | 0.52 | 0.27 | 0.02 | 0.14 | 0.19 |
| 1 | 63266720 | rs10493328 | 0.671 | 0.898 | 0.415 | -0.901 | -0.004 | 0.007 | 0.007 | 0 | 0.14 | 0.13 | 0.23 | 0.02 | 0.14 | 0.19 |
| 1 | 63268989 | rs17316337 | 1.82 | 0.896 | 0.285 | -0.921 | -0.023 | 0.047 | 0.047 | 2.013 | 0.13 | 0.02 | 0.07 | 0.04 | 0.12 | 0.07 |
| 1 | 63274538 | rs11208035 | 0.474 | 0.975 | 0.419 | -0.909 | 0.028 | 0.04 | 0.04 | 0 | 0.51 | 0.63 | 0.52 | 0.67 | 0.44 | 0.39 |
| 1 | 63277459 | rs11582138 | 1.069 | 0.888 | -0.007 | -0.932 | -0.028 | 0.012 | 0.012 | 0.011 | 0.17 | 0.04 | 0.18 | 0.14 | 0.22 | 0.16 |
| 1 | 63283005 | rs7539616 | 1.602 | 0.83 | 0.24 | -0.954 | 0.043 | -0.067 | -0.067 | 0 | 0.31 | 0.13 | 0.29 | 0.53 | 0.23 | 0.24 |
| 1 | 63288932 | rs7513520 | 0.879 | 1.088 | 0.952 | -0.715 | 0.027 | 0.007 | 0.007 | 0 | 0.51 | 0.55 | 0.5 | 0.57 | 0.44 | 0.38 |
| 1 | 63290128 | rs7519205 | 0.985 | 1.109 | 0.78 | -1.233 | 0.04 | 0.046 | 0.046 | 0 | 0.48 | 0.19 | 0.48 | 0.66 | 0.44 | 0.39 |
| 1 | 63296271 | rs6679582 | -0.142 | 1.176 | 0.798 | -1.258 | -0.004 | 0 | 0 | 0 | 0.64 | 0.63 | 0.73 | 0.68 | 0.58 | 0.59 |
| 1 | 63303845 | rs10493327 | 0.894 | 1.176 | 0.6 | -1.259 | 0.004 | 0.006 | 0.006 | 0 | 0.51 | 0.46 | 0.5 | 0.67 | 0.44 | 0.39 |
| 1 | 63311836 | rs11208047 | -0.279 | 1.195 | 0.452 | -1.245 | -0.021 | 0 | 0 | 0 | 0.65 | 0.69 | 0.73 | 0.68 | 0.58 | 0.59 |
| 1 | 63330217 | rs3211074 | 0.719 | 1.241 | -0.033 | -1.115 | 0.001 | 0.001 | 0.001 | 0 | 0.5 | 0.45 | 0.49 | 0.66 | 0.44 | 0.39 |
| 1 | 63333210 | rs2803245 | -1.01 | 1.377 | -1.115 | -1.09 | -0.049 | -0.046 | -0.046 | 0 | 0.8 | 0.93 | 0.83 | 0.72 | 0.7 | 0.66 |
| 1 | 63336271 | rs11208050 | 0.407 | 0.895 | -1.06 | -0.944 | 0.04 | 0.057 | 0.057 | 0 | 0.48 | 0.18 | 0.47 | 0.67 | 0.44 | 0.39 |
| 1 | 63358816 | rs7512480 | 0.462 | 0.712 | -1.102 | -0.992 | -0.019 | 0.041 | 0.041 | 0 | 0.44 | 0.17 | 0.51 | 0.69 | 0.5 | 0.39 |
| 1 | 63361221 | rs12751505 | 0.642 | 0.303 | -0.378 | -1.476 | 0.004 | -0.006 | -0.006 | 3.461 | 0.67 | 0.28 | 0.73 | 0.97 | 0.74 | 0.79 |
| 1 | 63362324 | rs12137912 | 0.714 | 0.159 | -0.643 | -1.598 | -0.001 | 0.033 | 0.033 | 0 | 0.07 | 0 | 0.12 | 0 | 0.05 | 0.05 |
| 1 | 63363624 | rs2780875 | -0.295 | 0.322 | -0.169 | -1.376 | 0.001 | 0.006 | 0.006 | 0 | 0.88 | 0.88 | 0.87 | 1 | 0.92 | 0.9 |
| 1 | 63368676 | rs2803242 | -0.747 | 0.34 | 0.231 | -0.875 | -0.006 | -0.075 | -0.075 | 0.033 | 0.22 | 0.63 | 0.14 | 0.03 | 0.18 | 0.11 |
| 1 | 63369333 | rs2803243 | -0.778 | 0.35 | 0.529 | -0.871 | -0.015 | -0.121 | -0.121 | 0.739 | 0.23 | 0.54 | 0.13 | 0.03 | 0.18 | 0.11 |
| 1 | 63370965 | rs17123995 | -0.824 | 0.936 | 0.288 | -0.18 | 0 | -0.024 | -0.024 | 1.053 | 0.05 | 0.09 | 0.05 | 0 | 0.05 | 0.03 |
| 1 | 63384292 | rs7556152 | -1.184 | 1.023 | 0.317 | 0.353 | 0.064 | 0.166 | 0.166 | 0 | 0.05 | 0.2 | 0.05 | 0.17 | 0.08 | 0.25 |
| 1 | 63392665 | rs11208061 | -0.378 | 0.881 | 0.238 | -0.865 | -0.002 | 0.083 | 0.083 | 0 | 0.32 | 0.22 | 0.35 | 0 | 0.3 | 0.21 |
| 1 | 63396290 | rs12024707 | -0.135 | 0.928 | 0.596 | -1.324 | 0.015 | 0.026 | 0.026 | 0 | 0.14 | 0.36 | 0.13 | 0.13 | 0.13 | 0.25 |
| 1 | 63396342 | rs12134779 | 0.117 | 1.006 | 0.434 | -1.463 | 0.004 | -0.013 | -0.013 | 0 | 0.3 | 0.3 | 0.34 | 0 | 0.26 | 0.21 |
| 1 | 63401632 | rs2803234 | -0.01 | 0.721 | -0.16 | -1.626 | 0 | 0.043 | 0.043 | 0 | 0.92 | 0.98 | 0.95 | 1 | 0.92 | 0.96 |
| 1 | 63401945 | rs7532163 | -0.598 | 0.746 | 0.047 | -1.595 | -0.007 | -0.071 | -0.071 | 0 | 0.41 | 0.55 | 0.44 | 0.1 | 0.37 | 0.43 |
| 1 | 63402478 | rs1360679 | 0.867 | 0.785 | -0.142 | -1.481 | -0.001 | -0.015 | -0.015 | 3.363 | 0.09 | 0.06 | 0.1 | 0 | 0.11 | 0.03 |
| 1 | 63402923 | rs11208065 | 0.471 | 0.978 | -0.393 | -1.606 | -0.036 | 0.079 | 0.079 | 0 | 0.22 | 0.12 | 0.24 | 0.04 | 0.27 | 0.11 |
| 1 | 63403020 | rs17124071 | 0.208 | 0.099 | -0.915 | -1.036 | 0.028 | -0.027 | -0.027 | 0 | 0.39 | 0.25 | 0.32 | 0.43 | 0.31 | 0.25 |
| 1 | 63404689 | rs10889392 | 0.285 | -0.099 | -0.211 | -1.192 | -0.012 | 0.045 | 0.045 | 0 | 0.28 | 0.4 | 0.27 | 0.42 | 0.22 | 0.24 |
| 1 | 63404702 | rs2803237 | -0.299 | -0.121 | 0.096 | -1.421 | 0.007 | 0.012 | 0.012 | 0 | 0.41 | 0.59 | 0.5 | 0.88 | 0.34 | 0.58 |
| 1 | 63405189 | rs2780881 | -0.158 | -0.141 | 0.248 | -1.479 | -0.017 | 0.048 | 0.048 | 0 | 0.28 | 0.41 | 0.34 | 0.46 | 0.24 | 0.35 |

| | | | | | | | | | | | | | | | | |
|---|---|---|---|---|---|---|---|---|---|---|---|---|---|---|---|---|
| 1 | 63406451 | rs1572109 | -0.981 | -0.062 | -0.071 | -1.468 | -0.053 | 0.16 | 0.16 | 0 | 0.59 | 0.84 | 0.63 | 0.89 | 0.48 | 0.61 |
| 1 | 63409807 | rs11208069 | 0.322 | -0.082 | 0.287 | -1.403 | -0.015 | 0.142 | 0.142 | 0 | 0.15 | 0.27 | 0.23 | 0.46 | 0.13 | 0.35 |
| 1 | 63412545 | rs11208070 | -0.09 | -0.055 | 0.156 | -1.38 | 0.002 | -0.002 | -0.002 | 0.78 | 0.42 | 0.44 | 0.39 | 0.42 | 0.35 | 0.26 |
| 1 | 63414401 | rs2065585 | -0.099 | -0.042 | 0.102 | -1.375 | -0.005 | 0.001 | 0.001 | 0 | 0.42 | 0.49 | 0.39 | 0.42 | 0.35 | 0.26 |
| 1 | 63415962 | rs12136978 | -0.867 | 0.573 | -0.263 | -1.034 | 0 | -0.003 | -0.003 | 0.019 | 0.39 | 0.63 | 0.37 | 0.51 | 0.38 | 0.48 |
| 1 | 63417567 | rs1333731 | -0.478 | 0.635 | -0.387 | -0.504 | -0.001 | 0 | 0 | 0 | 0.23 | 0.73 | 0.22 | 0.42 | 0.21 | 0.35 |
| 1 | 63431570 | rs12138360 | -0.345 | 0.444 | -0.801 | -0.384 | -0.001 | -0.006 | -0.006 | 0.078 | 0.3 | 0.3 | 0.38 | 0.54 | 0.28 | 0.34 |
| 1 | 63433037 | rs12121455 | 0.207 | 0.453 | -1.269 | -0.332 | -0.071 | -0.139 | -0.139 | 0 | 0.24 | 0.06 | 0.35 | 0.51 | 0.26 | 0.25 |
| 1 | 63436850 | rs6682421 | 0.207 | 0.467 | -1.269 | -0.282 | -0.074 | -0.143 | -0.143 | 1.154 | 0.24 | 0.06 | 0.35 | 0.47 | 0.27 | 0.25 |
| 1 | 63439838 | rs1888174 | -0.699 | 0.095 | -1.286 | -0.613 | 0.012 | 0.012 | 0.012 | 0 | 0.09 | 0.04 | 0.04 | 0.03 | 0.06 | 0.1 |
| 1 | 63446754 | rs6662502 | -0.698 | 0.083 | -1.22 | -0.456 | -0.002 | -0.036 | -0.036 | 0.018 | 0.35 | 0.21 | 0.39 | 0.54 | 0.34 | 0.35 |
| 1 | 63452076 | rs11208080 | 0.494 | 0.525 | -1.208 | -0.117 | -0.083 | -0.147 | -0.147 | 0 | 0.21 | 0.04 | 0.34 | 0.51 | 0.27 | 0.19 |
| 1 | 63452363 | rs6683386 | 0.22 | 0.498 | -0.982 | -0.13 | -0.007 | -0.014 | -0.014 | 0 | 0.23 | 0.27 | 0.37 | 0.52 | 0.27 | 0.22 |
| 1 | 63452419 | rs6696216 | -0.042 | 0.234 | -0.869 | -0.268 | 0.051 | 0.042 | 0.042 | 0 | 0.51 | 0.36 | 0.63 | 0.61 | 0.59 | 0.53 |
| 1 | 63452789 | rs6696619 | -0.381 | 0.531 | -0.837 | -1.05 | 0.018 | 0.058 | 0.058 | 0 | 0.59 | 0.76 | 0.69 | 0.91 | 0.65 | 0.77 |
| 1 | 63454731 | rs12119857 | -0.124 | 0.987 | -1.694 | -1.185 | -0.024 | 0.189 | 0.189 | 0 | 0.39 | 0.13 | 0.29 | 0.07 | 0.4 | 0.29 |
| 1 | 63457259 | rs4915627 | -0.125 | 0.805 | -1.69 | -1.423 | -0.014 | 0.073 | 0.073 | 0 | 0.36 | 0.23 | 0.46 | 0.83 | 0.39 | 0.55 |
| 1 | 63457362 | rs4915629 | -0.125 | 0.805 | -1.69 | -1.423 | -0.014 | 0.073 | 0.073 | 0 | 0.36 | 0.23 | 0.45 | 0.83 | 0.39 | 0.55 |
| 1 | 63458845 | rs17124198 | -0.965 | 0.104 | -0.544 | -1.062 | 0.041 | 0.182 | 0.182 | 0.01 | 0.06 | 0.35 | 0.09 | 0.12 | 0.02 | 0.02 |
| 1 | 63460850 | rs10889394 | -0.098 | 0.578 | -1.865 | -1.55 | -0.073 | 0.215 | 0.215 | 1.686 | 0.55 | 0.86 | 0.54 | 0.85 | 0.53 | 0.63 |
| 1 | 63461381 | rs11208082 | -0.514 | 0.454 | -1.358 | -0.506 | 0.001 | 0.022 | 0.022 | 0 | 0.19 | 0.26 | 0.25 | 0.63 | 0.23 | 0.39 |
| 1 | 63467957 | rs12130613 | -0.228 | 0.49 | -1.341 | -1.28 | -0.012 | 0.13 | 0.13 | 0.113 | 0.53 | 0.28 | 0.58 | 0.19 | 0.49 | 0.54 |
| 1 | 63470519 | rs2150030 | -0.162 | 0.373 | -0.801 | -1.144 | -0.001 | -0.014 | -0.014 | 1.623 | 0.48 | 0.43 | 0.43 | 0.8 | 0.54 | 0.64 |
| 1 | 63472670 | rs12131561 | -1.847 | 0.284 | -0.858 | -1.125 | -0.001 | 0.006 | 0.006 | 6.445 | 0.78 | 0.76 | 0.87 | 0.84 | 0.81 | 0.9 |
| 1 | 63472802 | rs12742218 | 0.308 | 0.173 | -0.87 | -1.162 | 0.014 | 0.121 | 0.121 | 0.001 | 0.5 | 0.32 | 0.53 | 0.04 | 0.46 | 0.31 |
| 1 | 63474236 | rs17124275 | 1.032 | 0.07 | -1.316 | -1.518 | -0.041 | 0.114 | 0.114 | 0 | 0.24 | 0.09 | 0.42 | 0.03 | 0.28 | 0.13 |
| 1 | 63475829 | rs4915864 | -2.122 | 0.16 | -1.11 | -1.303 | 0 | -0.001 | -0.001 | 0.319 | 0.78 | 0.74 | 0.87 | 0.84 | 0.81 | 0.9 |
| 1 | 63478358 | rs7541201 | 0.161 | 0.261 | -0.887 | -0.963 | 0.005 | 0.018 | 0.018 | 0 | 0.17 | 0.21 | 0.28 | 0.63 | 0.27 | 0.35 |
| 1 | 63480512 | rs7514406 | -0.169 | 0.156 | -0.828 | -0.874 | -0.014 | 0.053 | 0.053 | 0 | 0.05 | 0.14 | 0.06 | 0.15 | 0.02 | 0.22 |
| 1 | 63482751 | rs17124332 | 0.756 | -0.093 | -0.933 | -1.208 | 0.014 | 0.047 | 0.047 | 1.635 | 0.35 | 0.24 | 0.19 | 0.15 | 0.3 | 0.34 |
| 1 | 63483169 | rs11208089 | 1.422 | 0.014 | -0.996 | -1.147 | -0.027 | -0.017 | -0.017 | 0.171 | 0.11 | 0.05 | 0.21 | 0.49 | 0.24 | 0.13 |
| 1 | 63483556 | rs12566149 | 1.395 | 0.013 | -0.956 | -1.147 | -0.007 | 0.03 | 0.03 | 0 | 0.12 | 0.1 | 0.22 | 0.49 | 0.24 | 0.13 |
| 1 | 63484182 | rs7543262 | 0.745 | -0.144 | -1.202 | -1.406 | -0.114 | 0.206 | 0.206 | 0 | 0.34 | 0.03 | 0.45 | 0.03 | 0.32 | 0.21 |
| 1 | 63484216 | rs1411390 | -0.569 | -0.218 | -1.039 | -1.425 | -0.029 | 0.11 | 0.11 | 0 | 0.82 | 0.56 | 0.88 | 0.67 | 0.86 | 0.69 |
| 1 | 63484437 | rs12119690 | 0.517 | -0.127 | -1.166 | -1.421 | -0.019 | 0.005 | 0.005 | 0 | 0.14 | 0.2 | 0.09 | 0.15 | 0.11 | 0.29 |
| 1 | 63484941 | rs12142862 | 0.375 | 0.262 | -0.997 | -1.162 | -0.002 | 0.134 | 0.134 | 0.186 | 0.44 | 0.22 | 0.49 | 0.03 | 0.41 | 0.28 |
| 1 | 63486692 | rs713311 | 1.256 | 0.641 | -1.157 | -0.813 | 0.013 | 0.061 | 0.061 | 0 | 0.1 | 0.23 | 0.06 | 0.16 | 0.08 | 0.23 |
| 1 | 63487473 | rs11208093 | -0.83 | 0.734 | -1.372 | -0.787 | -0.033 | -0.055 | -0.055 | 0 | 0.66 | 0.78 | 0.67 | 0.51 | 0.57 | 0.81 |
| 1 | 63488228 | rs4915866 | -0.662 | 1.346 | -0.85 | 0.232 | -0.007 | 0.029 | 0.029 | 0 | 0.88 | 0.9 | 0.78 | 0.51 | 0.76 | 0.86 |
| 1 | 63493897 | rs4915868 | 0.469 | 1.318 | -1.134 | -0.08 | -0.07 | 0.078 | 0.078 | 0 | 0.19 | 0.02 | 0.38 | 0.03 | 0.22 | 0.1 |
| 1 | 63495975 | rs1330221 | 0.734 | 1.062 | -0.07 | 0.721 | 0.043 | 0.077 | 0.077 | 0.101 | 0.79 | 0.68 | 0.71 | 0.36 | 0.67 | 0.62 |
| 1 | 63496207 | rs12091913 | -0.689 | 1.207 | -0.609 | 0.903 | -0.015 | 0.053 | 0.053 | 0 | 0.07 | 0.19 | 0.05 | 0.17 | 0.05 | 0.07 |
| 1 | 63499652 | rs6588005 | 0.375 | 1.606 | -0.958 | 1.007 | 0 | 0.038 | 0.038 | 0 | 0.87 | 0.88 | 0.77 | 0.51 | 0.74 | 0.84 |
| 1 | 63501851 | rs6664042 | -0.414 | 1.506 | -1.223 | -0.046 | -0.027 | 0.046 | 0.046 | 0 | 0.31 | 0.17 | 0.25 | 0.16 | 0.35 | 0.18 |
| 1 | 63502074 | rs6677129 | -0.983 | 0.764 | -0.798 | -0.978 | -0.032 | -0.131 | -0.131 | 0.01 | 0.63 | 0.52 | 0.82 | 0.98 | 0.76 | 0.72 |
| 1 | 63503606 | rs6694848 | -0.306 | 0.67 | -1.012 | -1.265 | -0.011 | 0.019 | 0.019 | 0.98 | 0.39 | 0.31 | 0.61 | 0.84 | 0.48 | 0.61 |
| 1 | 63507395 | rs1874930 | -0.272 | 0.511 | -0.627 | -0.771 | 0 | 0 | 0 | 0.365 | 0.33 | 0.34 | 0.48 | 0.28 | 0.34 | 0.41 |
| 1 | 63508630 | rs11208103 | 1.006 | 0.587 | -0.611 | -0.796 | 0.01 | 0.018 | 0.018 | 0.098 | 0.18 | 0.08 | 0.12 | 0 | 0.15 | 0.14 |
| 1 | 63513116 | rs1027520 | -0.38 | 0.541 | -0.131 | -0.818 | 0.001 | -0.002 | -0.002 | 0.009 | 0.34 | 0.38 | 0.49 | 0.28 | 0.34 | 0.41 |
| 1 | 63517230 | rs881939 | 1.022 | 0.541 | -0.173 | -0.819 | 0.011 | 0.026 | 0.026 | 0.131 | 0.18 | 0.09 | 0.12 | 0 | 0.15 | 0.13 |
| 1 | 63517581 | rs1391813 | 0.72 | 0.1 | -0.088 | -0.724 | 0.004 | -0.005 | -0.005 | 0 | 0.53 | 0.18 | 0.37 | 0.59 | 0.57 | 0.43 |
| 1 | 63521755 | rs11208115 | -0.004 | 0.1 | 0.493 | -0.724 | 0 | -0.004 | -0.004 | 0 | 0.34 | 0.44 | 0.49 | 0.28 | 0.34 | 0.41 |
| 1 | 63521891 | rs6696559 | 0.951 | 0.148 | 0.575 | -0.689 | 0.023 | 0.056 | 0.056 | 0 | 0.07 | 0.2 | 0.11 | 0.12 | 0.07 | 0.14 |
| 1 | 63522943 | rs1124625 | 0.388 | 0.131 | 0.157 | -0.812 | -0.083 | -0.078 | -0.078 | 0 | 0.33 | 0.11 | 0.32 | 0.59 | 0.44 | 0.37 |
| 1 | 63524019 | rs11208116 | 1.344 | 0.16 | 0.247 | -0.79 | 0.003 | 0.015 | 0.015 | 0 | 0.14 | 0.14 | 0.35 | 0.28 | 0.19 | 0.28 |
| 1 | 63525756 | rs1888176 | -0.097 | 0.176 | 0.077 | -0.696 | 0 | 0 | 0 | 0 | 0.38 | 0.57 | 0.52 | 0.26 | 0.36 | 0.42 |
| 1 | 63527538 | rs17318040 | 2.112 | 0.071 | -0.177 | -0.645 | 0.043 | 0.058 | 0.058 | 2.74 | 0.2 | 0.07 | 0.05 | 0 | 0.14 | 0.06 |
| 1 | 63528577 | rs6684405 | 0.647 | 0.313 | -0.067 | -0.437 | 0.002 | 0.012 | 0.012 | 0 | 0.07 | 0.15 | 0.1 | 0.12 | 0.07 | 0.14 |
| 1 | 63529371 | rs1888177 | 1.007 | 0.586 | -0.184 | -0.27 | -0.007 | 0.054 | 0.054 | 0 | 0.07 | 0.04 | 0.13 | 0.43 | 0.16 | 0.22 |
| 1 | 63532094 | rs17124486 | 0.757 | 0.484 | -0.447 | -0.843 | -0.102 | 0.02 | 0.02 | 0 | 0.27 | 0.03 | 0.18 | 0.2 | 0.3 | 0.17 |
| 1 | 63532307 | rs17318082 | 1.32 | -0.024 | 0.103 | -1.026 | 0.014 | 0.037 | 0.037 | 0 | 0.13 | 0.15 | 0.35 | 0.25 | 0.19 | 0.26 |
| 1 | 63533935 | rs7537459 | -0.78 | -0.218 | 0.481 | -1.183 | -0.026 | -0.098 | -0.098 | 0 | 0.76 | 0.71 | 0.85 | 1 | 0.85 | 0.85 |
| 1 | 63535086 | rs4915873 | -0.626 | -0.217 | 0.438 | -1.183 | -0.055 | -0.166 | -0.166 | 0 | 0.75 | 0.56 | 0.83 | 1 | 0.85 | 0.85 |
| 1 | 63536077 | rs10889400 | -0.714 | -0.164 | 0.332 | -1.201 | -0.016 | -0.116 | -0.116 | 0.03 | 0.55 | 0.51 | 0.78 | 1 | 0.71 | 0.79 |
| 1 | 63536838 | rs12036206 | 0.746 | 0.244 | 0.012 | -0.846 | 0.009 | 0.006 | 0.006 | 0 | 0.34 | 0.23 | 0.41 | 0.25 | 0.33 | 0.33 |
| 1 | 63537603 | rs6659893 | -0.954 | 0.214 | 0.002 | -0.963 | -0.004 | -0.044 | -0.044 | 0 | 0.82 | 0.82 | 0.87 | 1 | 0.86 | 0.88 |
| 1 | 63540667 | rs10889401 | -0.292 | -0.07 | -0.095 | -1.241 | -0.023 | -0.142 | -0.142 | 2.659 | 0.59 | 0.45 | 0.76 | 0.98 | 0.66 | 0.68 |
| 1 | 63541253 | rs6679559 | -0.536 | 0.354 | 0.025 | -0.458 | -0.002 | 0.011 | 0.011 | 0 | 0.3 | 0.33 | 0.32 | 0.63 | 0.36 | 0.47 |
| 1 | 63545178 | rs11808494 | -0.229 | 1.1 | -0.215 | -0.16 | 0.004 | 0.005 | 0.005 | 0.532 | 0.2 | 0.2 | 0.37 | 0.29 | 0.27 | 0.4 |

| | | | | | | | | | | | | | | | | |
|---|---|---|---|---|---|---|---|---|---|---|---|---|---|---|---|---|
| 1 | 63545249 | rs10749737 | -0.88 | 0.835 | -0.404 | -0.923 | 0.023 | 0.038 | 0.038 | 0.037 | 0.67 | 0.78 | 0.84 | 1 | 0.7 | 0.82 |
| 1 | 63546715 | rs17318187 | 0.561 | 1.272 | -0.043 | -0.493 | 0.017 | -0.039 | -0.039 | 0 | 0.1 | 0.12 | 0.1 | 0 | 0.18 | 0.04 |
| 1 | 63547673 | rs17124579 | 0.202 | 1.089 | 0.231 | -0.413 | 0.029 | -0.028 | -0.028 | 0.057 | 0.06 | 0.2 | 0.05 | 0 | 0.07 | 0.03 |
| 1 | 63549230 | rs1391812 | 0.067 | 1.353 | 0.154 | -0.236 | -0.004 | -0.004 | -0.004 | 0.693 | 0.74 | 0.78 | 0.75 | 0.7 | 0.66 | 0.59 |
| 1 | 63549282 | rs953654 | 0.839 | 0.809 | -0.002 | -0.674 | -0.018 | 0.019 | 0.019 | 0.278 | 0.42 | 0.33 | 0.59 | 0.69 | 0.49 | 0.47 |
| 1 | 63550500 | rs10889402 | -0.193 | 0.268 | 0.312 | 0.134 | -0.013 | 0.004 | 0.004 | 0.342 | 0.33 | 0.34 | 0.2 | 0.43 | 0.2 | 0.25 |
| 1 | 63553556 | rs12041139 | -1.258 | 1.284 | 0.679 | 0.763 | 0.053 | 0.209 | 0.209 | 0.038 | 0.14 | 0.24 | 0.24 | 0.47 | 0.19 | 0.21 |
| 1 | 63553810 | rs13373927 | -0.588 | 1.022 | 0.33 | -0.22 | 0.012 | 0.035 | 0.035 | 0 | 0.49 | 0.57 | 0.34 | 0.12 | 0.38 | 0.28 |
| 1 | 63554295 | rs6663920 | -0.988 | 1.254 | 0.94 | 0.129 | 0.037 | 0.07 | 0.07 | 0 | 0.09 | 0.36 | 0.1 | 0.11 | 0.07 | 0.15 |
| 1 | 63554499 | rs17124604 | -0.766 | 1.853 | 0.669 | -0.244 | 0.013 | -0.03 | -0.03 | 0 | 0.11 | 0.15 | 0.1 | 0.05 | 0.14 | 0.2 |
| 1 | 63561813 | rs1498934 | -0.755 | 1.203 | 0.239 | -0.398 | -0.005 | 0.005 | 0.005 | 0 | 0.7 | 0.84 | 0.7 | 0.76 | 0.75 | 0.81 |
| 1 | 63561950 | rs10889403 | 0.934 | 1.396 | 0.411 | -0.459 | 0.024 | 0.015 | 0.015 | 0 | 0.44 | 0.28 | 0.49 | 0.65 | 0.44 | 0.38 |
| 1 | 63564069 | rs6677463 | -0.499 | 1.19 | 0.727 | -0.485 | 0 | -0.002 | -0.002 | 0 | 0.21 | 0.21 | 0.17 | 0.1 | 0.28 | 0.31 |
| 1 | 63567217 | rs11208127 | -1.256 | 1.196 | 0.837 | -0.452 | 0.003 | 0 | 0 | 0 | 0.73 | 0.72 | 0.7 | 0.75 | 0.74 | 0.81 |
| 1 | 63567656 | rs1498941 | -0.871 | 1.274 | 0.316 | -0.776 | -0.039 | 0.051 | 0.051 | 0 | 0.32 | 0.22 | 0.26 | 0.1 | 0.43 | 0.35 |
| 1 | 63570551 | rs1391819 | 1.229 | 1.091 | 0.212 | -0.616 | 0.029 | -0.007 | -0.007 | 0.069 | 0.37 | 0.27 | 0.42 | 0.63 | 0.29 | 0.35 |
| 1 | 63575255 | rs1498936 | -0.158 | 1.304 | 0.217 | -0.377 | 0.02 | -0.001 | -0.001 | 0 | 0.53 | 0.69 | 0.53 | 0.55 | 0.47 | 0.58 |
| 1 | 63575415 | rs17390595 | 0.197 | 1.48 | 0 | -0.413 | -0.042 | 0.034 | 0.034 | 0 | 0.18 | 0.06 | 0.13 | 0.08 | 0.24 | 0.21 |
| 1 | 63578064 | rs11208131 | -0.033 | 1.465 | 0.243 | -0.3 | -0.004 | -0.001 | -0.001 | 2.377 | 0.54 | 0.44 | 0.54 | 0.55 | 0.54 | 0.51 |
| 1 | 63580025 | rs1498930 | -1.121 | 1.354 | -0.02 | -0.103 | -0.012 | -0.015 | -0.015 | 0.018 | 0.72 | 0.8 | 0.66 | 0.63 | 0.71 | 0.79 |
| 1 | 63582114 | rs10493335 | 0.138 | 1.037 | 0.091 | -0.203 | 0.022 | 0.005 | 0.005 | 0 | 0.46 | 0.44 | 0.45 | 0.57 | 0.4 | 0.52 |
| 1 | 63586468 | rs7525585 | -0.792 | 1.186 | 0.023 | -0.343 | -0.01 | 0.018 | 0.018 | 0 | 0.12 | 0.08 | 0.1 | 0.01 | 0.13 | 0.22 |
| 1 | 63588287 | rs17390657 | -0.166 | 0.919 | -0.023 | -0.411 | 0 | -0.013 | -0.013 | 0 | 0.08 | 0.1 | 0.08 | 0.01 | 0.09 | 0.12 |
| 1 | 63590222 | rs924337 | 0.85 | 0.937 | 0.054 | -0.637 | 0.046 | 0.056 | 0.056 | 0 | 0.37 | 0.27 | 0.27 | 0.01 | 0.26 | 0.17 |
| 1 | 63594427 | rs17124674 | -0.507 | 1.041 | -0.012 | -0.585 | -0.002 | -0.012 | -0.012 | 0.408 | 0.12 | 0.13 | 0.1 | 0.01 | 0.13 | 0.22 |
| 1 | 63596457 | rs1556201 | -0.598 | 1.252 | 0.129 | -0.487 | -0.02 | 0.007 | 0.007 | 0 | 0.55 | 0.33 | 0.52 | 0.56 | 0.54 | 0.63 |
| 1 | 63597995 | rs11208137 | -0.734 | 1.232 | 0.49 | -0.518 | -0.007 | 0.001 | 0.001 | 0 | 0.57 | 0.62 | 0.54 | 0.56 | 0.54 | 0.63 |
| 1 | 63599492 | rs17124687 | 0.986 | 0.99 | 0.157 | -1.011 | 0.001 | -0.012 | -0.012 | 0 | 0.06 | 0.1 | 0.07 | 0 | 0.08 | 0.02 |
| 1 | 63602362 | rs11583639 | 0.314 | 0.736 | 0.066 | -0.965 | 0.005 | -0.055 | -0.055 | 0 | 0.41 | 0.38 | 0.3 | 0.01 | 0.3 | 0.27 |
| 1 | 63603161 | rs6692500 | -1.554 | 0.962 | -0.302 | -0.69 | -0.005 | 0.003 | 0.003 | 0 | 0.75 | 0.83 | 0.72 | 0.76 | 0.76 | 0.84 |
| 1 | 63612281 | rs11576443 | 0.662 | 0.142 | -1.012 | -1.041 | 0.062 | 0.104 | 0.104 | 0 | 0.31 | 0.13 | 0.13 | 0.01 | 0.2 | 0.14 |
| 1 | 63617566 | rs17318933 | 0.901 | -0.053 | -1.224 | -1.083 | 0.055 | 0.119 | 0.119 | 0.375 | 0.26 | 0.07 | 0.08 | 0.01 | 0.15 | 0.16 |
| 1 | 63619559 | rs7535047 | 0.447 | 0.386 | -0.905 | -0.466 | -0.003 | -0.009 | -0.009 | 0 | 0.21 | 0.19 | 0.24 | 0.28 | 0.23 | 0.24 |
| 1 | 63621743 | rs1353799 | -0.115 | 0.35 | -0.48 | -0.383 | 0.004 | 0.007 | 0.007 | 0 | 0.29 | 0.44 | 0.31 | 0.33 | 0.31 | 0.27 |
| 1 | 63626108 | rs17124726 | -0.339 | 0.408 | -0.501 | 0.055 | 0.006 | -0.005 | -0.005 | 0 | 0.05 | 0.16 | 0.03 | 0.01 | 0.02 | 0.09 |
| 1 | 63626202 | rs3887428 | -0.498 | 0.434 | -1.266 | -0.06 | -0.021 | 0.032 | 0.032 | 0.016 | 0.57 | 0.77 | 0.49 | 0.35 | 0.52 | 0.59 |
| 1 | 63628648 | rs4269797 | -0.688 | 1.08 | -0.791 | 0.904 | -0.048 | -0.055 | -0.055 | 0 | 0.09 | 0.01 | 0.29 | 0.74 | 0.18 | 0.32 |
| 1 | 63632518 | rs11208143 | 0.004 | 0.564 | -0.284 | 1.174 | -0.012 | 0.076 | 0.076 | 0 | 0.85 | 0.74 | 0.88 | 0.68 | 0.87 | 0.81 |
| 1 | 63632591 | rs10493341 | 1.061 | 0.522 | -0.954 | 0.651 | -0.048 | 0.064 | 0.064 | 0 | 0.26 | 0.05 | 0.28 | 0.18 | 0.29 | 0.19 |
| 1 | 63634135 | rs6588010 | -0.318 | 0.453 | -0.683 | 0.735 | -0.016 | 0.084 | 0.084 | 0 | 0.84 | 0.7 | 0.88 | 0.68 | 0.87 | 0.81 |
| 1 | 63637312 | rs4915878 | 0.104 | 0.69 | -0.577 | 0.618 | 0.011 | 0.063 | 0.063 | 0 | 0.81 | 0.68 | 0.82 | 0.67 | 0.81 | 0.79 |
| 1 | 63637911 | rs4915880 | 0.695 | 0.618 | -0.147 | 0.728 | 0.003 | 0.003 | 0.003 | 0 | 0.41 | 0.33 | 0.33 | 0.34 | 0.37 | 0.34 |
| 1 | 63638714 | rs6588015 | 0.01 | 0.786 | -0.075 | 0.897 | 0.012 | 0.067 | 0.067 | 0 | 0.81 | 0.67 | 0.82 | 0.66 | 0.81 | 0.79 |
| 1 | 63638791 | rs6588016 | -0.236 | 0.53 | -0.137 | 0.8 | -0.003 | 0.014 | 0.014 | 0 | 0.54 | 0.43 | 0.54 | 0.36 | 0.52 | 0.68 |
| 1 | 63643251 | rs7515372 | 0.335 | 0.421 | -0.135 | 0.919 | -0.028 | 0.003 | 0.003 | 0 | 0.51 | 0.15 | 0.46 | 0.43 | 0.48 | 0.53 |
| 1 | 63645215 | rs11208148 | -0.492 | 0.444 | 0.369 | 0.841 | 0.008 | 0.072 | 0.072 | 0 | 0.22 | 0.29 | 0.35 | 0.46 | 0.25 | 0.37 |
| 1 | 63650851 | rs7515650 | 0.988 | 0.446 | -0.535 | 0.672 | -0.033 | 0.078 | 0.078 | 0 | 0.38 | 0.1 | 0.38 | 0.3 | 0.38 | 0.31 |
| 1 | 63654902 | rs2031885 | 1.275 | 0.687 | -0.388 | 0.795 | -0.015 | 0.047 | 0.047 | 0 | 0.24 | 0.14 | 0.17 | 0.13 | 0.27 | 0.11 |
| 1 | 63655914 | rs4253945 | 0.354 | 0.725 | -0.239 | 0.661 | -0.013 | 0.023 | 0.023 | 0 | 0.52 | 0.24 | 0.46 | 0.4 | 0.49 | 0.53 |
| 1 | 63658137 | rs4915639 | 0.168 | 0.637 | 0.201 | 0.589 | 0.017 | -0.022 | -0.022 | 0 | 0.79 | 0.49 | 0.71 | 0.83 | 0.8 | 0.68 |
| 1 | 63659847 | rs11208154 | 0.22 | 0.624 | -0.43 | 0.786 | -0.004 | -0.015 | -0.015 | 0 | 0.11 | 0.06 | 0.07 | 0.14 | 0.11 | 0.19 |
| 1 | 63667713 | rs6689698 | 0.053 | 0.621 | -0.569 | 0.451 | -0.02 | -0.003 | -0.003 | 0 | 0.33 | 0.16 | 0.22 | 0.25 | 0.35 | 0.29 |
| 1 | 63668754 | rs6693289 | 0.875 | 0.554 | -0.835 | 0.624 | -0.032 | 0.022 | 0.022 | 0 | 0.42 | 0.11 | 0.4 | 0.39 | 0.43 | 0.36 |
| 1 | 63674545 | rs7514965 | -0.311 | 0.662 | -0.075 | 0.605 | 0.005 | 0.008 | 0.008 | 0 | 0.16 | 0.21 | 0.12 | 0.13 | 0.17 | 0.23 |
| 1 | 63674914 | rs1328589 | 0.964 | 0.644 | 0.361 | 0.567 | 0.015 | 0.031 | 0.031 | 0 | 0.69 | 0.55 | 0.59 | 0.51 | 0.68 | 0.46 |
| 1 | 63684460 | rs6675635 | -0.276 | 0.644 | 0.423 | 0.602 | 0.007 | 0.062 | 0.062 | 0 | 0.85 | 0.72 | 0.71 | 0.64 | 0.84 | 0.69 |
| 1 | 63687520 | rs17319248 | 0.908 | 0.653 | 0.316 | 0.599 | -0.027 | -0.042 | -0.042 | 0 | 0.09 | 0.02 | 0.07 | 0.13 | 0.11 | 0.19 |
| 1 | 63688381 | rs7552970 | 1.316 | 0.611 | 0.307 | 0.521 | 0 | 0.051 | 0.051 | 0 | 0.24 | 0.14 | 0.16 | 0.12 | 0.24 | 0.1 |
| 1 | 63689246 | rs1410581 | -0.252 | 0.616 | 0.77 | 0.549 | 0.017 | 0.109 | 0.109 | 0 | 0.84 | 0.67 | 0.7 | 0.64 | 0.84 | 0.69 |
| 1 | 63689707 | rs7556604 | 0.55 | 0.379 | 0.621 | 0.64 | -0.006 | -0.012 | -0.012 | 0 | 0.1 | 0.09 | 0.07 | 0.13 | 0.11 | 0.19 |
| 1 | 63694517 | rs997791 | -0.874 | 0.393 | 0.745 | 0.643 | 0.004 | 0.03 | 0.03 | 0 | 0.21 | 0.29 | 0.21 | 0.29 | 0.19 | 0.42 |
| 1 | 63696679 | rs6661668 | -0.435 | 0.356 | 0.735 | 0.542 | -0.001 | -0.01 | -0.01 | 0.016 | 0.63 | 0.53 | 0.62 | 0.68 | 0.62 | 0.78 |
| 1 | 63716758 | rs621058 | -0.467 | 0.402 | 0.5 | 0.544 | 0 | -0.002 | -0.002 | 7.047 | 0.64 | 0.42 | 0.61 | 0.68 | 0.62 | 0.79 |
| 1 | 63724097 | rs13374496 | 0.218 | 0.558 | 0.605 | 0.553 | 0.01 | -0.037 | -0.037 | 0.957 | 0.07 | 0.18 | 0.06 | 0 | 0.06 | 0.04 |
| 1 | 63727240 | rs7554136 | 0.416 | 0.606 | 0.684 | 0.576 | 0.014 | -0.031 | -0.031 | 0 | 0.07 | 0.14 | 0.05 | 0.01 | 0.06 | 0.04 |
| 1 | 63744393 | rs2031402 | -1.267 | 0.493 | -0.144 | 0.32 | 0.004 | -0.032 | -0.032 | 0 | 0.38 | 0.85 | 0.3 | 0.34 | 0.31 | 0.56 |
| 1 | 63753736 | rs4915891 | -1.056 | 0.608 | -0.024 | 0.375 | -0.001 | 0.003 | 0.003 | 0 | 0.34 | 0.6 | 0.28 | 0.34 | 0.31 | 0.57 |
| 1 | 63773214 | rs12024843 | 0.661 | 0.549 | -0.474 | 0.581 | -0.016 | -0.056 | -0.056 | 0 | 0.09 | 0.01 | 0.05 | 0.17 | 0.07 | 0.18 |
| 1 | 63774201 | rs11208184 | 0.473 | 0.361 | -0.646 | 0.645 | 0.006 | -0.023 | -0.023 | 0 | 0.08 | 0.04 | 0.17 | 0.19 | 0.08 | 0.09 |

| | | | | | | | | | | | | | | | | |
|---|---|---|---|---|---|---|---|---|---|---|---|---|---|---|---|---|
| 1 | 63775056 | rs2367270 | -1.094 | 0.324 | -0.524 | 0.316 | -0.003 | -0.011 | -0.011 | 0 | 0.68 | 0.71 | 0.63 | 0.53 | 0.65 | 0.61 |
| 1 | 63777021 | rs6701595 | 0.679 | 0.422 | -0.472 | 0.148 | -0.032 | 0.089 | 0.089 | 0 | 0.36 | 0.17 | 0.29 | 0.13 | 0.4 | 0.21 |
| 1 | 63780407 | rs6666222 | -0.32 | 0.43 | -0.071 | 0.19 | 0.005 | 0.011 | 0.011 | 0 | 0.57 | 0.38 | 0.53 | 0.62 | 0.59 | 0.63 |
| 1 | 63783472 | rs11208186 | -0.122 | 0.136 | -0.672 | -0.215 | -0.032 | -0.047 | -0.047 | 0.915 | 0.71 | 0.39 | 0.81 | 0.83 | 0.8 | 0.69 |
| 1 | 63785389 | rs2235948 | 0.208 | 0.031 | -0.832 | 0.031 | 0.002 | 0.017 | 0.017 | 0 | 0.47 | 0.3 | 0.42 | 0.6 | 0.43 | 0.59 |
| 1 | 63785711 | rs2235950 | -0.086 | 0.808 | -0.592 | 0.261 | 0.029 | 0.02 | 0.02 | 0 | 0.15 | 0.2 | 0.17 | 0.17 | 0.16 | 0.13 |
| 1 | 63790503 | rs6588025 | -0.024 | 0.77 | -0.532 | 0.234 | -0.007 | -0.019 | -0.019 | 0 | 0.23 | 0.37 | 0.18 | 0.16 | 0.18 | 0.32 |
| 1 | 63793155 | rs10749738 | 0.133 | 1.3 | -0.853 | 0.554 | -0.005 | -0.003 | -0.003 | 0 | 0.77 | 0.76 | 0.63 | 0.67 | 0.68 | 0.75 |
| 1 | 63798761 | rs10489494 | -1.08 | 0.85 | -0.633 | 0.325 | -0.008 | -0.009 | -0.009 | 1.716 | 0.13 | 0.22 | 0.1 | 0.14 | 0.12 | 0.14 |
| 1 | 63799525 | rs12123368 | -0.177 | 0.541 | -0.857 | 0.164 | 0.004 | 0.005 | 0.005 | 0 | 0.14 | 0.1 | 0.08 | 0.08 | 0.15 | 0.15 |
| 1 | 63805122 | rs932748 | -0.441 | 1.035 | -0.933 | -0.097 | 0.005 | -0.02 | -0.02 | 1.157 | 0.93 | 0.91 | 0.89 | 1 | 0.88 | 0.95 |
| 1 | 63805129 | rs932747 | 0.274 | 0.893 | -1.527 | -0.321 | -0.03 | -0.029 | -0.029 | 0.189 | 0.14 | 0.02 | 0.13 | 0.16 | 0.17 | 0.17 |
| 1 | 63818393 | rs6675284 | -0.906 | 1.601 | -1.948 | -0.33 | -0.028 | 0.025 | 0.025 | 0 | 0.38 | 0.22 | 0.31 | 0.35 | 0.43 | 0.46 |
| 1 | 63820276 | rs12407262 | 1.325 | 0.578 | -1.63 | -0.952 | 0.007 | 0.007 | 0.007 | 0 | 0.26 | 0.19 | 0.16 | 0.18 | 0.13 | 0.2 |
| 1 | 63821329 | rs7522426 | 0.571 | 1.335 | -1.694 | -0.619 | 0.01 | 0.013 | 0.013 | 0 | 0.37 | 0.26 | 0.22 | 0.3 | 0.3 | 0.29 |
| 1 | 63832640 | rs12119056 | -0.004 | 1.287 | -1.145 | -0.566 | -0.003 | 0.008 | 0.008 | 0 | 0.41 | 0.38 | 0.52 | 0.64 | 0.44 | 0.48 |
| 1 | 63835901 | rs10889417 | -1.662 | 1.209 | -1.206 | -0.694 | -0.025 | 0.024 | 0.024 | 0 | 0.89 | 0.95 | 0.84 | 0.97 | 0.79 | 0.94 |
| 1 | 63842972 | rs11208199 | 0.427 | 0.912 | -0.902 | -0.619 | -0.002 | -0.007 | -0.007 | 0 | 0.35 | 0.4 | 0.41 | 0.63 | 0.3 | 0.43 |
| 1 | 63844736 | rs4915899 | 1.34 | 0.791 | -1.031 | -0.777 | 0.074 | 0.082 | 0.082 | 0 | 0.3 | 0.07 | 0.17 | 0.18 | 0.18 | 0.23 |
| 1 | 63845218 | rs10749739 | -0.411 | 1.02 | -0.83 | -0.727 | -0.028 | 0.02 | 0.02 | 0 | 0.77 | 0.85 | 0.66 | 0.85 | 0.57 | 0.84 |
| 1 | 63856892 | rs11208204 | 1.109 | 0.387 | -0.495 | -0.626 | 0.038 | 0.035 | 0.035 | 0 | 0.27 | 0.19 | 0.16 | 0.18 | 0.15 | 0.18 |
| 1 | 63865055 | rs2886919 | 0.877 | 0.673 | -0.505 | -0.436 | -0.001 | 0.002 | 0.002 | 0 | 0.37 | 0.36 | 0.47 | 0.48 | 0.33 | 0.48 |
| 1 | 63871758 | rs2273122 | 1.182 | 0.612 | -0.474 | -0.456 | 0.028 | 0.025 | 0.025 | 0.009 | 0.28 | 0.22 | 0.17 | 0.19 | 0.15 | 0.18 |
| 1 | 63871881 | rs1012872 | 0.912 | 0.65 | -0.466 | -0.46 | -0.001 | 0.002 | 0.002 | 0 | 0.37 | 0.36 | 0.47 | 0.48 | 0.33 | 0.48 |
| 1 | 63872871 | rs6588028 | 0.199 | 0.624 | -0.538 | -0.472 | -0.041 | -0.015 | -0.015 | 0 | 0.67 | 0.84 | 0.66 | 0.67 | 0.47 | 0.65 |
| 1 | 63874182 | rs10789139 | 0.75 | 0.674 | -0.54 | -0.442 | -0.005 | 0.009 | 0.009 | 0 | 0.38 | 0.55 | 0.49 | 0.48 | 0.33 | 0.48 |
| 1 | 63874916 | rs6657738 | 0.463 | 0.871 | -0.714 | -0.406 | -0.035 | -0.033 | -0.033 | 0 | 0.16 | 0.07 | 0.1 | 0.19 | 0.21 | 0.25 |
| 1 | 63881934 | rs11208207 | 1.517 | 0.878 | -0.734 | -0.38 | -0.053 | -0.061 | -0.061 | 0.369 | 0.1 | 0.01 | 0.05 | 0.15 | 0.14 | 0.13 |
| 1 | 63887514 | rs9436228 | -1.919 | 0.89 | -0.663 | -0.367 | -0.067 | 0.026 | 0.026 | 0 | 0.89 | 0.99 | 0.81 | 0.97 | 0.78 | 0.95 |
| 1 | 63900901 | rs10493345 | 1.002 | -0.11 | 0.113 | -0.356 | -0.002 | 0.001 | 0.001 | 0.02 | 0.28 | 0.3 | 0.17 | 0.19 | 0.15 | 0.18 |
| 1 | 63908980 | rs6671446 | 0.912 | -0.512 | 0.299 | -0.328 | -0.063 | -0.052 | -0.052 | 1.349 | 0.28 | 0.42 | 0.18 | 0.19 | 0.14 | 0.18 |
| 1 | 63918388 | rs1057815 | -0.926 | -0.566 | 0.492 | -0.352 | -0.052 | -0.043 | -0.043 | 0 | 0.69 | 0.37 | 0.8 | 0.81 | 0.85 | 0.82 |
| 1 | 63956190 | rs6588043 | 0.826 | -0.5 | 0.435 | -0.377 | 0.008 | -0.026 | -0.026 | 0.066 | 0.36 | 0.26 | 0.47 | 0.48 | 0.33 | 0.48 |
| 1 | 63991267 | rs217478 | 0.817 | -0.489 | 0.122 | -0.373 | 0.002 | -0.007 | -0.007 | 0 | 0.37 | 0.3 | 0.47 | 0.48 | 0.33 | 0.48 |
| 1 | 63999868 | rs6657480 | 0.794 | -0.529 | 0.043 | -0.386 | -0.077 | -0.063 | -0.063 | 0 | 0.31 | 0.58 | 0.2 | 0.19 | 0.15 | 0.19 |
| 1 | 64002856 | rs217452 | 1.606 | -0.716 | 0.051 | -0.193 | 0.002 | -0.03 | -0.03 | 0 | 0.17 | 0.09 | 0.18 | 0.38 | 0.16 | 0.2 |
| 1 | 64015952 | rs34598062 | 0.767 | -0.159 | 0.435 | -0.01 | 0.074 | 0.057 | 0.057 | 0.011 | 0.26 | 0.07 | 0.16 | 0.19 | 0.14 | 0.17 |
| 1 | 64017830 | rs217483 | -0.842 | -0.038 | 0.414 | -0.015 | -0.021 | 0.005 | 0.005 | 0.091 | 0.83 | 0.91 | 0.77 | 0.84 | 0.68 | 0.9 |
| 1 | 64021096 | rs2273367 | 0.195 | -1.274 | 0.566 | 0.252 | 0.074 | 0.057 | 0.057 | 0.018 | 0.26 | 0.07 | 0.16 | 0.19 | 0.14 | 0.17 |
| 1 | 64038107 | rs6588050 | 0.07 | -1.272 | 0.442 | 0.253 | -0.004 | -0.002 | -0.002 | 2.038 | 0.27 | 0.27 | 0.17 | 0.19 | 0.14 | 0.17 |
| 1 | 64046766 | rs11800361 | 0.07 | -1.272 | 0.442 | 0.253 | -0.004 | -0.002 | -0.002 | 0 | 0.27 | 0.27 | 0.17 | 0.19 | 0.14 | 0.17 |
| 1 | 64056411 | rs11208250 | -0.256 | -0.751 | -1.435 | 0.601 | -0.076 | -0.07 | -0.07 | 0.415 | 0.16 | 0.01 | 0.1 | 0.18 | 0.21 | 0.25 |
| 1 | 64075768 | rs17125177 | -0.153 | -1.014 | -1.119 | 0.445 | 0.073 | 0.05 | 0.05 | 0.325 | 0.25 | 0.07 | 0.15 | 0.19 | 0.12 | 0.18 |
| 1 | 64078224 | rs855300 | -0.021 | -0.562 | -0.817 | 0.655 | 0.041 | 0.038 | 0.038 | 0.039 | 0.21 | 0.07 | 0.15 | 0.15 | 0.12 | 0.16 |
| 1 | 64079677 | rs2269266 | -0.905 | 0.119 | -0.201 | 1.134 | 0.011 | -0.001 | -0.001 | 0.332 | 0.26 | 0.65 | 0.18 | 0.24 | 0.29 | 0.28 |
| 1 | 64085337 | rs2269260 | -0.017 | -0.046 | -0.253 | 1.528 | -0.018 | -0.019 | -0.019 | 0 | 0.1 | 0.07 | 0.06 | 0.12 | 0.16 | 0.13 |
| 1 | 64086901 | rs2301059 | -0.894 | -0.764 | 0.337 | 1.579 | -0.001 | 0.151 | 0.151 | 0.12 | 0.1 | 0.36 | 0.13 | 0.16 | 0.05 | 0.11 |
| 1 | 64087444 | rs3790861 | -0.759 | -0.224 | -0.663 | 1.098 | 0.023 | -0.01 | -0.01 | 0 | 0.14 | 0.07 | 0.13 | 0.22 | 0.08 | 0.15 |
| 1 | 64094780 | rs10489612 | -1.338 | 0.237 | -0.276 | 0.908 | 0.008 | -0.009 | -0.009 | 0 | 0.13 | 0.22 | 0.12 | 0.15 | 0.12 | 0.13 |
| 1 | 64096094 | rs7535245 | -0.735 | -0.668 | -0.518 | 0.937 | 0.03 | 0.018 | 0.018 | 0 | 0.4 | 0.29 | 0.17 | 0.3 | 0.25 | 0.3 |
| 1 | 64098065 | rs2269252 | -0.514 | -0.999 | -0.746 | 1.357 | 0.031 | 0.017 | 0.017 | 0 | 0.36 | 0.19 | 0.27 | 0.27 | 0.28 | 0.28 |
| 1 | 64098910 | rs855307 | -1.357 | -0.927 | -0.913 | 1.029 | 0.058 | 0.11 | 0.11 | 0 | 0.52 | 0.78 | 0.46 | 0.67 | 0.41 | 0.41 |
| 1 | 64102114 | rs2269248 | -0.922 | -0.66 | -0.545 | 1.207 | 0.023 | 0.025 | 0.025 | 0 | 0.42 | 0.67 | 0.41 | 0.62 | 0.37 | 0.32 |
| 1 | 64102590 | rs17301901 | -0.623 | -0.089 | -0.256 | 1.281 | -0.019 | 0.025 | 0.025 | 0 | 0.05 | 0.01 | 0.07 | 0 | 0.11 | 0.02 |
| 1 | 64105775 | rs2284997 | -1.366 | -0.287 | -0.837 | 1.345 | -0.069 | -0.019 | -0.019 | 0 | 0.56 | 0.93 | 0.68 | 0.6 | 0.52 | 0.67 |
| 1 | 64105799 | rs2284996 | -0.97 | -0.281 | -0.526 | 1.661 | 0.012 | -0.04 | -0.04 | 0 | 0.31 | 0.2 | 0.21 | 0.34 | 0.27 | 0.44 |
| 1 | 64109264 | rs2269240 | -0.698 | -0.174 | 0.075 | 1.765 | -0.002 | -0.001 | -0.001 | 0 | 0.15 | 0.18 | 0.4 | 0.21 | 0.23 | 0.16 |
| 1 | 64117913 | rs2269237 | -0.614 | -0.302 | -0.055 | 1.977 | 0.014 | -0.043 | -0.043 | 0.538 | 0.11 | 0.05 | 0.11 | 0.3 | 0.09 | 0.17 |
| 1 | 64118212 | rs2269236 | -2.681 | -0.409 | 0.661 | 1.854 | -0.001 | -0.003 | -0.003 | 0 | 0.4 | 0.42 | 0.6 | 0.56 | 0.47 | 0.55 |
| 1 | 64118915 | rs2269235 | -1.096 | -0.628 | 1.706 | 1.689 | 0.017 | 0.009 | 0.009 | 0 | 0.17 | 0.39 | 0.41 | 0.21 | 0.22 | 0.17 |
| 1 | 64123868 | rs6671606 | -1.932 | -0.537 | 2.153 | 1.522 | -0.004 | 0.021 | 0.021 | 0 | 0.24 | 0.44 | 0.39 | 0.29 | 0.27 | 0.2 |
| 1 | 64125439 | rs4643 | -0.76 | -0.42 | 2.289 | 1.579 | -0.026 | -0.031 | -0.031 | 0.354 | 0.13 | 0.06 | 0.33 | 0.22 | 0.23 | 0.16 |
| 1 | 64125544 | rs10909 | -2.077 | -0.467 | 2.52 | 1.482 | -0.011 | -0.066 | -0.066 | 0 | 0.13 | 0.33 | 0.08 | 0.02 | 0.14 | 0.11 |
| 1 | 64127004 | rs855350 | -2.839 | -0.472 | 2.384 | 1.495 | 0.001 | 0.006 | 0.006 | 0.158 | 0.3 | 0.71 | 0.44 | 0.24 | 0.38 | 0.27 |
| 1 | 64128209 | rs2013012 | -1.885 | -0.393 | 1.988 | 1.512 | -0.006 | -0.001 | -0.001 | 4.017 | 0.21 | 0.17 | 0.37 | 0.23 | 0.3 | 0.21 |
| 1 | 64129393 | rs855332 | -2.331 | -0.415 | 2.17 | 1.585 | -0.01 | -0.027 | -0.027 | 0 | 0.13 | 0.35 | 0.08 | 0.07 | 0.15 | 0.09 |
| 1 | 64140572 | rs1740402 | -1.824 | -0.41 | 1.833 | 1.76 | 0.001 | -0.002 | -0.002 | 0 | 0.12 | 0.13 | 0.08 | 0.2 | 0.12 | 0.17 |
| 1 | 64149699 | rs12127151 | -1.03 | -0.244 | 1.817 | 1.77 | -0.02 | -0.01 | -0.01 | 0.513 | 0.19 | 0.09 | 0.36 | 0.21 | 0.26 | 0.14 |
| 1 | 64153652 | rs2658399 | -3.477 | -0.555 | 2.124 | 1.884 | 0.027 | -0.019 | -0.019 | 0 | 0.44 | 0.73 | 0.55 | 0.54 | 0.47 | 0.37 |

| | | | | | | | | | | | | | | | | |
|---|---|---|---|---|---|---|---|---|---|---|---|---|---|---|---|---|
| 1 | 64159344 | rs4915644 | -2.388 | -0.543 | 2.624 | 1.84 | -0.017 | 0.054 | 0.054 | 0 | 0.18 | 0.38 | 0.15 | 0.34 | 0.19 | 0.22 |
| 1 | 64161699 | rs17125348 | -1.062 | -0.5 | 2.47 | 1.666 | -0.003 | -0.016 | -0.016 | 0 | 0.07 | 0.26 | 0.05 | 0 | 0.03 | 0.02 |
| 1 | 64162806 | rs4915645 | -1.972 | -0.479 | 2.551 | 1.699 | -0.025 | 0.022 | 0.022 | 1.178 | 0.13 | 0.53 | 0.12 | 0.14 | 0.11 | 0.07 |
| 1 | 64163143 | rs4915646 | -1.549 | -0.448 | 1.925 | 1.722 | -0.022 | 0.008 | 0.008 | 0.276 | 0.12 | 0.28 | 0.1 | 0.14 | 0.1 | 0.07 |
| 1 | 64163348 | rs2174596 | -2.173 | -0.37 | 1.317 | 1.766 | -0.013 | -0.062 | -0.062 | 0 | 0.29 | 0.12 | 0.39 | 0.4 | 0.35 | 0.3 |
| 1 | 64165243 | rs7414489 | -2.704 | -0.363 | 1.654 | 1.478 | 0.006 | -0.006 | -0.006 | 0 | 0.37 | 0.66 | 0.49 | 0.43 | 0.4 | 0.3 |
| 1 | 64165490 | rs12145639 | -1.314 | -0.151 | 1.531 | 1.46 | -0.029 | -0.026 | -0.026 | 0.265 | 0.21 | 0.1 | 0.36 | 0.24 | 0.26 | 0.19 |
| 1 | 64165545 | rs10493350 | -2.052 | -0.184 | 1.813 | 1.575 | -0.041 | 0.024 | 0.024 | 0 | 0.17 | 0.51 | 0.14 | 0.2 | 0.16 | 0.11 |
| 1 | 64166213 | rs2819157 | -1.438 | -0.122 | 1.523 | 1.532 | 0 | 0.005 | 0.005 | 0 | 0.23 | 0.21 | 0.43 | 0.36 | 0.33 | 0.19 |
| 1 | 64168032 | rs1354719 | -1.457 | -0.27 | 1.919 | 0.735 | -0.017 | -0.024 | -0.024 | 0 | 0.13 | 0.15 | 0.06 | 0.07 | 0.1 | 0.1 |
| 1 | 64173484 | rs1506399 | -1.583 | -0.132 | 2.902 | 1.404 | 0.053 | 0.601 | 0.601 | 0 | 0.06 | 0.31 | 0.1 | 0.49 | 0.04 | 0.11 |
| 1 | 64173753 | rs937805 | 2.452 | 0.122 | 2.745 | 0.641 | 0.011 | -0.033 | -0.033 | 0.021 | 0.67 | 0.19 | 0.72 | 0.61 | 0.7 | 0.69 |
| 1 | 64174386 | rs2819159 | -1.353 | -0.078 | 2.829 | 0.565 | 0.005 | 0.011 | 0.011 | 0 | 0.18 | 0.14 | 0.12 | 0.02 | 0.17 | 0.13 |
| 1 | 64177299 | rs4915914 | 2.458 | -0.101 | 3.06 | 0.593 | -0.002 | 0 | 0 | 0 | 0.67 | 0.32 | 0.71 | 0.62 | 0.72 | 0.68 |
| 1 | 64179312 | rs1848024 | 2.328 | -0.089 | 3.069 | 0.597 | 0 | -0.001 | -0.001 | 0 | 0.68 | 0.31 | 0.73 | 0.61 | 0.73 | 0.7 |
| 1 | 64181014 | rs10458570 | 2.492 | -0.098 | 2.986 | 0.555 | 0.015 | -0.011 | -0.011 | 0 | 0.67 | 0.23 | 0.71 | 0.62 | 0.72 | 0.69 |
| 1 | 64189638 | rs4915649 | 2.478 | 0.123 | 2.824 | 0.536 | -0.001 | 0.001 | 0.001 | 0 | 0.65 | 0.36 | 0.71 | 0.62 | 0.69 | 0.65 |
| 1 | 64189713 | rs2819168 | -2.769 | 0.123 | 2.85 | 0.536 | 0.002 | 0.013 | 0.013 | 0 | 0.33 | 0.54 | 0.27 | 0.38 | 0.31 | 0.35 |
| 1 | 64189938 | rs4915650 | 2.254 | 0.223 | 2.583 | -0.689 | 0.008 | -0.139 | -0.139 | 0.369 | 0.7 | 0.51 | 0.81 | 0.97 | 0.72 | 0.77 |
| 1 | 64193197 | rs13374611 | -1.918 | 0.168 | 2.34 | -0.482 | 0.01 | -0.039 | -0.039 | 1.221 | 0.1 | 0.25 | 0.08 | 0.03 | 0.08 | 0.14 |
| 1 | 64195612 | rs17320271 | -1.707 | 0.086 | 2.25 | -0.336 | -0.003 | 0.039 | 0.039 | 0 | 0.15 | 0.29 | 0.08 | 0.21 | 0.11 | 0.25 |
| 1 | 64209711 | rs1463942 | 2.522 | -0.037 | 2.259 | -0.486 | 0.003 | 0.01 | 0.01 | 0 | 0.67 | 0.29 | 0.76 | 0.78 | 0.72 | 0.65 |
| 1 | 64212298 | rs17097089 | -1.706 | -0.034 | 2.32 | -0.485 | 0.001 | 0.083 | 0.083 | 0 | 0.16 | 0.49 | 0.1 | 0.21 | 0.1 | 0.24 |
| 1 | 64217123 | rs1701530 | -2.784 | -0.044 | 2.301 | -0.49 | -0.001 | -0.004 | -0.004 | 0 | 0.32 | 0.66 | 0.23 | 0.22 | 0.28 | 0.35 |
| 1 | 64221188 | rs17125501 | -1.563 | -0.011 | 2.236 | -0.483 | 0.002 | 0.059 | 0.059 | 0 | 0.14 | 0.33 | 0.08 | 0.2 | 0.1 | 0.25 |
| 1 | 64231541 | rs2819130 | 1.11 | 0.06 | 1.39 | -0.608 | 0.001 | -0.039 | -0.039 | 0 | 0.84 | 0.8 | 0.85 | 0.99 | 0.82 | 0.89 |
| 1 | 64238724 | rs589465 | -2.885 | -0.379 | 1.619 | -0.425 | 0.006 | 0.005 | 0.005 | 0.051 | 0.29 | 0.68 | 0.2 | 0.19 | 0.22 | 0.35 |
| 1 | 64240929 | rs620019 | -0.791 | -0.356 | 1.624 | -0.416 | 0.003 | 0.024 | 0.024 | 0.686 | 0.1 | 0.05 | 0.08 | 0.01 | 0.1 | 0.1 |
| 1 | 64245352 | rs583192 | -2.937 | -0.356 | 1.6 | -0.415 | 0.023 | 0.021 | 0.021 | 0.031 | 0.3 | 0.78 | 0.2 | 0.19 | 0.22 | 0.35 |
| 1 | 64247390 | rs619535 | -2.548 | -0.253 | 1.508 | -0.358 | -0.012 | -0.009 | -0.009 | 0 | 0.27 | 0.59 | 0.19 | 0.19 | 0.22 | 0.35 |
| 1 | 64249422 | rs648965 | -0.884 | -0.12 | 1.149 | -0.326 | -0.004 | -0.026 | -0.026 | 0 | 0.1 | 0.13 | 0.09 | 0.01 | 0.1 | 0.1 |
| 1 | 64256018 | rs659406 | -1.634 | -0.08 | 0.574 | -0.252 | -0.007 | -0.006 | -0.006 | 0 | 0.26 | 0.64 | 0.19 | 0.19 | 0.21 | 0.34 |
| 1 | 64262559 | rs2220894 | -1.157 | -0.172 | 0.282 | -0.3 | 0.002 | 0.016 | 0.016 | 0 | 0.16 | 0.14 | 0.13 | 0.07 | 0.15 | 0.14 |
| 1 | 64265539 | rs645558 | -1.865 | -0.741 | 0.59 | -0.393 | 0.016 | 0.004 | 0.004 | 0 | 0.29 | 0.74 | 0.19 | 0.24 | 0.19 | 0.35 |
| 1 | 64267606 | rs640407 | -1.764 | -0.711 | 1.236 | -0.479 | -0.012 | -0.005 | -0.005 | 5.517 | 0.28 | 0.64 | 0.19 | 0.23 | 0.2 | 0.35 |
| 1 | 64274356 | rs645945 | -1.856 | -0.602 | 1.572 | -0.276 | -0.021 | 0.151 | 0.151 | 1.988 | 0.07 | 0.64 | 0.09 | 0.19 | 0.02 | 0.2 |
| 1 | 64291940 | rs683574 | -1.785 | -0.441 | 1.248 | -1.11 | 0.006 | 0.016 | 0.016 | 0 | 0.29 | 0.74 | 0.2 | 0.08 | 0.21 | 0.34 |
| 1 | 64295971 | rs685458 | -1.701 | -0.436 | 1.566 | -1.107 | -0.006 | -0.042 | -0.042 | 0 | 0.28 | 0.64 | 0.19 | 0.08 | 0.21 | 0.34 |
| 1 | 64309093 | rs4915931 | -0.231 | -0.174 | 0.057 | -1.916 | -0.007 | 0.093 | 0.093 | 0 | 0.27 | 0.13 | 0.17 | 0.04 | 0.26 | 0.19 |
| 1 | 64309302 | rs583102 | -1.408 | -0.126 | 0.196 | -2.03 | 0.006 | 0.158 | 0.158 | 0 | 0.39 | 0.85 | 0.32 | 0.14 | 0.33 | 0.43 |
| 1 | 64312043 | rs677806 | -1.311 | -0.303 | 0.331 | -2.026 | 0.019 | 0.115 | 0.115 | 0 | 0.38 | 0.79 | 0.31 | 0.14 | 0.31 | 0.43 |
| 1 | 64312919 | rs663458 | -1.311 | -0.303 | 0.333 | -2.026 | 0.019 | 0.117 | 0.117 | 0 | 0.38 | 0.79 | 0.31 | 0.14 | 0.31 | 0.43 |
| 1 | 64313498 | rs2132161 | -1.649 | -0.404 | 0.554 | -2.118 | 0.006 | 0.028 | 0.028 | 0 | 0.5 | 0.45 | 0.32 | 0.11 | 0.42 | 0.42 |
| 1 | 64313696 | rs649608 | -1.398 | -0.416 | 0.691 | -1.895 | -0.016 | -0.001 | -0.001 | 0 | 0.11 | 0.63 | 0.14 | 0.1 | 0.05 | 0.23 |
| 1 | 64320736 | rs11208306 | -0.581 | -0.346 | 0.507 | -1.741 | -0.002 | 0.087 | 0.087 | 0 | 0.28 | 0.16 | 0.17 | 0.04 | 0.26 | 0.2 |
| 1 | 64339801 | rs7513180 | -0.699 | -0.811 | 0.497 | -1.403 | -0.037 | -0.128 | -0.128 | 0.163 | 0.27 | 0.42 | 0.14 | 0.03 | 0.18 | 0.18 |
| 1 | 64340452 | rs11208308 | -0.939 | -0.536 | -0.095 | -1.377 | 0.02 | 0.032 | 0.032 | 0 | 0.44 | 0.6 | 0.24 | 0.1 | 0.35 | 0.31 |
| 1 | 64348148 | rs662986 | 0.845 | -0.638 | 0.844 | -0.875 | -0.044 | -0.07 | -0.07 | 0.267 | 0.88 | 0.67 | 0.93 | 1 | 0.94 | 0.93 |
| 1 | 64350084 | rs637420 | 0.89 | -0.704 | 0.923 | -0.972 | -0.031 | -0.06 | -0.06 | 0 | 0.88 | 0.84 | 0.94 | 1 | 0.93 | 0.93 |
| 1 | 64352117 | rs855850 | 1.19 | -0.721 | 0.953 | -0.98 | -0.041 | -0.074 | -0.074 | 6.681 | 0.87 | 0.74 | 0.93 | 1 | 0.92 | 0.93 |
| 1 | 64353473 | rs855849 | 2.026 | -0.798 | 0.922 | -1.419 | 0.041 | 0.08 | 0.08 | 0 | 0.73 | 0.14 | 0.75 | 0.96 | 0.77 | 0.81 |
| 1 | 64354536 | rs855845 | 1.276 | -0.824 | 1.948 | -1.412 | -0.059 | -0.1 | -0.1 | 0 | 0.85 | 0.55 | 0.9 | 0.99 | 0.91 | 0.91 |
| 1 | 64357060 | rs855835 | 0.514 | -0.817 | 1.912 | -1.408 | -0.03 | -0.06 | -0.06 | 0 | 0.88 | 0.83 | 0.93 | 1 | 0.93 | 0.93 |
| 1 | 64361960 | rs855823 | 1.311 | -0.847 | 2.605 | -1.425 | -0.066 | -0.112 | -0.112 | 0 | 0.84 | 0.59 | 0.9 | 0.99 | 0.91 | 0.91 |
| 1 | 64374159 | rs17358977 | -0.711 | -0.754 | 2.341 | -1.386 | -0.004 | 0.007 | 0.007 | 0 | 0.09 | 0.06 | 0.1 | 0.02 | 0.12 | 0.08 |
| 1 | 64381917 | rs705529 | 1.186 | -0.581 | 2.754 | -1.522 | -0.021 | -0.085 | -0.085 | 0.047 | 0.85 | 0.69 | 0.89 | 0.99 | 0.88 | 0.89 |
| 1 | 64383389 | rs3861946 | 2.142 | -0.238 | 1.517 | -2.148 | -0.101 | 0.241 | 0.241 | 0.508 | 0.51 | 0.04 | 0.45 | 0.82 | 0.52 | 0.56 |
| 1 | 64412138 | rs6660751 | -2.06 | -0.218 | 2.209 | -2.066 | -0.003 | -0.028 | -0.028 | 0.019 | 0.26 | 0.65 | 0.22 | 0.05 | 0.22 | 0.17 |
| 1 | 64417555 | rs10889456 | -1.884 | -0.237 | 2.257 | -2.065 | -0.027 | -0.108 | -0.108 | 0.417 | 0.24 | 0.5 | 0.2 | 0.05 | 0.22 | 0.17 |
| 1 | 64435108 | rs1530887 | 2.055 | -0.691 | 1.393 | -1.825 | -0.129 | -0.076 | -0.076 | 0 | 0.38 | 0.03 | 0.39 | 0.65 | 0.4 | 0.33 |
| 1 | 64442169 | rs285339 | 1.183 | -0.617 | 1.695 | -1.768 | -0.02 | -0.085 | -0.085 | 0.031 | 0.76 | 0.62 | 0.8 | 0.95 | 0.78 | 0.83 |
| 1 | 64443641 | rs285342 | 1.282 | -0.617 | 1.663 | -1.767 | -0.027 | -0.108 | -0.108 | 0.378 | 0.76 | 0.49 | 0.79 | 0.95 | 0.78 | 0.83 |
| 1 | 64459519 | rs285364 | 1.578 | -0.644 | 1.445 | -1.801 | 0.028 | 0.055 | 0.055 | 0 | 0.72 | 0.19 | 0.77 | 0.95 | 0.78 | 0.81 |
| 1 | 64462582 | rs285370 | 1.451 | -0.649 | 1.506 | -1.805 | -0.014 | -0.053 | -0.053 | 0 | 0.73 | 0.35 | 0.78 | 0.95 | 0.77 | 0.83 |
| 1 | 64470166 | rs285386 | 1.316 | -0.622 | 1.63 | -1.793 | -0.03 | -0.114 | -0.114 | 0 | 0.75 | 0.52 | 0.79 | 0.95 | 0.78 | 0.83 |
| 1 | 64484485 | rs834485 | -0.362 | -0.817 | 0.907 | -1.411 | 0.006 | 0.011 | 0.011 | 0 | 0.39 | 0.19 | 0.34 | 0.3 | 0.34 | 0.49 |
| 1 | 64486207 | rs834487 | -0.584 | -0.893 | 0.981 | -1.433 | -0.023 | -0.063 | -0.063 | 0 | 0.89 | 0.79 | 0.93 | 1 | 0.92 | 0.93 |
| 1 | 64490893 | rs1341511 | 0.958 | -0.42 | 0.39 | -1.299 | -0.014 | 0.011 | 0.011 | 0 | 0.45 | 0.24 | 0.52 | 0.63 | 0.53 | 0.4 |
| 1 | 64498297 | rs1746927 | 1.584 | -0.165 | 0.857 | -1.2 | 0.023 | 0.071 | 0.071 | 0.511 | 0.72 | 0.19 | 0.8 | 0.86 | 0.76 | 0.75 |

| | | | | | | | | | | | | | | | | |
|---|---|---|---|---|---|---|---|---|---|---|---|---|---|---|---|---|
| 1 | 64508126 | rs705532 | 1.181 | -0.196 | 1.257 | -1.192 | -0.018 | -0.066 | -0.066 | 0 | 0.76 | 0.41 | 0.84 | 0.92 | 0.8 | 0.78 |
| 1 | 64515379 | rs1772626 | 0.75 | -0.233 | 0.988 | -1.138 | -0.034 | -0.11 | -0.11 | 0 | 0.8 | 0.55 | 0.86 | 0.94 | 0.83 | 0.8 |
| 1 | 64519223 | rs3844055 | 1.038 | -0.035 | 0.619 | -1.164 | -0.02 | -0.08 | -0.08 | 0.065 | 0.66 | 0.5 | 0.75 | 0.87 | 0.67 | 0.78 |
| 1 | 64520155 | rs7540568 | -0.561 | 0.689 | 0.81 | -1.1 | 0.063 | -0.026 | -0.026 | 0 | 0.06 | 0.27 | 0.08 | 0.01 | 0.08 | 0.07 |
| 1 | 64525961 | rs863087 | 0.145 | 0.68 | 0.599 | -1.197 | 0 | 0.012 | 0.012 | 0 | 0.84 | 0.89 | 0.85 | 0.92 | 0.8 | 0.85 |
| 1 | 64527865 | rs1149262 | 1.235 | 0.498 | 0.876 | -1.273 | 0 | -0.014 | -0.014 | 0 | 0.74 | 0.28 | 0.75 | 0.91 | 0.72 | 0.75 |
| 1 | 64531564 | rs6676244 | -0.802 | 0.57 | 0.878 | -1.143 | 0.097 | -0.02 | -0.02 | 0.092 | 0.08 | 0.39 | 0.09 | 0.01 | 0.08 | 0.07 |
| 1 | 64534922 | rs1260773 | -1.722 | 0.744 | -0.099 | -0.795 | -0.033 | 0.082 | 0.082 | 0.392 | 0.39 | 0.83 | 0.4 | 0.24 | 0.4 | 0.37 |
| 1 | 64537743 | rs11580998 | 0.336 | -0.484 | -1.374 | -0.633 | 0.024 | -0.072 | -0.072 | 1.546 | 0.19 | 0 | 0.06 | 0.23 | 0.11 | 0.22 |
| 1 | 64538961 | rs1747924 | 0.126 | -0.714 | -1.214 | -1.463 | -0.02 | -0.106 | -0.106 | 0 | 0.69 | 0.43 | 0.79 | 0.97 | 0.78 | 0.85 |
| 1 | 64540133 | rs1772618 | 0.585 | -1.08 | -2.178 | -1.486 | 0.064 | -0.088 | -0.088 | 1.963 | 0.29 | 0.03 | 0.12 | 0.3 | 0.2 | 0.21 |
| 1 | 64542800 | rs2360720 | 1.137 | -0.335 | -2.503 | -1.703 | -0.083 | -0.116 | -0.116 | 0 | 0.31 | 0.07 | 0.54 | 0.6 | 0.36 | 0.48 |
| 1 | 64545124 | rs834343 | -0.454 | -0.179 | -2.253 | -1.822 | 0.006 | -0.042 | -0.042 | 0 | 0.79 | 0.26 | 0.8 | 0.98 | 0.75 | 0.86 |
| 1 | 64549091 | rs834350 | 0.591 | -0.238 | -2.381 | -1.911 | -0.087 | 0.203 | 0.203 | 0.622 | 0.6 | 0.1 | 0.66 | 0.92 | 0.58 | 0.71 |
| 1 | 64550630 | rs6687610 | 0.917 | -0.938 | -2.199 | -1.724 | 0.001 | 0.005 | 0.005 | 0.216 | 0.19 | 0.16 | 0.13 | 0.06 | 0.18 | 0.15 |
| 1 | 64551422 | rs1619058 | -1.12 | -0.823 | -1.436 | -1.704 | 0.041 | -0.08 | -0.08 | 0 | 0.87 | 0.61 | 0.9 | 1 | 0.85 | 0.93 |
| 1 | 64553040 | rs11208349 | 0.553 | -0.834 | -1.357 | -1.71 | 0.024 | -0.092 | -0.092 | 0 | 0.14 | 0.51 | 0.12 | 0 | 0.15 | 0.07 |
| 1 | 64554013 | rs11208351 | 0.75 | -0.725 | -1.722 | -1.484 | -0.016 | -0.016 | -0.016 | 0 | 0.38 | 0.3 | 0.62 | 0.61 | 0.46 | 0.54 |
| 1 | 64555255 | rs7533927 | -0.546 | -0.219 | -0.977 | -1.533 | -0.003 | -0.042 | -0.042 | 0 | 0.32 | 0.61 | 0.25 | 0.06 | 0.32 | 0.21 |
| 1 | 64555464 | rs7533960 | -0.508 | 0.06 | -1.057 | -1.432 | 0.004 | -0.078 | -0.078 | 0 | 0.29 | 0.53 | 0.2 | 0.06 | 0.29 | 0.2 |
| 1 | 64560089 | rs1073712 | -0.122 | 0.101 | -1.781 | -0.444 | -0.011 | -0.106 | -0.106 | 0.499 | 0.27 | 0.08 | 0.38 | 0.55 | 0.31 | 0.33 |
| 1 | 64566396 | rs2806541 | 0.491 | -0.198 | -2.022 | -0.874 | -0.062 | -0.037 | -0.037 | 0 | 0.4 | 0.13 | 0.33 | 0.38 | 0.45 | 0.43 |
| 1 | 64567779 | rs2806545 | 0.438 | -0.438 | -2.156 | -0.974 | -0.078 | -0.041 | -0.041 | 0.61 | 0.41 | 0.1 | 0.32 | 0.37 | 0.46 | 0.44 |
| 1 | 64573067 | rs7521771 | 0.438 | -0.176 | -1.254 | -1.131 | 0.004 | -0.056 | -0.056 | 0 | 0.13 | 0.21 | 0.07 | 0 | 0.14 | 0.05 |
| 1 | 64573161 | rs12036326 | 0.825 | 0.216 | -1.62 | -1.214 | -0.02 | 0.023 | 0.023 | 0 | 0.16 | 0.07 | 0.12 | 0.08 | 0.17 | 0.13 |
| 1 | 64577770 | rs2806535 | 0.702 | -0.412 | -1.164 | -1.588 | -0.001 | -0.003 | -0.003 | 0 | 0.4 | 0.4 | 0.39 | 0.41 | 0.4 | 0.49 |
| 1 | 64579157 | rs2806537 | 0.709 | -0.225 | -0.745 | -1.985 | 0.009 | -0.137 | -0.137 | 0.872 | 0.66 | 0.56 | 0.79 | 0.96 | 0.69 | 0.77 |
| 1 | 64579350 | rs6677577 | 1.09 | -0.102 | -0.644 | -2.038 | 0.006 | -0.061 | -0.061 | 0 | 0.12 | 0.19 | 0.08 | 0.01 | 0.1 | 0.08 |
| 1 | 64582357 | rs17357054 | -0.868 | 0.143 | -1.192 | -1.916 | 0.006 | -0.068 | -0.068 | 1.428 | 0.31 | 0.34 | 0.2 | 0.04 | 0.3 | 0.18 |
| 1 | 64590297 | rs12127830 | 0.966 | 0.379 | -2.011 | -1.644 | -0.001 | 0.001 | 0.001 | 0 | 0.15 | 0.08 | 0.09 | 0.1 | 0.12 | 0.17 |
| 1 | 64590697 | rs1408417 | -0.193 | 0.511 | -1.693 | -1.567 | -0.069 | 0.078 | 0.078 | 0 | 0.17 | 0.03 | 0.11 | 0.03 | 0.2 | 0.1 |
| 1 | 64592838 | rs10489904 | 0.778 | 0.415 | -1.185 | -1.108 | -0.024 | 0.036 | 0.036 | 0 | 0.09 | 0.02 | 0.07 | 0.01 | 0.1 | 0.08 |
| 1 | 64601845 | rs7528814 | -0.579 | 0.189 | -0.972 | -1.088 | -0.004 | -0.028 | -0.028 | 0 | 0.24 | 0.66 | 0.17 | 0.04 | 0.18 | 0.12 |
| 1 | 64603898 | rs10399612 | 0.067 | 0.558 | -1.464 | -0.911 | -0.001 | 0.021 | 0.021 | 0 | 0.19 | 0.11 | 0.12 | 0.04 | 0.17 | 0.12 |
| 1 | 64612756 | rs2806542 | 0.041 | 1.117 | -1.715 | -0.803 | -0.106 | 0.138 | 0.138 | 0 | 0.21 | 0.02 | 0.2 | 0.01 | 0.31 | 0.17 |
| 1 | 64614011 | rs6588083 | -0.455 | 0.401 | -0.804 | 0.414 | -0.016 | 0.109 | 0.109 | 0.06 | 0.2 | 0.33 | 0.37 | 0.56 | 0.21 | 0.3 |
| 1 | 64615775 | rs1535872 | -0.75 | 0.266 | -0.912 | -0.001 | -0.004 | 0.008 | 0.008 | 0.251 | 0.36 | 0.55 | 0.49 | 0.6 | 0.36 | 0.38 |
| 1 | 64622069 | rs4619008 | -0.182 | 0.411 | -1.378 | 0.343 | -0.013 | -0.029 | -0.029 | 0 | 0.26 | 0.18 | 0.31 | 0.36 | 0.31 | 0.32 |
| 1 | 64624403 | rs4565754 | -0.685 | 0.333 | -1.528 | 0.25 | -0.01 | -0.05 | -0.05 | 0 | 0.36 | 0.22 | 0.36 | 0.41 | 0.4 | 0.39 |
| 1 | 64632885 | rs4423051 | 0.371 | 0.762 | -2.148 | -0.471 | -0.026 | 0.075 | 0.075 | 0 | 0.25 | 0.13 | 0.23 | 0.06 | 0.31 | 0.24 |
| 1 | 64634163 | rs7515058 | 1.549 | 0.266 | -2.608 | -0.708 | -0.042 | 0.059 | 0.059 | 0 | 0.16 | 0.01 | 0.1 | 0.06 | 0.17 | 0.09 |
| 1 | 64643277 | rs7527017 | 0.488 | 0.567 | -2.487 | -0.262 | -0.13 | -0.083 | -0.083 | 0 | 0.25 | 0.04 | 0.3 | 0.31 | 0.35 | 0.29 |
| 1 | 64647608 | rs11208377 | -0.443 | 0.052 | -1.658 | -0.397 | -0.004 | 0.002 | 0.002 | 0.543 | 0.58 | 0.57 | 0.56 | 0.42 | 0.66 | 0.57 |
| 1 | 64656538 | rs4644531 | -0.536 | -0.034 | -1.295 | -0.44 | 0.023 | -0.016 | -0.016 | 0 | 0.6 | 0.76 | 0.59 | 0.39 | 0.66 | 0.57 |
| 1 | 64656971 | rs11208380 | 0.71 | -0.382 | -0.516 | -0.265 | -0.045 | -0.001 | -0.001 | 0 | 0.18 | 0.35 | 0.09 | 0.1 | 0.1 | 0.1 |
| 1 | 64657942 | rs4304629 | -0.066 | -0.359 | -0.781 | -0.474 | 0.007 | 0.011 | 0.011 | 0 | 0.52 | 0.71 | 0.44 | 0.38 | 0.53 | 0.42 |
| 1 | 64658597 | rs4266926 | -0.035 | -0.239 | -0.773 | -0.439 | 0.013 | 0.021 | 0.021 | 0 | 0.51 | 0.71 | 0.44 | 0.38 | 0.51 | 0.41 |
| 1 | 64658676 | rs4454578 | 1.033 | -0.357 | -0.467 | -0.444 | -0.024 | -0.001 | -0.001 | 0.124 | 0.17 | 0.27 | 0.08 | 0.1 | 0.1 | 0.09 |
| 1 | 64665046 | rs7545208 | 0.22 | -0.352 | -0.491 | -0.514 | 0.039 | 0.043 | 0.043 | 0.6 | 0.5 | 0.36 | 0.53 | 0.29 | 0.63 | 0.48 |
| 1 | 64674428 | rs2132417 | 0.369 | -0.305 | -0.528 | -0.338 | 0.002 | -0.002 | -0.002 | 0 | 0.47 | 0.34 | 0.41 | 0.41 | 0.5 | 0.39 |
| 1 | 64677055 | rs2806532 | -0.596 | -0.624 | -0.536 | -0.279 | -0.007 | 0.006 | 0.006 | 0.323 | 0.68 | 0.7 | 0.64 | 0.4 | 0.75 | 0.57 |
| 1 | 64679749 | rs825175 | -0.943 | 0.344 | -0.403 | -0.05 | -0.055 | 0.001 | 0.001 | 0 | 0.69 | 0.85 | 0.65 | 0.7 | 0.59 | 0.7 |
| 1 | 64684038 | rs7526950 | 0.802 | 0.177 | -0.246 | -0.028 | -0.009 | -0.002 | -0.002 | 0 | 0.18 | 0.13 | 0.19 | 0.01 | 0.22 | 0.19 |
| 1 | 64693901 | rs17126246 | 0.825 | -0.062 | -0.045 | 0.548 | 0.013 | 0.144 | 0.144 | 0 | 0.09 | 0.15 | 0.26 | 0.41 | 0.11 | 0.17 |
| 1 | 64694167 | rs17126247 | 0.825 | -0.062 | -0.045 | 0.548 | 0.013 | 0.144 | 0.144 | 0 | 0.09 | 0.15 | 0.26 | 0.41 | 0.11 | 0.17 |
| 1 | 64696077 | rs17126249 | 2.222 | 0.25 | -0.157 | 0.516 | -0.022 | -0.02 | -0.02 | 0 | 0.06 | 0 | 0.04 | 0.09 | 0.07 | 0.14 |
| 1 | 64698411 | rs2029868 | 0.495 | -0.39 | 0.003 | 0.355 | 0.073 | 0.026 | 0.026 | 0 | 0.39 | 0.14 | 0.17 | 0.29 | 0.3 | 0.33 |
| 1 | 64699602 | rs825184 | -0.581 | -0.232 | 0.295 | 0.434 | 0.012 | -0.013 | -0.013 | 2.101 | 0.6 | 0.55 | 0.38 | 0.32 | 0.54 | 0.53 |
| 1 | 64703324 | rs11580076 | 0.568 | -0.319 | -0.232 | 0.411 | -0.004 | 0.035 | 0.035 | 0 | 0.16 | 0.16 | 0.25 | 0.39 | 0.14 | 0.24 |
| 1 | 64709891 | rs10789152 | 0.751 | -0.376 | -0.196 | 0.356 | 0.051 | -0.037 | -0.037 | 0 | 0.33 | 0.17 | 0.18 | 0.33 | 0.27 | 0.27 |
| 1 | 64725787 | rs705540 | -0.151 | -0.217 | -0.221 | 0.31 | 0.007 | 0.048 | 0.048 | 0 | 0.52 | 0.23 | 0.35 | 0.34 | 0.5 | 0.45 |
| 1 | 64727239 | rs11588217 | 0.471 | 0.31 | -0.019 | 0.619 | -0.001 | 0.059 | 0.059 | 0 | 0.14 | 0.16 | 0.24 | 0.37 | 0.14 | 0.21 |
| 1 | 64737015 | rs705551 | -0.432 | 0.112 | 0.223 | 0.558 | 0.019 | 0.088 | 0.088 | 0 | 0.21 | 0.56 | 0.31 | 0.37 | 0.21 | 0.24 |
| 1 | 64746781 | rs1983299 | 0.596 | -0.152 | 0.102 | 0.505 | 0.032 | -0.003 | -0.003 | 0 | 0.36 | 0.28 | 0.23 | 0.31 | 0.23 | 0.27 |
| 1 | 64747636 | rs825241 | 0.207 | -0.114 | 0.274 | 0.496 | -0.007 | -0.001 | -0.001 | 0.124 | 0.39 | 0.43 | 0.27 | 0.31 | 0.3 | 0.27 |
| 1 | 64749809 | rs825239 | -1.832 | 0.087 | 0.032 | 0.472 | -0.161 | -0.075 | -0.075 | 5.288 | 0.73 | 0.98 | 0.66 | 0.69 | 0.62 | 0.67 |
| 1 | 64752871 | rs705559 | -1.642 | 0.033 | 0.559 | 0.39 | 0.12 | 0.007 | 0.007 | 2.665 | 0.07 | 0.56 | 0.08 | 0 | 0.07 | 0.01 |
| 1 | 64753640 | rs17126275 | 0.784 | -0.006 | 0.602 | 0.408 | -0.001 | 0.042 | 0.042 | 0 | 0.15 | 0.15 | 0.25 | 0.37 | 0.15 | 0.24 |
| 1 | 64755943 | rs12726057 | 0.167 | -0.019 | 0.412 | 0.409 | 0.099 | 0.083 | 0.083 | 0 | 0.5 | 0.15 | 0.32 | 0.32 | 0.4 | 0.42 |

| | | | | | | | | | | | | | | | | |
|---|---|---|---|---|---|---|---|---|---|---|---|---|---|---|---|---|
| 1 | 64756763 | rs12727354 | 0.705 | -0.006 | 0.438 | 0.444 | 0.005 | 0.119 | 0.119 | 0 | 0.15 | 0.26 | 0.26 | 0.37 | 0.15 | 0.24 |
| 1 | 64761515 | rs17266284 | 0.626 | -0.167 | 0.556 | 0.216 | 0.088 | -0.017 | -0.017 | 0 | 0.34 | 0.12 | 0.21 | 0.31 | 0.22 | 0.26 |
| 1 | 64762226 | rs825213 | -0.013 | -0.137 | 0.694 | 0.187 | 0.022 | -0.115 | -0.115 | 0 | 0.21 | 0.41 | 0.17 | 0.01 | 0.25 | 0.17 |
| 1 | 64762259 | rs6683502 | 0.496 | -0.15 | 0.667 | 0.259 | 0.015 | 0 | 0 | 0 | 0.36 | 0.3 | 0.23 | 0.31 | 0.22 | 0.26 |
| 1 | 64784574 | rs957576 | -0.606 | -0.162 | 0.331 | 0.4 | -0.002 | -0.001 | -0.001 | 0 | 0.55 | 0.57 | 0.39 | 0.37 | 0.41 | 0.47 |
| 1 | 64785386 | rs12408613 | 0.16 | -0.295 | 0.35 | 0.494 | -0.037 | -0.004 | -0.004 | 0 | 0.38 | 0.46 | 0.25 | 0.31 | 0.23 | 0.3 |
| 1 | 64790514 | rs11208420 | 0.477 | -0.228 | 0.107 | 0.548 | 0.089 | -0.029 | -0.029 | 0 | 0.35 | 0.09 | 0.21 | 0.32 | 0.23 | 0.31 |
| 1 | 64791688 | rs4915972 | 0.439 | -0.121 | 0.386 | 0.711 | 0.02 | 0.157 | 0.157 | 0 | 0.15 | 0.38 | 0.23 | 0.33 | 0.14 | 0.23 |
| 1 | 64794749 | rs10889476 | 0.579 | 0.055 | 0.309 | 0.633 | -0.04 | 0.043 | 0.043 | 0.161 | 0.44 | 0.08 | 0.48 | 0.35 | 0.55 | 0.45 |
| 1 | 64800809 | rs1390475 | 0.494 | 0.076 | 0.657 | 0.644 | 0.013 | 0.046 | 0.046 | 0.003 | 0.46 | 0.34 | 0.5 | 0.35 | 0.55 | 0.44 |
| 1 | 64804300 | rs4915665 | -0.941 | 0.195 | 0.391 | 0.584 | -0.009 | -0.006 | -0.006 | 0.19 | 0.33 | 0.26 | 0.39 | 0.4 | 0.39 | 0.39 |
| 1 | 64804890 | rs10889478 | 1.174 | 0.108 | 0.525 | 0.595 | -0.001 | 0 | 0 | 0 | 0.32 | 0.34 | 0.4 | 0.29 | 0.45 | 0.3 |
| 1 | 64805503 | rs1908635 | 0.304 | 0.172 | 0.487 | 0.604 | -0.001 | -0.017 | -0.017 | 0.438 | 0.18 | 0.18 | 0.16 | 0.06 | 0.18 | 0.15 |
| 1 | 64806201 | rs17126413 | 0.545 | 0.251 | 0.011 | 0.722 | 0.001 | 0.046 | 0.046 | 0 | 0.14 | 0.16 | 0.22 | 0.34 | 0.15 | 0.23 |
| 1 | 64806671 | rs10889479 | 0.344 | 0.252 | 0.003 | 0.713 | 0 | -0.007 | -0.007 | 2.149 | 0.69 | 0.73 | 0.63 | 0.6 | 0.68 | 0.62 |
| 1 | 64825150 | rs2047284 | -0.661 | 0.522 | -0.093 | 0.431 | -0.006 | 0.05 | 0.05 | 0.047 | 0.83 | 0.92 | 0.85 | 0.94 | 0.83 | 0.83 |
| 1 | 64827193 | rs11208433 | 0.665 | 0.701 | 0.308 | 0.62 | 0.05 | -0.004 | -0.004 | 0 | 0.3 | 0.45 | 0.37 | 0.31 | 0.38 | 0.3 |
| 1 | 64834710 | rs1514966 | 0.444 | -0.231 | 0.267 | 0.379 | 0.066 | 0.026 | 0.026 | 0 | 0.49 | 0.38 | 0.53 | 0.64 | 0.64 | 0.6 |
| 1 | 64837143 | rs11208437 | -0.509 | -0.586 | 0.298 | 0.213 | -0.024 | 0.005 | 0.005 | 0 | 0.82 | 0.72 | 0.76 | 0.84 | 0.86 | 0.8 |
| 1 | 64837796 | rs7523530 | 1.419 | 0.078 | -0.544 | 0.227 | -0.171 | -0.182 | -0.182 | 0.44 | 0.38 | 0.04 | 0.42 | 0.51 | 0.49 | 0.43 |
| 1 | 64841469 | rs7349133 | 0.656 | -0.375 | -0.197 | 0.195 | 0.128 | 0.036 | 0.036 | 0 | 0.49 | 0.27 | 0.52 | 0.63 | 0.64 | 0.6 |
| 1 | 64845122 | rs6677170 | 0.788 | -0.332 | -0.197 | 0.217 | 0.13 | 0.043 | 0.043 | 0 | 0.5 | 0.3 | 0.52 | 0.63 | 0.64 | 0.6 |
| 1 | 64862973 | rs6695900 | -0.047 | -0.249 | -0.348 | 0.166 | 0.059 | 0.024 | 0.024 | 0 | 0.31 | 0.16 | 0.21 | 0.21 | 0.22 | 0.2 |
| 1 | 64868352 | rs11208446 | -1.068 | 0.365 | 0.144 | 0.212 | 0.062 | -0.011 | -0.011 | 0 | 0.1 | 0.31 | 0.07 | 0.08 | 0.12 | 0.15 |
| 1 | 64868506 | rs6703218 | 0.304 | 0.264 | 0.155 | 0.215 | 0.013 | -0.017 | -0.017 | 0 | 0.25 | 0.16 | 0.23 | 0.25 | 0.24 | 0.17 |
| 1 | 64870457 | rs12407601 | -0.383 | 0.304 | 0.17 | 0.128 | 0.064 | -0.027 | -0.027 | 0 | 0.05 | 0.17 | 0.04 | 0 | 0.09 | 0.01 |
| 1 | 64871720 | rs1155542 | 1.553 | -0.289 | -0.214 | 0.16 | 0.015 | 0.011 | 0.011 | 0.157 | 0.45 | 0.27 | 0.53 | 0.45 | 0.55 | 0.43 |
| 1 | 64871767 | rs1155541 | -0.717 | -0.175 | -0.1 | 0.223 | 0.014 | -0.024 | -0.024 | 0 | 0.86 | 0.95 | 0.86 | 0.83 | 0.89 | 0.81 |
| 1 | 64872227 | rs1567933 | 0.604 | -0.167 | 0.453 | 0.226 | -0.035 | 0.003 | 0.003 | 0 | 0.77 | 0.68 | 0.81 | 0.79 | 0.84 | 0.75 |
| 1 | 64892397 | rs11208448 | 0.226 | 0.026 | 0.281 | 0.408 | 0.005 | -0.007 | -0.007 | 0 | 0.24 | 0.18 | 0.24 | 0.23 | 0.25 | 0.18 |
| 1 | 64894096 | rs9988591 | -0.101 | 0.177 | 0.784 | 0.529 | -0.007 | 0.006 | 0.006 | 0 | 0.86 | 0.83 | 0.86 | 0.83 | 0.89 | 0.81 |
| 1 | 64913611 | rs6588094 | -0.507 | 0.575 | 0.819 | 0.256 | 0.017 | 0.046 | 0.046 | 0 | 0.08 | 0.21 | 0.07 | 0.11 | 0.08 | 0.05 |
| 1 | 64914030 | rs305533 | -0.239 | 0.423 | 0.494 | -0.103 | 0.005 | 0.032 | 0.032 | 0 | 0.65 | 0.73 | 0.59 | 0.87 | 0.63 | 0.74 |
| 1 | 64914437 | rs305535 | 0.298 | 0.577 | 0.694 | -0.002 | 0.003 | -0.004 | -0.004 | 0.635 | 0.11 | 0.21 | 0.07 | 0.07 | 0.11 | 0.23 |
| 1 | 64914487 | rs305536 | 0.298 | 0.577 | 0.694 | -0.002 | 0.003 | -0.004 | -0.004 | 0 | 0.11 | 0.2 | 0.07 | 0.07 | 0.11 | 0.23 |
| 1 | 64918776 | rs305544 | 0.541 | 0.581 | 0.315 | 0 | 0 | -0.001 | -0.001 | 0 | 0.11 | 0.11 | 0.06 | 0.07 | 0.11 | 0.23 |
| 1 | 64922435 | rs12139788 | -0.514 | 0.815 | -0.096 | -0.126 | 0.075 | 0.023 | 0.023 | 0 | 0.12 | 0.28 | 0.11 | 0.11 | 0.17 | 0.06 |
| 1 | 64923930 | rs11208456 | 1.389 | 0.485 | -1.049 | -0.074 | -0.012 | -0.077 | -0.077 | 0.553 | 0.37 | 0.12 | 0.4 | 0.67 | 0.34 | 0.41 |
| 1 | 64925123 | rs305549 | 0.372 | 0.115 | -0.563 | -0.067 | 0.016 | 0.077 | 0.077 | 0 | 0.63 | 0.81 | 0.6 | 0.86 | 0.63 | 0.73 |
| 1 | 64933188 | rs2484245 | -1.351 | -0.667 | 0.432 | 0.136 | 0.005 | -0.046 | -0.046 | 0 | 0.26 | 0.46 | 0.17 | 0.13 | 0.22 | 0.26 |
| 1 | 64941609 | rs7517824 | 2.003 | -1.296 | 0.355 | 0.03 | 0.054 | 0.082 | 0.082 | 0.722 | 0.56 | 0.19 | 0.69 | 0.74 | 0.69 | 0.68 |
| 1 | 64951754 | rs305540 | 0.443 | -1.624 | 1.32 | 0.102 | -0.046 | -0.039 | -0.039 | 0 | 0.85 | 0.73 | 0.9 | 0.94 | 0.93 | 0.9 |
| 1 | 64972899 | rs305576 | -0.947 | -1.645 | 1.396 | -0.122 | -0.003 | 0.005 | 0.005 | 0 | 0.1 | 0.11 | 0.1 | 0.17 | 0.08 | 0.1 |
| 1 | 64974601 | rs1932028 | 2.247 | -1.738 | 1.506 | -0.183 | 0.038 | 0.035 | 0.035 | 1.369 | 0.69 | 0.23 | 0.78 | 0.81 | 0.83 | 0.76 |
| 1 | 64976086 | rs305578 | -2.316 | -1.752 | 1.801 | -0.185 | -0.049 | 0.016 | 0.016 | 0.417 | 0.16 | 0.52 | 0.15 | 0.17 | 0.09 | 0.13 |
| 1 | 64981488 | rs764457 | 1.689 | -1.794 | 1.653 | -0.145 | -0.049 | -0.036 | -0.036 | 0 | 0.75 | 0.62 | 0.84 | 0.81 | 0.85 | 0.79 |
| 1 | 64985855 | rs90268 | -1.072 | -1.645 | 1.115 | -0.057 | -0.012 | 0.031 | 0.031 | 0 | 0.1 | 0.18 | 0.09 | 0.17 | 0.07 | 0.1 |
| 1 | 64990577 | rs305574 | 2.259 | -1.767 | 0.471 | -0.142 | 0.045 | 0.08 | 0.08 | 0.891 | 0.61 | 0.14 | 0.73 | 0.78 | 0.72 | 0.7 |
| 1 | 64992561 | rs305575 | 2.642 | -1.816 | 0.331 | 0.064 | 0.089 | 0.048 | 0.048 | 1.006 | 0.53 | 0.12 | 0.66 | 0.7 | 0.67 | 0.67 |
| 1 | 64994828 | rs646443 | 0.548 | -1.298 | 0.846 | -0.248 | -0.001 | -0.001 | -0.001 | 0 | 0.83 | 0.78 | 0.92 | 0.82 | 0.87 | 0.87 |
| 1 | 64997130 | rs596504 | -2.064 | -2.319 | 0.918 | -0.299 | -0.045 | -0.07 | -0.07 | 0 | 0.14 | 0.64 | 0.12 | 0.01 | 0.07 | 0.06 |
| 1 | 65003469 | rs6588096 | 1.729 | -2.085 | 0.744 | -0.108 | -0.06 | -0.026 | -0.026 | 0 | 0.73 | 0.49 | 0.85 | 0.8 | 0.86 | 0.85 |
| 1 | 65004354 | rs7556367 | 1.588 | -2.086 | 0.731 | -0.106 | -0.064 | -0.019 | -0.019 | 1.396 | 0.76 | 0.49 | 0.88 | 0.8 | 0.88 | 0.87 |
| 1 | 65005619 | rs307334 | 0.964 | -1.984 | 0.18 | -0.064 | -0.04 | -0.004 | -0.004 | 0 | 0.79 | 0.66 | 0.91 | 0.8 | 0.9 | 0.87 |
| 1 | 65008206 | rs635154 | 1.525 | -2.303 | -0.189 | 0.283 | 0.065 | -0.002 | -0.002 | 0 | 0.64 | 0.22 | 0.8 | 0.69 | 0.75 | 0.74 |
| 1 | 65010606 | rs17126643 | -1.196 | -1.417 | 0.441 | 0.377 | 0.068 | 0.015 | 0.015 | 0 | 0.07 | 0.34 | 0.06 | 0 | 0.05 | 0.01 |
| 1 | 65013586 | rs6681578 | 1.506 | -1.776 | -0.762 | 0.482 | 0.038 | -0.045 | -0.045 | 0 | 0.69 | 0.09 | 0.79 | 0.72 | 0.74 | 0.67 |
| 1 | 65015252 | rs17126653 | 0.244 | -1.91 | -0.559 | 0.505 | 0.013 | -0.004 | -0.004 | 0 | 0.05 | 0.02 | 0.02 | 0.08 | 0.03 | 0.13 |
| 1 | 65016399 | rs11808959 | -1.349 | -2.053 | -0.621 | 0.409 | 0.006 | -0.003 | -0.003 | 0.677 | 0.12 | 0.11 | 0.11 | 0.11 | 0.1 | 0.09 |
| 1 | 65016681 | rs12408795 | 2.25 | -2.612 | -0.618 | 0.19 | 0.063 | 0.023 | 0.023 | 0.443 | 0.61 | 0.23 | 0.68 | 0.69 | 0.73 | 0.57 |
| 1 | 65022445 | rs7512417 | 2.039 | -2.604 | -0.191 | 0.413 | 0.048 | 0.004 | 0.004 | 2.15 | 0.67 | 0.24 | 0.73 | 0.69 | 0.79 | 0.62 |
| 1 | 65029454 | rs12118994 | -1.441 | -2.661 | -0.183 | 0.382 | 0.005 | -0.003 | -0.003 | 0 | 0.12 | 0.13 | 0.11 | 0.11 | 0.09 | 0.08 |
| 1 | 65031000 | rs6660732 | 0.262 | -2.271 | 0.474 | 0.184 | -0.018 | -0.079 | -0.079 | 0 | 0.88 | 0.56 | 0.92 | 0.99 | 0.93 | 0.85 |
| 1 | 65031202 | rs11208467 | 0.96 | -2.184 | -0.052 | 0.243 | -0.035 | -0.014 | -0.014 | 0 | 0.78 | 0.38 | 0.84 | 0.81 | 0.88 | 0.69 |
| 1 | 65036321 | rs4915671 | -0.888 | -1.621 | -0.466 | 0.264 | 0.009 | 0.008 | 0.008 | 0 | 0.06 | 0.07 | 0.04 | 0 | 0.05 | 0.03 |
| 1 | 65041861 | rs6686231 | 1.236 | -2.432 | -0.469 | 0.508 | 0.079 | 0.04 | 0.04 | 0.649 | 0.55 | 0.28 | 0.47 | 0.33 | 0.73 | 0.37 |
| 1 | 65043551 | rs1436750 | -0.532 | -1.942 | -0.206 | 0.344 | 0.008 | 0.006 | 0.006 | 0 | 0.06 | 0.06 | 0.04 | 0 | 0.05 | 0.03 |
| 1 | 65043855 | rs1469876 | -0.87 | -2.347 | 0.179 | 0.644 | -0.003 | -0.002 | -0.002 | 0 | 0.17 | 0.18 | 0.22 | 0.13 | 0.14 | 0.18 |
| 1 | 65046494 | rs4915988 | -1.815 | -2.598 | 0.413 | 0.2 | 0.013 | 0.015 | 0.015 | 0 | 0.34 | 0.31 | 0.31 | 0.14 | 0.25 | 0.25 |

| | | | | | | | | | | | | | | | | |
|---|---|---|---|---|---|---|---|---|---|---|---|---|---|---|---|---|
| 1 | 65049950 | rs12068675 | -1.718 | -2.426 | 0.95 | 0.542 | -0.007 | -0.002 | -0.002 | 0 | 0.19 | 0.22 | 0.16 | 0.16 | 0.13 | 0.16 |
| 1 | 65068763 | rs10158570 | -2.253 | -1.885 | 1.411 | 0.54 | -0.021 | -0.027 | -0.027 | 0.416 | 0.3 | 0.48 | 0.34 | 0.22 | 0.24 | 0.24 |
| 1 | 65074235 | rs17126762 | -0.236 | -2.495 | 1.764 | 0.43 | 0.005 | 0 | 0 | 0 | 0.06 | 0.06 | 0.04 | 0 | 0.04 | 0.03 |
| 1 | 65074489 | rs7533098 | 2.59 | -2.53 | 1.99 | 0.411 | 0.014 | 0.013 | 0.013 | 0.558 | 0.63 | 0.34 | 0.62 | 0.8 | 0.77 | 0.76 |
| 1 | 65079551 | rs17126766 | -0.754 | -2.35 | 1.906 | 0.481 | -0.009 | -0.012 | -0.012 | 1.227 | 0.11 | 0.17 | 0.19 | 0.06 | 0.09 | 0.05 |
| 1 | 65085348 | rs11208484 | -1.569 | -2.381 | 1.81 | 0.453 | -0.009 | -0.012 | -0.012 | 0.111 | 0.15 | 0.17 | 0.19 | 0.06 | 0.12 | 0.1 |
| 1 | 65089766 | rs1436756 | 2.133 | -2.147 | 1.723 | 0.663 | -0.045 | -0.022 | -0.022 | 0 | 0.68 | 0.55 | 0.66 | 0.73 | 0.77 | 0.73 |
| 1 | 65093542 | rs12022915 | -1.424 | -2.272 | 1.611 | 0.818 | -0.02 | 0.004 | 0.004 | 0 | 0.17 | 0.24 | 0.14 | 0.21 | 0.09 | 0.15 |
| 1 | 65093974 | rs12023725 | 1.68 | -2.124 | 1.675 | 0.704 | -0.048 | -0.066 | -0.066 | 0 | 0.8 | 0.64 | 0.77 | 0.94 | 0.87 | 0.91 |
| 1 | 65097263 | rs1436753 | 1.309 | -2.035 | 1.673 | 0.697 | -0.01 | -0.055 | -0.055 | 0 | 0.9 | 0.67 | 0.93 | 1 | 0.94 | 0.97 |
| 1 | 65098136 | rs1075473 | -0.548 | -1.858 | 1.209 | 0.847 | 0.016 | 0.01 | 0.01 | 0.317 | 0.11 | 0.01 | 0.16 | 0.06 | 0.08 | 0.07 |
| 1 | 65100865 | rs751666 | 2.084 | -0.936 | 1.835 | 1.395 | -0.008 | 0.002 | 0.002 | 0 | 0.74 | 0.35 | 0.77 | 0.71 | 0.83 | 0.81 |
| 1 | 65103510 | rs7552434 | -1.965 | -0.824 | 1.622 | 1.674 | -0.003 | 0.014 | 0.014 | 0 | 0.18 | 0.25 | 0.14 | 0.29 | 0.12 | 0.17 |
| 1 | 65115319 | rs1561168 | -2.238 | -1.508 | 1.666 | 2.186 | -0.052 | 0.068 | 0.068 | 0 | 0.22 | 0.37 | 0.28 | 0.63 | 0.08 | 0.39 |
| 1 | 65116332 | rs10493362 | -2.501 | -1.537 | 1.726 | 2.172 | -0.079 | 0.089 | 0.089 | 0 | 0.24 | 0.47 | 0.29 | 0.63 | 0.08 | 0.41 |
| 1 | 65118909 | rs12742304 | 0.023 | -1.376 | 1.578 | 2.175 | -0.027 | -0.014 | -0.014 | 0 | 0.08 | 0 | 0.17 | 0.08 | 0.1 | 0.05 |
| 1 | 65125167 | rs12565594 | -2.332 | -1.378 | 1.63 | 2.218 | -0.067 | 0.083 | 0.083 | 0 | 0.23 | 0.43 | 0.29 | 0.63 | 0.08 | 0.39 |
| 1 | 65135068 | rs11208499 | -2.431 | -1.366 | 1.668 | 1.966 | -0.078 | 0.042 | 0.042 | 0 | 0.24 | 0.47 | 0.3 | 0.72 | 0.09 | 0.39 |
| 1 | 65141995 | rs1926306 | -2.863 | -1.337 | 1.703 | 1.943 | -0.002 | -0.001 | -0.001 | 0 | 0.33 | 0.67 | 0.48 | 0.79 | 0.19 | 0.44 |
| 1 | 65143441 | rs2499823 | -2.863 | -1.337 | 1.7 | 1.943 | -0.001 | 0 | 0 | 0.168 | 0.33 | 0.67 | 0.48 | 0.79 | 0.19 | 0.44 |
| 1 | 65146290 | rs12044266 | -1.862 | -1.236 | 1.372 | 2.291 | -0.005 | 0.078 | 0.078 | 0 | 0.11 | 0.16 | 0.21 | 0.52 | 0.08 | 0.27 |
| 1 | 65147002 | rs3828064 | -1.189 | -1.212 | 1.201 | 2.183 | -0.016 | 0.011 | 0.011 | 0 | 0.11 | 0.18 | 0.09 | 0.19 | 0.05 | 0.1 |
| 1 | 65148312 | rs12081048 | -1.533 | -0.865 | 1.002 | 2.242 | -0.001 | 0.397 | 0.397 | 0.081 | 0.07 | 0.28 | 0.18 | 0.45 | 0.02 | 0.2 |
| 1 | 65148949 | rs12757683 | -1.544 | -0.606 | 0.165 | 1.179 | -0.003 | 0 | 0 | 0 | 0.08 | 0.09 | 0.04 | 0.06 | 0.06 | 0.11 |
| 1 | 65153085 | rs1926301 | -1.268 | -0.954 | 0.12 | 0.711 | -0.001 | -0.001 | -0.001 | 0 | 0.38 | 0.58 | 0.46 | 0.74 | 0.24 | 0.44 |
| 1 | 65154595 | rs12089806 | -0.843 | -1.266 | 0.525 | 0.851 | 0.011 | 0.022 | 0.022 | 0 | 0.1 | 0.38 | 0.09 | 0.06 | 0.04 | 0.04 |
| 1 | 65155315 | rs17126884 | -0.514 | -1.279 | 0.939 | 0.845 | -0.006 | -0.003 | -0.003 | 0.106 | 0.08 | 0.12 | 0.07 | 0.06 | 0.04 | 0.04 |
| 1 | 65155765 | rs10889495 | -0.928 | -1.306 | 0.834 | 1.152 | 0.023 | -0.053 | -0.053 | 1.284 | 0.17 | 0.09 | 0.32 | 0.6 | 0.15 | 0.28 |
| 1 | 65156862 | rs17126889 | -0.269 | -1.305 | 0.825 | 1.139 | -0.001 | -0.001 | -0.001 | 3.236 | 0.11 | 0.11 | 0.06 | 0.05 | 0.09 | 0.12 |
| 1 | 65158724 | rs1926303 | -1.192 | -0.521 | 0.348 | 0.645 | -0.041 | 0.042 | 0.042 | 0 | 0.92 | 0.99 | 0.93 | 1 | 0.87 | 0.97 |
| 1 | 65160569 | rs11208508 | -0.86 | -0.848 | 0.799 | 0.912 | 0.004 | 0.014 | 0.014 | 0 | 0.1 | 0.31 | 0.08 | 0.06 | 0.05 | 0.05 |
| 1 | 65161230 | rs1926304 | -1.738 | -0.836 | 0.627 | 0.872 | -0.043 | 0.035 | 0.035 | 0 | 0.26 | 0.42 | 0.41 | 0.66 | 0.19 | 0.29 |
| 1 | 65161633 | rs2148323 | -1.744 | -0.846 | 0.622 | 0.889 | -0.044 | 0.037 | 0.037 | 0 | 0.26 | 0.42 | 0.41 | 0.66 | 0.19 | 0.31 |
| 1 | 65164851 | rs7539994 | -2.229 | -0.925 | 0.028 | 0.413 | 0.042 | 0.04 | 0.04 | 0.25 | 0.38 | 0.77 | 0.6 | 0.72 | 0.27 | 0.33 |
| 1 | 65164880 | rs12402072 | -0.92 | -0.34 | 0.121 | 0.532 | -0.011 | 0.009 | 0.009 | 0 | 0.07 | 0.03 | 0.16 | 0 | 0.06 | 0.05 |
| 1 | 65164955 | rs17126906 | -1.167 | -0.44 | 0.503 | 0.909 | 0.026 | 0.097 | 0.097 | 0 | 0.05 | 0.22 | 0.06 | 0.13 | 0.04 | 0.06 |
| 1 | 65170405 | rs11208512 | -2.038 | -0.03 | 0.112 | 0.352 | 0 | 0.018 | 0.018 | 0 | 0.4 | 0.66 | 0.61 | 0.78 | 0.37 | 0.47 |
| 1 | 65180463 | rs11799558 | -1.744 | 0.173 | 0.583 | 1.541 | 0.001 | 0.085 | 0.085 | 0.414 | 0.18 | 0.34 | 0.25 | 0.6 | 0.18 | 0.23 |
| 1 | 65189604 | rs10493364 | -1.789 | -0.603 | 1.152 | 1.711 | -0.043 | 0.045 | 0.045 | 0.19 | 0.21 | 0.47 | 0.27 | 0.26 | 0.16 | 0.18 |
| 1 | 65189751 | rs3920616 | -2.583 | -0.395 | 1.468 | 2.359 | -0.006 | 0.026 | 0.026 | 0.116 | 0.27 | 0.64 | 0.43 | 0.47 | 0.26 | 0.28 |
| 1 | 65190713 | rs10493365 | -1.7 | -0.445 | 1.447 | 2.304 | -0.047 | 0.042 | 0.042 | 0.239 | 0.21 | 0.47 | 0.27 | 0.25 | 0.16 | 0.18 |
| 1 | 65192187 | rs12073135 | -0.95 | -0.515 | 2.023 | 2.275 | -0.046 | 0.128 | 0.128 | 0 | 0.11 | 0.31 | 0.18 | 0.25 | 0.03 | 0.16 |
| 1 | 65200056 | rs2375473 | -2.738 | -0.58 | 2.351 | 2.189 | -0.003 | 0.022 | 0.022 | 1.761 | 0.21 | 0.69 | 0.28 | 0.28 | 0.23 | 0.15 |
| 1 | 65206148 | rs10493367 | -2.125 | -0.531 | 2.185 | 2.251 | 0.017 | 0.159 | 0.159 | 0 | 0.07 | 0.18 | 0.15 | 0.28 | 0.1 | 0.13 |
| 1 | 65221842 | rs6676160 | -0.818 | 0.177 | 2.101 | 2.13 | 0.012 | 0.062 | 0.062 | 0 | 0.1 | 0.2 | 0.21 | 0.25 | 0.11 | 0.16 |
| 1 | 65247394 | rs2478151 | -2.488 | 0.109 | 2.354 | 2.115 | 0.009 | 0.043 | 0.043 | 0.155 | 0.23 | 0.49 | 0.31 | 0.32 | 0.24 | 0.19 |
| 1 | 65250503 | rs10493372 | -0.028 | 0.13 | 1.945 | 2.051 | -0.013 | 0.032 | 0.032 | 0 | 0.07 | 0.02 | 0.06 | 0 | 0.11 | 0.01 |
| 1 | 65254893 | rs12128391 | -1.23 | 0.089 | 1.914 | 2.009 | -0.018 | 0.047 | 0.047 | 0 | 0.1 | 0.03 | 0.19 | 0.01 | 0.14 | 0.06 |
| 1 | 65294331 | rs11208527 | -1.949 | 0.103 | 1.998 | 2.13 | 0 | 0.007 | 0.007 | 0 | 0.13 | 0.15 | 0.22 | 0.22 | 0.21 | 0.09 |
| 1 | 65297240 | rs1048007 | -2.475 | -0.609 | 1.009 | 1.408 | -0.002 | -0.029 | -0.029 | 0.053 | 0.46 | 0.91 | 0.54 | 0.64 | 0.39 | 0.35 |
| 1 | 65302413 | rs2780816 | -2.385 | -0.667 | 1.374 | 1.531 | 0.015 | 0.005 | 0.005 | 0.181 | 0.31 | 0.69 | 0.32 | 0.29 | 0.24 | 0.19 |
| 1 | 65305877 | rs310244 | -1.514 | -0.693 | 1.681 | 1.454 | 0.005 | -0.018 | -0.018 | 0 | 0.22 | 0.18 | 0.15 | 0 | 0.22 | 0.11 |
| 1 | 65309387 | rs4916005 | -1.76 | -0.727 | 2.01 | 1.545 | -0.02 | 0.021 | 0.021 | 0 | 0.21 | 0.31 | 0.24 | 0.3 | 0.13 | 0.21 |
| 1 | 65313945 | rs2780898 | -3.149 | -0.757 | 2.05 | 1.791 | -0.003 | 0.041 | 0.041 | 1.492 | 0.22 | 0.66 | 0.39 | 0.51 | 0.26 | 0.17 |
| 1 | 65315298 | rs310219 | -2.683 | -1.346 | 2.089 | 1.61 | 0.005 | 0.002 | 0.002 | 0.204 | 0.3 | 0.66 | 0.32 | 0.3 | 0.25 | 0.22 |
| 1 | 65316675 | rs310216 | -2.66 | -1.346 | 2.104 | 1.61 | 0.002 | 0.001 | 0.001 | 0.131 | 0.3 | 0.63 | 0.32 | 0.3 | 0.25 | 0.22 |
| 1 | 65318507 | rs4916008 | -0.657 | -1.34 | 2.096 | 1.61 | -0.007 | 0.001 | 0.001 | 0 | 0.1 | 0.09 | 0.2 | 0.32 | 0.12 | 0.15 |
| 1 | 65318514 | rs2780900 | -0.325 | -1.331 | 2.058 | 1.595 | 0.01 | -0.047 | -0.047 | 0.106 | 0.08 | 0.22 | 0.07 | 0 | 0.11 | 0.01 |
| 1 | 65325970 | rs310224 | -3.068 | -1.331 | 1.979 | 1.611 | 0.009 | 0.006 | 0.006 | 0.01 | 0.34 | 0.64 | 0.45 | 0.31 | 0.29 | 0.28 |
| 1 | 65330682 | rs2256298 | -3.075 | -1.366 | 1.964 | 1.574 | -0.008 | 0.003 | 0.003 | 0.698 | 0.25 | 0.62 | 0.4 | 0.31 | 0.23 | 0.21 |
| 1 | 65331897 | rs2780886 | -3.754 | -1.375 | 1.884 | 1.59 | -0.017 | -0.026 | -0.026 | 0.079 | 0.44 | 0.87 | 0.65 | 0.62 | 0.41 | 0.43 |
| 1 | 65341567 | rs3790541 | -0.761 | -1.329 | 1.982 | 1.616 | 0.021 | 0.09 | 0.09 | 0 | 0.1 | 0.2 | 0.2 | 0.32 | 0.12 | 0.15 |
| 1 | 65343066 | rs11208534 | -1.659 | -1.359 | 2.047 | 1.614 | -0.013 | 0.015 | 0.015 | 0 | 0.18 | 0.24 | 0.22 | 0.3 | 0.11 | 0.2 |
| 1 | 65345616 | rs310209 | -2.757 | -1.344 | 1.857 | 1.616 | -0.028 | -0.006 | -0.006 | 0 | 0.3 | 0.47 | 0.41 | 0.31 | 0.26 | 0.24 |
| 1 | 65349544 | rs310202 | -2.868 | -1.32 | 1.731 | 1.785 | 0.001 | -0.007 | -0.007 | 0.041 | 0.3 | 0.69 | 0.42 | 0.6 | 0.22 | 0.35 |
| 1 | 65368042 | rs12743599 | 2.722 | -1.212 | 1.371 | 1.801 | -0.025 | -0.072 | -0.072 | 0.729 | 0.63 | 0.13 | 0.42 | 0.4 | 0.72 | 0.61 |
| 1 | 65381861 | rs12563017 | -1.468 | -1.131 | 1.255 | 1.659 | 0.002 | -0.013 | -0.013 | 0.233 | 0.16 | 0.13 | 0.18 | 0.28 | 0.11 | 0.21 |
| 1 | 65382792 | rs7528403 | 2.363 | -1.6 | 2.01 | 2.122 | 0.036 | 0.201 | 0.201 | 0 | 0.81 | 0.44 | 0.64 | 0.45 | 0.81 | 0.79 |
| 1 | 65383002 | rs7539178 | -1.97 | -1.578 | 2.076 | 2.152 | 0.023 | 0.155 | 0.155 | 0 | 0.14 | 0.33 | 0.21 | 0.55 | 0.16 | 0.19 |

| | | | | | | | | | | | | | | | | |
|---|---|---|---|---|---|---|---|---|---|---|---|---|---|---|---|---|
| 1 | 65405070 | rs1353595 | -2.182 | -1.537 | 2.234 | 2.263 | 0.049 | 0.219 | 0.219 | 0 | 0.12 | 0.34 | 0.21 | 0.54 | 0.15 | 0.16 |
| 1 | 65408096 | rs7531799 | -2.536 | -1.566 | 2.398 | 2.246 | 0.067 | 0.283 | 0.283 | 0.029 | 0.14 | 0.46 | 0.22 | 0.55 | 0.16 | 0.16 |
| 1 | 65408513 | rs17127168 | -0.543 | -1.926 | 2.041 | 2.049 | 0.001 | 0.017 | 0.017 | 0 | 0.09 | 0.07 | 0.1 | 0.27 | 0.02 | 0.14 |
| 1 | 65412137 | rs17127169 | -1.422 | -2.09 | 2.022 | 1.865 | 0.002 | -0.016 | -0.016 | 0 | 0.12 | 0.1 | 0.06 | 0 | 0.08 | 0.11 |
| 1 | 65421058 | rs4916014 | 3.58 | -2.301 | 1.873 | 2.11 | 0.03 | 0.046 | 0.046 | 0 | 0.61 | 0.13 | 0.57 | 0.29 | 0.72 | 0.59 |
| 1 | 65427476 | rs4915675 | 2.891 | -2.308 | 1.745 | 2.093 | 0.008 | -0.042 | -0.042 | 0 | 0.69 | 0.2 | 0.68 | 0.56 | 0.74 | 0.71 |
| 1 | 65439566 | rs7545743 | -2.882 | -2.6 | 1.928 | 2.137 | -0.086 | 0.012 | 0.012 | 0 | 0.29 | 0.49 | 0.32 | 0.28 | 0.11 | 0.26 |
| 1 | 65442015 | rs10749749 | -2.696 | -2.609 | 1.924 | 2.133 | -0.082 | 0.029 | 0.029 | 0 | 0.26 | 0.49 | 0.32 | 0.28 | 0.11 | 0.25 |
| 1 | 65445542 | rs11579758 | 2.955 | -2.559 | 1.263 | 2.017 | 0.056 | 0.004 | 0.004 | 0 | 0.63 | 0.12 | 0.6 | 0.32 | 0.76 | 0.64 |
| 1 | 65459206 | rs11208566 | -1.664 | -2.432 | 1.576 | 2.005 | -0.005 | -0.036 | -0.036 | 0.036 | 0.07 | 0.22 | 0.15 | 0 | 0.03 | 0.03 |
| 1 | 65464855 | rs2993612 | -2.916 | -2.499 | 1.496 | 2.312 | -0.02 | 0.003 | 0.003 | 0 | 0.28 | 0.63 | 0.31 | 0.6 | 0.13 | 0.3 |
| 1 | 65472323 | rs555038 | -2.526 | -2.449 | 1.381 | 2.232 | -0.101 | 0.05 | 0.05 | 0 | 0.23 | 0.41 | 0.3 | 0.29 | 0.09 | 0.22 |
| 1 | 65492194 | rs535857 | -2.55 | -2.728 | 1.109 | 2.095 | -0.092 | 0.041 | 0.041 | 0 | 0.29 | 0.39 | 0.33 | 0.38 | 0.1 | 0.25 |
| 1 | 65503789 | rs3014974 | -2.051 | -2.531 | 1.217 | 2.089 | -0.013 | 0.009 | 0.009 | 0 | 0.19 | 0.17 | 0.24 | 0.31 | 0.08 | 0.13 |
| 1 | 65506483 | rs3014975 | -2.902 | -2.511 | 1.315 | 2.168 | -0.062 | 0.053 | 0.053 | 0.111 | 0.33 | 0.47 | 0.44 | 0.42 | 0.17 | 0.23 |
| 1 | 65508424 | rs11208574 | -2.404 | -2.531 | 1.408 | 2.026 | -0.023 | -0.024 | -0.024 | 0.262 | 0.23 | 0.28 | 0.34 | 0.11 | 0.15 | 0.13 |
| 1 | 65512968 | rs11208576 | -2.273 | -2.48 | 1.368 | 2.132 | -0.048 | 0.05 | 0.05 | 0.043 | 0.19 | 0.27 | 0.26 | 0.31 | 0.08 | 0.13 |
| 1 | 65513522 | rs528645 | -1.072 | -2.539 | 1.178 | 2.061 | -0.014 | 0.046 | 0.046 | 0 | 0.11 | 0.11 | 0.09 | 0.31 | 0.04 | 0.09 |
| 1 | 65514604 | rs1338516 | 2.669 | -2.404 | 1.134 | 1.927 | -0.016 | 0.011 | 0.011 | 0 | 0.62 | 0.5 | 0.5 | 0.59 | 0.73 | 0.74 |
| 1 | 65515533 | rs478665 | -0.904 | -2.394 | 1.113 | 1.945 | 0.021 | -0.015 | -0.015 | 0 | 0.1 | 0.06 | 0.09 | 0.31 | 0.04 | 0.09 |
| 1 | 65515753 | rs17127247 | -0.446 | -2.048 | 0.859 | 2.057 | -0.011 | -0.006 | -0.006 | 0 | 0.06 | 0.02 | 0.03 | 0.2 | 0.11 | 0.08 |
| 1 | 65516055 | rs6588109 | 2.215 | -2.057 | 0.884 | 2.047 | -0.016 | 0.011 | 0.011 | 0 | 0.62 | 0.49 | 0.5 | 0.59 | 0.73 | 0.75 |
| 1 | 65526176 | rs1011210 | 2.663 | -2.141 | 0.256 | 1.527 | 0.044 | 0.073 | 0.073 | 0.236 | 0.51 | 0.14 | 0.45 | 0.4 | 0.61 | 0.64 |
| 1 | 65527407 | rs555146 | 1.22 | -1.517 | 0.598 | 1.614 | 0.018 | -0.011 | -0.011 | 0 | 0.85 | 0.77 | 0.79 | 0.89 | 0.86 | 0.95 |
| 1 | 65530114 | rs2201689 | 2.196 | -1.469 | -0.192 | 1.477 | -0.077 | 0.086 | 0.086 | 0 | 0.49 | 0.03 | 0.4 | 0.31 | 0.54 | 0.56 |
| 1 | 65534491 | rs11208579 | -2.382 | -1.519 | 0.197 | 1.456 | 0.077 | 0.055 | 0.055 | 0.052 | 0.43 | 0.83 | 0.5 | 0.69 | 0.3 | 0.37 |
| 1 | 65543802 | rs12089641 | -2.075 | -1.6 | 0.257 | 1.733 | 0.112 | -0.07 | -0.07 | 0 | 0.34 | 0.83 | 0.39 | 0.44 | 0.11 | 0.25 |
| 1 | 65546924 | rs6673673 | -1.197 | -1.634 | 0.523 | 1.788 | -0.022 | -0.005 | -0.005 | 0 | 0.19 | 0.22 | 0.27 | 0.15 | 0.07 | 0.05 |
| 1 | 65552976 | rs12023280 | -2.914 | -1.633 | 0.474 | 1.829 | 0.094 | 0.064 | 0.064 | 0.369 | 0.44 | 0.91 | 0.5 | 0.68 | 0.3 | 0.36 |
| 1 | 65553193 | rs2375675 | -1.192 | -1.633 | 0.552 | 1.759 | -0.069 | 0.096 | 0.096 | 0.476 | 0.15 | 0.6 | 0.12 | 0.29 | 0.04 | 0.19 |
| 1 | 65553909 | rs6698882 | -0.991 | -1.513 | 0.597 | 1.612 | -0.018 | -0.002 | -0.002 | 0.315 | 0.18 | 0.21 | 0.27 | 0.15 | 0.07 | 0.05 |
| 1 | 65557432 | rs6663598 | -1.891 | -1.49 | 1.056 | 1.576 | -0.031 | -0.047 | -0.047 | 0 | 0.28 | 0.43 | 0.36 | 0.15 | 0.18 | 0.11 |
| 1 | 65557876 | rs6699671 | -1.534 | -1.417 | 1.507 | 2.013 | -0.026 | -0.006 | -0.006 | 0 | 0.19 | 0.27 | 0.29 | 0.15 | 0.09 | 0.05 |
| 1 | 65560015 | rs7513840 | 2.778 | -1.44 | 0.721 | 2.033 | -0.01 | 0.062 | 0.062 | 0.166 | 0.55 | 0.05 | 0.45 | 0.33 | 0.62 | 0.69 |
| 1 | 65570028 | rs12729349 | -1.312 | -1.353 | 1.067 | 1.897 | -0.03 | 0.001 | 0.001 | 0.212 | 0.18 | 0.25 | 0.27 | 0.16 | 0.07 | 0.05 |
| 1 | 65571312 | rs17127323 | -1.168 | -1.281 | 0.8 | 2.093 | -0.023 | -0.035 | -0.035 | 0 | 0.1 | 0.06 | 0.12 | 0.22 | 0.19 | 0.13 |
| 1 | 65573794 | rs7524607 | 1.68 | -1.383 | 1.26 | 1.761 | -0.033 | 0.043 | 0.043 | 2.551 | 0.74 | 0.46 | 0.64 | 0.66 | 0.81 | 0.83 |
| 1 | 65581487 | rs10889520 | 1.244 | -1.511 | 1.028 | 1.674 | -0.016 | 0.015 | 0.015 | 1.272 | 0.75 | 0.65 | 0.66 | 0.67 | 0.83 | 0.81 |
| 1 | 65583858 | rs11208591 | 2.522 | -1.618 | 0.058 | 1.752 | 0.028 | 0.01 | 0.01 | 1.341 | 0.53 | 0.04 | 0.43 | 0.36 | 0.64 | 0.64 |
| 1 | 65584570 | rs12566370 | -0.737 | -1.446 | 0.202 | 1.727 | 0.001 | 0.061 | 0.061 | 0 | 0.16 | 0.32 | 0.18 | 0.3 | 0.15 | 0.16 |
| 1 | 65586815 | rs6696301 | 1.243 | -1.313 | 0.263 | 1.275 | -0.004 | 0.009 | 0.009 | 0.944 | 0.81 | 0.87 | 0.67 | 0.85 | 0.81 | 0.91 |
| 1 | 65593407 | rs17127359 | -1.506 | -1.419 | 0.624 | 1.256 | -0.059 | 0.078 | 0.078 | 0.555 | 0.15 | 0.49 | 0.08 | 0.18 | 0.08 | 0.13 |
| 1 | 65607538 | rs10047101 | -1.29 | -1.264 | 0.156 | 0.916 | -0.016 | -0.022 | -0.022 | 0.54 | 0.08 | 0.15 | 0.06 | 0.03 | 0.02 | 0.02 |
| 1 | 65611633 | rs2375754 | -2.111 | -2.585 | 0.067 | 1.319 | 0.083 | -0.013 | -0.013 | 0 | 0.34 | 0.78 | 0.38 | 0.38 | 0.09 | 0.16 |
| 1 | 65619728 | rs10749750 | -2.122 | -1.873 | -0.644 | 1.3 | 0.053 | -0.003 | -0.003 | 0 | 0.48 | 0.91 | 0.52 | 0.65 | 0.32 | 0.25 |
| 1 | 65619880 | rs10789171 | -0.868 | -2.075 | 0.632 | 1.41 | 0.021 | 0.158 | 0.158 | 0 | 0.23 | 0.46 | 0.18 | 0.55 | 0.24 | 0.16 |
| 1 | 65625752 | rs12401887 | -1.26 | -2.217 | 0.891 | 1.247 | -0.033 | -0.021 | -0.021 | 0 | 0.15 | 0.22 | 0.3 | 0.1 | 0.05 | 0.08 |
| 1 | 65638930 | rs6699614 | -2.823 | -2.375 | 0.559 | 1.378 | 0.053 | 0.101 | 0.101 | 0.056 | 0.49 | 0.97 | 0.54 | 0.73 | 0.32 | 0.31 |
| 1 | 65644186 | rs12030227 | -2.266 | -2.355 | 1.083 | 1.369 | -0.013 | -0.01 | -0.01 | 0.659 | 0.4 | 0.5 | 0.47 | 0.69 | 0.3 | 0.3 |
| 1 | 65648380 | rs2375597 | -1.407 | -2.359 | 1.286 | 1.511 | -0.006 | 0.079 | 0.079 | 0 | 0.27 | 0.53 | 0.21 | 0.54 | 0.25 | 0.19 |
| 1 | 65653425 | rs7549901 | -1.445 | -2.359 | 1.34 | 1.51 | -0.005 | 0.041 | 0.041 | 0 | 0.28 | 0.57 | 0.21 | 0.55 | 0.25 | 0.19 |
| 1 | 65658412 | rs6677316 | -1.266 | -2.357 | 1.266 | 1.512 | -0.003 | 0.1 | 0.1 | 0 | 0.27 | 0.44 | 0.2 | 0.55 | 0.25 | 0.19 |
| 1 | 65661665 | rs4511159 | -2.497 | -2.298 | 0.919 | 1.285 | 0.077 | 0.064 | 0.064 | 0 | 0.46 | 0.82 | 0.51 | 0.69 | 0.31 | 0.29 |
| 1 | 65668191 | rs4916030 | -2.13 | -2.069 | 1.33 | 1.267 | -0.004 | -0.004 | -0.004 | 0 | 0.39 | 0.42 | 0.49 | 0.68 | 0.32 | 0.37 |
| 1 | 65674836 | rs7526688 | -2.002 | -2.023 | 1.18 | 1.317 | 0.004 | 0.004 | 0.004 | 0 | 0.38 | 0.33 | 0.48 | 0.7 | 0.32 | 0.37 |
| 1 | 65677667 | rs12119363 | 0.61 | -2.056 | 0.879 | 1.534 | -0.052 | -0.093 | -0.093 | 0 | 0.13 | 0.01 | 0.12 | 0.39 | 0.17 | 0.07 |
| 1 | 65677933 | rs10789176 | -1.936 | -1.942 | 1.032 | 0.658 | -0.016 | -0.013 | -0.013 | 0 | 0.39 | 0.47 | 0.49 | 0.69 | 0.32 | 0.37 |
| 1 | 65681543 | rs4916031 | -1.554 | -1.416 | 0.645 | 0.496 | -0.003 | -0.003 | -0.003 | 0 | 0.34 | 0.34 | 0.47 | 0.72 | 0.28 | 0.49 |
| 1 | 65687556 | rs1889925 | -0.856 | -1.276 | 0.748 | 0.604 | -0.01 | 0.011 | 0.011 | 0 | 0.25 | 0.34 | 0.33 | 0.72 | 0.2 | 0.43 |
| 1 | 65688461 | rs6677191 | -0.898 | -1.216 | 0.519 | 0.62 | -0.023 | 0.005 | 0.005 | 0 | 0.27 | 0.42 | 0.36 | 0.72 | 0.21 | 0.43 |
| 1 | 65694612 | rs17127415 | -1.437 | -0.61 | 0.317 | 0.322 | -0.012 | -0.085 | -0.085 | 0 | 0.16 | 0.27 | 0.18 | 0 | 0.11 | 0.1 |
| 1 | 65699203 | rs10465850 | 0.087 | -0.209 | -0.16 | 0.425 | -0.001 | 0.011 | 0.011 | 0 | 0.43 | 0.38 | 0.31 | 0.33 | 0.41 | 0.33 |
| 1 | 65701787 | rs11581070 | 0.193 | -0.591 | 0.581 | 0.199 | 0.002 | -0.011 | -0.011 | 0 | 0.42 | 0.4 | 0.31 | 0.33 | 0.41 | 0.31 |
| 1 | 65703484 | rs12032616 | -0.078 | -0.515 | 0.167 | 0.666 | -0.001 | -0.002 | -0.002 | 0 | 0.2 | 0.21 | 0.23 | 0.29 | 0.25 | 0.24 |
| 1 | 65708834 | rs7539419 | -0.891 | -0.741 | -0.058 | 0.838 | -0.042 | 0.006 | 0.006 | 0 | 0.74 | 0.93 | 0.81 | 0.83 | 0.72 | 0.6 |
| 1 | 65710726 | rs1889929 | -0.228 | -0.776 | 0.239 | 1.036 | 0 | -0.017 | -0.017 | 0 | 0.67 | 0.69 | 0.55 | 0.59 | 0.68 | 0.57 |
| 1 | 65717710 | rs11208615 | -0.54 | -0.761 | 0.304 | 1.034 | -0.014 | 0.058 | 0.058 | 0 | 0.05 | 0.14 | 0.24 | 0.22 | 0.04 | 0.02 |
| 1 | 65723851 | rs11208616 | 0.892 | -0.753 | 0.223 | 0.621 | 0.001 | 0.001 | 0.001 | 0 | 0.49 | 0.4 | 0.44 | 0.57 | 0.47 | 0.5 |
| 1 | 65726516 | rs724685 | 1.371 | -0.839 | 0.224 | 0.847 | 0.015 | -0.021 | -0.021 | 0.036 | 0.78 | 0.52 | 0.62 | 0.79 | 0.74 | 0.91 |

| | | | | | | | | | | | | | | | |
|---|---|---|---|---|---|---|---|---|---|---|---|---|---|---|---|
| 1 | 65730862 | rs1749971 | 1.589 | -0.845 | 0.205 | 0.86 | 0.001 | -0.012 | -0.012 | 0 | 0.77 | 0.42 | 0.62 | 0.77 | 0.74 | 0.9 |
| 1 | 65732242 | rs2477786 | 0.794 | -0.863 | 0.129 | 0.822 | -0.078 | -0.064 | -0.064 | 0.258 | 0.28 | 0.02 | 0.17 | 0.21 | 0.27 | 0.4 |
| 1 | 65736258 | rs1749972 | 1.223 | -0.877 | 0.196 | 0.738 | 0.028 | -0.119 | -0.119 | 0 | 0.8 | 0.51 | 0.84 | 0.98 | 0.77 | 0.92 |
| 1 | 65748733 | rs12024972 | 0.533 | -1.164 | 0.355 | 0.808 | -0.018 | 0.005 | 0.005 | 0 | 0.11 | 0.12 | 0.07 | 0.18 | 0.04 | 0.19 |
| 1 | 65753072 | rs7511810 | 1.642 | -0.998 | 0.309 | 0.794 | -0.029 | 0.03 | 0.03 | 0 | 0.37 | 0.23 | 0.29 | 0.3 | 0.42 | 0.31 |
| 1 | 65762640 | rs2985902 | -0.567 | -0.637 | -0.007 | 0.731 | -0.143 | -0.009 | -0.009 | 0 | 0.67 | 0.97 | 0.75 | 0.79 | 0.58 | 0.59 |
| 1 | 65765184 | rs10399621 | 0.281 | -0.642 | 0 | 0.728 | -0.012 | 0.039 | 0.039 | 0.104 | 0.07 | 0.02 | 0.04 | 0 | 0.08 | 0.02 |
| 1 | 65772315 | rs10789179 | 0.254 | -0.648 | 0.35 | 0.69 | -0.012 | 0.034 | 0.034 | 0 | 0.58 | 0.71 | 0.69 | 0.79 | 0.5 | 0.56 |
| 1 | 65776702 | rs6677968 | 1.603 | -0.741 | 0.637 | 1.018 | 0.007 | -0.005 | -0.005 | 0.009 | 0.55 | 0.41 | 0.59 | 0.57 | 0.59 | 0.47 |
| 1 | 65782290 | rs4147109 | -0.537 | -0.731 | 0.56 | 1.052 | 0.015 | 0.001 | 0.001 | 0 | 0.78 | 0.74 | 0.78 | 0.83 | 0.71 | 0.69 |
| 1 | 65783231 | rs10493374 | -0.386 | -0.847 | 0.031 | 1.175 | -0.045 | 0.027 | 0.027 | 0 | 0.16 | 0.25 | 0.16 | 0.26 | 0.05 | 0.23 |
| 1 | 65803765 | rs1359533 | -0.061 | -0.78 | -0.358 | 1.239 | -0.067 | -0.093 | -0.093 | 0 | 0.72 | 0.88 | 0.54 | 0.61 | 0.62 | 0.61 |
| 1 | 65805144 | rs7528114 | 0.403 | -0.819 | -0.314 | 0.839 | -0.032 | -0.031 | -0.031 | 0 | 0.67 | 0.78 | 0.4 | 0.42 | 0.57 | 0.53 |
| 1 | 65805554 | rs6588132 | -0.499 | -0.714 | -0.403 | 0.859 | -0.13 | -0.044 | -0.044 | 0 | 0.75 | 0.98 | 0.76 | 0.83 | 0.63 | 0.65 |
| 1 | 65807412 | rs12086738 | 0.21 | -0.746 | -0.389 | 0.812 | -0.081 | -0.077 | -0.077 | 0.945 | 0.69 | 0.87 | 0.43 | 0.42 | 0.58 | 0.53 |
| 1 | 65808281 | rs2375699 | 1.488 | -0.749 | -0.287 | 0.527 | 0.005 | 0.014 | 0.014 | 0.025 | 0.51 | 0.53 | 0.37 | 0.34 | 0.54 | 0.37 |
| 1 | 65816567 | rs10789181 | -0.059 | 0.003 | -1.076 | 0.055 | -0.104 | -0.007 | -0.007 | 0.361 | 0.73 | 0.94 | 0.75 | 0.83 | 0.62 | 0.66 |
| 1 | 65830608 | rs1570868 | 0.641 | -0.146 | -0.614 | 0.1 | 0.001 | -0.006 | -0.006 | 0 | 0.61 | 0.46 | 0.52 | 0.29 | 0.57 | 0.6 |
| 1 | 65830905 | rs1570869 | -0.066 | -0.157 | -0.909 | 0.032 | -0.069 | 0.022 | 0.022 | 0 | 0.67 | 0.87 | 0.65 | 0.81 | 0.58 | 0.7 |
| 1 | 65838229 | rs7537733 | -0.114 | -0.08 | -0.906 | -0.05 | -0.051 | 0.024 | 0.024 | 0 | 0.67 | 0.85 | 0.62 | 0.81 | 0.59 | 0.69 |
| 1 | 65839065 | rs6588140 | -1.13 | -0.633 | -0.888 | -0.208 | -0.075 | 0.079 | 0.079 | 0 | 0.21 | 0.62 | 0.24 | 0.63 | 0.08 | 0.3 |
| 1 | 65851759 | rs10889546 | -0.159 | -0.413 | -0.867 | -0.261 | -0.042 | 0.027 | 0.027 | 0 | 0.66 | 0.82 | 0.61 | 0.79 | 0.58 | 0.68 |
| 1 | 65854556 | rs6588143 | -0.178 | -0.41 | -0.864 | -0.258 | -0.046 | 0.026 | 0.026 | 0 | 0.66 | 0.82 | 0.61 | 0.79 | 0.57 | 0.68 |
| 1 | 65858088 | rs6588144 | 0.332 | -0.507 | -1.06 | -0.31 | -0.013 | 0.045 | 0.045 | 0 | 0.58 | 0.38 | 0.46 | 0.21 | 0.55 | 0.52 |
| 1 | 65861798 | rs11208645 | 0.87 | -0.655 | -1.122 | -0.39 | 0.005 | 0.161 | 0.161 | 0 | 0.49 | 0.28 | 0.43 | 0.19 | 0.53 | 0.39 |
| 1 | 65867519 | rs4915691 | -1.082 | -1.249 | -0.912 | -0.09 | -0.095 | 0.145 | 0.145 | 0 | 0.17 | 0.56 | 0.18 | 0.6 | 0.05 | 0.3 |
| 1 | 65869274 | rs4655762 | 0.634 | -0.688 | -1.418 | -0.729 | 0.016 | 0.01 | 0.01 | 0 | 0.42 | 0.55 | 0.32 | 0.74 | 0.34 | 0.39 |
| 1 | 65873872 | rs17127601 | -0.212 | -0.302 | -1.6 | -0.788 | 0.007 | -0.014 | -0.014 | 0 | 0.14 | 0.34 | 0.24 | 0.08 | 0.12 | 0.14 |
| 1 | 65874037 | rs3818513 | -0.299 | -0.097 | -2.114 | -0.98 | -0.093 | 0.125 | 0.125 | 0 | 0.61 | 0.93 | 0.57 | 0.84 | 0.47 | 0.64 |
| 1 | 65879252 | rs10493377 | 0.192 | 0.376 | -1.292 | -1.139 | 0 | -0.022 | -0.022 | 0 | 0.52 | 0.5 | 0.47 | 0.87 | 0.44 | 0.56 |
| 1 | 65885357 | rs3806318 | 0.913 | 0.321 | -1.708 | -1.181 | 0.025 | 0.082 | 0.082 | 0 | 0.33 | 0.03 | 0.16 | 0.14 | 0.23 | 0.37 |
| 1 | 65887013 | rs12145690 | 0.555 | 0.542 | -0.911 | -1.256 | 0.011 | 0.067 | 0.067 | 0 | 0.51 | 0.39 | 0.46 | 0.87 | 0.44 | 0.57 |
| 1 | 65888854 | rs9436297 | -0.787 | 1.017 | -0.729 | -0.544 | 0.01 | -0.042 | -0.042 | 0.178 | 0.12 | 0.17 | 0.1 | 0 | 0.14 | 0.09 |
| 1 | 65894160 | rs12077210 | -0.927 | -0.27 | 0.072 | -0.418 | -0.044 | -0.033 | -0.033 | 0.112 | 0.11 | 0.31 | 0.07 | 0.04 | 0.04 | 0.1 |
| 1 | 65894342 | rs3790433 | -0.696 | -1.839 | -0.757 | -1.422 | 0.17 | 0.19 | 0.19 | 1.31 | 0.48 | 0.77 | 0.33 | 0.16 | 0.23 | 0.42 |
| 1 | 65895130 | rs7534511 | -0.485 | -0.624 | -1.54 | -2.031 | -0.024 | 0.003 | 0.003 | 0.181 | 0.78 | 0.8 | 0.71 | 0.19 | 0.67 | 0.66 |
| 1 | 65895927 | rs9436301 | 0.806 | -0.598 | -1.382 | -2.142 | -0.017 | -0.032 | -0.032 | 0 | 0.38 | 0.39 | 0.27 | 0.12 | 0.23 | 0.34 |
| 1 | 65896308 | rs12065099 | 0.329 | -0.677 | -1.384 | -2.161 | -0.056 | -0.044 | -0.044 | 0.651 | 0.12 | 0.38 | 0.07 | 0.04 | 0.04 | 0.1 |
| 1 | 65896612 | rs11208648 | 0.38 | -0.675 | -1.383 | -2.16 | -0.053 | -0.042 | -0.042 | 0 | 0.12 | 0.38 | 0.07 | 0.04 | 0.04 | 0.1 |
| 1 | 65897747 | rs1887285 | 0.963 | -1.684 | -1.554 | -2.788 | 0.019 | 0.007 | 0.007 | 0 | 0.17 | 0.09 | 0.08 | 0 | 0.09 | 0.07 |
| 1 | 65906762 | rs970467 | 1.452 | -1.647 | -1.437 | -2.718 | -0.011 | -0.009 | -0.009 | 0.397 | 0.19 | 0.24 | 0.17 | 0.12 | 0.1 | 0.24 |
| 1 | 65908473 | rs9436746 | 0.397 | -1.979 | -1.681 | -3.079 | 0 | 0.043 | 0.043 | 0 | 0.59 | 0.34 | 0.58 | 0.14 | 0.59 | 0.58 |
| 1 | 65911672 | rs9436748 | 1.536 | -1.895 | -1.532 | -3.274 | -0.066 | 0.1 | 0.1 | 0 | 0.31 | 0.16 | 0.38 | 0.03 | 0.44 | 0.22 |
| 1 | 65925349 | rs6669354 | 1.232 | -1.654 | -1.488 | -3.193 | 0.013 | 0.017 | 0.017 | 0.003 | 0.19 | 0.14 | 0.16 | 0.11 | 0.12 | 0.24 |
| 1 | 65935494 | rs6588147 | -0.414 | -1.181 | -1.254 | -2.919 | -0.038 | 0.007 | 0.007 | 1.324 | 0.82 | 0.86 | 0.69 | 0.16 | 0.65 | 0.74 |
| 1 | 65946506 | rs2025803 | 0.894 | -1.425 | -1.54 | -3.21 | -0.176 | 0.231 | 0.231 | 0 | 0.35 | 0.02 | 0.38 | 0.03 | 0.47 | 0.27 |
| 1 | 65950340 | rs7513047 | 0.942 | -1.461 | -1.539 | -3.209 | -0.184 | 0.226 | 0.226 | 0 | 0.34 | 0.02 | 0.38 | 0.03 | 0.48 | 0.27 |
| 1 | 65952781 | rs11808888 | 0.712 | -2.119 | -0.549 | -3.514 | -0.087 | -0.105 | -0.105 | 0 | 0.33 | 0.54 | 0.23 | 0.06 | 0.13 | 0.28 |
| 1 | 65955725 | rs17127673 | 0.929 | -2.173 | -0.228 | -3.539 | -0.06 | -0.076 | -0.076 | 0.025 | 0.31 | 0.34 | 0.22 | 0.06 | 0.13 | 0.28 |
| 1 | 65957141 | rs11579567 | -0.047 | -2.256 | 0.078 | -3.385 | -0.025 | -0.014 | -0.014 | 2.437 | 0.16 | 0.21 | 0.07 | 0.07 | 0.06 | 0.18 |
| 1 | 65963016 | rs6694528 | 0.775 | -2.214 | -0.049 | -3.372 | -0.085 | -0.104 | -0.104 | 1.785 | 0.32 | 0.44 | 0.23 | 0.06 | 0.13 | 0.27 |
| 1 | 65966839 | rs1327115 | 0.758 | -2.086 | -0.028 | -3.315 | 0.02 | -0.074 | -0.074 | 0.017 | 0.66 | 0.58 | 0.62 | 0.09 | 0.6 | 0.54 |
| 1 | 65978917 | rs2767485 | 0.658 | -2.157 | 0.29 | -3.262 | 0.016 | 0.003 | 0.003 | 0 | 0.35 | 0.28 | 0.25 | 0.06 | 0.16 | 0.27 |
| 1 | 65979280 | rs11208659 | -0.805 | -2.336 | 0.46 | -3.37 | -0.09 | -0.071 | -0.071 | 0.076 | 0.2 | 0.4 | 0.11 | 0.07 | 0.08 | 0.22 |
| 1 | 65983158 | rs1475397 | -0.738 | -2.309 | 0.39 | -3.378 | 0.098 | 0.106 | 0.106 | 0 | 0.54 | 0.69 | 0.35 | 0.13 | 0.24 | 0.47 |
| 1 | 65986804 | rs1782755 | 0.561 | -2.196 | 0.365 | -3.353 | -0.064 | -0.104 | -0.104 | 0.009 | 0.37 | 0.48 | 0.27 | 0.06 | 0.17 | 0.27 |
| 1 | 65988146 | rs1171278 | 0.561 | -2.196 | 0.365 | -3.353 | -0.064 | -0.104 | -0.104 | 4.047 | 0.37 | 0.48 | 0.27 | 0.06 | 0.17 | 0.27 |
| 1 | 65998467 | rs10749753 | 0.788 | -1.319 | 0.097 | -3.198 | 0 | -0.047 | -0.047 | 0 | 0.35 | 0.3 | 0.42 | 0.05 | 0.48 | 0.27 |
| 1 | 66002660 | rs12140910 | -0.523 | -1.172 | 0.329 | -2.805 | 0.055 | 0.042 | 0.042 | 0 | 0.15 | 0.05 | 0.06 | 0.06 | 0.06 | 0.29 |
| 1 | 66043019 | rs3790426 | -0.56 | -0.071 | 0.277 | -2.936 | 0.007 | -0.041 | -0.041 | 0 | 0.2 | 0.26 | 0.14 | 0.07 | 0.23 | 0.3 |
| 1 | 66046117 | rs10493380 | 1.187 | -0.082 | -0.187 | -3.257 | 0.016 | 0.062 | 0.062 | 0 | 0.19 | 0.05 | 0.29 | 0.04 | 0.17 | 0.12 |
| 1 | 66047072 | rs12059300 | -0.545 | 0.261 | 0.165 | -2.95 | 0.002 | -0.022 | -0.022 | 0 | 0.2 | 0.21 | 0.14 | 0.07 | 0.23 | 0.31 |
| 1 | 66048622 | rs1938489 | -0.572 | 0.26 | 0.178 | -2.95 | 0.007 | -0.038 | -0.038 | 0.613 | 0.2 | 0.25 | 0.14 | 0.07 | 0.23 | 0.31 |
| 1 | 66058801 | rs4655537 | -1.105 | 0.308 | 0.165 | -3.06 | 0.011 | 0.034 | 0.034 | 0.369 | 0.54 | 0.61 | 0.63 | 0.92 | 0.64 | 0.67 |
| 1 | 66073361 | rs4606347 | 0.61 | -0.194 | 0.352 | -2.784 | 0.006 | -0.005 | -0.005 | 1.934 | 0.24 | 0.2 | 0.19 | 0.06 | 0.15 | 0.11 |
| 1 | 66075136 | rs11801408 | -0.202 | -0.305 | 0.609 | -2.801 | -0.065 | -0.091 | -0.091 | 0.203 | 0.32 | 0.44 | 0.22 | 0.1 | 0.17 | 0.36 |
| 1 | 66077624 | rs6678033 | -0.634 | 0.063 | 0.62 | -1.978 | -0.002 | -0.026 | -0.026 | 0 | 0.38 | 0.56 | 0.5 | 0.87 | 0.37 | 0.47 |
| 1 | 66089582 | rs4567312 | 0.575 | 0.092 | -0.077 | -1.875 | -0.028 | -0.018 | -0.018 | 0 | 0.09 | 0.16 | 0.02 | 0.04 | 0.02 | 0.25 |
| 1 | 66099013 | rs17415296 | 0.806 | 0.049 | -0.29 | -1.94 | 0.049 | 0.114 | 0.114 | 0 | 0.23 | 0.03 | 0.18 | 0.06 | 0.16 | 0.11 |

| | | | | | | | | | | | | | | | | |
|---|---|---|---|---|---|---|---|---|---|---|---|---|---|---|---|---|
| 1 | 66102257 | rs1805096 | -0.651 | 0.142 | -0.131 | -1.938 | -0.009 | -0.064 | -0.064 | 0.422 | 0.37 | 0.45 | 0.48 | 0.87 | 0.38 | 0.48 |
| 1 | 66105944 | rs1892534 | -0.701 | 0.11 | -0.148 | -1.974 | -0.01 | -0.065 | -0.065 | 0.299 | 0.38 | 0.45 | 0.5 | 0.87 | 0.38 | 0.48 |
| 1 | 66125318 | rs11812068 | -0.131 | 0.154 | -0.087 | -1.944 | -0.006 | -0.018 | -0.018 | 0 | 0.32 | 0.31 | 0.21 | 0.1 | 0.18 | 0.36 |
| 1 | 66130398 | rs4655772 | -1.259 | 0.306 | -0.55 | -2.07 | 0 | -0.021 | -0.021 | 0 | 0.49 | 0.51 | 0.59 | 0.9 | 0.57 | 0.59 |
| 1 | 66138575 | rs11807752 | 1.073 | 0.259 | -0.564 | -2.068 | 0.049 | 0.114 | 0.114 | 0 | 0.23 | 0.03 | 0.18 | 0.06 | 0.16 | 0.11 |
| 1 | 66143044 | rs4655783 | -1.176 | 0.194 | -0.509 | -2.071 | 0 | -0.021 | -0.021 | 0.153 | 0.49 | 0.51 | 0.59 | 0.9 | 0.57 | 0.6 |
| 1 | 66153933 | rs12022410 | -1.176 | 0.194 | -0.509 | -2.071 | 0 | -0.021 | -0.021 | 0 | 0.49 | 0.51 | 0.59 | 0.9 | 0.58 | 0.6 |
| 1 | 66156021 | rs2211651 | -0.814 | -0.051 | -0.134 | -1.486 | -0.009 | -0.062 | -0.062 | 0 | 0.37 | 0.44 | 0.48 | 0.86 | 0.38 | 0.48 |
| 1 | 66161461 | rs4420065 | 0.563 | -0.035 | -0.34 | -1.478 | -0.009 | -0.057 | -0.057 | 0 | 0.62 | 0.58 | 0.52 | 0.14 | 0.62 | 0.53 |
| 1 | 66165034 | rs11805519 | 0.511 | -0.155 | -0.33 | -1.413 | -0.017 | -0.008 | -0.008 | 0.653 | 0.08 | 0.08 | 0.01 | 0.04 | 0.02 | 0.25 |
| 1 | 66166499 | rs7524581 | 0.216 | 0.382 | -0.448 | -1.018 | 0.003 | -0.022 | -0.022 | 0 | 0.71 | 0.66 | 0.53 | 0.18 | 0.65 | 0.77 |
| 1 | 66170362 | rs17407594 | 0.142 | -0.625 | -1.287 | -1.212 | 0.067 | 0.096 | 0.096 | 0 | 0.23 | 0.03 | 0.17 | 0.09 | 0.15 | 0.11 |
| 1 | 66171001 | rs11208722 | -0.183 | -0.514 | -1.299 | -1.697 | -0.036 | 0.129 | 0.129 | 0 | 0.67 | 0.89 | 0.72 | 0.98 | 0.67 | 0.68 |
| 1 | 66171970 | rs12121677 | 0.06 | -0.827 | -1.181 | -1.57 | 0.019 | 0.028 | 0.028 | 0 | 0.2 | 0.14 | 0.21 | 0.1 | 0.13 | 0.23 |
| 1 | 66175395 | rs4278348 | 0.996 | -1.365 | -0.743 | -1.671 | -0.037 | 0.115 | 0.115 | 0 | 0.42 | 0.17 | 0.48 | 0.8 | 0.55 | 0.41 |
| 1 | 66179064 | rs17416384 | 1.047 | -1.387 | -0.749 | -1.682 | -0.041 | 0.115 | 0.115 | 0 | 0.42 | 0.15 | 0.48 | 0.8 | 0.55 | 0.41 |
| 1 | 66179203 | rs11208724 | 0.187 | -1.449 | 0.195 | -1.711 | -0.073 | -0.056 | -0.056 | 0 | 0.73 | 0.45 | 0.76 | 0.83 | 0.86 | 0.65 |
| 1 | 66185403 | rs7539471 | 0.368 | -1.389 | 0.221 | -1.683 | -0.023 | 0.053 | 0.053 | 0 | 0.29 | 0.15 | 0.27 | 0.03 | 0.32 | 0.24 |
| 1 | 66191075 | rs4291477 | 0.906 | -1.283 | 0.205 | -1.406 | -0.042 | 0.112 | 0.112 | 0 | 0.41 | 0.16 | 0.48 | 0.79 | 0.54 | 0.42 |
| 1 | 66191777 | rs7416472 | 0.906 | -1.283 | 0.205 | -1.385 | -0.042 | 0.11 | 0.11 | 0 | 0.41 | 0.15 | 0.48 | 0.79 | 0.55 | 0.42 |
| 1 | 66194959 | rs12035209 | 0.911 | -1.282 | 0.206 | -1.363 | -0.042 | 0.107 | 0.107 | 0 | 0.41 | 0.15 | 0.48 | 0.79 | 0.55 | 0.42 |
| 1 | 66197680 | rs4655587 | 0.096 | -0.498 | 0.408 | -1.08 | -0.002 | -0.002 | -0.002 | 0.043 | 0.77 | 0.71 | 0.84 | 0.91 | 0.72 | 0.77 |
| 1 | 66202755 | rs10789198 | 1.016 | -0.617 | 0.391 | -0.511 | 0.017 | 0.041 | 0.041 | 0 | 0.5 | 0.34 | 0.61 | 0.67 | 0.59 | 0.49 |
| 1 | 66205540 | rs11208726 | -0.081 | -0.565 | -0.099 | -0.449 | 0.002 | -0.008 | -0.008 | 0 | 0.78 | 0.71 | 0.84 | 0.91 | 0.73 | 0.76 |
| 1 | 66211916 | rs11809779 | -0.527 | -0.985 | -0.26 | 0.066 | 0.006 | 0.001 | 0.001 | 0.964 | 0.26 | 0.23 | 0.21 | 0.25 | 0.13 | 0.27 |
| 1 | 66217215 | rs6656519 | -0.9 | -0.712 | -0.483 | 0.288 | 0.031 | 0 | 0 | 0.163 | 0.31 | 0.16 | 0.26 | 0.3 | 0.22 | 0.3 |
| 1 | 66217735 | rs12086495 | -1.092 | -1.17 | -0.343 | 1.139 | -0.046 | 0.01 | 0.01 | 1.773 | 0.33 | 0.47 | 0.16 | 0.34 | 0.19 | 0.28 |
| 1 | 66221643 | rs4633264 | -0.761 | -1.161 | -0.19 | 1.085 | -0.007 | 0.097 | 0.097 | 0 | 0.15 | 0.52 | 0.1 | 0.22 | 0.09 | 0.17 |
| 1 | 66222161 | rs12742722 | 0.168 | -1.142 | -0.252 | 1.083 | 0.018 | 0.019 | 0.019 | 0.726 | 0.19 | 0.02 | 0.08 | 0.12 | 0.12 | 0.14 |
| 1 | 66222381 | rs12742800 | 0.168 | -1.142 | -0.252 | 1.083 | 0.018 | 0.019 | 0.019 | 0.726 | 0.19 | 0.02 | 0.08 | 0.12 | 0.12 | 0.14 |
| 1 | 66228578 | rs17408644 | -0.648 | -0.673 | -0.486 | 1.118 | -0.004 | -0.099 | -0.099 | 0.527 | 0.27 | 0.08 | 0.29 | 0.54 | 0.33 | 0.32 |
| 1 | 66231308 | rs12401401 | -0.635 | -0.673 | -0.486 | 1.118 | -0.004 | -0.099 | -0.099 | 0 | 0.27 | 0.08 | 0.29 | 0.54 | 0.33 | 0.31 |
| 1 | 66234139 | rs4313369 | -1.309 | -0.734 | -0.487 | 1.082 | -0.006 | 0.017 | 0.017 | 0.86 | 0.2 | 0.7 | 0.24 | 0.38 | 0.1 | 0.26 |
| 1 | 66236207 | rs11808116 | -1.951 | -0.733 | -0.523 | 1.081 | 0.037 | 0.273 | 0.273 | 1.327 | 0.47 | 0.81 | 0.53 | 0.92 | 0.43 | 0.57 |
| 1 | 66238379 | rs12046844 | 0.017 | -0.724 | -0.524 | 1.093 | 0.002 | 0.069 | 0.069 | 0 | 0.08 | 0.06 | 0.21 | 0.42 | 0.21 | 0.18 |
| 1 | 66239487 | rs11208742 | 0.017 | -0.724 | -0.524 | 1.093 | 0.002 | 0.069 | 0.069 | 0 | 0.08 | 0.06 | 0.21 | 0.42 | 0.21 | 0.18 |
| 1 | 66242820 | rs11208743 | -0.599 | -0.726 | -0.524 | 1.092 | -0.005 | -0.102 | -0.102 | 8.623 | 0.27 | 0.08 | 0.29 | 0.54 | 0.33 | 0.31 |
| 1 | 66251418 | rs12121489 | -0.066 | -0.646 | -0.481 | 1.115 | 0.001 | 0.067 | 0.067 | 0 | 0.08 | 0.06 | 0.2 | 0.42 | 0.21 | 0.18 |
| 1 | 66257416 | rs9326069 | -1.517 | -0.557 | -0.431 | 0.805 | 0.037 | 0.273 | 0.273 | 1.327 | 0.47 | 0.81 | 0.53 | 0.92 | 0.43 | 0.58 |
| 1 | 66269066 | rs11208755 | -0.765 | -0.63 | 0.113 | 1.104 | 0.008 | 0.131 | 0.131 | 0 | 0.14 | 0.45 | 0.09 | 0.21 | 0.09 | 0.16 |
| 1 | 66269311 | rs11208756 | -0.812 | -0.6 | 0.041 | 1.105 | 0.005 | 0.115 | 0.115 | 0 | 0.14 | 0.48 | 0.09 | 0.21 | 0.09 | 0.16 |
| 1 | 66279125 | rs4384209 | 0.647 | -1.058 | -0.114 | 1.044 | -0.04 | 0.004 | 0.004 | 0.058 | 0.67 | 0.52 | 0.61 | 0.35 | 0.79 | 0.6 |
| 1 | 66292706 | rs11576180 | -0.25 | -1.621 | -0.167 | 1.719 | 0.022 | 0.034 | 0.034 | 0 | 0.16 | 0.05 | 0.08 | 0.08 | 0.12 | 0.15 |
| 1 | 66297732 | rs7514810 | 0.182 | -1.628 | -0.134 | 2.068 | -0.006 | 0.132 | 0.132 | 0 | 0.93 | 0.9 | 0.86 | 0.65 | 0.95 | 0.83 |
| 1 | 66298161 | rs7517475 | 1.475 | -1.668 | 0.018 | 2.047 | -0.053 | 0.021 | 0.021 | 0 | 0.69 | 0.54 | 0.6 | 0.35 | 0.81 | 0.6 |
| 1 | 66308651 | rs10493388 | -0.12 | -1.619 | -0.048 | 2.148 | 0.002 | 0.055 | 0.055 | 0 | 0.06 | 0.03 | 0.13 | 0.33 | 0.05 | 0.17 |
| 1 | 66310865 | rs10493389 | -0.223 | -1.744 | -0.1 | 2.033 | 0.025 | 0.031 | 0.031 | 0 | 0.16 | 0.07 | 0.09 | 0.08 | 0.12 | 0.14 |
| 1 | 66311760 | rs12125534 | -0.534 | -1.764 | 0.105 | 2.052 | 0.012 | 0.002 | 0.002 | 0 | 0.17 | 0.12 | 0.15 | 0.16 | 0.13 | 0.2 |
| 1 | 66314610 | rs6681607 | 1.096 | -1.811 | 0.503 | 2.043 | -0.028 | 0.179 | 0.179 | 0 | 0.89 | 0.77 | 0.77 | 0.53 | 0.94 | 0.8 |
| 1 | 66315450 | rs6684621 | 1.902 | -1.27 | -0.299 | 1.399 | -0.001 | -0.005 | -0.005 | 0 | 0.54 | 0.49 | 0.5 | 0.32 | 0.58 | 0.39 |
| 1 | 66324394 | rs12751825 | 0.22 | -1.157 | -0.316 | 1.402 | -0.097 | 0.07 | 0.07 | 0 | 0.16 | 0.02 | 0.1 | 0.05 | 0.23 | 0.21 |
| 1 | 66326917 | rs2801581 | 1.307 | -1.126 | -0.279 | 1.424 | 0.009 | 0.029 | 0.029 | 0.005 | 0.71 | 0.62 | 0.65 | 0.48 | 0.71 | 0.59 |
| 1 | 66332895 | rs2186124 | -0.707 | -1.168 | -0.776 | 1.722 | 0.02 | -0.05 | -0.05 | 0 | 0.2 | 0.12 | 0.22 | 0.35 | 0.18 | 0.32 |
| 1 | 66335318 | rs6588169 | -0.102 | -1.454 | -1.392 | 0.906 | -0.049 | 0.106 | 0.106 | 0 | 0.22 | 0.04 | 0.08 | 0.04 | 0.18 | 0.2 |
| 1 | 66339048 | rs1316871 | 2.138 | -1.818 | -1.603 | 0.862 | -0.013 | -0.001 | -0.001 | 0 | 0.42 | 0.6 | 0.45 | 0.35 | 0.51 | 0.24 |
| 1 | 66341764 | rs12760605 | -0.391 | -1.882 | -1.979 | 0.939 | -0.011 | 0.035 | 0.035 | 0 | 0.27 | 0.06 | 0.15 | 0.17 | 0.18 | 0.27 |
| 1 | 66352755 | rs4655804 | -0.8 | -1.659 | -1.774 | 1.035 | 0.02 | -0.083 | -0.083 | 0 | 0.24 | 0.02 | 0.21 | 0.4 | 0.19 | 0.42 |
| 1 | 66362971 | rs12744291 | -0.76 | -1.682 | -1.878 | 0.976 | -0.041 | -0.087 | -0.087 | 0.012 | 0.34 | 0.1 | 0.35 | 0.51 | 0.29 | 0.5 |
| 1 | 66371682 | rs11578009 | -0.835 | -1.751 | -1.901 | 0.846 | 0.001 | 0.04 | 0.04 | 0 | 0.19 | 0.07 | 0.17 | 0.09 | 0.19 | 0.19 |
| 1 | 66372089 | rs11208765 | -2.07 | -1.782 | -1.917 | 0.877 | -0.029 | -0.029 | -0.029 | 0 | 0.54 | 0.2 | 0.53 | 0.6 | 0.48 | 0.69 |
| 1 | 66374004 | rs4329483 | -2.167 | -1.798 | -1.799 | 0.854 | -0.012 | -0.014 | -0.014 | 0.999 | 0.55 | 0.34 | 0.55 | 0.6 | 0.48 | 0.68 |
| 1 | 66380342 | rs17128123 | 0.349 | -1.709 | -1.968 | 0.95 | -0.007 | 0.006 | 0.006 | 0.065 | 0.05 | 0.02 | 0.13 | 0.37 | 0.04 | 0.26 |
| 1 | 66387487 | rs17451403 | 0.016 | -1.678 | -1.957 | 0.378 | -0.08 | 0.142 | 0.142 | 0 | 0.28 | 0.02 | 0.2 | 0.05 | 0.24 | 0.23 |
| 1 | 66388153 | rs17451451 | -1.076 | -1.285 | -1.596 | 0.579 | 0.013 | 0.039 | 0.039 | 0 | 0.19 | 0.06 | 0.16 | 0.09 | 0.17 | 0.18 |
| 1 | 66389097 | rs4655809 | -1.245 | -1.621 | -1.582 | 0.703 | -0.003 | -0.009 | -0.009 | 0 | 0.41 | 0.28 | 0.39 | 0.47 | 0.35 | 0.47 |
| 1 | 66392405 | rs2310752 | -1.808 | -1.602 | -1.6 | 0.624 | 0 | 0.002 | 0.002 | 0.001 | 0.49 | 0.34 | 0.46 | 0.54 | 0.43 | 0.58 |
| 1 | 66399635 | rs2997091 | -0.098 | -2.18 | -1.297 | 0.716 | -0.009 | 0.273 | 0.273 | 0.949 | 0.91 | 0.47 | 0.89 | 0.74 | 0.96 | 0.83 |
| 1 | 66399895 | rs4255357 | 2.113 | -2.166 | -1.412 | 0.61 | -0.074 | 0.135 | 0.135 | 0 | 0.43 | 0.12 | 0.42 | 0.19 | 0.53 | 0.24 |
| 1 | 66402424 | rs12058296 | -0.712 | -2.218 | -1.427 | 0.63 | -0.027 | 0.146 | 0.146 | 0 | 0.36 | 0.19 | 0.33 | 0.13 | 0.34 | 0.26 |

| | | | | | | | | | | | | | | | | |
|---|---|---|---|---|---|---|---|---|---|---|---|---|---|---|---|---|
| 1 | 66404726 | rs11208772 | -0.231 | -2.334 | -1.521 | 0.534 | 0.033 | 0.04 | 0.04 | 0 | 0.18 | 0.14 | 0.2 | 0.09 | 0.09 | 0.08 |
| 1 | 66410541 | rs11208775 | 2.369 | -2.378 | -1.542 | 0.546 | -0.077 | 0.135 | 0.135 | 0 | 0.43 | 0.12 | 0.42 | 0.19 | 0.53 | 0.24 |
| 1 | 66411870 | rs11208777 | -0.284 | -2.241 | -1.522 | 0.508 | 0.034 | 0.04 | 0.04 | 0 | 0.18 | 0.13 | 0.19 | 0.09 | 0.09 | 0.08 |
| 1 | 66416309 | rs1937458 | -1.228 | -2.257 | -1.521 | 0.579 | -0.044 | -0.036 | -0.036 | 0.014 | 0.39 | 0.18 | 0.38 | 0.45 | 0.36 | 0.43 |
| 1 | 66416939 | rs1937455 | -1.766 | -2.291 | -1.329 | 0.603 | -0.002 | 0.009 | 0.009 | 0.377 | 0.48 | 0.33 | 0.46 | 0.53 | 0.43 | 0.58 |
| 1 | 66426100 | rs10789205 | 0.927 | -2.279 | -1.019 | 0.583 | -0.005 | -0.004 | -0.004 | 2.863 | 0.61 | 0.7 | 0.61 | 0.55 | 0.64 | 0.56 |
| 1 | 66430203 | rs12096433 | 0.14 | -2.323 | -1.119 | 0.493 | 0.042 | 0.07 | 0.07 | 0 | 0.14 | 0.05 | 0.05 | 0.03 | 0.07 | 0.06 |
| 1 | 66434743 | rs7519259 | 2.213 | -2.394 | -1.063 | 0.538 | -0.069 | 0.114 | 0.114 | 0.403 | 0.43 | 0.12 | 0.42 | 0.22 | 0.53 | 0.24 |
| 1 | 66436675 | rs17452121 | 0.062 | -2.044 | -1.174 | 0.395 | 0.031 | 0.075 | 0.075 | 0 | 0.15 | 0.05 | 0.06 | 0.03 | 0.09 | 0.06 |
| 1 | 66440553 | rs12085651 | 0.088 | -2.129 | -1.148 | 0.499 | 0.042 | 0.07 | 0.07 | 0 | 0.14 | 0.05 | 0.05 | 0.03 | 0.07 | 0.06 |
| 1 | 66446787 | rs1937457 | -0.175 | -2.081 | -1.087 | 0.514 | -0.023 | -0.029 | -0.029 | 0 | 0.24 | 0.18 | 0.34 | 0.43 | 0.29 | 0.39 |
| 1 | 66457244 | rs2253745 | -0.03 | -2.083 | -1.057 | 0.513 | 0.01 | 0.015 | 0.015 | 2.66 | 0.75 | 0.72 | 0.65 | 0.57 | 0.71 | 0.62 |
| 1 | 66470379 | rs7534143 | 1.834 | -2.06 | -0.339 | 0.171 | -0.019 | 0.066 | 0.066 | 0.16 | 0.45 | 0.29 | 0.45 | 0.22 | 0.54 | 0.25 |
| 1 | 66478644 | rs1937451 | 1.298 | -1.892 | -0.434 | 0.107 | -0.035 | -0.049 | -0.049 | 0 | 0.74 | 0.62 | 0.72 | 0.84 | 0.84 | 0.77 |
| 1 | 66481523 | rs2455020 | -0.933 | -1.374 | -0.519 | 0.464 | 0 | 0 | 0 | 0 | 0.1 | 0.09 | 0.21 | 0.13 | 0.1 | 0.17 |
| 1 | 66496108 | rs12757542 | 0.082 | -1.079 | -0.622 | 0.048 | -0.035 | 0.103 | 0.103 | 0.099 | 0.28 | 0.1 | 0.16 | 0.02 | 0.28 | 0.18 |
| 1 | 66496139 | rs2455032 | 0.748 | -1.339 | 0.094 | 0.416 | 0.002 | -0.014 | -0.014 | 0.401 | 0.58 | 0.51 | 0.69 | 0.34 | 0.63 | 0.47 |
| 1 | 66499557 | rs2503220 | 0.105 | -1.72 | 0.008 | 0.219 | -0.002 | 0 | 0 | 4.582 | 0.11 | 0.16 | 0.21 | 0.13 | 0.09 | 0.22 |
| 1 | 66509014 | rs2228693 | 1.194 | -1.755 | 0.128 | 0.608 | 0 | 0.002 | 0.002 | 0 | 0.51 | 0.44 | 0.54 | 0.57 | 0.55 | 0.47 |
| 1 | 66520998 | rs2455012 | 0.034 | -1.69 | 0.184 | 0.622 | -0.004 | 0.015 | 0.015 | 0 | 0.94 | 0.97 | 0.97 | 0.97 | 0.95 | 0.97 |
| 1 | 66524757 | rs6588184 | 0.416 | -1.623 | 0.232 | 0.689 | 0 | 0.001 | 0.001 | 0 | 0.83 | 0.81 | 0.76 | 0.85 | 0.86 | 0.71 |
| 1 | 66538408 | rs7552924 | 1.159 | -1.633 | -0.201 | 0.477 | -0.001 | 0.048 | 0.048 | 0 | 0.47 | 0.36 | 0.48 | 0.21 | 0.54 | 0.26 |
| 1 | 66539456 | rs1392817 | 0.875 | -1.145 | -0.318 | 0.3 | 0 | 0.001 | 0.001 | 0 | 0.49 | 0.44 | 0.54 | 0.56 | 0.54 | 0.42 |
| 1 | 66547009 | rs6588186 | 0.896 | -0.985 | 0.058 | 0.952 | -0.001 | 0.149 | 0.149 | 0 | 0.79 | 0.54 | 0.7 | 0.58 | 0.84 | 0.59 |
| 1 | 66555792 | rs12406476 | -0.834 | -0.618 | -0.251 | 0.771 | -0.009 | 0.001 | 0.001 | 0.159 | 0.64 | 0.69 | 0.52 | 0.64 | 0.48 | 0.72 |
| 1 | 66558759 | rs4655595 | -0.5 | -1.347 | -0.249 | 0.795 | 0.014 | -0.01 | -0.01 | 0 | 0.21 | 0.18 | 0.14 | 0.3 | 0.09 | 0.38 |
| 1 | 66559173 | rs10158178 | 0.38 | -1.314 | -0.134 | 0.682 | -0.006 | -0.007 | -0.007 | 2.305 | 0.43 | 0.5 | 0.37 | 0.35 | 0.39 | 0.34 |
| 1 | 66580943 | rs650440 | 0.602 | -1.434 | 0.329 | 0.71 | 0.002 | -0.001 | -0.001 | 0 | 0.4 | 0.4 | 0.57 | 0.65 | 0.53 | 0.44 |
| 1 | 66585962 | rs17421832 | 0.434 | -0.48 | 0.108 | 0.655 | -0.005 | 0 | 0 | 0 | 0.07 | 0.06 | 0.05 | 0.11 | 0.14 | 0.07 |
| 1 | 66586760 | rs10157331 | -0.254 | -0.643 | 0.355 | 0.632 | 0.085 | 0.132 | 0.132 | 0.756 | 0.05 | 0.4 | 0.15 | 0.04 | 0.02 | 0.02 |
| 1 | 66587376 | rs12745871 | 1.234 | -0.772 | -0.113 | 0.298 | -0.075 | 0.002 | 0.002 | 0.258 | 0.33 | 0.07 | 0.21 | 0.21 | 0.34 | 0.25 |
| 1 | 66588029 | rs17421942 | -0.748 | -0.72 | -0.273 | 0.063 | -0.014 | 0.084 | 0.084 | 0 | 0.23 | 0.07 | 0.11 | 0.11 | 0.21 | 0.3 |
| 1 | 66589748 | rs539322 | 1.255 | -0.254 | -0.101 | 0.325 | -0.013 | 0.037 | 0.037 | 0.575 | 0.31 | 0.2 | 0.36 | 0.17 | 0.37 | 0.24 |
| 1 | 66598763 | rs563573 | 1.236 | 0.039 | -0.157 | 0.289 | -0.003 | 0 | 0 | 0.001 | 0.24 | 0.2 | 0.27 | 0.13 | 0.32 | 0.18 |
| 1 | 66603875 | rs596662 | 1.39 | -0.156 | -0.306 | 0.326 | -0.034 | 0.043 | 0.043 | 0.528 | 0.36 | 0.07 | 0.22 | 0.22 | 0.34 | 0.3 |
| 1 | 66609154 | rs2503174 | -0.225 | -0.23 | -0.049 | 0.391 | 0.001 | -0.023 | -0.023 | 0 | 0.67 | 0.79 | 0.68 | 0.42 | 0.68 | 0.5 |
| 1 | 66615238 | rs1604269 | 0.89 | -0.218 | 0.003 | 0.395 | 0.009 | 0.006 | 0.006 | 0.126 | 0.61 | 0.3 | 0.5 | 0.35 | 0.67 | 0.48 |
| 1 | 66615971 | rs2455035 | 0.898 | -0.225 | -0.05 | 0.392 | 0.021 | 0.011 | 0.011 | 0.116 | 0.61 | 0.22 | 0.49 | 0.35 | 0.67 | 0.48 |
| 1 | 66622634 | rs1577844 | 1.218 | 0.106 | -0.604 | 0.475 | -0.036 | 0.058 | 0.058 | 0 | 0.25 | 0.11 | 0.28 | 0.13 | 0.33 | 0.18 |
| 1 | 66627732 | rs2485381 | -0.6 | 0.145 | -0.59 | 0.865 | -0.016 | 0.023 | 0.023 | 0.306 | 0.73 | 0.71 | 0.73 | 0.59 | 0.83 | 0.56 |
| 1 | 66634921 | rs11208807 | 0.177 | 0.073 | -0.452 | 0.426 | -0.001 | 0.002 | 0.002 | 0 | 0.33 | 0.3 | 0.32 | 0.58 | 0.31 | 0.5 |
| 1 | 66643562 | rs693257 | -0.509 | 0.623 | -0.559 | 0.652 | -0.04 | 0.058 | 0.058 | 0 | 0.76 | 0.89 | 0.73 | 0.87 | 0.67 | 0.82 |
| 1 | 66644337 | rs496713 | -0.093 | 0.6 | -0.459 | 0.883 | -0.01 | -0.012 | -0.012 | 0.472 | 0.73 | 0.74 | 0.67 | 0.65 | 0.67 | 0.81 |
| 1 | 66646184 | rs12135034 | 0.038 | -0.585 | -0.325 | -0.297 | 0.008 | 0.043 | 0.043 | 0 | 0.25 | 0.1 | 0.15 | 0.13 | 0.22 | 0.16 |
| 1 | 66663239 | rs11208816 | 0.992 | -0.454 | 0.25 | -0.244 | -0.007 | 0 | 0 | 0 | 0.56 | 0.31 | 0.62 | 0.57 | 0.52 | 0.64 |
| 1 | 66675699 | rs12045279 | -0.014 | -0.522 | 0.639 | -0.246 | -0.003 | -0.002 | -0.002 | 0 | 0.09 | 0.08 | 0.04 | 0.14 | 0.03 | 0.08 |
| 1 | 66682087 | rs1416312 | 1.096 | -0.532 | 0.325 | -0.431 | -0.009 | 0.045 | 0.045 | 0 | 0.55 | 0.26 | 0.63 | 0.71 | 0.53 | 0.72 |
| 1 | 66683452 | rs544361 | 0.016 | -0.111 | -0.275 | -0.257 | 0.105 | 0.043 | 0.043 | 0 | 0.07 | 0.13 | 0.16 | 0.14 | 0.22 | 0.08 |
| 1 | 66689109 | rs17423910 | 0.192 | -0.807 | -0.574 | -0.231 | 0.008 | 0.04 | 0.04 | 0 | 0.25 | 0.09 | 0.15 | 0.14 | 0.22 | 0.16 |
| 1 | 66691885 | rs952635 | -0.74 | -1.446 | -0.584 | -0.413 | -0.002 | 0.001 | 0.001 | 0 | 0.62 | 0.66 | 0.48 | 0.43 | 0.72 | 0.41 |
| 1 | 66697753 | rs12731764 | 1.209 | -0.929 | -0.662 | -0.759 | -0.088 | 0.101 | 0.101 | 0 | 0.17 | 0.02 | 0.14 | 0 | 0.27 | 0.07 |
| 1 | 66700542 | rs6588191 | -1.491 | -1.975 | 0.226 | 0.004 | -0.083 | 0.153 | 0.153 | 0.629 | 0.16 | 0.61 | 0.16 | 0.37 | 0.06 | 0.13 |
| 1 | 66706933 | rs6704282 | -1.418 | -1.939 | 0.346 | 0.007 | -0.076 | 0.213 | 0.213 | 0.579 | 0.15 | 0.54 | 0.15 | 0.37 | 0.06 | 0.13 |
| 1 | 66714584 | rs6664618 | 0.814 | -2.172 | 0.045 | -0.184 | -0.097 | 0.102 | 0.102 | 0 | 0.4 | 0.04 | 0.28 | 0.17 | 0.57 | 0.27 |
| 1 | 66716207 | rs12043611 | -1.461 | -1.862 | 0.561 | 0.232 | -0.077 | 0.155 | 0.155 | 1.017 | 0.16 | 0.63 | 0.16 | 0.37 | 0.06 | 0.14 |
| 1 | 66724745 | rs6668312 | -0.876 | -1.838 | 0.583 | 0.132 | -0.046 | -0.039 | -0.039 | 0 | 0.11 | 0.49 | 0.08 | 0.02 | 0.03 | 0 |
| 1 | 66730713 | rs1321172 | 1.019 | -1.857 | 0.426 | 0.041 | -0.049 | 0.079 | 0.079 | 0 | 0.36 | 0.14 | 0.28 | 0.16 | 0.54 | 0.29 |
| 1 | 66735601 | rs6700403 | 0.119 | -1.782 | 0.494 | 0.1 | 0.023 | -0.002 | -0.002 | 2.984 | 0.46 | 0.32 | 0.38 | 0.51 | 0.6 | 0.43 |
| 1 | 66737060 | rs543589 | -0.091 | -1.428 | 0.559 | 0.374 | 0.037 | 0 | 0 | 2.099 | 0.53 | 0.73 | 0.43 | 0.52 | 0.62 | 0.49 |
| 1 | 66745352 | rs12060491 | 0.567 | -1.509 | 0.175 | 0.426 | 0.001 | -0.002 | -0.002 | 0 | 0.24 | 0.19 | 0.34 | 0.41 | 0.17 | 0.37 |
| 1 | 66752207 | rs529601 | -1.396 | -1.512 | -0.039 | 0.385 | -0.02 | 0.066 | 0.066 | 0 | 0.78 | 0.94 | 0.78 | 0.94 | 0.79 | 0.86 |
| 1 | 66756717 | rs519044 | 0.261 | -0.471 | 0.498 | 0.613 | -0.032 | 0.105 | 0.105 | 0 | 0.39 | 0.15 | 0.28 | 0.16 | 0.56 | 0.36 |
| 1 | 66757975 | rs486136 | 0.005 | -0.646 | 0.543 | 0.62 | 0 | -0.002 | -0.002 | 0 | 0.42 | 0.46 | 0.35 | 0.19 | 0.57 | 0.38 |
| 1 | 66766222 | rs524770 | -0.862 | -0.516 | -0.351 | 0.686 | 0.03 | 0.001 | 0.001 | 0 | 0.54 | 0.73 | 0.44 | 0.53 | 0.63 | 0.49 |
| 1 | 66766741 | rs499170 | -1.419 | -0.347 | 0.403 | 1.283 | -0.049 | 0.231 | 0.231 | 0.681 | 0.14 | 0.49 | 0.11 | 0.36 | 0.06 | 0.1 |
| 1 | 66776137 | rs6689595 | 0.456 | 0.651 | -1.268 | -0.135 | -0.034 | 0.103 | 0.103 | 0 | 0.57 | 0.22 | 0.58 | 0.8 | 0.53 | 0.52 |
| 1 | 66784679 | rs1556805 | 0.512 | 0.586 | -1.206 | -0.138 | -0.071 | 0.147 | 0.147 | 0.016 | 0.57 | 0.1 | 0.56 | 0.81 | 0.54 | 0.52 |
| 1 | 66786000 | rs6667596 | -0.316 | 0.303 | -0.027 | -0.551 | -0.048 | -0.113 | -0.113 | 0 | 0.17 | 0.41 | 0.06 | 0.01 | 0.07 | 0.19 |
| 1 | 66790248 | rs1040715 | -0.24 | 0.237 | 0.112 | -0.575 | -0.084 | 0.038 | 0.038 | 0 | 0.79 | 0.92 | 0.72 | 0.9 | 0.62 | 0.76 |

| | | | | | | | | | | | | | | | | |
|---|---|---|---|---|---|---|---|---|---|---|---|---|---|---|---|---|
| 1 | 66796248 | rs910693 | 0.446 | 0.22 | 0.143 | -0.594 | 0.043 | -0.02 | -0.02 | 0 | 0.74 | 0.63 | 0.64 | 0.82 | 0.61 | 0.71 |
| 1 | 66801495 | rs6696880 | -0.431 | -0.378 | 1.191 | -0.743 | -0.022 | -0.067 | -0.067 | 0 | 0.15 | 0.23 | 0.05 | 0.01 | 0.07 | 0.22 |
| 1 | 66801552 | rs518476 | 1.414 | -0.189 | 1.539 | -0.872 | -0.001 | 0.001 | 0.001 | 0.087 | 0.61 | 0.39 | 0.68 | 0.82 | 0.62 | 0.56 |
| 1 | 66802591 | rs693826 | 1.018 | -0.155 | 1.354 | -1.07 | -0.019 | 0.112 | 0.112 | 0 | 0.66 | 0.85 | 0.74 | 0.9 | 0.67 | 0.57 |
| 1 | 66802772 | rs1890197 | 1.625 | -0.328 | 1.98 | -1.055 | 0.001 | 0.006 | 0.006 | 0 | 0.58 | 0.38 | 0.64 | 0.81 | 0.61 | 0.45 |
| 1 | 66808852 | rs12093599 | -1.838 | -0.248 | 1.832 | -1.006 | 0.047 | 0.138 | 0.138 | 1.554 | 0.06 | 0.43 | 0.08 | 0.09 | 0.01 | 0.12 |
| 1 | 66815071 | rs1321177 | 0.548 | -0.326 | 1.793 | -1.099 | 0.019 | -0.029 | -0.029 | 0.846 | 0.77 | 0.68 | 0.71 | 0.89 | 0.69 | 0.73 |
| 1 | 66819623 | rs598961 | 1.398 | -0.992 | 0.681 | -1.287 | -0.007 | 0.006 | 0.006 | 0 | 0.44 | 0.37 | 0.59 | 0.67 | 0.49 | 0.48 |
| 1 | 66823134 | rs2144719 | 0.321 | -0.481 | 0.035 | -1.697 | -0.129 | 0.074 | 0.074 | 0 | 0.73 | 0.94 | 0.71 | 0.91 | 0.61 | 0.81 |
| 1 | 66825944 | rs7525321 | -0.328 | 0.063 | 0.038 | -2.006 | -0.065 | 0.055 | 0.055 | 0 | 0.87 | 0.95 | 0.78 | 0.95 | 0.74 | 0.91 |
| 1 | 66830107 | rs783037 | 1.367 | 0.121 | 0.064 | -2.12 | -0.019 | -0.019 | -0.019 | 0 | 0.05 | 0 | 0.04 | 0.06 | 0.06 | 0.11 |
| 1 | 66831370 | rs783036 | -1.85 | 0.067 | 0.289 | -2.44 | -0.019 | 0.244 | 0.244 | 0 | 0.5 | 0.79 | 0.45 | 0.15 | 0.49 | 0.59 |
| 1 | 66838733 | rs783067 | 0.08 | 0.452 | 0.206 | -2.084 | -0.066 | 0.047 | 0.047 | 0 | 0.13 | 0.05 | 0.22 | 0.06 | 0.27 | 0.11 |
| 1 | 66851147 | rs11208844 | -0.927 | 0.179 | 0.703 | -2.172 | -0.015 | -0.083 | -0.083 | 0 | 0.24 | 0.37 | 0.14 | 0.04 | 0.16 | 0.39 |
| 1 | 66853485 | rs12128881 | 0.98 | 0.171 | 0.262 | -2.073 | -0.018 | 0.044 | 0.044 | 0 | 0.08 | 0.01 | 0.05 | 0 | 0.1 | 0.02 |
| 1 | 66857024 | rs1999856 | -0.419 | 0.396 | 0.22 | -2.095 | -0.055 | 0.05 | 0.05 | 0 | 0.14 | 0.05 | 0.19 | 0.05 | 0.22 | 0.08 |
| 1 | 66862492 | rs945969 | 0.726 | 0.331 | 0.58 | -2.059 | -0.016 | 0.06 | 0.06 | 0 | 0.1 | 0.01 | 0.05 | 0 | 0.11 | 0.02 |
| 1 | 66866041 | rs11208847 | -0.991 | 0.54 | 0.961 | -2.123 | -0.015 | -0.139 | -0.139 | 0 | 0.29 | 0.49 | 0.16 | 0.07 | 0.22 | 0.38 |
| 1 | 66873659 | rs1194487 | 0.073 | 0.473 | 0.562 | -2.117 | -0.05 | 0.003 | 0.003 | 0 | 0.12 | 0.05 | 0.17 | 0.09 | 0.19 | 0.12 |
| 1 | 66887451 | rs12093028 | -0.697 | 0.338 | 0.893 | -2.141 | -0.033 | -0.119 | -0.119 | 0 | 0.27 | 0.45 | 0.14 | 0.07 | 0.17 | 0.36 |
| 1 | 66905141 | rs12140647 | -0.538 | 0.331 | 1.083 | -2.174 | -0.012 | -0.062 | -0.062 | 0 | 0.26 | 0.32 | 0.13 | 0.07 | 0.17 | 0.36 |
| 1 | 66915125 | rs783435 | 1.301 | 0.546 | 0.97 | -2.198 | 0.007 | 0.074 | 0.074 | 1.831 | 0.62 | 0.23 | 0.75 | 0.91 | 0.68 | 0.58 |
| 1 | 66917166 | rs1819626 | 0.896 | 0.64 | 0.743 | -1.935 | -0.026 | -0.04 | -0.04 | 0 | 0.07 | 0 | 0.13 | 0.12 | 0.1 | 0.14 |
| 1 | 66920451 | rs699267 | -0.731 | 0.631 | 0.781 | -1.967 | -0.002 | -0.01 | -0.01 | 0 | 0.92 | 0.9 | 0.93 | 1 | 0.9 | 0.97 |
| 1 | 66920654 | rs10889612 | -0.616 | 0.509 | 0.817 | -1.956 | -0.029 | -0.112 | -0.112 | 0.069 | 0.27 | 0.43 | 0.14 | 0.09 | 0.17 | 0.35 |
| 1 | 66920877 | rs10493405 | 0.568 | 0.503 | 0.421 | -1.969 | -0.014 | 0.039 | 0.039 | 0 | 0.1 | 0.01 | 0.04 | 0.06 | 0.06 | 0.21 |
| 1 | 66921372 | rs12116697 | 0.187 | 0.645 | -0.406 | -2.43 | -0.001 | 0.207 | 0.207 | 0 | 0.48 | 0.23 | 0.33 | 0.11 | 0.46 | 0.49 |
| 1 | 66926566 | rs7539015 | -0.278 | 0.163 | 0.419 | -1.59 | -0.01 | -0.057 | -0.057 | 0.079 | 0.25 | 0.31 | 0.13 | 0.09 | 0.17 | 0.36 |
| 1 | 66929805 | rs10889617 | -0.278 | 0.185 | 0.241 | -1.581 | -0.003 | -0.033 | -0.033 | 0 | 0.25 | 0.26 | 0.12 | 0.09 | 0.17 | 0.36 |
| 1 | 66930964 | rs699260 | 0.987 | 0.204 | -0.593 | -1.333 | -0.004 | 0.126 | 0.126 | 0.797 | 0.55 | 0.23 | 0.6 | 0.79 | 0.58 | 0.44 |
| 1 | 66931240 | rs6684192 | 0.085 | 0.294 | -1.298 | -1.287 | 0.015 | 0.042 | 0.042 | 0 | 0.09 | 0.19 | 0.16 | 0.12 | 0.1 | 0.14 |
| 1 | 66934039 | rs783329 | 0.584 | 0.352 | -2.153 | -1.07 | -0.052 | 0.188 | 0.188 | 1.355 | 0.53 | 0.08 | 0.59 | 0.77 | 0.57 | 0.44 |
| 1 | 66939920 | rs1832420 | -0.657 | 0.308 | -1.24 | -0.394 | 0 | -0.008 | -0.008 | 0 | 0.11 | 0.1 | 0.12 | 0.07 | 0.06 | 0.27 |
| 1 | 66940704 | rs17128938 | -0.959 | 0.028 | -1.336 | -0.218 | 0.015 | 0.006 | 0.006 | 0 | 0.12 | 0.08 | 0.05 | 0.14 | 0.05 | 0.24 |
| 1 | 66941200 | rs699252 | 0.186 | 0.482 | -1.842 | -0.36 | -0.062 | 0.003 | 0.003 | 0.009 | 0.28 | 0.15 | 0.39 | 0.25 | 0.5 | 0.38 |
| 1 | 66941630 | rs783322 | 0.125 | 0.422 | -1.18 | -0.388 | -0.001 | 0 | 0 | 0 | 0.28 | 0.25 | 0.4 | 0.25 | 0.5 | 0.38 |
| 1 | 66941782 | rs12140300 | 1.496 | -0.865 | -1.86 | -0.117 | 0.055 | -0.028 | -0.028 | 3.069 | 0.22 | 0.09 | 0.09 | 0.17 | 0.11 | 0.11 |
| 1 | 66943781 | rs6694790 | 0.538 | -0.837 | -1.845 | -0.103 | 0.015 | 0.006 | 0.006 | 0 | 0.12 | 0.08 | 0.05 | 0.14 | 0.05 | 0.23 |
| 1 | 66945358 | rs814146 | 0.173 | -1.241 | -1.566 | -0.445 | 0.002 | 0.002 | 0.002 | 0.331 | 0.42 | 0.36 | 0.61 | 0.67 | 0.64 | 0.49 |
| 1 | 66953482 | rs4567260 | -0.282 | -1.379 | -1.636 | -0.686 | 0 | 0.027 | 0.027 | 0 | 0.8 | 0.84 | 0.87 | 0.86 | 0.84 | 0.74 |
| 1 | 66955878 | rs11208878 | -0.451 | -1.389 | -1.265 | -0.734 | 0.022 | 0.013 | 0.013 | 1.084 | 0.57 | 0.68 | 0.62 | 0.41 | 0.65 | 0.76 |
| 1 | 66956685 | rs9651195 | 0.351 | -0.82 | -1.545 | -0.819 | -0.007 | -0.005 | -0.005 | 0 | 0.38 | 0.59 | 0.4 | 0.06 | 0.44 | 0.29 |
| 1 | 66957337 | rs1025993 | 1.068 | -0.846 | -1.538 | -0.766 | 0.042 | -0.02 | -0.02 | 0 | 0.35 | 0.21 | 0.3 | 0.59 | 0.24 | 0.22 |
| 1 | 66958986 | rs12143938 | 2.149 | -0.927 | -1.587 | -0.778 | 0.052 | -0.027 | -0.027 | 0 | 0.21 | 0.09 | 0.08 | 0.17 | 0.11 | 0.11 |
| 1 | 66959434 | rs1338211 | -0.626 | -0.949 | -1.505 | -0.703 | -0.013 | -0.016 | -0.016 | 0 | 0.54 | 0.46 | 0.67 | 0.79 | 0.69 | 0.73 |
| 1 | 66961122 | rs961378 | -0.002 | -0.843 | -1.471 | -0.673 | -0.004 | -0.001 | -0.001 | 2.406 | 0.62 | 0.44 | 0.54 | 0.58 | 0.66 | 0.72 |
| 1 | 66962193 | rs1415975 | -1.97 | -0.809 | -1.525 | -0.669 | -0.01 | 0.008 | 0.008 | 0 | 0.92 | 0.92 | 0.92 | 1 | 0.89 | 0.98 |
| 1 | 66962906 | rs1338187 | 0.508 | -0.883 | -1.497 | -0.698 | 0.044 | -0.02 | -0.02 | 0.104 | 0.4 | 0.2 | 0.31 | 0.56 | 0.31 | 0.57 |
| 1 | 66965475 | rs765685 | 0.428 | -0.739 | -1.338 | -0.584 | -0.037 | -0.015 | -0.015 | 0 | 0.34 | 0.2 | 0.44 | 0.43 | 0.48 | 0.27 |
| 1 | 66967395 | rs17432003 | 1.607 | -0.714 | -1.429 | -0.599 | -0.045 | 0.047 | 0.047 | 0 | 0.2 | 0.1 | 0.23 | 0.04 | 0.35 | 0.14 |
| 1 | 66969013 | rs7555470 | 1.358 | -0.887 | -0.98 | -0.152 | 0.001 | -0.034 | -0.034 | 2.122 | 0.14 | 0.06 | 0.21 | 0.39 | 0.14 | 0.13 |
| 1 | 66969954 | rs2121047 | 0.968 | -0.945 | -0.104 | -0.413 | -0.013 | -0.051 | -0.051 | 0 | 0.15 | 0.21 | 0.15 | 0.01 | 0.09 | 0.14 |
| 1 | 66970057 | rs1545346 | -0.215 | -0.389 | -0.2 | 0.142 | -0.012 | -0.059 | -0.059 | 0.18 | 0.19 | 0.08 | 0.22 | 0.38 | 0.2 | 0.47 |
| 1 | 66972478 | rs6674096 | -0.347 | -0.363 | -0.227 | 0.155 | -0.002 | 0.028 | 0.028 | 1.885 | 0.21 | 0.27 | 0.24 | 0.38 | 0.2 | 0.47 |
| 1 | 66986715 | rs7533806 | 0.657 | -0.71 | -0.294 | 0.515 | 0.009 | -0.038 | -0.038 | 0.243 | 0.16 | 0.07 | 0.13 | 0.32 | 0.12 | 0.13 |
| 1 | 66988401 | rs12093875 | 0.457 | -0.94 | -0.289 | -0.031 | -0.004 | -0.007 | -0.007 | 0 | 0.26 | 0.3 | 0.22 | 0.22 | 0.24 | 0.18 |
| 1 | 66988477 | rs2121052 | 0.892 | -0.514 | -0.309 | -0.636 | 0.016 | -0.07 | -0.07 | 0.676 | 0.37 | 0.14 | 0.41 | 0.46 | 0.3 | 0.64 |
| 1 | 66988499 | rs2121053 | 1.57 | -0.732 | -0.476 | -0.78 | 0.057 | -0.105 | -0.105 | 0 | 0.26 | 0.06 | 0.32 | 0.42 | 0.16 | 0.48 |
| 1 | 66988869 | rs11208897 | -0.135 | -0.736 | -0.174 | -0.793 | -0.001 | 0.021 | 0.021 | 0 | 0.3 | 0.53 | 0.26 | 0.36 | 0.26 | 0.3 |
| 1 | 66990824 | rs6693644 | -0.644 | -0.772 | -0.131 | -0.772 | -0.012 | 0 | 0 | 0 | 0.66 | 0.71 | 0.66 | 0.68 | 0.54 | 0.82 |
| 1 | 66991390 | rs12135213 | 1.568 | -1.034 | -0.196 | -0.762 | 0.057 | -0.105 | -0.105 | 1.456 | 0.26 | 0.06 | 0.25 | 0.42 | 0.16 | 0.48 |
| 1 | 66991699 | rs10493407 | 0.275 | -1.03 | -0.184 | -0.776 | 0.063 | 0.061 | 0.061 | 6.391 | 0.43 | 0.24 | 0.46 | 0.74 | 0.28 | 0.61 |
| 1 | 66992613 | rs1338203 | 1.685 | -0.793 | 0.186 | -0.716 | 0 | 0.006 | 0.006 | 0.217 | 0.16 | 0.17 | 0.14 | 0.32 | 0.12 | 0.13 |
| 1 | 66992833 | rs12043594 | 1.764 | -0.782 | 0.053 | -0.711 | 0.009 | -0.041 | -0.041 | 0 | 0.16 | 0.07 | 0.13 | 0.32 | 0.12 | 0.13 |
| 1 | 66998436 | rs7532400 | 1.663 | -0.658 | 0.291 | -0.691 | 0.002 | -0.046 | -0.046 | 0 | 0.16 | 0.07 | 0.18 | 0.33 | 0.14 | 0.15 |
| 1 | 67002765 | rs4655616 | -0.569 | -0.535 | 0.387 | -0.777 | 0.012 | -0.001 | -0.001 | 0 | 0.64 | 0.59 | 0.6 | 0.67 | 0.49 | 0.8 |
| 1 | 67005072 | rs1338184 | -1.546 | -0.355 | 0.424 | -0.887 | 0.053 | -0.098 | -0.098 | 0 | 0.8 | 0.68 | 0.78 | 1 | 0.65 | 0.95 |
| 1 | 67005503 | rs1415974 | -1.53 | -0.357 | 0.424 | -0.888 | 0.052 | -0.098 | -0.098 | 0 | 0.8 | 0.68 | 0.78 | 1 | 0.65 | 0.95 |
| 1 | 67007389 | rs12136922 | -0.553 | -0.396 | 0.519 | -0.954 | 0.041 | -0.091 | -0.091 | 0.63 | 0.69 | 0.59 | 0.69 | 0.96 | 0.51 | 0.81 |

| | | | | | | | | | | | | | | | | |
|---|---|---|---|---|---|---|---|---|---|---|---|---|---|---|---|---|
| 1 | 67007679 | rs17486706 | 1.04 | -0.384 | 0.112 | -0.884 | -0.003 | 0.002 | 0.002 | 0 | 0.12 | 0.09 | 0.09 | 0.04 | 0.14 | 0.14 |
| 1 | 67010654 | rs4655617 | -0.13 | -0.626 | -0.022 | -0.97 | 0.011 | 0.139 | 0.139 | 4.758 | 0.53 | 0.26 | 0.55 | 0.82 | 0.41 | 0.73 |
| 1 | 67011019 | rs1856316 | 1.278 | -0.693 | -0.097 | -1.005 | -0.017 | 0.02 | 0.02 | 0 | 0.12 | 0.07 | 0.06 | 0.04 | 0.14 | 0.14 |
| 1 | 67011986 | rs6662771 | 1.643 | -1.693 | 0.051 | -1.011 | -0.017 | -0.044 | -0.044 | 0 | 0.11 | 0.17 | 0.12 | 0.02 | 0.07 | 0.09 |
| 1 | 67012188 | rs2068381 | 0.115 | -1.612 | -0.123 | -0.98 | -0.022 | -0.038 | -0.038 | 0 | 0.28 | 0.39 | 0.23 | 0.18 | 0.23 | 0.22 |
| 1 | 67013055 | rs11208905 | 1.193 | -1.461 | -0.818 | -0.404 | 0.059 | -0.057 | -0.057 | 0 | 0.24 | 0.12 | 0.22 | 0.41 | 0.11 | 0.44 |
| 1 | 67015272 | rs12048885 | 1.427 | -1.452 | -0.833 | -0.413 | -0.002 | -0.035 | -0.035 | 0 | 0.16 | 0.09 | 0.27 | 0.36 | 0.16 | 0.15 |
| 1 | 67015972 | rs1856317 | 0.389 | -1.019 | -0.623 | -0.743 | 0.068 | 0.082 | 0.082 | 6.764 | 0.41 | 0.21 | 0.49 | 0.78 | 0.27 | 0.59 |
| 1 | 67022204 | rs1832417 | 0.888 | -0.516 | -0.322 | -0.227 | -0.001 | -0.003 | -0.003 | 0 | 0.14 | 0.12 | 0.09 | 0.2 | 0.15 | 0.16 |
| 1 | 67030601 | rs17129100 | 1.041 | -0.517 | -0.324 | -0.241 | -0.001 | 0 | 0 | 0.433 | 0.14 | 0.11 | 0.09 | 0.12 | 0.15 | 0.15 |
| 1 | 67030831 | rs1338189 | -0.015 | -0.878 | 0.226 | -0.7 | -0.016 | -0.04 | -0.04 | 0 | 0.23 | 0.36 | 0.19 | 0.12 | 0.15 | 0.2 |
| 1 | 67037786 | rs963141 | -0.715 | -0.91 | 0.437 | -0.706 | 0.001 | 0.002 | 0.002 | 3.336 | 0.37 | 0.6 | 0.29 | 0.24 | 0.3 | 0.35 |
| 1 | 67043120 | rs920987 | -0.654 | -0.905 | 0.456 | -0.708 | -0.008 | -0.025 | -0.025 | 0 | 0.37 | 0.53 | 0.28 | 0.24 | 0.3 | 0.35 |
| 1 | 67047442 | rs1445573 | 0.288 | -1.596 | 0.205 | -0.715 | -0.013 | 0.004 | 0.004 | 0 | 0.38 | 0.38 | 0.3 | 0.59 | 0.27 | 0.67 |
| 1 | 67052736 | rs2166307 | 0.91 | -1.537 | 0.254 | -0.73 | 0.001 | -0.001 | -0.001 | 2.657 | 0.17 | 0.35 | 0.11 | 0.14 | 0.15 | 0.25 |
| 1 | 67062126 | rs10749763 | -1.584 | -1.632 | 0.038 | -1.078 | -0.098 | 0.092 | 0.092 | 0 | 0.76 | 0.91 | 0.77 | 1 | 0.56 | 0.91 |
| 1 | 67066555 | rs6588207 | -1.21 | -1.695 | 0.414 | -1.109 | 0.037 | -0.103 | -0.103 | 0 | 0.74 | 0.65 | 0.75 | 1 | 0.56 | 0.91 |
| 1 | 67075312 | rs11208920 | -0.336 | -1.812 | 0.334 | -1.058 | 0.01 | 0.012 | 0.012 | 0.07 | 0.5 | 0.42 | 0.54 | 0.54 | 0.44 | 0.39 |
| 1 | 67080226 | rs11208922 | -0.509 | -1.878 | 0.145 | -1.082 | 0 | -0.017 | -0.017 | 0.061 | 0.61 | 0.59 | 0.66 | 0.88 | 0.41 | 0.76 |
| 1 | 67082018 | rs12141816 | 0.962 | -1.998 | -0.062 | -1.128 | 0.001 | 0.041 | 0.041 | 0.452 | 0.19 | 0.08 | 0.17 | 0.1 | 0.15 | 0.1 |
| 1 | 67083647 | rs11208924 | 1.865 | -2.019 | -0.118 | -1.109 | -0.01 | -0.1 | -0.1 | 0 | 0.15 | 0.01 | 0.26 | 0.31 | 0.14 | 0.14 |
| 1 | 67085295 | rs4462119 | 1.964 | -1.938 | -0.047 | -1.102 | 0.043 | -0.082 | -0.082 | 0 | 0.21 | 0.07 | 0.2 | 0.45 | 0.12 | 0.42 |
| 1 | 67089561 | rs11208927 | -0.31 | -1.941 | 0.133 | -1.11 | -0.017 | 0.01 | 0.01 | 0 | 0.39 | 0.51 | 0.45 | 0.43 | 0.29 | 0.33 |
| 1 | 67092248 | rs5008753 | -0.542 | -1.901 | 0.072 | -1.197 | -0.003 | 0.022 | 0.022 | 0.017 | 0.61 | 0.64 | 0.66 | 0.88 | 0.41 | 0.76 |
| 1 | 67100309 | rs6696812 | -0.775 | -1.859 | 0.061 | -1.073 | 0.003 | 0.002 | 0.002 | 0.353 | 0.54 | 0.75 | 0.56 | 0.5 | 0.44 | 0.48 |
| 1 | 67109335 | rs7526812 | 0.855 | -1.444 | 0.278 | -1.115 | -0.037 | 0.103 | 0.103 | 0.007 | 0.2 | 0.43 | 0.29 | 0.27 | 0.14 | 0.23 |
| 1 | 67113050 | rs6689755 | 0.125 | -1.448 | 0.001 | -1.187 | 0.019 | 0.075 | 0.075 | 0.262 | 0.55 | 0.65 | 0.58 | 0.9 | 0.41 | 0.81 |
| 1 | 67115084 | rs1925341 | 0.13 | -1.251 | 0.047 | -1.055 | -0.005 | 0.03 | 0.03 | 0 | 0.3 | 0.6 | 0.38 | 0.4 | 0.29 | 0.38 |
| 1 | 67123057 | rs6681460 | -0.873 | -0.912 | 0.137 | -1.433 | -0.003 | -0.045 | -0.045 | 0 | 0.74 | 0.74 | 0.75 | 1 | 0.56 | 0.91 |
| 1 | 67130530 | rs12407460 | 1.486 | -0.931 | -0.071 | -1.357 | -0.003 | 0.001 | 0.001 | 0 | 0.1 | 0.07 | 0.05 | 0.1 | 0.07 | 0.08 |
| 1 | 67134326 | rs6692318 | 0.718 | -1.05 | -0.213 | -1.473 | -0.006 | 0.014 | 0.014 | 0 | 0.14 | 0.07 | 0.07 | 0.1 | 0.11 | 0.14 |
| 1 | 67135100 | rs7536634 | 1.749 | -0.86 | 0.15 | -1.402 | 0.013 | 0.01 | 0.01 | 0 | 0.1 | 0.15 | 0.09 | 0.13 | 0.15 | 0.15 |
| 1 | 67140723 | rs7519747 | -0.858 | -0.687 | 0.238 | -1.591 | -0.033 | 0.003 | 0.003 | 0 | 0.75 | 0.8 | 0.76 | 1 | 0.56 | 0.9 |
| 1 | 67146817 | rs10789214 | -0.972 | -0.852 | 0.528 | -1.478 | -0.062 | 0.044 | 0.044 | 0 | 0.76 | 0.84 | 0.76 | 1 | 0.56 | 0.91 |
| 1 | 67151185 | rs10789215 | -0.895 | -0.791 | 0.296 | -1.45 | -0.039 | 0.011 | 0.011 | 0 | 0.75 | 0.81 | 0.76 | 1 | 0.57 | 0.91 |
| 1 | 67159658 | rs11208949 | 0.645 | -1.015 | 0.253 | -1.13 | 0.003 | -0.019 | -0.019 | 0.586 | 0.17 | 0.09 | 0.26 | 0.29 | 0.15 | 0.14 |
| 1 | 67160602 | rs1325266 | 0.433 | -1.125 | 0.591 | -1.198 | 0.004 | 0.009 | 0.009 | 0.044 | 0.55 | 0.44 | 0.46 | 0.71 | 0.41 | 0.68 |
| 1 | 67160814 | rs1325267 | 1.738 | -1.547 | 0.307 | -1.212 | 0.049 | -0.07 | -0.07 | 0.885 | 0.21 | 0.1 | 0.19 | 0.49 | 0.09 | 0.34 |
| 1 | 67168606 | rs12117649 | 0.184 | -1.352 | 0.332 | -1.187 | 0.012 | 0.026 | 0.026 | 0.046 | 0.55 | 0.36 | 0.45 | 0.71 | 0.41 | 0.68 |
| 1 | 67177270 | rs602513 | -0.651 | -1.315 | 0.37 | -1.233 | 0.052 | -0.134 | -0.134 | 0 | 0.71 | 0.42 | 0.71 | 1 | 0.56 | 0.82 |
| 1 | 67182636 | rs566057 | 0.927 | -1.364 | 0.313 | -1.208 | 0.002 | -0.036 | -0.036 | 0 | 0.16 | 0.07 | 0.26 | 0.29 | 0.15 | 0.14 |
| 1 | 67188829 | rs7515848 | 1.732 | 0.016 | 0.458 | -1.63 | 0.006 | -0.04 | -0.04 | 0.744 | 0.14 | 0.15 | 0.13 | 0 | 0.19 | 0.11 |
| 1 | 67191084 | rs536410 | -0.155 | 0.023 | 0.184 | -1.62 | 0.028 | -0.101 | -0.101 | 1.102 | 0.3 | 0.63 | 0.29 | 0 | 0.44 | 0.19 |
| 1 | 67195353 | rs2483704 | 0.179 | 0.053 | 0.141 | -1.582 | 0.064 | -0.133 | -0.133 | 0 | 0.27 | 0.41 | 0.27 | 0 | 0.42 | 0.19 |
| 1 | 67196277 | rs9988481 | -0.912 | -0.185 | -0.134 | -1.547 | 0.058 | 0.042 | 0.042 | 0 | 0.63 | 0.82 | 0.55 | 0.24 | 0.75 | 0.52 |
| 1 | 67203574 | rs693956 | -0.889 | 0.171 | -0.045 | -1.376 | 0.057 | 0.048 | 0.048 | 0 | 0.6 | 0.78 | 0.54 | 0.25 | 0.73 | 0.51 |
| 1 | 67209088 | rs604737 | 0.665 | 0.327 | -0.038 | -1.408 | -0.022 | 0.007 | 0.007 | 0 | 0.14 | 0.09 | 0.13 | 0 | 0.25 | 0.08 |
| 1 | 67209818 | rs657808 | -0.697 | -0.555 | -0.043 | -1.16 | 0.049 | 0.037 | 0.037 | 0 | 0.6 | 0.73 | 0.53 | 0.26 | 0.73 | 0.49 |
| 1 | 67216697 | rs2916 | 1.013 | -0.204 | -0.182 | -1.121 | 0.081 | -0.105 | -0.105 | 2.499 | 0.22 | 0.37 | 0.24 | 0 | 0.41 | 0.13 |
| 1 | 67221387 | rs2274987 | 0.238 | -0.655 | -0.653 | -0.895 | 0 | 0.009 | 0.009 | 0.221 | 0.32 | 0.25 | 0.26 | 0.26 | 0.31 | 0.32 |
| 1 | 67222507 | rs502376 | -1.281 | -0.883 | -0.781 | -0.347 | 0.002 | 0.027 | 0.027 | 0.475 | 0.91 | 0.94 | 0.77 | 0.66 | 0.92 | 0.89 |
| 1 | 67222697 | rs504169 | -0.221 | -1.167 | -0.835 | -0.978 | 0.118 | 0.086 | 0.086 | 0 | 0.53 | 0.74 | 0.53 | 0.26 | 0.75 | 0.45 |
| 1 | 67231916 | rs1391547 | -0.975 | 0.504 | -0.265 | -0.375 | -0.01 | 0.024 | 0.024 | 0 | 0.43 | 0.63 | 0.37 | 0.25 | 0.47 | 0.38 |
| 1 | 67235320 | rs12727422 | -0.035 | 0.691 | -0.164 | -0.432 | -0.013 | -0.025 | -0.025 | 0.446 | 0.23 | 0.32 | 0.13 | 0.15 | 0.18 | 0.21 |
| 1 | 67236723 | rs10493416 | 0.782 | 0.683 | -0.459 | -0.435 | -0.002 | 0.016 | 0.016 | 0 | 0.17 | 0.09 | 0.1 | 0.15 | 0.15 | 0.18 |
| 1 | 67239462 | rs7515248 | 0.543 | 0.704 | -0.467 | -0.443 | -0.005 | 0.022 | 0.022 | 0 | 0.19 | 0.09 | 0.12 | 0.15 | 0.17 | 0.18 |
| 1 | 67242571 | rs471093 | 1.166 | 0.807 | -0.509 | -0.011 | 0.022 | 0.059 | 0.059 | 0 | 0.47 | 0.22 | 0.39 | 0.35 | 0.44 | 0.51 |
| 1 | 67243024 | rs2133173 | -0.436 | 1.689 | -0.191 | 0.717 | 0.032 | -0.018 | -0.018 | 0 | 0.18 | 0.21 | 0.22 | 0.1 | 0.29 | 0.17 |
| 1 | 67245658 | rs562910 | 0.371 | 0.563 | -0.531 | 1.133 | -0.003 | 0.131 | 0.131 | 0.071 | 0.9 | 0.84 | 0.75 | 0.6 | 0.91 | 0.89 |
| 1 | 67256670 | rs1391546 | 0.497 | 0.564 | -0.779 | -0.266 | 0 | -0.046 | -0.046 | 0 | 0.75 | 0.76 | 0.83 | 0.98 | 0.77 | 0.79 |
| 1 | 67257871 | rs11208957 | 0.102 | -0.145 | -1.02 | -0.466 | 0.01 | 0.038 | 0.038 | 0 | 0.46 | 0.26 | 0.4 | 0.32 | 0.52 | 0.54 |
| 1 | 67259284 | rs474409 | -0.868 | -0.093 | -1.016 | -0.634 | 0.006 | -0.061 | -0.061 | 0.124 | 0.84 | 0.8 | 0.89 | 0.99 | 0.83 | 0.9 |
| 1 | 67261066 | rs12408354 | 0.803 | 0.462 | -1.094 | -0.951 | 0.006 | -0.085 | -0.085 | 0 | 0.3 | 0.26 | 0.2 | 0 | 0.34 | 0.13 |
| 1 | 67261079 | rs7533363 | 0.726 | 0.041 | -0.734 | -0.625 | 0.009 | 0.001 | 0.001 | 0 | 0.35 | 0.35 | 0.49 | 0.68 | 0.32 | 0.36 |
| 1 | 67262748 | rs559809 | 1.056 | -0.54 | -1.27 | -0.351 | 0.002 | -0.061 | -0.061 | 0.015 | 0.2 | 0.15 | 0.24 | 0.43 | 0.19 | 0.14 |
| 1 | 67268953 | rs12080201 | 0.911 | -0.665 | -1.029 | -0.48 | 0.024 | -0.017 | -0.017 | 0.557 | 0.14 | 0.06 | 0.22 | 0.24 | 0.13 | 0.22 |
| 1 | 67280515 | rs7550905 | 0.839 | -0.801 | -1.166 | -0.51 | 0.037 | -0.031 | -0.031 | 2.586 | 0.13 | 0.04 | 0.2 | 0.25 | 0.12 | 0.2 |
| 1 | 67283439 | rs17495916 | -0.413 | -0.488 | -1.005 | -1.061 | -0.002 | 0.03 | 0.03 | 0.063 | 0.49 | 0.38 | 0.45 | 0.37 | 0.52 | 0.65 |
| 1 | 67287302 | rs562372 | 0.953 | -0.647 | -0.702 | -1.058 | -0.005 | -0.004 | -0.004 | 0 | 0.32 | 0.28 | 0.39 | 0.57 | 0.33 | 0.23 |

| | | | | | | | | | | | | | | | | |
|---|---|---|---|---|---|---|---|---|---|---|---|---|---|---|---|---|
| 1 | 67297122 | rs7548267 | -0.376 | -0.647 | -0.593 | -0.844 | -0.004 | 0.016 | 0.016 | 0.027 | 0.41 | 0.32 | 0.38 | 0.35 | 0.44 | 0.54 |
| 1 | 67301096 | rs3738168 | 0.842 | -2.216 | -0.516 | -1.196 | 0.02 | 0.025 | 0.025 | 0 | 0.17 | 0.06 | 0.07 | 0.09 | 0.11 | 0.19 |
| 1 | 67306054 | rs11208966 | 0.212 | -2.544 | -0.176 | -0.642 | -0.003 | 0.007 | 0.007 | 0.018 | 0.25 | 0.23 | 0.26 | 0.55 | 0.17 | 0.39 |
| 1 | 67306616 | rs12030322 | -0.742 | -2.751 | -0.04 | -0.759 | 0.043 | 0.022 | 0.022 | 0 | 0.51 | 0.54 | 0.39 | 0.66 | 0.33 | 0.61 |
| 1 | 67312720 | rs476232 | 0.4 | -1.76 | 0.079 | 0.142 | -0.024 | 0.049 | 0.049 | 0 | 0.74 | 0.69 | 0.74 | 0.45 | 0.82 | 0.61 |
| 1 | 67314581 | rs2454320 | 0.387 | -1.743 | 0.044 | 0.15 | -0.006 | 0.013 | 0.013 | 0.016 | 0.74 | 0.73 | 0.74 | 0.45 | 0.82 | 0.61 |
| 1 | 67320816 | rs1104705 | -0.591 | -1.737 | 0.034 | 0.153 | -0.002 | 0.005 | 0.005 | 0.076 | 0.25 | 0.23 | 0.26 | 0.55 | 0.18 | 0.39 |
| 1 | 67337285 | rs1280338 | -0.569 | -0.807 | 0.652 | -0.831 | -0.01 | -0.101 | -0.101 | 0.108 | 0.18 | 0.3 | 0.09 | 0 | 0.14 | 0.06 |
| 1 | 67352960 | rs1925400 | -0.783 | -0.698 | 0.013 | -0.75 | 0.001 | -0.035 | -0.035 | 2.221 | 0.2 | 0.19 | 0.1 | 0 | 0.14 | 0.06 |
| 1 | 67372726 | rs2985809 | -1.271 | -0.488 | 0.661 | -0.221 | -0.061 | -0.049 | -0.049 | 1.986 | 0.17 | 0.57 | 0.11 | 0.04 | 0.08 | 0.11 |
| 1 | 67379284 | rs3008848 | 0.711 | -0.049 | 0.489 | -0.353 | -0.01 | 0.015 | 0.015 | 1.505 | 0.79 | 0.84 | 0.83 | 0.88 | 0.73 | 0.89 |
| 1 | 67386204 | rs3008855 | 0.699 | -0.041 | 0.492 | -0.35 | -0.01 | 0.014 | 0.014 | 1.606 | 0.8 | 0.83 | 0.83 | 0.88 | 0.73 | 0.89 |
| 1 | 67436446 | rs4553158 | -0.189 | -0.088 | 1.358 | -0.419 | -0.048 | -0.011 | -0.011 | 0 | 0.11 | 0.01 | 0.09 | 0.07 | 0.18 | 0.1 |
| 1 | 67442275 | rs11208979 | -1.526 | -0.096 | 1.39 | -0.421 | -0.013 | 0.019 | 0.019 | 1.637 | 0.2 | 0.16 | 0.16 | 0.12 | 0.26 | 0.11 |
| 1 | 67452322 | rs17497828 | -1.516 | -0.094 | 1.415 | -0.424 | -0.018 | 0.024 | 0.024 | 1.11 | 0.2 | 0.12 | 0.16 | 0.12 | 0.26 | 0.11 |
| 1 | 67457199 | rs17497947 | -0.671 | -0.076 | 1.338 | -0.427 | -0.018 | 0.005 | 0.005 | 2.463 | 0.13 | 0.07 | 0.12 | 0.12 | 0.21 | 0.1 |
| 1 | 67458463 | rs6656890 | -0.146 | -0.079 | 1.338 | -0.429 | -0.009 | 0.021 | 0.021 | 0.615 | 0.07 | 0.03 | 0.04 | 0 | 0.06 | 0.01 |
| 1 | 67460791 | rs4486425 | -2.878 | -0.119 | 1.355 | -0.45 | 0.006 | 0.036 | 0.036 | 2.724 | 0.35 | 0.72 | 0.27 | 0.16 | 0.34 | 0.22 |
| 1 | 67473378 | rs17129595 | -0.958 | -0.123 | 1.274 | -0.442 | -0.047 | -0.038 | -0.038 | 0.554 | 0.15 | 0.52 | 0.1 | 0.04 | 0.08 | 0.11 |
| 1 | 67494591 | rs2815361 | 1.251 | -0.116 | 1.22 | -0.449 | -0.009 | 0.012 | 0.012 | 1.672 | 0.8 | 0.84 | 0.84 | 0.88 | 0.74 | 0.91 |
| 1 | 67496425 | rs2755251 | 2.2 | -0.089 | 1.052 | -0.435 | 0 | 0 | 0 | 2.399 | 0.66 | 0.35 | 0.73 | 0.84 | 0.66 | 0.8 |
| 1 | 67509698 | rs2815378 | 0.794 | -0.072 | 0.59 | 0.005 | 0.001 | -0.048 | -0.048 | 0.436 | 0.73 | 0.86 | 0.73 | 0.64 | 0.7 | 0.79 |
| 1 | 67512903 | rs1024229 | 1.352 | -0.097 | 0.63 | 0.002 | 0.001 | -0.027 | -0.027 | 1.013 | 0.66 | 0.77 | 0.69 | 0.64 | 0.64 | 0.78 |
| 1 | 67512920 | rs1024230 | 1.352 | -0.097 | 0.63 | 0.002 | 0.001 | -0.027 | -0.027 | 1.589 | 0.66 | 0.77 | 0.69 | 0.64 | 0.64 | 0.78 |
| 1 | 67517022 | rs2755259 | 1.398 | -0.045 | 0.765 | 0.008 | 0 | -0.011 | -0.011 | 1.112 | 0.64 | 0.7 | 0.6 | 0.64 | 0.62 | 0.77 |
| 1 | 67518534 | rs10157422 | -0.213 | -0.106 | 0.901 | 0.002 | 0.009 | -0.054 | -0.054 | 0.001 | 0.1 | 0.27 | 0.06 | 0 | 0.06 | 0.07 |
| 1 | 67526029 | rs12061545 | -0.874 | -0.119 | 0.783 | 0.02 | -0.007 | -0.108 | -0.108 | 0 | 0.17 | 0.35 | 0.11 | 0 | 0.11 | 0.08 |
| 1 | 67526409 | rs11811977 | -0.626 | -0.205 | 0.928 | 0.03 | 0.019 | 0.059 | 0.059 | 0 | 0.06 | 0.24 | 0.08 | 0.05 | 0.04 | 0.05 |
| 1 | 67528006 | rs12097741 | -1.131 | -0.137 | 0.078 | 0.075 | -0.021 | -0.066 | -0.066 | 0.041 | 0.22 | 0.59 | 0.19 | 0.05 | 0.15 | 0.13 |
| 1 | 67529089 | rs2815347 | -0.469 | -0.116 | -0.47 | 0.07 | -0.002 | 0.056 | 0.056 | 0.58 | 0.84 | 0.96 | 0.94 | 0.96 | 0.83 | 0.98 |
| 1 | 67529243 | rs2755263 | 1.315 | 0.155 | -0.466 | 0.236 | 0.013 | 0.039 | 0.039 | 0.556 | 0.52 | 0.2 | 0.53 | 0.63 | 0.56 | 0.72 |
| 1 | 67539356 | rs983111 | 0.802 | 0.243 | 0.033 | 0.178 | -0.008 | -0.041 | -0.041 | 0 | 0.68 | 0.55 | 0.74 | 0.85 | 0.69 | 0.81 |
| 1 | 67543884 | rs6701163 | -1.066 | 0.144 | 0.209 | 0.041 | -0.013 | 0.014 | 0.014 | 0 | 0.16 | 0.28 | 0.17 | 0.12 | 0.13 | 0.17 |
| 1 | 67551209 | rs10789220 | 1.957 | 0.094 | 0.672 | -0.113 | 0.012 | 0.041 | 0.041 | 0.476 | 0.6 | 0.29 | 0.67 | 0.85 | 0.63 | 0.79 |
| 1 | 67553765 | rs6658195 | -0.08 | 0.118 | 0.288 | -0.068 | -0.004 | -0.007 | -0.007 | 0.211 | 0.1 | 0.02 | 0.09 | 0.06 | 0.08 | 0.12 |
| 1 | 67558169 | rs6687611 | -0.609 | 0.128 | 0.309 | -0.081 | -0.04 | 0.078 | 0.078 | 3.154 | 0.86 | 0.99 | 0.92 | 0.97 | 0.81 | 0.98 |
| 1 | 67558275 | rs11806560 | -0.564 | 0.12 | 0.319 | -0.083 | 0.029 | 0.088 | 0.088 | 0 | 0.06 | 0.26 | 0.09 | 0.06 | 0.05 | 0.06 |
| 1 | 67560956 | rs11208997 | -0.656 | 0.13 | 0.285 | -0.085 | -0.04 | 0.077 | 0.077 | 3.135 | 0.86 | 0.99 | 0.92 | 0.97 | 0.81 | 0.98 |
| 1 | 67568824 | rs1885276 | -0.544 | 0.134 | 0.326 | -0.083 | -0.009 | 0.013 | 0.013 | 3.292 | 0.85 | 0.92 | 0.91 | 0.97 | 0.81 | 0.98 |
| 1 | 67573730 | rs10749771 | 0.233 | 0.152 | 0.183 | -0.039 | 0.009 | -0.045 | -0.045 | 0 | 0.79 | 0.66 | 0.83 | 0.91 | 0.76 | 0.92 |
| 1 | 67580350 | rs7520634 | 1.64 | 0.105 | 0.281 | 0.13 | -0.008 | -0.034 | -0.034 | 0 | 0.61 | 0.47 | 0.69 | 0.79 | 0.63 | 0.79 |
| 1 | 67583298 | rs17129664 | -1.058 | 0.125 | -0.337 | 0.915 | -0.001 | -0.001 | -0.001 | 0.007 | 0.13 | 0.18 | 0.1 | 0.12 | 0.08 | 0.12 |
| 1 | 67592114 | rs6670134 | -0.375 | 0.472 | -0.898 | 1.108 | -0.016 | -0.07 | -0.07 | 0 | 0.17 | 0.08 | 0.28 | 0.31 | 0.19 | 0.2 |
| 1 | 67597119 | rs3762318 | 0.449 | 0.446 | -0.607 | 0.701 | 0.003 | 0.009 | 0.009 | 1.556 | 0.77 | 0.78 | 0.84 | 0.9 | 0.81 | 0.94 |
| 1 | 67599657 | rs4655679 | -0.565 | 0.644 | -0.719 | 1.008 | -0.023 | -0.042 | -0.042 | 0.788 | 0.22 | 0.16 | 0.48 | 0.3 | 0.27 | 0.3 |
| 1 | 67601115 | rs10489631 | 1.65 | 0.38 | 0.323 | 0.907 | 0 | -0.001 | -0.001 | 0 | 0.51 | 0.36 | 0.33 | 0.62 | 0.53 | 0.63 |
| 1 | 67601132 | rs11209003 | -0.489 | 0.439 | 0.227 | 0.976 | -0.02 | -0.036 | -0.036 | 0.047 | 0.22 | 0.17 | 0.49 | 0.29 | 0.27 | 0.3 |
| 1 | 67605134 | rs10789224 | -0.794 | 0.296 | 0.683 | 0.896 | 0 | 0.001 | 0.001 | 0 | 0.26 | 0.29 | 0.46 | 0.29 | 0.28 | 0.31 |
| 1 | 67611613 | rs4655683 | -1.334 | 0.284 | 0.912 | 0.806 | 0 | 0.025 | 0.025 | 0 | 0.31 | 0.43 | 0.53 | 0.38 | 0.31 | 0.34 |
| 1 | 67633812 | rs1884444 | 1.646 | 0.143 | 1.053 | 0.64 | 0 | 0.007 | 0.007 | 0.795 | 0.49 | 0.47 | 0.37 | 0.64 | 0.52 | 0.62 |
| 1 | 67636652 | rs10889665 | -0.34 | 0.127 | 1.129 | 0.632 | -0.008 | -0.075 | -0.075 | 0 | 0.79 | 0.68 | 0.89 | 1 | 0.84 | 0.97 |
| 1 | 67651733 | rs10127763 | -0.415 | 0.129 | 1.026 | 0.632 | 0.005 | 0.051 | 0.051 | 2.013 | 0.8 | 0.86 | 0.9 | 1 | 0.84 | 0.97 |
| 1 | 67652984 | rs6588248 | 1.798 | 0.121 | 1.132 | 0.635 | -0.002 | 0.014 | 0.014 | 0.823 | 0.48 | 0.3 | 0.36 | 0.64 | 0.53 | 0.62 |
| 1 | 67653010 | rs2064689 | -0.729 | 0.122 | 1.077 | 0.69 | -0.008 | -0.016 | -0.016 | 0.139 | 0.26 | 0.21 | 0.49 | 0.32 | 0.29 | 0.31 |
| 1 | 67654110 | rs1321157 | 1.776 | 0.122 | 1.112 | 0.69 | -0.001 | 0.005 | 0.005 | 0.878 | 0.48 | 0.35 | 0.36 | 0.64 | 0.53 | 0.62 |
| 1 | 67655954 | rs6688383 | -0.749 | 0.149 | 0.842 | 0.646 | -0.005 | -0.011 | -0.011 | 0 | 0.26 | 0.22 | 0.5 | 0.32 | 0.29 | 0.31 |
| 1 | 67659896 | rs4655690 | -0.376 | 0.112 | 0.835 | 0.515 | 0.001 | -0.024 | -0.024 | 0.007 | 0.8 | 0.78 | 0.9 | 1 | 0.84 | 0.97 |
| 1 | 67662622 | rs10489630 | -1.365 | 0.615 | 0.63 | 0.847 | 0.001 | 0.002 | 0.002 | 0 | 0.33 | 0.41 | 0.57 | 0.39 | 0.37 | 0.37 |
| 1 | 67667291 | rs11209018 | 1.486 | 0.564 | 0.555 | 0.833 | -0.001 | 0.002 | 0.002 | 1.224 | 0.46 | 0.29 | 0.33 | 0.6 | 0.47 | 0.6 |
| 1 | 67670213 | rs1004819 | 1.411 | 1.31 | 0.308 | 0.55 | 0.027 | -0.004 | -0.004 | 0 | 0.45 | 0.3 | 0.22 | 0.59 | 0.3 | 0.56 |
| 1 | 67670916 | rs2902440 | -0.532 | 1.769 | 0.286 | 0.446 | -0.077 | -0.065 | -0.065 | 0.011 | 0.23 | 0.17 | 0.52 | 0.37 | 0.42 | 0.33 |
| 1 | 67672765 | rs7539625 | 1.651 | 1.761 | 0.312 | 0.588 | 0.027 | -0.008 | -0.008 | 0 | 0.45 | 0.3 | 0.17 | 0.54 | 0.3 | 0.56 |
| 1 | 67675516 | rs11805303 | 1.658 | 1.789 | 0.309 | 0.616 | 0.026 | -0.006 | -0.006 | 0 | 0.45 | 0.3 | 0.17 | 0.54 | 0.3 | 0.55 |
| 1 | 67676922 | rs790631 | 0.387 | 1.783 | 0.239 | 0.552 | 0.01 | 0.071 | 0.071 | 1.361 | 0.7 | 0.81 | 0.76 | 0.97 | 0.73 | 0.89 |
| 1 | 67681669 | rs7517847 | -0.622 | 2.044 | 0.097 | 0.735 | -0.075 | -0.074 | -0.074 | 0.013 | 0.23 | 0.16 | 0.57 | 0.42 | 0.42 | 0.33 |
| 1 | 67685116 | rs2863212 | -0.759 | 1.798 | 0.344 | 0.301 | -0.007 | -0.058 | -0.058 | 0 | 0.87 | 0.78 | 0.86 | 0.99 | 0.87 | 0.98 |
| 1 | 67685387 | rs7530511 | -0.408 | 1.662 | 0.316 | 0.194 | -0.009 | -0.07 | -0.07 | 0 | 0.85 | 0.76 | 0.85 | 0.99 | 0.86 | 0.95 |
| 1 | 67685598 | rs11465802 | 2.062 | 1.491 | 0.173 | 0.151 | 0.085 | 0.066 | 0.066 | 0 | 0.44 | 0.14 | 0.24 | 0.73 | 0.29 | 0.59 |
| 1 | 67688349 | rs10489629 | -1.24 | 1.489 | 0.226 | 0.121 | 0.024 | -0.01 | -0.01 | 0 | 0.35 | 0.58 | 0.55 | 0.26 | 0.45 | 0.3 |

| | | | | | | | | | | | | | | | | |
|---|---|---|---|---|---|---|---|---|---|---|---|---|---|---|---|---|
| 1 | 67689036 | rs7528924 | 0.091 | 1.487 | 0.228 | 0.093 | 0.003 | -0.08 | -0.08 | 0 | 0.79 | 0.72 | 0.79 | 0.99 | 0.75 | 0.89 |
| 1 | 67694202 | rs2201841 | 2.055 | 1.49 | 0.078 | 0.145 | 0.071 | 0.069 | 0.069 | 0 | 0.44 | 0.14 | 0.24 | 0.73 | 0.3 | 0.6 |
| 1 | 67697271 | rs12070470 | 0.892 | 1.47 | 0.053 | 0.149 | -0.007 | 0.003 | 0.003 | 0 | 0.06 | 0.05 | 0.05 | 0 | 0.11 | 0.06 |
| 1 | 67701765 | rs12030948 | 1.959 | 1.575 | -0.007 | 0.039 | 0.076 | 0.11 | 0.11 | 0 | 0.51 | 0.18 | 0.3 | 0.73 | 0.36 | 0.61 |
| 1 | 67702526 | rs11465804 | 0.326 | 1.545 | -0.002 | -0.011 | 0.003 | 0.019 | 0.019 | 0 | 0.07 | 0.04 | 0.06 | 0 | 0.07 | 0.01 |
| 1 | 67704863 | rs12751814 | 0.869 | 2.212 | 0.159 | -0.038 | 0.009 | -0.09 | -0.09 | 0 | 0.66 | 0.58 | 0.69 | 0.94 | 0.6 | 0.8 |
| 1 | 67708670 | rs6682033 | -0.539 | 2.245 | 0.386 | -0.023 | 0.039 | -0.104 | -0.104 | 0 | 0.21 | 0.44 | 0.18 | 0.05 | 0.25 | 0.15 |
| 1 | 67719129 | rs1343151 | -1.577 | 2.224 | 0.227 | -0.031 | -0.006 | -0.009 | -0.009 | 0 | 0.3 | 0.73 | 0.28 | 0.05 | 0.32 | 0.16 |
| 1 | 67720867 | rs6693831 | 0.104 | 2.305 | 0.085 | 0.037 | -0.041 | -0.042 | -0.042 | 0 | 0.8 | 0.94 | 0.57 | 0.79 | 0.73 | 0.82 |
| 1 | 67721097 | rs11465817 | 3.153 | 1.859 | -0.212 | 0.108 | -0.003 | -0.165 | -0.165 | 0 | 0.37 | 0.04 | 0.2 | 0.57 | 0.29 | 0.47 |
| 1 | 67722216 | rs10889675 | 0.251 | 2.339 | 0.047 | 0.265 | -0.037 | -0.021 | -0.021 | 0.015 | 0.06 | 0.01 | 0.3 | 0.2 | 0.12 | 0.14 |
| 1 | 67722567 | rs10889676 | 2.37 | 2.251 | 0.083 | 0.274 | 0.084 | 0.074 | 0.074 | 0 | 0.44 | 0.13 | 0.25 | 0.73 | 0.3 | 0.6 |
| 1 | 67725120 | rs10889677 | 2.37 | 2.297 | 0.083 | 0.269 | 0.084 | 0.07 | 0.07 | 0 | 0.44 | 0.13 | 0.25 | 0.73 | 0.3 | 0.6 |
| 1 | 67727555 | rs2000252 | -0.457 | 2.514 | 0.358 | 0.045 | 0.03 | -0.087 | -0.087 | 3.825 | 0.21 | 0.41 | 0.18 | 0.05 | 0.25 | 0.15 |
| 1 | 67730628 | rs10789230 | 2.002 | 2.128 | -0.294 | 0.07 | 0.087 | 0.172 | 0.172 | 0 | 0.48 | 0.16 | 0.32 | 0.79 | 0.34 | 0.61 |
| 1 | 67737775 | rs11209030 | -0.269 | 2.246 | 0.003 | 0.126 | 0.024 | 0 | 0 | 1.83 | 0.22 | 0.37 | 0.31 | 0.23 | 0.25 | 0.26 |
| 1 | 67740092 | rs11209032 | 1.507 | 1.309 | 0.164 | 0.717 | 0.057 | -0.022 | -0.022 | 0.283 | 0.44 | 0.2 | 0.23 | 0.5 | 0.33 | 0.52 |
| 1 | 67740835 | rs12568764 | 0.503 | 1.351 | 0.12 | 0.731 | -0.016 | 0.023 | 0.023 | 0 | 0.05 | 0.01 | 0.22 | 0.26 | 0.07 | 0.12 |
| 1 | 67744601 | rs6660226 | -0.555 | 1.346 | 0.097 | 0.644 | -0.004 | 0.003 | 0.003 | 0 | 0.38 | 0.41 | 0.41 | 0.24 | 0.45 | 0.34 |
| 1 | 67747415 | rs12119179 | 1.131 | 0.844 | -0.483 | 0.311 | 0.056 | -0.035 | -0.035 | 0 | 0.43 | 0.2 | 0.23 | 0.51 | 0.34 | 0.52 |
| 1 | 67753508 | rs1495965 | -1.066 | 1.138 | -0.563 | 0.78 | 0 | -0.002 | -0.002 | 0 | 0.44 | 0.58 | 0.65 | 0.48 | 0.52 | 0.46 |
| 1 | 67769006 | rs12131065 | 0.818 | 0.804 | -0.278 | 0.963 | -0.003 | 0.018 | 0.018 | 0 | 0.7 | 0.35 | 0.69 | 0.6 | 0.76 | 0.68 |
| 1 | 67770799 | rs11209045 | -1.271 | 0.165 | -0.338 | 0.078 | 0.005 | 0.031 | 0.031 | 0 | 0.45 | 0.56 | 0.45 | 0.25 | 0.38 | 0.34 |
| 1 | 67772011 | rs3762315 | -0.268 | -0.015 | -0.053 | 0.148 | 0.013 | 0.015 | 0.015 | 0 | 0.15 | 0.3 | 0.21 | 0.19 | 0.14 | 0.16 |
| 1 | 67773351 | rs12142823 | 0.081 | 0.294 | 0.473 | 0.312 | 0.029 | 0.03 | 0.03 | 0 | 0.11 | 0.14 | 0.2 | 0.19 | 0.16 | 0.17 |
| 1 | 67777021 | rs3790558 | 1.56 | 0.267 | 0.376 | 0.541 | -0.076 | -0.023 | -0.023 | 0 | 0.47 | 0.14 | 0.42 | 0.53 | 0.56 | 0.52 |
| 1 | 67779676 | rs10489627 | -1.389 | -0.083 | 1.098 | 0.512 | 0.007 | 0.078 | 0.078 | 0 | 0.42 | 0.66 | 0.45 | 0.26 | 0.33 | 0.33 |
| 1 | 67781117 | rs2066446 | -0.895 | 0.08 | 1.376 | 0.618 | 0.008 | 0.001 | 0.001 | 0 | 0.2 | 0.24 | 0.28 | 0.25 | 0.19 | 0.24 |
| 1 | 67786902 | rs4655538 | 0.251 | -0.055 | 1.317 | 0.615 | -0.016 | 0.077 | 0.077 | 0 | 0.93 | 0.73 | 0.91 | 0.84 | 0.98 | 0.91 |
| 1 | 67787715 | rs2201584 | -0.012 | 0.005 | 1.532 | 0.842 | -0.023 | -0.034 | -0.034 | 0 | 0.11 | 0.04 | 0.2 | 0.34 | 0.16 | 0.18 |
| 1 | 67791736 | rs11209050 | 1.268 | -0.262 | 1.619 | 0.485 | -0.003 | -0.003 | -0.003 | 0.05 | 0.66 | 0.3 | 0.73 | 0.75 | 0.81 | 0.79 |
| 1 | 67793171 | rs10489626 | -0.77 | -0.235 | 1.425 | 0.491 | 0.005 | 0.017 | 0.017 | 0 | 0.24 | 0.21 | 0.16 | 0.06 | 0.16 | 0.12 |
| 1 | 67795319 | rs1495963 | 0.247 | -0.099 | 1.46 | 0.709 | -0.015 | 0.11 | 0.11 | 0 | 0.93 | 0.72 | 0.9 | 0.81 | 0.98 | 0.91 |
| 1 | 67800018 | rs6693065 | 1.278 | -0.082 | 1.5 | 0.962 | 0.005 | 0.089 | 0.089 | 0 | 0.77 | 0.43 | 0.64 | 0.4 | 0.76 | 0.72 |
| 1 | 67811356 | rs3790565 | 1.451 | -0.193 | 1.309 | 0.776 | 0 | 0.001 | 0.001 | 1.065 | 0.74 | 0.26 | 0.75 | 0.8 | 0.8 | 0.84 |
| 1 | 67814440 | rs3790566 | 1.866 | -0.408 | 1.381 | 0.681 | 0.035 | 0.041 | 0.041 | 0 | 0.62 | 0.21 | 0.68 | 0.75 | 0.76 | 0.77 |
| 1 | 67821447 | rs6664757 | -1.167 | -0.22 | 1.668 | 0.835 | -0.016 | 0.074 | 0.074 | 0 | 0.1 | 0.31 | 0.15 | 0.19 | 0.07 | 0.11 |
| 1 | 67833501 | rs2252596 | -0.327 | -0.211 | 1.6 | 0.842 | -0.021 | 0.049 | 0.049 | 0 | 0.07 | 0.19 | 0.09 | 0.19 | 0.02 | 0.1 |
| 1 | 67837158 | rs10493423 | -0.286 | -0.321 | 1.567 | 0.846 | -0.02 | 0.051 | 0.051 | 0 | 0.07 | 0.2 | 0.09 | 0.19 | 0.02 | 0.1 |
| 1 | 67846156 | rs881087 | 0.901 | -0.465 | 1.482 | 0.727 | 0.046 | 0.048 | 0.048 | 0 | 0.72 | 0.52 | 0.68 | 0.59 | 0.7 | 0.74 |
| 1 | 67854602 | rs7536192 | -1.242 | -0.481 | 1.448 | 0.72 | -0.023 | 0.069 | 0.069 | 0.003 | 0.11 | 0.31 | 0.14 | 0.2 | 0.06 | 0.1 |
| 1 | 67855142 | rs11209063 | -1.355 | -0.498 | 1.42 | 0.706 | -0.025 | 0.003 | 0.003 | 0 | 0.23 | 0.32 | 0.23 | 0.24 | 0.13 | 0.18 |
| 1 | 67860627 | rs7555183 | 1.292 | -0.425 | 1.286 | 0.585 | 0.001 | 0.005 | 0.005 | 0 | 0.65 | 0.33 | 0.59 | 0.39 | 0.69 | 0.65 |
| 1 | 67861520 | rs2229546 | 1.174 | -0.431 | 1.181 | 0.597 | 0.008 | 0.015 | 0.015 | 0 | 0.66 | 0.51 | 0.59 | 0.38 | 0.68 | 0.65 |
| 1 | 67875414 | rs4655707 | 1.662 | -0.326 | 0.921 | 0.622 | 0.007 | 0.054 | 0.054 | 0 | 0.5 | 0.16 | 0.47 | 0.34 | 0.59 | 0.56 |
| 1 | 67890526 | rs6702742 | 1.629 | -0.314 | 0.927 | 0.617 | 0.012 | 0.058 | 0.058 | 0 | 0.5 | 0.19 | 0.47 | 0.34 | 0.58 | 0.56 |
| 1 | 67896672 | rs3762314 | 1.784 | -0.259 | 0.942 | 1.02 | 0.016 | 0.089 | 0.089 | 0 | 0.52 | 0.2 | 0.44 | 0.27 | 0.61 | 0.53 |
| 1 | 67908644 | rs11209072 | -1.192 | -0.332 | 0.914 | 0.677 | 0.034 | -0.035 | -0.035 | 0 | 0.33 | 0.23 | 0.37 | 0.66 | 0.23 | 0.4 |
| 1 | 67909782 | rs10889687 | -0.432 | -0.233 | 0.611 | 0.688 | 0.017 | -0.046 | -0.046 | 0.828 | 0.88 | 0.8 | 0.88 | 0.99 | 0.84 | 0.94 |
| 1 | 67912259 | rs12116575 | 1.741 | -0.307 | 0.197 | 0.779 | 0.023 | 0.048 | 0.048 | 0 | 0.51 | 0.15 | 0.48 | 0.33 | 0.61 | 0.53 |
| 1 | 67916910 | rs10889689 | -1.056 | -0.623 | 0.478 | 0.894 | -0.082 | 0.018 | 0.018 | 0.385 | 0.24 | 0.48 | 0.2 | 0.26 | 0.09 | 0.2 |
| 1 | 67917914 | rs12138671 | 1.236 | -0.565 | -0.884 | 0.593 | -0.05 | -0.037 | -0.037 | 0 | 0.45 | 0.12 | 0.48 | 0.43 | 0.56 | 0.54 |
| 1 | 67919086 | rs1572845 | -0.096 | -0.639 | -0.51 | 0.953 | 0.011 | -0.01 | -0.01 | 0 | 0.16 | 0.09 | 0.12 | 0.19 | 0.08 | 0.11 |
| 1 | 67926980 | rs4633262 | 1.461 | -0.793 | -0.376 | 0.973 | 0.047 | -0.014 | -0.014 | 0 | 0.58 | 0.23 | 0.6 | 0.38 | 0.75 | 0.54 |
| 1 | 67927786 | rs10458577 | -0.171 | -0.55 | -0.566 | 1.069 | 0.009 | -0.012 | -0.012 | 0 | 0.17 | 0.09 | 0.13 | 0.19 | 0.08 | 0.11 |
| 1 | 67935310 | rs6657603 | 0.795 | -0.452 | -0.093 | 1.041 | -0.046 | 0.043 | 0.043 | 0 | 0.73 | 0.44 | 0.76 | 0.57 | 0.83 | 0.65 |
| 1 | 67941167 | rs1977785 | 0.825 | 0.214 | -0.673 | 0.787 | 0.027 | 0.054 | 0.054 | 0.017 | 0.51 | 0.21 | 0.38 | 0.34 | 0.6 | 0.58 |
| 1 | 67944943 | rs4655715 | 0.64 | -0.303 | 0.27 | 0.879 | -0.014 | 0.051 | 0.051 | 0 | 0.9 | 0.72 | 0.94 | 0.88 | 0.97 | 0.9 |
| 1 | 67945005 | rs4655716 | 0.845 | -0.406 | 0.662 | 1.225 | 0.007 | -0.005 | -0.005 | 0.05 | 0.66 | 0.28 | 0.74 | 0.63 | 0.83 | 0.78 |
| 1 | 67951218 | rs6695089 | 0.491 | 0.344 | 0.393 | 0.752 | -0.038 | 0.046 | 0.046 | 0 | 0.1 | 0.06 | 0.19 | 0 | 0.17 | 0.07 |
| 1 | 67953681 | rs7554654 | 1.064 | 0.235 | 0.884 | 0.754 | 0.009 | -0.035 | -0.035 | 0 | 0.78 | 0.57 | 0.73 | 0.88 | 0.79 | 0.83 |
| 1 | 67959033 | rs12730759 | 0.219 | 0.608 | 1 | 0.759 | 0.002 | -0.02 | -0.02 | 0 | 0.11 | 0.11 | 0.19 | 0 | 0.17 | 0.07 |
| 1 | 67968725 | rs9285766 | 1.398 | 0.666 | 1.307 | 0.953 | -0.016 | 0.029 | 0.029 | 0.238 | 0.54 | 0.24 | 0.42 | 0.37 | 0.55 | 0.66 |
| 1 | 67972372 | rs12034578 | 0.282 | 0.632 | 0.784 | 1.1 | -0.002 | 0.003 | 0.003 | 0.609 | 0.23 | 0.31 | 0.19 | 0.26 | 0.16 | 0.12 |
| 1 | 67972435 | rs12037907 | 0.569 | 0.681 | 0.588 | 1.09 | -0.009 | 0.006 | 0.006 | 0 | 0.08 | 0.04 | 0.17 | 0.38 | 0.11 | 0.16 |
| 1 | 67978985 | rs10789235 | -0.293 | 0.461 | 0.814 | 0.544 | -0.026 | -0.15 | -0.15 | 0 | 0.77 | 0.66 | 0.9 | 1 | 0.86 | 0.98 |
| 1 | 67980297 | rs7538875 | 0.934 | 0.519 | 0.816 | 0.622 | -0.001 | 0.011 | 0.011 | 0.739 | 0.64 | 0.28 | 0.6 | 0.74 | 0.67 | 0.82 |
| 1 | 67982128 | rs11209099 | -0.111 | 0.364 | 0.2 | 0.992 | -0.004 | 0.008 | 0.008 | 0.117 | 0.23 | 0.33 | 0.19 | 0.26 | 0.16 | 0.12 |
| 1 | 67989691 | rs11209101 | -0.757 | -0.078 | -0.184 | 0.299 | 0.041 | 0.05 | 0.05 | 0 | 0.45 | 0.7 | 0.31 | 0.26 | 0.29 | 0.39 |

| | | | | | | | | | | | | | | | | |
|---|---|---|---|---|---|---|---|---|---|---|---|---|---|---|---|---|
| 1 | 67990154 | rs12134055 | -0.095 | 0.549 | -0.654 | 0.823 | -0.066 | -0.086 | -0.086 | 0.64 | 0.19 | 0.09 | 0.38 | 0.43 | 0.29 | 0.21 |
| 1 | 67993397 | rs17130073 | -0.337 | 0.58 | -0.604 | -0.161 | -0.002 | 0 | 0 | 0 | 0.32 | 0.32 | 0.28 | 0.33 | 0.32 | 0.32 |
| 1 | 67996114 | rs6692344 | 0.081 | 0.251 | -0.346 | -0.564 | -0.01 | -0.125 | -0.125 | 0 | 0.16 | 0.45 | 0.1 | 0 | 0.1 | 0.04 |
| 1 | 67998133 | rs7539855 | -1.039 | 0.239 | 0.075 | -0.588 | -0.001 | -0.015 | -0.015 | 0.943 | 0.45 | 0.48 | 0.36 | 0.33 | 0.43 | 0.37 |
| 1 | 68001373 | rs4233318 | 1.157 | 0.371 | 0.152 | -0.511 | -0.007 | 0.003 | 0.003 | 0 | 0.6 | 0.26 | 0.52 | 0.53 | 0.58 | 0.46 |
| 1 | 68005643 | rs4655561 | 0.275 | 0.504 | 0.418 | -0.515 | 0.006 | -0.052 | -0.052 | 0.123 | 0.73 | 0.64 | 0.72 | 0.83 | 0.72 | 0.79 |
| 1 | 68010127 | rs10493425 | -0.295 | 0.589 | 0.822 | -0.342 | -0.001 | 0.188 | 0.188 | 0.217 | 0.11 | 0.37 | 0.17 | 0.3 | 0.1 | 0.28 |
| 1 | 68012254 | rs12120145 | -0.252 | 0.679 | 0.436 | -0.526 | 0.004 | -0.012 | -0.012 | 0.429 | 0.2 | 0.25 | 0.26 | 0.17 | 0.25 | 0.21 |
| 1 | 68018455 | rs11209112 | -0.569 | 0.372 | 0.597 | -0.35 | 0.005 | 0.099 | 0.099 | 0 | 0.12 | 0.29 | 0.19 | 0.29 | 0.15 | 0.32 |
| 1 | 68021103 | rs1340592 | -1.015 | 0.204 | 0.073 | -0.91 | 0.004 | 0.004 | 0.004 | 0.8 | 0.35 | 0.4 | 0.29 | 0.26 | 0.27 | 0.31 |
| 1 | 68021906 | rs7530157 | -1.492 | 0.189 | 0.22 | -0.94 | 0.031 | 0.048 | 0.048 | 0.532 | 0.4 | 0.74 | 0.36 | 0.26 | 0.33 | 0.32 |
| 1 | 68024209 | rs11209115 | 1.177 | 0.15 | 0.254 | -0.931 | 0.048 | 0.075 | 0.075 | 0.528 | 0.6 | 0.18 | 0.64 | 0.74 | 0.69 | 0.68 |
| 1 | 68025638 | rs4655733 | -0.52 | 0.197 | -0.046 | -0.726 | -0.01 | -0.003 | -0.003 | 0 | 0.27 | 0.4 | 0.27 | 0.26 | 0.24 | 0.31 |
| 1 | 68027219 | rs4655735 | -0.424 | 0.184 | -0.032 | -0.779 | -0.015 | -0.016 | -0.016 | 0.191 | 0.25 | 0.39 | 0.22 | 0.18 | 0.18 | 0.25 |
| 1 | 68031411 | rs4336822 | -0.154 | 0.881 | -0.676 | -0.791 | 0.039 | -0.009 | -0.009 | 0 | 0.2 | 0.45 | 0.31 | 0.15 | 0.23 | 0.21 |
| 1 | 68032769 | rs4081356 | 0.939 | 0.857 | -1.845 | -0.517 | -0.05 | -0.054 | -0.054 | 0 | 0.47 | 0.07 | 0.43 | 0.47 | 0.46 | 0.4 |
| 1 | 68038313 | rs12085093 | 0.633 | 1.038 | -1.633 | -0.416 | -0.056 | -0.062 | -0.062 | 0 | 0.55 | 0.16 | 0.47 | 0.53 | 0.52 | 0.45 |
| 1 | 68040209 | rs787522 | -0.064 | 1.043 | -1.249 | -0.604 | 0.017 | -0.037 | -0.037 | 0 | 0.23 | 0.48 | 0.33 | 0.15 | 0.26 | 0.23 |
| 1 | 68041561 | rs968544 | 1.438 | 0.981 | -1.985 | -0.703 | -0.017 | 0.036 | 0.036 | 0 | 0.15 | 0.06 | 0.13 | 0.06 | 0.2 | 0.08 |
| 1 | 68043745 | rs699235 | -0.162 | 0.953 | -2.068 | -0.761 | 0.004 | -0.022 | -0.022 | 0 | 0.28 | 0.62 | 0.36 | 0.15 | 0.28 | 0.23 |
| 1 | 68046274 | rs9919329 | 1.584 | 0.961 | -2.16 | -0.747 | 0.012 | -0.072 | -0.072 | 0 | 0.36 | 0.11 | 0.32 | 0.47 | 0.31 | 0.36 |
| 1 | 68046309 | rs787533 | 0.087 | 0.861 | -1.968 | -0.47 | -0.01 | -0.01 | -0.01 | 0 | 0.64 | 0.74 | 0.68 | 0.62 | 0.59 | 0.59 |
| 1 | 68046354 | rs699233 | -0.835 | 0.715 | -1.849 | -0.13 | -0.003 | -0.008 | -0.008 | 0 | 0.79 | 0.83 | 0.82 | 0.68 | 0.79 | 0.66 |
| 1 | 68050630 | rs787538 | -0.671 | 0.246 | -2.366 | -0.485 | -0.038 | 0.11 | 0.11 | 0 | 0.4 | 0.87 | 0.46 | 0.2 | 0.41 | 0.32 |
| 1 | 68051868 | rs699232 | -0.624 | 0.44 | -2.205 | -0.218 | -0.062 | -0.076 | -0.076 | 0 | 0.76 | 0.95 | 0.78 | 0.67 | 0.7 | 0.63 |
| 1 | 68055306 | rs787520 | -0.153 | -0.438 | -0.971 | -0.61 | -0.007 | -0.1 | -0.1 | 0.428 | 0.2 | 0.6 | 0.24 | 0.02 | 0.17 | 0.08 |
| 1 | 68055347 | rs11585171 | -0.35 | -0.621 | -1.387 | -0.37 | -0.008 | -0.064 | -0.064 | 0 | 0.22 | 0.15 | 0.33 | 0.54 | 0.28 | 0.27 |
| 1 | 68062148 | rs12120849 | 0.498 | -0.716 | -1.469 | -0.578 | 0.018 | 0.02 | 0.02 | 0 | 0.17 | 0.12 | 0.06 | 0 | 0.09 | 0.01 |
| 1 | 68063786 | rs787507 | -0.235 | -0.686 | -1.422 | -0.492 | -0.004 | -0.014 | -0.014 | 0 | 0.23 | 0.68 | 0.32 | 0.14 | 0.19 | 0.18 |
| 1 | 68065339 | rs787508 | -0.282 | -0.691 | -1.424 | -0.495 | -0.004 | -0.013 | -0.013 | 0 | 0.23 | 0.68 | 0.33 | 0.14 | 0.2 | 0.18 |
| 1 | 68069073 | rs2418455 | 0.067 | -0.748 | -1.452 | -0.467 | -0.002 | -0.031 | -0.031 | 0 | 0.22 | 0.15 | 0.26 | 0.44 | 0.26 | 0.17 |
| 1 | 68070775 | rs805112 | -1.467 | -0.744 | -1.474 | -0.491 | 0.006 | -0.035 | -0.035 | 5.01 | 0.47 | 0.84 | 0.58 | 0.57 | 0.45 | 0.35 |
| 1 | 68071100 | rs787498 | -0.999 | -0.964 | -1.119 | -0.57 | 0.003 | 0.012 | 0.012 | 2.431 | 0.28 | 0.73 | 0.34 | 0.14 | 0.22 | 0.18 |
| 1 | 68071196 | rs9970835 | 1.019 | -1.257 | -0.984 | -0.43 | 0.008 | -0.005 | -0.005 | 0 | 0.12 | 0.1 | 0.05 | 0 | 0.07 | 0.01 |
| 1 | 68072927 | rs787501 | -1.291 | -1.368 | -0.391 | -0.406 | -0.049 | -0.052 | -0.052 | 0 | 0.16 | 0.57 | 0.1 | 0.06 | 0.07 | 0.11 |
| 1 | 68079175 | rs17130134 | -0.767 | -1.535 | -0.51 | -0.309 | 0.023 | -0.018 | -0.018 | 0 | 0.32 | 0.24 | 0.29 | 0.38 | 0.27 | 0.13 |
| 1 | 68083054 | rs787496 | -0.772 | -1.158 | -0.002 | -0.226 | -0.015 | -0.084 | -0.084 | 0.041 | 0.24 | 0.54 | 0.26 | 0.1 | 0.19 | 0.19 |
| 1 | 68084290 | rs12069919 | -1.681 | -1.122 | 0.056 | -0.136 | 0.004 | -0.001 | -0.001 | 0.004 | 0.45 | 0.58 | 0.5 | 0.49 | 0.39 | 0.33 |
| 1 | 68085586 | rs787494 | -0.714 | -0.69 | -0.332 | 0.118 | -0.008 | 0.015 | 0.015 | 0 | 0.25 | 0.34 | 0.26 | 0.39 | 0.21 | 0.12 |
| 1 | 68090386 | rs787486 | -0.158 | -0.052 | -0.062 | 0.252 | 0.008 | -0.028 | -0.028 | 0 | 0.16 | 0.28 | 0.2 | 0.08 | 0.18 | 0.14 |
| 1 | 68095207 | rs787481 | -1.096 | -0.667 | 0.017 | 0.814 | -0.017 | 0.033 | 0.033 | 0 | 0.33 | 0.56 | 0.35 | 0.48 | 0.25 | 0.24 |
| 1 | 68095872 | rs787480 | -1.376 | -0.835 | 0.088 | 0.781 | -0.012 | 0.046 | 0.046 | 0 | 0.24 | 0.62 | 0.29 | 0.46 | 0.19 | 0.23 |
| 1 | 68101543 | rs12132307 | -0.79 | -0.692 | -0.025 | 0.734 | -0.003 | -0.007 | -0.007 | 0 | 0.19 | 0.26 | 0.1 | 0.12 | 0.15 | 0.08 |
| 1 | 68105186 | rs12129053 | -0.208 | -0.721 | 0.041 | 0.632 | 0.002 | -0.022 | -0.022 | 0 | 0.15 | 0.16 | 0.06 | 0 | 0.1 | 0.05 |
| 1 | 68105453 | rs344924 | -1.062 | -0.812 | 0.536 | 1.653 | 0.098 | 0.531 | 0.531 | 0.536 | 0.06 | 0.44 | 0.12 | 0.32 | 0.05 | 0.19 |
| 1 | 68106505 | rs11209133 | -1.317 | -0.902 | -0.165 | 1.573 | 0.01 | 0.003 | 0.003 | 0 | 0.34 | 0.3 | 0.28 | 0.36 | 0.28 | 0.17 |
| 1 | 68109096 | rs344923 | -1.629 | -0.918 | -0.755 | 1.318 | 0.037 | -0.006 | -0.006 | 0 | 0.4 | 0.89 | 0.42 | 0.65 | 0.33 | 0.36 |
| 1 | 68110795 | rs687328 | -1.342 | -1.456 | -1.135 | 1.115 | 0.089 | -0.011 | -0.011 | 0 | 0.41 | 0.9 | 0.4 | 0.34 | 0.3 | 0.33 |
| 1 | 68111728 | rs12565229 | 0.469 | -2.149 | -0.922 | 0.305 | 0.035 | -0.011 | -0.011 | 0.095 | 0.58 | 0.36 | 0.53 | 0.47 | 0.67 | 0.64 |
| 1 | 68114809 | rs344922 | 0.664 | -0.926 | -0.259 | 0.21 | -0.003 | -0.007 | -0.007 | 0 | 0.45 | 0.4 | 0.53 | 0.48 | 0.49 | 0.48 |
| 1 | 68115186 | rs173180 | -0.862 | -1.139 | 0.239 | 0.003 | -0.01 | -0.068 | -0.068 | 0 | 0.38 | 0.44 | 0.33 | 0.15 | 0.32 | 0.29 |
| 1 | 68115401 | rs4655568 | 1.146 | -1.184 | 0.567 | 0.147 | -0.017 | -0.015 | -0.015 | 0 | 0.39 | 0.32 | 0.47 | 0.52 | 0.49 | 0.55 |
| 1 | 68117203 | rs12134425 | 0.304 | -1.502 | 0.653 | 0.127 | 0.001 | -0.005 | -0.005 | 0 | 0.21 | 0.17 | 0.12 | 0.08 | 0.18 | 0.14 |
| 1 | 68126096 | rs182943 | -0.002 | -1.676 | 0.842 | 0.293 | -0.035 | 0.098 | 0.098 | 0 | 0.72 | 0.51 | 0.71 | 0.52 | 0.77 | 0.74 |
| 1 | 68128130 | rs12748843 | 1.548 | -1.717 | 0.614 | 0.15 | -0.085 | -0.068 | -0.068 | 1.426 | 0.38 | 0.08 | 0.45 | 0.39 | 0.51 | 0.56 |
| 1 | 68128639 | rs11209138 | 1.548 | -1.717 | 0.614 | 0.15 | -0.085 | -0.068 | -0.068 | 0 | 0.38 | 0.08 | 0.45 | 0.39 | 0.51 | 0.56 |
| 1 | 68133074 | rs344929 | 0.823 | -1.751 | 0.942 | 0.129 | -0.019 | -0.02 | -0.02 | 0 | 0.12 | 0.12 | 0.13 | 0.06 | 0.06 | 0.04 |
| 1 | 68135838 | rs344931 | -0.79 | -1.786 | 1.22 | 0.191 | -0.018 | 0.059 | 0.059 | 0 | 0.78 | 0.63 | 0.8 | 0.63 | 0.85 | 0.84 |
| 1 | 68137613 | rs344934 | -1.191 | -1.772 | 1.214 | 0.166 | -0.008 | -0.012 | -0.012 | 0 | 0.4 | 0.55 | 0.35 | 0.24 | 0.33 | 0.28 |
| 1 | 68142619 | rs344916 | -1.324 | -1.781 | 1.417 | 0.172 | -0.023 | -0.011 | -0.011 | 1.378 | 0.27 | 0.62 | 0.3 | 0.23 | 0.23 | 0.25 |
| 1 | 68143540 | rs12097345 | 0.359 | -1.719 | 1.399 | 0.212 | -0.007 | 0.045 | 0.045 | 0 | 0.06 | 0.16 | 0.08 | 0.1 | 0.07 | 0.09 |
| 1 | 68147143 | rs488778 | -0.991 | -1.645 | 1.328 | 0.065 | 0 | 0.002 | 0.002 | 0 | 0.8 | 0.8 | 0.81 | 0.63 | 0.85 | 0.84 |
| 1 | 68147774 | rs624790 | -0.469 | -1.528 | 1.235 | -0.12 | -0.02 | -0.042 | -0.042 | 0 | 0.19 | 0.31 | 0.19 | 0.13 | 0.15 | 0.15 |
| 1 | 68152363 | rs3783468 | 1.233 | -1.664 | 0.774 | -0.066 | -0.012 | -0.036 | -0.036 | 0 | 0.42 | 0.15 | 0.5 | 0.44 | 0.53 | 0.6 |
| 1 | 68154693 | rs685724 | 0.171 | -1.771 | 1.218 | -0.106 | -0.005 | 0.002 | 0.002 | 0 | 0.6 | 0.41 | 0.68 | 0.56 | 0.68 | 0.73 |
| 1 | 68158037 | rs11583718 | 0.696 | -1.779 | 1.536 | -0.17 | 0.02 | 0.004 | 0.004 | 0.235 | 0.49 | 0.43 | 0.55 | 0.48 | 0.61 | 0.69 |
| 1 | 68158951 | rs7546055 | -0.43 | -1.535 | 1.641 | -0.193 | -0.027 | -0.1 | -0.1 | 0 | 0.26 | 0.54 | 0.19 | 0.09 | 0.26 | 0.15 |
| 1 | 68161789 | rs511041 | 0.646 | -1.446 | 1.485 | -0.149 | 0.024 | -0.004 | -0.004 | 0 | 0.54 | 0.35 | 0.64 | 0.49 | 0.59 | 0.65 |
| 1 | 68164498 | rs663612 | 0.665 | -1.513 | 1.155 | -0.2 | -0.012 | -0.018 | -0.018 | 0 | 0.13 | 0.13 | 0.14 | 0.07 | 0.07 | 0.05 |

| | | | | | | | | | | | | | | | | |
|---|---|---|---|---|---|---|---|---|---|---|---|---|---|---|---|---|
| 1 | 68167491 | rs7419 | 1.318 | -1.58 | 1.09 | -0.242 | -0.066 | -0.048 | -0.048 | 1.807 | 0.4 | 0.13 | 0.49 | 0.43 | 0.51 | 0.59 |
| 1 | 68167935 | rs506731 | 0.361 | -1.532 | 1.294 | -0.277 | -0.03 | -0.041 | -0.041 | 0 | 0.14 | 0.22 | 0.14 | 0.06 | 0.07 | 0.06 |
| 1 | 68195741 | rs7528997 | 1.492 | -1.62 | 1.022 | -0.289 | -0.07 | -0.037 | -0.037 | 0 | 0.37 | 0.16 | 0.49 | 0.41 | 0.51 | 0.59 |
| 1 | 68199090 | rs12085997 | 0.574 | -1.492 | 1.225 | -0.15 | -0.003 | -0.005 | -0.005 | 0.251 | 0.13 | 0.11 | 0.13 | 0.07 | 0.06 | 0.06 |
| 1 | 68208946 | rs557986 | -1.365 | -1.515 | 1.039 | -0.134 | -0.023 | -0.124 | -0.124 | 0.167 | 0.81 | 0.97 | 0.84 | 0.58 | 0.84 | 0.82 |
| 1 | 68215424 | rs3766270 | -1.096 | -1.099 | 1.687 | -0.037 | -0.012 | -0.051 | -0.051 | 0 | 0.22 | 0.3 | 0.19 | 0.13 | 0.16 | 0.12 |
| 1 | 68223227 | rs12121525 | 0.174 | -0.799 | 1.253 | 0.084 | 0.014 | -0.006 | -0.006 | 0.023 | 0.05 | 0.09 | 0.04 | 0.06 | 0.09 | 0.05 |
| 1 | 68231285 | rs7542492 | -0.387 | -1.15 | 1.424 | 0.084 | -0.051 | -0.039 | -0.039 | 0 | 0.17 | 0.4 | 0.15 | 0.07 | 0.05 | 0.07 |
| 1 | 68231736 | rs7542954 | -0.078 | -1.147 | 1.342 | 0.086 | -0.032 | -0.022 | -0.022 | 0 | 0.14 | 0.32 | 0.14 | 0.07 | 0.05 | 0.06 |
| 1 | 68235077 | rs1033106 | -1.107 | -1.441 | 0.949 | -0.031 | -0.052 | -0.047 | -0.047 | 0 | 0.23 | 0.55 | 0.19 | 0.12 | 0.13 | 0.15 |
| 1 | 68238855 | rs577305 | -1.091 | -1.575 | 0.654 | 0.03 | 0.024 | -0.068 | -0.068 | 0 | 0.69 | 0.87 | 0.78 | 0.61 | 0.76 | 0.84 |
| 1 | 68241665 | rs17130263 | 0.065 | -0.934 | 0.514 | 0.218 | -0.002 | -0.006 | -0.006 | 0.037 | 0.07 | 0.13 | 0.02 | 0.05 | 0.08 | 0.08 |
| 1 | 68243397 | rs1949133 | -1.27 | 0.611 | -0.484 | 0.153 | -0.014 | 0 | 0 | 0 | 0.78 | 0.86 | 0.69 | 0.76 | 0.66 | 0.7 |
| 1 | 68247614 | rs2802462 | -0.066 | -0.167 | -0.342 | 0.223 | 0.023 | -0.002 | -0.002 | 2.128 | 0.55 | 0.68 | 0.42 | 0.51 | 0.38 | 0.32 |
| 1 | 68260778 | rs1276303 | 0.143 | 0.25 | -0.092 | 0.326 | 0.006 | 0.014 | 0.014 | 0 | 0.73 | 0.69 | 0.71 | 0.68 | 0.72 | 0.64 |
| 1 | 68277375 | rs17474654 | 0.432 | 0.034 | -0.163 | 0.251 | -0.001 | 0.003 | 0.003 | 0 | 0.06 | 0.04 | 0.03 | 0.03 | 0.1 | 0.08 |
| 1 | 68278542 | rs17130309 | 0.538 | -0.222 | 0.182 | 0.023 | 0.025 | -0.005 | -0.005 | 0.779 | 0.44 | 0.4 | 0.38 | 0.44 | 0.32 | 0.2 |
| 1 | 68278722 | rs12091105 | 0.538 | -0.222 | 0.182 | 0.023 | 0.025 | -0.005 | -0.005 | 0 | 0.44 | 0.4 | 0.38 | 0.44 | 0.32 | 0.2 |
| 1 | 68279010 | rs17130310 | 0.497 | -0.209 | 0.354 | -0.061 | -0.001 | -0.007 | -0.007 | 0.716 | 0.06 | 0.08 | 0.04 | 0.03 | 0.1 | 0.09 |
| 1 | 68282901 | rs6671283 | -0.131 | -0.594 | 0.211 | -0.241 | -0.036 | -0.026 | -0.026 | 0 | 0.1 | 0.31 | 0.04 | 0.02 | 0.03 | 0.06 |
| 1 | 68286941 | rs9633418 | 0.581 | -0.331 | 0.38 | -0.013 | 0.012 | -0.001 | -0.001 | 0 | 0.43 | 0.42 | 0.38 | 0.41 | 0.33 | 0.21 |
| 1 | 68290946 | rs2246675 | 1.315 | -0.281 | 0.365 | -0.13 | 0 | 0 | 0 | 0 | 0.25 | 0.24 | 0.3 | 0.22 | 0.32 | 0.35 |
| 1 | 68291883 | rs11209162 | 0.985 | -0.313 | 0.562 | -0.124 | 0.042 | -0.007 | -0.007 | 2.273 | 0.39 | 0.24 | 0.35 | 0.4 | 0.31 | 0.2 |
| 1 | 68299552 | rs7540541 | 0.126 | -0.293 | 0.554 | -0.119 | 0.001 | 0.003 | 0.003 | 0 | 0.7 | 0.74 | 0.69 | 0.62 | 0.65 | 0.57 |
| 1 | 68299797 | rs6588283 | 0.311 | -0.287 | 0.474 | -0.116 | 0.014 | 0.027 | 0.027 | 0 | 0.67 | 0.52 | 0.66 | 0.62 | 0.64 | 0.56 |
| 1 | 68313038 | rs6671298 | 0.804 | -0.355 | 0.505 | -0.125 | 0.037 | 0.004 | 0.004 | 0 | 0.42 | 0.3 | 0.36 | 0.4 | 0.31 | 0.2 |
| 1 | 68316422 | rs6588286 | 1.231 | -0.243 | 0.401 | -0.097 | -0.017 | 0.008 | 0.008 | 0 | 0.26 | 0.17 | 0.3 | 0.23 | 0.33 | 0.38 |
| 1 | 68334436 | rs1938445 | 0.941 | -0.363 | 0.619 | -0.038 | 0.028 | -0.004 | -0.004 | 0 | 0.38 | 0.28 | 0.36 | 0.39 | 0.31 | 0.19 |
| 1 | 68335624 | rs1938446 | 0.686 | -0.214 | 0.631 | 0.034 | 0.011 | 0.002 | 0.002 | 0 | 0.42 | 0.39 | 0.37 | 0.39 | 0.31 | 0.19 |
| 1 | 68343642 | rs12144887 | -0.178 | -0.059 | 0.599 | 0.135 | -0.009 | -0.016 | -0.016 | 0 | 0.75 | 0.78 | 0.7 | 0.63 | 0.68 | 0.61 |
| 1 | 68358177 | rs7520899 | 0.6 | -0.322 | 0.417 | 0.03 | 0.01 | 0.005 | 0.005 | 0 | 0.45 | 0.41 | 0.29 | 0.36 | 0.32 | 0.22 |
| 1 | 68364218 | rs1938451 | -0.41 | -0.16 | -0.377 | 0.213 | -0.015 | -0.031 | -0.031 | 0 | 0.77 | 0.84 | 0.6 | 0.59 | 0.7 | 0.66 |
| 1 | 68366925 | rs7552577 | -0.111 | -0.152 | -0.366 | 0.131 | -0.017 | -0.031 | -0.031 | 0 | 0.74 | 0.79 | 0.58 | 0.55 | 0.63 | 0.59 |
| 1 | 68373757 | rs11583096 | 0.721 | -0.365 | -0.277 | -0.24 | -0.043 | 0.047 | 0.047 | 0 | 0.28 | 0.09 | 0.27 | 0.19 | 0.32 | 0.38 |
| 1 | 68384394 | rs2186238 | -0.124 | -0.595 | -0.236 | -0.202 | 0.003 | 0.002 | 0.002 | 0 | 0.86 | 0.87 | 0.93 | 0.92 | 0.92 | 0.87 |
| 1 | 68384979 | rs2186237 | 0.177 | -0.879 | 0.07 | -0.271 | 0.004 | 0.021 | 0.021 | 0 | 0.52 | 0.54 | 0.6 | 0.72 | 0.59 | 0.46 |
| 1 | 68389426 | rs1938432 | 0.791 | -1.756 | -0.034 | 0.098 | 0.039 | 0.001 | 0.001 | 0 | 0.34 | 0.2 | 0.13 | 0.36 | 0.2 | 0.16 |
| 1 | 68393522 | rs7520193 | 1.008 | -1.913 | 0.245 | 0.041 | 0.002 | -0.001 | -0.001 | 1.38 | 0.31 | 0.28 | 0.14 | 0.36 | 0.2 | 0.16 |
| 1 | 68395041 | rs17097357 | 0.563 | -2.206 | 0.782 | -0.124 | -0.033 | -0.028 | -0.028 | 0 | 0.12 | 0.21 | 0.05 | 0.04 | 0.02 | 0.06 |
| 1 | 68400546 | rs10493429 | 0.685 | -1.86 | 0.221 | -0.122 | -0.11 | -0.087 | -0.087 | 0 | 0.22 | 0.09 | 0.47 | 0.34 | 0.44 | 0.35 |
| 1 | 68401005 | rs6665420 | 0.231 | -2.11 | 0.458 | -0.169 | 0.019 | 0.008 | 0.008 | 2.612 | 0.46 | 0.61 | 0.19 | 0.43 | 0.23 | 0.22 |
| 1 | 68405111 | rs6688063 | 0.149 | -1.941 | 0.155 | -0.165 | 0.001 | 0 | 0 | 0.089 | 0.56 | 0.41 | 0.63 | 0.72 | 0.68 | 0.53 |
| 1 | 68406484 | rs1938429 | -0.974 | -1.652 | -0.239 | -0.071 | -0.005 | 0.02 | 0.02 | 0 | 0.69 | 0.8 | 0.69 | 0.79 | 0.71 | 0.58 |
| 1 | 68407584 | rs1938426 | 0.588 | -1.378 | -0.372 | 0.078 | -0.134 | -0.101 | -0.101 | 3.435 | 0.21 | 0.07 | 0.49 | 0.34 | 0.48 | 0.36 |
| 1 | 68411675 | rs10493430 | 0.373 | -1.473 | -0.306 | 0.068 | -0.053 | -0.04 | -0.04 | 0.49 | 0.24 | 0.18 | 0.5 | 0.37 | 0.48 | 0.36 |
| 1 | 68416023 | rs11209181 | 0.084 | -2.316 | -0.059 | 0.02 | -0.013 | -0.001 | -0.001 | 0 | 0.42 | 0.57 | 0.18 | 0.45 | 0.22 | 0.21 |
| 1 | 68421069 | rs11209184 | -0.395 | -1.89 | -0.399 | -0.093 | -0.022 | 0.03 | 0.03 | 1.055 | 0.65 | 0.77 | 0.59 | 0.78 | 0.61 | 0.57 |
| 1 | 68423721 | rs12029652 | 0.854 | -1.943 | -0.589 | -0.014 | -0.151 | -0.137 | -0.137 | 0 | 0.24 | 0.02 | 0.42 | 0.33 | 0.42 | 0.36 |
| 1 | 68441949 | rs1926298 | -0.339 | -1.945 | -0.003 | -0.314 | -0.079 | -0.043 | -0.043 | 0 | 0.17 | 0.6 | 0.09 | 0.08 | 0.03 | 0.06 |
| 1 | 68446291 | rs1926284 | 0.446 | -1.897 | -0.768 | -0.322 | -0.023 | 0.015 | 0.015 | 0.41 | 0.3 | 0.22 | 0.39 | 0.21 | 0.38 | 0.4 |
| 1 | 68446944 | rs12130147 | 0.529 | -1.866 | -0.764 | -0.301 | -0.044 | 0.025 | 0.025 | 1.07 | 0.29 | 0.15 | 0.39 | 0.2 | 0.38 | 0.41 |
| 1 | 68447267 | rs6662448 | -0.966 | -1.94 | -0.83 | -0.328 | -0.057 | 0.12 | 0.12 | 4.983 | 0.51 | 0.92 | 0.51 | 0.29 | 0.42 | 0.47 |
| 1 | 68448998 | rs1926280 | -0.517 | -1.869 | -0.724 | -0.397 | -0.05 | -0.032 | -0.032 | 0 | 0.21 | 0.7 | 0.12 | 0.08 | 0.04 | 0.07 |
| 1 | 68452388 | rs9326078 | 0.022 | -0.691 | -0.465 | 0.167 | -0.027 | 0.007 | 0.007 | 0.917 | 0.3 | 0.2 | 0.39 | 0.23 | 0.39 | 0.42 |
| 1 | 68458532 | rs12040310 | 0.637 | -1.51 | -0.064 | 0.302 | -0.089 | 0.005 | 0.005 | 0 | 0.28 | 0.42 | 0.11 | 0.28 | 0.15 | 0.21 |
| 1 | 68459693 | rs1926276 | 0.911 | -1.525 | 0.085 | 0.294 | -0.022 | 0.001 | 0.001 | 0 | 0.27 | 0.27 | 0.09 | 0.28 | 0.15 | 0.2 |
| 1 | 68460368 | rs10789253 | -0.077 | -1.02 | -0.627 | 0.391 | -0.002 | 0.005 | 0.005 | 0 | 0.54 | 0.6 | 0.4 | 0.59 | 0.51 | 0.48 |
| 1 | 68468718 | rs10889737 | -0.379 | -0.975 | -0.647 | 0.423 | -0.014 | -0.001 | -0.001 | 0 | 0.61 | 0.44 | 0.82 | 0.68 | 0.78 | 0.76 |
| 1 | 68475641 | rs4655768 | -0.697 | 0.511 | -0.608 | 0.59 | -0.008 | -0.013 | -0.013 | 0 | 0.89 | 0.88 | 0.93 | 0.96 | 0.93 | 0.97 |
| 1 | 68475955 | rs2209620 | -0.659 | 1.151 | -0.769 | 0.928 | -0.079 | -0.04 | -0.04 | 0 | 0.32 | 0.11 | 0.49 | 0.37 | 0.41 | 0.5 |
| 1 | 68480671 | rs11209195 | -0.86 | 1.388 | -0.113 | 1.017 | 0.032 | 0.028 | 0.028 | 0 | 0.24 | 0.32 | 0.44 | 0.47 | 0.39 | 0.45 |
| 1 | 68492521 | rs10889741 | -1.106 | -0.067 | 0.672 | 0.194 | -0.055 | -0.093 | -0.093 | 0 | 0.14 | 0.62 | 0.09 | 0 | 0.05 | 0.01 |
| 1 | 68507394 | rs12076293 | -1.218 | -0.106 | 0.719 | 0.175 | -0.055 | -0.088 | -0.088 | 0 | 0.14 | 0.67 | 0.09 | 0 | 0.05 | 0.01 |
| 1 | 68508995 | rs7550608 | -1.218 | -0.106 | 0.719 | 0.175 | -0.055 | -0.088 | -0.088 | 0.178 | 0.14 | 0.67 | 0.09 | 0 | 0.05 | 0.01 |
| 1 | 68512171 | rs11801053 | -0.645 | -0.113 | 0.943 | 0.172 | -0.037 | -0.069 | -0.069 | 0 | 0.11 | 0.28 | 0.05 | 0 | 0.04 | 0.01 |
| 1 | 68513484 | rs3790688 | 1.639 | -0.094 | -0.19 | 0.46 | 0.05 | -0.088 | -0.088 | 0.091 | 0.61 | 0.11 | 0.55 | 0.53 | 0.69 | 0.56 |
| 1 | 68531336 | rs269354 | -1.092 | 0.145 | 0.265 | -0.019 | -0.039 | -0.07 | -0.07 | 0 | 0.16 | 0.72 | 0.11 | 0 | 0.06 | 0.01 |
| 1 | 68538268 | rs669923 | -1.226 | 0.278 | 0.391 | -0.031 | -0.045 | -0.135 | -0.135 | 0 | 0.19 | 0.56 | 0.14 | 0 | 0.11 | 0.02 |
| 1 | 68549482 | rs269332 | 0.189 | 0.622 | -0.522 | 0.515 | -0.007 | -0.022 | -0.022 | 0.003 | 0.84 | 0.9 | 0.72 | 0.4 | 0.8 | 0.56 |

| | | | | | | | | | | | | | | | | |
|---|---|---|---|---|---|---|---|---|---|---|---|---|---|---|---|---|
| 1 | 68551909 | rs198866 | 0.232 | 0.624 | -0.507 | 0.511 | 0.002 | 0.024 | 0.024 | 0.422 | 0.84 | 0.85 | 0.72 | 0.41 | 0.8 | 0.56 |
| 1 | 68566576 | rs7526959 | -0.914 | 0.271 | -0.716 | -0.281 | -0.002 | -0.024 | -0.024 | 2.803 | 0.17 | 0.19 | 0.1 | 0.02 | 0.12 | 0.04 |
| 1 | 68566993 | rs1033973 | 0.649 | 0.605 | -1.665 | -0.923 | -0.089 | 0.173 | 0.173 | 0 | 0.39 | 0.09 | 0.39 | 0.15 | 0.46 | 0.28 |
| 1 | 68567299 | rs11209214 | 0.161 | 0.798 | -1.455 | -1.17 | -0.017 | 0.046 | 0.046 | 0.005 | 0.61 | 0.26 | 0.62 | 0.88 | 0.56 | 0.67 |
| 1 | 68567418 | rs9436797 | 0.271 | 0.714 | -1.325 | -0.847 | -0.009 | -0.029 | -0.029 | 1.049 | 0.21 | 0.25 | 0.18 | 0.11 | 0.17 | 0.15 |
| 1 | 68567848 | rs1033975 | -0.84 | 1.731 | -1.651 | -0.198 | -0.092 | -0.013 | -0.013 | 0 | 0.2 | 0.03 | 0.3 | 0.18 | 0.3 | 0.18 |
| 1 | 68568921 | rs11805150 | -0.568 | 1.418 | -1.029 | -0.168 | 0.01 | -0.009 | -0.009 | 0 | 0.18 | 0.22 | 0.32 | 0.13 | 0.25 | 0.18 |
| 1 | 68569771 | rs9436367 | -0.211 | 0.982 | -1.28 | -0.553 | -0.003 | 0.035 | 0.035 | 0 | 0.1 | 0.04 | 0.04 | 0.01 | 0.1 | 0.06 |
| 1 | 68571220 | rs6671127 | -0.659 | 1.492 | -1.332 | -0.01 | 0.004 | 0.002 | 0.002 | 0.003 | 0.24 | 0.25 | 0.38 | 0.34 | 0.34 | 0.31 |
| 1 | 68571431 | rs11209216 | -1.317 | 0.931 | -0.951 | -0.017 | -0.009 | 0.006 | 0.006 | 0 | 0.42 | 0.4 | 0.44 | 0.41 | 0.46 | 0.42 |
| 1 | 68579397 | rs269352 | -0.169 | 0.882 | -0.774 | 0.738 | 0.002 | 0 | 0 | 0 | 0.31 | 0.29 | 0.45 | 0.34 | 0.39 | 0.35 |
| 1 | 68579833 | rs12726865 | 1.341 | 0.37 | -1.231 | -0.107 | -0.053 | 0.039 | 0.039 | 0.621 | 0.34 | 0.04 | 0.33 | 0.17 | 0.34 | 0.27 |
| 1 | 68581085 | rs12134024 | 0.729 | 0.097 | -0.666 | -0.1 | -0.001 | -0.002 | -0.002 | 0.281 | 0.46 | 0.44 | 0.42 | 0.2 | 0.47 | 0.33 |
| 1 | 68581702 | rs2566760 | 0.34 | 0.412 | -0.613 | -0.174 | -0.005 | -0.022 | -0.022 | 0 | 0.57 | 0.43 | 0.55 | 0.18 | 0.57 | 0.4 |
| 1 | 68581840 | rs17130484 | 0.99 | 0.181 | -0.356 | -0.302 | -0.008 | -0.045 | -0.045 | 0 | 0.08 | 0.19 | 0.04 | 0 | 0.07 | 0.04 |
| 1 | 68581861 | rs2566761 | 0.313 | 0.476 | -0.397 | -0.32 | -0.001 | -0.014 | -0.014 | 0 | 0.55 | 0.5 | 0.53 | 0.18 | 0.5 | 0.36 |
| 1 | 68581947 | rs2435841 | 0.851 | 0.573 | -0.654 | 0.256 | -0.016 | 0.049 | 0.049 | 0.363 | 0.05 | 0.02 | 0.06 | 0.36 | 0.12 | 0.24 |
| 1 | 68582457 | rs2566762 | 0.095 | 0.409 | -0.468 | -0.044 | -0.007 | -0.02 | -0.02 | 0 | 0.07 | 0.11 | 0.02 | 0.02 | 0.04 | 0.03 |
| 1 | 68582582 | rs2988019 | 0.484 | 0.333 | -0.558 | 0.035 | -0.004 | 0.005 | 0.005 | 0 | 0.3 | 0.23 | 0.38 | 0.74 | 0.37 | 0.55 |
| 1 | 68584745 | rs14130 | 0.485 | 0.515 | -0.686 | 0.05 | -0.018 | -0.018 | -0.018 | 0 | 0.31 | 0.2 | 0.41 | 0.4 | 0.35 | 0.35 |
| 1 | 68585667 | rs2271 | 1.029 | 0.231 | -0.644 | 0.104 | 0.017 | 0.002 | 0.002 | 0 | 0.08 | 0.07 | 0.03 | 0.07 | 0.05 | 0.05 |
| 1 | 68586620 | rs12046920 | 1.018 | 0.332 | -0.596 | 0.123 | 0.001 | -0.006 | -0.006 | 0 | 0.25 | 0.21 | 0.34 | 0.38 | 0.26 | 0.28 |
| 1 | 68587458 | rs12740674 | 1.09 | 0.257 | -0.815 | -0.215 | -0.093 | 0.147 | 0.147 | 0.247 | 0.39 | 0.09 | 0.37 | 0.13 | 0.4 | 0.31 |
| 1 | 68589935 | rs10889748 | 0.618 | 0.259 | -0.255 | 0.108 | -0.001 | 0.017 | 0.017 | 0 | 0.7 | 0.42 | 0.72 | 0.6 | 0.67 | 0.64 |
| 1 | 68591104 | rs1046852 | 0.662 | 0.213 | 0.11 | 0.097 | -0.002 | 0.041 | 0.041 | 0 | 0.7 | 0.5 | 0.73 | 0.6 | 0.67 | 0.64 |
| 1 | 68591173 | rs1046835 | 0.662 | 0.213 | 0.109 | 0.097 | -0.002 | 0.042 | 0.042 | 0 | 0.7 | 0.5 | 0.73 | 0.6 | 0.67 | 0.64 |
| 1 | 68591253 | rs7062 | 0.66 | 0.071 | -0.179 | -0.135 | -0.008 | 0.003 | 0.003 | 0.001 | 0.37 | 0.4 | 0.39 | 0.45 | 0.32 | 0.33 |
| 1 | 68591405 | rs269350 | 0.861 | -0.076 | 0.02 | -0.152 | -0.003 | 0.006 | 0.006 | 0 | 0.64 | 0.39 | 0.71 | 0.53 | 0.65 | 0.6 |
| 1 | 68597121 | rs7552304 | 0.206 | -0.087 | 0.437 | -0.175 | 0.01 | 0.045 | 0.045 | 0.145 | 0.2 | 0.26 | 0.22 | 0.38 | 0.28 | 0.33 |
| 1 | 68597672 | rs4655576 | 0 | -0.143 | 0.376 | -0.204 | 0.036 | 0.011 | 0.011 | 2.227 | 0.53 | 0.37 | 0.56 | 0.54 | 0.63 | 0.64 |
| 1 | 68599427 | rs6658452 | 2.156 | 0.135 | 0.208 | -0.255 | -0.029 | 0.032 | 0.032 | 0.348 | 0.27 | 0.1 | 0.28 | 0.16 | 0.25 | 0.23 |
| 1 | 68599886 | rs2566783 | 1.256 | 0.01 | 0.329 | -0.25 | 0.015 | -0.013 | -0.013 | 0 | 0.6 | 0.32 | 0.68 | 0.51 | 0.63 | 0.59 |
| 1 | 68599901 | rs2772283 | 0.85 | -0.001 | 0.567 | -0.253 | -0.014 | 0.035 | 0.035 | 0 | 0.65 | 0.49 | 0.71 | 0.51 | 0.66 | 0.6 |
| 1 | 68600278 | rs12062874 | 1.179 | 0.009 | 0.444 | -0.231 | 0.008 | 0.005 | 0.005 | 0 | 0.09 | 0.08 | 0.03 | 0.06 | 0.02 | 0.04 |
| 1 | 68602735 | rs2566784 | 1.566 | 0.067 | 0.419 | -0.218 | 0.008 | -0.014 | -0.014 | 0.024 | 0.27 | 0.22 | 0.34 | 0.36 | 0.28 | 0.28 |
| 1 | 68603586 | rs983034 | 1.314 | 0.205 | 0.23 | -0.191 | -0.079 | 0.076 | 0.076 | 0.342 | 0.33 | 0.11 | 0.34 | 0.14 | 0.35 | 0.32 |
| 1 | 68608003 | rs1816317 | 0.889 | 0.31 | 0.38 | 0.275 | -0.008 | 0.018 | 0.018 | 0 | 0.65 | 0.54 | 0.72 | 0.54 | 0.66 | 0.62 |
| 1 | 68609491 | rs2052959 | 0.788 | -0.081 | 0.146 | 0.007 | -0.001 | 0.004 | 0.004 | 0 | 0.34 | 0.31 | 0.33 | 0.2 | 0.29 | 0.26 |
| 1 | 68611243 | rs1556157 | 1.041 | -0.593 | 0.222 | 0.173 | -0.063 | 0.07 | 0.07 | 0 | 0.35 | 0.17 | 0.36 | 0.14 | 0.38 | 0.3 |
| 1 | 68613933 | rs2303949 | 0.584 | -0.607 | 0.347 | 0.337 | 0 | -0.003 | -0.003 | 0.274 | 0.16 | 0.16 | 0.1 | 0.37 | 0.19 | 0.29 |
| 1 | 68614320 | rs17130529 | 0.932 | -0.636 | 0.246 | 0.299 | 0.001 | 0.001 | 0.001 | 0 | 0.09 | 0.08 | 0.05 | 0.06 | 0.05 | 0.05 |
| 1 | 68617627 | rs1430758 | -0.09 | -0.66 | 0.084 | 0.339 | -0.012 | 0.005 | 0.005 | 0 | 0.19 | 0.29 | 0.11 | 0.27 | 0.19 | 0.28 |
| 1 | 68619511 | rs1430759 | 1.024 | -0.584 | 0.127 | 0.143 | -0.004 | -0.036 | -0.036 | 0.301 | 0.33 | 0.19 | 0.45 | 0.4 | 0.37 | 0.36 |
| 1 | 68620762 | rs3762371 | 1.146 | -0.555 | 0.096 | 0.133 | -0.063 | 0.066 | 0.066 | 0 | 0.34 | 0.17 | 0.37 | 0.16 | 0.38 | 0.3 |
| 1 | 68622994 | rs12733835 | 1.16 | -0.548 | 0.099 | 0.122 | -0.063 | 0.068 | 0.068 | 0 | 0.34 | 0.17 | 0.36 | 0.14 | 0.38 | 0.3 |
| 1 | 68624878 | rs3748705 | 0.512 | -0.126 | 0.004 | 0.449 | 0.003 | -0.022 | -0.022 | 0 | 0.33 | 0.24 | 0.45 | 0.41 | 0.36 | 0.35 |
| 1 | 68628540 | rs2820489 | -0.138 | -0.009 | -0.387 | 0.574 | -0.01 | 0.057 | 0.057 | 0.03 | 0.18 | 0.29 | 0.1 | 0.38 | 0.19 | 0.29 |
| 1 | 68629530 | rs930145 | -0.069 | 0.019 | -0.373 | 0.588 | -0.008 | 0.061 | 0.061 | 0.01 | 0.18 | 0.28 | 0.1 | 0.38 | 0.19 | 0.29 |
| 1 | 68630667 | rs2419343 | 0.393 | 0.06 | -0.448 | 0.544 | 0.004 | -0.02 | -0.02 | 0 | 0.34 | 0.24 | 0.46 | 0.41 | 0.37 | 0.36 |
| 1 | 68631502 | rs11209225 | 0.45 | 0.063 | -0.618 | 0.556 | 0.002 | -0.035 | -0.035 | 0 | 0.34 | 0.18 | 0.45 | 0.41 | 0.37 | 0.36 |
| 1 | 68631522 | rs2820492 | -0.045 | 0.062 | -0.551 | 0.562 | -0.008 | 0.087 | 0.087 | 0 | 0.18 | 0.33 | 0.1 | 0.36 | 0.19 | 0.28 |
| 1 | 68631792 | rs2566759 | -0.216 | 0.194 | -0.618 | 0.082 | -0.001 | -0.011 | -0.011 | 0 | 0.52 | 0.5 | 0.55 | 0.79 | 0.56 | 0.65 |
| 1 | 68633924 | rs1367448 | -0.761 | -0.109 | -1.055 | 0.441 | 0.011 | -0.015 | -0.015 | 0.541 | 0.4 | 0.35 | 0.52 | 0.51 | 0.34 | 0.5 |
| 1 | 68634030 | rs1367447 | 1.365 | -0.186 | -1.289 | 0.375 | 0.007 | -0.016 | -0.016 | 0 | 0.19 | 0.11 | 0.13 | 0.27 | 0.21 | 0.25 |
| 1 | 68634144 | rs2566757 | 0.085 | -0.482 | -1.505 | -0.178 | 0.01 | 0.047 | 0.047 | 0 | 0.8 | 0.85 | 0.84 | 0.94 | 0.84 | 0.89 |
| 1 | 68634196 | rs11587907 | 0.75 | -0.422 | -1.368 | -0.278 | 0.002 | 0.005 | 0.005 | 0 | 0.13 | 0.08 | 0.05 | 0 | 0.09 | 0.12 |
| 1 | 68634257 | rs1864569 | 0.308 | -0.377 | -1.225 | 0.181 | -0.007 | 0.005 | 0.005 | 0.218 | 0.46 | 0.65 | 0.33 | 0.35 | 0.46 | 0.39 |
| 1 | 68634642 | rs12132335 | -0.352 | -0.572 | -0.773 | 0.413 | 0.012 | -0.013 | -0.013 | 0 | 0.19 | 0.11 | 0.1 | 0.14 | 0.15 | 0.1 |
| 1 | 68635390 | rs2566755 | 1.784 | -0.549 | -0.904 | 0.426 | 0.006 | -0.005 | -0.005 | 3.468 | 0.22 | 0.15 | 0.15 | 0.25 | 0.22 | 0.28 |
| 1 | 68636580 | rs2772304 | 0.926 | -0.547 | -0.712 | 0.389 | 0 | 0.001 | 0.001 | 0.518 | 0.28 | 0.26 | 0.26 | 0.32 | 0.29 | 0.31 |
| 1 | 68638451 | rs4233320 | 1.712 | -0.673 | -0.517 | 0.398 | 0.031 | -0.02 | -0.02 | 0 | 0.29 | 0.36 | 0.35 | 0.2 | 0.37 | 0.26 |
| 1 | 68638729 | rs12037394 | 1.239 | -0.697 | -0.699 | 0.361 | 0.019 | 0.022 | 0.022 | 0 | 0.11 | 0.06 | 0.04 | 0.03 | 0.08 | 0.07 |
| 1 | 68641238 | rs2566754 | -0.004 | -0.704 | -0.698 | 0.328 | -0.002 | 0.053 | 0.053 | 0 | 0.43 | 0.28 | 0.32 | 0.33 | 0.43 | 0.39 |
| 1 | 68642569 | rs12034358 | 0.155 | -0.713 | -0.712 | 0.327 | 0.006 | 0.034 | 0.034 | 0 | 0.17 | 0.12 | 0.15 | 0.08 | 0.15 | 0.09 |
| 1 | 68643091 | rs944082 | -1.46 | -0.502 | -0.278 | 0.553 | 0.006 | 0.005 | 0.005 | 0 | 0.68 | 0.67 | 0.44 | 0.47 | 0.56 | 0.54 |
| 1 | 68643147 | rs17130546 | 0.812 | -0.173 | -0.399 | 0.65 | 0.005 | -0.003 | -0.003 | 0.009 | 0.06 | 0.08 | 0.11 | 0.06 | 0.07 | 0.04 |
| 1 | 68645765 | rs17130550 | 0.812 | -0.157 | -0.459 | 0.641 | 0.006 | -0.004 | -0.004 | 0 | 0.06 | 0.08 | 0.11 | 0.06 | 0.07 | 0.04 |
| 1 | 68645917 | rs729884 | -0.836 | -0.131 | -0.668 | 0.654 | 0.005 | -0.007 | -0.007 | 0.867 | 0.93 | 0.88 | 0.88 | 0.94 | 0.93 | 0.96 |
| 1 | 68648026 | rs2772300 | 1.09 | -0.39 | -0.749 | 0.854 | -0.008 | 0.006 | 0.006 | 0.199 | 0.26 | 0.16 | 0.16 | 0.24 | 0.28 | 0.28 |

| | | | | | | | | | | | | | | | | |
|---|---|---|---|---|---|---|---|---|---|---|---|---|---|---|---|---|
| 1 | 68651948 | rs891526 | -0.048 | -0.889 | -0.674 | 0.926 | -0.013 | 0.005 | 0.005 | 0 | 0.24 | 0.25 | 0.28 | 0.35 | 0.19 | 0.35 |
| 1 | 68658961 | rs2820500 | -0.815 | -0.419 | -0.969 | 1.199 | 0.001 | -0.002 | -0.002 | 0 | 0.39 | 0.4 | 0.34 | 0.43 | 0.34 | 0.33 |
| 1 | 68660092 | rs1036067 | -0.324 | -0.378 | -0.904 | 0.999 | -0.032 | 0.055 | 0.055 | 0 | 0.13 | 0.28 | 0.09 | 0.14 | 0.09 | 0.06 |
| 1 | 68663847 | rs17130567 | 0.026 | 0.128 | -1.281 | 0.619 | -0.003 | -0.003 | -0.003 | 1.508 | 0.26 | 0.11 | 0.15 | 0.26 | 0.25 | 0.27 |
| 1 | 68665023 | rs7554551 | -0.213 | 0.094 | -1.227 | 0.546 | -0.029 | 0.054 | 0.054 | 0.44 | 0.32 | 0.16 | 0.34 | 0.23 | 0.32 | 0.35 |
| 1 | 68686391 | rs10493436 | -1.316 | -1.253 | 0.192 | 1.547 | -0.021 | -0.006 | -0.006 | 0.812 | 0.27 | 0.33 | 0.4 | 0.8 | 0.18 | 0.49 |
| 1 | 68687697 | rs10493438 | -0.647 | -1.39 | 0.309 | 1.938 | -0.005 | 0.015 | 0.015 | 0 | 0.16 | 0.15 | 0.33 | 0.43 | 0.1 | 0.26 |
| 1 | 68688069 | rs10493439 | -0.639 | -1.372 | 0.276 | 1.943 | -0.011 | 0.095 | 0.095 | 0 | 0.11 | 0.18 | 0.08 | 0.37 | 0.08 | 0.23 |
| 1 | 68689997 | rs17130595 | -0.474 | -1.377 | 0.269 | 1.946 | -0.004 | 0.067 | 0.067 | 0 | 0.1 | 0.15 | 0.08 | 0.37 | 0.08 | 0.23 |
| 1 | 68692642 | rs1430753 | 0.333 | -1.765 | 0.513 | 1.391 | -0.022 | -0.056 | -0.056 | 0 | 0.75 | 0.59 | 0.78 | 0.9 | 0.81 | 0.86 |
| 1 | 68692744 | rs12563946 | -0.345 | -1.319 | 0.303 | 1.722 | 0.004 | -0.003 | -0.003 | 0 | 0.15 | 0.12 | 0.33 | 0.43 | 0.11 | 0.26 |
| 1 | 68695148 | rs2772280 | -0.938 | -1.25 | 0.357 | 1.773 | 0.002 | 0.165 | 0.165 | 0 | 0.13 | 0.28 | 0.07 | 0.38 | 0.12 | 0.28 |
| 1 | 68712358 | rs1360180 | -0.216 | -1.208 | 0.241 | 1.43 | -0.003 | 0.009 | 0.009 | 0 | 0.15 | 0.06 | 0.16 | 0.08 | 0.13 | 0.14 |
| 1 | 68713347 | rs17130627 | -0.963 | -1.103 | 0.443 | 1.848 | -0.013 | 0.012 | 0.012 | 0 | 0.22 | 0.26 | 0.18 | 0.4 | 0.12 | 0.29 |
| 1 | 68714135 | rs1360181 | -1.217 | -0.964 | 0.365 | 1.888 | -0.03 | 0.072 | 0.072 | 0 | 0.24 | 0.38 | 0.21 | 0.43 | 0.21 | 0.35 |
| 1 | 68719016 | rs6686875 | 1.059 | -0.996 | 0.29 | 1.296 | -0.022 | -0.011 | -0.011 | 0 | 0.72 | 0.58 | 0.57 | 0.19 | 0.77 | 0.51 |
| 1 | 68725907 | rs11209240 | -0.583 | -0.752 | 0.54 | 1.372 | 0.005 | 0.01 | 0.01 | 0 | 0.2 | 0.12 | 0.17 | 0.08 | 0.16 | 0.13 |
| 1 | 68728518 | rs7554674 | -1.238 | -0.955 | 0.797 | 1.554 | -0.002 | -0.008 | -0.008 | 0 | 0.27 | 0.25 | 0.35 | 0.36 | 0.27 | 0.32 |
| 1 | 68729652 | rs12045284 | -1.06 | -1.121 | 0.667 | 1.458 | 0.027 | 0.027 | 0.027 | 0 | 0.27 | 0.16 | 0.2 | 0.12 | 0.19 | 0.19 |
| 1 | 68732819 | rs12065581 | -0.514 | -1.039 | 0.907 | 1.383 | 0.017 | 0.193 | 0.193 | 1.374 | 0.14 | 0.3 | 0.29 | 0.52 | 0.17 | 0.26 |
| 1 | 68733478 | rs885098 | 1.13 | -1.49 | 0.395 | 1.122 | -0.001 | 0 | 0 | 0 | 0.59 | 0.57 | 0.61 | 0.6 | 0.7 | 0.62 |
| 1 | 68735304 | rs12730489 | 1.705 | -1.277 | -0.129 | 0.702 | -0.021 | 0.242 | 0.242 | 0 | 0.46 | 0.11 | 0.3 | 0.12 | 0.53 | 0.4 |
| 1 | 68738278 | rs1360182 | 1.098 | -1.371 | 0.05 | 1.29 | 0.045 | -0.023 | -0.023 | 0.157 | 0.57 | 0.25 | 0.48 | 0.48 | 0.65 | 0.58 |
| 1 | 68739203 | rs10889756 | 1.013 | -1.565 | 0.654 | 1.098 | -0.017 | -0.019 | -0.019 | 0 | 0.64 | 0.43 | 0.56 | 0.76 | 0.71 | 0.72 |
| 1 | 68741905 | rs7529683 | 1.152 | -1.502 | 0.693 | 1.636 | -0.055 | 0.093 | 0.093 | 0 | 0.76 | 0.35 | 0.78 | 0.52 | 0.89 | 0.76 |
| 1 | 68746507 | rs10889757 | 0.396 | -0.764 | 0.584 | 1.635 | 0.005 | 0.014 | 0.014 | 0.022 | 0.92 | 0.85 | 0.73 | 0.84 | 0.89 | 0.81 |
| 1 | 68750782 | rs7546334 | -0.41 | -0.79 | 0.388 | 1.418 | 0.004 | 0 | 0 | 0 | 0.14 | 0.15 | 0.21 | 0.12 | 0.16 | 0.09 |
| 1 | 68757334 | rs3923451 | 0.584 | -1.238 | 0.147 | 1.163 | 0.001 | 0.004 | 0.004 | 0 | 0.63 | 0.36 | 0.38 | 0.57 | 0.59 | 0.57 |
| 1 | 68764274 | rs7541094 | 1.545 | -2.102 | -0.649 | 0.673 | -0.111 | 0.254 | 0.254 | 0 | 0.42 | 0.04 | 0.29 | 0.08 | 0.52 | 0.37 |
| 1 | 68765106 | rs12039098 | -0.185 | -2.126 | -0.139 | 0.97 | -0.107 | 0.207 | 0.207 | 0 | 0.19 | 0.46 | 0.08 | 0.51 | 0.05 | 0.17 |
| 1 | 68773120 | rs9436372 | -0.198 | -1.714 | -0.218 | 1.023 | 0.002 | 0.007 | 0.007 | 0.123 | 0.09 | 0.11 | 0.09 | 0.18 | 0.1 | 0.19 |
| 1 | 68779124 | rs4259701 | 0.304 | -1.753 | -0.142 | 0.923 | -0.003 | -0.007 | -0.007 | 0 | 0.67 | 0.67 | 0.42 | 0.59 | 0.62 | 0.54 |
| 1 | 68792113 | rs12124380 | 0.157 | -0.608 | -0.632 | 0.641 | 0.001 | -0.001 | -0.001 | 0 | 0.6 | 0.42 | 0.4 | 0.68 | 0.54 | 0.68 |
| 1 | 68797702 | rs12139131 | 0.861 | -0.535 | -1.36 | 0.59 | -0.032 | -0.033 | -0.033 | 0 | 0.06 | 0 | 0.09 | 0.19 | 0.09 | 0.13 |
| 1 | 68803646 | rs4926354 | -1 | -0.688 | -1.129 | 0.43 | -0.012 | 0.008 | 0.008 | 0 | 0.77 | 0.82 | 0.53 | 0.8 | 0.74 | 0.75 |
| 1 | 68804187 | rs4926355 | 0.038 | 0.341 | -1.51 | 0.196 | -0.003 | 0.002 | 0.002 | 1.447 | 0.43 | 0.29 | 0.37 | 0.39 | 0.55 | 0.42 |
| 1 | 68816800 | rs4926381 | -0.281 | 0.727 | -1.815 | 0.719 | -0.066 | -0.022 | -0.022 | 0 | 0.43 | 0.14 | 0.33 | 0.39 | 0.49 | 0.52 |
| 1 | 68821539 | rs9436812 | 0.114 | 1.277 | -1.197 | 1.312 | -0.057 | -0.063 | -0.063 | 0 | 0.29 | 0.13 | 0.26 | 0.39 | 0.38 | 0.38 |
| 1 | 68826316 | rs10889765 | 0.677 | 1.488 | -0.438 | 1.908 | -0.002 | 0.005 | 0.005 | 0 | 0.82 | 0.78 | 0.84 | 0.51 | 0.8 | 0.69 |
| 1 | 68842068 | rs1952124 | -0.845 | 1.574 | 0.653 | 1.857 | -0.001 | 0.005 | 0.005 | 0 | 0.22 | 0.22 | 0.41 | 0.4 | 0.28 | 0.25 |
| 1 | 68842414 | rs747253 | -1.097 | 1.574 | 0.669 | 1.856 | 0.024 | 0.082 | 0.082 | 0 | 0.23 | 0.41 | 0.43 | 0.4 | 0.28 | 0.25 |
| 1 | 68842785 | rs747252 | -0.818 | 1.58 | 0.637 | 1.859 | -0.001 | 0.005 | 0.005 | 0 | 0.22 | 0.19 | 0.41 | 0.4 | 0.28 | 0.25 |
| 1 | 68845203 | rs11209273 | -0.764 | 1.582 | 0.623 | 1.861 | -0.012 | -0.024 | -0.024 | 0.293 | 0.21 | 0.16 | 0.41 | 0.4 | 0.28 | 0.25 |
| 1 | 68845611 | rs4926338 | -0.714 | 1.576 | 0.645 | 1.858 | -0.031 | -0.062 | -0.062 | 0 | 0.21 | 0.09 | 0.4 | 0.4 | 0.28 | 0.25 |
| 1 | 68847248 | rs9436816 | -0.971 | 1.419 | 1.071 | 1.757 | -0.006 | 0.195 | 0.195 | 0 | 0.16 | 0.35 | 0.23 | 0.45 | 0.1 | 0.29 |
| 1 | 68849687 | rs3118378 | -1.14 | 1.45 | 1.304 | 1.847 | 0.046 | 0.078 | 0.078 | 0 | 0.25 | 0.44 | 0.61 | 0.53 | 0.31 | 0.27 |
| 1 | 68851568 | rs7515572 | -1.407 | 1.108 | 1.575 | 2.157 | -0.03 | 0.108 | 0.108 | 0.017 | 0.14 | 0.34 | 0.05 | 0.33 | 0.06 | 0.28 |
| 1 | 68858479 | rs3125923 | -1.583 | 1.154 | 1.419 | 2.04 | -0.014 | 0.19 | 0.19 | 0 | 0.2 | 0.49 | 0.3 | 0.46 | 0.14 | 0.38 |
| 1 | 68875372 | rs1925952 | -1.855 | 1.986 | 1.578 | 2.099 | 0.034 | 0.085 | 0.085 | 0.344 | 0.21 | 0.45 | 0.28 | 0.38 | 0.22 | 0.39 |
| 1 | 68876172 | rs3125880 | -1.755 | 1.984 | 1.595 | 2.114 | 0.031 | 0.077 | 0.077 | 0.293 | 0.2 | 0.41 | 0.28 | 0.38 | 0.22 | 0.39 |
| 1 | 68881919 | rs3118405 | -1.304 | 1.998 | 1.546 | 2.178 | 0.023 | 0.011 | 0.011 | 0 | 0.14 | 0.26 | 0.25 | 0.16 | 0.17 | 0.2 |
| 1 | 68884722 | rs10493443 | -0.841 | 2.254 | 0.964 | 2.487 | 0.014 | 0.161 | 0.161 | 0 | 0.1 | 0.19 | 0.17 | 0.36 | 0.13 | 0.21 |
| 1 | 68886499 | rs12403707 | 0.204 | 2.26 | 0.845 | 2.477 | 0.012 | 0.007 | 0.007 | 0 | 0.08 | 0.08 | 0.23 | 0.13 | 0.15 | 0.1 |
| 1 | 68893754 | rs4264030 | -0.914 | 1.824 | 0.521 | 2.501 | 0.009 | 0.133 | 0.133 | 0 | 0.09 | 0.18 | 0.16 | 0.36 | 0.13 | 0.2 |
| 1 | 68909113 | rs17130694 | -0.558 | 1.694 | -0.003 | 2.121 | 0.01 | 0.075 | 0.075 | 0 | 0.07 | 0.18 | 0.03 | 0.23 | 0.05 | 0.19 |
| 1 | 68910999 | rs3790472 | 1.051 | 1.839 | -0.678 | 1.456 | -0.027 | 0 | 0 | 0 | 0.58 | 0.22 | 0.44 | 0.38 | 0.45 | 0.49 |
| 1 | 68918895 | rs436070 | -1.062 | 2.278 | -0.513 | 1.881 | -0.008 | -0.005 | -0.005 | 0 | 0.13 | 0.14 | 0.14 | 0.32 | 0.21 | 0.27 |
| 1 | 68922352 | rs10518297 | -0.073 | 2.051 | -0.539 | 1.502 | 0.006 | 0.024 | 0.024 | 0.127 | 0.16 | 0.07 | 0.07 | 0.1 | 0.16 | 0.07 |
| 1 | 68927513 | rs435066 | -1.052 | 2.587 | -0.01 | 1.936 | 0.028 | 0.137 | 0.137 | 0.002 | 0.15 | 0.23 | 0.28 | 0.57 | 0.18 | 0.42 |
| 1 | 68929478 | rs4926340 | 0.389 | 2.482 | 0.629 | 1.945 | 0.019 | 0.049 | 0.049 | 0 | 0.05 | 0.2 | 0.05 | 0.07 | 0.05 | 0.04 |
| 1 | 68931366 | rs10889769 | 0.143 | 2.967 | 0.449 | 1.73 | -0.006 | -0.009 | -0.009 | 0 | 0.05 | 0.04 | 0.15 | 0.08 | 0.12 | 0.07 |
| 1 | 68936519 | rs12080547 | -0.523 | 2.668 | 0.952 | 1.099 | -0.007 | -0.01 | -0.01 | 0 | 0.21 | 0.27 | 0.12 | 0.17 | 0.21 | 0.13 |
| 1 | 68990012 | rs2147317 | 1.296 | 2.894 | 1.065 | 1.116 | -0.001 | 0 | 0 | 0 | 0.73 | 0.77 | 0.52 | 0.22 | 0.64 | 0.45 |
| 1 | 68993460 | rs17473413 | 0.245 | 2.783 | 0.707 | 1.149 | -0.004 | 0.02 | 0.02 | 0.74 | 0.13 | 0.04 | 0.07 | 0.1 | 0.16 | 0.09 |
| 1 | 68994335 | rs1159579 | 0.082 | 2.869 | 0.681 | 1.396 | -0.009 | 0.043 | 0.043 | 0.04 | 0.93 | 0.97 | 0.8 | 0.4 | 0.88 | 0.67 |
| 1 | 69009194 | rs12756859 | -0.972 | 2.72 | 1.304 | 0.884 | 0.028 | 0.021 | 0.021 | 0.063 | 0.21 | 0.38 | 0.25 | 0.24 | 0.28 | 0.17 |
| 1 | 69009933 | rs10889779 | 0.317 | 2.642 | 1.04 | 1.008 | -0.011 | -0.005 | -0.005 | 0 | 0.05 | 0.03 | 0.14 | 0.14 | 0.12 | 0.08 |
| 1 | 69045273 | rs12406361 | 0.471 | 2.644 | 1.204 | 0.68 | 0 | -0.001 | -0.001 | 0.019 | 0.74 | 0.78 | 0.57 | 0.32 | 0.72 | 0.54 |
| 1 | 69047142 | rs12132065 | -1.112 | 3 | 1.545 | 0.389 | 0.033 | -0.068 | -0.068 | 0 | 0.12 | 0.41 | 0.14 | 0.01 | 0.11 | 0.05 |

| | | | | | | | | | | | | | | | | |
|---|---|---|---|---|---|---|---|---|---|---|---|---|---|---|---|---|
| 1 | 69053937 | rs2064375 | 2.363 | 3.151 | 1.548 | 0.221 | 0.051 | 0.007 | 0.007 | 1.036 | 0.73 | 0.37 | 0.65 | 0.72 | 0.64 | 0.7 |
| 1 | 69066072 | rs2255093 | 1.609 | 3.141 | 1.609 | 0.227 | 0.064 | 0.063 | 0.063 | 0.91 | 0.84 | 0.66 | 0.77 | 0.73 | 0.75 | 0.75 |
| 1 | 69070320 | rs2507208 | -0.34 | 3.271 | 1.617 | 0.254 | -0.011 | -0.016 | -0.016 | 0 | 0.25 | 0.15 | 0.42 | 0.68 | 0.28 | 0.46 |
| 1 | 69070945 | rs2507209 | 0.781 | 2.618 | 1.989 | 0.29 | 0.019 | 0.011 | 0.011 | 0.022 | 0.92 | 0.89 | 0.81 | 0.83 | 0.81 | 0.83 |
| 1 | 69083551 | rs12123584 | -0.756 | 2.543 | 2.226 | -0.336 | 0.038 | -0.026 | -0.026 | 1.061 | 0.06 | 0.2 | 0.07 | 0 | 0.06 | 0.01 |
| 1 | 69087317 | rs12080219 | -0.666 | 2.521 | 2.233 | -0.342 | 0.009 | -0.023 | -0.023 | 0.747 | 0.07 | 0.23 | 0.06 | 0.01 | 0.05 | 0.04 |
| 1 | 69088662 | rs11209332 | -0.849 | 2.531 | 2.237 | -0.346 | 0.044 | -0.023 | -0.023 | 0 | 0.06 | 0.25 | 0.07 | 0 | 0.06 | 0.01 |
| 1 | 69088934 | rs12083070 | -0.667 | 2.342 | 1.833 | -0.342 | 0.009 | -0.023 | -0.023 | 0 | 0.07 | 0.23 | 0.06 | 0.01 | 0.05 | 0.04 |
| 1 | 69099017 | rs11209333 | -3.186 | 2.263 | 1.685 | -0.438 | 0.009 | 0.043 | 0.043 | 2.648 | 0.21 | 0.84 | 0.21 | 0.11 | 0.18 | 0.11 |
| 1 | 69118705 | rs2206779 | 0.735 | 2.318 | 1.701 | -0.412 | 0.02 | -0.001 | -0.001 | 0 | 0.91 | 0.91 | 0.81 | 0.93 | 0.8 | 0.84 |
| 1 | 69121134 | rs12091311 | -1.162 | 2.2 | 1.772 | -0.41 | 0.018 | -0.008 | -0.008 | 0 | 0.08 | 0.34 | 0.09 | 0.02 | 0.06 | 0.04 |
| 1 | 69133160 | rs1738013 | -0.702 | 2.201 | 1.65 | -0.408 | 0.002 | -0.016 | -0.016 | 0 | 0.07 | 0.22 | 0.06 | 0.02 | 0.05 | 0.04 |
| 1 | 69133840 | rs10889786 | 3.074 | 2.23 | 1.347 | -0.397 | -0.108 | 0.058 | 0.058 | 1.847 | 0.7 | 0.09 | 0.6 | 0.82 | 0.63 | 0.72 |
| 1 | 69136972 | rs872850 | 2.19 | 0.614 | 1.473 | -0.708 | -0.036 | -0.06 | -0.06 | 0 | 0.78 | 0.41 | 0.87 | 0.89 | 0.85 | 0.81 |
| 1 | 69146910 | rs9436845 | -1.915 | 0.63 | 1.32 | -0.693 | -0.021 | -0.026 | -0.026 | 0 | 0.15 | 0.43 | 0.11 | 0.1 | 0.11 | 0.1 |
| 1 | 69155787 | rs17130785 | -0.702 | 0.653 | 0.84 | -0.674 | -0.002 | -0.001 | -0.001 | 0 | 0.09 | 0.09 | 0.05 | 0.1 | 0.09 | 0.1 |
| 1 | 69156133 | rs1492075 | -0.702 | 0.588 | 0.84 | -0.703 | -0.002 | -0.001 | -0.001 | 0 | 0.09 | 0.09 | 0.05 | 0.1 | 0.09 | 0.1 |
| 1 | 69158190 | rs854285 | -2.218 | 0.619 | 0.501 | -0.683 | 0.032 | 0.072 | 0.072 | 1.196 | 0.27 | 0.82 | 0.19 | 0.12 | 0.19 | 0.2 |
| 1 | 69162800 | rs860534 | -1.37 | 0.837 | 0.953 | -0.29 | 0.042 | 0.015 | 0.015 | 1.959 | 0.05 | 0.25 | 0.06 | 0 | 0.02 | 0 |
| 1 | 69174297 | rs1018418 | 0.6 | 0.258 | -0.109 | -0.761 | 0.046 | 0.086 | 0.086 | 2.383 | 0.58 | 0.24 | 0.59 | 0.81 | 0.68 | 0.73 |
| 1 | 69184038 | rs7541833 | -0.122 | 0.134 | 0.472 | -0.879 | -0.018 | -0.023 | -0.023 | 0.37 | 0.31 | 0.34 | 0.14 | 0.15 | 0.21 | 0.2 |
| 1 | 69184214 | rs854286 | -0.061 | 0.233 | 0.253 | -0.822 | 0.021 | -0.012 | -0.012 | 1.769 | 0.92 | 0.86 | 0.76 | 0.96 | 0.89 | 0.94 |
| 1 | 69217209 | rs4140577 | 0.867 | 0.244 | -0.226 | -0.592 | -0.044 | -0.07 | -0.07 | 0 | 0.11 | 0 | 0.19 | 0.52 | 0.12 | 0.32 |
| 1 | 69237446 | rs17395202 | 1.712 | -0.205 | -0.286 | -1.579 | 0.029 | 0.036 | 0.036 | 0.41 | 0.14 | 0.01 | 0.05 | 0.05 | 0.08 | 0.05 |
| 1 | 69239442 | rs6692084 | 1.083 | 0.004 | 0.12 | -1.496 | -0.006 | -0.054 | -0.054 | 0 | 0.74 | 0.5 | 0.66 | 0.86 | 0.77 | 0.79 |
| 1 | 69243474 | rs854273 | 0.549 | -0.014 | 0.151 | -1.52 | 0.011 | -0.004 | -0.004 | 0 | 0.84 | 0.75 | 0.72 | 0.86 | 0.81 | 0.81 |
| 1 | 69243544 | rs10493449 | 0.908 | -0.015 | 0.122 | -1.518 | -0.013 | -0.024 | -0.024 | 0.451 | 0.07 | 0.01 | 0.03 | 0.1 | 0.08 | 0.12 |
| 1 | 69254610 | rs12082973 | 0.091 | -0.007 | 0.169 | -1.517 | 0.021 | -0.024 | -0.024 | 0 | 0.09 | 0.19 | 0.25 | 0.04 | 0.1 | 0.06 |
| 1 | 69267189 | rs1492062 | 0.549 | -0.027 | 0.126 | -1.523 | -0.011 | -0.017 | -0.017 | 0 | 0.83 | 0.71 | 0.91 | 0.9 | 0.87 | 0.85 |
| 1 | 69278274 | rs12143405 | 2.081 | 0.001 | -0.161 | -1.809 | -0.061 | 0.11 | 0.11 | 0.182 | 0.27 | 0.02 | 0.15 | 0.06 | 0.3 | 0.16 |
| 1 | 69279982 | rs3981199 | -0.159 | 0.596 | 0.841 | -0.904 | -0.015 | -0.052 | -0.052 | 0 | 0.91 | 0.75 | 0.95 | 1 | 0.95 | 0.97 |
| 1 | 69290631 | rs1492074 | 1.51 | 0.634 | 0.857 | -0.892 | -0.008 | -0.104 | -0.104 | 3.321 | 0.8 | 0.52 | 0.69 | 0.96 | 0.85 | 0.9 |
| 1 | 69323222 | rs7414744 | -0.142 | 0.806 | 0.207 | -0.685 | -0.001 | -0.003 | -0.003 | 0 | 0.08 | 0.07 | 0.26 | 0.03 | 0.1 | 0.06 |
| 1 | 69334834 | rs6674177 | 0.497 | 0.603 | 0.568 | -0.723 | 0.008 | -0.063 | -0.063 | 0 | 0.87 | 0.52 | 0.72 | 0.97 | 0.89 | 0.94 |
| 1 | 69347850 | rs11209362 | -0.532 | 0.608 | 0.569 | -0.729 | -0.001 | -0.052 | -0.052 | 0 | 0.13 | 0.33 | 0.26 | 0.03 | 0.11 | 0.06 |
| 1 | 69354742 | rs9436417 | -0.476 | 0.614 | 0.523 | -0.726 | 0.001 | -0.05 | -0.05 | 0 | 0.12 | 0.31 | 0.26 | 0.02 | 0.11 | 0.06 |
| 1 | 69385766 | rs4558025 | 0.616 | 0.833 | 0.114 | -0.52 | -0.057 | -0.094 | -0.094 | 0 | 0.13 | 0.01 | 0.24 | 0.65 | 0.15 | 0.28 |
| 1 | 69415205 | rs12068594 | 0.827 | 0.101 | 0.552 | -0.981 | -0.029 | -0.064 | -0.064 | 0 | 0.17 | 0.29 | 0.09 | 0.04 | 0.1 | 0.09 |
| 1 | 69419136 | rs13375505 | -0.241 | 0.272 | 0.545 | -0.87 | 0.006 | -0.048 | -0.048 | 0 | 0.11 | 0.3 | 0.26 | 0.02 | 0.1 | 0.07 |
| 1 | 69431263 | rs12030971 | 0.247 | 0.569 | -0.545 | -0.975 | -0.127 | -0.026 | -0.026 | 0 | 0.28 | 0.06 | 0.42 | 0.76 | 0.4 | 0.46 |
| 1 | 69432044 | rs1022013 | 0.507 | 0.352 | 0.559 | -0.448 | 0.012 | -0.065 | -0.065 | 0.188 | 0.87 | 0.54 | 0.73 | 0.98 | 0.9 | 0.94 |
| 1 | 69435479 | rs12097254 | -0.098 | 0.251 | 0.051 | -0.104 | -0.025 | -0.063 | -0.063 | 0 | 0.17 | 0.28 | 0.09 | 0.04 | 0.1 | 0.1 |
| 1 | 69442664 | rs1507853 | -1.082 | 1.077 | -0.791 | 0.82 | 0.001 | 0.018 | 0.018 | 0 | 0.07 | 0.12 | 0.03 | 0.16 | 0.09 | 0.12 |
| 1 | 69442881 | rs11577703 | -0.567 | 1.752 | -1.13 | 0.991 | -0.051 | -0.038 | -0.038 | 0 | 0.1 | 0.01 | 0.1 | 0.17 | 0.17 | 0.19 |
| 1 | 69445925 | rs4650310 | 0.064 | 1.099 | -1.026 | 0.625 | 0.016 | 0.064 | 0.064 | 3.296 | 0.38 | 0.27 | 0.33 | 0.15 | 0.36 | 0.27 |
| 1 | 69446281 | rs6588344 | -0.947 | 1.401 | -0.539 | 1.297 | 0.006 | 0.071 | 0.071 | 0 | 0.05 | 0.12 | 0.06 | 0.18 | 0.06 | 0.13 |
| 1 | 69449431 | rs12562973 | -0.72 | 2.114 | -1.053 | 2.003 | -0.103 | -0.111 | -0.111 | 0 | 0.14 | 0.01 | 0.25 | 0.42 | 0.3 | 0.28 |
| 1 | 69482704 | rs7524839 | 0.348 | 1.278 | -0.438 | 1.489 | 0.038 | -0.001 | -0.001 | 1.467 | 0.58 | 0.28 | 0.59 | 0.56 | 0.67 | 0.6 |
| 1 | 69515706 | rs6691636 | 1.065 | 1.09 | -0.274 | 1.558 | -0.006 | -0.001 | -0.001 | 0 | 0.61 | 0.3 | 0.64 | 0.39 | 0.58 | 0.47 |
| 1 | 69516639 | rs2842040 | 0.8 | 1.074 | 0.238 | 1.513 | 0.006 | 0.002 | 0.002 | 0.21 | 0.64 | 0.53 | 0.67 | 0.39 | 0.6 | 0.47 |
| 1 | 69520596 | rs1507860 | 1.447 | 0.984 | 0.06 | 1.371 | 0.012 | 0.041 | 0.041 | 0.304 | 0.45 | 0.32 | 0.39 | 0.25 | 0.4 | 0.36 |
| 1 | 69550796 | rs2661460 | 0.566 | 0.959 | 0.186 | 1.964 | 0 | 0.011 | 0.011 | 0 | 0.72 | 0.4 | 0.72 | 0.71 | 0.72 | 0.74 |
| 1 | 69551411 | rs2842053 | -0.721 | 1.135 | 0.152 | 2.092 | -0.002 | 0.009 | 0.009 | 0 | 0.1 | 0.09 | 0.09 | 0.34 | 0.1 | 0.28 |
| 1 | 69554417 | rs2456910 | -1.014 | 1.027 | 0.404 | 2.017 | -0.001 | -0.011 | -0.011 | 0.442 | 0.28 | 0.3 | 0.36 | 0.48 | 0.29 | 0.42 |
| 1 | 69569143 | rs6671705 | 1.853 | 1.213 | 0.357 | 2.134 | 0.011 | 0.02 | 0.02 | 0.037 | 0.47 | 0.3 | 0.42 | 0.37 | 0.41 | 0.42 |
| 1 | 69579252 | rs4147141 | 1.513 | 1.169 | 0.403 | 2.147 | -0.004 | -0.011 | -0.011 | 2.026 | 0.62 | 0.3 | 0.66 | 0.49 | 0.6 | 0.54 |
| 1 | 69588193 | rs955093 | -0.294 | 1.116 | 0.626 | 2.11 | -0.04 | 0.001 | 0.001 | 0.114 | 0.81 | 0.93 | 0.82 | 0.84 | 0.73 | 0.83 |
| 1 | 69597419 | rs4650112 | 1.43 | 1.102 | 0.646 | 2.124 | 0.009 | 0.044 | 0.044 | 0 | 0.65 | 0.51 | 0.7 | 0.5 | 0.63 | 0.56 |
| 1 | 69612469 | rs17130868 | -0.764 | 0.901 | 0.626 | 2.198 | 0.012 | -0.012 | -0.012 | 0.375 | 0.07 | 0.03 | 0.03 | 0.1 | 0.02 | 0.05 |
| 1 | 69619701 | rs1577611 | 0.268 | 0.987 | 0.743 | 2.075 | 0.026 | 0.034 | 0.034 | 0 | 0.77 | 0.61 | 0.66 | 0.63 | 0.7 | 0.64 |
| 1 | 69624644 | rs1613058 | 1.432 | 0.986 | 0.893 | 2.079 | 0.084 | 0.175 | 0.175 | 0.252 | 0.89 | 0.48 | 0.89 | 0.75 | 0.89 | 0.76 |
| 1 | 69627731 | rs2784828 | -0.402 | 0.989 | 0.895 | 2.05 | 0.004 | -0.015 | -0.015 | 0.401 | 0.82 | 0.75 | 0.71 | 0.87 | 0.77 | 0.86 |
| 1 | 69632907 | rs11810068 | -1.058 | 0.781 | 1.012 | 2.165 | 0.031 | 0.115 | 0.115 | 0 | 0.08 | 0.26 | 0.07 | 0.24 | 0.08 | 0.22 |
| 1 | 69634134 | rs1414663 | 1.752 | 0.815 | 0.725 | 2.435 | -0.001 | 0.229 | 0.229 | 0 | 0.83 | 0.55 | 0.86 | 0.48 | 0.87 | 0.65 |
| 1 | 69644108 | rs2784830 | -0.153 | 0.486 | 0.344 | 1.991 | -0.003 | -0.053 | -0.053 | 0 | 0.18 | 0.08 | 0.21 | 0.35 | 0.18 | 0.16 |
| 1 | 69646463 | rs1360931 | -1.859 | 0.623 | 0.316 | 1.864 | 0 | 0 | 0 | 0 | 0.48 | 0.51 | 0.55 | 0.59 | 0.5 | 0.46 |
| 1 | 69648655 | rs2250760 | -1.689 | 0.589 | 0.305 | 1.933 | -0.01 | 0.032 | 0.032 | 0 | 0.34 | 0.57 | 0.37 | 0.53 | 0.33 | 0.31 |
| 1 | 69649328 | rs12135476 | 0.246 | 0.604 | 0.195 | 1.589 | -0.033 | -0.043 | -0.043 | 0 | 0.11 | 0 | 0.11 | 0.18 | 0.15 | 0.14 |
| 1 | 69659840 | rs10493454 | -0.053 | 0.632 | 0.326 | 1.603 | -0.001 | 0.001 | 0.001 | 0 | 0.13 | 0.16 | 0.13 | 0.19 | 0.16 | 0.14 |

| | | | | | | | | | | | | | | | | |
|---|---|---|---|---|---|---|---|---|---|---|---|---|---|---|---|---|
| 1 | 69662082 | rs1796938 | -1.338 | 0.596 | 0.289 | 1.681 | -0.004 | 0.052 | 0.052 | 0.472 | 0.33 | 0.56 | 0.32 | 0.47 | 0.34 | 0.33 |
| 1 | 69664700 | rs1763926 | -1.653 | 0.51 | 0.18 | 1.614 | -0.002 | 0 | 0 | 0.014 | 0.47 | 0.57 | 0.46 | 0.53 | 0.48 | 0.47 |
| 1 | 69665956 | rs1572421 | -0.206 | 0.529 | 0.19 | 1.628 | -0.003 | 0.049 | 0.049 | 0 | 0.19 | 0.37 | 0.18 | 0.28 | 0.18 | 0.19 |
| 1 | 69680062 | rs11209403 | -0.088 | 0.516 | 0.186 | 1.629 | -0.001 | 0.011 | 0.011 | 0 | 0.19 | 0.28 | 0.18 | 0.27 | 0.19 | 0.19 |
| 1 | 69683138 | rs12036450 | -0.43 | 0.456 | -0.027 | 1.272 | 0.005 | -0.016 | -0.016 | 0.011 | 0.34 | 0.28 | 0.33 | 0.34 | 0.33 | 0.34 |
| 1 | 69684599 | rs11209405 | -0.321 | 0.516 | -0.297 | 1.288 | 0 | -0.002 | -0.002 | 0.823 | 0.33 | 0.37 | 0.33 | 0.34 | 0.33 | 0.33 |
| 1 | 69684997 | rs4650135 | -0.259 | 0.539 | -0.283 | 1.305 | 0.022 | 0.111 | 0.111 | 0.022 | 0.16 | 0.37 | 0.15 | 0.33 | 0.18 | 0.23 |
| 1 | 69685100 | rs4650137 | -0.618 | 0.566 | -1.15 | 0.635 | 0.001 | 0.08 | 0.08 | 0 | 0.5 | 0.74 | 0.48 | 0.66 | 0.51 | 0.57 |
| 1 | 69692232 | rs2248186 | 0.42 | 0.669 | -1.149 | 0.425 | -0.011 | 0.028 | 0.028 | 0 | 0.89 | 0.97 | 0.95 | 0.96 | 0.9 | 0.9 |
| 1 | 69695698 | rs2248556 | -0.54 | 0.576 | -0.182 | 0.461 | 0.041 | 0.074 | 0.074 | 0 | 0.09 | 0.33 | 0.1 | 0.13 | 0.07 | 0.15 |
| 1 | 69695923 | rs2248562 | 0.525 | 0.418 | 0.025 | 0.158 | 0.001 | 0.007 | 0.007 | 0 | 0.88 | 0.89 | 0.94 | 0.96 | 0.9 | 0.89 |
| 1 | 69711165 | rs2246312 | 0.668 | -0.228 | -0.286 | 0.051 | -0.02 | -0.003 | -0.003 | 0 | 0.65 | 0.54 | 0.81 | 0.69 | 0.72 | 0.73 |
| 1 | 69713109 | rs2420826 | 0.351 | -0.339 | 0.034 | -0.053 | -0.011 | 0.011 | 0.011 | 0 | 0.22 | 0.27 | 0.13 | 0.27 | 0.17 | 0.16 |
| 1 | 69715220 | rs2249487 | 0.791 | -0.324 | -0.032 | -0.052 | -0.02 | -0.003 | -0.003 | 0.083 | 0.65 | 0.54 | 0.81 | 0.69 | 0.72 | 0.73 |
| 1 | 69723765 | rs2791211 | 0.746 | -0.435 | -0.007 | -0.084 | -0.021 | -0.002 | -0.002 | 0 | 0.64 | 0.56 | 0.81 | 0.68 | 0.72 | 0.71 |
| 1 | 69724482 | rs2791209 | 0.756 | -0.435 | -0.026 | -0.084 | -0.024 | -0.002 | -0.002 | 0 | 0.64 | 0.53 | 0.81 | 0.68 | 0.72 | 0.7 |
| 1 | 69727964 | rs2815432 | -0.073 | -0.384 | -0.188 | -0.054 | 0.001 | -0.001 | -0.001 | 0 | 0.87 | 0.88 | 0.94 | 0.96 | 0.9 | 0.89 |
| 1 | 69728843 | rs4146494 | 0.986 | -0.334 | -0.317 | 0.057 | 0.044 | 0.043 | 0.043 | 0.686 | 0.56 | 0.2 | 0.71 | 0.68 | 0.66 | 0.58 |
| 1 | 69736009 | rs4650166 | 1.052 | -0.202 | -0.972 | -0.288 | -0.078 | -0.052 | -0.052 | 0.254 | 0.35 | 0.06 | 0.58 | 0.64 | 0.42 | 0.44 |
| 1 | 69743750 | rs12123555 | -0.516 | 0.074 | -0.342 | -0.491 | 0.002 | 0.004 | 0.004 | 0 | 0.41 | 0.54 | 0.24 | 0.3 | 0.38 | 0.28 |
| 1 | 69746933 | rs1824448 | -0.459 | 0.08 | -0.213 | -0.487 | 0 | 0 | 0 | 0 | 0.39 | 0.38 | 0.23 | 0.3 | 0.38 | 0.28 |
| 1 | 69779437 | rs720233 | -0.07 | -0.001 | -0.973 | -0.3 | -0.012 | 0.004 | 0.004 | 0 | 0.32 | 0.53 | 0.17 | 0.33 | 0.27 | 0.26 |
| 1 | 69787484 | rs2815468 | -0.256 | -0.23 | -1.565 | -0.815 | -0.018 | 0.091 | 0.091 | 0 | 0.68 | 0.79 | 0.7 | 0.95 | 0.66 | 0.74 |
| 1 | 69789580 | rs2815466 | 0.36 | 0.136 | -1.498 | -0.246 | -0.022 | 0.001 | 0.001 | 0 | 0.37 | 0.21 | 0.53 | 0.66 | 0.39 | 0.48 |
| 1 | 69808978 | rs2612242 | -0.159 | 0.314 | -1.593 | -0.765 | -0.052 | 0.11 | 0.11 | 0 | 0.77 | 0.91 | 0.71 | 0.99 | 0.7 | 0.81 |
| 1 | 69815440 | rs12404688 | -0.18 | -0.614 | -0.858 | -1.252 | -0.066 | -0.073 | -0.073 | 0 | 0.17 | 0.34 | 0.06 | 0.07 | 0.06 | 0.11 |
| 1 | 69829687 | rs288941 | -0.721 | -0.626 | -0.935 | -1.295 | -0.083 | 0.146 | 0.146 | 0 | 0.78 | 0.96 | 0.75 | 1 | 0.7 | 0.81 |
| 1 | 69830063 | rs2765864 | -0.151 | -0.656 | -0.789 | -1.295 | -0.063 | -0.07 | -0.07 | 0 | 0.17 | 0.32 | 0.06 | 0.07 | 0.05 | 0.1 |
| 1 | 69847591 | rs12043869 | 0.732 | -0.277 | -1.673 | -0.704 | 0.006 | -0.093 | -0.093 | 0.217 | 0.26 | 0.03 | 0.22 | 0.65 | 0.22 | 0.41 |
| 1 | 69851618 | rs548825 | 0.2 | -0.131 | -0.569 | -0.755 | 0.015 | 0.032 | 0.032 | 0.631 | 0.59 | 0.34 | 0.62 | 0.92 | 0.64 | 0.7 |
| 1 | 69865203 | rs7513164 | 0.247 | -0.105 | -0.67 | -0.743 | 0.025 | 0.085 | 0.085 | 0.592 | 0.58 | 0.28 | 0.62 | 0.92 | 0.64 | 0.7 |
| 1 | 69872197 | rs4650000 | 0.209 | -0.099 | -0.627 | -0.74 | 0.019 | 0.052 | 0.052 | 0.409 | 0.59 | 0.33 | 0.62 | 0.92 | 0.64 | 0.7 |
| 1 | 69877882 | rs1578356 | -0.027 | -0.154 | -0.039 | -0.773 | -0.004 | -0.092 | -0.092 | 0.035 | 0.63 | 0.48 | 0.65 | 0.92 | 0.64 | 0.7 |
| 1 | 69879802 | rs10518315 | 1.194 | -0.144 | -0.208 | -0.014 | -0.025 | 0.056 | 0.056 | 0.597 | 0.05 | 0.01 | 0.17 | 0.46 | 0.08 | 0.25 |
| 1 | 69885329 | rs868402 | 0.582 | 0.468 | -0.347 | -0.183 | -0.002 | 0.056 | 0.056 | 0.113 | 0.52 | 0.41 | 0.58 | 0.87 | 0.5 | 0.63 |
| 1 | 69886629 | rs1926257 | -0.362 | 0.469 | -0.335 | -0.183 | -0.004 | -0.092 | -0.092 | 0.478 | 0.63 | 0.48 | 0.65 | 0.92 | 0.64 | 0.7 |
| 1 | 69886839 | rs1926258 | -0.362 | 0.469 | -0.335 | -0.183 | -0.004 | -0.092 | -0.092 | 0 | 0.63 | 0.48 | 0.65 | 0.92 | 0.64 | 0.7 |
| 1 | 69889691 | rs11209454 | 0.998 | 0.431 | -0.109 | -0.165 | 0.014 | 0.016 | 0.016 | 0 | 0.45 | 0.31 | 0.4 | 0.41 | 0.42 | 0.38 |
| 1 | 69891657 | rs11209455 | -0.384 | 0.443 | -0.293 | -0.47 | -0.001 | -0.046 | -0.046 | 0 | 0.64 | 0.63 | 0.65 | 0.92 | 0.65 | 0.71 |
| 1 | 69895080 | rs12135013 | 0.342 | 0.899 | -0.326 | 0.121 | -0.012 | 0.01 | 0.01 | 0 | 0.12 | 0.07 | 0.07 | 0.06 | 0.14 | 0.08 |
| 1 | 69900116 | rs1926264 | 0.589 | 0.584 | -0.031 | 0.397 | 0.006 | 0.06 | 0.06 | 0 | 0.71 | 0.52 | 0.6 | 0.48 | 0.69 | 0.52 |
| 1 | 69901557 | rs1998846 | 0.479 | 0.767 | 0.077 | 0.667 | 0 | 0.028 | 0.028 | 0 | 0.71 | 0.55 | 0.67 | 0.63 | 0.71 | 0.57 |
| 1 | 69904379 | rs4650278 | -1.309 | 0.776 | -0.305 | 0.401 | -0.053 | 0.094 | 0.094 | 0 | 0.85 | 0.98 | 0.87 | 0.99 | 0.8 | 0.81 |
| 1 | 69904826 | rs4650001 | 0.718 | 0.203 | -0.183 | 0.13 | -0.011 | -0.049 | -0.049 | 0.51 | 0.64 | 0.62 | 0.78 | 0.84 | 0.69 | 0.66 |
| 1 | 69906307 | rs12085918 | 0.059 | 0.006 | -0.215 | 0.316 | 0.012 | 0.009 | 0.009 | 0.023 | 0.19 | 0.12 | 0.07 | 0.14 | 0.11 | 0.14 |
| 1 | 69909410 | rs9326089 | -1.086 | 0.013 | -0.418 | 0.22 | 0.054 | 0.099 | 0.099 | 0 | 0.49 | 0.76 | 0.44 | 0.76 | 0.39 | 0.52 |
| 1 | 69909474 | rs12042471 | 1.342 | 0.013 | -0.657 | 0.467 | 0.002 | -0.007 | -0.007 | 0 | 0.2 | 0.25 | 0.11 | 0.33 | 0.16 | 0.15 |
| 1 | 69909512 | rs12088026 | 0.094 | 0.15 | -0.811 | 0.384 | 0.012 | 0.009 | 0.009 | 0.012 | 0.19 | 0.12 | 0.07 | 0.14 | 0.11 | 0.14 |
| 1 | 69914861 | rs6424486 | 0.38 | 0.715 | -1.057 | 0.406 | 0.004 | -0.006 | -0.006 | 4.15 | 0.33 | 0.27 | 0.38 | 0.27 | 0.32 | 0.26 |
| 1 | 69915596 | rs17495923 | 0.638 | 1.008 | -1.093 | 0.642 | -0.012 | -0.026 | -0.026 | 0 | 0.1 | 0.03 | 0.13 | 0.15 | 0.1 | 0.12 |
| 1 | 69920333 | rs1926268 | -0.263 | 1.474 | -1.134 | 0.392 | -0.006 | 0.005 | 0.005 | 0 | 0.79 | 0.81 | 0.73 | 0.82 | 0.69 | 0.67 |
| 1 | 69920839 | rs11209464 | 0.859 | 1.235 | -0.866 | 0.322 | -0.003 | 0.006 | 0.006 | 0 | 0.58 | 0.6 | 0.53 | 0.65 | 0.55 | 0.46 |
| 1 | 69925811 | rs12062616 | -0.072 | 1.721 | -1.062 | 0.443 | -0.008 | -0.009 | -0.009 | 0 | 0.24 | 0.19 | 0.31 | 0.4 | 0.34 | 0.4 |
| 1 | 69933450 | rs2025721 | 1.077 | 1.6 | -0.659 | 0.485 | -0.007 | -0.002 | -0.002 | 0 | 0.46 | 0.51 | 0.42 | 0.44 | 0.37 | 0.28 |
| 1 | 69934056 | rs3928272 | 0.155 | 1.644 | -0.709 | 0.3 | -0.024 | 0.037 | 0.037 | 0 | 0.6 | 0.81 | 0.68 | 0.62 | 0.54 | 0.48 |
| 1 | 69935119 | rs11209468 | 0.589 | 1.513 | -0.648 | 0.327 | 0.004 | -0.01 | -0.01 | 0 | 0.57 | 0.53 | 0.59 | 0.61 | 0.49 | 0.39 |
| 1 | 69938888 | rs12728917 | 1.1 | 1.398 | -0.862 | 0.345 | -0.003 | -0.007 | -0.007 | 0 | 0.16 | 0.1 | 0.12 | 0.26 | 0.18 | 0.25 |
| 1 | 69943001 | rs1120087 | 0.402 | 1.411 | -0.88 | 0.034 | -0.005 | 0.056 | 0.056 | 0 | 0.56 | 0.63 | 0.54 | 0.75 | 0.53 | 0.49 |
| 1 | 69944198 | rs12028493 | -0.668 | 1.389 | -0.986 | -0.004 | 0.002 | 0.025 | 0.025 | 0 | 0.76 | 0.83 | 0.75 | 0.83 | 0.74 | 0.63 |
| 1 | 69948309 | rs12563277 | 2.051 | 1.085 | -0.94 | -0.125 | -0.004 | 0.023 | 0.023 | 0 | 0.22 | 0.24 | 0.31 | 0.4 | 0.18 | 0.17 |
| 1 | 69951220 | rs1341771 | -0.625 | 1.097 | -1.037 | -0.226 | 0.003 | 0.013 | 0.013 | 0 | 0.75 | 0.77 | 0.75 | 0.84 | 0.74 | 0.63 |
| 1 | 69951760 | rs10789295 | -0.918 | 1.141 | -1.196 | -0.205 | -0.004 | 0.031 | 0.031 | 0 | 0.77 | 0.89 | 0.75 | 0.84 | 0.74 | 0.63 |
| 1 | 69965403 | rs11209479 | 0.539 | 1.116 | -0.685 | 0.154 | -0.005 | 0.012 | 0.012 | 0 | 0.2 | 0.13 | 0.2 | 0.1 | 0.23 | 0.15 |
| 1 | 69969838 | rs12048011 | 0.614 | 1.462 | -0.166 | 0.324 | 0.005 | -0.018 | -0.018 | 0 | 0.14 | 0.15 | 0.13 | 0.1 | 0.12 | 0.11 |
| 1 | 69972693 | rs4259589 | 0.146 | 1.467 | -0.112 | 0.39 | -0.008 | 0.008 | 0.008 | 0.093 | 0.31 | 0.36 | 0.33 | 0.37 | 0.28 | 0.27 |
| 1 | 69973079 | rs11209481 | 2.488 | 1.516 | -0.482 | 0.405 | -0.073 | -0.084 | -0.084 | 0 | 0.3 | 0.03 | 0.21 | 0.28 | 0.32 | 0.16 |
| 1 | 69975660 | rs10889833 | 0.429 | 1.145 | -0.268 | 0.347 | 0 | 0.001 | 0.001 | 0.002 | 0.3 | 0.28 | 0.34 | 0.38 | 0.33 | 0.29 |
| 1 | 69983647 | rs6424550 | 0.665 | 1.135 | -0.178 | 0.179 | -0.002 | 0.011 | 0.011 | 0 | 0.66 | 0.69 | 0.66 | 0.77 | 0.67 | 0.49 |
| 1 | 69993512 | rs4589058 | 1.896 | 0.965 | -0.132 | 0.332 | 0.003 | -0.019 | -0.019 | 0 | 0.34 | 0.17 | 0.29 | 0.38 | 0.32 | 0.21 |

| | | | | | | | | | | | | | | | | |
|---|---|---|---|---|---|---|---|---|---|---|---|---|---|---|---|---|
| 1 | 70010325 | rs10889836 | -0.108 | 0.781 | -0.259 | 0.242 | 0.004 | 0.024 | 0.024 | 0 | 0.28 | 0.3 | 0.25 | 0.47 | 0.29 | 0.31 |
| 1 | 70012554 | rs17395792 | 0.973 | 0.822 | -0.301 | 0.196 | -0.041 | 0.06 | 0.06 | 0.379 | 0.2 | 0.07 | 0.34 | 0.06 | 0.27 | 0.28 |
| 1 | 70019811 | rs1341788 | 0.306 | 0.995 | -0.261 | 0.311 | 0.001 | 0.047 | 0.047 | 0.01 | 0.19 | 0.22 | 0.13 | 0.38 | 0.2 | 0.28 |
| 1 | 70029922 | rs4650295 | -1.754 | 0.951 | -0.114 | 0.35 | 0.001 | -0.006 | -0.006 | 0.099 | 0.64 | 0.68 | 0.69 | 0.62 | 0.67 | 0.79 |
| 1 | 70033296 | rs912774 | -0.269 | 0.98 | -0.199 | 0.526 | 0.003 | 0.018 | 0.018 | 0 | 0.29 | 0.3 | 0.26 | 0.38 | 0.28 | 0.3 |
| 1 | 70038818 | rs11209493 | 0.879 | 0.809 | -0.271 | 0.205 | 0.007 | -0.005 | -0.005 | 0 | 0.44 | 0.26 | 0.31 | 0.36 | 0.43 | 0.38 |
| 1 | 70042477 | rs12084695 | 1.519 | 0.582 | -0.373 | 0.021 | 0.002 | -0.066 | -0.066 | 0 | 0.34 | 0.14 | 0.21 | 0.36 | 0.31 | 0.19 |
| 1 | 70047731 | rs1245060 | 1.114 | 0.521 | -0.422 | -0.25 | -0.025 | 0.012 | 0.012 | 0 | 0.49 | 0.2 | 0.43 | 0.37 | 0.48 | 0.37 |
| 1 | 70048269 | rs6664878 | 1.431 | 0.616 | -0.321 | -0.26 | 0.001 | 0.007 | 0.007 | 0 | 0.1 | 0.05 | 0.07 | 0 | 0.09 | 0.09 |
| 1 | 70057783 | rs10489549 | -0.608 | 0.648 | 0.289 | -0.245 | 0.008 | -0.023 | -0.023 | 0 | 0.08 | 0.23 | 0.03 | 0.01 | 0.05 | 0.05 |
| 1 | 70063259 | rs4650296 | 0.012 | 0.911 | 0.081 | -0.017 | 0.027 | 0.091 | 0.091 | 0 | 0.1 | 0.21 | 0.2 | 0.39 | 0.18 | 0.21 |
| 1 | 70074131 | rs1749504 | -0.629 | 0.729 | 0.119 | -0.284 | 0.003 | -0.004 | -0.004 | 0.413 | 0.36 | 0.67 | 0.35 | 0.04 | 0.33 | 0.33 |
| 1 | 70084699 | rs12036330 | -0.365 | 0.916 | 0.158 | 0.115 | 0.001 | 0.01 | 0.01 | 0 | 0.29 | 0.35 | 0.37 | 0.6 | 0.31 | 0.43 |
| 1 | 70084797 | rs11809230 | 0.419 | 1.031 | 0.096 | 0.27 | 0.04 | 0.132 | 0.132 | 0.023 | 0.07 | 0.2 | 0.14 | 0.36 | 0.14 | 0.19 |
| 1 | 70085078 | rs11209501 | -0.753 | 0.9 | 0.548 | -0.009 | 0.02 | -0.01 | -0.01 | 0.01 | 0.06 | 0.2 | 0.04 | 0.01 | 0.05 | 0.04 |
| 1 | 70097033 | rs1773349 | -0.685 | 0.912 | 0.021 | -0.151 | -0.021 | 0 | 0 | 0 | 0.46 | 0.28 | 0.57 | 0.62 | 0.49 | 0.63 |
| 1 | 70097256 | rs6662257 | -0.171 | 0.941 | 0.069 | -0.08 | -0.004 | -0.019 | -0.019 | 0 | 0.29 | 0.25 | 0.36 | 0.6 | 0.31 | 0.45 |
| 1 | 70097708 | rs2066296 | 1.777 | 0.878 | -0.11 | 0.016 | 0 | -0.044 | -0.044 | 0.569 | 0.33 | 0.16 | 0.22 | 0.35 | 0.31 | 0.19 |
| 1 | 70116825 | rs7515290 | 0.514 | 0.66 | 0.324 | 0.019 | 0.003 | -0.001 | -0.001 | 0 | 0.09 | 0.18 | 0.13 | 0.09 | 0.07 | 0.12 |
| 1 | 70117404 | rs1024268 | -0.162 | 0.76 | 0.254 | -0.093 | -0.009 | -0.022 | -0.022 | 0 | 0.36 | 0.27 | 0.27 | 0.5 | 0.4 | 0.47 |
| 1 | 70124742 | rs227114 | 0.826 | 0.764 | 0.268 | -0.169 | -0.002 | 0.006 | 0.006 | 0 | 0.19 | 0.14 | 0.24 | 0.03 | 0.18 | 0.19 |
| 1 | 70124891 | rs227115 | -2.32 | 0.71 | -0.046 | -0.117 | 0.003 | -0.048 | -0.048 | 0 | 0.69 | 0.82 | 0.73 | 0.62 | 0.7 | 0.82 |
| 1 | 70126607 | rs800930 | 0.575 | 0.693 | 0.213 | 0.114 | 0.01 | 0.005 | 0.005 | 0 | 0.09 | 0.2 | 0.13 | 0.09 | 0.07 | 0.12 |
| 1 | 70143124 | rs12118129 | 0.084 | 0.652 | -0.072 | 0.139 | 0.002 | 0.021 | 0.021 | 0 | 0.25 | 0.17 | 0.14 | 0.23 | 0.24 | 0.29 |
| 1 | 70151123 | rs1417438 | -1.569 | 0.694 | 0.023 | 0.224 | -0.008 | -0.01 | -0.01 | 1.091 | 0.52 | 0.7 | 0.48 | 0.56 | 0.52 | 0.64 |
| 1 | 70154480 | rs17325299 | 0.643 | 0.953 | -0.362 | 0.153 | -0.002 | 0.018 | 0.018 | 0 | 0.19 | 0.11 | 0.11 | 0.02 | 0.17 | 0.11 |
| 1 | 70155099 | rs17131017 | -0.46 | 1.173 | 0.315 | 1.086 | 0.009 | 0.05 | 0.05 | 0 | 0.2 | 0.26 | 0.25 | 0.36 | 0.22 | 0.3 |
| 1 | 70160178 | rs1417436 | -1.022 | 1.202 | -0.198 | 0.811 | 0.009 | 0.011 | 0.011 | 0 | 0.37 | 0.54 | 0.5 | 0.46 | 0.45 | 0.56 |
| 1 | 70164686 | rs1740900 | -1.255 | 1.154 | -0.341 | 1.022 | -0.001 | -0.004 | -0.004 | 0 | 0.43 | 0.47 | 0.53 | 0.31 | 0.41 | 0.5 |
| 1 | 70176245 | rs1740909 | -1.905 | 1.031 | -1.55 | 0.891 | 0.006 | 0.021 | 0.021 | 0 | 0.53 | 0.84 | 0.7 | 0.62 | 0.6 | 0.74 |
| 1 | 70176931 | rs10489555 | -0.379 | 0.911 | -1.515 | 0.819 | -0.001 | -0.027 | -0.027 | 0.243 | 0.08 | 0.03 | 0.08 | 0.26 | 0.1 | 0.21 |
| 1 | 70177233 | rs6687106 | -0.571 | 0.877 | -1.314 | 0.39 | -0.042 | 0.03 | 0.03 | 0 | 0.24 | 0.07 | 0.32 | 0.14 | 0.27 | 0.33 |
| 1 | 70186036 | rs1340773 | -0.516 | 0.885 | -1.309 | 0.239 | -0.037 | 0.05 | 0.05 | 0 | 0.25 | 0.07 | 0.32 | 0.15 | 0.27 | 0.33 |
| 1 | 70207079 | rs10889850 | 0.428 | 0.952 | -0.692 | 0.226 | -0.01 | 0.029 | 0.029 | 0.957 | 0.7 | 0.77 | 0.79 | 0.91 | 0.66 | 0.79 |
| 1 | 70207668 | rs17397693 | 1.717 | 0.821 | -1.412 | 0.326 | -0.104 | 0.042 | 0.042 | 0.13 | 0.36 | 0.02 | 0.44 | 0.25 | 0.36 | 0.32 |
| 1 | 70219936 | rs1340769 | 1.624 | 0.983 | -0.08 | 0.741 | -0.024 | 0.027 | 0.027 | 0.57 | 0.52 | 0.21 | 0.57 | 0.6 | 0.5 | 0.47 |
| 1 | 70221562 | rs11209536 | 0.144 | 1.285 | -0.218 | 0.439 | -0.033 | 0.009 | 0.009 | 0.01 | 0.14 | 0.06 | 0.13 | 0.07 | 0.24 | 0.11 |
| 1 | 70223922 | rs6424593 | 1.162 | 1.284 | 0.032 | 0.502 | -0.009 | 0.018 | 0.018 | 0.342 | 0.62 | 0.31 | 0.64 | 0.77 | 0.57 | 0.7 |
| 1 | 70225340 | rs7523521 | -0.669 | 0.783 | 0.486 | 0.56 | -0.007 | -0.009 | -0.009 | 0.077 | 0.17 | 0.29 | 0.2 | 0.14 | 0.17 | 0.15 |
| 1 | 70229665 | rs17097452 | -0.836 | 1.451 | -0.469 | 0.846 | -0.013 | -0.014 | -0.014 | 0 | 0.08 | 0 | 0.07 | 0.08 | 0.07 | 0.14 |
| 1 | 70244251 | rs12037989 | -1.298 | 1.979 | 0.109 | 0.967 | 0.065 | 0.019 | 0.019 | 0 | 0.14 | 0.3 | 0.25 | 0.22 | 0.23 | 0.09 |
| 1 | 70248115 | rs7520521 | -0.478 | 1.443 | -0.357 | 0.936 | 0.009 | 0.004 | 0.004 | 0 | 0.47 | 0.63 | 0.45 | 0.6 | 0.43 | 0.41 |
| 1 | 70263007 | rs6681847 | -0.591 | 1.553 | -0.698 | 0.967 | -0.011 | 0.007 | 0.007 | 0 | 0.53 | 0.67 | 0.51 | 0.6 | 0.5 | 0.42 |
| 1 | 70272063 | rs12028771 | -1.336 | 1.841 | -0.61 | 1.952 | 0.005 | 0.069 | 0.069 | 0 | 0.09 | 0.1 | 0.27 | 0.51 | 0.16 | 0.2 |
| 1 | 70279152 | rs12027206 | -1.332 | 1.806 | -0.668 | 2.35 | -0.007 | 0.081 | 0.081 | 0 | 0.06 | 0.03 | 0.1 | 0.51 | 0.11 | 0.17 |
| 1 | 70279860 | rs1913268 | 0.095 | 1.49 | -0.749 | 1.692 | -0.002 | -0.007 | -0.007 | 0 | 0.63 | 0.56 | 0.64 | 0.77 | 0.67 | 0.58 |
| 1 | 70282875 | rs11578884 | 0.504 | 0.771 | -1.363 | 0.995 | -0.086 | -0.083 | -0.083 | 0 | 0.36 | 0.06 | 0.43 | 0.53 | 0.48 | 0.31 |
| 1 | 70297993 | rs11209552 | -0.149 | 0.288 | -0.994 | 0.686 | 0.004 | 0.004 | 0.004 | 0 | 0.53 | 0.37 | 0.53 | 0.54 | 0.55 | 0.34 |
| 1 | 70343674 | rs4623650 | -0.14 | 0.312 | -1.099 | 0.534 | 0.003 | 0.012 | 0.012 | 0 | 0.46 | 0.41 | 0.51 | 0.68 | 0.56 | 0.48 |
| 1 | 70363601 | rs6663915 | 0.293 | 0.232 | -1.094 | 0.482 | -0.039 | -0.023 | -0.023 | 0 | 0.4 | 0.21 | 0.44 | 0.54 | 0.48 | 0.34 |
| 1 | 70363647 | rs6424609 | -0.25 | 0.195 | -1.278 | 0.309 | 0.02 | 0.03 | 0.03 | 0 | 0.5 | 0.56 | 0.57 | 0.69 | 0.62 | 0.5 |
| 1 | 70364313 | rs6681560 | -0.483 | 0.193 | -1.245 | 0.339 | 0.113 | 0.13 | 0.13 | 0.04 | 0.07 | 0.35 | 0.09 | 0.14 | 0.08 | 0.15 |
| 1 | 70365414 | rs10889862 | -0.483 | 0.193 | -1.217 | 0.339 | 0.111 | 0.128 | 0.128 | 0.04 | 0.07 | 0.35 | 0.09 | 0.14 | 0.08 | 0.15 |
| 1 | 70405236 | rs2025896 | 0.559 | 0.205 | -1.356 | 0.29 | -0.068 | -0.044 | -0.044 | 1.017 | 0.4 | 0.12 | 0.43 | 0.54 | 0.48 | 0.34 |
| 1 | 70414873 | rs1244791 | -1.178 | -0.131 | -1.64 | -0.122 | -0.057 | 0.032 | 0.032 | 0.891 | 0.79 | 0.97 | 0.7 | 0.92 | 0.76 | 0.82 |
| 1 | 70450024 | rs12048343 | -1.053 | 0.324 | -0.706 | 0.167 | 0.154 | 0.109 | 0.109 | 0.745 | 0.07 | 0.45 | 0.07 | 0.08 | 0.09 | 0.16 |
| 1 | 70454240 | rs12034677 | 0.06 | 0.235 | -1.101 | 0.653 | 0.056 | 0.051 | 0.051 | 0.017 | 0.49 | 0.25 | 0.53 | 0.65 | 0.59 | 0.54 |
| 1 | 70499063 | rs17131138 | 1.254 | 0.984 | -0.664 | 0.393 | 0 | -0.012 | -0.012 | 0 | 0.07 | 0.06 | 0.06 | 0.02 | 0.13 | 0.05 |
| 1 | 70499373 | rs6674401 | 0.693 | 1.133 | -1.11 | 0.494 | -0.021 | -0.071 | -0.071 | 0.344 | 0.31 | 0.15 | 0.39 | 0.52 | 0.37 | 0.3 |
| 1 | 70500117 | rs12025814 | -0.553 | 1.489 | -0.816 | 0.738 | -0.001 | 0.001 | 0.001 | 0 | 0.05 | 0.08 | 0.1 | 0.1 | 0.1 | 0.13 |
| 1 | 70501140 | rs17097511 | -0.564 | 1.397 | -0.772 | 0.736 | 0.037 | 0.001 | 0.001 | 0 | 0.14 | 0.25 | 0.17 | 0.14 | 0.23 | 0.2 |
| 1 | 70521914 | rs631167 | 0.148 | 1.526 | -0.98 | 1.319 | -0.035 | -0.041 | -0.041 | 0 | 0.84 | 0.89 | 0.76 | 0.69 | 0.72 | 0.76 |
| 1 | 70524076 | rs4650019 | 0.06 | 0.845 | -0.576 | 1.211 | -0.046 | 0.004 | 0.004 | 0 | 0.18 | 0.38 | 0.13 | 0.17 | 0.08 | 0.22 |
| 1 | 70541127 | rs616511 | 1.235 | 0.848 | -0.569 | 1.204 | 0.007 | 0.041 | 0.041 | 0 | 0.65 | 0.51 | 0.63 | 0.53 | 0.64 | 0.54 |
| 1 | 70543349 | rs10493471 | 0.06 | 0.848 | -0.57 | 1.217 | -0.046 | 0.002 | 0.002 | 0 | 0.18 | 0.38 | 0.13 | 0.16 | 0.08 | 0.22 |
| 1 | 70558599 | rs12033206 | 1.277 | 0.986 | -1.483 | 0.691 | -0.12 | -0.048 | -0.048 | 0 | 0.3 | 0.02 | 0.31 | 0.32 | 0.36 | 0.31 |
| 1 | 70565386 | rs826900 | -0.773 | 1.011 | -1.032 | 0.746 | 0.006 | 0.011 | 0.011 | 0.002 | 0.84 | 0.8 | 0.75 | 0.69 | 0.72 | 0.76 |
| 1 | 70565511 | rs583035 | 0.674 | 0.982 | -1.035 | 0.665 | 0.004 | 0.026 | 0.026 | 0 | 0.65 | 0.43 | 0.62 | 0.53 | 0.64 | 0.54 |
| 1 | 70571471 | rs587562 | 0.958 | 1.089 | -1.022 | 0.566 | 0.002 | 0.004 | 0.004 | 0 | 0.62 | 0.59 | 0.63 | 0.53 | 0.61 | 0.55 |

| | | | | | | | | | | | | | | | | |
|---|---|---|---|---|---|---|---|---|---|---|---|---|---|---|---|---|
| 1 | 70579097 | rs17131212 | 0.904 | 0.915 | -1.003 | 0.612 | 0.025 | 0 | 0 | 0 | 0.16 | 0.08 | 0.08 | 0.14 | 0.08 | 0.2 |
| 1 | 70580058 | rs17131215 | 0.08 | 0.913 | -0.967 | 0.606 | -0.033 | -0.017 | -0.017 | 2.692 | 0.2 | 0.28 | 0.1 | 0.14 | 0.09 | 0.23 |
| 1 | 70583508 | rs12026267 | 1.798 | 0.962 | -1.362 | 0.496 | -0.096 | -0.075 | -0.075 | 0 | 0.27 | 0.02 | 0.29 | 0.31 | 0.34 | 0.29 |
| 1 | 70587360 | rs641909 | -0.292 | 1.258 | -0.821 | 0.718 | -0.014 | -0.016 | -0.016 | 0 | 0.82 | 0.84 | 0.73 | 0.63 | 0.71 | 0.74 |
| 1 | 70594571 | rs7547666 | -0.713 | 1.252 | -0.873 | 0.716 | -0.049 | -0.066 | -0.066 | 4.441 | 0.83 | 0.92 | 0.75 | 0.63 | 0.73 | 0.75 |
| 1 | 70596535 | rs6424382 | 1.373 | 1.047 | -0.565 | 0.572 | 0 | 0.006 | 0.006 | 0 | 0.61 | 0.58 | 0.63 | 0.53 | 0.62 | 0.51 |
| 1 | 70609129 | rs522320 | 0.548 | 1.078 | -0.679 | 0.816 | 0.012 | -0.021 | -0.021 | 0 | 0.74 | 0.8 | 0.74 | 0.63 | 0.82 | 0.59 |
| 1 | 70615141 | rs3767214 | 0.03 | 1.738 | -0.616 | 0.846 | 0.002 | 0.063 | 0.063 | 0 | 0.07 | 0.07 | 0.17 | 0.28 | 0.08 | 0.18 |
| 1 | 70620329 | rs587567 | -0.747 | 1.706 | -0.684 | 0.828 | 0.006 | -0.017 | -0.017 | 0 | 0.24 | 0.2 | 0.26 | 0.41 | 0.18 | 0.38 |
| 1 | 70646431 | rs6664480 | -1.035 | 1.836 | -0.618 | 0.779 | -0.004 | 0.007 | 0.007 | 0 | 0.26 | 0.26 | 0.27 | 0.41 | 0.18 | 0.42 |
| 1 | 70651979 | rs3767210 | 0.811 | 1.837 | -0.613 | 0.778 | 0.016 | 0.012 | 0.012 | 0 | 0.17 | 0.12 | 0.09 | 0.12 | 0.1 | 0.2 |
| 1 | 70669613 | rs3753817 | -0.184 | 1.835 | -0.61 | 0.736 | 0.002 | 0.069 | 0.069 | 0 | 0.07 | 0.08 | 0.17 | 0.29 | 0.08 | 0.18 |
| 1 | 70670312 | rs6696040 | 0.866 | 1.829 | -0.631 | 0.728 | 0.024 | 0.024 | 0.024 | 0 | 0.17 | 0.1 | 0.09 | 0.09 | 0.1 | 0.2 |
| 1 | 70679891 | rs17131241 | -0.991 | 1.822 | -0.643 | 0.726 | -0.004 | 0.007 | 0.007 | 0 | 0.26 | 0.25 | 0.26 | 0.38 | 0.18 | 0.41 |
| 1 | 70682550 | rs11581894 | 1.952 | 1.773 | -1.117 | 0.61 | -0.066 | 0.018 | 0.018 | 0.07 | 0.16 | 0.01 | 0.24 | 0.08 | 0.2 | 0.19 |
| 1 | 70705315 | rs10489927 | -0.264 | 1.018 | -0.876 | 0.773 | -0.004 | 0.027 | 0.027 | 0 | 0.07 | 0.04 | 0.17 | 0.29 | 0.08 | 0.18 |
| 1 | 70715884 | rs10489928 | -0.604 | 1.033 | -0.832 | 0.78 | 0.001 | 0.087 | 0.087 | 0 | 0.09 | 0.13 | 0.17 | 0.29 | 0.08 | 0.21 |
| 1 | 70731885 | rs269276 | -0.16 | 1.148 | -0.413 | 0.528 | 0.028 | 0.04 | 0.04 | 0.396 | 0.3 | 0.43 | 0.43 | 0.36 | 0.33 | 0.43 |
| 1 | 70744507 | rs12565719 | 0.884 | 0.485 | -1.319 | 0.186 | 0.018 | -0.031 | -0.031 | 0.123 | 0.24 | 0.06 | 0.11 | 0.26 | 0.2 | 0.13 |
| 1 | 70753318 | rs270491 | 1.564 | 1.053 | -0.984 | 0.104 | -0.047 | 0.06 | 0.06 | 0.303 | 0.17 | 0.06 | 0.13 | 0.07 | 0.22 | 0.18 |
| 1 | 70762970 | rs6686181 | -0.001 | 0.963 | -0.369 | 0.715 | 0.05 | 0.214 | 0.214 | 0 | 0.06 | 0.16 | 0.22 | 0.33 | 0.1 | 0.19 |
| 1 | 70764703 | rs11209600 | 0.187 | 0.962 | -0.344 | 0.714 | 0.024 | 0.142 | 0.142 | 0 | 0.06 | 0.08 | 0.21 | 0.33 | 0.1 | 0.19 |
| 1 | 70765659 | rs12135469 | 0.836 | 0.666 | -0.808 | 0.353 | -0.001 | 0.041 | 0.041 | 0 | 0.16 | 0.03 | 0.09 | 0.06 | 0.11 | 0.2 |
| 1 | 70780947 | rs12048356 | 1.913 | 0.88 | -1.002 | 0.206 | -0.042 | 0.055 | 0.055 | 0 | 0.18 | 0.04 | 0.13 | 0.08 | 0.21 | 0.17 |
| 1 | 70808932 | rs6424385 | -1.313 | 0.07 | 0.106 | 0.261 | 0.017 | 0.039 | 0.039 | 0 | 0.51 | 0.56 | 0.43 | 0.66 | 0.41 | 0.59 |
| 1 | 70816563 | rs10889868 | -0.945 | 0.269 | 0.447 | 0.437 | -0.001 | 0.059 | 0.059 | 0 | 0.25 | 0.35 | 0.32 | 0.38 | 0.2 | 0.39 |
| 1 | 70818576 | rs12036256 | -0.772 | 0.405 | 0.598 | 0.68 | 0.047 | 0.222 | 0.222 | 0 | 0.08 | 0.23 | 0.22 | 0.32 | 0.1 | 0.19 |
| 1 | 70821307 | rs4650041 | -0.228 | 0.434 | 0.368 | 0.71 | 0.023 | 0.138 | 0.138 | 0 | 0.06 | 0.09 | 0.21 | 0.32 | 0.1 | 0.19 |
| 1 | 70823238 | rs12140479 | 0.103 | 0.164 | 0.21 | 0.403 | 0.004 | 0.047 | 0.047 | 0 | 0.28 | 0.13 | 0.16 | 0.14 | 0.26 | 0.32 |
| 1 | 70825758 | rs998523 | 0.326 | 0.21 | 0.21 | 0.472 | 0.01 | 0.096 | 0.096 | 0 | 0.06 | 0.06 | 0.2 | 0.31 | 0.1 | 0.19 |
| 1 | 70828536 | rs4620474 | -0.872 | 0.09 | 0.351 | 0.354 | 0 | -0.007 | -0.007 | 0 | 0.37 | 0.32 | 0.36 | 0.47 | 0.36 | 0.51 |
| 1 | 70831841 | rs7539544 | 0.031 | 0.17 | 0.288 | 0.457 | 0.005 | 0.085 | 0.085 | 0 | 0.06 | 0.06 | 0.2 | 0.31 | 0.09 | 0.19 |
| 1 | 70858435 | rs7523188 | -0.499 | -0.094 | 0.402 | 0.267 | -0.003 | 0.001 | 0.001 | 0.12 | 0.33 | 0.31 | 0.32 | 0.53 | 0.22 | 0.42 |
| 1 | 70869116 | rs664051 | -0.982 | -0.159 | 0.319 | 0.107 | 0 | -0.001 | -0.001 | 0 | 0.82 | 0.82 | 0.9 | 0.88 | 0.8 | 0.84 |
| 1 | 70875779 | rs648743 | 0.925 | -0.061 | 0.364 | 0.15 | -0.001 | -0.004 | -0.004 | 0 | 0.49 | 0.53 | 0.58 | 0.31 | 0.58 | 0.41 |
| 1 | 70877743 | rs12030275 | 0.541 | 0.031 | 0.004 | 0.353 | 0.005 | 0.109 | 0.109 | 0 | 0.05 | 0.05 | 0.2 | 0.35 | 0.09 | 0.2 |
| 1 | 70877779 | rs487773 | 0.96 | -0.123 | 0.257 | 0.056 | 0.029 | 0.077 | 0.077 | 0 | 0.48 | 0.32 | 0.56 | 0.31 | 0.58 | 0.41 |
| 1 | 70877964 | rs10889869 | 1.908 | 0.182 | -0.016 | 0.135 | -0.02 | 0.017 | 0.017 | 0 | 0.05 | 0 | 0.07 | 0.05 | 0.06 | 0.1 |
| 1 | 70880717 | rs681475 | 0.018 | 0.009 | 0.419 | 0.24 | 0.025 | 0.075 | 0.075 | 0.076 | 0.73 | 0.59 | 0.66 | 0.54 | 0.65 | 0.62 |
| 1 | 70883840 | rs1145920 | -0.434 | 0.09 | 0.372 | 0.518 | -0.003 | 0.002 | 0.002 | 0 | 0.76 | 0.78 | 0.88 | 0.83 | 0.72 | 0.81 |
| 1 | 70887099 | rs672203 | 0.718 | 0.063 | -0.491 | 0.125 | 0.001 | 0.001 | 0.001 | 2.707 | 0.41 | 0.34 | 0.24 | 0.24 | 0.36 | 0.26 |
| 1 | 70896165 | rs17131305 | 1.076 | 0.344 | -0.744 | 0.23 | 0.002 | 0.105 | 0.105 | 0 | 0.05 | 0.07 | 0.2 | 0.3 | 0.08 | 0.19 |
| 1 | 70897183 | rs663649 | 1.165 | 0.443 | -0.994 | 0.011 | -0.002 | -0.019 | -0.019 | 0 | 0.28 | 0.23 | 0.19 | 0.05 | 0.29 | 0.19 |
| 1 | 70904800 | rs1021737 | 1.11 | 0.577 | -1.537 | 0.424 | -0.052 | -0.031 | -0.031 | 0 | 0.2 | 0.06 | 0.37 | 0.22 | 0.26 | 0.21 |
| 1 | 70905674 | rs6693082 | 1.342 | 0.537 | -1.603 | 0.472 | -0.09 | -0.067 | -0.067 | 0.034 | 0.18 | 0.01 | 0.37 | 0.23 | 0.26 | 0.23 |
| 1 | 70910065 | rs12561944 | 0.716 | -0.099 | -0.983 | 0.535 | 0.033 | -0.012 | -0.012 | 0 | 0.21 | 0.13 | 0.07 | 0.21 | 0.09 | 0.2 |
| 1 | 70913726 | rs555671 | -0.76 | -0.42 | -0.609 | 0.536 | 0.003 | -0.058 | -0.058 | 0 | 0.1 | 0.22 | 0.05 | 0 | 0.07 | 0.07 |
| 1 | 70920737 | rs12562375 | 0.946 | -0.501 | -1.537 | 0.659 | 0.031 | -0.02 | -0.02 | 0 | 0.18 | 0.09 | 0.03 | 0.21 | 0.05 | 0.16 |
| 1 | 70924020 | rs4650051 | 0.71 | 0.137 | -1.634 | 0.44 | -0.037 | 0.087 | 0.087 | 0.348 | 0.26 | 0.14 | 0.21 | 0.11 | 0.31 | 0.19 |
| 1 | 70928786 | rs4650052 | -0.107 | 0.922 | -1.162 | 1.013 | 0.006 | 0.003 | 0.003 | 0 | 0.09 | 0.1 | 0.27 | 0.14 | 0.14 | 0.17 |
| 1 | 70930788 | rs6424388 | -0.558 | 0.742 | -1.22 | 0.893 | -0.01 | 0.016 | 0.016 | 0 | 0.55 | 0.32 | 0.59 | 0.63 | 0.56 | 0.54 |
| 1 | 70934259 | rs8179319 | 0.078 | 0.588 | -1.376 | 0.85 | 0.05 | 0.063 | 0.063 | 0 | 0.45 | 0.22 | 0.27 | 0.29 | 0.4 | 0.34 |
| 1 | 70944520 | rs17131315 | -0.915 | 0.748 | -0.99 | 1.293 | -0.014 | -0.007 | -0.007 | 0 | 0.11 | 0.08 | 0.32 | 0.35 | 0.19 | 0.21 |
| 1 | 70945884 | rs9988559 | -0.478 | -0.263 | -0.568 | -0.113 | -0.016 | -0.036 | -0.036 | 0.01 | 0.19 | 0.28 | 0.14 | 0.11 | 0.16 | 0.19 |
| 1 | 70947395 | rs12128717 | -0.319 | -0.48 | -0.838 | -0.269 | -0.009 | 0.036 | 0.036 | 0 | 0.09 | 0.01 | 0.13 | 0.02 | 0.08 | 0.1 |
| 1 | 70947746 | rs17131317 | 0.143 | -0.607 | -0.197 | -0.545 | 0.029 | -0.017 | -0.017 | 0 | 0.07 | 0.29 | 0.05 | 0.01 | 0.06 | 0.08 |
| 1 | 70948010 | rs527148 | 1.012 | -1.056 | -0.564 | -0.831 | -0.023 | 0.005 | 0.005 | 0.034 | 0.36 | 0.24 | 0.43 | 0.35 | 0.43 | 0.35 |
| 1 | 70948788 | rs10493474 | 0.976 | -1.324 | -0.84 | -0.785 | 0.008 | 0.01 | 0.01 | 0 | 0.21 | 0.17 | 0.19 | 0.13 | 0.18 | 0.14 |
| 1 | 70952314 | rs2807912 | 0.445 | -1.508 | -1.073 | -0.735 | -0.018 | -0.109 | -0.109 | 0 | 0.23 | 0.04 | 0.15 | 0.47 | 0.25 | 0.33 |
| 1 | 70954299 | rs2807911 | -0.954 | -1.447 | -0.841 | -0.882 | -0.001 | 0.047 | 0.047 | 0.004 | 0.54 | 0.77 | 0.44 | 0.64 | 0.48 | 0.54 |
| 1 | 70957413 | rs2651248 | -0.631 | -0.841 | -0.792 | -0.519 | -0.008 | 0.04 | 0.04 | 0 | 0.32 | 0.45 | 0.22 | 0.51 | 0.31 | 0.39 |
| 1 | 70963368 | rs1333680 | -0.844 | -0.715 | -0.806 | -0.117 | -0.004 | -0.019 | -0.019 | 0 | 0.17 | 0.29 | 0.18 | 0.12 | 0.14 | 0.18 |
| 1 | 70966453 | rs655938 | 0.584 | -0.924 | -1.332 | -0.206 | -0.011 | 0.017 | 0.017 | 0 | 0.44 | 0.22 | 0.47 | 0.38 | 0.5 | 0.47 |
| 1 | 70968414 | rs17131320 | 1.276 | -0.968 | -1.398 | -0.059 | -0.014 | -0.035 | -0.035 | 0 | 0.08 | 0 | 0.1 | 0.4 | 0.1 | 0.16 |
| 1 | 70974117 | rs682717 | 0.166 | -0.638 | -1.35 | -0.108 | 0.053 | 0.033 | 0.033 | 0 | 0.51 | 0.26 | 0.58 | 0.41 | 0.57 | 0.61 |
| 1 | 70981747 | rs10518338 | 0.225 | -0.409 | -1.336 | 0.285 | -0.013 | 0.022 | 0.022 | 0 | 0.07 | 0.02 | 0.13 | 0.03 | 0.07 | 0.15 |
| 1 | 70983975 | rs625578 | -0.173 | -0.477 | -1.19 | 0.412 | -0.004 | 0.013 | 0.013 | 0 | 0.75 | 0.72 | 0.82 | 0.55 | 0.77 | 0.74 |
| 1 | 70986547 | rs10889879 | -0.075 | 0.041 | -1.779 | 0.46 | 0.007 | 0.018 | 0.018 | 0 | 0.48 | 0.19 | 0.44 | 0.33 | 0.57 | 0.37 |
| 1 | 70990002 | rs615367 | 0.505 | -0.329 | -1.337 | 0.351 | 0.026 | 0.031 | 0.031 | 0 | 0.84 | 0.87 | 0.94 | 0.92 | 0.87 | 0.77 |

| | | | | | | | | | | | | | | | | |
|---|---|---|---|---|---|---|---|---|---|---|---|---|---|---|---|---|
| 1 | 70990438 | rs617211 | 1.16 | -0.835 | -0.935 | 1.362 | -0.008 | 0.001 | 0.001 | 0 | 0.77 | 0.73 | 0.73 | 0.53 | 0.85 | 0.56 |
| 1 | 70990955 | rs629807 | 1.243 | -1.179 | -0.143 | 1.008 | -0.039 | -0.037 | -0.037 | 2.326 | 0.75 | 0.6 | 0.67 | 0.87 | 0.83 | 0.72 |
| 1 | 70992015 | rs580789 | 0.247 | -0.591 | -0.074 | 1.024 | -0.002 | 0.006 | 0.006 | 0.015 | 0.46 | 0.31 | 0.37 | 0.53 | 0.44 | 0.46 |
| 1 | 70993695 | rs2202044 | 0.384 | -0.434 | 0.256 | 0.962 | -0.001 | 0.033 | 0.033 | 0.656 | 0.16 | 0.29 | 0.17 | 0.26 | 0.12 | 0.16 |
| 1 | 70995562 | rs2651244 | 0.297 | -0.5 | 0.306 | 0.705 | -0.046 | 0.121 | 0.121 | 0 | 0.33 | 0.09 | 0.22 | 0.14 | 0.4 | 0.3 |
| 1 | 71001920 | rs2651236 | -0.416 | -0.039 | 0.37 | 0.347 | -0.014 | 0.111 | 0.111 | 0 | 0.45 | 0.3 | 0.32 | 0.18 | 0.52 | 0.39 |
| 1 | 71003226 | rs12075963 | -1.054 | -0.009 | 0.527 | 0.356 | 0.008 | -0.023 | -0.023 | 0 | 0.15 | 0.27 | 0.32 | 0.08 | 0.16 | 0.14 |
| 1 | 71003750 | rs517735 | 0.016 | -0.411 | 0.867 | 0.987 | -0.004 | 0.042 | 0.042 | 0 | 0.23 | 0.26 | 0.2 | 0.37 | 0.19 | 0.17 |
| 1 | 71007926 | rs9424945 | -1.124 | -0.491 | 0.897 | 0.811 | -0.005 | 0.015 | 0.015 | 0 | 0.66 | 0.76 | 0.74 | 0.84 | 0.58 | 0.69 |
| 1 | 71010207 | rs582435 | -1.11 | -0.534 | 1.198 | 0.947 | 0.005 | -0.018 | -0.018 | 0 | 0.34 | 0.53 | 0.3 | 0.3 | 0.3 | 0.25 |
| 1 | 71018986 | rs4233083 | -0.1 | -0.534 | 1.171 | 0.964 | 0.001 | 0.001 | 0.001 | 0.019 | 0.45 | 0.51 | 0.55 | 0.55 | 0.4 | 0.54 |
| 1 | 71019658 | rs12410281 | 1.793 | -0.645 | 1.055 | 1.213 | -0.044 | -0.016 | -0.016 | 0 | 0.25 | 0.02 | 0.21 | 0.19 | 0.24 | 0.2 |
| 1 | 71022016 | rs12035687 | -1.881 | -0.627 | 1.145 | 1.199 | 0.009 | 0.307 | 0.307 | 0.203 | 0.07 | 0.3 | 0.23 | 0.35 | 0.05 | 0.19 |
| 1 | 71023684 | rs10889881 | 0.836 | -0.626 | 1.168 | 1.207 | 0.002 | -0.003 | -0.003 | 0.667 | 0.33 | 0.4 | 0.46 | 0.54 | 0.29 | 0.39 |
| 1 | 71028815 | rs3121983 | -0.479 | -0.633 | 0.997 | 1.198 | -0.003 | 0.009 | 0.009 | 0 | 0.78 | 0.75 | 0.81 | 0.71 | 0.83 | 0.84 |
| 1 | 71034911 | rs12088046 | 0.279 | -0.374 | 0.581 | 1.234 | -0.047 | 0.063 | 0.063 | 0.043 | 0.38 | 0.15 | 0.5 | 0.26 | 0.41 | 0.34 |
| 1 | 71035504 | rs12117246 | 0.254 | -0.434 | 0.313 | 0.9 | -0.042 | 0.043 | 0.043 | 0 | 0.08 | 0.01 | 0.07 | 0 | 0.11 | 0.09 |
| 1 | 71038867 | rs4130191 | 0.5 | -0.688 | 0.534 | 1.109 | 0.016 | -0.023 | -0.023 | 0 | 0.33 | 0.32 | 0.45 | 0.54 | 0.3 | 0.43 |
| 1 | 71044680 | rs11209643 | -0.997 | -0.434 | 0.41 | 0.836 | 0.008 | -0.022 | -0.022 | 0 | 0.17 | 0.13 | 0.24 | 0.72 | 0.13 | 0.45 |
| 1 | 71045066 | rs7522196 | 0.41 | -0.554 | 0.545 | 0.349 | 0.011 | 0.068 | 0.068 | 0 | 0.45 | 0.32 | 0.39 | 0.17 | 0.54 | 0.4 |
| 1 | 71046578 | rs991302 | 0.814 | -0.483 | 0.427 | 0.23 | -0.024 | -0.012 | -0.012 | 0 | 0.86 | 0.47 | 0.71 | 0.89 | 0.89 | 0.92 |
| 1 | 71060083 | rs17131358 | -0.681 | 0.04 | -0.164 | 0.394 | -0.008 | -0.005 | -0.005 | 0 | 0.12 | 0.17 | 0.27 | 0.11 | 0.11 | 0.08 |
| 1 | 71067296 | rs6689757 | 1.097 | 0.34 | -0.844 | 0.426 | -0.108 | 0.09 | 0.09 | 0 | 0.32 | 0.04 | 0.26 | 0.16 | 0.39 | 0.28 |
| 1 | 71070312 | rs17131360 | 0.166 | 0.526 | -0.443 | 1.163 | -0.015 | -0.04 | -0.04 | 2.529 | 0.1 | 0.02 | 0.13 | 0.46 | 0.11 | 0.32 |
| 1 | 71084638 | rs1932065 | 1.294 | 0.494 | 0.12 | 0.752 | 0.039 | 0.023 | 0.023 | 0.294 | 0.81 | 0.55 | 0.63 | 0.77 | 0.78 | 0.87 |
| 1 | 71085728 | rs1007630 | 0.726 | 0.175 | -0.685 | 0.55 | -0.152 | 0.098 | 0.098 | 0.125 | 0.33 | 0.04 | 0.32 | 0.16 | 0.46 | 0.32 |
| 1 | 71095174 | rs12087314 | 0.409 | -0.268 | -0.111 | 1.044 | 0 | -0.018 | -0.018 | 0.433 | 0.1 | 0.09 | 0.05 | 0 | 0.06 | 0.01 |
| 1 | 71098437 | rs2421794 | 0.302 | -0.314 | 0.427 | 1.25 | 0.004 | 0.197 | 0.197 | 0 | 0.86 | 0.63 | 0.84 | 0.64 | 0.88 | 0.64 |
| 1 | 71100213 | rs2152119 | 0.714 | -0.318 | 0.215 | 0.994 | -0.094 | 0.131 | 0.131 | 0.223 | 0.39 | 0.05 | 0.4 | 0.19 | 0.52 | 0.36 |
| 1 | 71100726 | rs1932048 | 0.093 | -0.317 | 0.204 | 0.986 | 0.007 | 0.001 | 0.001 | 1.054 | 0.47 | 0.58 | 0.43 | 0.45 | 0.36 | 0.28 |
| 1 | 71103392 | rs6686879 | 0.579 | -0.36 | -0.373 | 0.791 | 0.042 | 0.09 | 0.09 | 3.447 | 0.4 | 0.23 | 0.3 | 0.19 | 0.34 | 0.16 |
| 1 | 71111456 | rs1335228 | 0.635 | -0.079 | -0.87 | 0.753 | -0.12 | 0.127 | 0.127 | 0.39 | 0.38 | 0.03 | 0.4 | 0.19 | 0.52 | 0.36 |
| 1 | 71116437 | rs2065717 | -0.095 | -0.138 | -0.562 | 0.89 | 0.095 | 0.014 | 0.014 | 2.336 | 0.5 | 0.21 | 0.56 | 0.55 | 0.63 | 0.72 |
| 1 | 71116689 | rs17131373 | -0.287 | 0.106 | -0.333 | 1.187 | -0.001 | 0.029 | 0.029 | 0 | 0.11 | 0.09 | 0.14 | 0.4 | 0.11 | 0.37 |
| 1 | 71119659 | rs17131376 | 0.246 | -0.115 | -0.272 | 0.685 | 0.012 | 0.012 | 0.012 | 0 | 0.11 | 0.06 | 0.05 | 0 | 0.08 | 0.01 |
| 1 | 71142202 | rs2782763 | -0.753 | -0.038 | 0.847 | 1.223 | -0.005 | 0.13 | 0.13 | 0 | 0.16 | 0.37 | 0.32 | 0.36 | 0.13 | 0.34 |
| 1 | 71144279 | rs11209673 | 1.748 | -0.164 | 0.175 | 1.045 | -0.024 | -0.005 | -0.005 | 0.038 | 0.24 | 0.03 | 0.09 | 0.17 | 0.24 | 0.09 |
| 1 | 71153714 | rs2820526 | -0.525 | -0.148 | 0.328 | 1.152 | -0.002 | 0.094 | 0.094 | 0 | 0.15 | 0.32 | 0.3 | 0.29 | 0.12 | 0.21 |
| 1 | 71154117 | rs1334997 | 1.063 | 0.505 | -0.209 | 1.398 | -0.026 | -0.023 | -0.023 | 0 | 0.31 | 0.18 | 0.23 | 0.35 | 0.35 | 0.22 |
| 1 | 71154417 | rs2152134 | -0.526 | 0.549 | -0.203 | 1.25 | -0.009 | -0.009 | -0.009 | 1.81 | 0.16 | 0.13 | 0.35 | 0.29 | 0.21 | 0.25 |
| 1 | 71157661 | rs12731606 | 0.027 | 0.317 | -0.257 | 0.203 | 0.012 | -0.002 | -0.002 | 0 | 0.2 | 0.15 | 0.09 | 0 | 0.15 | 0.02 |
| 1 | 71162976 | rs2820550 | -0.716 | 0.466 | -0.069 | 0.511 | 0.017 | 0.018 | 0.018 | 0 | 0.95 | 0.92 | 0.92 | 0.83 | 0.9 | 0.89 |
| 1 | 71163908 | rs2820551 | -0.269 | 0.39 | -0.307 | 0.301 | -0.012 | 0.094 | 0.094 | 0 | 0.44 | 0.29 | 0.36 | 0.17 | 0.46 | 0.27 |
| 1 | 71166263 | rs1889600 | 1.42 | 0.477 | -0.344 | 0.339 | -0.041 | 0.011 | 0.011 | 0.614 | 0.24 | 0.14 | 0.27 | 0.18 | 0.31 | 0.25 |
| 1 | 71170183 | rs2152126 | 1.584 | 0.238 | 0.086 | 0.242 | -0.009 | 0.043 | 0.043 | 0.055 | 0.31 | 0.17 | 0.16 | 0.21 | 0.33 | 0.3 |
| 1 | 71178475 | rs1855648 | -0.23 | 0.11 | 0.615 | 0.116 | 0.023 | 0.171 | 0.171 | 5.968 | 0.5 | 0.22 | 0.71 | 0.78 | 0.55 | 0.6 |
| 1 | 71180389 | rs2782792 | 0.861 | 0.147 | 0.689 | 0.173 | -0.014 | -0.058 | -0.058 | 4.03 | 0.14 | 0.2 | 0.07 | 0 | 0.08 | 0.01 |
| 1 | 71197126 | rs1932063 | 0.493 | 0.259 | 0.948 | 0.167 | 0.001 | 0.006 | 0.006 | 4.841 | 0.43 | 0.43 | 0.23 | 0.18 | 0.42 | 0.2 |
| 1 | 71198580 | rs7523542 | 1.77 | 0.274 | 0.868 | 0.177 | -0.026 | 0.055 | 0.055 | 0.147 | 0.29 | 0.14 | 0.16 | 0.18 | 0.33 | 0.18 |
| 1 | 71200411 | rs2184471 | 1.745 | 0.275 | 0.865 | 0.177 | -0.01 | 0.041 | 0.041 | 0.091 | 0.29 | 0.19 | 0.17 | 0.18 | 0.33 | 0.18 |
| 1 | 71215084 | rs1591385 | 0.89 | 0.269 | 0.886 | 0.498 | -0.015 | -0.071 | -0.071 | 0 | 0.14 | 0.22 | 0.07 | 0 | 0.09 | 0.01 |
| 1 | 71220683 | rs12033260 | 0.926 | 0.329 | 0.707 | 0.479 | -0.033 | 0.053 | 0.053 | 0.406 | 0.32 | 0.15 | 0.3 | 0.17 | 0.37 | 0.25 |
| 1 | 71229637 | rs11209689 | 0.162 | 0.305 | 0.925 | 0.516 | -0.011 | 0.085 | 0.085 | 0 | 0.21 | 0.32 | 0.38 | 0.38 | 0.16 | 0.31 |
| 1 | 71230547 | rs998550 | 0.097 | 0.387 | 1.03 | 0.596 | 0 | 0.019 | 0.019 | 0 | 0.52 | 0.71 | 0.57 | 0.6 | 0.48 | 0.63 |
| 1 | 71240621 | rs12080803 | 0.699 | 0.335 | 1.039 | 0.583 | -0.006 | -0.004 | -0.004 | 0 | 0.46 | 0.48 | 0.48 | 0.39 | 0.39 | 0.38 |
| 1 | 71245703 | rs12046339 | 1.007 | 0.356 | 0.784 | 0.568 | -0.058 | 0.057 | 0.057 | 0.445 | 0.32 | 0.07 | 0.29 | 0.18 | 0.37 | 0.25 |
| 1 | 71250352 | rs10889889 | 0.835 | 0.307 | 0.917 | 0.515 | 0.023 | 0.008 | 0.008 | 0 | 0.44 | 0.22 | 0.46 | 0.39 | 0.39 | 0.37 |
| 1 | 71258914 | rs2260575 | -1.928 | 0.284 | 0.769 | 0.429 | 0 | -0.022 | -0.022 | 3.058 | 0.89 | 0.93 | 0.95 | 1 | 0.91 | 0.98 |
| 1 | 71271871 | rs2257598 | 0.403 | 0.341 | 0.769 | 0.543 | -0.027 | -0.028 | -0.028 | 4.163 | 0.36 | 0.18 | 0.36 | 0.4 | 0.41 | 0.34 |
| 1 | 71274425 | rs2257759 | 0.448 | 0.408 | 0.78 | 0.445 | 0.042 | -0.029 | -0.029 | 0 | 0.07 | 0.12 | 0.11 | 0.01 | 0.13 | 0.12 |
| 1 | 71275174 | rs2257772 | -0.647 | 0.606 | 0.813 | 0.723 | -0.01 | 0.042 | 0.042 | 0.701 | 0.47 | 0.76 | 0.47 | 0.74 | 0.49 | 0.7 |
| 1 | 71286947 | rs4603079 | 1.101 | 0.433 | 0.687 | 0.689 | -0.032 | 0.057 | 0.057 | 0.055 | 0.31 | 0.11 | 0.29 | 0.17 | 0.36 | 0.25 |
| 1 | 71288661 | rs11209700 | 2.356 | 0.345 | 0.6 | 0.496 | 0.017 | 0.024 | 0.024 | 4.797 | 0.24 | 0.18 | 0.1 | 0 | 0.21 | 0.04 |
| 1 | 71301892 | rs7534636 | 0.326 | 0.449 | 0.851 | 0.863 | -0.001 | -0.025 | -0.025 | 0 | 0.69 | 0.84 | 0.56 | 0.61 | 0.71 | 0.66 |
| 1 | 71306749 | rs1327461 | 0.25 | 0.621 | 0.732 | 0.728 | 0.094 | -0.027 | -0.027 | 0 | 0.08 | 0.21 | 0.11 | 0.01 | 0.12 | 0.11 |
| 1 | 71314813 | rs7556177 | -1.575 | 0.488 | 0.466 | 0.654 | 0.003 | -0.007 | -0.007 | 0.4 | 0.89 | 0.94 | 0.95 | 1 | 0.91 | 0.98 |
| 1 | 71316861 | rs12065547 | 0.167 | 0.459 | 0.454 | 0.789 | 0.001 | 0.039 | 0.039 | 0.182 | 0.21 | 0.31 | 0.4 | 0.4 | 0.2 | 0.34 |
| 1 | 71321026 | rs6424405 | -0.941 | 0.433 | 0.417 | 0.837 | -0.013 | 0.02 | 0.02 | 0 | 0.86 | 0.75 | 0.91 | 0.81 | 0.89 | 0.97 |
| 1 | 71321740 | rs6664305 | 0.548 | 0.47 | 0.364 | 0.897 | -0.007 | -0.03 | -0.03 | 0 | 0.19 | 0.11 | 0.38 | 0.27 | 0.19 | 0.24 |

| | | | | | | | | | | | | | | | | |
|---|---|---|---|---|---|---|---|---|---|---|---|---|---|---|---|---|
| 1 | 71325044 | rs1327460 | -0.283 | 0.424 | 0.397 | 0.714 | 0 | -0.002 | -0.002 | 2.84 | 0.55 | 0.59 | 0.64 | 0.83 | 0.55 | 0.73 |
| 1 | 71325299 | rs6656853 | 1.214 | 0.269 | 0.453 | 0.75 | 0.001 | -0.027 | -0.027 | 0 | 0.33 | 0.34 | 0.21 | 0.14 | 0.35 | 0.24 |
| 1 | 71325340 | rs6672081 | 2.133 | 0.277 | 0.366 | 0.733 | 0.012 | 0.064 | 0.064 | 4.967 | 0.25 | 0.07 | 0.1 | 0.13 | 0.22 | 0.13 |
| 1 | 71327092 | rs7530345 | -0.147 | 0.635 | 0.274 | 0.617 | 0.001 | -0.006 | -0.006 | 0 | 0.55 | 0.59 | 0.66 | 0.83 | 0.58 | 0.74 |
| 1 | 71327362 | rs7540868 | -0.147 | 0.635 | 0.274 | 0.617 | 0.001 | -0.006 | -0.006 | 0 | 0.55 | 0.59 | 0.66 | 0.83 | 0.58 | 0.74 |
| 1 | 71327463 | rs7530738 | 0.902 | 0.68 | 0.262 | 0.697 | -0.079 | -0.098 | -0.098 | 0 | 0.18 | 0.01 | 0.39 | 0.27 | 0.21 | 0.26 |
| 1 | 71328975 | rs6685546 | 2.921 | 0.406 | -0.064 | 0.528 | -0.031 | 0.132 | 0.132 | 5.087 | 0.2 | 0.01 | 0.08 | 0 | 0.19 | 0.04 |
| 1 | 71331220 | rs12119442 | 0.053 | 0.495 | 0.055 | 0.588 | 0.114 | -0.021 | -0.021 | 0.06 | 0.08 | 0.28 | 0.11 | 0.01 | 0.12 | 0.12 |
| 1 | 71331430 | rs5702 | 0.759 | 0.493 | 0.049 | 0.596 | -0.016 | -0.03 | -0.03 | 0 | 0.19 | 0.12 | 0.39 | 0.27 | 0.21 | 0.26 |
| 1 | 71331498 | rs1409986 | -2.658 | 0.427 | -0.112 | 0.531 | -0.02 | 0.039 | 0.039 | 0 | 0.93 | 0.99 | 0.97 | 1 | 0.92 | 0.98 |
| 1 | 71332220 | rs12135331 | -0.004 | 0.611 | 0.008 | 0.574 | 0.114 | -0.02 | -0.02 | 0 | 0.08 | 0.28 | 0.11 | 0.01 | 0.12 | 0.12 |
| 1 | 71334870 | rs1536537 | 0.87 | 0.652 | -0.257 | 0.581 | -0.022 | 0.038 | 0.038 | 0.232 | 0.34 | 0.21 | 0.37 | 0.18 | 0.42 | 0.28 |
| 1 | 71337193 | rs1325949 | 0.863 | 0.545 | -0.387 | 0.731 | -0.077 | -0.092 | -0.092 | 0 | 0.18 | 0.01 | 0.39 | 0.26 | 0.21 | 0.24 |
| 1 | 71337925 | rs1327449 | 2.456 | 0.625 | -0.724 | 0.417 | -0.057 | 0.14 | 0.14 | 3.675 | 0.21 | 0.01 | 0.08 | 0 | 0.23 | 0.04 |
| 1 | 71338397 | rs1536261 | 1.036 | 0.208 | -0.295 | 0.7 | 0.011 | 0.021 | 0.021 | 0 | 0.56 | 0.35 | 0.46 | 0.33 | 0.62 | 0.41 |
| 1 | 71339973 | rs7543182 | 0.835 | 0.356 | -0.295 | 0.932 | -0.077 | -0.088 | -0.088 | 0 | 0.18 | 0.01 | 0.39 | 0.25 | 0.21 | 0.24 |
| 1 | 71347417 | rs17131476 | -0.847 | 0.273 | 0.109 | 0.985 | 0.031 | 0.245 | 0.245 | 0.513 | 0.07 | 0.36 | 0.04 | 0.23 | 0.03 | 0.32 |
| 1 | 71351785 | rs11209707 | -1.272 | 0.267 | 0.111 | 0.985 | 0.007 | 0.446 | 0.446 | 0.485 | 0.09 | 0.44 | 0.09 | 0.42 | 0.05 | 0.33 |
| 1 | 71355201 | rs6424409 | -0.703 | 0.315 | -0.212 | 0.674 | -0.029 | 0.182 | 0.182 | 5.691 | 0.55 | 0.78 | 0.77 | 0.86 | 0.54 | 0.81 |
| 1 | 71358190 | rs4998697 | -0.726 | 0.288 | 0.053 | 0.723 | -0.03 | 0.168 | 0.168 | 4.47 | 0.56 | 0.75 | 0.77 | 0.86 | 0.54 | 0.8 |
| 1 | 71360150 | rs17131481 | 0.854 | 0.24 | -0.143 | 0.751 | -0.071 | -0.089 | -0.089 | 0.132 | 0.19 | 0.01 | 0.39 | 0.25 | 0.21 | 0.22 |
| 1 | 71360342 | rs35864143 | 0.697 | 0.233 | -0.106 | 0.747 | -0.024 | -0.04 | -0.04 | 0.486 | 0.19 | 0.1 | 0.4 | 0.25 | 0.21 | 0.22 |
| 1 | 71360468 | rs17131483 | 0.703 | 0.236 | -0.104 | 0.749 | -0.03 | -0.047 | -0.047 | 0.007 | 0.19 | 0.08 | 0.4 | 0.25 | 0.21 | 0.22 |
| 1 | 71360580 | rs12039590 | 1.444 | 0.197 | 0.015 | 0.772 | -0.004 | 0.021 | 0.021 | 0.36 | 0.26 | 0.12 | 0.27 | 0.19 | 0.28 | 0.24 |
| 1 | 71360623 | rs12046641 | 0.956 | 0.197 | 0.076 | 0.771 | 0.002 | -0.006 | -0.006 | 0.169 | 0.3 | 0.24 | 0.28 | 0.41 | 0.29 | 0.49 |
| 1 | 71360672 | rs12032742 | 0.969 | 0.197 | 0.072 | 0.771 | 0.001 | -0.005 | -0.005 | 0 | 0.3 | 0.24 | 0.28 | 0.41 | 0.29 | 0.49 |
| 1 | 71361326 | rs35702222 | 0.79 | 0.204 | 0.063 | 0.781 | -0.07 | -0.089 | -0.089 | 0 | 0.19 | 0.01 | 0.39 | 0.25 | 0.22 | 0.22 |
| 1 | 71362587 | rs6687859 | 0.686 | 0.182 | 0.122 | 0.789 | -0.008 | -0.017 | -0.017 | 0 | 0.2 | 0.16 | 0.39 | 0.25 | 0.21 | 0.22 |
| 1 | 71362978 | rs17131484 | 0.853 | 0.182 | 0.073 | 0.793 | -0.055 | -0.076 | -0.076 | 0 | 0.19 | 0.02 | 0.38 | 0.25 | 0.21 | 0.22 |
| 1 | 71363236 | rs11209709 | 0.904 | 0.184 | 0.113 | 0.809 | 0.001 | -0.005 | -0.005 | 0.022 | 0.3 | 0.24 | 0.28 | 0.41 | 0.29 | 0.49 |
| 1 | 71363699 | rs34999609 | -1.047 | 0.186 | 0.131 | 0.812 | 0.026 | 0.46 | 0.46 | 0.257 | 0.08 | 0.38 | 0.08 | 0.43 | 0.04 | 0.34 |
| 1 | 71363749 | rs35368675 | -1.047 | 0.186 | 0.14 | 0.812 | 0.026 | 0.46 | 0.46 | 0.257 | 0.08 | 0.38 | 0.08 | 0.43 | 0.04 | 0.34 |
| 1 | 71363881 | rs1409166 | 0.253 | 0.18 | 0.125 | 0.804 | -0.008 | 0.008 | 0.008 | 0.654 | 0.35 | 0.5 | 0.36 | 0.62 | 0.33 | 0.58 |
| 1 | 71364105 | rs1409165 | -0.057 | 0.099 | 0.006 | 0.672 | -0.055 | 0.084 | 0.084 | 0 | 0.76 | 0.47 | 0.86 | 0.57 | 0.84 | 0.64 |
| 1 | 71365018 | rs11209710 | 0.481 | 0.336 | 0 | 0.608 | -0.008 | 0.013 | 0.013 | 0.755 | 0.34 | 0.48 | 0.35 | 0.62 | 0.33 | 0.57 |
| 1 | 71365772 | rs1409989 | 0.93 | 0.405 | 0.04 | 0.698 | 0.001 | -0.004 | -0.004 | 0 | 0.3 | 0.25 | 0.28 | 0.4 | 0.29 | 0.5 |
| 1 | 71367193 | rs942978 | 0.942 | 0.404 | 0.058 | 0.658 | 0.004 | -0.013 | -0.013 | 0 | 0.3 | 0.22 | 0.28 | 0.39 | 0.29 | 0.5 |
| 1 | 71367464 | rs942977 | 0.996 | 0.417 | -0.042 | 0.665 | 0.001 | -0.004 | -0.004 | 0 | 0.3 | 0.25 | 0.28 | 0.39 | 0.29 | 0.5 |
| 1 | 71368281 | rs10889896 | 0.996 | 0.417 | -0.042 | 0.665 | 0.001 | -0.004 | -0.004 | 0 | 0.3 | 0.25 | 0.28 | 0.39 | 0.29 | 0.5 |
| 1 | 71368758 | rs1409162 | -0.247 | 0.491 | -0.158 | 0.376 | -0.001 | 0.001 | 0.001 | 5.563 | 0.51 | 0.5 | 0.69 | 0.66 | 0.5 | 0.71 |
| 1 | 71369827 | rs4420040 | 0.542 | 0.805 | -0.523 | 0.548 | 0.011 | -0.02 | -0.02 | 0 | 0.35 | 0.4 | 0.28 | 0.24 | 0.39 | 0.2 |
| 1 | 71371420 | rs2182325 | 0.126 | 0.615 | -1.016 | 0.702 | -0.002 | -0.024 | -0.024 | 0 | 0.64 | 0.71 | 0.47 | 0.61 | 0.67 | 0.68 |
| 1 | 71373738 | rs875727 | 0.275 | 0.501 | -0.5 | 1.004 | -0.006 | 0.094 | 0.094 | 0 | 0.12 | 0.13 | 0.16 | 0.46 | 0.1 | 0.27 |
| 1 | 71375275 | rs12123324 | 0.312 | 0.321 | -0.701 | 0.555 | -0.004 | -0.021 | -0.021 | 0 | 0.22 | 0.15 | 0.27 | 0.3 | 0.21 | 0.2 |
| 1 | 71375363 | rs11209715 | 0.539 | 0.922 | -0.194 | 0.796 | -0.002 | -0.005 | -0.005 | 0 | 0.55 | 0.5 | 0.39 | 0.29 | 0.38 | 0.24 |
| 1 | 71379180 | rs7539384 | -0.685 | 1.268 | -0.135 | 0.7 | -0.004 | 0.002 | 0.002 | 0.166 | 0.72 | 0.72 | 0.64 | 0.79 | 0.55 | 0.62 |
| 1 | 71384680 | rs6424410 | 0.452 | 1.019 | -0.536 | 0.613 | -0.002 | -0.005 | -0.005 | 0.026 | 0.56 | 0.5 | 0.39 | 0.31 | 0.39 | 0.24 |
| 1 | 71391646 | rs602383 | 0.103 | 1.515 | -0.798 | 0.394 | -0.004 | 0.002 | 0.002 | 0 | 0.27 | 0.27 | 0.33 | 0.21 | 0.4 | 0.33 |
| 1 | 71394683 | rs578096 | 0.019 | 1.827 | -1.062 | 0.454 | -0.042 | -0.01 | -0.01 | 0.873 | 0.92 | 1 | 0.91 | 0.94 | 0.86 | 0.85 |
| 1 | 71398588 | rs12067140 | 0.182 | 1.196 | -0.821 | -0.3 | -0.007 | -0.054 | -0.054 | 0 | 0.18 | 0.22 | 0.06 | 0 | 0.12 | 0.02 |
| 1 | 71398902 | rs12097924 | 0.505 | 0.717 | -0.934 | -0.45 | -0.021 | -0.023 | -0.023 | 0.464 | 0.33 | 0.47 | 0.21 | 0.22 | 0.23 | 0.15 |
| 1 | 71399414 | rs625617 | 0.582 | 0.677 | -0.636 | -0.477 | -0.029 | 0.061 | 0.061 | 2.951 | 0.44 | 0.16 | 0.64 | 0.73 | 0.55 | 0.68 |
| 1 | 71406691 | rs475468 | 1.728 | 0.916 | -1.024 | -0.248 | -0.097 | -0.016 | -0.016 | 0 | 0.2 | 0.02 | 0.24 | 0.19 | 0.34 | 0.19 |
| 1 | 71413324 | rs622721 | 1.305 | 0.734 | -0.772 | 0.54 | -0.025 | -0.024 | -0.024 | 0.034 | 0.3 | 0.14 | 0.34 | 0.44 | 0.41 | 0.48 |
| 1 | 71413440 | rs1409984 | 1.284 | 0.722 | -0.79 | 0.562 | -0.03 | -0.095 | -0.095 | 0 | 0.14 | 0.01 | 0.3 | 0.29 | 0.14 | 0.2 |
| 1 | 71414863 | rs650194 | -0.286 | 0.719 | -0.659 | 0.244 | -0.002 | -0.006 | -0.006 | 0 | 0.62 | 0.38 | 0.7 | 0.73 | 0.67 | 0.7 |
| 1 | 71415135 | rs651265 | -1.008 | 0.707 | -0.377 | 0.211 | 0.045 | 0.015 | 0.015 | 0 | 0.94 | 0.79 | 0.91 | 0.94 | 0.89 | 0.85 |
| 1 | 71415597 | rs7541936 | 0.991 | 0.603 | -0.088 | 0.089 | -0.013 | -0.072 | -0.072 | 0 | 0.18 | 0.28 | 0.07 | 0 | 0.12 | 0.02 |
| 1 | 71421564 | rs661000 | -0.008 | 0.511 | 0.088 | 0.084 | 0 | -0.02 | -0.02 | 0 | 0.6 | 0.49 | 0.7 | 0.73 | 0.63 | 0.7 |
| 1 | 71426260 | rs977214 | 1.543 | 0.501 | 0.1 | 0.037 | -0.026 | -0.086 | -0.086 | 0.053 | 0.14 | 0.3 | 0.06 | 0 | 0.08 | 0.02 |
| 1 | 71426569 | rs573688 | -0.308 | 0.506 | 0.153 | 0.035 | 0.027 | -0.016 | -0.016 | 0 | 0.9 | 0.81 | 0.9 | 0.96 | 0.86 | 0.85 |
| 1 | 71427842 | rs6665776 | 1.656 | 0.502 | 0.15 | 0.036 | -0.023 | -0.081 | -0.081 | 0 | 0.14 | 0.29 | 0.06 | 0 | 0.08 | 0.02 |
| 1 | 71432984 | rs626398 | -0.348 | 0.532 | 0.173 | 0.069 | 0.029 | -0.018 | -0.018 | 0 | 0.9 | 0.77 | 0.89 | 0.96 | 0.86 | 0.85 |
| 1 | 71436256 | rs2300161 | 0.64 | 0.499 | 0.155 | 0.111 | 0 | 0 | 0 | 0 | 0.3 | 0.29 | 0.19 | 0.21 | 0.23 | 0.15 |
| 1 | 71436747 | rs5697 | 0.617 | 0.502 | 0.109 | 0.113 | -0.004 | -0.005 | -0.005 | 0 | 0.31 | 0.33 | 0.19 | 0.21 | 0.23 | 0.15 |
| 1 | 71439528 | rs17090700 | 0.676 | 0.504 | 0.098 | 0.114 | -0.002 | -0.002 | -0.002 | 0.641 | 0.3 | 0.3 | 0.19 | 0.21 | 0.23 | 0.15 |
| 1 | 71439590 | rs11587601 | 0.666 | 0.496 | 0.095 | 0.11 | -0.002 | -0.002 | -0.002 | 0 | 0.3 | 0.3 | 0.19 | 0.21 | 0.23 | 0.15 |
| 1 | 71440217 | rs2072947 | 0.444 | 0.602 | 0.03 | 0.112 | -0.029 | 0.075 | 0.075 | 2.339 | 0.45 | 0.17 | 0.64 | 0.74 | 0.55 | 0.68 |
| 1 | 71442206 | rs479934 | -0.794 | 0.499 | 0.197 | -0.183 | 0.001 | -0.08 | -0.08 | 0 | 0.77 | 0.72 | 0.84 | 0.98 | 0.79 | 0.84 |

| | | | | | | | | | | | | | | | | |
|---|---|---|---|---|---|---|---|---|---|---|---|---|---|---|---|---|
| 1 | 71443399 | rs726764 | -0.971 | 0.483 | 0.292 | -0.191 | -0.002 | -0.011 | -0.011 | 0.338 | 0.78 | 0.82 | 0.85 | 0.97 | 0.79 | 0.84 |
| 1 | 71449130 | rs2206343 | 0.101 | 0.506 | 0.378 | 0.151 | 0 | 0.008 | 0.008 | 0 | 0.63 | 0.6 | 0.47 | 0.53 | 0.65 | 0.64 |
| 1 | 71454352 | rs2300164 | 0.738 | 0.623 | 0.321 | 0.152 | -0.001 | -0.001 | -0.001 | 0.887 | 0.26 | 0.24 | 0.18 | 0.24 | 0.21 | 0.14 |
| 1 | 71458162 | rs2300167 | 1.19 | 0.657 | 0.138 | 0.138 | -0.023 | 0.023 | 0.023 | 0 | 0.31 | 0.16 | 0.28 | 0.29 | 0.41 | 0.49 |
| 1 | 71465292 | rs2268055 | 1.212 | 0.689 | 0.329 | 0.101 | -0.016 | 0.011 | 0.011 | 0.043 | 0.31 | 0.2 | 0.28 | 0.3 | 0.43 | 0.5 |
| 1 | 71473791 | rs2300168 | 1.031 | 0.569 | 0.366 | 0.107 | 0.014 | -0.064 | -0.064 | 0 | 0.19 | 0.12 | 0.37 | 0.44 | 0.16 | 0.2 |
| 1 | 71474782 | rs5693 | 0.505 | 0.494 | 0.534 | 0.04 | -0.01 | -0.01 | -0.01 | 0.218 | 0.27 | 0.47 | 0.2 | 0.23 | 0.23 | 0.14 |
| 1 | 71475063 | rs5691 | 1.62 | 0.503 | 0.406 | 0.095 | 0.001 | -0.021 | -0.021 | 0 | 0.17 | 0.14 | 0.07 | 0 | 0.09 | 0.02 |
| 1 | 71476872 | rs5682 | 1.391 | 0.613 | 0.204 | 0.189 | -0.038 | 0.017 | 0.017 | 0.073 | 0.3 | 0.14 | 0.27 | 0.3 | 0.43 | 0.5 |
| 1 | 71477539 | rs5675 | 1.217 | 0.6 | 0.178 | 0.204 | -0.018 | -0.151 | -0.151 | 0.074 | 0.18 | 0.01 | 0.36 | 0.44 | 0.16 | 0.2 |
| 1 | 71486202 | rs997997 | 1.203 | 0.871 | 0.38 | 0.25 | -0.03 | 0.017 | 0.017 | 0.061 | 0.3 | 0.15 | 0.27 | 0.3 | 0.43 | 0.5 |
| 1 | 71494365 | rs8179390 | 0.801 | 0.578 | 0.393 | 0.194 | -0.03 | 0.037 | 0.037 | 0.756 | 0.37 | 0.21 | 0.31 | 0.3 | 0.51 | 0.55 |
| 1 | 71499286 | rs2300179 | 0.212 | 0.46 | 0.764 | 0.334 | 0.041 | 0.22 | 0.22 | 0 | 0.05 | 0.22 | 0.08 | 0.24 | 0.07 | 0.1 |
| 1 | 71503105 | rs2817864 | 0.781 | 0.504 | 0.592 | -0.186 | -0.026 | 0.031 | 0.031 | 0.09 | 0.37 | 0.26 | 0.3 | 0.29 | 0.47 | 0.49 |
| 1 | 71509107 | rs10889906 | 0.226 | 0.247 | 0.613 | -0.474 | 0.02 | 0.036 | 0.036 | 4.955 | 0.6 | 0.25 | 0.72 | 0.97 | 0.7 | 0.82 |
| 1 | 71510419 | rs2050065 | -0.405 | 0.184 | 0.898 | -0.554 | -0.007 | -0.132 | -0.132 | 0.367 | 0.77 | 0.53 | 0.82 | 0.97 | 0.81 | 0.85 |
| 1 | 71512160 | rs2817867 | 0.881 | 0.148 | 0.824 | -0.513 | -0.024 | -0.111 | -0.111 | 0.623 | 0.18 | 0.4 | 0.11 | 0 | 0.1 | 0.03 |
| 1 | 71527548 | rs4520371 | 1.236 | 0.671 | 0.776 | -0.148 | -0.073 | 0.007 | 0.007 | 0.575 | 0.34 | 0.09 | 0.36 | 0.38 | 0.45 | 0.48 |
| 1 | 71544839 | rs10128001 | 1.268 | 0.595 | 1.175 | 0.295 | -0.023 | -0.088 | -0.088 | 0.255 | 0.16 | 0.28 | 0.08 | 0 | 0.09 | 0.01 |
| 1 | 71545970 | rs12035210 | 0.621 | 0.599 | 1.072 | 0.326 | 0.007 | -0.022 | -0.022 | 0.115 | 0.26 | 0.14 | 0.37 | 0.59 | 0.26 | 0.34 |
| 1 | 71548960 | rs17090804 | 1.037 | 0.59 | 1.185 | -0.235 | -0.015 | -0.075 | -0.075 | 1.607 | 0.17 | 0.28 | 0.09 | 0 | 0.11 | 0.03 |
| 1 | 71556167 | rs7531139 | 1.254 | 0.592 | 1.018 | -0.18 | 0 | -0.174 | -0.174 | 0 | 0.21 | 0.01 | 0.36 | 0.41 | 0.16 | 0.23 |
| 1 | 71560350 | rs12409826 | 1.218 | 0.555 | 1.141 | -0.282 | 0.01 | 0.003 | 0.003 | 1.103 | 0.17 | 0.11 | 0.09 | 0 | 0.1 | 0.03 |
| 1 | 71620851 | rs12142055 | 1.654 | 0.742 | 0.865 | -0.477 | -0.01 | 0.007 | 0.007 | 0 | 0.05 | 0.03 | 0.09 | 0.03 | 0.1 | 0.18 |
| 1 | 71632687 | rs1418449 | -0.323 | 0.72 | 0.946 | -0.486 | -0.008 | 0.031 | 0.031 | 2.142 | 0.69 | 0.82 | 0.47 | 0.26 | 0.63 | 0.48 |
| 1 | 71654508 | rs17090906 | 1.587 | 0.384 | 1.11 | -0.517 | -0.013 | -0.066 | -0.066 | 1.868 | 0.15 | 0.2 | 0.07 | 0 | 0.09 | 0.01 |
| 1 | 71658888 | rs2796200 | 0.856 | 0.404 | 1.221 | -0.56 | -0.038 | -0.11 | -0.11 | 0 | 0.18 | 0.41 | 0.1 | 0 | 0.08 | 0.02 |
| 1 | 71676720 | rs2639336 | -0.362 | 0.422 | 1.215 | -0.504 | 0.034 | 0.085 | 0.085 | 0.07 | 0.9 | 0.86 | 0.83 | 0.67 | 0.8 | 0.7 |
| 1 | 71694861 | rs2796188 | 0.452 | 0.264 | 1.155 | -0.72 | 0 | -0.007 | -0.007 | 0.624 | 0.52 | 0.5 | 0.39 | 0.26 | 0.53 | 0.46 |
| 1 | 71695806 | rs1891417 | 1.038 | -0.056 | 1.045 | -0.691 | -0.022 | -0.075 | -0.075 | 1.654 | 0.17 | 0.28 | 0.09 | 0 | 0.08 | 0.02 |
| 1 | 71696940 | rs10493483 | 1.523 | 0.064 | 0.466 | -0.571 | -0.013 | -0.066 | -0.066 | 0 | 0.15 | 0.21 | 0.07 | 0 | 0.09 | 0.01 |
| 1 | 71712223 | rs12131265 | 1.073 | 0.447 | 0.216 | -0.434 | -0.079 | 0.068 | 0.068 | 0.568 | 0.34 | 0.1 | 0.29 | 0.26 | 0.46 | 0.44 |
| 1 | 71728900 | rs11209757 | -0.461 | 0.294 | 0.156 | -0.466 | 0 | -0.002 | -0.002 | 0 | 0.42 | 0.41 | 0.61 | 0.75 | 0.43 | 0.54 |
| 1 | 71734612 | rs6673749 | 1.342 | 0.285 | 0.229 | -0.312 | -0.008 | -0.067 | -0.067 | 0.006 | 0.12 | 0.24 | 0.07 | 0 | 0.07 | 0.01 |
| 1 | 71740028 | rs962231 | 0.673 | 0.2 | 0.114 | -0.458 | 0.008 | 0.072 | 0.072 | 0.087 | 0.24 | 0.09 | 0.15 | 0.05 | 0.2 | 0.2 |
| 1 | 71740325 | rs7527741 | 0.582 | 0.218 | 0.085 | -0.42 | 0.005 | -0.002 | -0.002 | 0 | 0.24 | 0.11 | 0.45 | 0.7 | 0.27 | 0.36 |
| 1 | 71750361 | rs12749022 | 0.48 | 0.171 | 0.241 | -0.431 | 0.033 | 0.063 | 0.063 | 0.844 | 0.55 | 0.17 | 0.68 | 0.66 | 0.64 | 0.66 |
| 1 | 71762203 | rs6695355 | 0.445 | 0.228 | 0.055 | -0.379 | 0.005 | 0.001 | 0.001 | 0.176 | 0.24 | 0.19 | 0.46 | 0.7 | 0.27 | 0.36 |
| 1 | 71769465 | rs12048727 | -0.198 | 0.177 | 0.027 | -0.357 | -0.008 | -0.019 | -0.019 | 0 | 0.37 | 0.43 | 0.53 | 0.7 | 0.33 | 0.37 |
| 1 | 71777548 | rs17091107 | 1.255 | 0.326 | -0.55 | -0.189 | -0.005 | -0.161 | -0.161 | 0 | 0.19 | 0.01 | 0.38 | 0.41 | 0.18 | 0.22 |
| 1 | 71781585 | rs1506452 | 1.035 | 0.446 | -0.419 | -0.983 | -0.027 | 0.034 | 0.034 | 0.188 | 0.36 | 0.22 | 0.3 | 0.25 | 0.47 | 0.44 |
| 1 | 71799403 | rs2174602 | -0.389 | 0.477 | -0.543 | -0.242 | 0.008 | 0.019 | 0.019 | 0 | 0.36 | 0.3 | 0.52 | 0.7 | 0.33 | 0.37 |
| 1 | 71823940 | rs11209773 | 0.376 | 0.544 | -1.356 | 0.34 | -0.033 | -0.128 | -0.128 | 0.058 | 0.23 | 0.02 | 0.37 | 0.6 | 0.19 | 0.23 |
| 1 | 71830161 | rs2163502 | 0.663 | 0.535 | -1.8 | -0.434 | -0.099 | 0.094 | 0.094 | 0.837 | 0.37 | 0.09 | 0.35 | 0.24 | 0.54 | 0.51 |
| 1 | 71840498 | rs1517754 | 0.113 | 0.141 | -0.863 | -0.581 | 0.009 | 0.021 | 0.021 | 5.49 | 0.34 | 0.29 | 0.18 | 0.13 | 0.25 | 0.16 |
| 1 | 71849037 | rs12127469 | 0.749 | 0.402 | -0.941 | -0.533 | -0.149 | 0.085 | 0.085 | 1.11 | 0.36 | 0.04 | 0.35 | 0.24 | 0.54 | 0.51 |
| 1 | 71850820 | rs2221513 | 1.474 | 0.825 | -1.02 | -0.775 | -0.029 | 0.05 | 0.05 | 0 | 0.11 | 0.02 | 0.09 | 0 | 0.17 | 0.03 |
| 1 | 71876406 | rs12405821 | 1.251 | 0.477 | -0.547 | -0.836 | 0.002 | -0.009 | -0.009 | 0.982 | 0.1 | 0.12 | 0.07 | 0 | 0.06 | 0.02 |
| 1 | 71880016 | rs1506457 | -0.256 | 0.42 | -0.377 | -0.803 | 0.035 | 0.063 | 0.063 | 0.055 | 0.5 | 0.66 | 0.54 | 0.38 | 0.61 | 0.64 |
| 1 | 71902018 | rs7519228 | 1.142 | 0.405 | -0.456 | -0.901 | 0.001 | 0.027 | 0.027 | 0.339 | 0.2 | 0.1 | 0.1 | 0 | 0.17 | 0.07 |
| 1 | 71905723 | rs17091221 | 1.152 | 0.518 | -0.506 | -0.755 | -0.005 | 0.013 | 0.013 | 0.161 | 0.1 | 0.19 | 0.08 | 0.12 | 0.07 | 0.11 |
| 1 | 71912912 | rs12123319 | -0.692 | 0.606 | -0.566 | -0.497 | -0.01 | 0.016 | 0.016 | 0.005 | 0.74 | 0.62 | 0.66 | 0.4 | 0.79 | 0.79 |
| 1 | 71920314 | rs7550923 | 0.534 | 0.607 | -0.656 | -0.54 | -0.039 | 0.085 | 0.085 | 0.048 | 0.41 | 0.18 | 0.44 | 0.25 | 0.54 | 0.52 |
| 1 | 71924061 | rs12126700 | 0.787 | 0.581 | -0.971 | -0.509 | -0.102 | 0.061 | 0.061 | 0.083 | 0.34 | 0.03 | 0.36 | 0.24 | 0.4 | 0.46 |
| 1 | 71929289 | rs977145 | -0.774 | 0.427 | -0.758 | -0.408 | 0.007 | -0.021 | -0.021 | 0.155 | 0.75 | 0.74 | 0.67 | 0.4 | 0.78 | 0.79 |
| 1 | 71934229 | rs12135145 | -0.681 | 0.426 | -0.757 | -0.447 | 0.005 | -0.01 | -0.01 | 0.031 | 0.74 | 0.69 | 0.67 | 0.38 | 0.78 | 0.72 |
| 1 | 71939755 | rs17562024 | 0.459 | 0.83 | -1.325 | -0.519 | -0.098 | 0.074 | 0.074 | 0.242 | 0.38 | 0.05 | 0.4 | 0.27 | 0.46 | 0.52 |
| 1 | 71949903 | rs17091269 | 0.068 | 0.801 | -0.314 | 0.899 | -0.004 | -0.046 | -0.046 | 0 | 0.22 | 0.1 | 0.4 | 0.4 | 0.22 | 0.22 |
| 1 | 71960291 | rs357221 | 0.064 | 0.834 | -0.897 | 0.584 | -0.022 | -0.083 | -0.083 | 0 | 0.73 | 0.91 | 0.51 | 0.38 | 0.73 | 0.64 |
| 1 | 71963931 | rs6687400 | -0.356 | 0.924 | -0.168 | 0.317 | 0.006 | -0.005 | -0.005 | 0.68 | 0.07 | 0.1 | 0.04 | 0.04 | 0.09 | 0.08 |
| 1 | 71969514 | rs357200 | 0.695 | 0.915 | -0.071 | 0.426 | -0.001 | -0.004 | -0.004 | 0 | 0.65 | 0.71 | 0.46 | 0.34 | 0.64 | 0.57 |
| 1 | 71987496 | rs12754625 | -1.12 | 0.863 | 0.064 | 0.382 | 0.004 | -0.011 | -0.011 | 0 | 0.09 | 0.13 | 0.08 | 0.05 | 0.1 | 0.11 |
| 1 | 71996090 | rs357237 | 0.939 | 0.579 | 0.706 | 0.131 | -0.004 | -0.002 | -0.002 | 0 | 0.6 | 0.47 | 0.39 | 0.34 | 0.63 | 0.54 |
| 1 | 72007798 | rs17091348 | 0.043 | -0.117 | 0.515 | 0.121 | 0.029 | -0.078 | -0.078 | 0 | 0.31 | 0.11 | 0.48 | 0.61 | 0.25 | 0.34 |
| 1 | 72012933 | rs357233 | 0.349 | -0.103 | 0.498 | 0.128 | -0.025 | 0.011 | 0.011 | 0 | 0.63 | 0.56 | 0.45 | 0.39 | 0.74 | 0.64 |
| 1 | 72025716 | rs357210 | 0.403 | 0.223 | 0.24 | 0.075 | 0.005 | 0.001 | 0.001 | 0 | 0.62 | 0.64 | 0.44 | 0.33 | 0.67 | 0.58 |
| 1 | 72034921 | rs6424431 | 0.538 | 0.039 | 0.657 | 0.008 | -0.012 | -0.013 | -0.013 | 1.332 | 0.61 | 0.54 | 0.45 | 0.27 | 0.69 | 0.61 |
| 1 | 72038497 | rs1335223 | 0.542 | 0.04 | 0.687 | 0.009 | -0.005 | -0.006 | -0.006 | 0 | 0.61 | 0.58 | 0.45 | 0.27 | 0.69 | 0.61 |
| 1 | 72038923 | rs9919188 | -0.89 | 0.006 | 0.712 | 0.119 | -0.02 | 0.002 | 0.002 | 0 | 0.07 | 0.12 | 0.18 | 0.08 | 0.02 | 0.1 |

| | | | | | | | | | | | | | | | | |
|---|---|---|---|---|---|---|---|---|---|---|---|---|---|---|---|---|
| 1 | 72058405 | rs1334005 | 0.199 | 0.015 | 1.078 | 0.461 | 0.037 | 0.168 | 0.168 | 0.366 | 0.95 | 0.66 | 0.93 | 0.85 | 1 | 0.9 |
| 1 | 72061784 | rs2007130 | 0.761 | -0.022 | 1.134 | 0.443 | 0.032 | 0.171 | 0.171 | 1.206 | 0.93 | 0.56 | 0.91 | 0.85 | 1 | 0.9 |
| 1 | 72066623 | rs11209800 | -0.32 | -0.169 | 0.853 | 0.295 | -0.032 | -0.079 | -0.079 | 0.088 | 0.34 | 0.05 | 0.49 | 0.61 | 0.31 | 0.3 |
| 1 | 72070767 | rs12757229 | 0.586 | -0.129 | 0.884 | 0.246 | -0.015 | -0.017 | -0.017 | 0.181 | 0.59 | 0.47 | 0.42 | 0.26 | 0.69 | 0.6 |
| 1 | 72077549 | rs928615 | 0.948 | -0.274 | 0.799 | 0.15 | 0.053 | 0.064 | 0.064 | 0 | 0.54 | 0.26 | 0.4 | 0.26 | 0.68 | 0.58 |
| 1 | 72080604 | rs1762732 | -0.715 | -0.219 | 0.189 | 0.335 | 0.008 | 0 | 0 | 0.191 | 0.42 | 0.39 | 0.45 | 0.6 | 0.36 | 0.42 |
| 1 | 72090075 | rs6696052 | -0.579 | -0.083 | 0.067 | 0.599 | 0.032 | 0.008 | 0.008 | 0 | 0.42 | 0.24 | 0.54 | 0.63 | 0.35 | 0.41 |
| 1 | 72104447 | rs1041676 | 0.268 | -0.033 | -0.131 | 0.531 | -0.041 | -0.165 | -0.165 | 0 | 0.31 | 0.02 | 0.34 | 0.45 | 0.29 | 0.22 |
| 1 | 72121585 | rs954299 | 0.638 | 0.12 | -0.004 | 0.416 | -0.002 | 0.009 | 0.009 | 0 | 0.48 | 0.53 | 0.37 | 0.37 | 0.5 | 0.5 |
| 1 | 72126388 | rs2422014 | 0.424 | 0.093 | -0.184 | 0.26 | -0.004 | -0.002 | -0.002 | 0 | 0.49 | 0.64 | 0.41 | 0.46 | 0.51 | 0.59 |
| 1 | 72136346 | rs10889924 | 0.188 | -0.09 | -0.163 | 0.392 | -0.004 | -0.012 | -0.012 | 0 | 0.55 | 0.68 | 0.56 | 0.52 | 0.55 | 0.64 |
| 1 | 72150052 | rs1591382 | 1.073 | -0.059 | 0.112 | 0.313 | 0.043 | 0.08 | 0.08 | 0 | 0.93 | 0.72 | 0.91 | 0.89 | 0.92 | 0.89 |
| 1 | 72206799 | rs4147264 | -0.134 | -0.211 | -0.037 | 0.339 | -0.002 | 0.005 | 0.005 | 0 | 0.41 | 0.26 | 0.54 | 0.6 | 0.38 | 0.34 |
| 1 | 72208917 | rs17091704 | -0.295 | -0.242 | 0.117 | 0.356 | 0.004 | 0.08 | 0.08 | 0 | 0.05 | 0.16 | 0.07 | 0.15 | 0.05 | 0.13 |
| 1 | 72217553 | rs10789324 | -0.716 | -0.255 | 0.086 | 0.314 | -0.002 | -0.016 | -0.016 | 0 | 0.47 | 0.54 | 0.63 | 0.75 | 0.43 | 0.47 |
| 1 | 72222487 | rs12740031 | -0.098 | -0.267 | -0.007 | 0.343 | 0 | -0.01 | -0.01 | 0.157 | 0.4 | 0.22 | 0.46 | 0.55 | 0.37 | 0.29 |
| 1 | 72234070 | rs1599179 | 1.275 | -0.323 | -0.202 | 0.167 | -0.016 | 0.024 | 0.024 | 0 | 0.82 | 0.42 | 0.83 | 0.82 | 0.85 | 0.87 |
| 1 | 72239513 | rs4649946 | 0.285 | -0.312 | -0.101 | -0.173 | 0.015 | -0.085 | -0.085 | 1.117 | 0.88 | 0.56 | 0.9 | 1 | 0.9 | 0.96 |
| 1 | 72266725 | rs1580232 | 0.333 | -0.374 | -0.414 | -0.217 | -0.003 | -0.088 | -0.088 | 0 | 0.87 | 0.59 | 0.91 | 1 | 0.92 | 0.96 |
| 1 | 72267554 | rs1158573 | 1.177 | -0.295 | -1.42 | -0.158 | 0.017 | 0.007 | 0.007 | 1.084 | 0.78 | 0.15 | 0.73 | 0.81 | 0.86 | 0.79 |
| 1 | 72271264 | rs531071 | 1.186 | -0.475 | -1.823 | -0.182 | -0.06 | 0.083 | 0.083 | 0 | 0.38 | 0.13 | 0.28 | 0.24 | 0.47 | 0.47 |
| 1 | 72271300 | rs12031844 | 0.125 | -0.326 | -1.644 | 0.147 | -0.096 | -0.127 | -0.127 | 3.486 | 0.34 | 0.02 | 0.44 | 0.58 | 0.36 | 0.25 |
| 1 | 72282482 | rs12144209 | 0.539 | -0.597 | -1.046 | -0.115 | -0.018 | 0.063 | 0.063 | 0 | 0.39 | 0.24 | 0.38 | 0.22 | 0.42 | 0.4 |
| 1 | 72284206 | rs6424449 | 0.026 | -1.017 | -1.01 | 0.148 | -0.028 | 0.019 | 0.019 | 0 | 0.55 | 0.82 | 0.67 | 0.41 | 0.54 | 0.59 |
| 1 | 72288021 | rs1814054 | 0.31 | -1.144 | -0.836 | 0.22 | -0.001 | -0.134 | -0.134 | 0 | 0.32 | 0.05 | 0.15 | 0.5 | 0.26 | 0.29 |
| 1 | 72289671 | rs546741 | 0.157 | -0.889 | 0.261 | 0.225 | 0.022 | 0.013 | 0.013 | 0 | 0.52 | 0.64 | 0.71 | 0.38 | 0.67 | 0.56 |
| 1 | 72299344 | rs6683448 | 0.304 | -0.869 | 0.36 | 0.228 | 0.019 | 0.017 | 0.017 | 0.346 | 0.47 | 0.44 | 0.69 | 0.34 | 0.65 | 0.55 |
| 1 | 72309985 | rs513086 | 1.126 | -0.159 | 0.877 | 0.561 | 0.036 | -0.016 | -0.016 | 0 | 0.9 | 0.83 | 0.92 | 0.92 | 0.86 | 0.86 |
| 1 | 72331815 | rs591540 | -1.136 | -0.518 | 0.631 | 0.529 | -0.023 | 0.102 | 0.102 | 0.497 | 0.52 | 0.81 | 0.64 | 0.71 | 0.5 | 0.5 |
| 1 | 72336913 | rs17091873 | -1.256 | -0.156 | 0.308 | 0.539 | 0.032 | -0.011 | -0.011 | 0 | 0.09 | 0.16 | 0.1 | 0.08 | 0.15 | 0.14 |
| 1 | 72346331 | rs629079 | -0.166 | -0.097 | 0.284 | 0.399 | -0.022 | 0.001 | 0.001 | 0.2 | 0.38 | 0.47 | 0.26 | 0.61 | 0.3 | 0.35 |
| 1 | 72352237 | rs648556 | -0.089 | 0.004 | -0.369 | 0.448 | 0.015 | -0.003 | -0.003 | 1.454 | 0.38 | 0.34 | 0.25 | 0.61 | 0.3 | 0.35 |
| 1 | 72370307 | rs1765607 | -0.315 | 0.118 | -0.54 | 0.697 | -0.001 | 0.002 | 0.002 | 0 | 0.46 | 0.46 | 0.34 | 0.61 | 0.43 | 0.38 |
| 1 | 72381596 | rs601452 | -0.377 | 0.079 | -1.181 | 0.696 | 0.028 | -0.008 | -0.008 | 0.462 | 0.48 | 0.75 | 0.51 | 0.55 | 0.39 | 0.47 |
| 1 | 72384596 | rs1426173 | -0.533 | -0.321 | -1.119 | 0.5 | -0.017 | 0.029 | 0.029 | 0 | 0.59 | 0.79 | 0.63 | 0.63 | 0.4 | 0.59 |
| 1 | 72385590 | rs7520086 | 0.72 | -1.137 | -1.44 | 0.474 | 0.005 | -0.1 | -0.1 | 0.401 | 0.21 | 0.06 | 0.09 | 0.42 | 0.13 | 0.25 |
| 1 | 72388052 | rs2114214 | -1.185 | -1.147 | -1.445 | 0.513 | -0.017 | 0.044 | 0.044 | 0 | 0.59 | 0.77 | 0.63 | 0.64 | 0.4 | 0.59 |
| 1 | 72391457 | rs7527501 | 0.198 | -1.581 | -0.839 | 0.279 | 0.003 | 0.004 | 0.004 | 0.283 | 0.21 | 0.23 | 0.43 | 0.11 | 0.17 | 0.15 |
| 1 | 72398723 | rs3856029 | 0.155 | -1.588 | -0.73 | 0.276 | 0 | -0.001 | -0.001 | 0 | 0.21 | 0.25 | 0.43 | 0.11 | 0.17 | 0.15 |
| 1 | 72402030 | rs2768395 | -1.547 | -1.594 | -0.554 | 0.275 | -0.013 | 0.04 | 0.04 | 0 | 0.59 | 0.76 | 0.63 | 0.64 | 0.4 | 0.59 |
| 1 | 72402880 | rs2256913 | 0.275 | -1.595 | -0.556 | 0.378 | -0.031 | -0.031 | -0.031 | 1.471 | 0.16 | 0.45 | 0.11 | 0.12 | 0.1 | 0.19 |
| 1 | 72404727 | rs2114213 | -0.531 | -1.586 | -0.562 | 0.379 | -0.012 | 0.005 | 0.005 | 0 | 0.37 | 0.51 | 0.2 | 0.53 | 0.23 | 0.44 |
| 1 | 72407409 | rs7544722 | -1.488 | -1.613 | 0.204 | 0.356 | -0.006 | 0.033 | 0.033 | 0 | 0.58 | 0.7 | 0.62 | 0.64 | 0.4 | 0.59 |
| 1 | 72421519 | rs2768394 | -0.861 | -1.733 | 0.915 | 0.248 | -0.037 | -0.074 | -0.074 | 0 | 0.33 | 0.49 | 0.22 | 0.15 | 0.22 | 0.24 |
| 1 | 72424117 | rs7513441 | -0.783 | -1.73 | 0.924 | 0.25 | -0.021 | -0.046 | -0.046 | 0 | 0.33 | 0.41 | 0.21 | 0.15 | 0.22 | 0.24 |
| 1 | 72428834 | rs12024769 | -1.316 | -1.736 | 0.931 | 0.249 | -0.006 | -0.004 | -0.004 | 0 | 0.54 | 0.55 | 0.31 | 0.57 | 0.35 | 0.49 |
| 1 | 72433346 | rs1365208 | 0.73 | -1.769 | 0.863 | 0.201 | 0.018 | 0.049 | 0.049 | 1.185 | 0.16 | 0.05 | 0.11 | 0.03 | 0.12 | 0.05 |
| 1 | 72436403 | rs1016126 | -1.673 | -1.757 | 0.853 | 0.227 | -0.016 | 0.046 | 0.046 | 0 | 0.59 | 0.79 | 0.63 | 0.65 | 0.4 | 0.59 |
| 1 | 72436448 | rs17092004 | 0.508 | -1.717 | 0.655 | 0.227 | 0.016 | 0.024 | 0.024 | 1.06 | 0.17 | 0.11 | 0.12 | 0.03 | 0.12 | 0.05 |
| 1 | 72442018 | rs10518394 | 0.451 | -1.645 | 0.633 | 0.301 | 0.002 | 0.002 | 0.002 | 0 | 0.15 | 0.14 | 0.09 | 0.12 | 0.1 | 0.19 |
| 1 | 72444128 | rs11209878 | 0.918 | -1.599 | 0.519 | 0.347 | 0.003 | -0.106 | -0.106 | 0 | 0.21 | 0.04 | 0.09 | 0.42 | 0.13 | 0.25 |
| 1 | 72455331 | rs11802650 | -0.004 | -1.872 | 0.835 | 0.103 | -0.023 | -0.025 | -0.025 | 0.074 | 0.18 | 0.27 | 0.1 | 0.12 | 0.11 | 0.2 |
| 1 | 72458685 | rs6661979 | 0.344 | -1.922 | 1.129 | 0.139 | 0 | -0.001 | -0.001 | 0.019 | 0.26 | 0.24 | 0.33 | 0.5 | 0.18 | 0.35 |
| 1 | 72458866 | rs2029155 | -0.665 | -1.992 | 1.164 | 0.092 | 0.023 | 0.01 | 0.01 | 0 | 0.43 | 0.32 | 0.44 | 0.53 | 0.3 | 0.4 |
| 1 | 72461042 | rs1937186 | -0.605 | -2.001 | 1.247 | 0.09 | -0.026 | -0.007 | -0.007 | 0 | 0.22 | 0.4 | 0.36 | 0.2 | 0.16 | 0.29 |
| 1 | 72469148 | rs1954600 | -0.681 | -2 | 1.108 | 0.09 | -0.032 | -0.009 | -0.009 | 0 | 0.22 | 0.44 | 0.36 | 0.2 | 0.16 | 0.29 |
| 1 | 72478099 | rs12565657 | 0.417 | -1.739 | 1.013 | 0.185 | -0.002 | 0.003 | 0.003 | 0 | 0.07 | 0.12 | 0.06 | 0.1 | 0.06 | 0.12 |
| 1 | 72478194 | rs3102916 | -0.959 | -1.754 | 1.121 | 0.188 | 0.04 | 0.112 | 0.112 | 0 | 0.12 | 0.5 | 0.34 | 0.18 | 0.11 | 0.21 |
| 1 | 72480008 | rs12407054 | 0.457 | -1.884 | 0.942 | 0.096 | 0.036 | 0.049 | 0.049 | 1.85 | 0.17 | 0.09 | 0.1 | 0.03 | 0.1 | 0.03 |
| 1 | 72488815 | rs3102906 | -1.2 | -1.861 | 1.019 | 0.112 | -0.024 | -0.02 | -0.02 | 0 | 0.29 | 0.65 | 0.45 | 0.22 | 0.21 | 0.24 |
| 1 | 72492583 | rs12023418 | 0.457 | -1.903 | 0.859 | 0.146 | 0.015 | -0.154 | -0.154 | 0.294 | 0.32 | 0.03 | 0.1 | 0.44 | 0.18 | 0.35 |
| 1 | 72495217 | rs12403024 | 0.49 | -1.888 | 0.878 | 0.139 | 0.021 | 0.02 | 0.02 | 2.027 | 0.17 | 0.11 | 0.11 | 0.03 | 0.1 | 0.03 |
| 1 | 72499082 | rs3102918 | -1.544 | -1.889 | 0.871 | 0.141 | -0.015 | 0.031 | 0.031 | 0.1 | 0.62 | 0.75 | 0.56 | 0.65 | 0.4 | 0.6 |
| 1 | 72499912 | rs3128557 | -1.492 | -1.868 | 0.734 | 0.151 | -0.005 | 0.013 | 0.013 | 0.078 | 0.61 | 0.7 | 0.55 | 0.66 | 0.4 | 0.6 |
| 1 | 72505250 | rs11803665 | 0.158 | -1.474 | 1.091 | -0.033 | -0.002 | 0.004 | 0.004 | 0 | 0.07 | 0.13 | 0.06 | 0.1 | 0.06 | 0.12 |
| 1 | 72505345 | rs1486110 | 0.25 | -1.466 | 0.716 | -0.077 | 0.034 | 0.049 | 0.049 | 2.347 | 0.16 | 0.08 | 0.1 | 0.03 | 0.1 | 0.03 |
| 1 | 72508296 | rs1486090 | -1.245 | -0.66 | 0.877 | 0.386 | 0.024 | 0.087 | 0.087 | 0 | 0.12 | 0.58 | 0.35 | 0.18 | 0.11 | 0.21 |
| 1 | 72509490 | rs1486087 | -0.112 | -0.383 | 0.228 | -0.2 | 0.002 | -0.045 | -0.045 | 0 | 0.39 | 0.45 | 0.45 | 0.13 | 0.58 | 0.39 |
| 1 | 72516287 | rs1026997 | -0.392 | -0.213 | -0.016 | -0.979 | -0.011 | 0.038 | 0.038 | 0 | 0.44 | 0.4 | 0.53 | 0.13 | 0.49 | 0.33 |

| | | | | | | | | | | | | | | | | |
|---|---|---|---|---|---|---|---|---|---|---|---|---|---|---|---|---|
| 1 | 72517928 | rs975480 | -0.767 | -0.355 | 0.572 | -0.072 | 0.013 | 0.021 | 0.021 | 0 | 0.53 | 0.7 | 0.59 | 0.41 | 0.56 | 0.52 |
| 1 | 72520941 | rs1486084 | 0.575 | -0.421 | 0.699 | -0.089 | 0.01 | 0.025 | 0.025 | 0.033 | 0.46 | 0.19 | 0.4 | 0.59 | 0.44 | 0.48 |
| 1 | 72531925 | rs2821267 | 0.246 | -0.408 | 1.204 | -0.025 | 0.017 | 0.151 | 0.151 | 0.11 | 0.08 | 0.18 | 0.06 | 0.28 | 0.07 | 0.18 |
| 1 | 72532506 | rs2200485 | -0.375 | -0.421 | 1.264 | -0.031 | 0.023 | 0.188 | 0.188 | 0.797 | 0.09 | 0.28 | 0.06 | 0.28 | 0.07 | 0.18 |
| 1 | 72535536 | rs12024008 | 0.375 | -0.435 | 1.188 | -0.022 | -0.002 | 0.069 | 0.069 | 0.215 | 0.07 | 0.09 | 0.04 | 0.28 | 0.05 | 0.18 |
| 1 | 72549836 | rs988421 | -0.051 | -0.514 | 1.064 | -0.183 | -0.024 | 0.089 | 0.089 | 0 | 0.43 | 0.36 | 0.53 | 0.13 | 0.49 | 0.34 |
| 1 | 72553991 | rs12091740 | 0.01 | -0.542 | 1.021 | -0.197 | -0.04 | 0.134 | 0.134 | 0 | 0.42 | 0.27 | 0.51 | 0.13 | 0.49 | 0.34 |
| 1 | 72555807 | rs2821257 | -0.494 | -0.826 | 0.901 | -0.088 | -0.037 | 0.071 | 0.071 | 0 | 0.86 | 0.78 | 0.95 | 0.67 | 0.97 | 0.89 |
| 1 | 72569309 | rs10493493 | 0.521 | 0.093 | 0.501 | -0.145 | -0.04 | 0.029 | 0.029 | 0.346 | 0.22 | 0.07 | 0.22 | 0.17 | 0.27 | 0.32 |
| 1 | 72571364 | rs1486091 | -1.252 | -0.118 | 0.578 | -0.046 | 0.012 | 0.008 | 0.008 | 0.102 | 0.5 | 0.73 | 0.44 | 0.51 | 0.51 | 0.47 |
| 1 | 72575560 | rs2821248 | 0.762 | -0.178 | 0.652 | -0.098 | 0.016 | -0.036 | -0.036 | 0 | 0.11 | 0.18 | 0.1 | 0.01 | 0.19 | 0.03 |
| 1 | 72581966 | rs2630397 | -0.513 | -0.274 | 0.766 | -0.058 | -0.011 | 0.016 | 0.016 | 0.027 | 0.27 | 0.45 | 0.19 | 0.34 | 0.23 | 0.17 |
| 1 | 72590193 | rs2821239 | -1.277 | -0.262 | 0.773 | -0.109 | 0.004 | 0.009 | 0.009 | 0 | 0.49 | 0.64 | 0.43 | 0.5 | 0.5 | 0.5 |
| 1 | 72595125 | rs2821236 | -0.434 | -0.206 | 0.492 | 0.318 | -0.006 | 0.01 | 0.01 | 0 | 0.27 | 0.35 | 0.18 | 0.34 | 0.23 | 0.17 |
| 1 | 72615118 | rs11587434 | 1.406 | -0.544 | -0.557 | -0.496 | -0.034 | 0.088 | 0.088 | 0.016 | 0.35 | 0.02 | 0.26 | 0.14 | 0.28 | 0.27 |
| 1 | 72628347 | rs12755450 | 1.179 | 0.125 | -0.042 | -0.616 | -0.049 | 0.049 | 0.049 | 0 | 0.09 | 0.01 | 0.08 | 0 | 0.19 | 0.03 |
| 1 | 72629795 | rs2821296 | 0.524 | 0.024 | 0.551 | -0.691 | 0.005 | -0.018 | -0.018 | 0 | 0.64 | 0.38 | 0.64 | 0.82 | 0.52 | 0.66 |
| 1 | 72654730 | rs2298152 | 0.64 | -0.002 | 0.523 | -0.669 | -0.04 | 0.03 | 0.03 | 0.346 | 0.21 | 0.04 | 0.22 | 0.16 | 0.26 | 0.31 |
| 1 | 72664938 | rs1342874 | 0.733 | 0.036 | 0.728 | -0.633 | -0.037 | 0.027 | 0.027 | 0.61 | 0.2 | 0.06 | 0.22 | 0.16 | 0.26 | 0.31 |
| 1 | 72701100 | rs2797543 | -0.79 | -0.225 | 0.919 | -0.756 | -0.013 | 0.013 | 0.013 | 0 | 0.36 | 0.71 | 0.37 | 0.18 | 0.48 | 0.34 |
| 1 | 72703860 | rs1194269 | -0.666 | -0.157 | 0.662 | -0.723 | 0.035 | -0.052 | -0.052 | 8.566 | 0.34 | 0.46 | 0.35 | 0.18 | 0.48 | 0.34 |
| 1 | 72723863 | rs2000228 | 0.13 | -0.177 | 0.017 | -1.309 | -0.084 | 0.054 | 0.054 | 0.067 | 0.26 | 0.02 | 0.26 | 0.12 | 0.3 | 0.34 |
| 1 | 72729142 | rs1620977 | -1.953 | -0.534 | -0.278 | -1.672 | -0.09 | 0.056 | 0.056 | 0 | 0.7 | 0.99 | 0.74 | 0.87 | 0.71 | 0.81 |
| 1 | 72735458 | rs12137231 | -0.532 | -0.788 | 0.005 | -2.157 | -0.151 | 0.136 | 0.136 | 0.52 | 0.34 | 0.04 | 0.33 | 0.12 | 0.48 | 0.4 |
| 1 | 72741382 | rs11584542 | 0.228 | -0.827 | 0.022 | -2.139 | -0.059 | 0.069 | 0.069 | 0.064 | 0.26 | 0.03 | 0.26 | 0.12 | 0.3 | 0.34 |
| 1 | 72744144 | rs3101340 | -0.955 | -0.829 | 0.235 | -2.111 | 0.019 | -0.072 | -0.072 | 0.15 | 0.39 | 0.5 | 0.37 | 0.14 | 0.48 | 0.4 |
| 1 | 72744996 | rs1870676 | 0.784 | -0.867 | 0.255 | -2.122 | -0.011 | 0.033 | 0.033 | 0 | 0.59 | 0.3 | 0.61 | 0.86 | 0.52 | 0.6 |
| 1 | 72751185 | rs3101336 | 0.791 | -1.158 | 0.664 | -1.963 | 0.031 | -0.107 | -0.107 | 0.025 | 0.71 | 0.53 | 0.7 | 0.93 | 0.64 | 0.62 |
| 1 | 72765116 | rs2568958 | 0.791 | -1.158 | 0.664 | -1.963 | 0.031 | -0.107 | -0.107 | 0 | 0.71 | 0.53 | 0.7 | 0.93 | 0.64 | 0.62 |
| 1 | 72812440 | rs2815752 | 0.777 | -1.159 | 0.664 | -1.964 | 0.032 | -0.107 | -0.107 | 0 | 0.71 | 0.53 | 0.7 | 0.93 | 0.64 | 0.62 |
| 1 | 72814783 | rs2815749 | -0.645 | -1.1 | 0.718 | -1.952 | 0.104 | -0.056 | -0.056 | 0.816 | 0.89 | 0.63 | 0.9 | 0.98 | 0.82 | 0.94 |
| 1 | 72818226 | rs17526561 | 0.36 | -1.153 | 0.615 | -1.988 | -0.004 | 0.029 | 0.029 | 0 | 0.16 | 0.08 | 0.18 | 0.05 | 0.17 | 0.31 |
| 1 | 72819612 | rs2012697 | 0.976 | -1.157 | 0.645 | -1.991 | 0.028 | -0.098 | -0.098 | 9.627 | 0.71 | 0.51 | 0.71 | 0.93 | 0.64 | 0.62 |
| 1 | 72826172 | rs12143898 | 0.004 | -1.154 | 0.309 | -1.984 | -0.005 | 0.038 | 0.038 | 0 | 0.18 | 0.09 | 0.19 | 0.05 | 0.18 | 0.32 |
| 1 | 72828221 | rs10889947 | 1.051 | -1.122 | 0.367 | -2 | -0.002 | -0.011 | -0.011 | 0 | 0.69 | 0.28 | 0.68 | 0.93 | 0.63 | 0.62 |
| 1 | 72829530 | rs11581403 | 0.299 | -1.146 | -0.544 | -1.933 | -0.004 | 0.029 | 0.029 | 0 | 0.16 | 0.08 | 0.18 | 0.05 | 0.17 | 0.31 |
| 1 | 72834503 | rs2613507 | -0.901 | -1.106 | -0.482 | -1.919 | 0.053 | -0.05 | -0.05 | 0 | 0.9 | 0.79 | 0.91 | 0.98 | 0.82 | 0.94 |
| 1 | 72837239 | rs7531118 | 1.478 | -1.381 | -0.893 | -1.929 | -0.055 | 0.096 | 0.096 | 0.535 | 0.46 | 0.05 | 0.38 | 0.26 | 0.55 | 0.29 |
| 1 | 72838529 | rs1993709 | -0.326 | 0.103 | 0.32 | -1.891 | 0.033 | -0.038 | -0.038 | 0 | 0.91 | 0.87 | 0.92 | 0.98 | 0.82 | 0.94 |
| 1 | 72843946 | rs12120564 | -0.259 | 0.026 | 0.097 | -1.826 | -0.008 | 0.034 | 0.034 | 0 | 0.16 | 0.07 | 0.18 | 0.05 | 0.17 | 0.31 |
| 1 | 72846190 | rs1599339 | -0.116 | -0.221 | 0.38 | -1.757 | 0.039 | -0.051 | -0.051 | 0 | 0.88 | 0.8 | 0.91 | 0.98 | 0.81 | 0.93 |
| 1 | 72861847 | rs12123766 | -1.044 | -0.709 | 0.02 | -2.076 | 0.058 | 0.092 | 0.092 | 0 | 0.24 | 0.09 | 0.21 | 0.06 | 0.16 | 0.26 |
| 1 | 72866574 | rs1599334 | 2.109 | -0.779 | 0.467 | -2.218 | -0.008 | -0.077 | -0.077 | 0 | 0.57 | 0.46 | 0.65 | 0.92 | 0.61 | 0.66 |
| 1 | 72866847 | rs2815767 | 1.816 | -0.77 | 0.497 | -2.214 | 0.004 | -0.012 | -0.012 | 0 | 0.6 | 0.62 | 0.68 | 0.92 | 0.62 | 0.66 |
| 1 | 72876346 | rs290702 | 1.524 | -0.781 | 0.531 | -2.219 | 0.008 | 0.034 | 0.034 | 0.117 | 0.63 | 0.73 | 0.7 | 0.92 | 0.65 | 0.67 |
| 1 | 72884712 | rs10789339 | 2.084 | -0.78 | 0.562 | -2.219 | -0.006 | -0.064 | -0.064 | 0.527 | 0.57 | 0.49 | 0.65 | 0.92 | 0.61 | 0.66 |
| 1 | 72885281 | rs2590942 | 0.132 | -0.764 | 0.555 | -2.416 | 0.017 | -0.033 | -0.033 | 0.068 | 0.12 | 0.12 | 0.09 | 0.02 | 0.19 | 0.07 |
| 1 | 72913499 | rs7535539 | 2.047 | -0.767 | 0.51 | -2.403 | -0.011 | -0.115 | -0.115 | 0 | 0.57 | 0.48 | 0.65 | 0.96 | 0.61 | 0.66 |
| 1 | 72940273 | rs10789340 | 1.76 | -0.797 | 0.342 | -2.419 | -0.002 | -0.132 | -0.132 | 0 | 0.65 | 0.47 | 0.69 | 0.96 | 0.67 | 0.67 |
| 1 | 72947342 | rs12033462 | 1.647 | -0.68 | 0.283 | -2.356 | -0.007 | -0.093 | -0.093 | 1.869 | 0.61 | 0.55 | 0.67 | 0.96 | 0.64 | 0.66 |
| 1 | 72953251 | rs782207 | -1.713 | -0.606 | -0.035 | -2.239 | 0 | -0.022 | -0.022 | 0 | 0.37 | 0.39 | 0.31 | 0.08 | 0.35 | 0.33 |
| 1 | 72956037 | rs812462 | -1.574 | 0.002 | 0.583 | -1.63 | -0.036 | -0.054 | -0.054 | 0 | 0.2 | 0.34 | 0.1 | 0.06 | 0.11 | 0.25 |
| 1 | 72960011 | rs1445591 | -1.702 | 0.826 | 0.074 | -1.638 | -0.038 | 0.074 | 0.074 | 0 | 0.22 | 0.08 | 0.18 | 0.03 | 0.25 | 0.25 |
| 1 | 72960531 | rs7551627 | -1.34 | 0.421 | 0.694 | -1.235 | -0.002 | -0.015 | -0.015 | 0.471 | 0.14 | 0.15 | 0.16 | 0.02 | 0.19 | 0.13 |
| 1 | 72975877 | rs782228 | -2.189 | 0.205 | 1.671 | -1.302 | 0.016 | -0.122 | -0.122 | 0.141 | 0.2 | 0.42 | 0.2 | 0.02 | 0.21 | 0.13 |
| 1 | 72987229 | rs12060746 | -2.101 | 0.097 | 1.887 | -1.201 | -0.016 | 0.022 | 0.022 | 0.116 | 0.08 | 0.24 | 0.03 | 0.05 | 0.02 | 0.01 |
| 1 | 72987960 | rs782230 | -2.607 | 0.138 | 1.969 | -1.264 | 0.001 | -0.07 | -0.07 | 0.76 | 0.25 | 0.59 | 0.3 | 0.08 | 0.25 | 0.14 |
| 1 | 72991433 | rs1775367 | -1.811 | 0.2 | 1.986 | -1.139 | 0.023 | -0.048 | -0.048 | 0 | 0.18 | 0.35 | 0.28 | 0.08 | 0.23 | 0.13 |
| 1 | 72992856 | rs1775370 | -1.563 | 0.213 | 1.939 | -1.133 | 0.024 | -0.037 | -0.037 | 0.669 | 0.17 | 0.31 | 0.28 | 0.08 | 0.23 | 0.13 |
| 1 | 73011443 | rs2797092 | -1.432 | 0.194 | 2.18 | -1.158 | 0.058 | -0.108 | -0.108 | 0.433 | 0.17 | 0.35 | 0.21 | 0.02 | 0.23 | 0.1 |
| 1 | 73012228 | rs1445583 | -1.357 | 0.196 | 2.169 | -1.157 | 0.046 | -0.095 | -0.095 | 0 | 0.16 | 0.31 | 0.21 | 0.02 | 0.23 | 0.1 |
| 1 | 73035775 | rs2244695 | -2.068 | 0.117 | 2.303 | -1.13 | 0.01 | -0.062 | -0.062 | 0 | 0.22 | 0.53 | 0.3 | 0.08 | 0.24 | 0.14 |
| 1 | 73040590 | rs6424477 | -1.512 | 0.219 | 2.104 | -1.304 | -0.004 | -0.009 | -0.009 | 0 | 0.18 | 0.17 | 0.19 | 0.02 | 0.23 | 0.13 |
| 1 | 73075423 | rs4498754 | 0.269 | -0.672 | 0.522 | -1.032 | 0.081 | 0.081 | 0.081 | 0.057 | 0.61 | 0.16 | 0.62 | 0.21 | 0.76 | 0.49 |
| 1 | 73079152 | rs11209984 | 0 | -0.828 | 1.512 | -1.074 | -0.008 | -0.006 | -0.006 | 0.002 | 0.33 | 0.35 | 0.32 | 0.77 | 0.23 | 0.5 |
| 1 | 73082539 | rs4620479 | 0.381 | -0.82 | 1.429 | -1.071 | 0.076 | 0.071 | 0.071 | 0.038 | 0.61 | 0.17 | 0.62 | 0.22 | 0.76 | 0.49 |
| 1 | 73100055 | rs11209988 | -0.037 | -0.769 | 1.815 | -1.103 | -0.021 | -0.021 | -0.021 | 0.109 | 0.33 | 0.39 | 0.31 | 0.78 | 0.25 | 0.51 |
| 1 | 73102010 | rs7526444 | -0.136 | -0.775 | 1.812 | -1.106 | -0.021 | -0.021 | -0.021 | 0 | 0.33 | 0.4 | 0.31 | 0.78 | 0.25 | 0.51 |
| 1 | 73103734 | rs10889968 | -0.142 | -0.779 | 1.839 | -1.108 | -0.036 | -0.036 | -0.036 | 0 | 0.66 | 0.48 | 0.68 | 0.22 | 0.75 | 0.49 |

| | | | | | | | | | | | | | | | | |
|---|---|---|---|---|---|---|---|---|---|---|---|---|---|---|---|---|
| 1 | 73104160 | rs12024784 | -0.013 | -0.781 | 2.078 | -1.109 | -0.008 | -0.008 | -0.008 | 0 | 0.33 | 0.35 | 0.31 | 0.78 | 0.25 | 0.51 |
| 1 | 73107472 | rs4111519 | -0.047 | -0.781 | 2.096 | -1.109 | -0.026 | -0.026 | -0.026 | 0 | 0.33 | 0.41 | 0.31 | 0.78 | 0.25 | 0.51 |
| 1 | 73111887 | rs12030081 | 0.002 | -0.791 | 2.003 | -1.114 | -0.009 | -0.009 | -0.009 | 0 | 0.32 | 0.35 | 0.31 | 0.78 | 0.25 | 0.51 |
| 1 | 73116491 | rs4650146 | 0.401 | -0.792 | 1.704 | -1.118 | 0.083 | 0.089 | 0.089 | 0.713 | 0.61 | 0.15 | 0.63 | 0.22 | 0.74 | 0.48 |
| 1 | 73164809 | rs6424487 | -0.029 | -0.738 | 1.821 | -1.087 | -0.036 | -0.035 | -0.035 | 0 | 0.33 | 0.46 | 0.31 | 0.78 | 0.25 | 0.51 |
| 1 | 73203025 | rs2422280 | -0.258 | -0.681 | 2.062 | -1.204 | -0.013 | -0.018 | -0.018 | 0 | 0.67 | 0.6 | 0.68 | 0.22 | 0.73 | 0.48 |
| 1 | 73204521 | rs7542606 | -0.295 | -0.692 | 2.207 | -1.207 | -0.009 | -0.012 | -0.012 | 2.864 | 0.67 | 0.62 | 0.68 | 0.22 | 0.73 | 0.48 |
| 1 | 73214975 | rs12749370 | 0.099 | -0.712 | 2.209 | -1.221 | 0.01 | 0.012 | 0.012 | 0.915 | 0.63 | 0.33 | 0.64 | 0.22 | 0.72 | 0.49 |
| 1 | 73217509 | rs12133234 | 0.216 | -0.713 | 2.207 | -1.221 | 0.015 | 0.018 | 0.018 | 0.44 | 0.62 | 0.32 | 0.64 | 0.22 | 0.72 | 0.49 |
| 1 | 73218976 | rs4269729 | -0.298 | -0.714 | 2.22 | -1.222 | -0.011 | -0.016 | -0.016 | 0.051 | 0.67 | 0.6 | 0.68 | 0.22 | 0.72 | 0.5 |
| 1 | 73221921 | rs11210049 | 0.214 | -0.71 | 2.176 | -1.22 | 0.015 | 0.018 | 0.018 | 0.44 | 0.62 | 0.32 | 0.64 | 0.22 | 0.72 | 0.49 |
| 1 | 73227348 | rs12145482 | 0.926 | -0.674 | 1.502 | -1.749 | -0.072 | 0.191 | 0.191 | 0.03 | 0.45 | 0.03 | 0.39 | 0.17 | 0.56 | 0.4 |
| 1 | 73230347 | rs11210055 | 0.581 | -0.654 | 1.599 | -1.75 | 0.03 | 0.112 | 0.112 | 0 | 0.5 | 0.32 | 0.43 | 0.17 | 0.56 | 0.4 |
| 1 | 73231948 | rs2841188 | 0.266 | -0.528 | 1.653 | -1.729 | 0.014 | -0.082 | -0.082 | 0 | 0.82 | 0.65 | 0.75 | 0.95 | 0.83 | 0.9 |
| 1 | 73234829 | rs2422286 | -0.311 | -0.561 | 1.587 | -1.729 | 0 | -0.022 | -0.022 | 0 | 0.17 | 0.16 | 0.24 | 0.05 | 0.16 | 0.1 |
| 1 | 73237902 | rs1525986 | 0.923 | -0.642 | 1.465 | -1.775 | -0.076 | 0.196 | 0.196 | 0 | 0.45 | 0.03 | 0.39 | 0.17 | 0.56 | 0.39 |
| 1 | 73239170 | rs2204499 | 0.781 | -0.666 | 1.462 | -1.496 | -0.001 | -0.056 | -0.056 | 0 | 0.28 | 0.02 | 0.27 | 0.68 | 0.25 | 0.46 |
| 1 | 73251013 | rs17615698 | -0.224 | -0.573 | 1.339 | -1.532 | -0.002 | -0.002 | -0.002 | 0 | 0.17 | 0.12 | 0.23 | 0.05 | 0.16 | 0.1 |
| 1 | 73259276 | rs2259784 | 0.09 | -0.491 | 1.319 | -1.345 | -0.008 | -0.011 | -0.011 | 0 | 0.33 | 0.39 | 0.32 | 0.78 | 0.27 | 0.51 |
| 1 | 73286806 | rs11210072 | 0.507 | -0.328 | 0.648 | -1.742 | -0.028 | 0.037 | 0.037 | 0 | 0.36 | 0.27 | 0.38 | 0.13 | 0.47 | 0.31 |
| 1 | 73305593 | rs11210074 | 0.405 | -0.532 | 0.413 | -1.273 | 0.061 | 0 | 0 | 0.051 | 0.38 | 0.11 | 0.24 | 0.65 | 0.3 | 0.49 |
| 1 | 73313762 | rs1917881 | -0.4 | -0.58 | 1.062 | -1.552 | 0.004 | 0.045 | 0.045 | 0 | 0.58 | 0.69 | 0.41 | 0.71 | 0.43 | 0.57 |
| 1 | 73317559 | rs1405062 | 1.45 | -0.751 | 1.276 | -1.555 | 0.022 | 0.111 | 0.111 | 0.833 | 0.93 | 0.65 | 0.89 | 0.87 | 0.96 | 0.96 |
| 1 | 73318565 | rs10889994 | -0.042 | -0.778 | 1.61 | -1.572 | 0.018 | 0.07 | 0.07 | 0 | 0.54 | 0.31 | 0.37 | 0.71 | 0.43 | 0.57 |
| 1 | 73327954 | rs2692869 | 1.034 | -0.57 | 1.615 | -1.333 | 0.006 | 0.084 | 0.084 | 0.088 | 0.94 | 0.83 | 0.92 | 0.87 | 0.96 | 0.96 |
| 1 | 73390986 | rs6660036 | 1.812 | -0.064 | 1.869 | -1.411 | 0.041 | 0.093 | 0.093 | 0.057 | 0.92 | 0.58 | 0.89 | 0.89 | 0.96 | 0.96 |
| 1 | 73401018 | rs4262488 | 2.007 | -0.05 | 1.966 | -1.404 | 0.065 | 0.122 | 0.122 | 0.649 | 0.93 | 0.56 | 0.89 | 0.89 | 0.96 | 0.96 |
| 1 | 73401801 | rs4434807 | 1.838 | -0.049 | 1.973 | -1.404 | 0.06 | 0.114 | 0.114 | 0.277 | 0.93 | 0.59 | 0.89 | 0.88 | 0.96 | 0.96 |
| 1 | 73408437 | rs7536364 | -0.064 | -0.237 | 1.718 | -1.657 | -0.005 | -0.022 | -0.022 | 0 | 0.55 | 0.47 | 0.38 | 0.73 | 0.43 | 0.58 |
| 1 | 73408983 | rs6424507 | -0.092 | -0.246 | 1.8 | -1.661 | 0.019 | 0.064 | 0.064 | 0 | 0.54 | 0.31 | 0.37 | 0.73 | 0.43 | 0.58 |
| 1 | 73489496 | rs12119161 | 0.565 | -0.278 | 1.243 | -1.73 | -0.075 | 0.141 | 0.141 | 0 | 0.38 | 0.06 | 0.54 | 0.15 | 0.56 | 0.39 |
| 1 | 73492303 | rs12734738 | 0.919 | -0.509 | 1.243 | -1.887 | 0.08 | 0.051 | 0.051 | 0.359 | 0.35 | 0.09 | 0.22 | 0.72 | 0.26 | 0.48 |
| 1 | 73497953 | rs11210105 | 0.453 | -0.154 | 1.119 | -1.621 | -0.074 | 0.144 | 0.144 | 0 | 0.38 | 0.06 | 0.54 | 0.15 | 0.55 | 0.39 |
| 1 | 73552927 | rs7512691 | 0.121 | -0.165 | 1.229 | -1.616 | 0.01 | 0.077 | 0.077 | 0 | 0.42 | 0.23 | 0.62 | 0.26 | 0.58 | 0.43 |
| 1 | 73573941 | rs12033354 | 0.098 | -0.147 | 1.242 | -1.636 | 0.034 | 0.079 | 0.079 | 0 | 0.53 | 0.3 | 0.33 | 0.73 | 0.4 | 0.57 |
| 1 | 73579237 | rs4603080 | 0.333 | -0.034 | 0.703 | -1.666 | -0.046 | 0.146 | 0.146 | 0 | 0.39 | 0.12 | 0.55 | 0.15 | 0.56 | 0.4 |
| 1 | 73613560 | rs4452995 | 0.427 | -0.029 | 0.637 | -1.657 | -0.068 | 0.14 | 0.14 | 0 | 0.38 | 0.07 | 0.55 | 0.15 | 0.56 | 0.4 |
| 1 | 73625777 | rs7523688 | -0.307 | -0.034 | 0.816 | -1.656 | 0.001 | 0.04 | 0.04 | 0 | 0.58 | 0.65 | 0.43 | 0.84 | 0.44 | 0.6 |
| 1 | 73652122 | rs4650199 | -0.018 | -0.233 | 1.04 | -1.693 | 0.009 | 0.048 | 0.048 | 0 | 0.46 | 0.33 | 0.62 | 0.25 | 0.55 | 0.45 |
| 1 | 73658061 | rs11210178 | 0.831 | -0.259 | 1.053 | -1.756 | 0.008 | -0.01 | -0.01 | 0 | 0.93 | 0.7 | 0.96 | 1 | 0.98 | 0.99 |
| 1 | 73662169 | rs10890018 | 0.218 | -0.352 | 1.228 | -1.806 | -0.006 | 0.079 | 0.079 | 0 | 0.43 | 0.23 | 0.59 | 0.24 | 0.53 | 0.43 |
| 1 | 73664563 | rs7522520 | 0.228 | -0.352 | 1.262 | -1.807 | -0.009 | 0.091 | 0.091 | 0 | 0.43 | 0.21 | 0.59 | 0.24 | 0.53 | 0.43 |
| 1 | 73665844 | rs4650201 | 0.209 | -0.352 | 1.257 | -1.806 | -0.005 | 0.079 | 0.079 | 0 | 0.43 | 0.24 | 0.59 | 0.24 | 0.53 | 0.43 |
| 1 | 73669845 | rs6692884 | 0.118 | -0.315 | 1.194 | -1.782 | 0.006 | 0.015 | 0.015 | 0 | 0.51 | 0.46 | 0.36 | 0.76 | 0.45 | 0.55 |
| 1 | 73714714 | rs12030954 | 1.011 | -0.604 | 0.388 | -1.896 | 0 | -0.011 | -0.011 | 0 | 0.31 | 0.02 | 0.23 | 0.74 | 0.29 | 0.47 |
| 1 | 73736562 | rs4571923 | 0.119 | -0.122 | 0.385 | -1.828 | -0.064 | 0.137 | 0.137 | 0 | 0.43 | 0.11 | 0.57 | 0.2 | 0.52 | 0.43 |
| 1 | 73744843 | rs6684341 | 0.338 | -0.255 | 1.243 | -1.902 | 0.009 | 0.059 | 0.059 | 0 | 0.48 | 0.34 | 0.36 | 0.79 | 0.46 | 0.54 |
| 1 | 73766431 | rs11210201 | -0.208 | 0.206 | 0.901 | -2.002 | 0.007 | 0.07 | 0.07 | 0 | 0.44 | 0.33 | 0.58 | 0.21 | 0.42 | 0.33 |
| 1 | 73793499 | rs1885246 | 0.164 | 0.189 | 1.037 | -2.01 | 0.008 | 0.031 | 0.031 | 0 | 0.53 | 0.54 | 0.41 | 0.79 | 0.58 | 0.67 |
| 1 | 73816564 | rs1160682 | -0.521 | 0.181 | 0.889 | -2.018 | -0.04 | 0.099 | 0.099 | 0 | 0.61 | 0.89 | 0.46 | 0.8 | 0.6 | 0.68 |
| 1 | 73845734 | rs10732842 | -0.096 | 0.844 | 0.522 | -2.358 | 0 | -0.001 | -0.001 | 0 | 0.83 | 0.82 | 0.71 | 0.89 | 0.82 | 0.89 |
| 1 | 73851687 | rs6671130 | 0.753 | 0.544 | 0.557 | -2.404 | 0 | -0.006 | -0.006 | 0.314 | 0.49 | 0.48 | 0.38 | 0.78 | 0.56 | 0.66 |
| 1 | 73882478 | rs1475064 | 0.798 | 0.549 | 0.554 | -2.323 | 0.007 | 0.024 | 0.024 | 0 | 0.49 | 0.41 | 0.38 | 0.77 | 0.56 | 0.66 |
| 1 | 73882880 | rs1475065 | 0.52 | 0.796 | 0.506 | -2.485 | 0.026 | 0.066 | 0.066 | 0 | 0.56 | 0.72 | 0.44 | 0.78 | 0.64 | 0.67 |
| 1 | 73910055 | rs4113050 | 1.368 | 0.84 | 0.546 | -2.483 | 0.004 | 0.016 | 0.016 | 0 | 0.47 | 0.41 | 0.35 | 0.78 | 0.44 | 0.55 |
| 1 | 73920352 | rs501899 | 0.821 | 0.861 | 0.522 | -2.483 | -0.008 | 0.036 | 0.036 | 0 | 0.55 | 0.66 | 0.4 | 0.79 | 0.54 | 0.56 |
| 1 | 73932886 | rs12140865 | 0.71 | 0.864 | 0.551 | -2.485 | 0.004 | 0.006 | 0.006 | 0 | 0.57 | 0.55 | 0.42 | 0.79 | 0.66 | 0.68 |
| 1 | 73935251 | rs11210259 | 1.327 | 0.875 | 0.53 | -2.484 | 0.008 | 0.051 | 0.051 | 0 | 0.47 | 0.35 | 0.35 | 0.78 | 0.45 | 0.55 |
| 1 | 73938246 | rs17570390 | 1.103 | 1.012 | 0.047 | -2.513 | -0.048 | 0.042 | 0.042 | 0.038 | 0.12 | 0.02 | 0.11 | 0.06 | 0.15 | 0.07 |
| 1 | 73945480 | rs1501483 | 0.134 | 0.94 | 0.119 | -2.517 | 0.002 | -0.001 | -0.001 | 0.008 | 0.14 | 0.16 | 0.28 | 0.11 | 0.18 | 0.12 |
| 1 | 73951515 | rs9425120 | 0.73 | 0.89 | 0.097 | -2.514 | 0.004 | 0.046 | 0.046 | 0 | 0.56 | 0.66 | 0.41 | 0.78 | 0.58 | 0.67 |
| 1 | 73974388 | rs12742409 | 1.251 | 0.825 | 0.042 | -2.364 | 0.008 | 0.049 | 0.049 | 0 | 0.47 | 0.35 | 0.35 | 0.78 | 0.45 | 0.52 |
| 1 | 73981238 | rs4361946 | -0.732 | 0.821 | -0.19 | -2.324 | 0.035 | 0.094 | 0.094 | 0.012 | 0.74 | 0.91 | 0.73 | 0.9 | 0.83 | 0.8 |
| 1 | 73982084 | rs4384177 | -0.745 | 0.822 | -0.19 | -2.324 | 0.034 | 0.094 | 0.094 | 0.02 | 0.75 | 0.91 | 0.72 | 0.9 | 0.83 | 0.8 |
| 1 | 73993804 | rs10890042 | 0.406 | 0.897 | -0.438 | -2.365 | 0.001 | 0.122 | 0.122 | 2.387 | 0.64 | 0.82 | 0.61 | 0.84 | 0.64 | 0.82 |
| 1 | 73996208 | rs1501488 | 0.124 | 0.655 | 0.07 | -1.977 | 0.005 | 0.002 | 0.002 | 0.122 | 0.77 | 0.78 | 0.73 | 0.89 | 0.85 | 0.91 |
| 1 | 74000649 | rs10736420 | -0.052 | 0.554 | 0.036 | -1.966 | 0.045 | 0.052 | 0.052 | 0.292 | 0.78 | 0.87 | 0.75 | 0.89 | 0.86 | 0.91 |
| 1 | 74017178 | rs2062664 | 0.892 | 0.636 | 0.116 | -1.969 | 0 | -0.091 | -0.091 | 0 | 0.67 | 0.56 | 0.67 | 0.89 | 0.66 | 0.77 |
| 1 | 74022034 | rs6669337 | 1.489 | 0.567 | 0.08 | -1.91 | -0.006 | 0.045 | 0.045 | 0.061 | 0.54 | 0.34 | 0.4 | 0.78 | 0.5 | 0.63 |

| | | | | | | | | | | | | | | | | |
|---|---|---|---|---|---|---|---|---|---|---|---|---|---|---|---|---|
| 1 | 74031554 | rs7524751 | 0.658 | 0.693 | 0.106 | -1.619 | 0.036 | -0.019 | -0.019 | 0 | 0.05 | 0.15 | 0.04 | 0.01 | 0.08 | 0.01 |
| 1 | 74039936 | rs7512635 | -0.569 | 0.521 | -0.1 | -1.7 | 0.042 | 0.047 | 0.047 | 1.296 | 0.82 | 0.94 | 0.78 | 0.9 | 0.88 | 0.92 |
| 1 | 74041079 | rs1329989 | -0.427 | 0.506 | 0.079 | -1.707 | 0.01 | 0.01 | 0.01 | 0.857 | 0.81 | 0.85 | 0.77 | 0.9 | 0.88 | 0.92 |
| 1 | 74041576 | rs12025606 | 1.363 | 0.695 | 0.262 | -1.807 | 0.01 | -0.031 | -0.031 | 0 | 0.6 | 0.48 | 0.44 | 0.78 | 0.53 | 0.64 |
| 1 | 74042455 | rs7542794 | 1.447 | 0.669 | 0.304 | -1.819 | 0 | -0.007 | -0.007 | 2.267 | 0.59 | 0.41 | 0.43 | 0.78 | 0.53 | 0.64 |
| 1 | 74042517 | rs10890050 | 0.651 | 0.771 | 0.348 | -1.739 | 0.004 | 0.003 | 0.003 | 0 | 0.11 | 0.16 | 0.26 | 0.14 | 0.15 | 0.14 |
| 1 | 74043705 | rs12133927 | 0.88 | 0.733 | 0.361 | -1.78 | 0.036 | -0.018 | -0.018 | 0 | 0.05 | 0.16 | 0.04 | 0.01 | 0.08 | 0.01 |
| 1 | 74048649 | rs4649988 | -0.24 | 0.738 | 0.133 | -1.757 | 0.026 | -0.006 | -0.006 | 0.557 | 0.16 | 0.21 | 0.29 | 0.11 | 0.28 | 0.26 |
| 1 | 74055240 | rs7546041 | 1.128 | 0.715 | 0.297 | -1.769 | 0.006 | -0.037 | -0.037 | 0.888 | 0.63 | 0.49 | 0.46 | 0.79 | 0.61 | 0.66 |
| 1 | 74057686 | rs6700402 | -0.438 | 0.697 | 0.227 | -1.75 | 0.01 | 0.01 | 0.01 | 0.857 | 0.81 | 0.85 | 0.77 | 0.9 | 0.88 | 0.92 |
| 1 | 74083035 | rs2341263 | -0.562 | 0.883 | 0.313 | -1.637 | 0.06 | -0.014 | -0.014 | 0 | 0.17 | 0.32 | 0.3 | 0.11 | 0.28 | 0.26 |
| 1 | 74092495 | rs1576429 | 0.215 | 0.672 | 0.158 | -1.376 | 0.008 | 0.01 | 0.01 | 0.24 | 0.8 | 0.83 | 0.78 | 0.88 | 0.84 | 0.91 |
| 1 | 74106106 | rs6680131 | 1.247 | 0.531 | 0.289 | -1.327 | 0.003 | -0.029 | -0.029 | 0.062 | 0.66 | 0.48 | 0.5 | 0.77 | 0.64 | 0.65 |
| 1 | 74109213 | rs7553862 | -0.671 | 0.885 | 0.263 | -1.189 | -0.016 | -0.014 | -0.014 | 0.349 | 0.89 | 0.93 | 0.75 | 0.89 | 0.85 | 0.86 |
| 1 | 74110629 | rs10157788 | 0.113 | 0.891 | 0.366 | -1.186 | 0.009 | 0.006 | 0.006 | 0 | 0.86 | 0.67 | 0.73 | 0.89 | 0.84 | 0.86 |
| 1 | 74112883 | rs4606267 | -0.419 | 0.892 | 0.383 | -1.186 | 0.002 | 0.001 | 0.001 | 0 | 0.88 | 0.84 | 0.74 | 0.89 | 0.85 | 0.86 |
| 1 | 74113031 | rs4367755 | 0.056 | 0.901 | 0.325 | -1.181 | 0.009 | 0.006 | 0.006 | 0 | 0.86 | 0.67 | 0.73 | 0.89 | 0.84 | 0.86 |
| 1 | 74123576 | rs1497848 | 0.619 | 0.929 | 0.327 | -1.189 | 0.053 | -0.016 | -0.016 | 1.074 | 0.82 | 0.62 | 0.69 | 0.88 | 0.72 | 0.74 |
| 1 | 74129905 | rs9425123 | 0.616 | 0.902 | 0.203 | -0.645 | 0.053 | 0.033 | 0.033 | 0 | 0.82 | 0.62 | 0.69 | 0.77 | 0.72 | 0.73 |
| 1 | 74138765 | rs4617355 | -0.095 | 0.259 | -0.459 | -1.205 | 0.014 | 0.049 | 0.049 | 0 | 0.48 | 0.39 | 0.45 | 0.8 | 0.44 | 0.68 |
| 1 | 74139105 | rs11210325 | 1.045 | 0.159 | -0.762 | -1.191 | 0.028 | 0.001 | 0.001 | 0.333 | 0.31 | 0.15 | 0.25 | 0.7 | 0.25 | 0.48 |
| 1 | 74139340 | rs9425012 | 0.34 | 0.233 | -0.779 | -1.102 | -0.029 | 0.022 | 0.022 | 0.005 | 0.15 | 0.04 | 0.19 | 0.1 | 0.19 | 0.19 |
| 1 | 74145078 | rs9425014 | -0.209 | 0.194 | -0.725 | -1.087 | 0.011 | 0.068 | 0.068 | 0 | 0.52 | 0.36 | 0.46 | 0.77 | 0.49 | 0.67 |
| 1 | 74145595 | rs4431793 | -0.103 | 0.223 | -1.624 | -1.032 | -0.021 | 0.18 | 0.18 | 0.363 | 0.49 | 0.08 | 0.4 | 0.76 | 0.47 | 0.65 |
| 1 | 74151316 | rs2880289 | 0.404 | 0.548 | -1.19 | -0.429 | -0.015 | -0.073 | -0.073 | 0 | 0.3 | 0.02 | 0.22 | 0.67 | 0.27 | 0.39 |
| 1 | 74163209 | rs1497841 | -0.841 | 1.143 | -0.28 | -0.151 | 0.066 | 0.024 | 0.024 | 0.37 | 0.24 | 0.48 | 0.42 | 0.23 | 0.28 | 0.28 |
| 1 | 74166048 | rs4291455 | -0.877 | 0.863 | -0.628 | -0.473 | -0.002 | -0.035 | -0.035 | 0 | 0.57 | 0.56 | 0.58 | 0.82 | 0.59 | 0.75 |
| 1 | 74166959 | rs4650224 | 0.599 | 1.088 | -0.652 | -0.397 | -0.042 | -0.081 | -0.081 | 0 | 0.16 | 0.03 | 0.34 | 0.26 | 0.16 | 0.16 |
| 1 | 74167281 | rs9425020 | -0.401 | 1.112 | -0.273 | -0.412 | 0.148 | 0.003 | 0.003 | 1.272 | 0.07 | 0.34 | 0.06 | 0 | 0.1 | 0.11 |
| 1 | 74169283 | rs9425128 | 0.007 | 1.04 | -0.431 | -0.399 | -0.001 | 0.002 | 0.002 | 0 | 0.41 | 0.54 | 0.27 | 0.6 | 0.43 | 0.61 |
| 1 | 74170011 | rs1909067 | -0.435 | 1.045 | -0.396 | -0.376 | 0.056 | 0.038 | 0.038 | 0.416 | 0.24 | 0.49 | 0.41 | 0.26 | 0.27 | 0.27 |
| 1 | 74170796 | rs12145301 | 0.642 | 1.064 | -0.654 | -0.352 | 0.018 | -0.078 | -0.078 | 0 | 0.34 | 0.09 | 0.18 | 0.55 | 0.32 | 0.48 |
| 1 | 74174778 | rs1497840 | -0.532 | 1.054 | -0.552 | -0.392 | 0.018 | 0.007 | 0.007 | 0.692 | 0.25 | 0.65 | 0.43 | 0.26 | 0.28 | 0.29 |
| 1 | 74175290 | rs4630083 | -0.875 | 0.359 | -0.507 | -0.406 | 0.003 | 0.041 | 0.041 | 1.895 | 0.58 | 0.67 | 0.59 | 0.82 | 0.59 | 0.75 |
| 1 | 74179179 | rs6692022 | 0.292 | 0.463 | -0.963 | -0.183 | -0.102 | -0.125 | -0.125 | 0 | 0.35 | 0.04 | 0.21 | 0.57 | 0.42 | 0.56 |
| 1 | 74183738 | rs7515492 | -1.038 | 0.17 | 0.284 | 0.143 | -0.017 | 0.156 | 0.156 | 0.587 | 0.2 | 0.54 | 0.37 | 0.42 | 0.17 | 0.25 |
| 1 | 74184409 | rs2341264 | 0.998 | 0.067 | -0.247 | -0.527 | 0.003 | 0.133 | 0.133 | 0.118 | 0.39 | 0.2 | 0.38 | 0.04 | 0.4 | 0.17 |
| 1 | 74193940 | rs12750968 | -0.977 | -0.148 | 0.258 | -0.274 | -0.002 | 0.097 | 0.097 | 0 | 0.92 | 0.76 | 0.85 | 0.78 | 0.93 | 0.91 |
| 1 | 74200925 | rs10789377 | 0.992 | -0.105 | -0.142 | -0.419 | -0.101 | -0.173 | -0.173 | 0 | 0.29 | 0.02 | 0.16 | 0.49 | 0.34 | 0.51 |
| 1 | 74204559 | rs11210335 | 0.85 | -0.017 | -0.038 | -0.37 | -0.019 | -0.04 | -0.04 | 0 | 0.08 | 0.01 | 0.17 | 0.2 | 0.09 | 0.14 |
| 1 | 74220796 | rs11210343 | 0.056 | -0.248 | 0.132 | -0.912 | -0.005 | 0.122 | 0.122 | 0.122 | 0.49 | 0.31 | 0.49 | 0.08 | 0.5 | 0.24 |
| 1 | 74223811 | rs10081995 | 1.397 | -0.188 | -0.409 | -0.988 | -0.002 | 0.009 | 0.009 | 0 | 0.09 | 0.03 | 0.04 | 0 | 0.07 | 0.05 |
| 1 | 74225534 | rs2342208 | 0.613 | -0.278 | -0.383 | -0.851 | -0.046 | 0.21 | 0.21 | 0.167 | 0.38 | 0.11 | 0.41 | 0.05 | 0.43 | 0.17 |
| 1 | 74228366 | rs6680989 | 0.484 | 0.077 | 0.346 | -0.003 | -0.009 | 0.098 | 0.098 | 0.173 | 0.9 | 0.71 | 0.85 | 0.78 | 0.93 | 0.91 |
| 1 | 74236278 | rs1532480 | -0.224 | -0.058 | 0.123 | -0.796 | 0 | -0.033 | -0.033 | 0.935 | 0.49 | 0.52 | 0.61 | 0.88 | 0.55 | 0.74 |
| 1 | 74236657 | rs2132999 | 0.212 | -0.296 | 0.663 | -0.6 | -0.011 | 0.023 | 0.023 | 0 | 0.42 | 0.34 | 0.51 | 0.7 | 0.47 | 0.64 |
| 1 | 74240214 | rs17094362 | 0.342 | -0.403 | 0.816 | -1.155 | 0 | -0.057 | -0.057 | 0 | 0.14 | 0.18 | 0.05 | 0 | 0.1 | 0.02 |
| 1 | 74241248 | rs7537401 | 0.448 | -0.522 | 0.502 | -0.924 | -0.003 | 0.094 | 0.094 | 0 | 0.12 | 0.3 | 0.24 | 0.24 | 0.12 | 0.22 |
| 1 | 74245076 | rs17576753 | 1.531 | -0.557 | 0.512 | -1.256 | -0.001 | -0.006 | -0.006 | 2.455 | 0.09 | 0.07 | 0.05 | 0 | 0.06 | 0.04 |
| 1 | 74249035 | rs1874731 | 0.735 | -0.578 | 0.497 | -1.248 | -0.031 | 0.055 | 0.055 | 0.009 | 0.41 | 0.23 | 0.46 | 0.76 | 0.47 | 0.63 |
| 1 | 74251125 | rs4147207 | 0.859 | -0.528 | -0.028 | -1.104 | 0.014 | 0.014 | 0.014 | 0 | 0.3 | 0.19 | 0.36 | 0.23 | 0.28 | 0.27 |
| 1 | 74258672 | rs2342213 | 0.722 | -0.635 | -0.309 | -1.126 | 0.006 | 0.018 | 0.018 | 0 | 0.33 | 0.1 | 0.33 | 0.26 | 0.32 | 0.3 |
| 1 | 74260853 | rs1353671 | 1.096 | -0.691 | 0.155 | -1.247 | 0.004 | -0.067 | -0.067 | 0.018 | 0.11 | 0.24 | 0.07 | 0 | 0.06 | 0.06 |
| 1 | 74264128 | rs1340430 | 0.03 | -0.755 | 0.177 | -1.182 | 0.011 | 0.026 | 0.026 | 0 | 0.44 | 0.35 | 0.4 | 0.25 | 0.38 | 0.36 |
| 1 | 74264228 | rs1340429 | 0.619 | -0.584 | 0.297 | -1.147 | -0.002 | 0.003 | 0.003 | 0 | 0.39 | 0.38 | 0.51 | 0.74 | 0.43 | 0.6 |
| 1 | 74271482 | rs1340443 | 0.041 | -0.524 | 0.211 | -1.023 | 0.032 | 0.09 | 0.09 | 0.011 | 0.55 | 0.21 | 0.56 | 0.71 | 0.63 | 0.67 |
| 1 | 74274122 | rs10493524 | 0.861 | -0.564 | 0.152 | -1.077 | -0.019 | 0.036 | 0.036 | 0.542 | 0.31 | 0.1 | 0.31 | 0.22 | 0.31 | 0.27 |
| 1 | 74277734 | rs12039842 | 0.891 | -0.598 | 0.025 | -0.948 | 0.007 | -0.063 | -0.063 | 0 | 0.11 | 0 | 0.1 | 0.18 | 0.07 | 0.2 |
| 1 | 74279635 | rs826369 | 0.885 | -0.554 | 0.291 | -0.977 | -0.001 | 0.036 | 0.036 | 0.621 | 0.31 | 0.16 | 0.32 | 0.22 | 0.31 | 0.27 |
| 1 | 74284212 | rs12238977 | 1.126 | -0.52 | 0.344 | -0.969 | -0.009 | -0.059 | -0.059 | 0 | 0.1 | 0.22 | 0.06 | 0 | 0.05 | 0.03 |
| 1 | 74286123 | rs12409558 | 1.143 | -0.236 | -0.317 | -0.978 | -0.024 | 0.02 | 0.02 | 0 | 0.09 | 0.03 | 0.06 | 0 | 0.12 | 0.02 |
| 1 | 74286408 | rs10890079 | 0.159 | -0.351 | -0.028 | -0.7 | 0.033 | 0.098 | 0.098 | 0.018 | 0.55 | 0.19 | 0.55 | 0.71 | 0.63 | 0.67 |
| 1 | 74298228 | rs1948615 | -0.085 | -0.16 | -0.098 | -0.753 | -0.01 | 0.053 | 0.053 | 0 | 0.34 | 0.24 | 0.37 | 0.2 | 0.37 | 0.29 |
| 1 | 74298254 | rs2016423 | -0.132 | -0.071 | -0.353 | -0.719 | 0.002 | 0 | 0 | 0 | 0.78 | 0.79 | 0.75 | 0.8 | 0.73 | 0.72 |
| 1 | 74299658 | rs12401808 | 1.181 | 0.206 | -0.92 | -0.606 | -0.044 | 0.005 | 0.005 | 1.291 | 0.23 | 0.08 | 0.29 | 0.2 | 0.26 | 0.24 |
| 1 | 74321238 | rs1454610 | 0.861 | 0.377 | -1.014 | -0.337 | 0.018 | -0.01 | -0.01 | 0 | 0.42 | 0.24 | 0.34 | 0.54 | 0.35 | 0.44 |
| 1 | 74323287 | rs12036352 | -0.818 | 0.461 | -0.367 | -0.094 | -0.068 | -0.05 | -0.05 | 0 | 0.19 | 0.03 | 0.23 | 0.2 | 0.26 | 0.26 |
| 1 | 74323399 | rs12046593 | -0.805 | 0.373 | -0.363 | -0.089 | -0.063 | -0.05 | -0.05 | 0 | 0.19 | 0.03 | 0.23 | 0.2 | 0.25 | 0.26 |
| 1 | 74326154 | rs958820 | 0.66 | 0.61 | -0.258 | 0.126 | 0.009 | 0.004 | 0.004 | 0.164 | 0.75 | 0.73 | 0.72 | 0.75 | 0.72 | 0.72 |

| | | | | | | | | | | | | | | | | |
|---|---|---|---|---|---|---|---|---|---|---|---|---|---|---|---|---|
| 1 | 74327108 | rs7520000 | 0.139 | 0.191 | -0.355 | -0.124 | 0.001 | 0.001 | 0.001 | 0.438 | 0.41 | 0.38 | 0.4 | 0.64 | 0.31 | 0.41 |
| 1 | 74332387 | rs6685128 | -0.135 | -0.182 | -0.198 | -0.429 | -0.017 | -0.037 | -0.037 | 0 | 0.11 | 0.14 | 0.03 | 0 | 0.06 | 0.01 |
| 1 | 74333708 | rs11210357 | -0.498 | -0.093 | -0.172 | -0.547 | -0.001 | 0 | 0 | 0 | 0.11 | 0.12 | 0.14 | 0.08 | 0.11 | 0.19 |
| 1 | 74338530 | rs6672527 | 0.674 | 0.136 | -0.206 | -1.102 | -0.035 | 0.052 | 0.052 | 0 | 0.26 | 0.16 | 0.22 | 0.06 | 0.36 | 0.17 |
| 1 | 74339054 | rs1979217 | -0.682 | 0.119 | 0.063 | -1.123 | 0.018 | 0.002 | 0.002 | 0.022 | 0.13 | 0.28 | 0.09 | 0.11 | 0.12 | 0.22 |
| 1 | 74344914 | rs17094539 | 0.612 | 0.092 | 0.008 | -0.938 | -0.015 | -0.03 | -0.03 | 0.167 | 0.1 | 0.12 | 0.03 | 0 | 0.05 | 0.01 |
| 1 | 74345462 | rs1551391 | 0.932 | 0.035 | -0.287 | -0.961 | -0.077 | 0.234 | 0.234 | 0.509 | 0.48 | 0.05 | 0.63 | 0.83 | 0.47 | 0.61 |
| 1 | 74345867 | rs2014651 | 1.175 | 0.027 | -0.287 | -0.968 | -0.054 | 0.094 | 0.094 | 0.618 | 0.45 | 0.04 | 0.51 | 0.72 | 0.41 | 0.46 |
| 1 | 74350248 | rs11210364 | 0.162 | 0.032 | -0.155 | -1.045 | 0 | 0.019 | 0.019 | 0.049 | 0.61 | 0.33 | 0.71 | 0.94 | 0.59 | 0.83 |
| 1 | 74354806 | rs6700377 | -0.563 | 0.292 | 0 | -0.726 | 0.008 | -0.002 | -0.002 | 0 | 0.11 | 0.17 | 0.08 | 0.1 | 0.12 | 0.22 |
| 1 | 74365646 | rs1340434 | -0.449 | -0.198 | 0.311 | -0.842 | -0.005 | 0.011 | 0.011 | 0 | 0.64 | 0.65 | 0.68 | 0.88 | 0.6 | 0.79 |
| 1 | 74369034 | rs17094586 | -0.614 | -0.431 | 0.405 | -0.827 | -0.048 | -0.019 | -0.019 | 0 | 0.15 | 0.46 | 0.2 | 0.11 | 0.1 | 0.16 |
| 1 | 74385848 | rs1340425 | -0.597 | -0.469 | 0.453 | -0.8 | -0.047 | -0.018 | -0.018 | 0 | 0.14 | 0.46 | 0.2 | 0.11 | 0.1 | 0.16 |
| 1 | 74396896 | rs2733262 | 0.311 | -0.298 | -0.117 | -0.828 | 0.004 | 0.162 | 0.162 | 0.02 | 0.5 | 0.2 | 0.49 | 0.78 | 0.5 | 0.65 |
| 1 | 74402318 | rs10890092 | 0.348 | -0.301 | -0.13 | -0.829 | -0.005 | 0.168 | 0.168 | 0.056 | 0.49 | 0.17 | 0.49 | 0.78 | 0.5 | 0.63 |
| 1 | 74417742 | rs2603541 | 0.329 | -0.255 | -0.386 | -0.805 | -0.009 | 0.174 | 0.174 | 0.034 | 0.5 | 0.15 | 0.48 | 0.77 | 0.5 | 0.63 |
| 1 | 74445584 | rs10890101 | -0.492 | -0.256 | -0.108 | -0.814 | 0.003 | 0.032 | 0.032 | 0 | 0.47 | 0.64 | 0.37 | 0.17 | 0.5 | 0.39 |
| 1 | 74452323 | rs9662322 | 0.282 | -0.285 | 0.086 | -0.892 | 0.012 | -0.054 | -0.054 | 0 | 0.35 | 0.42 | 0.29 | 0.09 | 0.41 | 0.19 |
| 1 | 74454701 | rs11210389 | -0.039 | -0.058 | -0.915 | -0.752 | 0 | 0.107 | 0.107 | 0 | 0.54 | 0.27 | 0.62 | 0.83 | 0.54 | 0.78 |
| 1 | 74455745 | rs11806015 | -0.277 | -0.008 | -1.493 | -0.726 | -0.012 | 0.011 | 0.011 | 0 | 0.11 | 0.06 | 0.07 | 0.07 | 0.09 | 0.2 |
| 1 | 74461828 | rs4556311 | 0.595 | 0.009 | -1.399 | -0.538 | -0.051 | 0.04 | 0.04 | 1.062 | 0.42 | 0.07 | 0.54 | 0.77 | 0.45 | 0.57 |
| 1 | 74464107 | rs11210394 | 0.763 | -0.086 | -1.71 | -0.569 | -0.055 | -0.075 | -0.075 | 0.683 | 0.36 | 0.03 | 0.27 | 0.73 | 0.35 | 0.38 |
| 1 | 74464625 | rs12039894 | 0.62 | 0.498 | -1.755 | -1.021 | -0.126 | 0.043 | 0.043 | 0.157 | 0.26 | 0.02 | 0.28 | 0.09 | 0.32 | 0.21 |
| 1 | 74465302 | rs17588031 | -0.801 | 0.455 | -0.669 | -0.789 | 0 | -0.032 | -0.032 | 0.009 | 0.18 | 0.23 | 0.2 | 0.01 | 0.21 | 0.18 |
| 1 | 74470568 | rs10890102 | 0.865 | 0.354 | 0.332 | -0.621 | 0.012 | -0.035 | -0.035 | 0 | 0.89 | 0.48 | 0.89 | 0.96 | 0.91 | 0.84 |
| 1 | 74501477 | rs1384883 | 0.138 | 0.118 | 0.586 | -1.134 | 0.002 | 0.153 | 0.153 | 0.7 | 0.5 | 0.22 | 0.47 | 0.84 | 0.51 | 0.67 |
| 1 | 74513945 | rs6424572 | 0.187 | 0.08 | 0.694 | -1.152 | 0.022 | -0.03 | -0.03 | 1.094 | 0.89 | 0.53 | 0.89 | 0.96 | 0.91 | 0.89 |
| 1 | 74518464 | rs4650243 | 0.916 | 0.006 | 0.376 | -1.331 | -0.06 | 0.042 | 0.042 | 0.681 | 0.38 | 0.06 | 0.33 | 0.79 | 0.38 | 0.42 |
| 1 | 74535130 | rs1483793 | 0.274 | 0.04 | 0.434 | -1.268 | -0.022 | 0.145 | 0.145 | 0.229 | 0.47 | 0.16 | 0.41 | 0.8 | 0.48 | 0.52 |
| 1 | 74539128 | rs12742029 | 0.533 | 0.015 | 0.495 | -1.297 | -0.011 | -0.056 | -0.056 | 1.549 | 0.88 | 0.39 | 0.88 | 0.96 | 0.91 | 0.89 |
| 1 | 74540820 | rs17094802 | -0.136 | 0.011 | 1.047 | -1.299 | 0.006 | -0.03 | -0.03 | 0 | 0.09 | 0.25 | 0.09 | 0.04 | 0.09 | 0.11 |
| 1 | 74547515 | rs11210405 | -0.03 | 0.03 | 0.776 | -1.409 | 0.009 | 0.141 | 0.141 | 0.654 | 0.53 | 0.2 | 0.5 | 0.84 | 0.57 | 0.67 |
| 1 | 74560411 | rs10789387 | -0.045 | 0.037 | 0.894 | -1.302 | 0.004 | 0.106 | 0.106 | 0 | 0.51 | 0.3 | 0.47 | 0.84 | 0.51 | 0.67 |
| 1 | 74560603 | rs11210407 | 0.68 | 0.026 | 0.97 | -1.313 | -0.011 | 0.065 | 0.065 | 0 | 0.38 | 0.2 | 0.41 | 0.11 | 0.4 | 0.22 |
| 1 | 74561386 | rs4650246 | 0.358 | 0.042 | 0.945 | -1.285 | 0.02 | -0.033 | -0.033 | 0 | 0.89 | 0.51 | 0.89 | 0.96 | 0.91 | 0.89 |
| 1 | 74568667 | rs982792 | 0.358 | 0.042 | 0.945 | -1.285 | 0.02 | -0.033 | -0.033 | 0.432 | 0.89 | 0.51 | 0.89 | 0.96 | 0.91 | 0.89 |
| 1 | 74592761 | rs1601153 | 0.06 | 0.044 | 0.896 | -1.298 | 0.006 | 0.045 | 0.045 | 0 | 0.5 | 0.41 | 0.42 | 0.79 | 0.48 | 0.53 |
| 1 | 74627358 | rs1483790 | 0.968 | -0.006 | 0.896 | -1.345 | -0.033 | 0.121 | 0.121 | 0 | 0.35 | 0.12 | 0.34 | 0.08 | 0.39 | 0.16 |
| 1 | 74644454 | rs1333032 | 0.271 | 0.076 | 1.008 | -1.277 | 0.019 | -0.027 | -0.027 | 0.223 | 0.9 | 0.62 | 0.9 | 0.96 | 0.91 | 0.88 |
| 1 | 74644516 | rs11210427 | 0.787 | -0.045 | 0.878 | -1.32 | -0.03 | 0.087 | 0.087 | 0 | 0.37 | 0.14 | 0.43 | 0.12 | 0.41 | 0.18 |
| 1 | 74646058 | rs17591292 | 0.211 | -0.063 | 0.882 | -1.355 | -0.002 | -0.029 | -0.029 | 0 | 0.09 | 0.09 | 0.07 | 0 | 0.09 | 0.1 |
| 1 | 74648182 | rs1047127 | 0.211 | -0.033 | 0.882 | -1.355 | -0.001 | -0.029 | -0.029 | 0 | 0.09 | 0.09 | 0.07 | 0 | 0.09 | 0.11 |
| 1 | 74663709 | rs11806946 | 0.314 | -0.05 | 0.856 | -1.131 | -0.017 | 0.091 | 0.091 | 0.196 | 0.47 | 0.15 | 0.39 | 0.74 | 0.47 | 0.55 |
| 1 | 74665493 | rs792310 | 0.326 | -0.095 | 0.842 | -1.152 | -0.02 | 0.081 | 0.081 | 0.182 | 0.46 | 0.16 | 0.39 | 0.74 | 0.46 | 0.55 |
| 1 | 74670172 | rs12046751 | 0.808 | -0.1 | 0.846 | -1.157 | -0.026 | 0.095 | 0.095 | 0 | 0.38 | 0.14 | 0.43 | 0.12 | 0.41 | 0.18 |
| 1 | 74672323 | rs489941 | 0.307 | -0.087 | 0.769 | -1.135 | -0.022 | 0.083 | 0.083 | 0.182 | 0.46 | 0.15 | 0.39 | 0.74 | 0.46 | 0.55 |
| 1 | 74678285 | rs485929 | 0.986 | -0.066 | 0.475 | -1.106 | -0.061 | 0.001 | 0.001 | 0.681 | 0.37 | 0.06 | 0.32 | 0.74 | 0.37 | 0.45 |
| 1 | 74696439 | rs17094973 | -1.118 | 0.039 | 0.747 | -0.946 | 0.044 | 0.083 | 0.083 | 0.463 | 0.06 | 0.31 | 0.08 | 0.1 | 0.04 | 0.15 |
| 1 | 74699099 | rs3753183 | 1.057 | 0.035 | 0.092 | -0.797 | -0.004 | -0.02 | -0.02 | 0.549 | 0.06 | 0.02 | 0.03 | 0.11 | 0.05 | 0.07 |
| 1 | 74699912 | rs518769 | 0.182 | 0.63 | 0.21 | -0.949 | 0.054 | -0.052 | -0.052 | 0 | 0.88 | 0.62 | 0.88 | 0.96 | 0.86 | 0.85 |
| 1 | 74704513 | rs11210443 | 1.16 | 0.517 | -0.985 | -0.94 | -0.115 | 0.153 | 0.153 | 0 | 0.34 | 0.03 | 0.34 | 0.07 | 0.4 | 0.13 |
| 1 | 74710450 | rs481387 | -0.029 | 0.406 | -0.579 | -0.681 | 0.016 | 0.068 | 0.068 | 0 | 0.56 | 0.84 | 0.63 | 0.26 | 0.59 | 0.48 |
| 1 | 74712181 | rs2034453 | 1.118 | 0.805 | 0.472 | -0.252 | 0.01 | 0.019 | 0.019 | 0 | 0.89 | 0.8 | 0.91 | 0.89 | 0.88 | 0.88 |
| 1 | 74730853 | rs6700161 | -1.01 | 1.495 | 0.723 | -0.269 | -0.022 | -0.012 | -0.012 | 1.34 | 0.06 | 0.03 | 0.12 | 0.07 | 0.15 | 0.17 |
| 1 | 74734443 | rs10493540 | -1.992 | 1.262 | 1.369 | -0.359 | -0.012 | -0.019 | -0.019 | 0 | 0.13 | 0.23 | 0.09 | 0.06 | 0.07 | 0.06 |
| 1 | 74737081 | rs514012 | -2.155 | 1.425 | 1.498 | -0.304 | 0.028 | 0.008 | 0.008 | 0 | 0.09 | 0.14 | 0.15 | 0.14 | 0.18 | 0.31 |
| 1 | 74743034 | rs473834 | -1.831 | 1.445 | 1.487 | -0.288 | 0.015 | 0.004 | 0.004 | 0 | 0.08 | 0.12 | 0.14 | 0.14 | 0.15 | 0.3 |
| 1 | 74756402 | rs11210445 | 2.488 | 1.276 | 2.236 | -0.392 | 0.079 | 0.011 | 0.011 | 0 | 0.85 | 0.43 | 0.81 | 0.86 | 0.81 | 0.68 |
| 1 | 74762198 | rs6684409 | 3.372 | 1.259 | 1.899 | -0.399 | -0.014 | 0.017 | 0.017 | 0.025 | 0.72 | 0.19 | 0.71 | 0.8 | 0.74 | 0.62 |
| 1 | 74767844 | rs477920 | -2.092 | 1.38 | 1.761 | -0.326 | 0.046 | 0.019 | 0.019 | 0 | 0.08 | 0.14 | 0.12 | 0.14 | 0.17 | 0.29 |
| 1 | 74777775 | rs17095121 | -1.176 | 1.154 | 1.684 | -0.327 | 0.001 | 0.002 | 0.002 | 0.063 | 0.1 | 0.06 | 0.07 | 0.06 | 0.07 | 0.07 |
| 1 | 74822937 | rs1527683 | -2.223 | 1.405 | 2.174 | -0.334 | 0.024 | -0.011 | -0.011 | 7.614 | 0.15 | 0.18 | 0.16 | 0.09 | 0.21 | 0.17 |
| 1 | 74825037 | rs1417888 | -1.178 | 1.321 | 1.619 | -0.309 | 0.021 | 0.015 | 0.015 | 0 | 0.06 | 0.12 | 0.06 | 0.11 | 0.09 | 0.18 |
| 1 | 74857855 | rs6669186 | -0.117 | 0.858 | 1.284 | -0.282 | -0.017 | 0.008 | 0.008 | 0 | 0.69 | 0.47 | 0.57 | 0.29 | 0.72 | 0.59 |
| 1 | 74858230 | rs9988473 | 1.721 | 1.069 | 1.846 | -0.926 | 0.026 | 0.018 | 0.018 | 0.506 | 0.89 | 0.67 | 0.86 | 0.89 | 0.9 | 0.87 |
| 1 | 74858724 | rs1569092 | -2.134 | 1.295 | 1.886 | -1.076 | 0.048 | 0.001 | 0.001 | 0.182 | 0.11 | 0.24 | 0.09 | 0.11 | 0.15 | 0.14 |
| 1 | 74858887 | rs1011542 | 1.736 | 1.26 | 1.938 | -1.069 | 0.022 | -0.015 | -0.015 | 0 | 0.89 | 0.7 | 0.87 | 0.92 | 0.9 | 0.87 |
| 1 | 74861767 | rs6693806 | 3.184 | 1.065 | 1.437 | -1.171 | -0.056 | 0.058 | 0.058 | 1.692 | 0.71 | 0.1 | 0.74 | 0.81 | 0.74 | 0.73 |
| 1 | 74872533 | rs17095265 | -1.94 | 1.101 | 1.987 | -0.915 | 0.011 | 0.018 | 0.018 | 0.05 | 0.08 | 0.22 | 0.04 | 0.07 | 0.06 | 0.03 |

| | | | | | | | | | | | | | | | | |
|---|---|---|---|---|---|---|---|---|---|---|---|---|---|---|---|---|
| 1 | 74878253 | rs699844 | -2.567 | 1.152 | 2.163 | -0.771 | -0.01 | 0.039 | 0.039 | 0 | 0.11 | 0.27 | 0.07 | 0.18 | 0.07 | 0.11 |
| 1 | 74879364 | rs12028841 | 0.946 | 1.078 | 0.976 | -1.355 | -0.01 | 0.075 | 0.075 | 0.018 | 0.29 | 0.02 | 0.15 | 0.1 | 0.3 | 0.21 |
| 1 | 74880175 | rs699846 | 0.543 | 1.41 | 1.755 | -1.264 | 0.053 | -0.022 | -0.022 | 0 | 0.93 | 0.89 | 0.92 | 0.96 | 0.89 | 0.9 |
| 1 | 74885754 | rs2203369 | 3.033 | 1.469 | 1.881 | -1.319 | -0.013 | 0.01 | 0.01 | 1.277 | 0.76 | 0.2 | 0.75 | 0.79 | 0.76 | 0.77 |
| 1 | 74896816 | rs1341573 | 2.575 | 1.687 | 1.699 | -1.437 | -0.083 | 0.065 | 0.065 | 0.501 | 0.68 | 0.12 | 0.7 | 0.79 | 0.67 | 0.7 |
| 1 | 74899573 | rs792884 | 2.464 | 1.319 | 2.384 | -0.996 | 0.081 | 0.012 | 0.012 | 0.014 | 0.86 | 0.68 | 0.86 | 0.86 | 0.82 | 0.84 |
| 1 | 74904353 | rs17095322 | -0.851 | 0.915 | 1.611 | -0.832 | 0.003 | 0.006 | 0.006 | 0 | 0.05 | 0.07 | 0.05 | 0.11 | 0.07 | 0.06 |
| 1 | 74907002 | rs1022915 | 3.295 | 0.818 | 1.728 | -0.884 | -0.001 | 0 | 0 | 0.635 | 0.8 | 0.26 | 0.83 | 0.79 | 0.82 | 0.79 |
| 1 | 74907316 | rs1325604 | 2.677 | 1.016 | 1.407 | -0.83 | 0.112 | 0.02 | 0.02 | 0.182 | 0.86 | 0.58 | 0.85 | 0.86 | 0.82 | 0.84 |
| 1 | 74918027 | rs274596 | -1.679 | 0.665 | 0.736 | -0.679 | 0.021 | 0.042 | 0.042 | 0 | 0.07 | 0.21 | 0.06 | 0.11 | 0.07 | 0.06 |
| 1 | 74926794 | rs640493 | -0.172 | 0.149 | -0.006 | -1.175 | -0.021 | 0.191 | 0.191 | 0.029 | 0.61 | 0.9 | 0.79 | 0.92 | 0.61 | 0.61 |
| 1 | 74934996 | rs612695 | -0.156 | 0.16 | 1.128 | -0.834 | -0.001 | 0.004 | 0.004 | 0 | 0.63 | 0.69 | 0.7 | 0.82 | 0.58 | 0.59 |
| 1 | 74937749 | rs3765674 | 0.279 | 0.103 | 1.449 | -0.84 | -0.005 | -0.014 | -0.014 | 0 | 0.58 | 0.48 | 0.67 | 0.69 | 0.58 | 0.53 |
| 1 | 74939570 | rs507181 | 0.444 | 0.058 | 1.608 | -0.862 | 0.005 | 0.011 | 0.011 | 0 | 0.56 | 0.43 | 0.66 | 0.69 | 0.58 | 0.53 |
| 1 | 74941293 | rs649058 | -0.788 | 0.065 | 1.449 | -0.89 | -0.065 | 0.114 | 0.114 | 0 | 0.69 | 0.92 | 0.73 | 0.87 | 0.6 | 0.62 |
| 1 | 74943465 | rs274587 | 0.644 | 0.161 | 1.671 | -0.735 | 0.027 | 0.095 | 0.095 | 0 | 0.48 | 0.14 | 0.6 | 0.67 | 0.55 | 0.46 |
| 1 | 74948186 | rs9326113 | -1.634 | -0.056 | 1.685 | -0.637 | -0.004 | 0.037 | 0.037 | 0 | 0.07 | 0.26 | 0.03 | 0.05 | 0.02 | 0.07 |
| 1 | 74961817 | rs9730586 | -0.153 | 0.157 | 1.178 | -0.642 | -0.01 | 0.059 | 0.059 | 0 | 0.59 | 0.79 | 0.67 | 0.72 | 0.56 | 0.53 |
| 1 | 74975017 | rs17095482 | -1.786 | -0.495 | 1.791 | -0.357 | 0.009 | 0.046 | 0.046 | 0 | 0.06 | 0.3 | 0.03 | 0.05 | 0.02 | 0.07 |
| 1 | 74977425 | rs953567 | -0.24 | -0.333 | 1.551 | -0.586 | 0.007 | 0.046 | 0.046 | 0 | 0.52 | 0.41 | 0.54 | 0.82 | 0.41 | 0.49 |
| 1 | 74978977 | rs11210470 | -2.101 | -0.34 | 1.388 | -0.493 | -0.052 | 0.014 | 0.014 | 0 | 0.1 | 0.33 | 0.05 | 0.08 | 0.03 | 0.07 |
| 1 | 74983721 | rs17095501 | -1.423 | -0.306 | 1.428 | -0.477 | -0.02 | 0.049 | 0.049 | 0 | 0.07 | 0.27 | 0.04 | 0.07 | 0.03 | 0.07 |
| 1 | 74983835 | rs10493544 | 0.43 | -0.283 | 1.196 | -0.503 | -0.075 | 0.224 | 0.224 | 0 | 0.41 | 0.11 | 0.39 | 0.08 | 0.55 | 0.4 |
| 1 | 74991644 | rs1514175 | -0.235 | -0.21 | 1.349 | -0.347 | 0.005 | 0.077 | 0.077 | 0 | 0.49 | 0.31 | 0.41 | 0.23 | 0.57 | 0.48 |
| 1 | 75002667 | rs3845345 | -0.139 | 0.182 | 0.242 | -0.206 | -0.005 | 0.068 | 0.068 | 0 | 0.56 | 0.7 | 0.58 | 0.75 | 0.42 | 0.51 |
| 1 | 75004611 | rs6604872 | 0.347 | 0.187 | 0.237 | -0.204 | -0.086 | 0.173 | 0.173 | 0 | 0.4 | 0.11 | 0.38 | 0.15 | 0.55 | 0.41 |
| 1 | 75005238 | rs7553158 | 0.314 | 0.201 | 0.215 | -0.197 | -0.053 | 0.138 | 0.138 | 0 | 0.41 | 0.19 | 0.38 | 0.15 | 0.55 | 0.41 |
| 1 | 75008008 | rs6604871 | -0.158 | 0.218 | 0.034 | -0.189 | -0.001 | 0.081 | 0.081 | 0 | 0.54 | 0.68 | 0.57 | 0.75 | 0.41 | 0.51 |
| 1 | 75010536 | rs6694355 | 0.947 | 0.438 | -0.092 | 0.188 | -0.005 | 0.023 | 0.023 | 0 | 0.95 | 0.88 | 0.96 | 0.89 | 0.96 | 0.92 |
| 1 | 75014195 | rs7541513 | -0.244 | 0.541 | -0.141 | 0.102 | 0.01 | 0.057 | 0.057 | 0 | 0.5 | 0.67 | 0.59 | 0.67 | 0.43 | 0.5 |
| 1 | 75016147 | rs4287129 | -1.235 | 0.453 | 0.746 | 0.358 | 0.012 | 0.034 | 0.034 | 0 | 0.16 | 0.4 | 0.16 | 0.25 | 0.15 | 0.26 |
| 1 | 75018421 | rs10789398 | 0.829 | 0.351 | 0.599 | -0.152 | -0.004 | 0.145 | 0.145 | 0.002 | 0.48 | 0.27 | 0.45 | 0.13 | 0.5 | 0.38 |
| 1 | 75019436 | rs1443372 | -0.042 | 0.359 | 0.942 | 0.148 | 0.001 | 0.003 | 0.003 | 0 | 0.63 | 0.59 | 0.66 | 0.41 | 0.64 | 0.64 |
| 1 | 75035240 | rs12728662 | -0.113 | 0.349 | 1.168 | 0.121 | -0.044 | -0.051 | -0.051 | 1.663 | 0.35 | 0.06 | 0.32 | 0.59 | 0.36 | 0.37 |
| 1 | 75038228 | rs11580409 | -0.091 | 0.34 | 1.566 | 0.137 | -0.004 | -0.009 | -0.009 | 0.018 | 0.36 | 0.25 | 0.33 | 0.58 | 0.36 | 0.39 |
| 1 | 75045729 | rs10890136 | 1.022 | 0.101 | 1.899 | -0.006 | -0.013 | 0.09 | 0.09 | 0.19 | 0.45 | 0.24 | 0.42 | 0.13 | 0.49 | 0.35 |
| 1 | 75050303 | rs9661135 | -0.199 | 0.008 | 1.888 | 0.031 | -0.016 | -0.034 | -0.034 | 0 | 0.09 | 0.05 | 0.15 | 0.23 | 0.1 | 0.1 |
| 1 | 75051340 | rs696688 | -2.051 | -0.005 | 2.002 | 0.024 | -0.02 | 0.047 | 0.047 | 1.628 | 0.19 | 0.62 | 0.23 | 0.29 | 0.14 | 0.27 |
| 1 | 75051422 | rs9661462 | 1.18 | -0.041 | 1.802 | 0 | -0.036 | 0.151 | 0.151 | 0.134 | 0.44 | 0.16 | 0.45 | 0.13 | 0.5 | 0.35 |
| 1 | 75055419 | rs2305549 | -0.127 | 0.043 | 1.743 | -0.052 | -0.015 | -0.033 | -0.033 | 0.458 | 0.09 | 0.05 | 0.15 | 0.23 | 0.1 | 0.11 |
| 1 | 75060313 | rs696694 | -1.888 | 0.035 | 2.034 | -0.056 | -0.011 | 0.072 | 0.072 | 0.928 | 0.18 | 0.53 | 0.22 | 0.28 | 0.14 | 0.27 |
| 1 | 75065441 | rs696698 | -0.383 | 0.048 | 1.908 | -0.069 | -0.007 | -0.002 | -0.002 | 0.629 | 0.06 | 0.14 | 0.05 | 0.06 | 0.04 | 0.16 |
| 1 | 75071003 | rs10890137 | 1.078 | 0.066 | 1.744 | -0.098 | -0.023 | 0.153 | 0.153 | 0.042 | 0.46 | 0.21 | 0.46 | 0.12 | 0.5 | 0.35 |
| 1 | 75072038 | rs696702 | -0.787 | 0.121 | 1.444 | -0.021 | 0 | -0.017 | -0.017 | 0 | 0.14 | 0.17 | 0.19 | 0.28 | 0.14 | 0.27 |
| 1 | 75086665 | rs10493546 | 0.777 | 0.091 | 1.51 | -0.113 | -0.001 | -0.039 | -0.039 | 0 | 0.48 | 0.47 | 0.48 | 0.11 | 0.5 | 0.35 |
| 1 | 75089136 | rs699856 | -0.286 | 0.107 | 1.118 | 0.039 | -0.021 | -0.018 | -0.018 | 2.206 | 0.64 | 0.87 | 0.68 | 0.38 | 0.64 | 0.6 |
| 1 | 75095878 | rs686329 | -0.42 | 0.043 | 1.065 | 0.17 | -0.03 | -0.044 | -0.044 | 0 | 0.69 | 0.86 | 0.69 | 0.42 | 0.66 | 0.65 |
| 1 | 75117824 | rs699857 | -0.343 | 0.124 | 0.99 | 0.126 | -0.027 | -0.05 | -0.05 | 0 | 0.71 | 0.88 | 0.71 | 0.39 | 0.68 | 0.65 |
| 1 | 75118172 | rs730645 | -0.709 | 0.078 | 1.04 | 0.075 | -0.01 | 0.019 | 0.019 | 0 | 0.2 | 0.29 | 0.21 | 0.28 | 0.16 | 0.31 |
| 1 | 75121016 | rs564725 | -0.314 | 0.179 | 0.27 | 0.14 | -0.002 | -0.005 | -0.005 | 0.151 | 0.7 | 0.72 | 0.69 | 0.39 | 0.68 | 0.65 |
| 1 | 75130289 | rs12239912 | 0.555 | 0.533 | -0.301 | -0.282 | 0.005 | 0.09 | 0.09 | 1.38 | 0.5 | 0.34 | 0.49 | 0.17 | 0.53 | 0.31 |
| 1 | 75140299 | rs10789400 | -0.421 | 0.624 | -0.952 | -0.662 | -0.004 | 0.166 | 0.166 | 0.303 | 0.51 | 0.29 | 0.57 | 0.89 | 0.46 | 0.7 |
| 1 | 75147840 | rs17095739 | -0.14 | 0.43 | -1.275 | -0.137 | -0.028 | -0.124 | -0.124 | 0 | 0.24 | 0.02 | 0.28 | 0.61 | 0.16 | 0.39 |
| 1 | 75151932 | rs10493547 | 0.138 | 0.398 | -1.192 | -0.238 | 0.002 | -0.01 | -0.01 | 8.643 | 0.22 | 0.13 | 0.25 | 0.27 | 0.26 | 0.29 |
| 1 | 75152186 | rs10518473 | 0.559 | 0.437 | -1.204 | -0.583 | -0.001 | -0.001 | -0.001 | 0 | 0.08 | 0.08 | 0.02 | 0.08 | 0.1 | 0.11 |
| 1 | 75155271 | rs696672 | -1.931 | 0.201 | -1.243 | -0.798 | -0.084 | 0.205 | 0.205 | 4.923 | 0.63 | 0.9 | 0.64 | 0.96 | 0.57 | 0.81 |
| 1 | 75155657 | rs575533 | -0.807 | 0.127 | -0.178 | -0.662 | -0.013 | -0.004 | -0.004 | 1.522 | 0.35 | 0.43 | 0.34 | 0.69 | 0.26 | 0.51 |
| 1 | 75157273 | rs699801 | -0.967 | -0.027 | -0.844 | -0.625 | 0.023 | 0.089 | 0.089 | 2.561 | 0.51 | 0.66 | 0.55 | 0.82 | 0.44 | 0.64 |
| 1 | 75158118 | rs12061533 | 0.409 | -0.152 | -1.024 | -0.305 | 0.001 | -0.096 | -0.096 | 0 | 0.21 | 0.1 | 0.21 | 0.47 | 0.14 | 0.36 |
| 1 | 75159349 | rs17095774 | 0.517 | -0.173 | -1.154 | -0.303 | -0.027 | -0.15 | -0.15 | 0 | 0.2 | 0.01 | 0.2 | 0.47 | 0.14 | 0.36 |
| 1 | 75160724 | rs1340985 | -0.851 | 0.041 | -0.938 | -0.543 | 0.03 | 0.109 | 0.109 | 2.71 | 0.51 | 0.72 | 0.55 | 0.82 | 0.44 | 0.64 |
| 1 | 75164678 | rs728547 | 0.42 | 0.03 | -1.094 | -0.526 | -0.002 | -0.005 | -0.005 | 0 | 0.08 | 0.04 | 0.04 | 0.08 | 0.09 | 0.12 |
| 1 | 75170143 | rs6662846 | 1.012 | -0.197 | -1.864 | -1.223 | -0.126 | 0.172 | 0.172 | 0.219 | 0.37 | 0.08 | 0.38 | 0.11 | 0.46 | 0.22 |
| 1 | 75171751 | rs17459 | 0.335 | -0.604 | -1.161 | -0.948 | -0.009 | -0.014 | -0.014 | 0 | 0.6 | 0.51 | 0.72 | 0.69 | 0.61 | 0.56 |
| 1 | 75173051 | rs4650281 | 0.335 | -0.627 | -1.171 | -0.969 | -0.01 | -0.014 | -0.014 | 0 | 0.6 | 0.51 | 0.72 | 0.69 | 0.61 | 0.56 |
| 1 | 75175188 | rs2032968 | 1.303 | -0.528 | -1.439 | -1.605 | -0.066 | 0.119 | 0.119 | 0 | 0.34 | 0.14 | 0.39 | 0.01 | 0.48 | 0.22 |
| 1 | 75175262 | rs3753190 | -0.669 | -0.417 | -1.757 | -1.686 | 0.021 | -0.016 | -0.016 | 0.007 | 0.33 | 0.14 | 0.19 | 0.35 | 0.24 | 0.37 |
| 1 | 75175664 | rs2032969 | 0.187 | -0.584 | -0.718 | -1.513 | -0.007 | 0 | 0 | 0 | 0.25 | 0.25 | 0.32 | 0.69 | 0.12 | 0.42 |
| 1 | 75175886 | rs3819946 | -0.37 | -0.635 | -0.034 | -1.676 | -0.067 | -0.015 | -0.015 | 0 | 0.29 | 0.53 | 0.34 | 0.77 | 0.14 | 0.47 |

| | | | | | | | | | | | | | | | | |
|---|---|---|---|---|---|---|---|---|---|---|---|---|---|---|---|---|
| 1 | 75178360 | rs7522428 | -0.328 | -0.127 | 0.018 | -1.822 | 0.001 | 0.003 | 0.003 | 0 | 0.75 | 0.85 | 0.8 | 0.77 | 0.75 | 0.71 |
| 1 | 75178713 | rs13375787 | 1.402 | -0.314 | -0.015 | -1.802 | -0.028 | 0.024 | 0.024 | 0 | 0.17 | 0.18 | 0.28 | 0.64 | 0.05 | 0.32 |
| 1 | 75200486 | rs277374 | 0.144 | -0.131 | 0.051 | -1.808 | -0.006 | -0.002 | -0.002 | 2.632 | 0.24 | 0.11 | 0.2 | 0.23 | 0.25 | 0.29 |
| 1 | 75214441 | rs1133891 | 0.423 | -0.476 | -0.154 | -1.57 | -0.009 | 0 | 0 | 0 | 0.25 | 0.25 | 0.32 | 0.71 | 0.13 | 0.42 |
| 1 | 75215689 | rs28378934 | 1.258 | -0.339 | -0.448 | -1.498 | 0.009 | 0 | 0 | 0 | 0.11 | 0 | 0.04 | 0.09 | 0.07 | 0.2 |
| 1 | 75230606 | rs277368 | 0.037 | -0.28 | -0.362 | -1.499 | 0 | 0.003 | 0.003 | 0 | 0.25 | 0.14 | 0.2 | 0.23 | 0.25 | 0.29 |
| 1 | 75235082 | rs277370 | -1.552 | -0.155 | -0.279 | -1.516 | 0.002 | -0.017 | -0.017 | 0.018 | 0.94 | 0.95 | 0.95 | 1 | 0.94 | 0.97 |
| 1 | 75235317 | rs3931020 | -0.013 | -0.406 | -0.262 | -1.37 | -0.002 | 0.018 | 0.018 | 0 | 0.68 | 0.78 | 0.75 | 0.77 | 0.69 | 0.68 |
| 1 | 75241971 | rs12125889 | 1.707 | -0.383 | -0.486 | -1.445 | -0.118 | 0.195 | 0.195 | 0.011 | 0.37 | 0.08 | 0.39 | 0.07 | 0.51 | 0.2 |
| 1 | 75247617 | rs17618340 | 1.694 | -0.446 | -0.492 | -1.442 | -0.118 | 0.2 | 0.2 | 0 | 0.37 | 0.08 | 0.4 | 0.07 | 0.51 | 0.21 |
| 1 | 75252764 | rs7547190 | 0.187 | -0.652 | -0.144 | -1.328 | -0.007 | -0.001 | -0.001 | 0 | 0.26 | 0.25 | 0.3 | 0.75 | 0.14 | 0.47 |
| 1 | 75256135 | rs11163464 | 1.227 | -0.259 | -0.629 | -1.145 | 0.001 | 0.003 | 0.003 | 0 | 0.08 | 0.07 | 0.02 | 0.07 | 0.07 | 0.1 |
| 1 | 75257881 | rs2185695 | 0.302 | 0.195 | -0.756 | -1.145 | -0.004 | 0.015 | 0.015 | 0 | 0.68 | 0.78 | 0.75 | 0.76 | 0.68 | 0.68 |
| 1 | 75275272 | rs277384 | 0.174 | 0.629 | -1.087 | -1.258 | 0.013 | -0.01 | -0.01 | 0 | 0.22 | 0.3 | 0.24 | 0.14 | 0.3 | 0.13 |
| 1 | 75275749 | rs12738201 | -0.102 | -0.062 | -1.152 | -1.176 | -0.005 | -0.007 | -0.007 | 0 | 0.59 | 0.48 | 0.62 | 0.13 | 0.77 | 0.34 |
| 1 | 75277426 | rs12120636 | 0.527 | 0.79 | -1.671 | -1.126 | -0.072 | 0.17 | 0.17 | 0 | 0.32 | 0.09 | 0.28 | 0.01 | 0.38 | 0.11 |
| 1 | 75278236 | rs2134837 | -0.479 | -0.058 | -0.653 | -0.108 | -0.054 | 0.084 | 0.084 | 0 | 0.27 | 0.43 | 0.24 | 0.45 | 0.15 | 0.37 |
| 1 | 75283643 | rs277395 | -0.548 | 1.117 | -0.746 | -1.137 | -0.028 | 0.012 | 0.012 | 0 | 0.19 | 0.14 | 0.29 | 0.13 | 0.26 | 0.39 |
| 1 | 75290369 | rs277357 | -0.775 | 1.26 | -0.386 | -1.705 | 0 | -0.017 | -0.017 | 0.401 | 0.45 | 0.46 | 0.42 | 0.19 | 0.46 | 0.49 |
| 1 | 75300318 | rs277367 | -0.936 | 0.922 | -0.353 | -2.105 | 0.001 | 0.01 | 0.01 | 0 | 0.57 | 0.59 | 0.55 | 0.2 | 0.57 | 0.56 |
| 1 | 75300917 | rs277366 | -1.146 | 0.799 | -0.062 | -2.184 | -0.008 | 0.095 | 0.095 | 0 | 0.59 | 0.82 | 0.56 | 0.2 | 0.57 | 0.56 |
| 1 | 75309013 | rs277351 | 0.197 | 0.99 | 0.023 | -2.015 | 0.016 | -0.052 | -0.052 | 0 | 0.14 | 0.24 | 0.16 | 0.02 | 0.18 | 0.11 |
| 1 | 75311357 | rs1327080 | 1.053 | 0.814 | 0.007 | -1.995 | -0.006 | -0.028 | -0.028 | 0 | 0.11 | 0.14 | 0.13 | 0.01 | 0.11 | 0.07 |
| 1 | 75357094 | rs7367789 | -0.149 | 0.746 | -0.127 | -2.138 | 0.007 | 0.048 | 0.048 | 0 | 0.56 | 0.7 | 0.62 | 0.82 | 0.6 | 0.52 |
| 1 | 75362429 | rs6593547 | 0.418 | 0.8 | -0.292 | -2.376 | 0.001 | 0 | 0 | 0.908 | 0.15 | 0.15 | 0.27 | 0.8 | 0.19 | 0.33 |
| 1 | 75373403 | rs1413994 | -1.385 | 0.76 | -0.4 | -2.508 | -0.005 | -0.047 | -0.047 | 0.276 | 0.71 | 0.72 | 0.87 | 0.94 | 0.78 | 0.87 |
| 1 | 75380989 | rs12569094 | 0.44 | 0.865 | -0.596 | -2.347 | -0.029 | -0.023 | -0.023 | 0 | 0.14 | 0.06 | 0.26 | 0.8 | 0.19 | 0.33 |
| 1 | 75381179 | rs4949857 | -0.454 | 0.807 | -0.519 | -2.386 | -0.018 | 0.013 | 0.013 | 0 | 0.25 | 0.16 | 0.39 | 0.81 | 0.3 | 0.39 |
| 1 | 75403587 | rs10782637 | -1.494 | 0.777 | -0.425 | -2.427 | -0.003 | -0.089 | -0.089 | 0.054 | 0.71 | 0.64 | 0.79 | 0.99 | 0.7 | 0.88 |
| 1 | 75404117 | rs1413991 | 0.379 | 0.927 | -1.026 | -2.3 | -0.029 | -0.023 | -0.023 | 0 | 0.14 | 0.07 | 0.27 | 0.8 | 0.19 | 0.33 |
| 1 | 75442826 | rs7515377 | 1.253 | 0.803 | -1.025 | -2.379 | 0.002 | -0.001 | -0.001 | 0 | 0.11 | 0.09 | 0.13 | 0.01 | 0.11 | 0.07 |
| 1 | 75449592 | rs3845358 | -1.597 | 0.857 | -0.805 | -2.46 | 0 | -0.025 | -0.025 | 0 | 0.72 | 0.73 | 0.79 | 0.99 | 0.69 | 0.87 |
| 1 | 75462461 | rs3912426 | -1.53 | 0.854 | -0.804 | -2.461 | -0.002 | -0.072 | -0.072 | 0 | 0.71 | 0.62 | 0.78 | 0.99 | 0.69 | 0.87 |
| 1 | 75464506 | rs12033089 | 0.434 | 0.962 | -0.787 | -2.315 | -0.046 | -0.038 | -0.038 | 0.028 | 0.14 | 0.04 | 0.28 | 0.8 | 0.19 | 0.31 |
| 1 | 75464592 | rs3856036 | 0.993 | 0.812 | -0.726 | -2.36 | -0.026 | -0.118 | -0.118 | 0.223 | 0.3 | 0.44 | 0.14 | 0.06 | 0.21 | 0.14 |
| 1 | 75470087 | rs12129281 | 1.334 | 1.021 | -0.95 | -2.164 | -0.043 | -0.017 | -0.017 | 0 | 0.1 | 0.01 | 0.2 | 0.08 | 0.13 | 0.25 |
| 1 | 75471898 | rs12081831 | 0.894 | 0.909 | -0.847 | -2.385 | -0.009 | -0.036 | -0.036 | 1.746 | 0.11 | 0.18 | 0.12 | 0.02 | 0.1 | 0.06 |
| 1 | 75472141 | rs11807616 | 0.906 | 0.799 | -0.631 | -2.564 | -0.022 | -0.088 | -0.088 | 0 | 0.33 | 0.42 | 0.14 | 0.07 | 0.22 | 0.14 |
| 1 | 75473978 | rs795576 | 0.538 | 0.638 | -0.714 | -2.552 | -0.004 | 0.003 | 0.003 | 0.327 | 0.17 | 0.1 | 0.19 | 0.15 | 0.18 | 0.36 |
| 1 | 75486214 | rs12089365 | 1.218 | 0.67 | -0.599 | -2.586 | -0.012 | -0.073 | -0.073 | 0.168 | 0.3 | 0.33 | 0.14 | 0.05 | 0.22 | 0.15 |
| 1 | 75491039 | rs7523491 | 0.624 | 0.821 | -0.668 | -2.697 | 0.005 | 0.003 | 0.003 | 0 | 0.15 | 0.13 | 0.34 | 0.78 | 0.18 | 0.32 |
| 1 | 75502014 | rs1581586 | -0.7 | 0.746 | -0.715 | -2.756 | -0.045 | 0.157 | 0.157 | 0.386 | 0.59 | 0.82 | 0.61 | 0.85 | 0.51 | 0.53 |
| 1 | 75520600 | rs12028349 | 1.122 | 0.764 | -0.744 | -2.703 | -0.039 | -0.067 | -0.067 | 0 | 0.12 | 0.01 | 0.31 | 0.78 | 0.12 | 0.29 |
| 1 | 75544969 | rs7520936 | 1.104 | 0.766 | -0.721 | -2.72 | 0 | 0.004 | 0.004 | 0 | 0.12 | 0.09 | 0.32 | 0.8 | 0.12 | 0.29 |
| 1 | 75549806 | rs7541725 | 1.066 | 0.296 | -0.567 | -2.985 | -0.028 | -0.12 | -0.12 | 0 | 0.32 | 0.46 | 0.15 | 0.05 | 0.22 | 0.15 |
| 1 | 75563291 | rs6593566 | 0.1 | 0.272 | -0.664 | -3.004 | 0.044 | 0.111 | 0.111 | 0.132 | 0.43 | 0.73 | 0.27 | 0.06 | 0.33 | 0.21 |
| 1 | 75565047 | rs169453 | 0.583 | 0.441 | -0.681 | -2.916 | -0.005 | 0.005 | 0.005 | 0.411 | 0.17 | 0.08 | 0.19 | 0.13 | 0.18 | 0.36 |
| 1 | 75585157 | rs280888 | -1.273 | 0.943 | -0.596 | -2.424 | 0.005 | 0.005 | 0.005 | 0 | 0.9 | 0.94 | 0.89 | 0.98 | 0.9 | 0.95 |
| 1 | 75593875 | rs11162567 | 0.483 | 0.845 | -0.473 | -2.376 | 0.002 | -0.002 | -0.002 | 0 | 0.47 | 0.42 | 0.32 | 0.19 | 0.39 | 0.49 |
| 1 | 75595151 | rs11162571 | -0.917 | 0.618 | -0.784 | -1.776 | 0.005 | 0.004 | 0.004 | 0 | 0.9 | 0.95 | 0.89 | 0.98 | 0.9 | 0.94 |
| 1 | 75598900 | rs1526505 | 0.346 | 0.72 | -0.832 | -1.815 | -0.001 | -0.001 | -0.001 | 1.407 | 0.13 | 0.1 | 0.34 | 0.79 | 0.18 | 0.3 |
| 1 | 75609049 | rs12041465 | 0.393 | 0.722 | -0.831 | -1.814 | 0 | 0 | 0 | 0 | 0.13 | 0.1 | 0.33 | 0.8 | 0.17 | 0.29 |
| 1 | 75611510 | rs3925749 | -0.776 | 0.694 | -0.793 | -1.834 | -0.004 | -0.022 | -0.022 | 2.229 | 0.89 | 0.89 | 0.87 | 0.98 | 0.89 | 0.92 |
| 1 | 75613097 | rs1526512 | -0.972 | 0.706 | -0.853 | -1.821 | 0.008 | 0.012 | 0.012 | 0 | 0.9 | 0.96 | 0.88 | 0.98 | 0.9 | 0.93 |
| 1 | 75613259 | rs1526513 | 0.138 | 0.774 | -0.835 | -1.77 | 0 | -0.001 | -0.001 | 0 | 0.76 | 0.8 | 0.54 | 0.19 | 0.72 | 0.63 |
| 1 | 75622616 | rs941032 | 0.021 | 0.155 | -0.716 | -2.024 | -0.001 | -0.011 | -0.011 | 0.034 | 0.46 | 0.48 | 0.3 | 0.21 | 0.39 | 0.55 |
| 1 | 75636971 | rs17565565 | 0.982 | 0.093 | -1.115 | -2.23 | -0.016 | 0.078 | 0.078 | 0.173 | 0.15 | 0.02 | 0.1 | 0 | 0.17 | 0.1 |
| 1 | 75637919 | rs6593568 | -0.458 | -0.332 | -0.839 | -2.51 | -0.001 | -0.042 | -0.042 | 0.019 | 0.28 | 0.31 | 0.23 | 0.01 | 0.23 | 0.19 |
| 1 | 75660269 | rs11162729 | 0.379 | -0.302 | -0.785 | -2.477 | -0.038 | -0.065 | -0.065 | 0 | 0.13 | 0.29 | 0.13 | 0.01 | 0.06 | 0.1 |
| 1 | 75665476 | rs11162738 | 0.507 | -0.668 | -0.969 | -2.83 | -0.026 | 0.046 | 0.046 | 0 | 0.32 | 0.19 | 0.47 | 0.84 | 0.4 | 0.42 |
| 1 | 75671215 | rs12132601 | 1.537 | -0.69 | -1.009 | -2.851 | -0.023 | 0.084 | 0.084 | 0.247 | 0.15 | 0.02 | 0.1 | 0 | 0.17 | 0.1 |
| 1 | 75672376 | rs1948874 | 0.515 | -0.685 | -1.001 | -2.853 | -0.026 | 0.043 | 0.043 | 0 | 0.32 | 0.18 | 0.47 | 0.83 | 0.4 | 0.42 |
| 1 | 75681511 | rs595009 | -1.383 | -0.652 | -0.85 | -2.796 | 0.005 | 0.011 | 0.011 | 0.015 | 0.83 | 0.88 | 0.89 | 1 | 0.83 | 0.9 |
| 1 | 75684529 | rs610159 | -1.531 | -0.789 | -0.237 | -2.875 | 0.004 | 0.041 | 0.041 | 0.102 | 0.84 | 0.94 | 0.89 | 1 | 0.83 | 0.9 |
| 1 | 75686746 | rs627494 | 0.466 | -0.769 | -0.064 | -2.809 | -0.001 | 0.009 | 0.009 | 0 | 0.42 | 0.66 | 0.31 | 0.16 | 0.38 | 0.39 |
| 1 | 75692192 | rs611899 | 0.591 | -0.744 | -0.078 | -2.797 | 0.001 | -0.029 | -0.029 | 0.066 | 0.41 | 0.54 | 0.3 | 0.15 | 0.37 | 0.39 |
| 1 | 75699771 | rs10493565 | -0.012 | -1.022 | -0.266 | -3.417 | -0.004 | -0.05 | -0.05 | 0 | 0.37 | 0.37 | 0.23 | 0.05 | 0.27 | 0.21 |
| 1 | 75701824 | rs1505262 | -0.012 | -0.977 | -0.266 | -3.417 | -0.003 | -0.05 | -0.05 | 0 | 0.37 | 0.37 | 0.23 | 0.05 | 0.27 | 0.21 |
| 1 | 75702674 | rs17096508 | 1.27 | -1.054 | -0.344 | -3.438 | -0.003 | 0.059 | 0.059 | 0 | 0.23 | 0.12 | 0.08 | 0.04 | 0.2 | 0.13 |

| | | | | | | | | | | | | | | | | |
|---|---|---|---|---|---|---|---|---|---|---|---|---|---|---|---|---|
| 1 | 75703209 | rs616827 | 0.069 | -1.452 | -0.683 | -3.745 | 0.002 | -0.03 | -0.03 | 0 | 0.62 | 0.64 | 0.74 | 0.95 | 0.71 | 0.8 |
| 1 | 75714554 | rs649352 | -0.063 | -1.49 | -0.453 | -3.767 | 0.002 | -0.023 | -0.023 | 0 | 0.66 | 0.68 | 0.78 | 0.95 | 0.73 | 0.81 |
| 1 | 75716925 | rs588098 | 0.063 | -1.499 | -0.403 | -3.771 | -0.008 | -0.065 | -0.065 | 2.677 | 0.64 | 0.54 | 0.76 | 0.95 | 0.73 | 0.81 |
| 1 | 75722527 | rs604745 | -0.647 | -1.466 | -0.353 | -3.749 | 0 | -0.019 | -0.019 | 0 | 0.75 | 0.73 | 0.9 | 0.96 | 0.8 | 0.87 |
| 1 | 75740332 | rs1144297 | 1.37 | -1.377 | -0.377 | -3.744 | -0.05 | 0.073 | 0.073 | 0 | 0.3 | 0.09 | 0.48 | 0.87 | 0.41 | 0.4 |
| 1 | 75742038 | rs601788 | 0.421 | -1.47 | -0.365 | -3.84 | -0.005 | -0.057 | -0.057 | 0 | 0.37 | 0.36 | 0.3 | 0.09 | 0.32 | 0.35 |
| 1 | 75754161 | rs625357 | -0.922 | -1.183 | -0.3 | -3.458 | 0.023 | -0.074 | -0.074 | 0 | 0.81 | 0.64 | 0.9 | 0.97 | 0.8 | 0.88 |
| 1 | 75760951 | rs639866 | 0.14 | -1.148 | -0.374 | -3.449 | -0.004 | -0.085 | -0.085 | 0 | 0.7 | 0.58 | 0.78 | 0.96 | 0.74 | 0.81 |
| 1 | 75763491 | rs671919 | -1.174 | -0.813 | -0.696 | -3.17 | -0.01 | 0.011 | 0.011 | 1.042 | 0.83 | 0.86 | 0.87 | 0.92 | 0.76 | 0.79 |
| 1 | 75768066 | rs17096647 | 0.359 | -1.158 | -0.867 | -3.258 | 0.027 | 0.11 | 0.11 | 0 | 0.24 | 0.09 | 0.19 | 0.01 | 0.2 | 0.16 |
| 1 | 75770625 | rs1249675 | -0.224 | -1.382 | -0.626 | -3.473 | -0.002 | 0.018 | 0.018 | 0 | 0.57 | 0.62 | 0.66 | 0.84 | 0.58 | 0.48 |
| 1 | 75779404 | rs12127347 | 0.488 | -1.459 | -0.549 | -3.384 | 0.017 | 0.016 | 0.016 | 0 | 0.24 | 0.18 | 0.2 | 0.02 | 0.2 | 0.16 |
| 1 | 75779988 | rs1249667 | 0.454 | -1.606 | -0.204 | -3.473 | -0.065 | -0.112 | -0.112 | 0 | 0.16 | 0.4 | 0.15 | 0.01 | 0.07 | 0.13 |
| 1 | 75780028 | rs10493570 | 0.785 | -1.722 | -0.418 | -3.494 | 0.051 | 0.087 | 0.087 | 5.077 | 0.22 | 0.1 | 0.09 | 0 | 0.14 | 0.09 |
| 1 | 75785718 | rs945795 | 1.126 | -1.565 | 0.172 | -3.199 | -0.076 | -0.085 | -0.085 | 0.217 | 0.13 | 0.43 | 0.04 | 0 | 0 | 0 |
| 1 | 75787413 | rs12030510 | 0.961 | -0.998 | 0.195 | -3.092 | -0.035 | 0.036 | 0.036 | 0 | 0.29 | 0.11 | 0.44 | 0.82 | 0.38 | 0.32 |
| 1 | 75794538 | rs2657206 | 0.456 | -0.957 | 0.188 | -3.089 | 0.005 | -0.02 | -0.02 | 0.679 | 0.33 | 0.57 | 0.48 | 0.82 | 0.37 | 0.32 |
| 1 | 75811152 | rs17097955 | 0.2 | -1.15 | -0.101 | -2.941 | -0.001 | -0.016 | -0.016 | 2.728 | 0.16 | 0.21 | 0.13 | 0.01 | 0.07 | 0.13 |
| 1 | 75819102 | rs1249837 | 0.729 | -0.706 | -0.325 | -3.092 | -0.026 | 0.075 | 0.075 | 0 | 0.31 | 0.16 | 0.31 | 0.1 | 0.36 | 0.36 |
| 1 | 75830438 | rs1249849 | -1.079 | -0.732 | -0.272 | -3.4 | -0.076 | 0.146 | 0.146 | 0 | 0.63 | 0.86 | 0.64 | 0.97 | 0.53 | 0.58 |
| 1 | 75834454 | rs1249810 | -0.334 | -0.999 | -0.348 | -3.603 | 0.029 | 0.19 | 0.19 | 0 | 0.47 | 0.77 | 0.38 | 0.96 | 0.42 | 0.46 |
| 1 | 75839145 | rs1249808 | -0.414 | -1.015 | -0.334 | -3.595 | 0.036 | 0.198 | 0.198 | 0 | 0.48 | 0.77 | 0.39 | 0.96 | 0.42 | 0.45 |
| 1 | 75840172 | rs12411159 | 1.19 | -1.012 | -0.347 | -3.585 | 0.019 | 0.088 | 0.088 | 0.092 | 0.16 | 0.02 | 0.25 | 0.02 | 0.12 | 0.13 |
| 1 | 75862526 | rs1249813 | -0.368 | -1.029 | -0.1 | -3.599 | 0.027 | 0.107 | 0.107 | 0 | 0.48 | 0.67 | 0.38 | 0.96 | 0.42 | 0.45 |
| 1 | 75864719 | rs1249780 | -1.254 | -0.971 | -0.223 | -3.517 | -0.08 | 0.131 | 0.131 | 0 | 0.65 | 0.87 | 0.64 | 0.97 | 0.54 | 0.6 |
| 1 | 75876192 | rs1249794 | -0.383 | -0.937 | 0.074 | -3.141 | -0.009 | -0.02 | -0.02 | 5.666 | 0.42 | 0.44 | 0.39 | 0.15 | 0.33 | 0.34 |
| 1 | 75886843 | rs1249783 | 0.599 | -0.819 | 0.123 | -3.032 | -0.009 | -0.013 | -0.013 | 0 | 0.27 | 0.38 | 0.14 | 0.15 | 0.22 | 0.22 |
| 1 | 75893080 | rs17096860 | 0.761 | -0.813 | 0.074 | -3.029 | 0.001 | 0.002 | 0.002 | 0 | 0.26 | 0.23 | 0.13 | 0.15 | 0.22 | 0.22 |
| 1 | 75901137 | rs17096877 | 0.793 | -0.673 | -0.222 | -3.018 | 0.007 | 0.014 | 0.014 | 0 | 0.25 | 0.17 | 0.12 | 0.15 | 0.22 | 0.22 |
| 1 | 75906421 | rs2040079 | 1.301 | -0.684 | -0.329 | -3.036 | 0.009 | 0.002 | 0.002 | 0.121 | 0.15 | 0.07 | 0.25 | 0.02 | 0.11 | 0.13 |
| 1 | 75912000 | rs2799050 | -1.059 | -0.592 | -0.349 | -3.037 | -0.011 | -0.02 | -0.02 | 0 | 0.61 | 0.66 | 0.61 | 0.97 | 0.53 | 0.57 |
| 1 | 75917304 | rs6676696 | 0.299 | -0.298 | -0.443 | -2.401 | 0.003 | 0.005 | 0.005 | 0 | 0.26 | 0.27 | 0.13 | 0.15 | 0.22 | 0.22 |
| 1 | 75937429 | rs10874305 | 0.382 | 0.18 | -0.351 | -2.002 | 0.007 | 0.006 | 0.006 | 0 | 0.21 | 0.25 | 0.24 | 0.8 | 0.24 | 0.27 |
| 1 | 75938556 | rs1889036 | 0.494 | 0.201 | -0.269 | -2.018 | 0.005 | 0.001 | 0.001 | 0 | 0.24 | 0.24 | 0.25 | 0.79 | 0.27 | 0.34 |
| 1 | 75941966 | rs11163461 | -0.098 | -0.494 | 0.041 | -2.381 | -0.026 | -0.021 | -0.021 | 0 | 0.25 | 0.43 | 0.12 | 0.16 | 0.17 | 0.2 |
| 1 | 75959487 | rs10874314 | -0.848 | -0.615 | -0.218 | -2.723 | -0.03 | 0.148 | 0.148 | 0 | 0.57 | 0.8 | 0.43 | 0.94 | 0.48 | 0.58 |
| 1 | 75966345 | rs12738260 | 0.817 | -0.65 | -0.101 | -2.734 | -0.002 | 0.011 | 0.011 | 0 | 0.29 | 0.18 | 0.29 | 0.79 | 0.3 | 0.34 |
| 1 | 75967778 | rs12401888 | 1.834 | -0.444 | -0.523 | -2.943 | -0.02 | 0.085 | 0.085 | 0.318 | 0.14 | 0.02 | 0.27 | 0.02 | 0.16 | 0.13 |
| 1 | 75970795 | rs13373826 | 0.91 | -0.55 | -0.309 | -3.002 | 0.005 | -0.025 | -0.025 | 0.113 | 0.13 | 0.2 | 0.05 | 0.04 | 0.14 | 0.1 |
| 1 | 75974819 | rs6658825 | 1.016 | -0.278 | -0.665 | -2.802 | -0.028 | 0.035 | 0.035 | 0 | 0.28 | 0.18 | 0.3 | 0.04 | 0.36 | 0.29 |
| 1 | 75982426 | rs1571983 | -0.607 | -0.426 | -0.405 | -2.272 | -0.016 | -0.041 | -0.041 | 0.017 | 0.39 | 0.56 | 0.4 | 0.18 | 0.33 | 0.33 |
| 1 | 75998695 | rs184159 | -0.158 | -0.628 | -0.759 | -2.2 | 0.025 | 0.043 | 0.043 | 2.055 | 0.27 | 0.23 | 0.37 | 0.05 | 0.23 | 0.23 |
| 1 | 76015460 | rs1443845 | 0.809 | -0.643 | -0.536 | -2.257 | 0 | -0.019 | -0.019 | 0 | 0.17 | 0.13 | 0.31 | 0.02 | 0.18 | 0.18 |
| 1 | 76023350 | rs6670304 | 0.513 | -0.806 | -0.377 | -2.79 | -0.021 | 0.022 | 0.022 | 0 | 0.37 | 0.27 | 0.32 | 0.17 | 0.44 | 0.37 |
| 1 | 76023860 | rs7518629 | 0.931 | -0.931 | -0.435 | -2.904 | -0.02 | 0.015 | 0.015 | 0 | 0.3 | 0.21 | 0.26 | 0.05 | 0.39 | 0.29 |
| 1 | 76024221 | rs12139331 | 1.147 | -1.084 | -0.542 | -2.973 | -0.168 | 0.122 | 0.122 | 0 | 0.28 | 0.03 | 0.25 | 0.05 | 0.38 | 0.29 |
| 1 | 76025003 | rs211699 | 0.79 | -1.129 | 0.06 | -2.991 | 0.005 | 0.013 | 0.013 | 0.069 | 0.36 | 0.3 | 0.29 | 0.78 | 0.33 | 0.39 |
| 1 | 76032904 | rs211762 | 0.339 | -0.69 | 0.245 | -2.779 | -0.032 | -0.045 | -0.045 | 0.134 | 0.67 | 0.52 | 0.55 | 0.83 | 0.72 | 0.7 |
| 1 | 76037238 | rs211757 | 0.998 | -1 | 0.106 | -2.902 | 0.001 | -0.001 | -0.001 | 1.521 | 0.27 | 0.28 | 0.29 | 0.78 | 0.33 | 0.39 |
| 1 | 76038320 | rs11578532 | 1.356 | -0.977 | -0.195 | -2.894 | 0.001 | -0.001 | -0.001 | 0.223 | 0.17 | 0.09 | 0.3 | 0.02 | 0.19 | 0.17 |
| 1 | 76041010 | rs4949673 | -0.288 | -1.084 | -0.197 | -2.955 | -0.012 | 0.159 | 0.159 | 0 | 0.48 | 0.25 | 0.67 | 0.92 | 0.56 | 0.65 |
| 1 | 76041667 | rs10489587 | 1.264 | -0.486 | -0.56 | -2.97 | -0.1 | -0.069 | -0.069 | 0 | 0.25 | 0.02 | 0.26 | 0.77 | 0.33 | 0.39 |
| 1 | 76055098 | rs396106 | 0.392 | -0.552 | 0.094 | -2.587 | -0.021 | -0.039 | -0.039 | 0 | 0.67 | 0.55 | 0.57 | 0.82 | 0.67 | 0.69 |
| 1 | 76067894 | rs1332791 | 0.765 | -0.272 | -0.05 | -2.445 | 0.004 | -0.011 | -0.011 | 0 | 0.06 | 0.04 | 0.05 | 0.12 | 0.05 | 0.09 |
| 1 | 76068279 | rs6662665 | 0.872 | -0.254 | 0.005 | -2.593 | -0.001 | -0.022 | -0.022 | 0 | 0.17 | 0.13 | 0.33 | 0.02 | 0.24 | 0.17 |
| 1 | 76080162 | rs6593584 | -0.413 | -0.022 | 0.26 | -2.625 | 0 | -0.015 | -0.015 | 0 | 0.62 | 0.67 | 0.73 | 0.94 | 0.62 | 0.69 |
| 1 | 76085120 | rs11163827 | 0.77 | 0.295 | 0.128 | -2.458 | -0.002 | 0.017 | 0.017 | 0 | 0.3 | 0.22 | 0.23 | 0.83 | 0.29 | 0.46 |
| 1 | 76088392 | rs6675067 | 0.825 | 0.302 | 0.095 | -2.455 | -0.004 | 0.02 | 0.02 | 0 | 0.29 | 0.18 | 0.22 | 0.83 | 0.29 | 0.46 |
| 1 | 76091876 | rs4141464 | 0.968 | 0.313 | 0.04 | -2.446 | -0.011 | 0.021 | 0.021 | 0 | 0.28 | 0.13 | 0.22 | 0.83 | 0.29 | 0.44 |
| 1 | 76092166 | rs1332783 | 0.966 | 0.316 | 0.042 | -2.445 | -0.011 | 0.021 | 0.021 | 0 | 0.28 | 0.13 | 0.22 | 0.83 | 0.29 | 0.44 |
| 1 | 76092180 | rs17097106 | 0.906 | 0.304 | 0.025 | -2.483 | -0.001 | -0.018 | -0.018 | 0 | 0.06 | 0.04 | 0.06 | 0.09 | 0.07 | 0.07 |
| 1 | 76092192 | rs4949870 | 0.761 | 0.214 | 0.348 | -2.545 | -0.005 | 0.001 | 0.001 | 0 | 0.31 | 0.33 | 0.27 | 0.04 | 0.34 | 0.24 |
| 1 | 76094234 | rs11163860 | 0.629 | -0.101 | 0.298 | -1.998 | -0.011 | 0.021 | 0.021 | 0 | 0.28 | 0.13 | 0.22 | 0.83 | 0.29 | 0.44 |
| 1 | 76106675 | rs211718 | -0.847 | 0.503 | 0.626 | -1.988 | -0.016 | 0.058 | 0.058 | 0.056 | 0.75 | 0.84 | 0.69 | 0.98 | 0.7 | 0.8 |
| 1 | 76108055 | rs12745456 | 1.061 | 0.394 | 0.364 | -1.513 | -0.038 | -0.115 | -0.115 | 0 | 0.29 | 0.02 | 0.2 | 0.7 | 0.24 | 0.47 |
| 1 | 76119597 | rs814885 | 0.358 | 0.602 | 0.7 | -1.427 | -0.002 | 0.055 | 0.055 | 0 | 0.07 | 0.08 | 0.18 | 0.25 | 0.06 | 0.07 |
| 1 | 76120300 | rs440826 | -0.237 | 0.63 | 1.155 | -1.533 | 0.014 | 0.019 | 0.019 | 0 | 0.47 | 0.63 | 0.49 | 0.95 | 0.38 | 0.59 |
| 1 | 76129223 | rs12129523 | 0.991 | 0.636 | 1.141 | -1.522 | -0.004 | -0.021 | -0.021 | 0.016 | 0.28 | 0.21 | 0.21 | 0.02 | 0.32 | 0.21 |
| 1 | 76132070 | rs11163924 | 0.887 | 0.661 | 1.081 | -1.513 | -0.016 | 0.061 | 0.061 | 0.055 | 0.25 | 0.16 | 0.25 | 0.02 | 0.3 | 0.2 |

| | | | | | | | | | | | | | | | | |
|---|---|---|---|---|---|---|---|---|---|---|---|---|---|---|---|---|
| 1 | 76134765 | rs1535740 | 0.727 | 0.628 | 1.413 | -1.495 | -0.007 | -0.062 | -0.062 | 0 | 0.36 | 0.06 | 0.25 | 0.7 | 0.3 | 0.51 |
| 1 | 76154430 | rs7518038 | 0.771 | 0.688 | 1.051 | -1.503 | -0.019 | 0.064 | 0.064 | 0.075 | 0.25 | 0.15 | 0.31 | 0.02 | 0.31 | 0.2 |
| 1 | 76178440 | rs11164042 | 0.966 | 0.7 | 1.05 | -1.525 | -0.004 | -0.021 | -0.021 | 0 | 0.28 | 0.21 | 0.21 | 0.03 | 0.32 | 0.21 |
| 1 | 76185308 | rs7534754 | 0.855 | 0.741 | 0.878 | -1.501 | -0.018 | 0.069 | 0.069 | 0.05 | 0.25 | 0.14 | 0.3 | 0.03 | 0.31 | 0.2 |
| 1 | 76189445 | rs1251079 | -0.185 | 0.731 | 1.03 | -1.457 | 0 | 0 | 0 | 0 | 0.67 | 0.7 | 0.5 | 0.73 | 0.62 | 0.72 |
| 1 | 76196602 | rs1250880 | -0.57 | 0.713 | 0.928 | -1.476 | -0.003 | 0 | 0 | 1.756 | 0.66 | 0.73 | 0.62 | 0.72 | 0.63 | 0.72 |
| 1 | 76210289 | rs11161510 | 0.814 | 0.84 | 1.051 | -1.486 | -0.018 | 0.068 | 0.068 | 0.046 | 0.25 | 0.14 | 0.25 | 0.02 | 0.3 | 0.2 |
| 1 | 76213024 | rs1146581 | 0.128 | 0.732 | 0.907 | -1.496 | 0.009 | -0.101 | -0.101 | 0 | 0.29 | 0.6 | 0.33 | 0.02 | 0.32 | 0.2 |
| 1 | 76235084 | rs6664254 | 0.939 | 0.83 | 0.692 | -1.464 | -0.014 | 0.028 | 0.028 | 0.409 | 0.27 | 0.17 | 0.2 | 0.02 | 0.31 | 0.21 |
| 1 | 76244382 | rs11161602 | 0.893 | 0.692 | 0.208 | -1.214 | -0.032 | -0.126 | -0.126 | 0 | 0.32 | 0.01 | 0.21 | 0.69 | 0.24 | 0.48 |
| 1 | 76252011 | rs1146634 | -0.052 | 0.722 | 0.597 | -1.295 | 0.027 | 0.148 | 0.148 | 0 | 0.55 | 0.79 | 0.53 | 0.04 | 0.61 | 0.41 |
| 1 | 76255228 | rs11161620 | 0.724 | 0.795 | 0.706 | -1.336 | -0.011 | 0.079 | 0.079 | 0.448 | 0.25 | 0.11 | 0.22 | 0.02 | 0.28 | 0.2 |
| 1 | 76265712 | rs1146647 | -0.781 | 0.898 | 0.838 | -1.303 | -0.002 | 0.112 | 0.112 | 0 | 0.94 | 0.92 | 0.81 | 0.74 | 0.94 | 0.96 |
| 1 | 76269460 | rs5745325 | 0.909 | 0.933 | 1.177 | -1.463 | 0.003 | -0.073 | -0.073 | 0 | 0.28 | 0.3 | 0.2 | 0.02 | 0.31 | 0.2 |
| 1 | 76273805 | rs5745343 | 0.447 | 0.948 | 1.275 | -1.417 | 0.077 | 0.36 | 0.36 | 0 | 0.92 | 0.61 | 0.76 | 0.74 | 0.92 | 0.95 |
| 1 | 76275100 | rs1144333 | -0.201 | 0.953 | 1.433 | -1.413 | 0.029 | 0.214 | 0.214 | 0 | 0.93 | 0.81 | 0.77 | 0.74 | 0.92 | 0.95 |
| 1 | 76284395 | rs12024645 | 0.922 | 0.912 | 1.176 | -1.478 | -0.047 | -0.12 | -0.12 | 0.026 | 0.34 | 0.01 | 0.22 | 0.69 | 0.27 | 0.49 |
| 1 | 76320441 | rs1565717 | -0.936 | 0.968 | 1.341 | -1.476 | 0 | -0.092 | -0.092 | 0 | 0.75 | 0.65 | 0.77 | 0.98 | 0.74 | 0.8 |
| 1 | 76331320 | rs2172507 | -0.949 | 0.968 | 1.39 | -1.478 | 0 | -0.062 | -0.062 | 0 | 0.75 | 0.68 | 0.77 | 0.98 | 0.74 | 0.8 |
| 1 | 76342180 | rs12164500 | 0.158 | 0.97 | 1.329 | -1.504 | 0.001 | 0.192 | 0.192 | 0 | 0.43 | 0.19 | 0.47 | 0.96 | 0.38 | 0.55 |
| 1 | 76346712 | rs5745464 | 0.83 | 0.961 | 1.277 | -1.457 | -0.036 | -0.093 | -0.093 | 0 | 0.37 | 0.02 | 0.23 | 0.7 | 0.3 | 0.5 |
| 1 | 76358591 | rs1001160 | 0.887 | 0.982 | 1.443 | -1.443 | 0.009 | -0.113 | -0.113 | 0 | 0.28 | 0.34 | 0.21 | 0.02 | 0.31 | 0.2 |
| 1 | 76364825 | rs6687882 | 0.095 | 1.014 | 1.419 | -1.439 | 0.015 | 0.004 | 0.004 | 2.465 | 0.35 | 0.52 | 0.49 | 0.28 | 0.4 | 0.25 |
| 1 | 76367722 | rs11161848 | 0.973 | 0.991 | 1.423 | -1.531 | 0.001 | -0.058 | -0.058 | 1.092 | 0.27 | 0.28 | 0.2 | 0.02 | 0.31 | 0.19 |
| 1 | 76383297 | rs10782582 | -0.024 | 0.978 | 1.137 | -1.351 | -0.043 | 0.013 | 0.013 | 2.525 | 0.6 | 0.18 | 0.47 | 0.72 | 0.55 | 0.71 |
| 1 | 76397383 | rs2028212 | 0.792 | 0.959 | 1.039 | -1.369 | -0.034 | -0.077 | -0.077 | 0 | 0.37 | 0.03 | 0.26 | 0.71 | 0.3 | 0.5 |
| 1 | 76397972 | rs3795251 | 0.581 | 0.959 | 1.192 | -1.383 | 0.004 | -0.001 | -0.001 | 0 | 0.38 | 0.19 | 0.3 | 0.71 | 0.32 | 0.5 |
| 1 | 76398305 | rs6593520 | -0.007 | 0.978 | 1.201 | -1.451 | 0.01 | 0.132 | 0.132 | 0 | 0.47 | 0.28 | 0.51 | 0.96 | 0.44 | 0.59 |
| 1 | 76400261 | rs12090849 | 0.863 | 0.968 | 1.22 | -1.443 | -0.001 | -0.018 | -0.018 | 0 | 0.23 | 0.22 | 0.22 | 0.01 | 0.25 | 0.21 |
| 1 | 76411549 | rs1770882 | 0.801 | 0.967 | 1.046 | -1.364 | 0.009 | -0.106 | -0.106 | 0 | 0.28 | 0.34 | 0.21 | 0.02 | 0.31 | 0.19 |
| 1 | 76420745 | rs1251588 | -0.002 | 0.296 | 1.041 | -1.152 | -0.004 | 0.219 | 0.219 | 0 | 0.86 | 0.48 | 0.68 | 0.75 | 0.88 | 0.89 |
| 1 | 76438011 | rs12566984 | 0.874 | 0.336 | 0.53 | -1.34 | 0.003 | -0.096 | -0.096 | 0.287 | 0.33 | 0.01 | 0.24 | 0.71 | 0.25 | 0.41 |
| 1 | 76453857 | rs1251567 | 0.838 | 0.59 | -0.409 | -1.314 | -0.063 | 0.109 | 0.109 | 3.057 | 0.3 | 0.08 | 0.25 | 0.02 | 0.37 | 0.26 |
| 1 | 76456329 | rs1770878 | 0.363 | 0.74 | 0.257 | -0.577 | 0.027 | 0.179 | 0.179 | 0.239 | 0.86 | 0.43 | 0.66 | 0.75 | 0.85 | 0.89 |
| 1 | 76469334 | rs1251531 | 0.21 | 0.881 | -0.379 | -0.436 | 0 | 0.008 | 0.008 | 1.117 | 0.63 | 0.21 | 0.49 | 0.73 | 0.62 | 0.67 |
| 1 | 76485133 | rs2289241 | 0.141 | 0.364 | -1.457 | -1.144 | 0.006 | 0.027 | 0.027 | 0 | 0.33 | 0.04 | 0.27 | 0.79 | 0.3 | 0.38 |
| 1 | 76485783 | rs6703034 | 0.02 | 0.073 | -1.029 | -1.904 | -0.02 | -0.021 | -0.021 | 2.509 | 0.79 | 0.34 | 0.66 | 0.86 | 0.82 | 0.84 |
| 1 | 76486908 | rs6593523 | 0.262 | 0.636 | -1.504 | -2.468 | -0.029 | 0.255 | 0.255 | 0.06 | 0.5 | 0.08 | 0.43 | 0.85 | 0.48 | 0.63 |
| 1 | 76487624 | rs1251525 | -0.096 | 0.408 | -1.121 | -2.487 | 0.024 | 0.023 | 0.023 | 0.228 | 0.85 | 0.41 | 0.75 | 0.87 | 0.82 | 0.85 |
| 1 | 76497033 | rs1370714 | 0.215 | 0.093 | -0.304 | -2.936 | 0.011 | 0.046 | 0.046 | 0 | 0.45 | 0.58 | 0.46 | 0.9 | 0.47 | 0.52 |
| 1 | 76499313 | rs11162070 | 1.538 | -0.286 | -0.642 | -3.199 | -0.009 | 0.061 | 0.061 | 0.811 | 0.26 | 0.05 | 0.25 | 0.88 | 0.29 | 0.37 |
| 1 | 76500705 | rs17599859 | -0.139 | -0.287 | -0.508 | -3.202 | 0 | -0.021 | -0.021 | 0 | 0.44 | 0.38 | 0.36 | 0.9 | 0.47 | 0.51 |
| 1 | 76501151 | rs17097968 | -0.017 | -0.288 | -0.513 | -3.202 | -0.003 | -0.078 | -0.078 | 0.921 | 0.17 | 0.27 | 0.12 | 0.02 | 0.18 | 0.14 |
| 1 | 76503195 | rs10493579 | 0.778 | -0.281 | -0.64 | -3.252 | -0.1 | 0.21 | 0.21 | 0 | 0.34 | 0.04 | 0.24 | 0.04 | 0.35 | 0.21 |
| 1 | 76504146 | rs1251484 | -1.104 | -0.186 | -0.393 | -3.034 | 0.002 | -0.035 | -0.035 | 0.002 | 0.79 | 0.66 | 0.7 | 0.94 | 0.82 | 0.73 |
| 1 | 76509560 | rs7529565 | 1.271 | -0.187 | -0.665 | -3.035 | 0.006 | 0.052 | 0.052 | 0.131 | 0.27 | 0.12 | 0.24 | 0.88 | 0.29 | 0.37 |
| 1 | 76513759 | rs1013629 | 0.171 | -0.134 | -0.645 | -3.067 | 0.013 | 0.082 | 0.082 | 0 | 0.5 | 0.33 | 0.51 | 0.93 | 0.47 | 0.65 |
| 1 | 76518693 | rs1251490 | -0.073 | -0.145 | -0.216 | -3.063 | 0 | -0.042 | -0.042 | 0 | 0.53 | 0.46 | 0.62 | 0.95 | 0.47 | 0.64 |
| 1 | 76530882 | rs1251474 | -0.243 | -0.075 | -0.389 | -3.03 | -0.053 | 0.138 | 0.138 | 1.009 | 0.3 | 0.07 | 0.25 | 0.05 | 0.3 | 0.34 |
| 1 | 76535434 | rs1146658 | 0.033 | -0.104 | -0.209 | -3.038 | 0.006 | -0.063 | -0.063 | 0.008 | 0.12 | 0.21 | 0.1 | 0 | 0.12 | 0.06 |
| 1 | 76536649 | rs1251467 | -0.244 | 0.072 | -0.214 | -2.98 | 0.006 | -0.118 | -0.118 | 0 | 0.34 | 0.5 | 0.4 | 0.07 | 0.31 | 0.36 |
| 1 | 76549233 | rs11803060 | -0.588 | -0.114 | 0.132 | -3.012 | 0.028 | 0.019 | 0.019 | 0 | 0.07 | 0.32 | 0.03 | 0.02 | 0.02 | 0.09 |
| 1 | 76550721 | rs1251578 | 0.348 | 0.26 | -0.22 | -3.398 | 0.004 | -0.006 | -0.006 | 0.053 | 0.31 | 0.28 | 0.31 | 0.05 | 0.25 | 0.14 |
| 1 | 76552475 | rs1144346 | -0.829 | 0.503 | -0.374 | -3.089 | -0.001 | -0.008 | -0.008 | 0 | 0.67 | 0.66 | 0.63 | 0.13 | 0.59 | 0.59 |
| 1 | 76556292 | rs289692 | -1.166 | 0.814 | -0.196 | -2.61 | 0.037 | -0.011 | -0.011 | 0.259 | 0.11 | 0.21 | 0.22 | 0.06 | 0.19 | 0.14 |
| 1 | 76557784 | rs1251575 | -1.538 | 0.721 | -0.308 | -2.915 | 0.001 | -0.008 | -0.008 | 0.065 | 0.28 | 0.63 | 0.35 | 0.15 | 0.31 | 0.26 |
| 1 | 76559451 | rs50564 | -1.416 | 0.728 | -0.171 | -2.914 | 0.014 | -0.036 | -0.036 | 0 | 0.27 | 0.56 | 0.34 | 0.15 | 0.31 | 0.26 |
| 1 | 76568506 | rs10873866 | 0.008 | 0.677 | -0.605 | -3.203 | -0.1 | 0.214 | 0.214 | 0.43 | 0.43 | 0.09 | 0.44 | 0.15 | 0.38 | 0.44 |
| 1 | 76569151 | rs289679 | -1.278 | 0.474 | -0.599 | -3.283 | 0.011 | 0.048 | 0.048 | 0.322 | 0.61 | 0.67 | 0.69 | 0.21 | 0.57 | 0.58 |
| 1 | 76570926 | rs10493583 | 0.088 | 0.508 | -0.638 | -3.298 | -0.097 | 0.217 | 0.217 | 0.441 | 0.43 | 0.09 | 0.43 | 0.14 | 0.36 | 0.45 |
| 1 | 76573769 | rs409245 | -0.626 | 0.59 | -0.353 | -3.167 | 0.023 | -0.05 | -0.05 | 0.012 | 0.17 | 0.46 | 0.25 | 0.06 | 0.19 | 0.14 |
| 1 | 76585643 | rs17098119 | -0.162 | 0.587 | -0.27 | -3.186 | 0.03 | -0.058 | -0.058 | 0.577 | 0.14 | 0.41 | 0.22 | 0.03 | 0.14 | 0.07 |
| 1 | 76586272 | rs427589 | -1.185 | 0.696 | -0.4 | -3.297 | 0.006 | 0.047 | 0.047 | 0.377 | 0.61 | 0.67 | 0.69 | 0.21 | 0.55 | 0.58 |
| 1 | 76588080 | rs17609927 | 0.469 | 1.057 | -0.81 | -3.045 | -0.023 | 0.022 | 0.022 | 0 | 0.05 | 0.02 | 0.09 | 0 | 0.13 | 0.05 |
| 1 | 76589102 | rs11162084 | 0.73 | 0.907 | -0.797 | -3.033 | -0.031 | 0.105 | 0.105 | 1.508 | 0.29 | 0.12 | 0.35 | 0.06 | 0.24 | 0.34 |
| 1 | 76598209 | rs289697 | 0.143 | 0.663 | -0.825 | -3.133 | 0.008 | -0.053 | -0.053 | 0 | 0.39 | 0.53 | 0.38 | 0.83 | 0.49 | 0.42 |
| 1 | 76598267 | rs6702844 | 1.169 | 0.846 | -1.145 | -3.292 | -0.104 | 0.127 | 0.127 | 4.534 | 0.29 | 0.01 | 0.33 | 0.06 | 0.27 | 0.38 |
| 1 | 76601214 | rs11162086 | 1.159 | 0.825 | -0.367 | -2.797 | 0 | 0.004 | 0.004 | 1.121 | 0.27 | 0.2 | 0.25 | 0.79 | 0.31 | 0.3 |
| 1 | 76602125 | rs289701 | -1.187 | 1.097 | -0.621 | -2.437 | 0.032 | -0.039 | -0.039 | 0 | 0.12 | 0.3 | 0.13 | 0.04 | 0.17 | 0.11 |

| | | | | | | | | | | | | | | | | |
|---|---|---|---|---|---|---|---|---|---|---|---|---|---|---|---|---|
| 1 | 76602605 | rs915403 | -1.126 | 1.118 | -0.957 | -2.428 | 0.017 | -0.028 | -0.028 | 0.09 | 0.12 | 0.25 | 0.13 | 0.04 | 0.17 | 0.11 |
| 1 | 76608018 | rs209256 | -0.006 | 0.881 | -1.534 | -2.591 | -0.024 | 0.272 | 0.272 | 1.41 | 0.49 | 0.09 | 0.47 | 0.16 | 0.49 | 0.56 |
| 1 | 76608072 | rs7548049 | 0.473 | 0.661 | -0.468 | -2.408 | -0.023 | -0.039 | -0.039 | 0.096 | 0.28 | 0.47 | 0.29 | 0.8 | 0.26 | 0.3 |
| 1 | 76608777 | rs184095 | 0.816 | 1.157 | -0.295 | -2.025 | -0.009 | 0.014 | 0.014 | 0 | 0.68 | 0.31 | 0.7 | 0.86 | 0.59 | 0.73 |
| 1 | 76608949 | rs10493584 | 0.551 | 0.812 | -0.051 | -2.566 | 0.014 | 0.072 | 0.072 | 2.286 | 0.56 | 0.3 | 0.68 | 0.89 | 0.62 | 0.73 |
| 1 | 76610257 | rs4245647 | 0.682 | 0.76 | 0.863 | -2.55 | 0.025 | 0.162 | 0.162 | 2.609 | 0.54 | 0.17 | 0.68 | 0.89 | 0.62 | 0.72 |
| 1 | 76610397 | rs418644 | 0.975 | 0.733 | 0.917 | -2.584 | 0.019 | 0.029 | 0.029 | 4.001 | 0.24 | 0.17 | 0.11 | 0.02 | 0.12 | 0.08 |
| 1 | 76612729 | rs12057667 | -0.925 | 0.899 | 0.981 | -2.524 | 0.041 | 0 | 0 | 0.372 | 0.1 | 0.27 | 0.1 | 0.06 | 0.13 | 0.11 |
| 1 | 76617485 | rs571399 | 0.749 | 0.853 | 0.987 | -2.647 | 0.006 | -0.069 | -0.069 | 0.077 | 0.78 | 0.33 | 0.79 | 0.96 | 0.75 | 0.84 |
| 1 | 76617993 | rs4949623 | 0.073 | 0.654 | 1.062 | -2.65 | -0.021 | -0.087 | -0.087 | 0 | 0.62 | 0.42 | 0.79 | 0.95 | 0.74 | 0.83 |
| 1 | 76620313 | rs2783658 | 0.257 | 0.63 | 0.831 | -2.441 | -0.003 | -0.03 | -0.03 | 0 | 0.12 | 0.26 | 0.1 | 0.03 | 0.13 | 0.1 |
| 1 | 76624862 | rs11162089 | 0.034 | 0.95 | 0.9 | -2.397 | 0.033 | -0.034 | -0.034 | 5.719 | 0.07 | 0.16 | 0.09 | 0 | 0.12 | 0.1 |
| 1 | 76626260 | rs11162090 | 0.303 | 0.766 | 0.994 | -2.638 | 0.006 | -0.001 | -0.001 | 0.351 | 0.62 | 0.27 | 0.75 | 0.93 | 0.7 | 0.79 |
| 1 | 76626550 | rs402394 | 0.904 | 0.665 | 0.862 | -2.637 | 0.005 | 0.107 | 0.107 | 4.067 | 0.25 | 0.05 | 0.11 | 0.07 | 0.17 | 0.11 |
| 1 | 76628659 | rs1414406 | -0.514 | 0.709 | 0.948 | -2.615 | -0.026 | -0.085 | -0.085 | 0 | 0.7 | 0.5 | 0.85 | 0.93 | 0.82 | 0.89 |
| 1 | 76629673 | rs7513539 | 0.923 | 1.221 | -0.078 | -2.837 | -0.072 | 0.02 | 0.02 | 0 | 0.14 | 0.01 | 0.28 | 0.07 | 0.19 | 0.31 |
| 1 | 76637090 | rs7556363 | 0.785 | 0.794 | 0.464 | -2.69 | 0.008 | 0.219 | 0.219 | 0.835 | 0.53 | 0.14 | 0.7 | 0.88 | 0.6 | 0.71 |
| 1 | 76652957 | rs4949697 | 0.839 | 0.816 | 0.466 | -2.732 | 0.017 | 0.197 | 0.197 | 0.681 | 0.52 | 0.17 | 0.69 | 0.88 | 0.59 | 0.71 |
| 1 | 76658460 | rs923246 | 0.41 | 0.768 | 0.814 | -2.723 | 0.047 | -0.041 | -0.041 | 4.979 | 0.91 | 0.63 | 0.89 | 1 | 0.88 | 0.91 |
| 1 | 76658494 | rs923245 | 1.08 | 0.761 | 0.9 | -2.776 | 0.031 | -0.105 | -0.105 | 0 | 0.81 | 0.51 | 0.83 | 0.95 | 0.78 | 0.82 |
| 1 | 76659892 | rs10493585 | 0.304 | 0.788 | 0.555 | -2.696 | 0.003 | -0.018 | -0.018 | 0 | 0.09 | 0.13 | 0.06 | 0.05 | 0.09 | 0.09 |
| 1 | 76664973 | rs17098324 | 0.954 | 0.752 | 0.385 | -2.53 | -0.027 | -0.032 | -0.032 | 0.25 | 0.11 | 0.01 | 0.24 | 0.09 | 0.15 | 0.09 |
| 1 | 76665498 | rs1999734 | 0.114 | 0.842 | 0.759 | -2.631 | -0.005 | -0.063 | -0.063 | 0 | 0.63 | 0.4 | 0.77 | 0.93 | 0.7 | 0.81 |
| 1 | 76668470 | rs17098341 | -0.319 | 0.731 | 0.427 | -2.351 | 0.001 | -0.024 | -0.024 | 0 | 0.11 | 0.15 | 0.07 | 0.05 | 0.1 | 0.09 |
| 1 | 76680525 | rs517036 | 0.319 | 0.503 | 0.055 | -2.505 | 0.016 | 0.04 | 0.04 | 5.326 | 0.27 | 0.21 | 0.13 | 0.02 | 0.19 | 0.05 |
| 1 | 76685608 | rs551877 | -0.177 | 0.357 | 0.133 | -2.451 | -0.045 | -0.052 | -0.052 | 0 | 0.91 | 0.73 | 0.96 | 1 | 0.98 | 0.96 |
| 1 | 76685705 | rs12035017 | 0.999 | 0.788 | -0.667 | -2.338 | -0.017 | 0.234 | 0.234 | 0 | 0.5 | 0.11 | 0.67 | 0.84 | 0.52 | 0.67 |
| 1 | 76690002 | rs1124171 | 0.593 | 0.298 | -0.346 | -2.045 | 0.038 | 0.152 | 0.152 | 2.523 | 0.59 | 0.15 | 0.77 | 0.89 | 0.7 | 0.86 |
| 1 | 76692832 | rs479650 | 0.154 | 0.262 | -0.396 | -2.175 | 0.008 | 0.096 | 0.096 | 1.664 | 0.26 | 0.1 | 0.12 | 0.07 | 0.2 | 0.1 |
| 1 | 76695328 | rs17098418 | 0.427 | 0.115 | 0.103 | -2.227 | -0.031 | -0.033 | -0.033 | 0.189 | 0.06 | 0.17 | 0.01 | 0 | 0.01 | 0.03 |
| 1 | 76697767 | rs573657 | -0.86 | 0.141 | 0.632 | -2.254 | 0.004 | -0.024 | -0.024 | 0 | 0.38 | 0.67 | 0.18 | 0.07 | 0.23 | 0.13 |
| 1 | 76699911 | rs512082 | -1.15 | 0.216 | 0.634 | -2.307 | 0.028 | 0.14 | 0.14 | 1.776 | 0.43 | 0.77 | 0.22 | 0.14 | 0.31 | 0.16 |
| 1 | 76700571 | rs490570 | -0.941 | 0.236 | 0.63 | -2.301 | 0.018 | 0.072 | 0.072 | 1.094 | 0.41 | 0.71 | 0.21 | 0.14 | 0.31 | 0.16 |
| 1 | 76714019 | rs472536 | -1.091 | 0.462 | -0.523 | -2.443 | 0.004 | 0.056 | 0.056 | 0.265 | 0.45 | 0.69 | 0.27 | 0.09 | 0.4 | 0.19 |
| 1 | 76717966 | rs572581 | -0.325 | 0.428 | -0.527 | -2.513 | 0.001 | -0.066 | -0.066 | 0 | 0.28 | 0.67 | 0.18 | 0.05 | 0.23 | 0.09 |
| 1 | 76719571 | rs536811 | 0.495 | 0.592 | -0.775 | -2.418 | -0.025 | 0.05 | 0.05 | 0 | 0.2 | 0.11 | 0.13 | 0.05 | 0.21 | 0.08 |
| 1 | 76721164 | rs501426 | -1.218 | 0.475 | -0.891 | -2.474 | -0.002 | 0.286 | 0.286 | 0.62 | 0.48 | 0.84 | 0.28 | 0.08 | 0.4 | 0.22 |
| 1 | 76724652 | rs572200 | 0.512 | 0.362 | -0.81 | -2.261 | 0.004 | -0.016 | -0.016 | 0 | 0.2 | 0.22 | 0.13 | 0.08 | 0.2 | 0.06 |
| 1 | 76725259 | rs544858 | -0.565 | 0.517 | -1.366 | -1.932 | 0.002 | 0.238 | 0.238 | 0.186 | 0.47 | 0.83 | 0.31 | 0.11 | 0.41 | 0.24 |
| 1 | 76729651 | rs12069137 | 0.637 | 0.615 | -1.473 | -1.593 | -0.006 | 0.05 | 0.05 | 0.092 | 0.25 | 0.12 | 0.18 | 0.19 | 0.26 | 0.32 |
| 1 | 76729911 | rs11162104 | 0.135 | 0.362 | -1.759 | -1.873 | -0.007 | -0.005 | -0.005 | 0.391 | 0.23 | 0.09 | 0.48 | 0.69 | 0.23 | 0.3 |
| 1 | 76731907 | rs6696780 | 0.469 | -0.298 | -0.935 | -2.334 | -0.069 | -0.101 | -0.101 | 0.384 | 0.3 | 0.46 | 0.11 | 0.09 | 0.17 | 0.11 |
| 1 | 76732617 | rs4949624 | -1.066 | 0.201 | -0.885 | -2.124 | -0.015 | 0.052 | 0.052 | 0.365 | 0.72 | 0.85 | 0.79 | 0.8 | 0.68 | 0.61 |
| 1 | 76739856 | rs6695042 | 0.82 | -0.63 | 0.327 | -0.929 | -0.054 | 0.129 | 0.129 | 2.088 | 0.15 | 0.4 | 0.15 | 0.33 | 0.04 | 0.09 |
| 1 | 76746406 | rs12131613 | 0.486 | -0.35 | 0.152 | -1.547 | -0.063 | 0.063 | 0.063 | 0 | 0.24 | 0.1 | 0.37 | 0.02 | 0.39 | 0.21 |
| 1 | 76746909 | rs17098538 | 1.543 | -0.44 | 0.166 | -1.48 | -0.03 | 0.056 | 0.056 | 2.043 | 0.15 | 0.26 | 0.16 | 0.33 | 0.06 | 0.09 |
| 1 | 76749527 | rs11162113 | 1.055 | -0.829 | 0.375 | -1.625 | -0.001 | 0.034 | 0.034 | 0 | 0.21 | 0.21 | 0.31 | 0.5 | 0.19 | 0.29 |
| 1 | 76750203 | rs6662992 | -1.761 | -1.04 | 0.33 | -1.732 | -0.007 | 0.012 | 0.012 | 0 | 0.82 | 0.89 | 0.92 | 0.85 | 0.8 | 0.73 |
| 1 | 76750428 | rs4607942 | 0.06 | -1.059 | 0.322 | -1.753 | -0.007 | 0.003 | 0.003 | 0 | 0.39 | 0.36 | 0.53 | 0.35 | 0.45 | 0.3 |
| 1 | 76752116 | rs1537783 | 0.043 | -1.155 | 0.289 | -1.76 | 0.011 | -0.012 | -0.012 | 0.222 | 0.41 | 0.53 | 0.54 | 0.35 | 0.46 | 0.3 |
| 1 | 76754957 | rs9437407 | 1.813 | -1.122 | 0.074 | -1.7 | -0.031 | 0.041 | 0.041 | 1.972 | 0.15 | 0.26 | 0.16 | 0.29 | 0.06 | 0.09 |
| 1 | 76755492 | rs4949626 | -1.678 | -0.887 | -0.087 | -1.761 | -0.002 | 0.005 | 0.005 | 0.011 | 0.82 | 0.87 | 0.88 | 0.85 | 0.81 | 0.73 |
| 1 | 76756343 | rs3862899 | 0.635 | -0.689 | -0.196 | -1.64 | -0.07 | 0.067 | 0.067 | 0 | 0.25 | 0.1 | 0.37 | 0.06 | 0.41 | 0.21 |
| 1 | 76768447 | rs2893415 | 0.12 | -0.885 | -0.456 | -1.402 | -0.013 | 0.025 | 0.025 | 0 | 0.41 | 0.33 | 0.52 | 0.22 | 0.45 | 0.3 |
| 1 | 76770639 | rs12754064 | 1.125 | -0.563 | -0.566 | -1.137 | -0.022 | 0.012 | 0.012 | 0 | 0.18 | 0.08 | 0.12 | 0.17 | 0.22 | 0.33 |
| 1 | 76771114 | rs6701166 | -0.527 | -1.123 | -0.595 | -1.225 | 0.035 | 0.019 | 0.019 | 0.178 | 0.66 | 0.74 | 0.79 | 0.73 | 0.8 | 0.81 |
| 1 | 76772328 | rs10873876 | -0.398 | -1.115 | -0.364 | -1.111 | -0.047 | -0.017 | -0.017 | 0 | 0.64 | 0.52 | 0.78 | 0.71 | 0.8 | 0.81 |
| 1 | 76774201 | rs3884182 | 0.29 | -0.718 | -0.658 | -0.91 | -0.002 | -0.016 | -0.016 | 0.281 | 0.24 | 0.18 | 0.36 | 0.57 | 0.22 | 0.37 |
| 1 | 76774380 | rs2893416 | -0.988 | -0.898 | -0.68 | -0.824 | 0.002 | -0.004 | -0.004 | 0 | 0.79 | 0.81 | 0.78 | 0.74 | 0.89 | 0.85 |
| 1 | 76777065 | rs6681765 | 0.164 | -0.359 | -0.777 | -0.896 | 0.031 | -0.045 | -0.045 | 0 | 0.33 | 0.4 | 0.39 | 0.17 | 0.46 | 0.26 |
| 1 | 76779877 | rs10873878 | -0.271 | -0.61 | -0.831 | 0.311 | -0.003 | 0.071 | 0.071 | 0.127 | 0.14 | 0.18 | 0.3 | 0.41 | 0.12 | 0.3 |
| 1 | 76781576 | rs3904686 | 0.868 | -0.746 | -1.164 | -0.228 | 0.037 | 0.065 | 0.065 | 0.015 | 0.17 | 0.07 | 0.04 | 0 | 0.14 | 0.08 |
| 1 | 76782834 | rs12046510 | -0.357 | -0.827 | -0.896 | 0.235 | -0.009 | 0.006 | 0.006 | 0 | 0.3 | 0.31 | 0.33 | 0.38 | 0.23 | 0.34 |
| 1 | 76784636 | rs2392030 | -0.379 | -0.19 | -1.09 | -0.056 | 0.002 | 0.003 | 0.003 | 1.344 | 0.52 | 0.44 | 0.65 | 0.59 | 0.52 | 0.68 |
| 1 | 76785319 | rs11162121 | -0.293 | -0.076 | -1.115 | -1.143 | -0.02 | 0.138 | 0.138 | 0 | 0.54 | 0.7 | 0.5 | 0.16 | 0.51 | 0.32 |
| 1 | 76785983 | rs12039112 | 0.455 | -0.093 | -1.24 | -1.179 | -0.069 | 0.168 | 0.168 | 0 | 0.41 | 0.12 | 0.46 | 0.82 | 0.46 | 0.65 |
| 1 | 76786084 | rs9437421 | 0.505 | -0.099 | -1.243 | -1.182 | -0.07 | 0.167 | 0.167 | 0 | 0.41 | 0.12 | 0.46 | 0.82 | 0.46 | 0.65 |
| 1 | 76788220 | rs9437136 | -0.575 | -0.239 | -0.747 | -1.236 | -0.015 | -0.067 | -0.067 | 0.263 | 0.71 | 0.3 | 0.87 | 0.98 | 0.83 | 0.88 |
| 1 | 76796785 | rs10493591 | 1.114 | -0.122 | -0.736 | -1.191 | -0.057 | -0.152 | -0.152 | 0.5 | 0.24 | 0.41 | 0.09 | 0 | 0.15 | 0.09 |

| | | | | | | | | | | | | | | | | |
|---|---|---|---|---|---|---|---|---|---|---|---|---|---|---|---|---|
| 1 | 76801058 | rs6669101 | 0.638 | 0.637 | -1.062 | -1.209 | -0.023 | 0.018 | 0.018 | 0 | 0.09 | 0.04 | 0.08 | 0 | 0.12 | 0.08 |
| 1 | 76809963 | rs12069012 | 0.628 | 0.256 | -0.921 | -1.319 | -0.046 | -0.15 | -0.15 | 4.27 | 0.31 | 0.53 | 0.13 | 0.02 | 0.19 | 0.13 |
| 1 | 76810983 | rs6685141 | 0.482 | 0.483 | -1.408 | -1.095 | -0.012 | -0.022 | -0.022 | 0 | 0.38 | 0.1 | 0.35 | 0.67 | 0.3 | 0.58 |
| 1 | 76812062 | rs17672913 | 1.256 | 0.979 | -1.433 | -0.867 | -0.037 | 0.024 | 0.024 | 0.122 | 0.17 | 0.01 | 0.35 | 0.14 | 0.24 | 0.14 |
| 1 | 76815789 | rs12122625 | 1.39 | 0.977 | -1.239 | -0.785 | 0 | 0 | 0 | 0.085 | 0.1 | 0.11 | 0.15 | 0.16 | 0.26 | 0.14 |
| 1 | 76820435 | rs942699 | 1.364 | 1.034 | -1.24 | -0.778 | -0.004 | -0.004 | -0.004 | 0.023 | 0.1 | 0.09 | 0.15 | 0.16 | 0.26 | 0.14 |
| 1 | 76822376 | rs1325287 | 0.555 | 0.914 | -1.185 | -0.889 | 0.006 | 0.004 | 0.004 | 0.949 | 0.63 | 0.58 | 0.33 | 0.63 | 0.42 | 0.65 |
| 1 | 76824240 | rs4638174 | 0.161 | 0.384 | -0.752 | -0.963 | 0.019 | 0.017 | 0.017 | 0 | 0.52 | 0.37 | 0.63 | 0.34 | 0.71 | 0.4 |
| 1 | 76832860 | rs12071740 | 1.304 | 0.554 | -0.89 | -0.838 | -0.044 | 0.017 | 0.017 | 0.237 | 0.16 | 0.03 | 0.38 | 0.16 | 0.25 | 0.14 |
| 1 | 76833594 | rs12032409 | 1.064 | 0.277 | -0.943 | -0.957 | 0.036 | -0.035 | -0.035 | 0 | 0.3 | 0.2 | 0.17 | 0.61 | 0.15 | 0.47 |
| 1 | 76834705 | rs12128122 | -0.412 | 0.29 | -0.768 | -1.003 | -0.023 | -0.001 | -0.001 | 0 | 0.61 | 0.47 | 0.69 | 0.36 | 0.79 | 0.47 |
| 1 | 76837237 | rs12239582 | 1.228 | 0.3 | -1.048 | -1.143 | -0.107 | -0.003 | -0.003 | 0 | 0.33 | 0.06 | 0.21 | 0.16 | 0.43 | 0.24 |
| 1 | 76838919 | rs11162133 | 1.528 | 0.346 | -0.928 | -1.037 | -0.04 | 0.023 | 0.023 | 0.145 | 0.16 | 0.03 | 0.36 | 0.15 | 0.24 | 0.13 |
| 1 | 76840373 | rs7339997 | 0.811 | 0.23 | -0.557 | -0.944 | -0.036 | 0.024 | 0.024 | 0.155 | 0.33 | 0.5 | 0.2 | 0.61 | 0.17 | 0.47 |
| 1 | 76852740 | rs1925359 | -0.17 | 0.611 | -0.724 | -0.82 | -0.007 | 0.008 | 0.008 | 0.394 | 0.24 | 0.23 | 0.49 | 0.21 | 0.31 | 0.19 |
| 1 | 76862648 | rs2209458 | 0.83 | 0.547 | -0.904 | -0.789 | -0.033 | 0.044 | 0.044 | 0 | 0.36 | 0.2 | 0.24 | 0.16 | 0.46 | 0.25 |
| 1 | 76864069 | rs4949715 | -0.628 | 0.776 | -0.327 | -0.552 | 0.021 | -0.037 | -0.037 | 0.035 | 0.81 | 0.6 | 0.6 | 0.84 | 0.76 | 0.87 |
| 1 | 76865258 | rs4949716 | 0.052 | 1.071 | -0.379 | -0.428 | 0.134 | 0.091 | 0.091 | 0 | 0.81 | 0.52 | 0.65 | 0.8 | 0.67 | 0.79 |
| 1 | 76865351 | rs10782620 | 0.666 | 0.476 | -0.696 | -0.713 | -0.002 | -0.018 | -0.018 | 0 | 0.43 | 0.53 | 0.46 | 0.23 | 0.47 | 0.29 |
| 1 | 76865591 | rs12096348 | 1.005 | 0.988 | -0.76 | -0.22 | -0.01 | -0.001 | -0.001 | 0 | 0.14 | 0.06 | 0.24 | 0.18 | 0.21 | 0.19 |
| 1 | 76867853 | rs6667721 | 0.16 | 0.993 | -0.877 | -0.279 | -0.061 | 0.004 | 0.004 | 0 | 0.13 | 0.01 | 0.21 | 0.06 | 0.14 | 0.08 |
| 1 | 76870939 | rs10218698 | 0.55 | 0.288 | -0.603 | -0.259 | 0.011 | -0.016 | -0.016 | 0 | 0.58 | 0.66 | 0.19 | 0.54 | 0.35 | 0.56 |
| 1 | 76876498 | rs2147791 | 0.145 | 0.085 | -0.832 | -0.194 | -0.002 | -0.014 | -0.014 | 0.853 | 0.55 | 0.5 | 0.41 | 0.32 | 0.47 | 0.37 |
| 1 | 76880630 | rs10128020 | -0.827 | 0.085 | -0.554 | 0.401 | 0.069 | 0.048 | 0.048 | 0 | 0.81 | 0.6 | 0.56 | 0.74 | 0.69 | 0.79 |
| 1 | 76883631 | rs11162150 | 1.538 | -0.043 | -1.108 | 0.78 | 0.021 | -0.009 | -0.009 | 0 | 0.15 | 0.02 | 0.09 | 0.15 | 0.09 | 0.37 |
| 1 | 76890999 | rs7529530 | 0.991 | 0.141 | -1.272 | 0.358 | -0.039 | -0.046 | -0.046 | 0 | 0.18 | 0.03 | 0.26 | 0.27 | 0.24 | 0.21 |
| 1 | 76891796 | rs6593537 | -1.196 | -0.024 | -1.1 | 0.361 | -0.041 | -0.035 | -0.035 | 0 | 0.91 | 0.98 | 0.96 | 0.88 | 0.89 | 0.94 |
| 1 | 76899464 | rs4949718 | -0.334 | -0.369 | -0.95 | -0.033 | -0.001 | -0.014 | -0.014 | 0.913 | 0.42 | 0.27 | 0.69 | 0.56 | 0.59 | 0.44 |
| 1 | 76901499 | rs4478861 | -0.644 | -0.258 | -1.238 | 0.055 | -0.002 | 0.039 | 0.039 | 0 | 0.73 | 0.64 | 0.84 | 0.58 | 0.8 | 0.55 |
| 1 | 76906960 | rs17629475 | 1.45 | -0.25 | -2.134 | -0.259 | 0.062 | 0.132 | 0.132 | 0.024 | 0.27 | 0.06 | 0.1 | 0 | 0.18 | 0.11 |
| 1 | 76907906 | rs12742405 | 0.402 | 0.665 | -1.71 | 0.517 | -0.037 | -0.059 | -0.059 | 0.211 | 0.19 | 0.03 | 0.24 | 0.26 | 0.25 | 0.22 |
| 1 | 76910399 | rs7546379 | -0.454 | 1.192 | -1.103 | 0.454 | -0.049 | -0.044 | -0.044 | 0 | 0.83 | 0.91 | 0.66 | 0.69 | 0.68 | 0.79 |
| 1 | 76927674 | rs7413797 | 0.536 | 0.376 | -0.891 | -0.058 | -0.013 | -0.069 | -0.069 | 0.514 | 0.52 | 0.08 | 0.63 | 0.49 | 0.63 | 0.5 |
| 1 | 76931051 | rs7544496 | -0.864 | 1.036 | 0.818 | -0.406 | 0.028 | -0.006 | -0.006 | 0 | 0.13 | 0.25 | 0.12 | 0.07 | 0.16 | 0.03 |
| 1 | 76934790 | rs6691403 | -1.437 | 1.023 | 1.002 | -0.41 | 0.038 | -0.034 | -0.034 | 0.141 | 0.19 | 0.51 | 0.15 | 0.08 | 0.18 | 0.05 |
| 1 | 76945339 | rs12075814 | -0.817 | 0.728 | 0.681 | -0.391 | 0.014 | 0.01 | 0.01 | 0 | 0.13 | 0.34 | 0.12 | 0.07 | 0.14 | 0.05 |
| 1 | 76947447 | rs11162165 | -1.343 | 0.737 | 0.687 | -0.37 | 0.008 | 0.022 | 0.022 | 0 | 0.17 | 0.53 | 0.19 | 0.11 | 0.17 | 0.06 |
| 1 | 76957469 | rs6685154 | -1.504 | 0.821 | 0.146 | -0.458 | -0.005 | 0.028 | 0.028 | 1.305 | 0.26 | 0.9 | 0.25 | 0.11 | 0.23 | 0.08 |
| 1 | 76961960 | rs10873889 | 1.224 | 0.846 | 0.049 | -0.564 | -0.014 | 0.064 | 0.064 | 0.323 | 0.68 | 0.09 | 0.73 | 0.89 | 0.71 | 0.89 |
| 1 | 76965618 | rs1436085 | 0.469 | 1.035 | 0.18 | -0.305 | -0.001 | 0.004 | 0.004 | 0 | 0.81 | 0.73 | 0.74 | 0.8 | 0.79 | 0.9 |
| 1 | 76974109 | rs315093 | 0.494 | 1.059 | -2.282 | -0.509 | -0.065 | 0.145 | 0.145 | 0 | 0.55 | 0.07 | 0.52 | 0.77 | 0.54 | 0.79 |
| 1 | 76975019 | rs387831 | 0.478 | 0.939 | -2.153 | -0.468 | 0.018 | 0.128 | 0.128 | 0 | 0.53 | 0.16 | 0.55 | 0.75 | 0.54 | 0.71 |
| 1 | 76975059 | rs315091 | 0.191 | 0.954 | -2.285 | -0.53 | -0.03 | 0.1 | 0.1 | 1.651 | 0.64 | 0.07 | 0.59 | 0.82 | 0.66 | 0.83 |
| 1 | 76975437 | rs1436087 | -0.964 | 1.276 | -1.234 | -0.35 | 0.005 | -0.04 | -0.04 | 0.1 | 0.23 | 0.36 | 0.42 | 0.15 | 0.23 | 0.2 |
| 1 | 76975717 | rs10873890 | -0.655 | 1.434 | -1.649 | 0.516 | -0.043 | -0.067 | -0.067 | 1.81 | 0.29 | 0.18 | 0.38 | 0.45 | 0.39 | 0.5 |
| 1 | 76976522 | rs4949632 | -0.602 | 1.04 | -0.991 | 0.622 | 0.008 | 0.011 | 0.011 | 0.695 | 0.3 | 0.43 | 0.39 | 0.33 | 0.3 | 0.28 |
| 1 | 76979428 | rs188789 | 0.41 | 0.777 | -0.425 | 0.494 | 0.008 | 0.01 | 0.01 | 5.682 | 0.68 | 0.64 | 0.58 | 0.66 | 0.64 | 0.67 |
| 1 | 76980646 | rs12749279 | -0.197 | 0.255 | -0.787 | 0.644 | 0.008 | -0.027 | -0.027 | 0 | 0.12 | 0.05 | 0.07 | 0.22 | 0.1 | 0.23 |
| 1 | 76985313 | rs423480 | -1.418 | -0.027 | -0.312 | 0.635 | -0.004 | 0.044 | 0.044 | 3.223 | 0.49 | 0.72 | 0.57 | 0.61 | 0.47 | 0.59 |
| 1 | 76988637 | rs315084 | 0.256 | -0.202 | -0.173 | 0.31 | -0.018 | -0.065 | -0.065 | 0.041 | 0.85 | 0.8 | 0.88 | 1 | 0.87 | 0.94 |
| 1 | 76989264 | rs315087 | -1.11 | -0.436 | -0.19 | 0.245 | -0.003 | -0.016 | -0.016 | 0.438 | 0.4 | 0.72 | 0.51 | 0.4 | 0.36 | 0.37 |
| 1 | 76997071 | rs315081 | 0.635 | -1.349 | -0.052 | -0.672 | -0.018 | -0.088 | -0.088 | 0 | 0.19 | 0.33 | 0.11 | 0.01 | 0.15 | 0.08 |
| 1 | 77000914 | rs315079 | 0.337 | -1.481 | 0.004 | -0.68 | -0.036 | -0.124 | -0.124 | 0 | 0.2 | 0.45 | 0.11 | 0.01 | 0.13 | 0.06 |
| 1 | 77011016 | rs315066 | -0.707 | -1.709 | -0.028 | -0.638 | -0.004 | 0.015 | 0.015 | 0 | 0.12 | 0.16 | 0.2 | 0.25 | 0.12 | 0.21 |
| 1 | 77012251 | rs315063 | -0.309 | -1.689 | -0.114 | -0.633 | -0.041 | -0.079 | -0.079 | 1.333 | 0.76 | 0.39 | 0.83 | 0.95 | 0.87 | 0.92 |
| 1 | 77015113 | rs172714 | -0.866 | -1.846 | -0.06 | -0.511 | -0.013 | 0.004 | 0.004 | 0.046 | 0.17 | 0.21 | 0.2 | 0.22 | 0.16 | 0.27 |
| 1 | 77017425 | rs12025413 | 0.93 | -2.112 | -0.059 | -0.597 | 0.06 | 0.046 | 0.046 | 0.053 | 0.59 | 0.27 | 0.66 | 0.72 | 0.73 | 0.7 |
| 1 | 77018278 | rs366469 | -0.8 | -2.066 | 0.252 | -0.548 | -0.002 | -0.001 | -0.001 | 0 | 0.15 | 0.19 | 0.16 | 0.21 | 0.1 | 0.18 |
| 1 | 77019986 | rs17099155 | 0.744 | -2.51 | 0.661 | -1.005 | -0.012 | -0.056 | -0.056 | 0 | 0.22 | 0.25 | 0.11 | 0.01 | 0.18 | 0.1 |
| 1 | 77023273 | rs315049 | -1.31 | -2.819 | 1.072 | -1.046 | 0.057 | 0.058 | 0.058 | 0.092 | 0.44 | 0.67 | 0.35 | 0.27 | 0.28 | 0.29 |
| 1 | 77031816 | rs7544201 | 1.126 | -2.696 | 1.062 | -0.994 | 0.002 | -0.016 | -0.016 | 0.511 | 0.19 | 0.17 | 0.1 | 0.01 | 0.17 | 0.09 |
| 1 | 77033959 | rs315014 | -0.734 | -2.529 | 1.161 | -0.833 | -0.054 | 0.035 | 0.035 | 0 | 0.13 | 0.32 | 0.12 | 0.2 | 0.04 | 0.12 |
| 1 | 77034771 | rs315015 | 0.288 | -2.413 | 1.163 | -0.786 | -0.01 | 0.1 | 0.1 | 0 | 0.05 | 0.19 | 0.1 | 0.2 | 0.01 | 0.08 |
| 1 | 77039732 | rs315025 | -0.025 | -2.249 | 0.692 | -0.822 | 0.005 | 0.001 | 0.001 | 0 | 0.83 | 0.88 | 0.88 | 0.85 | 0.86 | 0.79 |
| 1 | 77040188 | rs436806 | -0.039 | -2.247 | 0.831 | -0.698 | -0.011 | 0.143 | 0.143 | 0 | 0.06 | 0.22 | 0.15 | 0.25 | 0.01 | 0.1 |
| 1 | 77041553 | rs11162180 | 0.983 | -2.826 | 0.821 | -1.204 | -0.001 | -0.006 | -0.006 | 0 | 0.4 | 0.22 | 0.49 | 0.58 | 0.54 | 0.49 |
| 1 | 77041694 | rs7513970 | -0.232 | -2.634 | 0.998 | -1.14 | 0.033 | 0.036 | 0.036 | 0 | 0.41 | 0.63 | 0.38 | 0.26 | 0.32 | 0.29 |
| 1 | 77042204 | rs369258 | 0.598 | -2.7 | 1.126 | -1.169 | -0.033 | -0.025 | -0.025 | 0 | 0.32 | 0.43 | 0.3 | 0.26 | 0.25 | 0.23 |
| 1 | 77042252 | rs315028 | 0.211 | -2.657 | 1.114 | -1.136 | -0.011 | 0.142 | 0.142 | 0 | 0.06 | 0.21 | 0.15 | 0.25 | 0.01 | 0.1 |

| | | | | | | | | | | | | | | | | |
|---|---|---|---|---|---|---|---|---|---|---|---|---|---|---|---|---|
| 1 | 77048617 | rs12117865 | 1.375 | -2.651 | 0.969 | -1.201 | 0.005 | -0.022 | -0.022 | 0 | 0.26 | 0.22 | 0.15 | 0.01 | 0.23 | 0.13 |
| 1 | 77051174 | rs315034 | -1.14 | -2.533 | 0.843 | -1.088 | -0.045 | 0.06 | 0.06 | 0 | 0.15 | 0.36 | 0.18 | 0.23 | 0.08 | 0.15 |
| 1 | 77054590 | rs315039 | -1.452 | -2.509 | 0.719 | -1.177 | -0.046 | 0.025 | 0.025 | 0 | 0.18 | 0.37 | 0.27 | 0.23 | 0.11 | 0.19 |
| 1 | 77061507 | rs7521965 | 0.41 | -2.166 | 0.291 | -0.895 | -0.012 | 0.127 | 0.127 | 0 | 0.06 | 0.2 | 0.1 | 0.22 | 0.01 | 0.1 |
| 1 | 77069228 | rs2574775 | 1.227 | -2.133 | -0.101 | -1.517 | 0.024 | 0.053 | 0.053 | 0 | 0.26 | 0.12 | 0.14 | 0.01 | 0.23 | 0.13 |
| 1 | 77070851 | rs563885 | 0.541 | -2.048 | -0.186 | -1.508 | 0.026 | 0.096 | 0.096 | 0 | 0.45 | 0.3 | 0.62 | 0.82 | 0.54 | 0.6 |
| 1 | 77072458 | rs2647389 | -0.396 | -1.092 | -0.168 | -1.042 | -0.001 | -0.001 | -0.001 | 0 | 0.15 | 0.15 | 0.13 | 0.15 | 0.14 | 0.22 |
| 1 | 77073771 | rs341028 | -0.059 | -0.944 | -0.125 | -0.997 | -0.001 | -0.001 | -0.001 | 0 | 0.86 | 0.85 | 0.88 | 0.85 | 0.87 | 0.8 |
| 1 | 77075248 | rs163603 | -0.756 | -1.026 | 0.14 | -1.045 | -0.031 | -0.002 | -0.002 | 0 | 0.16 | 0.37 | 0.14 | 0.15 | 0.09 | 0.18 |
| 1 | 77075591 | rs665178 | 0.134 | -1.306 | -0.74 | -1.445 | 0.075 | 0.212 | 0.212 | 0 | 0.49 | 0.21 | 0.7 | 0.8 | 0.6 | 0.63 |
| 1 | 77076004 | rs12354033 | -0.393 | -0.361 | -0.427 | -1.236 | 0.002 | -0.025 | -0.025 | 0.728 | 0.08 | 0.11 | 0.08 | 0 | 0.1 | 0.07 |
| 1 | 77081102 | rs341024 | -0.273 | -0.467 | 0.091 | -1.302 | 0.084 | 0.007 | 0.007 | 0 | 0.07 | 0.38 | 0.06 | 0 | 0.06 | 0.02 |
| 1 | 77084763 | rs645124 | 0.439 | 0.469 | -0.954 | -0.802 | -0.065 | -0.12 | -0.12 | 0 | 0.16 | 0.01 | 0.08 | 0.39 | 0.2 | 0.3 |
| 1 | 77085067 | rs11162183 | 1.639 | 0.692 | -1.44 | -1.041 | 0.017 | -0.018 | -0.018 | 0.236 | 0.38 | 0.1 | 0.36 | 0.37 | 0.37 | 0.31 |
| 1 | 77091045 | rs12129945 | 2.019 | -0.057 | -1.25 | -1.521 | 0.053 | 0.138 | 0.138 | 0 | 0.29 | 0.08 | 0.14 | 0 | 0.24 | 0.14 |
| 1 | 77093180 | rs1184626 | -1.406 | 0.479 | -0.366 | -0.763 | 0.006 | 0.039 | 0.039 | 0 | 0.42 | 0.64 | 0.52 | 0.24 | 0.4 | 0.38 |
| 1 | 77093530 | rs627873 | 0.799 | 0.445 | -0.165 | -0.763 | 0.099 | 0.025 | 0.025 | 0 | 0.05 | 0.32 | 0.05 | 0 | 0.06 | 0.02 |
| 1 | 77093829 | rs1173686 | -0.262 | 0.481 | -0.23 | -0.853 | -0.001 | -0.001 | -0.001 | 0.037 | 0.26 | 0.2 | 0.19 | 0.22 | 0.22 | 0.26 |
| 1 | 77094826 | rs676269 | -1.112 | 0.438 | -0.273 | -0.885 | -0.002 | -0.004 | -0.004 | 0.001 | 0.3 | 0.62 | 0.24 | 0.22 | 0.27 | 0.28 |
| 1 | 77095355 | rs12403583 | -0.121 | 0.381 | -0.523 | -0.892 | -0.037 | 0.06 | 0.06 | 0 | 0.12 | 0.01 | 0.28 | 0.02 | 0.14 | 0.1 |
| 1 | 77098729 | rs484299 | -1.948 | 0.295 | -0.465 | -0.889 | 0.003 | 0.036 | 0.036 | 0 | 0.42 | 0.62 | 0.51 | 0.24 | 0.41 | 0.38 |
| 1 | 77099238 | rs10518531 | 0.883 | 0.406 | -0.473 | -0.785 | -0.041 | 0.03 | 0.03 | 0 | 0.08 | 0 | 0.26 | 0.02 | 0.13 | 0.09 |
| 1 | 77102862 | rs341016 | 0.138 | 0.017 | -0.256 | -0.611 | -0.031 | -0.04 | -0.04 | 0.114 | 0.18 | 0.23 | 0.11 | 0.05 | 0.12 | 0.08 |
| 1 | 77104773 | rs2349044 | 0.498 | 0.123 | -0.42 | -0.535 | -0.013 | -0.01 | -0.01 | 0 | 0.05 | 0.04 | 0.07 | 0.17 | 0.08 | 0.17 |
| 1 | 77109465 | rs1925415 | -1.24 | 0.074 | -0.358 | -0.594 | -0.002 | -0.002 | -0.002 | 0 | 0.31 | 0.61 | 0.24 | 0.22 | 0.26 | 0.28 |
| 1 | 77112051 | rs544606 | -1.27 | 0.093 | -0.352 | -0.589 | -0.001 | -0.002 | -0.002 | 0 | 0.31 | 0.61 | 0.24 | 0.23 | 0.27 | 0.28 |
| 1 | 77113373 | rs491513 | -1.274 | 0.091 | -0.274 | -0.596 | -0.016 | -0.024 | -0.024 | 0.109 | 0.32 | 0.47 | 0.23 | 0.22 | 0.27 | 0.27 |
| 1 | 77124311 | rs4949737 | -0.927 | -0.007 | -0.249 | -0.689 | -0.003 | -0.068 | -0.068 | 0.014 | 0.14 | 0.23 | 0.12 | 0 | 0.14 | 0.08 |
| 1 | 77124408 | rs614117 | -0.887 | 0.31 | -0.36 | -0.689 | -0.013 | 0.02 | 0.02 | 0 | 0.94 | 0.97 | 0.96 | 1 | 0.94 | 0.93 |
| 1 | 77126373 | rs551585 | 0.924 | 0.152 | -0.175 | -0.663 | -0.05 | -0.035 | -0.035 | 0 | 0.15 | 0.18 | 0.06 | 0.05 | 0.06 | 0.04 |
| 1 | 77126434 | rs7543819 | 0.902 | 0.242 | -0.233 | -0.727 | -0.002 | -0.024 | -0.024 | 0.018 | 0.11 | 0.16 | 0.27 | 0.03 | 0.12 | 0.09 |
| 1 | 77127235 | rs645776 | -0.459 | 0.658 | -0.529 | -0.481 | -0.03 | -0.021 | -0.021 | 0.663 | 0.32 | 0.48 | 0.39 | 0.25 | 0.26 | 0.3 |
| 1 | 77128618 | rs4606356 | -0.803 | 0.708 | -0.44 | -0.389 | 0.011 | 0.008 | 0.008 | 0 | 0.37 | 0.6 | 0.43 | 0.09 | 0.3 | 0.19 |
| 1 | 77128655 | rs1886689 | -0.844 | 0.716 | -0.865 | 0.01 | -0.046 | -0.103 | -0.103 | 0 | 0.86 | 0.99 | 0.92 | 0.63 | 0.85 | 0.69 |
| 1 | 77132763 | rs11162187 | -0.493 | 0.648 | -0.35 | -0.202 | 0.021 | -0.05 | -0.05 | 0 | 0.1 | 0.19 | 0.1 | 0 | 0.12 | 0.06 |
| 1 | 77138723 | rs6665401 | 0.478 | 0.629 | -0.293 | -0.074 | 0.039 | -0.017 | -0.017 | 0 | 0.05 | 0.18 | 0.05 | 0 | 0.06 | 0.02 |
| 1 | 77138904 | rs689423 | -0.598 | 0.497 | -1.069 | 0.035 | 0.025 | 0.046 | 0.046 | 0 | 0.42 | 0.69 | 0.49 | 0.26 | 0.37 | 0.35 |
| 1 | 77142824 | rs505867 | -0.366 | -0.235 | -1.092 | 0.017 | 0.027 | -0.03 | -0.03 | 0 | 0.58 | 0.79 | 0.8 | 0.57 | 0.68 | 0.58 |
| 1 | 77146173 | rs10873899 | -0.049 | -0.362 | -0.411 | -0.171 | -0.001 | 0 | 0 | 0 | 0.44 | 0.5 | 0.23 | 0.43 | 0.32 | 0.38 |
| 1 | 77149207 | rs507630 | 0.639 | -0.556 | -0.101 | -0.331 | -0.02 | -0.022 | -0.022 | 0 | 0.4 | 0.24 | 0.69 | 0.51 | 0.56 | 0.37 |
| 1 | 77149849 | rs1498365 | 0.273 | -0.488 | 0.114 | -0.454 | 0.009 | -0.006 | -0.006 | 0 | 0.9 | 0.85 | 0.92 | 0.94 | 0.89 | 0.77 |
| 1 | 77154226 | rs10430054 | 0.776 | -0.665 | 0.107 | -0.25 | 0.009 | 0.001 | 0.001 | 2.373 | 0.37 | 0.37 | 0.19 | 0.33 | 0.27 | 0.33 |
| 1 | 77154319 | rs638072 | -0.806 | -0.629 | 0.099 | -0.139 | -0.007 | -0.005 | -0.005 | 0.008 | 0.94 | 0.96 | 0.96 | 0.87 | 0.92 | 0.79 |
| 1 | 77161524 | rs924296 | -0.657 | -0.655 | 0.173 | -0.151 | 0.042 | 0.06 | 0.06 | 0.296 | 0.94 | 0.81 | 0.95 | 0.87 | 0.91 | 0.79 |
| 1 | 77162485 | rs562580 | -0.39 | -0.692 | 0.068 | -0.169 | -0.011 | -0.012 | -0.012 | 0.8 | 0.93 | 0.96 | 0.96 | 0.87 | 0.91 | 0.79 |
| 1 | 77165502 | rs2764657 | -0.782 | -0.624 | 0.084 | -0.134 | -0.009 | -0.007 | -0.007 | 0 | 0.94 | 0.96 | 0.96 | 0.86 | 0.91 | 0.79 |
| 1 | 77168568 | rs9324157 | 0.928 | -0.742 | 0.022 | -0.229 | 0.041 | -0.01 | -0.01 | 0 | 0.35 | 0.17 | 0.17 | 0.33 | 0.27 | 0.33 |
| 1 | 77170017 | rs489723 | 1.657 | -0.745 | 0.421 | 0.054 | -0.005 | 0.014 | 0.014 | 0 | 0.68 | 0.34 | 0.79 | 0.54 | 0.74 | 0.54 |
| 1 | 77171575 | rs584208 | 0.141 | -0.646 | -0.024 | 0.098 | 0.001 | 0.001 | 0.001 | 1.979 | 0.89 | 0.86 | 0.92 | 0.87 | 0.88 | 0.77 |
| 1 | 77179690 | rs664576 | 1.698 | -0.641 | 0.003 | 0.097 | -0.012 | 0.08 | 0.08 | 0.053 | 0.7 | 0.53 | 0.81 | 0.54 | 0.74 | 0.55 |
| 1 | 77187344 | rs7552569 | 0.456 | -0.858 | -0.195 | 0.088 | 0.001 | 0.027 | 0.027 | 0.319 | 0.09 | 0.1 | 0.07 | 0.33 | 0.1 | 0.2 |
| 1 | 77187595 | rs1874806 | 0.619 | -0.861 | -0.179 | 0.083 | -0.009 | -0.043 | -0.043 | 0 | 0.08 | 0.18 | 0.04 | 0 | 0.04 | 0.03 |
| 1 | 77188392 | rs7534158 | -0.303 | -0.81 | -0.332 | 0.13 | -0.019 | 0.076 | 0.076 | 0 | 0.17 | 0.28 | 0.11 | 0.33 | 0.14 | 0.22 |
| 1 | 77190010 | rs649328 | -1.055 | -1.027 | -1.141 | -0.267 | 0.046 | -0.014 | -0.014 | 0 | 0.34 | 0.81 | 0.22 | 0.34 | 0.22 | 0.27 |
| 1 | 77190350 | rs647890 | -0.504 | -1.295 | -1.476 | -0.908 | 0.005 | 0.063 | 0.063 | 0.512 | 0.57 | 0.79 | 0.3 | 0.3 | 0.38 | 0.4 |
| 1 | 77195164 | rs12732852 | 1.377 | -2.172 | -1.487 | -1.508 | 0.03 | 0.021 | 0.021 | 0.36 | 0.17 | 0.08 | 0.03 | 0 | 0.07 | 0.06 |
| 1 | 77196032 | rs619463 | 0.202 | -1.909 | -1.745 | -1.905 | -0.051 | 0.032 | 0.032 | 1.449 | 0.34 | 0.17 | 0.45 | 0.76 | 0.49 | 0.51 |
| 1 | 77196594 | rs651752 | -0.523 | -2.122 | -1.291 | -2.055 | 0.033 | 0.101 | 0.101 | 0.021 | 0.49 | 0.61 | 0.52 | 0.98 | 0.57 | 0.68 |
| 1 | 77210385 | rs2173399 | -0.911 | -1.553 | -1.42 | -1.784 | -0.069 | -0.02 | -0.02 | 0 | 0.73 | 0.86 | 0.63 | 0.31 | 0.58 | 0.58 |
| 1 | 77214364 | rs1566250 | 0.102 | -1.113 | -0.805 | -1.201 | -0.044 | 0.053 | 0.053 | 0.241 | 0.15 | 0.06 | 0.34 | 0.02 | 0.3 | 0.23 |
| 1 | 77217180 | rs962668 | -0.45 | -1.558 | -0.032 | -0.563 | -0.025 | 0.103 | 0.103 | 0 | 0.14 | 0.36 | 0.09 | 0.23 | 0.08 | 0.19 |
| 1 | 77221362 | rs2221083 | -0.899 | -1.573 | -0.164 | -0.798 | -0.07 | -0.117 | -0.117 | 0.235 | 0.82 | 0.67 | 0.92 | 1 | 0.93 | 0.94 |
| 1 | 77223778 | rs924297 | 0.357 | -1.226 | -0.342 | -0.816 | 0.008 | -0.002 | -0.002 | 0 | 0.07 | 0.1 | 0.28 | 0.02 | 0.19 | 0.2 |
| 1 | 77224007 | rs17376644 | 0.551 | -1.227 | -0.374 | -0.816 | -0.043 | -0.006 | -0.006 | 0 | 0.07 | 0.01 | 0.27 | 0.02 | 0.19 | 0.2 |
| 1 | 77226258 | rs4255423 | 0.982 | -1.078 | -0.729 | -0.969 | -0.043 | 0.03 | 0.03 | 0 | 0.44 | 0.2 | 0.46 | 0.67 | 0.47 | 0.48 |
| 1 | 77233774 | rs12410542 | 1.004 | -1.778 | -1.018 | -1.061 | 0.063 | 0.027 | 0.027 | 0 | 0.47 | 0.11 | 0.5 | 0.67 | 0.59 | 0.5 |
| 1 | 77238731 | rs7530899 | -0.193 | -1.479 | -0.745 | -0.882 | -0.001 | 0.002 | 0.002 | 0 | 0.16 | 0.16 | 0.29 | 0.1 | 0.2 | 0.24 |
| 1 | 77240543 | rs10873910 | 1.119 | -1.518 | -0.761 | -0.897 | 0.076 | 0.05 | 0.05 | 0 | 0.47 | 0.18 | 0.5 | 0.67 | 0.57 | 0.5 |
| 1 | 77242042 | rs11162205 | 0.116 | -1.364 | -0.297 | -0.593 | -0.061 | -0.022 | -0.022 | 0 | 0.68 | 0.57 | 0.83 | 0.77 | 0.8 | 0.75 |

| | | | | | | | | | | | | | | | | |
|---|---|---|---|---|---|---|---|---|---|---|---|---|---|---|---|---|
| 1 | 77254137 | rs11162211 | -0.289 | -0.672 | -0.999 | -0.333 | 0.015 | -0.014 | -0.014 | 0 | 0.19 | 0.12 | 0.12 | 0.22 | 0.16 | 0.25 |
| 1 | 77256336 | rs4506508 | -0.077 | -0.107 | -0.897 | 0.114 | -0.026 | -0.011 | -0.011 | 0.512 | 0.64 | 0.47 | 0.82 | 0.77 | 0.79 | 0.7 |
| 1 | 77259107 | rs11577464 | -0.977 | 1.867 | -1.019 | 0.704 | -0.034 | 0.004 | 0.004 | 0 | 0.21 | 0.15 | 0.39 | 0.19 | 0.39 | 0.27 |
| 1 | 77263227 | rs12118675 | -0.926 | 2.023 | -1.043 | 0.922 | -0.03 | -0.011 | -0.011 | 0 | 0.19 | 0.14 | 0.47 | 0.24 | 0.39 | 0.28 |
| 1 | 77265060 | rs11578253 | -0.411 | 1.902 | -0.743 | 1.116 | 0.027 | 0.032 | 0.032 | 0 | 0.34 | 0.56 | 0.27 | 0.51 | 0.4 | 0.42 |
| 1 | 77266676 | rs7534641 | -0.451 | 0.939 | -0.49 | 1.256 | -0.012 | 0.016 | 0.016 | 0 | 0.18 | 0.25 | 0.11 | 0.22 | 0.13 | 0.16 |
| 1 | 77280831 | rs4077186 | 0.065 | 0.736 | -0.997 | 0.576 | -0.006 | 0.053 | 0.053 | 0 | 0.11 | 0.02 | 0.06 | 0 | 0.08 | 0.06 |
| 1 | 77284346 | rs11162217 | 0.731 | -0.416 | 0.165 | 0.642 | 0.029 | -0.016 | -0.016 | 0 | 0.7 | 0.22 | 0.71 | 0.62 | 0.82 | 0.77 |
| 1 | 77287528 | rs4949749 | 0.089 | -0.417 | 0.24 | 0.612 | 0.006 | -0.027 | -0.027 | 0 | 0.08 | 0.37 | 0.07 | 0 | 0.04 | 0.04 |
| 1 | 77291022 | rs12126169 | 0.983 | -0.22 | -0.241 | 0.875 | 0.033 | 0.163 | 0.163 | 0 | 0.48 | 0.16 | 0.5 | 0.24 | 0.46 | 0.43 |
| 1 | 77291188 | rs10873913 | 0.251 | -0.274 | -0.123 | 0.905 | -0.042 | -0.036 | -0.036 | 0.462 | 0.88 | 0.48 | 0.82 | 0.96 | 0.95 | 0.94 |
| 1 | 77295109 | rs4268394 | 0.622 | -0.291 | -0.241 | 0.94 | 0.03 | -0.016 | -0.016 | 0 | 0.7 | 0.22 | 0.71 | 0.61 | 0.82 | 0.77 |
| 1 | 77297392 | rs2055496 | 0.054 | -0.301 | 0.137 | 0.93 | -0.036 | -0.025 | -0.025 | 0.388 | 0.89 | 0.57 | 0.83 | 0.96 | 0.96 | 0.94 |
| 1 | 77299392 | rs1316659 | 0.326 | -0.263 | 0.154 | 0.945 | -0.021 | 0.07 | 0.07 | 0.162 | 0.19 | 0.34 | 0.12 | 0.35 | 0.14 | 0.17 |
| 1 | 77316849 | rs13376256 | 0.444 | -0.264 | 0.21 | 0.874 | -0.009 | -0.034 | -0.034 | 0 | 0.08 | 0.3 | 0.06 | 0 | 0.03 | 0.05 |
| 1 | 77323124 | rs9662183 | 0.82 | -0.244 | 0.11 | 0.952 | 0.023 | -0.013 | -0.013 | 0 | 0.7 | 0.23 | 0.71 | 0.62 | 0.82 | 0.77 |
| 1 | 77323384 | rs9660892 | 1.254 | -0.184 | -0.12 | 0.897 | -0.131 | -0.125 | -0.125 | 0 | 0.21 | 0.03 | 0.21 | 0.38 | 0.36 | 0.34 |
| 1 | 77324640 | rs1004878 | 0.133 | -0.257 | 0.144 | 0.652 | -0.026 | -0.01 | -0.01 | 0 | 0.9 | 0.56 | 0.82 | 0.96 | 0.96 | 0.94 |
| 1 | 77325124 | rs17099662 | 0.589 | -0.204 | 0.003 | 0.605 | -0.01 | -0.031 | -0.031 | 0.038 | 0.06 | 0.19 | 0.05 | 0 | 0.03 | 0.05 |
| 1 | 77325990 | rs1004454 | 0.726 | -0.186 | -0.287 | 0.709 | 0.017 | 0.175 | 0.175 | 0 | 0.48 | 0.12 | 0.49 | 0.24 | 0.46 | 0.43 |
| 1 | 77328297 | rs11162231 | -0.202 | -0.695 | -0.344 | 0.52 | 0.009 | -0.054 | -0.054 | 0 | 0.24 | 0.12 | 0.13 | 0.47 | 0.22 | 0.18 |
| 1 | 77331308 | rs1403243 | 0.259 | -0.959 | 0.144 | 0.226 | -0.004 | -0.027 | -0.027 | 0.773 | 0.05 | 0.22 | 0.03 | 0 | 0.03 | 0 |
| 1 | 77332627 | rs1522618 | 0.253 | -0.957 | 0.156 | 0.227 | -0.003 | -0.026 | -0.026 | 0 | 0.05 | 0.22 | 0.03 | 0 | 0.03 | 0 |
| 1 | 77370924 | rs2350527 | 0.212 | -0.962 | 0.173 | 0.224 | 0.001 | -0.024 | -0.024 | 2.724 | 0.05 | 0.25 | 0.03 | 0 | 0.03 | 0 |
| 1 | 77387631 | rs1766285 | 0.42 | -3.2 | -0.457 | -0.165 | 0.027 | 0.012 | 0.012 | 0 | 0.51 | 0.75 | 0.48 | 0.46 | 0.57 | 0.62 |
| 1 | 77396048 | rs17368577 | -1.473 | -3.492 | -0.56 | -0.303 | -0.004 | -0.024 | -0.024 | 0.208 | 0.93 | 0.92 | 0.96 | 1 | 0.94 | 0.95 |
| 1 | 77396166 | rs17368584 | 1.933 | -3.569 | -0.957 | -0.185 | -0.158 | -0.039 | -0.039 | 0 | 0.29 | 0.05 | 0.33 | 0.25 | 0.45 | 0.33 |
| 1 | 77397124 | rs3113982 | 0.427 | -3.558 | -0.721 | -0.108 | 0.016 | 0.011 | 0.011 | 0 | 0.52 | 0.66 | 0.46 | 0.43 | 0.59 | 0.62 |
| 1 | 77405284 | rs1252593 | 0.522 | -3.482 | -0.927 | -0.071 | 0.001 | 0.001 | 0.001 | 0 | 0.51 | 0.59 | 0.45 | 0.43 | 0.59 | 0.64 |
| 1 | 77405884 | rs10873923 | 0.753 | -2.694 | -0.962 | 0.653 | -0.003 | -0.014 | -0.014 | 0 | 0.07 | 0 | 0.05 | 0.42 | 0.05 | 0.1 |
| 1 | 77412583 | rs12124645 | 0.198 | -2.59 | -0.555 | 0.08 | -0.006 | -0.001 | -0.001 | 0.458 | 0.15 | 0.15 | 0.08 | 0.2 | 0.1 | 0.29 |
| 1 | 77425159 | rs858600 | -0.819 | -2.495 | -0.528 | -0.141 | 0.021 | 0.034 | 0.034 | 0 | 0.47 | 0.31 | 0.58 | 0.35 | 0.44 | 0.45 |
| 1 | 77437423 | rs12567699 | 0.193 | -2.107 | -0.466 | 0.117 | -0.001 | 0.001 | 0.001 | 0 | 0.09 | 0.05 | 0.05 | 0.13 | 0.11 | 0.09 |
| 1 | 77444267 | rs11162245 | 1.576 | -2.345 | -0.66 | -0.306 | -0.08 | -0.036 | -0.036 | 0 | 0.35 | 0.11 | 0.33 | 0.38 | 0.46 | 0.36 |
| 1 | 77453593 | rs2143582 | 0.865 | -2.52 | -0.129 | -0.337 | 0.038 | 0.049 | 0.049 | 0 | 0.5 | 0.4 | 0.43 | 0.73 | 0.64 | 0.53 |
| 1 | 77456303 | rs1933393 | 1.734 | -2.438 | -0.366 | -0.279 | -0.119 | -0.061 | -0.061 | 0 | 0.35 | 0.07 | 0.32 | 0.38 | 0.46 | 0.35 |
| 1 | 77461735 | rs10873925 | -0.511 | -2.691 | -0.39 | -0.463 | 0.001 | -0.004 | -0.004 | 0 | 0.29 | 0.22 | 0.21 | 0.35 | 0.29 | 0.19 |
| 1 | 77462662 | rs4949764 | -0.462 | -2.656 | -0.169 | -0.842 | -0.001 | 0 | 0 | 0 | 0.92 | 0.93 | 0.69 | 0.94 | 0.9 | 0.87 |
| 1 | 77464831 | rs1933399 | -0.048 | -2.834 | 0.169 | -0.925 | -0.074 | -0.011 | -0.011 | 4.778 | 0.21 | 0.34 | 0.12 | 0.21 | 0.12 | 0.31 |
| 1 | 77473081 | rs12065122 | 0.643 | -2.807 | 0.136 | -0.918 | -0.042 | 0.007 | 0.007 | 0.089 | 0.71 | 0.58 | 0.57 | 0.73 | 0.78 | 0.56 |
| 1 | 77477215 | rs10782639 | -0.086 | -2.48 | -0.108 | -0.616 | 0.009 | 0.01 | 0.01 | 0 | 0.17 | 0.25 | 0.13 | 0.26 | 0.24 | 0.19 |
| 1 | 77477794 | rs6693427 | -0.338 | -2.55 | -0.48 | -0.596 | -0.079 | -0.024 | -0.024 | 0 | 0.26 | 0.32 | 0.2 | 0.21 | 0.09 | 0.32 |
| 1 | 77480445 | rs199667 | -1.417 | -1.732 | -1.346 | -0.365 | -0.119 | -0.12 | -0.12 | 0 | 0.62 | 0.96 | 0.68 | 0.54 | 0.5 | 0.67 |
| 1 | 77482187 | rs199687 | -1.382 | -1.151 | -0.435 | -0.31 | -0.007 | 0.001 | 0.001 | 0.034 | 0.12 | 0.15 | 0.26 | 0.06 | 0.18 | 0.15 |
| 1 | 77487417 | rs199679 | -1.467 | -1.799 | -0.131 | -0.336 | -0.014 | -0.033 | -0.033 | 0 | 0.32 | 0.42 | 0.34 | 0.09 | 0.25 | 0.19 |
| 1 | 77492689 | rs1502524 | -1.398 | -1.811 | -0.075 | -0.338 | -0.052 | -0.044 | -0.044 | 0 | 0.23 | 0.49 | 0.26 | 0.09 | 0.14 | 0.18 |
| 1 | 77502479 | rs199691 | -1.134 | -1.446 | 0.244 | -0.147 | -0.023 | -0.113 | -0.113 | 6.813 | 0.27 | 0.45 | 0.16 | 0.03 | 0.2 | 0.06 |
| 1 | 77505413 | rs199696 | -1.991 | -1.64 | 0.165 | -0.03 | -0.028 | -0.01 | -0.01 | 0 | 0.39 | 0.56 | 0.51 | 0.35 | 0.31 | 0.44 |
| 1 | 77507127 | rs12084236 | -1.106 | -1.624 | 0.124 | -0.022 | 0.007 | -0.012 | -0.012 | 0.496 | 0.26 | 0.23 | 0.46 | 0.32 | 0.25 | 0.39 |
| 1 | 77509517 | rs2281305 | -0.098 | -1.607 | -0.014 | -0.001 | 0.004 | 0.003 | 0.003 | 1.887 | 0.11 | 0.1 | 0.35 | 0.32 | 0.11 | 0.37 |
| 1 | 77514455 | rs199707 | -1.915 | -1.605 | -0.062 | 0.005 | -0.024 | -0.009 | -0.009 | 0 | 0.39 | 0.57 | 0.51 | 0.35 | 0.31 | 0.44 |
| 1 | 77531223 | rs11162262 | -0.169 | -1.522 | -0.108 | 0.047 | -0.002 | 0.019 | 0.019 | 1.519 | 0.12 | 0.13 | 0.35 | 0.32 | 0.1 | 0.37 |
| 1 | 77531780 | rs199724 | -1.465 | -1.273 | -0.334 | 0.136 | -0.018 | -0.007 | -0.007 | 0 | 0.39 | 0.58 | 0.5 | 0.35 | 0.31 | 0.42 |
| 1 | 77546091 | rs169215 | -0.8 | -1.029 | -0.398 | 0.671 | -0.041 | 0.023 | 0.023 | 0.432 | 0.24 | 0.4 | 0.39 | 0.31 | 0.15 | 0.39 |
| 1 | 77547831 | rs199675 | -0.229 | -0.638 | -0.869 | 0.624 | 0.04 | 0.007 | 0.007 | 0.014 | 0.51 | 0.64 | 0.34 | 0.47 | 0.38 | 0.44 |
| 1 | 77548386 | rs7516609 | 0.267 | 0.76 | -0.919 | 0.677 | -0.007 | -0.009 | -0.009 | 0 | 0.74 | 0.76 | 0.78 | 0.48 | 0.67 | 0.48 |
| 1 | 77548518 | rs191827 | 0.051 | 1.029 | -1.464 | -0.218 | -0.014 | 0.177 | 0.177 | 0 | 0.54 | 0.8 | 0.48 | 0.8 | 0.47 | 0.55 |
| 1 | 77548843 | rs2223697 | -0.07 | 0.833 | -1.141 | -0.213 | 0 | -0.031 | -0.031 | 0 | 0.1 | 0.22 | 0.23 | 0.03 | 0.09 | 0.09 |
| 1 | 77550431 | rs199710 | -0.27 | 0.653 | -1.124 | -0.247 | -0.035 | -0.027 | -0.027 | 0 | 0.72 | 0.84 | 0.66 | 0.59 | 0.62 | 0.71 |
| 1 | 77550946 | rs6688757 | 0.69 | 0.285 | -0.751 | -0.213 | -0.021 | 0.04 | 0.04 | 0 | 0.08 | 0.17 | 0.05 | 0.16 | 0.05 | 0.26 |
| 1 | 77551537 | rs12744696 | -0.606 | 0.229 | -0.637 | -0.25 | -0.047 | 0.023 | 0.023 | 0 | 0.18 | 0.39 | 0.25 | 0.19 | 0.11 | 0.33 |
| 1 | 77552639 | rs199712 | 1.322 | 0.227 | -0.52 | -0.413 | -0.016 | 0.025 | 0.025 | 0 | 0.26 | 0.3 | 0.19 | 0.39 | 0.16 | 0.21 |
| 1 | 77553461 | rs199714 | 1.434 | 0.203 | -0.43 | -0.426 | -0.001 | 0.003 | 0.003 | 0 | 0.25 | 0.25 | 0.19 | 0.39 | 0.16 | 0.21 |
| 1 | 77554775 | rs17099885 | 1.694 | 0.226 | -0.481 | -0.407 | 0.013 | -0.02 | -0.02 | 0 | 0.05 | 0.01 | 0.02 | 0.16 | 0.04 | 0.26 |
| 1 | 77558358 | rs2036444 | 1.04 | 0.091 | -0.495 | -0.443 | -0.065 | -0.057 | -0.057 | 1.292 | 0.13 | 0.01 | 0.14 | 0.21 | 0.17 | 0.13 |
| 1 | 77569489 | rs4949766 | -0.572 | -0.526 | -0.075 | -0.409 | 0.07 | 0.025 | 0.025 | 0 | 0.45 | 0.72 | 0.44 | 0.59 | 0.27 | 0.54 |
| 1 | 77569545 | rs7550325 | 1.328 | -0.399 | -0.129 | -0.41 | -0.074 | 0.155 | 0.155 | 0 | 0.26 | 0.04 | 0.2 | 0 | 0.33 | 0.15 |
| 1 | 77586190 | rs7522727 | 0.406 | -0.39 | 0.03 | -0.268 | -0.017 | -0.025 | -0.025 | 0 | 0.11 | 0.25 | 0.2 | 0.04 | 0.06 | 0.07 |
| 1 | 77592351 | rs6683950 | 1.229 | -0.453 | 0.059 | -0.255 | 0.002 | -0.004 | -0.004 | 0 | 0.25 | 0.2 | 0.18 | 0.39 | 0.16 | 0.22 |

| | | | | | | | | | | | | | | | | |
|---|---|---|---|---|---|---|---|---|---|---|---|---|---|---|---|---|
| 1 | 77594164 | rs12760157 | -0.057 | 0.276 | 0.481 | 0.801 | -0.015 | -0.023 | -0.023 | 0.076 | 0.1 | 0.22 | 0.2 | 0.04 | 0.06 | 0.07 |
| 1 | 77594574 | rs2689674 | -1.379 | 0.267 | 0.497 | 0.807 | -0.049 | 0.023 | 0.023 | 0 | 0.18 | 0.41 | 0.25 | 0.19 | 0.11 | 0.33 |
| 1 | 77607160 | rs858447 | -1.595 | 0.266 | 0.484 | 0.807 | -0.063 | 0.022 | 0.022 | 0 | 0.19 | 0.47 | 0.25 | 0.19 | 0.11 | 0.33 |
| 1 | 77640166 | rs1098125 | -0.957 | 0.808 | 0.492 | 0.37 | -0.026 | -0.042 | -0.042 | 0.806 | 0.13 | 0.4 | 0.23 | 0.04 | 0.07 | 0.08 |
| 1 | 77644806 | rs1829913 | -0.25 | 0.822 | 0.273 | 0.375 | -0.015 | -0.023 | -0.023 | 0 | 0.1 | 0.23 | 0.2 | 0.04 | 0.06 | 0.07 |
| 1 | 77669114 | rs1028737 | -0.288 | 0.989 | -0.473 | 0.425 | -0.039 | -0.027 | -0.027 | 0 | 0.29 | 0.16 | 0.35 | 0.41 | 0.4 | 0.3 |
| 1 | 77685042 | rs12723684 | -1.348 | 0.417 | 0.167 | 0.317 | -0.033 | -0.052 | -0.052 | 1.756 | 0.14 | 0.46 | 0.23 | 0.04 | 0.07 | 0.08 |
| 1 | 77695854 | rs17100051 | -1.076 | 1.497 | -0.4 | 0.44 | -0.001 | -0.003 | -0.003 | 0 | 0.11 | 0.1 | 0.18 | 0.04 | 0.11 | 0.13 |
| 1 | 77695947 | rs4949781 | -0.064 | 1.364 | -1.31 | 0.418 | -0.034 | -0.029 | -0.029 | 0 | 0.56 | 0.23 | 0.54 | 0.42 | 0.51 | 0.52 |
| 1 | 77696605 | rs11162293 | -0.528 | 1.629 | -1.108 | 0.483 | -0.004 | -0.003 | -0.003 | 0 | 0.86 | 0.88 | 0.84 | 0.8 | 0.78 | 0.9 |
| 1 | 77696822 | rs10518440 | 1.369 | 1.485 | -0.886 | 0.467 | -0.004 | -0.002 | -0.002 | 0 | 0.08 | 0.06 | 0.12 | 0.21 | 0.16 | 0.11 |
| 1 | 77697582 | rs12131286 | 1.06 | 0.803 | -1.631 | -0.467 | -0.007 | 0.163 | 0.163 | 0 | 0.4 | 0.17 | 0.36 | 0.04 | 0.38 | 0.32 |
| 1 | 77700928 | rs11162297 | 0.473 | 0.858 | -0.818 | -0.31 | -0.016 | -0.013 | -0.013 | 0 | 0.57 | 0.39 | 0.56 | 0.41 | 0.5 | 0.51 |
| 1 | 77705365 | rs12040629 | 1.776 | 0.853 | -0.831 | -0.295 | -0.004 | -0.003 | -0.003 | 0.004 | 0.08 | 0.06 | 0.12 | 0.21 | 0.15 | 0.11 |
| 1 | 77705820 | rs1357570 | 0.318 | 0.824 | -0.714 | -0.257 | -0.022 | 0.021 | 0.021 | 0.062 | 0.29 | 0.46 | 0.27 | 0.38 | 0.28 | 0.38 |
| 1 | 77707391 | rs2689672 | 0.414 | 0.796 | -0.139 | -0.27 | -0.008 | 0.01 | 0.01 | 0.075 | 0.27 | 0.3 | 0.26 | 0.38 | 0.28 | 0.38 |
| 1 | 77718917 | rs1253220 | -1.115 | 0.902 | -0.269 | -0.234 | -0.006 | -0.005 | -0.005 | 0 | 0.86 | 0.9 | 0.84 | 0.8 | 0.78 | 0.9 |
| 1 | 77726109 | rs17100075 | 1.615 | 0.524 | -0.525 | -0.351 | 0.005 | -0.033 | -0.033 | 1.075 | 0.06 | 0.01 | 0.06 | 0.17 | 0.04 | 0.26 |
| 1 | 77726241 | rs10493596 | 0.726 | 0.524 | 0.025 | -0.499 | -0.002 | -0.008 | -0.008 | 0 | 0.21 | 0.22 | 0.19 | 0.21 | 0.24 | 0.12 |
| 1 | 77731287 | rs1988302 | 1.027 | 0.891 | 0.124 | -0.447 | -0.014 | -0.012 | -0.012 | 0.014 | 0.12 | 0.1 | 0.16 | 0.2 | 0.21 | 0.11 |
| 1 | 77733420 | rs12119637 | 1.834 | 0.601 | -0.412 | -1.075 | 0.019 | 0.14 | 0.14 | 0 | 0.26 | 0.06 | 0.11 | 0 | 0.2 | 0.16 |
| 1 | 77734562 | rs6692536 | 0.198 | 0.554 | -0.263 | -1.086 | 0.002 | -0.004 | -0.004 | 0.03 | 0.31 | 0.25 | 0.41 | 0.43 | 0.34 | 0.45 |
| 1 | 77734634 | rs10518445 | 0.528 | 0.677 | -0.056 | -0.969 | -0.01 | 0 | 0 | 1.902 | 0.12 | 0.06 | 0.1 | 0 | 0.13 | 0.1 |
| 1 | 77737217 | rs940283 | -1.926 | 0.407 | 0.272 | -0.886 | -0.013 | 0.11 | 0.11 | 0 | 0.84 | 0.77 | 0.8 | 0.63 | 0.88 | 0.81 |
| 1 | 77747248 | rs1851736 | 1.145 | 0.626 | 0.263 | -0.931 | -0.045 | -0.028 | -0.028 | 0 | 0.13 | 0.04 | 0.16 | 0.19 | 0.21 | 0.12 |
| 1 | 77748664 | rs1619903 | -0.674 | -0.169 | 0.313 | -1.315 | 0.002 | -0.022 | -0.022 | 0 | 0.45 | 0.55 | 0.45 | 0.72 | 0.46 | 0.58 |
| 1 | 77748786 | rs2689680 | -0.894 | -0.246 | 0.429 | -1.37 | 0.003 | -0.002 | -0.002 | 0 | 0.52 | 0.51 | 0.5 | 0.81 | 0.5 | 0.67 |
| 1 | 77749322 | rs7534033 | 1.638 | -0.267 | 0.369 | -1.26 | -0.007 | 0.011 | 0.011 | 0 | 0.1 | 0.17 | 0.07 | 0.21 | 0.06 | 0.25 |
| 1 | 77749527 | rs1779173 | -0.079 | -0.391 | 0.601 | -1.436 | -0.001 | -0.001 | -0.001 | 0.176 | 0.61 | 0.6 | 0.63 | 0.28 | 0.62 | 0.47 |
| 1 | 77751193 | rs10747353 | 0.284 | -0.494 | 0.666 | -1.604 | -0.002 | 0.008 | 0.008 | 0 | 0.33 | 0.23 | 0.44 | 0.2 | 0.41 | 0.27 |
| 1 | 77752622 | rs11162308 | 2.638 | -0.925 | 0.623 | -1.732 | -0.018 | -0.022 | -0.022 | 0 | 0.06 | 0.08 | 0.03 | 0 | 0.01 | 0 |
| 1 | 77754360 | rs1167201 | -0.968 | -0.717 | 0.516 | -1.704 | -0.006 | -0.009 | -0.009 | 0 | 0.72 | 0.77 | 0.71 | 0.49 | 0.69 | 0.72 |
| 1 | 77754853 | rs10782645 | -0.48 | -0.387 | 0.28 | -1.927 | -0.007 | 0.026 | 0.026 | 0 | 0.46 | 0.31 | 0.55 | 0.35 | 0.53 | 0.38 |
| 1 | 77756532 | rs11806344 | -0.229 | -0.411 | 0.326 | -1.927 | -0.006 | 0.015 | 0.015 | 0 | 0.41 | 0.31 | 0.53 | 0.27 | 0.49 | 0.36 |
| 1 | 77761956 | rs17100241 | 1.779 | -0.39 | 0.207 | -1.873 | 0.002 | -0.005 | -0.005 | 0.897 | 0.07 | 0.05 | 0.07 | 0.07 | 0.04 | 0.09 |
| 1 | 77765794 | rs1721165 | -2.114 | -0.406 | 0.3 | -1.891 | -0.019 | 0.035 | 0.035 | 0 | 0.9 | 0.76 | 0.91 | 0.79 | 0.94 | 0.75 |
| 1 | 77766847 | rs10782646 | 1.042 | -0.406 | 0.318 | -1.954 | -0.001 | 0.009 | 0.009 | 0 | 0.28 | 0.2 | 0.38 | 0.2 | 0.36 | 0.21 |
| 1 | 77768016 | rs1616948 | -0.262 | -0.293 | 0.257 | -1.914 | 0.002 | 0.002 | 0.002 | 0.049 | 0.78 | 0.71 | 0.81 | 0.65 | 0.74 | 0.66 |
| 1 | 77773679 | rs1779177 | -2.114 | -0.309 | 0.248 | -1.896 | -0.019 | 0.035 | 0.035 | 0 | 0.9 | 0.76 | 0.91 | 0.79 | 0.94 | 0.75 |
| 1 | 77775853 | rs9439799 | 0.701 | -0.333 | 0.279 | -1.927 | -0.002 | -0.003 | -0.003 | 0.795 | 0.34 | 0.34 | 0.2 | 0.14 | 0.34 | 0.19 |
| 1 | 77777898 | rs12141929 | 2.48 | -0.403 | 0.09 | -1.913 | 0.019 | 0.123 | 0.123 | 0 | 0.2 | 0.03 | 0.08 | 0.01 | 0.13 | 0.11 |
| 1 | 77778608 | rs1993266 | 1.435 | -0.273 | 0.185 | -1.849 | -0.002 | -0.008 | -0.008 | 0 | 0.07 | 0.04 | 0.07 | 0 | 0.09 | 0.06 |
| 1 | 77779836 | rs1167198 | -1.909 | -0.319 | 0.261 | -1.8 | -0.023 | 0.038 | 0.038 | 0 | 0.89 | 0.73 | 0.91 | 0.79 | 0.94 | 0.75 |
| 1 | 77781567 | rs12144794 | 0.493 | -0.335 | 0.101 | -1.801 | 0.029 | 0.003 | 0.003 | 0 | 0.49 | 0.26 | 0.41 | 0.45 | 0.37 | 0.45 |
| 1 | 77782761 | rs7514177 | 0.51 | -0.486 | -0.571 | -1.923 | 0.027 | -0.003 | -0.003 | 0 | 0.47 | 0.26 | 0.41 | 0.45 | 0.37 | 0.45 |
| 1 | 77784762 | rs12751884 | 2.056 | -0.305 | -0.801 | -1.94 | -0.038 | -0.032 | -0.032 | 4.228 | 0.12 | 0.02 | 0.29 | 0.2 | 0.23 | 0.19 |
| 1 | 77785561 | rs7549462 | 2.031 | -0.307 | -0.771 | -1.941 | -0.029 | -0.024 | -0.024 | 0.068 | 0.12 | 0.02 | 0.29 | 0.2 | 0.23 | 0.19 |
| 1 | 77791025 | rs2602951 | 0.731 | -0.261 | -0.679 | -1.94 | -0.056 | -0.007 | -0.007 | 0 | 0.13 | 0.03 | 0.1 | 0.13 | 0.21 | 0.09 |
| 1 | 77802376 | rs2815327 | 0.846 | -0.299 | -0.687 | -1.958 | -0.054 | -0.008 | -0.008 | 0 | 0.13 | 0.03 | 0.1 | 0.13 | 0.21 | 0.09 |
| 1 | 77802930 | rs12402184 | 1.531 | -0.415 | -0.734 | -2.051 | 0.002 | 0 | 0 | 0 | 0.22 | 0.18 | 0.25 | 0.45 | 0.15 | 0.28 |
| 1 | 77804242 | rs17378915 | 1.584 | 0.442 | -0.772 | -2.003 | -0.052 | -0.019 | -0.019 | 0.248 | 0.08 | 0 | 0.08 | 0.08 | 0.15 | 0.04 |
| 1 | 77808677 | rs11162319 | 2.015 | 0.24 | -0.696 | -2.031 | 0.003 | -0.004 | -0.004 | 0 | 0.1 | 0.18 | 0.08 | 0.22 | 0.06 | 0.25 |
| 1 | 77809846 | rs6696968 | 0.434 | 0.266 | -0.555 | -2.085 | 0.035 | 0.001 | 0.001 | 0 | 0.69 | 0.49 | 0.72 | 0.65 | 0.64 | 0.61 |
| 1 | 77812803 | rs17379082 | 1.744 | 0.177 | -0.966 | -2.151 | 0.038 | 0.095 | 0.095 | 0 | 0.26 | 0.07 | 0.15 | 0.08 | 0.17 | 0.21 |
| 1 | 77815566 | rs1010196 | 0.858 | 0.287 | -0.731 | -2.096 | -0.002 | 0.01 | 0.01 | 0 | 0.28 | 0.19 | 0.38 | 0.2 | 0.36 | 0.21 |
| 1 | 77818143 | rs6672375 | 2.08 | 0.301 | -0.665 | -2.075 | -0.01 | -0.06 | -0.06 | 0.121 | 0.16 | 0.18 | 0.1 | 0 | 0.13 | 0.02 |
| 1 | 77819743 | rs1377733 | 0.232 | 0.48 | -0.587 | -1.931 | 0.084 | -0.035 | -0.035 | 0 | 0.79 | 0.61 | 0.8 | 0.87 | 0.7 | 0.85 |
| 1 | 77820127 | rs12030598 | 2.72 | 0.482 | -0.698 | -1.805 | -0.031 | -0.035 | -0.035 | 0 | 0.08 | 0.01 | 0.14 | 0.18 | 0.17 | 0.12 |
| 1 | 77822519 | rs4949653 | 0.61 | -0.147 | -1.003 | -1.675 | -0.012 | -0.023 | -0.023 | 0.13 | 0.16 | 0.08 | 0.14 | 0.22 | 0.15 | 0.31 |
| 1 | 77823028 | rs1377735 | 2.048 | -0.232 | -0.784 | -1.528 | 0.001 | 0.01 | 0.01 | 0 | 0.15 | 0.11 | 0.18 | 0.37 | 0.11 | 0.19 |
| 1 | 77828152 | rs2054020 | 0.7 | -0.19 | -0.773 | -1.244 | 0.017 | 0.024 | 0.024 | 0.009 | 0.5 | 0.37 | 0.63 | 0.58 | 0.57 | 0.56 |
| 1 | 77828321 | rs12565526 | 0.358 | -0.469 | -1.022 | -1.425 | 0.025 | 0.085 | 0.085 | 0 | 0.5 | 0.32 | 0.65 | 0.83 | 0.56 | 0.74 |
| 1 | 77828665 | rs12565558 | 1.107 | -0.284 | -1.138 | -1.326 | -0.08 | -0.09 | -0.09 | 0 | 0.2 | 0.07 | 0.28 | 0.43 | 0.35 | 0.38 |
| 1 | 77829709 | rs11162324 | 1.323 | -0.538 | -0.845 | -1.551 | 0.008 | -0.024 | -0.024 | 2.469 | 0.25 | 0.21 | 0.15 | 0 | 0.21 | 0.04 |
| 1 | 77831799 | rs17100458 | 1.87 | -0.556 | -0.783 | -1.586 | 0.007 | -0.055 | -0.055 | 0.252 | 0.13 | 0.23 | 0.1 | 0 | 0.12 | 0.12 |
| 1 | 77832250 | rs2815307 | 1.533 | -0.584 | -0.894 | -1.584 | 0.004 | -0.028 | -0.028 | 0 | 0.12 | 0.02 | 0.08 | 0.17 | 0.11 | 0.11 |
| 1 | 77836489 | rs11162326 | 0.02 | -0.541 | -0.798 | -1.523 | -0.003 | -0.031 | -0.031 | 0 | 0.74 | 0.57 | 0.8 | 0.83 | 0.77 | 0.78 |
| 1 | 77839480 | rs17100475 | 1.924 | -0.777 | -0.707 | -1.451 | -0.002 | 0.005 | 0.005 | 0.02 | 0.29 | 0.25 | 0.37 | 0.4 | 0.21 | 0.36 |
| 1 | 77840100 | rs10873933 | 1.143 | -0.686 | -0.777 | -1.637 | -0.009 | -0.073 | -0.073 | 4.872 | 0.25 | 0.25 | 0.16 | 0 | 0.21 | 0.04 |

| | | | | | | | | | | | | | | | | |
|---|---|---|---|---|---|---|---|---|---|---|---|---|---|---|---|---|
| 1 | 77844602 | rs7546131 | -0.838 | -0.703 | -0.811 | -1.429 | 0.003 | 0.001 | 0.001 | 0 | 0.61 | 0.57 | 0.69 | 0.4 | 0.74 | 0.58 |
| 1 | 77850800 | rs11162331 | 1.707 | -0.726 | -0.781 | -1.191 | 0.029 | -0.044 | -0.044 | 0 | 0.31 | 0.19 | 0.23 | 0.41 | 0.2 | 0.23 |
| 1 | 77854238 | rs3844165 | 3.207 | -0.207 | -0.938 | -1.306 | -0.052 | -0.032 | -0.032 | 0 | 0.11 | 0.02 | 0.21 | 0.19 | 0.29 | 0.19 |
| 1 | 77857683 | rs4949800 | -0.963 | -0.376 | -0.779 | -1.522 | -0.002 | -0.014 | -0.014 | 0 | 0.71 | 0.72 | 0.79 | 0.84 | 0.73 | 0.85 |
| 1 | 77863068 | rs17379910 | 2.421 | -0.451 | -0.652 | -1.571 | -0.008 | -0.016 | -0.016 | 0.168 | 0.07 | 0.16 | 0.16 | 0.03 | 0.05 | 0.2 |
| 1 | 77864321 | rs17100513 | 1.524 | -0.367 | -0.826 | -1.578 | -0.002 | -0.03 | -0.03 | 0 | 0.1 | 0.01 | 0.07 | 0.16 | 0.09 | 0.1 |
| 1 | 77865140 | rs6663047 | 0.512 | -0.433 | -0.766 | -1.6 | -0.01 | -0.011 | -0.011 | 0.162 | 0.47 | 0.53 | 0.48 | 0.64 | 0.32 | 0.54 |
| 1 | 77865771 | rs11162334 | 2.329 | -0.312 | -0.76 | -1.464 | -0.002 | -0.054 | -0.054 | 0 | 0.19 | 0.18 | 0.13 | 0 | 0.18 | 0.06 |
| 1 | 77867129 | rs17100521 | 2.546 | -0.256 | -0.967 | -1.121 | 0.011 | -0.025 | -0.025 | 0 | 0.2 | 0.11 | 0.17 | 0.41 | 0.12 | 0.2 |
| 1 | 77867923 | rs17098287 | 1.295 | 0.233 | -1.069 | -0.988 | 0.001 | -0.033 | -0.033 | 0 | 0.12 | 0.12 | 0.1 | 0.01 | 0.12 | 0.12 |
| 1 | 77870133 | rs17380036 | 1.003 | 0.376 | -1.278 | -0.623 | -0.109 | -0.087 | -0.087 | 0 | 0.21 | 0.03 | 0.25 | 0.34 | 0.37 | 0.27 |
| 1 | 77870149 | rs17100524 | 0.532 | -1.139 | -0.957 | -0.637 | 0.008 | 0.024 | 0.024 | 0.147 | 0.16 | 0.12 | 0.11 | 0.1 | 0.12 | 0.14 |
| 1 | 77872142 | rs2882289 | 1.468 | -1.291 | -0.249 | -0.288 | 0.008 | -0.002 | -0.002 | 0 | 0.06 | 0.19 | 0.17 | 0.03 | 0.05 | 0.11 |
| 1 | 77876353 | rs4949802 | 1.478 | -1.533 | -0.201 | -0.206 | -0.003 | 0.022 | 0.022 | 0 | 0.23 | 0.24 | 0.28 | 0.38 | 0.19 | 0.24 |
| 1 | 77880168 | rs7512893 | 0.89 | -1.803 | -0.31 | -0.375 | 0.009 | 0.016 | 0.016 | 0 | 0.21 | 0.21 | 0.15 | 0.11 | 0.15 | 0.12 |
| 1 | 77880188 | rs7512897 | -0.807 | -1.541 | -0.222 | -0.207 | -0.001 | 0.022 | 0.022 | 0 | 0.83 | 0.64 | 0.78 | 0.78 | 0.84 | 0.75 |
| 1 | 77880369 | rs12022440 | 1.697 | -1.675 | -0.724 | -0.232 | -0.163 | 0.076 | 0.076 | 1.356 | 0.31 | 0.04 | 0.33 | 0.14 | 0.48 | 0.34 |
| 1 | 77881769 | rs1498404 | 0.976 | -1.682 | -0.681 | -0.225 | -0.002 | 0.001 | 0.001 | 0 | 0.08 | 0.11 | 0.07 | 0.11 | 0.05 | 0.09 |
| 1 | 77884712 | rs1498407 | 1.303 | -1.4 | -0.592 | -0.106 | -0.02 | 0.023 | 0.023 | 0.305 | 0.08 | 0.16 | 0.02 | 0.15 | 0.03 | 0.05 |
| 1 | 77885282 | rs9633478 | -1.649 | -1.047 | -0.342 | 0.048 | 0.002 | -0.005 | -0.005 | 2.216 | 0.89 | 0.89 | 0.96 | 0.81 | 0.9 | 0.87 |
| 1 | 77886339 | rs3104461 | -1.636 | -0.717 | -0.422 | -0.128 | 0.007 | -0.005 | -0.005 | 0 | 0.94 | 0.85 | 0.83 | 0.97 | 0.95 | 0.88 |
| 1 | 77886499 | rs10493606 | 0.517 | -0.751 | -0.701 | -0.159 | -0.039 | 0.046 | 0.046 | 0.338 | 0.39 | 0.19 | 0.35 | 0.29 | 0.51 | 0.39 |
| 1 | 77889505 | rs2647514 | -0.695 | -0.371 | -0.169 | 0.322 | -0.002 | -0.013 | -0.013 | 0.536 | 0.94 | 0.87 | 0.84 | 0.97 | 0.95 | 0.88 |
| 1 | 77892274 | rs2647519 | -0.805 | -0.363 | -0.241 | 0.326 | -0.004 | -0.013 | -0.013 | 0 | 0.94 | 0.89 | 0.84 | 0.97 | 0.95 | 0.88 |
| 1 | 77895517 | rs2647522 | 0.141 | -0.522 | -0.295 | 0.304 | -0.004 | -0.01 | -0.01 | 0 | 0.73 | 0.73 | 0.68 | 0.83 | 0.78 | 0.72 |
| 1 | 77895705 | rs1498401 | 0.438 | -0.874 | 0.066 | 0.631 | -0.028 | 0.118 | 0.118 | 0.26 | 0.76 | 0.52 | 0.77 | 0.55 | 0.81 | 0.62 |
| 1 | 77896080 | rs6675781 | -0.335 | -1.002 | 0.163 | 0.586 | -0.035 | -0.005 | -0.005 | 0 | 0.3 | 0.38 | 0.28 | 0.25 | 0.18 | 0.37 |
| 1 | 77896355 | rs12033923 | 0.09 | -0.827 | -0.345 | 0.662 | 0.007 | -0.042 | -0.042 | 0 | 0.1 | 0.03 | 0.12 | 0.23 | 0.06 | 0.16 |
| 1 | 77896934 | rs7537574 | 1.411 | -0.465 | -0.462 | 0.455 | -0.042 | -0.034 | -0.034 | 0 | 0.28 | 0.2 | 0.35 | 0.39 | 0.46 | 0.26 |
| 1 | 77897688 | rs4949655 | 0.297 | -0.541 | -0.79 | 0.307 | -0.001 | 0.004 | 0.004 | 0 | 0.14 | 0.22 | 0.14 | 0.19 | 0.13 | 0.26 |
| 1 | 77897713 | rs11162341 | 1.148 | -0.721 | -0.986 | 0.179 | 0.001 | 0.008 | 0.008 | 0.555 | 0.18 | 0.05 | 0.07 | 0.15 | 0.13 | 0.14 |
| 1 | 77902981 | rs10873941 | 1.252 | -0.681 | -1.159 | 0.147 | 0.024 | 0.063 | 0.063 | 0 | 0.23 | 0.12 | 0.26 | 0.04 | 0.18 | 0.18 |
| 1 | 77903611 | rs12034899 | -0.188 | -0.114 | -0.636 | 0.952 | 0.009 | -0.016 | -0.016 | 0.138 | 0.15 | 0.11 | 0.16 | 0.23 | 0.11 | 0.16 |
| 1 | 77904451 | rs12407481 | -1.295 | 0.084 | -0.56 | 0.828 | 0.015 | 0.004 | 0.004 | 0 | 0.48 | 0.4 | 0.37 | 0.57 | 0.36 | 0.56 |
| 1 | 77908985 | rs6658302 | -1.153 | 0.293 | -0.211 | 1.138 | -0.004 | 0.01 | 0.01 | 0 | 0.29 | 0.35 | 0.3 | 0.44 | 0.23 | 0.42 |
| 1 | 77914159 | rs2647506 | -1.112 | 1.166 | 0.321 | 1.254 | 0.006 | 0.007 | 0.007 | 0 | 0.18 | 0.23 | 0.22 | 0.26 | 0.24 | 0.43 |
| 1 | 77918347 | rs12044354 | -1.332 | 1.124 | 0.141 | 1.591 | -0.032 | -0.068 | -0.068 | 0 | 0.2 | 0.11 | 0.23 | 0.4 | 0.27 | 0.49 |
| 1 | 77932095 | rs6704141 | 1.147 | 0.641 | 0.563 | 1.445 | 0.003 | 0.076 | 0.076 | 0 | 0.75 | 0.61 | 0.75 | 0.53 | 0.74 | 0.54 |
| 1 | 77939418 | rs2647503 | 1.729 | 1.162 | 0.978 | 1.105 | 0.026 | 0.018 | 0.018 | 0 | 0.64 | 0.55 | 0.5 | 0.42 | 0.57 | 0.38 |
| 1 | 77946687 | rs2245189 | -0.37 | 1.085 | 1.189 | 1.197 | -0.045 | 0.06 | 0.06 | 0.423 | 0.91 | 0.79 | 0.97 | 0.79 | 0.98 | 0.95 |
| 1 | 77951460 | rs7522356 | 2.133 | 0.672 | 1.205 | 0.767 | 0.033 | 0.008 | 0.008 | 0 | 0.49 | 0.3 | 0.46 | 0.43 | 0.48 | 0.3 |
| 1 | 77952669 | rs1909203 | -0.101 | 0.81 | 1.522 | 0.569 | 0.078 | -0.068 | -0.068 | 0 | 0.11 | 0.22 | 0.25 | 0 | 0.19 | 0.13 |
| 1 | 77952756 | rs1909202 | 0.386 | 0.803 | 1.661 | 0.565 | 0.123 | -0.101 | -0.101 | 1.946 | 0.86 | 0.58 | 0.71 | 1 | 0.8 | 0.86 |
| 1 | 77954804 | rs9324162 | 1.118 | 0.942 | 1.517 | 0.792 | -0.017 | -0.018 | -0.018 | 0 | 0.06 | 0.02 | 0.08 | 0.15 | 0.17 | 0.08 |
| 1 | 77955556 | rs2647497 | 0.514 | 0.882 | 1.572 | 0.756 | 0.108 | -0.044 | -0.044 | 0 | 0.08 | 0.23 | 0.24 | 0 | 0.16 | 0.11 |
| 1 | 77955577 | rs6680677 | 1.02 | 0.831 | 1.417 | 0.805 | 0.009 | -0.01 | -0.01 | 0 | 0.78 | 0.45 | 0.69 | 0.78 | 0.79 | 0.83 |
| 1 | 77958712 | rs17380544 | -1.423 | 1.562 | 1.469 | 0.844 | 0.016 | 0.013 | 0.013 | 0 | 0.27 | 0.36 | 0.26 | 0.36 | 0.3 | 0.47 |
| 1 | 77959674 | rs12568528 | 0.504 | 1.574 | 1.117 | 0.941 | -0.001 | 0.04 | 0.04 | 0 | 0.06 | 0.08 | 0.06 | 0.22 | 0.05 | 0.06 |
| 1 | 77961074 | rs11162355 | 1.523 | 1.525 | 1.165 | 0.729 | 0 | 0.001 | 0.001 | 0 | 0.67 | 0.36 | 0.66 | 0.4 | 0.66 | 0.45 |
| 1 | 77964269 | rs7514041 | 1.636 | 1.527 | 1.05 | 0.73 | -0.006 | -0.014 | -0.014 | 0.617 | 0.66 | 0.31 | 0.66 | 0.4 | 0.66 | 0.45 |
| 1 | 77971047 | rs3104467 | -1.277 | 1.555 | 0.984 | 0.6 | 0.005 | 0.007 | 0.007 | 0 | 0.39 | 0.44 | 0.44 | 0.46 | 0.34 | 0.62 |
| 1 | 77976117 | rs17380796 | -0.24 | 1.68 | 0.557 | 0.695 | 0.03 | 0.005 | 0.005 | 0 | 0.12 | 0.2 | 0.14 | 0.14 | 0.21 | 0.11 |
| 1 | 77986516 | rs10873947 | 1.597 | 0.635 | -0.054 | 0.521 | 0.038 | -0.013 | -0.013 | 0 | 0.43 | 0.22 | 0.39 | 0.4 | 0.43 | 0.26 |
| 1 | 77988371 | rs17100672 | -1.017 | 0.636 | 0.089 | 0.631 | 0.002 | 0.006 | 0.006 | 0 | 0.15 | 0.17 | 0.13 | 0.23 | 0.17 | 0.28 |
| 1 | 77992695 | rs2803136 | 0.107 | -1.074 | -0.049 | -0.24 | -0.031 | -0.02 | -0.02 | 0 | 0.64 | 0.53 | 0.82 | 0.74 | 0.72 | 0.63 |
| 1 | 77994269 | rs3767035 | 0.755 | -0.828 | -0.567 | -0.21 | -0.023 | -0.031 | -0.031 | 0 | 0.06 | 0 | 0.05 | 0.15 | 0.06 | 0.1 |
| 1 | 78006340 | rs6695989 | 0.213 | -0.961 | -0.199 | -0.481 | 0.051 | 0.047 | 0.047 | 0 | 0.6 | 0.24 | 0.78 | 0.74 | 0.7 | 0.52 |
| 1 | 78017571 | rs4949659 | 0.22 | -0.962 | -0.204 | -0.477 | 0.053 | 0.047 | 0.047 | 0 | 0.6 | 0.23 | 0.77 | 0.73 | 0.7 | 0.5 |
| 1 | 78022217 | rs1565851 | 0.349 | -1.137 | -0.063 | -0.492 | 0.061 | 0.038 | 0.038 | 0 | 0.6 | 0.29 | 0.79 | 0.74 | 0.73 | 0.51 |
| 1 | 78023173 | rs7536433 | 2.514 | -1.399 | -0.83 | -0.775 | 0.003 | 0.032 | 0.032 | 0 | 0.17 | 0.19 | 0.17 | 0.42 | 0.22 | 0.06 |
| 1 | 78024602 | rs10873950 | -0.414 | -1.644 | -0.491 | -0.883 | -0.047 | -0.016 | -0.016 | 0 | 0.65 | 0.56 | 0.83 | 0.74 | 0.76 | 0.63 |
| 1 | 78025454 | rs4949811 | -0.496 | -1.647 | -0.492 | -0.885 | -0.047 | -0.016 | -0.016 | 1.841 | 0.65 | 0.56 | 0.83 | 0.74 | 0.76 | 0.63 |
| 1 | 78025551 | rs3509 | -0.136 | -1.644 | -0.329 | -0.879 | 0.007 | 0.005 | 0.005 | 0 | 0.62 | 0.39 | 0.8 | 0.74 | 0.74 | 0.52 |
| 1 | 78027725 | rs4949813 | -0.354 | -1.644 | -0.272 | -0.878 | -0.017 | -0.005 | -0.005 | 0.069 | 0.64 | 0.42 | 0.83 | 0.74 | 0.76 | 0.63 |
| 1 | 78048331 | rs17381664 | 1.36 | -1.567 | -0.831 | -1.268 | -0.092 | 0.095 | 0.095 | 0 | 0.19 | 0.04 | 0.45 | 0 | 0.33 | 0.24 |
| 1 | 78075766 | rs10873951 | 0.066 | -2.288 | -0.996 | -1.422 | -0.006 | -0.004 | -0.004 | 0.839 | 0.57 | 0.44 | 0.78 | 0.73 | 0.7 | 0.57 |
| 1 | 78083771 | rs7513619 | -0.967 | -2.476 | -0.804 | -1.52 | 0.049 | 0.004 | 0.004 | 0 | 0.7 | 0.81 | 0.85 | 0.73 | 0.76 | 0.6 |
| 1 | 78130243 | rs278853 | -2.746 | -2.57 | -1.141 | -1.601 | -0.026 | -0.058 | -0.058 | 0 | 0.78 | 0.84 | 0.8 | 0.58 | 0.72 | 0.92 |
| 1 | 78141164 | rs17100916 | 1.375 | -2.834 | -0.91 | -1.438 | 0.014 | -0.021 | -0.021 | 0 | 0.12 | 0.04 | 0.1 | 0.17 | 0.07 | 0.13 |

| | | | | | | | | | | | | | | | | |
|---|---|---|---|---|---|---|---|---|---|---|---|---|---|---|---|---|
| 1 | 78168758 | rs12021502 | 2.536 | -2.937 | -0.979 | -1.38 | -0.01 | -0.033 | -0.033 | 0 | 0.06 | 0 | 0.03 | 0.12 | 0.07 | 0.07 |
| 1 | 78183861 | rs11162378 | -0.813 | -2.929 | -0.953 | -1.39 | 0.113 | 0.06 | 0.06 | 0 | 0.48 | 0.74 | 0.26 | 0.41 | 0.33 | 0.56 |
| 1 | 78184325 | rs12142671 | -0.274 | -2.957 | -0.72 | -1.415 | -0.018 | -0.011 | -0.011 | 1.705 | 0.4 | 0.44 | 0.2 | 0.27 | 0.27 | 0.52 |
| 1 | 78198322 | rs12030080 | 1.837 | -3.005 | -0.79 | -1.414 | 0.007 | -0.029 | -0.029 | 0.438 | 0.11 | 0.02 | 0.09 | 0.17 | 0.07 | 0.13 |
| 1 | 78201745 | rs2296227 | -0.852 | -2.934 | -0.74 | -1.405 | 0.113 | 0.055 | 0.055 | 0 | 0.48 | 0.75 | 0.26 | 0.41 | 0.33 | 0.55 |
| 1 | 78209205 | rs12128041 | 0.851 | -2.893 | -0.393 | -1.379 | 0.014 | 0.004 | 0.004 | 0 | 0.31 | 0.29 | 0.15 | 0.27 | 0.24 | 0.39 |
| 1 | 78242638 | rs2147318 | 0.041 | -2.645 | -0.268 | -1.256 | -0.016 | -0.012 | -0.012 | 0 | 0.41 | 0.44 | 0.2 | 0.26 | 0.27 | 0.51 |
| 1 | 78243547 | rs12142441 | -0.647 | -2.608 | -0.775 | -1.205 | 0.112 | 0.058 | 0.058 | 0 | 0.48 | 0.74 | 0.26 | 0.4 | 0.33 | 0.55 |
| 1 | 78254258 | rs1166588 | 0.627 | -2.649 | -0.799 | -1.279 | 0.024 | 0.018 | 0.018 | 0 | 0.34 | 0.29 | 0.15 | 0.23 | 0.23 | 0.39 |
| 1 | 78256947 | rs1166587 | 0.404 | -2.653 | -0.873 | -1.281 | 0.037 | 0.028 | 0.028 | 0 | 0.35 | 0.26 | 0.15 | 0.22 | 0.21 | 0.34 |
| 1 | 78264259 | rs12122686 | 2.247 | -2.515 | -1.671 | -1.247 | 0.062 | 0.049 | 0.049 | 0 | 0.21 | 0.06 | 0.09 | 0.12 | 0.13 | 0.28 |
| 1 | 78266417 | rs1539739 | -0.137 | -1.578 | -1.281 | -0.757 | 0.11 | 0.078 | 0.078 | 0 | 0.5 | 0.72 | 0.27 | 0.38 | 0.33 | 0.57 |
| 1 | 78276977 | rs4121165 | 2.786 | -1.439 | -1.286 | -0.701 | 0 | 0.021 | 0.021 | 0 | 0.16 | 0.18 | 0.16 | 0.42 | 0.22 | 0.06 |
| 1 | 78281002 | rs11162386 | -0.293 | -2.013 | -0.844 | -0.229 | 0.015 | 0.009 | 0.009 | 0 | 0.57 | 0.43 | 0.79 | 0.62 | 0.72 | 0.47 |
| 1 | 78284211 | rs10873955 | -0.28 | -1.999 | -0.99 | -0.223 | 0.026 | 0.014 | 0.014 | 0 | 0.57 | 0.41 | 0.79 | 0.62 | 0.72 | 0.47 |
| 1 | 78288315 | rs10518561 | 0.23 | -0.648 | -0.592 | 0.258 | 0.006 | -0.024 | -0.024 | 0.087 | 0.11 | 0.04 | 0.09 | 0.2 | 0.07 | 0.12 |
| 1 | 78289045 | rs6604877 | 0.603 | -0.279 | -0.349 | 0.011 | -0.002 | -0.027 | -0.027 | 0 | 0.05 | 0.12 | 0.05 | 0 | 0.04 | 0.02 |
| 1 | 78305371 | rs1539737 | 0.289 | -0.316 | -0.482 | 0.161 | -0.002 | 0.001 | 0.001 | 0 | 0.91 | 0.88 | 0.8 | 0.9 | 0.91 | 0.86 |
| 1 | 78305783 | rs6604878 | 1.768 | -0.633 | 0.251 | 0.019 | -0.013 | -0.002 | -0.002 | 0 | 0.76 | 0.36 | 0.68 | 0.75 | 0.8 | 0.71 |
| 1 | 78313494 | rs10873956 | 0.134 | -0.509 | 0.099 | 0.046 | -0.001 | 0.001 | 0.001 | 0 | 0.91 | 0.88 | 0.8 | 0.9 | 0.9 | 0.86 |
| 1 | 78359029 | rs1166706 | 0.164 | -0.83 | 0.25 | -0.036 | 0 | -0.001 | -0.001 | 0.039 | 0.9 | 0.87 | 0.8 | 0.9 | 0.9 | 0.85 |
| 1 | 78366894 | rs1166704 | 0.164 | -0.787 | 0.25 | -0.016 | 0 | -0.001 | -0.001 | 1.79 | 0.9 | 0.87 | 0.8 | 0.9 | 0.9 | 0.85 |
| 1 | 78392446 | rs1166698 | 1.942 | -0.791 | -0.495 | 0.272 | -0.065 | -0.117 | -0.117 | 0.198 | 0.15 | 0.01 | 0.12 | 0.38 | 0.21 | 0.07 |
| 1 | 78396626 | rs9659938 | -0.152 | -0.767 | -0.425 | 0.306 | -0.002 | 0 | 0 | 0 | 0.91 | 0.87 | 0.8 | 0.91 | 0.92 | 0.86 |
| 1 | 78400540 | rs1780050 | 0.91 | -1.644 | -0.62 | -0.34 | 0.043 | -0.005 | -0.005 | 0 | 0.49 | 0.3 | 0.27 | 0.52 | 0.45 | 0.38 |
| 1 | 78401207 | rs17101082 | 0.392 | -1.69 | -0.087 | -0.347 | 0.03 | 0.081 | 0.081 | 0 | 0.05 | 0.16 | 0.04 | 0.14 | 0.05 | 0.13 |
| 1 | 78418776 | rs11807240 | 0.583 | -1.696 | -0.101 | -0.368 | -0.002 | 0 | 0 | 0 | 0.09 | 0.13 | 0.2 | 0.09 | 0.08 | 0.14 |
| 1 | 78442764 | rs12046966 | 1.233 | -1.482 | -0.009 | -0.65 | 0 | 0.011 | 0.011 | 0 | 0.1 | 0.07 | 0.1 | 0.25 | 0.07 | 0.14 |
| 1 | 78469660 | rs11162405 | 1.36 | -1.558 | -0.02 | -0.657 | 0.042 | -0.007 | -0.007 | 0 | 0.49 | 0.3 | 0.27 | 0.52 | 0.45 | 0.39 |
| 1 | 78475199 | rs6424648 | 0.252 | -1.437 | 0.105 | -0.584 | 0.002 | 0.055 | 0.055 | 0 | 0.86 | 0.71 | 0.76 | 0.77 | 0.88 | 0.73 |
| 1 | 78481640 | rs1039863 | 1.067 | -1.433 | 0.017 | -0.582 | -0.015 | -0.004 | -0.004 | 0 | 0.77 | 0.43 | 0.69 | 0.77 | 0.82 | 0.7 |
| 1 | 78492221 | rs2136682 | -1.042 | -1.463 | -0.115 | -0.51 | 0 | -0.002 | -0.002 | 0.182 | 0.92 | 0.93 | 0.81 | 0.91 | 0.93 | 0.87 |
| 1 | 78501215 | rs6424659 | 1.879 | -1.696 | -0.224 | -0.709 | -0.011 | -0.024 | -0.024 | 0.028 | 0.2 | 0.16 | 0.17 | 0.38 | 0.29 | 0.09 |
| 1 | 78502520 | rs12411130 | 1.564 | -1.677 | -0.276 | -1.084 | -0.068 | 0.057 | 0.057 | 0 | 0.18 | 0.06 | 0.34 | 0 | 0.32 | 0.18 |
| 1 | 78507648 | rs10735751 | 1.024 | -2.102 | 0.084 | -0.892 | -0.02 | -0.001 | -0.001 | 0 | 0.77 | 0.41 | 0.71 | 0.76 | 0.83 | 0.7 |
| 1 | 78510544 | rs6675105 | 0.924 | -2.003 | 0.002 | -0.788 | -0.014 | 0.025 | 0.025 | 0 | 0.82 | 0.64 | 0.73 | 0.76 | 0.85 | 0.71 |
| 1 | 78513624 | rs12140633 | 1.72 | -2.022 | -0.269 | -1.013 | 0.08 | 0.155 | 0.155 | 0 | 0.28 | 0.03 | 0.1 | 0.04 | 0.16 | 0.2 |
| 1 | 78515147 | rs12123953 | 1.632 | -1.953 | -0.236 | -0.994 | 0.074 | 0.081 | 0.081 | 0 | 0.28 | 0.14 | 0.1 | 0.14 | 0.16 | 0.28 |
| 1 | 78518668 | rs1855704 | -0.524 | -1.085 | 0.378 | -0.456 | 0.001 | 0.014 | 0.014 | 0 | 0.92 | 0.84 | 0.8 | 0.91 | 0.92 | 0.86 |
| 1 | 78527876 | rs638335 | 0.052 | -1.08 | 0.288 | -0.451 | -0.004 | -0.006 | -0.006 | 0 | 0.88 | 0.79 | 0.77 | 0.91 | 0.89 | 0.84 |
| 1 | 78540733 | rs4245660 | 0.651 | -1.19 | -0.067 | -0.28 | 0.009 | 0.002 | 0.002 | 0 | 0.58 | 0.25 | 0.57 | 0.4 | 0.6 | 0.63 |
| 1 | 78557608 | rs630640 | 0.819 | -1.142 | -0.103 | -0.084 | -0.003 | -0.043 | -0.043 | 3.123 | 0.23 | 0.16 | 0.16 | 0.5 | 0.26 | 0.1 |
| 1 | 78558419 | rs3790592 | 0.819 | -1.142 | -0.103 | -0.084 | -0.003 | -0.043 | -0.043 | 3.333 | 0.23 | 0.16 | 0.16 | 0.5 | 0.26 | 0.1 |
| 1 | 78563540 | rs633580 | -0.253 | -1 | 0.16 | -0.53 | -0.01 | -0.007 | -0.007 | 0 | 0.91 | 0.95 | 0.76 | 0.89 | 0.89 | 0.82 |
| 1 | 78566234 | rs812097 | 0.851 | -1.147 | 0.039 | -0.489 | 0.007 | -0.026 | -0.026 | 0 | 0.24 | 0.15 | 0.16 | 0.62 | 0.23 | 0.15 |
| 1 | 78574129 | rs6684364 | -0.212 | -1.251 | 0.571 | -0.545 | -0.015 | 0.006 | 0.006 | 0.43 | 0.33 | 0.51 | 0.22 | 0.63 | 0.27 | 0.19 |
| 1 | 78578177 | rs6666954 | -0.506 | -1.281 | 0.804 | -0.729 | 0.002 | -0.001 | -0.001 | 0 | 0.36 | 0.65 | 0.29 | 0.63 | 0.29 | 0.26 |
| 1 | 78585086 | rs540742 | 0.846 | -1.353 | 0.81 | -0.731 | 0.001 | -0.003 | -0.003 | 0 | 0.24 | 0.22 | 0.18 | 0.66 | 0.23 | 0.24 |
| 1 | 78589598 | rs1022103 | 2.047 | -2.079 | 0.152 | -1.261 | 0.053 | 0.038 | 0.038 | 0 | 0.25 | 0.16 | 0.12 | 0.17 | 0.16 | 0.27 |
| 1 | 78590027 | rs7542998 | 0.657 | -1.632 | 0.253 | -1.123 | -0.031 | -0.036 | -0.036 | 0 | 0.09 | 0.02 | 0.21 | 0.14 | 0.12 | 0.17 |
| 1 | 78590204 | rs573952 | 0.152 | -1.641 | 0.346 | -1.126 | 0.044 | 0.033 | 0.033 | 0.122 | 0.85 | 0.62 | 0.74 | 0.86 | 0.84 | 0.81 |
| 1 | 78590255 | rs12758374 | 1.237 | -1.726 | 0.273 | -1.253 | 0.024 | 0.188 | 0.188 | 0.146 | 0.51 | 0.23 | 0.52 | 0.17 | 0.58 | 0.53 |
| 1 | 78603383 | rs661136 | 1.139 | -1.621 | 0.272 | -1.203 | 0.008 | -0.123 | -0.123 | 0 | 0.78 | 0.56 | 0.71 | 0.95 | 0.83 | 0.73 |
| 1 | 78604614 | rs554298 | 0.921 | -1.494 | 0.262 | -0.889 | 0.001 | 0.003 | 0.003 | 0 | 0.24 | 0.21 | 0.19 | 0.76 | 0.24 | 0.23 |
| 1 | 78607268 | rs586300 | 2.619 | -1.587 | 0.111 | -0.731 | -0.039 | -0.125 | -0.125 | 0.011 | 0.15 | 0.01 | 0.14 | 0.47 | 0.19 | 0.09 |
| 1 | 78611730 | rs624766 | 0.897 | -1.517 | 0.327 | -1.659 | -0.004 | -0.011 | -0.011 | 0.139 | 0.24 | 0.28 | 0.2 | 0.77 | 0.25 | 0.23 |
| 1 | 78629908 | rs7555504 | -0.053 | -1.523 | 0.397 | -1.64 | -0.014 | -0.053 | -0.053 | 0 | 0.65 | 0.51 | 0.62 | 0.21 | 0.66 | 0.64 |
| 1 | 78632090 | rs12184300 | 1.53 | -1.7 | 0.229 | -1.845 | 0.045 | 0.08 | 0.08 | 0 | 0.3 | 0.14 | 0.16 | 0.17 | 0.21 | 0.33 |
| 1 | 78633857 | rs17101426 | 0.397 | -1.391 | 1.169 | -0.961 | 0 | -0.035 | -0.035 | 0.154 | 0.07 | 0.13 | 0.16 | 0 | 0.07 | 0.11 |
| 1 | 78634053 | rs11799753 | -0.538 | -1.375 | 1.087 | -0.946 | -0.014 | -0.06 | -0.06 | 0 | 0.16 | 0.29 | 0.21 | 0.02 | 0.13 | 0.17 |
| 1 | 78639244 | rs689374 | -1.622 | -1.392 | 1.078 | -0.982 | 0.043 | 0.158 | 0.158 | 0.161 | 0.45 | 0.78 | 0.47 | 0.83 | 0.38 | 0.49 |
| 1 | 78643674 | rs17101475 | -0.986 | -1.305 | 1.007 | -0.926 | -0.011 | -0.084 | -0.084 | 0 | 0.18 | 0.35 | 0.26 | 0.02 | 0.15 | 0.24 |
| 1 | 78666990 | rs683097 | 0.968 | -1.233 | 0.609 | -0.752 | -0.001 | 0 | 0 | 0 | 0.19 | 0.2 | 0.17 | 0.77 | 0.21 | 0.2 |
| 1 | 78691660 | rs669764 | -1.118 | -1.206 | 0.511 | -0.729 | 0.016 | 0.029 | 0.029 | 0.281 | 0.42 | 0.64 | 0.41 | 0.67 | 0.36 | 0.38 |
| 1 | 78696522 | rs2181453 | 0.261 | -0.994 | 0.736 | -0.666 | -0.009 | -0.016 | -0.016 | 0 | 0.65 | 0.49 | 0.74 | 0.37 | 0.68 | 0.69 |
| 1 | 78699130 | rs4650383 | -0.168 | -1.003 | 0.546 | -0.683 | -0.009 | -0.004 | -0.004 | 0 | 0.05 | 0.07 | 0.06 | 0.05 | 0.03 | 0.13 |
| 1 | 78699394 | rs942100 | 0.806 | -0.967 | 0.865 | -0.658 | 0.001 | 0.015 | 0.015 | 0.06 | 0.2 | 0.22 | 0.18 | 0.61 | 0.22 | 0.17 |
| 1 | 78703232 | rs11162435 | 0.307 | -0.972 | 0.653 | -0.514 | 0.002 | 0 | 0 | 0.55 | 0.64 | 0.39 | 0.76 | 0.41 | 0.68 | 0.68 |
| 1 | 78707881 | rs4650639 | -0.625 | -0.712 | 0.548 | -0.743 | 0.005 | 0.024 | 0.024 | 1.11 | 0.77 | 0.44 | 0.77 | 0.38 | 0.76 | 0.77 |

| | | | | | | | | | | | | | | | | |
|---|---|---|---|---|---|---|---|---|---|---|---|---|---|---|---|---|
| 1 | 78708393 | rs12409525 | -0.264 | -0.612 | 0.079 | -0.509 | 0 | -0.004 | -0.004 | 0 | 0.1 | 0.1 | 0.19 | 0.25 | 0.1 | 0.29 |
| 1 | 78714497 | rs644224 | 0.749 | -0.612 | 0 | -0.514 | -0.028 | -0.008 | -0.008 | 0.282 | 0.71 | 0.46 | 0.69 | 0.68 | 0.78 | 0.53 |
| 1 | 78716854 | rs585677 | 1.066 | 0.257 | 0.109 | 0.317 | 0.006 | 0.034 | 0.034 | 0 | 0.16 | 0.19 | 0.19 | 0.61 | 0.21 | 0.16 |
| 1 | 78740026 | rs1417102 | -0.212 | -0.05 | 0.529 | -0.153 | -0.008 | 0.002 | 0.002 | 0 | 0.27 | 0.33 | 0.28 | 0.67 | 0.28 | 0.23 |
| 1 | 78742185 | rs6685021 | 1.013 | 0.081 | 0.015 | -0.945 | -0.007 | 0.022 | 0.022 | 0 | 0.21 | 0.14 | 0.13 | 0.05 | 0.18 | 0.26 |
| 1 | 78743308 | rs1952363 | 0.287 | 0.023 | 0.337 | -0.942 | -0.015 | 0.005 | 0.005 | 0 | 0.27 | 0.39 | 0.3 | 0.67 | 0.28 | 0.23 |
| 1 | 78749908 | rs6424762 | 0.763 | -0.312 | -0.302 | -1.603 | 0.018 | 0.121 | 0.121 | 0 | 0.46 | 0.18 | 0.44 | 0.24 | 0.52 | 0.46 |
| 1 | 78752369 | rs7544813 | 0.351 | -0.503 | -0.324 | -1.732 | 0.029 | 0.052 | 0.052 | 0 | 0.53 | 0.37 | 0.47 | 0.28 | 0.56 | 0.47 |
| 1 | 78756602 | rs12144203 | 0.749 | -0.471 | -0.328 | -1.581 | -0.026 | 0.035 | 0.035 | 0 | 0.32 | 0.23 | 0.34 | 0.23 | 0.38 | 0.22 |
| 1 | 78760605 | rs11162451 | -0.393 | -0.585 | -0.193 | -1.605 | -0.041 | -0.131 | -0.131 | 0 | 0.78 | 0.55 | 0.74 | 0.95 | 0.83 | 0.71 |
| 1 | 78762998 | rs2882802 | 0.287 | -0.6 | -0.202 | -1.62 | -0.045 | -0.129 | -0.129 | 0 | 0.72 | 0.5 | 0.72 | 0.91 | 0.79 | 0.69 |
| 1 | 78766380 | rs720887 | -0.032 | -0.409 | 0.064 | -1.364 | -0.005 | 0.015 | 0.015 | 0 | 0.58 | 0.39 | 0.47 | 0.73 | 0.54 | 0.58 |
| 1 | 78772046 | rs11162456 | 1.239 | -0.258 | -0.423 | -1.273 | -0.057 | 0.039 | 0.039 | 0 | 0.27 | 0.11 | 0.31 | 0.2 | 0.33 | 0.2 |
| 1 | 78775182 | rs1328452 | 0.634 | -0.729 | -0.164 | -0.3 | -0.017 | 0.038 | 0.038 | 0 | 0.21 | 0.29 | 0.15 | 0.44 | 0.15 | 0.05 |
| 1 | 78781923 | rs11162463 | 0.634 | -0.729 | -0.164 | -0.321 | -0.017 | 0.037 | 0.037 | 0 | 0.21 | 0.29 | 0.15 | 0.46 | 0.15 | 0.05 |
| 1 | 78790397 | rs1328449 | 1.949 | -0.732 | -0.171 | -0.299 | -0.001 | 0.036 | 0.036 | 0 | 0.13 | 0.14 | 0.12 | 0.42 | 0.1 | 0.03 |
| 1 | 78790740 | rs2352037 | -0.267 | -0.741 | 0.079 | -0.316 | -0.037 | 0.039 | 0.039 | 0 | 0.28 | 0.41 | 0.34 | 0.53 | 0.22 | 0.28 |
| 1 | 78797784 | rs1328441 | -0.695 | -0.793 | 0.008 | -0.627 | -0.002 | 0.001 | 0.001 | 0 | 0.58 | 0.6 | 0.5 | 0.58 | 0.47 | 0.52 |
| 1 | 78800715 | rs1340161 | 1.047 | -0.685 | -0.37 | -0.636 | -0.004 | -0.002 | -0.002 | 0 | 0.07 | 0.1 | 0.18 | 0.07 | 0.07 | 0.23 |
| 1 | 78800843 | rs12063386 | 1.029 | -0.698 | -0.4 | -0.648 | -0.011 | 0.049 | 0.049 | 0 | 0.3 | 0.19 | 0.16 | 0.05 | 0.25 | 0.24 |
| 1 | 78801456 | rs11804627 | 0.607 | -0.387 | -0.562 | -0.271 | -0.005 | -0.002 | -0.002 | 0 | 0.07 | 0.12 | 0.18 | 0.07 | 0.07 | 0.23 |
| 1 | 78812335 | rs6657253 | 0.605 | -0.077 | -0.897 | -0.313 | -0.052 | 0.022 | 0.022 | 0 | 0.31 | 0.13 | 0.37 | 0.31 | 0.37 | 0.3 |
| 1 | 78819945 | rs2352039 | 0.377 | -0.513 | -0.164 | 0.127 | -0.035 | 0.09 | 0.09 | 0 | 0.21 | 0.35 | 0.31 | 0.49 | 0.18 | 0.27 |
| 1 | 78821497 | rs10873978 | -0.301 | 0.049 | -0.572 | 0.184 | -0.042 | -0.061 | -0.061 | 0 | 0.29 | 0.15 | 0.48 | 0.42 | 0.38 | 0.4 |
| 1 | 78825912 | rs7533564 | 0.287 | -0.175 | 0.168 | 0.188 | -0.021 | -0.037 | -0.037 | 0.36 | 0.93 | 0.76 | 0.98 | 1 | 0.98 | 0.89 |
| 1 | 78830851 | rs1328447 | -0.499 | 0.329 | 0.453 | 0.222 | 0.062 | 0.08 | 0.08 | 0.009 | 0.17 | 0.28 | 0.29 | 0.31 | 0.28 | 0.23 |
| 1 | 78831791 | rs12132664 | 0.365 | 0.34 | 0.429 | 0.227 | -0.011 | -0.007 | -0.007 | 0 | 0.13 | 0.08 | 0.27 | 0.31 | 0.26 | 0.23 |
| 1 | 78873062 | rs538855 | 0.459 | 0.275 | 0.591 | 0.201 | 0.105 | 0.134 | 0.134 | 0 | 0.82 | 0.57 | 0.7 | 0.69 | 0.72 | 0.77 |
| 1 | 78894362 | rs11162488 | 0.402 | 0.323 | 0.4 | 0.306 | -0.047 | -0.049 | -0.049 | 0 | 0.12 | 0.04 | 0.26 | 0.31 | 0.27 | 0.23 |
| 1 | 78894934 | rs674345 | 0.154 | -0.013 | 0.23 | 0.068 | 0.016 | -0.057 | -0.057 | 1.586 | 0.38 | 0.2 | 0.37 | 0.46 | 0.37 | 0.27 |
| 1 | 78896492 | rs518341 | 1.963 | 0.018 | 0.132 | 0.375 | 0.003 | 0.003 | 0.003 | 0 | 0.1 | 0.06 | 0.06 | 0.33 | 0.1 | 0.03 |
| 1 | 78921860 | rs10782665 | -0.071 | -0.225 | -0.374 | 0.014 | 0.001 | 0 | 0 | 0 | 0.35 | 0.35 | 0.37 | 0.34 | 0.39 | 0.45 |
| 1 | 78934451 | rs6678699 | 0.418 | -0.231 | -0.645 | 0.016 | -0.019 | -0.012 | -0.012 | 0 | 0.31 | 0.25 | 0.36 | 0.34 | 0.39 | 0.34 |
| 1 | 78936411 | rs10873985 | -0.409 | -0.286 | -0.843 | 0.004 | 0.001 | -0.009 | -0.009 | 0 | 0.39 | 0.51 | 0.4 | 0.34 | 0.4 | 0.46 |
| 1 | 78937828 | rs6692239 | 0.122 | 0.312 | -0.995 | 0.476 | -0.001 | -0.001 | -0.001 | 0.168 | 0.6 | 0.62 | 0.67 | 0.42 | 0.6 | 0.54 |
| 1 | 78938777 | rs1412414 | -0.133 | 0.491 | -1.024 | 0.712 | -0.001 | 0 | 0 | 0 | 0.63 | 0.63 | 0.67 | 0.47 | 0.61 | 0.67 |
| 1 | 78942270 | rs4650547 | -1.164 | 0.761 | -1.337 | 1.549 | -0.016 | -0.053 | -0.053 | 0 | 0.91 | 0.99 | 0.95 | 0.69 | 0.9 | 0.97 |
| 1 | 78946879 | rs1543675 | 0.07 | 0.737 | -1.182 | 1.15 | -0.05 | -0.045 | -0.045 | 0 | 0.26 | 0.12 | 0.33 | 0.32 | 0.33 | 0.23 |
| 1 | 78957550 | rs3766354 | 0.578 | 0.199 | -0.307 | 1.098 | -0.003 | 0.015 | 0.015 | 0 | 0.1 | 0.12 | 0.18 | 0.2 | 0.1 | 0.19 |
| 1 | 78963305 | rs1322935 | -0.253 | 0.78 | 0.109 | 0.728 | 0.004 | -0.013 | -0.013 | 0 | 0.18 | 0.23 | 0.11 | 0.07 | 0.22 | 0.15 |
| 1 | 78963437 | rs1322934 | -0.377 | 0.78 | 0.083 | 0.728 | 0.017 | -0.034 | -0.034 | 0.594 | 0.18 | 0.28 | 0.12 | 0.08 | 0.22 | 0.15 |
| 1 | 78976570 | rs6701594 | -0.352 | 0.758 | -0.311 | 0.987 | -0.045 | -0.066 | -0.066 | 0 | 0.71 | 0.92 | 0.75 | 0.63 | 0.69 | 0.65 |
| 1 | 78977715 | rs6700362 | -0.362 | 0.675 | 0.393 | 0.724 | 0.019 | -0.036 | -0.036 | 0 | 0.18 | 0.28 | 0.12 | 0.08 | 0.22 | 0.15 |
| 1 | 78987302 | rs672561 | -0.619 | 0.572 | 0.476 | 0.683 | 0.019 | -0.053 | -0.053 | 0.213 | 0.2 | 0.34 | 0.13 | 0.08 | 0.22 | 0.16 |
| 1 | 78989850 | rs520171 | -0.31 | 0.277 | 0.534 | 1.066 | -0.006 | 0.002 | 0.002 | 0 | 0.78 | 0.74 | 0.65 | 0.27 | 0.82 | 0.69 |
| 1 | 78998759 | rs33966768 | -0.861 | 0.658 | 0.716 | 1.167 | -0.027 | 0.056 | 0.056 | 0.061 | 0.18 | 0.08 | 0.11 | 0.06 | 0.2 | 0.15 |
| 1 | 78999088 | rs501078 | -2.073 | 0.474 | 1.176 | 1.618 | 0 | -0.011 | -0.011 | 6.044 | 0.3 | 0.7 | 0.29 | 0.59 | 0.26 | 0.43 |
| 1 | 78999269 | rs3766335 | -0.758 | 0.48 | 1.107 | 1.591 | -0.023 | 0.056 | 0.056 | 0.061 | 0.18 | 0.08 | 0.11 | 0.06 | 0.19 | 0.15 |
| 1 | 78999367 | rs686262 | -1.753 | 0.511 | 1.188 | 1.595 | -0.001 | 0.003 | 0.003 | 0.543 | 0.3 | 0.66 | 0.27 | 0.56 | 0.26 | 0.37 |
| 1 | 79003258 | rs898 | 0.169 | 0.646 | 0.65 | 1.028 | -0.001 | -0.01 | -0.01 | 0.007 | 0.13 | 0.15 | 0.1 | 0.01 | 0.17 | 0.12 |
| 1 | 79003355 | rs899 | 0.013 | 0.656 | 0.489 | 0.809 | -0.003 | -0.006 | -0.006 | 0 | 0.17 | 0.17 | 0.12 | 0.01 | 0.18 | 0.14 |
| 1 | 79005798 | rs12065196 | 0.013 | 0.656 | 0.489 | 0.809 | -0.003 | -0.006 | -0.006 | 0 | 0.17 | 0.17 | 0.12 | 0.01 | 0.18 | 0.14 |
| 1 | 79010670 | rs17101862 | -0.005 | 0.659 | 0.368 | 1 | 0 | 0.013 | 0.013 | 0 | 0.07 | 0.08 | 0.09 | 0.21 | 0.05 | 0.19 |
| 1 | 79019357 | rs514514 | -1.164 | 0.623 | -0.206 | 0.656 | 0.004 | -0.068 | -0.068 | 0.096 | 0.63 | 0.95 | 0.58 | 0.64 | 0.65 | 0.73 |
| 1 | 79043381 | rs2184229 | 0.785 | 0.431 | 1.225 | 0.741 | 0.039 | 0.225 | 0.225 | 0 | 0.9 | 0.54 | 0.88 | 0.74 | 0.95 | 0.74 |
| 1 | 79051869 | rs1023253 | 1.347 | 0.473 | 0.858 | 0.734 | -0.001 | -0.001 | -0.001 | 1.459 | 0.72 | 0.31 | 0.77 | 0.7 | 0.76 | 0.61 |
| 1 | 79052935 | rs1023254 | 1.349 | 0.475 | 0.728 | 0.735 | -0.003 | -0.002 | -0.002 | 0 | 0.71 | 0.33 | 0.77 | 0.7 | 0.76 | 0.61 |
| 1 | 79056787 | rs984107 | -1.31 | 0.569 | 0.532 | 1.498 | 0.029 | 0.177 | 0.177 | 0 | 0.08 | 0.28 | 0.1 | 0.26 | 0.05 | 0.25 |
| 1 | 79067549 | rs1413358 | -0.219 | 0.048 | -0.982 | 0.866 | 0.002 | -0.013 | -0.013 | 0 | 0.27 | 0.04 | 0.27 | 0.25 | 0.24 | 0.23 |
| 1 | 79079566 | rs11162519 | -0.211 | 0.225 | -0.992 | 0.872 | 0.002 | -0.003 | -0.003 | 0 | 0.27 | 0.04 | 0.27 | 0.25 | 0.25 | 0.23 |
| 1 | 79082772 | rs273238 | -0.902 | 0.043 | -0.225 | 1.287 | -0.007 | -0.016 | -0.016 | 0.461 | 0.4 | 0.45 | 0.36 | 0.75 | 0.34 | 0.41 |
| 1 | 79084734 | rs12405276 | 0.909 | -0.575 | -0.33 | 0.33 | -0.027 | 0.057 | 0.057 | 0.231 | 0.37 | 0.18 | 0.52 | 0.22 | 0.51 | 0.52 |
| 1 | 79090654 | rs1333969 | 0.981 | -0.605 | -0.449 | 0.356 | 0.001 | -0.018 | -0.018 | 0 | 0.26 | 0.03 | 0.26 | 0.25 | 0.24 | 0.23 |
| 1 | 79093818 | rs273259 | -0.278 | -0.654 | -0.071 | 0.33 | -0.001 | -0.005 | -0.005 | 0.127 | 0.39 | 0.36 | 0.35 | 0.75 | 0.34 | 0.41 |
| 1 | 79095581 | rs987495 | 0.935 | -0.663 | -0.23 | 0.796 | 0.013 | 0.012 | 0.012 | 0.946 | 0.26 | 0.12 | 0.27 | 0.25 | 0.25 | 0.23 |
| 1 | 79102726 | rs1981071 | 0.281 | -0.608 | -0.126 | 0.826 | 0.003 | 0.074 | 0.074 | 0 | 0.06 | 0.05 | 0.05 | 0.41 | 0.05 | 0.08 |
| 1 | 79109615 | rs166642 | -0.831 | -0.602 | -0.043 | 0.229 | -0.034 | -0.102 | -0.102 | 0.158 | 0.78 | 0.63 | 0.89 | 0.98 | 0.87 | 0.94 |
| 1 | 79110518 | rs4650590 | -0.198 | -0.395 | -0.275 | 1.048 | 0.015 | 0.004 | 0.004 | 0.968 | 0.37 | 0.2 | 0.37 | 0.67 | 0.36 | 0.41 |
| 1 | 79111142 | rs1333972 | 0.013 | 0.526 | -0.445 | 0.73 | -0.019 | -0.011 | -0.011 | 0 | 0.07 | 0.03 | 0.08 | 0.09 | 0.09 | 0.1 |

| | | | | | | | | | | | | | | | | |
|---|---|---|---|---|---|---|---|---|---|---|---|---|---|---|---|---|
| 1 | 79111419 | rs2148686 | -0.857 | 0.127 | -0.082 | 0.474 | -0.005 | -0.012 | -0.012 | 0 | 0.14 | 0.24 | 0.1 | 0.12 | 0.11 | 0.12 |
| 1 | 79111862 | rs695172 | 0.195 | 0.085 | -0.069 | 0.489 | 0 | 0 | 0 | 0.04 | 0.35 | 0.32 | 0.31 | 0.27 | 0.39 | 0.3 |
| 1 | 79115306 | rs273250 | -0.686 | 0.051 | -0.346 | 0.462 | -0.028 | -0.02 | -0.02 | 0 | 0.7 | 0.89 | 0.81 | 0.71 | 0.66 | 0.71 |
| 1 | 79123094 | rs7514391 | -0.229 | 0.013 | -0.047 | 0.061 | 0.002 | 0.003 | 0.003 | 0 | 0.66 | 0.66 | 0.42 | 0.46 | 0.58 | 0.6 |
| 1 | 79131805 | rs6669498 | 0.478 | -0.138 | -0.101 | -0.063 | -0.004 | -0.004 | -0.004 | 0.664 | 0.45 | 0.52 | 0.32 | 0.43 | 0.41 | 0.45 |
| 1 | 79132244 | rs1107320 | 1.521 | -0.134 | -0.247 | -0.06 | -0.024 | -0.018 | -0.018 | 0 | 0.29 | 0.12 | 0.19 | 0.3 | 0.34 | 0.3 |
| 1 | 79133774 | rs1333971 | 0.505 | -0.157 | 0.034 | -0.34 | -0.015 | -0.01 | -0.01 | 0 | 0.13 | 0.29 | 0.12 | 0.13 | 0.07 | 0.14 |
| 1 | 79143192 | rs6691836 | 0.628 | -0.146 | -0.1 | -0.333 | -0.008 | -0.005 | -0.005 | 0 | 0.13 | 0.18 | 0.12 | 0.13 | 0.07 | 0.14 |
| 1 | 79144319 | rs11162526 | 0.495 | -0.193 | -0.228 | -0.433 | 0.005 | 0.04 | 0.04 | 0 | 0.21 | 0.13 | 0.1 | 0.03 | 0.17 | 0.15 |
| 1 | 79155460 | rs17102085 | 0.783 | -0.407 | -0.23 | -0.405 | -0.001 | 0 | 0 | 0 | 0.12 | 0.14 | 0.11 | 0.13 | 0.07 | 0.14 |
| 1 | 79160070 | rs2148689 | 0.89 | -0.465 | -0.672 | -0.593 | -0.14 | 0.09 | 0.09 | 0 | 0.23 | 0.02 | 0.43 | 0.07 | 0.37 | 0.16 |
| 1 | 79167440 | rs7526675 | 0.372 | -0.676 | -0.295 | -0.451 | 0.015 | 0.033 | 0.033 | 0 | 0.21 | 0.14 | 0.1 | 0.03 | 0.15 | 0.15 |
| 1 | 79171286 | rs1926902 | 1.301 | -0.656 | -0.254 | -0.267 | -0.001 | 0.036 | 0.036 | 0.069 | 0.31 | 0.17 | 0.17 | 0.18 | 0.32 | 0.32 |
| 1 | 79174955 | rs12403117 | 1.293 | -0.663 | -0.257 | -0.27 | -0.001 | 0.036 | 0.036 | 0.057 | 0.31 | 0.17 | 0.17 | 0.18 | 0.32 | 0.32 |
| 1 | 79181290 | rs1780748 | -0.277 | -0.738 | 0.114 | -0.321 | -0.002 | 0.005 | 0.005 | 0 | 0.46 | 0.46 | 0.68 | 0.78 | 0.46 | 0.53 |
| 1 | 79182561 | rs1780746 | -0.566 | -0.732 | -0.143 | -0.42 | 0.013 | 0.017 | 0.017 | 0 | 0.82 | 0.87 | 0.89 | 0.97 | 0.84 | 0.88 |
| 1 | 79184105 | rs17102100 | 0.827 | -0.609 | -0.142 | -0.158 | -0.001 | -0.008 | -0.008 | 0.283 | 0.36 | 0.2 | 0.3 | 0.63 | 0.4 | 0.44 |
| 1 | 79186639 | rs1780719 | -0.909 | -0.949 | -0.046 | -1.032 | 0.008 | -0.001 | -0.001 | 0 | 0.85 | 0.9 | 0.9 | 0.98 | 0.85 | 0.91 |
| 1 | 79188388 | rs1751520 | 0.458 | -0.846 | 0.1 | -1.259 | -0.017 | 0.042 | 0.042 | 0.04 | 0.6 | 0.69 | 0.43 | 0.82 | 0.51 | 0.61 |
| 1 | 79195469 | rs4551552 | 2.028 | -1.069 | 0.03 | -1.235 | -0.012 | 0.017 | 0.017 | 0 | 0.28 | 0.17 | 0.17 | 0.22 | 0.32 | 0.3 |
| 1 | 79201589 | rs1751526 | 0.373 | -0.899 | 0.43 | -1.165 | -0.011 | 0.024 | 0.024 | 0.223 | 0.6 | 0.65 | 0.43 | 0.82 | 0.51 | 0.61 |
| 1 | 79202159 | rs6424789 | 1.39 | -1.038 | 0.26 | -1.24 | -0.009 | 0.027 | 0.027 | 1.284 | 0.18 | 0.1 | 0.12 | 0 | 0.14 | 0.05 |
| 1 | 79202300 | rs1751527 | 0.672 | -1.064 | 0.368 | -1.221 | -0.045 | 0.107 | 0.107 | 0 | 0.1 | 0.19 | 0.13 | 0.6 | 0.04 | 0.26 |
| 1 | 79206735 | rs1780732 | 0.478 | -0.718 | 0.316 | -1.072 | 0.01 | -0.057 | -0.057 | 0.08 | 0.59 | 0.52 | 0.42 | 0.82 | 0.51 | 0.61 |
| 1 | 79211929 | rs1033999 | -0.959 | -0.719 | 0.285 | -1.156 | 0.006 | -0.006 | -0.006 | 0 | 0.85 | 0.9 | 0.9 | 0.98 | 0.85 | 0.91 |
| 1 | 79214489 | rs1751529 | -1.038 | -0.721 | 0.287 | -1.158 | -0.002 | -0.03 | -0.03 | 0 | 0.85 | 0.86 | 0.9 | 0.98 | 0.85 | 0.91 |
| 1 | 79215138 | rs12135932 | 0.271 | -0.699 | 0.201 | -0.939 | 0.029 | -0.009 | -0.009 | 0 | 0.26 | 0.38 | 0.49 | 0.24 | 0.34 | 0.3 |
| 1 | 79215225 | rs1625911 | -1.062 | -0.718 | 0.055 | -0.954 | 0.009 | 0.067 | 0.067 | 0 | 0.55 | 0.7 | 0.73 | 0.76 | 0.53 | 0.61 |
| 1 | 79215916 | rs1780742 | -1.083 | -0.6 | -0.559 | -0.905 | -0.008 | -0.037 | -0.037 | 0 | 0.9 | 0.85 | 0.92 | 0.98 | 0.89 | 0.96 |
| 1 | 79217345 | rs1926896 | 0.502 | -0.449 | -0.567 | -0.759 | -0.035 | 0.001 | 0.001 | 0.719 | 0.25 | 0.17 | 0.47 | 0.24 | 0.34 | 0.3 |
| 1 | 79227956 | rs1937020 | 0.89 | -1.154 | -0.289 | -0.24 | -0.003 | 0 | 0 | 0.381 | 0.56 | 0.54 | 0.65 | 0.46 | 0.67 | 0.6 |
| 1 | 79233493 | rs7521977 | 0.064 | -0.885 | -0.524 | -0.029 | 0 | 0.001 | 0.001 | 0.517 | 0.72 | 0.73 | 0.76 | 0.48 | 0.82 | 0.69 |
| 1 | 79246505 | rs10873996 | 0.193 | -0.192 | -0.714 | -0.266 | 0.01 | -0.01 | -0.01 | 0 | 0.81 | 0.89 | 0.84 | 0.77 | 0.81 | 0.83 |
| 1 | 79255347 | rs1937009 | -0.157 | -0.154 | -0.106 | -0.729 | 0.003 | -0.135 | -0.135 | 0 | 0.2 | 0.51 | 0.14 | 0 | 0.15 | 0.06 |
| 1 | 79263805 | rs10873998 | -0.127 | 0.164 | -0.805 | -0.559 | -0.011 | 0.065 | 0.065 | 0 | 0.46 | 0.29 | 0.64 | 0.72 | 0.52 | 0.58 |
| 1 | 79264728 | rs1937011 | 0.38 | 0.189 | -0.517 | -0.577 | 0.011 | -0.026 | -0.026 | 0.88 | 0.37 | 0.44 | 0.27 | 0.28 | 0.41 | 0.3 |
| 1 | 79272660 | rs12034709 | 0.375 | 0.348 | -0.681 | -0.482 | -0.002 | 0.002 | 0.002 | 0.034 | 0.07 | 0.06 | 0.03 | 0.03 | 0.05 | 0.02 |
| 1 | 79274028 | rs6676466 | 0.275 | 0.19 | -0.647 | -0.458 | 0.009 | 0.011 | 0.011 | 0 | 0.54 | 0.32 | 0.43 | 0.44 | 0.58 | 0.46 |
| 1 | 79274420 | rs1937006 | 0.18 | -0.03 | -1.015 | -0.193 | 0 | -0.007 | -0.007 | 2.09 | 0.67 | 0.32 | 0.61 | 0.84 | 0.71 | 0.72 |
| 1 | 79274644 | rs12406640 | -0.435 | -0.126 | -1.384 | -0.425 | -0.044 | -0.089 | -0.089 | 0 | 0.32 | 0.05 | 0.36 | 0.61 | 0.33 | 0.43 |
| 1 | 79278264 | rs6659264 | 0.833 | -0.215 | -1.279 | -0.486 | 0.027 | 0.039 | 0.039 | 0 | 0.49 | 0.22 | 0.41 | 0.44 | 0.57 | 0.45 |
| 1 | 79278918 | rs1937029 | 2.088 | -0.15 | -1.177 | -0.43 | -0.019 | 0.027 | 0.027 | 0 | 0.3 | 0.16 | 0.23 | 0.23 | 0.37 | 0.27 |
| 1 | 79282661 | rs1937019 | -0.714 | -0.147 | -0.399 | -0.282 | 0.012 | -0.048 | -0.048 | 0.144 | 0.77 | 0.59 | 0.67 | 0.87 | 0.76 | 0.76 |
| 1 | 79293815 | rs7522086 | 0.204 | -0.126 | -0.302 | -0.284 | -0.007 | 0.015 | 0.015 | 0 | 0.63 | 0.52 | 0.48 | 0.48 | 0.63 | 0.49 |
| 1 | 79299411 | rs6695185 | 0.751 | 0.021 | -0.189 | 0.071 | 0.001 | 0.005 | 0.005 | 0 | 0.6 | 0.42 | 0.6 | 0.5 | 0.62 | 0.62 |
| 1 | 79321679 | rs12033775 | 0.803 | -0.104 | -0.025 | 0.318 | -0.004 | -0.085 | -0.085 | 0.757 | 0.13 | 0.01 | 0.14 | 0.26 | 0.13 | 0.2 |
| 1 | 79330525 | rs2171714 | 2.045 | 0.174 | -0.108 | 0.059 | -0.031 | -0.052 | -0.052 | 0 | 0.33 | 0.1 | 0.25 | 0.38 | 0.38 | 0.34 |
| 1 | 79336164 | rs3012079 | 0.044 | 0.344 | 0.267 | 0.022 | 0.003 | -0.013 | -0.013 | 0 | 0.76 | 0.68 | 0.77 | 0.76 | 0.77 | 0.83 |
| 1 | 79336607 | rs2996691 | 0.488 | 0.326 | 0.291 | 0.011 | -0.002 | -0.042 | -0.042 | 0 | 0.73 | 0.48 | 0.75 | 0.76 | 0.77 | 0.81 |
| 1 | 79337233 | rs2996692 | 1.496 | 0.483 | 0.246 | 0.138 | -0.001 | -0.009 | -0.009 | 0 | 0.05 | 0.04 | 0.04 | 0 | 0.05 | 0.03 |
| 1 | 79348506 | rs931413 | -0.614 | 0.479 | 0.25 | 0.094 | -0.009 | 0.018 | 0.018 | 0 | 0.23 | 0.37 | 0.22 | 0.26 | 0.2 | 0.28 |
| 1 | 79350364 | rs12037077 | -0.51 | 0.645 | -0.037 | 0.037 | -0.012 | 0.005 | 0.005 | 0 | 0.23 | 0.18 | 0.23 | 0.26 | 0.26 | 0.19 |
| 1 | 79353399 | rs10518588 | 1.748 | 0.705 | -0.195 | -0.22 | -0.024 | -0.029 | -0.029 | 0.048 | 0.06 | 0 | 0.22 | 0.09 | 0.1 | 0.14 |
| 1 | 79353655 | rs1261980 | 0.259 | 0.668 | -0.057 | 0.027 | 0 | 0.005 | 0.005 | 0 | 0.78 | 0.75 | 0.82 | 0.65 | 0.78 | 0.8 |
| 1 | 79377992 | rs971064 | 0.8 | 1.08 | -0.257 | 0.078 | -0.012 | -0.088 | -0.088 | 0 | 0.14 | 0.01 | 0.15 | 0.25 | 0.15 | 0.24 |
| 1 | 79379411 | rs1948209 | -0.29 | 1.238 | -0.043 | 0.047 | 0.026 | -0.005 | -0.005 | 0 | 0.11 | 0.23 | 0.14 | 0.08 | 0.11 | 0.15 |
| 1 | 79379776 | rs10158521 | -0.754 | 1.415 | -0.067 | -0.001 | 0.016 | 0.027 | 0.027 | 0 | 0.23 | 0.35 | 0.2 | 0.35 | 0.23 | 0.17 |
| 1 | 79380466 | rs7552978 | 2.498 | 1.28 | -0.105 | -0.175 | 0.005 | 0.019 | 0.019 | 0.829 | 0.34 | 0.29 | 0.21 | 0.23 | 0.32 | 0.2 |
| 1 | 79382009 | rs1489400 | -0.357 | 1.301 | -0.219 | -0.157 | 0.001 | -0.038 | -0.038 | 0 | 0.34 | 0.13 | 0.47 | 0.35 | 0.35 | 0.45 |
| 1 | 79382535 | rs977915 | -1.367 | 1.464 | -0.141 | -0.128 | 0.013 | 0.042 | 0.042 | 0 | 0.49 | 0.27 | 0.59 | 0.35 | 0.54 | 0.53 |
| 1 | 79392756 | rs12754818 | -2.196 | 1.445 | -0.054 | -0.125 | -0.009 | -0.028 | -0.028 | 0 | 0.61 | 0.49 | 0.68 | 0.7 | 0.62 | 0.66 |
| 1 | 79393984 | rs12036315 | 0.917 | 1.488 | -0.063 | -0.104 | 0.011 | 0.177 | 0.177 | 0 | 0.06 | 0.12 | 0.07 | 0.35 | 0.05 | 0.09 |
| 1 | 79394126 | rs12750112 | 0.857 | 1.493 | -0.077 | -0.126 | -0.006 | 0.027 | 0.027 | 0 | 0.19 | 0.08 | 0.31 | 0.1 | 0.2 | 0.22 |
| 1 | 79395424 | rs2352613 | -2.018 | 1.432 | -0.015 | -0.075 | -0.004 | -0.012 | -0.012 | 0 | 0.6 | 0.44 | 0.68 | 0.7 | 0.62 | 0.66 |
| 1 | 79396655 | rs6424618 | -1.337 | 1.435 | -0.055 | -0.087 | 0.014 | 0.041 | 0.041 | 0 | 0.49 | 0.3 | 0.59 | 0.35 | 0.54 | 0.53 |
| 1 | 79411390 | rs2035726 | -0.134 | 1.439 | -0.068 | -0.025 | 0 | 0.062 | 0.062 | 0 | 0.11 | 0.14 | 0.1 | 0.35 | 0.08 | 0.14 |
| 1 | 79411828 | rs2035724 | -0.134 | 1.439 | -0.068 | -0.025 | 0 | 0.062 | 0.062 | 0 | 0.11 | 0.14 | 0.1 | 0.35 | 0.08 | 0.14 |
| 1 | 79411888 | rs723898 | -1.281 | 1.446 | -0.058 | -0.092 | 0.011 | 0.029 | 0.029 | 0 | 0.5 | 0.35 | 0.59 | 0.36 | 0.54 | 0.53 |
| 1 | 79419229 | rs1489391 | -2.475 | 1.466 | -0.069 | -0.104 | -0.004 | -0.02 | -0.02 | 0 | 0.65 | 0.6 | 0.77 | 0.79 | 0.68 | 0.78 |

| | | | | | | | | | | | | | | | | |
|---|---|---|---|---|---|---|---|---|---|---|---|---|---|---|---|---|
| 1 | 79419439 | rs17413911 | 0.649 | 1.547 | -0.093 | -0.135 | 0 | 0.013 | 0.013 | 0 | 0.2 | 0.16 | 0.31 | 0.1 | 0.2 | 0.22 |
| 1 | 79430351 | rs6673093 | -1.962 | 1.493 | -0.12 | -0.113 | -0.008 | -0.03 | -0.03 | 0 | 0.6 | 0.54 | 0.69 | 0.72 | 0.62 | 0.66 |
| 1 | 79438696 | rs17414106 | 0.404 | 1.644 | -0.326 | 0.125 | -0.011 | 0.014 | 0.014 | 0 | 0.1 | 0.07 | 0.07 | 0 | 0.14 | 0.05 |
| 1 | 79440766 | rs11162582 | 0.606 | 1.392 | -0.252 | 0.393 | -0.001 | 0.079 | 0.079 | 0 | 0.06 | 0.05 | 0.07 | 0.36 | 0.05 | 0.09 |
| 1 | 79451015 | rs7527803 | -0.673 | 1.369 | -0.158 | 0.525 | 0 | 0.114 | 0.114 | 0 | 0.11 | 0.21 | 0.1 | 0.37 | 0.08 | 0.14 |
| 1 | 79453248 | rs17102586 | 0.719 | 1.366 | -0.183 | 0.482 | -0.001 | -0.012 | -0.012 | 0 | 0.05 | 0.09 | 0.03 | 0 | 0.04 | 0.05 |
| 1 | 79453395 | rs17414877 | 0.511 | 1.041 | -0.545 | 0.202 | 0.006 | 0.036 | 0.036 | 0 | 0.2 | 0.08 | 0.29 | 0.11 | 0.17 | 0.21 |
| 1 | 79454383 | rs11584172 | 0.61 | 0.99 | -0.258 | 0.487 | 0.009 | 0.197 | 0.197 | 0 | 0.06 | 0.11 | 0.07 | 0.37 | 0.05 | 0.09 |
| 1 | 79456112 | rs12134068 | 0.163 | 1.056 | -0.275 | 0.195 | 0.011 | -0.003 | -0.003 | 0 | 0.08 | 0.1 | 0.14 | 0.06 | 0.13 | 0.13 |
| 1 | 79458347 | rs6685355 | -1.562 | 0.887 | -0.242 | 0.089 | -0.004 | 0.005 | 0.005 | 0 | 0.61 | 0.46 | 0.71 | 0.41 | 0.64 | 0.68 |
| 1 | 79471959 | rs1571991 | 0.672 | 0.952 | -0.086 | -0.013 | -0.005 | 0.07 | 0.07 | 0.167 | 0.78 | 0.5 | 0.72 | 0.7 | 0.75 | 0.65 |
| 1 | 79482417 | rs2065486 | -0.121 | 0.937 | -0.022 | -0.019 | -0.001 | 0.006 | 0.006 | 0 | 0.14 | 0.14 | 0.14 | 0.24 | 0.16 | 0.21 |
| 1 | 79484287 | rs12406049 | 2.595 | 0.859 | -0.122 | 0.075 | 0.005 | 0.022 | 0.022 | 0.172 | 0.31 | 0.26 | 0.21 | 0.22 | 0.31 | 0.2 |
| 1 | 79484716 | rs2065485 | -1.019 | 0.933 | 0.062 | 0.203 | 0.012 | -0.008 | -0.008 | 0 | 0.94 | 0.89 | 0.88 | 0.95 | 0.9 | 0.87 |
| 1 | 79487080 | rs6424626 | 0.438 | 0.896 | 0.017 | 0.201 | -0.002 | 0.05 | 0.05 | 0 | 0.8 | 0.66 | 0.73 | 0.71 | 0.75 | 0.65 |
| 1 | 79488882 | rs1571989 | 0.354 | 0.863 | -0.434 | 0.234 | -0.015 | 0.095 | 0.095 | 0.21 | 0.82 | 0.48 | 0.75 | 0.71 | 0.81 | 0.67 |
| 1 | 79490302 | rs6691975 | -0.769 | 0.944 | -0.883 | 0.284 | -0.034 | -0.037 | -0.037 | 3.442 | 0.46 | 0.16 | 0.5 | 0.49 | 0.44 | 0.44 |
| 1 | 79491206 | rs6677175 | -0.041 | 0.962 | -0.661 | 0.132 | -0.035 | 0.067 | 0.067 | 0 | 0.25 | 0.09 | 0.35 | 0.11 | 0.25 | 0.24 |
| 1 | 79493936 | rs11807825 | -0.945 | 0.939 | 0.173 | 0.612 | -0.018 | 0.113 | 0.113 | 0.415 | 0.17 | 0.42 | 0.17 | 0.25 | 0.16 | 0.23 |
| 1 | 79501129 | rs1831704 | -0.598 | 0.697 | 0.409 | 0.649 | -0.026 | -0.006 | -0.006 | 0 | 0.3 | 0.45 | 0.31 | 0.34 | 0.25 | 0.16 |
| 1 | 79503456 | rs12043235 | 2.025 | 0.968 | 0.076 | 0.765 | -0.026 | -0.031 | -0.031 | 0 | 0.36 | 0.11 | 0.28 | 0.36 | 0.36 | 0.37 |
| 1 | 79507828 | rs1332819 | -0.399 | 1.075 | 0.369 | 0.723 | -0.01 | -0.016 | -0.016 | 0 | 0.81 | 0.64 | 0.73 | 0.84 | 0.82 | 0.71 |
| 1 | 79508151 | rs1332817 | 1.262 | 1.14 | 0.117 | 0.745 | -0.079 | -0.054 | -0.054 | 0.318 | 0.13 | 0.01 | 0.14 | 0.15 | 0.22 | 0.19 |
| 1 | 79509109 | rs6687262 | 1.221 | 0.986 | 0.195 | 0.721 | 0.062 | 0.028 | 0.028 | 0.079 | 0.5 | 0.16 | 0.42 | 0.51 | 0.58 | 0.55 |
| 1 | 79515310 | rs17098468 | 0.416 | 0.95 | 0.269 | 0.697 | 0.001 | 0.005 | 0.005 | 0.015 | 0.18 | 0.1 | 0.25 | 0.16 | 0.17 | 0.29 |
| 1 | 79519427 | rs10493621 | 0.111 | 0.857 | 0.558 | 0.684 | 0.019 | -0.011 | -0.011 | 0 | 0.27 | 0.11 | 0.27 | 0.33 | 0.24 | 0.15 |
| 1 | 79519555 | rs17102675 | -0.29 | 0.879 | 0.61 | 0.687 | -0.014 | -0.005 | -0.005 | 0.958 | 0.3 | 0.39 | 0.3 | 0.33 | 0.24 | 0.15 |
| 1 | 79539394 | rs1906281 | -1.243 | 0.909 | 0.564 | 0.71 | 0.006 | -0.003 | -0.003 | 0.827 | 0.38 | 0.59 | 0.53 | 0.32 | 0.47 | 0.54 |
| 1 | 79543571 | rs1332829 | 2.315 | 0.957 | 0.286 | 0.703 | -0.017 | -0.037 | -0.037 | 0 | 0.36 | 0.12 | 0.27 | 0.37 | 0.34 | 0.34 |
| 1 | 79549699 | rs1571993 | -0.841 | 0.971 | 0.343 | 0.621 | 0.01 | -0.007 | -0.007 | 0 | 0.36 | 0.56 | 0.43 | 0.3 | 0.4 | 0.51 |
| 1 | 79550278 | rs12760876 | 2.352 | 0.965 | 0.334 | 0.63 | -0.022 | -0.049 | -0.049 | 0 | 0.35 | 0.1 | 0.27 | 0.37 | 0.34 | 0.34 |
| 1 | 79550458 | rs1157230 | -0.496 | 0.95 | 0.378 | 0.632 | 0.017 | 0.002 | 0.002 | 0 | 0.74 | 0.81 | 0.8 | 0.69 | 0.81 | 0.88 |
| 1 | 79552777 | rs10874010 | 0.098 | 1.122 | 0.5 | 0.649 | 0.05 | 0.012 | 0.012 | 0 | 0.18 | 0.43 | 0.17 | 0.14 | 0.23 | 0.22 |
| 1 | 79556037 | rs7531214 | 0.348 | 1.004 | 0.045 | 0.523 | -0.001 | 0 | 0 | 0 | 0.18 | 0.16 | 0.26 | 0.15 | 0.18 | 0.29 |
| 1 | 79566969 | rs6424633 | 1.863 | 1.003 | -0.225 | 0.475 | -0.007 | -0.038 | -0.038 | 0.173 | 0.37 | 0.1 | 0.26 | 0.34 | 0.32 | 0.38 |
| 1 | 79571617 | rs12097617 | -1.142 | 1.341 | -0.38 | 0.589 | 0.003 | -0.001 | -0.001 | 0.002 | 0.47 | 0.4 | 0.56 | 0.57 | 0.46 | 0.43 |
| 1 | 79574878 | rs901680 | -1.403 | 1.462 | -0.667 | 0.593 | -0.004 | 0.003 | 0.003 | 0 | 0.87 | 0.86 | 0.85 | 0.92 | 0.78 | 0.81 |
| 1 | 79580281 | rs10874016 | 0.084 | 0.814 | -0.66 | 0.817 | -0.005 | -0.007 | -0.007 | 2.256 | 0.18 | 0.13 | 0.26 | 0.26 | 0.21 | 0.26 |
| 1 | 79580392 | rs11162632 | -1.037 | 0.928 | -0.683 | 0.543 | 0.005 | -0.003 | -0.003 | 0.095 | 0.46 | 0.36 | 0.56 | 0.59 | 0.46 | 0.44 |
| 1 | 79581567 | rs11589546 | -0.003 | 0.847 | -0.94 | 0.352 | -0.015 | 0.065 | 0.065 | 0 | 0.13 | 0.04 | 0.11 | 0.02 | 0.14 | 0.02 |
| 1 | 79582937 | rs11162634 | 0.579 | 0.703 | -0.512 | 0.767 | -0.008 | 0.002 | 0.002 | 0.137 | 0.67 | 0.65 | 0.69 | 0.63 | 0.73 | 0.76 |
| 1 | 79583385 | rs17408443 | -0.271 | 1.593 | -1.038 | 0.731 | -0.062 | -0.066 | -0.066 | 0.74 | 0.18 | 0.06 | 0.24 | 0.32 | 0.29 | 0.21 |
| 1 | 79587385 | rs17408603 | 1.132 | 1.846 | -1.068 | 0.745 | -0.053 | -0.024 | -0.024 | 0.058 | 0.1 | 0.01 | 0.08 | 0.12 | 0.17 | 0.1 |
| 1 | 79589903 | rs1545971 | 0.76 | 1.878 | -0.012 | 0.373 | 0.008 | 0 | 0 | 0 | 0.76 | 0.35 | 0.72 | 0.72 | 0.73 | 0.73 |
| 1 | 79590786 | rs17102806 | 0.597 | 1.94 | -0.13 | 0.409 | -0.02 | -0.035 | -0.035 | 0.061 | 0.08 | 0.03 | 0.15 | 0.19 | 0.12 | 0.11 |
| 1 | 79593296 | rs17102812 | 0.079 | 1.875 | 0.215 | 0.372 | 0.017 | -0.032 | -0.032 | 0 | 0.14 | 0.27 | 0.1 | 0.08 | 0.15 | 0.18 |
| 1 | 79594157 | rs12063711 | 0.917 | 1.806 | 0.115 | 0.376 | -0.001 | -0.001 | -0.001 | 0 | 0.12 | 0.11 | 0.08 | 0.08 | 0.12 | 0.02 |
| 1 | 79594565 | rs17102815 | 0.397 | 2.088 | 0.365 | 0.696 | 0 | 0 | 0 | 0 | 0.15 | 0.06 | 0.14 | 0.16 | 0.11 | 0.17 |
| 1 | 79595722 | rs6690294 | 1.464 | 2.218 | 0.72 | 0.637 | 0.004 | -0.008 | -0.008 | 0.262 | 0.73 | 0.33 | 0.58 | 0.73 | 0.69 | 0.65 |
| 1 | 79595821 | rs17102821 | -0.022 | 2.219 | 0.708 | 0.617 | -0.013 | -0.024 | -0.024 | 0 | 0.09 | 0.06 | 0.16 | 0.19 | 0.13 | 0.12 |
| 1 | 79614989 | rs1555385 | -1.633 | 2.234 | 0.833 | 0.57 | 0.004 | -0.004 | -0.004 | 0 | 0.26 | 0.31 | 0.4 | 0.27 | 0.32 | 0.37 |
| 1 | 79621939 | rs390842 | -1.747 | 2.042 | 0.585 | 0.603 | 0.008 | -0.015 | -0.015 | 0 | 0.27 | 0.62 | 0.42 | 0.27 | 0.3 | 0.37 |
| 1 | 79641217 | rs12027976 | -0.471 | 1.319 | 0.613 | 0.738 | 0.035 | 0.008 | 0.008 | 0 | 0.11 | 0.36 | 0.17 | 0.09 | 0.13 | 0.12 |
| 1 | 79646081 | rs228564 | -1.163 | 1.368 | 0.075 | 0.772 | -0.054 | -0.02 | -0.02 | 2.781 | 0.9 | 0.99 | 0.92 | 0.88 | 0.83 | 0.91 |
| 1 | 79646219 | rs228563 | -1.744 | 1.22 | 0.168 | 0.746 | -0.015 | -0.008 | -0.008 | 0.247 | 0.23 | 0.65 | 0.4 | 0.23 | 0.2 | 0.33 |
| 1 | 79653433 | rs10489655 | -0.597 | 1.026 | 0.022 | 0.807 | -0.017 | 0.001 | 0.001 | 0 | 0.12 | 0.17 | 0.22 | 0.14 | 0.07 | 0.2 |
| 1 | 79658861 | rs5008836 | 0.724 | 1.268 | -0.388 | 0.809 | 0 | -0.001 | -0.001 | 0.005 | 0.67 | 0.34 | 0.52 | 0.64 | 0.63 | 0.58 |
| 1 | 79673001 | rs10489653 | -0.508 | 1.279 | -0.649 | 1.273 | -0.041 | -0.079 | -0.079 | 1.058 | 0.19 | 0.06 | 0.19 | 0.4 | 0.21 | 0.36 |
| 1 | 79677027 | rs228513 | -1.766 | 0.79 | -1.493 | 0.832 | -0.097 | 0.175 | 0.175 | 0.093 | 0.53 | 0.95 | 0.52 | 0.82 | 0.47 | 0.73 |
| 1 | 79682535 | rs7529406 | -0.032 | 0.639 | -1.442 | 0.938 | -0.015 | -0.011 | -0.011 | 0.037 | 0.15 | 0.09 | 0.08 | 0.14 | 0.13 | 0.16 |
| 1 | 79687429 | rs228493 | 0.155 | 0.539 | -0.306 | 0.651 | -0.01 | -0.05 | -0.05 | 0.669 | 0.09 | 0.26 | 0.08 | 0 | 0.06 | 0.05 |
| 1 | 79710545 | rs417181 | -0.948 | 0.516 | 0.193 | 0.802 | -0.003 | 0.003 | 0.003 | 0 | 0.26 | 0.32 | 0.28 | 0.37 | 0.2 | 0.29 |
| 1 | 79728427 | rs7520601 | 0.203 | 0.546 | 0.228 | 0.852 | -0.039 | -0.076 | -0.076 | 0 | 0.19 | 0.08 | 0.19 | 0.4 | 0.22 | 0.35 |
| 1 | 79729175 | rs10158555 | -0.517 | 0.507 | 1.009 | 0.95 | 0.073 | 0.106 | 0.106 | 1.309 | 0.07 | 0.42 | 0.04 | 0.04 | 0.05 | 0.08 |
| 1 | 79741744 | rs7555545 | -0.802 | 0.514 | 1.084 | 0.903 | 0 | 0.008 | 0.008 | 0 | 0.33 | 0.58 | 0.29 | 0.43 | 0.32 | 0.43 |
| 1 | 79753199 | rs10874030 | -1.555 | 0.451 | 0.656 | 0.481 | -0.101 | 0.159 | 0.159 | 0.043 | 0.53 | 0.95 | 0.53 | 0.8 | 0.47 | 0.73 |
| 1 | 79770776 | rs1412139 | -0.954 | 0.455 | 0.852 | 0.486 | 0 | -0.016 | -0.016 | 0 | 0.46 | 0.5 | 0.48 | 0.77 | 0.41 | 0.65 |
| 1 | 79775411 | rs1412136 | -1.319 | 0.464 | 0.547 | 0.487 | -0.032 | 0.173 | 0.173 | 0.087 | 0.51 | 0.85 | 0.51 | 0.81 | 0.44 | 0.72 |
| 1 | 79780054 | rs11162685 | -1.319 | 0.464 | 0.547 | 0.487 | -0.032 | 0.173 | 0.173 | 0.087 | 0.51 | 0.85 | 0.51 | 0.81 | 0.44 | 0.72 |

| | | | | | | | | | | | | | | | | |
|---|---|---|---|---|---|---|---|---|---|---|---|---|---|---|---|---|
| 1 | 79794719 | rs10732727 | -1.532 | 0.442 | 0.36 | 0.491 | -0.103 | 0.155 | 0.155 | 0.011 | 0.53 | 0.95 | 0.52 | 0.81 | 0.47 | 0.73 |
| 1 | 79799912 | rs1331707 | -1.323 | 0.457 | 0.45 | 0.525 | -0.036 | 0.139 | 0.139 | 0.002 | 0.5 | 0.88 | 0.51 | 0.77 | 0.44 | 0.65 |
| 1 | 79826759 | rs6666584 | 0.842 | 0.538 | 0.745 | 0.415 | 0.004 | 0.006 | 0.006 | 1.132 | 0.51 | 0.53 | 0.29 | 0.22 | 0.5 | 0.33 |
| 1 | 79831147 | rs4414002 | 0.88 | 0.492 | 0.935 | 0.483 | -0.003 | -0.016 | -0.016 | 0.1 | 0.51 | 0.49 | 0.29 | 0.19 | 0.48 | 0.29 |
| 1 | 79851608 | rs3000112 | -1.485 | 0.455 | 1.209 | 0.528 | 0.005 | 0.02 | 0.02 | 0.229 | 0.07 | 0.28 | 0.04 | 0.03 | 0.06 | 0.09 |
| 1 | 79853766 | rs3015050 | 0.558 | 0.427 | 1.214 | 0.467 | 0.005 | 0.016 | 0.016 | 0.338 | 0.54 | 0.58 | 0.3 | 0.23 | 0.5 | 0.37 |
| 1 | 79856086 | rs3000156 | -1.692 | 0.437 | 1.137 | 0.48 | 0.005 | 0.02 | 0.02 | 0.842 | 0.08 | 0.31 | 0.04 | 0.03 | 0.06 | 0.09 |
| 1 | 79860397 | rs3015077 | -1.637 | 0.446 | 1.113 | 0.508 | 0.002 | 0.021 | 0.021 | 0.657 | 0.08 | 0.3 | 0.04 | 0.04 | 0.06 | 0.09 |
| 1 | 79869873 | rs3015098 | 0.955 | 0.411 | 1.263 | 0.474 | 0.04 | 0.093 | 0.093 | 0.005 | 0.48 | 0.26 | 0.27 | 0.22 | 0.46 | 0.35 |
| 1 | 79887916 | rs3000182 | 1.28 | 0.435 | 0.963 | 0.478 | 0.037 | 0.135 | 0.135 | 0.232 | 0.45 | 0.14 | 0.25 | 0.19 | 0.44 | 0.28 |
| 1 | 79896187 | rs3015119 | -0.052 | 0.723 | 0.954 | 1.064 | -0.003 | -0.014 | -0.014 | 7.36 | 0.67 | 0.74 | 0.57 | 0.45 | 0.68 | 0.56 |
| 1 | 79904383 | rs3000110 | 0.711 | 0.472 | 1.071 | 0.861 | 0.003 | 0.006 | 0.006 | 0.177 | 0.53 | 0.54 | 0.3 | 0.22 | 0.5 | 0.36 |
| 1 | 79917078 | rs3015006 | 0.741 | 0.485 | 1.001 | 0.648 | -0.002 | -0.005 | -0.005 | 0 | 0.53 | 0.46 | 0.3 | 0.22 | 0.5 | 0.36 |
| 1 | 79942754 | rs3000154 | 0.898 | 0.504 | 0.892 | 0.66 | 0.015 | 0.03 | 0.03 | 0 | 0.49 | 0.41 | 0.28 | 0.23 | 0.47 | 0.35 |
| 1 | 79970061 | rs2934871 | 0.799 | 0.501 | 0.853 | 0.624 | 0.005 | 0.016 | 0.016 | 0 | 0.52 | 0.41 | 0.29 | 0.23 | 0.5 | 0.4 |
| 1 | 79984533 | rs17453273 | 1.479 | 0.236 | 0.054 | 0.197 | -0.085 | 0.062 | 0.062 | 0 | 0.21 | 0.02 | 0.12 | 0.07 | 0.27 | 0.13 |
| 1 | 79997655 | rs2172794 | 0.933 | 0.786 | 0.839 | 0.961 | 0.008 | 0.012 | 0.012 | 0 | 0.54 | 0.19 | 0.52 | 0.35 | 0.61 | 0.43 |
| 1 | 80004566 | rs1353287 | 1.301 | 0.806 | 1.056 | 0.95 | 0.033 | 0.055 | 0.055 | 0 | 0.93 | 0.69 | 0.96 | 0.96 | 0.94 | 0.91 |
| 1 | 80014353 | rs958261 | 0.649 | 0.822 | 1.09 | 1.031 | 0.002 | 0.002 | 0.002 | 0 | 0.59 | 0.36 | 0.55 | 0.35 | 0.65 | 0.44 |
| 1 | 80025255 | rs1587076 | 0.366 | 0.91 | 1.084 | 1.071 | 0.089 | 0.106 | 0.106 | 0.023 | 0.89 | 0.7 | 0.72 | 0.79 | 0.82 | 0.81 |
| 1 | 80026228 | rs1603130 | -0.586 | 0.854 | 1.012 | 1.041 | 0 | -0.003 | -0.003 | 0 | 0.38 | 0.36 | 0.43 | 0.61 | 0.33 | 0.49 |
| 1 | 80034684 | rs2016809 | 1.165 | 0.821 | 0.908 | 0.842 | 0.015 | 0.042 | 0.042 | 0.033 | 0.51 | 0.41 | 0.29 | 0.18 | 0.49 | 0.32 |
| 1 | 80040353 | rs2201453 | 0.961 | 0.837 | 0.88 | 0.905 | 0.001 | 0.047 | 0.047 | 0 | 0.53 | 0.3 | 0.53 | 0.32 | 0.58 | 0.41 |
| 1 | 80052033 | rs6424661 | 0.676 | 0.785 | 0.857 | 0.885 | 0 | 0.004 | 0.004 | 0 | 0.58 | 0.35 | 0.55 | 0.36 | 0.62 | 0.47 |
| 1 | 80054305 | rs11162749 | -0.308 | 0.915 | 0.836 | 0.91 | 0.02 | 0 | 0 | 0 | 0.11 | 0.19 | 0.28 | 0.12 | 0.17 | 0.11 |
| 1 | 80058984 | rs12083183 | -0.731 | 0.911 | 0.839 | 0.907 | 0.02 | 0.012 | 0.012 | 0 | 0.13 | 0.22 | 0.29 | 0.16 | 0.19 | 0.17 |
| 1 | 80059329 | rs17454292 | -0.4 | 1.31 | 0.878 | 0.807 | 0.015 | -0.017 | -0.017 | 0 | 0.07 | 0.06 | 0.04 | 0.01 | 0.12 | 0.12 |
| 1 | 80066259 | rs11586640 | -0.671 | 1.355 | 0.81 | 0.859 | 0.012 | 0.008 | 0.008 | 0 | 0.13 | 0.19 | 0.29 | 0.17 | 0.19 | 0.17 |
| 1 | 80069082 | rs10465721 | -0.491 | 1.241 | 1.025 | 0.841 | 0.009 | -0.012 | -0.012 | 0 | 0.34 | 0.27 | 0.4 | 0.63 | 0.26 | 0.41 |
| 1 | 80069451 | rs10465729 | 0.732 | 1.17 | 0.583 | 1.109 | -0.042 | -0.075 | -0.075 | 0 | 0.16 | 0.01 | 0.19 | 0.24 | 0.16 | 0.19 |
| 1 | 80070748 | rs1382968 | -2.045 | 1.255 | 1.015 | 0.568 | 0.033 | 0.092 | 0.092 | 0 | 0.55 | 0.76 | 0.73 | 0.81 | 0.56 | 0.7 |
| 1 | 80072305 | rs1339784 | -1.217 | 1.265 | 1.013 | 0.572 | -0.002 | 0.007 | 0.007 | 0 | 0.47 | 0.47 | 0.68 | 0.79 | 0.45 | 0.58 |
| 1 | 80075820 | rs1479737 | -1.597 | 0.923 | 0.472 | 0.658 | 0.036 | 0.122 | 0.122 | 0 | 0.55 | 0.82 | 0.74 | 0.81 | 0.56 | 0.7 |
| 1 | 80084647 | rs7534346 | -1.425 | 0.973 | 0.5 | 0.731 | 0.021 | 0.042 | 0.042 | 0 | 0.52 | 0.62 | 0.72 | 0.77 | 0.54 | 0.65 |
| 1 | 80098006 | rs12138392 | -1.205 | 1.308 | 0.868 | 0.895 | 0.05 | -0.004 | -0.004 | 0 | 0.15 | 0.38 | 0.3 | 0.13 | 0.21 | 0.16 |
| 1 | 80098823 | rs10782691 | -1.35 | 1.297 | 0.702 | 0.893 | 0.065 | 0.024 | 0.024 | 0 | 0.18 | 0.52 | 0.31 | 0.17 | 0.23 | 0.2 |
| 1 | 80102675 | rs10874051 | -0.814 | 1.116 | 0.67 | 1.07 | 0.019 | 0.027 | 0.027 | 0.085 | 0.19 | 0.36 | 0.14 | 0.24 | 0.18 | 0.31 |
| 1 | 80115793 | rs10493653 | -0.504 | 1.163 | 0.776 | 1.059 | 0 | 0 | 0 | 0 | 0.17 | 0.15 | 0.13 | 0.24 | 0.19 | 0.31 |
| 1 | 80116138 | rs11162772 | 0.546 | 0.941 | 0.445 | 0.338 | -0.001 | -0.014 | -0.014 | 0 | 0.68 | 0.6 | 0.45 | 0.28 | 0.69 | 0.66 |
| 1 | 80126241 | rs6662701 | -0.301 | 0.937 | 0.901 | 1.099 | 0.016 | 0.11 | 0.11 | 0 | 0.92 | 0.81 | 0.88 | 0.8 | 0.94 | 0.94 |
| 1 | 80128969 | rs12028524 | 1.155 | 0.886 | 0.389 | 1.113 | -0.005 | -0.02 | -0.02 | 0 | 0.06 | 0 | 0.03 | 0.05 | 0.05 | 0.08 |
| 1 | 80135445 | rs11162780 | -0.716 | 0.826 | 0.443 | 0.962 | -0.002 | 0.027 | 0.027 | 0.022 | 0.23 | 0.3 | 0.42 | 0.47 | 0.21 | 0.22 |
| 1 | 80139113 | rs2000021 | 1.129 | 0.78 | 0.584 | 0.981 | 0.005 | 0.025 | 0.025 | 0.042 | 0.66 | 0.51 | 0.7 | 0.49 | 0.69 | 0.47 |
| 1 | 80145482 | rs2892636 | 0.567 | 0.757 | 1.001 | 0.991 | 0.036 | 0.005 | 0.005 | 0.772 | 0.89 | 0.59 | 0.93 | 0.92 | 0.93 | 0.77 |
| 1 | 80147123 | rs6681417 | 0.537 | 0.764 | 1.004 | 0.994 | 0.04 | 0.008 | 0.008 | 1.029 | 0.89 | 0.59 | 0.92 | 0.92 | 0.92 | 0.77 |
| 1 | 80148857 | rs11162785 | 1.158 | 0.394 | 0.887 | 1.036 | 0 | -0.006 | -0.006 | 0 | 0.68 | 0.36 | 0.52 | 0.43 | 0.72 | 0.56 |
| 1 | 80150181 | rs12076080 | -0.399 | 0.474 | 0.957 | 1.388 | 0 | -0.001 | -0.001 | 0.368 | 0.2 | 0.23 | 0.37 | 0.2 | 0.21 | 0.14 |
| 1 | 80155982 | rs989580 | -1.473 | 0.508 | 1.099 | 1.413 | -0.003 | -0.001 | -0.001 | 0.13 | 0.35 | 0.39 | 0.51 | 0.69 | 0.35 | 0.43 |
| 1 | 80157156 | rs6689140 | -1.9 | 0.479 | 1.139 | 1.292 | -0.001 | -0.02 | -0.02 | 0.431 | 0.43 | 0.47 | 0.62 | 0.88 | 0.42 | 0.51 |
| 1 | 80159221 | rs1524174 | -0.029 | 0.451 | 1.198 | 1.257 | 0.019 | 0.002 | 0.002 | 0.454 | 0.92 | 0.72 | 0.94 | 0.95 | 0.94 | 0.85 |
| 1 | 80171368 | rs7539480 | 0.978 | 0.498 | 1.11 | 1.368 | 0.004 | 0.005 | 0.005 | 0 | 0.12 | 0.18 | 0.09 | 0.19 | 0.12 | 0.21 |
| 1 | 80172400 | rs12144677 | 0.443 | 0.487 | 0.963 | 1.419 | -0.034 | -0.009 | -0.009 | 0 | 0.17 | 0.05 | 0.35 | 0.23 | 0.19 | 0.11 |
| 1 | 80176903 | rs2388919 | 1.253 | 0.476 | 1.235 | 1.566 | 0.002 | 0.039 | 0.039 | 0.005 | 0.68 | 0.51 | 0.73 | 0.56 | 0.73 | 0.53 |
| 1 | 80196238 | rs6699172 | 1.543 | 0.685 | 0.723 | 0.894 | 0.005 | 0.153 | 0.153 | 0 | 0.51 | 0.24 | 0.57 | 0.25 | 0.5 | 0.34 |
| 1 | 80197792 | rs6665734 | -0.295 | 0.473 | 1.127 | 1.14 | 0 | 0.042 | 0.042 | 0 | 0.13 | 0.11 | 0.17 | 0.48 | 0.11 | 0.18 |
| 1 | 80203708 | rs1404064 | 0.799 | 0.499 | 1.06 | 1.043 | 0.007 | 0.285 | 0.285 | 2.584 | 0.89 | 0.46 | 0.87 | 0.72 | 0.93 | 0.78 |
| 1 | 80217533 | rs2037617 | -0.095 | 0.592 | 0.634 | 0.432 | 0.022 | 0.01 | 0.01 | 0.705 | 0.92 | 0.74 | 0.94 | 0.95 | 0.94 | 0.85 |
| 1 | 80220979 | rs1464732 | -0.032 | 0.591 | 0.633 | 0.431 | 0.026 | 0.013 | 0.013 | 0.745 | 0.92 | 0.71 | 0.94 | 0.95 | 0.94 | 0.85 |
| 1 | 80225837 | rs4460589 | 1.933 | 0.519 | -0.167 | 0.187 | -0.031 | 0.193 | 0.193 | 0 | 0.33 | 0.01 | 0.22 | 0.04 | 0.33 | 0.23 |
| 1 | 80226884 | rs11162813 | 1.345 | 0.865 | 0.06 | 0.458 | -0.002 | -0.002 | -0.002 | 0 | 0.13 | 0.15 | 0.09 | 0.19 | 0.11 | 0.19 |
| 1 | 80226910 | rs7517325 | -0.147 | 0.791 | 0.456 | 0.59 | 0.001 | -0.011 | -0.011 | 0.165 | 0.13 | 0.39 | 0.07 | 0.13 | 0.09 | 0.12 |
| 1 | 80233281 | rs3927739 | -0.211 | 0.97 | 0.593 | 0.839 | -0.045 | -0.042 | -0.042 | 0 | 0.32 | 0.13 | 0.54 | 0.58 | 0.39 | 0.33 |
| 1 | 80234546 | rs940401 | 1.145 | 0.868 | 0.695 | 0.831 | 0.002 | -0.004 | -0.004 | 0 | 0.51 | 0.46 | 0.32 | 0.23 | 0.46 | 0.43 |
| 1 | 80235819 | rs12022561 | 0.304 | 0.82 | 0.18 | 1.064 | -0.005 | -0.05 | -0.05 | 2.775 | 0.17 | 0.09 | 0.45 | 0.55 | 0.17 | 0.21 |
| 1 | 80238627 | rs952309 | 0.402 | 0.964 | 0.573 | 0.531 | 0.001 | -0.023 | -0.023 | 0 | 0.68 | 0.43 | 0.62 | 0.22 | 0.74 | 0.58 |
| 1 | 80239398 | rs5008948 | 0.23 | 0.933 | 0.654 | 1.009 | 0.006 | 0.017 | 0.017 | 0 | 0.7 | 0.48 | 0.7 | 0.39 | 0.75 | 0.6 |
| 1 | 80241873 | rs12033756 | 0.758 | 0.958 | 0.802 | 1.371 | 0.001 | -0.01 | -0.01 | 1.765 | 0.14 | 0.09 | 0.14 | 0.19 | 0.11 | 0.1 |
| 1 | 80242598 | rs12023389 | -0.911 | 0.78 | 0.878 | 1.131 | -0.003 | -0.031 | -0.031 | 0 | 0.19 | 0.36 | 0.3 | 0.16 | 0.2 | 0.23 |
| 1 | 80248034 | rs10782700 | 0.534 | 0.795 | 0.92 | 1.179 | 0.107 | 0.345 | 0.345 | 0.91 | 0.95 | 0.68 | 0.88 | 0.82 | 0.98 | 0.93 |

| | | | | | | | | | | | | | | | | |
|---|---|---|---|---|---|---|---|---|---|---|---|---|---|---|---|---|
| 1 | 80248531 | rs7515884 | -0.667 | 0.777 | 1.007 | 1.122 | 0 | 0.006 | 0.006 | 0 | 0.17 | 0.13 | 0.28 | 0.16 | 0.2 | 0.23 |
| 1 | 80251768 | rs17458029 | 1.372 | 0.537 | 0.866 | 1.061 | -0.064 | 0.069 | 0.069 | 1.017 | 0.14 | 0.01 | 0.08 | 0.03 | 0.17 | 0.11 |
| 1 | 80255867 | rs1949779 | 1.066 | 0.481 | 0.909 | 1.068 | -0.001 | -0.008 | -0.008 | 0.398 | 0.13 | 0.1 | 0.13 | 0.19 | 0.1 | 0.07 |
| 1 | 80258075 | rs1568764 | -0.946 | 0.485 | 1.036 | 1.406 | 0.049 | 0.248 | 0.248 | 0.391 | 0.83 | 0.57 | 0.8 | 0.61 | 0.86 | 0.74 |
| 1 | 80265367 | rs9659877 | 0.213 | 0.447 | 1.202 | 1.327 | 0.005 | 0.002 | 0.002 | 0 | 0.1 | 0.07 | 0.04 | 0.13 | 0.06 | 0.14 |
| 1 | 80269886 | rs1880268 | -0.02 | 0.497 | 1.337 | 1.29 | 0.019 | -0.003 | -0.003 | 0 | 0.7 | 0.47 | 0.78 | 0.71 | 0.68 | 0.65 |
| 1 | 80270261 | rs10874076 | 1.838 | 0.495 | 1.148 | 1.353 | 0.001 | 0.009 | 0.009 | 0 | 0.12 | 0.13 | 0.09 | 0.24 | 0.13 | 0.19 |
| 1 | 80276868 | rs11162826 | -0.826 | 0.487 | 1.237 | 1.366 | 0.039 | 0.225 | 0.225 | 0.363 | 0.83 | 0.48 | 0.79 | 0.59 | 0.86 | 0.74 |
| 1 | 80278733 | rs12744944 | 1.668 | -0.157 | 0.943 | 1.088 | -0.092 | 0.283 | 0.283 | 0 | 0.4 | 0.02 | 0.45 | 0.03 | 0.48 | 0.29 |
| 1 | 80286411 | rs17425287 | 0.428 | -0.051 | 0.949 | 1.376 | -0.008 | 0.018 | 0.018 | 0 | 0.57 | 0.39 | 0.61 | 0.31 | 0.58 | 0.52 |
| 1 | 80289838 | rs11162830 | 0.492 | -0.18 | 0.869 | 1.347 | 0.003 | 0.067 | 0.067 | 0 | 0.55 | 0.33 | 0.52 | 0.2 | 0.56 | 0.49 |
| 1 | 80300618 | rs7552100 | 1.613 | -0.034 | 0.694 | 1.361 | -0.037 | -0.044 | -0.044 | 0 | 0.2 | 0.13 | 0.26 | 0.42 | 0.28 | 0.3 |
| 1 | 80305984 | rs12025436 | 0.767 | -0.07 | 0.753 | 1.226 | 0.001 | -0.015 | -0.015 | 0.056 | 0.13 | 0.1 | 0.14 | 0.2 | 0.1 | 0.07 |
| 1 | 80311188 | rs11162839 | 1.572 | 0.034 | 0.726 | 1.216 | -0.017 | -0.002 | -0.002 | 0 | 0.2 | 0.17 | 0.23 | 0.23 | 0.28 | 0.23 |
| 1 | 80312051 | rs12059430 | -0.141 | -1.056 | 0.097 | 0.497 | 0.021 | 0.039 | 0.039 | 0 | 0.5 | 0.59 | 0.51 | 0.34 | 0.51 | 0.45 |
| 1 | 80313191 | rs17103834 | 1.371 | -1.138 | 0.303 | 0.486 | -0.014 | -0.035 | -0.035 | 1.033 | 0.07 | 0.17 | 0.05 | 0.02 | 0.04 | 0.08 |
| 1 | 80313712 | rs10782705 | -0.761 | -1.418 | 0.151 | 0.479 | 0.012 | -0.017 | -0.017 | 0.69 | 0.55 | 0.72 | 0.48 | 0.59 | 0.51 | 0.58 |
| 1 | 80316225 | rs11162840 | 2.109 | -1.231 | 0.103 | 0.572 | 0 | -0.001 | -0.001 | 0 | 0.2 | 0.21 | 0.23 | 0.23 | 0.29 | 0.22 |
| 1 | 80316802 | rs12037755 | 1.322 | -1.223 | -0.247 | 0.623 | -0.001 | -0.025 | -0.025 | 1.06 | 0.13 | 0.07 | 0.13 | 0.2 | 0.1 | 0.07 |
| 1 | 80319975 | rs10874092 | -1.294 | -1.065 | 0.011 | 0.142 | -0.006 | 0.006 | 0.006 | 0 | 0.7 | 0.77 | 0.57 | 0.79 | 0.61 | 0.79 |
| 1 | 80322585 | rs10874093 | 1.713 | -1.064 | -0.364 | 0.152 | 0.011 | 0 | 0 | 0.298 | 0.22 | 0.27 | 0.23 | 0.23 | 0.29 | 0.23 |
| 1 | 80329280 | rs10874095 | -0.954 | -0.901 | -0.884 | 0.327 | -0.002 | 0.002 | 0.002 | 0 | 0.7 | 0.72 | 0.57 | 0.78 | 0.6 | 0.79 |
| 1 | 80331672 | rs7521332 | -1.126 | -0.834 | -1.058 | 0.169 | -0.002 | -0.01 | -0.01 | 0.091 | 0.71 | 0.77 | 0.58 | 0.94 | 0.6 | 0.8 |
| 1 | 80338715 | rs1370722 | -0.138 | -1.447 | -1.333 | 0.788 | 0.023 | -0.103 | -0.103 | 0.46 | 0.27 | 0.07 | 0.22 | 0.39 | 0.19 | 0.3 |
| 1 | 80341071 | rs4576621 | 0.694 | -1.311 | -1.219 | 0.5 | 0.049 | 0.017 | 0.017 | 0 | 0.59 | 0.27 | 0.66 | 0.4 | 0.69 | 0.53 |
| 1 | 80351989 | rs11162854 | 1.099 | -2.247 | -0.163 | 0.39 | -0.008 | 0 | 0 | 0.108 | 0.66 | 0.37 | 0.71 | 0.4 | 0.83 | 0.48 |
| 1 | 80369902 | rs912759 | 0.054 | -2.381 | 0.217 | 0.101 | 0.005 | 0.013 | 0.013 | 0 | 0.51 | 0.42 | 0.5 | 0.08 | 0.55 | 0.44 |
| 1 | 80376439 | rs17104038 | 0.89 | -2.57 | 0.548 | 0.203 | -0.062 | -0.059 | -0.059 | 0 | 0.36 | 0.14 | 0.39 | 0.48 | 0.42 | 0.29 |
| 1 | 80377618 | rs7547365 | 0.611 | -2.59 | 0.708 | 0.224 | -0.104 | 0.146 | 0.146 | 0 | 0.76 | 0.54 | 0.76 | 0.44 | 0.91 | 0.65 |
| 1 | 80386626 | rs1340675 | 0.537 | -2.526 | 0.81 | 0.164 | 0.001 | -0.026 | -0.026 | 0 | 0.09 | 0.15 | 0.04 | 0.03 | 0.07 | 0.19 |
| 1 | 80403167 | rs159665 | 0.018 | -2.546 | 0.884 | 0.203 | -0.042 | 0.066 | 0.066 | 0 | 0.89 | 0.69 | 0.95 | 0.79 | 0.97 | 0.85 |
| 1 | 80416562 | rs12085600 | 0.954 | -2.551 | 0.809 | 0.042 | 0.015 | 0.129 | 0.129 | 0 | 0.41 | 0.23 | 0.45 | 0.05 | 0.47 | 0.24 |
| 1 | 80416987 | rs7524563 | 1.153 | -2.955 | 0.672 | -0.075 | -0.087 | 0.041 | 0.041 | 0 | 0.68 | 0.45 | 0.72 | 0.41 | 0.87 | 0.6 |
| 1 | 80417950 | rs1325201 | 0.594 | -2.894 | 0.465 | -0.228 | -0.009 | 0.083 | 0.083 | 0 | 0.43 | 0.28 | 0.45 | 0.05 | 0.51 | 0.38 |
| 1 | 80439944 | rs17104166 | 0.741 | -2.837 | 0.283 | -0.201 | -0.001 | -0.018 | -0.018 | 0 | 0.09 | 0.12 | 0.04 | 0.03 | 0.07 | 0.19 |
| 1 | 80440298 | rs1325198 | 0.725 | -2.77 | -0.089 | -0.215 | 0.005 | 0.206 | 0.206 | 0 | 0.41 | 0.16 | 0.44 | 0.05 | 0.47 | 0.24 |
| 1 | 80441456 | rs997753 | 1.084 | -2.803 | -0.179 | -0.192 | 0.032 | 0.022 | 0.022 | 0.405 | 0.07 | 0.02 | 0.03 | 0.03 | 0.04 | 0.05 |
| 1 | 80442475 | rs10493634 | 2.178 | -2.667 | -0.131 | -0.115 | -0.083 | -0.068 | -0.068 | 0 | 0.24 | 0.1 | 0.26 | 0.36 | 0.35 | 0.22 |
| 1 | 80443109 | rs7416326 | 0.997 | -2.355 | -0.077 | -0.276 | -0.039 | 0.162 | 0.162 | 0 | 0.34 | 0.13 | 0.42 | 0.02 | 0.44 | 0.19 |
| 1 | 80450993 | rs17104197 | 1.185 | -2.757 | 0.5 | 0.352 | 0.003 | 0 | 0 | 0 | 0.11 | 0.06 | 0.14 | 0.12 | 0.07 | 0.09 |
| 1 | 80451812 | rs11162883 | 1.094 | -3.092 | 0.332 | -0.014 | 0.007 | 0.052 | 0.052 | 0 | 0.25 | 0.19 | 0.14 | 0.02 | 0.22 | 0.15 |
| 1 | 80452152 | rs10874105 | -1.569 | -3.387 | 0.569 | 0.147 | 0.01 | -0.022 | -0.022 | 0 | 0.6 | 0.79 | 0.44 | 0.63 | 0.42 | 0.69 |
| 1 | 80452633 | rs12038057 | 0.473 | -3.397 | 0.848 | 0.221 | -0.004 | 0.001 | 0.001 | 0 | 0.09 | 0.11 | 0.04 | 0.12 | 0.06 | 0.22 |
| 1 | 80454235 | rs1010470 | -0.836 | -3.408 | 1.299 | 0.307 | 0 | 0 | 0 | 0.148 | 0.5 | 0.51 | 0.37 | 0.54 | 0.37 | 0.63 |
| 1 | 80454564 | rs10493633 | -0.615 | -3.408 | 1.345 | 0.21 | -0.007 | -0.001 | -0.001 | 0 | 0.48 | 0.5 | 0.37 | 0.52 | 0.36 | 0.59 |
| 1 | 80455734 | rs17104213 | -1.442 | -3.337 | 1.106 | 0.207 | 0.003 | -0.006 | -0.006 | 0 | 0.36 | 0.6 | 0.3 | 0.61 | 0.2 | 0.54 |
| 1 | 80460168 | rs12085038 | 0.194 | -3.23 | 1.126 | 0.253 | -0.006 | -0.002 | -0.002 | 0 | 0.39 | 0.4 | 0.33 | 0.4 | 0.3 | 0.37 |
| 1 | 80463537 | rs1041705 | -1.72 | -3.189 | 1.02 | 0.281 | -0.007 | -0.024 | -0.024 | 0.002 | 0.63 | 0.8 | 0.44 | 0.65 | 0.43 | 0.72 |
| 1 | 80477880 | rs10493631 | -0.049 | -3.132 | 1.029 | 0.194 | -0.008 | -0.003 | -0.003 | 0 | 0.39 | 0.4 | 0.34 | 0.39 | 0.3 | 0.38 |
| 1 | 80484171 | rs4650447 | 1.754 | -3.049 | 0.995 | 0.328 | -0.038 | -0.037 | -0.037 | 0 | 0.36 | 0.12 | 0.54 | 0.36 | 0.56 | 0.31 |
| 1 | 80484625 | rs1340674 | -1.101 | -3.089 | 1.355 | 0.339 | 0.015 | 0.009 | 0.009 | 0 | 0.58 | 0.68 | 0.4 | 0.37 | 0.41 | 0.56 |
| 1 | 80498496 | rs6677996 | -0.739 | -3.177 | 1.509 | 0.305 | -0.058 | 0.019 | 0.019 | 0 | 0.21 | 0.34 | 0.11 | 0.25 | 0.11 | 0.3 |
| 1 | 80503523 | rs2184613 | -1.135 | -3.128 | 1.449 | 0.321 | 0.017 | 0.01 | 0.01 | 0 | 0.58 | 0.73 | 0.41 | 0.37 | 0.42 | 0.56 |
| 1 | 80508854 | rs2152391 | 1.754 | -3.019 | 1.406 | 0.345 | -0.026 | -0.028 | -0.028 | 3.054 | 0.36 | 0.14 | 0.54 | 0.37 | 0.56 | 0.31 |
| 1 | 80512853 | rs10493657 | 1.466 | -3.096 | 1.47 | 0.299 | -0.005 | 0 | 0 | 0 | 0.11 | 0.08 | 0.15 | 0.11 | 0.07 | 0.11 |
| 1 | 80514544 | rs6424682 | 1.664 | -3.089 | 1.506 | 0.293 | -0.011 | -0.012 | -0.012 | 0 | 0.37 | 0.24 | 0.55 | 0.37 | 0.56 | 0.31 |
| 1 | 80514706 | rs6699236 | -1.032 | -3.086 | 1.562 | 0.293 | 0.003 | 0.002 | 0.002 | 0 | 0.71 | 0.68 | 0.81 | 0.72 | 0.76 | 0.75 |
| 1 | 80517224 | rs6657785 | 1.741 | -3.014 | 1.397 | 0.17 | -0.012 | -0.007 | -0.007 | 1.711 | 0.34 | 0.27 | 0.51 | 0.38 | 0.49 | 0.3 |
| 1 | 80518186 | rs2388958 | 1.715 | -3.126 | 1.542 | 0.319 | -0.009 | -0.016 | -0.016 | 0 | 0.37 | 0.2 | 0.56 | 0.38 | 0.56 | 0.31 |
| 1 | 80519839 | rs10874110 | -1.186 | -3.206 | 1.471 | 0.298 | 0.009 | -0.001 | -0.001 | 0 | 0.73 | 0.82 | 0.83 | 0.72 | 0.76 | 0.75 |
| 1 | 80522918 | rs12038492 | -1.212 | -3.214 | 1.668 | 0.303 | -0.029 | 0.001 | 0.001 | 0 | 0.33 | 0.33 | 0.24 | 0.34 | 0.18 | 0.41 |
| 1 | 80523256 | rs1999726 | -1.202 | -3.205 | 1.707 | 0.307 | -0.029 | 0.001 | 0.001 | 0 | 0.33 | 0.33 | 0.24 | 0.34 | 0.18 | 0.41 |
| 1 | 80523448 | rs1999727 | -0.916 | -3.158 | 1.621 | 0.308 | -0.005 | -0.004 | -0.004 | 0 | 0.7 | 0.61 | 0.8 | 0.72 | 0.74 | 0.72 |
| 1 | 80524090 | rs12069558 | -0.832 | -2.995 | 1.355 | 0.363 | -0.019 | 0.001 | 0.001 | 0 | 0.21 | 0.23 | 0.1 | 0.23 | 0.12 | 0.3 |
| 1 | 80529386 | rs1029200 | -0.828 | -2.999 | 1.356 | 0.395 | -0.014 | 0.002 | 0.002 | 0 | 0.21 | 0.19 | 0.1 | 0.25 | 0.12 | 0.29 |
| 1 | 80534444 | rs12140474 | 1.566 | -2.798 | 1.273 | 0.373 | -0.008 | 0 | 0 | 0.632 | 0.36 | 0.26 | 0.55 | 0.35 | 0.56 | 0.3 |
| 1 | 80536798 | rs1572347 | 1.016 | -2.719 | 0.797 | 0.643 | 0.008 | -0.003 | -0.003 | 0 | 0.12 | 0.08 | 0.15 | 0.13 | 0.06 | 0.12 |
| 1 | 80542078 | rs6424687 | -0.118 | -2.825 | 0.61 | 0.665 | -0.036 | 0.051 | 0.051 | 0 | 0.83 | 0.78 | 0.91 | 0.62 | 0.92 | 0.81 |
| 1 | 80547435 | rs7545328 | 1.126 | -2.673 | 0.118 | 0.444 | 0.132 | 0.034 | 0.034 | 0 | 0.55 | 0.25 | 0.66 | 0.36 | 0.77 | 0.42 |

| | | | | | | | | | | | | | | | | |
|---|---|---|---|---|---|---|---|---|---|---|---|---|---|---|---|---|
| 1 | 80563300 | rs4618922 | 0.252 | -2.127 | 0.728 | 0.711 | -0.001 | 0.146 | 0.146 | 0 | 0.94 | 0.81 | 0.96 | 0.84 | 0.97 | 0.9 |
| 1 | 80563570 | rs7548641 | 1.194 | -2.22 | 0.389 | 0.321 | -0.008 | 0.069 | 0.069 | 0 | 0.2 | 0.09 | 0.11 | 0.01 | 0.2 | 0.11 |
| 1 | 80567175 | rs4454493 | 0.325 | -2.356 | 0.543 | 0.289 | 0.006 | 0.065 | 0.065 | 0 | 0.27 | 0.18 | 0.14 | 0.04 | 0.25 | 0.27 |
| 1 | 80581276 | rs6424691 | -0.163 | -2.347 | 0.566 | 0.363 | -0.012 | 0.059 | 0.059 | 0 | 0.95 | 0.9 | 0.96 | 0.84 | 0.97 | 0.9 |
| 1 | 80584740 | rs11162927 | 0.43 | -2.64 | 1.006 | 0.446 | -0.037 | 0.074 | 0.074 | 0 | 0.15 | 0.28 | 0.11 | 0.31 | 0.08 | 0.18 |
| 1 | 80586000 | rs4418540 | 0.905 | -2.619 | 0.97 | 0.427 | 0.079 | -0.012 | -0.012 | 0 | 0.51 | 0.12 | 0.68 | 0.47 | 0.62 | 0.44 |
| 1 | 80586993 | rs4384179 | 0.869 | -2.615 | 1.093 | 0.429 | 0.073 | 0.005 | 0.005 | 0 | 0.51 | 0.2 | 0.69 | 0.47 | 0.62 | 0.44 |
| 1 | 80587807 | rs4650452 | -0.072 | -1.961 | 1.131 | 0.552 | 0.014 | 0.194 | 0.194 | 0 | 0.95 | 0.79 | 0.95 | 0.83 | 0.97 | 0.91 |
| 1 | 80589739 | rs12407709 | 0.205 | -1.889 | 0.56 | 0.494 | 0.003 | -0.005 | -0.005 | 0 | 0.28 | 0.24 | 0.25 | 0.43 | 0.16 | 0.34 |
| 1 | 80613801 | rs12407162 | 0.79 | -1.396 | 0.363 | 0.52 | -0.098 | -0.037 | -0.037 | 0 | 0.38 | 0.06 | 0.53 | 0.36 | 0.55 | 0.25 |
| 1 | 80631231 | rs4330881 | 0.779 | -1.425 | 0.366 | 0.524 | -0.093 | -0.037 | -0.037 | 0 | 0.38 | 0.06 | 0.53 | 0.36 | 0.55 | 0.25 |
| 1 | 80638947 | rs10874118 | -0.365 | -1.48 | 0.812 | 0.564 | 0.02 | -0.012 | -0.012 | 0 | 0.56 | 0.7 | 0.41 | 0.48 | 0.4 | 0.64 |
| 1 | 80641525 | rs4503307 | 0.494 | -1.485 | 1.054 | 0.818 | 0.016 | 0.143 | 0.143 | 0 | 0.94 | 0.76 | 0.95 | 0.84 | 0.96 | 0.89 |
| 1 | 80644813 | rs7521092 | 0.023 | -1.625 | 1.052 | 0.707 | 0.021 | -0.017 | -0.017 | 0 | 0.33 | 0.28 | 0.28 | 0.48 | 0.2 | 0.49 |
| 1 | 80656114 | rs4310401 | -0.112 | -0.284 | 0.783 | 0.522 | -0.007 | 0.008 | 0.008 | 0.013 | 0.31 | 0.3 | 0.26 | 0.16 | 0.42 | 0.26 |
| 1 | 80659055 | rs10493661 | -0.028 | -0.322 | 1.051 | 0.529 | 0.017 | -0.019 | -0.019 | 0 | 0.67 | 0.53 | 0.73 | 0.82 | 0.57 | 0.71 |
| 1 | 80659876 | rs4375210 | 1.116 | -0.26 | 0.623 | 0.609 | -0.046 | -0.043 | -0.043 | 0.454 | 0.29 | 0.09 | 0.52 | 0.4 | 0.34 | 0.21 |
| 1 | 80662898 | rs10782722 | 0.079 | -0.771 | 0.73 | 0.699 | -0.001 | 0 | 0 | 1.246 | 0.38 | 0.36 | 0.2 | 0.43 | 0.22 | 0.5 |
| 1 | 80675529 | rs3001446 | -0.602 | -0.018 | -0.257 | 0.743 | -0.051 | 0.009 | 0.009 | 0 | 0.67 | 0.83 | 0.37 | 0.69 | 0.5 | 0.73 |
| 1 | 80683370 | rs1343931 | 0.096 | -0.702 | 0.186 | 1.078 | -0.017 | 0.014 | 0.014 | 0 | 0.68 | 0.54 | 0.77 | 0.67 | 0.68 | 0.78 |
| 1 | 80683606 | rs952603 | 0.479 | -0.791 | 0.159 | 1.11 | 0.009 | 0.005 | 0.005 | 0 | 0.35 | 0.32 | 0.2 | 0.32 | 0.24 | 0.5 |
| 1 | 80685710 | rs1687846 | -0.245 | -0.794 | 0.286 | 1.104 | -0.011 | 0.014 | 0.014 | 0 | 0.69 | 0.6 | 0.77 | 0.67 | 0.69 | 0.82 |
| 1 | 80689338 | rs2797654 | -0.451 | -0.835 | 0.319 | 1.192 | -0.008 | -0.036 | -0.036 | 0 | 0.72 | 0.65 | 0.79 | 0.9 | 0.71 | 0.84 |
| 1 | 80694377 | rs2211432 | -0.24 | -0.835 | 0.344 | 1.233 | -0.019 | -0.033 | -0.033 | 0 | 0.7 | 0.57 | 0.79 | 0.85 | 0.7 | 0.77 |
| 1 | 80700025 | rs1687833 | -0.482 | -0.823 | 0.286 | 1.239 | -0.008 | -0.015 | -0.015 | 0 | 0.72 | 0.65 | 0.79 | 0.85 | 0.71 | 0.82 |
| 1 | 80702191 | rs1776322 | 0.919 | -0.811 | -0.163 | 1.244 | -0.057 | 0.015 | 0.015 | 0 | 0.33 | 0.16 | 0.56 | 0.3 | 0.45 | 0.25 |
| 1 | 80703306 | rs1776325 | 0.034 | -0.804 | -0.144 | 1.309 | -0.022 | 0.038 | 0.038 | 0 | 0.67 | 0.48 | 0.76 | 0.61 | 0.69 | 0.79 |
| 1 | 80717516 | rs11162962 | 0.502 | -0.801 | -0.136 | 1.314 | 0.025 | 0.017 | 0.017 | 0 | 0.34 | 0.24 | 0.19 | 0.31 | 0.24 | 0.5 |
| 1 | 80722041 | rs1539486 | 0.54 | -0.245 | -0.381 | 1.227 | -0.037 | 0.011 | 0.011 | 0 | 0.33 | 0.2 | 0.57 | 0.31 | 0.45 | 0.29 |
| 1 | 80735096 | rs7529438 | -0.626 | -0.731 | -0.431 | 1.717 | -0.002 | 0.075 | 0.075 | 0 | 0.07 | 0.1 | 0.04 | 0.35 | 0.04 | 0.07 |
| 1 | 80737948 | rs1539484 | 0.506 | -0.313 | -0.589 | 1.607 | -0.005 | -0.008 | -0.008 | 0 | 0.07 | 0.07 | 0.22 | 0.1 | 0.09 | 0.06 |
| 1 | 80739543 | rs1012455 | 0.755 | -0.495 | -0.408 | 1.458 | -0.032 | 0.017 | 0.017 | 0 | 0.32 | 0.24 | 0.38 | 0.28 | 0.44 | 0.29 |
| 1 | 80741675 | rs1954338 | -0.633 | -1.035 | -0.109 | 1.315 | -0.03 | -0.03 | -0.03 | 0 | 0.65 | 0.56 | 0.72 | 0.74 | 0.71 | 0.53 |
| 1 | 80741898 | rs1776286 | 1.175 | -0.996 | -0.239 | 1.345 | -0.031 | 0.018 | 0.018 | 0 | 0.32 | 0.22 | 0.38 | 0.24 | 0.44 | 0.29 |
| 1 | 80744080 | rs12144659 | 0.433 | -1.017 | 0.148 | 1.316 | 0.018 | 0.019 | 0.019 | 0 | 0.34 | 0.27 | 0.27 | 0.26 | 0.29 | 0.47 |
| 1 | 80749361 | rs1776290 | -0.213 | -0.14 | 0.268 | 1.536 | 0.003 | 0.016 | 0.016 | 0 | 0.09 | 0.19 | 0.23 | 0.1 | 0.09 | 0.11 |
| 1 | 80758728 | rs1416467 | -0.058 | -0.148 | 0.208 | 1.551 | -0.006 | 0.008 | 0.008 | 0 | 0.38 | 0.28 | 0.59 | 0.34 | 0.54 | 0.35 |
| 1 | 80760473 | rs7412523 | 0.376 | -0.223 | 0.143 | 1.444 | 0.004 | 0.008 | 0.008 | 0 | 0.07 | 0.11 | 0.15 | 0.12 | 0.14 | 0.18 |
| 1 | 80761197 | rs11588728 | 0.669 | -0.179 | 0.079 | 1.462 | -0.006 | -0.01 | -0.01 | 0 | 0.07 | 0.08 | 0.22 | 0.1 | 0.09 | 0.07 |
| 1 | 80768813 | rs12033709 | 0.404 | -0.106 | -0.609 | 1.17 | 0.006 | 0.009 | 0.009 | 0 | 0.07 | 0.12 | 0.15 | 0.11 | 0.13 | 0.18 |
| 1 | 80770605 | rs1937791 | -0.597 | -0.087 | -0.779 | 1.147 | -0.006 | -0.005 | -0.005 | 0 | 0.44 | 0.53 | 0.64 | 0.61 | 0.57 | 0.41 |
| 1 | 80781284 | rs1781761 | 0.141 | 0.408 | -1.667 | 0.626 | -0.189 | -0.033 | -0.033 | 0 | 0.31 | 0.03 | 0.37 | 0.29 | 0.46 | 0.28 |
| 1 | 80782268 | rs11576485 | -0.814 | -0.371 | -1.099 | 0.356 | 0.012 | 0.011 | 0.011 | 0 | 0.13 | 0.1 | 0.03 | 0.1 | 0.07 | 0.15 |
| 1 | 80782665 | rs2389012 | -0.778 | -0.827 | -0.564 | -0.352 | -0.02 | -0.096 | -0.096 | 0 | 0.29 | 0.54 | 0.18 | 0.06 | 0.25 | 0.29 |
| 1 | 80782806 | rs2389011 | -0.13 | -0.929 | 0.004 | -0.456 | -0.006 | -0.101 | -0.101 | 0 | 0.16 | 0.38 | 0.08 | 0 | 0.12 | 0.11 |
| 1 | 80788599 | rs1781776 | 0.936 | -0.503 | -0.052 | -0.008 | -0.025 | -0.027 | -0.027 | 0 | 0.29 | 0.21 | 0.55 | 0.45 | 0.4 | 0.35 |
| 1 | 80791787 | rs2154298 | 0.134 | -0.595 | 0.244 | 0.171 | 0.032 | 0.046 | 0.046 | 0.724 | 0.44 | 0.28 | 0.2 | 0.25 | 0.29 | 0.38 |
| 1 | 80793118 | rs12065553 | 0.04 | -0.598 | 0.463 | 0.239 | 0.02 | 0.023 | 0.023 | 0.218 | 0.45 | 0.35 | 0.21 | 0.28 | 0.3 | 0.37 |
| 1 | 80805845 | rs2389024 | 0.156 | -0.642 | 0.572 | 0.35 | 0.014 | 0.02 | 0.02 | 0.63 | 0.44 | 0.38 | 0.21 | 0.1 | 0.3 | 0.35 |
| 1 | 80809034 | rs10493662 | 0.222 | -0.624 | 0.51 | 0.351 | 0.03 | 0.09 | 0.09 | 1.045 | 0.43 | 0.27 | 0.2 | 0.1 | 0.3 | 0.35 |
| 1 | 80816962 | rs17436212 | 0.984 | -0.595 | 0.563 | 0.343 | -0.004 | -0.008 | -0.008 | 0 | 0.1 | 0.11 | 0.09 | 0.07 | 0.12 | 0.17 |
| 1 | 80818530 | rs12118987 | 0.15 | -0.615 | 0.575 | 0.381 | -0.005 | -0.005 | -0.005 | 2.4 | 0.46 | 0.48 | 0.22 | 0.28 | 0.31 | 0.39 |
| 1 | 80821220 | rs11162975 | -0.14 | -0.535 | 0.671 | 0.493 | 0 | 0 | 0 | 0 | 0.49 | 0.52 | 0.25 | 0.35 | 0.36 | 0.44 |
| 1 | 80823037 | rs3014987 | -0.122 | -0.55 | 0.6 | 0.449 | -0.019 | 0.018 | 0.018 | 0.34 | 0.84 | 0.76 | 0.85 | 0.82 | 0.83 | 0.81 |
| 1 | 80824657 | rs12240077 | 1.229 | -0.797 | 0.518 | 0.264 | -0.015 | 0.086 | 0.086 | 0.492 | 0.26 | 0.06 | 0.12 | 0 | 0.24 | 0.14 |
| 1 | 80825083 | rs10158964 | 0.729 | 0.189 | 1.269 | 1.09 | 0.014 | 0.014 | 0.014 | 0.167 | 0.46 | 0.39 | 0.23 | 0.3 | 0.32 | 0.42 |
| 1 | 80835085 | rs1687812 | -0.651 | 0.146 | 1.251 | 1.165 | 0.009 | 0.114 | 0.114 | 0.913 | 0.94 | 0.81 | 0.93 | 0.89 | 0.95 | 0.99 |
| 1 | 80838880 | rs2997195 | 1.01 | 0.199 | 0.453 | 1.044 | -0.03 | -0.022 | -0.022 | 0.134 | 0.28 | 0.21 | 0.53 | 0.43 | 0.39 | 0.33 |
| 1 | 80849656 | rs6696084 | -0.211 | 0.529 | -0.099 | 0.535 | -0.002 | -0.009 | -0.009 | 0.321 | 0.54 | 0.5 | 0.28 | 0.32 | 0.41 | 0.44 |
| 1 | 80867324 | rs10518645 | -0.023 | 1.01 | -0.46 | 1.066 | -0.018 | 0.022 | 0.022 | 0 | 0.07 | 0.01 | 0.04 | 0.02 | 0.08 | 0.02 |
| 1 | 80876340 | rs17390593 | -1.38 | 0.731 | -0.051 | 1.577 | 0.011 | 0.1 | 0.1 | 0 | 0.16 | 0.33 | 0.16 | 0.32 | 0.2 | 0.13 |
| 1 | 80877080 | rs3909721 | 0.939 | 0.728 | -0.037 | 1.442 | 0.024 | 0.038 | 0.038 | 0 | 0.88 | 0.71 | 0.88 | 0.87 | 0.83 | 0.88 |
| 1 | 80879897 | rs10518644 | -1.598 | 0.364 | -0.175 | 1.413 | 0.001 | -0.007 | -0.007 | 0 | 0.18 | 0.15 | 0.13 | 0.25 | 0.17 | 0.14 |
| 1 | 80881727 | rs4650458 | 0.365 | 0.475 | 0.154 | 1.15 | 0.033 | 0.049 | 0.049 | 0 | 0.28 | 0.39 | 0.6 | 0.52 | 0.46 | 0.36 |
| 1 | 80892408 | rs1427239 | 1.405 | -0.059 | 1.062 | 1.388 | -0.03 | 0.052 | 0.052 | 5.08 | 0.83 | 0.27 | 0.83 | 0.67 | 0.89 | 0.85 |
| 1 | 80912432 | rs4607823 | 1.41 | -0.031 | 1.202 | 1.38 | -0.026 | 0.053 | 0.053 | 5.453 | 0.83 | 0.27 | 0.83 | 0.71 | 0.88 | 0.84 |
| 1 | 80929598 | rs1365750 | 1.617 | -0.505 | 1.136 | 0.95 | -0.014 | 0.007 | 0.007 | 2.079 | 0.73 | 0.31 | 0.8 | 0.7 | 0.81 | 0.78 |
| 1 | 80932212 | rs12565086 | -0.342 | -1.057 | 0.384 | 0.119 | -0.033 | 0.111 | 0.111 | 1.739 | 0.32 | 0.1 | 0.19 | 0.12 | 0.3 | 0.13 |
| 1 | 80933266 | rs12087192 | -1.118 | -1.09 | 1.232 | 0.441 | 0.022 | 0.037 | 0.037 | 0.001 | 0.07 | 0.32 | 0.08 | 0.05 | 0.04 | 0.1 |

| | | | | | | | | | | | | | | | | |
|---|---|---|---|---|---|---|---|---|---|---|---|---|---|---|---|---|
| 1 | 80944604 | rs1155549 | 1.552 | -1.099 | 1.075 | 0.478 | 0.011 | 0.059 | 0.059 | 1.96 | 0.9 | 0.5 | 0.89 | 0.93 | 0.95 | 0.89 |
| 1 | 80953659 | rs4650467 | 1.357 | -1.049 | 1.171 | 0.508 | -0.016 | 0.12 | 0.12 | 0.071 | 0.86 | 0.53 | 0.86 | 0.83 | 0.91 | 0.86 |
| 1 | 80954735 | rs10493669 | 1.383 | -1.071 | 1.151 | 0.485 | -0.019 | 0.104 | 0.104 | 0 | 0.86 | 0.6 | 0.87 | 0.83 | 0.91 | 0.86 |
| 1 | 80955429 | rs10493668 | 1.63 | -1.307 | 1.207 | 0.672 | -0.036 | 0.038 | 0.038 | 1.698 | 0.73 | 0.4 | 0.74 | 0.59 | 0.82 | 0.76 |
| 1 | 80956319 | rs718003 | 1.755 | -1.439 | 1.081 | 0.653 | 0.027 | -0.019 | -0.019 | 0 | 0.67 | 0.23 | 0.71 | 0.45 | 0.8 | 0.69 |
| 1 | 80964490 | rs10493666 | -1.067 | -1.24 | 1.514 | 0.62 | 0.022 | 0.032 | 0.032 | 0.022 | 0.07 | 0.26 | 0.06 | 0.04 | 0.03 | 0.09 |
| 1 | 80970978 | rs184812 | 0.475 | -1.685 | 1.084 | 0.059 | 0.016 | 0.043 | 0.043 | 0 | 0.6 | 0.67 | 0.77 | 0.77 | 0.68 | 0.74 |
| 1 | 80977160 | rs1593679 | -0.042 | -1.851 | 1.049 | 0.022 | 0.005 | -0.006 | -0.006 | 0 | 0.16 | 0.2 | 0.05 | 0.24 | 0.09 | 0.17 |
| 1 | 80977963 | rs356287 | -1.838 | -1.963 | 1.21 | 0.091 | 0.004 | 0.011 | 0.011 | 1.252 | 0.56 | 0.78 | 0.42 | 0.56 | 0.4 | 0.39 |
| 1 | 80982632 | rs242294 | -0.934 | -1.837 | 1.312 | 0.207 | -0.011 | 0.002 | 0.002 | 0.315 | 0.22 | 0.3 | 0.13 | 0.24 | 0.15 | 0.22 |
| 1 | 80985935 | rs17104916 | -0.014 | -2.014 | 1.623 | 0.289 | -0.019 | 0.215 | 0.215 | 0.17 | 0.1 | 0.31 | 0.16 | 0.29 | 0.05 | 0.06 |
| 1 | 80989370 | rs12031162 | 0.099 | -2.013 | 1.734 | 0.288 | -0.021 | 0.165 | 0.165 | 0.012 | 0.1 | 0.23 | 0.15 | 0.29 | 0.05 | 0.05 |
| 1 | 80993368 | rs356295 | -2.222 | -2.069 | 1.737 | 0.248 | -0.009 | 0.001 | 0.001 | 2.107 | 0.35 | 0.65 | 0.27 | 0.56 | 0.2 | 0.34 |
| 1 | 80996970 | rs356298 | -1.969 | -2.233 | 0.987 | -0.13 | -0.088 | -0.032 | -0.032 | 0 | 0.66 | 0.9 | 0.43 | 0.6 | 0.46 | 0.59 |
| 1 | 80997745 | rs12070796 | -1.093 | -1.911 | 1.299 | 0.495 | -0.025 | 0.008 | 0.008 | 0.016 | 0.22 | 0.37 | 0.09 | 0.24 | 0.12 | 0.24 |
| 1 | 81004543 | rs11163045 | 0.635 | -1.703 | 0.88 | 0.731 | -0.011 | 0.061 | 0.061 | 0 | 0.08 | 0.11 | 0.11 | 0.24 | 0.05 | 0.04 |
| 1 | 81009476 | rs11163048 | -0.578 | -1.837 | 0.758 | 0.676 | 0.001 | -0.003 | -0.003 | 0 | 0.18 | 0.23 | 0.06 | 0.27 | 0.09 | 0.18 |
| 1 | 81024433 | rs12060807 | -0.56 | -1.83 | 0.633 | 0.663 | -0.017 | 0.012 | 0.012 | 0 | 0.18 | 0.25 | 0.06 | 0.26 | 0.09 | 0.18 |
| 1 | 81027980 | rs2389057 | -1.119 | -1.774 | 0.89 | 0.891 | -0.063 | 0.128 | 0.128 | 0.513 | 0.22 | 0.45 | 0.18 | 0.46 | 0.11 | 0.16 |
| 1 | 81028590 | rs10874140 | -0.164 | -1.779 | 0.724 | 0.929 | -0.028 | 0.035 | 0.035 | 0 | 0.13 | 0.22 | 0.05 | 0.22 | 0.07 | 0.13 |
| 1 | 81029179 | rs17105040 | -0.324 | -1.832 | 0.796 | 0.907 | -0.012 | 0.015 | 0.015 | 0.025 | 0.09 | 0.21 | 0.08 | 0.07 | 0.06 | 0.1 |
| 1 | 81038969 | rs11163055 | -1.957 | -1.86 | 0.815 | 0.905 | -0.006 | -0.002 | -0.002 | 0 | 0.31 | 0.72 | 0.27 | 0.53 | 0.17 | 0.26 |
| 1 | 81042755 | rs10874143 | -0.124 | -1.815 | 0.748 | 0.917 | -0.03 | 0.037 | 0.037 | 0.001 | 0.13 | 0.22 | 0.05 | 0.22 | 0.07 | 0.13 |
| 1 | 81048650 | rs17399290 | 1.18 | -1.506 | -0.102 | 0.339 | -0.065 | 0.05 | 0.05 | 5.842 | 0.16 | 0.01 | 0.23 | 0.06 | 0.17 | 0.06 |
| 1 | 81052058 | rs10874144 | -0.814 | -1.774 | 0.57 | 0.914 | -0.077 | 0.134 | 0.134 | 0.869 | 0.23 | 0.48 | 0.18 | 0.45 | 0.11 | 0.16 |
| 1 | 81056073 | rs17105114 | -0.958 | -1.092 | 0.068 | 0.805 | -0.013 | 0.02 | 0.02 | 0.049 | 0.09 | 0.21 | 0.08 | 0.09 | 0.06 | 0.1 |
| 1 | 81059253 | rs10874145 | -1.355 | -1.658 | -0.377 | 0.861 | 0.039 | -0.02 | -0.02 | 0 | 0.33 | 0.8 | 0.27 | 0.55 | 0.18 | 0.26 |
| 1 | 81063642 | rs1853332 | -1.134 | -2.17 | 0.043 | 0.856 | -0.034 | 0.004 | 0.004 | 0.03 | 0.31 | 0.58 | 0.17 | 0.34 | 0.2 | 0.37 |
| 1 | 81070850 | rs1279686 | -1.147 | -2.073 | 0.561 | 0.989 | -0.034 | -0.014 | -0.014 | 0 | 0.29 | 0.51 | 0.16 | 0.27 | 0.21 | 0.36 |
| 1 | 81071433 | rs1279688 | -1.644 | -2.31 | 0.385 | 0.766 | 0.055 | 0.012 | 0.012 | 0 | 0.39 | 0.74 | 0.21 | 0.38 | 0.24 | 0.38 |
| 1 | 81071933 | rs6666532 | -0.345 | -2.375 | 0.5 | 0.668 | -0.04 | 0.027 | 0.027 | 0.252 | 0.18 | 0.34 | 0.1 | 0.24 | 0.11 | 0.25 |
| 1 | 81085999 | rs6658764 | 1.662 | -1.448 | -0.667 | 0.7 | -0.115 | -0.081 | -0.081 | 2.512 | 0.25 | 0.02 | 0.31 | 0.31 | 0.42 | 0.3 |
| 1 | 81087175 | rs7552365 | -1.027 | -1.869 | 0.057 | 0.777 | 0.013 | 0.007 | 0.007 | 0 | 0.36 | 0.62 | 0.2 | 0.32 | 0.22 | 0.34 |
| 1 | 81090382 | rs2800878 | -0.839 | -1.882 | 0.466 | 0.775 | -0.033 | -0.016 | -0.016 | 0 | 0.33 | 0.45 | 0.17 | 0.29 | 0.22 | 0.31 |
| 1 | 81113456 | rs11163072 | -0.801 | -1.816 | 0.92 | 1.305 | -0.037 | 0.104 | 0.104 | 0.206 | 0.23 | 0.5 | 0.19 | 0.39 | 0.13 | 0.22 |
| 1 | 81114489 | rs10493673 | 0.582 | -2.663 | 0.869 | 1.049 | -0.006 | 0 | 0 | 10.604 | 0.18 | 0.16 | 0.08 | 0.26 | 0.11 | 0.19 |
| 1 | 81116640 | rs17105199 | 0.589 | -2.659 | 0.958 | 0.792 | -0.002 | -0.062 | -0.062 | 0 | 0.14 | 0.22 | 0.08 | 0.04 | 0.1 | 0.16 |
| 1 | 81120474 | rs17105223 | -1.097 | -2.702 | 0.813 | 1.094 | -0.016 | 0.007 | 0.007 | 0.133 | 0.31 | 0.62 | 0.22 | 0.54 | 0.17 | 0.27 |
| 1 | 81130054 | rs1279690 | -1.785 | -3.352 | 0.477 | 0.84 | 0.106 | 0.018 | 0.018 | 0.004 | 0.53 | 0.86 | 0.33 | 0.59 | 0.3 | 0.43 |
| 1 | 81133616 | rs11163073 | 0.043 | -3.195 | 0.923 | 0.986 | -0.009 | 0.2 | 0.2 | 0 | 0.11 | 0.27 | 0.11 | 0.4 | 0.05 | 0.1 |
| 1 | 81137243 | rs17105288 | 0.314 | -3.295 | 1.076 | 0.88 | -0.059 | -0.021 | -0.021 | 0 | 0.21 | 0.32 | 0.1 | 0.15 | 0.12 | 0.18 |
| 1 | 81137567 | rs12035797 | 0.076 | -3.239 | 1.051 | 0.925 | -0.008 | 0.189 | 0.189 | 0 | 0.11 | 0.27 | 0.11 | 0.39 | 0.05 | 0.1 |
| 1 | 81140382 | rs6688321 | 2.107 | -3.202 | 0.959 | 0.883 | 0.003 | 0.026 | 0.026 | 1.806 | 0.06 | 0.03 | 0.06 | 0.25 | 0.02 | 0.06 |
| 1 | 81143088 | rs17105337 | -0.911 | -3.259 | 1.001 | 0.845 | -0.03 | 0.019 | 0.019 | 0.165 | 0.3 | 0.57 | 0.18 | 0.41 | 0.17 | 0.27 |
| 1 | 81143121 | rs10493675 | -0.825 | -3.252 | 1.075 | 0.849 | -0.063 | 0.062 | 0.062 | 0 | 0.29 | 0.44 | 0.17 | 0.41 | 0.17 | 0.26 |
| 1 | 81145258 | rs1936934 | -0.559 | -3.225 | 1.089 | 0.86 | -0.04 | 0.045 | 0.045 | 0 | 0.27 | 0.38 | 0.16 | 0.41 | 0.17 | 0.26 |
| 1 | 81154232 | rs12023504 | 0.647 | -2.747 | 1.062 | 0.402 | -0.001 | 0.083 | 0.083 | 0 | 0.08 | 0.13 | 0.07 | 0.26 | 0.05 | 0.07 |
| 1 | 81155350 | rs17105403 | 0.443 | -2.752 | 0.932 | -0.409 | -0.009 | -0.045 | -0.045 | 0 | 0.2 | 0.21 | 0.11 | 0.03 | 0.12 | 0.15 |
| 1 | 81156849 | rs841671 | -0.337 | -1.765 | -0.01 | -0.438 | 0.006 | -0.03 | -0.03 | 0 | 0.75 | 0.62 | 0.81 | 0.84 | 0.78 | 0.75 |
| 1 | 81159163 | rs841668 | -0.812 | -1.59 | -0.608 | -0.914 | -0.002 | -0.016 | -0.016 | 0 | 0.81 | 0.83 | 0.87 | 0.98 | 0.82 | 0.83 |
| 1 | 81160673 | rs11163087 | -0.235 | -1.956 | -0.807 | -1.052 | 0.036 | 0.024 | 0.024 | 0 | 0.48 | 0.32 | 0.32 | 0.61 | 0.39 | 0.3 |
| 1 | 81161337 | rs1832331 | 0.524 | -2.776 | -0.603 | -1.506 | 0.017 | 0.002 | 0.002 | 0 | 0.27 | 0.21 | 0.11 | 0.04 | 0.1 | 0.28 |
| 1 | 81163114 | rs11163088 | 1.26 | -3.171 | -0.884 | -1.677 | 0.01 | -0.028 | -0.028 | 0 | 0.21 | 0.18 | 0.15 | 0.57 | 0.14 | 0.19 |
| 1 | 81164242 | rs1337642 | -0.491 | -3.171 | -0.548 | -1.794 | 0 | -0.027 | -0.027 | 0 | 0.45 | 0.51 | 0.22 | 0.09 | 0.21 | 0.37 |
| 1 | 81176850 | rs1591678 | 1.912 | -3.088 | -0.408 | -1.729 | -0.029 | -0.038 | -0.038 | 0 | 0.15 | 0.17 | 0.08 | 0.03 | 0.09 | 0.07 |
| 1 | 81177390 | rs10874154 | -0.111 | -3.067 | -0.448 | -1.707 | 0.02 | 0.012 | 0.012 | 0 | 0.45 | 0.58 | 0.35 | 0.64 | 0.33 | 0.29 |
| 1 | 81177525 | rs841666 | -1.319 | -2.755 | -0.439 | -1.569 | -0.037 | -0.019 | -0.019 | 0 | 0.72 | 0.79 | 0.43 | 0.68 | 0.44 | 0.57 |
| 1 | 81181741 | rs11163090 | -1.295 | -2.652 | -0.486 | -1.603 | 0.004 | 0.001 | 0.001 | 0.76 | 0.72 | 0.72 | 0.43 | 0.66 | 0.45 | 0.55 |
| 1 | 81188002 | rs2152856 | 0.28 | -2.935 | -0.693 | -1.791 | 0.071 | 0.072 | 0.072 | 0 | 0.33 | 0.15 | 0.09 | 0.05 | 0.11 | 0.31 |
| 1 | 81192795 | rs17105462 | 1.98 | -2.529 | -0.696 | -1.525 | 0.011 | 0.006 | 0.006 | 0 | 0.13 | 0.1 | 0.08 | 0.03 | 0.1 | 0.06 |
| 1 | 81195339 | rs1337638 | -0.063 | -2.58 | -0.504 | -1.501 | -0.002 | -0.001 | -0.001 | 0 | 0.4 | 0.45 | 0.34 | 0.61 | 0.35 | 0.29 |
| 1 | 81202219 | rs12566647 | 0.854 | -2.326 | -0.594 | -1.479 | 0.002 | -0.039 | -0.039 | 0 | 0.25 | 0.17 | 0.23 | 0.57 | 0.24 | 0.21 |
| 1 | 81202427 | rs4414082 | 0.078 | -2.573 | -0.863 | -1.646 | 0.057 | 0.072 | 0.072 | 0 | 0.35 | 0.2 | 0.14 | 0.08 | 0.19 | 0.36 |
| 1 | 81203878 | rs7512682 | 0.219 | -1.876 | -0.7 | -1.145 | -0.007 | 0.035 | 0.035 | 0 | 0.29 | 0.42 | 0.27 | 0.58 | 0.25 | 0.23 |
| 1 | 81207701 | rs1765454 | -0.418 | -1.075 | -0.934 | -0.756 | -0.005 | -0.017 | -0.017 | 0 | 0.22 | 0.19 | 0.07 | 0.05 | 0.08 | 0.3 |
| 1 | 81209339 | rs1539362 | 0.526 | -0.058 | -0.983 | -0.372 | 0 | 0.01 | 0.01 | 5.202 | 0.62 | 0.75 | 0.47 | 0.65 | 0.41 | 0.32 |
| 1 | 81209906 | rs2892671 | -0.713 | 0.14 | -0.905 | -0.921 | -0.001 | -0.023 | -0.023 | 0.299 | 0.24 | 0.27 | 0.12 | 0.05 | 0.24 | 0.13 |
| 1 | 81212467 | rs6692988 | -1.499 | -0.148 | -1.108 | -0.356 | 0.086 | 0.059 | 0.059 | 2.215 | 0.58 | 0.75 | 0.79 | 0.76 | 0.76 | 0.81 |
| 1 | 81212945 | rs1577879 | 0.719 | 1.511 | -0.883 | 0.274 | 0.016 | 0.035 | 0.035 | 0.052 | 0.1 | 0.11 | 0.27 | 0.28 | 0.19 | 0.2 |

| | | | | | | | | | | | | | | | | |
|---|---|---|---|---|---|---|---|---|---|---|---|---|---|---|---|---|
| 1 | 81213348 | rs7541278 | -0.152 | 1.761 | -0.996 | 0.238 | -0.023 | -0.022 | -0.022 | 0 | 0.16 | 0.12 | 0.39 | 0.29 | 0.31 | 0.27 |
| 1 | 81222277 | rs6660382 | -0.518 | 1.607 | -0.488 | 0.2 | 0.068 | 0.055 | 0.055 | 0 | 0.18 | 0.33 | 0.34 | 0.28 | 0.32 | 0.23 |
| 1 | 81223511 | rs7553881 | -1.87 | 1.678 | -0.651 | 0.23 | 0.013 | 0.001 | 0.001 | 0.924 | 0.46 | 0.7 | 0.56 | 0.44 | 0.56 | 0.68 |
| 1 | 81227145 | rs11163106 | -1.17 | 2.49 | -0.3 | 0.886 | -0.062 | -0.038 | -0.038 | 0 | 0.22 | 0.13 | 0.43 | 0.29 | 0.44 | 0.3 |
| 1 | 81228366 | rs12129785 | -0.62 | 2.366 | -0.13 | 1.024 | -0.011 | -0.012 | -0.012 | 0 | 0.16 | 0.14 | 0.31 | 0.28 | 0.32 | 0.23 |
| 1 | 81230013 | rs12758138 | -0.603 | 2.331 | -0.159 | 0.954 | -0.016 | -0.016 | -0.016 | 0 | 0.16 | 0.15 | 0.31 | 0.29 | 0.32 | 0.23 |
| 1 | 81232434 | rs12120655 | 0.107 | 1.84 | 0.073 | 0.879 | 0.001 | 0.035 | 0.035 | 0 | 0.18 | 0.22 | 0.21 | 0.39 | 0.17 | 0.26 |
| 1 | 81272301 | rs10782741 | -0.704 | 1.668 | 0.2 | 0.621 | 0.019 | 0 | 0 | 0 | 0.36 | 0.48 | 0.61 | 0.67 | 0.49 | 0.51 |
| 1 | 81282234 | rs10874175 | -1.048 | 1.519 | 0.998 | 0.414 | -0.026 | -0.064 | -0.064 | 0 | 0.3 | 0.59 | 0.21 | 0.13 | 0.27 | 0.31 |
| 1 | 81287825 | rs1338138 | -1.184 | 1.53 | 0.876 | 0.415 | -0.02 | -0.055 | -0.055 | 0 | 0.3 | 0.48 | 0.2 | 0.13 | 0.27 | 0.31 |
| 1 | 81294721 | rs1361166 | 0.345 | 1.407 | 0.593 | 0.414 | 0.006 | 0.006 | 0.006 | 0 | 0.16 | 0.18 | 0.09 | 0.03 | 0.12 | 0.12 |
| 1 | 81300642 | rs11163137 | -0.547 | 1.436 | 0.55 | 0.57 | -0.011 | 0.011 | 0.011 | 0 | 0.05 | 0.09 | 0.03 | 0.1 | 0.02 | 0.15 |
| 1 | 81315043 | rs17105538 | 0.191 | 1.528 | 0.635 | 0.23 | -0.007 | -0.025 | -0.025 | 0.016 | 0.17 | 0.24 | 0.1 | 0.03 | 0.12 | 0.13 |
| 1 | 81316333 | rs12075367 | 0.157 | 1.526 | 0.643 | 0.229 | -0.008 | -0.026 | -0.026 | 0 | 0.17 | 0.25 | 0.1 | 0.03 | 0.12 | 0.12 |
| 1 | 81316894 | rs10874177 | -1.385 | 1.615 | 0.879 | 0.184 | -0.014 | -0.061 | -0.061 | 0 | 0.29 | 0.43 | 0.19 | 0.1 | 0.27 | 0.35 |
| 1 | 81317902 | rs1338145 | -0.272 | 2.107 | 0.6 | 0.587 | 0.006 | 0.067 | 0.067 | 3.798 | 0.92 | 0.93 | 0.8 | 0.65 | 0.86 | 0.81 |
| 1 | 81322436 | rs2152986 | 1.379 | 1.864 | 0.632 | 0.448 | 0.051 | 0.07 | 0.07 | 2.206 | 0.72 | 0.5 | 0.57 | 0.57 | 0.58 | 0.49 |
| 1 | 81329765 | rs4650353 | 0.753 | 1.625 | 0.399 | 0.664 | 0.031 | 0.055 | 0.055 | 1.905 | 0.81 | 0.74 | 0.64 | 0.63 | 0.68 | 0.68 |
| 1 | 81336283 | rs17105542 | -0.266 | 0.423 | -0.012 | 0.112 | -0.005 | -0.009 | -0.009 | 0.129 | 0.18 | 0.23 | 0.1 | 0.07 | 0.12 | 0.24 |
| 1 | 81342496 | rs4459149 | 0.359 | 1.569 | -0.596 | 0.642 | -0.008 | -0.002 | -0.002 | 0 | 0.83 | 0.85 | 0.82 | 0.47 | 0.74 | 0.72 |
| 1 | 81342872 | rs12091192 | -0.017 | 1.381 | -1.032 | 0.334 | -0.002 | 0.026 | 0.026 | 0 | 0.26 | 0.17 | 0.25 | 0.06 | 0.25 | 0.3 |
| 1 | 81345264 | rs6659760 | -1.065 | 1.26 | -1.068 | 0.352 | -0.005 | 0.103 | 0.103 | 0.265 | 0.55 | 0.75 | 0.54 | 0.85 | 0.5 | 0.64 |
| 1 | 81352249 | rs4650491 | -0.754 | 0.688 | -1.316 | 0.364 | 0.015 | 0.038 | 0.038 | 0 | 0.34 | 0.15 | 0.38 | 0.32 | 0.29 | 0.37 |
| 1 | 81355884 | rs1953745 | 0.458 | 0.952 | -0.641 | 0.694 | 0.007 | 0.021 | 0.021 | 0 | 0.84 | 0.74 | 0.88 | 0.75 | 0.84 | 0.8 |
| 1 | 81364776 | rs1415933 | 0.569 | 0.797 | -1.537 | -0.859 | -0.029 | 0.185 | 0.185 | 0.061 | 0.42 | 0.19 | 0.41 | 0.05 | 0.59 | 0.24 |
| 1 | 81369485 | rs11581930 | 0.532 | -0.101 | -1.653 | -0.66 | 0.027 | 0 | 0 | 1.381 | 0.13 | 0.07 | 0.03 | 0.08 | 0.08 | 0.11 |
| 1 | 81378292 | rs1338128 | 0.995 | -0.671 | -1.638 | -1.076 | 0.03 | 0.14 | 0.14 | 0 | 0.53 | 0.3 | 0.5 | 0.06 | 0.63 | 0.32 |
| 1 | 81384148 | rs1338127 | 1.482 | -0.864 | -1.398 | -0.971 | 0.01 | 0.171 | 0.171 | 0.03 | 0.43 | 0.22 | 0.46 | 0.05 | 0.62 | 0.26 |
| 1 | 81386389 | rs1933329 | -0.833 | -1.033 | -0.699 | -0.568 | 0 | 0.013 | 0.013 | 0 | 0.83 | 0.8 | 0.87 | 0.69 | 0.85 | 0.78 |
| 1 | 81386925 | rs11163153 | 1.472 | -1.078 | -0.581 | -0.432 | -0.009 | 0.016 | 0.016 | 0.013 | 0.09 | 0.03 | 0.25 | 0.45 | 0.1 | 0.11 |
| 1 | 81392951 | rs1338137 | 0.345 | -1.224 | 0.45 | -0.591 | -0.002 | -0.022 | -0.022 | 0.193 | 0.72 | 0.6 | 0.57 | 0.19 | 0.75 | 0.63 |
| 1 | 81394942 | rs6680798 | 0.801 | -1.234 | 0.454 | -0.628 | 0.001 | -0.026 | -0.026 | 0.498 | 0.6 | 0.61 | 0.59 | 0.1 | 0.68 | 0.57 |
| 1 | 81400663 | rs11588414 | -0.209 | -1.245 | 0.438 | -0.48 | 0.015 | 0.009 | 0.009 | 0.396 | 0.24 | 0.2 | 0.06 | 0.15 | 0.09 | 0.16 |
| 1 | 81401098 | rs10874182 | -0.84 | -1.239 | 0.493 | -0.487 | -0.055 | -0.032 | -0.032 | 0.017 | 0.29 | 0.39 | 0.1 | 0.15 | 0.14 | 0.37 |
| 1 | 81411025 | rs4307601 | 0.13 | -1.092 | 0.326 | -0.398 | 0 | 0 | 0 | 0 | 0.74 | 0.73 | 0.62 | 0.25 | 0.76 | 0.67 |
| 1 | 81420940 | rs12409530 | 1.933 | -0.815 | -0.398 | -1.153 | -0.09 | 0.096 | 0.096 | 0 | 0.22 | 0.05 | 0.29 | 0.05 | 0.36 | 0.16 |
| 1 | 81427862 | rs11163165 | 1.06 | -1.306 | 0.052 | -0.696 | -0.045 | -0.016 | -0.016 | 0 | 0.11 | 0.17 | 0.08 | 0.06 | 0.02 | 0.1 |
| 1 | 81430056 | rs7515605 | -0.88 | -1.165 | 0.21 | -0.554 | -0.018 | 0.008 | 0.008 | 0 | 0.3 | 0.34 | 0.25 | 0.58 | 0.19 | 0.42 |
| 1 | 81430792 | rs11803328 | 0.939 | -1.238 | -0.067 | -0.565 | -0.006 | 0.003 | 0.003 | 0.455 | 0.1 | 0.11 | 0.03 | 0.13 | 0.06 | 0.1 |
| 1 | 81431917 | rs4492663 | 1.432 | -1.327 | 0.021 | -0.545 | 0.035 | -0.014 | -0.014 | 0 | 0.54 | 0.35 | 0.66 | 0.46 | 0.68 | 0.38 |
| 1 | 81432224 | rs6424729 | 0.06 | -1.154 | 0.109 | -0.414 | 0.005 | 0.008 | 0.008 | 0 | 0.75 | 0.72 | 0.61 | 0.28 | 0.73 | 0.67 |
| 1 | 81436522 | rs6694777 | 0.364 | -1.212 | -0.154 | -0.309 | -0.007 | -0.044 | -0.044 | 0 | 0.71 | 0.83 | 0.77 | 0.54 | 0.72 | 0.66 |
| 1 | 81445360 | rs4433444 | 0.999 | -1.279 | -0.248 | -0.209 | 0.021 | -0.072 | -0.072 | 0.246 | 0.11 | 0 | 0.17 | 0.42 | 0.07 | 0.13 |
| 1 | 81447213 | rs4077887 | 0.767 | -1.367 | -0.005 | -0.362 | -0.047 | 0.003 | 0.003 | 0 | 0.1 | 0.16 | 0.07 | 0.07 | 0.02 | 0.1 |
| 1 | 81448130 | rs4650503 | 1.902 | -1.241 | -0.065 | -0.526 | 0.019 | 0.134 | 0.134 | 0 | 0.45 | 0.28 | 0.47 | 0.05 | 0.58 | 0.26 |
| 1 | 81448546 | rs4350241 | 1.898 | -1.203 | -0.069 | -0.53 | 0.017 | 0.135 | 0.135 | 0 | 0.45 | 0.28 | 0.47 | 0.05 | 0.58 | 0.26 |
| 1 | 81450244 | rs1863618 | 1.914 | -1.2 | -0.067 | -0.528 | 0.016 | 0.133 | 0.133 | 0 | 0.45 | 0.28 | 0.47 | 0.05 | 0.58 | 0.26 |
| 1 | 81450625 | rs12039604 | 0.652 | -0.975 | -0.159 | -0.263 | -0.014 | -0.013 | -0.013 | 0 | 0.16 | 0.1 | 0.2 | 0.33 | 0.22 | 0.23 |
| 1 | 81452973 | rs10747381 | -0.695 | -1.516 | 0.101 | -0.368 | -0.099 | 0.128 | 0.128 | 0 | 0.24 | 0.4 | 0.28 | 0.49 | 0.12 | 0.39 |
| 1 | 81452990 | rs10874192 | 0.09 | -1.361 | -0.147 | -0.22 | -0.032 | 0.054 | 0.054 | 0 | 0.17 | 0.17 | 0.19 | 0.45 | 0.06 | 0.17 |
| 1 | 81453585 | rs1426787 | 0.506 | -0.718 | -0.522 | -0.24 | -0.011 | -0.044 | -0.044 | 0 | 0.71 | 0.83 | 0.77 | 0.54 | 0.71 | 0.66 |
| 1 | 81453818 | rs1365523 | 0.147 | -0.815 | -0.442 | -0.172 | -0.003 | -0.032 | -0.032 | 0 | 0.76 | 0.83 | 0.79 | 0.55 | 0.76 | 0.67 |
| 1 | 81455877 | rs10782744 | -0.486 | -1.102 | -0.19 | -0.309 | -0.007 | 0.001 | 0.001 | 1.644 | 0.16 | 0.2 | 0.24 | 0.82 | 0.09 | 0.27 |
| 1 | 81457046 | rs996812 | 0.179 | -1.076 | -0.488 | -0.871 | -0.004 | 0.201 | 0.201 | 0.065 | 0.44 | 0.14 | 0.29 | 0.05 | 0.44 | 0.51 |
| 1 | 81457639 | rs11163179 | -0.303 | -1.032 | -0.353 | -0.848 | -0.007 | 0.034 | 0.034 | 0 | 0.56 | 0.34 | 0.52 | 0.86 | 0.51 | 0.76 |
| 1 | 81462249 | rs11163181 | 0.742 | -0.867 | -0.645 | 0.082 | 0 | 0.058 | 0.058 | 0 | 0.09 | 0.08 | 0.17 | 0.45 | 0.06 | 0.14 |
| 1 | 81469162 | rs11163182 | 0.395 | -0.738 | -1.219 | -1.018 | -0.053 | 0.143 | 0.143 | 0.086 | 0.37 | 0.16 | 0.28 | 0.04 | 0.45 | 0.18 |
| 1 | 81470086 | rs2042749 | 0.035 | -0.619 | -1.067 | -0.452 | -0.029 | -0.004 | -0.004 | 0 | 0.42 | 0.16 | 0.45 | 0.36 | 0.52 | 0.51 |
| 1 | 81471586 | rs12723074 | -1.071 | -0.33 | -0.505 | 0.073 | -0.013 | -0.068 | -0.068 | 2.068 | 0.58 | 0.53 | 0.77 | 0.95 | 0.62 | 0.77 |
| 1 | 81474889 | rs12043929 | -0.919 | -0.005 | -0.369 | 1.432 | -0.013 | -0.043 | -0.043 | 0 | 0.18 | 0.08 | 0.25 | 0.3 | 0.13 | 0.38 |
| 1 | 81477059 | rs1896176 | -0.025 | 0.001 | -0.381 | 1.445 | -0.005 | -0.008 | -0.008 | 0 | 0.13 | 0.14 | 0.06 | 0.06 | 0.06 | 0.07 |
| 1 | 81478037 | rs11163183 | -0.923 | 0.17 | -0.305 | 1.158 | 0.005 | 0.04 | 0.04 | 0 | 0.26 | 0.16 | 0.35 | 0.88 | 0.19 | 0.52 |
| 1 | 81482945 | rs12041774 | -0.785 | 0.969 | -0.06 | 1.893 | -0.03 | 0.028 | 0.028 | 0.449 | 0.05 | 0.01 | 0.19 | 0.3 | 0.07 | 0.3 |
| 1 | 81489207 | rs7531507 | 0.606 | 0.636 | -0.352 | 0.711 | 0.003 | -0.035 | -0.035 | 0 | 0.93 | 0.87 | 0.87 | 1 | 0.94 | 0.97 |
| 1 | 81489305 | rs7541874 | -0.047 | 0.273 | -0.974 | 0.656 | 0.029 | 0.001 | 0.001 | 0 | 0.41 | 0.28 | 0.39 | 0.32 | 0.42 | 0.37 |
| 1 | 81491004 | rs11163189 | -0.378 | 0.484 | -1.077 | 0.484 | -0.002 | -0.002 | -0.002 | 0 | 0.39 | 0.27 | 0.46 | 0.41 | 0.52 | 0.44 |
| 1 | 81491983 | rs2162393 | 0.355 | 0.545 | -1.922 | -0.624 | -0.058 | 0.195 | 0.195 | 0 | 0.33 | 0.04 | 0.31 | 0.06 | 0.35 | 0.4 |
| 1 | 81507666 | rs10782745 | 0.059 | 0.215 | -1.584 | -0.475 | 0.008 | 0.15 | 0.15 | 0 | 0.48 | 0.26 | 0.48 | 0.79 | 0.55 | 0.48 |
| 1 | 81514784 | rs1890547 | -0.591 | -0.066 | -0.315 | -0.561 | 0.015 | 0.133 | 0.133 | 0 | 0.9 | 0.53 | 0.93 | 0.85 | 0.94 | 0.89 |

| | | | | | | | | | | | | | | | | |
|---|---|---|---|---|---|---|---|---|---|---|---|---|---|---|---|---|
| 1 | 81516862 | rs1426782 | -0.656 | -0.119 | 0.478 | -0.609 | 0 | -0.03 | -0.03 | 1.167 | 0.61 | 0.61 | 0.54 | 0.89 | 0.61 | 0.53 |
| 1 | 81518782 | rs1426783 | 0.233 | 0.116 | -0.172 | -0.245 | -0.058 | 0.007 | 0.007 | 0 | 0.45 | 0.18 | 0.46 | 0.66 | 0.52 | 0.4 |
| 1 | 81528413 | rs17105841 | 1.031 | 0.087 | -0.819 | -0.248 | -0.006 | 0.002 | 0.002 | 0.362 | 0.16 | 0.02 | 0.12 | 0.1 | 0.14 | 0.19 |
| 1 | 81529677 | rs12041018 | 1.556 | 1.239 | -0.743 | -0.024 | -0.076 | -0.089 | -0.089 | 0 | 0.12 | 0.01 | 0.22 | 0.45 | 0.25 | 0.13 |
| 1 | 81530739 | rs11583080 | 0.203 | 1.077 | -0.176 | -0.24 | 0.001 | -0.002 | -0.002 | 0 | 0.16 | 0.13 | 0.14 | 0.2 | 0.16 | 0.13 |
| 1 | 81536949 | rs10782747 | 0.751 | 1.025 | -0.345 | -0.349 | -0.002 | 0.015 | 0.015 | 0.409 | 0.74 | 0.29 | 0.78 | 0.59 | 0.79 | 0.74 |
| 1 | 81541384 | rs1426786 | 1.52 | 1.026 | -0.968 | -0.705 | -0.072 | 0.202 | 0.202 | 0.003 | 0.31 | 0.02 | 0.29 | 0.02 | 0.31 | 0.37 |
| 1 | 81560055 | rs11163193 | 1.354 | 0.768 | -0.781 | -0.631 | -0.068 | 0.152 | 0.152 | 0 | 0.32 | 0.06 | 0.31 | 0.06 | 0.34 | 0.38 |
| 1 | 81563249 | rs575528 | 0.185 | 0.72 | -0.062 | -0.557 | -0.007 | 0.033 | 0.033 | 0 | 0.46 | 0.61 | 0.49 | 0.22 | 0.44 | 0.51 |
| 1 | 81566167 | rs630115 | -0.63 | 0.73 | -0.103 | -0.473 | -0.001 | -0.061 | -0.061 | 2.185 | 0.61 | 0.85 | 0.63 | 0.47 | 0.58 | 0.65 |
| 1 | 81567853 | rs6424736 | 0.828 | 0.685 | 0.284 | -0.988 | -0.017 | 0.09 | 0.09 | 0 | 0.43 | 0.25 | 0.45 | 0.1 | 0.42 | 0.42 |
| 1 | 81572438 | rs2993749 | 0.665 | 0.601 | 0.376 | -0.775 | 0.011 | -0.016 | -0.016 | 0 | 0.05 | 0.11 | 0.1 | 0.02 | 0.06 | 0.03 |
| 1 | 81584524 | rs1195702 | -0.154 | 0.756 | 0.378 | -0.795 | 0.022 | 0.026 | 0.026 | 0 | 0.56 | 0.75 | 0.52 | 0.37 | 0.54 | 0.61 |
| 1 | 81586357 | rs12128534 | 1.013 | 0.839 | 0.744 | -0.877 | -0.001 | -0.048 | -0.048 | 0 | 0.42 | 0.55 | 0.37 | 0.08 | 0.36 | 0.43 |
| 1 | 81587463 | rs583007 | -0.76 | 0.818 | 0.695 | -0.893 | 0.021 | -0.024 | -0.024 | 0 | 0.62 | 0.88 | 0.64 | 0.39 | 0.6 | 0.64 |
| 1 | 81591135 | rs1768400 | 1.172 | 0.779 | 1.016 | -0.949 | 0 | -0.06 | -0.06 | 0 | 0.41 | 0.4 | 0.36 | 0.06 | 0.36 | 0.38 |
| 1 | 81592966 | rs1752212 | -0.266 | 0.775 | 1.017 | -0.93 | 0.023 | 0.026 | 0.026 | 0 | 0.55 | 0.7 | 0.52 | 0.37 | 0.53 | 0.61 |
| 1 | 81593406 | rs17406922 | 0.751 | 0.789 | 0.769 | -0.916 | -0.003 | 0 | 0 | 0 | 0.13 | 0.11 | 0.14 | 0.28 | 0.18 | 0.18 |
| 1 | 81594126 | rs1768395 | -0.237 | 0.756 | 0.639 | -0.901 | 0.022 | 0.028 | 0.028 | 0 | 0.56 | 0.8 | 0.53 | 0.37 | 0.54 | 0.61 |
| 1 | 81595980 | rs596101 | -0.735 | 0.797 | 0.439 | -0.894 | -0.018 | -0.061 | -0.061 | 0.191 | 0.62 | 0.94 | 0.63 | 0.39 | 0.59 | 0.63 |
| 1 | 81599140 | rs1146413 | 1.121 | 0.755 | 0.868 | -0.916 | -0.005 | 0.018 | 0.018 | 0 | 0.39 | 0.3 | 0.34 | 0.07 | 0.35 | 0.37 |
| 1 | 81603437 | rs1146418 | 0.384 | 0.717 | 0.811 | -0.847 | 0.002 | 0.003 | 0.003 | 0 | 0.5 | 0.49 | 0.49 | 0.3 | 0.5 | 0.49 |
| 1 | 81609182 | rs17106024 | -0.19 | 0.374 | 0.407 | -0.844 | -0.011 | -0.012 | -0.012 | 0.68 | 0.37 | 0.61 | 0.31 | 0.19 | 0.26 | 0.35 |
| 1 | 81610300 | rs17106044 | 1.472 | 0.626 | 0.357 | -0.766 | -0.006 | -0.005 | -0.005 | 0.005 | 0.09 | 0.06 | 0.13 | 0.24 | 0.14 | 0.12 |
| 1 | 81614826 | rs1327712 | -0.669 | 0.608 | 0.333 | -0.998 | 0.021 | 0.026 | 0.026 | 0.135 | 0.41 | 0.71 | 0.33 | 0.21 | 0.29 | 0.38 |
| 1 | 81615013 | rs1195689 | -1.341 | 0.612 | 0.296 | -1.009 | 0.029 | 0.037 | 0.037 | 0.29 | 0.5 | 0.85 | 0.47 | 0.45 | 0.43 | 0.49 |
| 1 | 81617971 | rs1195685 | -1.991 | 0.703 | 0.111 | -1.077 | -0.062 | -0.131 | -0.131 | 0 | 0.84 | 0.99 | 0.78 | 0.49 | 0.75 | 0.84 |
| 1 | 81617981 | rs12066876 | -1.368 | 0.718 | 0.238 | -1.033 | 0.041 | 0.027 | 0.027 | 0.364 | 0.49 | 0.77 | 0.45 | 0.45 | 0.41 | 0.49 |
| 1 | 81622201 | rs10493685 | 1.602 | 0.655 | 0.429 | -1.022 | -0.006 | -0.006 | -0.006 | 0 | 0.09 | 0.06 | 0.13 | 0.24 | 0.14 | 0.12 |
| 1 | 81623348 | rs11163201 | -0.75 | 0.74 | 0.381 | -1.094 | 0.016 | 0.025 | 0.025 | 0.011 | 0.42 | 0.69 | 0.33 | 0.21 | 0.31 | 0.39 |
| 1 | 81625321 | rs3850865 | -0.618 | 0.643 | 0.551 | -1.045 | -0.007 | -0.014 | -0.014 | 0 | 0.39 | 0.61 | 0.32 | 0.19 | 0.29 | 0.33 |
| 1 | 81630069 | rs3908551 | -1.073 | 0.628 | 0.568 | -1.034 | 0.096 | 0.142 | 0.142 | 3.239 | 0.06 | 0.46 | 0.06 | 0.06 | 0.04 | 0.07 |
| 1 | 81633378 | rs3908548 | -0.718 | 0.57 | 0.49 | -1.089 | 0.013 | 0.041 | 0.041 | 0 | 0.43 | 0.29 | 0.4 | 0.37 | 0.39 | 0.39 |
| 1 | 81633403 | rs3862251 | 0.053 | 0.585 | 0.151 | -1.072 | 0.035 | 0.065 | 0.065 | 0 | 0.34 | 0.23 | 0.27 | 0.13 | 0.25 | 0.27 |
| 1 | 81639173 | rs1831109 | -1.27 | 0.67 | -0.217 | -1.047 | 0.01 | 0.027 | 0.027 | 0.033 | 0.52 | 0.77 | 0.46 | 0.44 | 0.45 | 0.54 |
| 1 | 81641330 | rs3887577 | 0.314 | 0.752 | -0.11 | -1.106 | 0.001 | 0.001 | 0.001 | 0 | 0.09 | 0.11 | 0.13 | 0.06 | 0.08 | 0.12 |
| 1 | 81647246 | rs4650362 | -1.055 | 0.259 | -0.178 | -1.264 | 0.021 | 0.046 | 0.046 | 0.042 | 0.52 | 0.71 | 0.45 | 0.39 | 0.41 | 0.49 |
| 1 | 81655249 | rs6664265 | 0.259 | 0.43 | -0.102 | -1.138 | -0.005 | -0.009 | -0.009 | 0 | 0.09 | 0.16 | 0.07 | 0.04 | 0.06 | 0.07 |
| 1 | 81655674 | rs11809541 | 1.555 | 0.63 | -0.375 | -1.196 | -0.022 | 0.043 | 0.043 | 0 | 0.28 | 0.12 | 0.3 | 0.05 | 0.31 | 0.28 |
| 1 | 81656725 | rs4120877 | -1.16 | -0.111 | 0.205 | -1.074 | -0.005 | 0.011 | 0.011 | 3.686 | 0.26 | 0.65 | 0.29 | 0.38 | 0.2 | 0.25 |
| 1 | 81656935 | rs6679342 | 0.693 | -0.156 | 0.594 | -1.072 | -0.026 | 0.018 | 0.018 | 0 | 0.88 | 0.7 | 0.87 | 0.82 | 0.94 | 0.86 |
| 1 | 81659678 | rs4259706 | 0.067 | 0.117 | 0.16 | -1.055 | -0.006 | -0.005 | -0.005 | 0 | 0.09 | 0.07 | 0.12 | 0.21 | 0.14 | 0.15 |
| 1 | 81661538 | rs12565210 | 0.437 | -0.612 | -0.077 | -2.014 | -0.005 | -0.006 | -0.006 | 0 | 0.61 | 0.39 | 0.74 | 0.75 | 0.74 | 0.64 |
| 1 | 81668315 | rs2225966 | 0.792 | -0.793 | -0.064 | -2 | 0.036 | 0.021 | 0.021 | 0.005 | 0.53 | 0.3 | 0.59 | 0.51 | 0.65 | 0.48 |
| 1 | 81670050 | rs11163210 | 1.179 | -0.604 | -0.571 | -1.934 | -0.003 | 0.003 | 0.003 | 0 | 0.16 | 0.16 | 0.24 | 0.48 | 0.24 | 0.13 |
| 1 | 81671841 | rs17106149 | 1.404 | -0.585 | -0.657 | -2.448 | -0.017 | -0.029 | -0.029 | 0 | 0.08 | 0.03 | 0.14 | 0.23 | 0.09 | 0.16 |
| 1 | 81672013 | rs5016898 | 1.145 | -0.521 | -0.645 | -2.646 | -0.05 | 0.133 | 0.133 | 0.085 | 0.36 | 0.1 | 0.34 | 0.03 | 0.4 | 0.35 |
| 1 | 81674497 | rs17131753 | -0.651 | -0.848 | 0.002 | -2.298 | -0.027 | -0.051 | -0.051 | 1.103 | 0.13 | 0.27 | 0.07 | 0.03 | 0.05 | 0.05 |
| 1 | 81677398 | rs7553222 | 0.507 | -0.667 | -0.107 | -2.267 | -0.004 | -0.004 | -0.004 | 0.141 | 0.18 | 0.2 | 0.26 | 0.68 | 0.3 | 0.23 |
| 1 | 81678254 | rs1195866 | 0.069 | -1.038 | 0.338 | -2.168 | -0.058 | -0.048 | -0.048 | 0 | 0.26 | 0.44 | 0.29 | 0.25 | 0.17 | 0.29 |
| 1 | 81692968 | rs17106228 | 1.192 | -1.027 | 0.274 | -2.665 | 0.023 | 0.016 | 0.016 | 0.999 | 0.15 | 0.11 | 0.13 | 0.01 | 0.07 | 0.13 |
| 1 | 81694861 | rs4639809 | 0.661 | -0.829 | 0.262 | -2.882 | -0.039 | 0.045 | 0.045 | 0.179 | 0.25 | 0.14 | 0.39 | 0.92 | 0.37 | 0.35 |
| 1 | 81698720 | rs1195854 | -1.056 | -0.979 | 0.433 | -3.054 | -0.021 | 0.166 | 0.166 | 0.213 | 0.55 | 0.81 | 0.6 | 0.97 | 0.5 | 0.57 |
| 1 | 81702178 | rs11163216 | -1.026 | -1.003 | 0.526 | -3.052 | -0.008 | 0.136 | 0.136 | 0.254 | 0.54 | 0.79 | 0.6 | 0.97 | 0.48 | 0.57 |
| 1 | 81704476 | rs17106261 | 0.214 | -1.15 | 0.662 | -3.11 | 0.034 | 0.04 | 0.04 | 0 | 0.23 | 0.16 | 0.16 | 0.04 | 0.12 | 0.18 |
| 1 | 81707228 | rs1536571 | -1.042 | -1.138 | 0.852 | -3.136 | 0.002 | 0.034 | 0.034 | 0 | 0.54 | 0.7 | 0.58 | 0.97 | 0.48 | 0.57 |
| 1 | 81708846 | rs12070922 | 0.477 | -1.01 | 0.633 | -2.879 | 0.022 | 0.01 | 0.01 | 0.309 | 0.2 | 0.17 | 0.15 | 0.01 | 0.11 | 0.15 |
| 1 | 81717415 | rs17450523 | 0.919 | -0.603 | 0.303 | -3.08 | -0.039 | 0.119 | 0.119 | 0.114 | 0.25 | 0.1 | 0.29 | 0.03 | 0.26 | 0.23 |
| 1 | 81721216 | rs2148916 | -0.865 | -0.683 | 0.59 | -3.226 | -0.029 | 0.076 | 0.076 | 0.073 | 0.72 | 0.88 | 0.85 | 0.97 | 0.69 | 0.74 |
| 1 | 81721241 | rs4650520 | -0.125 | -0.686 | 0.772 | -3.227 | -0.002 | -0.076 | -0.076 | 0.208 | 0.54 | 0.58 | 0.67 | 0.96 | 0.61 | 0.61 |
| 1 | 81721596 | rs12069796 | 1.008 | -0.665 | 0.324 | -3.229 | -0.038 | 0.116 | 0.116 | 0.085 | 0.25 | 0.1 | 0.29 | 0.03 | 0.26 | 0.22 |
| 1 | 81725122 | rs713333 | 0.113 | -0.554 | 0.441 | -3.111 | 0.014 | 0.246 | 0.246 | 5.657 | 0.5 | 0.2 | 0.64 | 0.89 | 0.58 | 0.57 |
| 1 | 81739593 | rs1557020 | 0.83 | -0.548 | 0.33 | -3.155 | -0.022 | 0.063 | 0.063 | 0.101 | 0.28 | 0.17 | 0.37 | 0.93 | 0.35 | 0.4 |
| 1 | 81741715 | rs2031773 | -0.265 | -0.54 | 0.325 | -3.112 | 0.013 | 0.072 | 0.072 | 0.554 | 0.57 | 0.7 | 0.7 | 0.96 | 0.62 | 0.63 |
| 1 | 81746732 | rs7551904 | -0.963 | -0.5 | 0.086 | -3.092 | -0.059 | 0.133 | 0.133 | 0.057 | 0.72 | 0.94 | 0.85 | 0.97 | 0.69 | 0.75 |
| 1 | 81748612 | rs6666131 | 0.425 | -0.382 | 0.321 | -2.982 | 0.005 | -0.083 | -0.083 | 0 | 0.34 | 0.45 | 0.4 | 0.93 | 0.38 | 0.4 |
| 1 | 81749474 | rs17417493 | 0.745 | -0.194 | -0.362 | -2.817 | -0.01 | 0.036 | 0.036 | 0 | 0.25 | 0.15 | 0.3 | 0.03 | 0.26 | 0.22 |
| 1 | 81753803 | rs4650527 | -1.77 | -0.073 | -1.004 | -2.731 | -0.027 | 0.06 | 0.06 | 0.002 | 0.82 | 0.93 | 0.9 | 0.97 | 0.8 | 0.81 |
| 1 | 81754170 | rs11163236 | 0.67 | -0.118 | -1.107 | -2.786 | -0.028 | 0.076 | 0.076 | 0 | 0.31 | 0.14 | 0.36 | 0.86 | 0.37 | 0.4 |

| | | | | | | | | | | | | | | | | |
|---|---|---|---|---|---|---|---|---|---|---|---|---|---|---|---|---|
| 1 | 81755444 | rs1327609 | 0.057 | -0.162 | -0.905 | -2.82 | -0.004 | -0.086 | -0.086 | 0 | 0.46 | 0.48 | 0.49 | 0.05 | 0.38 | 0.36 |
| 1 | 81756972 | rs10735763 | -1.623 | 0.094 | -1.209 | -2.563 | -0.041 | 0.089 | 0.089 | 0 | 0.82 | 0.97 | 0.9 | 0.97 | 0.79 | 0.8 |
| 1 | 81757936 | rs7525213 | -0.498 | 0.046 | -1.242 | -2.498 | -0.009 | 0.018 | 0.018 | 0 | 0.6 | 0.68 | 0.6 | 0.84 | 0.54 | 0.55 |
| 1 | 81759170 | rs12026119 | 1.349 | 0.072 | -1.519 | -2.433 | -0.044 | -0.06 | -0.06 | 0 | 0.23 | 0.05 | 0.28 | 0.75 | 0.28 | 0.31 |
| 1 | 81759831 | rs1327611 | -0.031 | -0.165 | -1.331 | -2.471 | -0.029 | -0.076 | -0.076 | 0 | 0.35 | 0.56 | 0.31 | 0.1 | 0.25 | 0.23 |
| 1 | 81761082 | rs17451493 | 0.594 | 0.512 | -1.634 | -1.702 | -0.074 | 0.008 | 0.008 | 0 | 0.19 | 0.01 | 0.26 | 0.08 | 0.22 | 0.24 |
| 1 | 81766935 | rs6683338 | -0.421 | 0.701 | -1.047 | -1.713 | 0.002 | -0.078 | -0.078 | 0 | 0.64 | 0.48 | 0.75 | 0.88 | 0.62 | 0.67 |
| 1 | 81766989 | rs10493690 | -0.396 | 0.529 | -1.443 | -1.199 | -0.008 | 0.144 | 0.144 | 0 | 0.34 | 0.12 | 0.42 | 0.08 | 0.33 | 0.36 |
| 1 | 81768971 | rs10158280 | -0.903 | 0.585 | -0.247 | -0.655 | 0.023 | 0 | 0 | 0.003 | 0.07 | 0.2 | 0.08 | 0.02 | 0.05 | 0.07 |
| 1 | 81769938 | rs12059808 | -1.084 | 0.518 | 0.051 | -0.7 | 0.032 | 0.01 | 0.01 | 0 | 0.07 | 0.26 | 0.07 | 0.02 | 0.04 | 0.07 |
| 1 | 81770151 | rs17106362 | -1.63 | -0.023 | 0.764 | -1.104 | -0.029 | -0.079 | -0.079 | 0 | 0.15 | 0.52 | 0.13 | 0.02 | 0.1 | 0.1 |
| 1 | 81770597 | rs1998981 | 0.099 | -0.107 | 0.231 | -1.224 | 0.001 | 0.002 | 0.002 | 0 | 0.27 | 0.23 | 0.27 | 0.23 | 0.31 | 0.3 |
| 1 | 81770827 | rs4650365 | -1.538 | 0.252 | -0.198 | -1.73 | -0.048 | 0.091 | 0.091 | 0.011 | 0.54 | 0.85 | 0.53 | 0.26 | 0.56 | 0.46 |
| 1 | 81774068 | rs11163245 | 0.088 | 0.889 | -0.557 | -2.089 | -0.047 | 0.047 | 0.047 | 0 | 0.37 | 0.22 | 0.46 | 0.22 | 0.5 | 0.33 |
| 1 | 81774898 | rs7553397 | -0.675 | 0.74 | -0.216 | -1.731 | 0.008 | 0.01 | 0.01 | 0 | 0.52 | 0.46 | 0.64 | 0.35 | 0.63 | 0.47 |
| 1 | 81778651 | rs12137941 | -0.202 | 0.634 | 0.034 | -1.871 | -0.015 | -0.054 | -0.054 | 0 | 0.13 | 0.27 | 0.12 | 0.02 | 0.1 | 0.09 |
| 1 | 81780540 | rs17452799 | -0.035 | 0.629 | 0.203 | -1.994 | -0.017 | 0.01 | 0.01 | 0.056 | 0.39 | 0.34 | 0.5 | 0.34 | 0.5 | 0.33 |
| 1 | 81791213 | rs4650366 | -0.719 | 0.541 | 0.165 | -1.995 | 0.007 | 0.038 | 0.038 | 2.219 | 0.49 | 0.7 | 0.59 | 0.39 | 0.55 | 0.51 |
| 1 | 81795252 | rs10493689 | 0.575 | 0.551 | 0.229 | -1.997 | 0.033 | 0.003 | 0.003 | 0 | 0.07 | 0.27 | 0.08 | 0.02 | 0.05 | 0.07 |
| 1 | 81796081 | rs4362025 | 1.159 | 0.396 | 0.147 | -2.145 | -0.01 | 0.016 | 0.016 | 0 | 0.21 | 0.13 | 0.22 | 0.15 | 0.3 | 0.11 |
| 1 | 81800625 | rs17453329 | 0.481 | 0.364 | -0.124 | -2.242 | 0.001 | -0.034 | -0.034 | 0 | 0.14 | 0.15 | 0.13 | 0.01 | 0.13 | 0.06 |
| 1 | 81803379 | rs11163254 | 0.269 | 0.428 | -0.127 | -2.162 | 0.005 | -0.047 | -0.047 | 0.366 | 0.3 | 0.44 | 0.31 | 0.18 | 0.35 | 0.19 |
| 1 | 81804154 | rs1327600 | 0.139 | 0.482 | -0.193 | -2.069 | -0.004 | -0.001 | -0.001 | 0.616 | 0.18 | 0.19 | 0.28 | 0.18 | 0.2 | 0.22 |
| 1 | 81813866 | rs12731404 | -0.375 | 0.27 | -0.192 | -1.973 | -0.001 | 0 | 0 | 0 | 0.46 | 0.55 | 0.57 | 0.36 | 0.55 | 0.4 |
| 1 | 81820712 | rs7534535 | 1.364 | 0.069 | -0.621 | -1.879 | 0.008 | -0.037 | -0.037 | 0 | 0.32 | 0.13 | 0.24 | 0.62 | 0.27 | 0.45 |
| 1 | 81824004 | rs12093730 | 0.635 | 0.268 | 0.246 | -1.811 | 0.005 | -0.045 | -0.045 | 0 | 0.11 | 0.16 | 0.15 | 0.02 | 0.12 | 0.03 |
| 1 | 81825125 | rs10874220 | 1.324 | 0.514 | 0.14 | -1.876 | -0.008 | 0.08 | 0.08 | 4.376 | 0.57 | 0.21 | 0.52 | 0.74 | 0.6 | 0.63 |
| 1 | 81827957 | rs4477328 | -0.785 | 0.506 | 0.748 | -1.887 | 0.033 | 0 | 0 | 0 | 0.94 | 0.77 | 0.94 | 0.98 | 0.95 | 0.94 |
| 1 | 81828029 | rs4512695 | 1.352 | 0.502 | 0.554 | -1.903 | -0.009 | 0.079 | 0.079 | 4.063 | 0.57 | 0.21 | 0.53 | 0.74 | 0.6 | 0.63 |
| 1 | 81829161 | rs10874223 | 1.052 | 0.557 | 0.713 | -1.562 | 0.014 | -0.004 | -0.004 | 2.641 | 0.75 | 0.4 | 0.72 | 0.79 | 0.76 | 0.72 |
| 1 | 81831533 | rs4526653 | 0.785 | 0.594 | 0.413 | -1.491 | 0.03 | 0.017 | 0.017 | 1.438 | 0.79 | 0.59 | 0.75 | 0.79 | 0.8 | 0.75 |
| 1 | 81846293 | rs6695272 | 0.241 | 0.078 | -0.155 | -1.667 | -0.002 | 0.004 | 0.004 | 2.646 | 0.45 | 0.43 | 0.52 | 0.72 | 0.47 | 0.66 |
| 1 | 81846350 | rs6695372 | 0.18 | 0.368 | -0.718 | -1.894 | -0.026 | 0.044 | 0.044 | 0 | 0.28 | 0.18 | 0.3 | 0.07 | 0.4 | 0.12 |
| 1 | 81852813 | rs7539302 | 0.132 | 0.621 | -0.82 | -1.312 | -0.032 | 0.027 | 0.027 | 5.371 | 0.33 | 0.19 | 0.35 | 0.77 | 0.35 | 0.55 |
| 1 | 81866112 | rs4561073 | -0.352 | 0.413 | -0.39 | -1.008 | -0.006 | -0.067 | -0.067 | 0 | 0.69 | 0.6 | 0.81 | 0.93 | 0.71 | 0.83 |
| 1 | 81870185 | rs3890825 | -0.612 | 0.54 | -0.434 | -0.511 | -0.001 | 0 | 0 | 0 | 0.17 | 0.26 | 0.18 | 0.21 | 0.17 | 0.17 |
| 1 | 81870764 | rs7546194 | 0.185 | 0.183 | -0.458 | -1.428 | -0.003 | -0.048 | -0.048 | 0 | 0.4 | 0.47 | 0.41 | 0.04 | 0.5 | 0.34 |
| 1 | 81872593 | rs10874238 | 0.017 | -0.316 | -0.815 | -1.154 | -0.016 | 0.004 | 0.004 | 0 | 0.64 | 0.48 | 0.78 | 0.53 | 0.74 | 0.47 |
| 1 | 81873838 | rs4650540 | -0.165 | -0.306 | -0.6 | -1.326 | 0.003 | -0.001 | -0.001 | 0 | 0.39 | 0.52 | 0.55 | 0.52 | 0.4 | 0.35 |
| 1 | 81873869 | rs11808193 | 1.651 | -0.535 | -0.475 | -1.465 | 0 | 0.003 | 0.003 | 0 | 0.18 | 0.12 | 0.12 | 0.17 | 0.17 | 0.28 |
| 1 | 81876723 | rs11163281 | 0.444 | -0.526 | -0.41 | -1.454 | -0.003 | -0.004 | -0.004 | 0.59 | 0.36 | 0.35 | 0.5 | 0.47 | 0.39 | 0.32 |
| 1 | 81878665 | rs4233110 | 0.52 | -0.461 | -0.463 | -1.617 | 0.009 | -0.004 | -0.004 | 0 | 0.6 | 0.37 | 0.73 | 0.48 | 0.68 | 0.46 |
| 1 | 81879287 | rs12760194 | 1.917 | -0.379 | -0.512 | -1.707 | -0.062 | 0.14 | 0.14 | 0 | 0.24 | 0.03 | 0.23 | 0.01 | 0.29 | 0.12 |
| 1 | 81881212 | rs6681688 | 0.589 | -0.495 | -0.407 | -1.721 | -0.007 | 0.081 | 0.081 | 0 | 0.47 | 0.29 | 0.42 | 0.21 | 0.51 | 0.45 |
| 1 | 81885380 | rs12722932 | 1.897 | -0.387 | -0.735 | -1.684 | -0.081 | 0.169 | 0.169 | 0 | 0.24 | 0.01 | 0.24 | 0.01 | 0.29 | 0.13 |
| 1 | 81890814 | rs4362026 | 0.699 | -0.528 | -0.146 | -1.426 | -0.001 | 0.023 | 0.023 | 0 | 0.48 | 0.42 | 0.43 | 0.21 | 0.51 | 0.45 |
| 1 | 81891826 | rs4272665 | -0.093 | -0.76 | -0.62 | -0.773 | -0.022 | 0.018 | 0.018 | 0 | 0.83 | 0.78 | 0.93 | 0.71 | 0.91 | 0.81 |
| 1 | 81891851 | rs10874242 | 0.934 | -0.317 | -0.783 | -0.535 | -0.016 | -0.083 | -0.083 | 0 | 0.2 | 0.07 | 0.36 | 0.4 | 0.23 | 0.11 |
| 1 | 81892985 | rs11163290 | -0.367 | -0.531 | -0.405 | -1.085 | -0.01 | -0.043 | -0.043 | 1.693 | 0.8 | 0.7 | 0.89 | 0.93 | 0.83 | 0.89 |
| 1 | 81895097 | rs11163291 | -0.301 | -0.73 | -0.806 | -1.242 | 0.021 | 0.023 | 0.023 | 0 | 0.38 | 0.33 | 0.19 | 0.23 | 0.23 | 0.43 |
| 1 | 81896746 | rs10782760 | -1.592 | 0.092 | -1.079 | -1.296 | -0.042 | 0.045 | 0.045 | 0 | 0.82 | 0.87 | 0.77 | 1 | 0.68 | 0.89 |
| 1 | 81897255 | rs11163293 | 0.638 | -0.705 | -0.618 | -0.926 | 0.012 | 0.051 | 0.051 | 0.084 | 0.18 | 0.07 | 0.06 | 0.06 | 0.14 | 0.15 |
| 1 | 81897388 | rs12410914 | -0.133 | -0.601 | -0.023 | -0.407 | 0 | -0.002 | -0.002 | 0 | 0.26 | 0.27 | 0.38 | 0.73 | 0.24 | 0.38 |
| 1 | 81900158 | rs4233112 | 0.934 | -0.803 | 1.19 | 0.157 | 0.045 | -0.012 | -0.012 | 0.022 | 0.93 | 0.74 | 0.95 | 0.99 | 0.94 | 0.97 |
| 1 | 81901137 | rs4520454 | -0.685 | -1.011 | 1.121 | 0.548 | 0 | 0.006 | 0.006 | 0 | 0.18 | 0.19 | 0.2 | 0.58 | 0.15 | 0.22 |
| 1 | 81908617 | rs7543033 | 0.705 | -1.013 | 1.388 | 0.576 | -0.008 | 0.042 | 0.042 | 0.097 | 0.81 | 0.73 | 0.79 | 0.41 | 0.84 | 0.79 |
| 1 | 81911370 | rs4390217 | 0.885 | -1.415 | 1.57 | 0.122 | -0.012 | -0.027 | -0.027 | 0.436 | 0.63 | 0.59 | 0.69 | 0.74 | 0.71 | 0.56 |
| 1 | 81911567 | rs12118952 | -0.376 | -1.413 | 1.461 | 0.125 | -0.001 | 0.033 | 0.033 | 0 | 0.13 | 0.16 | 0.17 | 0.59 | 0.12 | 0.23 |
| 1 | 81911989 | rs11163296 | 1.813 | -1.409 | 1.469 | -0.002 | 0 | 0.151 | 0.151 | 0 | 0.46 | 0.24 | 0.49 | 0.1 | 0.57 | 0.31 |
| 1 | 81922516 | rs6701355 | -0.298 | -1.47 | 1.21 | 0.265 | 0.004 | -0.002 | -0.002 | 0 | 0.07 | 0.08 | 0.14 | 0.03 | 0.1 | 0.12 |
| 1 | 81924106 | rs4420124 | 1.657 | -1.66 | 1.184 | 0.156 | 0 | 0.008 | 0.008 | 0.059 | 0.63 | 0.36 | 0.51 | 0.32 | 0.64 | 0.54 |
| 1 | 81929838 | rs6677307 | 1.311 | -1.763 | 1.61 | 0.353 | 0.044 | -0.005 | -0.005 | 0 | 0.9 | 0.67 | 0.87 | 0.96 | 0.91 | 0.84 |
| 1 | 81930778 | rs4456135 | 1.973 | -2.038 | 1.168 | 0.318 | 0.083 | 0.162 | 0.162 | 0 | 0.51 | 0.25 | 0.5 | 0.09 | 0.69 | 0.35 |
| 1 | 81938646 | rs4650370 | 0.982 | -1.849 | 1.569 | 1.605 | -0.055 | 0.038 | 0.038 | 0.554 | 0.73 | 0.58 | 0.75 | 0.32 | 0.86 | 0.58 |
| 1 | 81941270 | rs11586668 | -0.595 | -1.159 | 1.642 | 1.425 | -0.001 | -0.015 | -0.015 | 0.258 | 0.16 | 0.16 | 0.21 | 0.23 | 0.15 | 0.18 |
| 1 | 81943906 | rs11163302 | -0.485 | -1.256 | 1.674 | 1.383 | -0.048 | 0.031 | 0.031 | 0 | 0.17 | 0.35 | 0.09 | 0.3 | 0.07 | 0.26 |
| 1 | 81946851 | rs12728389 | -0.454 | -1.173 | 1.537 | 1.371 | -0.002 | -0.013 | -0.013 | 0 | 0.15 | 0.13 | 0.22 | 0.17 | 0.16 | 0.19 |
| 1 | 81947398 | rs11163306 | 1.456 | -1.135 | 1.75 | 1.489 | 0.002 | 0.045 | 0.045 | 0 | 0.75 | 0.51 | 0.65 | 0.69 | 0.76 | 0.67 |
| 1 | 81947930 | rs10874250 | 1.062 | -0.983 | 1.46 | 1.352 | 0.045 | 0.103 | 0.103 | 0 | 0.9 | 0.53 | 0.86 | 0.87 | 0.92 | 0.87 |

| | | | | | | | | | | | | | | | | |
|---|---|---|---|---|---|---|---|---|---|---|---|---|---|---|---|---|
| 1 | 81949079 | rs12142040 | 1.755 | -1.029 | 0.906 | 1.31 | 0.037 | -0.006 | -0.006 | 0.312 | 0.58 | 0.24 | 0.56 | 0.4 | 0.69 | 0.4 |
| 1 | 81951500 | rs4650559 | 1.49 | -0.431 | 0.9 | 0.936 | 0.045 | -0.04 | -0.04 | 0 | 0.9 | 0.53 | 0.88 | 1 | 0.94 | 0.93 |
| 1 | 81969284 | rs1108500 | -0.574 | -0.235 | -0.075 | 1.184 | -0.001 | -0.002 | -0.002 | 0 | 0.13 | 0.16 | 0.19 | 0.19 | 0.16 | 0.2 |
| 1 | 81971886 | rs2296234 | -2.024 | -0.823 | -0.583 | 1.218 | 0.042 | -0.064 | -0.064 | 0.648 | 0.41 | 0.89 | 0.46 | 0.57 | 0.3 | 0.52 |
| 1 | 81972032 | rs6670348 | -0.116 | -0.96 | -0.032 | 1.289 | -0.006 | 0.006 | 0.006 | 0.811 | 0.13 | 0.14 | 0.11 | 0.25 | 0.04 | 0.13 |
| 1 | 81974432 | rs11163320 | -0.239 | -0.848 | -0.08 | 1.274 | 0.003 | -0.004 | -0.004 | 0 | 0.14 | 0.13 | 0.12 | 0.25 | 0.05 | 0.14 |
| 1 | 81983176 | rs1535880 | -1.779 | -0.836 | -0.349 | 1.243 | 0.053 | -0.058 | -0.058 | 0 | 0.37 | 0.87 | 0.4 | 0.54 | 0.25 | 0.46 |
| 1 | 81983650 | rs1535881 | -0.564 | -0.824 | 0.034 | 1.198 | 0.017 | 0.042 | 0.042 | 0 | 0.14 | 0.27 | 0.2 | 0.21 | 0.16 | 0.22 |
| 1 | 81984158 | rs1408426 | -1.595 | -0.791 | -0.749 | 1.117 | 0.034 | -0.036 | -0.036 | 0 | 0.36 | 0.74 | 0.38 | 0.51 | 0.24 | 0.44 |
| 1 | 81987161 | rs11586072 | -0.744 | 0.359 | -1.052 | 0.877 | -0.021 | -0.038 | -0.038 | 0 | 0.12 | 0.06 | 0.18 | 0.19 | 0.15 | 0.2 |
| 1 | 81995870 | rs11806028 | -0.181 | 0.206 | -1.455 | -0.121 | 0.01 | 0.13 | 0.13 | 0.115 | 0.66 | 0.88 | 0.72 | 0.89 | 0.7 | 0.81 |
| 1 | 81996353 | rs17454592 | 0.518 | 1.002 | -1.911 | -0.286 | -0.04 | 0.005 | 0.005 | 0 | 0.08 | 0.02 | 0.07 | 0.07 | 0.12 | 0.15 |
| 1 | 81998027 | rs17106647 | 0.266 | 0.493 | -2.342 | 0.359 | 0.015 | -0.058 | -0.058 | 2.344 | 0.09 | 0 | 0.16 | 0.23 | 0.05 | 0.18 |
| 1 | 81999759 | rs17106650 | 1.232 | 1.145 | -2.211 | 0.484 | -0.014 | -0.009 | -0.009 | 0 | 0.08 | 0.03 | 0.12 | 0.08 | 0.12 | 0.06 |
| 1 | 82001286 | rs17106655 | 0.048 | 1.103 | -1.109 | 0.405 | 0.054 | -0.03 | -0.03 | 0 | 0.11 | 0.27 | 0.12 | 0.08 | 0.16 | 0.17 |
| 1 | 82009424 | rs7527341 | 0.569 | 0.827 | -0.767 | 0.327 | -0.022 | -0.065 | -0.065 | 0.186 | 0.34 | 0.18 | 0.28 | 0.5 | 0.37 | 0.43 |
| 1 | 82010446 | rs10157210 | -0.371 | 0.842 | -0.462 | 0.308 | 0.027 | -0.008 | -0.008 | 0.65 | 0.18 | 0.26 | 0.23 | 0.16 | 0.28 | 0.22 |
| 1 | 82015752 | rs7544613 | -0.48 | 0.695 | -0.355 | 0.172 | 0.026 | 0.024 | 0.024 | 0 | 0.51 | 0.41 | 0.51 | 0.66 | 0.65 | 0.66 |
| 1 | 82016757 | rs17421240 | 0.506 | 0.947 | -0.491 | 0.183 | 0.006 | -0.005 | -0.005 | 0.566 | 0.08 | 0.09 | 0.07 | 0.07 | 0.13 | 0.15 |
| 1 | 82019285 | rs4313439 | 0.442 | 0.424 | -0.766 | 0.296 | -0.047 | 0.041 | 0.041 | 0 | 0.17 | 0.03 | 0.11 | 0.06 | 0.18 | 0.12 |
| 1 | 82024405 | rs11163332 | -1.117 | -0.478 | 0.087 | 0.606 | -0.063 | 0.055 | 0.055 | 0 | 0.22 | 0.55 | 0.3 | 0.3 | 0.11 | 0.18 |
| 1 | 82026353 | rs12022306 | -1.117 | -0.578 | 0.154 | 0.618 | -0.073 | 0.051 | 0.051 | 0 | 0.22 | 0.58 | 0.29 | 0.33 | 0.09 | 0.18 |
| 1 | 82028246 | rs12145665 | -0.624 | -0.466 | 0.595 | 0.469 | -0.007 | -0.075 | -0.075 | 0.07 | 0.13 | 0.27 | 0.07 | 0 | 0.1 | 0.03 |
| 1 | 82029505 | rs7532624 | -1.559 | -0.534 | 0.241 | 0.496 | -0.056 | 0.033 | 0.033 | 0 | 0.25 | 0.62 | 0.32 | 0.37 | 0.12 | 0.25 |
| 1 | 82045070 | rs12030604 | -0.631 | -0.464 | 0.016 | 0.603 | -0.019 | 0.06 | 0.06 | 0 | 0.08 | 0.08 | 0.2 | 0.36 | 0.03 | 0.26 |
| 1 | 82048650 | rs4426018 | -1.24 | -0.483 | 0.055 | 0.457 | 0.024 | -0.032 | -0.032 | 0 | 0.19 | 0.11 | 0.25 | 0.33 | 0.13 | 0.3 |
| 1 | 82052342 | rs17106740 | -0.599 | -0.768 | 0.606 | 0.387 | -0.025 | 0.035 | 0.035 | 0 | 0.14 | 0.21 | 0.22 | 0.37 | 0.05 | 0.25 |
| 1 | 82053387 | rs10493695 | -0.484 | -0.346 | 0.257 | 0.061 | -0.015 | -0.003 | -0.003 | 0 | 0.7 | 0.81 | 0.77 | 0.69 | 0.66 | 0.67 |
| 1 | 82054524 | rs10874255 | -0.334 | -0.457 | -0.153 | -0.356 | 0 | -0.002 | -0.002 | 0 | 0.52 | 0.54 | 0.62 | 0.67 | 0.51 | 0.62 |
| 1 | 82055936 | rs1359309 | 1.674 | -0.364 | -0.194 | -0.415 | -0.001 | 0.002 | 0.002 | 0 | 0.19 | 0.2 | 0.23 | 0.08 | 0.24 | 0.22 |
| 1 | 82056068 | rs1359308 | 0.556 | -0.405 | -0.328 | -0.687 | -0.002 | -0.006 | -0.006 | 0 | 0.32 | 0.27 | 0.36 | 0.3 | 0.39 | 0.38 |
| 1 | 82057528 | rs531940 | -2.3 | -0.602 | -0.47 | -0.909 | 0.018 | 0.009 | 0.009 | 0.499 | 0.85 | 0.92 | 0.95 | 0.93 | 0.95 | 0.92 |
| 1 | 82059198 | rs526674 | -1.604 | -0.107 | -0.443 | -0.987 | 0.004 | -0.01 | -0.01 | 0 | 0.79 | 0.91 | 0.88 | 0.8 | 0.82 | 0.81 |
| 1 | 82061064 | rs1924558 | 0.237 | -0.199 | -0.397 | -0.968 | 0.001 | 0.001 | 0.001 | 0 | 0.37 | 0.38 | 0.39 | 0.38 | 0.26 | 0.3 |
| 1 | 82062764 | rs17106776 | 1.791 | -0.196 | -0.433 | -0.83 | 0.004 | 0.043 | 0.043 | 0 | 0.17 | 0.1 | 0.14 | 0.01 | 0.14 | 0.06 |
| 1 | 82064696 | rs12717771 | 0.889 | -0.158 | -0.418 | -0.861 | 0 | -0.003 | -0.003 | 0 | 0.32 | 0.27 | 0.34 | 0.31 | 0.38 | 0.38 |
| 1 | 82065813 | rs17106784 | 1.394 | -0.226 | -0.383 | -0.868 | -0.006 | -0.048 | -0.048 | 0 | 0.19 | 0.25 | 0.16 | 0.01 | 0.15 | 0.06 |
| 1 | 82066625 | rs4317851 | -0.567 | -0.21 | -0.432 | -0.909 | 0.011 | 0.002 | 0.002 | 0 | 0.7 | 0.66 | 0.77 | 0.69 | 0.7 | 0.66 |
| 1 | 82066699 | rs4417125 | -0.18 | 0.046 | -0.338 | -0.768 | 0.002 | -0.01 | -0.01 | 0 | 0.54 | 0.5 | 0.62 | 0.68 | 0.55 | 0.6 |
| 1 | 82075444 | rs2389163 | -1.36 | 0.319 | -0.782 | -0.658 | 0.017 | -0.001 | -0.001 | 0 | 0.92 | 0.93 | 0.9 | 0.89 | 0.82 | 0.88 |
| 1 | 82078790 | rs1924651 | -1.312 | -0.081 | -0.736 | -0.624 | -0.004 | -0.003 | -0.003 | 0 | 0.73 | 0.81 | 0.78 | 0.69 | 0.7 | 0.66 |
| 1 | 82087412 | rs7523717 | -0.14 | -0.211 | -0.827 | -0.587 | -0.003 | -0.005 | -0.005 | 0.415 | 0.41 | 0.5 | 0.42 | 0.39 | 0.32 | 0.28 |
| 1 | 82096075 | rs17106847 | 0.007 | 0.012 | -0.757 | 0.158 | 0.002 | -0.003 | -0.003 | 0 | 0.23 | 0.24 | 0.27 | 0.37 | 0.18 | 0.23 |
| 1 | 82096227 | rs17456745 | -0.714 | 0.35 | -0.828 | 0.222 | 0 | 0.009 | 0.009 | 0 | 0.3 | 0.29 | 0.34 | 0.3 | 0.36 | 0.35 |
| 1 | 82102924 | rs10493696 | -0.314 | 0.592 | -0.957 | 0.484 | 0.003 | 0.016 | 0.016 | 0 | 0.24 | 0.25 | 0.3 | 0.29 | 0.31 | 0.28 |
| 1 | 82117297 | rs2389165 | -0.702 | 0.107 | -1.209 | -0.225 | 0.014 | 0.01 | 0.01 | 0 | 0.21 | 0.16 | 0.09 | 0.01 | 0.13 | 0.18 |
| 1 | 82126526 | rs571361 | -0.269 | 1.155 | -0.89 | -0.504 | 0.023 | -0.041 | -0.041 | 0 | 0.27 | 0.5 | 0.43 | 0.21 | 0.29 | 0.25 |
| 1 | 82127429 | rs9324178 | 0.124 | 0.876 | -2.003 | -0.981 | -0.029 | 0.104 | 0.104 | 0 | 0.34 | 0.01 | 0.13 | 0.12 | 0.3 | 0.32 |
| 1 | 82130153 | rs9728976 | 0.609 | 1.072 | -2.021 | -1.047 | 0.006 | 0 | 0 | 0 | 0.18 | 0.22 | 0.15 | 0.15 | 0.21 | 0.26 |
| 1 | 82132298 | rs561278 | -0.145 | 1.001 | -2.026 | -1.042 | -0.024 | 0.027 | 0.027 | 0 | 0.43 | 0.69 | 0.54 | 0.36 | 0.48 | 0.45 |
| 1 | 82138740 | rs627084 | -1.188 | 1.056 | -1.973 | -0.925 | 0 | 0.01 | 0.01 | 0 | 0.78 | 0.74 | 0.68 | 0.46 | 0.78 | 0.78 |
| 1 | 82143805 | rs489438 | 0.472 | 0.845 | -1.313 | -1.241 | 0.027 | -0.013 | -0.013 | 0 | 0.3 | 0.38 | 0.45 | 0.27 | 0.34 | 0.29 |
| 1 | 82146102 | rs17423764 | 1.293 | 1.02 | -1.788 | -1.825 | 0.002 | 0.002 | 0.002 | 0 | 0.12 | 0.09 | 0.09 | 0.08 | 0.13 | 0.16 |
| 1 | 82147200 | rs10493701 | 1.936 | 0.68 | -1.413 | -2.007 | -0.004 | -0.002 | -0.002 | 0 | 0.11 | 0.07 | 0.3 | 0.03 | 0.16 | 0.1 |
| 1 | 82148070 | rs11163349 | 0.391 | 0.215 | -1.374 | -2.069 | 0.036 | 0.115 | 0.115 | 0 | 0.36 | 0.2 | 0.2 | 0.1 | 0.32 | 0.36 |
| 1 | 82156682 | rs645997 | 0.263 | 0.201 | -1.096 | -2.057 | 0.031 | -0.023 | -0.023 | 0.012 | 0.29 | 0.4 | 0.41 | 0.24 | 0.32 | 0.24 |
| 1 | 82162001 | rs660851 | -0.801 | 0.131 | -1.143 | -1.924 | 0 | 0 | 0 | 0 | 0.67 | 0.71 | 0.65 | 0.33 | 0.67 | 0.65 |
| 1 | 82173048 | rs654498 | -0.396 | 0.069 | -0.524 | -1.99 | 0.009 | 0.012 | 0.012 | 0 | 0.58 | 0.63 | 0.64 | 0.31 | 0.66 | 0.62 |
| 1 | 82173308 | rs540287 | 0.264 | -0.066 | -0.733 | -2.363 | 0.031 | -0.028 | -0.028 | 0 | 0.31 | 0.4 | 0.44 | 0.23 | 0.4 | 0.24 |
| 1 | 82175317 | rs12732009 | 0.303 | -0.298 | -0.615 | -2.191 | 0.013 | 0.042 | 0.042 | 0 | 0.25 | 0.18 | 0.16 | 0.07 | 0.23 | 0.33 |
| 1 | 82175982 | rs483233 | -0.474 | -0.279 | -0.787 | -2.541 | 0.004 | 0.001 | 0.001 | 1.177 | 0.67 | 0.69 | 0.73 | 0.31 | 0.78 | 0.66 |
| 1 | 82178052 | rs9662349 | 0.945 | 0.02 | -0.846 | -2.525 | 0.01 | -0.004 | -0.004 | 0 | 0.29 | 0.3 | 0.48 | 0.23 | 0.43 | 0.25 |
| 1 | 82181580 | rs9728534 | 0.554 | 0.169 | -1.081 | -2.601 | -0.003 | 0 | 0 | 0 | 0.18 | 0.14 | 0.13 | 0.1 | 0.22 | 0.31 |
| 1 | 82181963 | rs9660481 | 1.527 | 0.124 | -0.871 | -2.666 | -0.017 | 0.013 | 0.013 | 0 | 0.25 | 0.18 | 0.44 | 0.15 | 0.35 | 0.21 |
| 1 | 82183055 | rs9285611 | 0.726 | 0.166 | -0.885 | -2.631 | -0.001 | -0.001 | -0.001 | 0 | 0.18 | 0.14 | 0.09 | 0.1 | 0.19 | 0.25 |
| 1 | 82184463 | rs9724784 | -1.087 | -0.166 | -0.993 | -2.617 | 0.009 | -0.066 | -0.066 | 0.126 | 0.78 | 0.91 | 0.76 | 0.38 | 0.8 | 0.8 |
| 1 | 82186939 | rs9659032 | -0.919 | 0.079 | -0.416 | -2.217 | 0.006 | -0.009 | -0.009 | 0.826 | 0.81 | 0.8 | 0.78 | 0.39 | 0.88 | 0.82 |
| 1 | 82193885 | rs12731373 | 1.403 | 0.129 | -0.451 | -2.251 | -0.043 | 0.079 | 0.079 | 0.111 | 0.37 | 0.18 | 0.48 | 0.16 | 0.45 | 0.24 |
| 1 | 82194294 | rs11163354 | 1.227 | 0.111 | -0.582 | -2.092 | -0.012 | -0.011 | -0.011 | 0.004 | 0.12 | 0.19 | 0.1 | 0.05 | 0.11 | 0.16 |

| | | | | | | | | | | | | | | | | |
|---|---|---|---|---|---|---|---|---|---|---|---|---|---|---|---|---|
| 1 | 82195760 | rs9324183 | -0.527 | 0.104 | -0.617 | -2.123 | -0.001 | 0.005 | 0.005 | 0.003 | 0.32 | 0.34 | 0.24 | 0.15 | 0.33 | 0.47 |
| 1 | 82198945 | rs2039829 | 1.591 | 0.284 | -0.689 | -1.919 | -0.009 | -0.016 | -0.016 | 0 | 0.09 | 0.11 | 0.02 | 0 | 0.01 | 0.01 |
| 1 | 82199372 | rs9659808 | -1.071 | 1.282 | -0.591 | -1.44 | 0.001 | -0.017 | -0.017 | 0.023 | 0.31 | 0.32 | 0.19 | 0.18 | 0.32 | 0.42 |
| 1 | 82203562 | rs12090448 | 0.906 | 0.553 | -0.878 | -0.884 | 0 | 0.002 | 0.002 | 0.026 | 0.08 | 0.05 | 0.08 | 0.06 | 0.07 | 0.03 |
| 1 | 82204738 | rs12737523 | 0.265 | 0.555 | -0.664 | -0.601 | 0 | 0.018 | 0.018 | 0 | 0.12 | 0.07 | 0.05 | 0.05 | 0.11 | 0.17 |
| 1 | 82215080 | rs9729825 | -0.494 | 0.673 | -0.65 | -0.505 | -0.005 | -0.015 | -0.015 | 0 | 0.24 | 0.26 | 0.12 | 0.13 | 0.2 | 0.3 |
| 1 | 82219539 | rs9782905 | -0.916 | 1.753 | -0.89 | 0.254 | -0.041 | -0.003 | -0.003 | 3.691 | 0.15 | 0.07 | 0.2 | 0.16 | 0.21 | 0.18 |
| 1 | 82228427 | rs10874266 | -1.021 | 1.577 | -0.223 | 0.146 | -0.012 | -0.017 | -0.017 | 0.196 | 0.23 | 0.42 | 0.31 | 0.77 | 0.23 | 0.31 |
| 1 | 82232757 | rs12726534 | 0.436 | 1.639 | -0.228 | -0.015 | 0 | -0.006 | -0.006 | 0 | 0.06 | 0.03 | 0.05 | 0.03 | 0.08 | 0.15 |
| 1 | 82237323 | rs11163360 | -0.425 | 1.522 | 0.121 | 0.618 | -0.001 | 0.015 | 0.015 | 0 | 0.15 | 0.18 | 0.13 | 0.24 | 0.16 | 0.17 |
| 1 | 82242203 | rs9728563 | 0.377 | 1.622 | 0.347 | 0.601 | -0.007 | 0.18 | 0.18 | 0 | 0.07 | 0.14 | 0.18 | 0.53 | 0.07 | 0.14 |
| 1 | 82243083 | rs12072054 | -0.853 | 1.54 | 0.818 | 0.099 | 0.065 | -0.038 | -0.038 | 0.086 | 0.13 | 0.31 | 0.18 | 0.09 | 0.17 | 0.14 |
| 1 | 82244575 | rs11163371 | -0.562 | 1.658 | 0.812 | 0.067 | 0.058 | -0.039 | -0.039 | 0.007 | 0.08 | 0.18 | 0.15 | 0.02 | 0.13 | 0.06 |
| 1 | 82246289 | rs1498210 | -0.709 | 1.649 | 0.766 | 0.064 | 0.064 | -0.038 | -0.038 | 0 | 0.09 | 0.23 | 0.15 | 0.02 | 0.13 | 0.1 |
| 1 | 82265905 | rs3748770 | -1.098 | 1.059 | 0.374 | 0.093 | 0.003 | -0.001 | -0.001 | 0.093 | 0.28 | 0.26 | 0.19 | 0.24 | 0.26 | 0.35 |
| 1 | 82276586 | rs696726 | -0.296 | 1.645 | 0.455 | 0.032 | 0.045 | -0.005 | -0.005 | 0 | 0.06 | 0.13 | 0.21 | 0.03 | 0.09 | 0.05 |
| 1 | 82299039 | rs709702 | -0.585 | 1.615 | 1.248 | -0.077 | 0.049 | -0.003 | -0.003 | 0 | 0.07 | 0.14 | 0.23 | 0.07 | 0.12 | 0.05 |
| 1 | 82304384 | rs3790952 | -0.284 | 1.485 | 1.14 | -0.051 | 0.007 | -0.027 | -0.027 | 0 | 0.17 | 0.07 | 0.12 | 0.22 | 0.17 | 0.16 |
| 1 | 82318968 | rs12144010 | -0.72 | 1.375 | 1.168 | -0.393 | 0.001 | -0.042 | -0.042 | 0 | 0.11 | 0.19 | 0.08 | 0.03 | 0.11 | 0.18 |
| 1 | 82328415 | rs3790932 | -0.701 | 2.11 | 0.68 | -0.829 | -0.027 | 0.083 | 0.083 | 0 | 0.31 | 0.15 | 0.45 | 0.86 | 0.41 | 0.54 |
| 1 | 82330750 | rs1546086 | 1.356 | 1.894 | 1.288 | -0.474 | 0.142 | -0.023 | -0.023 | 0.032 | 0.9 | 0.62 | 0.74 | 0.95 | 0.86 | 0.95 |
| 1 | 82340753 | rs406304 | 0.525 | 1.946 | 0.865 | -0.449 | 0.071 | -0.024 | -0.024 | 0 | 0.93 | 0.81 | 0.77 | 0.95 | 0.88 | 0.96 |
| 1 | 82348717 | rs367881 | -0.018 | 1.482 | 0.523 | -0.358 | -0.022 | -0.008 | -0.008 | 0 | 0.91 | 0.98 | 0.95 | 0.94 | 0.9 | 0.82 |
| 1 | 82364673 | rs2026596 | -0.543 | 0.067 | 0.725 | -0.591 | 0.022 | -0.019 | -0.019 | 0.078 | 0.24 | 0.2 | 0.32 | 0.62 | 0.17 | 0.37 |
| 1 | 82368812 | rs654654 | 1.127 | 0.437 | 0.671 | -0.842 | -0.004 | 0.021 | 0.021 | 0 | 0.59 | 0.62 | 0.41 | 0.32 | 0.56 | 0.48 |
| 1 | 82379446 | rs284227 | -0.056 | 0.813 | 0.728 | -0.684 | 0.002 | -0.005 | -0.005 | 0.006 | 0.84 | 0.83 | 0.73 | 0.94 | 0.73 | 0.9 |
| 1 | 82383979 | rs12032052 | -0.407 | 0.767 | 0.508 | -0.711 | 0.001 | -0.095 | -0.095 | 0.008 | 0.13 | 0.02 | 0.28 | 0.62 | 0.12 | 0.34 |
| 1 | 82387711 | rs12758555 | 1.619 | 0.786 | 0.144 | -1.27 | 0.029 | 0.152 | 0.152 | 0 | 0.3 | 0.07 | 0.15 | 0.02 | 0.21 | 0.13 |
| 1 | 82388902 | rs993099 | 0.546 | 0.819 | 0.635 | -1.186 | -0.004 | 0.019 | 0.019 | 0 | 0.36 | 0.29 | 0.24 | 0.26 | 0.39 | 0.37 |
| 1 | 82389289 | rs2201163 | 0.803 | 0.813 | 0.61 | -1.194 | -0.002 | 0.038 | 0.038 | 0 | 0.33 | 0.17 | 0.19 | 0.17 | 0.33 | 0.28 |
| 1 | 82393655 | rs12079242 | 0.812 | 0.82 | 0.613 | -1.191 | -0.002 | 0.037 | 0.037 | 0 | 0.33 | 0.17 | 0.19 | 0.17 | 0.33 | 0.28 |
| 1 | 82398752 | rs11163394 | 0.359 | 0.881 | 0.531 | -1.257 | -0.007 | 0.012 | 0.012 | 0 | 0.38 | 0.32 | 0.26 | 0.26 | 0.42 | 0.38 |
| 1 | 82411874 | rs2172285 | 0.321 | 0.86 | 0.644 | -1.248 | -0.005 | 0.008 | 0.008 | 0 | 0.38 | 0.34 | 0.27 | 0.26 | 0.42 | 0.37 |
| 1 | 82413263 | rs11163396 | -0.401 | 0.787 | 0.62 | -1.224 | 0.001 | 0.006 | 0.006 | 0 | 0.15 | 0.17 | 0.31 | 0.62 | 0.13 | 0.37 |
| 1 | 82432618 | rs2296699 | 1.246 | 0.784 | 0.434 | -1.086 | -0.006 | -0.008 | -0.008 | 3.654 | 0.07 | 0.01 | 0.26 | 0.57 | 0.08 | 0.23 |
| 1 | 82436281 | rs1327022 | 0.211 | 0.774 | 0.542 | -1.326 | -0.017 | -0.015 | -0.015 | 0 | 0.46 | 0.48 | 0.47 | 0.68 | 0.31 | 0.53 |
| 1 | 82438148 | rs9438724 | 0.213 | 1.016 | 0.377 | -1.27 | -0.001 | -0.002 | -0.002 | 0 | 0.16 | 0.16 | 0.27 | 0.06 | 0.27 | 0.1 |
| 1 | 82447191 | rs12130650 | 1.725 | 0.757 | 0.138 | -1.379 | 0.058 | 0.161 | 0.161 | 0 | 0.28 | 0.02 | 0.13 | 0.02 | 0.16 | 0.11 |
| 1 | 82450194 | rs1154816 | -0.325 | 0.733 | 0.495 | -1.069 | -0.005 | 0.064 | 0.064 | 0 | 0.17 | 0.22 | 0.32 | 0.59 | 0.17 | 0.37 |
| 1 | 82454122 | rs17431604 | 1.123 | 1.015 | 0.529 | -1.2 | 0.006 | 0.083 | 0.083 | 0.191 | 0.32 | 0.13 | 0.18 | 0.16 | 0.29 | 0.21 |
| 1 | 82462304 | rs7412827 | 0.206 | 1.043 | 0.606 | -1.238 | 0 | -0.051 | -0.051 | 0 | 0.45 | 0.4 | 0.41 | 0.07 | 0.45 | 0.23 |
| 1 | 82463594 | rs1926840 | 1.608 | 0.817 | 0.451 | -1.154 | 0.054 | 0.092 | 0.092 | 0 | 0.29 | 0.09 | 0.13 | 0.02 | 0.17 | 0.12 |
| 1 | 82484131 | rs6696194 | -0.894 | 0.477 | 0.739 | -0.728 | -0.016 | 0.041 | 0.041 | 0 | 0.22 | 0.31 | 0.34 | 0.65 | 0.2 | 0.4 |
| 1 | 82492195 | rs12145656 | 0.432 | 0.677 | 0.845 | -0.832 | 0.013 | -0.01 | -0.01 | 0.005 | 0.14 | 0.18 | 0.25 | 0.06 | 0.28 | 0.1 |
| 1 | 82495835 | rs12027404 | -0.305 | 0.513 | 0.68 | -0.888 | -0.007 | 0.065 | 0.065 | 0 | 0.54 | 0.35 | 0.57 | 0.82 | 0.55 | 0.69 |
| 1 | 82503980 | rs12125527 | 1.429 | 0.463 | 0.643 | -0.899 | 0.035 | 0.038 | 0.038 | 0 | 0.29 | 0.17 | 0.13 | 0.09 | 0.15 | 0.13 |
| 1 | 82506257 | rs12135377 | 1.429 | 0.463 | 0.643 | -0.897 | 0.035 | 0.039 | 0.039 | 0 | 0.29 | 0.17 | 0.13 | 0.09 | 0.15 | 0.13 |
| 1 | 82508628 | rs10782773 | -0.389 | 0.496 | 0.672 | -0.907 | -0.001 | -0.009 | -0.009 | 0.289 | 0.55 | 0.43 | 0.58 | 0.86 | 0.57 | 0.76 |
| 1 | 82509065 | rs4970663 | 0.158 | 0.812 | 0.682 | -0.846 | 0.106 | -0.064 | -0.064 | 0.281 | 0.15 | 0.31 | 0.28 | 0.06 | 0.28 | 0.1 |
| 1 | 82517491 | rs7515935 | 0.87 | 0.754 | 0.549 | -0.866 | -0.002 | -0.013 | -0.013 | 0 | 0.34 | 0.34 | 0.22 | 0.12 | 0.27 | 0.27 |
| 1 | 82517645 | rs11163417 | -0.018 | 0.655 | 0.252 | -0.819 | 0.03 | 0.133 | 0.133 | 0 | 0.5 | 0.33 | 0.5 | 0.82 | 0.44 | 0.63 |
| 1 | 82539588 | rs11163421 | -0.253 | 0.68 | 0.166 | -0.636 | 0.002 | -0.03 | -0.03 | 0.368 | 0.16 | 0.12 | 0.3 | 0.58 | 0.17 | 0.37 |
| 1 | 82541810 | rs4609472 | 0.853 | 0.606 | -0.098 | -0.698 | 0.018 | 0.047 | 0.047 | 0.134 | 0.34 | 0.21 | 0.18 | 0.22 | 0.28 | 0.27 |
| 1 | 82542281 | rs6598985 | -1.319 | 0.929 | 0.076 | -0.645 | 0.023 | 0.086 | 0.086 | 0 | 0.7 | 0.88 | 0.82 | 0.9 | 0.77 | 0.84 |
| 1 | 82552067 | rs4970666 | -0.304 | 0.963 | 0.162 | -0.546 | 0.001 | 0.001 | 0.001 | 0 | 0.55 | 0.55 | 0.74 | 0.68 | 0.63 | 0.49 |
| 1 | 82561559 | rs962249 | -0.033 | 0.468 | 0.071 | -0.165 | -0.007 | -0.014 | -0.014 | 0.023 | 0.55 | 0.56 | 0.51 | 0.34 | 0.6 | 0.51 |
| 1 | 82562877 | rs10874286 | 0.145 | 0.49 | -0.091 | -0.105 | -0.009 | -0.007 | -0.007 | 0 | 0.41 | 0.41 | 0.47 | 0.63 | 0.34 | 0.38 |
| 1 | 82566764 | rs12083586 | 0.191 | 0.489 | -0.266 | -0.041 | 0.008 | 0.017 | 0.017 | 0 | 0.36 | 0.31 | 0.3 | 0.29 | 0.34 | 0.29 |
| 1 | 82571186 | rs10493708 | 0.078 | 0.765 | -0.368 | -0.069 | -0.007 | 0.085 | 0.085 | 0 | 0.38 | 0.17 | 0.33 | 0.26 | 0.39 | 0.31 |
| 1 | 82575142 | rs6687277 | -0.227 | 0.952 | -0.459 | -0.048 | -0.02 | 0.111 | 0.111 | 0 | 0.44 | 0.18 | 0.43 | 0.28 | 0.47 | 0.38 |
| 1 | 82575538 | rs4970672 | 1.433 | 0.878 | 0.925 | 0.449 | 0.057 | 0.115 | 0.115 | 3.292 | 0.94 | 0.46 | 0.96 | 0.95 | 0.99 | 0.94 |
| 1 | 82583859 | rs4970648 | -0.497 | 1.467 | 0.256 | 0.953 | 0.001 | 0.026 | 0.026 | 1.155 | 0.21 | 0.33 | 0.33 | 0.45 | 0.24 | 0.43 |
| 1 | 82595241 | rs4970675 | -0.43 | 1.47 | 0.237 | 0.954 | 0 | 0.014 | 0.014 | 1.589 | 0.21 | 0.3 | 0.32 | 0.45 | 0.24 | 0.43 |
| 1 | 82603634 | rs12407762 | -0.229 | 1.651 | 0.67 | 0.67 | 0.047 | -0.006 | -0.006 | 0 | 0.07 | 0.2 | 0.12 | 0.02 | 0.11 | 0.1 |
| 1 | 82605135 | rs11163433 | -1.239 | 1.676 | 0.51 | 0.828 | 0.043 | 0.099 | 0.099 | 2.918 | 0.28 | 0.56 | 0.44 | 0.44 | 0.35 | 0.52 |
| 1 | 82612212 | rs6681181 | -0.469 | 1.415 | -0.168 | 0.954 | -0.017 | -0.056 | -0.056 | 0.038 | 0.2 | 0.13 | 0.29 | 0.41 | 0.24 | 0.43 |
| 1 | 82653168 | rs7514492 | 0.473 | 1.493 | -0.352 | 0.871 | -0.016 | -0.045 | -0.045 | 2.214 | 0.78 | 0.81 | 0.67 | 0.58 | 0.72 | 0.57 |
| 1 | 82675112 | rs4970650 | -0.398 | 1.433 | -0.485 | 0.905 | -0.034 | -0.095 | -0.095 | 0 | 0.19 | 0.08 | 0.29 | 0.42 | 0.24 | 0.44 |
| 1 | 82686415 | rs695978 | 0.554 | 1.396 | -0.263 | 0.828 | -0.002 | -0.005 | -0.005 | 0.043 | 0.74 | 0.66 | 0.64 | 0.58 | 0.7 | 0.56 |

| | | | | | | | | | | | | | | | | |
|---|---|---|---|---|---|---|---|---|---|---|---|---|---|---|---|---|
| 1 | 82687054 | rs1505550 | 1.059 | 1.413 | 0.223 | 0.596 | 0.009 | 0.017 | 0.017 | 0.403 | 0.65 | 0.38 | 0.5 | 0.57 | 0.57 | 0.46 |
| 1 | 82689732 | rs10493711 | 0.63 | 1.452 | 0.289 | 0.629 | 0.031 | 0.065 | 0.065 | 0.625 | 0.72 | 0.6 | 0.57 | 0.58 | 0.66 | 0.5 |
| 1 | 82690006 | rs10493712 | 1.291 | 1.829 | -0.13 | 0.18 | -0.002 | -0.002 | -0.002 | 0.23 | 0.58 | 0.35 | 0.43 | 0.58 | 0.48 | 0.43 |
| 1 | 82690730 | rs12123603 | 1.315 | 1.504 | -1.083 | -0.118 | -0.034 | -0.018 | -0.018 | 0 | 0.17 | 0.01 | 0.1 | 0.18 | 0.14 | 0.12 |
| 1 | 82694699 | rs1144349 | -0.467 | 1.351 | -0.272 | 0.565 | 0.018 | 0.005 | 0.005 | 2.771 | 0.27 | 0.37 | 0.38 | 0.25 | 0.35 | 0.5 |
| 1 | 82696502 | rs10747391 | 0.743 | 1.348 | -0.166 | 0.58 | 0.044 | -0.003 | -0.003 | 5.809 | 0.67 | 0.46 | 0.55 | 0.72 | 0.57 | 0.49 |
| 1 | 82708274 | rs4106233 | 0.384 | 1.374 | -0.262 | 0.633 | -0.083 | -0.072 | -0.072 | 0 | 0.19 | 0.02 | 0.21 | 0.23 | 0.24 | 0.4 |
| 1 | 82712113 | rs796800 | -0.534 | 1.359 | -0.129 | 0.623 | 0.004 | 0.001 | 0.001 | 3.08 | 0.27 | 0.32 | 0.38 | 0.26 | 0.35 | 0.49 |
| 1 | 82714297 | rs17108037 | 1.337 | 1.038 | -0.289 | 0.467 | 0.005 | 0.006 | 0.006 | 0.504 | 0.12 | 0.08 | 0.16 | 0.14 | 0.09 | 0.07 |
| 1 | 82723394 | rs785605 | -0.56 | 1.305 | -0.201 | 0.558 | 0.029 | 0.006 | 0.006 | 3.189 | 0.28 | 0.42 | 0.38 | 0.26 | 0.36 | 0.49 |
| 1 | 82744600 | rs785612 | 0.714 | 1.336 | -0.217 | 0.479 | 0.024 | -0.008 | -0.008 | 0 | 0.67 | 0.51 | 0.56 | 0.73 | 0.59 | 0.49 |
| 1 | 82748577 | rs785627 | 0.619 | 1.398 | -0.193 | 0.476 | 0.053 | -0.023 | -0.023 | 0.022 | 0.87 | 0.64 | 0.77 | 0.94 | 0.82 | 0.89 |
| 1 | 82751201 | rs785631 | -0.317 | 1.416 | -0.197 | 0.514 | 0.054 | 0.016 | 0.016 | 0.013 | 0.92 | 0.71 | 0.83 | 0.96 | 0.89 | 0.9 |
| 1 | 82762033 | rs17436254 | 0.897 | 1.316 | -0.176 | 0.546 | -0.001 | -0.01 | -0.01 | 0 | 0.05 | 0.08 | 0.06 | 0 | 0.06 | 0.01 |
| 1 | 82762700 | rs785584 | 0.385 | 1.334 | -0.012 | 0.547 | 0.031 | 0.009 | 0.009 | 0 | 0.06 | 0.21 | 0.09 | 0.05 | 0.07 | 0.02 |
| 1 | 82764942 | rs1770679 | 0.163 | 1.304 | -0.025 | 0.524 | 0.015 | -0.004 | -0.004 | 0 | 0.73 | 0.64 | 0.64 | 0.77 | 0.66 | 0.51 |
| 1 | 82765052 | rs1770678 | 0.453 | 1.305 | -0.031 | 0.531 | 0.018 | 0.001 | 0.001 | 0 | 0.06 | 0.19 | 0.09 | 0.05 | 0.07 | 0.02 |
| 1 | 82777956 | rs709741 | 0.505 | 1.184 | -0.166 | 0.573 | 0.026 | -0.009 | -0.009 | 0 | 0.67 | 0.52 | 0.56 | 0.72 | 0.59 | 0.49 |
| 1 | 82787818 | rs1857814 | 0.434 | 1.242 | -0.256 | 0.58 | 0.03 | -0.021 | -0.021 | 0 | 0.68 | 0.5 | 0.57 | 0.76 | 0.6 | 0.5 |
| 1 | 82797633 | rs786077 | 0.374 | 1.256 | -0.402 | 0.557 | 0.002 | -0.002 | -0.002 | 0 | 0.68 | 0.62 | 0.57 | 0.76 | 0.61 | 0.5 |
| 1 | 82800233 | rs785643 | 0.765 | 0.719 | -1.147 | 0.868 | -0.008 | 0.012 | 0.012 | 0 | 0.19 | 0.24 | 0.18 | 0.25 | 0.17 | 0.12 |
| 1 | 82801007 | rs785639 | -0.096 | 0.969 | -1.245 | 0.737 | -0.001 | -0.037 | -0.037 | 0 | 0.36 | 0.26 | 0.37 | 0.46 | 0.37 | 0.51 |
| 1 | 82805566 | rs12408229 | -0.515 | 1.397 | -1.491 | 0.561 | -0.049 | 0.003 | 0.003 | 0 | 0.1 | 0.03 | 0.21 | 0.06 | 0.2 | 0.12 |
| 1 | 82809419 | rs7549211 | 0.694 | 0.675 | -1.397 | 0.58 | 0.017 | -0.007 | -0.007 | 0 | 0.44 | 0.34 | 0.42 | 0.52 | 0.39 | 0.39 |
| 1 | 82812305 | rs2047965 | -0.099 | 0.459 | -1.734 | 0.288 | 0.007 | 0.028 | 0.028 | 0.622 | 0.42 | 0.32 | 0.44 | 0.29 | 0.38 | 0.36 |
| 1 | 82817239 | rs950485 | -0.573 | 0.587 | -2.088 | 0.619 | -0.081 | -0.073 | -0.073 | 0 | 0.43 | 0.11 | 0.32 | 0.45 | 0.43 | 0.5 |
| 1 | 82818466 | rs1948885 | 1.067 | 0.72 | -2.12 | 0.638 | 0.015 | -0.008 | -0.008 | 0 | 0.09 | 0.16 | 0.18 | 0.04 | 0.16 | 0.15 |
| 1 | 82826465 | rs1144283 | -1.178 | 0.614 | -2.819 | 0.658 | 0.036 | 0.013 | 0.013 | 0 | 0.55 | 0.78 | 0.57 | 0.5 | 0.6 | 0.63 |
| 1 | 82829468 | rs17108253 | 1.476 | 0.78 | -3.288 | 1.139 | -0.003 | -0.055 | -0.055 | 0.759 | 0.14 | 0.01 | 0.09 | 0.16 | 0.09 | 0.13 |
| 1 | 82829953 | rs17437521 | 1.471 | 0.949 | -3.234 | 1.119 | -0.093 | -0.078 | -0.078 | 0.112 | 0.17 | 0.01 | 0.15 | 0.24 | 0.23 | 0.27 |
| 1 | 82833216 | rs12403706 | 1.469 | 0.304 | -2.695 | 0.897 | 0.007 | -0.004 | -0.004 | 0 | 0.11 | 0.02 | 0.13 | 0.12 | 0.08 | 0.06 |
| 1 | 82835418 | rs11579502 | -0.008 | 0.686 | -2.227 | 1.013 | 0.016 | -0.078 | -0.078 | 0 | 0.19 | 0.02 | 0.1 | 0.23 | 0.14 | 0.19 |
| 1 | 82836012 | rs11581340 | -0.468 | 1.35 | -1.612 | 0.749 | 0.015 | 0.001 | 0.001 | 0 | 0.09 | 0.16 | 0.18 | 0.07 | 0.16 | 0.13 |
| 1 | 82837212 | rs4295913 | 0.206 | 0.947 | -2.786 | 0.529 | -0.066 | -0.115 | -0.115 | 0.398 | 0.46 | 0.03 | 0.42 | 0.59 | 0.4 | 0.47 |
| 1 | 82843604 | rs1281614 | 0.446 | 0.988 | -2.42 | 0.412 | 0.002 | -0.005 | -0.005 | 0 | 0.1 | 0.11 | 0.06 | 0.15 | 0.09 | 0.11 |
| 1 | 82844732 | rs1856736 | 0.446 | 0.988 | -2.424 | 0.531 | 0.003 | -0.008 | -0.008 | 0 | 0.1 | 0.11 | 0.11 | 0.2 | 0.09 | 0.12 |
| 1 | 82848122 | rs11163467 | 0.924 | 0.951 | -2.786 | 0.223 | 0.031 | -0.055 | -0.055 | 0 | 0.39 | 0.09 | 0.4 | 0.41 | 0.33 | 0.42 |
| 1 | 82848779 | rs1281603 | -1.684 | 1.196 | -2.867 | 0.128 | -0.026 | 0.018 | 0.018 | 0 | 0.9 | 0.98 | 0.93 | 0.93 | 0.88 | 0.91 |
| 1 | 82850151 | rs946831 | -0.072 | 1.119 | -2.401 | 0.107 | 0.002 | -0.002 | -0.002 | 0.331 | 0.19 | 0.28 | 0.13 | 0.18 | 0.21 | 0.16 |
| 1 | 82852017 | rs12405458 | 1.371 | 0.908 | -2.964 | -0.082 | -0.028 | -0.111 | -0.111 | 0.054 | 0.16 | 0.01 | 0.3 | 0.35 | 0.16 | 0.29 |
| 1 | 82860892 | rs4970693 | 0.587 | 1.406 | -2.177 | 0.09 | 0.004 | 0.007 | 0.007 | 0 | 0.11 | 0.09 | 0.13 | 0.09 | 0.11 | 0.08 |
| 1 | 82865819 | rs1281595 | 0.11 | 1.596 | -2.19 | -0.703 | -0.003 | 0.008 | 0.008 | 0.053 | 0.14 | 0.11 | 0.07 | 0 | 0.12 | 0.08 |
| 1 | 82869015 | rs11163472 | 0.705 | 1.812 | -2.216 | -0.603 | -0.004 | 0.041 | 0.041 | 0 | 0.51 | 0.19 | 0.47 | 0.36 | 0.49 | 0.46 |
| 1 | 82872034 | rs17491330 | 0.885 | 1.351 | -1.819 | -0.408 | 0.001 | -0.014 | -0.014 | 0 | 0.06 | 0.06 | 0.05 | 0 | 0.07 | 0 |
| 1 | 82880659 | rs12043749 | 0.647 | 1.168 | -1.794 | -0.097 | -0.006 | 0.004 | 0.004 | 0 | 0.42 | 0.1 | 0.39 | 0.36 | 0.4 | 0.4 |
| 1 | 82883239 | rs1281597 | -0.806 | 0.866 | -0.9 | -0.758 | -0.003 | -0.078 | -0.078 | 0 | 0.36 | 0.55 | 0.29 | 0.13 | 0.35 | 0.24 |
| 1 | 82883310 | rs1281598 | -0.65 | 1 | -0.922 | 0.043 | 0.003 | -0.034 | -0.034 | 0 | 0.07 | 0.16 | 0.07 | 0 | 0.06 | 0 |
| 1 | 82884779 | rs12140871 | -0.803 | 1.116 | -1.453 | 0.166 | -0.059 | -0.013 | -0.013 | 0 | 0.17 | 0.01 | 0.18 | 0.13 | 0.18 | 0.16 |
| 1 | 82893001 | rs13373897 | -0.764 | 0.75 | -0.601 | 0.225 | 0.01 | 0.049 | 0.049 | 0 | 0.05 | 0.24 | 0.05 | 0.08 | 0.03 | 0.08 |
| 1 | 82896475 | rs1281610 | -0.067 | 0.613 | -1.118 | -0.285 | 0.004 | 0.072 | 0.072 | 0 | 0.4 | 0.21 | 0.22 | 0.23 | 0.35 | 0.26 |
| 1 | 82899747 | rs1986902 | -0.057 | 0.619 | -0.98 | -0.316 | 0.009 | 0.064 | 0.064 | 0 | 0.4 | 0.24 | 0.21 | 0.22 | 0.34 | 0.25 |
| 1 | 82901264 | rs10437024 | -1.154 | 0.722 | -0.133 | -0.272 | 0.048 | 0.048 | 0.048 | 0 | 0.09 | 0.41 | 0.07 | 0.11 | 0.06 | 0.11 |
| 1 | 82906724 | rs4550100 | 0.472 | 0.268 | -0.783 | -0.708 | -0.057 | -0.017 | -0.017 | 0.169 | 0.27 | 0.03 | 0.19 | 0.23 | 0.27 | 0.2 |
| 1 | 82913726 | rs11163476 | 0.249 | 0.939 | -0.776 | -0.584 | -0.049 | 0.01 | 0.01 | 0 | 0.18 | 0.04 | 0.36 | 0.15 | 0.28 | 0.13 |
| 1 | 82934401 | rs3863840 | 0.144 | 0.459 | 0.417 | -0.864 | 0.034 | 0.054 | 0.054 | 0.18 | 0.43 | 0.23 | 0.21 | 0.26 | 0.31 | 0.24 |
| 1 | 82935587 | rs11163480 | -0.85 | 0.675 | 0.505 | -0.71 | 0.074 | 0.012 | 0.012 | 0.015 | 0.09 | 0.37 | 0.09 | 0.05 | 0.1 | 0.1 |
| 1 | 82939537 | rs11163482 | -0.467 | 0.498 | 0.771 | -0.661 | 0.008 | 0.009 | 0.009 | 1.253 | 0.65 | 0.42 | 0.62 | 0.42 | 0.61 | 0.48 |
| 1 | 82944571 | rs10874312 | -1.355 | 0.445 | 0.25 | -0.655 | -0.024 | -0.046 | -0.046 | 0 | 0.74 | 0.81 | 0.68 | 0.47 | 0.67 | 0.58 |
| 1 | 82949643 | rs2026598 | 0.512 | 0.398 | -0.023 | -0.723 | 0.053 | 0.067 | 0.067 | 0.107 | 0.41 | 0.14 | 0.18 | 0.26 | 0.3 | 0.27 |
| 1 | 82956657 | rs12749816 | 0.436 | 0.291 | -0.053 | -0.775 | 0.039 | 0.049 | 0.049 | 0 | 0.39 | 0.21 | 0.18 | 0.26 | 0.3 | 0.26 |
| 1 | 82960385 | rs2225083 | 0.054 | 0.459 | 0.099 | -0.702 | -0.039 | 0.019 | 0.019 | 0.016 | 0.21 | 0.09 | 0.39 | 0.15 | 0.29 | 0.13 |
| 1 | 82964864 | rs7523716 | -0.99 | 0.36 | 0.443 | -0.709 | 0 | -0.012 | -0.012 | 0.035 | 0.33 | 0.63 | 0.48 | 0.22 | 0.35 | 0.31 |
| 1 | 82965451 | rs719793 | -1.043 | 0.322 | 0.37 | -0.728 | -0.001 | 0.012 | 0.012 | 0.121 | 0.33 | 0.72 | 0.49 | 0.22 | 0.35 | 0.32 |
| 1 | 82965883 | rs12067308 | -1.896 | 0.09 | 1.009 | -0.664 | -0.018 | -0.016 | -0.016 | 1.025 | 0.07 | 0.66 | 0.06 | 0.02 | 0.01 | 0.09 |
| 1 | 82969372 | rs10489862 | -0.628 | -0.191 | 0.001 | -0.525 | 0.011 | 0.01 | 0.01 | 0 | 0.53 | 0.62 | 0.43 | 0.61 | 0.57 | 0.53 |
| 1 | 82969770 | rs12734849 | -0.263 | -0.283 | 0.131 | -0.606 | 0.002 | 0.001 | 0.001 | 0 | 0.44 | 0.36 | 0.39 | 0.59 | 0.51 | 0.42 |
| 1 | 82986888 | rs17576309 | 0.201 | -0.318 | 0.4 | -0.562 | -0.024 | -0.037 | -0.037 | 0.625 | 0.07 | 0.11 | 0.02 | 0 | 0.05 | 0.06 |
| 1 | 82987838 | rs2209696 | -0.112 | 0.119 | -0.353 | -0.508 | -0.047 | -0.051 | -0.051 | 0.029 | 0.6 | 0.81 | 0.67 | 0.51 | 0.56 | 0.61 |
| 1 | 83014374 | rs318405 | 0.794 | 0.114 | -0.145 | -0.328 | -0.001 | -0.002 | -0.002 | 0.237 | 0.45 | 0.41 | 0.55 | 0.35 | 0.45 | 0.47 |

| | | | | | | | | | | | | | | | | |
|---|---|---|---|---|---|---|---|---|---|---|---|---|---|---|---|---|
| 1 | 83019476 | rs318422 | 0.223 | 0.323 | -0.953 | -0.265 | -0.044 | -0.06 | -0.06 | 0 | 0.58 | 0.83 | 0.66 | 0.46 | 0.56 | 0.61 |
| 1 | 83022595 | rs318419 | 0.88 | 0.33 | -1.066 | -0.261 | 0.016 | 0.039 | 0.039 | 0 | 0.48 | 0.75 | 0.58 | 0.38 | 0.45 | 0.52 |
| 1 | 83029025 | rs114723 | 0.143 | 0.383 | -1.118 | -0.204 | -0.095 | -0.106 | -0.106 | 3.068 | 0.61 | 0.92 | 0.68 | 0.49 | 0.56 | 0.66 |
| 1 | 83030504 | rs1690546 | 0.109 | 0.409 | -1.166 | -0.199 | -0.107 | -0.122 | -0.122 | 0 | 0.61 | 0.92 | 0.68 | 0.49 | 0.56 | 0.66 |
| 1 | 83035362 | rs6604916 | -0.021 | 0.389 | -1.003 | -0.107 | -0.037 | -0.051 | -0.051 | 0 | 0.64 | 0.82 | 0.72 | 0.55 | 0.64 | 0.65 |
| 1 | 83036341 | rs11579295 | 0.611 | 0.36 | -1.106 | -0.131 | 0.001 | -0.001 | -0.001 | 0.07 | 0.11 | 0.11 | 0.09 | 0.1 | 0.11 | 0.14 |
| 1 | 83037587 | rs10747399 | -1.281 | 0.371 | -1.476 | -0.296 | -0.099 | -0.129 | -0.129 | 0 | 0.79 | 0.96 | 0.76 | 0.59 | 0.74 | 0.79 |
| 1 | 83037900 | rs9661164 | 0.134 | 0.366 | -1.456 | -0.288 | -0.079 | -0.091 | -0.091 | 0.024 | 0.67 | 0.93 | 0.73 | 0.58 | 0.64 | 0.7 |
| 1 | 83052634 | rs4363415 | -1.205 | -0.06 | -1.011 | -0.352 | 0.019 | 0.028 | 0.028 | 0 | 0.26 | 0.21 | 0.12 | 0.14 | 0.2 | 0.28 |
| 1 | 83054799 | rs10874322 | -0.135 | -0.005 | -0.823 | -0.265 | -0.006 | -0.003 | -0.003 | 0 | 0.14 | 0.2 | 0.1 | 0.14 | 0.11 | 0.2 |
| 1 | 83057803 | rs954572 | 0.675 | 0.178 | -0.872 | -0.501 | -0.004 | 0 | 0 | 0.184 | 0.45 | 0.44 | 0.57 | 0.42 | 0.43 | 0.49 |
| 1 | 83058379 | rs12757364 | 0.933 | -0.226 | -1.357 | -0.665 | 0.004 | -0.008 | -0.008 | 0 | 0.28 | 0.2 | 0.39 | 0.31 | 0.27 | 0.35 |
| 1 | 83058812 | rs1768739 | -0.855 | 0.149 | -1.683 | -0.516 | -0.106 | -0.071 | -0.071 | 1.58 | 0.71 | 0.86 | 0.71 | 0.59 | 0.62 | 0.73 |
| 1 | 83063455 | rs11163513 | -0.052 | -0.117 | -1.404 | -0.598 | 0.005 | -0.008 | -0.008 | 0.017 | 0.34 | 0.28 | 0.28 | 0.41 | 0.3 | 0.32 |
| 1 | 83063712 | rs10493719 | 1.191 | -0.408 | -1.85 | -0.834 | -0.004 | 0.02 | 0.02 | 0 | 0.15 | 0.01 | 0.1 | 0.08 | 0.12 | 0.16 |
| 1 | 83063846 | rs7552548 | 0.091 | -0.427 | -1.487 | -0.811 | -0.01 | 0.01 | 0.01 | 0 | 0.36 | 0.49 | 0.3 | 0.49 | 0.3 | 0.4 |
| 1 | 83066525 | rs6604915 | -0.54 | -0.428 | -1.361 | -0.703 | 0.006 | 0.002 | 0.002 | 0 | 0.49 | 0.53 | 0.39 | 0.58 | 0.42 | 0.52 |
| 1 | 83068682 | rs1493982 | -1.572 | -0.316 | -1.189 | -0.815 | 0.023 | -0.035 | -0.035 | 0 | 0.87 | 0.7 | 0.9 | 0.93 | 0.89 | 0.94 |
| 1 | 83075377 | rs12563352 | 0.079 | -0.295 | -1.643 | -0.757 | -0.005 | 0.039 | 0.039 | 0 | 0.46 | 0.35 | 0.61 | 0.73 | 0.54 | 0.54 |
| 1 | 83076962 | rs2389318 | -0.145 | -0.309 | -1.647 | -0.711 | -0.001 | 0.032 | 0.032 | 0 | 0.49 | 0.43 | 0.63 | 0.8 | 0.54 | 0.64 |
| 1 | 83077523 | rs950269 | 1.408 | -0.392 | -1.775 | -0.418 | -0.013 | -0.033 | -0.033 | 0 | 0.23 | 0.09 | 0.19 | 0.32 | 0.24 | 0.17 |
| 1 | 83078971 | rs10493720 | -0.099 | -0.254 | -1.668 | -0.633 | -0.008 | 0.057 | 0.057 | 0 | 0.46 | 0.31 | 0.61 | 0.73 | 0.54 | 0.58 |
| 1 | 83083251 | rs1493977 | 0.045 | -0.199 | -1.261 | -0.297 | 0.003 | 0.02 | 0.02 | 0.049 | 0.25 | 0.34 | 0.39 | 0.32 | 0.31 | 0.32 |
| 1 | 83094954 | rs923877 | 0.578 | -0.711 | -1.492 | -0.553 | 0.009 | 0.005 | 0.005 | 0 | 0.16 | 0.03 | 0.1 | 0.1 | 0.12 | 0.18 |
| 1 | 83096252 | rs11163523 | 0.361 | -0.492 | -1.252 | -0.511 | -0.007 | 0.017 | 0.017 | 0 | 0.31 | 0.38 | 0.33 | 0.41 | 0.27 | 0.3 |
| 1 | 83096613 | rs11578309 | 0.5 | -0.674 | -1.197 | -0.441 | 0.031 | 0.052 | 0.052 | 0 | 0.19 | 0.04 | 0.09 | 0.07 | 0.14 | 0.16 |
| 1 | 83097971 | rs6660986 | 0.633 | -0.641 | -0.998 | -0.476 | -0.015 | -0.059 | -0.059 | 0 | 0.11 | 0.21 | 0.12 | 0 | 0.08 | 0.06 |
| 1 | 83098959 | rs6604921 | -0.575 | -0.729 | -0.972 | -0.611 | 0.009 | 0.019 | 0.019 | 0 | 0.61 | 0.55 | 0.47 | 0.56 | 0.47 | 0.56 |
| 1 | 83099614 | rs10518668 | 0.855 | -0.596 | -1.024 | -0.522 | -0.012 | -0.054 | -0.054 | 1.557 | 0.1 | 0.19 | 0.12 | 0 | 0.08 | 0.06 |
| 1 | 83101655 | rs17493817 | 1.07 | -0.432 | -1.202 | -0.575 | -0.088 | -0.086 | -0.086 | 0 | 0.27 | 0.1 | 0.43 | 0.37 | 0.44 | 0.38 |
| 1 | 83106539 | rs11163533 | 1.471 | -0.684 | -0.857 | -0.546 | -0.004 | -0.05 | -0.05 | 0 | 0.29 | 0.13 | 0.24 | 0.42 | 0.25 | 0.23 |
| 1 | 83106653 | rs12070417 | 0.843 | -0.723 | -0.721 | -0.607 | -0.012 | -0.06 | -0.06 | 0 | 0.1 | 0.22 | 0.12 | 0 | 0.08 | 0.06 |
| 1 | 83107700 | rs4662154 | 0.574 | -0.929 | -0.714 | -0.591 | 0.001 | -0.001 | -0.001 | 0.8 | 0.38 | 0.35 | 0.3 | 0.42 | 0.31 | 0.28 |
| 1 | 83110679 | rs1546125 | 1.486 | -0.914 | -0.751 | -0.585 | -0.003 | -0.053 | -0.053 | 0 | 0.29 | 0.12 | 0.25 | 0.42 | 0.25 | 0.23 |
| 1 | 83111268 | rs1577991 | 0.885 | -0.892 | -0.702 | -0.582 | -0.023 | -0.056 | -0.056 | 0 | 0.1 | 0.22 | 0.05 | 0 | 0.06 | 0.05 |
| 1 | 83112606 | rs409599 | -0.128 | -0.971 | -0.432 | -0.674 | 0.017 | 0.007 | 0.007 | 0.021 | 0.51 | 0.46 | 0.62 | 0.52 | 0.62 | 0.66 |
| 1 | 83113252 | rs4662155 | 1.589 | -0.991 | -0.426 | -0.688 | -0.004 | -0.055 | -0.055 | 0 | 0.29 | 0.13 | 0.25 | 0.42 | 0.25 | 0.23 |
| 1 | 83118044 | rs2787985 | -0.021 | -0.959 | -0.396 | -0.832 | 0.032 | -0.001 | -0.001 | 0 | 0.5 | 0.39 | 0.61 | 0.45 | 0.62 | 0.55 |
| 1 | 83119056 | rs1766628 | -1.676 | -0.935 | -0.313 | -0.722 | -0.001 | -0.028 | -0.028 | 2.305 | 0.89 | 0.81 | 0.92 | 0.94 | 0.92 | 0.94 |
| 1 | 83120757 | rs1766629 | -0.599 | -0.804 | -0.862 | -0.643 | -0.039 | -0.11 | -0.11 | 1.41 | 0.8 | 0.59 | 0.87 | 0.94 | 0.87 | 0.88 |
| 1 | 83120822 | rs365222 | 0.574 | -0.52 | -1.116 | -0.513 | -0.018 | -0.013 | -0.013 | 0 | 0.3 | 0.26 | 0.51 | 0.38 | 0.48 | 0.4 |
| 1 | 83127763 | rs319656 | 0.315 | -1.139 | -0.574 | -0.703 | 0.022 | 0.023 | 0.023 | 0 | 0.15 | 0.11 | 0.04 | 0.08 | 0.06 | 0.11 |
| 1 | 83128812 | rs319661 | 0.238 | -0.47 | -0.161 | -0.438 | -0.036 | -0.056 | -0.056 | 0.899 | 0.81 | 0.58 | 0.87 | 0.91 | 0.88 | 0.89 |
| 1 | 83131659 | rs3097974 | -0.838 | 0.546 | -0.224 | -0.002 | 0.053 | 0.045 | 0.045 | 0.498 | 0.29 | 0.47 | 0.52 | 0.35 | 0.4 | 0.43 |
| 1 | 83132400 | rs1433761 | -1.335 | 0.204 | -0.45 | -0.401 | 0.003 | -0.001 | -0.001 | 0 | 0.4 | 0.52 | 0.55 | 0.36 | 0.45 | 0.49 |
| 1 | 83135055 | rs3845616 | 0.077 | 0.263 | -0.625 | -0.397 | 0.01 | 0.034 | 0.034 | 0 | 0.14 | 0.07 | 0.06 | 0.01 | 0.09 | 0.11 |
| 1 | 83135671 | rs4662158 | -0.54 | 0.741 | -0.027 | 0 | 0.015 | 0.008 | 0.008 | 0.593 | 0.88 | 0.69 | 0.89 | 0.92 | 0.92 | 0.91 |
| 1 | 83136135 | rs17108829 | 0.635 | 0.809 | -0.267 | 0.056 | -0.032 | -0.033 | -0.033 | 0 | 0.12 | 0.07 | 0.38 | 0.24 | 0.28 | 0.26 |
| 1 | 83136590 | rs10874326 | 0.429 | 0.68 | -0.135 | 0.048 | 0.029 | 0.018 | 0.018 | 0 | 0.76 | 0.68 | 0.64 | 0.71 | 0.64 | 0.65 |
| 1 | 83137699 | rs319671 | -0.864 | -0.013 | -0.34 | -0.082 | -0.001 | 0.001 | 0.001 | 0 | 0.66 | 0.67 | 0.72 | 0.52 | 0.72 | 0.76 |
| 1 | 83139387 | rs374773 | 0.408 | -0.054 | -0.158 | -0.266 | 0.005 | 0.001 | 0.001 | 0.246 | 0.38 | 0.44 | 0.43 | 0.37 | 0.51 | 0.56 |
| 1 | 83144687 | rs984697 | 0.816 | 0.029 | -0.041 | -0.241 | 0.01 | 0.006 | 0.006 | 0.028 | 0.16 | 0.21 | 0.28 | 0.24 | 0.31 | 0.32 |
| 1 | 83147308 | rs12057639 | -0.651 | -0.146 | -0.012 | -0.323 | 0.004 | -0.003 | -0.003 | 0 | 0.54 | 0.57 | 0.49 | 0.48 | 0.61 | 0.69 |
| 1 | 83149729 | rs393717 | 0.579 | -0.207 | 0.078 | -0.368 | 0.014 | 0.033 | 0.033 | 0 | 0.36 | 0.21 | 0.19 | 0.21 | 0.3 | 0.34 |
| 1 | 83151734 | rs2892744 | 0.949 | -0.196 | 0.148 | -0.367 | 0.028 | 0.019 | 0.019 | 0.036 | 0.16 | 0.24 | 0.29 | 0.24 | 0.31 | 0.32 |
| 1 | 83156170 | rs403191 | 1.996 | -0.233 | 0.097 | -0.399 | 0.004 | 0.018 | 0.018 | 0 | 0.21 | 0.08 | 0.13 | 0.13 | 0.2 | 0.24 |
| 1 | 83162893 | rs17443254 | 0.927 | -0.42 | 0.028 | -0.505 | 0.006 | 0.025 | 0.025 | 0 | 0.14 | 0.06 | 0.23 | 0.04 | 0.12 | 0.07 |
| 1 | 83163132 | rs17108916 | 0.756 | -0.406 | 0.055 | -0.579 | 0.003 | 0.005 | 0.005 | 0.396 | 0.15 | 0.1 | 0.06 | 0.08 | 0.1 | 0.09 |
| 1 | 83166637 | rs2641944 | -0.617 | -0.422 | 0.092 | -0.575 | 0.004 | -0.003 | -0.003 | 0.503 | 0.53 | 0.57 | 0.49 | 0.46 | 0.61 | 0.69 |
| 1 | 83171984 | rs11585110 | 0.546 | -0.343 | 0.14 | -0.564 | 0.048 | 0.038 | 0.038 | 0 | 0.18 | 0.32 | 0.3 | 0.27 | 0.31 | 0.35 |
| 1 | 83171994 | rs10493722 | 0.921 | -0.391 | 0.093 | -0.601 | 0.006 | 0.018 | 0.018 | 0 | 0.14 | 0.09 | 0.21 | 0.03 | 0.11 | 0.07 |
| 1 | 83172112 | rs10493723 | -0.19 | -0.437 | 0.102 | -0.63 | -0.03 | -0.019 | -0.019 | 1.931 | 0.28 | 0.19 | 0.32 | 0.31 | 0.38 | 0.34 |
| 1 | 83172671 | rs393735 | -0.3 | -0.446 | 0.137 | -0.692 | 0.013 | 0.004 | 0.004 | 0.608 | 0.51 | 0.43 | 0.62 | 0.43 | 0.62 | 0.67 |
| 1 | 83172684 | rs389112 | 1.766 | -0.463 | 0.183 | -0.758 | -0.003 | -0.006 | -0.006 | 0 | 0.22 | 0.24 | 0.15 | 0.13 | 0.2 | 0.25 |
| 1 | 83173015 | rs406110 | 0.591 | -0.425 | 0.108 | -0.731 | 0.015 | 0.055 | 0.055 | 0 | 0.35 | 0.18 | 0.33 | 0.16 | 0.32 | 0.32 |
| 1 | 83177365 | rs12032817 | -0.36 | -0.453 | 0.02 | -0.738 | 0.035 | 0.114 | 0.114 | 0 | 0.49 | 0.25 | 0.39 | 0.24 | 0.41 | 0.4 |
| 1 | 83181179 | rs12089552 | 0.685 | -0.416 | 0.022 | -0.71 | 0.002 | 0.005 | 0.005 | 0.369 | 0.15 | 0.12 | 0.06 | 0.07 | 0.1 | 0.09 |
| 1 | 83184129 | rs12744723 | 1.372 | -0.354 | -0.06 | -0.612 | -0.078 | -0.069 | -0.069 | 0.155 | 0.14 | 0.03 | 0.25 | 0.23 | 0.29 | 0.27 |
| 1 | 83193770 | rs9660208 | -1.232 | -0.433 | 0.131 | -0.651 | -0.011 | 0.031 | 0.031 | 0 | 0.65 | 0.47 | 0.67 | 0.51 | 0.7 | 0.69 |

| | | | | | | | | | | | | | | | | |
|---|---|---|---|---|---|---|---|---|---|---|---|---|---|---|---|---|
| 1 | 83196421 | rs1338361 | 1.395 | -0.388 | 0.08 | -0.632 | -0.078 | -0.069 | -0.069 | 0 | 0.14 | 0.03 | 0.25 | 0.23 | 0.29 | 0.27 |
| 1 | 83202936 | rs11588508 | 1.395 | -0.388 | 0.08 | -0.632 | -0.078 | -0.069 | -0.069 | 0.011 | 0.14 | 0.03 | 0.25 | 0.24 | 0.29 | 0.27 |
| 1 | 83204106 | rs2060361 | 0.946 | -0.476 | 0.146 | -0.675 | 0.007 | 0.033 | 0.033 | 0 | 0.14 | 0.05 | 0.2 | 0.03 | 0.11 | 0.07 |
| 1 | 83205734 | rs320922 | -1.592 | -0.467 | 0.19 | -0.641 | -0.001 | 0.005 | 0.005 | 0 | 0.68 | 0.69 | 0.7 | 0.51 | 0.72 | 0.76 |
| 1 | 83213903 | rs114808 | -1.214 | -0.462 | 0.16 | -0.66 | -0.009 | 0.025 | 0.025 | 0 | 0.65 | 0.58 | 0.68 | 0.51 | 0.7 | 0.69 |
| 1 | 83220434 | rs17498581 | 0.686 | -0.417 | 0.153 | -0.634 | 0.002 | 0.004 | 0.004 | 0.387 | 0.15 | 0.12 | 0.06 | 0.07 | 0.1 | 0.09 |
| 1 | 83231866 | rs6429835 | -0.392 | -0.391 | 0.081 | -0.598 | 0.005 | 0.002 | 0.002 | 0 | 0.3 | 0.32 | 0.34 | 0.34 | 0.38 | 0.38 |
| 1 | 83234010 | rs7555550 | -0.441 | -0.36 | 0.043 | -0.557 | 0.005 | -0.003 | -0.003 | 0 | 0.54 | 0.61 | 0.49 | 0.47 | 0.61 | 0.69 |
| 1 | 83237484 | rs320901 | -1.611 | -0.356 | 0.036 | -0.555 | 0 | -0.012 | -0.012 | 0 | 0.69 | 0.73 | 0.71 | 0.51 | 0.72 | 0.76 |
| 1 | 83245723 | rs7531603 | 0.542 | -0.104 | -0.259 | -0.47 | 0.014 | 0.044 | 0.044 | 0 | 0.35 | 0.15 | 0.18 | 0.21 | 0.3 | 0.32 |
| 1 | 83249040 | rs7541474 | 0.532 | -0.103 | -0.258 | -0.474 | 0.015 | 0.041 | 0.041 | 0 | 0.35 | 0.15 | 0.18 | 0.24 | 0.3 | 0.34 |
| 1 | 83249252 | rs7528291 | -0.388 | -0.112 | -0.149 | -0.385 | 0.001 | -0.002 | -0.002 | 0 | 0.54 | 0.42 | 0.48 | 0.48 | 0.61 | 0.69 |
| 1 | 83251624 | rs2078370 | 0.553 | -0.374 | -0.431 | -0.494 | 0.028 | 0.038 | 0.038 | 0 | 0.33 | 0.15 | 0.18 | 0.27 | 0.27 | 0.31 |
| 1 | 83257232 | rs2094335 | -1.06 | 0.049 | -0.158 | -0.202 | 0.008 | 0.075 | 0.075 | 0 | 0.86 | 0.81 | 0.78 | 0.58 | 0.8 | 0.8 |
| 1 | 83260379 | rs4662165 | 0.686 | -0.835 | -0.312 | -1.501 | 0.059 | 0.057 | 0.057 | 2.086 | 0.23 | 0.1 | 0.06 | 0 | 0.1 | 0.11 |
| 1 | 83263047 | rs6685504 | 0.301 | 0.47 | 0.502 | -0.816 | 0.009 | 0.001 | 0.001 | 0.155 | 0.07 | 0.11 | 0.04 | 0.05 | 0.06 | 0.03 |
| 1 | 83265008 | rs7554166 | -0.391 | 0.436 | 0.688 | -0.844 | 0.044 | 0.021 | 0.021 | 0 | 0.09 | 0.41 | 0.09 | 0.07 | 0.07 | 0.14 |
| 1 | 83267760 | rs17506918 | -0.089 | 0.42 | 0.834 | -0.716 | -0.005 | -0.014 | -0.014 | 0 | 0.11 | 0.1 | 0.17 | 0.07 | 0.08 | 0.12 |
| 1 | 83279558 | rs493742 | 0.26 | 1.221 | -0.196 | -0.366 | 0.016 | -0.015 | -0.015 | 0 | 0.67 | 0.63 | 0.44 | 0.22 | 0.53 | 0.5 |
| 1 | 83297866 | rs11163582 | 1.044 | 1.245 | 0.526 | 0.626 | 0.132 | 0.273 | 0.273 | 0 | 0.93 | 0.64 | 0.88 | 0.75 | 0.88 | 0.89 |
| 1 | 83298665 | rs10782797 | 1.563 | 1.194 | 0.764 | 0.524 | 0.151 | 0.261 | 0.261 | 0.005 | 0.92 | 0.49 | 0.86 | 0.76 | 0.87 | 0.88 |
| 1 | 83306575 | rs12132310 | 0.411 | 1.387 | 0.072 | -0.14 | -0.04 | 0.09 | 0.09 | 0.094 | 0.26 | 0.09 | 0.2 | 0.03 | 0.3 | 0.29 |
| 1 | 83311835 | rs11163585 | 0.214 | 1.311 | 0.154 | -0.158 | -0.024 | 0.071 | 0.071 | 0 | 0.28 | 0.19 | 0.21 | 0.04 | 0.29 | 0.3 |
| 1 | 83312460 | rs1409799 | 0.309 | 1.219 | 0.244 | -0.15 | 0.008 | -0.02 | -0.02 | 1.13 | 0.63 | 0.41 | 0.67 | 0.73 | 0.59 | 0.59 |
| 1 | 83331118 | rs7532665 | 0.337 | 1.22 | 0.217 | -0.137 | 0.015 | -0.03 | -0.03 | 0 | 0.63 | 0.47 | 0.67 | 0.72 | 0.57 | 0.58 |
| 1 | 83339078 | rs988424 | 0.917 | 1.209 | 0.337 | -0.103 | 0.076 | 0.16 | 0.16 | 0.166 | 0.91 | 0.65 | 0.88 | 0.76 | 0.86 | 0.89 |
| 1 | 83340310 | rs876339 | 0.917 | 1.209 | 0.337 | -0.103 | 0.076 | 0.16 | 0.16 | 0.166 | 0.91 | 0.64 | 0.88 | 0.76 | 0.86 | 0.89 |
| 1 | 83354737 | rs501760 | 0.108 | 1.217 | 0.246 | -0.206 | 0.004 | -0.145 | -0.145 | 0 | 0.3 | 0.45 | 0.25 | 0.04 | 0.28 | 0.28 |
| 1 | 83364553 | rs11163602 | 0.218 | 1.23 | 0.349 | -0.027 | 0 | 0.001 | 0.001 | 0.827 | 0.64 | 0.35 | 0.69 | 0.65 | 0.65 | 0.61 |
| 1 | 83395322 | rs1144267 | -0.117 | 1.337 | -0.477 | 0.054 | -0.036 | -0.007 | -0.007 | 0.299 | 0.32 | 0.12 | 0.24 | 0.35 | 0.34 | 0.37 |
| 1 | 83405844 | rs7411897 | -0.202 | 1.317 | -0.496 | 0.044 | -0.009 | 0.008 | 0.008 | 0.436 | 0.32 | 0.22 | 0.25 | 0.35 | 0.34 | 0.37 |
| 1 | 83447063 | rs10782802 | -0.257 | 1.171 | -0.509 | 0.087 | -0.009 | 0.008 | 0.008 | 0.472 | 0.33 | 0.21 | 0.25 | 0.35 | 0.34 | 0.39 |
| 1 | 83448642 | rs10493734 | -0.264 | 1.169 | -0.496 | 0.086 | -0.004 | 0.008 | 0.008 | 0.487 | 0.33 | 0.28 | 0.26 | 0.35 | 0.35 | 0.39 |
| 1 | 83448667 | rs10874347 | 0.325 | 1.27 | -0.925 | -0.096 | -0.036 | 0.124 | 0.124 | 0.016 | 0.27 | 0.07 | 0.2 | 0.04 | 0.28 | 0.26 |
| 1 | 83451691 | rs2289445 | 0.106 | 0.964 | -1.103 | 0.489 | -0.036 | -0.012 | -0.012 | 0 | 0.14 | 0.02 | 0.2 | 0.1 | 0.18 | 0.14 |
| 1 | 83461181 | rs1440445 | -0.042 | 0.747 | -0.91 | 0.48 | 0.028 | 0.04 | 0.04 | 3.06 | 0.44 | 0.19 | 0.55 | 0.58 | 0.39 | 0.45 |
| 1 | 83465396 | rs6659184 | -0.84 | 1.057 | -0.717 | 1.48 | 0.005 | 0.05 | 0.05 | 0 | 0.14 | 0.25 | 0.13 | 0.3 | 0.16 | 0.19 |
| 1 | 83468851 | rs17109504 | -0.693 | 1.074 | -1.261 | 1.625 | -0.021 | -0.064 | -0.064 | 0 | 0.15 | 0.03 | 0.22 | 0.64 | 0.21 | 0.27 |
| 1 | 83469683 | rs1930276 | 0.583 | 1.015 | -0.761 | 1.949 | 0.017 | 0.137 | 0.137 | 0.174 | 0.94 | 0.83 | 0.81 | 0.73 | 0.93 | 0.92 |
| 1 | 83471955 | rs4450014 | -0.837 | 0.796 | -2.084 | 1.269 | -0.07 | 0.047 | 0.047 | 0.033 | 0.39 | 0.07 | 0.47 | 0.64 | 0.43 | 0.54 |
| 1 | 83474843 | rs2184006 | -0.933 | 0.625 | -2.028 | 0.759 | -0.004 | 0.166 | 0.166 | 0.008 | 0.5 | 0.22 | 0.53 | 0.65 | 0.5 | 0.55 |
| 1 | 83477213 | rs6664786 | -0.93 | 0.6 | -1.96 | 0.747 | -0.002 | 0.163 | 0.163 | 0 | 0.5 | 0.22 | 0.53 | 0.65 | 0.5 | 0.55 |
| 1 | 83479747 | rs12125199 | 0.989 | 0.838 | -2.389 | 0.098 | -0.022 | 0.08 | 0.08 | 0 | 0.2 | 0.05 | 0.08 | 0.06 | 0.15 | 0.1 |
| 1 | 83482870 | rs12732417 | -0.256 | 0.753 | -2.342 | 0.64 | -0.022 | -0.085 | -0.085 | 0 | 0.34 | 0.17 | 0.46 | 0.4 | 0.37 | 0.32 |
| 1 | 83483836 | rs17530979 | 1.183 | 1.164 | -2.53 | 0.317 | -0.033 | 0.032 | 0.032 | 0 | 0.08 | 0.03 | 0.12 | 0.01 | 0.21 | 0.07 |
| 1 | 83484494 | rs17531000 | 1.75 | 1.05 | -2.581 | 0.26 | 0 | 0.034 | 0.034 | 0.009 | 0.09 | 0.02 | 0.05 | 0 | 0.08 | 0.01 |
| 1 | 83487372 | rs6698372 | 0.223 | 0.627 | -2.006 | 0.01 | -0.009 | -0.046 | -0.046 | 0.968 | 0.42 | 0.16 | 0.52 | 0.4 | 0.43 | 0.56 |
| 1 | 83487728 | rs11802036 | 1.969 | 0.431 | -1.97 | -0.016 | 0.003 | -0.014 | -0.014 | 0 | 0.17 | 0.12 | 0.12 | 0 | 0.13 | 0.25 |
| 1 | 83489217 | rs10874351 | 0.666 | 0.614 | -2.104 | 0.034 | -0.058 | -0.155 | -0.155 | 0 | 0.23 | 0.03 | 0.39 | 0.39 | 0.27 | 0.3 |
| 1 | 83489466 | rs944726 | 0.918 | 0.531 | -2.146 | -0.057 | -0.044 | -0.023 | -0.023 | 0 | 0.35 | 0.08 | 0.19 | 0.31 | 0.29 | 0.33 |
| 1 | 83490161 | rs17465658 | 0.232 | 0.791 | -1.731 | 0.027 | -0.006 | 0.009 | 0.009 | 0.163 | 0.2 | 0.16 | 0.31 | 0.11 | 0.21 | 0.13 |
| 1 | 83490493 | rs17109599 | 0.641 | 0.783 | -1.637 | 0.036 | 0.032 | -0.005 | -0.005 | 0 | 0.34 | 0.16 | 0.28 | 0.29 | 0.28 | 0.44 |
| 1 | 83491590 | rs17531245 | 1.239 | 0.933 | -1.53 | -0.093 | -0.002 | -0.004 | -0.004 | 1.895 | 0.09 | 0.1 | 0.14 | 0.01 | 0.2 | 0.08 |
| 1 | 83491910 | rs1930272 | -0.273 | 0.741 | -1.666 | -0.178 | 0.003 | -0.009 | -0.009 | 1.336 | 0.49 | 0.19 | 0.56 | 0.4 | 0.48 | 0.57 |
| 1 | 83493060 | rs11163640 | -0.124 | 0.733 | -1.702 | -0.194 | -0.061 | -0.149 | -0.149 | 0.43 | 0.31 | 0.06 | 0.44 | 0.4 | 0.34 | 0.31 |
| 1 | 83500773 | rs7530348 | 2.113 | 0.724 | -1.644 | -0.206 | 0.006 | -0.002 | -0.002 | 0 | 0.17 | 0.1 | 0.12 | 0 | 0.13 | 0.26 |
| 1 | 83520031 | rs12132477 | 1.893 | 0.748 | -1.387 | -0.22 | -0.017 | 0.06 | 0.06 | 0 | 0.21 | 0.07 | 0.09 | 0.05 | 0.17 | 0.09 |
| 1 | 83526102 | rs1338500 | -0.11 | 0.677 | -1.424 | -0.194 | -0.062 | -0.143 | -0.143 | 0 | 0.3 | 0.06 | 0.44 | 0.41 | 0.33 | 0.31 |
| 1 | 83528240 | rs2022015 | 0.56 | 0.686 | -0.465 | 0.048 | 0.085 | -0.056 | -0.056 | 3.047 | 0.11 | 0.31 | 0.16 | 0.01 | 0.21 | 0.08 |
| 1 | 83535481 | rs323971 | -0.76 | 0.646 | -0.583 | 0.405 | -0.008 | 0.002 | 0.002 | 0 | 0.49 | 0.82 | 0.64 | 0.5 | 0.55 | 0.51 |
| 1 | 83535877 | rs323973 | -1.425 | 0.579 | -0.641 | 0.382 | 0.021 | -0.039 | -0.039 | 0 | 0.8 | 0.92 | 0.87 | 0.76 | 0.84 | 0.74 |
| 1 | 83541990 | rs17344386 | 1.195 | 0.648 | -0.556 | 0.356 | -0.002 | -0.005 | -0.005 | 0 | 0.09 | 0.1 | 0.13 | 0.01 | 0.2 | 0.08 |
| 1 | 83548341 | rs6676907 | 0.106 | 0.572 | -0.466 | 0.371 | 0.013 | 0.034 | 0.034 | 0.422 | 0.21 | 0.33 | 0.32 | 0.21 | 0.2 | 0.14 |
| 1 | 83552647 | rs2389394 | 1.673 | 0.515 | -0.511 | 0.361 | 0.004 | 0 | 0 | 0 | 0.2 | 0.16 | 0.18 | 0.28 | 0.14 | 0.28 |
| 1 | 83553657 | rs2022623 | -1.267 | 0.507 | -0.462 | 0.372 | -0.017 | 0.014 | 0.014 | 0 | 0.79 | 0.7 | 0.86 | 0.76 | 0.84 | 0.74 |
| 1 | 83562338 | rs12127333 | 1.37 | 0.579 | -0.522 | 0.347 | -0.038 | 0.003 | 0.003 | 0 | 0.3 | 0.1 | 0.23 | 0.25 | 0.3 | 0.23 |
| 1 | 83567501 | rs12405566 | 0.464 | 0.548 | -0.394 | 0.346 | -0.01 | -0.017 | -0.017 | 1.093 | 0.18 | 0.12 | 0.3 | 0.21 | 0.2 | 0.14 |
| 1 | 83570271 | rs984779 | 0.355 | 0.546 | -0.324 | 0.357 | 0.009 | -0.008 | -0.008 | 0 | 0.58 | 0.38 | 0.54 | 0.55 | 0.64 | 0.6 |
| 1 | 83572881 | rs537877 | 0.317 | 0.551 | -0.321 | 0.36 | 0.008 | -0.008 | -0.008 | 0.044 | 0.58 | 0.37 | 0.54 | 0.55 | 0.64 | 0.6 |

| | | | | | | | | | | | | | | | | |
|---|---|---|---|---|---|---|---|---|---|---|---|---|---|---|---|---|
| 1 | 83574925 | rs804068 | 0.585 | 0.57 | -0.364 | 0.345 | -0.065 | 0.048 | 0.048 | 0 | 0.38 | 0.21 | 0.36 | 0.27 | 0.5 | 0.32 |
| 1 | 83585592 | rs6682943 | -0.688 | 0.836 | 0.207 | 0.399 | -0.001 | -0.001 | -0.001 | 0 | 0.47 | 0.59 | 0.62 | 0.51 | 0.54 | 0.51 |
| 1 | 83586063 | rs6693586 | 1.063 | 0.897 | 0.159 | 0.413 | -0.001 | -0.005 | -0.005 | 0 | 0.08 | 0.1 | 0.13 | 0.01 | 0.2 | 0.08 |
| 1 | 83595041 | rs12754211 | 0.319 | 0.855 | -0.122 | 0.456 | -0.014 | -0.028 | -0.028 | 0 | 0.18 | 0.1 | 0.29 | 0.22 | 0.2 | 0.14 |
| 1 | 83596112 | rs10487982 | 0.178 | 0.853 | -0.083 | 0.455 | 0.01 | 0.029 | 0.029 | 0.547 | 0.18 | 0.28 | 0.31 | 0.22 | 0.2 | 0.14 |
| 1 | 83597301 | rs6664860 | -0.279 | 0.834 | -0.292 | 0.444 | -0.003 | 0.01 | 0.01 | 0 | 0.38 | 0.41 | 0.49 | 0.5 | 0.34 | 0.42 |
| 1 | 83597421 | rs17348539 | 1.536 | 0.575 | -0.664 | -0.015 | 0.001 | 0.037 | 0.037 | 0.177 | 0.1 | 0.02 | 0.05 | 0 | 0.09 | 0.01 |
| 1 | 83633428 | rs12039790 | -0.253 | 0.683 | -0.361 | 0.292 | 0.001 | 0.019 | 0.019 | 0.069 | 0.68 | 0.29 | 0.65 | 0.71 | 0.66 | 0.63 |
| 1 | 83701382 | rs11163695 | -0.048 | 0.385 | 0.419 | 0.75 | -0.007 | 0.002 | 0.002 | 0 | 0.33 | 0.42 | 0.19 | 0.35 | 0.27 | 0.28 |
| 1 | 83701728 | rs12403322 | 0.919 | 0.736 | -0.22 | 0.577 | -0.049 | 0.004 | 0.004 | 0 | 0.25 | 0.09 | 0.4 | 0.22 | 0.34 | 0.26 |
| 1 | 83712453 | rs12033542 | 0.297 | 0.867 | -0.372 | 0.851 | -0.022 | -0.046 | -0.046 | 0 | 0.08 | 0.01 | 0.15 | 0.2 | 0.1 | 0.16 |
| 1 | 83724335 | rs3904791 | 0.37 | 0.789 | -0.143 | 0.869 | 0.003 | -0.004 | -0.004 | 0.186 | 0.33 | 0.37 | 0.18 | 0.47 | 0.27 | 0.29 |
| 1 | 83772596 | rs1856149 | 0.767 | 0.84 | 0.229 | 0.945 | -0.01 | 0.013 | 0.013 | 0 | 0.29 | 0.37 | 0.16 | 0.44 | 0.21 | 0.22 |
| 1 | 83785316 | rs3861689 | 0.956 | 0.828 | 0.133 | 0.975 | 0.011 | -0.012 | -0.012 | 0.066 | 0.28 | 0.21 | 0.15 | 0.38 | 0.21 | 0.22 |
| 1 | 83789268 | rs11163718 | 1.302 | 0.903 | 0.405 | 0.355 | 0.02 | -0.101 | -0.101 | 0.482 | 0.14 | 0.43 | 0.09 | 0 | 0.12 | 0.02 |
| 1 | 83801769 | rs17131801 | 0.113 | 0.885 | 0.195 | 0.936 | -0.011 | -0.015 | -0.015 | 0.548 | 0.08 | 0.05 | 0.15 | 0.25 | 0.1 | 0.17 |
| 1 | 83808282 | rs1856150 | 1.022 | 0.863 | 0.478 | -0.055 | 0.008 | -0.03 | -0.03 | 0 | 0.2 | 0.32 | 0.18 | 0.17 | 0.18 | 0.28 |
| 1 | 83826710 | rs10782808 | 0.093 | 0.94 | -0.369 | -0.057 | -0.018 | 0.075 | 0.075 | 0 | 0.56 | 0.72 | 0.47 | 0.84 | 0.49 | 0.65 |
| 1 | 83834902 | rs12408136 | -0.274 | 0.905 | -0.831 | -0.04 | -0.009 | 0.067 | 0.067 | 0 | 0.42 | 0.27 | 0.49 | 0.21 | 0.48 | 0.31 |
| 1 | 83841946 | rs4403648 | -0.674 | 1.106 | -0.428 | 0.039 | -0.002 | 0.001 | 0.001 | 0 | 0.16 | 0.12 | 0.08 | 0.08 | 0.2 | 0.15 |
| 1 | 83847564 | rs12080397 | -0.095 | 1.065 | -0.151 | -0.066 | 0.02 | -0.028 | -0.028 | 0 | 0.19 | 0.34 | 0.16 | 0.15 | 0.19 | 0.15 |
| 1 | 83863396 | rs12139544 | 0.042 | 1.116 | -0.488 | -0.042 | 0.007 | -0.012 | -0.012 | 0 | 0.18 | 0.27 | 0.16 | 0.15 | 0.19 | 0.15 |
| 1 | 83897416 | rs12038205 | 0.415 | 0.278 | -0.827 | 0.589 | 0.003 | 0.003 | 0.003 | 0 | 0.1 | 0.05 | 0.22 | 0.46 | 0.08 | 0.28 |
| 1 | 83905111 | rs4907141 | -0.193 | -0.148 | -1.134 | -0.257 | 0.033 | 0.073 | 0.073 | 0.72 | 0.3 | 0.19 | 0.11 | 0.1 | 0.19 | 0.16 |
| 1 | 83905323 | rs35835790 | -0.337 | -0.323 | -0.717 | -0.32 | -0.027 | -0.059 | -0.059 | 0 | 0.15 | 0.18 | 0.07 | 0 | 0.07 | 0.11 |
| 1 | 83929243 | rs17130754 | -0.285 | -0.03 | -0.789 | -0.342 | -0.004 | -0.042 | -0.042 | 0.096 | 0.15 | 0.16 | 0.07 | 0 | 0.11 | 0.09 |
| 1 | 83933256 | rs3972749 | 0.708 | -0.166 | -0.985 | -0.212 | -0.004 | -0.044 | -0.044 | 0 | 0.67 | 0.81 | 0.66 | 0.53 | 0.68 | 0.59 |
| 1 | 83934440 | rs11163752 | 1.233 | 0.025 | -0.948 | -0.425 | -0.045 | 0.004 | 0.004 | 0 | 0.3 | 0.18 | 0.37 | 0.29 | 0.42 | 0.32 |
| 1 | 83938094 | rs10081983 | 1.345 | 0.32 | -1.01 | -0.485 | -0.027 | 0 | 0 | 0 | 0.27 | 0.16 | 0.29 | 0.29 | 0.35 | 0.3 |
| 1 | 83956770 | rs12731248 | 0.139 | -0.266 | -0.836 | -0.731 | 0.058 | 0.081 | 0.081 | 0 | 0.34 | 0.23 | 0.14 | 0.1 | 0.2 | 0.2 |
| 1 | 83958040 | rs12082801 | 1.241 | -0.247 | -0.438 | -0.717 | 0.037 | 0.071 | 0.071 | 0 | 0.12 | 0.32 | 0.2 | 0.15 | 0.17 | 0.12 |
| 1 | 83958331 | rs12723007 | 0.141 | -0.271 | -0.55 | -0.723 | 0.053 | 0.071 | 0.071 | 0 | 0.35 | 0.26 | 0.14 | 0.1 | 0.2 | 0.23 |
| 1 | 83959631 | rs17503362 | 1.9 | -0.143 | -0.767 | -0.635 | -0.103 | 0.005 | 0.005 | 0 | 0.26 | 0.04 | 0.28 | 0.24 | 0.34 | 0.26 |
| 1 | 83960539 | rs3850435 | -1.192 | -0.115 | -0.652 | -0.569 | -0.006 | -0.019 | -0.019 | 0 | 0.84 | 0.9 | 0.7 | 0.54 | 0.82 | 0.7 |
| 1 | 83961744 | rs7535867 | 0.988 | -0.42 | -0.731 | -0.766 | 0.031 | 0.04 | 0.04 | 0.008 | 0.25 | 0.19 | 0.08 | 0.09 | 0.15 | 0.11 |
| 1 | 83961895 | rs7528730 | 0.337 | -0.502 | -0.587 | -0.817 | 0.013 | 0 | 0 | 0 | 0.61 | 0.52 | 0.43 | 0.39 | 0.53 | 0.53 |
| 1 | 83964127 | rs6702689 | -0.239 | -0.645 | -0.37 | -1.107 | -0.026 | -0.018 | -0.018 | 0.283 | 0.35 | 0.52 | 0.27 | 0.25 | 0.29 | 0.23 |
| 1 | 83969716 | rs12076078 | -0.605 | -0.585 | -0.035 | -1.144 | -0.005 | -0.028 | -0.028 | 0 | 0.42 | 0.45 | 0.19 | 0.1 | 0.26 | 0.26 |
| 1 | 83971065 | rs6660757 | 0.42 | -0.639 | -0.123 | -1.173 | 0.019 | 0.043 | 0.043 | 0 | 0.54 | 0.34 | 0.67 | 0.74 | 0.61 | 0.68 |
| 1 | 83971236 | rs6670454 | -0.009 | -0.155 | -0.192 | -0.804 | -0.01 | -0.011 | -0.011 | 0 | 0.8 | 0.67 | 0.76 | 0.85 | 0.8 | 0.84 |
| 1 | 83977091 | rs4907142 | 0.109 | -0.195 | -0.161 | -0.646 | -0.003 | -0.03 | -0.03 | 0 | 0.85 | 0.77 | 0.92 | 0.97 | 0.89 | 0.89 |
| 1 | 83978214 | rs7537824 | 0.26 | 0.107 | -0.549 | -0.952 | -0.021 | 0.019 | 0.019 | 0 | 0.38 | 0.3 | 0.34 | 0.27 | 0.49 | 0.34 |
| 1 | 83978738 | rs12134784 | 1.026 | -0.102 | -0.755 | -0.859 | -0.029 | -0.069 | -0.069 | 0 | 0.12 | 0.01 | 0.11 | 0.2 | 0.15 | 0.12 |
| 1 | 83983252 | rs7553619 | -0.375 | -0.378 | -0.416 | -1.247 | 0.012 | 0.012 | 0.012 | 0 | 0.63 | 0.67 | 0.76 | 0.81 | 0.75 | 0.73 |
| 1 | 83987643 | rs12568728 | 0.337 | -0.055 | -0.418 | -1.11 | -0.002 | -0.001 | -0.001 | 0 | 0.28 | 0.24 | 0.46 | 0.74 | 0.32 | 0.44 |
| 1 | 83990906 | rs11163758 | 0.311 | 0.24 | -0.527 | -1.242 | 0.018 | -0.064 | -0.064 | 0 | 0.35 | 0.33 | 0.29 | 0.07 | 0.45 | 0.29 |
| 1 | 83991748 | rs12132108 | 0.066 | 0.095 | -0.52 | -0.902 | -0.003 | -0.005 | -0.005 | 0.023 | 0.14 | 0.13 | 0.12 | 0.13 | 0.16 | 0.12 |
| 1 | 83992961 | rs12133985 | 0.462 | 0.327 | -0.693 | -0.758 | -0.052 | -0.047 | -0.047 | 0.218 | 0.13 | 0.01 | 0.11 | 0.19 | 0.17 | 0.13 |
| 1 | 83993633 | rs17131049 | 0.204 | 0.088 | -0.391 | -0.839 | 0 | -0.033 | -0.033 | 0 | 0.1 | 0.09 | 0.03 | 0 | 0.07 | 0.08 |
| 1 | 84000353 | rs17131303 | 0.422 | -0.239 | -0.655 | -0.423 | 0.009 | -0.041 | -0.041 | 1.678 | 0.15 | 0.09 | 0.35 | 0.56 | 0.15 | 0.31 |
| 1 | 84002236 | rs4907144 | 0.249 | -0.02 | -0.58 | -1.375 | 0.001 | -0.031 | -0.031 | 0.157 | 0.44 | 0.53 | 0.38 | 0.2 | 0.52 | 0.4 |
| 1 | 84009549 | rs10499825 | 1.487 | 0.151 | -0.911 | -1.227 | -0.028 | -0.019 | -0.019 | 0.304 | 0.07 | 0.01 | 0.04 | 0.1 | 0.07 | 0.05 |
| 1 | 84012000 | rs977889 | -0.855 | -0.544 | -0.656 | -1.622 | -0.063 | -0.111 | -0.111 | 0.083 | 0.83 | 0.57 | 0.91 | 0.99 | 0.93 | 0.89 |
| 1 | 84016320 | rs1038148 | 1.576 | -0.346 | -1.613 | -1.825 | -0.089 | 0.019 | 0.019 | 0 | 0.3 | 0.12 | 0.27 | 0.22 | 0.4 | 0.3 |
| 1 | 84021429 | rs2291115 | 0.486 | -0.734 | -1.402 | -1.687 | -0.002 | -0.016 | -0.016 | 0 | 0.16 | 0.13 | 0.38 | 0.66 | 0.21 | 0.37 |
| 1 | 84023383 | rs4907149 | -0.288 | -0.854 | -1.33 | -1.841 | -0.022 | 0.039 | 0.039 | 2.088 | 0.36 | 0.23 | 0.53 | 0.75 | 0.41 | 0.46 |
| 1 | 84024341 | rs4142781 | 1.107 | -0.783 | -1.309 | -1.805 | -0.076 | 0.084 | 0.084 | 0 | 0.39 | 0.18 | 0.29 | 0.23 | 0.49 | 0.36 |
| 1 | 84025460 | rs12025304 | -0.65 | -0.623 | -1.053 | -1.694 | -0.001 | 0.05 | 0.05 | 0.115 | 0.46 | 0.36 | 0.62 | 0.77 | 0.47 | 0.54 |
| 1 | 84026447 | rs7518527 | 0.541 | -0.465 | -0.933 | -1.808 | -0.002 | 0.016 | 0.016 | 3.585 | 0.48 | 0.39 | 0.34 | 0.04 | 0.5 | 0.4 |
| 1 | 84027396 | rs12037871 | 1.337 | -0.346 | -1.157 | -1.269 | -0.014 | -0.039 | -0.039 | 0 | 0.09 | 0.02 | 0.34 | 0.53 | 0.12 | 0.33 |
| 1 | 84028076 | rs11585440 | 1.179 | -0.509 | -0.976 | -1.82 | -0.028 | 0.076 | 0.076 | 0.085 | 0.37 | 0.23 | 0.24 | 0.01 | 0.42 | 0.32 |
| 1 | 84028851 | rs7522657 | 1.868 | -0.302 | -0.697 | -1.373 | -0.014 | -0.032 | -0.032 | 0.908 | 0.11 | 0.16 | 0.1 | 0.02 | 0.07 | 0.07 |
| 1 | 84030634 | rs12035630 | 0.446 | -0.415 | -0.618 | -1.318 | 0.002 | -0.006 | -0.006 | 0 | 0.14 | 0.13 | 0.37 | 0.68 | 0.15 | 0.4 |
| 1 | 84032486 | rs4465227 | 0.982 | -0.351 | -0.221 | -1.347 | 0 | -0.014 | -0.014 | 0 | 0.14 | 0.17 | 0.11 | 0.03 | 0.11 | 0.12 |
| 1 | 84032625 | rs4243779 | -0.84 | -0.535 | 0.285 | -1.666 | -0.003 | 0.009 | 0.009 | 1.239 | 0.42 | 0.62 | 0.36 | 0.31 | 0.42 | 0.25 |
| 1 | 84034548 | rs6704084 | 0.447 | -0.387 | 0.388 | -1.67 | 0.055 | 0.112 | 0.112 | 1.01 | 0.15 | 0.37 | 0.15 | 0.27 | 0.16 | 0.1 |
| 1 | 84038198 | rs12033001 | 0.794 | -0.341 | 0.617 | -1.613 | 0.037 | 0.07 | 0.07 | 0.171 | 0.14 | 0.26 | 0.14 | 0.27 | 0.16 | 0.1 |
| 1 | 84039480 | rs12037319 | 1.011 | -0.32 | 0.372 | -1.603 | 0.004 | 0.011 | 0.011 | 0 | 0.13 | 0.15 | 0.13 | 0.27 | 0.16 | 0.1 |
| 1 | 84040528 | rs4243780 | -0.957 | -0.473 | -0.046 | -1.689 | 0.004 | 0.041 | 0.041 | 0 | 0.49 | 0.61 | 0.37 | 0.3 | 0.48 | 0.3 |

| | | | | | | | | | | | | | | | | |
|---|---|---|---|---|---|---|---|---|---|---|---|---|---|---|---|---|
| 1 | 84057173 | rs6661191 | -0.248 | -1.182 | 0.086 | -2.162 | 0.003 | -0.039 | -0.039 | 0 | 0.73 | 0.71 | 0.85 | 0.99 | 0.83 | 0.86 |
| 1 | 84058458 | rs12738873 | 1.929 | -0.873 | -0.508 | -2.523 | -0.078 | 0.156 | 0.156 | 0.076 | 0.32 | 0.09 | 0.22 | 0.05 | 0.38 | 0.3 |
| 1 | 84061515 | rs4370778 | -0.824 | -1.026 | -0.354 | -2.53 | 0.021 | -0.001 | -0.001 | 0.343 | 0.74 | 0.81 | 0.86 | 1 | 0.83 | 0.91 |
| 1 | 84062434 | rs4128486 | 1.69 | -0.748 | -0.224 | -2.491 | -0.026 | 0.034 | 0.034 | 0 | 0.32 | 0.19 | 0.22 | 0.01 | 0.38 | 0.28 |
| 1 | 84062884 | rs4907154 | 0.231 | -0.992 | -0.189 | -2.51 | 0.039 | 0.044 | 0.044 | 1.515 | 0.57 | 0.36 | 0.72 | 0.84 | 0.74 | 0.74 |
| 1 | 84062998 | rs11808968 | 1.756 | -0.811 | -0.515 | -2.501 | -0.08 | 0.16 | 0.16 | 0.093 | 0.32 | 0.09 | 0.22 | 0.05 | 0.38 | 0.3 |
| 1 | 84076139 | rs6673005 | -1.738 | -1.467 | -0.198 | -2.043 | -0.035 | 0.022 | 0.022 | 10.872 | 0.88 | 0.73 | 0.92 | 0.84 | 0.94 | 0.86 |
| 1 | 84080290 | rs4907156 | -0.845 | -1.596 | -0.118 | -2.106 | -0.067 | 0.004 | 0.004 | 6.154 | 0.84 | 0.63 | 0.91 | 0.84 | 0.94 | 0.8 |
| 1 | 84084512 | rs12410537 | 1.931 | -1.44 | -0.68 | -2.289 | -0.109 | 0.168 | 0.168 | 0 | 0.31 | 0.07 | 0.21 | 0.04 | 0.39 | 0.25 |
| 1 | 84091642 | rs12145760 | 1.587 | -1.276 | -0.51 | -2.134 | 0.002 | 0.048 | 0.048 | 0 | 0.15 | 0.06 | 0.11 | 0 | 0.15 | 0.02 |
| 1 | 84093357 | rs4347229 | -1.796 | -1.35 | -0.187 | -2.126 | -0.037 | 0.037 | 0.037 | 10.964 | 0.87 | 0.63 | 0.91 | 0.84 | 0.94 | 0.86 |
| 1 | 84094411 | rs12563303 | -0.652 | -1.335 | -0.56 | -1.946 | -0.02 | 0.041 | 0.041 | 0.273 | 0.4 | 0.32 | 0.54 | 0.75 | 0.5 | 0.44 |
| 1 | 84101123 | rs11163792 | 0.285 | -1.166 | -0.588 | -1.847 | -0.009 | -0.002 | -0.002 | 0.167 | 0.24 | 0.26 | 0.43 | 0.75 | 0.34 | 0.41 |
| 1 | 84101280 | rs4631716 | 1.389 | -1.324 | -0.849 | -1.95 | -0.044 | 0.166 | 0.166 | 0.034 | 0.44 | 0.2 | 0.36 | 0.09 | 0.44 | 0.37 |
| 1 | 84103977 | rs4907162 | 1.401 | -0.871 | -0.988 | -1.441 | -0.068 | -0.015 | -0.015 | 0 | 0.13 | 0.02 | 0.11 | 0.11 | 0.19 | 0.06 |
| 1 | 84105145 | rs7419078 | 1.421 | -0.847 | -0.89 | -1.358 | -0.031 | -0.058 | -0.058 | 0 | 0.1 | 0.02 | 0.3 | 0.63 | 0.15 | 0.32 |
| 1 | 84107543 | rs8179363 | 0.273 | -0.912 | -0.654 | -1.585 | -0.009 | -0.002 | -0.002 | 1.301 | 0.24 | 0.24 | 0.34 | 0.75 | 0.34 | 0.41 |
| 1 | 84109056 | rs12132624 | 1.633 | -1.052 | -0.684 | -1.696 | 0.003 | -0.006 | -0.006 | 0 | 0.14 | 0.07 | 0.1 | 0 | 0.14 | 0.02 |
| 1 | 84109270 | rs10128052 | -0.591 | -1.316 | -0.434 | -1.805 | -0.037 | -0.04 | -0.04 | 0 | 0.74 | 0.67 | 0.84 | 0.84 | 0.82 | 0.84 |
| 1 | 84111016 | rs11163794 | -0.348 | -1.272 | -0.757 | -1.827 | -0.022 | 0.039 | 0.039 | 0.343 | 0.38 | 0.28 | 0.43 | 0.75 | 0.48 | 0.43 |
| 1 | 84112571 | rs12408481 | 2.222 | -1.468 | -1.163 | -2.04 | -0.16 | 0.172 | 0.172 | 0 | 0.28 | 0.02 | 0.2 | 0.04 | 0.37 | 0.24 |
| 1 | 84113159 | rs4539154 | -1.197 | -1.437 | -0.918 | -1.974 | -0.009 | 0.038 | 0.038 | 0.04 | 0.57 | 0.67 | 0.72 | 0.81 | 0.57 | 0.62 |
| 1 | 84116142 | rs10782813 | -1.994 | -1.71 | -0.33 | -1.386 | -0.037 | 0.021 | 0.021 | 10.86 | 0.9 | 0.81 | 0.94 | 0.84 | 0.96 | 0.87 |
| 1 | 84117891 | rs11163798 | 1.022 | -1.841 | -0.34 | -1.494 | 0.045 | 0.044 | 0.044 | 0.744 | 0.56 | 0.38 | 0.54 | 0.77 | 0.73 | 0.66 |
| 1 | 84118933 | rs12145179 | 1.331 | -1.39 | -0.535 | -1.469 | -0.001 | 0.056 | 0.056 | 0 | 0.14 | 0.04 | 0.09 | 0 | 0.14 | 0.02 |
| 1 | 84119882 | rs12130451 | 0.151 | -1.404 | -0.444 | -1.476 | -0.013 | -0.103 | -0.103 | 0 | 0.19 | 0.28 | 0.12 | 0 | 0.16 | 0.09 |
| 1 | 84120354 | rs10158671 | -0.743 | -1.335 | -0.497 | -1.428 | -0.036 | -0.084 | -0.084 | 0.715 | 0.33 | 0.44 | 0.4 | 0.08 | 0.23 | 0.21 |
| 1 | 84121660 | rs4907172 | -0.54 | -1.063 | -0.455 | -1.373 | -0.023 | -0.086 | -0.086 | 0 | 0.34 | 0.44 | 0.43 | 0.07 | 0.26 | 0.15 |
| 1 | 84121781 | rs12141980 | 0.812 | -0.87 | -0.79 | -1.02 | 0.009 | 0.01 | 0.01 | 0 | 0.17 | 0.16 | 0.31 | 0.07 | 0.1 | 0.12 |
| 1 | 84123079 | rs10782814 | 1.37 | -0.599 | -1.218 | -0.572 | -0.104 | -0.121 | -0.121 | 9.897 | 0.37 | 0.09 | 0.42 | 0.53 | 0.48 | 0.5 |
| 1 | 84124122 | rs11590583 | 0.751 | -0.693 | -1.226 | -0.602 | 0.001 | -0.036 | -0.036 | 0 | 0.47 | 0.23 | 0.57 | 0.51 | 0.59 | 0.52 |
| 1 | 84126330 | rs4907175 | 0.092 | -1.148 | -0.918 | -0.608 | 0.009 | 0.002 | 0.002 | 0.957 | 0.67 | 0.74 | 0.84 | 0.68 | 0.8 | 0.76 |
| 1 | 84127328 | rs10874404 | 1.46 | -0.564 | -1.136 | -0.616 | -0.111 | -0.145 | -0.145 | 0 | 0.36 | 0.09 | 0.48 | 0.51 | 0.47 | 0.5 |
| 1 | 84130384 | rs12087224 | 1.084 | -0.719 | -0.553 | -0.687 | -0.017 | -0.052 | -0.052 | 0 | 0.17 | 0.33 | 0.09 | 0.06 | 0.09 | 0.09 |
| 1 | 84131789 | rs4243781 | -0.961 | -0.759 | -0.165 | -0.716 | -0.024 | -0.082 | -0.082 | 0 | 0.81 | 0.54 | 0.9 | 0.94 | 0.89 | 0.91 |
| 1 | 84134042 | rs11485345 | -0.201 | -0.822 | -0.166 | -0.705 | 0 | 0 | 0 | 0 | 0.28 | 0.25 | 0.18 | 0.38 | 0.34 | 0.29 |
| 1 | 84140500 | rs10747405 | -1.208 | -0.507 | -0.366 | -0.406 | -0.018 | -0.053 | -0.053 | 0 | 0.83 | 0.67 | 0.91 | 0.94 | 0.91 | 0.91 |
| 1 | 84146671 | rs12098061 | 1.538 | 0.019 | -0.874 | -0.014 | -0.023 | 0.045 | 0.045 | 0.036 | 0.08 | 0.01 | 0.06 | 0 | 0.09 | 0.01 |
| 1 | 84149799 | rs11163801 | 0.601 | 0.128 | -0.766 | 0.388 | -0.094 | -0.146 | -0.146 | 0.02 | 0.35 | 0.09 | 0.47 | 0.51 | 0.44 | 0.47 |
| 1 | 84161566 | rs6679802 | -0.689 | 0.23 | -0.722 | 0.421 | -0.019 | -0.015 | -0.015 | 0 | 0.66 | 0.75 | 0.55 | 0.61 | 0.56 | 0.53 |
| 1 | 84163667 | rs4287202 | 0.339 | -0.121 | -0.586 | 0.249 | 0.004 | 0.012 | 0.012 | 0.265 | 0.13 | 0.09 | 0.21 | 0.05 | 0.11 | 0.17 |
| 1 | 84168682 | rs11163805 | -0.627 | -0.417 | -0.253 | 0.439 | 0.009 | 0.008 | 0.008 | 0 | 0.39 | 0.36 | 0.3 | 0.27 | 0.25 | 0.38 |
| 1 | 84172192 | rs12031401 | 0.214 | -0.723 | 0.693 | 0.241 | -0.042 | -0.009 | -0.009 | 0 | 0.25 | 0.47 | 0.12 | 0.2 | 0.14 | 0.2 |
| 1 | 84172930 | rs7414950 | 1.522 | -0.206 | 0.02 | -0.072 | -0.006 | -0.005 | -0.005 | 0 | 0.16 | 0.15 | 0.15 | 0.16 | 0.21 | 0.18 |
| 1 | 84173558 | rs6667561 | -0.594 | -0.23 | -0.185 | 0 | -0.032 | 0 | 0 | 0 | 0.55 | 0.75 | 0.5 | 0.46 | 0.46 | 0.51 |
| 1 | 84181060 | rs6689132 | -1.25 | -0.26 | -0.167 | 0.023 | 0.011 | -0.042 | -0.042 | 0 | 0.69 | 0.8 | 0.65 | 0.55 | 0.66 | 0.61 |
| 1 | 84184059 | rs4243784 | -1.656 | -0.281 | -0.201 | 0.009 | -0.01 | -0.022 | -0.022 | 0 | 0.71 | 0.91 | 0.66 | 0.71 | 0.67 | 0.69 |
| 1 | 84187007 | rs9433610 | 0.553 | -0.307 | 0.098 | -0.264 | -0.006 | 0.014 | 0.014 | 0 | 0.29 | 0.28 | 0.35 | 0.07 | 0.3 | 0.29 |
| 1 | 84190263 | rs4636482 | 1.364 | -0.388 | 0.52 | -0.289 | 0 | -0.026 | -0.026 | 0 | 0.22 | 0.24 | 0.11 | 0.05 | 0.15 | 0.11 |
| 1 | 84196920 | rs9787364 | 0.677 | -0.495 | 0.368 | -0.053 | -0.036 | 0.029 | 0.029 | 0.336 | 0.26 | 0.44 | 0.22 | 0.38 | 0.17 | 0.21 |
| 1 | 84197066 | rs9787335 | 0.042 | -0.488 | 0.348 | -0.066 | -0.014 | 0.073 | 0.073 | 0 | 0.58 | 0.69 | 0.57 | 0.81 | 0.51 | 0.59 |
| 1 | 84201658 | rs10782817 | 1.267 | -0.434 | 0.117 | -0.014 | 0.001 | -0.007 | -0.007 | 0 | 0.31 | 0.27 | 0.35 | 0.44 | 0.34 | 0.38 |
| 1 | 84204198 | rs17577039 | 1.485 | -0.367 | -0.176 | -0.022 | -0.061 | -0.023 | -0.023 | 0.628 | 0.12 | 0.02 | 0.07 | 0.11 | 0.19 | 0.12 |
| 1 | 84216308 | rs6691243 | -0.236 | -0.434 | 0.004 | -0.052 | -0.001 | -0.005 | -0.005 | 0 | 0.4 | 0.62 | 0.3 | 0.49 | 0.36 | 0.35 |
| 1 | 84217159 | rs761401 | -0.693 | -0.379 | -0.204 | -0.159 | -0.014 | 0.068 | 0.068 | 0.034 | 0.7 | 0.82 | 0.65 | 0.93 | 0.69 | 0.73 |
| 1 | 84220313 | rs6576941 | -0.629 | -0.308 | -0.227 | -0.354 | -0.01 | 0.107 | 0.107 | 0.074 | 0.65 | 0.82 | 0.74 | 0.97 | 0.67 | 0.72 |
| 1 | 84220731 | rs10874417 | -1.502 | -0.326 | -0.285 | -0.333 | 0.029 | 0.056 | 0.056 | 0 | 0.79 | 0.91 | 0.89 | 0.98 | 0.87 | 0.83 |
| 1 | 84222134 | rs7547085 | 1.788 | -0.122 | -0.276 | -0.28 | -0.016 | 0.016 | 0.016 | 0 | 0.15 | 0.11 | 0.16 | 0 | 0.21 | 0.11 |
| 1 | 84222204 | rs7518150 | 2.022 | -0.09 | -0.354 | -0.252 | -0.033 | 0.041 | 0.041 | 0 | 0.14 | 0.07 | 0.15 | 0 | 0.2 | 0.1 |
| 1 | 84223416 | rs6682282 | -0.991 | -0.694 | -0.121 | 0.107 | 0.014 | -0.017 | -0.017 | 0 | 0.77 | 0.84 | 0.69 | 0.6 | 0.85 | 0.73 |
| 1 | 84225532 | rs7513510 | 0.672 | -0.322 | 0.002 | -0.158 | -0.005 | -0.066 | -0.066 | 0.148 | 0.17 | 0.29 | 0.09 | 0.05 | 0.12 | 0.11 |
| 1 | 84232835 | rs4285721 | 0.94 | -0.289 | -0.62 | 0.028 | -0.004 | -0.015 | -0.015 | 0.126 | 0.3 | 0.13 | 0.36 | 0.29 | 0.32 | 0.32 |
| 1 | 84234973 | rs17129321 | 1.177 | -0.244 | -0.641 | -0.144 | -0.014 | -0.037 | -0.037 | 0 | 0.16 | 0.04 | 0.06 | 0.18 | 0.13 | 0.18 |
| 1 | 84235072 | rs12093748 | 0.418 | -0.325 | 0.263 | -0.202 | -0.014 | 0.005 | 0.005 | 0.911 | 0.2 | 0.41 | 0.1 | 0.18 | 0.14 | 0.21 |
| 1 | 84236405 | rs7524397 | 0.905 | -0.384 | 0.074 | -0.357 | -0.09 | 0.067 | 0.067 | 0 | 0.24 | 0.09 | 0.22 | 0.1 | 0.38 | 0.2 |
| 1 | 84236661 | rs10489502 | 0.5 | -0.351 | 0.278 | -0.248 | -0.019 | 0.002 | 0.002 | 1.063 | 0.2 | 0.43 | 0.11 | 0.18 | 0.14 | 0.21 |
| 1 | 84247430 | rs6576943 | -0.569 | -0.588 | 0.107 | -0.354 | 0.028 | 0.079 | 0.079 | 0 | 0.48 | 0.72 | 0.37 | 0.28 | 0.54 | 0.44 |
| 1 | 84254735 | rs7539409 | -1.863 | -0.76 | -0.297 | -0.51 | -0.012 | 0.089 | 0.089 | 0 | 0.8 | 0.99 | 0.91 | 0.95 | 0.86 | 0.86 |
| 1 | 84258609 | rs6699405 | -0.454 | -0.688 | -0.097 | -0.271 | 0.029 | 0.065 | 0.065 | 0 | 0.47 | 0.8 | 0.37 | 0.31 | 0.54 | 0.44 |

| | | | | | | | | | | | | | | | | |
|---|---|---|---|---|---|---|---|---|---|---|---|---|---|---|---|---|
| 1 | 84259660 | rs17520771 | 1.25 | -0.189 | -0.138 | -0.346 | 0 | 0 | 0 | 0 | 0.11 | 0.14 | 0.09 | 0.12 | 0.14 | 0.1 |
| 1 | 84260360 | rs12048434 | 0.408 | -0.237 | 0.106 | -0.381 | -0.051 | 0.045 | 0.045 | 0 | 0.33 | 0.17 | 0.23 | 0.19 | 0.39 | 0.31 |
| 1 | 84266117 | rs742779 | 0.4 | -0.233 | 0.016 | -0.384 | -0.061 | 0.05 | 0.05 | 0 | 0.33 | 0.16 | 0.24 | 0.19 | 0.39 | 0.32 |
| 1 | 84269299 | rs4140462 | 0.719 | -0.34 | 0.332 | -0.397 | -0.022 | 0.012 | 0.012 | 0 | 0.34 | 0.48 | 0.56 | 0.62 | 0.32 | 0.44 |
| 1 | 84272649 | rs12735437 | 1.523 | -0.293 | 0.172 | -0.388 | -0.027 | -0.022 | -0.022 | 0 | 0.1 | 0.05 | 0.07 | 0.12 | 0.14 | 0.1 |
| 1 | 84273341 | rs6576949 | 0.83 | -0.397 | 0.342 | -0.433 | 0.002 | -0.002 | -0.002 | 0 | 0.33 | 0.29 | 0.54 | 0.62 | 0.32 | 0.45 |
| 1 | 84274266 | rs1040576 | 0.173 | -0.094 | 0.011 | -0.304 | -0.027 | 0.031 | 0.031 | 0 | 0.34 | 0.28 | 0.26 | 0.19 | 0.39 | 0.32 |
| 1 | 84276046 | rs11802417 | 0.299 | 0.044 | -0.012 | -0.383 | -0.015 | 0.013 | 0.013 | 0 | 0.28 | 0.27 | 0.26 | 0.12 | 0.39 | 0.2 |
| 1 | 84279470 | rs11163847 | 1.004 | -0.014 | 0.098 | -0.277 | -0.002 | 0.001 | 0.001 | 0.644 | 0.18 | 0.29 | 0.09 | 0.18 | 0.14 | 0.22 |
| 1 | 84287873 | rs12083999 | 0.148 | 0.002 | -0.037 | -0.306 | -0.026 | -0.012 | -0.012 | 0 | 0.27 | 0.17 | 0.14 | 0.28 | 0.28 | 0.32 |
| 1 | 84289764 | rs3850438 | 0.161 | 0.002 | -0.038 | -0.306 | -0.036 | -0.02 | -0.02 | 0 | 0.27 | 0.15 | 0.14 | 0.28 | 0.28 | 0.32 |
| 1 | 84289948 | rs4628522 | 0.148 | 0.002 | -0.038 | -0.306 | -0.026 | -0.012 | -0.012 | 1.869 | 0.27 | 0.17 | 0.14 | 0.28 | 0.28 | 0.32 |
| 1 | 84293113 | rs12135529 | 1.309 | -0.223 | 0.008 | -0.373 | 0.007 | -0.037 | -0.037 | 0 | 0.28 | 0.13 | 0.53 | 0.61 | 0.32 | 0.42 |
| 1 | 84295861 | rs2179763 | -0.129 | -0.277 | -0.026 | -0.073 | 0.004 | 0.005 | 0.005 | 0 | 0.41 | 0.54 | 0.38 | 0.51 | 0.42 | 0.49 |
| 1 | 84296540 | rs970457 | -1.711 | -0.173 | 0.037 | -0.101 | -0.003 | -0.012 | -0.012 | 0 | 0.76 | 0.75 | 0.88 | 0.9 | 0.84 | 0.86 |
| 1 | 84296983 | rs3738575 | 1.118 | -0.092 | 0.032 | -0.041 | 0.038 | 0.079 | 0.079 | 0 | 0.2 | 0.35 | 0.29 | 0.33 | 0.24 | 0.23 |
| 1 | 84303037 | rs12033859 | 1.636 | -0.094 | -0.277 | -0.049 | -0.023 | -0.034 | -0.034 | 0 | 0.16 | 0.06 | 0.07 | 0.18 | 0.14 | 0.22 |
| 1 | 84304441 | rs1408477 | 0.088 | -0.135 | -0.201 | -0.052 | 0.001 | 0.015 | 0.015 | 0 | 0.48 | 0.53 | 0.73 | 0.61 | 0.55 | 0.56 |
| 1 | 84304638 | rs17129909 | 1.579 | -0.146 | -0.745 | -0.117 | -0.023 | -0.034 | -0.034 | 0 | 0.16 | 0.05 | 0.07 | 0.18 | 0.14 | 0.22 |
| 1 | 84310193 | rs12134669 | 1.381 | -0.053 | -0.654 | -0.04 | -0.005 | -0.012 | -0.012 | 0 | 0.27 | 0.12 | 0.44 | 0.29 | 0.31 | 0.31 |
| 1 | 84313764 | rs4907182 | -0.461 | 0.174 | -0.416 | 0.046 | 0.023 | 0.036 | 0.036 | 0 | 0.33 | 0.5 | 0.37 | 0.43 | 0.38 | 0.36 |
| 1 | 84319949 | rs2092596 | -0.708 | -0.042 | -0.874 | -0.366 | 0.024 | 0.074 | 0.074 | 0 | 0.71 | 0.87 | 0.87 | 0.9 | 0.8 | 0.87 |
| 1 | 84323701 | rs6576954 | 0.662 | 0.231 | -1.111 | -0.205 | -0.08 | -0.013 | -0.013 | 0 | 0.3 | 0.15 | 0.56 | 0.31 | 0.45 | 0.31 |
| 1 | 84332235 | rs726607 | -1.247 | 0.015 | -1.052 | -0.407 | 0.031 | 0.089 | 0.089 | 0 | 0.66 | 0.88 | 0.85 | 0.89 | 0.75 | 0.73 |
| 1 | 84370393 | rs1570691 | 0.089 | 0.287 | -0.656 | -0.521 | -0.023 | 0.026 | 0.026 | 0 | 0.38 | 0.25 | 0.52 | 0.28 | 0.55 | 0.36 |
| 1 | 84384825 | rs1324486 | 0.066 | 0.293 | -0.579 | -0.528 | -0.023 | 0.026 | 0.026 | 0 | 0.39 | 0.28 | 0.52 | 0.28 | 0.55 | 0.36 |
| 1 | 84416408 | rs12757632 | 0.347 | 0.339 | -0.712 | -0.473 | -0.071 | 0.033 | 0.033 | 0 | 0.34 | 0.19 | 0.49 | 0.27 | 0.52 | 0.34 |
| 1 | 84427052 | rs1536166 | 1.008 | 0.978 | -0.961 | -0.643 | -0.089 | -0.087 | -0.087 | 0 | 0.19 | 0.02 | 0.34 | 0.28 | 0.33 | 0.28 |
| 1 | 84434425 | rs12759527 | 0.133 | 0.942 | -0.918 | -0.688 | -0.088 | 0.019 | 0.019 | 0 | 0.34 | 0.14 | 0.49 | 0.28 | 0.52 | 0.34 |
| 1 | 84437311 | rs663136 | 0.192 | 0.623 | -0.561 | -0.842 | -0.028 | 0.087 | 0.087 | 0.909 | 0.25 | 0.5 | 0.31 | 0.62 | 0.18 | 0.37 |
| 1 | 84444766 | rs12042443 | 1.86 | 0.444 | -0.284 | -0.392 | -0.011 | 0.222 | 0.222 | 0 | 0.07 | 0.16 | 0.23 | 0.37 | 0.05 | 0.22 |
| 1 | 84451820 | rs10874426 | -0.207 | 0.512 | -0.735 | -0.524 | -0.007 | 0.007 | 0.007 | 0 | 0.42 | 0.34 | 0.53 | 0.43 | 0.57 | 0.43 |
| 1 | 84476650 | rs639666 | 0.088 | 0.754 | -0.651 | -0.827 | -0.019 | 0.03 | 0.03 | 0.058 | 0.32 | 0.48 | 0.37 | 0.61 | 0.28 | 0.39 |
| 1 | 84476779 | rs12403631 | 1.779 | 1.008 | -1.158 | -0.656 | -0.061 | -0.075 | -0.075 | 0.11 | 0.12 | 0.02 | 0.28 | 0.28 | 0.24 | 0.26 |
| 1 | 84480212 | rs12401314 | 0.805 | 0.547 | -1.059 | -0.608 | -0.101 | -0.033 | -0.033 | 0 | 0.26 | 0.07 | 0.42 | 0.28 | 0.42 | 0.32 |
| 1 | 84485512 | rs680905 | -0.994 | -0.366 | -0.842 | -0.722 | 0.039 | 0.113 | 0.113 | 0 | 0.65 | 0.87 | 0.79 | 0.9 | 0.76 | 0.73 |
| 1 | 84499003 | rs615352 | -0.978 | -0.391 | -0.77 | -0.776 | 0.029 | 0.062 | 0.062 | 0 | 0.64 | 0.8 | 0.78 | 0.9 | 0.75 | 0.72 |
| 1 | 84514988 | rs670564 | -1.027 | -0.385 | -0.869 | -0.762 | 0.035 | 0.089 | 0.089 | 0 | 0.64 | 0.83 | 0.78 | 0.9 | 0.75 | 0.72 |
| 1 | 84517261 | rs12119275 | 0.155 | 0.327 | -0.798 | -0.648 | -0.093 | -0.011 | -0.011 | 0 | 0.29 | 0.11 | 0.38 | 0.3 | 0.47 | 0.32 |
| 1 | 84528434 | rs3903924 | 0.116 | 0.38 | -0.711 | -0.647 | -0.06 | 0 | 0 | 1.908 | 0.29 | 0.15 | 0.39 | 0.3 | 0.49 | 0.32 |
| 1 | 84531117 | rs3861699 | 1.418 | 0.307 | -0.743 | -0.593 | -0.022 | -0.029 | -0.029 | 0 | 0.1 | 0.04 | 0.16 | 0.29 | 0.23 | 0.22 |
| 1 | 84539714 | rs11163897 | -0.14 | 0.011 | -0.851 | -0.958 | 0.015 | 0.018 | 0.018 | 0 | 0.47 | 0.39 | 0.52 | 0.4 | 0.63 | 0.5 |
| 1 | 84548284 | rs12563929 | 1.181 | -0.267 | -0.327 | -0.775 | -0.02 | -0.019 | -0.019 | 0 | 0.1 | 0 | 0.02 | 0.1 | 0.08 | 0.06 |
| 1 | 84550849 | rs6701486 | -0.91 | -0.475 | -0.019 | -0.913 | -0.031 | 0.042 | 0.042 | 0 | 0.67 | 0.5 | 0.56 | 0.5 | 0.76 | 0.61 |
| 1 | 84555908 | rs6703653 | 0.018 | -0.452 | -0.025 | -0.9 | 0.012 | 0.008 | 0.008 | 0 | 0.55 | 0.44 | 0.54 | 0.39 | 0.68 | 0.53 |
| 1 | 84560010 | rs10874428 | -0.447 | -0.438 | 0.05 | -0.885 | -0.005 | 0.01 | 0.01 | 0 | 0.44 | 0.4 | 0.38 | 0.39 | 0.53 | 0.36 |
| 1 | 84576203 | rs2812452 | 1.314 | -0.483 | 0.006 | -0.735 | -0.003 | 0.007 | 0.007 | 0 | 0.25 | 0.14 | 0.23 | 0.17 | 0.23 | 0.28 |
| 1 | 84580593 | rs1603000 | -1.044 | -0.497 | 0.117 | -0.741 | -0.028 | 0.038 | 0.038 | 0 | 0.68 | 0.56 | 0.61 | 0.56 | 0.77 | 0.64 |
| 1 | 84584894 | rs12057561 | -0.34 | -0.443 | 0.02 | -0.751 | -0.012 | 0.018 | 0.018 | 0 | 0.43 | 0.36 | 0.38 | 0.38 | 0.53 | 0.37 |
| 1 | 84590129 | rs1016379 | 1.503 | -0.418 | -0.033 | -0.746 | -0.011 | 0.039 | 0.039 | 0 | 0.23 | 0.11 | 0.18 | 0.11 | 0.23 | 0.24 |
| 1 | 84590315 | rs12723299 | -0.402 | -0.48 | 0.061 | -0.77 | -0.004 | 0.008 | 0.008 | 0 | 0.44 | 0.42 | 0.38 | 0.39 | 0.54 | 0.37 |
| 1 | 84590527 | rs2139931 | 1.543 | -0.453 | 0.011 | -0.758 | -0.013 | 0.042 | 0.042 | 0 | 0.23 | 0.1 | 0.18 | 0.11 | 0.23 | 0.24 |
| 1 | 84604937 | rs6576962 | -0.279 | -0.388 | 0.028 | -0.722 | -0.022 | 0.023 | 0.023 | 0 | 0.42 | 0.31 | 0.38 | 0.38 | 0.53 | 0.35 |
| 1 | 84613259 | rs12129674 | 0.457 | -0.295 | -0.088 | -0.699 | -0.002 | 0.011 | 0.011 | 0.041 | 0.15 | 0.09 | 0.07 | 0.1 | 0.13 | 0.07 |
| 1 | 84622513 | rs903263 | -0.005 | -0.319 | 0.22 | -0.74 | -0.003 | 0.024 | 0.024 | 0 | 0.45 | 0.37 | 0.38 | 0.38 | 0.54 | 0.37 |
| 1 | 84640834 | rs12132002 | 0.67 | -0.202 | -0.009 | -0.743 | -0.032 | -0.039 | -0.039 | 0 | 0.29 | 0.11 | 0.24 | 0.38 | 0.37 | 0.3 |
| 1 | 84658591 | rs7547892 | 0.883 | -0.774 | 0.698 | -0.04 | -0.03 | 0.03 | 0.03 | 0 | 0.43 | 0.26 | 0.36 | 0.38 | 0.5 | 0.33 |
| 1 | 84659129 | rs6661411 | 0.924 | -0.74 | 0.671 | -0.005 | 0.011 | -0.025 | -0.025 | 3.375 | 0.1 | 0.13 | 0.12 | 0 | 0.16 | 0.05 |
| 1 | 84667452 | rs603939 | 0.167 | -1.079 | 0.705 | 0.351 | -0.026 | 0.038 | 0.038 | 0 | 0.31 | 0.43 | 0.38 | 0.44 | 0.23 | 0.36 |
| 1 | 84682179 | rs970318 | 0.472 | -0.844 | 0.072 | 0.292 | -0.063 | 0.031 | 0.031 | 0 | 0.39 | 0.14 | 0.37 | 0.33 | 0.52 | 0.33 |
| 1 | 84700766 | rs11163923 | -0.225 | -0.568 | 1.209 | 1.139 | -0.017 | 0.147 | 0.147 | 0.071 | 0.09 | 0.18 | 0.22 | 0.34 | 0.04 | 0.22 |
| 1 | 84701251 | rs600674 | -0.203 | -0.706 | 0.959 | 0.863 | 0.004 | -0.004 | -0.004 | 0 | 0.77 | 0.84 | 0.81 | 0.78 | 0.79 | 0.71 |
| 1 | 84702474 | rs17131308 | -0.029 | -0.67 | 1.104 | 0.77 | 0.016 | -0.039 | -0.039 | 0 | 0.07 | 0.15 | 0.06 | 0 | 0.08 | 0.01 |
| 1 | 84705481 | rs787858 | 1.353 | -0.706 | 1.237 | 0.817 | 0.024 | 0 | 0 | 0 | 0.58 | 0.23 | 0.5 | 0.42 | 0.66 | 0.48 |
| 1 | 84720173 | rs605969 | -0.083 | -0.549 | 1.223 | 0.756 | 0.014 | -0.042 | -0.042 | 0 | 0.08 | 0.17 | 0.06 | 0 | 0.09 | 0.02 |
| 1 | 84720776 | rs4907047 | 0.696 | -0.595 | 1.333 | 0.741 | -0.003 | -0.03 | -0.03 | 0 | 0.67 | 0.47 | 0.72 | 0.77 | 0.71 | 0.7 |
| 1 | 84722102 | rs634904 | 1.073 | -0.767 | 0.57 | 1.181 | -0.003 | 0 | 0 | 0 | 0.65 | 0.39 | 0.56 | 0.41 | 0.75 | 0.49 |
| 1 | 84722166 | rs583720 | 0.106 | -0.46 | -0.692 | 1.328 | -0.001 | -0.004 | -0.004 | 0 | 0.77 | 0.86 | 0.81 | 0.78 | 0.79 | 0.73 |
| 1 | 84728697 | rs11163934 | -0.252 | -0.087 | -1.072 | 0.809 | -0.014 | 0.1 | 0.1 | 0.037 | 0.08 | 0.12 | 0.2 | 0.36 | 0.04 | 0.23 |

| | | | | | | | | | | | | | | | | |
|---|---|---|---|---|---|---|---|---|---|---|---|---|---|---|---|---|
| 1 | 84729134 | rs12047850 | -0.542 | 0.342 | -1.335 | -0.429 | -0.042 | 0.096 | 0.096 | 0.215 | 0.24 | 0.02 | 0.17 | 0.08 | 0.22 | 0.28 |
| 1 | 84732644 | rs316689 | 0.796 | 0.289 | -0.08 | -0.27 | -0.011 | -0.089 | -0.089 | 0 | 0.86 | 0.65 | 0.84 | 1 | 0.89 | 0.97 |
| 1 | 84732963 | rs7528615 | 0.322 | 0.734 | -1.251 | 0.047 | -0.09 | -0.09 | -0.09 | 0 | 0.3 | 0.03 | 0.32 | 0.58 | 0.31 | 0.51 |
| 1 | 84735710 | rs316686 | -0.779 | 0.993 | 0.371 | -0.452 | 0.011 | -0.114 | -0.114 | 0 | 0.18 | 0.33 | 0.18 | 0.01 | 0.18 | 0.12 |
| 1 | 84743949 | rs316672 | -1.87 | 1.205 | 1.008 | -0.33 | 0.005 | -0.055 | -0.055 | 0 | 0.32 | 0.58 | 0.38 | 0.12 | 0.34 | 0.19 |
| 1 | 84744277 | rs7538115 | -0.597 | 1.259 | 0.717 | -0.104 | -0.004 | -0.01 | -0.01 | 0 | 0.15 | 0.09 | 0.15 | 0.01 | 0.17 | 0.12 |
| 1 | 84745373 | rs1983077 | 1.652 | 1.105 | 0.782 | -0.166 | -0.003 | -0.067 | -0.067 | 0 | 0.7 | 0.46 | 0.73 | 0.88 | 0.72 | 0.84 |
| 1 | 84745573 | rs12120336 | -0.514 | 1.176 | -0.106 | 0.325 | -0.025 | -0.033 | -0.033 | 0.077 | 0.07 | 0.01 | 0.07 | 0.11 | 0.1 | 0.04 |
| 1 | 84746227 | rs626118 | -1.489 | 1.254 | 0.202 | 0.712 | 0.003 | 0.019 | 0.019 | 0 | 0.2 | 0.3 | 0.24 | 0.25 | 0.2 | 0.2 |
| 1 | 84746845 | rs680308 | 0.814 | 1.239 | -0.268 | 1.07 | 0.002 | -0.046 | -0.046 | 0 | 0.72 | 0.8 | 0.59 | 0.52 | 0.76 | 0.75 |
| 1 | 84751614 | rs316635 | -0.628 | 1.691 | -0.375 | 0.977 | -0.074 | -0.089 | -0.089 | 2.19 | 0.33 | 0.13 | 0.25 | 0.58 | 0.43 | 0.42 |
| 1 | 84754264 | rs316641 | 0.275 | 1.367 | -0.148 | 0.527 | -0.011 | -0.053 | -0.053 | 0 | 0.84 | 0.74 | 0.81 | 1 | 0.88 | 0.9 |
| 1 | 84759423 | rs11163939 | 1.368 | 1.24 | 1.192 | 0.662 | 0.025 | 0.068 | 0.068 | 0.129 | 0.47 | 0.27 | 0.28 | 0.25 | 0.4 | 0.38 |
| 1 | 84760218 | rs813056 | 1.013 | 1.19 | 1.401 | 0.685 | 0.007 | -0.104 | -0.104 | 1.281 | 0.82 | 0.6 | 0.77 | 1 | 0.82 | 0.88 |
| 1 | 84761174 | rs784833 | 1.18 | 0.949 | 1.371 | 0.712 | -0.013 | -0.001 | -0.001 | 0.829 | 0.81 | 0.53 | 0.57 | 0.83 | 0.83 | 0.85 |
| 1 | 84762953 | rs868612 | 0.567 | 0.991 | 0.869 | 0.64 | -0.006 | -0.05 | -0.05 | 0 | 0.84 | 0.77 | 0.81 | 1 | 0.87 | 0.9 |
| 1 | 84787312 | rs316662 | 2.311 | 0.713 | 1.549 | 1.078 | 0.012 | 0.174 | 0.174 | 0.146 | 0.93 | 0.64 | 0.8 | 0.85 | 0.98 | 0.92 |
| 1 | 84791239 | rs315555 | 1.289 | 0.661 | 1.568 | 0.838 | 0.003 | 0.017 | 0.017 | 0 | 0.92 | 0.88 | 0.82 | 0.86 | 0.92 | 0.9 |
| 1 | 84795283 | rs315552 | 2.402 | 0.654 | 1.533 | 0.835 | 0.007 | 0.062 | 0.062 | 0.052 | 0.88 | 0.57 | 0.77 | 0.85 | 0.91 | 0.85 |
| 1 | 84812664 | rs17131613 | -1.955 | 0.551 | 1.739 | 0.781 | -0.033 | -0.042 | -0.042 | 0 | 0.1 | 0.18 | 0.07 | 0.01 | 0.05 | 0.13 |
| 1 | 84815468 | rs17131617 | -0.537 | 0.554 | 1.649 | 0.772 | 0.011 | 0.013 | 0.013 | 0 | 0.06 | 0.02 | 0.02 | 0 | 0.04 | 0.07 |
| 1 | 84816587 | rs315540 | 2.156 | 0.55 | 1.724 | 0.781 | -0.046 | 0.042 | 0.042 | 0 | 0.88 | 0.7 | 0.78 | 0.85 | 0.94 | 0.85 |
| 1 | 84820409 | rs315535 | 2.098 | 0.523 | 1.781 | 0.767 | 0.017 | 0.067 | 0.067 | 0 | 0.88 | 0.58 | 0.76 | 0.85 | 0.91 | 0.84 |
| 1 | 84832120 | rs6669915 | -2.316 | 0.798 | 1.551 | 0.721 | 0.008 | -0.076 | -0.076 | 0 | 0.12 | 0.29 | 0.08 | 0 | 0.09 | 0.09 |
| 1 | 84835699 | rs17131635 | -0.101 | 1.081 | 0.523 | 0.756 | -0.029 | -0.059 | -0.059 | 0.657 | 0.3 | 0.16 | 0.29 | 0.59 | 0.35 | 0.41 |
| 1 | 84840651 | rs551156 | 1.205 | 1.102 | 0.566 | 0.539 | -0.013 | 0.121 | 0.121 | 0 | 0.53 | 0.26 | 0.42 | 0.17 | 0.51 | 0.41 |
| 1 | 84842764 | rs953796 | -1.502 | 0.918 | 0.865 | 0.447 | 0.014 | -0.05 | -0.05 | 0 | 0.13 | 0.14 | 0.09 | 0 | 0.14 | 0.12 |
| 1 | 84843465 | rs642774 | 1.453 | 0.78 | 0.99 | 0.698 | 0.008 | 0.046 | 0.046 | 0 | 0.86 | 0.45 | 0.76 | 0.82 | 0.9 | 0.8 |
| 1 | 84851201 | rs11163949 | -0.859 | 0.349 | 0.709 | 0.396 | -0.014 | -0.028 | -0.028 | 2.033 | 0.09 | 0.13 | 0.06 | 0.01 | 0.05 | 0.08 |
| 1 | 84852558 | rs7554646 | -0.318 | -0.145 | 0.998 | -0.069 | -0.025 | -0.056 | -0.056 | 0 | 0.24 | 0.35 | 0.2 | 0.12 | 0.2 | 0.27 |
| 1 | 84853611 | rs576888 | 0.083 | -0.263 | 1.398 | -0.154 | -0.032 | -0.07 | -0.07 | 4.905 | 0.75 | 0.6 | 0.78 | 0.88 | 0.8 | 0.73 |
| 1 | 84856328 | rs17131653 | -0.372 | -0.306 | 1.416 | -0.164 | -0.016 | -0.034 | -0.034 | 0 | 0.09 | 0.15 | 0.06 | 0.01 | 0.06 | 0.08 |
| 1 | 84856633 | rs12752696 | -0.066 | 0.065 | 0.476 | 0.251 | -0.037 | -0.071 | -0.071 | 0.053 | 0.24 | 0.11 | 0.31 | 0.42 | 0.28 | 0.32 |
| 1 | 84856784 | rs621869 | 1.567 | -0.255 | 0.34 | 0.093 | -0.036 | -0.034 | -0.034 | 0.147 | 0.38 | 0.26 | 0.39 | 0.44 | 0.43 | 0.32 |
| 1 | 84861518 | rs7523888 | 0.109 | -0.082 | 0.422 | -0.158 | -0.015 | -0.047 | -0.047 | 0.194 | 0.69 | 0.56 | 0.64 | 0.83 | 0.72 | 0.69 |
| 1 | 84866408 | rs646681 | 0.771 | 0.281 | 0.337 | -0.004 | 0.013 | -0.012 | -0.012 | 0.775 | 0.2 | 0.29 | 0.27 | 0.17 | 0.22 | 0.22 |
| 1 | 84867148 | rs1246386 | -0.837 | 0.265 | 0.234 | -0.076 | 0.001 | 0 | 0 | 0 | 0.43 | 0.42 | 0.56 | 0.57 | 0.54 | 0.5 |
| 1 | 84867610 | rs3754274 | 1.081 | 0.177 | 0.123 | 0.024 | -0.059 | -0.069 | -0.069 | 0.175 | 0.14 | 0.04 | 0.25 | 0.37 | 0.27 | 0.26 |
| 1 | 84871923 | rs507772 | -0.896 | -0.117 | 0.249 | -0.067 | 0 | 0.002 | 0.002 | 0 | 0.48 | 0.53 | 0.59 | 0.61 | 0.52 | 0.52 |
| 1 | 84878228 | rs7511984 | 1.191 | -0.137 | 0.299 | -0.107 | 0.005 | -0.052 | -0.052 | 0.972 | 0.17 | 0.2 | 0.09 | 0.01 | 0.19 | 0.15 |
| 1 | 84880380 | rs3768250 | -0.134 | -0.109 | 0.225 | -0.044 | -0.014 | -0.032 | -0.032 | 0.083 | 0.29 | 0.21 | 0.36 | 0.46 | 0.33 | 0.31 |
| 1 | 84896062 | rs7512262 | 0.898 | -0.097 | 0.297 | -0.033 | 0.007 | -0.011 | -0.011 | 0 | 0.18 | 0.27 | 0.22 | 0.15 | 0.19 | 0.21 |
| 1 | 84898321 | rs959363 | -0.094 | -0.101 | 0.144 | -0.039 | -0.02 | -0.047 | -0.047 | 0 | 0.28 | 0.17 | 0.36 | 0.46 | 0.33 | 0.31 |
| 1 | 84899282 | rs7514211 | -0.002 | -0.158 | 0.144 | -0.083 | -0.006 | -0.016 | -0.016 | 0 | 0.06 | 0.11 | 0.04 | 0.01 | 0.05 | 0.04 |
| 1 | 84915300 | rs6677436 | -0.113 | -0.158 | 0.022 | -0.081 | -0.008 | -0.043 | -0.043 | 1.756 | 0.26 | 0.43 | 0.27 | 0.16 | 0.25 | 0.25 |
| 1 | 84917707 | rs2066197 | 1.394 | -0.349 | -0.387 | -0.212 | 0.01 | 0.006 | 0.006 | 0 | 0.41 | 0.3 | 0.24 | 0.38 | 0.34 | 0.36 |
| 1 | 84920939 | rs1395415 | 0.682 | -0.437 | -0.65 | -0.123 | -0.002 | 0.011 | 0.011 | 0 | 0.57 | 0.28 | 0.43 | 0.38 | 0.59 | 0.46 |
| 1 | 84922553 | rs12729133 | -0.012 | -0.119 | -0.864 | 0.275 | -0.008 | -0.021 | -0.021 | 0.019 | 0.27 | 0.2 | 0.34 | 0.41 | 0.32 | 0.33 |
| 1 | 84926750 | rs11578928 | 0.83 | -0.043 | -1.09 | 0.007 | -0.032 | 0.017 | 0.017 | 0 | 0.44 | 0.18 | 0.26 | 0.36 | 0.45 | 0.35 |
| 1 | 84941577 | rs12031171 | 0.687 | -0.772 | -0.862 | -0.321 | -0.011 | -0.048 | -0.048 | 0 | 0.23 | 0.15 | 0.27 | 0.52 | 0.28 | 0.21 |
| 1 | 84944940 | rs11163977 | 0.276 | -0.907 | 0.226 | -0.497 | 0.034 | -0.008 | -0.008 | 0.007 | 0.05 | 0.21 | 0.03 | 0 | 0.03 | 0.05 |
| 1 | 84944989 | rs2292191 | 1.02 | -0.956 | -0.14 | -0.827 | 0.032 | 0.131 | 0.131 | 0 | 0.54 | 0.2 | 0.46 | 0.73 | 0.66 | 0.48 |
| 1 | 84947055 | rs4907076 | 0.454 | -1.214 | 0.372 | -1.041 | 0.001 | 0.001 | 0.001 | 0.003 | 0.17 | 0.18 | 0.24 | 0.21 | 0.16 | 0.32 |
| 1 | 84953415 | rs11163981 | -2.006 | -1.284 | 0.526 | -1.097 | 0.018 | 0.031 | 0.031 | 0 | 0.39 | 0.64 | 0.48 | 0.26 | 0.27 | 0.46 |
| 1 | 84955233 | rs11163983 | 0.721 | -1.182 | 0.604 | -1.062 | 0.034 | -0.008 | -0.008 | 0 | 0.05 | 0.21 | 0.03 | 0 | 0.03 | 0.05 |
| 1 | 84959734 | rs7524576 | -1.309 | -1.212 | 0.611 | -1.079 | -0.057 | -0.125 | -0.125 | 0 | 0.21 | 0.45 | 0.22 | 0.05 | 0.1 | 0.14 |
| 1 | 84961108 | rs7528067 | 0.067 | -1.242 | 0.66 | -1.094 | -0.038 | -0.05 | -0.05 | 0 | 0.1 | 0.14 | 0.08 | 0 | 0.03 | 0.06 |
| 1 | 84969019 | rs17365723 | 1.923 | -1.234 | 0.568 | -1.093 | 0.006 | -0.025 | -0.025 | 0 | 0.05 | 0.12 | 0.05 | 0 | 0.07 | 0.06 |
| 1 | 84977773 | rs10493754 | -1.907 | -1.223 | 0.525 | -1.04 | -0.062 | -0.122 | -0.122 | 0 | 0.23 | 0.45 | 0.24 | 0.06 | 0.11 | 0.14 |
| 1 | 84978432 | rs17110590 | 0.415 | -0.872 | 0.362 | -0.744 | -0.001 | -0.006 | -0.006 | 0 | 0.06 | 0.1 | 0.12 | 0.05 | 0.04 | 0.03 |
| 1 | 84992110 | rs186360 | -0.313 | -0.972 | 0.562 | -0.952 | -0.001 | 0.002 | 0.002 | 0.781 | 0.55 | 0.6 | 0.63 | 0.76 | 0.49 | 0.45 |
| 1 | 84996604 | rs6698596 | -1.681 | -1.3 | 1.628 | -0.746 | -0.054 | -0.063 | -0.063 | 0 | 0.15 | 0.53 | 0.21 | 0.06 | 0.06 | 0.08 |
| 1 | 85000267 | rs6669593 | -1.483 | -1.301 | 1.676 | -0.747 | -0.047 | -0.056 | -0.056 | 0 | 0.14 | 0.46 | 0.2 | 0.06 | 0.06 | 0.08 |
| 1 | 85003568 | rs6576713 | -2.087 | -1.385 | 1.58 | -0.824 | 0.03 | 0.121 | 0.121 | 0 | 0.45 | 0.76 | 0.4 | 0.26 | 0.35 | 0.49 |
| 1 | 85007613 | rs7524406 | -0.365 | -1.386 | 1.801 | -0.813 | -0.037 | -0.043 | -0.043 | 1.635 | 0.09 | 0.21 | 0.08 | 0.01 | 0.03 | 0.06 |
| 1 | 85009894 | rs12143652 | -0.442 | -1.372 | 1.786 | -0.852 | 0 | 0.002 | 0.002 | 0.22 | 0.26 | 0.23 | 0.16 | 0.2 | 0.21 | 0.36 |
| 1 | 85011862 | rs11164005 | -0.307 | -1.38 | 1.732 | -0.867 | -0.005 | 0.019 | 0.019 | 0.385 | 0.24 | 0.15 | 0.14 | 0.03 | 0.21 | 0.28 |
| 1 | 85014989 | rs17111283 | -0.28 | -1.368 | 1.751 | -0.847 | -0.034 | -0.048 | -0.048 | 0 | 0.09 | 0.14 | 0.08 | 0 | 0.03 | 0.06 |
| 1 | 85015189 | rs17111299 | -0.555 | -1.368 | 1.72 | -0.846 | -0.04 | -0.046 | -0.046 | 0 | 0.09 | 0.22 | 0.09 | 0.01 | 0.03 | 0.06 |

| | | | | | | | | | | | | | | | | |
|---|---|---|---|---|---|---|---|---|---|---|---|---|---|---|---|---|
| 1 | 85016112 | rs17111308 | -0.555 | -1.368 | 1.72 | -0.846 | -0.04 | -0.046 | -0.046 | 0 | 0.09 | 0.22 | 0.09 | 0.01 | 0.03 | 0.06 |
| 1 | 85016177 | rs1400777 | -2.128 | -1.379 | 1.702 | -0.858 | 0.026 | 0.119 | 0.119 | 0 | 0.45 | 0.76 | 0.41 | 0.26 | 0.35 | 0.49 |
| 1 | 85020649 | rs1043493 | -0.922 | -1.493 | 1.797 | -0.891 | -0.048 | -0.055 | -0.055 | 0 | 0.11 | 0.25 | 0.09 | 0.01 | 0.03 | 0.06 |
| 1 | 85029077 | rs15911 | -0.358 | -1.385 | 1.736 | -1.007 | -0.015 | 0.069 | 0.069 | 0.881 | 0.24 | 0.11 | 0.13 | 0.03 | 0.2 | 0.28 |
| 1 | 85030053 | rs7547026 | -0.967 | -1.332 | 1.725 | -0.942 | -0.045 | 0.048 | 0.048 | 0 | 0.1 | 0.27 | 0.09 | 0.16 | 0.03 | 0.13 |
| 1 | 85031630 | rs2994953 | -1.059 | -0.962 | 1.524 | -0.787 | -0.001 | 0.001 | 0.001 | 0.016 | 0.3 | 0.34 | 0.26 | 0.23 | 0.25 | 0.38 |
| 1 | 85038958 | rs6681870 | -1.097 | -1.069 | 1.698 | -0.609 | 0.026 | -0.001 | -0.001 | 0 | 0.08 | 0.49 | 0.11 | 0 | 0.02 | 0.06 |
| 1 | 85048641 | rs2945152 | -1.867 | -1.092 | 0.904 | -0.67 | 0.034 | 0.134 | 0.134 | 0 | 0.44 | 0.81 | 0.4 | 0.24 | 0.33 | 0.49 |
| 1 | 85062615 | rs2945145 | -0.898 | -1.058 | 1.225 | -0.718 | -0.014 | -0.096 | -0.096 | 1.526 | 0.32 | 0.54 | 0.28 | 0.08 | 0.24 | 0.31 |
| 1 | 85064272 | rs17112247 | -0.189 | -1.084 | 1.288 | -0.744 | 0 | -0.056 | -0.056 | 0 | 0.11 | 0.2 | 0.11 | 0.01 | 0.09 | 0.11 |
| 1 | 85065769 | rs2911597 | -0.757 | -1.136 | 1.222 | -0.737 | -0.009 | -0.08 | -0.08 | 3.093 | 0.31 | 0.44 | 0.26 | 0.08 | 0.24 | 0.31 |
| 1 | 85070877 | rs2911603 | -1.721 | -1.056 | 0.679 | -0.534 | 0.037 | 0.078 | 0.078 | 0 | 0.44 | 0.8 | 0.39 | 0.32 | 0.33 | 0.49 |
| 1 | 85079410 | rs2007454 | -1.627 | -0.981 | 0.594 | -0.505 | 0.023 | 0.047 | 0.047 | 0 | 0.43 | 0.69 | 0.37 | 0.32 | 0.33 | 0.49 |
| 1 | 85084120 | rs2945138 | -0.121 | -0.927 | 0.397 | -0.041 | -0.008 | 0.079 | 0.079 | 0 | 0.05 | 0.11 | 0.12 | 0.28 | 0.03 | 0.09 |
| 1 | 85089402 | rs11164023 | -0.548 | -0.833 | 0.45 | -0.147 | 0.001 | 0.005 | 0.005 | 0.194 | 0.26 | 0.24 | 0.29 | 0.11 | 0.21 | 0.28 |
| 1 | 85091557 | rs2945133 | -1.3 | -0.648 | 0.983 | 0.237 | -0.017 | 0.082 | 0.082 | 0 | 0.08 | 0.23 | 0.09 | 0.17 | 0.02 | 0.12 |
| 1 | 85092463 | rs2994931 | -1.576 | -0.638 | 1.153 | 0.271 | -0.002 | 0.165 | 0.165 | 0 | 0.09 | 0.31 | 0.11 | 0.25 | 0.02 | 0.14 |
| 1 | 85095457 | rs12127432 | -1.314 | -0.725 | 1.193 | 0.11 | -0.013 | 0.031 | 0.031 | 0 | 0.09 | 0.13 | 0.08 | 0.19 | 0.05 | 0.15 |
| 1 | 85096920 | rs6695249 | -0.611 | -1.414 | 1.118 | -0.601 | -0.007 | -0.02 | -0.02 | 0 | 0.28 | 0.26 | 0.26 | 0.05 | 0.16 | 0.21 |
| 1 | 85100180 | rs17367018 | 0.356 | -0.825 | 0.842 | -0.694 | 0.008 | -0.014 | -0.014 | 2.52 | 0.05 | 0.07 | 0.05 | 0 | 0.11 | 0.05 |
| 1 | 85104040 | rs12035493 | -0.786 | -1.386 | 0.983 | -1.005 | 0.041 | 0.068 | 0.068 | 0 | 0.35 | 0.22 | 0.3 | 0.05 | 0.25 | 0.3 |
| 1 | 85106040 | rs2943672 | 0.67 | -1.337 | 1.304 | -0.952 | -0.009 | -0.001 | -0.001 | 0 | 0.84 | 0.83 | 0.87 | 0.81 | 0.88 | 0.75 |
| 1 | 85110325 | rs3782 | 0.328 | -1.328 | 1.458 | -0.937 | 0 | 0.089 | 0.089 | 0 | 0.92 | 0.79 | 0.91 | 0.79 | 0.95 | 0.84 |
| 1 | 85111124 | rs2911613 | -0.535 | -1.478 | 1.7 | -0.838 | -0.008 | -0.024 | -0.024 | 0 | 0.3 | 0.36 | 0.32 | 0.09 | 0.23 | 0.23 |
| 1 | 85113203 | rs2994949 | 1.019 | -1.595 | 1.881 | -0.895 | -0.038 | -0.003 | -0.003 | 0 | 0.81 | 0.61 | 0.86 | 0.79 | 0.88 | 0.75 |
| 1 | 85113229 | rs1057746 | 1.663 | -1.564 | 1.708 | -0.884 | 0.075 | 0.103 | 0.103 | 0 | 0.52 | 0.25 | 0.53 | 0.7 | 0.65 | 0.52 |
| 1 | 85117113 | rs6681911 | 1.482 | -1.688 | 1.619 | -0.969 | 0.066 | 0.077 | 0.077 | 0 | 0.55 | 0.25 | 0.54 | 0.7 | 0.67 | 0.52 |
| 1 | 85126143 | rs2911562 | 0.602 | -1.647 | 1.728 | -0.923 | -0.037 | -0.014 | -0.014 | 1.183 | 0.66 | 0.63 | 0.63 | 0.72 | 0.79 | 0.62 |
| 1 | 85131405 | rs4907001 | 0.345 | -1.549 | 1.367 | -0.898 | -0.013 | -0.049 | -0.049 | 0 | 0.1 | 0.17 | 0.05 | 0 | 0.07 | 0.09 |
| 1 | 85137984 | rs12022113 | 1.659 | -1.57 | 1.155 | -0.842 | -0.013 | -0.012 | -0.012 | 0.123 | 0.77 | 0.46 | 0.81 | 0.78 | 0.82 | 0.75 |
| 1 | 85140555 | rs6687220 | -0.759 | -1.569 | 1.124 | -0.835 | -0.017 | 0.007 | 0.007 | 0 | 0.16 | 0.22 | 0.14 | 0.21 | 0.12 | 0.24 |
| 1 | 85143872 | rs1340523 | 1.069 | -1.661 | 1.18 | -0.789 | 0.013 | 0.021 | 0.021 | 0 | 0.59 | 0.36 | 0.63 | 0.83 | 0.7 | 0.66 |
| 1 | 85166561 | rs2994449 | 0.727 | -1.659 | 1.143 | -0.778 | -0.028 | 0.008 | 0.008 | 0 | 0.82 | 0.64 | 0.85 | 0.79 | 0.87 | 0.76 |
| 1 | 85167337 | rs11164043 | -0.297 | -1.422 | 1.363 | -0.676 | -0.031 | -0.034 | -0.034 | 0.089 | 0.26 | 0.29 | 0.28 | 0.08 | 0.16 | 0.22 |
| 1 | 85188231 | rs17114425 | -0.27 | -1.478 | 1.098 | -0.638 | -0.041 | -0.035 | -0.035 | 0.029 | 0.24 | 0.3 | 0.27 | 0.07 | 0.14 | 0.22 |
| 1 | 85193058 | rs1770573 | 0.948 | -1.37 | 1.121 | -0.636 | 0.047 | -0.074 | -0.074 | 0 | 0.88 | 0.62 | 0.86 | 1 | 0.89 | 0.9 |
| 1 | 85198264 | rs818526 | 1.085 | -1.347 | 1.092 | -0.628 | 0.043 | -0.051 | -0.051 | 0.908 | 0.89 | 0.62 | 0.87 | 0.98 | 0.86 | 0.9 |
| 1 | 85201228 | rs818524 | 1.407 | -1.432 | 0.693 | -0.647 | 0.051 | 0.023 | 0.023 | 0.238 | 0.61 | 0.16 | 0.56 | 0.7 | 0.71 | 0.59 |
| 1 | 85203780 | rs11805570 | 1.135 | -1.205 | 1.053 | -0.533 | 0.022 | -0.048 | -0.048 | 0.012 | 0.89 | 0.66 | 0.88 | 0.98 | 0.89 | 0.9 |
| 1 | 85205632 | rs9725395 | -1.04 | -1.177 | 1.024 | -0.528 | 0.063 | -0.014 | -0.014 | 0.533 | 0.05 | 0.25 | 0.07 | 0 | 0.09 | 0.02 |
| 1 | 85206477 | rs11161436 | 1.173 | -1.896 | 0.42 | -0.854 | 0.08 | 0.029 | 0.029 | 0 | 0.6 | 0.15 | 0.55 | 0.7 | 0.73 | 0.57 |
| 1 | 85206774 | rs12027987 | 0.821 | -1.495 | -0.253 | -0.924 | 0.004 | -0.008 | -0.008 | 1.115 | 0.28 | 0.22 | 0.23 | 0.36 | 0.23 | 0.32 |
| 1 | 85207526 | rs818521 | -0.128 | -1.431 | 0.019 | -0.902 | -0.038 | -0.034 | -0.034 | 0.021 | 0.26 | 0.3 | 0.28 | 0.07 | 0.13 | 0.25 |
| 1 | 85208096 | rs11161438 | 0.634 | -0.411 | -0.139 | -0.473 | 0.003 | -0.003 | -0.003 | 0.023 | 0.89 | 0.82 | 0.91 | 0.87 | 0.91 | 0.85 |
| 1 | 85210779 | rs1998826 | -0.042 | -0.145 | -0.514 | -0.947 | 0.007 | 0.056 | 0.056 | 0 | 0.58 | 0.72 | 0.74 | 0.85 | 0.63 | 0.65 |
| 1 | 85212026 | rs17114809 | 0.63 | 0.406 | -0.703 | -0.51 | -0.014 | -0.023 | -0.023 | 0.012 | 0.22 | 0.12 | 0.37 | 0.24 | 0.26 | 0.27 |
| 1 | 85212582 | rs11161441 | 0.419 | 0.385 | -0.411 | -0.369 | 0.001 | 0.009 | 0.009 | 0 | 0.36 | 0.6 | 0.36 | 0.59 | 0.36 | 0.37 |
| 1 | 85213900 | rs12085269 | 0.364 | 0.334 | -0.101 | -0.393 | 0.006 | 0.026 | 0.026 | 0 | 0.35 | 0.54 | 0.36 | 0.59 | 0.36 | 0.37 |
| 1 | 85213914 | rs9660406 | 0.349 | 0.319 | -0.034 | -0.216 | 0.003 | 0.02 | 0.02 | 0 | 0.35 | 0.58 | 0.35 | 0.55 | 0.36 | 0.35 |
| 1 | 85214543 | rs4471288 | -0.417 | -0.034 | -0.116 | -0.607 | -0.009 | -0.013 | -0.013 | 0 | 0.4 | 0.54 | 0.47 | 0.29 | 0.37 | 0.32 |
| 1 | 85215534 | rs11161442 | -0.443 | -0.048 | -0.194 | -0.614 | -0.006 | -0.01 | -0.01 | 0.128 | 0.41 | 0.55 | 0.48 | 0.3 | 0.37 | 0.33 |
| 1 | 85216685 | rs12744552 | 0.981 | 0.021 | -0.697 | -0.625 | -0.001 | -0.009 | -0.009 | 0 | 0.16 | 0.18 | 0.08 | 0.06 | 0.12 | 0.05 |
| 1 | 85217231 | rs11161445 | 1.094 | 0.058 | -0.889 | -0.518 | -0.036 | -0.053 | -0.053 | 0.05 | 0.21 | 0.06 | 0.36 | 0.24 | 0.25 | 0.27 |
| 1 | 85219337 | rs4907091 | 0.996 | 0.002 | -0.594 | -0.441 | -0.053 | -0.088 | -0.088 | 0 | 0.3 | 0.11 | 0.32 | 0.58 | 0.36 | 0.36 |
| 1 | 85219868 | rs12065881 | 0.005 | -0.121 | -0.449 | -0.483 | 0.007 | -0.001 | -0.001 | 0.287 | 0.33 | 0.25 | 0.42 | 0.3 | 0.32 | 0.29 |
| 1 | 85220837 | rs4907093 | 1.022 | -0.009 | -0.775 | -0.376 | -0.033 | -0.066 | -0.066 | 0 | 0.27 | 0.15 | 0.3 | 0.59 | 0.33 | 0.37 |
| 1 | 85223952 | rs1770569 | -0.906 | 0.025 | -0.455 | -0.702 | 0.002 | 0.012 | 0.012 | 0.018 | 0.66 | 0.7 | 0.77 | 0.88 | 0.72 | 0.68 |
| 1 | 85224368 | rs1750493 | -0.066 | 0.077 | -0.505 | -0.49 | 0.006 | 0.017 | 0.017 | 0.75 | 0.51 | 0.58 | 0.69 | 0.78 | 0.62 | 0.63 |
| 1 | 85227586 | rs1770566 | 1.168 | 0.506 | -0.82 | 0.18 | -0.042 | -0.083 | -0.083 | 0 | 0.18 | 0.04 | 0.35 | 0.33 | 0.27 | 0.21 |
| 1 | 85229561 | rs1750485 | -0.164 | 0.49 | -0.558 | -0.101 | 0.01 | 0.055 | 0.055 | 0.441 | 0.49 | 0.4 | 0.63 | 0.77 | 0.6 | 0.6 |
| 1 | 85230072 | rs10873653 | -0.278 | 0.536 | -0.376 | 0.081 | 0 | -0.001 | -0.001 | 2.34 | 0.43 | 0.46 | 0.35 | 0.51 | 0.43 | 0.45 |
| 1 | 85233989 | rs846845 | -0.504 | 0.738 | -0.394 | -0.06 | 0.043 | 0.073 | 0.073 | 0.931 | 0.73 | 0.54 | 0.69 | 0.56 | 0.66 | 0.53 |
| 1 | 85234227 | rs4448534 | 0.415 | 0.382 | -1.236 | -0.598 | 0.014 | -0.102 | -0.102 | 0.067 | 0.32 | 0.12 | 0.4 | 0.48 | 0.28 | 0.35 |
| 1 | 85235399 | rs9659298 | 0.104 | 0.63 | -0.863 | -0.635 | 0.011 | -0.025 | -0.025 | 0 | 0.06 | 0.12 | 0.12 | 0 | 0.1 | 0.05 |
| 1 | 85239564 | rs9701126 | 0.158 | 0.089 | -0.754 | -0.753 | -0.005 | -0.013 | -0.013 | 4.411 | 0.1 | 0.18 | 0.07 | 0.05 | 0.08 | 0.05 |
| 1 | 85242494 | rs1326004 | 0.062 | 0.149 | -0.715 | -0.778 | 0.052 | 0.051 | 0.051 | 0 | 0.34 | 0.4 | 0.39 | 0.48 | 0.46 | 0.53 |
| 1 | 85242787 | rs1326001 | -0.573 | -0.459 | -1.179 | -1.168 | 0.019 | -0.089 | -0.089 | 0 | 0.58 | 0.88 | 0.54 | 0.49 | 0.62 | 0.65 |
| 1 | 85246018 | rs11161452 | -0.271 | -0.568 | -1.029 | -1.229 | 0.032 | -0.009 | -0.009 | 0 | 0.53 | 0.57 | 0.5 | 0.48 | 0.61 | 0.64 |
| 1 | 85246297 | rs9724696 | -0.788 | -0.542 | -1.285 | -1.222 | 0.012 | -0.115 | -0.115 | 0.023 | 0.61 | 0.92 | 0.59 | 0.49 | 0.65 | 0.66 |

| | | | | | | | | | | | | | | | | |
|---|---|---|---|---|---|---|---|---|---|---|---|---|---|---|---|---|
| 1 | 85252003 | rs10782526 | 0.62 | -0.115 | -1.036 | -1.144 | 0.038 | 0.077 | 0.077 | 0 | 0.3 | 0.42 | 0.31 | 0.5 | 0.36 | 0.45 |
| 1 | 85252556 | rs10782527 | -0.264 | -0.127 | -1.148 | -1.207 | -0.033 | -0.02 | -0.02 | 0 | 0.39 | 0.73 | 0.43 | 0.51 | 0.47 | 0.55 |
| 1 | 85253349 | rs4907003 | -1.754 | -0.58 | -1.666 | -1.251 | -0.077 | -0.091 | -0.091 | 0 | 0.76 | 0.98 | 0.6 | 0.67 | 0.68 | 0.76 |
| 1 | 85260648 | rs4907096 | 0.485 | -0.359 | -1.385 | -1.127 | 0.012 | 0.028 | 0.028 | 0 | 0.34 | 0.52 | 0.37 | 0.5 | 0.39 | 0.47 |
| 1 | 85264974 | rs12564032 | -1.17 | -0.438 | -1.482 | -1.44 | 0.009 | -0.062 | -0.062 | 0.112 | 0.73 | 0.9 | 0.61 | 0.55 | 0.73 | 0.67 |
| 1 | 85268208 | rs10782528 | 0.106 | -0.52 | -1.361 | -1.502 | -0.014 | 0.012 | 0.012 | 0.085 | 0.4 | 0.63 | 0.35 | 0.64 | 0.44 | 0.57 |
| 1 | 85268243 | rs10782529 | -0.763 | -0.605 | -1.359 | -1.517 | 0.001 | 0.011 | 0.011 | 0.789 | 0.63 | 0.79 | 0.53 | 0.65 | 0.62 | 0.69 |
| 1 | 85271708 | rs11161458 | 0.933 | -0.519 | -1.072 | -1.393 | -0.022 | 0.001 | 0.001 | 0 | 0.07 | 0.14 | 0.02 | 0.1 | 0.03 | 0.09 |
| 1 | 85273362 | rs17115896 | 0.578 | -0.544 | -1.356 | -1.592 | 0.03 | 0.054 | 0.054 | 0.87 | 0.3 | 0.19 | 0.14 | 0 | 0.19 | 0.09 |
| 1 | 85275714 | rs2892844 | -1.713 | -0.387 | -1.429 | -1.33 | -0.028 | -0.026 | -0.026 | 0.35 | 0.77 | 0.9 | 0.66 | 0.68 | 0.7 | 0.71 |
| 1 | 85276650 | rs9729377 | -0.455 | -0.069 | -0.999 | -1.712 | 0 | -0.077 | -0.077 | 0.134 | 0.43 | 0.55 | 0.31 | 0 | 0.35 | 0.26 |
| 1 | 85278665 | rs11161462 | -1.828 | -0.056 | -0.895 | -1.4 | -0.003 | -0.005 | -0.005 | 0 | 0.77 | 0.79 | 0.7 | 0.75 | 0.71 | 0.75 |
| 1 | 85282020 | rs12024356 | 1.843 | -0.057 | -0.792 | -1.204 | -0.003 | 0.004 | 0.004 | 0 | 0.07 | 0.08 | 0.05 | 0.08 | 0.07 | 0.05 |
| 1 | 85285744 | rs4907007 | 0.874 | -0.068 | -0.695 | -1.306 | 0.009 | 0.044 | 0.044 | 0.052 | 0.26 | 0.35 | 0.28 | 0.54 | 0.34 | 0.39 |
| 1 | 85287327 | rs17377758 | 2.118 | -0.26 | -1.041 | -1.514 | 0.002 | 0.099 | 0.099 | 0 | 0.17 | 0.01 | 0.08 | 0 | 0.11 | 0.08 |
| 1 | 85289748 | rs9729034 | 0.505 | -0.189 | -0.834 | -1.572 | 0.041 | 0.033 | 0.033 | 0.137 | 0.32 | 0.41 | 0.39 | 0.53 | 0.43 | 0.5 |
| 1 | 85290409 | rs9728717 | 0.484 | -0.25 | -0.923 | -1.634 | 0.016 | -0.152 | -0.152 | 0 | 0.24 | 0.43 | 0.21 | 0 | 0.2 | 0.13 |
| 1 | 85290827 | rs1409213 | 1.507 | -0.196 | -1.243 | -1.403 | -0.01 | -0.021 | -0.021 | 0.005 | 0.21 | 0.16 | 0.25 | 0.54 | 0.31 | 0.38 |
| 1 | 85294434 | rs1359607 | 1.052 | -0.239 | -1.404 | -1.378 | 0.014 | 0.044 | 0.044 | 0 | 0.22 | 0.11 | 0.11 | 0 | 0.17 | 0.13 |
| 1 | 85298014 | rs12077089 | 0.636 | -0.005 | -1.106 | -0.88 | 0 | -0.04 | -0.04 | 4.967 | 0.11 | 0.02 | 0.14 | 0.2 | 0.08 | 0.14 |
| 1 | 85298111 | rs11161464 | -1.85 | 0.242 | -0.883 | -0.77 | -0.009 | -0.017 | -0.017 | 1.616 | 0.8 | 0.88 | 0.73 | 0.74 | 0.75 | 0.77 |
| 1 | 85308062 | rs10493756 | 2.18 | 0.471 | -0.994 | -0.431 | -0.043 | -0.074 | -0.074 | 0.205 | 0.14 | 0.04 | 0.17 | 0.34 | 0.23 | 0.32 |
| 1 | 85308524 | rs9728153 | 0.256 | 0.238 | -0.443 | -0.884 | 0.065 | -0.031 | -0.031 | 0 | 0.08 | 0.25 | 0.13 | 0 | 0.09 | 0.1 |
| 1 | 85312366 | rs1325272 | -1.743 | 0.137 | -1.108 | -0.779 | -0.011 | -0.008 | -0.008 | 0.017 | 0.82 | 0.88 | 0.79 | 0.77 | 0.77 | 0.76 |
| 1 | 85314611 | rs17369401 | 1.287 | 0.167 | -1.365 | -1.16 | -0.015 | 0.091 | 0.091 | 0 | 0.21 | 0.05 | 0.13 | 0 | 0.2 | 0.09 |
| 1 | 85314642 | rs4907102 | -0.003 | -0.025 | -1.039 | -1.183 | 0 | -0.003 | -0.003 | 0 | 0.48 | 0.54 | 0.42 | 0.56 | 0.5 | 0.51 |
| 1 | 85315353 | rs17116652 | 0.922 | -0.054 | -0.962 | -1.34 | -0.001 | -0.093 | -0.093 | 0 | 0.19 | 0.27 | 0.16 | 0 | 0.16 | 0.09 |
| 1 | 85316840 | rs4907103 | 0.413 | -0.088 | -1.119 | -0.859 | -0.001 | -0.001 | -0.001 | 0 | 0.46 | 0.46 | 0.41 | 0.48 | 0.48 | 0.49 |
| 1 | 85316955 | rs4907104 | -0.05 | -0.069 | -1.21 | -0.612 | -0.006 | 0.04 | 0.04 | 0 | 0.68 | 0.6 | 0.55 | 0.49 | 0.68 | 0.6 |
| 1 | 85318584 | rs10127852 | 0.007 | 0.397 | -1.099 | -0.514 | -0.005 | -0.028 | -0.028 | 0.167 | 0.1 | 0.12 | 0.04 | 0 | 0.07 | 0.08 |
| 1 | 85328096 | rs10873661 | -0.611 | 0.652 | -1.396 | -0.217 | 0.004 | 0.023 | 0.023 | 0 | 0.57 | 0.67 | 0.54 | 0.59 | 0.54 | 0.57 |
| 1 | 85332120 | rs1325273 | -0.499 | 1.044 | -1.533 | 0.437 | -0.059 | -0.137 | -0.137 | 0.073 | 0.79 | 0.95 | 0.79 | 0.53 | 0.74 | 0.64 |
| 1 | 85333796 | rs9730423 | -0.397 | 0.69 | -0.839 | -0.12 | 0.005 | -0.058 | -0.058 | 0 | 0.13 | 0.27 | 0.07 | 0.01 | 0.13 | 0.05 |
| 1 | 85335709 | rs7512497 | 0.966 | 0.436 | -1.787 | -0.177 | -0.012 | -0.002 | -0.002 | 0 | 0.39 | 0.09 | 0.32 | 0.36 | 0.41 | 0.48 |
| 1 | 85336115 | rs10493757 | -0.591 | 0.28 | -1.301 | -0.282 | 0.006 | 0.007 | 0.007 | 0.1 | 0.3 | 0.74 | 0.38 | 0.16 | 0.21 | 0.13 |
| 1 | 85340005 | rs4907105 | -0.157 | 0.771 | -0.858 | 0.17 | -0.018 | -0.026 | -0.026 | 0.973 | 0.73 | 0.79 | 0.72 | 0.49 | 0.66 | 0.67 |
| 1 | 85346224 | rs17116972 | -1.13 | 0.009 | -0.003 | -0.44 | -0.019 | -0.094 | -0.094 | 0 | 0.14 | 0.34 | 0.1 | 0 | 0.1 | 0.1 |
| 1 | 85346891 | rs942682 | -0.882 | 0.146 | -0.313 | -0.045 | -0.002 | 0.02 | 0.02 | 0 | 0.25 | 0.51 | 0.32 | 0.35 | 0.23 | 0.25 |
| 1 | 85347522 | rs1998720 | 0.82 | 0.011 | -1.779 | -0.621 | -0.057 | 0.033 | 0.033 | 0 | 0.35 | 0.02 | 0.19 | 0.2 | 0.3 | 0.26 |
| 1 | 85350658 | rs1886644 | 0.347 | 0.066 | -1.126 | -1.082 | 0 | -0.041 | -0.041 | 0.077 | 0.74 | 0.73 | 0.71 | 1 | 0.74 | 0.7 |
| 1 | 85354563 | rs1886647 | 0.936 | -0.209 | -1.261 | -0.445 | 0.005 | -0.002 | -0.002 | 0 | 0.35 | 0.25 | 0.41 | 0.36 | 0.32 | 0.26 |
| 1 | 85366164 | rs12023575 | 0.232 | 0.29 | -1.428 | -0.16 | -0.008 | -0.098 | -0.098 | 0.717 | 0.25 | 0.09 | 0.18 | 0.48 | 0.28 | 0.36 |
| 1 | 85371265 | rs4907110 | -0.304 | 0.833 | -1.112 | 0.025 | -0.011 | 0.036 | 0.036 | 0 | 0.35 | 0.55 | 0.27 | 0.48 | 0.35 | 0.4 |
| 1 | 85371391 | rs4907013 | 0.483 | 1.743 | -1.549 | -0.269 | -0.051 | 0.029 | 0.029 | 2.049 | 0.75 | 0.89 | 0.69 | 0.84 | 0.69 | 0.67 |
| 1 | 85382107 | rs1536120 | 0.556 | 1.601 | -1.339 | -0.176 | 0.008 | -0.011 | -0.011 | 0 | 0.69 | 0.65 | 0.65 | 0.84 | 0.63 | 0.65 |
| 1 | 85384524 | rs7511675 | 0.024 | 1.97 | -1.555 | 0.058 | 0.01 | -0.019 | -0.019 | 0.289 | 0.88 | 0.87 | 0.77 | 1 | 0.8 | 0.89 |
| 1 | 85391140 | rs3738572 | 0.187 | 1.127 | -1.915 | 0.337 | -0.032 | 0.037 | 0.037 | 0 | 0.6 | 0.76 | 0.41 | 0.3 | 0.58 | 0.45 |
| 1 | 85392324 | rs6681025 | 0.111 | 1.056 | -2.103 | 0.915 | -0.029 | -0.031 | -0.031 | 0 | 0.31 | 0.14 | 0.3 | 0.37 | 0.33 | 0.23 |
| 1 | 85392616 | rs12057637 | -0.423 | 0.183 | -2.101 | 0.857 | 0.057 | -0.008 | -0.008 | 0 | 0.39 | 0.23 | 0.49 | 0.6 | 0.29 | 0.37 |
| 1 | 85394258 | rs12145073 | -0.291 | 1.098 | -2.12 | 0.656 | -0.081 | 0.026 | 0.026 | 0.556 | 0.16 | 0.02 | 0.21 | 0.1 | 0.25 | 0.26 |
| 1 | 85396158 | rs607345 | -0.949 | 0.775 | -2.079 | 0.549 | 0.004 | 0.045 | 0.045 | 0 | 0.57 | 0.81 | 0.54 | 0.69 | 0.57 | 0.65 |
| 1 | 85400114 | rs1325277 | 0.473 | 0.642 | -1.528 | 0.916 | 0 | -0.005 | -0.005 | 0 | 0.23 | 0.17 | 0.18 | 0.31 | 0.23 | 0.3 |
| 1 | 85400182 | rs1325278 | -0.832 | 0.544 | -0.66 | 0.822 | 0 | 0.011 | 0.011 | 0.007 | 0.46 | 0.41 | 0.46 | 0.7 | 0.53 | 0.63 |
| 1 | 85402997 | rs634294 | -0.179 | 0.599 | -0.165 | 0.868 | 0.023 | 0.041 | 0.041 | 0 | 0.23 | 0.35 | 0.29 | 0.39 | 0.3 | 0.32 |
| 1 | 85407863 | rs17117919 | -0.323 | 0.114 | 0.339 | 0.641 | -0.025 | -0.067 | -0.067 | 0 | 0.11 | 0.31 | 0.07 | 0 | 0.04 | 0.02 |
| 1 | 85408324 | rs12402681 | 0.436 | 0.049 | -0.21 | 0.699 | -0.037 | -0.067 | -0.067 | 0 | 0.21 | 0.04 | 0.14 | 0.27 | 0.22 | 0.29 |
| 1 | 85409036 | rs639500 | -0.183 | 0.032 | -0.067 | 0.651 | -0.005 | 0.019 | 0.019 | 0 | 0.26 | 0.54 | 0.22 | 0.29 | 0.24 | 0.3 |
| 1 | 85413765 | rs570564 | -0.907 | 0.052 | -0.448 | 0.648 | 0.016 | 0.015 | 0.015 | 0 | 0.36 | 0.75 | 0.28 | 0.29 | 0.29 | 0.32 |
| 1 | 85416796 | rs12138367 | -0.351 | 0.527 | -0.749 | 0.721 | -0.015 | -0.001 | -0.001 | 0.024 | 0.26 | 0.15 | 0.15 | 0.23 | 0.26 | 0.28 |
| 1 | 85420044 | rs504504 | -0.718 | 0.386 | -0.452 | 0.861 | 0.027 | 0.041 | 0.041 | 0.009 | 0.3 | 0.45 | 0.3 | 0.41 | 0.33 | 0.33 |
| 1 | 85422061 | rs632545 | -1.38 | 0.335 | -0.594 | 0.838 | 0.001 | -0.003 | -0.003 | 0 | 0.43 | 0.55 | 0.54 | 0.66 | 0.49 | 0.52 |
| 1 | 85450920 | rs6691970 | -1.24 | 0.263 | -0.283 | 0.876 | -0.035 | 0.016 | 0.016 | 0 | 0.41 | 0.28 | 0.52 | 0.63 | 0.49 | 0.52 |
| 1 | 85456902 | rs7547224 | -0.042 | 0.241 | -0.322 | 0.901 | -0.033 | -0.028 | -0.028 | 15.045 | 0.07 | 0.04 | 0.23 | 0.26 | 0.14 | 0.16 |
| 1 | 85457066 | rs7533496 | -0.843 | 0.179 | -0.151 | 0.836 | -0.009 | -0.022 | -0.022 | 0 | 0.92 | 0.68 | 0.97 | 0.99 | 0.98 | 0.96 |
| 1 | 85467045 | rs10873679 | 1.301 | 0.216 | -0.146 | 0.922 | 0.003 | -0.004 | -0.004 | 0 | 0.57 | 0.52 | 0.48 | 0.3 | 0.51 | 0.47 |
| 1 | 85467130 | rs12119445 | 0.35 | 0.158 | -0.21 | 0.866 | -0.07 | -0.096 | -0.096 | 0 | 0.22 | 0.1 | 0.22 | 0.42 | 0.27 | 0.31 |
| 1 | 85481700 | rs12730329 | 0.559 | 0.333 | -0.235 | 1.255 | -0.059 | -0.02 | -0.02 | 0 | 0.2 | 0.09 | 0.21 | 0.24 | 0.24 | 0.29 |
| 1 | 85484366 | rs7522239 | 0.576 | 0.33 | -0.032 | 1.252 | -0.003 | 0.015 | 0.015 | 0 | 0.85 | 0.83 | 0.75 | 0.73 | 0.84 | 0.8 |
| 1 | 85484473 | rs10782537 | 1.567 | 0.329 | -0.009 | 1.252 | 0.003 | 0.097 | 0.097 | 0 | 0.78 | 0.66 | 0.69 | 0.54 | 0.75 | 0.76 |

| | | | | | | | | | | | | | | | | |
|---|---|---|---|---|---|---|---|---|---|---|---|---|---|---|---|---|
| 1 | 85485768 | rs10443172 | 1.207 | 0.329 | -0.009 | 1.252 | -0.007 | 0.01 | 0.01 | 0 | 0.79 | 0.75 | 0.71 | 0.73 | 0.79 | 0.79 |
| 1 | 85487520 | rs10518327 | 1.711 | 0.459 | -0.563 | 0.939 | -0.106 | -0.008 | -0.008 | 0.09 | 0.16 | 0.01 | 0.21 | 0.1 | 0.27 | 0.24 |
| 1 | 85488030 | rs2028304 | 0.451 | 0.515 | -0.024 | 1.442 | -0.003 | 0.014 | 0.014 | 0 | 0.85 | 0.83 | 0.75 | 0.73 | 0.84 | 0.8 |
| 1 | 85496484 | rs11161512 | -0.698 | 0.512 | -0.066 | 1.364 | 0.007 | 0.089 | 0.089 | 0 | 0.09 | 0.11 | 0.25 | 0.47 | 0.18 | 0.18 |
| 1 | 85497812 | rs10873682 | 1.469 | 0.533 | -0.126 | 1.32 | 0.002 | -0.004 | -0.004 | 0.013 | 0.56 | 0.44 | 0.47 | 0.31 | 0.51 | 0.47 |
| 1 | 85501110 | rs7545349 | 0.779 | 0.642 | -0.236 | 1.224 | -0.003 | -0.026 | -0.026 | 0 | 0.06 | 0.08 | 0.04 | 0 | 0.05 | 0.02 |
| 1 | 85519418 | rs12755226 | 0.322 | 0.798 | -0.506 | 1.329 | -0.02 | -0.001 | -0.001 | 0 | 0.21 | 0.18 | 0.19 | 0.24 | 0.24 | 0.27 |
| 1 | 85527701 | rs6681619 | 0.135 | 0.493 | -0.316 | 1.286 | -0.02 | 0.015 | 0.015 | 0 | 0.82 | 0.73 | 0.91 | 0.79 | 0.88 | 0.9 |
| 1 | 85527974 | rs4907128 | -0.175 | 0.502 | -0.3 | 1.287 | -0.02 | -0.053 | -0.053 | 0.022 | 0.84 | 0.8 | 0.93 | 0.99 | 0.91 | 0.93 |
| 1 | 85528006 | rs12034421 | -0.348 | 0.449 | -0.218 | 1.348 | -0.009 | -0.012 | -0.012 | 4.497 | 0.27 | 0.29 | 0.41 | 0.49 | 0.38 | 0.42 |
| 1 | 85529523 | rs12035525 | 1.463 | -0.519 | -0.421 | 1.244 | 0 | -0.061 | -0.061 | 0.085 | 0.1 | 0 | 0.09 | 0.19 | 0.06 | 0.19 |
| 1 | 85531711 | rs6695223 | 0.326 | -0.182 | 0.042 | 1.134 | -0.011 | 0.008 | 0.008 | 0 | 0.83 | 0.77 | 0.91 | 0.79 | 0.87 | 0.9 |
| 1 | 85537890 | rs7527292 | 0.595 | -0.239 | -0.138 | 1.169 | -0.037 | 0.044 | 0.044 | 0 | 0.81 | 0.56 | 0.88 | 0.71 | 0.87 | 0.88 |
| 1 | 85538947 | rs4907133 | 0.958 | -0.222 | -0.391 | 1.281 | -0.008 | 0.037 | 0.037 | 0 | 0.76 | 0.7 | 0.7 | 0.53 | 0.77 | 0.75 |
| 1 | 85539791 | rs1037741 | 0.78 | -0.462 | -0.537 | 1.036 | -0.029 | 0.019 | 0.019 | 0 | 0.68 | 0.6 | 0.67 | 0.57 | 0.77 | 0.68 |
| 1 | 85541164 | rs12079571 | 0.45 | -0.266 | -0.811 | 0.914 | -0.013 | -0.007 | -0.007 | 0 | 0.16 | 0.14 | 0.16 | 0.24 | 0.2 | 0.26 |
| 1 | 85541721 | rs7517848 | 1.475 | -0.361 | -0.848 | 0.861 | 0.005 | 0.002 | 0.002 | 0 | 0.52 | 0.56 | 0.52 | 0.47 | 0.58 | 0.49 |
| 1 | 85545248 | rs1030934 | 1.104 | -0.136 | -0.752 | 0.914 | 0 | -0.001 | -0.001 | 0.154 | 0.58 | 0.6 | 0.54 | 0.49 | 0.61 | 0.52 |
| 1 | 85545707 | rs1975365 | 0.993 | -0.156 | -0.762 | 0.904 | -0.001 | 0.001 | 0.001 | 0 | 0.61 | 0.6 | 0.54 | 0.49 | 0.61 | 0.52 |
| 1 | 85552682 | rs12722826 | 0.614 | 0.096 | -0.683 | 0.927 | -0.06 | 0.035 | 0.035 | 0.595 | 0.14 | 0.02 | 0.15 | 0.1 | 0.19 | 0.19 |
| 1 | 85552771 | rs12722966 | 0.473 | 0.067 | -0.649 | 0.912 | -0.013 | -0.002 | -0.002 | 0 | 0.16 | 0.14 | 0.16 | 0.18 | 0.2 | 0.25 |
| 1 | 85554402 | rs1542427 | 0.25 | -0.132 | -0.583 | 0.71 | -0.013 | -0.003 | -0.003 | 0 | 0.74 | 0.74 | 0.7 | 0.73 | 0.81 | 0.77 |
| 1 | 85556791 | rs1770691 | -0.835 | 0.183 | -0.567 | 0.599 | -0.001 | 0 | 0 | 0 | 0.45 | 0.55 | 0.48 | 0.73 | 0.43 | 0.53 |
| 1 | 85560264 | rs709764 | -0.055 | -0.376 | 0.086 | 0.062 | -0.014 | -0.079 | -0.079 | 0 | 0.12 | 0.36 | 0.12 | 0 | 0.12 | 0.02 |
| 1 | 85561954 | rs709767 | -0.477 | -0.019 | -0.587 | -0.044 | -0.006 | 0.068 | 0.068 | 0 | 0.53 | 0.66 | 0.44 | 0.26 | 0.5 | 0.44 |
| 1 | 85568972 | rs1026469 | 0.801 | -0.044 | -0.841 | 0.15 | -0.009 | -0.037 | -0.037 | 0 | 0.34 | 0.21 | 0.32 | 0.41 | 0.36 | 0.45 |
| 1 | 85569106 | rs709771 | -0.422 | -0.022 | -0.883 | 0.275 | -0.002 | -0.004 | -0.004 | 0.993 | 0.59 | 0.66 | 0.48 | 0.41 | 0.57 | 0.46 |
| 1 | 85574748 | rs709760 | -0.629 | 0.039 | -0.781 | 0.211 | -0.003 | -0.006 | -0.006 | 0 | 0.59 | 0.67 | 0.63 | 0.44 | 0.57 | 0.53 |
| 1 | 85578057 | rs817482 | -0.953 | -0.261 | 0.03 | 0.556 | -0.038 | -0.024 | -0.024 | 0 | 0.2 | 0.42 | 0.31 | 0.18 | 0.18 | 0.11 |
| 1 | 85582549 | rs2050842 | 0.435 | 0.116 | -0.206 | 0.432 | 0.003 | 0.007 | 0.007 | 0 | 0.61 | 0.58 | 0.66 | 0.58 | 0.55 | 0.64 |
| 1 | 85583985 | rs817472 | 0.995 | -0.235 | 0.518 | 0.682 | -0.012 | 0.008 | 0.008 | 0 | 0.9 | 0.7 | 0.92 | 0.93 | 0.95 | 0.93 |
| 1 | 85584046 | rs4556376 | 1.26 | -0.245 | 0.62 | 0.793 | -0.009 | 0.025 | 0.025 | 0.448 | 0.7 | 0.37 | 0.73 | 0.58 | 0.69 | 0.81 |
| 1 | 85585260 | rs6576750 | -0.669 | -0.265 | 0.675 | 0.806 | -0.016 | 0.001 | 0.001 | 0 | 0.08 | 0.2 | 0.07 | 0.07 | 0.05 | 0.07 |
| 1 | 85585700 | rs17099217 | 1.85 | -0.257 | 0.639 | 0.912 | -0.019 | -0.035 | -0.035 | 0 | 0.06 | 0 | 0.04 | 0.14 | 0.07 | 0.02 |
| 1 | 85586170 | rs6576752 | -0.165 | -0.254 | 0.658 | 0.914 | 0.01 | 0.083 | 0.083 | 0 | 0.85 | 0.59 | 0.85 | 0.8 | 0.81 | 0.9 |
| 1 | 85588015 | rs1026468 | 0.517 | -0.248 | 0.675 | 0.913 | 0.002 | 0.091 | 0.091 | 0 | 0.78 | 0.56 | 0.8 | 0.65 | 0.74 | 0.88 |
| 1 | 85589842 | rs17121745 | 1.852 | -0.252 | 0.651 | 0.914 | -0.019 | -0.035 | -0.035 | 0 | 0.06 | 0 | 0.04 | 0.14 | 0.07 | 0.02 |
| 1 | 85590246 | rs11161532 | 1.517 | -0.264 | 0.665 | 0.906 | -0.011 | 0.035 | 0.035 | 0.109 | 0.71 | 0.39 | 0.73 | 0.58 | 0.69 | 0.81 |
| 1 | 85594020 | rs709782 | 0.409 | -0.305 | 0.614 | 0.748 | -0.013 | -0.003 | -0.003 | 0.738 | 0.91 | 0.89 | 0.93 | 0.93 | 0.95 | 0.92 |
| 1 | 85596749 | rs817476 | -0.27 | -0.193 | 0.182 | 0.806 | -0.002 | 0.057 | 0.057 | 0 | 0.15 | 0.33 | 0.24 | 0.21 | 0.18 | 0.1 |
| 1 | 85602412 | rs1870327 | -1.1 | 0.046 | 0.124 | 0.811 | -0.006 | 0.06 | 0.06 | 0 | 0.23 | 0.45 | 0.31 | 0.31 | 0.25 | 0.17 |
| 1 | 85609928 | rs7532792 | 0.599 | -0.268 | -0.371 | -0.138 | 0.015 | 0.058 | 0.058 | 0 | 0.46 | 0.33 | 0.69 | 0.76 | 0.55 | 0.67 |
| 1 | 85610585 | rs7513769 | 1.242 | -0.358 | -0.169 | -0.167 | -0.005 | 0.011 | 0.011 | 0 | 0.09 | 0.14 | 0.06 | 0.17 | 0.06 | 0.02 |
| 1 | 85613172 | rs817395 | -1.031 | -0.386 | 0.123 | -0.186 | -0.004 | 0.026 | 0.026 | 0 | 0.91 | 0.78 | 0.94 | 0.82 | 0.94 | 0.98 |
| 1 | 85615649 | rs12750492 | 1.085 | -0.14 | -0.314 | -0.003 | -0.088 | -0.071 | -0.071 | 0 | 0.29 | 0.08 | 0.33 | 0.35 | 0.38 | 0.43 |
| 1 | 85617351 | rs817403 | 1.699 | -0.049 | 0.09 | -0.06 | -0.001 | -0.017 | -0.017 | 0 | 0.07 | 0.07 | 0.08 | 0 | 0.1 | 0.02 |
| 1 | 85625855 | rs817413 | -0.069 | -0.221 | 0.62 | -0.108 | -0.022 | -0.039 | -0.039 | 1.092 | 0.22 | 0.42 | 0.16 | 0.17 | 0.19 | 0.05 |
| 1 | 85637648 | rs817432 | -0.296 | -0.202 | 0.32 | -0.099 | -0.03 | -0.051 | -0.051 | 0 | 0.23 | 0.5 | 0.17 | 0.18 | 0.19 | 0.05 |
| 1 | 85647868 | rs17122989 | 1.036 | -0.183 | -0.063 | -0.094 | -0.006 | -0.027 | -0.027 | 0 | 0.06 | 0.18 | 0.03 | 0 | 0.02 | 0.01 |
| 1 | 85648905 | rs11161539 | 0.193 | -0.236 | -0.791 | 0.035 | 0.011 | -0.015 | -0.015 | 1.436 | 0.62 | 0.24 | 0.52 | 0.54 | 0.64 | 0.72 |
| 1 | 85653831 | rs11161543 | 0.798 | -0.426 | -0.774 | -0.096 | 0.013 | 0.029 | 0.029 | 0.123 | 0.29 | 0.15 | 0.16 | 0.18 | 0.25 | 0.28 |
| 1 | 85656680 | rs817452 | -0.595 | -0.237 | -0.235 | 0.055 | -0.023 | -0.035 | -0.035 | 0 | 0.23 | 0.6 | 0.18 | 0.18 | 0.19 | 0.05 |
| 1 | 85662851 | rs4907139 | 1.208 | 0.243 | -0.672 | 0.42 | -0.016 | -0.04 | -0.04 | 0.119 | 0.15 | 0.1 | 0.3 | 0.28 | 0.19 | 0.26 |
| 1 | 85679397 | rs709790 | 0.038 | -0.236 | -0.343 | -0.039 | -0.008 | -0.012 | -0.012 | 0 | 0.41 | 0.46 | 0.42 | 0.32 | 0.35 | 0.38 |
| 1 | 85680408 | rs11161548 | -0.417 | 0.323 | -0.704 | -0.21 | 0.006 | 0.004 | 0.004 | 0 | 0.42 | 0.2 | 0.36 | 0.6 | 0.39 | 0.5 |
| 1 | 85681267 | rs12117219 | 0.276 | 0.346 | -0.684 | 0.036 | 0.001 | -0.002 | -0.002 | 0 | 0.21 | 0.16 | 0.17 | 0.25 | 0.23 | 0.12 |
| 1 | 85681278 | rs10218661 | -0.77 | 0.365 | -0.409 | 0.026 | 0 | 0 | 0 | 0 | 0.54 | 0.48 | 0.5 | 0.59 | 0.57 | 0.53 |
| 1 | 85682261 | rs12065219 | -0.213 | 0.458 | -0.317 | -0.033 | -0.008 | -0.01 | -0.01 | 0 | 0.43 | 0.36 | 0.44 | 0.5 | 0.46 | 0.41 |
| 1 | 85684842 | rs10873694 | 0.785 | 0.41 | -0.026 | -0.055 | -0.024 | -0.035 | -0.035 | 0.082 | 0.31 | 0.15 | 0.32 | 0.35 | 0.33 | 0.38 |
| 1 | 85686552 | rs963208 | 0.794 | 0.455 | 0.092 | -0.034 | -0.004 | -0.005 | -0.005 | 1.561 | 0.31 | 0.26 | 0.33 | 0.35 | 0.33 | 0.37 |
| 1 | 85694671 | rs17123650 | 2.044 | 0.31 | -0.166 | -0.25 | -0.043 | -0.071 | -0.071 | 0.231 | 0.16 | 0.05 | 0.31 | 0.3 | 0.2 | 0.28 |
| 1 | 85696004 | rs6660237 | -0.358 | 0.237 | -0.097 | -0.315 | 0.001 | 0.01 | 0.01 | 0 | 0.33 | 0.61 | 0.48 | 0.43 | 0.38 | 0.37 |
| 1 | 85700596 | rs9432848 | -1.32 | 0.27 | -0.299 | -0.285 | 0.011 | 0.091 | 0.091 | 0 | 0.7 | 0.88 | 0.84 | 0.93 | 0.74 | 0.75 |
| 1 | 85701071 | rs11161557 | 0.661 | 0.536 | 0.362 | 0.292 | 0 | 0.01 | 0.01 | 0 | 0.08 | 0.07 | 0.07 | 0.12 | 0.08 | 0.07 |
| 1 | 85701567 | rs9433250 | 0.455 | 0.54 | 0.269 | 0.295 | 0.008 | 0.038 | 0.038 | 0 | 0.09 | 0.17 | 0.07 | 0.12 | 0.08 | 0.07 |
| 1 | 85704193 | rs7524465 | 0.184 | 0.538 | -0.27 | 0.176 | 0.015 | 0.076 | 0.076 | 0 | 0.26 | 0.52 | 0.4 | 0.43 | 0.28 | 0.36 |
| 1 | 85709133 | rs17382610 | 1.552 | 0.646 | -0.737 | 0.036 | -0.003 | 0.013 | 0.013 | 0 | 0.06 | 0.05 | 0.07 | 0 | 0.09 | 0.02 |
| 1 | 85712287 | rs3942955 | 0.508 | 0.507 | -1.063 | -0.086 | -0.012 | 0.121 | 0.121 | 0.107 | 0.48 | 0.23 | 0.38 | 0.23 | 0.5 | 0.37 |
| 1 | 85713223 | rs10782545 | -0.255 | 0.397 | -0.253 | 0.311 | -0.003 | 0.097 | 0.097 | 0 | 0.22 | 0.42 | 0.18 | 0.35 | 0.17 | 0.36 |

| | | | | | | | | | | | | | | | | |
|---|---|---|---|---|---|---|---|---|---|---|---|---|---|---|---|---|
| 1 | 85716807 | rs10782546 | -0.044 | 0.364 | 0.252 | 0.295 | -0.001 | 0.04 | 0.04 | 0 | 0.21 | 0.26 | 0.17 | 0.35 | 0.17 | 0.36 |
| 1 | 85719610 | rs4074156 | 1.475 | 0.279 | 0.318 | 0.427 | -0.002 | 0.002 | 0.002 | 0 | 0.09 | 0.06 | 0.09 | 0.21 | 0.09 | 0.19 |
| 1 | 85720007 | rs10158486 | 2.398 | 0.421 | -0.044 | 0.076 | 0.003 | 0.001 | 0.001 | 0 | 0.15 | 0.18 | 0.19 | 0.16 | 0.21 | 0.1 |
| 1 | 85720013 | rs750517 | -0.347 | 0.393 | -0.032 | 0.022 | 0.003 | 0.001 | 0.001 | 0 | 0.55 | 0.48 | 0.37 | 0.42 | 0.46 | 0.63 |
| 1 | 85721911 | rs11161570 | -0.989 | 0.305 | 0.114 | 0.079 | 0.007 | 0.006 | 0.006 | 0 | 0.62 | 0.6 | 0.41 | 0.54 | 0.49 | 0.65 |
| 1 | 85724581 | rs11161573 | 0.877 | 0.241 | 0.187 | 0.021 | 0.003 | -0.012 | -0.012 | 0 | 0.36 | 0.34 | 0.21 | 0.07 | 0.3 | 0.27 |
| 1 | 85726103 | rs17124345 | 0.199 | 0.357 | 0.373 | -0.012 | -0.002 | 0 | 0 | 0 | 0.19 | 0.19 | 0.17 | 0.21 | 0.18 | 0.21 |
| 1 | 85727044 | rs11161574 | -2.822 | 0.388 | 0.342 | 0.014 | 0.012 | 0.024 | 0.024 | 0 | 0.85 | 0.84 | 0.68 | 0.71 | 0.79 | 0.76 |
| 1 | 85727218 | rs10493763 | 2.508 | 0.381 | 0.287 | 0.047 | 0.001 | 0 | 0 | 2.047 | 0.15 | 0.15 | 0.19 | 0.16 | 0.21 | 0.1 |
| 1 | 85727335 | rs12143319 | 1.012 | 0.357 | 0.215 | 0.057 | 0.009 | -0.034 | -0.034 | 0 | 0.09 | 0.02 | 0.07 | 0.15 | 0.07 | 0.17 |
| 1 | 85729987 | rs2647394 | 0.242 | 0.373 | 0.262 | 0.043 | 0.017 | 0.045 | 0.045 | 0 | 0.44 | 0.34 | 0.28 | 0.22 | 0.37 | 0.44 |
| 1 | 85732941 | rs962409 | -0.076 | 0.37 | 0.208 | 0.185 | -0.001 | 0 | 0 | 0 | 0.52 | 0.37 | 0.36 | 0.44 | 0.45 | 0.63 |
| 1 | 85738308 | rs12134420 | 2.462 | 0.44 | 0.264 | 0.16 | 0.005 | 0.001 | 0.001 | 0 | 0.15 | 0.19 | 0.2 | 0.16 | 0.2 | 0.1 |
| 1 | 85738395 | rs11161581 | 1.468 | 1.041 | 0.223 | 0.014 | -0.009 | -0.009 | -0.009 | 0 | 0.1 | 0.07 | 0.11 | 0.21 | 0.15 | 0.2 |
| 1 | 85740024 | rs4949927 | 0.998 | 0.947 | 0.222 | -0.051 | 0.018 | 0.021 | 0.021 | 0 | 0.33 | 0.27 | 0.18 | 0.07 | 0.24 | 0.26 |
| 1 | 85743452 | rs11555520 | 1.793 | 0.632 | -0.489 | -0.37 | -0.064 | 0.162 | 0.162 | 1.558 | 0.24 | 0.01 | 0.11 | 0.01 | 0.24 | 0.11 |
| 1 | 85743741 | rs2735592 | -1.271 | 0.745 | -0.264 | 0.033 | 0.002 | 0.001 | 0.001 | 0 | 0.77 | 0.78 | 0.61 | 0.72 | 0.69 | 0.75 |
| 1 | 85744472 | rs2735591 | -1.271 | 0.745 | -0.264 | 0.033 | 0.002 | 0.001 | 0.001 | 0 | 0.77 | 0.78 | 0.61 | 0.72 | 0.69 | 0.75 |
| 1 | 85744577 | rs9887812 | 1.094 | 0.412 | -0.54 | 0.09 | -0.003 | -0.005 | -0.005 | 0 | 0.09 | 0.07 | 0.09 | 0.21 | 0.08 | 0.2 |
| 1 | 85755925 | rs6670921 | 0.933 | 0.2 | -0.667 | -0.225 | 0.055 | 0.119 | 0.119 | 0 | 0.32 | 0.12 | 0.17 | 0.08 | 0.24 | 0.22 |
| 1 | 85763899 | rs2030075 | -0.289 | 0.513 | -0.623 | -0.126 | 0.018 | 0.023 | 0.023 | 0 | 0.53 | 0.4 | 0.34 | 0.39 | 0.4 | 0.53 |
| 1 | 85765619 | rs12098220 | 1.433 | 0.698 | -0.856 | -0.05 | 0 | -0.005 | -0.005 | 0 | 0.08 | 0.05 | 0.09 | 0.19 | 0.06 | 0.22 |
| 1 | 85765798 | rs11587512 | 0.782 | 0.706 | -0.909 | -0.123 | 0.035 | 0.116 | 0.116 | 0 | 0.35 | 0.16 | 0.2 | 0.08 | 0.29 | 0.26 |
| 1 | 85768379 | rs17125439 | -0.28 | 0.812 | -0.707 | -0.034 | 0.013 | -0.004 | -0.004 | 0 | 0.25 | 0.21 | 0.23 | 0.29 | 0.22 | 0.3 |
| 1 | 85769032 | rs17125445 | 0.934 | 0.623 | -0.762 | -0.149 | 0.019 | 0.058 | 0.058 | 0 | 0.29 | 0.18 | 0.17 | 0.08 | 0.23 | 0.22 |
| 1 | 85770794 | rs7530433 | -0.326 | 1.085 | -0.466 | 0.679 | 0.088 | 0.188 | 0.188 | 0.776 | 0.95 | 0.84 | 0.79 | 0.72 | 0.89 | 0.98 |
| 1 | 85772009 | rs233100 | -0.335 | 0.997 | -0.725 | 0.476 | -0.003 | -0.004 | -0.004 | 0 | 0.42 | 0.32 | 0.39 | 0.55 | 0.45 | 0.34 |
| 1 | 85775186 | rs233104 | 0.261 | 0.368 | -0.471 | 0.63 | 0.002 | -0.007 | -0.007 | 0 | 0.12 | 0.08 | 0.07 | 0.22 | 0.08 | 0.25 |
| 1 | 85775865 | rs12122154 | 1.696 | 0.418 | -0.514 | 0.617 | 0.003 | -0.014 | -0.014 | 0.057 | 0.07 | 0.03 | 0.03 | 0.1 | 0.04 | 0.2 |
| 1 | 85785655 | rs233111 | 0.037 | 0.405 | -0.017 | 0.562 | -0.002 | 0.011 | 0.011 | 0.894 | 0.76 | 0.67 | 0.67 | 0.66 | 0.71 | 0.85 |
| 1 | 85785751 | rs233112 | 0.616 | 0.672 | -0.4 | 0.536 | -0.009 | -0.01 | -0.01 | 0 | 0.34 | 0.32 | 0.33 | 0.41 | 0.38 | 0.42 |
| 1 | 85786977 | rs233115 | 1.039 | 0.642 | -0.396 | 0.524 | -0.006 | -0.003 | -0.003 | 0 | 0.31 | 0.3 | 0.32 | 0.29 | 0.36 | 0.41 |
| 1 | 85787334 | rs1498374 | 2.569 | 0.792 | -0.524 | 0.604 | -0.065 | -0.048 | -0.048 | 0 | 0.13 | 0.02 | 0.19 | 0.16 | 0.2 | 0.1 |
| 1 | 85789270 | rs17590006 | 1.932 | 0.45 | -0.646 | 0.409 | -0.015 | 0.094 | 0.094 | 0 | 0.25 | 0.04 | 0.26 | 0.1 | 0.23 | 0.18 |
| 1 | 85795112 | rs233130 | 0.974 | 0.638 | -0.438 | 0.593 | -0.005 | -0.002 | -0.002 | 0 | 0.31 | 0.32 | 0.32 | 0.29 | 0.36 | 0.41 |
| 1 | 85807695 | rs233072 | 0.626 | 0.641 | -0.397 | 0.635 | -0.006 | -0.008 | -0.008 | 0.292 | 0.34 | 0.33 | 0.34 | 0.42 | 0.38 | 0.42 |
| 1 | 85837793 | rs233050 | 1.093 | 0.67 | -0.5 | 0.616 | -0.051 | -0.032 | -0.032 | 0 | 0.31 | 0.15 | 0.32 | 0.3 | 0.37 | 0.42 |
| 1 | 85838413 | rs233051 | -0.238 | 0.605 | -0.362 | 0.57 | -0.003 | 0.012 | 0.012 | 1.2 | 0.74 | 0.64 | 0.65 | 0.65 | 0.7 | 0.84 |
| 1 | 85851308 | rs761602 | -2.215 | 0.526 | -0.381 | 0.519 | 0 | 0 | 0 | 0 | 0.91 | 0.89 | 0.91 | 0.91 | 0.89 | 0.95 |
| 1 | 85853902 | rs11161606 | 0.364 | 0.508 | -0.358 | 0.532 | -0.013 | -0.02 | -0.02 | 0 | 0.4 | 0.35 | 0.35 | 0.43 | 0.42 | 0.62 |
| 1 | 85874495 | rs6657732 | -0.585 | 0.403 | -0.304 | 0.464 | 0.001 | -0.001 | -0.001 | 0 | 0.61 | 0.55 | 0.49 | 0.34 | 0.51 | 0.55 |
| 1 | 85875622 | rs6673833 | 1.395 | 0.379 | -0.106 | 0.387 | 0.004 | 0.028 | 0.028 | 0 | 0.3 | 0.18 | 0.28 | 0.11 | 0.27 | 0.22 |
| 1 | 85882905 | rs1975755 | -1.13 | 0.569 | -0.272 | 0.539 | 0 | 0 | 0 | 0 | 0.75 | 0.71 | 0.86 | 0.73 | 0.8 | 0.49 |
| 1 | 85889596 | rs7521189 | -0.249 | 0.561 | -0.19 | 0.41 | 0.001 | 0.007 | 0.007 | 0 | 0.55 | 0.53 | 0.47 | 0.21 | 0.47 | 0.35 |
| 1 | 85898042 | rs551140 | 1.002 | 0.503 | -0.18 | 0.393 | -0.004 | -0.013 | -0.013 | 0 | 0.1 | 0.11 | 0.07 | 0.01 | 0.08 | 0.08 |
| 1 | 85898330 | rs553257 | 0.298 | 0.486 | -0.143 | 0.372 | -0.008 | -0.018 | -0.018 | 0 | 0.19 | 0.24 | 0.17 | 0.1 | 0.19 | 0.13 |
| 1 | 85899128 | rs1523993 | 1.119 | 0.506 | -0.212 | 0.34 | 0.002 | -0.007 | -0.007 | 0.02 | 0.32 | 0.26 | 0.29 | 0.11 | 0.27 | 0.22 |
| 1 | 85902045 | rs593728 | -1.603 | 0.808 | -0.377 | 0.698 | 0.001 | 0.01 | 0.01 | 0 | 0.92 | 0.93 | 0.97 | 0.87 | 0.93 | 0.8 |
| 1 | 85904338 | rs539714 | -2.043 | 0.776 | -0.385 | 0.715 | 0.006 | 0.001 | 0.001 | 0 | 0.83 | 0.74 | 0.88 | 0.86 | 0.84 | 0.69 |
| 1 | 85906816 | rs11161615 | 0.959 | 0.835 | -0.507 | 0.807 | -0.027 | -0.083 | -0.083 | 0.013 | 0.23 | 0.09 | 0.24 | 0.41 | 0.26 | 0.33 |
| 1 | 85907625 | rs11161616 | 1.822 | 0.865 | -0.501 | 0.806 | -0.038 | -0.049 | -0.049 | 2.629 | 0.17 | 0.06 | 0.23 | 0.28 | 0.22 | 0.12 |
| 1 | 85915134 | rs648310 | 0.132 | 0.865 | -0.306 | 0.717 | 0.002 | 0.003 | 0.003 | 0 | 0.51 | 0.4 | 0.38 | 0.26 | 0.42 | 0.51 |
| 1 | 85915149 | rs659070 | -2.018 | 0.97 | -0.69 | 0.813 | 0.002 | 0 | 0 | 0 | 0.83 | 0.76 | 0.88 | 0.87 | 0.85 | 0.7 |
| 1 | 85915402 | rs11161618 | 1.147 | 0.762 | -0.92 | 0.652 | -0.095 | 0.113 | 0.113 | 0.506 | 0.42 | 0.06 | 0.47 | 0.24 | 0.45 | 0.28 |
| 1 | 85918101 | rs1146382 | 0.067 | 1.223 | -0.351 | 1.134 | -0.008 | 0.005 | 0.005 | 0 | 0.58 | 0.64 | 0.63 | 0.45 | 0.58 | 0.37 |
| 1 | 85918329 | rs1146383 | -1.134 | 1.269 | -0.217 | 1.16 | 0.006 | 0.018 | 0.018 | 0.325 | 0.81 | 0.71 | 0.86 | 0.74 | 0.84 | 0.66 |
| 1 | 85922661 | rs506733 | -0.323 | 1.118 | -0.661 | 1.119 | -0.041 | -0.061 | -0.061 | 0 | 0.7 | 0.86 | 0.63 | 0.59 | 0.65 | 0.65 |
| 1 | 85923920 | rs877041 | 1.153 | 1.092 | -0.215 | 0.892 | 0 | 0 | 0 | 0 | 0.35 | 0.3 | 0.43 | 0.3 | 0.37 | 0.22 |
| 1 | 85923999 | rs582145 | -1.029 | 1.092 | -0.111 | 0.938 | 0.001 | -0.004 | -0.004 | 0 | 0.61 | 0.47 | 0.54 | 0.59 | 0.61 | 0.76 |
| 1 | 85927463 | rs480414 | -0.042 | 1.188 | -0.12 | 1.007 | 0.004 | 0.002 | 0.002 | 0 | 0.36 | 0.38 | 0.27 | 0.28 | 0.3 | 0.45 |
| 1 | 85927879 | rs1241321 | 0.309 | 1.085 | -0.037 | 0.893 | 0.003 | 0.024 | 0.024 | 0 | 0.28 | 0.31 | 0.33 | 0.4 | 0.29 | 0.43 |
| 1 | 85929214 | rs527762 | -1.2 | 1.067 | -0.443 | 0.457 | -0.006 | -0.032 | -0.032 | 0 | 0.64 | 0.61 | 0.71 | 0.81 | 0.7 | 0.78 |
| 1 | 85933638 | rs1403956 | 0.682 | 1.14 | -0.983 | 0.623 | -0.021 | 0.118 | 0.118 | 0 | 0.47 | 0.21 | 0.46 | 0.24 | 0.51 | 0.32 |
| 1 | 85939275 | rs10782551 | 0.796 | 1.221 | -0.806 | 1.021 | 0 | 0.025 | 0.025 | 0 | 0.08 | 0.09 | 0.04 | 0.23 | 0.06 | 0.16 |
| 1 | 85939350 | rs7538364 | -0.941 | 1.983 | -0.686 | 0.749 | 0.006 | 0.018 | 0.018 | 0 | 0.5 | 0.59 | 0.5 | 0.36 | 0.53 | 0.55 |
| 1 | 85939476 | rs10158333 | 0.049 | 2.71 | -1.047 | 1.231 | -0.015 | -0.018 | -0.018 | 0 | 0.25 | 0.16 | 0.29 | 0.32 | 0.29 | 0.3 |
| 1 | 85940105 | rs1403955 | -0.244 | 1.693 | -1.255 | 0.725 | -0.008 | 0.001 | 0.001 | 0 | 0.37 | 0.38 | 0.39 | 0.31 | 0.41 | 0.51 |
| 1 | 85941403 | rs10493765 | -0.52 | 2.313 | -1.758 | 0.784 | -0.094 | -0.045 | -0.045 | 0 | 0.15 | 0.02 | 0.14 | 0.17 | 0.27 | 0.09 |
| 1 | 85941419 | rs7512147 | 0.26 | 2.196 | -2.463 | 1.203 | -0.058 | -0.119 | -0.119 | 0 | 0.21 | 0.05 | 0.2 | 0.34 | 0.28 | 0.22 |

| | | | | | | | | | | | | | | | | |
|---|---|---|---|---|---|---|---|---|---|---|---|---|---|---|---|---|
| 1 | 85942086 | rs483784 | 0.434 | 2.417 | -1.613 | 0.737 | 0.019 | -0.001 | -0.001 | 0 | 0.74 | 0.32 | 0.59 | 0.73 | 0.65 | 0.9 |
| 1 | 85942879 | rs512298 | 0.221 | 1.896 | -2.132 | 0.475 | -0.064 | 0.052 | 0.052 | 1.726 | 0.35 | 0.06 | 0.24 | 0.24 | 0.37 | 0.38 |
| 1 | 85945703 | rs4949907 | 0.045 | 1.841 | -1.469 | 0.484 | 0.013 | -0.018 | -0.018 | 0.123 | 0.17 | 0.22 | 0.18 | 0.14 | 0.2 | 0.11 |
| 1 | 85946120 | rs1146391 | -0.097 | 1.901 | -1.078 | 0.522 | 0.021 | -0.008 | -0.008 | 0.016 | 0.83 | 0.43 | 0.72 | 0.86 | 0.84 | 0.98 |
| 1 | 85947200 | rs1146394 | 0.139 | 1.914 | -1.305 | 0.371 | -0.016 | 0.062 | 0.062 | 0.904 | 0.4 | 0.2 | 0.29 | 0.29 | 0.41 | 0.46 |
| 1 | 85951597 | rs1146395 | -0.599 | 1.869 | -0.893 | 0.35 | 0.056 | 0.068 | 0.068 | 0.498 | 0.53 | 0.31 | 0.44 | 0.3 | 0.67 | 0.48 |
| 1 | 85954113 | rs562080 | -0.149 | 1.879 | -0.948 | 0.318 | -0.036 | 0.15 | 0.15 | 1.033 | 0.47 | 0.13 | 0.36 | 0.26 | 0.52 | 0.4 |
| 1 | 85955261 | rs6665825 | -0.369 | 2.112 | -0.826 | 0.547 | 0.074 | 0.022 | 0.022 | 0 | 0.06 | 0.18 | 0.08 | 0.08 | 0.15 | 0.12 |
| 1 | 85955709 | rs17099243 | 0.946 | 1.885 | -1.028 | 0.607 | 0.094 | -0.085 | -0.085 | 0 | 0.4 | 0.09 | 0.33 | 0.43 | 0.26 | 0.43 |
| 1 | 85960156 | rs1361714 | 0 | 1.955 | -0.651 | 0.521 | 0.011 | 0.022 | 0.022 | 0.308 | 0.54 | 0.64 | 0.63 | 0.6 | 0.43 | 0.5 |
| 1 | 85967977 | rs12060378 | 0.274 | 1.516 | -1.052 | 0.614 | 0.009 | 0.013 | 0.013 | 0 | 0.47 | 0.27 | 0.54 | 0.47 | 0.45 | 0.48 |
| 1 | 85971852 | rs17392306 | -0.768 | 2.066 | -0.939 | 1.023 | 0.005 | -0.001 | -0.001 | 0.014 | 0.1 | 0.08 | 0.31 | 0.13 | 0.21 | 0.17 |
| 1 | 85972024 | rs10493768 | 0.006 | 1.797 | -0.885 | 0.907 | -0.005 | -0.012 | -0.012 | 0 | 0.7 | 0.29 | 0.61 | 0.5 | 0.68 | 0.69 |
| 1 | 85972426 | rs10493769 | 0.966 | 1.431 | -0.94 | 0.542 | 0.094 | 0.05 | 0.05 | 0 | 0.49 | 0.23 | 0.33 | 0.52 | 0.34 | 0.49 |
| 1 | 85974030 | rs677691 | 1.347 | 2.091 | -0.847 | 0.947 | -0.043 | -0.037 | -0.037 | 0 | 0.63 | 0.19 | 0.32 | 0.59 | 0.56 | 0.7 |
| 1 | 85975076 | rs4420136 | -1.322 | 1.493 | -0.402 | 0.655 | -0.001 | -0.003 | -0.003 | 0.424 | 0.12 | 0.14 | 0.29 | 0.09 | 0.13 | 0.2 |
| 1 | 85995684 | rs683146 | 0.853 | 0.963 | -0.194 | 0.934 | -0.001 | 0.033 | 0.033 | 0 | 0.9 | 0.83 | 0.87 | 0.86 | 0.9 | 0.79 |
| 1 | 86008398 | rs17099679 | -0.634 | 0.526 | 0.096 | 0.826 | -0.034 | -0.015 | -0.015 | 0.067 | 0.19 | 0.35 | 0.17 | 0.17 | 0.13 | 0.1 |
| 1 | 86008536 | rs12734233 | -0.623 | 0.526 | 0.102 | 0.826 | -0.032 | -0.015 | -0.015 | 0 | 0.19 | 0.35 | 0.17 | 0.17 | 0.13 | 0.1 |
| 1 | 86008940 | rs11161635 | 0.409 | 0.864 | -0.816 | 0.788 | -0.042 | -0.025 | -0.025 | 0.176 | 0.17 | 0.05 | 0.1 | 0.21 | 0.21 | 0.21 |
| 1 | 86010248 | rs11161636 | 1.187 | 0.67 | -0.349 | 0.919 | 0.037 | 0.001 | 0.001 | 0 | 0.54 | 0.24 | 0.61 | 0.53 | 0.6 | 0.48 |
| 1 | 86011057 | rs1524004 | -0.255 | 1.912 | -0.012 | 0.964 | 0.001 | -0.001 | -0.001 | 0 | 0.19 | 0.22 | 0.16 | 0.23 | 0.3 | 0.23 |
| 1 | 86017278 | rs1880209 | 0.017 | 2.073 | -0.287 | 0.632 | -0.006 | 0.011 | 0.011 | 0 | 0.2 | 0.12 | 0.2 | 0.14 | 0.26 | 0.11 |
| 1 | 86017485 | rs1524001 | -0.319 | 2.564 | 0.132 | 1.122 | 0.045 | 0.078 | 0.078 | 0 | 0.95 | 0.85 | 0.85 | 0.87 | 0.92 | 0.95 |
| 1 | 86018324 | rs17127877 | -0.279 | 2.734 | 0.054 | 1.255 | -0.009 | -0.009 | -0.009 | 0 | 0.07 | 0.05 | 0.04 | 0.16 | 0.08 | 0.12 |
| 1 | 86019307 | rs17127878 | 0.237 | 2.365 | 0.374 | 1.061 | -0.006 | -0.006 | -0.006 | 0 | 0.14 | 0.32 | 0.28 | 0.13 | 0.13 | 0.17 |
| 1 | 86021169 | rs11161637 | -0.197 | 2.322 | 0.306 | 0.852 | -0.013 | 0.014 | 0.014 | 0 | 0.33 | 0.26 | 0.31 | 0.29 | 0.46 | 0.18 |
| 1 | 86021214 | rs17127881 | 0.254 | 2.124 | 0.37 | 0.948 | -0.009 | -0.009 | -0.009 | 0 | 0.07 | 0.05 | 0.04 | 0.16 | 0.08 | 0.12 |
| 1 | 86028530 | rs1378227 | -0.219 | 2.181 | -0.064 | 0.967 | -0.002 | -0.002 | -0.002 | 2.525 | 0.83 | 0.84 | 0.6 | 0.75 | 0.79 | 0.79 |
| 1 | 86034864 | rs10493770 | -0.709 | 2.145 | -0.201 | 0.963 | -0.015 | -0.017 | -0.017 | 0.047 | 0.11 | 0.05 | 0.06 | 0.19 | 0.13 | 0.13 |
| 1 | 86035513 | rs10493772 | 2.109 | 2.01 | -0.406 | 0.749 | 0.056 | -0.017 | -0.017 | 0 | 0.36 | 0.14 | 0.2 | 0.28 | 0.21 | 0.48 |
| 1 | 86036573 | rs1378228 | 1.054 | 2.048 | -0.34 | 0.765 | 0.048 | -0.015 | -0.015 | 0 | 0.47 | 0.23 | 0.27 | 0.47 | 0.34 | 0.6 |
| 1 | 86037931 | rs2297138 | 0.237 | 1.929 | -0.532 | 0.716 | -0.021 | 0.005 | 0.005 | 0 | 0.21 | 0.03 | 0.28 | 0.15 | 0.19 | 0.15 |
| 1 | 86038476 | rs6682848 | -0.665 | 2.066 | -0.374 | 0.803 | 0.004 | 0.032 | 0.032 | 0 | 0.93 | 0.9 | 0.83 | 0.82 | 0.92 | 0.93 |
| 1 | 86040054 | rs12086058 | -0.078 | 1.968 | -0.171 | 0.56 | 0.043 | -0.01 | -0.01 | 0 | 0.3 | 0.38 | 0.29 | 0.36 | 0.47 | 0.23 |
| 1 | 86045104 | rs3753795 | 1.311 | 1.298 | -0.286 | 0.601 | 0.054 | 0.022 | 0.022 | 0 | 0.4 | 0.26 | 0.23 | 0.27 | 0.26 | 0.51 |
| 1 | 86045888 | rs3753793 | 1.244 | 1.265 | -0.331 | 0.613 | 0.073 | 0.029 | 0.029 | 0 | 0.41 | 0.22 | 0.22 | 0.28 | 0.25 | 0.55 |
| 1 | 86046924 | rs2297141 | 1.189 | 1.561 | -0.095 | 0.606 | -0.003 | 0.002 | 0.002 | 0 | 0.56 | 0.44 | 0.48 | 0.42 | 0.44 | 0.58 |
| 1 | 86050700 | rs1329961 | 0.615 | 1.116 | -0.027 | 0.016 | 0.018 | 0.015 | 0.015 | 0 | 0.22 | 0.32 | 0.29 | 0.33 | 0.29 | 0.17 |
| 1 | 86050965 | rs1571549 | -1.39 | 1.062 | -0.178 | -0.038 | 0.006 | 0.048 | 0.048 | 0 | 0.5 | 0.73 | 0.47 | 0.46 | 0.51 | 0.41 |
| 1 | 86062222 | rs12072027 | 0.703 | 1.139 | -0.086 | -0.114 | 0.011 | 0.005 | 0.005 | 0 | 0.21 | 0.25 | 0.29 | 0.3 | 0.29 | 0.18 |
| 1 | 86064486 | rs7518036 | -0.05 | 0.865 | -0.285 | -0.159 | -0.014 | -0.033 | -0.033 | 0 | 0.06 | 0.16 | 0.04 | 0 | 0.05 | 0.04 |
| 1 | 86069153 | rs2389969 | 1.854 | 0.995 | -0.759 | -0.043 | 0.064 | -0.075 | -0.075 | 0 | 0.36 | 0.13 | 0.29 | 0.42 | 0.24 | 0.47 |
| 1 | 86072216 | rs10873710 | 1.33 | 0.659 | -0.922 | -0.263 | -0.049 | -0.062 | -0.062 | 0 | 0.39 | 0.13 | 0.46 | 0.46 | 0.39 | 0.48 |
| 1 | 86073045 | rs4949915 | 0.835 | 0.78 | -0.671 | -0.109 | -0.02 | -0.029 | -0.029 | 0 | 0.56 | 0.16 | 0.51 | 0.46 | 0.52 | 0.59 |
| 1 | 86074710 | rs17398175 | 1.763 | 0.91 | -0.586 | -0.102 | -0.023 | -0.014 | -0.014 | 0 | 0.16 | 0.08 | 0.27 | 0.24 | 0.27 | 0.09 |
| 1 | 86074975 | rs2026773 | 0.274 | 0.779 | -0.04 | -0.18 | -0.003 | -0.012 | -0.012 | 0 | 0.84 | 0.64 | 0.88 | 0.85 | 0.86 | 0.8 |
| 1 | 86076069 | rs2026774 | -0.206 | 0.739 | -0.007 | -0.192 | -0.008 | -0.03 | -0.03 | 0 | 0.88 | 0.69 | 0.89 | 0.93 | 0.89 | 0.91 |
| 1 | 86081149 | rs12066742 | 1.145 | 0.789 | 0.067 | -0.078 | -0.008 | -0.007 | -0.007 | 0.348 | 0.19 | 0.17 | 0.28 | 0.31 | 0.3 | 0.15 |
| 1 | 86081453 | rs12061951 | 1.231 | 0.793 | 0.024 | -0.076 | -0.02 | -0.016 | -0.016 | 0.502 | 0.18 | 0.12 | 0.28 | 0.31 | 0.3 | 0.14 |
| 1 | 86085545 | rs1999186 | -0.246 | 0.729 | 0.109 | -0.073 | 0.023 | 0.081 | 0.081 | 0 | 0.92 | 0.67 | 0.91 | 0.86 | 0.94 | 0.89 |
| 1 | 86087287 | rs12565129 | 0.931 | 0.649 | -0.092 | -0.131 | -0.002 | -0.043 | -0.043 | 0.014 | 0.1 | 0 | 0.06 | 0.15 | 0.07 | 0.12 |
| 1 | 86092872 | rs11161645 | 0.698 | 0.634 | -0.058 | -0.169 | -0.005 | -0.041 | -0.041 | 0 | 0.63 | 0.2 | 0.55 | 0.45 | 0.6 | 0.62 |
| 1 | 86095613 | rs1360240 | 0.918 | 0.501 | -0.022 | -0.094 | 0.006 | -0.029 | -0.029 | 0 | 0.1 | 0.02 | 0.06 | 0.15 | 0.07 | 0.13 |
| 1 | 86100264 | rs821395 | 2.249 | 0.321 | -0.178 | -0.334 | 0.037 | -0.067 | -0.067 | 0 | 0.36 | 0.14 | 0.19 | 0.39 | 0.25 | 0.51 |
| 1 | 86117583 | rs2069208 | -0.275 | 0.098 | 0.501 | 0.024 | -0.021 | -0.029 | -0.029 | 0 | 0.3 | 0.18 | 0.34 | 0.48 | 0.39 | 0.26 |
| 1 | 86118447 | rs2970548 | 0.301 | 0.019 | 0.801 | -0.033 | 0.043 | 0.068 | 0.068 | 0 | 0.93 | 0.55 | 0.91 | 0.95 | 0.97 | 0.93 |
| 1 | 86121096 | rs821402 | 0.68 | 0.043 | 0.712 | -0.117 | -0.038 | -0.078 | -0.078 | 0 | 0.78 | 0.59 | 0.74 | 0.92 | 0.84 | 0.84 |
| 1 | 86121895 | rs3767163 | 1.219 | 0.262 | 0.547 | -0.069 | -0.035 | -0.016 | -0.016 | 0 | 0.18 | 0.08 | 0.29 | 0.25 | 0.31 | 0.1 |
| 1 | 86125753 | rs10493773 | 1.248 | 0.28 | 0.556 | -0.064 | -0.035 | -0.014 | -0.014 | 0 | 0.18 | 0.08 | 0.29 | 0.25 | 0.31 | 0.1 |
| 1 | 86139089 | rs821378 | -0.3 | 0.141 | 0.625 | -0.15 | -0.02 | -0.043 | -0.043 | 4.597 | 0.86 | 0.71 | 0.88 | 0.95 | 0.89 | 0.92 |
| 1 | 86166943 | rs1764296 | 1.081 | 0.122 | 0.698 | -0.182 | -0.037 | -0.074 | -0.074 | 0.048 | 0.76 | 0.35 | 0.71 | 0.92 | 0.84 | 0.84 |
| 1 | 86172978 | rs952837 | 1.448 | 0.15 | 0.563 | 0.054 | -0.036 | -0.035 | -0.035 | 0 | 0.47 | 0.15 | 0.37 | 0.43 | 0.45 | 0.58 |
| 1 | 86173298 | rs1539444 | -1.275 | 0.171 | 0.661 | 0.06 | -0.004 | 0.005 | 0.005 | 0 | 0.47 | 0.36 | 0.53 | 0.51 | 0.52 | 0.36 |
| 1 | 86174563 | rs1764285 | 1.142 | 0.276 | 0.638 | 0.176 | 0 | 0 | 0 | 0 | 0.49 | 0.44 | 0.39 | 0.43 | 0.45 | 0.58 |
| 1 | 86175554 | rs3753786 | -1.024 | 0.327 | 0.34 | 0.186 | -0.019 | 0 | 0 | 0 | 0.46 | 0.23 | 0.53 | 0.51 | 0.52 | 0.35 |
| 1 | 86195047 | rs1047893 | -0.985 | 0.331 | 0.342 | 0.173 | -0.021 | 0.001 | 0.001 | 0 | 0.45 | 0.23 | 0.52 | 0.51 | 0.51 | 0.35 |
| 1 | 86233871 | rs392038 | 0.933 | -0.207 | 0.533 | 0.058 | -0.006 | 0 | 0 | 0 | 0.74 | 0.66 | 0.72 | 0.7 | 0.81 | 0.71 |
| 1 | 86245155 | rs313728 | 0.805 | -0.22 | 0.39 | 0.071 | -0.052 | -0.015 | -0.015 | 0 | 0.76 | 0.56 | 0.72 | 0.78 | 0.84 | 0.76 |

| | | | | | | | | | | | | | | | | |
|---|---|---|---|---|---|---|---|---|---|---|---|---|---|---|---|---|
| 1 | 86247318 | rs12563027 | -0.713 | -0.102 | -0.489 | 0.109 | 0.014 | -0.019 | -0.019 | 0 | 0.21 | 0.06 | 0.09 | 0.23 | 0.16 | 0.18 |
| 1 | 86249454 | rs17128307 | -0.593 | -0.217 | -0.408 | 0.028 | 0.021 | 0.009 | 0.009 | 0 | 0.19 | 0.14 | 0.18 | 0.17 | 0.12 | 0.17 |
| 1 | 86251836 | rs313710 | 0.427 | -0.179 | -1.004 | 0.001 | -0.001 | 0.019 | 0.019 | 0.849 | 0.5 | 0.11 | 0.34 | 0.57 | 0.4 | 0.53 |
| 1 | 86252025 | rs12746325 | 0.473 | -0.278 | -0.62 | -0.249 | -0.041 | 0.07 | 0.07 | 0 | 0.33 | 0.16 | 0.28 | 0.07 | 0.47 | 0.29 |
| 1 | 86256732 | rs313719 | 1.282 | -0.441 | -0.94 | -0.365 | -0.02 | -0.089 | -0.089 | 0.074 | 0.38 | 0.08 | 0.31 | 0.5 | 0.33 | 0.42 |
| 1 | 86258949 | rs3753785 | 0.615 | -0.239 | -0.081 | -0.065 | -0.016 | 0.042 | 0.042 | 0.006 | 0.07 | 0.14 | 0.15 | 0.18 | 0.04 | 0.03 |
| 1 | 86259699 | rs313733 | 0.455 | -0.248 | 0.525 | -0.172 | -0.003 | 0.204 | 0.204 | 0.194 | 0.85 | 0.41 | 0.63 | 0.64 | 0.87 | 0.81 |
| 1 | 86265259 | rs366187 | 0.745 | -0.269 | 0.957 | -0.238 | 0.021 | 0.029 | 0.029 | 0.504 | 0.51 | 0.31 | 0.37 | 0.57 | 0.4 | 0.52 |
| 1 | 86265967 | rs1977331 | 0.582 | -0.213 | 0.952 | -0.233 | 0.002 | 0.004 | 0.004 | 0.383 | 0.52 | 0.5 | 0.48 | 0.57 | 0.4 | 0.55 |
| 1 | 86268032 | rs12727784 | 0.37 | -0.064 | 0.841 | -0.475 | -0.052 | 0.095 | 0.095 | 0.019 | 0.33 | 0.14 | 0.28 | 0.07 | 0.47 | 0.29 |
| 1 | 86272858 | rs313740 | 0.658 | 0.011 | 0.997 | -0.084 | -0.002 | 0 | 0 | 0 | 0.56 | 0.48 | 0.41 | 0.59 | 0.44 | 0.56 |
| 1 | 86287580 | rs6693669 | 0.029 | 0.049 | 1.071 | -0.148 | 0.009 | -0.076 | -0.076 | 0 | 0.34 | 0.53 | 0.39 | 0.07 | 0.42 | 0.33 |
| 1 | 86301180 | rs11161686 | 0.38 | 0.056 | 1.033 | -0.159 | -0.003 | -0.017 | -0.017 | 0 | 0.31 | 0.22 | 0.25 | 0.07 | 0.42 | 0.26 |
| 1 | 86305704 | rs184339 | 1.522 | 0.199 | 0.526 | 0.037 | -0.049 | -0.034 | -0.034 | 1.5 | 0.09 | 0.01 | 0.14 | 0.11 | 0.15 | 0.12 |
| 1 | 86309622 | rs313702 | 0.885 | -0.209 | 0.834 | -0.114 | 0.018 | 0.026 | 0.026 | 0.205 | 0.5 | 0.21 | 0.38 | 0.59 | 0.41 | 0.55 |
| 1 | 86325998 | rs1904947 | 0.342 | -0.041 | 0.732 | -0.25 | 0.002 | 0.002 | 0.002 | 0 | 0.66 | 0.64 | 0.52 | 0.6 | 0.54 | 0.77 |
| 1 | 86326832 | rs12133414 | 1.651 | -0.058 | 0.718 | -0.314 | -0.001 | -0.012 | -0.012 | 0 | 0.12 | 0.12 | 0.1 | 0 | 0.13 | 0.15 |
| 1 | 86328120 | rs12759687 | 1.776 | 0.102 | 0.646 | -0.338 | 0.001 | -0.016 | -0.016 | 0 | 0.14 | 0.12 | 0.13 | 0.03 | 0.25 | 0.04 |
| 1 | 86333141 | rs313769 | 1.33 | 0.028 | 0.698 | -0.316 | 0.013 | 0.02 | 0.02 | 0.197 | 0.51 | 0.33 | 0.39 | 0.59 | 0.41 | 0.63 |
| 1 | 86345486 | rs1072508 | 0.268 | 0.164 | 0.611 | -0.145 | 0 | 0.017 | 0.017 | 0 | 0.8 | 0.76 | 0.66 | 0.63 | 0.79 | 0.82 |
| 1 | 86350554 | rs12755267 | 0.618 | 0.225 | 0.534 | -0.051 | 0.01 | -0.003 | -0.003 | 0 | 0.2 | 0.2 | 0.38 | 0.21 | 0.28 | 0.07 |
| 1 | 86352629 | rs11589719 | 1.269 | 0.233 | 0.57 | -0.038 | -0.005 | 0.032 | 0.032 | 0 | 0.05 | 0.06 | 0.24 | 0.18 | 0.03 | 0.03 |
| 1 | 86352664 | rs11589722 | 0.61 | 0.236 | 0.571 | -0.048 | 0.011 | -0.003 | -0.003 | 0 | 0.19 | 0.2 | 0.38 | 0.21 | 0.28 | 0.07 |
| 1 | 86364433 | rs12088838 | 1.051 | -0.423 | 0.064 | -0.125 | -0.012 | -0.041 | -0.041 | 0 | 0.35 | 0.22 | 0.5 | 0.53 | 0.32 | 0.31 |
| 1 | 86378429 | rs6703317 | 1.028 | -0.414 | 0.028 | -0.136 | -0.023 | -0.061 | -0.061 | 0 | 0.34 | 0.16 | 0.49 | 0.53 | 0.32 | 0.31 |
| 1 | 86380396 | rs12568614 | 1.641 | -0.603 | -0.008 | -0.242 | 0.044 | 0.004 | 0.004 | 0 | 0.21 | 0.1 | 0.14 | 0.23 | 0.13 | 0.25 |
| 1 | 86380987 | rs12136742 | 1.701 | -0.485 | -0.021 | -0.221 | -0.05 | -0.069 | -0.069 | 0 | 0.29 | 0.1 | 0.25 | 0.36 | 0.29 | 0.29 |
| 1 | 86381650 | rs12136958 | 1.886 | -0.34 | 0.009 | -0.267 | -0.037 | 0.024 | 0.024 | 0 | 0.1 | 0.01 | 0.08 | 0.03 | 0.14 | 0.11 |
| 1 | 86384082 | rs6702912 | -0.499 | -0.341 | 0.287 | -0.263 | 0.006 | 0.004 | 0.004 | 0.081 | 0.78 | 0.7 | 0.81 | 0.8 | 0.72 | 0.74 |
| 1 | 86412177 | rs11161711 | 0.341 | -0.367 | 0.235 | -0.275 | 0.001 | -0.007 | -0.007 | 0 | 0.72 | 0.56 | 0.77 | 0.8 | 0.7 | 0.67 |
| 1 | 86416725 | rs605060 | 0.706 | -0.375 | 0.236 | -0.329 | -0.021 | 0.058 | 0.058 | 0 | 0.57 | 0.29 | 0.63 | 0.76 | 0.45 | 0.63 |
| 1 | 86417699 | rs1698733 | -0.712 | -0.325 | 0.34 | -0.305 | -0.005 | 0.009 | 0.009 | 0.04 | 0.42 | 0.63 | 0.36 | 0.24 | 0.55 | 0.37 |
| 1 | 86424650 | rs6665006 | 2.056 | 0.122 | 0.332 | -0.029 | 0.016 | -0.033 | -0.033 | 0 | 0.14 | 0.14 | 0.13 | 0.04 | 0.25 | 0.04 |
| 1 | 86425382 | rs560876 | 0.033 | 0.1 | 0.315 | -0.016 | -0.004 | -0.002 | -0.002 | 0 | 0.22 | 0.23 | 0.17 | 0.19 | 0.28 | 0.26 |
| 1 | 86431623 | rs571691 | 1.197 | 0.022 | 0.3 | 0.04 | -0.008 | -0.016 | -0.016 | 0 | 0.11 | 0.11 | 0.09 | 0.17 | 0.13 | 0.15 |
| 1 | 86432364 | rs12751341 | 1.561 | -0.158 | 0.35 | -0.002 | 0.069 | -0.079 | -0.079 | 0.247 | 0.15 | 0.27 | 0.14 | 0.04 | 0.25 | 0.04 |
| 1 | 86435850 | rs7538391 | -0.149 | -0.909 | -0.259 | 0.416 | -0.014 | 0.002 | 0.002 | 0 | 0.3 | 0.28 | 0.4 | 0.41 | 0.16 | 0.34 |
| 1 | 86438327 | rs608460 | -0.748 | -0.435 | -0.764 | 0.238 | -0.004 | 0.009 | 0.009 | 0 | 0.65 | 0.56 | 0.7 | 0.64 | 0.7 | 0.64 |
| 1 | 86439443 | rs593146 | 0.147 | -0.469 | -0.921 | 0.282 | -0.001 | 0 | 0 | 0.071 | 0.43 | 0.42 | 0.56 | 0.42 | 0.58 | 0.39 |
| 1 | 86450400 | rs10493780 | -0.105 | -0.625 | -1.262 | 0.028 | 0.041 | 0.004 | 0.004 | 0 | 0.22 | 0.11 | 0.13 | 0.23 | 0.13 | 0.25 |
| 1 | 86457334 | rs4912453 | -0.315 | 0.17 | -1 | -0.127 | -0.004 | 0.011 | 0.011 | 0 | 0.42 | 0.37 | 0.44 | 0.61 | 0.49 | 0.58 |
| 1 | 86457375 | rs9988431 | 0.899 | -0.199 | -0.632 | -0.599 | -0.001 | -0.083 | -0.083 | 0 | 0.15 | 0.23 | 0.11 | 0 | 0.17 | 0.02 |
| 1 | 86459250 | rs12120406 | -0.053 | -0.07 | -0.69 | -0.432 | -0.004 | 0.011 | 0.011 | 0 | 0.42 | 0.37 | 0.44 | 0.6 | 0.49 | 0.58 |
| 1 | 86462901 | rs12404609 | 0.215 | 0.297 | -0.786 | -0.523 | -0.003 | -0.003 | -0.003 | 0.436 | 0.28 | 0.3 | 0.36 | 0.58 | 0.39 | 0.51 |
| 1 | 86463845 | rs7547149 | 0.215 | 0.297 | -0.786 | -0.523 | -0.003 | -0.003 | -0.003 | 0 | 0.28 | 0.3 | 0.36 | 0.58 | 0.39 | 0.51 |
| 1 | 86464981 | rs11161717 | 0.128 | 0.284 | -0.803 | -0.53 | 0.007 | 0.004 | 0.004 | 0 | 0.29 | 0.34 | 0.36 | 0.58 | 0.39 | 0.51 |
| 1 | 86465736 | rs17128651 | 0.099 | 0.286 | -0.657 | 0.005 | 0.002 | 0.33 | 0.33 | 0 | 0.06 | 0.21 | 0.08 | 0.45 | 0.04 | 0.17 |
| 1 | 86466085 | rs12040977 | 0.059 | 0.344 | -0.825 | -0.164 | 0.032 | 0.026 | 0.026 | 0 | 0.24 | 0.33 | 0.34 | 0.58 | 0.37 | 0.45 |
| 1 | 86468500 | rs585708 | -1.411 | 0.393 | -0.878 | -0.257 | -0.074 | 0.003 | 0.003 | 0 | 0.83 | 0.93 | 0.76 | 0.88 | 0.66 | 0.68 |
| 1 | 86469506 | rs9701002 | 0.516 | 0.046 | -0.9 | -0.55 | -0.001 | -0.005 | -0.005 | 0 | 0.58 | 0.62 | 0.64 | 0.87 | 0.56 | 0.55 |
| 1 | 86474964 | rs4912455 | 0.831 | -0.073 | -0.85 | -0.549 | 0.012 | 0.063 | 0.063 | 2.429 | 0.18 | 0.04 | 0.11 | 0.03 | 0.12 | 0.11 |
| 1 | 86477220 | rs6687775 | 0.482 | -0.027 | -0.594 | -0.483 | 0.002 | -0.005 | -0.005 | 0 | 0.06 | 0.23 | 0.03 | 0 | 0.02 | 0 |
| 1 | 86479771 | rs6679444 | 0.829 | -0.007 | -0.584 | -0.474 | 0.005 | -0.002 | -0.002 | 0 | 0.05 | 0.2 | 0.03 | 0 | 0.01 | 0 |
| 1 | 86481084 | rs618555 | -1.79 | 0.138 | -0.762 | -0.506 | -0.122 | 0.028 | 0.028 | 0.046 | 0.84 | 0.98 | 0.79 | 0.94 | 0.7 | 0.7 |
| 1 | 86483632 | rs10518362 | 0.695 | -0.25 | -0.715 | -0.534 | 0.013 | 0.038 | 0.038 | 1.971 | 0.18 | 0.04 | 0.12 | 0.07 | 0.12 | 0.14 |
| 1 | 86487914 | rs11161721 | 1.293 | -1.013 | -1.067 | -1.031 | 0.006 | 0.004 | 0.004 | 0 | 0.28 | 0.23 | 0.35 | 0.27 | 0.29 | 0.21 |
| 1 | 86488232 | rs641712 | -0.821 | -0.937 | -1.254 | -1.119 | 0.007 | -0.008 | -0.008 | 0 | 0.46 | 0.42 | 0.3 | 0.13 | 0.3 | 0.27 |
| 1 | 86490043 | rs565860 | 0.272 | -0.852 | -1.131 | -1.005 | -0.026 | -0.086 | -0.086 | 0 | 0.28 | 0.38 | 0.19 | 0.06 | 0.18 | 0.13 |
| 1 | 86512253 | rs17406755 | 1.113 | -0.674 | -1.237 | -1.7 | 0.006 | 0.091 | 0.091 | 0.82 | 0.21 | 0.08 | 0.14 | 0.07 | 0.17 | 0.15 |
| 1 | 86512834 | rs541395 | 0.742 | -0.669 | -1.247 | -1.771 | -0.016 | -0.101 | -0.101 | 0.031 | 0.17 | 0.3 | 0.13 | 0 | 0.18 | 0.03 |
| 1 | 86526084 | rs12057842 | 0.615 | -0.918 | -1.346 | -1.679 | 0.024 | 0.044 | 0.044 | 1.742 | 0.2 | 0.09 | 0.13 | 0.07 | 0.13 | 0.13 |
| 1 | 86527070 | rs689040 | 1.642 | -0.904 | -1.338 | -1.706 | 0 | -0.032 | -0.032 | 0 | 0.13 | 0.15 | 0.11 | 0 | 0.16 | 0.03 |
| 1 | 86527492 | rs6682503 | -0.248 | -0.93 | -1.346 | -1.712 | 0.018 | 0.05 | 0.05 | 1.015 | 0.33 | 0.22 | 0.23 | 0.07 | 0.29 | 0.16 |
| 1 | 86529390 | rs12742014 | 1.291 | -0.268 | -0.913 | -1.217 | 0 | -0.028 | -0.028 | 0 | 0.12 | 0.13 | 0.11 | 0 | 0.16 | 0.03 |
| 1 | 86547798 | rs4912461 | -0.253 | -0.167 | -0.897 | -1.082 | 0.002 | -0.028 | -0.028 | 0 | 0.44 | 0.4 | 0.3 | 0.11 | 0.37 | 0.26 |
| 1 | 86552269 | rs10782563 | -0.756 | 0.222 | -0.736 | -0.229 | 0.005 | -0.005 | -0.005 | 0.044 | 0.36 | 0.42 | 0.46 | 0.27 | 0.5 | 0.51 |
| 1 | 86552334 | rs12723978 | 0.272 | 0.348 | -0.879 | -0.328 | 0 | -0.027 | -0.027 | 0 | 0.07 | 0.07 | 0.03 | 0.01 | 0.05 | 0.07 |
| 1 | 86557724 | rs17407336 | -0.055 | 0.443 | -0.909 | -0.353 | 0 | 0.004 | 0.004 | 0 | 0.16 | 0.13 | 0.13 | 0.06 | 0.14 | 0.12 |
| 1 | 86558898 | rs12082787 | 0.884 | 0.229 | -1.143 | -0.559 | -0.004 | 0.032 | 0.032 | 0 | 0.07 | 0.02 | 0.06 | 0 | 0.07 | 0.02 |

| | | | | | | | | | | | | | | | | |
|---|---|---|---|---|---|---|---|---|---|---|---|---|---|---|---|---|
| 1 | 86562513 | rs997110 | 0.825 | 0.253 | -1.12 | -0.51 | -0.07 | 0.025 | 0.025 | 0 | 0.18 | 0.07 | 0.26 | 0.12 | 0.35 | 0.32 |
| 1 | 86563718 | rs1605416 | -0.853 | -0.705 | -0.777 | -0.022 | -0.047 | 0.063 | 0.063 | 0 | 0.85 | 0.74 | 0.94 | 0.64 | 0.94 | 0.84 |
| 1 | 86564339 | rs883201 | -0.81 | -0.103 | -1.111 | -0.002 | 0.007 | 0.017 | 0.017 | 0 | 0.25 | 0.2 | 0.24 | 0.13 | 0.22 | 0.23 |
| 1 | 86565367 | rs6672107 | 0.414 | 0.197 | -0.6 | 0.258 | -0.024 | 0.013 | 0.013 | 0 | 0.19 | 0.16 | 0.27 | 0.12 | 0.36 | 0.32 |
| 1 | 86566278 | rs12044349 | -0.654 | -0.116 | -0.633 | 0.385 | -0.009 | 0 | 0 | 0.008 | 0.36 | 0.3 | 0.43 | 0.31 | 0.52 | 0.53 |
| 1 | 86571562 | rs11161735 | 0.212 | -0.326 | -0.289 | 0.339 | 0.002 | 0.002 | 0.002 | 1.247 | 0.54 | 0.46 | 0.6 | 0.53 | 0.62 | 0.49 |
| 1 | 86572463 | rs1360903 | 1.116 | -0.269 | -0.509 | 0.501 | -0.041 | -0.044 | -0.044 | 0 | 0.36 | 0.19 | 0.46 | 0.38 | 0.46 | 0.25 |
| 1 | 86578951 | rs12075854 | 0.043 | -0.657 | 0.352 | 1.053 | -0.002 | 0.009 | 0.009 | 1.812 | 0.16 | 0.1 | 0.17 | 0.06 | 0.17 | 0.25 |
| 1 | 86579895 | rs12024954 | -0.61 | -0.687 | 0.453 | 1.006 | -0.001 | 0.004 | 0.004 | 0.279 | 0.19 | 0.13 | 0.18 | 0.06 | 0.19 | 0.27 |
| 1 | 86580585 | rs2390014 | 0.082 | -0.69 | 0.437 | 1.007 | -0.013 | 0.057 | 0.057 | 1.89 | 0.16 | 0.03 | 0.15 | 0.06 | 0.17 | 0.27 |
| 1 | 86584695 | rs6680321 | 1.165 | -0.752 | 0.846 | 1.435 | -0.001 | 0.002 | 0.002 | 0.299 | 0.73 | 0.73 | 0.73 | 0.62 | 0.74 | 0.66 |
| 1 | 86587793 | rs6690591 | -0.611 | -0.761 | 0.821 | 1.466 | 0.008 | 0.035 | 0.035 | 1.171 | 0.21 | 0.11 | 0.19 | 0.13 | 0.17 | 0.28 |
| 1 | 86590948 | rs11161746 | 0.152 | -0.753 | 0.82 | 1.459 | -0.006 | 0.033 | 0.033 | 1.848 | 0.16 | 0.08 | 0.16 | 0.06 | 0.17 | 0.25 |
| 1 | 86592526 | rs1507264 | -0.483 | -0.982 | 1.417 | 1.411 | 0.018 | -0.03 | -0.03 | 0 | 0.18 | 0.13 | 0.2 | 0.32 | 0.11 | 0.3 |
| 1 | 86593968 | rs1507263 | 1.415 | -0.976 | 1.484 | 1.31 | 0.001 | -0.004 | -0.004 | 0 | 0.63 | 0.65 | 0.56 | 0.37 | 0.66 | 0.39 |
| 1 | 86594301 | rs1507262 | 2.164 | -0.963 | 1.48 | 1.251 | 0.012 | 0.02 | 0.02 | 0 | 0.54 | 0.4 | 0.52 | 0.3 | 0.63 | 0.37 |
| 1 | 86603942 | rs7516054 | 1.434 | -1.27 | 1.406 | 1.495 | -0.038 | 0.044 | 0.044 | 0.101 | 0.73 | 0.52 | 0.7 | 0.36 | 0.82 | 0.63 |
| 1 | 86613540 | rs1889851 | 1.807 | -1.231 | 1.409 | 1.506 | -0.028 | 0.016 | 0.016 | 0.27 | 0.7 | 0.54 | 0.71 | 0.35 | 0.8 | 0.62 |
| 1 | 86616271 | rs11161755 | -0.048 | -1.002 | 1.115 | 1.589 | -0.001 | 0.021 | 0.021 | 0 | 0.06 | 0.04 | 0.13 | 0.23 | 0.07 | 0.07 |
| 1 | 86621160 | rs12080986 | -1.521 | -1.065 | 1.315 | 1.564 | -0.005 | 0.048 | 0.048 | 0.206 | 0.1 | 0.18 | 0.17 | 0.25 | 0.09 | 0.08 |
| 1 | 86621256 | rs12070992 | -1.392 | -1.053 | 1.21 | 1.568 | -0.001 | 0.015 | 0.015 | 0.221 | 0.1 | 0.1 | 0.16 | 0.25 | 0.09 | 0.08 |
| 1 | 86635245 | rs1112865 | 0.037 | -1.048 | 1.447 | 1.498 | 0.002 | 0.059 | 0.059 | 0.232 | 0.06 | 0.21 | 0.03 | 0.06 | 0.01 | 0.02 |
| 1 | 86636876 | rs1507295 | 2.061 | -1.277 | 1.358 | 1.454 | 0.024 | 0.026 | 0.026 | 0 | 0.53 | 0.35 | 0.57 | 0.31 | 0.66 | 0.38 |
| 1 | 86641674 | rs1507279 | 1.303 | -1.276 | 0.99 | 1.479 | 0.028 | -0.019 | -0.019 | 0 | 0.62 | 0.7 | 0.74 | 0.53 | 0.73 | 0.45 |
| 1 | 86651383 | rs1108395 | 2.277 | -1.319 | 1.104 | 1.448 | 0.053 | 0.06 | 0.06 | 0 | 0.52 | 0.28 | 0.57 | 0.31 | 0.67 | 0.38 |
| 1 | 86651482 | rs1108394 | 0.122 | -0.999 | 0.527 | 1.473 | 0.007 | 0.011 | 0.011 | 1.543 | 0.81 | 0.9 | 0.87 | 0.89 | 0.86 | 0.73 |
| 1 | 86655440 | rs7418327 | -0.511 | -0.798 | 0.397 | 1.648 | 0.013 | -0.023 | -0.023 | 0 | 0.14 | 0.12 | 0.1 | 0.31 | 0.1 | 0.16 |
| 1 | 86664081 | rs1507269 | -0.763 | -0.609 | 0.666 | 1.81 | -0.01 | 0.038 | 0.038 | 0 | 0.14 | 0.25 | 0.11 | 0.37 | 0.1 | 0.25 |
| 1 | 86664322 | rs7536402 | -0.345 | -0.469 | 0.588 | 1.804 | -0.001 | 0.023 | 0.023 | 0 | 0.06 | 0.06 | 0.13 | 0.23 | 0.07 | 0.06 |
| 1 | 86666893 | rs7531445 | -0.896 | -0.469 | 0.674 | 1.515 | -0.002 | 0.072 | 0.072 | 0 | 0.07 | 0.12 | 0.17 | 0.23 | 0.07 | 0.07 |
| 1 | 86674550 | rs6576818 | -0.377 | -0.499 | 0.646 | 1.371 | -0.005 | -0.013 | -0.013 | 0 | 0.07 | 0.07 | 0.03 | 0.01 | 0.03 | 0.04 |
| 1 | 86683345 | rs7411591 | -0.348 | -0.573 | 0.603 | 1.295 | 0.003 | 0.004 | 0.004 | 1.588 | 0.85 | 0.88 | 0.88 | 0.91 | 0.88 | 0.76 |
| 1 | 86684713 | rs1507290 | 0.039 | -0.563 | 0.657 | 1.365 | -0.002 | -0.003 | -0.003 | 0.66 | 0.82 | 0.91 | 0.8 | 0.81 | 0.83 | 0.73 |
| 1 | 86685846 | rs12037768 | 1.799 | -0.617 | 0.891 | 1.381 | -0.018 | -0.001 | -0.001 | 0 | 0.61 | 0.49 | 0.62 | 0.36 | 0.69 | 0.41 |
| 1 | 86692117 | rs1395713 | -1.068 | -0.352 | 0.57 | 1.189 | -0.002 | 0.006 | 0.006 | 0 | 0.18 | 0.24 | 0.13 | 0.38 | 0.14 | 0.28 |
| 1 | 86697145 | rs6576821 | -1.009 | -0.098 | 0.508 | 0.485 | -0.009 | -0.028 | -0.028 | 0 | 0.07 | 0.22 | 0.04 | 0.01 | 0.03 | 0.04 |
| 1 | 86714730 | rs6677603 | -1.253 | 0.127 | 0.605 | 0.63 | 0.019 | 0.029 | 0.029 | 0 | 0.21 | 0.54 | 0.27 | 0.26 | 0.23 | 0.18 |
| 1 | 86714784 | rs6576824 | -0.888 | -0.092 | 0.569 | 0.131 | -0.003 | -0.032 | -0.032 | 0.948 | 0.89 | 0.91 | 0.94 | 0.99 | 0.92 | 0.8 |
| 1 | 86715854 | rs4655926 | 1 | -0.546 | 0.756 | 0.268 | -0.062 | -0.016 | -0.016 | 0.018 | 0.79 | 0.63 | 0.88 | 0.84 | 0.91 | 0.65 |
| 1 | 86717874 | rs7519336 | 0.745 | -0.505 | 0.51 | 0.177 | -0.016 | -0.022 | -0.022 | 0 | 0.81 | 0.83 | 0.91 | 0.92 | 0.91 | 0.74 |
| 1 | 86720853 | rs6420973 | 1.208 | 0.644 | -0.275 | -0.499 | -0.036 | 0.039 | 0.039 | 0 | 0.65 | 0.84 | 0.49 | 0.8 | 0.61 | 0.6 |
| 1 | 86734119 | rs36002452 | 0.525 | 0.594 | -0.525 | -0.291 | -0.04 | -0.034 | -0.034 | 0.136 | 0.11 | 0.01 | 0.08 | 0.14 | 0.14 | 0.07 |
| 1 | 86734898 | rs10873770 | 0.285 | 0.676 | -0.206 | -0.405 | -0.075 | 0.096 | 0.096 | 0 | 0.79 | 0.94 | 0.68 | 1 | 0.73 | 0.85 |
| 1 | 86736880 | rs7546158 | 1.557 | 0.566 | 0.456 | -0.489 | 0.018 | -0.042 | -0.042 | 2.226 | 0.64 | 0.49 | 0.46 | 0.82 | 0.6 | 0.61 |
| 1 | 86737698 | rs11161788 | 0.577 | 0.636 | 0.583 | -0.427 | -0.011 | -0.018 | -0.018 | 0.881 | 0.13 | 0.16 | 0.08 | 0.07 | 0.08 | 0.22 |
| 1 | 86745177 | rs1998450 | 0.792 | 0.668 | 0.483 | -0.469 | -0.01 | -0.048 | -0.048 | 0 | 0.13 | 0.18 | 0.06 | 0.02 | 0.07 | 0.17 |
| 1 | 86745979 | rs7513039 | 0.19 | 0.56 | 0.429 | -0.589 | 0.01 | -0.048 | -0.048 | 1.289 | 0.82 | 0.8 | 0.68 | 1 | 0.74 | 0.84 |
| 1 | 86751115 | rs1407721 | -0.361 | 0.603 | 0.085 | -0.569 | -0.039 | 0.029 | 0.029 | 0 | 0.85 | 0.92 | 0.69 | 1 | 0.74 | 0.84 |
| 1 | 86751754 | rs7529124 | 1.065 | 0.484 | 0.052 | -0.386 | 0.027 | -0.047 | -0.047 | 0.497 | 0.64 | 0.52 | 0.46 | 0.81 | 0.6 | 0.58 |
| 1 | 86758589 | rs976332 | 0.483 | -0.063 | -0.979 | -0.085 | 0.059 | -0.034 | -0.034 | 0 | 0.42 | 0.22 | 0.28 | 0.44 | 0.32 | 0.54 |
| 1 | 86760937 | rs10158226 | 1.134 | 0.143 | -1.363 | -0.223 | -0.025 | -0.113 | -0.113 | 0.079 | 0.3 | 0.08 | 0.23 | 0.41 | 0.3 | 0.34 |
| 1 | 86769951 | rs1323029 | -0.198 | -0.297 | -0.51 | -0.638 | -0.025 | -0.003 | -0.003 | 0 | 0.33 | 0.49 | 0.3 | 0.31 | 0.26 | 0.32 |
| 1 | 86773046 | rs969396 | -0.446 | -0.295 | -0.461 | -0.896 | 0.001 | -0.001 | -0.001 | 0 | 0.76 | 0.78 | 0.63 | 0.8 | 0.65 | 0.81 |
| 1 | 86784804 | rs1323035 | 0.686 | -0.3 | -0.429 | -0.909 | 0.021 | -0.033 | -0.033 | 0.14 | 0.63 | 0.55 | 0.51 | 0.71 | 0.55 | 0.66 |
| 1 | 86787767 | rs17099731 | 0.426 | -0.4 | -0.652 | -0.899 | -0.003 | -0.026 | -0.026 | 0 | 0.07 | 0.08 | 0.05 | 0 | 0.08 | 0.04 |
| 1 | 86805880 | rs272487 | -0.071 | -0.532 | -0.318 | -0.952 | -0.013 | 0.015 | 0.015 | 0 | 0.27 | 0.36 | 0.31 | 0.36 | 0.21 | 0.26 |
| 1 | 86807418 | rs10873771 | 1.965 | -0.196 | -1.025 | -0.878 | -0.028 | -0.106 | -0.106 | 0 | 0.3 | 0.08 | 0.23 | 0.41 | 0.32 | 0.35 |
| 1 | 86807618 | rs272489 | -0.286 | 0.175 | -0.651 | -0.808 | -0.075 | 0.033 | 0.033 | 0 | 0.77 | 0.92 | 0.68 | 0.83 | 0.66 | 0.82 |
| 1 | 86820267 | rs1182098 | -0.218 | 0.161 | -0.369 | -0.809 | 0.013 | -0.008 | -0.008 | 0 | 0.76 | 0.78 | 0.67 | 0.83 | 0.66 | 0.8 |
| 1 | 86822231 | rs385437 | 1.008 | -0.087 | -0.167 | -1.067 | 0 | -0.005 | -0.005 | 0 | 0.14 | 0.11 | 0.12 | 0.03 | 0.17 | 0.18 |
| 1 | 86833653 | rs272504 | -0.173 | -0.31 | 0.048 | -0.97 | -0.01 | -0.007 | -0.007 | 0 | 0.35 | 0.43 | 0.32 | 0.29 | 0.3 | 0.32 |
| 1 | 86838158 | rs421105 | -0.468 | -0.27 | -0.056 | -0.946 | 0.013 | -0.008 | -0.008 | 3.713 | 0.76 | 0.78 | 0.67 | 0.83 | 0.66 | 0.8 |
| 1 | 86839385 | rs272508 | -1.596 | 0.226 | -0.213 | -1.029 | -0.029 | -0.002 | -0.002 | 0 | 0.92 | 0.97 | 0.77 | 0.92 | 0.84 | 0.91 |
| 1 | 86847955 | rs272516 | -0.602 | -0.55 | -0.087 | -1.032 | -0.022 | -0.02 | -0.02 | 0 | 0.4 | 0.54 | 0.4 | 0.29 | 0.33 | 0.34 |
| 1 | 86851420 | rs1407719 | 1.745 | -0.617 | -0.055 | -0.988 | 0.006 | -0.028 | -0.028 | 0 | 0.34 | 0.24 | 0.26 | 0.42 | 0.32 | 0.35 |
| 1 | 86864866 | rs1116108 | 1.617 | -0.649 | -0.067 | -0.993 | 0.012 | -0.031 | -0.031 | 0 | 0.36 | 0.23 | 0.28 | 0.53 | 0.32 | 0.44 |
| 1 | 86869753 | rs12131620 | 1.617 | -0.649 | -0.067 | -0.993 | 0.012 | -0.031 | -0.031 | 0 | 0.36 | 0.23 | 0.28 | 0.53 | 0.32 | 0.44 |
| 1 | 86872524 | rs11161812 | 0.096 | -0.634 | -0.151 | -1.016 | 0.012 | -0.008 | -0.008 | 0 | 0.24 | 0.22 | 0.33 | 0.17 | 0.35 | 0.22 |
| 1 | 86874082 | rs1999600 | 1.997 | -0.661 | -0.201 | -0.924 | -0.033 | -0.109 | -0.109 | 0.249 | 0.31 | 0.08 | 0.23 | 0.42 | 0.32 | 0.35 |

| | | | | | | | | | | | | | | | | |
|---|---|---|---|---|---|---|---|---|---|---|---|---|---|---|---|---|
| 1 | 86884539 | rs12072655 | 2.191 | -0.905 | -0.217 | -1.159 | -0.011 | -0.06 | -0.06 | 0 | 0.26 | 0.14 | 0.21 | 0.42 | 0.26 | 0.35 |
| 1 | 86889127 | rs12747430 | 0.923 | -0.877 | -0.254 | -1.122 | 0.014 | -0.025 | -0.025 | 0 | 0.07 | 0 | 0.04 | 0.11 | 0.06 | 0.1 |
| 1 | 86895953 | rs7415758 | 1.507 | -0.905 | -0.102 | -1.182 | 0 | -0.003 | -0.003 | 2.912 | 0.31 | 0.3 | 0.26 | 0.42 | 0.28 | 0.37 |
| 1 | 86896541 | rs3737672 | 1.222 | -0.932 | -0.097 | -1.213 | 0.005 | -0.015 | -0.015 | 0 | 0.34 | 0.27 | 0.27 | 0.53 | 0.32 | 0.45 |
| 1 | 86898409 | rs1413431 | 0.276 | -0.964 | -0.142 | -1.291 | -0.02 | 0.012 | 0.012 | 0 | 0.24 | 0.15 | 0.32 | 0.18 | 0.35 | 0.23 |
| 1 | 86898557 | rs1931363 | -0.122 | -1.022 | 0.187 | -1.258 | -0.024 | -0.015 | -0.015 | 0 | 0.63 | 0.46 | 0.66 | 0.71 | 0.7 | 0.74 |
| 1 | 86902967 | rs11161828 | -0.059 | -1.002 | 0.078 | -1.187 | -0.022 | -0.032 | -0.032 | 0 | 0.14 | 0.17 | 0.11 | 0.05 | 0.09 | 0.1 |
| 1 | 86903209 | rs1814203 | 0.9 | -0.942 | 0.074 | -1.206 | 0.002 | 0.001 | 0.001 | 0 | 0.17 | 0.12 | 0.29 | 0.14 | 0.29 | 0.16 |
| 1 | 86904325 | rs1889337 | -1.819 | -1.386 | 0.258 | -1.293 | 0.02 | 0.066 | 0.066 | 0 | 0.54 | 0.62 | 0.59 | 0.33 | 0.61 | 0.49 |
| 1 | 86904588 | rs11161830 | -0.067 | -1.189 | 0.274 | -1.136 | -0.022 | -0.032 | -0.032 | 0 | 0.14 | 0.17 | 0.11 | 0.05 | 0.09 | 0.1 |
| 1 | 86905563 | rs12092398 | 0.915 | -1.173 | 0.245 | -1.124 | -0.008 | -0.009 | -0.009 | 0.914 | 0.1 | 0.09 | 0.05 | 0.05 | 0.07 | 0.07 |
| 1 | 86909582 | rs1413426 | -0.078 | -1.232 | 0.212 | -1.133 | -0.006 | -0.01 | -0.01 | 0 | 0.14 | 0.14 | 0.11 | 0.06 | 0.09 | 0.11 |
| 1 | 86909703 | rs3820039 | 0.961 | -1.036 | 0.387 | -1.144 | 0.03 | -0.005 | -0.005 | 0 | 0.16 | 0.14 | 0.3 | 0.13 | 0.3 | 0.15 |
| 1 | 86910264 | rs2249296 | -1.434 | -1.204 | 0.334 | -1.099 | 0.012 | -0.014 | -0.014 | 0.318 | 0.8 | 0.81 | 0.81 | 0.72 | 0.87 | 0.87 |
| 1 | 86912451 | rs2791463 | 0.058 | -1.214 | 0.378 | -1.104 | 0.013 | 0.003 | 0.003 | 0 | 0.62 | 0.55 | 0.48 | 0.59 | 0.57 | 0.72 |
| 1 | 86914272 | rs3765987 | 1.211 | -1.283 | 0.018 | -1.007 | -0.001 | 0 | 0 | 0 | 0.07 | 0.07 | 0.06 | 0.11 | 0.06 | 0.11 |
| 1 | 86916856 | rs969788 | -0.614 | -1.359 | -0.07 | -1.053 | 0.024 | 0.012 | 0.012 | 0 | 0.21 | 0.13 | 0.15 | 0.17 | 0.15 | 0.22 |
| 1 | 86916948 | rs969789 | 1.045 | -1.286 | -0.084 | -1.003 | 0.018 | 0.017 | 0.017 | 0 | 0.1 | 0.04 | 0.04 | 0.06 | 0.07 | 0.07 |
| 1 | 86920904 | rs2390057 | 1.309 | -1.324 | -0.016 | -1.065 | 0.011 | -0.039 | -0.039 | 0.544 | 0.36 | 0.21 | 0.24 | 0.44 | 0.34 | 0.43 |
| 1 | 86925400 | rs4656108 | -0.055 | -1.376 | 0.205 | -1.045 | -0.022 | 0.005 | 0.005 | 0 | 0.71 | 0.56 | 0.68 | 0.7 | 0.77 | 0.74 |
| 1 | 86929452 | rs4001077 | -1.038 | -1.534 | 0.066 | -1.276 | -0.019 | -0.048 | -0.048 | 1.083 | 0.23 | 0.33 | 0.19 | 0.1 | 0.18 | 0.25 |
| 1 | 86931529 | rs9659676 | 2.292 | -1.591 | -0.021 | -1.321 | -0.004 | -0.046 | -0.046 | 0 | 0.24 | 0.16 | 0.18 | 0.39 | 0.22 | 0.34 |
| 1 | 86932265 | rs3753276 | -0.936 | -1.619 | 0.028 | -1.328 | -0.025 | -0.061 | -0.061 | 0.009 | 0.24 | 0.36 | 0.19 | 0.11 | 0.18 | 0.26 |
| 1 | 86934176 | rs2791519 | -0.823 | -1.482 | -0.019 | -1.197 | -0.001 | 0 | 0 | 0 | 0.59 | 0.57 | 0.7 | 0.51 | 0.64 | 0.46 |
| 1 | 86935036 | rs2753338 | -1.024 | -1.651 | -0.32 | -1.267 | 0.009 | -0.001 | -0.001 | 0 | 0.53 | 0.64 | 0.63 | 0.54 | 0.54 | 0.5 |
| 1 | 86935293 | rs2753345 | 1.626 | -1.754 | -0.342 | -1.231 | 0.018 | 0.034 | 0.034 | 1.157 | 0.17 | 0.1 | 0.11 | 0.08 | 0.16 | 0.12 |
| 1 | 86936247 | rs1358826 | -0.086 | -1.24 | -0.24 | -1.116 | -0.007 | 0.016 | 0.016 | 0 | 0.69 | 0.56 | 0.67 | 0.67 | 0.73 | 0.75 |
| 1 | 86936310 | rs5744306 | 0.966 | -1.19 | -0.488 | -1.064 | -0.011 | -0.003 | -0.003 | 2 | 0.11 | 0.12 | 0.05 | 0.09 | 0.05 | 0.13 |
| 1 | 86938706 | rs2791516 | -0.334 | -1.028 | -0.672 | -1.166 | 0.051 | -0.056 | -0.056 | 0 | 0.76 | 0.63 | 0.63 | 0.86 | 0.66 | 0.78 |
| 1 | 86939130 | rs2145412 | -0.267 | -1.295 | -0.627 | -1.104 | 0.007 | -0.019 | -0.019 | 1.108 | 0.92 | 0.67 | 0.93 | 0.99 | 0.95 | 0.95 |
| 1 | 86940715 | rs5744329 | 2.027 | -1.227 | -0.966 | -1.02 | -0.001 | -0.063 | -0.063 | 0.43 | 0.25 | 0.14 | 0.19 | 0.38 | 0.23 | 0.35 |
| 1 | 86940872 | rs2753384 | 1.298 | -2.114 | -0.537 | -0.812 | -0.007 | 0.047 | 0.047 | 0 | 0.15 | 0.18 | 0.19 | 0.32 | 0.12 | 0.13 |
| 1 | 86941488 | rs926065 | 0.478 | -1.841 | -0.686 | -1.028 | -0.021 | -0.08 | -0.08 | 0 | 0.75 | 0.64 | 0.82 | 0.9 | 0.8 | 0.75 |
| 1 | 86948169 | rs2791514 | 0.98 | -1.794 | -1.308 | -0.445 | 0.033 | 0.046 | 0.046 | 0.787 | 0.18 | 0.1 | 0.12 | 0.08 | 0.16 | 0.15 |
| 1 | 86948708 | rs2791512 | 0.404 | -1.42 | -0.851 | 0.034 | -0.057 | 0.007 | 0.007 | 0 | 0.65 | 0.53 | 0.71 | 0.63 | 0.78 | 0.63 |
| 1 | 86949930 | rs2791510 | 0.719 | -0.813 | -1.249 | 0.284 | 0.019 | 0.004 | 0.004 | 0 | 0.81 | 0.89 | 0.89 | 0.81 | 0.84 | 0.83 |
| 1 | 86950320 | rs2734700 | -1.121 | -0.735 | -1.333 | 0.671 | 0.007 | -0.022 | -0.022 | 0 | 0.22 | 0.17 | 0.39 | 0.66 | 0.25 | 0.38 |
| 1 | 86951806 | rs5744367 | -0.668 | -0.922 | -1.429 | 1.154 | -0.012 | -0.007 | -0.007 | 0 | 0.09 | 0.03 | 0.12 | 0.5 | 0.13 | 0.31 |
| 1 | 86952601 | rs2075630 | -0.157 | -0.94 | -1.526 | -0.066 | 0.016 | 0.043 | 0.043 | 0 | 0.56 | 0.36 | 0.77 | 0.76 | 0.66 | 0.71 |
| 1 | 86959014 | rs2753346 | -0.481 | -0.719 | -0.651 | -0.202 | 0.032 | -0.012 | -0.012 | 0.048 | 0.87 | 0.73 | 0.9 | 0.91 | 0.86 | 0.92 |
| 1 | 86959173 | rs2791494 | 0.308 | -0.888 | -0.655 | -0.362 | -0.004 | -0.032 | -0.032 | 0 | 0.81 | 0.78 | 0.88 | 0.91 | 0.82 | 0.9 |
| 1 | 86959424 | rs1321690 | 0.475 | -1.17 | -0.644 | -0.303 | -0.012 | 0.011 | 0.011 | 0 | 0.77 | 0.7 | 0.68 | 0.7 | 0.78 | 0.81 |
| 1 | 86961157 | rs1407141 | 0.434 | -1.1 | -0.767 | -0.249 | 0.013 | -0.014 | -0.014 | 0 | 0.1 | 0.05 | 0.22 | 0.21 | 0.08 | 0.11 |
| 1 | 86962331 | rs2791491 | 1.497 | -1.286 | -0.893 | -0.461 | -0.008 | 0.145 | 0.145 | 0 | 0.46 | 0.26 | 0.4 | 0.15 | 0.48 | 0.35 |
| 1 | 86965419 | rs1882753 | -0.197 | -1.259 | -0.766 | -0.306 | 0.001 | 0.032 | 0.032 | 0 | 0.23 | 0.28 | 0.23 | 0.54 | 0.26 | 0.43 |
| 1 | 86974115 | rs1321686 | 1.851 | -1.581 | -0.809 | -0.434 | -0.095 | 0.12 | 0.12 | 2.869 | 0.36 | 0.09 | 0.4 | 0.16 | 0.49 | 0.35 |
| 1 | 86981256 | rs10489693 | 0.804 | -1.656 | -0.791 | -0.422 | 0.013 | -0.014 | -0.014 | 0.152 | 0.1 | 0.05 | 0.22 | 0.21 | 0.08 | 0.11 |
| 1 | 86986357 | rs2791475 | -0.355 | -1.771 | -0.632 | -0.516 | -0.02 | -0.077 | -0.077 | 0 | 0.79 | 0.62 | 0.87 | 0.91 | 0.84 | 0.9 |
| 1 | 86986617 | rs13376522 | -1.036 | -1.74 | -0.77 | -0.512 | -0.001 | -0.005 | -0.005 | 0 | 0.42 | 0.44 | 0.45 | 0.75 | 0.35 | 0.56 |
| 1 | 86988622 | rs1547036 | -0.254 | -1.505 | -0.959 | -0.404 | -0.003 | 0.002 | 0.002 | 0.136 | 0.33 | 0.36 | 0.23 | 0.55 | 0.27 | 0.44 |
| 1 | 86988709 | rs2791473 | 1.049 | -1.206 | -1.019 | -0.335 | -0.014 | 0.081 | 0.081 | 0.742 | 0.46 | 0.14 | 0.63 | 0.36 | 0.57 | 0.46 |
| 1 | 86995816 | rs12750427 | 0.088 | -0.845 | -0.639 | 0.088 | 0.008 | 0.012 | 0.012 | 0 | 0.14 | 0.26 | 0.07 | 0.11 | 0.14 | 0.13 |
| 1 | 87000308 | rs1575178 | 1.221 | -0.661 | -1.434 | -0.304 | -0.09 | 0.078 | 0.078 | 0.215 | 0.35 | 0.11 | 0.54 | 0.17 | 0.46 | 0.39 |
| 1 | 87000484 | rs1575177 | 0.139 | -0.801 | -1.378 | -0.068 | -0.004 | -0.006 | -0.006 | 0 | 0.75 | 0.93 | 0.84 | 0.74 | 0.76 | 0.82 |
| 1 | 87005402 | rs1321681 | 0.035 | -0.794 | -0.371 | -0.043 | 0.033 | -0.043 | -0.043 | 0 | 0.18 | 0.47 | 0.13 | 0.12 | 0.17 | 0.16 |
| 1 | 87014985 | rs2390134 | 0.77 | -1.053 | -0.342 | -0.039 | -0.028 | 0.064 | 0.064 | 0.017 | 0.08 | 0.2 | 0.06 | 0.2 | 0.04 | 0.04 |
| 1 | 87015179 | rs974843 | -1.262 | -1.227 | -0.91 | -0.296 | 0.032 | 0.148 | 0.148 | 0.256 | 0.46 | 0.91 | 0.29 | 0.76 | 0.35 | 0.53 |
| 1 | 87020617 | rs10782575 | 1.362 | -0.884 | 0.303 | -0.172 | -0.001 | -0.056 | -0.056 | 0.708 | 0.61 | 0.58 | 0.78 | 0.93 | 0.63 | 0.77 |
| 1 | 87026284 | rs1321678 | -0.818 | -0.832 | 0.11 | -0.156 | -0.023 | 0.036 | 0.036 | 0 | 0.93 | 1 | 0.97 | 1 | 0.91 | 0.96 |
| 1 | 87028669 | rs772605 | 1.335 | -0.896 | 0.342 | -0.199 | 0.001 | -0.004 | -0.004 | 0.134 | 0.63 | 0.62 | 0.78 | 0.93 | 0.63 | 0.77 |
| 1 | 87028855 | rs772604 | 0.337 | -0.838 | 0.273 | -0.185 | -0.013 | 0.047 | 0.047 | 0 | 0.79 | 0.88 | 0.86 | 0.94 | 0.75 | 0.89 |
| 1 | 87032324 | rs772601 | 0.33 | -0.84 | 0.27 | -0.186 | -0.014 | 0.05 | 0.05 | 0 | 0.79 | 0.89 | 0.86 | 0.94 | 0.75 | 0.89 |
| 1 | 87039945 | rs772598 | 1.765 | -0.953 | 0.576 | -0.038 | -0.005 | -0.02 | -0.02 | 1.062 | 0.57 | 0.51 | 0.73 | 0.73 | 0.59 | 0.72 |
| 1 | 87044583 | rs772607 | 0.798 | -0.877 | 0.604 | 0.054 | 0.007 | 0.01 | 0.01 | 0.047 | 0.76 | 0.56 | 0.82 | 0.73 | 0.74 | 0.85 |
| 1 | 87047478 | rs17129354 | 0.275 | -0.857 | 0.408 | 0.032 | -0.003 | 0.058 | 0.058 | 0 | 0.14 | 0.06 | 0.06 | 0 | 0.13 | 0.12 |
| 1 | 87051201 | rs1953652 | -1.377 | -0.813 | 0.679 | 0.195 | 0.002 | 0.017 | 0.017 | 0.088 | 0.35 | 0.3 | 0.21 | 0.27 | 0.35 | 0.18 |
| 1 | 87054030 | rs10157106 | -2.977 | -0.054 | 1.363 | 0.776 | -0.005 | 0.127 | 0.127 | 0.089 | 0.44 | 0.85 | 0.51 | 0.79 | 0.45 | 0.46 |
| 1 | 87061970 | rs12758538 | -0.326 | 0.129 | 1.398 | 0.368 | -0.004 | 0.01 | 0.01 | 0 | 0.14 | 0.08 | 0.1 | 0.06 | 0.17 | 0.09 |
| 1 | 87062201 | rs6679741 | -2.35 | 0.112 | 1.625 | 0.653 | 0.003 | 0.016 | 0.016 | 0.296 | 0.35 | 0.47 | 0.42 | 0.42 | 0.38 | 0.2 |

| | | | | | | | | | | | | | | | | |
|---|---|---|---|---|---|---|---|---|---|---|---|---|---|---|---|---|
| 1 | 87085266 | rs6662699 | -0.413 | 0.114 | 1.604 | 0.642 | -0.001 | -0.002 | -0.002 | 0 | 0.14 | 0.13 | 0.1 | 0.06 | 0.17 | 0.09 |
| 1 | 87101649 | rs2292829 | -0.048 | 0.107 | 1.652 | 0.656 | -0.002 | -0.042 | -0.042 | 0 | 0.12 | 0.06 | 0.22 | 0.37 | 0.11 | 0.07 |
| 1 | 87106136 | rs538417 | -2.658 | 0.107 | 1.686 | 0.647 | -0.001 | -0.011 | -0.011 | 0 | 0.37 | 0.71 | 0.44 | 0.42 | 0.39 | 0.2 |
| 1 | 87115922 | rs535037 | -2.737 | 0.168 | 1.267 | 0.676 | 0.001 | 0.003 | 0.003 | 0.246 | 0.37 | 0.64 | 0.46 | 0.43 | 0.42 | 0.28 |
| 1 | 87117642 | rs1517902 | -0.411 | 0.185 | 0.626 | 0.768 | -0.004 | 0.01 | 0.01 | 0 | 0.14 | 0.08 | 0.1 | 0.06 | 0.17 | 0.09 |
| 1 | 87118715 | rs6686918 | -1.303 | 0.18 | 0.635 | 0.782 | 0.002 | -0.028 | -0.028 | 0 | 0.22 | 0.26 | 0.19 | 0.07 | 0.28 | 0.13 |
| 1 | 87160811 | rs10489891 | -0.411 | 0.144 | -0.324 | 1.227 | -0.002 | -0.042 | -0.042 | 0.074 | 0.12 | 0.06 | 0.22 | 0.36 | 0.11 | 0.07 |
| 1 | 87166276 | rs263444 | 0.509 | 0.23 | -0.064 | 0.413 | -0.004 | 0.01 | 0.01 | 0 | 0.86 | 0.91 | 0.9 | 0.94 | 0.83 | 0.91 |
| 1 | 87169408 | rs1058978 | -0.081 | 0.612 | 0.051 | 0.69 | 0.002 | 0.004 | 0.004 | 0 | 0.07 | 0.06 | 0.2 | 0.17 | 0.14 | 0.05 |
| 1 | 87188881 | rs263463 | -2.323 | 0.02 | 0.537 | 0.32 | 0.004 | 0.016 | 0.016 | 0 | 0.29 | 0.7 | 0.28 | 0.21 | 0.24 | 0.15 |
| 1 | 87209658 | rs263476 | -2.338 | 0.108 | 1.156 | 0.311 | 0.006 | 0.028 | 0.028 | 0 | 0.29 | 0.76 | 0.29 | 0.21 | 0.24 | 0.15 |
| 1 | 87211940 | rs10489890 | -0.499 | 0.099 | 1.897 | 0.28 | -0.004 | -0.031 | -0.031 | 1.493 | 0.07 | 0.21 | 0.05 | 0 | 0.04 | 0.03 |
| 1 | 87215608 | rs12097495 | -1.985 | 0.115 | 2.325 | 0.264 | 0.01 | -0.066 | -0.066 | 0.303 | 0.17 | 0.37 | 0.1 | 0.04 | 0.14 | 0.11 |
| 1 | 87221551 | rs263493 | -2.458 | 0.067 | 2.437 | 0.273 | -0.005 | -0.018 | -0.018 | 0 | 0.24 | 0.39 | 0.25 | 0.21 | 0.19 | 0.13 |
| 1 | 87221781 | rs4523496 | -0.171 | 0.051 | 2.357 | 0.265 | -0.036 | 0.003 | 0.003 | 0 | 0.07 | 0.01 | 0.02 | 0.04 | 0.09 | 0.09 |
| 1 | 87223004 | rs263492 | -1.607 | 0.095 | 2.517 | 0.272 | -0.01 | -0.005 | -0.005 | 0 | 0.16 | 0.4 | 0.23 | 0.17 | 0.12 | 0.07 |
| 1 | 87223622 | rs17129466 | -0.052 | 0.058 | 2.239 | 0.257 | 0.003 | -0.031 | -0.031 | 1.245 | 0.06 | 0.13 | 0.05 | 0 | 0.06 | 0.03 |
| 1 | 87239013 | rs263459 | -2.4 | 0.028 | 2.179 | 0.248 | -0.037 | -0.03 | -0.03 | 0 | 0.2 | 0.57 | 0.25 | 0.17 | 0.13 | 0.07 |
| 1 | 87240077 | rs263458 | -3.028 | 0.035 | 2.139 | 0.471 | -0.007 | -0.008 | -0.008 | 0 | 0.28 | 0.64 | 0.28 | 0.25 | 0.22 | 0.16 |
| 1 | 87242062 | rs166412 | -2.549 | 0.024 | 2.201 | 0.313 | -0.021 | -0.019 | -0.019 | 0 | 0.22 | 0.52 | 0.23 | 0.2 | 0.16 | 0.13 |
| 1 | 87245278 | rs691531 | -3.186 | 0.157 | 2.131 | 0.328 | 0.005 | 0.029 | 0.029 | 0 | 0.29 | 0.76 | 0.29 | 0.21 | 0.23 | 0.17 |
| 1 | 87254444 | rs690676 | -1.07 | 0.274 | 2.216 | 0.43 | -0.019 | 0.015 | 0.015 | 0 | 0.13 | 0.26 | 0.19 | 0.18 | 0.09 | 0.04 |
| 1 | 87259306 | rs12074463 | -0.968 | 0.247 | 2.238 | 0.365 | -0.014 | 0.04 | 0.04 | 0 | 0.13 | 0.44 | 0.19 | 0.17 | 0.07 | 0.04 |
| 1 | 87278542 | rs6693416 | -0.708 | 0.062 | 1.225 | 0.332 | -0.003 | 0.004 | 0.004 | 0 | 0.1 | 0.08 | 0.03 | 0.05 | 0.09 | 0.12 |
| 1 | 87293905 | rs4282790 | 2.392 | 0.002 | 1.178 | 0.377 | 0.005 | 0.009 | 0.009 | 0.02 | 0.7 | 0.29 | 0.71 | 0.78 | 0.78 | 0.79 |
| 1 | 87297849 | rs12142266 | 1.415 | -0.198 | 0.108 | 0.387 | -0.098 | -0.014 | -0.014 | 0 | 0.39 | 0.06 | 0.45 | 0.4 | 0.47 | 0.37 |
| 1 | 87300389 | rs17129530 | 1.123 | 0.282 | 0.647 | 0.348 | -0.001 | 0 | 0 | 0 | 0.07 | 0.07 | 0.14 | 0.12 | 0.05 | 0.02 |
| 1 | 87311182 | rs493341 | -2.168 | 0.579 | 1.021 | 0.121 | -0.002 | -0.045 | -0.045 | 0 | 0.21 | 0.66 | 0.17 | 0.05 | 0.17 | 0.17 |
| 1 | 87316418 | rs526985 | -2.276 | 0.593 | 1.057 | 0.048 | -0.001 | -0.064 | -0.064 | 0.01 | 0.2 | 0.65 | 0.17 | 0.04 | 0.17 | 0.17 |
| 1 | 87324075 | rs492745 | 0.104 | 0.525 | 1.26 | 0.045 | -0.004 | -0.018 | -0.018 | 0.863 | 0.06 | 0.09 | 0.05 | 0.01 | 0.05 | 0.03 |
| 1 | 87326018 | rs552483 | -2.349 | 0.762 | 1.439 | 0.014 | 0.022 | -0.087 | -0.087 | 0 | 0.2 | 0.56 | 0.17 | 0.04 | 0.21 | 0.17 |
| 1 | 87327273 | rs496114 | -0.986 | 0.743 | 1.791 | 0.012 | 0.011 | -0.025 | -0.025 | 0.477 | 0.08 | 0.31 | 0.06 | 0.01 | 0.05 | 0.03 |
| 1 | 87328308 | rs540049 | -2.614 | 0.812 | 1.864 | 0.005 | 0.025 | -0.101 | -0.101 | 0 | 0.2 | 0.5 | 0.16 | 0.04 | 0.2 | 0.15 |
| 1 | 87328839 | rs5845 | -2.57 | 0.78 | 1.891 | 0.007 | 0.009 | -0.064 | -0.064 | 0 | 0.2 | 0.66 | 0.18 | 0.04 | 0.21 | 0.17 |
| 1 | 87330141 | rs488181 | -2.601 | 0.784 | 1.953 | 0.011 | 0.022 | -0.101 | -0.101 | 0.042 | 0.2 | 0.5 | 0.16 | 0.04 | 0.2 | 0.15 |
| 1 | 87371762 | rs561104 | -0.979 | 0.558 | 1.585 | 0.025 | -0.058 | -0.087 | -0.087 | 0 | 0.72 | 0.87 | 0.62 | 0.44 | 0.62 | 0.65 |
| 1 | 87378627 | rs1407131 | -0.951 | 0.589 | 2.216 | 0.661 | 0.01 | -0.01 | -0.01 | 4.805 | 0.1 | 0.12 | 0.08 | 0.03 | 0.15 | 0.12 |
| 1 | 87379309 | rs9433110 | -1.141 | 0.573 | 2.256 | 0.651 | 0.011 | -0.025 | -0.025 | 0 | 0.08 | 0.31 | 0.06 | 0.01 | 0.05 | 0.03 |
| 1 | 87379986 | rs754001 | -0.869 | 0.574 | 2.257 | 0.651 | 0.002 | -0.002 | -0.002 | 0 | 0.09 | 0.1 | 0.08 | 0.03 | 0.15 | 0.12 |
| 1 | 87379996 | rs101028 | -2.84 | 0.573 | 2.204 | 0.651 | 0.02 | -0.097 | -0.097 | 1.302 | 0.2 | 0.46 | 0.16 | 0.04 | 0.2 | 0.15 |
| 1 | 87380631 | rs548649 | -0.238 | 0.513 | 2.296 | 0.661 | -0.006 | -0.023 | -0.023 | 0.117 | 0.06 | 0.11 | 0.05 | 0.01 | 0.05 | 0.03 |
| 1 | 87381454 | rs487006 | -2.674 | 0.533 | 2.254 | 0.68 | 0.019 | -0.093 | -0.093 | 0 | 0.19 | 0.44 | 0.16 | 0.04 | 0.19 | 0.15 |
| 1 | 87428302 | rs1970607 | 2.469 | 0.532 | 2.182 | 0.681 | 0.031 | -0.101 | -0.101 | 0.864 | 0.8 | 0.5 | 0.83 | 0.96 | 0.79 | 0.83 |
| 1 | 87477786 | rs4233338 | 0.681 | 0.527 | 2.033 | 0.72 | -0.006 | -0.035 | -0.035 | 0 | 0.93 | 0.78 | 0.94 | 0.99 | 0.95 | 0.97 |
| 1 | 87485139 | rs11584931 | -0.126 | 0.586 | 2.045 | 0.712 | -0.006 | 0.003 | 0.003 | 0 | 0.07 | 0.05 | 0.06 | 0.03 | 0.12 | 0.07 |
| 1 | 87505590 | rs10127488 | 2.645 | 0.54 | 2.312 | 0.674 | 0.007 | -0.063 | -0.063 | 0 | 0.8 | 0.34 | 0.82 | 0.96 | 0.79 | 0.84 |
| 1 | 87506127 | rs6696540 | 2.453 | 0.548 | 2.349 | 0.686 | 0.028 | -0.076 | -0.076 | 0 | 0.82 | 0.44 | 0.84 | 0.97 | 0.82 | 0.88 |
| 1 | 87511398 | rs7534656 | 2.22 | 0.551 | 2.381 | 0.687 | 0.05 | -0.077 | -0.077 | 0 | 0.83 | 0.55 | 0.85 | 0.97 | 0.82 | 0.88 |
| 1 | 87520970 | rs4610985 | 1.524 | 0.526 | 2.401 | 0.689 | 0.012 | -0.048 | -0.048 | 0 | 0.9 | 0.61 | 0.91 | 0.99 | 0.94 | 0.95 |
| 1 | 87521319 | rs12066353 | -0.489 | 0.573 | 2.342 | 0.696 | 0.044 | -0.01 | -0.01 | 0 | 0.07 | 0.13 | 0.07 | 0.03 | 0.13 | 0.09 |
| 1 | 87522312 | rs7529516 | 0.893 | 0.534 | 2.273 | 0.708 | 0.052 | -0.061 | -0.061 | 0.012 | 0.85 | 0.56 | 0.87 | 0.97 | 0.83 | 0.89 |
| 1 | 87528599 | rs1936280 | 0.577 | 0.546 | 2.163 | 0.714 | 0.026 | -0.038 | -0.038 | 0.464 | 0.87 | 0.73 | 0.88 | 0.97 | 0.83 | 0.89 |
| 1 | 87554556 | rs1778012 | 1.921 | 0.531 | 2.024 | 0.742 | 0.007 | -0.053 | -0.053 | 0 | 0.8 | 0.34 | 0.81 | 0.94 | 0.78 | 0.83 |
| 1 | 87554612 | rs1778013 | 0.473 | 0.529 | 2.074 | 0.741 | 0.026 | -0.027 | -0.027 | 0 | 0.87 | 0.75 | 0.88 | 0.94 | 0.83 | 0.89 |
| 1 | 87555358 | rs1419132 | 1.978 | 0.547 | 2.061 | 1.03 | 0.008 | -0.002 | -0.002 | 0.102 | 0.8 | 0.35 | 0.81 | 0.81 | 0.78 | 0.82 |
| 1 | 87570398 | rs1621247 | 2.032 | 0.511 | 2.022 | 0.596 | 0.003 | -0.049 | -0.049 | 0 | 0.79 | 0.31 | 0.81 | 0.97 | 0.78 | 0.83 |
| 1 | 87572817 | rs2764430 | 0.516 | 0.593 | 1.462 | 0.619 | 0.04 | -0.05 | -0.05 | 0.889 | 0.87 | 0.69 | 0.87 | 0.97 | 0.83 | 0.89 |
| 1 | 87576481 | rs1419130 | 0.734 | 0.631 | 1.073 | 0.434 | 0.019 | -0.075 | -0.075 | 0 | 0.83 | 0.64 | 0.84 | 0.97 | 0.8 | 0.85 |
| 1 | 87581491 | rs1199668 | 0.631 | 0.24 | -0.073 | 0.336 | 0.074 | -0.03 | -0.03 | 0.465 | 0.51 | 0.17 | 0.54 | 0.54 | 0.61 | 0.53 |
| 1 | 87584639 | rs1199672 | 1.419 | 1.814 | 1.071 | 0.023 | 0.018 | -0.114 | -0.114 | 0 | 0.76 | 0.44 | 0.78 | 0.97 | 0.72 | 0.8 |
| 1 | 87587849 | rs1199673 | -2.115 | 2.819 | 1.176 | -0.112 | 0.042 | -0.141 | -0.141 | 0 | 0.32 | 0.5 | 0.36 | 0.01 | 0.46 | 0.28 |
| 1 | 87595088 | rs1201783 | -2.081 | 1.964 | 1.632 | 0.434 | 0.003 | -0.035 | -0.035 | 0 | 0.28 | 0.41 | 0.25 | 0.16 | 0.3 | 0.31 |
| 1 | 87598117 | rs6686259 | 1.566 | 1.985 | 1.441 | 0.446 | 0.006 | -0.012 | -0.012 | 0 | 0.77 | 0.67 | 0.79 | 0.84 | 0.73 | 0.71 |
| 1 | 87609509 | rs12405406 | 0.19 | 2.403 | 1.299 | 0.374 | -0.012 | 0.01 | 0.01 | 0 | 0.05 | 0.03 | 0.14 | 0 | 0.11 | 0.06 |
| 1 | 87610246 | rs1886632 | 1.43 | 2.455 | 1.4 | 0.369 | 0.003 | -0.012 | -0.012 | 0 | 0.78 | 0.65 | 0.83 | 0.84 | 0.79 | 0.74 |
| 1 | 87611710 | rs719015 | 2.062 | 2.413 | 1.548 | 0.364 | 0.042 | -0.056 | -0.056 | 0.367 | 0.69 | 0.44 | 0.63 | 0.84 | 0.62 | 0.67 |
| 1 | 87613358 | rs7553864 | 2.168 | 2.243 | 1.533 | 0.399 | 0.021 | -0.032 | -0.032 | 0.165 | 0.7 | 0.56 | 0.64 | 0.85 | 0.63 | 0.69 |
| 1 | 87614191 | rs11161935 | 2.12 | 2.265 | 1.482 | 0.421 | 0.024 | -0.032 | -0.032 | 0.989 | 0.71 | 0.54 | 0.66 | 0.84 | 0.63 | 0.68 |
| 1 | 87614821 | rs1886634 | 1.722 | 2.263 | 1.437 | 0.428 | 0.002 | -0.019 | -0.019 | 0 | 0.76 | 0.6 | 0.78 | 0.84 | 0.74 | 0.75 |

| | | | | | | | | | | | | | | | | |
|---|---|---|---|---|---|---|---|---|---|---|---|---|---|---|---|---|
| 1 | 87615412 | rs12136695 | -0.812 | 2.301 | 1.308 | 0.434 | -0.009 | -0.032 | -0.032 | 0 | 0.14 | 0.05 | 0.11 | 0.15 | 0.13 | 0.2 |
| 1 | 87626529 | rs6576869 | 0.955 | 2.605 | 1.82 | 0.731 | 0.005 | -0.026 | -0.026 | 0 | 0.93 | 0.88 | 0.9 | 1 | 0.87 | 0.95 |
| 1 | 87628173 | rs10873821 | -2.086 | 2.69 | 2.063 | 1.102 | 0.032 | -0.006 | -0.006 | 0 | 0.21 | 0.6 | 0.38 | 0.19 | 0.22 | 0.2 |
| 1 | 87628834 | rs12092928 | -0.495 | 2.7 | 2.129 | 1.1 | 0.083 | -0.033 | -0.033 | 0 | 0.07 | 0.24 | 0.15 | 0 | 0.11 | 0.06 |
| 1 | 87628868 | rs11161939 | -1.253 | 2.692 | 2.053 | 1.097 | 0.003 | 0.027 | 0.027 | 0 | 0.14 | 0.24 | 0.22 | 0.19 | 0.11 | 0.14 |
| 1 | 87629493 | rs17129854 | -0.099 | 2.805 | 2.014 | 1.128 | 0.042 | -0.03 | -0.03 | 0 | 0.06 | 0.15 | 0.14 | 0 | 0.11 | 0.06 |
| 1 | 87632354 | rs4655841 | 1.772 | 2.819 | 1.632 | 1.131 | 0.115 | -0.111 | -0.111 | 0.066 | 0.85 | 0.51 | 0.75 | 1 | 0.76 | 0.89 |
| 1 | 87633929 | rs17129860 | -0.787 | 2.215 | 1.733 | 1.4 | 0.137 | -0.018 | -0.018 | 0.435 | 0.07 | 0.36 | 0.16 | 0 | 0.11 | 0.06 |
| 1 | 87638729 | rs6690136 | -1.16 | 2.047 | 1.015 | 1.569 | 0.01 | 0.041 | 0.041 | 0 | 0.13 | 0.23 | 0.22 | 0.19 | 0.11 | 0.12 |
| 1 | 87640058 | rs2390270 | -1.264 | 2.017 | 0.006 | 1.649 | 0.005 | 0.026 | 0.026 | 0.008 | 0.28 | 0.6 | 0.45 | 0.35 | 0.27 | 0.25 |
| 1 | 87641938 | rs7512999 | -1.021 | 1.995 | -0.601 | 1.589 | -0.033 | 0.008 | 0.008 | 0.035 | 0.39 | 0.72 | 0.53 | 0.31 | 0.44 | 0.27 |
| 1 | 87643452 | rs912764 | -1.021 | 2.011 | -0.626 | 1.597 | -0.032 | 0.008 | 0.008 | 0 | 0.39 | 0.73 | 0.53 | 0.31 | 0.44 | 0.27 |
| 1 | 87649865 | rs7556318 | -0.987 | 1.555 | -0.785 | 1.144 | -0.035 | -0.029 | -0.029 | 0 | 0.39 | 0.74 | 0.55 | 0.51 | 0.45 | 0.26 |
| 1 | 87663215 | rs12563224 | -0.803 | 1.599 | -0.287 | 1.382 | -0.006 | -0.023 | -0.023 | 0.407 | 0.22 | 0.17 | 0.28 | 0.36 | 0.24 | 0.38 |
| 1 | 87667705 | rs723006 | -1.38 | 1.865 | -0.204 | 1.502 | 0.001 | 0.013 | 0.013 | 0 | 0.14 | 0.15 | 0.16 | 0.25 | 0.15 | 0.18 |
| 1 | 87681151 | rs7554095 | -1.841 | 1.698 | 0.553 | 1.003 | 0.017 | 0.011 | 0.011 | 0 | 0.26 | 0.53 | 0.25 | 0.23 | 0.26 | 0.19 |
| 1 | 87684844 | rs6690180 | -0.738 | 1.663 | 1.264 | 0.892 | 0.043 | -0.072 | -0.072 | 0.466 | 0.12 | 0.41 | 0.1 | 0 | 0.11 | 0.02 |
| 1 | 87687466 | rs7542364 | -1.272 | 1.554 | 1.306 | 1.224 | 0.024 | 0.112 | 0.112 | 0 | 0.15 | 0.36 | 0.17 | 0.28 | 0.15 | 0.19 |
| 1 | 87688394 | rs10873824 | -0.292 | 1.431 | 0.254 | 1.293 | -0.085 | -0.134 | -0.134 | 0 | 0.81 | 0.98 | 0.75 | 0.6 | 0.75 | 0.78 |
| 1 | 87690073 | rs6576876 | -1.332 | 1.257 | 0.597 | 1.166 | 0.03 | 0.112 | 0.112 | 0 | 0.18 | 0.48 | 0.2 | 0.28 | 0.18 | 0.2 |
| 1 | 87691451 | rs7526555 | -1.362 | 1.266 | 0.377 | 1.159 | 0.019 | 0.08 | 0.08 | 0 | 0.18 | 0.52 | 0.2 | 0.26 | 0.18 | 0.21 |
| 1 | 87696098 | rs12409961 | 0.069 | 1.37 | -0.306 | 1.305 | -0.041 | -0.055 | -0.055 | 0.669 | 0.81 | 0.9 | 0.86 | 0.67 | 0.74 | 0.79 |
| 1 | 87698441 | rs12144566 | 0.923 | 0.608 | -1.005 | 0.492 | -0.048 | 0.115 | 0.115 | 0 | 0.41 | 0.22 | 0.46 | 0.21 | 0.44 | 0.25 |
| 1 | 87698626 | rs11801833 | -0.272 | 0.118 | -0.334 | 0.207 | -0.011 | -0.06 | -0.06 | 0 | 0.16 | 0.22 | 0.06 | 0 | 0.11 | 0.14 |
| 1 | 87702362 | rs6576877 | -1.64 | -0.26 | -0.188 | 0.389 | 0.032 | 0.082 | 0.082 | 0 | 0.49 | 0.65 | 0.33 | 0.24 | 0.37 | 0.4 |
| 1 | 87703049 | rs10782594 | -1.656 | -0.189 | -0.023 | 0.377 | 0.035 | 0.105 | 0.105 | 0 | 0.49 | 0.64 | 0.33 | 0.23 | 0.38 | 0.39 |
| 1 | 87705031 | rs6576878 | -1.077 | -0.172 | 0.198 | 0.571 | -0.019 | 0.063 | 0.063 | 0 | 0.29 | 0.49 | 0.28 | 0.44 | 0.22 | 0.33 |
| 1 | 87708655 | rs7540876 | -0.456 | -0.274 | 0.523 | 0.64 | 0.017 | 0.116 | 0.116 | 0 | 0.15 | 0.36 | 0.26 | 0.37 | 0.17 | 0.13 |
| 1 | 87710877 | rs10158460 | -1.028 | -0.403 | 0.557 | 0.601 | -0.011 | -0.013 | -0.013 | 0 | 0.4 | 0.45 | 0.22 | 0.28 | 0.31 | 0.39 |
| 1 | 87711892 | rs10747342 | -2.461 | -0.394 | 0.236 | 0.608 | -0.1 | 0.012 | 0.012 | 0.022 | 0.57 | 0.89 | 0.49 | 0.64 | 0.48 | 0.52 |
| 1 | 87719075 | rs6680623 | -0.096 | -0.411 | 0.83 | 0.544 | -0.002 | -0.003 | -0.003 | 0.369 | 0.19 | 0.18 | 0.07 | 0.07 | 0.13 | 0.19 |
| 1 | 87720853 | rs4655941 | 3.136 | -0.335 | 0.308 | 0.493 | -0.068 | -0.002 | -0.002 | 0 | 0.28 | 0.1 | 0.42 | 0.27 | 0.33 | 0.25 |
| 1 | 87722965 | rs1330864 | 0.278 | -0.294 | 0.589 | 0.564 | -0.004 | -0.017 | -0.017 | 0.168 | 0.7 | 0.7 | 0.64 | 0.55 | 0.64 | 0.63 |
| 1 | 87732429 | rs7544647 | 0.635 | -0.225 | 0.9 | 0.688 | 0.027 | 0.179 | 0.179 | 0 | 0.08 | 0.2 | 0.21 | 0.36 | 0.1 | 0.13 |
| 1 | 87739930 | rs12035326 | 0.303 | -0.229 | 0.932 | 0.654 | -0.007 | -0.009 | -0.009 | 0 | 0.19 | 0.09 | 0.13 | 0.21 | 0.18 | 0.2 |
| 1 | 87743047 | rs12407239 | 0.932 | -0.034 | 1.07 | 0.785 | 0.001 | 0.066 | 0.066 | 0.017 | 0.07 | 0.09 | 0.2 | 0.36 | 0.1 | 0.12 |
| 1 | 87744420 | rs6678638 | 0.654 | -0.078 | 0.991 | 0.743 | -0.006 | -0.007 | -0.007 | 0.347 | 0.16 | 0.11 | 0.13 | 0.22 | 0.18 | 0.2 |
| 1 | 87746317 | rs6576888 | -1.214 | -0.246 | 0.855 | 0.657 | -0.022 | 0.039 | 0.039 | 0.752 | 0.45 | 0.79 | 0.43 | 0.65 | 0.4 | 0.51 |
| 1 | 87746591 | rs4320762 | -1.214 | -0.246 | 0.878 | 0.657 | -0.013 | 0.044 | 0.044 | 0.752 | 0.45 | 0.78 | 0.43 | 0.65 | 0.4 | 0.51 |
| 1 | 87746998 | rs4556331 | -1.349 | -0.245 | 1.04 | 0.658 | 0.003 | 0.037 | 0.037 | 0.542 | 0.44 | 0.68 | 0.42 | 0.65 | 0.4 | 0.51 |
| 1 | 87750413 | rs4233323 | -1.539 | -0.264 | 0.889 | 0.584 | -0.042 | -0.003 | -0.003 | 0 | 0.86 | 0.98 | 0.9 | 0.92 | 0.81 | 0.77 |
| 1 | 87750853 | rs11161968 | -0.137 | 0.02 | 1.841 | 0.848 | 0.111 | 0.313 | 0.313 | 0.121 | 0.09 | 0.34 | 0.22 | 0.36 | 0.11 | 0.13 |
| 1 | 87752886 | rs6688319 | -2.103 | -0.003 | 1.155 | 0.496 | -0.041 | 0.05 | 0.05 | 0.79 | 0.53 | 0.84 | 0.49 | 0.65 | 0.49 | 0.52 |
| 1 | 87754684 | rs10873827 | -1.589 | 0 | 1.329 | 0.697 | -0.004 | 0.011 | 0.011 | 0.547 | 0.44 | 0.58 | 0.43 | 0.65 | 0.42 | 0.51 |
| 1 | 87757861 | rs11811308 | -1.5 | -0.55 | 1.636 | 0.508 | 0.032 | 0.019 | 0.019 | 0 | 0.17 | 0.45 | 0.21 | 0.19 | 0.15 | 0.12 |
| 1 | 87758358 | rs7512896 | -0.541 | -0.472 | 1.579 | 0.656 | 0.082 | 0.094 | 0.094 | 0 | 0.1 | 0.3 | 0.16 | 0.18 | 0.11 | 0.11 |
| 1 | 87760435 | rs6667726 | -0.843 | -0.835 | 0.948 | 0.299 | -0.026 | 0.001 | 0.001 | 0 | 0.13 | 0.19 | 0.06 | 0.11 | 0.05 | 0.15 |
| 1 | 87761387 | rs17129936 | -0.491 | -0.646 | 0.749 | 0.192 | 0.004 | -0.001 | -0.001 | 0 | 0.1 | 0.09 | 0.05 | 0.08 | 0.06 | 0.12 |
| 1 | 87761635 | rs4119928 | -1.428 | -0.674 | 0.984 | 0.077 | -0.001 | -0.015 | -0.015 | 0 | 0.17 | 0.41 | 0.18 | 0.15 | 0.13 | 0.12 |
| 1 | 87764202 | rs6691205 | -1.009 | -0.797 | 0.757 | 0.37 | -0.006 | 0.01 | 0.01 | 0 | 0.25 | 0.31 | 0.15 | 0.3 | 0.21 | 0.3 |
| 1 | 87769531 | rs11161973 | -1.446 | -0.395 | 0.741 | 0.353 | -0.008 | -0.012 | -0.012 | 0 | 0.57 | 0.75 | 0.45 | 0.55 | 0.54 | 0.58 |
| 1 | 87773431 | rs10873829 | 0.396 | -0.42 | 0.794 | 0.338 | 0.012 | -0.028 | -0.028 | 0.15 | 0.31 | 0.39 | 0.27 | 0.25 | 0.32 | 0.26 |
| 1 | 87773503 | rs17129945 | -1.108 | -0.623 | 0.972 | 1.26 | -0.001 | -0.015 | -0.015 | 0.14 | 0.18 | 0.15 | 0.12 | 0.3 | 0.19 | 0.28 |
| 1 | 87774994 | rs944367 | -1.437 | -0.621 | 0.949 | 1.263 | 0 | -0.003 | -0.003 | 0 | 0.2 | 0.21 | 0.13 | 0.3 | 0.2 | 0.3 |
| 1 | 87781157 | rs2144523 | -1.663 | -0.551 | 0.907 | 1.132 | -0.002 | 0.001 | 0.001 | 0 | 0.26 | 0.31 | 0.17 | 0.28 | 0.22 | 0.32 |
| 1 | 87790813 | rs4488005 | -1.535 | -0.393 | 0.383 | 1.255 | -0.022 | 0.09 | 0.09 | 0 | 0.26 | 0.49 | 0.23 | 0.45 | 0.2 | 0.32 |
| 1 | 87793532 | rs3806177 | -1.268 | -0.393 | -0.292 | 0.583 | -0.001 | 0 | 0 | 0 | 0.09 | 0.14 | 0.05 | 0.05 | 0.06 | 0.11 |
| 1 | 87803200 | rs10873830 | 0.78 | -0.791 | -0.979 | 0.154 | 0.069 | 0.062 | 0.062 | 1.45 | 0.68 | 0.17 | 0.72 | 0.81 | 0.79 | 0.77 |
| 1 | 87807876 | rs3766024 | -1.51 | -0.5 | -0.234 | 0.055 | -0.009 | -0.044 | -0.044 | 0.012 | 0.1 | 0.16 | 0.06 | 0.02 | 0.08 | 0.07 |
| 1 | 87808352 | rs3766022 | -1.298 | -0.146 | 0.961 | 0.381 | 0.13 | 0.202 | 0.202 | 1.265 | 0.07 | 0.51 | 0.15 | 0.14 | 0.07 | 0.09 |
| 1 | 87816796 | rs1336945 | -2.145 | -0.285 | 1.34 | 0.642 | -0.005 | 0.006 | 0.006 | 0.039 | 0.2 | 0.71 | 0.25 | 0.22 | 0.18 | 0.23 |
| 1 | 87826418 | rs7518015 | 0.171 | -1.025 | -0.045 | 0.519 | 0.015 | -0.043 | -0.043 | 0.571 | 0.8 | 0.98 | 0.78 | 0.77 | 0.86 | 0.83 |
| 1 | 87826745 | rs17456070 | 0.215 | -0.329 | -0.496 | -0.309 | -0.012 | 0.011 | 0.011 | 0 | 0.29 | 0.25 | 0.31 | 0.09 | 0.34 | 0.29 |
| 1 | 87826767 | rs7555691 | -1.158 | -0.517 | -0.745 | 0.187 | 0.001 | -0.035 | -0.035 | 0 | 0.09 | 0.05 | 0.08 | 0.22 | 0.09 | 0.12 |
| 1 | 87828033 | rs17129984 | -0.506 | -0.318 | -0.947 | 0.326 | -0.015 | -0.029 | -0.029 | 1.319 | 0.12 | 0.06 | 0.09 | 0.26 | 0.14 | 0.18 |
| 1 | 87833443 | rs11161991 | 0.303 | -0.658 | -1.275 | 0.608 | 0.001 | -0.015 | -0.015 | 0 | 0.14 | 0.06 | 0.1 | 0.12 | 0.13 | 0.08 |
| 1 | 87834259 | rs12736459 | 0.787 | -0.856 | -2.154 | 0.561 | 0.019 | 0.002 | 0.002 | 0 | 0.26 | 0.12 | 0.35 | 0.26 | 0.21 | 0.33 |
| 1 | 87834794 | rs17422436 | 2.403 | -0.514 | -2.217 | 0.397 | -0.067 | 0.094 | 0.094 | 0 | 0.29 | 0.12 | 0.34 | 0.09 | 0.34 | 0.26 |
| 1 | 87834877 | rs12067223 | 0.823 | -0.545 | -2.272 | 0.492 | 0.005 | -0.045 | -0.045 | 1.809 | 0.16 | 0.1 | 0.12 | 0.37 | 0.14 | 0.12 |

| | | | | | | | | | | | | | | | | |
|---|---|---|---|---|---|---|---|---|---|---|---|---|---|---|---|---|
| 1 | 87835674 | rs11161992 | -0.582 | -0.878 | -2.126 | 0.191 | 0.003 | -0.106 | -0.106 | 0.386 | 0.78 | 0.45 | 0.87 | 0.95 | 0.77 | 0.81 |
| 1 | 87838608 | rs11161993 | 0.151 | -0.627 | -2.255 | 0.4 | 0.008 | 0.017 | 0.017 | 0 | 0.65 | 0.37 | 0.8 | 0.86 | 0.7 | 0.76 |
| 1 | 87845112 | rs12402858 | 0.919 | -0.755 | -2.282 | 0.539 | 0.02 | -0.006 | -0.006 | 0.851 | 0.24 | 0.09 | 0.34 | 0.26 | 0.2 | 0.31 |
| 1 | 87849215 | rs746191 | 0.196 | -0.971 | -1.921 | 0.91 | -0.017 | -0.001 | -0.001 | 0 | 0.66 | 0.52 | 0.69 | 0.68 | 0.72 | 0.75 |
| 1 | 87850592 | rs1415173 | -0.257 | -0.819 | -1.854 | 0.366 | -0.027 | -0.066 | -0.066 | 0.216 | 0.7 | 0.53 | 0.81 | 0.9 | 0.76 | 0.83 |
| 1 | 87850703 | rs12046968 | 0.615 | 0.117 | -1.799 | 1.094 | -0.024 | -0.045 | -0.045 | 0 | 0.07 | 0.01 | 0.06 | 0.23 | 0.08 | 0.12 |
| 1 | 87862181 | rs10158426 | -1.583 | 0.97 | -2.109 | 0.24 | -0.126 | 0.096 | 0.096 | 0.119 | 0.71 | 0.98 | 0.67 | 0.92 | 0.61 | 0.75 |
| 1 | 87866857 | rs7554926 | 0.275 | 0.569 | -1.707 | 0.225 | 0.02 | 0.101 | 0.101 | 0 | 0.42 | 0.2 | 0.34 | 0.19 | 0.34 | 0.4 |
| 1 | 87867680 | rs12034213 | 0.495 | 0.57 | -1.422 | 0.851 | -0.006 | 0.07 | 0.07 | 0.016 | 0.1 | 0.16 | 0.05 | 0.37 | 0.08 | 0.08 |
| 1 | 87868540 | rs4655956 | -0.867 | 0.593 | -0.228 | 0.648 | -0.002 | 0.026 | 0.026 | 0 | 0.26 | 0.48 | 0.29 | 0.72 | 0.25 | 0.35 |
| 1 | 87869438 | rs11589492 | 0.062 | 0.567 | -0.235 | 0.602 | -0.005 | -0.017 | -0.017 | 0 | 0.45 | 0.46 | 0.37 | 0.2 | 0.34 | 0.4 |
| 1 | 87875198 | rs6697170 | 0.56 | 0.588 | -0.207 | 0.816 | -0.002 | 0.097 | 0.097 | 0 | 0.08 | 0.18 | 0.04 | 0.36 | 0.08 | 0.07 |
| 1 | 87875774 | rs1006598 | 0.8 | 0.569 | -0.068 | 0.781 | 0.007 | 0.083 | 0.083 | 0 | 0.17 | 0.31 | 0.25 | 0.35 | 0.17 | 0.25 |
| 1 | 87877226 | rs12758097 | 0.241 | 0.499 | 0.146 | 0.683 | 0.023 | 0.098 | 0.098 | 0.063 | 0.42 | 0.12 | 0.34 | 0.2 | 0.34 | 0.41 |
| 1 | 87877540 | rs1006625 | -0.442 | -0.381 | 1.029 | 1.306 | -0.01 | 0.073 | 0.073 | 0.733 | 0.21 | 0.63 | 0.28 | 0.35 | 0.17 | 0.26 |
| 1 | 87887040 | rs6695071 | -0.141 | -0.229 | 0.307 | 1.178 | 0 | 0.132 | 0.132 | 0.016 | 0.08 | 0.17 | 0.04 | 0.37 | 0.08 | 0.07 |
| 1 | 87897907 | rs1856047 | -0.565 | -0.461 | -0.757 | 0.118 | -0.111 | -0.125 | -0.125 | 0 | 0.55 | 0.97 | 0.53 | 0.52 | 0.47 | 0.45 |
| 1 | 87903024 | rs4295863 | -0.364 | -0.45 | -0.337 | 0.102 | -0.014 | -0.026 | -0.026 | 0 | 0.53 | 0.79 | 0.51 | 0.49 | 0.46 | 0.45 |
| 1 | 87908917 | rs6685678 | -0.255 | -0.529 | -0.129 | 0.037 | 0.001 | 0.006 | 0.006 | 0 | 0.52 | 0.7 | 0.5 | 0.48 | 0.46 | 0.45 |
| 1 | 87914880 | rs1831526 | 1.097 | -0.46 | -0.29 | 0.06 | -0.006 | 0.005 | 0.005 | 0.91 | 0.05 | 0.02 | 0.05 | 0.01 | 0.08 | 0.04 |
| 1 | 87915254 | rs12727441 | 0.323 | -0.598 | 0.307 | 0.034 | 0.007 | -0.003 | -0.003 | 0.555 | 0.43 | 0.33 | 0.43 | 0.47 | 0.38 | 0.41 |
| 1 | 87921397 | rs12565028 | 1.48 | -0.455 | -0.005 | 0.087 | 0.003 | 0.016 | 0.016 | 0 | 0.25 | 0.15 | 0.33 | 0.15 | 0.25 | 0.15 |
| 1 | 87925943 | rs12737419 | 0.917 | 0.15 | 0.041 | 0.254 | -0.021 | -0.031 | -0.031 | 0.089 | 0.1 | 0 | 0.1 | 0.09 | 0.1 | 0.05 |
| 1 | 87930889 | rs6683030 | -0.18 | 0.412 | 0.327 | 0.361 | -0.05 | -0.028 | -0.028 | 0 | 0.57 | 0.83 | 0.55 | 0.48 | 0.5 | 0.42 |
| 1 | 87932732 | rs2892987 | 0.643 | 0.536 | 0.188 | 0.396 | -0.012 | 0.016 | 0.016 | 0.633 | 0.05 | 0.02 | 0.05 | 0.01 | 0.08 | 0.04 |
| 1 | 87934293 | rs2065909 | 0.469 | 0.494 | 0.428 | 0.412 | 0.009 | 0.004 | 0.004 | 0 | 0.47 | 0.33 | 0.46 | 0.47 | 0.42 | 0.38 |
| 1 | 87954542 | rs10873850 | 0.603 | 0.654 | -0.17 | 0.743 | -0.083 | -0.095 | -0.095 | 0.108 | 0.13 | 0.01 | 0.18 | 0.35 | 0.28 | 0.23 |
| 1 | 87974961 | rs7349161 | -0.113 | -0.046 | -0.581 | 0.486 | -0.014 | -0.027 | -0.027 | 0 | 0.41 | 0.22 | 0.42 | 0.44 | 0.5 | 0.49 |
| 1 | 87979762 | rs10873853 | -0.365 | -1.112 | 0.044 | -0.328 | 0.009 | 0.016 | 0.016 | 0 | 0.41 | 0.62 | 0.34 | 0.1 | 0.32 | 0.36 |
| 1 | 87980505 | rs12141629 | 0.451 | -1.249 | 0.02 | -0.402 | -0.027 | 0.1 | 0.1 | 0 | 0.29 | 0.04 | 0.23 | 0.1 | 0.24 | 0.31 |
| 1 | 87982396 | rs1888728 | 0.463 | -1.25 | 0.019 | -0.426 | -0.027 | 0.104 | 0.104 | 0 | 0.29 | 0.04 | 0.23 | 0.1 | 0.24 | 0.31 |
| 1 | 87986477 | rs17130035 | -0.998 | -1.225 | 0.151 | -0.404 | 0.008 | -0.057 | -0.057 | 0 | 0.11 | 0.51 | 0.11 | 0.01 | 0.08 | 0.05 |
| 1 | 87988705 | rs10782601 | -0.195 | -1.446 | 0.112 | -0.113 | -0.005 | 0.002 | 0.002 | 0 | 0.41 | 0.39 | 0.33 | 0.45 | 0.29 | 0.43 |
| 1 | 87992241 | rs17130036 | -0.375 | -0.881 | -0.107 | 0.018 | -0.003 | -0.026 | -0.026 | 0 | 0.07 | 0.1 | 0.06 | 0.01 | 0.08 | 0.05 |
| 1 | 87998297 | rs10873855 | 0.804 | -1.128 | -0.412 | -0.058 | -0.006 | -0.039 | -0.039 | 0 | 0.43 | 0.2 | 0.57 | 0.47 | 0.61 | 0.45 |
| 1 | 88001013 | rs6657047 | 0.15 | -1.157 | -0.346 | -0.002 | 0.017 | 0.036 | 0.036 | 0.011 | 0.38 | 0.66 | 0.29 | 0.13 | 0.25 | 0.33 |
| 1 | 88005122 | rs7517567 | -0.449 | -0.866 | -0.255 | 0.142 | 0.005 | 0.004 | 0.004 | 0 | 0.88 | 0.9 | 0.91 | 0.61 | 0.93 | 0.82 |
| 1 | 88015080 | rs7355015 | -0.128 | -0.534 | -0.082 | 0.182 | 0.012 | 0.011 | 0.011 | 0 | 0.41 | 0.6 | 0.28 | 0.13 | 0.26 | 0.35 |
| 1 | 88025144 | rs6694829 | 0.183 | -0.312 | -1.214 | 0.347 | 0.014 | 0.019 | 0.019 | 0 | 0.36 | 0.26 | 0.25 | 0.13 | 0.24 | 0.35 |
| 1 | 88033668 | rs12046929 | -0.242 | 0.53 | -1.811 | 0.915 | -0.045 | -0.047 | -0.047 | 0.042 | 0.4 | 0.19 | 0.52 | 0.48 | 0.54 | 0.46 |
| 1 | 88035848 | rs7551430 | 0.712 | 0.384 | -1.564 | 1.579 | 0.005 | -0.039 | -0.039 | 0 | 0.82 | 0.9 | 0.63 | 0.49 | 0.83 | 0.62 |
| 1 | 88035857 | rs4655852 | -0.362 | 0.229 | -1.61 | 0.972 | 0.008 | 0.002 | 0.002 | 0 | 0.55 | 0.46 | 0.61 | 0.6 | 0.43 | 0.66 |
| 1 | 88036573 | rs12410191 | -1.017 | 0.16 | -2.072 | 1.595 | -0.004 | -0.079 | -0.079 | 2.787 | 0.2 | 0.04 | 0.32 | 0.31 | 0.17 | 0.25 |
| 1 | 88039284 | rs902296 | -0.816 | 0.192 | -2.084 | 1.77 | -0.006 | -0.092 | -0.092 | 0 | 0.17 | 0.03 | 0.31 | 0.32 | 0.14 | 0.21 |
| 1 | 88041999 | rs12139533 | 0.914 | -0.313 | -1.385 | 1.523 | 0.034 | -0.025 | -0.025 | 0 | 0.6 | 0.24 | 0.46 | 0.49 | 0.69 | 0.71 |
| 1 | 88043045 | rs12732334 | 1.15 | -0.129 | -1.181 | 1.455 | -0.011 | 0.123 | 0.123 | 0 | 0.48 | 0.19 | 0.33 | 0.3 | 0.53 | 0.38 |
| 1 | 88045534 | rs6681488 | 0.138 | 0.008 | -0.707 | 1.392 | -0.019 | 0.043 | 0.043 | 0 | 0.13 | 0.04 | 0.1 | 0.05 | 0.16 | 0.05 |
| 1 | 88052301 | rs10159232 | -0.872 | -0.149 | 0.388 | 2.476 | 0.074 | 0.475 | 0.475 | 1.15 | 0.07 | 0.48 | 0.18 | 0.36 | 0.02 | 0.07 |
| 1 | 88057291 | rs9433603 | 1.027 | 0.399 | -0.52 | 1.893 | -0.042 | -0.033 | -0.033 | 0 | 0.62 | 0.15 | 0.49 | 0.4 | 0.59 | 0.63 |
| 1 | 88065739 | rs17130067 | -0.058 | 0.619 | -0.116 | 2.359 | 0.009 | 0.045 | 0.045 | 0 | 0.1 | 0.16 | 0.31 | 0.27 | 0.17 | 0.21 |
| 1 | 88071255 | rs2390324 | -0.478 | 0.117 | -0.057 | 1.699 | -0.007 | -0.007 | -0.007 | 0 | 0.18 | 0.19 | 0.18 | 0.16 | 0.17 | 0.3 |
| 1 | 88072382 | rs6699989 | -0.695 | 0.058 | -0.089 | 1.271 | -0.008 | -0.014 | -0.014 | 0 | 0.81 | 0.79 | 0.9 | 0.91 | 0.85 | 0.74 |
| 1 | 88074688 | rs7513881 | 0.295 | -0.076 | -0.704 | 1.431 | -0.012 | -0.017 | -0.017 | 0 | 0.17 | 0.15 | 0.36 | 0.55 | 0.19 | 0.27 |
| 1 | 88077288 | rs6701916 | 1.738 | -0.516 | -1.058 | 1.046 | -0.098 | 0.199 | 0.199 | 1.71 | 0.39 | 0.06 | 0.28 | 0.13 | 0.47 | 0.15 |
| 1 | 88083782 | rs923545 | 0.453 | -0.409 | -0.465 | 1.014 | -0.025 | 0.01 | 0.01 | 0 | 0.07 | 0.19 | 0.03 | 0.1 | 0.01 | 0.07 |
| 1 | 88086894 | rs4655854 | -0.486 | -0.244 | -0.767 | 1.034 | -0.014 | -0.024 | -0.024 | 0 | 0.27 | 0.26 | 0.49 | 0.57 | 0.33 | 0.49 |
| 1 | 88090813 | rs12127587 | 0.795 | -0.089 | -1.016 | 0.431 | -0.019 | 0.055 | 0.055 | 0 | 0.13 | 0.05 | 0.09 | 0.02 | 0.16 | 0.23 |
| 1 | 88100939 | rs12090659 | 0.369 | -0.133 | -0.843 | 0.787 | -0.004 | 0.007 | 0.007 | 0 | 0.14 | 0.16 | 0.36 | 0.34 | 0.17 | 0.19 |
| 1 | 88102690 | rs17425377 | 0.786 | -0.568 | -0.818 | 0.398 | 0.005 | -0.007 | -0.007 | 0 | 0.1 | 0.06 | 0.01 | 0 | 0.03 | 0.09 |
| 1 | 88115934 | rs17130094 | 0.458 | -0.275 | -0.638 | 0.866 | 0 | 0.029 | 0.029 | 0 | 0.06 | 0.05 | 0.05 | 0.33 | 0.02 | 0.12 |
| 1 | 88117638 | rs1490168 | 0.916 | -0.016 | -0.434 | 0.786 | 0.001 | 0 | 0 | 0 | 0.58 | 0.42 | 0.38 | 0.55 | 0.52 | 0.42 |
| 1 | 88120252 | rs1352652 | -0.25 | 0.009 | -0.829 | 0.775 | -0.017 | -0.019 | -0.019 | 1.299 | 0.29 | 0.25 | 0.45 | 0.34 | 0.34 | 0.44 |
| 1 | 88126536 | rs1618898 | 0.45 | 0.007 | -1.131 | 0.106 | 0.019 | 0.037 | 0.037 | 0 | 0.41 | 0.26 | 0.26 | 0.22 | 0.33 | 0.28 |
| 1 | 88129038 | rs12132154 | 1.176 | -0.296 | -2.026 | -0.354 | -0.08 | 0.138 | 0.138 | 0 | 0.27 | 0.02 | 0.33 | 0.05 | 0.28 | 0.23 |
| 1 | 88129160 | rs1813398 | -0.853 | -0.514 | -1.543 | 0.318 | 0.011 | 0.021 | 0.021 | 0 | 0.19 | 0.08 | 0.11 | 0.14 | 0.15 | 0.14 |
| 1 | 88132380 | rs983332 | -1.534 | -0.788 | -1.17 | 0.614 | -0.003 | 0.001 | 0.001 | 0 | 0.28 | 0.32 | 0.17 | 0.32 | 0.19 | 0.18 |
| 1 | 88148076 | rs17426089 | -0.051 | -0.546 | -1.318 | 0.507 | 0.003 | 0.01 | 0.01 | 0 | 0.13 | 0.09 | 0.09 | 0 | 0.11 | 0.1 |
| 1 | 88149030 | rs10922746 | -1.235 | -0.621 | -1.191 | 0.88 | -0.013 | 0.059 | 0.059 | 0 | 0.12 | 0.2 | 0.07 | 0.32 | 0.07 | 0.08 |
| 1 | 88155673 | rs12136752 | -1.09 | -0.685 | -1.056 | 0.701 | 0.009 | -0.007 | -0.007 | 0 | 0.1 | 0.05 | 0.03 | 0.14 | 0.04 | 0.05 |

| | | | | | | | | | | | | | | | | |
|---|---|---|---|---|---|---|---|---|---|---|---|---|---|---|---|---|
| 1 | 88173396 | rs10518396 | -0.959 | -0.808 | -1.176 | 0.7 | 0.004 | -0.057 | -0.057 | 0.029 | 0.13 | 0.07 | 0.08 | 0.26 | 0.11 | 0.09 |
| 1 | 88176059 | rs10801869 | -1.016 | -0.836 | -1.176 | 0.695 | 0.014 | -0.029 | -0.029 | 0 | 0.15 | 0.1 | 0.09 | 0.25 | 0.12 | 0.1 |
| 1 | 88176255 | rs10801870 | 0.282 | -0.835 | -1.175 | 0.695 | -0.001 | 0 | 0 | 0 | 0.06 | 0.04 | 0.01 | 0.06 | 0.03 | 0.03 |
| 1 | 88176320 | rs10801871 | 0.39 | -0.835 | -1.175 | 0.695 | 0 | -0.001 | -0.001 | 0.04 | 0.05 | 0.03 | 0.01 | 0.06 | 0.03 | 0.03 |
| 1 | 88176714 | rs10518397 | -0.983 | -0.833 | -1.175 | 0.693 | 0.014 | -0.029 | -0.029 | 0 | 0.15 | 0.1 | 0.09 | 0.25 | 0.12 | 0.1 |
| 1 | 88177437 | rs10923036 | -1.708 | -0.827 | -1.137 | 0.652 | -0.016 | 0.03 | 0.03 | 0.532 | 0.2 | 0.28 | 0.12 | 0.32 | 0.14 | 0.12 |
| 1 | 88177836 | rs10754317 | -0.239 | -0.707 | -1.369 | 0.379 | 0.017 | 0.109 | 0.109 | 0.001 | 0.77 | 0.57 | 0.63 | 0.46 | 0.78 | 0.5 |
| 1 | 88177988 | rs7541276 | 1.983 | -0.66 | -1.828 | 0.131 | -0.102 | 0.306 | 0.306 | 0.028 | 0.43 | 0.09 | 0.42 | 0.06 | 0.55 | 0.28 |
| 1 | 88179317 | rs12121005 | -0.55 | -0.621 | -1.805 | 0.238 | 0.012 | -0.013 | -0.013 | 0 | 0.09 | 0.02 | 0.02 | 0.13 | 0.04 | 0.06 |
| 1 | 88182718 | rs10493801 | 1.605 | -0.749 | -1.75 | 0.252 | -0.014 | 0.297 | 0.297 | 0.11 | 0.48 | 0.1 | 0.44 | 0.13 | 0.59 | 0.32 |
| 1 | 88185216 | rs305464 | -0.494 | -0.757 | -1.711 | 0.779 | -0.005 | -0.007 | -0.007 | 0 | 0.85 | 0.87 | 0.82 | 0.78 | 0.83 | 0.85 |
| 1 | 88190037 | rs6662386 | 1.609 | -0.528 | -1.771 | 0.697 | -0.006 | 0.23 | 0.23 | 0.047 | 0.5 | 0.1 | 0.52 | 0.22 | 0.59 | 0.39 |
| 1 | 88192300 | rs10518398 | 0.123 | -0.802 | -1.322 | 1.002 | 0.007 | -0.037 | -0.037 | 0 | 0.11 | 0.05 | 0.07 | 0.19 | 0.1 | 0.07 |
| 1 | 88195672 | rs305457 | -0.43 | -0.627 | -0.961 | 1.077 | -0.007 | 0.054 | 0.054 | 0 | 0.2 | 0.58 | 0.13 | 0.32 | 0.13 | 0.35 |
| 1 | 88202160 | rs11587349 | 1.751 | -0.508 | -1.451 | 0.68 | -0.175 | 0.307 | 0.307 | 0.276 | 0.42 | 0.04 | 0.42 | 0.07 | 0.54 | 0.32 |
| 1 | 88202768 | rs305449 | -1.533 | -0.631 | -1.414 | 0.777 | 0.035 | 0.15 | 0.15 | 0.014 | 0.45 | 0.81 | 0.46 | 0.83 | 0.35 | 0.56 |
| 1 | 88208135 | rs305445 | 0.929 | -0.627 | -1.174 | 0.847 | 0.03 | 0.222 | 0.222 | 0 | 0.12 | 0.42 | 0.08 | 0.31 | 0.09 | 0.32 |
| 1 | 88209373 | rs17467793 | 1.671 | -0.486 | -1.666 | 0.975 | -0.147 | 0.31 | 0.31 | 0.198 | 0.42 | 0.05 | 0.42 | 0.07 | 0.54 | 0.32 |
| 1 | 88212437 | rs1024314 | 0.634 | -0.497 | -1.876 | 0.813 | -0.001 | -0.013 | -0.013 | 0 | 0.08 | 0.03 | 0.15 | 0.26 | 0.09 | 0.07 |
| 1 | 88213014 | rs12139208 | 0.956 | -0.483 | -1.886 | 0.456 | 0.005 | 0.077 | 0.077 | 0.433 | 0.58 | 0.22 | 0.52 | 0.27 | 0.67 | 0.4 |
| 1 | 88213827 | rs17130136 | 0.619 | -0.368 | -1.659 | 0.657 | -0.002 | 0.001 | 0.001 | 0 | 0.16 | 0.16 | 0.11 | 0.2 | 0.13 | 0.08 |
| 1 | 88220476 | rs305431 | -0.521 | 0.006 | -1.704 | 0.109 | -0.001 | 0.003 | 0.003 | 0 | 0.87 | 0.94 | 0.88 | 0.9 | 0.9 | 0.88 |
| 1 | 88231484 | rs2436985 | 0.185 | -0.276 | -1.521 | 0.371 | -0.005 | 0.059 | 0.059 | 0 | 0.11 | 0.2 | 0.2 | 0.28 | 0.09 | 0.11 |
| 1 | 88231787 | rs2503259 | 0.259 | -0.282 | -1.657 | 0.378 | -0.005 | 0.046 | 0.046 | 0 | 0.11 | 0.16 | 0.19 | 0.28 | 0.09 | 0.11 |
| 1 | 88232116 | rs305413 | -0.328 | -0.382 | -1.769 | 0.064 | -0.01 | 0.012 | 0.012 | 0 | 0.35 | 0.63 | 0.29 | 0.51 | 0.25 | 0.45 |
| 1 | 88234437 | rs12139718 | 0.243 | 0.068 | -2.27 | -0.909 | -0.039 | 0.202 | 0.202 | 0 | 0.43 | 0.16 | 0.39 | 0.11 | 0.54 | 0.34 |
| 1 | 88236589 | rs305404 | -0.293 | 0.334 | -1.945 | -0.241 | -0.071 | 0.117 | 0.117 | 0.487 | 0.59 | 0.87 | 0.6 | 0.78 | 0.53 | 0.68 |
| 1 | 88239608 | rs17130142 | -0.977 | -0.129 | -0.991 | -0.047 | -0.016 | -0.034 | -0.034 | 0 | 0.07 | 0.14 | 0.03 | 0 | 0.04 | 0.09 |
| 1 | 88239954 | rs6663225 | -0.905 | 0.122 | -1.275 | 0.528 | -0.023 | -0.047 | -0.047 | 0 | 0.13 | 0.05 | 0.22 | 0.28 | 0.18 | 0.19 |
| 1 | 88242201 | rs2503263 | 1.864 | -0.662 | -0.329 | 0.201 | -0.012 | 0.003 | 0.003 | 1.537 | 0.67 | 0.39 | 0.69 | 0.65 | 0.74 | 0.57 |
| 1 | 88242446 | rs2503264 | 1.678 | -0.614 | -0.905 | 0.143 | 0.001 | 0 | 0 | 0.014 | 0.38 | 0.35 | 0.47 | 0.4 | 0.42 | 0.22 |
| 1 | 88251957 | rs6694037 | 0.826 | -0.582 | 0.69 | 0.152 | 0.011 | -0.035 | -0.035 | 0 | 0.88 | 0.53 | 0.93 | 0.97 | 0.93 | 0.85 |
| 1 | 88252763 | rs17130163 | -0.267 | -0.651 | 0.493 | 0.092 | 0.013 | 0.018 | 0.018 | 0.077 | 0.28 | 0.09 | 0.21 | 0.25 | 0.27 | 0.39 |
| 1 | 88257998 | rs6679195 | 0.615 | -0.666 | 0.56 | 0.076 | -0.013 | -0.032 | -0.032 | 0 | 0.65 | 0.5 | 0.74 | 0.74 | 0.68 | 0.55 |
| 1 | 88260539 | rs4655988 | 1.661 | -0.423 | 1.382 | 0.287 | 0.006 | 0.018 | 0.018 | 0.878 | 0.76 | 0.4 | 0.71 | 0.7 | 0.76 | 0.65 |
| 1 | 88265202 | rs6656291 | -1.323 | -0.45 | 1.022 | 0.259 | -0.006 | -0.008 | -0.008 | 0 | 0.1 | 0.15 | 0.08 | 0.07 | 0.07 | 0.09 |
| 1 | 88269313 | rs12061052 | -1.167 | -0.366 | 0.909 | 0.405 | -0.032 | 0.008 | 0.008 | 0 | 0.08 | 0.14 | 0.1 | 0.08 | 0.04 | 0.11 |
| 1 | 88273790 | rs2503273 | -1.118 | 0.102 | 1.025 | 0.461 | 0.02 | 0.085 | 0.085 | 0 | 0.11 | 0.24 | 0.19 | 0.21 | 0.12 | 0.21 |
| 1 | 88279171 | rs2117353 | -1.665 | 0.148 | 1.117 | 0.099 | 0.009 | -0.037 | -0.037 | 0 | 0.14 | 0.59 | 0.14 | 0.07 | 0.12 | 0.15 |
| 1 | 88281761 | rs12750772 | -0.906 | 0.511 | 0.808 | 0.38 | 0.019 | 0.031 | 0.031 | 0 | 0.06 | 0.14 | 0.1 | 0.11 | 0.08 | 0.16 |
| 1 | 88282170 | rs12755273 | -0.798 | 0.52 | 0.253 | -0.107 | -0.03 | 0.027 | 0.027 | 0.853 | 0.27 | 0.13 | 0.23 | 0.18 | 0.34 | 0.39 |
| 1 | 88287371 | rs2491281 | -1.916 | -0.042 | 0.901 | -0.191 | -0.013 | -0.043 | -0.043 | 0 | 0.17 | 0.53 | 0.14 | 0.08 | 0.13 | 0.19 |
| 1 | 88287879 | rs2491282 | -1.741 | 0.074 | 0.974 | -0.199 | 0.023 | -0.047 | -0.047 | 0 | 0.12 | 0.53 | 0.13 | 0.04 | 0.1 | 0.14 |
| 1 | 88290369 | rs4655858 | -0.446 | 0.164 | 0.511 | -0.027 | -0.008 | -0.007 | -0.007 | 1.89 | 0.06 | 0.02 | 0.14 | 0.1 | 0.09 | 0.1 |
| 1 | 88313724 | rs2491302 | 0.775 | -0.196 | 1.747 | -0.106 | -0.001 | -0.002 | -0.002 | 0 | 0.72 | 0.71 | 0.62 | 0.68 | 0.7 | 0.58 |
| 1 | 88317228 | rs2491304 | 0.715 | -0.202 | 1.755 | -0.115 | 0 | -0.002 | -0.002 | 3.957 | 0.72 | 0.73 | 0.6 | 0.7 | 0.69 | 0.58 |
| 1 | 88321638 | rs2491307 | 0.73 | -0.218 | 1.747 | -0.081 | 0 | -0.001 | -0.001 | 0 | 0.72 | 0.73 | 0.6 | 0.68 | 0.69 | 0.58 |
| 1 | 88322831 | rs2503237 | -0.489 | -0.454 | 1.834 | -0.351 | 0.028 | 0.013 | 0.013 | 0.258 | 0.09 | 0.41 | 0.06 | 0.03 | 0.04 | 0.03 |
| 1 | 88327760 | rs6677939 | 1.231 | -0.443 | 1.666 | -0.344 | 0.006 | 0.014 | 0.014 | 0.783 | 0.64 | 0.32 | 0.54 | 0.66 | 0.65 | 0.54 |
| 1 | 88343418 | rs3008440 | -0.988 | -0.329 | 1.391 | -0.372 | -0.002 | -0.002 | -0.002 | 0 | 0.31 | 0.31 | 0.37 | 0.33 | 0.32 | 0.45 |
| 1 | 88346411 | rs1413533 | -2.043 | -0.487 | 0.236 | -0.385 | -0.142 | -0.119 | -0.119 | 0 | 0.59 | 0.97 | 0.55 | 0.45 | 0.53 | 0.74 |
| 1 | 88358924 | rs2503248 | -0.822 | -0.433 | 1.19 | -0.368 | 0.012 | 0.012 | 0.012 | 0.27 | 0.1 | 0.34 | 0.07 | 0.05 | 0.06 | 0.05 |
| 1 | 88363315 | rs2503254 | -0.797 | -0.484 | 1.385 | -0.394 | -0.009 | -0.008 | -0.008 | 2.566 | 0.1 | 0.24 | 0.06 | 0.05 | 0.06 | 0.06 |
| 1 | 88365049 | rs10922747 | 1.636 | -0.262 | 0.408 | -0.661 | -0.065 | -0.027 | -0.027 | 0 | 0.41 | 0.06 | 0.44 | 0.61 | 0.43 | 0.34 |
| 1 | 88371676 | rs945058 | -1.878 | 0.141 | 0.902 | -0.358 | -0.056 | 0.031 | 0.031 | 0 | 0.43 | 0.85 | 0.44 | 0.36 | 0.43 | 0.55 |
| 1 | 88372202 | rs17130207 | -1.175 | 0.389 | 1.641 | -0.204 | 0.055 | 0.067 | 0.067 | 1.419 | 0.05 | 0.26 | 0.03 | 0.05 | 0.04 | 0.04 |
| 1 | 88373874 | rs3008463 | -1.828 | 0.375 | 1.505 | -0.214 | 0.09 | 0.098 | 0.098 | 1.172 | 0.06 | 0.44 | 0.04 | 0.05 | 0.04 | 0.04 |
| 1 | 88380467 | rs1334334 | -1.146 | 0.463 | 1.061 | -0.256 | -0.007 | 0.003 | 0.003 | 0 | 0.21 | 0.14 | 0.29 | 0.22 | 0.23 | 0.26 |
| 1 | 88382412 | rs12141842 | -1.868 | 0.482 | 1.142 | -0.226 | -0.003 | 0.004 | 0.004 | 0 | 0.35 | 0.3 | 0.39 | 0.32 | 0.37 | 0.49 |
| 1 | 88392864 | rs10489944 | -1.294 | 0.349 | 1.455 | -0.018 | 0.005 | 0.004 | 0.004 | 0 | 0.12 | 0.15 | 0.34 | 0.21 | 0.15 | 0.12 |
| 1 | 88397483 | rs2503257 | -2.846 | 0.092 | 1.081 | -0.063 | -0.001 | 0 | 0 | 0.378 | 0.25 | 0.71 | 0.41 | 0.3 | 0.2 | 0.26 |
| 1 | 88399971 | rs17130221 | -0.159 | 0.425 | 0.256 | 0.146 | 0.03 | -0.032 | -0.032 | 0 | 0.06 | 0 | 0.01 | 0.09 | 0.02 | 0.11 |
| 1 | 88402963 | rs12741507 | -1.638 | 0.487 | 0.361 | 0.095 | -0.002 | 0.016 | 0.016 | 0 | 0.19 | 0.22 | 0.36 | 0.3 | 0.17 | 0.23 |
| 1 | 88410277 | rs1334339 | -1.441 | 0.64 | -0.342 | -0.261 | -0.009 | 0.044 | 0.044 | 0 | 0.39 | 0.74 | 0.33 | 0.27 | 0.41 | 0.51 |
| 1 | 88416590 | rs6691335 | -1.253 | 0.787 | 0.695 | -0.011 | 0.017 | 0.035 | 0.035 | 0 | 0.11 | 0.36 | 0.08 | 0.14 | 0.1 | 0.17 |
| 1 | 88423966 | rs2991716 | -1.056 | 0.899 | 0.37 | -0.193 | -0.004 | 0.006 | 0.006 | 0 | 0.47 | 0.39 | 0.46 | 0.41 | 0.52 | 0.6 |
| 1 | 88427851 | rs3008453 | -1.368 | 0.689 | 0.183 | -0.203 | -0.041 | 0 | 0 | 0.898 | 0.51 | 0.89 | 0.51 | 0.41 | 0.51 | 0.57 |
| 1 | 88428199 | rs2151842 | 0.017 | 0.87 | 0.115 | -0.262 | 0 | -0.002 | -0.002 | 0.781 | 0.18 | 0.17 | 0.16 | 0.12 | 0.27 | 0.26 |
| 1 | 88430037 | rs1413527 | -1.465 | 0.799 | 0.195 | -0.282 | 0 | 0.016 | 0.016 | 0.144 | 0.49 | 0.67 | 0.49 | 0.4 | 0.51 | 0.59 |

| | | | | | | | | | | | | | | | | |
|---|---|---|---|---|---|---|---|---|---|---|---|---|---|---|---|---|
| 1 | 88441066 | rs1334331 | -1.005 | 0.605 | -0.52 | -0.588 | 0.001 | 0.055 | 0.055 | 0 | 0.37 | 0.72 | 0.31 | 0.2 | 0.38 | 0.45 |
| 1 | 88445213 | rs1413529 | -1.147 | 0.545 | -0.515 | -0.799 | 0.003 | 0.054 | 0.054 | 0 | 0.39 | 0.71 | 0.32 | 0.17 | 0.39 | 0.48 |
| 1 | 88446605 | rs3008445 | -1.657 | 0.428 | -0.854 | -0.701 | -0.116 | 0.009 | 0.009 | 0.023 | 0.6 | 0.97 | 0.55 | 0.31 | 0.56 | 0.62 |
| 1 | 88447326 | rs17130231 | -0.602 | 0.14 | 0 | -0.443 | -0.008 | -0.025 | -0.025 | 0 | 0.09 | 0.01 | 0.02 | 0.1 | 0.07 | 0.15 |
| 1 | 88448150 | rs4655860 | -0.915 | 0.396 | 0.639 | -0.379 | -0.013 | -0.122 | -0.122 | 0 | 0.27 | 0.53 | 0.25 | 0.08 | 0.25 | 0.3 |
| 1 | 88453303 | rs3008465 | -0.948 | 0.519 | -0.299 | -0.293 | -0.008 | -0.121 | -0.121 | 0 | 0.28 | 0.49 | 0.26 | 0.09 | 0.26 | 0.31 |
| 1 | 88457516 | rs10489947 | -0.491 | 0.257 | -0.827 | 0.748 | 0 | 0.01 | 0.01 | 0 | 0.19 | 0.23 | 0.2 | 0.44 | 0.17 | 0.21 |
| 1 | 88457588 | rs2991701 | -1.291 | -0.523 | -1.078 | 0.145 | 0.052 | -0.052 | -0.052 | 0 | 0.37 | 0.76 | 0.31 | 0.48 | 0.29 | 0.35 |
| 1 | 88463421 | rs10922944 | 0.446 | 0.058 | -2.011 | -0.106 | -0.044 | -0.022 | -0.022 | 0 | 0.3 | 0.1 | 0.29 | 0.32 | 0.3 | 0.29 |
| 1 | 88465247 | rs1537560 | 0.19 | 0.093 | -1.89 | 0.066 | -0.001 | 0.013 | 0.013 | 0 | 0.2 | 0.25 | 0.23 | 0.53 | 0.2 | 0.22 |
| 1 | 88467667 | rs11801253 | -0.401 | -0.344 | -1.834 | -0.244 | -0.019 | 0.094 | 0.094 | 0 | 0.44 | 0.31 | 0.5 | 0.79 | 0.45 | 0.42 |
| 1 | 88478364 | rs2390389 | -0.299 | -0.376 | -1.673 | -0.363 | -0.042 | 0.008 | 0.008 | 0 | 0.3 | 0.14 | 0.31 | 0.24 | 0.32 | 0.28 |
| 1 | 88481128 | rs7525663 | -0.921 | -0.481 | -0.625 | -0.507 | 0.001 | 0.019 | 0.019 | 0 | 0.39 | 0.39 | 0.48 | 0.76 | 0.36 | 0.45 |
| 1 | 88481774 | rs10922987 | 1.767 | -0.713 | -0.757 | -1.122 | -0.052 | 0.077 | 0.077 | 0 | 0.32 | 0.13 | 0.36 | 0.19 | 0.4 | 0.27 |
| 1 | 88482229 | rs12759463 | -0.625 | -0.753 | -0.796 | -1.15 | -0.001 | 0.032 | 0.032 | 0 | 0.42 | 0.38 | 0.49 | 0.77 | 0.39 | 0.47 |
| 1 | 88482648 | rs12031344 | 1.829 | -0.739 | -0.824 | -1.111 | -0.098 | 0.111 | 0.111 | 0 | 0.32 | 0.06 | 0.35 | 0.14 | 0.38 | 0.25 |
| 1 | 88484600 | rs10922989 | 0.382 | -0.817 | -0.046 | -1.033 | -0.002 | 0.008 | 0.008 | 0 | 0.31 | 0.24 | 0.31 | 0.26 | 0.31 | 0.28 |
| 1 | 88494859 | rs6413942 | -0.38 | -0.638 | 0.122 | -0.996 | 0.001 | 0.008 | 0.008 | 0 | 0.85 | 0.89 | 0.91 | 0.97 | 0.86 | 0.81 |
| 1 | 88504370 | rs12143926 | 1.929 | -0.759 | 0.001 | -1.032 | -0.089 | 0.183 | 0.183 | 0.567 | 0.33 | 0.03 | 0.22 | 0.09 | 0.37 | 0.24 |
| 1 | 88508392 | rs7527342 | -0.014 | -0.764 | 0.054 | -0.756 | 0.002 | 0.012 | 0.012 | 0 | 0.83 | 0.88 | 0.86 | 0.92 | 0.83 | 0.8 |
| 1 | 88514365 | rs4655999 | 1.957 | -0.773 | -0.092 | -1.039 | -0.095 | 0.178 | 0.178 | 0.505 | 0.33 | 0.03 | 0.22 | 0.09 | 0.36 | 0.19 |
| 1 | 88516085 | rs12410457 | -0.133 | -0.836 | 0.494 | -0.942 | -0.011 | 0.017 | 0.017 | 0 | 0.38 | 0.26 | 0.45 | 0.33 | 0.4 | 0.43 |
| 1 | 88516267 | rs10923016 | -0.314 | -0.83 | 0.531 | -0.941 | -0.007 | 0.014 | 0.014 | 0 | 0.39 | 0.26 | 0.45 | 0.36 | 0.4 | 0.43 |
| 1 | 88521688 | rs6658841 | 0.239 | -0.8 | 0.7 | -1 | -0.006 | -0.077 | -0.077 | 5.663 | 0.82 | 0.55 | 0.84 | 0.92 | 0.84 | 0.75 |
| 1 | 88522282 | rs4656007 | -0.837 | -0.63 | 0.664 | -1.023 | 0 | 0.07 | 0.07 | 0 | 0.49 | 0.41 | 0.63 | 0.83 | 0.49 | 0.56 |
| 1 | 88524540 | rs4656008 | 1.892 | -0.511 | 0.491 | -1.01 | -0.023 | 0.144 | 0.144 | 0.325 | 0.33 | 0.12 | 0.22 | 0.09 | 0.34 | 0.18 |
| 1 | 88539988 | rs7522610 | 0.511 | -0.858 | 1.78 | -0.158 | 0.029 | -0.008 | -0.008 | 0 | 0.86 | 0.66 | 0.89 | 0.88 | 0.85 | 0.77 |
| 1 | 88545471 | rs4307538 | 0.081 | -0.855 | 1.441 | -0.192 | -0.026 | 0.007 | 0.007 | 0 | 0.05 | 0 | 0.04 | 0.02 | 0.07 | 0.16 |
| 1 | 88546460 | rs4656010 | -0.411 | -0.732 | 1.683 | 0.092 | -0.001 | 0.196 | 0.196 | 0 | 0.08 | 0.19 | 0.18 | 0.44 | 0.07 | 0.12 |
| 1 | 88547243 | rs12028716 | 1.687 | -0.774 | 1.795 | 0.055 | 0.016 | -0.026 | -0.026 | 0 | 0.71 | 0.26 | 0.67 | 0.41 | 0.76 | 0.62 |
| 1 | 88548801 | rs10737721 | -0.177 | -0.719 | 1.739 | -0.041 | -0.002 | -0.038 | -0.038 | 0.875 | 0.84 | 0.8 | 0.88 | 0.98 | 0.85 | 0.76 |
| 1 | 88555928 | rs4278350 | 0.206 | -0.721 | 1.737 | -0.046 | 0 | -0.048 | -0.048 | 0.231 | 0.82 | 0.78 | 0.88 | 0.98 | 0.81 | 0.75 |
| 1 | 88562823 | rs7516752 | 0.442 | -0.848 | 1.438 | -0.002 | -0.016 | -0.018 | -0.018 | 0.418 | 0.81 | 0.4 | 0.85 | 0.85 | 0.84 | 0.76 |
| 1 | 88578657 | rs12142415 | -1.347 | -0.492 | 0.527 | 0.762 | 0.009 | -0.043 | -0.043 | 0 | 0.25 | 0.18 | 0.26 | 0.47 | 0.22 | 0.4 |
| 1 | 88584102 | rs10801864 | 1.15 | -0.294 | 0.098 | 0.778 | 0.017 | 0.006 | 0.006 | 0 | 0.68 | 0.27 | 0.61 | 0.31 | 0.74 | 0.6 |
| 1 | 88593050 | rs6428632 | -0.384 | 0.029 | -0.257 | 0.858 | 0.013 | 0.033 | 0.033 | 0 | 0.36 | 0.23 | 0.26 | 0.23 | 0.35 | 0.43 |
| 1 | 88603134 | rs1932760 | -0.223 | 0.045 | -0.384 | 0.873 | -0.009 | 0.142 | 0.142 | 0 | 0.09 | 0.11 | 0.12 | 0.44 | 0.07 | 0.15 |
| 1 | 88604157 | rs4656013 | 0.484 | 0.24 | -0.456 | 0.159 | -0.003 | -0.004 | -0.004 | 0 | 0.55 | 0.48 | 0.61 | 0.38 | 0.6 | 0.44 |
| 1 | 88607247 | rs1932761 | 0.472 | -0.066 | -0.07 | -0.434 | -0.024 | -0.158 | -0.158 | 0 | 0.77 | 0.55 | 0.89 | 1 | 0.81 | 0.82 |
| 1 | 88610289 | rs1336578 | 1.071 | 0.351 | -0.366 | -0.07 | -0.001 | 0.012 | 0.012 | 0 | 0.62 | 0.37 | 0.69 | 0.69 | 0.64 | 0.58 |
| 1 | 88615308 | rs4347185 | 0.523 | 0.005 | -0.801 | -0.422 | 0.002 | -0.007 | -0.007 | 0.011 | 0.24 | 0.21 | 0.31 | 0.61 | 0.22 | 0.29 |
| 1 | 88617883 | rs10923030 | 0.311 | 0.141 | -1.009 | -0.49 | -0.01 | -0.011 | -0.011 | 0.02 | 0.17 | 0.16 | 0.18 | 0.29 | 0.25 | 0.22 |
| 1 | 88620172 | rs2390437 | -1.397 | 0.08 | -1.4 | -0.577 | 0.001 | 0.032 | 0.032 | 0 | 0.93 | 0.98 | 0.98 | 1 | 0.96 | 0.89 |
| 1 | 88621347 | rs2390439 | 1.418 | 0.141 | -1.245 | -0.406 | -0.05 | 0.178 | 0.178 | 0 | 0.37 | 0.11 | 0.35 | 0.09 | 0.42 | 0.29 |
| 1 | 88623666 | rs12086697 | 0.093 | 0.166 | -0.565 | -0.021 | -0.005 | -0.068 | -0.068 | 0 | 0.13 | 0.21 | 0.06 | 0 | 0.09 | 0.1 |
| 1 | 88623995 | rs10493808 | 0.081 | 0.33 | -0.882 | 0.49 | -0.083 | -0.088 | -0.088 | 3.899 | 0.15 | 0.01 | 0.23 | 0.3 | 0.24 | 0.22 |
| 1 | 88632421 | rs10923033 | 0.742 | 0.05 | -1.247 | -0.008 | -0.095 | 0.228 | 0.228 | 0.419 | 0.34 | 0.05 | 0.36 | 0.01 | 0.36 | 0.26 |
| 1 | 88633737 | rs2390441 | -0.336 | -0.157 | -0.585 | -0.112 | 0.001 | 0.034 | 0.034 | 0 | 0.57 | 0.38 | 0.57 | 0.82 | 0.59 | 0.6 |
| 1 | 88638931 | rs6694280 | 0.34 | -0.19 | -0.341 | 0.359 | -0.002 | 0.016 | 0.016 | 0 | 0.41 | 0.34 | 0.38 | 0.28 | 0.4 | 0.4 |
| 1 | 88641063 | rs6428637 | -0.213 | 0.137 | -0.892 | 0.345 | 0.04 | 0.013 | 0.013 | 1.337 | 0.53 | 0.74 | 0.35 | 0.59 | 0.41 | 0.53 |
| 1 | 88642627 | rs12731063 | 0.77 | 0.299 | -1.035 | 0.324 | -0.129 | 0.094 | 0.094 | 0.627 | 0.36 | 0.05 | 0.42 | 0.12 | 0.41 | 0.27 |
| 1 | 88643619 | rs4233331 | -0.589 | 0.401 | -0.901 | 0.47 | 0.006 | 0.019 | 0.019 | 0 | 0.19 | 0.3 | 0.25 | 0.35 | 0.26 | 0.31 |
| 1 | 88643972 | rs4656019 | -0.684 | 0.266 | -0.887 | 0.473 | 0.006 | 0.008 | 0.008 | 0 | 0.21 | 0.3 | 0.27 | 0.35 | 0.32 | 0.33 |
| 1 | 88645313 | rs12567486 | -0.095 | -0.646 | -0.156 | 0.73 | -0.011 | 0.007 | 0.007 | 0 | 0.41 | 0.43 | 0.31 | 0.52 | 0.32 | 0.41 |
| 1 | 88647092 | rs10923035 | -1.206 | -0.146 | -0.047 | 0.905 | -0.012 | -0.002 | -0.002 | 0.418 | 0.09 | 0.29 | 0.05 | 0.06 | 0.08 | 0.11 |
| 1 | 88657007 | rs12755168 | 0.775 | 0.013 | -0.889 | 0.888 | 0.014 | 0.068 | 0.068 | 0.295 | 0.54 | 0.16 | 0.46 | 0.36 | 0.57 | 0.54 |
| 1 | 88661096 | rs6699416 | -0.165 | -0.109 | -0.886 | 1.43 | 0.012 | -0.08 | -0.08 | 0 | 0.22 | 0.08 | 0.18 | 0.39 | 0.16 | 0.18 |
| 1 | 88663791 | rs10801875 | 0.863 | -0.512 | 0.539 | 1.377 | 0.023 | 0.039 | 0.039 | 0 | 0.78 | 0.52 | 0.69 | 0.74 | 0.73 | 0.72 |
| 1 | 88666005 | rs6698308 | -1.36 | -0.856 | 1.053 | 1.301 | 0.007 | 0.015 | 0.015 | 0.046 | 0.12 | 0.36 | 0.04 | 0.11 | 0.1 | 0.14 |
| 1 | 88668577 | rs1336573 | 1.923 | -1.055 | 0.753 | 1.295 | 0.051 | 0.118 | 0.118 | 0 | 0.51 | 0.16 | 0.48 | 0.3 | 0.55 | 0.51 |
| 1 | 88674545 | rs12403601 | -0.606 | -1.048 | 1.433 | 1.34 | 0.009 | 0.01 | 0.01 | 0 | 0.11 | 0.18 | 0.26 | 0.16 | 0.17 | 0.09 |
| 1 | 88681580 | rs10923044 | -1.214 | -1.067 | 1.67 | 1.364 | 0.015 | 0.018 | 0.018 | 0 | 0.13 | 0.28 | 0.29 | 0.16 | 0.18 | 0.09 |
| 1 | 88684970 | rs10801877 | -0.213 | -1.115 | 1.748 | 1.367 | 0.035 | -0.061 | -0.061 | 3.081 | 0.24 | 0.12 | 0.18 | 0.42 | 0.16 | 0.21 |
| 1 | 88685730 | rs1932758 | -0.498 | -0.956 | 1.86 | 1.327 | 0.077 | 0.094 | 0.094 | 0.016 | 0.07 | 0.33 | 0.03 | 0.11 | 0.07 | 0.13 |
| 1 | 88692175 | rs10493805 | -0.463 | -0.904 | 1.999 | 1.345 | 0.012 | 0.013 | 0.013 | 0 | 0.1 | 0.16 | 0.26 | 0.16 | 0.17 | 0.1 |
| 1 | 88697663 | rs6690435 | -0.483 | -0.923 | 2.07 | 1.38 | 0.08 | 0.097 | 0.097 | 0.306 | 0.07 | 0.32 | 0.03 | 0.11 | 0.08 | 0.14 |
| 1 | 88709451 | rs6428641 | 1.83 | -1.001 | 1.733 | 1.345 | 0.003 | 0.009 | 0.009 | 0 | 0.79 | 0.34 | 0.69 | 0.73 | 0.74 | 0.76 |
| 1 | 88710048 | rs6428642 | 1.728 | -1.092 | 1.644 | 1.619 | 0.016 | 0.005 | 0.005 | 0 | 0.59 | 0.32 | 0.53 | 0.37 | 0.6 | 0.57 |
| 1 | 88718155 | rs514070 | -1.192 | -1.098 | 1.817 | 1.588 | -0.038 | 0.053 | 0.053 | 0 | 0.3 | 0.45 | 0.21 | 0.46 | 0.22 | 0.32 |

| | | | | | | | | | | | | | | | | |
|---|---|---|---|---|---|---|---|---|---|---|---|---|---|---|---|---|
| 1 | 88732523 | rs527607 | -0.108 | -1.083 | 1.719 | 1.626 | 0.013 | -0.127 | -0.127 | 0.108 | 0.21 | 0.02 | 0.16 | 0.35 | 0.14 | 0.19 |
| 1 | 88736621 | rs525002 | 1.259 | -1.077 | 1.828 | 1.642 | 0.005 | -0.013 | -0.013 | 0.154 | 0.65 | 0.69 | 0.56 | 0.5 | 0.64 | 0.65 |
| 1 | 88738404 | rs472473 | -1.865 | -0.514 | 2.517 | 2.109 | 0.033 | 0.044 | 0.044 | 0.083 | 0.1 | 0.49 | 0.06 | 0.1 | 0.07 | 0.13 |
| 1 | 88739923 | rs17436482 | -1.858 | -0.467 | 2.387 | 2.404 | 0.008 | 0.022 | 0.022 | 0 | 0.13 | 0.03 | 0.11 | 0.06 | 0.1 | 0.12 |
| 1 | 88740566 | rs534914 | 0.966 | -0.396 | 2.363 | 2.4 | -0.015 | -0.005 | -0.005 | 0.092 | 0.88 | 0.93 | 0.75 | 0.89 | 0.82 | 0.86 |
| 1 | 88740865 | rs1295491 | -1.762 | -0.317 | 2.596 | 2.599 | 0.043 | 0.366 | 0.366 | 0 | 0.1 | 0.47 | 0.05 | 0.37 | 0.07 | 0.16 |
| 1 | 88757128 | rs6679982 | -1.738 | -0.325 | 2.596 | 2.488 | 0.045 | 0.061 | 0.061 | 0 | 0.09 | 0.39 | 0.05 | 0.12 | 0.07 | 0.13 |
| 1 | 88767199 | rs566685 | 2.502 | -0.314 | 2.324 | 2.479 | 0 | -0.015 | -0.015 | 0 | 0.7 | 0.3 | 0.58 | 0.46 | 0.68 | 0.62 |
| 1 | 88776307 | rs54397 | 2.54 | -0.335 | 2.319 | 2.431 | -0.009 | -0.012 | -0.012 | 0 | 0.72 | 0.38 | 0.67 | 0.77 | 0.7 | 0.71 |
| 1 | 88791432 | rs480054 | 2.159 | -0.125 | 1.985 | 2.552 | -0.003 | -0.012 | -0.012 | 0 | 0.75 | 0.78 | 0.63 | 0.52 | 0.71 | 0.62 |
| 1 | 88802012 | rs2179965 | -1.085 | -0.147 | 2.25 | 2.547 | 0.027 | 0.067 | 0.067 | 0 | 0.08 | 0.15 | 0.25 | 0.22 | 0.16 | 0.1 |
| 1 | 88802939 | rs4655872 | -0.552 | -0.151 | 2.291 | 2.562 | 0.001 | 0.09 | 0.09 | 0 | 0.07 | 0.07 | 0.08 | 0.31 | 0.04 | 0.09 |
| 1 | 88802972 | rs4656031 | -0.574 | -0.151 | 2.291 | 2.562 | -0.001 | 0.034 | 0.034 | 0 | 0.07 | 0.04 | 0.08 | 0.31 | 0.04 | 0.09 |
| 1 | 88803250 | rs12749687 | -1.101 | -0.088 | 2.316 | 2.581 | 0.027 | 0.067 | 0.067 | 0 | 0.08 | 0.16 | 0.25 | 0.22 | 0.16 | 0.09 |
| 1 | 88815650 | rs4656036 | 1.526 | -0.108 | 2.236 | 2.534 | 0.017 | 0.041 | 0.041 | 0 | 0.85 | 0.93 | 0.95 | 0.97 | 0.86 | 0.8 |
| 1 | 88821516 | rs766353 | 2.692 | -0.109 | 2.435 | 2.546 | 0.001 | 0.024 | 0.024 | 0 | 0.7 | 0.64 | 0.61 | 0.44 | 0.67 | 0.62 |
| 1 | 88823272 | rs926795 | 2.178 | -0.127 | 1.835 | 2.476 | 0.007 | -0.02 | -0.02 | 0 | 0.79 | 0.86 | 0.86 | 0.66 | 0.83 | 0.71 |
| 1 | 88825604 | rs2808112 | 0.902 | -0.093 | 1.782 | 1.944 | 0 | -0.009 | -0.009 | 0 | 0.89 | 0.9 | 0.95 | 0.97 | 0.89 | 0.81 |
| 1 | 88843632 | rs12141716 | 2.235 | -0.156 | 2.168 | 2.171 | 0.015 | 0.074 | 0.074 | 0 | 0.73 | 0.51 | 0.59 | 0.43 | 0.68 | 0.62 |
| 1 | 88844311 | rs2808111 | -1.867 | -0.052 | 2.061 | 2.26 | 0.066 | 0.254 | 0.254 | 0 | 0.14 | 0.37 | 0.36 | 0.53 | 0.18 | 0.18 |
| 1 | 88855885 | rs400666 | -1.877 | -0.035 | 1.87 | 2.256 | 0.038 | 0.196 | 0.196 | 0 | 0.14 | 0.29 | 0.37 | 0.47 | 0.18 | 0.17 |
| 1 | 88856081 | rs17130443 | -1.196 | -0.003 | 1.769 | 2.271 | -0.026 | -0.013 | -0.013 | 0 | 0.11 | 0.05 | 0.36 | 0.47 | 0.18 | 0.17 |
| 1 | 88860530 | rs17130444 | -1.168 | -0.096 | 2.105 | 1.996 | 0.008 | -0.045 | -0.045 | 0 | 0.12 | 0.44 | 0.05 | 0.04 | 0.1 | 0.14 |
| 1 | 88864082 | rs4656043 | 1.652 | -0.138 | 1.252 | 2.256 | 0.009 | -0.019 | -0.019 | 0 | 0.65 | 0.24 | 0.53 | 0.43 | 0.64 | 0.62 |
| 1 | 88868236 | rs1498914 | 2.127 | 1.148 | 1.864 | 3.819 | 0.148 | 0.423 | 0.423 | 0 | 0.93 | 0.75 | 0.88 | 0.54 | 0.86 | 0.86 |
| 1 | 88889925 | rs6696560 | 1.956 | 1.145 | 1.786 | 3.861 | 0.126 | 0.288 | 0.288 | 0 | 0.94 | 0.77 | 0.89 | 0.68 | 0.88 | 0.89 |
| 1 | 88906024 | rs10801891 | -2.242 | 1.806 | 1.883 | 3.957 | 0.084 | 0.035 | 0.035 | 0.066 | 0.12 | 0.2 | 0.35 | 0.26 | 0.25 | 0.18 |
| 1 | 88919469 | rs12723729 | -2.242 | 1.806 | 1.883 | 3.957 | 0.084 | 0.035 | 0.035 | 0.066 | 0.12 | 0.21 | 0.35 | 0.26 | 0.25 | 0.18 |
| 1 | 88941573 | rs1353796 | 2.085 | 1.739 | 1.882 | 3.341 | 0.123 | 0.06 | 0.06 | 0.347 | 0.87 | 0.68 | 0.64 | 0.74 | 0.73 | 0.82 |
| 1 | 88950775 | rs10923081 | -2.459 | 2.092 | 1.867 | 3.382 | 0.129 | 0.143 | 0.143 | 0 | 0.13 | 0.25 | 0.42 | 0.34 | 0.31 | 0.16 |
| 1 | 88961409 | rs306322 | -2.73 | 2.024 | 1.797 | 3.298 | 0.113 | 0.163 | 0.163 | 0 | 0.16 | 0.3 | 0.46 | 0.36 | 0.3 | 0.24 |
| 1 | 88966843 | rs306333 | -1.654 | 1.726 | 1.39 | 2.818 | 0.058 | 0.047 | 0.047 | 0.044 | 0.26 | 0.38 | 0.55 | 0.36 | 0.43 | 0.28 |
| 1 | 88975378 | rs6696199 | -1.553 | 1.694 | 1.145 | 2.831 | -0.03 | 0.016 | 0.016 | 0.112 | 0.25 | 0.18 | 0.49 | 0.15 | 0.41 | 0.26 |
| 1 | 88986066 | rs6674065 | 1.441 | 1.608 | 1.546 | 2.921 | 0.076 | 0.071 | 0.071 | 0.11 | 0.73 | 0.5 | 0.44 | 0.63 | 0.56 | 0.73 |
| 1 | 89000384 | rs10923094 | -0.629 | 1.571 | 0.991 | 3.499 | -0.058 | -0.114 | -0.114 | 0 | 0.17 | 0.06 | 0.38 | 0.52 | 0.28 | 0.15 |
| 1 | 89005795 | rs306321 | -0.645 | 1.548 | 0.024 | 1.794 | -0.004 | -0.011 | -0.011 | 0.215 | 0.31 | 0.34 | 0.2 | 0.2 | 0.29 | 0.35 |
| 1 | 89009265 | rs10923099 | -0.559 | 1.608 | -0.05 | 1.775 | -0.024 | -0.014 | -0.014 | 0 | 0.06 | 0.02 | 0.11 | 0.07 | 0.12 | 0.11 |
| 1 | 89012137 | rs6685860 | 0.294 | 1.285 | -0.033 | 1.706 | 0.017 | 0.031 | 0.031 | 0 | 0.22 | 0.12 | 0.07 | 0.07 | 0.16 | 0.17 |
| 1 | 89014943 | rs17473112 | 0.898 | 1.242 | -0.026 | 1.656 | 0.01 | 0.035 | 0.035 | 0.99 | 0.17 | 0.08 | 0.05 | 0.02 | 0.14 | 0.15 |
| 1 | 89016499 | rs10493815 | 0.59 | 1.429 | -0.113 | 1.903 | -0.064 | -0.05 | -0.05 | 0.034 | 0.14 | 0.01 | 0.33 | 0.16 | 0.25 | 0.12 |
| 1 | 89020534 | rs306298 | -1.017 | 1.079 | 0.105 | 1.478 | -0.006 | -0.014 | -0.014 | 0.189 | 0.3 | 0.36 | 0.19 | 0.2 | 0.27 | 0.36 |
| 1 | 89032203 | rs17433417 | 0.029 | 1.735 | 0.101 | 1.54 | 0 | 0 | 0 | 0 | 0.07 | 0.09 | 0.04 | 0.07 | 0.05 | 0.08 |
| 1 | 89033810 | rs17130519 | -1.15 | 1.716 | 0.095 | 1.458 | -0.016 | -0.018 | -0.018 | 0 | 0.09 | 0.03 | 0.11 | 0.11 | 0.12 | 0.2 |
| 1 | 89042433 | rs595339 | -1.362 | 1.761 | -0.071 | 1.163 | -0.011 | -0.032 | -0.032 | 0 | 0.38 | 0.21 | 0.59 | 0.5 | 0.52 | 0.41 |
| 1 | 89045338 | rs658980 | -0.889 | 1.748 | 0.333 | 1.077 | 0.007 | 0.021 | 0.021 | 0 | 0.31 | 0.34 | 0.47 | 0.48 | 0.39 | 0.25 |
| 1 | 89069023 | rs1760206 | 0.202 | 1.641 | 0.32 | 0.887 | -0.032 | 0.023 | 0.023 | 0 | 0.84 | 0.87 | 0.79 | 0.93 | 0.75 | 0.76 |
| 1 | 89071203 | rs1532158 | 1.615 | 1.621 | 0.311 | 0.887 | 0.001 | 0 | 0 | 0.906 | 0.53 | 0.54 | 0.32 | 0.46 | 0.37 | 0.51 |
| 1 | 89072924 | rs12024611 | -0.157 | 1.609 | 0.293 | 0.888 | -0.027 | -0.043 | -0.043 | 0 | 0.27 | 0.14 | 0.45 | 0.41 | 0.36 | 0.19 |
| 1 | 89090212 | rs10923117 | 1.888 | 1.545 | 0.362 | 0.854 | -0.005 | -0.003 | -0.003 | 0.12 | 0.49 | 0.49 | 0.3 | 0.37 | 0.35 | 0.5 |
| 1 | 89102071 | rs7520755 | -1.002 | 1.567 | 0.852 | 1.304 | -0.007 | -0.028 | -0.028 | 0 | 0.31 | 0.23 | 0.47 | 0.5 | 0.37 | 0.2 |
| 1 | 89107857 | rs12041174 | -0.979 | 1.565 | 0.88 | 1.305 | -0.014 | -0.049 | -0.049 | 0.035 | 0.31 | 0.2 | 0.47 | 0.5 | 0.37 | 0.2 |
| 1 | 89108528 | rs12043361 | -1.386 | 1.568 | 0.87 | 1.295 | -0.004 | -0.01 | -0.01 | 1.877 | 0.35 | 0.29 | 0.49 | 0.56 | 0.4 | 0.26 |
| 1 | 89115413 | rs574740 | -1.511 | 1.563 | 0.873 | 1.293 | 0.002 | 0.011 | 0.011 | 0 | 0.36 | 0.42 | 0.5 | 0.56 | 0.4 | 0.26 |
| 1 | 89116300 | rs7534720 | 0.246 | 1.874 | 0.921 | 1.368 | -0.051 | -0.061 | -0.061 | 0.58 | 0.2 | 0.09 | 0.43 | 0.35 | 0.32 | 0.15 |
| 1 | 89123443 | rs6699417 | 0.351 | 1.682 | 1.458 | 1.133 | 0.035 | 0.059 | 0.059 | 0.004 | 0.76 | 0.65 | 0.54 | 0.52 | 0.62 | 0.81 |
| 1 | 89133812 | rs10754252 | 0.232 | 1.771 | 1.664 | 1.148 | 0.007 | 0.014 | 0.014 | 0 | 0.77 | 0.71 | 0.54 | 0.54 | 0.62 | 0.81 |
| 1 | 89137459 | rs4457557 | 0.395 | 1.772 | 1.682 | 1.147 | 0.055 | 0.088 | 0.088 | 0.005 | 0.76 | 0.59 | 0.54 | 0.51 | 0.62 | 0.75 |
| 1 | 89143305 | rs6698181 | 0.642 | 1.734 | 1.651 | 1.061 | -0.074 | -0.072 | -0.072 | 0 | 0.17 | 0.08 | 0.42 | 0.35 | 0.35 | 0.17 |
| 1 | 89144053 | rs10922478 | 0.487 | 1.769 | 1.729 | 1.028 | 0.014 | 0.024 | 0.024 | 0 | 0.74 | 0.67 | 0.53 | 0.53 | 0.61 | 0.81 |
| 1 | 89146759 | rs1002437 | 1.041 | 1.674 | 2.092 | 1.014 | 0.009 | 0.009 | 0.009 | 0.099 | 0.7 | 0.36 | 0.49 | 0.47 | 0.56 | 0.73 |
| 1 | 89148793 | rs3806340 | 0.691 | 1.671 | 2.093 | 0.987 | 0.061 | 0.089 | 0.089 | 0 | 0.73 | 0.53 | 0.52 | 0.52 | 0.6 | 0.74 |
| 1 | 89164597 | rs17130580 | -0.669 | 1.518 | 2.034 | 0.975 | 0.003 | -0.004 | -0.004 | 0.901 | 0.08 | 0.08 | 0.04 | 0.12 | 0.04 | 0.03 |
| 1 | 89220472 | rs305217 | -0.64 | 1.567 | 2.151 | 0.993 | -0.011 | 0.037 | 0.037 | 0 | 0.08 | 0.18 | 0.05 | 0.15 | 0.04 | 0.03 |
| 1 | 89222420 | rs6704399 | 1.068 | 1.667 | 2.161 | 1.014 | 0.01 | 0.011 | 0.011 | 0.06 | 0.7 | 0.36 | 0.49 | 0.47 | 0.57 | 0.72 |
| 1 | 89225976 | rs430600 | -0.884 | 1.8 | 2.111 | 1.001 | -0.081 | -0.08 | -0.08 | 0 | 0.83 | 0.93 | 0.58 | 0.65 | 0.66 | 0.83 |
| 1 | 89265248 | rs786911 | -0.884 | 1.798 | 1.886 | 1.004 | -0.073 | -0.072 | -0.072 | 1.039 | 0.83 | 0.93 | 0.58 | 0.65 | 0.66 | 0.83 |
| 1 | 89286673 | rs786921 | -0.102 | 1.792 | 2.093 | 0.998 | 0.113 | 0.099 | 0.099 | 0.25 | 0.78 | 0.51 | 0.53 | 0.62 | 0.61 | 0.75 |
| 1 | 89302955 | rs12133012 | 0.205 | 0.946 | 1.895 | 0.499 | 0.024 | 0.088 | 0.088 | 0 | 0.42 | 0.23 | 0.25 | 0.18 | 0.34 | 0.37 |
| 1 | 89322809 | rs7534197 | -2.272 | 1.979 | 2.558 | 0.992 | 0.162 | 0.122 | 0.122 | 0.653 | 0.05 | 0.45 | 0.05 | 0.03 | 0.04 | 0.08 |

| | | | | | | | | | | | | | | | | |
|---|---|---|---|---|---|---|---|---|---|---|---|---|---|---|---|---|
| 1 | 89333557 | rs10493818 | 0.322 | 2.207 | 2.286 | 1.092 | -0.063 | -0.064 | -0.064 | 0 | 0.17 | 0.08 | 0.42 | 0.35 | 0.33 | 0.15 |
| 1 | 89342441 | rs4528095 | -0.057 | 1.852 | 2.368 | 1.061 | -0.006 | 0.021 | 0.021 | 0 | 0.94 | 0.87 | 0.96 | 0.85 | 0.96 | 0.97 |
| 1 | 89359789 | rs10922501 | -0.149 | 1.727 | 2.276 | 1.005 | 0.093 | 0.101 | 0.101 | 0.191 | 0.2 | 0.33 | 0.45 | 0.37 | 0.37 | 0.18 |
| 1 | 89362631 | rs4142614 | 0.085 | 1.628 | 2.583 | 1.3 | 0.088 | 0.091 | 0.091 | 0.286 | 0.79 | 0.46 | 0.54 | 0.62 | 0.64 | 0.82 |
| 1 | 89372589 | rs7524919 | -1.255 | 1.179 | 2.419 | 0.952 | -0.025 | -0.011 | -0.011 | 0 | 0.17 | 0.37 | 0.11 | 0.15 | 0.11 | 0.1 |
| 1 | 89384717 | rs7538497 | -1.258 | 1.184 | 2.301 | 0.955 | -0.024 | -0.012 | -0.012 | 0 | 0.17 | 0.34 | 0.11 | 0.16 | 0.11 | 0.1 |
| 1 | 89388911 | rs12072458 | -0.88 | 1.186 | 2.341 | 0.953 | -0.011 | -0.009 | -0.009 | 0.025 | 0.16 | 0.24 | 0.09 | 0.14 | 0.11 | 0.1 |
| 1 | 89388944 | rs7532151 | -1.791 | 1.26 | 2.426 | 1.04 | 0.029 | 0.025 | 0.025 | 0 | 0.36 | 0.49 | 0.53 | 0.48 | 0.48 | 0.25 |
| 1 | 89401613 | rs14490 | -1.23 | 1.281 | 2.143 | 0.968 | -0.034 | -0.018 | -0.018 | 0.03 | 0.17 | 0.42 | 0.12 | 0.14 | 0.11 | 0.05 |
| 1 | 89406541 | rs10801696 | 1.691 | 1.277 | 2.001 | 0.984 | -0.04 | -0.029 | -0.029 | 0.58 | 0.62 | 0.22 | 0.46 | 0.54 | 0.52 | 0.75 |
| 1 | 89415576 | rs1325923 | -1.682 | 1.314 | 1.951 | 1.002 | 0.006 | 0.004 | 0.004 | 0 | 0.35 | 0.4 | 0.52 | 0.46 | 0.48 | 0.25 |
| 1 | 89417695 | rs2765524 | 0.121 | 1.323 | 2.055 | 1.006 | 0.064 | 0.043 | 0.043 | 0.069 | 0.78 | 0.6 | 0.56 | 0.68 | 0.62 | 0.8 |
| 1 | 89417918 | rs11579178 | -0.025 | 1.322 | 1.978 | 1.006 | -0.052 | -0.044 | -0.044 | 0 | 0.2 | 0.11 | 0.43 | 0.34 | 0.38 | 0.21 |
| 1 | 89421899 | rs1325921 | -0.468 | 1.323 | 1.993 | 1.008 | 0.003 | 0.002 | 0.002 | 0.271 | 0.14 | 0.11 | 0.07 | 0.12 | 0.1 | 0.05 |
| 1 | 89426260 | rs2765527 | 0.984 | 1.015 | 2.15 | 0.915 | 0.015 | 0.011 | 0.011 | 0 | 0.63 | 0.37 | 0.46 | 0.54 | 0.52 | 0.75 |
| 1 | 89426902 | rs3738055 | -1.021 | 1.042 | 1.29 | 0.928 | -0.044 | -0.036 | -0.036 | 0.398 | 0.34 | 0.23 | 0.5 | 0.46 | 0.48 | 0.25 |
| 1 | 89457587 | rs6660788 | -1.264 | 0.582 | 0.958 | 1.403 | -0.007 | 0.129 | 0.129 | 0 | 0.19 | 0.37 | 0.31 | 0.38 | 0.12 | 0.22 |
| 1 | 89459475 | rs6428490 | -1.264 | 0.582 | 1.063 | 1.411 | -0.006 | 0.137 | 0.137 | 0 | 0.19 | 0.39 | 0.31 | 0.38 | 0.12 | 0.22 |
| 1 | 89461865 | rs17486391 | -0.547 | 0.481 | 1.836 | 1.575 | 0.025 | 0.011 | 0.011 | 0 | 0.09 | 0.04 | 0.05 | 0.07 | 0.03 | 0.09 |
| 1 | 89467787 | rs6428492 | -2.184 | 0.24 | 2.058 | 1.534 | 0.017 | -0.035 | -0.035 | 1.261 | 0.23 | 0.8 | 0.34 | 0.39 | 0.12 | 0.22 |
| 1 | 89473134 | rs2077694 | -1.678 | 0.213 | 2.164 | 1.526 | -0.018 | 0.132 | 0.132 | 0 | 0.19 | 0.39 | 0.31 | 0.38 | 0.12 | 0.22 |
| 1 | 89473175 | rs1409150 | -1.451 | 0.607 | 0.826 | 1.287 | -0.03 | 0.037 | 0.037 | 0 | 0.49 | 0.84 | 0.58 | 0.6 | 0.44 | 0.47 |
| 1 | 89479074 | rs4656078 | 1.603 | 0.385 | 0.471 | 1.202 | -0.006 | 0.052 | 0.052 | 0 | 0.41 | 0.18 | 0.29 | 0.31 | 0.44 | 0.49 |
| 1 | 89481583 | rs12121223 | 0.764 | 0.387 | 0.447 | 1.209 | -0.102 | -0.01 | -0.01 | 0 | 0.26 | 0.04 | 0.24 | 0.2 | 0.32 | 0.25 |
| 1 | 89486037 | rs7555039 | 1.148 | 0.284 | 0.486 | 1.175 | -0.02 | 0.063 | 0.063 | 0.045 | 0.71 | 0.6 | 0.58 | 0.52 | 0.77 | 0.74 |
| 1 | 89489915 | rs12142922 | 0.654 | 0.326 | 0.461 | 1.136 | -0.106 | -0.004 | -0.004 | 0 | 0.27 | 0.05 | 0.25 | 0.2 | 0.34 | 0.25 |
| 1 | 89490707 | rs6683705 | 1.368 | 0.354 | 0.519 | 1.165 | 0.015 | -0.027 | -0.027 | 0.021 | 0.68 | 0.27 | 0.56 | 0.52 | 0.77 | 0.74 |
| 1 | 89500792 | rs10922552 | -0.642 | 0.353 | 0.388 | 1.199 | 0 | 0.005 | 0.005 | 0 | 0.9 | 0.97 | 0.89 | 0.93 | 0.89 | 0.95 |
| 1 | 89503665 | rs4656083 | 1.344 | 0.342 | 0.345 | 1.037 | -0.015 | 0.042 | 0.042 | 0 | 0.4 | 0.15 | 0.3 | 0.31 | 0.43 | 0.49 |
| 1 | 89509499 | rs1074541 | 0.298 | 0.459 | 0.841 | 1.103 | -0.033 | 0.008 | 0.008 | 0.012 | 0.31 | 0.17 | 0.31 | 0.27 | 0.36 | 0.31 |
| 1 | 89517810 | rs6675866 | 0.34 | 0.683 | 1.118 | 1.156 | 0.012 | 0.067 | 0.067 | 0 | 0.81 | 0.74 | 0.63 | 0.58 | 0.8 | 0.83 |
| 1 | 89518623 | rs2624 | -0.409 | 0.73 | 1.028 | 1.045 | -0.03 | 0.02 | 0.02 | 0 | 0.67 | 0.83 | 0.68 | 0.74 | 0.62 | 0.68 |
| 1 | 89522512 | rs4376712 | 1.309 | 0.842 | 1.015 | 0.983 | -0.008 | 0.02 | 0.02 | 0 | 0.42 | 0.2 | 0.3 | 0.36 | 0.46 | 0.49 |
| 1 | 89529680 | rs10493822 | 0.336 | 0.75 | 1.508 | 1.087 | -0.002 | 0.002 | 0.002 | 0 | 0.3 | 0.29 | 0.29 | 0.12 | 0.3 | 0.3 |
| 1 | 89546803 | rs10922563 | 0.065 | 0.732 | 1.59 | 1.088 | 0 | -0.044 | -0.044 | 0.021 | 0.33 | 0.53 | 0.3 | 0.18 | 0.3 | 0.3 |
| 1 | 89547560 | rs10922565 | 1.384 | 0.729 | 1.57 | 1.093 | -0.011 | 0.007 | 0.007 | 0 | 0.42 | 0.2 | 0.31 | 0.38 | 0.46 | 0.5 |
| 1 | 89551004 | rs12740160 | 0.859 | 0.733 | 1.56 | 1.082 | 0.001 | -0.006 | -0.006 | 0 | 0.74 | 0.75 | 0.61 | 0.56 | 0.77 | 0.8 |
| 1 | 89553601 | rs1969962 | 0.826 | 0.726 | 1.729 | 1.079 | 0 | -0.019 | -0.019 | 0 | 0.75 | 0.8 | 0.62 | 0.56 | 0.77 | 0.8 |
| 1 | 89555600 | rs12408856 | 1.307 | 0.731 | 1.765 | 1.082 | -0.007 | 0.008 | 0.008 | 0 | 0.43 | 0.23 | 0.31 | 0.38 | 0.46 | 0.5 |
| 1 | 89555843 | rs2031834 | 0.223 | 0.688 | 1.725 | 1.095 | 0.003 | 0.024 | 0.024 | 0.077 | 0.82 | 0.81 | 0.63 | 0.57 | 0.8 | 0.85 |
| 1 | 89564885 | rs2031913 | -1.373 | 0.695 | 1.434 | 0.869 | -0.079 | 0.068 | 0.068 | 0 | 0.82 | 0.98 | 0.83 | 0.96 | 0.76 | 0.93 |
| 1 | 89575677 | rs2026043 | 0.039 | 0.816 | 1.786 | 0.953 | -0.003 | -0.032 | -0.032 | 0 | 0.61 | 0.46 | 0.68 | 0.77 | 0.64 | 0.64 |
| 1 | 89579615 | rs7517828 | 0.247 | 0.805 | 1.828 | 0.958 | -0.021 | -0.019 | -0.019 | 0.103 | 0.08 | 0.12 | 0.02 | 0.01 | 0.03 | 0.04 |
| 1 | 89584069 | rs6428503 | 1.254 | 0.818 | 1.761 | 1.014 | -0.039 | -0.024 | -0.024 | 0 | 0.42 | 0.13 | 0.3 | 0.4 | 0.46 | 0.5 |
| 1 | 89588380 | rs10922573 | 1.254 | 0.818 | 1.761 | 1.014 | -0.039 | -0.024 | -0.024 | 6.065 | 0.42 | 0.13 | 0.3 | 0.4 | 0.46 | 0.5 |
| 1 | 89589556 | rs10922575 | 0.12 | 0.773 | 1.827 | 0.968 | 0 | 0 | 0 | 0 | 0.6 | 0.4 | 0.67 | 0.78 | 0.64 | 0.64 |
| 1 | 89602884 | rs10158415 | -0.854 | 0.922 | 1.546 | 1.037 | -0.02 | 0.035 | 0.035 | 0 | 0.7 | 0.81 | 0.72 | 0.78 | 0.68 | 0.69 |
| 1 | 89619792 | rs4233336 | -0.67 | 0.913 | 1.7 | 1.037 | 0.003 | -0.015 | -0.015 | 0 | 0.68 | 0.63 | 0.71 | 0.78 | 0.67 | 0.69 |
| 1 | 89630721 | rs2893089 | -0.422 | 0.913 | 1.327 | 1.081 | 0.003 | 0.039 | 0.039 | 0 | 0.18 | 0.23 | 0.36 | 0.37 | 0.17 | 0.15 |
| 1 | 89632854 | rs616098 | -0.233 | 0.842 | 1.283 | 1.043 | -0.001 | 0.047 | 0.047 | 0 | 0.63 | 0.73 | 0.7 | 0.77 | 0.64 | 0.65 |
| 1 | 89637578 | rs676913 | -0.291 | 0.851 | 1.215 | 1.047 | -0.007 | 0.059 | 0.059 | 0 | 0.63 | 0.78 | 0.7 | 0.77 | 0.64 | 0.65 |
| 1 | 89643644 | rs615739 | 1.691 | 0.729 | 0.623 | 0.53 | -0.019 | 0.047 | 0.047 | 0 | 0.19 | 0.1 | 0.11 | 0.03 | 0.21 | 0.16 |
| 1 | 89647386 | rs627125 | -0.879 | 0.719 | 0.831 | 0.803 | -0.013 | 0.033 | 0.033 | 0 | 0.7 | 0.8 | 0.73 | 0.78 | 0.69 | 0.7 |
| 1 | 89649824 | rs674903 | 0.758 | 0.758 | 0.782 | 0.816 | 0 | -0.005 | -0.005 | 0 | 0.46 | 0.45 | 0.34 | 0.4 | 0.49 | 0.51 |
| 1 | 89655994 | rs1410871 | -0.027 | 0.78 | 0.457 | 0.728 | -0.013 | -0.017 | -0.017 | 0 | 0.09 | 0.32 | 0.04 | 0.01 | 0.03 | 0.04 |
| 1 | 89661228 | rs650448 | -0.145 | 0.969 | -0.111 | 0.838 | -0.002 | -0.014 | -0.014 | 3.912 | 0.61 | 0.41 | 0.69 | 0.79 | 0.66 | 0.66 |
| 1 | 89673299 | rs12138939 | 0.262 | 1.003 | -0.201 | 0.89 | -0.024 | -0.05 | -0.05 | 0 | 0.11 | 0.04 | 0.13 | 0.32 | 0.14 | 0.1 |
| 1 | 89685357 | rs10159070 | 0.17 | 0.658 | 0.145 | 0.188 | -0.001 | -0.012 | -0.012 | 0.422 | 0.61 | 0.41 | 0.69 | 0.8 | 0.67 | 0.66 |
| 1 | 89715467 | rs2077845 | 0.702 | 0.815 | 0.191 | 0.684 | 0.016 | 0.041 | 0.041 | 0 | 0.57 | 0.29 | 0.37 | 0.32 | 0.58 | 0.56 |
| 1 | 89721862 | rs611596 | 0.231 | 0.687 | 0.677 | 1.197 | 0.019 | 0.081 | 0.081 | 0 | 0.78 | 0.69 | 0.56 | 0.55 | 0.75 | 0.71 |
| 1 | 89730723 | rs9628688 | 1.123 | 0.102 | 0.226 | 1.312 | 0.012 | 0.036 | 0.036 | 0 | 0.57 | 0.33 | 0.37 | 0.32 | 0.57 | 0.54 |
| 1 | 89731959 | rs12729209 | 1.225 | 0.142 | 0.056 | 1.332 | 0.018 | 0.057 | 0.057 | 0 | 0.56 | 0.3 | 0.37 | 0.31 | 0.57 | 0.54 |
| 1 | 89733273 | rs6690597 | 1.062 | 0.413 | -0.109 | 1.439 | 0.002 | 0.008 | 0.008 | 0 | 0.59 | 0.45 | 0.38 | 0.33 | 0.57 | 0.54 |
| 1 | 89735314 | rs3806338 | -0.046 | 0.336 | -0.718 | 1.732 | 0.035 | -0.076 | -0.076 | 0 | 0.17 | 0.03 | 0.13 | 0.25 | 0.12 | 0.16 |
| 1 | 89746017 | rs687059 | 0.766 | 1.092 | 0.087 | 1.871 | 0.017 | 0.136 | 0.136 | 0 | 0.89 | 0.79 | 0.81 | 0.67 | 0.9 | 0.81 |
| 1 | 89748907 | rs612767 | 0.999 | 1.281 | -0.122 | 1.845 | 0.011 | 0.059 | 0.059 | 0 | 0.87 | 0.83 | 0.77 | 0.67 | 0.82 | 0.79 |
| 1 | 89759219 | rs10922605 | -0.497 | 0.906 | -0.509 | 1.312 | 0 | -0.064 | -0.064 | 0.253 | 0.33 | 0.18 | 0.26 | 0.51 | 0.33 | 0.55 |
| 1 | 89772057 | rs2893103 | -0.585 | 0.669 | 0.681 | 1.274 | -0.003 | 0.005 | 0.005 | 3.283 | 0.36 | 0.44 | 0.32 | 0.54 | 0.33 | 0.56 |
| 1 | 89776233 | rs10801723 | 0.056 | 1.201 | 0.584 | 0.817 | 0.012 | -0.014 | -0.014 | 1.629 | 0.24 | 0.26 | 0.23 | 0.1 | 0.37 | 0.25 |

| | | | | | | | | | | | | | | | | |
|---|---|---|---|---|---|---|---|---|---|---|---|---|---|---|---|---|
| 1 | 89779051 | rs10801724 | -0.446 | 1.013 | 0.311 | 0.622 | 0 | 0.033 | 0.033 | 0.323 | 0.9 | 0.92 | 0.91 | 0.64 | 0.9 | 0.88 |
| 1 | 89780262 | rs12122989 | 0.109 | 0.947 | 0.561 | 0.255 | 0.006 | 0.06 | 0.06 | 4.007 | 0.54 | 0.34 | 0.61 | 0.12 | 0.57 | 0.33 |
| 1 | 89783040 | rs6428514 | 0.439 | 0.933 | 0.438 | 0.699 | 0.011 | 0.018 | 0.018 | 0 | 0.92 | 0.85 | 0.94 | 0.9 | 0.91 | 0.93 |
| 1 | 89787286 | rs1117177 | -0.154 | 1.259 | 0.075 | 0.107 | 0.006 | -0.031 | -0.031 | 0 | 0.28 | 0.35 | 0.44 | 0.14 | 0.33 | 0.24 |
| 1 | 89788116 | rs12125854 | 1.146 | 0.914 | -0.547 | 0.192 | -0.005 | 0.011 | 0.011 | 0 | 0.25 | 0.12 | 0.15 | 0.26 | 0.23 | 0.34 |
| 1 | 89789301 | rs7530906 | 1.157 | 0.916 | -0.549 | 0.193 | -0.006 | 0.012 | 0.012 | 0 | 0.25 | 0.11 | 0.15 | 0.26 | 0.23 | 0.34 |
| 1 | 89799068 | rs4658132 | 0.348 | 0.982 | -0.454 | 0.26 | 0.01 | -0.009 | -0.009 | 0.269 | 0.64 | 0.45 | 0.5 | 0.76 | 0.58 | 0.69 |
| 1 | 89806926 | rs11585444 | 1.484 | 1.262 | -0.645 | 0.173 | -0.004 | 0.009 | 0.009 | 0 | 0.12 | 0.09 | 0.09 | 0.02 | 0.15 | 0.17 |
| 1 | 89808387 | rs10922610 | 0.058 | 1.461 | -0.464 | 0.37 | 0.025 | 0.096 | 0.096 | 0.076 | 0.89 | 0.8 | 0.8 | 0.41 | 0.84 | 0.81 |
| 1 | 89817783 | rs6672564 | -0.796 | 0.985 | -0.791 | 0.743 | -0.02 | -0.057 | -0.057 | 0 | 0.69 | 0.88 | 0.8 | 0.59 | 0.69 | 0.57 |
| 1 | 89825320 | rs12732279 | -0.363 | 0.842 | -0.899 | 0.375 | -0.027 | 0.037 | 0.037 | 0 | 0.28 | 0.21 | 0.4 | 0.05 | 0.4 | 0.29 |
| 1 | 89828716 | rs1414355 | -0.834 | 0.75 | -0.783 | 0.678 | -0.001 | -0.003 | -0.003 | 0.675 | 0.89 | 0.87 | 0.87 | 0.81 | 0.88 | 0.81 |
| 1 | 89832464 | rs1335647 | -0.393 | 0.565 | -0.515 | 0.429 | 0.001 | -0.035 | -0.035 | 0 | 0.34 | 0.36 | 0.4 | 0.13 | 0.38 | 0.26 |
| 1 | 89835926 | rs6681338 | -0.488 | 0.612 | -0.516 | 0.768 | 0.013 | 0.009 | 0.009 | 0 | 0.37 | 0.48 | 0.56 | 0.39 | 0.42 | 0.4 |
| 1 | 89840340 | rs4529696 | -0.243 | 0.693 | -0.239 | 0.569 | 0.002 | 0.005 | 0.005 | 0 | 0.56 | 0.55 | 0.69 | 0.68 | 0.62 | 0.67 |
| 1 | 89847372 | rs4658359 | -1.101 | 0.685 | -0.201 | 0.918 | -0.011 | -0.03 | -0.03 | 0 | 0.73 | 0.91 | 0.84 | 0.73 | 0.75 | 0.78 |
| 1 | 89847411 | rs4658360 | -1.09 | 0.698 | -0.195 | 0.924 | -0.01 | -0.03 | -0.03 | 0 | 0.72 | 0.91 | 0.84 | 0.73 | 0.75 | 0.78 |
| 1 | 89847444 | rs4658146 | -1.09 | 0.698 | -0.195 | 0.924 | -0.01 | -0.03 | -0.03 | 1.883 | 0.72 | 0.91 | 0.84 | 0.73 | 0.75 | 0.78 |
| 1 | 89873394 | rs4491050 | -0.335 | 1.473 | 0.609 | 1.703 | 0 | 0.039 | 0.039 | 0 | 0.93 | 0.92 | 0.92 | 0.73 | 0.89 | 0.85 |
| 1 | 89876211 | rs6682580 | 0.021 | 1.438 | 0.77 | 1.675 | -0.006 | 0.01 | 0.01 | 0.571 | 0.11 | 0.21 | 0.12 | 0.12 | 0.1 | 0.13 |
| 1 | 89877573 | rs10801736 | 0.17 | 1.431 | 0.771 | 1.672 | -0.002 | 0.057 | 0.057 | 0 | 0.91 | 0.91 | 0.87 | 0.62 | 0.89 | 0.81 |
| 1 | 89886229 | rs6698601 | 1.49 | 1.433 | 0.771 | 1.672 | 0.011 | 0.024 | 0.024 | 0.045 | 0.78 | 0.7 | 0.67 | 0.33 | 0.72 | 0.65 |
| 1 | 89888151 | rs4421571 | -0.367 | 1.4 | 0.59 | 1.508 | 0 | 0.039 | 0.039 | 0 | 0.93 | 0.92 | 0.92 | 0.73 | 0.89 | 0.85 |
| 1 | 89890973 | rs12098051 | 0.061 | 1.254 | 0.773 | 1.28 | -0.003 | 0.006 | 0.006 | 0.478 | 0.1 | 0.14 | 0.12 | 0.12 | 0.1 | 0.13 |
| 1 | 89898842 | rs10801739 | 0.203 | 1.314 | 0.753 | 1.172 | 0.005 | -0.053 | -0.053 | 0.286 | 0.09 | 0.2 | 0.07 | 0.02 | 0.1 | 0.09 |
| 1 | 89899019 | rs11577456 | -0.405 | 1.384 | 0.519 | 1.364 | 0.003 | 0.013 | 0.013 | 0.446 | 0.12 | 0.12 | 0.19 | 0.28 | 0.16 | 0.16 |
| 1 | 89899285 | rs10922643 | 0.098 | 1.338 | 0.691 | 1.256 | 0.011 | -0.058 | -0.058 | 0 | 0.1 | 0.24 | 0.08 | 0.02 | 0.1 | 0.09 |
| 1 | 89921575 | rs10801741 | -0.445 | 1.271 | 0.345 | 1.199 | -0.007 | -0.013 | -0.013 | 0 | 0.57 | 0.55 | 0.72 | 0.74 | 0.64 | 0.54 |
| 1 | 89928044 | rs7556550 | 1.079 | 1.199 | 0.311 | 1.172 | -0.063 | 0.096 | 0.096 | 0.185 | 0.18 | 0.03 | 0.33 | 0.02 | 0.26 | 0.2 |
| 1 | 89933828 | rs17130817 | 1.476 | 0.9 | 0.431 | 1.404 | 0.005 | 0.017 | 0.017 | 0.345 | 0.28 | 0.02 | 0.12 | 0.17 | 0.22 | 0.31 |
| 1 | 89935101 | rs6692501 | -0.469 | 1 | 0.646 | 1.446 | -0.005 | 0 | 0 | 0 | 0.58 | 0.56 | 0.65 | 0.55 | 0.62 | 0.53 |
| 1 | 89935489 | rs1459641 | 0.855 | 0.85 | 0.458 | 1.668 | -0.002 | 0.094 | 0.094 | 0 | 0.05 | 0.07 | 0.05 | 0.27 | 0.05 | 0.11 |
| 1 | 89970163 | rs10493826 | 0.033 | 1.407 | 0.187 | 1.309 | -0.065 | -0.082 | -0.082 | 0 | 0.13 | 0.02 | 0.21 | 0.44 | 0.21 | 0.2 |
| 1 | 89972062 | rs1585364 | 0.487 | 1.431 | 0.189 | 1.392 | -0.057 | -0.055 | -0.055 | 1.401 | 0.11 | 0.02 | 0.16 | 0.26 | 0.19 | 0.17 |
| 1 | 89976988 | rs359941 | -0.813 | 1.166 | 0.421 | 0.782 | 0.009 | 0.017 | 0.017 | 0 | 0.64 | 0.73 | 0.75 | 0.75 | 0.71 | 0.71 |
| 1 | 89982719 | rs359937 | 0.783 | 1.119 | 0.401 | 0.158 | 0.002 | -0.045 | -0.045 | 0 | 0.19 | 0.22 | 0.35 | 0.02 | 0.24 | 0.21 |
| 1 | 89989354 | rs359920 | -0.587 | 1.088 | 0.666 | 0.124 | 0.069 | 0.015 | 0.015 | 0 | 0.05 | 0.28 | 0.06 | 0.02 | 0.06 | 0.05 |
| 1 | 89995069 | rs716833 | 0.538 | 1.13 | 0.522 | 0.101 | -0.013 | -0.003 | -0.003 | 0 | 0.21 | 0.12 | 0.35 | 0.03 | 0.3 | 0.26 |
| 1 | 90005132 | rs9428043 | -0.045 | 1.239 | 0.68 | 0.264 | -0.001 | -0.015 | -0.015 | 0 | 0.48 | 0.44 | 0.51 | 0.2 | 0.52 | 0.55 |
| 1 | 90006887 | rs6696878 | 1.246 | 1.213 | 0.415 | 0.351 | -0.043 | 0.063 | 0.063 | 0 | 0.16 | 0.07 | 0.32 | 0.01 | 0.24 | 0.22 |
| 1 | 90019642 | rs359946 | -0.642 | 0.974 | 1.149 | 1.205 | -0.002 | 0.092 | 0.092 | 0 | 0.92 | 0.94 | 0.92 | 0.68 | 0.94 | 0.9 |
| 1 | 90022796 | rs922107 | 0.781 | 1.021 | 0.973 | 1.1 | -0.001 | 0.017 | 0.017 | 0 | 0.51 | 0.44 | 0.52 | 0.2 | 0.52 | 0.59 |
| 1 | 90023768 | rs6699344 | 0.799 | 0.991 | 0.965 | 1.099 | -0.001 | 0.024 | 0.024 | 0 | 0.51 | 0.4 | 0.52 | 0.2 | 0.52 | 0.59 |
| 1 | 90025519 | rs922106 | 1.242 | 0.956 | 0.48 | 1.124 | -0.033 | 0.137 | 0.137 | 0 | 0.43 | 0.12 | 0.46 | 0.19 | 0.46 | 0.55 |
| 1 | 90032940 | rs10922664 | 1.087 | 1.111 | 0.531 | 1.21 | -0.002 | 0.116 | 0.116 | 0 | 0.46 | 0.22 | 0.48 | 0.19 | 0.46 | 0.55 |
| 1 | 90048248 | rs12130207 | 0.579 | 1.192 | 0.382 | 1.196 | 0.021 | 0.08 | 0.08 | 0 | 0.28 | 0.16 | 0.19 | 0.04 | 0.22 | 0.14 |
| 1 | 90050831 | rs7516314 | 0.73 | 1.386 | 0.186 | 1.301 | -0.012 | 0.13 | 0.13 | 0 | 0.45 | 0.2 | 0.46 | 0.19 | 0.47 | 0.55 |
| 1 | 90053245 | rs12135419 | 0.555 | 1.637 | -0.063 | 1.424 | 0.005 | 0.063 | 0.063 | 0 | 0.48 | 0.39 | 0.49 | 0.19 | 0.47 | 0.55 |
| 1 | 90053449 | rs359943 | -0.078 | 1.427 | -0.545 | 1.311 | -0.008 | 0.003 | 0.003 | 0 | 0.6 | 0.59 | 0.69 | 0.62 | 0.66 | 0.71 |
| 1 | 90065471 | rs17130836 | -0.907 | 1.97 | -0.609 | 1.877 | -0.014 | -0.011 | -0.011 | 0 | 0.11 | 0.05 | 0.14 | 0.36 | 0.2 | 0.15 |
| 1 | 90066415 | rs6428558 | -1.525 | 2.433 | -0.028 | 2.167 | 0.09 | 0.216 | 0.216 | 0 | 0.13 | 0.25 | 0.23 | 0.49 | 0.27 | 0.2 |
| 1 | 90066806 | rs1215494 | -0.944 | 1.924 | 0.351 | 1.315 | 0.008 | -0.016 | -0.016 | 0 | 0.3 | 0.39 | 0.3 | 0.22 | 0.34 | 0.38 |
| 1 | 90067602 | rs6685584 | -1.368 | 1.441 | 0.529 | 0.597 | 0.003 | 0.042 | 0.042 | 1.245 | 0.49 | 0.64 | 0.57 | 0.26 | 0.48 | 0.52 |
| 1 | 90068275 | rs12129905 | -1.196 | 1.353 | 0.922 | 0.582 | 0 | 0.006 | 0.006 | 0.288 | 0.47 | 0.47 | 0.55 | 0.27 | 0.48 | 0.52 |
| 1 | 90069248 | rs10922670 | -0.062 | 1.317 | 0.744 | 0.442 | -0.007 | 0.115 | 0.115 | 0 | 0.28 | 0.12 | 0.16 | 0.06 | 0.28 | 0.21 |
| 1 | 90069640 | rs1885745 | -1.668 | 1.268 | 0.867 | 0.468 | 0.013 | -0.004 | -0.004 | 0 | 0.34 | 0.55 | 0.43 | 0.28 | 0.37 | 0.49 |
| 1 | 90072888 | rs1215547 | -1.266 | 1.206 | 1.141 | 0.601 | 0.044 | 0.081 | 0.081 | 0.041 | 0.15 | 0.58 | 0.35 | 0.2 | 0.19 | 0.26 |
| 1 | 90087511 | rs1215512 | -0.813 | 0.835 | 0.903 | 0.332 | 0.019 | 0.06 | 0.06 | 0 | 0.44 | 0.74 | 0.47 | 0.72 | 0.38 | 0.35 |
| 1 | 90112431 | rs10754283 | -0.84 | 1.027 | 1.273 | 0.184 | -0.001 | -0.005 | -0.005 | 0.243 | 0.47 | 0.55 | 0.69 | 0.77 | 0.51 | 0.43 |
| 1 | 90117833 | rs10493829 | -0.836 | 0.989 | 1.328 | 0.173 | 0.001 | -0.018 | -0.018 | 0 | 0.47 | 0.5 | 0.68 | 0.77 | 0.5 | 0.42 |
| 1 | 90121032 | rs1320563 | -1.032 | 0.985 | 1.327 | 0.172 | -0.006 | 0.103 | 0.103 | 0 | 0.48 | 0.71 | 0.7 | 0.77 | 0.5 | 0.42 |
| 1 | 90121751 | rs10801764 | -0.844 | 1.019 | 1.2 | 0.188 | -0.001 | -0.002 | -0.002 | 0 | 0.47 | 0.55 | 0.68 | 0.77 | 0.5 | 0.42 |
| 1 | 90125566 | rs2390794 | 0.33 | 1.057 | 0.939 | 1.002 | -0.008 | -0.037 | -0.037 | 0 | 0.27 | 0.21 | 0.34 | 0.35 | 0.29 | 0.26 |
| 1 | 90127075 | rs12406924 | -1.021 | 0.641 | 1.435 | 0.804 | 0.002 | -0.006 | -0.006 | 0.067 | 0.22 | 0.23 | 0.33 | 0.43 | 0.2 | 0.23 |
| 1 | 90132069 | rs10922690 | 0.245 | 0.871 | 1.937 | 0.754 | 0.002 | 0.012 | 0.012 | 0 | 0.3 | 0.39 | 0.37 | 0.36 | 0.3 | 0.27 |
| 1 | 90132582 | rs2185196 | -0.172 | 0.811 | 2.201 | 0.721 | 0.001 | -0.009 | -0.009 | 0.249 | 0.67 | 0.3 | 0.59 | 0.64 | 0.7 | 0.73 |
| 1 | 90134392 | rs6675409 | 0.365 | 0.813 | 2.264 | 0.722 | 0.005 | 0.024 | 0.024 | 0 | 0.3 | 0.44 | 0.37 | 0.36 | 0.3 | 0.27 |
| 1 | 90135433 | rs17441836 | 1.264 | 0.887 | 1.969 | 0.73 | -0.036 | 0.175 | 0.175 | 0.094 | 0.46 | 0.11 | 0.26 | 0.2 | 0.48 | 0.5 |
| 1 | 90143228 | rs4658309 | -0.203 | 0.891 | 2.154 | 0.789 | 0 | -0.002 | -0.002 | 0 | 0.67 | 0.36 | 0.61 | 0.64 | 0.7 | 0.73 |

| | | | | | | | | | | | | | | | | |
|---|---|---|---|---|---|---|---|---|---|---|---|---|---|---|---|---|
| 1 | 90145331 | rs17130852 | 0.493 | 0.953 | 1.808 | 1.007 | -0.024 | -0.05 | -0.05 | 0.339 | 0.09 | 0 | 0.05 | 0.37 | 0.09 | 0.1 |
| 1 | 90173698 | rs10801771 | -1.71 | 1.145 | 2.274 | 0.547 | 0.017 | 0.024 | 0.024 | 0 | 0.24 | 0.54 | 0.36 | 0.29 | 0.24 | 0.24 |
| 1 | 90179532 | rs12032393 | -0.57 | 1.156 | 2.117 | 0.587 | -0.026 | -0.06 | -0.06 | 5.005 | 0.16 | 0.03 | 0.21 | 0.25 | 0.15 | 0.16 |
| 1 | 90186466 | rs3261 | -1.463 | 1.172 | 2.134 | 0.43 | 0.014 | 0.082 | 0.082 | 0 | 0.53 | 0.7 | 0.73 | 0.82 | 0.55 | 0.53 |
| 1 | 90189439 | rs4271195 | -0.869 | 1.105 | 1.994 | 0.478 | -0.017 | -0.023 | -0.023 | 0.366 | 0.2 | 0.15 | 0.33 | 0.3 | 0.24 | 0.24 |
| 1 | 90196552 | rs2224652 | 0.293 | 1.26 | 2.19 | 0.49 | 0 | 0.012 | 0.012 | 0 | 0.32 | 0.33 | 0.38 | 0.52 | 0.32 | 0.31 |
| 1 | 90199458 | rs541122 | 1.195 | 1.249 | 2.192 | 0.437 | -0.008 | 0.18 | 0.18 | 0 | 0.45 | 0.13 | 0.26 | 0.18 | 0.45 | 0.46 |
| 1 | 90210821 | rs12727736 | -1.095 | 1.271 | 2.198 | 0.448 | 0.003 | -0.001 | -0.001 | 0 | 0.51 | 0.47 | 0.7 | 0.82 | 0.55 | 0.54 |
| 1 | 90223400 | rs519989 | 1.72 | 1.098 | 1.708 | 0.034 | -0.034 | 0.22 | 0.22 | 0 | 0.34 | 0.07 | 0.21 | 0.03 | 0.34 | 0.3 |
| 1 | 90227649 | rs473412 | 0.76 | 1.026 | 2.03 | 0.191 | 0.005 | 0.035 | 0.035 | 0 | 0.49 | 0.38 | 0.29 | 0.18 | 0.47 | 0.47 |
| 1 | 90227682 | rs10922710 | -0.456 | 0.976 | 1.936 | 0.192 | -0.01 | -0.028 | -0.028 | 0.59 | 0.19 | 0.15 | 0.32 | 0.31 | 0.21 | 0.23 |
| 1 | 90232180 | rs4658318 | 0.398 | 0.977 | 1.96 | 0.085 | -0.008 | -0.044 | -0.044 | 0 | 0.3 | 0.2 | 0.38 | 0.49 | 0.33 | 0.3 |
| 1 | 90233068 | rs483517 | -0.506 | 0.974 | 1.656 | 0.083 | 0.004 | 0.061 | 0.061 | 0 | 0.68 | 0.52 | 0.59 | 0.51 | 0.67 | 0.7 |
| 1 | 90238018 | rs12026201 | 1.209 | 1.049 | 1.147 | 0.193 | -0.023 | 0.042 | 0.042 | 0 | 0.06 | 0.01 | 0.15 | 0.45 | 0.1 | 0.19 |
| 1 | 90243771 | rs10737711 | -0.995 | 0.965 | 1.493 | -0.476 | 0.002 | -0.025 | -0.025 | 0.088 | 0.5 | 0.47 | 0.7 | 0.91 | 0.53 | 0.52 |
| 1 | 90244656 | rs6690748 | -0.493 | 1.019 | 1.547 | -0.551 | -0.002 | -0.002 | -0.002 | 0.011 | 0.2 | 0.17 | 0.33 | 0.42 | 0.2 | 0.22 |
| 1 | 90245167 | rs3120939 | 0.667 | 0.909 | 1.559 | -1.114 | 0.009 | 0.129 | 0.129 | 0 | 0.49 | 0.29 | 0.27 | 0.09 | 0.47 | 0.47 |
| 1 | 90248302 | rs4658138 | -0.856 | 0.885 | 1.168 | -1.105 | 0 | -0.041 | -0.041 | 0 | 0.51 | 0.52 | 0.71 | 0.91 | 0.53 | 0.53 |
| 1 | 90249829 | rs4658321 | -1.012 | 0.978 | 0.483 | -1.046 | -0.037 | 0.262 | 0.262 | 0 | 0.56 | 0.89 | 0.77 | 0.91 | 0.55 | 0.53 |
| 1 | 90254276 | rs12135351 | -0.908 | 0.941 | 0.581 | -0.978 | -0.043 | 0.272 | 0.272 | 0.01 | 0.52 | 0.86 | 0.63 | 0.91 | 0.48 | 0.49 |
| 1 | 90256113 | rs7535521 | -0.884 | 0.945 | 0.605 | -0.976 | -0.014 | 0.184 | 0.184 | 0.006 | 0.51 | 0.79 | 0.63 | 0.91 | 0.48 | 0.49 |
| 1 | 90259555 | rs11806231 | 0.067 | 1 | 0.593 | -0.915 | 0.017 | 0.163 | 0.163 | 0.764 | 0.11 | 0.25 | 0.27 | 0.6 | 0.12 | 0.23 |
| 1 | 90265414 | rs6684610 | -0.889 | 0.973 | 0.339 | -0.939 | -0.003 | 0.088 | 0.088 | 0 | 0.51 | 0.7 | 0.62 | 0.91 | 0.48 | 0.5 |
| 1 | 90265971 | rs1009964 | -0.172 | 0.965 | 0.206 | -0.971 | -0.002 | -0.002 | -0.002 | 0 | 0.37 | 0.38 | 0.5 | 0.6 | 0.38 | 0.34 |
| 1 | 90269272 | rs1078094 | 0.112 | 0.981 | -0.196 | -0.586 | -0.004 | 0.095 | 0.095 | 0 | 0.14 | 0.29 | 0.12 | 0.31 | 0.1 | 0.16 |
| 1 | 90272372 | rs6659997 | 0.218 | 0.955 | -0.958 | -1.453 | -0.001 | -0.066 | -0.066 | 0 | 0.27 | 0.3 | 0.25 | 0.01 | 0.27 | 0.1 |
| 1 | 90285190 | rs12565034 | -0.723 | 0.656 | -1.181 | -0.631 | -0.04 | 0.039 | 0.039 | 0.06 | 0.2 | 0.05 | 0.2 | 0.11 | 0.21 | 0.33 |
| 1 | 90289861 | rs4587551 | 0.259 | 0.47 | -0.715 | -0.084 | -0.025 | -0.009 | -0.009 | 0 | 0.07 | 0 | 0.03 | 0.07 | 0.06 | 0.16 |
| 1 | 90298305 | rs2186184 | -0.6 | -0.189 | -0.53 | -0.235 | 0.011 | 0.012 | 0.012 | 0 | 0.25 | 0.2 | 0.31 | 0.85 | 0.16 | 0.39 |
| 1 | 90304024 | rs4658335 | 0.088 | 0.425 | -0.82 | 0.041 | 0 | 0.003 | 0.003 | 1.734 | 0.76 | 0.23 | 0.7 | 0.82 | 0.76 | 0.78 |
| 1 | 90308101 | rs10922733 | -0.01 | 0.375 | -0.772 | 0.296 | 0 | 0 | 0 | 0 | 0.62 | 0.61 | 0.64 | 0.59 | 0.62 | 0.75 |
| 1 | 90322384 | rs10754295 | 0.295 | -0.264 | -0.584 | -0.198 | -0.038 | -0.049 | -0.049 | 2.399 | 0.63 | 0.5 | 0.67 | 0.82 | 0.75 | 0.64 |
| 1 | 90327555 | rs4658340 | -0.618 | 0.108 | -0.119 | -0.66 | -0.003 | -0.069 | -0.069 | 0 | 0.3 | 0.5 | 0.18 | 0.12 | 0.3 | 0.31 |
| 1 | 90336244 | rs12095496 | -0.114 | 0.061 | 0.685 | -0.838 | -0.001 | -0.03 | -0.03 | 0 | 0.18 | 0.19 | 0.1 | 0.02 | 0.17 | 0.14 |
| 1 | 90338129 | rs17130908 | 0.388 | 0.485 | 1.048 | -0.26 | 0.016 | 0.076 | 0.076 | 0.007 | 0.11 | 0.22 | 0.25 | 0.33 | 0.12 | 0.21 |
| 1 | 90339193 | rs2816853 | -1.125 | 0.481 | 0.916 | -0.961 | -0.008 | -0.071 | -0.071 | 0 | 0.3 | 0.49 | 0.18 | 0.12 | 0.3 | 0.31 |
| 1 | 90349347 | rs17498135 | 0.011 | 0.51 | 0.754 | -0.915 | -0.021 | -0.025 | -0.025 | 0 | 0.1 | 0.02 | 0.06 | 0.11 | 0.13 | 0.17 |
| 1 | 90356770 | rs4658342 | -0.051 | 0.481 | 0.888 | -0.895 | -0.018 | -0.029 | -0.029 | 0 | 0.11 | 0.02 | 0.1 | 0.12 | 0.13 | 0.17 |
| 1 | 90357651 | rs1938032 | 0.136 | 0.424 | 0.996 | -1.072 | -0.007 | 0.06 | 0.06 | 0 | 0.17 | 0.08 | 0.09 | 0.02 | 0.17 | 0.14 |
| 1 | 90362639 | rs17498595 | -0.835 | 0.511 | 0.464 | -1.609 | -0.015 | 0.12 | 0.12 | 0 | 0.3 | 0.12 | 0.17 | 0.02 | 0.29 | 0.19 |
| 1 | 90363941 | rs1344190 | -0.168 | 0.718 | 0.812 | -1.249 | 0.003 | -0.099 | -0.099 | 0 | 0.21 | 0.41 | 0.11 | 0.04 | 0.19 | 0.14 |
| 1 | 90370808 | rs1892431 | -0.127 | 0.769 | 0.165 | -1.085 | -0.03 | -0.007 | -0.007 | 0 | 0.12 | 0.01 | 0.07 | 0.09 | 0.15 | 0.18 |
| 1 | 90371776 | rs1344059 | -1.407 | 0.712 | 0.344 | -1.322 | -0.002 | -0.027 | -0.027 | 0 | 0.42 | 0.46 | 0.24 | 0.21 | 0.41 | 0.38 |
| 1 | 90371913 | rs2816850 | -0.216 | 0.516 | 0.668 | -1.158 | -0.016 | -0.117 | -0.117 | 0 | 0.21 | 0.4 | 0.14 | 0.02 | 0.18 | 0.15 |
| 1 | 90373067 | rs2816849 | -1.104 | 0.289 | 0.455 | -1.195 | -0.021 | -0.041 | -0.041 | 0 | 0.37 | 0.46 | 0.18 | 0.16 | 0.27 | 0.23 |
| 1 | 90373449 | rs17446011 | 0.014 | 0.889 | -0.221 | -1.442 | -0.033 | 0.125 | 0.125 | 0 | 0.22 | 0.03 | 0.12 | 0.02 | 0.2 | 0.15 |
| 1 | 90388187 | rs17505638 | -0.957 | 1.42 | 0.77 | -1.486 | -0.02 | 0.016 | 0.016 | 0 | 0.13 | 0.08 | 0.09 | 0 | 0.2 | 0.06 |
| 1 | 90423039 | rs2816893 | -1.231 | 0.699 | 0.898 | -0.999 | 0.034 | 0.066 | 0.066 | 0.298 | 0.06 | 0.32 | 0.05 | 0.08 | 0.04 | 0.07 |
| 1 | 90433725 | rs2464448 | -1.318 | 0.782 | 0.924 | -0.997 | 0.013 | 0.029 | 0.029 | 0.101 | 0.07 | 0.24 | 0.05 | 0.09 | 0.06 | 0.08 |
| 1 | 90455415 | rs2816866 | 0.955 | 1.047 | 0.939 | -1.675 | 0 | -0.01 | -0.01 | 0 | 0.53 | 0.52 | 0.68 | 0.83 | 0.5 | 0.58 |
| 1 | 90461348 | rs2802022 | -1.1 | 1.083 | 1.04 | -1.753 | 0.021 | -0.014 | -0.014 | 0.089 | 0.11 | 0.39 | 0.07 | 0.07 | 0.1 | 0.06 |
| 1 | 90473851 | rs7531015 | -0.313 | 1.306 | 1.044 | -1.895 | -0.074 | 0.069 | 0.069 | 0 | 0.22 | 0.05 | 0.14 | 0.08 | 0.26 | 0.26 |
| 1 | 90486412 | rs2816881 | 0.965 | 1.244 | 1.112 | -1.895 | -0.007 | 0.005 | 0.005 | 0.114 | 0.6 | 0.66 | 0.72 | 0.89 | 0.57 | 0.63 |
| 1 | 90492517 | rs10922743 | 1.088 | 1.254 | 0.836 | -1.89 | 0.005 | -0.045 | -0.045 | 1.255 | 0.58 | 0.47 | 0.71 | 0.89 | 0.57 | 0.63 |
| 1 | 90493026 | rs10922744 | 1.164 | 1.292 | 0.858 | -1.868 | 0.003 | -0.04 | -0.04 | 1.586 | 0.57 | 0.47 | 0.7 | 0.89 | 0.57 | 0.6 |
| 1 | 90498540 | rs954145 | 1.178 | 1.09 | 0.781 | -1.859 | -0.002 | 0.005 | 0.005 | 0.251 | 0.54 | 0.44 | 0.69 | 0.89 | 0.54 | 0.56 |
| 1 | 90501669 | rs2816855 | 0.435 | 0.878 | 0.532 | -0.892 | -0.053 | 0.065 | 0.065 | 0 | 0.79 | 0.95 | 0.87 | 0.92 | 0.77 | 0.76 |
| 1 | 90532229 | rs12134621 | -0.883 | 0.448 | 0.546 | -0.649 | -0.009 | -0.086 | -0.086 | 0 | 0.14 | 0.32 | 0.1 | 0 | 0.12 | 0.07 |
| 1 | 90532256 | rs10801783 | -0.499 | 0.566 | 0.666 | -1.448 | 0 | -0.044 | -0.044 | 0 | 0.17 | 0.18 | 0.1 | 0.01 | 0.14 | 0.16 |
| 1 | 90532766 | rs7542860 | -1.751 | 0.584 | 0.381 | -1.54 | -0.005 | 0.13 | 0.13 | 0 | 0.52 | 0.73 | 0.36 | 0.18 | 0.46 | 0.44 |
| 1 | 90535344 | rs6663620 | -1.011 | 0.412 | 0.396 | -1.571 | -0.009 | -0.042 | -0.042 | 0 | 0.33 | 0.49 | 0.26 | 0.17 | 0.32 | 0.28 |
| 1 | 90536797 | rs2703993 | -1.181 | 0.539 | 0.401 | -1.421 | 0 | 0.067 | 0.067 | 0.278 | 0.39 | 0.69 | 0.31 | 0.19 | 0.34 | 0.37 |
| 1 | 90538025 | rs2813744 | -1.13 | 0.858 | 0.252 | -0.916 | -0.014 | -0.009 | -0.009 | 0.134 | 0.31 | 0.75 | 0.33 | 0.32 | 0.3 | 0.26 |
| 1 | 90539764 | rs2813746 | 1.199 | 1.401 | -0.181 | -1.137 | 0.023 | 0.059 | 0.059 | 0.432 | 0.5 | 0.22 | 0.5 | 0.63 | 0.47 | 0.56 |
| 1 | 90539813 | rs2813747 | 0.162 | 1.472 | 0.612 | -1.325 | 0.002 | -0.033 | -0.033 | 0 | 0.75 | 0.75 | 0.79 | 0.93 | 0.72 | 0.81 |
| 1 | 90540501 | rs17130964 | 0.582 | 1.369 | 0.484 | -1.339 | 0.013 | -0.038 | -0.038 | 0 | 0.08 | 0.03 | 0.04 | 0.2 | 0.06 | 0.1 |
| 1 | 90540565 | rs2703999 | 1.288 | 1.386 | 0.919 | -1.327 | -0.005 | 0.032 | 0.032 | 0.108 | 0.55 | 0.32 | 0.6 | 0.73 | 0.55 | 0.62 |
| 1 | 90546731 | rs2390866 | 1.276 | 1.314 | 0.953 | -1.162 | 0.009 | -0.002 | -0.002 | 0.19 | 0.61 | 0.43 | 0.58 | 0.63 | 0.58 | 0.59 |
| 1 | 90553637 | rs12126957 | 0.124 | 1.348 | 0.546 | -1.19 | -0.028 | -0.032 | -0.032 | 0 | 0.06 | 0 | 0.09 | 0.1 | 0.09 | 0.07 |

| | | | | | | | | | | | | | | | | |
|---|---|---|---|---|---|---|---|---|---|---|---|---|---|---|---|---|
| 1 | 90556150 | rs17130969 | 0.99 | 1.128 | 0.014 | -1.413 | 0.011 | 0.034 | 0.034 | 0 | 0.15 | 0.05 | 0.14 | 0.04 | 0.11 | 0.11 |
| 1 | 90556980 | rs1329642 | -1.199 | 1.33 | 0.245 | -1.299 | -0.006 | 0.014 | 0.014 | 0 | 0.86 | 0.9 | 0.85 | 0.94 | 0.82 | 0.87 |
| 1 | 90559504 | rs2772337 | -0.166 | 1.303 | 0.36 | -1.293 | -0.013 | 0.005 | 0.005 | 0 | 0.8 | 0.88 | 0.76 | 0.85 | 0.73 | 0.8 |
| 1 | 90568410 | rs17508841 | 0.186 | 1.147 | 0.558 | -1.278 | 0.002 | -0.026 | -0.026 | 0.131 | 0.11 | 0.09 | 0.12 | 0 | 0.1 | 0.08 |
| 1 | 90574778 | rs10493836 | 0.23 | 1.268 | 0.912 | -1.025 | 0.003 | 0.02 | 0.02 | 0 | 0.06 | 0.08 | 0.1 | 0.11 | 0.08 | 0.1 |
| 1 | 90575147 | rs1329644 | 0.625 | 1.177 | 0.943 | -1.398 | -0.002 | 0.011 | 0.011 | 0 | 0.66 | 0.36 | 0.59 | 0.85 | 0.68 | 0.71 |
| 1 | 90577660 | rs10493835 | 0.357 | 1.196 | 0.925 | -1.372 | 0.042 | -0.065 | -0.065 | 0 | 0.11 | 0.26 | 0.14 | 0 | 0.1 | 0.07 |
| 1 | 90582562 | rs12409406 | 1.055 | 1.204 | -0.124 | -1.188 | 0.007 | 0.021 | 0.021 | 0.021 | 0.45 | 0.2 | 0.38 | 0.62 | 0.43 | 0.47 |
| 1 | 90583093 | rs12044528 | 1.76 | 1.115 | -0.547 | -0.868 | 0.023 | -0.048 | -0.048 | 0 | 0.23 | 0.05 | 0.13 | 0.2 | 0.19 | 0.11 |
| 1 | 90588306 | rs10922769 | -0.025 | 1.487 | 0.249 | -0.781 | -0.002 | -0.01 | -0.01 | 3.224 | 0.24 | 0.28 | 0.26 | 0.15 | 0.22 | 0.22 |
| 1 | 90589720 | rs6679194 | 0.529 | 1.514 | 0.207 | -0.656 | 0.012 | -0.045 | -0.045 | 0.032 | 0.66 | 0.53 | 0.59 | 0.85 | 0.66 | 0.71 |
| 1 | 90594990 | rs4456078 | 0.74 | 1.628 | -0.023 | -0.505 | 0.03 | -0.014 | -0.014 | 0 | 0.81 | 0.66 | 0.64 | 0.89 | 0.78 | 0.79 |
| 1 | 90595332 | rs10801786 | 0.283 | 1.485 | 0.09 | -0.405 | 0.034 | 0.001 | 0.001 | 0.267 | 0.84 | 0.67 | 0.79 | 0.89 | 0.84 | 0.84 |
| 1 | 90597198 | rs7552852 | 1.264 | 1.367 | -0.939 | -0.256 | 0.011 | -0.014 | -0.014 | 0.172 | 0.32 | 0.11 | 0.22 | 0.3 | 0.33 | 0.24 |
| 1 | 90598021 | rs12038587 | -1.27 | 1.81 | -0.354 | 0.154 | -0.004 | -0.006 | -0.006 | 0 | 0.4 | 0.36 | 0.53 | 0.55 | 0.41 | 0.55 |
| 1 | 90598884 | rs10922771 | -0.73 | 1.469 | -0.628 | 0.682 | -0.002 | -0.023 | -0.023 | 0 | 0.29 | 0.25 | 0.39 | 0.55 | 0.31 | 0.46 |
| 1 | 90600817 | rs7520257 | -0.92 | 1.589 | -0.392 | 0.282 | 0.018 | -0.014 | -0.014 | 0 | 0.31 | 0.46 | 0.44 | 0.74 | 0.33 | 0.51 |
| 1 | 90600934 | rs6428571 | -1.228 | 2.073 | 0.19 | -0.212 | 0.07 | -0.043 | -0.043 | 0.802 | 0.21 | 0.42 | 0.18 | 0.1 | 0.25 | 0.17 |
| 1 | 90601007 | rs6662919 | -1.246 | 1.796 | -0.293 | -1.01 | -0.008 | 0.003 | 0.003 | 0 | 0.3 | 0.26 | 0.24 | 0.06 | 0.35 | 0.21 |
| 1 | 90601121 | rs6428572 | -1.023 | 0.727 | -0.458 | -1.353 | -0.039 | -0.112 | -0.112 | 0 | 0.21 | 0.4 | 0.16 | 0.02 | 0.11 | 0.16 |
| 1 | 90601373 | rs4269759 | 0.113 | 0.971 | -1.712 | -1.287 | -0.084 | 0.096 | 0.096 | 0 | 0.6 | 0.86 | 0.54 | 0.21 | 0.58 | 0.47 |
| 1 | 90607718 | rs6428576 | -0.319 | 1.051 | -1.411 | -1.366 | -0.016 | -0.004 | -0.004 | 0 | 0.27 | 0.18 | 0.45 | 0.29 | 0.3 | 0.31 |
| 1 | 90612804 | rs12145344 | 0.956 | 0.85 | -2.265 | -1.875 | -0.093 | 0.137 | 0.137 | 0 | 0.21 | 0.01 | 0.13 | 0 | 0.32 | 0.09 |
| 1 | 90620142 | rs11800888 | 0.553 | -0.044 | -1.279 | -1.628 | 0.002 | -0.014 | -0.014 | 0 | 0.18 | 0.16 | 0.15 | 0.06 | 0.18 | 0.3 |
| 1 | 90624605 | rs10157978 | 0.63 | -0.259 | -0.693 | -1.715 | 0.009 | -0.006 | -0.006 | 0.166 | 0.29 | 0.29 | 0.22 | 0.64 | 0.17 | 0.26 |
| 1 | 90626011 | rs6679971 | 0.359 | -0.26 | -0.33 | -1.73 | 0.036 | 0.054 | 0.054 | 0 | 0.51 | 0.36 | 0.62 | 0.3 | 0.65 | 0.44 |
| 1 | 90626197 | rs3922968 | -0.181 | -0.405 | -0.157 | -1.77 | 0.001 | -0.096 | -0.096 | 0 | 0.76 | 0.52 | 0.83 | 0.94 | 0.81 | 0.68 |
| 1 | 90627000 | rs3922966 | 0.878 | -0.356 | -0.365 | -1.683 | 0.046 | -0.042 | -0.042 | 0 | 0.26 | 0.15 | 0.21 | 0.64 | 0.16 | 0.25 |
| 1 | 90631849 | rs4370755 | 0.512 | -0.252 | -0.38 | -1.637 | 0.024 | -0.014 | -0.014 | 0 | 0.29 | 0.23 | 0.23 | 0.64 | 0.18 | 0.26 |
| 1 | 90640945 | rs4388690 | -0.549 | -0.176 | -0.419 | -1.669 | 0.014 | 0.022 | 0.022 | 0 | 0.49 | 0.58 | 0.38 | 0.69 | 0.36 | 0.56 |
| 1 | 90642254 | rs6428580 | 0.125 | -0.464 | -0.457 | -1.18 | -0.011 | 0.003 | 0.003 | 0 | 0.3 | 0.37 | 0.23 | 0.64 | 0.18 | 0.26 |
| 1 | 90646872 | rs6675307 | -0.286 | -0.235 | -0.569 | -1.24 | -0.01 | -0.002 | -0.002 | 0 | 0.38 | 0.65 | 0.28 | 0.69 | 0.23 | 0.31 |
| 1 | 90647995 | rs12127618 | 0.172 | 0.741 | -1.383 | -1.478 | -0.1 | -0.055 | -0.055 | 0 | 0.2 | 0.08 | 0.45 | 0.26 | 0.34 | 0.25 |
| 1 | 90652038 | rs12036175 | -0.084 | 0.307 | -0.886 | -1.693 | -0.009 | -0.009 | -0.009 | 0.021 | 0.38 | 0.59 | 0.33 | 0.73 | 0.26 | 0.49 |
| 1 | 90652231 | rs10922802 | 0.463 | 0.247 | -0.775 | -1.474 | -0.055 | 0.015 | 0.015 | 0.007 | 0.31 | 0.48 | 0.28 | 0.69 | 0.16 | 0.28 |
| 1 | 90655967 | rs4330906 | -0.272 | 0.721 | -1.767 | -1.357 | -0.069 | -0.017 | -0.017 | 0 | 0.74 | 0.84 | 0.53 | 0.74 | 0.63 | 0.69 |
| 1 | 90657295 | rs10493837 | -0.066 | -0.101 | -1.496 | -0.97 | 0.054 | -0.04 | -0.04 | 0 | 0.32 | 0.12 | 0.23 | 0.68 | 0.18 | 0.28 |
| 1 | 90658051 | rs4415546 | 0.229 | 0.591 | -0.989 | -1.536 | 0 | -0.09 | -0.09 | 0 | 0.7 | 0.65 | 0.8 | 0.99 | 0.73 | 0.74 |
| 1 | 90659140 | rs4559484 | 0.476 | 0.254 | -1.571 | -0.566 | -0.017 | -0.03 | -0.03 | 0 | 0.26 | 0.04 | 0.24 | 0.3 | 0.24 | 0.31 |
| 1 | 90659565 | rs4626871 | -0.044 | 0.951 | -1.219 | -1.056 | 0.02 | -0.025 | -0.025 | 0.837 | 0.79 | 0.71 | 0.66 | 0.95 | 0.71 | 0.76 |
| 1 | 90660848 | rs6694119 | -0.013 | 0.896 | -1.366 | -0.996 | 0 | 0.088 | 0.088 | 0 | 0.31 | 0.12 | 0.29 | 0.12 | 0.26 | 0.25 |
| 1 | 90661711 | rs10922805 | -1.021 | 0.8 | -0.548 | -1.02 | 0.004 | 0.009 | 0.009 | 0 | 0.48 | 0.56 | 0.46 | 0.22 | 0.52 | 0.54 |
| 1 | 90664986 | rs17130997 | 1.392 | 0.573 | -0.514 | -0.987 | 0.01 | 0.013 | 0.013 | 0 | 0.31 | 0.16 | 0.37 | 0.77 | 0.29 | 0.36 |
| 1 | 90668105 | rs10754298 | -0.113 | 0.622 | -0.488 | -1.066 | -0.001 | -0.013 | -0.013 | 0 | 0.64 | 0.61 | 0.65 | 0.88 | 0.68 | 0.75 |
| 1 | 90671625 | rs7553296 | 1.615 | 0.38 | -0.432 | 0.219 | -0.002 | -0.028 | -0.028 | 0.418 | 0.4 | 0.12 | 0.34 | 0.61 | 0.36 | 0.4 |
| 1 | 90679734 | rs4495684 | -0.137 | 0.157 | -0.221 | 0.467 | 0.012 | 0.009 | 0.009 | 0 | 0.22 | 0.12 | 0.26 | 0.2 | 0.16 | 0.21 |
| 1 | 90686440 | rs12410130 | 0.154 | 0.077 | -0.12 | 0.427 | -0.003 | -0.007 | -0.007 | 0 | 0.25 | 0.13 | 0.27 | 0.21 | 0.26 | 0.23 |
| 1 | 90693880 | rs6428583 | -2.018 | 0.107 | 0.154 | 0.47 | 0.003 | -0.035 | -0.035 | 0 | 0.59 | 0.78 | 0.64 | 0.47 | 0.62 | 0.62 |
| 1 | 90721832 | rs4489539 | 0.605 | -0.252 | 0.333 | 0.058 | -0.003 | -0.004 | -0.004 | 0 | 0.13 | 0.09 | 0.03 | 0.22 | 0.03 | 0.11 |
| 1 | 90738800 | rs4233435 | -0.958 | 0.05 | 0.315 | 0.29 | -0.007 | 0.068 | 0.068 | 0 | 0.46 | 0.22 | 0.58 | 0.6 | 0.45 | 0.44 |
| 1 | 90739930 | rs4507991 | -2.136 | 0.063 | 0.486 | 0.26 | 0.004 | 0.039 | 0.039 | 0 | 0.59 | 0.76 | 0.68 | 0.64 | 0.62 | 0.62 |
| 1 | 90741571 | rs4417049 | -1.363 | 0.028 | 0.362 | 0.315 | 0.007 | 0.046 | 0.046 | 0 | 0.5 | 0.38 | 0.6 | 0.6 | 0.47 | 0.45 |
| 1 | 90750902 | rs7531149 | -1.074 | 0.105 | -0.027 | 0.361 | -0.008 | 0.07 | 0.07 | 0 | 0.46 | 0.22 | 0.58 | 0.6 | 0.46 | 0.47 |
| 1 | 90759618 | rs6689390 | -1.076 | 0.115 | -0.022 | 0.366 | -0.008 | 0.07 | 0.07 | 0 | 0.46 | 0.22 | 0.58 | 0.6 | 0.46 | 0.46 |
| 1 | 90761709 | rs12735216 | 0.711 | 0.412 | 0.03 | 0.557 | -0.037 | -0.016 | -0.016 | 0 | 0.08 | 0.02 | 0.11 | 0.05 | 0.14 | 0.13 |
| 1 | 90770566 | rs7517905 | -1.049 | 0.343 | 0.117 | 0.514 | -0.014 | -0.001 | -0.001 | 0 | 0.31 | 0.23 | 0.5 | 0.6 | 0.33 | 0.35 |
| 1 | 90776860 | rs12407088 | 0.749 | 0.442 | 0.064 | 0.568 | -0.036 | -0.017 | -0.017 | 0 | 0.08 | 0.03 | 0.11 | 0.04 | 0.14 | 0.13 |
| 1 | 90781761 | rs4446947 | -1.364 | 0.143 | 0.283 | 0.413 | 0.01 | 0.017 | 0.017 | 0 | 0.49 | 0.57 | 0.56 | 0.59 | 0.49 | 0.5 |
| 1 | 90783440 | rs7550206 | -0.159 | 0.084 | 0.326 | 0.514 | -0.004 | 0.004 | 0.004 | 0 | 0.23 | 0.16 | 0.3 | 0.21 | 0.22 | 0.21 |
| 1 | 90784635 | rs4565695 | 0.985 | 0.035 | 0.471 | 0.467 | 0.009 | -0.013 | -0.013 | 0 | 0.17 | 0.13 | 0.08 | 0 | 0.14 | 0.1 |
| 1 | 90785933 | rs7532538 | 0.679 | 0.089 | 0.516 | 0.53 | 0.05 | 0.297 | 0.297 | 0 | 0.08 | 0.28 | 0.18 | 0.38 | 0.12 | 0.19 |
| 1 | 90790141 | rs4453022 | -0.929 | 0.056 | 0.451 | 0.078 | 0.003 | 0.003 | 0.003 | 0 | 0.4 | 0.35 | 0.38 | 0.21 | 0.35 | 0.3 |
| 1 | 90791906 | rs7549113 | 0.891 | 0.25 | 0.298 | 0.48 | 0.009 | 0.01 | 0.01 | 0 | 0.13 | 0.24 | 0.11 | 0.2 | 0.16 | 0.21 |
| 1 | 90795388 | rs12093237 | -0.373 | 0.247 | -0.073 | 0.287 | -0.001 | 0.001 | 0.001 | 0 | 0.23 | 0.19 | 0.3 | 0.21 | 0.22 | 0.21 |
| 1 | 90801258 | rs7518543 | -1.718 | 0.421 | -0.12 | 0.148 | 0.015 | 0.101 | 0.101 | 0 | 0.6 | 0.79 | 0.78 | 0.88 | 0.66 | 0.67 |
| 1 | 90805127 | rs6674478 | -0.555 | 0.671 | 0.089 | 0.455 | 0.007 | -0.004 | -0.004 | 0.14 | 0.23 | 0.28 | 0.33 | 0.21 | 0.25 | 0.22 |
| 1 | 90813173 | rs12048345 | 0.472 | 0.209 | 0.189 | 0.717 | -0.003 | 0.137 | 0.137 | 0 | 0.06 | 0.16 | 0.04 | 0.26 | 0.04 | 0.09 |
| 1 | 90816109 | rs12141568 | -0.527 | 0.004 | 0.617 | 0.724 | -0.031 | 0.022 | 0.022 | 0 | 0.32 | 0.45 | 0.21 | 0.43 | 0.31 | 0.37 |
| 1 | 90818872 | rs10801803 | 0.166 | -0.14 | 0.542 | 0.528 | -0.003 | 0 | 0 | 0 | 0.18 | 0.19 | 0.09 | 0.26 | 0.16 | 0.17 |

| | | | | | | | | | | | | | | | | |
|---|---|---|---|---|---|---|---|---|---|---|---|---|---|---|---|---|
| 1 | 90819317 | rs10801804 | -1.179 | 0.039 | 0.684 | 0.668 | 0.008 | 0.019 | 0.019 | 0 | 0.32 | 0.42 | 0.34 | 0.55 | 0.35 | 0.48 |
| 1 | 90819447 | rs6689027 | -1.88 | -0.094 | 0.942 | 0.634 | 0.017 | 0.147 | 0.147 | 0 | 0.49 | 0.76 | 0.52 | 0.9 | 0.5 | 0.56 |
| 1 | 90820234 | rs9427994 | 0.852 | -0.092 | 1.195 | 0.689 | -0.026 | 0.061 | 0.061 | 0 | 0.14 | 0.2 | 0.17 | 0.35 | 0.13 | 0.08 |
| 1 | 90821175 | rs12022152 | -1.199 | 0.031 | 1.259 | 0.802 | 0.019 | 0.049 | 0.049 | 0 | 0.32 | 0.45 | 0.34 | 0.47 | 0.35 | 0.48 |
| 1 | 90824652 | rs7546709 | -0.723 | 0.063 | 1.199 | 0.506 | 0.012 | 0.004 | 0.004 | 0 | 0.27 | 0.35 | 0.32 | 0.3 | 0.34 | 0.38 |
| 1 | 90825173 | rs6428586 | -0.958 | 0.038 | 1.22 | 0.492 | 0.029 | 0.003 | 0.003 | 0 | 0.3 | 0.48 | 0.34 | 0.3 | 0.36 | 0.38 |
| 1 | 90825206 | rs6428588 | -2.387 | 0.245 | 0.895 | 0.121 | 0.014 | 0.131 | 0.131 | 0 | 0.62 | 0.81 | 0.78 | 0.91 | 0.68 | 0.67 |
| 1 | 90826932 | rs1396315 | -0.321 | -0.06 | 1.401 | 0.517 | -0.007 | 0.159 | 0.159 | 0 | 0.21 | 0.47 | 0.25 | 0.51 | 0.22 | 0.27 |
| 1 | 90829329 | rs2221660 | -0.285 | -0.052 | 1.416 | 0.41 | -0.001 | 0.166 | 0.166 | 0 | 0.21 | 0.49 | 0.25 | 0.53 | 0.22 | 0.27 |
| 1 | 90835677 | rs12127465 | 1.027 | -0.062 | 1.497 | 0.409 | -0.001 | 0 | 0 | 0 | 0.13 | 0.16 | 0.1 | 0.17 | 0.15 | 0.24 |
| 1 | 90836739 | rs9428050 | -2.329 | 0.061 | 1.383 | 0.292 | -0.011 | 0.101 | 0.101 | 0 | 0.58 | 0.75 | 0.73 | 0.91 | 0.61 | 0.66 |
| 1 | 90840268 | rs7522543 | 0.699 | 0.556 | 1.523 | 0.676 | -0.009 | 0.242 | 0.242 | 0 | 0.06 | 0.18 | 0.12 | 0.36 | 0.03 | 0.03 |
| 1 | 90854258 | rs17131031 | 0.503 | 0.638 | 1.366 | 0.586 | -0.016 | -0.016 | -0.016 | 0 | 0.08 | 0.07 | 0.2 | 0.16 | 0.13 | 0.13 |
| 1 | 90855529 | rs12752313 | 0.559 | 0.466 | 1.477 | 0.564 | 0.016 | -0.044 | -0.044 | 0 | 0.1 | 0.11 | 0.19 | 0 | 0.12 | 0.09 |
| 1 | 90860965 | rs10801810 | 0.183 | 0.378 | 1.485 | 0.821 | 0 | 0.1 | 0.1 | 0 | 0.8 | 0.63 | 0.77 | 0.47 | 0.78 | 0.72 |
| 1 | 90861006 | rs10754303 | 0.166 | 0.379 | 1.414 | 0.821 | -0.001 | 0.078 | 0.078 | 0 | 0.81 | 0.65 | 0.77 | 0.47 | 0.78 | 0.73 |
| 1 | 90872387 | rs2202787 | 0.087 | -0.112 | 1.26 | 0.674 | -0.023 | 0.004 | 0.004 | 0.052 | 0.18 | 0.22 | 0.11 | 0.26 | 0.09 | 0.16 |
| 1 | 90873828 | rs17131041 | 0.598 | -0.088 | 1.246 | 0.696 | -0.029 | 0.012 | 0.012 | 0 | 0.15 | 0.21 | 0.11 | 0.26 | 0.07 | 0.16 |
| 1 | 90909633 | rs12126321 | 0.684 | 0.281 | 1.124 | 0.78 | 0 | 0.001 | 0.001 | 0 | 0.14 | 0.17 | 0.11 | 0.17 | 0.18 | 0.24 |
| 1 | 90910074 | rs10922861 | 0.684 | 0.281 | 1.124 | 0.78 | 0 | 0.001 | 0.001 | 0 | 0.14 | 0.17 | 0.11 | 0.17 | 0.18 | 0.24 |
| 1 | 90916072 | rs1337576 | -0.738 | 0.161 | 1.205 | 0.708 | -0.024 | 0.002 | 0.002 | 0 | 0.34 | 0.43 | 0.23 | 0.41 | 0.28 | 0.43 |
| 1 | 90917395 | rs507821 | -1.554 | 0.43 | 0.724 | 0.269 | 0.007 | 0.034 | 0.034 | 0 | 0.45 | 0.65 | 0.55 | 0.93 | 0.44 | 0.57 |
| 1 | 90920022 | rs506368 | -0.955 | 0.341 | 0.756 | 0.314 | -0.007 | -0.005 | -0.005 | 0 | 0.37 | 0.61 | 0.34 | 0.77 | 0.31 | 0.44 |
| 1 | 90925451 | rs578509 | -1.252 | 0.396 | 0.639 | -0.289 | 0.016 | 0.071 | 0.071 | 0 | 0.46 | 0.71 | 0.55 | 0.93 | 0.43 | 0.57 |
| 1 | 90928816 | rs699539 | -1.548 | 0.351 | 0.308 | -0.424 | -0.019 | 0.133 | 0.133 | 0 | 0.56 | 0.78 | 0.74 | 0.94 | 0.54 | 0.65 |
| 1 | 90939247 | rs1337575 | 0.993 | 0.39 | 0.331 | -0.152 | -0.001 | -0.001 | -0.001 | 0 | 0.12 | 0.11 | 0.1 | 0.18 | 0.17 | 0.2 |
| 1 | 90940023 | rs699535 | 0.904 | 0.394 | 0.174 | -0.184 | -0.031 | 0.035 | 0.035 | 0 | 0.08 | 0.02 | 0.07 | 0 | 0.11 | 0.06 |
| 1 | 90943526 | rs7413520 | -0.703 | 0.246 | 0.835 | 0.074 | -0.014 | 0.001 | 0.001 | 0 | 0.34 | 0.46 | 0.46 | 0.38 | 0.24 | 0.37 |
| 1 | 90945085 | rs699531 | -1.988 | 0.323 | 0.613 | 0.019 | -0.015 | 0.126 | 0.126 | 0 | 0.59 | 0.8 | 0.76 | 0.93 | 0.57 | 0.67 |
| 1 | 90947969 | rs699529 | -1.035 | 0.248 | 0.478 | -0.188 | -0.007 | -0.004 | -0.004 | 0 | 0.41 | 0.54 | 0.4 | 0.6 | 0.41 | 0.41 |
| 1 | 90949962 | rs1591670 | -0.97 | 0.074 | 0.228 | 0.153 | 0 | -0.001 | -0.001 | 0 | 0.33 | 0.36 | 0.46 | 0.38 | 0.24 | 0.37 |
| 1 | 90953104 | rs12726749 | -0.438 | 0.364 | 0.231 | 0.311 | 0 | 0.02 | 0.02 | 0 | 0.18 | 0.21 | 0.38 | 0.34 | 0.18 | 0.3 |
| 1 | 90965565 | rs693260 | -0.611 | 0.313 | 0.318 | 0.038 | -0.017 | 0.086 | 0.086 | 0.294 | 0.23 | 0.46 | 0.26 | 0.43 | 0.2 | 0.13 |
| 1 | 90967306 | rs553314 | -2.407 | 0.352 | -0.496 | -0.486 | -0.119 | 0.161 | 0.161 | 0.482 | 0.67 | 0.98 | 0.8 | 0.94 | 0.64 | 0.74 |
| 1 | 90971259 | rs12065389 | 0.71 | 0.268 | -0.175 | -0.363 | 0 | -0.018 | -0.018 | 1.247 | 0.15 | 0.18 | 0.13 | 0.05 | 0.16 | 0.09 |
| 1 | 90973861 | rs527402 | -1.398 | 0.262 | -0.328 | -0.437 | -0.035 | 0.016 | 0.016 | 0.751 | 0.46 | 0.9 | 0.6 | 0.38 | 0.43 | 0.46 |
| 1 | 90980866 | rs699521 | -0.556 | 0.296 | 0.313 | -0.279 | -0.022 | 0.143 | 0.143 | 0.438 | 0.21 | 0.47 | 0.33 | 0.45 | 0.18 | 0.13 |
| 1 | 90994364 | rs954472 | 0.91 | 0.354 | 0.169 | -0.292 | -0.004 | 0.012 | 0.012 | 0 | 0.14 | 0.11 | 0.2 | 0.06 | 0.16 | 0.08 |
| 1 | 91012109 | rs10159380 | 1.717 | 0.332 | 0.412 | -0.47 | -0.006 | -0.031 | -0.031 | 0.189 | 0.1 | 0.11 | 0.05 | 0 | 0.09 | 0.07 |
| 1 | 91013805 | rs885141 | 0.906 | 0.418 | 0.372 | -0.409 | -0.004 | 0.012 | 0.012 | 0 | 0.14 | 0.11 | 0.2 | 0.06 | 0.16 | 0.08 |
| 1 | 91014198 | rs1336874 | 0.341 | 0.395 | 0.532 | -0.421 | 0.011 | -0.052 | -0.052 | 0 | 0.16 | 0.41 | 0.22 | 0.07 | 0.16 | 0.08 |
| 1 | 91018205 | rs17131084 | 1.385 | 0.329 | -0.02 | -0.439 | -0.037 | -0.041 | -0.041 | 0.637 | 0.13 | 0.01 | 0.09 | 0.16 | 0.17 | 0.21 |
| 1 | 91028707 | rs7546770 | -1.561 | 0.784 | -0.074 | -0.614 | -0.005 | 0.179 | 0.179 | 0.092 | 0.5 | 0.87 | 0.73 | 0.77 | 0.48 | 0.5 |
| 1 | 91045387 | rs12078288 | -0.61 | 0.737 | 0.313 | -0.526 | -0.009 | 0.032 | 0.032 | 0.02 | 0.33 | 0.59 | 0.43 | 0.45 | 0.31 | 0.2 |
| 1 | 91047423 | rs487440 | -2.702 | 1.369 | -0.33 | -0.717 | -0.019 | 0.131 | 0.131 | 0.545 | 0.7 | 0.99 | 0.9 | 0.94 | 0.78 | 0.77 |
| 1 | 91048510 | rs10922879 | 0.525 | 0.91 | 0.048 | -0.619 | -0.009 | -0.023 | -0.023 | 0 | 0.15 | 0.07 | 0.28 | 0.32 | 0.15 | 0.31 |
| 1 | 91049339 | rs12124829 | 1.686 | 0.943 | -0.101 | -0.583 | -0.038 | -0.041 | -0.041 | 0 | 0.13 | 0.01 | 0.09 | 0.16 | 0.17 | 0.2 |
| 1 | 91053707 | rs10922882 | -1.979 | 0.825 | 0.394 | -0.26 | -0.037 | 0.149 | 0.149 | 0.251 | 0.53 | 0.85 | 0.76 | 0.77 | 0.5 | 0.5 |
| 1 | 91061122 | rs6690040 | 0.027 | 0.867 | 0.043 | -0.181 | 0 | 0.005 | 0.005 | 0 | 0.18 | 0.18 | 0.3 | 0.32 | 0.18 | 0.32 |
| 1 | 91075849 | rs6667438 | -0.289 | 0.916 | -0.346 | -0.077 | 0 | 0 | 0 | 0.115 | 0.3 | 0.28 | 0.36 | 0.32 | 0.28 | 0.36 |
| 1 | 91077772 | rs1943448 | 0.503 | 1.235 | -1.153 | -0.65 | 0.009 | -0.017 | -0.017 | 2.308 | 0.16 | 0.19 | 0.25 | 0.07 | 0.2 | 0.09 |
| 1 | 91080115 | rs4537535 | -1.31 | 1.206 | -1.5 | -0.648 | -0.12 | 0.149 | 0.149 | 0.528 | 0.52 | 0.95 | 0.76 | 0.77 | 0.5 | 0.52 |
| 1 | 91081383 | rs6679215 | 0.096 | 1.401 | -0.544 | -0.355 | -0.006 | 0.007 | 0.007 | 0 | 0.15 | 0.13 | 0.25 | 0.07 | 0.2 | 0.09 |
| 1 | 91090125 | rs9287148 | -1.11 | 1.151 | -0.97 | -0.745 | -0.047 | 0.125 | 0.125 | 0.062 | 0.67 | 0.9 | 0.82 | 0.92 | 0.64 | 0.75 |
| 1 | 91096973 | rs12116789 | -1.321 | 1.061 | -1.043 | -0.651 | -0.1 | 0.115 | 0.115 | 0.082 | 0.69 | 0.97 | 0.8 | 0.91 | 0.69 | 0.75 |
| 1 | 91099790 | rs7511891 | 1.008 | 0.959 | -1.031 | -1.081 | -0.001 | 0.006 | 0.006 | 0 | 0.14 | 0.06 | 0.07 | 0 | 0.13 | 0.06 |
| 1 | 91099903 | rs4233429 | -1.853 | 1.02 | -1.34 | -1.237 | -0.107 | 0.125 | 0.125 | 0 | 0.73 | 0.98 | 0.85 | 0.94 | 0.73 | 0.77 |
| 1 | 91102694 | rs11581245 | 0.505 | 1.139 | -0.258 | -0.691 | 0.006 | -0.029 | -0.029 | 0.319 | 0.21 | 0.38 | 0.2 | 0.21 | 0.21 | 0.23 |
| 1 | 91103227 | rs2212940 | -1.143 | 1.189 | -0.518 | -0.017 | -0.005 | -0.024 | -0.024 | 0.571 | 0.33 | 0.23 | 0.54 | 0.5 | 0.39 | 0.3 |
| 1 | 91103885 | rs9787186 | 0.119 | 1.156 | -0.372 | 0.753 | -0.019 | -0.062 | -0.062 | 0.398 | 0.15 | 0.05 | 0.34 | 0.37 | 0.22 | 0.19 |
| 1 | 91107981 | rs2797293 | 0.488 | 0.865 | 0.162 | 0.298 | 0.001 | 0.074 | 0.074 | 0 | 0.05 | 0.14 | 0.13 | 0.13 | 0.05 | 0.05 |
| 1 | 91113178 | rs2970544 | -0.092 | 1.031 | -0.856 | -0.148 | 0.001 | -0.003 | -0.003 | 0 | 0.82 | 0.81 | 0.82 | 0.8 | 0.79 | 0.78 |
| 1 | 91120020 | rs1029197 | -0.452 | 0.06 | -0.596 | -1.206 | -0.006 | -0.073 | -0.073 | 0 | 0.2 | 0.36 | 0.24 | 0.06 | 0.23 | 0.16 |
| 1 | 91122149 | rs4658083 | -1.135 | -0.045 | -0.187 | -1.234 | -0.008 | -0.042 | -0.042 | 0 | 0.92 | 0.8 | 0.94 | 1 | 0.93 | 0.98 |
| 1 | 91136463 | rs12354259 | -0.2 | -0.056 | -0.165 | -1.657 | 0.004 | -0.061 | -0.061 | 0 | 0.4 | 0.5 | 0.34 | 0.13 | 0.4 | 0.37 |
| 1 | 91137596 | rs6680305 | 0.988 | -0.086 | -0.151 | -1.698 | -0.009 | 0.024 | 0.024 | 0 | 0.26 | 0.19 | 0.15 | 0.08 | 0.25 | 0.25 |
| 1 | 91141196 | rs12116456 | 1.235 | -0.163 | -0.158 | -1.714 | 0.002 | -0.002 | -0.002 | 0 | 0.06 | 0.07 | 0.16 | 0.06 | 0.11 | 0.1 |
| 1 | 91144055 | rs10922899 | 0.837 | -0.303 | -0.329 | -1.162 | -0.094 | -0.034 | -0.034 | 0.02 | 0.45 | 0.04 | 0.43 | 0.41 | 0.43 | 0.45 |
| 1 | 91149594 | rs6679774 | 0.286 | -0.142 | -0.129 | -0.943 | 0 | -0.007 | -0.007 | 0.063 | 0.12 | 0.14 | 0.18 | 0.06 | 0.15 | 0.13 |

| | | | | | | | | | | | | | | | | |
|---|---|---|---|---|---|---|---|---|---|---|---|---|---|---|---|---|
| 1 | 91154551 | rs6428597 | 0.999 | -0.083 | -0.283 | -0.937 | -0.008 | 0.022 | 0.022 | 0 | 0.26 | 0.19 | 0.15 | 0.08 | 0.25 | 0.25 |
| 1 | 91158104 | rs10493842 | -0.359 | -0.083 | -0.465 | -0.932 | 0.006 | 0.155 | 0.155 | 0 | 0.21 | 0.44 | 0.37 | 0.5 | 0.25 | 0.3 |
| 1 | 91165138 | rs7519925 | 0.798 | 0.074 | -1.009 | -1.122 | -0.002 | -0.006 | -0.006 | 0 | 0.26 | 0.22 | 0.14 | 0.07 | 0.25 | 0.25 |
| 1 | 91173988 | rs1335720 | 1.011 | 0.147 | -1.508 | -1.065 | -0.032 | 0.086 | 0.086 | 0 | 0.23 | 0.07 | 0.25 | 0.06 | 0.25 | 0.18 |
| 1 | 91175419 | rs12133004 | 1.108 | 0.434 | -1.099 | -0.629 | 0.001 | -0.002 | -0.002 | 0 | 0.06 | 0.06 | 0.16 | 0.05 | 0.11 | 0.09 |
| 1 | 91179699 | rs17478233 | 0.623 | 0.185 | -1.265 | -0.782 | -0.037 | 0.097 | 0.097 | 0 | 0.25 | 0.07 | 0.13 | 0.07 | 0.25 | 0.25 |
| 1 | 91189933 | rs12089815 | -0.238 | -0.701 | -0.933 | -0.363 | -0.029 | 0.048 | 0.048 | 0 | 0.55 | 0.76 | 0.53 | 0.63 | 0.5 | 0.6 |
| 1 | 91198676 | rs1342780 | 0.255 | -0.733 | -0.575 | -0.248 | 0 | 0.005 | 0.005 | 0.008 | 0.52 | 0.44 | 0.38 | 0.43 | 0.43 | 0.54 |
| 1 | 91199862 | rs10922910 | 0.949 | -0.725 | -1.161 | -0.597 | -0.059 | 0.008 | 0.008 | 0 | 0.25 | 0.02 | 0.22 | 0.19 | 0.22 | 0.3 |
| 1 | 91203929 | rs6689845 | -1.096 | -0.348 | -0.436 | -0.566 | -0.005 | -0.034 | -0.034 | 0.301 | 0.93 | 0.86 | 0.95 | 1 | 0.92 | 0.98 |
| 1 | 91206170 | rs10801825 | -0.321 | -0.025 | -0.792 | -0.148 | -0.015 | -0.001 | -0.001 | 0 | 0.36 | 0.27 | 0.46 | 0.38 | 0.4 | 0.4 |
| 1 | 91209986 | rs1526480 | -0.493 | -0.03 | -0.766 | -0.151 | 0.003 | -0.002 | -0.002 | 0.418 | 0.37 | 0.42 | 0.47 | 0.38 | 0.41 | 0.4 |
| 1 | 91212216 | rs6699397 | -0.394 | 0.054 | -1.514 | -0.111 | -0.033 | -0.01 | -0.01 | 3.862 | 0.35 | 0.18 | 0.45 | 0.38 | 0.4 | 0.4 |
| 1 | 91219359 | rs6656775 | -0.19 | 0.63 | -1.266 | 0.37 | -0.012 | -0.011 | -0.011 | 0 | 0.27 | 0.24 | 0.44 | 0.37 | 0.34 | 0.29 |
| 1 | 91223871 | rs3856224 | -0.6 | 0.716 | -1.192 | 0.368 | -0.007 | -0.007 | -0.007 | 0 | 0.3 | 0.29 | 0.44 | 0.38 | 0.35 | 0.3 |
| 1 | 91225594 | rs4658172 | -0.115 | 1.195 | -1.398 | 0.784 | -0.03 | -0.037 | -0.037 | 0 | 0.23 | 0.15 | 0.42 | 0.38 | 0.31 | 0.27 |
| 1 | 91226447 | rs6696987 | -0.583 | 0.431 | -0.839 | 1.069 | 0.002 | 0.037 | 0.037 | 0 | 0.07 | 0.12 | 0.11 | 0.18 | 0.07 | 0.06 |
| 1 | 91234966 | rs10922921 | 0.239 | 0.168 | -1.43 | 0.674 | 0.025 | 0.01 | 0.01 | 0.614 | 0.55 | 0.3 | 0.6 | 0.43 | 0.58 | 0.43 |
| 1 | 91235489 | rs12030148 | -0.739 | -0.607 | -0.69 | 0.578 | -0.01 | -0.037 | -0.037 | 1.316 | 0.23 | 0.39 | 0.2 | 0.17 | 0.2 | 0.28 |
| 1 | 91235929 | rs6428604 | 0.405 | -1.031 | -0.519 | 0.262 | -0.008 | -0.029 | -0.029 | 0 | 0.76 | 0.63 | 0.86 | 0.86 | 0.78 | 0.88 |
| 1 | 91242589 | rs860943 | -0.39 | -1.1 | -0.076 | 0.23 | 0.042 | 0.019 | 0.019 | 1.912 | 0.06 | 0.23 | 0.07 | 0.04 | 0.07 | 0.02 |
| 1 | 91243376 | rs3843305 | 1.02 | -1.656 | -0.928 | 0.13 | -0.108 | 0.054 | 0.054 | 0 | 0.34 | 0.14 | 0.44 | 0.25 | 0.45 | 0.36 |
| 1 | 91244137 | rs698275 | -0.772 | -1.861 | -0.908 | 0.02 | 0.011 | -0.017 | -0.017 | 1.187 | 0.1 | 0.25 | 0.08 | 0.05 | 0.09 | 0.04 |
| 1 | 91244177 | rs698274 | 0.682 | -1.866 | -0.912 | 0.022 | -0.012 | 0.022 | 0.022 | 0 | 0.41 | 0.35 | 0.48 | 0.29 | 0.47 | 0.4 |
| 1 | 91245109 | rs3856226 | -0.338 | -2.386 | -0.407 | -0.053 | 0.001 | 0.012 | 0.012 | 0 | 0.42 | 0.35 | 0.33 | 0.27 | 0.45 | 0.43 |
| 1 | 91251325 | rs3904678 | 0.568 | -2.524 | -0.43 | -0.071 | -0.006 | 0.009 | 0.009 | 0 | 0.26 | 0.19 | 0.27 | 0.15 | 0.33 | 0.33 |
| 1 | 91251381 | rs712026 | -0.643 | -2.589 | -0.373 | -0.091 | -0.002 | 0.017 | 0.017 | 0 | 0.45 | 0.54 | 0.36 | 0.28 | 0.49 | 0.44 |
| 1 | 91252863 | rs10922924 | 1.997 | -2.6 | -0.524 | 0.025 | -0.122 | 0.082 | 0.082 | 0 | 0.3 | 0.05 | 0.4 | 0.19 | 0.38 | 0.35 |
| 1 | 91255484 | rs1725375 | -0.844 | -2.561 | -0.182 | 0.076 | -0.039 | 0.097 | 0.097 | 0 | 0.83 | 0.73 | 0.81 | 0.51 | 0.92 | 0.83 |
| 1 | 91258572 | rs1725390 | 0.382 | -2.517 | -0.24 | -0.304 | -0.004 | -0.013 | -0.013 | 2.121 | 0.56 | 0.51 | 0.53 | 0.35 | 0.59 | 0.49 |
| 1 | 91260076 | rs1725384 | -0.489 | -2.632 | -0.214 | -0.367 | -0.024 | -0.055 | -0.055 | 0 | 0.11 | 0.38 | 0.07 | 0.02 | 0.06 | 0.03 |
| 1 | 91260316 | rs1775580 | 0.31 | -2.567 | -0.196 | -0.369 | -0.005 | -0.02 | -0.02 | 0.358 | 0.57 | 0.52 | 0.53 | 0.35 | 0.59 | 0.49 |
| 1 | 91264138 | rs2120802 | -0.765 | -2.387 | 0.028 | 0.434 | 0 | -0.002 | -0.002 | 0 | 0.39 | 0.58 | 0.43 | 0.41 | 0.39 | 0.37 |
| 1 | 91265088 | rs10922926 | 0.347 | -2.371 | -0.06 | 0.414 | -0.002 | -0.003 | -0.003 | 0.61 | 0.27 | 0.21 | 0.36 | 0.39 | 0.33 | 0.34 |
| 1 | 91265543 | rs1997155 | 1.02 | -2.441 | -0.502 | 0.37 | 0.009 | 0.147 | 0.147 | 0.076 | 0.5 | 0.08 | 0.47 | 0.33 | 0.55 | 0.46 |
| 1 | 91279915 | rs1335718 | 0.509 | -2.378 | -0.449 | 0.366 | 0.004 | -0.006 | -0.006 | 0.462 | 0.6 | 0.3 | 0.56 | 0.6 | 0.61 | 0.64 |
| 1 | 91293921 | rs1358156 | 0.495 | -2.104 | -0.356 | 0.374 | -0.009 | 0.075 | 0.075 | 0.017 | 0.12 | 0.21 | 0.14 | 0.26 | 0.11 | 0.17 |
| 1 | 91301807 | rs1766167 | -0.066 | -2.115 | -0.537 | 0.219 | -0.003 | 0.003 | 0.003 | 0 | 0.31 | 0.33 | 0.21 | 0.39 | 0.28 | 0.32 |
| 1 | 91304636 | rs1188937 | -0.745 | -2.094 | -0.42 | 0.251 | -0.001 | 0 | 0 | 0 | 0.35 | 0.62 | 0.39 | 0.4 | 0.33 | 0.34 |
| 1 | 91308396 | rs1188931 | -0.697 | -2.063 | -0.527 | 0.261 | -0.004 | 0.006 | 0.006 | 0 | 0.34 | 0.57 | 0.39 | 0.4 | 0.34 | 0.34 |
| 1 | 91311423 | rs1188929 | -0.372 | -1.893 | -1.059 | 0.29 | -0.015 | -0.04 | -0.04 | 0 | 0.1 | 0.27 | 0.07 | 0.02 | 0.07 | 0.03 |
| 1 | 91321687 | rs12737988 | 0.425 | -1.739 | -2.071 | 0.421 | -0.028 | -0.074 | -0.074 | 0.779 | 0.22 | 0.05 | 0.29 | 0.38 | 0.27 | 0.32 |
| 1 | 91325323 | rs12046172 | 0.929 | -1.393 | -1.862 | 0.06 | -0.098 | 0.084 | 0.084 | 0 | 0.34 | 0.12 | 0.4 | 0.21 | 0.39 | 0.33 |
| 1 | 91339437 | rs1616696 | -0.742 | -1.508 | -1.457 | 0.245 | 0.019 | 0.018 | 0.018 | 0 | 0.83 | 0.95 | 0.95 | 0.86 | 0.87 | 0.86 |
| 1 | 91352818 | rs17481838 | 0.571 | -1.528 | -1.86 | 0.048 | -0.053 | 0.021 | 0.021 | 0 | 0.14 | 0.01 | 0.14 | 0.08 | 0.16 | 0.21 |
| 1 | 91364568 | rs1188912 | -0.961 | -1.517 | -1.784 | 0.172 | 0 | -0.003 | -0.003 | 0 | 0.83 | 0.99 | 0.95 | 0.86 | 0.87 | 0.86 |
| 1 | 91380305 | rs1188951 | 0.604 | -1.535 | -1.764 | 0.246 | 0.022 | 0.004 | 0.004 | 0 | 0.18 | 0.06 | 0.05 | 0.16 | 0.14 | 0.15 |
| 1 | 91380440 | rs12726261 | 0.069 | -1.886 | -1.828 | -0.01 | -0.009 | 0.07 | 0.07 | 0 | 0.2 | 0.08 | 0.17 | 0.07 | 0.2 | 0.22 |
| 1 | 91383589 | rs2448020 | 0.769 | -1.883 | -1.831 | -0.008 | 0.022 | 0.004 | 0.004 | 0 | 0.18 | 0.06 | 0.05 | 0.16 | 0.14 | 0.15 |
| 1 | 91384826 | rs10801834 | 1.97 | -1.244 | -2.053 | -0.271 | -0.12 | 0.111 | 0.111 | 0 | 0.33 | 0.1 | 0.42 | 0.18 | 0.41 | 0.31 |
| 1 | 91393027 | rs12724992 | 0.282 | -1.242 | -1.787 | -0.239 | 0.001 | 0.05 | 0.05 | 0 | 0.21 | 0.11 | 0.18 | 0.07 | 0.2 | 0.24 |
| 1 | 91395394 | rs358695 | -0.614 | -1.302 | -0.922 | 0.378 | -0.004 | 0.116 | 0.116 | 0.23 | 0.84 | 0.77 | 0.76 | 0.63 | 0.82 | 0.84 |
| 1 | 91402075 | rs358694 | -1.959 | -1.319 | -1.367 | 0.466 | -0.017 | -0.011 | -0.011 | 0 | 0.94 | 1 | 0.85 | 0.7 | 0.89 | 0.91 |
| 1 | 91418026 | rs497891 | 1.27 | -1.226 | -1.118 | -0.328 | 0.011 | 0.004 | 0.004 | 0 | 0.18 | 0.16 | 0.05 | 0.16 | 0.14 | 0.15 |
| 1 | 91444192 | rs4658090 | 1.989 | -1.263 | -1.096 | -0.4 | -0.118 | 0.111 | 0.111 | 0 | 0.33 | 0.1 | 0.42 | 0.18 | 0.41 | 0.32 |
| 1 | 91444533 | rs358689 | 1.894 | -1.235 | -1.052 | -0.386 | -0.076 | 0.096 | 0.096 | 0 | 0.34 | 0.17 | 0.43 | 0.18 | 0.41 | 0.32 |
| 1 | 91456926 | rs358684 | -0.235 | -1.111 | -0.836 | -0.254 | -0.022 | 0.082 | 0.082 | 0.039 | 0.78 | 0.64 | 0.69 | 0.42 | 0.77 | 0.72 |
| 1 | 91460362 | rs643854 | 1.103 | -1.023 | -1.047 | -0.287 | 0.009 | 0.003 | 0.003 | 0 | 0.18 | 0.15 | 0.05 | 0.16 | 0.14 | 0.15 |
| 1 | 91471041 | rs356349 | -0.707 | -1.042 | -1.069 | -0.306 | 0.023 | 0.052 | 0.052 | 0 | 0.39 | 0.27 | 0.23 | 0.24 | 0.34 | 0.38 |
| 1 | 91475729 | rs17131254 | 1.137 | -1.027 | -1.286 | -0.284 | 0.017 | 0.005 | 0.005 | 0 | 0.18 | 0.1 | 0.05 | 0.16 | 0.14 | 0.15 |
| 1 | 91480823 | rs6428606 | 0.115 | -0.889 | -1.245 | -0.115 | -0.019 | 0.072 | 0.072 | 0 | 0.2 | 0.04 | 0.17 | 0.07 | 0.2 | 0.23 |
| 1 | 91504195 | rs355644 | 0.155 | -0.53 | -0.803 | 0.255 | -0.002 | 0.115 | 0.115 | 0.138 | 0.77 | 0.63 | 0.73 | 0.51 | 0.76 | 0.81 |
| 1 | 91513968 | rs354623 | -0.626 | -0.301 | -1.422 | -0.201 | 0.014 | 0.042 | 0.042 | 0 | 0.39 | 0.3 | 0.25 | 0.23 | 0.36 | 0.36 |
| 1 | 91517927 | rs164992 | -0.563 | -0.267 | -1.759 | -0.184 | 0.017 | 0.067 | 0.067 | 0 | 0.38 | 0.23 | 0.24 | 0.24 | 0.36 | 0.36 |
| 1 | 91519832 | rs164996 | -0.338 | 0.088 | -1.621 | 0.222 | -0.018 | -0.005 | -0.005 | 0 | 0.25 | 0.09 | 0.14 | 0.22 | 0.28 | 0.2 |
| 1 | 91524208 | rs4658186 | -0.694 | 0.542 | -1.711 | 0.055 | -0.125 | 0.041 | 0.041 | 0 | 0.66 | 0.91 | 0.54 | 0.73 | 0.57 | 0.48 |
| 1 | 91528186 | rs165277 | -0.74 | 0.074 | -0.724 | 0.089 | 0 | -0.002 | -0.002 | 0.258 | 0.18 | 0.14 | 0.1 | 0.09 | 0.15 | 0.09 |
| 1 | 91529589 | rs469846 | -0.431 | -0.304 | -0.756 | 0 | 0.01 | 0.01 | 0.01 | 0 | 0.11 | 0.07 | 0.19 | 0.07 | 0.07 | 0.16 |
| 1 | 91530305 | rs469772 | -1.225 | 0.046 | -0.418 | 0.459 | 0 | -0.001 | -0.001 | 0 | 0.21 | 0.19 | 0.12 | 0.15 | 0.19 | 0.09 |

| | | | | | | | | | | | | | | | | |
|---|---|---|---|---|---|---|---|---|---|---|---|---|---|---|---|---|
| 1 | 91536485 | rs164989 | 0.808 | -0.039 | -0.577 | 0.41 | -0.017 | 0.042 | 0.042 | 0 | 0.46 | 0.23 | 0.56 | 0.39 | 0.54 | 0.42 |
| 1 | 91541513 | rs346932 | -0.252 | -0.1 | 0.103 | 0.549 | -0.024 | -0.002 | -0.002 | 0 | 0.14 | 0.34 | 0.07 | 0.09 | 0.08 | 0.02 |
| 1 | 91550812 | rs164911 | -0.487 | -0.259 | 0.294 | 0.711 | -0.037 | 0.056 | 0.056 | 1.169 | 0.16 | 0.46 | 0.09 | 0.16 | 0.09 | 0.01 |
| 1 | 91550896 | rs12723509 | 0.648 | 0.018 | -0.302 | 1.114 | -0.044 | -0.056 | -0.056 | 0 | 0.1 | 0.01 | 0.14 | 0.2 | 0.14 | 0.23 |
| 1 | 91556690 | rs11582243 | 1.458 | 0.389 | -0.606 | 0.582 | -0.002 | 0 | 0 | 0 | 0.18 | 0.21 | 0.23 | 0.13 | 0.23 | 0.11 |
| 1 | 91558038 | rs972054 | -1.214 | -0.207 | -0.643 | 0.01 | -0.009 | -0.081 | -0.081 | 1.941 | 0.73 | 0.61 | 0.66 | 0.88 | 0.74 | 0.72 |
| 1 | 91558057 | rs10922953 | 0.252 | -0.28 | -0.844 | 0.231 | -0.006 | -0.026 | -0.026 | 0 | 0.33 | 0.14 | 0.17 | 0.34 | 0.29 | 0.34 |
| 1 | 91566457 | rs164983 | -0.416 | -0.516 | -0.158 | 0.334 | 0.008 | 0.01 | 0.01 | 0 | 0.66 | 0.77 | 0.64 | 0.71 | 0.67 | 0.7 |
| 1 | 91567210 | rs164893 | 1.071 | -0.498 | -0.049 | 0.407 | -0.015 | 0.027 | 0.027 | 0 | 0.14 | 0.16 | 0.14 | 0.25 | 0.09 | 0.21 |
| 1 | 91569689 | rs164898 | -0.599 | 0.072 | -0.264 | 0.224 | -0.005 | 0.061 | 0.061 | 0 | 0.72 | 0.85 | 0.67 | 0.87 | 0.69 | 0.71 |
| 1 | 91581942 | rs165291 | 0.425 | 0.16 | 0.007 | 0.552 | -0.007 | 0.006 | 0.006 | 0 | 0.44 | 0.5 | 0.41 | 0.47 | 0.37 | 0.38 |
| 1 | 91586000 | rs240504 | -0.071 | 0.357 | -0.361 | 0.402 | -0.048 | -0.005 | -0.005 | 0 | 0.39 | 0.2 | 0.37 | 0.41 | 0.46 | 0.49 |
| 1 | 91602551 | rs346994 | -0.44 | 0.167 | 0.045 | 0.699 | -0.022 | 0.098 | 0.098 | 0 | 0.78 | 0.48 | 0.65 | 0.64 | 0.8 | 0.88 |
| 1 | 91604295 | rs1507350 | 1.388 | 0.47 | -0.609 | 0.534 | -0.031 | 0.063 | 0.063 | 0.001 | 0.19 | 0.03 | 0.17 | 0.09 | 0.23 | 0.21 |
| 1 | 91605491 | rs347004 | 0.035 | 0.461 | -0.214 | 0.57 | -0.007 | 0.004 | 0.004 | 0 | 0.6 | 0.55 | 0.49 | 0.55 | 0.57 | 0.67 |
| 1 | 91618766 | rs347027 | 1.885 | 0.328 | -0.308 | 0.446 | -0.003 | 0.028 | 0.028 | 0 | 0.31 | 0.16 | 0.27 | 0.21 | 0.28 | 0.35 |
| 1 | 91618793 | rs347028 | 1.078 | 0.311 | -0.347 | 0.428 | -0.019 | 0.123 | 0.123 | 0.067 | 0.5 | 0.18 | 0.44 | 0.29 | 0.51 | 0.56 |
| 1 | 91619300 | rs982009 | 0.441 | 0.581 | -0.187 | 0.579 | 0.104 | 0.058 | 0.058 | 0.2 | 0.1 | 0.36 | 0.08 | 0.1 | 0.11 | 0.05 |
| 1 | 91620228 | rs613217 | 0.768 | 0.446 | -0.417 | 0.584 | 0.011 | 0.026 | 0.026 | 0 | 0.24 | 0.16 | 0.23 | 0.14 | 0.22 | 0.12 |
| 1 | 91621607 | rs969618 | -0.083 | 0.443 | -0.403 | 0.582 | 0.015 | 0.065 | 0.065 | 0 | 0.56 | 0.7 | 0.59 | 0.77 | 0.55 | 0.67 |
| 1 | 91622626 | rs10922966 | 1.176 | 0.955 | -0.01 | 1.036 | -0.037 | -0.072 | -0.072 | 0 | 0.13 | 0.01 | 0.14 | 0.24 | 0.14 | 0.26 |
| 1 | 91623097 | rs983040 | -0.843 | 0.964 | 0.02 | 1.007 | 0.005 | 0.019 | 0.019 | 0 | 0.26 | 0.65 | 0.34 | 0.57 | 0.27 | 0.32 |
| 1 | 91627098 | rs613176 | 0.791 | 0.964 | -0.253 | 0.971 | -0.031 | 0.041 | 0.041 | 0 | 0.55 | 0.29 | 0.49 | 0.35 | 0.51 | 0.47 |
| 1 | 91627379 | rs17482910 | 1.514 | 0.912 | -0.123 | 0.958 | -0.002 | 0.044 | 0.044 | 0 | 0.19 | 0.06 | 0.18 | 0.09 | 0.22 | 0.21 |
| 1 | 91628655 | rs404716 | -0.434 | 0.95 | -0.017 | 1.137 | 0.059 | 0.192 | 0.192 | 0.216 | 0.12 | 0.6 | 0.19 | 0.32 | 0.13 | 0.07 |
| 1 | 91631091 | rs384642 | -0.229 | 0.764 | -0.323 | 1.012 | 0.009 | 0.04 | 0.04 | 0.272 | 0.23 | 0.66 | 0.39 | 0.49 | 0.26 | 0.15 |
| 1 | 91633434 | rs620760 | -0.382 | 0.695 | 0.066 | 1.098 | 0.035 | 0.097 | 0.097 | 0 | 0.83 | 0.55 | 0.8 | 0.74 | 0.81 | 0.71 |
| 1 | 91647150 | rs1454328 | 0.769 | 0.758 | -0.341 | 1.068 | -0.004 | 0.019 | 0.019 | 0.002 | 0.22 | 0.29 | 0.27 | 0.32 | 0.24 | 0.22 |
| 1 | 91657763 | rs12040260 | 0.081 | 0.862 | -0.423 | 1.376 | -0.049 | -0.072 | -0.072 | 0 | 0.14 | 0.01 | 0.14 | 0.23 | 0.17 | 0.27 |
| 1 | 91658047 | rs584538 | 0.07 | 0.743 | 0.465 | 1.399 | 0.011 | 0.065 | 0.065 | 0 | 0.84 | 0.71 | 0.82 | 0.73 | 0.83 | 0.71 |
| 1 | 91658920 | rs648094 | -0.878 | 0.712 | -0.33 | 1.351 | -0.024 | -0.044 | -0.044 | 0 | 0.35 | 0.22 | 0.46 | 0.49 | 0.39 | 0.41 |
| 1 | 91659939 | rs10801844 | 1.063 | 0.556 | -0.204 | 1.044 | -0.001 | 0.013 | 0.013 | 0 | 0.57 | 0.41 | 0.48 | 0.33 | 0.56 | 0.57 |
| 1 | 91661385 | rs2089432 | -0.737 | 0.862 | -0.207 | 0.985 | -0.008 | -0.011 | -0.011 | 0 | 0.19 | 0.15 | 0.18 | 0.28 | 0.22 | 0.31 |
| 1 | 91670375 | rs721297 | 0.983 | 0.442 | -0.505 | 0.395 | -0.023 | 0.019 | 0.019 | 0 | 0.46 | 0.21 | 0.36 | 0.36 | 0.54 | 0.5 |
| 1 | 91675675 | rs674135 | -0.421 | 0.603 | 0.194 | 0.212 | -0.008 | 0.032 | 0.032 | 0 | 0.33 | 0.43 | 0.44 | 0.53 | 0.28 | 0.43 |
| 1 | 91695097 | rs11487636 | 0.181 | 1.088 | 0.086 | -0.175 | 0 | -0.001 | -0.001 | 0 | 0.22 | 0.24 | 0.26 | 0.18 | 0.28 | 0.18 |
| 1 | 91708513 | rs605146 | -0.604 | 0.953 | 0.35 | -0.346 | -0.012 | 0.042 | 0.042 | 0 | 0.34 | 0.47 | 0.48 | 0.53 | 0.3 | 0.41 |
| 1 | 91716979 | rs615020 | 1.416 | 0.984 | 0.203 | -0.21 | -0.008 | -0.005 | -0.005 | 0.644 | 0.15 | 0.14 | 0.14 | 0.01 | 0.19 | 0.12 |
| 1 | 91730294 | rs10801849 | 0.328 | 0.864 | 0.314 | -0.208 | -0.002 | 0.05 | 0.05 | 0 | 0.23 | 0.26 | 0.39 | 0.53 | 0.22 | 0.38 |
| 1 | 91730806 | rs681589 | 0.206 | 0.904 | 0.243 | -0.196 | 0.008 | -0.009 | -0.009 | 0 | 0.22 | 0.28 | 0.26 | 0.17 | 0.28 | 0.18 |
| 1 | 91742055 | rs281959 | 0.015 | 0.919 | 0.234 | -0.205 | 0.004 | -0.005 | -0.005 | 0 | 0.23 | 0.27 | 0.27 | 0.17 | 0.31 | 0.18 |
| 1 | 91747006 | rs10801851 | -1.061 | 0.98 | 0.156 | -0.272 | 0.003 | -0.016 | -0.016 | 0.14 | 0.43 | 0.68 | 0.54 | 0.53 | 0.4 | 0.45 |
| 1 | 91774858 | rs1158437 | 1.368 | 0.922 | 0.134 | -0.365 | -0.016 | 0.006 | 0.006 | 0 | 0.15 | 0.12 | 0.16 | 0.01 | 0.23 | 0.12 |
| 1 | 91781413 | rs281992 | -0.457 | 0.899 | 0.148 | -0.362 | 0.002 | -0.002 | -0.002 | 0 | 0.77 | 0.75 | 0.72 | 0.82 | 0.69 | 0.82 |
| 1 | 91782031 | rs11584478 | 1.212 | 0.96 | 0.176 | -0.333 | -0.015 | 0.005 | 0.005 | 0 | 0.16 | 0.12 | 0.16 | 0.01 | 0.23 | 0.12 |
| 1 | 91784088 | rs281997 | -0.448 | 0.967 | 0.172 | -0.316 | 0.004 | -0.003 | -0.003 | 0 | 0.77 | 0.75 | 0.72 | 0.81 | 0.69 | 0.82 |
| 1 | 91784887 | rs10493845 | 0.186 | 0.964 | 0.178 | -0.318 | 0.003 | -0.004 | -0.004 | 0 | 0.22 | 0.25 | 0.28 | 0.18 | 0.31 | 0.18 |
| 1 | 91785583 | rs282000 | -0.994 | 0.92 | 0.162 | -0.311 | 0.005 | -0.011 | -0.011 | 0 | 0.43 | 0.67 | 0.52 | 0.54 | 0.38 | 0.45 |
| 1 | 91786579 | rs282001 | 1.461 | 0.898 | 0.311 | -0.344 | -0.002 | -0.053 | -0.053 | 0 | 0.09 | 0.22 | 0.07 | 0 | 0.06 | 0.01 |
| 1 | 91792196 | rs2893151 | 0.608 | 0.883 | 0.2 | -0.378 | 0 | -0.008 | -0.008 | 0 | 0.22 | 0.2 | 0.39 | 0.54 | 0.22 | 0.38 |
| 1 | 91794248 | rs281973 | -0.408 | 0.903 | 0.269 | -0.369 | -0.008 | 0.044 | 0.044 | 0 | 0.32 | 0.5 | 0.48 | 0.54 | 0.28 | 0.39 |
| 1 | 91794494 | rs3893101 | 0.726 | 0.865 | -0.062 | -0.341 | -0.001 | -0.003 | -0.003 | 0.234 | 0.21 | 0.2 | 0.39 | 0.54 | 0.22 | 0.38 |
| 1 | 91802783 | rs12060327 | 1.239 | 0.942 | -0.389 | -0.319 | -0.014 | 0.002 | 0.002 | 0 | 0.16 | 0.13 | 0.16 | 0.01 | 0.24 | 0.12 |
| 1 | 91813455 | rs1034499 | 0.424 | 0.631 | -0.647 | 0.022 | 0 | 0.051 | 0.051 | 0 | 0.22 | 0.26 | 0.39 | 0.53 | 0.22 | 0.38 |
| 1 | 91821503 | rs282057 | -0.72 | 0.446 | -1.354 | 0.626 | 0.045 | -0.046 | -0.046 | 0 | 0.6 | 0.84 | 0.7 | 0.55 | 0.64 | 0.57 |
| 1 | 91831097 | rs12129878 | 0.822 | 0.471 | -1.513 | 0.542 | 0 | -0.032 | -0.032 | 1.544 | 0.11 | 0.15 | 0.05 | 0 | 0.09 | 0.06 |
| 1 | 91848784 | rs17131417 | 0.438 | 0.241 | -1.69 | 1.154 | 0 | 0.025 | 0.025 | 0.627 | 0.06 | 0.1 | 0.12 | 0.17 | 0.07 | 0.07 |
| 1 | 91849219 | rs17131418 | 0.123 | 0.245 | -1.834 | 0.815 | 0.022 | -0.025 | -0.025 | 0.19 | 0.34 | 0.03 | 0.18 | 0.27 | 0.29 | 0.36 |
| 1 | 91854383 | rs2099749 | 0.13 | 0.256 | -1.905 | 0.82 | 0.011 | -0.037 | -0.037 | 0 | 0.34 | 0.02 | 0.17 | 0.27 | 0.29 | 0.36 |
| 1 | 91872374 | rs11166001 | -0.886 | 0.887 | -0.416 | 1.292 | 0.011 | 0.038 | 0.038 | 0.231 | 0.22 | 0.34 | 0.26 | 0.35 | 0.25 | 0.46 |
| 1 | 91886640 | rs11166334 | -0.74 | 0.534 | -0.205 | 1.35 | -0.01 | 0.068 | 0.068 | 0.037 | 0.28 | 0.42 | 0.46 | 0.49 | 0.22 | 0.33 |
| 1 | 91887372 | rs17485896 | 0.343 | 0.395 | -0.91 | 0.93 | -0.01 | 0.043 | 0.043 | 0 | 0.38 | 0.15 | 0.19 | 0.23 | 0.39 | 0.37 |
| 1 | 91895432 | rs11166498 | 0.019 | 0.753 | -0.052 | 1.354 | 0.037 | 0.045 | 0.045 | 0 | 0.17 | 0.27 | 0.23 | 0.29 | 0.22 | 0.27 |
| 1 | 91900687 | rs7528478 | -0.22 | 0.54 | 0.084 | 1.259 | 0.005 | 0 | 0 | 0.01 | 0.54 | 0.46 | 0.42 | 0.51 | 0.6 | 0.64 |
| 1 | 91901292 | rs6694567 | -0.263 | 0.454 | 0.309 | 1.197 | -0.005 | 0.001 | 0.001 | 0 | 0.57 | 0.55 | 0.44 | 0.56 | 0.61 | 0.71 |
| 1 | 91903413 | rs519244 | 0.623 | 0.21 | 0.713 | 1.306 | 0.005 | 0.007 | 0.007 | 0 | 0.32 | 0.31 | 0.27 | 0.28 | 0.24 | 0.18 |
| 1 | 91917448 | rs3851270 | 0.086 | 0.208 | 0.972 | 1.169 | -0.006 | 0.001 | 0.001 | 0.041 | 0.57 | 0.52 | 0.44 | 0.56 | 0.62 | 0.71 |
| 1 | 91919058 | rs12060889 | 0.712 | 0.174 | 1.157 | 1.142 | -0.005 | -0.014 | -0.014 | 0.024 | 0.42 | 0.54 | 0.26 | 0.25 | 0.41 | 0.44 |
| 1 | 91919533 | rs7540260 | 1.392 | -0.021 | 1.032 | 1.015 | -0.017 | 0.024 | 0.024 | 0 | 0.34 | 0.18 | 0.17 | 0.22 | 0.37 | 0.32 |

| | | | | | | | | | | | | | | | | |
|---|---|---|---|---|---|---|---|---|---|---|---|---|---|---|---|---|
| 1 | 91919610 | rs17131455 | 0.025 | -0.009 | 1.044 | 1.019 | -0.013 | -0.025 | -0.025 | 0 | 0.06 | 0.09 | 0.03 | 0.03 | 0.02 | 0.11 |
| 1 | 91923254 | rs12058802 | 0.151 | -0.005 | 0.979 | 1.125 | 0.005 | 0.012 | 0.012 | 0 | 0.51 | 0.61 | 0.51 | 0.39 | 0.54 | 0.55 |
| 1 | 91930483 | rs12125038 | 0.932 | -0.103 | 0.918 | 1.109 | -0.018 | -0.029 | -0.029 | 0.234 | 0.3 | 0.05 | 0.26 | 0.31 | 0.24 | 0.18 |
| 1 | 91935632 | rs2135603 | 0.943 | -0.079 | 1.018 | 1.125 | -0.005 | -0.026 | -0.026 | 0 | 0.38 | 0.5 | 0.24 | 0.24 | 0.41 | 0.38 |
| 1 | 91936560 | rs495026 | -0.446 | 0.126 | 0.907 | 1.236 | -0.008 | -0.009 | -0.009 | 0 | 0.83 | 0.88 | 0.78 | 0.7 | 0.77 | 0.73 |
| 1 | 91937695 | rs505892 | 0.144 | 0.101 | 0.872 | 1.212 | 0.042 | 0.142 | 0.142 | 0 | 0.79 | 0.61 | 0.56 | 0.56 | 0.75 | 0.69 |
| 1 | 91952120 | rs10874743 | -1.286 | -0.199 | 0.603 | 1.085 | -0.011 | 0.016 | 0.016 | 0 | 0.17 | 0.48 | 0.33 | 0.16 | 0.16 | 0.23 |
| 1 | 91959859 | rs10518494 | 0.61 | -0.181 | -0.217 | 1.053 | -0.025 | -0.038 | -0.038 | 0.077 | 0.29 | 0.05 | 0.26 | 0.34 | 0.25 | 0.22 |
| 1 | 91965850 | rs13447450 | 1.029 | -0.016 | -0.511 | 0.479 | -0.114 | 0.177 | 0.177 | 0.114 | 0.31 | 0.03 | 0.2 | 0.05 | 0.39 | 0.2 |
| 1 | 91966406 | rs13447454 | 0.039 | 0.276 | -0.257 | 0.368 | 0.01 | -0.022 | -0.022 | 0 | 0.08 | 0.08 | 0.08 | 0 | 0.13 | 0.08 |
| 1 | 91966445 | rs13447455 | 1.074 | 0.225 | -0.345 | 0.4 | -0.092 | 0.174 | 0.174 | 0.1 | 0.31 | 0.06 | 0.21 | 0.05 | 0.39 | 0.21 |
| 1 | 91980447 | rs483532 | 0.394 | -0.412 | 0.315 | 0.771 | -0.003 | -0.002 | -0.002 | 0 | 0.77 | 0.78 | 0.78 | 0.76 | 0.79 | 0.72 |
| 1 | 91987034 | rs6688261 | 0.707 | -0.589 | 0.869 | 0.836 | -0.042 | 0.044 | 0.044 | 0 | 0.38 | 0.28 | 0.47 | 0.24 | 0.51 | 0.33 |
| 1 | 91988009 | rs10493850 | 0.702 | -0.614 | 1.488 | 0.877 | 0.017 | -0.027 | -0.027 | 0 | 0.36 | 0.3 | 0.29 | 0.49 | 0.27 | 0.35 |
| 1 | 91990487 | rs12125947 | 0.862 | -0.572 | 1.435 | 0.849 | -0.055 | 0.1 | 0.1 | 0 | 0.37 | 0.24 | 0.28 | 0.05 | 0.49 | 0.25 |
| 1 | 92002506 | rs11164909 | 0.081 | -0.586 | 1.448 | 0.588 | 0.009 | -0.014 | -0.014 | 0 | 0.18 | 0.09 | 0.19 | 0.25 | 0.18 | 0.27 |
| 1 | 92003919 | rs17501381 | 0.886 | -0.578 | 1.457 | 0.562 | -0.057 | 0.104 | 0.104 | 0 | 0.37 | 0.24 | 0.28 | 0.05 | 0.49 | 0.25 |
| 1 | 92011467 | rs745731 | -0.188 | -0.521 | 1.293 | 0.828 | -0.001 | 0 | 0 | 0 | 0.19 | 0.16 | 0.19 | 0.25 | 0.18 | 0.27 |
| 1 | 92014051 | rs963911 | 0.449 | -0.409 | 1.257 | 0.418 | 0.017 | -0.069 | -0.069 | 0.001 | 0.89 | 0.6 | 0.92 | 0.99 | 0.93 | 0.88 |
| 1 | 92016653 | rs1949009 | -1.143 | -0.385 | 0.581 | 0.8 | -0.036 | 0.063 | 0.063 | 0.018 | 0.3 | 0.52 | 0.42 | 0.53 | 0.25 | 0.4 |
| 1 | 92021705 | rs17501937 | 0.263 | -0.223 | 0.293 | 0.493 | 0.013 | 0.033 | 0.033 | 0 | 0.49 | 0.46 | 0.35 | 0.06 | 0.57 | 0.31 |
| 1 | 92022597 | rs6681073 | -0.409 | -0.189 | 0.187 | 0.908 | -0.014 | 0.016 | 0.016 | 0 | 0.67 | 0.59 | 0.68 | 0.61 | 0.76 | 0.67 |
| 1 | 92023391 | rs11164947 | 0.622 | 0.528 | -0.298 | 0.724 | -0.001 | 0.008 | 0.008 | 0 | 0.62 | 0.49 | 0.68 | 0.58 | 0.59 | 0.5 |
| 1 | 92024948 | rs11585319 | -0.17 | 0.623 | -0.496 | 0.756 | -0.01 | -0.034 | -0.034 | 0 | 0.18 | 0.12 | 0.24 | 0.28 | 0.22 | 0.32 |
| 1 | 92030967 | rs7536573 | 0.76 | 0.433 | -0.494 | -0.378 | -0.006 | 0.002 | 0.002 | 0 | 0.3 | 0.21 | 0.19 | 0.03 | 0.33 | 0.19 |
| 1 | 92031009 | rs11164953 | 0.32 | 0.232 | -0.246 | -0.243 | -0.001 | 0.023 | 0.023 | 0 | 0.18 | 0.21 | 0.23 | 0.38 | 0.17 | 0.37 |
| 1 | 92032179 | rs11164955 | 1.133 | 0.011 | -0.091 | -0.175 | 0.008 | 0.004 | 0.004 | 0 | 0.31 | 0.16 | 0.35 | 0.28 | 0.25 | 0.36 |
| 1 | 92032395 | rs10874792 | -0.529 | -0.008 | 0.202 | -0.534 | -0.009 | 0 | 0 | 0 | 0.57 | 0.63 | 0.71 | 0.97 | 0.48 | 0.77 |
| 1 | 92035123 | rs906634 | -0.964 | 0.138 | 0.37 | -0.291 | 0.002 | -0.03 | -0.03 | 0 | 0.87 | 0.84 | 0.9 | 1 | 0.8 | 0.96 |
| 1 | 92035303 | rs6689719 | 0.472 | -0.01 | 0.653 | -0.244 | 0.008 | 0.038 | 0.038 | 0 | 0.8 | 0.55 | 0.77 | 0.69 | 0.75 | 0.9 |
| 1 | 92037984 | rs925191 | -0.283 | -0.025 | 0.667 | -0.664 | 0.024 | -0.102 | -0.102 | 0 | 0.84 | 0.6 | 0.87 | 1 | 0.8 | 0.96 |
| 1 | 92038225 | rs925192 | -0.441 | -0.008 | 0.546 | -0.656 | 0.008 | -0.061 | -0.061 | 0 | 0.85 | 0.74 | 0.88 | 1 | 0.8 | 0.96 |
| 1 | 92040543 | rs936340 | -0.716 | -0.541 | 0.331 | -0.806 | -0.02 | 0.066 | 0.066 | 0 | 0.63 | 0.76 | 0.62 | 0.87 | 0.6 | 0.68 |
| 1 | 92041502 | rs4658231 | -0.402 | -0.823 | 0.567 | -0.913 | -0.015 | -0.023 | -0.023 | 0 | 0.29 | 0.34 | 0.35 | 0.83 | 0.25 | 0.5 |
| 1 | 92042198 | rs10493853 | 1.259 | -0.724 | 0.368 | -1.106 | -0.07 | 0.186 | 0.186 | 0.261 | 0.34 | 0.1 | 0.24 | 0.02 | 0.38 | 0.19 |
| 1 | 92042245 | rs11164971 | 1.055 | -0.792 | 0.355 | -0.635 | -0.004 | -0.072 | -0.072 | 0.151 | 0.18 | 0.05 | 0.2 | 0.31 | 0.16 | 0.33 |
| 1 | 92045585 | rs10874794 | 0.226 | -1.113 | 0.533 | -0.671 | -0.015 | 0.044 | 0.044 | 0 | 0.24 | 0.3 | 0.39 | 0.6 | 0.21 | 0.39 |
| 1 | 92048779 | rs4658234 | -1.429 | -0.807 | 0.15 | -0.657 | -0.01 | 0.004 | 0.004 | 0 | 0.82 | 0.86 | 0.89 | 1 | 0.77 | 0.95 |
| 1 | 92055995 | rs12059556 | -0.227 | -0.96 | 0.038 | -0.687 | 0.009 | 0.003 | 0.003 | 0 | 0.43 | 0.56 | 0.48 | 0.66 | 0.34 | 0.46 |
| 1 | 92058290 | rs6693314 | 1.45 | -0.71 | -0.35 | -0.546 | 0 | 0.001 | 0.001 | 0 | 0.16 | 0.09 | 0.2 | 0.15 | 0.13 | 0.25 |
| 1 | 92060264 | rs17503335 | 0.238 | -0.696 | -0.433 | -0.664 | 0 | -0.001 | -0.001 | 0.321 | 0.34 | 0.34 | 0.3 | 0.3 | 0.36 | 0.34 |
| 1 | 92061160 | rs17131499 | 0.591 | -0.907 | -0.281 | -0.575 | -0.022 | 0.147 | 0.147 | 0 | 0.07 | 0.13 | 0.13 | 0.47 | 0.05 | 0.12 |
| 1 | 92066310 | rs4658241 | 0.348 | -0.893 | -0.315 | -0.642 | -0.002 | -0.003 | -0.003 | 0 | 0.34 | 0.29 | 0.3 | 0.31 | 0.36 | 0.35 |
| 1 | 92066594 | rs6662147 | -2.185 | -0.902 | -0.406 | -0.896 | 0.01 | 0.012 | 0.012 | 0 | 0.82 | 0.85 | 0.92 | 0.93 | 0.87 | 0.93 |
| 1 | 92067746 | rs4658243 | -0.144 | -0.872 | -0.326 | -0.85 | -0.003 | -0.013 | -0.013 | 0.249 | 0.41 | 0.44 | 0.42 | 0.77 | 0.41 | 0.47 |
| 1 | 92068967 | rs1192404 | 1.223 | -0.594 | -0.206 | -0.732 | -0.007 | -0.01 | -0.01 | 0.313 | 0.19 | 0.33 | 0.29 | 0.14 | 0.14 | 0.27 |
| 1 | 92069188 | rs4658244 | 0.234 | -0.446 | -0.439 | -0.545 | -0.022 | -0.031 | -0.031 | 0 | 0.29 | 0.22 | 0.34 | 0.48 | 0.36 | 0.31 |
| 1 | 92076523 | rs1041159 | -0.202 | -0.4 | -0.159 | -0.625 | 0.008 | -0.052 | -0.052 | 0.496 | 0.64 | 0.56 | 0.61 | 0.86 | 0.61 | 0.65 |
| 1 | 92080059 | rs1192419 | -1.162 | -0.361 | -0.141 | -0.665 | -0.005 | -0.011 | -0.011 | 0 | 0.79 | 0.71 | 0.71 | 0.87 | 0.82 | 0.7 |
| 1 | 92082656 | rs4650270 | 0.618 | -0.356 | -0.624 | -0.542 | 0 | -0.001 | -0.001 | 1.701 | 0.53 | 0.47 | 0.51 | 0.46 | 0.51 | 0.53 |
| 1 | 92082962 | rs12034186 | -0.314 | -0.019 | -0.825 | 0.202 | -0.007 | -0.045 | -0.045 | 0.17 | 0.22 | 0.15 | 0.16 | 0.41 | 0.26 | 0.17 |
| 1 | 92087522 | rs4658104 | 0.981 | 0.207 | -1.055 | 0.179 | -0.03 | -0.091 | -0.091 | 0 | 0.13 | 0.01 | 0.17 | 0.35 | 0.11 | 0.27 |
| 1 | 92089259 | rs17569956 | 0.451 | 0.151 | -0.71 | -0.665 | -0.007 | -0.012 | -0.012 | 3.133 | 0.17 | 0.14 | 0.1 | 0 | 0.22 | 0.05 |
| 1 | 92095078 | rs11165102 | -0.422 | 0.082 | -0.192 | -0.313 | -0.029 | 0.001 | 0.001 | 0 | 0.26 | 0.46 | 0.28 | 0.77 | 0.17 | 0.44 |
| 1 | 92099741 | rs1887054 | -0.014 | 0.205 | -0.127 | -0.993 | 0.027 | -0.075 | -0.075 | 1.672 | 0.77 | 0.42 | 0.83 | 0.96 | 0.71 | 0.91 |
| 1 | 92101035 | rs11165115 | 1.48 | 0.295 | -1.286 | -1.691 | -0.101 | 0.091 | 0.091 | 0.311 | 0.22 | 0.02 | 0.21 | 0.03 | 0.34 | 0.25 |
| 1 | 92101761 | rs11576243 | -0.067 | -0.169 | -0.983 | -1.18 | 0.004 | -0.005 | -0.005 | 0 | 0.46 | 0.33 | 0.39 | 0.49 | 0.46 | 0.39 |
| 1 | 92105501 | rs2478172 | 0.272 | -0.947 | -0.41 | -0.904 | -0.004 | 0.004 | 0.004 | 0 | 0.64 | 0.41 | 0.71 | 0.6 | 0.67 | 0.83 |
| 1 | 92106637 | rs10874851 | 1.198 | -0.934 | -0.706 | -1.06 | -0.105 | 0.185 | 0.185 | 0 | 0.38 | 0.08 | 0.49 | 0.08 | 0.5 | 0.46 |
| 1 | 92107736 | rs2182424 | 1.626 | -0.963 | -0.509 | -0.739 | -0.028 | -0.105 | -0.105 | 0.098 | 0.16 | 0.02 | 0.17 | 0.42 | 0.13 | 0.31 |
| 1 | 92112890 | rs2182425 | 0.206 | -0.902 | -1.236 | -0.684 | -0.027 | 0.064 | 0.064 | 0 | 0.11 | 0.13 | 0.14 | 0.39 | 0.05 | 0.09 |
| 1 | 92113813 | rs1887055 | -0.77 | -0.949 | -1.273 | -0.985 | -0.033 | -0.071 | -0.071 | 1.767 | 0.39 | 0.48 | 0.37 | 0.89 | 0.23 | 0.46 |
| 1 | 92120137 | rs12410523 | 0.5 | -0.52 | -1.794 | -0.555 | 0.031 | -0.032 | -0.032 | 0.209 | 0.61 | 0.19 | 0.67 | 0.56 | 0.67 | 0.83 |
| 1 | 92121843 | rs1926271 | 0.013 | -0.14 | -1.697 | -1.352 | 0 | -0.009 | -0.009 | 0 | 0.69 | 0.3 | 0.74 | 0.94 | 0.68 | 0.89 |
| 1 | 92122248 | rs4459159 | 0.181 | -0.336 | -1.567 | -1.02 | -0.027 | -0.015 | -0.015 | 0 | 0.25 | 0.48 | 0.29 | 0.8 | 0.18 | 0.37 |
| 1 | 92122332 | rs2148322 | -0.317 | 0.04 | -1.395 | -0.949 | -0.007 | -0.083 | -0.083 | 0 | 0.78 | 0.55 | 0.64 | 0.92 | 0.8 | 0.79 |
| 1 | 92125282 | rs12144361 | 0.151 | 0.382 | -1.943 | -0.449 | -0.013 | 0.028 | 0.028 | 0.003 | 0.18 | 0.01 | 0.07 | 0.1 | 0.11 | 0.06 |
| 1 | 92129685 | rs4658257 | -0.356 | 0.784 | -1.44 | -0.246 | -0.005 | 0.011 | 0.011 | 1.182 | 0.64 | 0.39 | 0.52 | 0.89 | 0.58 | 0.66 |
| 1 | 92143234 | rs2810894 | -0.152 | 1.197 | -1.814 | 0.923 | -0.057 | -0.093 | -0.093 | 0 | 0.72 | 0.9 | 0.66 | 0.52 | 0.63 | 0.74 |

| | | | | | | | | | | | | | | | | |
|---|---|---|---|---|---|---|---|---|---|---|---|---|---|---|---|---|
| 1 | 92144970 | rs2810893 | 0.335 | 0.832 | -1.721 | 0.117 | -0.032 | 0.091 | 0.091 | 0 | 0.58 | 0.78 | 0.64 | 0.84 | 0.54 | 0.67 |
| 1 | 92146890 | rs17571088 | -0.008 | 0.93 | -1.583 | 0.181 | -0.007 | 0.029 | 0.029 | 0 | 0.18 | 0.08 | 0.1 | 0.1 | 0.17 | 0.1 |
| 1 | 92148013 | rs1804506 | -0.494 | 1.223 | -1.208 | 0.808 | 0.017 | 0.164 | 0.164 | 0 | 0.1 | 0.19 | 0.25 | 0.42 | 0.17 | 0.08 |
| 1 | 92148839 | rs1805117 | 0.27 | 1.056 | -1.274 | 0.159 | -0.01 | 0.025 | 0.025 | 0 | 0.16 | 0.06 | 0.09 | 0.11 | 0.16 | 0.1 |
| 1 | 92149096 | rs1805116 | 0.381 | 1.08 | -1.435 | 0.171 | -0.046 | 0.01 | 0.01 | 0 | 0.16 | 0.01 | 0.08 | 0.1 | 0.16 | 0.1 |
| 1 | 92151273 | rs17131522 | 0.445 | 0.573 | -1.526 | -0.21 | 0.008 | 0.028 | 0.028 | 0 | 0.21 | 0.06 | 0.11 | 0.14 | 0.18 | 0.35 |
| 1 | 92153974 | rs11165262 | 0.63 | 1.125 | -1.168 | -0.499 | -0.023 | 0.026 | 0.026 | 0 | 0.16 | 0.09 | 0.17 | 0.03 | 0.21 | 0.16 |
| 1 | 92154088 | rs2810891 | 1.07 | 0.973 | -1.081 | -0.42 | -0.001 | 0 | 0 | 0 | 0.13 | 0.11 | 0.13 | 0.24 | 0.12 | 0.29 |
| 1 | 92155337 | rs12082710 | -0.406 | 0.924 | -0.532 | -0.603 | -0.001 | -0.005 | -0.005 | 0 | 0.34 | 0.38 | 0.26 | 0.09 | 0.39 | 0.27 |
| 1 | 92158447 | rs2765887 | 0.82 | 0.715 | -0.419 | -1.008 | 0.034 | 0.115 | 0.115 | 0 | 0.49 | 0.32 | 0.48 | 0.29 | 0.39 | 0.59 |
| 1 | 92158571 | rs2810887 | -0.205 | 0.587 | -0.101 | -0.793 | 0.001 | 0.013 | 0.013 | 0 | 0.81 | 0.85 | 0.73 | 0.38 | 0.74 | 0.86 |
| 1 | 92158751 | rs3754021 | 1.106 | 0.636 | -0.273 | -0.795 | -0.024 | 0.033 | 0.033 | 0 | 0.16 | 0.07 | 0.08 | 0.06 | 0.19 | 0.12 |
| 1 | 92161991 | rs12141338 | 1.106 | 0.636 | -0.281 | -0.795 | -0.025 | 0.035 | 0.035 | 0 | 0.16 | 0.06 | 0.08 | 0.06 | 0.19 | 0.12 |
| 1 | 92162920 | rs1926261 | -0.263 | 0.564 | -0.077 | -0.654 | -0.007 | -0.029 | -0.029 | 0.097 | 0.82 | 0.91 | 0.75 | 0.38 | 0.75 | 0.86 |
| 1 | 92163174 | rs6693438 | 0.675 | 0.644 | 0.022 | -0.774 | 0.01 | -0.034 | -0.034 | 7.614 | 0.18 | 0.3 | 0.19 | 0.03 | 0.21 | 0.15 |
| 1 | 92166931 | rs3767576 | 0.374 | 0.662 | -0.166 | -0.141 | 0.005 | 0.045 | 0.045 | 0 | 0.11 | 0.15 | 0.23 | 0.36 | 0.19 | 0.09 |
| 1 | 92171267 | rs284876 | -0.768 | 0.393 | -0.176 | -0.45 | -0.01 | 0.014 | 0.014 | 0 | 0.43 | 0.4 | 0.46 | 0.68 | 0.5 | 0.51 |
| 1 | 92172608 | rs6660484 | -0.364 | 0.551 | -0.26 | -0.453 | -0.005 | -0.006 | -0.006 | 0 | 0.31 | 0.37 | 0.35 | 0.44 | 0.4 | 0.22 |
| 1 | 92173371 | rs3767569 | 2.304 | -0.34 | -0.473 | -0.704 | 0.015 | 0.085 | 0.085 | 0 | 0.21 | 0.07 | 0.27 | 0.03 | 0.15 | 0.26 |
| 1 | 92174260 | rs284878 | -1.281 | 0.255 | 0.033 | -0.206 | -0.001 | 0.029 | 0.029 | 0.741 | 0.92 | 0.94 | 0.97 | 0.74 | 0.94 | 0.95 |
| 1 | 92174415 | rs2038931 | 0.376 | 0.452 | -0.012 | -0.32 | -0.002 | 0.12 | 0.12 | 1.094 | 0.1 | 0.24 | 0.2 | 0.37 | 0.1 | 0.09 |
| 1 | 92176044 | rs12567680 | 0.937 | 0.388 | 0.021 | -0.442 | 0.025 | 0.005 | 0.005 | 1.006 | 0.39 | 0.28 | 0.49 | 0.41 | 0.31 | 0.38 |
| 1 | 92177062 | rs17512269 | 1.127 | 0.823 | -0.094 | -0.935 | -0.01 | 0.008 | 0.008 | 0 | 0.17 | 0.17 | 0.19 | 0.03 | 0.23 | 0.15 |
| 1 | 92177938 | rs1805113 | -0.132 | 0.878 | 0.062 | -0.957 | -0.033 | 0.045 | 0.045 | 0 | 0.35 | 0.31 | 0.29 | 0.09 | 0.43 | 0.28 |
| 1 | 92181670 | rs2029354 | 1.519 | 0.398 | -0.096 | -0.945 | 0.023 | 0.006 | 0.006 | 0 | 0.3 | 0.18 | 0.31 | 0.03 | 0.21 | 0.29 |
| 1 | 92184814 | rs7524066 | 0.013 | 0.763 | 0.197 | -0.801 | 0.031 | -0.068 | -0.068 | 0 | 0.22 | 0.41 | 0.16 | 0.06 | 0.29 | 0.13 |
| 1 | 92185185 | rs2279455 | 0.088 | 0.647 | 0.026 | -0.637 | 0.011 | -0.011 | -0.011 | 0 | 0.67 | 0.62 | 0.59 | 0.32 | 0.61 | 0.72 |
| 1 | 92190072 | rs12751148 | -0.321 | 0.692 | -0.465 | -0.589 | -0.018 | 0.018 | 0.018 | 0 | 0.33 | 0.24 | 0.25 | 0.27 | 0.4 | 0.43 |
| 1 | 92191378 | rs17131540 | 1.413 | 1.12 | -0.723 | 0.261 | -0.031 | -0.006 | -0.006 | 0 | 0.07 | 0 | 0.19 | 0.4 | 0.1 | 0.06 |
| 1 | 92191809 | rs17131541 | 0.998 | 0.883 | -0.494 | 0.171 | 0.001 | 0.028 | 0.028 | 0.306 | 0.07 | 0.04 | 0.03 | 0.26 | 0.06 | 0.05 |
| 1 | 92194322 | rs11165354 | 0.087 | 0.953 | -0.263 | 0.065 | -0.003 | -0.009 | -0.009 | 0 | 0.69 | 0.27 | 0.62 | 0.54 | 0.63 | 0.81 |
| 1 | 92195601 | rs10783002 | 0.705 | 0.725 | -0.435 | 0.001 | 0.039 | 0.001 | 0.001 | 0 | 0.53 | 0.19 | 0.46 | 0.51 | 0.4 | 0.66 |
| 1 | 92198162 | rs7517044 | -0.145 | 0.851 | 0.106 | 0.173 | 0.012 | 0.079 | 0.079 | 0 | 0.27 | 0.56 | 0.32 | 0.42 | 0.31 | 0.18 |
| 1 | 92199184 | rs11582136 | 1.177 | 0.723 | -0.089 | 0.134 | -0.012 | -0.038 | -0.038 | 0 | 0.13 | 0.04 | 0.12 | 0.26 | 0.11 | 0.28 |
| 1 | 92200218 | rs2306887 | -0.057 | 0.701 | 0 | 0.12 | 0.011 | 0.08 | 0.08 | 0 | 0.27 | 0.56 | 0.32 | 0.42 | 0.3 | 0.18 |
| 1 | 92200782 | rs12069176 | -0.12 | 0.684 | 0.156 | 0.136 | 0.007 | 0.078 | 0.078 | 0 | 0.73 | 0.49 | 0.68 | 0.58 | 0.7 | 0.82 |
| 1 | 92201002 | rs11165380 | -0.13 | 0.729 | 0.07 | 0.146 | 0.005 | 0.079 | 0.079 | 0 | 0.73 | 0.49 | 0.67 | 0.58 | 0.7 | 0.82 |
| 1 | 92201699 | rs12076549 | 0.496 | 0.832 | 0.017 | 0.128 | -0.001 | 0 | 0 | 0 | 0.59 | 0.41 | 0.56 | 0.33 | 0.58 | 0.53 |
| 1 | 92202467 | rs6677523 | -0.232 | 1.109 | 0.822 | 0.712 | 0.013 | 0.07 | 0.07 | 0.013 | 0.83 | 0.53 | 0.85 | 0.74 | 0.88 | 0.71 |
| 1 | 92203190 | rs11579207 | 0.263 | 1.122 | 0.96 | 0.718 | 0.001 | 0.018 | 0.018 | 0 | 0.15 | 0.25 | 0.13 | 0.26 | 0.12 | 0.29 |
| 1 | 92205197 | rs17513917 | -0.602 | 1.067 | 0.983 | 0.315 | 0.005 | 0.003 | 0.003 | 2.053 | 0.28 | 0.3 | 0.26 | 0.29 | 0.36 | 0.22 |
| 1 | 92209311 | rs2029356 | 0.391 | 0.99 | 0.007 | 0.536 | 0.008 | 0.022 | 0.022 | 0.875 | 0.71 | 0.57 | 0.78 | 0.71 | 0.73 | 0.78 |
| 1 | 92212703 | rs10493860 | -0.287 | 1.337 | -0.142 | 0.831 | 0.008 | 0.027 | 0.027 | 0 | 0.11 | 0.15 | 0.17 | 0.27 | 0.15 | 0.05 |
| 1 | 92213442 | rs284169 | -0.739 | 0.94 | -0.32 | 0.37 | 0.006 | 0.003 | 0.003 | 0 | 0.51 | 0.59 | 0.5 | 0.54 | 0.43 | 0.43 |
| 1 | 92214628 | rs284170 | -1.465 | 0.632 | -0.646 | 0.119 | -0.009 | -0.033 | -0.033 | 0.129 | 0.84 | 0.96 | 0.86 | 0.79 | 0.85 | 0.74 |
| 1 | 92215089 | rs12123436 | 1.705 | 0.863 | -0.911 | -0.202 | -0.052 | -0.059 | -0.059 | 0.327 | 0.36 | 0.11 | 0.41 | 0.42 | 0.42 | 0.44 |
| 1 | 92215725 | rs12404952 | 1.423 | 0.534 | -0.912 | -0.175 | 0 | 0.009 | 0.009 | 1.136 | 0.1 | 0.01 | 0.12 | 0.04 | 0.06 | 0.06 |
| 1 | 92221608 | rs284174 | 0.61 | 0.494 | -0.856 | -0.22 | 0.005 | -0.005 | -0.005 | 0 | 0.56 | 0.4 | 0.61 | 0.5 | 0.62 | 0.62 |
| 1 | 92223976 | rs284176 | -0.044 | 0.865 | -0.212 | 0.262 | 0 | 0.001 | 0.001 | 0 | 0.72 | 0.72 | 0.67 | 0.52 | 0.69 | 0.71 |
| 1 | 92224067 | rs3738441 | -0.678 | 0.84 | -0.084 | 0.074 | 0.008 | 0.045 | 0.045 | 0 | 0.22 | 0.36 | 0.34 | 0.65 | 0.26 | 0.19 |
| 1 | 92224698 | rs284177 | 0.965 | 0.783 | 0.095 | -0.469 | 0.003 | 0.073 | 0.073 | 0 | 0.43 | 0.28 | 0.34 | 0.05 | 0.45 | 0.41 |
| 1 | 92224815 | rs17131552 | -0.262 | 0.762 | 0.094 | -0.31 | 0.001 | 0.002 | 0.002 | 0.104 | 0.2 | 0.2 | 0.29 | 0.42 | 0.17 | 0.21 |
| 1 | 92227848 | rs284180 | 0.498 | 0.761 | -0.121 | -1.262 | 0.007 | 0.096 | 0.096 | 0 | 0.44 | 0.26 | 0.34 | 0.05 | 0.45 | 0.44 |
| 1 | 92236048 | rs284191 | 0.075 | 0.469 | -0.254 | -0.913 | 0.013 | 0.025 | 0.025 | 0 | 0.53 | 0.58 | 0.41 | 0.09 | 0.39 | 0.48 |
| 1 | 92238167 | rs284198 | 0.094 | 0.471 | -0.311 | -0.814 | 0.014 | 0.031 | 0.031 | 0 | 0.52 | 0.57 | 0.41 | 0.12 | 0.38 | 0.48 |
| 1 | 92240685 | rs4658265 | 0.633 | 0.25 | -0.243 | -1.048 | -0.001 | -0.05 | -0.05 | 0 | 0.34 | 0.35 | 0.26 | 0.06 | 0.31 | 0.4 |
| 1 | 92241033 | rs284202 | -0.774 | 0.564 | -0.515 | -0.367 | 0.012 | -0.021 | -0.021 | 0.428 | 0.4 | 0.69 | 0.38 | 0.53 | 0.36 | 0.39 |
| 1 | 92242568 | rs12090296 | 0.433 | 0.534 | -0.055 | -0.357 | -0.019 | 0.041 | 0.041 | 0 | 0.16 | 0.27 | 0.1 | 0.28 | 0.09 | 0.13 |
| 1 | 92244566 | rs6604053 | 0.877 | 0.382 | 0.045 | -0.736 | 0.041 | 0.078 | 0.078 | 0 | 0.42 | 0.3 | 0.3 | 0.05 | 0.31 | 0.35 |
| 1 | 92247654 | rs4658269 | 1.119 | 0.354 | -0.334 | -0.751 | 0.019 | 0.276 | 0.276 | 0 | 0.39 | 0.09 | 0.27 | 0.05 | 0.3 | 0.35 |
| 1 | 92248695 | rs7536947 | 0.511 | 0.416 | -0.061 | -0.547 | -0.024 | 0.06 | 0.06 | 0 | 0.17 | 0.34 | 0.1 | 0.28 | 0.09 | 0.13 |
| 1 | 92249817 | rs6683840 | 0.029 | 0.937 | -0.466 | -0.133 | -0.036 | 0.027 | 0.027 | 0 | 0.63 | 0.74 | 0.44 | 0.33 | 0.47 | 0.54 |
| 1 | 92255996 | rs284143 | -0.554 | 1.284 | -0.361 | 0.419 | 0 | 0 | 0 | 0 | 0.76 | 0.77 | 0.73 | 0.77 | 0.66 | 0.68 |
| 1 | 92257971 | rs1411276 | -0.36 | 0.887 | -0.38 | 0.062 | 0.002 | -0.011 | -0.011 | 0 | 0.3 | 0.51 | 0.21 | 0.31 | 0.24 | 0.22 |
| 1 | 92258103 | rs11799400 | 0.154 | 0.742 | -0.564 | -0.048 | -0.003 | 0.112 | 0.112 | 0 | 0.07 | 0.14 | 0.17 | 0.44 | 0.09 | 0.1 |
| 1 | 92258782 | rs4658270 | 1.44 | 0.698 | -0.694 | -0.245 | -0.016 | 0.244 | 0.244 | 0.047 | 0.34 | 0.03 | 0.33 | 0.03 | 0.3 | 0.33 |
| 1 | 92265144 | rs17574203 | 0.495 | 0.804 | -0.261 | -0.23 | 0.008 | -0.026 | -0.026 | 0.34 | 0.09 | 0.09 | 0.09 | 0 | 0.14 | 0.05 |
| 1 | 92269982 | rs284161 | 0.193 | 0.783 | -0.328 | 0.03 | -0.002 | 0.123 | 0.123 | 0 | 0.07 | 0.12 | 0.16 | 0.43 | 0.09 | 0.1 |
| 1 | 92274817 | rs284152 | -0.958 | 0.802 | -0.179 | -0.188 | 0.001 | 0.004 | 0.004 | 0 | 0.38 | 0.65 | 0.38 | 0.75 | 0.33 | 0.35 |

| | | | | | | | | | | | | | | | | |
|---|---|---|---|---|---|---|---|---|---|---|---|---|---|---|---|---|
| 1 | 92285875 | rs10874981 | -0.055 | 0.797 | -0.184 | 0.016 | 0.015 | 0.001 | 0.001 | 0 | 0.65 | 0.53 | 0.57 | 0.42 | 0.55 | 0.58 |
| 1 | 92292407 | rs901917 | -0.106 | 0.802 | -0.28 | 0.042 | -0.004 | 0.111 | 0.111 | 0 | 0.92 | 0.8 | 0.86 | 0.65 | 0.91 | 0.9 |
| 1 | 92297887 | rs11165578 | -1.032 | 0.758 | -0.251 | 0.008 | 0.032 | 0.011 | 0.011 | 0 | 0.58 | 0.75 | 0.53 | 0.62 | 0.61 | 0.56 |
| 1 | 92302369 | rs11165594 | 0.2 | 0.757 | 0.321 | 0.111 | 0.001 | 0.006 | 0.006 | 0 | 0.25 | 0.26 | 0.21 | 0.37 | 0.23 | 0.21 |
| 1 | 92305250 | rs2129972 | -0.274 | 0.977 | 0.507 | 0.552 | -0.004 | 0.163 | 0.163 | 0 | 0.08 | 0.21 | 0.14 | 0.35 | 0.09 | 0.11 |
| 1 | 92313638 | rs2634031 | -0.514 | 0.976 | 0.488 | 0.311 | 0.016 | -0.003 | -0.003 | 0 | 0.35 | 0.5 | 0.43 | 0.6 | 0.45 | 0.42 |
| 1 | 92316682 | rs2799523 | -0.482 | 0.613 | 0.4 | 0.688 | -0.008 | 0.103 | 0.103 | 0 | 0.08 | 0.16 | 0.13 | 0.37 | 0.08 | 0.13 |
| 1 | 92323809 | rs11576557 | 1.415 | 0.368 | 0.901 | 0.515 | -0.042 | 0.17 | 0.17 | 1.81 | 0.83 | 0.41 | 0.81 | 0.63 | 0.89 | 0.81 |
| 1 | 92327045 | rs1805110 | -0.763 | 0.381 | 0.912 | 0.526 | -0.012 | 0.112 | 0.112 | 0 | 0.09 | 0.18 | 0.14 | 0.37 | 0.08 | 0.15 |
| 1 | 92327126 | rs1805109 | -0.588 | 0.383 | 0.878 | 0.527 | -0.008 | 0.096 | 0.096 | 0 | 0.09 | 0.16 | 0.14 | 0.37 | 0.08 | 0.15 |
| 1 | 92327707 | rs17881430 | 0.761 | 0.786 | 0.715 | 0.255 | -0.067 | -0.051 | -0.051 | 6.499 | 0.22 | 0.07 | 0.24 | 0.24 | 0.32 | 0.27 |
| 1 | 92329059 | rs11165679 | 0.812 | 0.781 | 0.713 | 0.211 | -0.067 | -0.04 | -0.04 | 0 | 0.21 | 0.07 | 0.24 | 0.21 | 0.32 | 0.24 |
| 1 | 92334749 | rs10493856 | 0.672 | 0.644 | 0.747 | 0.256 | -0.004 | 0.007 | 0.007 | 0.181 | 0.61 | 0.31 | 0.56 | 0.4 | 0.59 | 0.5 |
| 1 | 92339976 | rs10875030 | -0.234 | 0.695 | 0.453 | 0.225 | 0 | 0 | 0 | 0.468 | 0.72 | 0.63 | 0.63 | 0.4 | 0.65 | 0.6 |
| 1 | 92358371 | rs6680614 | 0.286 | 0.448 | 0.45 | 0.02 | 0 | -0.003 | -0.003 | 0 | 0.52 | 0.46 | 0.51 | 0.39 | 0.49 | 0.43 |
| 1 | 92366174 | rs17576372 | 0.632 | 0.482 | 0.13 | 0.091 | -0.078 | 0.01 | 0.01 | 0.812 | 0.34 | 0.08 | 0.39 | 0.22 | 0.42 | 0.32 |
| 1 | 92378843 | rs2770186 | 0.007 | 0.829 | 0.578 | 0.083 | 0.001 | 0.002 | 0.002 | 0 | 0.54 | 0.49 | 0.46 | 0.39 | 0.59 | 0.44 |
| 1 | 92393752 | rs9943134 | -0.567 | 0.826 | 0.728 | 0.348 | 0.012 | 0.229 | 0.229 | 0.401 | 0.1 | 0.27 | 0.15 | 0.37 | 0.09 | 0.18 |
| 1 | 92396114 | rs7549306 | -0.68 | 0.997 | 0.799 | 0.345 | 0.025 | 0.246 | 0.246 | 0.202 | 0.1 | 0.3 | 0.16 | 0.38 | 0.1 | 0.2 |
| 1 | 92416821 | rs6666624 | 1.273 | 0.92 | 1.349 | -0.215 | 0 | 0.103 | 0.103 | 0.629 | 0.77 | 0.48 | 0.73 | 0.61 | 0.8 | 0.75 |
| 1 | 92425621 | rs9887935 | 0.805 | 0.738 | 0.919 | -0.802 | 0.006 | 0.016 | 0.016 | 0.032 | 0.52 | 0.33 | 0.48 | 0.58 | 0.57 | 0.53 |
| 1 | 92428495 | rs10783071 | -0.055 | 0.699 | 1.076 | -0.795 | -0.014 | 0.188 | 0.188 | 1.51 | 0.89 | 0.66 | 0.84 | 0.61 | 0.92 | 0.81 |
| 1 | 92433595 | rs7541712 | 0.26 | 0.707 | 1.008 | -0.791 | 0.002 | 0.033 | 0.033 | 0.188 | 0.09 | 0.15 | 0.13 | 0.39 | 0.08 | 0.19 |
| 1 | 92438578 | rs10783072 | 0.778 | 0.736 | 0.966 | -0.781 | 0.007 | 0.016 | 0.016 | 0.032 | 0.52 | 0.34 | 0.47 | 0.58 | 0.57 | 0.53 |
| 1 | 92444125 | rs4658284 | 0.967 | 0.705 | 0.865 | -0.755 | -0.005 | 0.09 | 0.09 | 0.773 | 0.76 | 0.45 | 0.72 | 0.61 | 0.79 | 0.74 |
| 1 | 92447668 | rs17576636 | 1.551 | 0.812 | 0.384 | -0.794 | -0.095 | -0.022 | -0.022 | 0 | 0.3 | 0.02 | 0.3 | 0.2 | 0.32 | 0.35 |
| 1 | 92453336 | rs10735794 | 0.285 | 0.745 | 0.732 | -0.669 | -0.002 | 0.106 | 0.106 | 0.517 | 0.81 | 0.59 | 0.78 | 0.61 | 0.85 | 0.79 |
| 1 | 92457843 | rs10747493 | 0.408 | 0.696 | 0.74 | -0.692 | -0.006 | 0.096 | 0.096 | 0.121 | 0.8 | 0.59 | 0.77 | 0.61 | 0.84 | 0.79 |
| 1 | 92462296 | rs1410583 | 0.739 | 0.799 | 0.57 | -0.639 | -0.008 | 0.083 | 0.083 | 0.528 | 0.75 | 0.45 | 0.72 | 0.6 | 0.79 | 0.74 |
| 1 | 92494310 | rs34042557 | 1.302 | 0.749 | 0.168 | -0.757 | -0.042 | 0.132 | 0.132 | 0 | 0.23 | 0.03 | 0.25 | 0.03 | 0.23 | 0.21 |
| 1 | 92502060 | rs1014415 | 1.162 | 0.78 | 0.255 | -0.778 | 0 | -0.014 | -0.014 | 0 | 0.23 | 0.16 | 0.26 | 0.02 | 0.24 | 0.21 |
| 1 | 92503338 | rs17131593 | 1.017 | 0.795 | 0.28 | -0.742 | -0.001 | -0.05 | -0.05 | 0.001 | 0.21 | 0.09 | 0.16 | 0.38 | 0.25 | 0.19 |
| 1 | 92504560 | rs6690705 | -0.803 | 0.883 | 0.642 | -0.654 | 0.012 | 0.118 | 0.118 | 0.46 | 0.23 | 0.48 | 0.26 | 0.42 | 0.21 | 0.26 |
| 1 | 92512379 | rs1073001 | -0.695 | 0.841 | 0.496 | -0.796 | -0.005 | 0.019 | 0.019 | 0.36 | 0.47 | 0.7 | 0.52 | 0.42 | 0.44 | 0.46 |
| 1 | 92530670 | rs9659484 | -1.566 | 0.775 | 0.765 | -1.426 | 0.026 | -0.012 | -0.012 | 0.006 | 0.08 | 0.4 | 0.08 | 0.01 | 0.04 | 0.04 |
| 1 | 92544692 | rs17131598 | 0.424 | 0.898 | 0.02 | -1.405 | -0.014 | -0.071 | -0.071 | 0 | 0.26 | 0.09 | 0.19 | 0.42 | 0.29 | 0.24 |
| 1 | 92558583 | rs563606 | -0.481 | 0.791 | 0.244 | -2.787 | -0.004 | -0.029 | -0.029 | 0.639 | 0.71 | 0.77 | 0.66 | 0.97 | 0.66 | 0.79 |
| 1 | 92561392 | rs11166141 | -0.16 | 0.678 | 0.451 | -2.822 | 0.002 | -0.151 | -0.151 | 4.663 | 0.67 | 0.52 | 0.63 | 0.97 | 0.64 | 0.76 |
| 1 | 92579206 | rs1226187 | -0.546 | 0.705 | 0.299 | -2.813 | -0.01 | 0.036 | 0.036 | 0 | 0.7 | 0.85 | 0.67 | 0.97 | 0.65 | 0.76 |
| 1 | 92591916 | rs3851272 | -0.401 | 0.727 | 0.388 | -2.795 | 0.001 | -0.087 | -0.087 | 0.206 | 0.69 | 0.65 | 0.64 | 0.97 | 0.63 | 0.74 |
| 1 | 92592905 | rs490800 | -0.731 | 0.76 | -0.026 | -2.764 | -0.055 | 0.144 | 0.144 | 0 | 0.73 | 0.96 | 0.68 | 0.97 | 0.65 | 0.76 |
| 1 | 92598258 | rs506375 | -0.814 | 0.697 | 0.124 | -2.74 | -0.045 | 0.133 | 0.133 | 0 | 0.73 | 0.93 | 0.68 | 0.97 | 0.66 | 0.78 |
| 1 | 92598554 | rs1157468 | 0.989 | 0.699 | -0.191 | -3.012 | -0.03 | 0.151 | 0.151 | 0 | 0.32 | 0.06 | 0.28 | 0.1 | 0.3 | 0.36 |
| 1 | 92604114 | rs575805 | -0.636 | 0.963 | 0.242 | -2.33 | -0.08 | 0.177 | 0.177 | 0 | 0.74 | 0.98 | 0.69 | 0.97 | 0.66 | 0.78 |
| 1 | 92608852 | rs563113 | -0.419 | 0.974 | 0.38 | -2.324 | -0.004 | -0.014 | -0.014 | 0 | 0.72 | 0.78 | 0.67 | 0.97 | 0.66 | 0.78 |
| 1 | 92677976 | rs3122323 | 0.633 | 1.058 | 0.432 | -2.436 | 0.006 | -0.095 | -0.095 | 0.398 | 0.58 | 0.49 | 0.49 | 0.95 | 0.46 | 0.72 |
| 1 | 92678105 | rs17578686 | 0.45 | 1.231 | 0.533 | -2.575 | -0.022 | 0.063 | 0.063 | 0 | 0.26 | 0.09 | 0.36 | 0.02 | 0.34 | 0.23 |
| 1 | 92682286 | rs35513239 | 0.45 | 1.231 | 0.488 | -2.575 | -0.055 | 0.145 | 0.145 | 0 | 0.26 | 0.05 | 0.35 | 0.02 | 0.34 | 0.23 |
| 1 | 92689936 | rs3131824 | -0.648 | 1.254 | 0.506 | -2.557 | -0.049 | 0.051 | 0.051 | 0 | 0.9 | 0.98 | 0.89 | 1 | 0.81 | 0.96 |
| 1 | 92708323 | rs11588153 | 0.534 | 1.196 | 0.988 | -2.611 | -0.075 | 0.171 | 0.171 | 0 | 0.26 | 0.03 | 0.34 | 0.02 | 0.33 | 0.23 |
| 1 | 92742349 | rs11577572 | 0.557 | 1.203 | 0.991 | -2.608 | -0.075 | 0.169 | 0.169 | 0 | 0.26 | 0.03 | 0.34 | 0.02 | 0.33 | 0.23 |
| 1 | 92756474 | rs3131814 | 0.739 | 1.14 | 1.046 | -2.592 | 0.005 | -0.111 | -0.111 | 0 | 0.58 | 0.46 | 0.49 | 0.97 | 0.46 | 0.72 |
| 1 | 92765102 | rs10875349 | 0.739 | 1.14 | 1.046 | -2.592 | 0.005 | -0.111 | -0.111 | 6.48 | 0.58 | 0.46 | 0.49 | 0.97 | 0.46 | 0.72 |
| 1 | 92808334 | rs7528377 | 1.226 | 0.864 | 0.759 | -2.974 | -0.005 | 0.204 | 0.204 | 0.63 | 0.4 | 0.09 | 0.4 | 0.87 | 0.33 | 0.47 |
| 1 | 92822870 | rs12240065 | 1.596 | 0.875 | 1.026 | -3.062 | 0.005 | 0.029 | 0.029 | 0 | 0.15 | 0.08 | 0.08 | 0.08 | 0.12 | 0.21 |
| 1 | 92831569 | rs2046616 | 1.531 | 0.859 | 0.762 | -3.167 | 0.022 | 0.155 | 0.155 | 0.768 | 0.35 | 0.09 | 0.28 | 0.87 | 0.27 | 0.41 |
| 1 | 92842268 | rs11164288 | 0.301 | 1.256 | 1.099 | -2.812 | -0.08 | 0.156 | 0.156 | 0 | 0.63 | 0.85 | 0.54 | 0.97 | 0.47 | 0.73 |
| 1 | 92846681 | rs6692060 | -0.658 | 1.334 | 1.01 | -2.829 | -0.049 | 0.053 | 0.053 | 0 | 0.9 | 0.98 | 0.89 | 1 | 0.81 | 0.96 |
| 1 | 92846960 | rs6668803 | 0.326 | 1.334 | 1.01 | -2.829 | -0.08 | 0.156 | 0.156 | 0 | 0.63 | 0.85 | 0.53 | 0.97 | 0.47 | 0.73 |
| 1 | 92865160 | rs7524120 | -0.823 | 1.381 | 0.9 | -2.819 | -0.073 | 0.167 | 0.167 | 0 | 0.75 | 0.97 | 0.67 | 0.98 | 0.67 | 0.77 |
| 1 | 92882554 | rs2391157 | -0.799 | 1.36 | 0.901 | -2.818 | -0.075 | 0.169 | 0.169 | 0 | 0.74 | 0.97 | 0.66 | 0.98 | 0.67 | 0.77 |
| 1 | 92890344 | rs2025607 | 0.118 | 1.318 | 1.207 | -2.504 | -0.049 | 0.053 | 0.053 | 0 | 0.1 | 0.03 | 0.11 | 0 | 0.19 | 0.04 |
| 1 | 92935411 | rs6662618 | 0.18 | 1.096 | 1.146 | -2.543 | -0.023 | -0.151 | -0.151 | 0 | 0.78 | 0.52 | 0.73 | 0.98 | 0.82 | 0.85 |
| 1 | 92944994 | rs4970702 | 0.673 | 0.96 | 0.901 | -2.206 | -0.021 | 0.056 | 0.056 | 1.505 | 0.72 | 0.83 | 0.75 | 0.96 | 0.63 | 0.87 |
| 1 | 92957826 | rs7540564 | 1.111 | 0.909 | 1.429 | -2.228 | 0.002 | -0.071 | -0.071 | 0 | 0.67 | 0.42 | 0.71 | 0.94 | 0.62 | 0.8 |
| 1 | 92970953 | rs962222 | -0.478 | 1.018 | 1.124 | -2.185 | -0.024 | 0.019 | 0.019 | 0 | 0.91 | 0.95 | 0.89 | 0.98 | 0.81 | 0.96 |
| 1 | 92972933 | rs4970698 | -0.581 | 1.017 | 1.125 | -2.186 | -0.044 | 0.024 | 0.024 | 0 | 0.91 | 0.97 | 0.9 | 0.98 | 0.81 | 0.96 |
| 1 | 92974595 | rs10782922 | -0.478 | 1.017 | 1.122 | -2.186 | -0.024 | 0.019 | 0.019 | 0 | 0.91 | 0.95 | 0.89 | 0.98 | 0.81 | 0.96 |
| 1 | 92975938 | rs1886684 | 1.022 | 0.956 | 1.403 | -2.252 | 0.01 | -0.101 | -0.101 | 0 | 0.68 | 0.57 | 0.72 | 0.94 | 0.62 | 0.8 |

| | | | | | | | | | | | | | | | | |
|---|---|---|---|---|---|---|---|---|---|---|---|---|---|---|---|---|
| 1 | 92977275 | rs1886682 | -0.51 | 0.989 | 1.234 | -2.238 | -0.024 | 0.019 | 0.019 | 0 | 0.91 | 0.95 | 0.89 | 0.97 | 0.81 | 0.96 |
| 1 | 92979354 | rs6603979 | -0.51 | 0.989 | 1.234 | -2.238 | -0.024 | 0.019 | 0.019 | 0 | 0.91 | 0.95 | 0.89 | 0.98 | 0.81 | 0.96 |
| 1 | 93009313 | rs11804321 | 0.012 | 1.051 | 1.392 | -2.273 | 0.008 | 0.053 | 0.053 | 3.003 | 0.19 | 0.06 | 0.13 | 0.02 | 0.18 | 0.1 |
| 1 | 93017221 | rs6603986 | 0.823 | 1.043 | 1.4 | -2.276 | -0.03 | 0.084 | 0.084 | 1.427 | 0.72 | 0.88 | 0.76 | 0.96 | 0.63 | 0.86 |
| 1 | 93034023 | rs1556562 | -0.268 | 1.072 | 1.403 | -2.243 | -0.024 | 0.019 | 0.019 | 0 | 0.91 | 0.95 | 0.89 | 0.98 | 0.81 | 0.96 |
| 1 | 93073228 | rs11808092 | -0.301 | 0.88 | 1.476 | -2.181 | -0.032 | 0.096 | 0.096 | 4.077 | 0.25 | 0.07 | 0.19 | 0.02 | 0.27 | 0.11 |
| 1 | 93090934 | rs3790451 | 1.368 | 0.578 | 1.411 | -2.195 | -0.035 | 0.135 | 0.135 | 0.112 | 0.43 | 0.13 | 0.51 | 0.77 | 0.44 | 0.55 |
| 1 | 93181013 | rs11800409 | -0.343 | 0.651 | 1.846 | -1.958 | -0.029 | 0.106 | 0.106 | 3.489 | 0.26 | 0.09 | 0.19 | 0.03 | 0.27 | 0.12 |
| 1 | 93218776 | rs1337107 | 1.107 | 0.65 | 1.955 | -1.958 | -0.001 | 0.003 | 0.003 | 2.321 | 0.65 | 0.34 | 0.69 | 0.93 | 0.62 | 0.78 |
| 1 | 93263147 | rs7513624 | 1.832 | 0.713 | 2.2 | -1.665 | 0.057 | 0.009 | 0.009 | 3.51 | 0.91 | 0.48 | 0.88 | 0.97 | 0.89 | 0.9 |
| 1 | 93269824 | rs4847221 | 1.001 | 0.754 | 2.101 | -1.628 | 0.049 | 0.018 | 0.018 | 0.204 | 0.93 | 0.71 | 0.9 | 0.97 | 0.89 | 0.93 |
| 1 | 93274450 | rs12738964 | -0.431 | 0.737 | 1.542 | -1.867 | -0.004 | 0.003 | 0.003 | 0.862 | 0.28 | 0.17 | 0.2 | 0.01 | 0.27 | 0.13 |
| 1 | 93289668 | rs10874742 | 0.826 | 0.754 | 1.497 | -1.802 | 0.033 | 0.005 | 0.005 | 0.009 | 0.93 | 0.73 | 0.9 | 0.97 | 0.9 | 0.91 |
| 1 | 93297180 | rs34244251 | -0.701 | 0.75 | 1.569 | -1.868 | -0.001 | -0.088 | -0.088 | 0.177 | 0.3 | 0.33 | 0.22 | 0.01 | 0.27 | 0.18 |
| 1 | 93303603 | rs6604026 | -0.599 | 0.786 | 1.378 | -1.84 | -0.001 | -0.047 | -0.047 | 0 | 0.29 | 0.27 | 0.23 | 0.01 | 0.28 | 0.14 |
| 1 | 93306317 | rs10874744 | 0.785 | 0.762 | 1.185 | -1.821 | 0 | -0.122 | -0.122 | 3.108 | 0.64 | 0.46 | 0.69 | 0.96 | 0.62 | 0.81 |
| 1 | 93324269 | rs2811602 | 0.669 | 0.78 | 1.133 | -1.573 | 0.019 | -0.002 | -0.002 | 0 | 0.94 | 0.81 | 0.93 | 0.97 | 0.9 | 0.94 |
| 1 | 93349046 | rs7536563 | 0.793 | 0.727 | 1.055 | -1.81 | 0 | -0.037 | -0.037 | 0 | 0.63 | 0.4 | 0.67 | 0.96 | 0.62 | 0.76 |
| 1 | 93352669 | rs10782946 | 0.765 | 0.722 | 1.114 | -1.812 | -0.001 | -0.065 | -0.065 | 0 | 0.63 | 0.42 | 0.67 | 0.96 | 0.62 | 0.76 |
| 1 | 93368309 | rs2255723 | 0.289 | 0.686 | 1.391 | -1.848 | -0.018 | 0.08 | 0.08 | 1.225 | 0.69 | 0.84 | 0.73 | 0.97 | 0.63 | 0.82 |
| 1 | 93385136 | rs9651257 | -0.964 | 0.679 | 1.656 | -1.849 | 0.001 | -0.083 | -0.083 | 0 | 0.34 | 0.37 | 0.29 | 0.04 | 0.37 | 0.19 |
| 1 | 93397300 | rs6698233 | -0.719 | 0.679 | 1.649 | -1.849 | -0.004 | -0.002 | -0.002 | 0 | 0.32 | 0.25 | 0.28 | 0.03 | 0.37 | 0.18 |
| 1 | 93399694 | rs10493863 | -0.937 | 0.682 | 1.559 | -1.848 | 0.003 | -0.095 | -0.095 | 0 | 0.33 | 0.38 | 0.29 | 0.04 | 0.37 | 0.19 |
| 1 | 93421930 | rs10874752 | 1.186 | 0.735 | 0.759 | -1.951 | 0 | 0.009 | 0.009 | 0 | 0.43 | 0.4 | 0.49 | 0.73 | 0.46 | 0.51 |
| 1 | 93439821 | rs6671268 | 1.19 | 0.899 | 0.735 | -2.009 | -0.001 | 0.01 | 0.01 | 0 | 0.43 | 0.4 | 0.49 | 0.75 | 0.46 | 0.51 |
| 1 | 93443043 | rs11164854 | 0.139 | 0.92 | 0.077 | -1.922 | -0.009 | 0.006 | 0.006 | 0 | 0.22 | 0.06 | 0.18 | 0.24 | 0.16 | 0.24 |
| 1 | 93465636 | rs557095 | -0.324 | 0.842 | 0.011 | -1.305 | 0.003 | -0.026 | -0.026 | 0 | 0.82 | 0.89 | 0.73 | 0.64 | 0.87 | 0.76 |
| 1 | 93466960 | rs11164855 | -1.133 | 0.622 | -0.013 | -1.353 | -0.006 | 0.06 | 0.06 | 0.361 | 0.51 | 0.27 | 0.51 | 0.39 | 0.49 | 0.49 |
| 1 | 93468179 | rs17381406 | -0.363 | 1.038 | -0.165 | -1.543 | 0.001 | 0.025 | 0.025 | 1.347 | 0.21 | 0.14 | 0.16 | 0 | 0.24 | 0.2 |
| 1 | 93470714 | rs10493864 | -0.004 | 1.118 | 0.172 | -1.489 | 0.007 | -0.041 | -0.041 | 0 | 0.11 | 0.17 | 0.12 | 0.01 | 0.11 | 0.09 |
| 1 | 93472100 | rs12043007 | -0.217 | 1.325 | -0.672 | -1.702 | -0.068 | -0.036 | -0.036 | 2.796 | 0.41 | 0.04 | 0.4 | 0.43 | 0.44 | 0.38 |
| 1 | 93473150 | rs17381426 | 0.578 | 1.32 | -0.521 | -1.716 | 0.004 | -0.044 | -0.044 | 0.575 | 0.1 | 0.18 | 0.12 | 0.01 | 0.1 | 0.05 |
| 1 | 93473744 | rs492054 | 1.363 | 1.27 | 0.21 | -1.86 | -0.002 | 0.004 | 0.004 | 0 | 0.39 | 0.32 | 0.35 | 0.6 | 0.37 | 0.47 |
| 1 | 93477292 | rs2928183 | 1.03 | 1.251 | 0.342 | -1.901 | -0.005 | 0.013 | 0.013 | 0 | 0.43 | 0.35 | 0.51 | 0.61 | 0.45 | 0.53 |
| 1 | 93478970 | rs17131682 | 0.276 | 1.042 | 0.646 | -1.465 | 0.009 | -0.056 | -0.056 | 0 | 0.1 | 0.21 | 0.12 | 0.01 | 0.1 | 0.05 |
| 1 | 93479560 | rs11164862 | 0.181 | 0.947 | 0.724 | -1.577 | 0.006 | -0.007 | -0.007 | 0.035 | 0.16 | 0.12 | 0.11 | 0 | 0.16 | 0.08 |
| 1 | 93481640 | rs10493865 | 0.483 | 1.031 | 0.675 | -1.528 | 0.005 | -0.053 | -0.053 | 0 | 0.1 | 0.21 | 0.11 | 0.01 | 0.09 | 0.04 |
| 1 | 93489113 | rs680285 | 0.919 | 0.936 | 0.769 | -1.363 | -0.014 | 0.005 | 0.005 | 0 | 0.68 | 0.6 | 0.73 | 0.62 | 0.73 | 0.69 |
| 1 | 93490650 | rs6688632 | -0.163 | 0.914 | 0.807 | -1.404 | -0.016 | -0.081 | -0.081 | 0 | 0.18 | 0.28 | 0.12 | 0 | 0.16 | 0.08 |
| 1 | 93495280 | rs12043577 | 0.255 | 0.896 | 0.717 | -1.332 | 0.001 | -0.015 | -0.015 | 0 | 0.13 | 0.09 | 0.15 | 0.38 | 0.11 | 0.23 |
| 1 | 93496487 | rs12023767 | -0.078 | 0.91 | 0.716 | -1.325 | -0.004 | 0.031 | 0.031 | 0 | 0.15 | 0.19 | 0.15 | 0.38 | 0.11 | 0.23 |
| 1 | 93502451 | rs1678778 | 1.371 | 1.067 | 0.522 | -1.271 | 0.002 | 0.008 | 0.008 | 0 | 0.55 | 0.39 | 0.54 | 0.61 | 0.56 | 0.59 |
| 1 | 93513896 | rs2265928 | 1.148 | 1.258 | 0.372 | -1.16 | 0.004 | 0.003 | 0.003 | 0 | 0.63 | 0.51 | 0.63 | 0.6 | 0.64 | 0.61 |
| 1 | 93515049 | rs563930 | 0.989 | 1.255 | 0.765 | -1.069 | 0.001 | 0.036 | 0.036 | 0 | 0.71 | 0.54 | 0.69 | 0.62 | 0.75 | 0.66 |
| 1 | 93518578 | rs544285 | -0.048 | 0.934 | 0.628 | -0.968 | -0.008 | 0.048 | 0.048 | 0 | 0.83 | 0.75 | 0.84 | 0.64 | 0.87 | 0.72 |
| 1 | 93523083 | rs2249590 | 1.283 | 0.95 | -0.251 | -1.463 | -0.021 | 0.052 | 0.052 | 0.189 | 0.49 | 0.19 | 0.52 | 0.6 | 0.47 | 0.55 |
| 1 | 93525902 | rs2785599 | 0.051 | 1.057 | -0.018 | -1.392 | 0.041 | -0.011 | -0.011 | 0.217 | 0.05 | 0.16 | 0.09 | 0.01 | 0.11 | 0.06 |
| 1 | 93530980 | rs549938 | -0.532 | 0.465 | 0.108 | -1.402 | -0.025 | 0.01 | 0.01 | 0 | 0.32 | 0.45 | 0.28 | 0.37 | 0.24 | 0.35 |
| 1 | 93531706 | rs575367 | -0.002 | 0.736 | -0.03 | -1.747 | 0.002 | -0.017 | -0.017 | 1.473 | 0.87 | 0.86 | 0.89 | 0.97 | 0.81 | 0.95 |
| 1 | 93544629 | rs546445 | 1.517 | 0.107 | -0.203 | -1.694 | 0.008 | 0.035 | 0.035 | 0.804 | 0.76 | 0.17 | 0.78 | 0.93 | 0.74 | 0.91 |
| 1 | 93553223 | rs2815424 | 0.156 | 0.088 | 0.345 | -1.713 | 0.002 | -0.032 | -0.032 | 0.376 | 0.88 | 0.41 | 0.88 | 0.94 | 0.92 | 0.93 |
| 1 | 93558818 | rs12038699 | 1.787 | -0.053 | -0.34 | -1.659 | -0.001 | 0.003 | 0.003 | 0 | 0.62 | 0.09 | 0.59 | 0.64 | 0.64 | 0.7 |
| 1 | 93581024 | rs2295423 | 1.787 | -0.023 | -0.206 | -1.653 | -0.005 | 0.003 | 0.003 | 0 | 0.62 | 0.09 | 0.59 | 0.64 | 0.63 | 0.7 |
| 1 | 93584606 | rs4847399 | 1.937 | -0.077 | -0.254 | -1.663 | -0.044 | -0.037 | -0.037 | 0 | 0.61 | 0.03 | 0.58 | 0.64 | 0.63 | 0.69 |
| 1 | 93596936 | rs1010476 | -1.112 | -0.075 | 0.38 | -1.658 | -0.009 | -0.064 | -0.064 | 3.712 | 0.88 | 0.97 | 0.82 | 0.7 | 0.91 | 0.79 |
| 1 | 93616202 | rs1572343 | 1.715 | -0.099 | 0.55 | -1.694 | 0.016 | 0.006 | 0.006 | 0 | 0.64 | 0.17 | 0.6 | 0.64 | 0.66 | 0.7 |
| 1 | 93617546 | rs546 | 0.277 | -0.122 | 0.642 | -1.779 | 0.001 | -0.063 | -0.063 | 0 | 0.86 | 0.79 | 0.89 | 1 | 0.81 | 0.98 |
| 1 | 93620393 | rs1060622 | 2.211 | -0.153 | 0.599 | -1.782 | -0.008 | 0.004 | 0.004 | 0 | 0.62 | 0.07 | 0.58 | 0.64 | 0.63 | 0.69 |
| 1 | 93626984 | rs2783489 | -0.428 | -0.163 | 0.701 | -1.775 | -0.008 | 0.031 | 0.031 | 0.23 | 0.19 | 0.31 | 0.25 | 0.34 | 0.15 | 0.28 |
| 1 | 93639101 | rs10874762 | 2.048 | -0.158 | 0.61 | -1.771 | 0.015 | 0.011 | 0.011 | 0 | 0.63 | 0.13 | 0.6 | 0.64 | 0.66 | 0.7 |
| 1 | 93645182 | rs1335679 | 2.04 | -0.159 | 0.623 | -1.772 | 0.013 | 0.009 | 0.009 | 0 | 0.63 | 0.13 | 0.6 | 0.64 | 0.66 | 0.7 |
| 1 | 93646418 | rs10874763 | 1.754 | -0.207 | -0.126 | -2.417 | -0.124 | 0.165 | 0.165 | 0.05 | 0.24 | 0.01 | 0.19 | 0 | 0.33 | 0.12 |
| 1 | 93646433 | rs2255722 | 0.895 | -0.217 | 1.014 | -1.105 | -0.002 | -0.051 | -0.051 | 0 | 0.1 | 0.03 | 0.16 | 0.29 | 0.07 | 0.21 |
| 1 | 93651547 | rs12118262 | 2.228 | -0.221 | 1.02 | -1.109 | 0.01 | 0.007 | 0.007 | 0 | 0.64 | 0.13 | 0.6 | 0.65 | 0.65 | 0.7 |
| 1 | 93666349 | rs7513705 | 1.919 | -0.215 | 1.086 | -1.107 | -0.008 | -0.008 | -0.008 | 0 | 0.67 | 0.47 | 0.63 | 0.66 | 0.66 | 0.7 |
| 1 | 93672688 | rs2783499 | -0.757 | -0.22 | 1.133 | -1.105 | 0 | 0 | 0 | 0.178 | 0.18 | 0.2 | 0.24 | 0.32 | 0.15 | 0.27 |
| 1 | 93680736 | rs581825 | 1.534 | -0.229 | 1.145 | -1.116 | -0.007 | -0.063 | -0.063 | 0 | 0.82 | 0.33 | 0.84 | 0.96 | 0.81 | 0.97 |
| 1 | 93704326 | rs519723 | -1.016 | -0.23 | 1.175 | -1.117 | 0.014 | -0.073 | -0.073 | 0 | 0.14 | 0.33 | 0.13 | 0.02 | 0.19 | 0.03 |
| 1 | 93716974 | rs797680 | -2.8 | -0.244 | 1.119 | -1.127 | -0.008 | -0.001 | -0.001 | 0.453 | 0.38 | 0.93 | 0.41 | 0.36 | 0.36 | 0.31 |

| | | | | | | | | | | | | | | | | |
|---|---|---|---|---|---|---|---|---|---|---|---|---|---|---|---|---|
| 1 | 93723476 | rs633789 | -1.791 | -0.171 | 1.175 | -1.074 | -0.006 | -0.063 | -0.063 | 0 | 0.17 | 0.67 | 0.16 | 0.04 | 0.2 | 0.03 |
| 1 | 93741025 | rs12085959 | -0.963 | -0.137 | 1.098 | -1.06 | 0.002 | -0.063 | -0.063 | 0 | 0.14 | 0.22 | 0.12 | 0 | 0.19 | 0.02 |
| 1 | 93770334 | rs11164896 | 0.712 | -0.132 | 1.036 | -1.036 | 0.003 | -0.046 | -0.046 | 0 | 0.1 | 0.04 | 0.16 | 0.29 | 0.07 | 0.21 |
| 1 | 93787867 | rs6541349 | -1.262 | -0.146 | 1.134 | -1.054 | -0.019 | 0.058 | 0.058 | 0.036 | 0.21 | 0.53 | 0.27 | 0.33 | 0.15 | 0.27 |
| 1 | 93795932 | rs4847409 | 1.526 | -0.119 | 1.236 | -1.043 | -0.007 | -0.072 | -0.072 | 0.752 | 0.82 | 0.34 | 0.84 | 0.96 | 0.81 | 0.97 |
| 1 | 93810649 | rs11164911 | 1.513 | -0.1 | 1.243 | -1.034 | -0.007 | -0.072 | -0.072 | 0 | 0.82 | 0.34 | 0.84 | 0.96 | 0.81 | 0.97 |
| 1 | 93810900 | rs1323628 | 0.581 | -0.101 | 1.324 | -1.038 | 0.003 | -0.063 | -0.063 | 1.956 | 0.86 | 0.78 | 0.88 | 1 | 0.81 | 0.98 |
| 1 | 93811569 | rs10782959 | 1.522 | -0.113 | 1.245 | -1.024 | -0.004 | 0.001 | 0.001 | 0 | 0.72 | 0.3 | 0.68 | 0.67 | 0.73 | 0.75 |
| 1 | 93816400 | rs12133576 | 2.372 | -0.167 | 1.219 | -1.092 | 0.018 | 0.026 | 0.026 | 0 | 0.61 | 0.17 | 0.58 | 0.64 | 0.64 | 0.65 |
| 1 | 93817946 | rs4847240 | 2.372 | -0.167 | 1.219 | -1.092 | 0.018 | 0.026 | 0.026 | 0 | 0.61 | 0.17 | 0.58 | 0.64 | 0.64 | 0.65 |
| 1 | 93827300 | rs17131741 | -0.261 | -0.014 | 0.906 | -1.128 | -0.002 | -0.042 | -0.042 | 0 | 0.11 | 0.17 | 0.1 | 0.03 | 0.1 | 0.1 |
| 1 | 93863897 | rs1323626 | 0.602 | 0.091 | 0.847 | -1.087 | 0.002 | -0.053 | -0.053 | 0.238 | 0.86 | 0.82 | 0.89 | 1 | 0.81 | 0.98 |
| 1 | 93865391 | rs12138486 | 1.253 | -0.407 | 0.914 | -0.959 | 0.027 | -0.007 | -0.007 | 0.542 | 0.69 | 0.18 | 0.67 | 0.64 | 0.8 | 0.63 |
| 1 | 93875371 | rs2162276 | 1.016 | -0.377 | 1.268 | -0.928 | -0.022 | 0.002 | 0.002 | 0 | 0.73 | 0.61 | 0.72 | 0.71 | 0.83 | 0.65 |
| 1 | 93897624 | rs7339890 | -0.79 | -0.234 | 1.513 | -1.004 | -0.001 | -0.051 | -0.051 | 0.371 | 0.14 | 0.42 | 0.12 | 0.06 | 0.09 | 0.09 |
| 1 | 93900247 | rs2162277 | -0.381 | -0.35 | 1.544 | -0.886 | -0.002 | 0.018 | 0.018 | 0 | 0.87 | 0.83 | 0.83 | 0.68 | 0.9 | 0.72 |
| 1 | 93907020 | rs2493645 | -0.636 | -0.274 | 1.441 | -1.023 | 0 | -0.032 | -0.032 | 1.406 | 0.13 | 0.13 | 0.15 | 0 | 0.2 | 0.02 |
| 1 | 93909887 | rs2391271 | 2.12 | -0.485 | 0.975 | -1.191 | -0.142 | -0.027 | -0.027 | 0 | 0.42 | 0.02 | 0.37 | 0.6 | 0.49 | 0.49 |
| 1 | 93925862 | rs7543847 | 1.42 | -0.511 | 1.734 | -0.574 | -0.008 | 0.003 | 0.003 | 0 | 0.72 | 0.61 | 0.66 | 0.67 | 0.69 | 0.68 |
| 1 | 93941449 | rs17109997 | -0.278 | -0.535 | 1.691 | -0.595 | -0.009 | -0.037 | -0.037 | 0 | 0.12 | 0.18 | 0.1 | 0.04 | 0.08 | 0.11 |
| 1 | 93942442 | rs4847428 | 2.433 | -0.501 | 1.23 | -0.567 | 0.025 | 0.034 | 0.034 | 0 | 0.56 | 0.07 | 0.53 | 0.61 | 0.61 | 0.57 |
| 1 | 93953301 | rs11164939 | 0.503 | -0.413 | 1.269 | -0.514 | -0.005 | -0.056 | -0.056 | 0.633 | 0.12 | 0.04 | 0.16 | 0.33 | 0.09 | 0.29 |
| 1 | 93997477 | rs3767966 | 0.532 | -0.479 | 0.292 | -0.521 | -0.003 | -0.055 | -0.055 | 0 | 0.12 | 0.04 | 0.16 | 0.33 | 0.08 | 0.28 |
| 1 | 94002813 | rs237438 | 1.356 | -0.325 | 0.308 | -0.451 | 0.028 | 0.034 | 0.034 | 0 | 0.61 | 0.21 | 0.57 | 0.63 | 0.65 | 0.58 |
| 1 | 94012287 | rs237425 | 0.952 | 0.362 | 0.853 | -0.317 | 0.009 | -0.063 | -0.063 | 1.627 | 0.87 | 0.83 | 0.84 | 1 | 0.79 | 0.98 |
| 1 | 94026861 | rs17110063 | -0.764 | 0.459 | 0.652 | -0.168 | 0.03 | -0.038 | -0.038 | 0 | 0.1 | 0.18 | 0.14 | 0.01 | 0.19 | 0.04 |
| 1 | 94030534 | rs237435 | -0.427 | 0.319 | 0.18 | 0.037 | -0.002 | -0.065 | -0.065 | 0.053 | 0.76 | 0.88 | 0.67 | 0.62 | 0.76 | 0.56 |
| 1 | 94042207 | rs17110111 | -0.47 | 0.405 | 0.152 | -0.044 | 0.013 | -0.036 | -0.036 | 0 | 0.1 | 0.15 | 0.14 | 0 | 0.19 | 0.04 |
| 1 | 94044597 | rs236316 | 0.993 | 0.164 | 0.262 | 0.007 | -0.022 | -0.07 | -0.07 | 0.846 | 0.59 | 0.12 | 0.49 | 0.48 | 0.54 | 0.48 |
| 1 | 94049237 | rs3950119 | -1.048 | 0.459 | 0.881 | 0.221 | 0.04 | -0.074 | -0.074 | 0.129 | 0.12 | 0.34 | 0.14 | 0 | 0.19 | 0.04 |
| 1 | 94054382 | rs236319 | -0.101 | -0.055 | 0.571 | -0.034 | 0.035 | 0.021 | 0.021 | 0 | 0.53 | 0.68 | 0.51 | 0.63 | 0.62 | 0.46 |
| 1 | 94056279 | rs236322 | -2.185 | -0.03 | 1.118 | 0.031 | 0.01 | 0.041 | 0.041 | 1.303 | 0.23 | 0.83 | 0.27 | 0.14 | 0.23 | 0.1 |
| 1 | 94066635 | rs6677770 | -1.425 | 0.19 | 1.733 | 0.027 | 0.058 | -0.092 | -0.092 | 0.861 | 0.13 | 0.47 | 0.15 | 0 | 0.19 | 0.04 |
| 1 | 94081407 | rs236306 | -2.171 | -0.363 | 2.022 | 0.19 | -0.025 | -0.038 | -0.038 | 0 | 0.19 | 0.59 | 0.17 | 0.14 | 0.17 | 0.11 |
| 1 | 94083372 | rs6683296 | -0.628 | -0.358 | 2.131 | 0.188 | -0.003 | -0.036 | -0.036 | 0.114 | 0.09 | 0.14 | 0.11 | 0 | 0.13 | 0.03 |
| 1 | 94088363 | rs932523 | 1.414 | -0.657 | 1.48 | 0.459 | -0.001 | -0.107 | -0.107 | 1.308 | 0.47 | 0.07 | 0.37 | 0.44 | 0.57 | 0.45 |
| 1 | 94088650 | rs236312 | -2.563 | -0.265 | 2.92 | 0.579 | -0.036 | -0.063 | -0.063 | 0.245 | 0.2 | 0.53 | 0.15 | 0.13 | 0.17 | 0.09 |
| 1 | 94097441 | rs399649 | -0.918 | -0.266 | 2.708 | 0.659 | -0.004 | 0.024 | 0.024 | 0 | 0.07 | 0.16 | 0.03 | 0.13 | 0.04 | 0.08 |
| 1 | 94118883 | rs17110225 | -1.219 | -0.206 | 2.306 | 0.173 | -0.003 | -0.054 | -0.054 | 0.391 | 0.11 | 0.17 | 0.12 | 0 | 0.13 | 0.03 |
| 1 | 94132788 | rs236285 | -2.223 | -0.239 | 1.291 | 0.172 | -0.024 | -0.056 | -0.056 | 0 | 0.22 | 0.62 | 0.17 | 0.09 | 0.17 | 0.15 |
| 1 | 94143719 | rs1011903 | -1.763 | 0.047 | 0.847 | 0.476 | 0 | 0 | 0 | 0.075 | 0.18 | 0.75 | 0.17 | 0.14 | 0.17 | 0.13 |
| 1 | 94151589 | rs6702036 | -1.748 | 0.057 | 0.851 | 0.151 | 0 | -0.004 | -0.004 | 0 | 0.18 | 0.75 | 0.17 | 0.1 | 0.17 | 0.13 |
| 1 | 94155292 | rs12084853 | -1.729 | 0.037 | 0.717 | 0.174 | 0.002 | 0.003 | 0.003 | 2.122 | 0.18 | 0.76 | 0.17 | 0.1 | 0.17 | 0.13 |
| 1 | 94160787 | rs2893248 | 1.189 | 0.059 | 0.677 | 0.067 | 0.004 | -0.005 | -0.005 | 0 | 0.8 | 0.3 | 0.83 | 0.98 | 0.81 | 0.9 |
| 1 | 94175909 | rs12022530 | 0.334 | 0.265 | 0.456 | 0.801 | 0.002 | 0 | 0 | 0 | 0.6 | 0.42 | 0.43 | 0.52 | 0.49 | 0.43 |
| 1 | 94178912 | rs10493866 | -1.065 | -0.472 | 0.631 | -0.34 | 0.002 | 0.001 | 0.001 | 0 | 0.16 | 0.15 | 0.05 | 0.08 | 0.1 | 0.09 |
| 1 | 94185344 | rs4847183 | 0.838 | -0.403 | 0.774 | -0.701 | 0.004 | 0.04 | 0.04 | 0 | 0.48 | 0.42 | 0.42 | 0.23 | 0.5 | 0.32 |
| 1 | 94190110 | rs6541393 | 0.878 | -0.453 | 1.238 | -0.771 | -0.018 | -0.013 | -0.013 | 0 | 0.8 | 0.74 | 0.85 | 0.85 | 0.88 | 0.88 |
| 1 | 94194707 | rs6684007 | -0.013 | -0.297 | 1.178 | -0.634 | 0.001 | -0.002 | -0.002 | 0.007 | 0.32 | 0.31 | 0.33 | 0.62 | 0.38 | 0.56 |
| 1 | 94198856 | rs1467129 | 0.541 | -0.164 | 1.037 | -0.497 | -0.001 | 0 | 0 | 0 | 0.07 | 0.05 | 0.06 | 0.05 | 0.09 | 0.1 |
| 1 | 94199666 | rs17110370 | 0.254 | -0.031 | 1.032 | -0.68 | -0.008 | 0.001 | 0.001 | 0.616 | 0.08 | 0.06 | 0.11 | 0 | 0.18 | 0.04 |
| 1 | 94206162 | rs1108528 | -0.45 | -0.122 | 1.45 | -0.623 | 0.013 | 0.008 | 0.008 | 0.017 | 0.93 | 0.8 | 0.93 | 0.94 | 0.92 | 0.9 |
| 1 | 94210906 | rs12410101 | -0.271 | -0.406 | 1.612 | -0.45 | -0.026 | 0.031 | 0.031 | 0 | 0.13 | 0.18 | 0.17 | 0.19 | 0.06 | 0.06 |
| 1 | 94214272 | rs11164974 | 1.324 | -0.61 | 1.234 | -0.937 | -0.061 | 0.138 | 0.138 | 0 | 0.4 | 0.17 | 0.39 | 0.23 | 0.5 | 0.3 |
| 1 | 94214518 | rs11164976 | -0.374 | -0.629 | 1.374 | -0.912 | -0.048 | 0.075 | 0.075 | 0.248 | 0.14 | 0.32 | 0.18 | 0.2 | 0.06 | 0.07 |
| 1 | 94217804 | rs4350209 | -0.577 | -0.662 | 1.223 | -1.038 | -0.064 | -0.062 | -0.062 | 0 | 0.65 | 0.92 | 0.65 | 0.37 | 0.66 | 0.48 |
| 1 | 94224065 | rs4526623 | -0.56 | -0.72 | 1.486 | -0.835 | 0.025 | -0.022 | -0.022 | 0 | 0.12 | 0.43 | 0.11 | 0.05 | 0.12 | 0.12 |
| 1 | 94228900 | rs10158448 | -0.168 | -0.711 | 1.462 | -0.835 | -0.04 | 0.063 | 0.063 | 0.09 | 0.14 | 0.26 | 0.17 | 0.2 | 0.06 | 0.07 |
| 1 | 94241556 | rs2391313 | -1.577 | -0.795 | 1.353 | -0.897 | 0.009 | -0.004 | -0.004 | 0.181 | 0.26 | 0.68 | 0.28 | 0.26 | 0.18 | 0.19 |
| 1 | 94272119 | rs12406729 | -0.399 | -0.732 | 1.325 | -0.867 | -0.05 | 0.079 | 0.079 | 0.443 | 0.14 | 0.35 | 0.18 | 0.19 | 0.06 | 0.07 |
| 1 | 94291481 | rs11165012 | -1.5 | -0.73 | 1.042 | -0.879 | 0.028 | -0.01 | -0.01 | 0.243 | 0.26 | 0.75 | 0.28 | 0.25 | 0.18 | 0.19 |
| 1 | 94320233 | rs12565817 | 0.746 | -0.612 | 0.889 | -0.916 | -0.04 | -0.139 | -0.139 | 0.268 | 0.27 | 0.06 | 0.23 | 0.46 | 0.28 | 0.49 |
| 1 | 94332151 | rs10489603 | 0.839 | -0.632 | 0.87 | -0.918 | -0.08 | -0.185 | -0.185 | 0 | 0.26 | 0.02 | 0.24 | 0.46 | 0.28 | 0.5 |
| 1 | 94332263 | rs10489604 | 0.421 | -0.52 | 1.036 | -0.841 | -0.06 | -0.081 | -0.081 | 0.002 | 0.33 | 0.08 | 0.33 | 0.56 | 0.33 | 0.51 |
| 1 | 94336935 | rs12041975 | -0.229 | -0.541 | 1.267 | -0.859 | 0.001 | 0.032 | 0.032 | 0 | 0.41 | 0.27 | 0.39 | 0.62 | 0.4 | 0.61 |
| 1 | 94337048 | rs2143992 | -0.499 | -0.543 | 1.486 | -0.865 | -0.001 | -0.006 | -0.006 | 0 | 0.43 | 0.55 | 0.43 | 0.62 | 0.42 | 0.62 |
| 1 | 94340556 | rs4847264 | -0.499 | -0.543 | 1.486 | -0.865 | -0.001 | -0.006 | -0.006 | 0 | 0.43 | 0.55 | 0.43 | 0.62 | 0.42 | 0.62 |
| 1 | 94341137 | rs2273166 | -0.187 | -0.543 | 1.459 | -0.864 | 0.001 | 0.031 | 0.031 | 0 | 0.41 | 0.26 | 0.39 | 0.62 | 0.4 | 0.61 |
| 1 | 94342564 | rs3747965 | 0.447 | -0.537 | 1.395 | -0.858 | -0.06 | -0.082 | -0.082 | 0 | 0.33 | 0.08 | 0.33 | 0.56 | 0.33 | 0.51 |

| | | | | | | | | | | | | | | | | |
|---|---|---|---|---|---|---|---|---|---|---|---|---|---|---|---|---|
| 1 | 94343023 | rs2391322 | 0.223 | -0.537 | 1.433 | -0.858 | 0 | 0 | 0 | 0 | 0.36 | 0.38 | 0.37 | 0.56 | 0.35 | 0.53 |
| 1 | 94343233 | rs3789457 | -1.793 | -0.712 | 1.275 | -0.998 | -0.022 | 0.072 | 0.072 | 0 | 0.66 | 0.81 | 0.81 | 0.79 | 0.66 | 0.73 |
| 1 | 94344110 | rs4141737 | 0.547 | -0.705 | 1.224 | -0.98 | -0.057 | -0.078 | -0.078 | 0 | 0.33 | 0.09 | 0.33 | 0.56 | 0.33 | 0.51 |
| 1 | 94344716 | rs2235971 | 0.537 | -0.705 | 1.229 | -0.98 | -0.053 | -0.074 | -0.074 | 1.067 | 0.33 | 0.09 | 0.33 | 0.56 | 0.33 | 0.51 |
| 1 | 94345137 | rs2235970 | -1.141 | -0.719 | 1.368 | -1.001 | -0.006 | 0.026 | 0.026 | 0.111 | 0.58 | 0.37 | 0.72 | 0.73 | 0.59 | 0.63 |
| 1 | 94346327 | rs6687387 | -1.739 | -0.64 | 1.414 | -0.973 | 0.018 | 0.03 | 0.03 | 0.743 | 0.33 | 0.75 | 0.48 | 0.23 | 0.34 | 0.22 |
| 1 | 94352381 | rs10489605 | 1.019 | -0.646 | 1.472 | -0.976 | 0.002 | 0.001 | 0.001 | 0 | 0.07 | 0.05 | 0.05 | 0.06 | 0.07 | 0.1 |
| 1 | 94354948 | rs10874809 | -1.751 | -0.584 | 1.434 | -0.969 | 0.016 | 0.03 | 0.03 | 0.743 | 0.33 | 0.75 | 0.48 | 0.23 | 0.34 | 0.22 |
| 1 | 94360107 | rs769211 | -0.8 | -0.634 | 1.436 | -0.97 | -0.004 | -0.005 | -0.005 | 0 | 0.23 | 0.25 | 0.37 | 0.16 | 0.24 | 0.11 |
| 1 | 94361019 | rs11165031 | -0.941 | -0.63 | 1.428 | -0.964 | -0.011 | -0.013 | -0.013 | 0.987 | 0.24 | 0.32 | 0.38 | 0.16 | 0.24 | 0.11 |
| 1 | 94365334 | rs7517826 | -1.495 | -0.61 | 1.216 | -0.875 | 0.018 | 0.03 | 0.03 | 0.743 | 0.33 | 0.75 | 0.48 | 0.23 | 0.34 | 0.22 |
| 1 | 94367097 | rs7515191 | -1.495 | -0.61 | 1.216 | -0.875 | 0.018 | 0.03 | 0.03 | 0.743 | 0.33 | 0.75 | 0.48 | 0.23 | 0.34 | 0.22 |
| 1 | 94371909 | rs718873 | -0.361 | -0.44 | 1.367 | -0.705 | -0.011 | 0 | 0 | 0 | 0.17 | 0.21 | 0.31 | 0.16 | 0.15 | 0.09 |
| 1 | 94372878 | rs2301022 | -0.956 | -0.481 | 1.395 | -0.759 | -0.001 | -0.006 | -0.006 | 0 | 0.59 | 0.39 | 0.67 | 0.77 | 0.65 | 0.69 |
| 1 | 94374026 | rs3789453 | -1.583 | -0.428 | 1.164 | -0.731 | -0.023 | 0.05 | 0.05 | 0.038 | 0.67 | 0.78 | 0.81 | 0.77 | 0.67 | 0.73 |
| 1 | 94385591 | rs1473711 | -0.585 | -0.409 | 1.067 | -0.694 | -0.037 | 0.015 | 0.015 | 0.17 | 0.18 | 0.48 | 0.34 | 0.16 | 0.16 | 0.1 |
| 1 | 94388569 | rs743111 | -0.11 | -0.402 | 0.431 | -0.685 | -0.035 | 0.031 | 0.031 | 0 | 0.45 | 0.17 | 0.45 | 0.6 | 0.49 | 0.62 |
| 1 | 94394079 | rs6696758 | -0.369 | -0.395 | 0.411 | -0.697 | -0.022 | 0.003 | 0.003 | 0 | 0.17 | 0.25 | 0.31 | 0.16 | 0.15 | 0.1 |
| 1 | 94400045 | rs7520019 | -0.332 | -0.153 | -0.044 | -0.518 | -0.036 | 0.025 | 0.025 | 0 | 0.45 | 0.17 | 0.45 | 0.59 | 0.49 | 0.62 |
| 1 | 94417219 | rs4847269 | -1.406 | 0.039 | 0.287 | -0.163 | 0.021 | 0.077 | 0.077 | 1.31 | 0.49 | 0.97 | 0.6 | 0.3 | 0.4 | 0.29 |
| 1 | 94419799 | rs10874819 | -1.053 | 0.261 | 0.926 | 0.139 | 0.033 | 0.073 | 0.073 | 1.178 | 0.42 | 0.69 | 0.41 | 0.2 | 0.36 | 0.25 |
| 1 | 94441950 | rs743117 | -0.478 | 0.3 | 0.859 | -0.302 | -0.006 | -0.018 | -0.018 | 0.196 | 0.35 | 0.36 | 0.36 | 0.19 | 0.32 | 0.2 |
| 1 | 94442739 | rs743113 | -0.726 | 0.395 | 0.493 | -0.298 | -0.008 | -0.053 | -0.053 | 0.228 | 0.38 | 0.45 | 0.38 | 0.19 | 0.36 | 0.22 |
| 1 | 94450131 | rs1889547 | -0.894 | 0 | -0.296 | 0.155 | 0.014 | 0.05 | 0.05 | 0.071 | 0.44 | 0.68 | 0.43 | 0.21 | 0.41 | 0.29 |
| 1 | 94455420 | rs11165057 | -1.194 | -0.55 | 0.478 | 0.348 | -0.083 | -0.028 | -0.028 | 0 | 0.19 | 0.54 | 0.17 | 0.13 | 0.07 | 0.11 |
| 1 | 94455721 | rs11165058 | -1.134 | -0.672 | 1.532 | 0.272 | -0.049 | -0.015 | -0.015 | 3.874 | 0.17 | 0.31 | 0.16 | 0.13 | 0.06 | 0.09 |
| 1 | 94458421 | rs3747961 | -0.806 | -0.742 | 1.627 | 0.231 | -0.009 | -0.007 | -0.007 | 0 | 0.14 | 0.15 | 0.05 | 0.05 | 0.04 | 0.06 |
| 1 | 94461717 | rs6666652 | -1.103 | -0.79 | 2.809 | 0.19 | -0.077 | -0.097 | -0.097 | 0 | 0.15 | 0.52 | 0.08 | 0.03 | 0.05 | 0.07 |
| 1 | 94461754 | rs1800717 | -1.188 | -0.789 | 3.109 | 0.191 | -0.069 | -0.087 | -0.087 | 0 | 0.15 | 0.42 | 0.07 | 0.03 | 0.05 | 0.07 |
| 1 | 94462769 | rs17110736 | -1.447 | -0.768 | 3.17 | 0.272 | -0.053 | -0.017 | -0.017 | 1.57 | 0.16 | 0.35 | 0.07 | 0.11 | 0.06 | 0.08 |
| 1 | 94465132 | rs3789375 | -2.445 | -1.465 | 3.206 | -0.271 | 0.034 | 0.029 | 0.029 | 6.963 | 0.3 | 0.76 | 0.14 | 0.16 | 0.12 | 0.17 |
| 1 | 94465461 | rs4147871 | -0.815 | -1.502 | 3.336 | -0.243 | -0.049 | -0.045 | -0.045 | 0 | 0.16 | 0.31 | 0.06 | 0.07 | 0.06 | 0.08 |
| 1 | 94465677 | rs4147869 | -0.815 | -1.502 | 3.336 | -0.251 | -0.049 | -0.049 | -0.049 | 0 | 0.16 | 0.32 | 0.06 | 0.07 | 0.06 | 0.08 |
| 1 | 94466088 | rs12070273 | -0.815 | -1.502 | 3.336 | -0.251 | -0.049 | -0.049 | -0.049 | 0.052 | 0.16 | 0.32 | 0.06 | 0.07 | 0.06 | 0.08 |
| 1 | 94466154 | rs4147866 | -1.575 | -1.501 | 3.354 | -0.252 | -0.037 | -0.002 | -0.002 | 2.602 | 0.14 | 0.41 | 0.07 | 0.09 | 0.06 | 0.09 |
| 1 | 94467238 | rs4147864 | -0.863 | -1.435 | 3.189 | -0.231 | -0.049 | -0.049 | -0.049 | 0 | 0.16 | 0.32 | 0.06 | 0.07 | 0.06 | 0.08 |
| 1 | 94467407 | rs17110761 | -0.863 | -1.435 | 3.189 | -0.231 | -0.049 | -0.049 | -0.049 | 0.419 | 0.16 | 0.32 | 0.06 | 0.07 | 0.06 | 0.08 |
| 1 | 94469631 | rs7537325 | -1.723 | -0.205 | 2.969 | -0.433 | 0.059 | -0.016 | -0.016 | 0.202 | 0.16 | 0.39 | 0.22 | 0.09 | 0.24 | 0.1 |
| 1 | 94471075 | rs1762114 | 0.909 | -0.657 | 3.125 | -0.185 | -0.054 | -0.026 | -0.026 | 0.185 | 0.82 | 0.37 | 0.9 | 0.9 | 0.94 | 0.91 |
| 1 | 94471154 | rs4147863 | -2.139 | -0.574 | 3.095 | -0.279 | -0.001 | 0.002 | 0.002 | 0 | 0.13 | 0.15 | 0.13 | 0.12 | 0.18 | 0.11 |
| 1 | 94471519 | rs4147862 | -2.394 | -0.6 | 3.093 | -0.292 | -0.004 | 0.005 | 0.005 | 0 | 0.14 | 0.25 | 0.13 | 0.12 | 0.18 | 0.11 |
| 1 | 94471948 | rs4147861 | -0.492 | -0.586 | 3.106 | -0.294 | -0.001 | 0.006 | 0.006 | 0 | 0.07 | 0.07 | 0.03 | 0.08 | 0.06 | 0.08 |
| 1 | 94472489 | rs557026 | 0.401 | -0.272 | 3.006 | -0.412 | -0.021 | -0.064 | -0.064 | 0 | 0.83 | 0.75 | 0.92 | 0.98 | 0.89 | 0.96 |
| 1 | 94472520 | rs3789379 | -2.387 | -0.271 | 3 | -0.411 | -0.005 | 0.004 | 0.004 | 0 | 0.15 | 0.26 | 0.14 | 0.12 | 0.18 | 0.11 |
| 1 | 94472909 | rs7531001 | -2.387 | -0.271 | 3 | -0.381 | -0.005 | 0.005 | 0.005 | 0 | 0.15 | 0.26 | 0.17 | 0.12 | 0.18 | 0.11 |
| 1 | 94473845 | rs2275029 | -2.083 | -0.28 | 2.987 | -0.406 | -0.017 | -0.002 | -0.002 | 0 | 0.16 | 0.26 | 0.16 | 0.12 | 0.14 | 0.11 |
| 1 | 94473896 | rs2275031 | -2.191 | -0.284 | 2.988 | -0.405 | -0.012 | -0.004 | -0.004 | 0 | 0.17 | 0.27 | 0.18 | 0.12 | 0.18 | 0.11 |
| 1 | 94474020 | rs1191234 | 0.471 | -0.299 | 2.986 | -0.413 | -0.021 | -0.062 | -0.062 | 0 | 0.83 | 0.76 | 0.93 | 0.98 | 0.89 | 0.95 |
| 1 | 94474185 | rs2275032 | -2.204 | -0.278 | 2.95 | -0.386 | -0.012 | -0.004 | -0.004 | 0 | 0.17 | 0.26 | 0.18 | 0.12 | 0.18 | 0.11 |
| 1 | 94474328 | rs4147857 | -2.219 | -0.291 | 2.944 | -0.392 | -0.012 | -0.005 | -0.005 | 0 | 0.17 | 0.27 | 0.23 | 0.12 | 0.19 | 0.12 |
| 1 | 94474452 | rs4147856 | -2.219 | -0.291 | 2.944 | -0.392 | -0.012 | -0.005 | -0.005 | 0 | 0.17 | 0.27 | 0.23 | 0.12 | 0.19 | 0.12 |
| 1 | 94474872 | rs17110808 | -1.049 | -0.291 | 2.954 | -0.392 | -0.014 | 0.002 | 0.002 | 0.011 | 0.1 | 0.25 | 0.13 | 0.08 | 0.07 | 0.09 |
| 1 | 94474961 | rs3789387 | -1.957 | -0.292 | 3.021 | -0.329 | -0.007 | 0.074 | 0.074 | 0.749 | 0.13 | 0.54 | 0.17 | 0.17 | 0.08 | 0.15 |
| 1 | 94476035 | rs2065712 | -2.402 | -0.288 | 2.992 | -0.332 | -0.002 | -0.009 | -0.009 | 0 | 0.2 | 0.27 | 0.21 | 0.12 | 0.25 | 0.13 |
| 1 | 94477232 | rs565155 | -0.13 | -0.31 | 2.787 | -0.176 | -0.048 | -0.032 | -0.032 | 0.093 | 0.86 | 0.77 | 0.96 | 0.94 | 0.96 | 0.96 |
| 1 | 94477634 | rs11165062 | -1.54 | -0.314 | 2.744 | -0.173 | 0.001 | -0.017 | -0.017 | 0 | 0.14 | 0.25 | 0.11 | 0.08 | 0.13 | 0.1 |
| 1 | 94477893 | rs945067 | 1.476 | -0.311 | 2.686 | -0.161 | -0.039 | -0.008 | -0.008 | 0 | 0.75 | 0.38 | 0.81 | 0.82 | 0.83 | 0.87 |
| 1 | 94478293 | rs12085639 | -1.782 | -0.293 | 2.733 | -0.166 | 0.078 | 0.131 | 0.131 | 0 | 0.08 | 0.4 | 0.14 | 0.11 | 0.14 | 0.08 |
| 1 | 94478425 | rs486879 | 1.126 | -0.28 | 2.353 | -0.17 | -0.03 | -0.011 | -0.011 | 0.333 | 0.74 | 0.34 | 0.81 | 0.83 | 0.82 | 0.87 |
| 1 | 94478573 | rs567370 | 2.206 | -0.287 | 2.063 | -0.178 | 0.06 | 0.042 | 0.042 | 2.447 | 0.62 | 0.19 | 0.65 | 0.75 | 0.69 | 0.77 |
| 1 | 94478784 | rs4147851 | 2.245 | -0.312 | 1.171 | -0.202 | 0.059 | 0.044 | 0.044 | 2.416 | 0.61 | 0.19 | 0.65 | 0.75 | 0.69 | 0.77 |
| 1 | 94478847 | rs17391612 | -1.721 | 0.168 | 0.948 | -0.128 | -0.003 | 0.008 | 0.008 | 0 | 0.13 | 0.13 | 0.1 | 0.08 | 0.13 | 0.11 |
| 1 | 94479338 | rs3789391 | -0.917 | 0.088 | 0.847 | 0.367 | 0.005 | -0.026 | -0.026 | 1.73 | 0.09 | 0.05 | 0.06 | 0.17 | 0.07 | 0.1 |
| 1 | 94479468 | rs12049183 | -1.141 | 0.058 | 0.864 | 0.306 | 0.008 | -0.015 | -0.015 | 0 | 0.09 | 0.07 | 0.07 | 0.15 | 0.07 | 0.09 |
| 1 | 94480439 | rs2275034 | -0.42 | 0.338 | 0.438 | -0.83 | -0.015 | 0.017 | 0.017 | 0.908 | 0.66 | 0.71 | 0.47 | 0.24 | 0.6 | 0.46 |
| 1 | 94480529 | rs3818778 | -0.042 | 0.01 | 1.058 | -1.206 | 0.007 | -0.047 | -0.047 | 0 | 0.58 | 0.43 | 0.37 | 0.17 | 0.55 | 0.41 |
| 1 | 94481068 | rs914958 | 0.655 | -0.183 | 1.27 | -1.987 | 0.006 | -0.056 | -0.056 | 0 | 0.38 | 0.39 | 0.15 | 0.03 | 0.24 | 0.25 |
| 1 | 94481596 | rs915201 | 0.297 | -0.176 | 1.234 | -1.982 | -0.013 | -0.067 | -0.067 | 0 | 0.42 | 0.56 | 0.2 | 0.1 | 0.26 | 0.29 |
| 1 | 94481904 | rs915200 | 1.153 | -1.012 | 0.985 | -2.467 | -0.026 | -0.037 | -0.037 | 0.486 | 0.22 | 0.27 | 0.09 | 0.07 | 0.08 | 0.09 |

| | | | | | | | | | | | | | | | | |
|---|---|---|---|---|---|---|---|---|---|---|---|---|---|---|---|---|
| 1 | 94481929 | rs915199 | -0.156 | -1.095 | 0.746 | -2.515 | -0.02 | -0.084 | -0.084 | 0 | 0.38 | 0.55 | 0.19 | 0.1 | 0.25 | 0.28 |
| 1 | 94484705 | rs6681968 | -1.457 | -0.875 | 0.567 | -2.509 | -0.022 | 0.036 | 0.036 | 0 | 0.63 | 0.69 | 0.46 | 0.24 | 0.57 | 0.47 |
| 1 | 94485491 | rs17110850 | 0.244 | -0.502 | -0.304 | -2.253 | -0.014 | 0 | 0 | 0 | 0.12 | 0.05 | 0.07 | 0.09 | 0.14 | 0.09 |
| 1 | 94486406 | rs10493867 | -0.213 | -0.778 | -0.069 | -2.401 | 0.001 | -0.005 | -0.005 | 0 | 0.33 | 0.29 | 0.17 | 0.12 | 0.26 | 0.23 |
| 1 | 94486587 | rs4147848 | -0.854 | -0.772 | 0.289 | -2.496 | 0.007 | -0.052 | -0.052 | 0 | 0.31 | 0.46 | 0.21 | 0.16 | 0.31 | 0.28 |
| 1 | 94486667 | rs933073 | 0.343 | -0.734 | 0.201 | -2.315 | 0.041 | 0.022 | 0.022 | 0 | 0.06 | 0.15 | 0.04 | 0.08 | 0.11 | 0.04 |
| 1 | 94487354 | rs472908 | -0.292 | -0.116 | -0.506 | -1.747 | -0.017 | -0.012 | -0.012 | 0.146 | 0.61 | 0.76 | 0.57 | 0.46 | 0.56 | 0.53 |
| 1 | 94488326 | rs2282229 | 1.36 | -0.196 | 0.021 | -1.982 | -0.006 | -0.011 | -0.011 | 0 | 0.09 | 0.16 | 0.12 | 0.08 | 0.06 | 0.1 |
| 1 | 94488497 | rs1932014 | -0.883 | -0.271 | 0.047 | -2.089 | 0 | -0.005 | -0.005 | 0 | 0.59 | 0.56 | 0.44 | 0.26 | 0.57 | 0.47 |
| 1 | 94489975 | rs2151847 | -0.919 | -0.269 | 0.04 | -2.091 | 0.001 | -0.008 | -0.008 | 0 | 0.6 | 0.56 | 0.45 | 0.26 | 0.57 | 0.48 |
| 1 | 94491468 | rs11165065 | 0.187 | -0.27 | 0.082 | -2.135 | -0.001 | 0.001 | 0.001 | 0 | 0.27 | 0.2 | 0.27 | 0.11 | 0.31 | 0.24 |
| 1 | 94492773 | rs3945204 | -0.531 | -0.597 | 0.305 | -1.83 | -0.008 | 0.03 | 0.03 | 0 | 0.57 | 0.64 | 0.46 | 0.27 | 0.57 | 0.47 |
| 1 | 94493231 | rs17110885 | 1.865 | -0.763 | 0.23 | -1.906 | -0.002 | -0.031 | -0.031 | 1.565 | 0.19 | 0.17 | 0.05 | 0 | 0.06 | 0.06 |
| 1 | 94495407 | rs4147846 | -0.23 | -0.709 | 0.257 | -1.888 | 0.022 | -0.031 | -0.031 | 1.272 | 0.37 | 0.45 | 0.4 | 0.26 | 0.5 | 0.4 |
| 1 | 94495417 | rs4147845 | 0.031 | -0.706 | 0.172 | -1.887 | -0.008 | 0.008 | 0.008 | 0 | 0.36 | 0.27 | 0.38 | 0.26 | 0.5 | 0.4 |
| 1 | 94495487 | rs4147844 | -0.002 | -0.708 | 0.172 | -1.888 | -0.007 | 0.007 | 0.007 | 0 | 0.37 | 0.28 | 0.38 | 0.26 | 0.5 | 0.4 |
| 1 | 94496253 | rs2297671 | -0.431 | -0.76 | 0.148 | -1.949 | 0.009 | -0.021 | -0.021 | 0 | 0.39 | 0.4 | 0.39 | 0.26 | 0.5 | 0.4 |
| 1 | 94497133 | rs7543524 | 1.878 | -0.68 | 0.167 | -1.915 | -0.033 | -0.074 | -0.074 | 0 | 0.19 | 0.27 | 0.06 | 0 | 0.06 | 0.05 |
| 1 | 94497178 | rs4147841 | -0.805 | -0.626 | 0.107 | -1.911 | -0.008 | 0.022 | 0.022 | 0 | 0.6 | 0.64 | 0.44 | 0.25 | 0.57 | 0.46 |
| 1 | 94499110 | rs17110905 | 2.11 | -0.383 | 0.049 | -1.751 | 0.03 | 0.022 | 0.022 | 1.824 | 0.17 | 0.09 | 0.04 | 0 | 0.06 | 0.06 |
| 1 | 94499133 | rs3789393 | -0.531 | -0.337 | 0.019 | -1.738 | -0.002 | 0.005 | 0.005 | 0 | 0.56 | 0.54 | 0.43 | 0.25 | 0.57 | 0.48 |
| 1 | 94501594 | rs3789395 | -0.743 | 0.049 | -0.002 | -1.21 | 0.004 | -0.017 | -0.017 | 0 | 0.42 | 0.42 | 0.41 | 0.25 | 0.57 | 0.44 |
| 1 | 94501799 | rs1320502 | -0.703 | 0.194 | -0.49 | -1.144 | -0.009 | 0.018 | 0.018 | 0 | 0.4 | 0.29 | 0.4 | 0.25 | 0.57 | 0.42 |
| 1 | 94503197 | rs1889548 | 0.423 | 0.459 | -0.339 | -0.736 | 0.005 | -0.003 | -0.003 | 0.071 | 0.73 | 0.37 | 0.72 | 0.77 | 0.69 | 0.8 |
| 1 | 94504545 | rs11165069 | 0.092 | -0.113 | -0.458 | -0.953 | -0.003 | -0.031 | -0.031 | 0 | 0.33 | 0.29 | 0.31 | 0.08 | 0.18 | 0.32 |
| 1 | 94509226 | rs4147839 | 0.042 | 0.359 | -0.252 | -0.27 | -0.021 | 0.001 | 0.001 | 0 | 0.3 | 0.44 | 0.29 | 0.66 | 0.26 | 0.42 |
| 1 | 94510673 | rs2297633 | 0.04 | 1.81 | -0.756 | -0.556 | -0.057 | 0.009 | 0.009 | 3.552 | 0.14 | 0.07 | 0.21 | 0.16 | 0.36 | 0.08 |
| 1 | 94511717 | rs3789399 | -0.745 | 1.701 | -1.291 | -1.176 | -0.145 | 0.104 | 0.104 | 2.125 | 0.29 | 0.07 | 0.27 | 0.16 | 0.46 | 0.15 |
| 1 | 94512893 | rs544830 | -1.082 | 1.672 | -0.687 | -1.149 | -0.024 | 0.022 | 0.022 | 0.546 | 0.33 | 0.27 | 0.35 | 0.22 | 0.51 | 0.26 |
| 1 | 94516474 | rs4147836 | 0.785 | 1.319 | -0.905 | -1.578 | 0.005 | 0.001 | 0.001 | 0 | 0.24 | 0.19 | 0.23 | 0.74 | 0.24 | 0.41 |
| 1 | 94516985 | rs1191231 | -0.33 | 1.379 | -1.016 | -1.672 | -0.161 | 0.135 | 0.135 | 0.967 | 0.31 | 0.08 | 0.29 | 0.13 | 0.5 | 0.18 |
| 1 | 94523113 | rs497511 | -0.893 | 1.307 | -0.75 | -1.589 | -0.009 | 0.009 | 0.009 | 0.755 | 0.35 | 0.32 | 0.37 | 0.18 | 0.54 | 0.29 |
| 1 | 94524856 | rs549848 | 0.864 | 1.457 | -0.839 | -1.565 | -0.096 | -0.002 | -0.002 | 0 | 0.14 | 0.02 | 0.2 | 0.12 | 0.36 | 0.09 |
| 1 | 94525623 | rs521538 | -1.356 | 0.996 | -0.506 | -1.594 | -0.032 | -0.089 | -0.089 | 0.62 | 0.59 | 0.54 | 0.59 | 0.92 | 0.79 | 0.7 |
| 1 | 94528363 | rs4147833 | 0.854 | 1.113 | -0.601 | -1.435 | 0.001 | 0.005 | 0.005 | 0 | 0.23 | 0.18 | 0.28 | 0.78 | 0.24 | 0.43 |
| 1 | 94529743 | rs4847273 | 0.478 | 1.143 | -0.334 | -1.358 | 0 | 0 | 0 | 0 | 0.26 | 0.25 | 0.27 | 0.67 | 0.29 | 0.45 |
| 1 | 94530518 | rs1007347 | 0.063 | 1.09 | -0.136 | -1.37 | -0.004 | -0.016 | -0.016 | 0 | 0.28 | 0.47 | 0.3 | 0.7 | 0.29 | 0.51 |
| 1 | 94531013 | rs553608 | -1.601 | 0.885 | 0.17 | -1.572 | 0.062 | 0.09 | 0.09 | 0 | 0.61 | 0.79 | 0.59 | 0.85 | 0.8 | 0.7 |
| 1 | 94531324 | rs3789405 | 0.48 | 0.88 | 0.168 | -1.525 | 0 | 0.001 | 0.001 | 0 | 0.26 | 0.3 | 0.27 | 0.68 | 0.28 | 0.45 |
| 1 | 94531558 | rs10493868 | 2.143 | -0.258 | -0.282 | -2.131 | 0.055 | 0.055 | 0.055 | 1.514 | 0.18 | 0.04 | 0.04 | 0 | 0.06 | 0.05 |
| 1 | 94531606 | rs3789407 | 0.727 | 0.387 | -0.06 | -1.757 | -0.002 | 0.002 | 0.002 | 0 | 0.25 | 0.29 | 0.26 | 0.67 | 0.24 | 0.42 |
| 1 | 94532013 | rs4140392 | 0.884 | 0.412 | -0.097 | -1.745 | 0 | 0.001 | 0.001 | 0 | 0.24 | 0.24 | 0.26 | 0.67 | 0.24 | 0.42 |
| 1 | 94532562 | rs1191228 | -1.033 | 0.273 | -0.52 | -1.299 | 0.055 | 0.051 | 0.051 | 0 | 0.61 | 0.78 | 0.59 | 0.77 | 0.8 | 0.69 |
| 1 | 94533014 | rs1931575 | 0.613 | 1.24 | -0.531 | -0.721 | -0.007 | -0.025 | -0.025 | 0 | 0.22 | 0.23 | 0.19 | 0.6 | 0.25 | 0.42 |
| 1 | 94534354 | rs549114 | 0.515 | 1.583 | -0.78 | -0.745 | -0.041 | 0.003 | 0.003 | 0.216 | 0.15 | 0.05 | 0.21 | 0.18 | 0.34 | 0.12 |
| 1 | 94535174 | rs2151849 | 0.345 | 1.459 | -0.431 | -0.777 | -0.004 | 0.03 | 0.03 | 0 | 0.25 | 0.37 | 0.27 | 0.6 | 0.26 | 0.44 |
| 1 | 94535546 | rs3789411 | 0.611 | 1.588 | -1.031 | -0.77 | -0.041 | 0.004 | 0.004 | 0.132 | 0.15 | 0.05 | 0.21 | 0.17 | 0.34 | 0.12 |
| 1 | 94536067 | rs3789412 | 0.37 | 1.642 | -0.996 | -0.689 | 0.001 | -0.02 | -0.02 | 0.011 | 0.24 | 0.24 | 0.26 | 0.59 | 0.26 | 0.44 |
| 1 | 94537295 | rs12758774 | 0.53 | 1.463 | -1.395 | -1.098 | -0.046 | 0.107 | 0.107 | 0 | 0.19 | 0.04 | 0.08 | 0 | 0.18 | 0.11 |
| 1 | 94538011 | rs1761375 | -0.021 | 1.88 | -0.734 | -0.59 | -0.003 | 0.001 | 0.001 | 0 | 0.16 | 0.19 | 0.21 | 0.11 | 0.31 | 0.08 |
| 1 | 94542569 | rs492220 | 0.082 | 1.887 | -0.725 | -0.089 | -0.001 | -0.001 | -0.001 | 0 | 0.16 | 0.13 | 0.3 | 0.26 | 0.29 | 0.18 |
| 1 | 94542770 | rs3120133 | -0.771 | 1.467 | -0.312 | 0.511 | -0.018 | 0.028 | 0.028 | 0.191 | 0.85 | 0.66 | 0.88 | 0.87 | 0.92 | 0.83 |
| 1 | 94547889 | rs4147828 | 1.068 | 1.813 | -0.608 | 0.932 | -0.042 | -0.084 | -0.084 | 0 | 0.14 | 0.03 | 0.21 | 0.42 | 0.22 | 0.25 |
| 1 | 94549083 | rs574741 | -0.369 | 1.238 | -0.927 | 0.212 | 0.021 | 0.026 | 0.026 | 0 | 0.6 | 0.64 | 0.79 | 0.87 | 0.77 | 0.7 |
| 1 | 94550555 | rs546550 | 1.483 | 1.323 | -0.911 | 0.204 | -0.008 | -0.005 | -0.005 | 0 | 0.16 | 0.12 | 0.29 | 0.23 | 0.29 | 0.15 |
| 1 | 94551450 | rs17461953 | 1.95 | 1.244 | -0.907 | 0.124 | -0.019 | 0.101 | 0.101 | 0.126 | 0.23 | 0.05 | 0.16 | 0.06 | 0.2 | 0.14 |
| 1 | 94553866 | rs563429 | 1.532 | 1.315 | -0.853 | 0.257 | -0.007 | -0.004 | -0.004 | 0.601 | 0.15 | 0.12 | 0.28 | 0.23 | 0.29 | 0.14 |
| 1 | 94554453 | rs4847196 | 0.314 | 0.978 | -0.311 | 0.849 | 0.016 | 0.107 | 0.107 | 0 | 0.2 | 0.3 | 0.32 | 0.52 | 0.24 | 0.36 |
| 1 | 94556894 | rs1191238 | -1.113 | 0.736 | -0.729 | 0.69 | -0.011 | 0.033 | 0.033 | 0.198 | 0.85 | 0.67 | 0.88 | 0.85 | 0.91 | 0.84 |
| 1 | 94557357 | rs554931 | -0.238 | 0.988 | -0.728 | 0.834 | 0.021 | 0.025 | 0.025 | 0 | 0.6 | 0.65 | 0.79 | 0.84 | 0.75 | 0.69 |
| 1 | 94557434 | rs483904 | -0.238 | 0.988 | -0.728 | 0.834 | 0.021 | 0.025 | 0.025 | 0 | 0.6 | 0.65 | 0.79 | 0.84 | 0.75 | 0.69 |
| 1 | 94559715 | rs2068334 | 0.054 | 1.515 | -0.577 | 1.81 | 0 | 0.019 | 0.019 | 0 | 0.14 | 0.13 | 0.12 | 0.41 | 0.2 | 0.25 |
| 1 | 94560938 | rs4147825 | -1.469 | 1.341 | -0.601 | 1.357 | -0.001 | 0.007 | 0.007 | 0 | 0.39 | 0.35 | 0.35 | 0.69 | 0.37 | 0.61 |
| 1 | 94561272 | rs4147823 | -0.732 | 1.245 | -0.61 | 1.351 | 0.001 | 0.002 | 0.002 | 0 | 0.19 | 0.19 | 0.16 | 0.27 | 0.24 | 0.3 |
| 1 | 94562084 | rs4147820 | -0.467 | 1.268 | -0.56 | 1.396 | 0.002 | 0.005 | 0.005 | 0 | 0.17 | 0.19 | 0.14 | 0.26 | 0.23 | 0.27 |
| 1 | 94563916 | rs12088309 | -0.992 | 0.908 | -0.485 | 1.321 | -0.002 | -0.008 | -0.008 | 0.881 | 0.31 | 0.34 | 0.26 | 0.45 | 0.3 | 0.38 |
| 1 | 94565577 | rs3789421 | -0.244 | 0.896 | -0.291 | 1.315 | 0 | 0 | 0 | 0 | 0.17 | 0.11 | 0.1 | 0.17 | 0.17 | 0.17 |
| 1 | 94567223 | rs950283 | 0.466 | 0.717 | -0.599 | 1.102 | 0.009 | 0.03 | 0.03 | 0 | 0.44 | 0.17 | 0.36 | 0.37 | 0.37 | 0.33 |
| 1 | 94569504 | rs4147819 | 1.039 | 0.729 | -0.68 | 1.248 | -0.01 | -0.029 | -0.029 | 0 | 0.1 | 0.04 | 0.15 | 0.22 | 0.1 | 0.16 |

| | | | | | | | | | | | | | | | | |
|---|---|---|---|---|---|---|---|---|---|---|---|---|---|---|---|---|
| 1 | 94570016 | rs481931 | 0.432 | 0.431 | -0.995 | 1.193 | -0.069 | -0.067 | -0.067 | 0.734 | 0.29 | 0.1 | 0.43 | 0.38 | 0.38 | 0.4 |
| 1 | 94570218 | rs570926 | 0.214 | 0.376 | -0.278 | 1.219 | -0.002 | -0.002 | -0.002 | 0 | 0.31 | 0.38 | 0.46 | 0.43 | 0.38 | 0.43 |
| 1 | 94570234 | rs570878 | -0.819 | 0.352 | 0.037 | 1.049 | 0.002 | -0.006 | -0.006 | 0 | 0.44 | 0.49 | 0.62 | 0.66 | 0.5 | 0.65 |
| 1 | 94571420 | rs1211213 | 0.691 | 0.526 | -0.536 | 1.003 | -0.108 | -0.083 | -0.083 | 0 | 0.25 | 0.01 | 0.39 | 0.28 | 0.32 | 0.38 |
| 1 | 94574780 | rs4147816 | 0.339 | 0.52 | -0.363 | 0.888 | -0.052 | -0.06 | -0.06 | 0.118 | 0.29 | 0.11 | 0.43 | 0.38 | 0.38 | 0.4 |
| 1 | 94575043 | rs4147812 | 0.339 | 0.52 | -0.363 | 0.888 | -0.052 | -0.06 | -0.06 | 0.054 | 0.29 | 0.11 | 0.43 | 0.39 | 0.38 | 0.4 |
| 1 | 94575171 | rs3827712 | 0.339 | 0.52 | -0.363 | 0.888 | -0.052 | -0.06 | -0.06 | 0 | 0.29 | 0.11 | 0.43 | 0.38 | 0.38 | 0.4 |
| 1 | 94575308 | rs3789432 | 0.339 | 0.52 | -0.363 | 0.888 | -0.052 | -0.06 | -0.06 | 0 | 0.29 | 0.11 | 0.43 | 0.38 | 0.37 | 0.4 |
| 1 | 94575440 | rs3789433 | 0.168 | 0.347 | -0.008 | 0.782 | 0.011 | 0.048 | 0.048 | 0 | 0.75 | 0.51 | 0.81 | 0.65 | 0.73 | 0.75 |
| 1 | 94575978 | rs3789434 | 0.737 | 0.477 | 0.159 | 0.936 | 0.01 | 0.104 | 0.104 | 0.524 | 0.12 | 0.32 | 0.17 | 0.27 | 0.1 | 0.19 |
| 1 | 94576360 | rs3789435 | -0.503 | 0.466 | 0.094 | 0.881 | 0.001 | -0.003 | -0.003 | 0 | 0.41 | 0.46 | 0.61 | 0.65 | 0.47 | 0.59 |
| 1 | 94576664 | rs4147810 | 0.654 | 0.47 | 0.147 | 0.928 | 0.016 | 0.127 | 0.127 | 0.649 | 0.13 | 0.35 | 0.17 | 0.28 | 0.1 | 0.19 |
| 1 | 94576893 | rs2297635 | 0.654 | 0.47 | 0.147 | 0.944 | 0.016 | 0.121 | 0.121 | 0.649 | 0.13 | 0.35 | 0.15 | 0.26 | 0.1 | 0.19 |
| 1 | 94576968 | rs2297634 | -0.487 | 0.476 | 0.083 | 0.862 | 0.001 | -0.003 | -0.003 | 0 | 0.41 | 0.46 | 0.61 | 0.65 | 0.47 | 0.59 |
| 1 | 94577410 | rs1889405 | -0.058 | 0.544 | 0.539 | 1.019 | -0.004 | 0.001 | 0.001 | 0 | 0.19 | 0.13 | 0.12 | 0.2 | 0.22 | 0.19 |
| 1 | 94577423 | rs1889404 | -0.116 | 0.545 | 0.647 | 1.019 | 0 | 0 | 0 | 0 | 0.19 | 0.16 | 0.12 | 0.2 | 0.22 | 0.18 |
| 1 | 94577462 | rs3789438 | 0.386 | 0.569 | 0.834 | 1.241 | 0.011 | 0.103 | 0.103 | 0.425 | 0.12 | 0.31 | 0.17 | 0.27 | 0.1 | 0.19 |
| 1 | 94579053 | rs4147807 | -0.368 | 0.262 | 0.937 | 0.689 | -0.008 | -0.014 | -0.014 | 0 | 0.21 | 0.29 | 0.12 | 0.21 | 0.2 | 0.18 |
| 1 | 94579426 | rs3789439 | -0.136 | 0.269 | 1.041 | 0.599 | -0.001 | -0.009 | -0.009 | 0.378 | 0.2 | 0.19 | 0.12 | 0.2 | 0.22 | 0.16 |
| 1 | 94581125 | rs10782976 | -1.703 | 0.114 | 1.401 | 0.797 | 0.005 | 0.008 | 0.008 | 0.209 | 0.4 | 0.57 | 0.3 | 0.65 | 0.35 | 0.36 |
| 1 | 94581384 | rs3789441 | 1.593 | 0.149 | 1.121 | 0.386 | 0.006 | 0.012 | 0.012 | 0 | 0.36 | 0.29 | 0.29 | 0.15 | 0.3 | 0.25 |
| 1 | 94581456 | rs3789442 | 0.403 | 0.159 | 1.052 | 0.391 | -0.001 | -0.005 | -0.005 | 0 | 0.2 | 0.18 | 0.13 | 0.2 | 0.22 | 0.16 |
| 1 | 94581529 | rs3789443 | 0.403 | 0.159 | 1.085 | 0.391 | 0 | -0.001 | -0.001 | 0 | 0.2 | 0.17 | 0.12 | 0.2 | 0.22 | 0.16 |
| 1 | 94581540 | rs3789444 | 0.403 | 0.159 | 1.081 | 0.391 | -0.001 | -0.005 | -0.005 | 0 | 0.2 | 0.18 | 0.12 | 0.2 | 0.22 | 0.16 |
| 1 | 94581905 | rs7535005 | 1.977 | 0.138 | 0.937 | 0.365 | 0.013 | 0.116 | 0.116 | 0.06 | 0.32 | 0.1 | 0.24 | 0.01 | 0.29 | 0.22 |
| 1 | 94582249 | rs3789445 | -0.416 | 0.536 | 1.398 | 0.805 | -0.001 | 0.002 | 0.002 | 0 | 0.23 | 0.2 | 0.17 | 0.34 | 0.24 | 0.19 |
| 1 | 94584172 | rs4147802 | 0.842 | 0.545 | 1.297 | 0.937 | -0.002 | 0.059 | 0.059 | 0.062 | 0.1 | 0.11 | 0.09 | 0.26 | 0.08 | 0.15 |
| 1 | 94585009 | rs4147798 | 0.118 | 0.598 | 1.334 | 0.852 | 0.001 | -0.009 | -0.009 | 0.014 | 0.2 | 0.21 | 0.13 | 0.2 | 0.23 | 0.15 |
| 1 | 94587362 | rs3761906 | 0.394 | 0.562 | 1.713 | 0.967 | -0.007 | 0.125 | 0.125 | 0.14 | 0.12 | 0.25 | 0.11 | 0.29 | 0.09 | 0.16 |
| 1 | 94587687 | rs2151846 | -0.133 | 0.587 | 1.394 | 0.737 | 0.014 | 0.032 | 0.032 | 0 | 0.27 | 0.4 | 0.49 | 0.49 | 0.35 | 0.42 |
| 1 | 94588992 | rs1931572 | -0.326 | 0.852 | 1.072 | 0.751 | -0.014 | -0.012 | -0.012 | 0.914 | 0.75 | 0.7 | 0.57 | 0.65 | 0.68 | 0.62 |
| 1 | 94591481 | rs10874835 | 0.269 | 0.784 | 1.067 | 0.657 | 0.001 | -0.004 | -0.004 | 0.067 | 0.2 | 0.19 | 0.14 | 0.21 | 0.25 | 0.16 |
| 1 | 94592290 | rs1105123 | -1.989 | 0.79 | 0.64 | 0.486 | 0.027 | 0.165 | 0.165 | 0.11 | 0.62 | 0.88 | 0.75 | 0.99 | 0.7 | 0.74 |
| 1 | 94593399 | rs12071152 | 0.717 | 0.811 | 0.767 | 0.63 | 0.003 | 0.172 | 0.172 | 0.069 | 0.11 | 0.25 | 0.11 | 0.3 | 0.09 | 0.16 |
| 1 | 94594043 | rs11802196 | -0.05 | 0.88 | 0.72 | 0.764 | 0.02 | 0.027 | 0.027 | 0 | 0.26 | 0.45 | 0.45 | 0.38 | 0.34 | 0.4 |
| 1 | 94594080 | rs11165081 | -0.044 | 0.897 | 0.404 | 0.78 | 0.004 | 0.006 | 0.006 | 0 | 0.25 | 0.35 | 0.44 | 0.38 | 0.34 | 0.4 |
| 1 | 94596831 | rs6686599 | 0.202 | 1.016 | -0.083 | 0.749 | -0.037 | -0.035 | -0.035 | 0 | 0.22 | 0.17 | 0.41 | 0.35 | 0.31 | 0.37 |
| 1 | 94596867 | rs1931565 | -1.08 | 0.672 | -0.158 | 0.351 | 0.034 | 0.071 | 0.071 | 0 | 0.51 | 0.77 | 0.64 | 0.73 | 0.61 | 0.59 |
| 1 | 94607234 | rs11581939 | 1.109 | 0.73 | -0.703 | 0.093 | 0.038 | 0.165 | 0.165 | 0.037 | 0.42 | 0.15 | 0.26 | 0.15 | 0.39 | 0.25 |
| 1 | 94607607 | rs2022378 | -0.51 | 1.089 | -0.114 | 0.241 | -0.011 | -0.04 | -0.04 | 1.78 | 0.27 | 0.47 | 0.21 | 0.22 | 0.25 | 0.2 |
| 1 | 94607848 | rs4847286 | 0.025 | 1.183 | 0.019 | 0.393 | -0.004 | -0.001 | -0.001 | 0 | 0.69 | 0.71 | 0.49 | 0.36 | 0.64 | 0.45 |
| 1 | 94611300 | rs2774920 | -1.361 | 0.985 | 0.557 | 0.399 | 0.006 | 0.056 | 0.056 | 1.02 | 0.09 | 0.37 | 0.07 | 0.15 | 0.11 | 0.05 |
| 1 | 94629643 | rs12742802 | -0.464 | 0.946 | 0.833 | 0.22 | 0 | -0.019 | -0.019 | 0 | 0.23 | 0.21 | 0.17 | 0.13 | 0.27 | 0.18 |
| 1 | 94635028 | rs1411701 | -0.076 | 1.236 | 0.89 | 0.801 | 0.002 | 0.004 | 0.004 | 0 | 0.25 | 0.22 | 0.41 | 0.4 | 0.32 | 0.46 |
| 1 | 94636836 | rs10874840 | 0.217 | 1.239 | 0.901 | 0.86 | 0.006 | 0.005 | 0.005 | 0 | 0.22 | 0.2 | 0.37 | 0.32 | 0.3 | 0.43 |
| 1 | 94637058 | rs12752790 | 0.7 | 1.196 | 0.96 | 0.86 | -0.004 | -0.006 | -0.006 | 0.257 | 0.67 | 0.31 | 0.53 | 0.45 | 0.63 | 0.5 |
| 1 | 94638711 | rs1048866 | -0.113 | 1.171 | 0.928 | 0.874 | 0.001 | 0.004 | 0.004 | 0 | 0.25 | 0.22 | 0.41 | 0.4 | 0.31 | 0.46 |
| 1 | 94646514 | rs11577575 | -0.137 | 1.28 | 1.016 | 0.906 | -0.034 | 0.052 | 0.052 | 2.773 | 0.2 | 0.03 | 0.14 | 0.11 | 0.24 | 0.17 |
| 1 | 94667970 | rs1541098 | 1.301 | 1.768 | 0.997 | 0.93 | -0.077 | -0.082 | -0.082 | 0 | 0.12 | 0.01 | 0.28 | 0.31 | 0.2 | 0.16 |
| 1 | 94674726 | rs2274788 | -0.138 | 1.662 | 0.684 | 0.814 | -0.043 | -0.088 | -0.088 | 0 | 0.24 | 0.03 | 0.24 | 0.31 | 0.24 | 0.15 |
| 1 | 94685585 | rs3789689 | -0.597 | 1.663 | 1.046 | 0.867 | 0.051 | 0.084 | 0.084 | 0.698 | 0.06 | 0.21 | 0.11 | 0.15 | 0.09 | 0.05 |
| 1 | 94689027 | rs6541343 | 1.24 | 1.237 | 1.119 | 0.833 | 0.013 | 0.051 | 0.051 | 0 | 0.92 | 0.82 | 0.89 | 0.85 | 0.9 | 0.85 |
| 1 | 94689734 | rs12724116 | 0.24 | 1.233 | 0.966 | 0.664 | 0.003 | 0.098 | 0.098 | 0.954 | 0.18 | 0.02 | 0.13 | 0 | 0.15 | 0.1 |
| 1 | 94693145 | rs6673491 | 1.372 | 1.218 | 1.113 | 0.842 | 0.012 | 0.052 | 0.052 | 0 | 0.91 | 0.74 | 0.88 | 0.85 | 0.9 | 0.85 |
| 1 | 94721660 | rs12750249 | 0.897 | 1.327 | 0.933 | 0.974 | -0.09 | -0.092 | -0.092 | 0.524 | 0.14 | 0.02 | 0.3 | 0.3 | 0.23 | 0.17 |
| 1 | 94729088 | rs11165101 | -0.413 | 1.08 | 1.311 | 0.937 | -0.005 | -0.003 | -0.003 | 0.892 | 0.26 | 0.28 | 0.39 | 0.31 | 0.33 | 0.45 |
| 1 | 94737583 | rs2391472 | -0.646 | 1.067 | 1.445 | 0.955 | 0.008 | 0.036 | 0.036 | 0 | 0.06 | 0.11 | 0.1 | 0.15 | 0.1 | 0.15 |
| 1 | 94738846 | rs6697764 | -0.159 | 1.291 | 1.28 | 0.935 | -0.016 | -0.013 | -0.013 | 0 | 0.22 | 0.18 | 0.37 | 0.31 | 0.29 | 0.45 |
| 1 | 94752469 | rs11165110 | -0.321 | 1.087 | 1.319 | 0.976 | -0.024 | -0.015 | -0.015 | 0 | 0.24 | 0.18 | 0.39 | 0.31 | 0.33 | 0.45 |
| 1 | 94760885 | rs1330855 | -0.782 | 0.738 | 1.507 | 1.066 | -0.033 | 0.08 | 0.08 | 0.88 | 0.27 | 0.48 | 0.22 | 0.45 | 0.21 | 0.16 |
| 1 | 94786514 | rs16928 | -2.108 | 0.795 | 1.596 | 1.021 | 0.029 | 0.091 | 0.091 | 0.015 | 0.1 | 0.48 | 0.16 | 0.16 | 0.09 | 0.16 |
| 1 | 94788623 | rs17111408 | -1.013 | 0.797 | 1.659 | 1.022 | 0.042 | 0.092 | 0.092 | 0 | 0.07 | 0.19 | 0.11 | 0.15 | 0.09 | 0.14 |
| 1 | 94788768 | rs12027548 | -0.881 | 0.808 | 1.457 | 1.027 | 0.008 | 0.036 | 0.036 | 0 | 0.07 | 0.1 | 0.1 | 0.15 | 0.09 | 0.14 |
| 1 | 94792449 | rs11584317 | 0.036 | 0.812 | 1.373 | 1.198 | 0.016 | -0.05 | -0.05 | 0.784 | 0.2 | 0.12 | 0.16 | 0.33 | 0.16 | 0.12 |
| 1 | 94807102 | rs2065971 | 0.76 | 1.292 | 1.666 | 1.47 | 0.019 | 0.008 | 0.008 | 0 | 0.73 | 0.59 | 0.59 | 0.69 | 0.66 | 0.56 |
| 1 | 94850443 | rs2391467 | 1.599 | 1.312 | 1.471 | 1.553 | 0.022 | 0.019 | 0.019 | 0 | 0.65 | 0.38 | 0.46 | 0.53 | 0.56 | 0.4 |
| 1 | 94867056 | rs12037634 | -1.471 | 0.855 | 0.95 | 1.484 | -0.005 | 0.02 | 0.02 | 0 | 0.07 | 0.16 | 0.05 | 0.12 | 0.08 | 0.16 |
| 1 | 94870535 | rs11165135 | -1.612 | 1.176 | 0.955 | 1.735 | 0.009 | 0.013 | 0.013 | 0 | 0.29 | 0.31 | 0.48 | 0.47 | 0.4 | 0.48 |
| 1 | 94874521 | rs6681849 | -1.616 | 1.02 | 0.996 | 1.752 | 0.001 | 0.002 | 0.002 | 0 | 0.34 | 0.33 | 0.5 | 0.48 | 0.39 | 0.51 |

| | | | | | | | | | | | | | | | | |
|---|---|---|---|---|---|---|---|---|---|---|---|---|---|---|---|---|
| 1 | 94888021 | rs6660167 | 0.001 | 1.271 | 0.947 | 1.96 | -0.025 | -0.033 | -0.033 | 0 | 0.2 | 0.1 | 0.38 | 0.29 | 0.21 | 0.29 |
| 1 | 94893928 | rs12750904 | -0.169 | 1.169 | 0.949 | 1.972 | -0.04 | -0.04 | -0.04 | 0 | 0.22 | 0.09 | 0.39 | 0.29 | 0.26 | 0.3 |
| 1 | 94958378 | rs1749541 | 1.988 | 0.759 | 1.709 | 2.131 | -0.017 | 0.04 | 0.04 | 0 | 0.78 | 0.67 | 0.85 | 0.5 | 0.77 | 0.6 |
| 1 | 94978199 | rs3849301 | -0.341 | 0.792 | 1.467 | 2.169 | -0.015 | -0.039 | -0.039 | 0.029 | 0.07 | 0.01 | 0.05 | 0.17 | 0.11 | 0.19 |
| 1 | 94981253 | rs582798 | -0.146 | 0.791 | 1.566 | 2.101 | -0.003 | 0.015 | 0.015 | 0 | 0.07 | 0.13 | 0.05 | 0.14 | 0.07 | 0.11 |
| 1 | 94983203 | rs698951 | -2.226 | 0.79 | 1.457 | 2.035 | -0.026 | 0.039 | 0.039 | 0.957 | 0.25 | 0.59 | 0.19 | 0.51 | 0.23 | 0.4 |
| 1 | 94997288 | rs1324214 | 0.252 | 0.914 | 1.139 | 1.893 | -0.006 | -0.008 | -0.008 | 0 | 0.19 | 0.13 | 0.3 | 0.26 | 0.2 | 0.16 |
| 1 | 95002211 | rs1144300 | 1.878 | 0.674 | 1 | 1.571 | 0 | 0.002 | 0.002 | 0.235 | 0.51 | 0.52 | 0.39 | 0.18 | 0.48 | 0.34 |
| 1 | 95004084 | rs762485 | 1.86 | 0.783 | 0.644 | 1.578 | 0.009 | 0.152 | 0.152 | 0.072 | 0.49 | 0.21 | 0.37 | 0.18 | 0.48 | 0.34 |
| 1 | 95004410 | rs762484 | 0.249 | 0.831 | 0.225 | 1.977 | -0.015 | -0.005 | -0.005 | 0 | 0.21 | 0.12 | 0.3 | 0.2 | 0.21 | 0.19 |
| 1 | 95005220 | rs696619 | -1.023 | 0.923 | -0.15 | 1.4 | -0.025 | 0.01 | 0.01 | 0 | 0.37 | 0.13 | 0.47 | 0.7 | 0.39 | 0.52 |
| 1 | 95007918 | rs1361600 | -1.337 | 0.509 | -0.139 | 1.12 | -0.003 | 0.047 | 0.047 | 0.633 | 0.48 | 0.36 | 0.57 | 0.83 | 0.5 | 0.67 |
| 1 | 95017380 | rs337601 | -0.97 | 0.256 | -0.501 | 1.374 | 0.031 | -0.05 | -0.05 | 0 | 0.09 | 0 | 0.04 | 0.2 | 0.06 | 0.14 |
| 1 | 95025421 | rs2762509 | 1.102 | 0.524 | 0.179 | 1.247 | 0.008 | 0.072 | 0.072 | 0 | 0.82 | 0.54 | 0.8 | 0.68 | 0.83 | 0.72 |
| 1 | 95028989 | rs34365422 | -0.228 | 0.901 | 0.102 | 1.459 | 0.008 | 0.018 | 0.018 | 0.185 | 0.1 | 0.12 | 0.1 | 0.15 | 0.11 | 0.16 |
| 1 | 95050326 | rs6668501 | -1.034 | 1.052 | 0.7 | 1.532 | 0.015 | 0.024 | 0.024 | 0 | 0.15 | 0.21 | 0.17 | 0.21 | 0.18 | 0.25 |
| 1 | 95050728 | rs841675 | 0.902 | 1.082 | 0.397 | 1.299 | -0.002 | -0.003 | -0.003 | 0.412 | 0.61 | 0.37 | 0.57 | 0.41 | 0.52 | 0.49 |
| 1 | 95052267 | rs12758064 | 0.233 | 1.09 | 0.377 | 1.234 | 0.007 | 0.013 | 0.013 | 2.373 | 0.12 | 0.14 | 0.11 | 0.15 | 0.13 | 0.18 |
| 1 | 95052325 | rs1772905 | -0.371 | 1.089 | 0.602 | 1.241 | 0.011 | 0.004 | 0.004 | 0 | 0.1 | 0.23 | 0.09 | 0.16 | 0.1 | 0.12 |
| 1 | 95052476 | rs2022309 | 0.459 | 1.033 | 0.603 | 1.233 | -0.029 | -0.058 | -0.058 | 1.66 | 0.22 | 0.07 | 0.23 | 0.27 | 0.29 | 0.26 |
| 1 | 95053353 | rs12029080 | 0.459 | 1.037 | 0.493 | 1.22 | -0.025 | -0.052 | -0.052 | 0 | 0.22 | 0.07 | 0.23 | 0.27 | 0.3 | 0.26 |
| 1 | 95055132 | rs1772904 | -0.773 | 0.658 | 0.041 | 0.655 | 0.035 | 0.05 | 0.05 | 0 | 0.16 | 0.38 | 0.19 | 0.21 | 0.18 | 0.25 |
| 1 | 95058428 | rs841685 | -0.586 | 0.8 | -0.12 | 0.532 | 0.004 | 0.001 | 0.001 | 1.088 | 0.68 | 0.75 | 0.64 | 0.67 | 0.75 | 0.74 |
| 1 | 95061355 | rs1778214 | -0.234 | 0.669 | -0.08 | 0.803 | 0.089 | 0.194 | 0.194 | 0 | 0.08 | 0.39 | 0.13 | 0.2 | 0.08 | 0.14 |
| 1 | 95068097 | rs17402334 | 0.597 | 0.596 | -0.531 | 0.585 | -0.037 | -0.072 | -0.072 | 1.139 | 0.26 | 0.07 | 0.26 | 0.34 | 0.36 | 0.27 |
| 1 | 95068347 | rs1772895 | -1.083 | 0.434 | -0.493 | 0.449 | -0.008 | 0.066 | 0.066 | 0.004 | 0.49 | 0.62 | 0.5 | 0.66 | 0.56 | 0.6 |
| 1 | 95068921 | rs4847313 | 0.988 | 0.443 | -0.94 | 0.376 | -0.005 | 0.006 | 0.006 | 0.108 | 0.31 | 0.26 | 0.29 | 0.19 | 0.36 | 0.3 |
| 1 | 95068957 | rs4847314 | 0.435 | 0.273 | -0.511 | 0.838 | 0.001 | -0.006 | -0.006 | 1.576 | 0.46 | 0.45 | 0.41 | 0.34 | 0.47 | 0.49 |
| 1 | 95070041 | rs7542900 | -0.626 | 0.24 | -0.466 | 0.67 | -0.024 | 0.009 | 0.009 | 1.035 | 0.72 | 0.53 | 0.67 | 0.67 | 0.83 | 0.76 |
| 1 | 95070114 | rs11807669 | 0.265 | 0.638 | -0.733 | 0.325 | -0.017 | 0.068 | 0.068 | 0 | 0.4 | 0.24 | 0.31 | 0.16 | 0.4 | 0.39 |
| 1 | 95073156 | rs11165182 | -0.87 | 0.535 | -0.267 | 1.29 | -0.002 | 0.068 | 0.068 | 0 | 0.1 | 0.16 | 0.23 | 0.22 | 0.1 | 0.13 |
| 1 | 95074087 | rs1245487 | -1.052 | 0.59 | -0.522 | 1.417 | 0 | -0.009 | -0.009 | 0 | 0.31 | 0.67 | 0.44 | 0.51 | 0.29 | 0.44 |
| 1 | 95074230 | rs1245488 | 0.696 | 0.637 | -0.476 | 1.829 | 0.003 | 0.026 | 0.026 | 0 | 0.91 | 0.88 | 0.85 | 0.67 | 0.88 | 0.81 |
| 1 | 95074455 | rs1245489 | 0.01 | 0.598 | -0.908 | 1.455 | -0.014 | -0.036 | -0.036 | 0.118 | 0.65 | 0.77 | 0.63 | 0.51 | 0.61 | 0.69 |
| 1 | 95076153 | rs12042986 | 0.804 | 0.261 | -1.196 | 0.713 | -0.046 | 0.049 | 0.049 | 1.792 | 0.35 | 0.2 | 0.35 | 0.27 | 0.36 | 0.44 |
| 1 | 95078129 | rs4847317 | -0.672 | -0.385 | -0.893 | 0.519 | -0.002 | 0.016 | 0.016 | 0 | 0.49 | 0.55 | 0.55 | 0.69 | 0.55 | 0.46 |
| 1 | 95081445 | rs4474258 | -0.322 | -0.611 | -0.756 | 0.584 | -0.001 | 0 | 0 | 0.039 | 0.38 | 0.34 | 0.39 | 0.53 | 0.36 | 0.38 |
| 1 | 95081587 | rs17411335 | 2.134 | -0.604 | -0.742 | 0.546 | -0.087 | 0.13 | 0.13 | 1.539 | 0.25 | 0.08 | 0.27 | 0.03 | 0.3 | 0.29 |
| 1 | 95083836 | rs10874867 | 0.558 | -0.387 | -0.401 | 0.709 | 0 | 0 | 0 | 0.003 | 0.26 | 0.19 | 0.3 | 0.3 | 0.28 | 0.26 |
| 1 | 95085258 | rs10782983 | -0.338 | -0.394 | -0.618 | 0.664 | -0.001 | 0.009 | 0.009 | 0 | 0.36 | 0.35 | 0.39 | 0.54 | 0.34 | 0.38 |
| 1 | 95085927 | rs1539622 | -0.964 | -0.497 | -0.515 | 0.632 | -0.046 | 0.031 | 0.031 | 0.471 | 0.76 | 0.54 | 0.82 | 0.72 | 0.85 | 0.74 |
| 1 | 95086409 | rs2391383 | 1.238 | -0.378 | -0.619 | 0.688 | -0.006 | -0.044 | -0.044 | 0 | 0.09 | 0.05 | 0.06 | 0.23 | 0.06 | 0.12 |
| 1 | 95087398 | rs10874868 | 0.476 | -0.743 | -0.56 | 0.648 | 0.006 | 0.005 | 0.005 | 0 | 0.26 | 0.23 | 0.35 | 0.3 | 0.31 | 0.26 |
| 1 | 95087604 | rs10782984 | -1.231 | -0.659 | -0.398 | 0.174 | -0.029 | 0.022 | 0.022 | 0 | 0.78 | 0.7 | 0.84 | 0.73 | 0.86 | 0.74 |
| 1 | 95090677 | rs12738550 | 1.499 | -0.406 | -0.669 | 0.053 | -0.095 | 0.15 | 0.15 | 0.794 | 0.28 | 0.08 | 0.28 | 0.03 | 0.32 | 0.3 |
| 1 | 95093351 | rs12725602 | 0.574 | -0.449 | -0.665 | 0.069 | -0.052 | 0.094 | 0.094 | 0.11 | 0.4 | 0.15 | 0.43 | 0.18 | 0.52 | 0.33 |
| 1 | 95093889 | rs11576456 | 1.354 | -0.57 | -0.595 | 0.219 | -0.024 | -0.039 | -0.039 | 0.132 | 0.12 | 0.02 | 0.14 | 0.15 | 0.2 | 0.07 |
| 1 | 95099540 | rs2211318 | 0.589 | -0.521 | -0.553 | -0.015 | -0.037 | 0.075 | 0.075 | 0.263 | 0.39 | 0.23 | 0.41 | 0.13 | 0.49 | 0.34 |
| 1 | 95105307 | rs2794489 | -0.742 | -0.503 | -0.408 | 0.004 | 0.004 | 0.039 | 0.039 | 0 | 0.51 | 0.65 | 0.59 | 0.65 | 0.57 | 0.46 |
| 1 | 95107254 | rs2996284 | 0.793 | -0.47 | -0.222 | 0.12 | 0.002 | -0.009 | -0.009 | 0.256 | 0.15 | 0.25 | 0.16 | 0.09 | 0.2 | 0.09 |
| 1 | 95108162 | rs1341645 | 0.036 | -0.563 | -0.386 | -0.009 | -0.017 | 0.017 | 0.017 | 0 | 0.63 | 0.49 | 0.68 | 0.59 | 0.65 | 0.62 |
| 1 | 95110225 | rs1778196 | -1.182 | -0.597 | -0.53 | 0.22 | 0.014 | -0.02 | -0.02 | 1.425 | 0.78 | 0.82 | 0.85 | 0.7 | 0.85 | 0.71 |
| 1 | 95110447 | rs3885983 | -0.622 | -0.198 | -0.212 | 0.581 | -0.001 | -0.007 | -0.007 | 0 | 0.34 | 0.26 | 0.39 | 0.56 | 0.36 | 0.35 |
| 1 | 95113417 | rs1778193 | -0.126 | -0.116 | -0.115 | 0.253 | 0.001 | -0.038 | -0.038 | 0.671 | 0.11 | 0.27 | 0.09 | 0.07 | 0.08 | 0.08 |
| 1 | 95116072 | rs2153717 | -1.097 | -0.198 | -1.188 | 0.295 | 0.026 | -0.011 | -0.011 | 0.069 | 0.38 | 0.73 | 0.46 | 0.67 | 0.34 | 0.51 |
| 1 | 95116807 | rs1245481 | 0.457 | 0.561 | -1.389 | -0.161 | -0.063 | 0.162 | 0.162 | 0.565 | 0.35 | 0.12 | 0.32 | 0.03 | 0.38 | 0.29 |
| 1 | 95117157 | rs947386 | -0.861 | 0.188 | -0.632 | 0.739 | -0.005 | 0.068 | 0.068 | 0.573 | 0.23 | 0.42 | 0.41 | 0.66 | 0.2 | 0.36 |
| 1 | 95117704 | rs1245480 | 0.109 | 0.256 | -1.066 | 0.794 | 0.002 | 0.006 | 0.006 | 0 | 0.82 | 0.85 | 0.68 | 0.71 | 0.77 | 0.75 |
| 1 | 95117783 | rs11803470 | -0.87 | -0.461 | -0.673 | 0.887 | -0.039 | 0.095 | 0.095 | 0.342 | 0.14 | 0.47 | 0.14 | 0.28 | 0.11 | 0.13 |
| 1 | 95118391 | rs12725402 | 0.34 | 0.382 | -1.914 | 0.155 | -0.043 | 0.132 | 0.132 | 0 | 0.41 | 0.24 | 0.3 | 0.06 | 0.46 | 0.41 |
| 1 | 95120228 | rs11165191 | -0.061 | 0.158 | -2.011 | 0.74 | 0.025 | -0.022 | -0.022 | 0.908 | 0.35 | 0.13 | 0.26 | 0.32 | 0.33 | 0.36 |
| 1 | 95122518 | rs17111996 | -0.286 | 0.394 | -1.94 | 0.846 | -0.013 | -0.008 | -0.008 | 0 | 0.46 | 0.23 | 0.43 | 0.43 | 0.47 | 0.49 |
| 1 | 95127463 | rs11165192 | 0.72 | 0.221 | -1.137 | 1.16 | -0.005 | 0.077 | 0.077 | 0.027 | 0.47 | 0.27 | 0.37 | 0.28 | 0.51 | 0.42 |
| 1 | 95134903 | rs6668794 | 0.14 | 0.039 | -1.276 | 1.399 | -0.059 | -0.003 | -0.003 | 0 | 0.35 | 0.11 | 0.32 | 0.31 | 0.4 | 0.35 |
| 1 | 95138084 | rs841344 | 0.478 | -0.211 | 0.1 | 1.546 | -0.02 | 0.208 | 0.208 | 0 | 0.89 | 0.56 | 0.79 | 0.72 | 0.94 | 0.89 |
| 1 | 95140173 | rs6699528 | 1.113 | -0.223 | 0.152 | 1.542 | -0.041 | 0.129 | 0.129 | 0 | 0.87 | 0.44 | 0.78 | 0.71 | 0.94 | 0.88 |
| 1 | 95142547 | rs7513758 | -1.046 | -0.215 | 0.733 | 1.366 | 0.018 | 0.078 | 0.078 | 0 | 0.05 | 0.26 | 0.05 | 0.09 | 0.03 | 0.13 |
| 1 | 95142843 | rs841339 | 0.576 | -0.175 | 1.036 | 1.654 | 0.042 | 0.311 | 0.311 | 0 | 0.93 | 0.64 | 0.9 | 0.78 | 0.98 | 0.96 |
| 1 | 95144800 | rs3789696 | 1.282 | 0.113 | 0.186 | 1.221 | -0.023 | 0.074 | 0.074 | 0.346 | 0.42 | 0.03 | 0.38 | 0.22 | 0.42 | 0.41 |

| | | | | | | | | | | | | | | | | |
|---|---|---|---|---|---|---|---|---|---|---|---|---|---|---|---|---|
| 1 | 95145453 | rs12021681 | -0.374 | 0.484 | 0.604 | 1.219 | 0.011 | 0.102 | 0.102 | 0 | 0.11 | 0.16 | 0.13 | 0.32 | 0.1 | 0.19 |
| 1 | 95151009 | rs7522944 | 0.059 | 0.532 | 0.577 | 1.113 | -0.004 | 0.008 | 0.008 | 0 | 0.34 | 0.35 | 0.31 | 0.3 | 0.34 | 0.29 |
| 1 | 95151813 | rs841361 | 1.254 | 0.535 | 0.517 | 1.138 | 0 | 0.007 | 0.007 | 0 | 0.5 | 0.43 | 0.49 | 0.32 | 0.53 | 0.47 |
| 1 | 95154279 | rs12025248 | -0.059 | 0.584 | 0.28 | 1.175 | 0 | 0.029 | 0.029 | 0 | 0.11 | 0.09 | 0.12 | 0.32 | 0.11 | 0.19 |
| 1 | 95154785 | rs4240960 | 1.148 | 0.579 | 0.087 | 1.078 | -0.001 | 0.048 | 0.048 | 0 | 0.48 | 0.29 | 0.48 | 0.32 | 0.52 | 0.46 |
| 1 | 95160537 | rs10493876 | 0.192 | 0.58 | 0.208 | 1.052 | -0.014 | -0.004 | -0.004 | 0.106 | 0.05 | 0.08 | 0.07 | 0.06 | 0.03 | 0.05 |
| 1 | 95161617 | rs12120217 | 1.153 | 0.657 | -0.055 | 1.091 | -0.015 | 0.057 | 0.057 | 0 | 0.47 | 0.23 | 0.47 | 0.32 | 0.52 | 0.47 |
| 1 | 95166754 | rs841326 | 0.134 | 0.702 | -0.041 | 1.351 | -0.001 | 0.031 | 0.031 | 0 | 0.88 | 0.89 | 0.88 | 0.62 | 0.9 | 0.81 |
| 1 | 95169374 | rs2391387 | 1.154 | 0.68 | -0.127 | 0.915 | -0.014 | 0.042 | 0.042 | 0 | 0.47 | 0.24 | 0.48 | 0.32 | 0.52 | 0.46 |
| 1 | 95169671 | rs1892466 | -0.772 | 1.074 | -0.141 | 0.947 | -0.001 | 0 | 0 | 1.602 | 0.39 | 0.43 | 0.38 | 0.39 | 0.4 | 0.32 |
| 1 | 95170056 | rs1891873 | 1.066 | 0.815 | -0.253 | 0.627 | 0.014 | 0.008 | 0.008 | 0 | 0.54 | 0.39 | 0.54 | 0.38 | 0.6 | 0.55 |
| 1 | 95171891 | rs6541361 | 1.178 | 0.937 | -0.416 | 0.465 | -0.012 | 0.007 | 0.007 | 0 | 0.64 | 0.43 | 0.66 | 0.64 | 0.72 | 0.58 |
| 1 | 95172169 | rs2802112 | 0.664 | 1.058 | -0.3 | 0.48 | 0.012 | -0.066 | -0.066 | 0.046 | 0.16 | 0.41 | 0.16 | 0.02 | 0.15 | 0.13 |
| 1 | 95173343 | rs1027307 | 0.149 | 0.901 | -0.13 | 0.65 | 0.006 | -0.009 | -0.009 | 0 | 0.29 | 0.45 | 0.31 | 0.32 | 0.31 | 0.2 |
| 1 | 95177368 | rs6665058 | 0.036 | 0.66 | -0.024 | 0.025 | 0 | -0.002 | -0.002 | 0.386 | 0.59 | 0.43 | 0.59 | 0.37 | 0.6 | 0.58 |
| 1 | 95185274 | rs2640062 | -0.206 | 0.675 | -0.271 | -0.021 | 0.001 | -0.002 | -0.002 | 2.312 | 0.41 | 0.43 | 0.41 | 0.65 | 0.42 | 0.42 |
| 1 | 95189230 | rs2997380 | -0.201 | 0.091 | -0.263 | 0.032 | 0.017 | 0.019 | 0.019 | 0 | 0.51 | 0.44 | 0.51 | 0.66 | 0.61 | 0.5 |
| 1 | 95191213 | rs10874887 | 0.121 | -0.078 | -0.177 | 0.066 | 0.021 | 0.016 | 0.016 | 0.031 | 0.46 | 0.39 | 0.44 | 0.63 | 0.6 | 0.42 |
| 1 | 95195432 | rs3933012 | 0.508 | -0.125 | -0.42 | 0.08 | -0.002 | -0.001 | -0.001 | 0 | 0.41 | 0.4 | 0.45 | 0.34 | 0.33 | 0.49 |
| 1 | 95198251 | rs1782393 | 0.744 | 0.065 | -0.041 | 0.252 | -0.026 | -0.03 | -0.03 | 0.565 | 0.89 | 0.85 | 0.93 | 0.98 | 0.95 | 0.94 |
| 1 | 95202984 | rs11165231 | -0.526 | 0.197 | 0.004 | 0.343 | -0.006 | -0.004 | -0.004 | 0 | 0.31 | 0.36 | 0.23 | 0.19 | 0.24 | 0.32 |
| 1 | 95203212 | rs2933912 | 0.094 | 0.714 | -0.206 | 0.54 | -0.041 | -0.041 | -0.041 | 3.485 | 0.26 | 0.16 | 0.33 | 0.38 | 0.37 | 0.22 |
| 1 | 95207524 | rs10493875 | 0.023 | 0.319 | 0.004 | 0.467 | 0.021 | 0.016 | 0.016 | 0 | 0.3 | 0.27 | 0.22 | 0.18 | 0.23 | 0.32 |
| 1 | 95207633 | rs12036542 | 0.92 | 0.034 | -0.07 | 0.414 | 0.004 | 0 | 0 | 0 | 0.16 | 0.17 | 0.05 | 0.15 | 0.07 | 0.08 |
| 1 | 95220414 | rs1146268 | -0.494 | 0.565 | 0.454 | 0.517 | -0.015 | -0.021 | -0.021 | 0 | 0.77 | 0.82 | 0.67 | 0.61 | 0.67 | 0.8 |
| 1 | 95220538 | rs17406046 | 1.148 | 0.353 | 0.469 | 0.399 | -0.01 | 0.026 | 0.026 | 0.235 | 0.41 | 0.21 | 0.36 | 0.34 | 0.42 | 0.44 |
| 1 | 95222875 | rs12140262 | 1.196 | 0.33 | 0.249 | 0.388 | -0.053 | 0.041 | 0.041 | 0.77 | 0.4 | 0.1 | 0.35 | 0.34 | 0.42 | 0.43 |
| 1 | 95223872 | rs12141345 | 1.196 | 0.33 | 0.257 | 0.388 | -0.051 | 0.042 | 0.042 | 0.77 | 0.4 | 0.1 | 0.35 | 0.34 | 0.42 | 0.43 |
| 1 | 95235919 | rs7525706 | 0.502 | 0.347 | 0.988 | 0.551 | -0.076 | -0.006 | -0.006 | 0.043 | 0.17 | 0.37 | 0.07 | 0.15 | 0.07 | 0.1 |
| 1 | 95236282 | rs3860336 | 0.686 | 0.717 | 0.831 | 0.406 | 0.003 | 0.004 | 0.004 | 0.586 | 0.64 | 0.36 | 0.72 | 0.78 | 0.71 | 0.65 |
| 1 | 95239080 | rs17414905 | 0.526 | 0.748 | 0.831 | 0.417 | -0.073 | 0.018 | 0.018 | 0 | 0.14 | 0.01 | 0.12 | 0.03 | 0.21 | 0.23 |
| 1 | 95240361 | rs2391423 | -0.279 | 0.904 | 0.676 | 0.373 | -0.055 | -0.054 | -0.054 | 0 | 0.3 | 0.18 | 0.38 | 0.41 | 0.43 | 0.27 |
| 1 | 95240622 | rs6541369 | -0.177 | 0.816 | 1.036 | 0.975 | -0.075 | -0.014 | -0.014 | 0 | 0.8 | 0.4 | 0.91 | 0.81 | 0.93 | 0.88 |
| 1 | 95240822 | rs4847214 | -0.177 | 0.816 | 1.036 | 0.975 | -0.075 | -0.014 | -0.014 | 0.004 | 0.8 | 0.4 | 0.91 | 0.81 | 0.93 | 0.88 |
| 1 | 95249306 | rs12743267 | 0.337 | 0.929 | 0.987 | 0.907 | -0.025 | -0.034 | -0.034 | 0 | 0.16 | 0.08 | 0.29 | 0.43 | 0.26 | 0.16 |
| 1 | 95259174 | rs6678909 | -0.806 | 1.025 | 1.093 | 0.867 | 0.008 | 0.008 | 0.008 | 0 | 0.34 | 0.38 | 0.38 | 0.57 | 0.39 | 0.26 |
| 1 | 95259359 | rs3849302 | 1.571 | 1.027 | 1.067 | 0.873 | 0.01 | 0.092 | 0.092 | 0.478 | 0.46 | 0.22 | 0.48 | 0.32 | 0.43 | 0.48 |
| 1 | 95269752 | rs11590799 | 0.143 | 1.101 | 1.184 | 0.715 | 0.01 | -0.035 | -0.035 | 0 | 0.16 | 0.22 | 0.13 | 0.03 | 0.17 | 0.23 |
| 1 | 95270383 | rs6690134 | 1.447 | 1.166 | 1.236 | 0.976 | 0.015 | 0.076 | 0.076 | 0.245 | 0.47 | 0.25 | 0.48 | 0.36 | 0.43 | 0.5 |
| 1 | 95273964 | rs6686047 | 0.679 | 1.116 | 0.941 | 0.962 | -0.002 | 0 | 0 | 0.898 | 0.62 | 0.57 | 0.6 | 0.39 | 0.59 | 0.71 |
| 1 | 95276691 | rs3860340 | 0.839 | 1.14 | 0.872 | 0.974 | 0.004 | -0.003 | -0.003 | 0.384 | 0.61 | 0.47 | 0.59 | 0.39 | 0.59 | 0.71 |
| 1 | 95285752 | rs3827752 | 0.529 | 1.146 | 0.443 | 1.013 | 0.02 | -0.011 | -0.011 | 0.129 | 0.18 | 0.06 | 0.06 | 0.18 | 0.13 | 0.12 |
| 1 | 95294089 | rs2640065 | -0.629 | 1.197 | 0.386 | 0.575 | 0.012 | -0.021 | -0.021 | 0 | 0.18 | 0.25 | 0.14 | 0.1 | 0.21 | 0.28 |
| 1 | 95294370 | rs2640066 | -1.2 | 1.212 | 0.376 | 0.427 | 0.003 | 0.008 | 0.008 | 0.176 | 0.52 | 0.52 | 0.52 | 0.7 | 0.57 | 0.53 |
| 1 | 95297374 | rs915012 | -1.059 | 1.265 | 0.388 | 0.452 | 0.005 | 0.012 | 0.012 | 0 | 0.51 | 0.44 | 0.51 | 0.7 | 0.57 | 0.53 |
| 1 | 95298090 | rs6671906 | 0.143 | 1.297 | 0.153 | 0.704 | 0.005 | 0.03 | 0.03 | 0 | 0.14 | 0.16 | 0.11 | 0.19 | 0.15 | 0.23 |
| 1 | 95309134 | rs859111 | -1.012 | 1.168 | -0.374 | 0.759 | -0.003 | -0.014 | -0.014 | 0 | 0.51 | 0.46 | 0.55 | 0.74 | 0.57 | 0.52 |
| 1 | 95309772 | rs698964 | -0.303 | 1.193 | -0.619 | 0.725 | -0.004 | -0.015 | -0.015 | 3.888 | 0.67 | 0.7 | 0.61 | 0.48 | 0.64 | 0.76 |
| 1 | 95310618 | rs698963 | -0.969 | 1.214 | -0.376 | 0.917 | 0.021 | 0.034 | 0.034 | 0 | 0.18 | 0.29 | 0.14 | 0.18 | 0.21 | 0.28 |
| 1 | 95317464 | rs696620 | -0.233 | 1.047 | -1.454 | 0.599 | -0.073 | -0.022 | -0.022 | 0.32 | 0.38 | 0.16 | 0.46 | 0.59 | 0.51 | 0.64 |
| 1 | 95320536 | rs3849303 | 0.165 | -0.071 | -1.555 | -0.369 | 0.039 | 0.056 | 0.056 | 0 | 0.21 | 0.09 | 0.05 | 0.01 | 0.12 | 0.09 |
| 1 | 95323325 | rs7552892 | 0.362 | 0.103 | -0.954 | -0.504 | 0.012 | 0.005 | 0.005 | 0 | 0.11 | 0.18 | 0.14 | 0.11 | 0.16 | 0.12 |
| 1 | 95324784 | rs859101 | 0.92 | 0.082 | -1.231 | -0.416 | -0.067 | -0.012 | -0.012 | 0 | 0.37 | 0.14 | 0.33 | 0.63 | 0.44 | 0.58 |
| 1 | 95324990 | rs9432593 | -0.233 | -0.198 | -0.994 | -0.816 | 0.039 | 0.087 | 0.087 | 0 | 0.31 | 0.17 | 0.14 | 0.02 | 0.21 | 0.18 |
| 1 | 95327466 | rs9432596 | -0.234 | -0.196 | -1.041 | -0.85 | 0.055 | 0.119 | 0.119 | 0 | 0.29 | 0.13 | 0.12 | 0.02 | 0.17 | 0.17 |
| 1 | 95330372 | rs859098 | 0.366 | -0.286 | -0.542 | -0.932 | -0.036 | -0.147 | -0.147 | 0 | 0.65 | 0.48 | 0.81 | 0.98 | 0.78 | 0.81 |
| 1 | 95332624 | rs3849306 | -0.233 | -0.459 | -0.684 | -0.534 | 0.002 | 0 | 0 | 0.579 | 0.19 | 0.19 | 0.23 | 0.22 | 0.13 | 0.19 |
| 1 | 95332864 | rs12137650 | -0.048 | 0.122 | -0.464 | -0.823 | 0.004 | -0.003 | -0.003 | 0 | 0.22 | 0.22 | 0.34 | 0.15 | 0.32 | 0.15 |
| 1 | 95334106 | rs6665783 | -1.103 | 0.301 | -0.513 | -1.026 | 0.01 | 0.02 | 0.02 | 0 | 0.41 | 0.3 | 0.5 | 0.32 | 0.35 | 0.34 |
| 1 | 95335769 | rs735937 | 1.661 | 0.09 | -0.354 | -0.904 | -0.019 | -0.033 | -0.033 | 0.533 | 0.38 | 0.21 | 0.32 | 0.55 | 0.41 | 0.56 |
| 1 | 95337132 | rs17112592 | 2.084 | 0.125 | -0.992 | -0.3 | -0.003 | -0.019 | -0.019 | 0 | 0.21 | 0.05 | 0.12 | 0.2 | 0.22 | 0.22 |
| 1 | 95337418 | rs10493878 | 0.296 | 0.16 | -0.608 | -0.106 | 0 | -0.001 | -0.001 | 0.134 | 0.2 | 0.2 | 0.23 | 0.24 | 0.15 | 0.17 |
| 1 | 95338091 | rs7531174 | -0.539 | 0.153 | -0.63 | -0.115 | -0.01 | -0.008 | -0.008 | 0 | 0.28 | 0.29 | 0.27 | 0.24 | 0.19 | 0.26 |
| 1 | 95342165 | rs859088 | 0.736 | 0.179 | -0.016 | -0.057 | -0.021 | 0.004 | 0.004 | 0 | 0.68 | 0.53 | 0.57 | 0.67 | 0.73 | 0.74 |
| 1 | 95343052 | rs698953 | -0.311 | 0.231 | -0.601 | -0.091 | -0.001 | -0.002 | -0.002 | 0 | 0.8 | 0.8 | 0.71 | 0.74 | 0.77 | 0.88 |
| 1 | 95346659 | rs6700314 | 0.417 | 0.149 | -0.972 | -0.17 | -0.025 | -0.037 | -0.037 | 0.062 | 0.09 | 0.11 | 0.04 | 0 | 0.03 | 0.1 |
| 1 | 95346830 | rs859081 | -0.119 | 0.609 | -1.244 | 0.086 | -0.013 | -0.015 | -0.015 | 0 | 0.79 | 0.9 | 0.83 | 0.78 | 0.78 | 0.88 |
| 1 | 95349437 | rs859074 | -0.146 | 0.807 | -0.853 | 0.351 | -0.017 | -0.025 | -0.025 | 0.045 | 0.3 | 0.27 | 0.25 | 0.46 | 0.36 | 0.39 |
| 1 | 95351921 | rs859067 | 0.473 | 0.281 | -0.369 | 0.847 | -0.002 | 0.013 | 0.013 | 0 | 0.78 | 0.69 | 0.69 | 0.57 | 0.84 | 0.67 |

| | | | | | | | | | | | | | | | | |
|---|---|---|---|---|---|---|---|---|---|---|---|---|---|---|---|---|
| 1 | 95352030 | rs859065 | 0.912 | -0.315 | -0.138 | 0.502 | -0.018 | 0.018 | 0.018 | 0 | 0.68 | 0.58 | 0.64 | 0.56 | 0.79 | 0.6 |
| 1 | 95352756 | rs7550014 | -1.052 | -0.253 | 0.415 | 0.673 | -0.045 | 0.098 | 0.098 | 0 | 0.26 | 0.41 | 0.31 | 0.51 | 0.17 | 0.4 |
| 1 | 95358096 | rs859063 | 1.79 | -0.21 | 0.42 | 0.233 | 0.027 | 0.038 | 0.038 | 0 | 0.51 | 0.33 | 0.51 | 0.42 | 0.6 | 0.47 |
| 1 | 95358256 | rs3860360 | 1.021 | -0.28 | 0.401 | 0.292 | -0.023 | -0.041 | -0.041 | 0 | 0.1 | 0.11 | 0.04 | 0 | 0.05 | 0.07 |
| 1 | 95360309 | rs12030683 | 0.309 | -0.227 | 0.296 | 0.48 | -0.023 | -0.011 | -0.011 | 0 | 0.19 | 0.25 | 0.1 | 0.13 | 0.13 | 0.13 |
| 1 | 95363001 | rs3789699 | 1.441 | 0.199 | -0.063 | 0.698 | 0.002 | 0.011 | 0.011 | 0 | 0.1 | 0.08 | 0.07 | 0.06 | 0.1 | 0.05 |
| 1 | 95367416 | rs2275612 | 1.416 | 0.199 | -0.063 | 0.698 | 0.002 | 0.011 | 0.011 | 0 | 0.1 | 0.08 | 0.07 | 0.06 | 0.1 | 0.05 |
| 1 | 95367894 | rs859057 | -0.172 | 0.216 | -0.027 | 0.675 | -0.001 | 0.013 | 0.013 | 0 | 0.82 | 0.76 | 0.71 | 0.57 | 0.85 | 0.68 |
| 1 | 95372100 | rs7534566 | 0.797 | 0.4 | 0.102 | 0.294 | 0.016 | -0.037 | -0.037 | 0.246 | 0.11 | 0.19 | 0.1 | 0.02 | 0.12 | 0.09 |
| 1 | 95379506 | rs859046 | 1.659 | 0.355 | -0.162 | -0.062 | 0.001 | 0.016 | 0.016 | 0 | 0.3 | 0.22 | 0.26 | 0.17 | 0.32 | 0.32 |
| 1 | 95379758 | rs12067920 | 1.852 | 0.411 | -0.039 | -0.11 | 0.015 | -0.019 | -0.019 | 0 | 0.08 | 0.23 | 0.06 | 0.03 | 0.07 | 0.04 |
| 1 | 95380934 | rs859045 | 1.835 | 0.384 | 0.008 | -0.121 | 0 | 0.027 | 0.027 | 0 | 0.29 | 0.18 | 0.25 | 0.17 | 0.32 | 0.32 |
| 1 | 95381030 | rs859044 | 1.745 | 0.387 | 0.002 | -0.119 | 0.001 | 0.016 | 0.016 | 0 | 0.3 | 0.22 | 0.26 | 0.17 | 0.32 | 0.32 |
| 1 | 95381388 | rs17112757 | 0.718 | 0.537 | -0.14 | -0.166 | 0.008 | -0.041 | -0.041 | 2.33 | 0.12 | 0.23 | 0.09 | 0.04 | 0.11 | 0.07 |
| 1 | 95382491 | rs3860362 | 0.849 | 0.714 | -0.282 | 0.025 | 0.001 | 0.001 | 0.001 | 0 | 0.2 | 0.19 | 0.18 | 0.22 | 0.22 | 0.12 |
| 1 | 95384670 | rs859040 | -0.481 | 0.544 | -0.059 | 0.325 | 0.002 | 0.071 | 0.071 | 0 | 0.86 | 0.81 | 0.69 | 0.47 | 0.84 | 0.66 |
| 1 | 95384905 | rs17112770 | 0.886 | 0.572 | -0.146 | 0.501 | -0.001 | 0.084 | 0.084 | 0 | 0.06 | 0.06 | 0.22 | 0.4 | 0.05 | 0.27 |
| 1 | 95387208 | rs860873 | 1.421 | 0.215 | -0.311 | -0.072 | 0.01 | 0.153 | 0.153 | 0 | 0.47 | 0.27 | 0.35 | 0.16 | 0.44 | 0.39 |
| 1 | 95396932 | rs767015 | -1.871 | -0.107 | 0.311 | 0.413 | 0.002 | -0.003 | -0.003 | 0.02 | 0.53 | 0.67 | 0.43 | 0.47 | 0.48 | 0.33 |
| 1 | 95401199 | rs11807661 | 0.39 | -0.167 | 0.458 | 0.285 | 0.003 | -0.019 | -0.019 | 1.094 | 0.15 | 0.13 | 0.09 | 0 | 0.13 | 0.1 |
| 1 | 95401359 | rs11809155 | 1.156 | -0.173 | 0.465 | 0.293 | -0.008 | -0.021 | -0.021 | 0.017 | 0.09 | 0.1 | 0.03 | 0 | 0.05 | 0.06 |
| 1 | 95408492 | rs1146461 | -2.595 | -0.104 | 0.528 | 0.359 | -0.01 | 0.077 | 0.077 | 0 | 0.61 | 0.82 | 0.7 | 0.88 | 0.58 | 0.6 |
| 1 | 95408920 | rs1146460 | -2.098 | -0.075 | 0.397 | 0.346 | -0.004 | -0.007 | -0.007 | 0 | 0.56 | 0.68 | 0.46 | 0.47 | 0.53 | 0.37 |
| 1 | 95409858 | rs12137024 | 0.632 | -0.123 | 0.256 | 0.441 | -0.049 | -0.063 | -0.063 | 0 | 0.14 | 0.03 | 0.19 | 0.22 | 0.19 | 0.12 |
| 1 | 95412414 | rs2040048 | -1.664 | -0.176 | 0.41 | 0.482 | 0.006 | 0.04 | 0.04 | 0 | 0.43 | 0.24 | 0.38 | 0.31 | 0.43 | 0.33 |
| 1 | 95412967 | rs17407908 | 2.656 | -0.166 | 0.536 | 0.461 | 0.007 | -0.021 | -0.021 | 0 | 0.07 | 0.14 | 0.04 | 0.03 | 0.07 | 0.03 |
| 1 | 95417348 | rs1271952 | -2.173 | -0.233 | 0.379 | 0.473 | -0.006 | -0.01 | -0.01 | 0 | 0.56 | 0.69 | 0.46 | 0.48 | 0.53 | 0.37 |
| 1 | 95420286 | rs2148366 | 0.635 | -0.161 | 0.494 | 0.383 | 0.046 | -0.054 | -0.054 | 0 | 0.13 | 0.48 | 0.09 | 0.04 | 0.09 | 0.04 |
| 1 | 95421315 | rs6667676 | -0.699 | -0.111 | -0.064 | 0.486 | -0.024 | 0.018 | 0.018 | 0 | 0.3 | 0.16 | 0.28 | 0.23 | 0.34 | 0.22 |
| 1 | 95424222 | rs4847372 | 1.598 | -0.153 | -0.113 | 0.469 | 0.026 | -0.017 | -0.017 | 0 | 0.11 | 0.01 | 0.07 | 0.09 | 0.07 | 0.07 |
| 1 | 95424317 | rs2766002 | -1.066 | -0.155 | 0.273 | 0.468 | -0.012 | 0.036 | 0.036 | 0 | 0.23 | 0.38 | 0.2 | 0.34 | 0.18 | 0.2 |
| 1 | 95426247 | rs10493880 | -2.392 | -0.337 | -0.029 | 0.051 | -0.031 | 0.083 | 0.083 | 0 | 0.61 | 0.85 | 0.65 | 0.78 | 0.58 | 0.58 |
| 1 | 95426365 | rs2766003 | -1.645 | -0.384 | 0.058 | 0.058 | -0.023 | 0.053 | 0.053 | 0 | 0.42 | 0.79 | 0.52 | 0.69 | 0.43 | 0.47 |
| 1 | 95427608 | rs11810387 | 0.219 | -0.399 | 0.386 | -0.063 | -0.002 | -0.066 | -0.066 | 0 | 0.18 | 0.22 | 0.1 | 0 | 0.16 | 0.13 |
| 1 | 95427877 | rs2254418 | 0.011 | -0.339 | 0.759 | 0.569 | 0.074 | 0.151 | 0.151 | 1.389 | 0.06 | 0.32 | 0.25 | 0.13 | 0.05 | 0.05 |
| 1 | 95428051 | rs10493881 | 1.637 | -0.341 | 0.346 | 0.386 | 0.013 | -0.04 | -0.04 | 0.315 | 0.1 | 0.13 | 0.08 | 0 | 0.11 | 0.06 |
| 1 | 95428414 | rs2766005 | -2.026 | -0.445 | 0.255 | 0.438 | 0.03 | 0.106 | 0.106 | 0 | 0.48 | 0.72 | 0.53 | 0.88 | 0.44 | 0.51 |
| 1 | 95429026 | rs6541375 | -1.337 | -0.461 | 0.262 | 0.503 | -0.002 | -0.003 | -0.003 | 0 | 0.33 | 0.66 | 0.44 | 0.69 | 0.32 | 0.4 |
| 1 | 95429780 | rs11165281 | 0.8 | -0.43 | 0.05 | 0.589 | 0.029 | -0.009 | -0.009 | 0 | 0.18 | 0.06 | 0.12 | 0.19 | 0.15 | 0.12 |
| 1 | 95445468 | rs2797623 | -2.248 | -0.555 | 0.197 | 0.498 | -0.004 | 0.028 | 0.028 | 0 | 0.53 | 0.65 | 0.58 | 0.86 | 0.5 | 0.6 |
| 1 | 95446762 | rs7551774 | 0.774 | -0.409 | 0.107 | 0.933 | -0.026 | -0.055 | -0.055 | 0 | 0.14 | 0.07 | 0.2 | 0.5 | 0.19 | 0.28 |
| 1 | 95449033 | rs2298162 | -1.025 | -0.492 | 0.381 | 0.833 | 0.024 | -0.025 | -0.025 | 0 | 0.3 | 0.44 | 0.45 | 0.76 | 0.37 | 0.43 |
| 1 | 95451926 | rs2391391 | -1.109 | -0.496 | 0.395 | 0.839 | -0.002 | -0.007 | -0.007 | 0 | 0.29 | 0.37 | 0.34 | 0.26 | 0.27 | 0.25 |
| 1 | 95462101 | rs12036045 | 0.715 | -0.428 | 0.02 | 1.028 | -0.023 | -0.051 | -0.051 | 0 | 0.14 | 0.07 | 0.2 | 0.47 | 0.19 | 0.27 |
| 1 | 95470811 | rs7527765 | 1.011 | -0.538 | 0.119 | 0.889 | 0.014 | 0.01 | 0.01 | 0 | 0.14 | 0.07 | 0.1 | 0.1 | 0.09 | 0.1 |
| 1 | 95471456 | rs11165285 | 0.526 | -0.555 | 0.139 | 0.979 | -0.013 | -0.035 | -0.035 | 0 | 0.15 | 0.12 | 0.2 | 0.56 | 0.19 | 0.28 |
| 1 | 95478126 | rs12137333 | -0.289 | -0.742 | 0.242 | 0.86 | 0.013 | 0.013 | 0.013 | 0 | 0.22 | 0.17 | 0.12 | 0.1 | 0.13 | 0.17 |
| 1 | 95482668 | rs12751633 | 0.009 | -0.689 | 0.507 | 0.818 | 0.039 | 0.016 | 0.016 | 0.029 | 0.16 | 0.34 | 0.25 | 0.16 | 0.18 | 0.15 |
| 1 | 95486866 | rs4847376 | 1.861 | -0.72 | 0.475 | 0.794 | 0.036 | 0.005 | 0.005 | 0 | 0.11 | 0.02 | 0.06 | 0.1 | 0.06 | 0.08 |
| 1 | 95491261 | rs10465759 | 0.811 | -0.635 | 0.645 | 0.836 | 0.017 | -0.003 | -0.003 | 0.453 | 0.12 | 0.25 | 0.21 | 0.1 | 0.14 | 0.09 |
| 1 | 95493261 | rs4427415 | 0.692 | -0.634 | 0.517 | 0.856 | -0.007 | -0.015 | -0.015 | 1.583 | 0.15 | 0.12 | 0.21 | 0.56 | 0.19 | 0.28 |
| 1 | 95505324 | rs12022525 | 0.778 | -0.613 | 0.458 | 1.137 | -0.025 | -0.059 | -0.059 | 0 | 0.14 | 0.07 | 0.13 | 0.39 | 0.19 | 0.27 |
| 1 | 95515124 | rs6687388 | 2.748 | -0.627 | 0.575 | 0.589 | -0.036 | 0.117 | 0.117 | 0 | 0.38 | 0.13 | 0.36 | 0.19 | 0.42 | 0.38 |
| 1 | 95515637 | rs6675668 | 2.211 | -0.643 | 0.675 | 0.607 | -0.009 | 0.069 | 0.069 | 0 | 0.46 | 0.27 | 0.41 | 0.19 | 0.49 | 0.4 |
| 1 | 95524503 | rs6671842 | -0.252 | -0.424 | 0.537 | 0.784 | 0.039 | 0.016 | 0.016 | 0.098 | 0.16 | 0.37 | 0.25 | 0.16 | 0.18 | 0.15 |
| 1 | 95536715 | rs12755096 | -1.046 | -0.432 | 0.526 | 0.735 | 0.007 | -0.027 | -0.027 | 0 | 0.23 | 0.45 | 0.27 | 0.15 | 0.23 | 0.21 |
| 1 | 95546430 | rs7417186 | 0.044 | -0.465 | 0.3 | 0.662 | 0.019 | -0.015 | -0.015 | 0 | 0.14 | 0.32 | 0.24 | 0.11 | 0.16 | 0.11 |
| 1 | 95550654 | rs11801110 | 0.603 | -0.469 | 0.308 | 0.662 | 0.022 | 0.001 | 0.001 | 0.015 | 0.12 | 0.32 | 0.21 | 0.1 | 0.14 | 0.1 |
| 1 | 95551129 | rs4950059 | 1.067 | -0.488 | 0.22 | 0.627 | 0.003 | 0.004 | 0.004 | 0 | 0.53 | 0.43 | 0.59 | 0.29 | 0.58 | 0.5 |
| 1 | 95554849 | rs10874903 | 0.541 | -0.433 | 0.157 | 0.666 | -0.002 | -0.023 | -0.023 | 1.46 | 0.6 | 0.52 | 0.61 | 0.29 | 0.62 | 0.56 |
| 1 | 95567336 | rs6665763 | 0.999 | -0.267 | 0.005 | 1.104 | 0.029 | 0.04 | 0.04 | 0 | 0.53 | 0.27 | 0.58 | 0.35 | 0.59 | 0.59 |
| 1 | 95567854 | rs6680551 | 0.308 | -0.317 | 0.121 | 1.355 | 0.013 | 0.148 | 0.148 | 0.058 | 0.13 | 0.32 | 0.19 | 0.39 | 0.15 | 0.17 |
| 1 | 95567932 | rs6593673 | 1.033 | -0.274 | -0.074 | 1.347 | 0.029 | 0.04 | 0.04 | 0 | 0.53 | 0.27 | 0.58 | 0.35 | 0.59 | 0.59 |
| 1 | 95577297 | rs4950076 | 0.283 | 0.054 | -0.268 | 1.09 | -0.003 | -0.001 | -0.001 | 0 | 0.47 | 0.37 | 0.61 | 0.55 | 0.51 | 0.54 |
| 1 | 95586773 | rs1023330 | 1.05 | 0.184 | -0.822 | 0.5 | -0.037 | 0.117 | 0.117 | 0 | 0.35 | 0.1 | 0.42 | 0.15 | 0.36 | 0.34 |
| 1 | 95592609 | rs11165315 | -0.383 | -0.274 | 0.194 | 1.204 | -0.022 | 0.075 | 0.075 | 0 | 0.2 | 0.35 | 0.09 | 0.39 | 0.14 | 0.25 |
| 1 | 95598593 | rs10874905 | -0.812 | -0.011 | 0.417 | 1.42 | -0.002 | 0.282 | 0.282 | 0.825 | 0.13 | 0.44 | 0.08 | 0.44 | 0.09 | 0.17 |
| 1 | 95611717 | rs17113072 | 0.34 | 0.108 | -0.465 | 0.83 | -0.002 | 0 | 0 | 0 | 0.14 | 0.14 | 0.17 | 0.15 | 0.13 | 0.14 |
| 1 | 95614566 | rs10157254 | 0.234 | 0.567 | -0.764 | 0.567 | -0.007 | 0.004 | 0.004 | 0 | 0.12 | 0.11 | 0.16 | 0.07 | 0.18 | 0.13 |

| | | | | | | | | | | | | | | | | |
|---|---|---|---|---|---|---|---|---|---|---|---|---|---|---|---|---|
| 1 | 95619549 | rs3900596 | -0.537 | 0.542 | -0.756 | 0.626 | -0.004 | 0.001 | 0.001 | 0 | 0.19 | 0.15 | 0.27 | 0.16 | 0.24 | 0.25 |
| 1 | 95619863 | rs964333 | -1.331 | 0.521 | -1.322 | 0.132 | 0.007 | 0.205 | 0.205 | 0 | 0.52 | 0.83 | 0.56 | 0.83 | 0.49 | 0.56 |
| 1 | 95630461 | rs11165328 | -0.503 | 0.715 | -0.777 | 0.792 | -0.006 | -0.001 | -0.001 | 1.19 | 0.2 | 0.15 | 0.28 | 0.18 | 0.27 | 0.25 |
| 1 | 95634601 | rs11165331 | -0.477 | 0.683 | -0.431 | 1.066 | -0.006 | -0.002 | -0.002 | 0.267 | 0.19 | 0.15 | 0.28 | 0.18 | 0.27 | 0.25 |
| 1 | 95638667 | rs11165334 | -1.025 | 1.137 | -0.091 | 1.048 | 0.015 | 0.039 | 0.039 | 2.527 | 0.25 | 0.62 | 0.32 | 0.61 | 0.29 | 0.24 |
| 1 | 95650796 | rs1336435 | -0.698 | 1.185 | 0.356 | 1.057 | 0.052 | 0.184 | 0.184 | 1.292 | 0.19 | 0.49 | 0.27 | 0.61 | 0.21 | 0.22 |
| 1 | 95657959 | rs17113161 | 1.262 | 1.24 | 0.299 | 1.064 | -0.032 | -0.032 | -0.032 | 0 | 0.06 | 0.01 | 0.11 | 0.09 | 0.11 | 0.11 |
| 1 | 95658500 | rs10874907 | -0.668 | 1.145 | 0.396 | 1.081 | 0.054 | 0.184 | 0.184 | 1.115 | 0.19 | 0.49 | 0.27 | 0.61 | 0.21 | 0.23 |
| 1 | 95672229 | rs933107 | -0.632 | 1.06 | 0.488 | 0.925 | 0.02 | -0.047 | -0.047 | 0 | 0.91 | 0.75 | 0.94 | 1 | 0.92 | 0.99 |
| 1 | 95674341 | rs1556742 | -1.211 | 1.079 | 0.511 | 0.947 | 0.018 | 0.01 | 0.01 | 0 | 0.3 | 0.54 | 0.42 | 0.71 | 0.36 | 0.37 |
| 1 | 95690269 | rs2039614 | 0.298 | 0.949 | 0.651 | 0.886 | -0.001 | -0.058 | -0.058 | 0.013 | 0.79 | 0.62 | 0.78 | 0.9 | 0.76 | 0.84 |
| 1 | 95691579 | rs7555339 | 0.09 | 0.936 | 0.805 | 1.088 | 0.025 | 0.07 | 0.07 | 0.024 | 0.89 | 0.72 | 0.85 | 0.8 | 0.87 | 0.93 |
| 1 | 95694770 | rs11165348 | 0.759 | 0.932 | 0.778 | 1.067 | -0.005 | -0.014 | -0.014 | 10.76 | 0.14 | 0.22 | 0.17 | 0.1 | 0.15 | 0.14 |
| 1 | 95697490 | rs6666528 | 0.375 | 0.876 | 1.1 | 1.073 | 0.036 | 0.088 | 0.088 | 0.491 | 0.88 | 0.65 | 0.84 | 0.8 | 0.87 | 0.93 |
| 1 | 95697529 | rs6671200 | -1.211 | 0.923 | 1.11 | 1.093 | 0.015 | -0.039 | -0.039 | 0 | 0.93 | 0.83 | 0.95 | 1 | 0.92 | 0.99 |
| 1 | 95699184 | rs3753872 | 0.183 | 0.923 | 1.108 | 1.095 | 0.035 | 0.083 | 0.083 | 0.022 | 0.89 | 0.68 | 0.85 | 0.8 | 0.87 | 0.93 |
| 1 | 95703444 | rs1160318 | -0.877 | 0.998 | 0.979 | 1.198 | 0.036 | 0.109 | 0.109 | 0 | 0.22 | 0.37 | 0.28 | 0.46 | 0.28 | 0.28 |
| 1 | 95709039 | rs259359 | -1.115 | 1.079 | 1.069 | 1.033 | 0.015 | -0.039 | -0.039 | 0 | 0.93 | 0.82 | 0.95 | 1 | 0.92 | 0.99 |
| 1 | 95709074 | rs7528790 | -0.092 | 1.077 | 1.074 | 1.043 | 0.035 | 0.083 | 0.083 | 0.022 | 0.89 | 0.68 | 0.85 | 0.8 | 0.87 | 0.93 |
| 1 | 95709939 | rs2296308 | -0.092 | 1.079 | 1.034 | 1.044 | 0.035 | 0.083 | 0.083 | 0.022 | 0.89 | 0.68 | 0.85 | 0.8 | 0.87 | 0.93 |
| 1 | 95710329 | rs1062076 | 0.885 | 1.07 | 0.985 | 1.025 | 0 | 0.001 | 0.001 | 0 | 0.13 | 0.13 | 0.16 | 0.1 | 0.15 | 0.14 |
| 1 | 95716754 | rs4949946 | 0.774 | 1.065 | 0.846 | 1.058 | -0.003 | 0.015 | 0.015 | 0 | 0.75 | 0.46 | 0.68 | 0.71 | 0.72 | 0.79 |
| 1 | 95718941 | rs10783001 | 0.852 | 1.07 | 0.52 | 1.15 | 0.017 | 0.027 | 0.027 | 0 | 0.7 | 0.55 | 0.57 | 0.62 | 0.61 | 0.7 |
| 1 | 95741611 | rs11165361 | -0.918 | 0.817 | 0.353 | 1.415 | 0.01 | 0.026 | 0.026 | 0 | 0.21 | 0.32 | 0.25 | 0.27 | 0.24 | 0.28 |
| 1 | 95741812 | rs259332 | -1.679 | 0.669 | 0.298 | 1.325 | 0.007 | 0.005 | 0.005 | 0 | 0.41 | 0.51 | 0.4 | 0.53 | 0.46 | 0.46 |
| 1 | 95745335 | rs2039615 | -1.697 | 0.458 | 0.729 | 1.456 | -0.001 | 0.055 | 0.055 | 0 | 0.31 | 0.51 | 0.38 | 0.45 | 0.3 | 0.35 |
| 1 | 95745539 | rs259327 | 1.121 | -0.093 | 0.117 | 0.925 | -0.003 | 0.007 | 0.007 | 0 | 0.32 | 0.22 | 0.24 | 0.28 | 0.38 | 0.26 |
| 1 | 95746349 | rs259325 | 0.952 | -0.152 | 0.474 | 0.999 | 0.002 | 0.006 | 0.006 | 0 | 0.39 | 0.31 | 0.29 | 0.37 | 0.43 | 0.2 |
| 1 | 95746524 | rs11165363 | 0.216 | -0.43 | 0.086 | 1.059 | -0.087 | -0.044 | -0.044 | 0 | 0.36 | 0.11 | 0.49 | 0.36 | 0.42 | 0.44 |
| 1 | 95749439 | rs259323 | 1.459 | -0.418 | 0.368 | 1.239 | -0.006 | 0.002 | 0.002 | 0 | 0.66 | 0.4 | 0.58 | 0.63 | 0.74 | 0.52 |
| 1 | 95754014 | rs650611 | -1.993 | -0.363 | 0.51 | 1.244 | -0.004 | 0.038 | 0.038 | 0 | 0.65 | 0.86 | 0.79 | 0.72 | 0.59 | 0.85 |
| 1 | 95766707 | rs4949959 | -0.664 | 0.238 | 0.969 | 1.242 | -0.03 | 0.007 | 0.007 | 0.074 | 0.25 | 0.52 | 0.26 | 0.29 | 0.14 | 0.38 |
| 1 | 95766797 | rs4950044 | -0.001 | 0.23 | 0.882 | 1.225 | -0.008 | 0.002 | 0.002 | 0 | 0.19 | 0.35 | 0.16 | 0.21 | 0.13 | 0.32 |
| 1 | 95773106 | rs6683355 | -0.864 | 0.197 | 0.871 | 1.21 | -0.023 | 0.006 | 0.006 | 0.989 | 0.24 | 0.58 | 0.26 | 0.29 | 0.14 | 0.38 |
| 1 | 95776618 | rs2236389 | 0.347 | 0.231 | 0.524 | 1.204 | -0.001 | 0.009 | 0.009 | 0 | 0.17 | 0.32 | 0.16 | 0.21 | 0.13 | 0.32 |
| 1 | 95776788 | rs2236390 | -0.291 | 0.256 | -0.284 | 1.311 | 0.003 | -0.019 | -0.019 | 0.111 | 0.12 | 0.09 | 0.19 | 0.16 | 0.14 | 0.15 |
| 1 | 95781217 | rs4949960 | -0.383 | 0.285 | -0.401 | 1.256 | -0.081 | -0.091 | -0.091 | 0 | 0.38 | 0.12 | 0.52 | 0.43 | 0.44 | 0.47 |
| 1 | 95782533 | rs6593652 | -2.343 | 0.247 | -0.336 | 1.192 | -0.037 | 0.008 | 0.008 | 0 | 0.69 | 0.93 | 0.82 | 0.73 | 0.63 | 0.86 |
| 1 | 95782971 | rs17113362 | 0.272 | 0.261 | -0.025 | 1.381 | -0.009 | 0.016 | 0.016 | 0 | 0.05 | 0.09 | 0.03 | 0.08 | 0.03 | 0.06 |
| 1 | 95786229 | rs17113368 | 0.299 | 0.309 | -0.104 | 1.21 | -0.002 | -0.008 | -0.008 | 1.431 | 0.08 | 0.06 | 0.16 | 0.08 | 0.11 | 0.09 |
| 1 | 95789364 | rs17113374 | -0.155 | 0.253 | -0.04 | 1.174 | 0 | -0.002 | -0.002 | 0 | 0.1 | 0.11 | 0.17 | 0.08 | 0.12 | 0.09 |
| 1 | 95789603 | rs4604740 | 0.775 | 0.381 | -0.292 | 1.188 | -0.1 | -0.027 | -0.027 | 0.968 | 0.26 | 0.05 | 0.33 | 0.26 | 0.3 | 0.33 |
| 1 | 95791800 | rs912649 | -1.994 | 0.071 | 0.053 | 0.908 | 0.006 | 0.034 | 0.034 | 0 | 0.65 | 0.85 | 0.8 | 0.72 | 0.62 | 0.86 |
| 1 | 95798900 | rs912651 | -2.145 | 0.172 | 0.135 | 1.252 | -0.001 | 0.018 | 0.018 | 0 | 0.67 | 0.88 | 0.8 | 0.72 | 0.61 | 0.85 |
| 1 | 95800553 | rs11165383 | 0.024 | 0.244 | 0.313 | 1.366 | -0.074 | -0.058 | -0.058 | 0 | 0.35 | 0.15 | 0.39 | 0.44 | 0.4 | 0.45 |
| 1 | 95805002 | rs7547539 | -0.964 | 0.286 | 1.194 | 1.422 | -0.029 | -0.019 | -0.019 | 3.193 | 0.27 | 0.57 | 0.31 | 0.21 | 0.2 | 0.36 |
| 1 | 95810750 | rs4950046 | -1.063 | 0.295 | 0.97 | 1.421 | -0.018 | -0.01 | -0.01 | 0 | 0.27 | 0.65 | 0.33 | 0.22 | 0.2 | 0.35 |
| 1 | 95811910 | rs4523562 | 1.114 | 0.288 | 0.958 | 1.425 | 0.012 | 0.002 | 0.002 | 0.699 | 0.68 | 0.26 | 0.59 | 0.71 | 0.79 | 0.6 |
| 1 | 95817718 | rs4573557 | 0.479 | 0.286 | 0.991 | 1.424 | 0.006 | 0.005 | 0.005 | 0 | 0.08 | 0.14 | 0.17 | 0.08 | 0.12 | 0.09 |
| 1 | 95818057 | rs3814011 | 1.091 | 0.28 | 1.038 | 1.426 | -0.008 | -0.002 | -0.002 | 0.628 | 0.69 | 0.31 | 0.59 | 0.71 | 0.78 | 0.6 |
| 1 | 95825187 | rs4291548 | 0.455 | 0.341 | 0.811 | 1.491 | -0.017 | -0.02 | -0.02 | 0 | 0.6 | 0.68 | 0.52 | 0.51 | 0.6 | 0.52 |
| 1 | 95828652 | rs12562298 | -0.194 | 0.372 | 0.705 | 1.504 | -0.063 | -0.049 | -0.049 | 0 | 0.35 | 0.16 | 0.39 | 0.42 | 0.39 | 0.43 |
| 1 | 95829831 | rs17113446 | 0.148 | 0.405 | 0.711 | 1.474 | -0.035 | -0.01 | -0.01 | 0 | 0.17 | 0.38 | 0.15 | 0.14 | 0.08 | 0.27 |
| 1 | 95832611 | rs11165406 | 1.082 | 0.574 | -0.077 | 1.433 | 0.002 | 0.02 | 0.02 | 0 | 0.43 | 0.25 | 0.36 | 0.37 | 0.51 | 0.25 |
| 1 | 95834888 | rs7534563 | -0.205 | 0.597 | 0.31 | 1.428 | -0.014 | 0.018 | 0.018 | 0 | 0.84 | 0.74 | 0.85 | 0.76 | 0.92 | 0.79 |
| 1 | 95835124 | rs7531902 | 0.731 | 1.31 | 0.134 | 1.469 | -0.001 | 0.001 | 0.001 | 0 | 0.68 | 0.67 | 0.56 | 0.73 | 0.69 | 0.61 |
| 1 | 95837686 | rs7524217 | 0.942 | 1.119 | 0.85 | 1.786 | -0.022 | 0.033 | 0.033 | 0 | 0.69 | 0.46 | 0.7 | 0.6 | 0.79 | 0.55 |
| 1 | 95845474 | rs10489557 | -0.829 | 1.067 | 0.751 | 1.981 | -0.013 | 0.029 | 0.029 | 0 | 0.23 | 0.31 | 0.3 | 0.43 | 0.15 | 0.28 |
| 1 | 95850648 | rs2391684 | -1.722 | 1.065 | 0.788 | 1.98 | -0.01 | 0.009 | 0.009 | 0 | 0.38 | 0.57 | 0.46 | 0.56 | 0.3 | 0.55 |
| 1 | 95857362 | rs2064662 | -0.899 | 1.38 | 0.735 | 1.891 | 0.078 | -0.043 | -0.043 | 1.693 | 0.89 | 0.83 | 0.76 | 0.98 | 0.82 | 0.87 |
| 1 | 95861202 | rs11586346 | 1.079 | 1.214 | 0.499 | 1.987 | -0.009 | -0.039 | -0.039 | 0 | 0.07 | 0 | 0.16 | 0.15 | 0.08 | 0.09 |
| 1 | 95865533 | rs4428940 | 0.9 | 1.2 | 0.522 | 1.756 | -0.026 | -0.028 | -0.028 | 0.005 | 0.11 | 0.03 | 0.15 | 0.13 | 0.15 | 0.19 |
| 1 | 95867674 | rs10874921 | -0.078 | 1.065 | 0.607 | 1.039 | -0.021 | 0.027 | 0.027 | 0 | 0.17 | 0.28 | 0.14 | 0.27 | 0.07 | 0.19 |
| 1 | 95872794 | rs11165419 | 0.454 | 1.108 | -0.138 | 0.81 | 0.012 | -0.012 | -0.012 | 0.009 | 0.14 | 0.1 | 0.06 | 0.16 | 0.06 | 0.16 |
| 1 | 95877746 | rs11590987 | -0.677 | 1.11 | -0.177 | 0.836 | 0.023 | -0.019 | -0.019 | 0 | 0.22 | 0.17 | 0.18 | 0.29 | 0.14 | 0.25 |
| 1 | 95880135 | rs4259715 | -2.115 | 1.109 | -0.208 | 0.733 | 0.026 | 0.089 | 0.089 | 0 | 0.49 | 0.69 | 0.51 | 0.92 | 0.38 | 0.68 |
| 1 | 95881126 | rs12061105 | -1.383 | 1.108 | -0.112 | 0.974 | 0.013 | -0.019 | -0.019 | 0 | 0.31 | 0.29 | 0.21 | 0.43 | 0.22 | 0.38 |
| 1 | 95884729 | rs7547980 | -1.033 | 1.26 | -0.07 | 0.422 | 0.076 | -0.039 | -0.039 | 0 | 0.9 | 0.83 | 0.76 | 0.97 | 0.82 | 0.86 |
| 1 | 95893305 | rs7527710 | -0.785 | 0.67 | 0.161 | 0.68 | -0.007 | -0.034 | -0.034 | 0 | 0.2 | 0.34 | 0.07 | 0.07 | 0.1 | 0.23 |

| | | | | | | | | | | | | | | | | |
|---|---|---|---|---|---|---|---|---|---|---|---|---|---|---|---|---|
| 1 | 95896007 | rs2224153 | -0.594 | 0.841 | 0.127 | 0.752 | 0.008 | 0.063 | 0.063 | 0 | 0.25 | 0.36 | 0.39 | 0.61 | 0.26 | 0.39 |
| 1 | 95897424 | rs7526471 | -0.941 | 0.836 | -0.333 | 0.785 | 0.014 | 0.012 | 0.012 | 0 | 0.23 | 0.25 | 0.07 | 0.2 | 0.14 | 0.28 |
| 1 | 95902901 | rs2064663 | -1.064 | 0.873 | -0.413 | 0.704 | 0.001 | 0.012 | 0.012 | 0 | 0.3 | 0.55 | 0.45 | 0.64 | 0.33 | 0.4 |
| 1 | 95903597 | rs1474883 | -2.27 | 0.901 | -0.445 | 0.697 | -0.008 | 0.105 | 0.105 | 0 | 0.53 | 0.74 | 0.51 | 0.82 | 0.47 | 0.68 |
| 1 | 95917695 | rs17113575 | -0.685 | 0.89 | -0.388 | 0.695 | 0.007 | 0.008 | 0.008 | 0 | 0.22 | 0.24 | 0.07 | 0.18 | 0.14 | 0.29 |
| 1 | 95920263 | rs12568065 | -0.988 | 0.994 | -0.35 | 0.747 | 0.011 | 0.053 | 0.053 | 0 | 0.28 | 0.39 | 0.43 | 0.61 | 0.33 | 0.39 |
| 1 | 95929401 | rs6692957 | -1.893 | 1.127 | -0.784 | 0.746 | -0.012 | 0.135 | 0.135 | 0 | 0.52 | 0.77 | 0.51 | 0.82 | 0.46 | 0.65 |
| 1 | 95939767 | rs957186 | -1.777 | 1.135 | -0.79 | 0.748 | -0.001 | 0.095 | 0.095 | 0 | 0.51 | 0.66 | 0.51 | 0.79 | 0.46 | 0.65 |
| 1 | 95947234 | rs1415564 | 0.908 | 0.908 | -0.235 | 0.83 | -0.01 | 0.003 | 0.003 | 0.003 | 0.81 | 0.49 | 0.88 | 0.82 | 0.83 | 0.78 |
| 1 | 95947820 | rs12409850 | 0.458 | 1.146 | -0.376 | 0.79 | 0.044 | -0.028 | -0.028 | 0 | 0.09 | 0.15 | 0.13 | 0.02 | 0.13 | 0.12 |
| 1 | 95966161 | rs10874924 | 2.254 | 1.14 | -0.711 | 1.018 | -0.112 | -0.07 | -0.07 | 0.068 | 0.51 | 0.03 | 0.55 | 0.52 | 0.56 | 0.43 |
| 1 | 95967614 | rs1577864 | -0.679 | 1.286 | -0.44 | 0.846 | 0.033 | 0.017 | 0.017 | 1.305 | 0.16 | 0.22 | 0.28 | 0.23 | 0.22 | 0.2 |
| 1 | 95967757 | rs1577863 | -0.679 | 1.286 | -0.44 | 0.846 | 0.033 | 0.017 | 0.017 | 1.305 | 0.16 | 0.23 | 0.28 | 0.23 | 0.22 | 0.2 |
| 1 | 95973030 | rs874821 | -1.452 | 1.055 | -0.516 | 0.828 | -0.014 | 0.004 | 0.004 | 0 | 0.3 | 0.3 | 0.33 | 0.31 | 0.28 | 0.38 |
| 1 | 95977445 | rs1582123 | -0.268 | 0.67 | -0.609 | 0.998 | -0.01 | 0.123 | 0.123 | 0.783 | 0.88 | 0.57 | 0.94 | 0.81 | 0.94 | 0.84 |
| 1 | 95985066 | rs881713 | 0.248 | 0.732 | -0.699 | 1.002 | -0.015 | -0.035 | -0.035 | 0.131 | 0.08 | 0.02 | 0.16 | 0.21 | 0.08 | 0.08 |
| 1 | 95989978 | rs12566519 | 1.934 | 0.784 | -0.771 | 0.933 | -0.102 | -0.108 | -0.108 | 0.74 | 0.5 | 0.03 | 0.54 | 0.48 | 0.56 | 0.4 |
| 1 | 96002882 | rs10783011 | 0.914 | 0.759 | -0.682 | 0.926 | -0.032 | 0.117 | 0.117 | 0.967 | 0.78 | 0.5 | 0.78 | 0.6 | 0.85 | 0.75 |
| 1 | 96013073 | rs12354293 | -0.017 | 0.819 | -0.788 | 0.805 | 0.098 | -0.095 | -0.095 | 0.141 | 0.17 | 0.32 | 0.2 | 0.04 | 0.25 | 0.2 |
| 1 | 96018531 | rs17328231 | 1.379 | 0.437 | -1.686 | 0.901 | -0.072 | -0.089 | -0.089 | 1.359 | 0.6 | 0.1 | 0.57 | 0.52 | 0.62 | 0.53 |
| 1 | 96019588 | rs528629 | -2.062 | -0.387 | -0.767 | 1.496 | -0.01 | 0.018 | 0.018 | 0.417 | 0.31 | 0.41 | 0.24 | 0.5 | 0.18 | 0.38 |
| 1 | 96029510 | rs524151 | -1.377 | -0.374 | -0.76 | 1.522 | -0.03 | 0.027 | 0.027 | 0.339 | 0.22 | 0.39 | 0.08 | 0.28 | 0.1 | 0.32 |
| 1 | 96034482 | rs6593669 | 1.284 | -0.257 | -0.774 | 0.955 | -0.015 | 0.022 | 0.022 | 1.017 | 0.68 | 0.53 | 0.76 | 0.5 | 0.82 | 0.62 |
| 1 | 96035056 | rs13375091 | -0.512 | 0.402 | -1.17 | 0.373 | 0.011 | 0.014 | 0.014 | 0 | 0.1 | 0.07 | 0.02 | 0.05 | 0.03 | 0.14 |
| 1 | 96043511 | rs11165443 | -0.555 | 0.423 | -1.29 | 0.113 | 0.003 | 0.034 | 0.034 | 0 | 0.39 | 0.26 | 0.41 | 0.81 | 0.38 | 0.55 |
| 1 | 96044847 | rs12031667 | -0.102 | 0.286 | -1.507 | 1.121 | 0.004 | -0.006 | -0.006 | 0 | 0.2 | 0.18 | 0.16 | 0.34 | 0.17 | 0.23 |
| 1 | 96045069 | rs10783012 | -0.763 | 0.524 | -1.432 | 0.298 | -0.004 | 0.014 | 0.014 | 0 | 0.55 | 0.59 | 0.61 | 0.75 | 0.53 | 0.49 |
| 1 | 96049429 | rs7545959 | -0.775 | -0.121 | -1.657 | -0.581 | -0.001 | 0.038 | 0.038 | 0 | 0.53 | 0.38 | 0.52 | 0.87 | 0.46 | 0.58 |
| 1 | 96051831 | rs661595 | -0.337 | -0.126 | -1.245 | -0.794 | 0.024 | -0.062 | -0.062 | 1.149 | 0.67 | 0.53 | 0.65 | 0.88 | 0.6 | 0.64 |
| 1 | 96052746 | rs10489560 | 0.07 | -0.1 | -1.243 | -0.779 | -0.025 | -0.007 | -0.007 | 0.256 | 0.1 | 0.06 | 0.12 | 0.1 | 0.16 | 0.18 |
| 1 | 96056162 | rs670074 | 1.579 | -0.121 | -1.242 | -0.794 | 0 | -0.001 | -0.001 | 2.045 | 0.23 | 0.25 | 0.25 | 0.13 | 0.31 | 0.24 |
| 1 | 96056974 | rs684149 | 1.587 | -0.174 | -1.14 | -0.829 | 0.01 | -0.074 | -0.074 | 0 | 0.2 | 0.28 | 0.22 | 0 | 0.24 | 0.18 |
| 1 | 96058550 | rs10489562 | 0.515 | -0.186 | -1.241 | -0.669 | -0.009 | 0.002 | 0.002 | 0 | 0.31 | 0.26 | 0.2 | 0.36 | 0.18 | 0.31 |
| 1 | 96072754 | rs573117 | 0.442 | -0.124 | -1.279 | -0.599 | -0.016 | -0.041 | -0.041 | 0 | 0.75 | 0.8 | 0.66 | 0.5 | 0.73 | 0.74 |
| 1 | 96084768 | rs12047025 | 1.515 | -0.373 | -1.343 | -0.669 | 0.014 | 0.002 | 0.002 | 0 | 0.1 | 0.03 | 0.03 | 0.09 | 0.07 | 0.05 |
| 1 | 96088192 | rs621485 | 0.286 | -0.344 | -1.213 | -0.652 | 0 | 0 | 0 | 0 | 0.55 | 0.59 | 0.58 | 0.26 | 0.7 | 0.62 |
| 1 | 96098413 | rs880686 | 0.664 | -0.131 | -1.323 | -0.255 | -0.06 | 0.041 | 0.041 | 0.535 | 0.51 | 0.07 | 0.32 | 0.34 | 0.51 | 0.55 |
| 1 | 96098508 | rs937928 | 0.744 | 0.14 | -0.47 | 0.007 | 0.068 | -0.042 | -0.042 | 3.399 | 0.91 | 0.71 | 0.88 | 1 | 0.87 | 0.97 |
| 1 | 96106197 | rs588093 | -0.281 | -0.511 | -0.536 | 0.55 | -0.026 | 0.031 | 0.031 | 0 | 0.88 | 0.83 | 0.88 | 0.71 | 0.97 | 0.82 |
| 1 | 96115365 | rs3108719 | 0.208 | -0.281 | 0.06 | -0.041 | -0.004 | -0.004 | -0.004 | 0.321 | 0.66 | 0.36 | 0.64 | 0.72 | 0.71 | 0.65 |
| 1 | 96118261 | rs11587357 | 1.713 | -0.07 | -0.637 | -0.566 | -0.046 | 0.087 | 0.087 | 0 | 0.16 | 0.01 | 0.1 | 0.01 | 0.2 | 0.08 |
| 1 | 96118307 | rs17114028 | 0.711 | 0.273 | -0.528 | 0.007 | -0.024 | 0.011 | 0.011 | 0 | 0.07 | 0.02 | 0.21 | 0.37 | 0.1 | 0.07 |
| 1 | 96124935 | rs11165460 | -0.106 | 0.181 | -0.261 | -0.153 | 0.012 | 0.013 | 0.013 | 1.092 | 0.65 | 0.27 | 0.62 | 0.72 | 0.7 | 0.65 |
| 1 | 96125973 | rs10874931 | 0.303 | 0.208 | -0.26 | -0.193 | -0.004 | 0.015 | 0.015 | 1.209 | 0.59 | 0.25 | 0.41 | 0.35 | 0.61 | 0.58 |
| 1 | 96138526 | rs6703657 | 1.126 | 0.155 | -0.145 | -0.317 | 0.01 | -0.091 | -0.091 | 0 | 0.18 | 0.34 | 0.21 | 0.02 | 0.21 | 0.16 |
| 1 | 96142596 | rs1463973 | -0.031 | 0.06 | -1.022 | -0.239 | 0.019 | 0.009 | 0.009 | 2.054 | 0.67 | 0.18 | 0.61 | 0.7 | 0.71 | 0.66 |
| 1 | 96143185 | rs585463 | 1.461 | 0.041 | -0.985 | -0.266 | -0.003 | -0.003 | -0.003 | 0.015 | 0.16 | 0.19 | 0.2 | 0.03 | 0.2 | 0.16 |
| 1 | 96143770 | rs12023322 | 0.305 | 0.334 | -2.561 | 0.273 | -0.063 | -0.124 | -0.124 | 0.052 | 0.18 | 0.03 | 0.3 | 0.46 | 0.23 | 0.12 |
| 1 | 96145267 | rs12120304 | -0.043 | 0.336 | -2.158 | 0.253 | -0.078 | 0.118 | 0.118 | 0.58 | 0.48 | 0.08 | 0.3 | 0.25 | 0.49 | 0.53 |
| 1 | 96145968 | rs6693882 | -0.034 | 0.367 | -2.209 | 0.295 | -0.103 | 0.107 | 0.107 | 1.408 | 0.51 | 0.05 | 0.31 | 0.25 | 0.51 | 0.55 |
| 1 | 96148182 | rs12142219 | 0.617 | 0.353 | -2.089 | -0.019 | -0.075 | 0.069 | 0.069 | 0 | 0.25 | 0.01 | 0.19 | 0.11 | 0.24 | 0.26 |
| 1 | 96149941 | rs1931258 | 0.027 | 0.313 | -1.387 | 0.35 | 0.011 | 0.037 | 0.037 | 0.167 | 0.55 | 0.28 | 0.5 | 0.62 | 0.54 | 0.6 |
| 1 | 96150713 | rs11165466 | 0.424 | 0.687 | -0.818 | 0.122 | -0.006 | -0.006 | -0.006 | 0.186 | 0.63 | 0.39 | 0.45 | 0.25 | 0.67 | 0.58 |
| 1 | 96151539 | rs11803770 | -0.746 | 0.826 | -0.161 | 0.195 | 0.01 | 0.042 | 0.042 | 0 | 0.13 | 0.23 | 0.25 | 0.2 | 0.17 | 0.22 |
| 1 | 96152973 | rs7554994 | 1.021 | 0.475 | 0.401 | 0.238 | -0.001 | 0.114 | 0.114 | 1.738 | 0.9 | 0.55 | 0.9 | 0.84 | 0.96 | 0.84 |
| 1 | 96154744 | rs2700338 | -0.71 | 0.48 | 0.961 | 0.209 | 0.002 | 0.034 | 0.034 | 0 | 0.16 | 0.23 | 0.3 | 0.28 | 0.19 | 0.36 |
| 1 | 96158128 | rs2700335 | -0.446 | 0.862 | 0.178 | -0.634 | 0.002 | -0.004 | -0.004 | 0 | 0.62 | 0.48 | 0.62 | 0.3 | 0.63 | 0.67 |
| 1 | 96159819 | rs12751587 | 1.214 | 0.887 | -0.151 | -0.874 | 0.006 | 0.088 | 0.088 | 0 | 0.38 | 0.23 | 0.24 | 0.02 | 0.35 | 0.23 |
| 1 | 96163965 | rs4950073 | 0.472 | 0.857 | -0.076 | -0.695 | -0.004 | -0.005 | -0.005 | 0 | 0.37 | 0.33 | 0.38 | 0.62 | 0.39 | 0.34 |
| 1 | 96169760 | rs6593680 | 0.589 | 1.076 | -0.588 | -0.977 | -0.018 | 0.144 | 0.144 | 0 | 0.45 | 0.24 | 0.33 | 0.02 | 0.46 | 0.3 |
| 1 | 96174912 | rs1931262 | 0.638 | 1.01 | -0.509 | -0.96 | -0.032 | 0.102 | 0.102 | 0.302 | 0.43 | 0.24 | 0.35 | 0.02 | 0.48 | 0.24 |
| 1 | 96176563 | rs6678734 | 0.338 | 0.945 | -0.272 | -0.986 | -0.006 | -0.018 | -0.018 | 0.44 | 0.44 | 0.37 | 0.37 | 0.02 | 0.48 | 0.24 |
| 1 | 96186066 | rs1931254 | 0.685 | 0.727 | -0.147 | -0.316 | -0.002 | -0.013 | -0.013 | 0 | 0.8 | 0.83 | 0.69 | 0.7 | 0.78 | 0.59 |
| 1 | 96198292 | rs944972 | 0.504 | 0.721 | -0.471 | -0.287 | 0 | -0.008 | -0.008 | 0 | 0.82 | 0.84 | 0.7 | 0.69 | 0.8 | 0.6 |
| 1 | 96200418 | rs1931244 | 0.301 | 0.804 | -0.455 | -0.247 | -0.003 | -0.027 | -0.027 | 0 | 0.83 | 0.87 | 0.7 | 0.69 | 0.8 | 0.6 |
| 1 | 96214251 | rs10874942 | 0.828 | 0.75 | -0.37 | -0.251 | 0 | 0.004 | 0.004 | 0 | 0.8 | 0.78 | 0.68 | 0.69 | 0.79 | 0.63 |
| 1 | 96220903 | rs6677579 | 0.951 | 0.747 | -0.273 | -0.196 | -0.006 | 0.065 | 0.065 | 0 | 0.8 | 0.68 | 0.68 | 0.64 | 0.8 | 0.63 |
| 1 | 96233623 | rs6664676 | 0.122 | 0.754 | -0.687 | -0.258 | 0 | 0.016 | 0.016 | 0 | 0.84 | 0.82 | 0.7 | 0.69 | 0.82 | 0.64 |
| 1 | 96248794 | rs10783019 | 0.174 | 0.688 | -0.401 | -0.337 | 0.001 | -0.002 | -0.002 | 0 | 0.4 | 0.4 | 0.35 | 0.62 | 0.38 | 0.41 |

| | | | | | | | | | | | | | | | | |
|---|---|---|---|---|---|---|---|---|---|---|---|---|---|---|---|---|
| 1 | 96249028 | rs7538619 | -0.114 | 0.704 | -1.127 | -0.34 | 0.029 | 0 | 0 | 0 | 0.44 | 0.81 | 0.48 | 0.66 | 0.37 | 0.44 |
| 1 | 96255065 | rs9285624 | -0.189 | 0.679 | -0.901 | -0.205 | -0.001 | -0.003 | -0.003 | 1.827 | 0.95 | 0.96 | 0.92 | 0.91 | 0.94 | 0.88 |
| 1 | 96258121 | rs4245671 | 0.194 | 0.941 | -0.489 | -0.2 | 0.004 | 0.004 | 0.004 | 0 | 0.51 | 0.48 | 0.45 | 0.58 | 0.55 | 0.46 |
| 1 | 96259807 | rs11806525 | -0.136 | 0.893 | -0.47 | -0.413 | -0.003 | -0.006 | -0.006 | 0.929 | 0.2 | 0.16 | 0.28 | 0.29 | 0.21 | 0.24 |
| 1 | 96269870 | rs6657506 | 0.65 | 1.003 | -0.408 | -0.522 | -0.002 | 0.005 | 0.005 | 0 | 0.48 | 0.43 | 0.4 | 0.58 | 0.5 | 0.38 |
| 1 | 96271126 | rs4345843 | 0.701 | 0.619 | -0.225 | -0.542 | 0.015 | 0.003 | 0.003 | 0 | 0.5 | 0.21 | 0.44 | 0.58 | 0.57 | 0.46 |
| 1 | 96272062 | rs11589942 | 0.063 | 0.717 | -0.358 | -0.482 | -0.001 | 0.004 | 0.004 | 0 | 0.18 | 0.16 | 0.28 | 0.29 | 0.21 | 0.25 |
| 1 | 96273989 | rs10127720 | -1.294 | 0.44 | -0.145 | -0.624 | 0.017 | 0.057 | 0.057 | 0 | 0.76 | 0.9 | 0.77 | 0.88 | 0.79 | 0.75 |
| 1 | 96274009 | rs10874955 | -0.542 | 0.529 | -0.132 | -0.596 | -0.028 | -0.082 | -0.082 | 0 | 0.7 | 0.58 | 0.74 | 0.88 | 0.77 | 0.7 |
| 1 | 96279245 | rs4545389 | 1.101 | 0.721 | -0.047 | -0.775 | 0.007 | 0.006 | 0.006 | 0 | 0.27 | 0.2 | 0.19 | 0.04 | 0.23 | 0.24 |
| 1 | 96280456 | rs4472800 | 0.51 | 0.888 | -0.635 | -0.739 | -0.049 | 0.038 | 0.038 | 0 | 0.36 | 0.14 | 0.38 | 0.26 | 0.41 | 0.36 |
| 1 | 96281108 | rs4494187 | -0.915 | 0.895 | -0.438 | -0.504 | -0.004 | 0.031 | 0.031 | 0 | 0.94 | 0.91 | 0.92 | 0.88 | 0.94 | 0.89 |
| 1 | 96293105 | rs1418460 | -0.606 | 0.959 | -0.279 | -0.439 | -0.007 | 0.011 | 0.011 | 0 | 0.6 | 0.53 | 0.49 | 0.53 | 0.61 | 0.5 |
| 1 | 96293810 | rs17114164 | 1.088 | 1.005 | -0.344 | -0.31 | -0.013 | 0.037 | 0.037 | 0 | 0.06 | 0.03 | 0.22 | 0.31 | 0.09 | 0.14 |
| 1 | 96296426 | rs6593686 | 0.189 | 0.932 | -0.258 | -0.448 | -0.006 | 0.014 | 0.014 | 0 | 0.91 | 0.82 | 0.91 | 0.88 | 0.91 | 0.85 |
| 1 | 96296450 | rs6421760 | 0.202 | 0.997 | -0.327 | -0.496 | -0.002 | 0.001 | 0.001 | 0 | 0.89 | 0.87 | 0.9 | 0.88 | 0.88 | 0.8 |
| 1 | 96303910 | rs1935569 | 0.878 | 0.967 | -0.241 | -0.543 | -0.001 | -0.004 | -0.004 | 0.215 | 0.63 | 0.39 | 0.58 | 0.31 | 0.64 | 0.6 |
| 1 | 96305433 | rs4950085 | -1.338 | 0.964 | -0.198 | -0.533 | -0.002 | 0.009 | 0.009 | 0 | 0.95 | 0.95 | 0.92 | 0.88 | 0.94 | 0.89 |
| 1 | 96305610 | rs12122071 | 0.974 | 1.017 | -0.377 | -0.651 | -0.117 | -0.017 | -0.017 | 0 | 0.33 | 0.03 | 0.36 | 0.26 | 0.39 | 0.31 |
| 1 | 96306787 | rs2391710 | 0.069 | 0.962 | 0.086 | -0.571 | -0.002 | 0.001 | 0.001 | 0 | 0.71 | 0.68 | 0.82 | 0.61 | 0.74 | 0.74 |
| 1 | 96308556 | rs6670360 | 0.869 | 0.962 | 0.082 | -0.588 | 0 | -0.002 | -0.002 | 0 | 0.2 | 0.18 | 0.1 | 0.27 | 0.18 | 0.11 |
| 1 | 96309286 | rs2051029 | 1.003 | 0.959 | 0.106 | -0.598 | 0.001 | 0.006 | 0.006 | 0.531 | 0.62 | 0.32 | 0.57 | 0.31 | 0.64 | 0.6 |
| 1 | 96316702 | rs4950088 | 0.44 | 0.948 | 0.184 | -0.594 | -0.007 | -0.004 | -0.004 | 0.107 | 0.22 | 0.47 | 0.13 | 0.27 | 0.18 | 0.11 |
| 1 | 96319900 | rs6682623 | 1.107 | 0.919 | 0.295 | -0.607 | 0.002 | 0.032 | 0.032 | 0.748 | 0.61 | 0.19 | 0.56 | 0.29 | 0.64 | 0.6 |
| 1 | 96321437 | rs6679428 | 0.843 | 0.913 | 0.374 | -0.571 | 0.023 | 0.253 | 0.253 | 0.083 | 0.82 | 0.59 | 0.67 | 0.55 | 0.81 | 0.67 |
| 1 | 96322205 | rs6593688 | 0.683 | 0.887 | 0.113 | -0.594 | -0.06 | 0.024 | 0.024 | 0 | 0.34 | 0.11 | 0.38 | 0.25 | 0.4 | 0.33 |
| 1 | 96334478 | rs1891418 | 0.474 | 0.921 | 0.269 | -0.416 | -0.004 | 0.037 | 0.037 | 0 | 0.08 | 0.07 | 0.23 | 0.3 | 0.12 | 0.24 |
| 1 | 96351754 | rs12754240 | 2.071 | 0.79 | -0.272 | -0.644 | 0.011 | 0.131 | 0.131 | 0.031 | 0.26 | 0.05 | 0.17 | 0.04 | 0.22 | 0.18 |
| 1 | 96361709 | rs12035421 | 1.99 | 0.841 | -0.275 | -0.522 | -0.035 | -0.058 | -0.058 | 0.131 | 0.1 | 0 | 0.06 | 0.26 | 0.13 | 0.09 |
| 1 | 96386226 | rs12405938 | 1.084 | 1.035 | -0.582 | -0.789 | -0.047 | -0.067 | -0.067 | 0 | 0.2 | 0.02 | 0.23 | 0.26 | 0.25 | 0.25 |
| 1 | 96393689 | rs6593689 | 0.419 | 1.002 | 0.065 | -0.78 | -0.004 | 0.011 | 0.011 | 0 | 0.25 | 0.48 | 0.28 | 0.26 | 0.26 | 0.27 |
| 1 | 96399483 | rs4376787 | -0.491 | 1.031 | -0.01 | -0.705 | -0.017 | -0.035 | -0.035 | 0.05 | 0.74 | 0.81 | 0.55 | 0.56 | 0.67 | 0.6 |
| 1 | 96419709 | rs10783029 | 0.626 | 0.891 | -0.148 | -0.761 | -0.001 | -0.002 | -0.002 | 0 | 0.19 | 0.16 | 0.09 | 0.27 | 0.17 | 0.11 |
| 1 | 96426052 | rs6593693 | 0.877 | 0.941 | -0.077 | -0.819 | 0.003 | 0.023 | 0.023 | 0 | 0.52 | 0.55 | 0.45 | 0.28 | 0.49 | 0.45 |
| 1 | 96432494 | rs6663724 | 0.566 | 0.958 | -0.138 | -0.809 | -0.002 | 0.04 | 0.04 | 0 | 0.54 | 0.63 | 0.46 | 0.28 | 0.5 | 0.47 |
| 1 | 96442134 | rs10783030 | 0.623 | 1.042 | -0.202 | -0.773 | -0.003 | 0.005 | 0.005 | 0 | 0.23 | 0.34 | 0.25 | 0.24 | 0.25 | 0.24 |
| 1 | 96442314 | rs10874974 | 0.791 | 1.032 | -0.133 | -0.778 | -0.006 | -0.017 | -0.017 | 0 | 0.22 | 0.12 | 0.24 | 0.24 | 0.25 | 0.23 |
| 1 | 96443739 | rs11165529 | 2.044 | 1.079 | -0.214 | -0.796 | -0.016 | -0.039 | -0.039 | 0.105 | 0.1 | 0.03 | 0.05 | 0.27 | 0.12 | 0.09 |
| 1 | 96445333 | rs4316403 | -0.034 | 1.202 | -0.135 | -0.545 | -0.056 | -0.081 | -0.081 | 0 | 0.28 | 0.09 | 0.48 | 0.54 | 0.37 | 0.47 |
| 1 | 96451261 | rs12037176 | 1.279 | 1.111 | -0.581 | -0.652 | -0.045 | -0.06 | -0.06 | 0.15 | 0.18 | 0.01 | 0.21 | 0.24 | 0.22 | 0.18 |
| 1 | 96454876 | rs321271 | 1.807 | 0.995 | -0.521 | -0.95 | 0.015 | 0.063 | 0.063 | 0 | 0.31 | 0.16 | 0.17 | 0.04 | 0.27 | 0.27 |
| 1 | 96457002 | rs321270 | -1.843 | 1.115 | -0.565 | -0.933 | -0.031 | 0.006 | 0.006 | 0 | 0.83 | 0.93 | 0.77 | 0.9 | 0.8 | 0.84 |
| 1 | 96457465 | rs321269 | -0.667 | 1.072 | -0.365 | -0.909 | -0.025 | -0.04 | -0.04 | 0.003 | 0.74 | 0.82 | 0.51 | 0.55 | 0.67 | 0.59 |
| 1 | 96458324 | rs2125421 | 0.507 | 0.97 | -0.068 | -0.929 | -0.004 | 0.007 | 0.007 | 0 | 0.21 | 0.4 | 0.1 | 0.28 | 0.17 | 0.11 |
| 1 | 96463024 | rs321262 | 1.926 | 0.96 | -0.144 | -0.988 | 0.015 | 0.059 | 0.059 | 0 | 0.31 | 0.16 | 0.17 | 0.04 | 0.28 | 0.27 |
| 1 | 96463324 | rs321261 | -0.705 | 1.003 | -0.127 | -0.974 | -0.024 | -0.038 | -0.038 | 0 | 0.74 | 0.82 | 0.51 | 0.55 | 0.67 | 0.59 |
| 1 | 96464512 | rs4950091 | -0.361 | 0.996 | -0.076 | -0.985 | -0.004 | 0.004 | 0.004 | 0 | 0.41 | 0.53 | 0.32 | 0.51 | 0.39 | 0.29 |
| 1 | 96467373 | rs616243 | 0.577 | 1.014 | -0.144 | -0.977 | 0.005 | 0.034 | 0.034 | 0 | 0.52 | 0.56 | 0.27 | 0.32 | 0.45 | 0.38 |
| 1 | 96467524 | rs615413 | 1.956 | 1.043 | -0.179 | -0.967 | 0.015 | 0.06 | 0.06 | 0 | 0.31 | 0.16 | 0.17 | 0.04 | 0.28 | 0.27 |
| 1 | 96468117 | rs167677 | -0.509 | 1.266 | -0.142 | -0.694 | -0.047 | -0.075 | -0.075 | 0 | 0.75 | 0.86 | 0.51 | 0.55 | 0.67 | 0.58 |
| 1 | 96475548 | rs321244 | 0.544 | 0.99 | -0.055 | -0.884 | 0.006 | 0.037 | 0.037 | 0 | 0.52 | 0.59 | 0.28 | 0.32 | 0.44 | 0.36 |
| 1 | 96481556 | rs607674 | -1.392 | 0.93 | 0.035 | -0.884 | -0.005 | -0.021 | -0.021 | 0 | 0.55 | 0.51 | 0.76 | 0.71 | 0.59 | 0.67 |
| 1 | 96481572 | rs17114439 | 1.236 | 0.93 | -0.036 | -0.657 | 0.015 | -0.027 | -0.027 | 1.856 | 0.09 | 0.03 | 0.04 | 0.21 | 0.05 | 0.04 |
| 1 | 96484940 | rs635754 | -0.605 | 0.916 | 0.041 | -0.667 | -0.001 | 0.008 | 0.008 | 0 | 0.43 | 0.36 | 0.7 | 0.49 | 0.53 | 0.61 |
| 1 | 96486756 | rs1379971 | 0.318 | 1.359 | -0.058 | -0.604 | 0.016 | 0.085 | 0.085 | 0 | 0.09 | 0.07 | 0.27 | 0.37 | 0.18 | 0.26 |
| 1 | 96486881 | rs513981 | 1.738 | 0.61 | 0.336 | -0.739 | 0.001 | -0.055 | -0.055 | 2.796 | 0.36 | 0.34 | 0.18 | 0.03 | 0.3 | 0.29 |
| 1 | 96486967 | rs589482 | 1.738 | 0.61 | 0.336 | -0.739 | 0.001 | -0.055 | -0.055 | 0 | 0.36 | 0.34 | 0.18 | 0.03 | 0.3 | 0.29 |
| 1 | 96492489 | rs1146530 | -1.804 | 0.655 | 0.303 | -0.647 | 0.005 | 0.041 | 0.041 | 0 | 0.91 | 0.85 | 0.94 | 0.74 | 0.89 | 0.96 |
| 1 | 96496399 | rs161120 | -0.457 | 0.555 | 0.265 | -0.726 | -0.015 | 0.012 | 0.012 | 0 | 0.38 | 0.27 | 0.56 | 0.64 | 0.45 | 0.49 |
| 1 | 96499450 | rs161116 | 0.356 | 0.495 | 0.378 | -0.711 | 0.028 | 0.03 | 0.03 | 0 | 0.29 | 0.28 | 0.51 | 0.44 | 0.39 | 0.45 |
| 1 | 96500593 | rs161114 | 2.022 | 0.457 | 0.225 | -0.813 | 0.027 | 0.088 | 0.088 | 0 | 0.36 | 0.21 | 0.18 | 0.03 | 0.3 | 0.29 |
| 1 | 96504265 | rs991220 | 0.155 | 0.387 | 0.252 | -0.627 | -0.003 | 0.006 | 0.006 | 0 | 0.74 | 0.61 | 0.75 | 0.64 | 0.73 | 0.77 |
| 1 | 96506530 | rs1146535 | -0.129 | 0.418 | 0.159 | -0.61 | 0 | 0.006 | 0.006 | 0 | 0.76 | 0.65 | 0.75 | 0.65 | 0.74 | 0.8 |
| 1 | 96510342 | rs6682297 | 0.991 | 0.447 | 0.173 | -0.545 | -0.012 | 0.07 | 0.07 | 0 | 0.1 | 0.29 | 0.05 | 0.2 | 0.05 | 0.03 |
| 1 | 96510531 | rs1144306 | -1.812 | 0.498 | 0.075 | -0.516 | 0.003 | 0.041 | 0.041 | 0 | 0.91 | 0.85 | 0.94 | 0.71 | 0.89 | 0.96 |
| 1 | 96511272 | rs66512 | 0.719 | 0.553 | -0.025 | -0.6 | -0.036 | -0.045 | -0.045 | 0 | 0.25 | 0.14 | 0.49 | 0.41 | 0.34 | 0.42 |
| 1 | 96516252 | rs150892 | 0.894 | 0.428 | -0.167 | -0.694 | 0.024 | 0.224 | 0.224 | 0 | 0.5 | 0.25 | 0.39 | 0.09 | 0.49 | 0.46 |
| 1 | 96516578 | rs161107 | 3.022 | 0.136 | -0.529 | -0.988 | 0.015 | 0.147 | 0.147 | 0 | 0.25 | 0.03 | 0.12 | 0.01 | 0.19 | 0.18 |
| 1 | 96524674 | rs1144320 | -1.655 | 0.561 | -0.062 | -0.548 | 0.006 | 0.037 | 0.037 | 0 | 0.9 | 0.85 | 0.94 | 0.71 | 0.88 | 0.96 |

| | | | | | | | | | | | | | | | | |
|---|---|---|---|---|---|---|---|---|---|---|---|---|---|---|---|---|
| 1 | 96528940 | rs1146540 | 0.433 | 0.557 | -0.431 | -0.551 | -0.028 | 0.052 | 0.052 | 0.027 | 0.75 | 0.41 | 0.87 | 0.5 | 0.82 | 0.89 |
| 1 | 96537564 | rs223259 | 2.222 | 0.552 | -0.482 | -0.568 | 0.017 | 0.177 | 0.177 | 0 | 0.34 | 0.08 | 0.17 | 0.03 | 0.3 | 0.28 |
| 1 | 96539243 | rs1144325 | -2.007 | 0.598 | -0.451 | -0.399 | -0.012 | -0.021 | -0.021 | 0 | 0.92 | 0.96 | 0.95 | 0.71 | 0.89 | 0.96 |
| 1 | 96541099 | rs1144326 | 0.518 | 0.604 | -0.49 | -0.39 | -0.032 | 0.05 | 0.05 | 0.064 | 0.75 | 0.41 | 0.87 | 0.5 | 0.82 | 0.89 |
| 1 | 96544805 | rs6672740 | 0.461 | 0.73 | -0.533 | -0.282 | -0.025 | 0.038 | 0.038 | 0 | 0.12 | 0.29 | 0.05 | 0.2 | 0.06 | 0.03 |
| 1 | 96564857 | rs686101 | 0.774 | 0.972 | -1.506 | -0.356 | 0.001 | 0.004 | 0.004 | 0 | 0.22 | 0.16 | 0.39 | 0.33 | 0.32 | 0.32 |
| 1 | 96566226 | rs17114640 | 0.475 | 0.856 | -1.926 | -0.053 | 0.003 | -0.001 | -0.001 | 0 | 0.11 | 0.09 | 0.08 | 0.27 | 0.05 | 0.13 |
| 1 | 96581785 | rs12140320 | 0.418 | 1.305 | -2.081 | -0.533 | -0.045 | 0.014 | 0.014 | 0 | 0.13 | 0.01 | 0.23 | 0.07 | 0.17 | 0.18 |
| 1 | 96584567 | rs12741915 | 0.449 | 1.285 | -2.073 | -0.525 | -0.043 | 0.013 | 0.013 | 0 | 0.13 | 0.01 | 0.23 | 0.07 | 0.17 | 0.18 |
| 1 | 96587711 | rs11165545 | 1.194 | 1.102 | -1.606 | -0.298 | -0.007 | -0.008 | -0.008 | 0 | 0.08 | 0.05 | 0.05 | 0.29 | 0.11 | 0.04 |
| 1 | 96595179 | rs1222069 | 0.504 | 0.987 | -1.694 | -0.981 | 0.009 | 0.094 | 0.094 | 0 | 0.41 | 0.27 | 0.27 | 0.03 | 0.38 | 0.34 |
| 1 | 96597055 | rs1402807 | 0.529 | 0.681 | -1.732 | -0.322 | 0.047 | 0.145 | 0.145 | 0.185 | 0.48 | 0.15 | 0.27 | 0.34 | 0.43 | 0.28 |
| 1 | 96604591 | rs2391734 | 0.324 | 0.931 | -1.24 | 0.653 | -0.007 | -0.006 | -0.006 | 0 | 0.19 | 0.13 | 0.37 | 0.29 | 0.29 | 0.34 |
| 1 | 96604986 | rs10874979 | 0.739 | 0.982 | -1.233 | -0.428 | -0.006 | -0.021 | -0.021 | 0.754 | 0.17 | 0.14 | 0.22 | 0.01 | 0.24 | 0.23 |
| 1 | 96610271 | rs12567412 | 0.002 | 0.569 | -1.275 | -0.451 | -0.024 | 0.004 | 0.004 | 0.079 | 0.35 | 0.09 | 0.58 | 0.64 | 0.38 | 0.36 |
| 1 | 96627332 | rs17422841 | 0.28 | 0.523 | -1.351 | -0.549 | -0.045 | 0.191 | 0.191 | 0 | 0.32 | 0.03 | 0.29 | 0.07 | 0.32 | 0.17 |
| 1 | 96630199 | rs17422897 | 0.59 | 0.512 | -1.343 | -0.54 | -0.049 | 0.176 | 0.176 | 0.059 | 0.3 | 0.03 | 0.28 | 0.07 | 0.3 | 0.16 |
| 1 | 96634994 | rs1146557 | -0.576 | 0.325 | -0.4 | -0.605 | 0 | 0.001 | 0.001 | 0.178 | 0.42 | 0.54 | 0.63 | 0.63 | 0.4 | 0.46 |
| 1 | 96638203 | rs321559 | 0.822 | 0.333 | -0.455 | -0.835 | -0.004 | 0.07 | 0.07 | 0 | 0.47 | 0.33 | 0.3 | 0.35 | 0.52 | 0.44 |
| 1 | 96653787 | rs321563 | -0.154 | 0.672 | -0.592 | -0.468 | -0.002 | -0.017 | -0.017 | 0.028 | 0.89 | 0.84 | 0.92 | 0.98 | 0.91 | 0.89 |
| 1 | 96656005 | rs321565 | 1.07 | 0.663 | -0.344 | -0.403 | 0 | 0.05 | 0.05 | 0 | 0.83 | 0.72 | 0.62 | 0.42 | 0.84 | 0.7 |
| 1 | 96656207 | rs321567 | -0.4 | 0.719 | -0.381 | -0.593 | -0.002 | 0.009 | 0.009 | 0 | 0.9 | 0.91 | 0.93 | 0.98 | 0.91 | 0.89 |
| 1 | 96660592 | rs321576 | 0.445 | 0.367 | -0.575 | -0.811 | 0.005 | 0.066 | 0.066 | 0 | 0.48 | 0.69 | 0.32 | 0.39 | 0.52 | 0.52 |
| 1 | 96666138 | rs17114704 | -0.421 | 0.382 | -0.616 | -0.721 | 0 | 0.018 | 0.018 | 0 | 0.09 | 0.08 | 0.28 | 0.57 | 0.08 | 0.22 |
| 1 | 96669942 | rs321556 | 0.584 | 0.368 | -0.388 | -0.801 | 0 | 0.025 | 0.025 | 0 | 0.47 | 0.61 | 0.31 | 0.39 | 0.52 | 0.51 |
| 1 | 96670802 | rs321557 | -0.396 | 0.452 | -0.451 | -0.88 | 0.001 | 0.006 | 0.006 | 0 | 0.9 | 0.92 | 0.93 | 0.98 | 0.92 | 0.9 |
| 1 | 96671376 | rs321558 | -0.559 | 0.449 | -0.419 | -0.882 | -0.003 | -0.026 | -0.026 | 0 | 0.9 | 0.85 | 0.92 | 0.98 | 0.92 | 0.9 |
| 1 | 96673887 | rs17374320 | 0.576 | 0.388 | -0.413 | -1.017 | 0.006 | -0.001 | -0.001 | 0.049 | 0.34 | 0.23 | 0.3 | 0.01 | 0.33 | 0.16 |
| 1 | 96680000 | rs7539465 | -0.851 | 0.717 | -0.414 | -1.067 | -0.001 | 0.002 | 0.002 | 0 | 0.57 | 0.46 | 0.67 | 0.66 | 0.57 | 0.65 |
| 1 | 96683642 | rs12730665 | 1.134 | 0.653 | -0.199 | -1.136 | -0.033 | 0.205 | 0.205 | 0 | 0.32 | 0.04 | 0.28 | 0.01 | 0.32 | 0.16 |
| 1 | 96686237 | rs4367826 | -1.351 | 0.51 | 0.25 | -0.693 | 0.001 | -0.003 | -0.003 | 0 | 0.25 | 0.31 | 0.38 | 0.66 | 0.24 | 0.49 |
| 1 | 96688818 | rs321588 | 1.528 | 0.546 | 0.358 | -0.722 | 0.002 | -0.007 | -0.007 | 2.733 | 0.71 | 0.45 | 0.58 | 0.34 | 0.72 | 0.5 |
| 1 | 96694568 | rs1872733 | -1.313 | 0.528 | 0.335 | -0.702 | -0.002 | 0.002 | 0.002 | 0 | 0.25 | 0.36 | 0.38 | 0.65 | 0.23 | 0.5 |
| 1 | 96731590 | rs9660464 | 0.723 | 0.592 | 0.403 | -0.657 | 0.06 | 0.006 | 0.006 | 0.009 | 0.94 | 0.8 | 0.94 | 0.98 | 0.92 | 0.99 |
| 1 | 96756638 | rs1480666 | -1.057 | 0.758 | 0.303 | -0.706 | 0 | 0.015 | 0.015 | 0.002 | 0.16 | 0.2 | 0.35 | 0.62 | 0.16 | 0.39 |
| 1 | 96759047 | rs511274 | 0.8 | 0.722 | 0.398 | -0.762 | 0.072 | 0.022 | 0.022 | 0.936 | 0.93 | 0.66 | 0.94 | 0.98 | 0.93 | 0.99 |
| 1 | 96763525 | rs680954 | 1.406 | 0.533 | 0.385 | -1.221 | 0.028 | 0.029 | 0.029 | 0.169 | 0.24 | 0.13 | 0.09 | 0 | 0.14 | 0.19 |
| 1 | 96787405 | rs326677 | 0.843 | 1.293 | 0.506 | -0.359 | 0.021 | -0.064 | -0.064 | 0 | 0.82 | 0.54 | 0.88 | 0.95 | 0.83 | 0.86 |
| 1 | 96792430 | rs12564989 | -1.132 | 1.229 | 0.322 | -0.279 | -0.006 | -0.032 | -0.032 | 0.057 | 0.15 | 0.1 | 0.34 | 0.62 | 0.18 | 0.4 |
| 1 | 96795203 | rs12744985 | -1.201 | 1.231 | 0.306 | -0.278 | -0.012 | -0.045 | -0.045 | 0 | 0.15 | 0.08 | 0.34 | 0.62 | 0.18 | 0.4 |
| 1 | 96818456 | rs325953 | 1.886 | 1.236 | 0.373 | -0.318 | 0.006 | -0.022 | -0.022 | 0 | 0.67 | 0.45 | 0.53 | 0.33 | 0.66 | 0.46 |
| 1 | 96836370 | rs11165603 | -1.261 | 1.266 | 0.233 | -0.235 | -0.006 | -0.038 | -0.038 | 0 | 0.16 | 0.12 | 0.34 | 0.62 | 0.18 | 0.41 |
| 1 | 96851552 | rs326910 | -1.121 | 1.28 | 0.67 | -0.154 | -0.001 | 0.01 | 0.01 | 0.434 | 0.12 | 0.14 | 0.34 | 0.64 | 0.17 | 0.37 |
| 1 | 96852354 | rs326911 | 1.922 | 1.264 | 0.722 | -0.251 | 0.005 | -0.01 | -0.01 | 0 | 0.68 | 0.44 | 0.53 | 0.34 | 0.66 | 0.48 |
| 1 | 96854358 | rs12132961 | -0.86 | 1.396 | -0.191 | -0.058 | -0.018 | -0.039 | -0.039 | 0 | 0.11 | 0.04 | 0.3 | 0.64 | 0.14 | 0.35 |
| 1 | 96869211 | rs17115122 | -0.121 | 1.29 | 0.068 | -0.162 | 0 | -0.03 | -0.03 | 0 | 0.12 | 0.16 | 0.07 | 0 | 0.12 | 0.13 |
| 1 | 96874164 | rs547703 | 1.108 | 1.844 | 0.069 | 0.015 | 0.015 | 0.002 | 0.002 | 0.177 | 0.77 | 0.63 | 0.77 | 0.76 | 0.72 | 0.84 |
| 1 | 96875440 | rs12022566 | 0.345 | 1.799 | 0.063 | -0.007 | -0.013 | -0.013 | -0.013 | 0 | 0.12 | 0.09 | 0.16 | 0.23 | 0.22 | 0.12 |
| 1 | 96877548 | rs671135 | 0.238 | 1.388 | -0.013 | -0.014 | -0.005 | -0.055 | -0.055 | 0 | 0.9 | 0.8 | 0.92 | 1 | 0.9 | 0.93 |
| 1 | 96882331 | rs1570727 | -0.228 | 1.661 | -0.452 | 1.253 | -0.031 | 0.007 | 0.007 | 0 | 0.05 | 0 | 0.07 | 0.37 | 0.08 | 0.22 |
| 1 | 96882971 | rs10493885 | 0.438 | 1.862 | -0.582 | 1.188 | -0.053 | -0.058 | -0.058 | 0 | 0.12 | 0.02 | 0.15 | 0.24 | 0.21 | 0.11 |
| 1 | 96890636 | rs1980684 | -0.264 | 1.775 | -0.589 | 0.622 | -0.012 | 0.007 | 0.007 | 0 | 0.33 | 0.33 | 0.58 | 0.68 | 0.5 | 0.29 |
| 1 | 96891671 | rs510552 | -0.302 | 1.755 | -0.491 | 0.64 | -0.016 | 0.009 | 0.009 | 0 | 0.33 | 0.34 | 0.59 | 0.68 | 0.5 | 0.29 |
| 1 | 96892655 | rs1159909 | -0.61 | 1.606 | -0.39 | 0.43 | 0.002 | -0.005 | -0.005 | 1.789 | 0.91 | 0.92 | 0.93 | 0.99 | 0.92 | 0.93 |
| 1 | 96941158 | rs7512263 | 1.422 | 1.541 | -0.13 | 0.482 | 0.014 | 0.009 | 0.009 | 0 | 0.51 | 0.48 | 0.36 | 0.11 | 0.34 | 0.42 |
| 1 | 96941580 | rs2208901 | 0.913 | 1.673 | -0.151 | 0.733 | -0.012 | -0.001 | -0.001 | 0 | 0.13 | 0.1 | 0.15 | 0.41 | 0.22 | 0.11 |
| 1 | 96942406 | rs10489741 | -0.894 | 1.623 | -0.128 | 0.612 | -0.005 | 0.044 | 0.044 | 0 | 0.4 | 0.39 | 0.57 | 0.88 | 0.59 | 0.51 |
| 1 | 96951281 | rs6593602 | 0.901 | 1.598 | -0.36 | 0.661 | 0.035 | 0.171 | 0.171 | 0 | 0.56 | 0.82 | 0.38 | 0.14 | 0.34 | 0.42 |
| 1 | 96954561 | rs6677997 | 0.025 | 1.651 | -0.395 | 0.853 | -0.121 | -0.164 | -0.164 | 0 | 0.68 | 0.95 | 0.54 | 0.5 | 0.56 | 0.54 |
| 1 | 96961107 | rs11165656 | 1.01 | 1.645 | -0.35 | 0.856 | -0.034 | -0.04 | -0.04 | 0.891 | 0.12 | 0.05 | 0.16 | 0.35 | 0.22 | 0.11 |
| 1 | 96962047 | rs10875019 | -0.437 | 1.673 | -0.372 | 0.865 | -0.071 | -0.133 | -0.133 | 0 | 0.73 | 0.96 | 0.61 | 0.61 | 0.65 | 0.75 |
| 1 | 96975515 | rs6701031 | 0.503 | 1.628 | -0.326 | 0.757 | -0.05 | 0.089 | 0.089 | 0 | 0.61 | 0.82 | 0.45 | 0.25 | 0.43 | 0.64 |
| 1 | 96978961 | rs2391769 | 0.324 | 1.753 | -0.443 | 0.688 | -0.022 | -0.023 | -0.023 | 1.221 | 0.77 | 0.84 | 0.77 | 0.64 | 0.67 | 0.79 |
| 1 | 96980709 | rs2800424 | -0.555 | 1.782 | -0.451 | 0.676 | -0.014 | 0.043 | 0.043 | 0 | 0.89 | 0.97 | 0.92 | 1 | 0.88 | 0.9 |
| 1 | 96984344 | rs7541994 | 1.365 | 1.786 | -0.367 | 0.687 | -0.023 | -0.063 | -0.063 | 1.448 | 0.19 | 0.08 | 0.33 | 0.39 | 0.27 | 0.18 |
| 1 | 96985039 | rs2256459 | 1.042 | 1.713 | -0.379 | 0.96 | 0.006 | 0.041 | 0.041 | 0 | 0.13 | 0.13 | 0.22 | 0.42 | 0.21 | 0.11 |
| 1 | 96993449 | rs1967769 | 1.549 | 1.542 | -0.108 | 0.812 | 0.005 | -0.029 | -0.029 | 0 | 0.48 | 0.51 | 0.3 | 0.1 | 0.28 | 0.39 |
| 1 | 96993561 | rs11165660 | 0.637 | 1.558 | -0.154 | 0.861 | -0.122 | -0.176 | -0.176 | 0.401 | 0.24 | 0.02 | 0.34 | 0.48 | 0.36 | 0.39 |
| 1 | 96993603 | rs3912027 | 1.324 | 1.551 | -0.091 | 0.856 | 0.053 | 0.073 | 0.073 | 0 | 0.5 | 0.64 | 0.3 | 0.1 | 0.28 | 0.39 |

| | | | | | | | | | | | | | | | | |
|---|---|---|---|---|---|---|---|---|---|---|---|---|---|---|---|---|
| 1 | 96997655 | rs12751763 | 1.281 | 1.608 | -0.079 | 0.907 | -0.025 | -0.053 | -0.053 | 0.034 | 0.19 | 0.08 | 0.28 | 0.36 | 0.28 | 0.19 |
| 1 | 97001996 | rs6593611 | 0.63 | 1.422 | -0.012 | 0.737 | 0.008 | 0.055 | 0.055 | 0 | 0.55 | 0.69 | 0.38 | 0.22 | 0.37 | 0.6 |
| 1 | 97018734 | rs6677110 | 0.438 | 1.418 | -0.231 | 0.948 | -0.004 | 0.046 | 0.046 | 0 | 0.57 | 0.84 | 0.39 | 0.4 | 0.37 | 0.6 |
| 1 | 97023641 | rs1925833 | 0.831 | 0.963 | 0.215 | 0.788 | -0.002 | -0.002 | -0.002 | 0.346 | 0.49 | 0.52 | 0.3 | 0.4 | 0.28 | 0.45 |
| 1 | 97031756 | rs2477893 | 0.06 | 1.146 | 0.456 | 0.55 | -0.006 | -0.006 | -0.006 | 0 | 0.18 | 0.16 | 0.31 | 0.25 | 0.3 | 0.18 |
| 1 | 97055328 | rs9943084 | 0.347 | 0.802 | 0.564 | 0.421 | -0.007 | 0.08 | 0.08 | 1.144 | 0.27 | 0.55 | 0.32 | 0.49 | 0.26 | 0.41 |
| 1 | 97086056 | rs1613142 | -0.358 | 0.29 | -0.202 | -0.857 | 0.015 | 0.066 | 0.066 | 0.103 | 0.62 | 0.76 | 0.71 | 0.98 | 0.7 | 0.68 |
| 1 | 97093996 | rs6692249 | 1.901 | 0.051 | -0.161 | -0.938 | -0.027 | -0.032 | -0.032 | 0 | 0.06 | 0.18 | 0.02 | 0 | 0 | 0 |
| 1 | 97098359 | rs6693873 | 0.023 | 0.013 | 0.157 | -1.041 | -0.006 | -0.099 | -0.099 | 0 | 0.56 | 0.47 | 0.69 | 0.98 | 0.7 | 0.68 |
| 1 | 97100867 | rs969952 | 0.233 | -0.359 | -0.171 | -1.247 | -0.037 | 0.138 | 0.138 | 0 | 0.4 | 0.15 | 0.63 | 0.82 | 0.6 | 0.64 |
| 1 | 97100996 | rs17115571 | 2.012 | -0.328 | -0.115 | -1.211 | -0.008 | -0.006 | -0.006 | 0 | 0.18 | 0.2 | 0.06 | 0.17 | 0.12 | 0.1 |
| 1 | 97107005 | rs1325755 | -0.544 | -0.207 | -0.211 | -1.182 | -0.009 | -0.113 | -0.113 | 0 | 0.63 | 0.59 | 0.72 | 0.98 | 0.73 | 0.74 |
| 1 | 97110940 | rs2220948 | -0.201 | 0.012 | -0.383 | -0.776 | -0.002 | 0.004 | 0.004 | 0.154 | 0.41 | 0.56 | 0.41 | 0.58 | 0.39 | 0.25 |
| 1 | 97115049 | rs6673531 | -0.652 | -1.58 | -0.368 | -0.692 | -0.002 | 0.026 | 0.026 | 0 | 0.54 | 0.74 | 0.45 | 0.62 | 0.53 | 0.65 |
| 1 | 97146954 | rs1409057 | 0.124 | -1.559 | -0.486 | -0.713 | -0.03 | -0.043 | -0.043 | 0.276 | 0.31 | 0.53 | 0.15 | 0.22 | 0.21 | 0.16 |
| 1 | 97164343 | rs7544736 | 0.628 | -1.55 | -0.428 | -0.714 | -0.019 | -0.035 | -0.035 | 1.711 | 0.26 | 0.37 | 0.13 | 0.21 | 0.21 | 0.15 |
| 1 | 97174123 | rs2391798 | 0.411 | -1.572 | -0.496 | -0.67 | 0.002 | -0.02 | -0.02 | 0.585 | 0.28 | 0.29 | 0.32 | 0.41 | 0.33 | 0.5 |
| 1 | 97177176 | rs3908925 | -0.637 | -1.578 | -0.472 | -0.682 | 0.001 | 0.011 | 0.011 | 0 | 0.54 | 0.66 | 0.46 | 0.61 | 0.54 | 0.65 |
| 1 | 97178241 | rs10875029 | 0.115 | -1.591 | -0.471 | -0.768 | -0.035 | -0.053 | -0.053 | 0 | 0.31 | 0.54 | 0.15 | 0.21 | 0.21 | 0.15 |
| 1 | 97182039 | rs12087010 | 0.407 | -1.589 | -0.48 | -0.753 | 0.003 | -0.021 | -0.021 | 0 | 0.28 | 0.29 | 0.32 | 0.4 | 0.33 | 0.5 |
| 1 | 97187083 | rs3748785 | -1.405 | -1.917 | -1.004 | -1.186 | -0.071 | -0.076 | -0.076 | 0 | 0.73 | 0.91 | 0.53 | 0.62 | 0.69 | 0.82 |
| 1 | 97196678 | rs10875031 | 0.703 | -1.953 | -0.658 | -0.902 | -0.01 | -0.017 | -0.017 | 0.7 | 0.25 | 0.3 | 0.13 | 0.21 | 0.21 | 0.15 |
| 1 | 97198331 | rs10875032 | -0.548 | -1.854 | -1.092 | -0.808 | 0.002 | 0.014 | 0.014 | 0 | 0.54 | 0.66 | 0.55 | 0.63 | 0.54 | 0.65 |
| 1 | 97199355 | rs12563540 | 1.206 | -1.839 | -1.088 | -0.807 | -0.027 | -0.09 | -0.09 | 0 | 0.22 | 0.12 | 0.4 | 0.43 | 0.33 | 0.49 |
| 1 | 97258468 | rs6699242 | -0.166 | -1.733 | -0.65 | -1.046 | -0.037 | -0.054 | -0.054 | 0.015 | 0.31 | 0.54 | 0.15 | 0.21 | 0.21 | 0.16 |
| 1 | 97265692 | rs720090 | 1.695 | -1.749 | -0.643 | -1.068 | -0.002 | -0.003 | -0.003 | 0 | 0.07 | 0.04 | 0.05 | 0 | 0.09 | 0.05 |
| 1 | 97272456 | rs6699932 | -0.132 | -1.699 | -0.692 | -1.027 | -0.037 | -0.054 | -0.054 | 0 | 0.31 | 0.54 | 0.15 | 0.21 | 0.21 | 0.15 |
| 1 | 97286381 | rs12060538 | 1.339 | -1.647 | -1.108 | -0.843 | 0.009 | 0.032 | 0.032 | 0 | 0.16 | 0.11 | 0.09 | 0.07 | 0.12 | 0.1 |
| 1 | 97286510 | rs479424 | -0.742 | -1.656 | -1.535 | -0.586 | -0.078 | -0.117 | -0.117 | 0 | 0.63 | 0.92 | 0.43 | 0.59 | 0.44 | 0.71 |
| 1 | 97293023 | rs12024594 | 1.656 | -1.609 | -2.145 | -0.571 | -0.01 | -0.009 | -0.009 | 0 | 0.16 | 0.01 | 0.06 | 0.17 | 0.13 | 0.09 |
| 1 | 97296935 | rs12740489 | 1.751 | -0.833 | -1.872 | -0.359 | -0.075 | 0.103 | 0.103 | 0 | 0.16 | 0.01 | 0.18 | 0.01 | 0.27 | 0.1 |
| 1 | 97297245 | rs654848 | -0.728 | -0.217 | -1.193 | 0.355 | -0.059 | -0.089 | -0.089 | 0 | 0.8 | 0.94 | 0.61 | 0.57 | 0.7 | 0.86 |
| 1 | 97304571 | rs635578 | -0.189 | -0.501 | -0.862 | 0.578 | 0.002 | 0.006 | 0.006 | 0 | 0.09 | 0.09 | 0.2 | 0.32 | 0.09 | 0.47 |
| 1 | 97308704 | rs17115879 | 1.217 | -0.6 | -0.629 | 0.126 | -0.002 | -0.018 | -0.018 | 0 | 0.08 | 0.08 | 0.03 | 0 | 0.03 | 0.07 |
| 1 | 97312779 | rs6701736 | -0.762 | -0.806 | -0.109 | 0.306 | -0.033 | 0.135 | 0.135 | 0.007 | 0.16 | 0.4 | 0.32 | 0.35 | 0.1 | 0.45 |
| 1 | 97313098 | rs17115886 | 1.231 | -0.612 | 0.026 | 0.375 | -0.009 | -0.033 | -0.033 | 0 | 0.08 | 0.1 | 0.03 | 0 | 0.03 | 0.07 |
| 1 | 97316690 | rs11584143 | -0.508 | -0.571 | 0.023 | 0.436 | -0.009 | 0.153 | 0.153 | 0 | 0.14 | 0.39 | 0.24 | 0.33 | 0.09 | 0.43 |
| 1 | 97333968 | rs1519891 | 1.389 | -0.556 | -0.297 | 0.419 | 0.008 | 0.009 | 0.009 | 0 | 0.07 | 0.02 | 0.02 | 0 | 0.03 | 0.07 |
| 1 | 97349337 | rs722102 | 0.497 | -0.661 | -0.096 | 0.129 | -0.017 | -0.056 | -0.056 | 0 | 0.1 | 0.22 | 0.04 | 0 | 0.03 | 0.14 |
| 1 | 97365977 | rs2681194 | 0.519 | -0.898 | -0.422 | -0.183 | -0.025 | -0.025 | -0.025 | 0 | 0.37 | 0.25 | 0.38 | 0.45 | 0.42 | 0.31 |
| 1 | 97372940 | rs12717805 | 0.981 | -1.012 | -0.549 | -0.327 | -0.004 | -0.033 | -0.033 | 0 | 0.08 | 0.1 | 0.03 | 0 | 0.05 | 0.04 |
| 1 | 97377456 | rs1976805 | -0.688 | -1 | -0.532 | -0.307 | 0.005 | 0.003 | 0.003 | 0 | 0.62 | 0.62 | 0.62 | 0.5 | 0.58 | 0.69 |
| 1 | 97385644 | rs4950016 | 0.132 | -0.798 | -1.835 | -0.762 | -0.002 | 0.16 | 0.16 | 0 | 0.44 | 0.14 | 0.36 | 0.22 | 0.48 | 0.25 |
| 1 | 97409943 | rs11165740 | 0.51 | -0.234 | -1.891 | -0.114 | -0.02 | 0.191 | 0.191 | 0 | 0.45 | 0.07 | 0.34 | 0.21 | 0.45 | 0.3 |
| 1 | 97421239 | rs12565126 | -0.079 | 0.193 | -1.947 | 0.137 | -0.011 | 0.088 | 0.088 | 0 | 0.05 | 0.04 | 0.21 | 0.4 | 0.1 | 0.32 |
| 1 | 97428030 | rs12066233 | -0.096 | 0.212 | -1.919 | 0.066 | -0.013 | -0.002 | -0.002 | 0 | 0.09 | 0.36 | 0.04 | 0.04 | 0.01 | 0.05 |
| 1 | 97429246 | rs11165746 | 1.125 | 0.632 | -3.258 | 0.06 | -0.014 | -0.042 | -0.042 | 0 | 0.1 | 0.02 | 0.05 | 0.19 | 0.12 | 0.08 |
| 1 | 97431052 | rs995258 | -0.182 | 0.942 | -2.823 | 0.263 | -0.139 | -0.026 | -0.026 | 0 | 0.38 | 0.07 | 0.38 | 0.35 | 0.43 | 0.26 |
| 1 | 97435890 | rs7512858 | -0.482 | 1.556 | -1.696 | 1.428 | 0 | -0.004 | -0.004 | 0 | 0.07 | 0.03 | 0.03 | 0 | 0.06 | 0.05 |
| 1 | 97439491 | rs3002273 | -0.219 | 2.148 | -1.717 | 1.767 | -0.061 | -0.067 | -0.067 | 0 | 0.15 | 0.05 | 0.28 | 0.33 | 0.26 | 0.11 |
| 1 | 97457893 | rs11165753 | -0.7 | 1.337 | -1.251 | 1.672 | -0.02 | 0.048 | 0.048 | 0 | 0.06 | 0.03 | 0.28 | 0.41 | 0.1 | 0.32 |
| 1 | 97462677 | rs544746 | -0.503 | 1.206 | -1.238 | 0.867 | -0.013 | -0.014 | -0.014 | 12.085 | 0.39 | 0.31 | 0.38 | 0.53 | 0.42 | 0.36 |
| 1 | 97463800 | rs11165754 | -0.525 | 2.339 | -1.048 | 1.125 | 0.006 | 0.001 | 0.001 | 0 | 0.13 | 0.15 | 0.22 | 0.22 | 0.21 | 0.1 |
| 1 | 97469804 | rs290821 | 0.069 | 1.902 | -1.6 | 0.866 | 0.002 | 0.013 | 0.013 | 0 | 0.6 | 0.3 | 0.48 | 0.62 | 0.63 | 0.73 |
| 1 | 97473707 | rs290826 | 0.159 | 1.957 | -1.783 | 1.01 | 0 | 0.042 | 0.042 | 0 | 0.11 | 0.12 | 0.13 | 0.4 | 0.12 | 0.4 |
| 1 | 97488820 | rs11165761 | 0.812 | 2.117 | -0.668 | 0.598 | -0.007 | -0.131 | -0.131 | 0.939 | 0.22 | 0.35 | 0.16 | 0.01 | 0.22 | 0.12 |
| 1 | 97499599 | rs67530 | -0.32 | 2.048 | -0.698 | 0.623 | -0.012 | -0.019 | -0.019 | 4.92 | 0.22 | 0.2 | 0.29 | 0.53 | 0.3 | 0.5 |
| 1 | 97526732 | rs290857 | -0.491 | 1.789 | -0.084 | 0.086 | -0.008 | -0.09 | -0.09 | 1.889 | 0.74 | 0.37 | 0.8 | 0.99 | 0.76 | 0.89 |
| 1 | 97533597 | rs17100610 | 1.648 | 1.644 | -0.072 | 0.017 | 0 | -0.028 | -0.028 | 0.264 | 0.15 | 0.13 | 0.12 | 0.01 | 0.17 | 0.1 |
| 1 | 97535350 | rs290868 | -0.026 | 1.638 | -0.075 | 0.014 | -0.008 | -0.09 | -0.09 | 0 | 0.74 | 0.37 | 0.8 | 1 | 0.76 | 0.89 |
| 1 | 97536362 | rs290867 | -0.799 | 1.715 | 0.158 | 0.041 | -0.024 | -0.097 | -0.097 | 0.872 | 0.79 | 0.67 | 0.86 | 1 | 0.83 | 0.89 |
| 1 | 97540371 | rs290865 | -1.085 | 1.718 | 0.163 | 0.059 | -0.017 | -0.019 | -0.019 | 1.581 | 0.95 | 0.8 | 0.97 | 1 | 0.99 | 0.99 |
| 1 | 97542236 | rs158634 | -0.818 | 1.716 | 0.138 | 0.042 | -0.032 | -0.119 | -0.119 | 0 | 0.79 | 0.65 | 0.85 | 1 | 0.83 | 0.89 |
| 1 | 97542797 | rs290861 | -0.818 | 1.716 | 0.138 | 0.042 | -0.032 | -0.119 | -0.119 | 0 | 0.79 | 0.65 | 0.85 | 1 | 0.83 | 0.89 |
| 1 | 97543752 | rs291593 | -0.576 | 1.829 | 0.076 | 0.064 | -0.011 | -0.018 | -0.018 | 0 | 0.27 | 0.22 | 0.29 | 0.62 | 0.31 | 0.52 |
| 1 | 97543764 | rs291592 | -1.21 | 1.857 | 0.048 | -0.054 | -0.034 | 0.049 | 0.049 | 0 | 0.38 | 0.29 | 0.61 | 0.98 | 0.45 | 0.67 |
| 1 | 97544771 | rs290855 | -0.977 | 1.837 | 0.066 | -0.001 | -0.029 | -0.112 | -0.112 | 0 | 0.8 | 0.67 | 0.85 | 1 | 0.83 | 0.89 |
| 1 | 97547530 | rs1399285 | 1.689 | 1.801 | 0.068 | -0.001 | 0 | -0.031 | -0.031 | 0 | 0.15 | 0.13 | 0.12 | 0.01 | 0.17 | 0.1 |
| 1 | 97548597 | rs12060037 | 1.669 | 1.8 | 0.068 | -0.001 | 0 | -0.031 | -0.031 | 0 | 0.15 | 0.14 | 0.12 | 0.01 | 0.17 | 0.1 |

| | | | | | | | | | | | | | | | | |
|---|---|---|---|---|---|---|---|---|---|---|---|---|---|---|---|---|
| 1 | 97550191 | rs290852 | -0.909 | 1.797 | 0.081 | -0.003 | -0.033 | -0.115 | -0.115 | 0.572 | 0.79 | 0.66 | 0.86 | 1 | 0.83 | 0.89 |
| 1 | 97550461 | rs290851 | -1.205 | 1.798 | 0.077 | -0.002 | -0.019 | -0.021 | -0.021 | 0.003 | 0.95 | 0.81 | 0.97 | 1 | 0.99 | 0.99 |
| 1 | 97551437 | rs1040090 | 1.69 | 1.798 | 0.08 | -0.002 | -0.001 | -0.033 | -0.033 | 0 | 0.15 | 0.14 | 0.12 | 0.01 | 0.17 | 0.1 |
| 1 | 97552056 | rs290806 | -1.201 | 1.797 | 0.08 | 0.005 | -0.019 | -0.021 | -0.021 | 0 | 0.95 | 0.81 | 0.97 | 1 | 0.99 | 0.99 |
| 1 | 97552243 | rs290807 | -1.201 | 1.797 | 0.08 | 0.005 | -0.019 | -0.021 | -0.021 | 0 | 0.95 | 0.81 | 0.97 | 1 | 0.99 | 0.99 |
| 1 | 97553128 | rs17471125 | 2.066 | 1.568 | 0 | -0.526 | -0.031 | 0.047 | 0.047 | 0 | 0.08 | 0.01 | 0.1 | 0 | 0.16 | 0.04 |
| 1 | 97557073 | rs1514494 | -1.306 | 1.67 | 0.295 | -0.222 | -0.019 | -0.021 | -0.021 | 0.07 | 0.95 | 0.8 | 0.97 | 1 | 0.99 | 0.99 |
| 1 | 97557184 | rs17116398 | 1.742 | 1.668 | 0.284 | -0.232 | 0 | -0.031 | -0.031 | 0.343 | 0.15 | 0.14 | 0.12 | 0.01 | 0.17 | 0.1 |
| 1 | 97558162 | rs2056048 | -1.247 | 1.55 | 0.218 | -0.322 | -0.03 | 0.048 | 0.048 | 0 | 0.39 | 0.28 | 0.61 | 0.98 | 0.45 | 0.67 |
| 1 | 97558368 | rs12062102 | 1.816 | 1.545 | 0.21 | -0.324 | 0 | -0.031 | -0.031 | 0.336 | 0.15 | 0.14 | 0.12 | 0.01 | 0.17 | 0.1 |
| 1 | 97559723 | rs10449721 | 1.652 | 1.542 | 0.33 | -0.324 | -0.001 | -0.087 | -0.087 | 0.128 | 0.4 | 0.42 | 0.25 | 0.02 | 0.38 | 0.22 |
| 1 | 97565233 | rs11165781 | 1.805 | 1.543 | 0.306 | -0.313 | 0 | -0.031 | -0.031 | 0 | 0.15 | 0.14 | 0.12 | 0.01 | 0.17 | 0.1 |
| 1 | 97569544 | rs11165783 | -0.381 | 1.553 | 0.297 | -0.196 | -0.01 | -0.016 | -0.016 | 0 | 0.27 | 0.22 | 0.29 | 0.63 | 0.31 | 0.52 |
| 1 | 97570107 | rs7524812 | -0.943 | 1.508 | 0.322 | -0.267 | -0.015 | -0.018 | -0.018 | 0 | 0.94 | 0.75 | 0.97 | 1 | 0.99 | 0.99 |
| 1 | 97571799 | rs993568 | -1.23 | 1.532 | 0.29 | -0.27 | -0.028 | 0.039 | 0.039 | 0 | 0.39 | 0.29 | 0.61 | 0.98 | 0.45 | 0.67 |
| 1 | 97574944 | rs6593630 | -2.073 | 1.527 | 0.223 | -0.262 | 0 | -0.031 | -0.031 | 1.667 | 0.85 | 0.87 | 0.88 | 0.99 | 0.83 | 0.9 |
| 1 | 97576922 | rs1399291 | -1.221 | 1.546 | 0.183 | -0.259 | -0.049 | 0.109 | 0.109 | 0 | 0.39 | 0.23 | 0.61 | 0.98 | 0.45 | 0.67 |
| 1 | 97580323 | rs2137975 | 1.482 | 1.507 | 0.096 | -0.162 | 0.004 | -0.021 | -0.021 | 0 | 0.4 | 0.32 | 0.25 | 0.02 | 0.38 | 0.22 |
| 1 | 97583471 | rs7525018 | -0.995 | 1.455 | 0.075 | -0.038 | 0 | 0.001 | 0.001 | 0 | 0.32 | 0.38 | 0.32 | 0.63 | 0.32 | 0.52 |
| 1 | 97584032 | rs12130586 | 0.97 | 1.65 | 0.066 | -0.023 | -0.044 | 0.011 | 0.011 | 0 | 0.12 | 0.05 | 0.16 | 0.13 | 0.19 | 0.08 |
| 1 | 97584161 | rs6681958 | 1.517 | 1.522 | 0.192 | -0.312 | 0.013 | 0.061 | 0.061 | 0 | 0.42 | 0.28 | 0.23 | 0.04 | 0.4 | 0.25 |
| 1 | 97584646 | rs970337 | 0.286 | 1.949 | 0.348 | -0.004 | -0.019 | 0.004 | 0.004 | 0 | 0.64 | 0.69 | 0.43 | 0.34 | 0.55 | 0.39 |
| 1 | 97585937 | rs9437663 | -1.7 | 2.013 | 0.23 | 0.025 | 0 | 0.003 | 0.003 | 0 | 0.83 | 0.88 | 0.72 | 0.95 | 0.81 | 0.86 |
| 1 | 97588830 | rs10493890 | 1.331 | 1.699 | 0.141 | 0.061 | 0.039 | 0.076 | 0.076 | 0 | 0.4 | 0.25 | 0.22 | 0.04 | 0.37 | 0.24 |
| 1 | 97588913 | rs10875045 | -0.792 | 1.982 | 1.106 | 1.047 | -0.015 | -0.022 | -0.022 | 0 | 0.17 | 0.11 | 0.18 | 0.3 | 0.17 | 0.15 |
| 1 | 97594994 | rs10875047 | -0.792 | 1.982 | 1.106 | 1.047 | -0.015 | -0.022 | -0.022 | 1.642 | 0.17 | 0.1 | 0.18 | 0.3 | 0.17 | 0.15 |
| 1 | 97600039 | rs12047910 | -0.395 | 1.954 | 1.078 | 1.122 | -0.023 | -0.042 | -0.042 | 0 | 0.15 | 0.03 | 0.15 | 0.24 | 0.13 | 0.14 |
| 1 | 97602726 | rs1709409 | -1.325 | 1.904 | -0.535 | 1.098 | 0 | 0.002 | 0.002 | 0 | 0.22 | 0.23 | 0.19 | 0.3 | 0.18 | 0.15 |
| 1 | 97602898 | rs11165789 | -0.684 | 1.92 | -0.9 | 1.106 | -0.042 | -0.056 | -0.056 | 0 | 0.17 | 0.05 | 0.17 | 0.3 | 0.17 | 0.14 |
| 1 | 97602994 | rs1760217 | -1.305 | 1.92 | -0.898 | 1.106 | -0.005 | -0.012 | -0.012 | 0 | 0.22 | 0.19 | 0.18 | 0.3 | 0.18 | 0.15 |
| 1 | 97603679 | rs12039249 | -0.79 | 2.316 | -1.391 | 1.296 | -0.042 | -0.056 | -0.056 | 2.33 | 0.17 | 0.05 | 0.17 | 0.3 | 0.17 | 0.14 |
| 1 | 97604531 | rs11799399 | -0.79 | 2.281 | -1.39 | 1.29 | -0.036 | -0.052 | -0.052 | 0.351 | 0.17 | 0.05 | 0.17 | 0.29 | 0.16 | 0.14 |
| 1 | 97606480 | rs12023630 | -0.495 | 1.902 | -1.338 | 1.366 | -0.017 | -0.057 | -0.057 | 0.005 | 0.15 | 0.05 | 0.16 | 0.3 | 0.14 | 0.14 |
| 1 | 97606764 | rs7552825 | -1.238 | 2.346 | -1.865 | 1.278 | -0.062 | -0.072 | -0.072 | 0 | 0.28 | 0.11 | 0.34 | 0.41 | 0.35 | 0.16 |
| 1 | 97607220 | rs6667550 | -1.364 | 2.078 | -1.655 | 1.853 | -0.004 | 0.004 | 0.004 | 0 | 0.12 | 0.14 | 0.18 | 0.35 | 0.21 | 0.27 |
| 1 | 97607837 | rs17702702 | -0.378 | 2.24 | -2.019 | 1.145 | -0.054 | -0.093 | -0.093 | 0 | 0.16 | 0.01 | 0.25 | 0.29 | 0.17 | 0.13 |
| 1 | 97608892 | rs828052 | 0.145 | 1.811 | -2.977 | 0.006 | -0.085 | 0.2 | 0.2 | 0 | 0.64 | 0.86 | 0.66 | 0.99 | 0.5 | 0.84 |
| 1 | 97609326 | rs828053 | 0.224 | 1.597 | -1.951 | -0.075 | -0.009 | 0.024 | 0.024 | 0 | 0.17 | 0.22 | 0.25 | 0.28 | 0.15 | 0.12 |
| 1 | 97610026 | rs828054 | 0.056 | 1.613 | -1.976 | -0.102 | -0.087 | 0.202 | 0.202 | 0 | 0.63 | 0.86 | 0.66 | 1 | 0.49 | 0.84 |
| 1 | 97611515 | rs1760215 | -0.622 | 1.536 | -2.147 | -0.262 | -0.001 | 0.057 | 0.057 | 0 | 0.9 | 0.99 | 0.96 | 1 | 0.93 | 0.96 |
| 1 | 97613855 | rs17432093 | 1.883 | 0.61 | -2.045 | -0.658 | -0.01 | 0.122 | 0.122 | 0 | 0.26 | 0.04 | 0.14 | 0.05 | 0.23 | 0.23 |
| 1 | 97615098 | rs6593632 | -1.075 | 0.711 | -2.097 | -0.709 | -0.129 | 0.21 | 0.21 | 0 | 0.7 | 0.97 | 0.72 | 1 | 0.61 | 0.85 |
| 1 | 97615250 | rs4390222 | 0.645 | 0.707 | -2.063 | -0.694 | -0.008 | 0.02 | 0.02 | 0 | 0.17 | 0.21 | 0.25 | 0.28 | 0.15 | 0.12 |
| 1 | 97616058 | rs2039449 | 0.167 | 0.609 | -1.28 | -0.741 | 0.011 | 0.073 | 0.073 | 0 | 0.24 | 0.47 | 0.29 | 0.66 | 0.23 | 0.49 |
| 1 | 97617789 | rs1514495 | 0.167 | 0.609 | -1.28 | -0.739 | 0.011 | 0.07 | 0.07 | 0 | 0.24 | 0.47 | 0.29 | 0.67 | 0.23 | 0.49 |
| 1 | 97617831 | rs1514497 | 0.135 | 0.588 | -1.29 | -0.765 | 0.004 | 0.063 | 0.063 | 0 | 0.25 | 0.47 | 0.3 | 0.66 | 0.24 | 0.5 |
| 1 | 97621204 | rs11165792 | 0.969 | 0.603 | -0.751 | -0.569 | -0.001 | -0.064 | -0.064 | 0 | 0.16 | 0.05 | 0.24 | 0.28 | 0.15 | 0.12 |
| 1 | 97621437 | rs11165793 | 0.588 | 0.597 | -0.736 | -0.572 | -0.01 | 0.028 | 0.028 | 0 | 0.17 | 0.23 | 0.25 | 0.28 | 0.15 | 0.12 |
| 1 | 97621606 | rs11165794 | 0.625 | 0.6 | -0.736 | -0.571 | -0.008 | 0.02 | 0.02 | 0 | 0.17 | 0.21 | 0.25 | 0.28 | 0.15 | 0.12 |
| 1 | 97622914 | rs17116506 | 1.724 | 0.581 | -0.649 | -0.58 | 0.011 | 0.078 | 0.078 | 0 | 0.27 | 0.12 | 0.15 | 0.05 | 0.23 | 0.23 |
| 1 | 97623853 | rs10875050 | 0.908 | 0.605 | -0.584 | -0.542 | 0.005 | -0.045 | -0.045 | 0 | 0.16 | 0.06 | 0.24 | 0.28 | 0.15 | 0.12 |
| 1 | 97625198 | rs10875051 | 1.098 | 0.604 | -0.612 | -0.543 | -0.019 | -0.087 | -0.087 | 4.863 | 0.15 | 0.03 | 0.24 | 0.28 | 0.15 | 0.12 |
| 1 | 97632298 | rs828059 | 0.62 | 0.622 | -0.623 | -0.561 | -0.008 | 0.02 | 0.02 | 0 | 0.17 | 0.22 | 0.25 | 0.28 | 0.15 | 0.12 |
| 1 | 97635212 | rs6593634 | 0.084 | 0.628 | -0.601 | -0.552 | 0.01 | 0.069 | 0.069 | 0 | 0.24 | 0.46 | 0.29 | 0.67 | 0.23 | 0.49 |
| 1 | 97638127 | rs12134860 | 0.588 | 0.51 | -0.628 | -0.41 | -0.008 | 0.02 | 0.02 | 0 | 0.17 | 0.21 | 0.26 | 0.28 | 0.15 | 0.12 |
| 1 | 97638241 | rs12124552 | 0.569 | 0.51 | -0.624 | -0.41 | -0.01 | 0.027 | 0.027 | 0 | 0.17 | 0.23 | 0.26 | 0.28 | 0.15 | 0.12 |
| 1 | 97638550 | rs11165798 | -0.474 | 0.511 | -0.637 | -0.426 | -0.002 | 0.002 | 0.002 | 0 | 0.23 | 0.26 | 0.3 | 0.28 | 0.26 | 0.14 |
| 1 | 97640563 | rs914959 | -1.153 | 0.529 | -0.661 | -0.44 | -0.129 | 0.209 | 0.209 | 0 | 0.7 | 0.98 | 0.72 | 1 | 0.61 | 0.85 |
| 1 | 97640673 | rs11165800 | 0.559 | 0.522 | -0.576 | -0.351 | -0.008 | 0.02 | 0.02 | 0 | 0.17 | 0.21 | 0.25 | 0.28 | 0.15 | 0.12 |
| 1 | 97641178 | rs12759852 | 0.559 | 0.522 | -0.576 | -0.351 | -0.008 | 0.02 | 0.02 | 0 | 0.17 | 0.21 | 0.25 | 0.28 | 0.15 | 0.12 |
| 1 | 97641771 | rs12035560 | 1.457 | 0.545 | -0.31 | -0.319 | -0.004 | -0.003 | -0.003 | 0 | 0.13 | 0.09 | 0.13 | 0.4 | 0.09 | 0.31 |
| 1 | 97644724 | rs6593636 | -1.21 | 0.586 | -0.434 | -0.473 | -0.089 | 0.173 | 0.173 | 0 | 0.7 | 0.92 | 0.71 | 1 | 0.61 | 0.85 |
| 1 | 97645277 | rs6593637 | -1.159 | 0.598 | -0.467 | -0.493 | -0.089 | 0.171 | 0.171 | 0 | 0.71 | 0.91 | 0.72 | 1 | 0.62 | 0.85 |
| 1 | 97645313 | rs6683351 | -1.214 | 0.595 | -0.405 | -0.448 | -0.058 | 0.121 | 0.121 | 0 | 0.7 | 0.88 | 0.71 | 1 | 0.61 | 0.85 |
| 1 | 97645607 | rs12041436 | 0.868 | 0.53 | -0.397 | -0.315 | -0.003 | -0.066 | -0.066 | 0.002 | 0.16 | 0.05 | 0.25 | 0.27 | 0.15 | 0.12 |
| 1 | 97645979 | rs12740796 | 1.014 | 0.535 | -0.443 | -0.313 | -0.019 | -0.086 | -0.086 | 0 | 0.15 | 0.03 | 0.25 | 0.27 | 0.15 | 0.12 |
| 1 | 97647338 | rs7529372 | -0.091 | 0.5 | -0.151 | -0.299 | 0.027 | 0.208 | 0.208 | 0 | 0.12 | 0.41 | 0.16 | 0.27 | 0.14 | 0.18 |
| 1 | 97649058 | rs6681752 | -1.302 | 0.552 | -0.196 | -0.436 | -0.078 | 0.179 | 0.179 | 0 | 0.7 | 0.91 | 0.72 | 1 | 0.62 | 0.85 |
| 1 | 97649209 | rs6656660 | -0.05 | 0.556 | -0.204 | -0.42 | 0.027 | 0.208 | 0.208 | 0 | 0.12 | 0.41 | 0.16 | 0.27 | 0.14 | 0.18 |

| | | | | | | | | | | | | | | | | |
|---|---|---|---|---|---|---|---|---|---|---|---|---|---|---|---|---|
| 1 | 97650998 | rs6703589 | 1.909 | 0.528 | -0.372 | -0.479 | -0.014 | 0.127 | 0.127 | 0 | 0.26 | 0.04 | 0.14 | 0.05 | 0.23 | 0.23 |
| 1 | 97651567 | rs11165804 | 1.123 | 0.531 | -0.417 | -0.462 | -0.019 | -0.086 | -0.086 | 0 | 0.15 | 0.03 | 0.25 | 0.27 | 0.15 | 0.12 |
| 1 | 97653070 | rs11587873 | 1.91 | 0.534 | -0.408 | -0.48 | -0.009 | 0.123 | 0.123 | 0 | 0.26 | 0.04 | 0.14 | 0.05 | 0.23 | 0.23 |
| 1 | 97657086 | rs3762412 | -1.247 | 0.569 | -0.4 | -0.474 | -0.113 | 0.209 | 0.209 | 0 | 0.7 | 0.97 | 0.73 | 1 | 0.62 | 0.85 |
| 1 | 97660692 | rs1413228 | 1.024 | 0.598 | -0.256 | -0.46 | 0.055 | -0.016 | -0.016 | 0 | 0.07 | 0.27 | 0.06 | 0 | 0.12 | 0.02 |
| 1 | 97663530 | rs7513333 | -1.253 | 0.61 | -1.004 | -0.42 | -0.114 | 0.207 | 0.207 | 0 | 0.71 | 0.97 | 0.73 | 1 | 0.62 | 0.85 |
| 1 | 97668900 | rs8179310 | 0.878 | 0.596 | -0.733 | -0.114 | -0.02 | -0.085 | -0.085 | 0.725 | 0.15 | 0.03 | 0.24 | 0.27 | 0.15 | 0.12 |
| 1 | 97670325 | rs6593639 | -0.205 | 0.788 | -0.42 | 0.049 | -0.001 | 0.01 | 0.01 | 0 | 0.53 | 0.53 | 0.56 | 0.59 | 0.52 | 0.52 |
| 1 | 97672838 | rs4128473 | -0.527 | 0.969 | 0.046 | 0.35 | 0.034 | 0.218 | 0.218 | 0 | 0.12 | 0.43 | 0.17 | 0.27 | 0.14 | 0.2 |
| 1 | 97676260 | rs7537668 | -0.964 | 1.318 | -0.676 | -0.181 | -0.062 | 0.182 | 0.182 | 0 | 0.71 | 0.96 | 0.74 | 1 | 0.66 | 0.86 |
| 1 | 97676519 | rs3897933 | 0.102 | 1.022 | -0.347 | -0.293 | -0.003 | 0 | 0 | 0.27 | 0.25 | 0.28 | 0.33 | 0.24 | 0.29 | 0.1 |
| 1 | 97676941 | rs10783056 | 0.17 | 1.013 | -0.363 | -0.298 | -0.029 | -0.003 | -0.003 | 0.932 | 0.24 | 0.14 | 0.33 | 0.24 | 0.29 | 0.1 |
| 1 | 97680316 | rs12120068 | 1.818 | 0.755 | -0.307 | -0.474 | 0.025 | 0.102 | 0.102 | 0 | 0.21 | 0.02 | 0.07 | 0.04 | 0.13 | 0.21 |
| 1 | 97680720 | rs2027054 | 0.299 | 0.752 | -0.309 | -0.478 | -0.029 | -0.003 | -0.003 | 0 | 0.25 | 0.14 | 0.33 | 0.24 | 0.3 | 0.1 |
| 1 | 97681342 | rs1333727 | -0.307 | 0.759 | 0.291 | -0.264 | 0.006 | 0.042 | 0.042 | 0.008 | 0.31 | 0.57 | 0.46 | 0.53 | 0.32 | 0.28 |
| 1 | 97682026 | rs6697132 | -1.758 | 0.773 | 0.265 | -0.289 | -0.092 | 0.214 | 0.214 | 0 | 0.7 | 0.97 | 0.73 | 1 | 0.64 | 0.86 |
| 1 | 97682376 | rs12046655 | 0.766 | 0.717 | 0.481 | -0.332 | 0.007 | -0.016 | -0.016 | 0 | 0.31 | 0.23 | 0.22 | 0.46 | 0.22 | 0.56 |
| 1 | 97683103 | rs12116905 | 1.918 | 0.726 | 0.481 | -0.328 | 0.025 | 0.112 | 0.112 | 0 | 0.21 | 0.01 | 0.07 | 0.04 | 0.13 | 0.21 |
| 1 | 97683997 | rs10875055 | -0.981 | 0.736 | 0.503 | -0.319 | -0.026 | 0.116 | 0.116 | 0.183 | 0.61 | 0.8 | 0.68 | 1 | 0.54 | 0.84 |
| 1 | 97684328 | rs4511173 | -0.266 | 0.739 | 0.511 | -0.306 | 0.006 | 0.042 | 0.042 | 0.008 | 0.31 | 0.58 | 0.46 | 0.53 | 0.32 | 0.28 |
| 1 | 97685486 | rs2039447 | -0.266 | 0.74 | 0.503 | -0.306 | 0.006 | 0.042 | 0.042 | 0.008 | 0.31 | 0.58 | 0.46 | 0.53 | 0.32 | 0.28 |
| 1 | 97685992 | rs1413234 | 1.367 | 0.749 | 0.437 | -0.317 | 0.004 | -0.04 | -0.04 | 0 | 0.08 | 0.16 | 0.05 | 0 | 0.1 | 0.02 |
| 1 | 97686036 | rs1413235 | -1.711 | 0.752 | 0.302 | -0.302 | -0.074 | 0.2 | 0.2 | 0 | 0.69 | 0.96 | 0.73 | 1 | 0.64 | 0.86 |
| 1 | 97688831 | rs4415625 | 1.507 | 0.842 | 0.208 | -0.365 | -0.035 | -0.061 | -0.061 | 0 | 0.12 | 0.04 | 0.23 | 0.24 | 0.13 | 0.08 |
| 1 | 97691005 | rs885622 | -1.846 | 0.829 | 0.204 | -0.504 | -0.093 | 0.212 | 0.212 | 0 | 0.7 | 0.97 | 0.73 | 1 | 0.64 | 0.86 |
| 1 | 97693323 | rs11165810 | 0.343 | 0.816 | 0.38 | -0.418 | 0.069 | 0.405 | 0.405 | 0 | 0.06 | 0.3 | 0.12 | 0.3 | 0.02 | 0.19 |
| 1 | 97694442 | rs827497 | 0.105 | 0.887 | -0.369 | -0.471 | -0.031 | -0.018 | -0.018 | 0 | 0.24 | 0.14 | 0.33 | 0.28 | 0.29 | 0.1 |
| 1 | 97696711 | rs866129 | 0.05 | 0.883 | -0.379 | -0.472 | -0.015 | -0.009 | -0.009 | 0.167 | 0.26 | 0.24 | 0.34 | 0.29 | 0.3 | 0.1 |
| 1 | 97699757 | rs1520658 | 1.424 | 0.929 | -0.482 | -0.436 | -0.035 | -0.068 | -0.068 | 0 | 0.12 | 0.04 | 0.24 | 0.29 | 0.13 | 0.08 |
| 1 | 97700589 | rs12137711 | 1.683 | 0.671 | -0.24 | -0.175 | 0.026 | 0.108 | 0.108 | 0 | 0.21 | 0.01 | 0.07 | 0.04 | 0.13 | 0.21 |
| 1 | 97703325 | rs533365 | -1.535 | 0.731 | -0.246 | -0.237 | -0.077 | 0.199 | 0.199 | 0 | 0.7 | 0.96 | 0.73 | 1 | 0.64 | 0.86 |
| 1 | 97703552 | rs17116602 | 1.025 | 0.799 | -0.091 | -0.001 | 0.004 | 0.066 | 0.066 | 0 | 0.09 | 0.12 | 0.12 | 0.36 | 0.09 | 0.31 |
| 1 | 97704798 | rs865242 | 0.856 | 0.677 | -0.146 | -0.007 | -0.022 | -0.013 | -0.013 | 1.013 | 0.25 | 0.16 | 0.33 | 0.28 | 0.29 | 0.11 |
| 1 | 97706237 | rs12133176 | 1.456 | 0.651 | -0.334 | -0.008 | 0.027 | 0.04 | 0.04 | 0 | 0.22 | 0.1 | 0.11 | 0.08 | 0.14 | 0.23 |
| 1 | 97708511 | rs12125127 | 1.411 | 0.651 | -0.334 | -0.008 | 0.028 | 0.042 | 0.042 | 0 | 0.23 | 0.11 | 0.11 | 0.08 | 0.14 | 0.23 |
| 1 | 97709053 | rs12124488 | 0.668 | 0.654 | -0.334 | -0.018 | 0.025 | 0.06 | 0.06 | 0 | 0.29 | 0.15 | 0.16 | 0.08 | 0.24 | 0.24 |
| 1 | 97711452 | rs592609 | -0.862 | 0.686 | -0.247 | -0.102 | 0.001 | -0.014 | -0.014 | 0 | 0.63 | 0.57 | 0.61 | 0.7 | 0.61 | 0.65 |
| 1 | 97711561 | rs12758854 | -0.099 | 0.7 | -0.334 | -0.033 | 0.006 | -0.007 | -0.007 | 0 | 0.38 | 0.33 | 0.28 | 0.43 | 0.32 | 0.55 |
| 1 | 97714665 | rs10875057 | 0.69 | 0.643 | -0.179 | 0.274 | -0.005 | 0.002 | 0.002 | 0 | 0.32 | 0.29 | 0.39 | 0.28 | 0.4 | 0.12 |
| 1 | 97714892 | rs10875058 | 0.501 | 0.697 | -0.44 | 0.3 | 0.012 | -0.018 | -0.018 | 0 | 0.33 | 0.5 | 0.4 | 0.27 | 0.4 | 0.12 |
| 1 | 97714918 | rs12128564 | 1.375 | 0.681 | -0.772 | 0.286 | 0.016 | 0.091 | 0.091 | 0 | 0.21 | 0.01 | 0.11 | 0.05 | 0.14 | 0.23 |
| 1 | 97716141 | rs12564297 | -0.204 | 0.922 | -0.382 | 0.514 | 0.021 | 0.219 | 0.219 | 0 | 0.07 | 0.23 | 0.12 | 0.3 | 0.04 | 0.21 |
| 1 | 97716167 | rs1760212 | 0.398 | 0.974 | -0.356 | 0.532 | 0.012 | -0.012 | -0.012 | 0 | 0.32 | 0.46 | 0.4 | 0.28 | 0.4 | 0.12 |
| 1 | 97717279 | rs4950025 | 0.031 | 1.003 | -0.342 | 0.523 | 0.029 | 0.238 | 0.238 | 0.094 | 0.07 | 0.22 | 0.12 | 0.3 | 0.04 | 0.21 |
| 1 | 97717987 | rs1520663 | 0.392 | 1.029 | -0.331 | 0.444 | 0.012 | -0.013 | -0.013 | 0 | 0.32 | 0.46 | 0.4 | 0.28 | 0.4 | 0.12 |
| 1 | 97718075 | rs1520664 | 0.392 | 1.029 | -0.33 | 0.444 | 0.014 | -0.014 | -0.014 | 0 | 0.32 | 0.47 | 0.4 | 0.28 | 0.4 | 0.12 |
| 1 | 97719934 | rs12735722 | 0.082 | 1.041 | -0.496 | 0.48 | 0.029 | 0.259 | 0.259 | 0 | 0.07 | 0.22 | 0.12 | 0.35 | 0.04 | 0.2 |
| 1 | 97720465 | rs576189 | 0.179 | 1.253 | -0.913 | -0.452 | 0.012 | -0.011 | -0.011 | 1.582 | 0.33 | 0.47 | 0.4 | 0.28 | 0.4 | 0.12 |
| 1 | 97721392 | rs495257 | 0.174 | 1.325 | -0.955 | -0.462 | 0.019 | -0.009 | -0.009 | 0 | 0.33 | 0.47 | 0.4 | 0.28 | 0.42 | 0.12 |
| 1 | 97721488 | rs496179 | -0.109 | 1.231 | -1.382 | -0.531 | -0.007 | 0.085 | 0.085 | 0 | 0.7 | 0.88 | 0.73 | 1 | 0.66 | 0.86 |
| 1 | 97725679 | rs11165823 | -0.988 | 1.384 | -0.845 | 0.423 | 0.026 | 0.019 | 0.019 | 0 | 0.06 | 0.09 | 0.13 | 0.1 | 0.11 | 0.1 |
| 1 | 97727083 | rs693680 | 0.078 | 2.252 | -1.672 | 1.061 | -0.097 | -0.095 | -0.095 | 0 | 0.81 | 0.97 | 0.6 | 0.67 | 0.69 | 0.67 |
| 1 | 97728497 | rs827500 | 0.081 | 2.244 | -1.677 | 1.056 | -0.097 | -0.095 | -0.095 | 0 | 0.81 | 0.97 | 0.6 | 0.67 | 0.69 | 0.67 |
| 1 | 97730626 | rs10783057 | 0.522 | 1.294 | -2.378 | 0.463 | -0.007 | 0.019 | 0.019 | 0 | 0.3 | 0.21 | 0.31 | 0.25 | 0.37 | 0.19 |
| 1 | 97731402 | rs499009 | -0.773 | 1.29 | -2.384 | 0.462 | -0.097 | -0.095 | -0.095 | 0 | 0.81 | 0.97 | 0.6 | 0.67 | 0.69 | 0.67 |
| 1 | 97732025 | rs7518848 | -0.773 | 1.29 | -2.384 | 0.462 | -0.097 | -0.095 | -0.095 | 3.512 | 0.81 | 0.97 | 0.6 | 0.67 | 0.69 | 0.67 |
| 1 | 97733932 | rs12048612 | 0.342 | 1.056 | -1.684 | 0.342 | 0.034 | 0.013 | 0.013 | 0 | 0.49 | 0.57 | 0.28 | 0.42 | 0.32 | 0.48 |
| 1 | 97737237 | rs7527272 | 0.879 | -0.304 | -0.672 | 0.209 | 0.006 | 0.006 | 0.006 | 0 | 0.29 | 0.21 | 0.15 | 0.26 | 0.24 | 0.26 |
| 1 | 97737704 | rs644428 | -1.4 | -0.263 | -0.685 | 0.224 | -0.092 | -0.091 | -0.091 | 0 | 0.81 | 0.97 | 0.6 | 0.67 | 0.69 | 0.67 |
| 1 | 97738354 | rs507170 | -1.357 | -0.263 | -0.578 | 0.224 | -0.045 | -0.045 | -0.045 | 0 | 0.81 | 0.92 | 0.6 | 0.67 | 0.69 | 0.67 |
| 1 | 97738860 | rs628959 | -1.357 | -0.263 | -0.578 | 0.224 | -0.045 | -0.045 | -0.045 | 0 | 0.81 | 0.92 | 0.6 | 0.67 | 0.69 | 0.67 |
| 1 | 97739908 | rs11165827 | 1.214 | -0.285 | -0.568 | 0.211 | -0.007 | 0.019 | 0.019 | 0 | 0.3 | 0.21 | 0.31 | 0.25 | 0.37 | 0.19 |
| 1 | 97741123 | rs7539889 | 1.685 | -0.288 | -0.576 | 0.206 | 0.004 | 0.056 | 0.056 | 0.087 | 0.27 | 0.15 | 0.2 | 0.18 | 0.29 | 0.14 |
| 1 | 97743805 | rs641805 | 0.302 | -0.388 | -0.511 | 0.341 | 0.014 | 0.037 | 0.037 | 0 | 0.46 | 0.41 | 0.37 | 0.29 | 0.42 | 0.28 |
| 1 | 97744048 | rs10875061 | 1.373 | -0.388 | -0.54 | 0.335 | -0.008 | 0.019 | 0.019 | 0 | 0.3 | 0.21 | 0.31 | 0.25 | 0.37 | 0.19 |
| 1 | 97746924 | rs10875062 | 1.347 | -0.384 | -0.559 | 0.383 | -0.003 | 0.016 | 0.016 | 0 | 0.3 | 0.21 | 0.31 | 0.26 | 0.35 | 0.18 |
| 1 | 97747589 | rs12029106 | 1.031 | -0.453 | -0.298 | 0.353 | 0.007 | 0.009 | 0.009 | 0 | 0.29 | 0.16 | 0.16 | 0.26 | 0.25 | 0.26 |
| 1 | 97749248 | rs667565 | 0.311 | -0.513 | -0.354 | 0.331 | 0.008 | 0.016 | 0.016 | 0 | 0.46 | 0.45 | 0.37 | 0.3 | 0.41 | 0.27 |
| 1 | 97749380 | rs526645 | -1.46 | -0.511 | -0.359 | 0.328 | -0.099 | -0.093 | -0.093 | 0 | 0.81 | 0.97 | 0.6 | 0.69 | 0.68 | 0.67 |

| | | | | | | | | | | | | | | | | |
|---|---|---|---|---|---|---|---|---|---|---|---|---|---|---|---|---|
| 1 | 97750838 | rs6593642 | -0.555 | -0.517 | -0.348 | 0.315 | 0.008 | 0.016 | 0.016 | 0 | 0.54 | 0.55 | 0.63 | 0.7 | 0.59 | 0.73 |
| 1 | 97750974 | rs12024830 | 1.191 | -0.537 | -0.392 | 0.296 | 0 | 0.021 | 0.021 | 0 | 0.32 | 0.21 | 0.31 | 0.26 | 0.35 | 0.18 |
| 1 | 97760661 | rs4950026 | -1.364 | -0.507 | -0.313 | 0.306 | -0.001 | -0.015 | -0.015 | 0 | 0.68 | 0.58 | 0.68 | 0.74 | 0.64 | 0.82 |
| 1 | 97761972 | rs10783058 | 1.397 | -0.497 | -0.45 | 0.312 | -0.003 | 0.015 | 0.015 | 0 | 0.3 | 0.21 | 0.31 | 0.26 | 0.35 | 0.18 |
| 1 | 97764073 | rs4264037 | 0.957 | -0.43 | -0.485 | 0.472 | 0.007 | -0.002 | -0.002 | 0 | 0.29 | 0.16 | 0.16 | 0.28 | 0.25 | 0.26 |
| 1 | 97771947 | rs7556439 | 1.128 | -0.393 | -0.515 | 0.49 | -0.099 | -0.093 | -0.093 | 0 | 0.19 | 0.03 | 0.4 | 0.31 | 0.32 | 0.33 |
| 1 | 97778724 | rs10875064 | 0.946 | -0.395 | -0.471 | 0.47 | 0.002 | 0.018 | 0.018 | 0 | 0.34 | 0.25 | 0.33 | 0.26 | 0.37 | 0.2 |
| 1 | 97779535 | rs12030332 | 0.886 | -0.315 | -0.514 | 0.497 | 0.007 | -0.002 | -0.002 | 0 | 0.29 | 0.17 | 0.15 | 0.28 | 0.25 | 0.26 |
| 1 | 97781039 | rs1609519 | 0.714 | 0.138 | -0.364 | 0.767 | -0.105 | -0.095 | -0.095 | 0 | 0.2 | 0.03 | 0.39 | 0.32 | 0.31 | 0.34 |
| 1 | 97793775 | rs10875069 | 0.781 | -0.786 | -0.176 | 0.236 | -0.001 | -0.008 | -0.008 | 0 | 0.32 | 0.33 | 0.32 | 0.19 | 0.33 | 0.13 |
| 1 | 97797970 | rs12038518 | 0.516 | -0.335 | -0.308 | 1.04 | -0.012 | -0.039 | -0.039 | 0 | 0.13 | 0.03 | 0.16 | 0.26 | 0.15 | 0.26 |
| 1 | 97799545 | rs7555294 | -0.584 | -0.589 | 0.112 | 0.871 | 0.008 | -0.003 | -0.003 | 0.608 | 0.24 | 0.17 | 0.22 | 0.29 | 0.19 | 0.33 |
| 1 | 97803594 | rs9783115 | 0.138 | -0.669 | 0.651 | 0.828 | -0.001 | 0.029 | 0.029 | 0 | 0.15 | 0.24 | 0.17 | 0.26 | 0.15 | 0.25 |
| 1 | 97803724 | rs9782950 | -1.613 | -0.771 | 0.98 | 1.08 | 0.009 | -0.018 | -0.018 | 0 | 0.37 | 0.65 | 0.38 | 0.47 | 0.3 | 0.44 |
| 1 | 97807335 | rs1879375 | -0.742 | -0.881 | 1.093 | 1.016 | -0.027 | -0.009 | -0.009 | 0 | 0.27 | 0.47 | 0.23 | 0.28 | 0.19 | 0.32 |
| 1 | 97809250 | rs12044858 | 1.415 | -0.99 | 0.425 | 0.943 | 0.006 | -0.047 | -0.047 | 0.595 | 0.16 | 0.11 | 0.29 | 0.41 | 0.13 | 0.21 |
| 1 | 97810607 | rs12046771 | -0.263 | -0.859 | 0.789 | 0.915 | -0.003 | 0 | 0 | 0.206 | 0.25 | 0.21 | 0.22 | 0.3 | 0.19 | 0.36 |
| 1 | 97814678 | rs10875071 | -0.351 | -0.862 | 0.815 | 0.914 | -0.011 | 0 | 0 | 0 | 0.26 | 0.24 | 0.22 | 0.3 | 0.19 | 0.36 |
| 1 | 97819405 | rs11165845 | 0.996 | -0.767 | 0.803 | 1.203 | 0.013 | -0.034 | -0.034 | 0 | 0.65 | 0.33 | 0.63 | 0.49 | 0.72 | 0.54 |
| 1 | 97821081 | rs4950028 | 0.517 | -0.787 | 0.831 | 1.209 | -0.001 | 0.019 | 0.019 | 0 | 0.12 | 0.11 | 0.15 | 0.2 | 0.09 | 0.12 |
| 1 | 97821302 | rs6679536 | 1.059 | -0.781 | 0.777 | 1.188 | 0.015 | -0.036 | -0.036 | 0 | 0.65 | 0.31 | 0.63 | 0.49 | 0.72 | 0.54 |
| 1 | 97830546 | rs12564011 | 0.41 | -0.704 | 0.778 | 0.977 | -0.001 | 0.02 | 0.02 | 0 | 0.12 | 0.11 | 0.15 | 0.2 | 0.09 | 0.13 |
| 1 | 97833918 | rs1879372 | 0.463 | -0.647 | 0.664 | 0.854 | -0.011 | -0.034 | -0.034 | 0 | 0.13 | 0.14 | 0.06 | 0.03 | 0.05 | 0.09 |
| 1 | 97835163 | rs7515904 | 1.144 | -0.693 | 0.057 | 0.721 | 0.002 | -0.025 | -0.025 | 0 | 0.44 | 0.31 | 0.57 | 0.51 | 0.46 | 0.28 |
| 1 | 97837761 | rs10875077 | -1.183 | -0.701 | 0.137 | 0.716 | 0.004 | -0.002 | -0.002 | 0.776 | 0.52 | 0.37 | 0.4 | 0.49 | 0.54 | 0.72 |
| 1 | 97838887 | rs11582179 | 1.262 | -0.658 | 0.082 | 0.44 | -0.027 | 0.031 | 0.031 | 0.32 | 0.17 | 0.11 | 0.13 | 0.04 | 0.26 | 0.19 |
| 1 | 97839016 | rs1890138 | -0.625 | -0.663 | -0.238 | 0.454 | -0.003 | 0.008 | 0.008 | 0 | 0.29 | 0.25 | 0.25 | 0.39 | 0.24 | 0.47 |
| 1 | 97841850 | rs12040045 | 0.869 | -0.667 | -0.244 | 0.466 | 0 | -0.004 | -0.004 | 0 | 0.14 | 0.09 | 0.09 | 0.19 | 0.15 | 0.32 |
| 1 | 97841879 | rs10875079 | 1.22 | -0.642 | -0.205 | 0.48 | -0.001 | -0.012 | -0.012 | 0 | 0.44 | 0.31 | 0.57 | 0.53 | 0.46 | 0.28 |
| 1 | 97844979 | rs34085458 | 0.466 | -0.623 | -0.254 | 0.49 | -0.106 | -0.075 | -0.075 | 0 | 0.22 | 0.03 | 0.23 | 0.22 | 0.31 | 0.24 |
| 1 | 97845053 | rs1415683 | -0.408 | -0.556 | -0.154 | 0.565 | -0.003 | 0.007 | 0.007 | 0 | 0.29 | 0.25 | 0.25 | 0.4 | 0.24 | 0.47 |
| 1 | 97849296 | rs10875080 | 0.268 | -0.547 | -0.149 | -0.203 | -0.022 | 0.044 | 0.044 | 0 | 0.23 | 0.17 | 0.15 | 0.08 | 0.3 | 0.26 |
| 1 | 97850697 | rs1112314 | 0.018 | -0.553 | -0.328 | -0.186 | 0.006 | 0.012 | 0.012 | 0 | 0.26 | 0.16 | 0.14 | 0.22 | 0.2 | 0.41 |
| 1 | 97852932 | rs6677502 | 0.018 | -0.553 | -0.328 | -0.186 | 0.006 | 0.012 | 0.012 | 0 | 0.26 | 0.16 | 0.14 | 0.22 | 0.2 | 0.42 |
| 1 | 97853443 | rs2152878 | 1.192 | -0.536 | -0.353 | -0.198 | -0.088 | 0.073 | 0.073 | 0.214 | 0.17 | 0.02 | 0.12 | 0.04 | 0.26 | 0.19 |
| 1 | 97853619 | rs6665429 | 1.964 | -0.647 | -0.401 | -0.319 | 0.002 | -0.019 | -0.019 | 0.002 | 0.31 | 0.3 | 0.21 | 0.04 | 0.31 | 0.11 |
| 1 | 97854625 | rs12040388 | -0.226 | -0.378 | -0.076 | 0.138 | 0.006 | 0.012 | 0.012 | 0 | 0.26 | 0.16 | 0.14 | 0.22 | 0.2 | 0.41 |
| 1 | 97855607 | rs4495747 | -0.226 | -0.374 | -0.043 | 0.181 | 0.006 | 0.012 | 0.012 | 0 | 0.26 | 0.16 | 0.14 | 0.22 | 0.2 | 0.42 |
| 1 | 97855821 | rs4503384 | -0.226 | -0.373 | -0.048 | 0.182 | 0.006 | 0.012 | 0.012 | 0 | 0.26 | 0.16 | 0.14 | 0.22 | 0.2 | 0.41 |
| 1 | 97856487 | rs7541618 | -0.677 | -0.374 | -0.046 | 0.184 | -0.006 | 0.015 | 0.015 | 0 | 0.29 | 0.27 | 0.25 | 0.4 | 0.23 | 0.47 |
| 1 | 97856946 | rs12021567 | -0.224 | -0.37 | -0.093 | 0.184 | 0.006 | 0.012 | 0.012 | 0 | 0.26 | 0.16 | 0.14 | 0.22 | 0.2 | 0.41 |
| 1 | 97857061 | rs12040763 | -0.224 | -0.37 | -0.093 | 0.184 | 0.006 | 0.012 | 0.012 | 0 | 0.26 | 0.16 | 0.14 | 0.22 | 0.2 | 0.41 |
| 1 | 97857145 | rs12043125 | -0.224 | -0.37 | -0.093 | 0.188 | 0.006 | 0.011 | 0.011 | 0 | 0.26 | 0.16 | 0.14 | 0.22 | 0.2 | 0.41 |
| 1 | 97857761 | rs955850 | 1.384 | -0.373 | -0.088 | 0.152 | 0 | -0.025 | -0.025 | 1.198 | 0.44 | 0.31 | 0.57 | 0.5 | 0.46 | 0.29 |
| 1 | 97860321 | rs7540201 | -0.764 | -0.271 | -0.272 | 0.366 | 0 | -0.001 | -0.001 | 0 | 0.29 | 0.24 | 0.25 | 0.4 | 0.23 | 0.47 |
| 1 | 97862237 | rs12566907 | -0.776 | -0.271 | -0.275 | 0.366 | -0.003 | 0.007 | 0.007 | 0 | 0.29 | 0.26 | 0.25 | 0.4 | 0.23 | 0.46 |
| 1 | 97862828 | rs12040699 | -0.392 | -0.274 | -0.393 | 0.234 | 0.006 | 0.012 | 0.012 | 0 | 0.26 | 0.16 | 0.14 | 0.22 | 0.2 | 0.4 |
| 1 | 97862883 | rs4428925 | -0.392 | -0.274 | -0.393 | 0.238 | 0.006 | 0.012 | 0.012 | 0 | 0.26 | 0.16 | 0.14 | 0.22 | 0.2 | 0.4 |
| 1 | 97863281 | rs4445510 | -0.384 | -0.277 | -0.394 | 0.229 | 0.006 | 0.011 | 0.011 | 0 | 0.26 | 0.16 | 0.14 | 0.22 | 0.2 | 0.4 |
| 1 | 97866719 | rs12047072 | -0.385 | -0.274 | -0.392 | 0.231 | 0.006 | 0.012 | 0.012 | 0 | 0.26 | 0.16 | 0.14 | 0.22 | 0.2 | 0.41 |
| 1 | 97867198 | rs1577897 | 0.329 | -0.289 | -0.388 | 0.345 | -0.007 | 0.025 | 0.025 | 0 | 0.15 | 0.17 | 0.16 | 0.2 | 0.09 | 0.15 |
| 1 | 97867433 | rs7512213 | 0.626 | -0.302 | -0.314 | 0.293 | -0.02 | -0.029 | -0.029 | 0.187 | 0.13 | 0.02 | 0.07 | 0.19 | 0.15 | 0.32 |
| 1 | 97867713 | rs7548189 | -0.518 | -0.314 | -0.309 | 0.284 | 0.006 | 0.012 | 0.012 | 0 | 0.26 | 0.16 | 0.14 | 0.22 | 0.2 | 0.41 |
| 1 | 97869199 | rs12725266 | -0.294 | -0.253 | -0.526 | 0.122 | -0.125 | 0.035 | 0.035 | 0 | 0.3 | 0.04 | 0.2 | 0.23 | 0.41 | 0.49 |
| 1 | 97869853 | rs12047625 | -0.567 | -0.202 | -0.287 | 0.315 | 0.006 | 0.012 | 0.012 | 0 | 0.26 | 0.16 | 0.14 | 0.22 | 0.2 | 0.4 |
| 1 | 97870258 | rs11165861 | -0.038 | -0.203 | -0.249 | 0.366 | -0.001 | -0.008 | -0.008 | 0 | 0.16 | 0.11 | 0.19 | 0.38 | 0.18 | 0.36 |
| 1 | 97870315 | rs12022771 | -0.593 | -0.212 | -0.255 | 0.354 | 0.007 | 0.012 | 0.012 | 0 | 0.26 | 0.16 | 0.14 | 0.22 | 0.2 | 0.4 |
| 1 | 97870432 | rs12022799 | 0.521 | -0.189 | -0.254 | 0.372 | -0.019 | -0.029 | -0.029 | 0.392 | 0.13 | 0.02 | 0.07 | 0.19 | 0.15 | 0.31 |
| 1 | 97870591 | rs12030174 | 0.546 | -0.192 | -0.255 | 0.371 | -0.019 | -0.029 | -0.029 | 0 | 0.13 | 0.02 | 0.07 | 0.19 | 0.15 | 0.31 |
| 1 | 97870876 | rs12048663 | 0.523 | -0.188 | -0.253 | 0.373 | -0.019 | -0.029 | -0.029 | 0.079 | 0.13 | 0.02 | 0.07 | 0.19 | 0.15 | 0.32 |
| 1 | 97872422 | rs11165863 | -0.999 | -0.095 | -0.384 | 0.164 | -0.002 | -0.009 | -0.009 | 1.239 | 0.3 | 0.25 | 0.17 | 0.22 | 0.23 | 0.43 |
| 1 | 97872655 | rs4300257 | -0.657 | -0.245 | -0.33 | 0.3 | 0.007 | 0.013 | 0.013 | 0 | 0.27 | 0.16 | 0.14 | 0.22 | 0.2 | 0.42 |
| 1 | 97873007 | rs4434871 | 0.277 | 0.059 | -0.203 | 0.681 | -0.019 | -0.029 | -0.029 | 0.431 | 0.13 | 0.02 | 0.07 | 0.19 | 0.15 | 0.32 |
| 1 | 97876372 | rs945879 | -0.542 | -0.04 | 0.037 | 0.648 | 0.003 | -0.001 | -0.001 | 0 | 0.2 | 0.16 | 0.11 | 0.26 | 0.19 | 0.38 |
| 1 | 97878068 | rs11165867 | 0.097 | 0.115 | 0.057 | 0.662 | -0.001 | -0.018 | -0.018 | 0 | 0.14 | 0.1 | 0.17 | 0.37 | 0.16 | 0.33 |
| 1 | 97880883 | rs12024396 | 0.833 | 0.132 | -0.719 | 0.6 | -0.013 | -0.018 | -0.018 | 0.04 | 0.42 | 0.32 | 0.57 | 0.53 | 0.47 | 0.28 |
| 1 | 97885040 | rs11165868 | -0.422 | 0.39 | -0.453 | 1.46 | -0.001 | -0.018 | -0.018 | 0 | 0.14 | 0.1 | 0.17 | 0.37 | 0.16 | 0.33 |
| 1 | 97885249 | rs10875085 | -0.815 | 0.378 | -0.418 | 1.423 | 0 | -0.006 | -0.006 | 0 | 0.17 | 0.15 | 0.18 | 0.38 | 0.18 | 0.35 |
| 1 | 97886171 | rs4949952 | -0.201 | 0.381 | -0.009 | 0.752 | -0.005 | -0.015 | -0.015 | 0 | 0.53 | 0.59 | 0.74 | 0.8 | 0.64 | 0.62 |

| | | | | | | | | | | | | | | | | |
|---|---|---|---|---|---|---|---|---|---|---|---|---|---|---|---|---|
| 1 | 97890347 | rs4950033 | -0.254 | 0.879 | 0.032 | 1.506 | 0.001 | 0.012 | 0.012 | 0 | 0.52 | 0.64 | 0.67 | 0.77 | 0.56 | 0.59 |
| 1 | 97891161 | rs12123121 | 0.992 | 1.159 | -0.261 | 1.228 | 0.008 | 0.033 | 0.033 | 0.082 | 0.25 | 0.17 | 0.15 | 0.05 | 0.25 | 0.1 |
| 1 | 97892858 | rs1556798 | -0.923 | 0.894 | -0.298 | 1.23 | -0.036 | -0.032 | -0.032 | 0 | 0.79 | 0.65 | 0.85 | 0.88 | 0.85 | 0.8 |
| 1 | 97895762 | rs11165870 | 0.004 | 0.942 | -0.437 | 1.38 | -0.005 | 0.069 | 0.069 | 0 | 0.44 | 0.34 | 0.31 | 0.23 | 0.48 | 0.4 |
| 1 | 97896489 | rs4353133 | -1.457 | 0.935 | -0.417 | 1.412 | -0.031 | -0.011 | -0.011 | 0.036 | 0.81 | 0.69 | 0.86 | 0.86 | 0.87 | 0.83 |
| 1 | 97896865 | rs12126093 | 1.308 | 1.033 | -0.449 | 1.347 | 0.005 | 0.028 | 0.028 | 0 | 0.24 | 0.15 | 0.16 | 0.06 | 0.29 | 0.09 |
| 1 | 97898232 | rs7517433 | -0.998 | 1.052 | -0.382 | 1.404 | -0.039 | -0.04 | -0.04 | 0.018 | 0.79 | 0.62 | 0.84 | 0.88 | 0.85 | 0.81 |
| 1 | 97898560 | rs10449723 | -1.454 | 1.053 | -0.385 | 1.386 | -0.031 | -0.024 | -0.024 | 0 | 0.81 | 0.69 | 0.86 | 0.89 | 0.87 | 0.83 |
| 1 | 97903111 | rs7516433 | 0.768 | 1.165 | -0.405 | 1.486 | -0.008 | 0.001 | 0.001 | 0 | 0.35 | 0.28 | 0.53 | 0.66 | 0.36 | 0.41 |
| 1 | 97903583 | rs12031561 | 1.065 | 1.152 | -0.416 | 1.474 | -0.011 | -0.016 | -0.016 | 0 | 0.33 | 0.2 | 0.47 | 0.55 | 0.35 | 0.38 |
| 1 | 97904780 | rs4949953 | -0.067 | 1.118 | -0.431 | 1.445 | -0.005 | 0.063 | 0.063 | 0 | 0.43 | 0.35 | 0.31 | 0.23 | 0.48 | 0.4 |
| 1 | 97907894 | rs10783063 | -1.334 | 1.129 | -0.402 | 1.719 | -0.033 | -0.015 | -0.015 | 0.954 | 0.81 | 0.68 | 0.86 | 0.87 | 0.86 | 0.83 |
| 1 | 97908878 | rs1415681 | -0.8 | 1.149 | -0.397 | 0.946 | -0.041 | -0.042 | -0.042 | 0.342 | 0.78 | 0.62 | 0.85 | 0.88 | 0.84 | 0.79 |
| 1 | 97912335 | rs4612663 | -1.233 | 1.112 | -0.383 | 0.89 | -0.034 | -0.016 | -0.016 | 0 | 0.81 | 0.69 | 0.86 | 0.87 | 0.86 | 0.81 |
| 1 | 97914806 | rs10747482 | -1.215 | 1.111 | -0.384 | 0.89 | -0.035 | -0.017 | -0.017 | 0 | 0.81 | 0.68 | 0.86 | 0.87 | 0.86 | 0.81 |
| 1 | 97915213 | rs11165875 | 0.922 | 1.137 | -0.398 | 0.878 | -0.008 | 0.001 | 0.001 | 0 | 0.35 | 0.26 | 0.54 | 0.66 | 0.37 | 0.4 |
| 1 | 97918306 | rs3897854 | 0.022 | 0.714 | -0.233 | 0.275 | -0.007 | 0.064 | 0.064 | 0 | 0.43 | 0.34 | 0.31 | 0.23 | 0.48 | 0.38 |
| 1 | 97920956 | rs12354219 | -0.142 | 0.736 | -0.203 | 0.264 | -0.006 | 0.09 | 0.09 | 0 | 0.44 | 0.33 | 0.31 | 0.23 | 0.48 | 0.41 |
| 1 | 97921434 | rs6593649 | -1.255 | 0.753 | -0.357 | 0.291 | -0.038 | -0.015 | -0.015 | 0 | 0.81 | 0.65 | 0.85 | 0.86 | 0.87 | 0.83 |
| 1 | 97925053 | rs7523116 | -1.224 | 0.849 | -0.365 | 0.335 | -0.034 | -0.012 | -0.012 | 0 | 0.81 | 0.69 | 0.86 | 0.85 | 0.86 | 0.81 |
| 1 | 97926839 | rs7550959 | 0.065 | 0.863 | -0.437 | 0.329 | -0.007 | 0.063 | 0.063 | 0 | 0.42 | 0.33 | 0.31 | 0.22 | 0.48 | 0.38 |
| 1 | 97938684 | rs10875091 | -1.093 | 0.807 | -0.34 | 0.241 | -0.039 | -0.016 | -0.016 | 1.45 | 0.81 | 0.65 | 0.86 | 0.86 | 0.86 | 0.83 |
| 1 | 97940231 | rs11811436 | 0.904 | 0.846 | -0.399 | 0.205 | -0.004 | 0.002 | 0.002 | 0 | 0.16 | 0.21 | 0.09 | 0.07 | 0.19 | 0.12 |
| 1 | 97944341 | rs12759539 | 1.133 | 0.592 | -0.492 | 0.23 | 0.004 | 0.01 | 0.01 | 0 | 0.23 | 0.16 | 0.15 | 0.05 | 0.27 | 0.08 |
| 1 | 97946840 | rs11165880 | -0.414 | 0.742 | -0.037 | 0.706 | -0.037 | -0.015 | -0.015 | 0.009 | 0.8 | 0.67 | 0.86 | 0.84 | 0.86 | 0.81 |
| 1 | 97947883 | rs11165881 | 0.602 | 0.782 | -0.352 | 0.753 | 0 | 0.001 | 0.001 | 0 | 0.58 | 0.37 | 0.71 | 0.58 | 0.59 | 0.51 |
| 1 | 97949375 | rs4495745 | -0.017 | 0.148 | -0.124 | 0.698 | -0.043 | -0.014 | -0.014 | 0 | 0.81 | 0.7 | 0.86 | 0.84 | 0.88 | 0.82 |
| 1 | 97949431 | rs4451581 | 0.06 | 0.337 | -0.092 | 0.718 | -0.026 | -0.013 | -0.013 | 0 | 0.8 | 0.67 | 0.84 | 0.84 | 0.84 | 0.81 |
| 1 | 97955273 | rs952501 | -0.575 | 0.751 | -0.199 | 1.14 | -0.001 | 0.005 | 0.005 | 0 | 0.13 | 0.13 | 0.27 | 0.27 | 0.19 | 0.08 |
| 1 | 97958108 | rs2811188 | -0.976 | 0.604 | -0.325 | 0.75 | 0.004 | 0.013 | 0.013 | 0 | 0.55 | 0.6 | 0.61 | 0.73 | 0.6 | 0.58 |
| 1 | 97959482 | rs2786779 | 0.251 | 0.605 | -0.326 | 0.751 | -0.001 | 0.005 | 0.005 | 0 | 0.13 | 0.14 | 0.27 | 0.27 | 0.19 | 0.08 |
| 1 | 97961206 | rs12062552 | 0.979 | 0.806 | -0.371 | 0.467 | -0.008 | 0 | 0 | 0 | 0.06 | 0.04 | 0.07 | 0.01 | 0.12 | 0.1 |
| 1 | 97962038 | rs2811187 | 0.414 | 0.805 | -0.371 | 0.467 | -0.001 | 0.005 | 0.005 | 0 | 0.13 | 0.13 | 0.27 | 0.27 | 0.19 | 0.08 |
| 1 | 97962979 | rs2786778 | 0.414 | 0.721 | -0.095 | 0.459 | -0.001 | 0.005 | 0.005 | 0 | 0.13 | 0.13 | 0.27 | 0.27 | 0.19 | 0.08 |
| 1 | 97963035 | rs2786776 | -1.327 | 0.718 | -0.124 | 0.367 | 0.003 | 0.01 | 0.01 | 0 | 0.57 | 0.63 | 0.62 | 0.74 | 0.64 | 0.62 |
| 1 | 97966205 | rs4520446 | -0.669 | 0.765 | -0.124 | 0.367 | 0.001 | 0.011 | 0.011 | 0 | 0.48 | 0.56 | 0.54 | 0.72 | 0.49 | 0.48 |
| 1 | 97967983 | rs10493892 | 1.04 | 0.741 | -0.125 | 0.367 | -0.008 | 0 | 0 | 0 | 0.06 | 0.04 | 0.07 | 0.01 | 0.12 | 0.1 |
| 1 | 97968615 | rs2786782 | -1.318 | 0.739 | -0.126 | 0.367 | 0.001 | 0.004 | 0.004 | 0 | 0.57 | 0.61 | 0.62 | 0.73 | 0.64 | 0.62 |
| 1 | 97971215 | rs11165886 | 0.389 | 0.701 | 0.168 | 0.381 | 0.007 | -0.021 | -0.021 | 0 | 0.34 | 0.24 | 0.27 | 0.45 | 0.3 | 0.4 |
| 1 | 97971252 | rs2811183 | 0.531 | 0.693 | 0.166 | 0.368 | -0.001 | 0.005 | 0.005 | 0 | 0.13 | 0.13 | 0.27 | 0.27 | 0.19 | 0.08 |
| 1 | 97971440 | rs2811182 | -1.231 | 0.693 | 0.155 | 0.357 | 0.004 | 0.014 | 0.014 | 0 | 0.56 | 0.63 | 0.62 | 0.73 | 0.62 | 0.61 |
| 1 | 97972686 | rs10875094 | 0.506 | 0.663 | 0.118 | 0.352 | 0.006 | -0.02 | -0.02 | 0 | 0.34 | 0.24 | 0.27 | 0.44 | 0.3 | 0.4 |
| 1 | 97973252 | rs17116806 | 0.592 | 0.688 | 0.411 | 0.363 | -0.003 | -0.006 | -0.006 | 0 | 0.13 | 0.1 | 0.26 | 0.27 | 0.19 | 0.08 |
| 1 | 97973891 | rs2786780 | 0.508 | 0.687 | 0.398 | 0.365 | 0 | 0.006 | 0.006 | 0 | 0.13 | 0.13 | 0.26 | 0.27 | 0.19 | 0.08 |
| 1 | 97974234 | rs12087910 | 1.033 | 0.723 | 0.386 | 0.353 | -0.007 | 0 | 0 | 0 | 0.06 | 0.04 | 0.07 | 0.01 | 0.12 | 0.1 |
| 1 | 97974659 | rs11165887 | 0.435 | 0.684 | 0.402 | 0.348 | 0.006 | -0.02 | -0.02 | 0.014 | 0.34 | 0.24 | 0.27 | 0.44 | 0.3 | 0.4 |
| 1 | 97975461 | rs11165888 | 1.046 | 0.697 | 0.429 | 0.347 | -0.005 | -0.003 | -0.003 | 0.193 | 0.06 | 0.04 | 0.07 | 0.01 | 0.12 | 0.1 |
| 1 | 97975700 | rs868184 | -1.315 | 0.613 | 0.712 | 0.308 | 0.007 | 0.036 | 0.036 | 0 | 0.57 | 0.7 | 0.62 | 0.73 | 0.62 | 0.61 |
| 1 | 97979726 | rs7545340 | 0.43 | 0.67 | 0.536 | 0.339 | 0.006 | -0.02 | -0.02 | 0.17 | 0.34 | 0.24 | 0.27 | 0.44 | 0.3 | 0.4 |
| 1 | 97981395 | rs1801159 | 0.479 | 0.67 | 0.563 | 0.355 | 0.001 | 0.014 | 0.014 | 0 | 0.13 | 0.15 | 0.27 | 0.27 | 0.19 | 0.08 |
| 1 | 97985966 | rs2105129 | -1.185 | 0.663 | 0.548 | 0.208 | 0.004 | 0.014 | 0.014 | 0 | 0.56 | 0.63 | 0.63 | 0.73 | 0.63 | 0.61 |
| 1 | 97989751 | rs12061124 | 1.03 | 0.673 | 0.495 | 0.217 | -0.008 | 0 | 0 | 0 | 0.06 | 0.04 | 0.07 | 0.01 | 0.12 | 0.1 |
| 1 | 97990575 | rs2811174 | -1.26 | 0.697 | 0.142 | 0.219 | 0.003 | 0.011 | 0.011 | 0 | 0.57 | 0.63 | 0.69 | 0.74 | 0.63 | 0.62 |
| 1 | 97991219 | rs2786770 | -1.325 | 0.674 | 0.15 | 0.226 | 0 | 0.001 | 0.001 | 0 | 0.57 | 0.62 | 0.69 | 0.73 | 0.64 | 0.62 |
| 1 | 97991606 | rs17368837 | 1.06 | 0.712 | 0.151 | 0.23 | -0.007 | 0 | 0 | 0 | 0.06 | 0.04 | 0.07 | 0.01 | 0.12 | 0.1 |
| 1 | 97992257 | rs2811173 | -0.655 | 0.669 | 0.162 | 0.228 | 0 | 0.001 | 0.001 | 0 | 0.48 | 0.54 | 0.62 | 0.71 | 0.51 | 0.52 |
| 1 | 97992465 | rs2786512 | -1.178 | 0.379 | 0.206 | 0.292 | 0.002 | 0.005 | 0.005 | 0 | 0.56 | 0.62 | 0.69 | 0.73 | 0.63 | 0.62 |
| 1 | 97999687 | rs1812389 | 0.948 | -0.447 | 0.213 | -0.007 | -0.006 | -0.017 | -0.017 | 0.971 | 0.17 | 0.19 | 0.07 | 0.03 | 0.07 | 0.06 |
| 1 | 98000861 | rs2786519 | -0.601 | -0.547 | 0.172 | 0.104 | -0.034 | -0.006 | -0.006 | 0 | 0.29 | 0.5 | 0.36 | 0.24 | 0.23 | 0.09 |
| 1 | 98000954 | rs12729863 | 0.039 | -0.478 | -0.386 | 0.129 | 0 | 0.007 | 0.007 | 0 | 0.13 | 0.13 | 0.28 | 0.23 | 0.17 | 0.08 |
| 1 | 98001389 | rs12063030 | -0.465 | -0.478 | -0.389 | 0.084 | -0.007 | -0.005 | -0.005 | 0 | 0.28 | 0.32 | 0.34 | 0.22 | 0.23 | 0.09 |
| 1 | 98001608 | rs12089969 | 0.65 | -0.374 | -0.51 | -0.169 | -0.006 | -0.017 | -0.017 | 0.15 | 0.17 | 0.18 | 0.07 | 0.03 | 0.07 | 0.06 |
| 1 | 98006222 | rs12756513 | -0.156 | -0.122 | -0.992 | 0.308 | 0.01 | -0.052 | -0.052 | 0 | 0.27 | 0.13 | 0.19 | 0.42 | 0.29 | 0.14 |
| 1 | 98007685 | rs2811169 | -0.728 | 0.363 | -0.677 | 0.032 | -0.006 | -0.004 | -0.004 | 0 | 0.08 | 0.08 | 0.07 | 0 | 0.13 | 0.09 |
| 1 | 98010949 | rs17368985 | -0.127 | -0.123 | -1.065 | -0.253 | 0.034 | 0.08 | 0.08 | 0 | 0.15 | 0.01 | 0.07 | 0.04 | 0.11 | 0.04 |
| 1 | 98013638 | rs7530858 | -0.174 | -0.207 | -0.529 | -0.29 | -0.003 | -0.008 | -0.008 | 0.999 | 0.16 | 0.19 | 0.08 | 0.06 | 0.12 | 0.05 |
| 1 | 98017090 | rs2786529 | -1.52 | -0.163 | -0.432 | -0.327 | -0.004 | -0.054 | -0.054 | 0 | 0.25 | 0.37 | 0.15 | 0.06 | 0.23 | 0.14 |
| 1 | 98017233 | rs7540983 | -0.178 | -0.108 | -0.28 | -0.301 | -0.005 | -0.01 | -0.01 | 0.312 | 0.16 | 0.19 | 0.08 | 0.06 | 0.11 | 0.05 |
| 1 | 98020955 | rs17100624 | 0.34 | -0.299 | -0.432 | -0.159 | 0.012 | -0.033 | -0.033 | 0.014 | 0.15 | 0.07 | 0.34 | 0.66 | 0.12 | 0.38 |

| | | | | | | | | | | | | | | | | |
|---|---|---|---|---|---|---|---|---|---|---|---|---|---|---|---|---|
| 1 | 98023377 | rs17116920 | -0.471 | 0.132 | 0.007 | -0.329 | -0.005 | -0.009 | -0.009 | 5.555 | 0.16 | 0.2 | 0.08 | 0.06 | 0.11 | 0.05 |
| 1 | 98026230 | rs12066229 | -0.432 | 0.129 | 0 | -0.339 | -0.007 | -0.012 | -0.012 | 0 | 0.16 | 0.21 | 0.07 | 0.06 | 0.11 | 0.05 |
| 1 | 98026801 | rs2811219 | -1.354 | 0.097 | -0.008 | -0.303 | -0.003 | -0.053 | -0.053 | 0 | 0.24 | 0.37 | 0.15 | 0.06 | 0.23 | 0.14 |
| 1 | 98035253 | rs11165895 | 0.853 | -0.43 | 0.685 | -0.32 | -0.049 | -0.129 | -0.129 | 0.642 | 0.77 | 0.49 | 0.86 | 0.96 | 0.84 | 0.9 |
| 1 | 98036832 | rs2811215 | -0.618 | -0.471 | 0.871 | -0.345 | 0.005 | -0.02 | -0.02 | 0 | 0.18 | 0.18 | 0.09 | 0.02 | 0.15 | 0.07 |
| 1 | 98036858 | rs2811214 | -0.302 | -0.482 | 0.826 | -0.349 | 0.014 | 0.004 | 0.004 | 0 | 0.16 | 0.13 | 0.09 | 0.02 | 0.13 | 0.05 |
| 1 | 98038179 | rs4949958 | 1.176 | -0.53 | 0.906 | -0.373 | -0.054 | -0.124 | -0.124 | 0 | 0.77 | 0.52 | 0.86 | 0.96 | 0.84 | 0.9 |
| 1 | 98040871 | rs34961061 | 0.682 | -0.469 | 1.109 | -0.62 | 0 | 0 | 0 | 0 | 0.14 | 0.13 | 0.11 | 0.1 | 0.15 | 0.11 |
| 1 | 98042712 | rs10875099 | 0.904 | -0.469 | 1.15 | -0.639 | -0.052 | -0.126 | -0.126 | 0 | 0.77 | 0.5 | 0.86 | 0.96 | 0.84 | 0.9 |
| 1 | 98044939 | rs6663357 | 0.889 | -0.47 | 1.151 | -0.638 | -0.055 | -0.126 | -0.126 | 0.105 | 0.77 | 0.5 | 0.86 | 0.96 | 0.84 | 0.9 |
| 1 | 98046492 | rs2811211 | 1.559 | -0.42 | 1.175 | -0.592 | 0.029 | 0.026 | 0.026 | 0.326 | 0.93 | 0.63 | 0.95 | 0.98 | 0.97 | 0.96 |
| 1 | 98056463 | rs7537165 | 0.884 | -0.505 | 1.384 | -0.646 | -0.054 | -0.127 | -0.127 | 0 | 0.76 | 0.5 | 0.86 | 0.96 | 0.84 | 0.89 |
| 1 | 98056575 | rs11165897 | -0.061 | -0.504 | 1.486 | -0.646 | 0.014 | 0.001 | 0.001 | 0 | 0.17 | 0.13 | 0.09 | 0.02 | 0.13 | 0.06 |
| 1 | 98057024 | rs2786490 | -0.175 | -0.504 | 1.488 | -0.646 | -0.009 | -0.05 | -0.05 | 0 | 0.17 | 0.2 | 0.09 | 0.02 | 0.13 | 0.06 |
| 1 | 98057444 | rs2811203 | -0.448 | -0.504 | 1.489 | -0.646 | -0.005 | -0.048 | -0.048 | 0.037 | 0.19 | 0.22 | 0.1 | 0.02 | 0.16 | 0.08 |
| 1 | 98064363 | rs10783070 | 0.877 | -0.51 | 1.488 | -0.658 | -0.055 | -0.124 | -0.124 | 0.078 | 0.77 | 0.51 | 0.86 | 0.96 | 0.84 | 0.89 |
| 1 | 98067147 | rs2786498 | -0.451 | -0.496 | 1.425 | -0.654 | -0.005 | -0.048 | -0.048 | 0.037 | 0.19 | 0.22 | 0.1 | 0.02 | 0.16 | 0.08 |
| 1 | 98067668 | rs2811200 | -0.178 | -0.497 | 1.425 | -0.653 | -0.01 | -0.05 | -0.05 | 0.267 | 0.17 | 0.2 | 0.09 | 0.02 | 0.13 | 0.06 |
| 1 | 98067896 | rs2811199 | -0.57 | -0.503 | 1.42 | -0.658 | -0.004 | -0.045 | -0.045 | 0.004 | 0.19 | 0.22 | 0.1 | 0.02 | 0.16 | 0.08 |
| 1 | 98068801 | rs2811198 | -0.181 | -0.48 | 1.432 | -0.646 | -0.009 | -0.05 | -0.05 | 0 | 0.17 | 0.2 | 0.09 | 0.02 | 0.13 | 0.06 |
| 1 | 98071426 | rs2065943 | 0.824 | -0.489 | 1.447 | -0.648 | -0.049 | -0.12 | -0.12 | 0 | 0.77 | 0.52 | 0.86 | 0.96 | 0.84 | 0.89 |
| 1 | 98073410 | rs12144462 | 1.559 | -0.676 | 1.09 | -0.238 | -0.001 | -0.06 | -0.06 | 0 | 0.1 | 0.02 | 0.05 | 0.24 | 0.08 | 0.1 |
| 1 | 98075724 | rs2786500 | -0.258 | -0.656 | 1.157 | -0.247 | -0.005 | -0.048 | -0.048 | 0 | 0.19 | 0.22 | 0.1 | 0.02 | 0.16 | 0.08 |
| 1 | 98075725 | rs2786501 | 0.017 | -0.682 | 1.157 | -0.246 | -0.01 | -0.05 | -0.05 | 0 | 0.17 | 0.2 | 0.09 | 0.02 | 0.13 | 0.06 |
| 1 | 98075762 | rs7554746 | 0.832 | -0.34 | 1.516 | -0.511 | 0.036 | -0.034 | -0.034 | 0.038 | 0.08 | 0.14 | 0.15 | 0.01 | 0.1 | 0.07 |
| 1 | 98078993 | rs2811196 | -0.078 | -0.504 | 1.338 | -0.43 | 0 | -0.037 | -0.037 | 0 | 0.19 | 0.2 | 0.1 | 0.02 | 0.16 | 0.08 |
| 1 | 98079228 | rs7533902 | 0.934 | -0.519 | 1.362 | -0.426 | -0.039 | -0.007 | -0.007 | 0 | 0.67 | 0.49 | 0.81 | 0.72 | 0.75 | 0.8 |
| 1 | 98079674 | rs12033263 | 0.829 | -0.329 | 1.486 | -0.834 | 0 | 0.001 | 0.001 | 0 | 0.15 | 0.13 | 0.12 | 0.13 | 0.15 | 0.11 |
| 1 | 98080308 | rs2786505 | 0.005 | -0.321 | 0.979 | -0.839 | -0.004 | -0.039 | -0.039 | 0 | 0.17 | 0.18 | 0.09 | 0.02 | 0.13 | 0.06 |
| 1 | 98080718 | rs2811195 | -0.231 | -0.319 | 0.983 | -0.831 | -0.003 | -0.042 | -0.042 | 0.375 | 0.19 | 0.21 | 0.1 | 0.02 | 0.15 | 0.08 |
| 1 | 98085570 | rs17369442 | 0.915 | 0.173 | 0.525 | -0.961 | -0.038 | 0.038 | 0.038 | 0.137 | 0.07 | 0.01 | 0.05 | 0 | 0.14 | 0.11 |
| 1 | 98087725 | rs2811193 | 0.563 | -0.409 | 0.864 | -0.751 | -0.056 | -0.124 | -0.124 | 0 | 0.76 | 0.51 | 0.86 | 0.96 | 0.84 | 0.89 |
| 1 | 98095024 | rs1337522 | 0.457 | -0.392 | 0.505 | -0.695 | -0.043 | -0.101 | -0.101 | 0 | 0.77 | 0.55 | 0.86 | 0.96 | 0.84 | 0.89 |
| 1 | 98098879 | rs10518636 | 0.784 | 0.147 | -0.217 | -0.648 | -0.038 | -0.016 | -0.016 | 0 | 0.13 | 0.01 | 0.11 | 0.12 | 0.15 | 0.11 |
| 1 | 98105438 | rs6663670 | -0.56 | 0.034 | 0.359 | -0.785 | -0.01 | -0.065 | -0.065 | 0 | 0.19 | 0.2 | 0.1 | 0.02 | 0.15 | 0.09 |
| 1 | 98106812 | rs7512910 | -0.522 | 0.025 | 0.371 | -0.789 | 0.007 | -0.016 | -0.016 | 0 | 0.19 | 0.14 | 0.09 | 0.02 | 0.15 | 0.09 |
| 1 | 98109958 | rs10518637 | 0.823 | 0.07 | 0.473 | -0.806 | 0.002 | -0.021 | -0.021 | 0 | 0.05 | 0.06 | 0.04 | 0 | 0.06 | 0.01 |
| 1 | 98118320 | rs2786507 | 0.214 | 0.034 | 0.511 | -0.832 | 0.007 | -0.035 | -0.035 | 0 | 0.08 | 0.14 | 0.14 | 0.01 | 0.07 | 0.07 |
| 1 | 98126717 | rs13375516 | -0.768 | -0.055 | 0.027 | -0.711 | -0.014 | -0.076 | -0.076 | 0 | 0.2 | 0.23 | 0.09 | 0.02 | 0.15 | 0.1 |
| 1 | 98143266 | rs2151563 | 0.006 | -0.033 | -0.32 | -0.677 | -0.004 | -0.037 | -0.037 | 2.772 | 0.84 | 0.85 | 0.93 | 0.98 | 0.88 | 0.94 |
| 1 | 98143293 | rs2786509 | -0.075 | -0.028 | -0.331 | -0.675 | 0.009 | -0.003 | -0.003 | 0 | 0.85 | 0.91 | 0.93 | 0.98 | 0.88 | 0.94 |
| 1 | 98156892 | rs3790389 | -0.61 | -0.037 | -0.318 | -0.698 | 0.002 | -0.026 | -0.026 | 0 | 0.17 | 0.15 | 0.07 | 0.02 | 0.15 | 0.09 |
| 1 | 98159558 | rs10875102 | 1.094 | 0.155 | -0.455 | -0.521 | -0.025 | 0.089 | 0.089 | 0 | 0.49 | 0.16 | 0.57 | 0.69 | 0.49 | 0.54 |
| 1 | 98164768 | rs6668296 | -0.687 | 0.26 | -0.216 | -0.589 | -0.009 | -0.057 | -0.057 | 0 | 0.18 | 0.2 | 0.08 | 0.02 | 0.14 | 0.09 |
| 1 | 98165091 | rs2297595 | 0.091 | 0.281 | -0.338 | -0.573 | 0.009 | 0.08 | 0.08 | 0 | 0.14 | 0.03 | 0.06 | 0.02 | 0.12 | 0.06 |
| 1 | 98194845 | rs17378455 | 0.05 | 0.258 | -0.706 | -0.544 | 0.024 | 0.037 | 0.037 | 0 | 0.14 | 0.05 | 0.05 | 0.01 | 0.1 | 0.06 |
| 1 | 98198206 | rs4554755 | 1.484 | 0.113 | -1.089 | -0.385 | -0.096 | -0.138 | -0.138 | 0 | 0.31 | 0.02 | 0.25 | 0.42 | 0.31 | 0.31 |
| 1 | 98200719 | rs6683957 | 0.963 | 0.2 | 0.087 | -0.529 | -0.034 | -0.106 | -0.106 | 2.632 | 0.81 | 0.67 | 0.93 | 0.99 | 0.88 | 0.92 |
| 1 | 98203820 | rs17369919 | -0.514 | 0.312 | -0.905 | -0.367 | 0.024 | 0.038 | 0.038 | 0 | 0.14 | 0.06 | 0.05 | 0.01 | 0.1 | 0.06 |
| 1 | 98209110 | rs17378539 | -0.236 | 1.045 | -0.902 | -0.242 | -0.045 | 0.042 | 0.042 | 0.354 | 0.08 | 0.01 | 0.09 | 0 | 0.18 | 0.12 |
| 1 | 98230887 | rs17378671 | -0.896 | 0.501 | -0.072 | -0.647 | -0.012 | 0.045 | 0.045 | 0.135 | 0.17 | 0.07 | 0.12 | 0.02 | 0.21 | 0.17 |
| 1 | 98241291 | rs12739159 | -1.077 | 0.449 | -0.08 | -0.674 | -0.003 | 0.013 | 0.013 | 0.329 | 0.18 | 0.14 | 0.13 | 0.02 | 0.21 | 0.16 |
| 1 | 98244251 | rs12565244 | 0.677 | 0.446 | -0.046 | -0.676 | -0.004 | 0.006 | 0.006 | 0.12 | 0.82 | 0.87 | 0.88 | 0.98 | 0.79 | 0.84 |
| 1 | 98253405 | rs7527085 | 0.677 | 0.389 | -0.072 | -0.624 | -0.004 | 0.006 | 0.006 | 0.12 | 0.82 | 0.87 | 0.88 | 0.98 | 0.79 | 0.84 |
| 1 | 98253666 | rs10747485 | 0.693 | 0.38 | -0.076 | -0.628 | -0.004 | 0.006 | 0.006 | 0.066 | 0.82 | 0.87 | 0.88 | 0.98 | 0.79 | 0.84 |
| 1 | 98253738 | rs10747486 | 1.426 | 0.313 | 0.244 | -0.667 | -0.003 | -0.125 | -0.125 | 2.227 | 0.77 | 0.6 | 0.84 | 0.98 | 0.77 | 0.82 |
| 1 | 98254702 | rs10747487 | 0.898 | 0.316 | 0.232 | -0.663 | -0.001 | -0.056 | -0.056 | 0.001 | 0.8 | 0.76 | 0.87 | 0.98 | 0.79 | 0.83 |
| 1 | 98264578 | rs1333717 | 1.486 | 0.295 | 0.604 | -0.676 | -0.003 | -0.147 | -0.147 | 0 | 0.77 | 0.56 | 0.84 | 0.98 | 0.77 | 0.82 |
| 1 | 98265667 | rs10493894 | 0.639 | 0.297 | 0.552 | -0.675 | -0.001 | -0.027 | -0.027 | 0.026 | 0.82 | 0.82 | 0.87 | 0.98 | 0.79 | 0.84 |
| 1 | 98267427 | rs17379095 | -0.806 | 0.296 | 0.541 | -0.675 | -0.012 | 0.045 | 0.045 | 0.103 | 0.17 | 0.07 | 0.12 | 0.02 | 0.21 | 0.16 |
| 1 | 98268137 | rs6703307 | 0.573 | 0.296 | 0.541 | -0.675 | -0.004 | 0.005 | 0.005 | 0.113 | 0.82 | 0.87 | 0.88 | 0.98 | 0.79 | 0.84 |
| 1 | 98274132 | rs12568335 | 1.845 | 0.103 | 0.494 | -0.512 | 0.007 | 0.088 | 0.088 | 2.153 | 0.54 | 0.12 | 0.6 | 0.7 | 0.56 | 0.57 |
| 1 | 98279091 | rs10493896 | -0.688 | 0.327 | 0.438 | -0.559 | -0.015 | 0.047 | 0.047 | 0.068 | 0.18 | 0.07 | 0.12 | 0.02 | 0.22 | 0.16 |
| 1 | 98283965 | rs1931066 | 0.598 | 0.332 | 0.513 | -0.557 | 0.001 | -0.053 | -0.053 | 0.059 | 0.81 | 0.78 | 0.87 | 0.98 | 0.79 | 0.84 |
| 1 | 98284940 | rs1034215 | 0.952 | 0.321 | 0.526 | -0.562 | -0.003 | -0.112 | -0.112 | 0 | 0.79 | 0.64 | 0.86 | 0.98 | 0.79 | 0.83 |
| 1 | 98285320 | rs1034216 | 0.578 | 0.328 | 0.522 | -0.559 | 0.001 | -0.053 | -0.053 | 0.234 | 0.81 | 0.78 | 0.87 | 0.98 | 0.79 | 0.84 |
| 1 | 98285422 | rs1034217 | 0.371 | 0.329 | 0.515 | -0.558 | -0.004 | 0.005 | 0.005 | 0.113 | 0.82 | 0.87 | 0.88 | 0.98 | 0.79 | 0.84 |
| 1 | 98287474 | rs4329543 | 0.445 | 0.329 | 0.515 | -0.558 | -0.005 | 0.007 | 0.007 | 0.074 | 0.82 | 0.87 | 0.87 | 0.98 | 0.78 | 0.83 |
| 1 | 98288517 | rs1023245 | 0.445 | 0.392 | 0.515 | -0.558 | -0.006 | 0.007 | 0.007 | 0.074 | 0.82 | 0.87 | 0.87 | 0.98 | 0.78 | 0.83 |

| | | | | | | | | | | | | | | | | |
|---|---|---|---|---|---|---|---|---|---|---|---|---|---|---|---|---|
| 1 | 98290238 | rs12134028 | 2.066 | 0.317 | 0.33 | -0.621 | -0.013 | -0.005 | -0.005 | 3.873 | 0.06 | 0 | 0.04 | 0.04 | 0.05 | 0.01 |
| 1 | 98290577 | rs1974204 | 0.906 | 0.406 | 0.506 | -0.554 | -0.003 | -0.112 | -0.112 | 0 | 0.79 | 0.64 | 0.86 | 0.98 | 0.79 | 0.83 |
| 1 | 98290685 | rs1974203 | 1.278 | 0.401 | 0.509 | -0.555 | 0 | -0.133 | -0.133 | 0.02 | 0.76 | 0.59 | 0.84 | 0.98 | 0.76 | 0.82 |
| 1 | 98292937 | rs10157611 | 0.518 | 0.433 | 0.239 | -0.507 | -0.003 | -0.008 | -0.008 | 0 | 0.82 | 0.85 | 0.87 | 0.98 | 0.78 | 0.83 |
| 1 | 98299475 | rs10747488 | 1.23 | 0.184 | 0.311 | -0.38 | -0.024 | -0.075 | -0.075 | 6.566 | 0.77 | 0.61 | 0.77 | 0.91 | 0.77 | 0.73 |
| 1 | 98305394 | rs7526108 | 1.212 | 0.156 | 0.413 | -0.394 | -0.015 | -0.049 | -0.049 | 0 | 0.77 | 0.67 | 0.77 | 0.91 | 0.77 | 0.73 |
| 1 | 98316285 | rs4421623 | 1.232 | 0.13 | 0.417 | -0.392 | -0.028 | -0.075 | -0.075 | 0 | 0.77 | 0.6 | 0.77 | 0.91 | 0.78 | 0.73 |
| 1 | 98322379 | rs4379706 | 1.235 | 0.116 | 0.68 | -0.399 | -0.032 | -0.087 | -0.087 | 0 | 0.77 | 0.56 | 0.77 | 0.91 | 0.78 | 0.73 |
| 1 | 98327831 | rs4523551 | 1.183 | 0.114 | 0.72 | -0.4 | -0.024 | -0.069 | -0.069 | 0 | 0.77 | 0.63 | 0.78 | 0.91 | 0.78 | 0.73 |
| 1 | 98332868 | rs11590802 | 0.912 | 0.027 | 1 | -0.623 | -0.001 | -0.016 | -0.016 | 6.787 | 0.07 | 0.05 | 0.04 | 0 | 0.08 | 0.03 |
| 1 | 98334990 | rs11165921 | 1.022 | 0.028 | 0.961 | -0.615 | -0.024 | -0.069 | -0.069 | 0 | 0.78 | 0.63 | 0.78 | 0.91 | 0.79 | 0.73 |
| 1 | 98336329 | rs9661794 | 1.118 | 0.028 | 0.961 | -0.615 | -0.031 | -0.086 | -0.086 | 0 | 0.77 | 0.56 | 0.78 | 0.91 | 0.79 | 0.73 |
| 1 | 98336823 | rs11165922 | 2.183 | 0.236 | 1.017 | -0.432 | 0.014 | -0.002 | -0.002 | 0 | 0.57 | 0.09 | 0.62 | 0.62 | 0.59 | 0.48 |
| 1 | 98337283 | rs6677116 | 0.978 | 0.223 | 1.125 | -0.46 | -0.031 | -0.086 | -0.086 | 1.146 | 0.77 | 0.56 | 0.78 | 0.91 | 0.79 | 0.73 |
| 1 | 98341152 | rs10747491 | 0.978 | 0.249 | 1.125 | -0.462 | -0.031 | -0.086 | -0.086 | 0 | 0.77 | 0.56 | 0.78 | 0.91 | 0.78 | 0.73 |
| 1 | 98342417 | rs10747492 | 0.897 | 0.249 | 1.129 | -0.462 | -0.024 | -0.069 | -0.069 | 0 | 0.78 | 0.63 | 0.78 | 0.91 | 0.79 | 0.73 |
| 1 | 98345019 | rs7524038 | 0.897 | 0.273 | 1.224 | -0.462 | -0.024 | -0.069 | -0.069 | 0.635 | 0.78 | 0.63 | 0.78 | 0.91 | 0.79 | 0.73 |
| 1 | 98346177 | rs10875113 | 0.957 | 0.273 | 1.223 | -0.462 | -0.031 | -0.086 | -0.086 | 0 | 0.77 | 0.56 | 0.78 | 0.91 | 0.79 | 0.73 |
| 1 | 98348885 | rs1801265 | 0.979 | 0.273 | 1.235 | -0.462 | -0.031 | -0.086 | -0.086 | 0.065 | 0.77 | 0.56 | 0.78 | 0.91 | 0.79 | 0.73 |
| 1 | 98362040 | rs7418577 | 0.882 | 0.273 | 1.235 | -0.306 | -0.024 | -0.069 | -0.069 | 0 | 0.78 | 0.63 | 0.78 | 0.91 | 0.79 | 0.73 |
| 1 | 98365310 | rs10875118 | 0.957 | 0.273 | 1.234 | -0.306 | -0.031 | -0.086 | -0.086 | 0.371 | 0.77 | 0.56 | 0.78 | 0.91 | 0.79 | 0.73 |
| 1 | 98369744 | rs9662719 | 0.747 | 0.271 | 1.254 | -0.308 | -0.012 | -0.04 | -0.04 | 0 | 0.79 | 0.71 | 0.78 | 0.91 | 0.79 | 0.74 |
| 1 | 98371201 | rs6604094 | 0.753 | 0.233 | 1.685 | -0.163 | -0.003 | 0.028 | 0.028 | 0 | 0.05 | 0.16 | 0.04 | 0.08 | 0.04 | 0.07 |
| 1 | 98371449 | rs4394694 | 0.934 | 0.258 | 1.192 | -0.22 | -0.024 | -0.069 | -0.069 | 1.41 | 0.78 | 0.64 | 0.78 | 0.91 | 0.79 | 0.73 |
| 1 | 98376245 | rs9727976 | 0.787 | 0.544 | 1.241 | -0.284 | -0.005 | -0.034 | -0.034 | 0 | 0.78 | 0.72 | 0.78 | 0.91 | 0.78 | 0.72 |
| 1 | 98378786 | rs9662870 | 1.123 | 0.547 | 1.223 | -0.282 | -0.02 | -0.092 | -0.092 | 0.814 | 0.76 | 0.55 | 0.77 | 0.91 | 0.77 | 0.71 |
| 1 | 98382114 | rs6692580 | 1.137 | 0.547 | 1.221 | -0.282 | -0.02 | -0.092 | -0.092 | 0 | 0.76 | 0.56 | 0.77 | 0.91 | 0.77 | 0.71 |
| 1 | 98394864 | rs4970725 | 1.176 | 0.722 | 1.237 | -0.251 | 0.013 | -0.076 | -0.076 | 0.063 | 0.8 | 0.56 | 0.79 | 0.91 | 0.78 | 0.72 |
| 1 | 98395881 | rs4378243 | 1.046 | 0.69 | 1.109 | -0.282 | 0.037 | -0.018 | -0.018 | 0 | 0.89 | 0.75 | 0.9 | 0.93 | 0.84 | 0.81 |
| 1 | 98417967 | rs11165934 | 1.717 | 0.731 | 1.151 | -0.285 | 0.017 | -0.073 | -0.073 | 0.15 | 0.8 | 0.57 | 0.79 | 0.91 | 0.77 | 0.72 |
| 1 | 98423149 | rs9659380 | 0.845 | 0.734 | 1.137 | -0.283 | 0.029 | -0.01 | -0.01 | 0 | 0.91 | 0.82 | 0.9 | 0.93 | 0.84 | 0.82 |
| 1 | 98436232 | rs12067700 | -2.065 | 0.728 | 1.084 | -0.284 | 0.005 | -0.043 | -0.043 | 0 | 0.19 | 0.31 | 0.21 | 0.09 | 0.23 | 0.28 |
| 1 | 98461443 | rs9662365 | -1.231 | 0.644 | 0.281 | -0.667 | 0.004 | -0.001 | -0.001 | 0 | 0.44 | 0.51 | 0.57 | 0.41 | 0.48 | 0.38 |
| 1 | 98466786 | rs12138817 | 0.572 | 0.677 | -0.271 | -0.66 | -0.074 | 0.037 | 0.037 | 2.802 | 0.24 | 0.03 | 0.38 | 0.12 | 0.28 | 0.14 |
| 1 | 98475522 | rs2154402 | -0.964 | 0.778 | -0.226 | -0.581 | -0.032 | -0.017 | -0.017 | 0 | 0.95 | 0.99 | 0.94 | 0.93 | 0.88 | 0.86 |
| 1 | 98480212 | rs11165944 | 1.71 | 0.713 | -0.224 | -0.598 | 0.001 | -0.093 | -0.093 | 0 | 0.43 | 0.11 | 0.32 | 0.52 | 0.35 | 0.43 |
| 1 | 98481750 | rs6683447 | 2.26 | 0.739 | -0.28 | -0.652 | -0.032 | -0.103 | -0.103 | 0 | 0.37 | 0.08 | 0.32 | 0.56 | 0.31 | 0.39 |
| 1 | 98485192 | rs1938560 | 0.22 | 1.028 | 0.031 | -0.655 | 0.008 | -0.008 | -0.008 | 0 | 0.9 | 0.87 | 0.9 | 0.93 | 0.83 | 0.83 |
| 1 | 98488863 | rs9659996 | 0.22 | 1.028 | 0.031 | -0.655 | 0.008 | -0.008 | -0.008 | 0 | 0.9 | 0.87 | 0.9 | 0.93 | 0.83 | 0.83 |
| 1 | 98498441 | rs11804556 | 1.149 | 1.003 | 0.067 | -0.685 | -0.001 | -0.019 | -0.019 | 0 | 0.07 | 0.06 | 0.05 | 0 | 0.07 | 0.03 |
| 1 | 98502340 | rs1782810 | 1.454 | 1.079 | 0.037 | -0.529 | 0.008 | -0.056 | -0.056 | 0 | 0.82 | 0.67 | 0.89 | 0.93 | 0.79 | 0.78 |
| 1 | 98508258 | rs2802535 | 1.437 | 1.061 | 0.025 | -0.543 | 0.005 | -0.049 | -0.049 | 0 | 0.82 | 0.72 | 0.89 | 0.93 | 0.79 | 0.78 |
| 1 | 98528452 | rs2660299 | 1.507 | 1.211 | -0.173 | -0.538 | 0.005 | -0.045 | -0.045 | 0.197 | 0.82 | 0.7 | 0.88 | 0.93 | 0.78 | 0.76 |
| 1 | 98529378 | rs1198583 | 0.296 | 1.245 | -0.66 | -0.523 | -0.007 | 0.005 | 0.005 | 0 | 0.89 | 0.93 | 0.91 | 0.93 | 0.82 | 0.81 |
| 1 | 98530559 | rs1198581 | 1.348 | 1.251 | -0.665 | -0.518 | -0.001 | -0.003 | -0.003 | 0 | 0.82 | 0.87 | 0.89 | 0.93 | 0.78 | 0.76 |
| 1 | 98541677 | rs4950142 | 0.218 | 1.573 | -0.903 | -0.431 | -0.081 | -0.003 | -0.003 | 0.013 | 0.15 | 0.01 | 0.34 | 0.08 | 0.23 | 0.12 |
| 1 | 98550411 | rs1198589 | 0.552 | 1.254 | -0.475 | -0.228 | 0.002 | -0.002 | -0.002 | 0.026 | 0.89 | 0.9 | 0.91 | 0.93 | 0.82 | 0.81 |
| 1 | 98564736 | rs17117562 | 1.093 | 1.005 | -0.52 | -0.186 | 0.007 | -0.014 | -0.014 | 0.432 | 0.41 | 0.29 | 0.34 | 0.53 | 0.36 | 0.42 |
| 1 | 98565035 | rs1198586 | -0.25 | 1.333 | -0.053 | 0.118 | 0.006 | 0.048 | 0.048 | 0 | 0.25 | 0.49 | 0.41 | 0.39 | 0.28 | 0.27 |
| 1 | 98565234 | rs1198585 | -0.239 | 1.377 | -0.683 | -0.232 | -0.102 | -0.024 | -0.024 | 1.596 | 0.79 | 0.99 | 0.81 | 0.82 | 0.71 | 0.75 |
| 1 | 98572374 | rs11165950 | 0.087 | 1.205 | 0.007 | -0.726 | -0.001 | -0.007 | -0.007 | 1.027 | 0.09 | 0.2 | 0.28 | 0.06 | 0.1 | 0.07 |
| 1 | 98576274 | rs17117613 | 0.2 | 1.203 | 0.048 | -0.807 | -0.005 | -0.016 | -0.016 | 1.19 | 0.09 | 0.15 | 0.28 | 0.05 | 0.1 | 0.07 |
| 1 | 98579211 | rs6679864 | 0.841 | 1.152 | 0.426 | -0.632 | 0.013 | -0.007 | -0.007 | 0 | 0.62 | 0.3 | 0.49 | 0.55 | 0.62 | 0.61 |
| 1 | 98588692 | rs355375 | -0.252 | 1.147 | 0.456 | -0.626 | -0.011 | 0.028 | 0.028 | 0 | 0.28 | 0.46 | 0.23 | 0.4 | 0.28 | 0.33 |
| 1 | 98588885 | rs1473474 | 1.308 | 0.601 | -0.192 | -0.991 | 0.022 | 0.008 | 0.008 | 0 | 0.52 | 0.1 | 0.44 | 0.55 | 0.54 | 0.57 |
| 1 | 98592888 | rs12082064 | 0.571 | 0.148 | 0.023 | -0.93 | -0.008 | -0.057 | -0.057 | 0 | 0.11 | 0.22 | 0.04 | 0 | 0.07 | 0.03 |
| 1 | 98595853 | rs1487302 | 0.599 | -0.054 | -0.183 | -1.117 | 0 | -0.009 | -0.009 | 0 | 0.19 | 0.16 | 0.32 | 0.05 | 0.13 | 0.08 |
| 1 | 98603170 | rs1487289 | 0.732 | -0.047 | -0.216 | -1.114 | 0.021 | 0.034 | 0.034 | 6.691 | 0.18 | 0.11 | 0.32 | 0.05 | 0.14 | 0.08 |
| 1 | 98604137 | rs17381287 | -0.86 | 0.067 | -0.187 | -1.138 | 0.02 | -0.032 | -0.032 | 0.011 | 0.23 | 0.3 | 0.17 | 0.1 | 0.31 | 0.25 |
| 1 | 98605732 | rs4950107 | 1.188 | -0.038 | -0.233 | -1.057 | 0.023 | 0.003 | 0.003 | 0.223 | 0.48 | 0.15 | 0.59 | 0.51 | 0.4 | 0.42 |
| 1 | 98616396 | rs1507069 | -0.701 | 0.054 | -0.178 | -0.992 | 0.007 | 0.002 | 0.002 | 0.419 | 0.41 | 0.47 | 0.36 | 0.49 | 0.52 | 0.54 |
| 1 | 98622349 | rs6701393 | 0.533 | 0.085 | -0.462 | -0.978 | -0.007 | -0.002 | -0.002 | 0 | 0.58 | 0.37 | 0.63 | 0.51 | 0.48 | 0.46 |
| 1 | 98639563 | rs17117845 | 2.134 | -0.003 | -0.901 | -1.012 | -0.034 | -0.146 | -0.146 | 0.333 | 0.3 | 0.06 | 0.28 | 0.45 | 0.27 | 0.35 |
| 1 | 98644672 | rs12033249 | 0.874 | -0.093 | -0.76 | -1.241 | 0.019 | 0.036 | 0.036 | 8.233 | 0.17 | 0.08 | 0.31 | 0.05 | 0.13 | 0.08 |
| 1 | 98645213 | rs10493900 | -0.238 | -0.094 | -0.614 | -1.271 | 0.016 | 0.027 | 0.027 | 6.395 | 0.26 | 0.17 | 0.3 | 0.05 | 0.2 | 0.11 |
| 1 | 98647074 | rs1487298 | -1.411 | -0.027 | -0.894 | -1.313 | -0.017 | 0.053 | 0.053 | 2.66 | 0.39 | 0.25 | 0.48 | 0.16 | 0.43 | 0.31 |
| 1 | 98651987 | rs10783085 | -0.124 | -0.086 | -0.835 | -1.293 | 0.005 | -0.002 | -0.002 | 0 | 0.25 | 0.19 | 0.35 | 0.06 | 0.18 | 0.1 |
| 1 | 98652088 | rs822559 | 0.113 | 0.121 | -0.496 | -1.163 | -0.002 | 0.004 | 0.004 | 0 | 0.28 | 0.49 | 0.24 | 0.27 | 0.31 | 0.27 |
| 1 | 98654113 | rs10875133 | -1.288 | 0.067 | -0.703 | -1.3 | -0.016 | 0.089 | 0.089 | 0.688 | 0.44 | 0.27 | 0.5 | 0.16 | 0.47 | 0.32 |

| | | | | | | | | | | | | | | | | |
|---|---|---|---|---|---|---|---|---|---|---|---|---|---|---|---|---|
| 1 | 98657119 | rs2391908 | 2.012 | -0.015 | -0.753 | -1.29 | -0.024 | -0.115 | -0.115 | 0 | 0.31 | 0.05 | 0.31 | 0.45 | 0.3 | 0.37 |
| 1 | 98658035 | rs12734525 | -0.143 | -0.084 | -0.726 | -1.336 | 0.01 | 0.01 | 0.01 | 3.923 | 0.31 | 0.24 | 0.39 | 0.06 | 0.24 | 0.13 |
| 1 | 98659152 | rs4950097 | 0.954 | -0.019 | -0.738 | -1.26 | 0.017 | 0.052 | 0.052 | 5.255 | 0.2 | 0.09 | 0.34 | 0.05 | 0.15 | 0.07 |
| 1 | 98660622 | rs12567765 | -0.074 | 0.756 | -0.346 | -0.801 | -0.061 | 0.003 | 0.003 | 0.001 | 0.13 | 0.04 | 0.11 | 0.11 | 0.23 | 0.19 |
| 1 | 98662961 | rs698994 | -0.09 | 0.11 | 0.213 | -0.575 | 0.005 | 0.048 | 0.048 | 0 | 0.76 | 0.53 | 0.71 | 0.67 | 0.75 | 0.69 |
| 1 | 98666125 | rs11577139 | -0.693 | 0.406 | 0.182 | -0.314 | -0.019 | 0.039 | 0.039 | 0 | 0.38 | 0.27 | 0.54 | 0.18 | 0.43 | 0.32 |
| 1 | 98667546 | rs822555 | -1.305 | -0.121 | 0.417 | -0.002 | 0.001 | 0.001 | 0.001 | 0.243 | 0.59 | 0.44 | 0.69 | 0.57 | 0.66 | 0.6 |
| 1 | 98683104 | rs822617 | -0.835 | 0.078 | -0.418 | -0.219 | -0.025 | -0.074 | -0.074 | 0 | 0.17 | 0.64 | 0.1 | 0.05 | 0.09 | 0.09 |
| 1 | 98687633 | rs2031245 | 0.754 | 0.637 | -1.452 | 0.091 | -0.043 | -0.136 | -0.136 | 0 | 0.16 | 0.01 | 0.11 | 0.33 | 0.15 | 0.22 |
| 1 | 98689392 | rs12724493 | -1.209 | 1.08 | -0.515 | -0.559 | -0.062 | 0.096 | 0.096 | 0 | 0.37 | 0.16 | 0.52 | 0.19 | 0.49 | 0.33 |
| 1 | 98720270 | rs1155644 | -0.556 | 1.394 | -1.565 | -0.268 | -0.06 | 0.045 | 0.045 | 0 | 0.28 | 0.14 | 0.5 | 0.18 | 0.42 | 0.28 |
| 1 | 98727000 | rs11165975 | -0.843 | 0.671 | -1.064 | -0.038 | -0.002 | -0.027 | -0.027 | 0 | 0.1 | 0.09 | 0.03 | 0 | 0.06 | 0.06 |
| 1 | 98728204 | rs12134140 | -0.647 | 1.121 | -1.643 | -0.021 | -0.061 | 0.027 | 0.027 | 0 | 0.28 | 0.1 | 0.3 | 0.19 | 0.36 | 0.26 |
| 1 | 98730925 | rs822607 | -0.484 | -0.65 | -0.938 | 0.316 | -0.077 | 0.005 | 0.005 | 0 | 0.31 | 0.48 | 0.21 | 0.31 | 0.18 | 0.25 |
| 1 | 98741750 | rs6664132 | -0.5 | -1.381 | -0.747 | -0.036 | -0.004 | -0.006 | -0.006 | 0.452 | 0.22 | 0.18 | 0.16 | 0.31 | 0.1 | 0.23 |
| 1 | 98755006 | rs11165976 | 0.547 | -1.288 | -1.359 | -0.102 | -0.07 | -0.02 | -0.02 | 0.083 | 0.23 | 0.06 | 0.24 | 0.22 | 0.29 | 0.24 |
| 1 | 98757816 | rs11165979 | -0.074 | -1.34 | -0.759 | -0.063 | -0.014 | -0.001 | -0.001 | 0 | 0.23 | 0.21 | 0.16 | 0.31 | 0.1 | 0.23 |
| 1 | 98764432 | rs12120750 | -0.404 | -1.322 | -0.865 | -0.087 | -0.004 | 0.008 | 0.008 | 0 | 0.35 | 0.35 | 0.29 | 0.22 | 0.37 | 0.26 |
| 1 | 98774883 | rs1408081 | -1.553 | -1.303 | -0.504 | -0.186 | -0.054 | -0.049 | -0.049 | 0.099 | 0.6 | 0.84 | 0.48 | 0.54 | 0.51 | 0.51 |
| 1 | 98778347 | rs1075032 | -1.438 | -1.272 | -0.341 | -0.131 | -0.007 | -0.006 | -0.006 | 0 | 0.59 | 0.66 | 0.47 | 0.54 | 0.5 | 0.5 |
| 1 | 98783722 | rs11165990 | -0.162 | -1.224 | -0.608 | 0.194 | -0.053 | 0.052 | 0.052 | 0 | 0.19 | 0.32 | 0.16 | 0.31 | 0.13 | 0.24 |
| 1 | 98796509 | rs1323727 | -0.184 | -0.591 | -1.238 | 0.923 | 0.008 | -0.011 | -0.011 | 0 | 0.87 | 0.92 | 0.75 | 0.67 | 0.91 | 0.87 |
| 1 | 98799369 | rs11165996 | -0.429 | -0.149 | -1.388 | 1.005 | -0.054 | -0.061 | -0.061 | 0.082 | 0.29 | 0.13 | 0.28 | 0.34 | 0.36 | 0.28 |
| 1 | 98802463 | rs17118192 | -0.188 | -0.168 | -1.395 | 1.228 | -0.005 | -0.09 | -0.09 | 0 | 0.21 | 0.05 | 0.16 | 0.33 | 0.18 | 0.35 |
| 1 | 98803940 | rs10783088 | 0.476 | -0.445 | -1.559 | 0.726 | 0.003 | 0.009 | 0.009 | 0 | 0.61 | 0.77 | 0.39 | 0.34 | 0.62 | 0.62 |
| 1 | 98806421 | rs1028889 | 0.459 | 0.04 | -1.409 | -0.075 | -0.002 | -0.017 | -0.017 | 0 | 0.31 | 0.46 | 0.2 | 0.21 | 0.28 | 0.47 |
| 1 | 98808760 | rs1328307 | -0.409 | -0.139 | -1.258 | 0.108 | 0.007 | -0.02 | -0.02 | 0.868 | 0.7 | 0.74 | 0.56 | 0.43 | 0.75 | 0.7 |
| 1 | 98812484 | rs12048166 | 0.711 | -0.319 | -1.165 | -1.108 | 0.005 | -0.03 | -0.03 | 0 | 0.29 | 0.26 | 0.19 | 0.13 | 0.32 | 0.14 |
| 1 | 98814626 | rs1328310 | -0.764 | -0.659 | -1.151 | -1.21 | 0.015 | 0.033 | 0.033 | 0.088 | 0.21 | 0.11 | 0.09 | 0.01 | 0.16 | 0.17 |
| 1 | 98818587 | rs17386441 | -0.673 | -0.457 | -0.93 | -0.949 | -0.001 | 0.028 | 0.028 | 0 | 0.13 | 0.06 | 0.08 | 0.02 | 0.12 | 0.13 |
| 1 | 98819537 | rs1013024 | 0.279 | -0.831 | -0.571 | -1.571 | -0.002 | -0.071 | -0.071 | 0.199 | 0.33 | 0.42 | 0.21 | 0.08 | 0.31 | 0.41 |
| 1 | 98820796 | rs6697562 | -0.351 | -0.678 | -0.627 | -1.428 | -0.004 | 0.031 | 0.031 | 0 | 0.13 | 0.05 | 0.07 | 0.02 | 0.12 | 0.12 |
| 1 | 98829960 | rs10875144 | -0.855 | -0.779 | -0.365 | -1.536 | 0.022 | 0.153 | 0.153 | 0.583 | 0.46 | 0.79 | 0.3 | 0.1 | 0.4 | 0.49 |
| 1 | 98834146 | rs4542263 | 0.01 | -0.761 | -0.447 | -1.505 | 0 | 0.024 | 0.024 | 0 | 0.1 | 0.03 | 0.05 | 0.02 | 0.07 | 0.08 |
| 1 | 98838035 | rs11166002 | 0.374 | -1.268 | -0.029 | -1.275 | -0.003 | -0.081 | -0.081 | 0.073 | 0.3 | 0.42 | 0.2 | 0.08 | 0.29 | 0.41 |
| 1 | 98843371 | rs10875145 | 0.36 | -1.27 | 0.09 | -1.276 | -0.002 | -0.057 | -0.057 | 0 | 0.3 | 0.38 | 0.19 | 0.08 | 0.29 | 0.41 |
| 1 | 98846773 | rs7541400 | -0.913 | -1.302 | 0.07 | -1.295 | 0.022 | 0.156 | 0.156 | 0.574 | 0.46 | 0.77 | 0.3 | 0.11 | 0.39 | 0.49 |
| 1 | 98848697 | rs17118281 | -0.878 | -1.383 | 0.664 | -1.036 | -0.007 | -0.047 | -0.047 | 0 | 0.13 | 0.37 | 0.1 | 0.02 | 0.09 | 0.07 |
| 1 | 98855166 | rs728656 | 0.554 | -1.68 | 0.084 | -0.685 | 0.006 | 0.008 | 0.008 | 0.015 | 0.49 | 0.41 | 0.39 | 0.43 | 0.52 | 0.41 |
| 1 | 98856577 | rs7548836 | -1.338 | -1.525 | -0.269 | -0.592 | 0.003 | -0.078 | -0.078 | 0 | 0.85 | 0.93 | 0.63 | 0.51 | 0.84 | 0.85 |
| 1 | 98856950 | rs10875147 | 0.215 | -1.015 | -0.095 | -0.848 | 0.001 | -0.071 | -0.071 | 0 | 0.27 | 0.37 | 0.17 | 0.06 | 0.27 | 0.38 |
| 1 | 98859508 | rs10875148 | 0.509 | -1.009 | -0.258 | -0.799 | 0 | -0.024 | -0.024 | 0 | 0.24 | 0.27 | 0.17 | 0.06 | 0.25 | 0.32 |
| 1 | 98863760 | rs3908575 | -0.55 | -1.031 | -0.149 | -0.463 | 0.015 | 0.026 | 0.026 | 0.125 | 0.52 | 0.43 | 0.57 | 0.66 | 0.45 | 0.53 |
| 1 | 98866206 | rs10875150 | -1.411 | -1.053 | -0.107 | -0.527 | 0.028 | -0.016 | -0.016 | 0 | 0.62 | 0.57 | 0.64 | 0.7 | 0.55 | 0.59 |
| 1 | 98871410 | rs12058177 | 0.042 | -0.977 | -0.307 | -0.464 | 0.003 | 0.008 | 0.008 | 0 | 0.32 | 0.33 | 0.18 | 0.08 | 0.28 | 0.3 |
| 1 | 98873604 | rs7524325 | 0.201 | -1.003 | -0.671 | -0.458 | -0.01 | 0.034 | 0.034 | 0.56 | 0.12 | 0.01 | 0.05 | 0.04 | 0.1 | 0.08 |
| 1 | 98881389 | rs17387268 | -0.079 | -0.955 | -0.65 | -0.436 | 0 | 0.073 | 0.073 | 2.42 | 0.17 | 0.02 | 0.07 | 0.04 | 0.13 | 0.08 |
| 1 | 98883541 | rs10157812 | 1.414 | -0.98 | -0.587 | -0.439 | 0.019 | -0.007 | -0.007 | 0 | 0.36 | 0.51 | 0.36 | 0.3 | 0.45 | 0.39 |
| 1 | 98893908 | rs12030530 | -0.693 | -0.809 | -0.886 | -0.346 | 0.001 | 0.01 | 0.01 | 0 | 0.73 | 0.71 | 0.84 | 0.92 | 0.75 | 0.7 |
| 1 | 98896256 | rs1540839 | 1.156 | -0.789 | -0.99 | -0.332 | -0.001 | -0.148 | -0.148 | 0 | 0.19 | 0.02 | 0.39 | 0.58 | 0.18 | 0.23 |
| 1 | 98904871 | rs11166015 | 0.081 | -0.789 | -0.983 | -0.335 | 0.014 | 0.018 | 0.018 | 0.717 | 0.54 | 0.59 | 0.44 | 0.34 | 0.58 | 0.47 |
| 1 | 98905040 | rs12131979 | 0.938 | -0.454 | -0.742 | -0.036 | 0.013 | -0.125 | -0.125 | 0 | 0.19 | 0.03 | 0.39 | 0.58 | 0.18 | 0.23 |
| 1 | 98913185 | rs684637 | 0.627 | 0.078 | -0.385 | 0.222 | 0.003 | -0.001 | -0.001 | 0.353 | 0.37 | 0.56 | 0.37 | 0.3 | 0.45 | 0.39 |
| 1 | 98913843 | rs7530588 | 0.258 | -0.039 | -0.456 | 0.222 | 0.033 | -0.027 | -0.027 | 0 | 0.32 | 0.24 | 0.44 | 0.6 | 0.28 | 0.26 |
| 1 | 98914478 | rs316041 | -0.43 | 0.131 | -0.374 | 0.217 | -0.009 | 0.056 | 0.056 | 0 | 0.68 | 0.78 | 0.8 | 0.9 | 0.7 | 0.64 |
| 1 | 98916248 | rs11166016 | -0.569 | 0.005 | -0.08 | 1.101 | 0.035 | -0.025 | -0.025 | 0 | 0.33 | 0.24 | 0.44 | 0.62 | 0.28 | 0.26 |
| 1 | 98928271 | rs4950129 | 1.494 | 1.349 | 0.936 | 1.593 | 0.055 | 0.149 | 0.149 | 0.171 | 0.84 | 0.53 | 0.8 | 0.72 | 0.83 | 0.8 |
| 1 | 98928937 | rs770918 | 0.938 | 1.458 | 0.447 | 0.966 | -0.016 | 0.004 | 0.004 | 0 | 0.89 | 0.92 | 0.88 | 0.91 | 0.84 | 0.81 |
| 1 | 98937092 | rs648124 | 1.239 | 1.676 | 0.58 | 0.944 | -0.005 | 0.003 | 0.003 | 0.088 | 0.87 | 0.88 | 0.86 | 0.91 | 0.81 | 0.77 |
| 1 | 98938230 | rs2479667 | -0.965 | 1.551 | 1.245 | 0.897 | 0.009 | 0.013 | 0.013 | 0.013 | 0.14 | 0.21 | 0.13 | 0.16 | 0.15 | 0.13 |
| 1 | 98938977 | rs316055 | 0.958 | 1.678 | 0.985 | 0.996 | 0.003 | -0.001 | -0.001 | 0 | 0.88 | 0.85 | 0.73 | 0.86 | 0.84 | 0.78 |
| 1 | 98952064 | rs316052 | 1.246 | 1.451 | 1.216 | 0.684 | 0.039 | 0.022 | 0.022 | 0 | 0.86 | 0.68 | 0.71 | 0.84 | 0.84 | 0.77 |
| 1 | 98953664 | rs3853696 | 2.08 | 1.415 | 1.256 | 0.651 | 0.013 | -0.04 | -0.04 | 0 | 0.76 | 0.58 | 0.64 | 0.84 | 0.75 | 0.75 |
| 1 | 98954240 | rs316053 | 0.996 | 1.436 | 0.951 | 0.684 | 0.013 | 0.006 | 0.006 | 0 | 0.87 | 0.82 | 0.72 | 0.84 | 0.84 | 0.77 |
| 1 | 98957554 | rs520327 | 1.041 | 1.44 | 0.889 | 0.647 | 0.027 | 0.01 | 0.01 | 0 | 0.87 | 0.76 | 0.72 | 0.85 | 0.84 | 0.77 |
| 1 | 98961931 | rs167567 | -0.563 | 1.079 | 0.19 | 0.52 | -0.003 | -0.026 | -0.026 | 0 | 0.1 | 0.1 | 0.07 | 0 | 0.09 | 0.02 |
| 1 | 98964063 | rs571344 | 0.347 | 1.569 | -0.01 | 0.212 | -0.003 | -0.001 | -0.001 | 0 | 0.74 | 0.78 | 0.79 | 0.84 | 0.65 | 0.79 |
| 1 | 98964075 | rs571320 | 0.833 | 1.698 | 0.062 | 1.175 | 0.003 | -0.001 | -0.001 | 0 | 0.66 | 0.61 | 0.49 | 0.31 | 0.56 | 0.68 |
| 1 | 98964498 | rs591520 | -0.212 | 0.521 | -0.11 | 1 | -0.003 | 0.001 | 0.001 | 0 | 0.24 | 0.29 | 0.15 | 0.28 | 0.16 | 0.22 |

| | | | | | | | | | | | | | | | | |
|---|---|---|---|---|---|---|---|---|---|---|---|---|---|---|---|---|
| 1 | 98964541 | rs566731 | -0.248 | 0.704 | -0.121 | 0.999 | -0.022 | 0.028 | 0.028 | 0 | 0.72 | 0.8 | 0.75 | 0.84 | 0.61 | 0.72 |
| 1 | 98964760 | rs3867319 | 0.417 | 0.323 | -0.323 | 0.725 | 0.009 | -0.006 | -0.006 | 0.042 | 0.14 | 0.11 | 0.05 | 0 | 0.08 | 0.08 |
| 1 | 98965621 | rs1724950 | -0.406 | 0.559 | -0.182 | 0.694 | 0.002 | 0.004 | 0.004 | 0 | 0.76 | 0.7 | 0.78 | 0.71 | 0.68 | 0.79 |
| 1 | 98966328 | rs1655567 | -1.046 | 0.562 | -0.219 | 0.552 | -0.027 | 0.037 | 0.037 | 0 | 0.8 | 0.88 | 0.86 | 0.98 | 0.71 | 0.83 |
| 1 | 98983135 | rs1594867 | -0.694 | 0.552 | -0.108 | 0.546 | -0.004 | -0.025 | -0.025 | 0.033 | 0.78 | 0.79 | 0.85 | 0.98 | 0.71 | 0.83 |
| 1 | 98986884 | rs1121773 | -0.052 | 0.517 | -0.053 | 0.687 | 0 | 0.001 | 0.001 | 0 | 0.73 | 0.69 | 0.76 | 0.7 | 0.67 | 0.76 |
| 1 | 99008004 | rs2099414 | -1.056 | 0.482 | 0.553 | 0.489 | -0.037 | 0.058 | 0.058 | 0 | 0.8 | 0.91 | 0.85 | 0.98 | 0.71 | 0.83 |
| 1 | 99010292 | rs12123631 | 0.52 | 0.389 | 0.733 | 0.623 | 0.006 | 0.011 | 0.011 | 0 | 0.64 | 0.4 | 0.65 | 0.53 | 0.57 | 0.65 |
| 1 | 99024847 | rs7518684 | 0.514 | 0.374 | 0.848 | 0.616 | 0.01 | 0.02 | 0.02 | 0 | 0.64 | 0.45 | 0.65 | 0.53 | 0.57 | 0.65 |
| 1 | 99030859 | rs4387225 | -0.613 | 0.383 | 0.768 | 0.53 | -0.028 | 0.059 | 0.059 | 0 | 0.76 | 0.86 | 0.84 | 0.98 | 0.69 | 0.8 |
| 1 | 99037251 | rs995271 | -0.19 | 0.299 | 0.989 | 0.538 | 0.001 | 0.006 | 0.006 | 0 | 0.73 | 0.66 | 0.76 | 0.67 | 0.67 | 0.76 |
| 1 | 99039299 | rs1434367 | 0.496 | 0.264 | 0.867 | 0.756 | 0.007 | -0.01 | -0.01 | 0 | 0.63 | 0.42 | 0.58 | 0.35 | 0.56 | 0.63 |
| 1 | 99072344 | rs12069484 | 0.632 | -0.032 | 0.773 | 0.808 | -0.01 | 0.035 | 0.035 | 0 | 0.06 | 0.11 | 0.15 | 0.2 | 0.03 | 0.07 |
| 1 | 99090942 | rs12759704 | 1.436 | -0.226 | 0.231 | -0.189 | 0.01 | -0.003 | -0.003 | 0.06 | 0.26 | 0.18 | 0.26 | 0.01 | 0.24 | 0.29 |
| 1 | 99093976 | rs10783098 | -0.579 | -0.296 | 0.351 | -0.239 | 0.014 | 0.017 | 0.017 | 0 | 0.37 | 0.49 | 0.47 | 0.64 | 0.38 | 0.41 |
| 1 | 99097714 | rs12079791 | 0.22 | -0.267 | 0.4 | -0.162 | 0.009 | 0.014 | 0.014 | 0 | 0.28 | 0.36 | 0.29 | 0.42 | 0.27 | 0.3 |
| 1 | 99101223 | rs12088344 | -0.318 | -0.249 | 0.305 | -0.396 | 0.028 | 0.002 | 0.002 | 0 | 0.34 | 0.46 | 0.37 | 0.42 | 0.37 | 0.32 |
| 1 | 99108828 | rs12088202 | 0.643 | -0.286 | 0.241 | -0.422 | -0.011 | -0.002 | -0.002 | 0 | 0.25 | 0.2 | 0.27 | 0.33 | 0.25 | 0.26 |
| 1 | 99113106 | rs12731647 | 1.067 | -0.397 | 0.234 | -0.528 | 0.007 | -0.013 | -0.013 | 0.159 | 0.33 | 0.24 | 0.29 | 0.02 | 0.28 | 0.3 |
| 1 | 99132070 | rs4469760 | 0.282 | -0.414 | -0.129 | -0.413 | -0.019 | 0.014 | 0.014 | 0.099 | 0.31 | 0.18 | 0.28 | 0.32 | 0.28 | 0.34 |
| 1 | 99141448 | rs4540678 | 1.051 | -0.689 | 0.136 | -0.63 | 0.011 | 0.036 | 0.036 | 0 | 0.3 | 0.21 | 0.28 | 0.07 | 0.27 | 0.3 |
| 1 | 99160827 | rs12138303 | 1.555 | -0.739 | 0.159 | -0.882 | -0.053 | 0.103 | 0.103 | 0 | 0.22 | 0.04 | 0.14 | 0.01 | 0.28 | 0.18 |
| 1 | 99164434 | rs6577322 | 1.215 | -0.576 | 0.274 | -0.776 | 0.011 | 0.036 | 0.036 | 0 | 0.31 | 0.21 | 0.28 | 0.06 | 0.27 | 0.3 |
| 1 | 99166681 | rs4351664 | 0.832 | -0.542 | 0.606 | -0.58 | 0.073 | 0.149 | 0.149 | 0.554 | 0.94 | 0.63 | 0.96 | 0.91 | 0.96 | 0.96 |
| 1 | 99173681 | rs4545356 | -0.527 | -0.603 | 0.546 | -0.372 | -0.001 | -0.001 | -0.001 | 1.996 | 0.66 | 0.39 | 0.7 | 0.69 | 0.66 | 0.68 |
| 1 | 99176296 | rs10783100 | -0.425 | -0.622 | 0.524 | -0.485 | 0 | 0.001 | 0.001 | 0 | 0.49 | 0.49 | 0.53 | 0.52 | 0.52 | 0.47 |
| 1 | 99180910 | rs4545357 | 0.047 | -0.526 | 0.446 | -0.521 | 0 | -0.004 | -0.004 | 0 | 0.62 | 0.4 | 0.55 | 0.27 | 0.61 | 0.62 |
| 1 | 99184272 | rs6678214 | 1.048 | -0.57 | 0.462 | -0.586 | -0.001 | -0.028 | -0.028 | 0 | 0.32 | 0.3 | 0.27 | 0.06 | 0.26 | 0.3 |
| 1 | 99184798 | rs12725220 | 0.02 | -0.612 | 0.611 | -0.451 | 0.014 | -0.012 | -0.012 | 0 | 0.36 | 0.46 | 0.35 | 0.4 | 0.37 | 0.33 |
| 1 | 99185615 | rs12118149 | 0.832 | -0.612 | 0.571 | -0.431 | -0.021 | 0.04 | 0.04 | 0 | 0.3 | 0.15 | 0.28 | 0.2 | 0.35 | 0.32 |
| 1 | 99186118 | rs12137692 | 0.856 | -0.608 | 0.573 | -0.429 | -0.022 | 0.04 | 0.04 | 0 | 0.3 | 0.15 | 0.28 | 0.2 | 0.35 | 0.33 |
| 1 | 99199954 | rs6702126 | -0.047 | -0.606 | 0.592 | -0.422 | 0.004 | -0.021 | -0.021 | 0.602 | 0.62 | 0.45 | 0.55 | 0.27 | 0.61 | 0.62 |
| 1 | 99207147 | rs12122275 | 0.849 | -0.601 | 0.546 | -0.421 | -0.021 | 0.039 | 0.039 | 0 | 0.3 | 0.16 | 0.28 | 0.2 | 0.35 | 0.33 |
| 1 | 99209727 | rs992708 | -0.162 | -0.614 | 0.622 | -0.439 | 0.022 | 0.057 | 0.057 | 0.442 | 0.51 | 0.61 | 0.53 | 0.38 | 0.43 | 0.44 |
| 1 | 99216783 | rs11166104 | 0.531 | -0.62 | 0.762 | -0.438 | -0.004 | 0.015 | 0.015 | 0 | 0.18 | 0.22 | 0.25 | 0.31 | 0.17 | 0.15 |
| 1 | 99219716 | rs12025763 | -0.2 | -0.621 | 0.664 | -0.422 | -0.02 | 0.078 | 0.078 | 0 | 0.78 | 0.52 | 0.65 | 0.35 | 0.82 | 0.82 |
| 1 | 99220941 | rs11166108 | 0.18 | -0.628 | 0.66 | -0.435 | 0.014 | 0.102 | 0.102 | 0 | 0.47 | 0.26 | 0.38 | 0.29 | 0.56 | 0.52 |
| 1 | 99223397 | rs1482153 | -0.275 | -0.631 | 0.659 | -0.436 | -0.018 | 0.081 | 0.081 | 0 | 0.79 | 0.52 | 0.66 | 0.35 | 0.82 | 0.82 |
| 1 | 99223508 | rs12406053 | 1.3 | -0.64 | 0.664 | -0.441 | 0.01 | 0.025 | 0.025 | 0 | 0.31 | 0.23 | 0.27 | 0.06 | 0.26 | 0.29 |
| 1 | 99224250 | rs3813786 | 1.283 | -0.633 | 0.655 | -0.438 | 0.008 | 0.013 | 0.013 | 0 | 0.32 | 0.24 | 0.27 | 0.06 | 0.26 | 0.29 |
| 1 | 99225068 | rs4908025 | 1.62 | -0.596 | 0.54 | -0.42 | -0.023 | 0.033 | 0.033 | 2.416 | 0.17 | 0.1 | 0.1 | 0.08 | 0.2 | 0.19 |
| 1 | 99230383 | rs11166113 | 0.196 | -0.805 | 0.465 | -0.403 | 0.014 | 0.102 | 0.102 | 0 | 0.47 | 0.26 | 0.38 | 0.29 | 0.56 | 0.52 |
| 1 | 99242767 | rs9661188 | 0.249 | -0.4 | 0.597 | -0.258 | -0.006 | 0.03 | 0.03 | 0 | 0.19 | 0.28 | 0.25 | 0.32 | 0.17 | 0.15 |
| 1 | 99245025 | rs10875183 | -0.494 | 0.222 | 0.505 | -0.129 | -0.037 | 0.045 | 0.045 | 0 | 0.69 | 0.81 | 0.74 | 0.8 | 0.62 | 0.66 |
| 1 | 99245060 | rs10875184 | -0.494 | 0.222 | 0.505 | -0.129 | -0.037 | 0.045 | 0.045 | 0 | 0.69 | 0.81 | 0.74 | 0.79 | 0.62 | 0.66 |
| 1 | 99263348 | rs9728271 | 0.216 | -0.286 | 0.6 | -0.433 | 0.015 | -0.086 | -0.086 | 0.054 | 0.23 | 0.4 | 0.28 | 0.06 | 0.25 | 0.25 |
| 1 | 99289282 | rs2696358 | 0.555 | -0.3 | 0.833 | -0.073 | 0.002 | 0.002 | 0.002 | 1.061 | 0.43 | 0.41 | 0.46 | 0.62 | 0.37 | 0.35 |
| 1 | 99290291 | rs1384164 | 0.751 | -0.31 | 0.851 | -0.007 | 0.005 | 0.021 | 0.021 | 0.083 | 0.41 | 0.31 | 0.35 | 0.29 | 0.36 | 0.32 |
| 1 | 99302782 | rs1013468 | 0.964 | -0.205 | 0.755 | -0.138 | -0.036 | 0.05 | 0.05 | 0 | 0.25 | 0.14 | 0.18 | 0.01 | 0.33 | 0.19 |
| 1 | 99310398 | rs4908084 | -0.643 | -0.457 | 0.703 | -0.057 | 0.008 | -0.01 | -0.01 | 0 | 0.87 | 0.88 | 0.77 | 0.8 | 0.92 | 0.74 |
| 1 | 99320254 | rs12040415 | 0.573 | -0.447 | 0.697 | -0.066 | 0.012 | -0.015 | -0.015 | 0 | 0.13 | 0.09 | 0.22 | 0.2 | 0.08 | 0.26 |
| 1 | 99334672 | rs1482144 | -0.552 | -0.393 | 0.25 | 0.006 | -0.02 | 0.052 | 0.052 | 0.328 | 0.87 | 0.8 | 0.77 | 0.7 | 0.92 | 0.73 |
| 1 | 99348900 | rs1482162 | -0.59 | 0.22 | 0.084 | -0.067 | 0 | -0.001 | -0.001 | 0.35 | 0.45 | 0.56 | 0.41 | 0.51 | 0.54 | 0.41 |
| 1 | 99352644 | rs6698102 | 0.836 | 0.549 | -0.639 | -1.138 | -0.049 | 0.085 | 0.085 | 0 | 0.24 | 0.1 | 0.17 | 0.01 | 0.31 | 0.18 |
| 1 | 99354312 | rs2062105 | 0.825 | 0.553 | -0.638 | -1.136 | -0.049 | 0.086 | 0.086 | 0 | 0.24 | 0.1 | 0.17 | 0.01 | 0.31 | 0.18 |
| 1 | 99356942 | rs1351562 | 0.487 | 0.465 | -0.396 | -0.994 | -0.028 | 0.026 | 0.026 | 0 | 0.13 | 0.06 | 0.1 | 0.06 | 0.2 | 0.13 |
| 1 | 99358358 | rs984271 | 0.961 | 0.094 | -0.111 | -1.099 | 0 | 0.001 | 0.001 | 0.063 | 0.62 | 0.37 | 0.6 | 0.75 | 0.64 | 0.55 |
| 1 | 99362556 | rs7529505 | 0.508 | 0.341 | -0.326 | -0.948 | -0.003 | 0.002 | 0.002 | 0 | 0.13 | 0.13 | 0.1 | 0.06 | 0.2 | 0.1 |
| 1 | 99366952 | rs6675949 | -0.619 | 0.217 | -0.304 | -0.956 | 0.002 | -0.002 | -0.002 | 1.79 | 0.83 | 0.83 | 0.73 | 0.8 | 0.86 | 0.72 |
| 1 | 99373498 | rs2391986 | 0.963 | 0.251 | -0.63 | -1.385 | 0.008 | 0.057 | 0.057 | 0 | 0.54 | 0.32 | 0.54 | 0.76 | 0.54 | 0.57 |
| 1 | 99373609 | rs2046178 | 0.131 | 0.22 | -0.528 | -1.43 | -0.008 | -0.012 | -0.012 | 0 | 0.69 | 0.61 | 0.66 | 0.8 | 0.75 | 0.65 |
| 1 | 99374045 | rs7555367 | 0.903 | 0.341 | -0.508 | -1.265 | 0.031 | -0.04 | -0.04 | 0 | 0.13 | 0.23 | 0.11 | 0.04 | 0.19 | 0.07 |
| 1 | 99383364 | rs1329465 | 0.439 | 0.143 | -0.693 | -1.112 | -0.009 | 0.005 | 0.005 | 0.12 | 0.18 | 0.27 | 0.29 | 0.2 | 0.14 | 0.27 |
| 1 | 99398580 | rs411115 | -0.072 | 0.174 | -0.926 | -1.294 | -0.001 | 0.018 | 0.018 | 0.042 | 0.57 | 0.65 | 0.51 | 0.31 | 0.55 | 0.62 |
| 1 | 99402224 | rs309058 | 0.4 | 0.439 | -1.086 | -1.435 | -0.007 | 0.018 | 0.018 | 0.305 | 0.35 | 0.31 | 0.38 | 0.25 | 0.37 | 0.36 |
| 1 | 99403197 | rs309057 | -0.557 | -0.093 | -0.892 | -0.67 | 0 | -0.002 | -0.002 | 0 | 0.67 | 0.68 | 0.72 | 0.5 | 0.72 | 0.66 |
| 1 | 99410959 | rs1329464 | 0.813 | 0.433 | -0.847 | 0.018 | 0.01 | 0.031 | 0.031 | 0 | 0.08 | 0.09 | 0.18 | 0.22 | 0.1 | 0.1 |
| 1 | 99413725 | rs309073 | -0.932 | 0.608 | -1.484 | -0.271 | -0.06 | 0.158 | 0.158 | 0 | 0.69 | 0.94 | 0.7 | 0.95 | 0.62 | 0.78 |
| 1 | 99415713 | rs309080 | -0.028 | 0.698 | -1.35 | -0.207 | -0.028 | 0.09 | 0.09 | 0 | 0.56 | 0.74 | 0.49 | 0.72 | 0.49 | 0.67 |

| | | | | | | | | | | | | | | | | |
|---|---|---|---|---|---|---|---|---|---|---|---|---|---|---|---|---|
| 1 | 99416102 | rs309083 | -0.484 | 0.596 | -1.347 | -0.494 | -0.029 | 0.113 | 0.113 | 0 | 0.67 | 0.85 | 0.7 | 0.95 | 0.61 | 0.78 |
| 1 | 99418647 | rs309087 | -0.753 | 0.255 | -0.59 | 0.859 | -0.006 | 0.004 | 0.004 | 0.049 | 0.22 | 0.27 | 0.16 | 0.24 | 0.18 | 0.23 |
| 1 | 99425374 | rs11801576 | -0.997 | 0.405 | -0.494 | 1.354 | 0 | 0.041 | 0.041 | 0 | 0.2 | 0.25 | 0.19 | 0.43 | 0.23 | 0.21 |
| 1 | 99426616 | rs382051 | -0.584 | -0.01 | 0.174 | 0.885 | -0.005 | -0.046 | -0.046 | 0.023 | 0.08 | 0.19 | 0.05 | 0 | 0.08 | 0.05 |
| 1 | 99429789 | rs309052 | -0.538 | -0.336 | -1.067 | 0.258 | -0.011 | -0.11 | -0.11 | 0.041 | 0.71 | 0.93 | 0.75 | 0.62 | 0.74 | 0.58 |
| 1 | 99435366 | rs2149738 | 0.11 | -0.013 | -0.862 | 0.777 | -0.002 | 0.006 | 0.006 | 0 | 0.17 | 0.19 | 0.14 | 0.27 | 0.18 | 0.16 |
| 1 | 99441320 | rs10489922 | 0.598 | -0.08 | -0.887 | 0.682 | 0 | 0.011 | 0.011 | 0 | 0.08 | 0.05 | 0.08 | 0.01 | 0.12 | 0.07 |
| 1 | 99443516 | rs10875198 | 0.881 | -0.94 | -1.345 | 0.108 | -0.078 | 0.028 | 0.028 | 0 | 0.39 | 0.1 | 0.38 | 0.3 | 0.44 | 0.3 |
| 1 | 99453287 | rs10443196 | 1.665 | -0.956 | -0.94 | 0.091 | -0.095 | 0.138 | 0.138 | 0 | 0.31 | 0.09 | 0.33 | 0.04 | 0.38 | 0.22 |
| 1 | 99457570 | rs12042547 | 0.468 | -0.641 | -0.366 | 0.625 | -0.004 | 0.052 | 0.052 | 0 | 0.09 | 0.14 | 0.06 | 0.26 | 0.06 | 0.08 |
| 1 | 99472175 | rs6692269 | -1.502 | -0.606 | -0.3 | 0.659 | 0 | 0.022 | 0.022 | 0 | 0.2 | 0.22 | 0.19 | 0.44 | 0.24 | 0.19 |
| 1 | 99480614 | rs11166154 | 0.149 | -0.641 | -0.222 | 0.681 | -0.002 | 0.076 | 0.076 | 0 | 0.1 | 0.15 | 0.07 | 0.28 | 0.08 | 0.1 |
| 1 | 99495135 | rs1350176 | -1.452 | -0.663 | -0.226 | 0.658 | -0.001 | 0.01 | 0.01 | 0 | 0.19 | 0.19 | 0.19 | 0.45 | 0.24 | 0.19 |
| 1 | 99499591 | rs11166158 | 1.942 | -0.865 | 0.257 | 0.476 | 0.037 | -0.035 | -0.035 | 0 | 0.61 | 0.2 | 0.56 | 0.4 | 0.64 | 0.62 |
| 1 | 99518345 | rs897603 | -0.103 | -0.743 | 0.738 | 0.474 | -0.004 | -0.048 | -0.048 | 0 | 0.91 | 0.75 | 0.94 | 0.99 | 0.93 | 0.93 |
| 1 | 99519111 | rs7515995 | 0.776 | -0.783 | 0.818 | 0.483 | 0 | -0.009 | -0.009 | 0 | 0.83 | 0.89 | 0.85 | 0.7 | 0.8 | 0.85 |
| 1 | 99522014 | rs12046217 | -0.468 | -0.784 | 0.804 | 0.484 | 0 | -0.024 | -0.024 | 0.561 | 0.14 | 0.06 | 0.13 | 0.3 | 0.16 | 0.12 |
| 1 | 99524047 | rs10747502 | 1.008 | -0.822 | 1.04 | 0.463 | -0.013 | -0.062 | -0.062 | 1.013 | 0.89 | 0.67 | 0.94 | 1 | 0.93 | 0.93 |
| 1 | 99532176 | rs10158645 | 1.285 | -0.817 | 0.995 | 0.48 | -0.017 | 0.077 | 0.077 | 0 | 0.82 | 0.63 | 0.88 | 0.7 | 0.87 | 0.88 |
| 1 | 99536341 | rs4908271 | -0.489 | -0.746 | 0.812 | 0.684 | 0 | -0.005 | -0.005 | 0 | 0.91 | 0.91 | 0.96 | 0.93 | 0.93 | 0.95 |
| 1 | 99539892 | rs11166167 | 0.328 | -0.718 | 0.617 | 0.719 | -0.014 | -0.049 | -0.049 | 0 | 0.08 | 0.01 | 0.12 | 0.23 | 0.13 | 0.1 |
| 1 | 99542610 | rs2124611 | 1.326 | -0.953 | 0.694 | 0.449 | -0.001 | -0.001 | -0.001 | 0 | 0.75 | 0.65 | 0.84 | 0.78 | 0.78 | 0.87 |
| 1 | 99546157 | rs6577339 | 1.612 | -0.975 | 0.672 | 0.127 | -0.001 | 0.005 | 0.005 | 0 | 0.6 | 0.48 | 0.67 | 0.55 | 0.56 | 0.73 |
| 1 | 99549327 | rs1505552 | 0.805 | -0.917 | 0.896 | 0.051 | -0.005 | -0.067 | -0.067 | 0 | 0.79 | 0.64 | 0.9 | 0.93 | 0.83 | 0.88 |
| 1 | 99551140 | rs12411241 | 0.19 | -0.877 | 0.834 | 0.334 | -0.025 | 0.013 | 0.013 | 0.149 | 0.1 | 0.19 | 0.16 | 0.15 | 0.05 | 0.14 |
| 1 | 99563065 | rs1747985 | 1.28 | -0.136 | 1.123 | 0.605 | 0.016 | 0.009 | 0.009 | 0 | 0.8 | 0.7 | 0.6 | 0.73 | 0.67 | 0.68 |
| 1 | 99566473 | rs3850458 | 2.055 | -0.646 | 1.107 | 0.921 | 0 | -0.02 | -0.02 | 1.015 | 0.55 | 0.21 | 0.47 | 0.44 | 0.52 | 0.53 |
| 1 | 99571007 | rs11166179 | -1.699 | -0.592 | 1.71 | 0.857 | 0.024 | -0.026 | -0.026 | 0 | 0.11 | 0.28 | 0.15 | 0.06 | 0.12 | 0.2 |
| 1 | 99573733 | rs302815 | -1.757 | -0.636 | 1.888 | 0.812 | 0.015 | -0.091 | -0.091 | 0 | 0.18 | 0.44 | 0.18 | 0.06 | 0.19 | 0.25 |
| 1 | 99580912 | rs3896046 | -1.415 | -0.714 | 1.912 | 0.907 | 0.021 | -0.004 | -0.004 | 0.06 | 0.08 | 0.25 | 0.12 | 0.06 | 0.1 | 0.16 |
| 1 | 99586516 | rs3850463 | 1.646 | -0.773 | 1.973 | 0.929 | 0.017 | -0.006 | -0.006 | 0.201 | 0.73 | 0.5 | 0.59 | 0.73 | 0.64 | 0.67 |
| 1 | 99589379 | rs303386 | 1.39 | -0.959 | 1.957 | 1.029 | -0.011 | 0.097 | 0.097 | 0 | 0.72 | 0.52 | 0.65 | 0.51 | 0.69 | 0.77 |
| 1 | 99597328 | rs302748 | 1.067 | -1.026 | 1.87 | 1.081 | -0.026 | 0.086 | 0.086 | 0 | 0.74 | 0.47 | 0.62 | 0.47 | 0.75 | 0.78 |
| 1 | 99601609 | rs3889240 | -0.151 | -1.006 | 1.676 | 1.129 | -0.019 | 0.053 | 0.053 | 0 | 0.14 | 0.23 | 0.09 | 0.28 | 0.1 | 0.11 |
| 1 | 99601672 | rs7539323 | 0.231 | -0.948 | 1.482 | 0.435 | -0.013 | -0.043 | -0.043 | 0 | 0.1 | 0.03 | 0.28 | 0.26 | 0.15 | 0.11 |
| 1 | 99608295 | rs12075686 | -0.706 | -0.962 | 1.356 | 0.204 | -0.016 | 0.028 | 0.028 | 0 | 0.25 | 0.33 | 0.3 | 0.68 | 0.18 | 0.29 |
| 1 | 99610153 | rs863686 | -2.342 | -0.561 | 2.009 | 0.39 | -0.001 | -0.001 | -0.001 | 0.344 | 0.31 | 0.68 | 0.45 | 0.29 | 0.37 | 0.31 |
| 1 | 99618085 | rs302765 | -0.108 | -0.562 | 1.932 | 0.461 | -0.001 | 0.011 | 0.011 | 0 | 0.1 | 0.11 | 0.27 | 0.26 | 0.16 | 0.11 |
| 1 | 99636431 | rs683131 | -2.06 | -0.977 | 2.09 | 0.131 | -0.012 | -0.072 | -0.072 | 0 | 0.24 | 0.66 | 0.19 | 0.01 | 0.21 | 0.18 |
| 1 | 99636921 | rs302802 | -2.047 | -0.976 | 2.239 | 0.131 | -0.031 | -0.126 | -0.126 | 0.173 | 0.24 | 0.55 | 0.18 | 0.01 | 0.21 | 0.18 |
| 1 | 99641557 | rs819790 | -1.698 | -1.003 | 2.328 | 0.141 | 0.012 | 0.004 | 0.004 | 3.959 | 0.06 | 0.26 | 0.02 | 0 | 0.01 | 0.06 |
| 1 | 99641669 | rs638723 | -2.73 | -1.029 | 2.036 | 0.232 | 0.027 | 0.02 | 0.02 | 0 | 0.35 | 0.75 | 0.47 | 0.28 | 0.36 | 0.29 |
| 1 | 99646787 | rs17119881 | -2.442 | -0.543 | 2.162 | 0.408 | 0.026 | -0.051 | -0.051 | 0 | 0.09 | 0.26 | 0.12 | 0.01 | 0.13 | 0.11 |
| 1 | 99647421 | rs611829 | -0.791 | -0.612 | 2.429 | 0.328 | 0.002 | -0.062 | -0.062 | 0 | 0.11 | 0.28 | 0.07 | 0 | 0.09 | 0.08 |
| 1 | 99655620 | rs819785 | -1.916 | -0.608 | 2.359 | 0.415 | 0.009 | 0.039 | 0.039 | 0 | 0.2 | 0.35 | 0.4 | 0.28 | 0.29 | 0.22 |
| 1 | 99659671 | rs632284 | -1.984 | -0.573 | 2.153 | 0.439 | -0.018 | 0.008 | 0.008 | 0 | 0.25 | 0.52 | 0.37 | 0.28 | 0.26 | 0.24 |
| 1 | 99667737 | rs680017 | -2.714 | -0.561 | 2.004 | 0.438 | 0.001 | -0.008 | -0.008 | 0 | 0.29 | 0.53 | 0.46 | 0.28 | 0.38 | 0.27 |
| 1 | 99670492 | rs12097141 | -1.403 | -0.445 | 1.7 | 0.155 | 0.02 | -0.083 | -0.083 | 0.247 | 0.15 | 0.32 | 0.16 | 0 | 0.2 | 0.1 |
| 1 | 99671893 | rs590254 | -1.727 | -0.35 | 1.647 | 0.584 | -0.004 | 0.001 | 0.001 | 0 | 0.17 | 0.54 | 0.24 | 0.2 | 0.17 | 0.22 |
| 1 | 99682616 | rs6660498 | -1.371 | 0.39 | 1.727 | 0.887 | 0.003 | 0.071 | 0.071 | 0 | 0.21 | 0.36 | 0.34 | 0.42 | 0.24 | 0.24 |
| 1 | 99683656 | rs2392011 | -1.25 | 0.384 | 1.792 | 0.884 | 0.005 | 0.049 | 0.049 | 0 | 0.19 | 0.27 | 0.33 | 0.41 | 0.22 | 0.2 |
| 1 | 99691732 | rs7530667 | 1.923 | -0.16 | 1.663 | 0.353 | 0.014 | -0.004 | -0.004 | 0 | 0.6 | 0.32 | 0.52 | 0.58 | 0.61 | 0.67 |
| 1 | 99693159 | rs997484 | 1.995 | 0.043 | 1.677 | 0.33 | 0.01 | -0.004 | -0.004 | 0 | 0.59 | 0.27 | 0.52 | 0.58 | 0.6 | 0.67 |
| 1 | 99693744 | rs819746 | -0.313 | 0.006 | 1.463 | 0.23 | 0.02 | 0.045 | 0.045 | 0.102 | 0.86 | 0.98 | 0.93 | 1 | 0.89 | 0.92 |
| 1 | 99697631 | rs12035062 | 0.035 | 0.161 | 1.5 | 0.45 | -0.001 | 0.01 | 0.01 | 0 | 0.14 | 0.15 | 0.29 | 0.31 | 0.18 | 0.12 |
| 1 | 99698821 | rs7528845 | -1.397 | 0.236 | 1.438 | 0.37 | 0.014 | -0.023 | -0.023 | 0 | 0.06 | 0.14 | 0.06 | 0.01 | 0.07 | 0.03 |
| 1 | 99703278 | rs17120117 | -1.544 | 0.156 | 1.419 | 0.341 | 0.008 | -0.027 | -0.027 | 1.488 | 0.06 | 0.16 | 0.06 | 0.01 | 0.07 | 0.03 |
| 1 | 99705540 | rs1350156 | 1.554 | 0.173 | 0.959 | -0.006 | 0.017 | 0.068 | 0.068 | 0 | 0.52 | 0.38 | 0.5 | 0.68 | 0.53 | 0.55 |
| 1 | 99711388 | rs712915 | -2.485 | 0.092 | 1.766 | 0.23 | 0.041 | -0.016 | -0.016 | 0 | 0.09 | 0.43 | 0.06 | 0.02 | 0.05 | 0.07 |
| 1 | 99723764 | rs712904 | -1.653 | -0.032 | 1.547 | 0.649 | 0.007 | 0.109 | 0.109 | 0 | 0.08 | 0.24 | 0.07 | 0.16 | 0.03 | 0.11 |
| 1 | 99725220 | rs12083740 | -1.049 | -0.033 | 1.593 | 0.648 | -0.005 | 0.052 | 0.052 | 0 | 0.06 | 0.14 | 0.06 | 0.16 | 0.03 | 0.1 |
| 1 | 99725473 | rs712902 | -1.904 | 0.4 | 1.594 | 0.891 | 0 | -0.005 | -0.005 | 0 | 0.34 | 0.62 | 0.46 | 0.59 | 0.35 | 0.46 |
| 1 | 99733964 | rs1456169 | 2.276 | 0.387 | 1.308 | 0.76 | -0.01 | 0.02 | 0.02 | 0 | 0.53 | 0.29 | 0.43 | 0.41 | 0.5 | 0.49 |
| 1 | 99744294 | rs2198772 | -1.293 | 0.418 | 1.367 | 0.917 | 0.002 | 0.099 | 0.099 | 0 | 0.07 | 0.21 | 0.07 | 0.16 | 0.03 | 0.1 |
| 1 | 99748324 | rs712889 | -0.656 | 0.421 | 1.189 | 0.827 | 0.002 | 0.006 | 0.006 | 0 | 0.24 | 0.22 | 0.37 | 0.37 | 0.3 | 0.32 |
| 1 | 99751518 | rs712886 | -2.187 | 0.096 | 1.341 | 0.763 | 0.001 | -0.001 | -0.001 | 0.223 | 0.13 | 0.23 | 0.12 | 0.14 | 0.12 | 0.19 |
| 1 | 99754810 | rs12027179 | -0.262 | 0.263 | 0.92 | 0.67 | -0.081 | -0.064 | -0.064 | 0 | 0.24 | 0.04 | 0.35 | 0.28 | 0.29 | 0.25 |
| 1 | 99756179 | rs712883 | -1.292 | 0.252 | 1.26 | 0.514 | 0.007 | 0.01 | 0.01 | 0 | 0.34 | 0.42 | 0.45 | 0.43 | 0.41 | 0.42 |
| 1 | 99760324 | rs17120188 | 0.666 | 0.058 | 1.279 | 0.231 | -0.001 | -0.024 | -0.024 | 0 | 0.08 | 0.07 | 0.05 | 0.01 | 0.07 | 0.05 |

| | | | | | | | | | | | | | | | |
|---|---|---|---|---|---|---|---|---|---|---|---|---|---|---|---|
| 1 | 99762061 | rs2219257 | 1.39 | 0 | 1.238 | 0.121 | -0.001 | -0.016 | -0.016 | 0.772 | 0.08 | 0.05 | 0.03 | 0 | 0.06 | 0.02 |
| 1 | 99762659 | rs819908 | -1.981 | -0.16 | 1.229 | 0.233 | -0.017 | 0.022 | 0.022 | 0 | 0.48 | 0.73 | 0.56 | 0.59 | 0.51 | 0.46 |
| 1 | 99764226 | rs819910 | -1.633 | -0.286 | 1.365 | 0.362 | 0.001 | 0 | 0 | 0 | 0.4 | 0.61 | 0.52 | 0.61 | 0.46 | 0.44 |
| 1 | 99765872 | rs819914 | -1.444 | -0.258 | 1.13 | 0.384 | -0.004 | -0.002 | -0.002 | 0 | 0.37 | 0.39 | 0.51 | 0.61 | 0.46 | 0.43 |
| 1 | 99770075 | rs819920 | -0.293 | -0.271 | 1.174 | 0.361 | -0.001 | -0.045 | -0.045 | 0 | 0.1 | 0.22 | 0.07 | 0.02 | 0.09 | 0.09 |
| 1 | 99776122 | rs1925972 | 0.298 | -0.231 | 1.076 | 0.436 | -0.025 | -0.045 | -0.045 | 0 | 0.2 | 0.14 | 0.32 | 0.42 | 0.3 | 0.25 |
| 1 | 99776724 | rs819923 | -2.238 | -1.15 | 0.472 | 0.059 | -0.032 | 0.01 | 0.01 | 0.014 | 0.65 | 0.74 | 0.75 | 0.71 | 0.57 | 0.62 |
| 1 | 99780284 | rs712878 | -1.488 | -0.035 | 1.178 | 0.554 | 0.015 | 0.002 | 0.002 | 0 | 0.38 | 0.51 | 0.51 | 0.61 | 0.46 | 0.43 |
| 1 | 99782478 | rs7514675 | -2.103 | 0.15 | 1.086 | 0.525 | -0.008 | 0.013 | 0.013 | 0.233 | 0.2 | 0.34 | 0.18 | 0.25 | 0.13 | 0.19 |
| 1 | 99782657 | rs7555492 | -1.05 | 0.027 | 0.418 | 0.196 | 0.028 | 0.062 | 0.062 | 0.956 | 0.48 | 0.74 | 0.56 | 0.61 | 0.41 | 0.49 |
| 1 | 99783335 | rs7520594 | -1.128 | -0.031 | 0.54 | 0.108 | 0.006 | 0.058 | 0.058 | 0.502 | 0.52 | 0.72 | 0.63 | 0.64 | 0.47 | 0.49 |
| 1 | 99783758 | rs7518011 | -0.665 | -0.048 | 0.67 | 0.08 | 0.028 | 0.041 | 0.041 | 0.068 | 0.46 | 0.33 | 0.59 | 0.63 | 0.43 | 0.41 |
| 1 | 99783891 | rs2148154 | -1.194 | -0.035 | 0.501 | 0.09 | -0.003 | 0.064 | 0.064 | 0.487 | 0.53 | 0.75 | 0.63 | 0.64 | 0.47 | 0.49 |
| 1 | 99789514 | rs12025099 | -0.559 | 0.187 | 0.215 | 0.213 | 0.008 | 0.022 | 0.022 | 0 | 0.41 | 0.3 | 0.58 | 0.63 | 0.42 | 0.41 |
| 1 | 99800131 | rs4908314 | -0.584 | 0.491 | -0.256 | 0.478 | 0.003 | -0.007 | -0.007 | 0 | 0.43 | 0.44 | 0.54 | 0.44 | 0.42 | 0.38 |
| 1 | 99800604 | rs7525165 | -0.08 | 0.445 | -0.091 | 0.095 | 0.008 | -0.013 | -0.013 | 0 | 0.28 | 0.37 | 0.25 | 0.2 | 0.33 | 0.33 |
| 1 | 99800813 | rs17120238 | -0.798 | -0.576 | -0.272 | -0.401 | 0.001 | -0.008 | -0.008 | 0 | 0.3 | 0.27 | 0.2 | 0.09 | 0.23 | 0.22 |
| 1 | 99802789 | rs13374397 | -0.494 | -1.021 | 0.163 | -0.195 | -0.068 | -0.033 | -0.033 | 0 | 0.19 | 0.34 | 0.12 | 0.13 | 0.08 | 0.25 |
| 1 | 99809451 | rs10875217 | -0.485 | -1.083 | 0.405 | -0.225 | -0.084 | -0.043 | -0.043 | 0 | 0.2 | 0.39 | 0.13 | 0.13 | 0.08 | 0.27 |
| 1 | 99811487 | rs10493911 | -0.895 | -1.178 | 0.648 | 0.4 | -0.059 | 0.021 | 0.021 | 0.384 | 0.25 | 0.52 | 0.17 | 0.35 | 0.12 | 0.37 |
| 1 | 99811891 | rs12059105 | -1.548 | -1.81 | 0.59 | 0.766 | 0.017 | -0.016 | -0.016 | 0 | 0.33 | 0.7 | 0.35 | 0.51 | 0.15 | 0.47 |
| 1 | 99835240 | rs6682316 | -1.459 | -1.863 | 0.898 | 0.633 | 0.028 | -0.007 | -0.007 | 0.067 | 0.32 | 0.73 | 0.3 | 0.38 | 0.15 | 0.39 |
| 1 | 99852714 | rs1148347 | 1.621 | -1.816 | 0.146 | -0.186 | 0.005 | 0.257 | 0.257 | 0 | 0.43 | 0.08 | 0.42 | 0.06 | 0.6 | 0.28 |
| 1 | 99853467 | rs12409296 | -0.329 | -1.759 | 0.682 | 0.07 | 0.036 | 0.153 | 0.153 | 0 | 0.06 | 0.25 | 0.06 | 0.18 | 0.05 | 0.11 |
| 1 | 99864803 | rs12021787 | -1.422 | -2.036 | 0.534 | -0.025 | 0.059 | -0.01 | -0.01 | 0 | 0.35 | 0.76 | 0.31 | 0.41 | 0.15 | 0.42 |
| 1 | 99882119 | rs6577377 | -0.768 | -2.077 | 0.983 | -0.045 | -0.033 | -0.01 | -0.01 | 0 | 0.28 | 0.59 | 0.13 | 0.26 | 0.13 | 0.31 |
| 1 | 99907647 | rs7542970 | -1.434 | -1.982 | 0.764 | 0.069 | -0.017 | -0.018 | -0.018 | 0 | 0.94 | 0.99 | 0.91 | 0.93 | 0.94 | 0.95 |
| 1 | 99911058 | rs1419059 | -0.845 | -1.357 | 0.157 | -0.874 | -0.027 | -0.012 | -0.012 | 0 | 0.62 | 0.77 | 0.54 | 0.45 | 0.46 | 0.47 |
| 1 | 99913450 | rs6695717 | -1.014 | -1.318 | 0.099 | -0.717 | -0.033 | 0.095 | 0.095 | 0.664 | 0.12 | 0.25 | 0.07 | 0.28 | 0.06 | 0.15 |
| 1 | 99918960 | rs11166247 | -0.256 | -1.22 | -0.034 | -0.685 | -0.006 | 0.114 | 0.114 | 0 | 0.07 | 0.13 | 0.06 | 0.3 | 0.05 | 0.11 |
| 1 | 99922216 | rs10875223 | -1.364 | -1.317 | -0.252 | -0.842 | -0.05 | 0.111 | 0.111 | 0.011 | 0.14 | 0.39 | 0.08 | 0.29 | 0.06 | 0.18 |
| 1 | 99924542 | rs12141092 | 1.739 | -1.115 | -1.075 | -0.931 | -0.085 | 0.109 | 0.109 | 5.242 | 0.27 | 0.09 | 0.26 | 0.09 | 0.32 | 0.07 |
| 1 | 99926891 | rs1935720 | -1.103 | -1.483 | -0.927 | -0.954 | 0.005 | 0.001 | 0.001 | 0.282 | 0.35 | 0.63 | 0.23 | 0.35 | 0.14 | 0.43 |
| 1 | 99933338 | rs12144940 | 1.896 | -1.057 | -1.242 | -1.157 | -0.064 | 0.165 | 0.165 | 5.338 | 0.25 | 0.01 | 0.22 | 0.04 | 0.26 | 0.06 |
| 1 | 99936756 | rs713266 | 0.623 | -1.245 | -0.985 | -0.912 | 0.064 | -0.017 | -0.017 | 0 | 0.2 | 0.1 | 0.16 | 0.19 | 0.09 | 0.27 |
| 1 | 99946091 | rs4408174 | -0.809 | -1.365 | -0.802 | -0.976 | -0.001 | 0.006 | 0.006 | 0.155 | 0.32 | 0.7 | 0.2 | 0.19 | 0.1 | 0.35 |
| 1 | 99947106 | rs12123486 | 1.898 | -0.977 | -0.952 | -1.108 | -0.032 | 0.043 | 0.043 | 0 | 0.19 | 0.08 | 0.28 | 0.02 | 0.29 | 0.22 |
| 1 | 99949506 | rs17120518 | -0.638 | -1.143 | -0.713 | -0.872 | -0.015 | -0.001 | -0.001 | 0 | 0.3 | 0.68 | 0.2 | 0.19 | 0.1 | 0.34 |
| 1 | 99951162 | rs17120522 | 0.333 | -1.093 | -0.55 | -1.385 | 0.065 | 0.055 | 0.055 | 0.237 | 0.06 | 0.39 | 0.02 | 0 | 0 | 0.02 |
| 1 | 99951627 | rs17120526 | 1.284 | -1.059 | -1.36 | -1.353 | 0.034 | 0.042 | 0.042 | 0.301 | 0.16 | 0.01 | 0.05 | 0.06 | 0.08 | 0.18 |
| 1 | 99952060 | rs17120530 | -0.672 | -0.403 | -1.302 | -1.3 | 0.024 | 0.034 | 0.034 | 0 | 0.37 | 0.71 | 0.19 | 0.11 | 0.26 | 0.4 |
| 1 | 99952500 | rs12563719 | 0.596 | 0.2 | -1.16 | -0.393 | -0.021 | -0.032 | -0.032 | 0 | 0.18 | 0.1 | 0.34 | 0.69 | 0.23 | 0.3 |
| 1 | 99956689 | rs17403124 | 0.022 | 0.272 | -0.986 | -0.773 | 0.014 | -0.037 | -0.037 | 0.872 | 0.36 | 0.43 | 0.36 | 0.13 | 0.46 | 0.4 |
| 1 | 99956800 | rs9628986 | 0.427 | -0.276 | -1.619 | -0.832 | 0.015 | 0.149 | 0.149 | 0.245 | 0.38 | 0.02 | 0.25 | 0.2 | 0.28 | 0.25 |
| 1 | 99957207 | rs2392047 | 0.246 | -0.525 | -1.096 | -0.501 | 0.017 | 0.057 | 0.057 | 0 | 0.54 | 0.27 | 0.5 | 0.37 | 0.59 | 0.49 |
| 1 | 99960670 | rs10493914 | 0.169 | -0.645 | -0.712 | -0.177 | 0.013 | -0.036 | -0.036 | 0 | 0.64 | 0.87 | 0.52 | 0.42 | 0.66 | 0.52 |
| 1 | 99963251 | rs6577380 | 0.24 | -0.662 | -0.622 | -0.185 | 0.019 | -0.019 | -0.019 | 0.033 | 0.63 | 0.8 | 0.51 | 0.42 | 0.66 | 0.52 |
| 1 | 99966319 | rs12132420 | 0.587 | -0.339 | 0.498 | -0.186 | 0.01 | -0.046 | -0.046 | 2.15 | 0.08 | 0.24 | 0.05 | 0 | 0.09 | 0.05 |
| 1 | 99970499 | rs2027358 | 1.398 | -0.338 | 0.227 | 0.062 | 0.001 | -0.027 | -0.027 | 0 | 0.12 | 0.04 | 0.24 | 0.22 | 0.12 | 0.1 |
| 1 | 99970527 | rs10875226 | 0.861 | -0.402 | 0.352 | 0.005 | 0.012 | 0.008 | 0.008 | 0 | 0.55 | 0.34 | 0.44 | 0.42 | 0.58 | 0.57 |
| 1 | 99971142 | rs12083521 | 0.621 | -0.358 | 0.278 | 0.026 | 0.002 | 0.001 | 0.001 | 0 | 0.57 | 0.59 | 0.45 | 0.42 | 0.58 | 0.58 |
| 1 | 99994648 | rs12060602 | -0.762 | -0.047 | 0.314 | -0.249 | 0.002 | -0.004 | -0.004 | 0 | 0.35 | 0.64 | 0.33 | 0.38 | 0.31 | 0.33 |
| 1 | 100000827 | rs6678176 | -0.825 | -0.075 | 0.457 | -0.256 | -0.004 | 0.006 | 0.006 | 0 | 0.34 | 0.55 | 0.32 | 0.38 | 0.31 | 0.33 |
| 1 | 100005477 | rs12069019 | 0.71 | -0.119 | 0.345 | -0.28 | 0.001 | -0.009 | -0.009 | 0 | 0.2 | 0.11 | 0.21 | 0.32 | 0.16 | 0.22 |
| 1 | 100006117 | rs6686057 | 0.987 | -0.184 | -0.299 | -0.598 | -0.081 | 0.103 | 0.103 | 0 | 0.28 | 0.08 | 0.32 | 0.07 | 0.41 | 0.26 |
| 1 | 100018587 | rs11807493 | 0.014 | -0.386 | 0.466 | 0.147 | 0.02 | -0.053 | -0.053 | 0 | 0.09 | 0.33 | 0.05 | 0 | 0.09 | 0.06 |
| 1 | 100019269 | rs1339866 | -1.036 | -0.356 | 0.102 | 0.201 | 0.007 | -0.009 | -0.009 | 0 | 0.35 | 0.69 | 0.33 | 0.37 | 0.31 | 0.33 |
| 1 | 100026535 | rs11166274 | -0.488 | -0.305 | 0.021 | 0.527 | -0.006 | 0.037 | 0.037 | 0 | 0.25 | 0.32 | 0.27 | 0.37 | 0.22 | 0.28 |
| 1 | 100031574 | rs17120680 | -1.188 | -0.135 | -0.147 | 0.027 | -0.004 | -0.003 | -0.003 | 0 | 0.06 | 0.14 | 0.07 | 0.06 | 0.06 | 0.05 |
| 1 | 100040100 | rs1000792 | 0.454 | -0.288 | -0.215 | 0.377 | -0.001 | -0.005 | -0.005 | 0 | 0.16 | 0.12 | 0.24 | 0.18 | 0.16 | 0.15 |
| 1 | 100043789 | rs1339856 | -0.565 | -0.527 | -0.454 | -0.47 | 0.085 | 0.073 | 0.073 | 0 | 0.52 | 0.75 | 0.61 | 0.38 | 0.68 | 0.43 |
| 1 | 100046246 | rs6702619 | 0.726 | -0.408 | -0.538 | -0.815 | -0.091 | 0.079 | 0.079 | 0.775 | 0.26 | 0.08 | 0.26 | 0.07 | 0.48 | 0.25 |
| 1 | 100049785 | rs7543130 | 0.627 | -0.382 | -0.277 | -0.803 | 0.009 | -0.021 | -0.021 | 0 | 0.27 | 0.22 | 0.26 | 0.07 | 0.48 | 0.25 |
| 1 | 100051668 | rs2893418 | -0.303 | -0.486 | 0.147 | -0.551 | 0.002 | 0.002 | 0.002 | 0 | 0.48 | 0.49 | 0.57 | 0.32 | 0.64 | 0.4 |
| 1 | 100059035 | rs6702910 | 0.169 | -0.446 | -0.192 | -0.551 | 0.004 | 0.019 | 0.019 | 0 | 0.42 | 0.33 | 0.54 | 0.31 | 0.61 | 0.36 |
| 1 | 100059600 | rs10875238 | 0.188 | -0.344 | -0.187 | -0.545 | 0.002 | 0.018 | 0.018 | 0 | 0.42 | 0.33 | 0.54 | 0.31 | 0.59 | 0.36 |
| 1 | 100062417 | rs6672117 | -0.06 | -0.492 | -1.093 | -0.231 | 0.08 | 0.094 | 0.094 | 0 | 0.5 | 0.68 | 0.58 | 0.32 | 0.66 | 0.4 |
| 1 | 100063082 | rs2392039 | -0.161 | -0.475 | -1.154 | -0.217 | 0.086 | 0.07 | 0.07 | 0 | 0.52 | 0.74 | 0.61 | 0.38 | 0.7 | 0.43 |
| 1 | 100066515 | rs7518025 | 0.001 | 0.364 | -0.994 | -0.842 | -0.013 | 0.006 | 0.006 | 0 | 0.38 | 0.34 | 0.39 | 0.05 | 0.47 | 0.21 |

| | | | | | | | | | | | | | | | | |
|---|---|---|---|---|---|---|---|---|---|---|---|---|---|---|---|---|
| 1 | 100067902 | rs4908329 | -0.503 | 0.206 | -1.187 | -0.5 | 0.011 | 0.07 | 0.07 | 0.01 | 0.49 | 0.4 | 0.44 | 0.28 | 0.57 | 0.42 |
| 1 | 100069801 | rs12025301 | 0.254 | 0.255 | -1.381 | -0.456 | -0.01 | -0.036 | -0.036 | 0 | 0.11 | 0.03 | 0.07 | 0.22 | 0.14 | 0.21 |
| 1 | 100075695 | rs7554246 | 0.847 | -0.048 | -1.623 | -0.95 | -0.081 | 0.118 | 0.118 | 0 | 0.28 | 0.1 | 0.3 | 0.02 | 0.44 | 0.14 |
| 1 | 100079760 | rs6689789 | 0.147 | -0.206 | -0.968 | -0.972 | -0.018 | 0.033 | 0.033 | 0 | 0.38 | 0.29 | 0.39 | 0.06 | 0.47 | 0.21 |
| 1 | 100081117 | rs12729121 | 0.434 | -0.184 | -0.888 | -0.806 | -0.003 | -0.013 | -0.013 | 2.045 | 0.12 | 0.05 | 0.07 | 0.15 | 0.14 | 0.21 |
| 1 | 100085388 | rs835004 | 0.915 | -0.748 | -0.423 | -0.678 | -0.023 | -0.001 | -0.001 | 0 | 0.15 | 0.27 | 0.09 | 0.15 | 0.12 | 0.08 |
| 1 | 100086858 | rs17648010 | 1.005 | -0.813 | -0.129 | -0.217 | 0 | 0.005 | 0.005 | 0.028 | 0.3 | 0.25 | 0.44 | 0.53 | 0.24 | 0.37 |
| 1 | 100087162 | rs834979 | -0.438 | -0.718 | -0.167 | 0.033 | -0.006 | 0.002 | 0.002 | 0 | 0.26 | 0.31 | 0.17 | 0.3 | 0.25 | 0.29 |
| 1 | 100091852 | rs844330 | -0.195 | -0.377 | -0.181 | -0.145 | -0.006 | -0.001 | -0.001 | 0 | 0.21 | 0.31 | 0.21 | 0.21 | 0.18 | 0.22 |
| 1 | 100092765 | rs11804292 | 0.262 | -0.609 | -0.396 | -0.304 | 0.027 | 0.063 | 0.063 | 0 | 0.2 | 0.09 | 0.08 | 0.02 | 0.15 | 0.26 |
| 1 | 100097419 | rs834972 | -0.825 | -0.595 | -0.368 | 0.272 | 0.033 | 0.004 | 0.004 | 0.172 | 0.65 | 0.78 | 0.62 | 0.66 | 0.81 | 0.61 |
| 1 | 100101515 | rs17120799 | 0.513 | -0.336 | -0.535 | 0.46 | -0.008 | 0.02 | 0.02 | 0 | 0.09 | 0.05 | 0.06 | 0.38 | 0.13 | 0.09 |
| 1 | 100102788 | rs834963 | 0.678 | -0.414 | -0.281 | 0.308 | -0.003 | -0.034 | -0.034 | 0 | 0.2 | 0.23 | 0.09 | 0.02 | 0.15 | 0.25 |
| 1 | 100105385 | rs834961 | -0.397 | -0.276 | -0.225 | 0.338 | -0.003 | 0.008 | 0.008 | 0 | 0.29 | 0.34 | 0.24 | 0.4 | 0.28 | 0.31 |
| 1 | 100107260 | rs17405754 | 0.775 | -0.17 | -0.082 | -0.38 | -0.01 | 0.006 | 0.006 | 0 | 0.34 | 0.33 | 0.44 | 0.26 | 0.51 | 0.26 |
| 1 | 100107407 | rs835002 | 0.056 | -0.109 | 0.052 | -0.449 | -0.019 | -0.084 | -0.084 | 0.044 | 0.2 | 0.33 | 0.1 | 0.02 | 0.15 | 0.22 |
| 1 | 100111956 | rs834997 | -0.574 | 0.542 | -0.203 | -0.57 | 0.005 | -0.124 | -0.124 | 0.021 | 0.28 | 0.43 | 0.16 | 0.01 | 0.33 | 0.28 |
| 1 | 100114430 | rs7513463 | 0.717 | 0.134 | -0.578 | -0.01 | -0.021 | -0.022 | -0.022 | 0 | 0.31 | 0.23 | 0.39 | 0.38 | 0.41 | 0.37 |
| 1 | 100115032 | rs834994 | -0.431 | -0.669 | -0.019 | -0.066 | -0.044 | -0.004 | -0.004 | 0 | 0.83 | 0.61 | 0.93 | 0.86 | 0.92 | 0.83 |
| 1 | 100116253 | rs834993 | -0.189 | -0.707 | -0.287 | 0.16 | -0.001 | 0.004 | 0.004 | 0 | 0.45 | 0.62 | 0.58 | 0.58 | 0.43 | 0.39 |
| 1 | 100116394 | rs834992 | -0.722 | -0.695 | -0.264 | 0.064 | 0 | -0.002 | -0.002 | 0 | 0.63 | 0.68 | 0.68 | 0.69 | 0.6 | 0.58 |
| 1 | 100118418 | rs3766580 | 0.386 | -1.245 | 0.029 | 0.249 | -0.01 | 0.013 | 0.013 | 0.91 | 0.08 | 0.1 | 0.04 | 0.13 | 0.04 | 0.06 |
| 1 | 100127618 | rs3766582 | 0.543 | -1.184 | 0.215 | 0.259 | 0.035 | 0.134 | 0.134 | 0 | 0.06 | 0.22 | 0.21 | 0.15 | 0.06 | 0.07 |
| 1 | 100135336 | rs10783115 | -1.47 | -1.304 | -0.063 | 0.058 | 0.03 | 0.025 | 0.025 | 0 | 0.7 | 0.89 | 0.78 | 0.76 | 0.76 | 0.64 |
| 1 | 100139730 | rs7524870 | 1.555 | -1.16 | -0.076 | 0.031 | 0 | 0.021 | 0.021 | 0 | 0.17 | 0.1 | 0.1 | 0.11 | 0.16 | 0.19 |
| 1 | 100143458 | rs1981193 | 0.053 | -0.836 | 0.27 | 0.192 | -0.006 | -0.003 | -0.003 | 1.722 | 0.56 | 0.55 | 0.51 | 0.54 | 0.59 | 0.51 |
| 1 | 100145443 | rs10875241 | 0.045 | -0.757 | 0.388 | 0.342 | 0.044 | 0.148 | 0.148 | 0 | 0.07 | 0.25 | 0.21 | 0.16 | 0.06 | 0.07 |
| 1 | 100147287 | rs12122854 | 0.087 | -0.97 | 0.249 | 0.162 | -0.002 | -0.001 | -0.001 | 0 | 0.56 | 0.47 | 0.54 | 0.54 | 0.67 | 0.49 |
| 1 | 100148279 | rs12067797 | -0.163 | -0.969 | 0.407 | 0.25 | -0.013 | -0.038 | -0.038 | 0 | 0.07 | 0.18 | 0.02 | 0 | 0.02 | 0.05 |
| 1 | 100149952 | rs12136177 | 0.73 | -1.039 | 0.503 | 0.33 | 0.002 | 0.032 | 0.032 | 0 | 0.06 | 0.17 | 0.04 | 0.08 | 0.04 | 0.05 |
| 1 | 100152443 | rs1338576 | 0.778 | -1.184 | 0.578 | 0.187 | -0.006 | -0.018 | -0.018 | 0 | 0.43 | 0.35 | 0.48 | 0.47 | 0.53 | 0.43 |
| 1 | 100153422 | rs1416239 | 0.685 | -1.206 | 0.592 | 0.176 | -0.005 | -0.018 | -0.018 | 0 | 0.44 | 0.31 | 0.48 | 0.47 | 0.53 | 0.43 |
| 1 | 100153674 | rs4908004 | -1.677 | -1.269 | 0.577 | 0.202 | -0.032 | 0.011 | 0.011 | 0.879 | 0.18 | 0.53 | 0.14 | 0.16 | 0.1 | 0.17 |
| 1 | 100154055 | rs3737899 | 0.478 | -1.232 | 0.647 | 0.164 | 0.015 | 0.017 | 0.017 | 0.373 | 0.49 | 0.42 | 0.55 | 0.55 | 0.58 | 0.49 |
| 1 | 100154467 | rs6678252 | -0.53 | -1.197 | 0.634 | 0.157 | -0.006 | -0.036 | -0.036 | 0.305 | 0.08 | 0.23 | 0.03 | 0 | 0.02 | 0.05 |
| 1 | 100156054 | rs17120949 | 0.395 | -1.163 | 0.384 | 0.173 | -0.009 | -0.025 | -0.025 | 0 | 0.06 | 0.1 | 0.02 | 0 | 0.02 | 0.05 |
| 1 | 100156658 | rs3766588 | -0.743 | -1.161 | 0.324 | 0.183 | -0.024 | -0.012 | -0.012 | 0.058 | 0.63 | 0.95 | 0.63 | 0.63 | 0.64 | 0.6 |
| 1 | 100159418 | rs12130813 | -0.968 | -1.129 | 0.492 | 0.176 | -0.022 | -0.016 | -0.016 | 0 | 0.13 | 0.29 | 0.07 | 0.08 | 0.06 | 0.1 |
| 1 | 100162805 | rs2185132 | -0.762 | -1.125 | 0.033 | 0.223 | -0.029 | -0.039 | -0.039 | 0 | 0.63 | 0.95 | 0.63 | 0.61 | 0.63 | 0.59 |
| 1 | 100163271 | rs6679943 | -0.075 | -0.989 | 0.116 | 0.24 | 0.037 | 0.039 | 0.039 | 0.184 | 0.32 | 0.46 | 0.42 | 0.39 | 0.4 | 0.29 |
| 1 | 100166046 | rs12406373 | 1.473 | -0.852 | 0.062 | 0.237 | 0.034 | 0.037 | 0.037 | 0 | 0.2 | 0.27 | 0.35 | 0.32 | 0.34 | 0.19 |
| 1 | 100175698 | rs12139063 | 0.253 | -1.527 | -0.137 | -0.193 | -0.008 | 0.015 | 0.015 | 0.047 | 0.07 | 0.15 | 0.04 | 0.09 | 0.04 | 0.05 |
| 1 | 100181472 | rs2039725 | -0.894 | -1.884 | -0.845 | -0.384 | 0.071 | 0.076 | 0.076 | 0 | 0.51 | 0.93 | 0.31 | 0.35 | 0.32 | 0.54 |
| 1 | 100187428 | rs12094588 | -1.814 | -2.013 | -1.04 | -0.875 | 0.039 | 0.146 | 0.146 | 0.77 | 0.78 | 0.99 | 0.93 | 0.99 | 0.82 | 0.84 |
| 1 | 100190097 | rs7531793 | -0.268 | -1.792 | 0 | -0.118 | -0.016 | -0.006 | -0.006 | 0.146 | 0.4 | 0.51 | 0.34 | 0.34 | 0.29 | 0.48 |
| 1 | 100190843 | rs12127323 | -0.272 | -1.687 | -0.004 | -0.122 | 0.02 | 0.065 | 0.065 | 0.627 | 0.08 | 0.38 | 0.06 | 0.09 | 0.04 | 0.05 |
| 1 | 100194962 | rs7547543 | -1.106 | -1.61 | -0.873 | -0.081 | 0.051 | -0.049 | -0.049 | 1.369 | 0.51 | 0.95 | 0.5 | 0.44 | 0.32 | 0.53 |
| 1 | 100198059 | rs951125 | -1.238 | -1.942 | -0.442 | -0.572 | -0.008 | -0.017 | -0.017 | 10.527 | 0.26 | 0.65 | 0.23 | 0.11 | 0.1 | 0.29 |
| 1 | 100200184 | rs6670361 | 0.639 | -1.971 | -0.584 | -0.499 | 0.015 | -0.017 | -0.017 | 0 | 0.24 | 0.14 | 0.25 | 0.28 | 0.21 | 0.23 |
| 1 | 100203648 | rs7537296 | -1.511 | -2.091 | -0.667 | -0.578 | 0.073 | 0.03 | 0.03 | 0 | 0.51 | 0.94 | 0.49 | 0.39 | 0.31 | 0.52 |
| 1 | 100203693 | rs7514021 | -1.511 | -2.091 | -0.667 | -0.578 | 0.073 | 0.03 | 0.03 | 0 | 0.51 | 0.94 | 0.49 | 0.39 | 0.31 | 0.52 |
| 1 | 100203902 | rs10875245 | -0.286 | -2.243 | -0.067 | -0.663 | -0.031 | -0.067 | -0.067 | 0 | 0.13 | 0.21 | 0.03 | 0.02 | 0.04 | 0.18 |
| 1 | 100206801 | rs10875246 | -0.633 | -2.255 | -0.084 | -0.669 | -0.026 | -0.026 | -0.026 | 0 | 0.14 | 0.57 | 0.21 | 0.09 | 0.06 | 0.11 |
| 1 | 100210815 | rs11166322 | 0.764 | -2.239 | -0.178 | -0.66 | 0.017 | -0.027 | -0.027 | 0 | 0.24 | 0.11 | 0.25 | 0.28 | 0.21 | 0.23 |
| 1 | 100214487 | rs1416236 | -1.481 | -2.168 | -0.154 | -0.574 | 0.075 | 0.031 | 0.031 | 0 | 0.51 | 0.94 | 0.49 | 0.39 | 0.31 | 0.52 |
| 1 | 100225317 | rs2788517 | -1 | -2.109 | -0.114 | -0.535 | 0.075 | 0.031 | 0.031 | 0.336 | 0.45 | 0.79 | 0.48 | 0.39 | 0.3 | 0.52 |
| 1 | 100232074 | rs12127190 | 0.223 | -2.093 | -0.084 | -0.531 | 0.015 | 0.002 | 0.002 | 0.774 | 0.31 | 0.22 | 0.27 | 0.3 | 0.24 | 0.41 |
| 1 | 100234367 | rs4908018 | -0.614 | -2.124 | 0.041 | -0.532 | -0.016 | -0.015 | -0.015 | 0 | 0.14 | 0.42 | 0.2 | 0.09 | 0.06 | 0.11 |
| 1 | 100237542 | rs11166329 | 0.176 | -1.96 | -0.487 | -0.438 | 0.023 | -0.002 | -0.002 | 0 | 0.3 | 0.18 | 0.26 | 0.3 | 0.24 | 0.4 |
| 1 | 100250197 | rs931520 | 0.956 | -1.878 | -0.154 | -0.448 | -0.044 | -0.041 | -0.041 | 0.003 | 0.83 | 0.35 | 0.84 | 0.92 | 0.95 | 0.96 |
| 1 | 100252145 | rs12137801 | -0.818 | -2 | -0.013 | -0.576 | -0.043 | -0.084 | -0.084 | 0.03 | 0.19 | 0.5 | 0.14 | 0.1 | 0.08 | 0.22 |
| 1 | 100268991 | rs6689305 | -0.233 | -2.433 | -0.341 | -0.69 | -0.034 | -0.001 | -0.001 | 0.091 | 0.71 | 0.62 | 0.7 | 0.71 | 0.79 | 0.75 |
| 1 | 100271638 | rs12401503 | 1.444 | -2.459 | -0.607 | -0.673 | 0.046 | -0.011 | -0.011 | 0 | 0.46 | 0.08 | 0.51 | 0.54 | 0.66 | 0.46 |
| 1 | 100274670 | rs1389383 | 1.582 | -2.522 | -0.41 | -0.669 | 0.08 | 0.115 | 0.115 | 0.169 | 0.67 | 0.15 | 0.78 | 0.83 | 0.87 | 0.7 |
| 1 | 100274681 | rs1908150 | -0.804 | -2.439 | -0.287 | -0.62 | -0.037 | -0.06 | -0.06 | 0 | 0.21 | 0.51 | 0.14 | 0.16 | 0.11 | 0.23 |
| 1 | 100276218 | rs12121352 | -0.087 | -2.314 | -0.074 | -0.376 | 0.004 | 0.002 | 0.002 | 0.583 | 0.94 | 0.78 | 0.9 | 0.97 | 0.98 | 0.93 |
| 1 | 100278603 | rs6696562 | -0.221 | -2.038 | -0.472 | -0.575 | -0.002 | 0.003 | 0.003 | 0.165 | 0.65 | 0.68 | 0.61 | 0.64 | 0.65 | 0.65 |
| 1 | 100280436 | rs6663234 | 0.557 | -1.998 | -0.647 | -0.554 | -0.054 | -0.039 | -0.039 | 0.049 | 0.29 | 0.1 | 0.33 | 0.33 | 0.33 | 0.27 |
| 1 | 100286234 | rs4907896 | -0.209 | -2.047 | -0.671 | -0.538 | -0.023 | 0.011 | 0.011 | 0 | 0.4 | 0.12 | 0.39 | 0.36 | 0.38 | 0.36 |

| | | | | | | | | | | | | | | | | |
|---|---|---|---|---|---|---|---|---|---|---|---|---|---|---|---|---|
| 1 | 100292904 | rs10875268 | -1.031 | -2.12 | -0.391 | -0.801 | -0.053 | -0.146 | -0.146 | 0 | 0.76 | 0.65 | 0.92 | 1 | 0.85 | 0.86 |
| 1 | 100314969 | rs17121311 | 0.653 | -1.967 | -0.089 | -0.463 | -0.014 | 0.004 | 0.004 | 0 | 0.05 | 0.09 | 0.02 | 0.1 | 0.02 | 0.03 |
| 1 | 100316589 | rs2307130 | 0.545 | -2.119 | -0.078 | -0.54 | -0.041 | -0.022 | -0.022 | 0 | 0.33 | 0.22 | 0.47 | 0.52 | 0.46 | 0.54 |
| 1 | 100327026 | rs2307129 | 0.439 | -2.462 | 0.181 | -0.774 | -0.069 | -0.093 | -0.093 | 0 | 0.29 | 0.42 | 0.13 | 0.13 | 0.18 | 0.18 |
| 1 | 100331265 | rs2640911 | 0.959 | -2.471 | 0.075 | -0.617 | -0.02 | -0.019 | -0.019 | 0 | 0.31 | 0.27 | 0.43 | 0.47 | 0.44 | 0.46 |
| 1 | 100331481 | rs574592 | -1.172 | -2.578 | -0.027 | -0.67 | -0.104 | -0.062 | -0.062 | 1.758 | 0.71 | 0.93 | 0.7 | 0.68 | 0.68 | 0.75 |
| 1 | 100332985 | rs12756830 | 1.064 | -2.41 | 0.08 | -0.577 | -0.072 | -0.068 | -0.068 | 0.034 | 0.3 | 0.15 | 0.42 | 0.47 | 0.44 | 0.46 |
| 1 | 100340225 | rs634880 | -1.153 | -2.192 | -0.271 | -0.26 | -0.032 | -0.035 | -0.035 | 0 | 0.73 | 0.87 | 0.68 | 0.7 | 0.69 | 0.72 |
| 1 | 100342227 | rs3736295 | -0.261 | -2.422 | -0.105 | -0.092 | 0.005 | 0.018 | 0.018 | 0 | 0.45 | 0.42 | 0.61 | 0.68 | 0.53 | 0.61 |
| 1 | 100342845 | rs3820155 | 0.878 | -2.336 | -0.095 | 0.012 | -0.018 | -0.019 | -0.019 | 0 | 0.33 | 0.29 | 0.41 | 0.45 | 0.43 | 0.47 |
| 1 | 100349381 | rs17409324 | 0.995 | -2.324 | -0.229 | -0.03 | -0.08 | -0.083 | -0.083 | 0 | 0.32 | 0.17 | 0.41 | 0.46 | 0.43 | 0.47 |
| 1 | 100358103 | rs3753494 | 1.262 | -2.512 | 0.141 | -0.824 | 0.008 | -0.014 | -0.014 | 0.23 | 0.23 | 0.12 | 0.1 | 0.03 | 0.16 | 0.15 |
| 1 | 100359333 | rs620281 | -1.726 | -2.434 | 0.138 | -0.765 | -0.016 | 0.027 | 0.027 | 0 | 0.78 | 0.84 | 0.82 | 0.85 | 0.73 | 0.8 |
| 1 | 100373601 | rs1577541 | 1.224 | -1.964 | 0.164 | -0.539 | -0.037 | -0.025 | -0.025 | 0 | 0.27 | 0.21 | 0.52 | 0.55 | 0.42 | 0.32 |
| 1 | 100375347 | rs12039074 | 1.186 | -1.968 | 0.345 | -0.526 | 0.005 | 0.001 | 0.001 | 0.427 | 0.27 | 0.25 | 0.52 | 0.57 | 0.42 | 0.32 |
| 1 | 100376448 | rs2274570 | 0.394 | -2.067 | 0.516 | -0.599 | 0.007 | 0 | 0 | 0 | 0.36 | 0.35 | 0.57 | 0.57 | 0.5 | 0.38 |
| 1 | 100378651 | rs6577147 | -0.291 | -2.354 | 0.586 | -0.802 | 0.023 | 0.038 | 0.038 | 0 | 0.55 | 0.62 | 0.67 | 0.64 | 0.65 | 0.52 |
| 1 | 100384402 | rs7519706 | 1.415 | -2.24 | 0.592 | -0.742 | -0.026 | -0.018 | -0.018 | 0 | 0.27 | 0.22 | 0.52 | 0.55 | 0.42 | 0.32 |
| 1 | 100385309 | rs11166375 | 1.427 | -2.24 | 0.607 | -0.743 | -0.058 | -0.04 | -0.04 | 0 | 0.27 | 0.18 | 0.52 | 0.55 | 0.42 | 0.32 |
| 1 | 100388503 | rs1804809 | 1.507 | -2.234 | 0.551 | -0.732 | -0.057 | -0.04 | -0.04 | 0 | 0.26 | 0.17 | 0.52 | 0.55 | 0.42 | 0.32 |
| 1 | 100388748 | rs3766603 | 1.363 | -2.224 | 0.489 | -0.727 | 0.014 | 0.008 | 0.008 | 0.404 | 0.27 | 0.27 | 0.52 | 0.55 | 0.42 | 0.32 |
| 1 | 100399757 | rs6577150 | -0.856 | -2.347 | 0.41 | -0.784 | 0.003 | 0.009 | 0.009 | 0 | 0.63 | 0.6 | 0.75 | 0.68 | 0.68 | 0.55 |
| 1 | 100401893 | rs2025936 | 1.739 | -2.391 | 0.359 | -0.787 | 0.008 | 0.006 | 0.006 | 0 | 0.07 | 0.05 | 0.09 | 0.04 | 0.02 | 0.03 |
| 1 | 100403265 | rs12074648 | 1.486 | -2.386 | 0.459 | -0.791 | -0.03 | -0.022 | -0.022 | 0.023 | 0.27 | 0.22 | 0.53 | 0.55 | 0.44 | 0.32 |
| 1 | 100405465 | rs6703159 | -0.857 | -2.281 | 0.381 | -0.78 | 0.007 | 0.018 | 0.018 | 0 | 0.63 | 0.68 | 0.76 | 0.69 | 0.69 | 0.55 |
| 1 | 100410636 | rs2149193 | 0.393 | -2.273 | 0.406 | -0.774 | 0 | 0.001 | 0.001 | 0 | 0.47 | 0.4 | 0.62 | 0.62 | 0.6 | 0.47 |
| 1 | 100411287 | rs1541044 | 1.515 | -2.272 | 0.39 | -0.891 | -0.004 | -0.033 | -0.033 | 0 | 0.18 | 0.12 | 0.1 | 0.03 | 0.16 | 0.15 |
| 1 | 100414740 | rs1571351 | -0.654 | -2.215 | 0.368 | -0.832 | 0.003 | 0.005 | 0.005 | 0 | 0.62 | 0.59 | 0.76 | 0.68 | 0.69 | 0.55 |
| 1 | 100426978 | rs6701658 | -0.241 | -2.139 | 0.261 | -0.906 | -0.013 | -0.012 | -0.012 | 1.695 | 0.13 | 0.15 | 0.04 | 0.05 | 0.07 | 0.01 |
| 1 | 100428784 | rs11166382 | 0.746 | -2.083 | 0.444 | -0.958 | -0.009 | -0.049 | -0.049 | 0 | 0.23 | 0.2 | 0.13 | 0.04 | 0.19 | 0.16 |
| 1 | 100449797 | rs11166387 | 0.551 | -1.336 | 1.148 | -0.29 | -0.011 | -0.003 | -0.003 | 0 | 0.07 | 0.13 | 0.02 | 0.05 | 0.05 | 0.01 |
| 1 | 100458417 | rs12146139 | -1.753 | -1.434 | 1.257 | -0.39 | -0.015 | -0.109 | -0.109 | 6.477 | 0.17 | 0.34 | 0.09 | 0.03 | 0.1 | 0.09 |
| 1 | 100466279 | rs11166389 | -1.517 | -1.434 | 1.25 | -0.39 | -0.009 | -0.103 | -0.103 | 0 | 0.16 | 0.33 | 0.09 | 0.03 | 0.1 | 0.09 |
| 1 | 100482396 | rs575323 | 1.266 | -1.442 | 1.212 | -0.39 | 0.002 | -0.03 | -0.03 | 0 | 0.08 | 0.12 | 0.03 | 0.02 | 0.05 | 0.06 |
| 1 | 100488177 | rs482466 | -2.752 | -1.473 | 1.377 | -0.406 | -0.01 | -0.033 | -0.033 | 0.453 | 0.25 | 0.62 | 0.12 | 0.07 | 0.15 | 0.1 |
| 1 | 100489264 | rs561735 | 1.944 | -1.421 | 1.268 | -0.377 | 0.011 | -0.026 | -0.026 | 0 | 0.06 | 0.12 | 0.03 | 0.02 | 0.05 | 0.06 |
| 1 | 100492867 | rs473688 | -0.065 | -1.43 | 1.315 | -0.382 | 0.006 | -0.066 | -0.066 | 0.196 | 0.12 | 0.27 | 0.06 | 0.02 | 0.07 | 0.07 |
| 1 | 100497091 | rs510463 | 1.456 | -1.43 | 1.322 | -0.382 | 0.003 | -0.03 | -0.03 | 0 | 0.07 | 0.12 | 0.03 | 0.02 | 0.05 | 0.06 |
| 1 | 100502690 | rs12406605 | 1.728 | -1.275 | 1.19 | -0.314 | -0.128 | -0.107 | -0.107 | 0.155 | 0.22 | 0.05 | 0.56 | 0.58 | 0.38 | 0.31 |
| 1 | 100506415 | rs555778 | 0.423 | -1.313 | 1.252 | -0.31 | 0.02 | -0.054 | -0.054 | 0.565 | 0.11 | 0.27 | 0.06 | 0.02 | 0.07 | 0.06 |
| 1 | 100515497 | rs17121792 | 1.598 | -1.253 | 1.203 | -0.276 | -0.002 | 0.006 | 0.006 | 0.486 | 0.07 | 0.05 | 0.03 | 0.02 | 0.05 | 0.06 |
| 1 | 100533387 | rs577539 | 0.54 | -1.255 | 1.243 | -0.267 | -0.005 | 0.007 | 0.007 | 1.419 | 0.07 | 0.19 | 0.03 | 0.05 | 0.05 | 0.01 |
| 1 | 100542638 | rs6699449 | -0.482 | -1.262 | 1.294 | -0.289 | 0.022 | -0.063 | -0.063 | 0 | 0.12 | 0.28 | 0.07 | 0.03 | 0.09 | 0.09 |
| 1 | 100543962 | rs644835 | -0.827 | -1.26 | 1.347 | -0.283 | 0.002 | -0.05 | -0.05 | 0 | 0.16 | 0.42 | 0.07 | 0.07 | 0.12 | 0.08 |
| 1 | 100544651 | rs17121850 | 1.103 | -1.254 | 1.283 | -0.279 | 0.026 | -0.032 | -0.032 | 0 | 0.08 | 0.22 | 0.04 | 0.02 | 0.05 | 0.06 |
| 1 | 100547994 | rs5876 | 1.661 | -1.226 | 1.034 | -0.268 | -0.112 | -0.091 | -0.091 | 1.174 | 0.22 | 0.07 | 0.56 | 0.58 | 0.39 | 0.32 |
| 1 | 100557261 | rs10875279 | 1.134 | -1.228 | 1.117 | -0.272 | -0.006 | 0.005 | 0.005 | 2.088 | 0.07 | 0.15 | 0.03 | 0.05 | 0.05 | 0.01 |
| 1 | 100569440 | rs10875280 | 1.134 | -1.228 | 0.928 | -0.271 | -0.008 | -0.002 | -0.002 | 0 | 0.07 | 0.11 | 0.02 | 0.05 | 0.05 | 0.01 |
| 1 | 100572459 | rs17121889 | 0.988 | -1.245 | 0.898 | -0.291 | 0 | -0.002 | -0.002 | 0 | 0.08 | 0.08 | 0.03 | 0.02 | 0.05 | 0.06 |
| 1 | 100575933 | rs13375867 | 1.667 | -1.168 | 0.904 | -0.236 | -0.111 | -0.093 | -0.093 | 0 | 0.22 | 0.07 | 0.56 | 0.58 | 0.39 | 0.31 |
| 1 | 100579327 | rs2810422 | -1.556 | -1.122 | 0.948 | -0.229 | -0.005 | -0.024 | -0.024 | 0 | 0.94 | 0.92 | 0.98 | 1 | 0.95 | 0.99 |
| 1 | 100587146 | rs17418475 | 0.21 | -1.124 | 0.77 | -0.229 | 0.001 | 0.018 | 0.018 | 7.738 | 0.13 | 0.08 | 0.04 | 0.02 | 0.1 | 0.07 |
| 1 | 100592653 | rs663519 | 1.146 | -1.312 | 0.749 | -0.312 | 0.02 | 0.09 | 0.09 | 0 | 0.47 | 0.29 | 0.32 | 0.33 | 0.46 | 0.46 |
| 1 | 100598867 | rs687513 | 1.833 | -0.919 | 1.022 | -0.149 | -0.009 | -0.039 | -0.039 | 0.028 | 0.78 | 0.38 | 0.89 | 0.93 | 0.85 | 0.9 |
| 1 | 100603891 | rs561119 | 1.855 | -0.998 | 1.02 | -0.151 | -0.011 | -0.039 | -0.039 | 0 | 0.78 | 0.38 | 0.89 | 0.93 | 0.86 | 0.89 |
| 1 | 100609639 | rs6681721 | 1.518 | -0.995 | 1.104 | -0.152 | -0.001 | -0.007 | -0.007 | 0.066 | 0.07 | 0.08 | 0.03 | 0.02 | 0.05 | 0.06 |
| 1 | 100612675 | rs499479 | 0.662 | -1.006 | 0.973 | -0.141 | -0.002 | 0.054 | 0.054 | 0 | 0.56 | 0.31 | 0.34 | 0.35 | 0.47 | 0.59 |
| 1 | 100643910 | rs2810418 | 0.457 | -0.859 | 1.377 | -0.631 | -0.012 | -0.027 | -0.027 | 0.048 | 0.89 | 0.77 | 0.95 | 0.95 | 0.92 | 0.96 |
| 1 | 100644500 | rs11166411 | -0.372 | -0.848 | 1.322 | -0.62 | -0.003 | -0.027 | -0.027 | 0.056 | 0.05 | 0.15 | 0.03 | 0.01 | 0.03 | 0.02 |
| 1 | 100645909 | rs7515676 | 1.361 | -0.851 | 1.289 | -0.642 | -0.008 | -0.066 | -0.066 | 0 | 0.82 | 0.46 | 0.9 | 0.93 | 0.86 | 0.9 |
| 1 | 100650664 | rs2784172 | 1.169 | -0.85 | 1.288 | -0.64 | -0.001 | -0.075 | -0.075 | 0 | 0.83 | 0.54 | 0.91 | 0.93 | 0.87 | 0.9 |
| 1 | 100656265 | rs11166412 | 1.261 | -0.853 | 1.2 | -0.642 | -0.009 | -0.078 | -0.078 | 0 | 0.82 | 0.52 | 0.91 | 0.93 | 0.88 | 0.9 |
| 1 | 100656494 | rs11166413 | 1.521 | -0.831 | 0.524 | -0.194 | -0.135 | -0.05 | -0.05 | 0 | 0.22 | 0.04 | 0.43 | 0.27 | 0.39 | 0.31 |
| 1 | 100683627 | rs6577156 | 1.663 | -0.792 | 0.851 | -0.076 | -0.011 | -0.045 | -0.045 | 0.252 | 0.81 | 0.4 | 0.89 | 0.93 | 0.86 | 0.9 |
| 1 | 100693294 | rs6682782 | 1.44 | -0.705 | 0.763 | -0.046 | -0.015 | -0.075 | -0.075 | 0 | 0.82 | 0.56 | 0.9 | 0.93 | 0.87 | 0.9 |
| 1 | 100715454 | rs3806235 | 1.352 | -0.834 | 0.542 | -0.127 | -0.005 | -0.019 | -0.019 | 0.009 | 0.79 | 0.32 | 0.87 | 0.92 | 0.86 | 0.83 |
| 1 | 100715782 | rs3806237 | 2.225 | -1.149 | 0.382 | -0.033 | -0.012 | 0 | 0 | 3.44 | 0.07 | 0.07 | 0.01 | 0.11 | 0.01 | 0.01 |
| 1 | 100719130 | rs12060945 | 0.675 | -0.745 | 0.052 | -0.145 | 0.037 | 0.094 | 0.094 | 0 | 0.48 | 0.18 | 0.43 | 0.63 | 0.43 | 0.38 |
| 1 | 100728138 | rs3122126 | 1.13 | -0.575 | 1.135 | -0.098 | -0.02 | -0.058 | -0.058 | 0 | 0.83 | 0.65 | 0.89 | 0.92 | 0.87 | 0.84 |

| | | | | | | | | | | | | | | | | |
|---|---|---|---|---|---|---|---|---|---|---|---|---|---|---|---|---|
| 1 | 100731163 | rs3806239 | 1.394 | -0.577 | 0.82 | -0.096 | -0.005 | -0.017 | -0.017 | 0 | 0.8 | 0.31 | 0.87 | 0.92 | 0.86 | 0.84 |
| 1 | 100735903 | rs4579779 | 1.375 | -0.552 | 0.127 | -0.08 | -0.006 | -0.021 | -0.021 | 0 | 0.8 | 0.31 | 0.87 | 0.92 | 0.86 | 0.84 |
| 1 | 100742399 | rs10158777 | 0.875 | -0.284 | 0.121 | 0.082 | -0.002 | -0.048 | -0.048 | 0 | 0.85 | 0.5 | 0.88 | 0.93 | 0.87 | 0.88 |
| 1 | 100765696 | rs6577166 | -0.296 | -0.445 | -0.829 | 0.507 | 0.06 | 0.006 | 0.006 | 0 | 0.5 | 0.76 | 0.67 | 0.49 | 0.59 | 0.57 |
| 1 | 100770760 | rs11807256 | -0.466 | -0.026 | -0.366 | 0.825 | 0.022 | 0.059 | 0.059 | 0.014 | 0.05 | 0.13 | 0.24 | 0.13 | 0.09 | 0.02 |
| 1 | 100781361 | rs3119187 | 0.083 | -0.5 | 0.016 | 0.42 | -0.017 | -0.005 | -0.005 | 0 | 0.15 | 0.35 | 0.07 | 0.15 | 0.09 | 0.12 |
| 1 | 100787265 | rs540486 | -0.956 | -0.737 | -0.282 | 0.379 | 0.011 | 0.007 | 0.007 | 2.128 | 0.23 | 0.74 | 0.15 | 0.2 | 0.1 | 0.16 |
| 1 | 100795207 | rs864633 | -0.077 | -0.553 | -0.446 | 0.341 | -0.004 | -0.018 | -0.018 | 0 | 0.17 | 0.32 | 0.09 | 0.15 | 0.14 | 0.15 |
| 1 | 100796120 | rs478859 | -0.773 | 0.422 | -1.636 | 0.355 | -0.11 | -0.072 | -0.072 | 0 | 0.55 | 0.95 | 0.47 | 0.46 | 0.54 | 0.59 |
| 1 | 100796296 | rs599215 | -0.885 | 0.447 | -1.652 | 0.339 | -0.108 | -0.065 | -0.065 | 0.239 | 0.54 | 0.95 | 0.47 | 0.46 | 0.54 | 0.57 |
| 1 | 100798137 | rs7534479 | 1.247 | 0.399 | -1.524 | 0.405 | -0.114 | -0.075 | -0.075 | 0.43 | 0.21 | 0.06 | 0.27 | 0.27 | 0.37 | 0.21 |
| 1 | 100800905 | rs11166438 | -0.467 | -0.248 | -0.824 | 0.054 | 0.001 | 0.003 | 0.003 | 0 | 0.53 | 0.56 | 0.45 | 0.39 | 0.57 | 0.46 |
| 1 | 100806404 | rs6577168 | 0.129 | 0.098 | -0.877 | 0.012 | 0.004 | -0.009 | -0.009 | 0.087 | 0.92 | 0.88 | 0.92 | 0.96 | 0.91 | 0.89 |
| 1 | 100808363 | rs11166440 | 0.979 | -0.117 | -0.899 | 0.294 | 0.01 | -0.083 | -0.083 | 0 | 0.4 | 0.09 | 0.45 | 0.51 | 0.36 | 0.46 |
| 1 | 100811859 | rs7516666 | 0.395 | -0.099 | -0.34 | -0.006 | 0 | -0.038 | -0.038 | 0.243 | 0.71 | 0.6 | 0.79 | 0.84 | 0.71 | 0.79 |
| 1 | 100818728 | rs17420882 | 1.594 | -0.205 | -0.663 | 0.381 | -0.009 | -0.056 | -0.056 | 0.267 | 0.33 | 0.02 | 0.19 | 0.32 | 0.27 | 0.42 |
| 1 | 100819273 | rs2297170 | 0.206 | -0.097 | -0.345 | 0.182 | -0.001 | -0.009 | -0.009 | 0 | 0.22 | 0.23 | 0.15 | 0.13 | 0.2 | 0.11 |
| 1 | 100822776 | rs630646 | 1.358 | 0.004 | -0.474 | 0.306 | -0.024 | -0.024 | -0.024 | 0 | 0.27 | 0.21 | 0.32 | 0.34 | 0.34 | 0.31 |
| 1 | 100831103 | rs7551111 | 0.995 | 0.03 | -0.388 | 0.222 | -0.007 | 0.016 | 0.016 | 0 | 0.12 | 0.08 | 0.08 | 0.05 | 0.16 | 0.08 |
| 1 | 100841993 | rs17122338 | 0.238 | -0.011 | 0.034 | 0.156 | 0.032 | -0.068 | -0.068 | 0 | 0.15 | 0.48 | 0.11 | 0.05 | 0.17 | 0.08 |
| 1 | 100853755 | rs10875290 | 0.999 | -0.222 | -0.216 | 0.326 | -0.02 | -0.122 | -0.122 | 0 | 0.39 | 0.06 | 0.41 | 0.5 | 0.38 | 0.45 |
| 1 | 100860339 | rs17122383 | 1.73 | -0.275 | 0 | 0.196 | -0.039 | -0.012 | -0.012 | 0 | 0.1 | 0.21 | 0.05 | 0.09 | 0.02 | 0.03 |
| 1 | 100869821 | rs7531643 | -0.467 | -0.277 | -0.078 | 0.119 | -0.002 | -0.011 | -0.011 | 4.249 | 0.26 | 0.66 | 0.16 | 0.13 | 0.19 | 0.12 |
| 1 | 100880025 | rs473072 | -0.51 | -0.184 | -0.277 | 0.201 | -0.013 | -0.021 | -0.021 | 0.412 | 0.84 | 0.76 | 0.89 | 0.89 | 0.89 | 0.87 |
| 1 | 100883188 | rs12564838 | 0.886 | -0.373 | -0.871 | 0.484 | -0.029 | -0.128 | -0.128 | 1.839 | 0.39 | 0.06 | 0.46 | 0.53 | 0.37 | 0.46 |
| 1 | 100886970 | rs6577171 | -0.131 | -0.33 | -0.315 | 0.384 | -0.014 | -0.039 | -0.039 | 0.177 | 0.25 | 0.65 | 0.16 | 0.13 | 0.19 | 0.12 |
| 1 | 100902936 | rs491675 | -1.232 | -0.394 | -0.61 | 0.445 | -0.184 | -0.096 | -0.096 | 0 | 0.68 | 0.97 | 0.64 | 0.68 | 0.57 | 0.59 |
| 1 | 100914270 | rs11809079 | 0.985 | -0.453 | 0.239 | 0.528 | -0.062 | -0.013 | -0.013 | 0 | 0.12 | 0.45 | 0.04 | 0.08 | 0.02 | 0.02 |
| 1 | 100914659 | rs980800 | 0.585 | -0.339 | -0.17 | 0.904 | -0.014 | -0.009 | -0.009 | 3.304 | 0.08 | 0.03 | 0.2 | 0.25 | 0.12 | 0.05 |
| 1 | 100918124 | rs6703191 | -0.351 | -0.471 | 0.144 | 0.495 | -0.025 | -0.062 | -0.062 | 0.544 | 0.26 | 0.58 | 0.15 | 0.13 | 0.2 | 0.12 |
| 1 | 100919788 | rs10493922 | -0.351 | -0.471 | 0.136 | 0.495 | -0.026 | -0.063 | -0.063 | 0 | 0.26 | 0.58 | 0.15 | 0.13 | 0.2 | 0.12 |
| 1 | 100943100 | rs3767832 | 0.649 | -0.045 | -0.355 | 0.797 | -0.123 | -0.072 | -0.072 | 0 | 0.33 | 0.06 | 0.39 | 0.37 | 0.44 | 0.43 |
| 1 | 100949860 | rs2270694 | 1.023 | -0.329 | 0.628 | 0.631 | 0.008 | 0.032 | 0.032 | 0 | 0.14 | 0.06 | 0.1 | 0.06 | 0.14 | 0.09 |
| 1 | 100955532 | rs508020 | 0.58 | -0.235 | 0.768 | 0.69 | -0.019 | -0.011 | -0.011 | 0 | 0.36 | 0.25 | 0.42 | 0.38 | 0.49 | 0.46 |
| 1 | 100959325 | rs1566212 | -1.18 | -0.466 | 1.137 | 0.571 | -0.061 | -0.021 | -0.021 | 0 | 0.89 | 0.43 | 0.96 | 0.91 | 0.99 | 0.98 |
| 1 | 100962483 | rs17122671 | 3.248 | -0.489 | 1.458 | 0.565 | -0.02 | -0.008 | -0.008 | 0 | 0.06 | 0.06 | 0.01 | 0.07 | 0 | 0.02 |
| 1 | 100963041 | rs904295 | -1.042 | -0.487 | 1.437 | 0.565 | -0.061 | -0.021 | -0.021 | 0 | 0.89 | 0.43 | 0.96 | 0.91 | 0.99 | 0.98 |
| 1 | 100965683 | rs6680879 | -2.472 | -0.487 | 1.584 | 0.565 | -0.04 | -0.004 | -0.004 | 0 | 0.92 | 0.75 | 0.97 | 0.91 | 1 | 0.98 |
| 1 | 100967569 | rs6666801 | -1.069 | -0.482 | 1.24 | 0.569 | -0.061 | -0.021 | -0.021 | 0 | 0.89 | 0.43 | 0.96 | 0.91 | 0.99 | 0.98 |
| 1 | 100970346 | rs10875295 | -0.163 | -0.747 | 0.525 | 0.411 | -0.002 | 0.001 | 0.001 | 0.027 | 0.49 | 0.62 | 0.43 | 0.51 | 0.47 | 0.37 |
| 1 | 100984092 | rs10783134 | -1.273 | -0.67 | 1.239 | 0.298 | -0.049 | -0.002 | -0.002 | 0 | 0.9 | 0.46 | 0.96 | 0.92 | 1 | 0.99 |
| 1 | 100984439 | rs1063302 | -0.158 | -0.656 | 1.284 | 0.25 | 0.04 | -0.011 | -0.011 | 0 | 0.1 | 0.2 | 0.09 | 0.07 | 0.11 | 0.16 |
| 1 | 100984519 | rs7516109 | -1.283 | -0.712 | 1.294 | 0.307 | -0.033 | 0.02 | 0.02 | 3.62 | 0.9 | 0.56 | 0.97 | 0.92 | 1 | 0.98 |
| 1 | 100988821 | rs7554943 | -1.726 | -0.701 | 1.321 | 0.314 | -0.026 | 0.022 | 0.022 | 2.655 | 0.91 | 0.65 | 0.97 | 0.92 | 1 | 0.99 |
| 1 | 100991649 | rs11166470 | 0.693 | -0.804 | 0.928 | 0.222 | 0.002 | 0.002 | 0.002 | 3.4 | 0.68 | 0.38 | 0.77 | 0.77 | 0.74 | 0.8 |
| 1 | 100991834 | rs7529831 | 0.765 | -0.167 | 0.464 | -0.194 | -0.006 | 0.158 | 0.158 | 0.806 | 0.54 | 0.2 | 0.64 | 0.77 | 0.54 | 0.75 |
| 1 | 100996074 | rs6577175 | -0.082 | -1.172 | 1.247 | 0.472 | -0.001 | -0.019 | -0.019 | 0 | 0.83 | 0.62 | 0.87 | 0.84 | 0.88 | 0.88 |
| 1 | 100997680 | rs7555025 | 1.659 | -1.145 | 0.606 | 0.716 | -0.084 | -0.076 | -0.076 | 0.263 | 0.26 | 0.1 | 0.32 | 0.35 | 0.39 | 0.3 |
| 1 | 101002833 | rs6577178 | 1.098 | -1.971 | 0.605 | 0.335 | 0.011 | 0.034 | 0.034 | 0.001 | 0.14 | 0.05 | 0.09 | 0.07 | 0.14 | 0.08 |
| 1 | 101003659 | rs2036212 | -2.448 | -2.031 | 0.931 | 0.322 | -0.03 | 0.019 | 0.019 | 1.559 | 0.93 | 0.78 | 0.97 | 0.92 | 1 | 0.99 |
| 1 | 101003762 | rs2809823 | -0.109 | -2.02 | 0.662 | 0.59 | -0.017 | 0.076 | 0.076 | 0 | 0.55 | 0.21 | 0.64 | 0.68 | 0.49 | 0.59 |
| 1 | 101006577 | rs2030048 | 0.605 | -2.059 | 0.578 | 0.635 | -0.02 | -0.031 | -0.031 | 0 | 0.4 | 0.1 | 0.54 | 0.6 | 0.36 | 0.51 |
| 1 | 101007036 | rs2809817 | 1.376 | -2.307 | 0.18 | 0.534 | -0.018 | 0.045 | 0.045 | 0 | 0.26 | 0.01 | 0.2 | 0.11 | 0.2 | 0.33 |
| 1 | 101010015 | rs11166473 | 0.102 | -1.614 | 1.364 | 0.717 | -0.027 | 0 | 0 | 4.277 | 0.83 | 0.4 | 0.87 | 0.8 | 0.91 | 0.9 |
| 1 | 101010077 | rs10875304 | 0.102 | -1.614 | 1.364 | 0.717 | -0.027 | 0 | 0 | 4.277 | 0.83 | 0.4 | 0.87 | 0.8 | 0.91 | 0.9 |
| 1 | 101019326 | rs12565245 | 0.95 | -1.633 | 1.18 | 0.712 | -0.02 | -0.027 | -0.027 | 0 | 0.4 | 0.1 | 0.54 | 0.6 | 0.36 | 0.51 |
| 1 | 101022112 | rs11166475 | -0.711 | -1.275 | 1.713 | 0.815 | 0.029 | -0.036 | -0.036 | 0 | 0.65 | 0.53 | 0.73 | 0.76 | 0.58 | 0.69 |
| 1 | 101024217 | rs2783671 | -2.826 | -1.29 | 1.842 | 0.842 | -0.026 | 0.057 | 0.057 | 1.335 | 0.94 | 0.88 | 0.98 | 0.88 | 1 | 0.99 |
| 1 | 101028907 | rs1414414 | -0.909 | -1.092 | 1.721 | 0.866 | 0.029 | -0.035 | -0.035 | 0 | 0.65 | 0.54 | 0.73 | 0.75 | 0.57 | 0.68 |
| 1 | 101029715 | rs721385 | -0.446 | -0.874 | 1.737 | 0.851 | 0.031 | 0 | 0 | 0.173 | 0.09 | 0.21 | 0.09 | 0.08 | 0.09 | 0.09 |
| 1 | 101033032 | rs1335760 | 1.234 | -1.249 | 1.96 | 0.74 | 0.015 | 0.014 | 0.014 | 0 | 0.15 | 0.09 | 0.1 | 0.07 | 0.12 | 0.09 |
| 1 | 101041054 | rs2809789 | -0.658 | -1.154 | 1.498 | 0.738 | -0.009 | 0.009 | 0.009 | 0 | 0.63 | 0.73 | 0.74 | 0.75 | 0.54 | 0.65 |
| 1 | 101042717 | rs12023028 | 0.793 | -1.12 | 1.499 | 0.766 | -0.04 | 0.034 | 0.034 | 0.227 | 0.34 | 0.16 | 0.25 | 0.25 | 0.43 | 0.33 |
| 1 | 101045438 | rs10875317 | 0.793 | -1.12 | 1.499 | 0.766 | -0.04 | 0.034 | 0.034 | 0.227 | 0.34 | 0.16 | 0.25 | 0.25 | 0.43 | 0.33 |
| 1 | 101047693 | rs2050476 | 1.287 | -1.175 | 1.423 | 0.395 | 0.015 | 0.014 | 0.014 | 0 | 0.15 | 0.09 | 0.1 | 0.07 | 0.12 | 0.09 |
| 1 | 101051656 | rs11166486 | -0.078 | -1.089 | 1.357 | 0.72 | 0.001 | 0.001 | 0.001 | 0 | 0.43 | 0.54 | 0.4 | 0.54 | 0.52 | 0.48 |
| 1 | 101051786 | rs11166487 | -0.106 | -1.092 | 1.336 | 0.718 | -0.001 | -0.001 | -0.001 | 0 | 0.43 | 0.57 | 0.41 | 0.54 | 0.52 | 0.48 |
| 1 | 101055002 | rs12728424 | 0.736 | -1.028 | 0.941 | 0.765 | -0.077 | -0.092 | -0.092 | 0.989 | 0.34 | 0.1 | 0.3 | 0.47 | 0.44 | 0.38 |
| 1 | 101062146 | rs12058729 | 1.144 | -1.09 | 1.084 | 0.416 | 0.003 | 0.003 | 0.003 | 0 | 0.16 | 0.16 | 0.11 | 0.07 | 0.13 | 0.09 |

| | | | | | | | | | | | | | | | | |
|---|---|---|---|---|---|---|---|---|---|---|---|---|---|---|---|---|
| 1 | 101073216 | rs17123016 | -0.298 | -1.055 | 1.366 | 0.417 | 0.044 | 0.008 | 0.008 | 0 | 0.07 | 0.3 | 0.1 | 0.07 | 0.08 | 0.09 |
| 1 | 101077323 | rs7415343 | 0.094 | -1.125 | 0.775 | 0.374 | -0.023 | -0.035 | -0.035 | 0.355 | 0.74 | 0.34 | 0.78 | 0.86 | 0.8 | 0.82 |
| 1 | 101078467 | rs1577489 | -0.705 | -1.103 | 0.717 | 0.406 | -0.065 | -0.061 | -0.061 | 0.209 | 0.8 | 0.48 | 0.87 | 0.92 | 0.87 | 0.91 |
| 1 | 101084273 | rs945329 | 0.521 | -0.851 | 0.29 | 0.64 | -0.019 | -0.012 | -0.012 | 0 | 0.33 | 0.24 | 0.29 | 0.38 | 0.44 | 0.39 |
| 1 | 101084542 | rs1815519 | -0.011 | -0.985 | 0.253 | 0.701 | -0.002 | 0.131 | 0.131 | 1.642 | 0.86 | 0.49 | 0.91 | 0.81 | 0.9 | 0.96 |
| 1 | 101085094 | rs11583008 | 0.137 | -0.943 | -0.042 | 0.684 | 0.002 | 0.023 | 0.023 | 0.02 | 0.53 | 0.34 | 0.62 | 0.42 | 0.46 | 0.57 |
| 1 | 101092955 | rs12135243 | 1.329 | -1.002 | -0.333 | 0.78 | -0.168 | 0.009 | 0.009 | 0.194 | 0.25 | 0.02 | 0.2 | 0.14 | 0.43 | 0.33 |
| 1 | 101094662 | rs17412243 | 0.848 | -0.968 | -0.286 | 0.321 | -0.001 | 0 | 0 | 0 | 0.07 | 0.06 | 0.09 | 0.04 | 0.11 | 0.04 |
| 1 | 101107748 | rs2484535 | -0.065 | -1.044 | -0.197 | 0.316 | -0.006 | 0.008 | 0.008 | 0 | 0.43 | 0.4 | 0.36 | 0.58 | 0.52 | 0.42 |
| 1 | 101111615 | rs3844028 | 1.592 | -1.122 | -0.457 | 0.268 | -0.009 | -0.012 | -0.012 | 0 | 0.28 | 0.03 | 0.24 | 0.25 | 0.21 | 0.38 |
| 1 | 101123278 | rs12049233 | 1.744 | -0.823 | -0.281 | 0.433 | -0.022 | -0.025 | -0.025 | 0.007 | 0.06 | 0.01 | 0.05 | 0.2 | 0.08 | 0.03 |
| 1 | 101123737 | rs871151 | 0.075 | -0.711 | -0.067 | -0.721 | 0.005 | 0.053 | 0.053 | 0 | 0.52 | 0.26 | 0.61 | 0.42 | 0.46 | 0.58 |
| 1 | 101168121 | rs1591474 | 1.145 | -0.479 | -0.263 | -0.735 | -0.143 | 0.07 | 0.07 | 0.055 | 0.24 | 0.02 | 0.19 | 0.08 | 0.4 | 0.26 |
| 1 | 101181576 | rs4584412 | 0.698 | -0.776 | 0.561 | -0.49 | 0.005 | 0.006 | 0.006 | 0 | 0.2 | 0.17 | 0.29 | 0.14 | 0.15 | 0.15 |
| 1 | 101183396 | rs1409419 | -0.984 | -0.783 | 0.556 | -0.495 | -0.007 | 0.011 | 0.011 | 0 | 0.58 | 0.61 | 0.65 | 0.43 | 0.48 | 0.58 |
| 1 | 101183825 | rs1041163 | 0.667 | -0.699 | 0.225 | -0.454 | 0 | -0.001 | -0.001 | 0.219 | 0.2 | 0.18 | 0.3 | 0.14 | 0.16 | 0.15 |
| 1 | 101187219 | rs3176860 | -0.342 | -0.694 | -0.443 | -0.425 | 0.014 | 0.017 | 0.017 | 0 | 0.5 | 0.52 | 0.56 | 0.38 | 0.38 | 0.54 |
| 1 | 101189410 | rs3917009 | -0.575 | 0.002 | -0.487 | -0.092 | -0.003 | 0.002 | 0.002 | 0 | 0.07 | 0.06 | 0.07 | 0.04 | 0.1 | 0.04 |
| 1 | 101190173 | rs2392221 | -0.216 | -0.329 | -0.506 | 0.067 | 0.025 | 0.032 | 0.032 | 0.054 | 0.19 | 0.06 | 0.29 | 0.13 | 0.14 | 0.15 |
| 1 | 101195659 | rs3917012 | 0.362 | -0.09 | -0.937 | 0.168 | 0.036 | 0.056 | 0.056 | 1.047 | 0.62 | 0.23 | 0.54 | 0.78 | 0.71 | 0.77 |
| 1 | 101198708 | rs3181088 | -0.411 | 1.033 | -0.866 | 0.683 | -0.052 | -0.031 | -0.031 | 0 | 0.09 | 0.01 | 0.09 | 0.1 | 0.17 | 0.15 |
| 1 | 101199886 | rs3176874 | -0.143 | -0.254 | -1.332 | 0.896 | 0.044 | -0.048 | -0.048 | 0 | 0.25 | 0.05 | 0.16 | 0.23 | 0.14 | 0.21 |
| 1 | 101200608 | rs3176876 | -1.035 | -0.298 | -0.127 | 1.086 | -0.005 | -0.001 | -0.001 | 0 | 0.47 | 0.51 | 0.48 | 0.38 | 0.33 | 0.5 |
| 1 | 101202222 | rs3917018 | 0.895 | -0.368 | 0.091 | 1.118 | 0.065 | 0.019 | 0.019 | 0 | 0.52 | 0.26 | 0.51 | 0.49 | 0.67 | 0.51 |
| 1 | 101202354 | rs3917019 | 0.623 | -0.057 | 0.419 | 0.855 | -0.012 | 0.003 | 0.003 | 0 | 0.59 | 0.52 | 0.57 | 0.62 | 0.7 | 0.68 |
| 1 | 101203698 | rs3176878 | -0.655 | 0.76 | 0.695 | 0.756 | -0.001 | -0.011 | -0.011 | 0 | 0.14 | 0.17 | 0.24 | 0.02 | 0.15 | 0.08 |
| 1 | 101213249 | rs6660837 | -0.782 | 1.219 | 0.467 | 1.064 | 0.031 | 0.009 | 0.009 | 0.084 | 0.2 | 0.4 | 0.32 | 0.24 | 0.24 | 0.37 |
| 1 | 101215995 | rs4908093 | -1.648 | 1.325 | 0.985 | 1.761 | 0.018 | 0.025 | 0.025 | 0.147 | 0.35 | 0.54 | 0.53 | 0.58 | 0.42 | 0.49 |
| 1 | 101222807 | rs1028819 | -1.152 | 1.456 | 1.134 | 2.074 | 0.06 | 0.017 | 0.017 | 0.498 | 0.27 | 0.47 | 0.42 | 0.33 | 0.39 | 0.4 |
| 1 | 101224146 | rs6680442 | -0.367 | 1.436 | 1.198 | 2.079 | 0.035 | 0.008 | 0.008 | 0.032 | 0.2 | 0.35 | 0.32 | 0.24 | 0.24 | 0.36 |
| 1 | 101224461 | rs17414954 | -0.139 | 1.435 | 1.188 | 2.032 | -0.014 | -0.009 | -0.009 | 0 | 0.07 | 0.05 | 0.09 | 0.08 | 0.15 | 0.03 |
| 1 | 101230852 | rs1887101 | -1.785 | 1.354 | 1.578 | 2.039 | 0.018 | 0.022 | 0.022 | 0.066 | 0.34 | 0.48 | 0.52 | 0.58 | 0.43 | 0.48 |
| 1 | 101231300 | rs10875335 | -1.94 | 1.312 | 1.774 | 2.019 | 0.013 | 0.016 | 0.016 | 0.152 | 0.35 | 0.58 | 0.53 | 0.58 | 0.43 | 0.5 |
| 1 | 101236137 | rs11586136 | 0.648 | 1.107 | 1.145 | 1.518 | -0.002 | 0.025 | 0.025 | 0 | 0.16 | 0.08 | 0.09 | 0.08 | 0.16 | 0.06 |
| 1 | 101242593 | rs12567000 | 0.479 | 1.203 | 1.067 | 1.604 | -0.022 | -0.017 | -0.017 | 0.006 | 0.08 | 0.04 | 0.27 | 0.15 | 0.19 | 0.2 |
| 1 | 101255052 | rs4517368 | -1.109 | 1.122 | 1.215 | 1.504 | 0.003 | 0.061 | 0.061 | 0 | 0.27 | 0.38 | 0.43 | 0.5 | 0.28 | 0.4 |
| 1 | 101255080 | rs6577209 | -1.329 | 1.088 | 1.183 | 1.488 | -0.001 | 0.093 | 0.093 | 0 | 0.28 | 0.52 | 0.44 | 0.51 | 0.28 | 0.42 |
| 1 | 101266356 | rs12035757 | 1.444 | 1.077 | 1.018 | 1.474 | 0.002 | 0.001 | 0.001 | 0 | 0.64 | 0.32 | 0.47 | 0.4 | 0.56 | 0.55 |
| 1 | 101272079 | rs12746227 | 0.87 | 1.215 | 0.966 | 1.416 | -0.001 | 0.026 | 0.026 | 0 | 0.15 | 0.04 | 0.07 | 0.09 | 0.15 | 0.06 |
| 1 | 101278386 | rs4908102 | 1.77 | 1.154 | 1.048 | 1.385 | 0.013 | 0.074 | 0.074 | 0 | 0.48 | 0.21 | 0.39 | 0.32 | 0.42 | 0.49 |
| 1 | 101280234 | rs11583221 | 0.429 | 1.141 | 1.084 | 1.374 | -0.004 | -0.011 | -0.011 | 0 | 0.17 | 0.18 | 0.08 | 0.09 | 0.15 | 0.07 |
| 1 | 101284027 | rs961912 | 1.742 | 1.144 | 1.041 | 1.38 | 0.013 | 0.074 | 0.074 | 0 | 0.48 | 0.21 | 0.39 | 0.32 | 0.42 | 0.49 |
| 1 | 101285547 | rs4907926 | 1.702 | 1.141 | 1.052 | 1.379 | 0.016 | 0.073 | 0.073 | 0 | 0.49 | 0.22 | 0.39 | 0.32 | 0.42 | 0.49 |
| 1 | 101299216 | rs1571046 | 0.783 | 1.287 | 1.133 | 1.443 | 0.003 | 0.096 | 0.096 | 0 | 0.73 | 0.52 | 0.56 | 0.49 | 0.73 | 0.61 |
| 1 | 101322866 | rs7541397 | -0.658 | 1.384 | 1.208 | 1.377 | 0.003 | -0.008 | -0.008 | 0.113 | 0.25 | 0.27 | 0.17 | 0.12 | 0.31 | 0.11 |
| 1 | 101328083 | rs10875346 | 0.755 | 1.318 | 1.588 | 1.462 | -0.003 | 0.107 | 0.107 | 0 | 0.74 | 0.55 | 0.56 | 0.45 | 0.74 | 0.62 |
| 1 | 101338324 | rs8888 | 1.287 | 1.313 | 1.605 | 1.668 | 0.002 | 0.001 | 0.001 | 0 | 0.58 | 0.39 | 0.48 | 0.38 | 0.59 | 0.59 |
| 1 | 101338482 | rs3188491 | 1.583 | 1.313 | 1.56 | 1.669 | 0.028 | 0.004 | 0.004 | 0 | 0.55 | 0.23 | 0.47 | 0.4 | 0.59 | 0.57 |
| 1 | 101342650 | rs10493937 | 1.454 | 0.671 | 0.476 | 1.137 | -0.035 | 0.076 | 0.076 | 0 | 0.42 | 0.21 | 0.42 | 0.21 | 0.48 | 0.36 |
| 1 | 101348100 | rs7543296 | 0.566 | 0.85 | 0.556 | 1.151 | -0.005 | -0.011 | -0.011 | 0 | 0.59 | 0.54 | 0.51 | 0.26 | 0.62 | 0.43 |
| 1 | 101350306 | rs11809572 | 1.332 | 0.843 | 0.529 | 1.145 | 0.003 | 0.009 | 0.009 | 0.173 | 0.16 | 0.12 | 0.07 | 0.04 | 0.14 | 0.06 |
| 1 | 101351661 | rs10218565 | 1.984 | 0.88 | 0.497 | 1.145 | -0.001 | 0.054 | 0.054 | 0 | 0.34 | 0.21 | 0.33 | 0.14 | 0.31 | 0.31 |
| 1 | 101360215 | rs3737580 | 1.339 | 0.854 | 0.442 | 1.19 | -0.006 | -0.057 | -0.057 | 0 | 0.15 | 0.01 | 0.05 | 0.19 | 0.11 | 0.19 |
| 1 | 101361178 | rs17123491 | -0.056 | 0.801 | 0.559 | 1.536 | 0.075 | -0.124 | -0.124 | 0 | 0.29 | 0.05 | 0.18 | 0.56 | 0.18 | 0.39 |
| 1 | 101362371 | rs1074739 | 0.728 | 0.559 | 0.451 | 1.516 | -0.054 | -0.042 | -0.042 | 0 | 0.09 | 0.01 | 0.09 | 0.1 | 0.17 | 0.11 |
| 1 | 101377845 | rs10493940 | 0.728 | 0.559 | 0.451 | 1.516 | -0.054 | -0.042 | -0.042 | 1.805 | 0.09 | 0.01 | 0.09 | 0.1 | 0.17 | 0.12 |
| 1 | 101378647 | rs4908107 | -1.582 | 0.844 | 0.646 | 1.591 | 0.023 | 0.006 | 0.006 | 0 | 0.93 | 0.91 | 0.78 | 0.89 | 0.85 | 0.85 |
| 1 | 101386897 | rs17123521 | 1.74 | 0.673 | 0.769 | 1.683 | 0.028 | -0.024 | -0.024 | 0 | 0.06 | 0 | 0.12 | 0.28 | 0.04 | 0.05 |
| 1 | 101396743 | rs11579723 | 1.566 | 0.86 | 0.879 | 1.318 | 0.005 | 0.022 | 0.022 | 0.265 | 0.16 | 0.11 | 0.07 | 0.04 | 0.14 | 0.05 |
| 1 | 101407519 | rs11581062 | -0.328 | 0.85 | 0.899 | 1.315 | 0.001 | -0.005 | -0.005 | 2.047 | 0.28 | 0.32 | 0.18 | 0.15 | 0.31 | 0.19 |
| 1 | 101441775 | rs3087816 | 1.498 | 1.017 | 0.89 | 1.353 | -0.041 | -0.025 | -0.025 | 0 | 0.07 | 0.01 | 0.07 | 0.07 | 0.15 | 0.04 |
| 1 | 101442211 | rs17123572 | 1.427 | 0.969 | 0.967 | 1.325 | -0.004 | -0.015 | -0.015 | 0 | 0.16 | 0.18 | 0.07 | 0.05 | 0.13 | 0.06 |
| 1 | 101442367 | rs17449022 | 0.676 | 0.958 | 0.962 | 1.33 | -0.044 | -0.037 | -0.037 | 0 | 0.08 | 0.03 | 0.08 | 0.1 | 0.15 | 0.09 |
| 1 | 101443701 | rs17100939 | -0.191 | 0.955 | 0.974 | 1.329 | 0.006 | -0.011 | -0.011 | 0 | 0.26 | 0.36 | 0.17 | 0.15 | 0.28 | 0.16 |
| 1 | 101445821 | rs10493941 | 1.412 | 0.952 | 0.965 | 1.327 | -0.004 | -0.015 | -0.015 | 2.682 | 0.16 | 0.18 | 0.07 | 0.05 | 0.14 | 0.06 |
| 1 | 101451262 | rs7518380 | -0.169 | 0.953 | 0.951 | 1.328 | -0.017 | -0.021 | -0.021 | 0 | 0.26 | 0.33 | 0.11 | 0.11 | 0.15 | 0.09 |
| 1 | 101466054 | rs11583043 | -0.194 | 0.971 | 0.833 | 1.338 | 0.006 | -0.01 | -0.01 | 0 | 0.26 | 0.36 | 0.18 | 0.15 | 0.29 | 0.15 |
| 1 | 101470217 | rs6692186 | 1.495 | 1.004 | 0.737 | 1.324 | -0.041 | -0.025 | -0.025 | 0.839 | 0.07 | 0.01 | 0.07 | 0.07 | 0.15 | 0.03 |
| 1 | 101494276 | rs6677329 | 1.592 | 0.994 | 0.617 | 1.033 | 0.002 | 0.005 | 0.005 | 0.526 | 0.15 | 0.13 | 0.07 | 0.05 | 0.14 | 0.06 |

| | | | | | | | | | | | | | | | | |
|---|---|---|---|---|---|---|---|---|---|---|---|---|---|---|---|---|
| 1 | 101504079 | rs6693186 | 1.592 | 0.985 | 0.617 | 1.033 | 0.001 | 0.005 | 0.005 | 0.526 | 0.15 | 0.13 | 0.07 | 0.05 | 0.14 | 0.06 |
| 1 | 101513708 | rs11166538 | 0.98 | 1.153 | 0.612 | 1.159 | 0.038 | 0.02 | 0.02 | 0 | 0.08 | 0.12 | 0.15 | 0.18 | 0.19 | 0.19 |
| 1 | 101532657 | rs3861736 | 0.18 | 0.399 | 0.998 | 0.895 | -0.007 | -0.037 | -0.037 | 0 | 0.27 | 0.41 | 0.15 | 0.15 | 0.24 | 0.12 |
| 1 | 101540999 | rs17450029 | 2.141 | 0.707 | 0.546 | 0.42 | -0.007 | 0.024 | 0.024 | 0 | 0.21 | 0.14 | 0.39 | 0.12 | 0.23 | 0.2 |
| 1 | 101543389 | rs4409674 | 2.112 | 0.761 | 0.609 | 0.496 | 0.014 | 0.027 | 0.027 | 2.376 | 0.17 | 0.07 | 0.07 | 0.06 | 0.12 | 0.03 |
| 1 | 101549316 | rs950038 | 0.886 | 1.089 | 0.594 | 0.706 | 0.011 | 0.009 | 0.009 | 0.427 | 0.12 | 0.13 | 0.2 | 0.22 | 0.24 | 0.24 |
| 1 | 101550980 | rs1335746 | 2.013 | 0.941 | 0.444 | 0.69 | -0.051 | -0.021 | -0.021 | 2.592 | 0.14 | 0.01 | 0.05 | 0.14 | 0.14 | 0.14 |
| 1 | 101551926 | rs17410015 | 1.89 | 1.186 | 0.746 | 0.503 | -0.029 | -0.005 | -0.005 | 0.968 | 0.06 | 0 | 0.05 | 0.04 | 0.09 | 0.02 |
| 1 | 101564585 | rs7554744 | -0.032 | 0.905 | 0.738 | 0.415 | -0.024 | 0.109 | 0.109 | 0.639 | 0.48 | 0.18 | 0.41 | 0.74 | 0.5 | 0.69 |
| 1 | 101583663 | rs7528842 | 0.442 | 0.506 | 1.081 | 0.427 | 0 | -0.037 | -0.037 | 0 | 0.27 | 0.4 | 0.16 | 0.14 | 0.25 | 0.11 |
| 1 | 101584241 | rs658861 | -0.307 | 0.494 | 1.036 | 0.421 | -0.002 | -0.01 | -0.01 | 0 | 0.71 | 0.34 | 0.82 | 0.86 | 0.75 | 0.89 |
| 1 | 101586819 | rs10875350 | 0.71 | 0.466 | 0.682 | 0.483 | 0.031 | -0.109 | -0.109 | 0.037 | 0.35 | 0.07 | 0.22 | 0.52 | 0.27 | 0.46 |
| 1 | 101588654 | rs12727382 | 0.829 | 0.583 | 1.388 | 0.274 | -0.012 | -0.014 | -0.014 | 0 | 0.19 | 0.25 | 0.09 | 0.1 | 0.13 | 0.07 |
| 1 | 101589082 | rs12746449 | 0.829 | 0.583 | 1.388 | 0.348 | -0.012 | -0.009 | -0.009 | 0.431 | 0.19 | 0.25 | 0.15 | 0.11 | 0.13 | 0.07 |
| 1 | 101589175 | rs12727971 | -0.138 | 0.603 | 1.081 | 0.218 | -0.002 | -0.039 | -0.039 | 0 | 0.27 | 0.41 | 0.15 | 0.14 | 0.24 | 0.11 |
| 1 | 101597229 | rs612422 | 0.111 | 0.636 | 0.781 | 0.226 | -0.005 | -0.029 | -0.029 | 0 | 0.72 | 0.46 | 0.84 | 0.87 | 0.76 | 0.9 |
| 1 | 101604116 | rs6658561 | -1.557 | 0.684 | 1.039 | 0.104 | 0.03 | -0.039 | -0.039 | 0.465 | 0.12 | 0.58 | 0.11 | 0.04 | 0.11 | 0.06 |
| 1 | 101605109 | rs17450806 | 0.98 | 0.73 | 0.975 | 0.126 | 0.006 | -0.001 | -0.001 | 1.212 | 0.06 | 0.05 | 0.05 | 0.04 | 0.1 | 0.03 |
| 1 | 101606903 | rs1983967 | -0.7 | 0.74 | 1.306 | 0.131 | 0.076 | -0.004 | -0.004 | 0 | 0.1 | 0.35 | 0.08 | 0.04 | 0.11 | 0.05 |
| 1 | 101616430 | rs7552136 | -0.799 | 0.765 | 1.098 | 0.143 | 0.091 | -0.003 | -0.003 | 0 | 0.11 | 0.47 | 0.08 | 0.04 | 0.11 | 0.02 |
| 1 | 101620729 | rs17417367 | 1.188 | 0.661 | -0.253 | 0.043 | -0.007 | 0.076 | 0.076 | 3.88 | 0.17 | 0.01 | 0.07 | 0.05 | 0.13 | 0.04 |
| 1 | 101622960 | rs17123757 | 1.152 | 0.645 | -0.37 | -0.814 | 0 | -0.02 | -0.02 | 0 | 0.13 | 0.14 | 0.34 | 0.01 | 0.15 | 0.11 |
| 1 | 101624468 | rs625146 | 0.18 | 1.212 | -0.662 | -0.838 | 0.01 | 0.001 | 0.001 | 0.349 | 0.19 | 0.25 | 0.21 | 0.23 | 0.29 | 0.28 |
| 1 | 101630828 | rs499531 | 0.195 | 1.213 | -0.708 | -0.837 | -0.002 | -0.001 | -0.001 | 0.128 | 0.19 | 0.22 | 0.21 | 0.23 | 0.29 | 0.28 |
| 1 | 101638999 | rs7367938 | 0.95 | 0.301 | -0.942 | -1.118 | -0.011 | 0.024 | 0.024 | 0 | 0.24 | 0.27 | 0.23 | 0.57 | 0.2 | 0.38 |
| 1 | 101639648 | rs12041118 | 1.595 | 0.534 | -1.511 | -0.751 | 0.008 | 0.03 | 0.03 | 0 | 0.17 | 0.05 | 0.09 | 0.11 | 0.13 | 0.12 |
| 1 | 101644898 | rs12090198 | 1.83 | 0.807 | -1.141 | -0.492 | 0.018 | 0.052 | 0.052 | 2.458 | 0.17 | 0.11 | 0.07 | 0.03 | 0.12 | 0.03 |
| 1 | 101655670 | rs10783155 | -0.254 | 1.314 | -0.512 | 0.274 | -0.01 | -0.003 | -0.003 | 1.037 | 0.81 | 0.79 | 0.81 | 0.76 | 0.72 | 0.72 |
| 1 | 101660122 | rs6676334 | 0.248 | 1.016 | -0.174 | 0.779 | 0.007 | 0.007 | 0.007 | 0 | 0.59 | 0.57 | 0.63 | 0.57 | 0.53 | 0.53 |
| 1 | 101669840 | rs1571498 | 0.61 | 1.526 | -0.206 | 0.501 | -0.002 | 0.008 | 0.008 | 0.544 | 0.25 | 0.18 | 0.21 | 0.06 | 0.24 | 0.43 |
| 1 | 101669928 | rs1041312 | 0.69 | 1.375 | 0.768 | 1.962 | 0.018 | 0.057 | 0.057 | 0 | 0.68 | 0.46 | 0.52 | 0.55 | 0.63 | 0.7 |
| 1 | 101675844 | rs12407211 | -0.46 | 1.841 | 0.281 | 2.198 | 0.029 | 0.037 | 0.037 | 0.179 | 0.21 | 0.36 | 0.44 | 0.36 | 0.28 | 0.2 |
| 1 | 101678531 | rs12408201 | -0.783 | 1.388 | -0.178 | 2.088 | 0.027 | 0.221 | 0.221 | 0 | 0.07 | 0.21 | 0.11 | 0.26 | 0.07 | 0.05 |
| 1 | 101683903 | rs4341378 | -0.132 | 0.864 | -0.855 | 0.265 | 0.001 | -0.01 | -0.01 | 0 | 0.73 | 0.79 | 0.84 | 0.92 | 0.78 | 0.93 |
| 1 | 101684269 | rs17123870 | 0.254 | 1.127 | -0.831 | -0.158 | 0.006 | -0.029 | -0.029 | 0 | 0.27 | 0.29 | 0.44 | 0.14 | 0.31 | 0.46 |
| 1 | 101684415 | rs6676516 | -0.575 | 0.956 | -0.918 | -0.25 | 0.001 | -0.009 | -0.009 | 0.093 | 0.54 | 0.56 | 0.72 | 0.81 | 0.56 | 0.73 |
| 1 | 101690270 | rs6673771 | 0.033 | 0.541 | -1.022 | 0.824 | -0.002 | -0.001 | -0.001 | 0 | 0.33 | 0.29 | 0.58 | 0.37 | 0.33 | 0.55 |
| 1 | 101698388 | rs12746702 | 0.149 | 0.498 | -1.232 | 1.654 | 0.012 | -0.016 | -0.016 | 0 | 0.08 | 0.03 | 0.03 | 0.29 | 0.05 | 0.04 |
| 1 | 101701432 | rs12407294 | -0.274 | 0.48 | -1.482 | 1.653 | -0.043 | -0.039 | -0.039 | 0 | 0.31 | 0.14 | 0.3 | 0.39 | 0.36 | 0.32 |
| 1 | 101702825 | rs3753194 | -0.059 | 0.024 | -1.388 | 1.784 | 0.011 | -0.002 | -0.002 | 0 | 0.1 | 0.06 | 0.03 | 0.31 | 0.05 | 0.04 |
| 1 | 101704496 | rs3737578 | -0.296 | -0.553 | -1.631 | 1.866 | 0.013 | -0.016 | -0.016 | 0 | 0.13 | 0.09 | 0.04 | 0.27 | 0.06 | 0.08 |
| 1 | 101704532 | rs3737577 | 0.826 | -0.342 | -2.669 | 1.082 | -0.104 | 0.062 | 0.062 | 0 | 0.27 | 0.02 | 0.17 | 0.13 | 0.34 | 0.12 |
| 1 | 101708752 | rs12409786 | 0.044 | -0.405 | -2.221 | 1.365 | 0.005 | -0.003 | -0.003 | 0 | 0.14 | 0.11 | 0.04 | 0.28 | 0.05 | 0.1 |
| 1 | 101711783 | rs2031959 | -1.252 | -0.377 | -1.503 | 1.307 | -0.011 | -0.005 | -0.005 | 0 | 0.28 | 0.65 | 0.23 | 0.72 | 0.21 | 0.28 |
| 1 | 101715646 | rs12044675 | 0.284 | -0.335 | -1.461 | 1.374 | 0.019 | -0.025 | -0.025 | 0 | 0.14 | 0.08 | 0.04 | 0.28 | 0.05 | 0.1 |
| 1 | 101722623 | rs9433755 | 0.295 | -0.181 | -0.786 | 1.142 | -0.023 | 0.042 | 0.042 | 0 | 0.81 | 0.74 | 0.91 | 0.7 | 0.88 | 0.78 |
| 1 | 101724951 | rs7532171 | -0.189 | 0.089 | -0.279 | 1.24 | 0.011 | 0.164 | 0.164 | 0 | 0.07 | 0.15 | 0.12 | 0.43 | 0.12 | 0.06 |
| 1 | 101725605 | rs9434153 | -0.042 | 0.102 | -0.304 | 1.262 | 0.007 | 0.135 | 0.135 | 0 | 0.07 | 0.13 | 0.11 | 0.42 | 0.12 | 0.06 |
| 1 | 101726526 | rs2038363 | 0.57 | -0.069 | 0.037 | 0.768 | 0.078 | 0.025 | 0.025 | 0 | 0.93 | 0.67 | 0.95 | 0.97 | 0.94 | 0.88 |
| 1 | 101731461 | rs9434158 | 0.468 | 0.055 | 0.124 | 0.574 | 0.002 | -0.037 | -0.037 | 0 | 0.14 | 0.34 | 0.09 | 0.07 | 0.09 | 0.16 |
| 1 | 101732641 | rs6693660 | 0.934 | 0.066 | -0.218 | 0.397 | 0.01 | -0.004 | -0.004 | 0 | 0.54 | 0.34 | 0.73 | 0.48 | 0.65 | 0.54 |
| 1 | 101732914 | rs12118086 | 1.44 | 0.237 | -0.05 | 0.271 | -0.05 | -0.002 | -0.002 | 0 | 0.09 | 0.01 | 0.25 | 0.07 | 0.19 | 0.07 |
| 1 | 101733105 | rs9433759 | 0.003 | 0.152 | 0.051 | 0.245 | 0.007 | -0.012 | -0.012 | 0 | 0.68 | 0.7 | 0.83 | 0.56 | 0.76 | 0.7 |
| 1 | 101733328 | rs12760559 | 1.285 | 0.169 | -0.154 | 0.231 | 0.013 | -0.038 | -0.038 | 0 | 0.14 | 0.36 | 0.1 | 0.08 | 0.1 | 0.16 |
| 1 | 101733861 | rs11166573 | 1.828 | 0.27 | -0.855 | 0.334 | -0.041 | -0.006 | -0.006 | 0 | 0.39 | 0.06 | 0.3 | 0.32 | 0.39 | 0.43 |
| 1 | 101736356 | rs2038364 | 1.786 | 0.379 | -0.744 | 0.469 | 0.021 | -0.074 | -0.074 | 0.377 | 0.13 | 0.03 | 0.03 | 0.27 | 0.06 | 0.1 |
| 1 | 101740406 | rs11166576 | -0.564 | 0.421 | -0.4 | 0.468 | -0.001 | -0.026 | -0.026 | 0 | 0.28 | 0.22 | 0.44 | 0.4 | 0.32 | 0.2 |
| 1 | 101743821 | rs7519801 | 1.156 | 0.422 | -0.274 | 0.387 | -0.011 | -0.009 | -0.009 | 0 | 0.59 | 0.66 | 0.44 | 0.51 | 0.49 | 0.64 |
| 1 | 101744332 | rs6692828 | 1.543 | 0.376 | -0.199 | 0.402 | 0 | -0.002 | -0.002 | 0.53 | 0.54 | 0.56 | 0.43 | 0.39 | 0.49 | 0.59 |
| 1 | 101744821 | rs17124020 | 0.646 | 0.439 | -0.416 | 0.428 | -0.061 | -0.036 | -0.036 | 0 | 0.14 | 0.02 | 0.39 | 0.14 | 0.25 | 0.1 |
| 1 | 101747767 | rs6577233 | 0.882 | 0.51 | -0.194 | 0.303 | -0.009 | -0.008 | -0.008 | 0 | 0.61 | 0.67 | 0.44 | 0.51 | 0.5 | 0.64 |
| 1 | 101749974 | rs910633 | 0.934 | 0.657 | -0.176 | 0.11 | -0.003 | 0.002 | 0.002 | 0 | 0.57 | 0.63 | 0.44 | 0.39 | 0.49 | 0.59 |
| 1 | 101751884 | rs7520706 | -0.341 | 0.716 | -0.553 | 0.23 | -0.011 | 0.01 | 0.01 | 0 | 0.88 | 0.91 | 0.87 | 0.91 | 0.81 | 0.85 |
| 1 | 101753101 | rs1922987 | 0.989 | 0.593 | -0.444 | 0.23 | 0.018 | 0.022 | 0.022 | 0 | 0.44 | 0.3 | 0.36 | 0.35 | 0.39 | 0.43 |
| 1 | 101756557 | rs7516960 | 0.654 | 0.51 | -0.584 | 0.222 | 0.002 | 0.001 | 0.001 | 0 | 0.47 | 0.32 | 0.4 | 0.46 | 0.45 | 0.4 |
| 1 | 101757026 | rs878485 | -0.207 | 0.603 | -0.804 | 0.151 | -0.001 | -0.006 | -0.006 | 0.446 | 0.65 | 0.66 | 0.49 | 0.51 | 0.58 | 0.65 |
| 1 | 101757864 | rs10875366 | 1.132 | 0.736 | -1.442 | 0.255 | -0.098 | -0.094 | -0.094 | 0.007 | 0.38 | 0.02 | 0.31 | 0.36 | 0.36 | 0.36 |
| 1 | 101760889 | rs2144555 | -0.564 | 1.087 | -0.802 | -0.266 | 0.027 | 0.006 | 0.006 | 0 | 0.27 | 0.41 | 0.51 | 0.32 | 0.37 | 0.25 |
| 1 | 101762118 | rs10875368 | 1.184 | 1.012 | -1.411 | -0.125 | 0.016 | 0.064 | 0.064 | 0 | 0.35 | 0.04 | 0.24 | 0.2 | 0.28 | 0.36 |

| | | | | | | | | | | | | | | | | |
|---|---|---|---|---|---|---|---|---|---|---|---|---|---|---|---|---|
| 1 | 101767274 | rs12041851 | 1.035 | 1.082 | -1.293 | -0.1 | -0.01 | 0.05 | 0.05 | 0.444 | 0.34 | 0.04 | 0.23 | 0.21 | 0.28 | 0.34 |
| 1 | 101768868 | rs1033634 | -0.861 | 1.28 | -0.474 | -0.247 | 0.017 | -0.005 | -0.005 | 0 | 0.34 | 0.6 | 0.58 | 0.32 | 0.49 | 0.29 |
| 1 | 101770101 | rs17124135 | 0.985 | 1.384 | -0.08 | -0.266 | 0.006 | -0.002 | -0.002 | 1.661 | 0.09 | 0.09 | 0.09 | 0.11 | 0.15 | 0.11 |
| 1 | 101772190 | rs6683369 | -1.139 | 1.414 | -0.185 | -0.266 | 0.018 | -0.005 | -0.005 | 0.328 | 0.33 | 0.59 | 0.58 | 0.32 | 0.49 | 0.29 |
| 1 | 101772220 | rs12137590 | -0.137 | 1.415 | -0.542 | -0.239 | -0.006 | 0 | 0 | 0 | 0.22 | 0.25 | 0.47 | 0.2 | 0.33 | 0.18 |
| 1 | 101773056 | rs11166584 | 0.552 | 1.239 | -0.526 | -0.287 | 0.007 | -0.008 | -0.008 | 0.4 | 0.12 | 0.16 | 0.27 | 0.1 | 0.18 | 0.09 |
| 1 | 101774930 | rs4907932 | 1.352 | 0.995 | -1.131 | -0.434 | -0.035 | 0.034 | 0.034 | 0.646 | 0.29 | 0.03 | 0.19 | 0.21 | 0.26 | 0.26 |
| 1 | 101778236 | rs2010845 | -0.71 | 0.716 | -0.72 | -0.364 | -0.005 | -0.015 | -0.015 | 0 | 0.64 | 0.78 | 0.75 | 0.65 | 0.65 | 0.74 |
| 1 | 101784340 | rs9433784 | 0.858 | 0.761 | -0.3 | -0.472 | 0.01 | -0.024 | -0.024 | 0.825 | 0.13 | 0.28 | 0.11 | 0.1 | 0.12 | 0.2 |
| 1 | 101786421 | rs17420712 | 1.415 | 0.85 | -0.305 | -0.445 | -0.011 | -0.005 | -0.005 | 0.161 | 0.11 | 0.07 | 0.26 | 0.12 | 0.18 | 0.08 |
| 1 | 101789562 | rs6577237 | -1.571 | 0.243 | -0.338 | -0.434 | 0.043 | -0.079 | -0.079 | 0 | 0.76 | 0.94 | 0.88 | 0.66 | 0.87 | 0.81 |
| 1 | 101795029 | rs4908127 | -0.029 | 0.279 | -0.073 | -0.341 | 0.002 | 0.02 | 0.02 | 0 | 0.43 | 0.35 | 0.32 | 0.35 | 0.4 | 0.51 |
| 1 | 101801342 | rs4907933 | 1.574 | 0.263 | 0.079 | -0.376 | -0.057 | -0.038 | -0.038 | 0 | 0.25 | 0.02 | 0.18 | 0.25 | 0.24 | 0.22 |
| 1 | 101803470 | rs4908130 | -0.956 | 0.23 | 0.15 | -0.358 | 0.012 | -0.026 | -0.026 | 0.054 | 0.87 | 0.72 | 0.88 | 0.9 | 0.88 | 0.8 |
| 1 | 101815604 | rs4907938 | 0.38 | 0.206 | -0.037 | -0.363 | -0.004 | -0.004 | -0.004 | 0 | 0.34 | 0.44 | 0.36 | 0.32 | 0.33 | 0.35 |
| 1 | 101826927 | rs7516360 | -0.479 | 0.245 | -0.07 | -0.339 | -0.001 | 0 | 0 | 0.135 | 0.62 | 0.63 | 0.7 | 0.65 | 0.64 | 0.58 |
| 1 | 101829924 | rs17124314 | 2.278 | 0.317 | -0.273 | -0.434 | 0.007 | -0.025 | -0.025 | 0 | 0.06 | 0 | 0.02 | 0.11 | 0.03 | 0.11 |
| 1 | 101830625 | rs12143342 | -0.805 | 0.312 | -0.262 | -0.457 | 0.01 | -0.031 | -0.031 | 0.875 | 0.86 | 0.71 | 0.89 | 0.9 | 0.88 | 0.79 |
| 1 | 101830858 | rs11164177 | 0.23 | 0.344 | -0.948 | -0.44 | 0.008 | 0.047 | 0.047 | 0 | 0.51 | 0.25 | 0.51 | 0.58 | 0.54 | 0.45 |
| 1 | 101833964 | rs6577240 | 1.651 | 0.652 | -0.769 | -0.459 | -0.002 | -0.019 | -0.019 | 0 | 0.06 | 0.07 | 0.03 | 0 | 0.05 | 0.09 |
| 1 | 101836640 | rs7541500 | 0.214 | 0.689 | -0.905 | -0.303 | -0.014 | -0.01 | -0.01 | 0 | 0.46 | 0.26 | 0.49 | 0.46 | 0.51 | 0.34 |
| 1 | 101838064 | rs7537249 | 0.007 | 0.997 | -0.64 | 0.139 | 0.002 | -0.02 | -0.02 | 3.185 | 0.87 | 0.81 | 0.89 | 0.9 | 0.88 | 0.79 |
| 1 | 101843149 | rs7553108 | 0.171 | 1.166 | -1.009 | 0.139 | -0.003 | 0.003 | 0.003 | 0 | 0.87 | 0.9 | 0.92 | 0.92 | 0.85 | 0.78 |
| 1 | 101847845 | rs11164183 | -0.163 | 0.721 | -0.941 | 0.2 | 0 | -0.001 | -0.001 | 0.56 | 0.43 | 0.48 | 0.48 | 0.29 | 0.41 | 0.53 |
| 1 | 101848523 | rs10874482 | -0.144 | 0.744 | -0.935 | 0.212 | 0 | -0.004 | -0.004 | 0 | 0.43 | 0.5 | 0.48 | 0.28 | 0.41 | 0.53 |
| 1 | 101848719 | rs6577244 | -0.099 | 0.932 | -0.966 | 0.417 | 0.002 | -0.001 | -0.001 | 0 | 0.65 | 0.67 | 0.62 | 0.65 | 0.58 | 0.79 |
| 1 | 101849423 | rs1491643 | 0.015 | 0.583 | -1.318 | 0.658 | 0.04 | 0.034 | 0.034 | 0.541 | 0.5 | 0.29 | 0.66 | 0.61 | 0.63 | 0.46 |
| 1 | 101849830 | rs6577245 | -0.21 | 0.681 | -1.18 | 0.822 | 0.003 | -0.002 | -0.002 | 0 | 0.45 | 0.28 | 0.45 | 0.52 | 0.47 | 0.58 |
| 1 | 101850130 | rs6577246 | 0.325 | 0.665 | -1.26 | 0.768 | 0.01 | -0.007 | -0.007 | 0 | 0.2 | 0.23 | 0.31 | 0.15 | 0.28 | 0.22 |
| 1 | 101851025 | rs12127134 | 0.593 | 0.37 | -1.142 | 0.963 | 0.004 | -0.01 | -0.01 | 0 | 0.12 | 0.09 | 0.26 | 0.3 | 0.1 | 0.14 |
| 1 | 101851347 | rs1388452 | 0.496 | 0.16 | -0.433 | 0.768 | 0.063 | 0.081 | 0.081 | 0.545 | 0.06 | 0.33 | 0.06 | 0.07 | 0.05 | 0.02 |
| 1 | 101854661 | rs7529061 | -0.401 | 0.053 | -0.892 | 0.449 | -0.007 | 0.095 | 0.095 | 0.148 | 0.28 | 0.48 | 0.41 | 0.5 | 0.23 | 0.4 |
| 1 | 101857117 | rs7549900 | 0.429 | -0.239 | -1.282 | -0.196 | -0.039 | 0.036 | 0.036 | 0 | 0.36 | 0.23 | 0.29 | 0.26 | 0.44 | 0.27 |
| 1 | 101862103 | rs7520882 | 0.344 | -0.476 | -0.82 | -0.459 | 0.014 | -0.038 | -0.038 | 0.029 | 0.27 | 0.35 | 0.36 | 0.09 | 0.37 | 0.16 |
| 1 | 101864215 | rs12403551 | 0.349 | -0.873 | -0.741 | -0.339 | -0.017 | 0.057 | 0.057 | 0 | 0.46 | 0.29 | 0.51 | 0.33 | 0.47 | 0.51 |
| 1 | 101865012 | rs12041998 | 2.171 | -1.099 | -1.096 | -0.467 | 0.007 | -0.077 | -0.077 | 0.12 | 0.14 | 0.01 | 0.1 | 0.24 | 0.09 | 0.16 |
| 1 | 101868793 | rs10493947 | 1.081 | -1.019 | -0.84 | -0.417 | -0.021 | 0.044 | 0.044 | 0 | 0.33 | 0.19 | 0.26 | 0.18 | 0.38 | 0.25 |
| 1 | 101868913 | rs1491642 | -0.382 | -0.885 | -0.541 | -0.36 | 0.006 | -0.024 | -0.024 | 0 | 0.39 | 0.43 | 0.61 | 0.19 | 0.46 | 0.39 |
| 1 | 101871324 | rs12727858 | 1.477 | -0.452 | -1.06 | -0.537 | -0.039 | 0.05 | 0.05 | 0 | 0.15 | 0.04 | 0.13 | 0 | 0.25 | 0.1 |
| 1 | 101872865 | rs1826316 | -0.798 | -0.571 | -0.802 | -0.466 | 0 | 0 | 0 | 0 | 0.86 | 0.87 | 0.56 | 0.88 | 0.85 | 0.74 |
| 1 | 101873070 | rs1826317 | 0.406 | -1.061 | -0.558 | -0.093 | 0.001 | 0.002 | 0.002 | 0 | 0.5 | 0.44 | 0.33 | 0.48 | 0.49 | 0.37 |
| 1 | 101876646 | rs1602548 | 1.108 | -0.908 | -0.779 | -0.163 | 0.011 | -0.019 | -0.019 | 0 | 0.15 | 0.08 | 0.06 | 0.29 | 0.1 | 0.12 |
| 1 | 101885368 | rs4908143 | 1.321 | -0.815 | -0.781 | -0.381 | 0.007 | -0.017 | -0.017 | 0 | 0.14 | 0.08 | 0.06 | 0.29 | 0.1 | 0.12 |
| 1 | 101894763 | rs17421741 | 0.71 | -0.619 | -0.685 | -0.596 | -0.001 | 0.03 | 0.03 | 0 | 0.4 | 0.31 | 0.29 | 0.25 | 0.4 | 0.3 |
| 1 | 101895945 | rs12401392 | 1.588 | -0.466 | -0.722 | -0.55 | -0.008 | 0.025 | 0.025 | 0 | 0.09 | 0.04 | 0.22 | 0.02 | 0.08 | 0.13 |
| 1 | 101899294 | rs12058460 | 0.67 | -0.569 | -0.44 | -0.459 | -0.015 | -0.036 | -0.036 | 0 | 0.07 | 0.14 | 0.04 | 0.01 | 0.02 | 0.05 |
| 1 | 101913162 | rs12031265 | 0.695 | -0.676 | -0.7 | -0.308 | 0.007 | -0.013 | -0.013 | 0 | 0.13 | 0.13 | 0.08 | 0.26 | 0.09 | 0.17 |
| 1 | 101918376 | rs6703277 | 1.453 | -0.65 | -0.624 | -0.585 | -0.002 | -0.028 | -0.028 | 0 | 0.17 | 0.17 | 0.13 | 0 | 0.25 | 0.07 |
| 1 | 101919802 | rs12118351 | 1.66 | -0.7 | -0.748 | -0.436 | 0.007 | -0.024 | -0.024 | 0 | 0.13 | 0.07 | 0.06 | 0.29 | 0.1 | 0.12 |
| 1 | 101920805 | rs11164201 | -0.023 | -0.733 | -0.728 | -0.607 | 0 | 0 | 0 | 1.132 | 0.31 | 0.33 | 0.21 | 0.26 | 0.34 | 0.24 |
| 1 | 101922351 | rs12048997 | 1.153 | -0.599 | -0.646 | -0.438 | 0 | 0.009 | 0.009 | 0.474 | 0.12 | 0.14 | 0.07 | 0.26 | 0.09 | 0.15 |
| 1 | 101924306 | rs12069565 | -0.087 | -0.603 | -0.696 | -0.528 | -0.004 | 0.009 | 0.009 | 0 | 0.2 | 0.26 | 0.29 | 0.28 | 0.15 | 0.31 |
| 1 | 101927038 | rs12405027 | 1.733 | -0.528 | -0.833 | -0.52 | -0.003 | 0.004 | 0.004 | 0.027 | 0.06 | 0.04 | 0.21 | 0.02 | 0.05 | 0.11 |
| 1 | 101928396 | rs1199711 | 0.333 | -0.61 | -0.774 | -0.529 | -0.002 | 0 | 0 | 0 | 0.66 | 0.68 | 0.65 | 0.41 | 0.75 | 0.6 |
| 1 | 101932014 | rs1473510 | -0.322 | -0.516 | -0.825 | -0.469 | 0 | 0.007 | 0.007 | 0 | 0.81 | 0.82 | 0.69 | 0.63 | 0.85 | 0.64 |
| 1 | 101933692 | rs1777876 | 0.19 | -0.595 | -0.894 | -0.559 | -0.001 | -0.009 | -0.009 | 2.76 | 0.61 | 0.75 | 0.56 | 0.63 | 0.6 | 0.57 |
| 1 | 101937714 | rs12406997 | 2.106 | -0.8 | -1.222 | -0.786 | -0.006 | -0.012 | -0.012 | 0 | 0.16 | 0.01 | 0.08 | 0.15 | 0.1 | 0.2 |
| 1 | 101938477 | rs1948368 | 0.257 | -0.811 | -1.034 | -0.805 | -0.001 | 0.04 | 0.04 | 0 | 0.48 | 0.63 | 0.5 | 0.32 | 0.5 | 0.46 |
| 1 | 101938561 | rs1948369 | -1.146 | -0.872 | -1.084 | -0.836 | -0.028 | 0.062 | 0.062 | 0 | 0.75 | 0.89 | 0.64 | 0.89 | 0.69 | 0.75 |
| 1 | 101939689 | rs1361812 | -0.235 | -0.889 | -1.08 | -0.804 | -0.002 | -0.014 | -0.014 | 0 | 0.61 | 0.74 | 0.57 | 0.58 | 0.59 | 0.64 |
| 1 | 101940474 | rs10493950 | 1.921 | -0.885 | -1.063 | -0.799 | 0.003 | -0.003 | -0.003 | 0 | 0.13 | 0.08 | 0.06 | 0.31 | 0.1 | 0.11 |
| 1 | 101940809 | rs1777894 | 0.305 | -0.858 | -1.046 | -0.797 | -0.001 | 0.04 | 0.04 | 0 | 0.48 | 0.64 | 0.5 | 0.32 | 0.5 | 0.46 |
| 1 | 101943194 | rs1388456 | 1.422 | -0.844 | -1.087 | -0.749 | 0.002 | 0.001 | 0.001 | 0.263 | 0.12 | 0.11 | 0.06 | 0.26 | 0.09 | 0.15 |
| 1 | 101943331 | rs1612840 | 0.776 | -0.564 | -0.808 | -0.856 | 0 | -0.019 | -0.019 | 0.001 | 0.42 | 0.47 | 0.31 | 0.29 | 0.43 | 0.37 |
| 1 | 101945171 | rs1777879 | 0.124 | -0.566 | -0.88 | -0.839 | -0.002 | -0.003 | -0.003 | 0 | 0.55 | 0.62 | 0.37 | 0.55 | 0.51 | 0.52 |
| 1 | 101945654 | rs3131386 | 0.747 | -0.538 | -0.856 | -0.846 | -0.003 | 0.045 | 0.045 | 0.578 | 0.19 | 0.23 | 0.25 | 0.34 | 0.18 | 0.2 |
| 1 | 101947713 | rs1813917 | 0.138 | -0.428 | -0.826 | -0.785 | -0.001 | 0 | 0 | 0.091 | 0.38 | 0.34 | 0.38 | 0.34 | 0.43 | 0.27 |
| 1 | 101950270 | rs1776904 | 0.753 | -0.409 | -0.784 | -0.705 | -0.002 | 0.007 | 0.007 | 0 | 0.41 | 0.4 | 0.3 | 0.33 | 0.43 | 0.37 |
| 1 | 101951693 | rs1777871 | 1.158 | -0.504 | -0.677 | -0.773 | -0.019 | 0.053 | 0.053 | 0 | 0.37 | 0.25 | 0.28 | 0.21 | 0.41 | 0.3 |

| | | | | | | | | | | | | | | | | |
|---|---|---|---|---|---|---|---|---|---|---|---|---|---|---|---|---|
| 1 | 101962952 | rs6668197 | 0.639 | -0.437 | -0.745 | -0.731 | -0.009 | -0.025 | -0.025 | 0 | 0.33 | 0.16 | 0.19 | 0.38 | 0.35 | 0.21 |
| 1 | 101965251 | rs12123743 | 1.338 | -0.195 | -0.766 | -0.685 | -0.001 | -0.001 | -0.001 | 0 | 0.13 | 0.11 | 0.1 | 0.26 | 0.15 | 0.16 |
| 1 | 101973475 | rs450404 | 0.48 | -0.59 | -0.636 | -0.647 | 0 | 0 | 0 | 0.346 | 0.54 | 0.5 | 0.38 | 0.49 | 0.53 | 0.45 |
| 1 | 101973613 | rs263908 | 0.193 | -0.582 | -0.659 | -0.642 | -0.001 | -0.001 | -0.001 | 0 | 0.54 | 0.56 | 0.38 | 0.48 | 0.51 | 0.49 |
| 1 | 101975151 | rs263912 | -0.81 | -0.536 | -0.632 | -0.614 | 0.022 | 0.018 | 0.018 | 0.386 | 0.87 | 0.73 | 0.57 | 0.86 | 0.87 | 0.71 |
| 1 | 101976237 | rs3844029 | 1.443 | -0.523 | -0.734 | -0.623 | 0.002 | 0.004 | 0.004 | 0.226 | 0.12 | 0.11 | 0.06 | 0.26 | 0.09 | 0.15 |
| 1 | 101979504 | rs11584636 | 1.64 | -0.298 | -0.848 | -0.828 | -0.026 | 0.068 | 0.068 | 0.261 | 0.19 | 0.08 | 0.13 | 0 | 0.25 | 0.05 |
| 1 | 101991399 | rs1343364 | 1.914 | -0.329 | -0.828 | -0.87 | 0.005 | -0.019 | -0.019 | 0 | 0.22 | 0.25 | 0.19 | 0.07 | 0.31 | 0.12 |
| 1 | 101992222 | rs1688582 | -0.328 | 0.048 | -0.39 | -0.146 | 0.004 | -0.01 | -0.01 | 0 | 0.79 | 0.81 | 0.71 | 0.62 | 0.84 | 0.69 |
| 1 | 101993635 | rs991010 | 0.141 | 0.188 | -0.272 | -0.047 | 0.007 | -0.02 | -0.02 | 0 | 0.25 | 0.28 | 0.34 | 0.14 | 0.33 | 0.2 |
| 1 | 101994207 | rs12137486 | 1.242 | 0.189 | -0.277 | -0.048 | 0.008 | -0.014 | -0.014 | 0.049 | 0.13 | 0.08 | 0.06 | 0.27 | 0.1 | 0.11 |
| 1 | 101995341 | rs1418931 | -0.197 | 0.764 | -0.189 | -0.902 | 0.006 | -0.021 | -0.021 | 0 | 0.26 | 0.28 | 0.35 | 0.14 | 0.34 | 0.22 |
| 1 | 102005028 | rs1201780 | -0.335 | 0.706 | -0.455 | -0.701 | 0.021 | 0.028 | 0.028 | 0 | 0.81 | 0.91 | 0.75 | 0.89 | 0.89 | 0.82 |
| 1 | 102009914 | rs1146714 | 0.827 | 0.781 | -0.28 | -0.683 | -0.003 | -0.002 | -0.002 | 0 | 0.73 | 0.72 | 0.53 | 0.75 | 0.81 | 0.62 |
| 1 | 102011828 | rs1688581 | -0.33 | 0.774 | -0.288 | -0.685 | 0.02 | 0.026 | 0.026 | 0 | 0.8 | 0.89 | 0.75 | 0.88 | 0.89 | 0.82 |
| 1 | 102011935 | rs1777888 | 0.837 | 0.782 | -0.277 | -0.683 | -0.003 | -0.002 | -0.002 | 0 | 0.73 | 0.72 | 0.53 | 0.75 | 0.81 | 0.62 |
| 1 | 102012725 | rs1776934 | -0.917 | 0.764 | -0.313 | -0.691 | 0.01 | 0.015 | 0.015 | 0 | 0.84 | 0.92 | 0.75 | 0.88 | 0.9 | 0.83 |
| 1 | 102012977 | rs1343093 | 0.808 | 0.784 | -0.291 | -0.683 | -0.002 | -0.002 | -0.002 | 0 | 0.73 | 0.73 | 0.53 | 0.75 | 0.81 | 0.61 |
| 1 | 102013468 | rs12044991 | 0.774 | 0.773 | -0.269 | -0.648 | -0.002 | 0.059 | 0.059 | 0.559 | 0.16 | 0.2 | 0.14 | 0.58 | 0.16 | 0.29 |
| 1 | 102016094 | rs1146717 | -0.373 | 0.594 | -0.538 | -0.847 | 0.02 | 0.029 | 0.029 | 0 | 0.8 | 0.89 | 0.75 | 0.89 | 0.89 | 0.82 |
| 1 | 102017100 | rs4626910 | 0.279 | 0.712 | -0.409 | -0.8 | 0.013 | 0.043 | 0.043 | 1.391 | 0.22 | 0.3 | 0.34 | 0.62 | 0.24 | 0.39 |
| 1 | 102024942 | rs1484308 | 0.242 | 0.6 | -0.779 | -0.961 | -0.011 | 0.128 | 0.128 | 0 | 0.64 | 0.85 | 0.61 | 0.89 | 0.64 | 0.78 |
| 1 | 102033539 | rs4908165 | 1.064 | 0.52 | -0.482 | -0.672 | -0.004 | 0.056 | 0.056 | 0 | 0.39 | 0.3 | 0.25 | 0.17 | 0.4 | 0.29 |
| 1 | 102034837 | rs17458619 | 1.012 | 0.53 | -0.787 | -0.636 | -0.001 | -0.004 | -0.004 | 0 | 0.39 | 0.4 | 0.27 | 0.17 | 0.4 | 0.32 |
| 1 | 102039242 | rs12057522 | -0.421 | 0.662 | -1.164 | -0.193 | 0.004 | 0.019 | 0.019 | 0 | 0.17 | 0.11 | 0.26 | 0.11 | 0.13 | 0.18 |
| 1 | 102039804 | rs12743305 | -0.517 | 0.873 | -0.81 | -0.133 | -0.002 | 0 | 0 | 0 | 0.18 | 0.14 | 0.15 | 0.08 | 0.24 | 0.11 |
| 1 | 102040519 | rs7541801 | -0.489 | 0.699 | -1.07 | 0.225 | -0.042 | 0.005 | 0.005 | 0 | 0.38 | 0.19 | 0.43 | 0.36 | 0.46 | 0.37 |
| 1 | 102044634 | rs12142390 | 1.41 | 0.076 | 0.705 | 0.296 | 0.053 | 0.09 | 0.09 | 6.505 | 0.95 | 0.43 | 0.96 | 0.96 | 1 | 0.86 |
| 1 | 102048431 | rs6685257 | -0.472 | -0.069 | 0.45 | 0.58 | -0.001 | -0.019 | -0.019 | 1.068 | 0.14 | 0.13 | 0.26 | 0.32 | 0.18 | 0.22 |
| 1 | 102051020 | rs1601229 | 1.748 | -0.469 | 1.053 | 0.325 | -0.016 | 0.009 | 0.009 | 0 | 0.72 | 0.31 | 0.66 | 0.63 | 0.78 | 0.56 |
| 1 | 102061180 | rs6660392 | -0.603 | -0.385 | 1.432 | 0.388 | 0.002 | -0.007 | -0.007 | 0 | 0.17 | 0.15 | 0.28 | 0.32 | 0.18 | 0.23 |
| 1 | 102069696 | rs10493958 | -1.896 | -0.356 | 1.782 | 0.16 | 0.011 | -0.007 | -0.007 | 0.446 | 0.09 | 0.36 | 0.04 | 0.04 | 0.05 | 0.18 |
| 1 | 102080843 | rs1484315 | -1.898 | -0.382 | 1.853 | 0.125 | -0.013 | -0.057 | -0.057 | 0 | 0.11 | 0.38 | 0.04 | 0.03 | 0.06 | 0.09 |
| 1 | 102082236 | rs10782840 | 2.189 | -0.419 | 1.858 | 0.146 | 0.04 | -0.035 | -0.035 | 0.855 | 0.68 | 0.11 | 0.64 | 0.62 | 0.73 | 0.52 |
| 1 | 102085498 | rs12080747 | -2.076 | -0.393 | 2.074 | 0.128 | -0.016 | -0.069 | -0.069 | 0 | 0.12 | 0.53 | 0.05 | 0.03 | 0.06 | 0.09 |
| 1 | 102087717 | rs6672980 | -2.45 | -0.398 | 2.11 | 0.13 | -0.031 | -0.066 | -0.066 | 0 | 0.15 | 0.59 | 0.08 | 0.04 | 0.09 | 0.25 |
| 1 | 102088142 | rs6662473 | -1.36 | -0.36 | 1.994 | 0.149 | -0.01 | -0.049 | -0.049 | 0 | 0.1 | 0.29 | 0.04 | 0.03 | 0.06 | 0.09 |
| 1 | 102090174 | rs4908170 | 0.975 | -0.347 | 1.906 | 0.2 | -0.004 | 0.021 | 0.021 | 0 | 0.8 | 0.75 | 0.69 | 0.65 | 0.79 | 0.61 |
| 1 | 102095912 | rs4907947 | 0.263 | -0.341 | 1.813 | 0.204 | -0.001 | 0.006 | 0.006 | 0 | 0.83 | 0.8 | 0.72 | 0.66 | 0.82 | 0.77 |
| 1 | 102101223 | rs1552290 | 2.473 | -0.599 | 1.889 | -0.1 | -0.05 | -0.087 | -0.087 | 0 | 0.82 | 0.58 | 0.91 | 0.96 | 0.91 | 0.74 |
| 1 | 102108081 | rs880513 | 2.079 | -0.571 | 1.596 | 0.391 | 0.051 | -0.079 | -0.079 | 0.139 | 0.69 | 0.14 | 0.62 | 0.54 | 0.76 | 0.57 |
| 1 | 102110713 | rs7514091 | -2.259 | -0.494 | 2.062 | 0.394 | -0.011 | -0.015 | -0.015 | 0 | 0.1 | 0.4 | 0.06 | 0.04 | 0.04 | 0.16 |
| 1 | 102113247 | rs10874504 | -2.888 | -0.509 | 2.16 | 0.386 | -0.061 | -0.068 | -0.068 | 1.905 | 0.15 | 0.62 | 0.08 | 0.05 | 0.05 | 0.2 |
| 1 | 102120197 | rs11800181 | -2.046 | -0.746 | 2.088 | 0.293 | -0.027 | -0.059 | -0.059 | 0 | 0.11 | 0.33 | 0.05 | 0 | 0.04 | 0.06 |
| 1 | 102127452 | rs9803762 | -2.625 | -0.772 | 2.139 | 0.357 | -0.014 | -0.019 | -0.019 | 0 | 0.11 | 0.47 | 0.07 | 0.05 | 0.04 | 0.19 |
| 1 | 102129604 | rs1385126 | -1.442 | -0.761 | 2.209 | 0.359 | -0.003 | -0.033 | -0.033 | 0.5 | 0.08 | 0.27 | 0.05 | 0 | 0.04 | 0.06 |
| 1 | 102147556 | rs2086852 | -1.393 | -0.762 | 1.391 | 0.358 | 0.002 | -0.029 | -0.029 | 0 | 0.08 | 0.29 | 0.05 | 0 | 0.04 | 0.06 |
| 1 | 102155106 | rs1351839 | -0.096 | -0.782 | 0.362 | 0.442 | 0 | 0.03 | 0.03 | 0 | 0.86 | 0.85 | 0.7 | 0.59 | 0.8 | 0.78 |
| 1 | 102159783 | rs9803851 | -1.758 | -0.771 | 0.508 | 0.047 | -0.067 | -0.061 | -0.061 | 0.015 | 0.12 | 0.6 | 0.05 | 0.01 | 0.01 | 0.1 |
| 1 | 102161329 | rs12755682 | 0.129 | -0.112 | 0.082 | 0.043 | -0.119 | -0.019 | -0.019 | 0.287 | 0.23 | 0.02 | 0.2 | 0.14 | 0.32 | 0.09 |
| 1 | 102166616 | rs4907951 | -1.676 | -0.43 | 0.757 | 0.281 | -0.038 | -0.042 | -0.042 | 0 | 0.12 | 0.62 | 0.07 | 0.04 | 0.04 | 0.2 |
| 1 | 102176419 | rs7529743 | -1.821 | -0.387 | 0.684 | 0.258 | -0.028 | -0.059 | -0.059 | 0.662 | 0.14 | 0.6 | 0.08 | 0.04 | 0.08 | 0.22 |
| 1 | 102184478 | rs11164278 | 0.231 | 0.054 | -0.289 | 0.433 | -0.062 | 0.028 | 0.028 | 0 | 0.34 | 0.18 | 0.47 | 0.23 | 0.46 | 0.42 |
| 1 | 102189472 | rs2289811 | 0.7 | -0.254 | -0.103 | 0.577 | -0.02 | -0.032 | -0.032 | 0 | 0.09 | 0.01 | 0.16 | 0.08 | 0.09 | 0.1 |
| 1 | 102192809 | rs11164280 | 0.564 | -0.338 | 0.088 | 0.95 | -0.001 | 0.102 | 0.102 | 0 | 0.06 | 0.07 | 0.05 | 0.41 | 0.05 | 0.11 |
| 1 | 102193273 | rs6664594 | -1.31 | -0.664 | 0.162 | 0.728 | -0.02 | -0.028 | -0.028 | 0.377 | 0.88 | 0.74 | 0.94 | 0.93 | 0.93 | 0.85 |
| 1 | 102193300 | rs6702059 | 0.012 | -0.933 | 0.447 | 0.534 | 0.001 | -0.002 | -0.002 | 0 | 0.24 | 0.24 | 0.27 | 0.23 | 0.21 | 0.17 |
| 1 | 102194625 | rs17426819 | 1.397 | -1.118 | 0.423 | 0.492 | 0.01 | 0.045 | 0.045 | 0 | 0.23 | 0.11 | 0.12 | 0.02 | 0.21 | 0.11 |
| 1 | 102197513 | rs10874506 | -0.022 | -0.922 | 0.421 | 0.472 | 0.002 | -0.011 | -0.011 | 0 | 0.38 | 0.32 | 0.36 | 0.5 | 0.38 | 0.33 |
| 1 | 102197790 | rs11164282 | -1.058 | -0.246 | 0.566 | 1.002 | 0.004 | 0.071 | 0.071 | 0 | 0.15 | 0.21 | 0.24 | 0.48 | 0.17 | 0.23 |
| 1 | 102200830 | rs4908177 | -1.039 | -0.246 | 0.496 | 1.002 | 0 | 0.029 | 0.029 | 0 | 0.15 | 0.16 | 0.24 | 0.48 | 0.17 | 0.23 |
| 1 | 102209045 | rs2197445 | 0.371 | -0.181 | 0.477 | 0.915 | -0.049 | 0.019 | 0.019 | 0 | 0.34 | 0.22 | 0.48 | 0.24 | 0.46 | 0.44 |
| 1 | 102211705 | rs1445222 | -1.202 | -0.037 | 0.756 | 0.484 | 0.014 | 0.017 | 0.017 | 0 | 0.66 | 0.73 | 0.83 | 0.91 | 0.72 | 0.76 |
| 1 | 102223473 | rs12145515 | -0.159 | 0.057 | 0.715 | 0.554 | 0 | -0.001 | -0.001 | 1.834 | 0.08 | 0.08 | 0.19 | 0.07 | 0.12 | 0.12 |
| 1 | 102228731 | rs959163 | -0.173 | 0.004 | 0.862 | 0.572 | -0.001 | 0.001 | 0.001 | 0 | 0.22 | 0.22 | 0.27 | 0.23 | 0.2 | 0.17 |
| 1 | 102229165 | rs2339135 | -1.221 | -0.012 | 0.854 | 0.509 | 0.032 | 0.078 | 0.078 | 0 | 0.65 | 0.77 | 0.84 | 0.91 | 0.73 | 0.76 |
| 1 | 102229920 | rs7553875 | -0.471 | 0.03 | 0.937 | 0.584 | -0.021 | 0.018 | 0.018 | 0 | 0.24 | 0.36 | 0.28 | 0.22 | 0.2 | 0.17 |
| 1 | 102236122 | rs11164296 | -0.681 | 0.067 | 0.794 | 0.791 | 0.003 | 0.113 | 0.113 | 0 | 0.16 | 0.4 | 0.22 | 0.28 | 0.13 | 0.13 |
| 1 | 102237818 | rs1327583 | -1.25 | 0.034 | 0.764 | 0.768 | -0.006 | 0.024 | 0.024 | 0 | 0.32 | 0.53 | 0.47 | 0.51 | 0.32 | 0.3 |

| | | | | | | | | | | | | | | | | |
|---|---|---|---|---|---|---|---|---|---|---|---|---|---|---|---|---|
| 1 | 102239205 | rs1470438 | 1.132 | 0.073 | 0.327 | 0.74 | -0.074 | -0.024 | -0.024 | 0.409 | 0.3 | 0.1 | 0.34 | 0.36 | 0.37 | 0.38 |
| 1 | 102246949 | rs17125417 | 0.656 | -0.383 | 0.233 | 0.547 | -0.019 | -0.039 | -0.039 | 0 | 0.08 | 0.14 | 0.02 | 0 | 0.02 | 0.01 |
| 1 | 102249468 | rs12726478 | -0.244 | -0.302 | -0.079 | 0.609 | 0.033 | 0.041 | 0.041 | 0 | 0.5 | 0.36 | 0.41 | 0.32 | 0.45 | 0.4 |
| 1 | 102253545 | rs1470436 | -1.07 | -0.261 | -0.559 | 0.565 | -0.016 | -0.005 | -0.005 | 0 | 0.64 | 0.7 | 0.47 | 0.57 | 0.53 | 0.53 |
| 1 | 102254704 | rs17468763 | 0.12 | 0.047 | -1.361 | 0.919 | 0.03 | 0.078 | 0.078 | 0 | 0.35 | 0.22 | 0.17 | 0.12 | 0.28 | 0.23 |
| 1 | 102263909 | rs10493976 | -0.916 | 0.623 | -1.156 | 0.897 | -0.005 | -0.038 | -0.038 | 0.146 | 0.15 | 0.07 | 0.23 | 0.22 | 0.19 | 0.17 |
| 1 | 102271774 | rs4907955 | -0.124 | 0.425 | -0.995 | 0.047 | 0.006 | 0.015 | 0.015 | 0 | 0.46 | 0.59 | 0.42 | 0.65 | 0.58 | 0.32 |
| 1 | 102272312 | rs10874518 | -0.298 | -0.076 | -1.231 | -0.316 | 0.015 | 0.024 | 0.024 | 0 | 0.56 | 0.65 | 0.59 | 0.66 | 0.64 | 0.45 |
| 1 | 102274276 | rs1854169 | -0.947 | -0.524 | -1.668 | -0.764 | 0.024 | 0.112 | 0.112 | 0 | 0.74 | 0.91 | 0.8 | 0.93 | 0.8 | 0.73 |
| 1 | 102275057 | rs724479 | 0 | -0.357 | -1.406 | -0.628 | -0.001 | -0.014 | -0.014 | 0.499 | 0.17 | 0.13 | 0.19 | 0.27 | 0.16 | 0.28 |
| 1 | 102275190 | rs724480 | 0.464 | 0.185 | -1.258 | -0.495 | 0 | 0.007 | 0.007 | 0 | 0.46 | 0.57 | 0.4 | 0.65 | 0.56 | 0.31 |
| 1 | 102279573 | rs10493975 | -0.084 | 0.447 | -1.525 | -0.179 | -0.019 | -0.024 | -0.024 | 0 | 0.05 | 0.01 | 0.06 | 0.23 | 0.07 | 0.09 |
| 1 | 102283539 | rs4908185 | 0.558 | 0.833 | -1.19 | -0.613 | -0.017 | -0.002 | -0.002 | 0.336 | 0.3 | 0.23 | 0.34 | 0.65 | 0.39 | 0.25 |
| 1 | 102284779 | rs6702193 | -0.2 | 0.289 | -0.859 | -0.819 | 0.011 | 0.013 | 0.013 | 0 | 0.48 | 0.44 | 0.45 | 0.83 | 0.61 | 0.35 |
| 1 | 102284828 | rs6577274 | -0.336 | -0.4 | -0.779 | -1.176 | -0.012 | -0.01 | -0.01 | 0 | 0.66 | 0.59 | 0.65 | 0.69 | 0.7 | 0.53 |
| 1 | 102285043 | rs6694095 | -1.064 | -0.651 | -0.229 | -0.369 | -0.037 | 0.152 | 0.152 | 0 | 0.17 | 0.41 | 0.21 | 0.39 | 0.11 | 0.25 |
| 1 | 102285285 | rs7547160 | -0.701 | 0.448 | -0.347 | 0.053 | 0.052 | 0.069 | 0.069 | 0 | 0.29 | 0.53 | 0.41 | 0.45 | 0.46 | 0.49 |
| 1 | 102285489 | rs1445225 | -0.414 | 0.617 | -0.495 | 0.205 | 0 | 0 | 0 | 0.072 | 0.44 | 0.39 | 0.47 | 0.63 | 0.55 | 0.52 |
| 1 | 102289185 | rs11164306 | -0.514 | 1.051 | -0.24 | -0.061 | 0.006 | -0.014 | -0.014 | 0 | 0.32 | 0.56 | 0.36 | 0.29 | 0.34 | 0.24 |
| 1 | 102289362 | rs10747410 | -1.056 | 1.22 | -0.533 | -0.1 | 0.022 | 0.004 | 0.004 | 0 | 0.57 | 0.68 | 0.61 | 0.43 | 0.66 | 0.46 |
| 1 | 102292110 | rs17470099 | 1.171 | 1.268 | -0.663 | -0.163 | -0.031 | 0.109 | 0.109 | 0.632 | 0.18 | 0.01 | 0.09 | 0.01 | 0.18 | 0.11 |
| 1 | 102292567 | rs4908187 | 0.8 | 1.339 | -0.239 | -0.073 | 0.001 | -0.006 | -0.006 | 0 | 0.26 | 0.2 | 0.24 | 0.14 | 0.32 | 0.22 |
| 1 | 102294507 | rs6661054 | -0.679 | 0.392 | -0.554 | 0.013 | 0.029 | 0.007 | 0.007 | 0 | 0.57 | 0.69 | 0.6 | 0.43 | 0.66 | 0.46 |
| 1 | 102296279 | rs10493973 | 0.022 | 0.521 | -0.275 | -0.184 | -0.006 | -0.077 | -0.077 | 0.087 | 0.2 | 0.29 | 0.11 | 0.01 | 0.18 | 0.1 |
| 1 | 102297845 | rs2039943 | -0.21 | 0.607 | -0.438 | 0.431 | 0 | 0.001 | 0.001 | 0 | 0.45 | 0.46 | 0.61 | 0.64 | 0.5 | 0.48 |
| 1 | 102297957 | rs2039944 | -0.027 | 0.621 | -0.554 | 0.493 | -0.016 | -0.011 | -0.011 | 0 | 0.42 | 0.29 | 0.61 | 0.57 | 0.5 | 0.46 |
| 1 | 102298765 | rs6656847 | 0.648 | -0.201 | -0.798 | 0.299 | 0.024 | 0.023 | 0.023 | 0.083 | 0.25 | 0.16 | 0.08 | 0.12 | 0.2 | 0.22 |
| 1 | 102304094 | rs9324336 | 0.344 | 0.983 | -0.007 | 1.299 | 0.002 | -0.001 | -0.001 | 0 | 0.31 | 0.33 | 0.47 | 0.31 | 0.32 | 0.33 |
| 1 | 102304152 | rs769163 | -0.308 | 1.031 | -0.867 | 1.04 | -0.012 | -0.013 | -0.013 | 0.005 | 0.64 | 0.29 | 0.48 | 0.54 | 0.66 | 0.63 |
| 1 | 102306723 | rs11164308 | -0.164 | 1.248 | -0.954 | 1.45 | -0.066 | -0.102 | -0.102 | 0 | 0.15 | 0.03 | 0.29 | 0.41 | 0.26 | 0.28 |
| 1 | 102308337 | rs11805492 | -1.193 | 0.572 | -0.111 | 1.556 | -0.021 | 0.201 | 0.201 | 0.156 | 0.09 | 0.21 | 0.13 | 0.6 | 0.03 | 0.19 |
| 1 | 102312703 | rs4907956 | -0.076 | 1.052 | -1.261 | 0.064 | -0.16 | 0.203 | 0.203 | 0 | 0.29 | 0.03 | 0.22 | 0.02 | 0.39 | 0.19 |
| 1 | 102312848 | rs10874521 | 0.327 | 0.809 | -1.318 | -0.605 | -0.007 | 0.019 | 0.019 | 0 | 0.22 | 0.14 | 0.12 | 0 | 0.29 | 0.09 |
| 1 | 102315214 | rs1415105 | -0.97 | 0.663 | -0.23 | -0.008 | -0.01 | 0.161 | 0.161 | 1.507 | 0.14 | 0.58 | 0.14 | 0.31 | 0.14 | 0.12 |
| 1 | 102326159 | rs2050572 | -0.694 | 0.557 | 0.344 | -0.119 | -0.008 | 0.015 | 0.015 | 0 | 0.43 | 0.67 | 0.38 | 0.33 | 0.54 | 0.31 |
| 1 | 102341819 | rs12035099 | -0.377 | 0.252 | 0.777 | 0.174 | -0.002 | 0.008 | 0.008 | 0 | 0.25 | 0.23 | 0.19 | 0.33 | 0.3 | 0.18 |
| 1 | 102344571 | rs7528218 | -1.431 | 0.737 | 0.787 | 0.397 | 0.007 | 0.017 | 0.017 | 0.03 | 0.55 | 0.73 | 0.41 | 0.56 | 0.58 | 0.4 |
| 1 | 102349186 | rs1010590 | -1.022 | 1.121 | 0.612 | 0.586 | 0.002 | 0.009 | 0.009 | 0 | 0.48 | 0.38 | 0.37 | 0.34 | 0.56 | 0.31 |
| 1 | 102350747 | rs12117918 | -0.417 | 0.971 | 0.411 | 0.679 | -0.002 | 0.011 | 0.011 | 0 | 0.25 | 0.24 | 0.18 | 0.33 | 0.3 | 0.18 |
| 1 | 102357104 | rs11800637 | -0.168 | 1.036 | 0.432 | 0.456 | 0.003 | 0.122 | 0.122 | 0 | 0.06 | 0.13 | 0.03 | 0.23 | 0.05 | 0.1 |
| 1 | 102357987 | rs17125667 | -0.188 | 0.971 | 0.424 | 0.521 | 0 | 0.021 | 0.021 | 0 | 0.23 | 0.23 | 0.2 | 0.33 | 0.29 | 0.18 |
| 1 | 102359002 | rs3748653 | -1.236 | 1.214 | 0.41 | 0.429 | -0.001 | -0.002 | -0.002 | 0.168 | 0.51 | 0.45 | 0.39 | 0.58 | 0.59 | 0.41 |
| 1 | 102359362 | rs17432141 | 0.291 | 1.465 | 0.579 | 0.921 | -0.072 | 0.135 | 0.135 | 0 | 0.21 | 0.02 | 0.16 | 0.02 | 0.25 | 0.14 |
| 1 | 102363318 | rs17432246 | -0.571 | 1.468 | 0.61 | 0.922 | -0.01 | 0.071 | 0.071 | 0 | 0.28 | 0.15 | 0.19 | 0.02 | 0.27 | 0.15 |
| 1 | 102366140 | rs11802288 | -1.481 | 1.087 | 0.758 | 1.088 | 0.018 | 0.061 | 0.061 | 0 | 0.08 | 0.15 | 0.05 | 0.23 | 0.13 | 0.12 |
| 1 | 102367162 | rs943556 | -0.956 | 1.146 | 0.809 | 1.121 | 0.019 | -0.002 | -0.002 | 0 | 0.37 | 0.5 | 0.36 | 0.59 | 0.44 | 0.35 |
| 1 | 102371735 | rs1360998 | -0.966 | 1.155 | 0.799 | 1.125 | 0.017 | -0.002 | -0.002 | 0 | 0.37 | 0.51 | 0.36 | 0.59 | 0.44 | 0.35 |
| 1 | 102376258 | rs2000150 | -0.862 | 1.165 | 0.634 | 1.13 | 0.01 | -0.001 | -0.001 | 0 | 0.37 | 0.43 | 0.35 | 0.59 | 0.45 | 0.35 |
| 1 | 102378636 | rs7534442 | -0.211 | 0.463 | 0.361 | 0.592 | -0.002 | -0.002 | -0.002 | 1.12 | 0.29 | 0.26 | 0.3 | 0.36 | 0.31 | 0.22 |
| 1 | 102386104 | rs12060073 | -0.465 | 0.733 | -0.185 | 0.819 | 0.002 | 0.018 | 0.018 | 0 | 0.35 | 0.48 | 0.33 | 0.36 | 0.33 | 0.24 |
| 1 | 102389733 | rs11164337 | -0.443 | 0.195 | -0.305 | 0.872 | -0.025 | 0.007 | 0.007 | 0 | 0.37 | 0.48 | 0.49 | 0.64 | 0.3 | 0.43 |
| 1 | 102393844 | rs1336806 | 0.342 | 0.592 | -0.739 | 0.757 | -0.009 | -0.047 | -0.047 | 0 | 0.4 | 0.21 | 0.35 | 0.45 | 0.4 | 0.48 |
| 1 | 102394835 | rs1981159 | -0.004 | 0.283 | -0.097 | 0.764 | 0 | 0 | 0 | 0.885 | 0.51 | 0.57 | 0.48 | 0.63 | 0.46 | 0.65 |
| 1 | 102402745 | rs11587732 | 0.315 | 0.377 | -0.163 | 0.604 | -0.003 | 0 | 0 | 1.054 | 0.08 | 0.08 | 0.1 | 0.15 | 0.04 | 0.16 |
| 1 | 102404247 | rs755181 | 0.605 | 0.342 | -0.159 | 0.624 | 0 | -0.001 | -0.001 | 14.508 | 0.41 | 0.44 | 0.37 | 0.49 | 0.41 | 0.47 |
| 1 | 102410515 | rs1932850 | 0.073 | -0.086 | 0.067 | 0.444 | -0.002 | 0 | 0 | 0 | 0.08 | 0.07 | 0.1 | 0.15 | 0.04 | 0.16 |
| 1 | 102417093 | rs10782864 | 0.12 | 0.493 | -0.111 | 0.548 | 0.008 | 0.019 | 0.019 | 0.186 | 0.49 | 0.48 | 0.45 | 0.58 | 0.58 | 0.53 |
| 1 | 102421714 | rs10493966 | -0.434 | 0.569 | -0.581 | 0.088 | 0.022 | 0.081 | 0.081 | 0.06 | 0.48 | 0.65 | 0.42 | 0.28 | 0.42 | 0.35 |
| 1 | 102424197 | rs12035347 | 0.448 | 0.81 | -0.33 | -0.22 | 0.01 | -0.049 | -0.049 | 0 | 0.78 | 0.74 | 0.75 | 0.97 | 0.72 | 0.85 |
| 1 | 102426168 | rs17487709 | 0.173 | 0.619 | -0.17 | -0.315 | 0.012 | -0.011 | -0.011 | 0 | 0.31 | 0.28 | 0.24 | 0.21 | 0.35 | 0.28 |
| 1 | 102427608 | rs12737750 | 0.004 | -0.285 | -0.2 | -0.123 | 0.007 | 0.028 | 0.028 | 0 | 0.46 | 0.43 | 0.48 | 0.28 | 0.49 | 0.35 |
| 1 | 102428709 | rs4908203 | 0.76 | -0.349 | -0.208 | -0.006 | 0.007 | -0.019 | -0.019 | 0.149 | 0.22 | 0.22 | 0.27 | 0.49 | 0.17 | 0.32 |
| 1 | 102435169 | rs17125886 | 2.301 | -0.465 | -0.336 | -0.161 | 0.005 | -0.001 | -0.001 | 0 | 0.07 | 0.03 | 0.05 | 0 | 0.03 | 0.01 |
| 1 | 102437203 | rs951897 | 0.858 | -0.418 | -0.047 | -0.182 | 0.032 | -0.014 | -0.014 | 0 | 0.09 | 0.18 | 0.18 | 0.02 | 0.2 | 0.1 |
| 1 | 102443692 | rs2339204 | -0.951 | -0.42 | -0.011 | -0.183 | 0.064 | -0.02 | -0.02 | 0.564 | 0.91 | 0.76 | 0.82 | 0.98 | 0.8 | 0.9 |
| 1 | 102447116 | rs1415102 | 0.341 | -0.617 | -0.15 | -0.219 | 0.066 | 0.059 | 0.059 | 0 | 0.36 | 0.13 | 0.29 | 0.25 | 0.28 | 0.26 |
| 1 | 102469386 | rs6657751 | -0.171 | -0.851 | 0.25 | 0.476 | 0.008 | -0.003 | -0.003 | 0.019 | 0.56 | 0.38 | 0.55 | 0.6 | 0.44 | 0.47 |
| 1 | 102471839 | rs1415094 | 1.085 | -0.839 | 0.194 | 0.509 | -0.008 | 0.031 | 0.031 | 1.375 | 0.2 | 0.26 | 0.26 | 0.35 | 0.16 | 0.22 |
| 1 | 102491704 | rs10874533 | 0.271 | -0.6 | 0.474 | 0.207 | -0.001 | 0.009 | 0.009 | 0.021 | 0.39 | 0.33 | 0.4 | 0.22 | 0.55 | 0.4 |

| | | | | | | | | | | | | | | | | |
|---|---|---|---|---|---|---|---|---|---|---|---|---|---|---|---|---|
| 1 | 102511803 | rs10874535 | 0.147 | -0.875 | 0.232 | 0.414 | -0.007 | 0.002 | 0.002 | 3.226 | 0.3 | 0.33 | 0.22 | 0.4 | 0.22 | 0.33 |
| 1 | 102512873 | rs10449717 | -0.459 | -0.452 | 0.073 | 0.33 | 0.015 | -0.036 | -0.036 | 1.517 | 0.84 | 0.75 | 0.78 | 0.97 | 0.77 | 0.9 |
| 1 | 102540084 | rs10493980 | 0.75 | -0.587 | -0.018 | 0.479 | 0.004 | 0.001 | 0.001 | 0.029 | 0.52 | 0.29 | 0.54 | 0.51 | 0.53 | 0.51 |
| 1 | 102546011 | rs10782868 | -0.055 | -0.651 | -0.078 | 0.283 | -0.021 | 0.025 | 0.025 | 0 | 0.85 | 0.9 | 0.83 | 0.98 | 0.77 | 0.87 |
| 1 | 102549180 | rs12049277 | 0.885 | -0.834 | 0.298 | 0.396 | 0.006 | 0.004 | 0.004 | 1.037 | 0.23 | 0.21 | 0.19 | 0.25 | 0.16 | 0.22 |
| 1 | 102554563 | rs11583493 | 0.602 | -0.7 | 0.117 | 0.386 | -0.049 | 0.042 | 0.042 | 0 | 0.09 | 0.03 | 0.16 | 0.02 | 0.21 | 0.1 |
| 1 | 102562640 | rs6684718 | 0.603 | -0.807 | 0.176 | 0.456 | 0 | 0 | 0 | 0 | 0.58 | 0.53 | 0.53 | 0.47 | 0.51 | 0.61 |
| 1 | 102564075 | rs4609450 | -1.116 | -0.904 | 0.115 | 0.522 | 0.008 | -0.017 | -0.017 | 0.184 | 0.77 | 0.8 | 0.74 | 0.66 | 0.77 | 0.85 |
| 1 | 102565669 | rs6661510 | -1.185 | -0.89 | 0.079 | 0.53 | 0.011 | -0.035 | -0.035 | 0 | 0.77 | 0.88 | 0.75 | 0.66 | 0.77 | 0.85 |
| 1 | 102566131 | rs6670467 | -1.185 | -0.89 | 0.079 | 0.524 | 0.011 | -0.033 | -0.033 | 0 | 0.77 | 0.88 | 0.75 | 0.66 | 0.77 | 0.85 |
| 1 | 102572944 | rs12088003 | -0.331 | -0.767 | 0.155 | 0.582 | -0.01 | -0.008 | -0.008 | 0 | 0.17 | 0.14 | 0.2 | 0.2 | 0.26 | 0.24 |
| 1 | 102587593 | rs12563410 | 0.765 | -1.089 | 0.327 | 0.522 | -0.025 | 0.016 | 0.016 | 0.918 | 0.25 | 0.4 | 0.25 | 0.3 | 0.16 | 0.29 |
| 1 | 102601950 | rs7527051 | 1.343 | -0.825 | 0.062 | 0.451 | -0.03 | 0.05 | 0.05 | 0 | 0.32 | 0.13 | 0.28 | 0.18 | 0.35 | 0.32 |
| 1 | 102608577 | rs2022858 | 0.323 | -0.91 | 0.375 | 0.631 | 0 | 0.012 | 0.012 | 0.126 | 0.59 | 0.47 | 0.57 | 0.5 | 0.59 | 0.5 |
| 1 | 102616451 | rs9729206 | 0.858 | -0.862 | 0.36 | 0.535 | -0.015 | -0.004 | -0.004 | 0 | 0.39 | 0.28 | 0.44 | 0.38 | 0.46 | 0.48 |
| 1 | 102618195 | rs2000398 | 0.033 | -0.959 | 0.607 | 0.476 | 0.004 | 0.012 | 0.012 | 0 | 0.49 | 0.61 | 0.49 | 0.38 | 0.5 | 0.53 |
| 1 | 102620560 | rs12088697 | 0.76 | -0.921 | 0.572 | 0.482 | -0.014 | -0.029 | -0.029 | 0 | 0.07 | 0.13 | 0.02 | 0 | 0.03 | 0.04 |
| 1 | 102628005 | rs6679483 | 0.137 | -1.09 | 0.328 | 0.402 | 0.001 | 0.002 | 0.002 | 0 | 0.52 | 0.43 | 0.49 | 0.38 | 0.54 | 0.54 |
| 1 | 102629663 | rs11164405 | 1.38 | -0.969 | 0.122 | 0.475 | 0.008 | -0.032 | -0.032 | 0 | 0.21 | 0.1 | 0.23 | 0.33 | 0.19 | 0.17 |
| 1 | 102632935 | rs1938350 | 0.577 | -0.546 | 0.05 | 0.433 | -0.017 | -0.011 | -0.011 | 0.17 | 0.06 | 0.03 | 0.15 | 0.18 | 0.11 | 0.16 |
| 1 | 102639332 | rs12140539 | 1.205 | -0.605 | 0.106 | 0.133 | -0.005 | 0.007 | 0.007 | 0 | 0.33 | 0.24 | 0.3 | 0.19 | 0.35 | 0.33 |
| 1 | 102647938 | rs3008763 | -0.627 | -0.673 | 0.24 | 0.373 | 0.024 | 0.005 | 0.005 | 0 | 0.8 | 0.71 | 0.76 | 0.84 | 0.72 | 0.76 |
| 1 | 102654557 | rs1938368 | 0.508 | -0.717 | 0.164 | 0.83 | 0.012 | 0.046 | 0.046 | 0 | 0.06 | 0.11 | 0.15 | 0.24 | 0.1 | 0.16 |
| 1 | 102667388 | rs1938364 | 0.6 | -1.191 | -0.192 | 0.567 | 0.053 | -0.045 | -0.045 | 0 | 0.32 | 0.21 | 0.28 | 0.47 | 0.22 | 0.21 |
| 1 | 102673499 | rs4301653 | 0.24 | -1.43 | -0.198 | 0.406 | 0.017 | -0.011 | -0.011 | 0 | 0.34 | 0.31 | 0.28 | 0.47 | 0.21 | 0.23 |
| 1 | 102681580 | rs7530562 | 1.618 | -1.395 | -0.039 | 0.367 | -0.024 | 0.058 | 0.058 | 0 | 0.34 | 0.14 | 0.29 | 0.14 | 0.35 | 0.34 |
| 1 | 102684721 | rs12082028 | 1.632 | -1.345 | 0.058 | 0.377 | -0.013 | -0.029 | -0.029 | 0 | 0.07 | 0.09 | 0.03 | 0 | 0.04 | 0.02 |
| 1 | 102686309 | rs11164417 | 0.125 | -1.338 | 0.021 | 0.418 | 0.006 | -0.017 | -0.017 | 0.243 | 0.36 | 0.33 | 0.4 | 0.58 | 0.3 | 0.33 |
| 1 | 102688814 | rs2096129 | 1.39 | -1.356 | -0.083 | 0.425 | 0.02 | -0.037 | -0.037 | 0 | 0.26 | 0.19 | 0.26 | 0.47 | 0.18 | 0.2 |
| 1 | 102691723 | rs12132269 | 0.489 | -1.111 | -0.096 | 0.258 | 0.002 | 0.034 | 0.034 | 0 | 0.41 | 0.27 | 0.33 | 0.14 | 0.39 | 0.35 |
| 1 | 102694690 | rs12139212 | 0.418 | -1.137 | -0.137 | 0.249 | 0.001 | 0.029 | 0.029 | 0 | 0.42 | 0.28 | 0.33 | 0.14 | 0.4 | 0.36 |
| 1 | 102701409 | rs10874564 | 0.449 | -1.049 | -0.177 | -0.059 | 0 | 0.034 | 0.034 | 0 | 0.42 | 0.27 | 0.33 | 0.14 | 0.4 | 0.36 |
| 1 | 102704421 | rs12125348 | 0.94 | -0.866 | -0.576 | -0.042 | -0.012 | 0.051 | 0.051 | 0 | 0.35 | 0.18 | 0.29 | 0.14 | 0.35 | 0.34 |
| 1 | 102705824 | rs17492399 | 0.979 | -0.406 | -0.798 | -0.411 | 0.006 | 0.014 | 0.014 | 0 | 0.08 | 0.04 | 0.12 | 0.04 | 0.05 | 0.04 |
| 1 | 102706588 | rs2375897 | -0.554 | -0.226 | -0.544 | -0.275 | 0.024 | -0.002 | -0.002 | 0 | 0.81 | 0.78 | 0.76 | 0.85 | 0.69 | 0.71 |
| 1 | 102711925 | rs7533173 | 0.54 | -0.269 | -0.69 | -0.435 | -0.01 | 0.048 | 0.048 | 0 | 0.35 | 0.2 | 0.41 | 0.16 | 0.35 | 0.24 |
| 1 | 102714517 | rs6577309 | -0.708 | -0.266 | -0.27 | -0.458 | 0.029 | -0.029 | -0.029 | 0 | 0.74 | 0.6 | 0.83 | 0.85 | 0.67 | 0.66 |
| 1 | 102717010 | rs952484 | 0.752 | -0.497 | -0.325 | -0.593 | -0.017 | 0.089 | 0.089 | 0 | 0.39 | 0.18 | 0.48 | 0.16 | 0.4 | 0.27 |
| 1 | 102726399 | rs17126268 | 0.38 | -0.157 | -0.233 | -0.447 | -0.005 | -0.037 | -0.037 | 0 | 0.08 | 0.11 | 0.05 | 0 | 0.07 | 0.06 |
| 1 | 102729275 | rs12135261 | 0.344 | -0.16 | -0.235 | -0.449 | -0.005 | -0.037 | -0.037 | 0 | 0.08 | 0.11 | 0.05 | 0 | 0.07 | 0.06 |
| 1 | 102734224 | rs12124015 | 0.524 | -0.156 | -0.205 | -0.447 | -0.001 | -0.019 | -0.019 | 0.016 | 0.08 | 0.06 | 0.05 | 0 | 0.07 | 0.06 |
| 1 | 102734951 | rs12120258 | 0.395 | -0.153 | -0.248 | -0.446 | -0.005 | -0.037 | -0.037 | 0 | 0.08 | 0.12 | 0.05 | 0 | 0.07 | 0.06 |
| 1 | 102742475 | rs2889234 | 1.496 | -0.187 | -0.694 | -0.116 | -0.032 | -0.096 | -0.096 | 0 | 0.2 | 0.05 | 0.27 | 0.46 | 0.19 | 0.21 |
| 1 | 102743350 | rs10158366 | -0.779 | -0.119 | 0.304 | 0.27 | 0.008 | -0.041 | -0.041 | 0 | 0.33 | 0.41 | 0.2 | 0.15 | 0.38 | 0.4 |
| 1 | 102759741 | rs12039496 | 1.442 | -1.332 | 0.525 | 0.437 | 0 | -0.001 | -0.001 | 0 | 0.63 | 0.31 | 0.52 | 0.65 | 0.68 | 0.71 |
| 1 | 102760382 | rs12407943 | 1.868 | -1.394 | 0.058 | 0.467 | 0.046 | -0.03 | -0.03 | 0 | 0.49 | 0.09 | 0.46 | 0.57 | 0.6 | 0.45 |
| 1 | 102791263 | rs1935273 | -0.488 | -1.408 | 0.259 | 0.373 | -0.001 | 0.031 | 0.031 | 0 | 0.22 | 0.15 | 0.37 | 0.1 | 0.24 | 0.11 |
| 1 | 102799057 | rs2792007 | -0.076 | -1.439 | 0.245 | 0.357 | -0.015 | 0.029 | 0.029 | 0 | 0.81 | 0.89 | 0.65 | 0.9 | 0.79 | 0.89 |
| 1 | 102816541 | rs12061042 | 1.126 | -1.451 | 0.547 | 0.378 | -0.006 | -0.014 | -0.014 | 0 | 0.71 | 0.59 | 0.6 | 0.88 | 0.74 | 0.84 |
| 1 | 102826835 | rs7411744 | 1.717 | -1.467 | 0.543 | 0.408 | -0.009 | -0.001 | -0.001 | 0 | 0.63 | 0.36 | 0.52 | 0.65 | 0.68 | 0.71 |
| 1 | 102827023 | rs1341910 | 0.137 | -1.459 | 0.593 | 0.407 | 0.022 | 0.028 | 0.028 | 0 | 0.08 | 0.35 | 0.06 | 0.02 | 0.05 | 0.09 |
| 1 | 102828823 | rs2791997 | 0.916 | -1.459 | 0.461 | 0.261 | 0.01 | 0.04 | 0.04 | 0 | 0.73 | 0.78 | 0.61 | 0.9 | 0.74 | 0.89 |
| 1 | 102843599 | rs1591960 | -0.354 | -1.434 | 0.039 | 0.411 | -0.033 | -0.023 | -0.023 | 0 | 0.88 | 0.55 | 0.91 | 0.97 | 0.93 | 0.9 |
| 1 | 102843725 | rs12402460 | 1.965 | -1.527 | -0.191 | 0.441 | 0.058 | -0.044 | -0.044 | 0 | 0.49 | 0.09 | 0.48 | 0.51 | 0.61 | 0.46 |
| 1 | 102846144 | rs12119077 | -0.962 | -0.652 | 0.221 | 0.388 | -0.01 | -0.026 | -0.026 | 0 | 0.19 | 0.14 | 0.36 | 0.27 | 0.22 | 0.19 |
| 1 | 102846147 | rs11164492 | 0.716 | -0.63 | 0.241 | 0.372 | -0.003 | -0.01 | -0.01 | 1.318 | 0.05 | 0.07 | 0.02 | 0 | 0.02 | 0.01 |
| 1 | 102862791 | rs12043531 | -1.01 | -0.519 | 0.095 | 0.46 | -0.01 | -0.028 | -0.028 | 0 | 0.19 | 0.14 | 0.36 | 0.27 | 0.22 | 0.19 |
| 1 | 102864381 | rs12038472 | 1.497 | -0.609 | 0.14 | 0.406 | -0.026 | -0.06 | -0.06 | 0 | 0.64 | 0.45 | 0.55 | 0.88 | 0.69 | 0.79 |
| 1 | 102869841 | rs10747427 | 0.469 | -0.539 | -0.116 | 0.395 | -0.083 | -0.075 | -0.075 | 0 | 0.81 | 0.48 | 0.89 | 0.97 | 0.91 | 0.9 |
| 1 | 102883271 | rs1417226 | 0.921 | -0.191 | -0.376 | 0.533 | -0.009 | -0.002 | -0.002 | 0 | 0.62 | 0.33 | 0.53 | 0.7 | 0.69 | 0.71 |
| 1 | 102885930 | rs12045655 | 0.906 | -0.173 | -0.368 | 0.542 | -0.009 | -0.002 | -0.002 | 0 | 0.62 | 0.33 | 0.53 | 0.7 | 0.69 | 0.71 |
| 1 | 102899784 | rs1591959 | -0.874 | 0.282 | -0.479 | 0.601 | 0.008 | 0.001 | 0.001 | 0.113 | 0.08 | 0.09 | 0.11 | 0.09 | 0.13 | 0.06 |
| 1 | 102916914 | rs2376037 | 0.789 | -0.584 | -0.869 | 0.135 | 0.051 | 0.048 | 0.048 | 0.035 | 0.67 | 0.22 | 0.81 | 0.86 | 0.8 | 0.75 |
| 1 | 102922025 | rs12060427 | -0.429 | -0.752 | 0.395 | 0.072 | 0 | 0.016 | 0.016 | 0 | 0.07 | 0.31 | 0.04 | 0.02 | 0.02 | 0.11 |
| 1 | 102934974 | rs3916220 | 1.945 | -1.065 | 0.09 | 0.41 | -0.037 | -0.009 | -0.009 | 0 | 0.46 | 0.1 | 0.5 | 0.45 | 0.54 | 0.48 |
| 1 | 102938426 | rs7550150 | 0.678 | -1.162 | 0.433 | 0.362 | 0.041 | -0.016 | -0.016 | 0 | 0.08 | 0.27 | 0.03 | 0.02 | 0.06 | 0.05 |
| 1 | 102939690 | rs12139288 | 0.382 | -1.201 | 0.216 | 0.439 | -0.002 | -0.034 | -0.034 | 0 | 0.13 | 0.07 | 0.17 | 0.34 | 0.12 | 0.19 |
| 1 | 102940956 | rs17495930 | 0.068 | -1.189 | 0.244 | 0.415 | 0.003 | -0.001 | -0.001 | 0 | 0.07 | 0.06 | 0.15 | 0.08 | 0.13 | 0.08 |
| 1 | 102955343 | rs1934710 | -1.538 | -1.269 | 0.265 | 0.413 | -0.013 | -0.062 | -0.062 | 1.05 | 0.25 | 0.14 | 0.33 | 0.44 | 0.27 | 0.36 |

| | | | | | | | | | | | | | | | | |
|---|---|---|---|---|---|---|---|---|---|---|---|---|---|---|---|---|
| 1 | 102957393 | rs11164526 | 0.64 | -1.265 | 0.245 | 0.403 | 0.005 | -0.016 | -0.016 | 0 | 0.11 | 0.07 | 0.16 | 0.34 | 0.09 | 0.17 |
| 1 | 102959259 | rs10874621 | -2.372 | -1.4 | 0.655 | 0.28 | 0.031 | -0.019 | -0.019 | 0.334 | 0.43 | 0.19 | 0.44 | 0.53 | 0.4 | 0.47 |
| 1 | 102961031 | rs1340392 | -1.232 | -1.716 | 1.088 | 0.213 | -0.032 | -0.04 | -0.04 | 0 | 0.27 | 0.38 | 0.15 | 0.11 | 0.19 | 0.15 |
| 1 | 102967150 | rs12752401 | -0.394 | -1.737 | 1.109 | 0.201 | 0.037 | 0.024 | 0.024 | 0 | 0.14 | 0.03 | 0.07 | 0.06 | 0.09 | 0.07 |
| 1 | 102968036 | rs1934709 | 0.887 | -1.756 | 1.158 | 0.073 | -0.013 | -0.023 | -0.023 | 0 | 0.81 | 0.29 | 0.91 | 0.95 | 0.91 | 0.83 |
| 1 | 102969517 | rs1934704 | 2.615 | -1.815 | 0.899 | 0.31 | 0.023 | 0.025 | 0.025 | 0 | 0.47 | 0.07 | 0.63 | 0.43 | 0.61 | 0.49 |
| 1 | 102995218 | rs12035823 | 0.863 | -1.836 | 1.185 | 0.204 | 0.009 | 0.025 | 0.025 | 0 | 0.09 | 0.08 | 0.11 | 0.39 | 0.04 | 0.19 |
| 1 | 102998264 | rs950061 | 1.868 | -1.894 | 1.343 | 0.112 | 0.018 | -0.008 | -0.008 | 0 | 0.59 | 0.39 | 0.72 | 0.48 | 0.72 | 0.63 |
| 1 | 103010867 | rs2889309 | 2.822 | -1.902 | 1.315 | 0.107 | 0.022 | -0.003 | -0.003 | 0 | 0.47 | 0.07 | 0.63 | 0.43 | 0.61 | 0.49 |
| 1 | 103012046 | rs7538167 | -0.018 | -1.859 | 1.512 | 0.112 | 0.039 | -0.016 | -0.016 | 0 | 0.07 | 0.17 | 0.07 | 0.02 | 0.09 | 0.04 |
| 1 | 103012510 | rs11164544 | 1.866 | -1.891 | 1.545 | 0.089 | 0.019 | -0.009 | -0.009 | 0 | 0.59 | 0.39 | 0.72 | 0.48 | 0.72 | 0.63 |
| 1 | 103045655 | rs17126738 | 0.279 | -1.728 | 1.167 | -0.661 | 0.015 | -0.023 | -0.023 | 0.095 | 0.1 | 0.05 | 0.06 | 0.28 | 0.06 | 0.11 |
| 1 | 103047426 | rs4907976 | 2.854 | -1.761 | 1.063 | -0.791 | 0.006 | -0.012 | -0.012 | 0 | 0.47 | 0.1 | 0.72 | 0.59 | 0.59 | 0.59 |
| 1 | 103062191 | rs12723721 | -1.956 | -1.7 | 1.012 | -0.791 | 0.027 | -0.03 | -0.03 | 1.534 | 0.3 | 0.15 | 0.16 | 0.34 | 0.26 | 0.21 |
| 1 | 103071512 | rs1878510 | 0.764 | -1.683 | 1.029 | -0.79 | -0.003 | 0.018 | 0.018 | 0 | 0.05 | 0.03 | 0.03 | 0.01 | 0.06 | 0.04 |
| 1 | 103080187 | rs11164556 | -0.755 | -1.691 | 1.027 | -0.793 | -0.006 | -0.03 | -0.03 | 0 | 0.15 | 0.09 | 0.1 | 0.29 | 0.16 | 0.14 |
| 1 | 103081423 | rs1113640 | -2.561 | -1.685 | 1.034 | -0.786 | 0.028 | -0.01 | -0.01 | 0.357 | 0.35 | 0.17 | 0.19 | 0.37 | 0.32 | 0.25 |
| 1 | 103087672 | rs12726148 | -0.53 | -1.545 | 1.058 | -0.809 | 0.04 | 0.042 | 0.042 | 0 | 0.15 | 0.05 | 0.06 | 0.06 | 0.1 | 0.07 |
| 1 | 103093419 | rs932175 | -0.164 | -1.25 | 1.848 | -0.705 | 0.046 | -0.05 | -0.05 | 0 | 0.88 | 0.55 | 0.95 | 0.98 | 0.92 | 0.94 |
| 1 | 103103131 | rs12119078 | 0.116 | -1.106 | 1.496 | -0.671 | 0.033 | -0.02 | -0.02 | 0 | 0.05 | 0.12 | 0.05 | 0.01 | 0.1 | 0.03 |
| 1 | 103106047 | rs7526984 | -0.589 | -1.116 | 1.523 | -0.676 | 0.107 | 0.003 | 0.003 | 0.915 | 0.06 | 0.33 | 0.07 | 0.01 | 0.1 | 0.03 |
| 1 | 103145383 | rs1451033 | 0.966 | -0.844 | 1.53 | -0.571 | 0.087 | 0.001 | 0.001 | 0 | 0.92 | 0.66 | 0.92 | 0.98 | 0.87 | 0.91 |
| 1 | 103152697 | rs7550400 | -0.836 | -1.223 | 1.341 | -0.636 | 0.032 | -0.046 | -0.046 | 0 | 0.17 | 0.06 | 0.08 | 0.29 | 0.11 | 0.1 |
| 1 | 103155605 | rs10782899 | 0.468 | -1.17 | 1.63 | -0.61 | 0.035 | 0.037 | 0.037 | 0.018 | 0.93 | 0.6 | 0.94 | 0.98 | 0.97 | 0.94 |
| 1 | 103171943 | rs11164585 | -1.234 | -1.231 | 1.871 | -0.608 | -0.054 | 0.147 | 0.147 | 0 | 0.17 | 0.52 | 0.11 | 0.3 | 0.08 | 0.12 |
| 1 | 103171983 | rs10493982 | -0.914 | -1.314 | 1.805 | -0.825 | -0.002 | -0.019 | -0.019 | 0 | 0.23 | 0.2 | 0.09 | 0.07 | 0.18 | 0.14 |
| 1 | 103176592 | rs12030173 | 0.404 | -1.25 | 1.333 | -0.774 | -0.003 | 0.022 | 0.022 | 0.09 | 0.09 | 0.09 | 0.05 | 0.28 | 0.05 | 0.06 |
| 1 | 103188273 | rs6699996 | 2.358 | -1.265 | 1.079 | -0.835 | 0.033 | 0.034 | 0.034 | 0 | 0.52 | 0.23 | 0.74 | 0.61 | 0.6 | 0.62 |
| 1 | 103203107 | rs1376359 | 1.892 | -1.305 | 1.233 | -0.678 | 0.071 | 0.002 | 0.002 | 0 | 0.58 | 0.24 | 0.79 | 0.62 | 0.73 | 0.67 |
| 1 | 103206371 | rs1597582 | 1.971 | -1.522 | 1.128 | -0.857 | 0.019 | 0.006 | 0.006 | 0 | 0.58 | 0.23 | 0.76 | 0.65 | 0.66 | 0.64 |
| 1 | 103226221 | rs579692 | -1.437 | -1.63 | 1.805 | -0.773 | -0.008 | -0.032 | -0.032 | 0.09 | 0.27 | 0.26 | 0.11 | 0.07 | 0.21 | 0.12 |
| 1 | 103230040 | rs478830 | -1.452 | -1.547 | 1.759 | -1.051 | -0.003 | 0.041 | 0.041 | 0 | 0.25 | 0.37 | 0.14 | 0.31 | 0.2 | 0.36 |
| 1 | 103231028 | rs618072 | 1.057 | -1.51 | 1.852 | -1.039 | -0.033 | -0.098 | -0.098 | 0 | 0.73 | 0.56 | 0.87 | 0.93 | 0.79 | 0.88 |
| 1 | 103238392 | rs606756 | 0.46 | -1.552 | 2.142 | -1.06 | -0.001 | -0.013 | -0.013 | 0 | 0.8 | 0.8 | 0.91 | 0.93 | 0.82 | 0.9 |
| 1 | 103250122 | rs7528798 | -0.71 | -1.568 | 2.124 | -1.058 | 0 | 0.005 | 0.005 | 0.036 | 0.17 | 0.16 | 0.1 | 0.3 | 0.17 | 0.3 |
| 1 | 103250731 | rs582076 | 1.535 | -1.6 | 2.183 | -1.076 | -0.035 | -0.08 | -0.08 | 0.929 | 0.67 | 0.55 | 0.86 | 0.91 | 0.76 | 0.82 |
| 1 | 103250988 | rs678426 | -1.266 | -1.612 | 2.044 | -1.1 | -0.03 | 0.039 | 0.039 | 0 | 0.93 | 0.99 | 0.98 | 1 | 0.92 | 0.93 |
| 1 | 103251006 | rs10782904 | 2.779 | -1.6 | 2.064 | -1.085 | 0.022 | 0.021 | 0.021 | 0 | 0.49 | 0.23 | 0.74 | 0.62 | 0.59 | 0.52 |
| 1 | 103255303 | rs1451029 | 2.804 | -1.598 | 1.932 | -1.085 | 0.016 | 0.014 | 0.014 | 0 | 0.49 | 0.18 | 0.74 | 0.62 | 0.59 | 0.52 |
| 1 | 103277571 | rs12086644 | -0.32 | -1.615 | 1.689 | -1.01 | -0.003 | 0.048 | 0.048 | 0 | 0.1 | 0.14 | 0.07 | 0.28 | 0.08 | 0.27 |
| 1 | 103280472 | rs1597584 | 1.485 | -1.578 | 1.553 | -1.273 | -0.02 | 0.01 | 0.01 | 0 | 0.06 | 0.02 | 0.02 | 0.02 | 0.09 | 0.03 |
| 1 | 103289931 | rs621734 | 0.925 | -1.645 | 1.657 | -1.307 | 0.009 | 0.022 | 0.022 | 0 | 0.77 | 0.84 | 0.91 | 0.93 | 0.81 | 0.9 |
| 1 | 103305021 | rs1241202 | -2.239 | -1.686 | 1.783 | -1.252 | -0.023 | 0.003 | 0.003 | 0 | 0.32 | 0.38 | 0.17 | 0.33 | 0.22 | 0.37 |
| 1 | 103322312 | rs11164621 | 0.558 | -1.612 | 1.826 | -1.25 | 0.049 | -0.01 | -0.01 | 0 | 0.08 | 0.21 | 0.04 | 0.02 | 0.09 | 0.03 |
| 1 | 103323618 | rs1241178 | -1.108 | -1.617 | 1.623 | -1.258 | 0.002 | -0.018 | -0.018 | 0.288 | 0.92 | 0.92 | 0.98 | 0.99 | 0.92 | 0.93 |
| 1 | 103325774 | rs12404334 | 2.764 | -1.684 | 1.524 | -1.283 | 0.031 | 0.054 | 0.054 | 0 | 0.49 | 0.18 | 0.74 | 0.65 | 0.59 | 0.52 |
| 1 | 103326762 | rs1031821 | -1.666 | -1.58 | 1.654 | -1.201 | 0.018 | -0.014 | -0.014 | 0 | 0.25 | 0.21 | 0.14 | 0.31 | 0.19 | 0.31 |
| 1 | 103342392 | rs9659030 | -0.731 | -1.501 | 0.93 | -1.434 | 0.004 | -0.008 | -0.008 | 0.245 | 0.21 | 0.17 | 0.23 | 0.25 | 0.18 | 0.32 |
| 1 | 103348100 | rs1241182 | -0.655 | -1.537 | 0.944 | -1.446 | -0.007 | -0.005 | -0.005 | 0.358 | 0.15 | 0.18 | 0.07 | 0.07 | 0.11 | 0.04 |
| 1 | 103352451 | rs2229783 | 2.65 | -1.331 | 1.094 | -1.274 | -0.025 | 0.008 | 0.008 | 0 | 0.53 | 0.05 | 0.75 | 0.66 | 0.61 | 0.57 |
| 1 | 103354138 | rs1676486 | 0.654 | -1.201 | 1.234 | -1.185 | -0.002 | -0.006 | -0.006 | 0 | 0.81 | 0.8 | 0.89 | 0.75 | 0.83 | 0.66 |
| 1 | 103354428 | rs1763347 | 1.248 | -1.189 | 1.089 | -1.179 | 0.005 | 0 | 0 | 0 | 0.74 | 0.26 | 0.84 | 0.75 | 0.8 | 0.65 |
| 1 | 103355765 | rs1241163 | 1.02 | -1.159 | 1.263 | -1.162 | -0.004 | 0.017 | 0.017 | 0.057 | 0.77 | 0.44 | 0.85 | 0.75 | 0.81 | 0.66 |
| 1 | 103362638 | rs1676495 | -0.573 | -1.148 | 0.639 | -1.39 | -0.008 | -0.006 | -0.006 | 0 | 0.12 | 0.17 | 0.05 | 0.07 | 0.08 | 0.04 |
| 1 | 103371652 | rs12145872 | -0.299 | -1.057 | 0.427 | -1.397 | -0.01 | 0.048 | 0.048 | 1.524 | 0.13 | 0.04 | 0.04 | 0.02 | 0.13 | 0.05 |
| 1 | 103372655 | rs1676504 | 0.567 | -1.09 | 0.521 | -1.397 | -0.027 | -0.036 | -0.036 | 0.226 | 0.07 | 0.14 | 0.02 | 0 | 0.02 | 0.01 |
| 1 | 103377968 | rs11164636 | 1.194 | -1.079 | 0.152 | -1.448 | 0.025 | 0.101 | 0.101 | 0.025 | 0.69 | 0.2 | 0.86 | 0.98 | 0.77 | 0.87 |
| 1 | 103379918 | rs3753841 | 2.007 | -0.963 | -0.068 | -1.325 | -0.045 | 0.048 | 0.048 | 0.011 | 0.55 | 0.05 | 0.76 | 0.7 | 0.61 | 0.59 |
| 1 | 103393457 | rs12138977 | 1.678 | -0.996 | -0.012 | -1.326 | -0.009 | 0.092 | 0.092 | 0.62 | 0.59 | 0.06 | 0.78 | 0.77 | 0.67 | 0.64 |
| 1 | 103402184 | rs1120519 | 0.582 | -1.152 | 0.695 | -1.754 | -0.021 | -0.093 | -0.093 | 0.753 | 0.82 | 0.35 | 0.91 | 0.99 | 0.89 | 0.92 |
| 1 | 103404384 | rs2126643 | 1.506 | -1.374 | 0.803 | -1.994 | -0.014 | -0.042 | -0.042 | 0 | 0.57 | 0.49 | 0.7 | 0.88 | 0.65 | 0.75 |
| 1 | 103405892 | rs17127270 | -0.106 | -1.411 | 0.834 | -2.009 | 0.013 | -0.032 | -0.032 | 0 | 0.15 | 0.2 | 0.06 | 0.03 | 0.18 | 0.11 |
| 1 | 103407999 | rs2622865 | 1.792 | -1.45 | 1.178 | -2.028 | 0.007 | 0.03 | 0.03 | 0 | 0.49 | 0.42 | 0.67 | 0.87 | 0.57 | 0.68 |
| 1 | 103409100 | rs12046389 | 2.04 | -1.448 | 1.207 | -2.027 | -0.012 | 0.191 | 0.191 | 0 | 0.47 | 0.16 | 0.65 | 0.87 | 0.57 | 0.68 |
| 1 | 103409558 | rs2616009 | -0.52 | -1.314 | 1.253 | -1.965 | -0.009 | -0.022 | -0.022 | 0 | 0.93 | 0.69 | 0.96 | 1 | 0.98 | 0.99 |
| 1 | 103410124 | rs12084902 | -0.169 | -1.199 | 0.905 | -1.914 | 0.008 | -0.024 | -0.024 | 0.175 | 0.15 | 0.17 | 0.06 | 0.03 | 0.18 | 0.11 |
| 1 | 103419938 | rs12743365 | -0.122 | -1.347 | 0.648 | -1.92 | 0.002 | 0.049 | 0.049 | 0 | 0.19 | 0.05 | 0.19 | 0.07 | 0.16 | 0.12 |
| 1 | 103420053 | rs2615996 | -0.103 | -1.538 | 1.04 | -2.012 | -0.033 | -0.043 | -0.043 | 0 | 0.08 | 0.2 | 0.02 | 0 | 0.02 | 0.02 |
| 1 | 103427381 | rs1012282 | -1.522 | -1.482 | 1.203 | -2.054 | -0.012 | -0.048 | -0.048 | 0 | 0.33 | 0.4 | 0.26 | 0.09 | 0.26 | 0.15 |

| | | | | | | | | | | | | | | | | |
|---|---|---|---|---|---|---|---|---|---|---|---|---|---|---|---|---|
| 1 | 103438795 | rs12118365 | -0.755 | -1.407 | 1.131 | -2.052 | -0.002 | 0.022 | 0.022 | 0 | 0.25 | 0.2 | 0.23 | 0.09 | 0.24 | 0.14 |
| 1 | 103441814 | rs2061705 | 2.622 | -1.567 | 0.988 | -2.131 | -0.14 | 0.052 | 0.052 | 0 | 0.37 | 0.09 | 0.59 | 0.8 | 0.49 | 0.46 |
| 1 | 103444497 | rs1903787 | -1.883 | -1.375 | 1.823 | -1.975 | -0.016 | -0.059 | -0.059 | 0 | 0.33 | 0.56 | 0.27 | 0.09 | 0.26 | 0.15 |
| 1 | 103446620 | rs12124588 | -0.358 | -1.436 | 1.457 | -1.89 | 0.002 | 0.049 | 0.049 | 0 | 0.19 | 0.05 | 0.19 | 0.07 | 0.16 | 0.12 |
| 1 | 103455217 | rs1463034 | 1.812 | -1.443 | 1.72 | -1.831 | -0.087 | -0.064 | -0.064 | 0.265 | 0.8 | 0.49 | 0.92 | 0.93 | 0.91 | 0.92 |
| 1 | 103462386 | rs2622868 | -0.009 | -1.317 | 1.8 | -1.766 | -0.006 | -0.02 | -0.02 | 0 | 0.92 | 0.67 | 0.96 | 1 | 0.97 | 0.99 |
| 1 | 103463976 | rs2622870 | 0.882 | -1.32 | 1.797 | -1.767 | -0.033 | 0.006 | 0.006 | 0 | 0.88 | 0.61 | 0.94 | 0.93 | 0.94 | 0.94 |
| 1 | 103467615 | rs2929162 | -0.216 | -1.264 | 1.502 | -1.743 | -0.024 | -0.035 | -0.035 | 0 | 0.92 | 0.76 | 0.96 | 1 | 0.97 | 0.99 |
| 1 | 103496186 | rs12738133 | -0.655 | -1.139 | 1.12 | -1.759 | -0.007 | -0.003 | -0.003 | 0 | 0.1 | 0.12 | 0.04 | 0.07 | 0.07 | 0.07 |
| 1 | 103496620 | rs7523441 | -0.472 | -1.182 | 1.214 | -1.782 | -0.027 | -0.037 | -0.037 | 0.012 | 0.93 | 0.83 | 0.97 | 1 | 0.97 | 0.99 |
| 1 | 103499896 | rs11164658 | 1.588 | -1.291 | 1.31 | -1.909 | -0.044 | -0.089 | -0.089 | 1.113 | 0.66 | 0.49 | 0.74 | 0.93 | 0.77 | 0.86 |
| 1 | 103508275 | rs12123031 | -1.347 | -1.253 | 1.138 | -1.89 | -0.008 | -0.026 | -0.026 | 3.084 | 0.26 | 0.28 | 0.23 | 0.07 | 0.2 | 0.13 |
| 1 | 103544927 | rs1932986 | 0.41 | -0.741 | 0.72 | -1.644 | -0.012 | -0.067 | -0.067 | 0.012 | 0.78 | 0.78 | 0.8 | 1 | 0.83 | 0.93 |
| 1 | 103548497 | rs11164663 | 0.514 | -0.366 | 0.375 | -1.437 | 0.009 | -0.006 | -0.006 | 0 | 0.06 | 0.12 | 0.04 | 0.02 | 0.08 | 0.02 |
| 1 | 103551498 | rs4908290 | 1.41 | -0.499 | 0.114 | -1.899 | 0.009 | 0.229 | 0.229 | 0 | 0.57 | 0.19 | 0.67 | 0.97 | 0.63 | 0.83 |
| 1 | 103552935 | rs1415363 | 0.349 | -0.672 | 0.429 | -1.98 | -0.011 | 0.074 | 0.074 | 0 | 0.75 | 0.88 | 0.78 | 0.98 | 0.75 | 0.85 |
| 1 | 103554167 | rs11164669 | 2.012 | -0.654 | 0.313 | -1.983 | -0.006 | 0.267 | 0.267 | 0 | 0.53 | 0.11 | 0.64 | 0.91 | 0.57 | 0.75 |
| 1 | 103554293 | rs1337195 | -1.146 | -0.663 | 1.452 | -2.055 | 0.005 | -0.086 | -0.086 | 0 | 0.33 | 0.34 | 0.28 | 0.02 | 0.33 | 0.16 |
| 1 | 103556959 | rs6577353 | -0.173 | -0.509 | 1.948 | -1.806 | -0.03 | -0.043 | -0.043 | 0.331 | 0.08 | 0.16 | 0.04 | 0 | 0.04 | 0.01 |
| 1 | 103570432 | rs10493985 | -0.878 | -0.517 | 2.332 | -1.809 | -0.016 | -0.038 | -0.038 | 0 | 0.1 | 0.42 | 0.06 | 0 | 0.04 | 0.01 |
| 1 | 103578334 | rs7543626 | 0.899 | -0.172 | 2.218 | -2.242 | 0.029 | -0.111 | -0.111 | 0.016 | 0.84 | 0.46 | 0.83 | 0.99 | 0.82 | 0.97 |
| 1 | 103584570 | rs11802357 | -0.945 | -0.48 | 1.829 | -2.656 | -0.002 | -0.056 | -0.056 | 0 | 0.36 | 0.38 | 0.27 | 0.07 | 0.32 | 0.25 |
| 1 | 103603452 | rs11164689 | 0.892 | -0.431 | 1.889 | -2.7 | -0.008 | -0.036 | -0.036 | 1.171 | 0.07 | 0.18 | 0.06 | 0 | 0.04 | 0 |
| 1 | 103603691 | rs7544745 | -1.568 | -0.437 | 1.903 | -2.713 | 0.005 | -0.003 | -0.003 | 0 | 0.43 | 0.59 | 0.34 | 0.07 | 0.36 | 0.25 |
| 1 | 103612069 | rs6692914 | 2.062 | -0.47 | 2.051 | -2.607 | -0.074 | 0.212 | 0.212 | 0 | 0.43 | 0.13 | 0.55 | 0.9 | 0.48 | 0.71 |
| 1 | 103612149 | rs12092400 | -1.086 | -0.339 | 2.169 | -2.646 | -0.037 | -0.117 | -0.117 | 0 | 0.19 | 0.33 | 0.13 | 0 | 0.13 | 0.03 |
| 1 | 103612995 | rs7546530 | -0.948 | -0.408 | 2.195 | -2.619 | -0.046 | -0.115 | -0.115 | 0 | 0.18 | 0.33 | 0.13 | 0 | 0.11 | 0.02 |
| 1 | 103617675 | rs11586335 | 0.282 | -0.41 | 2.217 | -2.62 | -0.013 | -0.04 | -0.04 | 0 | 0.11 | 0.12 | 0.07 | 0 | 0.07 | 0.02 |
| 1 | 103621727 | rs12131191 | -0.388 | -0.402 | 2.052 | -2.677 | -0.018 | 0.134 | 0.134 | 0 | 0.29 | 0.09 | 0.23 | 0.02 | 0.27 | 0.17 |
| 1 | 103623133 | rs7550030 | -2.033 | -0.332 | 2.178 | -2.577 | 0.009 | 0.142 | 0.142 | 0 | 0.54 | 0.71 | 0.4 | 0.07 | 0.43 | 0.27 |
| 1 | 103662258 | rs1528202 | -0.782 | -0.349 | 2.027 | -2.704 | 0.016 | 0.03 | 0.03 | 0 | 0.26 | 0.21 | 0.21 | 0.06 | 0.18 | 0.16 |
| 1 | 103669167 | rs2889352 | -2.374 | -0.369 | 2.198 | -2.727 | 0.008 | 0.137 | 0.137 | 0 | 0.54 | 0.71 | 0.4 | 0.07 | 0.44 | 0.27 |
| 1 | 103671410 | rs12130012 | -0.658 | -0.366 | 2.085 | -2.726 | 0.019 | 0.088 | 0.088 | 0 | 0.24 | 0.1 | 0.2 | 0.06 | 0.18 | 0.17 |
| 1 | 103671460 | rs6683683 | -0.936 | -0.363 | 2.116 | -2.722 | -0.004 | -0.08 | -0.08 | 0.478 | 0.27 | 0.32 | 0.18 | 0.02 | 0.26 | 0.1 |
| 1 | 103685695 | rs17127592 | -0.937 | -0.097 | 2.244 | -2.251 | 0.013 | 0.03 | 0.03 | 0 | 0.24 | 0.16 | 0.2 | 0.06 | 0.18 | 0.17 |
| 1 | 103703266 | rs12046628 | 1.352 | -0.125 | 2.264 | -2.31 | 0.005 | 0.02 | 0.02 | 0 | 0.68 | 0.27 | 0.71 | 0.91 | 0.72 | 0.78 |
| 1 | 103715386 | rs6689743 | 1.216 | -0.111 | 2.268 | -2.305 | -0.008 | -0.077 | -0.077 | 0.288 | 0.69 | 0.49 | 0.74 | 0.91 | 0.72 | 0.79 |
| 1 | 103725629 | rs1406130 | 0.896 | -0.136 | 2.271 | -2.306 | -0.014 | -0.109 | -0.109 | 0.324 | 0.74 | 0.49 | 0.76 | 0.97 | 0.77 | 0.86 |
| 1 | 103728254 | rs10493998 | 0.029 | -0.085 | 1.952 | -2.258 | 0.003 | -0.02 | -0.02 | 0 | 0.18 | 0.14 | 0.18 | 0 | 0.12 | 0.07 |
| 1 | 103730700 | rs4617426 | 1.215 | -0.098 | 1.821 | -2.204 | 0.006 | 0.022 | 0.022 | 0 | 0.69 | 0.27 | 0.71 | 0.91 | 0.73 | 0.8 |
| 1 | 103741344 | rs7528387 | -1.101 | -0.169 | 1.496 | -2.297 | -0.008 | 0.055 | 0.055 | 0 | 0.36 | 0.26 | 0.3 | 0.05 | 0.37 | 0.18 |
| 1 | 103744171 | rs4907988 | 1.31 | -0.172 | 1.908 | -1.62 | 0.021 | 0.027 | 0.027 | 0 | 0.9 | 0.45 | 0.94 | 0.94 | 0.93 | 0.89 |
| 1 | 103750676 | rs2376362 | 1.14 | -0.135 | 1.453 | -1.608 | 0.044 | 0.045 | 0.045 | 0 | 0.91 | 0.61 | 0.96 | 0.94 | 0.93 | 0.89 |
| 1 | 103757778 | rs2222748 | -0.721 | 0.297 | 0.086 | -1.93 | 0.003 | 0.058 | 0.058 | 0 | 0.25 | 0.13 | 0.25 | 0.05 | 0.2 | 0.16 |
| 1 | 103761463 | rs17127731 | -0.913 | -0.613 | -0.508 | -2.173 | 0.028 | 0.067 | 0.067 | 0 | 0.44 | 0.34 | 0.38 | 0.23 | 0.35 | 0.31 |
| 1 | 103763252 | rs4907992 | 0.651 | -0.633 | -0.497 | -2.232 | 0.012 | 0.013 | 0.013 | 0 | 0.22 | 0.19 | 0.15 | 0.18 | 0.19 | 0.17 |
| 1 | 103764640 | rs1406135 | -0.239 | -0.373 | -0.556 | -2.134 | 0.006 | 0.037 | 0.037 | 0 | 0.21 | 0.15 | 0.23 | 0.05 | 0.16 | 0.14 |
| 1 | 103916755 | rs10785777 | 0.299 | 0.788 | -1.276 | -0.964 | -0.05 | -0.071 | -0.071 | 0 | 0.72 | 0.89 | 0.62 | 0.4 | 0.65 | 0.61 |
| 1 | 104017778 | rs6696797 | 0.278 | 0.76 | -1.025 | -0.926 | -0.033 | -0.034 | -0.034 | 0 | 0.72 | 0.81 | 0.56 | 0.38 | 0.65 | 0.6 |
| 1 | 104019631 | rs7524694 | 1.23 | 1.038 | -0.737 | -0.677 | 0.021 | 0.032 | 0.032 | 0 | 0.07 | 0.18 | 0.1 | 0.12 | 0.1 | 0.11 |
| 1 | 104030621 | rs10158225 | -0.047 | 1.146 | -0.909 | -0.593 | -0.054 | -0.055 | -0.055 | 0.308 | 0.73 | 0.88 | 0.56 | 0.37 | 0.65 | 0.6 |
| 1 | 104094998 | rs1060309 | -0.189 | 1.168 | -0.902 | -0.614 | -0.069 | -0.069 | -0.069 | 0 | 0.74 | 0.9 | 0.55 | 0.37 | 0.64 | 0.65 |
| 1 | 104096692 | rs11184430 | -0.004 | 1.143 | -0.608 | -0.597 | -0.016 | -0.017 | -0.017 | 0 | 0.72 | 0.75 | 0.54 | 0.37 | 0.64 | 0.65 |
| 1 | 104103057 | rs2050592 | 1.634 | 0.627 | -0.869 | -1.143 | 0.019 | 0.018 | 0.018 | 0 | 0.18 | 0.09 | 0.06 | 0 | 0.11 | 0.09 |
| 1 | 104113158 | rs12132911 | 1.347 | 0.922 | -0.956 | -1.092 | -0.037 | 0.006 | 0.006 | 0 | 0.17 | 0.05 | 0.13 | 0.13 | 0.19 | 0.32 |
| 1 | 104122556 | rs12085609 | -0.883 | 0.934 | -0.977 | -1.076 | -0.098 | -0.1 | -0.1 | 4.612 | 0.74 | 0.93 | 0.55 | 0.37 | 0.64 | 0.65 |
| 1 | 104123531 | rs17467099 | 1.386 | 0.935 | -0.964 | -1.078 | -0.031 | 0.009 | 0.009 | 0 | 0.17 | 0.06 | 0.13 | 0.13 | 0.19 | 0.32 |
| 1 | 104123662 | rs4847151 | -0.839 | 1.029 | -0.978 | -1.03 | -0.069 | -0.071 | -0.071 | 0.005 | 0.74 | 0.9 | 0.55 | 0.37 | 0.65 | 0.66 |
| 1 | 104125556 | rs11184556 | 1.176 | 1.172 | -1.208 | -1.328 | -0.025 | -0.152 | -0.152 | 0 | 0.36 | 0.41 | 0.18 | 0 | 0.3 | 0.19 |
| 1 | 104128618 | rs6661612 | -0.355 | 1.672 | -1.504 | -0.442 | -0.016 | -0.017 | -0.017 | 0 | 0.72 | 0.74 | 0.54 | 0.37 | 0.64 | 0.65 |
| 1 | 104309132 | rs2646249 | -0.503 | 1.843 | -1.523 | -0.236 | -0.064 | -0.063 | -0.063 | 0 | 0.75 | 0.89 | 0.57 | 0.37 | 0.65 | 0.67 |
| 1 | 104324819 | rs1930212 | 1.012 | 1.857 | -1.556 | -0.261 | -0.03 | -0.061 | -0.061 | 5.99 | 0.18 | 0.03 | 0.25 | 0.24 | 0.19 | 0.15 |
| 1 | 104329877 | rs1566154 | 0.896 | 2.168 | -0.823 | 1.311 | -0.021 | -0.063 | -0.063 | 0.159 | 0.17 | 0.06 | 0.31 | 0.35 | 0.16 | 0.23 |
| 1 | 104357400 | rs10881479 | -0.074 | 1.616 | -0.681 | 0.601 | -0.012 | -0.011 | -0.011 | 0 | 0.73 | 0.73 | 0.55 | 0.37 | 0.65 | 0.67 |
| 1 | 104365377 | rs1930180 | -0.272 | 1.715 | -1.283 | 0.657 | -0.064 | -0.066 | -0.066 | 0 | 0.75 | 0.89 | 0.57 | 0.37 | 0.65 | 0.67 |
| 1 | 104375470 | rs11185202 | -0.747 | 1.183 | -0.222 | 0.487 | 0.003 | 0.013 | 0.013 | 1.437 | 0.43 | 0.62 | 0.41 | 0.2 | 0.41 | 0.27 |
| 1 | 104376795 | rs11185216 | -0.653 | 1.19 | 0.048 | 0.48 | -0.003 | -0.03 | -0.03 | 0 | 0.42 | 0.51 | 0.4 | 0.18 | 0.41 | 0.27 |
| 1 | 104383330 | rs12133475 | 0.714 | 1.242 | -0.344 | 0.36 | -0.052 | -0.009 | -0.009 | 0 | 0.16 | 0.02 | 0.21 | 0.12 | 0.18 | 0.16 |
| 1 | 104383975 | rs1930199 | -0.92 | 1.173 | -0.023 | 0.336 | -0.002 | -0.028 | -0.028 | 0 | 0.41 | 0.51 | 0.4 | 0.18 | 0.41 | 0.26 |

| | | | | | | | | | | | | | | | | |
|---|---|---|---|---|---|---|---|---|---|---|---|---|---|---|---|---|
| 1 | 104401489 | rs12136807 | -1.021 | 1.242 | 0.369 | 1.087 | 0.001 | 0 | 0 | 0.149 | 0.42 | 0.51 | 0.53 | 0.39 | 0.48 | 0.32 |
| 1 | 104408821 | rs1930192 | -1.032 | 1.241 | 0.369 | 1.087 | 0.002 | 0.001 | 0.001 | 0.167 | 0.42 | 0.55 | 0.53 | 0.39 | 0.48 | 0.32 |
| 1 | 104436329 | rs7531107 | -0.925 | 1.168 | 0.391 | 1.026 | -0.008 | -0.009 | -0.009 | 0 | 0.39 | 0.37 | 0.5 | 0.39 | 0.44 | 0.31 |
| 1 | 104445316 | rs1930208 | 1.361 | 1.134 | -0.486 | 1.057 | -0.041 | 0.012 | 0.012 | 0 | 0.27 | 0.06 | 0.15 | 0.23 | 0.23 | 0.41 |
| 1 | 104447410 | rs12122420 | 0.365 | 1.472 | 0.466 | 1.789 | -0.064 | -0.011 | -0.011 | 0 | 0.16 | 0.02 | 0.21 | 0.13 | 0.19 | 0.15 |
| 1 | 104470622 | rs1475190 | 1.055 | 1.441 | 1.436 | 1.481 | 0.008 | 0.051 | 0.051 | 0.968 | 0.58 | 0.3 | 0.62 | 0.74 | 0.59 | 0.61 |
| 1 | 104471490 | rs4847149 | 1.037 | 1.428 | 1.524 | 1.475 | 0.001 | 0.005 | 0.005 | 0.576 | 0.58 | 0.4 | 0.62 | 0.74 | 0.59 | 0.61 |
| 1 | 104480469 | rs12731458 | -0.916 | 1.584 | 1.502 | 1.576 | 0.002 | 0.029 | 0.029 | 0.088 | 0.37 | 0.31 | 0.34 | 0.25 | 0.36 | 0.38 |
| 1 | 104483838 | rs3849298 | -0.748 | 1.526 | 1.688 | 1.461 | 0.039 | 0.007 | 0.007 | 0 | 0.06 | 0.26 | 0.03 | 0 | 0.05 | 0.01 |
| 1 | 104484407 | rs551284 | -0.93 | 1.617 | 1.257 | 1.691 | -0.015 | 0.065 | 0.065 | 0.15 | 0.36 | 0.2 | 0.34 | 0.23 | 0.36 | 0.38 |
| 1 | 104498943 | rs1576516 | -1.289 | 1.85 | 1.746 | 2.225 | -0.006 | -0.009 | -0.009 | 0 | 0.39 | 0.32 | 0.56 | 0.48 | 0.41 | 0.31 |
| 1 | 104502333 | rs11185345 | -1.658 | 1.834 | 1.847 | 2.177 | -0.022 | 0.099 | 0.099 | 0.371 | 0.46 | 0.74 | 0.7 | 0.73 | 0.5 | 0.41 |
| 1 | 104505231 | rs6694128 | 1.241 | 1.859 | 1.718 | 2.217 | -0.003 | 0.007 | 0.007 | 0.218 | 0.57 | 0.42 | 0.35 | 0.28 | 0.54 | 0.63 |
| 1 | 104506202 | rs6700491 | -1.251 | 1.929 | 1.144 | 2.118 | -0.014 | -0.021 | -0.021 | 0.033 | 0.39 | 0.27 | 0.56 | 0.48 | 0.41 | 0.3 |
| 1 | 104509012 | rs12725332 | 0.235 | 2.12 | 0.58 | 2.037 | -0.067 | -0.083 | -0.083 | 5.059 | 0.16 | 0.01 | 0.21 | 0.22 | 0.2 | 0.12 |
| 1 | 104524676 | rs17474256 | 0.437 | 1.755 | 0.628 | 1.745 | -0.005 | -0.034 | -0.034 | 0 | 0.13 | 0.01 | 0.19 | 0.13 | 0.12 | 0.08 |
| 1 | 104524702 | rs2019634 | -1.124 | 1.67 | 1.283 | 1.765 | -0.005 | -0.008 | -0.008 | 0 | 0.38 | 0.33 | 0.61 | 0.47 | 0.42 | 0.3 |
| 1 | 104529736 | rs2208963 | 0.918 | 1.556 | 1.667 | 1.236 | 0.002 | -0.007 | -0.007 | 0.007 | 0.59 | 0.44 | 0.33 | 0.29 | 0.52 | 0.61 |
| 1 | 104545721 | rs12741722 | -0.759 | 1.464 | 1.468 | 1.195 | -0.015 | -0.028 | -0.028 | 0.177 | 0.05 | 0.15 | 0.04 | 0 | 0.03 | 0.04 |
| 1 | 104554482 | rs1590902 | -1.214 | 1.492 | 1.581 | 1.201 | -0.01 | 0.01 | 0.01 | 0 | 0.39 | 0.33 | 0.64 | 0.71 | 0.48 | 0.39 |
| 1 | 104561488 | rs11185362 | -1.395 | 1.59 | 1.24 | 1.265 | 0.008 | -0.023 | -0.023 | 2.099 | 0.43 | 0.49 | 0.68 | 0.71 | 0.49 | 0.39 |
| 1 | 104573697 | rs12759387 | -0.656 | 1.648 | 0.5 | 1.215 | -0.061 | 0.063 | 0.063 | 3.921 | 0.35 | 0.09 | 0.22 | 0.2 | 0.36 | 0.13 |
| 1 | 104576359 | rs9435629 | 1.44 | 1.517 | 0.826 | 1.228 | -0.005 | 0.13 | 0.13 | 1.76 | 0.49 | 0.23 | 0.27 | 0.24 | 0.47 | 0.57 |
| 1 | 104591008 | rs12736718 | -0.997 | 1.425 | 1.221 | 1.274 | 0.031 | 0.002 | 0.002 | 0 | 0.07 | 0.31 | 0.06 | 0.01 | 0.05 | 0.08 |
| 1 | 104598941 | rs3934285 | -0.591 | 1.56 | 0.814 | 1.252 | -0.068 | 0.051 | 0.051 | 1.901 | 0.32 | 0.09 | 0.22 | 0.2 | 0.33 | 0.11 |
| 1 | 104603845 | rs11185369 | -1.022 | 1.515 | 1.845 | 2.049 | 0.06 | 0.302 | 0.302 | 0 | 0.08 | 0.21 | 0.37 | 0.5 | 0.13 | 0.21 |
| 1 | 104638728 | rs10494007 | -2.127 | 1.088 | 1.725 | 1.247 | -0.002 | -0.017 | -0.017 | 0 | 0.1 | 0.53 | 0.16 | 0.04 | 0.07 | 0.11 |
| 1 | 104690071 | rs11185384 | -1.751 | 1.556 | 1.533 | 1.433 | 0 | -0.026 | -0.026 | 0 | 0.27 | 0.58 | 0.31 | 0.23 | 0.3 | 0.2 |
| 1 | 104693341 | rs12754151 | -1.565 | 1.481 | 1.677 | 1.424 | -0.007 | -0.001 | -0.001 | 0 | 0.08 | 0.36 | 0.13 | 0.04 | 0.04 | 0.1 |
| 1 | 104696000 | rs11577888 | -1.837 | 1.465 | 1.567 | 1.435 | -0.004 | -0.022 | -0.022 | 0 | 0.26 | 0.58 | 0.3 | 0.23 | 0.29 | 0.19 |
| 1 | 104697678 | rs10157087 | -0.036 | 1.44 | 1.961 | 1.794 | 0.043 | 0.164 | 0.164 | 0 | 0.05 | 0.13 | 0.34 | 0.24 | 0.07 | 0.1 |
| 1 | 104727130 | rs17020664 | -1.98 | 1.293 | 1.736 | 1.5 | -0.014 | 0.037 | 0.037 | 0.183 | 0.09 | 0.46 | 0.12 | 0.07 | 0.03 | 0.09 |
| 1 | 104736011 | rs7527726 | -1.618 | 1.664 | 1.516 | 1.811 | 0.033 | 0.064 | 0.064 | 0 | 0.25 | 0.45 | 0.53 | 0.59 | 0.35 | 0.29 |
| 1 | 104747846 | rs11185389 | -1.028 | 1.66 | 0.882 | 1.113 | 0 | -0.001 | -0.001 | 0.003 | 0.44 | 0.53 | 0.62 | 0.6 | 0.52 | 0.39 |
| 1 | 104751861 | rs11185392 | -0.369 | 1.691 | 0.758 | 1.286 | -0.057 | -0.019 | -0.019 | 0 | 0.37 | 0.13 | 0.27 | 0.37 | 0.46 | 0.31 |
| 1 | 104752001 | rs11185393 | -1.382 | 1.728 | 0.571 | 1.292 | 0.022 | 0.077 | 0.077 | 0 | 0.49 | 0.71 | 0.71 | 0.68 | 0.57 | 0.48 |
| 1 | 104766179 | rs1928329 | 1.358 | 1.524 | 0.781 | 1.505 | 0.075 | 0.222 | 0.222 | 0.192 | 0.9 | 0.67 | 0.57 | 0.69 | 0.89 | 0.83 |
| 1 | 104770475 | rs1928327 | 0.489 | 1.333 | -0.075 | 0.696 | -0.002 | -0.017 | -0.017 | 1.033 | 0.59 | 0.31 | 0.59 | 0.53 | 0.58 | 0.6 |
| 1 | 104789235 | rs2317080 | 0.401 | 1.173 | -0.102 | 0.737 | -0.029 | -0.025 | -0.025 | 0.54 | 0.63 | 0.28 | 0.66 | 0.53 | 0.54 | 0.59 |
| 1 | 104792257 | rs11185408 | -0.87 | 1.098 | -0.033 | 0.777 | -0.036 | -0.031 | -0.031 | 0 | 0.41 | 0.75 | 0.43 | 0.59 | 0.51 | 0.47 |
| 1 | 104798765 | rs12406661 | 0.809 | 1.059 | -0.088 | 0.716 | -0.067 | -0.069 | -0.069 | 0 | 0.57 | 0.11 | 0.55 | 0.42 | 0.47 | 0.53 |
| 1 | 104800294 | rs12086216 | 0.346 | 0.9 | 0.337 | 0.581 | -0.005 | 0.009 | 0.009 | 0 | 0.05 | 0.12 | 0.1 | 0.13 | 0.06 | 0.06 |
| 1 | 104801950 | rs11185412 | 0.879 | 0.852 | 1.385 | 0.558 | 0.021 | 0.071 | 0.071 | 0.716 | 0.92 | 0.6 | 0.88 | 0.88 | 0.94 | 0.94 |
| 1 | 104807341 | rs1497659 | -0.478 | 0.93 | 1.552 | 0.575 | 0.001 | 0 | 0 | 0 | 0.34 | 0.34 | 0.32 | 0.47 | 0.49 | 0.41 |
| 1 | 104818575 | rs7526560 | 0.365 | 0.964 | 1.567 | 0.619 | -0.001 | -0.001 | -0.001 | 0 | 0.63 | 0.39 | 0.66 | 0.54 | 0.51 | 0.6 |
| 1 | 104832285 | rs6667386 | 0.559 | 0.962 | 1.503 | 0.618 | -0.024 | -0.017 | -0.017 | 0 | 0.62 | 0.29 | 0.65 | 0.53 | 0.51 | 0.59 |
| 1 | 104837082 | rs7521048 | 0.381 | 0.991 | 1.532 | 0.623 | 0.008 | 0.005 | 0.005 | 0 | 0.63 | 0.41 | 0.66 | 0.54 | 0.51 | 0.6 |
| 1 | 104851904 | rs12034300 | -0.162 | 0.89 | 1.462 | 0.318 | 0.003 | 0.005 | 0.005 | 0 | 0.27 | 0.29 | 0.49 | 0.51 | 0.38 | 0.28 |
| 1 | 104851938 | rs12035919 | -0.854 | 0.991 | 1.365 | 0.27 | 0.034 | 0.004 | 0.004 | 0 | 0.35 | 0.59 | 0.53 | 0.64 | 0.48 | 0.47 |
| 1 | 104853075 | rs17479455 | 0.215 | 0.305 | 0.537 | -0.105 | 0.025 | 0.075 | 0.075 | 0 | 0.14 | 0 | 0.03 | 0.02 | 0.1 | 0.17 |
| 1 | 104853997 | rs2055235 | 1.07 | 0.476 | 0.754 | -0.124 | 0.028 | 0.033 | 0.033 | 0.358 | 0.45 | 0.28 | 0.39 | 0.34 | 0.38 | 0.32 |
| 1 | 104870982 | rs11809638 | -0.641 | 0.559 | 0.739 | -0.375 | 0.013 | 0.009 | 0.009 | 0 | 0.38 | 0.48 | 0.42 | 0.46 | 0.44 | 0.3 |
| 1 | 104872140 | rs6700638 | -0.701 | 0.55 | 0.829 | -0.38 | 0.011 | 0.008 | 0.008 | 0 | 0.39 | 0.52 | 0.42 | 0.46 | 0.44 | 0.3 |
| 1 | 104886613 | rs6664343 | -0.099 | 0.547 | 0.942 | -0.384 | 0 | -0.015 | -0.015 | 0.311 | 0.32 | 0.3 | 0.25 | 0.15 | 0.4 | 0.27 |
| 1 | 104886713 | rs11811323 | 0.187 | 0.548 | 0.903 | -0.369 | 0.005 | 0.158 | 0.158 | 0 | 0.06 | 0.18 | 0.18 | 0.32 | 0.04 | 0.03 |
| 1 | 104887259 | rs10785866 | -2.046 | 0.484 | 0.752 | -0.605 | 0.015 | 0.144 | 0.144 | 0 | 0.65 | 0.87 | 0.81 | 1 | 0.7 | 0.76 |
| 1 | 104889158 | rs9325372 | 1.277 | 0.548 | 0.672 | -0.536 | -0.044 | 0.075 | 0.075 | 0.005 | 0.2 | 0.07 | 0.16 | 0 | 0.25 | 0.03 |
| 1 | 104890559 | rs17021432 | -0.455 | 0.527 | 0.736 | -0.383 | -0.001 | 0.049 | 0.049 | 0 | 0.26 | 0.37 | 0.38 | 0.54 | 0.27 | 0.46 |
| 1 | 104892674 | rs1497671 | -0.505 | -0.011 | 0.892 | 0.004 | 0.003 | 0.007 | 0.007 | 0 | 0.49 | 0.31 | 0.48 | 0.45 | 0.54 | 0.5 |
| 1 | 104892901 | rs17021437 | 0.315 | 0.034 | 0.613 | 0.11 | -0.001 | 0.056 | 0.056 | 0 | 0.06 | 0.04 | 0.25 | 0.31 | 0.04 | 0.04 |
| 1 | 104893789 | rs9726970 | 0.444 | 0.082 | 1.021 | 0.022 | 0.016 | 0.027 | 0.027 | 0 | 0.34 | 0.4 | 0.44 | 0.4 | 0.4 | 0.31 |
| 1 | 104893852 | rs17021444 | 0.28 | 0.265 | 0.932 | 0.134 | -0.001 | -0.001 | -0.001 | 0.043 | 0.1 | 0.13 | 0.08 | 0.21 | 0.15 | 0.22 |
| 1 | 104896161 | rs4409662 | -1.751 | 0.18 | 0.94 | -0.008 | 0.002 | 0.087 | 0.087 | 0 | 0.69 | 0.87 | 0.84 | 0.99 | 0.73 | 0.78 |
| 1 | 104898703 | rs11185445 | -1.399 | 0.168 | 1.184 | -0.015 | -0.012 | -0.042 | -0.042 | 0 | 0.3 | 0.57 | 0.32 | 0.23 | 0.28 | 0.38 |
| 1 | 104901525 | rs17021478 | 0.089 | 0.167 | 1.269 | 0.04 | -0.01 | -0.02 | -0.02 | 0 | 0.16 | 0.23 | 0.21 | 0.03 | 0.1 | 0.15 |
| 1 | 104902046 | rs12070214 | -0.69 | 0.18 | 1.204 | 0.044 | 0.036 | -0.004 | -0.004 | 0 | 0.14 | 0.34 | 0.12 | 0.2 | 0.18 | 0.24 |
| 1 | 104902248 | rs12064032 | -0.69 | 0.18 | 1.204 | 0.044 | 0.036 | -0.004 | -0.004 | 0 | 0.14 | 0.34 | 0.12 | 0.2 | 0.18 | 0.24 |
| 1 | 104907462 | rs12723426 | -1.786 | 0.183 | 0.859 | 0.059 | 0.003 | -0.011 | -0.011 | 0 | 0.34 | 0.66 | 0.47 | 0.58 | 0.32 | 0.46 |
| 1 | 104910603 | rs17021491 | -0.299 | 0.2 | 0.76 | 0.069 | -0.004 | 0.013 | 0.013 | 0 | 0.2 | 0.09 | 0.12 | 0.23 | 0.24 | 0.4 |

| | | | | | | | | | | | | | | | | |
|---|---|---|---|---|---|---|---|---|---|---|---|---|---|---|---|---|
| 1 | 104911326 | rs315154 | 0.687 | 0.275 | 0.747 | 0.184 | -0.033 | -0.052 | -0.052 | 3.057 | 0.36 | 0.21 | 0.37 | 0.41 | 0.41 | 0.32 |
| 1 | 104930486 | rs12057598 | -1.435 | 0.429 | 0.973 | 0.737 | -0.014 | 0.046 | 0.046 | 0 | 0.23 | 0.59 | 0.39 | 0.35 | 0.17 | 0.18 |
| 1 | 104945065 | rs10881154 | -1.085 | 0.781 | 0.988 | 1.075 | 0.099 | 0.428 | 0.428 | 0 | 0.07 | 0.33 | 0.19 | 0.33 | 0.07 | 0.03 |
| 1 | 104957563 | rs1448396 | 0.542 | 0.694 | -0.164 | 0.773 | 0.006 | -0.014 | -0.014 | 0 | 0.35 | 0.43 | 0.37 | 0.25 | 0.38 | 0.19 |
| 1 | 104959536 | rs2132960 | 0.429 | 0.691 | -0.171 | 0.895 | 0.005 | 0.011 | 0.011 | 0.142 | 0.39 | 0.49 | 0.43 | 0.45 | 0.41 | 0.3 |
| 1 | 104959857 | rs12757818 | -0.637 | 0.778 | -0.823 | 1.017 | -0.013 | 0.023 | 0.023 | 0 | 0.2 | 0.09 | 0.12 | 0.16 | 0.23 | 0.39 |
| 1 | 104975728 | rs2594664 | 0.619 | 0.503 | -0.773 | 0.635 | 0.003 | 0.002 | 0.002 | 0 | 0.43 | 0.5 | 0.47 | 0.45 | 0.46 | 0.35 |
| 1 | 104978179 | rs11184169 | 0.889 | 0.517 | -0.765 | 0.59 | -0.01 | 0.137 | 0.137 | 0.154 | 0.33 | 0.12 | 0.26 | 0.01 | 0.28 | 0.24 |
| 1 | 104978403 | rs12745577 | -0.639 | 0.644 | -0.966 | 0.726 | -0.016 | 0.013 | 0.013 | 0 | 0.2 | 0.08 | 0.12 | 0.16 | 0.23 | 0.38 |
| 1 | 104980964 | rs12036496 | -0.107 | 0.611 | -0.394 | 1.289 | -0.001 | 0 | 0 | 0.544 | 0.51 | 0.52 | 0.54 | 0.54 | 0.59 | 0.59 |
| 1 | 104986030 | rs10881159 | -1.179 | 0.602 | -0.935 | 1.406 | 0.003 | 0.001 | 0.001 | 0 | 0.25 | 0.17 | 0.17 | 0.25 | 0.21 | 0.25 |
| 1 | 104986052 | rs11184173 | -1.392 | 0.262 | -0.725 | 1.89 | -0.003 | 0.023 | 0.023 | 0 | 0.17 | 0.18 | 0.12 | 0.41 | 0.15 | 0.3 |
| 1 | 104987296 | rs12744616 | -0.782 | 0.489 | -0.771 | 1.284 | -0.012 | 0 | 0 | 4.092 | 0.09 | 0.04 | 0.12 | 0.12 | 0.13 | 0.17 |
| 1 | 104987416 | rs7552867 | -0.594 | -0.534 | -0.987 | 1.225 | 0.043 | -0.034 | -0.034 | 0 | 0.27 | 0.19 | 0.2 | 0.6 | 0.17 | 0.31 |
| 1 | 104987484 | rs10881160 | -0.214 | -0.525 | -0.952 | 1.384 | 0.02 | -0.023 | -0.023 | 0.133 | 0.1 | 0.06 | 0.04 | 0.54 | 0.08 | 0.16 |
| 1 | 104988625 | rs12038343 | 0.029 | -0.534 | -0.956 | 1.938 | 0.026 | -0.023 | -0.023 | 1.063 | 0.09 | 0.05 | 0.03 | 0.33 | 0.05 | 0.14 |
| 1 | 104989780 | rs12041707 | -0.314 | -0.567 | -0.954 | 1.922 | -0.02 | 0.054 | 0.054 | 0 | 0.1 | 0.15 | 0.03 | 0.33 | 0.05 | 0.15 |
| 1 | 104993773 | rs10785700 | 1.386 | -1.147 | -0.564 | 0.984 | 0.001 | 0.07 | 0.07 | 0.041 | 0.43 | 0.28 | 0.35 | 0.09 | 0.55 | 0.23 |
| 1 | 104997184 | rs1597178 | 0.688 | -1.222 | 0.087 | 1.066 | 0.015 | 0.029 | 0.029 | 0 | 0.56 | 0.65 | 0.42 | 0.63 | 0.63 | 0.39 |
| 1 | 104997471 | rs7411648 | 1.228 | -1.252 | 0.184 | 1.184 | 0.004 | 0.005 | 0.005 | 0 | 0.46 | 0.54 | 0.39 | 0.3 | 0.58 | 0.24 |
| 1 | 105006519 | rs1115851 | -1.005 | -1.244 | 0.22 | 1.163 | 0.015 | 0.025 | 0.025 | 0 | 0.45 | 0.39 | 0.59 | 0.37 | 0.37 | 0.6 |
| 1 | 105011378 | rs9724701 | -1.607 | -1.19 | 0.494 | 1.174 | 0.004 | 0.007 | 0.007 | 0 | 0.55 | 0.56 | 0.62 | 0.7 | 0.42 | 0.76 |
| 1 | 105013220 | rs12023545 | 0.511 | -1.184 | 0.414 | 1.224 | 0.018 | -0.01 | -0.01 | 0 | 0.09 | 0.05 | 0.02 | 0.33 | 0.05 | 0.15 |
| 1 | 105026270 | rs10458520 | -1.179 | -1.274 | 0.594 | 1.134 | 0.022 | 0.023 | 0.023 | 0 | 0.47 | 0.62 | 0.53 | 0.8 | 0.32 | 0.6 |
| 1 | 105032798 | rs12030464 | 0.612 | -1.262 | 0.336 | 1.167 | 0.025 | -0.026 | -0.026 | 0 | 0.09 | 0.03 | 0.02 | 0.33 | 0.04 | 0.14 |
| 1 | 105040651 | rs316977 | -1.084 | -1.258 | 0.451 | 1.16 | 0.047 | 0.003 | 0.003 | 0 | 0.44 | 0.37 | 0.5 | 0.59 | 0.29 | 0.57 |
| 1 | 105041495 | rs11184189 | 0.659 | -1.126 | -0.006 | 0.289 | -0.009 | 0.034 | 0.034 | 0 | 0.31 | 0.19 | 0.28 | 0.12 | 0.3 | 0.27 |
| 1 | 105057389 | rs12079720 | 1.959 | -1.302 | -0.13 | 0.167 | -0.001 | 0.026 | 0.026 | 0.406 | 0.18 | 0.07 | 0.15 | 0 | 0.15 | 0.08 |
| 1 | 105057555 | rs373489 | -1.615 | -1.32 | -0.033 | 0.291 | 0.08 | 0.073 | 0.073 | 0 | 0.75 | 0.58 | 0.77 | 0.69 | 0.59 | 0.85 |
| 1 | 105057760 | rs316973 | -0.404 | -1.336 | -0.065 | 0.279 | 0.034 | -0.003 | -0.003 | 0.274 | 0.45 | 0.41 | 0.49 | 0.57 | 0.29 | 0.58 |
| 1 | 105060691 | rs2218672 | 1.127 | -1.625 | 0.096 | 0.14 | 0.013 | 0.022 | 0.022 | 0 | 0.17 | 0.12 | 0.2 | 0.04 | 0.12 | 0.17 |
| 1 | 105061457 | rs316970 | -2.017 | -1.502 | 0.253 | 0.3 | 0.052 | -0.037 | -0.037 | 0.233 | 0.78 | 0.71 | 0.8 | 0.91 | 0.62 | 0.87 |
| 1 | 105063644 | rs12042136 | 1.267 | -1.578 | -0.069 | 0.395 | 0.024 | -0.027 | -0.027 | 0 | 0.08 | 0.02 | 0.02 | 0.33 | 0.04 | 0.14 |
| 1 | 105067105 | rs316960 | -0.635 | -1.611 | -0.088 | 0.288 | 0.033 | -0.011 | -0.011 | 0.621 | 0.4 | 0.35 | 0.42 | 0.46 | 0.23 | 0.44 |
| 1 | 105076810 | rs316951 | -0.092 | -1.594 | -0.122 | 0.152 | 0 | 0.002 | 0.002 | 0 | 0.33 | 0.35 | 0.44 | 0.25 | 0.2 | 0.3 |
| 1 | 105076879 | rs316950 | 1.32 | -1.616 | -0.147 | 0.101 | -0.005 | -0.009 | -0.009 | 0 | 0.17 | 0.19 | 0.22 | 0.08 | 0.11 | 0.15 |
| 1 | 105077882 | rs2034388 | 0.914 | -1.527 | -0.325 | 0.148 | 0.012 | 0.012 | 0.012 | 0 | 0.16 | 0.13 | 0.17 | 0.05 | 0.08 | 0.15 |
| 1 | 105086051 | rs6695172 | -0.789 | -1.527 | -0.283 | 0.105 | -0.011 | 0.013 | 0.013 | 0 | 0.61 | 0.32 | 0.52 | 0.71 | 0.51 | 0.71 |
| 1 | 105088785 | rs12026360 | 0.171 | -1.511 | -0.358 | 0.181 | -0.01 | -0.001 | -0.001 | 0 | 0.42 | 0.17 | 0.32 | 0.45 | 0.38 | 0.44 |
| 1 | 105099912 | rs11184204 | 1.348 | -1.498 | -0.36 | 0.161 | -0.005 | -0.005 | -0.005 | 1.613 | 0.25 | 0.12 | 0.21 | 0.22 | 0.24 | 0.23 |
| 1 | 105111413 | rs1597181 | -0.307 | -1.602 | -0.313 | -0.066 | 0.05 | 0.056 | 0.056 | 0 | 0.49 | 0.28 | 0.41 | 0.6 | 0.37 | 0.49 |
| 1 | 105111535 | rs12172730 | 0.268 | -1.389 | -0.214 | 0.135 | 0.005 | 0.012 | 0.012 | 0 | 0.12 | 0.09 | 0.05 | 0.55 | 0.08 | 0.26 |
| 1 | 105128348 | rs1118861 | -0.581 | -1.412 | 0.04 | -0.098 | -0.001 | -0.005 | -0.005 | 0.889 | 0.62 | 0.41 | 0.56 | 0.84 | 0.51 | 0.73 |
| 1 | 105138007 | rs869583 | 1.534 | -1.137 | 0.027 | -0.057 | 0.078 | -0.032 | -0.032 | 0 | 0.2 | 0.3 | 0.2 | 0.08 | 0.38 | 0.13 |
| 1 | 105138454 | rs12060689 | 1.277 | -1.326 | -0.552 | -0.077 | -0.019 | 0.077 | 0.077 | 0.116 | 0.22 | 0.07 | 0.19 | 0.01 | 0.21 | 0.11 |
| 1 | 105140299 | rs12568560 | 1.094 | -1.319 | -0.539 | -0.043 | -0.008 | 0.039 | 0.039 | 0.397 | 0.23 | 0.13 | 0.23 | 0.14 | 0.21 | 0.12 |
| 1 | 105144179 | rs4847178 | -0.097 | -1.377 | -0.647 | 0.065 | 0.049 | -0.015 | -0.015 | 0.004 | 0.53 | 0.68 | 0.53 | 0.46 | 0.64 | 0.62 |
| 1 | 105174866 | rs6703314 | 0.219 | -1.352 | -0.512 | -0.619 | -0.013 | -0.044 | -0.044 | 0 | 0.68 | 0.63 | 0.67 | 0.9 | 0.73 | 0.62 |
| 1 | 105175991 | rs12404332 | -0.005 | -0.972 | -0.219 | -0.341 | -0.003 | -0.011 | -0.011 | 0 | 0.14 | 0.12 | 0.12 | 0.06 | 0.14 | 0.24 |
| 1 | 105180089 | rs12044370 | 0.754 | -0.184 | -0.262 | -0.219 | -0.001 | 0 | 0 | 0 | 0.15 | 0.15 | 0.16 | 0.25 | 0.19 | 0.23 |
| 1 | 105181524 | rs11184238 | 1.661 | -0.125 | -0.118 | 0.015 | 0.016 | 0.037 | 0.037 | 0 | 0.2 | 0.23 | 0.18 | 0.31 | 0.29 | 0.15 |
| 1 | 105181816 | rs7537242 | 0.243 | -0.345 | -0.034 | -0.014 | 0.014 | 0.008 | 0.008 | 0 | 0.36 | 0.4 | 0.34 | 0.56 | 0.48 | 0.38 |
| 1 | 105184308 | rs12036108 | 0.961 | -0.43 | 0.018 | 0.059 | 0.021 | -0.023 | -0.023 | 0 | 0.09 | 0.03 | 0.03 | 0.33 | 0.05 | 0.14 |
| 1 | 105184759 | rs1028120 | 0.259 | 0.002 | 0.337 | 0.295 | 0.029 | -0.011 | -0.011 | 0.173 | 0.07 | 0.21 | 0.05 | 0.02 | 0.09 | 0.12 |
| 1 | 105189711 | rs10494020 | 0.583 | 0.254 | -0.078 | 0.529 | -0.01 | -0.017 | -0.017 | 0.512 | 0.29 | 0.23 | 0.28 | 0.35 | 0.37 | 0.27 |
| 1 | 105190881 | rs958515 | 0.104 | -0.365 | -0.099 | 0.423 | 0 | 0.002 | 0.002 | 0 | 0.44 | 0.4 | 0.43 | 0.59 | 0.54 | 0.49 |
| 1 | 105191391 | rs872987 | -0.119 | -0.245 | -0.129 | 0.501 | -0.006 | -0.02 | -0.02 | 0 | 0.91 | 0.85 | 0.83 | 0.96 | 0.92 | 0.93 |
| 1 | 105199870 | rs11184244 | -0.926 | -0.848 | -0.167 | 1.188 | -0.039 | 0.08 | 0.08 | 0 | 0.2 | 0.32 | 0.18 | 0.51 | 0.08 | 0.28 |
| 1 | 105219763 | rs7532749 | 1.077 | -1.883 | -0.282 | 0.887 | -0.023 | -0.006 | -0.006 | 2.325 | 0.63 | 0.58 | 0.64 | 0.23 | 0.82 | 0.46 |
| 1 | 105227914 | rs7543757 | 1.201 | -1.97 | -0.094 | -0.294 | -0.012 | 0.073 | 0.073 | 4.421 | 0.37 | 0.13 | 0.34 | 0.19 | 0.58 | 0.36 |
| 1 | 105230023 | rs12076721 | 0.744 | -1.876 | -0.015 | -0.273 | -0.01 | -0.118 | -0.118 | 0 | 0.26 | 0.45 | 0.3 | 0.03 | 0.23 | 0.1 |
| 1 | 105231358 | rs7546370 | 1.061 | -1.922 | 0.158 | -0.293 | 0.008 | 0.031 | 0.031 | 1.771 | 0.21 | 0.13 | 0.14 | 0.03 | 0.19 | 0.07 |
| 1 | 105234637 | rs10881189 | -0.089 | -2.159 | 0.289 | 0.019 | -0.055 | 0.185 | 0.185 | 0 | 0.22 | 0.53 | 0.32 | 0.5 | 0.13 | 0.24 |
| 1 | 105246855 | rs11184268 | -0.166 | -1.298 | 0.224 | -0.155 | 0 | 0.011 | 0.011 | 0 | 0.77 | 0.62 | 0.64 | 0.71 | 0.78 | 0.64 |
| 1 | 105254006 | rs6676838 | 1.275 | -1.473 | 0.038 | -0.277 | -0.039 | 0.066 | 0.066 | 2.545 | 0.19 | 0.07 | 0.15 | 0.03 | 0.24 | 0.05 |
| 1 | 105254921 | rs4272631 | -0.991 | -1.508 | 0.085 | -0.237 | 0.001 | -0.003 | -0.003 | 0.092 | 0.81 | 0.86 | 0.81 | 0.71 | 0.9 | 0.67 |
| 1 | 105256299 | rs12037801 | 0.268 | -1.591 | 0.217 | -0.158 | -0.06 | 0.12 | 0.12 | 0 | 0.2 | 0.41 | 0.18 | 0.52 | 0.08 | 0.22 |
| 1 | 105259884 | rs1904982 | 1.295 | -1.513 | 0.166 | -0.213 | -0.041 | 0.097 | 0.097 | 3.469 | 0.39 | 0.14 | 0.32 | 0.17 | 0.53 | 0.37 |
| 1 | 105270235 | rs10881193 | -0.412 | -1.492 | 0.328 | -0.099 | -0.035 | 0.018 | 0.018 | 0 | 0.78 | 0.63 | 0.65 | 0.71 | 0.85 | 0.64 |

| | | | | | | | | | | | | | | | | |
|---|---|---|---|---|---|---|---|---|---|---|---|---|---|---|---|---|
| 1 | 105277365 | rs7525816 | 0.429 | -1.623 | 0.323 | -0.069 | -0.029 | 0.063 | 0.063 | 0 | 0.19 | 0.31 | 0.17 | 0.52 | 0.08 | 0.22 |
| 1 | 105286219 | rs5000688 | 0.578 | -1.625 | 0.332 | -0.069 | -0.024 | 0.052 | 0.052 | 0 | 0.18 | 0.27 | 0.17 | 0.52 | 0.08 | 0.19 |
| 1 | 105286754 | rs2317613 | -0.723 | -1.547 | 0.36 | -0.122 | -0.001 | -0.003 | -0.003 | 0 | 0.82 | 0.84 | 0.87 | 0.87 | 0.83 | 0.73 |
| 1 | 105292212 | rs11579151 | -0.073 | -1.683 | 0.271 | -0.162 | 0.01 | -0.007 | -0.007 | 1.098 | 0.36 | 0.39 | 0.3 | 0.54 | 0.25 | 0.25 |
| 1 | 105293381 | rs7535918 | -0.159 | -1.071 | 0.401 | -0.107 | -0.009 | 0.021 | 0.021 | 0 | 0.19 | 0.28 | 0.3 | 0.28 | 0.14 | 0.3 |
| 1 | 105294293 | rs10399652 | -1.294 | -1.056 | 0.277 | -0.256 | 0.001 | 0 | 0 | 0 | 0.81 | 0.82 | 0.87 | 0.97 | 0.83 | 0.95 |
| 1 | 105295245 | rs11577870 | 1.155 | -1.041 | 0.258 | -0.247 | 0.002 | 0.027 | 0.027 | 2.775 | 0.18 | 0.12 | 0.13 | 0.03 | 0.18 | 0.05 |
| 1 | 105300481 | rs4539145 | -0.374 | -0.858 | 0.964 | 0.083 | 0.004 | 0.017 | 0.017 | 0 | 0.14 | 0.18 | 0.22 | 0.19 | 0.14 | 0.21 |
| 1 | 105306516 | rs4620554 | -1.152 | -0.988 | 1.484 | 0.02 | -0.005 | 0.029 | 0.029 | 1.64 | 0.18 | 0.52 | 0.25 | 0.19 | 0.14 | 0.22 |
| 1 | 105312623 | rs11184300 | 1.066 | -0.713 | 1.006 | -0.179 | 0.015 | 0.102 | 0.102 | 0.28 | 0.5 | 0.16 | 0.39 | 0.31 | 0.57 | 0.53 |
| 1 | 105341180 | rs4483421 | 0.735 | -0.873 | 1.073 | -0.242 | -0.018 | 0.001 | 0.001 | 2.352 | 0.8 | 0.35 | 0.74 | 0.81 | 0.86 | 0.83 |
| 1 | 105365300 | rs6583061 | 0.237 | -0.755 | 0.925 | -0.134 | -0.009 | -0.061 | -0.061 | 0 | 0.3 | 0.2 | 0.35 | 0.49 | 0.3 | 0.31 |
| 1 | 105370881 | rs11184316 | -1.494 | -0.788 | 1.548 | 0.052 | 0 | 0.019 | 0.019 | 0.474 | 0.2 | 0.52 | 0.26 | 0.19 | 0.16 | 0.18 |
| 1 | 105374759 | rs11802067 | -1.518 | -0.788 | 1.492 | 0.052 | -0.008 | 0.007 | 0.007 | 0.685 | 0.2 | 0.57 | 0.26 | 0.19 | 0.16 | 0.18 |
| 1 | 105380072 | rs12036702 | -0.946 | -0.783 | 1.297 | 0.061 | -0.004 | 0.012 | 0.012 | 0 | 0.16 | 0.25 | 0.23 | 0.19 | 0.14 | 0.18 |
| 1 | 105395113 | rs9803714 | -0.569 | -0.579 | 0.774 | 0.152 | -0.004 | 0.01 | 0.01 | 0 | 0.15 | 0.23 | 0.26 | 0.16 | 0.16 | 0.16 |
| 1 | 105401997 | rs7522559 | -0.537 | -0.596 | 0.818 | 0.165 | -0.002 | 0.006 | 0.006 | 0 | 0.15 | 0.21 | 0.25 | 0.16 | 0.16 | 0.15 |
| 1 | 105403089 | rs12092586 | -1.092 | -0.651 | 0.747 | 0.146 | -0.014 | 0.005 | 0.005 | 0 | 0.2 | 0.45 | 0.28 | 0.17 | 0.18 | 0.16 |
| 1 | 105424155 | rs7411085 | -0.18 | -0.801 | -0.667 | 0.134 | -0.002 | -0.016 | -0.016 | 0.009 | 0.13 | 0.01 | 0.03 | 0.1 | 0.1 | 0.22 |
| 1 | 105424848 | rs12741933 | 0.211 | -0.695 | -0.525 | 0.727 | 0.009 | -0.031 | -0.031 | 0.23 | 0.64 | 0.27 | 0.64 | 0.51 | 0.72 | 0.77 |
| 1 | 105429577 | rs7542735 | 0.236 | -0.704 | -0.545 | 0.723 | 0.011 | -0.042 | -0.042 | 0.182 | 0.64 | 0.22 | 0.64 | 0.51 | 0.72 | 0.77 |
| 1 | 105447842 | rs2122688 | 0.426 | -0.962 | -0.48 | 0.794 | -0.033 | -0.122 | -0.122 | 1.132 | 0.22 | 0.02 | 0.25 | 0.44 | 0.21 | 0.19 |
| 1 | 105448030 | rs12126615 | 1.291 | -1.322 | -0.348 | 0.555 | 0.045 | 0.004 | 0.004 | 0 | 0.52 | 0.28 | 0.42 | 0.39 | 0.64 | 0.56 |
| 1 | 105449139 | rs1376368 | 1.574 | -1.542 | -0.433 | 0.46 | 0.026 | 0.023 | 0.023 | 0 | 0.47 | 0.3 | 0.38 | 0.31 | 0.61 | 0.37 |
| 1 | 105451694 | rs1157148 | 0.168 | -1.501 | 0.03 | 0.443 | -0.029 | -0.021 | -0.021 | 0 | 0.86 | 0.6 | 0.93 | 0.91 | 0.93 | 0.81 |
| 1 | 105455168 | rs17015247 | 0.645 | -1.416 | -0.197 | 0.5 | 0.006 | -0.001 | -0.001 | 0.054 | 0.11 | 0.02 | 0.27 | 0.12 | 0.1 | 0.16 |
| 1 | 105455404 | rs12405334 | 0.645 | -1.416 | -0.197 | 0.5 | 0.006 | -0.001 | -0.001 | 1.073 | 0.11 | 0.02 | 0.27 | 0.12 | 0.1 | 0.16 |
| 1 | 105465012 | rs11184355 | 0.782 | -1.41 | -0.088 | 0.621 | -0.007 | -0.076 | -0.076 | 0 | 0.23 | 0.07 | 0.25 | 0.46 | 0.2 | 0.24 |
| 1 | 105469987 | rs2084670 | 0.48 | -1.421 | 0.283 | 0.47 | -0.034 | -0.022 | -0.022 | 0 | 0.89 | 0.71 | 0.95 | 0.94 | 0.95 | 0.83 |
| 1 | 105477683 | rs9787106 | 0.15 | -1.222 | 0.104 | 0.474 | 0.016 | 0.012 | 0.012 | 0 | 0.13 | 0.07 | 0.28 | 0.13 | 0.11 | 0.17 |
| 1 | 105484805 | rs6657845 | 0.346 | -1.323 | 0.316 | 0.552 | -0.001 | -0.006 | -0.006 | 0 | 0.27 | 0.2 | 0.28 | 0.49 | 0.24 | 0.28 |
| 1 | 105486240 | rs17015278 | -0.651 | -1.27 | 0.308 | 0.552 | 0.008 | -0.008 | -0.008 | 0 | 0.41 | 0.28 | 0.56 | 0.61 | 0.35 | 0.45 |
| 1 | 105504651 | rs2167300 | 1.336 | -1.319 | 0.386 | 0.517 | 0.038 | 0.038 | 0.038 | 0 | 0.49 | 0.3 | 0.38 | 0.33 | 0.61 | 0.4 |
| 1 | 105507896 | rs6695731 | 1.345 | -1.341 | 0.384 | 0.518 | 0.041 | 0.039 | 0.039 | 0 | 0.49 | 0.29 | 0.38 | 0.33 | 0.61 | 0.4 |
| 1 | 105516008 | rs1451055 | 0.386 | -1.394 | 0.429 | 0.388 | -0.014 | -0.003 | -0.003 | 0.008 | 0.9 | 0.57 | 0.95 | 0.94 | 0.96 | 0.85 |
| 1 | 105518136 | rs896199 | 0.82 | -1.392 | 0.386 | 0.404 | -0.042 | -0.034 | -0.034 | 0.112 | 0.77 | 0.59 | 0.67 | 0.82 | 0.84 | 0.69 |
| 1 | 105535421 | rs12718026 | 1.351 | -1.486 | 0.312 | 0.559 | 0.028 | 0.017 | 0.017 | 0 | 0.48 | 0.36 | 0.41 | 0.36 | 0.61 | 0.38 |
| 1 | 105538026 | rs2878086 | 0.728 | -1.593 | 0.379 | 0.552 | -0.032 | -0.028 | -0.028 | 0 | 0.77 | 0.65 | 0.67 | 0.83 | 0.84 | 0.67 |
| 1 | 105547584 | rs6699420 | 0.588 | -1.287 | 0.369 | 0.681 | 0 | -0.004 | -0.004 | 0 | 0.23 | 0.16 | 0.23 | 0.44 | 0.2 | 0.25 |
| 1 | 105554364 | rs12145330 | 0.265 | -1.409 | 0.388 | 0.621 | -0.008 | 0.028 | 0.028 | 0 | 0.66 | 0.54 | 0.71 | 0.5 | 0.75 | 0.71 |
| 1 | 105561345 | rs2923300 | 0.807 | -1.391 | 0.257 | 0.605 | -0.004 | -0.018 | -0.018 | 0 | 0.09 | 0.16 | 0.05 | 0.02 | 0.05 | 0.04 |
| 1 | 105569132 | rs3007632 | -0.569 | -1.438 | 0.28 | 0.73 | 0.006 | -0.017 | -0.017 | 0 | 0.33 | 0.29 | 0.23 | 0.47 | 0.25 | 0.3 |
| 1 | 105583017 | rs9659111 | -0.795 | -1.058 | 0.608 | 0.343 | -0.002 | -0.011 | -0.011 | 0 | 0.91 | 0.87 | 0.95 | 0.98 | 0.95 | 0.96 |
| 1 | 105588516 | rs10881240 | 1.661 | -1.091 | 0.595 | 0.3 | -0.025 | -0.023 | -0.023 | 0 | 0.76 | 0.65 | 0.71 | 0.86 | 0.84 | 0.81 |
| 1 | 105601544 | rs12567808 | -0.374 | -1.216 | 0.185 | 0.202 | 0.018 | -0.008 | -0.008 | 0.144 | 0.15 | 0.03 | 0.23 | 0.12 | 0.12 | 0.16 |
| 1 | 105607364 | rs2034524 | 1.637 | -1.053 | 0.498 | 0.295 | -0.037 | -0.035 | -0.035 | 0 | 0.76 | 0.62 | 0.71 | 0.86 | 0.84 | 0.81 |
| 1 | 105610279 | rs10881242 | 0.535 | -0.822 | 0.191 | 0.48 | -0.001 | 0 | 0 | 0 | 0.85 | 0.85 | 0.75 | 0.88 | 0.87 | 0.86 |
| 1 | 105629942 | rs7414261 | -0.324 | -0.864 | 0.207 | 0.634 | 0 | -0.003 | -0.003 | 0 | 0.76 | 0.73 | 0.78 | 0.79 | 0.77 | 0.75 |
| 1 | 105643103 | rs12030832 | 0.734 | 0.365 | 0.173 | 1.117 | -0.007 | -0.007 | -0.007 | 0 | 0.58 | 0.36 | 0.55 | 0.52 | 0.53 | 0.55 |
| 1 | 105663975 | rs10785729 | -0.66 | -0.239 | 0.489 | 0.539 | -0.019 | -0.067 | -0.067 | 0 | 0.61 | 0.54 | 0.68 | 0.85 | 0.71 | 0.76 |
| 1 | 105666096 | rs11184443 | -0.856 | -0.262 | 0.721 | 0.508 | 0.009 | 0.015 | 0.015 | 1.563 | 0.63 | 0.69 | 0.68 | 0.85 | 0.71 | 0.79 |
| 1 | 105666613 | rs10881260 | -0.027 | -0.313 | 0.279 | 0.827 | -0.003 | 0.003 | 0.003 | 0.094 | 0.43 | 0.38 | 0.58 | 0.37 | 0.56 | 0.55 |
| 1 | 105670343 | rs7554044 | -0.952 | -0.042 | 0.479 | 0.756 | 0.014 | 0.085 | 0.085 | 5.249 | 0.56 | 0.71 | 0.66 | 0.86 | 0.66 | 0.74 |
| 1 | 105673424 | rs2820355 | -1.012 | -0.064 | 0.644 | 0.824 | 0.002 | -0.006 | -0.006 | 0.345 | 0.58 | 0.6 | 0.66 | 0.87 | 0.67 | 0.78 |
| 1 | 105674169 | rs4847097 | 0.048 | -0.156 | 0.591 | 1.087 | -0.089 | -0.016 | -0.016 | 0 | 0.39 | 0.13 | 0.54 | 0.36 | 0.53 | 0.52 |
| 1 | 105674536 | rs6687976 | 0.099 | -0.14 | 0.903 | 1.104 | 0.003 | 0.069 | 0.069 | 0 | 0.12 | 0.27 | 0.08 | 0.16 | 0.11 | 0.12 |
| 1 | 105677615 | rs4545390 | -1.124 | -0.134 | 0.825 | 1.002 | 0 | -0.017 | -0.017 | 0.441 | 0.58 | 0.6 | 0.66 | 0.87 | 0.67 | 0.77 |
| 1 | 105680379 | rs10881264 | -0.372 | -0.109 | 0.553 | 1.086 | -0.007 | 0.044 | 0.044 | 5.305 | 0.46 | 0.32 | 0.6 | 0.72 | 0.56 | 0.65 |
| 1 | 105681907 | rs4421638 | -0.752 | -0.093 | 0.592 | 1.632 | -0.015 | 0.092 | 0.092 | 0 | 0.06 | 0.14 | 0.09 | 0.23 | 0.02 | 0.18 |
| 1 | 105682011 | rs4240949 | -0.892 | -0.155 | 0.646 | 1.462 | 0.032 | 0.112 | 0.112 | 6.8 | 0.55 | 0.34 | 0.67 | 0.87 | 0.69 | 0.79 |
| 1 | 105688084 | rs12758649 | -0.33 | -0.178 | 0.593 | 1.622 | 0.016 | 0.025 | 0.025 | 7.274 | 0.47 | 0.19 | 0.57 | 0.33 | 0.64 | 0.53 |
| 1 | 105698163 | rs5027215 | -1.077 | -0.134 | 0.752 | 1.237 | 0.029 | 0.101 | 0.101 | 5.349 | 0.58 | 0.77 | 0.68 | 0.88 | 0.68 | 0.78 |
| 1 | 105700935 | rs4325188 | -0.579 | -0.081 | 1.037 | 1.311 | -0.002 | -0.002 | -0.002 | 0 | 0.52 | 0.51 | 0.65 | 0.57 | 0.66 | 0.68 |
| 1 | 105703377 | rs10785733 | -1.073 | -0.056 | 0.571 | 1.148 | 0.015 | 0.11 | 0.11 | 5.153 | 0.58 | 0.8 | 0.67 | 0.88 | 0.66 | 0.77 |
| 1 | 105704285 | rs4403670 | -1.258 | -0.506 | 0.661 | 1.17 | 0.028 | 0.097 | 0.097 | 3.167 | 0.6 | 0.76 | 0.72 | 0.85 | 0.68 | 0.8 |
| 1 | 105707220 | rs10399884 | 0.159 | -0.278 | 0.735 | 1.224 | 0.001 | 0.007 | 0.007 | 0 | 0.1 | 0.15 | 0.06 | 0.13 | 0.08 | 0.15 |
| 1 | 105717631 | rs4847034 | -1.106 | -0.412 | 0.705 | 1.303 | 0.017 | -0.008 | -0.008 | 0 | 0.57 | 0.66 | 0.57 | 0.56 | 0.67 | 0.64 |
| 1 | 105720186 | rs6676097 | 0.058 | -0.564 | 0.826 | 1.088 | -0.006 | 0.009 | 0.009 | 1.668 | 0.4 | 0.38 | 0.47 | 0.19 | 0.54 | 0.39 |
| 1 | 105725458 | rs6667576 | -0.934 | -0.612 | 0.709 | 1.136 | -0.004 | 0.039 | 0.039 | 0 | 0.06 | 0.07 | 0.03 | 0.26 | 0.05 | 0.1 |

| | | | | | | | | | | | | | | | | |
|---|---|---|---|---|---|---|---|---|---|---|---|---|---|---|---|---|
| 1 | 105727098 | rs7528406 | -1.444 | -0.735 | 0.712 | 0.981 | 0.014 | 0.065 | 0.065 | 4.916 | 0.59 | 0.7 | 0.71 | 0.84 | 0.65 | 0.76 |
| 1 | 105728264 | rs11184462 | -1.088 | -0.757 | 0.857 | 1.053 | 0.002 | 0.003 | 0.003 | 4.132 | 0.54 | 0.52 | 0.68 | 0.58 | 0.64 | 0.67 |
| 1 | 105739503 | rs4384283 | -0.061 | -0.812 | 0.856 | 1.218 | -0.007 | 0.087 | 0.087 | 0 | 0.11 | 0.14 | 0.19 | 0.4 | 0.07 | 0.25 |
| 1 | 105741329 | rs11184474 | 0.038 | -0.786 | 0.855 | 1.023 | 0.001 | -0.01 | -0.01 | 0 | 0.38 | 0.36 | 0.47 | 0.17 | 0.53 | 0.36 |
| 1 | 105741765 | rs6583129 | -1.308 | -0.825 | 0.917 | 1.028 | -0.032 | 0.045 | 0.045 | 0 | 0.21 | 0.35 | 0.24 | 0.67 | 0.12 | 0.38 |
| 1 | 105743667 | rs4240951 | -1.588 | -0.793 | 0.797 | 0.974 | 0.013 | 0.071 | 0.071 | 4.619 | 0.59 | 0.71 | 0.71 | 0.84 | 0.65 | 0.74 |
| 1 | 105748581 | rs6700732 | -0.597 | -0.656 | 0.785 | 1.057 | 0 | -0.002 | -0.002 | 0 | 0.45 | 0.56 | 0.5 | 0.44 | 0.54 | 0.46 |
| 1 | 105756050 | rs2842237 | -0.525 | -0.672 | 0.797 | 1.036 | 0.001 | -0.005 | -0.005 | 0 | 0.44 | 0.53 | 0.5 | 0.43 | 0.54 | 0.46 |
| 1 | 105782314 | rs12565217 | -1.202 | -0.622 | 0.748 | 1.106 | -0.009 | 0.116 | 0.116 | 1.617 | 0.05 | 0.23 | 0.03 | 0.22 | 0.01 | 0.1 |
| 1 | 105785646 | rs12759058 | 1.555 | -0.87 | 0.639 | 0.976 | -0.014 | 0.114 | 0.114 | 2.964 | 0.37 | 0.13 | 0.26 | 0.17 | 0.34 | 0.25 |
| 1 | 105799204 | rs2809325 | -0.49 | -0.823 | 0.702 | 1.097 | -0.005 | 0.024 | 0.024 | 0 | 0.18 | 0.18 | 0.23 | 0.41 | 0.12 | 0.29 |
| 1 | 105802587 | rs2809312 | -0.721 | -0.827 | 0.708 | 1.095 | -0.009 | 0.03 | 0.03 | 0 | 0.2 | 0.23 | 0.23 | 0.41 | 0.12 | 0.29 |
| 1 | 105805227 | rs2842217 | -0.31 | -0.844 | 0.728 | 1.021 | -0.01 | 0.014 | 0.014 | 2.034 | 0.62 | 0.7 | 0.53 | 0.79 | 0.46 | 0.65 |
| 1 | 105812373 | rs7549655 | 1.103 | -1.203 | 0.992 | 0.974 | -0.013 | 0.052 | 0.052 | 2.451 | 0.43 | 0.37 | 0.46 | 0.15 | 0.46 | 0.29 |
| 1 | 105817094 | rs12096770 | 0.88 | -1.202 | 1.105 | 0.949 | -0.011 | 0.053 | 0.053 | 3.558 | 0.45 | 0.43 | 0.46 | 0.15 | 0.46 | 0.27 |
| 1 | 105821542 | rs17015613 | -0.685 | -1.383 | 1.317 | 1.151 | -0.042 | 0.045 | 0.045 | 0 | 0.18 | 0.26 | 0.22 | 0.66 | 0.09 | 0.33 |
| 1 | 105825561 | rs2653976 | -1.585 | -1.458 | 1.692 | 1.09 | -0.066 | 0.024 | 0.024 | 0 | 0.26 | 0.52 | 0.27 | 0.67 | 0.13 | 0.38 |
| 1 | 105830385 | rs12022556 | 0.214 | -1.357 | 1.349 | 1.23 | 0.005 | 0.02 | 0.02 | 0 | 0.13 | 0.11 | 0.2 | 0.45 | 0.08 | 0.22 |
| 1 | 105835847 | rs17418172 | 0.495 | -1.38 | 1.199 | 1.418 | 0.022 | -0.035 | -0.035 | 0 | 0.12 | 0.03 | 0.07 | 0.13 | 0.08 | 0.16 |
| 1 | 105836241 | rs6583142 | -0.371 | -1.249 | 1.344 | 1.205 | -0.009 | 0.021 | 0.021 | 1.718 | 0.67 | 0.75 | 0.71 | 0.81 | 0.59 | 0.65 |
| 1 | 105841681 | rs12082510 | 0.281 | -1.142 | 1.11 | 1.27 | -0.002 | -0.032 | -0.032 | 0 | 0.06 | 0.14 | 0.04 | 0 | 0.04 | 0.06 |
| 1 | 105845218 | rs2455650 | 0.302 | -1.147 | 1.054 | 1.267 | -0.001 | -0.021 | -0.021 | 0 | 0.06 | 0.12 | 0.03 | 0 | 0.04 | 0.05 |
| 1 | 105846616 | rs11184521 | -0.292 | -1.227 | 1.195 | 1.515 | -0.019 | 0.064 | 0.064 | 0 | 0.18 | 0.26 | 0.24 | 0.41 | 0.14 | 0.24 |
| 1 | 105856143 | rs2491642 | 0.153 | -1.214 | 1.259 | 1.362 | 0.001 | -0.005 | -0.005 | 0 | 0.59 | 0.47 | 0.7 | 0.6 | 0.57 | 0.5 |
| 1 | 105868193 | rs2491615 | 0.888 | -0.946 | 1.276 | 1.569 | -0.012 | -0.017 | -0.017 | 0.114 | 0.41 | 0.32 | 0.57 | 0.43 | 0.44 | 0.33 |
| 1 | 105909821 | rs12126523 | -0.611 | -0.988 | 1.531 | 1.572 | 0.003 | 0.033 | 0.033 | 3.601 | 0.52 | 0.28 | 0.39 | 0.36 | 0.56 | 0.55 |
| 1 | 105928580 | rs11800189 | 0.944 | -0.966 | 1.406 | 1.554 | -0.04 | 0.119 | 0.119 | 1.902 | 0.4 | 0.19 | 0.44 | 0.17 | 0.43 | 0.29 |
| 1 | 105941966 | rs2502916 | 0.808 | -0.954 | 1.759 | 1.699 | 0.002 | -0.037 | -0.037 | 0 | 0.42 | 0.46 | 0.45 | 0.17 | 0.43 | 0.29 |
| 1 | 105949895 | rs4847116 | -0.94 | -0.955 | 1.868 | 1.705 | 0.004 | 0.005 | 0.005 | 1.914 | 0.58 | 0.51 | 0.41 | 0.57 | 0.56 | 0.65 |
| 1 | 105952798 | rs6673429 | -0.96 | -0.958 | 1.897 | 1.704 | 0.002 | 0.002 | 0.002 | 2.447 | 0.58 | 0.53 | 0.42 | 0.57 | 0.56 | 0.65 |
| 1 | 105953996 | rs12026105 | -0.815 | -0.958 | 1.889 | 1.703 | -0.003 | -0.003 | -0.003 | 0 | 0.57 | 0.39 | 0.41 | 0.57 | 0.56 | 0.65 |
| 1 | 105954220 | rs4356105 | 0.911 | -0.959 | 1.811 | 1.61 | -0.022 | 0.084 | 0.084 | 1.864 | 0.4 | 0.23 | 0.44 | 0.17 | 0.43 | 0.29 |
| 1 | 105957249 | rs987243 | 0.93 | -0.994 | 1.782 | 1.596 | -0.006 | 0.018 | 0.018 | 2.581 | 0.41 | 0.37 | 0.45 | 0.17 | 0.43 | 0.29 |
| 1 | 105960628 | rs1913978 | -0.874 | -1 | 1.749 | 1.602 | 0.003 | -0.007 | -0.007 | 0 | 0.57 | 0.5 | 0.41 | 0.37 | 0.56 | 0.56 |
| 1 | 105974470 | rs6698864 | 0.996 | -0.966 | 1.261 | 1.256 | -0.023 | 0.081 | 0.081 | 2.432 | 0.4 | 0.25 | 0.45 | 0.17 | 0.43 | 0.29 |
| 1 | 105984113 | rs12749064 | 1.187 | -1.318 | 0.947 | 1.071 | -0.054 | 0.078 | 0.078 | 0 | 0.35 | 0.12 | 0.43 | 0.18 | 0.44 | 0.28 |
| 1 | 105993658 | rs6583163 | 1.11 | -1.765 | 1.004 | 1.049 | -0.046 | 0.111 | 0.111 | 0 | 0.37 | 0.12 | 0.48 | 0.17 | 0.54 | 0.3 |
| 1 | 105998384 | rs11184568 | 0.995 | -1.94 | 0.707 | 0.896 | -0.005 | 0.131 | 0.131 | 0 | 0.41 | 0.15 | 0.5 | 0.17 | 0.59 | 0.41 |
| 1 | 105999499 | rs1356211 | -0.185 | -2.213 | 0.723 | 0.856 | -0.023 | 0.129 | 0.129 | 0 | 0.77 | 0.55 | 0.71 | 0.55 | 0.82 | 0.81 |
| 1 | 105999870 | rs10881300 | -0.185 | -2.213 | 0.723 | 0.856 | -0.023 | 0.129 | 0.129 | 0 | 0.77 | 0.55 | 0.71 | 0.55 | 0.82 | 0.81 |
| 1 | 106005333 | rs12143797 | 1.007 | -2.221 | 0.775 | 0.733 | 0.015 | 0.081 | 0.081 | 0 | 0.44 | 0.31 | 0.52 | 0.17 | 0.59 | 0.41 |
| 1 | 106006480 | rs12023475 | -0.426 | -2.045 | 0.747 | 0.92 | 0.044 | -0.004 | -0.004 | 0 | 0.36 | 0.18 | 0.22 | 0.4 | 0.28 | 0.49 |
| 1 | 106007141 | rs2501603 | -0.675 | -1.946 | 0.624 | 1.018 | 0 | 0.099 | 0.099 | 0 | 0.82 | 0.72 | 0.74 | 0.57 | 0.83 | 0.8 |
| 1 | 106009370 | rs2502869 | 1.262 | -1.944 | 0.53 | 0.888 | -0.006 | 0.135 | 0.135 | 0.123 | 0.43 | 0.23 | 0.49 | 0.16 | 0.54 | 0.3 |
| 1 | 106010189 | rs10881302 | 0.763 | -2.134 | 0.726 | 0.849 | -0.027 | -0.006 | -0.006 | 0.035 | 0.62 | 0.51 | 0.75 | 0.6 | 0.7 | 0.5 |
| 1 | 106011788 | rs1516161 | -0.652 | -2.085 | 0.613 | 0.881 | 0.005 | 0.002 | 0.002 | 0 | 0.33 | 0.32 | 0.22 | 0.36 | 0.25 | 0.41 |
| 1 | 106025908 | rs11184578 | -0.141 | -2.124 | -0.115 | 1.016 | -0.018 | 0.107 | 0.107 | 0.007 | 0.17 | 0.27 | 0.19 | 0.47 | 0.13 | 0.39 |
| 1 | 106026464 | rs7415139 | -0.083 | -2.297 | -0.124 | 0.933 | -0.038 | 0.012 | 0.012 | 0 | 0.25 | 0.4 | 0.2 | 0.23 | 0.19 | 0.18 |
| 1 | 106027747 | rs1524108 | -1.838 | -2.421 | -0.683 | 0.704 | 0.007 | 0.221 | 0.221 | 0.058 | 0.58 | 0.87 | 0.58 | 0.98 | 0.44 | 0.69 |
| 1 | 106037417 | rs11184581 | -1.091 | -2.412 | -0.656 | 0.754 | 0.029 | 0.028 | 0.028 | 0 | 0.41 | 0.66 | 0.38 | 0.7 | 0.31 | 0.56 |
| 1 | 106037595 | rs1543251 | 0.492 | -2.445 | -0.668 | 0.734 | 0.007 | -0.012 | -0.012 | 0 | 0.15 | 0.1 | 0.19 | 0.28 | 0.13 | 0.14 |
| 1 | 106037628 | rs1710809 | -1.74 | -1.94 | 0.112 | 1.528 | 0.066 | 0.064 | 0.064 | 0.104 | 0.43 | 0.78 | 0.39 | 0.7 | 0.31 | 0.56 |
| 1 | 106040066 | rs6695016 | -0.522 | -1.509 | -0.612 | 1.795 | -0.012 | 0.059 | 0.059 | 0 | 0.17 | 0.23 | 0.19 | 0.47 | 0.12 | 0.38 |
| 1 | 106052011 | rs2255422 | -0.789 | -1.563 | -0.618 | 1.797 | -0.032 | 0.111 | 0.111 | 0.015 | 0.18 | 0.35 | 0.19 | 0.47 | 0.12 | 0.38 |
| 1 | 106054105 | rs11586079 | -0.046 | -1.519 | -1.019 | 1.834 | -0.022 | -0.093 | -0.093 | 0.656 | 0.2 | 0.01 | 0.16 | 0.23 | 0.19 | 0.17 |
| 1 | 106078007 | rs1606971 | -0.674 | -0.88 | -0.578 | 1.541 | -0.007 | 0.045 | 0.045 | 0 | 0.16 | 0.18 | 0.18 | 0.47 | 0.12 | 0.37 |
| 1 | 106085976 | rs34761229 | 0.709 | -0.87 | -1.689 | 0.662 | 0.139 | 0.097 | 0.097 | 0 | 0.51 | 0.12 | 0.6 | 0.34 | 0.67 | 0.46 |
| 1 | 106086980 | rs12120660 | 0.475 | -1.589 | -0.646 | 1.338 | -0.079 | 0.061 | 0.061 | 0.987 | 0.79 | 0.43 | 0.82 | 0.62 | 0.92 | 0.74 |
| 1 | 106089142 | rs2341709 | -0.174 | -0.786 | -1.134 | 1.049 | -0.022 | 0.038 | 0.038 | 0.904 | 0.45 | 0.21 | 0.5 | 0.51 | 0.52 | 0.32 |
| 1 | 106097579 | rs12040852 | -0.226 | -0.529 | -1.324 | 1.091 | -0.074 | -0.048 | -0.048 | 0 | 0.4 | 0.06 | 0.45 | 0.49 | 0.45 | 0.27 |
| 1 | 106100248 | rs777082 | 0.369 | -0.948 | -0.119 | 1.074 | -0.04 | 0.145 | 0.145 | 0 | 0.9 | 0.66 | 0.96 | 0.72 | 0.98 | 0.79 |
| 1 | 106100758 | rs12753622 | -0.074 | -0.965 | 0.037 | 0.928 | 0.004 | 0.034 | 0.034 | 1.369 | 0.42 | 0.19 | 0.47 | 0.54 | 0.45 | 0.33 |
| 1 | 106102016 | rs1710746 | 0.037 | -0.966 | 0.23 | 0.921 | -0.035 | 0.138 | 0.138 | 0 | 0.91 | 0.73 | 0.96 | 0.72 | 0.98 | 0.79 |
| 1 | 106123365 | rs6421299 | 0.563 | -0.959 | 0.311 | 0.83 | 0.005 | 0.003 | 0.003 | 0.797 | 0.48 | 0.45 | 0.48 | 0.18 | 0.53 | 0.46 |
| 1 | 106128123 | rs7539636 | 0.369 | -0.945 | 0.343 | 0.833 | 0.033 | 0.11 | 0.11 | 1.101 | 0.51 | 0.7 | 0.49 | 0.18 | 0.53 | 0.49 |
| 1 | 106130218 | rs12024670 | 0.055 | -0.88 | 0.288 | 0.972 | -0.018 | -0.02 | -0.02 | 0 | 0.41 | 0.08 | 0.38 | 0.54 | 0.42 | 0.33 |
| 1 | 106135739 | rs2140544 | 0.531 | 0.208 | 0.362 | 0.887 | 0.001 | 0.007 | 0.007 | 1.55 | 0.47 | 0.58 | 0.29 | 0.18 | 0.43 | 0.43 |
| 1 | 106144662 | rs7512214 | -0.444 | 0.258 | 0.473 | 0.95 | 0.022 | 0.047 | 0.047 | 0 | 0.91 | 0.87 | 0.68 | 0.73 | 0.86 | 0.8 |
| 1 | 106152813 | rs12563805 | 0.601 | 0.13 | 0.802 | 0.747 | 0 | 0.002 | 0.002 | 1.088 | 0.46 | 0.46 | 0.28 | 0.17 | 0.43 | 0.45 |

| | | | | | | | | | | | | | | | | |
|---|---|---|---|---|---|---|---|---|---|---|---|---|---|---|---|---|
| 1 | 106164864 | rs7550122 | 0.239 | 0.078 | -0.081 | 1.33 | -0.003 | -0.028 | -0.028 | 0 | 0.36 | 0.1 | 0.27 | 0.3 | 0.37 | 0.26 |
| 1 | 106179205 | rs4553248 | -0.265 | 0.36 | 0.384 | 1.523 | 0.019 | 0.043 | 0.043 | 0 | 0.1 | 0.15 | 0.3 | 0.28 | 0.15 | 0.21 |
| 1 | 106189791 | rs10785768 | -0.739 | -0.074 | 0.726 | 1.307 | -0.001 | -0.001 | -0.001 | 0 | 0.46 | 0.53 | 0.38 | 0.49 | 0.49 | 0.44 |
| 1 | 106194840 | rs1548337 | -0.323 | -0.163 | 0.929 | 0.902 | -0.028 | 0.052 | 0.052 | 0.047 | 0.38 | 0.16 | 0.32 | 0.24 | 0.43 | 0.35 |
| 1 | 106195966 | rs12743318 | 0.005 | -0.229 | 0.667 | 0.656 | -0.026 | 0.014 | 0.014 | 0 | 0.26 | 0.14 | 0.26 | 0.19 | 0.36 | 0.24 |
| 1 | 106198567 | rs1361464 | -1.353 | -0.537 | 0.922 | 0.415 | 0.005 | 0.044 | 0.044 | 0 | 0.47 | 0.8 | 0.46 | 0.34 | 0.46 | 0.36 |
| 1 | 106210496 | rs11184668 | -1.183 | -0.493 | 0.841 | 0.38 | -0.001 | -0.002 | -0.002 | 0 | 0.45 | 0.53 | 0.43 | 0.34 | 0.46 | 0.36 |
| 1 | 106215029 | rs17424754 | 0.752 | -0.239 | 0.431 | 0.362 | -0.086 | 0.019 | 0.019 | 0.308 | 0.19 | 0.02 | 0.18 | 0.12 | 0.29 | 0.17 |
| 1 | 106221506 | rs11184671 | -0.986 | -0.258 | 0.652 | 0.408 | -0.025 | 0.014 | 0.014 | 0.09 | 0.41 | 0.19 | 0.4 | 0.34 | 0.46 | 0.36 |
| 1 | 106224822 | rs2840187 | -0.986 | -0.258 | 0.652 | 0.408 | -0.025 | 0.014 | 0.014 | 0.09 | 0.41 | 0.19 | 0.4 | 0.34 | 0.46 | 0.36 |
| 1 | 106228100 | rs790995 | -1.4 | -0.24 | 0.753 | 0.419 | -0.001 | 0.001 | 0.001 | 0 | 0.45 | 0.43 | 0.42 | 0.34 | 0.46 | 0.36 |
| 1 | 106243672 | rs12131617 | -0.219 | -0.392 | 0.361 | 0.409 | -0.004 | 0.028 | 0.028 | 0 | 0.28 | 0.14 | 0.23 | 0.18 | 0.34 | 0.24 |
| 1 | 106249145 | rs790970 | -0.373 | -0.431 | 0.637 | 0.416 | -0.029 | 0.003 | 0.003 | 0 | 0.17 | 0.33 | 0.17 | 0.15 | 0.08 | 0.13 |
| 1 | 106263540 | rs790951 | -1.477 | -0.452 | 0.435 | 0.408 | 0.027 | 0.03 | 0.03 | 0 | 0.46 | 0.7 | 0.43 | 0.35 | 0.42 | 0.37 |
| 1 | 106273120 | rs17425570 | 2.169 | -0.369 | -0.248 | 0.243 | -0.104 | 0.077 | 0.077 | 0.251 | 0.28 | 0.02 | 0.19 | 0.14 | 0.33 | 0.22 |
| 1 | 106275064 | rs12143543 | -0.229 | -0.43 | 0.603 | 0.428 | -0.003 | 0.021 | 0.021 | 0.043 | 0.28 | 0.15 | 0.24 | 0.18 | 0.33 | 0.25 |
| 1 | 106279726 | rs791007 | -0.265 | -0.376 | 0.498 | 0.267 | -0.02 | 0.011 | 0.011 | 0 | 0.15 | 0.27 | 0.17 | 0.17 | 0.08 | 0.14 |
| 1 | 106286316 | rs791004 | -1.578 | -0.298 | -0.033 | 0.251 | 0.011 | 0.018 | 0.018 | 0 | 0.44 | 0.68 | 0.38 | 0.35 | 0.42 | 0.38 |
| 1 | 106287469 | rs11184683 | 0.369 | -0.2 | -0.3 | 0.523 | 0.002 | -0.032 | -0.032 | 0.165 | 0.11 | 0.04 | 0.15 | 0.18 | 0.08 | 0.12 |
| 1 | 106298276 | rs1856784 | 0.213 | -0.218 | -0.395 | 0.488 | -0.024 | 0.009 | 0.009 | 0.099 | 0.21 | 0.05 | 0.12 | 0.15 | 0.26 | 0.19 |
| 1 | 106303166 | rs791001 | 0.856 | -0.329 | 0.292 | 0.426 | -0.006 | 0.004 | 0.004 | 0 | 0.63 | 0.52 | 0.68 | 0.62 | 0.63 | 0.67 |
| 1 | 106306104 | rs1528685 | -0.489 | -0.34 | 0.023 | 0.567 | -0.055 | -0.123 | -0.123 | 0 | 0.3 | 0.03 | 0.25 | 0.4 | 0.34 | 0.31 |
| 1 | 106306182 | rs791002 | 0.968 | -0.405 | 0.559 | 0.529 | 0.004 | -0.006 | -0.006 | 0 | 0.62 | 0.31 | 0.67 | 0.55 | 0.64 | 0.58 |
| 1 | 106314780 | rs1163530 | 0.665 | -0.309 | 0.716 | 0.539 | -0.004 | 0.007 | 0.007 | 0 | 0.64 | 0.38 | 0.68 | 0.55 | 0.64 | 0.58 |
| 1 | 106360040 | rs1163482 | 0.803 | -0.362 | 0.317 | 0.474 | -0.019 | 0.022 | 0.022 | 0 | 0.64 | 0.44 | 0.7 | 0.55 | 0.64 | 0.57 |
| 1 | 106368717 | rs12033818 | -0.025 | -0.089 | -0.325 | 0.578 | -0.016 | -0.062 | -0.062 | 0.367 | 0.1 | 0.02 | 0.13 | 0.23 | 0.09 | 0.11 |
| 1 | 106384407 | rs7532790 | 0.655 | 0.031 | -0.902 | -0.564 | -0.027 | 0.085 | 0.085 | 0.327 | 0.32 | 0.14 | 0.21 | 0.02 | 0.38 | 0.31 |
| 1 | 106386899 | rs1528688 | 0.475 | -0.308 | -1.016 | -0.7 | 0.018 | 0.09 | 0.09 | 0.922 | 0.39 | 0.13 | 0.23 | 0.22 | 0.35 | 0.35 |
| 1 | 106394002 | rs12727940 | 0.668 | -0.43 | -0.815 | -0.838 | 0.011 | 0.054 | 0.054 | 0.991 | 0.36 | 0.22 | 0.21 | 0.22 | 0.33 | 0.28 |
| 1 | 106394977 | rs17426528 | 0.365 | -0.477 | -0.526 | -0.861 | 0.01 | 0.031 | 0.031 | 1.06 | 0.38 | 0.31 | 0.22 | 0.22 | 0.33 | 0.29 |
| 1 | 106412429 | rs790615 | 0.366 | -0.514 | -0.047 | -0.639 | 0 | 0 | 0 | 0.452 | 0.48 | 0.44 | 0.61 | 0.54 | 0.55 | 0.54 |
| 1 | 106419876 | rs12406882 | 0.545 | -0.476 | -0.045 | -0.564 | 0.004 | -0.018 | -0.018 | 0 | 0.13 | 0.1 | 0.16 | 0.24 | 0.13 | 0.17 |
| 1 | 106435371 | rs11184709 | 0.548 | -0.477 | -0.062 | -0.564 | 0.004 | -0.02 | -0.02 | 0 | 0.13 | 0.08 | 0.16 | 0.24 | 0.13 | 0.18 |
| 1 | 106436919 | rs17467366 | 1.503 | -0.663 | -0.461 | -0.811 | -0.042 | 0.012 | 0.012 | 0.552 | 0.22 | 0.03 | 0.14 | 0.16 | 0.24 | 0.19 |
| 1 | 106438336 | rs17016501 | 1.393 | -0.423 | -0.093 | -0.677 | 0.006 | 0.015 | 0.015 | 0.08 | 0.13 | 0.07 | 0.06 | 0.06 | 0.08 | 0.09 |
| 1 | 106449970 | rs940426 | 1.215 | 0.138 | 0.277 | -0.501 | 0.023 | 0.102 | 0.102 | 0 | 0.12 | 0.16 | 0.38 | 0.53 | 0.16 | 0.22 |
| 1 | 106453318 | rs1380289 | 0.319 | -0.419 | 0.139 | -0.433 | 0.002 | 0.001 | 0.001 | 0.05 | 0.48 | 0.43 | 0.61 | 0.54 | 0.55 | 0.51 |
| 1 | 106454673 | rs12407398 | -0.344 | -0.42 | -0.218 | -0.496 | 0.034 | 0.029 | 0.029 | 0.177 | 0.48 | 0.34 | 0.33 | 0.39 | 0.4 | 0.42 |
| 1 | 106461662 | rs3907134 | 0.051 | -0.103 | -0.245 | -0.633 | 0.003 | -0.004 | -0.004 | 0.828 | 0.56 | 0.42 | 0.49 | 0.41 | 0.47 | 0.53 |
| 1 | 106474028 | rs329095 | -0.865 | -0.12 | -0.081 | -0.541 | 0 | -0.001 | -0.001 | 0 | 0.72 | 0.73 | 0.7 | 0.82 | 0.59 | 0.77 |
| 1 | 106477681 | rs11184717 | 0.709 | -0.07 | -0.036 | -0.399 | 0 | 0.005 | 0.005 | 0.11 | 0.15 | 0.09 | 0.21 | 0.39 | 0.14 | 0.16 |
| 1 | 106508563 | rs3850613 | 0.575 | -0.201 | -0.657 | -0.82 | -0.01 | -0.043 | -0.043 | 0 | 0.35 | 0.13 | 0.41 | 0.43 | 0.37 | 0.45 |
| 1 | 106508601 | rs17427795 | 2.125 | -0.227 | -0.544 | -0.868 | 0.014 | 0.016 | 0.016 | 0.417 | 0.09 | 0.03 | 0.03 | 0 | 0.02 | 0.01 |
| 1 | 106516755 | rs3912311 | -0.845 | 0.031 | 0.017 | -0.685 | 0.008 | 0.035 | 0.035 | 0 | 0.5 | 0.62 | 0.63 | 0.82 | 0.46 | 0.69 |
| 1 | 106518710 | rs2342893 | -0.882 | 0.123 | 0.017 | -0.687 | 0 | 0.033 | 0.033 | 0 | 0.51 | 0.62 | 0.64 | 0.82 | 0.52 | 0.7 |
| 1 | 106529404 | rs7536668 | 0.836 | 0.195 | -0.1 | -0.486 | -0.009 | 0.076 | 0.076 | 1.602 | 0.14 | 0.25 | 0.2 | 0.39 | 0.1 | 0.19 |
| 1 | 106534683 | rs12117288 | -0.746 | 0.185 | -0.101 | -0.526 | 0.005 | 0.037 | 0.037 | 0.005 | 0.47 | 0.38 | 0.61 | 0.82 | 0.45 | 0.62 |
| 1 | 106540583 | rs11184726 | 0.374 | 0.266 | -0.176 | -0.411 | -0.008 | -0.03 | -0.03 | 0 | 0.37 | 0.13 | 0.42 | 0.43 | 0.4 | 0.47 |
| 1 | 106568613 | rs12404525 | 1.031 | 0.741 | -0.223 | -0.325 | -0.053 | -0.095 | -0.095 | 0 | 0.17 | 0.02 | 0.31 | 0.37 | 0.2 | 0.17 |
| 1 | 106577155 | rs480255 | 2.056 | 0.731 | 0.082 | -0.456 | 0.002 | -0.035 | -0.035 | 0 | 0.08 | 0.09 | 0.02 | 0 | 0.08 | 0.05 |
| 1 | 106577814 | rs12076092 | 0.327 | 0.524 | -0.178 | -0.041 | 0.025 | 0.023 | 0.023 | 0 | 0.16 | 0.11 | 0.07 | 0.07 | 0.1 | 0.22 |
| 1 | 106577890 | rs7517606 | 0.835 | 0.886 | -0.079 | -0.136 | 0.036 | -0.047 | -0.047 | 0 | 0.1 | 0.24 | 0.17 | 0 | 0.13 | 0.03 |
| 1 | 106578086 | rs12130096 | -1.511 | 0.324 | -0.357 | 0.002 | 0.003 | 0.039 | 0.039 | 0 | 0.52 | 0.39 | 0.62 | 0.81 | 0.5 | 0.66 |
| 1 | 106589618 | rs12407915 | -0.368 | 0.432 | 0.268 | 0.191 | -0.007 | -0.02 | -0.02 | 0 | 0.17 | 0.12 | 0.32 | 0.37 | 0.21 | 0.16 |
| 1 | 106595812 | rs12090319 | -1.562 | -0.041 | 0.281 | 0.206 | 0.012 | -0.01 | -0.01 | 1.405 | 0.31 | 0.22 | 0.38 | 0.43 | 0.28 | 0.38 |
| 1 | 106607211 | rs12058715 | -0.512 | 0.056 | 0.021 | 0.159 | -0.007 | -0.005 | -0.005 | 0 | 0.14 | 0.13 | 0.07 | 0.08 | 0.1 | 0.15 |
| 1 | 106609805 | rs10494035 | -0.7 | 0.744 | 0.002 | 0.466 | 0 | -0.023 | -0.023 | 0.632 | 0.18 | 0.12 | 0.31 | 0.36 | 0.18 | 0.18 |
| 1 | 106617673 | rs11184747 | -0.865 | 1.069 | 0.517 | 0.451 | 0.052 | -0.063 | -0.063 | 1.47 | 0.11 | 0.26 | 0.2 | 0 | 0.16 | 0.07 |
| 1 | 106622689 | rs6582953 | -0.636 | 0.69 | 0.415 | 0.515 | 0.1 | -0.061 | -0.061 | 1.492 | 0.11 | 0.3 | 0.21 | 0 | 0.19 | 0.07 |
| 1 | 106624365 | rs12411062 | 0.529 | -0.677 | -0.176 | 1.158 | 0.004 | 0.005 | 0.005 | 3.216 | 0.07 | 0.07 | 0.19 | 0.3 | 0.04 | 0.06 |
| 1 | 106630863 | rs994712 | -0.877 | -0.676 | -0.161 | 0.973 | -0.018 | 0.021 | 0.021 | 0.148 | 0.21 | 0.26 | 0.27 | 0.41 | 0.13 | 0.19 |
| 1 | 106639555 | rs6700635 | -0.882 | -0.723 | -0.156 | 0.797 | -0.019 | 0.014 | 0.014 | 0.158 | 0.21 | 0.26 | 0.27 | 0.39 | 0.13 | 0.25 |
| 1 | 106650029 | rs12046537 | -1.027 | 0.188 | 0.05 | 0.365 | -0.003 | 0.008 | 0.008 | 0 | 0.08 | 0.2 | 0.12 | 0.09 | 0.07 | 0.09 |
| 1 | 106650268 | rs587675 | 1.971 | -0.187 | 0.52 | 0.005 | -0.04 | -0.069 | -0.069 | 0 | 0.81 | 0.43 | 0.83 | 0.91 | 0.9 | 0.77 |
| 1 | 106652260 | rs617243 | 0.631 | -0.218 | 0.734 | -0.009 | -0.006 | -0.045 | -0.045 | 0.18 | 0.92 | 0.81 | 0.97 | 1 | 0.96 | 0.99 |
| 1 | 106653622 | rs1889463 | -0.169 | 0.079 | -0.053 | 0.06 | -0.067 | -0.117 | -0.117 | 0 | 0.24 | 0.05 | 0.37 | 0.57 | 0.34 | 0.33 |
| 1 | 106655045 | rs12035891 | -1.537 | -0.111 | 0.438 | 0.14 | -0.014 | 0 | 0 | 0.001 | 0.11 | 0.37 | 0.13 | 0.09 | 0.07 | 0.22 |
| 1 | 106678764 | rs11184769 | -0.879 | -0.154 | 0.317 | 0.135 | -0.005 | -0.03 | -0.03 | 0 | 0.08 | 0.12 | 0.03 | 0 | 0.04 | 0.01 |
| 1 | 106678806 | rs11590544 | -1.07 | -0.165 | 0.403 | 0.133 | -0.008 | 0.009 | 0.009 | 0 | 0.09 | 0.26 | 0.12 | 0.09 | 0.06 | 0.08 |

| | | | | | | | | | | | | | | | | |
|---|---|---|---|---|---|---|---|---|---|---|---|---|---|---|---|---|
| 1 | 106679169 | rs11184770 | -2.346 | -0.165 | 0.318 | 0.132 | -0.043 | -0.071 | -0.071 | 0 | 0.18 | 0.45 | 0.16 | 0.09 | 0.1 | 0.23 |
| 1 | 106687949 | rs17016826 | -1.17 | 0.042 | -0.027 | 0.139 | -0.002 | -0.05 | -0.05 | 0.095 | 0.09 | 0.23 | 0.03 | 0 | 0.04 | 0.14 |
| 1 | 106695375 | rs12026453 | -0.906 | 0.33 | -0.687 | 0.332 | -0.007 | -0.002 | -0.002 | 0 | 0.08 | 0.13 | 0.23 | 0.07 | 0.07 | 0.11 |
| 1 | 106705605 | rs10158160 | -0.957 | 0.336 | -0.686 | 0.283 | -0.007 | -0.008 | -0.008 | 0 | 0.09 | 0.13 | 0.23 | 0.06 | 0.07 | 0.11 |
| 1 | 106721791 | rs7528765 | -1.273 | 0.299 | -0.684 | 0.265 | -0.014 | -0.016 | -0.016 | 0.183 | 0.1 | 0.26 | 0.24 | 0.06 | 0.08 | 0.11 |
| 1 | 106728999 | rs12021804 | -1.225 | 0.316 | -0.679 | 0.273 | -0.012 | -0.015 | -0.015 | 0 | 0.1 | 0.29 | 0.25 | 0.06 | 0.08 | 0.11 |
| 1 | 106740758 | rs4480388 | 0.006 | 0.229 | -1.449 | 0.432 | -0.02 | -0.129 | -0.129 | 0 | 0.79 | 0.99 | 0.77 | 0.43 | 0.82 | 0.83 |
| 1 | 106750803 | rs11576730 | -1.535 | 0.855 | -0.549 | 0.304 | -0.008 | -0.011 | -0.011 | 0 | 0.11 | 0.32 | 0.24 | 0.06 | 0.07 | 0.1 |
| 1 | 106752234 | rs12023696 | -0.813 | 1.623 | -1.419 | 0.216 | -0.011 | 0.01 | 0.01 | 0.823 | 0.24 | 0.2 | 0.44 | 0.19 | 0.33 | 0.21 |
| 1 | 106767908 | rs7534309 | -1.241 | 2.042 | -1.678 | 0.761 | -0.052 | -0.055 | -0.055 | 0.373 | 0.14 | 0.05 | 0.31 | 0.3 | 0.22 | 0.16 |
| 1 | 106769130 | rs7551463 | 0.057 | 1.673 | -1.964 | -0.794 | 0.006 | 0.016 | 0.016 | 0.037 | 0.31 | 0.23 | 0.18 | 0.01 | 0.28 | 0.3 |
| 1 | 106770076 | rs6702428 | 0.569 | 2.287 | -2.373 | -0.455 | -0.05 | -0.051 | -0.051 | 0 | 0.76 | 0.88 | 0.62 | 0.62 | 0.66 | 0.77 |
| 1 | 106770110 | rs6687440 | -0.292 | 1.293 | -2.981 | -1.382 | 0.022 | 0.251 | 0.251 | 0.736 | 0.42 | 0.89 | 0.26 | 0.01 | 0.36 | 0.3 |
| 1 | 106774931 | rs1334126 | 0.189 | 1.992 | -2.552 | -0.572 | -0.029 | 0.012 | 0.012 | 0 | 0.06 | 0.02 | 0.27 | 0.34 | 0.1 | 0.13 |
| 1 | 106775048 | rs1334127 | 1.034 | 1.944 | -1.15 | -0.8 | 0.11 | 0.25 | 0.25 | 0.338 | 0.87 | 0.6 | 0.69 | 0.66 | 0.85 | 0.82 |
| 1 | 106781612 | rs9435409 | 0.62 | 1.584 | -1.061 | -1.339 | 0.013 | 0 | 0 | 0 | 0.5 | 0.38 | 0.43 | 0.48 | 0.47 | 0.58 |
| 1 | 106787317 | rs4503378 | 0.526 | 1.563 | -1.06 | -1.578 | 0.002 | 0.002 | 0.002 | 0.684 | 0.69 | 0.36 | 0.59 | 0.66 | 0.66 | 0.78 |
| 1 | 106792422 | rs12568304 | 0.613 | 1.127 | -1.118 | -1.64 | -0.043 | 0 | 0 | 0 | 0.19 | 0.05 | 0.16 | 0.17 | 0.18 | 0.2 |
| 1 | 106799250 | rs12090269 | -0.651 | 0.836 | -0.254 | -1.622 | 0.067 | 0.015 | 0.015 | 0.245 | 0.07 | 0.32 | 0.03 | 0.01 | 0.06 | 0.04 |
| 1 | 106802357 | rs7524060 | 0.852 | 0.814 | -0.053 | -1.721 | 0.003 | -0.072 | -0.072 | 0 | 0.14 | 0.25 | 0.12 | 0 | 0.17 | 0.01 |
| 1 | 106807178 | rs12046841 | 1.344 | 0.804 | 0.287 | -1.784 | 0.097 | -0.011 | -0.011 | 0 | 0.44 | 0.21 | 0.35 | 0.39 | 0.27 | 0.45 |
| 1 | 106807581 | rs12094362 | -0.366 | 0.827 | 0.315 | -1.78 | 0.067 | 0.01 | 0.01 | 0.245 | 0.07 | 0.32 | 0.03 | 0 | 0.06 | 0.04 |
| 1 | 106807983 | rs7533503 | 1.137 | 0.841 | 0.226 | -1.776 | 0.001 | -0.056 | -0.056 | 0 | 0.14 | 0.23 | 0.11 | 0 | 0.18 | 0.01 |
| 1 | 106810236 | rs4431871 | 0.226 | 0.844 | 0.053 | -1.775 | -0.028 | -0.023 | -0.023 | 0.021 | 0.64 | 0.77 | 0.49 | 0.39 | 0.48 | 0.48 |
| 1 | 106810917 | rs7519669 | 0.228 | 0.853 | 0.126 | -1.776 | 0.012 | -0.137 | -0.137 | 0 | 0.19 | 0.49 | 0.14 | 0 | 0.21 | 0.03 |
| 1 | 106815064 | rs1537531 | 0.432 | 0.966 | -0.151 | -1.638 | -0.026 | -0.02 | -0.02 | 0.04 | 0.63 | 0.75 | 0.49 | 0.39 | 0.48 | 0.48 |
| 1 | 106819798 | rs4304612 | 0.275 | 1.009 | -0.36 | -1.611 | -0.061 | -0.048 | -0.048 | 0 | 0.66 | 0.84 | 0.5 | 0.39 | 0.5 | 0.5 |
| 1 | 106824204 | rs1360707 | 0.275 | 1.009 | -0.382 | -1.611 | -0.063 | -0.05 | -0.05 | 0 | 0.66 | 0.84 | 0.5 | 0.39 | 0.5 | 0.5 |
| 1 | 106834169 | rs11184840 | 0.289 | 0.337 | -0.397 | -2.28 | 0.015 | -0.025 | -0.025 | 0.1 | 0.43 | 0.66 | 0.27 | 0.01 | 0.34 | 0.3 |
| 1 | 106838528 | rs4111473 | 0.599 | 0.846 | -0.788 | -1.985 | -0.089 | -0.025 | -0.025 | 3.08 | 0.18 | 0.02 | 0.19 | 0.18 | 0.27 | 0.22 |
| 1 | 106843482 | rs6694017 | 1.152 | 0.497 | -0.187 | -2.053 | 0.004 | -0.098 | -0.098 | 1.269 | 0.15 | 0.32 | 0.13 | 0 | 0.18 | 0.02 |
| 1 | 106848878 | rs1928718 | 0.294 | 0.33 | -0.166 | -2.351 | -0.01 | 0.087 | 0.087 | 0 | 0.46 | 0.34 | 0.34 | 0.18 | 0.45 | 0.5 |
| 1 | 106849846 | rs11184847 | -0.942 | 0.297 | -0.142 | -2.405 | -0.032 | 0.082 | 0.082 | 0 | 0.67 | 0.89 | 0.49 | 0.19 | 0.66 | 0.54 |
| 1 | 106860057 | rs1888181 | 0.312 | 0.296 | -0.273 | -2.369 | -0.01 | 0.084 | 0.084 | 0 | 0.46 | 0.34 | 0.34 | 0.18 | 0.45 | 0.5 |
| 1 | 106866722 | rs12059925 | 0.312 | 0.296 | -0.283 | -2.364 | -0.01 | 0.085 | 0.085 | 0 | 0.46 | 0.34 | 0.34 | 0.18 | 0.45 | 0.5 |
| 1 | 106867772 | rs11809005 | -0.261 | 0.843 | 0.253 | -1.717 | -0.002 | -0.145 | -0.145 | 0 | 0.21 | 0.5 | 0.14 | 0 | 0.22 | 0.04 |
| 1 | 106870121 | rs12734803 | 1.004 | 0.853 | -0.373 | -1.652 | 0 | -0.037 | -0.037 | 0 | 0.15 | 0.18 | 0.12 | 0 | 0.19 | 0.02 |
| 1 | 106881800 | rs6670501 | -0.656 | 0.883 | -0.328 | -1.547 | 0.004 | -0.01 | -0.01 | 0 | 0.29 | 0.59 | 0.23 | 0.18 | 0.32 | 0.28 |
| 1 | 106885146 | rs11184867 | -0.98 | 0.799 | -0.624 | -1.632 | -0.038 | 0.113 | 0.113 | 0.207 | 0.43 | 0.72 | 0.34 | 0.18 | 0.51 | 0.29 |
| 1 | 106887181 | rs11184870 | 0.018 | 0.883 | 0.149 | -1.682 | 0.027 | -0.015 | -0.015 | 0 | 0.22 | 0.29 | 0.21 | 0.17 | 0.29 | 0.25 |
| 1 | 106896377 | rs4915122 | -1.178 | 0.773 | 0.421 | -1.807 | -0.011 | 0.031 | 0.031 | 0 | 0.4 | 0.65 | 0.33 | 0.18 | 0.47 | 0.23 |
| 1 | 106896765 | rs11579468 | 0.517 | 0.713 | 0.635 | -1.708 | -0.009 | -0.111 | -0.111 | 0 | 0.18 | 0.38 | 0.12 | 0 | 0.19 | 0.03 |
| 1 | 106897428 | rs7552126 | 0.129 | 0.799 | 0.571 | -1.553 | 0.009 | -0.004 | -0.004 | 0 | 0.21 | 0.22 | 0.2 | 0.17 | 0.27 | 0.21 |
| 1 | 106911367 | rs1411398 | -0.042 | 0.842 | 0.529 | -1.613 | 0.024 | -0.007 | -0.007 | 0 | 0.19 | 0.27 | 0.2 | 0.17 | 0.25 | 0.2 |
| 1 | 106925187 | rs12736194 | -0.018 | 0.805 | 0.253 | -1.62 | -0.071 | -0.005 | -0.005 | 0 | 0.19 | 0.03 | 0.19 | 0.17 | 0.25 | 0.18 |
| 1 | 106933363 | rs17473350 | 1.506 | -0.05 | -0.032 | -2.454 | 0.031 | 0.07 | 0.07 | 1.068 | 0.32 | 0.21 | 0.15 | 0.01 | 0.21 | 0.36 |
| 1 | 106933779 | rs1536989 | 0.592 | 0.185 | 0.089 | -2.447 | -0.018 | 0.05 | 0.05 | 0 | 0.61 | 0.35 | 0.66 | 0.83 | 0.54 | 0.79 |
| 1 | 106937801 | rs11184892 | 0.338 | 0.721 | 0.201 | -2.404 | 0.017 | -0.055 | -0.055 | 0 | 0.84 | 0.4 | 0.82 | 0.94 | 0.85 | 0.85 |
| 1 | 106943366 | rs17486419 | 2.234 | 0.452 | -0.439 | -2.762 | -0.044 | 0.047 | 0.047 | 0 | 0.08 | 0.01 | 0.1 | 0.01 | 0.13 | 0.23 |
| 1 | 106945362 | rs11184894 | 0.1 | 0.523 | -0.226 | -2.636 | 0.018 | -0.049 | -0.049 | 0 | 0.85 | 0.39 | 0.84 | 0.94 | 0.87 | 0.86 |
| 1 | 106948755 | rs1831384 | 0.157 | 0.284 | -0.554 | -2.735 | 0.008 | 0.359 | 0.359 | 0.136 | 0.5 | 0.12 | 0.69 | 0.93 | 0.65 | 0.49 |
| 1 | 106949216 | rs11184896 | 0.868 | 0.326 | -0.617 | -2.621 | -0.145 | 0.222 | 0.222 | 0.453 | 0.35 | 0.05 | 0.58 | 0.93 | 0.47 | 0.48 |
| 1 | 106953168 | rs11184897 | 1.69 | -0.029 | -0.829 | -1.645 | -0.034 | -0.156 | -0.156 | 0.191 | 0.2 | 0.01 | 0.23 | 0.48 | 0.19 | 0.19 |
| 1 | 106967467 | rs2182149 | 0.514 | 0.186 | -0.475 | -2.029 | -0.008 | 0.036 | 0.036 | 0 | 0.35 | 0.29 | 0.56 | 0.86 | 0.38 | 0.39 |
| 1 | 106967482 | rs2182150 | -0.977 | 0.224 | -0.811 | -2.178 | 0.025 | 0.072 | 0.072 | 0.009 | 0.62 | 0.73 | 0.82 | 0.92 | 0.7 | 0.53 |
| 1 | 106972853 | rs7514699 | -0.14 | 0.388 | -0.495 | -1.848 | 0.012 | -0.133 | -0.133 | 0 | 0.28 | 0.52 | 0.21 | 0.04 | 0.32 | 0.14 |
| 1 | 106975048 | rs4915125 | 0.337 | 0.311 | -1.062 | -0.802 | -0.041 | 0.062 | 0.062 | 0.71 | 0.46 | 0.16 | 0.63 | 0.56 | 0.5 | 0.39 |
| 1 | 106979036 | rs10158153 | 0.222 | 0.038 | -0.316 | -0.646 | 0.019 | 0.031 | 0.031 | 0 | 0.2 | 0.33 | 0.19 | 0.29 | 0.21 | 0.28 |
| 1 | 106982478 | rs6660032 | 1.126 | 0.026 | -0.362 | -0.706 | -0.008 | -0.034 | -0.034 | 0 | 0.1 | 0.12 | 0.02 | 0 | 0.05 | 0.06 |
| 1 | 106983341 | rs1925429 | -0.8 | 0.051 | -0.435 | -0.668 | -0.031 | -0.012 | -0.012 | 3.518 | 0.54 | 0.85 | 0.36 | 0.51 | 0.42 | 0.57 |
| 1 | 106984123 | rs12040644 | 1.786 | 0.36 | -0.093 | -0.387 | -0.006 | 0.005 | 0.005 | 0 | 0.09 | 0.1 | 0.16 | 0.29 | 0.15 | 0.23 |
| 1 | 106994923 | rs956282 | 1.786 | 0.419 | -0.219 | -0.35 | 0.003 | 0.019 | 0.019 | 0 | 0.09 | 0.12 | 0.16 | 0.29 | 0.15 | 0.22 |
| 1 | 106995026 | rs725703 | -0.264 | 0.351 | -0.176 | -0.502 | 0.022 | -0.001 | -0.001 | 0 | 0.29 | 0.56 | 0.28 | 0.32 | 0.3 | 0.33 |
| 1 | 106995427 | rs7417032 | 1.219 | 0.388 | -0.068 | -0.446 | 0.078 | 0.152 | 0.152 | 0 | 0.11 | 0.34 | 0.18 | 0.29 | 0.16 | 0.22 |
| 1 | 106999448 | rs12032785 | -1.079 | -0.039 | -0.616 | -0.593 | -0.042 | -0.027 | -0.027 | 0 | 0.54 | 0.88 | 0.36 | 0.51 | 0.43 | 0.55 |
| 1 | 107041174 | rs6677772 | -1.669 | 0.406 | -0.688 | -0.223 | -0.081 | -0.065 | -0.065 | 0.788 | 0.88 | 0.98 | 0.66 | 0.77 | 0.71 | 0.75 |
| 1 | 107045474 | rs5008684 | 0.466 | -0.563 | -0.074 | -0.84 | -0.029 | -0.015 | -0.015 | 1.296 | 0.25 | 0.27 | 0.08 | 0.18 | 0.12 | 0.22 |
| 1 | 107046847 | rs12752606 | 1.285 | -0.534 | -0.132 | -0.77 | -0.002 | -0.015 | -0.015 | 0 | 0.07 | 0.08 | 0.05 | 0 | 0.09 | 0.01 |
| 1 | 107053627 | rs726669 | -1.28 | -0.432 | 0.031 | -0.68 | -0.022 | -0.01 | -0.01 | 0 | 0.63 | 0.72 | 0.42 | 0.5 | 0.54 | 0.55 |

| | | | | | | | | | | | | | | | | |
|---|---|---|---|---|---|---|---|---|---|---|---|---|---|---|---|---|
| 1 | 107055711 | rs10494043 | -0.643 | -0.315 | -0.083 | -0.741 | 0.001 | -0.014 | -0.014 | 0 | 0.53 | 0.47 | 0.26 | 0.21 | 0.43 | 0.33 |
| 1 | 107076801 | rs1925427 | 0.867 | 0.033 | 0.153 | -0.067 | 0.026 | 0.159 | 0.159 | 0 | 0.1 | 0.35 | 0.18 | 0.29 | 0.1 | 0.21 |
| 1 | 107090200 | rs17434983 | 0.454 | -0.112 | -0.18 | -0.382 | -0.005 | 0.064 | 0.064 | 0.043 | 0.21 | 0.08 | 0.1 | 0.02 | 0.17 | 0.11 |
| 1 | 107092445 | rs12034371 | 1.063 | -0.133 | -0.086 | -0.248 | 0 | 0.032 | 0.032 | 0 | 0.08 | 0.13 | 0.15 | 0.28 | 0.1 | 0.19 |
| 1 | 107107254 | rs12567271 | 1.246 | -0.428 | -0.345 | -0.436 | -0.077 | -0.032 | -0.032 | 0.1 | 0.42 | 0.07 | 0.63 | 0.54 | 0.56 | 0.49 |
| 1 | 107107290 | rs4601615 | -0.154 | -0.471 | -0.005 | -0.419 | -0.002 | -0.003 | -0.003 | 0 | 0.3 | 0.41 | 0.25 | 0.3 | 0.26 | 0.28 |
| 1 | 107109564 | rs12040716 | -0.116 | -0.472 | 0.106 | -0.419 | -0.005 | -0.007 | -0.007 | 0.011 | 0.3 | 0.48 | 0.26 | 0.3 | 0.26 | 0.28 |
| 1 | 107113504 | rs4915129 | -0.237 | -0.495 | 0.106 | -0.41 | -0.004 | -0.008 | -0.008 | 0.219 | 0.31 | 0.47 | 0.25 | 0.3 | 0.27 | 0.3 |
| 1 | 107116802 | rs7530540 | -0.674 | -0.598 | 0.185 | -0.432 | -0.022 | -0.039 | -0.039 | 0.008 | 0.75 | 0.68 | 0.9 | 0.83 | 0.82 | 0.77 |
| 1 | 107125001 | rs4545364 | -0.235 | -0.382 | 0.068 | -0.336 | -0.001 | -0.001 | -0.001 | 0 | 0.3 | 0.39 | 0.24 | 0.3 | 0.27 | 0.3 |
| 1 | 107146735 | rs10881425 | -0.304 | -0.321 | -0.208 | -0.326 | -0.006 | -0.01 | -0.01 | 0 | 0.31 | 0.42 | 0.23 | 0.29 | 0.27 | 0.3 |
| 1 | 107157877 | rs11184938 | -0.298 | -0.423 | -0.585 | -0.308 | -0.004 | -0.005 | -0.005 | 0.695 | 0.31 | 0.39 | 0.23 | 0.27 | 0.27 | 0.29 |
| 1 | 107160841 | rs7513071 | 0.801 | -0.267 | -1.517 | -0.048 | -0.053 | -0.011 | -0.011 | 0 | 0.41 | 0.11 | 0.64 | 0.57 | 0.56 | 0.48 |
| 1 | 107162694 | rs6697704 | 0.63 | -0.142 | -1.482 | 0.062 | -0.026 | -0.085 | -0.085 | 0 | 0.49 | 0.03 | 0.55 | 0.51 | 0.6 | 0.5 |
| 1 | 107163365 | rs12076959 | 0.679 | 0.366 | -0.376 | 0.682 | -0.001 | 0.019 | 0.019 | 0 | 0.07 | 0.08 | 0.12 | 0.24 | 0.09 | 0.19 |
| 1 | 107180296 | rs12023595 | -0.678 | 0.465 | 0.039 | 0.356 | 0.021 | 0.018 | 0.018 | 0 | 0.37 | 0.4 | 0.54 | 0.51 | 0.45 | 0.53 |
| 1 | 107182165 | rs4388724 | 1.002 | 0.234 | 0.208 | 0.284 | 0.035 | 0.04 | 0.04 | 0.054 | 0.46 | 0.38 | 0.22 | 0.2 | 0.34 | 0.29 |
| 1 | 107195059 | rs10494048 | -0.064 | -0.057 | 0.519 | 0.148 | 0.013 | 0.012 | 0.012 | 1.673 | 0.3 | 0.32 | 0.51 | 0.5 | 0.43 | 0.44 |
| 1 | 107198806 | rs11184955 | -0.09 | -0.08 | 0.568 | 0.137 | -0.002 | -0.002 | -0.002 | 0.125 | 0.29 | 0.28 | 0.51 | 0.5 | 0.42 | 0.44 |
| 1 | 107200476 | rs6681791 | 0.428 | -0.269 | 0.833 | 0.192 | -0.029 | -0.016 | -0.016 | 0 | 0.72 | 0.51 | 0.71 | 0.7 | 0.77 | 0.74 |
| 1 | 107205384 | rs7555950 | 0.525 | -0.272 | 0.827 | 0.191 | -0.028 | -0.018 | -0.018 | 0 | 0.71 | 0.49 | 0.71 | 0.7 | 0.77 | 0.74 |
| 1 | 107218527 | rs10881441 | 0.514 | -0.271 | 0.832 | 0.191 | -0.03 | -0.018 | -0.018 | 0.296 | 0.71 | 0.51 | 0.71 | 0.7 | 0.77 | 0.74 |
| 1 | 107218775 | rs12030878 | 1.592 | -0.12 | 0.777 | 0.38 | -0.021 | -0.021 | -0.021 | 0 | 0.06 | 0.02 | 0.09 | 0.22 | 0.09 | 0.14 |
| 1 | 107218922 | rs7545951 | 0.118 | 0.031 | 0.854 | 0.308 | -0.017 | 0.035 | 0.035 | 0 | 0.16 | 0.26 | 0.2 | 0.29 | 0.11 | 0.21 |
| 1 | 107219644 | rs10881442 | -0.907 | 0.024 | 0.806 | 0.287 | 0.001 | -0.056 | -0.056 | 0.063 | 0.87 | 0.77 | 0.91 | 0.99 | 0.87 | 0.94 |
| 1 | 107228575 | rs4494161 | -0.956 | 0.071 | 0.776 | 0.31 | 0.008 | -0.067 | -0.067 | 0.957 | 0.87 | 0.72 | 0.9 | 0.99 | 0.88 | 0.94 |
| 1 | 107230730 | rs4462181 | -0.812 | 0.064 | 0.821 | 0.306 | 0.012 | -0.078 | -0.078 | 0 | 0.87 | 0.65 | 0.9 | 0.99 | 0.88 | 0.94 |
| 1 | 107232816 | rs4336881 | -0.966 | 0.077 | 0.813 | 0.312 | 0.008 | -0.067 | -0.067 | 0.032 | 0.87 | 0.73 | 0.9 | 0.99 | 0.88 | 0.94 |
| 1 | 107235329 | rs12034671 | 1.223 | 0.07 | 0.815 | 0.341 | 0.013 | 0.096 | 0.096 | 0 | 0.07 | 0.13 | 0.1 | 0.22 | 0.09 | 0.14 |
| 1 | 107243425 | rs12138646 | 0.967 | 0.038 | 0.92 | 0.005 | -0.008 | 0.02 | 0.02 | 0 | 0.11 | 0.08 | 0.08 | 0.01 | 0.12 | 0.06 |
| 1 | 107244711 | rs11589101 | 0.862 | 0.013 | 0.964 | 0.016 | -0.026 | -0.015 | -0.015 | 0 | 0.71 | 0.48 | 0.71 | 0.7 | 0.78 | 0.74 |
| 1 | 107256227 | rs7340024 | -0.838 | 0.146 | 0.934 | 0.033 | 0.008 | -0.064 | -0.064 | 0 | 0.87 | 0.73 | 0.9 | 0.99 | 0.88 | 0.95 |
| 1 | 107265058 | rs1533670 | -0.291 | 0.292 | 0.982 | 0.012 | 0.044 | -0.093 | -0.093 | 0.24 | 0.85 | 0.6 | 0.86 | 0.96 | 0.82 | 0.88 |
| 1 | 107267115 | rs1401830 | 0.934 | 0.142 | 1.043 | -0.041 | -0.012 | -0.01 | -0.01 | 0 | 0.7 | 0.42 | 0.69 | 0.7 | 0.75 | 0.74 |
| 1 | 107277107 | rs12060755 | -0.499 | 0.19 | 1.008 | -0.078 | 0.022 | -0.097 | -0.097 | 3.64 | 0.86 | 0.54 | 0.89 | 0.99 | 0.88 | 0.94 |
| 1 | 107294677 | rs12085613 | -0.114 | 0.112 | 0.749 | -0.014 | -0.006 | 0.006 | 0.006 | 0 | 0.17 | 0.18 | 0.2 | 0.29 | 0.13 | 0.21 |
| 1 | 107294919 | rs7552695 | 0.717 | 0.132 | 0.781 | -0.033 | -0.005 | -0.004 | -0.004 | 0 | 0.69 | 0.4 | 0.69 | 0.7 | 0.75 | 0.74 |
| 1 | 107298541 | rs12133448 | 1.258 | 0.258 | 0.541 | -0.081 | 0.033 | -0.02 | -0.02 | 0 | 0.45 | 0.27 | 0.28 | 0.43 | 0.42 | 0.43 |
| 1 | 107313610 | rs11184985 | -0.815 | 0.339 | 0.17 | -0.037 | -0.008 | -0.007 | -0.007 | 0 | 0.29 | 0.34 | 0.29 | 0.3 | 0.25 | 0.26 |
| 1 | 107316651 | rs10430083 | 0.23 | 0.55 | -0.123 | 0 | -0.085 | -0.006 | -0.006 | 2.476 | 0.25 | 0.09 | 0.42 | 0.27 | 0.35 | 0.32 |
| 1 | 107332554 | rs17017723 | 0.056 | -0.017 | 0.123 | 0.134 | 0.01 | -0.032 | -0.032 | 0 | 0.15 | 0.06 | 0.19 | 0.29 | 0.13 | 0.2 |
| 1 | 107343141 | rs6671394 | 0.906 | 0.134 | 0.374 | -0.071 | 0.002 | 0.002 | 0.002 | 0 | 0.66 | 0.39 | 0.66 | 0.69 | 0.7 | 0.7 |
| 1 | 107344299 | rs1519887 | -0.295 | -0.038 | 0.556 | -0.014 | 0.006 | -0.012 | -0.012 | 0 | 0.84 | 0.88 | 0.8 | 0.7 | 0.87 | 0.79 |
| 1 | 107376140 | rs12080495 | 1.229 | -0.652 | 0.225 | -0.21 | 0.027 | -0.003 | -0.003 | 0 | 0.33 | 0.17 | 0.22 | 0.23 | 0.29 | 0.28 |
| 1 | 107387839 | rs6678521 | 0.867 | -0.431 | 0.178 | 0.127 | 0.027 | -0.017 | -0.017 | 0 | 0.44 | 0.31 | 0.26 | 0.42 | 0.39 | 0.42 |
| 1 | 107388843 | rs10881450 | 0.142 | -0.184 | 0.579 | 0.405 | -0.033 | 0.057 | 0.057 | 0 | 0.84 | 0.68 | 0.8 | 0.7 | 0.9 | 0.8 |
| 1 | 107404931 | rs12024651 | -1.075 | -0.174 | 0.122 | 0.117 | -0.03 | 0.007 | 0.007 | 0 | 0.2 | 0.3 | 0.09 | 0.2 | 0.13 | 0.23 |
| 1 | 107418424 | rs12128098 | -1.499 | -0.183 | -0.006 | 0.027 | 0.003 | 0.003 | 0.003 | 0 | 0.33 | 0.32 | 0.18 | 0.19 | 0.28 | 0.32 |
| 1 | 107430479 | rs12137571 | -0.894 | -0.086 | -0.185 | 0.01 | -0.023 | -0.094 | -0.094 | 0 | 0.24 | 0.33 | 0.12 | 0.01 | 0.16 | 0.17 |
| 1 | 107431974 | rs1519878 | -1.14 | -0.021 | -0.453 | 0.108 | -0.026 | -0.04 | -0.04 | 0 | 0.31 | 0.46 | 0.18 | 0.2 | 0.26 | 0.32 |
| 1 | 107432576 | rs12726652 | -1.121 | -0.07 | -0.148 | 0.613 | -0.003 | 0.105 | 0.105 | 0 | 0.21 | 0.5 | 0.15 | 0.42 | 0.21 | 0.29 |
| 1 | 107433085 | rs6679315 | -0.388 | -0.586 | -1.274 | 0.189 | 0.063 | 0.012 | 0.012 | 1.066 | 0.41 | 0.16 | 0.26 | 0.31 | 0.3 | 0.42 |
| 1 | 107440872 | rs9651163 | 0.616 | -0.692 | -0.429 | -0.301 | -0.012 | -0.035 | -0.035 | 0 | 0.71 | 0.68 | 0.81 | 0.88 | 0.74 | 0.61 |
| 1 | 107444101 | rs12406116 | 1.173 | -0.884 | -0.707 | -0.151 | 0.006 | -0.06 | -0.06 | 0.983 | 0.12 | 0.03 | 0.12 | 0.22 | 0.07 | 0.13 |
| 1 | 107448618 | rs6677003 | -0.126 | -0.975 | -0.648 | -0.229 | -0.005 | -0.062 | -0.062 | 0 | 0.19 | 0.25 | 0.14 | 0.03 | 0.2 | 0.19 |
| 1 | 107450588 | rs12022772 | 0.837 | -0.967 | -0.733 | -0.22 | 0.015 | -0.005 | -0.005 | 7.14 | 0.09 | 0.04 | 0.05 | 0.1 | 0.07 | 0.21 |
| 1 | 107450890 | rs4914934 | -1.365 | -0.992 | -0.489 | -0.227 | -0.008 | -0.003 | -0.003 | 0 | 0.43 | 0.43 | 0.32 | 0.34 | 0.34 | 0.54 |
| 1 | 107453603 | rs17017931 | 0.975 | -0.95 | -0.604 | -0.087 | 0.025 | -0.021 | -0.021 | 0 | 0.13 | 0.02 | 0.04 | 0.1 | 0.05 | 0.1 |
| 1 | 107455290 | rs4304614 | 1.327 | -1.162 | -0.295 | -0.323 | 0.052 | 0.028 | 0.028 | 0 | 0.52 | 0.36 | 0.66 | 0.65 | 0.66 | 0.44 |
| 1 | 107465208 | rs7553476 | 0.91 | -0.645 | 0.205 | -0.812 | -0.013 | -0.034 | -0.034 | 1.953 | 0.66 | 0.45 | 0.79 | 0.87 | 0.73 | 0.6 |
| 1 | 107480854 | rs4244120 | 0.656 | -0.214 | 0.04 | -0.624 | 0.072 | 0.014 | 0.014 | 2.259 | 0.91 | 0.51 | 0.92 | 0.95 | 0.92 | 0.91 |
| 1 | 107481837 | rs11185019 | 0.656 | -0.214 | 0.053 | -0.624 | 0.07 | 0.012 | 0.012 | 2.259 | 0.91 | 0.52 | 0.92 | 0.95 | 0.92 | 0.91 |
| 1 | 107484628 | rs9435418 | -0.537 | -0.228 | -0.296 | -0.558 | -0.006 | -0.031 | -0.031 | 0 | 0.26 | 0.3 | 0.19 | 0.13 | 0.24 | 0.38 |
| 1 | 107485489 | rs9435419 | -0.541 | -0.231 | -0.297 | -0.559 | -0.006 | -0.031 | -0.031 | 0 | 0.26 | 0.3 | 0.19 | 0.13 | 0.24 | 0.38 |
| 1 | 107488772 | rs6704278 | -0.932 | 0.361 | -0.014 | -0.724 | 0.057 | -0.09 | -0.09 | 0 | 0.17 | 0.45 | 0.15 | 0.03 | 0.17 | 0.17 |
| 1 | 107488850 | rs11185023 | 0.339 | -0.262 | -0.32 | -1.125 | -0.018 | -0.036 | -0.036 | 0 | 0.15 | 0.22 | 0.07 | 0.06 | 0.1 | 0.08 |
| 1 | 107489399 | rs11185024 | -0.724 | -0.475 | -0.695 | -1.46 | -0.029 | -0.061 | -0.061 | 0 | 0.26 | 0.38 | 0.15 | 0.1 | 0.18 | 0.18 |
| 1 | 107489766 | rs6699202 | -0.633 | -0.346 | -0.656 | -1.532 | -0.009 | -0.104 | -0.104 | 0.423 | 0.19 | 0.47 | 0.13 | 0.03 | 0.13 | 0.16 |
| 1 | 107489844 | rs6684256 | -0.647 | -0.187 | -0.69 | -1.37 | -0.008 | -0.036 | -0.036 | 0.149 | 0.86 | 0.78 | 0.94 | 0.95 | 0.89 | 0.92 |

| | | | | | | | | | | | | | | | | |
|---|---|---|---|---|---|---|---|---|---|---|---|---|---|---|---|---|
| 1 | 107494264 | rs4641328 | -0.235 | 0.66 | -0.783 | -1.056 | 0.053 | -0.018 | -0.018 | 0 | 0.3 | 0.46 | 0.26 | 0.26 | 0.41 | 0.37 |
| 1 | 107497665 | rs6582999 | -0.521 | 0.38 | -0.751 | -0.833 | -0.002 | -0.033 | -0.033 | 0 | 0.86 | 0.79 | 0.93 | 0.95 | 0.87 | 0.93 |
| 1 | 107501181 | rs17495607 | -0.281 | 0.21 | -1.006 | -0.581 | -0.001 | 0.052 | 0.052 | 0 | 0.1 | 0.01 | 0.05 | 0.02 | 0.07 | 0.1 |
| 1 | 107504932 | rs6583000 | -1.02 | 0.579 | -0.862 | -0.48 | -0.003 | 0.017 | 0.017 | 0 | 0.15 | 0.11 | 0.07 | 0.15 | 0.15 | 0.21 |
| 1 | 107506014 | rs9435334 | 0.82 | 0.545 | -0.761 | -0.551 | -0.01 | 0.044 | 0.044 | 2.437 | 0.62 | 0.28 | 0.72 | 0.68 | 0.62 | 0.62 |
| 1 | 107516208 | rs17443102 | 0.646 | 0.503 | 0.13 | -0.554 | 0.019 | -0.016 | -0.016 | 0.01 | 0.2 | 0.27 | 0.17 | 0.17 | 0.22 | 0.16 |
| 1 | 107525991 | rs9435337 | -0.47 | 0.449 | 0.479 | -0.356 | 0.011 | -0.017 | -0.017 | 7.239 | 0.77 | 0.53 | 0.76 | 0.77 | 0.77 | 0.8 |
| 1 | 107526857 | rs9435429 | 1.065 | 0.356 | 0.443 | -0.423 | -0.031 | 0.055 | 0.055 | 2.533 | 0.6 | 0.16 | 0.66 | 0.63 | 0.62 | 0.58 |
| 1 | 107527020 | rs9435430 | -0.743 | 0.394 | 0.487 | -0.447 | 0.02 | -0.036 | -0.036 | 1.108 | 0.78 | 0.63 | 0.82 | 0.83 | 0.77 | 0.84 |
| 1 | 107527544 | rs4118326 | -0.743 | 0.394 | 0.487 | -0.447 | 0.02 | -0.036 | -0.036 | 0 | 0.78 | 0.63 | 0.82 | 0.83 | 0.77 | 0.84 |
| 1 | 107528432 | rs12093106 | -0.493 | 0.377 | 0.137 | -0.386 | 0.001 | -0.011 | -0.011 | 0.001 | 0.15 | 0.25 | 0.09 | 0.15 | 0.15 | 0.22 |
| 1 | 107538505 | rs2141551 | 1.006 | 0.093 | -0.771 | -0.1 | -0.07 | -0.143 | -0.143 | 0 | 0.33 | 0.07 | 0.52 | 0.42 | 0.35 | 0.27 |
| 1 | 107543055 | rs11185037 | 1.219 | 0.261 | -0.733 | -0.244 | -0.019 | 0.016 | 0.016 | 0.529 | 0.19 | 0.07 | 0.15 | 0.16 | 0.22 | 0.2 |
| 1 | 107544176 | rs3861909 | -0.693 | 0.098 | -0.645 | -0.238 | -0.012 | 0.01 | 0.01 | 0 | 0.7 | 0.8 | 0.79 | 0.7 | 0.68 | 0.68 |
| 1 | 107545615 | rs12738156 | 1.273 | 0.337 | -1.083 | 0.244 | -0.13 | 0.028 | 0.028 | 0.673 | 0.32 | 0.06 | 0.32 | 0.18 | 0.38 | 0.28 |
| 1 | 107547919 | rs10494056 | 0.906 | 0.061 | -0.303 | 0.002 | -0.022 | 0.019 | 0.019 | 0.823 | 0.2 | 0.08 | 0.15 | 0.16 | 0.22 | 0.2 |
| 1 | 107548745 | rs2863991 | 0.688 | 0.13 | -0.077 | 0.191 | -0.012 | 0.003 | 0.003 | 0.016 | 0.42 | 0.31 | 0.6 | 0.54 | 0.45 | 0.48 |
| 1 | 107553330 | rs6685716 | 0.58 | 0.201 | 0.123 | 0.225 | 0 | 0 | 0 | 0 | 0.43 | 0.49 | 0.6 | 0.54 | 0.45 | 0.48 |
| 1 | 107558304 | rs12032059 | 0.101 | 0.249 | -0.149 | 0.175 | -0.001 | -0.003 | -0.003 | 0 | 0.23 | 0.3 | 0.17 | 0.16 | 0.23 | 0.2 |
| 1 | 107561546 | rs10881455 | -0.007 | 0.175 | -0.18 | 0.169 | 0 | 0.001 | 0.001 | 2.207 | 0.24 | 0.28 | 0.16 | 0.16 | 0.21 | 0.22 |
| 1 | 107572997 | rs3108680 | -0.363 | 0.162 | 0.156 | 0.142 | -0.005 | -0.027 | -0.027 | 0 | 0.37 | 0.52 | 0.23 | 0.3 | 0.33 | 0.35 |
| 1 | 107576399 | rs10494057 | 0.727 | 0.226 | 0.179 | 0.171 | -0.002 | 0.011 | 0.011 | 0.648 | 0.2 | 0.15 | 0.15 | 0.16 | 0.21 | 0.2 |
| 1 | 107577832 | rs4118325 | 0.501 | 0.194 | 0.466 | 0.156 | -0.001 | -0.006 | -0.006 | 0 | 0.22 | 0.26 | 0.15 | 0.16 | 0.21 | 0.22 |
| 1 | 107579295 | rs17018185 | 0.497 | 0.177 | 0.581 | 0.123 | 0.002 | -0.022 | -0.022 | 0 | 0.22 | 0.28 | 0.19 | 0.16 | 0.23 | 0.22 |
| 1 | 107591693 | rs7543089 | 0.676 | 0.187 | 0.598 | 0.126 | 0.003 | -0.025 | -0.025 | 0 | 0.21 | 0.28 | 0.17 | 0.15 | 0.21 | 0.2 |
| 1 | 107596964 | rs11185057 | 0.739 | 0.188 | 0.488 | 0.126 | -0.002 | 0.013 | 0.013 | 0.648 | 0.2 | 0.16 | 0.15 | 0.15 | 0.21 | 0.2 |
| 1 | 107599002 | rs1762482 | -0.124 | 0.033 | 0.868 | 0.027 | -0.001 | -0.001 | -0.001 | 0 | 0.66 | 0.65 | 0.78 | 0.7 | 0.68 | 0.65 |
| 1 | 107599918 | rs2232016 | 0.921 | 0.014 | 0.531 | -0.116 | -0.074 | -0.015 | -0.015 | 0.04 | 0.19 | 0.01 | 0.09 | 0.14 | 0.19 | 0.22 |
| 1 | 107600869 | rs3791185 | 1.172 | 0.023 | 0.67 | -0.064 | -0.061 | -0.006 | -0.006 | 0.533 | 0.18 | 0.01 | 0.14 | 0.15 | 0.21 | 0.2 |
| 1 | 107610540 | rs10785814 | 0.829 | 0.019 | 0.778 | -0.075 | 0.001 | -0.007 | -0.007 | 1.359 | 0.2 | 0.2 | 0.15 | 0.15 | 0.21 | 0.2 |
| 1 | 107617707 | rs1730859 | -0.301 | 0.004 | 0.36 | -0.092 | -0.074 | 0.001 | 0.001 | 0 | 0.7 | 0.96 | 0.81 | 0.71 | 0.68 | 0.68 |
| 1 | 107623224 | rs6684571 | 0.815 | 0.06 | 0.367 | -0.017 | -0.002 | 0.012 | 0.012 | 0.582 | 0.2 | 0.15 | 0.15 | 0.15 | 0.21 | 0.21 |
| 1 | 107625615 | rs6668299 | 0.867 | 0 | 0.307 | -0.019 | -0.016 | 0.018 | 0.018 | 0.536 | 0.43 | 0.23 | 0.61 | 0.55 | 0.47 | 0.43 |
| 1 | 107634116 | rs1762495 | -0.445 | 0.238 | 1.059 | 0 | -0.002 | -0.049 | -0.049 | 0 | 0.79 | 0.77 | 0.85 | 0.97 | 0.79 | 0.79 |
| 1 | 107637882 | rs4537589 | -0.564 | 0.174 | 1.007 | 0.442 | -0.003 | 0.038 | 0.038 | 0 | 0.15 | 0.18 | 0.33 | 0.38 | 0.15 | 0.26 |
| 1 | 107642605 | rs10494059 | -0.563 | 0.15 | 1.149 | 0.431 | -0.002 | 0.031 | 0.031 | 0.017 | 0.15 | 0.17 | 0.32 | 0.38 | 0.15 | 0.26 |
| 1 | 107645245 | rs1881822 | 1.413 | 0.073 | 1.118 | 0.367 | 0.015 | -0.099 | -0.099 | 0.241 | 0.6 | 0.09 | 0.48 | 0.58 | 0.64 | 0.53 |
| 1 | 107646623 | rs1572282 | -0.666 | 0.058 | 1.37 | 0.358 | -0.002 | -0.058 | -0.058 | 0 | 0.8 | 0.72 | 0.84 | 0.96 | 0.8 | 0.8 |
| 1 | 107648423 | rs1632771 | -0.759 | 0.062 | 1.466 | 0.36 | 0 | 0.018 | 0.018 | 1.359 | 0.81 | 0.87 | 0.86 | 0.96 | 0.8 | 0.8 |
| 1 | 107656356 | rs1762506 | -1.709 | 0.12 | 1.748 | 0.367 | 0.029 | -0.021 | -0.021 | 0 | 0.39 | 0.72 | 0.51 | 0.42 | 0.35 | 0.48 |
| 1 | 107658862 | rs1335051 | 0.262 | 0.241 | 1.408 | 0.436 | -0.002 | 0.026 | 0.026 | 0.008 | 0.85 | 0.84 | 0.67 | 0.62 | 0.84 | 0.73 |
| 1 | 107660600 | rs1572281 | -0.04 | 0.242 | 1.361 | 0.269 | 0 | 0.001 | 0.001 | 0 | 0.87 | 0.87 | 0.81 | 0.83 | 0.84 | 0.76 |
| 1 | 107666980 | rs6656996 | 0.695 | 0.056 | 1.351 | -0.036 | -0.004 | 0.033 | 0.033 | 1.103 | 0.19 | 0.1 | 0.14 | 0.05 | 0.2 | 0.2 |
| 1 | 107676684 | rs6583007 | -0.871 | 0.091 | 1.445 | -0.226 | -0.015 | -0.127 | -0.127 | 0 | 0.29 | 0.5 | 0.22 | 0.04 | 0.28 | 0.27 |
| 1 | 107678734 | rs11185063 | -0.892 | 0.09 | 1.445 | -0.226 | -0.016 | -0.127 | -0.127 | 0 | 0.29 | 0.5 | 0.22 | 0.04 | 0.28 | 0.26 |
| 1 | 107682004 | rs6698801 | -0.902 | 0.09 | 1.503 | -0.225 | -0.015 | -0.123 | -0.123 | 0 | 0.29 | 0.46 | 0.23 | 0.05 | 0.28 | 0.26 |
| 1 | 107689172 | rs11185064 | -1.938 | 0.242 | 1.216 | -0.036 | 0.028 | 0.056 | 0.056 | 0.069 | 0.44 | 0.75 | 0.35 | 0.31 | 0.41 | 0.5 |
| 1 | 107697660 | rs17505643 | -1.051 | 0.344 | 0.749 | 0.029 | 0.005 | -0.005 | -0.005 | 0.234 | 0.22 | 0.13 | 0.19 | 0.27 | 0.23 | 0.32 |
| 1 | 107701106 | rs12743478 | 1.118 | 0.25 | 0.746 | 0.235 | -0.024 | 0.036 | 0.036 | 3.461 | 0.14 | 0.02 | 0.11 | 0.04 | 0.16 | 0.16 |
| 1 | 107703873 | rs4364907 | -1.782 | 0.52 | 0.966 | 0.229 | 0.001 | 0.004 | 0.004 | 0.247 | 0.24 | 0.47 | 0.21 | 0.27 | 0.26 | 0.3 |
| 1 | 107704952 | rs7513206 | -1.295 | 0.571 | 0.384 | 0.254 | 0.008 | 0.01 | 0.01 | 0.313 | 0.21 | 0.31 | 0.19 | 0.27 | 0.26 | 0.3 |
| 1 | 107706141 | rs7553752 | -1.638 | 0.424 | -0.673 | 0.194 | -0.027 | 0.086 | 0.086 | 0.725 | 0.46 | 0.89 | 0.37 | 0.31 | 0.45 | 0.53 |
| 1 | 107707343 | rs4462177 | -0.499 | 0.78 | -0.28 | 0.748 | -0.008 | -0.046 | -0.046 | 0.114 | 0.11 | 0.04 | 0.23 | 0.3 | 0.13 | 0.12 |
| 1 | 107707384 | rs10082001 | -1.163 | 0.838 | 0.313 | 0.522 | 0.017 | 0.026 | 0.026 | 0 | 0.29 | 0.37 | 0.43 | 0.4 | 0.36 | 0.43 |
| 1 | 107709720 | rs9628993 | 0.312 | 0.441 | 0.176 | 0.033 | -0.021 | -0.009 | -0.009 | 1.202 | 0.74 | 0.5 | 0.68 | 0.8 | 0.79 | 0.71 |
| 1 | 107710378 | rs10494061 | -0.709 | 1.201 | -0.088 | 0.467 | -0.109 | -0.083 | -0.083 | 0 | 0.33 | 0.11 | 0.45 | 0.57 | 0.48 | 0.47 |
| 1 | 107714478 | rs6659943 | 0.276 | 0.779 | 0.972 | 0.393 | 0.01 | 0.063 | 0.063 | 0 | 0.87 | 0.72 | 0.8 | 0.8 | 0.87 | 0.8 |
| 1 | 107714954 | rs4287207 | 0.568 | 0.731 | 0.923 | 0.392 | 0.016 | 0.059 | 0.059 | 0 | 0.84 | 0.61 | 0.76 | 0.8 | 0.84 | 0.76 |
| 1 | 107716642 | rs4615866 | 2.013 | 0.782 | 0.409 | 0.402 | -0.016 | -0.012 | -0.012 | 0 | 0.63 | 0.08 | 0.51 | 0.66 | 0.65 | 0.45 |
| 1 | 107726492 | rs10881456 | 1.573 | 0.938 | 1.361 | 0.715 | 0.003 | 0.052 | 0.052 | 0.358 | 0.82 | 0.53 | 0.76 | 0.8 | 0.84 | 0.77 |
| 1 | 107729118 | rs12025396 | 2.462 | 0.843 | 1.557 | 0.366 | -0.002 | -0.075 | -0.075 | 3.948 | 0.86 | 0.36 | 0.9 | 0.99 | 0.9 | 0.95 |
| 1 | 107743984 | rs4915025 | 2.224 | 0.807 | 2.232 | 0.455 | 0.008 | 0.02 | 0.02 | 4.174 | 0.94 | 0.63 | 0.96 | 0.99 | 1 | 0.98 |
| 1 | 107748904 | rs4915029 | 2.734 | 0.773 | 2.571 | 0.428 | -0.001 | -0.04 | -0.04 | 0 | 0.89 | 0.4 | 0.92 | 0.99 | 0.95 | 0.96 |
| 1 | 107749627 | rs6679858 | -2.353 | 0.854 | 2.293 | 0.459 | 0.013 | -0.029 | -0.029 | 2.163 | 0.07 | 0.26 | 0.05 | 0 | 0.04 | 0.04 |
| 1 | 107775378 | rs5012609 | 3.133 | 0.911 | 2.378 | 0.63 | -0.001 | -0.018 | -0.018 | 0.694 | 0.86 | 0.3 | 0.89 | 0.91 | 0.88 | 0.94 |
| 1 | 107776461 | rs11185072 | -1.621 | 0.924 | 2.658 | 0.637 | 0.128 | 0.157 | 0.157 | 0.206 | 0.06 | 0.43 | 0.06 | 0.08 | 0.05 | 0.02 |
| 1 | 107786234 | rs11185074 | 3.037 | 0.91 | 2.405 | 0.648 | 0.002 | -0.014 | -0.014 | 0 | 0.86 | 0.33 | 0.89 | 0.91 | 0.88 | 0.94 |
| 1 | 107814895 | rs3125677 | -2.403 | 0.842 | 2.304 | 0.625 | 0.036 | -0.01 | -0.01 | 0 | 0.09 | 0.56 | 0.07 | 0.01 | 0.05 | 0.04 |
| 1 | 107816751 | rs4244126 | -1.241 | 1.109 | 2.046 | 1.368 | 0.005 | 0.065 | 0.065 | 0 | 0.13 | 0.15 | 0.3 | 0.4 | 0.14 | 0.35 |

| | | | | | | | | | | | | | | | | |
|---|---|---|---|---|---|---|---|---|---|---|---|---|---|---|---|---|
| 1 | 107817550 | rs12133394 | -2.323 | 1.1 | 1.975 | 1.255 | 0.011 | -0.041 | -0.041 | 5.845 | 0.11 | 0.3 | 0.06 | 0.02 | 0.1 | 0.08 |
| 1 | 107822185 | rs3125673 | -2.073 | 1.133 | 0.202 | 0.368 | -0.063 | 0.176 | 0.176 | 0 | 0.65 | 0.92 | 0.82 | 0.9 | 0.64 | 0.81 |
| 1 | 107823943 | rs7517024 | -0.151 | 1.036 | 0.335 | 0.478 | -0.004 | -0.05 | -0.05 | 0 | 0.4 | 0.17 | 0.42 | 0.48 | 0.4 | 0.38 |
| 1 | 107825866 | rs3123376 | -0.938 | 1.028 | -0.109 | 0.64 | -0.003 | -0.006 | -0.006 | 0 | 0.51 | 0.7 | 0.51 | 0.51 | 0.5 | 0.46 |
| 1 | 107837159 | rs4604720 | 1.121 | 1.742 | -0.871 | 0.774 | -0.085 | 0.111 | 0.111 | 0 | 0.46 | 0.07 | 0.41 | 0.23 | 0.51 | 0.39 |
| 1 | 107837591 | rs12404866 | -0.584 | 2.114 | -0.88 | 0.753 | -0.028 | -0.006 | -0.006 | 0 | 0.13 | 0.07 | 0.24 | 0.13 | 0.19 | 0.21 |
| 1 | 107842708 | rs11185082 | -0.551 | 1.543 | 0.023 | 1.022 | 0.003 | -0.004 | -0.004 | 0 | 0.43 | 0.37 | 0.42 | 0.49 | 0.41 | 0.36 |
| 1 | 107846098 | rs11185083 | -1.18 | 1.321 | 1.04 | 1.138 | 0.029 | 0.137 | 0.137 | 0 | 0.06 | 0.2 | 0.13 | 0.2 | 0.07 | 0.2 |
| 1 | 107848612 | rs12566028 | -1.212 | 1.281 | 1.039 | 0.854 | 0.011 | 0.057 | 0.057 | 0.616 | 0.22 | 0.31 | 0.33 | 0.34 | 0.2 | 0.39 |
| 1 | 107849040 | rs4244127 | -0.143 | 1.294 | 0.772 | 0.698 | -0.001 | 0.014 | 0.014 | 0 | 0.1 | 0.07 | 0.06 | 0.08 | 0.09 | 0.08 |
| 1 | 107855013 | rs17509160 | 0.554 | 0.928 | 0.443 | -0.156 | 0.024 | -0.012 | -0.012 | 0 | 0.13 | 0.01 | 0.21 | 0.1 | 0.07 | 0.21 |
| 1 | 107857623 | rs4244129 | -0.384 | 1.786 | 1.128 | 0.245 | -0.001 | 0.017 | 0.017 | 0 | 0.1 | 0.06 | 0.06 | 0.06 | 0.09 | 0.08 |
| 1 | 107862053 | rs7542677 | -0.36 | 1.641 | 1.191 | -0.017 | -0.001 | 0.017 | 0.017 | 0 | 0.1 | 0.06 | 0.06 | 0.07 | 0.09 | 0.08 |
| 1 | 107863638 | rs11185087 | 1.19 | 1.568 | 1.783 | -0.051 | -0.008 | -0.067 | -0.067 | 0 | 0.85 | 0.57 | 0.9 | 0.92 | 0.91 | 0.92 |
| 1 | 107865978 | rs7411507 | 2.223 | 1.728 | 1.152 | -0.075 | -0.037 | -0.059 | -0.059 | 0 | 0.58 | 0.12 | 0.54 | 0.58 | 0.67 | 0.51 |
| 1 | 107868724 | rs11185090 | -1.865 | 1.545 | 1.677 | 0.132 | 0.015 | 0.047 | 0.047 | 0.085 | 0.27 | 0.38 | 0.36 | 0.34 | 0.24 | 0.4 |
| 1 | 107872446 | rs6583021 | 0.951 | 1.652 | 0.76 | 0.132 | -0.015 | 0.121 | 0.121 | 6.197 | 0.82 | 0.43 | 0.75 | 0.66 | 0.91 | 0.79 |
| 1 | 107872936 | rs7555412 | 1.184 | 1.562 | -0.02 | 0.022 | -0.014 | -0.052 | -0.052 | 0.504 | 0.7 | 0.27 | 0.61 | 0.6 | 0.74 | 0.58 |
| 1 | 107883043 | rs12066815 | -0.871 | 0.938 | 0.159 | -1.266 | -0.01 | -0.053 | -0.053 | 0 | 0.3 | 0.33 | 0.3 | 0.15 | 0.21 | 0.21 |
| 1 | 107889648 | rs7555385 | 0.978 | 1.572 | -0.256 | -1.289 | 0.001 | -0.027 | -0.027 | 0 | 0.59 | 0.51 | 0.56 | 0.72 | 0.64 | 0.51 |
| 1 | 107895076 | rs12029417 | -0.211 | 1.192 | -0.754 | -1.199 | 0.023 | 0.019 | 0.019 | 5.15 | 0.2 | 0.13 | 0.22 | 0.14 | 0.06 | 0.15 |
| 1 | 107898606 | rs17509795 | -0.805 | 3.103 | -0.448 | -0.584 | -0.004 | 0.001 | 0.001 | 0 | 0.09 | 0.07 | 0.07 | 0.01 | 0.18 | 0.08 |
| 1 | 107909878 | rs6664221 | -1.177 | 1.798 | -0.301 | -0.124 | 0.033 | -0.055 | -0.055 | 0 | 0.14 | 0.18 | 0.09 | 0.02 | 0.16 | 0.06 |
| 1 | 107919366 | rs6662274 | -0.738 | 1.291 | -0.589 | -0.032 | 0 | -0.011 | -0.011 | 5.465 | 0.34 | 0.4 | 0.45 | 0.78 | 0.34 | 0.48 |
| 1 | 107919409 | rs6685989 | 0.473 | -0.096 | -1.006 | -0.072 | 0.028 | 0.006 | 0.006 | 1.041 | 0.8 | 0.82 | 0.91 | 0.86 | 0.9 | 0.92 |
| 1 | 107919425 | rs6677537 | 0.347 | 0.356 | -0.916 | 0.176 | -0.001 | 0.005 | 0.005 | 0 | 0.66 | 0.43 | 0.5 | 0.62 | 0.68 | 0.75 |
| 1 | 107923094 | rs12724090 | -0.604 | 1.185 | -1.171 | 0.55 | -0.054 | 0.011 | 0.011 | 0.541 | 0.12 | 0.01 | 0.09 | 0.05 | 0.18 | 0.08 |
| 1 | 107925625 | rs7541885 | 0.33 | 1.592 | -1.016 | 0.881 | 0.008 | 0.024 | 0.024 | 0.037 | 0.79 | 0.4 | 0.71 | 0.76 | 0.78 | 0.72 |
| 1 | 107927163 | rs12048014 | 0.313 | 2.162 | -0.842 | 0.984 | 0.047 | 0.057 | 0.057 | 0 | 0.79 | 0.45 | 0.69 | 0.67 | 0.73 | 0.71 |
| 1 | 107927932 | rs17463719 | -0.424 | 2.133 | -1.363 | 0.968 | -0.012 | -0.003 | -0.003 | 0.583 | 0.13 | 0.09 | 0.1 | 0.13 | 0.18 | 0.1 |
| 1 | 107930365 | rs6681988 | 1.035 | 1.867 | -1.534 | 0.904 | -0.071 | -0.059 | -0.059 | 0.044 | 0.63 | 0.19 | 0.53 | 0.55 | 0.55 | 0.6 |
| 1 | 107931160 | rs746603 | -0.685 | 1.5 | -1.855 | 0.599 | -0.116 | -0.071 | -0.071 | 0 | 0.26 | 0.02 | 0.36 | 0.24 | 0.31 | 0.34 |
| 1 | 107934646 | rs10494069 | -0.235 | 1.026 | -1.106 | 0.506 | -0.011 | 0.042 | 0.042 | 0 | 0.46 | 0.32 | 0.51 | 0.34 | 0.5 | 0.43 |
| 1 | 107935133 | rs17014199 | -0.302 | 0.92 | -0.456 | 0.432 | -0.002 | 0.026 | 0.026 | 0 | 0.47 | 0.41 | 0.52 | 0.34 | 0.5 | 0.43 |
| 1 | 107936393 | rs2587895 | -0.43 | 0.802 | 0.056 | 1.032 | 0 | 0 | 0 | 0.427 | 0.48 | 0.51 | 0.54 | 0.52 | 0.5 | 0.48 |
| 1 | 107941708 | rs17018870 | 0.966 | 0.692 | 0.078 | 1.186 | -0.002 | -0.012 | -0.012 | 0.211 | 0.15 | 0.12 | 0.11 | 0.22 | 0.16 | 0.2 |
| 1 | 107946802 | rs920835 | 1.013 | 0.72 | 0.031 | 0.751 | -0.033 | -0.034 | -0.034 | 0.296 | 0.25 | 0.1 | 0.27 | 0.28 | 0.3 | 0.26 |
| 1 | 107953826 | rs10748505 | 0.502 | 0.565 | 0.601 | 0.699 | -0.001 | -0.004 | -0.004 | 0 | 0.49 | 0.25 | 0.45 | 0.48 | 0.5 | 0.47 |
| 1 | 107958969 | rs12078289 | -0.355 | 0.506 | 0.792 | 0.604 | -0.005 | 0.017 | 0.017 | 0 | 0.43 | 0.36 | 0.51 | 0.3 | 0.47 | 0.39 |
| 1 | 107960860 | rs17018982 | 0.768 | 0.481 | 0.786 | 0.561 | -0.003 | 0.023 | 0.023 | 0 | 0.17 | 0.13 | 0.22 | 0.02 | 0.17 | 0.12 |
| 1 | 107962328 | rs11185111 | 1.223 | 0.527 | 0.745 | 0.546 | -0.01 | -0.037 | -0.037 | 0 | 0.26 | 0.17 | 0.28 | 0.45 | 0.3 | 0.32 |
| 1 | 107964274 | rs11185112 | 0.798 | 0.463 | 0.96 | 0.5 | -0.009 | -0.011 | -0.011 | 0 | 0.48 | 0.22 | 0.44 | 0.48 | 0.5 | 0.47 |
| 1 | 107966962 | rs583169 | 0.538 | 0.414 | 1.073 | 0.503 | -0.007 | -0.006 | -0.006 | 0 | 0.79 | 0.51 | 0.75 | 0.8 | 0.81 | 0.79 |
| 1 | 107981316 | rs1444038 | -0.321 | 0.431 | 0.889 | 0.54 | 0 | -0.001 | -0.001 | 6.431 | 0.17 | 0.2 | 0.22 | 0.22 | 0.17 | 0.17 |
| 1 | 107982992 | rs11185115 | 1.209 | 0.451 | 0.934 | 0.651 | -0.007 | -0.009 | -0.009 | 0.142 | 0.25 | 0.21 | 0.29 | 0.28 | 0.31 | 0.26 |
| 1 | 107990065 | rs640554 | 0.586 | 0.485 | 0.566 | 0.209 | 0.006 | -0.01 | -0.01 | 0 | 0.44 | 0.33 | 0.43 | 0.5 | 0.4 | 0.47 |
| 1 | 107994248 | rs17532072 | 1.368 | 0.521 | 0.474 | 0.183 | -0.015 | 0.031 | 0.031 | 0 | 0.1 | 0.03 | 0.04 | 0 | 0.11 | 0.06 |
| 1 | 107995842 | rs12117762 | 0.971 | 0.48 | 0.469 | 0.257 | -0.001 | -0.026 | -0.026 | 0 | 0.39 | 0.2 | 0.41 | 0.46 | 0.39 | 0.42 |
| 1 | 107997106 | rs628117 | -0.409 | 0.575 | 0.507 | 0.226 | 0.011 | -0.012 | -0.012 | 0.01 | 0.62 | 0.54 | 0.64 | 0.7 | 0.55 | 0.67 |
| 1 | 108001448 | rs521721 | 0.308 | 0.416 | 0.432 | 0.282 | 0.009 | -0.005 | -0.005 | 0 | 0.45 | 0.36 | 0.29 | 0.5 | 0.38 | 0.52 |
| 1 | 108010639 | rs10881468 | 1.092 | 0.39 | 0.007 | 0.366 | 0.008 | -0.029 | -0.029 | 0.005 | 0.35 | 0.12 | 0.26 | 0.37 | 0.32 | 0.43 |
| 1 | 108011220 | rs12354155 | 1.164 | 0.649 | -0.072 | 0.235 | -0.043 | 0.059 | 0.059 | 0 | 0.1 | 0.01 | 0.06 | 0 | 0.14 | 0.07 |
| 1 | 108014380 | rs7522375 | 0.199 | 0.484 | 0.35 | 0.342 | -0.001 | -0.001 | -0.001 | 0.216 | 0.53 | 0.42 | 0.61 | 0.43 | 0.5 | 0.55 |
| 1 | 108016292 | rs3813218 | 0.179 | 0.528 | -0.053 | 0.356 | -0.036 | 0.025 | 0.025 | 0 | 0.41 | 0.21 | 0.34 | 0.32 | 0.53 | 0.36 |
| 1 | 108016331 | rs3813219 | -1.181 | 0.516 | 0.418 | 0.241 | 0.001 | -0.054 | -0.054 | 0.03 | 0.2 | 0.2 | 0.09 | 0.07 | 0.18 | 0.16 |
| 1 | 108017559 | rs621222 | -0.976 | 0.626 | 0.312 | 0.688 | 0.008 | 0.01 | 0.01 | 0 | 0.26 | 0.39 | 0.43 | 0.26 | 0.28 | 0.2 |
| 1 | 108018552 | rs606373 | 0.273 | 0.469 | -0.211 | 0.503 | -0.001 | 0.003 | 0.003 | 0.127 | 0.59 | 0.47 | 0.64 | 0.6 | 0.56 | 0.56 |
| 1 | 108018964 | rs594165 | 0.88 | -0.021 | 0.43 | 0.361 | -0.03 | -0.02 | -0.02 | 0 | 0.79 | 0.62 | 0.66 | 0.89 | 0.84 | 0.82 |
| 1 | 108020197 | rs507203 | 0.038 | -0.039 | 0.453 | 0.304 | -0.029 | -0.049 | -0.049 | 0 | 0.91 | 0.8 | 0.96 | 1 | 0.97 | 0.96 |
| 1 | 108020853 | rs512719 | 0.006 | -0.056 | 0.408 | 0.295 | -0.032 | -0.058 | -0.058 | 1.893 | 0.9 | 0.7 | 0.95 | 1 | 0.97 | 0.96 |
| 1 | 108023589 | rs4915045 | 0.123 | 0.043 | -0.096 | 0.594 | 0.006 | -0.026 | -0.026 | 0 | 0.11 | 0.06 | 0.09 | 0.17 | 0.08 | 0.07 |
| 1 | 108024868 | rs597332 | 0.133 | 0.734 | -0.283 | -0.041 | -0.037 | -0.027 | -0.027 | 0 | 0.34 | 0.24 | 0.37 | 0.57 | 0.39 | 0.33 |
| 1 | 108028555 | rs17019212 | -0.091 | 0.504 | -0.864 | 0.385 | 0.002 | 0.019 | 0.019 | 0 | 0.12 | 0.17 | 0.31 | 0.11 | 0.12 | 0.14 |
| 1 | 108031468 | rs17019224 | -0.245 | 0 | -1.231 | 0.377 | -0.003 | -0.031 | -0.031 | 0 | 0.06 | 0.15 | 0.03 | 0 | 0.06 | 0.08 |
| 1 | 108037997 | rs12756322 | 0.349 | 0.062 | -1.75 | 0.479 | 0.015 | 0.001 | 0.001 | 1.4 | 0.33 | 0.04 | 0.17 | 0.31 | 0.27 | 0.31 |
| 1 | 108040596 | rs1006799 | -0.761 | 0.622 | -0.917 | 0.778 | 0.001 | 0.009 | 0.009 | 0 | 0.22 | 0.21 | 0.42 | 0.29 | 0.24 | 0.19 |
| 1 | 108045438 | rs12089394 | 0.532 | 0.436 | -1.513 | 0.323 | -0.029 | 0.09 | 0.09 | 0 | 0.37 | 0.22 | 0.32 | 0.19 | 0.44 | 0.33 |
| 1 | 108055112 | rs12092544 | 0.477 | 0.312 | -1.365 | 0.653 | -0.016 | 0.028 | 0.028 | 0 | 0.12 | 0.17 | 0.26 | 0.17 | 0.08 | 0.16 |
| 1 | 108061647 | rs11185125 | 0.453 | 0.859 | -1.886 | 0.714 | -0.018 | 0.052 | 0.052 | 0 | 0.35 | 0.23 | 0.31 | 0.19 | 0.41 | 0.31 |

| | | | | | | | | | | | | | | | | |
|---|---|---|---|---|---|---|---|---|---|---|---|---|---|---|---|---|
| 1 | 108067232 | rs1887631 | -0.432 | 1.941 | -1.792 | 1.553 | -0.08 | -0.059 | -0.059 | 0.108 | 0.14 | 0.01 | 0.18 | 0.17 | 0.2 | 0.1 |
| 1 | 108070435 | rs11185129 | -0.619 | 1.303 | -1.726 | 1.136 | -0.082 | -0.074 | -0.074 | 0 | 0.3 | 0.12 | 0.29 | 0.56 | 0.38 | 0.33 |
| 1 | 108074305 | rs525007 | -0.5 | 1.048 | -0.926 | 1.227 | 0.014 | 0.069 | 0.069 | 0 | 0.29 | 0.45 | 0.28 | 0.49 | 0.32 | 0.24 |
| 1 | 108074445 | rs12406309 | 0.183 | 0.99 | -1.752 | 1.199 | -0.029 | -0.098 | -0.098 | 0 | 0.17 | 0.04 | 0.22 | 0.31 | 0.19 | 0.18 |
| 1 | 108075536 | rs569454 | -0.056 | 0.843 | -1.976 | 0.791 | -0.02 | -0.061 | -0.061 | 0.141 | 0.7 | 0.88 | 0.7 | 0.68 | 0.69 | 0.61 |
| 1 | 108077102 | rs12739227 | 0.389 | 1.161 | -1.728 | 1.262 | 0.012 | 0.037 | 0.037 | 0 | 0.09 | 0.12 | 0.13 | 0.28 | 0.17 | 0.14 |
| 1 | 108078183 | rs11185131 | 0.552 | 0.792 | -1.034 | 1.17 | 0.008 | -0.002 | -0.002 | 0.348 | 0.55 | 0.63 | 0.63 | 0.53 | 0.58 | 0.52 |
| 1 | 108079073 | rs651844 | -0.36 | 0.758 | -0.959 | 1.135 | 0.023 | 0.076 | 0.076 | 0 | 0.25 | 0.42 | 0.21 | 0.42 | 0.28 | 0.25 |
| 1 | 108079945 | rs666003 | 0.478 | 0.757 | -0.978 | 1.102 | -0.026 | 0.015 | 0.015 | 0 | 0.59 | 0.68 | 0.56 | 0.4 | 0.52 | 0.48 |
| 1 | 108083870 | rs633869 | 1.236 | 0.63 | -0.391 | 1.038 | 0.003 | 0.029 | 0.029 | 0.478 | 0.15 | 0.32 | 0.09 | 0.14 | 0.12 | 0.11 |
| 1 | 108086798 | rs531284 | -0.381 | 0.606 | -0.906 | 0.806 | -0.022 | -0.074 | -0.074 | 0.019 | 0.69 | 0.88 | 0.69 | 0.68 | 0.69 | 0.6 |
| 1 | 108090429 | rs10785826 | 0.446 | 0.547 | -0.607 | 0.295 | -0.002 | -0.019 | -0.019 | 2.87 | 0.45 | 0.47 | 0.49 | 0.26 | 0.41 | 0.37 |
| 1 | 108095617 | rs10881471 | -0.423 | 0.797 | -0.707 | 0.414 | -0.001 | -0.001 | -0.001 | 0 | 0.16 | 0.13 | 0.21 | 0.23 | 0.15 | 0.29 |
| 1 | 108113702 | rs12410842 | -1.552 | 0.242 | 0.058 | -0.032 | -0.028 | -0.071 | -0.071 | 0 | 0.87 | 0.77 | 0.95 | 1 | 0.94 | 0.94 |
| 1 | 108113865 | rs8676 | 0.974 | 0.167 | -0.373 | 0.596 | -0.004 | 0.004 | 0.004 | 0.339 | 0.59 | 0.6 | 0.48 | 0.43 | 0.64 | 0.46 |
| 1 | 108115145 | rs2769668 | -1.882 | 0.117 | -0.386 | 0.471 | -0.03 | -0.07 | -0.07 | 0 | 0.88 | 0.78 | 0.95 | 1 | 0.94 | 0.94 |
| 1 | 108116169 | rs7549255 | 0.829 | 0.136 | -0.665 | -0.026 | -0.004 | 0.001 | 0.001 | 0.241 | 0.13 | 0.1 | 0.12 | 0 | 0.14 | 0.13 |
| 1 | 108124480 | rs17477236 | 1.927 | 0.346 | -1.178 | -0.453 | -0.095 | 0.236 | 0.236 | 0 | 0.32 | 0.03 | 0.21 | 0.02 | 0.35 | 0.16 |
| 1 | 108127895 | rs1328199 | -0.088 | 0.284 | -1.15 | -0.28 | -0.024 | -0.012 | -0.012 | 0 | 0.19 | 0.09 | 0.24 | 0.28 | 0.21 | 0.32 |
| 1 | 108135527 | rs7554525 | 1.286 | 0.306 | -0.741 | -0.094 | -0.009 | -0.079 | -0.079 | 0 | 0.14 | 0.06 | 0.25 | 0.35 | 0.15 | 0.19 |
| 1 | 108137016 | rs7537311 | -0.838 | 0.29 | -0.577 | -0.199 | -0.013 | 0.077 | 0.077 | 0 | 0.43 | 0.26 | 0.4 | 0.29 | 0.4 | 0.51 |
| 1 | 108139975 | rs3790676 | -0.314 | 0.302 | -0.392 | -0.101 | 0.009 | 0.153 | 0.153 | 0 | 0.8 | 0.56 | 0.71 | 0.58 | 0.8 | 0.75 |
| 1 | 108142912 | rs12729735 | 1.027 | 0.331 | -0.568 | -0.144 | -0.004 | -0.012 | -0.012 | 0 | 0.13 | 0.04 | 0.2 | 0.17 | 0.12 | 0.26 |
| 1 | 108143671 | rs3828085 | 1.514 | 0.407 | -0.568 | -0.2 | -0.007 | -0.001 | -0.001 | 0 | 0.39 | 0.25 | 0.32 | 0.36 | 0.43 | 0.29 |
| 1 | 108148686 | rs9804074 | -0.395 | 0.57 | -0.463 | -0.307 | -0.005 | -0.003 | -0.003 | 0 | 0.05 | 0.05 | 0.04 | 0.12 | 0.09 | 0.06 |
| 1 | 108163649 | rs12049132 | 0.997 | 0.472 | -0.403 | 0.093 | -0.003 | 0.009 | 0.009 | 0 | 0.14 | 0.18 | 0.2 | 0.19 | 0.14 | 0.15 |
| 1 | 108172396 | rs6583034 | 0.69 | 0.492 | -0.361 | 0.087 | 0.106 | 0.041 | 0.041 | 0.116 | 0.94 | 0.66 | 0.95 | 0.95 | 0.95 | 0.95 |
| 1 | 108173375 | rs12408984 | 0.833 | 0.489 | -0.648 | 0.238 | -0.003 | -0.027 | -0.027 | 1.28 | 0.12 | 0.04 | 0.16 | 0.2 | 0.11 | 0.25 |
| 1 | 108181947 | rs6693140 | -0.213 | 0.5 | -0.537 | 0.274 | 0.011 | 0.049 | 0.049 | 0.386 | 0.81 | 0.52 | 0.77 | 0.76 | 0.81 | 0.8 |
| 1 | 108185309 | rs12410676 | 0.823 | 0.38 | -1.051 | 0.325 | -0.004 | -0.027 | -0.027 | 0 | 0.12 | 0.04 | 0.16 | 0.21 | 0.11 | 0.23 |
| 1 | 108185717 | rs4550085 | 0.787 | 0.25 | -0.97 | 0.178 | 0.004 | -0.009 | -0.009 | 0 | 0.15 | 0.15 | 0.19 | 0.18 | 0.14 | 0.17 |
| 1 | 108188858 | rs17019602 | 0.469 | 0.04 | -1.101 | 0.01 | 0.026 | 0.011 | 0.011 | 0 | 0.26 | 0.17 | 0.23 | 0.19 | 0.2 | 0.22 |
| 1 | 108190581 | rs4915063 | 0.121 | 0.254 | -1.671 | -0.706 | -0.051 | -0.029 | -0.029 | 0 | 0.3 | 0.14 | 0.44 | 0.63 | 0.44 | 0.49 |
| 1 | 108195332 | rs11185156 | 1.926 | 0.131 | -1.784 | -0.824 | -0.026 | 0.032 | 0.032 | 0.297 | 0.23 | 0.06 | 0.16 | 0.18 | 0.21 | 0.13 |
| 1 | 108202710 | rs7543435 | -0.271 | 0.066 | -1.711 | -0.858 | -0.034 | 0.013 | 0.013 | 0.379 | 0.59 | 0.81 | 0.52 | 0.37 | 0.51 | 0.46 |
| 1 | 108203956 | rs12130964 | 1.703 | 0.386 | -1.553 | -0.613 | 0 | 0.03 | 0.03 | 0.385 | 0.23 | 0.05 | 0.15 | 0.18 | 0.19 | 0.12 |
| 1 | 108208282 | rs4378232 | -0.251 | 0.273 | -1.292 | -0.558 | -0.04 | -0.032 | -0.032 | 0 | 0.71 | 0.84 | 0.57 | 0.39 | 0.57 | 0.55 |
| 1 | 108212980 | rs4639792 | 1.171 | 0.394 | -1.157 | -0.4 | -0.097 | -0.133 | -0.133 | 0 | 0.17 | 0.02 | 0.24 | 0.46 | 0.31 | 0.25 |
| 1 | 108213078 | rs12140390 | 1.881 | 0.149 | -1.082 | -0.678 | -0.007 | 0.001 | 0.001 | 0.483 | 0.22 | 0.05 | 0.16 | 0.2 | 0.18 | 0.13 |
| 1 | 108214333 | rs6680436 | 0.342 | 0.108 | -1.104 | -0.776 | 0.003 | -0.13 | -0.13 | 1.288 | 0.21 | 0.4 | 0.16 | 0.01 | 0.18 | 0.15 |
| 1 | 108215310 | rs17019693 | 1.303 | -0.153 | -1.268 | -0.486 | 0.033 | 0.021 | 0.021 | 0 | 0.25 | 0.12 | 0.23 | 0.15 | 0.2 | 0.28 |
| 1 | 108218276 | rs2504469 | 0.451 | -0.006 | -1.331 | -0.454 | 0.037 | 0.038 | 0.038 | 0.011 | 0.49 | 0.35 | 0.4 | 0.35 | 0.4 | 0.42 |
| 1 | 108227361 | rs2504458 | 0.176 | 0.538 | -1.082 | -0.321 | 0.005 | 0.027 | 0.027 | 0 | 0.38 | 0.34 | 0.33 | 0.16 | 0.35 | 0.39 |
| 1 | 108235719 | rs7530592 | 0.36 | 0.99 | -0.641 | 0.16 | 0.035 | 0.004 | 0.004 | 0.003 | 0.66 | 0.58 | 0.51 | 0.37 | 0.53 | 0.52 |
| 1 | 108238093 | rs2494057 | -0.84 | 0.575 | -0.366 | 0.375 | 0.022 | 0.053 | 0.053 | 0 | 0.91 | 0.79 | 0.78 | 0.82 | 0.91 | 0.84 |
| 1 | 108239924 | rs12039907 | 1.498 | 0.246 | -0.945 | 0.123 | 0.015 | -0.023 | -0.023 | 0.005 | 0.23 | 0.03 | 0.14 | 0.17 | 0.16 | 0.27 |
| 1 | 108241166 | rs12090234 | 0.614 | 0.335 | 0.068 | 0.228 | 0.042 | 0.024 | 0.024 | 1.777 | 0.05 | 0.28 | 0.03 | 0 | 0.02 | 0.01 |
| 1 | 108243851 | rs12043814 | -0.045 | 0.553 | -0.338 | 0.556 | 0.012 | 0 | 0 | 0.01 | 0.57 | 0.36 | 0.55 | 0.52 | 0.66 | 0.64 |
| 1 | 108245410 | rs2494066 | -0.119 | 0.305 | -0.222 | 0.465 | -0.011 | -0.012 | -0.012 | 0.027 | 0.38 | 0.56 | 0.26 | 0.3 | 0.27 | 0.2 |
| 1 | 108245615 | rs2494067 | -0.33 | 0.634 | -0.329 | 0.927 | -0.003 | -0.001 | -0.001 | 0 | 0.46 | 0.5 | 0.47 | 0.59 | 0.39 | 0.4 |
| 1 | 108245664 | rs17019729 | 0.109 | 0.82 | -0.856 | 1.162 | -0.019 | -0.022 | -0.022 | 0.107 | 0.08 | 0.03 | 0.22 | 0.3 | 0.12 | 0.19 |
| 1 | 108246069 | rs10218617 | -0.397 | 0.458 | -0.498 | 0.625 | -0.006 | -0.004 | -0.004 | 0 | 0.44 | 0.51 | 0.28 | 0.38 | 0.32 | 0.3 |
| 1 | 108246265 | rs11185168 | -0.022 | 0.973 | -0.456 | 0.489 | -0.002 | -0.011 | -0.011 | 0 | 0.4 | 0.42 | 0.26 | 0.3 | 0.36 | 0.2 |
| 1 | 108247048 | rs2494070 | -0.289 | 0.819 | -0.35 | 0.511 | 0.001 | -0.018 | -0.018 | 0 | 0.42 | 0.48 | 0.29 | 0.35 | 0.4 | 0.2 |
| 1 | 108251323 | rs12403629 | -0.085 | 0.599 | -0.535 | 1.068 | -0.001 | 0.015 | 0.015 | 0 | 0.14 | 0.15 | 0.4 | 0.44 | 0.15 | 0.33 |
| 1 | 108258265 | rs10494073 | 1.055 | 0.302 | -0.747 | 1.051 | 0.015 | -0.031 | -0.031 | 0 | 0.12 | 0.03 | 0.21 | 0.17 | 0.08 | 0.15 |
| 1 | 108265699 | rs2494043 | -0.353 | -0.276 | -0.844 | 0.665 | -0.007 | -0.019 | -0.019 | 0.35 | 0.9 | 0.86 | 0.94 | 1 | 0.92 | 0.91 |
| 1 | 108268630 | rs4503376 | -0.545 | -0.116 | -0.542 | 0.579 | -0.018 | -0.009 | -0.009 | 0 | 0.64 | 0.58 | 0.72 | 0.69 | 0.72 | 0.8 |
| 1 | 108281720 | rs7519428 | 2.697 | -0.048 | -1.066 | 0.218 | 0.054 | -0.004 | -0.004 | 0 | 0.26 | 0.07 | 0.14 | 0.18 | 0.17 | 0.27 |
| 1 | 108281880 | rs12405182 | 1.754 | -0.064 | -1.101 | 0.263 | 0.015 | -0.021 | -0.021 | 0 | 0.12 | 0.03 | 0.21 | 0.15 | 0.08 | 0.15 |
| 1 | 108285388 | rs17019855 | 1.988 | -0.07 | -0.959 | 0.266 | 0.022 | -0.047 | -0.047 | 0.611 | 0.31 | 0.21 | 0.33 | 0.43 | 0.24 | 0.43 |
| 1 | 108286191 | rs6662917 | 0.595 | -0.061 | -0.817 | 0.243 | -0.012 | -0.005 | -0.005 | 0 | 0.35 | 0.39 | 0.27 | 0.33 | 0.28 | 0.2 |
| 1 | 108286632 | rs11185172 | -0.905 | 0.022 | -1.048 | 0.055 | 0.004 | 0.007 | 0.007 | 0 | 0.52 | 0.57 | 0.66 | 0.73 | 0.44 | 0.52 |
| 1 | 108286971 | rs6583045 | -0.441 | 0.846 | -0.698 | -0.022 | 0.126 | -0.093 | -0.093 | 0 | 0.8 | 0.65 | 0.81 | 0.96 | 0.62 | 0.84 |
| 1 | 108288220 | rs17019880 | 0.718 | 0.27 | -0.969 | 0.114 | 0.015 | -0.019 | -0.019 | 0.639 | 0.12 | 0.03 | 0.2 | 0.15 | 0.08 | 0.15 |
| 1 | 108294496 | rs4915071 | 0.863 | -0.042 | -0.65 | 0.114 | -0.023 | 0.042 | 0.042 | 0 | 0.72 | 0.67 | 0.51 | 0.56 | 0.76 | 0.59 |
| 1 | 108296508 | rs4531312 | -0.812 | 0.374 | -0.902 | -0.121 | -0.001 | -0.004 | -0.004 | 0 | 0.44 | 0.52 | 0.5 | 0.68 | 0.58 | 0.52 |
| 1 | 108300141 | rs11808011 | -0.732 | 0.491 | -0.997 | -0.437 | -0.011 | -0.003 | -0.003 | 0 | 0.35 | 0.25 | 0.25 | 0.38 | 0.38 | 0.19 |
| 1 | 108302754 | rs17542229 | 0.503 | 0.513 | -0.81 | -0.566 | -0.014 | -0.038 | -0.038 | 0 | 0.07 | 0.15 | 0.05 | 0 | 0.06 | 0.05 |

| | | | | | | | | | | | | | | | | |
|---|---|---|---|---|---|---|---|---|---|---|---|---|---|---|---|---|
| 1 | 108302922 | rs12142335 | 1.192 | 0.605 | -0.68 | -0.577 | 0.045 | -0.036 | -0.036 | 0 | 0.13 | 0.26 | 0.12 | 0.03 | 0.25 | 0.14 |
| 1 | 108304878 | rs4415601 | -1.009 | -0.241 | -0.466 | -0.417 | -0.017 | -0.009 | -0.009 | 0 | 0.57 | 0.59 | 0.43 | 0.39 | 0.7 | 0.38 |
| 1 | 108310224 | rs7523266 | 0.398 | -0.198 | -0.489 | -0.487 | 0.007 | -0.008 | -0.008 | 0.239 | 0.22 | 0.19 | 0.16 | 0.33 | 0.19 | 0.15 |
| 1 | 108312656 | rs6690837 | -1.479 | -0.239 | -0.48 | -0.455 | -0.016 | -0.002 | -0.002 | 0 | 0.61 | 0.66 | 0.62 | 0.66 | 0.77 | 0.57 |
| 1 | 108313258 | rs6583048 | -1.709 | -0.224 | -0.464 | -0.451 | -0.004 | 0 | 0 | 0 | 0.63 | 0.69 | 0.62 | 0.68 | 0.78 | 0.57 |
| 1 | 108315137 | rs4462178 | 1.367 | -0.121 | -0.421 | -0.486 | -0.004 | -0.015 | -0.015 | 0.01 | 0.22 | 0.26 | 0.19 | 0.04 | 0.37 | 0.2 |
| 1 | 108316189 | rs6686061 | 0.357 | -0.174 | -0.44 | -0.431 | -0.003 | 0.002 | 0.002 | 0 | 0.22 | 0.24 | 0.17 | 0.33 | 0.19 | 0.15 |
| 1 | 108319079 | rs4914959 | -1.796 | -0.226 | -0.488 | -0.431 | -0.013 | -0.002 | -0.002 | 0.014 | 0.65 | 0.69 | 0.63 | 0.72 | 0.78 | 0.58 |
| 1 | 108322535 | rs7521681 | 1.404 | -0.24 | -0.473 | -0.441 | 0.005 | -0.03 | -0.03 | 0.131 | 0.16 | 0.13 | 0.14 | 0.33 | 0.15 | 0.12 |
| 1 | 108325221 | rs10881483 | -1.785 | -0.032 | -0.095 | -0.229 | 0.031 | 0.003 | 0.003 | 0 | 0.66 | 0.76 | 0.67 | 0.72 | 0.79 | 0.6 |
| 1 | 108329926 | rs11185178 | -1.609 | 0.001 | -0.094 | -0.385 | -0.009 | -0.002 | -0.002 | 2.749 | 0.65 | 0.69 | 0.64 | 0.75 | 0.79 | 0.59 |
| 1 | 108330285 | rs12116592 | -1.062 | 0.622 | -0.081 | -0.823 | -0.002 | -0.001 | -0.001 | 0 | 0.41 | 0.46 | 0.45 | 0.35 | 0.58 | 0.42 |
| 1 | 108331296 | rs11576720 | 0.776 | 0.785 | -0.367 | -0.715 | 0.005 | -0.026 | -0.026 | 0 | 0.11 | 0.1 | 0.07 | 0.01 | 0.13 | 0.03 |
| 1 | 108334434 | rs4915074 | -0.549 | 0.799 | -0.365 | -0.692 | 0.015 | 0.033 | 0.033 | 0.004 | 0.17 | 0.19 | 0.26 | 0.34 | 0.2 | 0.2 |
| 1 | 108339344 | rs17020067 | -0.554 | 0.619 | -0.395 | -0.824 | -0.003 | -0.021 | -0.021 | 3.809 | 0.35 | 0.36 | 0.26 | 0.08 | 0.51 | 0.28 |
| 1 | 108343087 | rs7537605 | -0.996 | 1.264 | -0.266 | -0.599 | 0.11 | 0.081 | 0.081 | 0 | 0.19 | 0.36 | 0.36 | 0.32 | 0.33 | 0.22 |
| 1 | 108353670 | rs17020104 | -0.764 | 0.097 | -0.035 | -0.272 | 0.017 | 0.176 | 0.176 | 0 | 0.08 | 0.21 | 0.25 | 0.31 | 0.08 | 0.15 |
| 1 | 108356000 | rs17485868 | -0.642 | -0.317 | -0.202 | -0.674 | 0 | -0.001 | -0.001 | 1.744 | 0.67 | 0.69 | 0.57 | 0.38 | 0.69 | 0.58 |
| 1 | 108356293 | rs4612651 | 0.893 | -0.308 | -0.336 | -0.795 | -0.046 | 0.04 | 0.04 | 0 | 0.21 | 0.15 | 0.18 | 0.04 | 0.37 | 0.2 |
| 1 | 108356820 | rs12126655 | 0.264 | -0.438 | -0.652 | -0.68 | 0.026 | 0.021 | 0.021 | 0 | 0.43 | 0.33 | 0.32 | 0.34 | 0.3 | 0.35 |
| 1 | 108379043 | rs4460667 | -0.015 | -0.206 | -0.607 | -0.463 | 0.008 | 0.002 | 0.002 | 0 | 0.49 | 0.49 | 0.53 | 0.47 | 0.35 | 0.49 |
| 1 | 108385791 | rs11185183 | 0.503 | 0.22 | -0.536 | -0.52 | 0.012 | 0.019 | 0.019 | 0 | 0.4 | 0.37 | 0.35 | 0.2 | 0.29 | 0.35 |
| 1 | 108386404 | rs1380445 | 0.419 | 0.232 | -0.627 | -0.515 | -0.007 | -0.029 | -0.029 | 0 | 0.41 | 0.46 | 0.35 | 0.2 | 0.29 | 0.35 |
| 1 | 108393409 | rs6583052 | -0.585 | 0.293 | -0.63 | -0.484 | 0.002 | -0.003 | -0.003 | 0.071 | 0.66 | 0.69 | 0.71 | 0.5 | 0.7 | 0.68 |
| 1 | 108402075 | rs10494080 | 1.26 | 0.061 | -0.913 | -0.422 | 0.017 | 0.121 | 0.121 | 0 | 0.32 | 0.08 | 0.25 | 0.11 | 0.24 | 0.32 |
| 1 | 108402103 | rs10494081 | -0.132 | 0.236 | -1.016 | -0.469 | -0.013 | 0.005 | 0.005 | 0 | 0.63 | 0.58 | 0.67 | 0.41 | 0.68 | 0.7 |
| 1 | 108406826 | rs6677889 | -0.329 | 0.189 | -1.075 | 0.352 | 0 | 0 | 0 | 0.219 | 0.43 | 0.47 | 0.52 | 0.47 | 0.34 | 0.52 |
| 1 | 108408983 | rs1999487 | -0.125 | -0.238 | -1.16 | 0.335 | 0.007 | 0.003 | 0.003 | 0 | 0.4 | 0.41 | 0.4 | 0.37 | 0.28 | 0.35 |
| 1 | 108414230 | rs1332684 | -0.58 | -0.217 | -1.428 | 0.446 | -0.024 | -0.036 | -0.036 | 0 | 0.07 | 0.02 | 0.04 | 0.22 | 0.08 | 0.13 |
| 1 | 108415339 | rs17020234 | -0.536 | -0.227 | -1.798 | 0.441 | -0.024 | -0.035 | -0.035 | 4.212 | 0.07 | 0.02 | 0.04 | 0.22 | 0.08 | 0.12 |
| 1 | 108417044 | rs7527438 | 0.249 | -0.651 | -2.062 | 0.24 | 0.023 | 0.017 | 0.017 | 1.363 | 0.1 | 0.02 | 0.04 | 0.03 | 0.05 | 0.03 |
| 1 | 108421058 | rs345288 | -0.723 | -1.281 | -2.613 | -0.035 | 0.104 | 0.101 | 0.101 | 0 | 0.67 | 0.94 | 0.85 | 0.84 | 0.83 | 0.85 |
| 1 | 108429934 | rs2035015 | -0.3 | -1.1 | -2.486 | -0.029 | 0.086 | 0.018 | 0.018 | 0 | 0.6 | 0.91 | 0.8 | 0.63 | 0.75 | 0.71 |
| 1 | 108430813 | rs17020274 | -0.161 | -0.107 | -1.448 | 0.342 | 0.023 | 0.016 | 0.016 | 1.295 | 0.1 | 0.02 | 0.04 | 0.03 | 0.05 | 0.03 |
| 1 | 108436557 | rs345298 | -0.947 | 0.638 | -1.52 | 0.301 | -0.027 | 0.175 | 0.175 | 1.925 | 0.54 | 0.89 | 0.63 | 0.81 | 0.53 | 0.74 |
| 1 | 108441642 | rs17020294 | 0.011 | 0.795 | -1.293 | 0.863 | -0.006 | -0.035 | -0.035 | 0 | 0.19 | 0.14 | 0.25 | 0.35 | 0.22 | 0.19 |
| 1 | 108448202 | rs10494083 | -0.572 | 0.961 | -0.183 | 0.958 | 0.021 | -0.018 | -0.018 | 0 | 0.26 | 0.42 | 0.23 | 0.23 | 0.26 | 0.32 |
| 1 | 108450627 | rs11185215 | -0.017 | 0.961 | 0.072 | 0.868 | -0.004 | -0.004 | -0.004 | 12.32 | 0.14 | 0.11 | 0.13 | 0.02 | 0.18 | 0.26 |
| 1 | 108459260 | rs7548905 | -1.248 | 0.88 | 0.264 | 0.99 | -0.034 | 0.071 | 0.071 | 0.545 | 0.18 | 0.43 | 0.13 | 0.3 | 0.13 | 0.2 |
| 1 | 108462844 | rs345314 | -0.084 | 0.886 | 0.494 | 0.953 | 0.052 | -0.067 | -0.067 | 0.044 | 0.85 | 0.73 | 0.85 | 0.98 | 0.82 | 0.74 |
| 1 | 108467386 | rs345306 | 2.035 | 0.874 | 0.162 | 1.015 | 0.005 | 0.099 | 0.099 | 0.966 | 0.5 | 0.17 | 0.48 | 0.33 | 0.49 | 0.36 |
| 1 | 108474641 | rs17020387 | -0.402 | 0.891 | 1.028 | 0.694 | -0.001 | 0.013 | 0.013 | 0 | 0.08 | 0.16 | 0.07 | 0.15 | 0.08 | 0.05 |
| 1 | 108478253 | rs10494086 | -1.07 | 0.886 | 0.953 | 0.685 | 0.022 | -0.028 | -0.028 | 0 | 0.18 | 0.32 | 0.16 | 0.08 | 0.18 | 0.27 |
| 1 | 108484117 | rs17020437 | 0 | 0.876 | 0.931 | 0.705 | 0.005 | -0.025 | -0.025 | 0 | 0.05 | 0 | 0.02 | 0.1 | 0.04 | 0.13 |
| 1 | 108484632 | rs1020812 | -1.755 | 0.883 | 0.949 | 0.7 | 0.006 | -0.005 | -0.005 | 2.16 | 0.23 | 0.33 | 0.19 | 0.18 | 0.23 | 0.4 |
| 1 | 108489708 | rs6696531 | -1.239 | 1.067 | 1.167 | 0.459 | 0.02 | -0.028 | -0.028 | 0 | 0.18 | 0.31 | 0.16 | 0.08 | 0.18 | 0.27 |
| 1 | 108498145 | rs345271 | 1.886 | 1.04 | 1.096 | 0.451 | -0.013 | 0.019 | 0.019 | 1.97 | 0.73 | 0.21 | 0.75 | 0.77 | 0.73 | 0.68 |
| 1 | 108499929 | rs6679376 | -1.297 | 0.915 | 1.345 | 0.455 | -0.013 | 0.047 | 0.047 | 0.419 | 0.09 | 0.31 | 0.08 | 0.09 | 0.05 | 0.05 |
| 1 | 108511501 | rs404634 | 1.547 | 0.991 | 1.347 | 0.452 | 0.009 | -0.038 | -0.038 | 0 | 0.77 | 0.45 | 0.78 | 0.88 | 0.79 | 0.68 |
| 1 | 108514072 | rs345261 | 1.511 | 1.046 | 0.916 | 0.417 | 0.02 | -0.038 | -0.038 | 0 | 0.76 | 0.4 | 0.78 | 0.83 | 0.76 | 0.66 |
| 1 | 108523670 | rs12121990 | 1.806 | 1.033 | -0.297 | 0.466 | -0.025 | 0.009 | 0.009 | 2.623 | 0.71 | 0.17 | 0.73 | 0.74 | 0.72 | 0.64 |
| 1 | 108524089 | rs12409813 | -0.407 | 1.461 | -0.379 | 0.155 | -0.029 | 0.026 | 0.026 | 0 | 0.11 | 0.06 | 0.13 | 0.02 | 0.18 | 0.26 |
| 1 | 108524270 | rs7543760 | 0.811 | 0.496 | -0.119 | 0.456 | 0.038 | 0.039 | 0.039 | 0 | 0.93 | 0.65 | 0.93 | 0.96 | 0.96 | 0.96 |
| 1 | 108525730 | rs11185224 | -0.441 | 0.901 | -1.831 | 0.463 | -0.089 | -0.03 | -0.03 | 0 | 0.36 | 0.06 | 0.55 | 0.66 | 0.4 | 0.4 |
| 1 | 108527281 | rs1350629 | -0.123 | 0.509 | -2.247 | -0.129 | -0.023 | 0.084 | 0.084 | 0 | 0.44 | 0.09 | 0.65 | 0.68 | 0.56 | 0.49 |
| 1 | 108530811 | rs12569255 | 0.167 | -0.211 | -1.545 | -0.722 | 0.05 | 0.06 | 0.06 | 0 | 0.4 | 0.26 | 0.2 | 0.06 | 0.24 | 0.33 |
| 1 | 108534241 | rs2000259 | 0.584 | 0.12 | -1.567 | -1.443 | 0.023 | 0.085 | 0.085 | 0.168 | 0.41 | 0.27 | 0.28 | 0.04 | 0.36 | 0.32 |
| 1 | 108536312 | rs3747945 | 0.546 | 0.222 | -1.552 | -1.464 | 0.012 | 0.075 | 0.075 | 0.076 | 0.42 | 0.28 | 0.3 | 0.04 | 0.38 | 0.33 |
| 1 | 108538338 | rs4915085 | -0.231 | 0.409 | -0.782 | -0.981 | -0.001 | 0 | 0 | 0.559 | 0.6 | 0.59 | 0.43 | 0.38 | 0.58 | 0.56 |
| 1 | 108538402 | rs12563016 | 0.084 | 0.337 | -0.401 | -0.903 | 0.005 | 0.006 | 0.006 | 0 | 0.14 | 0.13 | 0.12 | 0.26 | 0.18 | 0.22 |
| 1 | 108538626 | rs1219769 | -0.035 | 0.448 | -0.118 | 0.188 | -0.004 | -0.013 | -0.013 | 0.178 | 0.38 | 0.3 | 0.39 | 0.38 | 0.36 | 0.37 |
| 1 | 108549532 | rs12039210 | -0.018 | 0.605 | -0.185 | 0.051 | -0.001 | -0.007 | -0.007 | 0 | 0.07 | 0.05 | 0.05 | 0.09 | 0.07 | 0.04 |
| 1 | 108554161 | rs1777467 | 0.52 | 0.779 | 0.03 | 0.034 | 0.032 | 0.04 | 0.04 | 0 | 0.79 | 0.66 | 0.8 | 0.67 | 0.74 | 0.74 |
| 1 | 108557108 | rs12076264 | 0.668 | 0.793 | 0.01 | -0.043 | -0.001 | -0.008 | -0.008 | 0 | 0.57 | 0.42 | 0.41 | 0.28 | 0.57 | 0.55 |
| 1 | 108557705 | rs1781050 | -0.346 | 0.811 | -0.013 | -0.001 | 0.022 | 0.017 | 0.017 | 0 | 0.84 | 0.75 | 0.88 | 0.76 | 0.81 | 0.78 |
| 1 | 108568011 | rs1781067 | -0.667 | 0.791 | -0.068 | -0.005 | -0.004 | 0.031 | 0.031 | 0.288 | 0.41 | 0.36 | 0.58 | 0.73 | 0.43 | 0.45 |
| 1 | 108568620 | rs1418992 | 1.379 | 0.776 | -0.239 | -0.09 | 0.014 | 0.032 | 0.032 | 0.012 | 0.43 | 0.35 | 0.3 | 0.04 | 0.38 | 0.33 |
| 1 | 108576267 | rs9803640 | 0.336 | 0.628 | -0.618 | 0.126 | -0.03 | -0.038 | -0.038 | 0 | 0.26 | 0.12 | 0.29 | 0.27 | 0.26 | 0.28 |
| 1 | 108581278 | rs6583058 | -1.082 | 0.557 | -0.592 | 0.04 | -0.019 | -0.017 | -0.017 | 0 | 0.43 | 0.27 | 0.46 | 0.52 | 0.48 | 0.49 |

| | | | | | | | | | | | | | | | | |
|---|---|---|---|---|---|---|---|---|---|---|---|---|---|---|---|---|
| 1 | 108585993 | rs1777458 | 1.289 | 0.846 | -0.686 | -0.081 | 0.023 | 0.012 | 0.012 | 0 | 0.52 | 0.63 | 0.3 | 0.05 | 0.39 | 0.39 |
| 1 | 108586767 | rs1618973 | -0.225 | 0.884 | -0.782 | 0.026 | -0.048 | -0.009 | -0.009 | 0 | 0.7 | 0.87 | 0.51 | 0.37 | 0.64 | 0.63 |
| 1 | 108588372 | rs12569350 | 1.524 | 1.031 | -0.914 | 0.212 | -0.02 | -0.009 | -0.009 | 0 | 0.13 | 0.01 | 0.07 | 0.07 | 0.11 | 0.11 |
| 1 | 108591583 | rs1618693 | 0.612 | 0.987 | -0.863 | 0.152 | -0.048 | 0.055 | 0.055 | 0.041 | 0.65 | 0.83 | 0.46 | 0.3 | 0.58 | 0.6 |
| 1 | 108594594 | rs17020774 | 0.099 | 0.915 | -0.949 | 0.204 | 0.026 | 0.023 | 0.023 | 0 | 0.14 | 0.19 | 0.16 | 0.26 | 0.2 | 0.21 |
| 1 | 108596968 | rs1777454 | 1.664 | 1.15 | -0.82 | 0.038 | 0.014 | -0.021 | -0.021 | 0 | 0.5 | 0.53 | 0.3 | 0.04 | 0.41 | 0.38 |
| 1 | 108602821 | rs1418999 | 1.716 | 1.096 | -0.843 | 0.014 | 0.016 | -0.018 | -0.018 | 0 | 0.5 | 0.53 | 0.3 | 0.04 | 0.41 | 0.37 |
| 1 | 108606575 | rs17020833 | 1.291 | 1.11 | -1.078 | 0.076 | 0.004 | -0.009 | -0.009 | 0.074 | 0.07 | 0.02 | 0.05 | 0.07 | 0.05 | 0.03 |
| 1 | 108607571 | rs2995706 | 1.591 | 1.096 | -1.062 | 0.043 | 0.025 | 0.042 | 0.042 | 0.123 | 0.52 | 0.65 | 0.31 | 0.05 | 0.42 | 0.38 |
| 1 | 108607992 | rs12126230 | 0.541 | 1.328 | -1.101 | 0.19 | -0.032 | -0.001 | -0.001 | 0.041 | 0.06 | 0 | 0.16 | 0.34 | 0.1 | 0.09 |
| 1 | 108608347 | rs1619485 | 1.155 | 1.313 | -1.095 | 0.16 | -0.005 | 0.037 | 0.037 | 0 | 0.59 | 0.69 | 0.36 | 0.12 | 0.47 | 0.42 |
| 1 | 108609194 | rs7531042 | 0.538 | 1.324 | -1.037 | 0.235 | 0.002 | 0.006 | 0.006 | 0 | 0.12 | 0.1 | 0.13 | 0.26 | 0.16 | 0.2 |
| 1 | 108613888 | rs1343192 | 0.243 | 1.333 | -1.067 | 0.232 | -0.072 | -0.081 | -0.081 | 0 | 0.22 | 0.08 | 0.29 | 0.27 | 0.26 | 0.3 |
| 1 | 108614697 | rs1343207 | 0.21 | 1.308 | -1.036 | 0.227 | -0.001 | -0.002 | -0.002 | 0 | 0.81 | 0.84 | 0.81 | 0.67 | 0.79 | 0.77 |
| 1 | 108615524 | rs1781064 | 0.474 | 1.31 | -1.035 | 0.227 | 0.002 | 0.028 | 0.028 | 0 | 0.8 | 0.74 | 0.81 | 0.67 | 0.79 | 0.77 |
| 1 | 108616600 | rs10494087 | -0.705 | 0.734 | -1.595 | -0.049 | 0.042 | 0.186 | 0.186 | 0 | 0.53 | 0.22 | 0.68 | 0.88 | 0.6 | 0.61 |
| 1 | 108616858 | rs10494088 | 0.351 | 0.716 | -1.693 | 0.028 | -0.031 | -0.024 | -0.024 | 0 | 0.08 | 0.01 | 0.22 | 0.35 | 0.12 | 0.1 |
| 1 | 108621962 | rs12565741 | 1.608 | 0.591 | -1.632 | -0.268 | -0.021 | -0.003 | -0.003 | 0 | 0.13 | 0.01 | 0.07 | 0.06 | 0.11 | 0.11 |
| 1 | 108635284 | rs12034547 | 1.481 | 0.495 | -1.393 | -0.274 | 0.007 | 0.002 | 0.002 | 0.147 | 0.22 | 0.07 | 0.12 | 0.26 | 0.22 | 0.2 |
| 1 | 108640905 | rs2336126 | -0.863 | 0.695 | -1.355 | -0.217 | -0.052 | 0.055 | 0.055 | 0 | 0.36 | 0.21 | 0.6 | 0.69 | 0.43 | 0.43 |
| 1 | 108642664 | rs11185266 | -0.787 | 0.718 | -1.451 | -0.217 | -0.102 | 0.061 | 0.061 | 0.055 | 0.36 | 0.12 | 0.59 | 0.69 | 0.43 | 0.42 |
| 1 | 108646557 | rs4083872 | 1.585 | 0.567 | -1.383 | -0.254 | 0.006 | -0.002 | -0.002 | 0 | 0.22 | 0.07 | 0.18 | 0.28 | 0.22 | 0.19 |
| 1 | 108651122 | rs3904718 | 0.654 | 0.433 | -1.112 | -0.278 | -0.008 | -0.005 | -0.005 | 0 | 0.09 | 0.17 | 0.06 | 0.07 | 0.06 | 0.05 |
| 1 | 108652454 | rs12742113 | -0.045 | 0.551 | -1.019 | -0.061 | -0.02 | -0.005 | -0.005 | 0 | 0.44 | 0.28 | 0.52 | 0.56 | 0.49 | 0.5 |
| 1 | 108654328 | rs501273 | -0.729 | 0.488 | -0.989 | -0.096 | -0.013 | -0.009 | -0.009 | 0 | 0.89 | 0.8 | 0.94 | 0.93 | 0.94 | 0.96 |
| 1 | 108655288 | rs17020971 | 0.388 | 0.539 | -0.994 | -0.086 | -0.008 | -0.01 | -0.01 | 0.435 | 0.22 | 0.21 | 0.34 | 0.28 | 0.27 | 0.31 |
| 1 | 108656238 | rs614499 | 0.459 | 0.528 | -0.944 | -0.062 | 0 | 0 | 0 | 0 | 0.45 | 0.52 | 0.42 | 0.37 | 0.45 | 0.46 |
| 1 | 108657934 | rs515194 | 1.144 | 0.488 | -0.89 | -0.233 | -0.007 | -0.11 | -0.11 | 0.242 | 0.37 | 0.5 | 0.26 | 0.04 | 0.34 | 0.36 |
| 1 | 108660916 | rs661202 | 0.649 | 0.647 | -0.464 | 0.074 | -0.009 | 0.022 | 0.022 | 0 | 0.81 | 0.73 | 0.79 | 0.69 | 0.84 | 0.78 |
| 1 | 108661274 | rs1777448 | -0.18 | 0.661 | -0.495 | 0.238 | -0.005 | -0.002 | -0.002 | 0 | 0.6 | 0.57 | 0.63 | 0.6 | 0.64 | 0.64 |
| 1 | 108661569 | rs12116716 | 0.881 | 0.692 | -0.491 | 1.014 | -0.033 | -0.031 | -0.031 | 0.388 | 0.06 | 0.01 | 0.15 | 0.23 | 0.11 | 0.09 |
| 1 | 108661636 | rs1660419 | -0.513 | 0.597 | -0.451 | 0.958 | -0.019 | 0.006 | 0.006 | 0 | 0.65 | 0.57 | 0.68 | 0.6 | 0.71 | 0.76 |
| 1 | 108662219 | rs11185272 | 2.377 | 0.33 | -0.2 | 0.726 | 0.002 | 0.003 | 0.003 | 0 | 0.23 | 0.18 | 0.14 | 0.16 | 0.2 | 0.17 |
| 1 | 108663295 | rs1777450 | -1.793 | 0.47 | 0.126 | 1.072 | 0.004 | -0.012 | -0.012 | 0.175 | 0.77 | 0.86 | 0.76 | 0.66 | 0.78 | 0.82 |
| 1 | 108665402 | rs11185274 | -0.555 | 0.757 | 0.303 | 1.137 | 0.03 | 0.019 | 0.019 | 0 | 0.89 | 0.85 | 0.79 | 0.83 | 0.82 | 0.76 |
| 1 | 108665658 | rs11185276 | -1.244 | 0.511 | 0.191 | 0.974 | -0.008 | 0.011 | 0.011 | 0 | 0.29 | 0.22 | 0.31 | 0.23 | 0.29 | 0.38 |
| 1 | 108665931 | rs10881505 | 1.219 | 0.465 | 0.092 | 0.857 | -0.001 | 0.042 | 0.042 | 0 | 0.39 | 0.27 | 0.27 | 0.21 | 0.37 | 0.34 |
| 1 | 108673405 | rs2000260 | 0.226 | 0.92 | 0.495 | 1.08 | -0.032 | -0.049 | -0.049 | 0 | 0.33 | 0.18 | 0.47 | 0.52 | 0.42 | 0.44 |
| 1 | 108678361 | rs575371 | -0.557 | 0.367 | 0.865 | 0.505 | -0.034 | -0.086 | -0.086 | 0.31 | 0.13 | 0.47 | 0.06 | 0.01 | 0.06 | 0.1 |
| 1 | 108680234 | rs17513833 | 1.47 | 0.613 | 0.331 | 0.7 | 0.005 | 0.029 | 0.029 | 0.004 | 0.21 | 0.11 | 0.17 | 0.08 | 0.21 | 0.15 |
| 1 | 108681944 | rs12076095 | -0.208 | 0.711 | 0.482 | 0.918 | 0.013 | 0.068 | 0.068 | 0 | 0.07 | 0.16 | 0.17 | 0.22 | 0.1 | 0.1 |
| 1 | 108691782 | rs2336507 | 0.315 | 0.564 | -0.117 | 0.382 | 0.007 | 0.013 | 0.013 | 0 | 0.3 | 0.24 | 0.2 | 0.08 | 0.26 | 0.24 |
| 1 | 108692661 | rs12410160 | -0.3 | 1.086 | -0.17 | 0.939 | 0 | 0.012 | 0.012 | 0 | 0.08 | 0.13 | 0.24 | 0.15 | 0.09 | 0.2 |
| 1 | 108693336 | rs10494090 | 0.151 | 1.38 | -0.303 | 0.628 | -0.009 | 0.009 | 0.009 | 0 | 0.62 | 0.72 | 0.53 | 0.74 | 0.57 | 0.49 |
| 1 | 108694894 | rs591141 | 0.801 | 1.733 | -0.412 | 0.932 | -0.037 | -0.047 | -0.047 | 0.175 | 0.84 | 0.91 | 0.6 | 0.63 | 0.74 | 0.67 |
| 1 | 108696264 | rs10785835 | 0.691 | 1.633 | -0.311 | 0.922 | -0.005 | -0.006 | -0.006 | 0 | 0.57 | 0.67 | 0.39 | 0.55 | 0.46 | 0.39 |
| 1 | 108697224 | rs6703140 | 0.691 | 1.633 | -0.311 | 0.922 | -0.005 | -0.006 | -0.006 | 0 | 0.57 | 0.67 | 0.39 | 0.55 | 0.46 | 0.39 |
| 1 | 108698658 | rs829001 | 1.369 | 1.669 | -0.303 | 0.891 | 0 | -0.002 | -0.002 | 0 | 0.22 | 0.17 | 0.16 | 0.08 | 0.22 | 0.17 |
| 1 | 108699096 | rs1157390 | 0.722 | 1.68 | -0.3 | 0.912 | -0.005 | -0.006 | -0.006 | 0 | 0.57 | 0.67 | 0.39 | 0.54 | 0.46 | 0.39 |
| 1 | 108711268 | rs829004 | 1.218 | 1.701 | -0.307 | 0.935 | -0.023 | -0.022 | -0.022 | 0 | 0.79 | 0.82 | 0.55 | 0.62 | 0.68 | 0.56 |
| 1 | 108726389 | rs597999 | -0.114 | 1.315 | -0.332 | 0.543 | -0.03 | -0.03 | -0.03 | 0 | 0.91 | 0.97 | 0.74 | 0.82 | 0.84 | 0.76 |
| 1 | 108728484 | rs862493 | 1.113 | 0.909 | -0.218 | 0.234 | -0.001 | -0.004 | -0.004 | 0 | 0.22 | 0.18 | 0.17 | 0.08 | 0.22 | 0.18 |
| 1 | 108741065 | rs594397 | 0.786 | 0.979 | -0.037 | -0.171 | -0.009 | -0.125 | -0.125 | 8.702 | 0.27 | 0.34 | 0.13 | 0.01 | 0.25 | 0.17 |
| 1 | 108744934 | rs828999 | -1.091 | 1.207 | -0.192 | 0.078 | 0.013 | 0.002 | 0.002 | 0 | 0.49 | 0.59 | 0.63 | 0.5 | 0.55 | 0.65 |
| 1 | 108753833 | rs7538977 | -1.263 | 1.613 | 0.064 | 0.846 | 0.021 | 0.022 | 0.022 | 0 | 0.14 | 0.29 | 0.25 | 0.2 | 0.14 | 0.17 |
| 1 | 108754099 | rs7418288 | -0.409 | 1.768 | 0.04 | 0.825 | -0.034 | -0.028 | -0.028 | 0.066 | 0.09 | 0.02 | 0.25 | 0.29 | 0.17 | 0.32 |
| 1 | 108755434 | rs6702877 | -1.598 | 1.143 | 0.09 | 0.405 | -0.007 | -0.008 | -0.008 | 0 | 0.33 | 0.3 | 0.57 | 0.54 | 0.36 | 0.52 |
| 1 | 108756690 | rs6583070 | -1.808 | 1.133 | 0.044 | 0.4 | 0.018 | 0.02 | 0.02 | 0 | 0.34 | 0.45 | 0.58 | 0.54 | 0.36 | 0.52 |
| 1 | 108860573 | rs6670555 | 0.104 | 0.766 | 0.028 | 0.198 | 0.008 | -0.044 | -0.044 | 0.067 | 0.08 | 0.25 | 0.06 | 0 | 0.06 | 0.09 |
| 1 | 108870525 | rs11185315 | -1.18 | 0.781 | -0.178 | 0.203 | -0.033 | -0.056 | -0.056 | 0 | 0.31 | 0.12 | 0.53 | 0.59 | 0.31 | 0.46 |
| 1 | 108871245 | rs1417300 | -1.18 | 0.781 | -0.224 | 0.203 | -0.038 | -0.063 | -0.063 | 0 | 0.31 | 0.12 | 0.53 | 0.59 | 0.31 | 0.46 |
| 1 | 108876270 | rs11185318 | -1.929 | 0.793 | -0.27 | 0.216 | -0.001 | -0.002 | -0.002 | 0 | 0.4 | 0.52 | 0.59 | 0.59 | 0.37 | 0.55 |
| 1 | 108912413 | rs9660743 | 0.15 | -0.487 | 0.046 | -0.209 | 0 | -0.014 | -0.014 | 0 | 0.08 | 0.1 | 0.04 | 0 | 0.06 | 0.09 |
| 1 | 109023455 | rs12408504 | -1.558 | -0.571 | -0.083 | -0.131 | -0.047 | -0.02 | -0.02 | 0 | 0.3 | 0.5 | 0.33 | 0.28 | 0.2 | 0.2 |
| 1 | 109042271 | rs7512643 | -0.504 | -0.569 | -0.455 | -0.141 | -0.033 | -0.087 | -0.087 | 0.107 | 0.13 | 0.33 | 0.08 | 0 | 0.06 | 0.08 |
| 1 | 109047536 | rs10776805 | -0.694 | -0.018 | -1.226 | 0.411 | -0.009 | -0.077 | -0.077 | 0 | 0.16 | 0.01 | 0.23 | 0.28 | 0.14 | 0.13 |
| 1 | 109069856 | rs12036929 | -0.411 | 0.095 | -1.018 | 0.398 | 0.009 | -0.027 | -0.027 | 0 | 0.15 | 0.08 | 0.23 | 0.27 | 0.14 | 0.12 |
| 1 | 109070032 | rs2454327 | -0.344 | -0.004 | -1.461 | 0.114 | -0.02 | 0.072 | 0.072 | 1.236 | 0.74 | 0.92 | 0.85 | 0.85 | 0.75 | 0.82 |
| 1 | 109073893 | rs11101974 | -1.47 | 0.238 | -0.639 | 0.954 | 0.001 | -0.047 | -0.047 | 0 | 0.2 | 0.14 | 0.28 | 0.43 | 0.19 | 0.2 |

| | | | | | | | | | | | | | | | | |
|---|---|---|---|---|---|---|---|---|---|---|---|---|---|---|---|---|
| 1 | 109074350 | rs6604133 | -1.483 | 0.225 | -0.31 | 0.946 | -0.006 | 0.056 | 0.056 | 0 | 0.21 | 0.34 | 0.29 | 0.43 | 0.2 | 0.2 |
| 1 | 109075236 | rs12039852 | -1.287 | 0.33 | -0.062 | 0.744 | 0.052 | 0.097 | 0.097 | 0 | 0.06 | 0.24 | 0.05 | 0.14 | 0.05 | 0.08 |
| 1 | 109093791 | rs1566254 | -1.12 | -0.453 | 0.136 | 0.59 | -0.069 | 0.033 | 0.033 | 3.444 | 0.18 | 0.48 | 0.21 | 0.24 | 0.07 | 0.12 |
| 1 | 109096332 | rs2798184 | 0.391 | -0.367 | -0.386 | -0.173 | -0.014 | 0.005 | 0.005 | 0.342 | 0.37 | 0.23 | 0.43 | 0.6 | 0.44 | 0.46 |
| 1 | 109101851 | rs11102031 | 0.125 | -0.463 | -0.108 | -0.164 | 0.012 | -0.001 | -0.001 | 0 | 0.14 | 0.13 | 0.19 | 0.17 | 0.06 | 0.08 |
| 1 | 109108395 | rs17616564 | 0.992 | -0.332 | -0.303 | -0.698 | -0.002 | -0.014 | -0.014 | 0 | 0.11 | 0.1 | 0.06 | 0 | 0.09 | 0.1 |
| 1 | 109113463 | rs6683552 | -0.015 | -0.505 | 0.258 | -0.758 | -0.05 | -0.002 | -0.002 | 0 | 0.16 | 0.32 | 0.2 | 0.18 | 0.06 | 0.15 |
| 1 | 109116069 | rs2804253 | 0.307 | -0.508 | 0.297 | -0.755 | -0.089 | 0.021 | 0.021 | 4.351 | 0.81 | 0.48 | 0.79 | 0.76 | 0.94 | 0.81 |
| 1 | 109126399 | rs17024373 | 1.012 | -0.6 | 0.272 | -0.797 | -0.006 | -0.002 | -0.002 | 0.533 | 0.09 | 0.1 | 0.16 | 0.16 | 0.02 | 0.05 |
| 1 | 109131023 | rs970860 | 0.943 | -0.722 | 0.165 | -0.771 | -0.013 | 0.004 | 0.004 | 0.822 | 0.35 | 0.23 | 0.43 | 0.63 | 0.43 | 0.44 |
| 1 | 109133839 | rs10494092 | 0.988 | -0.92 | -0.525 | -0.942 | -0.1 | 0.019 | 0.019 | 0.275 | 0.25 | 0.02 | 0.21 | 0.15 | 0.31 | 0.23 |
| 1 | 109150720 | rs17025036 | 0.153 | -0.915 | -0.176 | -0.735 | 0.012 | -0.001 | -0.001 | 1.928 | 0.14 | 0.12 | 0.19 | 0.18 | 0.07 | 0.14 |
| 1 | 109170199 | rs10776714 | 0.186 | -0.858 | -0.219 | -0.742 | -0.076 | -0.007 | -0.007 | 0 | 0.82 | 0.53 | 0.79 | 0.82 | 0.93 | 0.86 |
| 1 | 109187595 | rs10494093 | 1.975 | -0.783 | -0.896 | -0.467 | -0.057 | -0.141 | -0.141 | 0 | 0.17 | 0.01 | 0.3 | 0.5 | 0.19 | 0.18 |
| 1 | 109188382 | rs2358530 | -0.489 | -0.964 | -0.35 | -0.591 | -0.07 | -0.014 | -0.014 | 0.089 | 0.19 | 0.41 | 0.21 | 0.17 | 0.07 | 0.14 |
| 1 | 109194895 | rs10157512 | -0.534 | -0.972 | -0.334 | -0.594 | -0.073 | -0.016 | -0.016 | 0.221 | 0.19 | 0.42 | 0.21 | 0.18 | 0.07 | 0.14 |
| 1 | 109208995 | rs6665861 | -0.014 | -0.865 | -0.567 | -0.532 | -0.071 | -0.015 | -0.015 | 0 | 0.82 | 0.61 | 0.79 | 0.83 | 0.93 | 0.86 |
| 1 | 109241669 | rs2297563 | 0.751 | 0.14 | -1.502 | -0.591 | -0.088 | -0.048 | -0.048 | 0 | 0.22 | 0.02 | 0.2 | 0.19 | 0.28 | 0.24 |
| 1 | 109258150 | rs1333124 | 0.401 | -0.124 | -1.269 | -1.078 | 0.004 | 0.126 | 0.126 | 1.027 | 0.56 | 0.2 | 0.66 | 0.81 | 0.61 | 0.64 |
| 1 | 109258918 | rs11102288 | 0.592 | -0.062 | -1.767 | -1.075 | -0.026 | -0.03 | -0.03 | 0 | 0.13 | 0.02 | 0.05 | 0.12 | 0.17 | 0.15 |
| 1 | 109260034 | rs914807 | -0.183 | -0.345 | -0.742 | -1.403 | -0.056 | -0.031 | -0.031 | 0 | 0.84 | 0.44 | 0.93 | 0.88 | 0.94 | 0.89 |
| 1 | 109260357 | rs4970804 | 0.415 | -0.181 | -0.737 | -1.512 | 0.011 | 0.079 | 0.079 | 0.188 | 0.6 | 0.24 | 0.68 | 0.82 | 0.64 | 0.69 |
| 1 | 109261509 | rs17553619 | 1.281 | -0.15 | -0.997 | -1.777 | -0.015 | 0.008 | 0.008 | 0 | 0.08 | 0 | 0.03 | 0.03 | 0.08 | 0.11 |
| 1 | 109261971 | rs2184083 | 0.814 | -0.02 | -0.722 | -1.616 | -0.013 | 0.023 | 0.023 | 0.159 | 0.23 | 0.2 | 0.25 | 0.07 | 0.3 | 0.2 |
| 1 | 109265563 | rs4970808 | 0.35 | -0.031 | -0.627 | -1.47 | 0.006 | 0.119 | 0.119 | 1.027 | 0.56 | 0.25 | 0.67 | 0.86 | 0.6 | 0.64 |
| 1 | 109266523 | rs2026485 | 0.13 | -0.029 | -0.498 | -1.369 | 0.009 | 0.083 | 0.083 | 0.079 | 0.61 | 0.25 | 0.69 | 0.86 | 0.64 | 0.69 |
| 1 | 109268441 | rs11582005 | -0.59 | 0.012 | 0.206 | -1.152 | -0.019 | -0.014 | -0.014 | 0 | 0.16 | 0.32 | 0.09 | 0.1 | 0.14 | 0.12 |
| 1 | 109268615 | rs3006870 | 0.126 | -0.059 | 0.74 | -1.074 | 0.026 | 0.018 | 0.018 | 0.794 | 0.05 | 0.33 | 0.02 | 0 | 0 | 0 |
| 1 | 109270402 | rs10494096 | -0.032 | 0.057 | -0.136 | -1.551 | -0.032 | 0.097 | 0.097 | 0 | 0.44 | 0.22 | 0.32 | 0.19 | 0.48 | 0.46 |
| 1 | 109271037 | rs17026815 | -0.378 | -0.023 | 0.018 | -1.596 | 0.005 | -0.07 | -0.07 | 0 | 0.2 | 0.35 | 0.11 | 0.03 | 0.21 | 0.19 |
| 1 | 109273610 | rs4970810 | -0.057 | 0.032 | -0.104 | -1.727 | 0.013 | 0.049 | 0.049 | 0 | 0.48 | 0.57 | 0.44 | 0.2 | 0.54 | 0.47 |
| 1 | 109288766 | rs1759487 | -0.724 | 0.022 | -0.041 | -1.616 | 0.061 | 0.062 | 0.062 | 0 | 0.57 | 0.7 | 0.5 | 0.22 | 0.71 | 0.58 |
| 1 | 109303307 | rs4970811 | -0.968 | -0.077 | -0.006 | -1.605 | 0.065 | 0.051 | 0.051 | 0.012 | 0.61 | 0.74 | 0.52 | 0.22 | 0.74 | 0.63 |
| 1 | 109308814 | rs1767017 | -0.094 | 0.163 | 0.122 | -1.72 | 0.003 | -0.04 | -0.04 | 0 | 0.19 | 0.24 | 0.11 | 0.03 | 0.23 | 0.18 |
| 1 | 109341772 | rs1759492 | -0.59 | 0.138 | 0.11 | -1.729 | 0.005 | -0.031 | -0.031 | 0 | 0.83 | 0.79 | 0.89 | 0.97 | 0.77 | 0.82 |
| 1 | 109342295 | rs11586564 | 0.321 | 0.045 | 0.344 | -1.765 | 0 | -0.006 | -0.006 | 0 | 0.4 | 0.39 | 0.4 | 0.2 | 0.51 | 0.45 |
| 1 | 109352915 | rs1149152 | 0.872 | 0.263 | 0.393 | -1.704 | 0.07 | 0.041 | 0.041 | 0 | 0.38 | 0.23 | 0.47 | 0.76 | 0.26 | 0.37 |
| 1 | 109356216 | rs12140541 | 0.66 | 0.287 | 0.307 | -1.698 | -0.015 | 0.013 | 0.013 | 0 | 0.37 | 0.27 | 0.32 | 0.21 | 0.43 | 0.45 |
| 1 | 109356260 | rs10494098 | -1.084 | 0.261 | 0.338 | -1.696 | 0.01 | 0.008 | 0.008 | 0 | 0.61 | 0.62 | 0.52 | 0.23 | 0.74 | 0.63 |
| 1 | 109356524 | rs11807463 | -1.026 | 0.695 | -0.226 | -1.681 | 0.016 | -0.079 | -0.079 | 0.029 | 0.25 | 0.34 | 0.2 | 0.03 | 0.31 | 0.18 |
| 1 | 109357456 | rs10494099 | 0.209 | 0.52 | -0.857 | -1.327 | -0.019 | 0.03 | 0.03 | 0.729 | 0.16 | 0.08 | 0.2 | 0.07 | 0.23 | 0.18 |
| 1 | 109358958 | rs6696787 | 0.085 | 0.539 | -0.845 | -1.309 | -0.014 | 0.028 | 0.028 | 0.844 | 0.17 | 0.11 | 0.2 | 0.08 | 0.23 | 0.19 |
| 1 | 109359502 | rs1277213 | -0.384 | 0.245 | -1.026 | -1.346 | 0.006 | 0.027 | 0.027 | 0 | 0.55 | 0.43 | 0.59 | 0.78 | 0.48 | 0.55 |
| 1 | 109363232 | rs1333130 | 0.338 | -0.107 | -1.243 | -0.916 | 0.051 | 0.021 | 0.021 | 0 | 0.38 | 0.26 | 0.39 | 0.7 | 0.26 | 0.36 |
| 1 | 109366286 | rs7522157 | -0.357 | 1.147 | -1.578 | -1.05 | -0.062 | 0.039 | 0.039 | 0.129 | 0.24 | 0.06 | 0.18 | 0.11 | 0.29 | 0.23 |
| 1 | 109366370 | rs7522260 | -0.431 | 1.008 | -1.179 | -0.886 | 0.002 | 0.001 | 0.001 | 0 | 0.63 | 0.32 | 0.51 | 0.36 | 0.66 | 0.58 |
| 1 | 109370514 | rs7518022 | -0.6 | 1.458 | -0.382 | -0.928 | 0.044 | -0.05 | -0.05 | 0 | 0.31 | 0.47 | 0.37 | 0.14 | 0.36 | 0.28 |
| 1 | 109374087 | rs12059541 | 0.013 | 1.079 | 0.279 | -0.775 | 0.026 | -0.005 | -0.005 | 0 | 0.1 | 0.28 | 0.11 | 0.11 | 0.11 | 0.09 |
| 1 | 109376539 | rs6683747 | 0.009 | 1.089 | 0.283 | -0.771 | 0.029 | -0.002 | -0.002 | 0 | 0.1 | 0.28 | 0.11 | 0.12 | 0.11 | 0.09 |
| 1 | 109384436 | rs10857972 | -1.614 | 1.056 | 0.682 | -0.729 | 0.006 | -0.042 | -0.042 | 0 | 0.24 | 0.53 | 0.2 | 0.21 | 0.25 | 0.22 |
| 1 | 109384849 | rs11582294 | 0.034 | 1.114 | 0.78 | -0.746 | -0.004 | -0.002 | -0.002 | 0 | 0.1 | 0.07 | 0.09 | 0.11 | 0.13 | 0.06 |
| 1 | 109385463 | rs12075148 | -0.265 | 1.06 | 1.109 | -0.758 | 0.008 | -0.023 | -0.023 | 0 | 0.1 | 0.21 | 0.09 | 0.1 | 0.11 | 0.12 |
| 1 | 109386174 | rs6704266 | -1.774 | 1.049 | 1.064 | -0.737 | 0.011 | -0.033 | -0.033 | 0 | 0.23 | 0.51 | 0.21 | 0.22 | 0.25 | 0.22 |
| 1 | 109386466 | rs1277014 | 0.431 | 1.034 | 1.438 | -0.993 | 0.024 | -0.03 | -0.03 | 0.169 | 0.93 | 0.77 | 0.97 | 0.99 | 0.93 | 0.97 |
| 1 | 109389021 | rs7542337 | -1.302 | 1.072 | 1.42 | -1.016 | -0.01 | -0.093 | -0.093 | 0 | 0.2 | 0.45 | 0.16 | 0.1 | 0.18 | 0.17 |
| 1 | 109391027 | rs1149144 | 1.041 | 1.164 | 1.424 | -0.944 | 0.062 | 0.045 | 0.045 | 0 | 0.9 | 0.69 | 0.9 | 0.86 | 0.87 | 0.92 |
| 1 | 109392715 | rs10776762 | -0.51 | 1.094 | 1.224 | -0.972 | 0.02 | -0.044 | -0.044 | 0.131 | 0.09 | 0.3 | 0.07 | 0 | 0.07 | 0.05 |
| 1 | 109392837 | rs1277203 | 1.997 | 1.37 | 0.868 | -0.816 | -0.014 | 0.022 | 0.022 | 0.632 | 0.69 | 0.24 | 0.72 | 0.71 | 0.66 | 0.7 |
| 1 | 109392901 | rs9440550 | 0.651 | 1.163 | -0.011 | -1.22 | -0.023 | 0.068 | 0.068 | 0 | 0.25 | 0.09 | 0.18 | 0.1 | 0.21 | 0.14 |
| 1 | 109393302 | rs12035019 | -1.118 | 0.839 | 0.564 | -0.746 | 0.002 | 0.019 | 0.019 | 0 | 0.06 | 0.07 | 0.09 | 0.15 | 0.07 | 0.08 |
| 1 | 109394125 | rs4970815 | -1.788 | 0.365 | 0.333 | -1.178 | 0.003 | 0.026 | 0.026 | 0.07 | 0.29 | 0.17 | 0.21 | 0.22 | 0.29 | 0.25 |
| 1 | 109394564 | rs17030016 | -0.807 | 0.279 | 0.467 | -1.234 | 0.002 | 0.001 | 0.001 | 0.045 | 0.1 | 0.11 | 0.07 | 0.04 | 0.05 | 0.1 |
| 1 | 109394978 | rs17563390 | -0.453 | 0.321 | 0.436 | -1.249 | 0.016 | 0.006 | 0.006 | 0 | 0.1 | 0.15 | 0.11 | 0.14 | 0.14 | 0.07 |
| 1 | 109395105 | rs1277207 | 1.364 | 0.372 | 0.783 | -1.406 | -0.001 | 0.036 | 0.036 | 0.046 | 0.81 | 0.88 | 0.88 | 0.94 | 0.79 | 0.85 |
| 1 | 109395584 | rs17621489 | 1.074 | 0.386 | 0.686 | -1.411 | -0.002 | -0.006 | -0.006 | 0 | 0.06 | 0.05 | 0.04 | 0 | 0.07 | 0.02 |
| 1 | 109397229 | rs12405585 | -1.257 | 0.278 | 0.861 | -1.452 | 0.008 | 0.01 | 0.01 | 0 | 0.17 | 0.13 | 0.1 | 0.04 | 0.14 | 0.13 |
| 1 | 109399543 | rs7519293 | -0.21 | 0.292 | 0.836 | -1.443 | 0.001 | -0.008 | -0.008 | 0.686 | 0.1 | 0.08 | 0.04 | 0 | 0.09 | 0.06 |
| 1 | 109399660 | rs7519409 | -3.212 | 0.227 | 1.229 | -1.511 | -0.021 | -0.136 | -0.136 | 0 | 0.33 | 0.55 | 0.21 | 0.05 | 0.3 | 0.25 |
| 1 | 109399710 | rs7551142 | -0.218 | 0.271 | 0.906 | -1.433 | 0.001 | -0.008 | -0.008 | 0 | 0.1 | 0.08 | 0.04 | 0 | 0.09 | 0.06 |

| | | | | | | | | | | | | | | | | |
|---|---|---|---|---|---|---|---|---|---|---|---|---|---|---|---|---|
| 1 | 109400835 | rs17621851 | -0.218 | 0.271 | 0.906 | -1.433 | 0.001 | -0.008 | -0.008 | 0 | 0.1 | 0.08 | 0.04 | 0 | 0.09 | 0.06 |
| 1 | 109404282 | rs11587670 | -0.719 | 0.301 | 0.879 | -1.35 | -0.001 | -0.01 | -0.01 | 0.422 | 0.13 | 0.12 | 0.09 | 0.04 | 0.13 | 0.09 |
| 1 | 109407730 | rs11585358 | 1.027 | 0.356 | 0.768 | -1.309 | -0.002 | -0.005 | -0.005 | 0.344 | 0.06 | 0.06 | 0.04 | 0 | 0.08 | 0.02 |
| 1 | 109416225 | rs12090453 | 2.45 | 0.467 | 0.282 | -1.171 | -0.026 | -0.073 | -0.073 | 1.731 | 0.37 | 0.1 | 0.44 | 0.54 | 0.39 | 0.55 |
| 1 | 109417679 | rs7523050 | -1.249 | 0.036 | 1.016 | -1.43 | -0.032 | -0.068 | -0.068 | 0.859 | 0.2 | 0.3 | 0.12 | 0.05 | 0.15 | 0.15 |
| 1 | 109426288 | rs7542414 | -2.439 | 0.071 | 1.176 | -1.446 | -0.008 | -0.101 | -0.101 | 0 | 0.31 | 0.44 | 0.2 | 0.05 | 0.29 | 0.26 |
| 1 | 109445706 | rs9662782 | 0.092 | 0.17 | 0.975 | -1.343 | -0.001 | -0.024 | -0.024 | 0.021 | 0.09 | 0.11 | 0.05 | 0.01 | 0.08 | 0.08 |
| 1 | 109453129 | rs11102632 | -0.869 | 0.205 | 0.849 | -1.32 | 0.004 | -0.044 | -0.044 | 0 | 0.13 | 0.19 | 0.08 | 0.02 | 0.15 | 0.14 |
| 1 | 109470126 | rs839860 | -2.637 | 0.089 | 0.885 | -1.392 | -0.005 | -0.061 | -0.061 | 0 | 0.33 | 0.62 | 0.21 | 0.06 | 0.3 | 0.26 |
| 1 | 109470185 | rs839859 | -2.637 | 0.089 | 0.885 | -1.392 | -0.005 | -0.061 | -0.061 | 0 | 0.33 | 0.62 | 0.21 | 0.06 | 0.3 | 0.26 |
| 1 | 109471634 | rs839854 | -2.102 | 0.23 | 0.837 | -1.487 | 0.004 | 0.015 | 0.015 | 0 | 0.49 | 0.6 | 0.45 | 0.37 | 0.42 | 0.36 |
| 1 | 109472921 | rs14209 | 0.39 | 0.048 | 0.705 | -1.396 | 0.008 | 0.019 | 0.019 | 0 | 0.09 | 0.01 | 0.06 | 0.03 | 0.05 | 0.08 |
| 1 | 109473017 | rs10607 | -1.251 | -0.015 | 0.865 | -1.5 | 0.004 | -0.148 | -0.148 | 2.629 | 0.22 | 0.47 | 0.14 | 0 | 0.22 | 0.1 |
| 1 | 109473826 | rs1664415 | -1.995 | 0.148 | 0.607 | -1.395 | -0.006 | -0.066 | -0.066 | 0.802 | 0.36 | 0.62 | 0.22 | 0.04 | 0.28 | 0.26 |
| 1 | 109474221 | rs1769740 | -2.002 | 0.144 | 0.56 | -1.397 | -0.004 | -0.059 | -0.059 | 0 | 0.36 | 0.62 | 0.22 | 0.04 | 0.28 | 0.26 |
| 1 | 109479978 | rs168107 | -2.002 | 0.157 | 0.587 | -1.387 | -0.004 | -0.059 | -0.059 | 0 | 0.36 | 0.62 | 0.21 | 0.04 | 0.28 | 0.25 |
| 1 | 109483767 | rs839549 | -0.7 | 0.154 | 0.491 | -1.18 | -0.002 | -0.002 | -0.002 | 0 | 0.31 | 0.31 | 0.33 | 0.4 | 0.33 | 0.31 |
| 1 | 109483975 | rs193139 | -0.753 | -0.012 | 0.553 | -1.36 | -0.045 | -0.101 | -0.101 | 0 | 0.23 | 0.39 | 0.14 | 0.02 | 0.15 | 0.11 |
| 1 | 109486196 | rs338466 | -2.2 | 0.165 | 0.366 | -1.19 | -0.026 | 0.02 | 0.02 | 0 | 0.6 | 0.76 | 0.51 | 0.4 | 0.52 | 0.43 |
| 1 | 109495875 | rs865151 | -1.544 | 0.308 | 0.146 | -1.1 | -0.007 | 0.068 | 0.068 | 0 | 0.49 | 0.24 | 0.46 | 0.41 | 0.48 | 0.4 |
| 1 | 109500439 | rs517013 | -1.823 | 0.386 | 0.156 | -1.172 | -0.002 | 0.007 | 0.007 | 0 | 0.57 | 0.63 | 0.53 | 0.4 | 0.54 | 0.43 |
| 1 | 109504912 | rs705271 | 0.063 | 0.141 | 0.078 | -1.365 | -0.003 | -0.001 | -0.001 | 0.006 | 0.1 | 0.1 | 0.08 | 0.12 | 0.09 | 0.1 |
| 1 | 109505689 | rs550743 | -0.161 | 0.155 | -0.081 | -1.289 | -0.021 | 0.001 | 0.001 | 0 | 0.8 | 0.87 | 0.73 | 0.78 | 0.73 | 0.8 |
| 1 | 109513274 | rs11803800 | 0.56 | -0.047 | 0.144 | -1.34 | -0.008 | -0.05 | -0.05 | 0.768 | 0.12 | 0.2 | 0.07 | 0 | 0.09 | 0.03 |
| 1 | 109515028 | rs12032662 | 2.216 | -0.094 | 0.242 | -1.448 | 0.012 | 0.037 | 0.037 | 0 | 0.44 | 0.3 | 0.45 | 0.64 | 0.42 | 0.55 |
| 1 | 109515735 | rs570812 | -1.659 | -0.076 | 0.271 | -1.46 | -0.006 | -0.041 | -0.041 | 0 | 0.37 | 0.57 | 0.28 | 0.14 | 0.32 | 0.25 |
| 1 | 109517583 | rs17031647 | 0.562 | -0.28 | 0.047 | -1.29 | -0.007 | -0.049 | -0.049 | 0 | 0.12 | 0.18 | 0.07 | 0 | 0.09 | 0.03 |
| 1 | 109519561 | rs12066799 | 1.787 | -0.285 | 0.061 | -1.295 | 0.011 | 0.042 | 0.042 | 0.025 | 0.43 | 0.24 | 0.45 | 0.64 | 0.42 | 0.55 |
| 1 | 109521317 | rs11102735 | 0.524 | -0.209 | 0.01 | -1.229 | -0.007 | -0.049 | -0.049 | 0 | 0.12 | 0.18 | 0.07 | 0 | 0.09 | 0.03 |
| 1 | 109524818 | rs1025996 | 0.524 | -0.209 | 0.01 | -1.229 | -0.007 | -0.049 | -0.049 | 1.085 | 0.12 | 0.18 | 0.07 | 0 | 0.09 | 0.03 |
| 1 | 109563656 | rs11102813 | 1.717 | -0.145 | -0.242 | -1.134 | 0.009 | 0.031 | 0.031 | 0 | 0.43 | 0.3 | 0.45 | 0.65 | 0.42 | 0.55 |
| 1 | 109569132 | rs11485106 | 0.024 | -0.147 | 0.089 | -1.04 | -0.007 | -0.046 | -0.046 | 0 | 0.12 | 0.18 | 0.07 | 0 | 0.09 | 0.03 |
| 1 | 109578450 | rs28564628 | -0.047 | -0.17 | 0.071 | -1.014 | -0.007 | -0.045 | -0.045 | 0.802 | 0.13 | 0.18 | 0.07 | 0 | 0.09 | 0.03 |
| 1 | 109591433 | rs28503409 | -1.005 | -0.01 | 0.205 | -0.645 | 0.007 | 0.019 | 0.019 | 0 | 0.26 | 0.35 | 0.32 | 0.35 | 0.29 | 0.14 |
| 1 | 109602029 | rs28577885 | 1.259 | -0.2 | 0.303 | -0.786 | 0.007 | 0.024 | 0.024 | 0 | 0.7 | 0.43 | 0.7 | 0.63 | 0.71 | 0.79 |
| 1 | 109605305 | rs8856 | 0.409 | -0.147 | -0.163 | -0.783 | -0.054 | -0.08 | -0.08 | 0 | 0.12 | 0.03 | 0.21 | 0.36 | 0.18 | 0.09 |
| 1 | 109605378 | rs10466 | -0.281 | -0.132 | -0.03 | -0.789 | -0.011 | -0.067 | -0.067 | 0.522 | 0.17 | 0.05 | 0.22 | 0.36 | 0.18 | 0.17 |
| 1 | 109610537 | rs28715911 | -1.327 | -0.119 | 0.259 | -0.809 | -0.01 | -0.017 | -0.017 | 0 | 0.26 | 0.22 | 0.26 | 0.38 | 0.29 | 0.21 |
| 1 | 109611326 | rs28634276 | 1.852 | -0.39 | -0.307 | -1.068 | 0.004 | 0.005 | 0.005 | 0 | 0.48 | 0.28 | 0.58 | 0.52 | 0.55 | 0.58 |
| 1 | 109612484 | rs28553535 | 0.429 | -0.248 | -0.353 | -1.072 | 0.003 | 0.013 | 0.013 | 0 | 0.14 | 0.02 | 0.07 | 0.1 | 0.07 | 0.14 |
| 1 | 109612806 | rs2358956 | -0.447 | -0.127 | 0.024 | -0.954 | -0.001 | -0.037 | -0.037 | 0 | 0.93 | 0.82 | 0.95 | 1 | 0.96 | 0.98 |
| 1 | 109613466 | rs1562225 | 1.468 | -0.176 | -0.506 | -0.858 | -0.009 | -0.019 | -0.019 | 0.307 | 0.45 | 0.24 | 0.56 | 0.49 | 0.53 | 0.49 |
| 1 | 109633256 | rs521734 | -1.781 | -0.872 | -0.841 | -0.986 | -0.101 | -0.137 | -0.137 | 0 | 0.6 | 0.95 | 0.52 | 0.48 | 0.4 | 0.53 |
| 1 | 109634341 | rs616378 | -1.2 | -0.215 | 0.397 | -1.623 | -0.009 | -0.041 | -0.041 | 0.228 | 0.22 | 0.27 | 0.09 | 0.03 | 0.17 | 0.22 |
| 1 | 109672215 | rs17038458 | -0.763 | -0.492 | 0.343 | -1.313 | -0.009 | -0.034 | -0.034 | 0.388 | 0.11 | 0.26 | 0.08 | 0.05 | 0.08 | 0.06 |
| 1 | 109674281 | rs10494103 | -0.209 | -0.408 | 0.536 | -1.436 | -0.031 | -0.09 | -0.09 | 1.025 | 0.15 | 0.28 | 0.05 | 0 | 0.06 | 0.07 |
| 1 | 109685273 | rs587727 | 1.461 | -0.341 | -0.006 | -1.595 | 0.017 | 0.084 | 0.084 | 0 | 0.54 | 0.31 | 0.63 | 0.92 | 0.65 | 0.61 |
| 1 | 109689314 | rs531842 | 0.394 | 0.123 | -0.44 | -1.15 | -0.005 | -0.018 | -0.018 | 0 | 0.85 | 0.79 | 0.78 | 0.94 | 0.85 | 0.84 |
| 1 | 109696056 | rs565751 | 0.677 | 0.309 | -0.673 | -1.047 | 0.004 | 0.003 | 0.003 | 0 | 0.66 | 0.7 | 0.71 | 0.93 | 0.72 | 0.72 |
| 1 | 109696968 | rs595635 | -1.035 | 0.32 | -0.67 | -1.042 | 0.004 | 0.002 | 0.002 | 0 | 0.34 | 0.31 | 0.29 | 0.07 | 0.28 | 0.28 |
| 1 | 109705022 | rs676385 | 0.502 | 0.101 | -0.581 | -0.14 | 0.006 | -0.007 | -0.007 | 0 | 0.64 | 0.34 | 0.61 | 0.57 | 0.7 | 0.57 |
| 1 | 109706393 | rs649539 | 0.925 | -0.357 | -0.397 | -0.62 | 0.011 | -0.003 | -0.003 | 0 | 0.52 | 0.45 | 0.55 | 0.51 | 0.6 | 0.48 |
| 1 | 109714403 | rs17014495 | -1.305 | -0.502 | -0.163 | -0.633 | 0.049 | 0.005 | 0.005 | 0.082 | 0.5 | 0.7 | 0.45 | 0.51 | 0.39 | 0.51 |
| 1 | 109723188 | rs2093933 | -0.525 | -0.832 | 0.174 | -0.899 | -0.001 | 0.088 | 0.088 | 0 | 0.22 | 0.32 | 0.28 | 0.51 | 0.21 | 0.4 |
| 1 | 109724188 | rs587036 | 0.568 | -0.886 | 0.355 | -1.174 | -0.018 | -0.092 | -0.092 | 1.407 | 0.74 | 0.69 | 0.86 | 1 | 0.83 | 0.9 |
| 1 | 109727832 | rs12239854 | -0.702 | -0.749 | 0.316 | -1.162 | 0 | -0.044 | -0.044 | 0.026 | 0.23 | 0.24 | 0.13 | 0 | 0.17 | 0.05 |
| 1 | 109732080 | rs603285 | 1.276 | -1.434 | 0.215 | -1.775 | 0.025 | -0.012 | -0.012 | 1.748 | 0.6 | 0.28 | 0.79 | 1 | 0.75 | 0.87 |
| 1 | 109735416 | rs659543 | 1.018 | -1.636 | 0.224 | -1.869 | -0.012 | -0.111 | -0.111 | 0 | 0.67 | 0.36 | 0.81 | 1 | 0.76 | 0.93 |
| 1 | 109737063 | rs2359246 | 1.349 | -1.672 | 0.241 | -1.777 | 0.01 | -0.028 | -0.028 | 0.312 | 0.63 | 0.3 | 0.8 | 0.97 | 0.76 | 0.89 |
| 1 | 109737132 | rs2359244 | 0.881 | -1.677 | 0.332 | -1.78 | -0.027 | -0.138 | -0.138 | 0.309 | 0.68 | 0.47 | 0.81 | 0.97 | 0.77 | 0.93 |
| 1 | 109737172 | rs2359243 | 1.358 | -1.675 | 0.325 | -1.779 | 0.009 | -0.029 | -0.029 | 0 | 0.63 | 0.3 | 0.8 | 0.97 | 0.76 | 0.89 |
| 1 | 109737802 | rs17014528 | 1.059 | -1.598 | 0.34 | -1.637 | -0.007 | -0.06 | -0.06 | 6.291 | 0.67 | 0.34 | 0.79 | 0.96 | 0.77 | 0.92 |
| 1 | 109739071 | rs2478762 | 0.93 | -1.6 | 0.33 | -1.638 | -0.026 | -0.123 | -0.123 | 1.188 | 0.68 | 0.46 | 0.8 | 0.96 | 0.76 | 0.91 |
| 1 | 109740548 | rs2296696 | -0.791 | -2.008 | 0.202 | -1.208 | -0.001 | 0.05 | 0.05 | 0 | 0.24 | 0.27 | 0.27 | 0.51 | 0.24 | 0.42 |
| 1 | 109742485 | rs913179 | 0.78 | -1.312 | 0.081 | -1.552 | -0.031 | -0.144 | -0.144 | 0 | 0.68 | 0.47 | 0.81 | 0.96 | 0.79 | 0.88 |
| 1 | 109743167 | rs17034354 | -0.236 | -1.432 | 0.006 | -1.612 | 0.026 | 0.034 | 0.034 | 0.184 | 0.19 | 0.15 | 0.09 | 0 | 0.11 | 0.08 |
| 1 | 109745618 | rs1052878 | 0.029 | -1.083 | 0.349 | -1.238 | 0.014 | -0.039 | -0.039 | 0 | 0.89 | 0.78 | 0.9 | 0.96 | 0.9 | 0.95 |
| 1 | 109747635 | rs17034539 | -0.552 | -1.236 | 0.259 | -1.155 | -0.006 | -0.036 | -0.036 | 0 | 0.2 | 0.26 | 0.1 | 0.08 | 0.18 | 0.29 |
| 1 | 109748547 | rs4970827 | 0.117 | -1.055 | -0.036 | -0.924 | 0.014 | -0.039 | -0.039 | 0 | 0.89 | 0.78 | 0.9 | 0.96 | 0.9 | 0.95 |

| | | | | | | | | | | | | | | | | |
|---|---|---|---|---|---|---|---|---|---|---|---|---|---|---|---|---|
| 1 | 109749071 | rs3197233 | 0.117 | -1.055 | -0.036 | -0.916 | 0.014 | -0.018 | -0.018 | 0 | 0.89 | 0.78 | 0.9 | 0.95 | 0.9 | 0.95 |
| 1 | 109752204 | rs604500 | -1.323 | -0.943 | -0.167 | -0.865 | 0.002 | -0.028 | -0.028 | 0 | 0.25 | 0.17 | 0.31 | 0.47 | 0.23 | 0.25 |
| 1 | 109768889 | rs3120625 | -1.417 | -0.428 | -0.287 | -0.858 | -0.016 | -0.033 | -0.033 | 3.622 | 0.25 | 0.17 | 0.37 | 0.46 | 0.33 | 0.3 |
| 1 | 109771063 | rs683182 | 0.939 | -0.473 | 0.197 | -1.096 | -0.008 | -0.026 | -0.026 | 0 | 0.71 | 0.39 | 0.81 | 0.89 | 0.79 | 0.65 |
| 1 | 109771701 | rs669697 | -1.227 | -0.255 | 0.191 | -0.908 | 0.005 | 0.018 | 0.018 | 0 | 0.17 | 0.2 | 0.31 | 0.41 | 0.27 | 0.25 |
| 1 | 109774026 | rs600711 | 0.02 | -0.371 | 0.323 | -0.894 | -0.012 | 0.116 | 0.116 | 0 | 0.9 | 0.85 | 0.78 | 0.55 | 0.92 | 0.79 |
| 1 | 109782037 | rs611905 | -1.361 | -0.243 | 0.277 | -0.971 | 0.006 | 0.02 | 0.02 | 0 | 0.17 | 0.19 | 0.31 | 0.41 | 0.27 | 0.25 |
| 1 | 109782190 | rs611060 | -1.745 | -0.381 | 0.489 | -1.159 | -0.007 | -0.005 | -0.005 | 0 | 0.35 | 0.63 | 0.28 | 0.08 | 0.41 | 0.42 |
| 1 | 109783205 | rs4623734 | -1.135 | -0.429 | 0.736 | -0.891 | 0.013 | 0.105 | 0.105 | 0 | 0.22 | 0.31 | 0.34 | 0.45 | 0.26 | 0.38 |
| 1 | 109783261 | rs11102964 | -1.851 | -0.691 | 0.856 | -0.979 | -0.034 | 0.041 | 0.041 | 0.171 | 0.25 | 0.34 | 0.27 | 0.4 | 0.18 | 0.35 |
| 1 | 109783555 | rs4246519 | 2.522 | -0.334 | 0.908 | -1.077 | -0.004 | -0.002 | -0.002 | 0 | 0.51 | 0.37 | 0.52 | 0.51 | 0.54 | 0.38 |
| 1 | 109784781 | rs586254 | -2.766 | -0.248 | 0.773 | -0.979 | 0.024 | -0.003 | -0.003 | 0 | 0.5 | 0.68 | 0.51 | 0.5 | 0.53 | 0.59 |
| 1 | 109788713 | rs17035443 | -1.249 | 0.31 | 1.225 | -0.614 | -0.006 | -0.033 | -0.033 | 0.516 | 0.2 | 0.22 | 0.14 | 0.08 | 0.2 | 0.29 |
| 1 | 109789795 | rs585362 | 1.371 | 0.735 | 1.295 | -0.302 | 0.077 | 0.261 | 0.261 | 0.342 | 0.91 | 0.72 | 0.76 | 0.59 | 0.88 | 0.79 |
| 1 | 109794252 | rs454107 | 0.444 | 0.583 | 1.103 | -0.645 | 0.023 | -0.021 | -0.021 | 0.216 | 0.93 | 0.72 | 0.96 | 1 | 0.97 | 0.94 |
| 1 | 109795608 | rs437444 | 0.195 | 0.592 | 1.02 | -0.641 | 0.019 | -0.022 | -0.022 | 3.614 | 0.93 | 0.76 | 0.97 | 1 | 0.97 | 0.94 |
| 1 | 109804646 | rs4970833 | 2.119 | 0.655 | 0.232 | -0.767 | -0.028 | -0.06 | -0.06 | 0 | 0.47 | 0.08 | 0.47 | 0.5 | 0.47 | 0.38 |
| 1 | 109806313 | rs653635 | -0.539 | 0.768 | 0.741 | -0.849 | 0.053 | -0.041 | -0.041 | 1.345 | 0.08 | 0.25 | 0.09 | 0 | 0.1 | 0.02 |
| 1 | 109807099 | rs6689614 | 2.231 | 0.913 | 0.2 | -0.695 | -0.028 | -0.061 | -0.061 | 0 | 0.47 | 0.08 | 0.47 | 0.5 | 0.48 | 0.37 |
| 1 | 109810544 | rs2281894 | -0.487 | 0.975 | 0.559 | -0.267 | -0.018 | -0.083 | -0.083 | 0 | 0.18 | 0.05 | 0.22 | 0.42 | 0.19 | 0.3 |
| 1 | 109810981 | rs17035630 | -0.013 | 0.846 | 0.473 | -0.248 | -0.001 | -0.115 | -0.115 | 1.556 | 0.15 | 0.01 | 0.19 | 0.42 | 0.12 | 0.29 |
| 1 | 109814880 | rs4970834 | -1.537 | 1.361 | 1.678 | -0.853 | 0.071 | -0.064 | -0.064 | 0 | 0.15 | 0.3 | 0.11 | 0.04 | 0.19 | 0.16 |
| 1 | 109817192 | rs7528419 | -1.834 | 1.372 | 1.53 | -0.895 | 0.017 | -0.047 | -0.047 | 2.451 | 0.19 | 0.27 | 0.2 | 0.04 | 0.21 | 0.26 |
| 1 | 109817838 | rs660240 | 1.464 | 1.345 | 1.547 | -0.908 | 0.021 | -0.067 | -0.067 | 0 | 0.8 | 0.61 | 0.79 | 0.96 | 0.8 | 0.74 |
| 1 | 109818270 | rs658435 | -0.581 | 1.507 | 1.658 | -0.988 | 0.003 | -0.016 | -0.016 | 0.03 | 0.07 | 0.07 | 0.07 | 0 | 0.11 | 0.02 |
| 1 | 109818530 | rs646776 | 1.487 | 1.456 | 1.642 | -0.939 | 0.024 | -0.069 | -0.069 | 3.434 | 0.8 | 0.6 | 0.78 | 0.95 | 0.79 | 0.74 |
| 1 | 109820919 | rs17035949 | -0.861 | 1.423 | 1.395 | -0.904 | 0.06 | -0.003 | -0.003 | 0.183 | 0.08 | 0.35 | 0.13 | 0.01 | 0.04 | 0.1 |
| 1 | 109822166 | rs599839 | 1.605 | 1.397 | 0.548 | -0.945 | -0.016 | 0.037 | 0.037 | 1.124 | 0.75 | 0.18 | 0.75 | 0.93 | 0.78 | 0.74 |
| 1 | 109822509 | rs14000 | -0.963 | 1.43 | 0.664 | -0.594 | 0.033 | -0.04 | -0.04 | 0.011 | 0.07 | 0.15 | 0.08 | 0 | 0.12 | 0.02 |
| 1 | 109832283 | rs655246 | 1.273 | 0.943 | -0.102 | -0.331 | -0.012 | -0.016 | -0.016 | 0 | 0.48 | 0.17 | 0.51 | 0.5 | 0.52 | 0.43 |
| 1 | 109834940 | rs17036094 | 0.394 | 0.819 | 0.294 | -0.375 | -0.023 | -0.032 | -0.032 | 0.454 | 0.06 | 0.12 | 0.01 | 0 | 0.01 | 0.04 |
| 1 | 109835757 | rs17645143 | -0.867 | 0.987 | -0.059 | -0.404 | -0.003 | -0.118 | -0.118 | 1.027 | 0.36 | 0.07 | 0.38 | 0.47 | 0.32 | 0.45 |
| 1 | 109838918 | rs629001 | 1.348 | 1.292 | 1.089 | -0.604 | 0.076 | -0.024 | -0.024 | 0.059 | 0.93 | 0.66 | 0.95 | 1 | 0.93 | 0.94 |
| 1 | 109846278 | rs407102 | 0.883 | 1.356 | 1.198 | -0.925 | 0.026 | -0.09 | -0.09 | 0 | 0.75 | 0.58 | 0.84 | 0.9 | 0.7 | 0.71 |
| 1 | 109851126 | rs413582 | 2.212 | 0.858 | 0.43 | -1.091 | -0.116 | 0.046 | 0.046 | 0.813 | 0.46 | 0.03 | 0.49 | 0.61 | 0.49 | 0.42 |
| 1 | 109856306 | rs464218 | 2.007 | 0.854 | 0.896 | -1.11 | -0.034 | 0.104 | 0.104 | 0.171 | 0.49 | 0.1 | 0.56 | 0.63 | 0.54 | 0.46 |
| 1 | 109871787 | rs10745352 | 0.956 | 0.969 | 1.312 | -1.064 | -0.001 | -0.072 | -0.072 | 0 | 0.72 | 0.36 | 0.83 | 0.95 | 0.71 | 0.72 |
| 1 | 109880721 | rs11102972 | -0.595 | 1.068 | 1.804 | -1.097 | 0.003 | -0.014 | -0.014 | 1.574 | 0.22 | 0.16 | 0.12 | 0.04 | 0.23 | 0.22 |
| 1 | 109897103 | rs11142 | 0.71 | 1.068 | 1.959 | -1.096 | 0.003 | -0.087 | -0.087 | 0 | 0.75 | 0.69 | 0.85 | 0.96 | 0.71 | 0.72 |
| 1 | 109923677 | rs12037569 | -0.292 | 0.655 | 1.871 | -1.155 | -0.007 | -0.049 | -0.049 | 0 | 0.15 | 0.05 | 0.25 | 0.31 | 0.16 | 0.21 |
| 1 | 109943893 | rs10858092 | 1.389 | 0.633 | 1.971 | -1.428 | -0.001 | -0.072 | -0.072 | 0 | 0.72 | 0.36 | 0.83 | 0.96 | 0.71 | 0.72 |
| 1 | 110010289 | rs10857787 | 1.209 | 0.6 | 1.966 | -1.438 | 0.007 | -0.153 | -0.153 | 3.47 | 0.73 | 0.53 | 0.84 | 0.96 | 0.69 | 0.71 |
| 1 | 110011701 | rs2152924 | 1.374 | 0.586 | 2.191 | -1.445 | 0 | -0.084 | -0.084 | 0.012 | 0.72 | 0.41 | 0.83 | 0.96 | 0.69 | 0.71 |
| 1 | 110014170 | rs12136063 | 1.39 | 0.582 | 2.142 | -1.446 | 0.003 | 0.016 | 0.016 | 0 | 0.7 | 0.27 | 0.82 | 0.96 | 0.69 | 0.71 |
| 1 | 110015732 | rs10494040 | 1.216 | 0.635 | 1.954 | -1.422 | 0.007 | -0.153 | -0.153 | 0 | 0.73 | 0.53 | 0.84 | 0.96 | 0.69 | 0.71 |
| 1 | 110030945 | rs1144593 | -1.26 | 0.764 | 1.485 | -1.407 | 0.005 | -0.096 | -0.096 | 0.118 | 0.26 | 0.31 | 0.15 | 0.04 | 0.31 | 0.22 |
| 1 | 110031188 | rs12049330 | -0.191 | 0.71 | 1.114 | -1.168 | -0.02 | -0.063 | -0.063 | 2.46 | 0.14 | 0.02 | 0.12 | 0.29 | 0.15 | 0.21 |
| 1 | 110035159 | rs11590351 | -1.207 | 0.765 | 1.231 | -1.492 | -0.003 | -0.047 | -0.047 | 0 | 0.25 | 0.25 | 0.13 | 0.04 | 0.26 | 0.24 |
| 1 | 110040310 | rs6666119 | -1.364 | 0.7 | 1.193 | -1.511 | 0.012 | -0.152 | -0.152 | 0.413 | 0.27 | 0.41 | 0.16 | 0.04 | 0.31 | 0.23 |
| 1 | 110040680 | rs2297797 | 2.311 | 0.851 | 0.547 | -1.534 | -0.119 | 0.083 | 0.083 | 0.462 | 0.43 | 0.08 | 0.63 | 0.66 | 0.47 | 0.42 |
| 1 | 110047333 | rs1009437 | -0.633 | 0.776 | 0.899 | -1.055 | 0.002 | -0.004 | -0.004 | 4.453 | 0.14 | 0.09 | 0.08 | 0.29 | 0.1 | 0.21 |
| 1 | 110048436 | rs3738772 | -1.315 | 0.947 | 0.953 | -1.059 | -0.007 | -0.014 | -0.014 | 0.322 | 0.28 | 0.34 | 0.18 | 0.29 | 0.28 | 0.26 |
| 1 | 110056126 | rs6677291 | -1.031 | 0.669 | 0.987 | -1.011 | -0.035 | -0.022 | -0.022 | 0 | 0.3 | 0.36 | 0.16 | 0.29 | 0.23 | 0.34 |
| 1 | 110061625 | rs534135 | -2.567 | 0.754 | 0.433 | -1.523 | -0.008 | 0.146 | 0.146 | 0 | 0.54 | 0.87 | 0.39 | 0.36 | 0.46 | 0.61 |
| 1 | 110063540 | rs6537803 | -0.991 | 0.731 | 0.657 | -1.535 | -0.023 | -0.024 | -0.024 | 0 | 0.32 | 0.57 | 0.18 | 0.28 | 0.23 | 0.35 |
| 1 | 110065906 | rs529773 | -2.614 | 0.727 | 0.593 | -1.542 | -0.003 | 0.132 | 0.132 | 0.13 | 0.55 | 0.84 | 0.39 | 0.36 | 0.46 | 0.62 |
| 1 | 110070554 | rs532623 | -1.619 | 1.477 | 0.944 | -0.951 | 0 | -0.026 | -0.026 | 0 | 0.22 | 0.28 | 0.21 | 0.09 | 0.22 | 0.27 |
| 1 | 110080809 | rs1434285 | -1.83 | 1.752 | 1.103 | -0.635 | -0.002 | 0.033 | 0.033 | 0 | 0.22 | 0.56 | 0.18 | 0.36 | 0.18 | 0.36 |
| 1 | 110084408 | rs12044778 | -1.222 | 1.727 | 1.348 | -0.619 | -0.001 | -0.003 | -0.003 | 0 | 0.15 | 0.12 | 0.13 | 0.36 | 0.16 | 0.3 |
| 1 | 110084827 | rs12023963 | -2.071 | 1.712 | 1.602 | -0.591 | -0.004 | 0.049 | 0.049 | 0 | 0.22 | 0.49 | 0.17 | 0.37 | 0.17 | 0.36 |
| 1 | 110108742 | rs6692804 | -2.2 | 1.67 | 1.868 | -0.644 | -0.011 | 0.034 | 0.034 | 0.14 | 0.23 | 0.52 | 0.18 | 0.37 | 0.18 | 0.37 |
| 1 | 110109039 | rs518076 | -0.837 | 1.666 | 1.754 | -0.77 | 0 | -0.027 | -0.027 | 0.528 | 0.09 | 0.14 | 0.09 | 0.02 | 0.08 | 0.13 |
| 1 | 110119732 | rs6537837 | -1.492 | 1.678 | 1.705 | -0.757 | -0.001 | 0.002 | 0.002 | 0.002 | 0.18 | 0.17 | 0.16 | 0.34 | 0.17 | 0.32 |
| 1 | 110132098 | rs541320 | -0.018 | 2.143 | 1.404 | -1.04 | -0.018 | 0.019 | 0.019 | 0 | 0.88 | 0.95 | 0.9 | 1 | 0.82 | 0.92 |
| 1 | 110135308 | rs7371 | -1.283 | 2.125 | 1.416 | -1.047 | 0 | -0.001 | -0.001 | 0 | 0.18 | 0.14 | 0.15 | 0.34 | 0.17 | 0.32 |
| 1 | 110136458 | rs2301230 | 3.257 | 2.12 | 1.409 | -1.08 | -0.038 | 0.106 | 0.106 | 0.792 | 0.54 | 0.17 | 0.6 | 0.64 | 0.53 | 0.41 |
| 1 | 110140977 | rs505303 | -2.964 | 2.62 | 1.274 | -1.217 | -0.01 | 0.041 | 0.041 | 3.114 | 0.37 | 0.84 | 0.34 | 0.37 | 0.33 | 0.55 |
| 1 | 110147098 | rs17024249 | -0.057 | 2.594 | 1.123 | -1.117 | -0.01 | 0.043 | 0.043 | 0.454 | 0.06 | 0.06 | 0.03 | 0.24 | 0.03 | 0.14 |
| 1 | 110148974 | rs1799875 | 3.26 | 2.624 | 0.823 | -1.192 | -0.037 | 0.131 | 0.131 | 2.087 | 0.49 | 0.06 | 0.54 | 0.63 | 0.46 | 0.36 |

| | | | | | | | | | | | | | | | | |
|---|---|---|---|---|---|---|---|---|---|---|---|---|---|---|---|---|
| 1 | 110151395 | rs3738766 | 0.293 | 2.629 | 0.885 | -1.124 | 0.004 | -0.029 | -0.029 | 0 | 0.06 | 0 | 0.03 | 0.24 | 0.03 | 0.14 |
| 1 | 110155524 | rs2304355 | -0.572 | 2.667 | 0.818 | -1.134 | -0.013 | 0.007 | 0.007 | 4.23 | 0.11 | 0.06 | 0.12 | 0.11 | 0.14 | 0.19 |
| 1 | 110156212 | rs4970768 | 2.663 | 2.481 | 0.994 | -1.108 | 0.025 | 0.117 | 0.117 | 1.604 | 0.51 | 0.23 | 0.56 | 0.63 | 0.46 | 0.36 |
| 1 | 110157364 | rs510619 | -1.061 | 2.311 | 1.068 | -0.849 | -0.002 | -0.017 | -0.017 | 0 | 0.13 | 0.15 | 0.14 | 0.02 | 0.14 | 0.17 |
| 1 | 110164657 | rs12046107 | -1.222 | 2.274 | 1.364 | -0.579 | -0.014 | 0.099 | 0.099 | 0.257 | 0.14 | 0.35 | 0.08 | 0.31 | 0.08 | 0.3 |
| 1 | 110169190 | rs865774 | -0.858 | 2.657 | 0.762 | -0.431 | -0.01 | 0.009 | 0.009 | 0 | 0.13 | 0.13 | 0.13 | 0.04 | 0.17 | 0.16 |
| 1 | 110169957 | rs524998 | 0.221 | 3.197 | 0.79 | -0.579 | -0.017 | 0.006 | 0.006 | 0.095 | 0.82 | 0.89 | 0.79 | 0.98 | 0.68 | 0.89 |
| 1 | 110173775 | rs568686 | -0.813 | 3.019 | 0.783 | -0.408 | -0.009 | 0.008 | 0.008 | 0 | 0.13 | 0.12 | 0.13 | 0.04 | 0.17 | 0.16 |
| 1 | 110175980 | rs7544426 | -1.12 | 2.482 | 1.037 | -0.025 | -0.013 | 0.213 | 0.213 | 2.065 | 0.15 | 0.42 | 0.08 | 0.39 | 0.08 | 0.3 |
| 1 | 110177848 | rs3895838 | -1.261 | 2.286 | 0.956 | -0.135 | -0.032 | 0.156 | 0.156 | 1.313 | 0.17 | 0.42 | 0.09 | 0.35 | 0.1 | 0.31 |
| 1 | 110186563 | rs12745189 | 2.174 | 2.581 | 0.257 | -0.589 | 0.002 | 0.006 | 0.006 | 2.695 | 0.55 | 0.43 | 0.62 | 0.64 | 0.45 | 0.4 |
| 1 | 110195228 | rs681991 | -1.798 | 2.8 | -0.052 | -0.646 | 0.032 | -0.004 | -0.004 | 0 | 0.33 | 0.44 | 0.29 | 0.33 | 0.41 | 0.45 |
| 1 | 110195864 | rs668532 | -1.809 | 2.797 | -0.053 | -0.647 | 0.032 | -0.004 | -0.004 | 0 | 0.33 | 0.44 | 0.29 | 0.33 | 0.41 | 0.45 |
| 1 | 110195944 | rs668413 | -1.798 | 2.803 | -0.051 | -0.645 | 0.032 | -0.004 | -0.004 | 0.288 | 0.33 | 0.44 | 0.29 | 0.33 | 0.41 | 0.45 |
| 1 | 110200360 | rs560018 | -0.671 | 2.916 | -0.572 | -0.809 | -0.042 | 0.037 | 0.037 | 0.257 | 0.18 | 0.09 | 0.23 | 0.05 | 0.33 | 0.12 |
| 1 | 110201809 | rs562996 | 0.882 | 2.11 | -0.358 | -0.725 | 0.003 | -0.013 | -0.013 | 0 | 0.87 | 0.82 | 0.9 | 0.97 | 0.86 | 0.85 |
| 1 | 110210671 | rs625456 | -0.733 | 1.967 | -0.449 | -1.219 | 0.071 | -0.038 | -0.038 | 0.012 | 0.07 | 0.19 | 0.08 | 0 | 0.15 | 0.05 |
| 1 | 110213458 | rs673151 | 0.237 | 1.254 | -0.915 | -1.037 | 0.006 | 0.031 | 0.031 | 0 | 0.92 | 0.98 | 0.95 | 1 | 0.94 | 0.91 |
| 1 | 110215178 | rs655315 | 0.75 | 1.221 | -1.137 | -0.968 | -0.048 | 0.114 | 0.114 | 5.074 | 0.52 | 0.12 | 0.61 | 0.67 | 0.49 | 0.46 |
| 1 | 110234286 | rs2239892 | -0.967 | 0.898 | -1.104 | -0.286 | -0.003 | 0.008 | 0.008 | 4.41 | 0.06 | 0.12 | 0.1 | 0.09 | 0.05 | 0.22 |
| 1 | 110253241 | rs3754446 | 0.095 | 0.831 | -2.25 | -0.764 | -0.051 | -0.021 | -0.021 | 0 | 0.38 | 0.02 | 0.47 | 0.69 | 0.34 | 0.37 |
| 1 | 110260742 | rs11807 | -0.872 | 1.504 | -2.097 | -1.09 | -0.004 | -0.003 | -0.003 | 0 | 0.12 | 0.11 | 0.16 | 0.16 | 0.18 | 0.21 |
| 1 | 110266754 | rs11101992 | -1.945 | 0.551 | -2.042 | -1.729 | 0.014 | 0.035 | 0.035 | 1.23 | 0.29 | 0.81 | 0.24 | 0.16 | 0.23 | 0.29 |
| 1 | 110268828 | rs4970774 | -0.616 | 0.3 | -1.604 | -2.122 | 0.003 | 0.023 | 0.023 | 0.042 | 0.52 | 0.55 | 0.52 | 0.71 | 0.45 | 0.5 |
| 1 | 110269286 | rs1854962 | 1.51 | 0.22 | -1.874 | -2.302 | -0.07 | -0.086 | -0.086 | 0 | 0.25 | 0.01 | 0.37 | 0.68 | 0.25 | 0.34 |
| 1 | 110273541 | rs12059276 | -0.46 | 0.183 | -1.679 | -2.32 | -0.027 | -0.112 | -0.112 | 0 | 0.15 | 0.45 | 0.07 | 0 | 0.08 | 0.04 |
| 1 | 110275744 | rs1927328 | 0.635 | 0.166 | -1.686 | -2.324 | -0.039 | 0.019 | 0.019 | 0.168 | 0.33 | 0.06 | 0.41 | 0.73 | 0.3 | 0.44 |
| 1 | 110275861 | rs1360513 | -0.833 | 0.181 | -1.672 | -2.313 | -0.001 | -0.014 | -0.014 | 0 | 0.52 | 0.48 | 0.52 | 0.74 | 0.45 | 0.51 |
| 1 | 110277949 | rs1109138 | -1.615 | 0.182 | -1.798 | -2.286 | -0.053 | 0.142 | 0.142 | 0 | 0.62 | 0.91 | 0.67 | 0.87 | 0.56 | 0.72 |
| 1 | 110279602 | rs2234696 | -0.464 | 0.18 | -1.538 | -2.282 | 0.01 | -0.049 | -0.049 | 0 | 0.09 | 0.33 | 0.07 | 0 | 0.07 | 0.08 |
| 1 | 110279701 | rs7483 | 0.717 | 0.223 | -1.993 | -2.259 | -0.05 | 0.008 | 0.008 | 0.013 | 0.32 | 0.06 | 0.4 | 0.73 | 0.3 | 0.42 |
| 1 | 110281941 | rs10735234 | -1.29 | 0.178 | -1.865 | -2.308 | -0.064 | 0.146 | 0.146 | 0 | 0.62 | 0.88 | 0.66 | 0.86 | 0.55 | 0.71 |
| 1 | 110283785 | rs4970777 | -1.261 | 0.333 | -1.894 | -2.316 | -0.067 | 0.146 | 0.146 | 0 | 0.61 | 0.91 | 0.66 | 0.86 | 0.55 | 0.71 |
| 1 | 110294577 | rs2274536 | -1.528 | 0.855 | -1.731 | -2.313 | -0.002 | -0.04 | -0.04 | 0.068 | 0.31 | 0.67 | 0.28 | 0.13 | 0.27 | 0.31 |
| 1 | 110295772 | rs1887547 | -1.162 | 0.654 | -1.754 | -2.237 | -0.014 | 0.159 | 0.159 | 0 | 0.6 | 0.91 | 0.68 | 0.86 | 0.58 | 0.74 |
| 1 | 110298107 | rs11591171 | 0.313 | 0.783 | -1.487 | -2.076 | 0.002 | -0.009 | -0.009 | 0.62 | 0.05 | 0.05 | 0.09 | 0.02 | 0.08 | 0.05 |
| 1 | 110299550 | rs2094469 | -1.337 | 0.76 | -1.287 | -1.892 | 0 | -0.03 | -0.03 | 0.288 | 0.25 | 0.31 | 0.21 | 0.09 | 0.22 | 0.33 |
| 1 | 110299691 | rs11102001 | -0.65 | 0.599 | -0.855 | -1.349 | 0.006 | -0.059 | -0.059 | 0 | 0.11 | 0.43 | 0.13 | 0.01 | 0.06 | 0.07 |
| 1 | 110300441 | rs3818562 | -0.178 | 1.049 | -1.977 | -0.903 | -0.076 | 0.122 | 0.122 | 0 | 0.42 | 0.16 | 0.44 | 0.79 | 0.47 | 0.6 |
| 1 | 110305188 | rs7533771 | -0.983 | 1.36 | -1.251 | -0.065 | -0.05 | -0.036 | -0.036 | 1.93 | 0.17 | 0.08 | 0.18 | 0.23 | 0.24 | 0.2 |
| 1 | 110307190 | rs3814310 | -1.003 | 1.192 | -1.099 | 0.033 | -0.022 | -0.029 | -0.029 | 0.909 | 0.16 | 0.1 | 0.15 | 0.23 | 0.21 | 0.21 |
| 1 | 110307263 | rs1109113 | -1.022 | 1.185 | -1.103 | 0.029 | -0.022 | -0.029 | -0.029 | 0 | 0.16 | 0.1 | 0.15 | 0.23 | 0.21 | 0.21 |
| 1 | 110307440 | rs943640 | -1.858 | 0.48 | -0.065 | -0.595 | -0.007 | -0.025 | -0.025 | 0.045 | 0.32 | 0.47 | 0.23 | 0.24 | 0.32 | 0.28 |
| 1 | 110313033 | rs12062976 | -1.351 | 0.477 | 1.07 | -0.37 | 0.021 | 0.041 | 0.041 | 0 | 0.19 | 0.34 | 0.18 | 0.27 | 0.22 | 0.21 |
| 1 | 110320104 | rs12126552 | -0.908 | 0.25 | 1.224 | -0.374 | -0.022 | -0.055 | -0.055 | 0 | 0.09 | 0.26 | 0.07 | 0 | 0.06 | 0.07 |
| 1 | 110320249 | rs7519527 | -1.173 | 0.178 | 1.282 | -0.243 | 0.037 | 0.064 | 0.064 | 0 | 0.19 | 0.47 | 0.19 | 0.27 | 0.22 | 0.2 |
| 1 | 110325532 | rs6681343 | -1.357 | -0.2 | 1.262 | -0.42 | -0.005 | 0.011 | 0.011 | 0.429 | 0.24 | 0.58 | 0.2 | 0.29 | 0.21 | 0.27 |
| 1 | 110329003 | rs11581614 | -1.785 | -0.093 | 0.882 | -0.424 | -0.035 | -0.118 | -0.118 | 0 | 0.25 | 0.56 | 0.14 | 0.02 | 0.16 | 0.16 |
| 1 | 110332816 | rs7524737 | -0.855 | 0.208 | -0.092 | 0.132 | -0.001 | -0.027 | -0.027 | 0 | 0.17 | 0.09 | 0.14 | 0.29 | 0.2 | 0.15 |
| 1 | 110334034 | rs11102006 | 1.071 | 0.204 | 0.437 | 0.158 | -0.011 | -0.019 | -0.019 | 0 | 0.79 | 0.39 | 0.87 | 0.87 | 0.84 | 0.71 |
| 1 | 110335680 | rs4970781 | 0.588 | -0.204 | -0.148 | 0.154 | 0.005 | -0.002 | -0.002 | 0 | 0.6 | 0.36 | 0.61 | 0.61 | 0.68 | 0.7 |
| 1 | 110338434 | rs485074 | -1.031 | -0.337 | -0.129 | -0.361 | -0.018 | -0.013 | -0.013 | 0.237 | 0.11 | 0.66 | 0.08 | 0.06 | 0.04 | 0.06 |
| 1 | 110343644 | rs12727662 | -0.553 | 0 | -1.224 | -0.798 | 0.02 | 0.142 | 0.142 | 0 | 0.28 | 0.07 | 0.2 | 0 | 0.25 | 0.19 |
| 1 | 110359882 | rs480419 | -0.39 | 0.608 | -0.935 | -1.024 | -0.018 | 0.019 | 0.019 | 5.84 | 0.69 | 0.81 | 0.37 | 0.12 | 0.55 | 0.54 |
| 1 | 110361682 | rs499345 | -0.006 | -0.071 | -0.561 | -1.891 | 0.003 | -0.059 | -0.059 | 2.527 | 0.42 | 0.41 | 0.22 | 0.02 | 0.28 | 0.4 |
| 1 | 110364275 | rs2938616 | -0.384 | -0.802 | -0.78 | -2.074 | 0.016 | 0.042 | 0.042 | 1.833 | 0.47 | 0.68 | 0.27 | 0.14 | 0.36 | 0.38 |
| 1 | 110366083 | rs484959 | -1.335 | -0.733 | -0.726 | -2.03 | -0.079 | 0.045 | 0.045 | 5.352 | 0.7 | 0.9 | 0.39 | 0.13 | 0.55 | 0.54 |
| 1 | 110366310 | rs483096 | -0.248 | -1.001 | -0.048 | -2.029 | -0.038 | -0.066 | -0.066 | 0.998 | 0.23 | 0.65 | 0.14 | 0.02 | 0.09 | 0.24 |
| 1 | 110368230 | rs521093 | -0.477 | -0.239 | 0.47 | -1.801 | 0.011 | -0.054 | -0.054 | 0 | 0.2 | 0.38 | 0.16 | 0.09 | 0.16 | 0.21 |
| 1 | 110369740 | rs6537880 | 0.865 | -0.212 | 0.006 | -1.295 | -0.035 | 0.003 | 0.003 | 0 | 0.19 | 0.12 | 0.13 | 0.15 | 0.25 | 0.13 |
| 1 | 110371904 | rs12563382 | -0.564 | -0.583 | 0.572 | -0.077 | -0.029 | 0.018 | 0.018 | 0 | 0.27 | 0.33 | 0.18 | 0.35 | 0.15 | 0.23 |
| 1 | 110372198 | rs523213 | -0.768 | -0.582 | 0.652 | -0.03 | -0.052 | 0.025 | 0.025 | 0 | 0.29 | 0.51 | 0.19 | 0.35 | 0.15 | 0.23 |
| 1 | 110375695 | rs525566 | -0.547 | 0.505 | 0.436 | 0.407 | 0.011 | 0.018 | 0.018 | 0.002 | 0.81 | 0.78 | 0.49 | 0.59 | 0.69 | 0.63 |
| 1 | 110379315 | rs484673 | -0.705 | 0.579 | -0.221 | 0.251 | -0.035 | -0.043 | -0.043 | 0.079 | 0.83 | 0.9 | 0.59 | 0.62 | 0.72 | 0.65 |
| 1 | 110408241 | rs10857800 | -1.018 | -0.204 | 0.851 | 0.418 | -0.018 | 0.005 | 0.005 | 0.19 | 0.16 | 0.66 | 0.17 | 0.2 | 0.08 | 0.12 |
| 1 | 110415626 | rs6695237 | 0.008 | 0.318 | 0.33 | -0.257 | 0.004 | 0.136 | 0.136 | 0 | 0.49 | 0.28 | 0.4 | 0.11 | 0.46 | 0.33 |
| 1 | 110415799 | rs12036949 | 0.525 | 0.688 | 0.797 | 0.346 | 0.042 | 0.061 | 0.061 | 0 | 0.05 | 0.15 | 0.03 | 0.1 | 0.06 | 0.06 |
| 1 | 110416867 | rs7522621 | 0.57 | 0.688 | 0.887 | 0.361 | 0.046 | 0.069 | 0.069 | 0 | 0.05 | 0.15 | 0.02 | 0.1 | 0.06 | 0.05 |
| 1 | 110418429 | rs11102019 | -0.094 | 0.595 | 0.86 | 0.359 | 0.018 | -0.012 | -0.012 | 0.222 | 0.65 | 0.49 | 0.71 | 0.31 | 0.59 | 0.64 |

| | | | | | | | | | | | | | | | | |
|---|---|---|---|---|---|---|---|---|---|---|---|---|---|---|---|---|
| 1 | 110420222 | rs7527452 | 0.263 | 0.844 | 1.166 | 0.728 | 0.14 | 0.275 | 0.275 | 0.012 | 0.92 | 0.58 | 0.94 | 0.77 | 0.92 | 0.98 |
| 1 | 110424529 | rs1021880 | 0.053 | 0.613 | 0.852 | 0.594 | 0.069 | 0.182 | 0.182 | 0.108 | 0.93 | 0.64 | 0.95 | 0.86 | 0.95 | 0.99 |
| 1 | 110426778 | rs453577 | -0.245 | 0.448 | -0.026 | 0.017 | -0.002 | -0.017 | -0.017 | 0 | 0.62 | 0.65 | 0.66 | 0.78 | 0.67 | 0.58 |
| 1 | 110431378 | rs433192 | -0.088 | 0.565 | -0.238 | 0.506 | -0.011 | -0.024 | -0.024 | 0 | 0.41 | 0.13 | 0.27 | 0.35 | 0.4 | 0.32 |
| 1 | 110436506 | rs333958 | 0.376 | 0.841 | 0.056 | 0.421 | 0.006 | 0.002 | 0.002 | 0 | 0.66 | 0.48 | 0.69 | 0.68 | 0.67 | 0.69 |
| 1 | 110439480 | rs333960 | -0.045 | 0.822 | 0.104 | -0.228 | -0.01 | -0.091 | -0.091 | 0 | 0.76 | 0.68 | 0.84 | 0.98 | 0.78 | 0.89 |
| 1 | 110439636 | rs333961 | 0.3 | 1.149 | -0.63 | -1.024 | 0.024 | -0.098 | -0.098 | 0 | 0.3 | 0.52 | 0.32 | 0.08 | 0.36 | 0.22 |
| 1 | 110441792 | rs333966 | -0.199 | 1.247 | -0.616 | -0.527 | -0.004 | -0.017 | -0.017 | 0 | 0.7 | 0.59 | 0.81 | 0.79 | 0.73 | 0.72 |
| 1 | 110450177 | rs1999713 | 0.524 | 0.959 | -0.331 | -0.298 | 0.028 | 0.019 | 0.019 | 0 | 0.42 | 0.31 | 0.51 | 0.68 | 0.35 | 0.55 |
| 1 | 110451043 | rs333949 | -0.548 | 0.954 | -0.13 | -0.214 | 0.003 | -0.002 | -0.002 | 0 | 0.75 | 0.79 | 0.83 | 0.98 | 0.76 | 0.88 |
| 1 | 110451190 | rs333950 | 0.52 | 1.294 | 0.049 | -0.423 | -0.001 | 0.001 | 0.001 | 0 | 0.62 | 0.59 | 0.66 | 0.72 | 0.58 | 0.65 |
| 1 | 110451690 | rs333951 | -0.306 | 1.087 | -0.266 | 0.34 | 0.001 | -0.001 | -0.001 | 0.348 | 0.73 | 0.8 | 0.72 | 0.71 | 0.72 | 0.85 |
| 1 | 110455433 | rs915357 | 0.849 | 0.685 | -0.192 | -0.461 | -0.013 | 0.012 | 0.012 | 1.03 | 0.38 | 0.29 | 0.31 | 0 | 0.47 | 0.24 |
| 1 | 110458234 | rs2275123 | -0.565 | 1.036 | -0.15 | 0.56 | 0.019 | 0.074 | 0.074 | 0 | 0.12 | 0.17 | 0.17 | 0.38 | 0.13 | 0.11 |
| 1 | 110460520 | rs333968 | 0.606 | 0.7 | 0.024 | 0.747 | -0.028 | 0.07 | 0.07 | 0.134 | 0.84 | 0.39 | 0.78 | 0.79 | 0.9 | 0.79 |
| 1 | 110466338 | rs333970 | 1.761 | 0.641 | -1.087 | 0.429 | -0.041 | -0.039 | -0.039 | 4.657 | 0.6 | 0.11 | 0.57 | 0.4 | 0.62 | 0.6 |
| 1 | 110470764 | rs333947 | 0.85 | 0.372 | -1.614 | 0.76 | -0.026 | -0.115 | -0.115 | 1.874 | 0.2 | 0.01 | 0.28 | 0.28 | 0.13 | 0.28 |
| 1 | 110471906 | rs3093037 | -0.178 | 1.176 | -0.877 | 0.616 | -0.027 | 0.04 | 0.04 | 0 | 0.14 | 0.07 | 0.11 | 0 | 0.18 | 0.08 |
| 1 | 110478519 | rs1018738 | -0.279 | 1.259 | -0.344 | 1.203 | 0.053 | 0.187 | 0.187 | 0 | 0.06 | 0.16 | 0.09 | 0.36 | 0.08 | 0.05 |
| 1 | 110478803 | rs6537885 | -0.467 | 0.845 | -0.95 | 1.12 | -0.018 | -0.063 | -0.063 | 0 | 0.77 | 0.97 | 0.74 | 0.71 | 0.8 | 0.56 |
| 1 | 110478911 | rs7555325 | 0.825 | 0.892 | -0.827 | 0.978 | -0.029 | 0.037 | 0.037 | 0 | 0.63 | 0.85 | 0.57 | 0.71 | 0.61 | 0.48 |
| 1 | 110480564 | rs7417586 | 0.195 | 1.023 | -0.741 | 0.524 | -0.013 | 0.013 | 0.013 | 0 | 0.14 | 0.1 | 0.17 | 0 | 0.19 | 0.08 |
| 1 | 110482335 | rs11589677 | 1.781 | 0.635 | -0.435 | 0.673 | -0.018 | 0.143 | 0.143 | 0 | 0.4 | 0.19 | 0.32 | 0.06 | 0.43 | 0.26 |
| 1 | 110483773 | rs7540934 | 1.092 | 0.711 | -0.033 | 0.721 | 0.022 | 0.109 | 0.109 | 0 | 0.48 | 0.37 | 0.34 | 0.06 | 0.44 | 0.26 |
| 1 | 110486839 | rs17025216 | -1.148 | 0.804 | -0.033 | 1.303 | 0.039 | 0.161 | 0.161 | 0 | 0.11 | 0.19 | 0.07 | 0.48 | 0.15 | 0.1 |
| 1 | 110490341 | rs17025223 | -0.664 | 0.821 | -0.255 | 1.091 | 0.023 | 0.093 | 0.093 | 0 | 0.09 | 0.17 | 0.05 | 0.34 | 0.1 | 0.06 |
| 1 | 110492809 | rs7365459 | 0.793 | 0.895 | -0.112 | 0.969 | -0.001 | -0.003 | -0.003 | 0 | 0.53 | 0.46 | 0.37 | 0.4 | 0.51 | 0.32 |
| 1 | 110499669 | rs390923 | -0.896 | 1.116 | 0.223 | 1.367 | 0.003 | 0.001 | 0.001 | 0 | 0.47 | 0.5 | 0.59 | 0.45 | 0.41 | 0.67 |
| 1 | 110501597 | rs11579145 | 1.32 | 1.297 | -0.212 | 1.431 | -0.021 | 0.072 | 0.072 | 0.284 | 0.45 | 0.22 | 0.37 | 0.23 | 0.52 | 0.28 |
| 1 | 110503433 | rs17025286 | -1.036 | 1.155 | 0.029 | 1.679 | 0 | 0.009 | 0.009 | 0 | 0.11 | 0.13 | 0.07 | 0.31 | 0.07 | 0.06 |
| 1 | 110517396 | rs11102043 | -0.3 | 1.433 | -0.143 | 1.518 | -0.019 | -0.062 | -0.062 | 0 | 0.2 | 0.06 | 0.32 | 0.34 | 0.17 | 0.48 |
| 1 | 110525630 | rs333079 | -0.644 | 1.726 | 0.769 | 1.551 | -0.023 | -0.009 | -0.009 | 0 | 0.28 | 0.21 | 0.29 | 0.28 | 0.33 | 0.22 |
| 1 | 110529484 | rs9429768 | -0.357 | 1.889 | 0.757 | 1.355 | -0.067 | -0.064 | -0.064 | 0 | 0.23 | 0.06 | 0.17 | 0.31 | 0.28 | 0.17 |
| 1 | 110533997 | rs10857808 | -1.054 | 1.689 | 1.135 | 1.051 | 0.085 | 0.062 | 0.062 | 0.759 | 0.05 | 0.4 | 0.04 | 0 | 0.02 | 0.05 |
| 1 | 110534462 | rs3754444 | -0.178 | 1.699 | 0.928 | 1.076 | -0.013 | -0.058 | -0.058 | 0 | 0.21 | 0.07 | 0.32 | 0.35 | 0.17 | 0.5 |
| 1 | 110538926 | rs17025313 | -0.193 | 1.698 | 0.928 | 1.074 | -0.013 | -0.056 | -0.056 | 0 | 0.21 | 0.07 | 0.32 | 0.34 | 0.16 | 0.5 |
| 1 | 110543024 | rs720917 | -0.213 | 1.693 | 0.987 | 1.049 | -0.002 | -0.031 | -0.031 | 0 | 0.21 | 0.13 | 0.32 | 0.34 | 0.16 | 0.49 |
| 1 | 110587956 | rs7545139 | -1.709 | 1.872 | 1.169 | 1.021 | 0.002 | 0.001 | 0.001 | 0 | 0.53 | 0.48 | 0.62 | 0.76 | 0.56 | 0.73 |
| 1 | 110595528 | rs10857809 | -0.205 | 1.736 | 1.094 | 1.034 | -0.001 | -0.005 | -0.005 | 0 | 0.26 | 0.22 | 0.34 | 0.34 | 0.21 | 0.51 |
| 1 | 110604747 | rs2360635 | -0.516 | 1.649 | 0.071 | 1.087 | -0.031 | -0.051 | -0.051 | 0 | 0.26 | 0.19 | 0.27 | 0.42 | 0.32 | 0.17 |
| 1 | 110606675 | rs904023 | -1.783 | 1.838 | 0.091 | 0.856 | 0.007 | 0.012 | 0.012 | 0.255 | 0.57 | 0.64 | 0.65 | 0.76 | 0.57 | 0.73 |
| 1 | 110609288 | rs3754443 | -1.997 | 1.852 | -0.345 | 0.861 | 0.033 | 0.06 | 0.06 | 0.508 | 0.57 | 0.77 | 0.66 | 0.77 | 0.58 | 0.78 |
| 1 | 110619665 | rs10127659 | -0.459 | 1.935 | -0.53 | 0.785 | -0.043 | -0.065 | -0.065 | 0 | 0.26 | 0.16 | 0.27 | 0.41 | 0.33 | 0.23 |
| 1 | 110624104 | rs1869596 | -1.439 | 1.441 | -0.632 | 0.015 | 0.025 | 0.058 | 0.058 | 0.666 | 0.56 | 0.73 | 0.66 | 0.78 | 0.57 | 0.77 |
| 1 | 110640235 | rs756134 | -0.412 | 1.082 | -0.673 | 0.127 | -0.075 | -0.098 | -0.098 | 0.072 | 0.31 | 0.08 | 0.46 | 0.48 | 0.38 | 0.54 |
| 1 | 110640433 | rs555095 | -1.245 | 1.178 | -0.527 | 0.125 | -0.001 | 0.062 | 0.062 | 2.339 | 0.18 | 0.7 | 0.18 | 0.35 | 0.11 | 0.19 |
| 1 | 110645327 | rs17025426 | -0.06 | 1.154 | -0.41 | 0.122 | -0.008 | -0.066 | -0.066 | 0.114 | 0.11 | 0.22 | 0.05 | 0.01 | 0.06 | 0.07 |
| 1 | 110646283 | rs673792 | -1.455 | 1.221 | -0.977 | 0.146 | 0.025 | 0.214 | 0.214 | 0.426 | 0.51 | 0.81 | 0.64 | 0.82 | 0.5 | 0.67 |
| 1 | 110650174 | rs4839136 | -0.507 | 1.517 | -1.091 | 0.635 | 0.005 | 0.053 | 0.053 | 0 | 0.12 | 0.13 | 0.12 | 0.35 | 0.1 | 0.12 |
| 1 | 110657532 | rs2168144 | -0.229 | 1.587 | -1.309 | 0.513 | -0.013 | -0.022 | -0.022 | 0 | 0.21 | 0.17 | 0.44 | 0.61 | 0.24 | 0.53 |
| 1 | 110657949 | rs17669930 | -0.422 | 1.591 | -1.316 | 0.62 | -0.013 | 0.048 | 0.048 | 0 | 0.17 | 0.08 | 0.15 | 0 | 0.24 | 0.11 |
| 1 | 110671320 | rs593644 | -0.492 | 1.024 | -1.235 | -0.255 | -0.001 | -0.012 | -0.012 | 0 | 0.49 | 0.46 | 0.64 | 0.79 | 0.5 | 0.68 |
| 1 | 110674137 | rs697387 | 0.582 | 0.82 | -1.735 | 0.276 | -0.003 | 0.003 | 0.003 | 0 | 0.71 | 0.92 | 0.76 | 0.75 | 0.69 | 0.81 |
| 1 | 110677921 | rs7518801 | -0.458 | 0.694 | -1.7 | -0.165 | 0 | 0.073 | 0.073 | 1.603 | 0.47 | 0.32 | 0.33 | 0.23 | 0.55 | 0.44 |
| 1 | 110678929 | rs17670272 | -0.445 | 1.209 | -1.752 | -0.219 | -0.023 | 0.065 | 0.065 | 1.366 | 0.4 | 0.19 | 0.29 | 0.23 | 0.49 | 0.4 |
| 1 | 110679596 | rs1010386 | -0.844 | 1.295 | -1.892 | -0.404 | -0.006 | 0.075 | 0.075 | 0.518 | 0.47 | 0.34 | 0.37 | 0.24 | 0.56 | 0.43 |
| 1 | 110680114 | rs924181 | -0.342 | 1.216 | -1.947 | -0.494 | -0.013 | 0.042 | 0.042 | 0.651 | 0.38 | 0.26 | 0.3 | 0.23 | 0.47 | 0.37 |
| 1 | 110684287 | rs10494113 | -0.808 | 1.116 | -1.939 | -0.491 | -0.015 | 0.062 | 0.062 | 1.731 | 0.45 | 0.34 | 0.36 | 0.24 | 0.56 | 0.43 |
| 1 | 110685011 | rs970977 | -0.532 | 1.106 | -1.94 | -0.499 | -0.011 | 0.062 | 0.062 | 1.783 | 0.41 | 0.24 | 0.29 | 0.23 | 0.49 | 0.4 |
| 1 | 110686937 | rs6679244 | -0.84 | 1.072 | -1.911 | -0.492 | -0.014 | 0.064 | 0.064 | 1.585 | 0.46 | 0.34 | 0.35 | 0.23 | 0.55 | 0.42 |
| 1 | 110690577 | rs975512 | -0.976 | 1.075 | -1.927 | -0.491 | -0.03 | 0.104 | 0.104 | 1.52 | 0.45 | 0.25 | 0.51 | 0.23 | 0.55 | 0.41 |
| 1 | 110698651 | rs2002384 | -0.707 | 1.204 | -1.218 | -0.408 | -0.032 | 0.051 | 0.051 | 1.961 | 0.38 | 0.28 | 0.34 | 0.21 | 0.52 | 0.38 |
| 1 | 110701071 | rs4838931 | -0.865 | 1.198 | -1.218 | -0.411 | -0.011 | 0.02 | 0.02 | 0.986 | 0.4 | 0.37 | 0.35 | 0.21 | 0.52 | 0.38 |
| 1 | 110703619 | rs11579877 | -0.671 | 1.293 | -1.376 | -0.366 | -0.028 | 0.039 | 0.039 | 1.063 | 0.38 | 0.3 | 0.34 | 0.21 | 0.53 | 0.38 |
| 1 | 110712154 | rs1010891 | -0.843 | 1.234 | -1.185 | -0.377 | 0.004 | -0.014 | -0.014 | 0 | 0.39 | 0.41 | 0.35 | 0.21 | 0.52 | 0.37 |
| 1 | 110713909 | rs7554444 | -0.279 | 1.287 | -1.358 | -0.357 | -0.136 | 0.037 | 0.037 | 2.461 | 0.33 | 0.03 | 0.26 | 0.21 | 0.47 | 0.35 |
| 1 | 110715768 | rs12116787 | -0.809 | 1.331 | -1.277 | -0.298 | 0.078 | 0.017 | 0.017 | 0.035 | 0.05 | 0.31 | 0.08 | 0 | 0.06 | 0.02 |
| 1 | 110716511 | rs12747833 | -0.477 | 1.601 | -2.134 | -0.026 | -0.118 | 0.011 | 0.011 | 1.449 | 0.34 | 0.04 | 0.26 | 0.25 | 0.45 | 0.35 |
| 1 | 110725640 | rs11584494 | -0.226 | 1.621 | -1.945 | 0.539 | -0.101 | -0.08 | -0.08 | 0 | 0.24 | 0.04 | 0.31 | 0.31 | 0.38 | 0.31 |

| | | | | | | | | | | | | | | | | |
|---|---|---|---|---|---|---|---|---|---|---|---|---|---|---|---|---|
| 1 | 110728459 | rs4839223 | -0.342 | 1.521 | -2.045 | 0.638 | -0.118 | -0.106 | -0.106 | 0.04 | 0.27 | 0.02 | 0.31 | 0.34 | 0.32 | 0.31 |
| 1 | 110729404 | rs2011688 | -0.408 | 0.561 | -1.651 | 0.993 | -0.002 | 0.018 | 0.018 | 0 | 0.11 | 0.14 | 0.09 | 0.22 | 0.08 | 0.34 |
| 1 | 110733674 | rs17025524 | -0.803 | 0.777 | -1.402 | 0.403 | 0.005 | -0.02 | -0.02 | 0 | 0.07 | 0.12 | 0.07 | 0.01 | 0.09 | 0.02 |
| 1 | 110733794 | rs621514 | -0.588 | 0.177 | -1.926 | 0.032 | 0.044 | 0.048 | 0.048 | 0.494 | 0.54 | 0.78 | 0.54 | 0.35 | 0.61 | 0.41 |
| 1 | 110738296 | rs2784140 | 0.448 | -0.001 | -2.21 | -0.13 | -0.184 | -0.053 | -0.053 | 0.024 | 0.33 | 0.04 | 0.38 | 0.3 | 0.45 | 0.37 |
| 1 | 110750277 | rs2003154 | -0.934 | -0.131 | -1.87 | -0.316 | -0.001 | -0.017 | -0.017 | 0.824 | 0.19 | 0.76 | 0.13 | 0.04 | 0.13 | 0.05 |
| 1 | 110753808 | rs11810910 | 0.435 | -0.432 | -2.26 | -0.29 | -0.175 | -0.062 | -0.062 | 0.238 | 0.35 | 0.05 | 0.39 | 0.31 | 0.45 | 0.37 |
| 1 | 110758375 | rs2784144 | -0.791 | -0.603 | -1.704 | -0.221 | 0.035 | 0.054 | 0.054 | 0 | 0.55 | 0.7 | 0.53 | 0.32 | 0.6 | 0.42 |
| 1 | 110759455 | rs2603594 | 0.06 | -0.455 | -1.261 | -0.379 | 0.005 | -0.015 | -0.015 | 0 | 0.42 | 0.46 | 0.43 | 0.31 | 0.5 | 0.38 |
| 1 | 110759796 | rs615204 | 0.396 | -0.441 | -1.576 | -0.233 | -0.059 | 0.052 | 0.052 | 0 | 0.39 | 0.14 | 0.38 | 0.24 | 0.47 | 0.38 |
| 1 | 110760189 | rs11102064 | -0.366 | -0.528 | -1.734 | -0.204 | -0.017 | -0.044 | -0.044 | 1.29 | 0.2 | 0.67 | 0.16 | 0.09 | 0.13 | 0.05 |
| 1 | 110760444 | rs628594 | -0.082 | -0.14 | -1.616 | 0.359 | 0.041 | 0.025 | 0.025 | 0 | 0.57 | 0.82 | 0.54 | 0.36 | 0.61 | 0.4 |
| 1 | 110769589 | rs2282262 | -0.43 | -0.313 | -1.321 | 0.686 | 0.009 | -0.016 | -0.016 | 0 | 0.33 | 0.27 | 0.42 | 0.5 | 0.29 | 0.54 |
| 1 | 110770223 | rs958798 | -0.826 | 0.331 | -1.207 | 0.22 | 0.01 | -0.018 | -0.018 | 0 | 0.13 | 0.2 | 0.13 | 0.06 | 0.2 | 0.09 |
| 1 | 110770293 | rs2077402 | -0.978 | 0.12 | -1.202 | 0.573 | -0.002 | 0 | 0 | 0.022 | 0.17 | 0.17 | 0.15 | 0.16 | 0.14 | 0.42 |
| 1 | 110774238 | rs615819 | 0.165 | 0.147 | -1.636 | 1.015 | -0.01 | -0.111 | -0.111 | 0.099 | 0.78 | 0.94 | 0.83 | 0.6 | 0.76 | 0.84 |
| 1 | 110775195 | rs17025562 | -0.601 | -0.146 | -1.123 | 0.856 | -0.024 | 0.002 | 0.002 | 0.09 | 0.09 | 0.16 | 0.08 | 0.11 | 0.01 | 0.1 |
| 1 | 110779084 | rs11102066 | -0.62 | -0.044 | -1.389 | 0.875 | -0.002 | -0.017 | -0.017 | 0 | 0.07 | 0.04 | 0.23 | 0.1 | 0.05 | 0.07 |
| 1 | 110780834 | rs2603590 | 0.205 | -0.15 | -1.442 | -0.256 | 0.002 | 0 | 0 | 0 | 0.43 | 0.36 | 0.35 | 0.23 | 0.37 | 0.49 |
| 1 | 110781469 | rs2181695 | 0.999 | -0.245 | -1.908 | -0.338 | -0.083 | -0.014 | -0.014 | 0 | 0.19 | 0.06 | 0.19 | 0.22 | 0.3 | 0.26 |
| 1 | 110784219 | rs814324 | -0.625 | -0.729 | -1.422 | -0.38 | -0.005 | 0.032 | 0.032 | 0 | 0.74 | 0.68 | 0.8 | 0.59 | 0.75 | 0.81 |
| 1 | 110788825 | rs814318 | -0.444 | -0.824 | -1.454 | -0.347 | 0.025 | -0.081 | -0.081 | 0.574 | 0.69 | 0.87 | 0.64 | 0.61 | 0.74 | 0.78 |
| 1 | 110789107 | rs814319 | 1.319 | -0.989 | -1.15 | -0.595 | -0.002 | 0.024 | 0.024 | 0 | 0.34 | 0.21 | 0.28 | 0.15 | 0.36 | 0.39 |
| 1 | 110790977 | rs2198612 | 0.813 | -1.012 | -1.043 | -0.535 | -0.091 | -0.05 | -0.05 | 0 | 0.2 | 0.08 | 0.25 | 0.32 | 0.36 | 0.29 |
| 1 | 110794827 | rs2224807 | -0.363 | -1.045 | -1.078 | -0.745 | 0.016 | -0.027 | -0.027 | 0.589 | 0.66 | 0.74 | 0.61 | 0.58 | 0.7 | 0.76 |
| 1 | 110795018 | rs2209158 | -0.736 | -0.936 | -0.946 | -0.57 | 0.022 | 0.009 | 0.009 | 0 | 0.33 | 0.53 | 0.35 | 0.43 | 0.37 | 0.38 |
| 1 | 110803935 | rs12146055 | 0.876 | -0.754 | -0.624 | -0.517 | -0.014 | -0.002 | -0.002 | 0.031 | 0.18 | 0.17 | 0.21 | 0.21 | 0.3 | 0.26 |
| 1 | 110814007 | rs1323723 | -0.048 | -0.806 | -0.721 | -0.762 | 0.001 | -0.001 | -0.001 | 0 | 0.52 | 0.46 | 0.65 | 0.51 | 0.53 | 0.59 |
| 1 | 110816152 | rs4839287 | -0.441 | -0.884 | -0.673 | -0.817 | 0.001 | 0.006 | 0.006 | 0 | 0.58 | 0.56 | 0.67 | 0.52 | 0.54 | 0.66 |
| 1 | 110816491 | rs3904626 | 0.482 | -0.945 | -0.63 | -0.846 | -0.003 | -0.002 | -0.002 | 3.426 | 0.35 | 0.34 | 0.46 | 0.3 | 0.24 | 0.4 |
| 1 | 110825766 | rs3916176 | 1.361 | -0.79 | -0.816 | -0.775 | -0.034 | -0.01 | -0.01 | 0 | 0.17 | 0.13 | 0.19 | 0.21 | 0.29 | 0.19 |
| 1 | 110830317 | rs17025584 | 1.573 | -0.847 | -0.836 | -0.774 | 0.004 | 0 | 0 | 0 | 0.06 | 0.05 | 0.01 | 0.01 | 0.01 | 0.07 |
| 1 | 110832827 | rs643841 | 1.187 | -0.699 | -0.681 | -0.748 | 0.005 | 0 | 0 | 0 | 0.18 | 0.25 | 0.19 | 0.21 | 0.29 | 0.19 |
| 1 | 110833150 | rs645177 | -0.397 | -0.77 | -0.703 | -0.96 | -0.002 | -0.007 | -0.007 | 0.175 | 0.59 | 0.72 | 0.69 | 0.53 | 0.54 | 0.66 |
| 1 | 110842415 | rs4112315 | 1.31 | -0.41 | -0.797 | -0.895 | -0.036 | 0.04 | 0.04 | 0 | 0.17 | 0.11 | 0.18 | 0.05 | 0.26 | 0.18 |
| 1 | 110846362 | rs497451 | -0.283 | -0.761 | -0.967 | -1.015 | 0.01 | 0.028 | 0.028 | 0.016 | 0.53 | 0.76 | 0.6 | 0.48 | 0.41 | 0.6 |
| 1 | 110847884 | rs666493 | 0.029 | -0.657 | -0.394 | -1.158 | -0.004 | 0.025 | 0.025 | 0 | 0.54 | 0.64 | 0.61 | 0.31 | 0.53 | 0.54 |
| 1 | 110850317 | rs617293 | 0.947 | -0.415 | -0.619 | -0.731 | -0.007 | -0.019 | -0.019 | 3.25 | 0.55 | 0.5 | 0.61 | 0.74 | 0.66 | 0.63 |
| 1 | 110851358 | rs566689 | -0.267 | -0.481 | 0.082 | -0.649 | 0.015 | -0.125 | -0.125 | 0 | 0.32 | 0.49 | 0.44 | 0.05 | 0.36 | 0.27 |
| 1 | 110851995 | rs12407097 | 0.246 | -0.492 | 0.143 | -0.486 | -0.002 | -0.027 | -0.027 | 0 | 0.12 | 0.16 | 0.22 | 0 | 0.09 | 0.1 |
| 1 | 110853284 | rs7544525 | -0.452 | -0.532 | 0.165 | -0.515 | 0.004 | -0.023 | -0.023 | 0.093 | 0.17 | 0.19 | 0.24 | 0.01 | 0.1 | 0.16 |
| 1 | 110861635 | rs4839325 | -1.536 | -0.575 | 0.24 | -0.344 | -0.009 | 0.018 | 0.018 | 0 | 0.32 | 0.58 | 0.49 | 0.46 | 0.25 | 0.37 |
| 1 | 110862068 | rs2201808 | 0.173 | -0.522 | 0.356 | -0.309 | 0.032 | 0.157 | 0.157 | 0 | 0.15 | 0.31 | 0.25 | 0.46 | 0.15 | 0.21 |
| 1 | 110863079 | rs6537656 | 0.048 | -0.478 | 0.161 | -0.228 | -0.017 | -0.065 | -0.065 | 0 | 0.13 | 0.23 | 0.23 | 0 | 0.09 | 0.1 |
| 1 | 110885736 | rs1572263 | -1.22 | -0.559 | -0.194 | -0.114 | -0.019 | -0.011 | -0.011 | 0 | 0.71 | 0.91 | 0.85 | 0.77 | 0.68 | 0.81 |
| 1 | 110886538 | rs4839342 | -0.2 | -0.668 | 0.255 | -0.272 | -0.018 | 0.017 | 0.017 | 0 | 0.34 | 0.33 | 0.37 | 0.27 | 0.41 | 0.41 |
| 1 | 110887075 | rs3768460 | -1.196 | -0.717 | 0.389 | -0.308 | -0.002 | -0.001 | -0.001 | 0 | 0.47 | 0.58 | 0.66 | 0.51 | 0.49 | 0.53 |
| 1 | 110892044 | rs6694203 | 1.478 | -0.607 | 0.264 | -0.272 | 0.005 | -0.017 | -0.017 | 0.435 | 0.09 | 0.14 | 0.05 | 0.05 | 0.07 | 0.08 |
| 1 | 110902305 | rs12733724 | -0.194 | -0.633 | 0.246 | -0.333 | -0.034 | 0.035 | 0.035 | 0 | 0.34 | 0.28 | 0.37 | 0.27 | 0.41 | 0.41 |
| 1 | 110913120 | rs722570 | -0.271 | -0.649 | 0.238 | -0.342 | -0.036 | 0.039 | 0.039 | 0 | 0.34 | 0.28 | 0.37 | 0.27 | 0.41 | 0.43 |
| 1 | 110922140 | rs2946571 | -0.087 | -0.728 | 0.263 | -0.276 | 0.079 | -0.015 | -0.015 | 0 | 0.43 | 0.72 | 0.5 | 0.51 | 0.28 | 0.44 |
| 1 | 110922797 | rs12403693 | 0.26 | -0.693 | 0.333 | -0.306 | -0.001 | -0.024 | -0.024 | 0.021 | 0.11 | 0.1 | 0.22 | 0 | 0.08 | 0.09 |
| 1 | 110922814 | rs12126890 | -1.603 | -0.756 | 0.193 | -0.236 | -0.013 | 0.003 | 0.003 | 0.016 | 0.73 | 0.93 | 0.88 | 0.79 | 0.72 | 0.81 |
| 1 | 110924353 | rs3738750 | 2.694 | -0.761 | 0.254 | -0.375 | 0.014 | -0.048 | -0.048 | 0.031 | 0.21 | 0.03 | 0.16 | 0.26 | 0.19 | 0.28 |
| 1 | 110929476 | rs1334882 | -1.395 | -0.725 | 0.531 | -0.398 | 0.009 | 0.004 | 0.004 | 0 | 0.5 | 0.62 | 0.67 | 0.51 | 0.52 | 0.54 |
| 1 | 110931112 | rs7556440 | -1.59 | -0.723 | 0.613 | -0.38 | 0.003 | -0.001 | -0.001 | 1.22 | 0.73 | 0.86 | 0.86 | 0.78 | 0.71 | 0.82 |
| 1 | 110936383 | rs12126959 | 0.597 | -0.6 | 0.555 | -0.42 | -0.001 | 0 | 0 | 0 | 0.15 | 0.15 | 0.17 | 0.22 | 0.17 | 0.19 |
| 1 | 110936557 | rs12127781 | 1.152 | -0.633 | 0.571 | -0.404 | 0.001 | -0.011 | -0.011 | 0.127 | 0.13 | 0.14 | 0.06 | 0.05 | 0.11 | 0.11 |
| 1 | 110949720 | rs6671673 | -1.413 | -0.616 | 0.682 | -0.414 | 0.017 | 0.008 | 0.008 | 0 | 0.5 | 0.65 | 0.68 | 0.52 | 0.52 | 0.53 |
| 1 | 110950277 | rs6698159 | 1.157 | -0.6 | 0.433 | -0.582 | 0.001 | -0.01 | -0.01 | 0.031 | 0.13 | 0.14 | 0.06 | 0.05 | 0.11 | 0.11 |
| 1 | 110950439 | rs6674775 | -0.272 | -0.698 | 0.508 | -0.498 | -0.027 | 0.045 | 0.045 | 0 | 0.37 | 0.31 | 0.38 | 0.28 | 0.45 | 0.44 |
| 1 | 110950462 | rs11102099 | 0.462 | -0.687 | 0.516 | -0.484 | -0.004 | -0.031 | -0.031 | 0.005 | 0.11 | 0.11 | 0.22 | 0 | 0.07 | 0.07 |
| 1 | 110955220 | rs4839011 | 0.132 | -0.614 | 0.246 | -0.429 | -0.044 | 0.036 | 0.036 | 0 | 0.32 | 0.25 | 0.36 | 0.27 | 0.44 | 0.37 |
| 1 | 110958034 | rs7546690 | -0.275 | -0.57 | 0.244 | -0.375 | -0.026 | 0.044 | 0.044 | 0 | 0.37 | 0.32 | 0.38 | 0.28 | 0.45 | 0.44 |
| 1 | 110960311 | rs6657333 | -1.443 | -0.672 | -0.282 | -0.529 | -0.005 | -0.002 | -0.002 | 0 | 0.73 | 0.9 | 0.86 | 0.79 | 0.71 | 0.81 |
| 1 | 110961722 | rs6537657 | -1.443 | -0.672 | -0.281 | -0.529 | -0.004 | -0.001 | -0.001 | 1.062 | 0.73 | 0.89 | 0.86 | 0.79 | 0.71 | 0.81 |
| 1 | 110968264 | rs4839387 | 0.753 | -0.555 | -0.139 | -0.484 | -0.005 | 0.001 | 0.001 | 1.619 | 0.32 | 0.3 | 0.54 | 0.68 | 0.29 | 0.39 |
| 1 | 110974248 | rs7515826 | -1.39 | -0.449 | -0.68 | -0.504 | 0.036 | 0.047 | 0.047 | 0 | 0.5 | 0.82 | 0.67 | 0.44 | 0.5 | 0.5 |
| 1 | 110978427 | rs12563285 | 0.683 | -0.221 | -0.731 | -0.368 | -0.02 | -0.033 | -0.033 | 0 | 0.09 | 0.04 | 0.33 | 0.33 | 0.1 | 0.11 |

| | | | | | | | | | | | | | | | | |
|---|---|---|---|---|---|---|---|---|---|---|---|---|---|---|---|---|
| 1 | 110978516 | rs12130170 | 0.838 | -0.216 | -0.516 | -0.647 | 0.004 | -0.031 | -0.031 | 0 | 0.15 | 0.19 | 0.1 | 0.05 | 0.14 | 0.15 |
| 1 | 110981370 | rs4839015 | -1.273 | 0.202 | -1.162 | -0.397 | -0.003 | -0.026 | -0.026 | 1.684 | 0.73 | 0.9 | 0.84 | 0.7 | 0.71 | 0.82 |
| 1 | 110985784 | rs1391281 | 1.483 | 0.33 | -0.994 | -0.377 | -0.051 | 0.05 | 0.05 | 0 | 0.16 | 0.08 | 0.19 | 0.05 | 0.26 | 0.16 |
| 1 | 110988874 | rs7534042 | -0.702 | 0.178 | -0.728 | -0.169 | -0.04 | 0.025 | 0.025 | 0.059 | 0.57 | 0.76 | 0.67 | 0.63 | 0.49 | 0.53 |
| 1 | 110990929 | rs11102109 | 0.398 | 0.102 | -0.397 | -0.217 | -0.003 | 0.004 | 0.004 | 0.442 | 0.35 | 0.33 | 0.55 | 0.58 | 0.34 | 0.35 |
| 1 | 110991324 | rs4564141 | 1.347 | 0.118 | -0.512 | -0.056 | -0.022 | -0.021 | -0.021 | 0.947 | 0.27 | 0.09 | 0.25 | 0.27 | 0.33 | 0.24 |
| 1 | 110993349 | rs1857512 | 0.228 | 0.082 | -0.472 | -0.12 | -0.003 | -0.102 | -0.102 | 0 | 0.22 | 0.43 | 0.11 | 0.06 | 0.15 | 0.18 |
| 1 | 110993990 | rs4839391 | 0.152 | 0.011 | -0.462 | 0.568 | -0.001 | 0.014 | 0.014 | 0.079 | 0.09 | 0.09 | 0.33 | 0.29 | 0.09 | 0.1 |
| 1 | 110996325 | rs3795828 | 0.708 | 0.013 | -0.489 | 0.56 | -0.022 | -0.021 | -0.021 | 0 | 0.27 | 0.09 | 0.22 | 0.27 | 0.33 | 0.24 |
| 1 | 110996950 | rs17628304 | 0.033 | -0.137 | -0.341 | 0.029 | -0.017 | -0.016 | -0.016 | 0.35 | 0.37 | 0.22 | 0.55 | 0.56 | 0.42 | 0.34 |
| 1 | 110998854 | rs7514102 | -0.199 | 0.077 | -0.52 | 0.152 | -0.001 | 0 | 0 | 0 | 0.4 | 0.37 | 0.57 | 0.58 | 0.42 | 0.34 |
| 1 | 111003201 | rs11102110 | -0.272 | -0.721 | -0.517 | 0.124 | -0.008 | -0.004 | -0.004 | 0 | 0.63 | 0.6 | 0.76 | 0.7 | 0.67 | 0.81 |
| 1 | 111006869 | rs6668000 | 0.036 | -0.715 | -0.342 | 0.308 | 0.007 | -0.005 | -0.005 | 0 | 0.58 | 0.58 | 0.5 | 0.54 | 0.6 | 0.67 |
| 1 | 111009031 | rs11102112 | 0.737 | -0.643 | -0.718 | -0.132 | -0.063 | 0.025 | 0.025 | 0 | 0.3 | 0.12 | 0.31 | 0.19 | 0.33 | 0.41 |
| 1 | 111009859 | rs12118420 | -0.209 | -1.4 | -0.386 | -0.126 | 0.028 | 0.031 | 0.031 | 0.037 | 0.54 | 0.34 | 0.52 | 0.35 | 0.58 | 0.61 |
| 1 | 111011847 | rs11102115 | 1.012 | -1.428 | -0.672 | -0.088 | -0.013 | -0.083 | -0.083 | 0 | 0.29 | 0.15 | 0.23 | 0.42 | 0.33 | 0.25 |
| 1 | 111014215 | rs2361309 | -1.157 | -1.418 | -0.535 | -0.087 | 0 | -0.036 | -0.036 | 0.437 | 0.71 | 0.75 | 0.76 | 0.58 | 0.68 | 0.75 |
| 1 | 111014420 | rs4472753 | -0.321 | -0.992 | -0.102 | -0.184 | 0.015 | 0.038 | 0.038 | 0.073 | 0.45 | 0.58 | 0.43 | 0.23 | 0.58 | 0.56 |
| 1 | 111016758 | rs7537693 | -0.516 | -0.76 | -0.195 | -0.067 | -0.016 | 0.015 | 0.015 | 0.03 | 0.65 | 0.62 | 0.72 | 0.77 | 0.56 | 0.51 |
| 1 | 111018598 | rs4839417 | 0.899 | -0.681 | -0.309 | 0.219 | 0.024 | -0.025 | -0.025 | 0 | 0.08 | 0.03 | 0.17 | 0.16 | 0.02 | 0.04 |
| 1 | 111018874 | rs17025859 | 0.141 | -0.243 | 0.282 | 0.411 | 0.008 | 0.094 | 0.094 | 0 | 0.06 | 0.16 | 0.2 | 0.19 | 0.03 | 0.08 |
| 1 | 111021728 | rs4839418 | -0.108 | -0.412 | 0.103 | 0.418 | 0.018 | 0.021 | 0.021 | 0 | 0.51 | 0.34 | 0.48 | 0.41 | 0.54 | 0.6 |
| 1 | 111022475 | rs6677767 | 0.235 | -0.279 | 0.027 | 0.03 | -0.002 | 0.001 | 0.001 | 0 | 0.42 | 0.42 | 0.29 | 0.43 | 0.43 | 0.34 |
| 1 | 111023490 | rs17680634 | 0.069 | -0.097 | -0.558 | -0.097 | 0.005 | -0.004 | -0.004 | 0 | 0.49 | 0.31 | 0.35 | 0.49 | 0.54 | 0.46 |
| 1 | 111026570 | rs12048054 | -0.145 | -0.073 | -1.265 | -0.287 | 0.049 | 0.197 | 0.197 | 0 | 0.52 | 0.82 | 0.67 | 0.82 | 0.44 | 0.64 |
| 1 | 111027035 | rs1857514 | -0.534 | 0.361 | -0.547 | 0.272 | -0.016 | 0.127 | 0.127 | 0 | 0.1 | 0.21 | 0.34 | 0.27 | 0.04 | 0.2 |
| 1 | 111027692 | rs12023354 | -0.654 | 0.889 | -1.085 | -0.467 | 0.027 | 0.073 | 0.073 | 0 | 0.05 | 0.22 | 0.04 | 0.12 | 0.02 | 0.07 |
| 1 | 111028588 | rs10776712 | 0.052 | 0.981 | -1.876 | -0.922 | -0.027 | 0.098 | 0.098 | 0 | 0.79 | 0.94 | 0.86 | 1 | 0.74 | 0.85 |
| 1 | 111029568 | rs6537662 | 0.052 | 0.981 | -1.876 | -0.922 | -0.027 | 0.098 | 0.098 | 0 | 0.79 | 0.94 | 0.86 | 1 | 0.74 | 0.85 |
| 1 | 111033629 | rs17629893 | -0.36 | 0.878 | -1.901 | -1.051 | 0.009 | 0.123 | 0.123 | 0 | 0.31 | 0.12 | 0.34 | 0.03 | 0.33 | 0.2 |
| 1 | 111033809 | rs4839431 | -0.582 | 0.99 | -2.227 | -1.127 | -0.017 | 0.046 | 0.046 | 0 | 0.85 | 0.95 | 0.88 | 0.93 | 0.83 | 0.8 |
| 1 | 111035174 | rs6537664 | 0.53 | 1.044 | -1.621 | -0.607 | -0.003 | 0 | 0 | 0.641 | 0.37 | 0.28 | 0.42 | 0.59 | 0.37 | 0.46 |
| 1 | 111036717 | rs17630160 | 0.416 | 0.834 | -1.679 | -1.184 | -0.003 | 0.043 | 0.043 | 0 | 0.24 | 0.17 | 0.15 | 0.01 | 0.31 | 0.14 |
| 1 | 111038840 | rs815324 | -0.638 | 0.959 | -1.243 | -1.051 | 0.002 | 0.012 | 0.012 | 0 | 0.43 | 0.46 | 0.43 | 0.37 | 0.45 | 0.33 |
| 1 | 111038969 | rs815325 | -0.144 | 0.896 | -1.099 | -0.919 | -0.006 | 0.003 | 0.003 | 0 | 0.19 | 0.34 | 0.17 | 0.27 | 0.14 | 0.19 |
| 1 | 111040055 | rs641227 | -0.376 | 0.581 | -0.925 | -1.139 | -0.009 | -0.018 | -0.018 | 0 | 0.4 | 0.51 | 0.39 | 0.29 | 0.42 | 0.21 |
| 1 | 111040235 | rs12031009 | 0.184 | 0.535 | -0.639 | -0.989 | -0.012 | -0.012 | -0.012 | 0 | 0.29 | 0.43 | 0.31 | 0.26 | 0.32 | 0.15 |
| 1 | 111040498 | rs17025931 | -0.776 | 0.539 | -0.204 | -0.156 | 0.044 | 0.092 | 0.092 | 0 | 0.1 | 0.3 | 0.08 | 0.2 | 0.14 | 0.05 |
| 1 | 111041925 | rs561606 | -0.271 | -0.151 | 0.204 | -0.183 | 0.001 | 0.025 | 0.025 | 0 | 0.54 | 0.72 | 0.5 | 0.42 | 0.52 | 0.43 |
| 1 | 111044227 | rs10857825 | 0.565 | -0.236 | 0.162 | -0.12 | -0.007 | 0.035 | 0.035 | 0 | 0.07 | 0.16 | 0.05 | 0.26 | 0.05 | 0.07 |
| 1 | 111049872 | rs1281183 | 0.786 | -0.586 | 0.135 | -0.371 | -0.085 | 0.158 | 0.158 | 0.003 | 0.37 | 0.05 | 0.39 | 0.15 | 0.46 | 0.34 |
| 1 | 111050158 | rs1281182 | 0.035 | -0.529 | 0.557 | -0.328 | 0.003 | 0.021 | 0.021 | 0 | 0.47 | 0.45 | 0.46 | 0.4 | 0.51 | 0.45 |
| 1 | 111051999 | rs7543509 | 0.122 | -0.335 | 0.357 | 0.554 | 0 | 0.036 | 0.036 | 0 | 0.07 | 0.07 | 0.12 | 0.22 | 0.07 | 0.14 |
| 1 | 111056389 | rs1281179 | -1.555 | -0.579 | 0.203 | 0.417 | 0.005 | -0.033 | -0.033 | 0 | 0.87 | 0.83 | 0.89 | 0.95 | 0.85 | 0.8 |
| 1 | 111058469 | rs1281177 | 0.708 | -0.728 | -0.271 | 0.598 | 0.024 | 0.019 | 0.019 | 0.079 | 0.56 | 0.32 | 0.66 | 0.33 | 0.66 | 0.47 |
| 1 | 111061011 | rs3748731 | 1.091 | -0.649 | -0.312 | 0.558 | -0.005 | -0.002 | -0.002 | 0 | 0.21 | 0.19 | 0.25 | 0.17 | 0.2 | 0.12 |
| 1 | 111061062 | rs1281174 | 0.237 | -1.057 | 0.471 | 1.31 | -0.021 | 0.019 | 0.019 | 0 | 0.69 | 0.51 | 0.78 | 0.59 | 0.73 | 0.59 |
| 1 | 111061653 | rs3768456 | -1.293 | -0.426 | 0.348 | 1.111 | -0.011 | 0.06 | 0.06 | 0.286 | 0.12 | 0.2 | 0.13 | 0.26 | 0.07 | 0.11 |
| 1 | 111065753 | rs7526730 | 0.305 | -0.416 | 0.206 | 0.092 | 0.006 | -0.022 | -0.022 | 0 | 0.12 | 0.12 | 0.14 | 0.05 | 0.14 | 0.22 |
| 1 | 111068272 | rs1106151 | -0.456 | -0.523 | -0.098 | 0.305 | 0.005 | -0.015 | -0.015 | 0 | 0.08 | 0.04 | 0.17 | 0.17 | 0.06 | 0.2 |
| 1 | 111069611 | rs17025978 | -0.188 | -0.653 | -0.235 | 0.341 | 0.002 | -0.017 | -0.017 | 0 | 0.12 | 0.05 | 0.17 | 0.16 | 0.09 | 0.17 |
| 1 | 111070760 | rs7554231 | -1.356 | -1.066 | 0.41 | 0.092 | -0.021 | -0.027 | -0.027 | 1.004 | 0.3 | 0.4 | 0.33 | 0.23 | 0.21 | 0.43 |
| 1 | 111071279 | rs1339700 | -1.54 | -0.791 | 0.715 | -0.169 | -0.006 | -0.014 | -0.014 | 0 | 0.35 | 0.55 | 0.36 | 0.24 | 0.26 | 0.44 |
| 1 | 111071958 | rs1769203 | -0.158 | -0.72 | 0.38 | -0.249 | -0.001 | -0.002 | -0.002 | 0 | 0.07 | 0.06 | 0.03 | 0.1 | 0.02 | 0.05 |
| 1 | 111077162 | rs7522150 | -0.65 | -0.403 | 0.487 | -0.354 | -0.008 | 0.001 | 0.001 | 0.371 | 0.3 | 0.24 | 0.35 | 0.25 | 0.31 | 0.37 |
| 1 | 111080468 | rs12748140 | -0.914 | -0.414 | 0.324 | -0.312 | 0.02 | -0.011 | -0.011 | 2.156 | 0.33 | 0.38 | 0.37 | 0.26 | 0.31 | 0.38 |
| 1 | 111082521 | rs6537666 | -1.068 | -0.454 | 0.308 | -0.343 | 0.001 | -0.02 | -0.02 | 0 | 0.35 | 0.4 | 0.36 | 0.26 | 0.29 | 0.35 |
| 1 | 111087058 | rs7526729 | -0.881 | 0.021 | -0.001 | -0.127 | 0.007 | -0.014 | -0.014 | 0 | 0.33 | 0.49 | 0.36 | 0.25 | 0.29 | 0.33 |
| 1 | 111087124 | rs2790372 | -0.888 | 0.182 | -0.516 | 0.116 | 0.004 | 0.01 | 0.01 | 0.146 | 0.09 | 0.06 | 0.04 | 0.04 | 0.06 | 0.09 |
| 1 | 111091584 | rs6676386 | -0.565 | 0.261 | -0.322 | 0.305 | -0.007 | 0.002 | 0.002 | 1.306 | 0.1 | 0.15 | 0.16 | 0.11 | 0.06 | 0.14 |
| 1 | 111092468 | rs11102129 | -0.013 | 0.965 | -0.879 | 0.668 | -0.054 | -0.063 | -0.063 | 0 | 0.1 | 0.01 | 0.22 | 0.64 | 0.18 | 0.16 |
| 1 | 111093653 | rs1769207 | -1.03 | 0.398 | -0.364 | -0.468 | 0.005 | 0.023 | 0.023 | 0 | 0.42 | 0.66 | 0.37 | 0.24 | 0.33 | 0.38 |
| 1 | 111098766 | rs903050 | -0.956 | 0.262 | 0.24 | -0.647 | -0.024 | 0.027 | 0.027 | 1.649 | 0.18 | 0.3 | 0.21 | 0.22 | 0.1 | 0.22 |
| 1 | 111100562 | rs6537668 | -0.472 | 0.082 | 0.582 | -0.713 | -0.003 | 0.011 | 0.011 | 0.42 | 0.27 | 0.24 | 0.18 | 0.12 | 0.25 | 0.21 |
| 1 | 111102214 | rs12407862 | -0.1 | 0.209 | 0.338 | -1.079 | 0 | 0.026 | 0.026 | 0.663 | 0.17 | 0.07 | 0.19 | 0.11 | 0.21 | 0.19 |
| 1 | 111103688 | rs6656346 | -0.343 | 0.322 | 0.347 | -1.123 | -0.009 | 0.024 | 0.024 | 0.801 | 0.21 | 0.11 | 0.24 | 0.11 | 0.27 | 0.25 |
| 1 | 111104733 | rs17632564 | 0.09 | 0.232 | 0.514 | -1.197 | 0.002 | -0.065 | -0.065 | 0 | 0.24 | 0.33 | 0.15 | 0.01 | 0.21 | 0.16 |
| 1 | 111107668 | rs12062889 | -1.287 | 0.176 | 0.46 | -1.225 | -0.001 | -0.022 | -0.022 | 0 | 0.46 | 0.54 | 0.41 | 0.11 | 0.48 | 0.4 |
| 1 | 111108738 | rs12744965 | 0.353 | 0.234 | 0.335 | -1.152 | -0.007 | 0.008 | 0.008 | 0.159 | 0.22 | 0.17 | 0.14 | 0.01 | 0.21 | 0.16 |

| | | | | | | | | | | | | | | | | |
|---|---|---|---|---|---|---|---|---|---|---|---|---|---|---|---|---|
| 1 | 111109137 | rs11578748 | -0.736 | -0.003 | 0.614 | -0.941 | 0.001 | -0.037 | -0.037 | 0 | 0.45 | 0.48 | 0.39 | 0.11 | 0.48 | 0.4 |
| 1 | 111113317 | rs6672114 | 1.188 | -0.084 | 0.83 | -0.327 | -0.009 | 0.137 | 0.137 | 0.543 | 0.92 | 0.64 | 0.95 | 0.87 | 0.97 | 0.95 |
| 1 | 111114154 | rs923826 | 0.512 | 0.308 | 0.35 | -0.948 | -0.009 | 0.009 | 0.009 | 0 | 0.63 | 0.36 | 0.58 | 0.9 | 0.54 | 0.6 |
| 1 | 111117814 | rs6675315 | -0.437 | -0.376 | -0.94 | -0.22 | 0.012 | 0.025 | 0.025 | 0.011 | 0.34 | 0.12 | 0.26 | 0.27 | 0.28 | 0.16 |
| 1 | 111121822 | rs923824 | 0.32 | -0.953 | -0.471 | 0.733 | -0.011 | -0.002 | -0.002 | 0 | 0.62 | 0.57 | 0.59 | 0.38 | 0.74 | 0.57 |
| 1 | 111125481 | rs3860223 | 0.1 | -0.934 | -0.336 | 0.867 | 0.004 | 0.026 | 0.026 | 0 | 0.75 | 0.79 | 0.73 | 0.82 | 0.79 | 0.77 |
| 1 | 111135098 | rs1890675 | -1.309 | -1.192 | 0.042 | 0.829 | 0.036 | 0.037 | 0.037 | 0 | 0.47 | 0.7 | 0.48 | 0.64 | 0.35 | 0.4 |
| 1 | 111137811 | rs4839069 | -1.024 | -1.394 | 0.457 | 0.805 | 0.011 | 0.004 | 0.004 | 0 | 0.29 | 0.21 | 0.54 | 0.71 | 0.26 | 0.34 |
| 1 | 111141682 | rs9782928 | -1.755 | -1.476 | 0.226 | 0.824 | 0.055 | 0.011 | 0.011 | 0.115 | 0.48 | 0.73 | 0.49 | 0.55 | 0.34 | 0.42 |
| 1 | 111144007 | rs12047887 | -0.911 | -0.796 | 0.21 | 0.94 | 0.002 | 0.019 | 0.019 | 0 | 0.25 | 0.21 | 0.22 | 0.19 | 0.27 | 0.2 |
| 1 | 111144661 | rs12407942 | 0.339 | -0.93 | 0.52 | 1.238 | -0.001 | 0.079 | 0.079 | 0 | 0.1 | 0.14 | 0.35 | 0.39 | 0.07 | 0.2 |
| 1 | 111148080 | rs3887820 | -0.431 | -0.725 | 0.667 | 1.048 | 0.049 | 0.323 | 0.323 | 0 | 0.13 | 0.47 | 0.25 | 0.35 | 0.09 | 0.2 |
| 1 | 111155595 | rs4839542 | -0.891 | -0.9 | 0.26 | 0.858 | 0.011 | 0.013 | 0.013 | 0 | 0.22 | 0.2 | 0.19 | 0.16 | 0.19 | 0.23 |
| 1 | 111161677 | rs3860225 | -0.888 | -1.036 | 0.327 | 1.16 | -0.001 | -0.001 | -0.001 | 0 | 0.22 | 0.23 | 0.16 | 0.33 | 0.14 | 0.23 |
| 1 | 111163693 | rs12132073 | 2.096 | -0.97 | 0.332 | 1.014 | 0.067 | 0.004 | 0.004 | 0 | 0.53 | 0.17 | 0.54 | 0.44 | 0.7 | 0.52 |
| 1 | 111166960 | rs2172514 | 0.593 | -0.738 | 0.157 | 0.975 | 0.008 | -0.009 | -0.009 | 0 | 0.78 | 0.8 | 0.83 | 0.67 | 0.84 | 0.76 |
| 1 | 111169375 | rs1416811 | -0.307 | -0.568 | 0.348 | 1.101 | -0.002 | 0.054 | 0.054 | 0 | 0.94 | 0.84 | 0.98 | 0.84 | 0.97 | 0.94 |
| 1 | 111171407 | rs17026186 | 0.302 | -0.555 | 0.622 | 1.274 | 0.021 | 0.099 | 0.099 | 0 | 0.1 | 0.2 | 0.27 | 0.23 | 0.08 | 0.23 |
| 1 | 111172413 | rs11102140 | 0.279 | -0.556 | 0.644 | 1.274 | 0.026 | 0.112 | 0.112 | 0 | 0.1 | 0.23 | 0.27 | 0.23 | 0.08 | 0.23 |
| 1 | 111173494 | rs2797531 | 1.404 | -0.677 | 0.22 | 0.766 | 0.012 | 0.005 | 0.005 | 0 | 0.73 | 0.78 | 0.87 | 0.75 | 0.83 | 0.79 |
| 1 | 111177237 | rs4839549 | -1.101 | -0.46 | 0.506 | 0.816 | -0.018 | -0.016 | -0.016 | 0 | 0.14 | 0.26 | 0.1 | 0.1 | 0.09 | 0.13 |
| 1 | 111177405 | rs2640503 | 0.604 | -0.385 | 0.523 | 1.001 | -0.004 | 0.074 | 0.074 | 0 | 0.92 | 0.76 | 0.91 | 0.81 | 0.96 | 0.93 |
| 1 | 111181086 | rs2640500 | 0.987 | 0.39 | -0.118 | 0.467 | 0.026 | 0.012 | 0.012 | 0 | 0.5 | 0.34 | 0.55 | 0.49 | 0.5 | 0.3 |
| 1 | 111183367 | rs11585237 | 0.823 | 1.341 | 0.154 | 0.543 | 0.033 | 0.007 | 0.007 | 0 | 0.08 | 0.14 | 0.08 | 0.1 | 0.14 | 0.17 |
| 1 | 111187885 | rs2640493 | 1.899 | 1.085 | 0.104 | 0.236 | 0.029 | -0.016 | -0.016 | 0 | 0.42 | 0.32 | 0.48 | 0.48 | 0.36 | 0.29 |
| 1 | 111189722 | rs2797534 | -0.48 | 0.999 | 0.471 | 0.263 | -0.012 | -0.021 | -0.021 | 0 | 0.82 | 0.86 | 0.79 | 0.59 | 0.79 | 0.64 |
| 1 | 111190726 | rs2797535 | 0.893 | 0.984 | 0.718 | 0.234 | -0.004 | 0.017 | 0.017 | 0 | 0.67 | 0.66 | 0.7 | 0.58 | 0.71 | 0.52 |
| 1 | 111192893 | rs17635235 | 0.182 | 1.024 | 0.733 | 0.192 | -0.002 | -0.027 | -0.027 | 0.453 | 0.1 | 0.1 | 0.06 | 0 | 0.09 | 0.03 |
| 1 | 111193595 | rs2797537 | 1.597 | 0.982 | 0.685 | 0.276 | -0.002 | -0.001 | -0.001 | 1.642 | 0.55 | 0.53 | 0.63 | 0.59 | 0.61 | 0.47 |
| 1 | 111202768 | rs7531120 | 0.681 | 1.345 | 0.323 | 0.134 | 0.01 | 0.009 | 0.009 | 0 | 0.08 | 0.15 | 0.06 | 0.1 | 0.08 | 0.17 |
| 1 | 111204185 | rs2797523 | 1.59 | 1.132 | 0.302 | 0.255 | -0.002 | 0 | 0 | 0 | 0.57 | 0.59 | 0.55 | 0.58 | 0.6 | 0.46 |
| 1 | 111204247 | rs17635693 | -0.805 | 1.263 | -0.106 | 0.236 | 0.006 | -0.014 | -0.014 | 0.049 | 0.15 | 0.12 | 0.08 | 0.01 | 0.08 | 0.13 |
| 1 | 111206036 | rs11102146 | 0.759 | 1.523 | -0.128 | 0.362 | 0.001 | -0.002 | -0.002 | 0 | 0.08 | 0.11 | 0.08 | 0.1 | 0.13 | 0.17 |
| 1 | 111211533 | rs1319782 | 0.779 | 1.258 | -0.09 | 0.395 | -0.001 | 0.01 | 0.01 | 0 | 0.69 | 0.67 | 0.63 | 0.56 | 0.71 | 0.55 |
| 1 | 111213248 | rs2640480 | 0.78 | 1.25 | -0.108 | 0.39 | -0.001 | 0.005 | 0.005 | 0 | 0.69 | 0.69 | 0.63 | 0.56 | 0.71 | 0.55 |
| 1 | 111215472 | rs1058184 | 0.751 | 1.304 | -0.081 | 0.417 | -0.001 | 0.008 | 0.008 | 0 | 0.7 | 0.68 | 0.63 | 0.56 | 0.71 | 0.55 |
| 1 | 111218920 | rs2821557 | 0.864 | 1.84 | -0.317 | 0.656 | -0.039 | -0.027 | -0.027 | 0 | 0.69 | 0.74 | 0.59 | 0.43 | 0.6 | 0.44 |
| 1 | 111219344 | rs3762379 | 0.144 | 1.831 | -0.357 | 0.637 | -0.025 | 0 | 0 | 0.278 | 0.11 | 0.09 | 0.1 | 0.1 | 0.16 | 0.19 |
| 1 | 111226895 | rs1158050 | 0.147 | 1.829 | -0.351 | 0.636 | -0.017 | 0 | 0 | 0.19 | 0.11 | 0.1 | 0.1 | 0.1 | 0.16 | 0.16 |
| 1 | 111228409 | rs2840380 | -0.186 | 1.847 | -0.385 | 0.669 | -0.023 | -0.056 | -0.056 | 0 | 0.8 | 0.88 | 0.69 | 0.53 | 0.76 | 0.63 |
| 1 | 111229552 | rs1418747 | 1.246 | 1.467 | -0.293 | 0.351 | -0.004 | -0.003 | -0.003 | 0 | 0.6 | 0.6 | 0.53 | 0.43 | 0.51 | 0.41 |
| 1 | 111234601 | rs17636084 | -0.729 | 1.344 | -0.214 | 0.219 | -0.001 | -0.025 | -0.025 | 0 | 0.09 | 0.07 | 0.05 | 0 | 0.09 | 0.03 |
| 1 | 111237685 | rs12061304 | -0.988 | 1.739 | -0.048 | 0.254 | 0.032 | -0.037 | -0.037 | 0 | 0.21 | 0.32 | 0.19 | 0.08 | 0.31 | 0.18 |
| 1 | 111250018 | rs2840382 | 0.013 | 1.708 | 0.206 | 0.803 | 0.001 | 0.075 | 0.075 | 0 | 0.91 | 0.81 | 0.79 | 0.79 | 0.93 | 0.78 |
| 1 | 111250512 | rs7548006 | -0.713 | 1.573 | 0.081 | 0.518 | -0.048 | 0.047 | 0.047 | 0.763 | 0.2 | 0.12 | 0.17 | 0.01 | 0.31 | 0.14 |
| 1 | 111250619 | rs2840383 | 1.879 | 1.095 | -0.22 | 0.557 | 0.043 | -0.035 | -0.035 | 0.444 | 0.38 | 0.21 | 0.27 | 0.43 | 0.32 | 0.31 |
| 1 | 111256867 | rs12073706 | 0.044 | 1.654 | 0.024 | 0.456 | -0.012 | -0.002 | -0.002 | 0.004 | 0.15 | 0.14 | 0.11 | 0.01 | 0.19 | 0.1 |
| 1 | 111257724 | rs343602 | 0.023 | 1.395 | 0.097 | 0.995 | -0.007 | 0.061 | 0.061 | 0 | 0.88 | 0.85 | 0.78 | 0.66 | 0.92 | 0.75 |
| 1 | 111262809 | rs4839557 | -0.954 | 1.305 | 0.093 | 0.797 | 0.001 | -0.029 | -0.029 | 2.981 | 0.39 | 0.49 | 0.24 | 0.25 | 0.41 | 0.26 |
| 1 | 111270605 | rs343609 | 1.742 | 1.466 | 0.03 | 0.609 | -0.012 | 0.016 | 0.016 | 3.81 | 0.42 | 0.26 | 0.48 | 0.37 | 0.42 | 0.42 |
| 1 | 111276629 | rs343614 | 1.231 | 1.387 | 0.175 | 0.612 | 0 | 0.003 | 0.003 | 1.032 | 0.52 | 0.44 | 0.54 | 0.37 | 0.52 | 0.53 |
| 1 | 111288171 | rs343597 | 0.478 | 1.312 | 0.151 | 0.765 | -0.005 | 0.019 | 0.019 | 0 | 0.21 | 0.23 | 0.12 | 0.32 | 0.22 | 0.13 |
| 1 | 111290022 | rs1994918 | -0.327 | 1.184 | 0.185 | 0.732 | -0.013 | 0.06 | 0.06 | 0 | 0.12 | 0.2 | 0.22 | 0.3 | 0.08 | 0.24 |
| 1 | 111290542 | rs1953099 | -0.063 | 1.187 | 0.164 | 0.733 | -0.005 | 0.028 | 0.028 | 0 | 0.11 | 0.14 | 0.22 | 0.3 | 0.08 | 0.24 |
| 1 | 111295674 | rs420033 | 2.346 | 1.605 | 0.143 | 0.716 | 0.02 | 0.129 | 0.129 | 6.119 | 0.34 | 0.03 | 0.25 | 0.13 | 0.28 | 0.21 |
| 1 | 111295948 | rs440156 | 0.417 | 1.785 | 0.47 | 0.751 | 0.002 | -0.061 | -0.061 | 0.194 | 0.75 | 0.7 | 0.82 | 0.99 | 0.71 | 0.8 |
| 1 | 111296652 | rs569542 | -0.822 | 1.792 | 0.438 | 0.751 | -0.001 | -0.024 | -0.024 | 2.073 | 0.86 | 0.86 | 0.88 | 0.99 | 0.81 | 0.9 |
| 1 | 111301623 | rs343791 | 1.699 | 1.629 | 0.412 | 0.818 | 0.012 | 0.04 | 0.04 | 4.283 | 0.45 | 0.24 | 0.48 | 0.34 | 0.39 | 0.42 |
| 1 | 111302518 | rs343792 | 0.95 | 1.636 | 0.438 | 0.83 | 0.026 | -0.005 | -0.005 | 0.123 | 0.68 | 0.51 | 0.61 | 0.66 | 0.62 | 0.58 |
| 1 | 111306687 | rs343795 | 0.561 | 1.61 | 0.319 | 0.808 | -0.003 | 0.023 | 0.023 | 0 | 0.1 | 0.06 | 0.06 | 0.01 | 0.1 | 0.11 |
| 1 | 111306729 | rs12118017 | 0.011 | 1.457 | 0.637 | 1.181 | -0.011 | 0.031 | 0.031 | 0 | 0.23 | 0.26 | 0.13 | 0.32 | 0.23 | 0.16 |
| 1 | 111309340 | rs482620 | 0.61 | 1.611 | 0.329 | 1.165 | 0.043 | 0.04 | 0.04 | 0.271 | 0.78 | 0.57 | 0.67 | 0.67 | 0.72 | 0.69 |
| 1 | 111311541 | rs574508 | 0.06 | 1.641 | -0.064 | 0.925 | 0.012 | -0.064 | -0.064 | 0 | 0.83 | 0.74 | 0.87 | 0.99 | 0.78 | 0.9 |
| 1 | 111318115 | rs560950 | -0.947 | 1.702 | -0.063 | 0.974 | 0.025 | -0.048 | -0.048 | 0 | 0.89 | 0.81 | 0.9 | 0.99 | 0.83 | 0.92 |
| 1 | 111319892 | rs488147 | 0.582 | 1.654 | 0.022 | 1.144 | -0.01 | -0.005 | -0.005 | 0.115 | 0.64 | 0.71 | 0.77 | 0.63 | 0.56 | 0.74 |
| 1 | 111324727 | rs11102159 | -1.107 | 1.63 | 0.132 | 1.108 | 0.024 | 0.094 | 0.094 | 0 | 0.23 | 0.41 | 0.46 | 0.55 | 0.24 | 0.47 |
| 1 | 111330531 | rs520056 | -1.232 | 1.741 | 0.147 | 0.771 | 0.015 | -0.035 | -0.035 | 8.299 | 0.9 | 0.86 | 0.91 | 0.99 | 0.83 | 0.92 |
| 1 | 111331000 | rs4839566 | -0.627 | 1.613 | 0.023 | 0.992 | 0.045 | 0.069 | 0.069 | 0 | 0.17 | 0.31 | 0.43 | 0.35 | 0.2 | 0.35 |
| 1 | 111334391 | rs724032 | -0.008 | 1.308 | -0.029 | 0.636 | 0.043 | 0.049 | 0.049 | 0 | 0.12 | 0.23 | 0.22 | 0.18 | 0.13 | 0.19 |

| | | | | | | | | | | | | | | | | |
|---|---|---|---|---|---|---|---|---|---|---|---|---|---|---|---|---|
| 1 | 111336774 | rs343815 | -0.092 | 0.614 | -0.771 | 0.201 | 0.005 | -0.006 | -0.006 | 0 | 0.22 | 0.14 | 0.14 | 0.25 | 0.25 | 0.13 |
| 1 | 111342490 | rs343819 | 1.23 | 0.629 | -1.089 | 0.175 | 0.029 | 0.066 | 0.066 | 2.209 | 0.37 | 0.23 | 0.29 | 0.1 | 0.29 | 0.26 |
| 1 | 111355935 | rs343764 | -0.763 | 0.886 | -0.984 | 1.025 | 0.004 | -0.006 | -0.006 | 1.545 | 0.22 | 0.12 | 0.14 | 0.25 | 0.25 | 0.13 |
| 1 | 111370372 | rs343769 | -0.163 | 1.123 | -0.956 | 1.203 | -0.01 | -0.028 | -0.028 | 0 | 0.17 | 0.07 | 0.33 | 0.3 | 0.26 | 0.1 |
| 1 | 111371644 | rs6655953 | -0.776 | 0.869 | -0.245 | 1.404 | 0.008 | 0.167 | 0.167 | 0 | 0.18 | 0.51 | 0.12 | 0.33 | 0.15 | 0.27 |
| 1 | 111389792 | rs17026542 | -0.376 | 0.535 | -0.93 | 0.476 | 0.013 | 0.024 | 0.024 | 0 | 0.1 | 0.04 | 0.02 | 0 | 0.05 | 0.07 |
| 1 | 111397230 | rs6698741 | -0.437 | 0.978 | -0.748 | 0.129 | 0.02 | -0.054 | -0.054 | 0 | 0.1 | 0.21 | 0.05 | 0.01 | 0.1 | 0.09 |
| 1 | 111398877 | rs578513 | -0.43 | 1.745 | -1.17 | 0.014 | -0.027 | 0.011 | 0.011 | 0 | 0.89 | 0.93 | 0.9 | 0.92 | 0.82 | 0.9 |
| 1 | 111403793 | rs12026259 | -0.094 | 1.897 | -1.421 | 0.729 | -0.052 | -0.054 | -0.054 | 0 | 0.09 | 0.01 | 0.14 | 0.28 | 0.16 | 0.28 |
| 1 | 111415841 | rs1494324 | 1.425 | 1.009 | -1.801 | 0.661 | 0.069 | -0.012 | -0.012 | 0.292 | 0.39 | 0.07 | 0.3 | 0.33 | 0.27 | 0.38 |
| 1 | 111417446 | rs931547 | 1.403 | 1.007 | -1.802 | 0.729 | 0.07 | 0.007 | 0.007 | 1.551 | 0.39 | 0.07 | 0.3 | 0.31 | 0.27 | 0.38 |
| 1 | 111418217 | rs1389722 | 0.349 | 0.977 | -1.041 | 0.58 | 0.032 | 0.076 | 0.076 | 0.313 | 0.84 | 0.42 | 0.8 | 0.72 | 0.85 | 0.74 |
| 1 | 111421427 | rs10857833 | 1.223 | 0.884 | -1.537 | 0.308 | -0.017 | 0.033 | 0.033 | 1.337 | 0.56 | 0.16 | 0.64 | 0.62 | 0.5 | 0.5 |
| 1 | 111421712 | rs10857834 | 0.773 | 0.459 | -1.832 | 0.368 | -0.014 | -0.041 | -0.041 | 0 | 0.17 | 0.09 | 0.35 | 0.32 | 0.23 | 0.12 |
| 1 | 111422188 | rs1494320 | 1.916 | 0.438 | -1.85 | 0.354 | 0.074 | 0.018 | 0.018 | 2.854 | 0.39 | 0.03 | 0.29 | 0.28 | 0.26 | 0.38 |
| 1 | 111422676 | rs6658287 | -0.232 | 0.588 | -1.327 | 0.344 | -0.01 | 0.004 | 0.004 | 0 | 0.23 | 0.16 | 0.15 | 0.08 | 0.33 | 0.2 |
| 1 | 111422817 | rs1494322 | 0.183 | 0.467 | -1.172 | 0.474 | 0.017 | 0.074 | 0.074 | 0.556 | 0.82 | 0.37 | 0.8 | 0.7 | 0.84 | 0.72 |
| 1 | 111430632 | rs2154185 | 1.574 | 0.423 | -1.313 | 0.341 | -0.014 | -0.005 | -0.005 | 0 | 0.52 | 0.15 | 0.61 | 0.43 | 0.5 | 0.43 |
| 1 | 111430758 | rs4839580 | 1.574 | 0.423 | -1.313 | 0.341 | -0.014 | -0.005 | -0.005 | 0 | 0.52 | 0.15 | 0.61 | 0.43 | 0.5 | 0.43 |
| 1 | 111435335 | rs3790722 | -0.02 | 0.426 | -0.581 | 0.372 | 0.021 | 0.094 | 0.094 | 0.723 | 0.83 | 0.39 | 0.81 | 0.69 | 0.85 | 0.72 |
| 1 | 111437129 | rs1343817 | 1.668 | 0.392 | -0.669 | 0.33 | -0.018 | -0.006 | -0.006 | 0 | 0.52 | 0.14 | 0.61 | 0.43 | 0.5 | 0.43 |
| 1 | 111439544 | rs10494124 | 0.426 | 0.425 | -0.635 | 0.425 | 0.009 | -0.04 | -0.04 | 0 | 0.11 | 0.04 | 0.08 | 0.18 | 0.08 | 0.11 |
| 1 | 111439902 | rs4839586 | 1.101 | 0.422 | -0.565 | 0.416 | -0.006 | 0.055 | 0.055 | 0.368 | 0.72 | 0.34 | 0.73 | 0.5 | 0.76 | 0.6 |
| 1 | 111441485 | rs11102175 | -0.062 | 0.469 | -0.514 | 0.244 | 0.028 | 0.109 | 0.109 | 0.514 | 0.83 | 0.42 | 0.82 | 0.69 | 0.85 | 0.72 |
| 1 | 111449999 | rs7521468 | 1.07 | 0.096 | -0.747 | -0.312 | -0.013 | 0.074 | 0.074 | 1.985 | 0.54 | 0.27 | 0.39 | 0.19 | 0.5 | 0.45 |
| 1 | 111450408 | rs1936942 | 0.343 | 0.096 | -0.516 | -0.172 | 0.016 | -0.01 | -0.01 | 1.49 | 0.66 | 0.4 | 0.48 | 0.35 | 0.59 | 0.56 |
| 1 | 111459294 | rs7547322 | 2.581 | -0.02 | -0.928 | -0.312 | 0.036 | 0.11 | 0.11 | 4.613 | 0.32 | 0.07 | 0.26 | 0.11 | 0.22 | 0.28 |
| 1 | 111459847 | rs1282023 | -1.008 | -0.054 | -0.712 | -0.323 | -0.01 | 0.026 | 0.026 | 0.097 | 0.41 | 0.71 | 0.57 | 0.67 | 0.45 | 0.46 |
| 1 | 111463624 | rs12567355 | 1.078 | 0.052 | -0.9 | -0.221 | 0.005 | -0.008 | -0.008 | 3.463 | 0.08 | 0.06 | 0.06 | 0.15 | 0.06 | 0.09 |
| 1 | 111463683 | rs12566475 | 1.153 | 0.053 | -0.909 | -0.22 | 0.009 | -0.02 | -0.02 | 2.773 | 0.08 | 0.03 | 0.06 | 0.15 | 0.06 | 0.09 |
| 1 | 111464435 | rs7533260 | 0.45 | 0.066 | -0.92 | -0.276 | -0.025 | 0.034 | 0.034 | 0 | 0.18 | 0.12 | 0.11 | 0.07 | 0.27 | 0.17 |
| 1 | 111465615 | rs12077952 | -0.779 | -0.049 | -0.609 | -0.331 | -0.029 | 0.024 | 0.024 | 0.242 | 0.37 | 0.2 | 0.52 | 0.66 | 0.45 | 0.46 |
| 1 | 111466053 | rs1712674 | 0.638 | -0.094 | -0.257 | -0.294 | -0.014 | 0.013 | 0.013 | 0 | 0.1 | 0.18 | 0.07 | 0.15 | 0.06 | 0.09 |
| 1 | 111468886 | rs1418425 | -1.421 | -0.107 | -0.258 | -0.315 | 0.041 | 0.046 | 0.046 | 1.639 | 0.55 | 0.34 | 0.63 | 0.73 | 0.72 | 0.63 |
| 1 | 111470050 | rs2885806 | 2.162 | -0.085 | -0.258 | -0.337 | -0.032 | -0.074 | -0.074 | 0.016 | 0.35 | 0.44 | 0.3 | 0.12 | 0.22 | 0.28 |
| 1 | 111485207 | rs11102184 | -0.029 | 0.078 | -0.262 | 0.013 | -0.005 | -0.028 | -0.028 | 0.844 | 0.25 | 0.14 | 0.38 | 0.35 | 0.29 | 0.17 |
| 1 | 111488573 | rs12145219 | 1.423 | 0.164 | -0.429 | 0.025 | -0.071 | -0.116 | -0.116 | 2.269 | 0.15 | 0.01 | 0.32 | 0.35 | 0.24 | 0.15 |
| 1 | 111494194 | rs2232041 | 0.209 | 0.229 | -0.252 | -0.281 | -0.013 | 0.013 | 0.013 | 0 | 0.18 | 0.16 | 0.11 | 0.06 | 0.27 | 0.17 |
| 1 | 111498111 | rs1418426 | 0.896 | -0.116 | -0.107 | -0.117 | 0.002 | 0.002 | 0.002 | 0.402 | 0.46 | 0.41 | 0.44 | 0.44 | 0.38 | 0.57 |
| 1 | 111524493 | rs6687647 | 0.176 | -0.064 | -0.163 | -0.276 | -0.026 | -0.099 | -0.099 | 0 | 0.84 | 0.63 | 0.92 | 1 | 0.87 | 0.92 |
| 1 | 111525736 | rs9660523 | 1.035 | -0.06 | -0.652 | -0.586 | 0 | -0.021 | -0.021 | 0 | 0.21 | 0.4 | 0.23 | 0.12 | 0.19 | 0.35 |
| 1 | 111528210 | rs17026705 | 0.505 | 0.311 | -0.612 | -0.06 | -0.055 | -0.067 | -0.067 | 0 | 0.1 | 0.01 | 0.15 | 0.33 | 0.15 | 0.2 |
| 1 | 111528570 | rs4838884 | 0.92 | -0.455 | -0.251 | -0.865 | -0.036 | -0.142 | -0.142 | 0 | 0.77 | 0.56 | 0.87 | 0.99 | 0.82 | 0.85 |
| 1 | 111543527 | rs7545868 | -0.44 | -0.35 | -0.37 | -0.672 | -0.015 | -0.064 | -0.064 | 0 | 0.21 | 0.27 | 0.13 | 0.01 | 0.16 | 0.12 |
| 1 | 111546531 | rs1149185 | 0.658 | 0.268 | -0.166 | -0.172 | 0.007 | 0.001 | 0.001 | 0.031 | 0.67 | 0.63 | 0.55 | 0.36 | 0.56 | 0.32 |
| 1 | 111546887 | rs1149186 | 1.736 | -0.191 | -0.079 | -0.583 | 0.036 | -0.003 | -0.003 | 2.211 | 0.37 | 0.14 | 0.37 | 0.35 | 0.3 | 0.15 |
| 1 | 111553602 | rs1282131 | 0.313 | -0.134 | 0.016 | -0.553 | -0.09 | -0.038 | -0.038 | 0 | 0.69 | 0.82 | 0.57 | 0.36 | 0.56 | 0.32 |
| 1 | 111557266 | rs1282288 | 0.025 | -0.1 | -0.05 | -0.355 | -0.1 | -0.097 | -0.097 | 0.575 | 0.74 | 0.85 | 0.61 | 0.45 | 0.59 | 0.42 |
| 1 | 111559848 | rs12735344 | 0.255 | -0.202 | 0.238 | -0.456 | 0.024 | 0.049 | 0.049 | 0 | 0.29 | 0.16 | 0.16 | 0 | 0.26 | 0.17 |
| 1 | 111567469 | rs1282334 | 0.152 | 0.367 | 0.505 | 0.088 | -0.096 | -0.083 | -0.083 | 0.027 | 0.75 | 0.87 | 0.62 | 0.52 | 0.59 | 0.43 |
| 1 | 111570320 | rs11102199 | 0.126 | 0.875 | 0.425 | 0.229 | -0.076 | -0.071 | -0.071 | 0.073 | 0.13 | 0.03 | 0.18 | 0.32 | 0.26 | 0.23 |
| 1 | 111571080 | rs1282333 | -0.147 | 0.875 | 0.451 | 0.229 | -0.047 | -0.044 | -0.044 | 0 | 0.86 | 0.92 | 0.82 | 0.68 | 0.73 | 0.76 |
| 1 | 111573680 | rs1282330 | 2.048 | 0.534 | 0.466 | 0.19 | 0.042 | 0.014 | 0.014 | 2.878 | 0.4 | 0.15 | 0.38 | 0.35 | 0.31 | 0.16 |
| 1 | 111575921 | rs1578790 | 1.283 | 0.579 | 0.444 | 0.078 | 0.01 | -0.001 | -0.001 | 0 | 0.67 | 0.6 | 0.55 | 0.32 | 0.56 | 0.33 |
| 1 | 111585494 | rs1782838 | 0.726 | 0.486 | 0.373 | 0.101 | -0.1 | -0.096 | -0.096 | 0 | 0.73 | 0.85 | 0.61 | 0.49 | 0.58 | 0.42 |
| 1 | 111586053 | rs1659659 | 1.858 | 0.58 | 0.454 | 0.08 | -0.014 | -0.013 | -0.013 | 0 | 0.4 | 0.44 | 0.4 | 0.32 | 0.32 | 0.14 |
| 1 | 111586123 | rs11583268 | 0.383 | 0.709 | 0.508 | 0.002 | 0.01 | -0.017 | -0.017 | 0 | 0.26 | 0.19 | 0.16 | 0 | 0.24 | 0.17 |
| 1 | 111607219 | rs1144599 | 1.815 | 0.922 | 0.647 | 0.171 | 0.027 | 0.029 | 0.029 | 0.884 | 0.42 | 0.24 | 0.39 | 0.32 | 0.34 | 0.16 |
| 1 | 111610738 | rs11102208 | -1.042 | 1.069 | 0.624 | 0.245 | -0.065 | -0.068 | -0.068 | 0.972 | 0.27 | 0.18 | 0.36 | 0.5 | 0.39 | 0.58 |
| 1 | 111622739 | rs1282129 | 0.977 | 0.921 | 0.657 | 0.008 | 0.024 | 0.025 | 0.025 | 0 | 0.55 | 0.37 | 0.58 | 0.72 | 0.62 | 0.41 |
| 1 | 111628733 | rs646734 | -0.277 | 0.927 | 0.776 | -0.015 | -0.011 | 0.052 | 0.052 | 0.215 | 0.17 | 0.36 | 0.22 | 0.21 | 0.13 | 0.41 |
| 1 | 111641712 | rs12752511 | 0.658 | 0.929 | 0.781 | -0.022 | 0.019 | 0.053 | 0.053 | 0 | 0.22 | 0.1 | 0.16 | 0 | 0.23 | 0.17 |
| 1 | 111642031 | rs11102212 | -0.261 | 1.023 | 0.724 | 0.013 | 0.034 | 0.036 | 0.036 | 0 | 0.14 | 0.21 | 0.19 | 0.36 | 0.27 | 0.24 |
| 1 | 111646770 | rs17026872 | 0.545 | 0.999 | 0.771 | 0.005 | 0.003 | -0.036 | -0.036 | 0 | 0.23 | 0.17 | 0.17 | 0 | 0.23 | 0.17 |
| 1 | 111660540 | rs11002 | -0.941 | 1.098 | 0.826 | 0.127 | -0.025 | 0.061 | 0.061 | 0.208 | 0.22 | 0.47 | 0.25 | 0.28 | 0.15 | 0.42 |
| 1 | 111661528 | rs325917 | -0.954 | 1.098 | 0.827 | 0.128 | -0.026 | 0.06 | 0.06 | 0.233 | 0.22 | 0.47 | 0.25 | 0.28 | 0.15 | 0.42 |
| 1 | 111664343 | rs325921 | -0.954 | 1.098 | 0.827 | 0.128 | -0.026 | 0.06 | 0.06 | 0.233 | 0.22 | 0.47 | 0.25 | 0.28 | 0.15 | 0.42 |
| 1 | 111671707 | rs10494126 | 0.061 | 1.14 | 0.71 | 0.063 | -0.006 | 0.001 | 0.001 | 0 | 0.15 | 0.28 | 0.19 | 0.12 | 0.12 | 0.33 |
| 1 | 111682118 | rs3762374 | -0.63 | 1.348 | 0.78 | 0.248 | 0.023 | 0.026 | 0.026 | 0 | 0.15 | 0.21 | 0.2 | 0.35 | 0.28 | 0.25 |

| | | | | | | | | | | | | | | | | |
|---|---|---|---|---|---|---|---|---|---|---|---|---|---|---|---|---|
| 1 | 111682307 | rs1043096 | 0.303 | 1.311 | 0.776 | 0.199 | -0.003 | 0.012 | 0.012 | 0.064 | 0.13 | 0.26 | 0.19 | 0.12 | 0.09 | 0.3 |
| 1 | 111686054 | rs817582 | -0.488 | 1.29 | 0.782 | 0.209 | -0.011 | 0.042 | 0.042 | 0.617 | 0.18 | 0.32 | 0.24 | 0.29 | 0.12 | 0.4 |
| 1 | 111700001 | rs12745827 | 0.621 | 1.701 | 0.026 | 0.256 | -0.048 | 0.005 | 0.005 | 0 | 0.08 | 0.01 | 0.04 | 0.04 | 0.15 | 0.11 |
| 1 | 111726213 | rs694180 | -1.973 | 0.565 | -0.287 | 0.008 | 0.034 | 0.042 | 0.042 | 1.067 | 0.58 | 0.85 | 0.8 | 0.65 | 0.64 | 0.82 |
| 1 | 111726288 | rs2275122 | 0.889 | 0.587 | -0.266 | -0.016 | -0.001 | -0.008 | -0.008 | 0 | 0.11 | 0.08 | 0.35 | 0.13 | 0.09 | 0.34 |
| 1 | 111726869 | rs3795823 | 0.384 | 0.621 | -0.41 | -0.134 | -0.088 | -0.086 | -0.086 | 0.022 | 0.13 | 0.03 | 0.18 | 0.33 | 0.28 | 0.18 |
| 1 | 111728610 | rs3818797 | 0.325 | 0.714 | -0.324 | -0.119 | 0.041 | 0.045 | 0.045 | 0 | 0.15 | 0.24 | 0.19 | 0.34 | 0.28 | 0.26 |
| 1 | 111729325 | rs626339 | -1.16 | 0.564 | -0.29 | -0.139 | 0.012 | 0.013 | 0.013 | 0 | 0.44 | 0.61 | 0.6 | 0.32 | 0.36 | 0.61 |
| 1 | 111730901 | rs608881 | 0.196 | 0.899 | -0.253 | -0.205 | 0.014 | 0.008 | 0.008 | 0 | 0.23 | 0.15 | 0.16 | 0 | 0.23 | 0.19 |
| 1 | 111731158 | rs2297703 | -0.085 | 1.512 | -0.325 | 0.254 | 0.047 | 0.05 | 0.05 | 0 | 0.15 | 0.25 | 0.21 | 0.34 | 0.3 | 0.25 |
| 1 | 111736389 | rs657801 | -1.627 | 0.708 | -0.422 | 0.178 | 0.036 | 0.032 | 0.032 | 2.171 | 0.6 | 0.85 | 0.8 | 0.66 | 0.67 | 0.85 |
| 1 | 111736672 | rs656843 | 0.258 | 1.229 | -0.194 | 0.015 | 0.008 | -0.02 | -0.02 | 0 | 0.21 | 0.18 | 0.14 | 0 | 0.17 | 0.19 |
| 1 | 111737398 | rs9970286 | -0.324 | 1.54 | -0.44 | 0.377 | 0.079 | 0.083 | 0.083 | 0 | 0.17 | 0.31 | 0.22 | 0.38 | 0.34 | 0.24 |
| 1 | 111737916 | rs630505 | -0.045 | 1.576 | -0.779 | 0.381 | -0.018 | -0.056 | -0.056 | 1.772 | 0.32 | 0.42 | 0.2 | 0.1 | 0.26 | 0.25 |
| 1 | 111738116 | rs629608 | -0.332 | 0.905 | -1.14 | 0.611 | 0.019 | -0.003 | -0.003 | 0 | 0.48 | 0.66 | 0.44 | 0.48 | 0.61 | 0.53 |
| 1 | 111740942 | rs610261 | -1.405 | 1.126 | -1.117 | 0.617 | -0.001 | -0.001 | -0.001 | 0.077 | 0.15 | 0.17 | 0.09 | 0.09 | 0.15 | 0.13 |
| 1 | 111742761 | rs639210 | 0.823 | 1.599 | -1.647 | 0.243 | -0.049 | 0.087 | 0.087 | 0.036 | 0.85 | 0.98 | 0.92 | 0.99 | 0.81 | 0.85 |
| 1 | 111746355 | rs2296381 | -0.753 | 1.596 | -1.82 | 1.261 | -0.017 | 0.002 | 0.002 | 0 | 0.05 | 0.01 | 0.13 | 0.35 | 0.09 | 0.14 |
| 1 | 111748088 | rs623400 | -1.315 | 2.095 | -2.02 | 1.386 | -0.019 | -0.062 | -0.062 | 0.293 | 0.21 | 0.14 | 0.24 | 0.47 | 0.3 | 0.35 |
| 1 | 111748128 | rs623775 | -0.649 | 2.072 | -2.017 | 1.177 | -0.026 | -0.025 | -0.025 | 0 | 0.15 | 0.06 | 0.11 | 0.13 | 0.2 | 0.2 |
| 1 | 111748325 | rs2477582 | 0.711 | 2.626 | -2.297 | 1.454 | -0.109 | -0.051 | -0.051 | 0 | 0.8 | 0.98 | 0.66 | 0.84 | 0.69 | 0.71 |
| 1 | 111754654 | rs7515583 | -0.633 | 2.331 | -2.147 | 1.542 | -0.026 | -0.024 | -0.024 | 0.049 | 0.15 | 0.06 | 0.1 | 0.13 | 0.2 | 0.21 |
| 1 | 111755828 | rs12749641 | -0.423 | 2.309 | -2.024 | 1.586 | -0.015 | -0.004 | -0.004 | 0 | 0.06 | 0.03 | 0.24 | 0.16 | 0.12 | 0.13 |
| 1 | 111758446 | rs684559 | -0.31 | 2.58 | -2.812 | 0.774 | -0.088 | 0.043 | 0.043 | 0 | 0.28 | 0.05 | 0.2 | 0.18 | 0.37 | 0.25 |
| 1 | 111760278 | rs654997 | -0.494 | 2.32 | -2.884 | 0.61 | 0.003 | 0.054 | 0.054 | 0.139 | 0.89 | 0.98 | 0.95 | 0.99 | 0.9 | 0.9 |
| 1 | 111762482 | rs2764546 | -0.133 | 2.429 | -2.902 | 0.603 | -0.065 | 0.04 | 0.04 | 0 | 0.27 | 0.05 | 0.17 | 0.18 | 0.34 | 0.24 |
| 1 | 111763153 | rs17014713 | -0.684 | 2.258 | -2.473 | 1.052 | 0.003 | 0.076 | 0.076 | 0 | 0.08 | 0.1 | 0.14 | 0.37 | 0.11 | 0.18 |
| 1 | 111764026 | rs2477583 | 0.011 | 2.222 | -2.272 | 0.935 | 0.001 | 0 | 0 | 0 | 0.92 | 0.94 | 0.75 | 0.91 | 0.88 | 0.85 |
| 1 | 111770423 | rs755467 | -1.785 | 1.462 | -1.8 | 0.583 | 0.007 | 0.006 | 0.006 | 0 | 0.61 | 0.67 | 0.68 | 0.67 | 0.7 | 0.74 |
| 1 | 111770591 | rs2147790 | 0.307 | 1.484 | -1.811 | 0.559 | -0.041 | -0.036 | -0.036 | 0 | 0.07 | 0 | 0.25 | 0.09 | 0.13 | 0.15 |
| 1 | 111773600 | rs2274232 | -0.053 | 1.456 | -0.325 | 0.568 | -0.002 | 0.078 | 0.078 | 0 | 0.05 | 0.05 | 0.13 | 0.36 | 0.09 | 0.18 |
| 1 | 111778468 | rs961364 | 1.9 | 1.236 | -0.103 | 0.494 | -0.015 | -0.005 | -0.005 | 0 | 0.36 | 0.08 | 0.29 | 0.32 | 0.3 | 0.26 |
| 1 | 111778824 | rs2251608 | -1.652 | 1.232 | 0.394 | 0.49 | -0.009 | -0.012 | -0.012 | 0 | 0.59 | 0.53 | 0.66 | 0.68 | 0.67 | 0.74 |
| 1 | 111780334 | rs2820087 | 0.616 | 1.208 | 0.315 | 0.404 | -0.059 | 0.013 | 0.013 | 0 | 0.25 | 0.05 | 0.17 | 0.2 | 0.33 | 0.24 |
| 1 | 111784816 | rs11102221 | -0.967 | 1.263 | 0.305 | 0.424 | -0.007 | -0.039 | -0.039 | 0 | 0.28 | 0.37 | 0.34 | 0.12 | 0.24 | 0.32 |
| 1 | 111785394 | rs3934922 | -1.605 | 1.261 | 0.31 | 0.435 | -0.002 | -0.003 | -0.003 | 0 | 0.58 | 0.44 | 0.64 | 0.67 | 0.67 | 0.74 |
| 1 | 111785467 | rs3934923 | 0.401 | 1.31 | 0.172 | 0.465 | -0.018 | 0.018 | 0.018 | 0 | 0.27 | 0.15 | 0.19 | 0.2 | 0.34 | 0.24 |
| 1 | 111788779 | rs3886706 | 1.473 | 1.337 | 0.18 | 0.49 | 0.009 | -0.035 | -0.035 | 0 | 0.62 | 0.22 | 0.48 | 0.52 | 0.63 | 0.49 |
| 1 | 111789004 | rs17027123 | -0.268 | 1.874 | 0.215 | 0.995 | -0.031 | 0.001 | 0.001 | 0 | 0.06 | 0.01 | 0.15 | 0.36 | 0.11 | 0.25 |
| 1 | 111789610 | rs17699274 | 0.596 | 1.647 | 0.619 | 0.42 | 0.009 | -0.002 | -0.002 | 0 | 0.64 | 0.62 | 0.54 | 0.7 | 0.54 | 0.6 |
| 1 | 111794733 | rs7514853 | 0.777 | 1.7 | 0.275 | 0.464 | 0.024 | -0.009 | -0.009 | 0 | 0.63 | 0.52 | 0.52 | 0.69 | 0.52 | 0.59 |
| 1 | 111795352 | rs1075772 | -1.113 | 1.458 | 0.388 | 0.599 | 0.041 | 0.028 | 0.028 | 0 | 0.11 | 0.25 | 0.29 | 0.1 | 0.15 | 0.16 |
| 1 | 111798711 | rs12082033 | -0.493 | 1.103 | 0.369 | 0.323 | 0.013 | -0.013 | -0.013 | 0.501 | 0.12 | 0.32 | 0.12 | 0.08 | 0.13 | 0.06 |
| 1 | 111799137 | rs17654501 | -0.175 | 1.123 | -0.115 | 0.024 | -0.017 | 0.029 | 0.029 | 0 | 0.07 | 0.02 | 0.06 | 0.01 | 0.1 | 0.12 |
| 1 | 111800472 | rs2820082 | 0.641 | 0.893 | 0.575 | 0.677 | -0.004 | 0.06 | 0.06 | 1.209 | 0.84 | 0.66 | 0.86 | 0.79 | 0.84 | 0.78 |
| 1 | 111806570 | rs2820061 | 0.66 | 0.861 | 0.727 | 0.784 | -0.014 | 0.048 | 0.048 | 0 | 0.95 | 0.87 | 0.97 | 0.9 | 0.97 | 0.85 |
| 1 | 111810174 | rs3122591 | 1.368 | 0.768 | 0.546 | 0.322 | -0.003 | -0.008 | -0.008 | 0 | 0.68 | 0.54 | 0.77 | 0.77 | 0.69 | 0.6 |
| 1 | 111812709 | rs3118208 | 0.638 | 1.051 | -0.052 | 0.567 | 0 | 0 | 0 | 0.094 | 0.5 | 0.46 | 0.51 | 0.47 | 0.42 | 0.43 |
| 1 | 111813951 | rs3122595 | 0.733 | 0.962 | -0.043 | 0.576 | -0.011 | -0.004 | -0.004 | 0 | 0.71 | 0.55 | 0.78 | 0.78 | 0.73 | 0.64 |
| 1 | 111818052 | rs6702380 | -0.325 | 0.938 | -0.036 | 0.636 | -0.001 | -0.006 | -0.006 | 0 | 0.23 | 0.27 | 0.17 | 0.13 | 0.23 | 0.21 |
| 1 | 111820644 | rs6703883 | -0.191 | 0.924 | -0.025 | 0.63 | 0 | 0 | 0 | 0 | 0.22 | 0.22 | 0.17 | 0.13 | 0.23 | 0.21 |
| 1 | 111821434 | rs12048213 | 1.493 | 0.791 | -0.537 | 0.071 | 0.001 | -0.104 | -0.104 | 0.297 | 0.34 | 0.41 | 0.16 | 0.08 | 0.28 | 0.23 |
| 1 | 111821777 | rs4838896 | 0.245 | 0.795 | -0.599 | 0.106 | -0.032 | 0.012 | 0.012 | 0 | 0.37 | 0.14 | 0.63 | 0.69 | 0.45 | 0.41 |
| 1 | 111822648 | rs12028469 | 0.602 | 0.818 | -0.642 | 0.15 | -0.012 | -0.078 | -0.078 | 0 | 0.16 | 0.03 | 0.36 | 0.39 | 0.17 | 0.22 |
| 1 | 111823173 | rs12047673 | 2.398 | 0.517 | -0.853 | -0.501 | 0.005 | 0.023 | 0.023 | 1.749 | 0.13 | 0.06 | 0.03 | 0.06 | 0.07 | 0.06 |
| 1 | 111823220 | rs4323713 | 2.654 | 0.59 | -0.818 | -0.426 | -0.012 | -0.025 | -0.025 | 0 | 0.21 | 0.11 | 0.27 | 0.3 | 0.28 | 0.19 |
| 1 | 111824209 | rs6537690 | 0.891 | 0.541 | -0.634 | -0.434 | 0 | -0.003 | -0.003 | 7.918 | 0.22 | 0.24 | 0.17 | 0.13 | 0.23 | 0.21 |
| 1 | 111825226 | rs6687390 | 0.14 | 0.528 | -0.624 | -0.429 | -0.009 | 0.001 | 0.001 | 0.321 | 0.27 | 0.38 | 0.2 | 0.23 | 0.26 | 0.36 |
| 1 | 111826609 | rs4575084 | -1.095 | 0.541 | -0.34 | -0.512 | 0.007 | -0.055 | -0.055 | 0.022 | 0.74 | 0.56 | 0.83 | 0.89 | 0.76 | 0.68 |
| 1 | 111827749 | rs11102233 | 0.195 | 0.695 | -0.62 | -0.415 | 0.009 | -0.019 | -0.019 | 0 | 0.3 | 0.17 | 0.4 | 0.45 | 0.25 | 0.28 |
| 1 | 111829797 | rs12401767 | 0.695 | 0.704 | -0.68 | -0.557 | -0.007 | 0.018 | 0.018 | 0 | 0.39 | 0.23 | 0.63 | 0.7 | 0.46 | 0.42 |
| 1 | 111832205 | rs3806448 | 0.429 | 0.562 | -0.497 | -0.644 | -0.004 | 0.065 | 0.065 | 0 | 0.49 | 0.32 | 0.64 | 0.79 | 0.49 | 0.47 |
| 1 | 111832291 | rs3806446 | 0.691 | 0.555 | -0.646 | -0.69 | -0.004 | 0.047 | 0.047 | 0 | 0.44 | 0.24 | 0.63 | 0.74 | 0.46 | 0.4 |
| 1 | 111833661 | rs4838897 | -1.434 | 0.436 | -0.379 | -0.992 | -0.002 | -0.027 | -0.027 | 0.282 | 0.79 | 0.83 | 0.85 | 0.98 | 0.76 | 0.83 |
| 1 | 111846416 | rs12563158 | 1.479 | 0.243 | -0.534 | -0.685 | -0.006 | -0.017 | -0.017 | 0 | 0.3 | 0.18 | 0.3 | 0.34 | 0.32 | 0.19 |
| 1 | 111852352 | rs7531210 | 0.053 | 0.138 | 0.123 | -1.117 | -0.008 | -0.063 | -0.063 | 3.653 | 0.14 | 0.21 | 0.08 | 0 | 0.11 | 0.1 |
| 1 | 111854207 | rs4839122 | -0.412 | -0.187 | 0.091 | -1.762 | 0.009 | -0.05 | -0.05 | 0.367 | 0.69 | 0.55 | 0.81 | 0.84 | 0.68 | 0.62 |
| 1 | 111857087 | rs2786151 | -1.291 | 0.13 | 0.104 | -0.837 | 0.004 | 0.013 | 0.013 | 0 | 0.58 | 0.66 | 0.35 | 0.45 | 0.56 | 0.67 |
| 1 | 111857154 | rs2786152 | -2.025 | 0.262 | -0.011 | -0.999 | 0.014 | -0.011 | -0.011 | 0 | 0.69 | 0.79 | 0.42 | 0.48 | 0.71 | 0.77 |

| | | | | | | | | | | | | | | | | |
|---|---|---|---|---|---|---|---|---|---|---|---|---|---|---|---|---|
| 1 | 111857208 | rs3818822 | 1.51 | 0.362 | 0.209 | -0.953 | 0.031 | 0.031 | 0.031 | 0.485 | 0.08 | 0.19 | 0.08 | 0.09 | 0.1 | 0.15 |
| 1 | 111857485 | rs2820072 | -0.78 | 0.339 | 0.133 | -0.972 | 0.001 | 0.001 | 0.001 | 0 | 0.55 | 0.58 | 0.34 | 0.48 | 0.6 | 0.67 |
| 1 | 111858444 | rs2786156 | -2.66 | 0.306 | 0.118 | -0.647 | 0.019 | -0.045 | -0.045 | 0.023 | 0.73 | 0.86 | 0.45 | 0.64 | 0.75 | 0.85 |
| 1 | 111859838 | rs12034576 | 0.852 | 0.367 | 0.161 | -1.03 | -0.04 | 0.053 | 0.053 | 0 | 0.34 | 0.1 | 0.16 | 0.36 | 0.35 | 0.43 |
| 1 | 111859912 | rs10494134 | 0.195 | 0.221 | 0.24 | -0.953 | -0.053 | 0.122 | 0.122 | 0 | 0.43 | 0.13 | 0.23 | 0.38 | 0.49 | 0.52 |
| 1 | 111861822 | rs17027410 | 1.868 | 0.226 | 0.355 | -0.951 | 0.002 | 0.003 | 0.003 | 0 | 0.13 | 0.09 | 0.07 | 0 | 0.11 | 0.1 |
| 1 | 111861841 | rs2275253 | -2.022 | 0.279 | 0.53 | -0.707 | 0.016 | -0.013 | -0.013 | 0 | 0.69 | 0.8 | 0.43 | 0.48 | 0.71 | 0.77 |
| 1 | 111861974 | rs2275254 | -0.561 | 0.286 | 0.602 | -0.713 | 0.012 | 0.038 | 0.038 | 0 | 0.58 | 0.32 | 0.31 | 0.38 | 0.6 | 0.61 |
| 1 | 111862074 | rs2820092 | 0.398 | 0.335 | 0.403 | -0.673 | 0.039 | 0.039 | 0.039 | 0.163 | 0.11 | 0.35 | 0.1 | 0.09 | 0.1 | 0.15 |
| 1 | 111862952 | rs2256721 | -1.767 | -0.224 | 0.155 | -0.54 | 0.014 | -0.012 | -0.012 | 0 | 0.69 | 0.79 | 0.42 | 0.48 | 0.71 | 0.77 |
| 1 | 111869487 | rs10776724 | -0.426 | 0.161 | -0.094 | -0.477 | -0.003 | -0.009 | -0.009 | 0.262 | 0.52 | 0.47 | 0.29 | 0.42 | 0.53 | 0.56 |
| 1 | 111869948 | rs12137697 | 1.778 | 0.141 | -0.157 | -0.596 | 0.004 | -0.03 | -0.03 | 1.671 | 0.11 | 0.16 | 0.08 | 0.02 | 0.13 | 0.14 |
| 1 | 111873172 | rs1832286 | -1.88 | 0.104 | -0.241 | -0.467 | 0.016 | -0.033 | -0.033 | 0 | 0.7 | 0.78 | 0.43 | 0.54 | 0.72 | 0.82 |
| 1 | 111875202 | rs2820075 | 0.613 | 0.091 | -0.391 | -0.649 | -0.048 | 0.064 | 0.064 | 0 | 0.37 | 0.15 | 0.19 | 0.38 | 0.4 | 0.44 |
| 1 | 111875706 | rs7532160 | 1.019 | 0.149 | -0.13 | -0.424 | -0.002 | -0.044 | -0.044 | 0 | 0.13 | 0.18 | 0.09 | 0 | 0.12 | 0.11 |
| 1 | 111884929 | rs11589320 | 1.019 | 0.149 | -0.13 | -0.424 | -0.002 | -0.044 | -0.044 | 0 | 0.13 | 0.18 | 0.09 | 0 | 0.12 | 0.11 |
| 1 | 111885460 | rs10776725 | -1.731 | 0.044 | -0.286 | -0.338 | 0.015 | -0.033 | -0.033 | 0 | 0.7 | 0.79 | 0.43 | 0.54 | 0.72 | 0.82 |
| 1 | 111891192 | rs2184884 | 1.193 | 0.137 | -0.088 | -0.362 | 0 | -0.001 | -0.001 | 0.163 | 0.13 | 0.11 | 0.08 | 0 | 0.12 | 0.1 |
| 1 | 111892905 | rs3820443 | -0.273 | 0.165 | -0.182 | -0.574 | -0.005 | -0.008 | -0.008 | 0 | 0.35 | 0.39 | 0.19 | 0.37 | 0.31 | 0.41 |
| 1 | 111893958 | rs1668 | 0.005 | 0.187 | -0.284 | -0.636 | 0.003 | 0.021 | 0.021 | 0 | 0.33 | 0.27 | 0.17 | 0.32 | 0.29 | 0.4 |
| 1 | 111895487 | rs1058530 | -1.003 | 0.317 | -0.386 | -0.256 | -0.001 | 0 | 0 | 0.491 | 0.47 | 0.53 | 0.25 | 0.58 | 0.4 | 0.64 |
| 1 | 111897335 | rs11102257 | -1.11 | 0.339 | -0.387 | -0.233 | 0.01 | -0.001 | -0.001 | 0 | 0.48 | 0.61 | 0.27 | 0.58 | 0.4 | 0.65 |
| 1 | 111899929 | rs1106189 | -0.344 | 0.229 | -0.099 | 0.02 | 0.018 | 0.056 | 0.056 | 0 | 0.22 | 0.47 | 0.16 | 0.26 | 0.23 | 0.3 |
| 1 | 111900279 | rs11102258 | 0.878 | 0.309 | -0.13 | -0.035 | 0.006 | -0.032 | -0.032 | 0 | 0.14 | 0.19 | 0.11 | 0.03 | 0.17 | 0.24 |
| 1 | 111904494 | rs6663292 | -0.632 | 0.095 | -0.291 | 0.25 | 0.002 | -0.003 | -0.003 | 0.012 | 0.61 | 0.64 | 0.36 | 0.64 | 0.63 | 0.73 |
| 1 | 111905989 | rs17027501 | 0.43 | 0.089 | 0.454 | 0.379 | 0.004 | 0.166 | 0.166 | 0 | 0.06 | 0.18 | 0.04 | 0.23 | 0.07 | 0.05 |
| 1 | 111906034 | rs4838908 | 0.034 | -0.28 | 0.415 | -0.243 | -0.022 | -0.049 | -0.049 | 0 | 0.82 | 0.66 | 0.88 | 0.95 | 0.86 | 0.86 |
| 1 | 111908432 | rs5011851 | -0.708 | 0.094 | 0.009 | -0.222 | 0.009 | -0.059 | -0.059 | 0.001 | 0.18 | 0.51 | 0.16 | 0.05 | 0.16 | 0.14 |
| 1 | 111908450 | rs5011850 | -0.68 | 0.098 | 0.055 | -0.221 | 0.012 | -0.059 | -0.059 | 0.325 | 0.18 | 0.49 | 0.16 | 0.05 | 0.16 | 0.14 |
| 1 | 111908713 | rs1361049 | 0.602 | 0.043 | -0.7 | -0.501 | 0.006 | 0.11 | 0.11 | 0 | 0.51 | 0.25 | 0.67 | 0.78 | 0.53 | 0.55 |
| 1 | 111909048 | rs7541956 | -0.364 | 0.63 | -0.142 | -0.371 | 0.033 | -0.07 | -0.07 | 5.438 | 0.13 | 0.33 | 0.13 | 0.01 | 0.13 | 0.1 |
| 1 | 111909604 | rs2065929 | -0.53 | 1.012 | -0.717 | 0.325 | -0.03 | -0.066 | -0.066 | 0 | 0.86 | 0.97 | 0.89 | 0.66 | 0.85 | 0.75 |
| 1 | 111909959 | rs10453833 | 0.784 | 0.689 | -0.782 | -0.072 | -0.005 | 0.045 | 0.045 | 0.426 | 0.27 | 0.13 | 0.12 | 0.14 | 0.27 | 0.18 |
| 1 | 111913523 | rs961558 | 0.19 | 0.506 | -0.648 | 0.222 | 0.017 | 0.012 | 0.012 | 0 | 0.33 | 0.73 | 0.22 | 0.23 | 0.26 | 0.3 |
| 1 | 111914431 | rs12122084 | 1.545 | 0.45 | -0.347 | 0.137 | 0.002 | -0.024 | -0.024 | 1.685 | 0.23 | 0.2 | 0.14 | 0.01 | 0.18 | 0.22 |
| 1 | 111916670 | rs11102263 | 0.911 | 1.093 | 0.452 | 0.89 | -0.003 | 0.001 | 0.001 | 0 | 0.44 | 0.48 | 0.66 | 0.44 | 0.46 | 0.4 |
| 1 | 111922854 | rs12035026 | 0.882 | 1.116 | 0.052 | 0.89 | -0.019 | -0.051 | -0.051 | 0 | 0.11 | 0.02 | 0.07 | 0.33 | 0.1 | 0.22 |
| 1 | 111924907 | rs7513936 | 1.427 | 1.128 | 0.037 | 0.898 | 0.005 | 0.021 | 0.021 | 0.118 | 0.39 | 0.22 | 0.63 | 0.28 | 0.42 | 0.36 |
| 1 | 111925349 | rs4839130 | 0.551 | 1.141 | 0.072 | 0.825 | 0.016 | 0.037 | 0.037 | 0 | 0.49 | 0.74 | 0.71 | 0.5 | 0.51 | 0.43 |
| 1 | 111926153 | rs6537699 | -0.213 | 1.123 | -0.192 | 0.581 | -0.076 | 0.16 | 0.16 | 0 | 0.62 | 0.91 | 0.8 | 0.83 | 0.61 | 0.67 |
| 1 | 111929295 | rs7521484 | 1.131 | 0.754 | -0.027 | 0.495 | 0.009 | 0.016 | 0.016 | 0.178 | 0.41 | 0.27 | 0.64 | 0.28 | 0.43 | 0.38 |
| 1 | 111930916 | rs12077338 | -0.36 | 0.806 | 0.331 | 0.589 | 0.017 | 0.157 | 0.157 | 0.012 | 0.06 | 0.32 | 0.03 | 0.15 | 0.02 | 0.04 |
| 1 | 111930990 | rs17027591 | 0.062 | 0.816 | 0.249 | 0.594 | 0.004 | 0.127 | 0.127 | 0 | 0.05 | 0.24 | 0.03 | 0.15 | 0.02 | 0.04 |
| 1 | 111931251 | rs7513731 | 0.847 | 0.923 | 0.02 | 0.563 | -0.008 | 0.004 | 0.004 | 0.156 | 0.42 | 0.49 | 0.61 | 0.42 | 0.43 | 0.39 |
| 1 | 111933151 | rs7545946 | 1.663 | 1.683 | 0.986 | 1.43 | 0.012 | 0.012 | 0.012 | 0.199 | 0.62 | 0.49 | 0.72 | 0.47 | 0.61 | 0.44 |
| 1 | 111934304 | rs4839132 | 1.626 | 1.641 | 0.965 | 1.571 | -0.022 | -0.005 | -0.005 | 0 | 0.64 | 0.75 | 0.74 | 0.3 | 0.6 | 0.41 |
| 1 | 111944073 | rs1264880 | 1.368 | 1.694 | 0.969 | 1.629 | -0.033 | -0.038 | -0.038 | 0 | 0.68 | 0.8 | 0.76 | 0.47 | 0.63 | 0.46 |
| 1 | 111946958 | rs1539581 | -2.169 | 1.686 | 1.27 | 1.61 | -0.01 | 0.094 | 0.094 | 1.265 | 0.11 | 0.55 | 0.07 | 0.17 | 0.05 | 0.16 |
| 1 | 111955195 | rs945750 | -0.653 | 1.79 | 1.136 | 1.725 | -0.007 | 0.062 | 0.062 | 0 | 0.11 | 0.14 | 0.08 | 0.38 | 0.09 | 0.25 |
| 1 | 111956473 | rs1777833 | -0.574 | 1.792 | 1.127 | 1.726 | -0.005 | 0.055 | 0.055 | 0 | 0.11 | 0.14 | 0.09 | 0.38 | 0.09 | 0.25 |
| 1 | 111957245 | rs9629 | -1.482 | 1.919 | 1.03 | 1.862 | -0.046 | -0.012 | -0.012 | 0 | 0.19 | 0.12 | 0.19 | 0.19 | 0.27 | 0.21 |
| 1 | 111957693 | rs2485319 | -3.065 | 1.881 | 1.143 | 1.889 | -0.028 | 0.077 | 0.077 | 2.135 | 0.43 | 0.76 | 0.36 | 0.77 | 0.44 | 0.71 |
| 1 | 111959983 | rs4838910 | -2.555 | 1.898 | 1.187 | 1.848 | 0.003 | 0.001 | 0.001 | 0.789 | 0.33 | 0.62 | 0.29 | 0.38 | 0.36 | 0.47 |
| 1 | 111962455 | rs1264881 | -2.244 | 1.891 | 1.416 | 1.828 | 0.022 | 0.022 | 0.022 | 0.251 | 0.35 | 0.44 | 0.31 | 0.59 | 0.38 | 0.55 |
| 1 | 111966291 | rs954739 | -1.055 | 1.848 | 1.45 | 2.035 | -0.053 | 0.027 | 0.027 | 0 | 0.18 | 0.06 | 0.13 | 0.14 | 0.24 | 0.21 |
| 1 | 111967008 | rs7554372 | -2.122 | 1.838 | 1.732 | 2.075 | 0.032 | 0.009 | 0.009 | 0.313 | 0.26 | 0.42 | 0.18 | 0.31 | 0.29 | 0.37 |
| 1 | 111968316 | rs1264888 | -2.938 | 1.849 | 1.728 | 2.057 | 0.005 | -0.014 | -0.014 | 0 | 0.38 | 0.56 | 0.31 | 0.74 | 0.4 | 0.62 |
| 1 | 111987035 | rs1264901 | -0.238 | 1.78 | 1.73 | 2.1 | -0.002 | 0.034 | 0.034 | 0 | 0.11 | 0.12 | 0.08 | 0.39 | 0.09 | 0.24 |
| 1 | 111991845 | rs1270853 | -2.879 | 1.785 | 1.782 | 1.563 | 0.006 | -0.015 | -0.015 | 1.199 | 0.38 | 0.53 | 0.31 | 0.74 | 0.39 | 0.62 |
| 1 | 111992095 | rs1264899 | -2.729 | 1.786 | 1.642 | 1.563 | 0.004 | -0.007 | -0.007 | 0.532 | 0.36 | 0.39 | 0.29 | 0.74 | 0.39 | 0.62 |
| 1 | 112000344 | rs12039707 | -2.271 | 1.853 | 1.171 | 0.722 | -0.002 | -0.001 | -0.001 | 0 | 0.24 | 0.24 | 0.16 | 0.3 | 0.29 | 0.37 |
| 1 | 112001579 | rs12040764 | -2.636 | 1.884 | 1.116 | 0.729 | 0.016 | -0.001 | -0.001 | 0 | 0.28 | 0.6 | 0.2 | 0.3 | 0.29 | 0.37 |
| 1 | 112008135 | rs10489794 | -1.392 | 1.835 | 0.576 | 0.517 | -0.053 | 0.03 | 0.03 | 0 | 0.18 | 0.06 | 0.13 | 0.14 | 0.24 | 0.21 |
| 1 | 112009195 | rs12045175 | -0.371 | 1.959 | 0.47 | 0.525 | -0.04 | 0.007 | 0.007 | 0 | 0.09 | 0.01 | 0.05 | 0.09 | 0.12 | 0.15 |
| 1 | 112018657 | rs6703267 | -0.714 | 1.766 | 0.622 | 0.656 | 0.005 | -0.01 | -0.01 | 0 | 0.2 | 0.2 | 0.14 | 0.14 | 0.24 | 0.21 |
| 1 | 112021029 | rs1054680 | -1.195 | 1.688 | -0.166 | 0.705 | 0.004 | -0.007 | -0.007 | 0 | 0.31 | 0.64 | 0.29 | 0.24 | 0.33 | 0.39 |
| 1 | 112026954 | rs1361135 | 0.69 | 1.876 | -0.023 | 1.014 | 0.035 | 0.03 | 0.03 | 0.123 | 0.72 | 0.59 | 0.58 | 0.6 | 0.59 | 0.65 |
| 1 | 112027033 | rs1361133 | 0.764 | 1.198 | -0.611 | 0.295 | -0.012 | 0.081 | 0.081 | 6.883 | 0.15 | 0.06 | 0.08 | 0 | 0.14 | 0.02 |
| 1 | 112029763 | rs3767602 | 0.03 | 1.038 | -0.011 | 0.81 | 0.014 | 0.171 | 0.171 | 0.13 | 0.06 | 0.16 | 0.03 | 0.25 | 0.06 | 0.16 |

| | | | | | | | | | | | | | | | | |
|---|---|---|---|---|---|---|---|---|---|---|---|---|---|---|---|---|
| 1 | 112032205 | rs7544118 | -0.673 | 0.642 | 0.598 | 0.507 | -0.023 | 0.005 | 0.005 | 4.31 | 0.28 | 0.46 | 0.14 | 0.25 | 0.21 | 0.18 |
| 1 | 112033983 | rs7547951 | 0.677 | 0.606 | 0.69 | 0.409 | -0.001 | -0.032 | -0.032 | 0 | 0.17 | 0.22 | 0.1 | 0 | 0.14 | 0.02 |
| 1 | 112042131 | rs3394 | -0.346 | 1.364 | 0.714 | 0.792 | 0.05 | 0.077 | 0.077 | 0.221 | 0.88 | 0.74 | 0.86 | 0.76 | 0.83 | 0.72 |
| 1 | 112042149 | rs3393 | 1.335 | 1.334 | 0.79 | 0.847 | -0.017 | 0.006 | 0.006 | 0.314 | 0.56 | 0.17 | 0.66 | 0.44 | 0.56 | 0.48 |
| 1 | 112042195 | rs1415792 | -0.183 | 1.11 | 0.751 | 1.031 | 0.05 | 0.077 | 0.077 | 0.221 | 0.88 | 0.74 | 0.86 | 0.76 | 0.83 | 0.72 |
| 1 | 112042236 | rs1415793 | -0.183 | 1.11 | 0.751 | 1.031 | 0.05 | 0.077 | 0.077 | 0.221 | 0.88 | 0.74 | 0.86 | 0.76 | 0.83 | 0.72 |
| 1 | 112042383 | rs923 | -0.183 | 1.11 | 0.751 | 1.031 | 0.05 | 0.077 | 0.077 | 0.221 | 0.88 | 0.74 | 0.86 | 0.76 | 0.83 | 0.72 |
| 1 | 112042632 | rs2229155 | -0.183 | 1.11 | 0.751 | 1.031 | 0.05 | 0.077 | 0.077 | 0.221 | 0.88 | 0.74 | 0.86 | 0.76 | 0.83 | 0.72 |
| 1 | 112046540 | rs1544224 | 0.574 | 1.104 | 0.671 | 1.002 | 0.05 | 0.093 | 0.093 | 0.005 | 0.84 | 0.62 | 0.8 | 0.67 | 0.76 | 0.66 |
| 1 | 112046557 | rs1544223 | 0.59 | 1.103 | 0.671 | 1.001 | 0.049 | 0.091 | 0.091 | 0.011 | 0.84 | 0.62 | 0.8 | 0.67 | 0.76 | 0.66 |
| 1 | 112048264 | rs2298191 | 1.257 | 0.62 | -0.295 | 0.493 | 0.037 | 0.042 | 0.042 | 0 | 0.37 | 0.11 | 0.45 | 0.33 | 0.31 | 0.35 |
| 1 | 112052105 | rs6537704 | 0.273 | 0.652 | 0.125 | 0.501 | -0.005 | -0.02 | -0.02 | 0 | 0.84 | 0.68 | 0.9 | 0.89 | 0.84 | 0.85 |
| 1 | 112052305 | rs17027867 | -1.066 | 0.668 | 0.171 | 0.524 | 0.009 | 0.183 | 0.183 | 2.195 | 0.19 | 0.55 | 0.19 | 0.44 | 0.19 | 0.37 |
| 1 | 112054366 | rs10857888 | 0.586 | 0.637 | 0.071 | 0.422 | -0.006 | -0.061 | -0.061 | 0 | 0.78 | 0.54 | 0.84 | 0.89 | 0.78 | 0.8 |
| 1 | 112056187 | rs11102292 | -0.295 | 0.895 | 0.306 | 0.199 | 0.073 | 0.383 | 0.383 | 1.224 | 0.11 | 0.38 | 0.13 | 0.44 | 0.14 | 0.34 |
| 1 | 112056416 | rs4839139 | 0.161 | 0.889 | -0.166 | -0.684 | -0.128 | 0.076 | 0.076 | 1.196 | 0.28 | 0.03 | 0.25 | 0.12 | 0.35 | 0.13 |
| 1 | 112061288 | rs4839141 | 0.147 | 0.727 | -0.207 | -0.699 | -0.115 | 0.098 | 0.098 | 0.694 | 0.31 | 0.06 | 0.25 | 0.12 | 0.35 | 0.16 |
| 1 | 112061479 | rs4839142 | -0.117 | 0.773 | -0.118 | -0.676 | -0.068 | 0.111 | 0.111 | 0.806 | 0.34 | 0.12 | 0.26 | 0.12 | 0.35 | 0.16 |
| 1 | 112063002 | rs10857892 | 1.105 | 0.123 | 0.758 | -0.17 | -0.005 | -0.076 | -0.076 | 1.582 | 0.88 | 0.53 | 0.89 | 0.99 | 0.9 | 0.94 |
| 1 | 112075948 | rs2786967 | -0.173 | 0.345 | 0.043 | -0.418 | -0.034 | 0.045 | 0.045 | 0.433 | 0.31 | 0.23 | 0.3 | 0.09 | 0.42 | 0.34 |
| 1 | 112076856 | rs12028677 | -0.027 | 0.015 | 0.36 | 0.427 | 0.01 | 0.172 | 0.172 | 0.015 | 0.11 | 0.19 | 0.07 | 0.48 | 0.13 | 0.12 |
| 1 | 112078745 | rs10489796 | -1.221 | -0.307 | 0.796 | 0.489 | -0.008 | 0.19 | 0.19 | 0 | 0.18 | 0.35 | 0.1 | 0.48 | 0.14 | 0.15 |
| 1 | 112079314 | rs10776732 | 0.02 | -0.187 | 0.487 | 0.57 | 0.011 | 0.177 | 0.177 | 0.015 | 0.11 | 0.19 | 0.07 | 0.48 | 0.13 | 0.12 |
| 1 | 112080120 | rs11102303 | 1.065 | -0.547 | -0.505 | 0.056 | 0.03 | 0 | 0 | 0 | 0.33 | 0.1 | 0.51 | 0.29 | 0.3 | 0.4 |
| 1 | 112081591 | rs3849171 | 1.078 | -0.581 | -0.813 | 0.059 | 0.016 | -0.03 | -0.03 | 0 | 0.32 | 0.06 | 0.48 | 0.28 | 0.29 | 0.48 |
| 1 | 112082365 | rs2364813 | 0.448 | -0.45 | -0.216 | 0.45 | -0.014 | 0.159 | 0.159 | 0.286 | 0.09 | 0.17 | 0.05 | 0.49 | 0.07 | 0.09 |
| 1 | 112082564 | rs2789535 | 0.426 | -0.446 | -0.341 | 0.296 | -0.042 | 0.114 | 0.114 | 2.43 | 0.55 | 0.2 | 0.6 | 0.27 | 0.5 | 0.46 |
| 1 | 112086030 | rs2185029 | -0.176 | -0.278 | 0 | -0.132 | 0.029 | -0.026 | -0.026 | 0 | 0.65 | 0.52 | 0.66 | 0.76 | 0.57 | 0.57 |
| 1 | 112093989 | rs7553961 | 0.919 | -0.244 | -0.32 | -0.14 | -0.029 | 0.095 | 0.095 | 1.951 | 0.15 | 0.01 | 0.07 | 0 | 0.13 | 0.02 |
| 1 | 112101405 | rs7533936 | -0.958 | 0.181 | 0.291 | 0.399 | 0.077 | 0.209 | 0.209 | 0.177 | 0.05 | 0.37 | 0.05 | 0.15 | 0.01 | 0.03 |
| 1 | 112102030 | rs1890549 | -0.105 | 0.343 | -0.402 | 0.359 | -0.098 | 0.109 | 0.109 | 0.174 | 0.29 | 0.12 | 0.28 | 0.08 | 0.42 | 0.36 |
| 1 | 112103432 | rs2800903 | -0.529 | 0.7 | -1.249 | 0.627 | -0.058 | -0.018 | -0.018 | 0.469 | 0.37 | 0.74 | 0.37 | 0.61 | 0.46 | 0.52 |
| 1 | 112106247 | rs2364815 | -0.346 | 0.798 | -1.494 | 0.22 | -0.02 | 0.063 | 0.063 | 0 | 0.28 | 0.12 | 0.3 | 0.19 | 0.32 | 0.28 |
| 1 | 112106444 | rs1591712 | -0.174 | 0.58 | -1.178 | -0.209 | -0.007 | -0.029 | -0.029 | 0 | 0.87 | 0.82 | 0.92 | 0.95 | 0.91 | 0.9 |
| 1 | 112107669 | rs10776733 | 0.838 | 0.535 | -0.129 | -0.381 | -0.002 | -0.038 | -0.038 | 0.061 | 0.57 | 0.51 | 0.61 | 0.76 | 0.58 | 0.63 |
| 1 | 112113959 | rs4838917 | -0.019 | 0.584 | -0.064 | -0.377 | -0.002 | 0.029 | 0.029 | 0 | 0.29 | 0.25 | 0.3 | 0.13 | 0.31 | 0.26 |
| 1 | 112119065 | rs11806788 | 1.379 | 1.205 | -0.776 | 1.015 | -0.013 | 0.062 | 0.062 | 0 | 0.39 | 0.01 | 0.44 | 0.21 | 0.29 | 0.42 |
| 1 | 112120953 | rs10857898 | 0.097 | 1.296 | -0.524 | 0.947 | -0.001 | -0.001 | -0.001 | 0 | 0.59 | 0.58 | 0.61 | 0.62 | 0.6 | 0.61 |
| 1 | 112123946 | rs11807250 | 0.828 | 1.019 | -0.503 | 0.956 | 0.057 | 0.002 | 0.002 | 0.421 | 0.47 | 0.17 | 0.49 | 0.55 | 0.35 | 0.46 |
| 1 | 112127718 | rs10857899 | 0.256 | 1.491 | -0.632 | 0.611 | -0.086 | 0.112 | 0.112 | 0 | 0.3 | 0.12 | 0.34 | 0.14 | 0.45 | 0.33 |
| 1 | 112128242 | rs7516113 | -0.741 | 1.043 | 0.3 | 1.01 | -0.01 | -0.006 | -0.006 | 0 | 0.15 | 0.55 | 0.08 | 0.12 | 0.07 | 0.14 |
| 1 | 112129621 | rs17028042 | -0.882 | 1.138 | -0.389 | 1.33 | 0.003 | -0.01 | -0.01 | 0 | 0.14 | 0.08 | 0.2 | 0.32 | 0.1 | 0.07 |
| 1 | 112129888 | rs1609633 | -0.45 | 1.057 | -0.284 | 1.387 | -0.003 | 0.022 | 0.022 | 0.719 | 0.12 | 0.11 | 0.2 | 0.23 | 0.07 | 0.06 |
| 1 | 112130427 | rs10494138 | -1.113 | 1.472 | -0.564 | 1.262 | -0.008 | -0.019 | -0.019 | 1.286 | 0.18 | 0.12 | 0.27 | 0.22 | 0.2 | 0.15 |
| 1 | 112130624 | rs2800880 | -1.034 | 1.774 | 0.311 | 0.785 | 0.049 | -0.1 | -0.1 | 0 | 0.16 | 0.43 | 0.13 | 0.04 | 0.2 | 0.17 |
| 1 | 112131164 | rs2247867 | -0.211 | 1.616 | -0.154 | 0.862 | -0.009 | -0.03 | -0.03 | 0 | 0.87 | 0.98 | 0.76 | 0.9 | 0.88 | 0.92 |
| 1 | 112132221 | rs12139261 | -0.184 | 1.588 | 0.314 | 0.641 | 0.028 | -0.034 | -0.034 | 0 | 0.07 | 0.18 | 0.08 | 0 | 0.07 | 0.06 |
| 1 | 112133910 | rs2800881 | 0.466 | 1.881 | 0.258 | 0.89 | -0.002 | -0.004 | -0.004 | 0 | 0.69 | 0.66 | 0.59 | 0.45 | 0.64 | 0.72 |
| 1 | 112135194 | rs1338078 | 1.325 | 2.47 | 0.346 | 0.875 | 0.008 | 0.009 | 0.009 | 0.094 | 0.64 | 0.4 | 0.6 | 0.57 | 0.57 | 0.64 |
| 1 | 112135958 | rs1890548 | -1.305 | 1.979 | 0.544 | 0.317 | 0.021 | 0.013 | 0.013 | 0 | 0.29 | 0.48 | 0.28 | 0.34 | 0.3 | 0.33 |
| 1 | 112138656 | rs9661153 | -0.177 | 2.005 | 0.291 | 0.234 | 0.027 | -0.018 | -0.018 | 0 | 0.13 | 0.25 | 0.07 | 0.05 | 0.13 | 0.14 |
| 1 | 112140475 | rs1780555 | 0.579 | 1.943 | 0.065 | 0.29 | -0.005 | -0.003 | -0.003 | 0 | 0.14 | 0.09 | 0.14 | 0.2 | 0.17 | 0.19 |
| 1 | 112141765 | rs7548973 | 0.465 | 1.783 | 0.174 | 0.219 | -0.001 | -0.008 | -0.008 | 0 | 0.07 | 0.07 | 0.04 | 0.04 | 0.08 | 0.1 |
| 1 | 112143035 | rs2800883 | -1.817 | 1.659 | 0.644 | 0.192 | 0.002 | -0.006 | -0.006 | 0 | 0.36 | 0.56 | 0.32 | 0.37 | 0.38 | 0.43 |
| 1 | 112147569 | rs2786991 | 1.849 | 1.985 | 0.56 | -0.052 | 0.028 | -0.006 | -0.006 | 0 | 0.49 | 0.29 | 0.37 | 0.47 | 0.42 | 0.5 |
| 1 | 112153539 | rs11102314 | 0.363 | 2.043 | 0.633 | -0.139 | 0.053 | -0.053 | -0.053 | 0.458 | 0.09 | 0.22 | 0.12 | 0 | 0.13 | 0.08 |
| 1 | 112156965 | rs4839154 | 0.296 | 2.096 | 0.512 | -0.09 | -0.053 | -0.032 | -0.032 | 0.029 | 0.18 | 0.06 | 0.17 | 0.24 | 0.23 | 0.22 |
| 1 | 112161635 | rs1359414 | -0.838 | 2.072 | 0.752 | -0.101 | 0.004 | -0.003 | -0.003 | 0 | 0.25 | 0.32 | 0.21 | 0.25 | 0.29 | 0.3 |
| 1 | 112165171 | rs508594 | 1.465 | 2.01 | 0.762 | -0.116 | 0 | 0.002 | 0.002 | 0 | 0.57 | 0.43 | 0.49 | 0.47 | 0.55 | 0.58 |
| 1 | 112186337 | rs472159 | -1.188 | 1.981 | 0.582 | -0.124 | -0.01 | -0.066 | -0.066 | 0 | 0.85 | 0.99 | 0.78 | 0.79 | 0.86 | 0.89 |
| 1 | 112188403 | rs1980767 | 0.368 | 1.981 | 0.743 | -0.149 | 0.001 | -0.036 | -0.036 | 0.527 | 0.07 | 0.2 | 0.04 | 0.01 | 0.06 | 0.08 |
| 1 | 112191408 | rs17665274 | 1.203 | 2.057 | 0.559 | -0.129 | -0.044 | -0.027 | -0.027 | 0 | 0.12 | 0.01 | 0.13 | 0.18 | 0.16 | 0.17 |
| 1 | 112198198 | rs2746444 | 1.516 | 2.132 | 0.62 | -0.15 | 0.001 | -0.005 | -0.005 | 0 | 0.05 | 0.04 | 0.04 | 0.03 | 0.08 | 0.1 |
| 1 | 112203185 | rs11576942 | 2.892 | 2.155 | 0.638 | -0.209 | 0.022 | -0.086 | -0.086 | 0 | 0.41 | 0.08 | 0.32 | 0.44 | 0.33 | 0.4 |
| 1 | 112203452 | rs10776734 | 1.777 | 2.215 | 0.601 | -0.156 | -0.006 | -0.005 | -0.005 | 0.076 | 0.06 | 0.05 | 0.05 | 0.06 | 0.07 | 0.06 |
| 1 | 112206429 | rs2477429 | -0.568 | 2.27 | 0.711 | -0.173 | 0.004 | -0.023 | -0.023 | 0 | 0.28 | 0.16 | 0.36 | 0.39 | 0.3 | 0.27 |
| 1 | 112216961 | rs7525578 | 1.783 | 2.284 | 0.709 | -0.169 | -0.006 | -0.005 | -0.005 | 0 | 0.06 | 0.05 | 0.05 | 0.06 | 0.08 | 0.06 |
| 1 | 112216978 | rs474722 | -1.431 | 2.245 | 0.871 | -0.178 | -0.012 | 0.009 | 0.009 | 0 | 0.37 | 0.51 | 0.46 | 0.47 | 0.37 | 0.36 |
| 1 | 112218156 | rs11811128 | -0.282 | 2.231 | 0.869 | -0.174 | 0.015 | 0.007 | 0.007 | 0.003 | 0.09 | 0.35 | 0.1 | 0.08 | 0.08 | 0.08 |

| | | | | | | | | | | | | | | | | |
|---|---|---|---|---|---|---|---|---|---|---|---|---|---|---|---|---|
| 1 | 112234084 | rs10489469 | 1.309 | 2.243 | 0.784 | -0.175 | -0.04 | -0.028 | -0.028 | 0 | 0.12 | 0.01 | 0.13 | 0.18 | 0.15 | 0.17 |
| 1 | 112241795 | rs558989 | 0.925 | 2.238 | 0.792 | -0.177 | 0.007 | -0.037 | -0.037 | 0.657 | 0.15 | 0.05 | 0.23 | 0.21 | 0.15 | 0.11 |
| 1 | 112241952 | rs7526355 | 2.899 | 2.28 | 0.879 | -0.207 | 0.038 | -0.049 | -0.049 | 0.771 | 0.42 | 0.16 | 0.33 | 0.44 | 0.34 | 0.4 |
| 1 | 112243947 | rs12751967 | -0.401 | 1.996 | 0.619 | 0.083 | 0.015 | 0.007 | 0.007 | 0.003 | 0.09 | 0.35 | 0.1 | 0.08 | 0.08 | 0.09 |
| 1 | 112245587 | rs7536042 | 0.327 | 1.995 | 0.577 | 0.082 | 0.012 | -0.032 | -0.032 | 1.926 | 0.07 | 0.22 | 0.03 | 0.01 | 0.07 | 0.08 |
| 1 | 112251902 | rs563083 | 0.818 | 2.008 | 0.205 | 0.089 | -0.01 | -0.066 | -0.066 | 0.667 | 0.15 | 0.01 | 0.23 | 0.21 | 0.15 | 0.11 |
| 1 | 112252461 | rs480578 | -1.534 | 2.036 | 0.231 | 0.09 | -0.012 | 0.009 | 0.009 | 0 | 0.37 | 0.51 | 0.46 | 0.47 | 0.37 | 0.36 |
| 1 | 112253114 | rs486982 | -1.531 | 2.023 | 0.219 | 0.084 | -0.008 | 0.006 | 0.006 | 0 | 0.37 | 0.53 | 0.46 | 0.47 | 0.37 | 0.36 |
| 1 | 112253212 | rs506991 | -2.25 | 2.035 | 0.17 | 0.083 | -0.002 | -0.001 | -0.001 | 0 | 0.49 | 0.62 | 0.55 | 0.56 | 0.53 | 0.52 |
| 1 | 112255389 | rs6573 | 0.989 | 2.034 | 0.053 | 0.102 | -0.039 | -0.027 | -0.027 | 1.196 | 0.12 | 0.01 | 0.13 | 0.18 | 0.15 | 0.17 |
| 1 | 112260747 | rs504885 | 0.17 | 2.079 | 0.159 | 0.06 | 0.011 | -0.002 | -0.002 | 0.109 | 0.11 | 0.14 | 0.09 | 0.09 | 0.15 | 0.16 |
| 1 | 112261533 | rs565522 | -1.683 | 2.11 | 0.213 | 0.094 | -0.003 | 0 | 0 | 0 | 0.42 | 0.46 | 0.51 | 0.51 | 0.45 | 0.46 |
| 1 | 112263682 | rs2076590 | 0.882 | 1.766 | 0.027 | -0.275 | 0.004 | 0.006 | 0.006 | 0 | 0.72 | 0.67 | 0.59 | 0.63 | 0.66 | 0.74 |
| 1 | 112264274 | rs2076591 | -1.195 | 1.556 | 0.075 | -0.228 | 0.032 | -0.017 | -0.017 | 1.103 | 0.2 | 0.46 | 0.18 | 0.17 | 0.23 | 0.24 |
| 1 | 112273026 | rs2076586 | 1.158 | 0.936 | -0.837 | -0.354 | -0.031 | -0.141 | -0.141 | 0 | 0.23 | 0.02 | 0.15 | 0.41 | 0.24 | 0.3 |
| 1 | 112276608 | rs11102329 | 0.602 | 1.098 | 0.096 | -1.081 | 0.003 | -0.038 | -0.038 | 0 | 0.13 | 0.14 | 0.07 | 0 | 0.12 | 0.07 |
| 1 | 112277597 | rs1000947 | 1.35 | 0.839 | -0.077 | -1.498 | -0.002 | -0.047 | -0.047 | 0 | 0.31 | 0.13 | 0.33 | 0.62 | 0.35 | 0.56 |
| 1 | 112280674 | rs197438 | -1.468 | 0.787 | 0.726 | -1.176 | -0.002 | 0.014 | 0.014 | 0.349 | 0.3 | 0.68 | 0.28 | 0.16 | 0.26 | 0.19 |
| 1 | 112283838 | rs3754026 | 0.858 | 0.78 | 0.653 | -1.259 | 0.001 | -0.029 | -0.029 | 0 | 0.13 | 0.05 | 0.22 | 0.18 | 0.15 | 0.12 |
| 1 | 112286309 | rs197394 | 0.644 | 0.79 | 0.87 | -1.267 | 0.014 | -0.076 | -0.076 | 0.136 | 0.14 | 0.24 | 0.08 | 0 | 0.12 | 0.07 |
| 1 | 112289118 | rs197372 | -1.798 | 0.766 | 0.939 | -1.278 | 0.014 | 0.025 | 0.025 | 1.294 | 0.39 | 0.71 | 0.47 | 0.31 | 0.34 | 0.25 |
| 1 | 112292778 | rs197381 | 0.623 | 0.815 | 1.122 | -1.255 | 0.016 | -0.084 | -0.084 | 4.225 | 0.14 | 0.3 | 0.08 | 0 | 0.11 | 0.07 |
| 1 | 112298582 | rs197392 | -0.68 | 0.829 | 0.545 | -1.244 | 0.021 | -0.088 | -0.088 | 0 | 0.23 | 0.57 | 0.15 | 0.04 | 0.2 | 0.13 |
| 1 | 112302219 | rs538779 | -0.148 | 0.686 | 0.481 | -1.223 | -0.007 | -0.019 | -0.019 | 0 | 0.19 | 0.11 | 0.26 | 0.21 | 0.22 | 0.17 |
| 1 | 112308953 | rs197412 | -1.975 | 0.678 | 0.432 | -1.226 | -0.007 | 0.03 | 0.03 | 1.112 | 0.44 | 0.72 | 0.51 | 0.34 | 0.42 | 0.31 |
| 1 | 112308972 | rs197413 | -1.997 | 0.68 | 0.417 | -1.225 | -0.01 | 0.032 | 0.032 | 1.206 | 0.44 | 0.74 | 0.51 | 0.34 | 0.42 | 0.31 |
| 1 | 112309123 | rs197414 | 0.148 | 0.716 | 0.34 | -1.202 | 0.023 | -0.131 | -0.131 | 0.393 | 0.17 | 0.43 | 0.1 | 0 | 0.13 | 0.07 |
| 1 | 112309331 | rs85276 | 0.148 | 0.716 | 0.34 | -1.202 | 0.023 | -0.131 | -0.131 | 0.244 | 0.17 | 0.43 | 0.1 | 0 | 0.13 | 0.07 |
| 1 | 112315450 | rs197419 | -1.729 | 0.676 | -0.591 | -1.088 | -0.025 | 0.031 | 0.031 | 1.567 | 0.45 | 0.82 | 0.52 | 0.34 | 0.42 | 0.31 |
| 1 | 112320800 | rs535317 | 0.376 | 0.572 | -0.343 | -0.687 | 0.002 | -0.03 | -0.03 | 0.172 | 0.13 | 0.05 | 0.22 | 0.18 | 0.15 | 0.12 |
| 1 | 112329551 | rs3738298 | 0.297 | 0.478 | 0.062 | -0.673 | -0.009 | 0.025 | 0.025 | 1.81 | 0.14 | 0.21 | 0.11 | 0.23 | 0.16 | 0.23 |
| 1 | 112333983 | rs755720 | 0.129 | 0.785 | 0.251 | -1.265 | -0.011 | 0.009 | 0.009 | 0 | 0.16 | 0.09 | 0.27 | 0.08 | 0.19 | 0.13 |
| 1 | 112334474 | rs1246194 | -0.842 | 0.627 | 0.09 | -1.689 | -0.011 | -0.076 | -0.076 | 0.022 | 0.69 | 0.53 | 0.69 | 0.84 | 0.73 | 0.84 |
| 1 | 112335252 | rs475620 | -1.207 | 0.618 | -0.024 | -1.167 | 0 | -0.004 | -0.004 | 0 | 0.26 | 0.67 | 0.45 | 0.2 | 0.24 | 0.15 |
| 1 | 112339880 | rs17722309 | 0.914 | 1.071 | -0.58 | -1.135 | -0.001 | -0.001 | -0.001 | 2.241 | 0.06 | 0.04 | 0.2 | 0.08 | 0.07 | 0.06 |
| 1 | 112342509 | rs4839168 | 0.299 | 0.939 | -0.53 | -1.086 | -0.006 | 0.006 | 0.006 | 0 | 0.16 | 0.1 | 0.26 | 0.08 | 0.19 | 0.13 |
| 1 | 112346109 | rs817977 | -0.047 | 0.731 | -0.59 | -1.08 | -0.003 | 0.001 | 0.001 | 0 | 0.23 | 0.15 | 0.3 | 0.07 | 0.24 | 0.15 |
| 1 | 112346989 | rs2788610 | 0.378 | 0.647 | -1.098 | -0.955 | 0.005 | 0.013 | 0.013 | 0.052 | 0.16 | 0.13 | 0.07 | 0.1 | 0.17 | 0.22 |
| 1 | 112347256 | rs672757 | -0.468 | 0.858 | -0.846 | -0.803 | 0 | -0.033 | -0.033 | 0 | 0.44 | 0.46 | 0.45 | 0.23 | 0.44 | 0.37 |
| 1 | 112347677 | rs2618036 | -0.562 | 0.84 | -1.232 | -0.714 | -0.006 | 0.045 | 0.045 | 0 | 0.79 | 0.84 | 0.84 | 0.96 | 0.79 | 0.84 |
| 1 | 112348342 | rs2618037 | 0.237 | 0.583 | -0.75 | 0.374 | -0.005 | 0.011 | 0.011 | 0.599 | 0.77 | 0.75 | 0.62 | 0.67 | 0.81 | 0.72 |
| 1 | 112348592 | rs2618038 | -1.329 | 0.809 | -1.405 | -0.361 | -0.092 | 0.241 | 0.241 | 0 | 0.62 | 0.93 | 0.76 | 0.93 | 0.61 | 0.79 |
| 1 | 112349568 | rs17028610 | 0.519 | 0.345 | -1.209 | -0.05 | -0.013 | 0.064 | 0.064 | 0.256 | 0.07 | 0.21 | 0.13 | 0.12 | 0.03 | 0.02 |
| 1 | 112354653 | rs612231 | -0.767 | 0.772 | -0.943 | 0.778 | -0.001 | -0.023 | -0.023 | 0 | 0.42 | 0.22 | 0.39 | 0.44 | 0.41 | 0.58 |
| 1 | 112356676 | rs641724 | -1.115 | 0.896 | -0.658 | 0.368 | -0.003 | 0.065 | 0.065 | 0 | 0.52 | 0.66 | 0.6 | 0.82 | 0.52 | 0.73 |
| 1 | 112356814 | rs2788611 | -1.144 | 0.904 | -0.832 | 0.378 | -0.011 | 0.111 | 0.111 | 0 | 0.52 | 0.7 | 0.6 | 0.82 | 0.52 | 0.72 |
| 1 | 112356899 | rs517680 | -0.103 | 0.786 | -0.769 | 0.948 | -0.023 | 0.041 | 0.041 | 0 | 0.16 | 0.01 | 0.07 | 0.1 | 0.17 | 0.15 |
| 1 | 112357144 | rs713268 | -0.477 | 0.697 | -0.493 | 1.079 | 0.001 | -0.003 | -0.003 | 0.164 | 0.18 | 0.18 | 0.08 | 0.21 | 0.19 | 0.21 |
| 1 | 112357730 | rs7556364 | 1.259 | 0.573 | -0.277 | 1.354 | 0.015 | -0.019 | -0.019 | 0 | 0.63 | 0.27 | 0.64 | 0.6 | 0.67 | 0.6 |
| 1 | 112362645 | rs2365669 | 1.559 | 0.328 | -0.452 | 1.305 | 0.018 | 0.06 | 0.06 | 0.003 | 0.6 | 0.09 | 0.54 | 0.33 | 0.65 | 0.57 |
| 1 | 112362762 | rs639570 | -0.6 | 0.274 | -0.522 | 1.229 | -0.033 | 0.084 | 0.084 | 0 | 0.86 | 0.99 | 0.92 | 1 | 0.84 | 0.9 |
| 1 | 112364573 | rs1471437 | 1.235 | 0.282 | 0.024 | 1.268 | 0.026 | 0.023 | 0.023 | 0.123 | 0.65 | 0.26 | 0.57 | 0.33 | 0.69 | 0.62 |
| 1 | 112365016 | rs2169815 | 1.655 | 0.401 | 0.162 | 1.259 | 0.026 | 0.056 | 0.056 | 0 | 0.61 | 0.19 | 0.54 | 0.34 | 0.65 | 0.58 |
| 1 | 112369354 | rs4839171 | 1.277 | 0.42 | 0.372 | 1.276 | 0.011 | 0.026 | 0.026 | 0.032 | 0.64 | 0.26 | 0.57 | 0.31 | 0.66 | 0.62 |
| 1 | 112370606 | rs1904200 | 0.761 | 0.449 | 0.495 | 1.583 | -0.001 | 0.009 | 0.009 | 0 | 0.74 | 0.68 | 0.67 | 0.43 | 0.71 | 0.66 |
| 1 | 112376121 | rs11102342 | 1.036 | 0.657 | 0.708 | 1.399 | 0.001 | 0.024 | 0.024 | 0 | 0.65 | 0.51 | 0.59 | 0.44 | 0.63 | 0.55 |
| 1 | 112376191 | rs661407 | 0.314 | 0.747 | 0.779 | 1.329 | 0.019 | -0.033 | -0.033 | 0 | 0.94 | 0.81 | 0.96 | 1 | 0.93 | 0.94 |
| 1 | 112384100 | rs4839175 | -0.547 | 0.694 | 0.571 | 1.786 | -0.011 | 0.063 | 0.063 | 0 | 0.05 | 0 | 0.23 | 0.46 | 0.09 | 0.2 |
| 1 | 112384332 | rs643559 | -0.136 | 0.675 | 0.731 | 1.671 | -0.005 | 0.03 | 0.03 | 0 | 0.2 | 0.12 | 0.13 | 0.08 | 0.21 | 0.2 |
| 1 | 112385547 | rs475508 | -1.387 | 0.385 | 1.423 | 1.787 | -0.008 | 0 | 0 | 0 | 0.33 | 0.45 | 0.41 | 0.66 | 0.36 | 0.44 |
| 1 | 112391809 | rs11102347 | -2.066 | 0.463 | 1.439 | 2.027 | 0.003 | 0.205 | 0.205 | 0 | 0.09 | 0.19 | 0.25 | 0.52 | 0.12 | 0.25 |
| 1 | 112395432 | rs817974 | 0.668 | 0.485 | 1.742 | 1.637 | 0.037 | 0.079 | 0.079 | 0 | 0.9 | 0.66 | 0.93 | 0.88 | 0.9 | 0.93 |
| 1 | 112403295 | rs652334 | -0.814 | 0.472 | 1.44 | 1.813 | 0.001 | 0.14 | 0.14 | 0 | 0.06 | 0.1 | 0.23 | 0.43 | 0.09 | 0.2 |
| 1 | 112404225 | rs637813 | -0.823 | 0.151 | 1.386 | 1.356 | 0.009 | 0.079 | 0.079 | 0 | 0.33 | 0.14 | 0.19 | 0 | 0.3 | 0.23 |
| 1 | 112410433 | rs639622 | -1.178 | 0.171 | 1.4 | 1.511 | 0.006 | 0.231 | 0.231 | 0 | 0.08 | 0.2 | 0.24 | 0.44 | 0.09 | 0.21 |
| 1 | 112413022 | rs597205 | -1.33 | 0.152 | 1.458 | 1.616 | 0.006 | 0.262 | 0.262 | 0.036 | 0.08 | 0.23 | 0.25 | 0.56 | 0.09 | 0.24 |
| 1 | 112414858 | rs12139985 | 1.503 | -0.045 | 1.05 | 1.328 | 0.026 | 0.062 | 0.062 | 0 | 0.53 | 0.23 | 0.53 | 0.35 | 0.54 | 0.45 |
| 1 | 112415201 | rs11102350 | 1.255 | -0.036 | 1.098 | 1.289 | 0.013 | 0.001 | 0.001 | 0 | 0.56 | 0.32 | 0.54 | 0.44 | 0.57 | 0.54 |
| 1 | 112421677 | rs7415980 | 1.361 | 0.572 | 0.661 | 0.811 | -0.001 | 0.035 | 0.035 | 0 | 0.51 | 0.36 | 0.48 | 0.38 | 0.44 | 0.43 |

| | | | | | | | | | | | | | | | | |
|---|---|---|---|---|---|---|---|---|---|---|---|---|---|---|---|---|
| 1 | 112425797 | rs17028939 | -0.506 | 0.798 | 0.689 | 1.184 | 0 | 0.011 | 0.011 | 0 | 0.09 | 0.1 | 0.05 | 0.17 | 0.09 | 0.06 |
| 1 | 112426851 | rs1373291 | -0.526 | 0.587 | 0.834 | 1.151 | 0.005 | 0.073 | 0.073 | 0 | 0.16 | 0.3 | 0.28 | 0.45 | 0.22 | 0.33 |
| 1 | 112428953 | rs1767283 | -0.604 | 0.441 | 0.895 | 0.775 | -0.002 | -0.016 | -0.016 | 0 | 0.78 | 0.81 | 0.84 | 1 | 0.75 | 0.87 |
| 1 | 112433666 | rs11102354 | 1.675 | 0.098 | -0.028 | 0.305 | -0.019 | -0.055 | -0.055 | 0 | 0.3 | 0.16 | 0.35 | 0.44 | 0.31 | 0.31 |
| 1 | 112433863 | rs605604 | -1.265 | 0.074 | 0.269 | 0.215 | -0.001 | -0.035 | -0.035 | 0 | 0.77 | 0.8 | 0.84 | 1 | 0.74 | 0.87 |
| 1 | 112434196 | rs17028969 | 0.669 | -0.028 | 0.498 | 0.261 | 0.005 | 0 | 0 | 1.554 | 0.23 | 0.21 | 0.16 | 0.04 | 0.17 | 0.21 |
| 1 | 112435325 | rs17028976 | 0.72 | 0.114 | 0.398 | 0.509 | -0.007 | -0.026 | -0.026 | 0 | 0.08 | 0.05 | 0.05 | 0.17 | 0.08 | 0.06 |
| 1 | 112436001 | rs6697691 | 0.193 | 0.104 | 0.585 | 0.594 | 0.003 | 0.04 | 0.04 | 0 | 0.16 | 0.25 | 0.27 | 0.36 | 0.18 | 0.29 |
| 1 | 112437235 | rs10776736 | 0.118 | 0.138 | 0.286 | 0.562 | -0.011 | -0.033 | -0.033 | 0 | 0.14 | 0.09 | 0.26 | 0.49 | 0.19 | 0.33 |
| 1 | 112437879 | rs12080989 | 0.542 | 0.018 | 0.387 | -0.04 | -0.002 | -0.022 | -0.022 | 0.532 | 0.22 | 0.28 | 0.17 | 0.04 | 0.16 | 0.21 |
| 1 | 112437907 | rs2587368 | -0.23 | -0.066 | 0.189 | -0.12 | -0.002 | -0.049 | -0.049 | 0 | 0.67 | 0.6 | 0.78 | 0.9 | 0.64 | 0.8 |
| 1 | 112438818 | rs894848 | -0.625 | -0.3 | -0.107 | -0.434 | 0.005 | 0.005 | 0.005 | 0 | 0.85 | 0.86 | 0.88 | 0.97 | 0.92 | 0.87 |
| 1 | 112439757 | rs608673 | -0.467 | -0.258 | -0.05 | -0.551 | 0.008 | -0.038 | -0.038 | 0.264 | 0.67 | 0.6 | 0.76 | 0.91 | 0.56 | 0.82 |
| 1 | 112439770 | rs7539683 | 1.679 | -0.267 | -0.452 | -0.142 | -0.013 | -0.027 | -0.027 | 0 | 0.31 | 0.12 | 0.3 | 0.37 | 0.26 | 0.28 |
| 1 | 112440057 | rs4838925 | 0.71 | -0.031 | -0.573 | 0.039 | -0.038 | -0.062 | -0.062 | 0.104 | 0.13 | 0.03 | 0.27 | 0.2 | 0.17 | 0.29 |
| 1 | 112442367 | rs2813866 | 0.108 | -0.539 | -0.701 | -0.181 | 0.006 | 0.054 | 0.054 | 0 | 0.53 | 0.7 | 0.5 | 0.71 | 0.41 | 0.53 |
| 1 | 112443281 | rs818109 | 0.068 | -0.838 | -0.766 | -0.217 | 0.012 | 0.01 | 0.01 | 0 | 0.21 | 0.18 | 0.15 | 0.04 | 0.13 | 0.18 |
| 1 | 112444441 | rs572473 | -0.121 | -0.693 | -0.777 | -0.207 | 0.024 | 0.035 | 0.035 | 0 | 0.25 | 0.18 | 0.18 | 0.04 | 0.18 | 0.2 |
| 1 | 112450636 | rs2995807 | -0.372 | -0.843 | -0.809 | -0.196 | 0.042 | 0.053 | 0.053 | 0 | 0.77 | 0.87 | 0.83 | 0.96 | 0.86 | 0.78 |
| 1 | 112451134 | rs1036959 | 0.22 | -0.685 | -0.818 | -0.125 | 0.002 | -0.017 | -0.017 | 1.096 | 0.12 | 0.11 | 0.26 | 0.19 | 0.14 | 0.27 |
| 1 | 112451681 | rs2034124 | -0.591 | -0.283 | -1.002 | -0.229 | -0.021 | 0 | 0 | 0.005 | 0.43 | 0.7 | 0.5 | 0.59 | 0.54 | 0.5 |
| 1 | 112461637 | rs9429608 | 1.196 | -0.278 | -0.914 | -0.512 | 0.001 | 0.145 | 0.145 | 0 | 0.06 | 0.12 | 0.05 | 0.32 | 0.08 | 0.1 |
| 1 | 112462984 | rs6682872 | 0.121 | -0.208 | -0.605 | -0.673 | 0.025 | -0.003 | -0.003 | 0.033 | 0.28 | 0.31 | 0.21 | 0.31 | 0.37 | 0.23 |
| 1 | 112463441 | rs4839187 | 0.223 | 0.35 | -1.317 | -0.471 | -0.021 | -0.041 | -0.041 | 0.916 | 0.14 | 0.06 | 0.27 | 0.19 | 0.16 | 0.28 |
| 1 | 112464376 | rs17029069 | 1.564 | 0.144 | -1.414 | -0.606 | -0.043 | -0.089 | -0.089 | 0 | 0.3 | 0.11 | 0.31 | 0.46 | 0.31 | 0.26 |
| 1 | 112467014 | rs10494143 | 1.547 | 0.232 | -1.352 | -0.592 | -0.023 | 0.021 | 0.021 | 0 | 0.05 | 0.02 | 0.04 | 0 | 0.09 | 0.02 |
| 1 | 112467126 | rs9429422 | -0.633 | 0.1 | -1.257 | -0.608 | 0 | 0.025 | 0.025 | 0 | 0.21 | 0.19 | 0.32 | 0.49 | 0.25 | 0.38 |
| 1 | 112467932 | rs12408551 | 1.279 | 0.1 | -1.514 | -0.443 | 0.015 | -0.039 | -0.039 | 0 | 0.1 | 0.05 | 0.22 | 0.19 | 0.08 | 0.27 |
| 1 | 112468814 | rs12119724 | 1.51 | 0.072 | -1.603 | -0.54 | -0.04 | -0.094 | -0.094 | 0.118 | 0.3 | 0.11 | 0.31 | 0.46 | 0.31 | 0.26 |
| 1 | 112470093 | rs11102355 | 1.186 | 0.176 | -1.329 | -0.257 | 0.02 | -0.038 | -0.038 | 0 | 0.09 | 0.04 | 0.22 | 0.19 | 0.08 | 0.27 |
| 1 | 112474363 | rs2000265 | -0.014 | 0.12 | -1.912 | -0.778 | -0.123 | -0.105 | -0.105 | 0 | 0.36 | 0.06 | 0.38 | 0.45 | 0.49 | 0.31 |
| 1 | 112474929 | rs12025303 | -0.428 | -0.305 | -0.894 | -0.9 | -0.046 | 0.068 | 0.068 | 0.004 | 0.84 | 0.5 | 0.72 | 0.82 | 0.89 | 0.75 |
| 1 | 112475549 | rs11102357 | -0.682 | -0.123 | -1.102 | -1.325 | -0.005 | 0.04 | 0.04 | 0 | 0.4 | 0.35 | 0.43 | 0.79 | 0.44 | 0.51 |
| 1 | 112475936 | rs1936061 | -0.092 | 0.02 | -1.531 | -1.254 | -0.018 | 0.013 | 0.013 | 0 | 0.41 | 0.1 | 0.52 | 0.63 | 0.43 | 0.53 |
| 1 | 112477114 | rs17726989 | 1.193 | 0.364 | -1.085 | -1.057 | 0.032 | -0.036 | -0.036 | 0 | 0.07 | 0.11 | 0.06 | 0 | 0.11 | 0.02 |
| 1 | 112478366 | rs9429610 | 0.959 | 0.019 | -1.081 | -1.106 | 0.01 | 0.013 | 0.013 | 0 | 0.43 | 0.36 | 0.33 | 0.37 | 0.34 | 0.42 |
| 1 | 112479331 | rs2995810 | -0.607 | -0.044 | -1.07 | -1.093 | -0.003 | -0.007 | -0.007 | 0.862 | 0.7 | 0.72 | 0.68 | 0.54 | 0.65 | 0.76 |
| 1 | 112480010 | rs2995811 | 0.48 | -0.122 | -1.401 | -1.117 | 0 | 0.004 | 0.004 | 0 | 0.61 | 0.36 | 0.45 | 0.37 | 0.57 | 0.53 |
| 1 | 112480576 | rs9429425 | 1.125 | -0.537 | -1.09 | -1.485 | -0.04 | -0.057 | -0.057 | 0 | 0.18 | 0.3 | 0.14 | 0.03 | 0.09 | 0.15 |
| 1 | 112480855 | rs9429612 | 0.922 | -0.13 | -0.813 | -1.16 | 0.008 | 0.037 | 0.037 | 0 | 0.41 | 0.32 | 0.33 | 0.06 | 0.33 | 0.28 |
| 1 | 112485709 | rs17727131 | 0.305 | -0.345 | -0.594 | -0.42 | -0.011 | 0.006 | 0.006 | 0 | 0.13 | 0.15 | 0.22 | 0.21 | 0.07 | 0.29 |
| 1 | 112486470 | rs17029132 | 0.04 | 0.369 | -0.335 | -0.975 | 0.006 | -0.066 | -0.066 | 0.978 | 0.17 | 0.23 | 0.11 | 0 | 0.18 | 0.07 |
| 1 | 112486533 | rs12741034 | -0.364 | 0.266 | -0.266 | -0.714 | 0.003 | -0.027 | -0.027 | 0 | 0.08 | 0.13 | 0.07 | 0.01 | 0.12 | 0.09 |
| 1 | 112487033 | rs12072776 | -0.712 | -0.445 | 0.08 | -0.872 | -0.067 | -0.103 | -0.103 | 0.826 | 0.21 | 0.44 | 0.1 | 0.06 | 0.11 | 0.1 |
| 1 | 112487834 | rs1808973 | -1.132 | 0.292 | -0.057 | -0.483 | 0.009 | 0.031 | 0.031 | 0 | 0.49 | 0.61 | 0.53 | 0.45 | 0.54 | 0.47 |
| 1 | 112488677 | rs1822567 | 0.305 | 0.295 | 0.053 | -0.658 | -0.005 | 0.022 | 0.022 | 0 | 0.2 | 0.23 | 0.32 | 0.44 | 0.17 | 0.34 |
| 1 | 112488820 | rs11578917 | 1.789 | 0.426 | -0.247 | -0.621 | -0.004 | 0.027 | 0.027 | 1.132 | 0.4 | 0.24 | 0.27 | 0.31 | 0.38 | 0.31 |
| 1 | 112489299 | rs969100 | -2.006 | 0.453 | -0.293 | -0.608 | 0.001 | 0.026 | 0.026 | 1.225 | 0.6 | 0.74 | 0.72 | 0.69 | 0.62 | 0.69 |
| 1 | 112489832 | rs9429614 | -0.754 | 0.774 | 0.096 | -0.761 | 0.032 | -0.156 | -0.156 | 3.673 | 0.29 | 0.47 | 0.25 | 0.01 | 0.38 | 0.13 |
| 1 | 112496701 | rs11102362 | -0.345 | 1.197 | 0.288 | 0.101 | 0.041 | 0.02 | 0.02 | 0 | 0.25 | 0.32 | 0.45 | 0.32 | 0.38 | 0.34 |
| 1 | 112497742 | rs6699612 | 0.303 | 0.589 | -0.132 | 0.213 | 0.001 | 0.007 | 0.007 | 0 | 0.1 | 0.11 | 0.17 | 0.26 | 0.09 | 0.2 |
| 1 | 112504704 | rs17676470 | 1.28 | 0.291 | -0.466 | -0.657 | 0.049 | 0.099 | 0.099 | 0.65 | 0.39 | 0.17 | 0.21 | 0.23 | 0.31 | 0.37 |
| 1 | 112505214 | rs4839193 | -0.703 | 0.493 | -0.25 | -0.471 | -0.002 | -0.037 | -0.037 | 0 | 0.84 | 0.89 | 0.8 | 0.73 | 0.84 | 0.83 |
| 1 | 112505378 | rs4838932 | -0.184 | 0.511 | 0.135 | -0.537 | 0 | 0.001 | 0.001 | 0 | 0.29 | 0.26 | 0.45 | 0.45 | 0.37 | 0.39 |
| 1 | 112505519 | rs7525808 | 0.913 | 0.512 | 0.26 | -0.541 | 0 | 0 | 0 | 0.295 | 0.53 | 0.43 | 0.33 | 0.28 | 0.47 | 0.44 |
| 1 | 112505551 | rs7548910 | 0.922 | 0.511 | 0.218 | -0.541 | 0.001 | 0.023 | 0.023 | 0 | 0.53 | 0.41 | 0.33 | 0.28 | 0.47 | 0.44 |
| 1 | 112505892 | rs716954 | -0.8 | 0.536 | 0.056 | -0.534 | -0.003 | -0.039 | -0.039 | 0 | 0.84 | 0.91 | 0.8 | 0.73 | 0.84 | 0.83 |
| 1 | 112506564 | rs3008538 | -0.467 | 0.373 | 0.286 | -0.415 | 0.019 | 0.034 | 0.034 | 0 | 0.3 | 0.39 | 0.46 | 0.45 | 0.37 | 0.39 |
| 1 | 112510932 | rs10776739 | -0.474 | 0.683 | 0.094 | -0.214 | -0.007 | -0.008 | -0.008 | 0 | 0.23 | 0.21 | 0.45 | 0.44 | 0.34 | 0.31 |
| 1 | 112511555 | rs12753051 | 1.151 | -0.556 | -0.237 | -1.336 | 0.014 | 0.146 | 0.146 | 0.29 | 0.34 | 0.16 | 0.16 | 0.03 | 0.29 | 0.22 |
| 1 | 112513203 | rs894846 | 0.036 | -0.139 | 0.311 | -1.058 | -0.007 | 0.078 | 0.078 | 0 | 0.18 | 0.27 | 0.39 | 0.56 | 0.16 | 0.42 |
| 1 | 112515593 | rs730022 | 1.694 | -0.2 | -0.035 | -1.359 | 0.016 | 0.13 | 0.13 | 0.47 | 0.31 | 0.14 | 0.16 | 0.01 | 0.25 | 0.21 |
| 1 | 112516275 | rs17676657 | 1.169 | 0.066 | 0.023 | -1.004 | -0.078 | -0.04 | -0.04 | 0 | 0.22 | 0.02 | 0.26 | 0.24 | 0.32 | 0.2 |
| 1 | 112516455 | rs3017627 | 0.331 | 0.041 | 0.181 | -1.02 | -0.005 | 0.008 | 0.008 | 0 | 0.12 | 0.07 | 0.08 | 0.09 | 0.12 | 0.09 |
| 1 | 112517914 | rs1373286 | -0.679 | 0.002 | 0.474 | -1.007 | -0.002 | 0.001 | 0.001 | 0 | 0.43 | 0.39 | 0.42 | 0.41 | 0.54 | 0.37 |
| 1 | 112523095 | rs3008527 | -0.905 | 0.167 | 0.684 | -0.74 | 0.009 | -0.002 | -0.002 | 0 | 0.2 | 0.24 | 0.15 | 0.18 | 0.22 | 0.17 |
| 1 | 112538239 | rs7513088 | 0.993 | 0.117 | 0.22 | -0.835 | 0.001 | -0.018 | -0.018 | 0 | 0.63 | 0.45 | 0.34 | 0.22 | 0.63 | 0.49 |
| 1 | 112538743 | rs6681719 | 0.386 | 0.104 | 0.202 | -0.722 | -0.027 | 0.062 | 0.062 | 0 | 0.17 | 0.24 | 0.49 | 0.59 | 0.12 | 0.35 |
| 1 | 112539410 | rs7528073 | 0.721 | 0.258 | -0.365 | -0.834 | -0.046 | 0.049 | 0.049 | 0 | 0.31 | 0.09 | 0.17 | 0.22 | 0.36 | 0.27 |

| | | | | | | | | | | | | | | | | |
|---|---|---|---|---|---|---|---|---|---|---|---|---|---|---|---|---|
| 1 | 112545414 | rs3008530 | -0.393 | 0.119 | 0.038 | -0.407 | -0.027 | 0.016 | 0.016 | 0 | 0.11 | 0.05 | 0.09 | 0.1 | 0.14 | 0.08 |
| 1 | 112547125 | rs3000752 | -0.652 | -0.188 | 0.331 | -0.269 | 0.003 | 0.03 | 0.03 | 0.053 | 0.15 | 0.27 | 0.18 | 0.18 | 0.14 | 0.13 |
| 1 | 112548118 | rs3000754 | 0.019 | -0.108 | -0.139 | -0.294 | -0.001 | 0.001 | 0.001 | 0 | 0.12 | 0.1 | 0.16 | 0.11 | 0.13 | 0.09 |
| 1 | 112555423 | rs12562249 | -0.034 | -0.35 | 0.205 | 0.35 | -0.021 | 0.075 | 0.075 | 0 | 0.16 | 0.22 | 0.41 | 0.52 | 0.11 | 0.29 |
| 1 | 112559150 | rs7555530 | 0.616 | -0.147 | -0.582 | -0.367 | -0.019 | 0.144 | 0.144 | 0 | 0.44 | 0.24 | 0.32 | 0.12 | 0.42 | 0.32 |
| 1 | 112560278 | rs7526235 | -0.233 | 0.398 | -0.654 | -0.123 | 0.01 | -0.017 | -0.017 | 0 | 0.38 | 0.46 | 0.25 | 0.29 | 0.46 | 0.35 |
| 1 | 112563850 | rs4839200 | 0.525 | -0.291 | -0.517 | 0.405 | -0.02 | 0.02 | 0.02 | 0 | 0.77 | 0.71 | 0.57 | 0.38 | 0.85 | 0.65 |
| 1 | 112572442 | rs269110 | 0.374 | -0.445 | 0.435 | -0.97 | 0.005 | 0.001 | 0.001 | 0.102 | 0.68 | 0.69 | 0.58 | 0.11 | 0.73 | 0.51 |
| 1 | 112572509 | rs7413702 | -0.3 | -0.387 | 0.182 | -1.471 | 0.001 | 0.023 | 0.023 | 0 | 0.12 | 0.07 | 0.07 | 0 | 0.1 | 0.07 |
| 1 | 112575961 | rs3860210 | 0.143 | -0.557 | 1.205 | -1.587 | 0.005 | -0.001 | -0.001 | 0 | 0.08 | 0.06 | 0.04 | 0 | 0.05 | 0.04 |
| 1 | 112579882 | rs4838936 | 0.193 | -0.3 | 1.087 | -2.576 | -0.024 | 0.114 | 0.114 | 0 | 0.36 | 0.2 | 0.26 | 0.08 | 0.36 | 0.31 |
| 1 | 112582366 | rs11102370 | 0.221 | -0.379 | 1.208 | -2.661 | 0.004 | -0.004 | -0.004 | 0 | 0.36 | 0.35 | 0.45 | 0.92 | 0.32 | 0.53 |
| 1 | 112584207 | rs1372861 | 1.161 | -0.274 | 1.086 | -2.683 | -0.04 | 0.095 | 0.095 | 1.082 | 0.24 | 0.07 | 0.24 | 0.01 | 0.31 | 0.17 |
| 1 | 112586954 | rs4839202 | 0.255 | -0.224 | 1.166 | -2.542 | -0.01 | 0.053 | 0.053 | 0 | 0.37 | 0.28 | 0.28 | 0.09 | 0.36 | 0.31 |
| 1 | 112588522 | rs3017628 | -0.747 | -0.562 | 1.205 | -2.664 | -0.004 | -0.169 | -0.169 | 3.59 | 0.69 | 0.52 | 0.71 | 0.99 | 0.68 | 0.83 |
| 1 | 112590049 | rs174892 | -1.293 | -0.498 | 0.913 | -2.634 | -0.004 | -0.021 | -0.021 | 0 | 0.74 | 0.82 | 0.75 | 0.99 | 0.69 | 0.83 |
| 1 | 112591009 | rs10857922 | 0.273 | -0.388 | 0.702 | -2.568 | 0.006 | 0.104 | 0.104 | 0.156 | 0.35 | 0.19 | 0.43 | 0.89 | 0.33 | 0.51 |
| 1 | 112595728 | rs1838300 | -0.36 | -0.385 | 1.012 | -2.411 | 0 | -0.026 | -0.026 | 0.084 | 0.39 | 0.61 | 0.33 | 0.11 | 0.37 | 0.31 |
| 1 | 112597532 | rs269079 | -0.468 | -0.451 | 0.912 | -2.442 | -0.002 | 0.087 | 0.087 | 0 | 0.4 | 0.72 | 0.34 | 0.11 | 0.37 | 0.31 |
| 1 | 112601956 | rs269088 | -0.289 | -0.35 | 1.278 | -2.237 | 0.033 | 0.035 | 0.035 | 1.04 | 0.05 | 0.33 | 0.03 | 0 | 0 | 0 |
| 1 | 112604622 | rs4839203 | 0.885 | -0.365 | 1.266 | -2.331 | -0.023 | 0.035 | 0.035 | 0 | 0.24 | 0.13 | 0.22 | 0.11 | 0.3 | 0.26 |
| 1 | 112607677 | rs10857925 | 0.525 | -0.388 | 1.213 | -2.342 | 0.022 | -0.066 | -0.066 | 0 | 0.27 | 0.41 | 0.25 | 0.11 | 0.3 | 0.26 |
| 1 | 112610318 | rs10776740 | -0.023 | -0.698 | 0.593 | -2.615 | 0.004 | 0.086 | 0.086 | 0 | 0.37 | 0.23 | 0.47 | 0.88 | 0.38 | 0.56 |
| 1 | 112613028 | rs269131 | -0.473 | -0.326 | 0.583 | -1.896 | -0.024 | 0.065 | 0.065 | 0 | 0.46 | 0.74 | 0.36 | 0.4 | 0.46 | 0.4 |
| 1 | 112617049 | rs12026840 | 0.963 | -0.242 | 0.509 | -1.825 | 0 | -0.009 | -0.009 | 0 | 0.14 | 0.09 | 0.09 | 0.27 | 0.13 | 0.13 |
| 1 | 112626237 | rs385768 | 0.597 | -0.26 | 0.625 | -1.858 | 0.023 | -0.058 | -0.058 | 0.224 | 0.29 | 0.51 | 0.26 | 0.13 | 0.32 | 0.27 |
| 1 | 112628860 | rs269101 | -0.533 | -0.304 | 0.306 | -2.035 | -0.007 | 0.127 | 0.127 | 0 | 0.59 | 0.85 | 0.68 | 0.72 | 0.59 | 0.69 |
| 1 | 112629056 | rs11801758 | 1.281 | -0.409 | 0.362 | -2.078 | -0.073 | 0.052 | 0.052 | 0 | 0.24 | 0.07 | 0.22 | 0.13 | 0.32 | 0.27 |
| 1 | 112632440 | rs12122815 | 1.231 | -0.409 | 0.38 | -2.078 | -0.059 | 0.05 | 0.05 | 0 | 0.25 | 0.08 | 0.22 | 0.13 | 0.32 | 0.27 |
| 1 | 112639174 | rs269094 | -1.498 | -0.397 | 0.295 | -2.128 | -0.027 | 0.153 | 0.153 | 0 | 0.74 | 0.94 | 0.76 | 0.99 | 0.74 | 0.83 |
| 1 | 112639472 | rs269093 | 0.46 | -0.028 | 0.354 | -2.166 | 0 | -0.007 | -0.007 | 0 | 0.31 | 0.65 | 0.26 | 0.13 | 0.29 | 0.27 |
| 1 | 112654994 | rs461311 | -1.179 | -0.066 | -0.105 | -1.873 | -0.012 | 0.128 | 0.128 | 0 | 0.74 | 0.92 | 0.77 | 0.99 | 0.75 | 0.83 |
| 1 | 112659363 | rs682489 | -0.312 | 0.08 | 0.272 | -1.364 | 0.003 | 0.028 | 0.028 | 0 | 0.41 | 0.7 | 0.51 | 0.62 | 0.4 | 0.49 |
| 1 | 112659834 | rs466951 | -1.519 | -0.025 | 0.068 | -1.58 | -0.085 | 0.219 | 0.219 | 0.009 | 0.59 | 0.93 | 0.62 | 0.87 | 0.54 | 0.58 |
| 1 | 112667758 | rs406486 | -0.66 | 0.019 | 0.413 | -0.973 | 0.007 | 0.009 | 0.009 | 0 | 0.42 | 0.76 | 0.52 | 0.57 | 0.39 | 0.49 |
| 1 | 112668457 | rs457071 | -0.796 | -0.076 | 0.78 | -1.104 | 0.048 | 0.043 | 0.043 | 1.288 | 0.06 | 0.45 | 0.05 | 0 | 0 | 0 |
| 1 | 112671286 | rs2083429 | 0.03 | -0.005 | 0.155 | -1.058 | -0.059 | -0.037 | -0.037 | 0 | 0.34 | 0.12 | 0.45 | 0.59 | 0.39 | 0.49 |
| 1 | 112677859 | rs12756458 | -0.346 | 0.225 | -0.371 | -0.362 | -0.001 | 0 | 0 | 0 | 0.39 | 0.36 | 0.49 | 0.57 | 0.39 | 0.49 |
| 1 | 112682312 | rs4425984 | -0.205 | 0.308 | -0.846 | -0.35 | -0.004 | -0.002 | -0.002 | 0.045 | 0.39 | 0.3 | 0.48 | 0.56 | 0.38 | 0.49 |
| 1 | 112686349 | rs17742164 | 0.408 | 0.56 | -1.323 | -0.22 | 0.004 | 0.004 | 0.004 | 0 | 0.2 | 0.21 | 0.14 | 0.27 | 0.2 | 0.12 |
| 1 | 112687359 | rs155648 | -0.967 | 0.671 | -2.3 | -0.595 | -0.024 | 0.195 | 0.195 | 0 | 0.59 | 0.91 | 0.66 | 0.85 | 0.57 | 0.6 |
| 1 | 112688311 | rs155645 | -0.166 | 0.819 | -1.563 | 0.564 | -0.055 | -0.091 | -0.091 | 0 | 0.3 | 0.13 | 0.41 | 0.43 | 0.37 | 0.47 |
| 1 | 112707208 | rs155669 | 0.126 | 0.123 | -0.898 | 0.026 | 0.002 | 0.024 | 0.024 | 0 | 0.69 | 0.8 | 0.7 | 0.84 | 0.68 | 0.74 |
| 1 | 112707866 | rs155672 | 0.376 | -0.051 | -0.473 | -0.059 | -0.002 | -0.012 | -0.012 | 0.264 | 0.67 | 0.67 | 0.69 | 0.84 | 0.67 | 0.73 |
| 1 | 112709285 | rs10857934 | 0.167 | -0.06 | -0.137 | 0.069 | -0.006 | -0.012 | -0.012 | 0 | 0.32 | 0.29 | 0.24 | 0.46 | 0.35 | 0.22 |
| 1 | 112711169 | rs149802 | -0.007 | -0.026 | 0.015 | -0.315 | -0.003 | -0.023 | -0.023 | 0 | 0.67 | 0.66 | 0.68 | 0.88 | 0.67 | 0.73 |
| 1 | 112712847 | rs10745326 | 0.14 | -0.171 | 0.428 | -0.388 | -0.017 | -0.08 | -0.08 | 0 | 0.66 | 0.52 | 0.67 | 0.88 | 0.67 | 0.73 |
| 1 | 112721915 | rs11102404 | 0.443 | -0.254 | 0.409 | -0.401 | 0.031 | 0.083 | 0.083 | 0.838 | 0.64 | 0.27 | 0.65 | 0.88 | 0.68 | 0.73 |
| 1 | 112723031 | rs4292960 | 0.504 | -0.276 | 0.405 | -0.413 | 0.04 | 0.135 | 0.135 | 1.065 | 0.63 | 0.16 | 0.63 | 0.88 | 0.68 | 0.73 |
| 1 | 112727458 | rs2196887 | 0.376 | -0.398 | 0.529 | -0.577 | -0.005 | -0.039 | -0.039 | 0 | 0.2 | 0.39 | 0.16 | 0.12 | 0.21 | 0.17 |
| 1 | 112730350 | rs2581414 | -0.657 | -0.354 | 0.455 | -0.583 | 0.01 | 0.015 | 0.015 | 0 | 0.87 | 0.93 | 0.83 | 1 | 0.89 | 0.9 |
| 1 | 112731101 | rs1856390 | 1.904 | -0.382 | 0.393 | -0.342 | -0.054 | -0.027 | -0.027 | 1.409 | 0.32 | 0.06 | 0.4 | 0.32 | 0.31 | 0.43 |
| 1 | 112736898 | rs12138691 | 0.459 | -0.169 | 0.799 | -0.151 | 0.001 | -0.022 | -0.022 | 0 | 0.19 | 0.28 | 0.16 | 0.12 | 0.2 | 0.16 |
| 1 | 112761639 | rs2788407 | -0.256 | -0.097 | 0.395 | -0.118 | 0.005 | 0.007 | 0.007 | 1.333 | 0.23 | 0.75 | 0.2 | 0.12 | 0.21 | 0.17 |
| 1 | 112773825 | rs6537730 | -1.009 | -0.071 | 0.57 | -0.072 | 0 | 0.001 | 0.001 | 0 | 0.52 | 0.51 | 0.4 | 0.67 | 0.57 | 0.47 |
| 1 | 112773948 | rs6537731 | -0.476 | -0.062 | 0.784 | 0.139 | -0.003 | -0.008 | -0.008 | 0.358 | 0.35 | 0.34 | 0.27 | 0.55 | 0.37 | 0.33 |
| 1 | 112774577 | rs961799 | 1.318 | -0.025 | 1.12 | 0.018 | 0.001 | -0.001 | -0.001 | 0 | 0.94 | 0.65 | 0.95 | 1 | 0.99 | 0.99 |
| 1 | 112778604 | rs12564809 | 0.191 | 0.107 | 0.402 | 0.117 | -0.05 | -0.104 | -0.104 | 0 | 0.26 | 0.08 | 0.19 | 0.54 | 0.29 | 0.29 |
| 1 | 112779668 | rs12033004 | -0.004 | 0.088 | 0.604 | 0.108 | -0.003 | -0.008 | -0.008 | 0 | 0.27 | 0.26 | 0.21 | 0.54 | 0.29 | 0.29 |
| 1 | 112780618 | rs2998359 | 0.952 | 0.151 | 0.394 | 0.043 | -0.008 | 0.088 | 0.088 | 0.028 | 0.48 | 0.15 | 0.57 | 0.33 | 0.45 | 0.55 |
| 1 | 112783743 | rs1890400 | -0.286 | 0.285 | 0.586 | 0.04 | 0.001 | -0.018 | -0.018 | 0.012 | 0.13 | 0.13 | 0.18 | 0 | 0.14 | 0.1 |
| 1 | 112785271 | rs2581392 | -0.252 | 0.495 | 0.67 | 0.242 | 0.006 | -0.032 | -0.032 | 0 | 0.79 | 0.69 | 0.86 | 0.88 | 0.77 | 0.85 |
| 1 | 112785697 | rs7551793 | -0.042 | 0.563 | 0.469 | 0.275 | 0.011 | -0.043 | -0.043 | 0 | 0.22 | 0.34 | 0.15 | 0.13 | 0.23 | 0.15 |
| 1 | 112787396 | rs7520740 | 0.156 | 0.63 | -0.065 | 0.307 | 0.003 | -0.015 | -0.015 | 0 | 0.2 | 0.19 | 0.14 | 0.12 | 0.23 | 0.15 |
| 1 | 112788376 | rs2027278 | -1.278 | 0.528 | -0.744 | 0.45 | 0.002 | 0.027 | 0.027 | 0.063 | 0.59 | 0.81 | 0.48 | 0.66 | 0.66 | 0.54 |
| 1 | 112788660 | rs893835 | -0.064 | 0.365 | -0.49 | 0.511 | 0.004 | -0.028 | -0.028 | 0.371 | 0.06 | 0.12 | 0.09 | 0 | 0.1 | 0.08 |
| 1 | 112789288 | rs7527003 | -0.081 | 0.273 | -0.691 | 0.442 | -0.003 | -0.009 | -0.009 | 0.133 | 0.25 | 0.26 | 0.13 | 0.14 | 0.22 | 0.17 |
| 1 | 112789402 | rs1078317 | 1.095 | 0.522 | -1.269 | 0.664 | -0.037 | 0.082 | 0.082 | 0.224 | 0.46 | 0.15 | 0.52 | 0.32 | 0.49 | 0.44 |
| 1 | 112789639 | rs10157998 | 0.322 | 0.59 | -1.344 | 0.863 | -0.026 | -0.037 | -0.037 | 0.04 | 0.06 | 0 | 0.01 | 0.2 | 0.07 | 0.09 |

| | | | | | | | | | | | | | | | | |
|---|---|---|---|---|---|---|---|---|---|---|---|---|---|---|---|---|
| 1 | 112791108 | rs7520729 | 0.522 | 0.262 | -1.148 | 0.422 | 0.006 | 0.007 | 0.007 | 0 | 0.55 | 0.33 | 0.61 | 0.41 | 0.57 | 0.49 |
| 1 | 112792984 | rs1561582 | 0.54 | 0.736 | -1.71 | 0.752 | -0.031 | 0.027 | 0.027 | 0.368 | 0.49 | 0.14 | 0.53 | 0.41 | 0.5 | 0.43 |
| 1 | 112793307 | rs11102441 | 0.54 | 0.736 | -1.709 | 0.752 | -0.026 | 0.03 | 0.03 | 0.368 | 0.49 | 0.15 | 0.53 | 0.41 | 0.5 | 0.44 |
| 1 | 112801723 | rs12026605 | 0.519 | 0.738 | -1.952 | 0.618 | -0.071 | 0.047 | 0.047 | 0.212 | 0.48 | 0.07 | 0.49 | 0.37 | 0.45 | 0.43 |
| 1 | 112804559 | rs11102446 | 0.652 | 0.058 | -1.019 | 0.653 | 0.028 | -0.007 | -0.007 | 0 | 0.56 | 0.32 | 0.61 | 0.45 | 0.62 | 0.49 |
| 1 | 112813678 | rs2998333 | -1.506 | -0.279 | 0.034 | -0.486 | -0.031 | -0.136 | -0.136 | 0 | 0.17 | 0.5 | 0.11 | 0 | 0.11 | 0.13 |
| 1 | 112816851 | rs6662678 | -1.181 | -0.523 | 0.243 | -0.433 | 0.019 | 0.069 | 0.069 | 0 | 0.36 | 0.23 | 0.19 | 0.18 | 0.31 | 0.33 |
| 1 | 112822938 | rs3128353 | 1.474 | -0.6 | 0.419 | -0.45 | 0.009 | -0.017 | -0.017 | 1.269 | 0.61 | 0.32 | 0.78 | 0.94 | 0.69 | 0.72 |
| 1 | 112823822 | rs4073808 | -0.725 | -0.48 | 0.197 | -0.394 | 0.004 | 0.019 | 0.019 | 0 | 0.3 | 0.24 | 0.18 | 0.18 | 0.28 | 0.29 |
| 1 | 112824160 | rs4073809 | -0.583 | -0.476 | 0.002 | -0.392 | -0.025 | 0.059 | 0.059 | 0 | 0.28 | 0.04 | 0.16 | 0.18 | 0.28 | 0.29 |
| 1 | 112827185 | rs9308252 | 0.771 | -0.387 | 0.06 | -0.353 | -0.011 | 0.019 | 0.019 | 0 | 0.07 | 0.02 | 0.06 | 0 | 0.08 | 0.09 |
| 1 | 112829454 | rs3121992 | 1.034 | -0.268 | 0.371 | -0.305 | 0.001 | -0.005 | -0.005 | 0.562 | 0.61 | 0.61 | 0.79 | 0.82 | 0.68 | 0.66 |
| 1 | 112847591 | rs7543986 | -0.328 | -0.096 | 0.091 | -0.204 | -0.014 | -0.04 | -0.04 | 0 | 0.07 | 0.2 | 0.02 | 0 | 0.03 | 0.04 |
| 1 | 112848144 | rs12127798 | 0.645 | -0.157 | 0.024 | -0.246 | 0.017 | 0.045 | 0.045 | 0 | 0.62 | 0.7 | 0.81 | 0.82 | 0.69 | 0.67 |
| 1 | 112858294 | rs3128373 | 0.85 | 0.082 | 0.016 | -0.301 | -0.001 | -0.023 | -0.023 | 0 | 0.57 | 0.54 | 0.77 | 0.82 | 0.63 | 0.65 |
| 1 | 112865946 | rs2153995 | 0.63 | -0.712 | -0.21 | -0.345 | 0.071 | 0.101 | 0.101 | 1.414 | 0.51 | 0.29 | 0.57 | 0.78 | 0.7 | 0.7 |
| 1 | 112866597 | rs11102457 | -0.299 | -0.719 | -0.368 | 0.193 | -0.001 | 0.002 | 0.002 | 0 | 0.14 | 0.16 | 0.12 | 0.24 | 0.16 | 0.2 |
| 1 | 112871321 | rs3005774 | 0.287 | -0.974 | 0.32 | -0.226 | 0.011 | 0.011 | 0.011 | 0.109 | 0.53 | 0.46 | 0.59 | 0.79 | 0.71 | 0.67 |
| 1 | 112872684 | rs11102458 | -0.039 | -0.99 | 0.157 | -0.104 | 0.003 | 0.001 | 0.001 | 0 | 0.16 | 0.12 | 0.11 | 0.18 | 0.17 | 0.21 |
| 1 | 112872875 | rs2995774 | -0.216 | -1.374 | 0.47 | -0.312 | 0.031 | 0.068 | 0.068 | 0 | 0.64 | 0.93 | 0.64 | 0.79 | 0.75 | 0.7 |
| 1 | 112874469 | rs6678502 | -1.836 | -0.707 | 1.445 | 0.204 | -0.025 | -0.073 | -0.073 | 2.155 | 0.13 | 0.65 | 0.07 | 0 | 0.07 | 0.04 |
| 1 | 112875033 | rs1343274 | -1.07 | -0.682 | 1.325 | 0.29 | -0.017 | -0.051 | -0.051 | 0.097 | 0.09 | 0.23 | 0.03 | 0 | 0.04 | 0.03 |
| 1 | 112879437 | rs6695540 | 0.114 | -0.594 | 1.347 | 0.827 | 0 | 0.088 | 0.088 | 0 | 0.09 | 0.21 | 0.24 | 0.24 | 0.08 | 0.12 |
| 1 | 112881454 | rs2995770 | -1.506 | -0.707 | 1.339 | 0.576 | 0.018 | 0.006 | 0.006 | 1.257 | 0.47 | 0.62 | 0.45 | 0.44 | 0.38 | 0.56 |
| 1 | 112886545 | rs2995765 | -1.711 | -0.744 | 1.377 | 0.594 | 0.048 | 0.015 | 0.015 | 0.704 | 0.49 | 0.67 | 0.46 | 0.46 | 0.38 | 0.57 |
| 1 | 112888622 | rs11102461 | -1.416 | -0.718 | 1.098 | 0.56 | 0.003 | 0.001 | 0.001 | 1.258 | 0.46 | 0.53 | 0.45 | 0.43 | 0.39 | 0.56 |
| 1 | 112897590 | rs1343275 | -1.436 | -0.776 | 0.875 | 0.685 | 0.007 | -0.005 | -0.005 | 2.71 | 0.4 | 0.66 | 0.43 | 0.48 | 0.31 | 0.49 |
| 1 | 112899446 | rs1936050 | -1.619 | -0.784 | 0.497 | 0.67 | 0.042 | -0.021 | -0.021 | 0 | 0.42 | 0.72 | 0.43 | 0.48 | 0.31 | 0.5 |
| 1 | 112901203 | rs11804546 | -0.452 | -0.875 | 0.516 | 0.739 | 0.007 | 0.008 | 0.008 | 0 | 0.22 | 0.13 | 0.29 | 0.18 | 0.17 | 0.15 |
| 1 | 112911324 | rs6537736 | -0.721 | -0.599 | 0.271 | 0.766 | -0.042 | 0.089 | 0.089 | 0.029 | 0.18 | 0.44 | 0.14 | 0.31 | 0.14 | 0.34 |
| 1 | 112911843 | rs2488778 | -1.29 | -0.007 | -0.878 | 0.681 | 0.023 | -0.023 | -0.023 | 0 | 0.46 | 0.79 | 0.51 | 0.49 | 0.42 | 0.51 |
| 1 | 112918545 | rs2995762 | -1.552 | -0.181 | -1.386 | 0.658 | 0.017 | -0.074 | -0.074 | 3.284 | 0.52 | 0.96 | 0.47 | 0.55 | 0.4 | 0.52 |
| 1 | 112920252 | rs1954369 | -0.781 | -0.261 | -0.259 | 0.378 | -0.02 | -0.148 | -0.148 | 0 | 0.21 | 0.52 | 0.1 | 0 | 0.16 | 0.05 |
| 1 | 112926756 | rs2492495 | 0.713 | 0.094 | -0.98 | 1.103 | 0.003 | -0.111 | -0.111 | 0 | 0.77 | 0.91 | 0.65 | 0.47 | 0.74 | 0.58 |
| 1 | 112927912 | rs2492497 | 0.711 | 0.366 | -1.016 | 1.334 | -0.03 | -0.128 | -0.128 | 0 | 0.77 | 0.95 | 0.67 | 0.54 | 0.74 | 0.58 |
| 1 | 112928036 | rs2999471 | -1.774 | 0.131 | -0.408 | 1.233 | 0.009 | -0.001 | -0.001 | 0 | 0.4 | 0.68 | 0.59 | 0.6 | 0.33 | 0.41 |
| 1 | 112931256 | rs2492501 | -0.865 | 0.509 | -0.289 | 1.631 | -0.012 | -0.027 | -0.027 | 0 | 0.17 | 0.11 | 0.15 | 0.43 | 0.23 | 0.29 |
| 1 | 112939750 | rs2488762 | -2.226 | -0.651 | -0.616 | 1.134 | 0.076 | -0.006 | -0.006 | 1.026 | 0.44 | 0.83 | 0.28 | 0.42 | 0.33 | 0.29 |
| 1 | 112940944 | rs2492506 | -2.182 | -0.58 | -0.248 | 1.089 | 0.053 | -0.023 | -0.023 | 0 | 0.43 | 0.76 | 0.44 | 0.55 | 0.34 | 0.29 |
| 1 | 112941134 | rs2492508 | -0.883 | -0.714 | 0.808 | 1.017 | 0.045 | 0.041 | 0.041 | 4.217 | 0.06 | 0.53 | 0.05 | 0 | 0 | 0 |
| 1 | 112948319 | rs11102472 | -1.511 | -0.746 | 1.796 | 1.001 | 0.018 | -0.05 | -0.05 | 0.06 | 0.36 | 0.18 | 0.39 | 0.55 | 0.34 | 0.29 |
| 1 | 112951422 | rs1777608 | -1.55 | -0.746 | 1.781 | 1.001 | 0.016 | -0.035 | -0.035 | 0.168 | 0.36 | 0.23 | 0.39 | 0.55 | 0.34 | 0.29 |
| 1 | 112954994 | rs962402 | -0.284 | -0.801 | 1.727 | 0.937 | 0.006 | 0.056 | 0.056 | 0.349 | 0.83 | 0.95 | 0.94 | 1 | 0.86 | 0.94 |
| 1 | 112955870 | rs1175633 | -0.245 | -0.802 | 1.726 | 0.936 | 0.006 | 0.057 | 0.057 | 0.314 | 0.83 | 0.95 | 0.94 | 1 | 0.86 | 0.94 |
| 1 | 112964687 | rs2492516 | 2.336 | -0.823 | 1.763 | 0.95 | 0.068 | -0.048 | -0.048 | 0.156 | 0.56 | 0.14 | 0.55 | 0.45 | 0.66 | 0.71 |
| 1 | 112965697 | rs2483359 | 2.111 | -0.821 | 2.037 | 0.95 | 0.026 | -0.015 | -0.015 | 0 | 0.58 | 0.32 | 0.57 | 0.45 | 0.66 | 0.71 |
| 1 | 112981412 | rs1175645 | -0.159 | -0.825 | 1.814 | 0.933 | 0.01 | 0.042 | 0.042 | 0.249 | 0.83 | 0.93 | 0.93 | 1 | 0.86 | 0.94 |
| 1 | 112981604 | rs2483345 | 2.425 | -0.823 | 1.701 | 0.94 | 0.052 | -0.1 | -0.1 | 0 | 0.55 | 0.05 | 0.55 | 0.45 | 0.66 | 0.71 |
| 1 | 112985270 | rs2492530 | 2.102 | -0.809 | 1.731 | 0.948 | 0.026 | -0.015 | -0.015 | 0 | 0.59 | 0.32 | 0.57 | 0.45 | 0.66 | 0.71 |
| 1 | 112991698 | rs2256883 | 2.102 | -0.809 | 1.731 | 0.948 | 0.026 | -0.015 | -0.015 | 0 | 0.59 | 0.32 | 0.57 | 0.45 | 0.66 | 0.71 |
| 1 | 112991862 | rs3818729 | -0.794 | -0.695 | 1.581 | 1.006 | 0.003 | -0.069 | -0.069 | 0 | 0.19 | 0.09 | 0.32 | 0.55 | 0.2 | 0.23 |
| 1 | 112999227 | rs1175641 | -0.183 | -0.616 | 1.351 | 1.042 | 0.006 | 0.057 | 0.057 | 0.314 | 0.83 | 0.96 | 0.94 | 1 | 0.86 | 0.94 |
| 1 | 113000565 | rs3795821 | -0.823 | -0.576 | 1.361 | 1.102 | 0.003 | -0.069 | -0.069 | 0 | 0.19 | 0.09 | 0.32 | 0.55 | 0.2 | 0.23 |
| 1 | 113006804 | rs11102479 | -0.823 | -0.515 | 0.77 | 1.139 | 0.003 | -0.069 | -0.069 | 0 | 0.19 | 0.09 | 0.32 | 0.56 | 0.2 | 0.23 |
| 1 | 113007539 | rs2798245 | -0.026 | -0.693 | 0.761 | 0.944 | 0.006 | 0.057 | 0.057 | 0.271 | 0.83 | 0.96 | 0.94 | 1 | 0.86 | 0.94 |
| 1 | 113020298 | rs10776751 | -0.175 | -0.637 | 0.495 | 1.656 | 0.004 | 0.042 | 0.042 | 0 | 0.08 | 0.07 | 0.27 | 0.49 | 0.08 | 0.15 |
| 1 | 113021218 | rs4644492 | 1.506 | -0.62 | 0.317 | 1.25 | 0.064 | -0.009 | -0.009 | 0 | 0.57 | 0.19 | 0.55 | 0.37 | 0.65 | 0.66 |
| 1 | 113023980 | rs1620668 | -0.808 | -0.405 | 0.721 | 1.392 | 0.005 | 0.067 | 0.067 | 0 | 0.14 | 0.18 | 0.22 | 0.38 | 0.16 | 0.15 |
| 1 | 113026557 | rs971792 | 2.114 | -0.361 | 0.466 | 1.078 | 0.035 | -0.025 | -0.025 | 0 | 0.51 | 0.15 | 0.49 | 0.42 | 0.56 | 0.64 |
| 1 | 113028105 | rs1777606 | -1.645 | -0.402 | 0.955 | 1.036 | -0.009 | 0.061 | 0.061 | 0.011 | 0.23 | 0.57 | 0.28 | 0.38 | 0.21 | 0.2 |
| 1 | 113028271 | rs974442 | 1.878 | -0.278 | 0.416 | 1.114 | 0.035 | -0.025 | -0.025 | 0 | 0.51 | 0.15 | 0.49 | 0.42 | 0.56 | 0.63 |
| 1 | 113038795 | rs351354 | -0.728 | -0.199 | 0.41 | 1.117 | -0.001 | -0.001 | -0.001 | 0 | 0.84 | 0.9 | 0.83 | 0.8 | 0.83 | 0.87 |
| 1 | 113040277 | rs351359 | 1.101 | -0.262 | 0.836 | 0.862 | -0.005 | 0.008 | 0.008 | 0 | 0.6 | 0.6 | 0.56 | 0.42 | 0.61 | 0.68 |
| 1 | 113040421 | rs351360 | 1.915 | -0.009 | 0.142 | 0.843 | -0.055 | 0.012 | 0.012 | 0 | 0.43 | 0.18 | 0.47 | 0.36 | 0.48 | 0.58 |
| 1 | 113046395 | rs351365 | 0.034 | 0.186 | 0.388 | 1.099 | -0.002 | 0.012 | 0.012 | 0 | 0.77 | 0.68 | 0.68 | 0.7 | 0.77 | 0.82 |
| 1 | 113049862 | rs351367 | 1.086 | 0.416 | -0.061 | 1.08 | -0.005 | 0.055 | 0.055 | 0 | 0.51 | 0.78 | 0.51 | 0.38 | 0.47 | 0.62 |
| 1 | 113054659 | rs351370 | 0.87 | 0.429 | 0.706 | 1.023 | -0.006 | -0.014 | -0.014 | 0 | 0.58 | 0.51 | 0.76 | 0.74 | 0.6 | 0.73 |
| 1 | 113056990 | rs3790609 | -0.589 | 0.384 | 0.382 | 1.027 | -0.011 | -0.018 | -0.018 | 0 | 0.17 | 0.05 | 0.25 | 0.14 | 0.18 | 0.15 |
| 1 | 113063771 | rs2273368 | -0.363 | 0.586 | 0.456 | 1.332 | -0.03 | -0.077 | -0.077 | 0 | 0.15 | 0.06 | 0.27 | 0.43 | 0.21 | 0.21 |

| | | | | | | | | | | | | | | | | |
|---|---|---|---|---|---|---|---|---|---|---|---|---|---|---|---|---|
| 1 | 113067792 | rs3790611 | -0.321 | 0.688 | 0.455 | 0.999 | 0.008 | 0.023 | 0.023 | 0 | 0.29 | 0.21 | 0.15 | 0.19 | 0.25 | 0.12 |
| 1 | 113072477 | rs2273369 | -0.251 | 0.685 | 0.466 | 0.998 | 0.009 | 0.034 | 0.034 | 0 | 0.29 | 0.15 | 0.13 | 0.19 | 0.25 | 0.12 |
| 1 | 113076756 | rs12728001 | -1.696 | 0.557 | 0.857 | 0.939 | 0.005 | 0.016 | 0.016 | 0 | 0.49 | 0.58 | 0.45 | 0.62 | 0.48 | 0.34 |
| 1 | 113083439 | rs10745330 | -1.646 | 0.56 | 0.948 | 0.941 | 0 | -0.001 | -0.001 | 0.583 | 0.49 | 0.53 | 0.44 | 0.62 | 0.48 | 0.34 |
| 1 | 113085175 | rs6537741 | 0.184 | 0.513 | 0.889 | 0.878 | 0.012 | 0.066 | 0.066 | 0 | 0.24 | 0.12 | 0.11 | 0.08 | 0.2 | 0.1 |
| 1 | 113085721 | rs4838959 | -2.05 | 0.472 | 1.021 | 0.917 | -0.104 | 0.05 | 0.05 | 0.38 | 0.58 | 0.92 | 0.67 | 0.68 | 0.53 | 0.37 |
| 1 | 113098534 | rs6658555 | 0.077 | 0.476 | 0.93 | 0.881 | 0.004 | 0.103 | 0.103 | 0 | 0.29 | 0.09 | 0.12 | 0.09 | 0.24 | 0.09 |
| 1 | 113106633 | rs12137269 | -0.067 | 0.486 | 0.906 | 0.912 | -0.032 | -0.083 | -0.083 | 0 | 0.15 | 0.07 | 0.28 | 0.48 | 0.23 | 0.21 |
| 1 | 113123734 | rs7554916 | -1.507 | 0.485 | 0.918 | 0.855 | 0 | 0 | 0 | 0 | 0.5 | 0.53 | 0.44 | 0.62 | 0.47 | 0.34 |
| 1 | 113129128 | rs6671645 | -0.534 | 0.485 | 0.919 | 0.859 | -0.018 | -0.038 | -0.038 | 0 | 0.34 | 0.45 | 0.16 | 0.19 | 0.25 | 0.13 |
| 1 | 113132393 | rs6666579 | -1.484 | 0.483 | 0.911 | 0.857 | 0 | 0 | 0 | 0.581 | 0.5 | 0.53 | 0.44 | 0.62 | 0.47 | 0.34 |
| 1 | 113140583 | rs12049572 | -0.099 | 0.558 | 0.268 | 0.915 | -0.032 | -0.079 | -0.079 | 0 | 0.15 | 0.06 | 0.28 | 0.43 | 0.23 | 0.21 |
| 1 | 113186778 | rs10745331 | -0.261 | 0.79 | 0.465 | 0.565 | 0.002 | -0.001 | -0.001 | 0 | 0.69 | 0.67 | 0.75 | 0.68 | 0.66 | 0.76 |
| 1 | 113202571 | rs12120956 | -0.094 | 0.302 | -0.303 | -0.058 | -0.022 | -0.06 | -0.06 | 0 | 0.15 | 0.08 | 0.28 | 0.44 | 0.24 | 0.21 |
| 1 | 113214097 | rs1238 | -0.19 | 0.317 | -0.288 | -0.061 | 0.009 | 0.034 | 0.034 | 0 | 0.7 | 0.84 | 0.87 | 0.81 | 0.76 | 0.88 |
| 1 | 113214381 | rs6537748 | 1.807 | 0.282 | -0.362 | -0.131 | -0.157 | 0.02 | 0.02 | 0.301 | 0.4 | 0.03 | 0.32 | 0.31 | 0.45 | 0.62 |
| 1 | 113216543 | rs2932538 | 0.098 | 0.872 | -0.559 | 0.198 | 0.009 | 0.037 | 0.037 | 0 | 0.7 | 0.84 | 0.87 | 0.81 | 0.76 | 0.88 |
| 1 | 113236681 | rs3748656 | -0.831 | 0.948 | -0.049 | 0.75 | -0.003 | 0.004 | 0.004 | 0 | 0.16 | 0.17 | 0.29 | 0.54 | 0.23 | 0.22 |
| 1 | 113242122 | rs1106287 | -0.931 | 0.947 | -0.045 | 0.749 | -0.002 | 0.007 | 0.007 | 0 | 0.16 | 0.19 | 0.29 | 0.54 | 0.23 | 0.22 |
| 1 | 113243048 | rs6679 | -0.834 | 0.986 | -0.104 | 0.747 | -0.003 | 0.005 | 0.005 | 0 | 0.16 | 0.17 | 0.29 | 0.54 | 0.23 | 0.22 |
| 1 | 113244433 | rs2279500 | -0.486 | 0.691 | -0.019 | 0.736 | 0.005 | 0.022 | 0.022 | 0 | 0.8 | 0.89 | 0.93 | 0.91 | 0.84 | 0.92 |
| 1 | 113248791 | rs12144044 | -0.321 | 0.737 | -0.437 | 0.629 | 0.012 | 0.053 | 0.053 | 0 | 0.3 | 0.21 | 0.14 | 0.09 | 0.26 | 0.16 |
| 1 | 113250888 | rs11102522 | -0.011 | 1.319 | -0.446 | 1.192 | -0.053 | -0.089 | -0.089 | 0 | 0.15 | 0.07 | 0.33 | 0.45 | 0.23 | 0.15 |
| 1 | 113263711 | rs4450019 | 0.155 | 0.777 | -0.706 | 0.142 | 0 | 0.001 | 0.001 | 0 | 0.48 | 0.47 | 0.51 | 0.32 | 0.47 | 0.63 |
| 1 | 113268225 | rs11102524 | -0.266 | 0.583 | -0.865 | -0.26 | 0.004 | 0.007 | 0.007 | 0.466 | 0.43 | 0.44 | 0.47 | 0.34 | 0.43 | 0.58 |
| 1 | 113276156 | rs11102525 | -0.151 | 0.521 | -0.687 | -0.566 | 0.002 | 0.014 | 0.014 | 0.214 | 0.54 | 0.68 | 0.56 | 0.35 | 0.52 | 0.58 |
| 1 | 113278651 | rs6537754 | 0.015 | 0.544 | -0.289 | -0.829 | 0 | 0.018 | 0.018 | 0.621 | 0.49 | 0.57 | 0.52 | 0.34 | 0.46 | 0.54 |
| 1 | 113284079 | rs12402272 | 0.938 | 0.616 | -0.077 | -1.262 | 0.002 | -0.022 | -0.022 | 0.47 | 0.31 | 0.2 | 0.39 | 0.34 | 0.28 | 0.36 |
| 1 | 113286780 | rs4534375 | -1.035 | 0.839 | -0.209 | -1.046 | 0.017 | 0.113 | 0.113 | 0.584 | 0.62 | 0.78 | 0.81 | 0.88 | 0.66 | 0.8 |
| 1 | 113289902 | rs4839260 | -1.359 | 1.03 | 0.152 | -0.824 | -0.03 | -0.053 | -0.053 | 0.105 | 0.28 | 0.16 | 0.39 | 0.54 | 0.37 | 0.44 |
| 1 | 113290694 | rs12033013 | 0.038 | 1.118 | 0.04 | -0.73 | 0.001 | 0.138 | 0.138 | 0 | 0.06 | 0.07 | 0.23 | 0.49 | 0.1 | 0.23 |
| 1 | 113292658 | rs4838964 | 0.993 | 0.786 | -0.248 | -1.102 | -0.003 | 0.013 | 0.013 | 0.582 | 0.44 | 0.26 | 0.45 | 0.35 | 0.42 | 0.4 |
| 1 | 113293449 | rs7553091 | 0.661 | 0.766 | -0.257 | -1.067 | -0.006 | 0.059 | 0.059 | 0 | 0.28 | 0.14 | 0.17 | 0.13 | 0.29 | 0.19 |
| 1 | 113297767 | rs6537756 | 0.844 | 0.776 | -0.194 | -1.071 | 0.008 | 0.051 | 0.051 | 1.994 | 0.72 | 0.44 | 0.61 | 0.48 | 0.69 | 0.59 |
| 1 | 113313563 | rs11590090 | 0.471 | 0.467 | -0.467 | -1.171 | -0.004 | 0.074 | 0.074 | 0 | 0.29 | 0.16 | 0.17 | 0.12 | 0.29 | 0.25 |
| 1 | 113331134 | rs4378202 | 1.115 | 0.409 | -0.465 | -0.77 | 0.004 | 0.013 | 0.013 | 1.852 | 0.4 | 0.18 | 0.41 | 0.29 | 0.36 | 0.31 |
| 1 | 113331258 | rs4385708 | -0.107 | 0.495 | 0.051 | -0.701 | 0.042 | 0.235 | 0.235 | 0.04 | 0.89 | 0.79 | 0.72 | 0.48 | 0.82 | 0.75 |
| 1 | 113334191 | rs10857976 | 1.049 | 0.441 | -0.699 | -0.783 | -0.003 | -0.071 | -0.071 | 0 | 0.39 | 0.15 | 0.41 | 0.43 | 0.36 | 0.35 |
| 1 | 113334373 | rs10857977 | 1.049 | 0.441 | -0.699 | -0.783 | -0.003 | -0.071 | -0.071 | 0 | 0.39 | 0.14 | 0.41 | 0.43 | 0.36 | 0.35 |
| 1 | 113336905 | rs6681499 | 0.012 | 0.585 | -0.523 | -0.988 | -0.012 | 0.061 | 0.061 | 0 | 0.38 | 0.31 | 0.26 | 0.12 | 0.4 | 0.34 |
| 1 | 113338649 | rs10857978 | 1.308 | 0.633 | -0.223 | -0.615 | -0.003 | 0.041 | 0.041 | 0.02 | 0.74 | 0.46 | 0.66 | 0.55 | 0.75 | 0.69 |
| 1 | 113344950 | rs7546286 | -0.212 | 1.042 | -0.657 | -0.835 | -0.043 | 0.043 | 0.043 | 0 | 0.24 | 0.07 | 0.19 | 0.14 | 0.28 | 0.26 |
| 1 | 113345279 | rs4838967 | 1.644 | 0.821 | -0.509 | -0.906 | -0.027 | 0.163 | 0.163 | 0.638 | 0.53 | 0.14 | 0.67 | 0.8 | 0.54 | 0.61 |
| 1 | 113346108 | rs7535706 | -1.289 | 1.154 | -0.794 | -1.049 | -0.075 | 0.161 | 0.161 | 0 | 0.4 | 0.14 | 0.25 | 0.14 | 0.46 | 0.31 |
| 1 | 113348570 | rs11804649 | -0.873 | 0.963 | -0.455 | -1.105 | -0.003 | 0.045 | 0.045 | 0 | 0.17 | 0.08 | 0.07 | 0 | 0.16 | 0.12 |
| 1 | 113350929 | rs4839265 | -1.053 | 0.949 | 0.189 | -0.882 | 0 | -0.021 | -0.021 | 0 | 0.09 | 0.13 | 0.04 | 0 | 0.08 | 0.05 |
| 1 | 113351374 | rs7520797 | -2.016 | 0.929 | 0.768 | -0.986 | -0.002 | -0.02 | -0.02 | 0.929 | 0.44 | 0.5 | 0.47 | 0.16 | 0.41 | 0.33 |
| 1 | 113384015 | rs7539948 | -0.961 | 0.527 | 1.752 | -0.87 | -0.005 | -0.05 | -0.05 | 0 | 0.12 | 0.25 | 0.07 | 0.02 | 0.08 | 0.09 |
| 1 | 113390844 | rs7527283 | -2.569 | 0.52 | 1.809 | -0.877 | -0.013 | -0.066 | -0.066 | 0 | 0.22 | 0.69 | 0.14 | 0.02 | 0.16 | 0.14 |
| 1 | 113391029 | rs12090523 | -1.612 | 0.57 | 1.585 | -0.584 | -0.008 | -0.067 | -0.067 | 0.151 | 0.14 | 0.46 | 0.09 | 0.02 | 0.08 | 0.09 |
| 1 | 113413940 | rs10399710 | -1.239 | 1.091 | 1.114 | -1.133 | 0.015 | -0.113 | -0.113 | 2.933 | 0.25 | 0.37 | 0.15 | 0 | 0.28 | 0.15 |
| 1 | 113416122 | rs2996294 | -1.455 | 1.23 | 0.814 | -1.008 | -0.027 | 0.023 | 0.023 | 0 | 0.14 | 0.09 | 0.1 | 0.01 | 0.22 | 0.09 |
| 1 | 113418834 | rs1216539 | 0.013 | 1.593 | 0.203 | -1.213 | -0.097 | 0.038 | 0.038 | 0 | 0.77 | 0.98 | 0.76 | 0.91 | 0.67 | 0.74 |
| 1 | 113424469 | rs2996295 | 0.764 | 1.288 | 0.359 | -0.843 | -0.034 | 0.026 | 0.026 | 0 | 0.62 | 0.86 | 0.55 | 0.74 | 0.56 | 0.55 |
| 1 | 113426308 | rs11102559 | 2.131 | 1.024 | 1.065 | -0.905 | 0.028 | 0.041 | 0.041 | 0 | 0.42 | 0.27 | 0.4 | 0.72 | 0.32 | 0.46 |
| 1 | 113431786 | rs1539624 | 1.124 | 0.896 | 1.375 | -0.964 | 0.008 | 0.027 | 0.027 | 0 | 0.55 | 0.65 | 0.53 | 0.72 | 0.55 | 0.55 |
| 1 | 113445253 | rs7545602 | -0.662 | 1.221 | 1.79 | -0.937 | -0.074 | 0.072 | 0.072 | 0 | 0.12 | 0.03 | 0.09 | 0.02 | 0.23 | 0.09 |
| 1 | 113448466 | rs9429709 | -1.675 | 1.155 | 2.093 | -0.969 | 0.075 | -0.115 | -0.115 | 0 | 0.17 | 0.55 | 0.13 | 0.02 | 0.23 | 0.09 |
| 1 | 113449186 | rs4838971 | -1.768 | 1.175 | 2.169 | -0.971 | 0.05 | -0.105 | -0.105 | 0 | 0.18 | 0.61 | 0.14 | 0.02 | 0.23 | 0.09 |
| 1 | 113449627 | rs12037621 | 2.269 | 1.086 | 2.076 | -0.557 | 0.033 | 0.019 | 0.019 | 0 | 0.42 | 0.23 | 0.4 | 0.64 | 0.32 | 0.49 |
| 1 | 113452233 | rs7550843 | -1.804 | 1.057 | 2.105 | 0.18 | 0.041 | 0.024 | 0.024 | 0.03 | 0.18 | 0.63 | 0.31 | 0.26 | 0.24 | 0.2 |
| 1 | 113452284 | rs12727968 | 0.156 | 1.051 | 1.983 | -0.477 | 0.011 | 0.069 | 0.069 | 0 | 0.14 | 0.01 | 0.08 | 0.02 | 0.13 | 0.11 |
| 1 | 113454424 | rs17493313 | -0.845 | 1.094 | 2.476 | -0.391 | 0.005 | -0.023 | -0.023 | 0.225 | 0.13 | 0.14 | 0.1 | 0.02 | 0.23 | 0.09 |
| 1 | 113454571 | rs9429505 | -1.658 | 1.095 | 2.459 | -0.39 | 0.052 | -0.105 | -0.105 | 0 | 0.18 | 0.61 | 0.14 | 0.02 | 0.24 | 0.09 |
| 1 | 113455099 | rs7169 | 0.829 | 1.094 | 2.461 | -0.371 | -0.036 | -0.019 | -0.019 | 1.609 | 0.62 | 0.91 | 0.55 | 0.66 | 0.56 | 0.58 |
| 1 | 113460917 | rs3849174 | -1.655 | 0.925 | 2.469 | -0.364 | 0.052 | -0.105 | -0.105 | 0 | 0.18 | 0.61 | 0.14 | 0.02 | 0.24 | 0.09 |
| 1 | 113463853 | rs7518984 | -1.825 | 0.867 | 2.443 | -0.392 | 0.043 | -0.108 | -0.108 | 0 | 0.19 | 0.6 | 0.14 | 0.02 | 0.24 | 0.09 |
| 1 | 113471400 | rs3789592 | 0.853 | 0.971 | 2.591 | -0.333 | -0.036 | -0.015 | -0.015 | 0.344 | 0.62 | 0.91 | 0.55 | 0.66 | 0.56 | 0.58 |
| 1 | 113492862 | rs4301628 | 2.467 | 0.988 | 2.634 | -0.333 | 0.034 | 0.009 | 0.009 | 0 | 0.42 | 0.15 | 0.39 | 0.64 | 0.31 | 0.49 |

| | | | | | | | | | | | | | | | | |
|---|---|---|---|---|---|---|---|---|---|---|---|---|---|---|---|---|
| 1 | 113492909 | rs7552903 | -1.672 | 1.015 | 2.527 | -0.362 | 0.052 | -0.106 | -0.106 | 0.015 | 0.18 | 0.61 | 0.14 | 0.02 | 0.24 | 0.09 |
| 1 | 113493337 | rs6537765 | 0.837 | 1.016 | 2.347 | -0.357 | -0.038 | -0.018 | -0.018 | 0 | 0.62 | 0.91 | 0.55 | 0.66 | 0.56 | 0.58 |
| 1 | 113496790 | rs4839272 | -0.6 | 1.063 | 2.025 | -0.366 | -0.075 | 0.074 | 0.074 | 0 | 0.12 | 0.03 | 0.1 | 0.02 | 0.23 | 0.09 |
| 1 | 113502552 | rs4839273 | 2.368 | 1.053 | 2.07 | -0.317 | 0.036 | 0.019 | 0.019 | 0 | 0.42 | 0.23 | 0.39 | 0.64 | 0.32 | 0.49 |
| 1 | 113504658 | rs7524567 | -1.695 | 1.025 | 2.046 | -0.426 | 0.046 | -0.102 | -0.102 | 0 | 0.18 | 0.62 | 0.14 | 0.02 | 0.24 | 0.09 |
| 1 | 113579103 | rs2798691 | 1.977 | 0.215 | 1.888 | -0.216 | 0.034 | 0.047 | 0.047 | 0 | 0.43 | 0.3 | 0.37 | 0.68 | 0.32 | 0.46 |
| 1 | 113597564 | rs1216766 | 1.518 | 0.285 | 1.569 | -0.204 | 0.054 | 0.067 | 0.067 | 0.074 | 0.49 | 0.67 | 0.41 | 0.68 | 0.34 | 0.46 |
| 1 | 113601445 | rs2798686 | 1.658 | 0.363 | 1.692 | -0.097 | 0.028 | 0.045 | 0.045 | 0 | 0.51 | 0.6 | 0.42 | 0.69 | 0.39 | 0.46 |
| 1 | 113629424 | rs11102582 | -1.684 | 0.462 | 1.394 | -0.06 | 0.007 | 0.058 | 0.058 | 0.082 | 0.45 | 0.11 | 0.54 | 0.32 | 0.6 | 0.52 |
| 1 | 113663927 | rs10494148 | -0.453 | 0.825 | 1.513 | 0.171 | 0.006 | 0.015 | 0.015 | 0 | 0.11 | 0.06 | 0.17 | 0.02 | 0.14 | 0.07 |
| 1 | 113668344 | rs10494150 | -0.463 | 0.825 | 1.513 | 0.171 | 0.006 | 0.016 | 0.016 | 0 | 0.12 | 0.06 | 0.17 | 0.02 | 0.13 | 0.07 |
| 1 | 113676486 | rs7543653 | -0.641 | 0.819 | 1.543 | 0.168 | -0.004 | -0.023 | -0.023 | 0.542 | 0.12 | 0.14 | 0.17 | 0.02 | 0.14 | 0.07 |
| 1 | 113711034 | rs6537777 | 0.982 | 0.846 | 1.493 | 0.189 | -0.016 | -0.075 | -0.075 | 0 | 0.85 | 0.64 | 0.81 | 0.96 | 0.86 | 0.85 |
| 1 | 113717127 | rs12133521 | -0.315 | 0.867 | 1.251 | 0.199 | 0.003 | 0.007 | 0.007 | 0 | 0.11 | 0.08 | 0.17 | 0.02 | 0.13 | 0.07 |
| 1 | 113717470 | rs1216870 | -0.143 | 1.378 | 1.02 | 0.242 | -0.123 | -0.053 | -0.053 | 0 | 0.19 | 0.04 | 0.24 | 0.23 | 0.36 | 0.24 |
| 1 | 113721397 | rs946218 | -1.912 | 0.402 | 1.784 | 0.018 | 0.001 | 0.018 | 0.018 | 0 | 0.17 | 0.14 | 0.21 | 0.05 | 0.19 | 0.19 |
| 1 | 113726980 | rs12137636 | -1.559 | 0.137 | 1.925 | 0.274 | 0.001 | -0.003 | -0.003 | 0 | 0.13 | 0.12 | 0.19 | 0.25 | 0.15 | 0.19 |
| 1 | 113728629 | rs1216878 | 1.576 | 0.117 | 2.117 | 0.265 | -0.017 | 0.047 | 0.047 | 0 | 0.84 | 0.64 | 0.8 | 0.75 | 0.85 | 0.81 |
| 1 | 113735593 | rs17497065 | -0.572 | 0.154 | 1.602 | 0.203 | -0.007 | 0.044 | 0.044 | 0 | 0.1 | 0.01 | 0.16 | 0.02 | 0.13 | 0.07 |
| 1 | 113738848 | rs1216890 | 1.606 | 0.122 | 1.768 | 0.245 | -0.031 | 0.04 | 0.04 | 0 | 0.83 | 0.5 | 0.79 | 0.75 | 0.85 | 0.81 |
| 1 | 113742121 | rs1766884 | 0.969 | 0.144 | 1.19 | 0.252 | -0.019 | 0.052 | 0.052 | 0 | 0.85 | 0.67 | 0.72 | 0.7 | 0.84 | 0.83 |
| 1 | 113751640 | rs1216899 | 1.133 | 0.096 | 0.456 | -0.293 | -0.028 | -0.039 | -0.039 | 0 | 0.82 | 0.33 | 0.7 | 0.9 | 0.86 | 0.88 |
| 1 | 113755018 | rs7555991 | -1.199 | 0.492 | 0.011 | -0.174 | 0 | 0 | 0 | 0 | 0.07 | 0.09 | 0.09 | 0.1 | 0.1 | 0.12 |
| 1 | 113759903 | rs7526767 | 0.16 | 0.469 | -0.342 | -2.153 | -0.003 | 0.01 | 0.01 | 0.234 | 0.4 | 0.3 | 0.39 | 0.11 | 0.4 | 0.33 |
| 1 | 113770426 | rs11102602 | 0.045 | 0.341 | -0.388 | -2.87 | 0.002 | -0.031 | -0.031 | 0 | 0.29 | 0.35 | 0.48 | 0.85 | 0.29 | 0.33 |
| 1 | 113771402 | rs6677928 | -0.632 | 0.159 | -0.49 | -3.27 | -0.004 | -0.101 | -0.101 | 0 | 0.4 | 0.48 | 0.54 | 0.93 | 0.34 | 0.42 |
| 1 | 113771607 | rs12129538 | 0.281 | 0.239 | -0.44 | -3.114 | 0.008 | -0.027 | -0.027 | 0 | 0.05 | 0.11 | 0.07 | 0 | 0.07 | 0.07 |
| 1 | 113772733 | rs6696679 | -1.06 | 0.013 | -0.455 | -3.327 | -0.015 | 0.041 | 0.041 | 0.94 | 0.79 | 0.92 | 0.78 | 0.98 | 0.73 | 0.82 |
| 1 | 113773981 | rs17014869 | 0.301 | -0.04 | -0.422 | -3.094 | -0.001 | -0.001 | -0.001 | 1.929 | 0.09 | 0.07 | 0.06 | 0.08 | 0.05 | 0.09 |
| 1 | 113775761 | rs6688859 | 0.85 | -0.085 | -0.622 | -3.157 | -0.004 | 0.067 | 0.067 | 0 | 0.31 | 0.18 | 0.2 | 0.08 | 0.35 | 0.39 |
| 1 | 113784170 | rs11102603 | 0.618 | -0.114 | -0.559 | -3.37 | 0.001 | -0.016 | -0.016 | 0 | 0.29 | 0.33 | 0.44 | 0.83 | 0.28 | 0.33 |
| 1 | 113785301 | rs773620 | 0.265 | 0.003 | -1.044 | -3.47 | 0.012 | 0.137 | 0.137 | 0 | 0.4 | 0.75 | 0.51 | 0.91 | 0.32 | 0.41 |
| 1 | 113789136 | rs1886503 | 0.397 | 0.171 | -1.129 | -3.336 | 0.005 | 0.124 | 0.124 | 0 | 0.36 | 0.21 | 0.26 | 0.07 | 0.37 | 0.43 |
| 1 | 113791338 | rs773618 | 0.099 | 0.171 | -0.705 | -2.997 | 0.001 | 0.015 | 0.015 | 0 | 0.39 | 0.64 | 0.52 | 0.93 | 0.36 | 0.36 |
| 1 | 113805583 | rs773610 | -0.214 | 0.547 | -0.555 | -2.448 | -0.001 | -0.032 | -0.032 | 0 | 0.84 | 0.79 | 0.84 | 0.97 | 0.85 | 0.8 |
| 1 | 113810389 | rs11811051 | 0.225 | 0.829 | -0.849 | -2.282 | 0.007 | -0.039 | -0.039 | 0 | 0.09 | 0.15 | 0.07 | 0 | 0.12 | 0.05 |
| 1 | 113810931 | rs12725447 | 0.539 | 1.178 | -1.462 | -2.467 | -0.037 | 0.058 | 0.058 | 0 | 0.2 | 0.07 | 0.17 | 0.08 | 0.28 | 0.35 |
| 1 | 113810964 | rs773609 | -0.181 | 0.572 | -0.965 | -1.862 | 0.014 | 0.09 | 0.09 | 0 | 0.69 | 0.85 | 0.76 | 0.97 | 0.74 | 0.74 |
| 1 | 113818120 | rs773573 | 0.29 | 0.601 | -0.725 | -1.683 | -0.006 | -0.1 | -0.1 | 1.964 | 0.57 | 0.51 | 0.67 | 0.96 | 0.61 | 0.66 |
| 1 | 113819412 | rs750179 | -0.572 | 0.568 | -0.522 | -0.604 | -0.01 | 0.007 | 0.007 | 0 | 0.31 | 0.5 | 0.25 | 0.31 | 0.25 | 0.25 |
| 1 | 113819478 | rs773568 | -0.394 | 0.675 | -1.017 | -0.829 | 0.006 | 0.013 | 0.013 | 1.121 | 0.57 | 0.62 | 0.66 | 0.83 | 0.61 | 0.53 |
| 1 | 113822100 | rs6677404 | -0.203 | 0.915 | -0.999 | -0.37 | -0.003 | -0.004 | -0.004 | 0 | 0.14 | 0.09 | 0.11 | 0.2 | 0.14 | 0.15 |
| 1 | 113822284 | rs7533863 | -0.129 | 0.209 | -1.289 | -1.025 | -0.007 | 0.03 | 0.03 | 0 | 0.54 | 0.62 | 0.66 | 0.66 | 0.54 | 0.55 |
| 1 | 113824297 | rs773564 | -1.08 | 0.363 | -1.981 | -0.769 | -0.054 | -0.08 | -0.08 | 0 | 0.77 | 0.93 | 0.48 | 0.4 | 0.63 | 0.57 |
| 1 | 113828836 | rs12063762 | 0.775 | 0.091 | -1.733 | -1.231 | -0.013 | -0.042 | -0.042 | 0 | 0.16 | 0.45 | 0.1 | 0.08 | 0.11 | 0.13 |
| 1 | 113830085 | rs773554 | -0.169 | -0.218 | -1.999 | -1.585 | 0.016 | 0.059 | 0.059 | 0.873 | 0.33 | 0.81 | 0.21 | 0.15 | 0.28 | 0.24 |
| 1 | 113831578 | rs773575 | -1.068 | -0.324 | -2.107 | -1.655 | -0.038 | -0.061 | -0.061 | 0 | 0.69 | 0.9 | 0.46 | 0.4 | 0.6 | 0.55 |
| 1 | 113834428 | rs699670 | -1.891 | -0.36 | -2.053 | -0.627 | -0.047 | -0.007 | -0.007 | 0 | 0.77 | 0.9 | 0.68 | 0.78 | 0.64 | 0.64 |
| 1 | 113837852 | rs12564854 | 1.121 | -0.379 | -2.32 | -0.844 | 0.002 | 0.032 | 0.032 | 0 | 0.13 | 0.04 | 0.09 | 0.06 | 0.12 | 0.14 |
| 1 | 113841736 | rs773586 | -0.558 | -0.266 | -2.129 | -0.707 | -0.034 | 0.039 | 0.039 | 1.296 | 0.55 | 0.87 | 0.38 | 0.34 | 0.48 | 0.4 |
| 1 | 113843501 | rs12740908 | 0.527 | -0.022 | -1.78 | -0.142 | -0.059 | -0.008 | -0.008 | 0.007 | 0.22 | 0.03 | 0.17 | 0.19 | 0.2 | 0.16 |
| 1 | 113848334 | rs10857988 | 0.313 | -0.156 | -1.098 | 0.167 | -0.018 | 0.008 | 0.008 | 0 | 0.17 | 0.45 | 0.11 | 0.12 | 0.12 | 0.12 |
| 1 | 113849470 | rs4839306 | 0.876 | 0.086 | -1.663 | 0.565 | 0.022 | -0.041 | -0.041 | 0 | 0.09 | 0 | 0.23 | 0.37 | 0.05 | 0.1 |
| 1 | 113865048 | rs1342653 | -0.552 | 0.311 | -1.82 | -0.299 | 0.04 | 0.144 | 0.144 | 0 | 0.49 | 0.24 | 0.49 | 0.75 | 0.37 | 0.35 |
| 1 | 113871058 | rs6689098 | -0.51 | 0.148 | -1.863 | -0.38 | 0.023 | 0.12 | 0.12 | 0 | 0.46 | 0.27 | 0.49 | 0.76 | 0.37 | 0.35 |
| 1 | 113876419 | rs4839312 | -0.007 | 0.037 | -2.256 | -0.192 | -0.07 | -0.019 | -0.019 | 0 | 0.29 | 0.02 | 0.21 | 0.25 | 0.25 | 0.21 |
| 1 | 113876786 | rs4838985 | -0.821 | -0.036 | -2.259 | -0.218 | -0.053 | 0.141 | 0.141 | 0 | 0.42 | 0.05 | 0.49 | 0.76 | 0.38 | 0.32 |
| 1 | 113890470 | rs10494155 | 0.962 | -0.007 | -2.225 | -0.183 | 0.002 | -0.06 | -0.06 | 0.006 | 0.13 | 0.03 | 0.27 | 0.5 | 0.12 | 0.11 |
| 1 | 113891038 | rs10494157 | 1.002 | 0.025 | -1.642 | -0.771 | 0.015 | -0.008 | -0.008 | 0 | 0.15 | 0.47 | 0.11 | 0.09 | 0.12 | 0.16 |
| 1 | 113891651 | rs4838986 | -0.698 | 0.179 | -1.817 | -0.589 | 0.012 | 0.14 | 0.14 | 0 | 0.45 | 0.21 | 0.49 | 0.76 | 0.38 | 0.32 |
| 1 | 113894627 | rs1935745 | 0.803 | 0.796 | -1.889 | -0.683 | 0.006 | 0.162 | 0.162 | 0 | 0.5 | 0.75 | 0.46 | 0.24 | 0.52 | 0.63 |
| 1 | 113898769 | rs1006610 | 0.678 | 0.782 | -1.768 | -0.716 | 0.006 | 0.094 | 0.094 | 0 | 0.49 | 0.66 | 0.45 | 0.24 | 0.52 | 0.63 |
| 1 | 113914023 | rs6664279 | -0.089 | 0.517 | -2.174 | -0.753 | 0.021 | -0.034 | -0.034 | 0 | 0.25 | 0.15 | 0.34 | 0.56 | 0.18 | 0.17 |
| 1 | 113918302 | rs4839318 | -1.004 | 0.737 | -2.11 | -0.98 | 0.026 | 0.14 | 0.14 | 0 | 0.45 | 0.23 | 0.5 | 0.79 | 0.38 | 0.33 |
| 1 | 113919499 | rs17013288 | -0.56 | 0.683 | -1.953 | -0.738 | -0.001 | -0.013 | -0.013 | 0 | 0.29 | 0.22 | 0.38 | 0.59 | 0.29 | 0.22 |
| 1 | 113925870 | rs10218568 | 0.792 | 1.084 | -1.363 | -0.956 | 0.017 | -0.033 | -0.033 | 0 | 0.15 | 0.23 | 0.12 | 0.04 | 0.18 | 0.11 |
| 1 | 113931805 | rs1112543 | 0.653 | 0.724 | -1.987 | -0.879 | -0.033 | -0.109 | -0.109 | 0 | 0.18 | 0.01 | 0.3 | 0.6 | 0.18 | 0.13 |
| 1 | 113934304 | rs1342650 | 0.908 | 0.826 | -2.248 | -1.131 | -0.051 | -0.022 | -0.022 | 1.246 | 0.14 | 0.04 | 0.17 | 0.18 | 0.2 | 0.16 |
| 1 | 113945989 | rs6688827 | 0.086 | 0.796 | -2.15 | -1.136 | -0.002 | -0.025 | -0.025 | 0 | 0.23 | 0.17 | 0.34 | 0.61 | 0.24 | 0.17 |

| | | | | | | | | | | | | | | | | |
|---|---|---|---|---|---|---|---|---|---|---|---|---|---|---|---|---|
| 1 | 113949453 | rs11102625 | 1.321 | 0.562 | -2.136 | -1.298 | -0.003 | 0.166 | 0.166 | 0 | 0.5 | 0.71 | 0.43 | 0.2 | 0.5 | 0.62 |
| 1 | 113951107 | rs12023641 | 0.841 | 0.62 | -2.204 | -1.105 | -0.033 | -0.109 | -0.109 | 0 | 0.18 | 0.01 | 0.3 | 0.6 | 0.18 | 0.13 |
| 1 | 113959886 | rs11102627 | 0.184 | 0.607 | -2.18 | -1.096 | -0.006 | -0.035 | -0.035 | 0 | 0.23 | 0.15 | 0.33 | 0.61 | 0.24 | 0.17 |
| 1 | 113962031 | rs7552954 | 1.172 | 0.518 | -2.232 | -1.373 | -0.021 | 0.199 | 0.199 | 0 | 0.51 | 0.79 | 0.44 | 0.2 | 0.5 | 0.62 |
| 1 | 113971051 | rs1936930 | 1.172 | 0.433 | -2.202 | -1.34 | -0.021 | 0.199 | 0.199 | 0 | 0.51 | 0.79 | 0.44 | 0.2 | 0.5 | 0.62 |
| 1 | 114014105 | rs10157946 | 1.196 | 0.436 | -2.244 | -1.339 | -0.026 | 0.216 | 0.216 | 0 | 0.51 | 0.8 | 0.44 | 0.2 | 0.5 | 0.62 |
| 1 | 114048588 | rs4839322 | -0.923 | 0.281 | -1.965 | -0.957 | 0.023 | 0.059 | 0.059 | 0 | 0.89 | 0.98 | 0.94 | 0.99 | 0.94 | 0.95 |
| 1 | 114075796 | rs10858000 | -0.172 | 0.279 | -2.321 | -0.978 | 0.014 | -0.035 | -0.035 | 0 | 0.28 | 0.07 | 0.41 | 0.64 | 0.25 | 0.21 |
| 1 | 114078755 | rs12065882 | 0.505 | 0.231 | -1.993 | -1.679 | -0.001 | -0.034 | -0.034 | 0 | 0.19 | 0.23 | 0.16 | 0.02 | 0.17 | 0.21 |
| 1 | 114080071 | rs2153977 | 0.043 | 0.401 | -2.045 | -1.696 | 0.004 | -0.002 | -0.002 | 2.071 | 0.31 | 0.13 | 0.45 | 0.65 | 0.31 | 0.25 |
| 1 | 114085723 | rs7554019 | 0.42 | 0.45 | -2.83 | -2.286 | -0.026 | 0.073 | 0.073 | 0.946 | 0.62 | 0.17 | 0.67 | 0.75 | 0.66 | 0.66 |
| 1 | 114088416 | rs10745339 | 0.776 | 0.408 | -2.86 | -2.284 | 0.014 | -0.032 | -0.032 | 0 | 0.28 | 0.07 | 0.41 | 0.65 | 0.25 | 0.22 |
| 1 | 114095750 | rs34258372 | 1.935 | 0.408 | -2.819 | -2.272 | -0.066 | 0.148 | 0.148 | 0 | 0.31 | 0.02 | 0.21 | 0.09 | 0.35 | 0.4 |
| 1 | 114105855 | rs1343630 | -0.07 | 0.436 | -2.418 | -2.339 | -0.001 | -0.018 | -0.018 | 5.869 | 0.72 | 0.27 | 0.78 | 0.92 | 0.8 | 0.76 |
| 1 | 114108252 | rs1343128 | 0.771 | 0.466 | -2.435 | -2.229 | 0.018 | -0.033 | -0.033 | 0 | 0.28 | 0.07 | 0.41 | 0.64 | 0.25 | 0.22 |
| 1 | 114129474 | rs1113523 | -0.846 | 0.467 | -2.253 | -2.258 | 0.02 | -0.001 | -0.001 | 0.975 | 0.7 | 0.79 | 0.58 | 0.36 | 0.75 | 0.78 |
| 1 | 114134025 | rs1235000 | -0.327 | 0.547 | -2.392 | -2.391 | -0.001 | -0.016 | -0.016 | 0 | 0.3 | 0.69 | 0.27 | 0.08 | 0.28 | 0.28 |
| 1 | 114139991 | rs1217225 | -1.066 | 0.592 | -2.464 | -2.434 | -0.019 | 0.032 | 0.032 | 0.885 | 0.4 | 0.69 | 0.36 | 0.27 | 0.42 | 0.37 |
| 1 | 114151606 | rs4838994 | 0.914 | 0.457 | -2.484 | -2.406 | -0.053 | -0.048 | -0.048 | 0 | 0.09 | 0 | 0.1 | 0.19 | 0.14 | 0.09 |
| 1 | 114157023 | rs3761931 | 2.437 | 0.396 | -2.571 | -2.472 | -0.062 | 0.141 | 0.141 | 0 | 0.29 | 0.01 | 0.2 | 0.09 | 0.33 | 0.4 |
| 1 | 114166561 | rs6669008 | 0.925 | 0.421 | -2.557 | -2.47 | -0.072 | 0.102 | 0.102 | 0 | 0.4 | 0.11 | 0.31 | 0.28 | 0.47 | 0.5 |
| 1 | 114187371 | rs1777237 | -0.965 | 0.471 | -2.597 | -2.494 | 0.013 | -0.02 | -0.02 | 0 | 0.71 | 0.86 | 0.57 | 0.36 | 0.74 | 0.77 |
| 1 | 114196495 | rs2359173 | 2.429 | 0.475 | -2.606 | -2.503 | -0.06 | 0.149 | 0.149 | 0 | 0.3 | 0.01 | 0.2 | 0.09 | 0.33 | 0.4 |
| 1 | 114197693 | rs7515189 | 1.636 | 0.344 | -2.724 | -2.602 | -0.041 | 0.095 | 0.095 | 0.131 | 0.15 | 0.01 | 0.13 | 0 | 0.17 | 0.13 |
| 1 | 114216449 | rs1080307 | 2.456 | 0.411 | -2.435 | -2.534 | -0.06 | 0.155 | 0.155 | 0 | 0.3 | 0.01 | 0.2 | 0.09 | 0.33 | 0.41 |
| 1 | 114217527 | rs17274634 | 2.259 | 0.406 | -2.399 | -2.536 | -0.003 | 0.118 | 0.118 | 0 | 0.31 | 0.09 | 0.21 | 0.09 | 0.33 | 0.41 |
| 1 | 114217705 | rs7522138 | -1.271 | 0.456 | -2.415 | -2.55 | -0.037 | 0.074 | 0.074 | 1.85 | 0.41 | 0.77 | 0.36 | 0.27 | 0.42 | 0.38 |
| 1 | 114242122 | rs3752954 | 2.605 | 0.688 | -2.757 | -1.882 | -0.092 | -0.02 | -0.02 | 1.403 | 0.23 | 0.01 | 0.21 | 0.25 | 0.35 | 0.39 |
| 1 | 114243899 | rs1111695 | 0.106 | 0.395 | -1.959 | -1.408 | -0.036 | 0.129 | 0.129 | 0 | 0.61 | 0.82 | 0.48 | 0.17 | 0.6 | 0.68 |
| 1 | 114246453 | rs1936398 | -0.987 | 0.367 | -1.941 | -1.176 | 0.013 | -0.027 | -0.027 | 0 | 0.71 | 0.84 | 0.57 | 0.4 | 0.74 | 0.77 |
| 1 | 114282287 | rs11102678 | -0.516 | 0.376 | -1.606 | -1.215 | -0.001 | -0.032 | -0.032 | 0 | 0.29 | 0.66 | 0.26 | 0.07 | 0.28 | 0.27 |
| 1 | 114287902 | rs10858017 | -0.909 | 0.362 | -1.643 | -1.204 | 0.016 | -0.032 | -0.032 | 0 | 0.7 | 0.83 | 0.57 | 0.4 | 0.74 | 0.77 |
| 1 | 114295438 | rs2797414 | 0.175 | 0.381 | -1.654 | -1.226 | -0.031 | 0.13 | 0.13 | 0 | 0.61 | 0.82 | 0.48 | 0.17 | 0.6 | 0.68 |
| 1 | 114299516 | rs2359167 | 1.519 | 0.199 | -1.762 | -1.269 | -0.052 | -0.124 | -0.124 | 0 | 0.22 | 0.01 | 0.33 | 0.59 | 0.24 | 0.18 |
| 1 | 114302966 | rs2797412 | 0.693 | 0.321 | -1.67 | -1.215 | 0.015 | -0.034 | -0.034 | 0 | 0.29 | 0.16 | 0.43 | 0.6 | 0.26 | 0.23 |
| 1 | 114309172 | rs1935836 | 0.229 | 0.328 | -1.59 | -1.236 | -0.001 | -0.031 | -0.031 | 0 | 0.71 | 0.35 | 0.74 | 0.93 | 0.72 | 0.73 |
| 1 | 114312623 | rs2040041 | -0.181 | 0.341 | -1.47 | -1.255 | -0.01 | -0.066 | -0.066 | 0.22 | 0.74 | 0.4 | 0.8 | 0.93 | 0.79 | 0.73 |
| 1 | 114315493 | rs12144309 | 1.165 | 0.315 | -1.052 | -1.168 | -0.004 | -0.056 | -0.056 | 0 | 0.18 | 0.23 | 0.14 | 0.02 | 0.15 | 0.23 |
| 1 | 114318604 | rs12117799 | 1.643 | 0.329 | -1.876 | -1.168 | -0.001 | 0.031 | 0.031 | 0 | 0.16 | 0.08 | 0.12 | 0 | 0.15 | 0.14 |
| 1 | 114318727 | rs3789598 | -0.579 | 0.565 | -2.351 | -1.071 | -0.008 | 0.074 | 0.074 | 0 | 0.41 | 0.28 | 0.54 | 0.83 | 0.4 | 0.32 |
| 1 | 114323318 | rs3789599 | 2.178 | 0.558 | -2.35 | -1.067 | -0.057 | 0.168 | 0.168 | 0 | 0.31 | 0.02 | 0.23 | 0.09 | 0.37 | 0.42 |
| 1 | 114325914 | rs2636008 | -1.001 | 0.381 | -2.38 | -0.874 | -0.01 | 0.037 | 0.037 | 0 | 0.38 | 0.31 | 0.51 | 0.81 | 0.37 | 0.32 |
| 1 | 114330202 | rs2797409 | 0.019 | 0.137 | -2.332 | -0.878 | 0.015 | -0.011 | -0.011 | 0.191 | 0.31 | 0.25 | 0.44 | 0.61 | 0.26 | 0.23 |
| 1 | 114334222 | rs3789600 | -1.168 | 0.144 | -2.332 | -0.885 | -0.018 | 0.06 | 0.06 | 0 | 0.42 | 0.33 | 0.58 | 0.83 | 0.45 | 0.32 |
| 1 | 114341332 | rs6703387 | 2.273 | -0.018 | -2.234 | -0.785 | -0.06 | 0.15 | 0.15 | 0 | 0.3 | 0.01 | 0.2 | 0.09 | 0.33 | 0.41 |
| 1 | 114343336 | rs1217380 | -0.383 | -0.021 | -2.239 | -0.783 | 0.008 | -0.054 | -0.054 | 0 | 0.71 | 0.92 | 0.57 | 0.39 | 0.74 | 0.77 |
| 1 | 114344083 | rs1217379 | 0.862 | -0.02 | -2.236 | -0.786 | -0.025 | 0.086 | 0.086 | 0 | 0.59 | 0.7 | 0.42 | 0.17 | 0.55 | 0.68 |
| 1 | 114346904 | rs1217375 | 0.495 | -0.021 | -2.235 | -0.786 | -0.002 | -0.022 | -0.022 | 0.049 | 0.29 | 0.66 | 0.22 | 0.07 | 0.23 | 0.27 |
| 1 | 114351706 | rs6537798 | 0.894 | 0.042 | -2.269 | -0.749 | -0.015 | 0.049 | 0.049 | 0 | 0.58 | 0.65 | 0.42 | 0.17 | 0.55 | 0.68 |
| 1 | 114354654 | rs3195954 | 1.16 | 0.063 | -2.416 | -0.735 | -0.063 | -0.058 | -0.058 | 0 | 0.1 | 0 | 0.13 | 0.23 | 0.19 | 0.1 |
| 1 | 114356663 | rs3811021 | 1.016 | 0.088 | -2.404 | -0.723 | -0.032 | -0.029 | -0.029 | 0 | 0.11 | 0.05 | 0.14 | 0.23 | 0.19 | 0.1 |
| 1 | 114363459 | rs2476599 | 0.313 | 0.064 | -2.415 | -0.737 | -0.001 | -0.019 | -0.019 | 0 | 0.3 | 0.66 | 0.22 | 0.07 | 0.23 | 0.27 |
| 1 | 114364476 | rs1217388 | -0.36 | 0.14 | -2.414 | -0.702 | 0.008 | -0.054 | -0.054 | 0 | 0.71 | 0.92 | 0.57 | 0.39 | 0.74 | 0.78 |
| 1 | 114365752 | rs1217389 | -0.304 | 0.137 | -2.416 | -0.703 | 0.01 | -0.052 | -0.052 | 3.854 | 0.7 | 0.91 | 0.57 | 0.39 | 0.74 | 0.78 |
| 1 | 114366434 | rs3789607 | 2.322 | 0.179 | -2.397 | -0.686 | -0.06 | 0.15 | 0.15 | 0 | 0.3 | 0.01 | 0.2 | 0.09 | 0.33 | 0.41 |
| 1 | 114369259 | rs2797416 | 1.025 | 0.458 | -2.096 | -0.383 | -0.025 | 0.086 | 0.086 | 0 | 0.59 | 0.7 | 0.42 | 0.17 | 0.55 | 0.68 |
| 1 | 114369734 | rs2476600 | 1.018 | 0.469 | -2.092 | -0.378 | -0.026 | 0.085 | 0.085 | 0 | 0.59 | 0.7 | 0.42 | 0.17 | 0.55 | 0.68 |
| 1 | 114374435 | rs1217395 | -0.17 | 0.52 | -2.39 | -0.345 | 0.007 | -0.054 | -0.054 | 0 | 0.71 | 0.92 | 0.57 | 0.39 | 0.74 | 0.78 |
| 1 | 114394463 | rs3765598 | 0.805 | 0.67 | -2.215 | -0.19 | -0.03 | -0.028 | -0.028 | 0.029 | 0.1 | 0.05 | 0.14 | 0.23 | 0.19 | 0.1 |
| 1 | 114417843 | rs6665194 | -1.263 | 0.19 | -2.219 | -0.663 | -0.007 | 0.057 | 0.057 | 0 | 0.41 | 0.29 | 0.58 | 0.82 | 0.43 | 0.31 |
| 1 | 114418205 | rs1217385 | 0.982 | 0.19 | -2.204 | -0.664 | -0.005 | 0.037 | 0.037 | 0 | 0.59 | 0.69 | 0.42 | 0.18 | 0.57 | 0.69 |
| 1 | 114420328 | rs12566340 | -0.019 | -0.147 | -2.408 | -0.516 | -0.021 | -0.095 | -0.095 | 0 | 0.28 | 0.03 | 0.41 | 0.61 | 0.25 | 0.22 |
| 1 | 114420462 | rs7529353 | -0.291 | -0.142 | -2.378 | -0.512 | 0.016 | -0.012 | -0.012 | 0 | 0.3 | 0.22 | 0.44 | 0.61 | 0.25 | 0.22 |
| 1 | 114421313 | rs11102691 | -0.051 | -0.055 | -2.678 | -0.441 | 0.004 | -0.067 | -0.067 | 0.085 | 0.28 | 0.07 | 0.41 | 0.61 | 0.25 | 0.21 |
| 1 | 114423653 | rs1217382 | -0.086 | 0.436 | -2.722 | -0.462 | 0.01 | 0.205 | 0.205 | 0.67 | 0.39 | 0.9 | 0.26 | 0.08 | 0.26 | 0.3 |
| 1 | 114429340 | rs1746860 | -0.152 | 0.396 | -2.657 | -0.466 | 0.031 | 0.148 | 0.148 | 0 | 0.39 | 0.8 | 0.25 | 0.08 | 0.27 | 0.31 |
| 1 | 114429461 | rs2358994 | 0.775 | 0.432 | -2.789 | -0.25 | -0.056 | -0.123 | -0.123 | 0 | 0.17 | 0.02 | 0.38 | 0.6 | 0.19 | 0.15 |
| 1 | 114430356 | rs1539438 | -2.022 | 0.482 | -2.832 | -0.296 | -0.066 | 0.133 | 0.133 | 0 | 0.68 | 0.99 | 0.76 | 0.86 | 0.65 | 0.55 |
| 1 | 114433659 | rs1217394 | -1.994 | 0.482 | -2.849 | -0.533 | -0.073 | 0.133 | 0.133 | 0 | 0.68 | 0.99 | 0.76 | 0.86 | 0.64 | 0.55 |

| | | | | | | | | | | | | | | | | |
|---|---|---|---|---|---|---|---|---|---|---|---|---|---|---|---|---|
| 1 | 114436482 | rs10776775 | -0.726 | 0.552 | -2.809 | -0.485 | -0.088 | 0.023 | 0.023 | 0 | 0.28 | 0.08 | 0.5 | 0.78 | 0.37 | 0.24 |
| 1 | 114436970 | rs10745340 | -0.771 | 0.544 | -2.811 | -0.488 | -0.087 | 0.025 | 0.025 | 0 | 0.28 | 0.08 | 0.5 | 0.78 | 0.37 | 0.24 |
| 1 | 114438951 | rs1217401 | -0.112 | 0.578 | -2.774 | -0.44 | 0.034 | 0.151 | 0.151 | 0 | 0.39 | 0.8 | 0.25 | 0.08 | 0.27 | 0.31 |
| 1 | 114443899 | rs17464525 | 0.54 | 0.913 | -2.614 | -0.179 | -0.025 | -0.032 | -0.032 | 0.219 | 0.11 | 0.06 | 0.11 | 0.18 | 0.18 | 0.1 |
| 1 | 114447565 | rs1217397 | 0.028 | 0.673 | -2.223 | -0.201 | 0.006 | 0.03 | 0.03 | 0 | 0.34 | 0.67 | 0.22 | 0.08 | 0.25 | 0.29 |
| 1 | 114449662 | rs3761936 | -0.36 | 0.986 | -2.193 | 0.759 | -0.057 | -0.124 | -0.124 | 4.189 | 0.17 | 0.02 | 0.38 | 0.59 | 0.19 | 0.15 |
| 1 | 114451307 | rs1217390 | -0.576 | 0.673 | -2.024 | 0.156 | 0.035 | 0.135 | 0.135 | 0 | 0.51 | 0.78 | 0.61 | 0.7 | 0.57 | 0.5 |
| 1 | 114455907 | rs6674384 | -1.034 | 0.626 | -1.291 | 0.476 | 0.005 | 0.025 | 0.025 | 0 | 0.2 | 0.34 | 0.22 | 0.24 | 0.25 | 0.15 |
| 1 | 114457769 | rs10858024 | 0.651 | 0.46 | -1.694 | -0.254 | 0 | -0.001 | -0.001 | 0.113 | 0.48 | 0.48 | 0.51 | 0.61 | 0.43 | 0.44 |
| 1 | 114466110 | rs11102703 | -0.701 | 0.445 | -1.651 | -0.247 | 0 | 0.027 | 0.027 | 0 | 0.22 | 0.43 | 0.23 | 0.25 | 0.25 | 0.14 |
| 1 | 114471189 | rs1000528 | -1.031 | 0.503 | -1.719 | -0.229 | -0.056 | 0.109 | 0.109 | 0.087 | 0.7 | 0.97 | 0.75 | 0.86 | 0.69 | 0.59 |
| 1 | 114475350 | rs17032011 | 1.247 | 0.763 | -1.82 | 0.019 | -0.032 | 0.035 | 0.035 | 1.67 | 0.06 | 0 | 0.19 | 0.51 | 0.08 | 0.06 |
| 1 | 114482520 | rs7519860 | -0.578 | 0.648 | -1.683 | -0.146 | 0.006 | 0.034 | 0.034 | 0 | 0.2 | 0.4 | 0.21 | 0.25 | 0.24 | 0.13 |
| 1 | 114489769 | rs4381184 | -1.146 | 0.697 | -1.773 | 0.079 | -0.056 | 0.055 | 0.055 | 0.042 | 0.7 | 0.97 | 0.65 | 0.82 | 0.69 | 0.59 |
| 1 | 114493583 | rs11102707 | -0.191 | 0.59 | -1.631 | 0.056 | 0.009 | 0.024 | 0.024 | 0 | 0.14 | 0.2 | 0.18 | 0.24 | 0.22 | 0.1 |
| 1 | 114501968 | rs11485101 | -0.361 | 0.372 | -1.009 | -0.066 | -0.008 | -0.031 | -0.031 | 0 | 0.12 | 0.33 | 0.06 | 0.03 | 0.08 | 0.08 |
| 1 | 114513488 | rs7522987 | -0.975 | 0.628 | -1.218 | 0.111 | -0.05 | 0.022 | 0.022 | 0 | 0.68 | 0.97 | 0.65 | 0.77 | 0.69 | 0.56 |
| 1 | 114515717 | rs2358996 | 0.652 | 0.768 | -1.123 | 0.121 | 0.005 | -0.001 | -0.001 | 0.258 | 0.49 | 0.55 | 0.44 | 0.52 | 0.43 | 0.46 |
| 1 | 114516330 | rs10108 | -0.289 | 0.482 | -1.086 | -0.054 | 0.01 | 0.024 | 0.024 | 0 | 0.15 | 0.22 | 0.18 | 0.24 | 0.23 | 0.1 |
| 1 | 114519351 | rs10732635 | 0.477 | 0.517 | -1.03 | 0.001 | -0.002 | 0.03 | 0.03 | 0.942 | 0.51 | 0.6 | 0.55 | 0.62 | 0.46 | 0.47 |
| 1 | 114520712 | rs10494163 | -0.824 | 0.482 | -0.931 | -0.048 | 0.008 | 0.03 | 0.03 | 0 | 0.16 | 0.26 | 0.2 | 0.25 | 0.23 | 0.11 |
| 1 | 114523465 | rs3789614 | 1.262 | 0.362 | -0.855 | -0.339 | -0.001 | -0.046 | -0.046 | 0 | 0.35 | 0.33 | 0.24 | 0.04 | 0.36 | 0.38 |
| 1 | 114524661 | rs7364 | -0.832 | 0.56 | -0.655 | 0.209 | 0.053 | 0.1 | 0.1 | 0 | 0.13 | 0.34 | 0.2 | 0.25 | 0.21 | 0.08 |
| 1 | 114527375 | rs17032048 | -0.537 | 0.445 | -0.689 | 0.346 | -0.003 | 0.039 | 0.039 | 0.702 | 0.14 | 0.16 | 0.19 | 0.47 | 0.1 | 0.08 |
| 1 | 114527420 | rs3006988 | 0.148 | 0.446 | -0.87 | 0.078 | -0.012 | -0.015 | -0.015 | 0.06 | 0.52 | 0.81 | 0.46 | 0.49 | 0.47 | 0.37 |
| 1 | 114529141 | rs924778 | -1.076 | 0.468 | -0.72 | 0.189 | -0.02 | 0.075 | 0.075 | 0.033 | 0.19 | 0.3 | 0.23 | 0.48 | 0.12 | 0.11 |
| 1 | 114533498 | rs1995483 | 0.118 | 0.518 | -0.829 | 0.221 | -0.002 | -0.01 | -0.01 | 0.248 | 0.23 | 0.16 | 0.22 | 0.31 | 0.29 | 0.16 |
| 1 | 114535293 | rs7520320 | -0.225 | 0.55 | -1.006 | 0.286 | 0.004 | -0.002 | -0.002 | 0 | 0.14 | 0.1 | 0.2 | 0.47 | 0.1 | 0.08 |
| 1 | 114539586 | rs3006977 | -1.051 | 0.609 | -0.988 | -0.068 | -0.05 | -0.005 | -0.005 | 0 | 0.77 | 0.96 | 0.68 | 0.8 | 0.77 | 0.58 |
| 1 | 114541498 | rs1503836 | -0.919 | 0.532 | -0.843 | -0.086 | 0 | 0 | 0 | 0 | 0.38 | 0.34 | 0.42 | 0.77 | 0.4 | 0.24 |
| 1 | 114541955 | rs12402202 | 0.992 | 0.799 | -0.639 | -0.142 | 0.001 | -0.135 | -0.135 | 0 | 0.37 | 0.49 | 0.26 | 0.02 | 0.38 | 0.38 |
| 1 | 114546528 | rs10494164 | -0.057 | 1.221 | -0.941 | 1.243 | -0.029 | -0.057 | -0.057 | 0 | 0.09 | 0 | 0.1 | 0.29 | 0.09 | 0.05 |
| 1 | 114546804 | rs2938323 | -0.116 | 1.24 | -0.759 | 1.187 | -0.046 | -0.072 | -0.072 | 0.365 | 0.74 | 0.94 | 0.59 | 0.68 | 0.75 | 0.57 |
| 1 | 114555118 | rs2938330 | -0.741 | 1.277 | 0.18 | 0.917 | 0.093 | 0.235 | 0.235 | 0 | 0.09 | 0.32 | 0.14 | 0.3 | 0.16 | 0.1 |
| 1 | 114555337 | rs4274061 | 0.566 | 0.732 | -0.208 | -0.367 | 0.01 | 0.022 | 0.022 | 0 | 0.51 | 0.36 | 0.3 | 0.13 | 0.45 | 0.37 |
| 1 | 114561433 | rs11582986 | -0.504 | 0.608 | -0.02 | 0.585 | -0.022 | -0.027 | -0.027 | 0.019 | 0.06 | 0 | 0.13 | 0.2 | 0.07 | 0.32 |
| 1 | 114577615 | rs4839003 | -0.47 | 0.653 | 0.771 | 0.453 | -0.038 | -0.031 | -0.031 | 0.179 | 0.07 | 0.01 | 0.13 | 0.31 | 0.14 | 0.34 |
| 1 | 114578835 | rs2938341 | 0.135 | 0.553 | 1.234 | 0.074 | 0.007 | -0.054 | -0.054 | 0 | 0.78 | 0.38 | 0.78 | 0.92 | 0.81 | 0.91 |
| 1 | 114585077 | rs2938337 | -0.244 | 0.504 | 1.27 | 0.063 | 0.016 | -0.037 | -0.037 | 0.666 | 0.81 | 0.67 | 0.8 | 0.92 | 0.81 | 0.9 |
| 1 | 114587447 | rs2938321 | 0.148 | 0.233 | 1.365 | 0.08 | -0.011 | -0.048 | -0.048 | 0 | 0.79 | 0.41 | 0.79 | 0.92 | 0.84 | 0.9 |
| 1 | 114587766 | rs11102720 | 1.082 | 0.048 | 1.34 | 0.098 | 0 | -0.012 | -0.012 | 0 | 0.67 | 0.29 | 0.65 | 0.56 | 0.69 | 0.53 |
| 1 | 114596480 | rs11102724 | 1.027 | 0.323 | 1.119 | 0.167 | 0.004 | 0.015 | 0.015 | 0.418 | 0.7 | 0.38 | 0.66 | 0.55 | 0.7 | 0.55 |
| 1 | 114598158 | rs17032172 | -0.458 | -0.275 | 0.663 | 0.084 | -0.055 | -0.096 | -0.096 | 0 | 0.23 | 0.31 | 0.11 | 0.03 | 0.1 | 0.09 |
| 1 | 114602971 | rs950779 | -0.097 | -0.161 | 0.447 | 0.145 | -0.035 | -0.068 | -0.068 | 0 | 0.2 | 0.24 | 0.1 | 0.02 | 0.09 | 0.07 |
| 1 | 114616228 | rs496267 | -1.524 | 1.27 | 0.708 | 1.148 | 0.054 | 0.186 | 0.186 | 0.061 | 0.1 | 0.22 | 0.14 | 0.36 | 0.13 | 0.35 |
| 1 | 114620453 | rs4240538 | -1.266 | 1.355 | -0.63 | 1.289 | 0 | 0.027 | 0.027 | 0.005 | 0.09 | 0.1 | 0.12 | 0.33 | 0.13 | 0.35 |
| 1 | 114625510 | rs12125801 | -0.855 | 1.467 | -0.892 | 1.222 | -0.034 | -0.015 | -0.015 | 0 | 0.08 | 0.03 | 0.16 | 0.37 | 0.13 | 0.36 |
| 1 | 114631980 | rs2432 | -0.981 | 1.596 | -1.414 | -0.015 | -0.114 | 0.167 | 0.167 | 0.187 | 0.34 | 0.15 | 0.51 | 0.91 | 0.46 | 0.61 |
| 1 | 114633098 | rs901146 | 0.096 | 1.542 | -1.061 | -0.07 | -0.003 | -0.024 | -0.024 | 0.024 | 0.11 | 0.12 | 0.05 | 0 | 0.06 | 0.03 |
| 1 | 114634228 | rs2279380 | -0.889 | 1.723 | -0.337 | 0.581 | 0.011 | 0.012 | 0.012 | 0 | 0.23 | 0.27 | 0.44 | 0.52 | 0.37 | 0.3 |
| 1 | 114634485 | rs1053135 | -0.93 | 1.667 | 0.335 | 0.558 | 0.036 | 0.039 | 0.039 | 0 | 0.23 | 0.34 | 0.36 | 0.52 | 0.37 | 0.29 |
| 1 | 114639939 | rs680714 | -0.764 | 1.425 | 0.604 | 0.846 | 0.004 | 0.009 | 0.009 | 0 | 0.14 | 0.18 | 0.15 | 0.12 | 0.14 | 0.16 |
| 1 | 114641685 | rs2774285 | -1.597 | 0.59 | -0.569 | -0.224 | 0.022 | 0.179 | 0.179 | 1.153 | 0.68 | 0.97 | 0.74 | 0.93 | 0.74 | 0.84 |
| 1 | 114642562 | rs6537810 | 0.958 | 0.622 | 0.224 | -0.188 | -0.001 | -0.083 | -0.083 | 0 | 0.24 | 0.28 | 0.19 | 0.01 | 0.26 | 0.21 |
| 1 | 114646763 | rs750906 | -0.029 | 0.591 | -0.01 | -0.017 | 0.002 | -0.05 | -0.05 | 0 | 0.38 | 0.47 | 0.34 | 0.14 | 0.4 | 0.37 |
| 1 | 114649247 | rs729300 | -0.754 | 1.088 | -0.47 | 0.402 | 0.042 | 0.052 | 0.052 | 1.68 | 0.56 | 0.76 | 0.68 | 0.62 | 0.61 | 0.67 |
| 1 | 114651076 | rs611514 | -0.391 | 0.988 | -0.351 | 0.77 | 0.008 | -0.002 | -0.002 | 0 | 0.4 | 0.73 | 0.46 | 0.61 | 0.41 | 0.58 |
| 1 | 114653036 | rs2629812 | -0.213 | 1.078 | -0.504 | 0.492 | 0.001 | 0 | 0 | 0 | 0.36 | 0.4 | 0.42 | 0.36 | 0.41 | 0.34 |
| 1 | 114653675 | rs676715 | -0.059 | 1.036 | -0.379 | 0.464 | -0.006 | -0.005 | -0.005 | 0 | 0.33 | 0.26 | 0.4 | 0.35 | 0.41 | 0.3 |
| 1 | 114655222 | rs519553 | -1.164 | 1.007 | -1.64 | 0.062 | -0.049 | -0.156 | -0.156 | 0.002 | 0.69 | 0.97 | 0.61 | 0.47 | 0.7 | 0.72 |
| 1 | 114659327 | rs552581 | -0.053 | 1.291 | -0.807 | -0.091 | -0.001 | 0.003 | 0.003 | 0.238 | 0.37 | 0.62 | 0.43 | 0.32 | 0.4 | 0.33 |
| 1 | 114659419 | rs588983 | -0.606 | 1.362 | -0.907 | -0.206 | 0.013 | 0.049 | 0.049 | 0 | 0.48 | 0.65 | 0.5 | 0.33 | 0.55 | 0.41 |
| 1 | 114662870 | rs479359 | 0.023 | 1.151 | -0.882 | 0.176 | 0.002 | -0.008 | -0.008 | 0.511 | 0.36 | 0.48 | 0.41 | 0.32 | 0.38 | 0.33 |
| 1 | 114663751 | rs2629810 | -0.095 | 1.338 | -1.243 | 0.117 | -0.011 | 0.023 | 0.023 | 0 | 0.72 | 0.84 | 0.74 | 0.81 | 0.68 | 0.84 |
| 1 | 114665586 | rs6683573 | -0.003 | 1.329 | -1.008 | -0.105 | -0.006 | 0.04 | 0.04 | 0.112 | 0.61 | 0.69 | 0.73 | 0.84 | 0.56 | 0.74 |
| 1 | 114668299 | rs7546417 | -0.354 | 2.008 | -0.368 | 0.698 | 0.02 | 0.011 | 0.011 | 0 | 0.21 | 0.29 | 0.31 | 0.27 | 0.25 | 0.25 |
| 1 | 114669174 | rs11581857 | -0.467 | 1.984 | -0.347 | 0.6 | 0.043 | 0.01 | 0.01 | 0 | 0.22 | 0.44 | 0.32 | 0.23 | 0.25 | 0.25 |
| 1 | 114670969 | rs2774307 | -0.961 | 2.113 | -0.871 | 0.524 | -0.022 | -0.003 | -0.003 | 2.618 | 0.87 | 0.94 | 0.87 | 0.9 | 0.76 | 0.94 |
| 1 | 114675629 | rs670495 | 0.032 | 1.374 | -1.408 | 0.511 | -0.012 | -0.004 | -0.004 | 0 | 0.47 | 0.21 | 0.57 | 0.57 | 0.48 | 0.65 |

| | | | | | | | | | | | | | | | | |
|---|---|---|---|---|---|---|---|---|---|---|---|---|---|---|---|---|
| 1 | 114676106 | rs581392 | -0.001 | 1.409 | -1.303 | 0.603 | -0.016 | 0.024 | 0.024 | 0 | 0.48 | 0.2 | 0.55 | 0.62 | 0.48 | 0.65 |
| 1 | 114676391 | rs684231 | -1.018 | 1.517 | -1.763 | 0.418 | -0.12 | -0.037 | -0.037 | 0 | 0.79 | 0.98 | 0.83 | 0.86 | 0.69 | 0.87 |
| 1 | 114676717 | rs476567 | 0.562 | 1.311 | -0.917 | 0.508 | -0.001 | 0.003 | 0.003 | 0 | 0.39 | 0.38 | 0.47 | 0.5 | 0.39 | 0.53 |
| 1 | 114677948 | rs529989 | 0.123 | 0.985 | -0.707 | 0.111 | 0 | 0.012 | 0.012 | 0 | 0.56 | 0.59 | 0.68 | 0.76 | 0.56 | 0.67 |
| 1 | 114677997 | rs11102738 | 1.174 | 1.09 | -0.797 | 0.101 | 0 | 0.048 | 0.048 | 0 | 0.2 | 0.09 | 0.11 | 0.05 | 0.18 | 0.1 |
| 1 | 114678786 | rs537478 | -0.33 | 1.871 | -0.59 | 0.479 | 0 | -0.002 | -0.002 | 0 | 0.44 | 0.48 | 0.43 | 0.48 | 0.48 | 0.36 |
| 1 | 114678857 | rs7527134 | 1.856 | 1.806 | -0.693 | 0.359 | 0.009 | 0.022 | 0.022 | 0 | 0.18 | 0.1 | 0.11 | 0 | 0.14 | 0.09 |
| 1 | 114679068 | rs901137 | 0.958 | 1.857 | -0.643 | 0.599 | -0.011 | 0.007 | 0.007 | 0 | 0.23 | 0.16 | 0.19 | 0.23 | 0.26 | 0.17 |
| 1 | 114680540 | rs17032441 | 0.651 | 1.682 | -0.795 | 0.675 | 0.001 | 0.003 | 0.003 | 4.015 | 0.2 | 0.22 | 0.16 | 0.26 | 0.21 | 0.16 |
| 1 | 114681932 | rs2629808 | -1.177 | 1.522 | -1.175 | 0.524 | -0.043 | -0.039 | -0.039 | 0.019 | 0.92 | 0.99 | 0.94 | 0.87 | 0.84 | 0.91 |
| 1 | 114682001 | rs734928 | 0.196 | 0.358 | -1.144 | -0.427 | -0.005 | -0.022 | -0.022 | 0 | 0.3 | 0.31 | 0.21 | 0.09 | 0.18 | 0.18 |
| 1 | 114684071 | rs4567275 | -0.882 | 0.751 | -1.269 | 0.397 | -0.073 | -0.11 | -0.11 | 0.922 | 0.84 | 0.97 | 0.74 | 0.44 | 0.72 | 0.68 |
| 1 | 114684419 | rs1119040 | 0.204 | 0.395 | -1.27 | 0.181 | 0 | -0.013 | -0.013 | 0 | 0.18 | 0.12 | 0.31 | 0.32 | 0.22 | 0.34 |
| 1 | 114687012 | rs2774298 | -0.354 | 0.249 | -0.62 | 0.016 | -0.023 | 0.139 | 0.139 | 0 | 0.13 | 0.37 | 0.23 | 0.3 | 0.12 | 0.28 |
| 1 | 114687316 | rs2629824 | -0.863 | 0.575 | -1.133 | -0.08 | -0.091 | -0.124 | -0.124 | 0.095 | 0.84 | 0.97 | 0.73 | 0.46 | 0.71 | 0.67 |
| 1 | 114688326 | rs17475914 | 1.261 | -0.044 | -0.809 | -0.938 | -0.067 | 0.097 | 0.097 | 2.497 | 0.16 | 0.01 | 0.11 | 0 | 0.19 | 0.05 |
| 1 | 114690985 | rs6687460 | 0.349 | -0.125 | -0.384 | -0.95 | 0.012 | -0.095 | -0.095 | 0 | 0.21 | 0.32 | 0.14 | 0 | 0.21 | 0.08 |
| 1 | 114697134 | rs2251405 | -0.56 | -1.339 | -0.296 | -0.683 | -0.042 | 0.054 | 0.054 | 0 | 0.31 | 0.45 | 0.4 | 0.53 | 0.24 | 0.52 |
| 1 | 114697867 | rs923124 | 0.544 | -1.235 | -0.5 | -0.596 | -0.038 | -0.097 | -0.097 | 0 | 0.13 | 0.02 | 0.32 | 0.52 | 0.19 | 0.43 |
| 1 | 114703427 | rs2629821 | -0.951 | -1.244 | -0.25 | -0.773 | -0.009 | 0.065 | 0.065 | 0.111 | 0.53 | 0.66 | 0.67 | 0.83 | 0.51 | 0.71 |
| 1 | 114706770 | rs2150203 | 0.51 | -1.074 | -0.346 | -0.662 | -0.003 | 0 | 0 | 0 | 0.14 | 0.11 | 0.33 | 0.53 | 0.19 | 0.43 |
| 1 | 114709094 | rs10081970 | 1.147 | -1.167 | -0.174 | -0.735 | -0.031 | -0.036 | -0.036 | 0.592 | 0.07 | 0.16 | 0.04 | 0 | 0.02 | 0.08 |
| 1 | 114709317 | rs497304 | 0.601 | -1.229 | -0.076 | -0.78 | 0.002 | 0.004 | 0.004 | 2.705 | 0.3 | 0.35 | 0.3 | 0.3 | 0.3 | 0.21 |
| 1 | 114713205 | rs480226 | 0.498 | -1.285 | -0.071 | -0.916 | -0.035 | -0.036 | -0.036 | 0 | 0.07 | 0.2 | 0.04 | 0 | 0.02 | 0.08 |
| 1 | 114713359 | rs17535387 | 0.838 | -1.448 | -0.593 | -1.166 | 0.027 | 0.064 | 0.064 | 0 | 0.2 | 0.05 | 0.09 | 0 | 0.12 | 0.09 |
| 1 | 114715584 | rs2774292 | 0.42 | -0.623 | 0.021 | -0.309 | -0.03 | -0.036 | -0.036 | 0 | 0.07 | 0.14 | 0.03 | 0 | 0.02 | 0.08 |
| 1 | 114720018 | rs631230 | -0.494 | -0.392 | -0.083 | -0.203 | -0.002 | -0.003 | -0.003 | 0 | 0.46 | 0.63 | 0.53 | 0.51 | 0.47 | 0.61 |
| 1 | 114721046 | rs6684831 | -1.263 | -0.827 | 0.513 | 0.404 | -0.036 | 0.145 | 0.145 | 0 | 0.17 | 0.42 | 0.15 | 0.42 | 0.13 | 0.44 |
| 1 | 114722644 | rs209878 | -1.168 | -0.494 | 0.1 | -0.428 | -0.015 | -0.072 | -0.072 | 1.828 | 0.12 | 0.52 | 0.06 | 0 | 0.03 | 0.03 |
| 1 | 114723933 | rs209876 | 0.268 | -0.119 | -0.971 | -0.229 | 0.009 | 0.037 | 0.037 | 2.045 | 0.82 | 0.9 | 0.91 | 1 | 0.86 | 0.96 |
| 1 | 114733635 | rs4839004 | 0.527 | 0.075 | -0.445 | 0.604 | -0.011 | 0.066 | 0.066 | 0.065 | 0.73 | 0.54 | 0.69 | 0.58 | 0.69 | 0.55 |
| 1 | 114735242 | rs1158366 | -0.402 | 0.531 | -0.987 | 0.805 | -0.013 | -0.018 | -0.018 | 0 | 0.26 | 0.16 | 0.48 | 0.36 | 0.26 | 0.32 |
| 1 | 114736559 | rs12046577 | 0.125 | 0.607 | -1.049 | 0.806 | 0.005 | -0.001 | -0.001 | 0.599 | 0.11 | 0.07 | 0.06 | 0.1 | 0.07 | 0.13 |
| 1 | 114741351 | rs430928 | 0.527 | -0.3 | -0.763 | 0.303 | -0.002 | -0.003 | -0.003 | 0.311 | 0.42 | 0.5 | 0.38 | 0.45 | 0.4 | 0.28 |
| 1 | 114742357 | rs378188 | 1.488 | -0.351 | -0.59 | 0.243 | 0.025 | 0.02 | 0.02 | 1.205 | 0.28 | 0.32 | 0.31 | 0.34 | 0.33 | 0.15 |
| 1 | 114743352 | rs1028574 | 0.739 | -0.727 | -0.803 | -0.247 | 0.003 | 0.01 | 0.01 | 0 | 0.12 | 0.12 | 0.08 | 0.01 | 0.14 | 0.11 |
| 1 | 114749024 | rs148210 | 0.875 | -0.842 | -0.7 | -0.11 | 0.01 | -0.006 | -0.006 | 0.15 | 0.42 | 0.46 | 0.39 | 0.37 | 0.47 | 0.26 |
| 1 | 114749081 | rs768566 | 0.777 | -0.868 | -0.865 | -0.142 | -0.044 | -0.039 | -0.039 | 1.441 | 0.14 | 0.03 | 0.31 | 0.25 | 0.19 | 0.18 |
| 1 | 114750579 | rs6671620 | -0.854 | -0.824 | -0.83 | -0.321 | 0 | 0.009 | 0.009 | 0 | 0.37 | 0.32 | 0.32 | 0.35 | 0.38 | 0.39 |
| 1 | 114750905 | rs6537816 | -0.471 | -1.155 | -0.5 | -0.134 | -0.045 | -0.001 | -0.001 | 0 | 0.74 | 0.55 | 0.79 | 0.76 | 0.78 | 0.67 |
| 1 | 114753010 | rs12029255 | -0.415 | -1.16 | -0.72 | -0.181 | 0.007 | -0.026 | -0.026 | 0 | 0.28 | 0.23 | 0.26 | 0.4 | 0.31 | 0.48 |
| 1 | 114753083 | rs1385665 | 1.812 | -1.073 | -0.681 | -0.254 | -0.015 | -0.019 | -0.019 | 0 | 0.35 | 0.19 | 0.32 | 0.38 | 0.39 | 0.25 |
| 1 | 114753905 | rs11586444 | 0.175 | -1.12 | -0.696 | -0.171 | 0.003 | 0.028 | 0.028 | 0 | 0.21 | 0.15 | 0.15 | 0.14 | 0.21 | 0.21 |
| 1 | 114754578 | rs12068395 | 0.787 | -1.023 | -0.352 | 0.047 | 0.001 | 0.088 | 0.088 | 0 | 0.08 | 0.18 | 0.12 | 0.26 | 0.12 | 0.27 |
| 1 | 114756957 | rs176662 | -1.373 | -1.274 | -0.196 | -0.198 | 0.011 | -0.006 | -0.006 | 3.256 | 0.84 | 0.89 | 0.92 | 1 | 0.91 | 0.94 |
| 1 | 114757562 | rs7553373 | 0.469 | -1.124 | -0.173 | 0.01 | -0.003 | 0.063 | 0.063 | 0 | 0.1 | 0.18 | 0.12 | 0.27 | 0.14 | 0.33 |
| 1 | 114762107 | rs1485594 | 1.186 | -1.257 | -0.289 | -0.231 | 0.007 | -0.006 | -0.006 | 0 | 0.53 | 0.28 | 0.6 | 0.5 | 0.57 | 0.46 |
| 1 | 114764538 | rs6679065 | 1.21 | -1.515 | -0.497 | -0.328 | -0.004 | -0.018 | -0.018 | 0 | 0.58 | 0.28 | 0.61 | 0.5 | 0.58 | 0.49 |
| 1 | 114766687 | rs10776781 | 0.872 | -1.508 | -0.545 | -0.414 | -0.013 | -0.027 | -0.027 | 0 | 0.6 | 0.47 | 0.72 | 0.74 | 0.73 | 0.74 |
| 1 | 114773981 | rs7550248 | -0.725 | -1.412 | -0.372 | -0.323 | -0.043 | -0.088 | -0.088 | 0 | 0.83 | 0.71 | 0.92 | 1 | 0.91 | 0.93 |
| 1 | 114776791 | rs11102757 | 1.596 | -1.181 | -0.751 | -0.111 | -0.015 | -0.002 | -0.002 | 0.831 | 0.38 | 0.22 | 0.49 | 0.37 | 0.43 | 0.28 |
| 1 | 114780835 | rs6670053 | 0.134 | -0.824 | -0.44 | 0.403 | 0.003 | -0.012 | -0.012 | 0.024 | 0.11 | 0.08 | 0.12 | 0.23 | 0.13 | 0.35 |
| 1 | 114795885 | rs416908 | -1.521 | -0.312 | -0.448 | 0.319 | 0 | -0.002 | -0.002 | 0 | 0.4 | 0.66 | 0.43 | 0.48 | 0.34 | 0.6 |
| 1 | 114797698 | rs416190 | -0.56 | -0.348 | -0.43 | 0.315 | -0.012 | -0.004 | -0.004 | 0.028 | 0.28 | 0.36 | 0.26 | 0.22 | 0.24 | 0.47 |
| 1 | 114799870 | rs12146102 | 1.035 | -0.238 | -0.469 | 0.343 | -0.004 | 0.001 | 0.001 | 0 | 0.09 | 0.05 | 0.14 | 0.26 | 0.1 | 0.13 |
| 1 | 114801953 | rs2220211 | 0.333 | -0.288 | -0.487 | 0.211 | -0.01 | -0.055 | -0.055 | 0.084 | 0.2 | 0.24 | 0.09 | 0 | 0.11 | 0.1 |
| 1 | 114803749 | rs12562994 | 0.828 | 0.132 | -0.772 | 0.715 | -0.026 | -0.037 | -0.037 | 2.678 | 0.07 | 0.01 | 0.11 | 0.21 | 0.13 | 0.36 |
| 1 | 114806793 | rs6537822 | -1.215 | 0.236 | -0.67 | 0.321 | 0.006 | 0.006 | 0.006 | 0 | 0.69 | 0.69 | 0.61 | 0.64 | 0.65 | 0.79 |
| 1 | 114807559 | rs11589933 | -0.496 | 0.1 | -0.994 | -0.258 | -0.005 | 0.048 | 0.048 | 0 | 0.29 | 0.17 | 0.2 | 0.16 | 0.3 | 0.19 |
| 1 | 114808631 | rs12030629 | -0.386 | 0.167 | -1.314 | -0.222 | -0.027 | 0.06 | 0.06 | 0 | 0.28 | 0.09 | 0.19 | 0.16 | 0.3 | 0.19 |
| 1 | 114818994 | rs6664266 | -0.545 | 0.222 | -0.607 | 0.209 | -0.017 | -0.004 | -0.004 | 0 | 0.64 | 0.74 | 0.37 | 0.58 | 0.55 | 0.7 |
| 1 | 114828360 | rs11804407 | -0.515 | -0.13 | 0.11 | -0.287 | -0.01 | -0.035 | -0.035 | 3.382 | 0.15 | 0.37 | 0.29 | 0.09 | 0.12 | 0.08 |
| 1 | 114833410 | rs11588687 | 1.053 | -0.399 | -0.147 | -0.85 | 0.004 | 0.005 | 0.005 | 0 | 0.69 | 0.3 | 0.61 | 0.91 | 0.76 | 0.85 |
| 1 | 114833941 | rs535288 | -0.566 | -0.378 | -0.047 | -0.849 | -0.019 | 0.035 | 0.035 | 0 | 0.88 | 0.96 | 0.93 | 1 | 0.89 | 0.97 |
| 1 | 114835090 | rs6695414 | 0.739 | -0.406 | 0.435 | -0.877 | -0.001 | -0.037 | -0.037 | 0 | 0.73 | 0.66 | 0.64 | 0.92 | 0.76 | 0.9 |
| 1 | 114837021 | rs10489667 | 1.564 | 0.003 | -0.312 | -0.489 | -0.04 | -0.056 | -0.056 | 0 | 0.18 | 0.09 | 0.22 | 0.38 | 0.28 | 0.28 |
| 1 | 114837975 | rs714569 | 0.935 | 0.119 | -0.22 | -1.266 | 0.005 | 0.019 | 0.019 | 0 | 0.61 | 0.34 | 0.59 | 0.73 | 0.68 | 0.65 |
| 1 | 114857644 | rs6689069 | 1.242 | 0.485 | -0.685 | -1.46 | -0.007 | -0.092 | -0.092 | 0 | 0.76 | 0.34 | 0.84 | 0.99 | 0.82 | 0.89 |
| 1 | 114861832 | rs634405 | -0.355 | 0.668 | -0.647 | -1.278 | 0.004 | -0.04 | -0.04 | 0.146 | 0.12 | 0.15 | 0.09 | 0 | 0.12 | 0.02 |

| | | | | | | | | | | | | | | | | |
|---|---|---|---|---|---|---|---|---|---|---|---|---|---|---|---|---|
| 1 | 114865343 | rs668800 | -0.759 | 0.103 | -1.262 | -1.179 | -0.005 | 0.072 | 0.072 | 0.42 | 0.47 | 0.67 | 0.33 | 0.15 | 0.47 | 0.33 |
| 1 | 114873168 | rs642746 | -1.155 | -0.447 | -0.139 | -0.442 | -0.044 | -0.097 | -0.097 | 0 | 0.15 | 0.62 | 0.07 | 0 | 0.06 | 0.07 |
| 1 | 114875359 | rs570419 | -0.392 | -0.887 | -0.67 | -0.409 | 0.022 | 0.017 | 0.017 | 0 | 0.32 | 0.3 | 0.19 | 0.19 | 0.19 | 0.32 |
| 1 | 114878783 | rs11102782 | -0.267 | -0.714 | -0.644 | -0.679 | -0.005 | -0.017 | -0.017 | 0 | 0.22 | 0.25 | 0.22 | 0.13 | 0.22 | 0.16 |
| 1 | 114880092 | rs4448517 | -0.437 | -0.78 | -0.401 | -0.699 | -0.017 | -0.043 | -0.043 | 0 | 0.24 | 0.34 | 0.22 | 0.13 | 0.22 | 0.16 |
| 1 | 114884179 | rs486755 | -0.093 | -0.593 | -0.661 | -0.253 | -0.002 | 0.008 | 0.008 | 0.144 | 0.4 | 0.43 | 0.48 | 0.54 | 0.42 | 0.27 |
| 1 | 114885239 | rs9429736 | 0.092 | -0.406 | -1.289 | 0.142 | 0.01 | -0.016 | -0.016 | 0.034 | 0.1 | 0.04 | 0.05 | 0.24 | 0.09 | 0.09 |
| 1 | 114885641 | rs662353 | -0.298 | 0.343 | -1.336 | -0.245 | -0.04 | 0.038 | 0.038 | 0.141 | 0.78 | 0.89 | 0.75 | 0.9 | 0.69 | 0.67 |
| 1 | 114888412 | rs1535035 | -0.111 | -1.019 | -0.813 | -0.021 | -0.017 | 0.001 | 0.001 | 0 | 0.45 | 0.51 | 0.33 | 0.5 | 0.25 | 0.45 |
| 1 | 114893146 | rs574339 | -0.356 | -0.876 | -1.194 | -0.154 | 0.071 | 0.03 | 0.03 | 0.34 | 0.49 | 0.69 | 0.47 | 0.56 | 0.29 | 0.48 |
| 1 | 114894805 | rs516610 | -0.455 | -0.053 | -1.229 | -0.419 | -0.024 | -0.001 | -0.001 | 0.062 | 0.64 | 0.78 | 0.39 | 0.31 | 0.44 | 0.63 |
| 1 | 114900678 | rs556765 | 0.055 | -0.261 | -1.261 | -0.72 | -0.011 | -0.115 | -0.115 | 0 | 0.42 | 0.51 | 0.24 | 0.02 | 0.32 | 0.29 |
| 1 | 114906992 | rs1989138 | -0.008 | -0.301 | -1.486 | 0.355 | 0.02 | -0.029 | -0.029 | 0 | 0.2 | 0.13 | 0.14 | 0.28 | 0.12 | 0.34 |
| 1 | 114920097 | rs12753188 | -0.131 | -0.096 | -1.946 | -0.438 | -0.025 | 0.093 | 0.093 | 0 | 0.18 | 0.02 | 0.11 | 0.03 | 0.18 | 0.15 |
| 1 | 114927201 | rs9429541 | -1.126 | -0.5 | -1.505 | -1.462 | -0.003 | -0.065 | -0.065 | 0 | 0.4 | 0.48 | 0.24 | 0.04 | 0.32 | 0.33 |
| 1 | 114941326 | rs3789615 | -1.219 | -1.103 | -1.025 | -1.049 | -0.017 | -0.02 | -0.02 | 0.206 | 0.56 | 0.82 | 0.43 | 0.43 | 0.44 | 0.76 |
| 1 | 114948281 | rs6537825 | -1.449 | -1.449 | -1.477 | -1.153 | 0.003 | -0.081 | -0.081 | 0 | 0.87 | 0.98 | 0.8 | 0.58 | 0.94 | 0.93 |
| 1 | 114948638 | rs6677898 | 1.678 | -1.414 | -1.333 | -1.326 | 0.002 | -0.017 | -0.017 | 0 | 0.07 | 0.07 | 0.03 | 0 | 0.06 | 0.05 |
| 1 | 114953420 | rs6694768 | -0.404 | -1.26 | -1.322 | -1.334 | -0.002 | -0.058 | -0.058 | 0.625 | 0.39 | 0.47 | 0.24 | 0.04 | 0.33 | 0.33 |
| 1 | 114959888 | rs7547836 | 0.079 | -1.326 | -1.313 | -1.366 | 0.004 | -0.002 | -0.002 | 0.695 | 0.31 | 0.29 | 0.17 | 0.04 | 0.25 | 0.26 |
| 1 | 114973429 | rs12083584 | -0.613 | -1.333 | -1.316 | -1.378 | -0.003 | -0.062 | -0.062 | 0 | 0.39 | 0.47 | 0.24 | 0.04 | 0.33 | 0.33 |
| 1 | 114973690 | rs11585926 | 0.539 | -1.351 | -1.314 | -1.388 | -0.002 | 0.055 | 0.055 | 0 | 0.27 | 0.19 | 0.15 | 0.03 | 0.24 | 0.2 |
| 1 | 114989211 | rs2143583 | -0.106 | -1.272 | -1.272 | -1.35 | -0.005 | -0.059 | -0.059 | 0 | 0.33 | 0.41 | 0.18 | 0.04 | 0.25 | 0.26 |
| 1 | 114996112 | rs11102794 | -0.659 | -1.285 | -1.274 | -1.344 | -0.002 | -0.061 | -0.061 | 0 | 0.39 | 0.47 | 0.24 | 0.04 | 0.32 | 0.33 |
| 1 | 115012189 | rs17593470 | 0.358 | -1.02 | -1.189 | -1.169 | -0.002 | -0.02 | -0.02 | 0 | 0.13 | 0.13 | 0.08 | 0.01 | 0.13 | 0.12 |
| 1 | 115052708 | rs3827735 | 1.202 | -1.071 | -1.077 | -0.908 | -0.016 | 0.157 | 0.157 | 0.075 | 0.17 | 0.35 | 0.19 | 0.38 | 0.12 | 0.44 |
| 1 | 115061584 | rs11102807 | -0.276 | -0.42 | -1.525 | -1.778 | -0.002 | 0.014 | 0.014 | 0 | 0.46 | 0.44 | 0.31 | 0.03 | 0.47 | 0.35 |
| 1 | 115064304 | rs12128188 | 0.942 | -0.557 | -1.704 | -1.939 | -0.008 | 0.032 | 0.032 | 2.514 | 0.18 | 0.07 | 0.11 | 0 | 0.18 | 0.07 |
| 1 | 115073780 | rs10858046 | -0.709 | 0.094 | -1.641 | -1.809 | 0.007 | 0.014 | 0.014 | 0.343 | 0.66 | 0.48 | 0.54 | 0.49 | 0.6 | 0.41 |
| 1 | 115073876 | rs10858047 | 1.288 | 0.268 | -1.85 | -1.719 | -0.043 | -0.062 | -0.062 | 0 | 0.11 | 0.02 | 0.22 | 0.46 | 0.14 | 0.07 |
| 1 | 115079658 | rs11587400 | 1.786 | 0.225 | -2.006 | -1.886 | -0.101 | 0.002 | 0.002 | 0 | 0.15 | 0.01 | 0.18 | 0.09 | 0.31 | 0.11 |
| 1 | 115108738 | rs12040830 | 1.275 | 0.057 | -1.943 | -1.736 | -0.043 | -0.066 | -0.066 | 0.311 | 0.11 | 0.01 | 0.22 | 0.48 | 0.13 | 0.07 |
| 1 | 115123747 | rs1564824 | 1.301 | 0.02 | -2.006 | -1.778 | -0.043 | -0.065 | -0.065 | 0 | 0.11 | 0.01 | 0.22 | 0.48 | 0.13 | 0.07 |
| 1 | 115128087 | rs222493 | -1.531 | 0.023 | -2.005 | -1.757 | -0.043 | -0.066 | -0.066 | 0 | 0.89 | 0.99 | 0.78 | 0.52 | 0.87 | 0.93 |
| 1 | 115129123 | rs6700833 | 1.179 | -0.549 | -2.197 | -2.342 | 0.016 | 0.043 | 0.043 | 0.331 | 0.33 | 0.2 | 0.18 | 0 | 0.3 | 0.16 |
| 1 | 115136771 | rs222495 | 2.282 | -0.769 | -1.277 | -2.476 | -0.006 | 0.07 | 0.07 | 0 | 0.13 | 0.21 | 0.27 | 0.39 | 0.1 | 0.44 |
| 1 | 115146488 | rs222499 | 0.812 | -0.796 | -1.328 | -2.519 | 0.029 | 0.057 | 0.057 | 0.046 | 0.31 | 0.49 | 0.46 | 0.48 | 0.39 | 0.54 |
| 1 | 115152814 | rs11102824 | 1.243 | -0.675 | -1.548 | -2.468 | 0.016 | 0.043 | 0.043 | 0.331 | 0.33 | 0.2 | 0.18 | 0 | 0.3 | 0.16 |
| 1 | 115163388 | rs11102825 | 1.85 | -0.744 | -1.666 | -2.48 | -0.058 | 0.184 | 0.184 | 0 | 0.27 | 0.02 | 0.15 | 0 | 0.29 | 0.16 |
| 1 | 115168097 | rs12136548 | 2.578 | -0.553 | -1.509 | -2.341 | -0.078 | 0.007 | 0.007 | 0 | 0.14 | 0.02 | 0.18 | 0.09 | 0.29 | 0.1 |
| 1 | 115169866 | rs178445 | -1.269 | -0.614 | -1.457 | -2.314 | -0.007 | 0 | 0 | 0.673 | 0.86 | 0.86 | 0.76 | 0.52 | 0.83 | 0.92 |
| 1 | 115174843 | rs9308267 | -1.952 | -0.611 | -1.467 | -2.297 | -0.036 | -0.056 | -0.056 | 0 | 0.89 | 0.98 | 0.78 | 0.52 | 0.87 | 0.93 |
| 1 | 115180672 | rs6537838 | 1.813 | -0.762 | -1.524 | -2.476 | 0 | 0.012 | 0.012 | 0 | 0.28 | 0.18 | 0.16 | 0 | 0.29 | 0.16 |
| 1 | 115185923 | rs10489524 | 1.101 | -0.611 | -1.395 | -2.381 | 0 | -0.014 | -0.014 | 0.088 | 0.17 | 0.2 | 0.09 | 0.02 | 0.12 | 0.1 |
| 1 | 115196166 | rs10858049 | 2.431 | -0.588 | -1.286 | -2.374 | -0.026 | 0.012 | 0.012 | 0 | 0.15 | 0.07 | 0.18 | 0.09 | 0.29 | 0.1 |
| 1 | 115200808 | rs6537841 | -1.883 | -0.438 | -1.112 | -2.159 | -0.043 | -0.065 | -0.065 | 0 | 0.89 | 0.99 | 0.78 | 0.52 | 0.87 | 0.93 |
| 1 | 115212572 | rs11803022 | 0.86 | -0.619 | -0.891 | -2.068 | 0.008 | 0.011 | 0.011 | 0 | 0.17 | 0.16 | 0.09 | 0.02 | 0.12 | 0.1 |
| 1 | 115218413 | rs2070986 | 1.977 | -0.498 | -0.741 | -1.889 | -0.006 | 0.08 | 0.08 | 0 | 0.13 | 0.21 | 0.27 | 0.39 | 0.1 | 0.43 |
| 1 | 115227493 | rs2268697 | 0.845 | -0.541 | -0.948 | -1.891 | 0.029 | 0.055 | 0.055 | 0 | 0.32 | 0.51 | 0.46 | 0.47 | 0.38 | 0.57 |
| 1 | 115227600 | rs2268698 | 0.199 | -0.66 | -1.035 | -1.676 | 0.037 | -0.085 | -0.085 | 0 | 0.2 | 0.07 | 0.22 | 0.51 | 0.1 | 0.14 |
| 1 | 115234632 | rs12038990 | 0.338 | -0.622 | -1.036 | -1.66 | 0.029 | -0.079 | -0.079 | 0 | 0.18 | 0.06 | 0.21 | 0.5 | 0.09 | 0.1 |
| 1 | 115235010 | rs2010899 | -0.701 | -0.503 | -1.23 | -2.068 | -0.031 | 0.102 | 0.102 | 0 | 0.6 | 0.79 | 0.75 | 1 | 0.55 | 0.81 |
| 1 | 115238982 | rs6667218 | -1.577 | -0.499 | -1.384 | -1.666 | -0.021 | -0.005 | -0.005 | 1.602 | 0.74 | 0.81 | 0.82 | 1 | 0.66 | 0.85 |
| 1 | 115239355 | rs11588144 | 0.193 | -0.678 | -1.312 | -1.593 | -0.005 | 0.008 | 0.008 | 0 | 0.51 | 0.71 | 0.58 | 0.49 | 0.56 | 0.76 |
| 1 | 115239816 | rs926938 | 0.597 | -0.689 | -1.196 | -1.599 | -0.001 | -0.002 | -0.002 | 0 | 0.44 | 0.54 | 0.55 | 0.48 | 0.51 | 0.67 |
| 1 | 115249843 | rs14804 | 1.072 | -0.655 | -1.19 | -1.675 | -0.025 | 0.093 | 0.093 | 0 | 0.26 | 0.12 | 0.15 | 0.02 | 0.25 | 0.23 |
| 1 | 115256669 | rs969273 | -1.495 | -0.618 | -1.301 | -1.719 | -0.133 | 0.171 | 0.171 | 0 | 0.75 | 0.98 | 0.82 | 1 | 0.67 | 0.85 |
| 1 | 115259599 | rs8453 | 1.162 | -0.337 | -1.113 | -1.498 | -0.036 | -0.027 | -0.027 | 0 | 0.06 | 0 | 0.11 | 0.08 | 0.16 | 0.07 |
| 1 | 115259768 | rs1065634 | 2.191 | -0.501 | -1.209 | -1.567 | -0.003 | 0.04 | 0.04 | 0 | 0.1 | 0.01 | 0.05 | 0.02 | 0.07 | 0.05 |
| 1 | 115263685 | rs10489525 | 1.33 | -0.74 | -0.715 | -1.534 | -0.007 | 0.023 | 0.023 | 0 | 0.29 | 0.36 | 0.38 | 0.38 | 0.29 | 0.53 |
| 1 | 115263791 | rs6692870 | -1.986 | -0.788 | -0.728 | -1.493 | -0.022 | -0.092 | -0.092 | 0 | 0.8 | 0.94 | 0.75 | 0.5 | 0.8 | 0.88 |
| 1 | 115269211 | rs2144428 | 0.729 | -0.716 | -0.578 | -1.442 | 0.01 | 0.023 | 0.023 | 0 | 0.36 | 0.41 | 0.5 | 0.46 | 0.47 | 0.61 |
| 1 | 115283197 | rs10858052 | 0.14 | -0.607 | -0.394 | -1.666 | 0.007 | 0.032 | 0.032 | 0 | 0.32 | 0.27 | 0.18 | 0.02 | 0.26 | 0.23 |
| 1 | 115287833 | rs12130524 | 1.812 | -0.47 | -0.405 | -1.603 | -0.068 | 0.123 | 0.123 | 0 | 0.2 | 0.01 | 0.11 | 0 | 0.2 | 0.12 |
| 1 | 115293527 | rs10489526 | 1.812 | -0.47 | -0.405 | -1.603 | -0.068 | 0.123 | 0.123 | 0 | 0.2 | 0.01 | 0.11 | 0 | 0.2 | 0.12 |
| 1 | 115300685 | rs6681671 | 1.11 | -0.361 | -0.231 | -1.441 | -0.004 | 0.002 | 0.002 | 0 | 0.26 | 0.24 | 0.16 | 0.02 | 0.25 | 0.23 |
| 1 | 115302879 | rs12140675 | 1.175 | -0.364 | -0.314 | -1.43 | -0.02 | 0.09 | 0.09 | 0 | 0.26 | 0.12 | 0.15 | 0.02 | 0.25 | 0.23 |
| 1 | 115309512 | rs2207701 | -0.247 | -0.338 | -0.109 | -1.426 | 0.005 | -0.028 | -0.028 | 0 | 0.44 | 0.45 | 0.25 | 0.04 | 0.33 | 0.28 |
| 1 | 115316158 | rs3827739 | -0.437 | -0.261 | 0.111 | -1.297 | 0.021 | -0.011 | -0.011 | 0 | 0.19 | 0.34 | 0.19 | 0.1 | 0.24 | 0.27 |

| | | | | | | | | | | | | | | | | |
|---|---|---|---|---|---|---|---|---|---|---|---|---|---|---|---|---|
| 1 | 115342465 | rs2377415 | -1.201 | -0.311 | -0.317 | -1.229 | -0.026 | 0.062 | 0.062 | 0 | 0.62 | 0.83 | 0.44 | 0.15 | 0.56 | 0.53 |
| 1 | 115343885 | rs12126634 | 1.519 | -0.428 | -0.348 | -1.306 | -0.015 | 0.097 | 0.097 | 0 | 0.24 | 0.08 | 0.12 | 0.04 | 0.23 | 0.16 |
| 1 | 115348476 | rs3126218 | -0.04 | -0.439 | 0.164 | -1.278 | 0.022 | 0.007 | 0.007 | 0 | 0.82 | 0.58 | 0.81 | 0.9 | 0.8 | 0.7 |
| 1 | 115352043 | rs3126219 | 0.526 | -0.501 | -0.212 | -1.323 | 0.003 | 0.135 | 0.135 | 0.017 | 0.49 | 0.22 | 0.62 | 0.87 | 0.51 | 0.52 |
| 1 | 115355974 | rs3905243 | 1.133 | -0.385 | 0.03 | -1.32 | -0.001 | 0.008 | 0.008 | 0 | 0.27 | 0.24 | 0.14 | 0.03 | 0.25 | 0.17 |
| 1 | 115357015 | rs951307 | 0.016 | -0.438 | 0.117 | -1.118 | 0.002 | -0.003 | -0.003 | 0 | 0.42 | 0.43 | 0.33 | 0.17 | 0.33 | 0.23 |
| 1 | 115362785 | rs3104447 | 1.452 | -0.289 | -0.281 | -1.011 | -0.002 | -0.003 | -0.003 | 1.837 | 0.19 | 0.15 | 0.25 | 0.5 | 0.21 | 0.1 |
| 1 | 115363594 | rs2798664 | 1.11 | -0.368 | 0.102 | -1.009 | -0.004 | -0.007 | -0.007 | 0 | 0.07 | 0.36 | 0.04 | 0 | 0.01 | 0 |
| 1 | 115372525 | rs3126216 | 0.824 | -0.392 | -0.14 | -1.077 | -0.018 | -0.032 | -0.032 | 0 | 0.43 | 0.2 | 0.37 | 0.54 | 0.41 | 0.27 |
| 1 | 115379614 | rs869990 | 0.226 | -0.354 | -0.042 | -1.067 | -0.006 | -0.017 | -0.017 | 0 | 0.38 | 0.31 | 0.49 | 0.44 | 0.47 | 0.66 |
| 1 | 115404099 | rs360617 | 0.299 | -0.371 | -0.513 | -1.156 | 0.024 | -0.012 | -0.012 | 0 | 0.52 | 0.24 | 0.44 | 0.54 | 0.52 | 0.33 |
| 1 | 115407873 | rs12741053 | 0.454 | -0.183 | -0.565 | -1.052 | -0.027 | 0.004 | 0.004 | 0 | 0.09 | 0 | 0.07 | 0.03 | 0.13 | 0.09 |
| 1 | 115437986 | rs6661511 | 1.935 | -0.195 | -0.47 | -1.18 | -0.001 | 0.062 | 0.062 | 0 | 0.24 | 0.1 | 0.11 | 0.05 | 0.2 | 0.16 |
| 1 | 115470852 | rs360589 | 1.906 | -0.18 | -0.494 | -1.19 | -0.02 | -0.047 | -0.047 | 0 | 0.17 | 0.08 | 0.24 | 0.48 | 0.2 | 0.1 |
| 1 | 115490918 | rs360675 | -0.032 | -0.26 | -0.481 | -1.331 | 0.001 | -0.023 | -0.023 | 0 | 0.57 | 0.27 | 0.46 | 0.54 | 0.53 | 0.33 |
| 1 | 115492398 | rs11102855 | 1.806 | -0.059 | -0.327 | -1.196 | 0.007 | 0.054 | 0.054 | 0 | 0.28 | 0.12 | 0.15 | 0 | 0.24 | 0.21 |
| 1 | 115496181 | rs360680 | -0.322 | -0.002 | 0.163 | -0.887 | 0.006 | 0.04 | 0.04 | 0 | 0.66 | 0.57 | 0.52 | 0.55 | 0.61 | 0.42 |
| 1 | 115498726 | rs6696121 | 1.03 | -0.008 | -0.08 | -0.894 | 0.008 | 0.071 | 0.071 | 0.267 | 0.14 | 0.03 | 0.07 | 0.02 | 0.08 | 0.06 |
| 1 | 115537933 | rs11084 | 0.119 | -0.007 | 0.006 | -0.908 | -0.002 | -0.013 | -0.013 | 4.607 | 0.62 | 0.33 | 0.51 | 0.54 | 0.57 | 0.38 |
| 1 | 115563866 | rs1575067 | -0.285 | -0.087 | 0.26 | -0.838 | 0.004 | 0.009 | 0.009 | 0 | 0.81 | 0.84 | 0.89 | 0.98 | 0.84 | 0.89 |
| 1 | 115567885 | rs1575070 | 1.029 | -0.076 | 0.284 | -0.843 | -0.015 | -0.001 | -0.001 | 0 | 0.47 | 0.26 | 0.41 | 0.53 | 0.46 | 0.32 |
| 1 | 115571916 | rs1321109 | 0.043 | -0.082 | 0.336 | -0.851 | 0 | 0.038 | 0.038 | 0 | 0.33 | 0.55 | 0.47 | 0.46 | 0.38 | 0.58 |
| 1 | 115572365 | rs1321108 | 1.002 | -0.058 | 0.301 | -0.834 | -0.014 | 0 | 0 | 0 | 0.47 | 0.27 | 0.41 | 0.53 | 0.46 | 0.32 |
| 1 | 115575715 | rs11102875 | 1.479 | 0.015 | 0.618 | -0.626 | 0.006 | 0.013 | 0.013 | 0 | 0.2 | 0.24 | 0.33 | 0.34 | 0.26 | 0.39 |
| 1 | 115580540 | rs1998008 | 0.923 | 0.052 | 0.397 | -0.822 | 0 | 0.002 | 0.002 | 0 | 0.5 | 0.44 | 0.41 | 0.53 | 0.45 | 0.32 |
| 1 | 115582623 | rs7552175 | 0.229 | 0.075 | 0.404 | -0.819 | 0.006 | 0.038 | 0.038 | 0 | 0.31 | 0.41 | 0.47 | 0.45 | 0.39 | 0.58 |
| 1 | 115590501 | rs10776793 | -0.04 | -0.25 | 0.402 | -0.823 | 0.004 | 0.049 | 0.049 | 0 | 0.33 | 0.46 | 0.5 | 0.46 | 0.41 | 0.58 |
| 1 | 115590998 | rs11102878 | 0.979 | -0.3 | 0.419 | -0.879 | -0.003 | 0 | 0 | 0 | 0.49 | 0.2 | 0.4 | 0.53 | 0.48 | 0.33 |
| 1 | 115591058 | rs1052480 | -0.089 | -0.144 | 0.403 | -0.747 | 0.004 | 0.049 | 0.049 | 0 | 0.33 | 0.46 | 0.5 | 0.46 | 0.41 | 0.58 |
| 1 | 115592820 | rs3738706 | -0.089 | -0.144 | 0.403 | -0.747 | 0.004 | 0.049 | 0.049 | 0 | 0.33 | 0.46 | 0.5 | 0.46 | 0.41 | 0.58 |
| 1 | 115597836 | rs12026930 | 0.283 | -0.156 | 0.302 | -0.72 | 0 | 0.003 | 0.003 | 0 | 0.3 | 0.29 | 0.45 | 0.45 | 0.37 | 0.56 |
| 1 | 115600137 | rs3738704 | 0.581 | -0.203 | 0.313 | -0.853 | -0.014 | -0.043 | -0.043 | 0 | 0.14 | 0.21 | 0.09 | 0.02 | 0.08 | 0.06 |
| 1 | 115602119 | rs6537850 | -0.637 | -0.191 | 0.354 | -0.899 | 0 | -0.016 | -0.016 | 0 | 0.93 | 0.92 | 0.96 | 1 | 0.9 | 0.96 |
| 1 | 115602194 | rs6537851 | 0.266 | -0.215 | 0.385 | -0.921 | -0.007 | -0.057 | -0.057 | 0 | 0.79 | 0.72 | 0.87 | 0.99 | 0.83 | 0.9 |
| 1 | 115603844 | rs4240539 | -0.516 | -0.055 | 0.379 | -0.85 | 0.005 | -0.033 | -0.033 | 0.586 | 0.93 | 0.88 | 0.95 | 1 | 0.9 | 0.96 |
| 1 | 115603902 | rs4478805 | 0.356 | -0.108 | 0.206 | -1.024 | -0.002 | 0 | 0 | 0.015 | 0.6 | 0.61 | 0.62 | 0.51 | 0.6 | 0.78 |
| 1 | 115604241 | rs7526323 | 0.778 | -0.154 | 0.204 | -1.084 | 0 | 0.051 | 0.051 | 0 | 0.27 | 0.4 | 0.42 | 0.47 | 0.31 | 0.57 |
| 1 | 115604798 | rs10776795 | 0.02 | -0.129 | 0.166 | -1.047 | -0.002 | -0.01 | -0.01 | 0 | 0.33 | 0.3 | 0.33 | 0.49 | 0.3 | 0.18 |
| 1 | 115605646 | rs4839407 | 0.045 | -0.117 | 0.172 | -1.042 | -0.003 | -0.01 | -0.01 | 0 | 0.33 | 0.3 | 0.33 | 0.49 | 0.3 | 0.19 |
| 1 | 115609203 | rs1321106 | -0.412 | -0.124 | 0.158 | -1.035 | 0.017 | 0.016 | 0.016 | 0 | 0.7 | 0.64 | 0.67 | 0.52 | 0.63 | 0.83 |
| 1 | 115610091 | rs717275 | 1.495 | -0.128 | 0.214 | -0.991 | 0.002 | -0.02 | -0.02 | 0 | 0.31 | 0.29 | 0.17 | 0.04 | 0.25 | 0.2 |
| 1 | 115612702 | rs11102883 | 0.372 | 0.016 | 0.055 | -0.853 | 0 | -0.005 | -0.005 | 0 | 0.31 | 0.3 | 0.42 | 0.48 | 0.34 | 0.61 |
| 1 | 115617406 | rs1286364 | 0.672 | -0.064 | 0.078 | -0.841 | -0.002 | -0.034 | -0.034 | 1.217 | 0.41 | 0.41 | 0.25 | 0.08 | 0.31 | 0.22 |
| 1 | 115617856 | rs1286366 | -0.788 | 0.127 | -0.167 | -0.632 | 0.014 | 0.03 | 0.03 | 0 | 0.76 | 0.68 | 0.74 | 0.57 | 0.75 | 0.84 |
| 1 | 115618238 | rs12742348 | -0.214 | -0.176 | -0.685 | -0.543 | 0.017 | -0.033 | -0.033 | 0.011 | 0.18 | 0.09 | 0.16 | 0.15 | 0.18 | 0.22 |
| 1 | 115618813 | rs4839411 | 0.558 | 0.135 | -0.482 | -0.586 | 0.015 | 0.02 | 0.02 | 0 | 0.22 | 0.27 | 0.28 | 0.44 | 0.25 | 0.12 |
| 1 | 115620485 | rs12740360 | 0.589 | 0.057 | -0.59 | -0.703 | -0.004 | 0.034 | 0.034 | 0 | 0.42 | 0.14 | 0.3 | 0.25 | 0.42 | 0.37 |
| 1 | 115620739 | rs12137569 | 1.009 | 0.027 | -0.577 | -0.688 | 0.007 | 0.016 | 0.016 | 0 | 0.21 | 0.23 | 0.25 | 0.42 | 0.21 | 0.11 |
| 1 | 115622283 | rs4475751 | -0.29 | -0.018 | -0.508 | -0.607 | -0.008 | -0.002 | -0.002 | 0 | 0.66 | 0.43 | 0.57 | 0.67 | 0.68 | 0.49 |
| 1 | 115625348 | rs2144549 | 0.629 | -0.155 | -0.755 | -0.634 | -0.016 | -0.085 | -0.085 | 0 | 0.14 | 0.01 | 0.11 | 0.19 | 0.16 | 0.22 |
| 1 | 115626071 | rs1406859 | 0.491 | -0.101 | -0.538 | -0.718 | 0.006 | 0.007 | 0.007 | 0 | 0.52 | 0.42 | 0.45 | 0.48 | 0.52 | 0.28 |
| 1 | 115638044 | rs4839413 | 0.173 | -0.081 | -0.513 | -0.77 | 0.008 | 0.01 | 0.01 | 0 | 0.3 | 0.32 | 0.31 | 0.44 | 0.28 | 0.18 |
| 1 | 115642818 | rs6691374 | 1.155 | -0.242 | -0.873 | -0.809 | -0.006 | 0.017 | 0.017 | 0 | 0.22 | 0.32 | 0.36 | 0.31 | 0.22 | 0.39 |
| 1 | 115642847 | rs1750391 | -1.486 | -0.064 | -1.905 | -0.864 | -0.006 | 0.095 | 0.095 | 0 | 0.7 | 0.86 | 0.82 | 0.94 | 0.7 | 0.8 |
| 1 | 115648756 | rs1852989 | 0.217 | 0.18 | -1.825 | -0.379 | 0.003 | -0.047 | -0.047 | 0.974 | 0.15 | 0.07 | 0.12 | 0.19 | 0.15 | 0.22 |
| 1 | 115651160 | rs17479518 | 1.109 | 0.483 | -1.41 | -0.155 | 0 | 0.001 | 0.001 | 0 | 0.08 | 0.07 | 0.05 | 0.02 | 0.07 | 0.07 |
| 1 | 115654143 | rs2180230 | 0.049 | 0.475 | -1.492 | -0.033 | -0.039 | -0.043 | -0.043 | 0.003 | 0.28 | 0.12 | 0.29 | 0.44 | 0.28 | 0.18 |
| 1 | 115654866 | rs1286201 | 0.233 | 0.22 | -1.547 | -0.446 | 0.025 | 0.113 | 0.113 | 0 | 0.48 | 0.33 | 0.35 | 0.2 | 0.44 | 0.39 |
| 1 | 115658497 | rs2076901 | 0.479 | 0.698 | -2.195 | -0.546 | -0.075 | -0.044 | -0.044 | 0.376 | 0.57 | 0.12 | 0.59 | 0.46 | 0.56 | 0.62 |
| 1 | 115659236 | rs6672575 | 1.154 | 0.351 | -2.408 | -1.07 | -0.016 | 0.049 | 0.049 | 0 | 0.23 | 0.13 | 0.17 | 0 | 0.24 | 0.11 |
| 1 | 115664311 | rs6679900 | 1.156 | 0.409 | -2.431 | -1.043 | -0.02 | 0.115 | 0.115 | 0 | 0.25 | 0.06 | 0.16 | 0 | 0.24 | 0.11 |
| 1 | 115664930 | rs987267 | 0.576 | 0.454 | -2.55 | -0.899 | -0.024 | -0.135 | -0.135 | 0 | 0.31 | 0.04 | 0.42 | 0.46 | 0.3 | 0.5 |
| 1 | 115666678 | rs10218480 | 0.422 | 0.524 | -0.749 | -0.349 | 0.044 | 0.063 | 0.063 | 0.048 | 0.08 | 0.49 | 0.08 | 0.09 | 0.05 | 0.16 |
| 1 | 115672664 | rs12127083 | 0.538 | 1.018 | -0.916 | -0.26 | 0.001 | 0.007 | 0.007 | 0 | 0.14 | 0.11 | 0.19 | 0.36 | 0.23 | 0.08 |
| 1 | 115674952 | rs910625 | 0.596 | 0.87 | -0.716 | -0.347 | 0.012 | 0.006 | 0.006 | 0 | 0.66 | 0.57 | 0.68 | 0.55 | 0.64 | 0.78 |
| 1 | 115676278 | rs6702696 | -0.683 | 1.054 | -0.183 | -0.427 | -0.002 | -0.017 | -0.017 | 0 | 0.14 | 0.19 | 0.12 | 0.08 | 0.13 | 0.11 |
| 1 | 115676999 | rs997672 | 0.379 | 1.459 | -0.43 | 0.025 | -0.046 | -0.043 | -0.043 | 1.586 | 0.11 | 0.04 | 0.22 | 0.41 | 0.25 | 0.07 |
| 1 | 115677183 | rs2078371 | -0.605 | 1.106 | 0.165 | -0.263 | 0.001 | -0.03 | -0.03 | 0.065 | 0.14 | 0.22 | 0.1 | 0.07 | 0.11 | 0.09 |
| 1 | 115681071 | rs12138359 | 0.275 | 1.06 | 0.004 | -0.251 | 0.002 | 0.013 | 0.013 | 0 | 0.11 | 0.01 | 0.07 | 0.07 | 0.05 | 0.08 |

| | | | | | | | | | | | | | | | | |
|---|---|---|---|---|---|---|---|---|---|---|---|---|---|---|---|---|
| 1 | 115683469 | rs1286238 | -0.877 | 0.903 | 0.448 | -0.291 | 0.054 | 0.044 | 0.044 | 0 | 0.21 | 0.34 | 0.29 | 0.39 | 0.33 | 0.08 |
| 1 | 115685684 | rs4367784 | -0.144 | 0.923 | 0.487 | -0.283 | 0.073 | 0.065 | 0.065 | 0 | 0.15 | 0.25 | 0.26 | 0.39 | 0.32 | 0.08 |
| 1 | 115686952 | rs1777710 | -1.535 | 0.272 | 0.091 | -1.051 | 0.004 | -0.034 | -0.034 | 0 | 0.58 | 0.64 | 0.71 | 0.97 | 0.59 | 0.73 |
| 1 | 115688400 | rs1627351 | -1.293 | 0.068 | 0.013 | -1.113 | 0.001 | 0.001 | 0.001 | 0 | 0.38 | 0.6 | 0.37 | 0.46 | 0.36 | 0.23 |
| 1 | 115688472 | rs1627157 | -0.922 | -0.279 | 0.192 | -1.33 | 0.002 | -0.016 | -0.016 | 0 | 0.91 | 0.74 | 0.91 | 0.97 | 0.92 | 0.87 |
| 1 | 115696756 | rs7517073 | 1.094 | -0.252 | 0.244 | -1.35 | 0.017 | 0.025 | 0.025 | 0 | 0.13 | 0.13 | 0.23 | 0.39 | 0.27 | 0.07 |
| 1 | 115697580 | rs2798476 | -0.197 | -0.585 | 0.199 | -1.338 | -0.016 | 0.013 | 0.013 | 0.315 | 0.31 | 0.41 | 0.41 | 0.56 | 0.21 | 0.52 |
| 1 | 115699246 | rs1286235 | 0.145 | -0.481 | 0.344 | -1.422 | -0.058 | -0.03 | -0.03 | 0 | 0.11 | 0.37 | 0.07 | 0.05 | 0.01 | 0.03 |
| 1 | 115700763 | rs1286233 | -0.506 | 0.049 | -0.128 | -1.286 | 0.003 | -0.02 | -0.02 | 0 | 0.91 | 0.74 | 0.91 | 0.97 | 0.92 | 0.87 |
| 1 | 115702326 | rs17480309 | 1.201 | 0.191 | -0.782 | -1.179 | 0.001 | 0.173 | 0.173 | 0 | 0.33 | 0.07 | 0.2 | 0 | 0.33 | 0.14 |
| 1 | 115712579 | rs17033324 | 0.261 | 0.277 | -0.397 | -0.251 | -0.001 | -0.027 | -0.027 | 0 | 0.05 | 0 | 0.07 | 0.11 | 0.04 | 0.17 |
| 1 | 115723695 | rs195216 | 0.456 | -0.116 | 0.109 | -0.325 | 0.015 | 0.009 | 0.009 | 0 | 0.55 | 0.38 | 0.48 | 0.42 | 0.72 | 0.25 |
| 1 | 115728581 | rs1146165 | -0.943 | -0.111 | 0.293 | -0.33 | -0.037 | 0.028 | 0.028 | 0.212 | 0.19 | 0.53 | 0.19 | 0.2 | 0.1 | 0.31 |
| 1 | 115728898 | rs1146166 | -0.943 | -0.111 | 0.296 | -0.33 | -0.037 | 0.027 | 0.027 | 0.212 | 0.19 | 0.54 | 0.19 | 0.2 | 0.09 | 0.31 |
| 1 | 115733494 | rs9429549 | -0.105 | 0.309 | -0.074 | -0.673 | -0.006 | 0.01 | 0.01 | 0.275 | 0.59 | 0.67 | 0.43 | 0.22 | 0.51 | 0.48 |
| 1 | 115734091 | rs195204 | 0.832 | 0.431 | -0.232 | -0.512 | 0.002 | -0.081 | -0.081 | 0 | 0.28 | 0.09 | 0.34 | 0.39 | 0.24 | 0.45 |
| 1 | 115737280 | rs6657296 | -0.829 | 0.376 | -0.067 | -0.629 | -0.017 | 0 | 0 | 0.421 | 0.2 | 0.71 | 0.2 | 0.2 | 0.09 | 0.3 |
| 1 | 115738343 | rs1076754 | 1.86 | 0.184 | -0.751 | -0.975 | -0.075 | 0.192 | 0.192 | 0.101 | 0.28 | 0.03 | 0.19 | 0 | 0.33 | 0.13 |
| 1 | 115753482 | rs11806316 | 1.607 | 0.252 | -0.747 | -0.987 | -0.11 | 0.202 | 0.202 | 0.037 | 0.29 | 0.04 | 0.21 | 0 | 0.38 | 0.14 |
| 1 | 115754404 | rs12070042 | 0.048 | 0.221 | -0.138 | -0.826 | -0.008 | 0.007 | 0.007 | 0 | 0.12 | 0.17 | 0.1 | 0.16 | 0.06 | 0.19 |
| 1 | 115759393 | rs1146179 | -1.396 | 0.216 | 0.023 | -0.859 | -0.002 | -0.001 | -0.001 | 0 | 0.22 | 0.64 | 0.23 | 0.21 | 0.17 | 0.33 |
| 1 | 115759830 | rs1040605 | -1.213 | 0.409 | 0.047 | -0.534 | 0.027 | 0.058 | 0.058 | 0 | 0.88 | 0.86 | 0.77 | 0.59 | 0.75 | 0.93 |
| 1 | 115767978 | rs1294387 | 1.233 | 0.104 | 0.298 | -0.608 | -0.017 | -0.024 | -0.024 | 0 | 0.07 | 0.29 | 0.05 | 0 | 0.01 | 0 |
| 1 | 115768909 | rs4839424 | 0.357 | 0.144 | 0.37 | -0.572 | 0.003 | 0.035 | 0.035 | 0 | 0.06 | 0.07 | 0.07 | 0.16 | 0.05 | 0.19 |
| 1 | 115771616 | rs2798464 | 0.147 | 0.033 | 0.351 | -0.924 | 0 | -0.054 | -0.054 | 0 | 0.52 | 0.52 | 0.36 | 0.05 | 0.51 | 0.28 |
| 1 | 115774074 | rs971723 | -0.543 | 0.041 | 0.233 | -0.667 | 0.036 | 0.028 | 0.028 | 0 | 0.49 | 0.37 | 0.49 | 0.58 | 0.35 | 0.78 |
| 1 | 115779865 | rs1298590 | -0.058 | 0.032 | 0.277 | -0.685 | -0.036 | -0.063 | -0.063 | 0 | 0.18 | 0.38 | 0.11 | 0.05 | 0.12 | 0.14 |
| 1 | 115781479 | rs1738845 | 0.139 | 0.219 | 0.072 | -0.845 | -0.001 | -0.055 | -0.055 | 0 | 0.52 | 0.56 | 0.34 | 0.05 | 0.52 | 0.3 |
| 1 | 115782754 | rs7542599 | -1.028 | 0.19 | -0.023 | -0.579 | 0.029 | 0.066 | 0.066 | 0 | 0.85 | 0.78 | 0.74 | 0.58 | 0.76 | 0.94 |
| 1 | 115784029 | rs4839426 | 0.9 | 0.084 | -0.415 | -0.552 | 0.027 | -0.126 | -0.126 | 0.306 | 0.28 | 0.04 | 0.36 | 0.53 | 0.21 | 0.6 |
| 1 | 115786408 | rs1985652 | 0.814 | 0.238 | -0.305 | -0.521 | -0.039 | -0.04 | -0.04 | 0 | 0.63 | 0.16 | 0.6 | 0.53 | 0.59 | 0.8 |
| 1 | 115796490 | rs11102901 | 1.111 | 0.254 | -0.286 | -0.405 | -0.039 | 0.113 | 0.113 | 0 | 0.38 | 0.13 | 0.29 | 0.16 | 0.41 | 0.35 |
| 1 | 115797624 | rs12033518 | 0.653 | 0.206 | -0.069 | -0.403 | -0.004 | -0.005 | -0.005 | 1.729 | 0.66 | 0.31 | 0.61 | 0.53 | 0.6 | 0.8 |
| 1 | 115799299 | rs12745217 | 0.64 | 0.145 | 0.016 | 0.033 | 0.026 | -0.034 | -0.034 | 0.255 | 0.28 | 0.17 | 0.32 | 0.37 | 0.19 | 0.45 |
| 1 | 115804914 | rs10858070 | 0.136 | 0.228 | 0.208 | -0.393 | 0.032 | 0.049 | 0.049 | 0.415 | 0.05 | 0.25 | 0.07 | 0.05 | 0.03 | 0.12 |
| 1 | 115805053 | rs10776796 | -0.019 | 0.404 | -0.155 | -0.426 | 0 | -0.002 | -0.002 | 0 | 0.57 | 0.59 | 0.43 | 0.21 | 0.58 | 0.49 |
| 1 | 115812379 | rs11102914 | 0.374 | -0.257 | -0.541 | 0.116 | 0.058 | -0.015 | -0.015 | 0.121 | 0.51 | 0.77 | 0.48 | 0.51 | 0.38 | 0.56 |
| 1 | 115818779 | rs2982742 | -0.017 | 0.043 | -0.707 | -0.045 | -0.096 | -0.091 | -0.091 | 0 | 0.41 | 0.1 | 0.47 | 0.47 | 0.53 | 0.42 |
| 1 | 115821085 | rs2068797 | -0.002 | 0.028 | -0.714 | -0.053 | -0.096 | -0.091 | -0.091 | 0.554 | 0.41 | 0.09 | 0.47 | 0.47 | 0.53 | 0.42 |
| 1 | 115823387 | rs7523086 | 0.467 | 0.21 | -0.671 | -0.073 | 0.06 | -0.003 | -0.003 | 0 | 0.49 | 0.68 | 0.46 | 0.5 | 0.36 | 0.55 |
| 1 | 115829313 | rs6330 | 0.261 | 0.347 | -0.932 | -0.369 | -0.04 | 0.048 | 0.048 | 0.682 | 0.3 | 0.16 | 0.32 | 0.16 | 0.46 | 0.19 |
| 1 | 115830626 | rs11466108 | 0.353 | 0.283 | -0.507 | 0.199 | 0 | -0.009 | -0.009 | 0.196 | 0.07 | 0.09 | 0.04 | 0.01 | 0.03 | 0.01 |
| 1 | 115833581 | rs2254527 | -0.83 | -0.022 | -1.018 | 0.526 | 0.013 | -0.061 | -0.061 | 0 | 0.72 | 0.91 | 0.78 | 0.65 | 0.77 | 0.69 |
| 1 | 115835500 | rs910330 | -0.232 | -0.156 | -0.545 | 0.679 | 0.015 | -0.023 | -0.023 | 0 | 0.64 | 0.72 | 0.69 | 0.56 | 0.72 | 0.65 |
| 1 | 115837709 | rs2239622 | -0.569 | -0.219 | -0.327 | 0.685 | 0.021 | -0.028 | -0.028 | 0 | 0.66 | 0.82 | 0.76 | 0.57 | 0.73 | 0.63 |
| 1 | 115837919 | rs2856813 | 0.547 | 0.093 | -0.245 | 0.554 | -0.044 | 0.099 | 0.099 | 0 | 0.49 | 0.15 | 0.45 | 0.36 | 0.56 | 0.55 |
| 1 | 115838282 | rs2856811 | 0.329 | 0.029 | 0.304 | 0.584 | -0.002 | -0.004 | -0.004 | 0 | 0.53 | 0.49 | 0.5 | 0.36 | 0.64 | 0.57 |
| 1 | 115841110 | rs11102920 | 0.308 | -0.528 | -0.268 | 0.081 | -0.057 | -0.005 | -0.005 | 0 | 0.25 | 0.43 | 0.24 | 0.25 | 0.16 | 0.15 |
| 1 | 115845046 | rs6686615 | 0.586 | -0.265 | -0.325 | 0.079 | -0.046 | 0.08 | 0.08 | 0 | 0.34 | 0.21 | 0.28 | 0.1 | 0.46 | 0.31 |
| 1 | 115846221 | rs12058927 | 0.29 | -1.386 | -0.213 | 0.124 | -0.069 | -0.015 | -0.015 | 0 | 0.28 | 0.47 | 0.26 | 0.26 | 0.17 | 0.17 |
| 1 | 115847295 | rs10776797 | -0.939 | -0.791 | -0.036 | 0.785 | 0.015 | -0.052 | -0.052 | 0.881 | 0.75 | 0.9 | 0.8 | 0.65 | 0.8 | 0.72 |
| 1 | 115847392 | rs10858072 | -0.372 | -1.036 | 0.238 | 0.927 | -0.002 | 0.13 | 0.13 | 0 | 0.1 | 0.21 | 0.23 | 0.3 | 0.09 | 0.22 |
| 1 | 115862254 | rs10776798 | -0.857 | -1.027 | -0.674 | 0.408 | 0.031 | -0.103 | -0.103 | 1.42 | 0.73 | 0.98 | 0.77 | 0.63 | 0.84 | 0.75 |
| 1 | 115862568 | rs7535026 | 0.002 | -1.142 | -0.127 | 0.348 | 0.019 | 0.013 | 0.013 | 0 | 0.59 | 0.3 | 0.56 | 0.33 | 0.7 | 0.58 |
| 1 | 115863900 | rs7530686 | 0.009 | -1.167 | 0.192 | 0.349 | -0.017 | -0.01 | -0.01 | 0 | 0.6 | 0.5 | 0.57 | 0.33 | 0.71 | 0.58 |
| 1 | 115869499 | rs4565713 | 0.139 | -1.059 | 0.45 | 0.352 | 0.018 | 0.015 | 0.015 | 0 | 0.57 | 0.35 | 0.54 | 0.32 | 0.69 | 0.57 |
| 1 | 115869850 | rs17033692 | -0.053 | -0.895 | 0.457 | 0.587 | -0.001 | 0.026 | 0.026 | 0 | 0.08 | 0.09 | 0.16 | 0.26 | 0.08 | 0.15 |
| 1 | 115873047 | rs10776799 | -0.612 | -0.985 | -0.262 | 0.378 | -0.018 | 0.005 | 0.005 | 0 | 0.68 | 0.36 | 0.72 | 0.63 | 0.8 | 0.73 |
| 1 | 115875645 | rs4332358 | -0.155 | -1.679 | -1.23 | 0.177 | 0.051 | -0.002 | -0.002 | 0 | 0.43 | 0.7 | 0.46 | 0.57 | 0.26 | 0.34 |
| 1 | 115876503 | rs11102929 | 0.257 | -1.249 | -1.281 | 0.132 | 0.001 | 0.019 | 0.019 | 0 | 0.22 | 0.31 | 0.27 | 0.31 | 0.19 | 0.23 |
| 1 | 115877624 | rs17033706 | 0.747 | -0.966 | -1.572 | -0.335 | -0.023 | 0.003 | 0.003 | 0 | 0.27 | 0.26 | 0.26 | 0.29 | 0.14 | 0.2 |
| 1 | 115878242 | rs6673867 | 0.054 | 0.469 | -2.014 | -0.711 | -0.163 | 0.084 | 0.084 | 0.918 | 0.71 | 0.97 | 0.72 | 0.9 | 0.49 | 0.67 |
| 1 | 115880132 | rs17540656 | -0.412 | 0.339 | -1.697 | -0.67 | -0.042 | 0.144 | 0.144 | 0 | 0.37 | 0.14 | 0.32 | 0.09 | 0.55 | 0.32 |
| 1 | 115882541 | rs10776801 | -0.64 | 0.119 | -0.497 | -0.493 | 0.037 | -0.025 | -0.025 | 0 | 0.1 | 0.37 | 0.08 | 0.02 | 0.04 | 0.03 |
| 1 | 115883203 | rs4418592 | -0.271 | -0.098 | -0.641 | -0.189 | -0.02 | -0.046 | -0.046 | 0.561 | 0.1 | 0.02 | 0.04 | 0.16 | 0.12 | 0.19 |
| 1 | 115886386 | rs10858077 | -0.156 | -0.158 | -0.479 | -0.218 | 0.013 | -0.007 | -0.007 | 0 | 0.08 | 0.34 | 0.05 | 0.02 | 0.02 | 0.02 |
| 1 | 115891156 | rs4539131 | -1.101 | -0.945 | -1.396 | -0.066 | 0.06 | -0.028 | -0.028 | 0.435 | 0.7 | 0.9 | 0.63 | 0.67 | 0.79 | 0.72 |
| 1 | 115894321 | rs11102933 | -0.357 | -0.476 | -1.376 | -0.146 | -0.004 | -0.012 | -0.012 | 0.664 | 0.57 | 0.76 | 0.56 | 0.54 | 0.44 | 0.53 |
| 1 | 115895741 | rs9787265 | 1.644 | -0.315 | -1.878 | 0.115 | -0.005 | -0.068 | -0.068 | 0 | 0.17 | 0.02 | 0.09 | 0.19 | 0.16 | 0.23 |

| | | | | | | | | | | | | | | | | |
|---|---|---|---|---|---|---|---|---|---|---|---|---|---|---|---|---|
| 1 | 115895838 | rs12081407 | 0.897 | 0.086 | -1.364 | 0.054 | 0.002 | 0 | 0 | 0 | 0.37 | 0.31 | 0.42 | 0.36 | 0.26 | 0.32 |
| 1 | 115898335 | rs7513294 | 0.841 | 0.415 | -1.822 | 0.146 | -0.136 | 0.023 | 0.023 | 0 | 0.22 | 0.02 | 0.32 | 0.15 | 0.35 | 0.17 |
| 1 | 115906457 | rs4144397 | -0.264 | 0.426 | -1.407 | 0.287 | 0.01 | 0.003 | 0.003 | 0.169 | 0.51 | 0.42 | 0.53 | 0.52 | 0.44 | 0.51 |
| 1 | 115907009 | rs11584234 | -1.001 | 1.335 | -1.382 | 0.317 | -0.048 | -0.041 | -0.041 | 0.008 | 0.13 | 0.02 | 0.11 | 0.16 | 0.18 | 0.19 |
| 1 | 115907557 | rs11102944 | -0.999 | 1.347 | -1.357 | 0.358 | -0.048 | -0.039 | -0.039 | 0.075 | 0.13 | 0.03 | 0.1 | 0.16 | 0.18 | 0.16 |
| 1 | 115910424 | rs7534855 | 1.315 | 1.877 | -1.165 | 0.386 | 0.015 | 0 | 0 | 0.072 | 0.71 | 0.71 | 0.61 | 0.72 | 0.53 | 0.7 |
| 1 | 115921098 | rs11102945 | -0.35 | 2.159 | -1.713 | 0.484 | -0.062 | -0.018 | -0.018 | 0 | 0.12 | 0.02 | 0.15 | 0.11 | 0.23 | 0.11 |
| 1 | 115921355 | rs7552722 | 1.06 | 1.942 | -1.609 | 0.466 | -0.045 | -0.009 | -0.009 | 0 | 0.77 | 0.87 | 0.77 | 0.76 | 0.63 | 0.76 |
| 1 | 115921680 | rs4391665 | 1.151 | 1.88 | -1.618 | 0.426 | -0.044 | -0.004 | -0.004 | 0 | 0.76 | 0.86 | 0.76 | 0.76 | 0.63 | 0.76 |
| 1 | 115943077 | rs556723 | -0.142 | 0.734 | -2.234 | 0.473 | 0.035 | -0.024 | -0.024 | 0 | 0.32 | 0.11 | 0.29 | 0.29 | 0.28 | 0.34 |
| 1 | 115969592 | rs17033822 | 0.512 | 0.699 | -2.305 | 0.561 | 0.018 | 0.004 | 0.004 | 0 | 0.26 | 0.02 | 0.25 | 0.18 | 0.21 | 0.27 |
| 1 | 115972614 | rs17033823 | 0.695 | 0.889 | -3.081 | 0.665 | -0.032 | -0.117 | -0.117 | 0 | 0.23 | 0.07 | 0.3 | 0.45 | 0.24 | 0.29 |
| 1 | 115973920 | rs1327126 | 0.156 | 1.037 | -2.097 | 0.186 | -0.005 | -0.012 | -0.012 | 0 | 0.4 | 0.41 | 0.41 | 0.32 | 0.41 | 0.44 |
| 1 | 115994787 | rs2148691 | 1.112 | 0.839 | -2.373 | 0.173 | 0.029 | -0.001 | -0.001 | 0.025 | 0.28 | 0.02 | 0.34 | 0.22 | 0.23 | 0.31 |
| 1 | 115998480 | rs17556126 | 1.679 | 0.973 | -2.363 | 0.646 | -0.048 | -0.066 | -0.066 | 0 | 0.15 | 0.04 | 0.19 | 0.32 | 0.19 | 0.2 |
| 1 | 116000680 | rs2334915 | -0.589 | 0.965 | -2.154 | 0.486 | 0.037 | 0.022 | 0.022 | 0 | 0.58 | 0.79 | 0.56 | 0.63 | 0.63 | 0.55 |
| 1 | 116000763 | rs553146 | 0.216 | 0.967 | -1.573 | 0.757 | -0.028 | -0.048 | -0.048 | 0 | 0.72 | 0.82 | 0.72 | 0.53 | 0.69 | 0.68 |
| 1 | 116004970 | rs1170050 | 0.284 | 0.151 | -1.041 | 0.267 | 0.003 | -0.002 | -0.002 | 0 | 0.87 | 0.87 | 0.94 | 0.85 | 0.9 | 0.88 |
| 1 | 116011300 | rs913203 | 0.275 | 0.088 | -0.83 | 0.294 | -0.027 | 0.008 | 0.008 | 0 | 0.86 | 0.73 | 0.93 | 0.85 | 0.91 | 0.88 |
| 1 | 116011901 | rs735882 | -0.386 | 0.75 | -0.889 | 0.093 | 0.007 | -0.047 | -0.047 | 0.444 | 0.22 | 0.35 | 0.21 | 0.09 | 0.23 | 0.15 |
| 1 | 116021452 | rs6696763 | -0.355 | 0.59 | -1.235 | 0.663 | -0.017 | -0.023 | -0.023 | 0 | 0.17 | 0.07 | 0.25 | 0.28 | 0.2 | 0.31 |
| 1 | 116021584 | rs12080720 | -0.007 | 0.232 | -0.952 | 0.528 | -0.002 | -0.005 | -0.005 | 0.355 | 0.61 | 0.65 | 0.53 | 0.63 | 0.56 | 0.63 |
| 1 | 116040016 | rs17033866 | -1.22 | -0.428 | -0.335 | 0.176 | -0.04 | -0.059 | -0.059 | 0.012 | 0.1 | 0.31 | 0.05 | 0.02 | 0.03 | 0.05 |
| 1 | 116050880 | rs28645661 | -1.125 | -0.42 | -0.286 | 0.18 | -0.032 | -0.049 | -0.049 | 0.347 | 0.09 | 0.27 | 0.05 | 0.02 | 0.03 | 0.05 |
| 1 | 116069039 | rs552279 | -0.667 | -0.959 | -1.294 | 0.132 | 0.058 | 0.028 | 0.028 | 0 | 0.53 | 0.91 | 0.41 | 0.38 | 0.34 | 0.6 |
| 1 | 116089793 | rs3010406 | -1.268 | -1.411 | 0.562 | -0.112 | -0.078 | -0.097 | -0.097 | 0.171 | 0.14 | 0.63 | 0.08 | 0.02 | 0.03 | 0.06 |
| 1 | 116093778 | rs17033934 | 0.058 | -1.932 | 0.571 | 0.172 | 0.047 | -0.049 | -0.049 | 0 | 0.41 | 0.16 | 0.34 | 0.5 | 0.32 | 0.6 |
| 1 | 116100276 | rs17033969 | -0.62 | -1.538 | 1.876 | 0.201 | -0.013 | -0.043 | -0.043 | 0 | 0.08 | 0.21 | 0.05 | 0.01 | 0.05 | 0.05 |
| 1 | 116102456 | rs4839038 | 1.674 | -1.552 | 1.974 | 0.194 | -0.045 | -0.076 | -0.076 | 0 | 0.85 | 0.26 | 0.9 | 0.98 | 0.95 | 0.94 |
| 1 | 116109499 | rs3010399 | 1.325 | -1.535 | 1.875 | 0.202 | -0.043 | -0.088 | -0.088 | 0 | 0.87 | 0.53 | 0.92 | 0.98 | 0.95 | 0.94 |
| 1 | 116128484 | rs6413949 | 1.008 | -1.52 | 1.882 | 0.229 | -0.029 | -0.068 | -0.068 | 0 | 0.88 | 0.53 | 0.92 | 0.98 | 0.95 | 0.94 |
| 1 | 116128774 | rs4403633 | -0.234 | -2.095 | 1.34 | 0.029 | 0.001 | -0.011 | -0.011 | 0.862 | 0.18 | 0.1 | 0.24 | 0.33 | 0.17 | 0.45 |
| 1 | 116136810 | rs3860222 | 1.552 | -2.165 | 1.28 | -0.006 | 0.055 | -0.039 | -0.039 | 6.51 | 0.66 | 0.07 | 0.66 | 0.65 | 0.78 | 0.49 |
| 1 | 116137190 | rs10802151 | -1.199 | -2.066 | 1.75 | 0.064 | -0.026 | -0.073 | -0.073 | 0 | 0.12 | 0.44 | 0.08 | 0.02 | 0.05 | 0.06 |
| 1 | 116138871 | rs4240546 | 1.788 | -2.097 | 1.737 | 0.153 | 0.017 | 0.017 | 0.017 | 1.657 | 0.79 | 0.2 | 0.86 | 0.9 | 0.88 | 0.9 |
| 1 | 116140239 | rs6658990 | -1.159 | -2.088 | 1.763 | 0.149 | -0.021 | -0.073 | -0.073 | 0 | 0.12 | 0.44 | 0.08 | 0.02 | 0.05 | 0.06 |
| 1 | 116146472 | rs4332360 | 1.867 | -2.127 | 1.697 | 0.171 | 0.004 | -0.002 | -0.002 | 1.283 | 0.71 | 0.29 | 0.69 | 0.66 | 0.78 | 0.51 |
| 1 | 116154831 | rs4839460 | 0.138 | -2.071 | 1.82 | 0.198 | -0.021 | -0.041 | -0.041 | 0.103 | 0.92 | 0.73 | 0.97 | 0.99 | 0.98 | 0.96 |
| 1 | 116155918 | rs10494178 | 0.239 | -2.058 | 1.192 | 0.292 | -0.006 | -0.018 | -0.018 | 0.704 | 0.09 | 0.06 | 0.19 | 0.26 | 0.09 | 0.4 |
| 1 | 116156655 | rs6672017 | 1.61 | -2.001 | 1.194 | 0.292 | 0.005 | -0.005 | -0.005 | 0 | 0.73 | 0.27 | 0.72 | 0.63 | 0.81 | 0.52 |
| 1 | 116157111 | rs4303074 | 0.537 | -1.974 | 1.231 | 0.272 | -0.055 | -0.077 | -0.077 | 0 | 0.88 | 0.39 | 0.94 | 0.98 | 0.98 | 0.96 |
| 1 | 116165222 | rs4483407 | 0.401 | -2.009 | 1.084 | 0.273 | -0.055 | -0.073 | -0.073 | 0 | 0.88 | 0.43 | 0.95 | 0.99 | 0.98 | 0.96 |
| 1 | 116165273 | rs4576661 | 0.157 | -1.977 | 1.024 | 0.294 | -0.001 | 0.002 | 0.002 | 0 | 0.06 | 0.06 | 0.03 | 0.09 | 0.07 | 0.04 |
| 1 | 116170427 | rs4240548 | 0.54 | -1.124 | 0.888 | 0.121 | 0.031 | -0.101 | -0.101 | 1.17 | 0.85 | 0.58 | 0.91 | 0.98 | 0.87 | 0.94 |
| 1 | 116170705 | rs4839463 | 1.376 | -1.127 | 0.352 | 0.156 | 0.005 | -0.005 | -0.005 | 1.973 | 0.7 | 0.13 | 0.69 | 0.71 | 0.78 | 0.55 |
| 1 | 116171383 | rs4603135 | 1.743 | -1.293 | 0.263 | 0.072 | 0.003 | -0.025 | -0.025 | 0 | 0.62 | 0.1 | 0.65 | 0.62 | 0.7 | 0.49 |
| 1 | 116188971 | rs3010380 | -1.674 | -1.034 | -0.05 | 0.18 | 0.003 | -0.004 | -0.004 | 0 | 0.3 | 0.87 | 0.31 | 0.29 | 0.22 | 0.45 |
| 1 | 116196994 | rs17559092 | -0.417 | -1.322 | 0.307 | 0.061 | 0 | 0.014 | 0.014 | 0 | 0.09 | 0.16 | 0.04 | 0.1 | 0.07 | 0.06 |
| 1 | 116197610 | rs3010382 | -1.534 | -1.178 | -0.802 | 0.436 | 0.045 | -0.037 | -0.037 | 0 | 0.5 | 0.9 | 0.49 | 0.47 | 0.42 | 0.72 |
| 1 | 116202014 | rs9428201 | 0.597 | -0.865 | -0.512 | 0.561 | 0.001 | -0.005 | -0.005 | 0.026 | 0.11 | 0.13 | 0.1 | 0.16 | 0.1 | 0.09 |
| 1 | 116202811 | rs6658884 | -0.194 | -0.635 | -0.357 | 0.233 | -0.002 | -0.014 | -0.014 | 0.356 | 0.1 | 0.15 | 0.07 | 0.03 | 0.08 | 0.39 |
| 1 | 116203864 | rs10801911 | -0.996 | -0.706 | -0.225 | 0.422 | -0.038 | -0.023 | -0.023 | 0 | 0.2 | 0.35 | 0.27 | 0.14 | 0.14 | 0.18 |
| 1 | 116204661 | rs10923142 | -0.464 | -0.936 | -1.443 | 0.304 | 0.039 | 0.031 | 0.031 | 0 | 0.26 | 0.08 | 0.12 | 0.25 | 0.18 | 0.19 |
| 1 | 116204807 | rs10923143 | 0.83 | 0.191 | -1.041 | 0.661 | 0 | 0.003 | 0.003 | 0 | 0.7 | 0.41 | 0.7 | 0.7 | 0.72 | 0.74 |
| 1 | 116205021 | rs7540389 | -1.142 | 0.888 | -0.909 | 1.028 | 0.003 | 0.009 | 0.009 | 0 | 0.33 | 0.35 | 0.37 | 0.55 | 0.31 | 0.43 |
| 1 | 116205848 | rs4428890 | -0.431 | 0.984 | -0.714 | 1.002 | 0.001 | -0.001 | -0.001 | 0.328 | 0.1 | 0.08 | 0.07 | 0.1 | 0.09 | 0.15 |
| 1 | 116206423 | rs4839469 | -0.366 | 1.148 | -0.935 | 1.123 | -0.018 | -0.01 | -0.01 | 1.389 | 0.13 | 0.04 | 0.22 | 0.12 | 0.16 | 0.09 |
| 1 | 116207111 | rs12752458 | -0.439 | 0.913 | -0.727 | 0.951 | -0.009 | 0.01 | 0.01 | 0 | 0.13 | 0.1 | 0.12 | 0.1 | 0.13 | 0.2 |
| 1 | 116207484 | rs12039617 | -1.476 | 0.195 | -0.28 | 1.376 | -0.015 | 0.011 | 0.011 | 0 | 0.29 | 0.37 | 0.28 | 0.37 | 0.19 | 0.3 |
| 1 | 116208597 | rs4440851 | -0.497 | 0.229 | -0.2 | 1.266 | 0.009 | -0.02 | -0.02 | 0 | 0.16 | 0.3 | 0.14 | 0.11 | 0.13 | 0.17 |
| 1 | 116209709 | rs4433392 | -1.189 | 0.013 | -0.427 | 1.026 | 0.006 | -0.005 | -0.005 | 0.02 | 0.62 | 0.62 | 0.65 | 0.45 | 0.67 | 0.69 |
| 1 | 116211763 | rs17500488 | 1.306 | -0.064 | -0.684 | 0.853 | 0.01 | -0.01 | -0.01 | 0.524 | 0.12 | 0.06 | 0.07 | 0.14 | 0.11 | 0.05 |
| 1 | 116214279 | rs10801913 | -2.176 | -0.071 | -0.219 | 0.905 | -0.022 | 0 | 0 | 0 | 0.69 | 0.49 | 0.76 | 0.7 | 0.69 | 0.77 |
| 1 | 116214846 | rs12137412 | 0.277 | 0.148 | -0.64 | 0.728 | -0.032 | 0.025 | 0.025 | 0.062 | 0.31 | 0.17 | 0.43 | 0.19 | 0.4 | 0.42 |
| 1 | 116219323 | rs10923156 | -0.313 | 0.278 | -0.613 | 1.196 | -0.039 | -0.083 | -0.083 | 0 | 0.2 | 0.07 | 0.29 | 0.39 | 0.23 | 0.31 |
| 1 | 116220451 | rs4571967 | -1.1 | 0.244 | -0.552 | 1.185 | -0.004 | -0.084 | -0.084 | 0 | 0.3 | 0.19 | 0.31 | 0.51 | 0.28 | 0.35 |
| 1 | 116220533 | rs4284265 | -1.108 | 0.196 | -0.551 | 1.127 | -0.013 | -0.084 | -0.084 | 0.274 | 0.3 | 0.19 | 0.32 | 0.52 | 0.3 | 0.35 |
| 1 | 116223259 | rs12568559 | -0.075 | 0.321 | -0.565 | 1.139 | -0.034 | -0.077 | -0.077 | 0 | 0.21 | 0.09 | 0.26 | 0.39 | 0.23 | 0.31 |
| 1 | 116223306 | rs6699583 | -1.774 | 0.351 | 0.015 | 0.977 | -0.001 | 0.003 | 0.003 | 0 | 0.7 | 0.73 | 0.77 | 0.73 | 0.67 | 0.79 |

| | | | | | | | | | | | | | | | | |
|---|---|---|---|---|---|---|---|---|---|---|---|---|---|---|---|---|
| 1 | 116226459 | rs12144356 | -1.748 | 0.344 | 0.108 | 0.955 | -0.002 | -0.009 | -0.009 | 0 | 0.69 | 0.6 | 0.76 | 0.72 | 0.67 | 0.78 |
| 1 | 116234313 | rs3811012 | -1.786 | 0.319 | -0.156 | 0.815 | -0.019 | 0.006 | 0.006 | 0 | 0.67 | 0.86 | 0.75 | 0.74 | 0.62 | 0.7 |
| 1 | 116234416 | rs3811011 | 0.249 | 0.304 | 0.049 | 0.687 | 0 | -0.005 | -0.005 | 0 | 0.26 | 0.27 | 0.3 | 0.2 | 0.26 | 0.33 |
| 1 | 116234505 | rs6700610 | -2.14 | 0.306 | 0.042 | 0.688 | -0.019 | 0.006 | 0.006 | 0 | 0.67 | 0.86 | 0.75 | 0.74 | 0.62 | 0.7 |
| 1 | 116235277 | rs3204955 | 0.176 | 0.324 | 0.06 | 0.668 | 0 | -0.005 | -0.005 | 0.078 | 0.26 | 0.27 | 0.3 | 0.2 | 0.26 | 0.33 |
| 1 | 116237023 | rs10923204 | 0.254 | 0.323 | 0.056 | 0.669 | 0 | -0.005 | -0.005 | 0 | 0.26 | 0.25 | 0.3 | 0.2 | 0.26 | 0.33 |
| 1 | 116239266 | rs17034230 | 0.051 | 0.359 | 0.25 | 0.728 | -0.005 | -0.008 | -0.008 | 0 | 0.16 | 0.25 | 0.17 | 0.12 | 0.16 | 0.14 |
| 1 | 116239368 | rs10494179 | -1.187 | 0.348 | 0.36 | 0.721 | -0.001 | -0.009 | -0.009 | 0 | 0.41 | 0.44 | 0.47 | 0.32 | 0.42 | 0.47 |
| 1 | 116239633 | rs4348723 | 0.545 | 0.322 | 0.407 | 0.855 | 0 | 0.007 | 0.007 | 0 | 0.25 | 0.3 | 0.27 | 0.37 | 0.2 | 0.23 |
| 1 | 116239751 | rs10801933 | -1.251 | 0.325 | 0.421 | 0.853 | -0.003 | -0.016 | -0.016 | 0 | 0.42 | 0.49 | 0.47 | 0.32 | 0.42 | 0.47 |
| 1 | 116240026 | rs6428677 | 0.56 | 0.347 | 0.355 | 0.864 | 0 | 0.006 | 0.006 | 0 | 0.24 | 0.3 | 0.27 | 0.36 | 0.2 | 0.23 |
| 1 | 116240212 | rs3811006 | 0.075 | 0.409 | 0.356 | 0.45 | -0.004 | -0.006 | -0.006 | 0 | 0.16 | 0.25 | 0.17 | 0.12 | 0.16 | 0.14 |
| 1 | 116241211 | rs6673301 | 0.411 | 0.414 | 0.344 | 0.445 | -0.002 | 0.005 | 0.005 | 0 | 0.25 | 0.21 | 0.3 | 0.2 | 0.26 | 0.33 |
| 1 | 116243380 | rs7521023 | -1.712 | 0.434 | 0.099 | 0.495 | 0.002 | 0.019 | 0.019 | 0 | 0.63 | 0.74 | 0.73 | 0.74 | 0.61 | 0.68 |
| 1 | 116243877 | rs7413162 | -0.993 | 0.355 | 0.093 | 0.476 | -0.003 | -0.012 | -0.012 | 0.016 | 0.41 | 0.46 | 0.46 | 0.32 | 0.41 | 0.47 |
| 1 | 116245241 | rs6664262 | -1.095 | 0.43 | -0.106 | 0.55 | -0.008 | 0.033 | 0.033 | 0 | 0.57 | 0.72 | 0.61 | 0.72 | 0.53 | 0.58 |
| 1 | 116247790 | rs3811003 | -0.625 | 0.265 | -0.028 | 0.476 | 0.007 | -0.009 | -0.009 | 0.061 | 0.41 | 0.61 | 0.35 | 0.52 | 0.3 | 0.27 |
| 1 | 116251072 | rs3923977 | -1.751 | 0.286 | -0.001 | 0.469 | -0.048 | 0.013 | 0.013 | 0 | 0.63 | 0.86 | 0.63 | 0.71 | 0.55 | 0.6 |
| 1 | 116251296 | rs3923975 | 0.704 | 0.304 | -0.061 | 0.517 | -0.014 | -0.008 | -0.008 | 0.014 | 0.18 | 0.12 | 0.27 | 0.21 | 0.23 | 0.31 |
| 1 | 116251879 | rs10923235 | 0.184 | 0.19 | 0.364 | 0.561 | 0.007 | -0.006 | -0.006 | 0 | 0.32 | 0.24 | 0.28 | 0.39 | 0.23 | 0.24 |
| 1 | 116253342 | rs4423004 | -1.377 | 0.313 | 0.55 | 0.485 | -0.01 | 0.012 | 0.012 | 0 | 0.61 | 0.72 | 0.62 | 0.7 | 0.55 | 0.61 |
| 1 | 116257891 | rs4275457 | -1.375 | 0.296 | 0.789 | 0.453 | 0.004 | -0.007 | -0.007 | 0 | 0.61 | 0.53 | 0.62 | 0.67 | 0.56 | 0.61 |
| 1 | 116259062 | rs2999465 | -1.556 | 0.296 | 0.795 | 0.466 | -0.009 | 0.005 | 0.005 | 0 | 0.62 | 0.71 | 0.64 | 0.67 | 0.57 | 0.61 |
| 1 | 116260604 | rs2997742 | -1.462 | 0.298 | 0.789 | 0.467 | -0.004 | 0.003 | 0.003 | 0 | 0.62 | 0.66 | 0.63 | 0.67 | 0.57 | 0.61 |
| 1 | 116268699 | rs6665970 | -1.399 | 0.303 | 0.692 | 0.462 | -0.002 | 0.002 | 0.002 | 0 | 0.61 | 0.44 | 0.62 | 0.67 | 0.56 | 0.62 |
| 1 | 116270234 | rs35672929 | 0.528 | 0.463 | 0.559 | 0.713 | -0.046 | -0.02 | -0.02 | 0.528 | 0.16 | 0.08 | 0.26 | 0.16 | 0.22 | 0.29 |
| 1 | 116274332 | rs4255385 | 0.92 | 0.532 | 0.731 | 0.799 | -0.006 | -0.012 | -0.012 | 0 | 0.09 | 0.09 | 0.17 | 0.16 | 0.11 | 0.27 |
| 1 | 116278503 | rs6677074 | -1.011 | 0.401 | 0.966 | 0.774 | 0 | -0.001 | -0.001 | 0 | 0.47 | 0.44 | 0.54 | 0.53 | 0.44 | 0.57 |
| 1 | 116283343 | rs9428083 | 1.403 | 0.225 | 1.092 | 0.793 | -0.011 | 0.011 | 0.011 | 2.276 | 0.8 | 0.28 | 0.76 | 0.74 | 0.87 | 0.71 |
| 1 | 116283526 | rs7538337 | 1.403 | 0.225 | 1.092 | 0.793 | -0.011 | 0.011 | 0.011 | 2.276 | 0.8 | 0.28 | 0.76 | 0.74 | 0.87 | 0.71 |
| 1 | 116284997 | rs7554304 | 1.319 | 0.206 | 1.271 | 0.749 | 0.007 | 0.089 | 0.089 | 2.313 | 0.93 | 0.75 | 0.95 | 0.9 | 0.97 | 0.98 |
| 1 | 116295007 | rs4839478 | 0.994 | 0.216 | 1.237 | 0.755 | 0.013 | 0.094 | 0.094 | 0.597 | 0.95 | 0.83 | 0.95 | 0.89 | 0.97 | 0.98 |
| 1 | 116297758 | rs4073778 | 1.201 | 0.358 | 0.305 | 0.623 | -0.058 | 0.072 | 0.072 | 0 | 0.44 | 0.11 | 0.42 | 0.31 | 0.58 | 0.41 |
| 1 | 116301565 | rs9428087 | 1.518 | 0.44 | 0.311 | 0.7 | -0.048 | 0.038 | 0.038 | 0 | 0.4 | 0.14 | 0.29 | 0.29 | 0.48 | 0.35 |
| 1 | 116302592 | rs4143097 | 1.586 | 0.438 | 0.299 | 0.699 | -0.062 | 0.032 | 0.032 | 0 | 0.39 | 0.12 | 0.28 | 0.29 | 0.48 | 0.35 |
| 1 | 116302923 | rs9428225 | 0.917 | 0.442 | 0.685 | 0.762 | 0.012 | 0.002 | 0.002 | 0 | 0.51 | 0.39 | 0.46 | 0.44 | 0.58 | 0.62 |
| 1 | 116310967 | rs4074536 | -0.687 | 0.152 | 0.96 | 0.938 | -0.019 | 0.009 | 0.009 | 0 | 0.43 | 0.49 | 0.37 | 0.52 | 0.29 | 0.3 |
| 1 | 116320833 | rs10923445 | -0.036 | 0.068 | 1.017 | 0.905 | 0.001 | -0.003 | -0.003 | 0 | 0.3 | 0.27 | 0.3 | 0.4 | 0.23 | 0.25 |
| 1 | 116324707 | rs11580099 | 0.488 | 0.074 | 0.864 | 0.99 | -0.012 | -0.046 | -0.046 | 0.69 | 0.1 | 0.05 | 0.18 | 0.26 | 0.11 | 0.28 |
| 1 | 116325537 | rs4839484 | 1.742 | 0.124 | 0.957 | 0.938 | -0.023 | 0.066 | 0.066 | 0 | 0.45 | 0.18 | 0.34 | 0.31 | 0.54 | 0.37 |
| 1 | 116329944 | rs9428232 | 0.412 | 0.044 | 0.971 | 1.154 | -0.003 | 0.162 | 0.162 | 0.308 | 0.85 | 0.48 | 0.78 | 0.74 | 0.89 | 0.72 |
| 1 | 116331806 | rs7418954 | 0.655 | -0.058 | 0.934 | 1.115 | -0.017 | 0.141 | 0.141 | 0.03 | 0.84 | 0.47 | 0.78 | 0.74 | 0.89 | 0.73 |
| 1 | 116333111 | rs12039739 | -0.116 | 0.299 | 0.84 | 1.059 | -0.001 | -0.005 | -0.005 | 0 | 0.31 | 0.27 | 0.31 | 0.4 | 0.28 | 0.23 |
| 1 | 116333581 | rs9428235 | -0.074 | 0.301 | 0.908 | 1.082 | -0.003 | 0.042 | 0.042 | 0 | 0.87 | 0.65 | 0.83 | 0.85 | 0.9 | 0.73 |
| 1 | 116333734 | rs6669078 | 0.943 | 0.299 | 0.879 | 1.101 | 0.002 | 0 | 0 | 0 | 0.56 | 0.35 | 0.52 | 0.45 | 0.62 | 0.48 |
| 1 | 116333773 | rs12040622 | -0.186 | 0.378 | 0.59 | 1.032 | -0.001 | -0.006 | -0.006 | 0 | 0.31 | 0.27 | 0.32 | 0.41 | 0.28 | 0.25 |
| 1 | 116334813 | rs4515784 | -0.178 | 0.389 | 0.588 | 1.037 | -0.001 | -0.006 | -0.006 | 0.142 | 0.31 | 0.27 | 0.31 | 0.41 | 0.28 | 0.25 |
| 1 | 116336876 | rs4072907 | 0.524 | 0.425 | 0.524 | 1.295 | 0.006 | 0.052 | 0.052 | 0 | 0.86 | 0.7 | 0.71 | 0.76 | 0.86 | 0.67 |
| 1 | 116337217 | rs10494185 | 0.135 | 0.45 | 0.213 | 1.267 | -0.012 | -0.032 | -0.032 | 0 | 0.11 | 0.07 | 0.16 | 0.18 | 0.13 | 0.26 |
| 1 | 116339269 | rs4483405 | 0.941 | 0.305 | 0.428 | 0.971 | 0.001 | 0.007 | 0.007 | 0 | 0.53 | 0.45 | 0.38 | 0.34 | 0.59 | 0.41 |
| 1 | 116340260 | rs7523715 | -0.668 | -0.242 | -0.05 | 0.959 | -0.073 | -0.06 | -0.06 | 0 | 0.46 | 0.1 | 0.4 | 0.53 | 0.43 | 0.51 |
| 1 | 116347890 | rs4453038 | 0.873 | -0.314 | 0.508 | 0.866 | 0.012 | 0 | 0 | 0 | 0.49 | 0.51 | 0.59 | 0.5 | 0.62 | 0.5 |
| 1 | 116354571 | rs950369 | 1.168 | -0.1 | 0.789 | 0.918 | -0.014 | 0.024 | 0.024 | 0.223 | 0.89 | 0.75 | 0.84 | 0.86 | 0.93 | 0.91 |
| 1 | 116357354 | rs2797180 | 1.199 | -0.13 | 0.661 | 0.861 | 0.001 | -0.006 | -0.006 | 0 | 0.44 | 0.47 | 0.47 | 0.38 | 0.57 | 0.42 |
| 1 | 116361981 | rs1342714 | 1.608 | -0.225 | 0.859 | 0.79 | -0.042 | 0.002 | 0.002 | 0 | 0.39 | 0.24 | 0.42 | 0.38 | 0.52 | 0.41 |
| 1 | 116362598 | rs2811585 | 1.183 | -0.279 | 0.72 | 0.823 | 0.001 | -0.007 | -0.007 | 0 | 0.44 | 0.46 | 0.47 | 0.38 | 0.57 | 0.42 |
| 1 | 116369290 | rs2811587 | 0.674 | -0.442 | 0.608 | 0.946 | -0.023 | -0.008 | -0.008 | 0 | 0.92 | 0.82 | 0.98 | 0.95 | 0.97 | 0.99 |
| 1 | 116377439 | rs11579104 | 1.02 | -0.373 | 0.265 | 1.221 | -0.027 | -0.051 | -0.051 | 0 | 0.1 | 0 | 0.16 | 0.13 | 0.11 | 0.13 |
| 1 | 116378399 | rs2797185 | -0.579 | -0.522 | 0.573 | 1.191 | 0 | 0.004 | 0.004 | 0 | 0.34 | 0.31 | 0.48 | 0.46 | 0.31 | 0.41 |
| 1 | 116380172 | rs10754377 | -1.118 | -0.774 | 0.577 | 1.041 | 0.005 | 0.028 | 0.028 | 0.208 | 0.48 | 0.29 | 0.53 | 0.59 | 0.4 | 0.53 |
| 1 | 116384580 | rs2797208 | -0.905 | -0.793 | 0.519 | 0.95 | -0.006 | 0.016 | 0.016 | 0.052 | 0.39 | 0.52 | 0.51 | 0.48 | 0.33 | 0.44 |
| 1 | 116391331 | rs10923633 | 0.874 | -0.777 | 0.033 | 0.89 | 0.015 | -0.011 | -0.011 | 0 | 0.16 | 0.08 | 0.06 | 0.13 | 0.09 | 0.12 |
| 1 | 116394349 | rs12033106 | 0.504 | -0.69 | -0.027 | 0.915 | -0.041 | -0.112 | -0.112 | 0.496 | 0.26 | 0.04 | 0.23 | 0.36 | 0.23 | 0.33 |
| 1 | 116396243 | rs4839496 | -0.609 | -0.732 | 0.191 | 0.562 | -0.003 | 0.006 | 0.006 | 0.022 | 0.38 | 0.44 | 0.5 | 0.47 | 0.33 | 0.43 |
| 1 | 116397910 | rs4548442 | 1.368 | -0.582 | -0.542 | 0.57 | -0.072 | 0.033 | 0.033 | 0 | 0.4 | 0.13 | 0.4 | 0.35 | 0.57 | 0.44 |
| 1 | 116400329 | rs2811578 | 0.102 | -0.394 | -0.346 | 0.933 | 0.012 | -0.035 | -0.035 | 0 | 0.3 | 0.19 | 0.41 | 0.51 | 0.2 | 0.4 |
| 1 | 116401394 | rs2096210 | -0.502 | 0.283 | 0.097 | 1.038 | -0.023 | -0.026 | -0.026 | 0 | 0.19 | 0.08 | 0.11 | 0.2 | 0.19 | 0.12 |
| 1 | 116403691 | rs2066396 | -0.53 | 0.313 | -0.047 | 0.982 | -0.024 | -0.02 | -0.02 | 0 | 0.19 | 0.08 | 0.1 | 0.19 | 0.19 | 0.13 |
| 1 | 116406102 | rs10754389 | 1.724 | -0.2 | 0.842 | 0.749 | 0.001 | -0.036 | -0.036 | 0 | 0.92 | 0.42 | 0.95 | 1 | 0.98 | 0.98 |

| | | | | | | | | | | | | | | | | |
|---|---|---|---|---|---|---|---|---|---|---|---|---|---|---|---|---|
| 1 | 116411950 | rs11586273 | 0.068 | -0.618 | 0.372 | 0.912 | 0.066 | -0.015 | -0.015 | 0.057 | 0.42 | 0.2 | 0.4 | 0.45 | 0.32 | 0.36 |
| 1 | 116412304 | rs10802063 | 0.464 | -0.84 | 0.874 | 0.832 | 0.025 | -0.015 | -0.015 | 0 | 0.36 | 0.24 | 0.39 | 0.45 | 0.28 | 0.33 |
| 1 | 116421457 | rs814987 | -0.839 | -0.992 | 0.724 | 0.852 | 0.054 | -0.032 | -0.032 | 3.26 | 0.66 | 0.83 | 0.6 | 0.59 | 0.73 | 0.67 |
| 1 | 116426939 | rs10754390 | -0.252 | -0.786 | 1.265 | 1.069 | -0.01 | 0.008 | 0.008 | 0 | 0.58 | 0.52 | 0.45 | 0.45 | 0.67 | 0.63 |
| 1 | 116429910 | rs1617980 | -0.365 | -0.949 | 1.008 | 0.902 | 0.092 | -0.04 | -0.04 | 1.357 | 0.58 | 0.83 | 0.59 | 0.54 | 0.7 | 0.66 |
| 1 | 116432480 | rs6656298 | -0.417 | -0.981 | 1.3 | 0.875 | 0.004 | -0.021 | -0.021 | 0 | 0.05 | 0.19 | 0.03 | 0.01 | 0.03 | 0.04 |
| 1 | 116433182 | rs1342722 | 0.188 | -0.981 | 1.312 | 0.891 | 0.063 | 0.017 | 0.017 | 0 | 0.52 | 0.33 | 0.44 | 0.42 | 0.66 | 0.64 |
| 1 | 116433936 | rs2797188 | -0.115 | -0.983 | 1.323 | 0.893 | 0.008 | -0.001 | -0.001 | 0.048 | 0.54 | 0.45 | 0.56 | 0.54 | 0.7 | 0.66 |
| 1 | 116436143 | rs2131440 | 1.263 | -0.846 | 0.924 | 0.797 | -0.114 | -0.024 | -0.024 | 0 | 0.35 | 0.06 | 0.35 | 0.33 | 0.53 | 0.48 |
| 1 | 116444094 | rs1498113 | -0.312 | -0.831 | 1.173 | 0.842 | 0.034 | -0.001 | -0.001 | 0.039 | 0.55 | 0.37 | 0.56 | 0.53 | 0.69 | 0.66 |
| 1 | 116451413 | rs958690 | -0.568 | -0.853 | 1.171 | 0.827 | 0.074 | -0.027 | -0.027 | 0 | 0.59 | 0.76 | 0.59 | 0.53 | 0.69 | 0.66 |
| 1 | 116454332 | rs12080102 | -0.941 | -0.83 | 1.256 | 0.704 | -0.039 | -0.072 | -0.072 | 0.619 | 0.12 | 0.42 | 0.04 | 0 | 0.04 | 0.03 |
| 1 | 116469435 | rs10923806 | 0.065 | -0.877 | 1.019 | 0.848 | 0.112 | -0.032 | -0.032 | 0 | 0.52 | 0.14 | 0.48 | 0.44 | 0.66 | 0.65 |
| 1 | 116478690 | rs12568384 | 1.342 | -0.853 | 1.04 | 0.868 | 0.023 | -0.144 | -0.144 | 0.534 | 0.32 | 0.02 | 0.32 | 0.44 | 0.26 | 0.3 |
| 1 | 116496141 | rs699765 | -1.607 | -0.77 | 1.27 | 0.772 | 0.118 | 0.143 | 0.143 | 1.164 | 0.05 | 0.34 | 0.14 | 0.13 | 0.05 | 0.02 |
| 1 | 116499471 | rs699767 | -0.543 | -0.93 | 1.235 | 0.826 | -0.018 | 0.003 | 0.003 | 0 | 0.59 | 0.57 | 0.64 | 0.53 | 0.71 | 0.67 |
| 1 | 116501174 | rs10923890 | 1.188 | -0.73 | 1.082 | 0.741 | -0.131 | -0.015 | -0.015 | 2.89 | 0.34 | 0.06 | 0.35 | 0.29 | 0.49 | 0.44 |
| 1 | 116502789 | rs1157378 | -0.802 | -0.489 | 1.431 | 1.041 | 0.001 | -0.006 | -0.006 | 0 | 0.94 | 0.94 | 0.98 | 1 | 0.97 | 0.97 |
| 1 | 116518319 | rs1281693 | -2.159 | -0.478 | 1.467 | 1.047 | 0.072 | 0.067 | 0.067 | 0.046 | 0.1 | 0.43 | 0.17 | 0.13 | 0.07 | 0.06 |
| 1 | 116521057 | rs924277 | 1.002 | -0.49 | 1.486 | 1.035 | -0.036 | -0.082 | -0.082 | 0 | 0.87 | 0.64 | 0.95 | 1 | 0.94 | 0.93 |
| 1 | 116521089 | rs924276 | 1.002 | -0.49 | 1.486 | 1.035 | -0.036 | -0.082 | -0.082 | 0.03 | 0.87 | 0.64 | 0.95 | 1 | 0.94 | 0.93 |
| 1 | 116525672 | rs12726299 | 0.457 | -0.514 | 1.436 | 1.092 | 0.05 | -0.01 | -0.01 | 0 | 0.49 | 0.14 | 0.48 | 0.42 | 0.6 | 0.6 |
| 1 | 116527068 | rs2488432 | 0.38 | -0.607 | 1.442 | 1.099 | -0.023 | 0.021 | 0.021 | 0 | 0.37 | 0.5 | 0.46 | 0.58 | 0.3 | 0.33 |
| 1 | 116528090 | rs7547928 | 0.52 | -0.598 | 1.414 | 1.105 | 0.108 | -0.002 | -0.002 | 2.114 | 0.5 | 0.14 | 0.49 | 0.42 | 0.64 | 0.61 |
| 1 | 116533220 | rs1281699 | -0.83 | -0.588 | 1.325 | 1.103 | -0.001 | -0.009 | -0.009 | 1.867 | 0.94 | 0.94 | 0.98 | 1 | 0.97 | 0.97 |
| 1 | 116534852 | rs4046190 | 0.513 | -0.583 | 1.242 | 1.115 | 0.108 | -0.002 | -0.002 | 0 | 0.5 | 0.14 | 0.48 | 0.42 | 0.64 | 0.61 |
| 1 | 116540966 | rs1775699 | -1.363 | -0.551 | 1.271 | 1.083 | -0.039 | -0.092 | -0.092 | 0.082 | 0.13 | 0.46 | 0.06 | 0 | 0.06 | 0.07 |
| 1 | 116551979 | rs2051060 | 0.274 | -0.573 | 1.195 | 1.121 | 0.075 | 0.009 | 0.009 | 0 | 0.52 | 0.24 | 0.5 | 0.42 | 0.64 | 0.61 |
| 1 | 116552570 | rs1775703 | -0.285 | -0.572 | 1.207 | 1.122 | -0.015 | 0 | 0 | 0 | 0.44 | 0.52 | 0.36 | 0.45 | 0.31 | 0.37 |
| 1 | 116553207 | rs1565694 | 0.381 | -0.569 | 1.104 | 1.124 | 0.083 | 0.014 | 0.014 | 0 | 0.51 | 0.24 | 0.5 | 0.42 | 0.64 | 0.61 |
| 1 | 116559153 | rs7535090 | 1.076 | -0.56 | 1.165 | 1.131 | 0.046 | -0.09 | -0.09 | 0 | 0.31 | 0.09 | 0.31 | 0.45 | 0.25 | 0.31 |
| 1 | 116566678 | rs10802146 | 0.321 | -0.508 | 1.138 | 1.152 | 0.081 | 0.012 | 0.012 | 0 | 0.51 | 0.24 | 0.5 | 0.42 | 0.64 | 0.61 |
| 1 | 116567296 | rs1775711 | -0.236 | -0.522 | 1.171 | 1.138 | -0.017 | 0 | 0 | 0 | 0.44 | 0.52 | 0.35 | 0.45 | 0.31 | 0.37 |
| 1 | 116568032 | rs17035127 | -0.102 | -0.25 | 0.938 | 1.162 | -0.025 | -0.028 | -0.028 | 0 | 0.05 | 0.01 | 0.04 | 0.15 | 0.07 | 0.09 |
| 1 | 116571981 | rs12562912 | -1.343 | -0.396 | 1.208 | 1.043 | 0.012 | 0.028 | 0.028 | 0 | 0.08 | 0.18 | 0.06 | 0.12 | 0.08 | 0.1 |
| 1 | 116574306 | rs6666101 | 2.776 | -0.489 | 1.097 | 0.997 | 0.002 | -0.008 | -0.008 | 0 | 0.73 | 0.09 | 0.73 | 0.75 | 0.82 | 0.81 |
| 1 | 116577110 | rs10923960 | -1.346 | -0.478 | 1.329 | 1.011 | 0.013 | 0.016 | 0.016 | 0 | 0.08 | 0.19 | 0.06 | 0.1 | 0.08 | 0.1 |
| 1 | 116579016 | rs17035172 | 0.634 | -0.497 | 1.433 | 0.965 | -0.017 | -0.028 | -0.028 | 0.019 | 0.06 | 0.1 | 0.02 | 0 | 0.03 | 0.03 |
| 1 | 116581479 | rs10923962 | 0.778 | -0.609 | 1.45 | 0.886 | -0.042 | -0.073 | -0.073 | 0 | 0.88 | 0.67 | 0.96 | 1 | 0.96 | 0.97 |
| 1 | 116584842 | rs1935829 | 2.298 | -0.61 | 1.481 | 0.894 | -0.016 | -0.023 | -0.023 | 0 | 0.8 | 0.32 | 0.79 | 0.87 | 0.89 | 0.91 |
| 1 | 116602245 | rs12566568 | 1.181 | -0.678 | 1.323 | 0.957 | 0.01 | -0.147 | -0.147 | 0.251 | 0.3 | 0.01 | 0.29 | 0.42 | 0.24 | 0.31 |
| 1 | 116603163 | rs1281750 | 1.267 | -0.321 | 2.45 | 0.903 | -0.042 | -0.073 | -0.073 | 0.109 | 0.88 | 0.67 | 0.96 | 1 | 0.96 | 0.97 |
| 1 | 116608903 | rs1281745 | 3.623 | -0.319 | 2.12 | 0.872 | 0.019 | 0.016 | 0.016 | 1.613 | 0.7 | 0.12 | 0.73 | 0.75 | 0.81 | 0.82 |
| 1 | 116609833 | rs3748741 | -1.965 | -0.319 | 2.132 | 0.872 | 0.01 | 0.024 | 0.024 | 0 | 0.09 | 0.19 | 0.06 | 0.12 | 0.08 | 0.09 |
| 1 | 116610227 | rs1281744 | 2.671 | -0.318 | 2.128 | 0.873 | -0.016 | -0.023 | -0.023 | 0 | 0.79 | 0.31 | 0.79 | 0.87 | 0.89 | 0.91 |
| 1 | 116618453 | rs10923970 | -1.328 | -0.262 | 1.819 | 0.861 | 0 | 0.003 | 0.003 | 0 | 0.07 | 0.07 | 0.05 | 0.11 | 0.07 | 0.09 |
| 1 | 116645757 | rs1146353 | -3.874 | -0.32 | 1.778 | 0.906 | 0.019 | -0.015 | -0.015 | 0 | 0.3 | 0.88 | 0.27 | 0.28 | 0.19 | 0.18 |
| 1 | 116649821 | rs2479373 | -1.011 | -0.285 | 1.863 | 0.938 | -0.033 | -0.051 | -0.051 | 0 | 0.09 | 0.2 | 0.03 | 0 | 0.03 | 0.03 |
| 1 | 116652963 | rs1887540 | -1.545 | -0.21 | 1.91 | 0.981 | -0.04 | -0.064 | -0.064 | 0 | 0.11 | 0.28 | 0.04 | 0 | 0.04 | 0.03 |
| 1 | 116653452 | rs2488435 | -1.621 | -0.211 | 1.857 | 0.977 | -0.04 | -0.071 | -0.071 | 0 | 0.11 | 0.31 | 0.05 | 0 | 0.04 | 0.03 |
| 1 | 116655492 | rs1146356 | -2.855 | -0.189 | 1.693 | 0.985 | -0.02 | -0.015 | -0.015 | 0.086 | 0.21 | 0.63 | 0.21 | 0.16 | 0.12 | 0.09 |
| 1 | 116657911 | rs1146359 | -2.345 | 0.029 | 1.767 | 1.108 | 0.063 | 0.103 | 0.103 | 0 | 0.11 | 0.45 | 0.18 | 0.16 | 0.09 | 0.06 |
| 1 | 116669877 | rs6668233 | -3.286 | 0.048 | 0.822 | 1.029 | 0.009 | -0.011 | -0.011 | 0.272 | 0.29 | 0.82 | 0.29 | 0.28 | 0.22 | 0.2 |
| 1 | 116671061 | rs10802157 | -3.235 | 0.052 | 0.71 | 1.031 | 0.001 | -0.028 | -0.028 | 0.717 | 0.3 | 0.9 | 0.3 | 0.28 | 0.22 | 0.2 |
| 1 | 116674418 | rs10923976 | -3.238 | 0.068 | 0.703 | 1.038 | 0 | -0.032 | -0.032 | 0 | 0.31 | 0.9 | 0.31 | 0.28 | 0.23 | 0.2 |
| 1 | 116676035 | rs10802159 | -3.238 | 0.068 | 0.703 | 1.038 | 0 | -0.032 | -0.032 | 0 | 0.31 | 0.9 | 0.31 | 0.28 | 0.23 | 0.2 |
| 1 | 116676548 | rs6670668 | -1.913 | 0.063 | 0.869 | 1.034 | -0.014 | -0.009 | -0.009 | 0 | 0.18 | 0.41 | 0.11 | 0.12 | 0.15 | 0.14 |
| 1 | 116682954 | rs10802160 | -2.64 | 0.101 | 0.036 | 1.016 | 0 | -0.041 | -0.041 | 0 | 0.31 | 0.9 | 0.31 | 0.31 | 0.23 | 0.2 |
| 1 | 116691120 | rs10802162 | -2.667 | 0.287 | 0.032 | 0.977 | -0.014 | -0.031 | -0.031 | 0 | 0.34 | 0.88 | 0.33 | 0.32 | 0.27 | 0.22 |
| 1 | 116694130 | rs17459496 | -1.679 | 0.784 | 0.302 | 1.226 | 0.001 | 0.003 | 0.003 | 0 | 0.2 | 0.24 | 0.26 | 0.28 | 0.23 | 0.17 |
| 1 | 116704771 | rs12093582 | -1.653 | 0.779 | 0.709 | 1.109 | 0.054 | 0.014 | 0.014 | 0.171 | 0.15 | 0.46 | 0.11 | 0.1 | 0.15 | 0.13 |
| 1 | 116706783 | rs4839515 | 0.111 | 0.199 | -0.576 | 1.287 | 0.032 | -0.143 | -0.143 | 0.5 | 0.35 | 0.05 | 0.38 | 0.59 | 0.27 | 0.37 |
| 1 | 116708713 | rs10754432 | -0.274 | 0.354 | 0.391 | 1.048 | -0.004 | -0.013 | -0.013 | 0 | 0.63 | 0.65 | 0.78 | 0.76 | 0.71 | 0.65 |
| 1 | 116709648 | rs6664070 | 0.454 | 0.557 | 0.915 | 1.21 | 0.001 | -0.001 | -0.001 | 0 | 0.47 | 0.53 | 0.55 | 0.43 | 0.52 | 0.47 |
| 1 | 116717522 | rs10489749 | -1.578 | 0.276 | 1.822 | 0.882 | 0.006 | -0.003 | -0.003 | 0.157 | 0.09 | 0.23 | 0.06 | 0.08 | 0.07 | 0.06 |
| 1 | 116717619 | rs6658845 | -0.193 | 0.587 | 1.454 | 0.836 | 0.009 | 0.004 | 0.004 | 0 | 0.58 | 0.63 | 0.42 | 0.59 | 0.63 | 0.65 |
| 1 | 116718023 | rs2208363 | 0.28 | 0.664 | 0.864 | 0.929 | 0.035 | -0.006 | -0.006 | 0.563 | 0.51 | 0.3 | 0.3 | 0.44 | 0.53 | 0.55 |
| 1 | 116727982 | rs10802166 | 0.059 | 0.267 | 0.08 | 0.158 | 0.029 | 0.101 | 0.101 | 0 | 0.41 | 0.2 | 0.37 | 0.18 | 0.37 | 0.37 |
| 1 | 116733218 | rs2208365 | -0.043 | 0.194 | -0.104 | 0.585 | -0.009 | -0.049 | -0.049 | 0.528 | 0.54 | 0.85 | 0.38 | 0.5 | 0.51 | 0.62 |

| | | | | | | | | | | | | | | | | |
|---|---|---|---|---|---|---|---|---|---|---|---|---|---|---|---|---|
| 1 | 116735572 | rs7546742 | -1.795 | -0.083 | 1.114 | 0.833 | -0.002 | 0.169 | 0.169 | 2.245 | 0.09 | 0.51 | 0.05 | 0.12 | 0.01 | 0.05 |
| 1 | 116735588 | rs7537977 | -1.795 | -0.083 | 1.114 | 0.833 | -0.002 | 0.169 | 0.169 | 2.245 | 0.09 | 0.51 | 0.05 | 0.12 | 0.01 | 0.05 |
| 1 | 116736247 | rs17035430 | -1.795 | -0.083 | 1.114 | 0.833 | -0.002 | 0.169 | 0.169 | 2.245 | 0.09 | 0.51 | 0.05 | 0.12 | 0.01 | 0.05 |
| 1 | 116740195 | rs1884989 | -1.795 | -0.083 | 1.114 | 0.833 | -0.002 | 0.169 | 0.169 | 2.245 | 0.09 | 0.5 | 0.05 | 0.12 | 0.01 | 0.05 |
| 1 | 116743996 | rs10923993 | -1.492 | -0.083 | 1.475 | 0.83 | 0.01 | 0.158 | 0.158 | 1.777 | 0.07 | 0.4 | 0.04 | 0.11 | 0.01 | 0.05 |
| 1 | 116764781 | rs7548144 | -2.646 | -0.273 | 1.349 | 0.731 | -0.056 | 0.027 | 0.027 | 2.734 | 0.11 | 0.72 | 0.06 | 0.12 | 0.01 | 0.06 |
| 1 | 116769215 | rs12120197 | -0.813 | -0.111 | 1.19 | 0.63 | 0.001 | -0.023 | -0.023 | 0.271 | 0.06 | 0.14 | 0.04 | 0.01 | 0.04 | 0.06 |
| 1 | 116771069 | rs10924002 | -2.214 | -0.214 | 1.289 | 0.783 | -0.029 | 0.257 | 0.257 | 1.284 | 0.16 | 0.47 | 0.32 | 0.44 | 0.11 | 0.11 |
| 1 | 116776537 | rs12125273 | -0.584 | -0.026 | 0.675 | 0.612 | 0.007 | -0.028 | -0.028 | 0 | 0.06 | 0.14 | 0.05 | 0 | 0.06 | 0.05 |
| 1 | 116795524 | rs2184483 | 0.184 | 0.764 | 0.239 | 0.579 | -0.005 | -0.002 | -0.002 | 0.042 | 0.86 | 0.87 | 0.86 | 0.83 | 0.75 | 0.72 |
| 1 | 116797899 | rs10924021 | 0.304 | 0.543 | 0.058 | 0.78 | 0.008 | 0.064 | 0.064 | 0 | 0.06 | 0.03 | 0.25 | 0.43 | 0.07 | 0.05 |
| 1 | 116800201 | rs4839524 | 0.861 | 0.273 | -0.456 | 0.137 | -0.047 | 0.183 | 0.183 | 0 | 0.33 | 0.03 | 0.33 | 0.08 | 0.34 | 0.28 |
| 1 | 116801420 | rs2184484 | 0.694 | 0.362 | 0.319 | 0.131 | 0.002 | 0.01 | 0.01 | 0 | 0.42 | 0.4 | 0.25 | 0.32 | 0.34 | 0.39 |
| 1 | 116803322 | rs929362 | 0.773 | 0.093 | -0.043 | -0.378 | 0.003 | 0.139 | 0.139 | 0 | 0.33 | 0.11 | 0.33 | 0.08 | 0.34 | 0.28 |
| 1 | 116804727 | rs6694434 | 0.368 | 0.165 | -0.178 | -0.485 | 0.008 | 0.001 | 0.001 | 0 | 0.43 | 0.07 | 0.57 | 0.63 | 0.41 | 0.35 |
| 1 | 116807231 | rs9428340 | 1.24 | 0.431 | 0.107 | -0.622 | -0.012 | 0.01 | 0.01 | 0.184 | 0.73 | 0.21 | 0.77 | 0.82 | 0.68 | 0.65 |
| 1 | 116808177 | rs4839061 | 0.491 | 0.306 | 0.214 | -0.685 | -0.017 | 0.007 | 0.007 | 0 | 0.77 | 0.82 | 0.82 | 0.83 | 0.69 | 0.64 |
| 1 | 116812245 | rs9428197 | 0.685 | 0.319 | -0.127 | -0.865 | 0.01 | 0.037 | 0.037 | 0 | 0.48 | 0.12 | 0.59 | 0.63 | 0.45 | 0.38 |
| 1 | 116812376 | rs4839066 | 0.734 | 0.245 | -0.069 | -0.895 | -0.025 | 0.032 | 0.032 | 0 | 0.28 | 0.08 | 0.22 | 0.21 | 0.3 | 0.33 |
| 1 | 116819325 | rs12089467 | 1.167 | 0.273 | -0.061 | -0.88 | -0.001 | 0.007 | 0.007 | 0 | 0.25 | 0.16 | 0.19 | 0.2 | 0.25 | 0.29 |
| 1 | 116821111 | rs2335957 | 0.914 | 0.293 | -0.067 | -0.856 | 0.001 | -0.004 | -0.004 | 0 | 0.74 | 0.27 | 0.79 | 0.85 | 0.71 | 0.65 |
| 1 | 116822125 | rs7542792 | -0.113 | 0.321 | 0.163 | -0.802 | 0.07 | 0 | 0 | 0 | 0.82 | 0.62 | 0.85 | 0.85 | 0.77 | 0.71 |
| 1 | 116834105 | rs975351 | 0.67 | 0.229 | -0.868 | -0.838 | -0.039 | -0.01 | -0.01 | 0.401 | 0.47 | 0.03 | 0.59 | 0.63 | 0.45 | 0.35 |
| 1 | 116838208 | rs10924049 | 0.797 | 0.272 | -0.487 | -1.229 | -0.07 | 0.091 | 0.091 | 0 | 0.24 | 0.02 | 0.29 | 0.09 | 0.31 | 0.23 |
| 1 | 116851932 | rs4839532 | 0.199 | 0.397 | -0.059 | -1.149 | 0.065 | -0.01 | -0.01 | 0 | 0.8 | 0.49 | 0.85 | 0.84 | 0.76 | 0.73 |
| 1 | 116859356 | rs4839534 | 0.454 | 0.42 | -0.35 | -1.128 | 0.017 | -0.002 | -0.002 | 0.515 | 0.78 | 0.31 | 0.83 | 0.81 | 0.74 | 0.69 |
| 1 | 116860957 | rs17553121 | 0.596 | -0.396 | -0.97 | -0.722 | -0.049 | 0.063 | 0.063 | 0.176 | 0.46 | 0.03 | 0.5 | 0.72 | 0.39 | 0.4 |
| 1 | 116873813 | rs6428892 | -1.74 | -0.576 | 0.663 | -0.315 | -0.052 | -0.071 | -0.071 | 0 | 0.13 | 0.7 | 0.09 | 0 | 0.03 | 0.05 |
| 1 | 116884348 | rs1544450 | -2.318 | -0.455 | 0.206 | -0.345 | 0.032 | 0.062 | 0.062 | 3.74 | 0.17 | 0.93 | 0.12 | 0.03 | 0.08 | 0.15 |
| 1 | 116886230 | rs6428895 | -1.375 | -0.438 | 1.007 | -0.321 | -0.02 | -0.051 | -0.051 | 0 | 0.11 | 0.47 | 0.06 | 0 | 0.04 | 0.05 |
| 1 | 116887455 | rs12087334 | -2.145 | -0.474 | 0.34 | -0.344 | 0.033 | 0.068 | 0.068 | 4.369 | 0.17 | 0.95 | 0.1 | 0.03 | 0.07 | 0.14 |
| 1 | 116887742 | rs6674304 | -2.182 | -0.544 | 0.338 | -0.188 | 0.04 | 0.021 | 0.021 | 4.52 | 0.17 | 0.95 | 0.16 | 0.1 | 0.07 | 0.13 |
| 1 | 116903246 | rs7552721 | -2.202 | -0.633 | 1.286 | -0.193 | -0.014 | -0.007 | -0.007 | 0.109 | 0.16 | 0.78 | 0.14 | 0.09 | 0.05 | 0.12 |
| 1 | 116906839 | rs850617 | 0.873 | -1.001 | 0.003 | -0.573 | -0.072 | 0.106 | 0.106 | 0.394 | 0.4 | 0.05 | 0.41 | 0.2 | 0.58 | 0.59 |
| 1 | 116907359 | rs7539947 | -1.82 | -1.085 | 0.957 | -0.469 | -0.043 | -0.022 | -0.022 | 0 | 0.16 | 0.68 | 0.13 | 0.09 | 0.06 | 0.12 |
| 1 | 116918385 | rs757578 | 0.089 | -1.264 | 0.636 | -0.674 | 0.014 | 0.056 | 0.056 | 0 | 0.15 | 0.02 | 0.05 | 0.03 | 0.07 | 0.12 |
| 1 | 116919519 | rs10924073 | -2.134 | -0.83 | 1.67 | -0.256 | -0.027 | -0.026 | -0.026 | 0.092 | 0.1 | 0.7 | 0.11 | 0.04 | 0.03 | 0.04 |
| 1 | 116923792 | rs6428896 | -2.241 | -0.847 | 2.871 | -0.265 | -0.027 | -0.03 | -0.03 | 0.471 | 0.11 | 0.71 | 0.12 | 0.04 | 0.03 | 0.05 |
| 1 | 116931628 | rs12079419 | -1.613 | -0.847 | 2.849 | -0.264 | 0.017 | 0.064 | 0.064 | 4.636 | 0.07 | 0.42 | 0.09 | 0.04 | 0 | 0.01 |
| 1 | 116935068 | rs10924081 | -2.838 | -0.855 | 2.508 | -0.268 | 0.034 | 0.035 | 0.035 | 9.033 | 0.13 | 0.93 | 0.14 | 0.04 | 0.03 | 0.05 |
| 1 | 116945161 | rs1998447 | -2.49 | -0.838 | 2.526 | -0.26 | -0.005 | -0.006 | -0.006 | 0 | 0.12 | 0.82 | 0.13 | 0.04 | 0.03 | 0.05 |
| 1 | 116957604 | rs879651 | -2.495 | -0.838 | 2.522 | -0.26 | -0.003 | -0.004 | -0.004 | 0 | 0.12 | 0.83 | 0.13 | 0.04 | 0.03 | 0.05 |
| 1 | 116991956 | rs12128828 | -1.957 | -0.675 | 1.584 | -0.137 | 0.007 | 0.028 | 0.028 | 3.789 | 0.09 | 0.65 | 0.06 | 0.07 | 0.03 | 0.05 |
| 1 | 117000748 | rs6691908 | 0.23 | -0.961 | 1.05 | -0.326 | -0.014 | -0.017 | -0.017 | 0 | 0.08 | 0.3 | 0.03 | 0.01 | 0.01 | 0.03 |
| 1 | 117002636 | rs850598 | -1.122 | -1.076 | -0.092 | -0.155 | 0.055 | 0.054 | 0.054 | 0.825 | 0.43 | 0.94 | 0.44 | 0.73 | 0.35 | 0.36 |
| 1 | 117003588 | rs744908 | -1.658 | -0.654 | 0.699 | 0.345 | -0.026 | 0.007 | 0.007 | 2.198 | 0.15 | 0.81 | 0.08 | 0.17 | 0.03 | 0.06 |
| 1 | 117004657 | rs10494189 | -1.121 | -0.173 | 0.592 | 0.583 | 0.06 | 0.237 | 0.237 | 0 | 0.05 | 0.36 | 0.03 | 0.15 | 0.02 | 0.04 |
| 1 | 117010232 | rs17575616 | 0.277 | -0.011 | -0.8 | -0.021 | -0.127 | -0.015 | -0.015 | 0 | 0.35 | 0.06 | 0.44 | 0.28 | 0.52 | 0.43 |
| 1 | 117013732 | rs1885800 | -0.479 | -0.475 | -0.052 | -0.346 | -0.01 | -0.028 | -0.028 | 1.338 | 0.35 | 0.64 | 0.22 | 0.13 | 0.16 | 0.27 |
| 1 | 117015499 | rs12406751 | 0.213 | -0.351 | -0.345 | -0.515 | -0.045 | 0.115 | 0.115 | 0 | 0.25 | 0.01 | 0.16 | 0.05 | 0.19 | 0.24 |
| 1 | 117016646 | rs6661803 | -0.174 | -0.338 | 0.392 | -0.434 | -0.015 | 0.019 | 0.019 | 1.192 | 0.1 | 0.49 | 0.04 | 0.07 | 0.01 | 0.04 |
| 1 | 117019614 | rs6671978 | -0.55 | -0.385 | 0.374 | -0.473 | -0.023 | -0.07 | -0.07 | 0 | 0.36 | 0.45 | 0.21 | 0.13 | 0.21 | 0.28 |
| 1 | 117023431 | rs1176389 | -0.332 | 0.044 | 0.3 | -0.266 | -0.011 | -0.068 | -0.068 | 0.68 | 0.31 | 0.55 | 0.23 | 0.13 | 0.24 | 0.28 |
| 1 | 117024491 | rs1176387 | -0.369 | 0.77 | -0.681 | -0.361 | -0.011 | -0.007 | -0.007 | 0 | 0.83 | 0.86 | 0.7 | 0.77 | 0.64 | 0.64 |
| 1 | 117025609 | rs4839541 | 0.71 | 0.829 | -0.642 | -0.288 | 0.029 | 0.031 | 0.031 | 0 | 0.53 | 0.34 | 0.4 | 0.62 | 0.42 | 0.38 |
| 1 | 117026156 | rs1176386 | 0.633 | 1.486 | -0.716 | 0.197 | 0.054 | 0.004 | 0.004 | 0.238 | 0.79 | 0.7 | 0.7 | 0.77 | 0.65 | 0.61 |
| 1 | 117027445 | rs1176385 | 0.199 | 1.218 | -1.787 | 0.425 | -0.046 | -0.033 | -0.033 | 0 | 0.85 | 0.97 | 0.78 | 0.78 | 0.75 | 0.64 |
| 1 | 117030982 | rs10924102 | -0.344 | -0.001 | -2.121 | 0.154 | 0.057 | 0.056 | 0.056 | 0.096 | 0.44 | 0.91 | 0.34 | 0.37 | 0.32 | 0.47 |
| 1 | 117036551 | rs10924103 | -0.949 | -0.873 | -1.446 | 0.063 | -0.006 | -0.001 | -0.001 | 0 | 0.3 | 0.71 | 0.39 | 0.68 | 0.21 | 0.4 |
| 1 | 117038040 | rs766429 | 0.231 | -1.621 | -1.421 | -0.812 | 0.071 | 0.167 | 0.167 | 2.159 | 0.32 | 0.04 | 0.25 | 0.05 | 0.22 | 0.2 |
| 1 | 117049676 | rs10754444 | -0.38 | -1.584 | -0.353 | -0.113 | -0.02 | 0.088 | 0.088 | 0 | 0.21 | 0.36 | 0.32 | 0.58 | 0.14 | 0.4 |
| 1 | 117053745 | rs10924106 | 0.021 | -1.618 | -0.211 | -0.121 | -0.001 | 0.025 | 0.025 | 0.036 | 0.16 | 0.14 | 0.29 | 0.56 | 0.12 | 0.36 |
| 1 | 117056730 | rs10802189 | -1.348 | -2.043 | -0.337 | -0.333 | 0.091 | 0.007 | 0.007 | 0.204 | 0.54 | 0.92 | 0.5 | 0.62 | 0.35 | 0.56 |
| 1 | 117062474 | rs10924108 | -0.505 | -1.003 | 0.544 | 0.529 | -0.001 | 0.027 | 0.027 | 0.006 | 0.16 | 0.15 | 0.29 | 0.56 | 0.12 | 0.36 |
| 1 | 117066907 | rs1414275 | -0.96 | -1.005 | 0.539 | 0.552 | -0.015 | 0.122 | 0.122 | 0 | 0.18 | 0.36 | 0.3 | 0.56 | 0.12 | 0.39 |
| 1 | 117080166 | rs6677309 | -1.635 | -1.004 | 0.685 | 0.56 | -0.036 | 0.095 | 0.095 | 0.249 | 0.23 | 0.59 | 0.34 | 0.59 | 0.14 | 0.41 |
| 1 | 117081810 | rs758518 | -0.65 | -0.699 | 0.028 | 0.709 | -0.004 | 0.047 | 0.047 | 0.011 | 0.17 | 0.18 | 0.29 | 0.56 | 0.12 | 0.37 |
| 1 | 117087779 | rs12044852 | -0.48 | -0.628 | -0.416 | 0.755 | 0 | -0.046 | -0.046 | 0 | 0.15 | 0.07 | 0.28 | 0.55 | 0.12 | 0.36 |
| 1 | 117104215 | rs2300747 | -1.045 | -0.797 | -0.215 | 0.657 | -0.011 | 0.036 | 0.036 | 0 | 0.2 | 0.27 | 0.33 | 0.59 | 0.14 | 0.39 |

| | | | | | | | | | | | | | | | | |
|---|---|---|---|---|---|---|---|---|---|---|---|---|---|---|---|---|
| 1 | 117115129 | rs17036001 | -0.834 | -0.956 | -0.018 | 0.614 | -0.013 | 0.007 | 0.007 | 0 | 0.07 | 0.12 | 0.11 | 0.08 | 0.05 | 0.09 |
| 1 | 117115868 | rs1414272 | -1.262 | -0.921 | -0.729 | 0.171 | 0.007 | 0.055 | 0.055 | 0.543 | 0.31 | 0.91 | 0.34 | 0.15 | 0.33 | 0.27 |
| 1 | 117121865 | rs12066192 | -0.796 | -0.889 | -0.286 | 0.147 | 0.002 | 0.075 | 0.075 | 2.483 | 0.26 | 0.92 | 0.23 | 0.08 | 0.28 | 0.19 |
| 1 | 117122130 | rs6703791 | -0.966 | -0.179 | 0.141 | 0.615 | 0.016 | 0.137 | 0.137 | 0 | 0.2 | 0.35 | 0.3 | 0.52 | 0.17 | 0.37 |
| 1 | 117128172 | rs570440 | -0.964 | 0.686 | -1.148 | -0.002 | -0.103 | 0.136 | 0.136 | 1.055 | 0.35 | 0.93 | 0.37 | 0.12 | 0.43 | 0.29 |
| 1 | 117164591 | rs12750589 | 0.78 | 0.629 | -1.106 | -0.108 | -0.028 | 0.078 | 0.078 | 0.082 | 0.58 | 0.24 | 0.51 | 0.79 | 0.47 | 0.69 |
| 1 | 117199884 | rs606604 | -0.751 | 0.184 | -1.616 | 0.611 | -0.005 | -0.121 | -0.121 | 0.056 | 0.78 | 0.97 | 0.75 | 0.49 | 0.81 | 0.86 |
| 1 | 117206336 | rs1981183 | -0.524 | 0.583 | -1.427 | 0.519 | -0.007 | -0.03 | -0.03 | 0 | 0.2 | 0.17 | 0.26 | 0.42 | 0.22 | 0.46 |
| 1 | 117212594 | rs599563 | -1.586 | 0.346 | -1.506 | 0.684 | -0.029 | -0.121 | -0.121 | 1.402 | 0.84 | 0.97 | 0.82 | 0.55 | 0.84 | 0.86 |
| 1 | 117227513 | rs668224 | 1.496 | 0.253 | -1.542 | -0.042 | -0.101 | 0.172 | 0.172 | 0.364 | 0.36 | 0.03 | 0.32 | 0.11 | 0.44 | 0.2 |
| 1 | 117232158 | rs699713 | -0.383 | 0.523 | -0.392 | 0.798 | -0.001 | 0 | 0 | 0 | 0.17 | 0.2 | 0.23 | 0.4 | 0.17 | 0.47 |
| 1 | 117234297 | rs699714 | -0.748 | 0.58 | -0.124 | 0.749 | 0.012 | 0.071 | 0.071 | 0 | 0.19 | 0.52 | 0.26 | 0.31 | 0.17 | 0.35 |
| 1 | 117236379 | rs696663 | -0.718 | 0.594 | -0.162 | 0.755 | 0.013 | 0.069 | 0.069 | 0 | 0.19 | 0.51 | 0.25 | 0.32 | 0.17 | 0.35 |
| 1 | 117243440 | rs12127189 | 0.326 | 0.715 | -0.997 | 1.074 | -0.026 | -0.08 | -0.08 | 0.014 | 0.19 | 0.05 | 0.19 | 0.36 | 0.21 | 0.21 |
| 1 | 117245849 | rs699718 | -0.805 | 0.498 | -0.339 | 1.194 | -0.033 | 0.107 | 0.107 | 0 | 0.21 | 0.39 | 0.3 | 0.48 | 0.11 | 0.43 |
| 1 | 117255079 | rs6428666 | 0.48 | 1.326 | -0.461 | 0.889 | -0.01 | -0.008 | -0.008 | 5.854 | 0.75 | 0.77 | 0.63 | 0.64 | 0.69 | 0.62 |
| 1 | 117255270 | rs6428667 | 1.149 | 1.315 | -0.421 | 0.7 | -0.001 | -0.008 | -0.008 | 0 | 0.54 | 0.47 | 0.42 | 0.2 | 0.47 | 0.37 |
| 1 | 117255312 | rs10923135 | 0.577 | 1.107 | -0.643 | 1.018 | -0.012 | -0.052 | -0.052 | 4.271 | 0.17 | 0.09 | 0.17 | 0.36 | 0.19 | 0.2 |
| 1 | 117255426 | rs1775762 | -0.695 | 0.429 | -0.409 | 0.966 | 0 | 0.008 | 0.008 | 0.08 | 0.14 | 0.15 | 0.27 | 0.33 | 0.13 | 0.29 |
| 1 | 117256697 | rs9887785 | 0.628 | 0.539 | -0.056 | 0.331 | 0.006 | 0.009 | 0.009 | 0 | 0.66 | 0.47 | 0.58 | 0.58 | 0.65 | 0.65 |
| 1 | 117262791 | rs958434 | 0.623 | -0.313 | 0.406 | -0.124 | 0.009 | -0.013 | -0.013 | 0 | 0.34 | 0.29 | 0.31 | 0.52 | 0.24 | 0.33 |
| 1 | 117263790 | rs624988 | -0.283 | -0.269 | 0.537 | -0.34 | 0.002 | 0.004 | 0.004 | 0 | 0.6 | 0.44 | 0.51 | 0.59 | 0.59 | 0.49 |
| 1 | 117263868 | rs12405671 | 1.278 | -0.388 | 0.411 | -0.43 | -0.03 | 0.033 | 0.033 | 0 | 0.18 | 0.08 | 0.19 | 0.04 | 0.26 | 0.23 |
| 1 | 117264106 | rs612865 | -1.6 | -0.427 | 0.564 | -0.437 | -0.006 | 0.046 | 0.046 | 0 | 0.81 | 0.76 | 0.71 | 0.63 | 0.84 | 0.71 |
| 1 | 117265154 | rs699727 | 1.463 | -0.64 | 0.26 | -0.665 | 0.01 | 0.009 | 0.009 | 0.2 | 0.28 | 0.2 | 0.12 | 0.19 | 0.13 | 0.26 |
| 1 | 117268911 | rs7529979 | 1.222 | -0.628 | 0.133 | -0.666 | 0.01 | 0.01 | 0.01 | 0 | 0.3 | 0.22 | 0.12 | 0.18 | 0.14 | 0.26 |
| 1 | 117271568 | rs519264 | 0.485 | -0.517 | 0.073 | -0.688 | -0.064 | 0.098 | 0.098 | 0 | 0.25 | 0.08 | 0.18 | 0.08 | 0.33 | 0.14 |
| 1 | 117273577 | rs614448 | -0.457 | -0.916 | 0.523 | -0.294 | -0.016 | -0.004 | -0.004 | 0 | 0.69 | 0.45 | 0.49 | 0.31 | 0.73 | 0.63 |
| 1 | 117275764 | rs10923138 | -0.161 | -0.638 | 0.257 | -0.332 | -0.024 | 0.05 | 0.05 | 1.072 | 0.37 | 0.24 | 0.32 | 0.12 | 0.52 | 0.37 |
| 1 | 117282727 | rs12408586 | 0.623 | -0.509 | 0.626 | 0.327 | -0.021 | 0.039 | 0.039 | 0 | 0.18 | 0.11 | 0.13 | 0.02 | 0.26 | 0.12 |
| 1 | 117287643 | rs7540836 | 0.702 | -0.545 | 0.587 | 0.537 | -0.018 | 0.05 | 0.05 | 0 | 0.42 | 0.18 | 0.39 | 0.26 | 0.41 | 0.45 |
| 1 | 117293763 | rs797999 | 0.181 | -0.691 | 0.406 | 1.106 | 0.005 | -0.015 | -0.015 | 0 | 0.73 | 0.79 | 0.66 | 0.61 | 0.77 | 0.67 |
| 1 | 117299420 | rs3136700 | -1.734 | -0.834 | 1.515 | 1.114 | -0.02 | 0.078 | 0.078 | 0.027 | 0.14 | 0.29 | 0.21 | 0.28 | 0.11 | 0.11 |
| 1 | 117300926 | rs798045 | -1.603 | -0.95 | 1.949 | 1.099 | -0.015 | 0.152 | 0.152 | 0 | 0.12 | 0.35 | 0.22 | 0.29 | 0.1 | 0.1 |
| 1 | 117302613 | rs798044 | -1.552 | -1.071 | 2.401 | 1.021 | -0.036 | 0.129 | 0.129 | 0.183 | 0.14 | 0.37 | 0.22 | 0.29 | 0.1 | 0.09 |
| 1 | 117307358 | rs798038 | -2.213 | -1.248 | 2.377 | 0.881 | -0.051 | 0.025 | 0.025 | 1.001 | 0.18 | 0.63 | 0.25 | 0.28 | 0.1 | 0.1 |
| 1 | 117308238 | rs798037 | -1.559 | -1.244 | 2.5 | 0.893 | -0.038 | 0.076 | 0.076 | 0.108 | 0.15 | 0.32 | 0.22 | 0.29 | 0.1 | 0.1 |
| 1 | 117312119 | rs798034 | -2.16 | -1.281 | 2.245 | 0.85 | -0.051 | 0.026 | 0.026 | 1.064 | 0.18 | 0.63 | 0.25 | 0.29 | 0.1 | 0.1 |
| 1 | 117326135 | rs798055 | -0.625 | -1.282 | 1.712 | 0.823 | -0.013 | 0.018 | 0.018 | 1.714 | 0.1 | 0.08 | 0.07 | 0.26 | 0.04 | 0.05 |
| 1 | 117333461 | rs798046 | -1.862 | -1.375 | 1.43 | 0.778 | -0.067 | 0.011 | 0.011 | 3.423 | 0.18 | 0.63 | 0.12 | 0.27 | 0.05 | 0.06 |
| 1 | 117337524 | rs798029 | -1.484 | -1.366 | 1.329 | 0.783 | -0.08 | 0.039 | 0.039 | 1.885 | 0.17 | 0.54 | 0.11 | 0.26 | 0.05 | 0.05 |
| 1 | 117337726 | rs798028 | -1.634 | -1.352 | 1.112 | 0.789 | -0.077 | 0.02 | 0.02 | 3.116 | 0.18 | 0.6 | 0.12 | 0.26 | 0.05 | 0.05 |
| 1 | 117339802 | rs798023 | -1.672 | -1.34 | 0.962 | 0.796 | -0.07 | 0.013 | 0.013 | 2.993 | 0.18 | 0.62 | 0.12 | 0.26 | 0.05 | 0.05 |
| 1 | 117361083 | rs1182218 | -1.183 | -1.353 | 0.619 | 0.495 | -0.034 | -0.019 | -0.019 | 0 | 0.3 | 0.63 | 0.29 | 0.28 | 0.16 | 0.21 |
| 1 | 117367190 | rs6428670 | 0.9 | -1.408 | 0.822 | 0.73 | -0.042 | -0.025 | -0.025 | 0 | 0.71 | 0.43 | 0.72 | 0.71 | 0.84 | 0.79 |
| 1 | 117373464 | rs2767333 | 0.3 | -1.593 | 1.449 | 0.6 | -0.031 | -0.06 | -0.06 | 0 | 0.79 | 0.74 | 0.91 | 0.97 | 0.89 | 0.87 |
| 1 | 117376448 | rs2767326 | 1.399 | -1.631 | 1.111 | 1.116 | -0.011 | 0.004 | 0.004 | 0.001 | 0.57 | 0.43 | 0.59 | 0.42 | 0.67 | 0.6 |
| 1 | 117384443 | rs7527627 | -1.849 | -1.683 | 1.374 | 0.824 | 0.02 | 0.015 | 0.015 | 4.347 | 0.07 | 0.52 | 0.04 | 0 | 0 | 0 |
| 1 | 117392487 | rs6663197 | -1.159 | -1.637 | 1.176 | 0.846 | 0.02 | 0.016 | 0.016 | 2.434 | 0.06 | 0.38 | 0.03 | 0 | 0 | 0 |
| 1 | 117392940 | rs10494190 | 0.016 | -1.551 | 0.767 | 1.108 | -0.016 | -0.05 | -0.05 | 0.089 | 0.15 | 0.03 | 0.14 | 0.23 | 0.18 | 0.21 |
| 1 | 117411383 | rs1013910 | -0.426 | -1.396 | 0.404 | 0.76 | 0.016 | 0.025 | 0.025 | 0 | 0.83 | 1 | 0.89 | 0.94 | 0.88 | 0.88 |
| 1 | 117435020 | rs12726592 | 0.663 | -0.802 | 0.11 | 0.477 | -0.084 | 0.032 | 0.032 | 1.037 | 0.35 | 0.02 | 0.42 | 0.23 | 0.36 | 0.45 |
| 1 | 117442421 | rs10923158 | -0.169 | -1.052 | 1.254 | 0.871 | -0.03 | 0.037 | 0.037 | 0 | 0.12 | 0.28 | 0.05 | 0.14 | 0.06 | 0.1 |
| 1 | 117445514 | rs2251401 | -0.857 | -1.195 | 1.665 | 0.804 | 0.03 | 0.027 | 0.027 | 4.115 | 0.06 | 0.37 | 0.03 | 0 | 0 | 0 |
| 1 | 117447049 | rs2767336 | -1.456 | -1.237 | 1.781 | 1.058 | 0.022 | 0.018 | 0.018 | 6.184 | 0.07 | 0.47 | 0.04 | 0 | 0 | 0 |
| 1 | 117448282 | rs2251108 | -1.353 | -1.289 | 1.607 | 1.04 | -0.034 | -0.003 | -0.003 | 0 | 0.17 | 0.76 | 0.09 | 0.15 | 0.06 | 0.09 |
| 1 | 117458375 | rs2254121 | -1.233 | -1.303 | 1.591 | 1.024 | -0.056 | 0 | 0 | 0 | 0.16 | 0.68 | 0.09 | 0.15 | 0.05 | 0.08 |
| 1 | 117460934 | rs7365057 | 0.601 | -1.274 | 1.511 | 1.038 | -0.001 | 0.002 | 0.002 | 0 | 0.08 | 0.08 | 0.04 | 0.15 | 0.05 | 0.05 |
| 1 | 117463537 | rs2182605 | -0.047 | -0.851 | 1.23 | 0.98 | 0 | 0.002 | 0.002 | 0 | 0.52 | 0.55 | 0.57 | 0.42 | 0.47 | 0.54 |
| 1 | 117471094 | rs2806866 | -1.407 | -0.8 | 1.356 | 0.612 | 0.03 | 0.026 | 0.026 | 4.014 | 0.06 | 0.45 | 0.03 | 0 | 0 | 0 |
| 1 | 117472999 | rs2148634 | 0.041 | -0.313 | 0.481 | 0.432 | 0.008 | 0.045 | 0.045 | 0 | 0.5 | 0.38 | 0.58 | 0.3 | 0.44 | 0.53 |
| 1 | 117473419 | rs17036591 | -0.841 | -1.001 | 0.176 | -0.748 | -0.037 | -0.104 | -0.104 | 4.667 | 0.24 | 0.37 | 0.25 | 0.09 | 0.19 | 0.21 |
| 1 | 117473588 | rs11583252 | -1.141 | -0.447 | 0.374 | -0.81 | -0.016 | -0.089 | -0.089 | 0 | 0.14 | 0.36 | 0.08 | 0 | 0.12 | 0.07 |
| 1 | 117474229 | rs2478766 | -0.115 | -0.607 | 0.401 | -0.543 | -0.031 | 0.023 | 0.023 | 0.045 | 0.14 | 0.38 | 0.07 | 0.14 | 0.07 | 0.07 |
| 1 | 117477572 | rs2246315 | -0.533 | -0.216 | 0.016 | -0.977 | -0.014 | 0.017 | 0.017 | 0 | 0.82 | 0.87 | 0.82 | 0.95 | 0.76 | 0.91 |
| 1 | 117479245 | rs2806880 | 0.89 | -0.254 | 0.133 | -0.973 | -0.023 | 0.019 | 0.019 | 0 | 0.12 | 0.27 | 0.06 | 0.14 | 0.07 | 0.07 |
| 1 | 117479752 | rs6692981 | -1.228 | -0.085 | -0.037 | -0.998 | 0 | 0.028 | 0.028 | 0 | 0.86 | 0.95 | 0.88 | 0.95 | 0.87 | 0.92 |
| 1 | 117482765 | rs4539125 | 1.027 | -0.102 | 0.114 | -1.006 | -0.001 | -0.007 | -0.007 | 0 | 0.66 | 0.36 | 0.73 | 0.81 | 0.67 | 0.78 |
| 1 | 117482818 | rs6700778 | -0.695 | -0.096 | 0.114 | -0.97 | 0.07 | -0.029 | -0.029 | 0 | 0.07 | 0.26 | 0.08 | 0 | 0.12 | 0.06 |

| | | | | | | | | | | | | | | | | |
|---|---|---|---|---|---|---|---|---|---|---|---|---|---|---|---|---|
| 1 | 117491123 | rs4433378 | 0.901 | 0.065 | -0.852 | -0.657 | -0.032 | 0.015 | 0.015 | 0.853 | 0.41 | 0.06 | 0.49 | 0.33 | 0.39 | 0.54 |
| 1 | 117494820 | rs4128518 | 0.825 | 0.168 | -0.318 | -0.114 | -0.002 | 0.015 | 0.015 | 0 | 0.66 | 0.43 | 0.7 | 0.39 | 0.67 | 0.67 |
| 1 | 117508304 | rs12024946 | -1.015 | 0.435 | -0.014 | 0.4 | -0.016 | 0.038 | 0.038 | 0 | 0.06 | 0.05 | 0.09 | 0.38 | 0.13 | 0.17 |
| 1 | 117510804 | rs10801917 | -1.542 | 0.498 | 0.09 | 0.15 | 0.013 | 0.044 | 0.044 | 0.136 | 0.29 | 0.51 | 0.27 | 0.43 | 0.31 | 0.34 |
| 1 | 117512505 | rs12096164 | -0.045 | 0.458 | 0.087 | 0.132 | 0.002 | 0 | 0 | 0 | 0.19 | 0.17 | 0.16 | 0.05 | 0.18 | 0.17 |
| 1 | 117514334 | rs6687760 | -1.844 | 0.542 | 0.345 | 0.379 | 0.114 | 0.35 | 0.35 | 0.23 | 0.1 | 0.35 | 0.12 | 0.46 | 0.14 | 0.17 |
| 1 | 117514675 | rs17520400 | 0.049 | 0.399 | -0.015 | 0.064 | 0.018 | 0.044 | 0.044 | 0 | 0.17 | 0.07 | 0.09 | 0.01 | 0.14 | 0.15 |
| 1 | 117524022 | rs12063648 | -1.255 | 0.65 | 0.163 | 0.26 | 0.014 | 0.053 | 0.053 | 0.18 | 0.27 | 0.52 | 0.27 | 0.45 | 0.28 | 0.32 |
| 1 | 117526055 | rs11576262 | 0.798 | 0.587 | -0.632 | 0.167 | -0.07 | -0.03 | -0.03 | 0 | 0.25 | 0.01 | 0.28 | 0.17 | 0.23 | 0.33 |
| 1 | 117530273 | rs3885 | -1.5 | 0.586 | -0.389 | 0.146 | -0.001 | -0.003 | -0.003 | 0.011 | 0.53 | 0.51 | 0.65 | 0.66 | 0.51 | 0.65 |
| 1 | 117530477 | rs10801923 | -1.505 | 0.585 | -0.388 | 0.146 | -0.002 | -0.004 | -0.004 | 1.247 | 0.53 | 0.51 | 0.65 | 0.66 | 0.51 | 0.65 |
| 1 | 117530507 | rs1127655 | -1.381 | 0.581 | -0.357 | 0.144 | 0.001 | 0.002 | 0.002 | 0 | 0.52 | 0.47 | 0.65 | 0.66 | 0.51 | 0.65 |
| 1 | 117531620 | rs13951 | -1.154 | 0.612 | -0.29 | 0.132 | -0.006 | 0.042 | 0.042 | 0 | 0.48 | 0.29 | 0.59 | 0.66 | 0.44 | 0.64 |
| 1 | 117531813 | rs3829881 | -1.517 | 0.607 | -0.283 | 0.132 | -0.001 | -0.002 | -0.002 | 0 | 0.53 | 0.52 | 0.65 | 0.66 | 0.51 | 0.65 |
| 1 | 117531911 | rs10923191 | -1.302 | 0.603 | -0.231 | 0.13 | 0.028 | 0.056 | 0.056 | 0 | 0.51 | 0.29 | 0.64 | 0.66 | 0.51 | 0.61 |
| 1 | 117532291 | rs8254 | 0.747 | 0.607 | -0.257 | 0.141 | -0.035 | -0.007 | -0.007 | 0.067 | 0.26 | 0.09 | 0.39 | 0.21 | 0.23 | 0.37 |
| 1 | 117532372 | rs11836 | -1.041 | 0.608 | -0.242 | 0.143 | 0.012 | 0.043 | 0.043 | 0.111 | 0.27 | 0.37 | 0.26 | 0.45 | 0.28 | 0.28 |
| 1 | 117532790 | rs1127215 | -0.801 | 0.623 | -0.269 | 0.148 | 0.005 | 0.044 | 0.044 | 0 | 0.46 | 0.34 | 0.54 | 0.26 | 0.39 | 0.53 |
| 1 | 117536802 | rs10923193 | 0.336 | 0.46 | -0.495 | 0.208 | -0.031 | -0.057 | -0.057 | 4.84 | 0.26 | 0.13 | 0.45 | 0.61 | 0.26 | 0.43 |
| 1 | 117539373 | rs4659051 | -0.054 | 0.518 | -0.244 | 0.268 | -0.006 | -0.015 | -0.015 | 0 | 0.3 | 0.26 | 0.47 | 0.61 | 0.27 | 0.47 |
| 1 | 117541054 | rs10801925 | 0.099 | 0.531 | -0.28 | 0.276 | -0.039 | -0.071 | -0.071 | 0 | 0.29 | 0.11 | 0.45 | 0.61 | 0.28 | 0.43 |
| 1 | 117544702 | rs10494192 | 0.514 | 0.749 | -0.303 | 0.382 | -0.079 | -0.047 | -0.047 | 0 | 0.23 | 0.01 | 0.38 | 0.21 | 0.23 | 0.32 |
| 1 | 117547790 | rs2806870 | 0.376 | 0.788 | -0.335 | 0.24 | 0 | -0.019 | -0.019 | 0 | 0.17 | 0.14 | 0.15 | 0.04 | 0.16 | 0.14 |
| 1 | 117550623 | rs2764850 | -1.537 | 0.888 | -0.742 | 0.326 | 0.009 | 0.018 | 0.018 | 0 | 0.52 | 0.62 | 0.69 | 0.66 | 0.51 | 0.64 |
| 1 | 117560818 | rs3736908 | 0.092 | 0.671 | -1.457 | 0.389 | -0.093 | -0.123 | -0.123 | 2.46 | 0.27 | 0.04 | 0.44 | 0.62 | 0.28 | 0.47 |
| 1 | 117560929 | rs3736907 | 0.38 | 0.48 | -1.476 | 0.335 | -0.08 | -0.044 | -0.044 | 0.506 | 0.24 | 0.01 | 0.39 | 0.21 | 0.23 | 0.33 |
| 1 | 117561863 | rs2806874 | -0.868 | 0.276 | -0.986 | -0.174 | 0.01 | 0.022 | 0.022 | 0 | 0.53 | 0.64 | 0.7 | 0.67 | 0.53 | 0.64 |
| 1 | 117571079 | rs17587765 | -0.258 | -0.389 | -1.133 | -0.575 | 0.016 | 0.101 | 0.101 | 0.034 | 0.24 | 0.05 | 0.14 | 0.05 | 0.2 | 0.25 |
| 1 | 117574023 | rs3767780 | -0.712 | -0.053 | -0.486 | -0.275 | 0.016 | 0.027 | 0.027 | 0 | 0.49 | 0.57 | 0.59 | 0.29 | 0.41 | 0.52 |
| 1 | 117582373 | rs12121509 | -1.25 | 0.22 | 0.259 | 0.411 | 0.069 | 0.12 | 0.12 | 0 | 0.13 | 0.51 | 0.15 | 0.18 | 0.14 | 0.19 |
| 1 | 117596441 | rs10923201 | -0.35 | -0.047 | -0.636 | -0.227 | -0.003 | 0.065 | 0.065 | 0 | 0.23 | 0.06 | 0.16 | 0.06 | 0.23 | 0.22 |
| 1 | 117599054 | rs2764862 | 1.106 | -0.918 | -0.065 | -0.696 | 0.019 | 0.04 | 0.04 | 0 | 0.6 | 0.34 | 0.73 | 0.84 | 0.66 | 0.71 |
| 1 | 117600045 | rs2477725 | 1.448 | -1.476 | -0.401 | -0.653 | 0.073 | 0.113 | 0.113 | 0 | 0.52 | 0.1 | 0.66 | 0.75 | 0.6 | 0.52 |
| 1 | 117601619 | rs3754109 | -0.07 | -1.46 | -0.284 | -0.944 | -0.021 | 0.012 | 0.012 | 0 | 0.08 | 0.19 | 0.07 | 0.09 | 0.05 | 0.19 |
| 1 | 117601896 | rs1004986 | 0.824 | -1.533 | 0.695 | -0.992 | -0.058 | -0.052 | -0.052 | 0 | 0.79 | 0.38 | 0.86 | 0.91 | 0.88 | 0.79 |
| 1 | 117602960 | rs2244501 | 0.836 | -1.542 | 0.69 | -0.973 | -0.058 | -0.049 | -0.049 | 0 | 0.79 | 0.38 | 0.86 | 0.9 | 0.88 | 0.79 |
| 1 | 117603156 | rs2274253 | -0.062 | -1.513 | 0.624 | -0.954 | -0.021 | 0.014 | 0.014 | 0 | 0.08 | 0.2 | 0.07 | 0.1 | 0.05 | 0.19 |
| 1 | 117603188 | rs2244406 | 0.468 | -1.434 | 0.544 | -1.018 | -0.021 | -0.106 | -0.106 | 0 | 0.64 | 0.51 | 0.77 | 0.94 | 0.69 | 0.75 |
| 1 | 117612298 | rs10754334 | 1.59 | -1.845 | -0.055 | -1.165 | 0.012 | 0.079 | 0.079 | 0.16 | 0.47 | 0.07 | 0.62 | 0.7 | 0.57 | 0.49 |
| 1 | 117615935 | rs1289675 | -0.766 | -1.876 | 0.613 | -1.193 | -0.017 | -0.055 | -0.055 | 0 | 0.9 | 0.77 | 0.95 | 1 | 0.93 | 0.98 |
| 1 | 117617705 | rs998532 | 1.173 | -1.885 | 0.124 | -1.194 | 0.035 | 0.151 | 0.151 | 0 | 0.54 | 0.12 | 0.67 | 0.77 | 0.59 | 0.6 |
| 1 | 117621281 | rs3767767 | 0.147 | -1.578 | 0.247 | -1.055 | -0.011 | 0.077 | 0.077 | 0.008 | 0.24 | 0.08 | 0.17 | 0.06 | 0.26 | 0.2 |
| 1 | 117628620 | rs2764854 | 0.384 | -1.535 | 0.339 | -0.787 | 0.02 | -0.008 | -0.008 | 0.295 | 0.06 | 0.03 | 0.05 | 0.06 | 0.02 | 0.11 |
| 1 | 117635905 | rs10494197 | -0.135 | -1.357 | 0.728 | -0.755 | -0.02 | 0.045 | 0.045 | 0 | 0.05 | 0.17 | 0.05 | 0.09 | 0.02 | 0.13 |
| 1 | 117636493 | rs2057594 | -0.429 | -1.359 | 0.658 | -1.193 | 0 | -0.004 | -0.004 | 0 | 0.27 | 0.22 | 0.18 | 0.13 | 0.25 | 0.17 |
| 1 | 117637634 | rs1289667 | 0.953 | -1.267 | 1.041 | -0.974 | -0.016 | -0.029 | -0.029 | 0.185 | 0.84 | 0.64 | 0.89 | 0.86 | 0.88 | 0.92 |
| 1 | 117638270 | rs1289666 | 0.761 | -1.644 | 0.65 | -1.233 | -0.014 | -0.008 | -0.008 | 0.014 | 0.77 | 0.68 | 0.82 | 0.77 | 0.84 | 0.77 |
| 1 | 117643482 | rs1289663 | -0.613 | -0.873 | 0.323 | -1.789 | -0.002 | 0.002 | 0.002 | 0 | 0.41 | 0.61 | 0.48 | 0.18 | 0.43 | 0.38 |
| 1 | 117645744 | rs9728670 | 1.881 | -0.774 | -0.176 | -2.066 | -0.086 | 0.199 | 0.199 | 0 | 0.32 | 0.02 | 0.27 | 0.05 | 0.35 | 0.27 |
| 1 | 117653380 | rs10923207 | 0.317 | -0.823 | 0.442 | -2.053 | -0.021 | 0.029 | 0.029 | 0 | 0.27 | 0.21 | 0.37 | 0.03 | 0.35 | 0.21 |
| 1 | 117654638 | rs12118664 | -0.546 | -0.867 | 0.896 | -2.07 | 0.006 | -0.059 | -0.059 | 0 | 0.35 | 0.44 | 0.39 | 0.11 | 0.38 | 0.3 |
| 1 | 117656088 | rs1289658 | -1.245 | -0.873 | 0.71 | -2.07 | -0.007 | 0.06 | 0.06 | 0 | 0.43 | 0.68 | 0.48 | 0.19 | 0.44 | 0.38 |
| 1 | 117657855 | rs2296445 | -0.74 | -0.956 | 0.635 | -2.074 | 0.015 | 0 | 0 | 0 | 0.07 | 0.03 | 0.01 | 0.08 | 0.02 | 0.09 |
| 1 | 117660608 | rs1289655 | -0.859 | -0.861 | 0.71 | -1.962 | -0.005 | 0.006 | 0.006 | 0 | 0.88 | 0.8 | 0.88 | 0.87 | 0.91 | 0.92 |
| 1 | 117660754 | rs749902 | 0.247 | -0.773 | 0.254 | -1.939 | -0.021 | 0.029 | 0.029 | 0 | 0.27 | 0.21 | 0.37 | 0.03 | 0.35 | 0.21 |
| 1 | 117663913 | rs3795655 | -0.4 | -0.785 | 0.257 | -1.934 | -0.011 | 0.001 | 0.001 | 0 | 0.72 | 0.76 | 0.63 | 0.97 | 0.64 | 0.79 |
| 1 | 117664046 | rs6657335 | -0.354 | -0.789 | 0.254 | -1.937 | -0.012 | 0.003 | 0.003 | 0 | 0.72 | 0.76 | 0.63 | 0.97 | 0.64 | 0.79 |
| 1 | 117674546 | rs1342736 | 0.583 | -0.622 | 0.347 | -1.764 | -0.006 | 0.036 | 0.036 | 0 | 0.6 | 0.35 | 0.7 | 0.81 | 0.58 | 0.6 |
| 1 | 117674749 | rs1342735 | 1.522 | -0.412 | -0.394 | -1.614 | -0.108 | 0.001 | 0.001 | 0 | 0.43 | 0.02 | 0.55 | 0.68 | 0.46 | 0.52 |
| 1 | 117675943 | rs4659126 | 0.39 | -0.281 | 0.772 | -1.326 | 0.002 | -0.04 | -0.04 | 4.657 | 0.66 | 0.66 | 0.79 | 0.89 | 0.65 | 0.7 |
| 1 | 117680515 | rs7546885 | -1.171 | -0.22 | 0.089 | -1.222 | -0.002 | 0.157 | 0.157 | 1.346 | 0.45 | 0.8 | 0.32 | 0.18 | 0.42 | 0.36 |
| 1 | 117685560 | rs1241 | 1.17 | -0.376 | 1.577 | -1.147 | 0 | -0.018 | -0.018 | 1.509 | 0.62 | 0.42 | 0.76 | 0.9 | 0.71 | 0.69 |
| 1 | 117690272 | rs10754339 | 2.093 | -0.366 | 2.037 | -1.018 | -0.025 | -0.031 | -0.031 | 0 | 0.85 | 0.29 | 0.87 | 0.92 | 0.92 | 0.85 |
| 1 | 117690758 | rs2358817 | -1.963 | -0.32 | 2.388 | -0.964 | -0.016 | -0.026 | -0.026 | 0.532 | 0.13 | 0.45 | 0.11 | 0.08 | 0.07 | 0.15 |
| 1 | 117704016 | rs7512111 | -1.694 | -0.293 | 2.364 | -0.979 | 0.009 | -0.05 | -0.05 | 0 | 0.1 | 0.37 | 0.1 | 0.02 | 0.07 | 0.09 |
| 1 | 117708477 | rs7524748 | -2.509 | -0.388 | 2.001 | -1.003 | -0.021 | -0.026 | -0.026 | 0 | 0.16 | 0.74 | 0.13 | 0.08 | 0.08 | 0.17 |
| 1 | 117708664 | rs1892410 | -0.493 | -0.271 | 1.49 | -0.935 | -0.006 | 0.015 | 0.015 | 0 | 0.06 | 0.02 | 0.07 | 0.02 | 0.07 | 0.1 |
| 1 | 117715026 | rs6657810 | 1.404 | -0.212 | 1.451 | -1.318 | -0.005 | -0.067 | -0.067 | 0 | 0.55 | 0.49 | 0.66 | 0.92 | 0.62 | 0.64 |
| 1 | 117718797 | rs6428686 | 0.324 | -0.238 | 1.084 | -1.32 | -0.024 | 0.199 | 0.199 | 0.205 | 0.68 | 0.97 | 0.77 | 0.99 | 0.7 | 0.84 |

| | | | | | | | | | | | | | | | | |
|---|---|---|---|---|---|---|---|---|---|---|---|---|---|---|---|---|
| 1 | 117723631 | rs2185815 | 0.658 | -0.237 | 1.509 | -1.393 | -0.032 | -0.156 | -0.156 | 1.729 | 0.68 | 0.5 | 0.83 | 1 | 0.76 | 0.83 |
| 1 | 117728652 | rs10047089 | 1.528 | 0.168 | 1.012 | -0.911 | 0.018 | 0.186 | 0.186 | 0.516 | 0.51 | 0.24 | 0.63 | 0.83 | 0.55 | 0.65 |
| 1 | 117730074 | rs12025144 | -0.558 | 0.763 | 1.229 | -0.489 | 0.064 | 0.102 | 0.102 | 0 | 0.05 | 0.14 | 0.11 | 0.15 | 0.1 | 0.08 |
| 1 | 117732171 | rs17036925 | -0.309 | 0.618 | 0.841 | -0.636 | 0.003 | -0.04 | -0.04 | 0 | 0.12 | 0.17 | 0.09 | 0.01 | 0.14 | 0.1 |
| 1 | 117737486 | rs4659190 | 0.34 | 0.184 | 0.548 | -0.74 | 0.005 | -0.039 | -0.039 | 0 | 0.71 | 0.76 | 0.86 | 1 | 0.77 | 0.82 |
| 1 | 117738637 | rs1935780 | -1.766 | 0.889 | 1.192 | 0.562 | 0.022 | 0.007 | 0.007 | 0 | 0.25 | 0.67 | 0.33 | 0.25 | 0.33 | 0.24 |
| 1 | 117749809 | rs6674646 | 0.949 | -0.19 | 1.795 | 1.143 | 0.12 | 0.346 | 0.346 | 0.119 | 0.95 | 0.59 | 0.89 | 0.88 | 0.97 | 0.97 |
| 1 | 117750686 | rs10923228 | -0.946 | -0.778 | 1.657 | 1.451 | -0.009 | 0.002 | 0.002 | 0 | 0.23 | 0.22 | 0.14 | 0.29 | 0.14 | 0.26 |
| 1 | 117752498 | rs10923229 | 1.793 | -0.015 | 2.425 | 1.832 | -0.007 | 0.025 | 0.025 | 0 | 0.81 | 0.45 | 0.76 | 0.86 | 0.84 | 0.84 |
| 1 | 117759325 | rs1892409 | -1.589 | -0.081 | 2.896 | 2.006 | 0.015 | 0.118 | 0.118 | 0 | 0.14 | 0.47 | 0.21 | 0.14 | 0.13 | 0.12 |
| 1 | 117767736 | rs12568103 | -0.267 | -0.077 | 2.428 | 1.989 | 0.001 | -0.005 | -0.005 | 2.017 | 0.08 | 0.04 | 0.11 | 0.01 | 0.1 | 0.08 |
| 1 | 117767918 | rs10923243 | -1.173 | -0.115 | 2.092 | 2.067 | -0.015 | 0.112 | 0.112 | 0.082 | 0.07 | 0.21 | 0.03 | 0.16 | 0.02 | 0.12 |
| 1 | 117773202 | rs6692758 | 1.179 | -0.064 | 1.145 | 1.767 | -0.021 | 0.051 | 0.051 | 0.082 | 0.8 | 0.32 | 0.76 | 0.7 | 0.85 | 0.78 |
| 1 | 117776034 | rs1418667 | 1.257 | -0.044 | 0.022 | 1.761 | 0.023 | -0.037 | -0.037 | 0.082 | 0.79 | 0.2 | 0.75 | 0.7 | 0.85 | 0.78 |
| 1 | 117789283 | rs1342738 | 0.985 | 0.498 | 0.422 | 1.792 | 0.012 | 0.109 | 0.109 | 0 | 0.87 | 0.73 | 0.87 | 0.62 | 0.83 | 0.7 |
| 1 | 117796932 | rs1418451 | -1.971 | 0.25 | 1.106 | 1.431 | 0.001 | 0.092 | 0.092 | 0.993 | 0.23 | 0.62 | 0.21 | 0.51 | 0.21 | 0.35 |
| 1 | 117803822 | rs1146276 | -2.012 | 0.143 | 1.611 | 1.334 | -0.046 | 0.008 | 0.008 | 7.106 | 0.13 | 0.66 | 0.09 | 0.14 | 0.03 | 0.08 |
| 1 | 117809503 | rs17037061 | -1.419 | 0.223 | 1.793 | 1.445 | 0.015 | 0.075 | 0.075 | 0 | 0.16 | 0.28 | 0.16 | 0.37 | 0.19 | 0.27 |
| 1 | 117812481 | rs4659042 | -0.007 | -0.807 | 0.586 | -0.013 | 0.021 | 0.044 | 0.044 | 0.077 | 0.72 | 0.81 | 0.74 | 0.91 | 0.85 | 0.87 |
| 1 | 117815984 | rs2137400 | 0.561 | -0.496 | 0.347 | -0.756 | -0.002 | -0.003 | -0.003 | 0 | 0.63 | 0.33 | 0.68 | 0.88 | 0.64 | 0.83 |
| 1 | 117816368 | rs12125871 | -0.007 | -0.588 | 0.525 | -0.896 | 0.015 | 0.044 | 0.044 | 0 | 0.25 | 0.16 | 0.28 | 0.12 | 0.16 | 0.2 |
| 1 | 117818054 | rs7536105 | -1.175 | 0.226 | 0.63 | -0.932 | -0.001 | -0.002 | -0.002 | 0 | 0.51 | 0.54 | 0.64 | 0.62 | 0.52 | 0.53 |
| 1 | 117818497 | rs10923260 | -0.506 | 0.158 | 0.682 | -0.949 | 0.013 | 0.039 | 0.039 | 0 | 0.34 | 0.27 | 0.35 | 0.11 | 0.25 | 0.21 |
| 1 | 117820117 | rs1146291 | 0.955 | -0.496 | 0.162 | -1.292 | 0.006 | -0.025 | -0.025 | 0 | 0.29 | 0.13 | 0.24 | 0.3 | 0.3 | 0.25 |
| 1 | 117826699 | rs12135026 | 0.779 | -0.554 | 0.148 | -1.634 | 0.002 | 0.004 | 0.004 | 0 | 0.23 | 0.23 | 0.27 | 0.11 | 0.14 | 0.2 |
| 1 | 117829261 | rs7554348 | 1.512 | -0.593 | 0.499 | -1.708 | -0.001 | -0.062 | -0.062 | 0.774 | 0.21 | 0.39 | 0.11 | 0.08 | 0.18 | 0.22 |
| 1 | 117833996 | rs12142453 | 0.786 | -0.349 | 0.408 | -1.598 | 0.01 | 0.033 | 0.033 | 0 | 0.22 | 0.16 | 0.26 | 0.11 | 0.15 | 0.2 |
| 1 | 117838939 | rs2280077 | 0.943 | -0.338 | 0.357 | -1.592 | 0.005 | 0.038 | 0.038 | 0 | 0.22 | 0.12 | 0.26 | 0.11 | 0.15 | 0.2 |
| 1 | 117844251 | rs1877465 | 1.177 | -0.253 | 0.568 | -1.589 | -0.004 | -0.042 | -0.042 | 0 | 0.24 | 0.29 | 0.11 | 0.08 | 0.19 | 0.23 |
| 1 | 117854418 | rs1289862 | -1.113 | -0.301 | 0.44 | -1.669 | 0.003 | 0.004 | 0.004 | 0 | 0.77 | 0.78 | 0.75 | 0.89 | 0.87 | 0.81 |
| 1 | 117863190 | rs17037131 | 1.402 | -0.145 | 0.43 | -1.708 | -0.021 | -0.045 | -0.045 | 0.51 | 0.22 | 0.29 | 0.08 | 0.08 | 0.12 | 0.21 |
| 1 | 117864775 | rs1289873 | -0.617 | -0.095 | 0.449 | -1.567 | -0.005 | -0.006 | -0.006 | 0 | 0.64 | 0.67 | 0.56 | 0.45 | 0.59 | 0.49 |
| 1 | 117866924 | rs7517425 | 1.191 | 0.115 | 0.747 | -1.446 | -0.046 | -0.083 | -0.083 | 0 | 0.24 | 0.41 | 0.09 | 0.08 | 0.12 | 0.21 |
| 1 | 117867339 | rs1707109 | 0.227 | 0.136 | 0.722 | -1.446 | 0 | -0.001 | -0.001 | 0.258 | 0.51 | 0.55 | 0.48 | 0.45 | 0.51 | 0.46 |
| 1 | 117872629 | rs1707113 | 0.559 | 0.128 | 0.66 | -1.477 | -0.005 | 0.002 | 0.002 | 0 | 0.44 | 0.44 | 0.41 | 0.45 | 0.37 | 0.41 |
| 1 | 117883834 | rs17037149 | 0.633 | 0.253 | 0.372 | -1.459 | -0.01 | -0.015 | -0.015 | 0 | 0.36 | 0.32 | 0.28 | 0.56 | 0.4 | 0.57 |
| 1 | 117884022 | rs11586858 | 1.27 | 0.258 | 0.261 | -1.476 | 0.006 | 0.004 | 0.004 | 0 | 0.1 | 0.07 | 0.08 | 0 | 0.09 | 0.02 |
| 1 | 117892669 | rs1289859 | -1.219 | 0.279 | 0.305 | -1.413 | -0.004 | -0.012 | -0.012 | 0.016 | 0.76 | 0.75 | 0.76 | 0.93 | 0.88 | 0.84 |
| 1 | 117892818 | rs6657965 | 1.17 | 0.451 | 0.115 | -1.267 | 0.023 | -0.03 | -0.03 | 0 | 0.07 | 0.12 | 0.07 | 0 | 0.14 | 0.06 |
| 1 | 117910682 | rs1146297 | -1.792 | 0.286 | -0.028 | -1.252 | 0.009 | -0.077 | -0.077 | 0 | 0.8 | 0.95 | 0.68 | 0.62 | 0.76 | 0.81 |
| 1 | 117913965 | rs1289952 | 0.23 | 0.22 | 0.227 | -1.427 | -0.001 | -0.043 | -0.043 | 0 | 0.32 | 0.39 | 0.31 | 0.07 | 0.26 | 0.22 |
| 1 | 117915907 | rs1146298 | -1.7 | 0.195 | 0.341 | -1.4 | -0.001 | 0.009 | 0.009 | 0 | 0.79 | 0.83 | 0.67 | 0.62 | 0.76 | 0.81 |
| 1 | 117927803 | rs1009549 | -1.847 | 0.207 | 0.337 | -1.396 | 0.008 | -0.032 | -0.032 | 1.293 | 0.79 | 0.89 | 0.67 | 0.63 | 0.76 | 0.81 |
| 1 | 117939774 | rs3767794 | 1.714 | 0.173 | 0.396 | -1.497 | -0.002 | -0.012 | -0.012 | 0 | 0.21 | 0.22 | 0.07 | 0.07 | 0.12 | 0.22 |
| 1 | 117941924 | rs1316011 | -1.932 | 0.186 | 0.4 | -1.486 | 0.002 | -0.004 | -0.004 | 0 | 0.79 | 0.87 | 0.67 | 0.62 | 0.76 | 0.81 |
| 1 | 117942862 | rs12131109 | 1.52 | 0.235 | 0.434 | -1.512 | 0.009 | -0.018 | -0.018 | 0.019 | 0.07 | 0.09 | 0.07 | 0 | 0.14 | 0.05 |
| 1 | 117944732 | rs1289941 | 0.955 | 0.138 | 0.384 | -1.407 | -0.001 | -0.008 | -0.008 | 0 | 0.24 | 0.26 | 0.25 | 0.07 | 0.12 | 0.17 |
| 1 | 117945220 | rs3767795 | 1.384 | 0.211 | 0.335 | -1.413 | -0.004 | 0.006 | 0.006 | 0 | 0.15 | 0.12 | 0.2 | 0.49 | 0.29 | 0.35 |
| 1 | 117983815 | rs10923297 | -0.551 | 0.191 | 0.384 | -1.411 | 0.003 | -0.002 | -0.002 | 0.48 | 0.55 | 0.6 | 0.43 | 0.56 | 0.63 | 0.65 |
| 1 | 118029601 | rs1290539 | -1.885 | -0.168 | 0.284 | -1.547 | 0.011 | -0.054 | -0.054 | 0.852 | 0.79 | 0.92 | 0.68 | 0.62 | 0.75 | 0.82 |
| 1 | 118030256 | rs1290541 | -1.735 | -0.182 | 0.3 | -1.553 | -0.006 | 0.027 | 0.027 | 0 | 0.78 | 0.79 | 0.67 | 0.62 | 0.75 | 0.82 |
| 1 | 118033110 | rs1475393 | 1.139 | -0.207 | 0.264 | -1.576 | 0.018 | 0.032 | 0.032 | 0 | 0.23 | 0.14 | 0.23 | 0.07 | 0.12 | 0.17 |
| 1 | 118038557 | rs1290544 | -0.691 | -0.188 | 0.238 | -1.595 | 0 | 0.001 | 0.001 | 0 | 0.63 | 0.6 | 0.72 | 0.45 | 0.6 | 0.42 |
| 1 | 118042757 | rs3767802 | 1.31 | -0.182 | 0.231 | -1.598 | 0.001 | 0.013 | 0.013 | 0 | 0.16 | 0.14 | 0.2 | 0.49 | 0.29 | 0.36 |
| 1 | 118043411 | rs3767803 | 1.142 | -0.177 | 0.206 | -1.599 | 0.016 | 0.028 | 0.028 | 0 | 0.23 | 0.16 | 0.23 | 0.07 | 0.12 | 0.17 |
| 1 | 118065841 | rs1290558 | 0.333 | -0.129 | -0.127 | -1.579 | 0.002 | -0.001 | -0.001 | 0 | 0.45 | 0.4 | 0.57 | 0.45 | 0.37 | 0.35 |
| 1 | 118072302 | rs10801963 | 0.762 | -0.133 | -0.124 | -1.579 | 0.001 | -0.042 | -0.042 | 0 | 0.18 | 0.2 | 0.15 | 0 | 0.23 | 0.08 |
| 1 | 118075647 | rs2767319 | -1.923 | -0.62 | -0.102 | -1.536 | 0 | -0.008 | -0.008 | 0 | 0.79 | 0.79 | 0.93 | 0.93 | 0.88 | 0.78 |
| 1 | 118078875 | rs2622366 | 0.195 | -0.641 | -0.149 | -1.53 | 0.001 | -0.001 | -0.001 | 0 | 0.45 | 0.4 | 0.54 | 0.45 | 0.3 | 0.34 |
| 1 | 118083588 | rs2767322 | 1.344 | -0.67 | -0.366 | -0.879 | 0.038 | -0.073 | -0.073 | 0 | 0.2 | 0.04 | 0.29 | 0.38 | 0.18 | 0.18 |
| 1 | 118085375 | rs2622359 | -1.459 | -0.602 | -0.458 | -0.861 | -0.002 | -0.014 | -0.014 | 0 | 0.79 | 0.77 | 0.92 | 0.92 | 0.88 | 0.78 |
| 1 | 118087384 | rs2767323 | 0.434 | 0.326 | -0.523 | 0.056 | -0.002 | 0.004 | 0.004 | 0 | 0.21 | 0.18 | 0.31 | 0.37 | 0.18 | 0.18 |
| 1 | 118087449 | rs3795657 | -0.511 | 1.499 | -0.468 | 0.622 | 0.004 | 0.017 | 0.017 | 0 | 0.16 | 0.18 | 0.2 | 0.49 | 0.28 | 0.37 |
| 1 | 118105554 | rs6684265 | 0.507 | -0.023 | -0.423 | -0.346 | -0.009 | -0.028 | -0.028 | 0 | 0.79 | 0.76 | 0.77 | 0.93 | 0.84 | 0.84 |
| 1 | 118106076 | rs6661142 | 0.854 | -0.288 | 0.039 | -0.527 | -0.019 | -0.05 | -0.05 | 0 | 0.7 | 0.56 | 0.82 | 0.82 | 0.76 | 0.75 |
| 1 | 118108859 | rs4659053 | 0.457 | -0.441 | 0.387 | -0.53 | -0.002 | -0.005 | -0.005 | 0 | 0.74 | 0.74 | 0.84 | 0.85 | 0.77 | 0.76 |
| 1 | 118110520 | rs11804412 | -0.105 | -0.785 | -0.393 | -0.603 | -0.013 | -0.01 | -0.01 | 0 | 0.13 | 0.17 | 0.09 | 0.09 | 0.1 | 0.13 |
| 1 | 118115243 | rs1963278 | 0.874 | -0.453 | -0.511 | -0.686 | 0.022 | 0.044 | 0.044 | 0 | 0.57 | 0.4 | 0.72 | 0.74 | 0.62 | 0.61 |
| 1 | 118116834 | rs1751353 | -1.278 | -0.118 | -0.174 | -0.379 | -0.021 | -0.038 | -0.038 | 0.632 | 0.18 | 0.41 | 0.18 | 0.15 | 0.12 | 0.16 |

| | | | | | | | | | | | | | | | | |
|---|---|---|---|---|---|---|---|---|---|---|---|---|---|---|---|---|
| 1 | 118116974 | rs1290600 | -1.309 | 0.744 | -0.718 | -0.067 | 0.009 | -0.009 | -0.009 | 0 | 0.37 | 0.39 | 0.59 | 0.32 | 0.43 | 0.31 |
| 1 | 118118636 | rs10923346 | -1.137 | 0.698 | -0.872 | 0.21 | 0 | 0.001 | 0.001 | 0 | 0.16 | 0.17 | 0.27 | 0.07 | 0.16 | 0.14 |
| 1 | 118118765 | rs10923347 | -0.792 | 0.399 | -0.782 | 0.305 | -0.006 | -0.009 | -0.009 | 0.199 | 0.13 | 0.17 | 0.21 | 0.07 | 0.1 | 0.13 |
| 1 | 118121634 | rs17037423 | -0.818 | 0.294 | -1.04 | 0.462 | 0.008 | 0.005 | 0.005 | 0 | 0.13 | 0.1 | 0.25 | 0.09 | 0.12 | 0.17 |
| 1 | 118128176 | rs320365 | 0.086 | -0.075 | -1.554 | 0.92 | 0.01 | -0.086 | -0.086 | 0 | 0.84 | 0.97 | 0.85 | 0.75 | 0.89 | 0.88 |
| 1 | 118133639 | rs320348 | 0.076 | 0.004 | -1.6 | 1.568 | 0.013 | -0.072 | -0.072 | 0.006 | 0.88 | 0.97 | 0.87 | 0.76 | 0.92 | 0.91 |
| 1 | 118136861 | rs6687902 | -0.703 | -0.374 | -1.161 | 1.35 | -0.009 | 0.032 | 0.032 | 0 | 0.15 | 0.24 | 0.17 | 0.23 | 0.15 | 0.18 |
| 1 | 118139031 | rs12037710 | -0.298 | -0.651 | -0.45 | 1.454 | -0.015 | 0.04 | 0.04 | 2.308 | 0.1 | 0.15 | 0.09 | 0.19 | 0.06 | 0.05 |
| 1 | 118139540 | rs17613339 | -0.099 | -1.054 | -1.089 | 1.483 | 0.003 | 0.008 | 0.008 | 0.394 | 0.15 | 0.02 | 0.2 | 0.11 | 0.13 | 0.19 |
| 1 | 118141764 | rs10923351 | -1.3 | -1.363 | -1.342 | 1.203 | 0.038 | -0.012 | -0.012 | 0 | 0.3 | 0.18 | 0.31 | 0.3 | 0.22 | 0.25 |
| 1 | 118143517 | rs320369 | 0.525 | 0.392 | -1.571 | 1.033 | -0.029 | 0.038 | 0.038 | 0 | 0.76 | 0.87 | 0.72 | 0.91 | 0.7 | 0.82 |
| 1 | 118144332 | rs1966228 | -1.217 | 0.275 | -1.074 | 1.083 | 0.005 | -0.01 | -0.01 | 0.019 | 0.29 | 0.32 | 0.33 | 0.46 | 0.27 | 0.32 |
| 1 | 118147675 | rs2576570 | -2.298 | 0.127 | -0.23 | 0.985 | 0.002 | 0.013 | 0.013 | 0 | 0.55 | 0.63 | 0.61 | 0.55 | 0.57 | 0.52 |
| 1 | 118155620 | rs3767812 | -1.046 | 0.069 | -0.141 | 0.906 | -0.018 | -0.011 | -0.011 | 1.925 | 0.31 | 0.34 | 0.31 | 0.24 | 0.23 | 0.25 |
| 1 | 118158079 | rs1775831 | -2.317 | 0.032 | 0.168 | 0.891 | 0 | -0.012 | -0.012 | 0.082 | 0.56 | 0.69 | 0.63 | 0.48 | 0.57 | 0.52 |
| 1 | 118159654 | rs1657817 | 0.673 | 0.046 | 0.194 | 0.906 | -0.003 | 0 | 0 | 2.666 | 0.1 | 0.13 | 0.08 | 0.12 | 0.06 | 0.05 |
| 1 | 118163457 | rs860395 | 0.418 | 0.032 | 0.283 | 0.898 | -0.02 | 0.023 | 0.023 | 1.396 | 0.11 | 0.26 | 0.09 | 0.12 | 0.06 | 0.05 |
| 1 | 118166750 | rs2066411 | -0.948 | 0.045 | 0.234 | 0.859 | 0.025 | 0.014 | 0.014 | 0 | 0.3 | 0.19 | 0.31 | 0.24 | 0.22 | 0.27 |
| 1 | 118166877 | rs10923360 | -0.66 | 0.076 | 0.244 | 0.866 | 0.023 | 0.01 | 0.01 | 0 | 0.23 | 0.29 | 0.3 | 0.24 | 0.34 | 0.26 |
| 1 | 118167199 | rs826403 | -1.28 | 0.041 | 0.2 | 0.849 | 0.006 | 0.001 | 0.001 | 0.002 | 0.27 | 0.65 | 0.32 | 0.24 | 0.34 | 0.28 |
| 1 | 118167653 | rs866111 | -0.337 | 0.032 | 0.184 | 0.867 | -0.006 | 0.214 | 0.214 | 0.036 | 0.14 | 0.45 | 0.11 | 0.27 | 0.11 | 0.12 |
| 1 | 118168826 | rs11581122 | -0.11 | 0.041 | -0.016 | 0.783 | -0.004 | 0.023 | 0.023 | 0.052 | 0.2 | 0.01 | 0.22 | 0.11 | 0.16 | 0.22 |
| 1 | 118169463 | rs2282456 | -0.452 | 0.128 | 0.193 | 0.756 | 0.009 | -0.016 | -0.016 | 0 | 0.22 | 0.2 | 0.28 | 0.09 | 0.29 | 0.2 |
| 1 | 118172538 | rs6696923 | -1.178 | 0.091 | 0.253 | 0.78 | 0.009 | -0.016 | -0.016 | 0.007 | 0.32 | 0.36 | 0.36 | 0.22 | 0.36 | 0.25 |
| 1 | 118174159 | rs6428705 | -0.308 | 0.076 | 0.299 | 0.833 | -0.005 | 0.016 | 0.016 | 0 | 0.18 | 0.23 | 0.24 | 0.26 | 0.18 | 0.27 |
| 1 | 118174222 | rs1808806 | -1.311 | 0.096 | 0.124 | 0.581 | -0.003 | 0.003 | 0.003 | 0 | 0.37 | 0.33 | 0.37 | 0.22 | 0.39 | 0.28 |
| 1 | 118178654 | rs11802277 | 1.599 | -0.24 | 0.044 | 0.71 | -0.002 | 0.002 | 0.002 | 0 | 0.37 | 0.32 | 0.29 | 0.36 | 0.38 | 0.4 |
| 1 | 118182628 | rs422484 | -1.609 | 0.027 | -0.073 | 0.468 | -0.009 | 0.005 | 0.005 | 0 | 0.62 | 0.52 | 0.63 | 0.59 | 0.67 | 0.58 |
| 1 | 118182792 | rs7522280 | 0.522 | -0.011 | -0.535 | 0.313 | -0.023 | -0.034 | -0.034 | 0 | 0.2 | 0.03 | 0.26 | 0.19 | 0.21 | 0.2 |
| 1 | 118185332 | rs319506 | -1.62 | -0.027 | -0.272 | 0.411 | 0.004 | 0.003 | 0.003 | 0 | 0.59 | 0.33 | 0.64 | 0.4 | 0.62 | 0.54 |
| 1 | 118189080 | rs10923367 | 0.832 | 0.248 | -0.469 | 0.345 | -0.054 | -0.061 | -0.061 | 0 | 0.19 | 0.01 | 0.33 | 0.21 | 0.23 | 0.2 |
| 1 | 118189519 | rs12725245 | 1.101 | 0.353 | -0.63 | 0.161 | 0.003 | 0.016 | 0.016 | 1.394 | 0.13 | 0.02 | 0.09 | 0.12 | 0.14 | 0.09 |
| 1 | 118189751 | rs12028922 | -1.066 | 0.204 | -0.406 | 0.253 | -0.018 | 0.027 | 0.027 | 0.309 | 0.36 | 0.25 | 0.3 | 0.31 | 0.39 | 0.34 |
| 1 | 118192953 | rs319501 | -2.487 | 0.136 | -0.171 | 0.219 | -0.002 | 0.003 | 0.003 | 0 | 0.63 | 0.58 | 0.68 | 0.56 | 0.66 | 0.6 |
| 1 | 118193962 | rs12746383 | -0.486 | 0.083 | -0.38 | 0.144 | -0.035 | 0.054 | 0.054 | 0.379 | 0.28 | 0.17 | 0.27 | 0.15 | 0.33 | 0.27 |
| 1 | 118195592 | rs17185310 | -0.367 | 0.083 | -0.341 | 0.156 | 0.013 | -0.043 | -0.043 | 0.166 | 0.32 | 0.14 | 0.39 | 0.39 | 0.33 | 0.33 |
| 1 | 118199767 | rs7524574 | -1.012 | 0.633 | -0.123 | 0.443 | 0.017 | -0.053 | -0.053 | 0 | 0.31 | 0.39 | 0.28 | 0.17 | 0.33 | 0.31 |
| 1 | 118208245 | rs11588056 | 0.937 | 0.59 | -0.324 | 0.497 | -0.057 | -0.064 | -0.064 | 0 | 0.19 | 0.01 | 0.32 | 0.21 | 0.22 | 0.2 |
| 1 | 118214719 | rs7520590 | -1.321 | 0.529 | -0.09 | 0.369 | 0.022 | -0.048 | -0.048 | 0.715 | 0.33 | 0.51 | 0.32 | 0.19 | 0.36 | 0.32 |
| 1 | 118215086 | rs7544119 | 0.641 | 0.523 | -0.14 | 0.457 | -0.028 | -0.089 | -0.089 | 0.048 | 0.21 | 0.04 | 0.35 | 0.38 | 0.26 | 0.26 |
| 1 | 118218745 | rs7516381 | -1.4 | 0.722 | -0.032 | 0.394 | 0.032 | -0.058 | -0.058 | 0 | 0.32 | 0.44 | 0.31 | 0.18 | 0.36 | 0.3 |
| 1 | 118219568 | rs12116919 | -0.928 | 0.733 | -0.095 | 0.392 | -0.036 | 0.048 | 0.048 | 0 | 0.28 | 0.18 | 0.27 | 0.16 | 0.33 | 0.28 |
| 1 | 118224210 | rs10923386 | -1.489 | 0.893 | -0.017 | 0.407 | 0.023 | -0.045 | -0.045 | 0 | 0.33 | 0.54 | 0.32 | 0.18 | 0.37 | 0.32 |
| 1 | 118224598 | rs1539441 | 0.122 | 0.802 | 0.041 | 0.628 | -0.006 | -0.001 | -0.001 | 1.262 | 0.08 | 0.04 | 0.07 | 0.11 | 0.11 | 0.08 |
| 1 | 118242864 | rs1891556 | -1.478 | 0.675 | -0.85 | 0.539 | 0.007 | 0.004 | 0.004 | 0.02 | 0.49 | 0.48 | 0.66 | 0.58 | 0.58 | 0.57 |
| 1 | 118244639 | rs12735195 | -0.522 | 0.929 | -1.078 | 0.508 | -0.035 | 0.036 | 0.036 | 0 | 0.26 | 0.17 | 0.27 | 0.18 | 0.32 | 0.29 |
| 1 | 118249041 | rs319469 | 1.025 | 0.635 | -1.183 | 0.475 | 0 | -0.001 | -0.001 | 0 | 0.56 | 0.52 | 0.42 | 0.42 | 0.55 | 0.43 |
| 1 | 118253295 | rs1812607 | 0.757 | 0.988 | -1.166 | 0.683 | -0.041 | -0.054 | -0.054 | 0 | 0.17 | 0.03 | 0.2 | 0.22 | 0.19 | 0.18 |
| 1 | 118255058 | rs2002747 | -0.399 | 1.119 | -1.442 | 0.367 | 0.006 | 0.011 | 0.011 | 0 | 0.51 | 0.42 | 0.61 | 0.61 | 0.55 | 0.58 |
| 1 | 118276152 | rs1419369 | 0.233 | 1.174 | -0.932 | 0.026 | 0 | 0.004 | 0.004 | 0 | 0.33 | 0.3 | 0.39 | 0.22 | 0.32 | 0.31 |
| 1 | 118276535 | rs17257575 | 0.215 | 1.209 | -0.974 | 0.032 | -0.001 | -0.013 | -0.013 | 0 | 0.33 | 0.37 | 0.39 | 0.22 | 0.32 | 0.31 |
| 1 | 118277252 | rs17257582 | 0.224 | 1.062 | -1.42 | 0.066 | -0.038 | 0.128 | 0.128 | 0.032 | 0.39 | 0.14 | 0.41 | 0.23 | 0.39 | 0.42 |
| 1 | 118278275 | rs2164685 | 0.384 | 1.183 | -1.158 | 0.485 | -0.053 | -0.02 | -0.02 | 0 | 0.41 | 0.15 | 0.44 | 0.48 | 0.47 | 0.55 |
| 1 | 118281822 | rs4659339 | 0.439 | 0.858 | -0.91 | 0.23 | -0.04 | 0.156 | 0.156 | 0 | 0.43 | 0.17 | 0.41 | 0.19 | 0.43 | 0.53 |
| 1 | 118282675 | rs17185373 | 0.613 | 0.835 | -0.917 | 0.221 | -0.078 | 0.171 | 0.171 | 0 | 0.42 | 0.1 | 0.4 | 0.19 | 0.44 | 0.53 |
| 1 | 118283062 | rs319487 | 0.131 | 0.707 | 0.089 | 0.538 | 0 | -0.002 | -0.002 | 0 | 0.48 | 0.45 | 0.43 | 0.49 | 0.47 | 0.61 |
| 1 | 118293093 | rs319464 | -0.544 | 0.75 | 0.47 | 0.655 | 0.037 | 0.009 | 0.009 | 0.391 | 0.88 | 0.73 | 0.86 | 0.91 | 0.85 | 0.93 |
| 1 | 118294476 | rs12145990 | 0.535 | 0.746 | 0.503 | 0.662 | 0.071 | 0.246 | 0.246 | 0.767 | 0.85 | 0.59 | 0.84 | 0.62 | 0.82 | 0.84 |
| 1 | 118296638 | rs12731636 | 0.167 | 0.587 | 0.463 | 0.624 | 0.003 | -0.002 | -0.002 | 0 | 0.41 | 0.4 | 0.43 | 0.43 | 0.38 | 0.35 |
| 1 | 118300750 | rs544173 | -0.214 | 0.453 | 0.463 | 0.552 | 0.027 | 0.043 | 0.043 | 0 | 0.46 | 0.65 | 0.46 | 0.73 | 0.41 | 0.4 |
| 1 | 118311046 | rs1775751 | 0.115 | 0.175 | -0.248 | 0.49 | -0.017 | -0.02 | -0.02 | 0 | 0.39 | 0.22 | 0.45 | 0.42 | 0.45 | 0.35 |
| 1 | 118319990 | rs3754123 | 0.08 | 0.273 | -0.239 | 0.539 | -0.029 | -0.033 | -0.033 | 0 | 0.4 | 0.18 | 0.45 | 0.42 | 0.45 | 0.35 |
| 1 | 118327587 | rs1780041 | -0.079 | 0.252 | -0.175 | 0.451 | -0.001 | -0.056 | -0.056 | 1.433 | 0.83 | 0.52 | 0.88 | 0.93 | 0.89 | 0.93 |
| 1 | 118328656 | rs2570345 | 0.23 | 0.255 | -0.181 | 0.477 | -0.018 | -0.02 | -0.02 | 0 | 0.4 | 0.2 | 0.45 | 0.43 | 0.45 | 0.36 |
| 1 | 118329870 | rs1775752 | -0.201 | 0.313 | -0.206 | 0.428 | 0.006 | -0.012 | -0.012 | 0 | 0.83 | 0.54 | 0.88 | 0.92 | 0.89 | 0.93 |
| 1 | 118337206 | rs10923424 | 1.248 | 0.316 | -0.292 | 0.473 | -0.056 | 0.147 | 0.147 | 0.111 | 0.38 | 0.08 | 0.4 | 0.18 | 0.41 | 0.48 |
| 1 | 118338453 | rs11805137 | 0.238 | 0.317 | -0.275 | 0.483 | -0.014 | -0.02 | -0.02 | 0 | 0.4 | 0.21 | 0.45 | 0.45 | 0.45 | 0.36 |
| 1 | 118338828 | rs12140339 | 1.065 | 0.35 | -0.362 | 0.538 | -0.052 | 0.145 | 0.145 | 0.065 | 0.39 | 0.09 | 0.4 | 0.18 | 0.41 | 0.48 |
| 1 | 118339349 | rs1655200 | 0.682 | 0.382 | -0.355 | 0.494 | -0.004 | -0.007 | -0.007 | 0 | 0.42 | 0.29 | 0.43 | 0.47 | 0.44 | 0.57 |

| | | | | | | | | | | | | | | | | |
|---|---|---|---|---|---|---|---|---|---|---|---|---|---|---|---|---|
| 1 | 118346219 | rs1342813 | -0.317 | 0.403 | -0.356 | 0.346 | -0.001 | 0.018 | 0.018 | 0 | 0.45 | 0.4 | 0.47 | 0.74 | 0.48 | 0.45 |
| 1 | 118348085 | rs10801996 | -0.151 | 0.389 | -0.402 | 0.35 | -0.007 | 0.041 | 0.041 | 0 | 0.43 | 0.3 | 0.46 | 0.74 | 0.49 | 0.45 |
| 1 | 118351638 | rs7524918 | -0.105 | 0.362 | -0.29 | 0.336 | -0.006 | 0.035 | 0.035 | 0 | 0.57 | 0.68 | 0.53 | 0.26 | 0.52 | 0.55 |
| 1 | 118354933 | rs1342816 | 0.413 | 0.307 | -0.125 | 0.554 | 0.02 | 0.318 | 0.318 | 0.492 | 0.06 | 0.28 | 0.03 | 0.3 | 0.03 | 0.09 |
| 1 | 118355412 | rs7412527 | 0.303 | 0.277 | -0.194 | 0.396 | 0.001 | -0.002 | -0.002 | 0.001 | 0.44 | 0.37 | 0.43 | 0.48 | 0.44 | 0.56 |
| 1 | 118369833 | rs720162 | 0.238 | 0.156 | -0.102 | 0.579 | -0.001 | 0.001 | 0.001 | 0 | 0.44 | 0.39 | 0.46 | 0.56 | 0.44 | 0.67 |
| 1 | 118377788 | rs6681593 | 0.902 | 0.133 | 0.015 | 0.665 | 0.006 | 0.262 | 0.262 | 1.139 | 0.05 | 0.19 | 0.03 | 0.3 | 0.03 | 0.09 |
| 1 | 118406208 | rs1371 | 0.943 | 0.069 | -0.426 | 0.257 | -0.011 | 0.076 | 0.076 | 0 | 0.14 | 0.03 | 0.07 | 0 | 0.14 | 0.06 |
| 1 | 118421109 | rs12058607 | -1.404 | 0.125 | 0.062 | 0.65 | -0.019 | 0.307 | 0.307 | 0.068 | 0.1 | 0.34 | 0.08 | 0.38 | 0.04 | 0.23 |
| 1 | 118439995 | rs10489592 | 1.35 | 0.373 | -0.284 | 0.362 | -0.04 | -0.039 | -0.039 | 0 | 0.09 | 0.01 | 0.08 | 0.13 | 0.15 | 0.11 |
| 1 | 118455754 | rs4233455 | 0.325 | 0.11 | -0.056 | 0.37 | -0.003 | 0.004 | 0.004 | 0 | 0.46 | 0.44 | 0.48 | 0.56 | 0.46 | 0.67 |
| 1 | 118474225 | rs1321665 | 0.261 | 0.167 | 0.035 | 0.43 | 0 | -0.001 | -0.001 | 0 | 0.46 | 0.5 | 0.48 | 0.53 | 0.46 | 0.67 |
| 1 | 118480591 | rs10923448 | 0.609 | 0.166 | -0.008 | 0.349 | -0.01 | -0.007 | -0.007 | 2.801 | 0.42 | 0.32 | 0.44 | 0.48 | 0.44 | 0.54 |
| 1 | 118510784 | rs1033728 | 0.364 | 0.23 | -0.002 | 0.313 | -0.002 | 0.003 | 0.003 | 0 | 0.46 | 0.42 | 0.47 | 0.56 | 0.46 | 0.67 |
| 1 | 118513484 | rs6683735 | 0.023 | 0.175 | -0.182 | 0.291 | -0.016 | -0.019 | -0.019 | 0.492 | 0.52 | 0.85 | 0.5 | 0.52 | 0.47 | 0.59 |
| 1 | 118520042 | rs1409055 | 0.109 | 0.33 | -0.336 | 0.062 | 0.015 | -0.002 | -0.002 | 0 | 0.49 | 0.38 | 0.5 | 0.5 | 0.52 | 0.68 |
| 1 | 118521350 | rs10923452 | 0.415 | 0.363 | -0.781 | -0.163 | -0.006 | 0.141 | 0.141 | 0 | 0.46 | 0.2 | 0.46 | 0.21 | 0.49 | 0.58 |
| 1 | 118524718 | rs6428716 | -0.877 | -0.076 | -1.13 | -0.348 | 0.016 | 0.005 | 0.005 | 0.132 | 0.49 | 0.79 | 0.5 | 0.52 | 0.46 | 0.43 |
| 1 | 118526592 | rs4347199 | -1.265 | -0.466 | -1.023 | -0.213 | 0.041 | -0.056 | -0.056 | 0 | 0.85 | 0.96 | 0.93 | 0.66 | 0.9 | 0.8 |
| 1 | 118536418 | rs2765283 | 0.825 | 0.232 | -0.788 | -0.271 | 0.006 | 0.007 | 0.007 | 0 | 0.07 | 0.06 | 0.06 | 0.07 | 0.07 | 0.11 |
| 1 | 118538078 | rs2802663 | -0.001 | 0.139 | -0.191 | -0.372 | 0.015 | 0.019 | 0.019 | 0.013 | 0.49 | 0.76 | 0.5 | 0.48 | 0.52 | 0.55 |
| 1 | 118575757 | rs6428727 | 0.09 | 0.016 | -0.271 | -0.4 | 0.014 | 0.015 | 0.015 | 0 | 0.51 | 0.71 | 0.5 | 0.48 | 0.52 | 0.56 |
| 1 | 118577714 | rs1409053 | 1.114 | 0.241 | 0.121 | -0.387 | 0.009 | -0.011 | -0.011 | 0 | 0.05 | 0.28 | 0.03 | 0 | 0.02 | 0 |
| 1 | 118583408 | rs10923477 | 1.018 | 0.262 | -0.238 | -0.369 | -0.066 | 0.081 | 0.081 | 0.517 | 0.34 | 0.16 | 0.41 | 0.16 | 0.43 | 0.37 |
| 1 | 118598950 | rs12124243 | 2.014 | 0.284 | -0.449 | -0.605 | -0.059 | 0.094 | 0.094 | 0.164 | 0.18 | 0.04 | 0.24 | 0.02 | 0.22 | 0.12 |
| 1 | 118617274 | rs1935993 | 0.696 | -0.89 | 0.427 | -0.071 | 0.002 | -0.033 | -0.033 | 0 | 0.44 | 0.6 | 0.46 | 0.17 | 0.49 | 0.38 |
| 1 | 118623055 | rs6428729 | 0.492 | -0.462 | 0.984 | 0.409 | 0.009 | -0.013 | -0.013 | 0 | 0.07 | 0.38 | 0.04 | 0 | 0.02 | 0 |
| 1 | 118624336 | rs12043257 | -0.295 | -0.444 | 0.925 | 0.518 | 0.008 | 0.083 | 0.083 | 0 | 0.05 | 0.09 | 0.04 | 0.29 | 0.06 | 0.19 |
| 1 | 118657396 | rs12065339 | 0.616 | -0.456 | 0.937 | 0.505 | 0.015 | -0.007 | -0.007 | 0 | 0.07 | 0.36 | 0.04 | 0 | 0.02 | 0 |
| 1 | 118678953 | rs17257729 | 1.167 | -0.557 | 0.514 | 0.296 | 0.038 | 0.184 | 0.184 | 0.1 | 0.48 | 0.21 | 0.5 | 0.17 | 0.59 | 0.44 |
| 1 | 118693217 | rs1935992 | 0.336 | -0.549 | 0.453 | 0.451 | -0.006 | -0.125 | -0.125 | 0.048 | 0.27 | 0.01 | 0.35 | 0.4 | 0.24 | 0.24 |
| 1 | 118702287 | rs2153978 | 0.558 | -0.435 | 0.698 | 0.417 | 0.006 | -0.01 | -0.01 | 0 | 0.25 | 0.2 | 0.35 | 0.44 | 0.23 | 0.25 |
| 1 | 118730197 | rs6703152 | 0.038 | -0.03 | 0.466 | 0.425 | 0.049 | 0.03 | 0.03 | 0.42 | 0.53 | 0.28 | 0.54 | 0.45 | 0.64 | 0.61 |
| 1 | 118731761 | rs10923507 | 0.038 | -0.03 | 0.399 | 0.425 | 0.057 | 0.033 | 0.033 | 0.42 | 0.53 | 0.25 | 0.54 | 0.45 | 0.64 | 0.61 |
| 1 | 118739301 | rs1782414 | 0.333 | -0.035 | 0.477 | 0.502 | 0.025 | 0.141 | 0.141 | 0 | 0.95 | 0.87 | 0.95 | 0.71 | 0.95 | 0.84 |
| 1 | 118749926 | rs7514644 | 0.45 | -0.095 | 0.403 | 0.425 | -0.004 | 0.23 | 0.23 | 0.119 | 0.47 | 0.09 | 0.49 | 0.17 | 0.59 | 0.44 |
| 1 | 118757034 | rs4411121 | -0.112 | 0.005 | 0.56 | 0.544 | 0 | 0 | 0 | 0.035 | 0.46 | 0.59 | 0.44 | 0.54 | 0.34 | 0.4 |
| 1 | 118758446 | rs10494207 | 0.552 | 0.136 | 0.058 | 0.675 | 0.03 | 0.167 | 0.167 | 0.031 | 0.94 | 0.84 | 0.94 | 0.72 | 0.94 | 0.84 |
| 1 | 118760171 | rs6667043 | 0.75 | 0.245 | -0.266 | 0.624 | -0.028 | 0.063 | 0.063 | 0.283 | 0.41 | 0.27 | 0.45 | 0.18 | 0.49 | 0.42 |
| 1 | 118766534 | rs6669864 | -0.048 | 0.254 | -0.256 | 0.655 | -0.005 | -0.003 | -0.003 | 0 | 0.43 | 0.49 | 0.44 | 0.56 | 0.37 | 0.4 |
| 1 | 118784987 | rs7553660 | 0.284 | 0.176 | -0.163 | 0.749 | -0.006 | 0.016 | 0.016 | 0 | 0.84 | 0.77 | 0.88 | 0.76 | 0.86 | 0.81 |
| 1 | 118787119 | rs1938270 | 0.222 | 0.172 | -0.112 | 0.738 | -0.006 | -0.013 | -0.013 | 0.037 | 0.11 | 0.06 | 0.1 | 0.25 | 0.14 | 0.17 |
| 1 | 118792822 | rs7515409 | -0.798 | 0.208 | -0.284 | 0.61 | 0.004 | -0.025 | -0.025 | 0 | 0.58 | 0.58 | 0.53 | 0.82 | 0.51 | 0.55 |
| 1 | 118796804 | rs1938263 | 0.438 | 0.185 | -0.395 | 0.936 | -0.016 | 0.007 | 0.007 | 0 | 0.36 | 0.49 | 0.4 | 0.45 | 0.29 | 0.28 |
| 1 | 118799439 | rs9661113 | -0.977 | 0.187 | -0.384 | 0.929 | 0.001 | -0.015 | -0.015 | 0 | 0.58 | 0.59 | 0.54 | 0.81 | 0.51 | 0.54 |
| 1 | 118815799 | rs12044199 | 0.457 | 0.154 | -0.57 | 0.963 | -0.022 | 0.012 | 0.012 | 0.14 | 0.36 | 0.5 | 0.4 | 0.45 | 0.29 | 0.28 |
| 1 | 118817482 | rs17038096 | 0.72 | 0.298 | -0.678 | 0.975 | -0.026 | -0.027 | -0.027 | 0 | 0.05 | 0.01 | 0.03 | 0.11 | 0.05 | 0.11 |
| 1 | 118821908 | rs7554803 | -0.897 | 0.238 | -0.601 | 0.569 | -0.002 | -0.005 | -0.005 | 0 | 0.58 | 0.6 | 0.54 | 0.81 | 0.5 | 0.54 |
| 1 | 118830947 | rs11588400 | -0.332 | 0.298 | -0.794 | 0.617 | -0.004 | -0.037 | -0.037 | 0 | 0.22 | 0.13 | 0.15 | 0.39 | 0.21 | 0.29 |
| 1 | 118835195 | rs12092268 | -0.218 | 0.625 | -0.646 | -0.023 | 0.001 | -0.019 | -0.019 | 1.382 | 0.57 | 0.59 | 0.54 | 0.82 | 0.49 | 0.54 |
| 1 | 118841141 | rs1938275 | -0.051 | 0.43 | -0.716 | 0.351 | -0.005 | -0.003 | -0.003 | 0 | 0.62 | 0.6 | 0.71 | 0.7 | 0.67 | 0.71 |
| 1 | 118845756 | rs6428738 | 0.214 | 0.852 | -0.837 | -0.29 | 0.005 | 0.031 | 0.031 | 0.032 | 0.44 | 0.62 | 0.56 | 0.77 | 0.39 | 0.58 |
| 1 | 118846667 | rs10923537 | 0.488 | 0.978 | -1.326 | -0.331 | -0.094 | 0.03 | 0.03 | 0.529 | 0.35 | 0.02 | 0.27 | 0.22 | 0.36 | 0.28 |
| 1 | 118852817 | rs9428102 | -0.844 | 1.419 | -1.45 | -0.141 | -0.025 | 0.014 | 0.014 | 0 | 0.16 | 0.07 | 0.11 | 0.11 | 0.24 | 0.24 |
| 1 | 118852975 | rs12731372 | 0.882 | 1.351 | -1.802 | 0.283 | -0.027 | -0.036 | -0.036 | 0.2 | 0.24 | 0.08 | 0.28 | 0.28 | 0.24 | 0.18 |
| 1 | 118855012 | rs12750838 | 0.094 | 1.485 | -1.826 | 0.289 | -0.033 | 0 | 0 | 0.047 | 0.1 | 0.02 | 0.09 | 0.06 | 0.14 | 0.15 |
| 1 | 118855587 | rs9428104 | -0.563 | 1.208 | -1.271 | 0.276 | 0 | -0.001 | -0.001 | 0 | 0.71 | 0.73 | 0.7 | 0.66 | 0.75 | 0.67 |
| 1 | 118858218 | rs9428107 | 0.858 | 1.218 | -1.27 | 0.57 | -0.001 | -0.001 | -0.001 | 0 | 0.07 | 0.05 | 0.13 | 0.12 | 0.1 | 0.1 |
| 1 | 118859049 | rs9428109 | -0.583 | 1.185 | -1.264 | 0.563 | -0.026 | -0.014 | -0.014 | 0.006 | 0.17 | 0.07 | 0.22 | 0.18 | 0.24 | 0.25 |
| 1 | 118862669 | rs17038164 | 0.35 | 1.162 | -1.241 | 0.568 | 0 | -0.001 | -0.001 | 0.051 | 0.29 | 0.28 | 0.3 | 0.34 | 0.25 | 0.33 |
| 1 | 118864602 | rs7536458 | 0.464 | 1.102 | -1.253 | 0.627 | -0.007 | -0.019 | -0.019 | 0.018 | 0.29 | 0.16 | 0.28 | 0.34 | 0.26 | 0.33 |
| 1 | 118864968 | rs4375253 | 1.009 | 1.087 | -1.114 | 0.608 | 0.004 | 0.034 | 0.034 | 0 | 0.51 | 0.57 | 0.32 | 0.29 | 0.48 | 0.32 |
| 1 | 118869933 | rs6690477 | 0.797 | 1.149 | -1.266 | 0.715 | -0.021 | -0.047 | -0.047 | 0 | 0.8 | 0.87 | 0.59 | 0.63 | 0.74 | 0.69 |
| 1 | 118874221 | rs17185617 | 1.285 | 1.114 | -0.927 | 0.486 | -0.005 | 0.043 | 0.043 | 0.109 | 0.42 | 0.19 | 0.24 | 0.29 | 0.42 | 0.31 |
| 1 | 118881957 | rs6689939 | -0.166 | 1.252 | -0.798 | 0.371 | -0.005 | -0.003 | -0.003 | 0 | 0.9 | 0.92 | 0.83 | 0.87 | 0.85 | 0.89 |
| 1 | 118884512 | rs2474946 | -0.338 | 0.842 | -0.411 | 0.574 | 0.008 | 0.01 | 0.01 | 0 | 0.72 | 0.65 | 0.62 | 0.62 | 0.7 | 0.66 |
| 1 | 118889621 | rs2474942 | -0.527 | 0.625 | -0.22 | 0.48 | -0.007 | 0.025 | 0.025 | 0 | 0.19 | 0.13 | 0.18 | 0.09 | 0.23 | 0.29 |
| 1 | 118889632 | rs6428742 | 1.613 | 0.638 | 0.126 | 0.643 | -0.001 | 0.019 | 0.019 | 0.102 | 0.7 | 0.4 | 0.51 | 0.6 | 0.68 | 0.63 |
| 1 | 118889815 | rs7535961 | 1.524 | 0.595 | 0.066 | 0.645 | -0.016 | 0.009 | 0.009 | 1.857 | 0.45 | 0.19 | 0.27 | 0.35 | 0.44 | 0.31 |

| | | | | | | | | | | | | | | | | |
|---|---|---|---|---|---|---|---|---|---|---|---|---|---|---|---|---|
| 1 | 118902978 | rs2474937 | -0.821 | 0.63 | 0.455 | 0.647 | -0.003 | -0.039 | -0.039 | 0.576 | 0.22 | 0.36 | 0.2 | 0.09 | 0.23 | 0.29 |
| 1 | 118903196 | rs10923556 | 1.483 | 0.65 | 0.196 | 0.686 | -0.001 | 0.016 | 0.016 | 1.488 | 0.46 | 0.25 | 0.27 | 0.35 | 0.44 | 0.32 |
| 1 | 118906721 | rs1885012 | -0.158 | 0.859 | 0.022 | 0.841 | 0.004 | 0.062 | 0.062 | 0 | 0.94 | 0.92 | 0.73 | 0.69 | 0.91 | 0.92 |
| 1 | 118936321 | rs10489732 | -0.094 | 0.677 | -0.444 | 0.093 | 0.002 | -0.001 | -0.001 | 0 | 0.47 | 0.45 | 0.28 | 0.45 | 0.47 | 0.56 |
| 1 | 118938522 | rs2145419 | -1.203 | 0.826 | -0.655 | 0.464 | 0.002 | -0.004 | -0.004 | 1.716 | 0.77 | 0.85 | 0.63 | 0.68 | 0.82 | 0.84 |
| 1 | 118953104 | rs17038273 | -0.015 | 0.939 | -0.855 | 0.502 | 0.001 | 0.012 | 0.012 | 0.237 | 0.09 | 0.11 | 0.17 | 0.25 | 0.13 | 0.24 |
| 1 | 118966067 | rs12093200 | -1.385 | 0.926 | -0.847 | 0.488 | 0.002 | -0.008 | -0.008 | 0 | 0.78 | 0.88 | 0.64 | 0.69 | 0.83 | 0.84 |
| 1 | 118971450 | rs17038298 | -0.669 | 1.406 | -0.558 | 0.785 | -0.001 | 0.006 | 0.006 | 0.018 | 0.1 | 0.11 | 0.19 | 0.27 | 0.16 | 0.24 |
| 1 | 118973742 | rs1229137 | -2.067 | 1.412 | -0.519 | 0.906 | 0.004 | 0.006 | 0.006 | 0.23 | 0.31 | 0.36 | 0.36 | 0.38 | 0.39 | 0.56 |
| 1 | 118975209 | rs1146349 | -1.024 | 1.429 | -0.868 | 0.421 | 0 | -0.001 | -0.001 | 0.431 | 0.19 | 0.16 | 0.16 | 0.1 | 0.22 | 0.28 |
| 1 | 118981665 | rs1146344 | -0.997 | 1.326 | -0.856 | 0.432 | 0 | -0.001 | -0.001 | 0 | 0.19 | 0.16 | 0.15 | 0.1 | 0.2 | 0.28 |
| 1 | 118986187 | rs10489731 | -0.054 | 1.785 | -1.102 | 0.683 | -0.012 | -0.012 | -0.012 | 0 | 0.06 | 0.03 | 0.22 | 0.14 | 0.19 | 0.08 |
| 1 | 118987765 | rs1146337 | -0.788 | 2.008 | -1.269 | 0.249 | -0.027 | 0.052 | 0.052 | 0 | 0.32 | 0.27 | 0.22 | 0.08 | 0.38 | 0.29 |
| 1 | 118987888 | rs1146336 | 1.414 | 1.391 | -1.757 | 0.747 | 0.036 | -0.019 | -0.019 | 0 | 0.32 | 0.17 | 0.16 | 0.29 | 0.2 | 0.24 |
| 1 | 118990640 | rs1408592 | -0.83 | 1.635 | -0.922 | 0.394 | -0.021 | 0.058 | 0.058 | 0 | 0.43 | 0.67 | 0.47 | 0.27 | 0.52 | 0.42 |
| 1 | 119007709 | rs1213983 | -0.503 | 1.528 | -0.683 | 0.908 | -0.005 | 0.009 | 0.009 | 0 | 0.14 | 0.12 | 0.1 | 0.07 | 0.15 | 0.2 |
| 1 | 119023640 | rs1229119 | -1.417 | 1.6 | -0.499 | 1.528 | -0.002 | -0.004 | -0.004 | 0 | 0.29 | 0.32 | 0.44 | 0.51 | 0.4 | 0.48 |
| 1 | 119029083 | rs17038321 | 0.218 | 1.716 | -0.665 | 1.908 | -0.029 | -0.015 | -0.015 | 0 | 0.05 | 0.01 | 0.15 | 0.25 | 0.1 | 0.07 |
| 1 | 119029958 | rs1324727 | -1.048 | 1.538 | 0.19 | 1.966 | 0.07 | 0.295 | 0.295 | 0 | 0.08 | 0.3 | 0.23 | 0.34 | 0.1 | 0.08 |
| 1 | 119034534 | rs12408596 | -1.119 | 1.526 | 0.219 | 2.219 | 0.007 | 0.018 | 0.018 | 0 | 0.09 | 0.2 | 0.33 | 0.19 | 0.21 | 0.13 |
| 1 | 119048181 | rs1213989 | -0.314 | 1.405 | 0.141 | 2.142 | 0.001 | 0.042 | 0.042 | 0.004 | 0.18 | 0.1 | 0.1 | 0.07 | 0.16 | 0.22 |
| 1 | 119051781 | rs17257862 | -0.801 | 1.355 | 0.079 | 2.137 | -0.028 | -0.032 | -0.032 | 0 | 0.08 | 0.08 | 0.32 | 0.18 | 0.21 | 0.12 |
| 1 | 119069110 | rs1830853 | 1.615 | 1.007 | 0.115 | 1.944 | 0.016 | -0.053 | -0.053 | 0 | 0.63 | 0.17 | 0.62 | 0.36 | 0.69 | 0.53 |
| 1 | 119071421 | rs17021936 | -0.256 | 0.862 | 0.88 | 1.919 | -0.009 | 0.096 | 0.096 | 0 | 0.08 | 0.16 | 0.05 | 0.23 | 0.04 | 0.17 |
| 1 | 119074219 | rs1213884 | -1.858 | 0.852 | 1.047 | 1.925 | 0.017 | 0.211 | 0.211 | 0 | 0.18 | 0.42 | 0.23 | 0.49 | 0.17 | 0.28 |
| 1 | 119081459 | rs1324720 | -0.87 | 0.939 | 0.995 | 2.001 | 0.014 | 0.063 | 0.063 | 0 | 0.09 | 0.16 | 0.37 | 0.28 | 0.21 | 0.13 |
| 1 | 119095273 | rs4659102 | 1.742 | 0.872 | 0.975 | 1.913 | 0.018 | -0.042 | -0.042 | 0.075 | 0.61 | 0.25 | 0.66 | 0.44 | 0.67 | 0.5 |
| 1 | 119102839 | rs17185776 | 0.323 | 1.183 | 0.948 | 1.92 | -0.032 | -0.008 | -0.008 | 0 | 0.06 | 0.01 | 0.09 | 0.06 | 0.17 | 0.07 |
| 1 | 119104373 | rs1343301 | -1.065 | 1.15 | 1.229 | 1.957 | 0.04 | 0.111 | 0.111 | 0 | 0.1 | 0.29 | 0.18 | 0.26 | 0.12 | 0.12 |
| 1 | 119106936 | rs10923615 | -1.038 | 1.105 | 1.236 | 1.641 | -0.007 | -0.027 | -0.027 | 0.002 | 0.26 | 0.39 | 0.18 | 0.16 | 0.22 | 0.29 |
| 1 | 119117167 | rs10802043 | -1.017 | 1.123 | 1.257 | 1.78 | 0.037 | 0.065 | 0.065 | 0.015 | 0.1 | 0.32 | 0.26 | 0.19 | 0.12 | 0.08 |
| 1 | 119129814 | rs2798445 | -1.265 | 1.214 | 1.265 | 1.834 | -0.016 | -0.021 | -0.021 | 0.083 | 0.28 | 0.19 | 0.34 | 0.56 | 0.37 | 0.49 |
| 1 | 119142209 | rs1911167 | 0.349 | 1.25 | 1.24 | 2.025 | -0.005 | -0.035 | -0.035 | 0 | 0.77 | 0.91 | 0.87 | 0.65 | 0.8 | 0.63 |
| 1 | 119146085 | rs9661038 | 0.257 | 1.359 | 0.587 | 1.364 | 0.021 | -0.053 | -0.053 | 0.178 | 0.31 | 0.4 | 0.42 | 0.16 | 0.36 | 0.12 |
| 1 | 119153514 | rs2884876 | -0.934 | 1.494 | 1.223 | 1.963 | 0.049 | 0.081 | 0.081 | 0.024 | 0.1 | 0.37 | 0.26 | 0.19 | 0.12 | 0.08 |
| 1 | 119157490 | rs1742856 | 0.946 | 1.537 | 0.928 | 2.003 | -0.021 | -0.027 | -0.027 | 0.042 | 0.72 | 0.83 | 0.66 | 0.44 | 0.62 | 0.51 |
| 1 | 119160266 | rs7542082 | -2.287 | 1.168 | 1.027 | 1.838 | 0.013 | 0.135 | 0.135 | 2.112 | 0.13 | 0.67 | 0.25 | 0.28 | 0.18 | 0.1 |
| 1 | 119168450 | rs1701970 | 0.585 | 1.419 | 0.918 | 1.798 | -0.024 | -0.04 | -0.04 | 0 | 0.73 | 0.85 | 0.77 | 0.55 | 0.69 | 0.61 |
| 1 | 119188329 | rs2360627 | -1.307 | 1.407 | 1.147 | 1.762 | 0.04 | 0.088 | 0.088 | 0 | 0.11 | 0.46 | 0.26 | 0.19 | 0.1 | 0.06 |
| 1 | 119189778 | rs10923640 | -0.111 | 1.449 | 1.08 | 1.843 | -0.063 | 0.014 | 0.014 | 0.415 | 0.2 | 0.07 | 0.19 | 0.15 | 0.25 | 0.22 |
| 1 | 119193242 | rs2145789 | 0.151 | 1.449 | 1.095 | 1.832 | -0.008 | -0.015 | -0.015 | 0 | 0.07 | 0.03 | 0.23 | 0.19 | 0.1 | 0.06 |
| 1 | 119202269 | rs1213850 | 1.075 | 1.453 | 1.168 | 1.91 | -0.005 | -0.004 | -0.004 | 0.952 | 0.71 | 0.7 | 0.64 | 0.42 | 0.65 | 0.56 |
| 1 | 119203789 | rs1213852 | 0.951 | 1.45 | 1.176 | 1.908 | 0.018 | 0.015 | 0.015 | 1.042 | 0.7 | 0.63 | 0.63 | 0.42 | 0.65 | 0.56 |
| 1 | 119209832 | rs1742852 | 0.668 | 1.443 | 1.168 | 1.941 | -0.006 | 0.137 | 0.137 | 0.088 | 0.91 | 0.77 | 0.95 | 0.71 | 0.95 | 0.83 |
| 1 | 119210359 | rs1701971 | 0.96 | 1.451 | 1.165 | 1.955 | 0.028 | 0.336 | 0.336 | 0.066 | 0.9 | 0.7 | 0.82 | 0.57 | 0.9 | 0.79 |
| 1 | 119217675 | rs12041292 | -0.497 | 1.49 | 1.156 | 1.901 | -0.032 | -0.029 | -0.029 | 0.466 | 0.23 | 0.21 | 0.32 | 0.29 | 0.3 | 0.27 |
| 1 | 119226884 | rs10158862 | -1.344 | 1.431 | 1.308 | 2.027 | -0.006 | -0.013 | -0.013 | 0.185 | 0.14 | 0.12 | 0.32 | 0.33 | 0.24 | 0.18 |
| 1 | 119234540 | rs2755116 | -0.413 | 1.431 | 1.031 | 2.003 | 0.009 | 0.173 | 0.173 | 0 | 0.07 | 0.14 | 0.05 | 0.29 | 0.05 | 0.17 |
| 1 | 119246087 | rs9428155 | -0.103 | 1.445 | 0.89 | 2.009 | 0 | 0.101 | 0.101 | 0.019 | 0.07 | 0.08 | 0.05 | 0.29 | 0.05 | 0.17 |
| 1 | 119246407 | rs9428157 | -0.103 | 1.445 | 0.89 | 2.022 | 0 | 0.098 | 0.098 | 0.019 | 0.07 | 0.08 | 0.05 | 0.29 | 0.05 | 0.17 |
| 1 | 119252219 | rs10158804 | -0.259 | 1.5 | 0.922 | 2.067 | 0 | 0.005 | 0.005 | 0 | 0.07 | 0.08 | 0.08 | 0.14 | 0.14 | 0.11 |
| 1 | 119255045 | rs2953032 | -0.037 | 1.505 | 0.881 | 2.068 | 0.002 | 0.11 | 0.11 | 0 | 0.06 | 0.08 | 0.04 | 0.29 | 0.05 | 0.17 |
| 1 | 119259262 | rs6692676 | -1.424 | 1.535 | 0.812 | 2.053 | 0.002 | 0.012 | 0.012 | 0 | 0.15 | 0.18 | 0.32 | 0.33 | 0.24 | 0.18 |
| 1 | 119277851 | rs1961171 | -1.357 | 1.649 | 1.102 | 1.83 | 0.025 | 0.039 | 0.039 | 0.214 | 0.28 | 0.41 | 0.41 | 0.44 | 0.43 | 0.34 |
| 1 | 119282172 | rs17022367 | -0.255 | 1.594 | 1.129 | 1.762 | 0.005 | 0.02 | 0.02 | 0 | 0.07 | 0.12 | 0.08 | 0.14 | 0.13 | 0.1 |
| 1 | 119289072 | rs10923662 | -0.707 | 1.555 | 0.993 | 1.679 | 0.001 | 0.008 | 0.008 | 0 | 0.09 | 0.12 | 0.24 | 0.17 | 0.11 | 0.08 |
| 1 | 119291382 | rs1570816 | -0.707 | 1.555 | 0.993 | 1.679 | 0.001 | 0.008 | 0.008 | 0 | 0.09 | 0.12 | 0.24 | 0.17 | 0.11 | 0.08 |
| 1 | 119292887 | rs17022390 | 0.02 | 1.563 | 0.747 | 1.775 | 0 | 0.071 | 0.071 | 0 | 0.06 | 0.06 | 0.04 | 0.29 | 0.05 | 0.17 |
| 1 | 119293640 | rs17185832 | -0.319 | 1.617 | 0.568 | 1.398 | -0.028 | 0.002 | 0.002 | 0.31 | 0.18 | 0.09 | 0.18 | 0.15 | 0.25 | 0.2 |
| 1 | 119300747 | rs2764473 | -0.192 | 1.564 | 0.674 | 1.482 | 0.015 | 0.001 | 0.001 | 0 | 0.14 | 0.11 | 0.16 | 0.16 | 0.19 | 0.14 |
| 1 | 119300941 | rs9787392 | -1.462 | 0.746 | 0.547 | 1.512 | -0.02 | 0.103 | 0.103 | 0 | 0.2 | 0.37 | 0.3 | 0.46 | 0.15 | 0.24 |
| 1 | 119316263 | rs10923673 | -1.31 | 0.841 | 0.619 | 1.596 | 0.005 | 0.192 | 0.192 | 0.056 | 0.14 | 0.35 | 0.28 | 0.46 | 0.13 | 0.18 |
| 1 | 119319805 | rs4659120 | 0.754 | 1.017 | 0.18 | 1.239 | 0.034 | 0.066 | 0.066 | 0.276 | 0.73 | 0.47 | 0.59 | 0.55 | 0.62 | 0.69 |
| 1 | 119372737 | rs17185860 | 2.053 | 0.149 | -0.824 | 0.216 | 0.056 | 0.112 | 0.112 | 3.068 | 0.26 | 0 | 0.08 | 0.07 | 0.14 | 0.14 |
| 1 | 119380513 | rs1779415 | -0.301 | 0.551 | 0.257 | 0.522 | 0.023 | 0.045 | 0.045 | 0.034 | 0.1 | 0.29 | 0.1 | 0.15 | 0.16 | 0.1 |
| 1 | 119380928 | rs1338442 | 0.255 | 0.559 | 0.373 | 0.62 | -0.027 | 0.006 | 0.006 | 0.309 | 0.18 | 0.07 | 0.18 | 0.16 | 0.22 | 0.17 |
| 1 | 119383818 | rs1766803 | -0.79 | 0.525 | 0.327 | 0.619 | 0.025 | 0.004 | 0.004 | 0.446 | 0.27 | 0.36 | 0.29 | 0.3 | 0.38 | 0.27 |
| 1 | 119392083 | rs868157 | 0.388 | 0.505 | 0.276 | 0.618 | 0.001 | 0.013 | 0.013 | 2.684 | 0.09 | 0.13 | 0.24 | 0.16 | 0.09 | 0.04 |
| 1 | 119394195 | rs12408957 | 0.676 | 0.517 | 0.035 | 0.624 | -0.002 | -0.005 | -0.005 | 0 | 0.08 | 0.05 | 0.24 | 0.16 | 0.09 | 0.04 |

| | | | | | | | | | | | | | | | | |
|---|---|---|---|---|---|---|---|---|---|---|---|---|---|---|---|---|
| 1 | 119409085 | rs17185873 | 0.967 | 0.664 | -0.373 | 0.215 | -0.105 | 0.145 | 0.145 | 0 | 0.23 | 0.03 | 0.23 | 0.01 | 0.32 | 0.19 |
| 1 | 119410215 | rs2361028 | -1.309 | 0.839 | 0.106 | 0.524 | -0.011 | 0.014 | 0.014 | 0.477 | 0.35 | 0.68 | 0.32 | 0.73 | 0.44 | 0.47 |
| 1 | 119411415 | rs12040674 | 0.418 | 0.727 | -0.02 | 0.932 | -0.017 | 0.009 | 0.009 | 0 | 0.06 | 0.03 | 0.07 | 0.25 | 0.14 | 0.1 |
| 1 | 119416488 | rs8179426 | 0.45 | 0.725 | 0.221 | 0.939 | -0.017 | 0.011 | 0.011 | 0 | 0.06 | 0.03 | 0.08 | 0.25 | 0.14 | 0.1 |
| 1 | 119421209 | rs6681289 | 0.353 | 0.813 | 0.047 | 1.192 | -0.056 | -0.096 | -0.096 | 0 | 0.14 | 0.01 | 0.11 | 0.32 | 0.14 | 0.26 |
| 1 | 119431014 | rs10923694 | -0.277 | 0.431 | 0.605 | 1.238 | 0.013 | 0.105 | 0.105 | 0 | 0.08 | 0.14 | 0.09 | 0.23 | 0.15 | 0.15 |
| 1 | 119434896 | rs1766783 | -1.628 | -0.264 | -0.181 | 0.382 | 0.024 | 0.159 | 0.159 | 0.212 | 0.71 | 1 | 0.88 | 0.94 | 0.81 | 0.83 |
| 1 | 119446001 | rs2282322 | -0.583 | 0.462 | -0.802 | 1.071 | -0.04 | -0.088 | -0.088 | 0 | 0.13 | 0.01 | 0.41 | 0.37 | 0.19 | 0.11 |
| 1 | 119448317 | rs946205 | -0.079 | 0.537 | -0.55 | 0.266 | 0 | 0.001 | 0.001 | 0.48 | 0.86 | 0.9 | 0.76 | 0.88 | 0.88 | 0.72 |
| 1 | 119452758 | rs1766787 | -0.186 | 0.396 | -0.66 | 0.434 | 0.002 | -0.004 | -0.004 | 0.274 | 0.85 | 0.9 | 0.76 | 0.87 | 0.88 | 0.74 |
| 1 | 119453304 | rs1779418 | 1.199 | -0.006 | -0.513 | 0.278 | -0.002 | -0.005 | -0.005 | 0 | 0.61 | 0.66 | 0.41 | 0.35 | 0.65 | 0.6 |
| 1 | 119453869 | rs1779416 | 1.169 | -0.014 | -0.367 | 0.295 | 0.003 | 0.011 | 0.011 | 0 | 0.62 | 0.72 | 0.41 | 0.35 | 0.64 | 0.6 |
| 1 | 119454212 | rs12033524 | 0.263 | -0.032 | -0.354 | 0.308 | -0.015 | -0.086 | -0.086 | 1.845 | 0.2 | 0.06 | 0.34 | 0.51 | 0.24 | 0.13 |
| 1 | 119458215 | rs1014098 | 0.177 | 0.428 | 0.001 | 0.116 | 0.025 | 0.008 | 0.008 | 0 | 0.07 | 0.12 | 0.11 | 0.06 | 0.16 | 0.11 |
| 1 | 119458234 | rs10923698 | 2.497 | -0.307 | -0.768 | -0.922 | 0.027 | 0.034 | 0.034 | 0 | 0.24 | 0.1 | 0.12 | 0.23 | 0.14 | 0.14 |
| 1 | 119460481 | rs1779429 | 1.616 | -0.489 | -0.698 | -1.091 | 0.032 | 0.039 | 0.039 | 0 | 0.3 | 0.19 | 0.14 | 0.25 | 0.17 | 0.2 |
| 1 | 119461396 | rs1779431 | 0.502 | -0.409 | -0.402 | -1.022 | 0.006 | 0.015 | 0.015 | 0.557 | 0.74 | 0.54 | 0.63 | 0.8 | 0.72 | 0.62 |
| 1 | 119468760 | rs1419286 | -0.28 | -0.386 | -0.544 | -1.138 | -0.002 | -0.017 | -0.017 | 0 | 0.63 | 0.6 | 0.57 | 0.8 | 0.64 | 0.51 |
| 1 | 119469188 | rs10494217 | 1.651 | -0.366 | -0.622 | -1.141 | -0.04 | -0.03 | -0.03 | 0 | 0.07 | 0.01 | 0.1 | 0.07 | 0.17 | 0.11 |
| 1 | 119471908 | rs2645290 | -1.231 | -0.413 | -0.481 | -1.163 | -0.015 | -0.024 | -0.024 | 0 | 0.71 | 0.68 | 0.68 | 0.87 | 0.8 | 0.62 |
| 1 | 119473877 | rs17185926 | 3.45 | 0.316 | -0.757 | -1.377 | -0.028 | 0.03 | 0.03 | 0 | 0.07 | 0.03 | 0.07 | 0 | 0.14 | 0.02 |
| 1 | 119478611 | rs1325934 | 2.554 | -0.128 | -0.453 | -1.22 | -0.008 | 0 | 0 | 0 | 0.14 | 0.15 | 0.06 | 0.23 | 0.06 | 0.05 |
| 1 | 119485771 | rs1325945 | -0.073 | -0.104 | -0.444 | -1.259 | 0.01 | 0.01 | 0.01 | 0.012 | 0.49 | 0.46 | 0.52 | 0.57 | 0.58 | 0.46 |
| 1 | 119489882 | rs10923703 | 0.331 | 0.02 | -0.69 | -1.308 | -0.05 | -0.071 | -0.071 | 0 | 0.22 | 0.06 | 0.42 | 0.55 | 0.36 | 0.23 |
| 1 | 119493411 | rs1750330 | 0.458 | -0.025 | -0.615 | -1.367 | 0 | -0.007 | -0.007 | 0.556 | 0.49 | 0.55 | 0.26 | 0.32 | 0.43 | 0.39 |
| 1 | 119495681 | rs6667873 | 1.554 | 0.274 | -0.804 | -1.13 | -0.01 | -0.043 | -0.043 | 0.063 | 0.15 | 0.06 | 0.31 | 0.48 | 0.2 | 0.12 |
| 1 | 119496579 | rs1155949 | 1.986 | 0.222 | -0.803 | -1.303 | -0.005 | -0.012 | -0.012 | 0 | 0.24 | 0.28 | 0.16 | 0.04 | 0.31 | 0.27 |
| 1 | 119500773 | rs985036 | 0.143 | 0.416 | -1.013 | -1.147 | -0.028 | -0.042 | -0.042 | 1.125 | 0.22 | 0.12 | 0.42 | 0.55 | 0.37 | 0.23 |
| 1 | 119504284 | rs984225 | 0.03 | 0.139 | -0.595 | -1.136 | 0.005 | 0.01 | 0.01 | 0 | 0.45 | 0.38 | 0.57 | 0.54 | 0.61 | 0.45 |
| 1 | 119512913 | rs10923714 | 1.42 | 0.19 | -0.596 | -1.124 | -0.02 | 0.024 | 0.024 | 0 | 0.29 | 0.27 | 0.23 | 0.07 | 0.41 | 0.31 |
| 1 | 119521631 | rs6680737 | 1.364 | 0.233 | -0.638 | -1.108 | -0.001 | -0.013 | -0.013 | 0 | 0.29 | 0.34 | 0.24 | 0.07 | 0.41 | 0.31 |
| 1 | 119525788 | rs12742627 | 2.552 | 0.025 | -0.528 | -0.907 | -0.001 | -0.001 | -0.001 | 0 | 0.12 | 0.08 | 0.05 | 0.22 | 0.06 | 0.05 |
| 1 | 119538433 | rs1409159 | -1.346 | 0.282 | -0.301 | -0.887 | 0.002 | 0.003 | 0.003 | 0 | 0.67 | 0.75 | 0.67 | 0.86 | 0.76 | 0.65 |
| 1 | 119538705 | rs716217 | -0.849 | 0.185 | -0.162 | -0.847 | -0.04 | -0.007 | -0.007 | 0 | 0.25 | 0.43 | 0.1 | 0.32 | 0.16 | 0.22 |
| 1 | 119540842 | rs7551082 | 0.935 | 0.183 | -0.179 | -0.799 | -0.043 | 0.046 | 0.046 | 0.463 | 0.1 | 0.27 | 0.03 | 0.22 | 0.02 | 0.07 |
| 1 | 119543486 | rs10923722 | 1.382 | 0.616 | -0.644 | -0.936 | -0.004 | -0.006 | -0.006 | 0 | 0.28 | 0.31 | 0.22 | 0.07 | 0.37 | 0.29 |
| 1 | 119546153 | rs12145407 | 1.038 | 0.674 | -0.88 | -0.719 | -0.016 | 0.02 | 0.02 | 0 | 0.12 | 0.04 | 0.06 | 0.1 | 0.13 | 0.14 |
| 1 | 119552284 | rs4501872 | 1.563 | 0.748 | -0.851 | -0.49 | -0.059 | -0.101 | -0.101 | 0 | 0.13 | 0.01 | 0.45 | 0.48 | 0.2 | 0.12 |
| 1 | 119555146 | rs2645303 | 0.357 | 0.771 | -0.516 | -0.725 | 0.008 | 0.011 | 0.011 | 0 | 0.4 | 0.51 | 0.6 | 0.51 | 0.45 | 0.35 |
| 1 | 119555759 | rs1325933 | -1.428 | 0.688 | -0.548 | -0.891 | 0.019 | 0.062 | 0.062 | 0 | 0.67 | 0.9 | 0.8 | 0.86 | 0.74 | 0.64 |
| 1 | 119572057 | rs2148150 | -1.428 | 0.673 | -0.53 | -0.887 | 0.022 | 0.06 | 0.06 | 0 | 0.67 | 0.89 | 0.8 | 0.86 | 0.75 | 0.65 |
| 1 | 119574289 | rs12030972 | 2.005 | 0.755 | -0.564 | -0.806 | -0.058 | -0.094 | -0.094 | 0 | 0.12 | 0.01 | 0.45 | 0.46 | 0.19 | 0.12 |
| 1 | 119574537 | rs12026409 | 2.005 | 0.755 | -0.427 | -0.806 | -0.058 | -0.094 | -0.094 | 0 | 0.12 | 0.01 | 0.45 | 0.46 | 0.19 | 0.12 |
| 1 | 119574587 | rs2645294 | 0.134 | 0.785 | -0.359 | -0.828 | 0 | -0.003 | -0.003 | 0 | 0.45 | 0.61 | 0.7 | 0.6 | 0.57 | 0.4 |
| 1 | 119575170 | rs1057991 | 1.725 | 0.75 | -0.243 | -0.858 | 0.002 | -0.094 | -0.094 | 0.417 | 0.27 | 0.46 | 0.15 | 0.1 | 0.26 | 0.25 |
| 1 | 119575232 | rs1057990 | 1.725 | 0.75 | -0.243 | -0.858 | 0.002 | -0.094 | -0.094 | 1.121 | 0.27 | 0.46 | 0.15 | 0.1 | 0.26 | 0.25 |
| 1 | 119580252 | rs12029198 | 2.75 | 0.744 | -0.316 | -0.881 | -0.004 | 0.039 | 0.039 | 0 | 0.07 | 0.08 | 0.03 | 0.22 | 0.05 | 0.05 |
| 1 | 119581900 | rs2765529 | 1.728 | 0.75 | -0.368 | -0.87 | 0.001 | -0.097 | -0.097 | 0.755 | 0.27 | 0.45 | 0.15 | 0.09 | 0.26 | 0.25 |
| 1 | 119583854 | rs1409160 | 1.012 | 0.745 | -0.371 | -0.874 | 0.011 | -0.08 | -0.08 | 0.724 | 0.33 | 0.6 | 0.24 | 0.14 | 0.37 | 0.28 |
| 1 | 119589343 | rs17023164 | 1.415 | 0.811 | -0.419 | -0.883 | -0.08 | -0.119 | -0.119 | 2.686 | 0.15 | 0.01 | 0.48 | 0.51 | 0.26 | 0.19 |
| 1 | 119593268 | rs17023171 | 1.641 | 0.715 | -0.316 | -0.841 | 0.001 | -0.099 | -0.099 | 0.009 | 0.27 | 0.46 | 0.15 | 0.09 | 0.26 | 0.25 |
| 1 | 119619448 | rs12045138 | 0.298 | 0.723 | -0.473 | -0.661 | 0 | 0.004 | 0.004 | 0 | 0.39 | 0.71 | 0.26 | 0.4 | 0.4 | 0.34 |
| 1 | 119621894 | rs7525871 | 2.149 | 0.659 | -0.338 | -0.694 | 0.004 | -0.022 | -0.022 | 0 | 0.24 | 0.32 | 0.13 | 0.08 | 0.26 | 0.23 |
| 1 | 119624163 | rs2885226 | 1.12 | 0.895 | -0.718 | -0.567 | 0.014 | -0.045 | -0.045 | 0 | 0.33 | 0.59 | 0.24 | 0.18 | 0.37 | 0.28 |
| 1 | 119627148 | rs12044562 | 2.778 | 0.891 | -0.715 | -0.602 | -0.009 | 0.099 | 0.099 | 0 | 0.06 | 0.13 | 0.02 | 0.21 | 0.03 | 0.05 |
| 1 | 119627856 | rs6658479 | 0.487 | 0.903 | -0.768 | -0.626 | 0 | 0.006 | 0.006 | 0 | 0.39 | 0.71 | 0.26 | 0.39 | 0.4 | 0.34 |
| 1 | 119629704 | rs10754391 | -1.337 | 0.96 | -0.859 | -0.719 | 0.015 | 0.035 | 0.035 | 0.047 | 0.66 | 0.79 | 0.8 | 0.89 | 0.75 | 0.63 |
| 1 | 119638553 | rs6701378 | 1.245 | 1.039 | -0.798 | -0.609 | 0.017 | -0.035 | -0.035 | 0 | 0.3 | 0.51 | 0.23 | 0.15 | 0.36 | 0.25 |
| 1 | 119650928 | rs2885227 | -0.319 | 1.032 | -0.735 | -0.617 | 0.006 | -0.004 | -0.004 | 1.056 | 0.27 | 0.32 | 0.57 | 0.77 | 0.34 | 0.25 |
| 1 | 119657057 | rs1341580 | -0.769 | 1.068 | -0.75 | -0.637 | 0.01 | -0.012 | -0.012 | 1.392 | 0.3 | 0.32 | 0.62 | 0.83 | 0.43 | 0.32 |
| 1 | 119668640 | rs10923754 | 1.802 | 1.008 | -0.73 | -0.697 | 0.002 | -0.058 | -0.058 | 0 | 0.27 | 0.41 | 0.15 | 0.06 | 0.28 | 0.27 |
| 1 | 119673093 | rs2498500 | 2.465 | 0.97 | -0.698 | -0.607 | -0.01 | 0.116 | 0.116 | 0 | 0.07 | 0.16 | 0.02 | 0.23 | 0.03 | 0.05 |
| 1 | 119673344 | rs10923755 | 0.571 | 1.02 | -0.78 | -0.702 | 0.001 | -0.028 | -0.028 | 0.337 | 0.39 | 0.46 | 0.22 | 0.11 | 0.41 | 0.39 |
| 1 | 119680296 | rs4387216 | 0.513 | 0.879 | -1.083 | -0.932 | -0.007 | 0.012 | 0.012 | 0 | 0.58 | 0.69 | 0.32 | 0.18 | 0.48 | 0.61 |
| 1 | 119687012 | rs6699136 | 0.071 | 0.996 | -0.883 | -0.776 | 0.006 | -0.005 | -0.005 | 0.165 | 0.72 | 0.59 | 0.43 | 0.22 | 0.67 | 0.83 |
| 1 | 119688490 | rs6428801 | -0.128 | 0.962 | -0.964 | -0.801 | 0.004 | 0.117 | 0.117 | 0.027 | 0.45 | 0.24 | 0.28 | 0.18 | 0.39 | 0.56 |
| 1 | 119705966 | rs7543720 | 0.117 | 0.929 | -1.001 | -0.847 | 0.033 | 0.159 | 0.159 | 0.005 | 0.42 | 0.15 | 0.24 | 0.11 | 0.3 | 0.48 |
| 1 | 119728605 | rs12756326 | 1.726 | 0.931 | -1.029 | -0.87 | -0.001 | 0.091 | 0.091 | 0 | 0.19 | 0.03 | 0.1 | 0 | 0.13 | 0.16 |
| 1 | 119729171 | rs6678854 | 1.564 | 0.838 | -0.937 | -0.759 | 0.001 | 0.004 | 0.004 | 0.004 | 0.21 | 0.18 | 0.15 | 0.11 | 0.17 | 0.29 |

| | | | | | | | | | | | | | | | | |
|---|---|---|---|---|---|---|---|---|---|---|---|---|---|---|---|---|
| 1 | 119732347 | rs10923767 | 1.715 | 0.878 | -0.967 | -0.68 | 0.002 | -0.043 | -0.043 | 0 | 0.28 | 0.41 | 0.16 | 0.09 | 0.28 | 0.27 |
| 1 | 119751002 | rs1891222 | -0.296 | 0.797 | -1.122 | -0.446 | 0 | 0.159 | 0.159 | 0.031 | 0.45 | 0.21 | 0.29 | 0.16 | 0.39 | 0.56 |
| 1 | 119759690 | rs1557102 | 1.214 | 0.942 | -1.018 | -0.287 | 0.003 | 0.023 | 0.023 | 0.016 | 0.2 | 0.12 | 0.14 | 0.11 | 0.17 | 0.28 |
| 1 | 119763540 | rs6701188 | -0.18 | 0.818 | -0.793 | -0.063 | 0.003 | -0.001 | -0.001 | 0 | 0.7 | 0.8 | 0.82 | 0.6 | 0.76 | 0.69 |
| 1 | 119766273 | rs10923775 | 1.255 | 1.256 | -0.542 | 0.046 | -0.004 | -0.041 | -0.041 | 0 | 0.3 | 0.39 | 0.15 | 0.06 | 0.27 | 0.27 |
| 1 | 119766683 | rs7524644 | -0.481 | 1.48 | -0.162 | 0.973 | 0.007 | 0.14 | 0.14 | 0.026 | 0.95 | 0.84 | 0.97 | 0.75 | 0.97 | 0.92 |
| 1 | 119770702 | rs12060730 | 1.129 | 1.535 | -0.268 | 0.884 | -0.001 | -0.015 | -0.015 | 0.439 | 0.53 | 0.6 | 0.33 | 0.14 | 0.47 | 0.53 |
| 1 | 119772663 | rs871665 | 1.135 | 1.535 | -0.271 | 0.885 | 0 | -0.031 | -0.031 | 0 | 0.53 | 0.59 | 0.33 | 0.14 | 0.47 | 0.53 |
| 1 | 119773596 | rs868675 | 1.163 | 1.389 | -0.338 | 0.91 | -0.001 | -0.017 | -0.017 | 0.43 | 0.52 | 0.52 | 0.32 | 0.14 | 0.48 | 0.54 |
| 1 | 119776984 | rs17023344 | 0.459 | 1.021 | -0.389 | 1.2 | -0.038 | -0.044 | -0.044 | 0.128 | 0.13 | 0.04 | 0.37 | 0.21 | 0.19 | 0.14 |
| 1 | 119781299 | rs4658997 | 0.389 | 0.486 | 0.239 | 1.266 | 0.037 | 0.099 | 0.099 | 0.295 | 0.94 | 0.72 | 0.96 | 0.87 | 0.97 | 0.92 |
| 1 | 119787542 | rs11579916 | -0.233 | 0.481 | -0.213 | 0.995 | -0.021 | -0.029 | -0.029 | 0 | 0.25 | 0.25 | 0.12 | 0.1 | 0.16 | 0.21 |
| 1 | 119789289 | rs12730226 | -0.184 | 1.412 | -0.752 | 0.782 | -0.021 | -0.025 | -0.025 | 0 | 0.05 | 0.01 | 0.1 | 0.09 | 0.17 | 0.05 |
| 1 | 119795133 | rs1341797 | 0.562 | 0.051 | -0.517 | 0.458 | -0.013 | -0.013 | -0.013 | 0 | 0.67 | 0.48 | 0.67 | 0.21 | 0.73 | 0.59 |
| 1 | 119799665 | rs10802085 | 0.188 | -0.434 | -0.654 | 0.303 | 0.008 | -0.033 | -0.033 | 0 | 0.45 | 0.25 | 0.57 | 0.46 | 0.6 | 0.39 |
| 1 | 119812129 | rs12757996 | -0.011 | -0.024 | -0.11 | -0.241 | 0.006 | 0.02 | 0.02 | 0 | 0.07 | 0.2 | 0.05 | 0.04 | 0.09 | 0.02 |
| 1 | 119823046 | rs1341802 | 0.042 | -0.53 | -0.383 | -0.499 | 0.006 | 0.019 | 0.019 | 0 | 0.61 | 0.34 | 0.73 | 0.71 | 0.65 | 0.72 |
| 1 | 119828542 | rs1539188 | 1.001 | -0.599 | -0.358 | -0.489 | -0.043 | 0.004 | 0.004 | 0.023 | 0.39 | 0.19 | 0.55 | 0.56 | 0.5 | 0.39 |
| 1 | 119828591 | rs17186108 | 0.019 | -0.028 | -0.646 | -0.576 | 0.019 | 0.019 | 0.019 | 0 | 0.22 | 0.13 | 0.18 | 0.15 | 0.14 | 0.33 |
| 1 | 119830226 | rs17186115 | 0.108 | 0.598 | -0.245 | -0.364 | -0.01 | 0.025 | 0.025 | 1.985 | 0.24 | 0.16 | 0.19 | 0.05 | 0.25 | 0.17 |
| 1 | 119836143 | rs17023386 | -1.526 | 0.617 | -0.038 | 0.441 | 0.001 | 0.008 | 0.008 | 0.549 | 0.08 | 0.12 | 0.12 | 0.18 | 0.09 | 0.13 |
| 1 | 119856801 | rs10923799 | 1.438 | 0.102 | 0.862 | 0.171 | -0.047 | -0.003 | -0.003 | 2.566 | 0.83 | 0.58 | 0.9 | 0.82 | 0.88 | 0.79 |
| 1 | 119858082 | rs2253988 | -1.351 | 0.091 | 0.913 | 0.166 | -0.029 | 0.033 | 0.033 | 2.946 | 0.07 | 0.21 | 0.02 | 0.16 | 0.01 | 0.12 |
| 1 | 119858685 | rs7526517 | -2.109 | 0.087 | 0.841 | 0.162 | -0.044 | -0.009 | -0.009 | 0 | 0.17 | 0.39 | 0.1 | 0.18 | 0.12 | 0.21 |
| 1 | 119859674 | rs838519 | -2.128 | 0.017 | 0.972 | 0.171 | -0.03 | 0.054 | 0.054 | 2.989 | 0.08 | 0.33 | 0.03 | 0.16 | 0.02 | 0.12 |
| 1 | 119860022 | rs838518 | -0.872 | 0.193 | 0.715 | -0.686 | -0.013 | 0.107 | 0.107 | 3.738 | 0.55 | 0.72 | 0.35 | 0.21 | 0.5 | 0.35 |
| 1 | 119863356 | rs947425 | -1.647 | 0.284 | 1.005 | 0.198 | -0.001 | 0.098 | 0.098 | 1.927 | 0.07 | 0.34 | 0.03 | 0.16 | 0.02 | 0.08 |
| 1 | 119864997 | rs6695549 | -0.582 | 0.415 | 0.539 | 0.203 | 0.001 | 0.003 | 0.003 | 0 | 0.09 | 0.04 | 0.06 | 0.02 | 0.1 | 0.08 |
| 1 | 119871977 | rs699771 | -0.988 | 0.351 | 0.603 | 0.457 | -0.022 | 0.039 | 0.039 | 0.046 | 0.05 | 0.19 | 0.02 | 0.16 | 0.01 | 0.13 |
| 1 | 119875730 | rs17023457 | -0.473 | 0.743 | 0.189 | 1.337 | -0.041 | 0.026 | 0.026 | 0 | 0.07 | 0.01 | 0.35 | 0.41 | 0.16 | 0.11 |
| 1 | 119876076 | rs2153738 | -1.871 | -0.029 | 0.662 | 0.472 | -0.021 | 0.07 | 0.07 | 2.74 | 0.08 | 0.39 | 0.04 | 0.18 | 0.02 | 0.13 |
| 1 | 119893928 | rs838534 | -1.103 | -0.142 | 0.134 | 0.077 | 0.002 | 0.002 | 0.002 | 0 | 0.26 | 0.27 | 0.1 | 0.23 | 0.17 | 0.26 |
| 1 | 119895221 | rs699774 | -1.054 | -0.166 | -0.487 | 0.175 | -0.059 | -0.045 | -0.045 | 0 | 0.61 | 0.84 | 0.44 | 0.51 | 0.45 | 0.62 |
| 1 | 119895261 | rs838533 | -0.842 | -0.498 | 0.166 | 0.347 | -0.023 | 0.065 | 0.065 | 0.014 | 0.06 | 0.2 | 0.06 | 0.24 | 0.02 | 0.14 |
| 1 | 119897499 | rs838530 | -1.376 | -0.793 | 1.117 | 0.133 | -0.06 | -0.022 | -0.022 | 0.001 | 0.2 | 0.61 | 0.14 | 0.25 | 0.06 | 0.21 |
| 1 | 119908763 | rs3820032 | -0.753 | -0.779 | 0.771 | -0.039 | -0.004 | 0.002 | 0.002 | 0.405 | 0.33 | 0.61 | 0.27 | 0.48 | 0.24 | 0.5 |
| 1 | 119909208 | rs3753266 | -0.05 | -1.112 | 0.668 | -0.272 | -0.018 | -0.027 | -0.027 | 0 | 0.14 | 0.2 | 0.08 | 0.03 | 0.06 | 0.08 |
| 1 | 119909514 | rs3753264 | -1.332 | -0.988 | 0.276 | -0.142 | 0.101 | -0.002 | -0.002 | 0 | 0.47 | 0.82 | 0.35 | 0.53 | 0.3 | 0.58 |
| 1 | 119913824 | rs6667203 | 0.324 | -0.804 | 0.63 | 0.094 | -0.007 | 0.01 | 0.01 | 0 | 0.2 | 0.25 | 0.13 | 0.33 | 0.18 | 0.27 |
| 1 | 119916426 | rs7514283 | -0.301 | -0.799 | 1.204 | 0.262 | 0.005 | -0.022 | -0.022 | 0 | 0.09 | 0.3 | 0.12 | 0.04 | 0.05 | 0.12 |
| 1 | 119917253 | rs3765953 | 1.249 | -0.383 | 0.768 | 0.078 | -0.117 | -0.112 | -0.112 | 3.859 | 0.35 | 0.08 | 0.52 | 0.45 | 0.52 | 0.34 |
| 1 | 119917573 | rs3765952 | -0.889 | -0.486 | 0.722 | 0.014 | -0.047 | 0.067 | 0.067 | 0 | 0.18 | 0.46 | 0.19 | 0.22 | 0.1 | 0.28 |
| 1 | 119925936 | rs2298028 | -0.175 | -0.504 | 0.458 | 0 | 0.008 | 0.045 | 0.045 | 4.504 | 0.55 | 0.71 | 0.73 | 0.67 | 0.62 | 0.62 |
| 1 | 119928193 | rs3207643 | 0.583 | 0.265 | -0.093 | 0.622 | -0.03 | -0.013 | -0.013 | 0 | 0.07 | 0.02 | 0.35 | 0.28 | 0.21 | 0.11 |
| 1 | 119936263 | rs3765949 | 0.831 | -0.526 | 0.147 | 0.325 | -0.04 | -0.04 | -0.04 | 0 | 0.38 | 0.26 | 0.54 | 0.45 | 0.53 | 0.35 |
| 1 | 119940918 | rs879332 | 1.099 | -0.468 | -0.296 | 0.352 | -0.124 | -0.121 | -0.121 | 2.316 | 0.35 | 0.08 | 0.52 | 0.45 | 0.53 | 0.35 |
| 1 | 119970688 | rs10923819 | -0.552 | -0.849 | -0.27 | 0.214 | -0.016 | 0.032 | 0.032 | 0.684 | 0.66 | 0.84 | 0.77 | 0.68 | 0.66 | 0.7 |
| 1 | 119977687 | rs12035458 | 0.481 | -1.034 | 0.106 | 0.256 | 0.002 | 0.005 | 0.005 | 0 | 0.08 | 0.07 | 0.05 | 0.18 | 0.04 | 0.22 |
| 1 | 119977806 | rs1417609 | -1.341 | -1.248 | 0.286 | 0.11 | -0.046 | -0.008 | -0.008 | 2.352 | 0.29 | 0.62 | 0.23 | 0.24 | 0.12 | 0.36 |
| 1 | 119990027 | rs7546652 | 1.388 | -1.196 | 0.247 | 0.15 | -0.094 | -0.083 | -0.083 | 3.659 | 0.34 | 0.1 | 0.53 | 0.43 | 0.54 | 0.35 |
| 1 | 119990185 | rs1538987 | -0.541 | -0.991 | 0.317 | -0.142 | -0.001 | 0.016 | 0.016 | 0.441 | 0.64 | 0.73 | 0.76 | 0.67 | 0.66 | 0.71 |
| 1 | 119991055 | rs12722964 | 0.645 | -0.935 | 0.153 | -0.093 | -0.011 | 0.047 | 0.047 | 7.648 | 0.44 | 0.26 | 0.6 | 0.6 | 0.58 | 0.57 |
| 1 | 119991429 | rs10923825 | -0.777 | -1.092 | 0.52 | 0.045 | -0.04 | -0.078 | -0.078 | 0.41 | 0.19 | 0.33 | 0.15 | 0.06 | 0.08 | 0.14 |
| 1 | 119993484 | rs6673366 | -1.345 | -1.054 | 0.803 | 0.589 | -0.048 | 0.137 | 0.137 | 1.598 | 0.12 | 0.46 | 0.11 | 0.23 | 0.03 | 0.22 |
| 1 | 119994095 | rs7528420 | 0.608 | -1.239 | 0.201 | 0.632 | -0.002 | -0.006 | -0.006 | 0 | 0.36 | 0.32 | 0.5 | 0.41 | 0.41 | 0.32 |
| 1 | 119999212 | rs6672903 | -0.397 | -1.479 | 0.365 | 0.569 | 0.037 | 0.016 | 0.016 | 0 | 0.48 | 0.89 | 0.59 | 0.58 | 0.44 | 0.5 |
| 1 | 120000021 | rs1361529 | 0.987 | -1.463 | 0.35 | 0.588 | -0.066 | -0.054 | -0.054 | 0.26 | 0.33 | 0.08 | 0.45 | 0.35 | 0.41 | 0.28 |
| 1 | 120000183 | rs10923830 | -1.423 | -1.365 | 0.635 | 0.662 | -0.05 | 0.033 | 0.033 | 3.562 | 0.14 | 0.75 | 0.14 | 0.22 | 0.03 | 0.22 |
| 1 | 120009049 | rs12406375 | -0.378 | -1.243 | 0.58 | 0.64 | -0.027 | 0.11 | 0.11 | 0.297 | 0.1 | 0.42 | 0.08 | 0.16 | 0.03 | 0.19 |
| 1 | 120011081 | rs12406851 | -0.794 | -1.055 | -0.367 | 0.438 | -0.055 | -0.051 | -0.051 | 0 | 0.67 | 0.93 | 0.79 | 0.64 | 0.64 | 0.71 |
| 1 | 120014607 | rs7547046 | -0.792 | -0.844 | 0.203 | -0.016 | -0.008 | -0.018 | -0.018 | 0 | 0.31 | 0.56 | 0.27 | 0.23 | 0.22 | 0.4 |
| 1 | 120017900 | rs2064901 | 0.228 | -1.175 | 0.067 | -0.255 | 0 | 0.024 | 0.024 | 0 | 0.6 | 0.31 | 0.7 | 0.82 | 0.63 | 0.71 |
| 1 | 120047647 | rs6661258 | -0.725 | -1.255 | 1.896 | -0.258 | 0.066 | 0.119 | 0.119 | 1.366 | 0.06 | 0.42 | 0.07 | 0.05 | 0.01 | 0.07 |
| 1 | 120049735 | rs932603 | -0.669 | -1.264 | 2.025 | -0.263 | 0.053 | 0.102 | 0.102 | 1.478 | 0.05 | 0.38 | 0.07 | 0.05 | 0.01 | 0.07 |
| 1 | 120051446 | rs3765945 | 0.327 | -1.528 | 1.702 | -0.601 | 0.008 | 0.057 | 0.057 | 0 | 0.58 | 0.31 | 0.7 | 0.86 | 0.63 | 0.77 |
| 1 | 120052043 | rs7519379 | -0.598 | -1.446 | 1.53 | -0.516 | 0.056 | 0.129 | 0.129 | 3.184 | 0.06 | 0.45 | 0.08 | 0.06 | 0.01 | 0.07 |
| 1 | 120062933 | rs12035199 | -1.252 | -1.505 | 0.783 | -0.842 | -0.05 | 0.053 | 0.053 | 0 | 0.79 | 0.95 | 0.83 | 0.93 | 0.72 | 0.9 |
| 1 | 120077584 | rs7553527 | 1.558 | -1.587 | 0.314 | -0.598 | -0.131 | -0.109 | -0.109 | 0.069 | 0.28 | 0.02 | 0.49 | 0.67 | 0.43 | 0.39 |
| 1 | 120083340 | rs6681580 | -0.078 | -1.497 | 0.382 | -0.618 | 0.003 | 0.013 | 0.013 | 0 | 0.61 | 0.41 | 0.72 | 0.84 | 0.65 | 0.78 |

| | | | | | | | | | | | | | | | | |
|---|---|---|---|---|---|---|---|---|---|---|---|---|---|---|---|---|
| 1 | 120085131 | rs9887805 | -0.096 | -1.493 | 0.368 | -0.616 | -0.001 | -0.012 | -0.012 | 0 | 0.61 | 0.45 | 0.72 | 0.84 | 0.66 | 0.78 |
| 1 | 120091434 | rs10802112 | -0.057 | -1.492 | 0.032 | -0.371 | -0.004 | 0.022 | 0.022 | 0 | 0.28 | 0.14 | 0.22 | 0.21 | 0.24 | 0.39 |
| 1 | 120110175 | rs4659204 | 1.047 | -1.22 | -1.353 | -0.225 | -0.053 | -0.11 | -0.11 | 0.406 | 0.31 | 0.08 | 0.46 | 0.57 | 0.37 | 0.34 |
| 1 | 120115384 | rs12134798 | 0.357 | -0.76 | -1.343 | -0.785 | -0.013 | 0.069 | 0.069 | 0 | 0.22 | 0.06 | 0.22 | 0.03 | 0.2 | 0.24 |
| 1 | 120123179 | rs6428833 | -0.542 | -0.735 | -1.432 | -0.84 | -0.067 | 0.081 | 0.081 | 0.533 | 0.66 | 0.89 | 0.78 | 0.85 | 0.57 | 0.79 |
| 1 | 120123642 | rs10754411 | -0.484 | -0.578 | -0.684 | -0.573 | -0.003 | -0.022 | -0.022 | 0.151 | 0.27 | 0.24 | 0.27 | 0.1 | 0.2 | 0.23 |
| 1 | 120129258 | rs346671 | -0.795 | 0.167 | -1.327 | -0.613 | -0.006 | 0.064 | 0.064 | 0.266 | 0.64 | 0.85 | 0.62 | 0.76 | 0.57 | 0.68 |
| 1 | 120131597 | rs12139088 | 0.936 | 0.389 | -1.172 | -0.815 | 0.008 | -0.014 | -0.014 | 0 | 0.07 | 0.05 | 0.07 | 0 | 0.09 | 0.03 |
| 1 | 120137445 | rs10494227 | 0.961 | 0.814 | -1.503 | -1.166 | -0.046 | 0.068 | 0.068 | 1.183 | 0.14 | 0.03 | 0.08 | 0.01 | 0.17 | 0.06 |
| 1 | 120138083 | rs1923025 | 1.125 | 0.808 | -1.096 | -0.991 | 0.003 | 0.023 | 0.023 | 0.168 | 0.37 | 0.21 | 0.43 | 0.27 | 0.45 | 0.38 |
| 1 | 120138151 | rs10923871 | 0.731 | 0.541 | -1.54 | -0.724 | 0.016 | -0.016 | -0.016 | 1.227 | 0.16 | 0.04 | 0.07 | 0.16 | 0.1 | 0.14 |
| 1 | 120138246 | rs11586489 | 1.227 | 0.659 | -1.216 | -0.542 | 0.02 | -0.025 | -0.025 | 0 | 0.07 | 0.08 | 0.07 | 0 | 0.09 | 0.03 |
| 1 | 120138931 | rs346683 | 0.716 | 0.412 | -0.738 | -0.578 | 0 | -0.01 | -0.01 | 0 | 0.21 | 0.17 | 0.32 | 0.56 | 0.18 | 0.4 |
| 1 | 120145883 | rs4659015 | -0.745 | 0.801 | -0.655 | -0.661 | -0.005 | 0.023 | 0.023 | 0 | 0.56 | 0.68 | 0.55 | 0.78 | 0.47 | 0.65 |
| 1 | 120150949 | rs10923875 | -0.616 | 0.958 | -0.641 | 0.156 | 0.055 | 0.086 | 0.086 | 0 | 0.26 | 0.42 | 0.43 | 0.55 | 0.28 | 0.34 |
| 1 | 120152082 | rs6689350 | -0.244 | 0.865 | -0.54 | -0.061 | 0.051 | -0.03 | -0.03 | 0.879 | 0.14 | 0.42 | 0.15 | 0.04 | 0.11 | 0.09 |
| 1 | 120162787 | rs2281493 | -0.834 | 0.765 | -1.651 | -0.12 | 0.026 | 0.108 | 0.108 | 0.312 | 0.61 | 0.84 | 0.65 | 0.77 | 0.58 | 0.67 |
| 1 | 120163675 | rs35922771 | 0.961 | 0.874 | -1.733 | 0.095 | -0.065 | -0.095 | -0.095 | 0 | 0.13 | 0.01 | 0.27 | 0.47 | 0.17 | 0.22 |
| 1 | 120164966 | rs7552885 | -0.449 | 0.224 | -1.27 | -0.691 | 0.023 | 0.11 | 0.11 | 0.308 | 0.61 | 0.85 | 0.65 | 0.77 | 0.58 | 0.67 |
| 1 | 120165187 | rs3795663 | -0.434 | 0.234 | -1.27 | -0.686 | 0.026 | 0.108 | 0.108 | 0.312 | 0.61 | 0.84 | 0.65 | 0.77 | 0.58 | 0.67 |
| 1 | 120168353 | rs4659221 | -0.46 | 0.219 | -1.263 | -0.663 | 0.012 | 0.085 | 0.085 | 0.071 | 0.62 | 0.84 | 0.67 | 0.77 | 0.6 | 0.66 |
| 1 | 120177454 | rs10754417 | -0.217 | 0.06 | -0.703 | -1.142 | 0 | -0.002 | -0.002 | 0 | 0.49 | 0.47 | 0.58 | 0.82 | 0.52 | 0.56 |
| 1 | 120180034 | rs347905 | 0.183 | 0.121 | -0.595 | -1.289 | -0.009 | -0.033 | -0.033 | 0 | 0.19 | 0.37 | 0.13 | 0.1 | 0.16 | 0.12 |
| 1 | 120182906 | rs838989 | -0.259 | -0.077 | -0.098 | -1.441 | 0.008 | 0.013 | 0.013 | 0 | 0.6 | 0.67 | 0.65 | 0.91 | 0.65 | 0.67 |
| 1 | 120188180 | rs347911 | 0.096 | -0.342 | -0.136 | -1.631 | 0.002 | -0.031 | -0.031 | 0.199 | 0.53 | 0.47 | 0.62 | 0.9 | 0.62 | 0.64 |
| 1 | 120193579 | rs372814 | -0.825 | -0.285 | -0.622 | -1.3 | -0.012 | 0.072 | 0.072 | 0 | 0.72 | 0.92 | 0.84 | 0.87 | 0.7 | 0.72 |
| 1 | 120195042 | rs454510 | -0.728 | -0.269 | -0.592 | -1.343 | -0.01 | 0.087 | 0.087 | 0 | 0.71 | 0.89 | 0.74 | 0.9 | 0.69 | 0.68 |
| 1 | 120197685 | rs496886 | 0.178 | 0.152 | -0.049 | -1.519 | 0.01 | 0.016 | 0.016 | 0 | 0.48 | 0.39 | 0.54 | 0.9 | 0.42 | 0.56 |
| 1 | 120199189 | rs380155 | 0.261 | -0.01 | -0.099 | -0.623 | 0.014 | -0.01 | -0.01 | 0 | 0.44 | 0.34 | 0.47 | 0.53 | 0.38 | 0.34 |
| 1 | 120200843 | rs481357 | -0.836 | -0.12 | -0.229 | -0.751 | 0.018 | -0.03 | -0.03 | 0.063 | 0.64 | 0.78 | 0.57 | 0.44 | 0.69 | 0.64 |
| 1 | 120208472 | rs539799 | 0.227 | 0.513 | 0.029 | -1.037 | -0.002 | 0.008 | 0.008 | 0 | 0.45 | 0.43 | 0.37 | 0.11 | 0.51 | 0.43 |
| 1 | 120216287 | rs12025390 | 0.403 | 0.148 | -0.481 | -1.317 | -0.013 | -0.017 | -0.017 | 0 | 0.15 | 0.17 | 0.07 | 0.03 | 0.09 | 0.13 |
| 1 | 120217414 | rs639216 | -0.968 | 0.515 | -0.715 | -1.268 | -0.089 | 0.1 | 0.1 | 0.71 | 0.71 | 0.86 | 0.72 | 0.92 | 0.58 | 0.7 |
| 1 | 120230079 | rs620741 | 1.273 | 0.187 | -0.699 | -1.044 | 0.027 | -0.032 | -0.032 | 0 | 0.3 | 0.14 | 0.37 | 0.59 | 0.23 | 0.34 |
| 1 | 120240114 | rs1853048 | 0.241 | 0.143 | -0.266 | -1.249 | 0.013 | 0.103 | 0.103 | 0.064 | 0.17 | 0.34 | 0.19 | 0.32 | 0.18 | 0.21 |
| 1 | 120241771 | rs2764886 | -1.169 | 0.085 | -0.307 | -1.437 | -0.085 | 0.118 | 0.118 | 0.005 | 0.67 | 0.85 | 0.69 | 0.92 | 0.56 | 0.67 |
| 1 | 120242537 | rs10802132 | -1.1 | 0.087 | -0.25 | -1.36 | 0.004 | 0.003 | 0.003 | 0.008 | 0.36 | 0.68 | 0.31 | 0.33 | 0.33 | 0.33 |
| 1 | 120247782 | rs12030925 | 0.21 | 0.082 | -0.313 | -1.34 | -0.012 | -0.083 | -0.083 | 0.291 | 0.2 | 0.34 | 0.12 | 0.01 | 0.15 | 0.11 |
| 1 | 120256713 | rs636101 | 1.471 | -0.014 | -0.492 | -1.29 | 0.028 | -0.045 | -0.045 | 0 | 0.29 | 0.1 | 0.37 | 0.59 | 0.23 | 0.38 |
| 1 | 120257576 | rs477992 | -1.156 | 0.763 | -0.436 | -1.45 | -0.001 | -0.013 | -0.013 | 3.751 | 0.7 | 0.63 | 0.73 | 0.93 | 0.68 | 0.76 |
| 1 | 120258970 | rs666930 | -0.429 | 0.9 | -0.454 | -1.461 | -0.013 | 0.031 | 0.031 | 1.516 | 0.6 | 0.58 | 0.63 | 0.9 | 0.54 | 0.72 |
| 1 | 120260969 | rs586650 | 0.806 | 0.631 | -0.83 | -1.454 | 0.048 | 0.006 | 0.006 | 0 | 0.44 | 0.13 | 0.46 | 0.6 | 0.38 | 0.49 |
| 1 | 120263018 | rs942835 | -0.927 | 0.57 | -0.076 | -0.964 | 0.01 | 0.088 | 0.088 | 2.577 | 0.18 | 0.51 | 0.17 | 0.3 | 0.16 | 0.24 |
| 1 | 120267544 | rs1886736 | -0.129 | 0.791 | -0.211 | -0.843 | 0.052 | 0.16 | 0.16 | 0.451 | 0.13 | 0.43 | 0.15 | 0.29 | 0.14 | 0.19 |
| 1 | 120272477 | rs523395 | -0.698 | 0.63 | -0.508 | -1.147 | -0.058 | 0.169 | 0.169 | 0.857 | 0.59 | 0.79 | 0.75 | 0.93 | 0.51 | 0.65 |
| 1 | 120272726 | rs3828090 | -0.809 | 1.064 | 0.035 | -0.263 | 0.053 | 0.061 | 0.061 | 0 | 0.2 | 0.35 | 0.33 | 0.31 | 0.27 | 0.24 |
| 1 | 120274455 | rs2246410 | 0.41 | 0.99 | -0.03 | -0.502 | 0.016 | -0.01 | -0.01 | 3.992 | 0.79 | 0.75 | 0.72 | 0.83 | 0.74 | 0.81 |
| 1 | 120282232 | rs17024167 | 1.19 | -0.628 | -0.283 | -1.172 | 0.027 | 0.026 | 0.026 | 0 | 0.19 | 0.17 | 0.06 | 0.04 | 0.11 | 0.16 |
| 1 | 120284988 | rs517237 | 0.856 | -0.075 | -0.51 | -0.89 | -0.029 | -0.022 | -0.022 | 0 | 0.53 | 0.12 | 0.46 | 0.57 | 0.49 | 0.53 |
| 1 | 120285546 | rs543703 | -0.444 | 0.21 | -0.151 | -0.897 | 0.049 | 0.05 | 0.05 | 0.564 | 0.83 | 0.77 | 0.75 | 0.62 | 0.67 | 0.8 |
| 1 | 120286863 | rs17258425 | 1.434 | 0.187 | -0.018 | -0.913 | 0.022 | 0.017 | 0.017 | 0 | 0.19 | 0.16 | 0.06 | 0.03 | 0.11 | 0.16 |
| 1 | 120288333 | rs522277 | -0.224 | 0.268 | 0.378 | -0.799 | 0.123 | 0.107 | 0.107 | 0.364 | 0.82 | 0.66 | 0.74 | 0.64 | 0.67 | 0.8 |
| 1 | 120292279 | rs532208 | 0.858 | 0.163 | 0.155 | -0.78 | 0.003 | -0.002 | -0.002 | 0 | 0.54 | 0.33 | 0.47 | 0.56 | 0.49 | 0.52 |
| 1 | 120292733 | rs536662 | 0.806 | 0.13 | -0.067 | -0.888 | -0.001 | 0.003 | 0.003 | 0 | 0.54 | 0.33 | 0.47 | 0.57 | 0.5 | 0.53 |
| 1 | 120293367 | rs667246 | -0.132 | 0.184 | -0.373 | -0.876 | 0.081 | 0.071 | 0.071 | 0.228 | 0.81 | 0.64 | 0.74 | 0.64 | 0.67 | 0.79 |
| 1 | 120302419 | rs12121219 | 0.831 | 0.698 | -0.337 | -0.254 | 0.015 | 0.006 | 0.006 | 0 | 0.18 | 0.16 | 0.06 | 0.03 | 0.09 | 0.16 |
| 1 | 120303322 | rs4659019 | 0.346 | 0.806 | -0.421 | -0.086 | 0.013 | 0 | 0 | 1.178 | 0.21 | 0.21 | 0.08 | 0.03 | 0.11 | 0.16 |
| 1 | 146930254 | rs7525162 | -0.336 | 1.807 | 1.796 | 1.095 | 0.024 | 0.055 | 0.055 | 0 | 0.1 | 0.23 | 0.16 | 0.17 | 0.14 | 0.07 |
| 1 | 146964025 | rs4132958 | 0.733 | 1.906 | 0.355 | 1.25 | 0.002 | 0.004 | 0.004 | 0 | 0.72 | 0.69 | 0.69 | 0.63 | 0.69 | 0.73 |
| 1 | 146975765 | rs612963 | -0.234 | 1.333 | -0.7 | 1.327 | -0.007 | -0.061 | -0.061 | 0.011 | 0.81 | 0.92 | 0.88 | 0.63 | 0.83 | 0.75 |
| 1 | 146980988 | rs596561 | -0.582 | 1.451 | -0.993 | 1.285 | -0.021 | -0.069 | -0.069 | 3.852 | 0.82 | 0.93 | 0.88 | 0.63 | 0.82 | 0.77 |
| 1 | 146981725 | rs17160305 | -0.366 | 1.535 | 0.218 | 1.433 | 0.104 | 0.141 | 0.141 | 0 | 0.08 | 0.25 | 0.19 | 0.17 | 0.16 | 0.08 |
| 1 | 147004811 | rs4950328 | 0.378 | 0.264 | 0.173 | 0.683 | -0.012 | -0.003 | -0.003 | 1.054 | 0.66 | 0.64 | 0.77 | 0.69 | 0.73 | 0.76 |
| 1 | 147011783 | rs3806218 | 0.884 | 0.362 | -0.251 | 0.558 | -0.005 | 0.043 | 0.043 | 0.56 | 0.54 | 0.12 | 0.7 | 0.68 | 0.62 | 0.73 |
| 1 | 147013362 | rs10900376 | 0.853 | 0.344 | -0.189 | 0.546 | 0.001 | 0.056 | 0.056 | 0.463 | 0.54 | 0.12 | 0.69 | 0.7 | 0.64 | 0.73 |
| 1 | 147018361 | rs1027424 | -1.243 | 0.261 | 0.368 | 0.46 | 0.062 | 0.125 | 0.125 | 0 | 0.09 | 0.37 | 0.09 | 0.19 | 0.09 | 0.04 |
| 1 | 147020456 | rs594206 | -0.183 | 0.313 | 0.284 | 0.449 | -0.004 | -0.036 | -0.036 | 0 | 0.31 | 0.31 | 0.17 | 0.06 | 0.21 | 0.18 |
| 1 | 147022697 | rs12081028 | -0.866 | 0.447 | 0.122 | 0.81 | 0.032 | 0.096 | 0.096 | 0.135 | 0.06 | 0.19 | 0.07 | 0.19 | 0.05 | 0.04 |
| 1 | 147030457 | rs1546161 | 0.063 | -0.07 | -1.388 | 0.239 | -0.016 | 0.064 | 0.064 | 0 | 0.22 | 0.01 | 0.36 | 0.11 | 0.21 | 0.22 |

| | | | | | | | | | | | | | | | | |
|---|---|---|---|---|---|---|---|---|---|---|---|---|---|---|---|---|
| 1 | 147033573 | rs4950461 | 0.699 | -0.275 | -1.072 | 0.134 | -0.016 | -0.019 | -0.019 | 0 | 0.85 | 0.25 | 0.89 | 0.94 | 0.92 | 0.96 |
| 1 | 147043542 | rs1015235 | 0.32 | -0.261 | -1.502 | -0.008 | -0.017 | -0.008 | -0.008 | 0.016 | 0.27 | 0.13 | 0.2 | 0.29 | 0.27 | 0.22 |
| 1 | 147048600 | rs2353544 | 0.543 | -0.289 | -1.537 | 0.037 | -0.007 | 0.014 | 0.014 | 0.761 | 0.57 | 0.1 | 0.69 | 0.65 | 0.65 | 0.74 |
| 1 | 147050595 | rs11240082 | -0.954 | -0.452 | -1.426 | -0.082 | -0.026 | 0.047 | 0.047 | 0 | 0.55 | 0.24 | 0.58 | 0.4 | 0.52 | 0.48 |
| 1 | 147051058 | rs11240083 | -0.478 | -0.577 | -0.955 | 0.316 | 0.002 | 0.031 | 0.031 | 0 | 0.26 | 0.16 | 0.38 | 0.19 | 0.26 | 0.24 |
| 1 | 147057224 | rs614502 | 0.826 | -0.95 | 0.066 | 0.778 | -0.047 | 0.13 | 0.13 | 0 | 0.9 | 0.73 | 0.75 | 0.67 | 0.97 | 0.9 |
| 1 | 147060091 | rs6695965 | 2.13 | -1.204 | -0.124 | 0.689 | 0.071 | -0.065 | -0.065 | 0 | 0.76 | 0.11 | 0.63 | 0.62 | 0.9 | 0.76 |
| 1 | 147060234 | rs1390507 | 1.616 | -1.106 | 0.763 | 0.665 | -0.042 | 0.164 | 0.164 | 0.139 | 0.82 | 0.48 | 0.67 | 0.62 | 0.9 | 0.83 |
| 1 | 147062571 | rs2132441 | 2.376 | -1.109 | 0.222 | 0.665 | 0.055 | -0.055 | -0.055 | 0 | 0.75 | 0.13 | 0.62 | 0.62 | 0.88 | 0.76 |
| 1 | 147077283 | rs688325 | 1.037 | -1.1 | 0.804 | 1.258 | -0.026 | 0.038 | 0.038 | 0 | 0.75 | 0.71 | 0.76 | 0.43 | 0.87 | 0.78 |
| 1 | 147082026 | rs12122799 | 0.066 | -1.56 | 0.757 | 0.476 | -0.054 | 0.012 | 0.012 | 0 | 0.25 | 0.47 | 0.33 | 0.31 | 0.16 | 0.19 |
| 1 | 147083744 | rs677977 | 0.705 | -1.321 | 0.104 | 0.331 | 0.002 | 0.014 | 0.014 | 0 | 0.35 | 0.25 | 0.32 | 0.28 | 0.31 | 0.4 |
| 1 | 147086504 | rs7512513 | 0.282 | -1.275 | 0.127 | 0.293 | -0.051 | 0 | 0 | 2.356 | 0.27 | 0.57 | 0.34 | 0.32 | 0.16 | 0.2 |
| 1 | 147091972 | rs3820129 | 1.57 | -1.027 | -0.094 | 0.309 | -0.151 | 0.079 | 0.079 | 0 | 0.24 | 0.02 | 0.21 | 0.1 | 0.41 | 0.23 |
| 1 | 147093887 | rs2236570 | 0.874 | -0.959 | 0.014 | 0.379 | -0.012 | 0.016 | 0.016 | 0 | 0.34 | 0.15 | 0.32 | 0.28 | 0.31 | 0.4 |
| 1 | 147096794 | rs867270 | -0.875 | -1.029 | 0.137 | 0.565 | -0.003 | 0.033 | 0.033 | 0.005 | 0.87 | 0.86 | 0.89 | 0.71 | 0.88 | 0.86 |
| 1 | 147097266 | rs946903 | 0.515 | -1.092 | 0.195 | 0.459 | -0.059 | 0.003 | 0.003 | 0 | 0.26 | 0.47 | 0.33 | 0.31 | 0.17 | 0.2 |
| 1 | 147097635 | rs3766512 | -0.503 | -0.636 | -0.172 | 0.656 | -0.005 | 0.047 | 0.047 | 0.006 | 0.86 | 0.83 | 0.88 | 0.7 | 0.88 | 0.82 |
| 1 | 147107123 | rs10157759 | 0.564 | -0.563 | -0.196 | 0.64 | 0 | -0.025 | -0.025 | 0 | 0.13 | 0.06 | 0.12 | 0.3 | 0.12 | 0.16 |
| 1 | 147107893 | rs1417279 | 0.31 | -0.63 | 0.342 | 0.57 | 0.001 | 0.153 | 0.153 | 0 | 0.82 | 0.54 | 0.85 | 0.7 | 0.86 | 0.8 |
| 1 | 147111830 | rs7539714 | -0.858 | -0.526 | 0.225 | 0.676 | -0.001 | 0.037 | 0.037 | 0.059 | 0.87 | 0.87 | 0.88 | 0.66 | 0.88 | 0.84 |
| 1 | 147113387 | rs10494247 | 0.052 | -0.595 | 0.094 | 0.301 | -0.001 | 0.001 | 0.001 | 0 | 0.48 | 0.49 | 0.53 | 0.42 | 0.54 | 0.39 |
| 1 | 147118181 | rs12045115 | -0.319 | -0.625 | 0.043 | 0.306 | 0.002 | 0.018 | 0.018 | 0 | 0.5 | 0.44 | 0.37 | 0.58 | 0.42 | 0.6 |
| 1 | 147119273 | rs1344 | 0.037 | -0.622 | -0.025 | 0.298 | 0 | 0.02 | 0.02 | 0 | 0.49 | 0.43 | 0.63 | 0.42 | 0.58 | 0.4 |
| 1 | 147122831 | rs946904 | 1.279 | -0.741 | -0.358 | 0.161 | -0.137 | 0.105 | 0.105 | 0 | 0.26 | 0.07 | 0.31 | 0.11 | 0.44 | 0.25 |
| 1 | 147127602 | rs885239 | 0.897 | -0.815 | -0.225 | 0.215 | -0.039 | 0.006 | 0.006 | 0 | 0.23 | 0.32 | 0.32 | 0.31 | 0.15 | 0.15 |
| 1 | 147130275 | rs3766509 | 0.106 | -0.301 | -0.426 | 0.278 | -0.006 | -0.005 | -0.005 | 0.021 | 0.6 | 0.66 | 0.56 | 0.59 | 0.44 | 0.61 |
| 1 | 147130661 | rs33965274 | 1.07 | 0.103 | -0.697 | 0.341 | -0.156 | 0.083 | 0.083 | 0 | 0.24 | 0.03 | 0.3 | 0.1 | 0.43 | 0.24 |
| 1 | 147137368 | rs3766506 | -0.238 | -0.094 | 0.349 | 0.945 | 0.001 | 0.124 | 0.124 | 0 | 0.15 | 0.28 | 0.14 | 0.29 | 0.12 | 0.18 |
| 1 | 147138809 | rs2185437 | -0.634 | -0.215 | 0.234 | 0.91 | 0.026 | 0.015 | 0.015 | 0.061 | 0.42 | 0.69 | 0.42 | 0.61 | 0.32 | 0.32 |
| 1 | 147161777 | rs11804045 | 0.113 | 0.605 | -0.54 | 0.349 | 0.001 | -0.02 | -0.02 | 0 | 0.08 | 0.09 | 0.06 | 0.01 | 0.08 | 0.06 |
| 1 | 147162445 | rs7545497 | -0.052 | 0.06 | -0.475 | 0.606 | -0.025 | -0.005 | -0.005 | 0 | 0.61 | 0.47 | 0.61 | 0.39 | 0.68 | 0.71 |
| 1 | 147166377 | rs1001193 | -0.226 | 1.87 | -0.489 | 0.791 | 0.015 | 0 | 0 | 0 | 0.08 | 0.08 | 0.12 | 0.08 | 0.25 | 0.03 |
| 1 | 147173505 | rs1342710 | 0.97 | 1.35 | -0.276 | 1.344 | 0.02 | 0.031 | 0.031 | 0 | 0.8 | 0.73 | 0.78 | 0.69 | 0.67 | 0.88 |
| 1 | 147191841 | rs12755965 | -1.228 | 0.957 | -0.695 | 1.276 | -0.011 | -0.045 | -0.045 | 0 | 0.32 | 0.17 | 0.35 | 0.43 | 0.39 | 0.23 |
| 1 | 147194005 | rs11577892 | -1.545 | 1.331 | -0.772 | 1.555 | -0.007 | 0.026 | 0.026 | 0 | 0.38 | 0.19 | 0.4 | 0.36 | 0.4 | 0.38 |
| 1 | 147210227 | rs999095 | -1.199 | 1.426 | 0.349 | 1.58 | -0.017 | 0.003 | 0.003 | 0.247 | 0.24 | 0.4 | 0.3 | 0.28 | 0.16 | 0.37 |
| 1 | 147211298 | rs1342700 | 1.965 | 1.52 | 0.348 | 1.552 | 0.012 | -0.01 | -0.01 | 0.525 | 0.62 | 0.52 | 0.56 | 0.6 | 0.56 | 0.6 |
| 1 | 147212431 | rs11240115 | -1.542 | 2.733 | 0.827 | 2.133 | 0 | 0.073 | 0.073 | 0.079 | 0.29 | 0.46 | 0.4 | 0.47 | 0.26 | 0.37 |
| 1 | 147216860 | rs6684174 | 2.236 | 3.811 | 1.797 | 2.902 | -0.006 | -0.001 | -0.001 | 2.883 | 0.61 | 0.39 | 0.48 | 0.41 | 0.52 | 0.58 |
| 1 | 147220326 | rs4031118 | 1.842 | 2.748 | 1.348 | 2.158 | 0.03 | -0.015 | -0.015 | 0.018 | 0.64 | 0.51 | 0.55 | 0.65 | 0.54 | 0.63 |
| 1 | 147221438 | rs7513213 | -1.064 | 2.791 | 1.497 | 2.173 | 0.006 | 0.002 | 0.002 | 0 | 0.25 | 0.19 | 0.3 | 0.25 | 0.21 | 0.32 |
| 1 | 147225584 | rs11240118 | 1.905 | 2.521 | 1.29 | 1.982 | 0.098 | 0.04 | 0.04 | 0.027 | 0.8 | 0.52 | 0.73 | 0.73 | 0.69 | 0.77 |
| 1 | 147228874 | rs11240121 | -1.543 | 2.494 | 1.294 | 2.02 | -0.019 | 0.047 | 0.047 | 0.155 | 0.32 | 0.54 | 0.38 | 0.45 | 0.23 | 0.38 |
| 1 | 147232374 | rs6593809 | -1.68 | 2.277 | 0.908 | 1.61 | -0.02 | 0.021 | 0.021 | 0 | 0.4 | 0.35 | 0.47 | 0.34 | 0.48 | 0.4 |
| 1 | 147232740 | rs10465885 | 1.782 | 2.291 | 0.86 | 1.533 | -0.003 | -0.004 | -0.004 | 0.362 | 0.57 | 0.64 | 0.47 | 0.46 | 0.5 | 0.56 |
| 1 | 147236233 | rs2047280 | 0.813 | 2.924 | 1.491 | 1.613 | 0.038 | 0.094 | 0.094 | 0.109 | 0.84 | 0.55 | 0.85 | 0.81 | 0.83 | 0.86 |
| 1 | 147241723 | rs6670327 | -1.749 | 2.97 | 1.145 | 1.547 | -0.012 | -0.007 | -0.007 | 0.073 | 0.27 | 0.21 | 0.31 | 0.37 | 0.45 | 0.29 |
| 1 | 150175887 | rs13375906 | -0.561 | 1.937 | 1.565 | 1.106 | -0.022 | -0.028 | -0.028 | 0 | 0.06 | 0.31 | 0.02 | 0 | 0 | 0 |
| 1 | 150204731 | rs12063159 | -0.574 | 2.105 | 1.627 | 1.114 | -0.012 | -0.019 | -0.019 | 0.544 | 0.06 | 0.33 | 0.02 | 0 | 0 | 0 |
| 1 | 150250636 | rs10157197 | -0.912 | 1.546 | 0.532 | 0.433 | 0.001 | 0.03 | 0.03 | 0 | 0.41 | 0.27 | 0.52 | 0.72 | 0.42 | 0.62 |
| 1 | 150257262 | rs942453 | -0.565 | 1.511 | 0.933 | 0.429 | 0.096 | -0.057 | -0.057 | 0.882 | 0.1 | 0.33 | 0.09 | 0 | 0.14 | 0.03 |
| 1 | 150260993 | rs10888578 | -1.113 | 1.402 | 0.863 | 0.456 | -0.001 | -0.003 | -0.003 | 0 | 0.42 | 0.4 | 0.52 | 0.72 | 0.41 | 0.63 |
| 1 | 150270209 | rs1776273 | -1.317 | 1.397 | 0.805 | 0.436 | 0.003 | 0.009 | 0.009 | 0 | 0.44 | 0.57 | 0.63 | 0.77 | 0.41 | 0.63 |
| 1 | 150276022 | rs509345 | -1.473 | 1.512 | 0.523 | 0.156 | 0.01 | 0.195 | 0.195 | 0.499 | 0.5 | 0.8 | 0.68 | 0.77 | 0.48 | 0.65 |
| 1 | 150292417 | rs1694379 | 0.36 | 1.794 | 0.599 | 0.359 | 0.002 | 0.024 | 0.024 | 0 | 0.6 | 0.72 | 0.5 | 0.28 | 0.59 | 0.37 |
| 1 | 150306125 | rs1260407 | 0.512 | 1.826 | 0.557 | 0.326 | -0.002 | -0.016 | -0.016 | 0 | 0.58 | 0.49 | 0.38 | 0.23 | 0.59 | 0.37 |
| 1 | 150371394 | rs11582094 | -0.538 | 1.094 | -0.366 | -0.344 | 0.033 | -0.073 | -0.073 | 0 | 0.13 | 0.36 | 0.09 | 0 | 0.13 | 0.04 |
| 1 | 150388318 | rs698915 | -0.898 | 0.919 | -0.474 | -0.309 | -0.004 | -0.003 | -0.003 | 0 | 0.76 | 0.81 | 0.61 | 0.75 | 0.74 | 0.52 |
| 1 | 150416913 | rs16836940 | -1.355 | 1.003 | -0.23 | -0.636 | 0.022 | -0.042 | -0.042 | 0 | 0.18 | 0.67 | 0.25 | 0.06 | 0.21 | 0.05 |
| 1 | 150466925 | rs12116970 | -0.347 | 0.978 | -0.387 | -0.612 | 0.073 | -0.025 | -0.025 | 0 | 0.15 | 0.53 | 0.23 | 0.06 | 0.21 | 0.05 |
| 1 | 150478623 | rs3754214 | -0.369 | 0.407 | -0.724 | -0.739 | 0.003 | 0.01 | 0.01 | 0 | 0.58 | 0.62 | 0.49 | 0.28 | 0.59 | 0.35 |
| 1 | 150482738 | rs11205387 | 0.654 | 0.407 | -1.598 | -0.6 | -0.001 | -0.017 | -0.017 | 0 | 0.27 | 0.18 | 0.31 | 0.29 | 0.26 | 0.25 |
| 1 | 150484987 | rs13294 | 0.494 | 0.629 | -1.502 | -0.421 | 0.006 | 0.125 | 0.125 | 0.516 | 0.44 | 0.17 | 0.27 | 0.24 | 0.38 | 0.3 |
| 1 | 150510660 | rs6693567 | -0.524 | 0.52 | -1.09 | 0.21 | -0.013 | -0.012 | -0.012 | 0 | 0.77 | 0.85 | 0.61 | 0.74 | 0.74 | 0.47 |
| 1 | 150515021 | rs4971010 | 0.14 | 0.374 | -0.789 | -0.06 | -0.001 | -0.008 | -0.008 | 0 | 0.61 | 0.57 | 0.37 | 0.24 | 0.59 | 0.32 |
| 1 | 150527354 | rs9659073 | -0.042 | 0.357 | -0.526 | 0.397 | -0.003 | 0.003 | 0.003 | 0.289 | 0.47 | 0.46 | 0.39 | 0.54 | 0.49 | 0.33 |
| 1 | 150528885 | rs12119250 | 0.607 | 0.579 | -0.542 | 0.391 | -0.024 | 0.033 | 0.033 | 0.003 | 0.06 | 0 | 0.04 | 0 | 0.08 | 0.02 |
| 1 | 150531008 | rs10888382 | 1.302 | 0.373 | 0.337 | 0.652 | 0.019 | 0.06 | 0.06 | 0 | 0.9 | 0.65 | 0.9 | 0.84 | 0.93 | 0.83 |

| | | | | | | | | | | | | | | | | |
|---|---|---|---|---|---|---|---|---|---|---|---|---|---|---|---|---|
| 1 | 150531360 | rs12097596 | -1.114 | 0.151 | 0.786 | 0.689 | 0.03 | 0.092 | 0.092 | 0 | 0.07 | 0.35 | 0.11 | 0.07 | 0.02 | 0.03 |
| 1 | 150542128 | rs6655975 | -0.88 | -0.437 | -0.354 | 0.833 | 0.004 | 0 | 0 | 0 | 0.62 | 0.72 | 0.56 | 0.68 | 0.65 | 0.45 |
| 1 | 150543686 | rs11204668 | 0.603 | -0.647 | -0.398 | 0.685 | -0.025 | 0.069 | 0.069 | 0.691 | 0.44 | 0.12 | 0.38 | 0.45 | 0.55 | 0.32 |
| 1 | 150552042 | rs16837903 | 2.277 | -0.446 | -0.59 | 0.51 | -0.048 | 0.02 | 0.02 | 0 | 0.09 | 0.01 | 0.11 | 0.02 | 0.15 | 0.05 |
| 1 | 150554876 | rs12034302 | 1.242 | -0.616 | -0.421 | 0.613 | 0.001 | -0.065 | -0.065 | 0 | 0.1 | 0 | 0.05 | 0.16 | 0.09 | 0.1 |
| 1 | 150564476 | rs6698495 | 0.692 | -0.576 | -0.383 | 0.552 | -0.044 | 0.045 | 0.045 | 0.458 | 0.4 | 0.12 | 0.38 | 0.43 | 0.53 | 0.3 |
| 1 | 150570985 | rs11204675 | 0.25 | -0.585 | -0.099 | 0.552 | 0.007 | 0.043 | 0.043 | 0.125 | 0.46 | 0.26 | 0.4 | 0.43 | 0.55 | 0.31 |
| 1 | 150584103 | rs4970966 | 1.937 | -0.031 | -0.62 | 0.535 | -0.073 | -0.032 | -0.032 | 0.433 | 0.16 | 0.01 | 0.24 | 0.27 | 0.24 | 0.16 |
| 1 | 150584834 | rs12045807 | 0.054 | -0.245 | -0.128 | 0.523 | -0.032 | 0.095 | 0.095 | 0 | 0.14 | 0.41 | 0.07 | 0.17 | 0.1 | 0.09 |
| 1 | 150596411 | rs7517 | 0.647 | -0.22 | -0.136 | 0.516 | -0.022 | 0.075 | 0.075 | 0 | 0.13 | 0.31 | 0.06 | 0.16 | 0.09 | 0.1 |
| 1 | 150608894 | rs11204684 | 0.391 | -0.262 | -0.709 | 0.38 | -0.048 | 0.053 | 0.053 | 0.434 | 0.41 | 0.11 | 0.39 | 0.41 | 0.53 | 0.31 |
| 1 | 150615094 | rs12402939 | 1.009 | -0.317 | -0.572 | 0.384 | -0.02 | 0.003 | 0.003 | 0 | 0.4 | 0.23 | 0.45 | 0.36 | 0.42 | 0.57 |
| 1 | 150618632 | rs2055975 | 0.016 | -0.405 | -0.719 | 0.306 | 0.003 | 0.006 | 0.006 | 0 | 0.64 | 0.35 | 0.74 | 0.86 | 0.72 | 0.77 |
| 1 | 150618961 | rs9733 | 0.091 | -0.004 | -0.929 | 0.581 | -0.023 | 0.021 | 0.021 | 0.104 | 0.41 | 0.1 | 0.65 | 0.73 | 0.58 | 0.54 |
| 1 | 150638572 | rs10788792 | 1.134 | -0.235 | -0.678 | 0.041 | -0.006 | 0.004 | 0.004 | 0 | 0.4 | 0.28 | 0.45 | 0.36 | 0.41 | 0.56 |
| 1 | 150657011 | rs11204695 | 0.264 | -0.321 | -0.982 | 0.034 | -0.098 | -0.023 | -0.023 | 0 | 0.38 | 0.09 | 0.45 | 0.46 | 0.47 | 0.3 |
| 1 | 150658971 | rs11204697 | 0.48 | -0.148 | -0.853 | 0.091 | -0.004 | 0.012 | 0.012 | 0 | 0.43 | 0.29 | 0.46 | 0.36 | 0.43 | 0.57 |
| 1 | 150659545 | rs11204698 | 0.766 | -0.164 | -0.86 | 0.085 | -0.005 | 0.006 | 0.006 | 0 | 0.41 | 0.28 | 0.45 | 0.36 | 0.41 | 0.56 |
| 1 | 150664077 | rs11803569 | 0.456 | -0.139 | -0.849 | 0.097 | -0.004 | 0.013 | 0.013 | 0 | 0.43 | 0.29 | 0.46 | 0.36 | 0.43 | 0.57 |
| 1 | 150669952 | rs2275236 | 0.473 | -0.13 | -0.845 | 0.102 | -0.004 | 0.012 | 0.012 | 0 | 0.43 | 0.29 | 0.46 | 0.36 | 0.43 | 0.57 |
| 1 | 150671630 | rs11204704 | 0.464 | -0.134 | -0.847 | 0.1 | -0.004 | 0.013 | 0.013 | 0 | 0.43 | 0.29 | 0.46 | 0.36 | 0.43 | 0.57 |
| 1 | 150677719 | rs11807450 | 0.791 | -0.139 | -0.822 | 0.098 | -0.014 | 0.002 | 0.002 | 0 | 0.39 | 0.23 | 0.45 | 0.36 | 0.41 | 0.56 |
| 1 | 150679033 | rs1336900 | 0.799 | -0.063 | -0.847 | 0.104 | -0.014 | 0.001 | 0.001 | 0 | 0.39 | 0.23 | 0.45 | 0.36 | 0.41 | 0.56 |
| 1 | 150679358 | rs11586422 | 0.534 | 0.089 | -0.83 | 0.104 | -0.004 | 0.012 | 0.012 | 0 | 0.43 | 0.29 | 0.46 | 0.36 | 0.43 | 0.57 |
| 1 | 150727394 | rs11576175 | 0.888 | -0.127 | -0.959 | 0.253 | -0.002 | -0.062 | -0.062 | 0 | 0.1 | 0 | 0.05 | 0.17 | 0.09 | 0.1 |
| 1 | 150729793 | rs12568757 | 0.379 | -0.197 | -1.237 | -0.013 | -0.067 | -0.02 | -0.02 | 0.158 | 0.39 | 0.1 | 0.46 | 0.47 | 0.51 | 0.31 |
| 1 | 150729983 | rs10888391 | 0.881 | -0.453 | -0.892 | -0.136 | 0.013 | 0.009 | 0.009 | 0 | 0.42 | 0.26 | 0.45 | 0.35 | 0.39 | 0.56 |
| 1 | 150742700 | rs11204725 | 0.493 | -0.467 | -0.917 | -0.135 | -0.077 | -0.027 | -0.027 | 0 | 0.38 | 0.09 | 0.46 | 0.41 | 0.51 | 0.31 |
| 1 | 150743492 | rs11204726 | 0.785 | -0.465 | -0.898 | -0.134 | 0.018 | 0.013 | 0.013 | 0 | 0.43 | 0.26 | 0.45 | 0.41 | 0.39 | 0.56 |
| 1 | 150749913 | rs10888394 | 0.785 | -0.465 | -0.898 | -0.134 | 0.018 | 0.013 | 0.013 | 0 | 0.43 | 0.26 | 0.45 | 0.41 | 0.39 | 0.56 |
| 1 | 150759979 | rs11204727 | 1.216 | -0.465 | -0.899 | -0.139 | -0.002 | -0.062 | -0.062 | 1.529 | 0.1 | 0 | 0.05 | 0.17 | 0.09 | 0.1 |
| 1 | 150760174 | rs12125650 | 0.453 | -0.463 | -0.898 | -0.122 | -0.076 | -0.029 | -0.029 | 0 | 0.38 | 0.09 | 0.46 | 0.42 | 0.51 | 0.31 |
| 1 | 150762171 | rs10888395 | 0.758 | -0.431 | -0.881 | -0.131 | 0.017 | 0.014 | 0.014 | 0 | 0.43 | 0.26 | 0.45 | 0.4 | 0.39 | 0.56 |
| 1 | 150766822 | rs4319334 | 0.547 | -0.409 | 0.074 | -0.132 | 0.008 | 0.007 | 0.007 | 0 | 0.47 | 0.52 | 0.46 | 0.4 | 0.39 | 0.56 |
| 1 | 150780582 | rs4379678 | 1.158 | -0.528 | -0.383 | -0.136 | -0.002 | -0.062 | -0.062 | 0 | 0.1 | 0 | 0.05 | 0.17 | 0.09 | 0.09 |
| 1 | 150783985 | rs11552229 | 1.002 | -0.626 | -0.426 | -0.265 | 0.014 | 0.007 | 0.007 | 0 | 0.41 | 0.26 | 0.44 | 0.4 | 0.37 | 0.56 |
| 1 | 150792616 | rs11589458 | 1.008 | -0.735 | -0.491 | -0.233 | 0.017 | 0.009 | 0.009 | 0 | 0.41 | 0.22 | 0.43 | 0.4 | 0.37 | 0.56 |
| 1 | 150799755 | rs1889740 | 0.584 | -0.129 | -0.557 | -0.25 | 0.005 | 0.004 | 0.004 | 0 | 0.46 | 0.52 | 0.46 | 0.4 | 0.39 | 0.57 |
| 1 | 150800655 | rs10305714 | 0.98 | -0.129 | -0.553 | -0.249 | 0.016 | 0.009 | 0.009 | 0 | 0.42 | 0.26 | 0.44 | 0.4 | 0.37 | 0.56 |
| 1 | 150801466 | rs3738483 | -0.182 | -0.163 | -0.483 | -0.256 | -0.035 | 0.056 | 0.056 | 0 | 0.85 | 0.64 | 0.92 | 0.82 | 0.91 | 0.88 |
| 1 | 150804429 | rs10305704 | 1.131 | -0.142 | -0.815 | -0.384 | 0.016 | 0.01 | 0.01 | 0 | 0.42 | 0.26 | 0.44 | 0.4 | 0.37 | 0.56 |
| 1 | 150809919 | rs3768016 | 1.143 | -0.088 | -0.895 | -0.414 | 0.016 | 0.009 | 0.009 | 0 | 0.42 | 0.26 | 0.44 | 0.4 | 0.37 | 0.56 |
| 1 | 150814174 | rs2864873 | 0.871 | -0.081 | -0.857 | -0.409 | 0.018 | 0.015 | 0.015 | 0 | 0.43 | 0.26 | 0.45 | 0.4 | 0.39 | 0.57 |
| 1 | 150815411 | rs3768013 | 0.714 | -0.073 | -0.936 | -0.39 | 0.005 | 0.005 | 0.005 | 0 | 0.46 | 0.52 | 0.47 | 0.4 | 0.4 | 0.57 |
| 1 | 150816886 | rs2134688 | -0.205 | -0.079 | -0.93 | -0.404 | -0.039 | 0.061 | 0.061 | 0 | 0.85 | 0.61 | 0.92 | 0.82 | 0.91 | 0.88 |
| 1 | 150821847 | rs7532045 | 0.694 | -0.066 | -0.928 | -0.408 | 0.006 | 0.005 | 0.005 | 0 | 0.46 | 0.52 | 0.46 | 0.4 | 0.39 | 0.57 |
| 1 | 150836439 | rs12406660 | 2.569 | -0.366 | -1.851 | -0.746 | -0.011 | -0.105 | -0.105 | 0 | 0.15 | 0.01 | 0.28 | 0.36 | 0.14 | 0.46 |
| 1 | 150847647 | rs11204737 | 0.321 | 0.339 | -1.314 | -0.435 | -0.074 | -0.028 | -0.028 | 0 | 0.38 | 0.09 | 0.46 | 0.42 | 0.51 | 0.31 |
| 1 | 150850035 | rs7517566 | -0.631 | 0.362 | -1.215 | -0.86 | 0.003 | -0.003 | -0.003 | 0.107 | 0.87 | 0.85 | 0.94 | 0.82 | 0.91 | 0.88 |
| 1 | 150854767 | rs6679701 | 1.308 | 0.417 | -1.28 | -0.92 | -0.006 | -0.061 | -0.061 | 0 | 0.1 | 0 | 0.05 | 0.16 | 0.09 | 0.08 |
| 1 | 150856153 | rs12410869 | 0.594 | 0.385 | -1.261 | -0.922 | 0.021 | -0.001 | -0.001 | 0 | 0.47 | 0.58 | 0.55 | 0.47 | 0.39 | 0.57 |
| 1 | 150860471 | rs7412746 | 0.173 | 0.389 | -1.269 | -0.924 | -0.066 | 0.028 | 0.028 | 0.098 | 0.39 | 0.09 | 0.38 | 0.37 | 0.52 | 0.29 |
| 1 | 150861960 | rs10399947 | 0.664 | 0.408 | -1.263 | -0.912 | 0.019 | -0.003 | -0.003 | 0 | 0.46 | 0.58 | 0.54 | 0.47 | 0.38 | 0.57 |
| 1 | 150884271 | rs533928 | -0.817 | 0.416 | -1.267 | -0.994 | -0.005 | 0.009 | 0.009 | 0 | 0.88 | 0.8 | 0.93 | 0.83 | 0.9 | 0.84 |
| 1 | 150884776 | rs2292166 | 2.114 | 0.707 | -1.429 | -1.074 | -0.084 | -0.056 | -0.056 | 0.12 | 0.16 | 0.02 | 0.25 | 0.27 | 0.26 | 0.21 |
| 1 | 150916657 | rs2271076 | 0.593 | 0.625 | -1.289 | -1.026 | 0.02 | 0.003 | 0.003 | 0 | 0.47 | 0.58 | 0.55 | 0.44 | 0.39 | 0.52 |
| 1 | 150923099 | rs17661357 | 2.422 | 0.686 | -1.352 | -1.079 | -0.022 | 0.057 | 0.057 | 0 | 0.1 | 0 | 0.05 | 0 | 0.09 | 0.02 |
| 1 | 150937661 | rs1546376 | 1.505 | 1.279 | -0.613 | -0.652 | 0.009 | -0.008 | -0.008 | 0 | 0.41 | 0.28 | 0.52 | 0.43 | 0.37 | 0.51 |
| 1 | 150938571 | rs8444 | 1.514 | 1.28 | -0.631 | -0.651 | 0.01 | -0.009 | -0.009 | 0 | 0.41 | 0.27 | 0.51 | 0.43 | 0.37 | 0.52 |
| 1 | 150940625 | rs267738 | 1.293 | 1.749 | -0.861 | -0.675 | -0.07 | 0.066 | 0.066 | 0.491 | 0.16 | 0.01 | 0.1 | 0.03 | 0.19 | 0.05 |
| 1 | 150951857 | rs3754211 | 0.365 | 1.499 | -0.799 | -0.615 | -0.014 | 0.021 | 0.021 | 0 | 0.59 | 0.71 | 0.63 | 0.59 | 0.48 | 0.61 |
| 1 | 150958836 | rs267733 | 1.109 | 1.278 | -0.876 | -0.491 | 0.007 | 0.058 | 0.058 | 0.583 | 0.17 | 0.04 | 0.07 | 0.03 | 0.14 | 0.05 |
| 1 | 150960350 | rs7532008 | 0.422 | 1.222 | -0.309 | -0.227 | -0.004 | -0.005 | -0.005 | 0.471 | 0.52 | 0.51 | 0.41 | 0.58 | 0.34 | 0.59 |
| 1 | 150960928 | rs12033744 | 0.717 | 1.439 | -0.771 | -0.138 | 0.005 | 0.013 | 0.013 | 0 | 0.1 | 0.18 | 0.16 | 0.27 | 0.13 | 0.14 |
| 1 | 150961254 | rs11204754 | -0.409 | 1.487 | -0.891 | -0.518 | -0.01 | 0.026 | 0.026 | 0 | 0.66 | 0.71 | 0.6 | 0.85 | 0.55 | 0.75 |
| 1 | 150961869 | rs1722784 | 0.485 | 1.546 | -1.431 | -0.49 | -0.074 | 0.092 | 0.092 | 0 | 0.32 | 0.09 | 0.39 | 0.15 | 0.44 | 0.25 |
| 1 | 150980281 | rs7548516 | 0.596 | 1.09 | -1.28 | -0.272 | 0.037 | -0.066 | -0.066 | 0 | 0.39 | 0.08 | 0.36 | 0.42 | 0.27 | 0.51 |
| 1 | 150990825 | rs3738480 | 0.643 | 2.391 | 0.108 | 0.42 | 0.107 | -0.012 | -0.012 | 0 | 0.88 | 0.72 | 0.63 | 0.88 | 0.76 | 0.87 |
| 1 | 150991221 | rs11204758 | -0.4 | 1.564 | -0.527 | 0.732 | -0.014 | -0.011 | -0.011 | 0.211 | 0.12 | 0.07 | 0.18 | 0.4 | 0.18 | 0.18 |

| | | | | | | | | | | | | | | | | |
|---|---|---|---|---|---|---|---|---|---|---|---|---|---|---|---|---|
| 1 | 150997723 | rs3820539 | -0.294 | 1.514 | -0.656 | 0.634 | -0.032 | -0.062 | -0.062 | 0.886 | 0.12 | 0.03 | 0.29 | 0.42 | 0.18 | 0.26 |
| 1 | 151001583 | rs6691701 | 1.482 | 1.138 | -0.385 | -0.186 | 0.011 | 0.039 | 0.039 | 2.397 | 0.75 | 0.2 | 0.77 | 0.87 | 0.8 | 0.79 |
| 1 | 151004003 | rs4970991 | 1.374 | 1.135 | -0.019 | -0.183 | -0.002 | -0.01 | -0.01 | 0.007 | 0.76 | 0.34 | 0.79 | 0.87 | 0.81 | 0.79 |
| 1 | 151008852 | rs7540258 | 0.674 | 1.163 | 0.385 | -0.123 | 0.008 | 0.004 | 0.004 | 0 | 0.92 | 0.83 | 0.86 | 0.9 | 0.91 | 0.9 |
| 1 | 151012060 | rs2091219 | 2.064 | 1.771 | 0.911 | -0.416 | -0.01 | 0.025 | 0.025 | 3.225 | 0.7 | 0.28 | 0.75 | 0.87 | 0.7 | 0.73 |
| 1 | 151015868 | rs12068365 | -0.146 | 1.8 | 1.014 | -0.411 | 0.042 | -0.029 | -0.029 | 0.855 | 0.06 | 0.15 | 0.04 | 0 | 0.07 | 0.01 |
| 1 | 151016396 | rs11204764 | -0.694 | 1.747 | 1.393 | -0.342 | -0.032 | -0.004 | -0.004 | 0.43 | 0.1 | 0.34 | 0.05 | 0.07 | 0.04 | 0.11 |
| 1 | 151018861 | rs6587552 | 2.173 | 1.714 | 1.409 | -0.484 | 0.011 | -0.064 | -0.064 | 0 | 0.76 | 0.45 | 0.79 | 0.88 | 0.77 | 0.73 |
| 1 | 151019180 | rs6587553 | 1.414 | 1.883 | 1.865 | -0.393 | 0.041 | -0.029 | -0.029 | 0.431 | 0.85 | 0.71 | 0.82 | 0.91 | 0.81 | 0.84 |
| 1 | 151021509 | rs6664387 | -0.135 | 1.851 | 1.727 | -0.361 | 0.043 | -0.028 | -0.028 | 0.207 | 0.06 | 0.14 | 0.03 | 0 | 0.07 | 0.01 |
| 1 | 151031667 | rs3806386 | 0.67 | 2.06 | 1.691 | -0.364 | 0.021 | -0.006 | -0.006 | 1.15 | 0.88 | 0.83 | 0.82 | 0.91 | 0.81 | 0.84 |
| 1 | 151035628 | rs6694357 | 0.196 | 1.574 | 1.278 | -0.722 | 0.008 | 0.017 | 0.017 | 0 | 0.08 | 0.04 | 0.05 | 0 | 0.08 | 0.03 |
| 1 | 151038135 | rs16832976 | 0.266 | 1.594 | 1.374 | -0.669 | 0.049 | -0.024 | -0.024 | 0 | 0.05 | 0.14 | 0.03 | 0 | 0.07 | 0.01 |
| 1 | 151039609 | rs3738481 | 0.47 | 1.599 | 1.362 | -0.675 | 0.006 | -0.003 | -0.003 | 0 | 0.88 | 0.84 | 0.82 | 0.9 | 0.82 | 0.84 |
| 1 | 151055670 | rs10788798 | 0.482 | 1.644 | 1.357 | -0.679 | 0.006 | -0.003 | -0.003 | 0 | 0.88 | 0.84 | 0.82 | 0.9 | 0.82 | 0.84 |
| 1 | 151062957 | rs11204774 | 0.276 | 1.591 | 1.431 | -0.686 | 0.047 | -0.025 | -0.025 | 1.257 | 0.05 | 0.13 | 0.03 | 0 | 0.07 | 0.01 |
| 1 | 151079317 | rs6676190 | 2.527 | 1.41 | 1.341 | -0.813 | -0.022 | 0.048 | 0.048 | 3.04 | 0.71 | 0.21 | 0.74 | 0.88 | 0.72 | 0.73 |
| 1 | 151091181 | rs4970998 | 1.028 | 1.511 | 1.272 | -0.743 | 0.042 | -0.024 | -0.024 | 1.004 | 0.86 | 0.7 | 0.81 | 0.9 | 0.82 | 0.84 |
| 1 | 151104078 | rs4971004 | 1.805 | 1.081 | 0.421 | -0.733 | -0.082 | -0.118 | -0.118 | 0 | 0.4 | 0.02 | 0.48 | 0.41 | 0.42 | 0.4 |
| 1 | 151104598 | rs4971005 | 1.362 | 1.374 | 1.531 | -0.738 | 0.057 | -0.027 | -0.027 | 0.126 | 0.86 | 0.63 | 0.8 | 0.9 | 0.82 | 0.84 |
| 1 | 151108137 | rs4971007 | 0.838 | 1.387 | 1.202 | -0.732 | 0.025 | -0.011 | -0.011 | 0 | 0.87 | 0.76 | 0.81 | 0.9 | 0.82 | 0.84 |
| 1 | 151115997 | rs11807526 | -1.443 | 1.422 | 1.255 | -0.761 | 0.095 | -0.012 | -0.012 | 0.915 | 0.11 | 0.37 | 0.13 | 0.06 | 0.14 | 0.12 |
| 1 | 151117010 | rs12089658 | -0.975 | 1.489 | 1.019 | -0.82 | 0.056 | -0.043 | -0.043 | 0.161 | 0.08 | 0.18 | 0.04 | 0 | 0.08 | 0.02 |
| 1 | 151122489 | rs7554936 | 2.4 | 1.153 | 0.408 | -0.936 | -0.102 | 0.145 | 0.145 | 3.53 | 0.63 | 0.03 | 0.68 | 0.86 | 0.67 | 0.78 |
| 1 | 151129254 | rs2067606 | -2.471 | 1.716 | 1.044 | -0.655 | 0.003 | -0.013 | -0.013 | 0.875 | 0.21 | 0.71 | 0.19 | 0.07 | 0.21 | 0.11 |
| 1 | 151135855 | rs12097169 | -2.081 | 1.69 | 0.998 | -0.492 | 0.013 | -0.079 | -0.079 | 0.221 | 0.13 | 0.68 | 0.08 | 0 | 0.08 | 0.02 |
| 1 | 151137144 | rs6587557 | 1.395 | 2.387 | 0.839 | -0.404 | 0.096 | -0.045 | -0.045 | 0.02 | 0.84 | 0.71 | 0.8 | 0.92 | 0.7 | 0.88 |
| 1 | 151143166 | rs6689566 | -1.075 | 2.063 | 0.668 | -0.243 | 0.053 | -0.034 | -0.034 | 0.553 | 0.06 | 0.13 | 0.04 | 0 | 0.11 | 0.01 |
| 1 | 151150857 | rs11204785 | 0.759 | 1.193 | 0.84 | -0.005 | -0.012 | -0.001 | -0.001 | 0 | 0.63 | 0.48 | 0.57 | 0.62 | 0.68 | 0.59 |
| 1 | 151173113 | rs4971013 | 0.509 | 1.09 | 1.264 | -0.022 | -0.005 | 0.004 | 0.004 | 1.018 | 0.67 | 0.67 | 0.61 | 0.61 | 0.7 | 0.6 |
| 1 | 151174772 | rs10749660 | 0.942 | 1.067 | 1.261 | -0.036 | -0.018 | 0 | 0 | 0.097 | 0.63 | 0.55 | 0.61 | 0.61 | 0.7 | 0.59 |
| 1 | 151193897 | rs1892196 | 0.93 | 1.036 | 1.205 | -0.055 | -0.013 | 0 | 0 | 0 | 0.63 | 0.58 | 0.61 | 0.61 | 0.7 | 0.59 |
| 1 | 151195764 | rs10788801 | -1.091 | 1.004 | 1.26 | 0.051 | -0.005 | 0.024 | 0.024 | 1.124 | 0.86 | 0.84 | 0.8 | 0.67 | 0.88 | 0.7 |
| 1 | 151199514 | rs11204787 | 0.994 | 1.159 | 0.862 | 0.086 | -0.063 | -0.102 | -0.102 | 0 | 0.13 | 0 | 0.23 | 0.46 | 0.21 | 0.29 |
| 1 | 151210126 | rs4971018 | -0.994 | 0.46 | 1.612 | 0.054 | -0.005 | 0.024 | 0.024 | 1.213 | 0.86 | 0.84 | 0.8 | 0.67 | 0.88 | 0.7 |
| 1 | 151233080 | rs4636463 | -0.754 | 0.386 | 1.604 | 0.039 | 0 | 0.011 | 0.011 | 0.57 | 0.84 | 0.84 | 0.77 | 0.67 | 0.83 | 0.67 |
| 1 | 151250421 | rs4971027 | 1.118 | 0.174 | 1.747 | -0.02 | 0.002 | 0 | 0 | 0 | 0.66 | 0.59 | 0.6 | 0.63 | 0.64 | 0.6 |
| 1 | 151254041 | rs3811406 | -0.712 | 0.134 | 1.72 | 0.044 | 0.005 | 0.013 | 0.013 | 0.615 | 0.83 | 0.82 | 0.75 | 0.68 | 0.77 | 0.67 |
| 1 | 151259043 | rs2298265 | -0.476 | 0.048 | 1.8 | 0.071 | 0.004 | 0.006 | 0.006 | 0 | 0.15 | 0.12 | 0.14 | 0.06 | 0.13 | 0.07 |
| 1 | 151259543 | rs3748545 | -0.413 | 0.048 | 1.823 | 0.071 | -0.001 | -0.009 | -0.009 | 0 | 0.16 | 0.14 | 0.14 | 0.06 | 0.13 | 0.07 |
| 1 | 151263782 | rs10788803 | 1.569 | 0.018 | 2.038 | 0.057 | -0.004 | -0.006 | -0.006 | 0 | 0.61 | 0.45 | 0.59 | 0.62 | 0.64 | 0.59 |
| 1 | 151271782 | rs10888409 | 1.408 | 0.022 | 1.986 | 0.059 | -0.003 | -0.005 | -0.005 | 0 | 0.63 | 0.56 | 0.6 | 0.62 | 0.64 | 0.59 |
| 1 | 151272451 | rs10888410 | 1.573 | -0.039 | 2.182 | 0.029 | 0.011 | 0.014 | 0.014 | 1.198 | 0.6 | 0.27 | 0.58 | 0.62 | 0.64 | 0.59 |
| 1 | 151279855 | rs1780586 | -0.355 | -0.04 | 2.187 | 0.029 | -0.002 | -0.011 | -0.011 | 0.18 | 0.16 | 0.15 | 0.14 | 0.06 | 0.13 | 0.07 |
| 1 | 151280995 | rs4971030 | 1.575 | -0.04 | 2.176 | 0.029 | 0.012 | 0.014 | 0.014 | 1.145 | 0.6 | 0.27 | 0.58 | 0.62 | 0.63 | 0.59 |
| 1 | 151288172 | rs1056847 | 1.566 | -0.029 | 2.176 | 0.163 | 0.01 | -0.002 | -0.002 | 0 | 0.6 | 0.27 | 0.57 | 0.58 | 0.63 | 0.59 |
| 1 | 151295475 | rs1752382 | -1.129 | 0.167 | 1.212 | 0.239 | 0.005 | 0.005 | 0.005 | 0 | 0.19 | 0.15 | 0.15 | 0.11 | 0.13 | 0.07 |
| 1 | 151313774 | rs7552906 | 1.276 | 0.658 | 1.189 | 0.265 | 0.002 | -0.005 | -0.005 | 0 | 0.65 | 0.61 | 0.61 | 0.64 | 0.63 | 0.64 |
| 1 | 151314379 | rs1752387 | -0.281 | 1.029 | 0.986 | 0.623 | 0.006 | 0.009 | 0.009 | 0.532 | 0.83 | 0.85 | 0.75 | 0.7 | 0.77 | 0.72 |
| 1 | 151325055 | rs2800946 | -0.259 | 0.972 | 0.745 | 0.642 | -0.003 | -0.004 | -0.004 | 0 | 0.83 | 0.87 | 0.75 | 0.71 | 0.77 | 0.73 |
| 1 | 151327376 | rs2769273 | 0.079 | 0.978 | 0.739 | 0.645 | 0 | 0.001 | 0.001 | 0 | 0.79 | 0.8 | 0.75 | 0.71 | 0.75 | 0.72 |
| 1 | 151344150 | rs2769265 | -1.331 | 0.615 | 1.262 | 0.877 | -0.006 | 0.021 | 0.021 | 0 | 0.21 | 0.25 | 0.41 | 0.27 | 0.16 | 0.2 |
| 1 | 151346333 | rs12741361 | -0.323 | 0.842 | 1.115 | 0.298 | -0.018 | 0.001 | 0.001 | 0 | 0.27 | 0.25 | 0.22 | 0 | 0.37 | 0.15 |
| 1 | 151347746 | rs1752380 | -1.729 | 0.753 | 1.751 | 0.289 | 0.003 | 0.052 | 0.052 | 0 | 0.47 | 0.64 | 0.62 | 0.29 | 0.52 | 0.32 |
| 1 | 151347820 | rs11806477 | -0.482 | 0.828 | 1.581 | 0.002 | 0.017 | -0.11 | -0.11 | 0.014 | 0.28 | 0.39 | 0.22 | 0.01 | 0.37 | 0.15 |
| 1 | 151349174 | rs17564336 | 0.121 | 0.837 | 1.183 | 0.136 | -0.043 | 0.055 | 0.055 | 0 | 0.21 | 0.13 | 0.15 | 0 | 0.32 | 0.13 |
| 1 | 151357333 | rs6684085 | 1.256 | -0.03 | 2.307 | 1.458 | -0.007 | 0.077 | 0.077 | 0 | 0.8 | 0.62 | 0.62 | 0.66 | 0.84 | 0.81 |
| 1 | 151374025 | rs4603 | -1.404 | -0.652 | 2.331 | 1.878 | 0.012 | -0.068 | -0.068 | 0 | 0.18 | 0.07 | 0.3 | 0.32 | 0.15 | 0.2 |
| 1 | 151377407 | rs1571294 | -1.09 | -0.701 | 2.754 | 1.861 | 0.046 | 0.572 | 0.572 | 1.397 | 0.05 | 0.27 | 0.22 | 0.46 | 0.02 | 0.13 |
| 1 | 151384711 | rs1887545 | 1.397 | -0.65 | 2.278 | 1.862 | 0.009 | -0.02 | -0.02 | 0.654 | 0.81 | 0.84 | 0.7 | 0.68 | 0.85 | 0.8 |
| 1 | 151386465 | rs6587572 | 1.397 | -0.65 | 2.278 | 1.862 | 0.009 | -0.02 | -0.02 | 0.088 | 0.81 | 0.84 | 0.7 | 0.68 | 0.85 | 0.8 |
| 1 | 151402045 | rs6587577 | 1.584 | -0.671 | 2.346 | 1.852 | -0.007 | 0.019 | 0.019 | 0 | 0.8 | 0.75 | 0.69 | 0.68 | 0.85 | 0.8 |
| 1 | 151403948 | rs4246527 | 1.452 | -0.614 | 1.78 | 1.922 | -0.005 | 0.02 | 0.02 | 0 | 0.8 | 0.75 | 0.63 | 0.63 | 0.85 | 0.79 |
| 1 | 151405792 | rs12025432 | -1.483 | -0.543 | 1.247 | 1.669 | 0.012 | -0.069 | -0.069 | 0 | 0.18 | 0.07 | 0.3 | 0.32 | 0.15 | 0.2 |
| 1 | 151406375 | rs7546151 | 1.409 | -0.541 | 1.244 | 1.67 | 0.009 | -0.02 | -0.02 | 0 | 0.81 | 0.84 | 0.7 | 0.68 | 0.85 | 0.8 |
| 1 | 151413180 | rs4291502 | -0.903 | -0.564 | 1.205 | 1.66 | -0.009 | 0.043 | 0.043 | 0 | 0.11 | 0.03 | 0.1 | 0.03 | 0.11 | 0.11 |
| 1 | 151428605 | rs10736380 | 1.242 | -0.511 | 1.246 | 1.642 | 0.01 | -0.021 | -0.021 | 0 | 0.81 | 0.84 | 0.7 | 0.68 | 0.85 | 0.8 |
| 1 | 151444333 | rs11204824 | 1.264 | -0.261 | 0.156 | 1.334 | 0.016 | -0.036 | -0.036 | 0 | 0.8 | 0.85 | 0.7 | 0.68 | 0.86 | 0.78 |
| 1 | 151488782 | rs7552458 | -0.989 | 0.06 | -0.115 | 0.931 | 0.001 | -0.004 | -0.004 | 0 | 0.18 | 0.17 | 0.41 | 0.78 | 0.17 | 0.35 |

| | | | | | | | | | | | | | | | | |
|---|---|---|---|---|---|---|---|---|---|---|---|---|---|---|---|---|
| 1 | 151504690 | rs7549296 | 0.195 | -0.08 | -0.546 | -0.003 | 0.004 | 0.051 | 0.051 | 0 | 0.37 | 0.25 | 0.25 | 0.1 | 0.36 | 0.37 |
| 1 | 151504778 | rs6673116 | 0.146 | 0.054 | -0.739 | -0.18 | 0.014 | -0.018 | -0.018 | 0 | 0.37 | 0.18 | 0.51 | 0.63 | 0.35 | 0.3 |
| 1 | 151510825 | rs1045298 | 0.971 | 0.013 | -1.059 | 0.221 | -0.054 | -0.06 | -0.06 | 0 | 0.12 | 0 | 0.19 | 0.16 | 0.16 | 0.11 |
| 1 | 151511268 | rs3811411 | 1.085 | -0.053 | -1.049 | 0.208 | 0.033 | -0.13 | -0.13 | 0.022 | 0.23 | 0.02 | 0.3 | 0.46 | 0.19 | 0.19 |
| 1 | 151512895 | rs3828054 | 0.836 | -0.09 | -1.291 | -0.043 | -0.011 | -0.026 | -0.026 | 0 | 0.11 | 0.16 | 0.1 | 0.05 | 0.11 | 0.11 |
| 1 | 151513513 | rs9651181 | 1.304 | -0.1 | -1.33 | -0.057 | -0.025 | 0.105 | 0.105 | 0 | 0.26 | 0.08 | 0.16 | 0.04 | 0.25 | 0.22 |
| 1 | 151514603 | rs2337360 | 0.705 | 0.176 | -1.116 | 0.227 | 0.002 | 0.045 | 0.045 | 0 | 0.37 | 0.25 | 0.25 | 0.1 | 0.36 | 0.33 |
| 1 | 151515188 | rs12121608 | 0.568 | 0.29 | -1.329 | 0.313 | -0.009 | -0.01 | -0.01 | 0 | 0.13 | 0.11 | 0.21 | 0.16 | 0.16 | 0.11 |
| 1 | 151515654 | rs10158855 | 0.126 | 0.473 | -0.897 | 0.024 | -0.001 | -0.033 | -0.033 | 0 | 0.39 | 0.34 | 0.26 | 0.1 | 0.36 | 0.33 |
| 1 | 151524275 | rs17640598 | -0.289 | 1.451 | -1.426 | 0.934 | -0.046 | -0.04 | -0.04 | 0 | 0.19 | 0.15 | 0.2 | 0.25 | 0.28 | 0.37 |
| 1 | 151538366 | rs3790506 | -0.544 | 0.795 | -1.32 | 1.048 | -0.02 | -0.043 | -0.043 | 0 | 0.24 | 0.09 | 0.36 | 0.25 | 0.26 | 0.37 |
| 1 | 151555741 | rs12749 | 0.672 | 0.668 | 0.018 | 1.353 | 0.026 | 0.087 | 0.087 | 0.103 | 0.86 | 0.65 | 0.75 | 0.79 | 0.8 | 0.89 |
| 1 | 151570241 | rs1539492 | 0.469 | 0.649 | -0.025 | 1.302 | 0.001 | -0.008 | -0.008 | 0.165 | 0.91 | 0.84 | 0.82 | 0.93 | 0.9 | 0.98 |
| 1 | 151579662 | rs1196360 | 1.642 | 0.351 | 0.924 | 1.272 | 0.059 | 0.19 | 0.19 | 0 | 0.92 | 0.62 | 0.9 | 0.84 | 0.9 | 0.87 |
| 1 | 151588735 | rs12122920 | -0.421 | 0.259 | -0.535 | 0.707 | -0.075 | 0.101 | 0.101 | 0.38 | 0.45 | 0.02 | 0.41 | 0.2 | 0.56 | 0.33 |
| 1 | 151588815 | rs1196357 | -0.712 | 0.148 | 0.435 | 0.723 | 0.039 | 0.016 | 0.016 | 0.037 | 0.51 | 0.32 | 0.49 | 0.37 | 0.67 | 0.46 |
| 1 | 151599267 | rs1196350 | -0.265 | 0.221 | 0.471 | 0.723 | -0.014 | 0.135 | 0.135 | 0.129 | 0.45 | 0.13 | 0.4 | 0.2 | 0.53 | 0.33 |
| 1 | 151600441 | rs1196351 | 0.8 | -0.208 | 0.792 | 1.595 | -0.027 | 0.227 | 0.227 | 0.272 | 0.93 | 0.72 | 0.88 | 0.68 | 1 | 0.7 |
| 1 | 151606405 | rs1196348 | 0.746 | -0.231 | 0.815 | 1.584 | -0.024 | 0.179 | 0.179 | 0 | 0.94 | 0.87 | 0.89 | 0.68 | 1 | 0.7 |
| 1 | 151608096 | rs6702842 | 0.892 | -0.45 | 1.372 | 1.49 | 0.014 | 0.042 | 0.042 | 0 | 0.4 | 0.31 | 0.39 | 0.29 | 0.33 | 0.24 |
| 1 | 151627113 | rs6701385 | 0.384 | -0.632 | 1.245 | 1.193 | -0.003 | 0.003 | 0.003 | 0 | 0.1 | 0.07 | 0.06 | 0.05 | 0.12 | 0.13 |
| 1 | 151629958 | rs6690942 | 1.724 | -0.904 | 0.965 | 0.979 | -0.006 | 0.01 | 0.01 | 0 | 0.25 | 0.01 | 0.28 | 0.24 | 0.18 | 0.12 |
| 1 | 151653312 | rs6687866 | 0.264 | -0.846 | 1.264 | 1.056 | 0.01 | 0.025 | 0.025 | 0 | 0.41 | 0.36 | 0.39 | 0.29 | 0.33 | 0.24 |
| 1 | 151662113 | rs2495395 | -1.683 | -0.861 | 1.32 | 1.092 | -0.024 | 0.28 | 0.28 | 1.407 | 0.08 | 0.32 | 0.12 | 0.34 | 0 | 0.3 |
| 1 | 151667718 | rs1308135 | -1.236 | -0.861 | 1.333 | 1.093 | -0.025 | 0.201 | 0.201 | 0.569 | 0.07 | 0.24 | 0.12 | 0.32 | 0 | 0.3 |
| 1 | 151668295 | rs1308137 | -0.334 | -0.791 | 1.308 | 1.088 | -0.001 | 0 | 0 | 0 | 0.48 | 0.6 | 0.52 | 0.61 | 0.34 | 0.55 |
| 1 | 151669555 | rs3204344 | 0.381 | -0.816 | 1.318 | 1.087 | 0.015 | 0.043 | 0.043 | 0 | 0.4 | 0.27 | 0.38 | 0.29 | 0.33 | 0.25 |
| 1 | 151669997 | rs1048694 | -0.453 | -0.764 | 1.155 | 1.051 | 0.037 | 0.015 | 0.015 | 0.045 | 0.49 | 0.68 | 0.51 | 0.63 | 0.34 | 0.55 |
| 1 | 151672658 | rs3466 | 1.551 | -1.204 | 0.598 | 0.763 | -0.194 | 0.067 | 0.067 | 0 | 0.31 | 0.02 | 0.21 | 0.1 | 0.45 | 0.27 |
| 1 | 151673350 | rs1308145 | -0.172 | -1.118 | 0.801 | 0.899 | -0.002 | -0.001 | -0.001 | 0 | 0.48 | 0.59 | 0.51 | 0.6 | 0.34 | 0.55 |
| 1 | 151675486 | rs3790508 | -0.165 | -1.044 | 0.673 | 0.862 | 0.012 | 0.059 | 0.059 | 0.093 | 0.05 | 0.06 | 0.08 | 0.17 | 0.14 | 0.15 |
| 1 | 151679441 | rs1308152 | -0.386 | -1.231 | 0.562 | 0.709 | 0.002 | 0.001 | 0.001 | 0.024 | 0.51 | 0.51 | 0.53 | 0.64 | 0.35 | 0.58 |
| 1 | 151684262 | rs3762302 | -1.144 | -1.174 | 0.296 | 0.831 | 0.008 | 0.016 | 0.016 | 0 | 0.09 | 0.06 | 0.14 | 0.36 | 0.02 | 0.32 |
| 1 | 151686240 | rs7537265 | -1.351 | -1.112 | 0.549 | 0.759 | 0.002 | 0.002 | 0.002 | 0 | 0.69 | 0.72 | 0.7 | 0.78 | 0.81 | 0.86 |
| 1 | 151691148 | rs868360 | 0.25 | -0.92 | 0.331 | 0.595 | -0.006 | 0.011 | 0.011 | 0.015 | 0.1 | 0.05 | 0.09 | 0.02 | 0.13 | 0.13 |
| 1 | 151696875 | rs1781409 | -0.649 | -1.058 | 0.471 | 0.898 | -0.017 | 0.213 | 0.213 | 0 | 0.06 | 0.15 | 0.11 | 0.35 | 0.01 | 0.22 |
| 1 | 151701345 | rs2298294 | 1.285 | -1.231 | 0.361 | 0.746 | 0.018 | 0.016 | 0.016 | 0.398 | 0.3 | 0.21 | 0.31 | 0.27 | 0.21 | 0.19 |
| 1 | 151705137 | rs2096402 | 0.847 | -0.907 | 0.006 | 0.77 | -0.092 | 0.042 | 0.042 | 0 | 0.41 | 0.06 | 0.29 | 0.22 | 0.52 | 0.35 |
| 1 | 151707765 | rs1781417 | -1.163 | -0.897 | 0.42 | 0.856 | -0.026 | 0.234 | 0.234 | 0.106 | 0.07 | 0.22 | 0.11 | 0.33 | 0 | 0.25 |
| 1 | 151725359 | rs10888430 | 0.279 | -1.1 | 0.328 | 0.567 | 0.006 | 0.008 | 0.008 | 0 | 0.41 | 0.62 | 0.53 | 0.41 | 0.35 | 0.27 |
| 1 | 151728756 | rs11204880 | -0.255 | -0.99 | 0.212 | 0.61 | 0.03 | 0.079 | 0.079 | 0 | 0.5 | 0.69 | 0.6 | 0.43 | 0.45 | 0.33 |
| 1 | 151733335 | rs8480 | -0.268 | -0.94 | 0.217 | 0.616 | 0.022 | 0.078 | 0.078 | 0 | 0.5 | 0.69 | 0.6 | 0.44 | 0.46 | 0.33 |
| 1 | 151735576 | rs7007 | -0.757 | -0.647 | 0.351 | 0.682 | 0.003 | 0 | 0 | 0 | 0.07 | 0.13 | 0.12 | 0.11 | 0.13 | 0.12 |
| 1 | 151744168 | rs3790514 | -0.849 | -0.602 | 0.584 | 0.667 | 0.027 | 0.015 | 0.015 | 0 | 0.08 | 0.2 | 0.14 | 0.12 | 0.14 | 0.11 |
| 1 | 151744928 | rs2495396 | 0.419 | -0.603 | 0.105 | 0.464 | -0.035 | 0.061 | 0.061 | 0 | 0.4 | 0.22 | 0.27 | 0.2 | 0.51 | 0.41 |
| 1 | 151746235 | rs1054475 | 0.419 | -0.603 | 0.105 | 0.464 | -0.035 | 0.061 | 0.061 | 0 | 0.4 | 0.22 | 0.27 | 0.2 | 0.51 | 0.41 |
| 1 | 151761865 | rs868867 | 0.333 | -0.63 | -0.002 | 0.409 | -0.047 | 0.035 | 0.035 | 0 | 0.37 | 0.24 | 0.26 | 0.2 | 0.5 | 0.4 |
| 1 | 151773167 | rs12409493 | 0.742 | -0.866 | 0.115 | 0.589 | 0.03 | 0.09 | 0.09 | 0.861 | 0.46 | 0.18 | 0.36 | 0.34 | 0.35 | 0.21 |
| 1 | 151778683 | rs9017 | -0.291 | -0.917 | 0.116 | 0.637 | -0.002 | -0.004 | -0.004 | 0 | 0.47 | 0.55 | 0.41 | 0.32 | 0.64 | 0.51 |
| 1 | 151778899 | rs9826 | 0.704 | -0.767 | -0.763 | 0.48 | 0.019 | 0.098 | 0.098 | 0.317 | 0.45 | 0.12 | 0.35 | 0.34 | 0.35 | 0.21 |
| 1 | 151782090 | rs12045886 | -1.176 | -0.227 | -1.245 | 0.231 | -0.002 | -0.003 | -0.003 | 0 | 0.3 | 0.27 | 0.32 | 0.47 | 0.31 | 0.44 |
| 1 | 151782130 | rs1521177 | -0.74 | 0.696 | -1.399 | -0.139 | 0.004 | 0.105 | 0.105 | 0 | 0.49 | 0.7 | 0.52 | 0.69 | 0.53 | 0.77 |
| 1 | 151783688 | rs939595 | -1.087 | 0.389 | -1.462 | -0.177 | 0.008 | 0.06 | 0.06 | 0.01 | 0.54 | 0.7 | 0.61 | 0.71 | 0.63 | 0.82 |
| 1 | 151783736 | rs2280471 | -0.505 | -0.376 | -1.123 | 0.153 | 0.001 | -0.001 | -0.001 | 0 | 0.07 | 0.1 | 0.09 | 0.15 | 0.09 | 0.07 |
| 1 | 151790858 | rs7540530 | -0.103 | -0.301 | -1.353 | 0.047 | -0.024 | -0.021 | -0.021 | 0 | 0.41 | 0.27 | 0.28 | 0.44 | 0.55 | 0.42 |
| 1 | 151792814 | rs949969 | 0.958 | -0.275 | -1.808 | 0.243 | -0.008 | -0.112 | -0.112 | 0.717 | 0.31 | 0.02 | 0.33 | 0.36 | 0.2 | 0.4 |
| 1 | 151801951 | rs4845606 | -0.434 | 0.352 | -0.679 | 0.114 | 0.01 | 0.02 | 0.02 | 0 | 0.08 | 0.14 | 0.23 | 0.15 | 0.12 | 0.14 |
| 1 | 151804907 | rs3811417 | -0.656 | 0.381 | -0.508 | 0.099 | 0.023 | 0.037 | 0.037 | 0 | 0.08 | 0.23 | 0.24 | 0.16 | 0.12 | 0.13 |
| 1 | 151805049 | rs3811418 | -0.255 | 0.398 | -0.86 | 0.106 | -0.006 | -0.01 | -0.01 | 2.132 | 0.07 | 0.04 | 0.23 | 0.16 | 0.12 | 0.13 |
| 1 | 151814405 | rs6587622 | -0.211 | -0.345 | -1.004 | -0.364 | -0.022 | -0.015 | -0.015 | 0 | 0.66 | 0.68 | 0.51 | 0.35 | 0.45 | 0.64 |
| 1 | 151816718 | rs7536659 | 0.569 | -1.316 | -1.021 | -0.629 | 0.055 | 0.036 | 0.036 | 0 | 0.45 | 0.24 | 0.21 | 0.29 | 0.3 | 0.42 |
| 1 | 151817998 | rs998454 | -0.816 | -1.675 | -0.748 | -0.077 | -0.041 | -0.016 | -0.016 | 0 | 0.67 | 0.61 | 0.73 | 0.73 | 0.82 | 0.55 |
| 1 | 151820324 | rs6587625 | -1.298 | -1.606 | -1.622 | -0.373 | 0.025 | -0.041 | -0.041 | 0.065 | 0.72 | 0.88 | 0.87 | 0.68 | 0.83 | 0.63 |
| 1 | 151824089 | rs2279503 | -0.264 | -1.521 | -1.619 | -0.403 | 0.07 | 0.006 | 0.006 | 0 | 0.54 | 0.76 | 0.57 | 0.66 | 0.7 | 0.42 |
| 1 | 151827813 | rs4845624 | 0.149 | -1.011 | -2.244 | -0.147 | 0.066 | -0.06 | -0.06 | 0.051 | 0.44 | 0.08 | 0.54 | 0.61 | 0.68 | 0.38 |
| 1 | 151832901 | rs12727170 | 0.401 | -0.34 | -1.97 | 0.172 | -0.007 | -0.108 | -0.108 | 0 | 0.29 | 0.03 | 0.14 | 0.32 | 0.22 | 0.5 |
| 1 | 151836824 | rs7529451 | 0.048 | -0.514 | -0.878 | 0.112 | -0.004 | -0.024 | -0.024 | 0.494 | 0.9 | 0.74 | 0.91 | 0.9 | 0.92 | 0.93 |
| 1 | 151844396 | rs1490187 | 0.269 | -0.604 | -0.107 | 0.081 | -0.006 | -0.037 | -0.037 | 0 | 0.9 | 0.73 | 0.9 | 0.94 | 0.92 | 0.94 |
| 1 | 151844650 | rs6684312 | 0.478 | -0.197 | 0.144 | -0.167 | 0.009 | -0.015 | -0.015 | 0 | 0.34 | 0.16 | 0.19 | 0.3 | 0.28 | 0.54 |

| | | | | | | | | | | | | | | | | |
|---|---|---|---|---|---|---|---|---|---|---|---|---|---|---|---|---|
| 1 | 151846059 | rs6587627 | -0.019 | -0.188 | 0.228 | -0.157 | -0.003 | -0.022 | -0.022 | 0 | 0.9 | 0.78 | 0.91 | 0.9 | 0.92 | 0.93 |
| 1 | 151846517 | rs13320 | 0.296 | -0.225 | 0.205 | -0.258 | 0.02 | 0.017 | 0.017 | 0 | 0.5 | 0.4 | 0.68 | 0.6 | 0.6 | 0.37 |
| 1 | 151855156 | rs6587629 | -0.465 | -0.147 | 0.159 | -0.267 | 0 | -0.005 | -0.005 | 0 | 0.91 | 0.9 | 0.91 | 0.95 | 0.92 | 0.94 |
| 1 | 151858888 | rs6685187 | 0.227 | -0.168 | 0.174 | -0.262 | -0.002 | -0.002 | -0.002 | 0 | 0.52 | 0.55 | 0.7 | 0.61 | 0.6 | 0.38 |
| 1 | 151861477 | rs1352654 | -0.04 | -0.115 | 0.21 | -0.309 | -0.003 | -0.022 | -0.022 | 0 | 0.9 | 0.77 | 0.91 | 0.91 | 0.92 | 0.93 |
| 1 | 151865439 | rs902314 | -0.135 | -0.115 | 0.258 | -0.309 | -0.005 | -0.022 | -0.022 | 0.559 | 0.9 | 0.81 | 0.91 | 0.91 | 0.92 | 0.93 |
| 1 | 151876367 | rs1873312 | -0.471 | -0.113 | 0.2 | -0.309 | -0.001 | -0.006 | -0.006 | 0.072 | 0.91 | 0.9 | 0.91 | 0.95 | 0.92 | 0.95 |
| 1 | 151882826 | rs11204915 | 0.458 | -0.125 | 0.284 | -0.228 | 0.019 | 0.027 | 0.027 | 0 | 0.5 | 0.41 | 0.68 | 0.63 | 0.6 | 0.37 |
| 1 | 151894887 | rs4845713 | 0.002 | -0.162 | 0.019 | -0.269 | 0.016 | 0.02 | 0.02 | 0 | 0.55 | 0.71 | 0.71 | 0.67 | 0.63 | 0.39 |
| 1 | 151895093 | rs10788819 | 0.806 | 0.165 | -0.212 | -0.003 | 0.064 | -0.007 | -0.007 | 0 | 0.32 | 0.17 | 0.47 | 0.35 | 0.22 | 0.26 |
| 1 | 151909453 | rs6703448 | 0.857 | 0.314 | -0.175 | -0.02 | 0.008 | 0.002 | 0.002 | 0 | 0.36 | 0.23 | 0.19 | 0.28 | 0.28 | 0.55 |
| 1 | 151915541 | rs6663064 | 0.935 | 0.319 | -0.187 | -0.011 | 0.014 | 0.005 | 0.005 | 0 | 0.35 | 0.19 | 0.19 | 0.26 | 0.28 | 0.53 |
| 1 | 151917115 | rs9943251 | 0.556 | 0.336 | -0.073 | -0.013 | 0.009 | 0.009 | 0.009 | 0 | 0.39 | 0.28 | 0.2 | 0.26 | 0.31 | 0.53 |
| 1 | 151917652 | rs2279502 | -0.217 | 0.656 | -0.025 | -0.047 | 0.002 | 0.003 | 0.003 | 0 | 0.52 | 0.56 | 0.71 | 0.65 | 0.61 | 0.41 |
| 1 | 151919584 | rs17583707 | -0.626 | 0.633 | -0.042 | -0.03 | 0.003 | -0.022 | -0.022 | 0.036 | 0.92 | 0.86 | 0.92 | 0.95 | 0.92 | 0.94 |
| 1 | 151922808 | rs2932586 | 0.008 | 0.606 | 0.089 | -0.048 | 0.019 | 0.027 | 0.027 | 0 | 0.5 | 0.42 | 0.7 | 0.65 | 0.62 | 0.38 |
| 1 | 151934447 | rs2999508 | -0.2 | 0.681 | -0.154 | -0.009 | -0.001 | -0.001 | -0.001 | 0 | 0.49 | 0.52 | 0.7 | 0.64 | 0.61 | 0.36 |
| 1 | 151938763 | rs7541099 | 0.612 | 0.737 | -0.146 | -0.015 | 0.005 | 0.006 | 0.006 | 0 | 0.41 | 0.34 | 0.22 | 0.27 | 0.32 | 0.57 |
| 1 | 151940403 | rs1552604 | 0.82 | 1.484 | 0.088 | 0.275 | -0.093 | -0.087 | -0.087 | 0 | 0.13 | 0.02 | 0.19 | 0.34 | 0.33 | 0.1 |
| 1 | 151942428 | rs1038746 | -0.093 | 0.682 | 0.108 | -0.209 | -0.001 | -0.009 | -0.009 | 0 | 0.91 | 0.9 | 0.92 | 0.95 | 0.92 | 0.94 |
| 1 | 151947568 | rs2999514 | -0.104 | 0.69 | 0.112 | -0.205 | -0.001 | -0.009 | -0.009 | 0 | 0.91 | 0.9 | 0.92 | 0.95 | 0.92 | 0.94 |
| 1 | 151952075 | rs3007704 | 0.148 | 0.69 | 0.112 | -0.204 | -0.002 | -0.012 | -0.012 | 0 | 0.91 | 0.85 | 0.92 | 0.91 | 0.92 | 0.94 |
| 1 | 151978039 | rs6587641 | 0.336 | 0.555 | 0.085 | -0.141 | -0.009 | -0.018 | -0.018 | 0 | 0.91 | 0.82 | 0.93 | 0.9 | 0.94 | 0.94 |
| 1 | 151988019 | rs959945 | 0.038 | 0.573 | 0.07 | -0.142 | -0.007 | -0.022 | -0.022 | 0.531 | 0.91 | 0.83 | 0.93 | 0.95 | 0.94 | 0.94 |
| 1 | 151999347 | rs3007671 | -0.646 | 0.493 | -0.125 | -0.614 | 0.026 | 0.093 | 0.093 | 0 | 0.38 | 0.72 | 0.22 | 0.11 | 0.3 | 0.19 |
| 1 | 152002663 | rs2999547 | -0.52 | 0.447 | 0.205 | -0.64 | 0.007 | -0.017 | -0.017 | 0 | 0.36 | 0.61 | 0.21 | 0.04 | 0.3 | 0.19 |
| 1 | 152004435 | rs12134184 | 0.173 | 0.413 | 0.675 | -0.693 | -0.012 | -0.132 | -0.132 | 0.052 | 0.28 | 0.34 | 0.15 | 0 | 0.26 | 0.13 |
| 1 | 152004804 | rs12123975 | 0.065 | 0.399 | 0.665 | -0.699 | -0.021 | -0.176 | -0.176 | 2.44 | 0.29 | 0.4 | 0.16 | 0 | 0.26 | 0.13 |
| 1 | 152007599 | rs3007676 | -1.185 | 0.47 | 0.579 | -0.58 | -0.005 | -0.015 | -0.015 | 0.087 | 0.09 | 0.28 | 0.06 | 0.11 | 0.03 | 0.07 |
| 1 | 152008309 | rs2999551 | 0.172 | 0.486 | 0.481 | -0.575 | -0.001 | -0.007 | -0.007 | 0 | 0.06 | 0.1 | 0.05 | 0.04 | 0.03 | 0.06 |
| 1 | 152016425 | rs3007680 | -1.313 | 1.106 | 0.1 | 0.203 | -0.025 | -0.024 | -0.024 | 0 | 0.85 | 0.9 | 0.82 | 0.69 | 0.68 | 0.89 |
| 1 | 152022989 | rs6680866 | 0.745 | 0.596 | -0.679 | 0.148 | -0.042 | -0.058 | -0.058 | 0 | 0.15 | 0.05 | 0.11 | 0.2 | 0.19 | 0.46 |
| 1 | 152028561 | rs4333848 | 1.4 | 0.266 | -0.791 | 0.033 | 0.054 | -0.06 | -0.06 | 0 | 0.32 | 0.01 | 0.45 | 0.31 | 0.19 | 0.26 |
| 1 | 152062750 | rs1496037 | 1.173 | 0.545 | -0.217 | -0.187 | 0.017 | -0.008 | -0.008 | 0.728 | 0.05 | 0.16 | 0.06 | 0.03 | 0.03 | 0.05 |
| 1 | 152062767 | rs17646946 | 1.109 | 0.568 | -0.339 | -0.461 | -0.018 | 0.096 | 0.096 | 0 | 0.23 | 0.11 | 0.11 | 0 | 0.24 | 0.1 |
| 1 | 152079989 | rs1131471 | -0.404 | -0.195 | 0.326 | 0.296 | -0.001 | -0.002 | -0.002 | 0 | 0.55 | 0.51 | 0.65 | 0.6 | 0.4 | 0.78 |
| 1 | 152088605 | rs1496041 | 0.005 | -0.213 | -0.099 | 0.295 | 0.046 | 0.008 | 0.008 | 0 | 0.48 | 0.28 | 0.59 | 0.49 | 0.37 | 0.72 |
| 1 | 152095083 | rs3104205 | 0.836 | -0.206 | -0.088 | 0.296 | 0.002 | -0.013 | -0.013 | 0 | 0.05 | 0.11 | 0.06 | 0.03 | 0.03 | 0.05 |
| 1 | 152106853 | rs11588437 | -0.076 | -0.125 | -0.294 | 0.322 | 0.049 | 0.015 | 0.015 | 0 | 0.49 | 0.3 | 0.58 | 0.5 | 0.37 | 0.72 |
| 1 | 152126467 | rs3001978 | -0.47 | -0.076 | -0.094 | 0.412 | 0 | 0.001 | 0.001 | 0 | 0.57 | 0.6 | 0.65 | 0.6 | 0.4 | 0.78 |
| 1 | 152142087 | rs924088 | 0.426 | -0.957 | -0.868 | 0.777 | 0.016 | -0.008 | -0.008 | 0.747 | 0.3 | 0.25 | 0.44 | 0.37 | 0.15 | 0.37 |
| 1 | 152142637 | rs17648141 | 0.193 | -0.399 | -1.186 | 1.129 | -0.019 | -0.068 | -0.068 | 0.828 | 0.12 | 0 | 0.09 | 0.15 | 0.11 | 0.19 |
| 1 | 152144260 | rs969359 | -0.76 | -0.82 | -1.301 | 0.85 | 0.066 | -0.086 | -0.086 | 0 | 0.68 | 0.91 | 0.56 | 0.57 | 0.81 | 0.49 |
| 1 | 152151950 | rs11204930 | -1.194 | -0.612 | 1.218 | 0.602 | 0.087 | 0.083 | 0.083 | 1.457 | 0.05 | 0.45 | 0.03 | 0 | 0 | 0 |
| 1 | 152160437 | rs10749668 | 1.16 | -0.743 | 1.226 | 0.551 | 0.028 | 0.029 | 0.029 | 1.857 | 0.94 | 0.59 | 0.95 | 0.98 | 0.98 | 0.94 |
| 1 | 152161735 | rs10788826 | 0.202 | -1.296 | 0.381 | 0.476 | 0.002 | 0.002 | 0.002 | 0 | 0.54 | 0.52 | 0.47 | 0.24 | 0.7 | 0.34 |
| 1 | 152162106 | rs12130219 | 1.164 | -0.831 | 0.241 | -0.055 | 0 | -0.025 | -0.025 | 0 | 0.25 | 0.24 | 0.27 | 0.04 | 0.24 | 0.12 |
| 1 | 152165878 | rs1552991 | -0.268 | -1.264 | 0.032 | -0.054 | -0.011 | 0.025 | 0.025 | 2.455 | 0.4 | 0.21 | 0.28 | 0.29 | 0.58 | 0.39 |
| 1 | 152183474 | rs4845741 | -0.772 | -1.215 | 0.052 | 0.174 | -0.062 | 0.016 | 0.016 | 0.143 | 0.65 | 0.47 | 0.55 | 0.33 | 0.83 | 0.51 |
| 1 | 152184704 | rs1496045 | -0.997 | -1.22 | 0.121 | 0.192 | -0.038 | 0.014 | 0.014 | 0.056 | 0.67 | 0.55 | 0.56 | 0.38 | 0.83 | 0.51 |
| 1 | 152184950 | rs4845423 | -0.975 | -1.22 | 0.112 | 0.191 | -0.048 | 0.016 | 0.016 | 0.043 | 0.66 | 0.53 | 0.56 | 0.33 | 0.83 | 0.51 |
| 1 | 152193851 | rs11204937 | -0.768 | -1.22 | 0.124 | 0.202 | -0.064 | 0.016 | 0.016 | 0.143 | 0.65 | 0.47 | 0.55 | 0.33 | 0.83 | 0.51 |
| 1 | 152200472 | rs10888470 | -0.898 | -1.224 | 0.186 | 0.233 | -0.059 | 0.022 | 0.022 | 0.058 | 0.66 | 0.44 | 0.55 | 0.39 | 0.83 | 0.51 |
| 1 | 152203906 | rs11204943 | -0.898 | -1.224 | 0.186 | 0.233 | -0.059 | 0.021 | 0.021 | 0.058 | 0.66 | 0.44 | 0.55 | 0.39 | 0.83 | 0.51 |
| 1 | 152218257 | rs11204947 | 0.919 | -1.211 | 0.236 | 0.245 | 0.02 | -0.014 | -0.014 | 0 | 0.32 | 0.26 | 0.43 | 0.61 | 0.17 | 0.49 |
| 1 | 152219679 | rs12030667 | 1.668 | -0.263 | -0.055 | 0.06 | -0.071 | -0.061 | -0.061 | 0.107 | 0.12 | 0.04 | 0.12 | 0.21 | 0.28 | 0.16 |
| 1 | 152246726 | rs12746239 | 1.264 | -0.584 | 0.066 | 0.045 | 0.038 | -0.033 | -0.033 | 0 | 0.3 | 0.2 | 0.42 | 0.61 | 0.17 | 0.49 |
| 1 | 152251714 | rs4534386 | -1.078 | -0.619 | 0.154 | 0.035 | -0.051 | 0.018 | 0.018 | 0.003 | 0.66 | 0.43 | 0.55 | 0.34 | 0.83 | 0.51 |
| 1 | 152259078 | rs11204971 | 2.146 | -0.625 | -0.063 | 0.03 | 0.013 | -0.156 | -0.156 | 0.736 | 0.24 | 0.01 | 0.39 | 0.58 | 0.14 | 0.43 |
| 1 | 152259742 | rs2152862 | 0.854 | -0.615 | -0.051 | 0.033 | -0.052 | 0.022 | 0.022 | 0.008 | 0.33 | 0.57 | 0.45 | 0.61 | 0.17 | 0.49 |
| 1 | 152273168 | rs2184951 | 1.506 | -0.589 | -0.046 | 0.046 | 0.059 | -0.091 | -0.091 | 0 | 0.28 | 0.07 | 0.41 | 0.61 | 0.17 | 0.49 |
| 1 | 152282267 | rs12405278 | 2.078 | -0.613 | -0.055 | 0.077 | 0.017 | -0.157 | -0.157 | 0 | 0.24 | 0.01 | 0.39 | 0.58 | 0.14 | 0.43 |
| 1 | 152286126 | rs11582620 | 1.202 | -0.625 | -0.217 | -0.059 | -0.007 | -0.017 | -0.017 | 0 | 0.12 | 0 | 0.08 | 0.1 | 0.11 | 0.17 |
| 1 | 152296863 | rs6587666 | 1.316 | -0.522 | 0.025 | 0.14 | 0.064 | -0.058 | -0.058 | 0 | 0.3 | 0.12 | 0.42 | 0.61 | 0.17 | 0.49 |
| 1 | 152301576 | rs1933064 | -0.187 | -0.39 | 0.123 | -0.021 | 0.027 | 0.192 | 0.192 | 0 | 0.5 | 0.84 | 0.59 | 0.87 | 0.46 | 0.65 |
| 1 | 152308030 | rs3126093 | -1.372 | -0.524 | 0.437 | 0.255 | 0.059 | 0.067 | 0.067 | 4.921 | 0.06 | 0.5 | 0.04 | 0.05 | 0 | 0 |
| 1 | 152312597 | rs2050631 | -0.848 | -0.387 | -0.141 | 0.248 | 0.004 | -0.059 | -0.059 | 0 | 0.65 | 0.91 | 0.54 | 0.42 | 0.69 | 0.51 |
| 1 | 152321344 | rs2065954 | 1.026 | -0.842 | 0.382 | 0.599 | 0.005 | -0.008 | -0.008 | 0 | 0.27 | 0.29 | 0.4 | 0.45 | 0.14 | 0.33 |
| 1 | 152322042 | rs11204982 | 0.807 | -0.882 | 0.639 | 0.582 | -0.04 | 0.052 | 0.052 | 0 | 0.28 | 0.4 | 0.42 | 0.5 | 0.14 | 0.33 |

| | | | | | | | | | | | | | | | | |
|---|---|---|---|---|---|---|---|---|---|---|---|---|---|---|---|---|
| 1 | 152329435 | rs2282303 | 0.932 | -0.873 | 0.667 | 0.587 | -0.027 | 0.038 | 0.038 | 0 | 0.28 | 0.36 | 0.42 | 0.5 | 0.14 | 0.33 |
| 1 | 152343893 | rs11204988 | 1.41 | -0.931 | 0.251 | 0.404 | -0.047 | -0.026 | -0.026 | 2.119 | 0.22 | 0.01 | 0.23 | 0.19 | 0.21 | 0.11 |
| 1 | 152347403 | rs12029633 | -0.007 | -0.792 | 1.011 | 0.539 | -0.061 | 0.039 | 0.039 | 0.245 | 0.34 | 0.49 | 0.45 | 0.51 | 0.17 | 0.38 |
| 1 | 152360184 | rs1923504 | -0.987 | -0.371 | 0.613 | 0.495 | -0.135 | -0.006 | -0.006 | 1.156 | 0.79 | 0.98 | 0.83 | 0.81 | 0.58 | 0.8 |
| 1 | 152362727 | rs7534742 | -0.169 | -0.828 | 1.813 | 0.839 | 0.021 | -0.024 | -0.024 | 0.494 | 0.3 | 0.19 | 0.41 | 0.51 | 0.17 | 0.37 |
| 1 | 152365929 | rs12407319 | 0.111 | -0.826 | 1.72 | 0.84 | 0.045 | -0.068 | -0.068 | 0 | 0.28 | 0.11 | 0.41 | 0.46 | 0.17 | 0.37 |
| 1 | 152372035 | rs7515448 | -0.669 | -0.73 | 1.719 | 0.87 | -0.001 | -0.003 | -0.003 | 0 | 0.34 | 0.64 | 0.45 | 0.5 | 0.17 | 0.34 |
| 1 | 152382062 | rs10888486 | 0.631 | -0.524 | 1.607 | 0.87 | 0.024 | -0.158 | -0.158 | 0 | 0.22 | 0.02 | 0.37 | 0.43 | 0.14 | 0.32 |
| 1 | 152382120 | rs3829868 | 0.484 | -0.526 | 1.637 | 0.869 | 0.042 | -0.117 | -0.117 | 0.08 | 0.23 | 0.06 | 0.37 | 0.43 | 0.14 | 0.32 |
| 1 | 152384759 | rs7521839 | 0.155 | -0.529 | 1.744 | 0.867 | -0.019 | 0.032 | 0.032 | 0 | 0.25 | 0.28 | 0.39 | 0.43 | 0.15 | 0.32 |
| 1 | 152400293 | rs726864 | -0.243 | -0.497 | 1.715 | 0.878 | -0.059 | 0.07 | 0.07 | 0.013 | 0.28 | 0.53 | 0.4 | 0.47 | 0.15 | 0.33 |
| 1 | 152400361 | rs726863 | 0.129 | -0.463 | 1.312 | 0.905 | -0.004 | 0.006 | 0.006 | 0 | 0.25 | 0.25 | 0.38 | 0.46 | 0.14 | 0.33 |
| 1 | 152411153 | rs1923501 | 0.049 | -0.282 | 1.335 | 0.024 | -0.009 | 0.001 | 0.001 | 1.089 | 0.27 | 0.33 | 0.29 | 0.22 | 0.17 | 0.24 |
| 1 | 152411813 | rs11579266 | 0.451 | -0.273 | 1.186 | 0.028 | 0.023 | -0.075 | -0.075 | 1.33 | 0.24 | 0.02 | 0.26 | 0.22 | 0.16 | 0.24 |
| 1 | 152412196 | rs2146118 | -0.238 | 0.055 | 1.206 | -0.16 | -0.003 | -0.001 | -0.001 | 0.81 | 0.89 | 0.93 | 0.89 | 0.97 | 0.82 | 0.95 |
| 1 | 152416494 | rs4845769 | 0.268 | 0.014 | 1.228 | -0.172 | 0.005 | 0.017 | 0.017 | 1.676 | 0.65 | 0.85 | 0.63 | 0.77 | 0.67 | 0.71 |
| 1 | 152416630 | rs11205003 | 0.657 | -0.024 | 1.513 | -0.186 | -0.013 | -0.016 | -0.016 | 0 | 0.61 | 0.45 | 0.6 | 0.73 | 0.66 | 0.71 |
| 1 | 152423423 | rs1923497 | -0.282 | 0.112 | 1.447 | -0.097 | -0.04 | -0.014 | -0.014 | 0 | 0.9 | 0.98 | 0.9 | 0.94 | 0.82 | 0.95 |
| 1 | 152430152 | rs6587673 | 1.235 | 0.14 | 1.532 | -0.143 | 0.031 | 0.026 | 0.026 | 0 | 0.52 | 0.28 | 0.52 | 0.56 | 0.57 | 0.52 |
| 1 | 152430680 | rs4845771 | 0.399 | 0.221 | 1.437 | -0.128 | 0.002 | 0.003 | 0.003 | 0.865 | 0.63 | 0.62 | 0.61 | 0.77 | 0.66 | 0.71 |
| 1 | 152430842 | rs6696556 | 0.545 | 0.398 | 0.854 | -0.029 | 0.038 | 0.006 | 0.006 | 1.384 | 0.24 | 0.08 | 0.26 | 0.16 | 0.16 | 0.24 |
| 1 | 152431931 | rs12089980 | -0.545 | 0.834 | 1.335 | 0.036 | -0.005 | -0.009 | -0.009 | 0 | 0.08 | 0.05 | 0.07 | 0.17 | 0.09 | 0.19 |
| 1 | 152443495 | rs2146115 | 0.343 | 1.017 | 0.175 | 0.106 | -0.001 | 0 | 0 | 0 | 0.89 | 0.9 | 0.88 | 0.94 | 0.82 | 0.95 |
| 1 | 152480654 | rs1415432 | 0.553 | 0.755 | 0.344 | 0.266 | -0.034 | -0.022 | -0.022 | 1.481 | 0.82 | 0.89 | 0.82 | 0.75 | 0.72 | 0.75 |
| 1 | 152482587 | rs11581947 | 0.745 | 0.199 | 0.714 | -0.139 | -0.001 | -0.001 | -0.001 | 0 | 0.61 | 0.42 | 0.54 | 0.61 | 0.62 | 0.6 |
| 1 | 152484129 | rs2105117 | 0.74 | 0.152 | 0.662 | -0.161 | 0.007 | 0.003 | 0.003 | 0 | 0.59 | 0.31 | 0.53 | 0.61 | 0.62 | 0.6 |
| 1 | 152485227 | rs3753451 | 0.687 | 0.16 | 0.673 | -0.158 | 0.005 | 0.003 | 0.003 | 0 | 0.59 | 0.35 | 0.54 | 0.65 | 0.62 | 0.6 |
| 1 | 152488428 | rs1053590 | 0.506 | 0.116 | 0.783 | -0.349 | 0 | 0.001 | 0.001 | 0 | 0.63 | 0.63 | 0.56 | 0.65 | 0.63 | 0.6 |
| 1 | 152489548 | rs488393 | 0.428 | 0.118 | 0.794 | -0.348 | 0 | 0.001 | 0.001 | 0 | 0.64 | 0.65 | 0.56 | 0.65 | 0.63 | 0.6 |
| 1 | 152507156 | rs4845443 | 1.008 | 0.005 | 0.206 | -0.43 | 0.009 | -0.012 | -0.012 | 0.417 | 0.57 | 0.14 | 0.52 | 0.61 | 0.62 | 0.6 |
| 1 | 152510348 | rs6662637 | 0.885 | 0.567 | -0.566 | -0.576 | -0.095 | 0.096 | 0.096 | 0.163 | 0.22 | 0.01 | 0.14 | 0.07 | 0.25 | 0.14 |
| 1 | 152513431 | rs474086 | 1.107 | 0.458 | 0.491 | 0.184 | -0.001 | 0.003 | 0.003 | 0 | 0.61 | 0.5 | 0.55 | 0.61 | 0.62 | 0.6 |
| 1 | 152514332 | rs549044 | 1.107 | 0.458 | 0.491 | 0.184 | -0.001 | 0.003 | 0.003 | 0 | 0.61 | 0.5 | 0.55 | 0.61 | 0.62 | 0.6 |
| 1 | 152518294 | rs533917 | 0.92 | 0.889 | -0.269 | 0.391 | 0.001 | 0.002 | 0.002 | 0 | 0.65 | 0.65 | 0.56 | 0.61 | 0.62 | 0.61 |
| 1 | 152526812 | rs7530609 | 0.821 | 1.168 | -0.265 | 0.523 | 0.009 | 0.014 | 0.014 | 0 | 0.67 | 0.59 | 0.54 | 0.61 | 0.62 | 0.63 |
| 1 | 152534954 | rs4240889 | 0.424 | 1.094 | -0.52 | -0.167 | 0.002 | -0.023 | -0.023 | 0 | 0.76 | 0.74 | 0.8 | 0.94 | 0.73 | 0.93 |
| 1 | 152537155 | rs4845792 | -0.169 | 0.998 | -0.772 | -0.106 | -0.001 | 0 | 0 | 0.586 | 0.52 | 0.56 | 0.33 | 0.26 | 0.56 | 0.42 |
| 1 | 152537954 | rs10888501 | -0.571 | 0.452 | -1.354 | 0.06 | 0.027 | -0.024 | -0.024 | 0 | 0.4 | 0.75 | 0.49 | 0.56 | 0.37 | 0.51 |
| 1 | 152539393 | rs17659389 | 0.144 | 1.552 | -1.423 | -0.143 | -0.015 | 0.015 | 0.015 | 0 | 0.09 | 0.04 | 0.1 | 0.03 | 0.25 | 0.13 |
| 1 | 152542229 | rs7516108 | -0.98 | 0.932 | -1.94 | -0.133 | -0.012 | 0.009 | 0.009 | 0 | 0.57 | 0.77 | 0.64 | 0.61 | 0.6 | 0.61 |
| 1 | 152546889 | rs12046030 | -0.535 | -0.42 | -1.413 | 0.157 | -0.033 | 0.001 | 0.001 | 0.095 | 0.14 | 0.22 | 0.11 | 0.13 | 0.03 | 0.07 |
| 1 | 152548480 | rs4845445 | 0.01 | -0.252 | -1.437 | -0.047 | -0.044 | 0.02 | 0.02 | 1.938 | 0.3 | 0.16 | 0.23 | 0.23 | 0.37 | 0.24 |
| 1 | 152551801 | rs11589128 | 0.772 | -0.33 | -1.385 | -0.282 | -0.111 | 0.118 | 0.118 | 2.603 | 0.23 | 0.01 | 0.16 | 0.03 | 0.3 | 0.15 |
| 1 | 152552461 | rs4845313 | 1.144 | -0.41 | -1.41 | 0.05 | -0.027 | -0.04 | -0.04 | 0 | 0.2 | 0.07 | 0.3 | 0.3 | 0.25 | 0.32 |
| 1 | 152590187 | rs6677595 | -0.956 | -0.609 | -1.151 | 0.015 | 0.002 | 0.005 | 0.005 | 0 | 0.59 | 0.37 | 0.62 | 0.64 | 0.64 | 0.62 |
| 1 | 152592184 | rs4845454 | -0.956 | -0.609 | -1.151 | 0.015 | 0.002 | 0.005 | 0.005 | 0 | 0.59 | 0.37 | 0.62 | 0.64 | 0.64 | 0.62 |
| 1 | 152595194 | rs12032251 | -0.757 | -0.359 | -0.311 | -0.036 | -0.001 | -0.003 | -0.003 | 2.541 | 0.41 | 0.49 | 0.31 | 0.33 | 0.39 | 0.3 |
| 1 | 152596414 | rs7527788 | -0.282 | -0.434 | -0.185 | -0.072 | -0.014 | -0.006 | -0.006 | 0 | 0.35 | 0.26 | 0.3 | 0.33 | 0.38 | 0.3 |
| 1 | 152598435 | rs4845457 | -0.499 | -0.432 | -0.202 | -0.071 | -0.002 | -0.001 | -0.001 | 0 | 0.37 | 0.4 | 0.31 | 0.33 | 0.39 | 0.3 |
| 1 | 152607331 | rs7545817 | -1.709 | -1.095 | -0.014 | 0.275 | -0.109 | -0.07 | -0.07 | 0 | 0.8 | 0.99 | 0.87 | 0.74 | 0.73 | 0.8 |
| 1 | 152612910 | rs1325508 | -1.248 | -1.36 | 0.953 | 0.025 | -0.053 | 0.115 | 0.115 | 0 | 0.18 | 0.44 | 0.16 | 0.3 | 0.09 | 0.14 |
| 1 | 152617781 | rs942826 | -0.75 | -1.35 | 0.867 | 0.031 | -0.026 | 0.124 | 0.124 | 0 | 0.14 | 0.35 | 0.15 | 0.3 | 0.09 | 0.14 |
| 1 | 152625148 | rs10888505 | -1.333 | -1.36 | 1.49 | 0.025 | -0.049 | 0.079 | 0.079 | 0 | 0.18 | 0.61 | 0.17 | 0.3 | 0.09 | 0.14 |
| 1 | 152632433 | rs1332497 | -1.317 | -1.352 | 1.493 | -0.006 | -0.048 | 0.075 | 0.075 | 0 | 0.18 | 0.61 | 0.17 | 0.29 | 0.09 | 0.14 |
| 1 | 152635201 | rs11205062 | -1.43 | -1.363 | 1.487 | -0.012 | -0.05 | 0.069 | 0.069 | 0 | 0.19 | 0.61 | 0.17 | 0.3 | 0.09 | 0.14 |
| 1 | 152644161 | rs11205066 | -0.416 | -1.356 | 1.64 | -0.008 | -0.018 | 0.136 | 0.136 | 0 | 0.13 | 0.3 | 0.16 | 0.29 | 0.09 | 0.14 |
| 1 | 152651029 | rs2014368 | -1.255 | -1.401 | 1.569 | -0.116 | 0.005 | 0.017 | 0.017 | 0.016 | 0.43 | 0.67 | 0.33 | 0.33 | 0.39 | 0.31 |
| 1 | 152654385 | rs11205073 | -1.255 | -1.401 | 1.569 | -0.116 | 0.005 | 0.017 | 0.017 | 0.016 | 0.43 | 0.67 | 0.33 | 0.33 | 0.39 | 0.31 |
| 1 | 152659194 | rs35748742 | 0.596 | -1.406 | 1.514 | -0.12 | -0.111 | 0.137 | 0.137 | 1.989 | 0.24 | 0.01 | 0.16 | 0.03 | 0.3 | 0.16 |
| 1 | 152670341 | rs4845317 | -1.33 | -1.409 | 1.721 | -0.121 | 0.009 | 0.029 | 0.029 | 0.008 | 0.44 | 0.72 | 0.33 | 0.33 | 0.39 | 0.3 |
| 1 | 152672005 | rs17661905 | 0.446 | -1.298 | 1.663 | -0.144 | -0.074 | 0.138 | 0.138 | 1.04 | 0.26 | 0.08 | 0.16 | 0.03 | 0.3 | 0.17 |
| 1 | 152672349 | rs16834343 | 0.571 | -1.297 | 1.678 | -0.068 | -0.004 | 0.047 | 0.047 | 0 | 0.08 | 0.18 | 0.05 | 0.17 | 0.08 | 0.08 |
| 1 | 152675538 | rs6693713 | -0.139 | -1.284 | 1.657 | -0.34 | 0.007 | -0.098 | -0.098 | 0.04 | 0.3 | 0.36 | 0.17 | 0.03 | 0.31 | 0.17 |
| 1 | 152680909 | rs11205080 | 0.255 | -1.208 | 1.361 | 0.021 | -0.009 | 0.018 | 0.018 | 0 | 0.09 | 0.19 | 0.05 | 0.13 | 0.07 | 0.1 |
| 1 | 152681835 | rs10888510 | -0.289 | -1.211 | 1.67 | 0.041 | -0.008 | 0.119 | 0.119 | 0.011 | 0.07 | 0.27 | 0.1 | 0.13 | 0.01 | 0.05 |
| 1 | 152682909 | rs11205083 | -1.224 | -1.239 | 1.458 | -0.087 | 0 | 0.02 | 0.02 | 0 | 0.45 | 0.61 | 0.35 | 0.29 | 0.43 | 0.32 |
| 1 | 152685025 | rs12754243 | 0.335 | -1.102 | 1.151 | -0.121 | -0.074 | 0.13 | 0.13 | 0.871 | 0.27 | 0.1 | 0.21 | 0.03 | 0.35 | 0.17 |
| 1 | 152690448 | rs1332498 | -0.664 | -1.587 | 1.491 | 0.056 | 0.001 | -0.042 | -0.042 | 2.003 | 0.37 | 0.54 | 0.38 | 0.17 | 0.36 | 0.23 |
| 1 | 152691438 | rs7534164 | -0.664 | -1.591 | 1.543 | 0.054 | 0.001 | -0.04 | -0.04 | 0 | 0.37 | 0.55 | 0.38 | 0.17 | 0.36 | 0.23 |

| | | | | | | | | | | | | | | | |
|---|---|---|---|---|---|---|---|---|---|---|---|---|---|---|---|
| 1 | 152692472 | rs873775 | -0.664 | -1.591 | 1.543 | 0.054 | 0.001 | -0.04 | -0.04 | 0 | 0.37 | 0.55 | 0.38 | 0.17 | 0.36 | 0.23 |
| 1 | 152692588 | rs944682 | -0.664 | -1.591 | 1.543 | 0.054 | 0.001 | -0.04 | -0.04 | 0.899 | 0.37 | 0.55 | 0.38 | 0.17 | 0.36 | 0.23 |
| 1 | 152692781 | rs3814350 | -0.664 | -1.591 | 1.543 | 0.054 | 0.001 | -0.04 | -0.04 | 0 | 0.37 | 0.55 | 0.38 | 0.17 | 0.36 | 0.23 |
| 1 | 152697338 | rs6700613 | -0.664 | -1.587 | 1.384 | 0.056 | 0.002 | -0.047 | -0.047 | 0 | 0.37 | 0.54 | 0.38 | 0.17 | 0.36 | 0.23 |
| 1 | 152701395 | rs10788847 | -0.141 | -1.531 | 1.297 | 0.021 | 0.01 | -0.109 | -0.109 | 1.405 | 0.32 | 0.41 | 0.23 | 0.03 | 0.35 | 0.16 |
| 1 | 152708619 | rs11205094 | -1.367 | -1.683 | 0.736 | 0.07 | 0.008 | 0.062 | 0.062 | 0 | 0.51 | 0.7 | 0.45 | 0.29 | 0.46 | 0.34 |
| 1 | 152709456 | rs4845473 | 0.583 | -1.104 | 0.089 | -0.352 | -0.077 | 0.127 | 0.127 | 0.556 | 0.26 | 0.08 | 0.2 | 0.03 | 0.34 | 0.15 |
| 1 | 152712747 | rs16834421 | 0.292 | -0.937 | 0.739 | 0.498 | -0.004 | 0.041 | 0.041 | 0 | 0.14 | 0.16 | 0.22 | 0.34 | 0.11 | 0.16 |
| 1 | 152713558 | rs12091871 | -0.464 | -1.099 | 1.37 | 0.438 | -0.014 | 0.03 | 0.03 | 0.189 | 0.06 | 0.26 | 0.02 | 0.04 | 0.01 | 0.07 |
| 1 | 152719318 | rs12023710 | -0.156 | -1.132 | 1.466 | 0.452 | -0.013 | 0.055 | 0.055 | 0 | 0.16 | 0.24 | 0.23 | 0.35 | 0.11 | 0.17 |
| 1 | 152724053 | rs1332506 | -1.28 | -1.434 | 1.743 | 0.024 | -0.002 | 0.002 | 0.002 | 0 | 0.49 | 0.6 | 0.49 | 0.42 | 0.52 | 0.42 |
| 1 | 152728472 | rs11578927 | 0.719 | -1.44 | 1.705 | 0.003 | -0.057 | 0.098 | 0.098 | 0 | 0.27 | 0.08 | 0.24 | 0.03 | 0.39 | 0.17 |
| 1 | 152729136 | rs6689049 | -1.275 | -1.447 | 1.812 | 0.014 | -0.001 | 0 | 0 | 0 | 0.48 | 0.55 | 0.49 | 0.42 | 0.52 | 0.42 |
| 1 | 152732174 | rs944683 | 0.611 | -1.365 | 1.788 | 0.104 | -0.002 | 0.034 | 0.034 | 0 | 0.13 | 0.14 | 0.21 | 0.35 | 0.11 | 0.16 |
| 1 | 152732567 | rs16834461 | -0.173 | -1.363 | 1.786 | 0.106 | -0.025 | 0.099 | 0.099 | 0 | 0.17 | 0.36 | 0.25 | 0.34 | 0.11 | 0.16 |
| 1 | 152734390 | rs2297487 | -0.173 | -1.363 | 1.786 | 0.106 | -0.025 | 0.099 | 0.099 | 0 | 0.17 | 0.36 | 0.25 | 0.34 | 0.11 | 0.16 |
| 1 | 152747126 | rs11205102 | -1.476 | -1.476 | 1.732 | -0.043 | -0.021 | -0.004 | -0.004 | 0 | 0.54 | 0.78 | 0.51 | 0.41 | 0.52 | 0.42 |
| 1 | 152747890 | rs4845322 | -1.474 | -1.48 | 1.744 | -0.046 | -0.02 | -0.003 | -0.003 | 0.04 | 0.54 | 0.77 | 0.51 | 0.41 | 0.52 | 0.42 |
| 1 | 152749891 | rs913998 | 0.728 | -1.491 | 1.833 | -0.056 | -0.006 | 0.029 | 0.029 | 0 | 0.16 | 0.18 | 0.22 | 0.34 | 0.11 | 0.17 |
| 1 | 152750506 | rs1888964 | 0.482 | -1.477 | 1.968 | -0.106 | -0.012 | 0.003 | 0.003 | 0 | 0.3 | 0.23 | 0.25 | 0.03 | 0.39 | 0.17 |
| 1 | 152753798 | rs2339381 | 0.692 | -1.468 | 1.956 | -0.102 | -0.037 | 0.064 | 0.064 | 0 | 0.28 | 0.1 | 0.24 | 0.03 | 0.39 | 0.17 |
| 1 | 152756538 | rs7549803 | -0.54 | -1.442 | 2.144 | 0.016 | -0.053 | 0.096 | 0.096 | 0 | 0.23 | 0.44 | 0.26 | 0.38 | 0.12 | 0.24 |
| 1 | 152756774 | rs950337 | -0.813 | -1.415 | 2.018 | 0.028 | -0.068 | 0.093 | 0.093 | 0 | 0.24 | 0.56 | 0.26 | 0.37 | 0.12 | 0.25 |
| 1 | 152757741 | rs11205105 | -0.433 | -1.268 | 1.8 | 0.04 | -0.026 | 0.064 | 0.064 | 0 | 0.22 | 0.35 | 0.26 | 0.37 | 0.14 | 0.25 |
| 1 | 152758519 | rs11576947 | -1.346 | -1.219 | 1.262 | 0.04 | -0.014 | 0.003 | 0.003 | 0 | 0.53 | 0.76 | 0.5 | 0.41 | 0.5 | 0.43 |
| 1 | 152759678 | rs1034109 | -1.312 | -1.177 | 1.282 | 0.085 | -0.011 | 0.005 | 0.005 | 0 | 0.53 | 0.76 | 0.51 | 0.41 | 0.5 | 0.42 |
| 1 | 152771837 | rs12568028 | 0.118 | -1.033 | 0.904 | 0.376 | -0.012 | 0.04 | 0.04 | 0 | 0.17 | 0.23 | 0.22 | 0.31 | 0.11 | 0.24 |
| 1 | 152773905 | rs6694602 | 0.118 | -1.033 | 0.904 | 0.376 | -0.012 | 0.04 | 0.04 | 0 | 0.17 | 0.23 | 0.22 | 0.31 | 0.11 | 0.24 |
| 1 | 152777574 | rs1048534 | 0.344 | -1 | 0.618 | 0.392 | 0 | 0.007 | 0.007 | 0 | 0.16 | 0.14 | 0.22 | 0.31 | 0.11 | 0.24 |
| 1 | 152778526 | rs6701216 | -0.706 | -1.22 | 0.705 | 0.308 | -0.005 | 0.008 | 0.008 | 0 | 0.23 | 0.29 | 0.25 | 0.33 | 0.14 | 0.32 |
| 1 | 152778558 | rs6701221 | -0.57 | -1.203 | 0.689 | 0.166 | -0.001 | -0.001 | -0.001 | 2.595 | 0.22 | 0.24 | 0.12 | 0.12 | 0.14 | 0.19 |
| 1 | 152778576 | rs4845488 | -0.95 | -1.042 | 0.634 | -0.011 | -0.001 | 0.018 | 0.018 | 0 | 0.51 | 0.44 | 0.34 | 0.15 | 0.49 | 0.35 |
| 1 | 152780139 | rs7411365 | 0.581 | -0.955 | 0.288 | 0.128 | -0.002 | -0.003 | -0.003 | 0 | 0.15 | 0.11 | 0.09 | 0.08 | 0.11 | 0.18 |
| 1 | 152783255 | rs7524281 | -0.216 | -1.022 | 0.34 | 0.26 | 0.002 | 0.006 | 0.006 | 0 | 0.37 | 0.64 | 0.38 | 0.29 | 0.36 | 0.3 |
| 1 | 152783560 | rs7536191 | -0.871 | -1.34 | -0.02 | 0.836 | 0.064 | -0.018 | -0.018 | 0 | 0.41 | 0.82 | 0.37 | 0.53 | 0.28 | 0.44 |
| 1 | 152783724 | rs12023196 | -0.979 | -1.58 | -0.169 | 0.625 | 0.088 | -0.021 | -0.021 | 0 | 0.43 | 0.87 | 0.35 | 0.37 | 0.28 | 0.43 |
| 1 | 152784255 | rs11804609 | -1.684 | -1.398 | 1.132 | 0.283 | -0.034 | -0.023 | -0.023 | 0 | 0.12 | 0.6 | 0.06 | 0.04 | 0.04 | 0.05 |
| 1 | 152801416 | rs11205114 | 1.006 | -1.039 | 0.434 | -0.022 | -0.057 | 0.04 | 0.04 | 0 | 0.25 | 0.01 | 0.17 | 0.12 | 0.28 | 0.18 |
| 1 | 152815273 | rs1411618 | 0.347 | -0.793 | 1.822 | 0.491 | -0.01 | -0.012 | -0.012 | 0 | 0.05 | 0.23 | 0.02 | 0 | 0 | 0 |
| 1 | 152816340 | rs16834611 | -0.972 | -0.797 | 2.022 | 0.488 | -0.006 | -0.014 | -0.014 | 0 | 0.08 | 0.43 | 0.04 | 0 | 0.01 | 0 |
| 1 | 152862466 | rs6587709 | -0.423 | -0.426 | 0.775 | -0.115 | 0.008 | -0.039 | -0.039 | 0 | 0.39 | 0.55 | 0.32 | 0.14 | 0.45 | 0.2 |
| 1 | 152872294 | rs11580565 | -0.023 | -0.426 | 0.034 | -0.156 | -0.129 | 0.162 | 0.162 | 0 | 0.34 | 0.06 | 0.24 | 0.1 | 0.44 | 0.21 |
| 1 | 152888178 | rs4845496 | -1.051 | -0.723 | 0.207 | 0.046 | -0.01 | 0.014 | 0.014 | 0 | 0.54 | 0.8 | 0.51 | 0.63 | 0.55 | 0.5 |
| 1 | 152897923 | rs12027303 | -0.102 | -0.825 | 1.112 | 0.084 | -0.038 | 0.266 | 0.266 | 0 | 0.16 | 0.43 | 0.26 | 0.48 | 0.1 | 0.27 |
| 1 | 152908222 | rs756303 | -0.765 | -1.102 | 1.088 | -0.41 | -0.002 | -0.007 | -0.007 | 0 | 0.47 | 0.48 | 0.5 | 0.64 | 0.57 | 0.51 |
| 1 | 152918556 | rs11576457 | -0.108 | -1.106 | 0.937 | -0.743 | -0.065 | 0.181 | 0.181 | 0 | 0.41 | 0.08 | 0.3 | 0.13 | 0.56 | 0.27 |
| 1 | 152922896 | rs2066004 | -0.995 | -1.139 | 0.861 | -0.655 | 0.01 | 0.047 | 0.047 | 0 | 0.48 | 0.6 | 0.55 | 0.7 | 0.58 | 0.51 |
| 1 | 152924650 | rs12734348 | -0.012 | -1.14 | 0.636 | -0.695 | -0.141 | 0.172 | 0.172 | 0 | 0.4 | 0.02 | 0.29 | 0.13 | 0.56 | 0.28 |
| 1 | 152926691 | rs10494281 | 1.234 | -0.84 | 0.686 | -0.874 | -0.021 | 0.061 | 0.061 | 0 | 0.12 | 0.02 | 0.08 | 0 | 0.14 | 0.05 |
| 1 | 152930426 | rs6690549 | -1.426 | -1.023 | 0.799 | -0.876 | -0.006 | 0.072 | 0.072 | 0 | 0.55 | 0.82 | 0.5 | 0.65 | 0.58 | 0.51 |
| 1 | 152931207 | rs7546648 | -0.315 | -0.947 | 0.804 | -0.814 | 0.007 | -0.012 | -0.012 | 0 | 0.4 | 0.53 | 0.4 | 0.62 | 0.44 | 0.46 |
| 1 | 152933960 | rs6587715 | -2.175 | -0.57 | 0.169 | -0.93 | -0.07 | 0.027 | 0.027 | 0 | 0.87 | 0.99 | 0.82 | 0.87 | 0.82 | 0.86 |
| 1 | 152937401 | rs4845501 | -1.196 | -0.6 | 1.164 | -0.651 | 0.001 | 0.002 | 0.002 | 0 | 0.52 | 0.58 | 0.49 | 0.65 | 0.58 | 0.51 |
| 1 | 152941345 | rs883810 | -0.786 | -0.645 | 1.265 | -0.66 | -0.009 | 0.035 | 0.035 | 0 | 0.44 | 0.29 | 0.45 | 0.61 | 0.55 | 0.48 |
| 1 | 152944594 | rs3170863 | -1.453 | -0.875 | 1.554 | -0.689 | 0 | 0.076 | 0.076 | 0 | 0.54 | 0.83 | 0.5 | 0.63 | 0.57 | 0.49 |
| 1 | 152952220 | rs16834835 | 1.317 | -0.682 | 1.764 | -0.557 | -0.015 | -0.026 | -0.026 | 0 | 0.07 | 0.25 | 0.03 | 0 | 0.01 | 0.01 |
| 1 | 152954759 | rs7541720 | 1.106 | -0.739 | 1.595 | -0.647 | -0.01 | -0.036 | -0.036 | 2.215 | 0.09 | 0.11 | 0.04 | 0 | 0.05 | 0.05 |
| 1 | 152957439 | rs1611759 | -1.088 | -0.796 | 1.569 | -0.744 | -0.002 | 0.09 | 0.09 | 0 | 0.49 | 0.79 | 0.47 | 0.63 | 0.54 | 0.51 |
| 1 | 152959939 | rs4845503 | -0.065 | -0.835 | 1.652 | -0.73 | -0.067 | -0.006 | -0.006 | 0 | 0.35 | 0.16 | 0.39 | 0.6 | 0.48 | 0.46 |
| 1 | 152963468 | rs1933383 | -0.965 | -0.782 | 1.409 | -0.701 | 0.003 | 0.062 | 0.062 | 0 | 0.48 | 0.72 | 0.46 | 0.63 | 0.53 | 0.51 |
| 1 | 152970233 | rs1413849 | -0.744 | -0.738 | 0.999 | -0.657 | -0.001 | 0.006 | 0.006 | 0 | 0.46 | 0.57 | 0.44 | 0.63 | 0.52 | 0.51 |
| 1 | 152973690 | rs1415959 | -0.819 | -0.751 | 1.137 | -0.684 | 0 | -0.005 | -0.005 | 0 | 0.45 | 0.51 | 0.43 | 0.63 | 0.53 | 0.51 |
| 1 | 152974119 | rs6671414 | -0.819 | -0.751 | 1.137 | -0.684 | 0 | -0.005 | -0.005 | 0 | 0.45 | 0.5 | 0.43 | 0.63 | 0.53 | 0.51 |
| 1 | 152974254 | rs28924721 | 0.358 | -0.83 | 0.895 | -0.603 | -0.181 | 0.101 | 0.101 | 0 | 0.31 | 0.02 | 0.24 | 0.12 | 0.47 | 0.24 |
| 1 | 152976474 | rs3737867 | -1.048 | -0.807 | 0.922 | -0.596 | 0.001 | 0.05 | 0.05 | 0 | 0.48 | 0.69 | 0.45 | 0.63 | 0.53 | 0.51 |
| 1 | 152984397 | rs6683514 | 0.284 | -0.734 | 0.6 | -0.603 | -0.18 | 0.105 | 0.105 | 0 | 0.32 | 0.02 | 0.24 | 0.12 | 0.47 | 0.24 |
| 1 | 152989321 | rs4363385 | -1.464 | -0.854 | 0.761 | -0.651 | 0.004 | 0.054 | 0.054 | 0 | 0.55 | 0.78 | 0.51 | 0.63 | 0.59 | 0.54 |
| 1 | 153003295 | rs1933382 | -0.749 | -0.824 | 1.262 | -0.565 | -0.005 | 0.013 | 0.013 | 0 | 0.45 | 0.62 | 0.44 | 0.62 | 0.51 | 0.5 |
| 1 | 153004809 | rs3181787 | -0.672 | -0.83 | 1.209 | -0.549 | -0.002 | 0.002 | 0.002 | 0 | 0.43 | 0.44 | 0.43 | 0.59 | 0.51 | 0.5 |

| | | | | | | | | | | | | | | | | |
|---|---|---|---|---|---|---|---|---|---|---|---|---|---|---|---|---|
| 1 | 153007504 | rs407448 | -0.886 | -0.825 | 1.23 | -0.546 | -0.014 | 0.053 | 0.053 | 0 | 0.46 | 0.71 | 0.44 | 0.63 | 0.51 | 0.5 |
| 1 | 153012240 | rs1048296 | -0.674 | -0.817 | 1.209 | -0.544 | -0.003 | 0.002 | 0.002 | 0 | 0.43 | 0.44 | 0.43 | 0.59 | 0.51 | 0.5 |
| 1 | 153014741 | rs409986 | -0.886 | -0.812 | 0.914 | -0.534 | -0.014 | 0.054 | 0.054 | 0 | 0.46 | 0.71 | 0.44 | 0.63 | 0.51 | 0.5 |
| 1 | 153023650 | rs12082627 | 0.358 | -0.785 | 0.704 | -0.562 | -0.102 | 0.105 | 0.105 | 0 | 0.31 | 0.13 | 0.24 | 0.12 | 0.45 | 0.24 |
| 1 | 153024634 | rs12046394 | -0.02 | -0.936 | 0.904 | -0.346 | -0.023 | 0.309 | 0.309 | 0 | 0.13 | 0.37 | 0.19 | 0.47 | 0.06 | 0.26 |
| 1 | 153029961 | rs447187 | 1.127 | -0.893 | 0.763 | -0.473 | -0.002 | -0.02 | -0.02 | 0 | 0.09 | 0.08 | 0.04 | 0 | 0.06 | 0.05 |
| 1 | 153030365 | rs6664380 | 0.339 | -0.779 | 0.345 | -0.605 | -0.138 | 0.116 | 0.116 | 0 | 0.31 | 0.07 | 0.22 | 0.11 | 0.44 | 0.24 |
| 1 | 153037463 | rs310098 | 0.168 | -0.911 | 0.419 | -0.62 | -0.027 | 0.284 | 0.284 | 0 | 0.13 | 0.37 | 0.19 | 0.48 | 0.06 | 0.27 |
| 1 | 153041903 | rs6673356 | -0.127 | -0.944 | 0.619 | -0.637 | 0.006 | -0.022 | -0.022 | 0 | 0.35 | 0.4 | 0.26 | 0.15 | 0.44 | 0.24 |
| 1 | 153042554 | rs310103 | -1.516 | -0.915 | 0.217 | -0.648 | 0.003 | 0.051 | 0.051 | 0 | 0.56 | 0.78 | 0.51 | 0.64 | 0.59 | 0.54 |
| 1 | 153044996 | rs6686526 | 0.185 | -0.567 | 0.081 | -0.451 | -0.073 | 0.095 | 0.095 | 0 | 0.31 | 0.16 | 0.23 | 0.12 | 0.44 | 0.24 |
| 1 | 153055879 | rs310109 | -0.727 | -0.553 | 0.192 | -0.422 | -0.007 | 0.023 | 0.023 | 0 | 0.45 | 0.64 | 0.44 | 0.63 | 0.51 | 0.5 |
| 1 | 153059801 | rs310117 | -0.19 | -0.983 | 0.549 | -0.036 | -0.038 | 0.29 | 0.29 | 0 | 0.14 | 0.41 | 0.19 | 0.49 | 0.06 | 0.27 |
| 1 | 153064002 | rs6664183 | -0.288 | -1.311 | 0.054 | -1.015 | -0.014 | 0.04 | 0.04 | 0 | 0.4 | 0.3 | 0.34 | 0.18 | 0.53 | 0.29 |
| 1 | 153065287 | rs3843257 | -0.177 | -2.163 | -0.013 | -0.892 | -0.009 | -0.001 | -0.001 | 0 | 0.31 | 0.61 | 0.38 | 0.31 | 0.23 | 0.26 |
| 1 | 153079072 | rs2018193 | 0.203 | -1.671 | 0.576 | -0.375 | -0.072 | 0.125 | 0.125 | 0.239 | 0.19 | 0.33 | 0.15 | 0.43 | 0.04 | 0.27 |
| 1 | 153079521 | rs310127 | 0.567 | -1.417 | 0.29 | -0.744 | 0.006 | 0.01 | 0.01 | 0 | 0.1 | 0.05 | 0.06 | 0 | 0.08 | 0.03 |
| 1 | 153084246 | rs383582 | 0.239 | -1.413 | 0.255 | -0.785 | 0.006 | 0.019 | 0.019 | 0 | 0.11 | 0.05 | 0.07 | 0 | 0.09 | 0.05 |
| 1 | 153087661 | rs1500941 | -0.309 | -1.432 | 0.626 | -0.678 | 0.057 | 0.036 | 0.036 | 0.281 | 0.3 | 0.38 | 0.35 | 0.5 | 0.43 | 0.47 |
| 1 | 153091081 | rs11586440 | 0.213 | -1.455 | 0.443 | -0.698 | -0.172 | 0.009 | 0.009 | 0.215 | 0.25 | 0.03 | 0.22 | 0.14 | 0.42 | 0.24 |
| 1 | 153099345 | rs381115 | -0.11 | -1.62 | 0.506 | -0.867 | 0.047 | 0.038 | 0.038 | 1.807 | 0.27 | 0.19 | 0.09 | 0 | 0.13 | 0.06 |
| 1 | 153101488 | rs428913 | -1.57 | -1.566 | 0.376 | -0.758 | 0.002 | 0.009 | 0.009 | 0 | 0.59 | 0.66 | 0.48 | 0.57 | 0.56 | 0.53 |
| 1 | 153119969 | rs533437 | -0.883 | -1.866 | 0.437 | -0.711 | 0.009 | -0.003 | -0.003 | 1.174 | 0.44 | 0.21 | 0.43 | 0.5 | 0.55 | 0.5 |
| 1 | 153122310 | rs510277 | -0.864 | -1.868 | 0.436 | -0.738 | 0.008 | 0.003 | 0.003 | 0.111 | 0.43 | 0.21 | 0.43 | 0.51 | 0.55 | 0.5 |
| 1 | 153122620 | rs513555 | -1.059 | -1.889 | 0.621 | -0.943 | -0.001 | -0.003 | -0.003 | 0 | 0.47 | 0.47 | 0.5 | 0.61 | 0.55 | 0.5 |
| 1 | 153135946 | rs475182 | -0.9 | -1.85 | 0.133 | -0.924 | 0.01 | 0.024 | 0.024 | 0 | 0.46 | 0.36 | 0.49 | 0.61 | 0.55 | 0.5 |
| 1 | 153136355 | rs12068238 | 0.926 | -1.488 | -0.66 | -2.1 | -0.026 | -0.052 | -0.052 | 0 | 0.17 | 0.19 | 0.03 | 0 | 0.05 | 0.03 |
| 1 | 153140532 | rs12065199 | 1.326 | -1.401 | -0.705 | -2.058 | -0.017 | -0.04 | -0.04 | 0.041 | 0.14 | 0.16 | 0.02 | 0 | 0.04 | 0.03 |
| 1 | 153142328 | rs11585767 | -0.168 | -1.121 | -0.8 | -1.951 | -0.185 | 0.039 | 0.039 | 0 | 0.31 | 0.02 | 0.26 | 0.16 | 0.47 | 0.25 |
| 1 | 153143048 | rs493631 | -0.515 | -1.254 | -0.128 | -1.926 | -0.086 | -0.173 | -0.173 | 0 | 0.31 | 0.49 | 0.11 | 0 | 0.13 | 0.08 |
| 1 | 153146946 | rs986760 | 0.441 | 0.039 | -0.83 | -1.483 | 0.01 | 0.021 | 0.021 | 0 | 0.3 | 0.17 | 0.12 | 0 | 0.25 | 0.08 |
| 1 | 153151622 | rs530590 | -1.025 | 0.604 | -0.097 | -0.098 | -0.047 | -0.038 | -0.038 | 0.565 | 0.15 | 0.51 | 0.31 | 0.08 | 0.08 | 0.03 |
| 1 | 153163377 | rs943139 | -0.77 | 1.793 | -1.077 | 0.199 | -0.084 | -0.029 | -0.029 | 4.056 | 0.13 | 0.01 | 0.14 | 0.09 | 0.23 | 0.12 |
| 1 | 153167865 | rs12143597 | -1.131 | 0.941 | -0.13 | -0.019 | 0.029 | -0.063 | -0.063 | 0 | 0.11 | 0.32 | 0.08 | 0 | 0.08 | 0.04 |
| 1 | 153169522 | rs1831238 | -0.482 | 1.143 | -0.753 | 0.309 | -0.016 | -0.02 | -0.02 | 0 | 0.36 | 0.24 | 0.4 | 0.56 | 0.49 | 0.57 |
| 1 | 153171183 | rs4845528 | 0.08 | 0.4 | -1.311 | -0.31 | 0.015 | 0.044 | 0.044 | 0 | 0.16 | 0.05 | 0.08 | 0 | 0.12 | 0.08 |
| 1 | 153176366 | rs11576296 | -0.241 | 1.252 | -0.945 | 0.085 | 0.032 | 0.052 | 0.052 | 0 | 0.72 | 0.62 | 0.71 | 0.49 | 0.64 | 0.49 |
| 1 | 153177852 | rs7534334 | 0.653 | 0.27 | -1.96 | -0.146 | -0.029 | 0.11 | 0.11 | 0 | 0.34 | 0.03 | 0.2 | 0.13 | 0.34 | 0.26 |
| 1 | 153179923 | rs12145115 | 0.116 | 0.987 | -1.937 | 0.922 | -0.092 | -0.052 | -0.052 | 0.342 | 0.77 | 0.92 | 0.7 | 0.72 | 0.51 | 0.68 |
| 1 | 153180097 | rs7555477 | 0.634 | -0.229 | -1.888 | 0.304 | -0.008 | 0.044 | 0.044 | 0.043 | 0.32 | 0.16 | 0.44 | 0.14 | 0.33 | 0.2 |
| 1 | 153182979 | rs11205222 | 0.688 | -0.33 | -1.789 | -0.075 | 0.008 | 0.04 | 0.04 | 3.293 | 0.25 | 0.18 | 0.13 | 0.05 | 0.22 | 0.08 |
| 1 | 153192214 | rs1360100 | -0.123 | 0.638 | -1.827 | -0.461 | -0.003 | 0.057 | 0.057 | 0 | 0.45 | 0.39 | 0.31 | 0.16 | 0.5 | 0.24 |
| 1 | 153200839 | rs737302 | 0.654 | 0.08 | -1.748 | -0.243 | 0.056 | 0.044 | 0.044 | 2.289 | 0.51 | 0.31 | 0.5 | 0.35 | 0.39 | 0.48 |
| 1 | 153200987 | rs12081567 | 0.972 | 0.23 | -1.758 | -0.331 | -0.016 | 0.012 | 0.012 | 0.708 | 0.17 | 0.13 | 0.15 | 0.07 | 0.26 | 0.16 |
| 1 | 153202019 | rs12723408 | 0.963 | 0.224 | -1.759 | -0.333 | -0.02 | 0.018 | 0.018 | 0.751 | 0.17 | 0.12 | 0.15 | 0.07 | 0.26 | 0.16 |
| 1 | 153202504 | rs1329099 | 2.058 | 0.05 | -1.75 | -0.141 | -0.001 | 0.052 | 0.052 | 3.948 | 0.25 | 0.09 | 0.13 | 0.11 | 0.22 | 0.08 |
| 1 | 153207274 | rs1410863 | -0.547 | -0.085 | -0.699 | -0.148 | 0.005 | 0.018 | 0.018 | 0 | 0.72 | 0.7 | 0.66 | 0.43 | 0.65 | 0.62 |
| 1 | 153207416 | rs4517302 | 0.724 | -0.374 | -0.784 | -0.206 | 0.017 | -0.003 | -0.003 | 1.793 | 0.26 | 0.27 | 0.37 | 0.26 | 0.16 | 0.39 |
| 1 | 153209588 | rs10888541 | 0.403 | -0.128 | -1.057 | -0.294 | 0.035 | -0.031 | -0.031 | 0 | 0.29 | 0.13 | 0.43 | 0.28 | 0.24 | 0.41 |
| 1 | 153210760 | rs12122588 | 2.011 | -0.217 | -0.926 | -0.205 | -0.007 | -0.024 | -0.024 | 0 | 0.28 | 0.3 | 0.13 | 0.09 | 0.22 | 0.09 |
| 1 | 153210808 | rs4845342 | 0.715 | -0.069 | -1.178 | 0.028 | 0.058 | -0.025 | -0.025 | 0 | 0.25 | 0.13 | 0.36 | 0.26 | 0.16 | 0.39 |
| 1 | 153214867 | rs7523883 | -0.551 | -0.036 | -0.877 | 0.176 | -0.001 | 0.013 | 0.013 | 0 | 0.68 | 0.6 | 0.66 | 0.53 | 0.66 | 0.67 |
| 1 | 153217977 | rs1926234 | 0.287 | -0.122 | -0.765 | 0.009 | 0.008 | -0.002 | -0.002 | 0 | 0.55 | 0.65 | 0.56 | 0.48 | 0.43 | 0.48 |
| 1 | 153220716 | rs10494296 | 1.767 | -0.175 | -0.77 | 0.013 | 0 | -0.031 | -0.031 | 0 | 0.26 | 0.38 | 0.15 | 0.13 | 0.23 | 0.06 |
| 1 | 153225886 | rs7518712 | 1.115 | -0.037 | -0.753 | -0.342 | 0.03 | -0.04 | -0.04 | 0 | 0.19 | 0.25 | 0.24 | 0.07 | 0.29 | 0.17 |
| 1 | 153226983 | rs873233 | 0.158 | -0.188 | -0.848 | -0.304 | 0.007 | -0.001 | -0.001 | 0 | 0.55 | 0.61 | 0.56 | 0.42 | 0.43 | 0.49 |
| 1 | 153227177 | rs873234 | 0.491 | -0.252 | -0.919 | -0.321 | 0.003 | 0 | 0 | 0 | 0.49 | 0.47 | 0.51 | 0.42 | 0.38 | 0.46 |
| 1 | 153230954 | rs11589914 | 1.835 | -0.261 | -0.894 | -0.335 | 0.001 | -0.08 | -0.08 | 0 | 0.25 | 0.39 | 0.15 | 0.07 | 0.23 | 0.06 |
| 1 | 153231876 | rs943967 | -1.428 | -0.136 | -0.749 | -0.12 | -0.005 | -0.053 | -0.053 | 0 | 0.74 | 0.84 | 0.81 | 0.48 | 0.72 | 0.66 |
| 1 | 153242088 | rs10888547 | 0.488 | 1.13 | -0.272 | -0.614 | 0.004 | -0.017 | -0.017 | 1.076 | 0.17 | 0.21 | 0.19 | 0.06 | 0.2 | 0.13 |
| 1 | 153245082 | rs821405 | -0.872 | 0.871 | -0.363 | -0.424 | -0.02 | -0.013 | -0.013 | 0 | 0.93 | 0.99 | 0.93 | 0.9 | 0.87 | 0.8 |
| 1 | 153245625 | rs10888548 | 0.086 | 0.916 | 0.004 | -0.839 | 0.011 | -0.034 | -0.034 | 0.24 | 0.35 | 0.37 | 0.39 | 0.13 | 0.5 | 0.22 |
| 1 | 153249072 | rs10888549 | -0.81 | 0.825 | 0.02 | -0.872 | 0.044 | 0.164 | 0.164 | 0 | 0.48 | 0.79 | 0.46 | 0.16 | 0.57 | 0.25 |
| 1 | 153253740 | rs4345763 | 0.776 | 0.547 | 0.209 | -0.848 | -0.019 | -0.088 | -0.088 | 0.944 | 0.13 | 0.4 | 0.06 | 0 | 0.07 | 0.02 |
| 1 | 153265527 | rs6587735 | -0.293 | 1.203 | -0.08 | -0.454 | 0.013 | -0.039 | -0.039 | 0 | 0.35 | 0.52 | 0.36 | 0.15 | 0.41 | 0.19 |
| 1 | 153271524 | rs821431 | 0.588 | 1.079 | 0.135 | 0.049 | -0.001 | -0.003 | -0.003 | 0 | 0.61 | 0.57 | 0.52 | 0.71 | 0.4 | 0.57 |
| 1 | 153275977 | rs821427 | 0.691 | 0.845 | 0.11 | -0.092 | 0.062 | 0.045 | 0.045 | 0.334 | 0.5 | 0.65 | 0.49 | 0.69 | 0.36 | 0.54 |
| 1 | 153277423 | rs843971 | 0.851 | 0.932 | 0.235 | -0.036 | 0.018 | 0.006 | 0.006 | 0.387 | 0.48 | 0.39 | 0.46 | 0.58 | 0.4 | 0.55 |
| 1 | 153283179 | rs3006473 | -0.164 | 1.327 | -0.304 | 0.02 | -0.005 | 0.002 | 0.002 | 0 | 0.31 | 0.34 | 0.37 | 0.19 | 0.47 | 0.27 |

| | | | | | | | | | | | | | | | | |
|---|---|---|---|---|---|---|---|---|---|---|---|---|---|---|---|---|
| 1 | 153283495 | rs2987760 | -0.164 | 1.348 | -0.304 | 0.02 | -0.004 | 0.002 | 0.002 | 0 | 0.31 | 0.34 | 0.37 | 0.19 | 0.47 | 0.27 |
| 1 | 153294653 | rs2916229 | -0.277 | 1.114 | -0.234 | 0.049 | -0.03 | 0.026 | 0.026 | 0 | 0.33 | 0.29 | 0.38 | 0.19 | 0.51 | 0.3 |
| 1 | 153296027 | rs821408 | 1.124 | 1.081 | -0.288 | 0.06 | -0.013 | -0.001 | -0.001 | 0 | 0.46 | 0.47 | 0.48 | 0.57 | 0.36 | 0.54 |
| 1 | 153302827 | rs2570440 | -0.536 | 1.122 | -0.279 | 0.08 | 0.002 | -0.003 | -0.003 | 0 | 0.79 | 0.84 | 0.87 | 0.76 | 0.87 | 0.85 |
| 1 | 153302887 | rs10888551 | -0.31 | 1.142 | -0.385 | 0.09 | -0.041 | 0.037 | 0.037 | 0 | 0.33 | 0.27 | 0.38 | 0.19 | 0.51 | 0.3 |
| 1 | 153303930 | rs1655310 | 1.045 | 1.155 | -0.389 | 0.093 | 0.019 | -0.001 | -0.001 | 0 | 0.45 | 0.37 | 0.47 | 0.57 | 0.36 | 0.54 |
| 1 | 153314153 | rs3006448 | 0.019 | 1.038 | -0.196 | 0.867 | -0.009 | -0.022 | -0.022 | 0.06 | 0.81 | 0.75 | 0.87 | 0.91 | 0.88 | 0.87 |
| 1 | 153320372 | rs3006458 | 0.448 | 1.151 | 0.069 | 1.27 | -0.019 | 0.027 | 0.027 | 0 | 0.78 | 0.7 | 0.86 | 0.74 | 0.87 | 0.85 |
| 1 | 153322574 | rs3753460 | -0.529 | 1.235 | 0.069 | 1.149 | 0.016 | 0.008 | 0.008 | 0 | 0.15 | 0.25 | 0.23 | 0.19 | 0.23 | 0.24 |
| 1 | 153325156 | rs11205275 | -0.585 | 0.459 | -0.415 | 0.887 | -0.032 | 0.017 | 0.017 | 0 | 0.2 | 0.3 | 0.12 | 0.16 | 0.08 | 0.13 |
| 1 | 153328347 | rs2916196 | 0.719 | 1.558 | -0.806 | 0.993 | -0.001 | 0.003 | 0.003 | 0 | 0.91 | 0.95 | 0.87 | 0.8 | 0.88 | 0.87 |
| 1 | 153328571 | rs6691934 | -0.807 | 1.466 | -0.786 | 0.487 | 0.007 | -0.002 | -0.002 | 0 | 0.12 | 0.17 | 0.18 | 0.12 | 0.12 | 0.21 |
| 1 | 153329071 | rs3014866 | -0.298 | 1.16 | -1.169 | 0.579 | 0.018 | 0.08 | 0.08 | 0 | 0.5 | 0.72 | 0.56 | 0.6 | 0.54 | 0.37 |
| 1 | 153335384 | rs1560832 | -0.738 | 1.373 | -0.492 | 0.717 | 0.009 | 0.012 | 0.012 | 0 | 0.31 | 0.31 | 0.4 | 0.37 | 0.31 | 0.23 |
| 1 | 153338751 | rs3014875 | -0.664 | 1.083 | -0.132 | 0.03 | 0.003 | -0.023 | -0.023 | 7.316 | 0.31 | 0.54 | 0.28 | 0.12 | 0.34 | 0.17 |
| 1 | 153344636 | rs3006475 | -1.844 | 0.552 | 0.502 | 0.459 | -0.008 | -0.001 | -0.001 | 0.273 | 0.11 | 0.21 | 0.12 | 0.09 | 0.11 | 0.1 |
| 1 | 153346263 | rs4772 | -1.536 | 0.549 | 0.558 | 0.458 | 0 | 0 | 0 | 0.138 | 0.1 | 0.11 | 0.11 | 0.09 | 0.11 | 0.1 |
| 1 | 153354406 | rs12035710 | -1.293 | 0.549 | 0.541 | 0.457 | -0.015 | -0.022 | -0.022 | 0 | 0.09 | 0.01 | 0.1 | 0.09 | 0.1 | 0.1 |
| 1 | 153356513 | rs3006483 | -1.703 | 0.549 | 0.57 | 0.458 | -0.004 | -0.001 | -0.001 | 0 | 0.11 | 0.15 | 0.11 | 0.09 | 0.11 | 0.1 |
| 1 | 153362507 | rs3006488 | -2.071 | 0.469 | 0.97 | 0.42 | -0.006 | 0.005 | 0.005 | 1.416 | 0.12 | 0.34 | 0.13 | 0.09 | 0.11 | 0.1 |
| 1 | 153367113 | rs4845346 | -2.038 | 0.471 | 1.018 | 0.421 | -0.006 | 0.005 | 0.005 | 1.529 | 0.12 | 0.33 | 0.13 | 0.09 | 0.11 | 0.11 |
| 1 | 153377503 | rs11580467 | -1.809 | 0.552 | 0.798 | 0.444 | -0.005 | 0 | 0 | 0.44 | 0.12 | 0.25 | 0.12 | 0.09 | 0.11 | 0.1 |
| 1 | 153393554 | rs3014818 | -2.139 | 0.221 | 1.241 | 0.535 | -0.035 | 0.058 | 0.058 | 0.029 | 0.15 | 0.48 | 0.17 | 0.14 | 0.09 | 0.08 |
| 1 | 153401588 | rs3014824 | -0.066 | 0.677 | 0.484 | 0.68 | -0.075 | -0.046 | -0.046 | 0.612 | 0.78 | 0.96 | 0.81 | 0.75 | 0.72 | 0.63 |
| 1 | 153404808 | rs16835308 | -1.004 | 0.056 | 0.719 | 0.685 | 0.007 | 0.088 | 0.088 | 0.021 | 0.08 | 0.18 | 0.13 | 0.14 | 0.06 | 0.08 |
| 1 | 153407572 | rs12043556 | -1.368 | -0.023 | 0.722 | 0.694 | -0.001 | 0.062 | 0.062 | 0.234 | 0.08 | 0.14 | 0.13 | 0.13 | 0.06 | 0.07 |
| 1 | 153408831 | rs2026604 | 1.53 | 0.185 | 1.065 | 0.624 | -0.021 | 0.015 | 0.015 | 0 | 0.63 | 0.51 | 0.71 | 0.61 | 0.74 | 0.59 |
| 1 | 153420475 | rs3006430 | 0.362 | 0.007 | 0.995 | 0.613 | 0.02 | -0.027 | -0.027 | 0 | 0.77 | 0.96 | 0.87 | 0.77 | 0.83 | 0.67 |
| 1 | 153433696 | rs11205297 | -1.501 | -0.926 | 0.872 | 0.35 | 0 | 0.073 | 0.073 | 1.065 | 0.1 | 0.2 | 0.14 | 0.16 | 0.09 | 0.08 |
| 1 | 153440689 | rs2986208 | 1.187 | -1.014 | 0.837 | 0.271 | -0.03 | 0.036 | 0.036 | 0 | 0.67 | 0.57 | 0.73 | 0.61 | 0.78 | 0.6 |
| 1 | 153441260 | rs1998844 | -0.223 | -0.963 | 0.752 | 0.296 | 0.008 | -0.036 | -0.036 | 0 | 0.8 | 0.96 | 0.89 | 0.77 | 0.86 | 0.68 |
| 1 | 153450505 | rs10888567 | 1.457 | -1.389 | 0.96 | 0.141 | -0.035 | 0.012 | 0.012 | 0 | 0.62 | 0.59 | 0.73 | 0.61 | 0.77 | 0.57 |
| 1 | 153456349 | rs12025638 | -1.695 | -0.898 | 0.968 | 0.182 | -0.005 | 0.014 | 0.014 | 1.036 | 0.11 | 0.12 | 0.14 | 0.16 | 0.09 | 0.13 |
| 1 | 153466704 | rs6672390 | 1.297 | -1.069 | 1.226 | 0.115 | -0.011 | -0.038 | -0.038 | 7.949 | 0.8 | 0.67 | 0.88 | 0.92 | 0.88 | 0.91 |
| 1 | 153468106 | rs16835366 | 1.607 | -1.231 | 1.546 | 0.199 | -0.037 | -0.048 | -0.048 | 0 | 0.77 | 0.5 | 0.87 | 0.9 | 0.88 | 0.91 |
| 1 | 153474243 | rs10494299 | -1.048 | -1.508 | 1.31 | 0.213 | 0.02 | -0.028 | -0.028 | 0 | 0.12 | 0 | 0.08 | 0.11 | 0.09 | 0.11 |
| 1 | 153492017 | rs4845554 | 1.617 | -1.194 | 1.782 | 0.135 | -0.011 | -0.031 | -0.031 | 0 | 0.81 | 0.49 | 0.88 | 0.9 | 0.9 | 0.93 |
| 1 | 153493800 | rs4272569 | 0.689 | -1.085 | 1.54 | 0.187 | 0.001 | -0.002 | -0.002 | 0 | 0.86 | 0.85 | 0.91 | 0.9 | 0.9 | 0.93 |
| 1 | 153495244 | rs6684970 | 1.355 | -1.066 | 1.516 | 0.196 | 0 | -0.015 | -0.015 | 0 | 0.83 | 0.63 | 0.89 | 0.9 | 0.9 | 0.93 |
| 1 | 153505013 | rs585215 | -1.186 | -1.153 | 1.408 | 0.225 | -0.001 | -0.016 | -0.016 | 0 | 0.15 | 0.29 | 0.11 | 0.07 | 0.09 | 0.08 |
| 1 | 153508996 | rs2480695 | -1.313 | -1.189 | 1.452 | 0.223 | -0.001 | -0.015 | -0.015 | 0 | 0.15 | 0.32 | 0.12 | 0.07 | 0.09 | 0.08 |
| 1 | 153509890 | rs3795393 | -0.915 | -1.624 | 1.166 | 0.719 | 0.027 | -0.067 | -0.067 | 0.455 | 0.1 | 0 | 0.13 | 0.24 | 0.08 | 0.12 |
| 1 | 153517172 | rs1050923 | -1.249 | -1.352 | 1.849 | 0.143 | -0.023 | -0.058 | -0.058 | 0 | 0.14 | 0.3 | 0.09 | 0.01 | 0.05 | 0.06 |
| 1 | 153517519 | rs11488669 | -1.7 | -1.351 | 1.736 | 0.143 | -0.046 | -0.095 | -0.095 | 0 | 0.17 | 0.48 | 0.11 | 0.01 | 0.05 | 0.06 |
| 1 | 153522350 | rs3762268 | -1.522 | -1.361 | 1.691 | 0.176 | -0.04 | -0.073 | -0.073 | 2.942 | 0.16 | 0.42 | 0.1 | 0.03 | 0.05 | 0.05 |
| 1 | 153533898 | rs8401 | -1.515 | -1.324 | 1.474 | 0.13 | -0.041 | -0.092 | -0.092 | 0 | 0.16 | 0.57 | 0.09 | 0.01 | 0.05 | 0.05 |
| 1 | 153534027 | rs1047325 | -1.247 | -1.239 | 1.402 | 0.171 | -0.02 | -0.086 | -0.086 | 0 | 0.14 | 0.45 | 0.08 | 0.01 | 0.05 | 0.05 |
| 1 | 153540573 | rs3762270 | -0.868 | -0.99 | 0.232 | 0.808 | 0.01 | -0.05 | -0.05 | 0.591 | 0.08 | 0 | 0.05 | 0.26 | 0.06 | 0.11 |
| 1 | 153557122 | rs28463744 | -1.267 | -0.887 | 0.444 | 0.226 | -0.01 | -0.014 | -0.014 | 1.732 | 0.11 | 0.14 | 0.07 | 0.05 | 0.03 | 0.05 |
| 1 | 153560040 | rs28857273 | -1.917 | -0.988 | 0.812 | 0.041 | -0.047 | -0.086 | -0.086 | 0 | 0.24 | 0.54 | 0.13 | 0.08 | 0.1 | 0.15 |
| 1 | 153583066 | rs11497739 | 0.038 | -0.312 | -0.23 | 0.041 | -0.002 | 0 | 0 | 0.021 | 0.66 | 0.64 | 0.68 | 0.68 | 0.51 | 0.62 |
| 1 | 153587308 | rs11548104 | -0.054 | -0.344 | -0.455 | 0.047 | -0.052 | -0.01 | -0.01 | 0 | 0.67 | 0.77 | 0.7 | 0.68 | 0.51 | 0.62 |
| 1 | 153588843 | rs4072425 | 0.731 | -1.285 | -1.115 | -0.598 | -0.074 | 0.019 | 0.019 | 0 | 0.45 | 0.08 | 0.6 | 0.67 | 0.46 | 0.47 |
| 1 | 153590254 | rs9330298 | 0.798 | -1.28 | -1.2 | -0.613 | -0.113 | -0.017 | -0.017 | 0 | 0.44 | 0.03 | 0.59 | 0.67 | 0.46 | 0.47 |
| 1 | 153595381 | rs28416529 | -0.461 | -1.305 | -1.063 | -0.628 | -0.054 | -0.01 | -0.01 | 0 | 0.67 | 0.77 | 0.7 | 0.68 | 0.51 | 0.62 |
| 1 | 153600778 | rs3806233 | 0.769 | -1.488 | -0.895 | -0.725 | -0.011 | 0.056 | 0.056 | 0 | 0.46 | 0.14 | 0.6 | 0.68 | 0.44 | 0.51 |
| 1 | 153604519 | rs3818632 | -0.161 | -1.366 | 0.144 | -0.91 | 0.026 | 0.001 | 0.001 | 0 | 0.66 | 0.56 | 0.7 | 0.69 | 0.52 | 0.67 |
| 1 | 153605909 | rs28380060 | -0.528 | -1.395 | 0.647 | -0.924 | -0.027 | -0.008 | -0.008 | 0 | 0.68 | 0.78 | 0.72 | 0.69 | 0.53 | 0.67 |
| 1 | 153606387 | rs1888115 | -0.305 | -1.489 | 0.565 | -0.969 | 0.035 | 0.006 | 0.006 | 0 | 0.67 | 0.56 | 0.7 | 0.69 | 0.53 | 0.67 |
| 1 | 153612053 | rs2274740 | -0.063 | -1.483 | 0.585 | -0.967 | 0.029 | 0.002 | 0.002 | 0 | 0.66 | 0.56 | 0.7 | 0.69 | 0.52 | 0.67 |
| 1 | 153612133 | rs2274739 | -0.063 | -1.483 | 0.585 | -0.967 | 0.029 | 0.002 | 0.002 | 0 | 0.66 | 0.56 | 0.7 | 0.69 | 0.52 | 0.67 |
| 1 | 153613545 | rs12564925 | -0.063 | -1.483 | 0.585 | -0.967 | 0.029 | 0.002 | 0.002 | 0 | 0.66 | 0.56 | 0.7 | 0.69 | 0.52 | 0.67 |
| 1 | 153613783 | rs10797056 | -0.225 | -1.485 | 0.623 | -0.967 | 0.024 | 0.003 | 0.003 | 0 | 0.67 | 0.61 | 0.71 | 0.69 | 0.53 | 0.67 |
| 1 | 153614712 | rs7549308 | 0.196 | -1.76 | 0.927 | -1.114 | 0.008 | -0.013 | -0.013 | 0 | 0.16 | 0.17 | 0.05 | 0 | 0.05 | 0.09 |
| 1 | 153618766 | rs7934 | -0.299 | -1.695 | 0.854 | -1.062 | 0.024 | 0.004 | 0.004 | 0 | 0.67 | 0.61 | 0.71 | 0.69 | 0.53 | 0.67 |
| 1 | 153629026 | rs4845563 | 1.505 | -1.331 | 0.774 | -1.337 | 0.001 | 0.036 | 0.036 | 0 | 0.45 | 0.23 | 0.61 | 0.69 | 0.44 | 0.51 |
| 1 | 153631071 | rs3806234 | 1.523 | -1.001 | 0.757 | -1.308 | 0.018 | 0.03 | 0.03 | 0 | 0.44 | 0.23 | 0.59 | 0.68 | 0.39 | 0.5 |
| 1 | 153634058 | rs7345 | -0.224 | -1.105 | 0.852 | -1.306 | -0.031 | -0.009 | -0.009 | 0 | 0.69 | 0.78 | 0.71 | 0.7 | 0.52 | 0.66 |
| 1 | 153635425 | rs3795396 | -0.129 | -1.103 | 1.036 | -1.305 | -0.003 | -0.001 | -0.001 | 0 | 0.68 | 0.68 | 0.71 | 0.68 | 0.52 | 0.66 |

| | | | | | | | | | | | | | | | | |
|---|---|---|---|---|---|---|---|---|---|---|---|---|---|---|---|---|
| 1 | 153637404 | rs6427671 | -0.405 | -0.317 | 0.661 | -1.302 | -0.08 | -0.054 | -0.054 | 0.047 | 0.88 | 0.99 | 0.9 | 0.84 | 0.71 | 0.84 |
| 1 | 153640827 | rs7536700 | -0.476 | -0.668 | 0.876 | -1.275 | -0.041 | -0.091 | -0.091 | 2.904 | 0.22 | 0.35 | 0.09 | 0.01 | 0.08 | 0.15 |
| 1 | 153642909 | rs11265625 | 1.807 | -0.62 | 0.678 | -1.221 | -0.004 | 0.031 | 0.031 | 0 | 0.44 | 0.2 | 0.61 | 0.69 | 0.44 | 0.51 |
| 1 | 153656650 | rs10082235 | -0.943 | -0.609 | 0.836 | -1.249 | -0.071 | -0.142 | -0.142 | 0.175 | 0.24 | 0.47 | 0.1 | 0.01 | 0.07 | 0.15 |
| 1 | 153660989 | rs7533987 | -0.286 | -0.375 | 0.391 | -1.492 | 0.002 | -0.027 | -0.027 | 0 | 0.2 | 0.24 | 0.06 | 0 | 0.07 | 0.09 |
| 1 | 153678097 | rs6427005 | 0.941 | -0.508 | 0.255 | -1.363 | 0.004 | -0.017 | -0.017 | 2.534 | 0.59 | 0.47 | 0.71 | 0.71 | 0.52 | 0.56 |
| 1 | 153690013 | rs7550138 | -1.094 | -0.624 | -0.282 | -0.628 | -0.018 | -0.018 | -0.018 | 0 | 0.3 | 0.25 | 0.26 | 0.53 | 0.45 | 0.46 |
| 1 | 153696665 | rs7552428 | -1.1 | -0.557 | -0.273 | -0.116 | -0.017 | 0.001 | 0.001 | 1.45 | 0.3 | 0.25 | 0.37 | 0.68 | 0.46 | 0.47 |
| 1 | 153704034 | rs4845568 | -1.9 | -0.754 | -0.187 | -0.217 | 0.004 | 0.001 | 0.001 | 0 | 0.93 | 0.94 | 0.95 | 1 | 0.95 | 0.96 |
| 1 | 153708633 | rs11264336 | -0.309 | -0.824 | 0.084 | -0.381 | -0.05 | -0.114 | -0.114 | 0 | 0.23 | 0.36 | 0.06 | 0 | 0.07 | 0.07 |
| 1 | 153714252 | rs12041194 | -0.628 | -0.514 | -0.067 | -0.353 | -0.016 | -0.019 | -0.019 | 0.055 | 0.24 | 0.19 | 0.23 | 0.53 | 0.39 | 0.42 |
| 1 | 153718746 | rs9426902 | -1.656 | -0.65 | -0.221 | -0.37 | -0.017 | -0.014 | -0.014 | 0.564 | 0.4 | 0.69 | 0.28 | 0.54 | 0.43 | 0.51 |
| 1 | 153720690 | rs6427268 | -2.742 | -0.68 | -0.403 | -0.462 | -0.023 | -0.002 | -0.002 | 0 | 0.55 | 0.83 | 0.34 | 0.56 | 0.53 | 0.55 |
| 1 | 153723037 | rs12097583 | -0.26 | -0.497 | -0.364 | -0.707 | 0.018 | 0.002 | 0.002 | 0.714 | 0.21 | 0.16 | 0.05 | 0 | 0.07 | 0.07 |
| 1 | 153743234 | rs6675978 | -1.299 | 0.187 | -0.446 | -0.386 | 0.017 | 0.001 | 0.001 | 0.088 | 0.26 | 0.3 | 0.36 | 0.7 | 0.43 | 0.45 |
| 1 | 153743317 | rs4345834 | 1.023 | 0.038 | -0.947 | -0.256 | 0.006 | 0.017 | 0.017 | 0 | 0.56 | 0.36 | 0.48 | 0.22 | 0.42 | 0.44 |
| 1 | 153753079 | rs12030242 | -0.307 | 0.319 | -1.207 | 0.504 | 0.008 | -0.061 | -0.061 | 2.26 | 0.14 | 0.07 | 0.12 | 0.47 | 0.19 | 0.21 |
| 1 | 153753840 | rs3922806 | 0.26 | 0.395 | -0.742 | -0.57 | -0.008 | -0.024 | -0.024 | 2.02 | 0.81 | 0.63 | 0.77 | 0.83 | 0.86 | 0.83 |
| 1 | 153756083 | rs6665637 | 0.23 | 1.222 | -0.925 | -1.07 | 0.017 | -0.059 | -0.059 | 0 | 0.23 | 0.23 | 0.18 | 0.02 | 0.27 | 0.12 |
| 1 | 153765399 | rs6690619 | -1.055 | 0.835 | -1.158 | -1.181 | 0.007 | 0.085 | 0.085 | 0 | 0.45 | 0.25 | 0.33 | 0.07 | 0.51 | 0.34 |
| 1 | 153773429 | rs16835600 | 0.199 | 0.678 | -0.894 | -0.991 | -0.006 | 0.035 | 0.035 | 0 | 0.07 | 0.17 | 0.15 | 0.17 | 0.03 | 0.19 |
| 1 | 153776666 | rs4341393 | -0.906 | 0.655 | -0.911 | -1.089 | 0.003 | 0.081 | 0.081 | 0 | 0.45 | 0.26 | 0.32 | 0.07 | 0.51 | 0.34 |
| 1 | 153777848 | rs1139620 | -0.906 | 0.675 | -0.911 | -1.08 | 0.002 | 0.082 | 0.082 | 0 | 0.45 | 0.26 | 0.32 | 0.08 | 0.51 | 0.34 |
| 1 | 153779412 | rs1127091 | -0.926 | 0.626 | -0.912 | -1.072 | 0.004 | 0.087 | 0.087 | 0 | 0.45 | 0.26 | 0.44 | 0.16 | 0.52 | 0.34 |
| 1 | 153780298 | rs9426938 | -0.909 | 0.637 | -0.909 | -1.067 | 0.003 | 0.081 | 0.081 | 0 | 0.45 | 0.26 | 0.32 | 0.07 | 0.51 | 0.34 |
| 1 | 153781888 | rs6700115 | -0.876 | 0.538 | -0.956 | -1.115 | 0.01 | 0.097 | 0.097 | 0 | 0.46 | 0.27 | 0.44 | 0.16 | 0.52 | 0.34 |
| 1 | 153805735 | rs11264446 | -0.267 | 0.628 | -0.537 | -0.986 | -0.005 | 0.074 | 0.074 | 0.035 | 0.08 | 0.3 | 0.16 | 0.17 | 0.03 | 0.18 |
| 1 | 153822673 | rs4612664 | 2.025 | 0.529 | -0.533 | -0.974 | 0.022 | 0.027 | 0.027 | 0 | 0.36 | 0.11 | 0.41 | 0.28 | 0.29 | 0.33 |
| 1 | 153849169 | rs12142130 | 0.915 | 1.34 | -0.726 | -1.138 | -0.052 | 0.047 | 0.047 | 0 | 0.11 | 0.02 | 0.13 | 0.02 | 0.26 | 0.17 |
| 1 | 153854380 | rs12043350 | 1.912 | 0.784 | -0.545 | -0.953 | 0.013 | 0.022 | 0.022 | 0 | 0.35 | 0.08 | 0.4 | 0.28 | 0.29 | 0.33 |
| 1 | 153889042 | rs17364880 | 0.855 | 0.695 | -0.635 | -1.047 | 0.015 | 0.054 | 0.054 | 0 | 0.15 | 0.01 | 0.08 | 0.05 | 0.11 | 0.13 |
| 1 | 153893023 | rs10494303 | 0.606 | 0.964 | -0.103 | -0.661 | -0.004 | -0.015 | -0.015 | 0 | 0.47 | 0.45 | 0.52 | 0.76 | 0.44 | 0.47 |
| 1 | 153901302 | rs3748848 | 1.591 | 0.962 | -0.171 | -0.605 | 0.016 | 0.021 | 0.021 | 0 | 0.35 | 0.1 | 0.41 | 0.28 | 0.29 | 0.33 |
| 1 | 153913684 | rs2297895 | 1.399 | 0.957 | -0.17 | -0.653 | -0.013 | -0.042 | -0.042 | 0 | 0.1 | 0.03 | 0.09 | 0.48 | 0.16 | 0.15 |
| 1 | 153913699 | rs1110165 | 1.641 | 0.962 | -0.144 | -0.658 | 0.019 | 0.023 | 0.023 | 0 | 0.35 | 0.12 | 0.41 | 0.28 | 0.29 | 0.33 |
| 1 | 153919756 | rs946682 | 1.552 | 0.962 | -0.147 | -0.658 | 0.016 | 0.021 | 0.021 | 0 | 0.35 | 0.1 | 0.41 | 0.28 | 0.29 | 0.33 |
| 1 | 153920286 | rs8450 | 1.552 | 0.962 | -0.147 | -0.658 | 0.016 | 0.021 | 0.021 | 0 | 0.35 | 0.1 | 0.41 | 0.28 | 0.29 | 0.33 |
| 1 | 153920564 | rs10559 | 1.552 | 0.962 | -0.147 | -0.658 | 0.016 | 0.021 | 0.021 | 0 | 0.35 | 0.1 | 0.41 | 0.28 | 0.29 | 0.33 |
| 1 | 153923276 | rs10127983 | 1.553 | 0.965 | -0.16 | -0.669 | 0.022 | 0.022 | 0.022 | 0 | 0.36 | 0.21 | 0.41 | 0.28 | 0.29 | 0.33 |
| 1 | 153927270 | rs10908558 | 1.524 | 0.963 | -0.256 | -0.657 | 0.015 | 0.02 | 0.02 | 0 | 0.35 | 0.09 | 0.4 | 0.28 | 0.29 | 0.33 |
| 1 | 153935543 | rs2072704 | 0.83 | 1.025 | -0.085 | -0.567 | -0.002 | 0.197 | 0.197 | 0 | 0.12 | 0.24 | 0.1 | 0.48 | 0.16 | 0.15 |
| 1 | 153939130 | rs11264736 | -0.105 | 0.951 | -0.173 | -0.743 | 0.007 | 0.087 | 0.087 | 0 | 0.55 | 0.74 | 0.68 | 0.93 | 0.48 | 0.66 |
| 1 | 153941514 | rs11264743 | 1.635 | 0.971 | -0.266 | -0.689 | 0.009 | 0.014 | 0.014 | 0 | 0.34 | 0.08 | 0.41 | 0.28 | 0.29 | 0.33 |
| 1 | 153943079 | rs12403956 | 1.48 | 0.994 | -0.22 | -0.678 | 0.015 | 0.016 | 0.016 | 0 | 0.35 | 0.23 | 0.42 | 0.28 | 0.29 | 0.33 |
| 1 | 153954674 | rs10908578 | 0.053 | 1.009 | -0.315 | -0.678 | 0.003 | 0.065 | 0.065 | 0 | 0.54 | 0.66 | 0.68 | 0.93 | 0.49 | 0.68 |
| 1 | 153973171 | rs11264824 | 0.767 | 0.804 | -0.312 | -0.467 | 0 | 0.155 | 0.155 | 0 | 0.12 | 0.17 | 0.1 | 0.48 | 0.16 | 0.15 |
| 1 | 153989774 | rs3008665 | 0.029 | 0.872 | -0.52 | -0.712 | 0.009 | 0.037 | 0.037 | 0 | 0.54 | 0.63 | 0.66 | 0.93 | 0.45 | 0.61 |
| 1 | 153992734 | rs12048137 | 1.564 | 0.697 | -0.731 | -0.644 | 0.001 | 0.015 | 0.015 | 0 | 0.32 | 0.17 | 0.41 | 0.28 | 0.28 | 0.33 |
| 1 | 153994647 | rs11264875 | 1.62 | 0.694 | -0.733 | -0.646 | -0.002 | 0.013 | 0.013 | 0 | 0.31 | 0.14 | 0.4 | 0.28 | 0.28 | 0.33 |
| 1 | 154012001 | rs10908627 | 1.637 | 0.676 | -0.793 | -0.646 | -0.016 | 0.008 | 0.008 | 0 | 0.31 | 0.11 | 0.4 | 0.28 | 0.28 | 0.33 |
| 1 | 154017501 | rs2841102 | 0.444 | 0.751 | -0.547 | -0.568 | 0.003 | 0.056 | 0.056 | 0 | 0.44 | 0.31 | 0.51 | 0.74 | 0.45 | 0.5 |
| 1 | 154020378 | rs6671009 | 0.943 | 1.137 | -0.265 | -0.511 | -0.022 | -0.042 | -0.042 | 0 | 0.11 | 0.04 | 0.1 | 0.45 | 0.17 | 0.17 |
| 1 | 154068196 | rs4459102 | -1.074 | 0.863 | -0.267 | -0.356 | -0.005 | 0.109 | 0.109 | 0 | 0.89 | 0.86 | 0.92 | 0.63 | 0.86 | 0.84 |
| 1 | 154091730 | rs6664278 | 1.437 | 0.774 | -0.884 | -0.47 | 0.006 | -0.06 | -0.06 | 0.339 | 0.38 | 0.11 | 0.49 | 0.44 | 0.34 | 0.37 |
| 1 | 154105016 | rs4845363 | 1.404 | 0.772 | -0.86 | -0.471 | 0.014 | -0.042 | -0.042 | 0 | 0.38 | 0.14 | 0.5 | 0.44 | 0.34 | 0.37 |
| 1 | 154116872 | rs12091348 | -0.078 | 0.694 | -0.694 | -0.696 | -0.025 | -0.028 | -0.028 | 0 | 0.12 | 0.39 | 0.1 | 0.08 | 0.04 | 0.18 |
| 1 | 154123523 | rs4521985 | 1.489 | 0.654 | -0.798 | -0.601 | -0.005 | -0.078 | -0.078 | 0 | 0.38 | 0.09 | 0.5 | 0.44 | 0.34 | 0.37 |
| 1 | 154137165 | rs12026177 | 0.855 | 0.688 | -0.535 | -0.571 | -0.003 | 0.209 | 0.209 | 0 | 0.11 | 0.25 | 0.1 | 0.33 | 0.14 | 0.14 |
| 1 | 154141908 | rs4845364 | -0.205 | 0.659 | -0.557 | -0.658 | -0.053 | 0.14 | 0.14 | 0 | 0.62 | 0.85 | 0.71 | 0.86 | 0.53 | 0.68 |
| 1 | 154149767 | rs11265238 | 0.711 | 0.374 | -0.541 | -0.552 | -0.004 | 0.223 | 0.223 | 0 | 0.12 | 0.27 | 0.1 | 0.33 | 0.14 | 0.13 |
| 1 | 154157090 | rs11265251 | 0.683 | 0.616 | -0.611 | -0.881 | 0.004 | 0 | 0 | 0 | 0.5 | 0.52 | 0.62 | 0.52 | 0.4 | 0.56 |
| 1 | 154164139 | rs3811454 | 2.106 | 0.235 | -0.955 | -0.991 | -0.024 | -0.026 | -0.026 | 0.066 | 0.11 | 0 | 0.11 | 0.13 | 0.1 | 0.07 |
| 1 | 154167116 | rs4072431 | 0.05 | 0.547 | -0.361 | -0.865 | -0.065 | 0.155 | 0.155 | 0 | 0.63 | 0.83 | 0.73 | 0.95 | 0.56 | 0.71 |
| 1 | 154179679 | rs6719 | 1.024 | 0.475 | -0.165 | -0.849 | 0.01 | -0.01 | -0.01 | 0 | 0.48 | 0.3 | 0.62 | 0.52 | 0.41 | 0.67 |
| 1 | 154187305 | rs10908751 | 0.033 | 0.484 | -0.191 | -0.856 | -0.064 | 0.159 | 0.159 | 0 | 0.63 | 0.82 | 0.74 | 0.95 | 0.57 | 0.75 |
| 1 | 154189963 | rs11265303 | 0.673 | 0.498 | -0.199 | -0.863 | 0.001 | 0.002 | 0.002 | 0 | 0.08 | 0.07 | 0.11 | 0.09 | 0.05 | 0.2 |
| 1 | 154194579 | rs3811453 | 1.487 | 0.492 | -0.205 | -0.804 | -0.023 | -0.018 | -0.018 | 0 | 0.08 | 0.01 | 0.07 | 0.42 | 0.14 | 0.07 |
| 1 | 154195753 | rs4393149 | 0.007 | 0.481 | -0.195 | -0.908 | -0.066 | 0.162 | 0.162 | 0 | 0.63 | 0.82 | 0.74 | 0.95 | 0.57 | 0.75 |
| 1 | 154197126 | rs11265310 | 0.653 | 0.552 | -0.253 | -0.893 | 0.001 | 0.002 | 0.002 | 0 | 0.08 | 0.07 | 0.11 | 0.09 | 0.05 | 0.19 |

| | | | | | | | | | | | | | | | | |
|---|---|---|---|---|---|---|---|---|---|---|---|---|---|---|---|---|
| 1 | 154209490 | rs2274993 | 1.444 | 0.521 | -0.476 | -0.876 | -0.021 | -0.012 | -0.012 | 0 | 0.08 | 0.01 | 0.08 | 0.43 | 0.15 | 0.07 |
| 1 | 154210210 | rs12026615 | 1.444 | 0.521 | -0.488 | -0.876 | -0.025 | -0.018 | -0.018 | 0 | 0.08 | 0.01 | 0.08 | 0.43 | 0.15 | 0.07 |
| 1 | 154210702 | rs3828076 | 1.674 | 0.387 | -0.409 | -0.812 | -0.067 | -0.119 | -0.119 | 0 | 0.38 | 0.03 | 0.5 | 0.43 | 0.37 | 0.48 |
| 1 | 154225344 | rs2340473 | 0.82 | 0.539 | -0.335 | -0.733 | 0.008 | -0.001 | -0.001 | 0 | 0.5 | 0.44 | 0.62 | 0.53 | 0.41 | 0.68 |
| 1 | 154238383 | rs1194596 | -1.687 | 1.628 | 0.239 | 0.074 | -0.015 | -0.019 | -0.019 | 0 | 0.39 | 0.66 | 0.35 | 0.48 | 0.55 | 0.32 |
| 1 | 154239283 | rs1212352 | -2.152 | 1.612 | 0.253 | 0.061 | 0.004 | -0.017 | -0.017 | 0 | 0.48 | 0.87 | 0.47 | 0.57 | 0.6 | 0.52 |
| 1 | 154243115 | rs1044013 | -1.775 | 2.016 | 0.414 | 0.294 | -0.015 | -0.018 | -0.018 | 0 | 0.38 | 0.66 | 0.35 | 0.48 | 0.55 | 0.3 |
| 1 | 154252860 | rs7521837 | -0.802 | 2.01 | -0.889 | -0.463 | -0.017 | 0.02 | 0.02 | 0 | 0.17 | 0.15 | 0.16 | 0.05 | 0.22 | 0.15 |
| 1 | 154270877 | rs3103309 | -1.272 | 1.494 | -0.74 | -0.52 | -0.001 | 0.008 | 0.008 | 0 | 0.33 | 0.62 | 0.3 | 0.15 | 0.35 | 0.35 |
| 1 | 154278799 | rs12724639 | -0.919 | 0.516 | -0.626 | -0.495 | 0.025 | 0.02 | 0.02 | 0 | 0.19 | 0.12 | 0.15 | 0.12 | 0.16 | 0.25 |
| 1 | 154279975 | rs11578377 | 0.01 | 0.397 | -0.161 | -0.582 | -0.004 | -0.008 | -0.008 | 0 | 0.17 | 0.17 | 0.13 | 0.12 | 0.12 | 0.12 |
| 1 | 154288680 | rs35387092 | 0.229 | 1.199 | -0.218 | -0.481 | 0.005 | 0.001 | 0.001 | 0 | 0.1 | 0.11 | 0.14 | 0.09 | 0.24 | 0.14 |
| 1 | 154296076 | rs1194610 | -0.631 | 1.01 | 0.659 | -0.514 | 0.004 | -0.045 | -0.045 | 0 | 0.24 | 0.5 | 0.27 | 0.15 | 0.23 | 0.15 |
| 1 | 154303976 | rs6702754 | -1.43 | 0.883 | 0.541 | -0.592 | 0 | 0.001 | 0.001 | 0 | 0.29 | 0.34 | 0.24 | 0.21 | 0.32 | 0.32 |
| 1 | 154307630 | rs11586379 | -1.369 | 0.829 | 0.584 | -0.491 | -0.005 | -0.005 | -0.005 | 0 | 0.38 | 0.33 | 0.31 | 0.25 | 0.29 | 0.31 |
| 1 | 154308641 | rs2988721 | 0.479 | 0.884 | 0.61 | -0.489 | 0.138 | 0.007 | 0.007 | 0 | 0.85 | 0.53 | 0.81 | 0.88 | 0.75 | 0.85 |
| 1 | 154310495 | rs767804 | 0.448 | 0.859 | 0.371 | -0.49 | 0.004 | 0.004 | 0.004 | 0 | 0.18 | 0.12 | 0.18 | 0.13 | 0.12 | 0.12 |
| 1 | 154317153 | rs1194587 | 2.181 | 0.826 | 0.313 | -0.498 | -0.053 | 0.048 | 0.048 | 0.01 | 0.47 | 0.12 | 0.49 | 0.62 | 0.45 | 0.53 |
| 1 | 154323772 | rs11589949 | -0.212 | 0.778 | 0.343 | -0.507 | 0.019 | 0.02 | 0.02 | 0 | 0.18 | 0.02 | 0.12 | 0.12 | 0.17 | 0.19 |
| 1 | 154325149 | rs2483712 | 2.169 | 0.9 | 0.335 | -0.534 | -0.045 | 0.053 | 0.053 | 0 | 0.47 | 0.12 | 0.49 | 0.63 | 0.44 | 0.52 |
| 1 | 154325302 | rs1194589 | 0.706 | 0.95 | 0.49 | -0.584 | -0.047 | -0.119 | -0.119 | 0 | 0.76 | 0.58 | 0.82 | 0.96 | 0.8 | 0.81 |
| 1 | 154326279 | rs4040747 | 0.351 | 1.174 | 0.453 | -0.397 | 0.004 | -0.001 | -0.001 | 0 | 0.18 | 0.13 | 0.18 | 0.2 | 0.13 | 0.14 |
| 1 | 154326366 | rs6688376 | -1.455 | 1.213 | 0.585 | -0.362 | 0.002 | -0.033 | -0.033 | 0 | 0.28 | 0.39 | 0.24 | 0.14 | 0.3 | 0.31 |
| 1 | 154331186 | rs4414034 | 0.576 | 1.254 | 0.601 | -0.285 | 0.024 | 0.001 | 0.001 | 0 | 0.09 | 0.14 | 0.11 | 0.1 | 0.15 | 0.13 |
| 1 | 154332870 | rs6696089 | -1.064 | 1.141 | 0.508 | -0.327 | 0 | -0.008 | -0.008 | 0 | 0.31 | 0.45 | 0.31 | 0.3 | 0.29 | 0.27 |
| 1 | 154340789 | rs12025518 | -0.642 | 1.218 | 0.422 | -0.499 | -0.014 | 0.017 | 0.017 | 0.182 | 0.18 | 0.16 | 0.17 | 0.09 | 0.25 | 0.2 |
| 1 | 154346416 | rs4406622 | -0.089 | 1.235 | 0.63 | -0.458 | 0.04 | -0.008 | -0.008 | 0.243 | 0.11 | 0.27 | 0.11 | 0.09 | 0.15 | 0.12 |
| 1 | 154346967 | rs6672627 | -0.407 | 1.21 | 0.619 | -0.47 | 0.033 | -0.013 | -0.013 | 0.242 | 0.12 | 0.27 | 0.11 | 0.09 | 0.15 | 0.12 |
| 1 | 154350589 | rs3811451 | -0.05 | 1.229 | 0.65 | -0.461 | 0.05 | -0.004 | -0.004 | 0 | 0.11 | 0.3 | 0.11 | 0.09 | 0.14 | 0.12 |
| 1 | 154356481 | rs10908835 | -1.167 | 1.145 | 0.644 | -0.527 | 0.017 | -0.025 | -0.025 | 0 | 0.32 | 0.47 | 0.36 | 0.28 | 0.39 | 0.3 |
| 1 | 154363985 | rs11582424 | -0.719 | 1.224 | 0.688 | -0.508 | 0.001 | -0.005 | -0.005 | 0 | 0.28 | 0.3 | 0.3 | 0.25 | 0.29 | 0.26 |
| 1 | 154368928 | rs952146 | -0.98 | 1.019 | 0.631 | -0.474 | 0.012 | -0.015 | -0.015 | 0.487 | 0.31 | 0.44 | 0.36 | 0.27 | 0.39 | 0.3 |
| 1 | 154369252 | rs17654071 | 1.663 | 0.924 | 0.227 | -0.542 | -0.01 | 0.117 | 0.117 | 0 | 0.5 | 0.17 | 0.49 | 0.67 | 0.47 | 0.53 |
| 1 | 154370938 | rs2054855 | -0.799 | 0.849 | 0.676 | -0.619 | -0.029 | -0.045 | -0.045 | 0 | 0.19 | 0.25 | 0.13 | 0.05 | 0.13 | 0.16 |
| 1 | 154376671 | rs3887104 | -0.572 | 0.948 | 0.122 | -0.76 | 0.003 | 0.031 | 0.031 | 0.056 | 0.19 | 0.12 | 0.12 | 0.06 | 0.16 | 0.15 |
| 1 | 154377898 | rs4845617 | -1.112 | 1.048 | 0.579 | 0.094 | 0.012 | 0.018 | 0.018 | 0 | 0.3 | 0.32 | 0.47 | 0.42 | 0.4 | 0.33 |
| 1 | 154380486 | rs6427641 | 1.16 | 1.156 | 0.027 | 0.145 | -0.013 | -0.026 | -0.026 | 0.146 | 0.61 | 0.32 | 0.49 | 0.53 | 0.57 | 0.65 |
| 1 | 154382049 | rs1386821 | -0.399 | 1.211 | -0.185 | -0.118 | -0.008 | 0.011 | 0.011 | 0.156 | 0.16 | 0.11 | 0.13 | 0.06 | 0.19 | 0.15 |
| 1 | 154400015 | rs4845618 | -0.092 | 1.023 | -0.17 | -0.035 | 0.006 | 0.012 | 0.012 | 2.155 | 0.6 | 0.47 | 0.66 | 0.44 | 0.54 | 0.48 |
| 1 | 154404406 | rs7553796 | -0.618 | 0.915 | 0.142 | -0.124 | -0.004 | -0.006 | -0.006 | 0 | 0.63 | 0.66 | 0.68 | 0.43 | 0.56 | 0.48 |
| 1 | 154407419 | rs7518199 | 0.338 | 0.859 | -0.075 | -0.469 | 0.016 | -0.005 | -0.005 | 0.011 | 0.41 | 0.19 | 0.55 | 0.33 | 0.37 | 0.3 |
| 1 | 154418879 | rs4537545 | -0.236 | 0.722 | 0.256 | -0.507 | 0.046 | 0.034 | 0.034 | 3.012 | 0.47 | 0.66 | 0.58 | 0.32 | 0.37 | 0.29 |
| 1 | 154422067 | rs4845625 | -0.575 | 0.767 | 0.15 | -0.495 | -0.024 | -0.029 | -0.029 | 0 | 0.62 | 0.71 | 0.68 | 0.43 | 0.55 | 0.42 |
| 1 | 154426264 | rs4129267 | 0.44 | 0.658 | 0.106 | -0.546 | -0.01 | -0.031 | -0.031 | 0 | 0.39 | 0.09 | 0.54 | 0.32 | 0.36 | 0.3 |
| 1 | 154430092 | rs11265618 | 0.711 | 0.413 | 0.556 | -0.583 | 0.001 | -0.043 | -0.043 | 0 | 0.2 | 0.34 | 0.13 | 0.1 | 0.19 | 0.19 |
| 1 | 154432420 | rs4329505 | 0.728 | 0.433 | 0.679 | -0.586 | 0.011 | -0.077 | -0.077 | 0 | 0.2 | 0.41 | 0.12 | 0.1 | 0.17 | 0.19 |
| 1 | 154436195 | rs4240872 | -0.506 | 0.464 | 0.551 | -0.638 | 0.002 | -0.069 | -0.069 | 0 | 0.75 | 0.43 | 0.8 | 0.81 | 0.72 | 0.78 |
| 1 | 154437896 | rs2229238 | -1.4 | 0.56 | 0.247 | -0.632 | -0.001 | 0 | 0 | 0 | 0.82 | 0.77 | 0.86 | 0.81 | 0.78 | 0.79 |
| 1 | 154438084 | rs7514452 | -1.387 | 0.611 | 0.259 | -0.627 | -0.001 | 0 | 0 | 0 | 0.82 | 0.77 | 0.86 | 0.81 | 0.78 | 0.78 |
| 1 | 154438880 | rs4072391 | -1.081 | 0.535 | 0.271 | -0.628 | 0.005 | -0.03 | -0.03 | 1.23 | 0.8 | 0.63 | 0.86 | 0.8 | 0.78 | 0.78 |
| 1 | 154451420 | rs11265622 | 0.161 | 0.464 | -0.004 | -0.812 | -0.029 | 0.159 | 0.159 | 0 | 0.58 | 0.12 | 0.71 | 0.74 | 0.59 | 0.58 |
| 1 | 154455949 | rs12568083 | 0.161 | 0.464 | -0.004 | -0.812 | -0.029 | 0.159 | 0.159 | 0 | 0.58 | 0.12 | 0.71 | 0.74 | 0.59 | 0.58 |
| 1 | 154457855 | rs4474240 | -1.553 | 0.405 | 0.083 | -0.85 | 0.02 | -0.061 | -0.061 | 0 | 0.81 | 0.6 | 0.88 | 0.91 | 0.82 | 0.81 |
| 1 | 154462360 | rs12044132 | 1.478 | 0.105 | -0.451 | -1.085 | 0.013 | -0.126 | -0.126 | 0.064 | 0.2 | 0.02 | 0.33 | 0.36 | 0.15 | 0.15 |
| 1 | 154468135 | rs10047079 | 1.053 | 0.708 | 0.196 | -1.579 | -0.008 | 0.025 | 0.025 | 0.117 | 0.19 | 0.14 | 0.15 | 0.17 | 0.23 | 0.23 |
| 1 | 154481624 | rs10752642 | -1.505 | 0.572 | 0.279 | -1.515 | 0.017 | -0.062 | -0.062 | 0 | 0.81 | 0.58 | 0.88 | 0.91 | 0.82 | 0.81 |
| 1 | 154491683 | rs6674171 | 0.962 | 0.153 | 0.318 | -1.376 | -0.008 | 0.025 | 0.025 | 0.153 | 0.19 | 0.14 | 0.15 | 0.17 | 0.23 | 0.23 |
| 1 | 154516477 | rs12750774 | 1.154 | 0.209 | 0.218 | -1.449 | -0.101 | 0.117 | 0.117 | 0 | 0.21 | 0.01 | 0.25 | 0.03 | 0.31 | 0.13 |
| 1 | 154516578 | rs3811448 | 0.4 | -0.283 | 0.828 | -0.588 | -0.007 | 0.025 | 0.025 | 0.113 | 0.19 | 0.14 | 0.15 | 0.17 | 0.23 | 0.23 |
| 1 | 154531910 | rs4074436 | 0.363 | -0.394 | 0.568 | -0.192 | 0.047 | 0.06 | 0.06 | 2.021 | 0.77 | 0.13 | 0.82 | 0.89 | 0.89 | 0.81 |
| 1 | 154538205 | rs4845652 | 0.584 | 0.024 | 0.597 | 0.053 | -0.005 | -0.005 | -0.005 | 0 | 0.15 | 0.13 | 0.12 | 0.12 | 0.1 | 0.17 |
| 1 | 154548721 | rs2072660 | 0.291 | 0.735 | 0.552 | 0.658 | 0.023 | 0.068 | 0.068 | 0.001 | 0.79 | 0.53 | 0.76 | 0.7 | 0.74 | 0.8 |
| 1 | 154554357 | rs11264222 | -0.745 | 0.512 | 0.684 | 0.629 | 0.002 | 0.006 | 0.006 | 0 | 0.28 | 0.29 | 0.21 | 0.33 | 0.33 | 0.23 |
| 1 | 154556266 | rs1127314 | 0.713 | 0.32 | 1.357 | 0.534 | 0.002 | -0.006 | -0.006 | 0 | 0.68 | 0.28 | 0.76 | 0.67 | 0.67 | 0.77 |
| 1 | 154556425 | rs1127313 | 0.849 | 0.127 | 0.392 | 0.145 | -0.037 | -0.047 | -0.047 | 0 | 0.42 | 0.21 | 0.48 | 0.52 | 0.47 | 0.38 |
| 1 | 154556663 | rs1127311 | 0.849 | 0.127 | 0.392 | 0.145 | -0.037 | -0.047 | -0.047 | 0 | 0.42 | 0.21 | 0.48 | 0.52 | 0.47 | 0.38 |
| 1 | 154561925 | rs1127309 | -0.078 | 0.093 | 0.554 | 0.126 | 0.007 | 0.02 | 0.02 | 0 | 0.28 | 0.29 | 0.21 | 0.32 | 0.32 | 0.23 |
| 1 | 154568665 | rs9426823 | 0.105 | 0.117 | 0.725 | 0.095 | 0 | 0.01 | 0.01 | 0 | 0.69 | 0.4 | 0.76 | 0.67 | 0.67 | 0.77 |
| 1 | 154568683 | rs9427097 | 1.586 | 0.418 | 0.025 | 0.24 | -0.053 | -0.05 | -0.05 | 0 | 0.16 | 0.01 | 0.14 | 0.23 | 0.18 | 0.24 |

| | | | | | | | | | | | | | | | | |
|---|---|---|---|---|---|---|---|---|---|---|---|---|---|---|---|---|
| 1 | 154573419 | rs3738032 | 0.597 | 0.332 | 0.276 | 0.234 | 0 | -0.002 | -0.002 | 0 | 0.23 | 0.21 | 0.18 | 0.13 | 0.26 | 0.18 |
| 1 | 154573967 | rs2229857 | -0.087 | 0.267 | 0.106 | 0.185 | -0.001 | 0.004 | 0.004 | 0 | 0.68 | 0.37 | 0.76 | 0.67 | 0.67 | 0.77 |
| 1 | 154582970 | rs6699825 | 0.469 | 0.315 | -0.134 | 0.244 | -0.017 | -0.024 | -0.024 | 1.372 | 0.41 | 0.21 | 0.48 | 0.51 | 0.47 | 0.38 |
| 1 | 154589628 | rs2335230 | 1.452 | 0.136 | -0.244 | 0.193 | -0.003 | 0.035 | 0.035 | 0 | 0.23 | 0.07 | 0.25 | 0.15 | 0.19 | 0.36 |
| 1 | 154594690 | rs9426830 | 0.444 | 0.323 | -0.35 | 0.266 | -0.014 | -0.019 | -0.019 | 0 | 0.42 | 0.21 | 0.5 | 0.5 | 0.47 | 0.38 |
| 1 | 154600998 | rs884617 | -0.167 | 0.29 | -0.711 | 0.174 | 0.003 | 0.002 | 0.002 | 0.032 | 0.63 | 0.61 | 0.75 | 0.66 | 0.61 | 0.72 |
| 1 | 154607914 | rs9427108 | 0.006 | 0.803 | -1.518 | 0.597 | -0.103 | -0.122 | -0.122 | 0 | 0.35 | 0.04 | 0.47 | 0.49 | 0.43 | 0.36 |
| 1 | 154626705 | rs9427114 | 0.08 | 0.782 | -1.521 | 0.593 | -0.125 | -0.119 | -0.119 | 0.193 | 0.35 | 0.04 | 0.48 | 0.49 | 0.45 | 0.37 |
| 1 | 154630498 | rs10796927 | 0.758 | 0.941 | -1.377 | 0.745 | -0.07 | -0.112 | -0.112 | 0 | 0.18 | 0.01 | 0.39 | 0.34 | 0.27 | 0.13 |
| 1 | 154633513 | rs1876304 | 1.321 | 0.762 | -1.949 | 0.462 | -0.079 | -0.108 | -0.108 | 0 | 0.21 | 0.02 | 0.21 | 0.32 | 0.23 | 0.2 |
| 1 | 154635448 | rs7534678 | 0.757 | 0.204 | -1.32 | -0.032 | -0.001 | -0.006 | -0.006 | 0 | 0.32 | 0.32 | 0.23 | 0.14 | 0.29 | 0.35 |
| 1 | 154637584 | rs9427118 | 1.19 | 0.165 | -1.76 | -0.046 | -0.067 | 0.017 | 0.017 | 2.659 | 0.2 | 0.01 | 0.15 | 0.18 | 0.21 | 0.3 |
| 1 | 154638826 | rs7556080 | -0.366 | 0.023 | -0.785 | -0.1 | 0.032 | 0.046 | 0.046 | 0.42 | 0.64 | 0.33 | 0.75 | 0.86 | 0.73 | 0.65 |
| 1 | 154639255 | rs7547072 | 0.001 | 0.313 | -0.474 | 0.089 | -0.03 | -0.03 | -0.03 | 0 | 0.42 | 0.29 | 0.53 | 0.52 | 0.47 | 0.5 |
| 1 | 154653704 | rs10737169 | 0.738 | 0.596 | 0.46 | -0.204 | 0.01 | -0.024 | -0.024 | 0 | 0.93 | 0.85 | 0.92 | 0.99 | 0.91 | 0.95 |
| 1 | 154659317 | rs954213 | 0.157 | 0.637 | 0.436 | -0.193 | 0.007 | -0.016 | -0.016 | 0 | 0.94 | 0.89 | 0.93 | 0.99 | 0.9 | 0.95 |
| 1 | 154669294 | rs2172706 | 2.089 | 0.586 | 0.836 | -0.294 | 0.002 | -0.005 | -0.005 | 0 | 0.8 | 0.21 | 0.77 | 0.89 | 0.74 | 0.87 |
| 1 | 154673755 | rs6696760 | -0.71 | 0.754 | 0.512 | -0.203 | 0.037 | 0.015 | 0.015 | 0 | 0.08 | 0.21 | 0.12 | 0.09 | 0.16 | 0.09 |
| 1 | 154678986 | rs7543897 | 0.625 | 0.687 | -0.018 | -0.182 | 0.008 | -0.024 | -0.024 | 0.006 | 0.92 | 0.83 | 0.92 | 0.98 | 0.9 | 0.96 |
| 1 | 154681950 | rs6695232 | -0.52 | 0.744 | -0.168 | -0.076 | -0.004 | 0 | 0 | 0 | 0.34 | 0.34 | 0.35 | 0.28 | 0.46 | 0.35 |
| 1 | 154691445 | rs10128027 | -0.074 | 0.45 | -0.999 | -0.074 | -0.001 | -0.021 | -0.021 | 0 | 0.63 | 0.41 | 0.7 | 0.8 | 0.65 | 0.71 |
| 1 | 154692088 | rs4845663 | -0.35 | 0.325 | -2.148 | 0.086 | -0.003 | 0.066 | 0.066 | 0 | 0.53 | 0.91 | 0.36 | 0.26 | 0.41 | 0.42 |
| 1 | 154692883 | rs7554577 | 0.043 | 0.731 | -1.612 | 0.077 | 0.017 | 0.078 | 0.078 | 0 | 0.51 | 0.65 | 0.4 | 0.28 | 0.46 | 0.4 |
| 1 | 154693513 | rs11264250 | 0.026 | 1.008 | -1.882 | 0.631 | -0.036 | -0.028 | -0.028 | 0.01 | 0.55 | 0.21 | 0.55 | 0.55 | 0.5 | 0.45 |
| 1 | 154698817 | rs3738028 | 0.12 | 1.078 | -1.616 | 0.716 | 0.02 | 0.029 | 0.029 | 0 | 0.54 | 0.62 | 0.39 | 0.41 | 0.56 | 0.44 |
| 1 | 154700649 | rs6662748 | 0.084 | 1.474 | -1.499 | 0.919 | 0 | 0.002 | 0.002 | 0 | 0.7 | 0.78 | 0.82 | 0.77 | 0.68 | 0.83 |
| 1 | 154701218 | rs10494301 | 0.097 | 1.714 | -1.496 | 0.888 | -0.005 | 0.007 | 0.007 | 0 | 0.71 | 0.77 | 0.8 | 0.82 | 0.65 | 0.83 |
| 1 | 154701406 | rs12402320 | -0.095 | 1.989 | -1.347 | 0.768 | 0.008 | 0 | 0 | 7.778 | 0.2 | 0.19 | 0.29 | 0.17 | 0.28 | 0.12 |
| 1 | 154703959 | rs2102354 | 0.298 | 1.41 | -1.571 | 1.059 | 0.012 | 0.003 | 0.003 | 0 | 0.47 | 0.57 | 0.5 | 0.43 | 0.35 | 0.62 |
| 1 | 154704595 | rs925042 | 0.147 | 1.355 | -1.256 | 0.79 | -0.004 | 0 | 0 | 0 | 0.55 | 0.58 | 0.6 | 0.52 | 0.52 | 0.69 |
| 1 | 154710756 | rs960182 | -0.451 | 1.573 | -1.505 | 0.612 | -0.035 | 0.012 | 0.012 | 0.789 | 0.33 | 0.16 | 0.55 | 0.25 | 0.41 | 0.4 |
| 1 | 154710904 | rs10908429 | -0.117 | 0.865 | -1.33 | 0.62 | 0.001 | -0.003 | -0.003 | 0.056 | 0.31 | 0.24 | 0.35 | 0.41 | 0.27 | 0.29 |
| 1 | 154711226 | rs6682000 | -0.673 | 0.933 | -0.99 | -0.438 | 0.003 | -0.053 | -0.053 | 0.175 | 0.35 | 0.49 | 0.27 | 0.09 | 0.45 | 0.29 |
| 1 | 154712137 | rs11264253 | -0.616 | 0.281 | -0.649 | 0.077 | -0.007 | 0.005 | 0.005 | 0 | 0.21 | 0.26 | 0.23 | 0.21 | 0.14 | 0.24 |
| 1 | 154712358 | rs6697935 | -1.182 | 0.09 | -0.615 | -0.531 | 0.025 | 0.043 | 0.043 | 0.672 | 0.53 | 0.69 | 0.71 | 0.79 | 0.54 | 0.65 |
| 1 | 154714006 | rs4999127 | -0.68 | -0.168 | -1.075 | -0.863 | 0.026 | 0.053 | 0.053 | 0.065 | 0.83 | 0.94 | 0.95 | 1 | 0.85 | 0.86 |
| 1 | 154716883 | rs10908431 | 0.159 | 0.012 | -0.821 | -0.755 | -0.071 | -0.06 | -0.06 | 0 | 0.27 | 0.21 | 0.46 | 0.57 | 0.36 | 0.41 |
| 1 | 154717908 | rs16836320 | 1.796 | -0.137 | -0.538 | -0.765 | 0.014 | 0.016 | 0.016 | 1.909 | 0.31 | 0.2 | 0.22 | 0.23 | 0.32 | 0.21 |
| 1 | 154718738 | rs906278 | -0.27 | -0.226 | -0.531 | -0.692 | 0.013 | -0.007 | -0.007 | 1.845 | 0.79 | 0.85 | 0.91 | 1 | 0.82 | 0.84 |
| 1 | 154719917 | rs906281 | 0.346 | -0.113 | -0.312 | -0.718 | -0.016 | -0.077 | -0.077 | 0.074 | 0.73 | 0.7 | 0.87 | 0.99 | 0.78 | 0.81 |
| 1 | 154721780 | rs6698628 | 1.126 | 0.584 | -0.279 | -0.213 | 0.018 | -0.01 | -0.01 | 0.438 | 0.05 | 0.15 | 0.04 | 0.01 | 0.04 | 0.03 |
| 1 | 154724003 | rs870031 | 0.725 | 0.741 | -0.715 | 0.141 | -0.025 | -0.059 | -0.059 | 0 | 0.1 | 0.01 | 0.17 | 0.2 | 0.09 | 0.19 |
| 1 | 154726242 | rs4845668 | -0.863 | 0.624 | -0.41 | -0.323 | 0.004 | 0.008 | 0.008 | 0 | 0.46 | 0.5 | 0.67 | 0.75 | 0.49 | 0.63 |
| 1 | 154729394 | rs10908435 | -0.835 | 1.28 | -0.279 | -0.152 | -0.054 | 0.066 | 0.066 | 0 | 0.34 | 0.19 | 0.28 | 0.18 | 0.41 | 0.35 |
| 1 | 154732308 | rs1553611 | -0.293 | 1.164 | 0.316 | 0.554 | 0.001 | 0.091 | 0.091 | 0 | 0.87 | 0.75 | 0.64 | 0.74 | 0.88 | 0.85 |
| 1 | 154732776 | rs6682291 | 0.017 | 1.702 | 0.638 | -0.262 | 0.041 | -0.015 | -0.015 | 0 | 0.05 | 0.15 | 0.06 | 0.01 | 0.08 | 0.04 |
| 1 | 154736240 | rs7513331 | 0.223 | 1.986 | 0.665 | -0.349 | 0.004 | -0.01 | -0.01 | 0 | 0.2 | 0.2 | 0.2 | 0.16 | 0.21 | 0.27 |
| 1 | 154737236 | rs7553671 | 0.181 | 1.987 | 0.658 | -0.356 | 0.004 | -0.01 | -0.01 | 0.431 | 0.21 | 0.2 | 0.2 | 0.16 | 0.21 | 0.27 |
| 1 | 154742575 | rs4845392 | -0.139 | 2.33 | 0.607 | -0.691 | 0.039 | -0.07 | -0.07 | 0 | 0.86 | 0.77 | 0.9 | 0.99 | 0.82 | 0.86 |
| 1 | 154744852 | rs1131820 | 0.561 | 2.237 | 0.748 | -0.787 | 0.059 | -0.105 | -0.105 | 0.107 | 0.8 | 0.59 | 0.84 | 0.98 | 0.74 | 0.81 |
| 1 | 154749047 | rs11808053 | 1.271 | 1.516 | 0.545 | -0.421 | 0.06 | 0.093 | 0.093 | 0.41 | 0.45 | 0.16 | 0.26 | 0.33 | 0.43 | 0.36 |
| 1 | 154750816 | rs7532286 | 0.881 | 1.499 | 0.556 | -0.378 | 0.022 | 0.075 | 0.075 | 0.757 | 0.53 | 0.68 | 0.34 | 0.34 | 0.5 | 0.4 |
| 1 | 154750989 | rs6426929 | 0.846 | 1.609 | 0.455 | -0.33 | -0.002 | -0.007 | -0.007 | 0.154 | 0.53 | 0.5 | 0.33 | 0.34 | 0.5 | 0.4 |
| 1 | 154753697 | rs1506982 | 0.115 | 1.23 | 0.167 | 0.037 | -0.001 | 0.031 | 0.031 | 0 | 0.1 | 0.09 | 0.09 | 0.25 | 0.09 | 0.11 |
| 1 | 154755126 | rs4845671 | -0.88 | 1.349 | 0.116 | -0.201 | 0 | -0.002 | -0.002 | 1.201 | 0.35 | 0.58 | 0.32 | 0.42 | 0.33 | 0.37 |
| 1 | 154755843 | rs11589855 | 1.165 | 1.357 | -0.247 | -0.401 | 0.009 | 0.024 | 0.024 | 0.566 | 0.43 | 0.24 | 0.3 | 0.37 | 0.46 | 0.36 |
| 1 | 154756589 | rs11264262 | -0.239 | 1.366 | -0.042 | -0.284 | 0.025 | 0.083 | 0.083 | 0 | 0.12 | 0.25 | 0.11 | 0.25 | 0.12 | 0.12 |
| 1 | 154758422 | rs883319 | 0.404 | 1.487 | -0.463 | -0.787 | -0.004 | 0.011 | 0.011 | 0 | 0.2 | 0.16 | 0.2 | 0.17 | 0.2 | 0.23 |
| 1 | 154760965 | rs11584635 | 1.947 | 1.595 | -0.482 | -0.736 | 0.02 | 0.075 | 0.075 | 3.009 | 0.33 | 0.14 | 0.21 | 0.13 | 0.35 | 0.35 |
| 1 | 154761265 | rs4079531 | 0.975 | 1.775 | -0.545 | -0.786 | -0.012 | 0.053 | 0.053 | 0 | 0.6 | 0.73 | 0.48 | 0.29 | 0.64 | 0.48 |
| 1 | 154764715 | rs12728678 | 0.184 | 1.584 | -0.638 | -0.429 | -0.004 | 0.006 | 0.006 | 0 | 0.18 | 0.17 | 0.2 | 0.16 | 0.2 | 0.12 |
| 1 | 154766100 | rs11577297 | 0.999 | 1.189 | -0.269 | 0.13 | -0.004 | -0.001 | -0.001 | 0 | 0.76 | 0.58 | 0.56 | 0.32 | 0.8 | 0.69 |
| 1 | 154769201 | rs953559 | 0.727 | 1.308 | -0.836 | 0.62 | -0.001 | 0.015 | 0.015 | 0 | 0.89 | 0.86 | 0.91 | 0.66 | 0.88 | 0.92 |
| 1 | 154769561 | rs4492614 | -1.027 | 1.263 | -0.881 | -0.434 | 0 | -0.007 | -0.007 | 0 | 0.14 | 0.11 | 0.09 | 0.01 | 0.12 | 0.14 |
| 1 | 154770139 | rs954785 | 1.389 | 1.595 | -0.537 | -0.653 | 0.007 | -0.067 | -0.067 | 0.174 | 0.81 | 0.73 | 0.86 | 0.99 | 0.79 | 0.89 |
| 1 | 154772089 | rs12737434 | 0.601 | 0.966 | -1.014 | -0.804 | -0.014 | 0.045 | 0.045 | 0 | 0.45 | 0.27 | 0.64 | 0.7 | 0.51 | 0.3 |
| 1 | 154774916 | rs6699284 | 0.949 | 0.694 | -1.352 | -1.179 | -0.011 | -0.019 | -0.019 | 0 | 0.16 | 0.07 | 0.11 | 0.2 | 0.17 | 0.32 |
| 1 | 154774958 | rs2135695 | 0.343 | 0.515 | -1.492 | -1.415 | -0.005 | -0.038 | -0.038 | 0 | 0.77 | 0.32 | 0.82 | 0.92 | 0.81 | 0.8 |
| 1 | 154776412 | rs6426944 | -0.019 | 0.415 | -0.847 | -1.496 | 0.002 | -0.004 | -0.004 | 0 | 0.74 | 0.58 | 0.8 | 0.76 | 0.74 | 0.59 |

| | | | | | | | | | | | | | | | | |
|---|---|---|---|---|---|---|---|---|---|---|---|---|---|---|---|---|
| 1 | 154777157 | rs1702176 | -1.402 | 0.445 | -1.169 | -1.665 | -0.014 | -0.067 | -0.067 | 0 | 0.31 | 0.57 | 0.2 | 0.08 | 0.23 | 0.29 |
| 1 | 154778678 | rs6664312 | 0.902 | 0.492 | -1.519 | -1.633 | -0.009 | 0.041 | 0.041 | 0 | 0.1 | 0.02 | 0.05 | 0 | 0.08 | 0.09 |
| 1 | 154779544 | rs2256209 | -0.053 | 0.517 | -1.366 | -1.552 | -0.032 | -0.044 | -0.044 | 0.011 | 0.14 | 0.5 | 0.1 | 0.05 | 0.08 | 0.12 |
| 1 | 154779685 | rs2798601 | -1.011 | 0.843 | -1.537 | -1.311 | -0.013 | -0.007 | -0.007 | 1.304 | 0.84 | 0.89 | 0.89 | 0.8 | 0.81 | 0.69 |
| 1 | 154783756 | rs6700814 | -0.905 | 0.5 | -1.388 | -1.202 | 0.008 | 0.018 | 0.018 | 0.076 | 0.38 | 0.65 | 0.24 | 0.09 | 0.29 | 0.35 |
| 1 | 154786686 | rs2335406 | -0.807 | 0.504 | -1.986 | -0.8 | -0.044 | 0.062 | 0.062 | 0 | 0.56 | 0.84 | 0.39 | 0.31 | 0.52 | 0.69 |
| 1 | 154787213 | rs1106235 | -0.454 | 0.076 | -1.961 | -0.879 | 0.034 | 0.061 | 0.061 | 0.568 | 0.26 | 0.1 | 0.2 | 0.13 | 0.17 | 0.33 |
| 1 | 154789875 | rs1218603 | -0.334 | -0.029 | -2.09 | -0.26 | 0.052 | -0.012 | -0.012 | 0 | 0.6 | 0.81 | 0.67 | 0.57 | 0.72 | 0.64 |
| 1 | 154791128 | rs10752609 | -0.528 | -0.171 | -2.655 | -0.537 | -0.03 | 0.174 | 0.174 | 0.248 | 0.44 | 0.83 | 0.69 | 0.8 | 0.47 | 0.39 |
| 1 | 154791676 | rs1218601 | 1.122 | -0.124 | -2.642 | -0.508 | -0.011 | 0.115 | 0.115 | 1.234 | 0.29 | 0.1 | 0.15 | 0.1 | 0.27 | 0.39 |
| 1 | 154793296 | rs4845673 | 1.445 | 0.376 | -2.44 | -0.09 | -0.019 | -0.038 | -0.038 | 0 | 0.16 | 0.09 | 0.31 | 0.36 | 0.26 | 0.18 |
| 1 | 154793763 | rs951240 | 1.218 | 0.36 | -2.456 | -0.263 | -0.028 | 0.106 | 0.106 | 1.561 | 0.3 | 0.07 | 0.16 | 0.14 | 0.27 | 0.4 |
| 1 | 154793798 | rs951241 | 0.298 | 0.692 | -1.002 | -0.029 | 0.116 | 0.095 | 0.095 | 0 | 0.08 | 0.3 | 0.11 | 0.1 | 0.12 | 0.04 |
| 1 | 154794897 | rs12029542 | -0.167 | 0.649 | -0.826 | -0.024 | 0.134 | 0.052 | 0.052 | 0 | 0.1 | 0.27 | 0.13 | 0.1 | 0.19 | 0.06 |
| 1 | 154795389 | rs1218598 | -1.072 | 0.489 | -1.091 | -0.335 | 0.007 | 0.071 | 0.071 | 0.023 | 0.75 | 0.87 | 0.87 | 0.94 | 0.78 | 0.8 |
| 1 | 154796520 | rs11264268 | 0.983 | 0.38 | -0.829 | 0.013 | 0.012 | 0.044 | 0.044 | 0 | 0.17 | 0.26 | 0.32 | 0.32 | 0.25 | 0.18 |
| 1 | 154796712 | rs6681392 | 0.112 | 0.182 | -0.976 | -0.096 | -0.043 | 0.063 | 0.063 | 0.373 | 0.26 | 0.62 | 0.34 | 0.49 | 0.12 | 0.19 |
| 1 | 154796895 | rs1984285 | 1.024 | 0.687 | -1.308 | 0.27 | -0.023 | -0.045 | -0.045 | 0 | 0.16 | 0.07 | 0.34 | 0.33 | 0.26 | 0.18 |
| 1 | 154798627 | rs1218553 | -0.558 | 0.79 | -1.228 | -0.506 | -0.032 | 0.07 | 0.07 | 0 | 0.73 | 0.89 | 0.82 | 0.93 | 0.67 | 0.77 |
| 1 | 154801173 | rs1218551 | -0.589 | 0.81 | -1.163 | -0.508 | -0.02 | 0.04 | 0.04 | 0 | 0.74 | 0.84 | 0.82 | 0.93 | 0.67 | 0.77 |
| 1 | 154801247 | rs1218550 | 0.367 | 0.354 | -0.94 | -0.597 | 0.013 | 0.071 | 0.071 | 1.341 | 0.35 | 0.19 | 0.2 | 0.17 | 0.3 | 0.42 |
| 1 | 154802139 | rs6665642 | 0.565 | 0.315 | -0.672 | -0.043 | -0.008 | -0.043 | -0.043 | 0 | 0.11 | 0.04 | 0.13 | 0.24 | 0.08 | 0.2 |
| 1 | 154803645 | rs1218549 | 0.111 | 0.374 | -0.201 | -1.717 | -0.002 | -0.033 | -0.033 | 3.523 | 0.65 | 0.62 | 0.78 | 0.96 | 0.65 | 0.72 |
| 1 | 154807935 | rs11584630 | -0.44 | 0.408 | -0.399 | -1.747 | -0.015 | 0.025 | 0.025 | 0 | 0.34 | 0.23 | 0.22 | 0.03 | 0.37 | 0.29 |
| 1 | 154808287 | rs4845394 | 0.791 | 0.481 | -0.458 | -1.461 | 0.017 | -0.007 | -0.007 | 1.511 | 0.45 | 0.25 | 0.46 | 0.36 | 0.46 | 0.39 |
| 1 | 154810030 | rs1218578 | 0.07 | 0.529 | -0.472 | -1.546 | 0.001 | 0.002 | 0.002 | 0 | 0.52 | 0.39 | 0.53 | 0.63 | 0.52 | 0.54 |
| 1 | 154811127 | rs11264274 | -0.573 | 1.042 | -0.029 | -2.527 | -0.049 | 0.079 | 0.079 | 0 | 0.23 | 0.12 | 0.17 | 0.03 | 0.31 | 0.24 |
| 1 | 154811420 | rs12139030 | -0.328 | 0.62 | 0.38 | -2.498 | -0.002 | -0.037 | -0.037 | 0 | 0.09 | 0.11 | 0.06 | 0.01 | 0.1 | 0.14 |
| 1 | 154811435 | rs12128882 | -1.44 | 0.674 | 0.8 | -2.587 | 0.022 | -0.071 | -0.071 | 0 | 0.16 | 0.33 | 0.11 | 0.04 | 0.2 | 0.2 |
| 1 | 154811677 | rs1218574 | 0.778 | 0.112 | 0.867 | -2.649 | -0.052 | -0.152 | -0.152 | 0 | 0.8 | 0.5 | 0.88 | 0.99 | 0.85 | 0.81 |
| 1 | 154814917 | rs6666891 | -0.573 | 0.608 | 0.815 | -2.711 | 0.067 | -0.131 | -0.131 | 0.241 | 0.26 | 0.39 | 0.19 | 0.03 | 0.34 | 0.21 |
| 1 | 154815257 | rs6426987 | -0.716 | 0.601 | 0.597 | -2.715 | 0.055 | -0.13 | -0.13 | 0 | 0.27 | 0.48 | 0.2 | 0.03 | 0.34 | 0.21 |
| 1 | 154817687 | rs1218565 | 1.088 | 0.562 | -0.651 | -2.501 | -0.051 | 0.17 | 0.17 | 0.043 | 0.57 | 0.25 | 0.7 | 0.92 | 0.52 | 0.58 |
| 1 | 154825270 | rs11264275 | 0.802 | 0.77 | -0.517 | -2.515 | -0.045 | 0.206 | 0.206 | 0 | 0.57 | 0.22 | 0.67 | 0.96 | 0.49 | 0.75 |
| 1 | 154828409 | rs4845396 | 0.727 | 0.755 | -0.415 | -2.536 | -0.038 | 0.17 | 0.17 | 0 | 0.58 | 0.24 | 0.68 | 0.96 | 0.49 | 0.75 |
| 1 | 154831925 | rs1218587 | -0.48 | 0.536 | 0.387 | -2.499 | -0.017 | -0.142 | -0.142 | 0 | 0.29 | 0.41 | 0.17 | 0.01 | 0.23 | 0.12 |
| 1 | 154832290 | rs1061122 | -0.313 | 0.524 | 0.474 | -2.512 | -0.012 | -0.138 | -0.138 | 0 | 0.27 | 0.46 | 0.16 | 0.02 | 0.23 | 0.11 |
| 1 | 154832661 | rs1218585 | -0.016 | 0.558 | 0.183 | -2.502 | -0.009 | 0.079 | 0.079 | 0.003 | 0.23 | 0.1 | 0.13 | 0.02 | 0.23 | 0.11 |
| 1 | 154837796 | rs7553195 | 0.881 | 0.036 | 0.655 | -2.555 | 0.004 | 0.008 | 0.008 | 0 | 0.52 | 0.42 | 0.64 | 0.89 | 0.41 | 0.6 |
| 1 | 154840516 | rs7531728 | 1.029 | -0.055 | 0.674 | -2.585 | -0.015 | 0.288 | 0.288 | 0 | 0.48 | 0.14 | 0.61 | 0.89 | 0.4 | 0.6 |
| 1 | 154843347 | rs2335407 | 1.025 | -0.05 | 0.762 | -2.583 | 0.021 | 0.1 | 0.1 | 0 | 0.5 | 0.33 | 0.62 | 0.89 | 0.4 | 0.6 |
| 1 | 154849590 | rs10908445 | 0.724 | -0.085 | 1.339 | -2.616 | 0 | -0.024 | -0.024 | 2.685 | 0.54 | 0.44 | 0.65 | 0.89 | 0.43 | 0.62 |
| 1 | 154896032 | rs2211498 | -1.447 | -0.346 | 1.166 | -2.757 | -0.044 | -0.088 | -0.088 | 0.041 | 0.15 | 0.65 | 0.09 | 0 | 0.06 | 0.07 |
| 1 | 154903047 | rs11264286 | -1.447 | -0.307 | 1.144 | -2.757 | -0.041 | -0.09 | -0.09 | 0 | 0.15 | 0.64 | 0.08 | 0 | 0.06 | 0.07 |
| 1 | 154904840 | rs1891805 | -1.447 | -0.307 | 1.144 | -2.757 | -0.041 | -0.09 | -0.09 | 0.511 | 0.15 | 0.64 | 0.08 | 0 | 0.06 | 0.07 |
| 1 | 154913021 | rs6699043 | 0.644 | -0.206 | 1.185 | -2.686 | -0.001 | 0.069 | 0.069 | 0 | 0.55 | 0.64 | 0.67 | 0.9 | 0.44 | 0.62 |
| 1 | 154917314 | rs705094 | 0.97 | -0.126 | 0.76 | -2.679 | -0.008 | 0.28 | 0.28 | 0 | 0.51 | 0.14 | 0.63 | 0.89 | 0.43 | 0.62 |
| 1 | 154919080 | rs2061690 | 1.068 | -0.145 | 0.722 | -2.67 | -0.005 | 0.291 | 0.291 | 0 | 0.5 | 0.15 | 0.63 | 0.9 | 0.43 | 0.62 |
| 1 | 154928707 | rs6695407 | -0.191 | -0.154 | 1.684 | -2.154 | 0.01 | 0.006 | 0.006 | 0.71 | 0.08 | 0.08 | 0.02 | 0 | 0.03 | 0.03 |
| 1 | 154934665 | rs7528548 | 0.208 | -0.199 | 1.685 | -2.157 | -0.001 | -0.008 | -0.008 | 0.475 | 0.06 | 0.09 | 0.02 | 0 | 0.02 | 0.05 |
| 1 | 154943881 | rs12075984 | -1.863 | -0.193 | 1.824 | -2.266 | -0.042 | -0.072 | -0.072 | 0 | 0.12 | 0.5 | 0.06 | 0 | 0.03 | 0.06 |
| 1 | 154944156 | rs12076073 | -1.863 | -0.193 | 1.824 | -2.266 | -0.042 | -0.072 | -0.072 | 0.374 | 0.12 | 0.5 | 0.06 | 0 | 0.03 | 0.06 |
| 1 | 154946517 | rs11264293 | -1.813 | -0.183 | 1.831 | -2.261 | -0.039 | -0.07 | -0.07 | 0 | 0.12 | 0.5 | 0.06 | 0 | 0.03 | 0.06 |
| 1 | 154955724 | rs10908449 | -1.75 | -0.156 | 1.841 | -2.248 | -0.036 | -0.066 | -0.066 | 0 | 0.12 | 0.5 | 0.06 | 0 | 0.03 | 0.06 |
| 1 | 154956662 | rs6696397 | -0.384 | -0.104 | 1.507 | -2.205 | -0.02 | -0.033 | -0.033 | 0 | 0.08 | 0.14 | 0.03 | 0 | 0.02 | 0.06 |
| 1 | 154959882 | rs7525580 | -1.192 | -0.155 | 1.531 | -2.205 | -0.024 | -0.044 | -0.044 | 0 | 0.1 | 0.4 | 0.05 | 0 | 0.02 | 0.06 |
| 1 | 154961458 | rs2878517 | -0.057 | -0.118 | 1.06 | -2.188 | 0.002 | -0.004 | -0.004 | 0 | 0.07 | 0.09 | 0.02 | 0 | 0.02 | 0.06 |
| 1 | 154961706 | rs2089248 | -0.741 | -0.115 | 1.071 | -2.187 | -0.021 | -0.039 | -0.039 | 0.374 | 0.09 | 0.31 | 0.04 | 0 | 0.02 | 0.06 |
| 1 | 154977929 | rs4845402 | 0.06 | -0.053 | -0.012 | -2.4 | 0.001 | -0.029 | -0.029 | 0 | 0.8 | 0.84 | 0.89 | 0.99 | 0.86 | 0.92 |
| 1 | 154981708 | rs4845700 | 0.94 | -0.29 | -0.086 | -2.276 | 0.064 | 0.079 | 0.079 | 3.46 | 0.68 | 0.23 | 0.73 | 0.9 | 0.82 | 0.89 |
| 1 | 154986091 | rs3753639 | -1.259 | 0.778 | -0.138 | -2.607 | -0.002 | -0.003 | -0.003 | 0 | 0.18 | 0.17 | 0.13 | 0.07 | 0.22 | 0.12 |
| 1 | 154986793 | rs3820225 | 1.041 | 1.342 | -0.137 | -3.065 | 0.029 | -0.103 | -0.103 | 0.934 | 0.64 | 0.49 | 0.69 | 0.91 | 0.5 | 0.74 |
| 1 | 154987526 | rs11264295 | 1.549 | 1.188 | -0.172 | -3.058 | -0.002 | -0.079 | -0.079 | 0 | 0.58 | 0.46 | 0.67 | 0.91 | 0.45 | 0.67 |
| 1 | 154989595 | rs6673081 | 1.353 | 1.412 | -0.199 | -3.093 | 0.078 | -0.109 | -0.109 | 1.394 | 0.72 | 0.48 | 0.75 | 0.91 | 0.6 | 0.73 |
| 1 | 154994750 | rs11264296 | 1.602 | 1.279 | 0.075 | -3.042 | 0.001 | -0.019 | -0.019 | 0.019 | 0.56 | 0.37 | 0.66 | 0.9 | 0.45 | 0.68 |
| 1 | 154995503 | rs2242193 | 1.483 | 1.292 | 0.087 | -3.034 | -0.002 | -0.078 | -0.078 | 0 | 0.58 | 0.46 | 0.67 | 0.91 | 0.45 | 0.68 |
| 1 | 155001929 | rs10908452 | 1.341 | 0.777 | -0.47 | -3.349 | -0.012 | -0.071 | -0.071 | 0 | 0.45 | 0.43 | 0.59 | 0.88 | 0.36 | 0.62 |
| 1 | 155003954 | rs11582128 | 0.413 | 0.796 | -0.597 | -3.376 | -0.01 | 0.113 | 0.113 | 4.423 | 0.18 | 0.02 | 0.1 | 0 | 0.16 | 0.06 |
| 1 | 155006091 | rs10908453 | 1.026 | 0.646 | -0.255 | -3.095 | 0.006 | -0.014 | -0.014 | 0.227 | 0.45 | 0.54 | 0.61 | 0.88 | 0.36 | 0.63 |

| | | | | | | | | | | | | | | | | |
|---|---|---|---|---|---|---|---|---|---|---|---|---|---|---|---|---|
| 1 | 155007293 | rs4845404 | 0.998 | 0.662 | -0.393 | -3.114 | -0.01 | -0.064 | -0.064 | 0 | 0.44 | 0.43 | 0.59 | 0.88 | 0.36 | 0.62 |
| 1 | 155007302 | rs4845405 | 0.99 | 0.649 | -0.39 | -3.111 | -0.01 | -0.064 | -0.064 | 0 | 0.44 | 0.43 | 0.59 | 0.88 | 0.36 | 0.62 |
| 1 | 155008610 | rs4845406 | 0.142 | 0.855 | -0.706 | -3.063 | -0.037 | 0.15 | 0.15 | 0.614 | 0.59 | 0.78 | 0.73 | 0.94 | 0.56 | 0.69 |
| 1 | 155019710 | rs11264300 | 0.627 | 0.369 | -0.472 | -2.738 | 0.005 | -0.02 | -0.02 | 0 | 0.45 | 0.39 | 0.59 | 0.88 | 0.36 | 0.62 |
| 1 | 155026942 | rs6427128 | -0.92 | 0.254 | -0.951 | -2.695 | -0.001 | -0.033 | -0.033 | 0 | 0.84 | 0.87 | 0.93 | 0.99 | 0.84 | 0.9 |
| 1 | 155029253 | rs3765087 | 0.386 | 0.347 | -1.405 | -2.768 | -0.008 | 0.048 | 0.048 | 0 | 0.17 | 0.02 | 0.11 | 0.08 | 0.13 | 0.13 |
| 1 | 155033572 | rs11264305 | 0.088 | 0.169 | -1.642 | -3.154 | 0.022 | 0.131 | 0.131 | 0.012 | 0.62 | 0.85 | 0.8 | 0.98 | 0.68 | 0.75 |
| 1 | 155035611 | rs3806256 | 0.566 | 0.551 | -1.45 | -3.724 | -0.008 | 0.031 | 0.031 | 0 | 0.42 | 0.34 | 0.64 | 0.92 | 0.49 | 0.61 |
| 1 | 155038756 | rs4045192 | 0.531 | 0.78 | -1.4 | -3.759 | -0.009 | 0.086 | 0.086 | 0 | 0.39 | 0.23 | 0.61 | 0.93 | 0.44 | 0.58 |
| 1 | 155042886 | rs35902694 | -0.325 | 0.095 | -0.87 | -3.813 | 0.004 | -0.023 | -0.023 | 0.597 | 0.56 | 0.62 | 0.75 | 0.93 | 0.63 | 0.71 |
| 1 | 155043325 | rs11264306 | 0.274 | 0.648 | -0.716 | -3.636 | -0.011 | 0.035 | 0.035 | 0 | 0.41 | 0.32 | 0.66 | 0.92 | 0.54 | 0.6 |
| 1 | 155047379 | rs1001848 | 0.225 | 0.663 | -0.719 | -3.629 | -0.01 | 0.034 | 0.034 | 0 | 0.42 | 0.32 | 0.66 | 0.92 | 0.54 | 0.59 |
| 1 | 155048369 | rs6427137 | -0.01 | 0.645 | -0.619 | -3.647 | 0.041 | -0.01 | -0.01 | 0 | 0.06 | 0.29 | 0.05 | 0 | 0.05 | 0.06 |
| 1 | 155049402 | rs3766916 | -0.15 | 0.548 | -0.738 | -3.708 | -0.003 | -0.066 | -0.066 | 0 | 0.47 | 0.51 | 0.7 | 0.93 | 0.6 | 0.66 |
| 1 | 155058663 | rs17723260 | 0.72 | 0.716 | -1.15 | -3.593 | -0.019 | 0.107 | 0.107 | 0 | 0.19 | 0.02 | 0.1 | 0.01 | 0.13 | 0.14 |
| 1 | 155059851 | rs2306124 | -0.105 | 1.045 | -0.874 | -3.301 | -0.016 | 0.017 | 0.017 | 0 | 0.37 | 0.29 | 0.64 | 0.92 | 0.52 | 0.52 |
| 1 | 155079477 | rs7367207 | -0.385 | 1.39 | -0.879 | -3.184 | -0.008 | -0.007 | -0.007 | 0 | 0.34 | 0.3 | 0.62 | 0.91 | 0.45 | 0.5 |
| 1 | 155105882 | rs4971066 | 0.311 | 1.462 | -1.366 | -2.458 | -0.025 | -0.041 | -0.041 | 0 | 0.14 | 0.01 | 0.47 | 0.78 | 0.15 | 0.36 |
| 1 | 155106550 | rs9297 | -0.818 | 1.589 | -1.153 | -2.653 | -0.024 | 0.038 | 0.038 | 0 | 0.32 | 0.24 | 0.59 | 0.8 | 0.41 | 0.49 |
| 1 | 155106697 | rs12904 | -0.846 | 1.578 | -1.103 | -2.658 | -0.021 | 0.033 | 0.033 | 0 | 0.33 | 0.25 | 0.6 | 0.8 | 0.41 | 0.49 |
| 1 | 155122783 | rs6427184 | -1.083 | 1.501 | -1.684 | -3.023 | -0.049 | 0.107 | 0.107 | 0 | 0.82 | 0.99 | 0.9 | 0.99 | 0.87 | 0.86 |
| 1 | 155131673 | rs4276913 | -0.713 | 1.606 | -1.677 | -3.188 | -0.037 | 0.071 | 0.071 | 0 | 0.33 | 0.2 | 0.59 | 0.8 | 0.4 | 0.49 |
| 1 | 155132836 | rs12028043 | -0.89 | 1.547 | -1.163 | -3.216 | -0.006 | 0.009 | 0.009 | 0 | 0.35 | 0.34 | 0.6 | 0.8 | 0.4 | 0.49 |
| 1 | 155135335 | rs4460629 | -1.487 | 1.518 | -0.604 | -3.416 | 0.002 | -0.03 | -0.03 | 0.716 | 0.43 | 0.65 | 0.67 | 0.79 | 0.5 | 0.53 |
| 1 | 155138450 | rs11264337 | -1.487 | 1.518 | -0.611 | -3.418 | 0.001 | -0.023 | -0.023 | 0 | 0.43 | 0.65 | 0.67 | 0.79 | 0.49 | 0.53 |
| 1 | 155138557 | rs11264338 | -1.487 | 1.518 | -0.611 | -3.418 | 0.001 | -0.023 | -0.023 | 0 | 0.43 | 0.65 | 0.67 | 0.79 | 0.49 | 0.53 |
| 1 | 155139509 | rs7556304 | -1.488 | 1.521 | -0.609 | -3.418 | 0.001 | -0.024 | -0.024 | 0 | 0.43 | 0.65 | 0.67 | 0.79 | 0.5 | 0.53 |
| 1 | 155140648 | rs11264339 | -1.482 | 1.533 | -0.605 | -3.413 | 0.001 | -0.024 | -0.024 | 0 | 0.43 | 0.65 | 0.66 | 0.79 | 0.49 | 0.53 |
| 1 | 155143946 | rs4971092 | -1.343 | 1.449 | -0.77 | -3.289 | -0.037 | 0.098 | 0.098 | 0.08 | 0.84 | 1 | 0.9 | 0.99 | 0.89 | 0.86 |
| 1 | 155149718 | rs3814316 | -0.862 | 1.6 | -0.302 | -3.274 | -0.02 | 0.032 | 0.032 | 0 | 0.29 | 0.22 | 0.57 | 0.76 | 0.36 | 0.43 |
| 1 | 155151493 | rs11264341 | -1.118 | 1.904 | -0.074 | -2.544 | -0.036 | 0.094 | 0.094 | 0 | 0.41 | 0.2 | 0.63 | 0.7 | 0.41 | 0.47 |
| 1 | 155162067 | rs4072037 | -1.974 | 1.591 | 0.385 | -2.668 | 0.001 | -0.009 | -0.009 | 0 | 0.51 | 0.6 | 0.71 | 0.76 | 0.55 | 0.55 |
| 1 | 155172379 | rs2066981 | -2.013 | 1.529 | 0.463 | -2.677 | 0.001 | -0.007 | -0.007 | 1.717 | 0.52 | 0.62 | 0.71 | 0.78 | 0.55 | 0.55 |
| 1 | 155172725 | rs35154152 | -1.262 | 1.44 | 0.49 | -2.683 | 0.024 | -0.046 | -0.046 | 0 | 0.1 | 0.26 | 0.07 | 0.02 | 0.13 | 0.09 |
| 1 | 155175892 | rs914615 | -1.936 | 1.372 | 0.358 | -2.667 | 0.001 | -0.007 | -0.007 | 0 | 0.52 | 0.62 | 0.71 | 0.78 | 0.54 | 0.55 |
| 1 | 155194980 | rs2049805 | -1.977 | 1.296 | 0.479 | -2.125 | 0.001 | -0.004 | -0.004 | 0 | 0.49 | 0.62 | 0.69 | 0.76 | 0.52 | 0.55 |
| 1 | 155201190 | rs4043 | -2.529 | 1.244 | 1.362 | -1.063 | 0.022 | -0.02 | -0.02 | 0.879 | 0.28 | 0.55 | 0.32 | 0.69 | 0.34 | 0.38 |
| 1 | 155201235 | rs1045253 | -2.529 | 1.244 | 1.36 | -1.063 | 0.022 | -0.02 | -0.02 | 1.108 | 0.28 | 0.55 | 0.32 | 0.69 | 0.34 | 0.38 |
| 1 | 155206341 | rs9628662 | -2.515 | 1.308 | 1.384 | -1.039 | 0.027 | -0.018 | -0.018 | 1.206 | 0.28 | 0.53 | 0.32 | 0.7 | 0.34 | 0.37 |
| 1 | 155214473 | rs12034326 | -2.512 | 1.318 | 1.385 | -1.042 | 0.028 | -0.018 | -0.018 | 0 | 0.28 | 0.54 | 0.32 | 0.69 | 0.34 | 0.37 |
| 1 | 155218446 | rs734074 | -2.595 | 0.598 | 1.507 | -0.993 | -0.003 | -0.019 | -0.019 | 0 | 0.27 | 0.63 | 0.3 | 0.7 | 0.27 | 0.36 |
| 1 | 155222483 | rs12730005 | -2.806 | 0.656 | 1.591 | -0.966 | -0.011 | -0.02 | -0.02 | 0.789 | 0.28 | 0.59 | 0.3 | 0.7 | 0.27 | 0.34 |
| 1 | 155232074 | rs1046188 | -2.325 | 0.686 | 1.636 | -0.918 | 0.006 | -0.012 | -0.012 | 0.795 | 0.26 | 0.54 | 0.32 | 0.69 | 0.28 | 0.36 |
| 1 | 155239624 | rs2361543 | -2.35 | 0.597 | 1.636 | -0.917 | 0.009 | -0.014 | -0.014 | 0 | 0.26 | 0.54 | 0.32 | 0.69 | 0.28 | 0.35 |
| 1 | 155247707 | rs12749306 | -1.965 | 0.545 | 1.71 | -0.939 | 0.01 | 0.001 | 0.001 | 0.39 | 0.22 | 0.3 | 0.31 | 0.69 | 0.28 | 0.33 |
| 1 | 155255837 | rs11264352 | -2.455 | 0.628 | 1.68 | -0.897 | 0.006 | -0.013 | -0.013 | 0 | 0.26 | 0.5 | 0.33 | 0.7 | 0.28 | 0.35 |
| 1 | 155260096 | rs932972 | -2.574 | 0.697 | 1.766 | -0.832 | 0.007 | -0.008 | -0.008 | 0 | 0.26 | 0.49 | 0.33 | 0.69 | 0.28 | 0.35 |
| 1 | 155260350 | rs1052177 | -2.57 | 0.698 | 1.766 | -0.832 | 0.007 | -0.008 | -0.008 | 0 | 0.26 | 0.49 | 0.33 | 0.69 | 0.28 | 0.35 |
| 1 | 155260383 | rs1052176 | -2.565 | 0.697 | 1.766 | -0.832 | 0.007 | -0.007 | -0.007 | 0 | 0.26 | 0.49 | 0.33 | 0.69 | 0.28 | 0.35 |
| 1 | 155279482 | rs2297480 | -1.875 | 0.636 | 1.621 | -0.785 | 0.016 | 0.007 | 0.007 | 0.183 | 0.2 | 0.28 | 0.31 | 0.69 | 0.26 | 0.31 |
| 1 | 155286258 | rs10908463 | -1.929 | 0.672 | 1.626 | -0.807 | 0 | -0.001 | -0.001 | 0.019 | 0.19 | 0.2 | 0.3 | 0.71 | 0.26 | 0.31 |
| 1 | 155289545 | rs11264361 | -1.929 | 0.672 | 1.637 | -0.807 | 0.006 | 0.002 | 0.002 | 0 | 0.19 | 0.21 | 0.3 | 0.71 | 0.26 | 0.31 |
| 1 | 155412037 | rs11264371 | -2.251 | 0.594 | 1.835 | -0.871 | 0.003 | -0.003 | -0.003 | 0 | 0.25 | 0.36 | 0.31 | 0.71 | 0.24 | 0.3 |
| 1 | 155424065 | rs11264375 | -2.35 | 0.585 | 1.834 | -0.871 | 0.008 | -0.006 | -0.006 | 0 | 0.26 | 0.36 | 0.32 | 0.71 | 0.28 | 0.31 |
| 1 | 155478897 | rs6688636 | 2.881 | 0.475 | 1.842 | -0.998 | -0.026 | 0.069 | 0.069 | 6.671 | 0.63 | 0.13 | 0.62 | 0.29 | 0.67 | 0.63 |
| 1 | 155505967 | rs12746592 | -2.105 | 0.586 | 1.853 | -0.895 | -0.031 | -0.002 | -0.002 | 0.833 | 0.24 | 0.13 | 0.31 | 0.71 | 0.28 | 0.31 |
| 1 | 155540660 | rs1556488 | -2.209 | 0.616 | 1.844 | -0.88 | 0.006 | -0.005 | -0.005 | 0 | 0.26 | 0.34 | 0.32 | 0.71 | 0.28 | 0.31 |
| 1 | 155563152 | rs11264387 | -2.039 | 0.634 | 1.573 | -0.862 | -0.048 | -0.014 | -0.014 | 0 | 0.24 | 0.1 | 0.3 | 0.69 | 0.28 | 0.32 |
| 1 | 155625602 | rs2175391 | 2.358 | 0.684 | 2.669 | -0.76 | 0.001 | -0.002 | -0.002 | 0 | 0.71 | 0.33 | 0.66 | 0.32 | 0.73 | 0.68 |
| 1 | 155630752 | rs3738590 | -2.213 | 0.691 | 2.585 | -0.756 | -0.028 | -0.015 | -0.015 | 0 | 0.25 | 0.16 | 0.3 | 0.68 | 0.27 | 0.32 |
| 1 | 155667102 | rs12035524 | -1.836 | 0.717 | 2.465 | -0.735 | -0.024 | -0.027 | -0.027 | 0 | 0.22 | 0.11 | 0.29 | 0.68 | 0.24 | 0.31 |
| 1 | 155677223 | rs7525057 | -1.575 | 0.72 | 2.512 | -0.731 | -0.001 | -0.037 | -0.037 | 0 | 0.07 | 0.14 | 0.04 | 0 | 0.06 | 0.03 |
| 1 | 155687232 | rs621181 | -3 | 0.68 | 2.801 | -0.75 | -0.01 | -0.075 | -0.075 | 0 | 0.12 | 0.56 | 0.07 | 0 | 0.06 | 0.06 |
| 1 | 155687277 | rs552778 | -3.051 | 0.679 | 2.801 | -0.751 | -0.012 | -0.077 | -0.077 | 2.739 | 0.12 | 0.56 | 0.07 | 0 | 0.06 | 0.06 |
| 1 | 155713650 | rs10908481 | -1.656 | 0.715 | 2.808 | -0.699 | -0.03 | -0.011 | -0.011 | 0 | 0.24 | 0.18 | 0.33 | 0.68 | 0.3 | 0.32 |
| 1 | 155725531 | rs11803678 | -2.445 | 0.68 | 2.847 | -0.727 | 0.032 | -0.041 | -0.041 | 0 | 0.1 | 0.42 | 0.06 | 0 | 0.06 | 0.03 |
| 1 | 155726290 | rs16837037 | -1.466 | 0.682 | 2.83 | -0.726 | -0.001 | -0.036 | -0.036 | 0.437 | 0.07 | 0.14 | 0.04 | 0 | 0.06 | 0.03 |
| 1 | 155762509 | rs4612618 | -1.069 | 0.682 | 2.819 | -0.726 | 0 | -0.002 | -0.002 | 3.1 | 0.07 | 0.07 | 0.04 | 0 | 0.06 | 0.03 |
| 1 | 155776497 | rs1094462 | 2.439 | 0.678 | 2.838 | -0.728 | -0.01 | -0.075 | -0.075 | 1.561 | 0.88 | 0.44 | 0.93 | 1 | 0.94 | 0.94 |

| | | | | | | | | | | | | | | | | |
|---|---|---|---|---|---|---|---|---|---|---|---|---|---|---|---|---|
| 1 | 155815572 | rs16837063 | -1.415 | 0.702 | 2.823 | -0.717 | 0 | -0.037 | -0.037 | 2.401 | 0.07 | 0.13 | 0.04 | 0 | 0.06 | 0.03 |
| 1 | 155822971 | rs822490 | 2.533 | 0.728 | 2.744 | -0.717 | -0.023 | 0.037 | 0.037 | 3.719 | 0.64 | 0.19 | 0.6 | 0.31 | 0.66 | 0.62 |
| 1 | 155829511 | rs3820594 | -1.701 | 0.695 | 1.727 | -0.568 | -0.027 | -0.011 | -0.011 | 0 | 0.23 | 0.17 | 0.32 | 0.69 | 0.29 | 0.32 |
| 1 | 155842568 | rs822516 | 1.327 | 0.491 | 2.17 | -0.89 | 0.045 | -0.036 | -0.036 | 0 | 0.92 | 0.68 | 0.95 | 1 | 0.92 | 0.94 |
| 1 | 155860091 | rs1749405 | 2.386 | -0.005 | 1.879 | -0.911 | 0.021 | 0.152 | 0.152 | 1.903 | 0.58 | 0.14 | 0.6 | 0.18 | 0.65 | 0.57 |
| 1 | 155868984 | rs1055184 | 1.508 | -0.071 | 2.292 | -0.935 | -0.04 | -0.043 | -0.043 | 0.066 | 0.82 | 0.43 | 0.91 | 0.94 | 0.89 | 0.88 |
| 1 | 155870416 | rs1749409 | 1.508 | -0.071 | 2.292 | -0.935 | -0.04 | -0.043 | -0.043 | 0 | 0.82 | 0.43 | 0.91 | 0.94 | 0.89 | 0.88 |
| 1 | 155873314 | rs2993207 | 2.687 | -0.087 | 2.277 | -0.961 | 0.024 | 0.155 | 0.155 | 1.939 | 0.58 | 0.14 | 0.6 | 0.18 | 0.66 | 0.57 |
| 1 | 155887772 | rs605025 | 0.888 | -0.094 | 2.401 | -0.964 | -0.021 | -0.096 | -0.096 | 0 | 0.85 | 0.66 | 0.92 | 1 | 0.9 | 0.9 |
| 1 | 155891045 | rs625658 | 2.643 | -0.071 | 2.123 | -0.953 | 0.023 | 0.154 | 0.154 | 1.909 | 0.58 | 0.14 | 0.6 | 0.18 | 0.66 | 0.57 |
| 1 | 155904392 | rs2275081 | 1.366 | -0.053 | 2.107 | -1.027 | -0.039 | -0.043 | -0.043 | 0.15 | 0.82 | 0.43 | 0.91 | 0.94 | 0.89 | 0.88 |
| 1 | 155910830 | rs1018730 | 1.357 | -0.056 | 2.105 | -1.028 | -0.039 | -0.043 | -0.043 | 0 | 0.82 | 0.43 | 0.91 | 0.94 | 0.89 | 0.88 |
| 1 | 155914988 | rs2016251 | -1.644 | 0.012 | 1.853 | -0.867 | -0.001 | -0.001 | -0.001 | 0.168 | 0.23 | 0.25 | 0.31 | 0.78 | 0.24 | 0.3 |
| 1 | 155922195 | rs12081057 | -0.631 | -0.024 | 1.702 | -1.286 | -0.002 | 0 | 0 | 0 | 0.12 | 0.09 | 0.06 | 0 | 0.11 | 0.05 |
| 1 | 155951502 | rs1889532 | -1.297 | 0.283 | 1.829 | -1.121 | 0.005 | -0.008 | -0.008 | 2.176 | 0.21 | 0.24 | 0.32 | 0.83 | 0.24 | 0.33 |
| 1 | 155978725 | rs915178 | -1.032 | 0.254 | 1.711 | -1.168 | 0.007 | -0.016 | -0.016 | 0 | 0.23 | 0.28 | 0.35 | 0.82 | 0.26 | 0.3 |
| 1 | 155983648 | rs17381047 | 1.994 | 0.152 | 1.409 | -1.333 | 0.018 | 0.168 | 0.168 | 0.554 | 0.55 | 0.21 | 0.53 | 0.15 | 0.61 | 0.56 |
| 1 | 155983710 | rs11577179 | 1.797 | 0.185 | 1.481 | -1.317 | 0.009 | 0.072 | 0.072 | 0.055 | 0.57 | 0.31 | 0.54 | 0.15 | 0.61 | 0.56 |
| 1 | 155983869 | rs17385169 | -0.645 | 0.11 | 1.142 | -1.308 | 0.006 | 0.05 | 0.05 | 0 | 0.15 | 0.07 | 0.09 | 0.03 | 0.13 | 0.12 |
| 1 | 155995080 | rs12032631 | 1.349 | 0.205 | 1.182 | -1.174 | 0.017 | 0.128 | 0.128 | 0.095 | 0.56 | 0.26 | 0.54 | 0.15 | 0.61 | 0.56 |
| 1 | 156024373 | rs3754293 | 1.304 | 0.409 | -0.604 | -1.122 | -0.001 | 0.255 | 0.255 | 1.027 | 0.54 | 0.12 | 0.59 | 0.16 | 0.57 | 0.59 |
| 1 | 156027550 | rs6427299 | -0.525 | 0.332 | -0.68 | -1.314 | -0.036 | 0.074 | 0.074 | 0 | 0.16 | 0.04 | 0.15 | 0 | 0.21 | 0.07 |
| 1 | 156032887 | rs12049195 | 1.374 | 0.188 | -0.223 | -1.511 | 0.009 | 0.102 | 0.102 | 0.013 | 0.55 | 0.27 | 0.57 | 0.13 | 0.56 | 0.53 |
| 1 | 156042915 | rs11264432 | 1.623 | 0.123 | -0.611 | -1.521 | 0.013 | 0.307 | 0.307 | 0.507 | 0.51 | 0.08 | 0.55 | 0.13 | 0.55 | 0.54 |
| 1 | 156045662 | rs10047112 | -1.449 | 0.297 | -0.41 | -1.402 | 0 | 0.059 | 0.059 | 0.116 | 0.38 | 0.76 | 0.4 | 0.78 | 0.39 | 0.42 |
| 1 | 156045973 | rs3738593 | -0.646 | 0.598 | -0.123 | -0.906 | -0.028 | 0.018 | 0.018 | 0.009 | 0.13 | 0.07 | 0.15 | 0.05 | 0.24 | 0.07 |
| 1 | 156063880 | rs10737170 | -0.044 | 0.366 | -0.058 | -0.841 | -0.001 | -0.004 | -0.004 | 0 | 0.89 | 0.86 | 0.93 | 0.9 | 0.89 | 0.93 |
| 1 | 156083468 | rs2485662 | 1.227 | -0.384 | 0.268 | -1.36 | -0.006 | -0.042 | -0.042 | 0 | 0.66 | 0.37 | 0.79 | 0.91 | 0.7 | 0.82 |
| 1 | 156088526 | rs584025 | -0.718 | -0.506 | 0.972 | -1.261 | -0.035 | -0.096 | -0.096 | 0.039 | 0.13 | 0.42 | 0.06 | 0 | 0.06 | 0.03 |
| 1 | 156093195 | rs693671 | -0.798 | -0.533 | 1.239 | -1.275 | -0.037 | -0.099 | -0.099 | 2.013 | 0.14 | 0.48 | 0.06 | 0 | 0.07 | 0.03 |
| 1 | 156094872 | rs2485676 | -0.359 | -0.535 | 1.429 | -1.275 | -0.024 | -0.077 | -0.077 | 0.508 | 0.12 | 0.32 | 0.06 | 0 | 0.06 | 0.03 |
| 1 | 156100739 | rs577492 | -1.293 | -0.585 | 1.06 | -1.311 | -0.046 | -0.088 | -0.088 | 0 | 0.19 | 0.66 | 0.09 | 0.02 | 0.09 | 0.11 |
| 1 | 156105337 | rs2485664 | -0.833 | -0.723 | 1.271 | -1.397 | -0.046 | -0.107 | -0.107 | 0 | 0.17 | 0.52 | 0.09 | 0.02 | 0.09 | 0.11 |
| 1 | 156106185 | rs505058 | -1.295 | -0.742 | 1.25 | -1.406 | -0.034 | -0.069 | -0.069 | 0 | 0.19 | 0.72 | 0.1 | 0.02 | 0.09 | 0.11 |
| 1 | 156106863 | rs553016 | -0.674 | -0.674 | 1.056 | -1.373 | -0.039 | -0.098 | -0.098 | 3.73 | 0.16 | 0.49 | 0.08 | 0.02 | 0.09 | 0.11 |
| 1 | 156107534 | rs4641 | 0.168 | -0.511 | -0.586 | -0.993 | -0.014 | -0.04 | -0.04 | 0.111 | 0.24 | 0.07 | 0.3 | 0.27 | 0.22 | 0.31 |
| 1 | 156111287 | rs536857 | -0.394 | -0.611 | -0.751 | -1.09 | -0.006 | -0.028 | -0.028 | 0.045 | 0.73 | 0.93 | 0.69 | 0.73 | 0.76 | 0.69 |
| 1 | 156116488 | rs1468772 | 0.699 | 0.021 | -1.053 | -1.313 | -0.1 | -0.003 | -0.003 | 0 | 0.24 | 0.01 | 0.3 | 0.18 | 0.31 | 0.3 |
| 1 | 156128337 | rs10908489 | 1.035 | -0.235 | -0.519 | -1.084 | -0.009 | 0.02 | 0.02 | 0 | 0.06 | 0.13 | 0.07 | 0.09 | 0.02 | 0.08 |
| 1 | 156129748 | rs11264448 | -0.456 | -0.24 | -0.373 | -1.207 | 0.003 | -0.019 | -0.019 | 0.074 | 0.38 | 0.49 | 0.41 | 0.28 | 0.4 | 0.43 |
| 1 | 156129968 | rs510441 | 0.698 | -0.091 | -0.63 | -1.519 | 0.009 | 0.039 | 0.039 | 0.304 | 0.25 | 0.13 | 0.18 | 0.02 | 0.24 | 0.12 |
| 1 | 156131617 | rs1984508 | -1.255 | -0.101 | -0.513 | -1.502 | -0.005 | 0.007 | 0.007 | 0 | 0.64 | 0.71 | 0.59 | 0.31 | 0.64 | 0.55 |
| 1 | 156133038 | rs7555253 | -0.641 | 0.151 | -0.326 | -1.042 | 0 | -0.014 | -0.014 | 0.484 | 0.39 | 0.58 | 0.42 | 0.29 | 0.41 | 0.44 |
| 1 | 156140932 | rs11264452 | -0.229 | 0.492 | -0.099 | -0.822 | 0.001 | 0.015 | 0.015 | 0.04 | 0.24 | 0.46 | 0.31 | 0.28 | 0.23 | 0.34 |
| 1 | 156146218 | rs12401573 | -0.363 | 0.38 | -0.362 | -1.186 | 0.004 | 0.014 | 0.014 | 0 | 0.54 | 0.39 | 0.52 | 0.23 | 0.6 | 0.44 |
| 1 | 156147326 | rs7695 | -0.158 | 0.382 | -0.005 | -1.438 | -0.001 | -0.087 | -0.087 | 0.726 | 0.37 | 0.36 | 0.28 | 0.02 | 0.39 | 0.21 |
| 1 | 156147558 | rs3738581 | -0.752 | 0.274 | 0.173 | -1.484 | 0.021 | 0.098 | 0.098 | 0 | 0.47 | 0.7 | 0.37 | 0.11 | 0.42 | 0.29 |
| 1 | 156153785 | rs11264455 | -0.189 | 0.006 | 0.854 | -0.648 | -0.01 | 0.012 | 0.012 | 0 | 0.07 | 0.18 | 0.08 | 0.08 | 0.03 | 0.11 |
| 1 | 156163695 | rs2842879 | -1.84 | -0.109 | 1.208 | -0.634 | -0.046 | -0.01 | -0.01 | 0 | 0.15 | 0.57 | 0.13 | 0.11 | 0.07 | 0.14 |
| 1 | 156163718 | rs2241111 | -0.315 | -0.122 | 1.34 | -0.64 | -0.005 | 0.006 | 0.006 | 0 | 0.08 | 0.14 | 0.08 | 0.11 | 0.05 | 0.13 |
| 1 | 156169610 | rs2072499 | -1.758 | -0.052 | 1.114 | -0.917 | -0.001 | 0.018 | 0.018 | 0 | 0.36 | 0.67 | 0.33 | 0.14 | 0.36 | 0.3 |
| 1 | 156173082 | rs2241109 | -0.001 | -0.01 | 0.957 | -0.902 | -0.032 | 0.04 | 0.04 | 0 | 0.2 | 0.11 | 0.19 | 0.02 | 0.28 | 0.15 |
| 1 | 156181530 | rs2495270 | -1.482 | -0.03 | 1 | -0.872 | -0.036 | 0 | 0 | 0 | 0.14 | 0.4 | 0.12 | 0.12 | 0.07 | 0.14 |
| 1 | 156184268 | rs1060604 | -1.283 | 0.479 | 0.238 | -0.64 | -0.017 | 0.032 | 0.032 | 0 | 0.29 | 0.22 | 0.28 | 0.14 | 0.35 | 0.29 |
| 1 | 156189286 | rs2758596 | -0.185 | 0.378 | 0.187 | -0.563 | -0.004 | 0.005 | 0.005 | 0 | 0.08 | 0.11 | 0.08 | 0.12 | 0.05 | 0.13 |
| 1 | 156202300 | rs2251615 | -0.181 | 0.37 | 0.184 | -0.545 | -0.007 | 0.007 | 0.007 | 0 | 0.08 | 0.15 | 0.09 | 0.12 | 0.05 | 0.13 |
| 1 | 156204047 | rs2251847 | -1.086 | 0.409 | 0.038 | -0.187 | 0.016 | -0.012 | -0.012 | 0 | 0.31 | 0.46 | 0.32 | 0.33 | 0.35 | 0.34 |
| 1 | 156206121 | rs1052067 | -0.311 | 0.479 | -0.448 | -0.166 | -0.02 | 0.006 | 0.006 | 0 | 0.19 | 0.11 | 0.22 | 0.21 | 0.28 | 0.19 |
| 1 | 156209857 | rs1543294 | 0.081 | -0.632 | -0.306 | -0.333 | 0.004 | 0.004 | 0.004 | 0 | 0.25 | 0.2 | 0.36 | 0.26 | 0.19 | 0.3 |
| 1 | 156210557 | rs12090516 | -0.104 | -0.665 | -0.32 | -0.349 | -0.004 | -0.003 | -0.003 | 0 | 0.27 | 0.23 | 0.37 | 0.26 | 0.19 | 0.3 |
| 1 | 156213257 | rs759330 | -1.111 | -0.431 | -0.122 | -0.126 | -0.012 | -0.006 | -0.006 | 0 | 0.78 | 0.83 | 0.82 | 0.75 | 0.69 | 0.76 |
| 1 | 156215562 | rs2075166 | 1.14 | -0.472 | -0.52 | -0.121 | -0.012 | 0.024 | 0.024 | 1.403 | 0.26 | 0.09 | 0.26 | 0.22 | 0.26 | 0.21 |
| 1 | 156217401 | rs12401854 | 1.412 | -0.639 | -0.597 | -0.161 | 0.003 | -0.002 | -0.002 | 0 | 0.17 | 0.1 | 0.3 | 0.26 | 0.13 | 0.27 |
| 1 | 156235120 | rs933489 | -0.058 | -0.295 | -0.469 | 0.213 | -0.001 | 0.003 | 0.003 | 0 | 0.54 | 0.39 | 0.65 | 0.48 | 0.46 | 0.52 |
| 1 | 156241728 | rs11264462 | 1.109 | -0.332 | -0.601 | 0.159 | -0.031 | 0.02 | 0.02 | 0 | 0.26 | 0.06 | 0.27 | 0.23 | 0.26 | 0.23 |
| 1 | 156242806 | rs11264463 | 0.637 | -0.389 | -0.404 | 0.122 | -0.003 | 0.021 | 0.021 | 0 | 0.3 | 0.24 | 0.29 | 0.23 | 0.28 | 0.23 |
| 1 | 156246728 | rs2277872 | 1.181 | -0.392 | -0.555 | 0.146 | 0.006 | 0.007 | 0.007 | 0 | 0.25 | 0.2 | 0.36 | 0.25 | 0.19 | 0.3 |
| 1 | 156247907 | rs2248273 | 1.357 | -0.422 | -0.555 | 0.175 | 0.001 | 0 | 0 | 0 | 0.21 | 0.19 | 0.18 | 0.26 | 0.3 | 0.24 |
| 1 | 156254358 | rs12023438 | 0.823 | -0.264 | -1.136 | 0.268 | -0.022 | 0.03 | 0.03 | 0 | 0.27 | 0.09 | 0.28 | 0.23 | 0.27 | 0.22 |

| | | | | | | | | | | | | | | | | |
|---|---|---|---|---|---|---|---|---|---|---|---|---|---|---|---|---|
| 1 | 156255456 | rs6684514 | 0.964 | -0.265 | -1.135 | 0.267 | -0.028 | 0.025 | 0.025 | 0 | 0.27 | 0.08 | 0.28 | 0.23 | 0.27 | 0.22 |
| 1 | 156258953 | rs12563631 | 0.925 | -0.264 | -1.135 | 0.268 | -0.024 | 0.027 | 0.027 | 0 | 0.27 | 0.09 | 0.28 | 0.23 | 0.27 | 0.22 |
| 1 | 156263097 | rs10908494 | 0.823 | -0.264 | -1.135 | 0.268 | -0.022 | 0.03 | 0.03 | 0 | 0.27 | 0.09 | 0.28 | 0.23 | 0.27 | 0.22 |
| 1 | 156263940 | rs10908495 | 0.823 | -0.264 | -1.135 | 0.268 | -0.022 | 0.03 | 0.03 | 0 | 0.27 | 0.09 | 0.28 | 0.23 | 0.27 | 0.22 |
| 1 | 156264000 | rs10908496 | 0.991 | -0.265 | -1.136 | 0.267 | -0.028 | 0.025 | 0.025 | 0 | 0.26 | 0.07 | 0.27 | 0.23 | 0.27 | 0.22 |
| 1 | 156273061 | rs11264467 | 0.42 | -0.318 | -1.141 | 0.273 | 0 | 0 | 0 | 0 | 0.31 | 0.28 | 0.33 | 0.3 | 0.27 | 0.25 |
| 1 | 156275281 | rs7522860 | 0.029 | -0.001 | -1.007 | 0.317 | -0.003 | 0.034 | 0.034 | 0 | 0.53 | 0.29 | 0.64 | 0.46 | 0.46 | 0.52 |
| 1 | 156278175 | rs10908498 | 0.7 | 0.09 | -1.103 | 0.304 | -0.022 | 0.03 | 0.03 | 0 | 0.27 | 0.09 | 0.28 | 0.23 | 0.27 | 0.22 |
| 1 | 156300082 | rs10908502 | 0.802 | 0.09 | -1.103 | 0.305 | -0.024 | 0.027 | 0.027 | 0 | 0.27 | 0.09 | 0.28 | 0.23 | 0.27 | 0.22 |
| 1 | 156302575 | rs12141236 | 0.802 | 0.09 | -1.103 | 0.305 | -0.024 | 0.027 | 0.027 | 0 | 0.27 | 0.09 | 0.28 | 0.23 | 0.27 | 0.22 |
| 1 | 156306918 | rs11264475 | 0.802 | 0.09 | -1.103 | 0.305 | -0.024 | 0.027 | 0.027 | 0 | 0.27 | 0.09 | 0.28 | 0.23 | 0.27 | 0.22 |
| 1 | 156311165 | rs12117903 | 0.69 | 0.102 | -1.097 | 0.31 | -0.021 | 0.032 | 0.032 | 0 | 0.28 | 0.09 | 0.28 | 0.23 | 0.27 | 0.22 |
| 1 | 156314104 | rs2025577 | 0.044 | 0.198 | -1.085 | 0.4 | -0.006 | 0.023 | 0.023 | 0 | 0.54 | 0.28 | 0.64 | 0.48 | 0.46 | 0.58 |
| 1 | 156314147 | rs12022657 | -0.003 | 0.51 | -1.24 | 0.697 | -0.032 | 0.018 | 0.018 | 0 | 0.26 | 0.06 | 0.27 | 0.23 | 0.27 | 0.22 |
| 1 | 156321154 | rs11264477 | 0.225 | 0.817 | -1.501 | 1.024 | -0.027 | -0.001 | -0.001 | 0.049 | 0.21 | 0.06 | 0.25 | 0.23 | 0.24 | 0.21 |
| 1 | 156333831 | rs2764407 | 0.417 | 0.402 | -0.858 | 1.467 | 0 | 0.001 | 0.001 | 0.286 | 0.79 | 0.81 | 0.84 | 0.72 | 0.77 | 0.82 |
| 1 | 156340618 | rs2985714 | 0.567 | 0.174 | -0.569 | 1.332 | -0.003 | -0.001 | -0.001 | 0.691 | 0.85 | 0.81 | 0.91 | 0.86 | 0.9 | 0.81 |
| 1 | 156341494 | rs7525133 | -0.932 | 0.644 | -0.563 | 1.192 | 0 | -0.001 | -0.001 | 0 | 0.06 | 0.07 | 0.05 | 0.02 | 0.07 | 0.03 |
| 1 | 156342917 | rs4641304 | 0.075 | 0.392 | -0.41 | 1.266 | 0 | -0.002 | -0.002 | 5.37 | 0.27 | 0.22 | 0.36 | 0.4 | 0.26 | 0.28 |
| 1 | 156347131 | rs2245623 | 1.645 | 0.345 | -0.229 | 0.641 | -0.038 | 0.122 | 0.122 | 0.022 | 0.39 | 0.19 | 0.28 | 0.1 | 0.46 | 0.35 |
| 1 | 156351699 | rs3748569 | -0.829 | 0.172 | 0.253 | 0.949 | 0.001 | 0.007 | 0.007 | 0 | 0.42 | 0.57 | 0.6 | 0.7 | 0.41 | 0.44 |
| 1 | 156351792 | rs3748568 | -0.831 | 0.156 | 0.331 | 0.794 | -0.002 | -0.011 | -0.011 | 0 | 0.42 | 0.54 | 0.55 | 0.75 | 0.4 | 0.44 |
| 1 | 156352396 | rs872120 | -0.235 | 0.146 | 0.505 | 0.892 | -0.008 | 0.004 | 0.004 | 0 | 0.31 | 0.39 | 0.39 | 0.35 | 0.3 | 0.3 |
| 1 | 156352974 | rs867921 | -1.052 | -0.047 | 0.393 | 0.785 | -0.01 | 0.012 | 0.012 | 0 | 0.31 | 0.61 | 0.29 | 0.54 | 0.24 | 0.33 |
| 1 | 156353041 | rs3790462 | -0.001 | -0.031 | 0.355 | 0.804 | -0.002 | 0 | 0 | 0 | 0.29 | 0.28 | 0.38 | 0.35 | 0.28 | 0.29 |
| 1 | 156353970 | rs6668419 | -1.251 | 0.07 | 0.737 | 1.275 | -0.024 | 0.033 | 0.033 | 0 | 0.28 | 0.42 | 0.27 | 0.53 | 0.21 | 0.32 |
| 1 | 156356480 | rs6686886 | 0.878 | -0.09 | 0.456 | 1.203 | -0.002 | 0.02 | 0.02 | 0 | 0.83 | 0.82 | 0.74 | 0.55 | 0.88 | 0.8 |
| 1 | 156364720 | rs7551218 | -1.713 | -0.136 | 0.375 | 0.454 | 0 | -0.073 | -0.073 | 0.224 | 0.57 | 0.53 | 0.66 | 0.9 | 0.46 | 0.56 |
| 1 | 156367438 | rs11590409 | -0.505 | -0.289 | 0.434 | 0.778 | 0.034 | 0.003 | 0.003 | 0 | 0.32 | 0.19 | 0.34 | 0.33 | 0.24 | 0.29 |
| 1 | 156369815 | rs11264482 | -0.371 | -0.295 | 0.121 | 0.784 | -0.018 | -0.049 | -0.049 | 0 | 0.31 | 0.03 | 0.38 | 0.32 | 0.28 | 0.32 |
| 1 | 156374256 | rs3431 | -0.413 | -1.013 | 0.324 | 1.267 | 0.014 | -0.035 | -0.035 | 0 | 0.24 | 0.12 | 0.35 | 0.32 | 0.22 | 0.26 |
| 1 | 156378425 | rs7551781 | -1.462 | -1.022 | 0.185 | 1.26 | 0.043 | 0.026 | 0.026 | 0 | 0.4 | 0.75 | 0.56 | 0.71 | 0.26 | 0.39 |
| 1 | 156381160 | rs12022607 | -0.006 | -0.445 | 0.076 | 1.94 | 0 | -0.048 | -0.048 | 0 | 0.08 | 0 | 0.05 | 0.16 | 0.05 | 0.1 |
| 1 | 156386787 | rs2296373 | -1.351 | -0.127 | 0.544 | 1.595 | -0.03 | 0.068 | 0.068 | 0 | 0.12 | 0.19 | 0.18 | 0.37 | 0.04 | 0.12 |
| 1 | 156394000 | rs7548323 | -1.76 | 0.516 | 0.665 | 1.544 | 0.005 | 0.09 | 0.09 | 0 | 0.17 | 0.31 | 0.15 | 0.36 | 0.18 | 0.14 |
| 1 | 156395744 | rs16837386 | -2.165 | 0.2 | 0.745 | 1.557 | -0.016 | 0.06 | 0.06 | 2.758 | 0.18 | 0.72 | 0.13 | 0.36 | 0.12 | 0.12 |
| 1 | 156397981 | rs7534014 | -1.796 | 0.28 | 1.314 | 1.203 | 0.035 | 0.091 | 0.091 | 3.761 | 0.05 | 0.34 | 0.02 | 0.03 | 0 | 0.01 |
| 1 | 156400060 | rs10494305 | -2.102 | 0.239 | 1.131 | 1.294 | -0.001 | 0.006 | 0.006 | 0.263 | 0.1 | 0.24 | 0.06 | 0.09 | 0.09 | 0.12 |
| 1 | 156405037 | rs2025579 | 0.461 | 0.723 | 0.631 | 1.676 | -0.02 | -0.038 | -0.038 | 0 | 0.89 | 0.96 | 0.75 | 0.8 | 0.86 | 0.81 |
| 1 | 156406853 | rs4414033 | 1.848 | 0.784 | 1.036 | 1.017 | -0.001 | 0 | 0 | 0 | 0.6 | 0.38 | 0.56 | 0.74 | 0.62 | 0.66 |
| 1 | 156407051 | rs6665039 | -1.748 | 0.496 | 1.684 | 0.795 | -0.006 | -0.042 | -0.042 | 0 | 0.3 | 0.57 | 0.19 | 0.08 | 0.25 | 0.2 |
| 1 | 156421654 | rs2477972 | 0.853 | 1.017 | 2.044 | 0.684 | -0.034 | -0.054 | -0.054 | 0 | 0.78 | 0.57 | 0.89 | 0.94 | 0.87 | 0.89 |
| 1 | 156425048 | rs16837408 | -0.713 | 1.238 | 2.007 | 0.914 | 0.01 | 0.043 | 0.043 | 0 | 0.11 | 0.19 | 0.27 | 0.19 | 0.15 | 0.2 |
| 1 | 156430749 | rs750439 | 2.293 | 1.187 | 2.143 | 0.735 | 0.003 | -0.01 | -0.01 | 0 | 0.65 | 0.24 | 0.59 | 0.75 | 0.7 | 0.68 |
| 1 | 156431341 | rs1171556 | 0.015 | 1.065 | 2.221 | 0.675 | -0.007 | 0.012 | 0.012 | 0 | 0.9 | 0.96 | 0.92 | 1 | 0.91 | 0.96 |
| 1 | 156434703 | rs1050316 | 2.524 | 1.126 | 2.187 | 0.734 | -0.01 | -0.01 | -0.01 | 0.155 | 0.67 | 0.24 | 0.59 | 0.76 | 0.67 | 0.71 |
| 1 | 156446242 | rs2274316 | 2.524 | 1.126 | 2.187 | 0.72 | -0.01 | -0.013 | -0.013 | 0 | 0.67 | 0.24 | 0.59 | 0.76 | 0.67 | 0.71 |
| 1 | 156450719 | rs1185700 | -1.578 | 1.097 | 2.353 | 0.703 | 0.002 | 0.012 | 0.012 | 0 | 0.17 | 0.18 | 0.3 | 0.25 | 0.24 | 0.24 |
| 1 | 156450740 | rs1925950 | 2.651 | 1.083 | 2.312 | 0.694 | -0.01 | -0.013 | -0.013 | 0 | 0.67 | 0.24 | 0.6 | 0.76 | 0.67 | 0.72 |
| 1 | 156454383 | rs1750304 | 1.183 | 1.124 | 2.353 | 0.702 | -0.006 | -0.084 | -0.084 | 0 | 0.86 | 0.68 | 0.91 | 1 | 0.92 | 0.95 |
| 1 | 156460511 | rs746527 | -1.496 | 1.144 | 2.096 | 0.744 | -0.006 | -0.006 | -0.006 | 0.465 | 0.16 | 0.16 | 0.29 | 0.24 | 0.24 | 0.21 |
| 1 | 156462848 | rs1925951 | -0.214 | 1.192 | 2.043 | 0.77 | 0.001 | 0.006 | 0.006 | 0 | 0.93 | 0.97 | 0.95 | 1 | 0.94 | 0.97 |
| 1 | 156472214 | rs1171551 | -1.282 | 1.19 | 2.063 | 0.811 | -0.018 | -0.022 | -0.022 | 0 | 0.14 | 0.09 | 0.29 | 0.24 | 0.22 | 0.22 |
| 1 | 156480324 | rs10159180 | -1.059 | 0.345 | 1.531 | 0.255 | -0.006 | 0.023 | 0.023 | 0.074 | 0.57 | 0.66 | 0.53 | 0.71 | 0.51 | 0.75 |
| 1 | 156486009 | rs6685228 | 1.47 | 0.382 | 1.621 | 0.163 | -0.027 | -0.079 | -0.079 | 0 | 0.76 | 0.43 | 0.86 | 0.97 | 0.85 | 0.86 |
| 1 | 156486061 | rs6674079 | 0.373 | 0.449 | 1.538 | 0.232 | -0.008 | -0.011 | -0.011 | 0 | 0.34 | 0.26 | 0.4 | 0.68 | 0.36 | 0.6 |
| 1 | 156494916 | rs947661 | 0.485 | 0.54 | 1.714 | 0.411 | 0.001 | 0.003 | 0.003 | 0 | 0.28 | 0.28 | 0.37 | 0.68 | 0.3 | 0.56 |
| 1 | 156499969 | rs1171566 | -1.199 | -0.534 | 1.825 | 0.843 | 0.06 | -0.027 | -0.027 | 0.167 | 0.47 | 0.84 | 0.44 | 0.6 | 0.33 | 0.69 |
| 1 | 156504579 | rs745596 | -0.275 | -0.667 | 2.059 | 0.832 | -0.009 | -0.035 | -0.035 | 0.345 | 0.1 | 0.16 | 0.05 | 0.01 | 0.07 | 0.07 |
| 1 | 156505061 | rs942965 | 0.598 | -0.658 | 1.97 | 0.914 | 0.001 | -0.001 | -0.001 | 0 | 0.26 | 0.25 | 0.32 | 0.57 | 0.19 | 0.54 |
| 1 | 156508477 | rs1177696 | -1.19 | -0.65 | 1.915 | 0.915 | 0.067 | -0.032 | -0.032 | 0 | 0.47 | 0.84 | 0.44 | 0.6 | 0.32 | 0.68 |
| 1 | 156513556 | rs1171576 | 0.421 | -0.661 | 2.09 | 0.945 | -0.027 | 0.096 | 0.096 | 0.052 | 0.27 | 0.46 | 0.34 | 0.57 | 0.19 | 0.54 |
| 1 | 156516796 | rs7533781 | -0.085 | -0.509 | 2.413 | 1.105 | -0.012 | 0.015 | 0.015 | 0.186 | 0.35 | 0.36 | 0.37 | 0.57 | 0.26 | 0.58 |
| 1 | 156518379 | rs744224 | -0.016 | -0.501 | 2.337 | 1.108 | 0.005 | -0.01 | -0.01 | 0 | 0.34 | 0.3 | 0.36 | 0.57 | 0.26 | 0.57 |
| 1 | 156520320 | rs1778818 | 0.003 | -0.541 | 2.163 | 1.201 | 0.023 | -0.021 | -0.021 | 0 | 0.34 | 0.18 | 0.31 | 0.52 | 0.26 | 0.58 |
| 1 | 156521284 | rs4661178 | -1.473 | -0.549 | 2.329 | 1.386 | -0.015 | 0.038 | 0.038 | 0 | 0.12 | 0.43 | 0.21 | 0.15 | 0.06 | 0.1 |
| 1 | 156524221 | rs6694580 | -0.389 | -0.463 | 1.831 | 1.285 | 0.017 | -0.011 | -0.011 | 0.028 | 0.4 | 0.23 | 0.36 | 0.54 | 0.31 | 0.6 |
| 1 | 156525023 | rs12760461 | 1.039 | -0.478 | 1.822 | 1.279 | 0.016 | -0.063 | -0.063 | 0 | 0.47 | 0.14 | 0.55 | 0.43 | 0.63 | 0.27 |
| 1 | 156526387 | rs11264498 | 1.039 | -0.478 | 1.822 | 1.279 | 0.016 | -0.063 | -0.063 | 0.005 | 0.47 | 0.14 | 0.55 | 0.43 | 0.63 | 0.27 |

| | | | | | | | | | | | | | | | | |
|---|---|---|---|---|---|---|---|---|---|---|---|---|---|---|---|---|
| 1 | 156529034 | rs12757164 | 1.039 | -0.478 | 1.822 | 1.283 | 0.016 | -0.056 | -0.056 | 0 | 0.47 | 0.14 | 0.55 | 0.42 | 0.63 | 0.27 |
| 1 | 156532344 | rs7548309 | 0.256 | -0.522 | 1.794 | 1.313 | 0 | 0 | 0 | 0.579 | 0.6 | 0.63 | 0.63 | 0.46 | 0.71 | 0.42 |
| 1 | 156550254 | rs10908509 | -0.179 | -0.807 | 1.535 | 1.612 | 0.019 | -0.017 | -0.017 | 0 | 0.35 | 0.23 | 0.34 | 0.61 | 0.24 | 0.59 |
| 1 | 156550968 | rs6679425 | -1.59 | -0.834 | 1.727 | 1.076 | -0.033 | -0.024 | -0.024 | 0.003 | 0.14 | 0.41 | 0.07 | 0.03 | 0.06 | 0.07 |
| 1 | 156551848 | rs6682716 | 0.996 | -1.15 | 1.056 | 1.467 | 0.141 | 0.012 | 0.012 | 0 | 0.51 | 0.08 | 0.46 | 0.29 | 0.7 | 0.34 |
| 1 | 156560502 | rs4661188 | 0.475 | -1.115 | 1.807 | 0.812 | 0.007 | 0 | 0 | 0 | 0.58 | 0.42 | 0.67 | 0.52 | 0.71 | 0.36 |
| 1 | 156560561 | rs4661189 | -0.046 | -1.103 | 1.404 | 0.82 | -0.018 | -0.003 | -0.003 | 0 | 0.92 | 0.86 | 0.96 | 0.97 | 0.96 | 0.92 |
| 1 | 156563813 | rs6668178 | 0.012 | -1.103 | 1.404 | 0.82 | -0.018 | -0.004 | -0.004 | 0 | 0.92 | 0.86 | 0.96 | 0.97 | 0.96 | 0.92 |
| 1 | 156564577 | rs4661047 | 0.076 | -1.067 | 1.057 | 0.832 | -0.014 | -0.003 | -0.003 | 0 | 0.08 | 0.1 | 0.04 | 0.03 | 0.04 | 0.08 |
| 1 | 156564606 | rs4661190 | 0.076 | -1.067 | 1.057 | 0.832 | -0.014 | -0.003 | -0.003 | 0.247 | 0.08 | 0.1 | 0.04 | 0.03 | 0.04 | 0.08 |
| 1 | 156564922 | rs3795733 | 0.953 | -1.078 | 1.055 | 0.819 | 0.119 | -0.014 | -0.014 | 0 | 0.52 | 0.16 | 0.62 | 0.49 | 0.7 | 0.34 |
| 1 | 156568252 | rs11264505 | -0.025 | -1.702 | 1.328 | 1.104 | -0.014 | 0.006 | 0.006 | 1.015 | 0.08 | 0.1 | 0.04 | 0.05 | 0.04 | 0.08 |
| 1 | 156572658 | rs12023410 | -0.016 | -1.689 | 0.738 | 1.088 | 0.03 | -0.018 | -0.018 | 0 | 0.31 | 0.11 | 0.28 | 0.43 | 0.24 | 0.56 |
| 1 | 156581534 | rs7536235 | -0.105 | -1.652 | 0.497 | 0.932 | 0.022 | 0.003 | 0.003 | 0 | 0.32 | 0.21 | 0.28 | 0.38 | 0.25 | 0.55 |
| 1 | 156588749 | rs2365714 | -0.11 | -1.625 | 0.292 | 0.885 | 0.016 | 0.002 | 0.002 | 0.004 | 0.32 | 0.25 | 0.28 | 0.38 | 0.25 | 0.55 |
| 1 | 156590696 | rs11264508 | -0.113 | -1.636 | 0.294 | 0.887 | 0.016 | 0.002 | 0.002 | 0.005 | 0.32 | 0.25 | 0.28 | 0.38 | 0.24 | 0.55 |
| 1 | 156594656 | rs12567958 | 0.983 | -1.556 | 0.217 | 0.906 | 0.105 | 0.008 | 0.008 | 0 | 0.54 | 0.19 | 0.63 | 0.56 | 0.71 | 0.37 |
| 1 | 156597408 | rs10908510 | 1.533 | -0.97 | 0.825 | 0.531 | -0.058 | -0.015 | -0.015 | 0.484 | 0.86 | 0.36 | 0.87 | 0.94 | 0.96 | 0.91 |
| 1 | 156603422 | rs4319322 | 0.813 | -0.665 | 0.201 | 0.602 | -0.016 | 0.024 | 0.024 | 0.898 | 0.89 | 0.6 | 0.94 | 0.95 | 0.96 | 0.92 |
| 1 | 156608394 | rs12408758 | -0.422 | -0.517 | -1.133 | 0.591 | 0.014 | -0.033 | -0.033 | 0.673 | 0.37 | 0.14 | 0.31 | 0.41 | 0.32 | 0.62 |
| 1 | 156610688 | rs1361744 | 0.803 | -0.844 | -0.423 | 0.204 | -0.042 | -0.066 | -0.066 | 0 | 0.88 | 0.4 | 0.87 | 0.99 | 0.94 | 0.99 |
| 1 | 156612932 | rs3795736 | -0.267 | -0.728 | -0.994 | 0.29 | -0.008 | -0.046 | -0.046 | 0 | 0.33 | 0.13 | 0.27 | 0.41 | 0.29 | 0.62 |
| 1 | 156617968 | rs2277873 | -0.258 | -1.307 | -0.821 | -0.466 | 0.004 | 0.032 | 0.032 | 0 | 0.41 | 0.24 | 0.35 | 0.35 | 0.37 | 0.52 |
| 1 | 156618768 | rs4661209 | -0.954 | -1.791 | -1.157 | -0.903 | -0.09 | -0.021 | -0.021 | 0 | 0.6 | 0.89 | 0.47 | 0.4 | 0.47 | 0.68 |
| 1 | 156622252 | rs1056695 | -0.007 | -1.623 | -0.643 | -0.819 | 0.001 | 0.039 | 0.039 | 0 | 0.43 | 0.26 | 0.37 | 0.35 | 0.39 | 0.53 |
| 1 | 156627996 | rs2298135 | -0.104 | -1.522 | -0.713 | -0.541 | 0.012 | -0.027 | -0.027 | 0 | 0.07 | 0.02 | 0.14 | 0.29 | 0.04 | 0.28 |
| 1 | 156629037 | rs11360 | -0.602 | -1.669 | -0.533 | -1.095 | 0.004 | 0.045 | 0.045 | 0 | 0.51 | 0.66 | 0.47 | 0.41 | 0.45 | 0.67 |
| 1 | 156634639 | rs4661210 | 0.195 | -1.635 | -0.461 | -1.133 | -0.079 | 0.198 | 0.198 | 0.004 | 0.37 | 0.1 | 0.29 | 0.13 | 0.41 | 0.34 |
| 1 | 156640156 | rs3748570 | -0.42 | -1.551 | 0.005 | -1.041 | -0.004 | -0.006 | -0.006 | 0 | 0.65 | 0.71 | 0.71 | 0.87 | 0.66 | 0.67 |
| 1 | 156640308 | rs3748571 | 0.828 | -0.984 | -1.409 | -1.425 | -0.085 | -0.071 | -0.071 | 0.165 | 0.26 | 0.02 | 0.29 | 0.29 | 0.3 | 0.31 |
| 1 | 156640503 | rs3828043 | 0.823 | -1.022 | -1.407 | -1.425 | -0.081 | -0.071 | -0.071 | 0 | 0.27 | 0.02 | 0.29 | 0.29 | 0.3 | 0.31 |
| 1 | 156640933 | rs2365718 | 0.129 | -0.844 | -1.095 | -1.459 | -0.006 | 0.014 | 0.014 | 0 | 0.05 | 0.22 | 0.04 | 0.01 | 0.01 | 0.04 |
| 1 | 156641537 | rs951781 | 0.724 | -0.641 | -1.903 | -1.383 | -0.104 | 0.16 | 0.16 | 0 | 0.32 | 0.04 | 0.27 | 0.12 | 0.34 | 0.33 |
| 1 | 156650909 | rs11264516 | 0.021 | -0.17 | -1.37 | -0.543 | -0.017 | -0.047 | -0.047 | 2.097 | 0.3 | 0.16 | 0.32 | 0.54 | 0.36 | 0.33 |
| 1 | 156659768 | rs7532302 | -0.259 | 0.469 | -1.239 | -0.324 | -0.01 | -0.004 | -0.004 | 0 | 0.71 | 0.76 | 0.61 | 0.63 | 0.67 | 0.7 |
| 1 | 156662372 | rs3935541 | 0.181 | 0.361 | -1.647 | -0.638 | 0.007 | 0.126 | 0.126 | 0 | 0.36 | 0.18 | 0.27 | 0.08 | 0.29 | 0.45 |
| 1 | 156662759 | rs12759184 | 0.055 | 0.527 | -0.76 | -0.092 | 0 | 0.016 | 0.016 | 0 | 0.31 | 0.35 | 0.3 | 0.56 | 0.32 | 0.21 |
| 1 | 156663217 | rs4492611 | -0.016 | -0.006 | -1.603 | -0.249 | 0.074 | 0.08 | 0.08 | 0 | 0.48 | 0.21 | 0.42 | 0.64 | 0.62 | 0.5 |
| 1 | 156663278 | rs4402117 | -0.878 | -0.284 | -0.895 | -0.86 | -0.014 | -0.046 | -0.046 | 0.209 | 0.35 | 0.59 | 0.27 | 0.09 | 0.24 | 0.31 |
| 1 | 156667399 | rs11264527 | 0.48 | -0.141 | -0.41 | -0.975 | -0.003 | -0.006 | -0.006 | 0 | 0.61 | 0.5 | 0.48 | 0.78 | 0.65 | 0.7 |
| 1 | 156670545 | rs4661213 | 1.465 | -0.791 | 0.403 | -0.864 | -0.001 | -0.023 | -0.023 | 0.006 | 0.93 | 0.74 | 0.97 | 1 | 0.97 | 0.96 |
| 1 | 156686005 | rs12140437 | -0.369 | -0.556 | -0.854 | -1.253 | -0.079 | 0.084 | 0.084 | 0 | 0.23 | 0.02 | 0.22 | 0.11 | 0.32 | 0.21 |
| 1 | 156688372 | rs6427327 | 0.305 | -0.986 | -0.106 | -1.4 | -0.013 | 0.001 | 0.001 | 0 | 0.65 | 0.43 | 0.44 | 0.72 | 0.72 | 0.55 |
| 1 | 156689481 | rs11811735 | 0.569 | -0.96 | -0.079 | -1.395 | -0.003 | -0.023 | -0.023 | 0.934 | 0.08 | 0.1 | 0.03 | 0 | 0.04 | 0.01 |
| 1 | 156695504 | rs16837622 | -0.279 | -1.274 | 0.539 | -1.427 | -0.023 | 0.078 | 0.078 | 0 | 0.12 | 0.28 | 0.28 | 0.18 | 0.06 | 0.12 |
| 1 | 156695537 | rs12023776 | 0.516 | -1.409 | 0.706 | -1.968 | -0.003 | 0.024 | 0.024 | 0 | 0.21 | 0.12 | 0.25 | 0.07 | 0.21 | 0.34 |
| 1 | 156698346 | rs10908517 | 0.631 | -1.436 | 0.737 | -1.983 | -0.007 | 0.023 | 0.023 | 0 | 0.19 | 0.1 | 0.2 | 0.06 | 0.21 | 0.17 |
| 1 | 156700651 | rs12145743 | 0.358 | -1.584 | 0.562 | -2.1 | -0.092 | 0.088 | 0.088 | 0 | 0.23 | 0.02 | 0.22 | 0.12 | 0.33 | 0.22 |
| 1 | 156700845 | rs7550297 | 2.331 | -1.623 | 0.903 | -2.142 | -0.019 | -0.013 | -0.013 | 0 | 0.31 | 0.15 | 0.17 | 0.68 | 0.33 | 0.26 |
| 1 | 156707697 | rs11581044 | 2.309 | -1.57 | 0.938 | -2.152 | -0.026 | -0.015 | -0.015 | 0.306 | 0.31 | 0.14 | 0.17 | 0.69 | 0.33 | 0.27 |
| 1 | 156711623 | rs3806417 | 0.332 | -1.509 | 1.078 | -2.798 | -0.043 | 0.087 | 0.087 | 0 | 0.24 | 0.07 | 0.22 | 0.13 | 0.33 | 0.26 |
| 1 | 156713558 | rs4399146 | 0.321 | -1.508 | 1.271 | -2.798 | -0.042 | 0.088 | 0.088 | 0 | 0.24 | 0.07 | 0.22 | 0.13 | 0.33 | 0.26 |
| 1 | 156720382 | rs12404207 | 0.949 | -1.535 | 1.469 | -2.815 | 0.001 | 0.004 | 0.004 | 0 | 0.51 | 0.5 | 0.5 | 0.79 | 0.43 | 0.39 |
| 1 | 156720550 | rs11264534 | 0.319 | -1.485 | 1.358 | -2.794 | -0.042 | 0.088 | 0.088 | 0 | 0.24 | 0.07 | 0.21 | 0.13 | 0.33 | 0.26 |
| 1 | 156725653 | rs11805654 | 0.868 | -1.69 | 1.714 | -2.263 | -0.052 | 0.017 | 0.017 | 0 | 0.16 | 0.03 | 0.24 | 0.06 | 0.21 | 0.15 |
| 1 | 156737969 | rs11264542 | 0.839 | -1.677 | 1.72 | -2.257 | -0.051 | 0.02 | 0.02 | 0 | 0.17 | 0.03 | 0.23 | 0.07 | 0.22 | 0.15 |
| 1 | 156738271 | rs16837644 | 0.288 | -1.571 | 1.903 | -2.213 | -0.011 | -0.048 | -0.048 | 0.081 | 0.09 | 0.19 | 0.04 | 0 | 0.04 | 0.01 |
| 1 | 156743173 | rs11264546 | 1.998 | -1.571 | 1.614 | -2.248 | -0.099 | 0.08 | 0.08 | 0 | 0.39 | 0.09 | 0.36 | 0.72 | 0.43 | 0.28 |
| 1 | 156746608 | rs1412310 | -0.381 | -1.562 | 1.65 | -2.25 | 0.001 | 0.058 | 0.058 | 0 | 0.31 | 0.22 | 0.3 | 0.08 | 0.28 | 0.36 |
| 1 | 156754465 | rs3765787 | -0.11 | -1.158 | 1.932 | -1.998 | 0.004 | -0.052 | -0.052 | 0 | 0.81 | 0.69 | 0.84 | 0.88 | 0.77 | 0.75 |
| 1 | 156765973 | rs3765783 | 1.072 | -1.17 | 1.956 | -2.004 | -0.016 | -0.059 | -0.059 | 0.043 | 0.71 | 0.46 | 0.68 | 0.81 | 0.7 | 0.64 |
| 1 | 156767253 | rs6701637 | -0.25 | -1.114 | 1.846 | -1.886 | -0.023 | -0.022 | -0.022 | 0 | 0.11 | 0.22 | 0.16 | 0.06 | 0.06 | 0.11 |
| 1 | 156774021 | rs16837672 | -0.261 | -1.114 | 1.845 | -1.892 | -0.023 | -0.025 | -0.025 | 0 | 0.11 | 0.23 | 0.16 | 0.06 | 0.06 | 0.11 |
| 1 | 156774029 | rs2644608 | -1.119 | -1.055 | 1.71 | -1.883 | -0.001 | -0.003 | -0.003 | 0 | 0.46 | 0.48 | 0.45 | 0.21 | 0.49 | 0.61 |
| 1 | 156785356 | rs2150906 | 0.611 | -1.06 | 1.605 | -1.355 | -0.019 | -0.013 | -0.013 | 0 | 0.88 | 0.6 | 0.83 | 0.94 | 0.93 | 0.9 |
| 1 | 156785617 | rs1800601 | 1.349 | -1.132 | 1.583 | -1.425 | -0.008 | -0.031 | -0.031 | 0 | 0.7 | 0.44 | 0.67 | 0.81 | 0.69 | 0.64 |
| 1 | 156792944 | rs12073087 | -1.166 | -0.921 | 1.664 | -1.203 | -0.007 | 0.004 | 0.004 | 0.012 | 0.11 | 0.46 | 0.16 | 0.05 | 0.07 | 0.06 |
| 1 | 156796697 | rs7522395 | -0.441 | -0.833 | 1.465 | -1.28 | -0.004 | 0.006 | 0.006 | 0 | 0.86 | 0.91 | 0.86 | 0.88 | 0.8 | 0.76 |
| 1 | 156796844 | rs6427332 | -0.441 | -0.833 | 1.465 | -1.28 | -0.004 | 0.006 | 0.006 | 0 | 0.86 | 0.91 | 0.86 | 0.88 | 0.8 | 0.76 |

| | | | | | | | | | | | | | | | | |
|---|---|---|---|---|---|---|---|---|---|---|---|---|---|---|---|---|
| 1 | 156797559 | rs11264565 | -0.68 | -0.763 | 1.729 | -1.223 | -0.002 | -0.001 | -0.001 | 0 | 0.09 | 0.26 | 0.15 | 0.03 | 0.07 | 0.06 |
| 1 | 156802657 | rs4661061 | 0.568 | -0.699 | 1.315 | -1.174 | 0 | 0.001 | 0.001 | 0 | 0.75 | 0.78 | 0.81 | 0.85 | 0.76 | 0.73 |
| 1 | 156804705 | rs16837709 | -1.084 | -0.72 | 1.331 | -1.167 | -0.005 | 0.011 | 0.011 | 0.418 | 0.11 | 0.53 | 0.16 | 0.04 | 0.07 | 0.05 |
| 1 | 156805697 | rs11812062 | 0.453 | -0.455 | 0.314 | -1.031 | 0 | -0.008 | -0.008 | 0.017 | 0.07 | 0.05 | 0.03 | 0 | 0.04 | 0.01 |
| 1 | 156805803 | rs7534418 | 0.53 | -0.39 | 0.159 | -0.978 | -0.001 | -0.006 | -0.006 | 1.699 | 0.67 | 0.43 | 0.65 | 0.8 | 0.69 | 0.68 |
| 1 | 156809206 | rs4661229 | -0.209 | 0.027 | -0.592 | -0.466 | -0.017 | 0.063 | 0.063 | 0 | 0.25 | 0.08 | 0.26 | 0.12 | 0.25 | 0.27 |
| 1 | 156813650 | rs10908521 | -1.156 | -0.072 | 0.569 | 0.012 | -0.009 | -0.025 | -0.025 | 0 | 0.27 | 0.4 | 0.24 | 0.2 | 0.28 | 0.29 |
| 1 | 156824801 | rs4661063 | -1.524 | -0.356 | 0.937 | 0.089 | -0.047 | -0.034 | -0.034 | 0 | 0.15 | 0.49 | 0.14 | 0.07 | 0.08 | 0.11 |
| 1 | 156830218 | rs10908523 | -1.941 | -0.217 | 1.195 | 0.244 | 0.02 | 0.057 | 0.057 | 0 | 0.05 | 0.27 | 0.06 | 0.06 | 0.02 | 0.02 |
| 1 | 156836838 | rs2274497 | -2.639 | -0.461 | 1.842 | 0.362 | 0.001 | 0.093 | 0.093 | 4.687 | 0.08 | 0.46 | 0.05 | 0.1 | 0.02 | 0.02 |
| 1 | 156836967 | rs11264577 | -2.637 | -0.466 | 2.068 | 0.386 | 0.007 | 0.092 | 0.092 | 3.716 | 0.08 | 0.42 | 0.04 | 0.08 | 0.02 | 0.02 |
| 1 | 156837416 | rs11264578 | -1.163 | -0.553 | 2.056 | 0.523 | -0.022 | 0.003 | 0.003 | 0 | 0.26 | 0.43 | 0.22 | 0.32 | 0.24 | 0.32 |
| 1 | 156838141 | rs1800879 | 0.271 | 0.009 | 1.414 | -0.395 | 0.047 | -0.113 | -0.113 | 0 | 0.68 | 0.61 | 0.64 | 0.96 | 0.6 | 0.76 |
| 1 | 156841674 | rs1410082 | -1.764 | -0.858 | 1.789 | -0.336 | -0.083 | -0.091 | -0.091 | 0 | 0.15 | 0.31 | 0.05 | 0 | 0.05 | 0.08 |
| 1 | 156846233 | rs6334 | 0.302 | -1.476 | 0.842 | 0.254 | 0.001 | -0.035 | -0.035 | 0 | 0.19 | 0.11 | 0.22 | 0.32 | 0.2 | 0.25 |
| 1 | 156848106 | rs2644604 | 1.153 | -2.113 | 0.903 | 0.103 | -0.042 | 0.032 | 0.032 | 0 | 0.77 | 0.69 | 0.8 | 0.35 | 0.87 | 0.53 |
| 1 | 156855497 | rs943551 | -0.578 | -1.852 | 0.807 | 0.621 | -0.003 | 0.075 | 0.075 | 0.049 | 0.1 | 0.12 | 0.16 | 0.53 | 0.08 | 0.36 |
| 1 | 156861168 | rs2644620 | 0.767 | -2.102 | 1.175 | 0.376 | -0.022 | 0.104 | 0.104 | 0 | 0.82 | 0.45 | 0.82 | 0.69 | 0.88 | 0.8 |
| 1 | 156863043 | rs3118644 | -0.177 | -2.011 | 1.411 | 0.42 | 0.005 | 0.105 | 0.105 | 0 | 0.88 | 0.8 | 0.84 | 0.69 | 0.88 | 0.82 |
| 1 | 156869714 | rs12041331 | -1.662 | -2.649 | 1.422 | 0.706 | -0.102 | 0.1 | 0.1 | 0.023 | 0.23 | 0.47 | 0.2 | 0.46 | 0.09 | 0.37 |
| 1 | 156873463 | rs11264579 | -2.472 | -2.943 | 1.448 | 0.723 | 0.055 | 0.037 | 0.037 | 0.838 | 0.37 | 0.75 | 0.36 | 0.8 | 0.2 | 0.6 |
| 1 | 156875764 | rs2644592 | 1.419 | -2.923 | 1.835 | 0.706 | -0.059 | 0.072 | 0.072 | 0 | 0.74 | 0.61 | 0.76 | 0.63 | 0.85 | 0.68 |
| 1 | 156879580 | rs3737224 | -0.469 | -2.611 | 1.646 | 0.976 | -0.002 | 0.053 | 0.053 | 0 | 0.13 | 0.16 | 0.1 | 0.31 | 0.1 | 0.19 |
| 1 | 156883493 | rs11264581 | -0.52 | -1.807 | 1.106 | 0.794 | -0.021 | -0.007 | -0.007 | 0 | 0.09 | 0.05 | 0.3 | 0.47 | 0.18 | 0.25 |
| 1 | 156883546 | rs12137505 | -1.308 | -0.909 | 0.451 | -0.061 | -0.012 | -0.019 | -0.019 | 0 | 0.27 | 0.26 | 0.56 | 0.59 | 0.39 | 0.55 |
| 1 | 156884365 | rs749256 | -3.11 | -0.888 | -0.476 | -1.101 | -0.048 | 0.136 | 0.136 | 0.199 | 0.77 | 0.97 | 0.86 | 1 | 0.73 | 0.96 |
| 1 | 156891610 | rs12143768 | 0.27 | -0.754 | 0.082 | -0.683 | 0.001 | 0.009 | 0.009 | 0 | 0.19 | 0.2 | 0.17 | 0.28 | 0.17 | 0.16 |
| 1 | 156892394 | rs822436 | -1.086 | -1.01 | 0.364 | -0.732 | -0.006 | 0.003 | 0.003 | 0.103 | 0.27 | 0.65 | 0.2 | 0.29 | 0.17 | 0.17 |
| 1 | 156898830 | rs822426 | -0.844 | -1.011 | 0.764 | -0.73 | -0.017 | 0.04 | 0.04 | 0 | 0.25 | 0.46 | 0.21 | 0.32 | 0.19 | 0.22 |
| 1 | 156901888 | rs822429 | -1.085 | -1.023 | 0.908 | -0.716 | -0.022 | 0.029 | 0.029 | 0 | 0.27 | 0.52 | 0.21 | 0.33 | 0.19 | 0.23 |
| 1 | 156902222 | rs12119908 | 0.868 | -1.035 | 0.889 | -0.718 | 0.006 | 0.024 | 0.024 | 0 | 0.17 | 0.19 | 0.16 | 0.29 | 0.17 | 0.17 |
| 1 | 156902259 | rs822430 | -2.952 | -1.111 | 0.78 | -0.838 | -0.086 | 0.168 | 0.168 | 0 | 0.57 | 0.89 | 0.71 | 0.85 | 0.53 | 0.74 |
| 1 | 156902281 | rs822431 | 0.074 | -1.12 | 0.801 | -0.818 | 0.001 | 0.061 | 0.061 | 0 | 0.22 | 0.45 | 0.3 | 0.29 | 0.18 | 0.15 |
| 1 | 156904157 | rs822434 | 2.058 | -1.074 | 0.845 | -0.8 | -0.003 | -0.001 | -0.001 | 0 | 0.05 | 0.25 | 0.03 | 0 | 0 | 0.01 |
| 1 | 156907081 | rs945508 | -2.938 | -1.092 | 0.576 | -0.84 | -0.091 | 0.159 | 0.159 | 0 | 0.58 | 0.89 | 0.72 | 0.85 | 0.55 | 0.75 |
| 1 | 156909695 | rs2275199 | 0.867 | -0.912 | 0.792 | -0.639 | 0.002 | 0.01 | 0.01 | 0 | 0.17 | 0.16 | 0.16 | 0.29 | 0.17 | 0.17 |
| 1 | 156918348 | rs6684725 | 0.89 | -1.01 | 0.921 | -0.703 | -0.017 | -0.015 | -0.015 | 0 | 0.07 | 0.29 | 0.02 | 0 | 0 | 0.01 |
| 1 | 156921220 | rs2275202 | 1.256 | -0.932 | 0.658 | -0.698 | 0.001 | -0.012 | -0.012 | 0 | 0.09 | 0.12 | 0.15 | 0.05 | 0.09 | 0.16 |
| 1 | 156927751 | rs12132684 | -0.067 | -0.869 | 0.677 | -0.593 | -0.004 | 0.019 | 0.019 | 0 | 0.23 | 0.28 | 0.19 | 0.33 | 0.2 | 0.23 |
| 1 | 156939067 | rs2275206 | 1.072 | -0.581 | 0.651 | -0.485 | 0.009 | 0.026 | 0.026 | 0 | 0.14 | 0.15 | 0.16 | 0.27 | 0.18 | 0.17 |
| 1 | 156942228 | rs1999754 | 0.775 | -0.578 | 0.702 | -0.479 | 0.01 | 0.093 | 0.093 | 0 | 0.1 | 0.13 | 0.32 | 0.45 | 0.2 | 0.25 |
| 1 | 156952054 | rs1572416 | -0.266 | -0.647 | 0.878 | -0.516 | 0 | 0.032 | 0.032 | 0 | 0.21 | 0.28 | 0.19 | 0.33 | 0.2 | 0.23 |
| 1 | 156964052 | rs7542311 | 0.027 | -0.611 | 0.953 | -0.512 | 0.025 | 0.164 | 0.164 | 0 | 0.12 | 0.24 | 0.33 | 0.45 | 0.2 | 0.26 |
| 1 | 156968568 | rs869724 | 0.921 | -0.563 | 0.673 | -0.499 | 0.016 | 0.044 | 0.044 | 0 | 0.15 | 0.19 | 0.16 | 0.29 | 0.18 | 0.17 |
| 1 | 156972485 | rs967072 | -0.039 | -0.693 | 0.111 | -0.627 | 0.013 | 0.016 | 0.016 | 0 | 0.2 | 0.19 | 0.17 | 0.06 | 0.14 | 0.25 |
| 1 | 157005154 | rs1750802 | 2.799 | -0.179 | -0.119 | -0.676 | 0.006 | 0.135 | 0.135 | 0 | 0.42 | 0.12 | 0.27 | 0.16 | 0.37 | 0.22 |
| 1 | 157020401 | rs822587 | -0.796 | 0.077 | -0.129 | -0.52 | 0.029 | 0.164 | 0.164 | 0 | 0.89 | 0.8 | 0.68 | 0.55 | 0.8 | 0.74 |
| 1 | 157021272 | rs12139883 | 1.004 | 0.083 | -0.15 | -0.517 | -0.014 | 0.023 | 0.023 | 0 | 0.09 | 0.06 | 0.32 | 0.45 | 0.2 | 0.26 |
| 1 | 157036322 | rs12747244 | 1.201 | -0.368 | -0.207 | -1.507 | 0.002 | -0.02 | -0.02 | 0.338 | 0.1 | 0.12 | 0.04 | 0 | 0.09 | 0.06 |
| 1 | 157037055 | rs1359647 | -0.253 | -0.411 | -0.144 | -1.213 | 0.027 | 0.058 | 0.058 | 0 | 0.19 | 0.08 | 0.17 | 0.06 | 0.14 | 0.25 |
| 1 | 157046852 | rs11264612 | 0.02 | -0.079 | -0.691 | -1.021 | 0.016 | 0 | 0 | 1.339 | 0.24 | 0.25 | 0.49 | 0.72 | 0.37 | 0.42 |
| 1 | 157049759 | rs4661082 | 0.915 | -0.658 | -1.221 | -0.765 | -0.024 | 0.004 | 0.004 | 0 | 0.08 | 0.01 | 0.14 | 0.07 | 0.09 | 0.15 |
| 1 | 157051801 | rs17412946 | -0.629 | -0.651 | -1.145 | -0.744 | 0.012 | 0.01 | 0.01 | 0 | 0.21 | 0.22 | 0.18 | 0.07 | 0.15 | 0.26 |
| 1 | 157061973 | rs1055106 | 0.707 | -0.525 | -1.912 | -0.714 | 0.035 | 0.104 | 0.104 | 0 | 0.45 | 0.21 | 0.28 | 0.24 | 0.39 | 0.26 |
| 1 | 157064929 | rs7544345 | 1.025 | -0.699 | -2.292 | -0.813 | -0.064 | 0.005 | 0.005 | 0 | 0.32 | 0.06 | 0.24 | 0.24 | 0.34 | 0.19 |
| 1 | 157065174 | rs1467662 | 0.198 | -0.693 | -1.909 | -0.905 | 0.014 | 0.025 | 0.025 | 0 | 0.19 | 0.13 | 0.18 | 0.06 | 0.16 | 0.3 |
| 1 | 157066747 | rs10796969 | -0.289 | -0.403 | -1.831 | -0.973 | -0.042 | 0.059 | 0.059 | 5.087 | 0.39 | 0.25 | 0.63 | 0.8 | 0.55 | 0.64 |
| 1 | 157066893 | rs17404670 | 0.49 | -0.089 | -1.711 | -0.547 | -0.126 | 0.114 | 0.114 | 0 | 0.33 | 0.01 | 0.23 | 0.13 | 0.34 | 0.14 |
| 1 | 157066950 | rs6427351 | -1.855 | 0.206 | 0.203 | 1.107 | 0.022 | 0.311 | 0.311 | 0 | 0.06 | 0.18 | 0.2 | 0.42 | 0.08 | 0.13 |
| 1 | 157077305 | rs7521028 | -1.423 | 0.117 | -0.039 | -0.119 | -0.027 | -0.037 | -0.037 | 0.973 | 0.17 | 0.38 | 0.2 | 0.13 | 0.12 | 0.1 |
| 1 | 157077577 | rs6689840 | -1.4 | 0.067 | -0.036 | -0.117 | -0.029 | -0.037 | -0.037 | 0 | 0.17 | 0.38 | 0.2 | 0.13 | 0.12 | 0.1 |
| 1 | 157079130 | rs1176529 | 0.118 | 0.264 | -0.67 | -0.18 | 0.003 | 0.027 | 0.027 | 0 | 0.54 | 0.39 | 0.52 | 0.35 | 0.56 | 0.43 |
| 1 | 157082017 | rs7519739 | -0.94 | -0.06 | -0.946 | -0.26 | -0.006 | -0.017 | -0.017 | 0.404 | 0.13 | 0.21 | 0.07 | 0.05 | 0.06 | 0.09 |
| 1 | 157083817 | rs1183449 | 0.778 | 0.222 | -1.389 | -0.164 | -0.07 | 0.079 | 0.079 | 0 | 0.39 | 0.09 | 0.37 | 0.22 | 0.44 | 0.4 |
| 1 | 157086131 | rs946577 | 0.336 | 0.421 | -1.168 | 0.002 | -0.017 | 0.122 | 0.122 | 0 | 0.47 | 0.21 | 0.41 | 0.27 | 0.47 | 0.48 |
| 1 | 157089530 | rs16838103 | -1.107 | 0.256 | -0.577 | 0.587 | -0.012 | -0.003 | -0.003 | 0 | 0.14 | 0.1 | 0.13 | 0.1 | 0.16 | 0.21 |
| 1 | 157095470 | rs2231856 | 0.163 | -0.106 | -0.9 | 0.982 | 0.005 | -0.12 | -0.12 | 0.645 | 0.15 | 0.01 | 0.29 | 0.56 | 0.11 | 0.33 |
| 1 | 157102453 | rs998842 | -2.55 | -0.8 | 0.101 | 0.674 | 0.035 | 0.051 | 0.051 | 0 | 0.45 | 0.72 | 0.57 | 0.78 | 0.3 | 0.61 |
| 1 | 157102939 | rs7521824 | 0.293 | -0.672 | -0.137 | 0.74 | 0.005 | -0.12 | -0.12 | 0 | 0.15 | 0.01 | 0.29 | 0.56 | 0.11 | 0.33 |

| | | | | | | | | | | | | | | | | |
|---|---|---|---|---|---|---|---|---|---|---|---|---|---|---|---|---|
| 1 | 157105790 | rs3737269 | -0.837 | -0.634 | -0.102 | 0.703 | -0.01 | -0.002 | -0.002 | 1.858 | 0.14 | 0.1 | 0.13 | 0.1 | 0.16 | 0.21 |
| 1 | 157109479 | rs6427352 | -2.179 | -0.804 | 0.543 | 0.717 | -0.021 | -0.034 | -0.034 | 0 | 0.4 | 0.52 | 0.55 | 0.74 | 0.28 | 0.59 |
| 1 | 157109624 | rs6427353 | -2.179 | -0.804 | 0.543 | 0.717 | -0.021 | -0.034 | -0.034 | 2.157 | 0.4 | 0.52 | 0.55 | 0.74 | 0.28 | 0.59 |
| 1 | 157118421 | rs10494309 | -1.703 | -0.726 | 0.534 | 0.775 | 0.009 | -0.001 | -0.001 | 0.146 | 0.23 | 0.25 | 0.24 | 0.2 | 0.17 | 0.26 |
| 1 | 157130719 | rs6657097 | -0.914 | -0.612 | 0.649 | 0.449 | 0.014 | -0.012 | -0.012 | 0 | 0.29 | 0.2 | 0.39 | 0.62 | 0.25 | 0.52 |
| 1 | 157133150 | rs6700498 | -1.107 | -0.463 | 0.871 | -0.179 | 0.015 | 0.017 | 0.017 | 0 | 0.37 | 0.34 | 0.53 | 0.84 | 0.27 | 0.6 |
| 1 | 157147109 | rs883328 | -0.968 | -0.229 | 0.42 | -0.149 | -0.005 | 0.03 | 0.03 | 0 | 0.37 | 0.3 | 0.56 | 0.84 | 0.41 | 0.61 |
| 1 | 157153876 | rs10908538 | -1.035 | -0.275 | 0.355 | 0.067 | -0.002 | -0.022 | -0.022 | 0.027 | 0.39 | 0.42 | 0.48 | 0.81 | 0.39 | 0.52 |
| 1 | 157153900 | rs10908539 | -1.026 | -0.183 | -0.056 | 0.062 | 0.001 | -0.005 | -0.005 | 0 | 0.42 | 0.66 | 0.5 | 0.83 | 0.39 | 0.55 |
| 1 | 157156851 | rs2275447 | -1.446 | -0.61 | -0.069 | 1.016 | -0.013 | 0.04 | 0.04 | 0 | 0.24 | 0.45 | 0.22 | 0.29 | 0.19 | 0.2 |
| 1 | 157161346 | rs6676790 | -0.734 | -0.447 | -0.313 | 1.033 | 0.002 | -0.001 | -0.001 | 0 | 0.05 | 0.06 | 0.03 | 0.02 | 0.07 | 0.03 |
| 1 | 157176781 | rs11264631 | -0.99 | -0.92 | -0.493 | 1.345 | 0.029 | -0.081 | -0.081 | 0 | 0.16 | 0.07 | 0.17 | 0.23 | 0.11 | 0.16 |
| 1 | 157193893 | rs10494310 | 0.223 | -1.304 | -0.456 | 0.616 | -0.002 | -0.02 | -0.02 | 0 | 0.44 | 0.5 | 0.42 | 0.28 | 0.52 | 0.34 |
| 1 | 157196178 | rs7515516 | -0.338 | -1.597 | -0.639 | 0.884 | 0.085 | -0.049 | -0.049 | 0 | 0.58 | 0.84 | 0.61 | 0.46 | 0.68 | 0.48 |
| 1 | 157197805 | rs6656933 | -0.759 | -1.488 | -0.273 | 0.856 | 0.003 | 0.016 | 0.016 | 0 | 0.12 | 0.12 | 0.17 | 0.18 | 0.15 | 0.15 |
| 1 | 157202651 | rs6673923 | -0.287 | -1.99 | -0.316 | 0.684 | 0.074 | -0.033 | -0.033 | 0.192 | 0.59 | 0.8 | 0.66 | 0.43 | 0.7 | 0.46 |
| 1 | 157203517 | rs10908544 | -0.637 | -1.657 | -0.293 | 0.557 | 0.01 | 0.018 | 0.018 | 0 | 0.1 | 0.16 | 0.17 | 0.15 | 0.15 | 0.12 |
| 1 | 157204332 | rs7524426 | -0.416 | -1.834 | -0.358 | 0.556 | 0.084 | -0.061 | -0.061 | 0 | 0.61 | 0.87 | 0.67 | 0.4 | 0.73 | 0.49 |
| 1 | 157206284 | rs1591785 | 0.594 | -1.835 | -0.075 | 0.438 | 0.006 | 0.025 | 0.025 | 0.324 | 0.46 | 0.55 | 0.48 | 0.26 | 0.55 | 0.37 |
| 1 | 157223575 | rs1416282 | -0.654 | -2.264 | -0.178 | 0.388 | 0.055 | -0.086 | -0.086 | 0 | 0.67 | 0.88 | 0.71 | 0.41 | 0.75 | 0.54 |
| 1 | 157236007 | rs10796976 | -0.297 | -2.138 | -0.122 | 0.399 | 0.002 | 0.011 | 0.011 | 0 | 0.1 | 0.13 | 0.17 | 0.18 | 0.15 | 0.16 |
| 1 | 157237374 | rs12120788 | 0.946 | -2.371 | 0.487 | 0.259 | 0.009 | 0.101 | 0.101 | 0.304 | 0.44 | 0.23 | 0.34 | 0.15 | 0.55 | 0.3 |
| 1 | 157239514 | rs11264659 | -1.154 | -1.983 | 0.475 | 0.484 | 0.012 | 0.034 | 0.034 | 0 | 0.13 | 0.22 | 0.19 | 0.18 | 0.16 | 0.17 |
| 1 | 157244089 | rs12079878 | -1.969 | -1.991 | 0.449 | 0.487 | 0.001 | 0.107 | 0.107 | 0.446 | 0.2 | 0.53 | 0.34 | 0.26 | 0.18 | 0.22 |
| 1 | 157252102 | rs1338685 | -1.489 | -2.12 | 0.269 | 0.574 | -0.018 | 0.056 | 0.056 | 0.228 | 0.16 | 0.43 | 0.17 | 0.16 | 0.13 | 0.18 |
| 1 | 157256043 | rs1535982 | -0.803 | -2.527 | -0.268 | 0.458 | 0.056 | -0.15 | -0.15 | 0 | 0.68 | 0.95 | 0.72 | 0.45 | 0.76 | 0.53 |
| 1 | 157256739 | rs6703082 | -0.862 | -2.513 | -0.126 | 0.224 | -0.001 | 0.062 | 0.062 | 0.161 | 0.13 | 0.36 | 0.15 | 0.15 | 0.11 | 0.16 |
| 1 | 157259119 | rs12130232 | -0.101 | -2.151 | -0.273 | -0.115 | 0.056 | 0.08 | 0.08 | 0 | 0.56 | 0.67 | 0.58 | 0.24 | 0.65 | 0.36 |
| 1 | 157263272 | rs6682568 | 0.282 | -1.727 | -0.701 | -0.173 | 0.022 | 0.053 | 0.053 | 0 | 0.41 | 0.29 | 0.38 | 0.16 | 0.59 | 0.27 |
| 1 | 157274427 | rs2209358 | -0.712 | -1.321 | -1.151 | 0.25 | 0.068 | 0.032 | 0.032 | 0 | 0.47 | 0.81 | 0.5 | 0.66 | 0.34 | 0.64 |
| 1 | 157275294 | rs16838411 | -0.359 | -1.52 | -0.129 | -0.118 | 0.051 | -0.086 | -0.086 | 0 | 0.14 | 0.32 | 0.12 | 0 | 0.19 | 0.04 |
| 1 | 157276277 | rs9427247 | -1.595 | -1.442 | 0.487 | 0.299 | 0 | -0.003 | -0.003 | 0 | 0.26 | 0.72 | 0.38 | 0.2 | 0.23 | 0.21 |
| 1 | 157276464 | rs9427248 | -1.758 | -1.493 | 0.581 | 0.274 | -0.002 | 0.01 | 0.01 | 0 | 0.27 | 0.76 | 0.38 | 0.2 | 0.24 | 0.21 |
| 1 | 157278567 | rs7418099 | -0.959 | -1.446 | 0.624 | 0.31 | -0.012 | 0.123 | 0.123 | 0 | 0.11 | 0.41 | 0.26 | 0.2 | 0.04 | 0.17 |
| 1 | 157279561 | rs991707 | 0.279 | -1.407 | 0.495 | 0.33 | -0.004 | 0.034 | 0.034 | 0 | 0.07 | 0.12 | 0.24 | 0.2 | 0.04 | 0.17 |
| 1 | 157280599 | rs7536111 | -1.467 | -1.401 | 0.577 | 0.312 | 0.028 | -0.018 | -0.018 | 0 | 0.24 | 0.53 | 0.36 | 0.2 | 0.23 | 0.21 |
| 1 | 157283479 | rs1338678 | -1.396 | -1.398 | 0.671 | 0.317 | 0.036 | -0.015 | -0.015 | 0 | 0.23 | 0.5 | 0.36 | 0.2 | 0.24 | 0.21 |
| 1 | 157296028 | rs9426948 | -0.926 | -1.367 | 0.546 | 0.348 | 0.046 | 0.005 | 0.005 | 0 | 0.2 | 0.42 | 0.35 | 0.21 | 0.23 | 0.25 |
| 1 | 157306264 | rs4661119 | -0.027 | -1.329 | 0.257 | 0.279 | 0.024 | -0.057 | -0.057 | 5.178 | 0.14 | 0.22 | 0.11 | 0 | 0.19 | 0.04 |
| 1 | 157319351 | rs7535100 | -1.243 | -1.626 | -0.205 | 0.476 | 0.004 | 0.004 | 0.004 | 0 | 0.45 | 0.55 | 0.51 | 0.69 | 0.35 | 0.64 |
| 1 | 157322310 | rs4661028 | 0.328 | -1.65 | -1.274 | 0.648 | 0.073 | -0.105 | -0.105 | 0.324 | 0.23 | 0.04 | 0.2 | 0.49 | 0.11 | 0.4 |
| 1 | 157323211 | rs1408554 | -1.337 | -1.131 | -1.443 | 0.717 | 0.039 | -0.052 | -0.052 | 4.378 | 0.47 | 0.9 | 0.43 | 0.61 | 0.33 | 0.59 |
| 1 | 157327571 | rs12239592 | 0.31 | -1.227 | -1.136 | 1.005 | 0.066 | -0.078 | -0.078 | 0 | 0.22 | 0.05 | 0.11 | 0.44 | 0.11 | 0.39 |
| 1 | 157330887 | rs873448 | -0.186 | -0.397 | -1.509 | 0.274 | -0.006 | 0.026 | 0.026 | 0.099 | 0.23 | 0.16 | 0.3 | 0.17 | 0.24 | 0.26 |
| 1 | 157331407 | rs2224901 | 0.968 | 0.143 | -2.162 | 0.327 | -0.118 | -0.107 | -0.107 | 3.454 | 0.32 | 0.06 | 0.49 | 0.31 | 0.47 | 0.27 |
| 1 | 157335891 | rs2147445 | 0.879 | 0.138 | -2.1 | 0.326 | -0.097 | -0.091 | -0.091 | 0 | 0.33 | 0.07 | 0.48 | 0.31 | 0.47 | 0.27 |
| 1 | 157338665 | rs11264701 | 0.571 | -0.462 | -1.17 | 0.447 | -0.053 | 0.029 | 0.029 | 0 | 0.22 | 0.31 | 0.1 | 0.39 | 0.09 | 0.35 |
| 1 | 157341948 | rs9427270 | 0.039 | 0.257 | -0.796 | -0.497 | 0.054 | -0.117 | -0.117 | 0 | 0.71 | 0.56 | 0.81 | 0.92 | 0.71 | 0.88 |
| 1 | 157346107 | rs12408914 | 0.345 | 0.166 | -1.071 | -0.36 | -0.048 | 0.011 | 0.011 | 0 | 0.22 | 0.07 | 0.29 | 0.2 | 0.28 | 0.31 |
| 1 | 157347856 | rs12410496 | 0.184 | 0.528 | -1.23 | -0.796 | -0.025 | 0.081 | 0.081 | 0 | 0.66 | 0.25 | 0.8 | 0.92 | 0.68 | 0.86 |
| 1 | 157348712 | rs4661142 | 1.142 | 0.373 | -1.5 | 0.12 | 0.044 | -0.078 | -0.078 | 1.051 | 0.17 | 0.08 | 0.15 | 0.4 | 0.1 | 0.31 |
| 1 | 157349170 | rs10908561 | -0.461 | 0.66 | -0.254 | -0.2 | 0.097 | 0.055 | 0.055 | 0 | 0.06 | 0.34 | 0.05 | 0.05 | 0.05 | 0.04 |
| 1 | 157354879 | rs2209354 | -1.103 | 0.507 | -0.166 | -0.389 | -0.027 | 0.031 | 0.031 | 0 | 0.79 | 0.86 | 0.87 | 0.97 | 0.75 | 0.93 |
| 1 | 157360338 | rs11264712 | -0.286 | 0.392 | 0.209 | -0.195 | 0.078 | 0.039 | 0.039 | 0 | 0.06 | 0.26 | 0.04 | 0.05 | 0.05 | 0.04 |
| 1 | 157361577 | rs12043295 | -1.054 | 0.485 | -0.174 | -0.439 | -0.032 | 0.045 | 0.045 | 0 | 0.78 | 0.86 | 0.85 | 0.97 | 0.72 | 0.92 |
| 1 | 157365378 | rs10465957 | -0.286 | -0.2 | 0.293 | 0.018 | -0.051 | 0.098 | 0.098 | 0 | 0.29 | 0.51 | 0.23 | 0.46 | 0.16 | 0.39 |
| 1 | 157367179 | rs12734925 | 0.591 | -0.07 | 0.318 | 0.011 | -0.003 | 0.001 | 0.001 | 0 | 0.49 | 0.32 | 0.62 | 0.51 | 0.55 | 0.52 |
| 1 | 157369202 | rs10908563 | 0.401 | 0.036 | 0.283 | 0.01 | 0.064 | 0.033 | 0.033 | 0 | 0.05 | 0.27 | 0.04 | 0.05 | 0.05 | 0.05 |
| 1 | 157372996 | rs7549840 | -1.154 | 0.018 | -0.108 | -0.142 | -0.044 | 0.066 | 0.066 | 0 | 0.77 | 0.88 | 0.84 | 0.97 | 0.71 | 0.92 |
| 1 | 157375078 | rs12073794 | 1.339 | -0.316 | -0.612 | -0.132 | -0.027 | 0.022 | 0.022 | 0 | 0.26 | 0.07 | 0.32 | 0.21 | 0.32 | 0.31 |
| 1 | 157378527 | rs4518899 | 0.289 | -0.542 | -0.497 | -0.008 | 0.002 | -0.005 | -0.005 | 0 | 0.1 | 0.11 | 0.08 | 0.14 | 0.07 | 0.07 |
| 1 | 157380954 | rs6681602 | 0.589 | -0.724 | -0.148 | 0.278 | -0.013 | 0.042 | 0.042 | 0 | 0.45 | 0.27 | 0.6 | 0.36 | 0.56 | 0.54 |
| 1 | 157381033 | rs4971112 | -0.015 | -0.792 | -0.338 | 0.333 | 0.014 | -0.014 | -0.014 | 0.573 | 0.41 | 0.64 | 0.45 | 0.48 | 0.35 | 0.49 |
| 1 | 157383945 | rs4246535 | 1.038 | -0.648 | -0.118 | 0.302 | 0 | -0.002 | -0.002 | 0 | 0.24 | 0.23 | 0.3 | 0.16 | 0.26 | 0.23 |
| 1 | 157384175 | rs4246536 | -0.061 | -0.499 | -0.502 | 0.422 | -0.011 | 0.009 | 0.009 | 0 | 0.65 | 0.7 | 0.75 | 0.69 | 0.64 | 0.8 |
| 1 | 157385496 | rs16838537 | 1.618 | -1.312 | -0.569 | 0.742 | 0.04 | -0.065 | -0.065 | 0 | 0.15 | 0.06 | 0.13 | 0.32 | 0.08 | 0.27 |
| 1 | 157386282 | rs9427273 | 0.34 | -1.199 | -0.185 | 0.364 | -0.002 | 0.002 | 0.002 | 0 | 0.26 | 0.41 | 0.35 | 0.25 | 0.27 | 0.21 |
| 1 | 157386331 | rs7416817 | -1.788 | -1.09 | -0.689 | -0.121 | -0.044 | 0.09 | 0.09 | 0 | 0.76 | 0.88 | 0.83 | 0.97 | 0.74 | 0.93 |
| 1 | 157386394 | rs7416823 | -0.971 | -1.122 | -0.639 | -0.005 | -0.031 | 0.042 | 0.042 | 0 | 0.69 | 0.82 | 0.8 | 0.8 | 0.68 | 0.88 |

| | | | | | | | | | | | | | | | | |
|---|---|---|---|---|---|---|---|---|---|---|---|---|---|---|---|---|
| 1 | 157386936 | rs3845580 | 0.338 | -1.255 | -0.182 | 0.137 | -0.004 | -0.003 | -0.003 | 0 | 0.27 | 0.53 | 0.36 | 0.25 | 0.27 | 0.2 |
| 1 | 157387799 | rs12748574 | 1.3 | -1.357 | -0.32 | 0.377 | 0.006 | -0.019 | -0.019 | 0 | 0.25 | 0.23 | 0.19 | 0.35 | 0.19 | 0.41 |
| 1 | 157388083 | rs9427274 | -0.188 | -1.366 | -0.3 | 0.362 | 0 | 0 | 0 | 1.045 | 0.53 | 0.56 | 0.38 | 0.54 | 0.45 | 0.48 |
| 1 | 157390112 | rs12745427 | 1.577 | -1.329 | -0.33 | 0.589 | -0.011 | -0.034 | -0.034 | 0 | 0.23 | 0.07 | 0.16 | 0.25 | 0.19 | 0.4 |
| 1 | 157393023 | rs7535020 | -0.233 | -1.093 | 0.2 | 0.183 | -0.004 | 0.012 | 0.012 | 0.361 | 0.7 | 0.42 | 0.62 | 0.59 | 0.73 | 0.79 |
| 1 | 157396835 | rs7524312 | 1.027 | -0.701 | -0.38 | -0.36 | -0.036 | 0.076 | 0.076 | 0 | 0.24 | 0.12 | 0.17 | 0.03 | 0.26 | 0.07 |
| 1 | 157402121 | rs7523656 | 0.838 | -0.938 | -0.452 | -0.188 | -0.014 | 0.03 | 0.03 | 0 | 0.31 | 0.2 | 0.37 | 0.07 | 0.34 | 0.07 |
| 1 | 157402470 | rs1970101 | -1.669 | -1.079 | -1.081 | 0.046 | 0.002 | -0.101 | -0.101 | 0 | 0.76 | 0.91 | 0.81 | 0.54 | 0.8 | 0.56 |
| 1 | 157406190 | rs4246538 | -0.092 | -1.048 | -1.052 | -0.157 | -0.03 | 0.015 | 0.015 | 0 | 0.39 | 0.21 | 0.4 | 0.33 | 0.46 | 0.49 |
| 1 | 157419719 | rs2225573 | 0.725 | -0.795 | -0.578 | -0.284 | -0.001 | -0.009 | -0.009 | 0.062 | 0.3 | 0.25 | 0.37 | 0.11 | 0.34 | 0.07 |
| 1 | 157426570 | rs12562052 | 1.311 | -0.834 | -0.952 | -0.03 | -0.049 | -0.087 | -0.087 | 0 | 0.25 | 0.07 | 0.23 | 0.34 | 0.26 | 0.43 |
| 1 | 157441651 | rs6675938 | -0.826 | -0.839 | -0.501 | -0.053 | -0.007 | 0.017 | 0.017 | 0 | 0.68 | 0.64 | 0.74 | 0.65 | 0.74 | 0.56 |
| 1 | 157442117 | rs11264737 | 0.261 | -1.02 | -0.698 | -0.321 | 0.013 | -0.024 | -0.024 | 0 | 0.28 | 0.3 | 0.22 | 0.42 | 0.22 | 0.34 |
| 1 | 157445012 | rs4971124 | 0.588 | -0.431 | -0.97 | -0.078 | -0.009 | -0.034 | -0.034 | 0.339 | 0.28 | 0.14 | 0.24 | 0.35 | 0.26 | 0.48 |
| 1 | 157449254 | rs950453 | -0.435 | -0.052 | -1.355 | 0.072 | -0.012 | 0.019 | 0.019 | 0 | 0.65 | 0.48 | 0.7 | 0.61 | 0.74 | 0.52 |
| 1 | 157452142 | rs1412329 | 0.042 | 0.33 | -1.219 | -0.012 | 0.007 | 0.007 | 0.007 | 0 | 0.39 | 0.45 | 0.46 | 0.47 | 0.45 | 0.3 |
| 1 | 157453883 | rs6427381 | 0.384 | 0.825 | -1.524 | 0.338 | 0.017 | -0.018 | -0.018 | 0 | 0.58 | 0.12 | 0.64 | 0.48 | 0.68 | 0.46 |
| 1 | 157463366 | rs6680515 | -0.658 | 1.644 | -0.586 | -0.065 | 0.017 | -0.031 | -0.031 | 0 | 0.17 | 0.29 | 0.27 | 0.11 | 0.18 | 0.14 |
| 1 | 157463460 | rs12565002 | -0.233 | 1.275 | -1.33 | 0.192 | -0.015 | -0.101 | -0.101 | 0 | 0.25 | 0.08 | 0.29 | 0.5 | 0.25 | 0.4 |
| 1 | 157467562 | rs7547177 | 0.005 | 1.627 | -0.395 | -0.002 | 0.062 | -0.035 | -0.035 | 0 | 0.84 | 0.62 | 0.73 | 0.88 | 0.77 | 0.87 |
| 1 | 157470168 | rs11264747 | 0.396 | 0.933 | -1.031 | 0.236 | 0 | 0 | 0 | 0.654 | 0.54 | 0.61 | 0.59 | 0.55 | 0.55 | 0.55 |
| 1 | 157474737 | rs7340113 | -0.616 | 1.056 | -1.469 | 0.293 | -0.002 | 0.001 | 0.001 | 0.097 | 0.46 | 0.29 | 0.39 | 0.45 | 0.46 | 0.45 |
| 1 | 157477996 | rs3856264 | -0.786 | 1.651 | -1.533 | 1.031 | -0.028 | -0.037 | -0.037 | 0 | 0.14 | 0.06 | 0.23 | 0.41 | 0.23 | 0.36 |
| 1 | 157482371 | rs1807765 | -0.748 | 0.92 | -2.034 | -0.004 | -0.019 | 0.09 | 0.09 | 0.351 | 0.21 | 0.04 | 0.11 | 0.07 | 0.19 | 0.16 |
| 1 | 157484453 | rs3845586 | 0.679 | -0.019 | -2.989 | -0.56 | 0.043 | 0.195 | 0.195 | 0.75 | 0.49 | 0.08 | 0.67 | 0.8 | 0.61 | 0.65 |
| 1 | 157484575 | rs3843307 | 0.51 | -0.431 | -1.865 | -0.683 | 0.008 | 0.013 | 0.013 | 0 | 0.59 | 0.36 | 0.72 | 0.78 | 0.68 | 0.67 |
| 1 | 157484698 | rs3900770 | 0.087 | -0.259 | -1.688 | -0.604 | -0.007 | -0.053 | -0.053 | 0 | 0.1 | 0.28 | 0.05 | 0 | 0.07 | 0.01 |
| 1 | 157485561 | rs3811035 | 0.905 | -1.146 | -2.298 | -0.461 | 0.039 | 0.035 | 0.035 | 0.676 | 0.54 | 0.09 | 0.65 | 0.66 | 0.63 | 0.68 |
| 1 | 157490710 | rs6696309 | 0.742 | -0.961 | -1.299 | -0.812 | -0.018 | 0.134 | 0.134 | 2.093 | 0.58 | 0.12 | 0.65 | 0.82 | 0.62 | 0.7 |
| 1 | 157491322 | rs6696922 | 0.579 | -1.138 | -0.781 | -0.875 | 0.009 | -0.017 | -0.017 | 0.449 | 0.84 | 0.8 | 0.8 | 0.88 | 0.8 | 0.92 |
| 1 | 157503805 | rs17416676 | 0.753 | -0.736 | -1.59 | -1.126 | -0.058 | 0.117 | 0.117 | 0.295 | 0.32 | 0.01 | 0.18 | 0.13 | 0.26 | 0.26 |
| 1 | 157508882 | rs6427384 | 0.841 | -0.876 | -0.244 | -0.933 | 0.008 | 0.028 | 0.028 | 0.703 | 0.84 | 0.32 | 0.81 | 0.79 | 0.85 | 0.85 |
| 1 | 157509025 | rs2012199 | 0.069 | -0.876 | -0.201 | -0.934 | 0.031 | -0.009 | -0.009 | 0 | 0.89 | 0.73 | 0.84 | 0.88 | 0.86 | 0.93 |
| 1 | 157510939 | rs1977588 | 0.89 | -0.873 | -0.307 | -0.932 | 0.008 | 0.028 | 0.028 | 0.703 | 0.84 | 0.32 | 0.81 | 0.79 | 0.85 | 0.85 |
| 1 | 157513569 | rs10494316 | 0.605 | -0.756 | -0.41 | -0.71 | 0.004 | 0.041 | 0.041 | 1.44 | 0.58 | 0.35 | 0.65 | 0.75 | 0.62 | 0.76 |
| 1 | 157514097 | rs6679793 | 0.319 | -1.217 | -0.507 | -1.24 | -0.021 | -0.049 | -0.049 | 0 | 0.76 | 0.41 | 0.73 | 0.91 | 0.82 | 0.76 |
| 1 | 157517127 | rs7531685 | 0.338 | -1.145 | -0.889 | -1.2 | 0.009 | 0.015 | 0.015 | 2.829 | 0.29 | 0.23 | 0.17 | 0.08 | 0.24 | 0.17 |
| 1 | 157537530 | rs2758684 | -1.246 | -1.113 | -1.057 | -1.17 | -0.039 | 0.094 | 0.094 | 1.618 | 0.55 | 0.92 | 0.45 | 0.29 | 0.43 | 0.41 |
| 1 | 157537967 | rs12049389 | 0.35 | -1.076 | -0.666 | -1.032 | 0.009 | 0.033 | 0.033 | 5.295 | 0.27 | 0.16 | 0.16 | 0.08 | 0.24 | 0.17 |
| 1 | 157539317 | rs1412676 | 0.25 | -1.068 | -0.653 | -1.028 | 0.006 | 0.013 | 0.013 | 3.573 | 0.27 | 0.21 | 0.17 | 0.08 | 0.24 | 0.17 |
| 1 | 157540104 | rs7523341 | -0.447 | -1.175 | -0.21 | -1.001 | -0.024 | -0.033 | -0.033 | 1.179 | 0.4 | 0.51 | 0.26 | 0.17 | 0.28 | 0.34 |
| 1 | 157541485 | rs12080640 | -0.52 | -1.082 | 0.214 | -0.974 | -0.016 | -0.045 | -0.045 | 0 | 0.09 | 0.25 | 0.04 | 0.01 | 0.04 | 0.01 |
| 1 | 157542162 | rs849815 | -0.028 | -1.241 | 0.583 | -1.247 | -0.009 | 0.026 | 0.026 | 0.825 | 0.33 | 0.2 | 0.3 | 0.24 | 0.35 | 0.27 |
| 1 | 157542216 | rs849814 | -0.723 | -1.241 | 0.593 | -1.248 | -0.003 | -0.024 | -0.024 | 0.205 | 0.41 | 0.48 | 0.34 | 0.24 | 0.39 | 0.28 |
| 1 | 157543746 | rs2758680 | -0.028 | -1.238 | 0.513 | -1.246 | -0.009 | 0.026 | 0.026 | 0.825 | 0.33 | 0.2 | 0.3 | 0.24 | 0.35 | 0.27 |
| 1 | 157543762 | rs2758679 | -1.255 | -1.232 | -0.111 | -1.432 | -0.009 | 0.043 | 0.043 | 0.048 | 0.75 | 0.87 | 0.88 | 0.86 | 0.77 | 0.79 |
| 1 | 157544850 | rs14335 | 1.336 | -1.293 | -0.165 | -1.459 | -0.096 | -0.106 | -0.106 | 0 | 0.3 | 0.03 | 0.51 | 0.62 | 0.38 | 0.51 |
| 1 | 157545231 | rs849827 | -0.72 | -1.287 | -0.12 | -1.437 | 0.015 | 0.12 | 0.12 | 0.111 | 0.45 | 0.79 | 0.36 | 0.25 | 0.39 | 0.28 |
| 1 | 157553680 | rs11591055 | 0.911 | -1.29 | -0.26 | -1.41 | -0.019 | 0.005 | 0.005 | 0 | 0.32 | 0.17 | 0.5 | 0.72 | 0.39 | 0.55 |
| 1 | 157556532 | rs2777963 | -1.207 | -1.234 | -0.11 | -1.449 | 0.003 | -0.043 | -0.043 | 0.864 | 0.74 | 0.67 | 0.85 | 0.9 | 0.78 | 0.84 |
| 1 | 157557701 | rs2777962 | 0.239 | -1.263 | -0.097 | -1.41 | 0.002 | 0.01 | 0.01 | 0 | 0.4 | 0.36 | 0.54 | 0.68 | 0.43 | 0.54 |
| 1 | 157569194 | rs2778003 | 0.65 | -0.535 | -0.3 | -1.119 | 0 | 0.009 | 0.009 | 0 | 0.34 | 0.26 | 0.51 | 0.71 | 0.39 | 0.55 |
| 1 | 157584301 | rs2208753 | 0.341 | -0.57 | -0.325 | -1.116 | -0.023 | 0.062 | 0.062 | 2.301 | 0.32 | 0.14 | 0.3 | 0.18 | 0.36 | 0.27 |
| 1 | 157586167 | rs2758644 | 0.575 | -0.642 | -0.289 | -0.999 | -0.002 | 0.01 | 0.01 | 0 | 0.35 | 0.27 | 0.52 | 0.72 | 0.41 | 0.55 |
| 1 | 157588430 | rs2777971 | 0.796 | -1.151 | -0.285 | -0.914 | -0.037 | -0.012 | -0.012 | 0 | 0.34 | 0.1 | 0.51 | 0.66 | 0.42 | 0.54 |
| 1 | 157591350 | rs2777976 | 0.507 | -1.064 | -0.288 | -0.91 | -0.004 | 0.017 | 0.017 | 0 | 0.37 | 0.22 | 0.52 | 0.71 | 0.43 | 0.56 |
| 1 | 157596390 | rs2777987 | -0.579 | -0.822 | -0.584 | -0.993 | -0.029 | 0.068 | 0.068 | 2.762 | 0.68 | 0.89 | 0.74 | 0.82 | 0.67 | 0.72 |
| 1 | 157596949 | rs10489675 | 0.637 | -0.861 | -0.383 | -1.024 | 0.002 | -0.048 | -0.048 | 0.032 | 0.28 | 0.4 | 0.19 | 0.11 | 0.23 | 0.16 |
| 1 | 157605501 | rs2758620 | 0.641 | -0.854 | -0.483 | -1.02 | -0.001 | -0.003 | -0.003 | 0 | 0.4 | 0.41 | 0.54 | 0.7 | 0.44 | 0.57 |
| 1 | 157607377 | rs2758688 | -0.306 | -0.633 | -1.025 | -0.752 | -0.022 | 0.064 | 0.064 | 2.667 | 0.68 | 0.88 | 0.74 | 0.82 | 0.67 | 0.72 |
| 1 | 157619889 | rs10489676 | -0.18 | -0.928 | -1.108 | -0.739 | 0.004 | 0.097 | 0.097 | 2.249 | 0.36 | 0.11 | 0.26 | 0.19 | 0.32 | 0.27 |
| 1 | 157637964 | rs7523492 | -1.659 | -1.153 | -1.112 | -0.394 | -0.122 | -0.168 | -0.168 | 1.283 | 0.77 | 0.99 | 0.75 | 0.44 | 0.62 | 0.53 |
| 1 | 157641489 | rs12125366 | 1.275 | -1.064 | -0.179 | -0.54 | 0.016 | 0.017 | 0.017 | 0.368 | 0.22 | 0.31 | 0.18 | 0.26 | 0.21 | 0.15 |
| 1 | 157641683 | rs10908583 | -1.071 | -1.134 | -0.148 | -0.57 | -0.01 | -0.021 | -0.021 | 0.411 | 0.64 | 0.75 | 0.47 | 0.5 | 0.55 | 0.45 |
| 1 | 157647789 | rs11264794 | -1.282 | -1.169 | -0.199 | -0.587 | -0.106 | -0.144 | -0.144 | 0 | 0.66 | 0.94 | 0.48 | 0.5 | 0.55 | 0.45 |
| 1 | 157648098 | rs6691569 | 0.134 | -1.155 | -0.061 | -0.666 | 0.018 | 0.057 | 0.057 | 2.6 | 0.35 | 0.19 | 0.22 | 0.18 | 0.29 | 0.22 |
| 1 | 157648155 | rs11805944 | -1.149 | -0.656 | 0.588 | -0.232 | 0.022 | 0.015 | 0.015 | 0 | 0.06 | 0.24 | 0.06 | 0.05 | 0.05 | 0.08 |
| 1 | 157657842 | rs7549100 | 0.552 | -0.57 | 0.284 | -0.176 | 0.029 | 0.012 | 0.012 | 0.501 | 0.25 | 0.47 | 0.2 | 0.26 | 0.24 | 0.15 |
| 1 | 157666644 | rs6681271 | -1.197 | -0.93 | -0.318 | -0.566 | -0.068 | 0 | 0 | 1.858 | 0.53 | 0.92 | 0.47 | 0.43 | 0.5 | 0.33 |

| | | | | | | | | | | | | | | | | |
|---|---|---|---|---|---|---|---|---|---|---|---|---|---|---|---|---|
| 1 | 157668390 | rs7522061 | -0.881 | -0.973 | 0.208 | -0.587 | 0.001 | 0.024 | 0.024 | 0.067 | 0.5 | 0.62 | 0.45 | 0.43 | 0.5 | 0.29 |
| 1 | 157670816 | rs7528684 | -1.179 | -0.915 | -0.266 | -0.544 | -0.074 | -0.017 | -0.017 | 0 | 0.53 | 0.94 | 0.46 | 0.44 | 0.48 | 0.28 |
| 1 | 157672304 | rs10158440 | -0.692 | -0.561 | 0.388 | -0.512 | 0.023 | 0.017 | 0.017 | 0 | 0.05 | 0.23 | 0.07 | 0.05 | 0.05 | 0.08 |
| 1 | 157679691 | rs17676303 | 0.27 | -0.815 | 0.009 | -0.611 | -0.019 | 0 | 0 | 0.035 | 0.21 | 0.06 | 0.19 | 0.21 | 0.23 | 0.13 |
| 1 | 157685739 | rs7524764 | 0.674 | -0.679 | -0.047 | -0.554 | -0.045 | -0.034 | -0.034 | 0.408 | 0.15 | 0.03 | 0.18 | 0.21 | 0.2 | 0.08 |
| 1 | 157695412 | rs1999793 | -0.419 | -0.648 | 0.302 | -0.584 | 0.008 | 0.003 | 0.003 | 0 | 0.18 | 0.2 | 0.23 | 0.26 | 0.23 | 0.26 |
| 1 | 157702017 | rs6688945 | 0.633 | -1.012 | 0.024 | -0.714 | -0.009 | 0.049 | 0.049 | 0.341 | 0.28 | 0.08 | 0.19 | 0.17 | 0.24 | 0.16 |
| 1 | 157718325 | rs3818813 | 0.708 | -0.878 | 0.521 | -0.524 | 0.026 | 0.023 | 0.023 | 0 | 0.15 | 0.25 | 0.2 | 0.25 | 0.2 | 0.07 |
| 1 | 157737504 | rs2873413 | -0.949 | -1.062 | 0.092 | -0.622 | -0.091 | -0.005 | -0.005 | 0 | 0.56 | 0.92 | 0.47 | 0.48 | 0.48 | 0.49 |
| 1 | 157744004 | rs12568320 | 0.541 | -0.821 | 0.282 | -0.545 | -0.003 | -0.002 | -0.002 | 0.024 | 0.18 | 0.12 | 0.19 | 0.24 | 0.2 | 0.16 |
| 1 | 157751356 | rs6677622 | 0.475 | -0.818 | 0.581 | -0.53 | 0.001 | 0 | 0 | 0 | 0.18 | 0.14 | 0.19 | 0.24 | 0.2 | 0.16 |
| 1 | 157754359 | rs12743184 | 1.07 | -0.838 | 0.486 | -0.566 | -0.006 | 0.025 | 0.025 | 0.041 | 0.26 | 0.13 | 0.2 | 0.19 | 0.24 | 0.14 |
| 1 | 157780892 | rs6689427 | -0.549 | -0.732 | 0.514 | -0.442 | 0 | 0.015 | 0.015 | 0 | 0.48 | 0.59 | 0.4 | 0.44 | 0.44 | 0.3 |
| 1 | 157794774 | rs12143912 | 1.037 | -0.724 | 0.104 | -0.434 | -0.002 | 0.023 | 0.023 | 0.106 | 0.26 | 0.15 | 0.2 | 0.19 | 0.24 | 0.15 |
| 1 | 157798000 | rs2765493 | 0.904 | -1.165 | -0.34 | -0.692 | -0.109 | -0.046 | -0.046 | 0 | 0.4 | 0.07 | 0.5 | 0.52 | 0.5 | 0.48 |
| 1 | 157799708 | rs16839239 | 0.481 | -1.161 | -0.081 | -0.639 | 0 | 0.004 | 0.004 | 0 | 0.2 | 0.11 | 0.17 | 0.24 | 0.18 | 0.16 |
| 1 | 157804648 | rs2765501 | 1.477 | -0.933 | -0.412 | -0.136 | -0.124 | -0.137 | -0.137 | 0.241 | 0.24 | 0.01 | 0.23 | 0.34 | 0.4 | 0.37 |
| 1 | 157809662 | rs10489683 | -0.16 | -0.726 | -0.396 | 0.315 | -0.006 | -0.013 | -0.013 | 0.122 | 0.29 | 0.36 | 0.25 | 0.29 | 0.24 | 0.37 |
| 1 | 157811175 | rs2765505 | 0.68 | -0.429 | -1.443 | 0.135 | -0.076 | -0.033 | -0.033 | 0.04 | 0.4 | 0.07 | 0.28 | 0.35 | 0.47 | 0.43 |
| 1 | 157811392 | rs2260040 | -0.518 | 0.167 | -0.536 | 0.127 | -0.003 | 0.003 | 0.003 | 0.08 | 0.74 | 0.79 | 0.75 | 0.81 | 0.72 | 0.86 |
| 1 | 157825776 | rs16839299 | -0.621 | -0.471 | -0.088 | 0.299 | -0.033 | 0.004 | 0.004 | 0 | 0.14 | 0.31 | 0.08 | 0.15 | 0.05 | 0.01 |
| 1 | 157826691 | rs17740850 | 0.199 | -1.022 | -0.302 | -0.044 | 0.012 | 0.015 | 0.015 | 0 | 0.13 | 0.05 | 0.23 | 0.06 | 0.07 | 0.23 |
| 1 | 157832074 | rs4514234 | -0.797 | -1.054 | -0.141 | -0.103 | -0.018 | -0.01 | -0.01 | 0.069 | 0.28 | 0.33 | 0.3 | 0.21 | 0.13 | 0.24 |
| 1 | 157840353 | rs6427405 | -1.042 | -0.129 | -0.314 | -0.529 | -0.034 | 0.029 | 0.029 | 0 | 0.81 | 0.89 | 0.77 | 0.89 | 0.69 | 0.83 |
| 1 | 157842192 | rs11582594 | 1.181 | -0.603 | -0.016 | 0.062 | -0.007 | -0.005 | -0.005 | 0.263 | 0.43 | 0.35 | 0.37 | 0.46 | 0.53 | 0.52 |
| 1 | 157842936 | rs12074997 | -0.155 | -0.533 | -0.002 | -0.175 | -0.039 | -0.022 | -0.022 | 0 | 0.29 | 0.41 | 0.31 | 0.22 | 0.13 | 0.23 |
| 1 | 157848162 | rs7520880 | -0.984 | -0.45 | -0.021 | -0.149 | -0.008 | 0.005 | 0.005 | 0 | 0.79 | 0.84 | 0.76 | 0.88 | 0.69 | 0.82 |
| 1 | 157864742 | rs4614251 | 0.338 | -0.499 | 0.01 | -0.165 | -0.004 | 0.013 | 0.013 | 0 | 0.54 | 0.58 | 0.48 | 0.67 | 0.56 | 0.6 |
| 1 | 157864913 | rs7524550 | 0.752 | -0.527 | -0.097 | -0.179 | -0.001 | 0.031 | 0.031 | 0 | 0.51 | 0.4 | 0.46 | 0.67 | 0.56 | 0.59 |
| 1 | 157871097 | rs11264845 | 0.103 | -0.741 | -0.248 | -0.172 | 0.015 | 0.023 | 0.023 | 0.225 | 0.42 | 0.75 | 0.35 | 0.53 | 0.39 | 0.36 |
| 1 | 157879145 | rs4450009 | 0.53 | -0.863 | -0.386 | -0.214 | 0.05 | 0.006 | 0.006 | 0 | 0.28 | 0.16 | 0.3 | 0.26 | 0.14 | 0.26 |
| 1 | 157879294 | rs7524095 | 0.224 | -0.347 | 0.39 | 0.598 | 0.003 | 0.003 | 0.003 | 0.019 | 0.34 | 0.47 | 0.26 | 0.3 | 0.35 | 0.32 |
| 1 | 157884976 | rs11264857 | -0.249 | -0.335 | 0.136 | 0.52 | 0 | 0.001 | 0.001 | 0 | 0.4 | 0.61 | 0.34 | 0.53 | 0.39 | 0.35 |
| 1 | 157891524 | rs4971165 | 0.623 | -0.119 | -0.187 | 0.627 | 0.004 | -0.005 | -0.005 | 0.182 | 0.29 | 0.25 | 0.17 | 0.29 | 0.22 | 0.33 |
| 1 | 157891565 | rs7514610 | -1.084 | -0.836 | 0.001 | 0.67 | 0.015 | 0.183 | 0.183 | 3.162 | 0.13 | 0.52 | 0.14 | 0.23 | 0.14 | 0.05 |
| 1 | 157891597 | rs4971166 | -0.355 | -0.329 | -0.756 | 0.747 | 0.069 | 0.026 | 0.026 | 0 | 0.43 | 0.76 | 0.32 | 0.53 | 0.33 | 0.42 |
| 1 | 157891709 | rs7524979 | -0.095 | -0.711 | -0.657 | 0.241 | 0.064 | 0.007 | 0.007 | 0.264 | 0.37 | 0.76 | 0.22 | 0.29 | 0.2 | 0.35 |
| 1 | 157907476 | rs6704322 | 0.197 | 0.916 | -0.107 | 0.059 | 0.043 | -0.017 | -0.017 | 0 | 0.14 | 0.19 | 0.15 | 0.09 | 0.32 | 0.26 |
| 1 | 157908444 | rs10494321 | -0.005 | 0.821 | 0.494 | 0.014 | 0.106 | -0.038 | -0.038 | 0 | 0.15 | 0.33 | 0.16 | 0.09 | 0.32 | 0.26 |
| 1 | 157909525 | rs1925035 | 0.736 | 0.252 | -0.079 | -0.047 | -0.029 | -0.083 | -0.083 | 0.058 | 0.36 | 0.13 | 0.53 | 0.57 | 0.38 | 0.4 |
| 1 | 157910302 | rs1925036 | 0.093 | -0.363 | 0.511 | 0.021 | 0.003 | 0.009 | 0.009 | 1.123 | 0.54 | 0.45 | 0.65 | 0.71 | 0.53 | 0.51 |
| 1 | 157914330 | rs4971169 | 1.036 | -0.262 | 0.157 | 0.058 | -0.012 | 0.009 | 0.009 | 0 | 0.13 | 0.11 | 0.15 | 0.09 | 0.32 | 0.26 |
| 1 | 157918761 | rs3887587 | -0.209 | -0.368 | 0.252 | 0.02 | -0.002 | 0.019 | 0.019 | 0.645 | 0.43 | 0.29 | 0.33 | 0.29 | 0.47 | 0.49 |
| 1 | 157921100 | rs1925041 | 0.522 | -1.341 | 0.403 | 0.322 | 0.079 | -0.022 | -0.022 | 0 | 0.29 | 0.08 | 0.18 | 0.2 | 0.16 | 0.23 |
| 1 | 157927955 | rs753345 | -0.192 | -1.411 | 0.447 | 0.306 | -0.055 | -0.01 | -0.01 | 0.666 | 0.66 | 0.59 | 0.79 | 0.8 | 0.84 | 0.77 |
| 1 | 157930021 | rs10494320 | 0.471 | -1.242 | -0.47 | 0.388 | 0.082 | -0.02 | -0.02 | 0 | 0.29 | 0.09 | 0.18 | 0.2 | 0.16 | 0.23 |
| 1 | 157934242 | rs720773 | -0.08 | -0.889 | -0.367 | 0.387 | -0.04 | -0.004 | -0.004 | 0 | 0.6 | 0.5 | 0.75 | 0.69 | 0.81 | 0.73 |
| 1 | 157944945 | rs1929843 | -0.67 | -0.39 | -0.834 | -0.128 | -0.017 | 0.09 | 0.09 | 0 | 0.61 | 0.77 | 0.64 | 0.82 | 0.4 | 0.45 |
| 1 | 157947235 | rs6696582 | 0.385 | -1.32 | -0.343 | 0.553 | -0.037 | 0.003 | 0.003 | 0 | 0.35 | 0.39 | 0.2 | 0.37 | 0.13 | 0.26 |
| 1 | 157955130 | rs7518837 | -0.868 | -1.246 | -0.468 | 0.609 | -0.016 | -0.027 | -0.027 | 0 | 0.29 | 0.12 | 0.4 | 0.32 | 0.29 | 0.17 |
| 1 | 157955804 | rs4246539 | 0.438 | -1.173 | -0.607 | 0.528 | -0.052 | -0.091 | -0.091 | 0.4 | 0.13 | 0.01 | 0.31 | 0.31 | 0.15 | 0.13 |
| 1 | 157956610 | rs1217058 | 1.107 | -0.545 | 0.284 | 0.795 | 0.03 | 0.141 | 0.141 | 2.655 | 0.86 | 0.46 | 0.9 | 0.72 | 0.86 | 0.95 |
| 1 | 157958669 | rs1217060 | 1.248 | -1.436 | 0.431 | 0.687 | -0.016 | 0.015 | 0.015 | 0 | 0.63 | 0.48 | 0.58 | 0.56 | 0.7 | 0.75 |
| 1 | 157961861 | rs3845588 | -0.28 | -1.227 | -0.162 | 0.496 | -0.027 | -0.036 | -0.036 | 0 | 0.08 | 0.14 | 0.02 | 0 | 0.03 | 0.05 |
| 1 | 157964856 | rs6656063 | -1.354 | -1.094 | -0.912 | 0.36 | 0.016 | 0.002 | 0.002 | 0 | 0.42 | 0.63 | 0.48 | 0.61 | 0.34 | 0.29 |
| 1 | 157975608 | rs6698985 | -1.199 | -0.861 | -1.257 | 0.053 | 0 | 0.046 | 0.046 | 0 | 0.52 | 0.64 | 0.6 | 0.73 | 0.52 | 0.39 |
| 1 | 157976503 | rs3856265 | -0.674 | -1.076 | -1.535 | 0.645 | 0.01 | -0.08 | -0.08 | 0.588 | 0.33 | 0.16 | 0.22 | 0.46 | 0.27 | 0.16 |
| 1 | 157976566 | rs2777819 | 0.463 | -0.534 | -1.281 | 0.638 | -0.027 | 0.003 | 0.003 | 0.012 | 0.33 | 0.21 | 0.33 | 0.26 | 0.42 | 0.56 |
| 1 | 157976896 | rs11585913 | -0.233 | -0.677 | -0.675 | 0.787 | -0.001 | -0.029 | -0.029 | 0 | 0.11 | 0.09 | 0.06 | 0 | 0.1 | 0.05 |
| 1 | 157977572 | rs11580742 | -0.127 | -0.713 | -0.685 | 0.808 | 0.001 | -0.047 | -0.047 | 0.137 | 0.11 | 0.14 | 0.06 | 0 | 0.1 | 0.05 |
| 1 | 157977753 | rs1888821 | -0.163 | -0.483 | -1.046 | 1.242 | -0.023 | -0.045 | -0.045 | 0 | 0.08 | 0 | 0.32 | 0.25 | 0.08 | 0.13 |
| 1 | 157982579 | rs1925034 | -1.285 | -0.743 | -0.689 | 0.531 | -0.028 | -0.028 | -0.028 | 0 | 0.34 | 0.44 | 0.22 | 0.23 | 0.26 | 0.13 |
| 1 | 157986247 | rs16839562 | -0.456 | -0.584 | -1.429 | 0.628 | -0.031 | -0.113 | -0.113 | 0 | 0.2 | 0.01 | 0.15 | 0.33 | 0.19 | 0.24 |
| 1 | 157988926 | rs7537784 | -1.681 | -0.954 | 0.054 | 0.191 | -0.044 | -0.038 | -0.038 | 0 | 0.18 | 0.3 | 0.12 | 0.14 | 0.12 | 0.28 |
| 1 | 157990700 | rs1925037 | 0.705 | -0.983 | 0.003 | 0.106 | -0.026 | -0.037 | -0.037 | 0 | 0.92 | 0.85 | 0.96 | 1 | 0.98 | 0.99 |
| 1 | 157997084 | rs12039337 | -0.378 | -0.942 | -0.033 | 0.144 | -0.002 | -0.004 | -0.004 | 0 | 0.06 | 0.03 | 0.04 | 0.12 | 0.06 | 0.25 |
| 1 | 157997676 | rs16839617 | -0.429 | -0.959 | -0.041 | 0.188 | -0.002 | -0.007 | -0.007 | 1.081 | 0.06 | 0.03 | 0.05 | 0.13 | 0.07 | 0.25 |
| 1 | 158000740 | rs1972216 | 2.137 | -1.009 | 0.013 | 0.064 | -0.078 | -0.075 | -0.075 | 0 | 0.78 | 0.54 | 0.87 | 0.88 | 0.88 | 0.72 |
| 1 | 158006301 | rs10430456 | -1.001 | -0.693 | -0.314 | 0.186 | 0.001 | -0.005 | -0.005 | 0.189 | 0.08 | 0.04 | 0.05 | 0.11 | 0.07 | 0.26 |

| | | | | | | | | | | | | | | | | |
|---|---|---|---|---|---|---|---|---|---|---|---|---|---|---|---|---|
| 1 | 158008056 | rs6688137 | -2.035 | -0.729 | 0.012 | 0.088 | -0.016 | -0.022 | -0.022 | 0 | 0.05 | 0.17 | 0.01 | 0 | 0.01 | 0 |
| 1 | 158009712 | rs6686246 | -1.105 | -0.813 | -0.119 | -0.567 | -0.021 | -0.082 | -0.082 | 0 | 0.2 | 0.29 | 0.1 | 0.01 | 0.15 | 0.17 |
| 1 | 158017250 | rs4971122 | 1.003 | -0.761 | -0.151 | -0.542 | -0.052 | -0.139 | -0.139 | 0 | 0.76 | 0.59 | 0.89 | 0.99 | 0.84 | 0.85 |
| 1 | 158023332 | rs12561890 | -0.587 | -0.042 | -0.83 | 0.196 | 0.007 | 0.007 | 0.007 | 0.02 | 0.11 | 0.07 | 0.04 | 0.07 | 0.09 | 0.09 |
| 1 | 158023515 | rs7527735 | 0.385 | -1.329 | -1.167 | -1.01 | 0.089 | 0.123 | 0.123 | 4.186 | 0.5 | 0.26 | 0.72 | 0.85 | 0.66 | 0.72 |
| 1 | 158024499 | rs10796995 | -0.349 | -1.564 | -1.097 | -1.07 | -0.01 | -0.03 | -0.03 | 0 | 0.64 | 0.37 | 0.82 | 0.92 | 0.83 | 0.82 |
| 1 | 158026124 | rs17454196 | -0.501 | -1.795 | -1.486 | -1.199 | 0.062 | 0.129 | 0.129 | 3.414 | 0.32 | 0.1 | 0.15 | 0.11 | 0.22 | 0.24 |
| 1 | 158026290 | rs877872 | 0.138 | -0.153 | -1.529 | -0.645 | 0.026 | 0.02 | 0.02 | 0 | 0.12 | 0.13 | 0.26 | 0.2 | 0.21 | 0.1 |
| 1 | 158026539 | rs12061312 | 0.054 | -0.744 | -0.763 | -0.979 | -0.045 | -0.084 | -0.084 | 0.03 | 0.24 | 0.4 | 0.14 | 0.09 | 0.16 | 0.12 |
| 1 | 158029021 | rs7552600 | 0.052 | 0.469 | -1.227 | -1.081 | -0.027 | 0.038 | 0.038 | 0 | 0.18 | 0.08 | 0.28 | 0.04 | 0.26 | 0.2 |
| 1 | 158031160 | rs1925032 | 0.293 | 0.003 | -1.301 | -0.823 | 0.005 | 0 | 0 | 0 | 0.73 | 0.25 | 0.75 | 0.74 | 0.78 | 0.8 |
| 1 | 158032447 | rs6677623 | -0.411 | 0.52 | -1.889 | -0.545 | -0.051 | -0.072 | -0.072 | 0.065 | 0.78 | 0.92 | 0.62 | 0.37 | 0.68 | 0.58 |
| 1 | 158034283 | rs4971191 | -1.339 | 0.339 | -1.005 | -0.725 | 0.001 | -0.05 | -0.05 | 2.825 | 0.9 | 0.83 | 0.89 | 1 | 0.89 | 0.94 |
| 1 | 158036373 | rs17420502 | 1.011 | 0.304 | -1.154 | -0.989 | -0.019 | 0.032 | 0.032 | 0 | 0.18 | 0.12 | 0.28 | 0.04 | 0.26 | 0.2 |
| 1 | 158036740 | rs1886389 | 0.983 | -0.343 | -1.371 | -1.428 | -0.041 | -0.001 | -0.001 | 0 | 0.39 | 0.14 | 0.4 | 0.62 | 0.42 | 0.49 |
| 1 | 158038589 | rs12075777 | 1.428 | -0.466 | -0.734 | -1.623 | -0.007 | -0.031 | -0.031 | 0 | 0.12 | 0.15 | 0.05 | 0.04 | 0.06 | 0.07 |
| 1 | 158039877 | rs7367687 | 1.957 | -0.415 | -0.66 | -1.596 | 0.002 | 0.01 | 0.01 | 0 | 0.1 | 0.09 | 0.04 | 0.03 | 0.06 | 0.05 |
| 1 | 158040193 | rs7547764 | 0.498 | -0.384 | -0.693 | -1.621 | -0.004 | 0.002 | 0.002 | 0.258 | 0.3 | 0.27 | 0.32 | 0.07 | 0.32 | 0.27 |
| 1 | 158042153 | rs10908615 | -1.213 | -0.093 | -0.416 | -1.412 | -0.032 | -0.089 | -0.089 | 2.349 | 0.8 | 0.68 | 0.9 | 0.97 | 0.9 | 0.9 |
| 1 | 158043445 | rs12033891 | 0.085 | 0.787 | -0.265 | -0.804 | 0.031 | 0.012 | 0.012 | 0 | 0.12 | 0.17 | 0.15 | 0.19 | 0.19 | 0.14 |
| 1 | 158045111 | rs11264898 | 0.985 | 0.746 | -0.305 | -0.849 | 0.004 | 0.005 | 0.005 | 0 | 0.08 | 0.11 | 0.09 | 0.15 | 0.08 | 0.1 |
| 1 | 158045749 | rs11264900 | -0.528 | 0.333 | -0.607 | -1.034 | -0.011 | -0.022 | -0.022 | 0 | 0.58 | 0.49 | 0.76 | 0.77 | 0.68 | 0.64 |
| 1 | 158049646 | rs11264902 | 1.194 | 0.578 | -0.575 | -0.704 | -0.02 | -0.08 | -0.08 | 3.467 | 0.24 | 0.1 | 0.43 | 0.5 | 0.27 | 0.33 |
| 1 | 158050627 | rs6697006 | 0.317 | -0.299 | -0.44 | -1.095 | -0.009 | 0.005 | 0.005 | 0 | 0.37 | 0.32 | 0.59 | 0.72 | 0.45 | 0.46 |
| 1 | 158052966 | rs6695790 | 0.201 | -0.19 | -0.802 | -1.725 | -0.004 | 0.004 | 0.004 | 0 | 0.38 | 0.37 | 0.6 | 0.78 | 0.5 | 0.54 |
| 1 | 158053024 | rs11802995 | 0.375 | -0.379 | -0.94 | -1.41 | -0.003 | 0.03 | 0.03 | 0 | 0.2 | 0.14 | 0.17 | 0.03 | 0.22 | 0.17 |
| 1 | 158054499 | rs11264908 | -1.018 | -0.725 | -0.194 | -0.877 | 0.006 | -0.041 | -0.041 | 0 | 0.15 | 0.49 | 0.1 | 0.08 | 0.11 | 0.17 |
| 1 | 158056651 | rs7364493 | 0.026 | -1.006 | -1.171 | -0.986 | 0.048 | 0.071 | 0.071 | 0 | 0.43 | 0.15 | 0.67 | 0.74 | 0.65 | 0.51 |
| 1 | 158057691 | rs3820676 | 0.079 | -1.302 | -1.056 | -1.295 | 0.029 | 0.052 | 0.052 | 0.373 | 0.42 | 0.68 | 0.33 | 0.26 | 0.32 | 0.26 |
| 1 | 158058109 | rs6427419 | 0.469 | -1.227 | -1.681 | -1.521 | -0.125 | -0.003 | -0.003 | 0.056 | 0.41 | 0.03 | 0.58 | 0.7 | 0.51 | 0.62 |
| 1 | 158060499 | rs3768541 | 0.555 | -1.152 | -1.566 | -1.499 | -0.01 | 0.015 | 0.015 | 0 | 0.12 | 0.04 | 0.06 | 0.04 | 0.16 | 0.11 |
| 1 | 158061043 | rs2274165 | 0.917 | -1.132 | -1.194 | -1.471 | 0.016 | -0.018 | -0.018 | 0 | 0.06 | 0.12 | 0.04 | 0.02 | 0.07 | 0.08 |
| 1 | 158061626 | rs912572 | -0.538 | -1.115 | -0.902 | -1.36 | -0.019 | -0.019 | -0.019 | 0 | 0.63 | 0.38 | 0.7 | 0.75 | 0.74 | 0.81 |
| 1 | 158065762 | rs12749808 | 1.369 | -0.443 | -1.732 | -0.862 | -0.129 | -0.157 | -0.157 | 0 | 0.19 | 0.01 | 0.38 | 0.48 | 0.35 | 0.26 |
| 1 | 158069099 | rs11897 | -0.428 | -0.691 | -1.036 | -0.745 | -0.013 | -0.013 | -0.013 | 0 | 0.66 | 0.58 | 0.72 | 0.75 | 0.74 | 0.81 |
| 1 | 158071131 | rs942540 | 0.593 | -0.677 | -1.054 | -0.812 | 0.06 | 0.162 | 0.162 | 3.747 | 0.49 | 0.18 | 0.72 | 0.78 | 0.6 | 0.56 |
| 1 | 158073254 | rs12088518 | -0.629 | 0.048 | -0.844 | -0.197 | 0.006 | 0.015 | 0.015 | 0 | 0.33 | 0.24 | 0.18 | 0.19 | 0.32 | 0.31 |
| 1 | 158085565 | rs1324659 | -0.825 | 0.215 | -0.489 | 0.277 | 0.003 | 0.005 | 0.005 | 0 | 0.17 | 0.26 | 0.29 | 0.16 | 0.15 | 0.25 |
| 1 | 158087841 | rs12071018 | -1.475 | -0.346 | -0.926 | 0.78 | 0.017 | -0.025 | -0.025 | 10.025 | 0.42 | 0.84 | 0.48 | 0.58 | 0.32 | 0.57 |
| 1 | 158088687 | rs12061698 | -0.135 | -0.502 | 0.006 | 1.248 | -0.01 | 0.022 | 0.022 | 0 | 0.17 | 0.22 | 0.13 | 0.27 | 0.13 | 0.35 |
| 1 | 158089212 | rs1998639 | -0.932 | -0.729 | 0.454 | 1.169 | 0.006 | 0 | 0 | 0 | 0.24 | 0.42 | 0.33 | 0.25 | 0.21 | 0.31 |
| 1 | 158100580 | rs10797003 | -0.126 | -0.566 | -0.365 | 0.363 | -0.001 | -0.001 | -0.001 | 0 | 0.61 | 0.6 | 0.73 | 0.63 | 0.66 | 0.55 |
| 1 | 158102110 | rs927663 | -0.111 | -0.561 | -0.362 | 0.365 | -0.001 | -0.001 | -0.001 | 0 | 0.61 | 0.6 | 0.73 | 0.63 | 0.66 | 0.55 |
| 1 | 158102520 | rs4656184 | 1.513 | -0.481 | -0.664 | 0.451 | -0.08 | -0.08 | -0.08 | 0.174 | 0.35 | 0.1 | 0.4 | 0.37 | 0.45 | 0.22 |
| 1 | 158108217 | rs10752623 | -0.179 | -0.609 | -0.205 | 0.367 | -0.002 | -0.016 | -0.016 | 0 | 0.62 | 0.63 | 0.71 | 0.83 | 0.64 | 0.59 |
| 1 | 158109813 | rs12042121 | 0.021 | -0.673 | -0.078 | 0.432 | 0.005 | 0.003 | 0.003 | 0 | 0.21 | 0.25 | 0.29 | 0.24 | 0.2 | 0.29 |
| 1 | 158118787 | rs7548010 | -0.165 | -0.703 | -0.058 | 0.643 | 0.002 | 0.058 | 0.058 | 0 | 0.22 | 0.33 | 0.3 | 0.45 | 0.21 | 0.33 |
| 1 | 158140205 | rs980937 | 1.411 | -0.635 | -0.295 | 0.389 | -0.024 | -0.029 | -0.029 | 0 | 0.37 | 0.23 | 0.4 | 0.38 | 0.43 | 0.23 |
| 1 | 158140884 | rs1324656 | 0.001 | -0.514 | -0.232 | 0.423 | -0.001 | 0 | 0 | 0 | 0.63 | 0.64 | 0.71 | 0.63 | 0.64 | 0.55 |
| 1 | 158146028 | rs3087210 | -0.144 | -0.783 | -0.578 | 0.709 | 0.008 | -0.022 | -0.022 | 0 | 0.15 | 0.1 | 0.11 | 0.45 | 0.1 | 0.33 |
| 1 | 158148054 | rs859003 | -0.477 | -0.868 | -0.62 | 0.659 | 0.014 | -0.045 | -0.045 | 0 | 0.17 | 0.11 | 0.11 | 0.45 | 0.11 | 0.36 |
| 1 | 158150056 | rs859008 | -0.271 | -0.902 | -0.595 | 0.647 | 0.011 | -0.051 | -0.051 | 0 | 0.16 | 0.08 | 0.11 | 0.45 | 0.11 | 0.36 |
| 1 | 158150119 | rs3754471 | 1.504 | -1.366 | -0.988 | 0.073 | -0.046 | 0.001 | 0.001 | 0 | 0.31 | 0.21 | 0.42 | 0.31 | 0.45 | 0.21 |
| 1 | 158153526 | rs692565 | 0.098 | -1.551 | -0.775 | -0.11 | 0.019 | 0.096 | 0.096 | 0 | 0.5 | 0.38 | 0.53 | 0.75 | 0.56 | 0.57 |
| 1 | 158159393 | rs371380 | 1.358 | -2.05 | -0.586 | -0.025 | -0.031 | 0.016 | 0.016 | 0 | 0.4 | 0.27 | 0.48 | 0.32 | 0.52 | 0.23 |
| 1 | 158178492 | rs691175 | 0.802 | -2.361 | 0.567 | 0.299 | -0.063 | 0.108 | 0.108 | 0 | 0.77 | 0.59 | 0.71 | 0.51 | 0.88 | 0.61 |
| 1 | 158179236 | rs384414 | 1.403 | -2.459 | 0.406 | 0.251 | -0.009 | -0.001 | -0.001 | 1.025 | 0.71 | 0.29 | 0.69 | 0.51 | 0.88 | 0.61 |
| 1 | 158183942 | rs691314 | 1.515 | -2.452 | 0.253 | 0.274 | 0.041 | -0.039 | -0.039 | 0 | 0.67 | 0.22 | 0.63 | 0.51 | 0.8 | 0.55 |
| 1 | 158215299 | rs432611 | 0.155 | -2.556 | 0.784 | 0.229 | -0.034 | -0.028 | -0.028 | 0 | 0.91 | 0.68 | 0.81 | 0.93 | 0.98 | 0.9 |
| 1 | 158219965 | rs3181029 | -0.525 | -2.728 | 0.431 | 0.193 | -0.007 | -0.004 | -0.004 | 0 | 0.17 | 0.21 | 0.06 | 0.16 | 0.1 | 0.18 |
| 1 | 158224059 | rs16840041 | 0.687 | -2.895 | 0.036 | 0.534 | 0.029 | -0.036 | -0.036 | 0 | 0.11 | 0.04 | 0.09 | 0.39 | 0.06 | 0.23 |
| 1 | 158224825 | rs411089 | -2.581 | -3.041 | 0.26 | 0.437 | 0.053 | 0.01 | 0.01 | 0.68 | 0.43 | 0.72 | 0.39 | 0.66 | 0.26 | 0.58 |
| 1 | 158224904 | rs2269714 | 0.68 | -2.924 | 0.088 | 0.41 | 0.029 | -0.037 | -0.037 | 0.223 | 0.11 | 0.04 | 0.09 | 0.39 | 0.06 | 0.26 |
| 1 | 158228392 | rs389293 | -1.604 | -2.713 | 0.609 | 0.455 | -0.03 | 0.01 | 0.01 | 0 | 0.3 | 0.53 | 0.41 | 0.35 | 0.21 | 0.32 |
| 1 | 158237062 | rs12407446 | -1.733 | -2.659 | 0.782 | 0.435 | -0.038 | 0.018 | 0.018 | 0 | 0.3 | 0.46 | 0.41 | 0.36 | 0.21 | 0.32 |
| 1 | 158257838 | rs10797006 | -2.045 | -2.477 | 0.816 | 0.273 | 0 | 0 | 0 | 0 | 0.33 | 0.68 | 0.43 | 0.36 | 0.21 | 0.35 |
| 1 | 158267853 | rs1923142 | -0.216 | -2.31 | 0.869 | 0.311 | -0.05 | -0.025 | -0.025 | 0.757 | 0.11 | 0.23 | 0.18 | 0.08 | 0.02 | 0.08 |
| 1 | 158293942 | rs10908647 | -0.801 | -2.24 | 0.563 | 0.306 | 0 | 0.009 | 0.009 | 0 | 0.13 | 0.18 | 0.19 | 0.26 | 0.11 | 0.22 |
| 1 | 158295009 | rs1321646 | 0.814 | -2.139 | 0.23 | 0.225 | -0.013 | -0.083 | -0.083 | 0 | 0.15 | 0.04 | 0.13 | 0.4 | 0.14 | 0.27 |

| | | | | | | | | | | | | | | | | |
|---|---|---|---|---|---|---|---|---|---|---|---|---|---|---|---|---|
| 1 | 158295629 | rs10797007 | -2.327 | -2.366 | 0.455 | 0.057 | 0.062 | 0.096 | 0.096 | 0.104 | 0.52 | 0.84 | 0.41 | 0.66 | 0.36 | 0.6 |
| 1 | 158297960 | rs16840096 | 0.754 | -2.497 | 0.668 | 0.213 | -0.012 | -0.077 | -0.077 | 0.656 | 0.15 | 0.05 | 0.13 | 0.4 | 0.14 | 0.27 |
| 1 | 158311985 | rs10158210 | -2.031 | -2.531 | 0.645 | 0.167 | 0.077 | 0.032 | 0.032 | 0.585 | 0.37 | 0.85 | 0.29 | 0.25 | 0.21 | 0.33 |
| 1 | 158315924 | rs2317955 | 0.912 | -2.508 | 0.613 | 0.192 | -0.011 | -0.078 | -0.078 | 0 | 0.15 | 0.05 | 0.13 | 0.41 | 0.14 | 0.27 |
| 1 | 158324425 | rs1065457 | -2.254 | -2.468 | 0.622 | -0.34 | 0.063 | 0.099 | 0.099 | 0.053 | 0.52 | 0.9 | 0.42 | 0.66 | 0.36 | 0.6 |
| 1 | 158330242 | rs12742463 | -0.881 | -2.38 | 1.37 | -1.1 | -0.074 | -0.161 | -0.161 | 1.098 | 0.21 | 0.43 | 0.07 | 0 | 0.1 | 0.09 |
| 1 | 158332459 | rs10908648 | -0.953 | -2.339 | 1.342 | -1.033 | -0.002 | 0.024 | 0.024 | 0 | 0.14 | 0.3 | 0.2 | 0.21 | 0.11 | 0.19 |
| 1 | 158334546 | rs12139388 | 1.509 | -2.057 | 1.021 | -1.096 | -0.002 | 0.045 | 0.045 | 0 | 0.11 | 0.02 | 0.05 | 0 | 0.07 | 0.03 |
| 1 | 158338434 | rs13376124 | -0.142 | -1.763 | 1.787 | -0.386 | -0.067 | -0.083 | -0.083 | 1.521 | 0.14 | 0.21 | 0.02 | 0 | 0.02 | 0.06 |
| 1 | 158338906 | rs11801558 | -1.268 | -1.748 | 1.294 | -0.381 | -0.065 | -0.149 | -0.149 | 0 | 0.2 | 0.39 | 0.06 | 0 | 0.09 | 0.09 |
| 1 | 158341273 | rs7547997 | -2.307 | -1.717 | 0.99 | -0.309 | -0.001 | 0 | 0 | 0.065 | 0.29 | 0.66 | 0.22 | 0.25 | 0.13 | 0.26 |
| 1 | 158349097 | rs12736117 | -1.478 | -1.596 | 0.762 | -0.822 | -0.069 | -0.143 | -0.143 | 0 | 0.22 | 0.51 | 0.07 | 0 | 0.09 | 0.11 |
| 1 | 158366125 | rs1321642 | -0.929 | -0.948 | -0.194 | -0.358 | 0.001 | 0.001 | 0.001 | 0 | 0.2 | 0.2 | 0.2 | 0.14 | 0.12 | 0.26 |
| 1 | 158368846 | rs6662382 | -1.991 | -1.354 | -0.081 | -0.755 | -0.008 | -0.01 | -0.01 | 0.194 | 0.43 | 0.42 | 0.31 | 0.23 | 0.31 | 0.53 |
| 1 | 158370948 | rs12402548 | -0.591 | -1.166 | -0.569 | -0.536 | 0.009 | 0.009 | 0.009 | 0 | 0.17 | 0.1 | 0.15 | 0.09 | 0.07 | 0.23 |
| 1 | 158372524 | rs17673153 | 0.004 | -0.603 | -0.598 | -0.187 | 0.009 | -0.032 | -0.032 | 0 | 0.13 | 0.02 | 0.08 | 0.14 | 0.11 | 0.21 |
| 1 | 158375606 | rs11578937 | -1.056 | -0.751 | -0.205 | -0.442 | -0.044 | -0.073 | -0.073 | 0.027 | 0.21 | 0.45 | 0.23 | 0.1 | 0.13 | 0.33 |
| 1 | 158375629 | rs1321640 | -1.385 | -0.561 | -0.24 | -0.444 | -0.003 | -0.005 | -0.005 | 0 | 0.42 | 0.54 | 0.37 | 0.22 | 0.37 | 0.59 |
| 1 | 158377882 | rs10489828 | -0.822 | -0.105 | 0.098 | -0.327 | -0.005 | -0.06 | -0.06 | 0 | 0.1 | 0.3 | 0.1 | 0.01 | 0.07 | 0.2 |
| 1 | 158380828 | rs4656283 | -1.212 | 0.094 | -0.304 | -0.885 | -0.003 | 0.001 | 0.001 | 0.586 | 0.36 | 0.34 | 0.32 | 0.13 | 0.39 | 0.49 |
| 1 | 158381815 | rs4656284 | -1.163 | 0.086 | -0.159 | -1.597 | -0.001 | -0.007 | -0.007 | 0 | 0.46 | 0.51 | 0.31 | 0.17 | 0.48 | 0.34 |
| 1 | 158387196 | rs10489829 | -0.484 | 0.102 | -0.042 | -1.788 | 0.057 | 0.006 | 0.006 | 0.074 | 0.21 | 0.25 | 0.21 | 0.17 | 0.36 | 0.26 |
| 1 | 158388434 | rs17627858 | -0.428 | 0.101 | 0.001 | -1.788 | 0.007 | 0 | 0 | 0 | 0.2 | 0.19 | 0.21 | 0.17 | 0.36 | 0.26 |
| 1 | 158390834 | rs10489830 | -0.842 | -0.077 | -0.064 | -1.856 | -0.029 | 0.019 | 0.019 | 0 | 0.32 | 0.22 | 0.27 | 0.17 | 0.46 | 0.3 |
| 1 | 158403819 | rs7519343 | 0.159 | -0.109 | 0.038 | -1.852 | -0.004 | -0.015 | -0.015 | 0 | 0.1 | 0.14 | 0.02 | 0 | 0.02 | 0.05 |
| 1 | 158415517 | rs6699757 | -1.261 | 0.032 | 0.027 | -1.798 | 0.024 | -0.025 | -0.025 | 0.536 | 0.35 | 0.37 | 0.28 | 0.17 | 0.46 | 0.3 |
| 1 | 158417540 | rs989203 | -0.323 | 0.59 | -0.204 | -1.825 | 0.02 | 0.009 | 0.009 | 0.022 | 0.16 | 0.18 | 0.18 | 0.17 | 0.28 | 0.23 |
| 1 | 158425271 | rs12127971 | -0.979 | 1.268 | -0.239 | -1.313 | -0.033 | 0.021 | 0.021 | 0 | 0.32 | 0.22 | 0.27 | 0.17 | 0.46 | 0.3 |
| 1 | 158434049 | rs9651033 | -0.806 | 0.58 | 0.538 | -1.403 | -0.073 | -0.178 | -0.178 | 0.001 | 0.28 | 0.46 | 0.1 | 0 | 0.12 | 0.08 |
| 1 | 158446532 | rs10489831 | -0.456 | 0.865 | 0.101 | -0.9 | -0.007 | -0.009 | -0.009 | 0 | 0.15 | 0.14 | 0.15 | 0.19 | 0.22 | 0.21 |
| 1 | 158450238 | rs3820678 | -0.449 | 0.862 | 0.095 | -0.901 | -0.008 | -0.01 | -0.01 | 0.107 | 0.15 | 0.14 | 0.15 | 0.19 | 0.22 | 0.21 |
| 1 | 158450314 | rs6679056 | -0.929 | 0.401 | -0.339 | -0.891 | -0.004 | -0.007 | -0.007 | 0.22 | 0.52 | 0.48 | 0.44 | 0.57 | 0.56 | 0.42 |
| 1 | 158450382 | rs1418843 | -0.578 | 0.471 | -0.324 | -0.977 | 0.008 | -0.002 | -0.002 | 0 | 0.38 | 0.38 | 0.38 | 0.57 | 0.46 | 0.38 |
| 1 | 158454356 | rs10489832 | 2.014 | 0.567 | -0.424 | -0.982 | -0.017 | -0.065 | -0.065 | 0 | 0.13 | 0.02 | 0.19 | 0.38 | 0.15 | 0.14 |
| 1 | 158459975 | rs950488 | 1.396 | 0.334 | -0.319 | -0.92 | -0.061 | -0.095 | -0.095 | 0 | 0.19 | 0.03 | 0.22 | 0.38 | 0.24 | 0.17 |
| 1 | 158468676 | rs1342958 | -0.537 | 0.255 | -0.021 | -0.928 | 0.001 | -0.007 | -0.007 | 0 | 0.46 | 0.47 | 0.4 | 0.57 | 0.47 | 0.42 |
| 1 | 158484285 | rs1418846 | -0.653 | 0.189 | 0.008 | -0.96 | -0.002 | 0.021 | 0.021 | 0 | 0.48 | 0.57 | 0.41 | 0.57 | 0.47 | 0.42 |
| 1 | 158490324 | rs10797020 | -1.219 | 0.228 | 0.026 | -0.853 | -0.003 | 0.003 | 0.003 | 0 | 0.63 | 0.68 | 0.47 | 0.57 | 0.58 | 0.46 |
| 1 | 158493726 | rs7548349 | -1.236 | 0.233 | 0.028 | -0.851 | -0.003 | 0.003 | 0.003 | 0 | 0.63 | 0.68 | 0.47 | 0.57 | 0.58 | 0.46 |
| 1 | 158495209 | rs6699473 | -1.236 | 0.233 | 0.028 | -0.851 | -0.003 | 0.003 | 0.003 | 0 | 0.63 | 0.68 | 0.47 | 0.57 | 0.58 | 0.46 |
| 1 | 158510273 | rs12121983 | -0.628 | 0.331 | -0.053 | -0.775 | -0.001 | 0.005 | 0.005 | 0 | 0.47 | 0.52 | 0.4 | 0.57 | 0.48 | 0.42 |
| 1 | 158518939 | rs9804152 | -1.249 | 0.19 | -0.535 | -0.74 | 0.001 | 0.001 | 0.001 | 0 | 0.82 | 0.78 | 0.67 | 0.79 | 0.75 | 0.6 |
| 1 | 158520737 | rs6697656 | -0.645 | 0.134 | -0.476 | -0.823 | -0.004 | 0.004 | 0.004 | 0.052 | 0.63 | 0.74 | 0.48 | 0.58 | 0.57 | 0.46 |
| 1 | 158523671 | rs10908684 | 0.668 | 0.413 | -0.357 | -0.617 | 0.004 | 0.011 | 0.011 | 0 | 0.13 | 0.17 | 0.14 | 0.21 | 0.21 | 0.21 |
| 1 | 158549264 | rs863360 | -0.239 | 0.131 | -0.196 | -0.902 | -0.003 | 0.015 | 0.015 | 0 | 0.47 | 0.57 | 0.41 | 0.59 | 0.47 | 0.41 |
| 1 | 158549492 | rs863362 | -0.239 | 0.131 | -0.196 | -0.902 | -0.003 | 0.015 | 0.015 | 0 | 0.47 | 0.57 | 0.41 | 0.59 | 0.47 | 0.41 |
| 1 | 158549511 | rs863363 | -0.277 | 0.135 | -0.273 | -0.901 | -0.005 | 0.029 | 0.029 | 0 | 0.47 | 0.63 | 0.42 | 0.59 | 0.47 | 0.41 |
| 1 | 158553332 | rs863368 | 0.308 | 0.632 | -0.079 | -0.529 | 0 | 0.004 | 0.004 | 0.629 | 0.5 | 0.45 | 0.45 | 0.49 | 0.42 | 0.25 |
| 1 | 158554142 | rs2479861 | 0.075 | 0.596 | 0.023 | -0.547 | -0.002 | 0.009 | 0.009 | 0.233 | 0.52 | 0.55 | 0.45 | 0.49 | 0.43 | 0.25 |
| 1 | 158564383 | rs857707 | -0.515 | 0.721 | -0.381 | -0.444 | -0.003 | 0.01 | 0.01 | 0.03 | 0.62 | 0.73 | 0.54 | 0.7 | 0.58 | 0.45 |
| 1 | 158565115 | rs1332688 | 1.061 | 0.605 | -0.49 | -0.351 | -0.01 | 0.004 | 0.004 | 0 | 0.11 | 0.05 | 0.1 | 0.08 | 0.13 | 0.21 |
| 1 | 158580069 | rs2479868 | 0.717 | 0.432 | -0.618 | -0.063 | -0.006 | -0.012 | -0.012 | 0.338 | 0.26 | 0.22 | 0.31 | 0.43 | 0.28 | 0.17 |
| 1 | 158580477 | rs12128171 | 1.13 | 0.1 | -0.561 | -0.172 | -0.008 | -0.003 | -0.003 | 0 | 0.24 | 0.2 | 0.21 | 0.2 | 0.29 | 0.25 |
| 1 | 158584091 | rs952094 | -0.101 | 0.11 | -0.617 | -0.167 | 0.015 | 0.018 | 0.018 | 0 | 0.51 | 0.42 | 0.52 | 0.63 | 0.56 | 0.42 |
| 1 | 158585230 | rs2518491 | 1.413 | 0.135 | -0.776 | -0.138 | -0.063 | -0.08 | -0.08 | 0 | 0.23 | 0.06 | 0.3 | 0.43 | 0.28 | 0.17 |
| 1 | 158600194 | rs12138682 | 1.214 | 0.132 | -0.626 | -0.139 | -0.01 | -0.005 | -0.005 | 0.229 | 0.23 | 0.19 | 0.21 | 0.21 | 0.29 | 0.25 |
| 1 | 158601055 | rs12042819 | 2.485 | -0.195 | -1.334 | -0.413 | -0.047 | -0.086 | -0.086 | 0 | 0.13 | 0.01 | 0.28 | 0.28 | 0.16 | 0.29 |
| 1 | 158604391 | rs16830483 | 1.869 | -0.768 | -0.538 | -0.501 | -0.024 | -0.038 | -0.038 | 0.782 | 0.1 | 0.14 | 0.03 | 0 | 0.02 | 0.04 |
| 1 | 158606449 | rs3738791 | 0.464 | -0.396 | -0.174 | -0.266 | -0.012 | -0.026 | -0.026 | 0 | 0.35 | 0.47 | 0.25 | 0.21 | 0.31 | 0.29 |
| 1 | 158607372 | rs857724 | -0.64 | -0.308 | -0.319 | -0.241 | -0.003 | -0.003 | -0.003 | 0 | 0.63 | 0.68 | 0.56 | 0.64 | 0.59 | 0.46 |
| 1 | 158607935 | rs857725 | 1.089 | -0.3 | -0.326 | -0.221 | -0.032 | -0.051 | -0.051 | 0 | 0.25 | 0.1 | 0.3 | 0.44 | 0.28 | 0.17 |
| 1 | 158615702 | rs857676 | -0.322 | 0.006 | -0.212 | -0.04 | -0.004 | -0.003 | -0.003 | 0 | 0.63 | 0.68 | 0.56 | 0.64 | 0.59 | 0.46 |
| 1 | 158618223 | rs857677 | -0.064 | -0.044 | -0.237 | 0.029 | 0 | 0 | 0 | 0 | 0.59 | 0.59 | 0.54 | 0.59 | 0.56 | 0.42 |
| 1 | 158618455 | rs2246434 | 1.042 | 0.392 | -0.458 | 0.31 | -0.029 | -0.04 | -0.04 | 0 | 0.2 | 0.11 | 0.28 | 0.4 | 0.25 | 0.15 |
| 1 | 158625789 | rs16840445 | -0.673 | 0.057 | 0.599 | 0.268 | -0.003 | -0.007 | -0.007 | 0 | 0.08 | 0.42 | 0.04 | 0 | 0 | 0 |
| 1 | 158626378 | rs857691 | -0.518 | -0.225 | 0.491 | -0.02 | 0.009 | -0.016 | -0.016 | 0 | 0.33 | 0.79 | 0.36 | 0.41 | 0.25 | 0.16 |
| 1 | 158627315 | rs16840450 | -0.296 | -0.074 | 0.389 | 0.063 | -0.007 | -0.01 | -0.01 | 0 | 0.08 | 0.38 | 0.03 | 0 | 0 | 0 |
| 1 | 158631171 | rs2518493 | 0.911 | 0.177 | -0.264 | 0.176 | -0.018 | -0.025 | -0.025 | 0 | 0.21 | 0.14 | 0.3 | 0.41 | 0.25 | 0.16 |
| 1 | 158635852 | rs857680 | -0.48 | 0.094 | -0.759 | 0.149 | -0.08 | -0.097 | -0.097 | 0 | 0.68 | 0.9 | 0.55 | 0.62 | 0.58 | 0.46 |

| | | | | | | | | | | | | | | | | |
|---|---|---|---|---|---|---|---|---|---|---|---|---|---|---|---|---|
| 1 | 158639601 | rs2298801 | 0.002 | 0.086 | -0.394 | 0.2 | -0.022 | -0.019 | -0.019 | 0 | 0.33 | 0.28 | 0.5 | 0.34 | 0.46 | 0.38 |
| 1 | 158641441 | rs857932 | -0.52 | -0.018 | 0.328 | 0.062 | 0.005 | -0.01 | -0.01 | 1.674 | 0.41 | 0.62 | 0.49 | 0.42 | 0.39 | 0.57 |
| 1 | 158647522 | rs703121 | 0.428 | -0.062 | 0.586 | 0.031 | 0.021 | -0.041 | -0.041 | 0.101 | 0.58 | 0.23 | 0.51 | 0.58 | 0.62 | 0.43 |
| 1 | 158649429 | rs703123 | -0.405 | -0.057 | 0.944 | 0.033 | -0.005 | 0.019 | 0.019 | 0 | 0.39 | 0.48 | 0.47 | 0.42 | 0.38 | 0.57 |
| 1 | 158656973 | rs857940 | 1.591 | -0.231 | 0.108 | -0.319 | -0.066 | -0.028 | -0.028 | 0 | 0.32 | 0.09 | 0.37 | 0.39 | 0.38 | 0.17 |
| 1 | 158662926 | rs2023405 | 0.523 | -0.274 | 0.149 | -0.326 | 0.008 | 0.031 | 0.031 | 0 | 0.26 | 0.16 | 0.14 | 0.19 | 0.24 | 0.26 |
| 1 | 158664972 | rs12023371 | 0.494 | -0.27 | 0.208 | -0.324 | 0.009 | 0.028 | 0.028 | 0 | 0.26 | 0.17 | 0.14 | 0.19 | 0.24 | 0.26 |
| 1 | 158668378 | rs751258 | 1.654 | -0.271 | 0.156 | -0.32 | -0.086 | -0.039 | -0.039 | 4.808 | 0.32 | 0.06 | 0.37 | 0.39 | 0.38 | 0.17 |
| 1 | 158668589 | rs751257 | 0.097 | -0.317 | 0.293 | -0.321 | 0.019 | -0.033 | -0.033 | 0 | 0.58 | 0.24 | 0.51 | 0.59 | 0.61 | 0.42 |
| 1 | 158684278 | rs16840667 | 1.833 | -0.306 | -0.18 | -0.194 | -0.011 | -0.057 | -0.057 | 0.009 | 0.11 | 0.01 | 0.21 | 0.24 | 0.09 | 0.23 |
| 1 | 158685036 | rs451851 | -0.438 | -0.103 | 0.245 | -0.636 | -0.003 | 0.009 | 0.009 | 0.189 | 0.61 | 0.62 | 0.57 | 0.6 | 0.62 | 0.43 |
| 1 | 158687224 | rs857703 | 1.474 | -0.007 | -0.276 | -0.723 | -0.062 | -0.049 | -0.049 | 0 | 0.33 | 0.09 | 0.39 | 0.45 | 0.39 | 0.17 |
| 1 | 158687896 | rs857705 | -0.176 | -0.699 | 0.321 | -0.715 | -0.003 | -0.018 | -0.018 | 0 | 0.25 | 0.29 | 0.16 | 0.15 | 0.22 | 0.25 |
| 1 | 158701420 | rs16840748 | 1.531 | -0.191 | 0.189 | -0.461 | -0.015 | -0.042 | -0.042 | 0 | 0.26 | 0.08 | 0.25 | 0.29 | 0.26 | 0.12 |
| 1 | 158704144 | rs1429854 | 0.114 | -0.248 | 0.15 | -0.496 | -0.039 | -0.019 | -0.019 | 3.933 | 0.27 | 0.16 | 0.28 | 0.28 | 0.32 | 0.3 |
| 1 | 158704573 | rs12076675 | 2.02 | -0.284 | 0.325 | -0.439 | 0.002 | 0.001 | 0.001 | 2.902 | 0.2 | 0.19 | 0.12 | 0.17 | 0.15 | 0.06 |
| 1 | 158706639 | rs16840809 | 3.121 | -0.275 | 0.325 | -0.43 | -0.03 | -0.032 | -0.032 | 0 | 0.07 | 0.17 | 0.02 | 0 | 0 | 0 |
| 1 | 158707690 | rs2435282 | -0.384 | -0.231 | 0.495 | -0.43 | 0 | 0.001 | 0.001 | 0.888 | 0.5 | 0.54 | 0.42 | 0.46 | 0.48 | 0.36 |
| 1 | 158708549 | rs381510 | -0.745 | -0.078 | 0.15 | -0.357 | -0.022 | -0.016 | -0.016 | 1.23 | 0.56 | 0.72 | 0.44 | 0.46 | 0.48 | 0.36 |
| 1 | 158720188 | rs454127 | 0.518 | -0.3 | 0.12 | -0.292 | 0.009 | 0.009 | 0.009 | 6.791 | 0.32 | 0.66 | 0.16 | 0.17 | 0.16 | 0.06 |
| 1 | 158720758 | rs10489843 | 0.301 | 0.518 | 0.178 | -0.018 | -0.001 | -0.003 | -0.003 | 0 | 0.08 | 0.06 | 0.11 | 0.15 | 0.15 | 0.11 |
| 1 | 158723868 | rs11586535 | -0.078 | 0.01 | -0.277 | -0.324 | -0.009 | 0.012 | 0.012 | 0 | 0.25 | 0.23 | 0.26 | 0.13 | 0.29 | 0.25 |
| 1 | 158724073 | rs12085542 | 2.301 | -0.606 | 0.214 | -0.243 | 0 | -0.004 | -0.004 | 0.58 | 0.07 | 0.29 | 0.04 | 0 | 0 | 0 |
| 1 | 158725080 | rs16841001 | 1.294 | -0.666 | -0.344 | -0.24 | 0.021 | -0.001 | -0.001 | 0 | 0.19 | 0.04 | 0.11 | 0.17 | 0.15 | 0.06 |
| 1 | 158726746 | rs11265061 | 1.384 | -0.328 | 0.375 | -0.144 | -0.018 | -0.021 | -0.021 | 0 | 0.08 | 0.33 | 0.04 | 0 | 0 | 0 |
| 1 | 158729346 | rs6680399 | -0.031 | 0.326 | -0.185 | 0.291 | 0.005 | 0.016 | 0.016 | 0 | 0.07 | 0.1 | 0.28 | 0.19 | 0.11 | 0.03 |
| 1 | 158729713 | rs11810781 | 0.081 | 0.567 | -0.04 | -0.568 | 0.006 | -0.11 | -0.11 | 0 | 0.23 | 0.33 | 0.14 | 0 | 0.25 | 0.16 |
| 1 | 158734560 | rs857819 | 0.217 | 0.338 | -0.538 | -0.428 | -0.007 | -0.002 | -0.002 | 0 | 0.33 | 0.3 | 0.62 | 0.61 | 0.47 | 0.27 |
| 1 | 158735691 | rs857825 | -0.794 | 0.23 | -0.76 | -0.416 | 0.045 | 0.02 | 0.02 | 0 | 0.58 | 0.78 | 0.78 | 0.62 | 0.72 | 0.43 |
| 1 | 158735740 | rs857826 | -0.794 | 0.23 | -0.76 | -0.416 | 0.045 | 0.02 | 0.02 | 0 | 0.58 | 0.78 | 0.78 | 0.62 | 0.72 | 0.43 |
| 1 | 158735892 | rs857827 | -0.794 | 0.23 | -0.76 | -0.416 | 0.045 | 0.02 | 0.02 | 0 | 0.58 | 0.78 | 0.78 | 0.62 | 0.72 | 0.43 |
| 1 | 158736445 | rs1864346 | 1.132 | 0.176 | -0.854 | -0.159 | -0.102 | -0.148 | -0.148 | 0 | 0.2 | 0.02 | 0.33 | 0.42 | 0.33 | 0.22 |
| 1 | 158739691 | rs6427448 | -0.498 | -0.182 | -0.177 | -0.197 | 0.025 | 0.042 | 0.042 | 0 | 0.22 | 0.41 | 0.38 | 0.34 | 0.23 | 0.16 |
| 1 | 158740662 | rs2859971 | -0.484 | -0.17 | -0.402 | -0.173 | 0.003 | 0.008 | 0.008 | 0 | 0.22 | 0.29 | 0.36 | 0.34 | 0.23 | 0.16 |
| 1 | 158741091 | rs1864348 | 0.589 | -0.607 | -0.901 | -0.137 | 0 | 0.019 | 0.019 | 0.003 | 0.45 | 0.31 | 0.44 | 0.57 | 0.47 | 0.28 |
| 1 | 158744090 | rs16841088 | 0.814 | -0.656 | -0.552 | 0.472 | -0.002 | -0.004 | -0.004 | 0 | 0.12 | 0.1 | 0.09 | 0.13 | 0.14 | 0.07 |
| 1 | 158744303 | rs2325696 | -0.054 | -1.112 | -0.355 | 0.539 | -0.017 | 0.016 | 0.016 | 0 | 0.08 | 0.11 | 0.18 | 0.15 | 0.03 | 0.02 |
| 1 | 158746336 | rs2518524 | -0.797 | -1.321 | -0.727 | 0.66 | 0.004 | -0.002 | -0.002 | 0 | 0.23 | 0.23 | 0.09 | 0.34 | 0.16 | 0.51 |
| 1 | 158746847 | rs16841094 | -0.092 | -1.403 | -0.756 | 0.552 | -0.017 | 0.004 | 0.004 | 0 | 0.13 | 0.23 | 0.05 | 0.19 | 0.07 | 0.34 |
| 1 | 158750550 | rs857838 | -1.866 | -1.383 | -1.323 | 0.26 | -0.05 | -0.045 | -0.045 | 0 | 0.67 | 0.82 | 0.64 | 0.63 | 0.56 | 0.72 |
| 1 | 158750786 | rs857840 | -0.678 | -1.456 | -0.916 | 0.553 | 0.012 | -0.019 | -0.019 | 0 | 0.22 | 0.11 | 0.1 | 0.35 | 0.16 | 0.5 |
| 1 | 158753406 | rs857843 | -1.073 | -1.185 | -1.155 | 0.815 | 0.012 | -0.037 | -0.037 | 0 | 0.33 | 0.21 | 0.26 | 0.52 | 0.28 | 0.59 |
| 1 | 158754813 | rs857847 | -1.228 | -1.166 | -1.115 | 0.542 | 0 | 0 | 0 | 0 | 0.35 | 0.35 | 0.27 | 0.47 | 0.28 | 0.6 |
| 1 | 158756275 | rs12075966 | 1.965 | -0.638 | -0.908 | 0.62 | -0.009 | 0.007 | 0.007 | 1.152 | 0.07 | 0.08 | 0.12 | 0.12 | 0.05 | 0.03 |
| 1 | 158756914 | rs857849 | -0.545 | -0.257 | -0.829 | 0.344 | -0.007 | -0.009 | -0.009 | 0 | 0.15 | 0.19 | 0.06 | 0.12 | 0.1 | 0.11 |
| 1 | 158763983 | rs857854 | -0.855 | 0.328 | -1.983 | 0.695 | -0.172 | -0.169 | -0.169 | 0.332 | 0.69 | 0.98 | 0.57 | 0.57 | 0.53 | 0.71 |
| 1 | 158772845 | rs919477 | -0.311 | -0.59 | -1.434 | 0.746 | 0.026 | -0.028 | -0.028 | 0 | 0.33 | 0.21 | 0.36 | 0.5 | 0.26 | 0.59 |
| 1 | 158775997 | rs857863 | -0.343 | -0.642 | -1.089 | 0.463 | -0.004 | -0.006 | -0.006 | 0 | 0.12 | 0.18 | 0.06 | 0.11 | 0.09 | 0.06 |
| 1 | 158778764 | rs7550055 | -0.516 | -0.989 | -0.504 | 0.659 | -0.025 | 0.01 | 0.01 | 0.078 | 0.38 | 0.45 | 0.44 | 0.56 | 0.26 | 0.61 |
| 1 | 158784017 | rs16841134 | 0.033 | -0.767 | -0.596 | 1.129 | -0.004 | -0.014 | -0.014 | 0.011 | 0.1 | 0.05 | 0.21 | 0.32 | 0.1 | 0.35 |
| 1 | 158784259 | rs857785 | -0.072 | -0.745 | -0.915 | 0.337 | -0.028 | -0.02 | -0.02 | 0 | 0.4 | 0.23 | 0.33 | 0.38 | 0.49 | 0.25 |
| 1 | 158785137 | rs857788 | -0.033 | -0.822 | -0.728 | 0.3 | -0.017 | -0.011 | -0.011 | 0.48 | 0.41 | 0.29 | 0.34 | 0.38 | 0.49 | 0.25 |
| 1 | 158788997 | rs857791 | 0.592 | -0.906 | -0.842 | 0.332 | -0.011 | -0.009 | -0.009 | 0 | 0.29 | 0.2 | 0.24 | 0.26 | 0.39 | 0.14 |
| 1 | 158803873 | rs7513873 | 1.109 | -0.841 | -0.608 | 0.03 | 0.001 | -0.005 | -0.005 | 0 | 0.15 | 0.14 | 0.29 | 0.13 | 0.18 | 0.1 |
| 1 | 158810312 | rs857869 | 0.098 | -1.051 | -0.107 | 0.011 | -0.015 | -0.003 | -0.003 | 0.91 | 0.4 | 0.26 | 0.46 | 0.57 | 0.49 | 0.48 |
| 1 | 158814834 | rs860781 | -0.525 | -1.07 | -0.129 | -0.004 | 0.013 | 0.043 | 0.043 | 0.137 | 0.52 | 0.38 | 0.55 | 0.7 | 0.58 | 0.59 |
| 1 | 158816589 | rs2188112 | -1.716 | -1.1 | -0.334 | -0.054 | 0.005 | 0.009 | 0.009 | 0 | 0.72 | 0.78 | 0.87 | 0.83 | 0.77 | 0.69 |
| 1 | 158817474 | rs2261107 | 1.379 | -0.87 | -0.806 | -0.002 | -0.037 | -0.052 | -0.052 | 0 | 0.23 | 0.09 | 0.2 | 0.26 | 0.33 | 0.12 |
| 1 | 158820153 | rs960893 | 0.849 | -0.422 | -0.684 | -0.185 | -0.02 | -0.015 | -0.015 | 0.01 | 0.1 | 0.03 | 0.09 | 0.12 | 0.1 | 0.11 |
| 1 | 158821200 | rs2106091 | 0.571 | -0.251 | -0.429 | -0.228 | -0.003 | 0.005 | 0.005 | 0.399 | 0.14 | 0.09 | 0.28 | 0.13 | 0.18 | 0.1 |
| 1 | 158832444 | rs2106092 | 0.776 | 0.038 | -0.769 | -0.052 | -0.05 | 0.008 | 0.008 | 2.106 | 0.26 | 0.07 | 0.2 | 0.18 | 0.33 | 0.17 |
| 1 | 158833750 | rs2188114 | 0.794 | -0.243 | -1.071 | 0.04 | 0.004 | -0.07 | -0.07 | 0 | 0.19 | 0.03 | 0.12 | 0.24 | 0.16 | 0.14 |
| 1 | 158841260 | rs2852707 | 0.771 | -0.216 | -0.339 | -0.036 | 0.002 | -0.013 | -0.013 | 0.175 | 0.27 | 0.35 | 0.2 | 0.18 | 0.31 | 0.17 |
| 1 | 158842020 | rs2518510 | 0.771 | -0.216 | -0.339 | -0.036 | 0.002 | -0.013 | -0.013 | 0 | 0.27 | 0.36 | 0.21 | 0.18 | 0.31 | 0.17 |
| 1 | 158842453 | rs2494034 | -1.307 | 0.028 | -0.281 | -0.159 | 0 | 0.022 | 0.022 | 0 | 0.71 | 0.77 | 0.85 | 0.96 | 0.74 | 0.7 |
| 1 | 158847720 | rs2852657 | -1.224 | 0.047 | -0.289 | -0.15 | 0 | -0.011 | -0.011 | 0 | 0.7 | 0.71 | 0.84 | 0.96 | 0.74 | 0.71 |
| 1 | 158852953 | rs1894042 | 0.03 | 0.357 | -0.043 | 0.347 | 0.002 | -0.039 | -0.039 | 0 | 0.32 | 0.47 | 0.25 | 0.18 | 0.35 | 0.18 |
| 1 | 158854489 | rs12036526 | -0.398 | 0.522 | -0.016 | 0.662 | 0.006 | 0.065 | 0.065 | 0.344 | 0.1 | 0.11 | 0.25 | 0.4 | 0.12 | 0.31 |
| 1 | 158871453 | rs12088342 | -0.484 | 0.546 | -0.267 | 0.663 | 0.008 | 0.081 | 0.081 | 0.231 | 0.1 | 0.15 | 0.26 | 0.4 | 0.12 | 0.31 |

| | | | | | | | | | | | | | | | | |
|---|---|---|---|---|---|---|---|---|---|---|---|---|---|---|---|---|
| 1 | 158875409 | rs12569226 | 0.891 | 0.664 | -0.811 | 0.334 | -0.032 | -0.003 | -0.003 | 0.263 | 0.22 | 0.06 | 0.18 | 0.17 | 0.28 | 0.17 |
| 1 | 158876921 | rs2518544 | 1.066 | 0.03 | -0.708 | -0.119 | 0 | -0.048 | -0.048 | 0.047 | 0.29 | 0.35 | 0.16 | 0.04 | 0.26 | 0.29 |
| 1 | 158884519 | rs860637 | 1.353 | -0.03 | -0.739 | -0.148 | -0.007 | 0.039 | 0.039 | 0 | 0.27 | 0.18 | 0.15 | 0.04 | 0.26 | 0.29 |
| 1 | 158887208 | rs2106096 | 0.514 | 0.077 | -0.775 | -0.032 | -0.007 | 0.044 | 0.044 | 0 | 0.07 | 0.07 | 0.22 | 0.41 | 0.1 | 0.31 |
| 1 | 158891391 | rs856099 | -0.787 | 0.242 | -0.617 | -0.743 | 0.009 | -0.034 | -0.034 | 0 | 0.73 | 0.63 | 0.78 | 0.88 | 0.69 | 0.83 |
| 1 | 158893432 | rs11805756 | 1.073 | 0.207 | -1 | -0.659 | -0.009 | -0.04 | -0.04 | 0.776 | 0.11 | 0.01 | 0.23 | 0.15 | 0.11 | 0.08 |
| 1 | 158899720 | rs856118 | 1.298 | 0.3 | -0.94 | -0.71 | -0.019 | 0.054 | 0.054 | 0 | 0.24 | 0.11 | 0.14 | 0.07 | 0.25 | 0.3 |
| 1 | 158899747 | rs856119 | 1.223 | 0.323 | -0.928 | -0.699 | -0.001 | -0.01 | -0.01 | 0 | 0.25 | 0.23 | 0.15 | 0.07 | 0.25 | 0.29 |
| 1 | 158901127 | rs16841336 | -0.321 | 0.716 | -0.895 | -0.104 | 0 | 0 | 0 | 0 | 0.19 | 0.17 | 0.28 | 0.17 | 0.18 | 0.1 |
| 1 | 158902276 | rs1476438 | -0.172 | 1.219 | -1.077 | -0.224 | -0.063 | 0.043 | 0.043 | 0.959 | 0.25 | 0.06 | 0.21 | 0.15 | 0.36 | 0.2 |
| 1 | 158902344 | rs6675945 | -0.416 | 1.331 | -0.773 | -0.173 | 0.03 | -0.055 | -0.055 | 0 | 0.21 | 0.37 | 0.19 | 0.1 | 0.24 | 0.17 |
| 1 | 158903252 | rs2820187 | 0.25 | 1.558 | -1.738 | 0.255 | -0.099 | -0.027 | -0.027 | 0 | 0.63 | 0.89 | 0.75 | 0.7 | 0.57 | 0.64 |
| 1 | 158904114 | rs1633277 | -1.1 | 1.64 | -0.764 | 0.173 | 0.007 | -0.006 | -0.006 | 0.382 | 0.18 | 0.21 | 0.19 | 0.13 | 0.29 | 0.18 |
| 1 | 158909164 | rs2325704 | 0.871 | 1.25 | -0.358 | 0.08 | 0.014 | -0.022 | -0.022 | 0 | 0.82 | 0.76 | 0.83 | 0.85 | 0.77 | 0.8 |
| 1 | 158915141 | rs856084 | -0.67 | 0.982 | -0.652 | 0.671 | -0.04 | -0.059 | -0.059 | 0 | 0.18 | 0.05 | 0.18 | 0.35 | 0.25 | 0.36 |
| 1 | 158915917 | rs861319 | -0.051 | -0.985 | -1.205 | -0.265 | 0.065 | 0.073 | 0.073 | 6.363 | 0.63 | 0.28 | 0.6 | 0.86 | 0.8 | 0.87 |
| 1 | 158918264 | rs7547305 | 0.035 | -1.321 | -0.171 | -0.559 | 0.008 | -0.033 | -0.033 | 0.367 | 0.08 | 0.12 | 0.07 | 0.01 | 0.14 | 0.03 |
| 1 | 158922287 | rs856090 | -0.132 | -1.639 | 0.338 | -0.712 | -0.061 | -0.17 | -0.17 | 0 | 0.77 | 0.55 | 0.9 | 0.99 | 0.86 | 0.97 |
| 1 | 158932555 | rs1101999 | -0.409 | -1.458 | 0.235 | -0.504 | 0.001 | -0.002 | -0.002 | 0.125 | 0.93 | 0.69 | 0.96 | 1 | 1 | 1 |
| 1 | 158941219 | rs1633292 | 0.112 | -1.487 | -1.061 | -0.839 | -0.026 | -0.026 | -0.026 | 0 | 0.32 | 0.18 | 0.36 | 0.41 | 0.34 | 0.5 |
| 1 | 158941634 | rs10908697 | 0.937 | -1.477 | -1.208 | -0.749 | -0.108 | -0.083 | -0.083 | 0.248 | 0.35 | 0.12 | 0.39 | 0.5 | 0.47 | 0.43 |
| 1 | 158944127 | rs1623601 | 0.739 | -1.804 | -0.41 | -0.791 | -0.034 | -0.004 | -0.004 | 0.019 | 0.74 | 0.37 | 0.85 | 0.73 | 0.85 | 0.76 |
| 1 | 158944584 | rs1615480 | 0.302 | -1.906 | 0.156 | -0.82 | -0.036 | 0.2 | 0.2 | 2.723 | 0.9 | 0.61 | 0.96 | 0.73 | 0.98 | 0.83 |
| 1 | 158956412 | rs1361567 | 1.119 | -2.454 | -0.427 | -1.198 | -0.05 | -0.017 | -0.017 | 0 | 0.39 | 0.21 | 0.54 | 0.55 | 0.48 | 0.44 |
| 1 | 158957703 | rs856142 | 1.104 | -2.718 | -0.472 | -1.355 | 0.028 | 0.013 | 0.013 | 0.247 | 0.64 | 0.3 | 0.72 | 0.7 | 0.79 | 0.69 |
| 1 | 158959450 | rs856146 | 1.056 | -2.492 | -0.329 | -1.24 | -0.043 | 0.055 | 0.055 | 0 | 0.25 | 0.08 | 0.18 | 0.14 | 0.31 | 0.25 |
| 1 | 158971086 | rs4657616 | 1.562 | -2.559 | -0.812 | -0.878 | -0.148 | -0.184 | -0.184 | 0 | 0.23 | 0.01 | 0.56 | 0.47 | 0.4 | 0.33 |
| 1 | 158972490 | rs4657618 | 0.614 | -2.65 | -0.225 | -0.847 | 0.007 | 0.062 | 0.062 | 0 | 0.49 | 0.29 | 0.66 | 0.63 | 0.51 | 0.49 |
| 1 | 158978596 | rs9887904 | 0.043 | -1.872 | -0.546 | -0.786 | -0.001 | -0.037 | -0.037 | 0 | 0.13 | 0.17 | 0.11 | 0 | 0.13 | 0.06 |
| 1 | 158979191 | rs1417806 | -0.391 | -2.327 | -0.527 | -0.726 | -0.001 | 0.009 | 0.009 | 0 | 0.69 | 0.76 | 0.77 | 0.84 | 0.7 | 0.76 |
| 1 | 158979950 | rs2276404 | -0.686 | -3.031 | 0.055 | 0.178 | -0.012 | 0.003 | 0.003 | 0.699 | 0.27 | 0.27 | 0.1 | 0.4 | 0.13 | 0.31 |
| 1 | 158983285 | rs856064 | 1.421 | -3.103 | -0.409 | 0.04 | 0.114 | 0.01 | 0.01 | 4.741 | 0.6 | 0.19 | 0.83 | 0.59 | 0.79 | 0.68 |
| 1 | 158985135 | rs1361565 | -0.46 | -3.06 | -0.33 | 0.006 | -0.004 | -0.032 | -0.032 | 0 | 0.1 | 0.11 | 0.08 | 0 | 0.09 | 0.06 |
| 1 | 158987941 | rs856046 | 1.827 | -3.242 | -0.112 | 0.21 | 0.158 | -0.114 | -0.114 | 0.183 | 0.59 | 0.04 | 0.79 | 0.5 | 0.81 | 0.66 |
| 1 | 158990412 | rs856049 | 1.396 | -3.278 | 0.011 | 0.205 | 0.152 | -0.028 | -0.028 | 0 | 0.62 | 0.15 | 0.81 | 0.53 | 0.82 | 0.68 |
| 1 | 158996895 | rs1101993 | 1.198 | -3.096 | 0.235 | 0.148 | 0.068 | 0.005 | 0.005 | 5.208 | 0.65 | 0.21 | 0.85 | 0.61 | 0.81 | 0.7 |
| 1 | 158998665 | rs12094741 | -1.039 | -3.127 | 0.697 | 0.134 | -0.038 | 0.007 | 0.007 | 1.953 | 0.29 | 0.61 | 0.13 | 0.39 | 0.16 | 0.29 |
| 1 | 159001521 | rs856054 | 2.135 | -3.198 | 0.276 | 0.141 | 0.156 | -0.06 | -0.06 | 0.965 | 0.6 | 0.08 | 0.81 | 0.53 | 0.81 | 0.69 |
| 1 | 159002222 | rs861318 | 0.873 | -3.195 | 0.949 | 0.152 | -0.004 | -0.001 | -0.001 | 0 | 0.71 | 0.32 | 0.87 | 0.6 | 0.84 | 0.71 |
| 1 | 159002377 | rs1057027 | 0.873 | -3.195 | 0.949 | 0.157 | -0.004 | -0.001 | -0.001 | 0 | 0.71 | 0.32 | 0.87 | 0.59 | 0.84 | 0.71 |
| 1 | 159005977 | rs1633266 | 1.061 | -3.285 | 1.568 | 0.052 | -0.072 | 0 | 0 | 5.368 | 0.7 | 0.49 | 0.89 | 0.59 | 0.88 | 0.69 |
| 1 | 159006846 | rs3754460 | 0.172 | -3.275 | 1.357 | 0.05 | -0.023 | 0.016 | 0.016 | 0.016 | 0.19 | 0.25 | 0.06 | 0.35 | 0.12 | 0.29 |
| 1 | 159012646 | rs2570916 | 2.165 | -3.32 | 1.243 | 0.011 | 0.156 | 0.005 | 0.005 | 5.72 | 0.61 | 0.08 | 0.83 | 0.59 | 0.83 | 0.7 |
| 1 | 159016491 | rs2793843 | 0.253 | -3.248 | 1.341 | 0.002 | -0.019 | -0.02 | -0.02 | 0 | 0.95 | 0.93 | 0.99 | 1 | 1 | 1 |
| 1 | 159021445 | rs3737522 | 0.127 | -3.259 | 1.444 | 0.005 | -0.033 | 0.025 | 0.025 | 0 | 0.19 | 0.28 | 0.06 | 0.35 | 0.12 | 0.28 |
| 1 | 159028378 | rs855865 | 1.164 | -3.27 | 1.392 | 0.01 | -0.048 | 0.004 | 0.004 | 0.806 | 0.71 | 0.62 | 0.88 | 0.64 | 0.84 | 0.71 |
| 1 | 159032255 | rs2793845 | -0.248 | -3.205 | 1.298 | 0.052 | -0.018 | 0.015 | 0.015 | 0 | 0.8 | 0.78 | 0.94 | 0.64 | 0.88 | 0.72 |
| 1 | 159046430 | rs2298803 | 0.57 | -3.242 | 1.018 | 0.04 | 0.05 | -0.049 | -0.049 | 0.092 | 0.16 | 0.07 | 0.03 | 0.36 | 0.04 | 0.27 |
| 1 | 159055656 | rs2518562 | 1.147 | -3.343 | 1.206 | -0.022 | -0.056 | -0.001 | -0.001 | 0 | 0.71 | 0.39 | 0.89 | 0.64 | 0.9 | 0.71 |
| 1 | 159060560 | rs2852720 | 1.206 | -3.248 | 0.739 | -0.091 | -0.051 | -0.067 | -0.067 | 0 | 0.85 | 0.38 | 0.89 | 0.92 | 0.94 | 0.96 |
| 1 | 159069211 | rs1894043 | 1.333 | -3.108 | -0.289 | 0.028 | 0.155 | -0.004 | -0.004 | 0.408 | 0.59 | 0.04 | 0.8 | 0.58 | 0.81 | 0.69 |
| 1 | 159086986 | rs2852727 | 1.047 | -2.798 | -0.308 | -0.031 | 0.006 | -0.068 | -0.068 | 2.372 | 0.39 | 0.09 | 0.71 | 0.52 | 0.63 | 0.4 |
| 1 | 159090256 | rs1002408 | 1.564 | -2.946 | 0.373 | -0.15 | -0.063 | 0.009 | 0.009 | 2.866 | 0.12 | 0.27 | 0.04 | 0.12 | 0 | 0.02 |
| 1 | 159093732 | rs12130875 | 0.311 | -2.675 | -0.351 | -0.524 | -0.009 | 0.021 | 0.021 | 0 | 0.11 | 0.04 | 0.09 | 0 | 0.17 | 0.03 |
| 1 | 159098713 | rs2814779 | -0.093 | -2.38 | -0.072 | -0.248 | -0.056 | -0.054 | -0.054 | 2.334 | 0.75 | 0.55 | 0.92 | 0.85 | 0.89 | 0.64 |
| 1 | 159103988 | rs863027 | 0.326 | -1.364 | -1.225 | 0.174 | -0.086 | -0.052 | -0.052 | 3.328 | 0.3 | 0.15 | 0.56 | 0.37 | 0.46 | 0.29 |
| 1 | 159116681 | rs863035 | 0.614 | -2.057 | -0.469 | -0.106 | 0.055 | 0.035 | 0.035 | 1.173 | 0.54 | 0.39 | 0.82 | 0.69 | 0.8 | 0.62 |
| 1 | 159117690 | rs3026935 | -0.44 | -0.919 | -0.631 | 0.149 | 0.015 | 0.007 | 0.007 | 0 | 0.18 | 0.1 | 0.05 | 0.14 | 0.08 | 0.33 |
| 1 | 159131926 | rs3026943 | -0.149 | 0.077 | -0.639 | -0.17 | -0.002 | -0.018 | -0.018 | 0 | 0.47 | 0.51 | 0.68 | 0.71 | 0.6 | 0.56 |
| 1 | 159132649 | rs12042284 | 0.066 | 0.424 | -0.755 | -0.041 | -0.004 | -0.001 | -0.001 | 0.311 | 0.06 | 0.05 | 0.1 | 0.06 | 0.1 | 0.01 |
| 1 | 159135282 | rs3026946 | 0.224 | 0.938 | -1.137 | 0.668 | -0.002 | -0.141 | -0.141 | 0 | 0.21 | 0.03 | 0.35 | 0.45 | 0.21 | 0.36 |
| 1 | 159135353 | rs2814774 | -1.025 | -0.176 | -0.528 | -0.134 | 0.011 | 0.07 | 0.07 | 0 | 0.5 | 0.66 | 0.62 | 0.64 | 0.53 | 0.54 |
| 1 | 159140376 | rs965359 | 0.038 | -0.459 | -0.21 | -0.303 | -0.005 | 0.006 | 0.006 | 0 | 0.09 | 0.06 | 0.12 | 0.06 | 0.12 | 0.02 |
| 1 | 159147452 | rs3026968 | 0.685 | -0.577 | -0.404 | -0.112 | -0.01 | -0.113 | -0.113 | 2.971 | 0.2 | 0.03 | 0.37 | 0.53 | 0.24 | 0.36 |
| 1 | 159148513 | rs1474747 | -1.499 | -1.162 | -0.413 | -0.502 | -0.002 | 0.053 | 0.053 | 0 | 0.6 | 0.71 | 0.73 | 0.73 | 0.63 | 0.54 |
| 1 | 159149515 | rs2852714 | -0.14 | -1.212 | -0.38 | -0.518 | 0.015 | -0.036 | -0.036 | 0 | 0.26 | 0.35 | 0.21 | 0.14 | 0.28 | 0.17 |
| 1 | 159153480 | rs2852716 | -1.219 | -1.257 | -0.491 | -0.518 | -0.019 | 0.093 | 0.093 | 0.007 | 0.53 | 0.8 | 0.61 | 0.67 | 0.51 | 0.52 |
| 1 | 159156285 | rs2281300 | 0.772 | -1.432 | -0.544 | -0.319 | -0.008 | -0.114 | -0.114 | 0 | 0.21 | 0.03 | 0.37 | 0.53 | 0.24 | 0.37 |
| 1 | 159160880 | rs862988 | -0.919 | -3.129 | -0.623 | -1.007 | 0.03 | 0.025 | 0.025 | 0.476 | 0.46 | 0.59 | 0.53 | 0.65 | 0.34 | 0.42 |

| | | | | | | | | | | | | | | | | |
|---|---|---|---|---|---|---|---|---|---|---|---|---|---|---|---|---|
| 1 | 159161279 | rs862989 | -0.036 | -3.218 | -0.418 | -0.525 | -0.049 | 0.044 | 0.044 | 0 | 0.32 | 0.53 | 0.4 | 0.46 | 0.23 | 0.35 |
| 1 | 159164364 | rs862991 | 0.737 | -3.095 | -0.78 | -1.321 | -0.035 | -0.095 | -0.095 | 0 | 0.59 | 0.5 | 0.69 | 0.93 | 0.73 | 0.76 |
| 1 | 159165610 | rs2814755 | 1.379 | -2.805 | -0.863 | -1.125 | -0.094 | -0.107 | -0.107 | 0.142 | 0.17 | 0.32 | 0.03 | 0 | 0.01 | 0.02 |
| 1 | 159167574 | rs2746044 | 1.379 | -2.805 | -0.872 | -1.125 | -0.095 | -0.108 | -0.108 | 0 | 0.17 | 0.32 | 0.03 | 0 | 0.01 | 0.02 |
| 1 | 159169641 | rs862999 | 0.065 | -2.105 | -1.578 | -0.872 | -0.002 | 0.062 | 0.062 | 0 | 0.79 | 0.91 | 0.72 | 0.93 | 0.74 | 0.79 |
| 1 | 159171603 | rs3845622 | 0.025 | -1.659 | -2.244 | -0.018 | -0.027 | -0.074 | -0.074 | 0 | 0.11 | 0.01 | 0.19 | 0.58 | 0.12 | 0.26 |
| 1 | 159174920 | rs863002 | 0.014 | 0.98 | -2.946 | -0.787 | -0.148 | 0.077 | 0.077 | 0.557 | 0.22 | 0.01 | 0.29 | 0.08 | 0.4 | 0.31 |
| 1 | 159175354 | rs12075 | -0.954 | 0.489 | -3.87 | -1.673 | -0.152 | 0.171 | 0.171 | 0.72 | 0.67 | 0.98 | 0.54 | 0.08 | 0.6 | 0.36 |
| 1 | 159177748 | rs863006 | -0.748 | -0.447 | -3.357 | -2.174 | 0.016 | 0.102 | 0.102 | 0 | 0.51 | 0.66 | 0.51 | 0.08 | 0.58 | 0.36 |
| 1 | 159183713 | rs3027048 | 0.318 | -1.21 | -3.762 | -1.358 | 0.006 | -0.129 | -0.129 | 0.434 | 0.37 | 0.07 | 0.42 | 0.47 | 0.36 | 0.4 |
| 1 | 159186231 | rs3027056 | 0.267 | -1.253 | -3.82 | -1.335 | 0.008 | -0.099 | -0.099 | 0.083 | 0.38 | 0.07 | 0.42 | 0.53 | 0.37 | 0.4 |
| 1 | 159196765 | rs10489849 | -0.091 | -1.711 | -3.621 | -1.788 | 0.08 | 0.081 | 0.081 | 3.706 | 0.42 | 0.66 | 0.28 | 0.02 | 0.19 | 0.12 |
| 1 | 159200020 | rs863013 | 1.611 | -0.886 | -3.69 | -1.438 | -0.114 | -0.047 | -0.047 | 0.316 | 0.16 | 0.01 | 0.24 | 0.18 | 0.36 | 0.25 |
| 1 | 159211064 | rs6660102 | -0.146 | -1.751 | -3.585 | -1.479 | 0.064 | 0.082 | 0.082 | 0.7 | 0.31 | 0.8 | 0.28 | 0.08 | 0.18 | 0.11 |
| 1 | 159215287 | rs12072644 | -0.166 | -1.744 | -3.587 | -1.482 | 0.062 | 0.084 | 0.084 | 0.802 | 0.31 | 0.8 | 0.28 | 0.08 | 0.18 | 0.11 |
| 1 | 159217581 | rs16841937 | 1.459 | -1.578 | -3.324 | -1.352 | -0.092 | -0.095 | -0.095 | 5.062 | 0.15 | 0.56 | 0.04 | 0 | 0 | 0 |
| 1 | 159218266 | rs3845624 | -1.291 | -1.402 | -3.736 | -1.326 | -0.079 | -0.139 | -0.139 | 0.003 | 0.63 | 0.96 | 0.52 | 0.53 | 0.42 | 0.45 |
| 1 | 159228378 | rs10908702 | -0.948 | -1.788 | -3.691 | -1.283 | 0.124 | 0.015 | 0.015 | 3.512 | 0.53 | 0.92 | 0.41 | 0.59 | 0.28 | 0.41 |
| 1 | 159236439 | rs2325927 | 1.672 | -1.75 | -2.992 | -1.257 | -0.074 | 0.006 | 0.006 | 5.49 | 0.12 | 0.49 | 0.03 | 0.06 | 0 | 0 |
| 1 | 159244893 | rs2511212 | -2.578 | -1.611 | -3.249 | -1.429 | -0.068 | -0.026 | -0.026 | 0.001 | 0.79 | 0.97 | 0.75 | 0.77 | 0.78 | 0.71 |
| 1 | 159248815 | rs10489853 | 1.682 | -1.511 | -2.927 | -1.242 | -0.071 | -0.075 | -0.075 | 0.195 | 0.12 | 0.49 | 0.03 | 0 | 0 | 0 |
| 1 | 159253672 | rs2494262 | -0.029 | -1.357 | -3.556 | -1.282 | -0.146 | -0.048 | -0.048 | 0.486 | 0.33 | 0.06 | 0.44 | 0.65 | 0.56 | 0.56 |
| 1 | 159258545 | rs2427837 | 1.48 | -0.943 | -3.549 | -1.286 | -0.08 | 0.08 | 0.08 | 0 | 0.17 | 0.05 | 0.14 | 0.04 | 0.3 | 0.25 |
| 1 | 159259680 | rs2247584 | 0.629 | -1.479 | -3.425 | -0.893 | -0.155 | 0.017 | 0.017 | 0.043 | 0.26 | 0.05 | 0.34 | 0.18 | 0.5 | 0.32 |
| 1 | 159266066 | rs2427825 | -2.233 | -1.615 | -3.511 | -0.782 | -0.079 | -0.016 | -0.016 | 0 | 0.77 | 0.97 | 0.74 | 0.78 | 0.76 | 0.73 |
| 1 | 159267255 | rs7548864 | 1.445 | -2.014 | -3.377 | -0.887 | -0.017 | 0.005 | 0.005 | 0.973 | 0.26 | 0.29 | 0.15 | 0.53 | 0.09 | 0.3 |
| 1 | 159272764 | rs12119226 | 1.994 | -2.046 | -3.451 | -0.83 | 0.059 | -0.076 | -0.076 | 0 | 0.22 | 0.08 | 0.14 | 0.5 | 0.09 | 0.16 |
| 1 | 159276153 | rs2252226 | -1.216 | -1.822 | -3.477 | -0.766 | -0.131 | 0.024 | 0.024 | 1.538 | 0.66 | 0.95 | 0.53 | 0.71 | 0.58 | 0.46 |
| 1 | 159277868 | rs7549785 | 1.797 | -1.982 | -3.146 | -0.683 | -0.032 | 0.027 | 0.027 | 0 | 0.09 | 0.02 | 0.2 | 0.02 | 0.16 | 0.1 |
| 1 | 159279994 | rs16842010 | 1.758 | -2.065 | -3.118 | -0.559 | 0.007 | -0.042 | -0.042 | 0.53 | 0.09 | 0 | 0.12 | 0.42 | 0.08 | 0.12 |
| 1 | 159282011 | rs2427829 | -1.593 | -2.23 | -3.054 | -0.993 | -0.053 | 0.099 | 0.099 | 0.611 | 0.79 | 0.97 | 0.76 | 0.95 | 0.75 | 0.61 |
| 1 | 159283746 | rs11265165 | 1.851 | -2.537 | -2.671 | -0.764 | 0.029 | -0.033 | -0.033 | 0.312 | 0.24 | 0.18 | 0.13 | 0.42 | 0.1 | 0.13 |
| 1 | 159292850 | rs4282816 | -0.286 | -2.452 | -2.599 | -1.278 | -0.022 | -0.058 | -0.058 | 0.801 | 0.66 | 0.48 | 0.72 | 0.89 | 0.75 | 0.6 |
| 1 | 159300876 | rs7544351 | 1.302 | -2.493 | -2.383 | -1.104 | -0.098 | -0.123 | -0.123 | 0 | 0.27 | 0.56 | 0.06 | 0 | 0.03 | 0.01 |
| 1 | 159319803 | rs10218647 | -1.149 | -1.75 | -3.036 | -1.048 | 0.012 | 0.12 | 0.12 | 0 | 0.66 | 0.9 | 0.77 | 0.89 | 0.72 | 0.6 |
| 1 | 159321675 | rs4443888 | -0.002 | -0.951 | -3.207 | -0.708 | -0.153 | -0.092 | -0.092 | 0 | 0.35 | 0.05 | 0.49 | 0.62 | 0.54 | 0.41 |
| 1 | 159325834 | rs11265177 | 1.824 | -0.671 | -3.015 | -0.674 | -0.064 | -0.07 | -0.07 | 2.954 | 0.13 | 0.02 | 0.22 | 0.26 | 0.18 | 0.19 |
| 1 | 159329046 | rs11265178 | 0.687 | -0.707 | -2.984 | -0.558 | -0.131 | -0.044 | -0.044 | 0.052 | 0.22 | 0.04 | 0.37 | 0.25 | 0.47 | 0.29 |
| 1 | 159336948 | rs10494322 | 1.98 | -0.116 | -2.943 | -0.477 | -0.052 | -0.057 | -0.057 | 0.106 | 0.1 | 0.01 | 0.22 | 0.2 | 0.19 | 0.06 |
| 1 | 159342439 | rs6687840 | -0.003 | -0.73 | -3.259 | -0.701 | 0.028 | -0.007 | -0.007 | 0 | 0.44 | 0.93 | 0.42 | 0.38 | 0.29 | 0.29 |
| 1 | 159374604 | rs10494323 | 1.918 | -0.707 | -3.292 | -0.793 | -0.03 | 0.036 | 0.036 | 0 | 0.09 | 0.03 | 0.16 | 0.03 | 0.13 | 0.12 |
| 1 | 159391401 | rs11265190 | 0.029 | -0.922 | -3.409 | -0.817 | 0.024 | 0.116 | 0.116 | 2.289 | 0.48 | 0.07 | 0.48 | 0.24 | 0.65 | 0.6 |
| 1 | 159408803 | rs10908720 | 0.181 | -1.194 | -2.929 | -1.004 | 0.002 | 0.095 | 0.095 | 3.396 | 0.47 | 0.06 | 0.47 | 0.24 | 0.65 | 0.6 |
| 1 | 159409857 | rs12048482 | -0.149 | -1.221 | -2.189 | -1.008 | 0.004 | 0.009 | 0.009 | 0.347 | 0.55 | 0.43 | 0.51 | 0.24 | 0.65 | 0.6 |
| 1 | 159410975 | rs4656236 | 2.091 | -0.855 | -2.302 | -1.267 | -0.079 | 0.066 | 0.066 | 0.508 | 0.13 | 0.01 | 0.15 | 0.02 | 0.27 | 0.22 |
| 1 | 159415977 | rs7552361 | -2.762 | -1.61 | -1.618 | -0.897 | -0.102 | -0.13 | -0.13 | 0.021 | 0.81 | 0.66 | 0.97 | 1 | 0.98 | 0.99 |
| 1 | 159418212 | rs11265196 | 0.397 | -1.367 | -1.933 | -0.754 | 0.049 | 0.125 | 0.125 | 1.148 | 0.49 | 0.1 | 0.48 | 0.28 | 0.65 | 0.62 |
| 1 | 159439913 | rs11265204 | 0.857 | -1.249 | -2.008 | -0.852 | -0.091 | 0.022 | 0.022 | 0 | 0.37 | 0.09 | 0.43 | 0.27 | 0.55 | 0.36 |
| 1 | 159446370 | rs4233351 | 0.295 | -1.301 | -1.661 | -0.601 | 0.077 | -0.04 | -0.04 | 0 | 0.55 | 0.19 | 0.65 | 0.49 | 0.7 | 0.75 |
| 1 | 159449126 | rs4584388 | 1.148 | -1.244 | -1.554 | -0.437 | -0.001 | 0.06 | 0.06 | 0 | 0.1 | 0.11 | 0.11 | 0.48 | 0.07 | 0.11 |
| 1 | 159461877 | rs10752629 | -2.866 | -1.26 | -1.278 | -0.707 | -0.068 | -0.091 | -0.091 | 0 | 0.83 | 0.73 | 0.97 | 1 | 0.98 | 0.99 |
| 1 | 159462381 | rs4325129 | 1.12 | -0.791 | -1.393 | -0.564 | -0.051 | -0.006 | -0.006 | 1.975 | 0.11 | 0.03 | 0.18 | 0.09 | 0.21 | 0.13 |
| 1 | 159478230 | rs11811256 | 0.703 | -1.178 | -1.761 | -0.682 | 0.019 | 0.129 | 0.129 | 0.728 | 0.5 | 0.16 | 0.46 | 0.28 | 0.6 | 0.62 |
| 1 | 159479005 | rs12047230 | 1.47 | -1.307 | -2.125 | -0.975 | -0.124 | -0.041 | -0.041 | 0 | 0.29 | 0.04 | 0.39 | 0.27 | 0.47 | 0.34 |
| 1 | 159479781 | rs7532282 | -0.323 | -1.068 | -1.816 | -1.181 | 0.014 | 0.074 | 0.074 | 0 | 0.26 | 0.44 | 0.32 | 0.45 | 0.28 | 0.24 |
| 1 | 159502248 | rs1374484 | 1.648 | -0.752 | -2.538 | -1.067 | -0.027 | 0.087 | 0.087 | 0.398 | 0.14 | 0.01 | 0.06 | 0.01 | 0.13 | 0.26 |
| 1 | 159505297 | rs4656837 | 0.287 | -0.613 | -2.254 | -0.727 | 0.049 | -0.022 | -0.022 | 0 | 0.56 | 0.29 | 0.66 | 0.49 | 0.69 | 0.74 |
| 1 | 159517379 | rs10494325 | 0.357 | 0.558 | -1.754 | -0.135 | 0 | 0.018 | 0.018 | 0 | 0.12 | 0.13 | 0.17 | 0.44 | 0.12 | 0.12 |
| 1 | 159517860 | rs6702567 | 0.251 | 0.872 | -1.687 | -0.281 | -0.045 | 0.045 | 0.045 | 0 | 0.1 | 0.03 | 0.15 | 0.02 | 0.19 | 0.14 |
| 1 | 159527480 | rs2275674 | -0.492 | -0.024 | -1.591 | -0.16 | 0.007 | -0.019 | -0.019 | 0.233 | 0.29 | 0.29 | 0.25 | 0.46 | 0.24 | 0.36 |
| 1 | 159531750 | rs6681315 | -0.835 | -0.208 | -1.609 | -0.864 | 0.004 | 0.068 | 0.068 | 0 | 0.39 | 0.25 | 0.43 | 0.81 | 0.36 | 0.46 |
| 1 | 159533068 | rs10908734 | -1.882 | -0.977 | -1.436 | -1.557 | -0.044 | -0.132 | -0.132 | 0.137 | 0.76 | 0.59 | 0.91 | 0.97 | 0.88 | 0.9 |
| 1 | 159535626 | rs12047264 | -0.874 | -1.048 | -1.449 | -1.735 | -0.04 | -0.096 | -0.096 | 0.048 | 0.68 | 0.45 | 0.88 | 0.94 | 0.84 | 0.8 |
| 1 | 159538241 | rs12022043 | 2.761 | -1.084 | -1.728 | -1.499 | -0.025 | 0.022 | 0.022 | 0 | 0.07 | 0.01 | 0.11 | 0.41 | 0.07 | 0.11 |
| 1 | 159557369 | rs3753867 | -0.336 | -1.319 | -1.784 | -1.733 | -0.022 | -0.032 | -0.032 | 0 | 0.25 | 0.45 | 0.36 | 0.8 | 0.19 | 0.21 |
| 1 | 159558258 | rs2808661 | -1.996 | -1.276 | -1.791 | -1.746 | -0.038 | 0.102 | 0.102 | 0.073 | 0.84 | 0.98 | 0.91 | 0.99 | 0.84 | 0.75 |
| 1 | 159559531 | rs1446963 | 1.191 | -1.272 | -1.691 | -1.662 | -0.037 | -0.138 | -0.138 | 0.038 | 0.28 | 0.41 | 0.1 | 0.02 | 0.14 | 0.13 |
| 1 | 159562766 | rs1446965 | 1.228 | -1.089 | -1.597 | -1.404 | -0.011 | 0.004 | 0.004 | 0 | 0.09 | 0.03 | 0.21 | 0.39 | 0.13 | 0.1 |
| 1 | 159563483 | rs1446966 | 0.508 | -1.213 | -1.643 | -1.886 | -0.008 | -0.041 | -0.041 | 0 | 0.58 | 0.51 | 0.54 | 0.2 | 0.64 | 0.54 |

| | | | | | | | | | | | | | | | | |
|---|---|---|---|---|---|---|---|---|---|---|---|---|---|---|---|---|
| 1 | 159567813 | rs1037143 | 0.623 | -1.142 | -1.7 | -2.101 | -0.005 | -0.049 | -0.049 | 0 | 0.55 | 0.5 | 0.53 | 0.16 | 0.62 | 0.48 |
| 1 | 159573450 | rs6695377 | -0.745 | -0.891 | -1.618 | -1.885 | -0.025 | -0.067 | -0.067 | 0 | 0.28 | 0.45 | 0.37 | 0.83 | 0.22 | 0.28 |
| 1 | 159581387 | rs2121479 | 1.252 | -0.354 | -2.27 | -1.667 | -0.093 | 0.073 | 0.073 | 0.111 | 0.25 | 0.09 | 0.43 | 0.13 | 0.48 | 0.35 |
| 1 | 159582055 | rs2794502 | -1.044 | -1.101 | -2.208 | -1.797 | -0.005 | 0.066 | 0.066 | 0.008 | 0.8 | 0.92 | 0.89 | 0.98 | 0.82 | 0.72 |
| 1 | 159582103 | rs10494331 | -0.003 | -1.066 | -2.196 | -1.548 | -0.003 | -0.006 | -0.006 | 0.259 | 0.18 | 0.17 | 0.18 | 0.44 | 0.18 | 0.21 |
| 1 | 159582218 | rs4233355 | 0.549 | -0.814 | -2.214 | -1.454 | -0.056 | -0.065 | -0.065 | 0 | 0.34 | 0.12 | 0.63 | 0.51 | 0.61 | 0.45 |
| 1 | 159583346 | rs1446972 | 0.392 | -1.008 | -1.839 | -2.428 | -0.007 | 0.045 | 0.045 | 0.019 | 0.39 | 0.25 | 0.71 | 0.9 | 0.58 | 0.51 |
| 1 | 159592915 | rs2794510 | -0.151 | -1.56 | -1.971 | -2.55 | 0.091 | 0.139 | 0.139 | 2.344 | 0.41 | 0.8 | 0.19 | 0.06 | 0.22 | 0.17 |
| 1 | 159595682 | rs983789 | 0.364 | -1.578 | -1.381 | -2.44 | -0.057 | -0.095 | -0.095 | 0 | 0.31 | 0.63 | 0.11 | 0.05 | 0.1 | 0.1 |
| 1 | 159599414 | rs984553 | -0.003 | -1.368 | -1.468 | -2.329 | 0.006 | -0.024 | -0.024 | 0 | 0.34 | 0.72 | 0.16 | 0.06 | 0.2 | 0.17 |
| 1 | 159612046 | rs2808656 | -1.514 | -1.049 | -1.431 | -1.962 | -0.002 | -0.001 | -0.001 | 0.014 | 0.86 | 0.91 | 0.92 | 0.98 | 0.85 | 0.78 |
| 1 | 159613798 | rs2808658 | -0.627 | -1.157 | -1.423 | -2.013 | 0.001 | -0.006 | -0.006 | 1.186 | 0.79 | 0.85 | 0.91 | 0.96 | 0.83 | 0.69 |
| 1 | 159616159 | rs2808660 | 1.935 | -0.237 | -1.355 | -1.993 | -0.015 | 0.009 | 0.009 | 0 | 0.12 | 0.1 | 0.15 | 0.02 | 0.27 | 0.11 |
| 1 | 159616629 | rs2592900 | 0.671 | -0.356 | -1.365 | -2.044 | -0.003 | -0.005 | -0.005 | 2.496 | 0.2 | 0.18 | 0.17 | 0.04 | 0.28 | 0.19 |
| 1 | 159628964 | rs12076827 | 2.261 | -1.064 | -0.832 | -2.097 | -0.1 | -0.072 | -0.072 | 0 | 0.16 | 0.38 | 0.03 | 0.02 | 0.01 | 0.07 |
| 1 | 159629021 | rs1891186 | 0.146 | -0.562 | -1.185 | -1.824 | -0.005 | -0.05 | -0.05 | 0 | 0.43 | 0.52 | 0.26 | 0.17 | 0.39 | 0.29 |
| 1 | 159629342 | rs1446975 | -0.094 | -0.424 | -1.612 | -1.759 | 0.032 | 0.112 | 0.112 | 0 | 0.48 | 0.73 | 0.28 | 0.17 | 0.39 | 0.29 |
| 1 | 159641029 | rs11589667 | 0.043 | 0.093 | -1.972 | -0.604 | -0.053 | -0.12 | -0.12 | 0 | 0.28 | 0.06 | 0.33 | 0.5 | 0.35 | 0.37 |
| 1 | 159646544 | rs2794515 | -0.613 | -0.976 | -1.859 | -2.194 | 0.001 | -0.012 | -0.012 | 0 | 0.75 | 0.83 | 0.75 | 0.96 | 0.75 | 0.64 |
| 1 | 159646924 | rs16842484 | 1.095 | -1.084 | -1.751 | -2.3 | 0.001 | -0.096 | -0.096 | 1.354 | 0.27 | 0.35 | 0.16 | 0.01 | 0.28 | 0.12 |
| 1 | 159647988 | rs12093699 | 1.052 | -1.423 | -1.722 | -2.13 | 0.001 | -0.035 | -0.035 | 0.898 | 0.3 | 0.33 | 0.32 | 0.05 | 0.33 | 0.2 |
| 1 | 159652939 | rs2592887 | -0.4 | -1.788 | -1.921 | -2.004 | -0.003 | -0.002 | -0.002 | 1.029 | 0.48 | 0.5 | 0.43 | 0.56 | 0.39 | 0.35 |
| 1 | 159667190 | rs2027469 | 1.467 | -1.659 | -1.984 | -1.971 | -0.057 | 0.014 | 0.014 | 0 | 0.09 | 0.01 | 0.1 | 0.04 | 0.19 | 0.12 |
| 1 | 159668195 | rs2369146 | -1.416 | -1.935 | -1.952 | -2.295 | -0.006 | -0.031 | -0.031 | 0 | 0.72 | 0.57 | 0.78 | 0.8 | 0.77 | 0.71 |
| 1 | 159673585 | rs1572970 | 0.043 | -2.022 | -2.066 | -2.325 | 0.072 | 0.068 | 0.068 | 0.019 | 0.53 | 0.28 | 0.73 | 0.64 | 0.7 | 0.61 |
| 1 | 159674933 | rs876537 | -0.077 | -1.958 | -2.074 | -2.302 | -0.028 | -0.058 | -0.058 | 0 | 0.33 | 0.17 | 0.39 | 0.56 | 0.36 | 0.35 |
| 1 | 159676011 | rs2808628 | 0.115 | -1.986 | -1.942 | -2.169 | -0.014 | -0.05 | -0.05 | 0 | 0.29 | 0.18 | 0.37 | 0.57 | 0.31 | 0.33 |
| 1 | 159678816 | rs2794520 | 0.016 | -1.988 | -1.943 | -2.172 | -0.013 | -0.051 | -0.051 | 0 | 0.29 | 0.18 | 0.38 | 0.57 | 0.31 | 0.33 |
| 1 | 159679636 | rs3093077 | 0.948 | -2.208 | -1.891 | -2.307 | -0.081 | -0.045 | -0.045 | 0.075 | 0.25 | 0.36 | 0.06 | 0.15 | 0.07 | 0.1 |
| 1 | 159680868 | rs2808630 | -2.424 | -2.341 | -1.854 | -2.296 | -0.006 | 0 | 0 | 0 | 0.81 | 0.85 | 0.8 | 0.8 | 0.72 | 0.68 |
| 1 | 159682233 | rs1205 | 0.183 | -2.322 | -1.849 | -2.286 | -0.017 | -0.059 | -0.059 | 0 | 0.3 | 0.17 | 0.37 | 0.57 | 0.31 | 0.33 |
| 1 | 159683091 | rs1130864 | 2.298 | -2.381 | -1.89 | -2.29 | -0.034 | 0.042 | 0.042 | 0 | 0.21 | 0.13 | 0.33 | 0.06 | 0.33 | 0.24 |
| 1 | 159685096 | rs2794521 | -2.495 | -2.361 | -1.885 | -2.285 | -0.014 | -0.001 | -0.001 | 0 | 0.81 | 0.87 | 0.8 | 0.8 | 0.71 | 0.68 |
| 1 | 159685136 | rs3093059 | 0.881 | -2.263 | -1.867 | -2.224 | -0.057 | -0.031 | -0.031 | 0.55 | 0.25 | 0.32 | 0.05 | 0.15 | 0.07 | 0.08 |
| 1 | 159686483 | rs3116636 | 2.576 | -2.211 | -1.862 | -2.208 | -0.048 | 0.045 | 0.045 | 0 | 0.19 | 0.08 | 0.32 | 0.06 | 0.33 | 0.25 |
| 1 | 159689323 | rs3122012 | 2.54 | -2.194 | -1.862 | -2.208 | -0.047 | 0.046 | 0.046 | 0 | 0.19 | 0.08 | 0.32 | 0.06 | 0.33 | 0.25 |
| 1 | 159690466 | rs12079772 | 3.426 | -2.316 | -1.849 | -2.219 | -0.069 | -0.071 | -0.071 | 0 | 0.12 | 0.26 | 0.01 | 0 | 0 | 0 |
| 1 | 159693722 | rs2211321 | -1.678 | -2.253 | -2.009 | -2.188 | 0.018 | -0.016 | -0.016 | 0 | 0.75 | 0.64 | 0.78 | 0.8 | 0.71 | 0.68 |
| 1 | 159694209 | rs2808635 | -1.678 | -2.256 | -2.011 | -2.19 | 0.018 | -0.016 | -0.016 | 0 | 0.75 | 0.64 | 0.78 | 0.8 | 0.71 | 0.68 |
| 1 | 159696858 | rs16842596 | 2.862 | -2.228 | -1.985 | -2.189 | -0.087 | -0.09 | -0.09 | 0 | 0.14 | 0.32 | 0.01 | 0 | 0 | 0 |
| 1 | 159696910 | rs3116653 | 2.587 | -2.173 | -2.037 | -2.176 | -0.088 | 0.054 | 0.054 | 0 | 0.19 | 0.04 | 0.32 | 0.06 | 0.33 | 0.25 |
| 1 | 159697475 | rs16842599 | 1.446 | -2.206 | -2.049 | -2.184 | -0.002 | 0 | 0 | 0 | 0.12 | 0.1 | 0.05 | 0.15 | 0.07 | 0.09 |
| 1 | 159698485 | rs3116651 | 2.593 | -2.182 | -2.069 | -2.175 | -0.088 | 0.054 | 0.054 | 0 | 0.19 | 0.04 | 0.32 | 0.06 | 0.33 | 0.25 |
| 1 | 159698549 | rs7553007 | 0.065 | -2.184 | -1.996 | -2.089 | -0.008 | -0.038 | -0.038 | 0 | 0.29 | 0.2 | 0.37 | 0.57 | 0.31 | 0.33 |
| 1 | 159700039 | rs11265260 | 1.422 | -2.563 | -1.589 | -1.841 | 0.005 | -0.002 | -0.002 | 0 | 0.11 | 0.07 | 0.05 | 0.15 | 0.07 | 0.09 |
| 1 | 159706560 | rs12081264 | 0.413 | -2.629 | -1.601 | -1.878 | -0.102 | -0.061 | -0.061 | 0 | 0.26 | 0.41 | 0.06 | 0.15 | 0.07 | 0.09 |
| 1 | 159710093 | rs12728740 | 1.692 | -1.912 | -2.358 | -1.612 | -0.118 | 0.07 | 0.07 | 0 | 0.21 | 0.02 | 0.35 | 0.05 | 0.35 | 0.29 |
| 1 | 159713556 | rs11265265 | -0.238 | -0.374 | -1.523 | -0.764 | -0.065 | 0.038 | 0.038 | 4.342 | 0.64 | 0.84 | 0.75 | 0.69 | 0.52 | 0.54 |
| 1 | 159714149 | rs4285692 | -0.218 | -0.374 | -1.514 | -0.764 | -0.054 | 0.035 | 0.035 | 4.408 | 0.64 | 0.82 | 0.74 | 0.69 | 0.52 | 0.54 |
| 1 | 159717162 | rs11588887 | 0.387 | -0.411 | -1.446 | -0.626 | -0.002 | -0.019 | -0.019 | 0 | 0.16 | 0.12 | 0.26 | 0.62 | 0.14 | 0.23 |
| 1 | 159723521 | rs4656849 | -0.708 | 0.451 | -0.95 | -0.718 | -0.035 | 0.021 | 0.021 | 1.091 | 0.76 | 0.86 | 0.8 | 0.82 | 0.62 | 0.66 |
| 1 | 159723619 | rs12094103 | 0.879 | 0.652 | -0.958 | -0.686 | -0.009 | -0.083 | -0.083 | 0.147 | 0.45 | 0.51 | 0.42 | 0.05 | 0.38 | 0.31 |
| 1 | 159726123 | rs10908741 | -0.639 | 0.714 | -0.868 | 0.799 | -0.015 | -0.019 | -0.019 | 0 | 0.22 | 0.15 | 0.3 | 0.75 | 0.24 | 0.34 |
| 1 | 159728127 | rs11265269 | -0.739 | 0.815 | -0.86 | 0.847 | -0.001 | -0.003 | -0.003 | 0 | 0.25 | 0.22 | 0.31 | 0.75 | 0.24 | 0.34 |
| 1 | 159732811 | rs10908742 | -0.764 | 0.841 | -0.822 | 0.758 | 0.001 | 0.006 | 0.006 | 0 | 0.27 | 0.24 | 0.36 | 0.77 | 0.24 | 0.36 |
| 1 | 159736139 | rs4604689 | -1.385 | 1.587 | -0.151 | 1.629 | 0.044 | 0.055 | 0.055 | 0 | 0.15 | 0.26 | 0.32 | 0.29 | 0.25 | 0.2 |
| 1 | 159736755 | rs4364879 | 0.065 | 1.782 | -0.366 | 1.772 | -0.008 | -0.011 | -0.011 | 0 | 0.07 | 0.05 | 0.25 | 0.12 | 0.17 | 0.12 |
| 1 | 159738719 | rs6427494 | -1.723 | 1.47 | -0.133 | 1.916 | -0.002 | 0 | 0 | 0.659 | 0.37 | 0.61 | 0.52 | 0.67 | 0.43 | 0.56 |
| 1 | 159741700 | rs4424502 | 0.077 | 1.469 | -0.319 | 1.842 | -0.008 | -0.007 | -0.007 | 0 | 0.05 | 0.03 | 0.07 | 0.17 | 0.09 | 0.08 |
| 1 | 159743845 | rs7519478 | -1.843 | 1.291 | -0.336 | 1.275 | -0.01 | 0.052 | 0.052 | 2.469 | 0.44 | 0.68 | 0.56 | 0.72 | 0.47 | 0.58 |
| 1 | 159746857 | rs6674721 | -0.503 | 1.231 | -0.264 | 1.041 | -0.024 | -0.002 | -0.002 | 0 | 0.09 | 0.17 | 0.06 | 0.07 | 0.04 | 0.02 |
| 1 | 159748269 | rs12032950 | 0.376 | 1.34 | -0.74 | 1.094 | 0.001 | -0.002 | -0.002 | 0 | 0.07 | 0.06 | 0.05 | 0.07 | 0.04 | 0.02 |
| 1 | 159749188 | rs3820099 | -1.264 | 1.494 | -0.926 | 0.904 | -0.008 | 0.087 | 0.087 | 1.9 | 0.45 | 0.71 | 0.62 | 0.78 | 0.47 | 0.58 |
| 1 | 159750629 | rs3806187 | -0.003 | 2.357 | -1.141 | 0.975 | -0.024 | -0.011 | -0.011 | 0 | 0.11 | 0.06 | 0.25 | 0.12 | 0.17 | 0.12 |
| 1 | 159753097 | rs4403628 | -0.413 | 1.77 | -0.971 | 1.018 | -0.001 | -0.002 | -0.002 | 2.142 | 0.21 | 0.17 | 0.39 | 0.36 | 0.35 | 0.29 |
| 1 | 159753183 | rs4233356 | -0.372 | 0.27 | -1.014 | 0.256 | 0.04 | 0.037 | 0.037 | 0 | 0.51 | 0.59 | 0.65 | 0.69 | 0.64 | 0.71 |
| 1 | 159759058 | rs6662918 | -0.721 | 0.291 | -1.005 | 0.117 | 0.006 | 0.034 | 0.034 | 0.056 | 0.07 | 0.02 | 0.05 | 0 | 0.05 | 0.02 |
| 1 | 159761433 | rs11584887 | -1.364 | 0.315 | -0.834 | 0.059 | 0.002 | -0.003 | -0.003 | 1.083 | 0.12 | 0.1 | 0.08 | 0.01 | 0.11 | 0.06 |
| 1 | 159762120 | rs16842662 | -1.731 | 0.294 | -0.4 | 0.255 | -0.005 | -0.008 | -0.008 | 0 | 0.12 | 0.18 | 0.13 | 0.08 | 0.11 | 0.13 |

| | | | | | | | | | | | | | | | | |
|---|---|---|---|---|---|---|---|---|---|---|---|---|---|---|---|---|
| 1 | 159765249 | rs4630124 | -0.274 | 0.076 | -1.409 | 0.306 | -0.122 | -0.058 | -0.058 | 0 | 0.36 | 0.09 | 0.29 | 0.39 | 0.44 | 0.42 |
| 1 | 159766709 | rs7512164 | -0.325 | -0.762 | -0.651 | 0.203 | -0.041 | -0.01 | -0.01 | 0 | 0.26 | 0.53 | 0.24 | 0.17 | 0.2 | 0.1 |
| 1 | 159769794 | rs12092557 | 0.508 | -0.93 | 0.167 | 0.239 | -0.022 | 0.004 | 0.004 | 0 | 0.1 | 0.16 | 0.11 | 0.08 | 0.03 | 0.03 |
| 1 | 159774408 | rs11265282 | -0.761 | -1.024 | 0.334 | 0.162 | -0.04 | -0.006 | -0.006 | 0 | 0.25 | 0.49 | 0.23 | 0.17 | 0.2 | 0.11 |
| 1 | 159778699 | rs12042065 | 0.351 | -0.917 | -0.015 | 0.24 | -0.112 | -0.053 | -0.053 | 0.026 | 0.37 | 0.09 | 0.3 | 0.37 | 0.44 | 0.41 |
| 1 | 159782181 | rs6656979 | -1.033 | -1.621 | 0.508 | 0.273 | -0.019 | -0.003 | -0.003 | 0 | 0.23 | 0.35 | 0.22 | 0.17 | 0.2 | 0.12 |
| 1 | 159785425 | rs4443889 | -1.158 | -1.901 | 0.274 | 0.141 | -0.036 | -0.021 | -0.021 | 0.916 | 0.3 | 0.46 | 0.23 | 0.17 | 0.2 | 0.12 |
| 1 | 159794514 | rs2494495 | -0.57 | -1.057 | -0.237 | 0.323 | -0.031 | -0.039 | -0.039 | 0 | 0.2 | 0.44 | 0.14 | 0.1 | 0.17 | 0.09 |
| 1 | 159795478 | rs12046006 | -0.532 | -0.791 | -1.359 | 0.574 | 0.006 | -0.06 | -0.06 | 0 | 0.16 | 0 | 0.11 | 0.18 | 0.1 | 0.15 |
| 1 | 159798306 | rs2494523 | -0.216 | 0.172 | -0.952 | 0.539 | -0.001 | 0 | 0 | 0 | 0.54 | 0.58 | 0.35 | 0.5 | 0.51 | 0.42 |
| 1 | 159798407 | rs17373453 | -0.912 | 0.473 | -1.281 | 0.788 | -0.024 | -0.031 | -0.031 | 1.07 | 0.05 | 0.01 | 0.12 | 0.12 | 0.07 | 0.25 |
| 1 | 159801074 | rs10158752 | -0.636 | -0.046 | -1.491 | 0.034 | 0.019 | 0.03 | 0.03 | 0.038 | 0.2 | 0.17 | 0.11 | 0.01 | 0.17 | 0.08 |
| 1 | 159801111 | rs7516146 | 0.371 | 0.396 | -2.652 | -0.836 | -0.153 | 0.111 | 0.111 | 0.853 | 0.25 | 0.02 | 0.47 | 0.06 | 0.43 | 0.27 |
| 1 | 159801155 | rs2501341 | 0.065 | -0.91 | -2.749 | -0.298 | -0.006 | 0.017 | 0.017 | 0 | 0.56 | 0.62 | 0.67 | 0.64 | 0.56 | 0.66 |
| 1 | 159802627 | rs2494480 | 0.693 | -1.909 | -1.953 | 0.397 | -0.07 | 0.096 | 0.096 | 0 | 0.24 | 0.48 | 0.15 | 0.52 | 0.11 | 0.25 |
| 1 | 159808286 | rs2501312 | -0.471 | -1.994 | -2.173 | 0.098 | 0.141 | 0.005 | 0.005 | 1.367 | 0.48 | 0.79 | 0.28 | 0.53 | 0.29 | 0.31 |
| 1 | 159809342 | rs16842686 | 0.553 | -1.577 | -2.036 | -0.041 | 0.008 | -0.01 | -0.01 | 0 | 0.2 | 0.25 | 0.11 | 0.01 | 0.14 | 0.06 |
| 1 | 159815642 | rs2501354 | 0.233 | -0.709 | -1.854 | 0.178 | 0.003 | 0.006 | 0.006 | 0 | 0.58 | 0.6 | 0.7 | 0.71 | 0.61 | 0.62 |
| 1 | 159815940 | rs2501353 | -1.058 | -0.379 | -2.047 | 0.344 | -0.043 | -0.039 | -0.039 | 0.131 | 0.82 | 0.95 | 0.85 | 0.82 | 0.8 | 0.77 |
| 1 | 159816313 | rs2501351 | -0.38 | -1.038 | -1.782 | 0.258 | 0.049 | 0.002 | 0.002 | 1.557 | 0.44 | 0.69 | 0.32 | 0.5 | 0.31 | 0.26 |
| 1 | 159818304 | rs2501346 | -0.495 | -0.702 | -2.058 | 0.214 | -0.111 | -0.019 | -0.019 | 0.164 | 0.68 | 0.9 | 0.69 | 0.76 | 0.57 | 0.7 |
| 1 | 159818513 | rs6659742 | 0.002 | -0.073 | -1.664 | -0.073 | 0 | 0.003 | 0.003 | 0 | 0.37 | 0.36 | 0.5 | 0.26 | 0.38 | 0.38 |
| 1 | 159819217 | rs5010834 | -0.177 | -0.283 | -1.655 | 0.101 | 0.027 | 0.065 | 0.065 | 0 | 0.5 | 0.3 | 0.47 | 0.71 | 0.54 | 0.51 |
| 1 | 159819491 | rs4133290 | 0.135 | 0.239 | -1.866 | 0.692 | -0.014 | -0.025 | -0.025 | 7.316 | 0.34 | 0.25 | 0.48 | 0.45 | 0.42 | 0.35 |
| 1 | 159820026 | rs2501331 | -1.809 | 0.48 | -1.882 | 0.1 | -0.033 | 0.024 | 0.024 | 0 | 0.9 | 0.99 | 0.9 | 0.95 | 0.85 | 0.92 |
| 1 | 159824205 | rs11807946 | 1.15 | 0.328 | -1.715 | 0.092 | -0.011 | 0.009 | 0.009 | 0 | 0.2 | 0.15 | 0.13 | 0.09 | 0.24 | 0.22 |
| 1 | 159829535 | rs2494506 | -0.055 | -0.247 | -1.322 | 0.098 | -0.008 | -0.016 | -0.016 | 0 | 0.62 | 0.56 | 0.77 | 0.79 | 0.7 | 0.68 |
| 1 | 159831406 | rs4074846 | 1.086 | -0.226 | -1.506 | 0.254 | -0.092 | -0.03 | -0.03 | 0.672 | 0.29 | 0.02 | 0.42 | 0.31 | 0.4 | 0.36 |
| 1 | 159833733 | rs12409375 | -0.213 | -0.414 | -1.287 | 0.388 | -0.04 | 0.203 | 0.203 | 0.573 | 0.11 | 0.37 | 0.19 | 0.29 | 0.03 | 0.08 |
| 1 | 159834428 | rs3795332 | -0.268 | -0.415 | -1.28 | 0.387 | -0.04 | 0.219 | 0.219 | 0.42 | 0.11 | 0.39 | 0.19 | 0.29 | 0.03 | 0.08 |
| 1 | 159835673 | rs2501332 | -0.184 | -0.394 | -1.304 | 0.318 | -0.012 | -0.024 | -0.024 | 0 | 0.62 | 0.55 | 0.76 | 0.79 | 0.69 | 0.68 |
| 1 | 159839971 | rs2494515 | -0.708 | -0.414 | -1.272 | 0.317 | 0.019 | 0.045 | 0.045 | 0 | 0.7 | 0.82 | 0.82 | 0.84 | 0.76 | 0.74 |
| 1 | 159841056 | rs11265288 | 0.162 | -0.451 | -1.202 | 0.296 | -0.007 | -0.021 | -0.021 | 0 | 0.61 | 0.57 | 0.77 | 0.84 | 0.73 | 0.73 |
| 1 | 159842103 | rs11265289 | 0.893 | -0.448 | -1.266 | 0.34 | 0.093 | 0.002 | 0.002 | 0.412 | 0.5 | 0.16 | 0.57 | 0.5 | 0.66 | 0.61 |
| 1 | 159844819 | rs7521729 | 1.742 | -0.3 | -1.312 | 0.273 | -0.033 | 0.078 | 0.078 | 0.919 | 0.13 | 0 | 0.1 | 0 | 0.13 | 0.06 |
| 1 | 159849306 | rs16842774 | 0.015 | -1.09 | -1.235 | 0.32 | -0.015 | -0.022 | -0.022 | 0 | 0.16 | 0.39 | 0.11 | 0.06 | 0.1 | 0.07 |
| 1 | 159852745 | rs7514472 | 1.664 | -1.065 | -1.215 | 0.355 | 0.004 | 0.003 | 0.003 | 0 | 0.16 | 0.15 | 0.06 | 0.11 | 0.1 | 0.18 |
| 1 | 159853139 | rs6671929 | 0.008 | -0.467 | -0.817 | 0.821 | 0.041 | 0.063 | 0.063 | 0.164 | 0.52 | 0.32 | 0.71 | 0.79 | 0.67 | 0.67 |
| 1 | 159853321 | rs11265291 | 0.295 | -0.445 | -0.986 | 0.821 | 0.095 | 0.003 | 0.003 | 0.351 | 0.5 | 0.16 | 0.57 | 0.5 | 0.66 | 0.61 |
| 1 | 159853664 | rs12566514 | 1.08 | -0.304 | -0.934 | -0.274 | 0.023 | 0.03 | 0.03 | 0.307 | 0.08 | 0.23 | 0.06 | 0.06 | 0.07 | 0.06 |
| 1 | 159854251 | rs16842789 | 1.08 | -0.304 | -0.934 | -0.274 | 0.023 | 0.03 | 0.03 | 0.307 | 0.08 | 0.23 | 0.06 | 0.06 | 0.07 | 0.06 |
| 1 | 159859392 | rs4656856 | 0.676 | -0.159 | -1.411 | -0.227 | -0.1 | -0.035 | -0.035 | 0 | 0.29 | 0.02 | 0.41 | 0.31 | 0.41 | 0.35 |
| 1 | 159860259 | rs2501320 | 0.928 | -0.039 | -1.19 | -0.765 | -0.055 | 0.044 | 0.044 | 0 | 0.24 | 0.06 | 0.19 | 0.22 | 0.36 | 0.15 |
| 1 | 159860917 | rs4233361 | -0.134 | -0.863 | -0.217 | -0.586 | 0.009 | 0.012 | 0.012 | 0 | 0.68 | 0.7 | 0.81 | 0.84 | 0.76 | 0.74 |
| 1 | 159862153 | rs10752631 | 1.056 | -0.838 | -0.215 | -0.574 | 0.084 | 0.145 | 0.145 | 0.387 | 0.52 | 0.17 | 0.7 | 0.79 | 0.67 | 0.67 |
| 1 | 159868546 | rs2501319 | -0.309 | -0.845 | -0.183 | -0.969 | 0.021 | 0.061 | 0.061 | 0 | 0.71 | 0.83 | 0.82 | 0.91 | 0.76 | 0.75 |
| 1 | 159871738 | rs2494485 | 0.412 | -0.833 | -0.207 | -0.965 | 0.012 | 0.004 | 0.004 | 0.207 | 0.06 | 0.31 | 0.03 | 0 | 0.01 | 0 |
| 1 | 159875153 | rs11265294 | 0.882 | -0.703 | -1.381 | -0.909 | 0.08 | 0.205 | 0.205 | 0.4 | 0.51 | 0.17 | 0.7 | 0.85 | 0.66 | 0.68 |
| 1 | 159878126 | rs2494481 | -0.995 | -0.18 | -0.813 | 0.229 | -0.032 | -0.035 | -0.035 | 0.65 | 0.18 | 0.62 | 0.11 | 0.05 | 0.09 | 0.07 |
| 1 | 159885500 | rs2789423 | -0.071 | 0.258 | -2.02 | 0.358 | -0.097 | -0.001 | -0.001 | 0 | 0.42 | 0.05 | 0.64 | 0.61 | 0.53 | 0.55 |
| 1 | 159886422 | rs12122337 | -0.456 | 0.071 | -1.848 | 0.182 | -0.001 | -0.002 | -0.002 | 0 | 0.09 | 0.13 | 0.07 | 0.04 | 0.07 | 0.03 |
| 1 | 159889686 | rs2252815 | 0.197 | -0.212 | -0.87 | -0.222 | 0.005 | 0.01 | 0.01 | 0 | 0.45 | 0.38 | 0.63 | 0.64 | 0.51 | 0.47 |
| 1 | 159890459 | rs11265297 | 0.371 | -0.862 | -0.843 | -0.481 | -0.067 | 0.029 | 0.029 | 0 | 0.34 | 0.16 | 0.26 | 0.31 | 0.41 | 0.38 |
| 1 | 159892780 | rs7551141 | 0.113 | -0.869 | -0.586 | 0.126 | -0.029 | -0.024 | -0.024 | 0.224 | 0.38 | 0.26 | 0.35 | 0.43 | 0.44 | 0.36 |
| 1 | 159895536 | rs7513326 | -0.072 | -1.234 | -0.492 | 0.138 | 0.002 | -0.021 | -0.021 | 0.854 | 0.42 | 0.29 | 0.39 | 0.45 | 0.5 | 0.36 |
| 1 | 159895979 | rs1934075 | 0.418 | -1.075 | -0.714 | -0.158 | -0.036 | -0.047 | -0.047 | 0 | 0.32 | 0.23 | 0.36 | 0.43 | 0.36 | 0.32 |
| 1 | 159897327 | rs2153208 | -1.225 | -1.452 | -0.326 | -0.365 | -0.047 | -0.076 | -0.076 | 0 | 0.17 | 0.4 | 0.09 | 0.06 | 0.11 | 0.08 |
| 1 | 159897559 | rs1319080 | 0.951 | -1.391 | -0.859 | 0.065 | -0.004 | -0.036 | -0.036 | 0 | 0.06 | 0 | 0.15 | 0.21 | 0.07 | 0.1 |
| 1 | 159898913 | rs7543462 | -1.179 | -1.46 | -0.496 | -0.375 | -0.018 | 0.038 | 0.038 | 0 | 0.62 | 0.76 | 0.64 | 0.72 | 0.55 | 0.64 |
| 1 | 159906156 | rs10908750 | 0.442 | -1.085 | -0.712 | 0.083 | -0.042 | -0.068 | -0.068 | 0.179 | 0.35 | 0.15 | 0.36 | 0.46 | 0.35 | 0.37 |
| 1 | 159908045 | rs4262538 | -0.57 | -0.635 | -0.213 | 0.048 | 0.023 | -0.03 | -0.03 | 0 | 0.08 | 0.29 | 0.05 | 0 | 0.05 | 0.03 |
| 1 | 159910337 | rs4313394 | -0.079 | -0.596 | -0.363 | 0.103 | -0.005 | 0.024 | 0.024 | 0 | 0.08 | 0.07 | 0.15 | 0.23 | 0.08 | 0.11 |
| 1 | 159911041 | rs16842852 | -0.931 | -0.445 | -0.284 | -0.391 | 0.015 | -0.04 | -0.04 | 0 | 0.09 | 0.34 | 0.06 | 0 | 0.07 | 0.03 |
| 1 | 159912346 | rs1320568 | 0.041 | -0.295 | -1.629 | -0.407 | -0.135 | -0.06 | -0.06 | 0 | 0.34 | 0.04 | 0.4 | 0.38 | 0.45 | 0.42 |
| 1 | 159913388 | rs1316275 | 0.21 | -0.378 | -1.467 | -1.722 | -0.024 | 0.063 | 0.063 | 0.193 | 0.41 | 0.28 | 0.58 | 0.85 | 0.52 | 0.62 |
| 1 | 159919986 | rs2820579 | 0.226 | -0.556 | -0.754 | -1.788 | 0.001 | -0.052 | -0.052 | 0 | 0.89 | 0.81 | 0.95 | 1 | 0.9 | 0.93 |
| 1 | 159920719 | rs16831153 | -0.003 | -1.119 | -0.448 | -2.228 | -0.009 | -0.025 | -0.025 | 0.017 | 0.24 | 0.32 | 0.24 | 0.13 | 0.22 | 0.14 |
| 1 | 159922369 | rs2840583 | 0.144 | -1.39 | -0.732 | -2.344 | -0.016 | -0.072 | -0.072 | 0 | 0.83 | 0.76 | 0.94 | 1 | 0.9 | 0.93 |
| 1 | 159922959 | rs2249707 | 1.439 | -1.468 | -0.58 | -2.487 | 0.062 | 0.165 | 0.165 | 0.29 | 0.49 | 0.24 | 0.65 | 0.84 | 0.63 | 0.7 |

| | | | | | | | | | | | | | | | |
|---|---|---|---|---|---|---|---|---|---|---|---|---|---|---|---|
| 1 | 159923111 | rs2789417 | 0.751 | -1.052 | -0.476 | -2.191 | -0.019 | -0.075 | -0.075 | 0 | 0.82 | 0.75 | 0.94 | 1 | 0.9 | 0.93 |
| 1 | 159928059 | rs2789426 | 0.798 | -0.947 | -0.816 | -2.123 | 0.016 | 0.033 | 0.033 | 0 | 0.8 | 0.89 | 0.93 | 0.99 | 0.87 | 0.86 |
| 1 | 159931395 | rs2840588 | 0.217 | -0.471 | -0.424 | -1.786 | 0.012 | -0.037 | -0.037 | 0 | 0.93 | 0.84 | 0.96 | 1 | 0.93 | 0.93 |
| 1 | 159932319 | rs7541727 | 0.074 | -1.018 | -0.561 | -2.287 | -0.001 | -0.033 | -0.033 | 0 | 0.19 | 0.13 | 0.08 | 0.02 | 0.16 | 0.1 |
| 1 | 159933136 | rs7521490 | 0.144 | -1.217 | -0.357 | -2.537 | -0.008 | -0.057 | -0.057 | 0 | 0.13 | 0.21 | 0.08 | 0 | 0.11 | 0.04 |
| 1 | 159935569 | rs2820565 | -0.374 | -1.343 | -0.315 | -2.55 | -0.031 | -0.071 | -0.071 | 0 | 0.87 | 0.75 | 0.94 | 1 | 0.92 | 0.96 |
| 1 | 159940000 | rs2840591 | 1.047 | -1.552 | -1.068 | -2.704 | 0.086 | 0.174 | 0.174 | 0.037 | 0.54 | 0.23 | 0.79 | 0.96 | 0.68 | 0.78 |
| 1 | 159941535 | rs4378200 | 0.051 | -1.525 | -1.383 | -2.678 | 0 | -0.036 | -0.036 | 0.533 | 0.23 | 0.19 | 0.11 | 0.02 | 0.19 | 0.1 |
| 1 | 159942456 | rs2789424 | 0.423 | -0.861 | -1.962 | -2.256 | 0.12 | 0.278 | 0.278 | 0.274 | 0.52 | 0.15 | 0.78 | 0.96 | 0.68 | 0.78 |
| 1 | 159954369 | rs1317712 | 0.047 | 0.397 | -2.059 | -0.611 | -0.062 | -0.133 | -0.133 | 0 | 0.27 | 0.04 | 0.27 | 0.55 | 0.33 | 0.47 |
| 1 | 159954994 | rs10908752 | -0.023 | 0.261 | -1.571 | -0.881 | -0.009 | -0.009 | -0.009 | 0 | 0.16 | 0.25 | 0.31 | 0.1 | 0.16 | 0.13 |
| 1 | 159955486 | rs2185212 | -0.662 | 1.199 | -1.778 | -1.117 | 0.014 | -0.014 | -0.014 | 0 | 0.22 | 0.27 | 0.37 | 0.11 | 0.29 | 0.24 |
| 1 | 159958368 | rs16831222 | 0.789 | 1.414 | -2.014 | -1.048 | -0.059 | -0.116 | -0.116 | 1.023 | 0.25 | 0.05 | 0.26 | 0.57 | 0.31 | 0.48 |
| 1 | 159959505 | rs2789428 | 0.794 | 0.6 | -1.823 | -1.363 | -0.005 | -0.006 | -0.006 | 0 | 0.22 | 0.25 | 0.26 | 0.33 | 0.27 | 0.22 |
| 1 | 159969008 | rs6676862 | -0.027 | 0.772 | -1.552 | -1.274 | -0.003 | -0.045 | -0.045 | 0 | 0.2 | 0.27 | 0.28 | 0.06 | 0.22 | 0.13 |
| 1 | 159973452 | rs7544170 | 0.835 | 1.006 | -1.667 | -1.211 | -0.033 | -0.014 | -0.014 | 0.399 | 0.05 | 0.01 | 0.12 | 0.03 | 0.13 | 0.11 |
| 1 | 159973615 | rs2494208 | -0.455 | 0.615 | -1.608 | -1.105 | -0.044 | -0.04 | -0.04 | 6.34 | 0.66 | 0.82 | 0.64 | 0.42 | 0.59 | 0.44 |
| 1 | 159974740 | rs11809878 | -0.567 | 0.768 | -1.273 | -0.987 | 0.008 | -0.019 | -0.019 | 0 | 0.25 | 0.29 | 0.39 | 0.09 | 0.35 | 0.24 |
| 1 | 159976751 | rs2185214 | -0.201 | 1.163 | -1.402 | 0.064 | -0.012 | -0.013 | -0.013 | 0.155 | 0.1 | 0.04 | 0.16 | 0.31 | 0.14 | 0.15 |
| 1 | 159977542 | rs2211107 | -0.268 | 0.387 | -1.182 | -0.253 | -0.002 | 0.003 | 0.003 | 0 | 0.47 | 0.59 | 0.44 | 0.61 | 0.46 | 0.63 |
| 1 | 159986521 | rs6669066 | 0.056 | -0.133 | -1.042 | -0.734 | -0.002 | -0.002 | -0.002 | 0 | 0.62 | 0.59 | 0.56 | 0.36 | 0.58 | 0.66 |
| 1 | 159987008 | rs11577096 | 1.153 | -0.456 | -1.002 | -0.975 | -0.011 | -0.036 | -0.036 | 0 | 0.2 | 0.23 | 0.25 | 0.08 | 0.19 | 0.12 |
| 1 | 159992782 | rs12239927 | 0.569 | -0.829 | -0.844 | -1.152 | -0.053 | -0.109 | -0.109 | 0 | 0.26 | 0.44 | 0.27 | 0.08 | 0.19 | 0.14 |
| 1 | 159992919 | rs12239930 | 3.054 | -0.491 | -1.264 | -0.938 | -0.019 | -0.021 | -0.021 | 0 | 0.06 | 0.21 | 0.02 | 0 | 0 | 0.01 |
| 1 | 159995633 | rs6659129 | 0.754 | -0.371 | -1.675 | -0.846 | -0.044 | 0.016 | 0.016 | 0.434 | 0.35 | 0.15 | 0.26 | 0.27 | 0.39 | 0.52 |
| 1 | 160006472 | rs16831300 | 1.153 | -0.332 | -1.732 | -0.682 | -0.015 | -0.053 | -0.053 | 0 | 0.1 | 0.01 | 0.15 | 0.31 | 0.11 | 0.13 |
| 1 | 160006809 | rs17367150 | 1.9 | -0.17 | -1.898 | -1.063 | -0.048 | -0.057 | -0.057 | 0 | 0.22 | 0.03 | 0.16 | 0.24 | 0.26 | 0.37 |
| 1 | 160009121 | rs1053074 | 0.979 | 0.112 | -1.476 | -0.911 | 0.04 | 0.052 | 0.052 | 0 | 0.48 | 0.66 | 0.43 | 0.33 | 0.42 | 0.5 |
| 1 | 160009419 | rs2486253 | 0.342 | 0.18 | -0.996 | -0.787 | -0.001 | 0.001 | 0.001 | 0.021 | 0.81 | 0.82 | 0.85 | 0.96 | 0.86 | 0.83 |
| 1 | 160013353 | rs2820553 | -0.348 | 0.956 | -0.946 | -0.303 | -0.019 | 0.002 | 0.002 | 0 | 0.92 | 0.94 | 0.87 | 0.96 | 0.87 | 0.92 |
| 1 | 160023053 | rs1186689 | -0.496 | -0.069 | -0.595 | -0.067 | 0.003 | -0.004 | -0.004 | 0 | 0.44 | 0.52 | 0.47 | 0.33 | 0.49 | 0.26 |
| 1 | 160029212 | rs1186685 | 1.375 | 0.068 | -1.101 | -0.023 | -0.113 | -0.096 | -0.096 | 3.599 | 0.15 | 0.01 | 0.24 | 0.24 | 0.33 | 0.18 |
| 1 | 160029456 | rs11265313 | 0.819 | -0.592 | -0.626 | -0.28 | 0.032 | 0.035 | 0.035 | 0 | 0.5 | 0.32 | 0.48 | 0.67 | 0.48 | 0.66 |
| 1 | 160031330 | rs4656873 | 0.351 | -0.5 | -0.728 | -0.112 | 0 | -0.002 | -0.002 | 0 | 0.14 | 0.12 | 0.27 | 0.24 | 0.13 | 0.19 |
| 1 | 160033502 | rs6690889 | -0.112 | -0.477 | 0.136 | -0.294 | 0.006 | -0.001 | -0.001 | 0 | 0.66 | 0.63 | 0.55 | 0.72 | 0.52 | 0.75 |
| 1 | 160045858 | rs1321650 | 0.23 | -0.477 | 0.282 | -0.234 | 0.008 | -0.006 | -0.006 | 0 | 0.6 | 0.48 | 0.54 | 0.72 | 0.53 | 0.75 |
| 1 | 160055932 | rs2737703 | -0.205 | -0.347 | 0.532 | -0.36 | 0.003 | -0.002 | -0.002 | 0.1 | 0.34 | 0.37 | 0.3 | 0.26 | 0.37 | 0.21 |
| 1 | 160058252 | rs2753268 | 1.313 | -0.289 | -0.144 | -0.357 | -0.087 | -0.085 | -0.085 | 0 | 0.16 | 0.02 | 0.23 | 0.27 | 0.31 | 0.17 |
| 1 | 160059748 | rs12402888 | 1.524 | -0.103 | -0.102 | -0.259 | -0.034 | -0.011 | -0.011 | 0.265 | 0.06 | 0.01 | 0.16 | 0.05 | 0.1 | 0.06 |
| 1 | 160060272 | rs11265318 | 0.114 | -0.219 | 0.208 | -0.323 | -0.007 | -0.002 | -0.002 | 0 | 0.33 | 0.25 | 0.29 | 0.27 | 0.37 | 0.2 |
| 1 | 160061934 | rs12408242 | 0.068 | -0.371 | -0.186 | -0.148 | -0.065 | -0.065 | -0.065 | 2.495 | 0.13 | 0.02 | 0.26 | 0.23 | 0.19 | 0.16 |
| 1 | 160064418 | rs2369406 | -0.935 | -0.45 | 0.253 | -0.189 | 0.001 | 0 | 0 | 0 | 0.48 | 0.51 | 0.56 | 0.5 | 0.56 | 0.38 |
| 1 | 160073904 | rs11265321 | 0.327 | -1.135 | 0.006 | -0.427 | -0.01 | 0.015 | 0.015 | 0.034 | 0.39 | 0.27 | 0.33 | 0.26 | 0.45 | 0.26 |
| 1 | 160075437 | rs10908753 | -0.034 | -1.22 | 0.077 | -0.435 | -0.003 | 0.022 | 0.022 | 0 | 0.45 | 0.29 | 0.49 | 0.32 | 0.55 | 0.32 |
| 1 | 160076224 | rs11265323 | -0.321 | -1.059 | -0.065 | -0.403 | -0.001 | 0 | 0 | 0.216 | 0.55 | 0.53 | 0.61 | 0.49 | 0.65 | 0.44 |
| 1 | 160077854 | rs6688363 | 0.583 | -0.867 | -0.612 | -0.247 | -0.001 | -0.03 | -0.03 | 0 | 0.17 | 0.11 | 0.3 | 0.35 | 0.16 | 0.2 |
| 1 | 160083012 | rs4656879 | -0.376 | -1.271 | -0.29 | -0.83 | -0.03 | -0.077 | -0.077 | 0 | 0.76 | 0.65 | 0.81 | 0.94 | 0.86 | 0.91 |
| 1 | 160084643 | rs3761685 | 0.789 | -1.25 | -0.474 | -0.572 | -0.034 | -0.041 | -0.041 | 0.079 | 0.07 | 0.01 | 0.11 | 0.14 | 0.14 | 0.09 |
| 1 | 160092121 | rs2753267 | -0.244 | -1.533 | -0.088 | -0.655 | -0.039 | -0.036 | -0.036 | 0 | 0.72 | 0.58 | 0.92 | 0.85 | 0.84 | 0.64 |
| 1 | 160093222 | rs2820581 | -1.363 | -1.198 | 0.132 | -0.357 | -0.028 | -0.03 | -0.03 | 0 | 0.94 | 0.8 | 0.98 | 1 | 1 | 0.99 |
| 1 | 160094644 | rs1023421 | 0.361 | -1.051 | -0.473 | -0.277 | 0.022 | 0.009 | 0.009 | 0.735 | 0.51 | 0.42 | 0.51 | 0.52 | 0.62 | 0.75 |
| 1 | 160097666 | rs6695366 | -0.015 | -0.272 | -0.668 | 0.294 | 0.002 | -0.048 | -0.048 | 0 | 0.82 | 0.94 | 0.72 | 0.66 | 0.84 | 0.85 |
| 1 | 160102256 | rs10797059 | 0.079 | -0.253 | -0.175 | 0.412 | -0.02 | 0.038 | 0.038 | 0 | 0.82 | 0.7 | 0.72 | 0.66 | 0.86 | 0.86 |
| 1 | 160107588 | rs10494336 | 0.164 | 0.125 | -0.591 | 0.335 | 0 | -0.001 | -0.001 | 0.634 | 0.16 | 0.17 | 0.07 | 0.15 | 0.12 | 0.28 |
| 1 | 160108803 | rs12410866 | 0.724 | 0.825 | -0.806 | 0.407 | -0.031 | -0.017 | -0.017 | 0 | 0.05 | 0 | 0.12 | 0.07 | 0.11 | 0.07 |
| 1 | 160112138 | rs2070704 | -0.068 | 0.845 | -0.511 | 0.199 | 0 | -0.005 | -0.005 | 0 | 0.2 | 0.25 | 0.12 | 0.15 | 0.21 | 0.3 |
| 1 | 160112741 | rs1046995 | -0.068 | 0.845 | -0.511 | 0.199 | 0 | -0.005 | -0.005 | 0 | 0.2 | 0.25 | 0.12 | 0.15 | 0.21 | 0.3 |
| 1 | 160114070 | rs12753283 | 0.272 | 0.843 | -0.735 | 0.274 | 0.002 | -0.014 | -0.014 | 0 | 0.17 | 0.11 | 0.28 | 0.35 | 0.17 | 0.14 |
| 1 | 160114343 | rs11585375 | 1.123 | 0.705 | -0.658 | 0.031 | 0.017 | 0.036 | 0.036 | 0 | 0.45 | 0.25 | 0.35 | 0.32 | 0.4 | 0.41 |
| 1 | 160120335 | rs16831441 | 0.403 | 0.947 | -0.436 | 0.328 | 0.001 | -0.002 | -0.002 | 0.515 | 0.18 | 0.19 | 0.12 | 0.14 | 0.21 | 0.31 |
| 1 | 160121488 | rs2369476 | 0.813 | 0.715 | -1.071 | 0.305 | -0.025 | -0.047 | -0.047 | 0 | 0.25 | 0.11 | 0.28 | 0.31 | 0.3 | 0.38 |
| 1 | 160121746 | rs746362 | 0.777 | 0.707 | -1.05 | 0.301 | -0.019 | -0.037 | -0.037 | 0 | 0.25 | 0.13 | 0.27 | 0.31 | 0.3 | 0.38 |
| 1 | 160123556 | rs6697199 | 0.085 | 0.677 | -1.045 | 0.309 | -0.011 | 0.031 | 0.031 | 0 | 0.58 | 0.65 | 0.51 | 0.27 | 0.51 | 0.48 |
| 1 | 160124875 | rs6427504 | -0.622 | 0.844 | -0.864 | 0.459 | -0.006 | -0.013 | -0.013 | 0 | 0.74 | 0.75 | 0.71 | 0.68 | 0.7 | 0.62 |
| 1 | 160126138 | rs927203 | 0.218 | 1.227 | -0.58 | 0.654 | 0.007 | 0.002 | 0.002 | 0 | 0.6 | 0.49 | 0.61 | 0.59 | 0.56 | 0.66 |
| 1 | 160128671 | rs4656888 | -0.081 | 0.924 | -0.966 | 0.152 | -0.08 | -0.037 | -0.037 | 0.106 | 0.36 | 0.14 | 0.5 | 0.4 | 0.49 | 0.3 |
| 1 | 160129495 | rs7551983 | -0.68 | 0.806 | -0.53 | -0.047 | -0.005 | 0.003 | 0.003 | 0 | 0.57 | 0.55 | 0.66 | 0.46 | 0.63 | 0.52 |
| 1 | 160129557 | rs7541968 | 1.044 | 0.927 | -0.823 | -0.063 | -0.12 | -0.101 | -0.101 | 0 | 0.22 | 0.01 | 0.28 | 0.33 | 0.34 | 0.26 |
| 1 | 160131429 | rs953104 | -0.364 | 0.471 | -0.156 | -0.077 | 0.005 | -0.061 | -0.061 | 0 | 0.86 | 0.77 | 0.91 | 1 | 0.87 | 0.9 |

| | | | | | | | | | | | | | | | | |
|---|---|---|---|---|---|---|---|---|---|---|---|---|---|---|---|---|
| 1 | 160134205 | rs11265338 | -0.052 | 0.444 | -0.602 | -0.02 | 0.017 | -0.001 | -0.001 | 4.199 | 0.55 | 0.36 | 0.68 | 0.54 | 0.64 | 0.53 |
| 1 | 160136208 | rs10797062 | -0.348 | 0.721 | -0.026 | 0.082 | 0.063 | 0.079 | 0.079 | 0.193 | 0.77 | 0.55 | 0.73 | 0.58 | 0.71 | 0.78 |
| 1 | 160137300 | rs4656889 | -0.625 | 0.568 | -0.535 | -0.011 | 0.004 | 0.006 | 0.006 | 0 | 0.48 | 0.7 | 0.61 | 0.53 | 0.53 | 0.48 |
| 1 | 160141268 | rs12058214 | -0.257 | 0.718 | -0.15 | -0.879 | -0.004 | -0.075 | -0.075 | 6.301 | 0.2 | 0.39 | 0.2 | 0.08 | 0.18 | 0.18 |
| 1 | 160141636 | rs1891265 | -0.376 | 0.637 | -0.912 | -0.695 | -0.004 | 0.048 | 0.048 | 2.844 | 0.54 | 0.63 | 0.41 | 0.14 | 0.52 | 0.56 |
| 1 | 160144895 | rs11265343 | -0.026 | 0.545 | -1.029 | -0.255 | 0 | -0.001 | -0.001 | 0 | 0.45 | 0.47 | 0.38 | 0.62 | 0.45 | 0.35 |
| 1 | 160145298 | rs911228 | -0.676 | 0.33 | -1.37 | -0.655 | -0.029 | 0.062 | 0.062 | 0 | 0.74 | 0.89 | 0.8 | 0.89 | 0.7 | 0.75 |
| 1 | 160146173 | rs881291 | 1.447 | 0.23 | -1.692 | -1.088 | -0.078 | -0.116 | -0.116 | 0.015 | 0.28 | 0.05 | 0.27 | 0.62 | 0.33 | 0.28 |
| 1 | 160146645 | rs16831518 | 0.58 | 0.342 | -1.301 | -0.92 | 0.004 | 0.027 | 0.027 | 0.297 | 0.16 | 0.34 | 0.35 | 0.17 | 0.16 | 0.24 |
| 1 | 160146790 | rs6693877 | -0.758 | 0.307 | -1.082 | -0.975 | 0.004 | 0.017 | 0.017 | 2.429 | 0.33 | 0.7 | 0.46 | 0.18 | 0.32 | 0.32 |
| 1 | 160148810 | rs11265346 | 0.07 | 0.304 | -0.827 | -1.007 | -0.004 | -0.093 | -0.093 | 0 | 0.17 | 0.36 | 0.11 | 0.01 | 0.16 | 0.08 |
| 1 | 160151450 | rs7519748 | -0.314 | 0.346 | -1.149 | -0.942 | -0.035 | 0.057 | 0.057 | 0 | 0.73 | 0.89 | 0.8 | 0.89 | 0.69 | 0.72 |
| 1 | 160152057 | rs1935306 | 0.805 | 0.299 | -0.901 | -0.924 | 0.001 | -0.015 | -0.015 | 0 | 0.57 | 0.56 | 0.66 | 0.78 | 0.54 | 0.62 |
| 1 | 160157029 | rs7524566 | 0.156 | 0.329 | -1.029 | -0.906 | -0.005 | -0.038 | -0.038 | 0 | 0.15 | 0.24 | 0.13 | 0.07 | 0.12 | 0.09 |
| 1 | 160157365 | rs12070422 | 0.299 | 0.389 | -1.067 | -0.762 | -0.002 | -0.004 | -0.004 | 0 | 0.59 | 0.63 | 0.71 | 0.85 | 0.56 | 0.64 |
| 1 | 160159090 | rs11265352 | 0.66 | 0.492 | -1.611 | -0.487 | 0.025 | 0.082 | 0.082 | 2.269 | 0.54 | 0.15 | 0.72 | 0.7 | 0.63 | 0.65 |
| 1 | 160159156 | rs659235 | -0.905 | 0.737 | -1.05 | -0.307 | 0.004 | -0.003 | -0.003 | 0 | 0.12 | 0.14 | 0.07 | 0.1 | 0.14 | 0.15 |
| 1 | 160162872 | rs617698 | 0.691 | 0.013 | -1.481 | 0.057 | -0.019 | -0.072 | -0.072 | 0 | 0.37 | 0.1 | 0.38 | 0.57 | 0.38 | 0.27 |
| 1 | 160163509 | rs3747623 | 0.721 | 0.034 | -1.586 | 0.06 | -0.051 | -0.126 | -0.126 | 0 | 0.35 | 0.04 | 0.36 | 0.57 | 0.37 | 0.27 |
| 1 | 160166235 | rs6697826 | -1.136 | -0.528 | -0.396 | -0.038 | -0.044 | -0.066 | -0.066 | 2.566 | 0.24 | 0.47 | 0.17 | 0.11 | 0.15 | 0.35 |
| 1 | 160173901 | rs2795070 | 1.452 | -0.524 | -0.563 | 0.016 | -0.038 | -0.068 | -0.068 | 0 | 0.38 | 0.09 | 0.4 | 0.55 | 0.43 | 0.26 |
| 1 | 160185262 | rs4656252 | -0.199 | -0.591 | -0.45 | 0.032 | 0.009 | -0.012 | -0.012 | 0 | 0.09 | 0.05 | 0.03 | 0.09 | 0.06 | 0.09 |
| 1 | 160189500 | rs1061511 | -1.753 | -0.686 | 0.549 | 0.008 | -0.036 | -0.065 | -0.065 | 0 | 0.62 | 0.91 | 0.6 | 0.45 | 0.57 | 0.74 |
| 1 | 160212700 | rs11265362 | 0.596 | -0.779 | 0.663 | -0.04 | -0.014 | -0.03 | -0.03 | 0 | 0.08 | 0 | 0.05 | 0.14 | 0.08 | 0.11 |
| 1 | 160214027 | rs2275700 | -1.802 | -0.724 | 0.739 | -0.076 | -0.043 | -0.064 | -0.064 | 0 | 0.62 | 0.91 | 0.6 | 0.45 | 0.57 | 0.73 |
| 1 | 160246485 | rs1557524 | -0.555 | -0.92 | 1.194 | -0.179 | -0.003 | 0.007 | 0.007 | 0.591 | 0.43 | 0.33 | 0.52 | 0.33 | 0.51 | 0.63 |
| 1 | 160263124 | rs1317944 | -1.988 | -0.879 | 0.817 | -0.17 | -0.04 | -0.061 | -0.061 | 1.884 | 0.62 | 0.91 | 0.6 | 0.44 | 0.57 | 0.73 |
| 1 | 160276709 | rs7522166 | -0.362 | -1.351 | 1.124 | -0.184 | -0.019 | 0.025 | 0.025 | 0.766 | 0.41 | 0.22 | 0.52 | 0.33 | 0.51 | 0.64 |
| 1 | 160303416 | rs1802778 | -1.775 | -1.266 | 0.77 | -0.176 | -0.045 | -0.048 | -0.048 | 0 | 0.59 | 0.91 | 0.59 | 0.44 | 0.55 | 0.72 |
| 1 | 160312265 | rs10752637 | -1.985 | -1.248 | 0.773 | -0.15 | -0.041 | -0.063 | -0.063 | 0 | 0.62 | 0.91 | 0.6 | 0.44 | 0.57 | 0.73 |
| 1 | 160312625 | rs1324738 | -1.831 | -1.074 | 1.095 | -0.148 | -0.065 | -0.042 | -0.042 | 1.006 | 0.17 | 0.44 | 0.06 | 0.09 | 0.06 | 0.09 |
| 1 | 160318940 | rs6669689 | -1.024 | -0.945 | 0.817 | 0.278 | -0.003 | -0.005 | -0.005 | 0 | 0.55 | 0.59 | 0.49 | 0.48 | 0.54 | 0.66 |
| 1 | 160318984 | rs16831846 | -2.196 | -0.343 | 1.342 | 0.048 | 0.004 | -0.027 | -0.027 | 0 | 0.06 | 0.16 | 0.07 | 0 | 0.06 | 0.01 |
| 1 | 160321065 | rs12239747 | -1.089 | -0.336 | 1.414 | 0.155 | 0.005 | 0.007 | 0.007 | 0 | 0.08 | 0.26 | 0.05 | 0.03 | 0.05 | 0.05 |
| 1 | 160321359 | rs6664438 | -0.051 | -0.34 | 1.227 | 0.154 | 0.002 | 0.001 | 0.001 | 0 | 0.05 | 0.04 | 0.03 | 0.03 | 0.05 | 0.05 |
| 1 | 160323353 | rs1124379 | -2.13 | -0.531 | 1.023 | 0.174 | -0.039 | -0.061 | -0.061 | 1.792 | 0.58 | 0.88 | 0.56 | 0.45 | 0.53 | 0.72 |
| 1 | 160324771 | rs6677637 | -0.144 | -0.524 | 1.087 | 0.181 | 0.001 | 0.001 | 0.001 | 0 | 0.06 | 0.06 | 0.03 | 0.03 | 0.05 | 0.05 |
| 1 | 160328804 | rs12724113 | 0.529 | -0.523 | 0.959 | 0.108 | -0.112 | 0.045 | 0.045 | 0.379 | 0.23 | 0.02 | 0.32 | 0.1 | 0.32 | 0.28 |
| 1 | 160329430 | rs11265374 | -1.353 | -0.557 | 1.18 | 0.125 | -0.009 | 0.006 | 0.006 | 0 | 0.1 | 0.17 | 0.05 | 0.11 | 0.07 | 0.1 |
| 1 | 160331019 | rs4656256 | -2.899 | -0.525 | 0.896 | 0.121 | 0.042 | -0.014 | -0.014 | 0 | 0.34 | 0.85 | 0.24 | 0.35 | 0.21 | 0.45 |
| 1 | 160341142 | rs10908778 | -0.962 | -0.534 | 1.228 | 0.146 | 0.007 | 0.039 | 0.039 | 0 | 0.17 | 0.3 | 0.16 | 0.27 | 0.14 | 0.35 |
| 1 | 160342015 | rs7521304 | -2.079 | -0.516 | 0.82 | 0.213 | -0.005 | -0.063 | -0.063 | 0 | 0.57 | 0.84 | 0.56 | 0.52 | 0.54 | 0.73 |
| 1 | 160342944 | rs11265378 | -0.799 | -0.429 | 1.09 | 0.366 | 0.003 | 0.014 | 0.014 | 0 | 0.15 | 0.15 | 0.15 | 0.27 | 0.14 | 0.35 |
| 1 | 160346304 | rs7542200 | -2.592 | -0.541 | 1.261 | 0.015 | -0.063 | 0.018 | 0.018 | 4.568 | 0.13 | 0.62 | 0.06 | 0.13 | 0.02 | 0.07 |
| 1 | 160347420 | rs12080243 | -2.641 | -0.542 | 1.08 | 0.015 | -0.062 | 0.004 | 0.004 | 5.132 | 0.14 | 0.66 | 0.06 | 0.13 | 0.02 | 0.09 |
| 1 | 160349098 | rs11265381 | -0.892 | -0.287 | 0.902 | -0.071 | 0.002 | -0.001 | -0.001 | 0 | 0.37 | 0.41 | 0.46 | 0.38 | 0.43 | 0.58 |
| 1 | 160349624 | rs12028921 | -1.807 | -0.2 | 0.984 | 0.044 | -0.014 | 0.055 | 0.055 | 2.62 | 0.06 | 0.24 | 0.04 | 0.13 | 0.03 | 0.11 |
| 1 | 160349901 | rs12403970 | -0.735 | -0.345 | 1.137 | -0.114 | -0.021 | 0.016 | 0.016 | 0.321 | 0.34 | 0.27 | 0.43 | 0.26 | 0.39 | 0.47 |
| 1 | 160364947 | rs12739519 | -0.455 | -0.163 | 1.12 | -0.117 | -0.019 | 0.012 | 0.012 | 0.792 | 0.31 | 0.23 | 0.37 | 0.25 | 0.35 | 0.46 |
| 1 | 160369372 | rs11582932 | -0.362 | -0.143 | 1.121 | -0.094 | -0.012 | 0.002 | 0.002 | 0.325 | 0.31 | 0.3 | 0.38 | 0.27 | 0.34 | 0.53 |
| 1 | 160372109 | rs7513382 | -1.974 | -0.379 | 0.705 | -0.467 | -0.128 | -0.015 | -0.015 | 0 | 0.56 | 0.94 | 0.55 | 0.39 | 0.5 | 0.69 |
| 1 | 160376060 | rs12090877 | -0.888 | -0.165 | 1.49 | -0.378 | -0.015 | -0.039 | -0.039 | 0 | 0.13 | 0.2 | 0.07 | 0.04 | 0.09 | 0.07 |
| 1 | 160377499 | rs2369608 | -3.195 | 0.248 | 2.122 | -0.02 | 0.004 | -0.022 | -0.022 | 0.182 | 0.11 | 0.46 | 0.09 | 0.02 | 0.06 | 0.1 |
| 1 | 160388141 | rs4656907 | -1.408 | 0.302 | 1.963 | -0.004 | 0.004 | 0.031 | 0.031 | 1.088 | 0.35 | 0.69 | 0.28 | 0.21 | 0.3 | 0.38 |
| 1 | 160391398 | rs6427518 | -1.855 | 0.121 | 1.832 | 0.062 | -0.021 | 0.046 | 0.046 | 0.195 | 0.47 | 0.8 | 0.45 | 0.39 | 0.48 | 0.67 |
| 1 | 160392393 | rs12085898 | -0.624 | 0.028 | 2.015 | 0.017 | 0.002 | -0.033 | -0.033 | 0 | 0.3 | 0.31 | 0.23 | 0.09 | 0.29 | 0.32 |
| 1 | 160394937 | rs17380127 | -0.142 | 0.054 | 1.808 | 0.028 | -0.009 | 0.025 | 0.025 | 0.005 | 0.26 | 0.17 | 0.21 | 0.08 | 0.28 | 0.31 |
| 1 | 160402539 | rs10494343 | -0.987 | 0.388 | 1.44 | 0.144 | -0.034 | -0.01 | -0.01 | 0 | 0.12 | 0.06 | 0.18 | 0.17 | 0.2 | 0.28 |
| 1 | 160405578 | rs1014649 | 0.473 | 0.332 | 1.478 | 0.115 | -0.013 | -0.015 | -0.015 | 0 | 0.05 | 0.21 | 0.02 | 0 | 0 | 0 |
| 1 | 160406125 | rs11265399 | -1.491 | 0.29 | 1.661 | 0.095 | -0.019 | -0.022 | -0.022 | 0 | 0.08 | 0.36 | 0.03 | 0 | 0 | 0 |
| 1 | 160407840 | rs10908780 | -0.443 | 0.245 | 1.701 | 0.228 | -0.003 | -0.034 | -0.034 | 0 | 0.29 | 0.49 | 0.19 | 0.21 | 0.25 | 0.27 |
| 1 | 160408563 | rs4656258 | -0.751 | 0.248 | 1.57 | 0.263 | -0.003 | -0.003 | -0.003 | 0 | 0.32 | 0.65 | 0.26 | 0.28 | 0.25 | 0.27 |
| 1 | 160409987 | rs4656260 | -1.868 | 0.346 | 0.981 | 0.389 | -0.078 | -0.037 | -0.037 | 0 | 0.5 | 0.92 | 0.44 | 0.45 | 0.46 | 0.55 |
| 1 | 160420036 | rs11585691 | -0.229 | 0.694 | 1.639 | 0.704 | 0.003 | -0.018 | -0.018 | 0 | 0.24 | 0.27 | 0.16 | 0.08 | 0.25 | 0.24 |
| 1 | 160428188 | rs1149392 | -0.354 | 0.799 | 0.699 | 0.691 | 0.012 | 0.01 | 0.01 | 0.015 | 0.26 | 0.44 | 0.23 | 0.28 | 0.25 | 0.27 |
| 1 | 160430580 | rs11265410 | -1.757 | 0.909 | 0.968 | 0.989 | 0.024 | 0.081 | 0.081 | 2.317 | 0.11 | 0.24 | 0.16 | 0.18 | 0.11 | 0.1 |
| 1 | 160432428 | rs11805036 | -1.267 | 1.121 | 0.429 | 0.86 | -0.01 | -0.005 | -0.005 | 0.446 | 0.15 | 0.14 | 0.28 | 0.18 | 0.19 | 0.1 |
| 1 | 160452218 | rs2369611 | 1.916 | 1.315 | 0.733 | 0.847 | 0.032 | 0.031 | 0.031 | 0 | 0.74 | 0.45 | 0.59 | 0.63 | 0.71 | 0.69 |
| 1 | 160459909 | rs12119876 | -1.036 | 1.526 | -0.702 | 1.055 | -0.018 | -0.011 | -0.011 | 0 | 0.15 | 0.13 | 0.27 | 0.19 | 0.19 | 0.1 |

| | | | | | | | | | | | | | | | | |
|---|---|---|---|---|---|---|---|---|---|---|---|---|---|---|---|---|
| 1 | 160464548 | rs678456 | 0.495 | 1.029 | -0.749 | 0.807 | -0.021 | -0.021 | -0.021 | 0 | 0.94 | 0.97 | 0.9 | 0.82 | 0.91 | 0.89 |
| 1 | 160472139 | rs17385337 | -0.166 | 0.388 | -1.204 | 0.016 | -0.023 | 0.053 | 0.053 | 0 | 0.32 | 0.12 | 0.27 | 0.21 | 0.33 | 0.18 |
| 1 | 160472340 | rs10494344 | -0.101 | -0.204 | -0.148 | 0.102 | -0.005 | 0.058 | 0.058 | 5.49 | 0.06 | 0.3 | 0.03 | 0.09 | 0.01 | 0.12 |
| 1 | 160473163 | rs6660233 | 0.523 | -1.616 | -0.448 | -0.537 | -0.033 | -0.007 | -0.007 | 0.043 | 0.35 | 0.15 | 0.46 | 0.6 | 0.55 | 0.48 |
| 1 | 160481662 | rs12401994 | 1.159 | -0.733 | -0.724 | -0.057 | -0.054 | -0.049 | -0.049 | 0 | 0.08 | 0.01 | 0.14 | 0.15 | 0.16 | 0.14 |
| 1 | 160489974 | rs3845627 | -0.74 | -2.01 | 0.362 | -0.713 | -0.095 | -0.062 | -0.062 | 0 | 0.77 | 0.63 | 0.78 | 0.91 | 0.92 | 0.78 |
| 1 | 160499961 | rs7528976 | 0.441 | -2.151 | 0.227 | -0.789 | -0.015 | 0.004 | 0.004 | 0 | 0.37 | 0.16 | 0.47 | 0.6 | 0.57 | 0.48 |
| 1 | 160507084 | rs7418241 | 1.256 | -2.324 | 0.246 | -0.834 | -0.048 | 0.104 | 0.104 | 0 | 0.29 | 0.09 | 0.19 | 0.14 | 0.31 | 0.17 |
| 1 | 160509461 | rs12080742 | -1.115 | -2.362 | 0.793 | -0.795 | -0.044 | 0.049 | 0.049 | 0.059 | 0.11 | 0.36 | 0.11 | 0.15 | 0.04 | 0.14 |
| 1 | 160512224 | rs12134393 | 0.268 | -2.37 | 0.819 | -1 | 0.035 | 0.032 | 0.032 | 0.03 | 0.67 | 0.26 | 0.66 | 0.82 | 0.88 | 0.75 |
| 1 | 160513183 | rs1503859 | -1.037 | -2.336 | 0.86 | -0.986 | -0.099 | -0.055 | -0.055 | 0.744 | 0.78 | 0.65 | 0.77 | 0.9 | 0.92 | 0.78 |
| 1 | 160516262 | rs11585738 | 1.243 | -2.384 | 0.735 | -1.004 | -0.044 | 0.086 | 0.086 | 0 | 0.28 | 0.1 | 0.19 | 0.14 | 0.31 | 0.17 |
| 1 | 160516308 | rs6427528 | -0.099 | -2.353 | 0.993 | -0.974 | -0.129 | -0.087 | -0.087 | 0 | 0.71 | 0.53 | 0.75 | 0.88 | 0.9 | 0.71 |
| 1 | 160517692 | rs1055880 | 0.625 | -2.293 | 0.888 | -0.984 | 0.003 | -0.013 | -0.013 | 1.507 | 0.29 | 0.37 | 0.27 | 0.21 | 0.32 | 0.17 |
| 1 | 160536561 | rs6670721 | 0.985 | -2.329 | 0.949 | -1.039 | -0.015 | 0.033 | 0.033 | 0 | 0.27 | 0.18 | 0.2 | 0.14 | 0.32 | 0.18 |
| 1 | 160541308 | rs1998651 | 0.466 | -2.596 | 1.336 | -1.178 | 0.007 | 0.058 | 0.058 | 0 | 0.41 | 0.17 | 0.47 | 0.68 | 0.57 | 0.56 |
| 1 | 160548247 | rs10908787 | -0.487 | -2.701 | 1.79 | -1.186 | -0.07 | -0.033 | -0.033 | 0 | 0.73 | 0.62 | 0.76 | 0.86 | 0.91 | 0.73 |
| 1 | 160549579 | rs977020 | -0.112 | -2.7 | 1.786 | -1.186 | -0.103 | -0.034 | -0.034 | 0 | 0.71 | 0.54 | 0.68 | 0.8 | 0.89 | 0.73 |
| 1 | 160551592 | rs6692382 | 0.385 | -2.643 | 1.714 | -1.19 | 0.007 | 0.039 | 0.039 | 0 | 0.42 | 0.24 | 0.47 | 0.67 | 0.56 | 0.55 |
| 1 | 160556144 | rs349250 | -1.267 | -2.829 | 1.7 | -1.3 | 0.061 | 0.029 | 0.029 | 0 | 0.81 | 0.91 | 0.79 | 0.88 | 0.92 | 0.75 |
| 1 | 160559549 | rs7543788 | -0.821 | -2.816 | 1.703 | -1.293 | -0.068 | -0.028 | -0.028 | 0 | 0.78 | 0.67 | 0.77 | 0.88 | 0.92 | 0.75 |
| 1 | 160578141 | rs5002947 | 0.746 | -2.761 | 1.303 | -1.293 | 0 | -0.008 | -0.008 | 0 | 0.31 | 0.32 | 0.29 | 0.17 | 0.34 | 0.19 |
| 1 | 160580015 | rs1061217 | 0.591 | -2.821 | 0.707 | -1.311 | 0.002 | 0.064 | 0.064 | 0 | 0.41 | 0.17 | 0.47 | 0.69 | 0.57 | 0.56 |
| 1 | 160580807 | rs349251 | -1.508 | -2.906 | 0.087 | -1.554 | 0.011 | 0.012 | 0.012 | 0 | 0.81 | 0.83 | 0.79 | 0.88 | 0.92 | 0.76 |
| 1 | 160581306 | rs1394079 | 1.06 | -2.568 | -0.422 | -1.593 | -0.186 | -0.064 | -0.064 | 0.187 | 0.27 | 0.02 | 0.37 | 0.68 | 0.42 | 0.48 |
| 1 | 160581899 | rs912582 | 1.183 | -1.452 | 0.221 | -1.471 | -0.071 | 0.079 | 0.079 | 0 | 0.23 | 0.07 | 0.18 | 0.11 | 0.3 | 0.14 |
| 1 | 160588595 | rs2753260 | -0.932 | -1.456 | 0.35 | -1.479 | -0.034 | 0.055 | 0.055 | 0.879 | 0.75 | 0.88 | 0.82 | 0.86 | 0.69 | 0.84 |
| 1 | 160590765 | rs164283 | 0.775 | -1.373 | 0.699 | -1.56 | -0.006 | 0.124 | 0.124 | 0.022 | 0.56 | 0.26 | 0.69 | 0.87 | 0.55 | 0.8 |
| 1 | 160591997 | rs16832283 | 0.085 | -1.316 | 0.785 | -1.474 | 0.014 | -0.105 | -0.105 | 0 | 0.29 | 0.53 | 0.21 | 0.12 | 0.31 | 0.14 |
| 1 | 160599010 | rs1503854 | -0.139 | -1.137 | 0.262 | -1.409 | -0.016 | 0.035 | 0.035 | 0 | 0.33 | 0.24 | 0.25 | 0.12 | 0.39 | 0.18 |
| 1 | 160600319 | rs6672302 | -0.534 | -0.753 | 0.18 | -1.248 | -0.031 | -0.136 | -0.136 | 0 | 0.22 | 0.38 | 0.12 | 0.01 | 0.16 | 0.09 |
| 1 | 160602248 | rs6704124 | -0.228 | -0.09 | -0.372 | 0.158 | -0.015 | -0.013 | -0.013 | 0 | 0.28 | 0.19 | 0.33 | 0.37 | 0.36 | 0.23 |
| 1 | 160612139 | rs3766364 | -0.743 | -0.267 | -0.367 | -0.165 | -0.009 | -0.039 | -0.039 | 0.153 | 0.14 | 0.21 | 0.08 | 0.05 | 0.09 | 0.15 |
| 1 | 160613181 | rs1889792 | -0.042 | 0.268 | -0.992 | 0.442 | 0.025 | 0.007 | 0.007 | 0 | 0.34 | 0.13 | 0.2 | 0.32 | 0.3 | 0.44 |
| 1 | 160615677 | rs10489635 | -0.158 | 0.465 | -0.104 | 0.701 | -0.012 | -0.005 | -0.005 | 1.361 | 0.06 | 0.11 | 0.02 | 0.01 | 0.03 | 0.05 |
| 1 | 160616703 | rs2295612 | 0.114 | 1.414 | -0.474 | 0.496 | -0.075 | 0.041 | 0.041 | 0 | 0.11 | 0.01 | 0.21 | 0.03 | 0.22 | 0.09 |
| 1 | 160626383 | rs4656923 | -1.601 | 1.352 | -0.061 | 1.309 | 0 | 0.068 | 0.068 | 0 | 0.48 | 0.67 | 0.68 | 0.75 | 0.52 | 0.44 |
| 1 | 160630143 | rs11265461 | -1.295 | 1.729 | 0.464 | 1.492 | 0.062 | 0.058 | 0.058 | 0 | 0.24 | 0.42 | 0.38 | 0.41 | 0.36 | 0.2 |
| 1 | 160634588 | rs6427540 | -0.38 | 0.735 | 0.772 | 2.041 | 0.027 | 0.135 | 0.135 | 0 | 0.11 | 0.26 | 0.15 | 0.37 | 0.14 | 0.11 |
| 1 | 160643111 | rs3795324 | -1.278 | 0.798 | 0.497 | 1.85 | 0.025 | 0.251 | 0.251 | 0 | 0.19 | 0.49 | 0.18 | 0.48 | 0.18 | 0.18 |
| 1 | 160648396 | rs1553456 | 0.596 | 0.485 | 0.049 | 1.485 | 0.003 | 0.014 | 0.014 | 0 | 0.83 | 0.77 | 0.75 | 0.73 | 0.79 | 0.82 |
| 1 | 160648481 | rs12096589 | -0.725 | 0.396 | -0.094 | 1.285 | -0.012 | 0.008 | 0.008 | 0 | 0.07 | 0.11 | 0.02 | 0.05 | 0.03 | 0.05 |
| 1 | 160648549 | rs1126644 | 0.603 | 0.615 | -0.373 | 1.385 | 0 | 0.012 | 0.012 | 0.006 | 0.83 | 0.8 | 0.75 | 0.63 | 0.79 | 0.74 |
| 1 | 160651457 | rs12753811 | -0.941 | -0.361 | -0.375 | 0.461 | -0.017 | -0.013 | -0.013 | 0 | 0.16 | 0.17 | 0.16 | 0.09 | 0.06 | 0.16 |
| 1 | 160651750 | rs10489636 | -1.145 | -0.571 | -1.27 | -0.148 | 0.008 | 0.059 | 0.059 | 0.003 | 0.3 | 0.06 | 0.31 | 0.17 | 0.24 | 0.23 |
| 1 | 160652476 | rs10489637 | -1.071 | -1.1 | -0.572 | -0.919 | -0.025 | -0.036 | -0.036 | 0 | 0.36 | 0.54 | 0.32 | 0.18 | 0.23 | 0.26 |
| 1 | 160654066 | rs10489638 | 0.131 | -1.153 | -0.322 | -0.92 | 0.031 | 0.013 | 0.013 | 0 | 0.11 | 0.04 | 0.12 | 0.11 | 0.04 | 0.11 |
| 1 | 160654786 | rs1980606 | -0.933 | -1.138 | -0.145 | -0.952 | -0.014 | -0.035 | -0.035 | 0.184 | 0.28 | 0.33 | 0.26 | 0.14 | 0.19 | 0.23 |
| 1 | 160670454 | rs507260 | 0.642 | -1.071 | -0.097 | -0.979 | -0.023 | -0.043 | -0.043 | 0 | 0.73 | 0.65 | 0.83 | 0.86 | 0.82 | 0.77 |
| 1 | 160673779 | rs978763 | -0.117 | -0.94 | 0.03 | -0.998 | 0.022 | -0.05 | -0.05 | 0 | 0.09 | 0.2 | 0.1 | 0.02 | 0.1 | 0.05 |
| 1 | 160674460 | rs11265472 | 0.152 | -0.764 | -0.129 | -1.651 | 0.007 | -0.071 | -0.071 | 4.569 | 0.18 | 0.31 | 0.14 | 0 | 0.2 | 0.08 |
| 1 | 160679527 | rs10489639 | 0.537 | -0.892 | -0.65 | -0.97 | 0.016 | 0.023 | 0.023 | 0 | 0.48 | 0.38 | 0.54 | 0.32 | 0.59 | 0.29 |
| 1 | 160679935 | rs10752639 | 0.174 | -1.222 | -0.483 | -1.128 | 0.002 | 0.003 | 0.003 | 0 | 0.58 | 0.62 | 0.57 | 0.33 | 0.64 | 0.37 |
| 1 | 160680439 | rs352683 | 0.4 | -1.203 | -0.6 | -1.144 | -0.039 | -0.082 | -0.082 | 6.315 | 0.63 | 0.48 | 0.81 | 0.87 | 0.75 | 0.74 |
| 1 | 160686494 | rs352692 | -0.069 | -0.236 | -1.159 | -0.173 | -0.036 | -0.1 | -0.1 | 0.924 | 0.33 | 0.09 | 0.55 | 0.49 | 0.36 | 0.34 |
| 1 | 160687799 | rs352694 | 0.008 | 0.25 | -0.868 | 0.322 | -0.016 | -0.038 | -0.038 | 3.574 | 0.31 | 0.2 | 0.51 | 0.45 | 0.35 | 0.35 |
| 1 | 160689145 | rs184772 | 0.526 | 0.466 | -0.14 | 0.419 | -0.014 | 0.056 | 0.056 | 0.018 | 0.94 | 0.77 | 0.96 | 0.88 | 0.97 | 0.95 |
| 1 | 160692544 | rs570901 | 0.21 | 1.111 | -0.509 | 0.289 | 0.001 | 0 | 0 | 0 | 0.75 | 0.7 | 0.77 | 0.76 | 0.65 | 0.78 |
| 1 | 160696304 | rs503832 | 0.277 | 1.045 | -0.455 | 0.298 | 0.005 | -0.001 | -0.001 | 0 | 0.75 | 0.71 | 0.77 | 0.77 | 0.66 | 0.78 |
| 1 | 160703965 | rs983494 | -0.322 | 1.466 | -0.476 | 0.089 | -0.015 | 0.006 | 0.006 | 0 | 0.16 | 0.19 | 0.1 | 0.11 | 0.21 | 0.09 |
| 1 | 160705619 | rs549471 | 0.195 | 1.367 | 0.098 | -0.335 | 0.012 | -0.097 | -0.097 | 0 | 0.85 | 0.71 | 0.86 | 1 | 0.83 | 0.94 |
| 1 | 160707779 | rs508744 | -0.583 | 0.759 | -0.477 | -0.386 | 0.033 | 0.08 | 0.08 | 0 | 0.43 | 0.31 | 0.6 | 0.76 | 0.58 | 0.54 |
| 1 | 160710243 | rs573702 | 1.116 | 0.403 | -0.557 | -0.286 | -0.002 | -0.003 | -0.003 | 0 | 0.61 | 0.38 | 0.69 | 0.85 | 0.53 | 0.76 |
| 1 | 160710745 | rs535622 | 1.13 | 0.702 | -0.816 | 0.11 | -0.137 | -0.162 | -0.162 | 0.272 | 0.21 | 0.01 | 0.48 | 0.58 | 0.36 | 0.32 |
| 1 | 160714724 | rs3766373 | -0.204 | 0.847 | -0.047 | 0.134 | -0.06 | 0.022 | 0.022 | 0 | 0.19 | 0.12 | 0.13 | 0.11 | 0.29 | 0.12 |
| 1 | 160722550 | rs489286 | 1.258 | 0.748 | 0.225 | 0.153 | 0.017 | -0.02 | -0.02 | 0 | 0.7 | 0.49 | 0.81 | 0.8 | 0.68 | 0.77 |
| 1 | 160723517 | rs518721 | 1.249 | 0.708 | 0.227 | 0.155 | 0.013 | -0.02 | -0.02 | 0 | 0.7 | 0.49 | 0.81 | 0.8 | 0.69 | 0.77 |
| 1 | 160726891 | rs1328111 | -0.344 | 0.872 | -0.137 | 0.12 | -0.034 | 0.014 | 0.014 | 0 | 0.2 | 0.17 | 0.13 | 0.15 | 0.3 | 0.12 |
| 1 | 160728209 | rs522947 | -0.272 | 0.709 | 0.044 | 0.139 | -0.02 | -0.004 | -0.004 | 0 | 0.94 | 0.89 | 0.98 | 0.97 | 0.98 | 0.94 |

| | | | | | | | | | | | | | | | | |
|---|---|---|---|---|---|---|---|---|---|---|---|---|---|---|---|---|
| 1 | 160732784 | rs564883 | 0.053 | 0.903 | 0.004 | 0.109 | -0.015 | 0.008 | 0.008 | 0 | 0.8 | 0.75 | 0.86 | 0.85 | 0.7 | 0.87 |
| 1 | 160733764 | rs11805079 | 0.386 | 1.101 | -0.216 | 0.324 | -0.043 | -0.023 | -0.023 | 0 | 0.08 | 0.01 | 0.11 | 0.14 | 0.15 | 0.04 |
| 1 | 160734233 | rs518242 | 0.1 | 0.874 | -0.005 | 0.217 | -0.012 | 0.007 | 0.007 | 0 | 0.78 | 0.73 | 0.86 | 0.84 | 0.7 | 0.88 |
| 1 | 160735814 | rs574491 | 0.81 | 0.696 | 0.534 | 0.259 | 0.013 | -0.011 | -0.011 | 0 | 0.7 | 0.52 | 0.83 | 0.77 | 0.67 | 0.81 |
| 1 | 160738146 | rs528095 | 0.832 | 0.706 | 0.733 | 0.263 | 0.021 | -0.019 | -0.019 | 0 | 0.7 | 0.46 | 0.82 | 0.77 | 0.67 | 0.81 |
| 1 | 160744961 | rs497490 | 1.088 | 0.436 | 0.719 | 0.304 | -0.016 | -0.002 | -0.002 | 0 | 0.81 | 0.57 | 0.85 | 0.81 | 0.83 | 0.89 |
| 1 | 160747615 | rs7556098 | -0.74 | 0.804 | 0.678 | 0.277 | -0.015 | 0.003 | 0.003 | 0 | 0.24 | 0.15 | 0.16 | 0.18 | 0.31 | 0.22 |
| 1 | 160764759 | rs11265485 | -1.178 | 0.275 | 0.467 | 0.545 | 0.011 | 0.005 | 0.005 | 0 | 0.56 | 0.55 | 0.69 | 0.63 | 0.72 | 0.52 |
| 1 | 160766925 | rs3904492 | 0.577 | 0.397 | 0.281 | 0.529 | -0.109 | -0.097 | -0.097 | 0.016 | 0.25 | 0.04 | 0.49 | 0.41 | 0.37 | 0.24 |
| 1 | 160770515 | rs6427545 | -1.201 | -0.017 | 0.752 | 0.453 | -0.022 | -0.01 | -0.01 | 0.126 | 0.56 | 0.46 | 0.68 | 0.63 | 0.72 | 0.56 |
| 1 | 160779495 | rs4656932 | 1.314 | 0.164 | 0.399 | 0.617 | -0.052 | -0.074 | -0.074 | 0 | 0.19 | 0.03 | 0.39 | 0.29 | 0.23 | 0.24 |
| 1 | 160786670 | rs560681 | 1.272 | 0.023 | 0.948 | 0.551 | 0.029 | 0.014 | 0.014 | 0 | 0.42 | 0.31 | 0.31 | 0.37 | 0.28 | 0.43 |
| 1 | 160788131 | rs3817407 | 0.263 | 0.148 | 1.319 | 0.792 | 0.002 | 0.041 | 0.041 | 0 | 0.26 | 0.42 | 0.44 | 0.34 | 0.24 | 0.35 |
| 1 | 160791411 | rs571841 | -0.136 | -0.931 | 0.31 | -0.046 | 0.013 | 0.084 | 0.084 | 0 | 0.5 | 0.69 | 0.66 | 0.7 | 0.45 | 0.65 |
| 1 | 160791880 | rs567306 | 0.043 | -0.925 | 0.273 | -0.043 | 0.014 | 0.093 | 0.093 | 0 | 0.5 | 0.68 | 0.67 | 0.7 | 0.45 | 0.65 |
| 1 | 160792659 | rs2779800 | 0.038 | -0.935 | 0.344 | -0.048 | 0.013 | 0.084 | 0.084 | 0 | 0.5 | 0.68 | 0.67 | 0.7 | 0.45 | 0.65 |
| 1 | 160793442 | rs474131 | 0.038 | -0.935 | 0.384 | -0.048 | 0.013 | 0.09 | 0.09 | 0 | 0.5 | 0.69 | 0.66 | 0.7 | 0.45 | 0.65 |
| 1 | 160793560 | rs509749 | -0.076 | -0.928 | 0.158 | -0.045 | 0.005 | 0.112 | 0.112 | 0 | 0.51 | 0.79 | 0.67 | 0.7 | 0.45 | 0.65 |
| 1 | 160800480 | rs485618 | -0.166 | -1.007 | 0.39 | -0.104 | -0.003 | 0.067 | 0.067 | 0 | 0.53 | 0.68 | 0.67 | 0.7 | 0.47 | 0.65 |
| 1 | 160800559 | rs3766377 | 0.026 | -1.068 | 0.569 | 0.035 | -0.006 | 0.029 | 0.029 | 0 | 0.3 | 0.5 | 0.45 | 0.32 | 0.26 | 0.32 |
| 1 | 160803802 | rs11265493 | 0.123 | -0.971 | 0.592 | 0.221 | 0.002 | -0.011 | -0.011 | 2.586 | 0.6 | 0.34 | 0.6 | 0.46 | 0.6 | 0.59 |
| 1 | 160807409 | rs3766378 | -0.647 | -0.954 | 0.77 | 0.288 | -0.039 | 0.034 | 0.034 | 0 | 0.72 | 0.6 | 0.77 | 0.61 | 0.81 | 0.69 |
| 1 | 160807715 | rs3766379 | -0.058 | -1.021 | 0.619 | 0.187 | 0 | 0.014 | 0.014 | 0.085 | 0.6 | 0.57 | 0.61 | 0.46 | 0.56 | 0.62 |
| 1 | 160809285 | rs6671710 | 0.594 | -1.217 | 0.3 | 0.165 | 0.028 | -0.011 | -0.011 | 0 | 0.52 | 0.15 | 0.57 | 0.41 | 0.55 | 0.47 |
| 1 | 160813743 | rs512525 | -0.838 | -1.219 | 0.295 | 0.169 | 0.033 | 0.003 | 0.003 | 0.184 | 0.48 | 0.78 | 0.42 | 0.6 | 0.45 | 0.49 |
| 1 | 160814420 | rs539771 | 1.651 | -1.437 | -0.064 | 0.485 | -0.033 | -0.053 | -0.053 | 0 | 0.08 | 0 | 0.08 | 0.21 | 0.09 | 0.26 |
| 1 | 160816942 | rs549565 | -0.327 | -1.548 | -0.032 | 0.256 | -0.021 | -0.055 | -0.055 | 0 | 0.37 | 0.25 | 0.37 | 0.55 | 0.44 | 0.37 |
| 1 | 160823363 | rs11265496 | -1.343 | -1.546 | 0.015 | 0.256 | 0.032 | 0.003 | 0.003 | 0.101 | 0.48 | 0.8 | 0.42 | 0.6 | 0.45 | 0.51 |
| 1 | 160823812 | rs11582719 | -1.336 | -1.533 | 0.018 | 0.259 | 0.035 | 0.002 | 0.002 | 0.101 | 0.48 | 0.8 | 0.42 | 0.59 | 0.45 | 0.51 |
| 1 | 160825253 | rs4656937 | -0.299 | -1.509 | 0.004 | 0.267 | -0.026 | -0.064 | -0.064 | 0 | 0.36 | 0.23 | 0.36 | 0.55 | 0.44 | 0.37 |
| 1 | 160828285 | rs11265498 | -0.948 | -1.399 | 0.101 | 0.479 | 0.057 | -0.01 | -0.01 | 0 | 0.35 | 0.81 | 0.27 | 0.4 | 0.2 | 0.39 |
| 1 | 160828602 | rs12038600 | -0.486 | -1.226 | 0.203 | 0.564 | -0.027 | 0.015 | 0.015 | 0 | 0.3 | 0.57 | 0.26 | 0.4 | 0.2 | 0.34 |
| 1 | 160831048 | rs4656942 | 2.097 | -0.955 | -0.36 | 0.433 | -0.053 | -0.068 | -0.068 | 2.376 | 0.16 | 0.03 | 0.4 | 0.29 | 0.25 | 0.25 |
| 1 | 160832646 | rs12036607 | -0.07 | -1.166 | 0.14 | 0.489 | -0.01 | 0.009 | 0.009 | 0 | 0.27 | 0.36 | 0.23 | 0.38 | 0.2 | 0.34 |
| 1 | 160832777 | rs12037395 | -0.24 | -1.152 | 0.105 | 0.495 | -0.026 | 0.019 | 0.019 | 0 | 0.28 | 0.46 | 0.24 | 0.39 | 0.2 | 0.31 |
| 1 | 160840298 | rs4656947 | 0.75 | -1.066 | -0.645 | 0.513 | -0.107 | -0.072 | -0.072 | 0 | 0.34 | 0.04 | 0.48 | 0.36 | 0.41 | 0.38 |
| 1 | 160844745 | rs952804 | 0.38 | -0.898 | 0.006 | 0.59 | -0.001 | -0.001 | -0.001 | 0 | 0.65 | 0.34 | 0.69 | 0.62 | 0.68 | 0.62 |
| 1 | 160845701 | rs11265506 | 0.193 | -0.999 | -0.002 | 0.386 | -0.012 | -0.014 | -0.014 | 0 | 0.19 | 0.19 | 0.23 | 0.31 | 0.23 | 0.28 |
| 1 | 160846284 | rs1333062 | 0.562 | -0.929 | -0.266 | 0.341 | 0.001 | 0.008 | 0.008 | 0 | 0.66 | 0.32 | 0.7 | 0.69 | 0.68 | 0.63 |
| 1 | 160847617 | rs4656956 | 0.109 | -0.95 | -0.205 | 0.33 | 0.007 | -0.01 | -0.01 | 0 | 0.7 | 0.47 | 0.72 | 0.7 | 0.68 | 0.7 |
| 1 | 160850936 | rs2236515 | 0.161 | -0.903 | -0.123 | 0.352 | 0.007 | -0.006 | -0.006 | 0 | 0.71 | 0.6 | 0.73 | 0.7 | 0.68 | 0.7 |
| 1 | 160854445 | rs2039415 | 0.191 | -0.788 | -0.101 | 0.455 | 0.016 | -0.032 | -0.032 | 2.646 | 0.71 | 0.54 | 0.73 | 0.74 | 0.68 | 0.71 |
| 1 | 160876923 | rs10908810 | 0.805 | -0.656 | -0.275 | 0.603 | -0.007 | -0.001 | -0.001 | 0 | 0.65 | 0.29 | 0.68 | 0.64 | 0.66 | 0.6 |
| 1 | 160878913 | rs11265519 | 0.356 | -0.513 | -0.199 | 0.583 | 0 | -0.001 | -0.001 | 0 | 0.69 | 0.37 | 0.7 | 0.69 | 0.66 | 0.67 |
| 1 | 160886294 | rs4656279 | 1.096 | -0.407 | -0.459 | 0.412 | -0.023 | 0.026 | 0.026 | 0 | 0.53 | 0.27 | 0.63 | 0.59 | 0.56 | 0.49 |
| 1 | 160892144 | rs11265524 | 0.709 | -0.049 | -0.324 | 0.589 | 0.005 | 0.047 | 0.047 | 1.794 | 0.14 | 0.21 | 0.09 | 0.18 | 0.1 | 0.21 |
| 1 | 160895847 | rs10908813 | -0.033 | 0.135 | -0.263 | 0.529 | 0.034 | -0.017 | -0.017 | 0.105 | 0.75 | 0.6 | 0.73 | 0.75 | 0.67 | 0.77 |
| 1 | 160899481 | rs2774277 | -1.586 | 0.15 | -0.584 | 0.432 | 0.001 | -0.025 | -0.025 | 0 | 0.91 | 0.88 | 0.92 | 0.98 | 0.89 | 0.98 |
| 1 | 160917902 | rs1556519 | 0.806 | 0.006 | -2.157 | 0.772 | -0.047 | -0.017 | -0.017 | 0 | 0.31 | 0.04 | 0.39 | 0.25 | 0.32 | 0.32 |
| 1 | 160921230 | rs4656971 | 0.467 | 0.287 | -1.58 | 0.773 | -0.028 | -0.006 | -0.006 | 0 | 0.55 | 0.24 | 0.63 | 0.56 | 0.57 | 0.52 |
| 1 | 160922195 | rs2252389 | 0.22 | 0.587 | -1.054 | 0.957 | 0.01 | 0.023 | 0.023 | 0.151 | 0.88 | 0.76 | 0.87 | 0.82 | 0.91 | 0.85 |
| 1 | 160922751 | rs4596920 | 0.278 | 0.386 | -2.194 | 0.849 | -0.039 | -0.118 | -0.118 | 4.668 | 0.3 | 0.05 | 0.4 | 0.37 | 0.34 | 0.29 |
| 1 | 160924819 | rs12730072 | 0.413 | 0.474 | -2.154 | 0.582 | -0.058 | 0.037 | 0.037 | 0 | 0.12 | 0.01 | 0.1 | 0.03 | 0.19 | 0.09 |
| 1 | 160927045 | rs2255852 | 1.007 | -0.096 | -1.774 | 0.773 | -0.005 | -0.009 | -0.009 | 0 | 0.56 | 0.57 | 0.5 | 0.33 | 0.59 | 0.45 |
| 1 | 160927294 | rs16832570 | 0.625 | -0.725 | -1.582 | 1.254 | 0.038 | -0.032 | -0.032 | 2.115 | 0.15 | 0.06 | 0.17 | 0.19 | 0.08 | 0.18 |
| 1 | 160927727 | rs1999248 | 0.282 | -0.18 | -1.177 | 1.25 | -0.004 | 0.155 | 0.155 | 0 | 0.84 | 0.66 | 0.77 | 0.55 | 0.85 | 0.73 |
| 1 | 160933065 | rs2483139 | 0.12 | -0.297 | -2.16 | 0.59 | 0.008 | 0.03 | 0.03 | 0 | 0.78 | 0.89 | 0.78 | 0.83 | 0.76 | 0.76 |
| 1 | 160933467 | rs2256633 | 0.192 | -0.229 | -2.616 | 0.444 | -0.033 | 0.013 | 0.013 | 0 | 0.56 | 0.78 | 0.49 | 0.63 | 0.45 | 0.61 |
| 1 | 160937964 | rs12403334 | 1.321 | -0.189 | -2.665 | 0.468 | -0.073 | -0.035 | -0.035 | 0 | 0.21 | 0.05 | 0.29 | 0.2 | 0.32 | 0.14 |
| 1 | 160950966 | rs2250174 | -0.301 | -0.35 | -2.696 | 0.392 | 0.006 | 0.029 | 0.029 | 0 | 0.78 | 0.9 | 0.79 | 0.83 | 0.77 | 0.76 |
| 1 | 160956745 | rs2990701 | 0.094 | -0.282 | -2.686 | 0.392 | -0.065 | 0.005 | 0.005 | 0 | 0.57 | 0.82 | 0.51 | 0.63 | 0.48 | 0.62 |
| 1 | 160959844 | rs12140181 | 0.983 | -0.603 | -3.161 | 0.116 | -0.139 | 0.062 | 0.062 | 0 | 0.24 | 0.01 | 0.17 | 0.14 | 0.3 | 0.28 |
| 1 | 160966844 | rs1062827 | 1.62 | -0.54 | -2.93 | 0.304 | -0.062 | -0.036 | -0.036 | 0 | 0.2 | 0.05 | 0.28 | 0.2 | 0.3 | 0.13 |
| 1 | 160966963 | rs1044478 | 1.62 | -0.54 | -2.93 | 0.304 | -0.062 | -0.036 | -0.036 | 0.171 | 0.2 | 0.05 | 0.28 | 0.2 | 0.3 | 0.13 |
| 1 | 160969930 | rs869985 | 1.62 | -0.54 | -2.93 | 0.314 | -0.062 | -0.034 | -0.034 | 0.657 | 0.2 | 0.05 | 0.28 | 0.2 | 0.3 | 0.13 |
| 1 | 160985708 | rs2039223 | -0.464 | -0.958 | -2.809 | 0.171 | 0.01 | 0.014 | 0.014 | 0 | 0.78 | 0.82 | 0.79 | 0.84 | 0.79 | 0.76 |
| 1 | 160989614 | rs2481084 | 1.49 | -0.655 | -2.804 | 0.263 | -0.015 | -0.007 | -0.007 | 0.026 | 0.21 | 0.15 | 0.28 | 0.2 | 0.29 | 0.13 |
| 1 | 160991821 | rs6695707 | 1.344 | -1.243 | -3.069 | 0.612 | 0.01 | -0.079 | -0.079 | 0 | 0.16 | 0 | 0.18 | 0.2 | 0.13 | 0.18 |
| 1 | 161003874 | rs11576837 | 0.49 | -1.316 | -2.856 | 0.493 | 0.112 | 0.071 | 0.071 | 0 | 0.48 | 0.15 | 0.46 | 0.33 | 0.64 | 0.45 |

| | | | | | | | | | | | | | | | | |
|---|---|---|---|---|---|---|---|---|---|---|---|---|---|---|---|---|
| 1 | 161006019 | rs11265544 | 1.108 | -1.203 | -2.965 | 0.384 | -0.14 | 0.085 | 0.085 | 0 | 0.26 | 0.03 | 0.19 | 0.14 | 0.34 | 0.29 |
| 1 | 161009523 | rs3737787 | 1.992 | -1.129 | -2.993 | 0.43 | -0.065 | -0.05 | -0.05 | 0 | 0.2 | 0.04 | 0.26 | 0.19 | 0.29 | 0.13 |
| 1 | 161010762 | rs2073658 | 1.992 | -1.129 | -2.993 | 0.43 | -0.065 | -0.05 | -0.05 | 0 | 0.2 | 0.04 | 0.27 | 0.19 | 0.29 | 0.13 |
| 1 | 161012760 | rs2073653 | 0.827 | -1.494 | -2.725 | 0.482 | -0.038 | 0.035 | 0.035 | 0 | 0.26 | 0.63 | 0.32 | 0.5 | 0.13 | 0.25 |
| 1 | 161014727 | rs2516837 | 0.444 | -1.225 | -2.622 | 0.51 | 0.105 | 0.066 | 0.066 | 0 | 0.48 | 0.19 | 0.47 | 0.33 | 0.63 | 0.46 |
| 1 | 161017556 | rs2774279 | 0.964 | -1.155 | -2.66 | 0.492 | -0.064 | 0.069 | 0.069 | 0 | 0.27 | 0.14 | 0.2 | 0.14 | 0.33 | 0.3 |
| 1 | 161027492 | rs10737178 | -1.14 | -1.206 | -2.715 | 0.474 | -0.006 | -0.038 | -0.038 | 0.722 | 0.9 | 1 | 0.91 | 0.85 | 0.93 | 0.84 |
| 1 | 161030806 | rs17384360 | 1.75 | -0.592 | -2.686 | 0.49 | -0.078 | -0.042 | -0.042 | 0 | 0.2 | 0.04 | 0.27 | 0.18 | 0.31 | 0.14 |
| 1 | 161037042 | rs10797092 | -0.075 | -1.116 | -2.446 | 0.91 | 0.014 | -0.045 | -0.045 | 0 | 0.8 | 0.89 | 0.77 | 0.67 | 0.8 | 0.78 |
| 1 | 161037857 | rs16832617 | 0.495 | -1.573 | -2.489 | 0.851 | -0.044 | 0.012 | 0.012 | 0 | 0.28 | 0.6 | 0.28 | 0.35 | 0.13 | 0.36 |
| 1 | 161037979 | rs12118313 | 0.543 | -0.591 | -2.295 | 0.789 | 0.028 | -0.039 | -0.039 | 0.886 | 0.76 | 0.86 | 0.77 | 0.67 | 0.78 | 0.74 |
| 1 | 161042063 | rs12116949 | 0.331 | 0.018 | -2.517 | 0.369 | -0.129 | 0.053 | 0.053 | 0 | 0.23 | 0.01 | 0.15 | 0.13 | 0.29 | 0.33 |
| 1 | 161043764 | rs4656978 | 0.826 | -0.261 | -2.123 | 0.525 | 0.006 | -0.006 | -0.006 | 0 | 0.74 | 0.74 | 0.76 | 0.66 | 0.76 | 0.71 |
| 1 | 161045128 | rs4559524 | -0.16 | -0.075 | -2.234 | 0.496 | 0 | -0.018 | -0.018 | 0 | 0.07 | 0.12 | 0.11 | 0.01 | 0.13 | 0.05 |
| 1 | 161051526 | rs11265549 | -0.795 | -0.06 | -2.318 | 0.677 | -0.004 | -0.051 | -0.051 | 0.035 | 0.9 | 0.99 | 0.88 | 0.68 | 0.91 | 0.82 |
| 1 | 161051558 | rs12401309 | -0.603 | -0.054 | -2.315 | 0.691 | 0.006 | -0.054 | -0.054 | 0 | 0.9 | 0.99 | 0.89 | 0.68 | 0.93 | 0.82 |
| 1 | 161053773 | rs3892376 | -0.167 | -0.197 | -2.392 | 0.591 | 0.023 | -0.094 | -0.094 | 0.021 | 0.83 | 0.99 | 0.87 | 0.68 | 0.9 | 0.76 |
| 1 | 161054054 | rs3892375 | 0.322 | 0.174 | -2.539 | 0.015 | -0.125 | 0.077 | 0.077 | 0 | 0.22 | 0.01 | 0.15 | 0.08 | 0.29 | 0.34 |
| 1 | 161056618 | rs10159127 | -0.401 | 0.393 | -2.367 | 0.106 | -0.044 | 0.087 | 0.087 | 0 | 0.27 | 0.07 | 0.13 | 0.09 | 0.26 | 0.29 |
| 1 | 161057909 | rs1411614 | -0.468 | 0.481 | -2.258 | 0.089 | -0.029 | 0.063 | 0.063 | 0 | 0.28 | 0.13 | 0.15 | 0.1 | 0.28 | 0.29 |
| 1 | 161062264 | rs12739089 | -1.116 | 0.498 | -2.247 | 0.341 | -0.03 | 0.02 | 0.02 | 0 | 0.38 | 0.27 | 0.34 | 0.32 | 0.49 | 0.47 |
| 1 | 161064008 | rs4656287 | 0.041 | 0.449 | -2.032 | 0.702 | -0.009 | -0.042 | -0.042 | 0 | 0.08 | 0.01 | 0.1 | 0.22 | 0.07 | 0.19 |
| 1 | 161066446 | rs4656288 | 0.041 | 0.449 | -2.032 | 0.702 | -0.009 | -0.042 | -0.042 | 0 | 0.08 | 0.01 | 0.11 | 0.22 | 0.07 | 0.19 |
| 1 | 161074601 | rs4233365 | -1.208 | 0.445 | -2.149 | 0.819 | -0.032 | -0.022 | -0.022 | 0 | 0.4 | 0.25 | 0.37 | 0.45 | 0.43 | 0.37 |
| 1 | 161084007 | rs11265554 | -1.217 | 0.425 | -2.153 | 0.809 | -0.032 | -0.022 | -0.022 | 0 | 0.4 | 0.24 | 0.37 | 0.45 | 0.43 | 0.37 |
| 1 | 161092761 | rs17389237 | 0.817 | 0.443 | -2.206 | 0.794 | 0.005 | 0.015 | 0.015 | 0 | 0.12 | 0.11 | 0.07 | 0.07 | 0.07 | 0.14 |
| 1 | 161094190 | rs1135783 | 0.791 | 0.443 | -2.206 | 0.794 | 0.005 | 0.015 | 0.015 | 0 | 0.12 | 0.11 | 0.07 | 0.07 | 0.07 | 0.14 |
| 1 | 161098128 | rs11265555 | -1.528 | 0.676 | -2.235 | 0.766 | -0.019 | -0.027 | -0.027 | 0.013 | 0.39 | 0.25 | 0.37 | 0.45 | 0.4 | 0.36 |
| 1 | 161111931 | rs10908824 | 0.225 | 0.685 | -2.263 | 0.788 | -0.091 | -0.118 | -0.118 | 0 | 0.2 | 0.03 | 0.29 | 0.38 | 0.29 | 0.18 |
| 1 | 161116022 | rs11589567 | 0.182 | 0.687 | -2.26 | 0.789 | -0.087 | -0.114 | -0.114 | 0.748 | 0.2 | 0.04 | 0.3 | 0.37 | 0.29 | 0.18 |
| 1 | 161122903 | rs4656988 | -1.536 | 0.7 | -2.239 | 0.797 | -0.022 | -0.03 | -0.03 | 0 | 0.39 | 0.24 | 0.38 | 0.45 | 0.4 | 0.36 |
| 1 | 161135559 | rs2502806 | -2.259 | 0.5 | -2.641 | 0.093 | -0.113 | 0.098 | 0.098 | 0 | 0.7 | 0.98 | 0.74 | 0.88 | 0.63 | 0.8 |
| 1 | 161136224 | rs2301286 | -1.361 | 0.843 | -2.174 | 0.394 | -0.015 | -0.02 | -0.02 | 0 | 0.41 | 0.25 | 0.38 | 0.45 | 0.4 | 0.36 |
| 1 | 161143312 | rs11579514 | -0.226 | 0.966 | -2.146 | 0.484 | -0.042 | -0.061 | -0.061 | 0 | 0.25 | 0.11 | 0.31 | 0.38 | 0.32 | 0.21 |
| 1 | 161145005 | rs3813620 | 0.742 | 0.501 | -1.932 | 0.488 | 0.007 | 0.016 | 0.016 | 0 | 0.12 | 0.1 | 0.07 | 0.07 | 0.07 | 0.13 |
| 1 | 161147771 | rs17392570 | 1.077 | 0.474 | -1.962 | 0.497 | 0.01 | 0.023 | 0.023 | 0 | 0.11 | 0.07 | 0.06 | 0.07 | 0.07 | 0.14 |
| 1 | 161155392 | rs4575098 | 0.549 | 0.791 | -2.104 | 0.738 | -0.064 | -0.106 | -0.106 | 0 | 0.17 | 0.03 | 0.27 | 0.35 | 0.24 | 0.11 |
| 1 | 161156033 | rs11585858 | 0.549 | 0.785 | -2.104 | 0.738 | -0.062 | -0.106 | -0.106 | 0 | 0.17 | 0.03 | 0.27 | 0.35 | 0.24 | 0.11 |
| 1 | 161159147 | rs4233366 | 0.093 | 0.364 | -1.848 | 0.374 | -0.007 | 0.015 | 0.015 | 0 | 0.33 | 0.41 | 0.37 | 0.45 | 0.27 | 0.48 |
| 1 | 161162173 | rs7512012 | -1.846 | 0.321 | -1.822 | 0.19 | 0.025 | -0.082 | -0.082 | 0 | 0.68 | 0.6 | 0.71 | 0.89 | 0.61 | 0.78 |
| 1 | 161163037 | rs4233367 | -1.829 | 0.336 | -1.837 | 0.188 | 0.021 | -0.078 | -0.078 | 0 | 0.68 | 0.59 | 0.71 | 0.89 | 0.61 | 0.78 |
| 1 | 161167698 | rs4589131 | -1.967 | 0.297 | -1.724 | 0.206 | 0.022 | -0.068 | -0.068 | 0.955 | 0.68 | 0.64 | 0.71 | 0.89 | 0.59 | 0.78 |
| 1 | 161172525 | rs10908826 | 1.353 | 0.198 | -1.802 | 0.298 | -0.011 | -0.081 | -0.081 | 0.003 | 0.13 | 0.04 | 0.23 | 0.35 | 0.14 | 0.09 |
| 1 | 161176137 | rs4656993 | -1.908 | 0.224 | -1.782 | 0.237 | 0.021 | -0.069 | -0.069 | 0.301 | 0.67 | 0.61 | 0.69 | 0.89 | 0.58 | 0.78 |
| 1 | 161178684 | rs3924264 | -0.294 | 0.254 | -1.802 | 0.228 | 0 | 0.005 | 0.005 | 0 | 0.39 | 0.44 | 0.4 | 0.46 | 0.38 | 0.52 |
| 1 | 161179877 | rs4656994 | -0.498 | 0.251 | -1.823 | 0.246 | 0.028 | -0.044 | -0.044 | 0 | 0.28 | 0.17 | 0.29 | 0.43 | 0.21 | 0.26 |
| 1 | 161182208 | rs11576415 | 1.212 | 0.317 | -1.881 | 0.207 | -0.037 | 0.022 | 0.022 | 0 | 0.06 | 0.01 | 0.03 | 0.01 | 0.11 | 0.1 |
| 1 | 161182672 | rs10797094 | -2.118 | 0.293 | -1.852 | 0.19 | 0.019 | -0.06 | -0.06 | 0.192 | 0.68 | 0.65 | 0.7 | 0.89 | 0.59 | 0.78 |
| 1 | 161183516 | rs1136207 | 1.32 | 0.385 | -1.986 | 0.342 | -0.011 | -0.082 | -0.082 | 0 | 0.13 | 0.03 | 0.23 | 0.36 | 0.14 | 0.09 |
| 1 | 161184097 | rs1136224 | 0.773 | 0.022 | -1.885 | 0.393 | -0.009 | -0.056 | -0.056 | 0.182 | 0.18 | 0.03 | 0.22 | 0.28 | 0.15 | 0.3 |
| 1 | 161184875 | rs11587213 | -0.325 | 0.295 | -1.653 | 0.3 | -0.015 | 0.046 | 0.046 | 0.036 | 0.19 | 0.1 | 0.1 | 0.09 | 0.18 | 0.27 |
| 1 | 161186303 | rs12094497 | 0.162 | 0.47 | -1.476 | 0.473 | -0.007 | 0.005 | 0.005 | 0 | 0.08 | 0.07 | 0.04 | 0.01 | 0.11 | 0.1 |
| 1 | 161187665 | rs2070902 | 0.01 | 0.094 | -1.354 | 0.736 | -0.012 | 0.014 | 0.014 | 0.017 | 0.31 | 0.41 | 0.35 | 0.4 | 0.26 | 0.4 |
| 1 | 161189713 | rs4489574 | -0.057 | 0.347 | -1.302 | 0.744 | -0.001 | 0.001 | 0.001 | 0 | 0.35 | 0.37 | 0.37 | 0.42 | 0.34 | 0.41 |
| 1 | 161193683 | rs5082 | -1.732 | 0.395 | -1.227 | 0.592 | -0.001 | -0.01 | -0.01 | 0 | 0.72 | 0.78 | 0.71 | 0.9 | 0.61 | 0.78 |
| 1 | 161197244 | rs4233368 | 0.533 | 0.056 | -1.147 | 0.74 | -0.015 | 0.023 | 0.023 | 0 | 0.31 | 0.45 | 0.37 | 0.36 | 0.26 | 0.39 |
| 1 | 161200487 | rs4073054 | -1.862 | 0.26 | -1.27 | 0.517 | 0 | -0.012 | -0.012 | 1.218 | 0.73 | 0.78 | 0.72 | 0.9 | 0.62 | 0.79 |
| 1 | 161200586 | rs2307418 | 0.778 | -0.077 | -1.895 | 0.529 | -0.034 | 0.013 | 0.013 | 0 | 0.15 | 0.01 | 0.08 | 0.09 | 0.15 | 0.11 |
| 1 | 161202556 | rs2307420 | 2.942 | -0.171 | -1.842 | 0.348 | 0.015 | 0.011 | 0.011 | 0 | 0.06 | 0.05 | 0.01 | 0 | 0.01 | 0.05 |
| 1 | 161202605 | rs2307424 | -0.779 | -0.08 | -1.699 | 0.625 | -0.026 | -0.094 | -0.094 | 0 | 0.32 | 0.11 | 0.33 | 0.52 | 0.35 | 0.45 |
| 1 | 161204538 | rs2501873 | -0.487 | 0.149 | -1.486 | 0.67 | 0.003 | -0.014 | -0.014 | 0 | 0.46 | 0.69 | 0.48 | 0.58 | 0.44 | 0.48 |
| 1 | 161209841 | rs2502805 | 1.082 | 0.079 | -1.283 | 0.643 | 0.002 | 0.019 | 0.019 | 0 | 0.27 | 0.35 | 0.35 | 0.35 | 0.27 | 0.29 |
| 1 | 161210729 | rs7530560 | -1.582 | -0.086 | -1.29 | 0.602 | -0.004 | -0.002 | -0.002 | 1.326 | 0.39 | 0.46 | 0.41 | 0.6 | 0.36 | 0.43 |
| 1 | 161211678 | rs12562860 | 0.214 | -0.096 | -1.244 | 0.626 | 0 | 0.002 | 0.002 | 0.103 | 0.21 | 0.21 | 0.29 | 0.37 | 0.19 | 0.25 |
| 1 | 161211909 | rs10157822 | -1.529 | -0.03 | -1.27 | 0.653 | -0.004 | -0.002 | -0.002 | 0.549 | 0.39 | 0.46 | 0.41 | 0.6 | 0.36 | 0.43 |
| 1 | 161212453 | rs11584174 | -0.266 | 0.217 | -1.034 | 0.969 | 0 | 0.001 | 0.001 | 0 | 0.14 | 0.13 | 0.11 | 0.23 | 0.17 | 0.18 |
| 1 | 161212569 | rs2501870 | 0.852 | 0.266 | -1.037 | 0.957 | -0.002 | 0.019 | 0.019 | 0 | 0.27 | 0.35 | 0.35 | 0.35 | 0.26 | 0.3 |
| 1 | 161215212 | rs12029543 | -0.928 | 0.378 | -1.081 | 1.111 | -0.001 | 0 | 0 | 0 | 0.15 | 0.13 | 0.11 | 0.23 | 0.17 | 0.18 |
| 1 | 161217657 | rs12041273 | -0.441 | 0.569 | -1.548 | 1.203 | -0.042 | -0.053 | -0.053 | 1.099 | 0.13 | 0.01 | 0.1 | 0.23 | 0.17 | 0.18 |

| | | | | | | | | | | | | | | | | |
|---|---|---|---|---|---|---|---|---|---|---|---|---|---|---|---|---|
| 1 | 161228800 | rs4078110 | -0.784 | 0.468 | -1.504 | 1.2 | 0.004 | -0.022 | -0.022 | 0.145 | 0.25 | 0.19 | 0.32 | 0.36 | 0.23 | 0.33 |
| 1 | 161228843 | rs16832733 | 0.214 | 0.679 | -1.592 | 1.127 | -0.031 | -0.036 | -0.036 | 0 | 0.08 | 0.01 | 0.08 | 0.27 | 0.12 | 0.1 |
| 1 | 161229651 | rs4078109 | -0.41 | 0.322 | -1.846 | 0.14 | -0.035 | 0.079 | 0.079 | 0 | 0.57 | 0.77 | 0.57 | 0.73 | 0.54 | 0.55 |
| 1 | 161230065 | rs2501865 | -0.617 | 0.027 | -1.836 | -0.125 | 0.014 | -0.008 | -0.008 | 0 | 0.28 | 0.18 | 0.21 | 0.27 | 0.26 | 0.28 |
| 1 | 161232772 | rs2501862 | -1.333 | 0.453 | -1.486 | -0.53 | -0.002 | 0.016 | 0.016 | 0 | 0.43 | 0.42 | 0.32 | 0.29 | 0.4 | 0.34 |
| 1 | 161234623 | rs11265577 | -0.509 | 0.042 | -1.424 | -0.912 | 0.006 | 0.006 | 0.006 | 0 | 0.28 | 0.22 | 0.19 | 0.03 | 0.29 | 0.2 |
| 1 | 161234796 | rs2501880 | 0.547 | 0.042 | -1.289 | -0.963 | -0.002 | 0.003 | 0.003 | 0 | 0.15 | 0.18 | 0.1 | 0 | 0.1 | 0.1 |
| 1 | 161234965 | rs2501879 | -1.96 | 0.123 | -1.288 | -1.106 | -0.004 | 0.087 | 0.087 | 0.056 | 0.73 | 0.92 | 0.82 | 0.96 | 0.73 | 0.79 |
| 1 | 161236636 | rs10494345 | 0.497 | -0.699 | -0.864 | -0.619 | 0.003 | 0.006 | 0.006 | 0.259 | 0.15 | 0.12 | 0.11 | 0.02 | 0.15 | 0.14 |
| 1 | 161239143 | rs2502812 | -0.255 | -0.713 | -0.543 | -0.073 | 0.001 | 0.001 | 0.001 | 0.206 | 0.38 | 0.62 | 0.54 | 0.5 | 0.41 | 0.36 |
| 1 | 161239471 | rs2501875 | -0.048 | -0.74 | -0.529 | -0.218 | 0.006 | 0.005 | 0.005 | 0 | 0.36 | 0.6 | 0.54 | 0.43 | 0.39 | 0.35 |
| 1 | 161249787 | rs4271232 | 0.052 | -0.735 | -0.649 | -0.245 | 0.005 | 0.004 | 0.004 | 0 | 0.33 | 0.55 | 0.55 | 0.66 | 0.34 | 0.44 |
| 1 | 161253067 | rs4657009 | -0.248 | -0.93 | -0.585 | -0.404 | 0.008 | -0.006 | -0.006 | 0 | 0.38 | 0.55 | 0.58 | 0.66 | 0.4 | 0.47 |
| 1 | 161253657 | rs4255401 | -0.248 | -0.93 | -0.607 | -0.404 | 0.006 | -0.004 | -0.004 | 0 | 0.38 | 0.55 | 0.58 | 0.66 | 0.4 | 0.47 |
| 1 | 161260215 | rs4436408 | 0.889 | -1.128 | -0.515 | -0.45 | -0.004 | 0.031 | 0.031 | 0 | 0.24 | 0.33 | 0.48 | 0.67 | 0.25 | 0.4 |
| 1 | 161261041 | rs11265584 | -0.422 | -0.855 | -0.418 | 0.027 | 0.01 | 0.016 | 0.016 | 0 | 0.15 | 0.25 | 0.15 | 0.22 | 0.14 | 0.2 |
| 1 | 161263380 | rs12093075 | -0.53 | -0.789 | -0.402 | 0.003 | 0.007 | 0.014 | 0.014 | 0.389 | 0.17 | 0.33 | 0.15 | 0.23 | 0.14 | 0.2 |
| 1 | 161264923 | rs12098224 | -0.515 | -0.837 | -0.395 | 0.01 | 0.006 | 0.015 | 0.015 | 0.357 | 0.17 | 0.33 | 0.16 | 0.23 | 0.14 | 0.2 |
| 1 | 161269157 | rs11265585 | -0.417 | -0.86 | -0.899 | -1.019 | -0.023 | 0.165 | 0.165 | 1.378 | 0.6 | 0.9 | 0.76 | 0.96 | 0.57 | 0.74 |
| 1 | 161270716 | rs6427588 | 0.637 | -0.19 | -0.389 | -0.532 | -0.009 | -0.01 | -0.01 | 0 | 0.43 | 0.5 | 0.58 | 0.67 | 0.4 | 0.51 |
| 1 | 161272441 | rs4657015 | 0.039 | -0.265 | -0.343 | -0.329 | 0.002 | 0.017 | 0.017 | 0 | 0.59 | 0.71 | 0.71 | 0.76 | 0.58 | 0.66 |
| 1 | 161274311 | rs7532602 | -1.206 | -0.745 | 0.366 | 0.317 | -0.002 | 0.004 | 0.004 | 0 | 0.21 | 0.41 | 0.14 | 0.14 | 0.14 | 0.23 |
| 1 | 161274905 | rs16832786 | -1.13 | -0.632 | 0.133 | -0.098 | -0.008 | -0.033 | -0.033 | 0 | 0.11 | 0.21 | 0.09 | 0.05 | 0.11 | 0.19 |
| 1 | 161280249 | rs3813630 | -0.902 | -0.826 | -0.012 | 0.271 | -0.001 | -0.002 | -0.002 | 0 | 0.25 | 0.24 | 0.13 | 0.36 | 0.12 | 0.37 |
| 1 | 161280463 | rs12064957 | -0.294 | -0.805 | -0.061 | 0.286 | 0.012 | 0.004 | 0.004 | 0 | 0.19 | 0.15 | 0.08 | 0.19 | 0.07 | 0.21 |
| 1 | 161284297 | rs11265589 | -0.107 | -0.765 | -0.123 | 0.323 | 0.036 | 0.006 | 0.006 | 0 | 0.17 | 0.09 | 0.07 | 0.15 | 0.07 | 0.15 |
| 1 | 161291412 | rs12069801 | -0.064 | -0.756 | -0.12 | 0.327 | 0.034 | 0.005 | 0.005 | 0 | 0.17 | 0.09 | 0.07 | 0.15 | 0.07 | 0.15 |
| 1 | 161312656 | rs11585941 | -0.47 | -0.818 | 0.484 | 0.211 | 0.011 | -0.017 | -0.017 | 0 | 0.09 | 0.24 | 0.09 | 0.04 | 0.11 | 0.14 |
| 1 | 161333191 | rs4600063 | -0.887 | -0.92 | 0.387 | 0.221 | -0.007 | 0.095 | 0.095 | 0.04 | 0.08 | 0.25 | 0.06 | 0.21 | 0.05 | 0.22 |
| 1 | 161333558 | rs3935401 | -0.887 | -0.92 | 0.387 | 0.221 | -0.007 | 0.095 | 0.095 | 0.04 | 0.08 | 0.25 | 0.06 | 0.21 | 0.05 | 0.22 |
| 1 | 161334658 | rs1133805 | -0.414 | -0.92 | 0.399 | 0.204 | -0.006 | 0.033 | 0.033 | 0 | 0.06 | 0.1 | 0.04 | 0.21 | 0.05 | 0.22 |
| 1 | 161354580 | rs16832911 | -0.153 | -0.882 | 0.45 | 0.201 | 0.007 | -0.039 | -0.039 | 0 | 0.08 | 0.13 | 0.08 | 0 | 0.12 | 0.13 |
| 1 | 161357471 | rs16832922 | -1.377 | -0.499 | -0.342 | 0.11 | 0.005 | 0.002 | 0.002 | 0 | 0.18 | 0.25 | 0.14 | 0.21 | 0.2 | 0.29 |
| 1 | 161360435 | rs4077631 | -0.599 | -0.383 | -0.288 | 0.172 | -0.007 | 0.099 | 0.099 | 0.087 | 0.08 | 0.25 | 0.06 | 0.21 | 0.05 | 0.22 |
| 1 | 161361979 | rs12744234 | 0.201 | -0.134 | -0.746 | 0.129 | -0.049 | 0.045 | 0.045 | 0 | 0.08 | 0 | 0.08 | 0 | 0.15 | 0.06 |
| 1 | 161370985 | rs12745664 | -0.956 | -0.273 | -0.448 | 0.281 | 0.041 | 0.063 | 0.063 | 0 | 0.48 | 0.74 | 0.32 | 0.36 | 0.39 | 0.58 |
| 1 | 161394500 | rs3922744 | -0.408 | -0.261 | -0.383 | 0.928 | 0 | 0.002 | 0.002 | 0 | 0.53 | 0.43 | 0.43 | 0.42 | 0.39 | 0.44 |
| 1 | 161398477 | rs10157070 | 0 | -1.045 | -0.675 | 0.079 | 0.008 | -0.011 | -0.011 | 1.678 | 0.18 | 0.14 | 0.09 | 0 | 0.12 | 0.07 |
| 1 | 161398984 | rs11265601 | -0.053 | -1.046 | -0.649 | 0.122 | 0.011 | 0.009 | 0.009 | 0 | 0.18 | 0.16 | 0.22 | 0.02 | 0.12 | 0.07 |
| 1 | 161406056 | rs9427374 | -0.781 | -1.375 | -0.531 | -0.311 | 0.002 | -0.029 | -0.029 | 0 | 0.32 | 0.32 | 0.3 | 0.02 | 0.25 | 0.12 |
| 1 | 161408556 | rs12125079 | 1.128 | -1.554 | -0.584 | -0.348 | 0.019 | 0.013 | 0.013 | 0 | 0.44 | 0.33 | 0.44 | 0.32 | 0.67 | 0.41 |
| 1 | 161410458 | rs12731669 | -1.021 | -1.195 | -0.584 | -0.302 | -0.05 | 0.08 | 0.08 | 0 | 0.87 | 1 | 0.94 | 1 | 0.87 | 0.94 |
| 1 | 161456186 | rs6698806 | 0.301 | -0.884 | -1.052 | -0.512 | -0.001 | 0.002 | 0.002 | 0 | 0.64 | 0.62 | 0.59 | 0.74 | 0.64 | 0.78 |
| 1 | 161462728 | rs4657039 | 0.671 | -1.359 | -1.012 | -0.582 | 0.002 | 0.02 | 0.02 | 0 | 0.45 | 0.35 | 0.57 | 0.22 | 0.61 | 0.44 |
| 1 | 161463814 | rs6696854 | -0.285 | -1.415 | -1.321 | -0.56 | 0 | -0.001 | -0.001 | 0 | 0.72 | 0.74 | 0.73 | 0.26 | 0.82 | 0.54 |
| 1 | 161472158 | rs10800309 | -0.253 | -1.131 | -1.49 | -0.694 | 0.003 | 0.003 | 0.003 | 0 | 0.67 | 0.74 | 0.52 | 0.25 | 0.7 | 0.53 |
| 1 | 161478708 | rs6671753 | 0.68 | -0.816 | -1.86 | -0.706 | -0.017 | 0.073 | 0.073 | 0 | 0.18 | 0.05 | 0.07 | 0 | 0.14 | 0.07 |
| 1 | 161479599 | rs7529425 | 0.711 | -0.806 | -1.866 | -0.701 | -0.019 | 0.076 | 0.076 | 0 | 0.18 | 0.05 | 0.07 | 0 | 0.14 | 0.07 |
| 1 | 161479745 | rs1801274 | 0.287 | -0.667 | -1.169 | -0.155 | 0.009 | -0.011 | -0.011 | 0.255 | 0.41 | 0.53 | 0.45 | 0.28 | 0.51 | 0.42 |
| 1 | 161490896 | rs511278 | -1.039 | 0.23 | -1.565 | -0.284 | -0.007 | 0.022 | 0.022 | 0.09 | 0.85 | 0.91 | 0.74 | 0.91 | 0.8 | 0.83 |
| 1 | 161492587 | rs12129787 | 0.116 | 0.872 | -1.947 | 0.211 | -0.022 | -0.026 | -0.026 | 0.062 | 0.27 | 0.22 | 0.4 | 0.37 | 0.35 | 0.35 |
| 1 | 161500130 | rs2099684 | -0.156 | 0.298 | -2.154 | 0.031 | -0.029 | -0.013 | -0.013 | 0 | 0.35 | 0.14 | 0.24 | 0.36 | 0.36 | 0.32 |
| 1 | 161646807 | rs11799952 | -0.384 | 0.104 | -1.651 | -0.754 | -0.004 | -0.043 | -0.043 | 0 | 0.17 | 0.21 | 0.1 | 0.02 | 0.12 | 0.19 |
| 1 | 161646824 | rs12118043 | 1.35 | 1.371 | -2.215 | -0.71 | -0.06 | -0.031 | -0.031 | 0 | 0.09 | 0.01 | 0.1 | 0.06 | 0.2 | 0.11 |
| 1 | 161653708 | rs11801845 | 0.432 | 0.767 | -1.657 | -0.857 | -0.027 | -0.034 | -0.034 | 0 | 0.09 | 0.14 | 0.04 | 0.02 | 0.02 | 0.08 |
| 1 | 161655342 | rs1832739 | -0.861 | 0.116 | -1.947 | -0.648 | -0.015 | -0.001 | -0.001 | 0 | 0.55 | 0.8 | 0.4 | 0.53 | 0.52 | 0.54 |
| 1 | 161660696 | rs6697139 | 1.694 | 0.312 | -2.438 | -0.954 | -0.064 | 0.125 | 0.125 | 0 | 0.19 | 0.02 | 0.17 | 0.01 | 0.22 | 0.21 |
| 1 | 161661489 | rs1340976 | -1.803 | 0.215 | -2.094 | -0.679 | -0.013 | -0.076 | -0.076 | 0.42 | 0.81 | 0.94 | 0.59 | 0.54 | 0.78 | 0.79 |
| 1 | 161662484 | rs4657090 | 0.238 | 0.355 | -1.089 | 0.169 | 0.037 | 0.015 | 0.015 | 0 | 0.2 | 0.29 | 0.21 | 0.26 | 0.28 | 0.27 |
| 1 | 161669427 | rs12078337 | 0.901 | -0.221 | -0.859 | 0.436 | 0 | 0.012 | 0.012 | 0 | 0.09 | 0.06 | 0.07 | 0.18 | 0.11 | 0.09 |
| 1 | 161673508 | rs6699818 | -0.56 | -0.546 | -0.544 | -0.011 | -0.058 | -0.039 | -0.039 | 0 | 0.16 | 0.33 | 0.06 | 0.08 | 0.03 | 0.13 |
| 1 | 161675580 | rs17413333 | -0.012 | 0.334 | -1.245 | 0.373 | -0.081 | -0.07 | -0.07 | 0.348 | 0.34 | 0.1 | 0.53 | 0.34 | 0.4 | 0.33 |
| 1 | 161676394 | rs1538971 | 0.46 | 0.132 | -0.541 | -0.07 | -0.016 | -0.042 | -0.042 | 0 | 0.62 | 0.5 | 0.5 | 0.83 | 0.69 | 0.55 |
| 1 | 161676557 | rs1538972 | -1.077 | 0.01 | -0.216 | -0.131 | -0.003 | -0.03 | -0.03 | 0 | 0.8 | 0.79 | 0.92 | 0.97 | 0.88 | 0.86 |
| 1 | 161676970 | rs1340978 | 0.615 | 0.039 | -0.181 | -0.144 | -0.005 | -0.017 | -0.017 | 0 | 0.62 | 0.6 | 0.51 | 0.83 | 0.69 | 0.55 |
| 1 | 161678254 | rs12135523 | 1.423 | 0.135 | -0.763 | 0.066 | -0.016 | 0.166 | 0.166 | 0.762 | 0.49 | 0.1 | 0.37 | 0.21 | 0.58 | 0.45 |
| 1 | 161679644 | rs7523762 | -0.345 | 0.643 | -0.523 | -0.038 | 0 | -0.062 | -0.062 | 0.477 | 0.76 | 0.68 | 0.86 | 0.97 | 0.77 | 0.71 |
| 1 | 161680258 | rs6427618 | 0.716 | 0.611 | -0.957 | 0.201 | 0.041 | 0.046 | 0.046 | 0 | 0.58 | 0.18 | 0.48 | 0.36 | 0.66 | 0.63 |
| 1 | 161680868 | rs1934908 | 1.43 | 0.605 | -0.971 | 0.21 | -0.106 | 0.121 | 0.121 | 0.113 | 0.49 | 0.05 | 0.35 | 0.24 | 0.52 | 0.45 |

| | | | | | | | | | | | | | | | | |
|---|---|---|---|---|---|---|---|---|---|---|---|---|---|---|---|---|
| 1 | 161681780 | rs2275603 | -0.716 | 0.506 | -0.945 | 0.18 | 0.021 | -0.023 | -0.023 | 0.073 | 0.3 | 0.76 | 0.46 | 0.57 | 0.21 | 0.28 |
| 1 | 161683136 | rs11746 | 1.662 | 0.577 | -1.102 | 0.167 | -0.116 | 0.123 | 0.123 | 0.142 | 0.49 | 0.05 | 0.34 | 0.24 | 0.51 | 0.44 |
| 1 | 161684569 | rs6683580 | -1.287 | 0.384 | -0.94 | 0.196 | 0.039 | -0.004 | -0.004 | 0 | 0.35 | 0.87 | 0.46 | 0.64 | 0.25 | 0.32 |
| 1 | 161684621 | rs2333749 | 0.342 | 0.398 | -0.556 | 0.262 | -0.001 | -0.004 | -0.004 | 0 | 0.7 | 0.67 | 0.54 | 0.78 | 0.69 | 0.63 |
| 1 | 161685591 | rs10917750 | 0.451 | 0.349 | -0.402 | 0.234 | -0.015 | -0.029 | -0.029 | 0 | 0.66 | 0.41 | 0.54 | 0.81 | 0.74 | 0.68 |
| 1 | 161685725 | rs12042893 | -1.203 | 0.296 | -0.436 | 0.311 | 0.029 | 0.003 | 0.003 | 1.089 | 0.34 | 0.82 | 0.47 | 0.64 | 0.25 | 0.32 |
| 1 | 161687220 | rs16856590 | -0.108 | 0.381 | -0.406 | 0.388 | 0.012 | 0.14 | 0.14 | 0 | 0.18 | 0.38 | 0.4 | 0.55 | 0.17 | 0.16 |
| 1 | 161691366 | rs1891018 | -0.932 | 0.258 | -0.566 | 0.271 | -0.044 | -0.112 | -0.112 | 0 | 0.79 | 0.63 | 0.91 | 0.96 | 0.89 | 0.82 |
| 1 | 161691433 | rs1891019 | 0.566 | 0.404 | -0.694 | 0.383 | -0.001 | -0.001 | -0.001 | 0 | 0.67 | 0.32 | 0.83 | 0.52 | 0.76 | 0.63 |
| 1 | 161692154 | rs11590932 | 1.672 | 0.544 | -0.89 | 0.215 | -0.077 | 0.1 | 0.1 | 0 | 0.24 | 0.01 | 0.2 | 0.04 | 0.29 | 0.29 |
| 1 | 161693003 | rs4657093 | 0.127 | 0.575 | -0.061 | 0.905 | -0.001 | 0.012 | 0.012 | 0 | 0.15 | 0.24 | 0.37 | 0.16 | 0.15 | 0.14 |
| 1 | 161694857 | rs12079477 | -0.51 | 0.801 | -0.648 | 1.087 | 0.1 | 0.338 | 0.338 | 0.62 | 0.07 | 0.23 | 0.14 | 0.5 | 0.13 | 0.07 |
| 1 | 161700827 | rs2840310 | -0.009 | 0.692 | -1.101 | 0.97 | -0.04 | 0.045 | 0.045 | 0.033 | 0.69 | 0.84 | 0.5 | 0.77 | 0.63 | 0.63 |
| 1 | 161701810 | rs10917809 | -0.254 | 0.423 | -1.342 | 0.913 | -0.003 | 0.011 | 0.011 | 1.61 | 0.18 | 0.74 | 0.1 | 0.42 | 0.06 | 0.15 |
| 1 | 161702763 | rs6427623 | 0.466 | 0.512 | -1.058 | 1.109 | -0.032 | 0.094 | 0.094 | 0.509 | 0.15 | 0.54 | 0.08 | 0.25 | 0.06 | 0.15 |
| 1 | 161702792 | rs11576107 | 0.996 | 0.572 | -2.076 | 1.114 | -0.031 | 0.045 | 0.045 | 0.079 | 0.52 | 0.1 | 0.4 | 0.35 | 0.58 | 0.48 |
| 1 | 161703525 | rs6699725 | -0.578 | 0.031 | -1.363 | 1.117 | 0.001 | -0.002 | -0.002 | 0 | 0.19 | 0.74 | 0.13 | 0.33 | 0.1 | 0.17 |
| 1 | 161709317 | rs17415059 | 0.408 | 0.802 | -2.077 | 0.438 | -0.092 | 0.133 | 0.133 | 0 | 0.33 | 0.12 | 0.27 | 0.05 | 0.45 | 0.34 |
| 1 | 161711791 | rs2066247 | -0.131 | 0.203 | -2.344 | 0.579 | 0.028 | 0.084 | 0.084 | 0.103 | 0.49 | 0.9 | 0.34 | 0.71 | 0.39 | 0.37 |
| 1 | 161712857 | rs1027702 | 0.063 | 0.094 | -2.324 | 0.599 | 0.048 | 0.061 | 0.061 | 0.13 | 0.46 | 0.89 | 0.32 | 0.71 | 0.34 | 0.35 |
| 1 | 161713221 | rs12121500 | 0.245 | -0.099 | -1.618 | 0.636 | -0.001 | -0.001 | -0.001 | 0 | 0.4 | 0.42 | 0.25 | 0.48 | 0.29 | 0.32 |
| 1 | 161714087 | rs12026445 | 1.127 | -0.17 | -1.691 | 0.563 | -0.001 | -0.016 | -0.016 | 0 | 0.06 | 0.01 | 0.34 | 0.25 | 0.07 | 0.11 |
| 1 | 161719215 | rs12121310 | 0.241 | -0.127 | -1.622 | 0.695 | -0.002 | -0.002 | -0.002 | 0.003 | 0.39 | 0.41 | 0.25 | 0.38 | 0.29 | 0.32 |
| 1 | 161721707 | rs1063178 | 0.216 | -0.113 | -1.615 | 0.595 | -0.001 | -0.001 | -0.001 | 0 | 0.4 | 0.41 | 0.25 | 0.48 | 0.29 | 0.32 |
| 1 | 161738238 | rs4657101 | 0.053 | -0.022 | -2.154 | 0.212 | 0 | -0.017 | -0.017 | 1.342 | 0.25 | 0.15 | 0.46 | 0.33 | 0.27 | 0.14 |
| 1 | 161741274 | rs905594 | 0.982 | -0.763 | -2.578 | 0.046 | 0.048 | 0.001 | 0.001 | 0.474 | 0.59 | 0.07 | 0.44 | 0.3 | 0.7 | 0.61 |
| 1 | 161748056 | rs2070151 | 0.593 | -0.789 | -2.571 | 0.044 | -0.016 | 0.044 | 0.044 | 0 | 0.11 | 0.16 | 0.36 | 0.33 | 0.07 | 0.09 |
| 1 | 161751741 | rs1058405 | 1.744 | -0.485 | -2.749 | -0.098 | -0.076 | -0.08 | -0.08 | 0.161 | 0.22 | 0.01 | 0.17 | 0.26 | 0.27 | 0.17 |
| 1 | 161753802 | rs2271013 | 1.082 | -0.499 | -2.715 | -0.079 | 0.016 | -0.018 | -0.018 | 0 | 0.1 | 0.04 | 0.35 | 0.33 | 0.07 | 0.09 |
| 1 | 161754834 | rs6685369 | -1.74 | -0.571 | -2.781 | -0.107 | -0.005 | -0.03 | -0.03 | 0 | 0.87 | 0.98 | 0.9 | 0.88 | 0.88 | 0.85 |
| 1 | 161780979 | rs2134697 | 0.377 | -0.592 | -2.687 | -0.128 | 0.015 | -0.016 | -0.016 | 0 | 0.7 | 0.28 | 0.8 | 0.63 | 0.78 | 0.7 |
| 1 | 161793612 | rs11581556 | 1.764 | -0.497 | -2.667 | -0.121 | -0.073 | -0.08 | -0.08 | 0 | 0.22 | 0.01 | 0.17 | 0.26 | 0.27 | 0.17 |
| 1 | 161794336 | rs10800032 | 0.407 | -0.486 | -2.599 | -0.149 | -0.009 | 0.014 | 0.014 | 1.663 | 0.16 | 0.7 | 0.1 | 0.24 | 0.1 | 0.15 |
| 1 | 161795896 | rs10918029 | 1.01 | -0.382 | -2.312 | -0.099 | 0.151 | 0.345 | 0.345 | 7.566 | 0.06 | 0.35 | 0.06 | 0.21 | 0.07 | 0.05 |
| 1 | 161813765 | rs17408556 | 1.078 | -0.454 | -2.648 | -0.293 | -0.149 | 0.227 | 0.227 | 0 | 0.36 | 0.05 | 0.27 | 0.04 | 0.43 | 0.44 |
| 1 | 161820904 | rs3767635 | 1.447 | -0.367 | -2.706 | -0.274 | -0.117 | 0.151 | 0.151 | 0.131 | 0.29 | 0.01 | 0.21 | 0.04 | 0.34 | 0.39 |
| 1 | 161831507 | rs4657115 | 0.137 | -0.553 | -2.426 | 0 | -0.02 | 0.043 | 0.043 | 0 | 0.13 | 0.19 | 0.37 | 0.33 | 0.08 | 0.1 |
| 1 | 161885352 | rs2341479 | -1.568 | -0.37 | -2.473 | 0.039 | 0.002 | -0.001 | -0.001 | 0 | 0.87 | 0.88 | 0.9 | 0.88 | 0.88 | 0.85 |
| 1 | 161897797 | rs12043145 | 0.124 | -0.548 | -2.266 | 0.021 | -0.079 | 0.061 | 0.061 | 0 | 0.21 | 0.5 | 0.41 | 0.37 | 0.1 | 0.22 |
| 1 | 161908338 | rs10737517 | 0.546 | -0.564 | -2.309 | 0.008 | -0.029 | 0.013 | 0.013 | 0 | 0.66 | 0.39 | 0.49 | 0.51 | 0.79 | 0.64 |
| 1 | 161924749 | rs2341471 | 1.007 | -0.501 | -2.622 | -0.057 | 0.021 | -0.01 | -0.01 | 0 | 0.34 | 0.36 | 0.26 | 0.25 | 0.42 | 0.41 |
| 1 | 161928327 | rs9482 | 0.619 | -0.583 | -2.576 | -0.101 | -0.037 | 0.013 | 0.013 | 0 | 0.64 | 0.42 | 0.49 | 0.51 | 0.78 | 0.63 |
| 1 | 161928771 | rs1136046 | 0.152 | -0.495 | -2.688 | 0.001 | -0.071 | 0.053 | 0.053 | 0 | 0.21 | 0.43 | 0.4 | 0.37 | 0.1 | 0.22 |
| 1 | 161930730 | rs2499853 | 0.567 | 0.14 | -3.373 | 0.084 | -0.043 | 0.029 | 0.029 | 0 | 0.29 | 0.08 | 0.22 | 0.21 | 0.35 | 0.22 |
| 1 | 161936788 | rs2490438 | 0.746 | 0.073 | -3.413 | 0.137 | -0.054 | 0.023 | 0.023 | 0 | 0.29 | 0.05 | 0.22 | 0.21 | 0.35 | 0.18 |
| 1 | 161937624 | rs2490436 | 0.592 | 0.049 | -3.405 | 0.168 | -0.059 | -0.053 | -0.053 | 0.008 | 0.31 | 0.05 | 0.24 | 0.34 | 0.38 | 0.24 |
| 1 | 161941219 | rs2490431 | 1.056 | -0.489 | -2.937 | 0.526 | 0.08 | -0.068 | -0.068 | 0.573 | 0.57 | 0.06 | 0.45 | 0.42 | 0.71 | 0.6 |
| 1 | 161943634 | rs2340725 | -1.196 | -0.44 | -2.792 | 0.67 | 0.053 | -0.096 | -0.096 | 3.735 | 0.34 | 0.92 | 0.5 | 0.58 | 0.23 | 0.34 |
| 1 | 161946727 | rs2340727 | -0.722 | -0.385 | -2.143 | 0.536 | 0.031 | -0.011 | -0.011 | 0 | 0.23 | 0.86 | 0.12 | 0.25 | 0.13 | 0.16 |
| 1 | 161950451 | rs10918342 | 0.773 | -0.196 | -2.489 | 0.482 | -0.028 | -0.086 | -0.086 | 0.05 | 0.61 | 0.05 | 0.74 | 0.63 | 0.68 | 0.64 |
| 1 | 161952747 | rs10918361 | -1.233 | -0.209 | -1.124 | 0.797 | 0.009 | 0.04 | 0.04 | 0 | 0.28 | 0.39 | 0.35 | 0.46 | 0.34 | 0.38 |
| 1 | 161955205 | rs7523243 | -0.824 | 0.028 | -0.725 | 0.954 | 0.071 | 0.029 | 0.029 | 0.059 | 0.12 | 0.24 | 0.12 | 0.18 | 0.22 | 0.22 |
| 1 | 161956804 | rs2499841 | -0.468 | -0.423 | 0.116 | 1.051 | 0.03 | -0.004 | -0.004 | 0 | 0.82 | 0.84 | 0.82 | 0.84 | 0.9 | 0.79 |
| 1 | 161957719 | rs10494362 | -1.331 | -0.136 | 0.201 | 1.131 | 0.167 | 0.18 | 0.18 | 0.898 | 0.14 | 0.45 | 0.19 | 0.32 | 0.24 | 0.21 |
| 1 | 161963790 | rs4657131 | 0.883 | -0.323 | 0.464 | 1.061 | 0.003 | 0.007 | 0.007 | 0.064 | 0.67 | 0.53 | 0.68 | 0.66 | 0.67 | 0.57 |
| 1 | 161964536 | rs2490420 | 0.583 | -0.321 | 0.388 | 1.057 | 0.003 | 0.007 | 0.007 | 0 | 0.69 | 0.59 | 0.69 | 0.66 | 0.67 | 0.61 |
| 1 | 161967287 | rs6691548 | 1.204 | -0.29 | -0.048 | 0.928 | 0.002 | -0.022 | -0.022 | 0.224 | 0.63 | 0.32 | 0.6 | 0.5 | 0.65 | 0.51 |
| 1 | 161968297 | rs12756866 | 1.023 | -0.265 | -0.034 | 0.916 | 0 | 0 | 0 | 0 | 0.64 | 0.4 | 0.6 | 0.51 | 0.64 | 0.51 |
| 1 | 161969322 | rs7527826 | 0.593 | 0.099 | -0.543 | 0.378 | -0.001 | -0.006 | -0.006 | 3.772 | 0.41 | 0.33 | 0.45 | 0.55 | 0.4 | 0.38 |
| 1 | 161969636 | rs2499834 | -0.099 | 0.066 | -0.476 | 1.104 | 0.031 | -0.028 | -0.028 | 0.199 | 0.8 | 0.88 | 0.76 | 0.81 | 0.88 | 0.79 |
| 1 | 161973976 | rs2490424 | -0.73 | -0.877 | -0.294 | 0.95 | -0.005 | 0.019 | 0.019 | 0.358 | 0.14 | 0.39 | 0.09 | 0.09 | 0.08 | 0.06 |
| 1 | 161974584 | rs2490422 | 0.392 | -1.121 | -1.24 | 1.175 | 0.031 | -0.033 | -0.033 | 0 | 0.74 | 0.8 | 0.73 | 0.63 | 0.82 | 0.64 |
| 1 | 161975281 | rs7522269 | 0.554 | -1.625 | -1.45 | 1.135 | 0.046 | -0.021 | -0.021 | 0.115 | 0.63 | 0.28 | 0.72 | 0.59 | 0.75 | 0.58 |
| 1 | 161975491 | rs7555648 | 0.215 | -1.703 | -1.365 | 1.104 | 0.01 | -0.012 | -0.012 | 0 | 0.62 | 0.76 | 0.61 | 0.6 | 0.59 | 0.47 |
| 1 | 161976234 | rs12022536 | 0.868 | -3.284 | -0.495 | 1.333 | -0.03 | 0.01 | 0.01 | 0 | 0.65 | 0.44 | 0.79 | 0.46 | 0.82 | 0.55 |
| 1 | 161976473 | rs12093822 | -0.976 | -3.548 | -0.222 | 1.489 | -0.025 | 0.006 | 0.006 | 0 | 0.15 | 0.23 | 0.07 | 0.2 | 0.09 | 0.15 |
| 1 | 161980422 | rs10218544 | -1.069 | -3.509 | -0.374 | 1.305 | 0.005 | -0.016 | -0.016 | 0 | 0.07 | 0.11 | 0.03 | 0.03 | 0.04 | 0.03 |
| 1 | 161980502 | rs12025136 | 2.12 | -4.334 | -0.817 | 1.157 | 0.164 | 0.055 | 0.055 | 0.008 | 0.5 | 0.09 | 0.71 | 0.41 | 0.72 | 0.47 |
| 1 | 161981845 | rs16838789 | 0.375 | -4.325 | -0.666 | 1.549 | 0.011 | -0.022 | -0.022 | 0 | 0.1 | 0.05 | 0.13 | 0.26 | 0.08 | 0.27 |

| | | | | | | | | | | | | | | | | |
|---|---|---|---|---|---|---|---|---|---|---|---|---|---|---|---|---|
| 1 | 161983089 | rs6676438 | 2.39 | -4.639 | -0.99 | 1.32 | 0.059 | 0.014 | 0.014 | 0.245 | 0.46 | 0.03 | 0.69 | 0.43 | 0.69 | 0.49 |
| 1 | 161984197 | rs12121431 | 0.08 | -4.619 | -0.374 | 1.382 | 0.005 | 0 | 0 | 8.186 | 0.16 | 0.15 | 0.04 | 0.14 | 0.07 | 0.03 |
| 1 | 161987385 | rs16839677 | 0.632 | -4.61 | -0.344 | 1.616 | 0.009 | -0.017 | -0.017 | 0 | 0.1 | 0.09 | 0.14 | 0.23 | 0.08 | 0.25 |
| 1 | 161988206 | rs7532680 | -2.325 | -4.779 | 0.533 | 1.547 | 0.105 | -0.025 | -0.025 | 0 | 0.42 | 0.85 | 0.25 | 0.4 | 0.21 | 0.36 |
| 1 | 161989354 | rs12562972 | 0.601 | -4.663 | 0.57 | 1.605 | 0.008 | -0.013 | -0.013 | 0 | 0.1 | 0.09 | 0.14 | 0.23 | 0.08 | 0.25 |
| 1 | 161990991 | rs4657134 | 0.127 | -4.389 | 1.101 | 1.317 | -0.001 | 0.001 | 0.001 | 0 | 0.86 | 0.9 | 0.94 | 0.82 | 0.9 | 0.83 |
| 1 | 161991193 | rs12126296 | -0.093 | -4.218 | 1.017 | 1.23 | 0.003 | 0 | 0 | 0 | 0.16 | 0.16 | 0.04 | 0.14 | 0.07 | 0.03 |
| 1 | 161993192 | rs12130792 | -0.093 | -4.218 | 1.018 | 1.23 | 0.002 | 0 | 0 | 0 | 0.16 | 0.16 | 0.04 | 0.14 | 0.07 | 0.03 |
| 1 | 162000398 | rs7551221 | 1.261 | -4.019 | 0.823 | 1.312 | -0.018 | -0.003 | -0.003 | 0 | 0.59 | 0.55 | 0.85 | 0.66 | 0.77 | 0.73 |
| 1 | 162000432 | rs7514121 | 1.261 | -4.019 | 0.823 | 1.312 | -0.018 | -0.003 | -0.003 | 0.733 | 0.59 | 0.55 | 0.85 | 0.66 | 0.77 | 0.73 |
| 1 | 162003846 | rs885092 | -1.317 | -4.289 | 0.991 | 1.193 | -0.007 | -0.002 | -0.002 | 0.274 | 0.29 | 0.28 | 0.12 | 0.2 | 0.17 | 0.24 |
| 1 | 162008034 | rs7539281 | -2.082 | -4.488 | 1.289 | 1.673 | 0.053 | -0.002 | -0.002 | 0 | 0.46 | 0.67 | 0.31 | 0.51 | 0.28 | 0.52 |
| 1 | 162008079 | rs7539367 | -0.626 | -4.245 | 1.488 | 1.881 | -0.026 | 0.128 | 0.128 | 0.481 | 0.15 | 0.39 | 0.18 | 0.31 | 0.11 | 0.26 |
| 1 | 162010787 | rs10158975 | -0.785 | -4.271 | 0.831 | 1.44 | 0.031 | 0.005 | 0.005 | 0 | 0.23 | 0.15 | 0.07 | 0.2 | 0.12 | 0.21 |
| 1 | 162011178 | rs7524328 | -0.324 | -3.359 | 1.089 | 1.578 | -0.003 | 0.003 | 0.003 | 0.139 | 0.12 | 0.11 | 0.15 | 0.2 | 0.1 | 0.26 |
| 1 | 162013475 | rs6679284 | -1.664 | -3.177 | 1.647 | 1.449 | 0.018 | 0.08 | 0.08 | 0.381 | 0.1 | 0.36 | 0.16 | 0.17 | 0.08 | 0.18 |
| 1 | 162014632 | rs2880058 | -1.898 | -4.023 | 1.29 | 1.393 | 0.093 | 0.085 | 0.085 | 0 | 0.55 | 0.79 | 0.43 | 0.71 | 0.35 | 0.71 |
| 1 | 162021058 | rs12087446 | -2.128 | -3.983 | 1.766 | 1.761 | -0.031 | 0.104 | 0.104 | 6.932 | 0.16 | 0.68 | 0.2 | 0.35 | 0.1 | 0.2 |
| 1 | 162024818 | rs6658860 | -1.085 | -3.92 | 1.809 | 1.673 | -0.003 | 0.057 | 0.057 | 0 | 0.11 | 0.32 | 0.16 | 0.18 | 0.08 | 0.2 |
| 1 | 162029291 | rs7550692 | -1.615 | -4.035 | 1.762 | 1.642 | 0.056 | 0.001 | 0.001 | 0 | 0.52 | 0.55 | 0.4 | 0.53 | 0.34 | 0.71 |
| 1 | 162030654 | rs10918593 | -1.831 | -4.103 | 1.651 | 1.661 | 0.093 | 0.091 | 0.091 | 0 | 0.55 | 0.89 | 0.44 | 0.71 | 0.35 | 0.72 |
| 1 | 162031465 | rs10918595 | -1.517 | -4.087 | 1.793 | 1.472 | 0.011 | 0.001 | 0.001 | 0 | 0.51 | 0.48 | 0.4 | 0.53 | 0.34 | 0.71 |
| 1 | 162033890 | rs12143842 | -0.701 | -4.054 | 1.031 | 1.531 | 0.097 | 0.016 | 0.016 | 0 | 0.39 | 0.12 | 0.23 | 0.34 | 0.26 | 0.44 |
| 1 | 162036479 | rs12567209 | -0.932 | -3.722 | 0.833 | 1.526 | -0.001 | 0.001 | 0.001 | 0 | 0.09 | 0.08 | 0.14 | 0.18 | 0.08 | 0.25 |
| 1 | 162037856 | rs10753746 | -1.157 | -3.879 | 1.076 | 1.213 | 0.009 | -0.003 | -0.003 | 0 | 0.46 | 0.45 | 0.35 | 0.53 | 0.3 | 0.69 |
| 1 | 162039394 | rs12044487 | -1.093 | -3.769 | 1.083 | 1.188 | 0.041 | -0.015 | -0.015 | 0 | 0.44 | 0.32 | 0.34 | 0.53 | 0.3 | 0.7 |
| 1 | 162040878 | rs16849113 | -2.438 | -3.66 | 1.331 | 1.232 | 0.012 | 0.221 | 0.221 | 5.74 | 0.1 | 0.54 | 0.09 | 0.2 | 0.06 | 0.07 |
| 1 | 162042313 | rs1123015 | 1.543 | -3.79 | 1.184 | 1.252 | 0.047 | 0.068 | 0.068 | 0 | 0.45 | 0.09 | 0.54 | 0.3 | 0.64 | 0.29 |
| 1 | 162043710 | rs7515045 | 0.789 | -3.76 | 1.264 | 1.242 | 0.01 | -0.001 | -0.001 | 0 | 0.54 | 0.64 | 0.63 | 0.49 | 0.7 | 0.36 |
| 1 | 162063023 | rs1337062 | 1.178 | -3.463 | 1.428 | 1.311 | 0.003 | -0.001 | -0.001 | 0 | 0.52 | 0.56 | 0.59 | 0.51 | 0.67 | 0.33 |
| 1 | 162064913 | rs4656350 | 1.178 | -3.446 | 1.647 | 1.352 | 0.003 | -0.001 | -0.001 | 0 | 0.52 | 0.56 | 0.59 | 0.51 | 0.67 | 0.33 |
| 1 | 162067299 | rs10918671 | 1.422 | -3.463 | 1.7 | 1.343 | 0.069 | 0.084 | 0.084 | 0 | 0.48 | 0.17 | 0.56 | 0.34 | 0.65 | 0.3 |
| 1 | 162084886 | rs1572498 | 1.216 | -3.43 | 1.104 | 1.32 | 0.055 | 0.064 | 0.064 | 0 | 0.49 | 0.32 | 0.56 | 0.34 | 0.64 | 0.3 |
| 1 | 162085685 | rs10494366 | 1.216 | -3.43 | 1.104 | 1.32 | 0.055 | 0.064 | 0.064 | 0 | 0.49 | 0.32 | 0.56 | 0.34 | 0.64 | 0.3 |
| 1 | 162092110 | rs10458392 | -0.436 | -3.272 | 0.608 | 1.314 | 0.028 | -0.133 | -0.133 | 0.687 | 0.32 | 0.04 | 0.29 | 0.54 | 0.23 | 0.63 |
| 1 | 162099301 | rs1572495 | -1.765 | -3.55 | 0.721 | 0.992 | -0.038 | 0.016 | 0.016 | 4.774 | 0.17 | 0.7 | 0.12 | 0.17 | 0.08 | 0.06 |
| 1 | 162104658 | rs1337072 | -1.821 | -3.551 | 0.723 | 0.985 | -0.033 | 0.016 | 0.016 | 3.871 | 0.18 | 0.67 | 0.12 | 0.17 | 0.08 | 0.06 |
| 1 | 162108119 | rs4657154 | 0.378 | -3.599 | 0.657 | 1 | 0.008 | -0.009 | -0.009 | 0 | 0.63 | 0.71 | 0.65 | 0.45 | 0.71 | 0.35 |
| 1 | 162112650 | rs16857019 | -1.162 | -3.413 | 0.599 | 1.079 | -0.006 | 0.043 | 0.043 | 0 | 0.13 | 0.3 | 0.09 | 0.18 | 0.08 | 0.06 |
| 1 | 162113887 | rs10918740 | 1.573 | -3.503 | 0.101 | 0.967 | -0.008 | 0.069 | 0.069 | 0.39 | 0.44 | 0.04 | 0.53 | 0.26 | 0.63 | 0.29 |
| 1 | 162118902 | rs945708 | -1.393 | -3.415 | 0.934 | 0.801 | 0.037 | 0.064 | 0.064 | 0 | 0.5 | 0.68 | 0.4 | 0.81 | 0.37 | 0.7 |
| 1 | 162118984 | rs10918762 | -0.111 | -3.639 | 0.936 | 0.809 | 0.013 | -0.022 | -0.022 | 0 | 0.27 | 0.25 | 0.28 | 0.63 | 0.23 | 0.57 |
| 1 | 162126444 | rs951121 | 1.293 | -3.287 | 1.267 | 0.744 | -0.031 | -0.011 | -0.011 | 0 | 0.76 | 0.47 | 0.88 | 0.83 | 0.86 | 0.88 |
| 1 | 162129334 | rs12069336 | -1.22 | -3.283 | 0.888 | 0.68 | -0.025 | -0.017 | -0.017 | 0 | 0.22 | 0.35 | 0.1 | 0.16 | 0.13 | 0.1 |
| 1 | 162133117 | rs12029454 | 0.243 | -3.432 | 0.712 | 1.175 | -0.011 | 0.023 | 0.023 | 0.258 | 0.2 | 0.31 | 0.23 | 0.37 | 0.18 | 0.35 |
| 1 | 162139692 | rs7546353 | -1.334 | -3.924 | 0.705 | 1.147 | -0.046 | 0.026 | 0.026 | 0 | 0.31 | 0.45 | 0.23 | 0.39 | 0.19 | 0.25 |
| 1 | 162148700 | rs4526593 | -2.059 | -3.173 | 0.568 | 0.865 | 0.024 | 0.068 | 0.068 | 0.168 | 0.55 | 0.8 | 0.52 | 0.77 | 0.41 | 0.64 |
| 1 | 162160920 | rs4657164 | -1.033 | -3.326 | 0.566 | 0.773 | 0.017 | 0.01 | 0.01 | 0 | 0.26 | 0.16 | 0.19 | 0.19 | 0.14 | 0.24 |
| 1 | 162161219 | rs10918840 | -1.39 | -3.375 | 0.724 | 0.823 | -0.022 | 0.006 | 0.006 | 0 | 0.43 | 0.5 | 0.37 | 0.55 | 0.29 | 0.57 |
| 1 | 162161940 | rs12087800 | -1.948 | -3.377 | 0.696 | 0.745 | 0.007 | 0 | 0 | 0 | 0.52 | 0.55 | 0.53 | 0.52 | 0.39 | 0.63 |
| 1 | 162162334 | rs12047527 | -1.443 | -3.354 | 0.749 | 0.788 | -0.017 | 0.005 | 0.005 | 0.033 | 0.43 | 0.48 | 0.42 | 0.53 | 0.33 | 0.6 |
| 1 | 162165433 | rs7513132 | -1.865 | -3.36 | 0.726 | 0.748 | 0.011 | 0.003 | 0.003 | 0.11 | 0.5 | 0.53 | 0.46 | 0.5 | 0.39 | 0.63 |
| 1 | 162165708 | rs6700395 | -1.897 | -3.359 | 0.857 | 0.75 | 0.042 | 0.009 | 0.009 | 0.112 | 0.51 | 0.62 | 0.47 | 0.5 | 0.39 | 0.63 |
| 1 | 162168958 | rs3897374 | 1.588 | -3.42 | 1.011 | 0.804 | 0.004 | 0.004 | 0.004 | 0 | 0.71 | 0.75 | 0.79 | 0.88 | 0.83 | 0.73 |
| 1 | 162173395 | rs12061249 | -0.478 | -3.471 | 0.959 | 0.8 | 0.028 | 0.02 | 0.02 | 0 | 0.2 | 0.1 | 0.13 | 0.11 | 0.1 | 0.2 |
| 1 | 162175972 | rs6676737 | -1.21 | -3.441 | 0.962 | 0.815 | 0.021 | 0.022 | 0.022 | 0 | 0.25 | 0.2 | 0.15 | 0.11 | 0.13 | 0.23 |
| 1 | 162180456 | rs12116744 | 0.409 | -2.977 | 0.803 | 0.791 | -0.007 | 0.024 | 0.024 | 0.674 | 0.23 | 0.32 | 0.29 | 0.38 | 0.26 | 0.37 |
| 1 | 162187512 | rs7416392 | 1.032 | -3.013 | 0.736 | 0.754 | 0.014 | -0.002 | -0.002 | 0 | 0.57 | 0.43 | 0.55 | 0.51 | 0.64 | 0.44 |
| 1 | 162191412 | rs6683968 | 1.57 | -2.882 | 0.664 | 1.04 | 0.053 | 0.014 | 0.014 | 0 | 0.51 | 0.34 | 0.52 | 0.51 | 0.6 | 0.4 |
| 1 | 162194576 | rs12734991 | 2.089 | -2.88 | 0.4 | 0.943 | -0.078 | 0.115 | 0.115 | 0.568 | 0.43 | 0.04 | 0.43 | 0.22 | 0.54 | 0.35 |
| 1 | 162201217 | rs8179274 | 1.404 | -2.876 | 0.633 | 0.91 | -0.02 | -0.032 | -0.032 | 0 | 0.7 | 0.66 | 0.77 | 0.88 | 0.82 | 0.73 |
| 1 | 162202488 | rs10918936 | 1.545 | -2.748 | 0.42 | 0.989 | 0.035 | 0.003 | 0.003 | 0.027 | 0.48 | 0.33 | 0.47 | 0.49 | 0.56 | 0.36 |
| 1 | 162210471 | rs4431825 | 1.964 | -3.146 | -0.479 | 0.282 | -0.131 | 0.054 | 0.054 | 0 | 0.35 | 0.04 | 0.36 | 0.19 | 0.52 | 0.35 |
| 1 | 162210610 | rs4657178 | -0.317 | -3.26 | 0.401 | 0.4 | -0.011 | 0.041 | 0.041 | 0.353 | 0.29 | 0.39 | 0.36 | 0.53 | 0.29 | 0.42 |
| 1 | 162213867 | rs10918963 | 0.936 | -3.279 | 0.268 | 0.4 | 0.004 | -0.029 | -0.029 | 0.042 | 0.21 | 0.13 | 0.3 | 0.52 | 0.25 | 0.37 |
| 1 | 162219739 | rs6677052 | -0.475 | -3.483 | 0.005 | 0.261 | 0.026 | -0.009 | -0.009 | 1.638 | 0.26 | 0.15 | 0.21 | 0.28 | 0.17 | 0.2 |
| 1 | 162224550 | rs16859092 | -0.019 | -3.129 | 0.023 | 0.042 | -0.001 | -0.02 | -0.02 | 0.82 | 0.07 | 0.13 | 0.05 | 0.01 | 0.04 | 0.05 |
| 1 | 162224586 | rs12742393 | 1.533 | -3.23 | -0.003 | -0.001 | -0.002 | 0.113 | 0.113 | 0 | 0.42 | 0.17 | 0.41 | 0.2 | 0.54 | 0.38 |
| 1 | 162224979 | rs10800386 | -1.17 | -3.105 | 0.014 | 0.127 | 0.025 | 0.039 | 0.039 | 0 | 0.5 | 0.62 | 0.54 | 0.8 | 0.42 | 0.57 |

| | | | | | | | | | | | | | | | | |
|---|---|---|---|---|---|---|---|---|---|---|---|---|---|---|---|---|
| 1 | 162230356 | rs4131505 | -1.13 | -3.035 | -0.205 | 0.172 | -0.001 | 0.004 | 0.004 | 0 | 0.29 | 0.26 | 0.29 | 0.39 | 0.25 | 0.36 |
| 1 | 162235437 | rs4584372 | -1.217 | -2.897 | -0.215 | 0.028 | -0.004 | 0.008 | 0.008 | 0 | 0.31 | 0.29 | 0.28 | 0.38 | 0.24 | 0.33 |
| 1 | 162236582 | rs4298709 | 1.916 | -3.283 | 0.017 | -0.121 | 0.007 | 0.11 | 0.11 | 0 | 0.46 | 0.28 | 0.34 | 0.17 | 0.53 | 0.33 |
| 1 | 162237655 | rs12410644 | -0.85 | -3.17 | -0.044 | -0.035 | -0.002 | 0.009 | 0.009 | 0 | 0.28 | 0.26 | 0.25 | 0.38 | 0.24 | 0.35 |
| 1 | 162239031 | rs10919025 | 0.718 | -2.989 | 0.329 | 0.105 | -0.016 | 0.032 | 0.032 | 0 | 0.17 | 0.23 | 0.32 | 0.43 | 0.18 | 0.3 |
| 1 | 162247296 | rs10157390 | -1.434 | -3.11 | 0.314 | 0.097 | -0.005 | -0.007 | -0.007 | 1.443 | 0.41 | 0.56 | 0.57 | 0.74 | 0.38 | 0.51 |
| 1 | 162247738 | rs4657180 | -0.24 | -3.354 | 0.225 | -0.028 | -0.037 | 0.004 | 0.004 | 1.799 | 0.77 | 0.64 | 0.89 | 0.77 | 0.9 | 0.8 |
| 1 | 162247779 | rs7528328 | -0.161 | -3.223 | 0.095 | -0.042 | -0.008 | 0.04 | 0.04 | 0 | 0.2 | 0.25 | 0.21 | 0.3 | 0.18 | 0.19 |
| 1 | 162248078 | rs11800645 | 0.714 | -3.248 | 0.108 | -0.122 | 0.014 | 0.042 | 0.042 | 0 | 0.13 | 0.05 | 0.05 | 0 | 0.09 | 0.07 |
| 1 | 162249697 | rs6659953 | -2.098 | -3.07 | 0.217 | -0.043 | -0.012 | 0.105 | 0.105 | 0.012 | 0.61 | 0.79 | 0.64 | 0.96 | 0.45 | 0.69 |
| 1 | 162250009 | rs6671271 | 0.531 | -2.748 | 0.388 | 0.267 | 0.034 | 0.257 | 0.257 | 0 | 0.07 | 0.19 | 0.16 | 0.29 | 0.09 | 0.11 |
| 1 | 162252414 | rs12027674 | -0.49 | -2.775 | -0.155 | 0.041 | 0.002 | -0.004 | -0.004 | 0.742 | 0.31 | 0.31 | 0.38 | 0.62 | 0.26 | 0.38 |
| 1 | 162252870 | rs10919060 | -0.723 | -2.747 | -0.153 | 0.042 | 0.007 | -0.009 | -0.009 | 0 | 0.33 | 0.32 | 0.4 | 0.62 | 0.29 | 0.4 |
| 1 | 162260715 | rs10800409 | 1.098 | -2.344 | -0.377 | 0.292 | 0.004 | -0.008 | -0.008 | 0 | 0.14 | 0.09 | 0.31 | 0.44 | 0.17 | 0.31 |
| 1 | 162263270 | rs10919096 | 1.801 | -2.606 | -0.854 | -0.016 | -0.071 | 0.085 | 0.085 | 0 | 0.39 | 0.1 | 0.32 | 0.19 | 0.53 | 0.31 |
| 1 | 162263754 | rs6677529 | -0.699 | -2.325 | -0.249 | 0.185 | -0.035 | 0.06 | 0.06 | 0.775 | 0.29 | 0.48 | 0.27 | 0.35 | 0.18 | 0.28 |
| 1 | 162267823 | rs10919117 | -0.472 | -0.956 | -0.168 | -0.113 | 0.003 | -0.006 | -0.006 | 0 | 0.55 | 0.39 | 0.45 | 0.4 | 0.52 | 0.47 |
| 1 | 162268123 | rs12128479 | 1.535 | -0.914 | -0.458 | 0.117 | 0.011 | -0.067 | -0.067 | 1.439 | 0.11 | 0.01 | 0.26 | 0.47 | 0.11 | 0.29 |
| 1 | 162274971 | rs2221644 | -0.725 | -0.869 | 0.351 | -0.024 | -0.001 | 0.011 | 0.011 | 0 | 0.59 | 0.51 | 0.46 | 0.4 | 0.54 | 0.47 |
| 1 | 162278552 | rs2819316 | 0.728 | -0.799 | 0.112 | -0.027 | -0.003 | -0.037 | -0.037 | 0.685 | 0.38 | 0.26 | 0.51 | 0.6 | 0.44 | 0.51 |
| 1 | 162281028 | rs2819322 | 0.198 | -0.836 | -0.127 | -0.187 | 0.01 | 0.008 | 0.008 | 0 | 0.58 | 0.54 | 0.61 | 0.6 | 0.54 | 0.61 |
| 1 | 162283667 | rs10919166 | 0.08 | -0.673 | -0.363 | -0.264 | -0.023 | -0.043 | -0.043 | 0 | 0.2 | 0.12 | 0.41 | 0.53 | 0.27 | 0.19 |
| 1 | 162284057 | rs2819328 | 0.575 | -0.586 | 0.07 | 0.073 | 0.108 | 0.141 | 0.141 | 0.232 | 0.95 | 0.67 | 0.96 | 0.93 | 0.97 | 0.9 |
| 1 | 162299197 | rs9661075 | -0.037 | -0.83 | -0.454 | -0.079 | 0.004 | -0.003 | -0.003 | 0 | 0.42 | 0.41 | 0.42 | 0.52 | 0.3 | 0.28 |
| 1 | 162299823 | rs347300 | 0.257 | -0.618 | -1.259 | -0.189 | 0.055 | 0.063 | 0.063 | 0 | 0.5 | 0.15 | 0.53 | 0.4 | 0.66 | 0.61 |
| 1 | 162302635 | rs347306 | 0.169 | -0.838 | -1.266 | -0.585 | 0.013 | 0.098 | 0.098 | 0 | 0.43 | 0.25 | 0.46 | 0.21 | 0.56 | 0.31 |
| 1 | 162303459 | rs347309 | -1.014 | -1.204 | -0.964 | -0.559 | -0.033 | 0.02 | 0.02 | 0 | 0.9 | 0.77 | 0.95 | 0.85 | 0.96 | 0.81 |
| 1 | 162303942 | rs347312 | -0.802 | -1.232 | -0.743 | -0.572 | -0.039 | 0.039 | 0.039 | 0 | 0.89 | 0.69 | 0.94 | 0.85 | 0.96 | 0.81 |
| 1 | 162306207 | rs10753789 | 0.249 | -1.37 | -0.752 | -0.687 | -0.01 | -0.005 | -0.005 | 0 | 0.42 | 0.39 | 0.46 | 0.64 | 0.37 | 0.53 |
| 1 | 162306295 | rs6700867 | 0.945 | -1.371 | -0.906 | -0.625 | -0.029 | 0.016 | 0.016 | 0 | 0.1 | 0.2 | 0.05 | 0.15 | 0.04 | 0.15 |
| 1 | 162308721 | rs1396225 | 1.28 | -1.219 | -1.335 | -0.832 | -0.145 | 0.014 | 0.014 | 0 | 0.28 | 0.03 | 0.36 | 0.16 | 0.42 | 0.2 |
| 1 | 162309279 | rs423125 | -0.871 | -1.441 | -0.836 | -0.33 | -0.041 | 0.032 | 0.032 | 0 | 0.89 | 0.7 | 0.94 | 0.85 | 0.96 | 0.81 |
| 1 | 162312753 | rs347282 | 0.479 | -0.996 | -1.098 | -0.412 | 0.015 | 0.093 | 0.093 | 0 | 0.44 | 0.27 | 0.46 | 0.21 | 0.56 | 0.3 |
| 1 | 162317513 | rs347273 | 0.879 | -0.28 | -0.525 | 0.206 | -0.046 | -0.034 | -0.034 | 0 | 0.83 | 0.72 | 0.9 | 0.91 | 0.86 | 0.76 |
| 1 | 162319329 | rs347271 | 0.303 | -0.256 | -0.733 | 0.209 | 0.01 | -0.021 | -0.021 | 0 | 0.93 | 0.76 | 0.97 | 1 | 0.96 | 0.94 |
| 1 | 162319524 | rs11577628 | -0.081 | 0.168 | -1.738 | 0.554 | 0.022 | -0.018 | -0.018 | 0 | 0.1 | 0.01 | 0.06 | 0.09 | 0.09 | 0.18 |
| 1 | 162321600 | rs6685342 | -0.264 | 0.698 | -1.807 | 0.513 | -0.015 | -0.039 | -0.039 | 0.146 | 0.4 | 0.2 | 0.53 | 0.54 | 0.42 | 0.34 |
| 1 | 162322955 | rs12403202 | -0.948 | 1.691 | -1.949 | 0.566 | -0.068 | -0.041 | -0.041 | 3.369 | 0.15 | 0.04 | 0.21 | 0.18 | 0.24 | 0.13 |
| 1 | 162324445 | rs449908 | 1.23 | 0.895 | -1.448 | 0.203 | 0.003 | 0 | 0 | 0 | 0.76 | 0.74 | 0.76 | 0.76 | 0.71 | 0.66 |
| 1 | 162331408 | rs12122048 | 0.399 | 0.91 | -1.429 | -0.41 | 0.043 | 0.066 | 0.066 | 0 | 0.47 | 0.34 | 0.28 | 0.19 | 0.35 | 0.4 |
| 1 | 162331621 | rs6681981 | 0.817 | 1.261 | -1.46 | -0.315 | -0.089 | -0.077 | -0.077 | 0.036 | 0.26 | 0.1 | 0.4 | 0.54 | 0.39 | 0.21 |
| 1 | 162333990 | rs1963645 | 1.049 | 1.15 | -1.489 | -0.316 | 0.037 | 0.059 | 0.059 | 0 | 0.46 | 0.32 | 0.28 | 0.19 | 0.35 | 0.36 |
| 1 | 162334100 | rs905720 | 0.815 | 1.15 | -1.458 | -0.319 | -0.006 | -0.023 | -0.023 | 0.283 | 0.72 | 0.43 | 0.68 | 0.73 | 0.74 | 0.57 |
| 1 | 162335256 | rs348624 | -0.321 | 1.035 | -0.837 | -0.327 | -0.045 | -0.042 | -0.042 | 2.04 | 0.19 | 0.43 | 0.22 | 0.12 | 0.12 | 0.29 |
| 1 | 162336539 | rs7551382 | 0.83 | 1.071 | -1.057 | -0.331 | -0.091 | -0.075 | -0.075 | 1.77 | 0.26 | 0.1 | 0.4 | 0.55 | 0.4 | 0.23 |
| 1 | 162338485 | rs164148 | 0.34 | 1.196 | -0.629 | -0.37 | -0.043 | -0.059 | -0.059 | 0.95 | 0.79 | 0.49 | 0.78 | 0.87 | 0.84 | 0.71 |
| 1 | 162338496 | rs1876986 | 0.897 | 1.261 | -0.691 | -0.411 | -0.103 | -0.092 | -0.092 | 0 | 0.24 | 0.06 | 0.39 | 0.55 | 0.37 | 0.22 |
| 1 | 162338581 | rs164149 | 0.483 | 1.149 | -0.682 | -0.4 | 0 | 0 | 0 | 0 | 0.55 | 0.43 | 0.39 | 0.32 | 0.47 | 0.46 |
| 1 | 162342203 | rs164154 | -0.223 | 1.099 | -0.698 | -0.308 | -0.01 | 0.018 | 0.018 | 0 | 0.34 | 0.18 | 0.5 | 0.72 | 0.55 | 0.38 |
| 1 | 162343388 | rs164158 | 0.755 | 0.222 | -1.252 | -0.487 | 0.014 | 0.014 | 0.014 | 0 | 0.33 | 0.29 | 0.21 | 0.17 | 0.24 | 0.26 |
| 1 | 162343774 | rs7512126 | 0.588 | 0.982 | -1.249 | -0.198 | -0.042 | -0.041 | -0.041 | 0.246 | 0.31 | 0.19 | 0.43 | 0.51 | 0.41 | 0.3 |
| 1 | 162344069 | rs13823 | 0.66 | 0.699 | -1.409 | -0.247 | 0.019 | -0.013 | -0.013 | 0 | 0.07 | 0 | 0.11 | 0.08 | 0.06 | 0.16 |
| 1 | 162344102 | rs2282397 | 0.572 | 1.063 | -1.199 | -0.135 | -0.041 | -0.048 | -0.048 | 0 | 0.31 | 0.16 | 0.42 | 0.51 | 0.38 | 0.31 |
| 1 | 162345091 | rs164181 | -0.149 | 1.205 | -0.656 | -0.081 | -0.029 | -0.03 | -0.03 | 0.906 | 0.94 | 0.79 | 0.98 | 1 | 1 | 0.99 |
| 1 | 162345294 | rs3737312 | -0.707 | 1.802 | -1.231 | 0.084 | -0.066 | -0.01 | -0.01 | 0 | 0.2 | 0.09 | 0.21 | 0.19 | 0.28 | 0.22 |
| 1 | 162350635 | rs12131795 | -1.111 | 1.562 | -1.241 | 0.217 | -0.033 | -0.009 | -0.009 | 2.111 | 0.19 | 0.05 | 0.2 | 0.2 | 0.26 | 0.3 |
| 1 | 162351586 | rs901662 | -1.08 | 1.455 | -0.979 | 0.214 | -0.018 | -0.007 | -0.007 | 0.428 | 0.17 | 0.12 | 0.18 | 0.19 | 0.21 | 0.3 |
| 1 | 162352111 | rs164189 | 1.704 | 1.475 | -0.48 | 0.477 | -0.014 | 0.038 | 0.038 | 0 | 0.57 | 0.4 | 0.4 | 0.26 | 0.51 | 0.37 |
| 1 | 162353726 | rs164417 | 1.322 | 1.446 | 0.002 | 1.072 | -0.011 | -0.012 | -0.012 | 0.387 | 0.66 | 0.76 | 0.57 | 0.44 | 0.62 | 0.53 |
| 1 | 162356140 | rs164177 | 1.19 | 0.81 | -0.855 | 0.341 | 0.093 | -0.011 | -0.011 | 0 | 0.44 | 0.2 | 0.31 | 0.42 | 0.31 | 0.25 |
| 1 | 162356487 | rs1047802 | 0.229 | 0.807 | -0.232 | 0.34 | 0.028 | 0.011 | 0.011 | 0 | 0.28 | 0.41 | 0.43 | 0.37 | 0.4 | 0.39 |
| 1 | 162357653 | rs164178 | 0.985 | 0.709 | 0.256 | 0.299 | 0.065 | 0.011 | 0.011 | 0.485 | 0.5 | 0.3 | 0.34 | 0.44 | 0.36 | 0.35 |
| 1 | 162358395 | rs1394320 | 0.435 | 0.772 | 0.224 | 0.406 | 0.003 | 0.07 | 0.07 | 0 | 0.11 | 0.15 | 0.2 | 0.35 | 0.11 | 0.12 |
| 1 | 162360008 | rs10919317 | -0.186 | 1.012 | 0.397 | 0.221 | -0.009 | 0.005 | 0.005 | 0 | 0.22 | 0.16 | 0.23 | 0.19 | 0.27 | 0.26 |
| 1 | 162360279 | rs164423 | 1.064 | 0.993 | 0.534 | 0.285 | 0.018 | 0.005 | 0.005 | 0.035 | 0.68 | 0.54 | 0.59 | 0.65 | 0.62 | 0.49 |
| 1 | 162363732 | rs2655732 | 0.199 | 0.856 | 0.158 | 0.211 | 0.012 | 0 | 0 | 0 | 0.65 | 0.39 | 0.54 | 0.63 | 0.57 | 0.49 |
| 1 | 162365659 | rs164126 | 1.084 | 0.792 | -0.163 | 0.211 | 0.067 | 0.025 | 0.025 | 1.052 | 0.47 | 0.21 | 0.45 | 0.59 | 0.38 | 0.38 |
| 1 | 162366067 | rs16861748 | -0.088 | 0.725 | -0.063 | 0.224 | -0.016 | -0.011 | -0.011 | 0 | 0.05 | 0.1 | 0.02 | 0.02 | 0.02 | 0.12 |
| 1 | 162367235 | rs164124 | -0.182 | 0.714 | -0.206 | 0.182 | 0 | -0.001 | -0.001 | 0 | 0.7 | 0.31 | 0.68 | 0.76 | 0.66 | 0.64 |

| | | | | | | | | | | | | | | | | |
|---|---|---|---|---|---|---|---|---|---|---|---|---|---|---|---|---|
| 1 | 162368357 | rs164122 | 0.942 | 0.583 | -0.297 | 0.236 | 0.086 | 0.03 | 0.03 | 0.751 | 0.48 | 0.18 | 0.44 | 0.58 | 0.37 | 0.38 |
| 1 | 162371870 | rs2632549 | 1.497 | 0.331 | -0.318 | 0.213 | 0.074 | -0.034 | -0.034 | 0 | 0.39 | 0.21 | 0.26 | 0.44 | 0.25 | 0.23 |
| 1 | 162374809 | rs351450 | 0.477 | 0.436 | 0.121 | 0.202 | 0.027 | 0.041 | 0.041 | 0 | 0.88 | 0.65 | 0.8 | 0.83 | 0.88 | 0.82 |
| 1 | 162374853 | rs12746024 | -0.284 | 0.562 | -0.173 | 0.008 | -0.046 | 0.022 | 0.022 | 0.95 | 0.25 | 0.07 | 0.24 | 0.18 | 0.3 | 0.28 |
| 1 | 162375916 | rs351453 | 1.398 | 0.449 | 0.029 | 0.053 | 0.032 | -0.012 | -0.012 | 0 | 0.42 | 0.32 | 0.28 | 0.46 | 0.25 | 0.32 |
| 1 | 162385511 | rs164128 | 0.712 | 0.487 | 0.053 | 0.054 | 0.001 | -0.001 | -0.001 | 0 | 0.65 | 0.66 | 0.56 | 0.67 | 0.58 | 0.55 |
| 1 | 162387704 | rs164410 | 1.793 | 0.481 | -0.08 | 0.05 | 0.034 | -0.021 | -0.021 | 0.155 | 0.37 | 0.25 | 0.27 | 0.42 | 0.23 | 0.2 |
| 1 | 162390085 | rs905717 | 0.61 | 0.849 | -0.248 | -0.166 | 0.03 | -0.011 | -0.011 | 0 | 0.23 | 0.34 | 0.28 | 0.21 | 0.33 | 0.23 |
| 1 | 162416145 | rs469834 | -1.33 | 1.63 | -0.425 | -0.193 | -0.021 | 0.015 | 0.015 | 0 | 0.2 | 0.12 | 0.28 | 0.16 | 0.31 | 0.24 |
| 1 | 162416532 | rs469809 | -0.224 | 1.761 | -0.513 | -0.163 | -0.037 | -0.021 | -0.021 | 0 | 0.09 | 0.03 | 0.21 | 0.16 | 0.2 | 0.19 |
| 1 | 162428275 | rs10800559 | 0.363 | 2.222 | -0.781 | 0.395 | -0.051 | -0.109 | -0.109 | 0 | 0.16 | 0.02 | 0.25 | 0.46 | 0.2 | 0.29 |
| 1 | 162431320 | rs7532188 | 0.477 | 2.036 | -0.457 | 0.291 | -0.002 | -0.006 | -0.006 | 0.189 | 0.05 | 0.03 | 0.06 | 0 | 0.09 | 0.03 |
| 1 | 162433012 | rs164160 | -1.528 | 1.687 | -0.067 | 0.193 | 0.003 | -0.027 | -0.027 | 0.002 | 0.3 | 0.59 | 0.32 | 0.19 | 0.33 | 0.39 |
| 1 | 162439692 | rs164170 | -1.022 | 1.504 | -0.228 | 0.146 | 0.003 | 0.018 | 0.018 | 0 | 0.46 | 0.62 | 0.53 | 0.36 | 0.56 | 0.61 |
| 1 | 162439902 | rs164172 | -1.035 | 1.499 | -0.232 | 0.286 | 0.004 | 0.01 | 0.01 | 0 | 0.46 | 0.62 | 0.54 | 0.44 | 0.56 | 0.61 |
| 1 | 162440776 | rs12238992 | -1.013 | 1.05 | -0.267 | -0.32 | 0.019 | -0.045 | -0.045 | 0.112 | 0.34 | 0.41 | 0.34 | 0.21 | 0.42 | 0.29 |
| 1 | 162443785 | rs10919539 | -0.389 | -1.317 | -0.978 | -0.651 | -0.015 | -0.015 | -0.015 | 0 | 0.32 | 0.35 | 0.19 | 0.13 | 0.19 | 0.41 |
| 1 | 162445136 | rs12757408 | 0.071 | -1.208 | -0.974 | 0.229 | -0.027 | -0.006 | -0.006 | 0.019 | 0.4 | 0.26 | 0.43 | 0.37 | 0.46 | 0.44 |
| 1 | 162450093 | rs12733615 | -0.499 | -1.228 | -0.889 | 0.113 | -0.015 | -0.009 | -0.009 | 0.005 | 0.11 | 0.54 | 0.1 | 0.08 | 0.06 | 0.09 |
| 1 | 162455569 | rs461172 | -0.166 | -1.001 | -1.275 | 0.45 | -0.008 | -0.004 | -0.004 | 0 | 0.45 | 0.72 | 0.49 | 0.49 | 0.47 | 0.45 |
| 1 | 162458412 | rs6673387 | -0.998 | -0.774 | -1.468 | 0.439 | -0.049 | 0.007 | 0.007 | 0 | 0.66 | 0.84 | 0.74 | 0.72 | 0.57 | 0.58 |
| 1 | 162462526 | rs11803344 | 0.423 | -0.755 | -1.127 | 0.594 | 0.004 | 0.004 | 0.004 | 0 | 0.37 | 0.39 | 0.43 | 0.48 | 0.45 | 0.42 |
| 1 | 162465904 | rs1827635 | 0.718 | -0.906 | -1.253 | 0.569 | 0.014 | -0.066 | -0.066 | 0 | 0.16 | 0.04 | 0.24 | 0.23 | 0.1 | 0.13 |
| 1 | 162466523 | rs10919548 | -0.72 | -0.945 | -1.249 | 0.55 | 0 | 0.013 | 0.013 | 0 | 0.55 | 0.43 | 0.68 | 0.67 | 0.55 | 0.55 |
| 1 | 162468913 | rs10494370 | 0.559 | -0.896 | -1.173 | 0.598 | 0.029 | -0.034 | -0.034 | 0 | 0.17 | 0.09 | 0.25 | 0.27 | 0.11 | 0.14 |
| 1 | 162469475 | rs10919550 | 0.126 | -0.571 | -0.816 | 0.676 | 0 | 0 | 0 | 0 | 0.39 | 0.36 | 0.43 | 0.43 | 0.44 | 0.44 |
| 1 | 162490218 | rs6427680 | -0.043 | -0.558 | -0.807 | 0.682 | 0.01 | 0.009 | 0.009 | 0 | 0.39 | 0.45 | 0.44 | 0.48 | 0.44 | 0.44 |
| 1 | 162491166 | rs2342715 | 0.82 | -0.618 | -0.841 | 0.757 | -0.08 | -0.028 | -0.028 | 0.862 | 0.31 | 0.07 | 0.37 | 0.27 | 0.4 | 0.33 |
| 1 | 162493009 | rs1062174 | 0.16 | -0.591 | -0.7 | 0.699 | -0.003 | -0.003 | -0.003 | 0 | 0.39 | 0.35 | 0.43 | 0.46 | 0.44 | 0.44 |
| 1 | 162493123 | rs2236581 | 0.064 | -0.588 | -0.713 | 0.697 | 0.007 | 0.007 | 0.007 | 0 | 0.39 | 0.44 | 0.44 | 0.48 | 0.44 | 0.44 |
| 1 | 162505496 | rs16863247 | 1.345 | -0.815 | -0.773 | 0.805 | -0.03 | 0.051 | 0.051 | 4.98 | 0.07 | 0.16 | 0.03 | 0.12 | 0.02 | 0.05 |
| 1 | 162506369 | rs6665354 | 0.227 | -0.695 | -0.905 | 0.784 | -0.014 | -0.01 | -0.01 | 0 | 0.39 | 0.29 | 0.43 | 0.43 | 0.45 | 0.44 |
| 1 | 162521816 | rs10753578 | -0.302 | -0.823 | -1.141 | 0.45 | 0.007 | -0.004 | -0.004 | 0 | 0.26 | 0.25 | 0.32 | 0.32 | 0.23 | 0.28 |
| 1 | 162528505 | rs6704428 | 0.096 | -0.445 | -1.341 | 0.585 | 0.023 | -0.029 | -0.029 | 0 | 0.15 | 0.05 | 0.21 | 0.23 | 0.09 | 0.17 |
| 1 | 162543172 | rs12130399 | 0.185 | 0.044 | -1.276 | 0.547 | -0.019 | -0.031 | -0.031 | 0.051 | 0.41 | 0.18 | 0.52 | 0.51 | 0.52 | 0.46 |
| 1 | 162551486 | rs12736865 | 1.106 | 0.704 | -1.549 | 0.626 | -0.023 | -0.024 | -0.024 | 0 | 0.23 | 0.13 | 0.37 | 0.31 | 0.31 | 0.24 |
| 1 | 162556810 | rs10917570 | 0.725 | -0.007 | -1.485 | 0.587 | 0.016 | -0.07 | -0.07 | 0.18 | 0.12 | 0 | 0.2 | 0.26 | 0.09 | 0.17 |
| 1 | 162562775 | rs1510267 | -1.349 | -0.138 | -0.899 | 0.113 | -0.015 | -0.014 | -0.014 | 0.039 | 0.66 | 0.59 | 0.85 | 0.8 | 0.76 | 0.72 |
| 1 | 162567225 | rs16863833 | -0.956 | 0.172 | -0.223 | 0.293 | -0.003 | -0.002 | -0.002 | 0 | 0.25 | 0.29 | 0.33 | 0.27 | 0.24 | 0.28 |
| 1 | 162571610 | rs1995874 | 1.053 | 0.934 | -0.342 | 0.17 | -0.002 | -0.01 | -0.01 | 0 | 0.08 | 0.04 | 0.11 | 0 | 0.15 | 0.1 |
| 1 | 162587423 | rs4657210 | 0.367 | 0.317 | -0.413 | 0.309 | 0.004 | -0.002 | -0.002 | 1.929 | 0.09 | 0.11 | 0.13 | 0.11 | 0.13 | 0.07 |
| 1 | 162590088 | rs12027707 | 0.662 | 0.382 | -0.961 | 0.183 | -0.093 | -0.076 | -0.076 | 1.019 | 0.3 | 0.04 | 0.29 | 0.3 | 0.38 | 0.35 |
| 1 | 162591307 | rs17431968 | 0.292 | 0.314 | -0.413 | 0.218 | -0.012 | -0.021 | -0.021 | 0 | 0.36 | 0.26 | 0.35 | 0.46 | 0.42 | 0.54 |
| 1 | 162592702 | rs16864157 | 0.094 | 0.312 | -0.366 | 0.189 | -0.004 | -0.004 | -0.004 | 1.157 | 0.41 | 0.37 | 0.38 | 0.52 | 0.47 | 0.68 |
| 1 | 162597545 | rs6702652 | 0.004 | 0.185 | -0.358 | 0.092 | -0.001 | 0 | 0 | 0 | 0.21 | 0.2 | 0.34 | 0.23 | 0.22 | 0.19 |
| 1 | 162600119 | rs7544972 | -0.607 | 0.177 | -0.228 | 0.079 | -0.014 | -0.022 | -0.022 | 0 | 0.62 | 0.54 | 0.72 | 0.76 | 0.7 | 0.87 |
| 1 | 162606728 | rs4656375 | 1.132 | -0.222 | -0.048 | 0.307 | 0.013 | 0.018 | 0.018 | 1.516 | 0.6 | 0.23 | 0.49 | 0.58 | 0.62 | 0.54 |
| 1 | 162613122 | rs7524537 | -0.288 | -0.236 | -0.145 | 0.141 | -0.019 | -0.015 | -0.015 | 0 | 0.2 | 0.09 | 0.32 | 0.21 | 0.22 | 0.2 |
| 1 | 162619362 | rs10494373 | 0.497 | -0.266 | 0.128 | 0.092 | 0 | 0.001 | 0.001 | 0 | 0.05 | 0.05 | 0.08 | 0.02 | 0.08 | 0.03 |
| 1 | 162619828 | rs10799854 | 0.601 | -0.312 | 0.836 | 0.085 | 0.008 | -0.002 | -0.002 | 0.117 | 0.9 | 0.74 | 0.92 | 0.89 | 0.92 | 0.76 |
| 1 | 162625756 | rs988383 | 0.701 | -0.332 | 1.004 | 0.092 | 0.006 | 0.011 | 0.011 | 0 | 0.89 | 0.69 | 0.88 | 0.88 | 0.89 | 0.85 |
| 1 | 162630894 | rs10917583 | 1.123 | -0.353 | 1.19 | 0.093 | 0.001 | 0.02 | 0.02 | 0.051 | 0.88 | 0.56 | 0.87 | 0.82 | 0.89 | 0.84 |
| 1 | 162635875 | rs12044481 | 0.771 | -0.6 | 0.434 | -0.057 | -0.002 | -0.031 | -0.031 | 0 | 0.24 | 0.04 | 0.18 | 0.25 | 0.21 | 0.07 |
| 1 | 162639774 | rs4657219 | 1.76 | -0.536 | 1.448 | -0.199 | -0.002 | -0.04 | -0.04 | 0.094 | 0.84 | 0.55 | 0.83 | 0.87 | 0.83 | 0.74 |
| 1 | 162640649 | rs4656376 | 2.12 | -0.59 | 1.395 | -0.205 | 0.018 | 0.03 | 0.03 | 1.89 | 0.6 | 0.18 | 0.49 | 0.6 | 0.63 | 0.58 |
| 1 | 162641516 | rs12116496 | -0.84 | -0.331 | 1.932 | 0.301 | 0 | 0.002 | 0.002 | 0 | 0.06 | 0.08 | 0.04 | 0.05 | 0.06 | 0.12 |
| 1 | 162642183 | rs4657221 | 2.391 | -0.399 | 1.783 | 0.284 | -0.016 | -0.029 | -0.029 | 0 | 0.8 | 0.31 | 0.8 | 0.82 | 0.83 | 0.73 |
| 1 | 162644167 | rs4656379 | 0.565 | 0.007 | 1.709 | 0.464 | 0.015 | 0.084 | 0.084 | 0 | 0.9 | 0.75 | 0.74 | 0.74 | 0.92 | 0.88 |
| 1 | 162644411 | rs10799862 | 2.18 | -0.024 | 2.013 | 0.451 | 0.015 | 0.056 | 0.056 | 0 | 0.77 | 0.57 | 0.6 | 0.6 | 0.75 | 0.67 |
| 1 | 162653274 | rs4657226 | 2.103 | -0.005 | 1.893 | 0.405 | -0.001 | 0.03 | 0.03 | 0.317 | 0.75 | 0.37 | 0.55 | 0.62 | 0.75 | 0.69 |
| 1 | 162661011 | rs7553831 | 1.818 | 0.144 | 1.675 | 0.474 | 0.015 | 0.048 | 0.048 | 0.046 | 0.79 | 0.5 | 0.57 | 0.75 | 0.78 | 0.77 |
| 1 | 162669775 | rs7520418 | 0.609 | 0.08 | 1.646 | 0.506 | 0.016 | 0.106 | 0.106 | 0 | 0.88 | 0.62 | 0.72 | 0.79 | 0.91 | 0.9 |
| 1 | 162670043 | rs16843630 | -0.928 | 0.122 | 1.635 | 0.574 | 0.023 | 0.025 | 0.025 | 0 | 0.07 | 0.11 | 0.14 | 0.15 | 0.14 | 0.14 |
| 1 | 162670541 | rs4477251 | 2.053 | 0.1 | 1.539 | 0.561 | 0 | 0.056 | 0.056 | 0.167 | 0.76 | 0.39 | 0.54 | 0.58 | 0.76 | 0.68 |
| 1 | 162672011 | rs17433710 | -0.719 | 0.128 | 1.252 | 0.625 | 0.012 | 0.015 | 0.015 | 0 | 0.06 | 0.07 | 0.14 | 0.2 | 0.13 | 0.15 |
| 1 | 162675040 | rs6682905 | 0.659 | 0.038 | 1.396 | 0.631 | -0.024 | 0.091 | 0.091 | 0 | 0.86 | 0.6 | 0.71 | 0.79 | 0.93 | 0.9 |
| 1 | 162681151 | rs4559477 | 0.897 | 0.028 | 1.543 | 0.583 | -0.016 | 0.142 | 0.142 | 0 | 0.84 | 0.56 | 0.66 | 0.65 | 0.89 | 0.81 |
| 1 | 162684359 | rs6695961 | 1.001 | 0.015 | 1.595 | 0.58 | -0.019 | 0.162 | 0.162 | 0 | 0.84 | 0.53 | 0.66 | 0.61 | 0.89 | 0.8 |
| 1 | 162684606 | rs6698508 | 0.507 | -0.263 | 0.996 | 0.08 | -0.005 | 0.098 | 0.098 | 0 | 0.16 | 0 | 0.08 | 0 | 0.11 | 0.02 |

| | | | | | | | | | | | | | | | | |
|---|---|---|---|---|---|---|---|---|---|---|---|---|---|---|---|---|
| 1 | 162690939 | rs7516390 | -0.534 | 0.04 | 1.796 | 0.52 | 0.071 | 0.001 | 0.001 | 0 | 0.06 | 0.2 | 0.1 | 0.02 | 0.11 | 0.05 |
| 1 | 162694593 | rs1039871 | 1.307 | -0.362 | 1.541 | 0.494 | -0.017 | 0.09 | 0.09 | 0 | 0.8 | 0.55 | 0.62 | 0.61 | 0.82 | 0.79 |
| 1 | 162713074 | rs2665496 | 0.353 | -0.543 | 1.613 | 0.532 | -0.034 | 0.027 | 0.027 | 0 | 0.84 | 0.68 | 0.72 | 0.79 | 0.91 | 0.9 |
| 1 | 162722216 | rs1704740 | 0.641 | -0.581 | 1.713 | 0.514 | -0.051 | 0.033 | 0.033 | 0 | 0.82 | 0.59 | 0.71 | 0.79 | 0.91 | 0.89 |
| 1 | 162732491 | rs2805025 | 1.449 | -0.506 | 1.21 | 0.422 | 0.031 | -0.036 | -0.036 | 0 | 0.71 | 0.2 | 0.55 | 0.58 | 0.77 | 0.67 |
| 1 | 162736463 | rs1000050 | 0.829 | -0.459 | 1.238 | 0.508 | -0.034 | 0.062 | 0.062 | 0 | 0.77 | 0.39 | 0.64 | 0.59 | 0.85 | 0.72 |
| 1 | 162739009 | rs6676809 | -0.889 | -0.464 | 1.197 | 0.503 | -0.034 | 0.1 | 0.1 | 0 | 0.22 | 0.48 | 0.35 | 0.37 | 0.15 | 0.27 |
| 1 | 162741387 | rs16844884 | -0.096 | -0.498 | 0.132 | 0.457 | -0.003 | -0.012 | -0.012 | 0 | 0.06 | 0.09 | 0.06 | 0.05 | 0.07 | 0.07 |
| 1 | 162748025 | rs1780007 | 1.135 | -0.437 | -0.391 | 0.475 | 0.045 | -0.05 | -0.05 | 0 | 0.7 | 0.2 | 0.57 | 0.58 | 0.78 | 0.66 |
| 1 | 162766476 | rs1780019 | 1.108 | -0.424 | -0.376 | 0.481 | 0.037 | -0.041 | -0.041 | 0.06 | 0.7 | 0.21 | 0.57 | 0.56 | 0.78 | 0.66 |
| 1 | 162770647 | rs11589262 | 0.094 | 0.732 | -0.771 | 0.07 | -0.031 | 0.01 | 0.01 | 0 | 0.25 | 0.15 | 0.27 | 0.22 | 0.35 | 0.09 |
| 1 | 162772985 | rs10917597 | 0.361 | 1.009 | -0.893 | 0.041 | -0.099 | -0.018 | -0.018 | 0.119 | 0.2 | 0.01 | 0.21 | 0.15 | 0.28 | 0.08 |
| 1 | 162776137 | rs4656381 | -0.003 | 0.908 | -0.795 | 0.099 | -0.022 | 0.009 | 0.009 | 0 | 0.25 | 0.15 | 0.25 | 0.22 | 0.32 | 0.09 |
| 1 | 162787989 | rs7522935 | 0.034 | 0.576 | -0.624 | -0.152 | 0.008 | 0.022 | 0.022 | 0 | 0.1 | 0.04 | 0.07 | 0.06 | 0.08 | 0.05 |
| 1 | 162793865 | rs1510324 | 0.265 | 0.6 | -0.388 | -0.122 | -0.009 | 0.007 | 0.007 | 0 | 0.26 | 0.19 | 0.28 | 0.22 | 0.32 | 0.1 |
| 1 | 162794500 | rs6673329 | 2.212 | 0.481 | 0.205 | -0.313 | -0.006 | -0.038 | -0.038 | 2.725 | 0.07 | 0.14 | 0.04 | 0 | 0.02 | 0.01 |
| 1 | 162797691 | rs2684874 | -0.73 | 0.376 | -0.201 | -0.146 | -0.001 | 0.009 | 0.009 | 0 | 0.76 | 0.67 | 0.66 | 0.59 | 0.75 | 0.75 |
| 1 | 162798636 | rs10917603 | 1.773 | 0.108 | -0.748 | -0.17 | -0.028 | -0.02 | -0.02 | 0 | 0.24 | 0.06 | 0.21 | 0.22 | 0.24 | 0.48 |
| 1 | 162800207 | rs12026860 | 1.773 | 0.108 | -0.748 | -0.163 | -0.028 | -0.031 | -0.031 | 0.188 | 0.24 | 0.06 | 0.21 | 0.23 | 0.24 | 0.48 |
| 1 | 162800426 | rs7354897 | 0.559 | 0.243 | -0.359 | 0.054 | -0.004 | -0.017 | -0.017 | 0 | 0.42 | 0.18 | 0.31 | 0.4 | 0.4 | 0.62 |
| 1 | 162802615 | rs2684879 | 0.424 | 0.398 | -0.356 | 0.212 | -0.009 | 0.016 | 0.016 | 0 | 0.26 | 0.19 | 0.28 | 0.15 | 0.32 | 0.09 |
| 1 | 162810754 | rs2684880 | 0.418 | 0.416 | -0.57 | 0.137 | -0.009 | 0.011 | 0.011 | 0 | 0.26 | 0.19 | 0.28 | 0.17 | 0.32 | 0.1 |
| 1 | 162813583 | rs6691351 | 0.02 | 0.31 | -0.468 | 0.166 | -0.002 | 0.004 | 0.004 | 0 | 0.59 | 0.59 | 0.49 | 0.5 | 0.59 | 0.7 |
| 1 | 162813651 | rs6679069 | 0.017 | 0.35 | -0.473 | 0.102 | -0.001 | 0.001 | 0.001 | 0 | 0.59 | 0.59 | 0.48 | 0.48 | 0.58 | 0.69 |
| 1 | 162816410 | rs1848860 | -0.065 | 0.293 | -0.709 | -0.194 | -0.001 | -0.003 | -0.003 | 0 | 0.17 | 0.2 | 0.11 | 0.03 | 0.17 | 0.07 |
| 1 | 162817494 | rs1779993 | -0.591 | 0.133 | -0.498 | 0.041 | -0.005 | 0.118 | 0.118 | 0 | 0.91 | 0.86 | 0.78 | 0.58 | 0.93 | 0.78 |
| 1 | 162819508 | rs16845797 | 0.725 | -0.152 | -0.5 | -0.112 | -0.033 | 0.054 | 0.054 | 0 | 0.09 | 0.19 | 0.07 | 0.2 | 0.03 | 0.13 |
| 1 | 162820067 | rs2880639 | 0.324 | 0.157 | -0.954 | -0.538 | -0.035 | 0.051 | 0.051 | 0 | 0.39 | 0.21 | 0.36 | 0.25 | 0.41 | 0.48 |
| 1 | 162821291 | rs1912453 | -0.714 | 0.184 | -0.907 | -0.675 | -0.006 | 0.01 | 0.01 | 0 | 0.62 | 0.52 | 0.55 | 0.46 | 0.63 | 0.66 |
| 1 | 162822540 | rs12134171 | 0.302 | 0.509 | -1.189 | -0.747 | -0.012 | 0.034 | 0.034 | 0 | 0.37 | 0.26 | 0.34 | 0.25 | 0.37 | 0.46 |
| 1 | 162825258 | rs12036026 | -0.24 | 0.652 | -0.939 | -0.959 | -0.001 | -0.016 | -0.016 | 0.122 | 0.17 | 0.23 | 0.11 | 0.03 | 0.18 | 0.06 |
| 1 | 162826331 | rs12132001 | 0.809 | 0.662 | -1.251 | -0.985 | -0.027 | 0.033 | 0.033 | 0 | 0.14 | 0.06 | 0.12 | 0 | 0.19 | 0.06 |
| 1 | 162827816 | rs1355285 | -0.4 | 0.315 | -1.032 | -0.799 | 0 | -0.021 | -0.021 | 0.348 | 0.17 | 0.2 | 0.1 | 0.03 | 0.17 | 0.04 |
| 1 | 162828060 | rs1355284 | -0.033 | 0.375 | -1.129 | -0.77 | -0.006 | 0.021 | 0.021 | 0 | 0.16 | 0.13 | 0.09 | 0.03 | 0.16 | 0.04 |
| 1 | 162829986 | rs1510313 | 0.022 | 0.354 | -1.222 | -0.766 | -0.01 | 0.045 | 0.045 | 0 | 0.15 | 0.11 | 0.09 | 0.03 | 0.16 | 0.04 |
| 1 | 162831317 | rs11588419 | -0.219 | -0.196 | -1.138 | -0.516 | 0.013 | 0.009 | 0.009 | 0 | 0.6 | 0.37 | 0.52 | 0.37 | 0.71 | 0.38 |
| 1 | 162831535 | rs12141395 | 1.546 | -0.127 | -1.306 | -0.506 | -0.043 | -0.02 | -0.02 | 0 | 0.2 | 0.05 | 0.22 | 0.21 | 0.22 | 0.43 |
| 1 | 162832406 | rs12024415 | 0.092 | -0.409 | -1.214 | -0.584 | 0 | 0.008 | 0.008 | 0 | 0.19 | 0.2 | 0.15 | 0.13 | 0.19 | 0.1 |
| 1 | 162835703 | rs12137443 | 0.57 | -0.339 | -1.329 | -0.585 | -0.014 | 0.037 | 0.037 | 0 | 0.17 | 0.09 | 0.14 | 0.09 | 0.19 | 0.07 |
| 1 | 162838142 | rs1779995 | 0.255 | -0.281 | -0.546 | -0.165 | -0.014 | 0.165 | 0.165 | 0 | 0.84 | 0.62 | 0.67 | 0.43 | 0.88 | 0.73 |
| 1 | 162840531 | rs2175256 | 0.706 | -0.263 | -1.407 | -0.741 | -0.129 | 0.24 | 0.24 | 0 | 0.36 | 0.05 | 0.25 | 0.05 | 0.45 | 0.22 |
| 1 | 162842202 | rs12047510 | 1.004 | -0.176 | -0.87 | 0.024 | -0.018 | -0.008 | -0.008 | 0.138 | 0.13 | 0.07 | 0.12 | 0.12 | 0.15 | 0.07 |
| 1 | 162843606 | rs2803881 | 1.115 | -0.254 | -0.488 | 0.064 | -0.015 | -0.031 | -0.031 | 0.159 | 0.64 | 0.43 | 0.51 | 0.24 | 0.7 | 0.58 |
| 1 | 162846170 | rs10917623 | -0.494 | -0.164 | -0.589 | 0.483 | -0.004 | 0.142 | 0.142 | 0 | 0.89 | 0.82 | 0.72 | 0.44 | 0.91 | 0.7 |
| 1 | 162864313 | rs4657235 | 0.526 | -0.285 | -0.412 | 0.418 | -0.026 | 0.15 | 0.15 | 0 | 0.83 | 0.62 | 0.67 | 0.41 | 0.88 | 0.69 |
| 1 | 162866629 | rs7523184 | -0.567 | -0.115 | -0.896 | -0.159 | -0.005 | 0.1 | 0.1 | 0 | 0.47 | 0.25 | 0.38 | 0.25 | 0.53 | 0.25 |
| 1 | 162866915 | rs1986860 | 1.609 | -0.296 | -1.08 | -0.192 | -0.098 | 0.121 | 0.121 | 0.817 | 0.35 | 0.02 | 0.27 | 0.16 | 0.38 | 0.39 |
| 1 | 162870633 | rs4540634 | 0.958 | -0.208 | -0.869 | 0.404 | -0.001 | -0.002 | -0.002 | 0 | 0.43 | 0.58 | 0.37 | 0.38 | 0.4 | 0.48 |
| 1 | 162871675 | rs11583881 | 1.562 | 0.15 | -1.166 | 0.302 | -0.045 | 0.069 | 0.069 | 0 | 0.13 | 0.02 | 0.12 | 0.03 | 0.17 | 0.05 |
| 1 | 162874971 | rs10799880 | -1.262 | -0.329 | -0.669 | 0.684 | -0.011 | 0.119 | 0.119 | 0 | 0.89 | 0.83 | 0.74 | 0.64 | 0.93 | 0.74 |
| 1 | 162877157 | rs12128758 | -0.537 | -0.259 | -1.357 | -0.327 | -0.005 | 0.014 | 0.014 | 0 | 0.39 | 0.36 | 0.3 | 0.14 | 0.42 | 0.23 |
| 1 | 162877621 | rs10917635 | 0.627 | -0.119 | -1.399 | -0.487 | -0.044 | 0.071 | 0.071 | 0 | 0.26 | 0.11 | 0.22 | 0.07 | 0.36 | 0.16 |
| 1 | 162878023 | rs10917636 | 0.424 | -0.122 | -1.405 | -0.489 | -0.008 | -0.012 | -0.012 | 0 | 0.27 | 0.23 | 0.23 | 0.07 | 0.36 | 0.16 |
| 1 | 162879819 | rs2880640 | 1.638 | -0.207 | -2.148 | -0.523 | -0.11 | 0.151 | 0.151 | 0.747 | 0.35 | 0.02 | 0.29 | 0.14 | 0.39 | 0.4 |
| 1 | 162881307 | rs10917640 | 0.169 | 0.031 | -1.919 | -0.514 | 0.004 | 0.004 | 0.004 | 0 | 0.66 | 0.33 | 0.52 | 0.34 | 0.74 | 0.5 |
| 1 | 162883337 | rs10799883 | -0.367 | -0.053 | -1.942 | -0.513 | -0.013 | 0.089 | 0.089 | 0 | 0.45 | 0.27 | 0.37 | 0.26 | 0.52 | 0.2 |
| 1 | 162883597 | rs10753595 | 1.163 | -0.037 | -1.967 | -0.545 | -0.041 | 0.064 | 0.064 | 0 | 0.26 | 0.12 | 0.22 | 0.07 | 0.36 | 0.12 |
| 1 | 162883668 | rs11576066 | 2.413 | -0.325 | -1.727 | -0.28 | 0.013 | 0.001 | 0.001 | 1.069 | 0.1 | 0.09 | 0.05 | 0.05 | 0.04 | 0.02 |
| 1 | 162884258 | rs10753596 | 0.373 | 0.01 | -1.675 | -0.354 | -0.051 | 0.071 | 0.071 | 0 | 0.36 | 0.19 | 0.32 | 0.2 | 0.48 | 0.18 |
| 1 | 162887095 | rs4276890 | 0.685 | 0.02 | -1.423 | 0.023 | 0.013 | 0.093 | 0.093 | 0.02 | 0.31 | 0.14 | 0.2 | 0.13 | 0.26 | 0.31 |
| 1 | 162890041 | rs4500290 | -0.508 | 0.037 | -0.74 | 0.977 | -0.009 | -0.021 | -0.021 | 0 | 0.42 | 0.71 | 0.37 | 0.42 | 0.44 | 0.21 |
| 1 | 162890135 | rs4281307 | 0.452 | 0.182 | -0.652 | 0.522 | -0.034 | 0.048 | 0.048 | 0 | 0.26 | 0.14 | 0.26 | 0.11 | 0.36 | 0.15 |
| 1 | 162890325 | rs2343504 | 0.34 | 0.161 | -0.568 | 0.513 | -0.021 | 0.035 | 0.035 | 0 | 0.27 | 0.17 | 0.26 | 0.11 | 0.36 | 0.15 |
| 1 | 162892049 | rs6700130 | -0.947 | -0.214 | -0.767 | 0.532 | 0.026 | 0 | 0 | 0.738 | 0.38 | 0.74 | 0.33 | 0.6 | 0.24 | 0.33 |
| 1 | 162893276 | rs6657722 | -0.941 | -0.216 | -0.767 | 0.531 | 0.025 | 0 | 0 | 0.774 | 0.38 | 0.74 | 0.33 | 0.6 | 0.24 | 0.33 |
| 1 | 162893500 | rs12093487 | 0.351 | -0.186 | -0.549 | 0.519 | -0.005 | -0.017 | -0.017 | 0.389 | 0.13 | 0.33 | 0.07 | 0.06 | 0.07 | 0.01 |
| 1 | 162894492 | rs10753597 | -1.221 | -0.13 | -0.714 | 0.38 | 0 | 0.018 | 0.018 | 0.211 | 0.61 | 0.66 | 0.56 | 0.71 | 0.6 | 0.49 |
| 1 | 162896345 | rs6427697 | -0.739 | -0.372 | -0.379 | 0.153 | -0.009 | 0.022 | 0.022 | 0.037 | 0.54 | 0.63 | 0.48 | 0.67 | 0.53 | 0.5 |
| 1 | 162896495 | rs6427699 | -0.456 | -0.323 | -0.323 | 0.56 | 0.005 | 0.002 | 0.002 | 0 | 0.5 | 0.54 | 0.43 | 0.48 | 0.48 | 0.4 |

| | | | | | | | | | | | | | | | | |
|---|---|---|---|---|---|---|---|---|---|---|---|---|---|---|---|---|
| 1 | 162896697 | rs2343508 | -1.744 | -0.35 | -0.389 | 0.54 | -0.035 | -0.016 | -0.016 | 2.031 | 0.65 | 0.88 | 0.55 | 0.68 | 0.6 | 0.51 |
| 1 | 162898732 | rs10917642 | 1.872 | -0.657 | -0.612 | 0.553 | 0.033 | -0.08 | -0.08 | 0 | 0.12 | 0.01 | 0.1 | 0.33 | 0.03 | 0.18 |
| 1 | 162901055 | rs4233395 | -1.951 | -0.57 | -0.434 | 0.464 | -0.034 | -0.016 | -0.016 | 0 | 0.65 | 0.88 | 0.55 | 0.68 | 0.6 | 0.51 |
| 1 | 162905537 | rs10399626 | 0.966 | -0.655 | -0.327 | 0.384 | -0.011 | 0.012 | 0.012 | 0 | 0.25 | 0.21 | 0.25 | 0.15 | 0.34 | 0.15 |
| 1 | 162910384 | rs6664865 | -0.961 | -0.912 | -0.22 | 0.496 | -0.035 | 0.042 | 0.042 | 0.217 | 0.31 | 0.57 | 0.25 | 0.52 | 0.19 | 0.35 |
| 1 | 162910683 | rs4492592 | 1.375 | -0.953 | -0.346 | 0.469 | 0.043 | -0.087 | -0.087 | 0 | 0.13 | 0.01 | 0.1 | 0.34 | 0.03 | 0.18 |
| 1 | 162915745 | rs12091621 | 1.043 | -0.865 | -0.039 | 0.258 | 0.003 | -0.077 | -0.077 | 0.019 | 0.12 | 0.28 | 0.07 | 0 | 0.11 | 0.07 |
| 1 | 162915993 | rs17490891 | 0.216 | -0.508 | -0.474 | 0.109 | -0.014 | 0.019 | 0.019 | 0 | 0.37 | 0.31 | 0.34 | 0.16 | 0.47 | 0.18 |
| 1 | 162918233 | rs12124940 | 1.141 | -0.059 | -0.524 | 0.296 | -0.028 | -0.014 | -0.014 | 0 | 0.06 | 0 | 0.05 | 0.08 | 0.08 | 0.06 |
| 1 | 162920421 | rs12021880 | 1.389 | -0.309 | -0.662 | 0.281 | 0.051 | -0.097 | -0.097 | 0 | 0.15 | 0 | 0.11 | 0.34 | 0.05 | 0.15 |
| 1 | 162920524 | rs10917649 | -0.203 | -0.364 | -0.424 | 0.202 | -0.041 | 0.038 | 0.038 | 0 | 0.29 | 0.36 | 0.2 | 0.44 | 0.15 | 0.26 |
| 1 | 162920811 | rs2343490 | -1.142 | -0.347 | -0.419 | 0.21 | -0.025 | 0.017 | 0.017 | 0 | 0.37 | 0.56 | 0.27 | 0.5 | 0.23 | 0.32 |
| 1 | 162927922 | rs10917653 | -0.861 | -0.584 | -0.682 | -0.023 | 0.009 | 0.014 | 0.014 | 0 | 0.55 | 0.39 | 0.66 | 0.68 | 0.55 | 0.39 |
| 1 | 162932150 | rs10799891 | 1.248 | -0.554 | -0.697 | 0.118 | 0.02 | -0.025 | -0.025 | 2.243 | 0.16 | 0.09 | 0.31 | 0.46 | 0.05 | 0.17 |
| 1 | 162932163 | rs6686926 | -1.483 | 0.021 | -0.46 | 0.132 | -0.006 | -0.006 | -0.006 | 0 | 0.7 | 0.83 | 0.76 | 0.75 | 0.66 | 0.49 |
| 1 | 162932822 | rs11585636 | 1.098 | 0.249 | -0.02 | -0.061 | -0.011 | 0.003 | 0.003 | 0 | 0.17 | 0.15 | 0.2 | 0.15 | 0.3 | 0.13 |
| 1 | 162934692 | rs6427703 | -0.27 | 0.121 | 0.103 | -0.058 | 0.031 | -0.073 | -0.073 | 0.141 | 0.3 | 0.49 | 0.28 | 0.15 | 0.4 | 0.19 |
| 1 | 162938179 | rs17438400 | 0.036 | -0.021 | -0.214 | 0.128 | -0.006 | 0.007 | 0.007 | 0 | 0.26 | 0.26 | 0.27 | 0.14 | 0.38 | 0.2 |
| 1 | 162938703 | rs2343278 | -1.147 | -0.379 | -0.289 | 0.233 | -0.02 | 0.004 | 0.004 | 0.146 | 0.69 | 0.8 | 0.76 | 0.74 | 0.64 | 0.5 |
| 1 | 162940693 | rs12126781 | -0.153 | -0.366 | 0.255 | 0.255 | 0.012 | -0.027 | -0.027 | 0.532 | 0.29 | 0.44 | 0.28 | 0.22 | 0.38 | 0.22 |
| 1 | 162945443 | rs11808024 | 1.02 | -0.371 | 0.266 | 0.218 | -0.047 | 0.075 | 0.075 | 0 | 0.22 | 0.09 | 0.2 | 0.09 | 0.34 | 0.13 |
| 1 | 162945455 | rs7339985 | -0.988 | -0.375 | 0.414 | 0.309 | 0.059 | 0.142 | 0.142 | 3.202 | 0.07 | 0.35 | 0.09 | 0.13 | 0.04 | 0.09 |
| 1 | 162949567 | rs16847360 | 0.582 | -0.482 | 0.439 | 0.702 | 0.003 | 0.004 | 0.004 | 0 | 0.31 | 0.23 | 0.39 | 0.29 | 0.21 | 0.15 |
| 1 | 162957811 | rs1507741 | 0.118 | -0.397 | 0.258 | 0.486 | 0.001 | -0.026 | -0.026 | 0 | 0.27 | 0.29 | 0.25 | 0.1 | 0.36 | 0.17 |
| 1 | 162958403 | rs2841966 | 0.227 | -0.513 | 0.045 | 0.571 | 0.011 | -0.007 | -0.007 | 1.395 | 0.46 | 0.37 | 0.56 | 0.52 | 0.33 | 0.39 |
| 1 | 162958937 | rs2661320 | -0.439 | 0.017 | 0.345 | 0.55 | 0.002 | 0.043 | 0.043 | 0.007 | 0.3 | 0.4 | 0.31 | 0.55 | 0.37 | 0.31 |
| 1 | 162959836 | rs2841979 | -0.213 | 0.205 | 0.522 | -0.018 | -0.006 | 0.015 | 0.015 | 0 | 0.74 | 0.78 | 0.73 | 0.79 | 0.64 | 0.83 |
| 1 | 162961718 | rs1507742 | 0.015 | 0.191 | 0.582 | -0.024 | 0.002 | -0.018 | -0.018 | 0 | 0.72 | 0.65 | 0.72 | 0.78 | 0.64 | 0.83 |
| 1 | 162969472 | rs1507744 | -0.075 | 0.138 | 0.371 | -0.071 | 0.003 | 0.001 | 0.001 | 0 | 0.45 | 0.38 | 0.53 | 0.61 | 0.36 | 0.33 |
| 1 | 162970153 | rs2842002 | -0.467 | 0.4 | 0.086 | -0.317 | -0.019 | 0.045 | 0.045 | 0 | 0.81 | 0.91 | 0.84 | 0.95 | 0.69 | 0.92 |
| 1 | 162980295 | rs2842003 | 0.934 | 0.4 | 0.087 | -0.159 | -0.001 | 0.034 | 0.034 | 0.429 | 0.56 | 0.37 | 0.65 | 0.72 | 0.53 | 0.74 |
| 1 | 162995355 | rs1507746 | 0.122 | 0.355 | -0.302 | -0.141 | -0.001 | -0.049 | -0.049 | 0 | 0.73 | 0.75 | 0.83 | 0.95 | 0.73 | 0.92 |
| 1 | 163009498 | rs2842006 | 0.368 | 0.346 | -0.232 | 0.367 | -0.001 | -0.002 | -0.002 | 0 | 0.54 | 0.53 | 0.48 | 0.33 | 0.57 | 0.64 |
| 1 | 163017585 | rs2661337 | -0.354 | 0.356 | -0.072 | 0.76 | 0 | -0.001 | -0.001 | 0.313 | 0.91 | 0.92 | 0.93 | 0.85 | 0.94 | 0.93 |
| 1 | 163018934 | rs2661339 | -0.211 | 0.328 | 0.04 | 0.747 | -0.006 | 0.02 | 0.02 | 0.373 | 0.9 | 0.84 | 0.93 | 0.85 | 0.94 | 0.93 |
| 1 | 163027520 | rs2842020 | 0.538 | 0.326 | 0.191 | 0.749 | -0.004 | 0.011 | 0.011 | 0.183 | 0.85 | 0.75 | 0.89 | 0.83 | 0.9 | 0.77 |
| 1 | 163031580 | rs2661275 | 0.294 | 0.382 | -0.065 | 0.503 | -0.004 | -0.055 | -0.055 | 0.904 | 0.84 | 0.64 | 0.88 | 0.93 | 0.9 | 0.81 |
| 1 | 163033511 | rs951437 | 0.186 | 0.598 | -0.643 | 0.39 | -0.02 | -0.045 | -0.045 | 0 | 0.37 | 0.19 | 0.5 | 0.49 | 0.47 | 0.28 |
| 1 | 163033537 | rs951438 | 0.186 | 0.617 | -0.643 | 0.39 | -0.018 | -0.045 | -0.045 | 0 | 0.37 | 0.19 | 0.5 | 0.49 | 0.47 | 0.28 |
| 1 | 163033691 | rs951439 | 0.593 | 0.578 | -0.653 | 0.398 | 0 | 0.001 | 0.001 | 0.144 | 0.47 | 0.48 | 0.39 | 0.43 | 0.43 | 0.52 |
| 1 | 163038613 | rs7515900 | 0.661 | 0.619 | -0.598 | 0.342 | 0 | 0 | 0 | 0 | 0.45 | 0.48 | 0.38 | 0.43 | 0.43 | 0.52 |
| 1 | 163039777 | rs2661319 | -0.423 | 0.918 | -1.338 | 0.761 | -0.021 | -0.04 | -0.04 | 0 | 0.39 | 0.2 | 0.51 | 0.49 | 0.49 | 0.28 |
| 1 | 163046351 | rs10759 | -0.887 | 1.027 | -1.5 | 1.139 | -0.021 | -0.06 | -0.06 | 0 | 0.23 | 0.15 | 0.29 | 0.42 | 0.27 | 0.34 |
| 1 | 163053289 | rs2063142 | -0.924 | 0.73 | -1.571 | 1.371 | -0.033 | -0.094 | -0.094 | 1.394 | 0.2 | 0.06 | 0.13 | 0.38 | 0.22 | 0.23 |
| 1 | 163054445 | rs12077304 | 0.099 | 0.396 | -1.304 | 0.747 | -0.005 | -0.05 | -0.05 | 1.382 | 0.11 | 0.18 | 0.09 | 0 | 0.11 | 0.02 |
| 1 | 163056062 | rs966999 | -1.846 | 0.145 | -0.746 | 0.528 | 0 | 0 | 0 | 0 | 0.45 | 0.57 | 0.52 | 0.49 | 0.45 | 0.48 |
| 1 | 163058194 | rs2841959 | -2.038 | 0.14 | -0.769 | 0.526 | 0 | 0.001 | 0.001 | 0 | 0.44 | 0.47 | 0.51 | 0.49 | 0.45 | 0.48 |
| 1 | 163064483 | rs2661303 | -1.194 | 0.128 | -0.788 | 0.521 | 0 | -0.012 | -0.012 | 0 | 0.31 | 0.27 | 0.32 | 0.41 | 0.3 | 0.41 |
| 1 | 163073502 | rs10799899 | -0.281 | 0.239 | -0.861 | 0.564 | 0 | -0.017 | -0.017 | 0 | 0.13 | 0.2 | 0.2 | 0.08 | 0.16 | 0.07 |
| 1 | 163080200 | rs2940249 | -1.847 | 0.139 | -0.822 | 0.546 | -0.002 | -0.004 | -0.004 | 0 | 0.4 | 0.38 | 0.47 | 0.5 | 0.44 | 0.45 |
| 1 | 163080661 | rs2841977 | -1.466 | 0.006 | -1.562 | -0.071 | -0.062 | 0.083 | 0.083 | 0 | 0.73 | 0.94 | 0.73 | 0.89 | 0.67 | 0.86 |
| 1 | 163090421 | rs6691456 | 0.43 | 0.027 | -1.451 | 0.029 | 0.023 | 0.015 | 0.015 | 0 | 0.07 | 0.18 | 0.15 | 0.08 | 0.08 | 0.05 |
| 1 | 163093022 | rs10917680 | 0.753 | -0.379 | -2.203 | 0.176 | -0.019 | -0.024 | -0.024 | 0.994 | 0.2 | 0.1 | 0.32 | 0.26 | 0.25 | 0.31 |
| 1 | 163093382 | rs2841989 | 1.294 | -0.19 | -2.286 | 0.349 | -0.018 | -0.026 | -0.026 | 0 | 0.12 | 0.06 | 0.11 | 0.32 | 0.16 | 0.18 |
| 1 | 163104742 | rs11585883 | 1.804 | -0.246 | -2.375 | 0.274 | 0.003 | 0.028 | 0.028 | 0 | 0.07 | 0.02 | 0.05 | 0 | 0.07 | 0.02 |
| 1 | 163104848 | rs1507735 | -1.689 | -0.297 | -2.379 | 0.202 | -0.009 | 0.013 | 0.013 | 0 | 0.71 | 0.74 | 0.71 | 0.89 | 0.65 | 0.84 |
| 1 | 163107107 | rs10494382 | -0.055 | -0.314 | -2.443 | 0.217 | -0.007 | 0.011 | 0.011 | 0.756 | 0.29 | 0.17 | 0.37 | 0.27 | 0.32 | 0.35 |
| 1 | 163111066 | rs2841993 | -0.044 | -0.371 | -2.302 | 0.367 | -0.019 | -0.062 | -0.062 | 0.122 | 0.21 | 0.13 | 0.25 | 0.36 | 0.23 | 0.31 |
| 1 | 163112204 | rs10917682 | 1.045 | -0.359 | -2.393 | -0.167 | -0.012 | 0.005 | 0.005 | 0 | 0.08 | 0 | 0.13 | 0.04 | 0.07 | 0.15 |
| 1 | 163113260 | rs1056515 | 0.187 | -0.096 | -2.623 | -0.048 | -0.016 | -0.054 | -0.054 | 0 | 0.36 | 0.22 | 0.52 | 0.46 | 0.42 | 0.43 |
| 1 | 163113878 | rs3806368 | 1.415 | -0.114 | -2.639 | -0.021 | -0.068 | 0.013 | 0.013 | 2.047 | 0.21 | 0.51 | 0.1 | 0.26 | 0.09 | 0.18 |
| 1 | 163114358 | rs3820487 | 1.415 | -0.114 | -2.639 | -0.021 | -0.068 | 0.013 | 0.013 | 2.047 | 0.21 | 0.51 | 0.1 | 0.26 | 0.09 | 0.18 |
| 1 | 163114605 | rs2815274 | -2.123 | -0.093 | -2.716 | -0.02 | -0.009 | 0.02 | 0.02 | 0 | 0.77 | 0.82 | 0.82 | 0.88 | 0.74 | 0.92 |
| 1 | 163115321 | rs3806366 | 1.328 | -0.101 | -2.714 | -0.018 | -0.06 | 0.005 | 0.005 | 2.338 | 0.22 | 0.56 | 0.1 | 0.26 | 0.09 | 0.18 |
| 1 | 163115431 | rs3806365 | 1.328 | -0.101 | -2.714 | -0.018 | -0.06 | 0.005 | 0.005 | 2.338 | 0.22 | 0.56 | 0.1 | 0.26 | 0.09 | 0.18 |
| 1 | 163116878 | rs15049 | 1.396 | -0.103 | -2.759 | -0.019 | -0.067 | 0.01 | 0.01 | 1.976 | 0.21 | 0.53 | 0.1 | 0.26 | 0.09 | 0.18 |
| 1 | 163120753 | rs12127281 | 2.06 | 0.027 | -3.603 | -0.014 | -0.089 | 0.064 | 0.064 | 0 | 0.21 | 0.02 | 0.17 | 0.09 | 0.26 | 0.08 |
| 1 | 163126417 | rs2255642 | -1.56 | 1.149 | -2.877 | 0.544 | 0.031 | 0.052 | 0.052 | 3.082 | 0.56 | 0.26 | 0.72 | 0.62 | 0.64 | 0.74 |
| 1 | 163132634 | rs7540909 | 0.466 | 1.149 | -2.945 | 0.536 | -0.043 | -0.099 | -0.099 | 0 | 0.26 | 0.11 | 0.47 | 0.45 | 0.34 | 0.38 |

| | | | | | | | | | | | | | | | | |
|---|---|---|---|---|---|---|---|---|---|---|---|---|---|---|---|---|
| 1 | 163132803 | rs1876451 | -0.239 | 1.171 | -2.922 | 0.587 | -0.041 | 0.004 | 0.004 | 0 | 0.21 | 0.06 | 0.2 | 0.17 | 0.22 | 0.31 |
| 1 | 163136535 | rs2815284 | -1.343 | 1.251 | -2.819 | 0.645 | 0.033 | 0.059 | 0.059 | 3.326 | 0.56 | 0.22 | 0.72 | 0.63 | 0.64 | 0.74 |
| 1 | 163137589 | rs2456899 | 0.215 | 1.785 | -2.529 | 1.231 | -0.024 | -0.004 | -0.004 | 0 | 0.12 | 0.05 | 0.07 | 0.13 | 0.16 | 0.16 |
| 1 | 163141491 | rs12041294 | -0.283 | 1.343 | -2.187 | 1.436 | -0.092 | -0.161 | -0.161 | 0 | 0.22 | 0.02 | 0.39 | 0.42 | 0.31 | 0.38 |
| 1 | 163142168 | rs10917690 | -0.515 | 1.029 | -1.991 | 1.298 | -0.018 | -0.088 | -0.088 | 0 | 0.25 | 0.09 | 0.3 | 0.43 | 0.31 | 0.37 |
| 1 | 163142555 | rs12035879 | -0.751 | 0.697 | -2.123 | 0.994 | 0.004 | -0.099 | -0.099 | 0.843 | 0.33 | 0.11 | 0.45 | 0.4 | 0.37 | 0.39 |
| 1 | 163145390 | rs10753605 | 0.136 | 0.743 | -2.066 | 0.374 | 0.004 | -0.004 | -0.004 | 1.274 | 0.32 | 0.13 | 0.45 | 0.63 | 0.31 | 0.3 |
| 1 | 163145696 | rs7534573 | 0.406 | 0.098 | -1.789 | -0.499 | -0.008 | -0.002 | -0.002 | 0 | 0.26 | 0.28 | 0.11 | 0.24 | 0.16 | 0.31 |
| 1 | 163148728 | rs10917695 | 1.21 | -0.094 | -1.857 | -0.614 | -0.014 | -0.024 | -0.024 | 0 | 0.28 | 0.1 | 0.43 | 0.63 | 0.3 | 0.3 |
| 1 | 163149325 | rs10917696 | 0.754 | 0.499 | -1.602 | -1.176 | -0.004 | 0.006 | 0.006 | 0 | 0.2 | 0.2 | 0.16 | 0.09 | 0.22 | 0.13 |
| 1 | 163150975 | rs7533913 | -0.278 | 0.379 | -1.32 | -1.193 | -0.002 | -0.007 | -0.007 | 0.016 | 0.46 | 0.49 | 0.27 | 0.33 | 0.37 | 0.43 |
| 1 | 163157126 | rs4657252 | -0.768 | 0.711 | -1.33 | -1.448 | 0.046 | -0.063 | -0.063 | 0 | 0.86 | 0.66 | 0.79 | 0.95 | 0.82 | 0.95 |
| 1 | 163158175 | rs12143973 | 0.9 | 0.642 | -1.365 | -1.472 | -0.019 | -0.006 | -0.006 | 0.149 | 0.26 | 0.32 | 0.11 | 0.23 | 0.16 | 0.29 |
| 1 | 163159496 | rs7355070 | 0.859 | 0.688 | -1.412 | -1.532 | -0.003 | 0.087 | 0.087 | 0 | 0.48 | 0.29 | 0.6 | 0.71 | 0.51 | 0.43 |
| 1 | 163159682 | rs10799902 | 0.086 | 0.679 | -1.416 | -1.536 | 0.026 | 0.042 | 0.042 | 0.057 | 0.59 | 0.3 | 0.67 | 0.72 | 0.67 | 0.66 |
| 1 | 163162441 | rs12138022 | -0.221 | 0.686 | -1.402 | -1.524 | 0.004 | 0.007 | 0.007 | 0 | 0.38 | 0.39 | 0.2 | 0.24 | 0.31 | 0.51 |
| 1 | 163162458 | rs12137214 | 0.957 | 0.457 | -1.175 | -1.388 | 0.028 | 0.001 | 0.001 | 0 | 0.23 | 0.13 | 0.1 | 0.23 | 0.15 | 0.27 |
| 1 | 163171868 | rs10218752 | 0.846 | 0.61 | -1.088 | -1.335 | -0.012 | 0.026 | 0.026 | 0 | 0.19 | 0.16 | 0.15 | 0.1 | 0.22 | 0.15 |
| 1 | 163174426 | rs2815271 | 0.148 | 0.556 | -0.903 | -1.104 | 0.034 | -0.071 | -0.071 | 0.537 | 0.15 | 0.38 | 0.21 | 0.05 | 0.18 | 0.05 |
| 1 | 163181934 | rs2036702 | 0.007 | 0.755 | -1.078 | -1.064 | 0.019 | -0.019 | -0.019 | 0 | 0.08 | 0.13 | 0.16 | 0.04 | 0.14 | 0.05 |
| 1 | 163183196 | rs2036700 | -0.477 | 0.791 | -1.193 | -1.038 | 0.012 | -0.002 | -0.002 | 0 | 0.1 | 0.13 | 0.21 | 0.1 | 0.18 | 0.09 |
| 1 | 163183944 | rs2662770 | -0.798 | 0.291 | -1.349 | -1.208 | 0.01 | 0.014 | 0.014 | 0 | 0.4 | 0.64 | 0.36 | 0.29 | 0.32 | 0.36 |
| 1 | 163188476 | rs1509019 | -0.458 | -0.044 | -1.685 | -1.34 | 0.009 | -0.005 | -0.005 | 0 | 0.35 | 0.43 | 0.42 | 0.34 | 0.44 | 0.34 |
| 1 | 163189943 | rs2662769 | -2.129 | -0.097 | -1.962 | -1.283 | -0.099 | -0.119 | -0.119 | 0 | 0.73 | 0.98 | 0.6 | 0.43 | 0.67 | 0.72 |
| 1 | 163192579 | rs10917702 | -0.991 | -0.123 | -1.866 | -1.379 | 0.005 | 0.008 | 0.008 | 0 | 0.47 | 0.49 | 0.5 | 0.33 | 0.59 | 0.57 |
| 1 | 163199332 | rs12130030 | 0.598 | -0.179 | -1.897 | -1.298 | -0.016 | -0.033 | -0.033 | 1.453 | 0.06 | 0.01 | 0.08 | 0.12 | 0.07 | 0.16 |
| 1 | 163199590 | rs12130821 | 0.598 | -0.179 | -1.897 | -1.298 | -0.016 | -0.033 | -0.033 | 0 | 0.06 | 0.01 | 0.08 | 0.12 | 0.07 | 0.17 |
| 1 | 163200062 | rs2457185 | -2.281 | 0.217 | -1.791 | -1.285 | -0.059 | -0.102 | -0.102 | 0 | 0.74 | 0.98 | 0.65 | 0.35 | 0.76 | 0.72 |
| 1 | 163201904 | rs12080108 | 0.988 | 0.19 | -1.521 | -1.432 | 0.016 | -0.042 | -0.042 | 0 | 0.19 | 0.34 | 0.16 | 0.1 | 0.2 | 0.14 |
| 1 | 163206986 | rs2999963 | 0.133 | 0.028 | -0.708 | -1.52 | 0.015 | 0.055 | 0.055 | 0 | 0.07 | 0.27 | 0.07 | 0.06 | 0.03 | 0.05 |
| 1 | 163212633 | rs11804244 | 0.03 | -0.05 | -0.877 | -1.674 | -0.03 | 0.034 | 0.034 | 0 | 0.16 | 0.09 | 0.2 | 0.09 | 0.24 | 0.17 |
| 1 | 163218439 | rs6664060 | 0.03 | -0.05 | -0.874 | -1.674 | -0.04 | 0.041 | 0.041 | 0 | 0.16 | 0.08 | 0.2 | 0.09 | 0.24 | 0.17 |
| 1 | 163220214 | rs10494392 | 1.24 | -0.038 | -0.848 | -1.68 | 0.018 | -0.04 | -0.04 | 0 | 0.19 | 0.35 | 0.16 | 0.1 | 0.2 | 0.14 |
| 1 | 163220797 | rs10494390 | 0.687 | -0.039 | -0.9 | -1.646 | -0.018 | 0.016 | 0.016 | 0 | 0.08 | 0.06 | 0.16 | 0.02 | 0.18 | 0.05 |
| 1 | 163228151 | rs937925 | 1.399 | -0.139 | -0.911 | -1.679 | 0.037 | 0.012 | 0.012 | 0 | 0.13 | 0.05 | 0.08 | 0.08 | 0.08 | 0.12 |
| 1 | 163241407 | rs7518042 | -1.575 | -0.104 | -0.78 | -1.695 | -0.004 | -0.013 | -0.013 | 0 | 0.63 | 0.63 | 0.54 | 0.27 | 0.67 | 0.68 |
| 1 | 163251311 | rs6679239 | -0.593 | -0.13 | -0.996 | -1.606 | 0.002 | 0.02 | 0.02 | 0 | 0.31 | 0.27 | 0.29 | 0.17 | 0.32 | 0.29 |
| 1 | 163257164 | rs2999863 | -0.343 | -0.167 | -0.958 | -1.623 | 0 | 0.002 | 0.002 | 0 | 0.72 | 0.31 | 0.74 | 0.83 | 0.74 | 0.8 |
| 1 | 163257303 | rs10494388 | -0.578 | -0.255 | -0.755 | -1.504 | -0.078 | 0.106 | 0.106 | 0 | 0.22 | 0.07 | 0.24 | 0.03 | 0.33 | 0.3 |
| 1 | 163260844 | rs16852030 | 0.924 | -0.306 | -0.742 | -1.511 | 0.006 | 0.005 | 0.005 | 0 | 0.14 | 0.13 | 0.1 | 0.08 | 0.08 | 0.12 |
| 1 | 163272272 | rs16852295 | 1.599 | -0.117 | -0.892 | -1.287 | -0.001 | -0.002 | -0.002 | 0 | 0.09 | 0.08 | 0.04 | 0.06 | 0.06 | 0.11 |
| 1 | 163273585 | rs11582919 | -1.072 | -0.032 | -0.84 | -1.196 | -0.003 | 0.109 | 0.109 | 0 | 0.45 | 0.28 | 0.38 | 0.17 | 0.47 | 0.54 |
| 1 | 163276396 | rs4656391 | -1.021 | 0.008 | -0.8 | -1.173 | -0.002 | 0.106 | 0.106 | 0 | 0.47 | 0.28 | 0.43 | 0.22 | 0.48 | 0.58 |
| 1 | 163289571 | rs2136241 | 0.322 | 0.079 | -0.364 | -0.615 | 0.001 | -0.001 | -0.001 | 0.668 | 0.56 | 0.51 | 0.52 | 0.53 | 0.54 | 0.53 |
| 1 | 163292040 | rs2292274 | 0.766 | 0.137 | -0.744 | -1.08 | 0.012 | 0.047 | 0.047 | 0 | 0.33 | 0.23 | 0.27 | 0.23 | 0.28 | 0.29 |
| 1 | 163292217 | rs2292275 | 1.617 | 0.255 | -0.893 | -1.165 | -0.074 | -0.079 | -0.079 | 0 | 0.29 | 0.01 | 0.3 | 0.6 | 0.33 | 0.26 |
| 1 | 163292362 | rs2292276 | -0.971 | 0.242 | -0.636 | -1.2 | -0.003 | -0.011 | -0.011 | 0 | 0.63 | 0.5 | 0.61 | 0.38 | 0.63 | 0.71 |
| 1 | 163295576 | rs3010373 | 1.517 | 0.238 | -0.667 | -1.265 | -0.002 | -0.002 | -0.002 | 0 | 0.3 | 0.19 | 0.32 | 0.6 | 0.33 | 0.26 |
| 1 | 163297001 | rs10917728 | -0.998 | 0.306 | -1.796 | -1.228 | -0.001 | -0.006 | -0.006 | 0 | 0.63 | 0.5 | 0.6 | 0.37 | 0.63 | 0.71 |
| 1 | 163297322 | rs16852612 | 1.175 | 0.409 | -1.842 | -1.261 | -0.007 | 0.013 | 0.013 | 0 | 0.06 | 0.04 | 0.14 | 0.02 | 0.18 | 0.04 |
| 1 | 163297381 | rs12691496 | 0.814 | 0.414 | -1.801 | -1.225 | 0.02 | -0.022 | -0.022 | 0 | 0.1 | 0.16 | 0.17 | 0.02 | 0.19 | 0.07 |
| 1 | 163310510 | rs1022058 | -0.368 | 0.4 | -1.806 | -1.219 | 0.006 | 0.088 | 0.088 | 0 | 0.53 | 0.32 | 0.44 | 0.35 | 0.44 | 0.64 |
| 1 | 163313262 | rs10799912 | 0.31 | 0.369 | -1.835 | -1.25 | 0.039 | 0.083 | 0.083 | 0 | 0.42 | 0.23 | 0.36 | 0.31 | 0.32 | 0.39 |
| 1 | 163313539 | rs11802875 | 0.827 | 0.398 | -1.811 | -1.234 | 0.021 | -0.025 | -0.025 | 0 | 0.1 | 0.17 | 0.17 | 0.02 | 0.19 | 0.07 |
| 1 | 163322033 | rs17363117 | -0.402 | 0.373 | -1.919 | -1.191 | 0.006 | 0.088 | 0.088 | 0 | 0.53 | 0.32 | 0.44 | 0.35 | 0.44 | 0.64 |
| 1 | 163328898 | rs16852991 | -0.76 | 0.951 | -2.341 | -0.836 | 0.006 | 0.042 | 0.042 | 0 | 0.22 | 0.11 | 0.22 | 0.13 | 0.19 | 0.37 |
| 1 | 163330805 | rs1396544 | -1.423 | 1.683 | -1.897 | -0.443 | 0.027 | -0.052 | -0.052 | 1.637 | 0.24 | 0.34 | 0.29 | 0.09 | 0.34 | 0.34 |
| 1 | 163331167 | rs10917735 | -0.358 | 1.745 | -1.897 | -0.475 | -0.003 | 0.011 | 0.011 | 0 | 0.13 | 0.1 | 0.12 | 0.06 | 0.14 | 0.27 |
| 1 | 163331900 | rs12064397 | -0.383 | 1.751 | -1.898 | -0.475 | -0.005 | 0.011 | 0.011 | 0 | 0.13 | 0.1 | 0.12 | 0.06 | 0.15 | 0.27 |
| 1 | 163332067 | rs12064483 | -0.383 | 1.697 | -1.898 | -0.475 | -0.004 | 0.011 | 0.011 | 0 | 0.13 | 0.1 | 0.12 | 0.06 | 0.15 | 0.27 |
| 1 | 163334558 | rs17363152 | -0.384 | 1.717 | -1.87 | -0.466 | -0.004 | 0.011 | 0.011 | 0 | 0.13 | 0.1 | 0.12 | 0.06 | 0.15 | 0.27 |
| 1 | 163338418 | rs16853144 | 0.029 | 1.68 | -1.711 | -0.232 | 0.026 | -0.025 | -0.025 | 1.94 | 0.1 | 0.15 | 0.17 | 0.02 | 0.19 | 0.07 |
| 1 | 163338854 | rs3010342 | 0.029 | 0.499 | -1.594 | 0.067 | 0.041 | 0.07 | 0.07 | 0 | 0.52 | 0.33 | 0.59 | 0.7 | 0.66 | 0.6 |
| 1 | 163344343 | rs17359123 | -1.042 | 1.233 | -1.583 | 0.907 | -0.005 | -0.008 | -0.008 | 0 | 0.15 | 0.14 | 0.23 | 0.28 | 0.18 | 0.09 |
| 1 | 163344782 | rs4475722 | -0.445 | 1.355 | -2.711 | -0.103 | -0.145 | 0.105 | 0.105 | 0 | 0.37 | 0.04 | 0.34 | 0.15 | 0.47 | 0.27 |
| 1 | 163345829 | rs10917739 | 0.273 | 1.076 | -3.35 | 0.021 | 0.102 | 0.011 | 0.011 | 0 | 0.5 | 0.07 | 0.34 | 0.64 | 0.39 | 0.6 |
| 1 | 163348444 | rs3013501 | -0.414 | 1.621 | -2.425 | -0.666 | 0.094 | -0.075 | -0.075 | 0 | 0.26 | 0.36 | 0.37 | 0.1 | 0.46 | 0.22 |
| 1 | 163351383 | rs3013495 | -0.995 | 2.162 | -2.512 | -0.918 | 0.013 | -0.075 | -0.075 | 0 | 0.35 | 0.42 | 0.42 | 0.12 | 0.54 | 0.31 |
| 1 | 163351718 | rs3010344 | -0.296 | 2.17 | -2.503 | -0.914 | 0.1 | -0.081 | -0.081 | 0.01 | 0.26 | 0.38 | 0.37 | 0.1 | 0.46 | 0.22 |

| | | | | | | | | | | | | | | | | |
|---|---|---|---|---|---|---|---|---|---|---|---|---|---|---|---|---|
| 1 | 163353790 | rs6685251 | 0.761 | 1.988 | -1.529 | -0.995 | 0.021 | -0.106 | -0.106 | 2.407 | 0.66 | 0.53 | 0.6 | 0.93 | 0.46 | 0.75 |
| 1 | 163359017 | rs3010348 | -0.948 | 2.008 | -1.456 | -0.986 | 0.008 | -0.052 | -0.052 | 0 | 0.33 | 0.32 | 0.39 | 0.07 | 0.54 | 0.25 |
| 1 | 163363776 | rs3013487 | 0.891 | 1.741 | -1.073 | -1.128 | 0.021 | -0.066 | -0.066 | 0 | 0.74 | 0.47 | 0.8 | 0.9 | 0.77 | 0.76 |
| 1 | 163364886 | rs12407429 | 0.2 | 1.735 | -1.063 | -1.124 | 0 | -0.045 | -0.045 | 0 | 0.14 | 0.35 | 0.09 | 0.04 | 0.08 | 0.12 |
| 1 | 163365747 | rs3013484 | -0.881 | 1.731 | -1.069 | -1.128 | 0.001 | -0.024 | -0.024 | 0 | 0.9 | 0.85 | 0.94 | 0.98 | 0.92 | 0.92 |
| 1 | 163366854 | rs10799916 | 0.101 | 1.726 | -1.054 | -1.102 | 0.004 | -0.044 | -0.044 | 0 | 0.14 | 0.41 | 0.09 | 0.05 | 0.08 | 0.12 |
| 1 | 163369704 | rs3010355 | 0.061 | 1.761 | -0.636 | -1.118 | 0.021 | -0.062 | -0.062 | 0 | 0.21 | 0.19 | 0.28 | 0.02 | 0.39 | 0.14 |
| 1 | 163375247 | rs12046142 | 1.68 | 1.609 | -0.779 | -1.095 | 0.098 | 0.246 | 0.246 | 0 | 0.53 | 0.17 | 0.51 | 0.88 | 0.38 | 0.63 |
| 1 | 163375960 | rs6675964 | 1.68 | 1.609 | -0.793 | -1.095 | 0.098 | 0.246 | 0.246 | 0 | 0.53 | 0.17 | 0.51 | 0.87 | 0.38 | 0.63 |
| 1 | 163382050 | rs4409627 | 1.485 | 1.772 | -0.724 | -1.067 | 0.008 | 0.009 | 0.009 | 0 | 0.56 | 0.39 | 0.39 | 0.65 | 0.43 | 0.59 |
| 1 | 163384831 | rs6691127 | 1.984 | 1.65 | -0.445 | -1.09 | 0.089 | 0.001 | 0.001 | 0 | 0.44 | 0.13 | 0.34 | 0.65 | 0.35 | 0.55 |
| 1 | 163385492 | rs12047638 | 1.374 | 1.63 | -0.342 | -0.985 | 0.005 | 0.005 | 0.005 | 4.299 | 0.09 | 0.09 | 0.18 | 0.23 | 0.04 | 0.08 |
| 1 | 163391538 | rs7546952 | -0.351 | 1.652 | -0.209 | -1.009 | -0.025 | 0.039 | 0.039 | 0.742 | 0.21 | 0.38 | 0.24 | 0.23 | 0.12 | 0.12 |
| 1 | 163399083 | rs12735706 | 1.394 | 1.746 | -0.456 | -1.147 | -0.037 | 0.066 | 0.066 | 0.101 | 0.55 | 0.14 | 0.43 | 0.72 | 0.48 | 0.66 |
| 1 | 163411338 | rs4657276 | -1.101 | 1.895 | -0.251 | -1.258 | -0.012 | -0.127 | -0.127 | 0.23 | 0.31 | 0.52 | 0.15 | 0.02 | 0.24 | 0.16 |
| 1 | 163414559 | rs4265409 | 1.511 | 1.924 | -0.269 | -1.302 | 0.005 | 0.232 | 0.232 | 0 | 0.56 | 0.19 | 0.58 | 0.97 | 0.44 | 0.74 |
| 1 | 163415714 | rs4274035 | -0.319 | 1.676 | -0.21 | -1.223 | -0.003 | -0.04 | -0.04 | 0 | 0.19 | 0.27 | 0.11 | 0.02 | 0.17 | 0.12 |
| 1 | 163416975 | rs4657277 | 0.062 | 1.41 | -0.271 | -0.991 | 0.037 | -0.044 | -0.044 | 0 | 0.15 | 0.32 | 0.11 | 0.04 | 0.14 | 0.07 |
| 1 | 163422204 | rs12036398 | 1.242 | 1.393 | -0.669 | -1.099 | 0.077 | 0.191 | 0.191 | 0 | 0.54 | 0.22 | 0.54 | 0.93 | 0.38 | 0.73 |
| 1 | 163422320 | rs10917756 | 0.272 | 1.363 | -0.456 | -1.111 | 0.007 | -0.056 | -0.056 | 0.265 | 0.69 | 0.61 | 0.66 | 0.97 | 0.52 | 0.8 |
| 1 | 163424565 | rs4469696 | 1.382 | 1.516 | -0.508 | -0.84 | 0.088 | 0.008 | 0.008 | 0.027 | 0.44 | 0.14 | 0.34 | 0.66 | 0.34 | 0.57 |
| 1 | 163440717 | rs4657284 | -0.221 | 1.673 | -0.122 | -0.814 | 0.072 | -0.047 | -0.047 | 0 | 0.15 | 0.34 | 0.12 | 0.04 | 0.15 | 0.07 |
| 1 | 163444418 | rs12143307 | 0.633 | 0.774 | -1.139 | -0.393 | 0.092 | 0.121 | 0.121 | 0 | 0.47 | 0.2 | 0.32 | 0.27 | 0.38 | 0.37 |
| 1 | 163444495 | rs7528059 | 0.612 | 0.878 | -1.151 | -0.329 | 0.065 | 0.131 | 0.131 | 0.022 | 0.5 | 0.25 | 0.34 | 0.29 | 0.46 | 0.38 |
| 1 | 163448078 | rs11590745 | -0.157 | 1.081 | -0.897 | -0.19 | -0.007 | 0.003 | 0.003 | 0 | 0.05 | 0.04 | 0.04 | 0.06 | 0.08 | 0.12 |
| 1 | 163452016 | rs2791150 | 1.073 | 0.09 | -0.96 | -0.402 | 0.025 | 0.021 | 0.021 | 0.446 | 0.33 | 0.22 | 0.19 | 0.24 | 0.25 | 0.18 |
| 1 | 163453047 | rs2820568 | -0.52 | 0.293 | -0.831 | -0.389 | 0.008 | -0.01 | -0.01 | 0 | 0.85 | 0.77 | 0.87 | 0.87 | 0.8 | 0.8 |
| 1 | 163456961 | rs2211183 | -0.007 | 0.58 | -0.671 | 0.083 | 0.003 | 0.018 | 0.018 | 0 | 0.63 | 0.5 | 0.64 | 0.47 | 0.66 | 0.45 |
| 1 | 163457348 | rs1890798 | 1.394 | 0.816 | -1.422 | 0.314 | -0.039 | -0.026 | -0.026 | 0 | 0.16 | 0.09 | 0.31 | 0.17 | 0.24 | 0.12 |
| 1 | 163466520 | rs1934511 | 0.905 | -0.139 | -1.282 | 0.107 | 0.049 | 0.038 | 0.038 | 0.559 | 0.3 | 0.11 | 0.18 | 0.24 | 0.25 | 0.18 |
| 1 | 163473738 | rs1934504 | 0.005 | 0.028 | -1.052 | 0.284 | 0.007 | 0.057 | 0.057 | 0 | 0.58 | 0.76 | 0.68 | 0.75 | 0.56 | 0.43 |
| 1 | 163478287 | rs4657294 | -0.244 | -0.279 | -0.659 | 0.215 | -0.007 | -0.001 | -0.001 | 0 | 0.43 | 0.52 | 0.4 | 0.46 | 0.38 | 0.23 |
| 1 | 163479468 | rs12128972 | -0.088 | -0.362 | -0.133 | 0.395 | 0.012 | 0.018 | 0.018 | 0 | 0.41 | 0.38 | 0.36 | 0.35 | 0.38 | 0.22 |
| 1 | 163480019 | rs10799925 | 0.639 | -0.447 | 0.049 | 0.371 | -0.027 | 0.025 | 0.025 | 0 | 0.31 | 0.38 | 0.3 | 0.46 | 0.23 | 0.22 |
| 1 | 163480551 | rs1934523 | -0.726 | -0.455 | 0.003 | 0.439 | -0.019 | -0.002 | -0.002 | 0.01 | 0.69 | 0.56 | 0.75 | 0.73 | 0.71 | 0.44 |
| 1 | 163482009 | rs12565748 | -0.701 | -0.41 | 0.116 | 0.466 | -0.023 | -0.003 | -0.003 | 0 | 0.68 | 0.51 | 0.76 | 0.74 | 0.71 | 0.45 |
| 1 | 163483089 | rs10917783 | 1.214 | -0.526 | -0.222 | 0.56 | 0.011 | -0.014 | -0.014 | 0 | 0.22 | 0.14 | 0.23 | 0.38 | 0.16 | 0.16 |
| 1 | 163487264 | rs12035232 | 1.192 | -0.529 | -0.222 | 0.558 | 0.011 | -0.015 | -0.015 | 0 | 0.23 | 0.14 | 0.23 | 0.38 | 0.16 | 0.16 |
| 1 | 163491892 | rs6695991 | -0.099 | -0.317 | -0.067 | 0.438 | 0.018 | 0.009 | 0.009 | 0 | 0.58 | 0.34 | 0.67 | 0.65 | 0.61 | 0.38 |
| 1 | 163492132 | rs6681222 | 1.508 | -0.207 | -0.514 | 0.402 | -0.083 | -0.035 | -0.035 | 0 | 0.23 | 0.05 | 0.33 | 0.27 | 0.3 | 0.2 |
| 1 | 163492245 | rs10917787 | 1.035 | -0.391 | -0.276 | 0.535 | 0 | -0.001 | -0.001 | 0 | 0.09 | 0.17 | 0.08 | 0.1 | 0.1 | 0.07 |
| 1 | 163494597 | rs7512954 | -0.904 | -0.392 | -0.605 | 0.561 | 0.008 | -0.015 | -0.015 | 0.854 | 0.69 | 0.73 | 0.78 | 0.63 | 0.71 | 0.42 |
| 1 | 163494784 | rs10799928 | 0.209 | -0.448 | -0.222 | 0.32 | 0.026 | 0.021 | 0.021 | 0.387 | 0.12 | 0.43 | 0.11 | 0.1 | 0.1 | 0.07 |
| 1 | 163502326 | rs3001083 | -1.197 | -0.266 | -1.203 | 0.393 | 0.024 | -0.01 | -0.01 | 0.733 | 0.78 | 0.86 | 0.77 | 0.73 | 0.84 | 0.87 |
| 1 | 163503181 | rs7553090 | -0.03 | 0.004 | -1.172 | 0.322 | 0.063 | -0.01 | -0.01 | 0 | 0.45 | 0.69 | 0.34 | 0.48 | 0.31 | 0.3 |
| 1 | 163505425 | rs2994232 | -0.305 | 0.594 | -0.796 | 0.367 | 0.003 | -0.037 | -0.037 | 0 | 0.88 | 0.82 | 0.89 | 1 | 0.84 | 0.98 |
| 1 | 163505727 | rs12091875 | -0.249 | 0.552 | -1.021 | 0.62 | 0.008 | 0.003 | 0.003 | 0 | 0.33 | 0.2 | 0.31 | 0.32 | 0.32 | 0.18 |
| 1 | 163505777 | rs1934494 | -0.072 | 1.255 | -0.557 | 0.911 | 0 | 0 | 0 | 0 | 0.56 | 0.52 | 0.41 | 0.57 | 0.48 | 0.55 |
| 1 | 163506077 | rs1934496 | -1.77 | 3.099 | -0.214 | 1.662 | -0.009 | -0.009 | -0.009 | 0.457 | 0.19 | 0.2 | 0.29 | 0.37 | 0.34 | 0.36 |
| 1 | 163506121 | rs1934497 | -1.744 | 3.5 | 0.298 | 1.292 | 0.144 | 0.06 | 0.06 | 0.005 | 0.23 | 0.46 | 0.26 | 0.34 | 0.36 | 0.37 |
| 1 | 163506187 | rs10494335 | -1.541 | 3.003 | -0.104 | 0.839 | 0.118 | 0.024 | 0.024 | 0 | 0.12 | 0.37 | 0.19 | 0.22 | 0.24 | 0.22 |
| 1 | 163506835 | rs2994234 | -0.189 | 2.164 | -0.949 | 0.397 | -0.044 | 0.044 | 0.044 | 0 | 0.2 | 0.08 | 0.26 | 0.1 | 0.3 | 0.13 |
| 1 | 163508344 | rs2820871 | 0.047 | 1.727 | -1.511 | 0.727 | -0.018 | -0.1 | -0.1 | 0 | 0.76 | 0.88 | 0.78 | 0.59 | 0.8 | 0.84 |
| 1 | 163511425 | rs6674167 | -0.042 | 1.202 | -0.886 | 0.037 | 0.011 | -0.036 | -0.036 | 0 | 0.72 | 0.61 | 0.69 | 0.85 | 0.66 | 0.7 |
| 1 | 163525477 | rs10917799 | -0.176 | 1.205 | -1.42 | 0.434 | -0.029 | -0.067 | -0.067 | 0 | 0.24 | 0.09 | 0.21 | 0.32 | 0.2 | 0.16 |
| 1 | 163535283 | rs2486834 | 0.18 | 1.165 | -1.167 | 0.226 | -0.011 | 0.025 | 0.025 | 0 | 0.25 | 0.15 | 0.23 | 0.13 | 0.3 | 0.11 |
| 1 | 163543441 | rs1934533 | -0.663 | 0.47 | -0.712 | 0.245 | 0.003 | -0.003 | -0.003 | 0 | 0.26 | 0.72 | 0.23 | 0.31 | 0.17 | 0.17 |
| 1 | 163552464 | rs12758012 | 0.816 | 0.494 | -1.145 | 0.299 | 0.003 | 0.026 | 0.026 | 0.379 | 0.11 | 0.01 | 0.09 | 0.06 | 0.09 | 0.03 |
| 1 | 163558749 | rs10917804 | -0.341 | 0.938 | -1.84 | 0.735 | -0.002 | -0.013 | -0.013 | 1.431 | 0.1 | 0.08 | 0.09 | 0.25 | 0.07 | 0.14 |
| 1 | 163569354 | rs7525601 | 0.502 | 0.958 | -2.042 | 0.675 | -0.062 | -0.114 | -0.114 | 0.094 | 0.37 | 0.03 | 0.31 | 0.46 | 0.41 | 0.51 |
| 1 | 163579635 | rs10799935 | 0.252 | 0.911 | -1.932 | 0.211 | 0.003 | 0.045 | 0.045 | 0 | 0.37 | 0.15 | 0.27 | 0.22 | 0.34 | 0.24 |
| 1 | 163579688 | rs10917812 | -0.584 | 0.544 | -1.908 | 0.397 | 0.012 | -0.034 | -0.034 | 1.208 | 0.25 | 0.18 | 0.39 | 0.65 | 0.23 | 0.46 |
| 1 | 163579955 | rs2791149 | -0.076 | 1.302 | -1.859 | 0.167 | -0.039 | 0.001 | 0.001 | 0 | 0.19 | 0.03 | 0.19 | 0.19 | 0.25 | 0.17 |
| 1 | 163582039 | rs10753622 | -1.145 | 1.251 | -0.915 | -0.054 | 0.005 | -0.036 | -0.036 | 0 | 0.39 | 0.38 | 0.47 | 0.84 | 0.48 | 0.63 |
| 1 | 163586969 | rs12123582 | -0.102 | 1.618 | -1.034 | 0.193 | -0.036 | -0.026 | -0.026 | 0 | 0.07 | 0 | 0.05 | 0.11 | 0.1 | 0.2 |
| 1 | 163587355 | rs10917813 | -0.413 | 1.623 | -1.068 | 0.647 | -0.107 | -0.143 | -0.143 | 0 | 0.18 | 0.01 | 0.28 | 0.54 | 0.3 | 0.49 |
| 1 | 163604000 | rs2345965 | -0.86 | 1.347 | -0.665 | 0.311 | -0.003 | 0 | 0 | 0.049 | 0.25 | 0.17 | 0.39 | 0.73 | 0.4 | 0.6 |
| 1 | 163627709 | rs10494395 | 0.027 | 1.262 | -0.593 | 0.273 | -0.005 | -0.001 | -0.001 | 0 | 0.14 | 0.14 | 0.08 | 0.11 | 0.09 | 0.04 |
| 1 | 163635058 | rs10799942 | -0.825 | 1.335 | -0.522 | 0.305 | -0.056 | -0.001 | -0.001 | 0 | 0.25 | 0.11 | 0.39 | 0.73 | 0.4 | 0.59 |

| | | | | | | | | | | | | | | | | |
|---|---|---|---|---|---|---|---|---|---|---|---|---|---|---|---|---|
| 1 | 163645519 | rs4535983 | 0.065 | 1.304 | -0.008 | 0.084 | 0.002 | 0.002 | 0.002 | 0.167 | 0.1 | 0.16 | 0.23 | 0.16 | 0.09 | 0.13 |
| 1 | 163674732 | rs12239576 | 0.03 | 1.319 | 0.042 | 0.096 | 0.003 | 0.003 | 0.003 | 0.218 | 0.11 | 0.19 | 0.23 | 0.18 | 0.09 | 0.13 |
| 1 | 163677950 | rs1289004 | 0.964 | 1.254 | -0.051 | 0.043 | -0.078 | 0.104 | 0.104 | 0 | 0.66 | 0.88 | 0.53 | 0.18 | 0.52 | 0.37 |
| 1 | 163697998 | rs10799944 | -1.03 | 1.236 | 0.019 | 0.059 | -0.001 | 0 | 0 | 0 | 0.29 | 0.25 | 0.41 | 0.72 | 0.41 | 0.6 |
| 1 | 163714218 | rs10753623 | -1.81 | 1.186 | 0.083 | 0.03 | 0.003 | -0.03 | -0.03 | 0.128 | 0.4 | 0.38 | 0.49 | 0.82 | 0.49 | 0.63 |
| 1 | 163715990 | rs1932360 | 0.704 | 1.196 | -0.17 | 0.075 | 0.01 | -0.036 | -0.036 | 0 | 0.1 | 0 | 0.07 | 0.11 | 0.07 | 0.03 |
| 1 | 163717662 | rs3904944 | -0.614 | 1.254 | -0.105 | 0.07 | -0.043 | -0.001 | -0.001 | 0 | 0.25 | 0.13 | 0.4 | 0.72 | 0.41 | 0.6 |
| 1 | 163731087 | rs4657302 | -0.614 | 1.254 | -0.105 | 0.07 | -0.043 | -0.001 | -0.001 | 0 | 0.25 | 0.13 | 0.4 | 0.72 | 0.41 | 0.6 |
| 1 | 163736925 | rs10737495 | -1.957 | 1.192 | -0.051 | -0.122 | 0.006 | -0.072 | -0.072 | 0.924 | 0.49 | 0.46 | 0.71 | 1 | 0.58 | 0.76 |
| 1 | 163738347 | rs10753629 | -1.707 | 1.21 | -0.273 | -0.061 | -0.036 | 0.183 | 0.183 | 0 | 0.43 | 0.16 | 0.5 | 0.82 | 0.52 | 0.65 |
| 1 | 163738950 | rs3845534 | -1.324 | 1.132 | -0.082 | -0.009 | -0.003 | -0.007 | -0.007 | 0.458 | 0.37 | 0.35 | 0.49 | 0.81 | 0.47 | 0.62 |
| 1 | 163766491 | rs1596025 | 0.295 | 0.793 | -0.044 | 0.31 | 0.006 | -0.002 | -0.002 | 0 | 0.16 | 0.08 | 0.26 | 0.2 | 0.13 | 0.15 |
| 1 | 163766672 | rs1837732 | -1.389 | 0.875 | 0.404 | 0.259 | 0 | -0.008 | -0.008 | 0.174 | 0.45 | 0.45 | 0.66 | 0.89 | 0.51 | 0.76 |
| 1 | 163768115 | rs885256 | -0.021 | 0.906 | 0.233 | 0.646 | -0.082 | -0.108 | -0.108 | 0.95 | 0.15 | 0.01 | 0.28 | 0.5 | 0.25 | 0.44 |
| 1 | 163772455 | rs13374982 | -1.166 | -0.023 | 1.012 | -0.413 | -0.009 | 0.035 | 0.035 | 2.609 | 0.09 | 0.42 | 0.07 | 0.05 | 0.03 | 0.01 |
| 1 | 163775335 | rs6671893 | 0.282 | 0.188 | 0.795 | -0.321 | -0.01 | -0.023 | -0.023 | 0.064 | 0.23 | 0.13 | 0.3 | 0.38 | 0.23 | 0.28 |
| 1 | 163783586 | rs915395 | -1.665 | -0.139 | 1.263 | 0.64 | 0 | 0 | 0 | 0 | 0.36 | 0.63 | 0.43 | 0.62 | 0.35 | 0.43 |
| 1 | 163795105 | rs7533878 | -0.915 | -0.07 | 1.17 | 0.658 | -0.004 | -0.016 | -0.016 | 0 | 0.27 | 0.24 | 0.36 | 0.57 | 0.32 | 0.4 |
| 1 | 163802280 | rs10917855 | -1.597 | -0.036 | 1.067 | 0.673 | -0.003 | 0.002 | 0.002 | 8.862 | 0.34 | 0.42 | 0.41 | 0.62 | 0.35 | 0.42 |
| 1 | 163805912 | rs12030349 | -1.558 | 0.315 | -0.436 | 0.842 | -0.001 | 0 | 0 | 6.856 | 0.34 | 0.35 | 0.4 | 0.62 | 0.35 | 0.42 |
| 1 | 163807040 | rs10917864 | -0.693 | 0.646 | -0.216 | 0.89 | -0.036 | -0.079 | -0.079 | 0 | 0.24 | 0.09 | 0.35 | 0.57 | 0.31 | 0.4 |
| 1 | 163811147 | rs1348128 | -1.089 | 0.78 | -0.275 | 0.925 | 0 | 0.02 | 0.02 | 2.109 | 0.3 | 0.36 | 0.37 | 0.56 | 0.32 | 0.38 |
| 1 | 163812795 | rs964860 | -0.837 | 1.025 | -1.021 | 1.043 | -0.004 | -0.019 | -0.019 | 0.011 | 0.28 | 0.23 | 0.36 | 0.56 | 0.32 | 0.38 |
| 1 | 163817001 | rs781940 | -0.153 | 0.972 | -1.194 | 0.677 | -0.004 | 0.064 | 0.064 | 0 | 0.5 | 0.74 | 0.57 | 0.63 | 0.5 | 0.47 |
| 1 | 163829349 | rs796694 | 0.036 | 0.712 | -1.333 | 0.9 | 0.013 | -0.011 | -0.011 | 0.12 | 0.15 | 0.1 | 0.1 | 0.2 | 0.12 | 0.17 |
| 1 | 163831730 | rs1339995 | 0.767 | 0.463 | -0.953 | 1.14 | 0.013 | -0.002 | -0.002 | 0 | 0.51 | 0.23 | 0.48 | 0.46 | 0.53 | 0.58 |
| 1 | 163836535 | rs781936 | -0.112 | 0.706 | -0.478 | 1.323 | 0.027 | 0.008 | 0.008 | 0 | 0.9 | 0.85 | 0.89 | 0.92 | 0.85 | 0.95 |
| 1 | 163836720 | rs884518 | 1.15 | 0.697 | -0.441 | 1.309 | 0.002 | -0.006 | -0.006 | 0 | 0.51 | 0.19 | 0.44 | 0.46 | 0.52 | 0.58 |
| 1 | 163837511 | rs811435 | 0.224 | 0.774 | 0.369 | 1.217 | 0.055 | -0.049 | -0.049 | 0 | 0.91 | 0.69 | 0.92 | 1 | 0.91 | 0.97 |
| 1 | 163841622 | rs767544 | 1.306 | 1.016 | -0.078 | 1.165 | -0.031 | 0.201 | 0.201 | 0 | 0.46 | 0.15 | 0.32 | 0.12 | 0.49 | 0.38 |
| 1 | 163842143 | rs11582225 | 0.732 | 0.73 | -0.102 | 0.966 | 0.011 | -0.051 | -0.051 | 0 | 0.18 | 0.03 | 0.26 | 0.26 | 0.17 | 0.19 |
| 1 | 163850104 | rs11577916 | 0.106 | 0.863 | -0.389 | 0.377 | 0.004 | 0.003 | 0.003 | 0.01 | 0.25 | 0.13 | 0.29 | 0.27 | 0.26 | 0.23 |
| 1 | 163850670 | rs12120922 | -0.319 | 0.551 | 0.206 | -0.402 | -0.03 | -0.117 | -0.117 | 0 | 0.24 | 0.38 | 0.13 | 0.01 | 0.15 | 0.23 |
| 1 | 163852034 | rs7553359 | 0.117 | 0.532 | 0.355 | -0.413 | -0.02 | -0.074 | -0.074 | 0.049 | 0.22 | 0.25 | 0.1 | 0 | 0.12 | 0.12 |
| 1 | 163857878 | rs2033966 | -0.718 | 0.365 | 0.707 | -0.169 | -0.065 | -0.03 | -0.03 | 0 | 0.29 | 0.55 | 0.21 | 0.24 | 0.15 | 0.21 |
| 1 | 163859805 | rs6661534 | 1.138 | 1.105 | 0.55 | -0.698 | 0.002 | -0.02 | -0.02 | 0.84 | 0.1 | 0.1 | 0.1 | 0 | 0.16 | 0.07 |
| 1 | 163864334 | rs1441178 | -1.164 | 1.426 | 0.103 | -0.864 | 0.019 | 0.028 | 0.028 | 0.941 | 0.56 | 0.74 | 0.65 | 0.4 | 0.62 | 0.51 |
| 1 | 163866023 | rs1816756 | -1.304 | 1.468 | 0.069 | -0.895 | 0.025 | 0.02 | 0.02 | 0.428 | 0.58 | 0.74 | 0.67 | 0.4 | 0.65 | 0.54 |
| 1 | 163883114 | rs7556434 | -0.06 | 1.687 | 0.001 | -0.899 | 0.005 | -0.035 | -0.035 | 0 | 0.27 | 0.27 | 0.28 | 0.03 | 0.45 | 0.22 |
| 1 | 163886496 | rs12139078 | 0.147 | 1.682 | -0.024 | -0.897 | -0.012 | -0.011 | -0.011 | 0 | 0.26 | 0.23 | 0.26 | 0.02 | 0.42 | 0.12 |
| 1 | 163894487 | rs10436885 | 0.292 | 1.535 | -0.337 | -0.829 | 0.01 | 0.005 | 0.005 | 0.708 | 0.24 | 0.24 | 0.18 | 0.24 | 0.15 | 0.21 |
| 1 | 163898055 | rs7554212 | 0.176 | 1.543 | -0.337 | -0.834 | 0.007 | 0.005 | 0.005 | 0.829 | 0.26 | 0.25 | 0.2 | 0.25 | 0.18 | 0.32 |
| 1 | 163902358 | rs7550282 | 0.19 | 1.544 | -0.358 | -0.848 | 0.009 | 0.007 | 0.007 | 0.839 | 0.26 | 0.24 | 0.2 | 0.25 | 0.18 | 0.32 |
| 1 | 163913755 | rs12129997 | 1.469 | 1.683 | -0.456 | -0.902 | -0.089 | 0.074 | 0.074 | 0 | 0.15 | 0.02 | 0.16 | 0.02 | 0.26 | 0.05 |
| 1 | 163914507 | rs6662025 | 0.874 | 1.207 | -0.279 | -0.846 | 0.03 | 0.037 | 0.037 | 1.297 | 0.2 | 0.09 | 0.08 | 0 | 0.12 | 0.12 |
| 1 | 163916669 | rs10494403 | -0.993 | 1.314 | -0.092 | -0.757 | 0.004 | 0.025 | 0.025 | 0.041 | 0.43 | 0.69 | 0.51 | 0.37 | 0.4 | 0.52 |
| 1 | 163929747 | rs12127998 | 0.514 | 1.23 | -0.146 | -0.757 | 0.036 | 0.013 | 0.013 | 0.617 | 0.24 | 0.12 | 0.17 | 0.24 | 0.15 | 0.21 |
| 1 | 163941245 | rs1858534 | 0.053 | 1.971 | 0.322 | -0.55 | 0.036 | -0.084 | -0.084 | 0 | 0.13 | 0.33 | 0.12 | 0 | 0.16 | 0.07 |
| 1 | 163943873 | rs12082367 | -0.979 | 1.898 | -0.273 | -0.524 | 0.056 | -0.046 | -0.046 | 0 | 0.2 | 0.59 | 0.34 | 0.13 | 0.25 | 0.31 |
| 1 | 163947700 | rs1444445 | -0.916 | 1.933 | -0.349 | -0.57 | 0.029 | -0.057 | -0.057 | 0 | 0.33 | 0.62 | 0.46 | 0.13 | 0.45 | 0.34 |
| 1 | 163947842 | rs1444444 | -0.812 | 1.864 | -0.541 | -0.194 | 0.025 | 0.026 | 0.026 | 0 | 0.05 | 0.12 | 0.22 | 0.1 | 0.08 | 0.23 |
| 1 | 163948127 | rs12143508 | 0.072 | 1.944 | -0.629 | -0.624 | -0.015 | 0.009 | 0.009 | 1.505 | 0.19 | 0.15 | 0.15 | 0 | 0.23 | 0.05 |
| 1 | 163948373 | rs1479640 | -1.101 | 1.771 | -0.261 | 0.042 | 0.002 | 0.003 | 0.003 | 2.412 | 0.39 | 0.52 | 0.5 | 0.56 | 0.46 | 0.31 |
| 1 | 163948473 | rs10494405 | 0.559 | 1.309 | -0.164 | -0.135 | 0.001 | -0.012 | -0.012 | 0 | 0.14 | 0.13 | 0.1 | 0 | 0.15 | 0.03 |
| 1 | 163950439 | rs1444439 | -0.031 | 0.921 | 0.022 | -0.126 | -0.008 | 0.003 | 0.003 | 3.024 | 0.1 | 0.18 | 0.17 | 0.1 | 0.06 | 0.08 |
| 1 | 163953111 | rs1122279 | -0.791 | 0.702 | 0.305 | -0.141 | 0.018 | 0.014 | 0.014 | 0 | 0.28 | 0.46 | 0.43 | 0.32 | 0.3 | 0.29 |
| 1 | 163957374 | rs424175 | -1.253 | 0.22 | 0.037 | 0.454 | 0.005 | 0.005 | 0.005 | 0 | 0.91 | 0.95 | 0.93 | 0.74 | 0.91 | 0.85 |
| 1 | 163957638 | rs17376111 | 1 | 0.286 | 0.036 | 0.367 | 0.016 | 0.005 | 0.005 | 0 | 0.17 | 0.2 | 0.25 | 0.22 | 0.24 | 0.21 |
| 1 | 163968414 | rs260913 | 2.326 | -0.085 | -0.715 | -0.209 | -0.135 | 0.112 | 0.112 | 0.446 | 0.28 | 0.02 | 0.24 | 0.04 | 0.36 | 0.22 |
| 1 | 163973084 | rs260905 | -0.972 | -0.472 | 0.345 | 0.534 | 0.003 | 0.001 | 0.001 | 0 | 0.83 | 0.85 | 0.85 | 0.85 | 0.9 | 0.72 |
| 1 | 164001513 | rs7536844 | 0.857 | -0.638 | 0.591 | 0.56 | -0.015 | -0.018 | -0.018 | 0 | 0.16 | 0.34 | 0.21 | 0.11 | 0.14 | 0.1 |
| 1 | 164003370 | rs2346420 | -1.248 | -0.692 | 0.693 | 0.708 | -0.013 | 0.018 | 0.018 | 0.459 | 0.33 | 0.39 | 0.28 | 0.46 | 0.25 | 0.44 |
| 1 | 164006598 | rs1289586 | -1.178 | -0.672 | 0.483 | 0.591 | 0.001 | -0.003 | -0.003 | 0 | 0.33 | 0.32 | 0.28 | 0.44 | 0.25 | 0.44 |
| 1 | 164021052 | rs12760871 | 1.247 | -0.588 | 0.315 | 0.608 | 0.005 | -0.003 | -0.003 | 0 | 0.08 | 0.03 | 0.07 | 0.27 | 0.09 | 0.15 |
| 1 | 164031874 | rs10917897 | 1.926 | -0.475 | -0.145 | 0.509 | 0.013 | 0.005 | 0.005 | 0.175 | 0.51 | 0.37 | 0.51 | 0.45 | 0.61 | 0.46 |
| 1 | 164045309 | rs12144726 | 1.431 | 0.009 | -0.022 | 0.758 | 0.003 | -0.041 | -0.041 | 0.006 | 0.12 | 0.21 | 0.1 | 0 | 0.14 | 0.03 |
| 1 | 164046161 | rs4085003 | 1.011 | -0.098 | 0.038 | 0.921 | -0.007 | 0.016 | 0.016 | 2.524 | 0.69 | 0.32 | 0.6 | 0.57 | 0.7 | 0.62 |
| 1 | 164059430 | rs16824651 | 1.425 | 0.038 | -0.049 | 0.662 | -0.001 | -0.018 | -0.018 | 0.349 | 0.12 | 0.18 | 0.1 | 0 | 0.14 | 0.03 |
| 1 | 164060002 | rs6672232 | 0.517 | 0.224 | 0.184 | 1.201 | -0.014 | 0.129 | 0.129 | 0.215 | 0.83 | 0.56 | 0.81 | 0.68 | 0.85 | 0.73 |

| | | | | | | | | | | | | | | | | |
|---|---|---|---|---|---|---|---|---|---|---|---|---|---|---|---|---|
| 1 | 164063336 | rs1577834 | 0.544 | 0.271 | -1 | 1.152 | 0.012 | -0.017 | -0.017 | 0.018 | 0.13 | 0.05 | 0.11 | 0.14 | 0.08 | 0.16 |
| 1 | 164067167 | rs1337357 | 1.242 | 0.571 | -0.598 | 1.355 | 0.001 | 0.009 | 0.009 | 1.192 | 0.79 | 0.27 | 0.65 | 0.69 | 0.76 | 0.7 |
| 1 | 164067832 | rs12757975 | 1.058 | 0.352 | -1.452 | 1.199 | -0.064 | 0.108 | 0.108 | 0.162 | 0.43 | 0.11 | 0.34 | 0.2 | 0.43 | 0.26 |
| 1 | 164071162 | rs4657327 | 0.149 | 0.319 | -1.229 | 1.435 | 0.009 | -0.003 | -0.003 | 0 | 0.1 | 0.09 | 0.21 | 0.41 | 0.09 | 0.25 |
| 1 | 164073721 | rs7554160 | 1.3 | 0.388 | -1.35 | 1.286 | -0.07 | 0.158 | 0.158 | 0.593 | 0.39 | 0.11 | 0.33 | 0.13 | 0.41 | 0.24 |
| 1 | 164075176 | rs7545877 | 1.198 | 0.329 | -1.393 | 1.458 | -0.065 | 0.047 | 0.047 | 0.283 | 0.42 | 0.11 | 0.44 | 0.27 | 0.42 | 0.28 |
| 1 | 164078275 | rs7533006 | 0.399 | 0.305 | -0.922 | 1.382 | -0.009 | 0.082 | 0.082 | 0.031 | 0.53 | 0.33 | 0.65 | 0.68 | 0.51 | 0.53 |
| 1 | 164082055 | rs4657329 | 1.088 | 0.41 | -0.741 | 0.829 | -0.034 | 0.065 | 0.065 | 0.29 | 0.42 | 0.2 | 0.46 | 0.26 | 0.42 | 0.27 |
| 1 | 164099722 | rs4657343 | -1.228 | 0.375 | -0.521 | 0.827 | -0.026 | 0.049 | 0.049 | 0.18 | 0.58 | 0.71 | 0.53 | 0.74 | 0.55 | 0.73 |
| 1 | 164100378 | rs10917931 | -1.19 | 0.382 | -0.442 | 0.83 | -0.016 | 0.036 | 0.036 | 0.089 | 0.58 | 0.68 | 0.52 | 0.74 | 0.55 | 0.73 |
| 1 | 164108382 | rs1415439 | 0.211 | 0.163 | -0.194 | 1.468 | -0.018 | 0.068 | 0.068 | 0 | 0.88 | 0.83 | 0.81 | 0.65 | 0.91 | 0.77 |
| 1 | 164109036 | rs10800004 | 0.893 | 0.061 | 0.276 | 1.437 | -0.05 | 0.176 | 0.176 | 0 | 0.86 | 0.6 | 0.8 | 0.69 | 0.91 | 0.77 |
| 1 | 164119921 | rs10737498 | 0.558 | 0.07 | 0.207 | 1.429 | -0.04 | 0.142 | 0.142 | 0 | 0.87 | 0.7 | 0.8 | 0.65 | 0.91 | 0.77 |
| 1 | 164120219 | rs4657349 | 2.1 | -0.018 | 0.407 | 1.328 | -0.013 | 0.021 | 0.021 | 0 | 0.75 | 0.32 | 0.72 | 0.61 | 0.82 | 0.62 |
| 1 | 164125695 | rs10494410 | 0.694 | 0.309 | -0.315 | 1.337 | -0.065 | -0.102 | -0.102 | 0 | 0.17 | 0.04 | 0.26 | 0.38 | 0.22 | 0.15 |
| 1 | 164132019 | rs10800007 | -0.864 | 0.181 | -0.374 | 0.789 | -0.017 | 0.053 | 0.053 | 0 | 0.47 | 0.24 | 0.47 | 0.69 | 0.49 | 0.59 |
| 1 | 164144117 | rs12037444 | 2.326 | 0.149 | -0.394 | 0.311 | -0.013 | 0.072 | 0.072 | 0 | 0.37 | 0.24 | 0.33 | 0.07 | 0.4 | 0.23 |
| 1 | 164149433 | rs1361065 | -0.263 | 0.153 | -0.532 | 0.635 | -0.052 | -0.063 | -0.063 | 0.268 | 0.38 | 0.13 | 0.4 | 0.56 | 0.41 | 0.37 |
| 1 | 164151402 | rs7524933 | 1.361 | 0.191 | -0.387 | 0.598 | 0.009 | 0.076 | 0.076 | 0 | 0.47 | 0.64 | 0.36 | 0.1 | 0.42 | 0.38 |
| 1 | 164153205 | rs2210960 | -0.525 | 0.062 | -0.119 | 0.95 | -0.005 | 0.019 | 0.019 | 0 | 0.13 | 0.16 | 0.17 | 0.33 | 0.09 | 0.25 |
| 1 | 164153868 | rs945792 | 0.884 | 0.193 | -0.126 | 0.85 | 0.001 | 0.016 | 0.016 | 0.102 | 0.61 | 0.66 | 0.46 | 0.11 | 0.59 | 0.45 |
| 1 | 164156236 | rs7518128 | 0.429 | 0.195 | -0.232 | 0.881 | -0.023 | -0.072 | -0.072 | 0 | 0.23 | 0.09 | 0.31 | 0.56 | 0.26 | 0.3 |
| 1 | 164159810 | rs11800877 | 0.393 | 0.174 | -0.213 | 0.914 | -0.001 | -0.001 | -0.001 | 0 | 0.09 | 0.09 | 0.06 | 0.16 | 0.07 | 0.22 |
| 1 | 164164032 | rs12124350 | 0.363 | 0.173 | -0.233 | 0.88 | -0.022 | -0.074 | -0.074 | 0 | 0.23 | 0.09 | 0.31 | 0.56 | 0.26 | 0.3 |
| 1 | 164173124 | rs1415434 | -1.143 | 0.337 | -0.175 | 0.946 | -0.012 | 0.02 | 0.02 | 0 | 0.14 | 0.19 | 0.17 | 0.33 | 0.09 | 0.25 |
| 1 | 164174572 | rs12047676 | -0.849 | 0.458 | -0.145 | 0.99 | -0.015 | 0.031 | 0.031 | 0 | 0.13 | 0.19 | 0.17 | 0.36 | 0.09 | 0.25 |
| 1 | 164178654 | rs6656557 | 0.823 | 0.422 | 0.294 | 0.854 | -0.003 | -0.058 | -0.058 | 1.109 | 0.63 | 0.48 | 0.47 | 0.12 | 0.62 | 0.46 |
| 1 | 164180730 | rs4258194 | -0.46 | 0.437 | 0.292 | 0.998 | -0.009 | 0.006 | 0.006 | 0 | 0.31 | 0.38 | 0.37 | 0.7 | 0.3 | 0.5 |
| 1 | 164184997 | rs2346884 | 0.83 | 0.424 | 0.249 | 1.042 | -0.028 | -0.073 | -0.073 | 2.262 | 0.19 | 0.07 | 0.3 | 0.51 | 0.24 | 0.29 |
| 1 | 164195785 | rs12076657 | -0.405 | 0.379 | 0.374 | 1.123 | -0.049 | 0.052 | 0.052 | 0 | 0.11 | 0.29 | 0.07 | 0.16 | 0.06 | 0.21 |
| 1 | 164197996 | rs16826591 | -0.378 | 0.744 | 0.669 | 1.54 | -0.008 | -0.002 | -0.002 | 0 | 0.1 | 0.18 | 0.04 | 0.06 | 0.06 | 0.15 |
| 1 | 164199038 | rs7542616 | -0.251 | 0.863 | 0.539 | 1.667 | -0.003 | 0.012 | 0.012 | 0 | 0.05 | 0.06 | 0.15 | 0.17 | 0.08 | 0.03 |
| 1 | 164212456 | rs2881506 | -0.541 | 0.408 | 0.606 | 1.332 | -0.001 | 0.01 | 0.01 | 0 | 0.28 | 0.22 | 0.43 | 0.88 | 0.28 | 0.51 |
| 1 | 164213686 | rs1556867 | 0.113 | 0.51 | 0.625 | 1.419 | -0.002 | -0.003 | -0.003 | 0.688 | 0.22 | 0.18 | 0.4 | 0.67 | 0.24 | 0.31 |
| 1 | 164216796 | rs7520925 | 0.239 | 0.037 | 0.567 | 1.392 | 0.008 | -0.001 | -0.001 | 0 | 0.06 | 0.03 | 0.08 | 0.21 | 0.03 | 0.2 |
| 1 | 164226252 | rs2881507 | 0.136 | 0.073 | 0.732 | 1.385 | 0.024 | 0.05 | 0.05 | 0 | 0.25 | 0.26 | 0.37 | 0.52 | 0.33 | 0.39 |
| 1 | 164227437 | rs6682660 | -0.408 | 0.132 | 0.415 | 0.808 | 0.036 | 0.012 | 0.012 | 0 | 0.3 | 0.45 | 0.42 | 0.67 | 0.35 | 0.42 |
| 1 | 164230383 | rs10917967 | -0.488 | 0.154 | 0.085 | 0.829 | 0.008 | 0 | 0 | 0 | 0.32 | 0.57 | 0.43 | 0.68 | 0.34 | 0.42 |
| 1 | 164242365 | rs1953819 | -1.313 | -0.007 | -0.607 | 0.477 | -0.095 | 0.198 | 0.198 | 0.093 | 0.64 | 0.98 | 0.69 | 0.93 | 0.62 | 0.79 |
| 1 | 164268177 | rs499492 | -1.35 | 0.137 | 0.027 | 0.793 | 0.041 | 0.166 | 0.166 | 0 | 0.62 | 0.86 | 0.68 | 0.93 | 0.68 | 0.63 |
| 1 | 164274152 | rs10800013 | 0.138 | 0.261 | 1.029 | 1.187 | 0.054 | 0.03 | 0.03 | 0 | 0.29 | 0.55 | 0.41 | 0.61 | 0.37 | 0.27 |
| 1 | 164280278 | rs535988 | -1.067 | 0.247 | 1.25 | 1.176 | -0.012 | -0.01 | -0.01 | 0 | 0.36 | 0.55 | 0.28 | 0.32 | 0.3 | 0.35 |
| 1 | 164284530 | rs531802 | -1.135 | 0.187 | 1.601 | 1.547 | -0.005 | 0.003 | 0.003 | 0.055 | 0.36 | 0.54 | 0.29 | 0.48 | 0.3 | 0.36 |
| 1 | 164288668 | rs6426865 | 0.578 | 0.292 | 1.636 | 1.584 | 0.006 | 0.01 | 0.01 | 0 | 0.25 | 0.32 | 0.37 | 0.46 | 0.37 | 0.26 |
| 1 | 164293400 | rs2130151 | -0.723 | 0.281 | 1.684 | 1.341 | 0.001 | 0.002 | 0.002 | 0.076 | 0.31 | 0.26 | 0.28 | 0.32 | 0.29 | 0.34 |
| 1 | 164301943 | rs555908 | 0.036 | 0.342 | 1.765 | 1.432 | 0.066 | 0.03 | 0.03 | 0 | 0.29 | 0.52 | 0.4 | 0.61 | 0.38 | 0.27 |
| 1 | 164311974 | rs499321 | -1.786 | 0.272 | 1.703 | 1.399 | 0.033 | 0.151 | 0.151 | 0 | 0.62 | 0.86 | 0.68 | 0.91 | 0.68 | 0.61 |
| 1 | 164312587 | rs555162 | 0.466 | 0.287 | 1.677 | 1.42 | -0.014 | -0.01 | -0.01 | 0 | 0.26 | 0.22 | 0.37 | 0.58 | 0.37 | 0.27 |
| 1 | 164317596 | rs12725288 | -0.847 | 0.294 | 1.843 | 1.428 | 0 | 0 | 0 | 0 | 0.33 | 0.29 | 0.27 | 0.33 | 0.29 | 0.34 |
| 1 | 164327833 | rs574944 | -1.929 | 0.192 | 1.826 | 1.371 | 0.003 | -0.019 | -0.019 | 0 | 0.66 | 0.68 | 0.68 | 0.93 | 0.69 | 0.79 |
| 1 | 164332649 | rs2077405 | -0.459 | 0.218 | 1.642 | 1.375 | -0.006 | 0.004 | 0.004 | 1.681 | 0.23 | 0.25 | 0.22 | 0.33 | 0.19 | 0.28 |
| 1 | 164336084 | rs10917994 | -0.364 | 0.23 | 1.578 | 1.393 | -0.039 | -0.04 | -0.04 | 1.295 | 0.38 | 0.16 | 0.4 | 0.5 | 0.5 | 0.5 |
| 1 | 164336549 | rs535740 | -1.728 | 0.324 | 1.656 | 1.262 | -0.022 | -0.053 | -0.053 | 0 | 0.63 | 0.48 | 0.66 | 0.85 | 0.68 | 0.78 |
| 1 | 164339179 | rs1338982 | -0.425 | 0.026 | 1.521 | 1.433 | -0.065 | -0.056 | -0.056 | 0.155 | 0.39 | 0.11 | 0.41 | 0.51 | 0.5 | 0.51 |
| 1 | 164348564 | rs10917995 | -1.317 | 0.061 | 1.636 | 1.481 | -0.02 | 0.015 | 0.015 | 0.047 | 0.34 | 0.46 | 0.31 | 0.43 | 0.28 | 0.33 |
| 1 | 164349646 | rs7541743 | -1.92 | 0.003 | 1.581 | 1.453 | 0.02 | 0.044 | 0.044 | 0 | 0.64 | 0.78 | 0.65 | 0.82 | 0.69 | 0.78 |
| 1 | 164349769 | rs2881588 | -0.944 | -0.023 | 1.602 | 1.448 | 0.001 | 0.001 | 0.001 | 0 | 0.3 | 0.27 | 0.27 | 0.32 | 0.26 | 0.31 |
| 1 | 164350468 | rs526421 | -2.036 | -0.022 | 1.34 | 1.429 | 0.027 | 0.106 | 0.106 | 0 | 0.65 | 0.82 | 0.69 | 0.92 | 0.69 | 0.78 |
| 1 | 164352952 | rs7528809 | -0.43 | -0.255 | 1.313 | 1.476 | -0.041 | -0.051 | -0.051 | 0 | 0.41 | 0.14 | 0.4 | 0.5 | 0.53 | 0.51 |
| 1 | 164354594 | rs931404 | -1.318 | -0.815 | 1.829 | 1.879 | -0.002 | -0.03 | -0.03 | 0.03 | 0.64 | 0.65 | 0.68 | 0.92 | 0.69 | 0.79 |
| 1 | 164360749 | rs1416436 | -0.522 | -0.81 | 1.678 | 1.907 | 0.076 | 0.02 | 0.02 | 0 | 0.54 | 0.27 | 0.48 | 0.55 | 0.64 | 0.61 |
| 1 | 164365846 | rs12043375 | -2.117 | -0.551 | 3.047 | 3.246 | -0.045 | 0.04 | 0.04 | 0 | 0.2 | 0.48 | 0.22 | 0.28 | 0.15 | 0.22 |
| 1 | 164385690 | rs10800018 | -1.94 | -0.472 | 2.448 | 3.222 | -0.01 | 0.173 | 0.173 | 0 | 0.08 | 0.19 | 0.24 | 0.66 | 0.06 | 0.24 |
| 1 | 164388854 | rs12122949 | -1.199 | -0.505 | 1.958 | 2.998 | 0.018 | 0.048 | 0.048 | 0 | 0.14 | 0.05 | 0.05 | 0 | 0.1 | 0.06 |
| 1 | 164392446 | rs12030918 | -1.705 | -0.445 | 1.95 | 3.323 | -0.007 | 0.134 | 0.134 | 0 | 0.07 | 0.16 | 0.24 | 0.66 | 0.06 | 0.22 |
| 1 | 164404171 | rs2788526 | -1.855 | -0.108 | 1.053 | 2.598 | 0.013 | 0.254 | 0.254 | 0 | 0.17 | 0.4 | 0.24 | 0.56 | 0.14 | 0.34 |
| 1 | 164409349 | rs1873137 | -1.777 | -0.013 | 0.464 | 2.354 | -0.002 | 0.009 | 0.009 | 0 | 0.24 | 0.23 | 0.3 | 0.43 | 0.21 | 0.35 |
| 1 | 164412847 | rs1289726 | -1.141 | 0.36 | 0.684 | 2.865 | 0.011 | 0.126 | 0.126 | 0 | 0.08 | 0.12 | 0.16 | 0.45 | 0.1 | 0.12 |
| 1 | 164414741 | rs1387353 | -1.922 | -0.168 | 1.393 | 2.912 | -0.012 | 0.357 | 0.357 | 0.897 | 0.09 | 0.47 | 0.21 | 0.33 | 0.04 | 0.17 |

| | | | | | | | | | | | | | | | | |
|---|---|---|---|---|---|---|---|---|---|---|---|---|---|---|---|---|
| 1 | 164421385 | rs11806933 | -1.765 | 0.173 | 0.474 | 2.753 | -0.004 | 0.157 | 0.157 | 0 | 0.07 | 0.16 | 0.08 | 0.41 | 0.08 | 0.2 |
| 1 | 164444233 | rs10753643 | 1.138 | -0.056 | 0.377 | 1.901 | -0.026 | 0.063 | 0.063 | 0.408 | 0.77 | 0.53 | 0.66 | 0.33 | 0.82 | 0.61 |
| 1 | 164446853 | rs1538371 | -0.085 | -0.194 | -0.844 | 0.237 | 0.04 | 0.172 | 0.172 | 0 | 0.57 | 0.84 | 0.74 | 0.88 | 0.65 | 0.68 |
| 1 | 164447438 | rs10494417 | -0.29 | 0.507 | -0.851 | 0.195 | -0.017 | -0.004 | -0.004 | 0 | 0.12 | 0.06 | 0.09 | 0.15 | 0.16 | 0.23 |
| 1 | 164447829 | rs1416261 | 0.302 | 0.317 | -0.702 | -0.206 | 0.002 | 0.012 | 0.012 | 0 | 0.39 | 0.27 | 0.38 | 0.18 | 0.41 | 0.24 |
| 1 | 164450357 | rs1556853 | -0.286 | 0.138 | -0.376 | -0.628 | -0.018 | 0.024 | 0.024 | 0 | 0.57 | 0.64 | 0.48 | 0.24 | 0.53 | 0.39 |
| 1 | 164456024 | rs7526828 | 0.711 | 0.459 | -0.091 | -0.486 | -0.004 | -0.003 | -0.003 | 0 | 0.11 | 0.08 | 0.07 | 0.14 | 0.12 | 0.23 |
| 1 | 164457615 | rs10800024 | -0.123 | 0.227 | 0.173 | -0.661 | 0.017 | 0.002 | 0.002 | 0 | 0.53 | 0.54 | 0.5 | 0.38 | 0.59 | 0.53 |
| 1 | 164467246 | rs16827858 | -0.007 | 0.403 | 0.153 | -1.392 | -0.001 | -0.013 | -0.013 | 0 | 0.14 | 0.18 | 0.08 | 0.1 | 0.13 | 0.24 |
| 1 | 164474258 | rs10918018 | 0.618 | 0.288 | -0.153 | -1.555 | 0 | -0.016 | -0.016 | 0 | 0.39 | 0.31 | 0.26 | 0.07 | 0.4 | 0.28 |
| 1 | 164475201 | rs17385225 | 0.076 | 0.286 | -0.603 | -0.945 | -0.003 | -0.014 | -0.014 | 2.165 | 0.06 | 0.1 | 0.13 | 0 | 0.04 | 0.03 |
| 1 | 164478274 | rs10800027 | -1.052 | 0.73 | -0.784 | -0.962 | 0 | -0.001 | -0.001 | 0.012 | 0.25 | 0.27 | 0.2 | 0.13 | 0.22 | 0.34 |
| 1 | 164479479 | rs6662422 | -0.099 | 0.468 | -0.931 | -1.418 | 0.005 | 0.035 | 0.035 | 0.038 | 0.56 | 0.65 | 0.45 | 0.14 | 0.58 | 0.46 |
| 1 | 164481240 | rs868376 | -0.824 | -0.161 | -0.755 | -1.427 | -0.019 | -0.057 | -0.057 | 0 | 0.39 | 0.55 | 0.19 | 0.13 | 0.31 | 0.44 |
| 1 | 164481676 | rs1954114 | 0.392 | -0.155 | -0.862 | -1.236 | -0.001 | -0.003 | -0.003 | 0.12 | 0.05 | 0.05 | 0.03 | 0 | 0.05 | 0.01 |
| 1 | 164489175 | rs954049 | -0.167 | -0.055 | -0.739 | -1.262 | 0.005 | -0.013 | -0.013 | 0.542 | 0.6 | 0.51 | 0.34 | 0.33 | 0.5 | 0.42 |
| 1 | 164489804 | rs12060820 | 0.746 | -0.583 | -0.841 | -1.786 | -0.026 | -0.021 | -0.021 | 0 | 0.14 | 0.2 | 0.04 | 0.12 | 0.08 | 0.06 |
| 1 | 164492154 | rs12402920 | 0.914 | -0.41 | -1.453 | -2.05 | -0.028 | 0.015 | 0.015 | 0 | 0.13 | 0.06 | 0.18 | 0.06 | 0.24 | 0.1 |
| 1 | 164496611 | rs946253 | -0.147 | -1.091 | -1.043 | -2.264 | -0.045 | -0.048 | -0.048 | 0 | 0.3 | 0.42 | 0.15 | 0.2 | 0.17 | 0.18 |
| 1 | 164496822 | rs12091822 | 0.86 | -1.2 | -0.859 | -2.315 | -0.015 | -0.017 | -0.017 | 0.458 | 0.06 | 0.23 | 0.03 | 0 | 0.01 | 0 |
| 1 | 164497415 | rs12410040 | 0.595 | -1.297 | -1.674 | -2.376 | -0.004 | 0.007 | 0.007 | 0 | 0.07 | 0.13 | 0.06 | 0.13 | 0.04 | 0.02 |
| 1 | 164497528 | rs6657417 | 1.072 | -1.206 | -1.772 | -2.436 | 0.029 | 0.044 | 0.044 | 10.152 | 0.17 | 0.05 | 0.07 | 0.07 | 0.12 | 0.15 |
| 1 | 164503366 | rs1340519 | -1.543 | -0.645 | -2.84 | -2.501 | -0.03 | 0.041 | 0.041 | 0 | 0.81 | 0.89 | 0.85 | 0.92 | 0.79 | 0.81 |
| 1 | 164503450 | rs12121616 | 0.767 | -0.165 | -2.761 | -2.148 | -0.047 | 0.091 | 0.091 | 0.714 | 0.25 | 0.04 | 0.24 | 0.06 | 0.32 | 0.13 |
| 1 | 164504803 | rs1572827 | 1.03 | -0.049 | -2.787 | -2.155 | -0.022 | 0.027 | 0.027 | 0 | 0.07 | 0 | 0.03 | 0.02 | 0.09 | 0.09 |
| 1 | 164506189 | rs1338634 | 1.03 | -0.049 | -2.787 | -2.155 | -0.022 | 0.027 | 0.027 | 0 | 0.07 | 0 | 0.03 | 0.02 | 0.09 | 0.09 |
| 1 | 164506508 | rs11590969 | 0.496 | 0.175 | -2.397 | -1.822 | -0.008 | 0.062 | 0.062 | 0.377 | 0.26 | 0.14 | 0.25 | 0.06 | 0.32 | 0.13 |
| 1 | 164509706 | rs10918034 | 1.244 | 0.212 | -2.622 | -1.812 | -0.087 | -0.102 | -0.102 | 1.314 | 0.16 | 0.02 | 0.41 | 0.64 | 0.22 | 0.42 |
| 1 | 164512021 | rs12121992 | 0.117 | -0.558 | -1.882 | -1.622 | -0.018 | 0.075 | 0.075 | 0.384 | 0.25 | 0.09 | 0.25 | 0.06 | 0.32 | 0.13 |
| 1 | 164518580 | rs12404863 | -0.825 | -0.442 | -1.115 | -0.837 | -0.002 | -0.001 | -0.001 | 0 | 0.11 | 0.14 | 0.07 | 0.13 | 0.05 | 0.02 |
| 1 | 164529666 | rs2275560 | -0.25 | -0.599 | -1.316 | -1.204 | 0.06 | 0.07 | 0.07 | 0 | 0.45 | 0.65 | 0.28 | 0.13 | 0.31 | 0.28 |
| 1 | 164530130 | rs3767374 | 0.534 | 0.489 | -1.001 | -0.105 | -0.01 | -0.005 | -0.005 | 0 | 0.76 | 0.32 | 0.74 | 0.8 | 0.8 | 0.88 |
| 1 | 164535555 | rs1340521 | 0.266 | 1.929 | -1.695 | -0.068 | -0.152 | 0.002 | 0.002 | 0.862 | 0.81 | 0.97 | 0.79 | 0.86 | 0.62 | 0.57 |
| 1 | 164559536 | rs1780335 | -0.035 | 1.041 | -1.253 | 0.116 | -0.001 | 0 | 0 | 0.376 | 0.53 | 0.51 | 0.65 | 0.62 | 0.52 | 0.6 |
| 1 | 164561328 | rs12119483 | -0.195 | 0.266 | -1.813 | -0.596 | -0.042 | 0.145 | 0.145 | 0 | 0.35 | 0.11 | 0.21 | 0.06 | 0.37 | 0.24 |
| 1 | 164564532 | rs10800038 | -0.439 | -0.225 | -0.703 | -0.695 | -0.001 | 0.011 | 0.011 | 0 | 0.39 | 0.65 | 0.25 | 0.08 | 0.37 | 0.27 |
| 1 | 164565029 | rs1770554 | -0.668 | -0.362 | -0.84 | -0.379 | 0.001 | -0.004 | -0.004 | 0.032 | 0.87 | 0.91 | 0.72 | 0.57 | 0.86 | 0.79 |
| 1 | 164565739 | rs10918050 | 0.085 | -0.37 | -0.245 | -0.424 | -0.011 | 0.055 | 0.055 | 0 | 0.36 | 0.22 | 0.22 | 0.06 | 0.37 | 0.24 |
| 1 | 164568606 | rs2792253 | 1.224 | -0.609 | 0.078 | -0.423 | -0.011 | -0.029 | -0.029 | 1.19 | 0.39 | 0.31 | 0.58 | 0.62 | 0.42 | 0.46 |
| 1 | 164568820 | rs2792254 | 0.376 | -0.595 | 0.113 | -0.338 | 0.007 | -0.022 | -0.022 | 9.932 | 0.78 | 0.85 | 0.83 | 0.7 | 0.8 | 0.73 |
| 1 | 164573864 | rs11581789 | 1.383 | -0.381 | -0.203 | -0.846 | -0.028 | 0.065 | 0.065 | 0 | 0.22 | 0.06 | 0.13 | 0.06 | 0.24 | 0.17 |
| 1 | 164578817 | rs2800787 | 0.819 | -0.397 | 0.023 | -0.604 | -0.011 | 0.018 | 0.018 | 0.385 | 0.45 | 0.37 | 0.46 | 0.5 | 0.49 | 0.48 |
| 1 | 164589641 | rs1770557 | -0.67 | -0.381 | 0.307 | -0.702 | 0 | -0.099 | -0.099 | 0.103 | 0.64 | 0.54 | 0.77 | 0.94 | 0.62 | 0.75 |
| 1 | 164592465 | rs10494418 | -0.69 | -0.25 | 0.047 | -0.675 | -0.004 | -0.049 | -0.049 | 0.712 | 0.13 | 0.19 | 0.09 | 0 | 0.13 | 0.11 |
| 1 | 164595489 | rs2996669 | -0.057 | -0.211 | -0.074 | -0.536 | 0 | -0.025 | -0.025 | 0 | 0.53 | 0.57 | 0.7 | 0.94 | 0.52 | 0.66 |
| 1 | 164601072 | rs1620861 | 0.285 | 0.193 | -0.676 | -0.339 | -0.027 | -0.003 | -0.003 | 2.809 | 0.42 | 0.29 | 0.45 | 0.53 | 0.48 | 0.48 |
| 1 | 164603526 | rs10159124 | -0.027 | 0.069 | -0.872 | -0.467 | 0.004 | -0.036 | -0.036 | 0 | 0.07 | 0.16 | 0.05 | 0 | 0.06 | 0.06 |
| 1 | 164603902 | rs12048281 | -0.91 | 0.028 | -1.512 | -0.025 | 0.007 | -0.011 | -0.011 | 0 | 0.12 | 0.1 | 0.09 | 0.25 | 0.12 | 0.14 |
| 1 | 164604122 | rs2792249 | 0.118 | 0.38 | -1.762 | -0.022 | -0.058 | 0.02 | 0.02 | 0.498 | 0.37 | 0.09 | 0.31 | 0.38 | 0.42 | 0.39 |
| 1 | 164604957 | rs1780337 | -0.076 | 0.162 | -1.13 | 0.083 | 0.028 | 0.05 | 0.05 | 4.044 | 0.51 | 0.33 | 0.44 | 0.33 | 0.58 | 0.57 |
| 1 | 164605447 | rs2996667 | 0.816 | -0.955 | -0.385 | 0.288 | -0.069 | -0.033 | -0.033 | 0 | 0.77 | 0.47 | 0.87 | 0.82 | 0.85 | 0.83 |
| 1 | 164607346 | rs1618566 | 0.599 | -0.589 | -1.21 | 0.155 | -0.013 | 0.027 | 0.027 | 3.003 | 0.34 | 0.13 | 0.27 | 0.24 | 0.35 | 0.38 |
| 1 | 164609582 | rs16833488 | -0.209 | 0.004 | -1.273 | 0.473 | -0.001 | -0.013 | -0.013 | 1.247 | 0.08 | 0.04 | 0.1 | 0.13 | 0.08 | 0.17 |
| 1 | 164619900 | rs6426874 | 0.3 | -0.221 | -0.737 | 0.347 | -0.009 | -0.012 | -0.012 | 0 | 0.41 | 0.23 | 0.55 | 0.55 | 0.53 | 0.34 |
| 1 | 164625696 | rs12047043 | -1.013 | -0.551 | -0.556 | 0.376 | 0.003 | 0.005 | 0.005 | 0 | 0.18 | 0.13 | 0.17 | 0.16 | 0.18 | 0.21 |
| 1 | 164627551 | rs4657367 | -0.921 | -0.505 | -0.537 | 0.397 | 0.001 | 0.003 | 0.003 | 0 | 0.17 | 0.13 | 0.17 | 0.16 | 0.18 | 0.21 |
| 1 | 164635818 | rs2635034 | 1.476 | -0.963 | 0.156 | 0.579 | -0.019 | 0.072 | 0.072 | 0.092 | 0.91 | 0.58 | 0.9 | 0.89 | 0.97 | 0.91 |
| 1 | 164636498 | rs2800775 | 1.382 | -1.308 | 0.058 | 0.458 | -0.064 | -0.013 | -0.013 | 0.853 | 0.7 | 0.4 | 0.79 | 0.78 | 0.84 | 0.81 |
| 1 | 164645738 | rs2635028 | 1.168 | -0.931 | -0.519 | 0.588 | 0.01 | 0.007 | 0.007 | 0 | 0.76 | 0.77 | 0.86 | 0.85 | 0.86 | 0.86 |
| 1 | 164647693 | rs1892174 | 0.402 | -0.942 | -0.534 | 0.541 | 0.007 | -0.015 | -0.015 | 0.347 | 0.83 | 0.84 | 0.94 | 0.99 | 0.94 | 0.94 |
| 1 | 164649075 | rs6426878 | -0.896 | -0.87 | -0.434 | 0.61 | 0.001 | 0.023 | 0.023 | 0 | 0.08 | 0.14 | 0.09 | 0.13 | 0.08 | 0.08 |
| 1 | 164649240 | rs2800804 | 0.769 | -0.858 | -0.669 | 0.362 | 0.003 | 0.003 | 0.003 | 0 | 0.79 | 0.79 | 0.92 | 0.92 | 0.92 | 0.87 |
| 1 | 164651159 | rs10800041 | -0.99 | -0.501 | -0.91 | 1.047 | 0.004 | -0.016 | -0.016 | 1.125 | 0.17 | 0.13 | 0.15 | 0.29 | 0.17 | 0.21 |
| 1 | 164652451 | rs16833830 | -1.067 | -0.983 | -1.307 | 1.012 | 0.008 | -0.035 | -0.035 | 0 | 0.13 | 0 | 0.07 | 0.11 | 0.07 | 0.12 |
| 1 | 164655441 | rs17387607 | 0.347 | 0.176 | -1.85 | 0.662 | -0.064 | 0.016 | 0.016 | 0 | 0.18 | 0.05 | 0.24 | 0.16 | 0.27 | 0.2 |
| 1 | 164655831 | rs1780361 | 0.273 | 0.171 | -1.433 | 0.801 | -0.004 | 0.005 | 0.005 | 0 | 0.92 | 0.88 | 0.94 | 0.92 | 0.93 | 0.85 |
| 1 | 164656976 | rs1552560 | -0.118 | -0.347 | -1.118 | 0.767 | -0.023 | 0.007 | 0.007 | 0.706 | 0.38 | 0.49 | 0.23 | 0.39 | 0.31 | 0.25 |
| 1 | 164658057 | rs10800043 | 0.465 | 0.314 | -1.449 | 1.045 | -0.017 | -0.037 | -0.037 | 0 | 0.21 | 0.12 | 0.31 | 0.33 | 0.25 | 0.16 |
| 1 | 164659590 | rs1686195 | 0.141 | 0.09 | -1.267 | 0.734 | 0.002 | -0.002 | -0.002 | 0.005 | 0.64 | 0.35 | 0.65 | 0.61 | 0.7 | 0.67 |

| | | | | | | | | | | | | | | | | |
|---|---|---|---|---|---|---|---|---|---|---|---|---|---|---|---|---|
| 1 | 164660037 | rs1617366 | 0.101 | -0.098 | -1.816 | 0.4 | 0.071 | 0.072 | 0.072 | 0.845 | 0.55 | 0.27 | 0.68 | 0.73 | 0.67 | 0.7 |
| 1 | 164666253 | rs2800798 | 0.15 | 0.529 | -1.644 | 1.02 | -0.004 | -0.007 | -0.007 | 0 | 0.22 | 0.24 | 0.18 | 0.45 | 0.23 | 0.4 |
| 1 | 164666940 | rs2171692 | 0.561 | 0.009 | -2.066 | 0.524 | -0.006 | 0.083 | 0.083 | 0.011 | 0.5 | 0.21 | 0.57 | 0.33 | 0.52 | 0.38 |
| 1 | 164668349 | rs17383948 | 1.543 | 0.069 | -2.441 | 0.472 | -0.082 | -0.065 | -0.065 | 0 | 0.13 | 0.01 | 0.31 | 0.19 | 0.25 | 0.14 |
| 1 | 164670350 | rs10918059 | 0.724 | -0.239 | -1.313 | 0.639 | 0.007 | 0.051 | 0.051 | 0 | 0.51 | 0.28 | 0.56 | 0.36 | 0.5 | 0.37 |
| 1 | 164671255 | rs977330 | 0.656 | -0.155 | -1.46 | 0.708 | -0.009 | -0.025 | -0.025 | 0 | 0.23 | 0.21 | 0.17 | 0.44 | 0.22 | 0.42 |
| 1 | 164672441 | rs12089801 | -0.107 | -0.145 | -0.961 | 0.551 | 0.013 | -0.01 | -0.01 | 0 | 0.21 | 0.28 | 0.22 | 0.13 | 0.27 | 0.11 |
| 1 | 164678924 | rs1844294 | 0.632 | -0.325 | -1.196 | 0.608 | -0.03 | 0.052 | 0.052 | 0.262 | 0.52 | 0.16 | 0.58 | 0.35 | 0.53 | 0.37 |
| 1 | 164682390 | rs1686172 | 0.632 | -0.594 | -1.145 | 0.988 | -0.032 | 0.011 | 0.011 | 0 | 0.47 | 0.17 | 0.54 | 0.42 | 0.48 | 0.33 |
| 1 | 164684722 | rs12062545 | -0.274 | -1.069 | -0.505 | 1.785 | -0.012 | 0.024 | 0.024 | 0 | 0.08 | 0.15 | 0.04 | 0.17 | 0.05 | 0.18 |
| 1 | 164689328 | rs2171691 | -1.53 | -1.194 | 0.259 | 1.597 | 0.014 | 0.023 | 0.023 | 0 | 0.46 | 0.59 | 0.35 | 0.71 | 0.34 | 0.76 |
| 1 | 164689615 | rs2130166 | -1.166 | -1.244 | 0.477 | 2.071 | -0.009 | 0.025 | 0.025 | 0.002 | 0.35 | 0.45 | 0.29 | 0.58 | 0.3 | 0.56 |
| 1 | 164695090 | rs1489330 | 0.191 | -1.268 | 0.455 | 2.223 | -0.02 | -0.013 | -0.013 | 0 | 0.2 | 0.11 | 0.16 | 0.28 | 0.24 | 0.36 |
| 1 | 164697172 | rs16834140 | 0.432 | -1.26 | 0.271 | 2.21 | 0.001 | 0.002 | 0.002 | 0.324 | 0.08 | 0.07 | 0.03 | 0.12 | 0.04 | 0.15 |
| 1 | 164699116 | rs4656420 | -2.192 | -1.267 | 0.393 | 2.263 | -0.007 | -0.003 | -0.003 | 0.118 | 0.47 | 0.54 | 0.33 | 0.51 | 0.34 | 0.75 |
| 1 | 164701316 | rs12031116 | -0.587 | -1.214 | 0.376 | 2.251 | 0.004 | -0.004 | -0.004 | 0 | 0.11 | 0.09 | 0.06 | 0.11 | 0.04 | 0.2 |
| 1 | 164702044 | rs7518582 | -0.832 | -1.09 | 0.337 | 2.171 | -0.002 | -0.003 | -0.003 | 0 | 0.27 | 0.39 | 0.17 | 0.28 | 0.24 | 0.47 |
| 1 | 164714422 | rs12082443 | 0.442 | -0.569 | -0.235 | 1.626 | -0.058 | -0.035 | -0.035 | 0 | 0.17 | 0.01 | 0.12 | 0.14 | 0.19 | 0.38 |
| 1 | 164714538 | rs16834286 | -1.122 | -0.689 | -0.039 | 1.703 | -0.029 | 0.043 | 0.043 | 0 | 0.14 | 0.36 | 0.07 | 0.22 | 0.08 | 0.24 |
| 1 | 164715576 | rs990715 | -0.456 | -0.682 | -0.054 | 1.703 | 0 | 0.002 | 0.002 | 1.319 | 0.23 | 0.27 | 0.15 | 0.31 | 0.22 | 0.5 |
| 1 | 164716168 | rs11583670 | -1.036 | -0.554 | -0.143 | 1.687 | -0.008 | 0.002 | 0.002 | 0 | 0.34 | 0.44 | 0.23 | 0.37 | 0.27 | 0.66 |
| 1 | 164716209 | rs6681005 | -1.046 | -0.402 | -0.169 | 1.45 | -0.016 | 0.006 | 0.006 | 0 | 0.41 | 0.51 | 0.31 | 0.51 | 0.3 | 0.72 |
| 1 | 164720061 | rs10918063 | -0.755 | -0.153 | -0.107 | 1.244 | -0.021 | 0.071 | 0.071 | 0 | 0.1 | 0.36 | 0.08 | 0.17 | 0.04 | 0.17 |
| 1 | 164723835 | rs7543038 | -0.821 | 0.24 | -0.947 | 1.06 | -0.003 | 0.001 | 0.001 | 0 | 0.44 | 0.56 | 0.3 | 0.51 | 0.37 | 0.75 |
| 1 | 164729434 | rs2635038 | 0.241 | 0.471 | -1.968 | 0.982 | -0.06 | 0.032 | 0.032 | 1.654 | 0.53 | 0.12 | 0.41 | 0.4 | 0.52 | 0.4 |
| 1 | 164730178 | rs2800795 | 0.493 | 0.366 | -2.163 | 0.992 | -0.021 | 0.149 | 0.149 | 2.31 | 0.47 | 0.08 | 0.38 | 0.27 | 0.47 | 0.29 |
| 1 | 164732727 | rs9660091 | 0.284 | 0.407 | -2.423 | 1.43 | -0.123 | -0.056 | -0.056 | 0.147 | 0.31 | 0.02 | 0.23 | 0.26 | 0.36 | 0.36 |
| 1 | 164733504 | rs2789443 | -0.116 | 0.384 | -1.582 | 1.637 | 0.023 | 0.18 | 0.18 | 3.239 | 0.93 | 0.38 | 0.76 | 0.79 | 0.97 | 0.98 |
| 1 | 164734457 | rs2789444 | 0.64 | 0.432 | -1.566 | 1.424 | -0.024 | 0.044 | 0.044 | 0.968 | 0.52 | 0.2 | 0.41 | 0.4 | 0.52 | 0.4 |
| 1 | 164735294 | rs16834573 | 0.232 | 0.455 | -1.063 | 1.443 | 0.003 | 0.013 | 0.013 | 0 | 0.07 | 0.12 | 0.07 | 0.12 | 0.08 | 0.17 |
| 1 | 164737549 | rs7521521 | 0.434 | 0.273 | -1.858 | 1.013 | 0.055 | 0.04 | 0.04 | 0.224 | 0.51 | 0.2 | 0.42 | 0.41 | 0.61 | 0.44 |
| 1 | 164739171 | rs6670655 | -0.177 | -0.112 | -0.951 | 1.013 | -0.024 | 0.005 | 0.005 | 0 | 0.72 | 0.61 | 0.58 | 0.73 | 0.76 | 0.87 |
| 1 | 164740099 | rs2171690 | -0.235 | 0.384 | -0.725 | 0.59 | -0.009 | 0.012 | 0.012 | 0 | 0.44 | 0.37 | 0.37 | 0.38 | 0.47 | 0.53 |
| 1 | 164740891 | rs12098218 | 1.203 | 0.549 | -1.151 | 0.713 | -0.038 | -0.064 | -0.064 | 0.014 | 0.13 | 0.01 | 0.12 | 0.17 | 0.12 | 0.13 |
| 1 | 164741588 | rs10918068 | -0.44 | 0.795 | -0.821 | 0.135 | -0.012 | 0.047 | 0.047 | 1.787 | 0.67 | 0.8 | 0.69 | 0.79 | 0.6 | 0.59 |
| 1 | 164743778 | rs12737465 | -0.388 | 0.916 | -0.821 | 0.131 | -0.018 | 0.056 | 0.056 | 1.45 | 0.66 | 0.8 | 0.68 | 0.79 | 0.58 | 0.6 |
| 1 | 164744179 | rs10800047 | 0.499 | 1.049 | -0.677 | 0.033 | -0.001 | 0.011 | 0.011 | 0.766 | 0.22 | 0.19 | 0.16 | 0.13 | 0.23 | 0.29 |
| 1 | 164744889 | rs10753646 | 0.141 | 0.853 | -0.113 | -0.027 | -0.006 | -0.055 | -0.055 | 1.311 | 0.24 | 0.37 | 0.18 | 0.13 | 0.23 | 0.3 |
| 1 | 164745464 | rs10753647 | 0.334 | 0.855 | -0.079 | -0.026 | -0.002 | -0.035 | -0.035 | 0 | 0.23 | 0.32 | 0.18 | 0.13 | 0.23 | 0.3 |
| 1 | 164747408 | rs10800048 | -0.296 | 0.78 | 0.222 | 0.158 | -0.01 | -0.033 | -0.033 | 3.855 | 0.29 | 0.58 | 0.22 | 0.25 | 0.26 | 0.36 |
| 1 | 164749002 | rs6686727 | -0.782 | 0.699 | -0.018 | 0.382 | 0.002 | 0.012 | 0.012 | 0 | 0.5 | 0.62 | 0.4 | 0.58 | 0.54 | 0.65 |
| 1 | 164756267 | rs1387387 | -1.605 | 0.558 | 0.257 | 0.294 | 0.04 | 0.094 | 0.094 | 0.481 | 0.05 | 0.28 | 0.02 | 0.1 | 0.03 | 0.05 |
| 1 | 164764500 | rs4657375 | 0.403 | 0.69 | -0.163 | 0.442 | -0.016 | -0.021 | -0.021 | 0 | 0.22 | 0.14 | 0.17 | 0.31 | 0.28 | 0.27 |
| 1 | 164766209 | rs902038 | 0.993 | 0.725 | -0.097 | 0.181 | 0.001 | 0.002 | 0.002 | 0.063 | 0.22 | 0.23 | 0.15 | 0.14 | 0.21 | 0.31 |
| 1 | 164771701 | rs729071 | -1.536 | 0.477 | -0.57 | -0.003 | -0.061 | -0.136 | -0.136 | 0 | 0.68 | 0.94 | 0.49 | 0.53 | 0.64 | 0.73 |
| 1 | 164772640 | rs3767371 | -0.663 | 0.414 | -0.073 | -0.255 | -0.001 | 0.005 | 0.005 | 0 | 0.47 | 0.58 | 0.33 | 0.47 | 0.49 | 0.63 |
| 1 | 164775247 | rs3767369 | 0.654 | 0.371 | 0.003 | -0.305 | -0.017 | -0.056 | -0.056 | 0 | 0.24 | 0.4 | 0.15 | 0.14 | 0.21 | 0.29 |
| 1 | 164781015 | rs10494422 | 2.515 | 0.724 | -0.176 | -0.338 | 0.013 | -0.021 | -0.021 | 0 | 0.05 | 0.06 | 0.05 | 0 | 0.09 | 0.01 |
| 1 | 164786245 | rs2185217 | -0.683 | 0.557 | -0.204 | -0.312 | -0.001 | 0.001 | 0.001 | 0 | 0.48 | 0.51 | 0.34 | 0.47 | 0.51 | 0.63 |
| 1 | 164786799 | rs956409 | 0.642 | 0.619 | -0.411 | -0.339 | -0.065 | -0.07 | -0.07 | 0 | 0.21 | 0.03 | 0.18 | 0.27 | 0.29 | 0.27 |
| 1 | 164788638 | rs3767365 | -0.925 | 0.669 | -0.303 | -0.305 | -0.001 | 0.016 | 0.016 | 0 | 0.49 | 0.66 | 0.35 | 0.47 | 0.51 | 0.61 |
| 1 | 164793178 | rs6698381 | -0.216 | 0.519 | -0.067 | -0.813 | -0.003 | -0.037 | -0.037 | 1.55 | 0.44 | 0.45 | 0.31 | 0.16 | 0.38 | 0.39 |
| 1 | 164793479 | rs1890536 | 0.045 | 0.477 | 0.068 | -0.484 | -0.022 | -0.049 | -0.049 | 0 | 0.28 | 0.58 | 0.17 | 0.18 | 0.21 | 0.36 |
| 1 | 164799811 | rs4657378 | -0.388 | 0.797 | 0.012 | -0.397 | -0.001 | -0.009 | -0.009 | 0 | 0.44 | 0.4 | 0.31 | 0.19 | 0.45 | 0.42 |
| 1 | 164800483 | rs9887944 | -0.524 | 0.022 | 0.618 | 0.259 | 0.007 | 0.159 | 0.159 | 0 | 0.11 | 0.27 | 0.07 | 0.27 | 0.09 | 0.18 |
| 1 | 164807125 | rs1107533 | -0.169 | 0.154 | 0.508 | 0.061 | -0.014 | -0.041 | -0.041 | 0 | 0.34 | 0.39 | 0.21 | 0.14 | 0.28 | 0.28 |
| 1 | 164807929 | rs1107531 | -0.118 | 0.157 | 0.511 | 0.064 | -0.01 | -0.033 | -0.033 | 0 | 0.34 | 0.36 | 0.21 | 0.14 | 0.28 | 0.28 |
| 1 | 164810580 | rs10800055 | -0.386 | 0.017 | 0.381 | -0.294 | -0.014 | -0.047 | -0.047 | 0 | 0.37 | 0.46 | 0.23 | 0.21 | 0.34 | 0.34 |
| 1 | 164812653 | rs11808475 | -0.196 | -0.086 | 0.315 | -0.381 | -0.018 | -0.051 | -0.051 | 0 | 0.35 | 0.41 | 0.21 | 0.14 | 0.28 | 0.27 |
| 1 | 164813307 | rs6693637 | -0.196 | -0.086 | 0.315 | -0.381 | -0.018 | -0.051 | -0.051 | 0 | 0.35 | 0.41 | 0.2 | 0.14 | 0.28 | 0.28 |
| 1 | 164815467 | rs12048214 | -0.398 | 0.457 | 0.443 | -0.026 | 0.003 | 0.012 | 0.012 | 0 | 0.53 | 0.57 | 0.35 | 0.15 | 0.44 | 0.45 |
| 1 | 164825891 | rs10918081 | -0.685 | 0.387 | 0.53 | 0.346 | -0.001 | 0.003 | 0.003 | 0 | 0.58 | 0.56 | 0.4 | 0.33 | 0.54 | 0.53 |
| 1 | 164829035 | rs1890535 | -0.409 | 0.381 | 0.77 | 0.356 | 0 | -0.01 | -0.01 | 0 | 0.56 | 0.46 | 0.39 | 0.33 | 0.53 | 0.52 |
| 1 | 164835685 | rs12095760 | -0.07 | 0.44 | 0.685 | 0.368 | 0.002 | 0.025 | 0.025 | 0 | 0.06 | 0.08 | 0.04 | 0.17 | 0.06 | 0.16 |
| 1 | 164836521 | rs10918082 | -0.098 | 0.378 | 0.855 | 0.065 | -0.005 | 0.064 | 0.064 | 0.015 | 0.51 | 0.34 | 0.37 | 0.23 | 0.48 | 0.41 |
| 1 | 164839391 | rs17390990 | 0.127 | -0.065 | 0.68 | -0.136 | -0.029 | 0.09 | 0.09 | 0 | 0.19 | 0.07 | 0.15 | 0 | 0.24 | 0.13 |
| 1 | 164848510 | rs2488915 | 0.3 | -0.174 | 0.917 | -0.001 | 0.025 | 0.05 | 0.05 | 0.354 | 0.89 | 0.6 | 0.94 | 0.85 | 0.93 | 0.83 |
| 1 | 164862175 | rs1213293 | 0.857 | -0.189 | 0.842 | -0.019 | 0.008 | -0.037 | -0.037 | 0 | 0.69 | 0.51 | 0.77 | 0.75 | 0.64 | 0.65 |
| 1 | 164862578 | rs12410158 | 0.646 | -0.181 | 0.689 | -0.076 | -0.017 | -0.021 | -0.021 | 0 | 0.31 | 0.34 | 0.22 | 0.2 | 0.19 | 0.24 |

| | | | | | | | | | | | | | | | | |
|---|---|---|---|---|---|---|---|---|---|---|---|---|---|---|---|---|
| 1 | 164864196 | rs10800056 | 0.784 | -0.119 | 0.885 | -0.079 | 0.003 | 0.004 | 0.004 | 0 | 0.3 | 0.26 | 0.22 | 0.24 | 0.21 | 0.29 |
| 1 | 164864941 | rs2488912 | 1.667 | -0.176 | 0.391 | -0.11 | -0.148 | -0.143 | -0.143 | 0 | 0.33 | 0.03 | 0.54 | 0.56 | 0.45 | 0.33 |
| 1 | 164873859 | rs1213285 | 0.707 | -0.25 | 0.896 | -0.086 | 0 | -0.037 | -0.037 | 0 | 0.73 | 0.64 | 0.8 | 0.87 | 0.71 | 0.74 |
| 1 | 164875964 | rs1541243 | 0.522 | -0.269 | 0.989 | -0.031 | -0.012 | -0.01 | -0.01 | 0 | 0.31 | 0.35 | 0.23 | 0.25 | 0.21 | 0.28 |
| 1 | 164879202 | rs1936078 | 1.947 | -0.277 | 0.758 | 0.03 | -0.147 | -0.192 | -0.192 | 0 | 0.32 | 0.03 | 0.53 | 0.5 | 0.43 | 0.32 |
| 1 | 164879437 | rs1936076 | 0.511 | -0.296 | 0.976 | -0.156 | -0.016 | 0.048 | 0.048 | 0 | 0.16 | 0.08 | 0.14 | 0 | 0.22 | 0.09 |
| 1 | 164882733 | rs4347190 | -0.671 | -0.268 | 1.011 | -0.131 | 0.003 | 0.017 | 0.017 | 0 | 0.49 | 0.43 | 0.36 | 0.25 | 0.44 | 0.43 |
| 1 | 164883887 | rs7538001 | -1.056 | -0.219 | 0.775 | -0.105 | 0 | 0.044 | 0.044 | 0 | 0.54 | 0.68 | 0.39 | 0.32 | 0.47 | 0.48 |
| 1 | 164892677 | rs4526594 | -0.063 | -0.27 | 0.7 | -0.131 | -0.002 | 0.019 | 0.019 | 0 | 0.19 | 0.12 | 0.14 | 0 | 0.22 | 0.1 |
| 1 | 164897721 | rs7553620 | 0.367 | -0.226 | 0.626 | 0.015 | 0.008 | 0.06 | 0.06 | 0.56 | 0.89 | 0.7 | 0.92 | 0.8 | 0.89 | 0.79 |
| 1 | 164899436 | rs7512483 | 0.189 | -0.246 | 0.664 | -0.166 | 0.006 | 0.02 | 0.02 | 0.152 | 0.89 | 0.7 | 0.94 | 0.87 | 0.93 | 0.83 |
| 1 | 164899933 | rs1936096 | -0.921 | -0.242 | 0.66 | -0.184 | 0.003 | 0.039 | 0.039 | 0 | 0.53 | 0.67 | 0.4 | 0.32 | 0.46 | 0.47 |
| 1 | 164922082 | rs1936090 | 1.423 | -0.124 | 0.426 | -0.152 | -0.106 | -0.096 | -0.096 | 0 | 0.35 | 0.05 | 0.51 | 0.57 | 0.47 | 0.31 |
| 1 | 164922847 | rs1419067 | 0.647 | -0.1 | 0.776 | -0.104 | 0.004 | 0.069 | 0.069 | 1.425 | 0.91 | 0.71 | 0.94 | 0.84 | 0.94 | 0.84 |
| 1 | 164923650 | rs1936093 | 1.548 | -0.128 | 0.482 | -0.08 | -0.109 | -0.133 | -0.133 | 0 | 0.36 | 0.05 | 0.52 | 0.5 | 0.48 | 0.31 |
| 1 | 164924633 | rs10918095 | 0.426 | -0.161 | 0.684 | -0.094 | 0.013 | -0.021 | -0.021 | 0 | 0.32 | 0.14 | 0.23 | 0.33 | 0.24 | 0.31 |
| 1 | 164927336 | rs7550721 | 0.146 | 0.042 | 0.749 | -0.341 | 0.023 | 0.003 | 0.003 | 3.369 | 0.93 | 0.62 | 0.96 | 1 | 0.98 | 0.99 |
| 1 | 164928753 | rs1891737 | 0.106 | -0.044 | 0.798 | -0.357 | 0.005 | -0.008 | -0.008 | 0.089 | 0.37 | 0.34 | 0.28 | 0.5 | 0.28 | 0.48 |
| 1 | 164932033 | rs4259623 | 0.787 | -0.119 | 0.963 | -0.234 | 0.001 | -0.014 | -0.014 | 0 | 0.92 | 0.57 | 0.95 | 1 | 0.98 | 0.99 |
| 1 | 164948279 | rs1343309 | 0.234 | -0.086 | 0.559 | -0.27 | 0.005 | -0.012 | -0.012 | 0 | 0.36 | 0.3 | 0.21 | 0.5 | 0.27 | 0.43 |
| 1 | 164948745 | rs10800064 | 1.393 | 0.083 | -0.112 | -0.271 | -0.108 | -0.133 | -0.133 | 0 | 0.37 | 0.05 | 0.58 | 0.49 | 0.48 | 0.37 |
| 1 | 164948817 | rs4657386 | -0.416 | -0.175 | 0.131 | -0.322 | 0.026 | 0.013 | 0.013 | 2.664 | 0.94 | 0.67 | 0.96 | 1 | 0.98 | 0.99 |
| 1 | 164951242 | rs4300198 | 0.217 | -0.013 | -1.108 | -0.228 | 0.088 | -0.027 | -0.027 | 0 | 0.57 | 0.15 | 0.74 | 0.54 | 0.75 | 0.63 |
| 1 | 164951316 | rs12726675 | -0.795 | 0.514 | -0.303 | -0.402 | -0.057 | 0.087 | 0.087 | 0 | 0.15 | 0.04 | 0.14 | 0 | 0.21 | 0.13 |
| 1 | 164958839 | rs1573181 | 1.321 | 0.4 | 0.229 | -0.332 | -0.001 | -0.039 | -0.039 | 0.316 | 0.76 | 0.33 | 0.79 | 0.95 | 0.75 | 0.8 |
| 1 | 164966447 | rs2211479 | 1.445 | 0.317 | 0.302 | -0.372 | 0.003 | 0.026 | 0.026 | 0.622 | 0.75 | 0.24 | 0.79 | 0.95 | 0.75 | 0.8 |
| 1 | 164967360 | rs4656423 | 1.53 | 0.311 | 0.485 | -0.375 | 0 | -0.005 | -0.005 | 0 | 0.75 | 0.28 | 0.79 | 0.95 | 0.75 | 0.8 |
| 1 | 164969521 | rs1343313 | 1.024 | 0.343 | 0.472 | -0.386 | -0.001 | -0.108 | -0.108 | 0.377 | 0.78 | 0.38 | 0.8 | 1 | 0.78 | 0.87 |
| 1 | 164979815 | rs1343315 | 1.221 | 0.129 | -0.005 | -0.241 | 0.002 | -0.001 | -0.001 | 0 | 0.8 | 0.27 | 0.84 | 0.95 | 0.83 | 0.9 |
| 1 | 164992228 | rs1936075 | 1.33 | 0.157 | -0.169 | -0.261 | 0.014 | 0.034 | 0.034 | 5.218 | 0.79 | 0.19 | 0.82 | 0.94 | 0.84 | 0.88 |
| 1 | 164995713 | rs4387163 | 1.239 | -0.35 | -1.364 | 0.048 | -0.026 | -0.132 | -0.132 | 1.655 | 0.52 | 0.02 | 0.33 | 0.51 | 0.63 | 0.46 |
| 1 | 164995796 | rs1936072 | -0.428 | -0.26 | -0.733 | 0.496 | -0.008 | 0.122 | 0.122 | 0 | 0.07 | 0.1 | 0.23 | 0.35 | 0.05 | 0.19 |
| 1 | 164999850 | rs7542261 | -0.566 | -0.38 | -0.728 | 0.17 | -0.016 | 0.04 | 0.04 | 0 | 0.26 | 0.15 | 0.28 | 0.05 | 0.28 | 0.24 |
| 1 | 165013535 | rs7524769 | -1.867 | -0.498 | -0.023 | 0.27 | 0.001 | -0.004 | -0.004 | 0 | 0.42 | 0.4 | 0.65 | 0.54 | 0.37 | 0.56 |
| 1 | 165013647 | rs11800195 | -0.996 | -0.459 | -0.119 | 0.711 | -0.001 | 0.051 | 0.051 | 0 | 0.08 | 0.11 | 0.34 | 0.38 | 0.07 | 0.2 |
| 1 | 165016712 | rs723821 | -1.658 | -0.369 | -0.153 | 0.555 | -0.018 | 0.075 | 0.075 | 1.076 | 0.17 | 0.27 | 0.39 | 0.49 | 0.11 | 0.35 |
| 1 | 165018128 | rs6659813 | 0.894 | 0.123 | -0.639 | -0.029 | -0.001 | -0.003 | -0.003 | 0 | 0.72 | 0.81 | 0.5 | 0.67 | 0.71 | 0.62 |
| 1 | 165018496 | rs12058573 | -0.661 | 1.153 | -0.424 | 0.035 | -0.056 | -0.028 | -0.028 | 0 | 0.16 | 0.06 | 0.44 | 0.27 | 0.27 | 0.33 |
| 1 | 165018803 | rs1343316 | -0.447 | 0.097 | -0.488 | -0.319 | -0.024 | -0.027 | -0.027 | 0 | 0.33 | 0.58 | 0.18 | 0.17 | 0.19 | 0.27 |
| 1 | 165020001 | rs1419074 | -0.094 | -0.066 | -1.103 | -0.433 | 0.001 | 0.002 | 0.002 | 0 | 0.73 | 0.29 | 0.84 | 0.84 | 0.88 | 0.81 |
| 1 | 165025758 | rs16838572 | 0.147 | 0.024 | -1.411 | -0.385 | 0.002 | 0.024 | 0.024 | 0 | 0.08 | 0.03 | 0.04 | 0.01 | 0.07 | 0.1 |
| 1 | 165027342 | rs6426893 | 0.318 | 0.166 | -1.369 | -0.315 | 0.067 | 0.076 | 0.076 | 0.266 | 0.63 | 0.15 | 0.79 | 0.83 | 0.81 | 0.73 |
| 1 | 165029535 | rs10800069 | -0.379 | 0.954 | -1.493 | 0.018 | -0.107 | -0.067 | -0.067 | 6.083 | 0.16 | 0.02 | 0.43 | 0.27 | 0.3 | 0.27 |
| 1 | 165032101 | rs12049436 | 0.774 | 0.493 | -1.451 | -0.157 | -0.007 | 0.022 | 0.022 | 0 | 0.5 | 0.19 | 0.43 | 0.56 | 0.56 | 0.43 |
| 1 | 165037370 | rs12048476 | 1.178 | 0.353 | -1.812 | -0.554 | -0.065 | -0.114 | -0.114 | 0.016 | 0.31 | 0.04 | 0.45 | 0.58 | 0.35 | 0.3 |
| 1 | 165038271 | rs12119600 | 0.242 | 0.48 | -1.786 | -0.589 | -0.002 | -0.004 | -0.004 | 0 | 0.05 | 0.03 | 0.04 | 0.01 | 0.09 | 0.03 |
| 1 | 165038576 | rs730006 | 0.211 | -0.031 | -1.123 | -1.003 | -0.007 | 0.018 | 0.018 | 0 | 0.61 | 0.67 | 0.64 | 0.69 | 0.59 | 0.53 |
| 1 | 165038938 | rs4495669 | 1.631 | 0.038 | -0.75 | -1.074 | -0.013 | -0.022 | -0.022 | 0 | 0.32 | 0.13 | 0.45 | 0.63 | 0.35 | 0.3 |
| 1 | 165046004 | rs7538054 | 0.847 | -0.054 | -0.364 | -1.083 | 0.007 | 0.038 | 0.038 | 0.196 | 0.48 | 0.37 | 0.56 | 0.63 | 0.47 | 0.33 |
| 1 | 165046596 | rs6663388 | -0.216 | -0.255 | -0.379 | -1.159 | -0.018 | 0.007 | 0.007 | 0.654 | 0.12 | 0.21 | 0.1 | 0.14 | 0.07 | 0.24 |
| 1 | 165047786 | rs2881832 | 0.88 | -0.176 | -0.584 | -1.093 | 0.003 | 0.015 | 0.015 | 0.153 | 0.49 | 0.46 | 0.57 | 0.61 | 0.47 | 0.33 |
| 1 | 165051974 | rs12564912 | 1.156 | 0.111 | -1.171 | -0.988 | -0.096 | 0.019 | 0.019 | 0.075 | 0.41 | 0.07 | 0.51 | 0.67 | 0.48 | 0.46 |
| 1 | 165062200 | rs12133696 | -0.27 | 0.531 | -0.638 | -1.2 | 0.004 | 0.051 | 0.051 | 0 | 0.29 | 0.21 | 0.2 | 0.17 | 0.27 | 0.31 |
| 1 | 165065993 | rs7545659 | 0.954 | 0.367 | -0.845 | -1.29 | -0.022 | 0.005 | 0.005 | 0.212 | 0.12 | 0.01 | 0.08 | 0.05 | 0.12 | 0.09 |
| 1 | 165071376 | rs2348108 | 1.375 | 0.23 | -0.732 | -1.19 | -0.084 | 0.032 | 0.032 | 0.116 | 0.44 | 0.07 | 0.5 | 0.65 | 0.48 | 0.47 |
| 1 | 165079632 | rs10494431 | 0.892 | 0.321 | -0.027 | -1.293 | 0.007 | -0.088 | -0.088 | 0 | 0.14 | 0.25 | 0.11 | 0 | 0.12 | 0.03 |
| 1 | 165081033 | rs923937 | 0.32 | 0.334 | -0.31 | -1.289 | -0.01 | 0.027 | 0.027 | 0 | 0.66 | 0.7 | 0.73 | 0.8 | 0.61 | 0.64 |
| 1 | 165081595 | rs6686369 | -0.002 | 0.152 | 0.116 | -0.967 | -0.002 | -0.042 | -0.042 | 0 | 0.17 | 0.41 | 0.12 | 0.07 | 0.12 | 0.11 |
| 1 | 165086665 | rs4657399 | 0.5 | 0.517 | -0.654 | -1.311 | -0.015 | 0.103 | 0.103 | 0.175 | 0.45 | 0.3 | 0.49 | 0.84 | 0.48 | 0.65 |
| 1 | 165087492 | rs2172712 | 0.557 | 0.611 | -0.092 | -1.029 | 0.002 | -0.009 | -0.009 | 2.093 | 0.41 | 0.42 | 0.4 | 0.69 | 0.45 | 0.52 |
| 1 | 165088116 | rs1494408 | 0.667 | 0.581 | 0.151 | -1.04 | -0.019 | 0.028 | 0.028 | 0 | 0.4 | 0.25 | 0.41 | 0.69 | 0.46 | 0.52 |
| 1 | 165091285 | rs9803673 | -0.083 | 0.619 | 0.134 | -1.4 | -0.002 | -0.012 | -0.012 | 0 | 0.55 | 0.55 | 0.53 | 0.78 | 0.65 | 0.7 |
| 1 | 165096196 | rs12092467 | 0.31 | 0.66 | -0.067 | -1.422 | 0.014 | -0.075 | -0.075 | 0 | 0.12 | 0.32 | 0.07 | 0 | 0.09 | 0.04 |
| 1 | 165102499 | rs6426897 | 2.283 | 0.951 | -0.91 | -1.45 | -0.05 | -0.144 | -0.144 | 0.005 | 0.24 | 0.01 | 0.31 | 0.58 | 0.25 | 0.34 |
| 1 | 165103961 | rs4657400 | 0.378 | -0.359 | -0.483 | -1.767 | -0.002 | 0.004 | 0.004 | 0 | 0.14 | 0.12 | 0.18 | 0.07 | 0.13 | 0.19 |
| 1 | 165104368 | rs903336 | -1.153 | -0.691 | -0.646 | -1.914 | -0.014 | 0.001 | 0.001 | 0 | 0.89 | 0.81 | 0.83 | 0.86 | 0.93 | 0.91 |
| 1 | 165104658 | rs1466734 | 1.708 | -0.508 | -1.431 | -1.895 | -0.088 | 0.007 | 0.007 | 0.063 | 0.38 | 0.1 | 0.4 | 0.69 | 0.46 | 0.51 |
| 1 | 165104676 | rs17397108 | 1.394 | -0.617 | -1.311 | -1.878 | 0.008 | 0.079 | 0.079 | 1.969 | 0.18 | 0.04 | 0.12 | 0 | 0.15 | 0.02 |
| 1 | 165107381 | rs1494405 | 1.148 | -0.349 | -1.231 | -1.65 | -0.055 | 0.09 | 0.09 | 0 | 0.42 | 0.13 | 0.45 | 0.75 | 0.57 | 0.6 |

| | | | | | | | | | | | | | | | | |
|---|---|---|---|---|---|---|---|---|---|---|---|---|---|---|---|---|
| 1 | 165111242 | rs4657404 | 0.818 | -0.535 | -0.921 | -1.622 | -0.007 | 0.076 | 0.076 | 0 | 0.46 | 0.32 | 0.62 | 0.81 | 0.53 | 0.59 |
| 1 | 165111608 | rs10918127 | -0.07 | -0.671 | -0.552 | -1.664 | 0.003 | 0.003 | 0.003 | 0 | 0.63 | 0.63 | 0.72 | 0.89 | 0.66 | 0.79 |
| 1 | 165112096 | rs1038980 | 1.083 | -0.66 | -0.823 | -1.654 | -0.014 | 0.032 | 0.032 | 0 | 0.17 | 0.07 | 0.16 | 0.04 | 0.23 | 0.1 |
| 1 | 165116198 | rs2050986 | 0.144 | -1.134 | -0.821 | -1.802 | -0.005 | 0.094 | 0.094 | 0 | 0.67 | 0.87 | 0.78 | 0.9 | 0.69 | 0.72 |
| 1 | 165116421 | rs7553072 | 0.693 | -1.412 | -0.359 | -1.95 | -0.051 | -0.064 | -0.064 | 0 | 0.19 | 0.36 | 0.1 | 0.05 | 0.1 | 0.1 |
| 1 | 165117237 | rs10494434 | -0.525 | -0.896 | -0.577 | -1.517 | -0.036 | 0.086 | 0.086 | 0 | 0.76 | 0.92 | 0.81 | 0.96 | 0.74 | 0.89 |
| 1 | 165120633 | rs4657406 | 0.524 | -0.855 | -0.524 | -1.397 | 0.011 | 0.007 | 0.007 | 0 | 0.09 | 0.04 | 0.03 | 0.06 | 0.04 | 0.16 |
| 1 | 165121286 | rs10737505 | 1.157 | -0.764 | -0.423 | -0.838 | 0.006 | 0.009 | 0.009 | 0 | 0.49 | 0.41 | 0.51 | 0.57 | 0.55 | 0.48 |
| 1 | 165124178 | rs10918129 | -0.539 | -0.529 | -0.253 | -0.592 | -0.009 | 0.039 | 0.039 | 0 | 0.16 | 0.31 | 0.22 | 0.28 | 0.14 | 0.19 |
| 1 | 165126948 | rs6660004 | -0.545 | -0.52 | -0.393 | -1.319 | -0.002 | -0.087 | -0.087 | 0.114 | 0.25 | 0.38 | 0.19 | 0.06 | 0.26 | 0.14 |
| 1 | 165130289 | rs6671169 | 0.1 | -1.066 | -1.041 | -2.096 | 0.033 | 0.04 | 0.04 | 0.275 | 0.21 | 0.09 | 0.08 | 0.01 | 0.11 | 0.04 |
| 1 | 165131295 | rs2348247 | 1.294 | -1.829 | -1.453 | -2.184 | -0.066 | -0.006 | -0.006 | 0 | 0.35 | 0.11 | 0.57 | 0.65 | 0.52 | 0.61 |
| 1 | 165131967 | rs6678019 | 1.312 | -1.842 | -1.454 | -2.189 | -0.077 | -0.012 | -0.012 | 0.009 | 0.34 | 0.09 | 0.57 | 0.65 | 0.52 | 0.61 |
| 1 | 165132051 | rs7532137 | -0.624 | -2.242 | -1.033 | -2.294 | -0.067 | 0.024 | 0.024 | 0.442 | 0.81 | 0.44 | 0.84 | 0.77 | 0.9 | 0.8 |
| 1 | 165132992 | rs903343 | 0.323 | -2.285 | -0.59 | -2.305 | -0.048 | -0.067 | -0.067 | 0 | 0.62 | 0.47 | 0.76 | 0.88 | 0.79 | 0.81 |
| 1 | 165133001 | rs4656429 | 0.709 | -2.207 | -0.581 | -2.281 | -0.004 | 0.053 | 0.053 | 0.285 | 0.25 | 0.12 | 0.17 | 0.11 | 0.25 | 0.15 |
| 1 | 165133691 | rs730890 | 2.379 | -2.272 | -0.705 | -2.353 | -0.012 | 0.071 | 0.071 | 0.603 | 0.16 | 0.03 | 0.11 | 0 | 0.17 | 0.02 |
| 1 | 165135937 | rs2055596 | 0.144 | -2.431 | -0.588 | -2.4 | -0.072 | -0.075 | -0.075 | 0 | 0.66 | 0.5 | 0.78 | 0.89 | 0.82 | 0.82 |
| 1 | 165137813 | rs2027573 | 1.163 | -2.324 | -0.797 | -2.35 | 0.016 | 0 | 0 | 0 | 0.17 | 0.12 | 0.06 | 0.01 | 0.08 | 0.02 |
| 1 | 165140955 | rs10753664 | 1.189 | -1.744 | -0.758 | -2.002 | -0.052 | -0.012 | -0.012 | 0 | 0.35 | 0.13 | 0.57 | 0.63 | 0.53 | 0.61 |
| 1 | 165157400 | rs4656434 | 0.534 | -1.666 | -0.777 | -2.033 | 0.024 | 0.12 | 0.12 | 2.906 | 0.44 | 0.18 | 0.61 | 0.75 | 0.59 | 0.73 |
| 1 | 165158705 | rs6426901 | -0.628 | -1.666 | -0.242 | -1.781 | -0.049 | -0.017 | -0.017 | 0 | 0.83 | 0.6 | 0.84 | 0.9 | 0.9 | 0.84 |
| 1 | 165166002 | rs12029375 | -0.155 | -1.667 | -0.287 | -1.811 | -0.001 | 0.002 | 0.002 | 0 | 0.15 | 0.22 | 0.16 | 0.22 | 0.12 | 0.18 |
| 1 | 165169159 | rs1819768 | 1.167 | -1.657 | -1.002 | -2.166 | -0.102 | -0.027 | -0.027 | 0.433 | 0.35 | 0.08 | 0.54 | 0.64 | 0.51 | 0.61 |
| 1 | 165171677 | rs17402900 | 0.981 | -1.619 | -0.913 | -2.251 | -0.001 | 0.181 | 0.181 | 3.999 | 0.31 | 0.06 | 0.18 | 0 | 0.27 | 0.06 |
| 1 | 165173549 | rs4657411 | 0.003 | -0.899 | -1.043 | -2.231 | 0.032 | 0.202 | 0.202 | 0 | 0.51 | 0.22 | 0.4 | 0.17 | 0.57 | 0.42 |
| 1 | 165174959 | rs16841013 | 2.021 | -0.616 | -0.952 | -1.99 | -0.08 | -0.151 | -0.151 | 0 | 0.22 | 0.04 | 0.38 | 0.48 | 0.26 | 0.35 |
| 1 | 165175507 | rs6671290 | 1.151 | -0.225 | -0.357 | -2.002 | -0.001 | 0.007 | 0.007 | 0 | 0.06 | 0.1 | 0.04 | 0.11 | 0.05 | 0.02 |
| 1 | 165176571 | rs12742667 | -0.122 | -0.095 | -0.338 | -2.028 | -0.023 | 0.011 | 0.011 | 2.32 | 0.2 | 0.16 | 0.22 | 0.17 | 0.3 | 0.36 |
| 1 | 165179874 | rs6703000 | 0.641 | -0.471 | 0.083 | -1.998 | -0.029 | -0.043 | -0.043 | 0.006 | 0.09 | 0.33 | 0.05 | 0 | 0.03 | 0 |
| 1 | 165180820 | rs6426903 | -0.15 | -0.384 | -0.437 | -2.151 | -0.034 | -0.169 | -0.169 | 2.837 | 0.7 | 0.58 | 0.83 | 1 | 0.77 | 0.87 |
| 1 | 165182355 | rs3738193 | -0.399 | 0.196 | -0.766 | -1.347 | -0.007 | -0.018 | -0.018 | 0 | 0.6 | 0.58 | 0.48 | 0.29 | 0.63 | 0.49 |
| 1 | 165185859 | rs6674957 | -0.4 | -0.057 | -0.941 | -0.898 | -0.001 | 0 | 0 | 0 | 0.14 | 0.13 | 0.08 | 0.24 | 0.11 | 0.25 |
| 1 | 165186490 | rs10737508 | -0.74 | 0.636 | -1.06 | -1.025 | 0.001 | 0.009 | 0.009 | 0 | 0.54 | 0.45 | 0.39 | 0.29 | 0.56 | 0.39 |
| 1 | 165190876 | rs6686960 | 0.369 | 0.6 | -1.254 | -0.354 | 0 | -0.015 | -0.015 | 0 | 0.81 | 0.8 | 0.86 | 0.95 | 0.82 | 0.94 |
| 1 | 165191702 | rs16841280 | -0.742 | 0.633 | -1.176 | 0.235 | 0.001 | -0.006 | -0.006 | 0 | 0.11 | 0.1 | 0.07 | 0.15 | 0.11 | 0.17 |
| 1 | 165193662 | rs12742947 | -1.006 | 0.864 | -1.299 | 0.093 | -0.007 | -0.005 | -0.005 | 0.168 | 0.1 | 0.11 | 0.17 | 0.12 | 0.12 | 0.11 |
| 1 | 165195664 | rs969099 | -0.784 | 0.897 | -1.545 | -0.467 | -0.03 | 0.028 | 0.028 | 0 | 0.2 | 0.12 | 0.23 | 0.13 | 0.26 | 0.13 |
| 1 | 165196400 | rs10494436 | 0.573 | -0.05 | -2.707 | -0.905 | -0.099 | -0.021 | -0.021 | 0.114 | 0.43 | 0.06 | 0.53 | 0.63 | 0.51 | 0.55 |
| 1 | 165199023 | rs10494438 | 1.277 | -0.63 | -1.632 | -1.16 | 0.032 | -0.019 | -0.019 | 1.032 | 0.13 | 0.24 | 0.12 | 0.08 | 0.17 | 0.07 |
| 1 | 165199343 | rs12126617 | -0.701 | -0.72 | -1.413 | -1.171 | 0.014 | -0.019 | -0.019 | 0 | 0.35 | 0.4 | 0.36 | 0.21 | 0.43 | 0.2 |
| 1 | 165207875 | rs6665694 | -0.761 | -1.494 | -0.773 | -1.128 | -0.022 | -0.02 | -0.02 | 0 | 0.37 | 0.55 | 0.33 | 0.29 | 0.32 | 0.35 |
| 1 | 165213822 | rs1394439 | 0.146 | -1.619 | -0.254 | -1.185 | -0.027 | -0.021 | -0.021 | 0 | 0.27 | 0.4 | 0.15 | 0.16 | 0.19 | 0.25 |
| 1 | 165214162 | rs1394438 | -1.363 | -1.375 | -0.831 | -1.632 | -0.027 | -0.04 | -0.04 | 0 | 0.68 | 0.87 | 0.54 | 0.37 | 0.64 | 0.52 |
| 1 | 165216783 | rs10494440 | 1.191 | -1.296 | -1.076 | -1.747 | -0.009 | 0.015 | 0.015 | 0 | 0.09 | 0.02 | 0.11 | 0.07 | 0.12 | 0.06 |
| 1 | 165217356 | rs1506946 | -0.47 | -1.248 | -0.989 | -1.707 | -0.004 | -0.01 | -0.01 | 0 | 0.45 | 0.5 | 0.43 | 0.36 | 0.44 | 0.42 |
| 1 | 165218204 | rs1891928 | 0.678 | -1.234 | -1.001 | -1.711 | 0.006 | -0.05 | -0.05 | 0.414 | 0.31 | 0.36 | 0.21 | 0.08 | 0.32 | 0.17 |
| 1 | 165218679 | rs9970062 | 0.804 | -1.194 | -1.094 | -1.691 | -0.001 | -0.015 | -0.015 | 0 | 0.3 | 0.29 | 0.2 | 0.08 | 0.32 | 0.17 |
| 1 | 165218792 | rs10753668 | 1.064 | -0.634 | -1.103 | -1.189 | -0.006 | -0.047 | -0.047 | 0.26 | 0.23 | 0.13 | 0.35 | 0.56 | 0.24 | 0.42 |
| 1 | 165226552 | rs7513950 | -0.682 | -0.675 | -0.522 | -1.044 | 0 | -0.02 | -0.02 | 1.648 | 0.08 | 0.13 | 0.04 | 0 | 0.07 | 0.02 |
| 1 | 165243506 | rs16841924 | -0.389 | -0.825 | -0.179 | -0.882 | -0.007 | -0.007 | -0.007 | 0.172 | 0.14 | 0.21 | 0.2 | 0.14 | 0.12 | 0.04 |
| 1 | 165244061 | rs7519089 | -0.381 | -0.865 | -0.102 | -0.97 | -0.001 | -0.048 | -0.048 | 0 | 0.24 | 0.31 | 0.27 | 0.14 | 0.26 | 0.05 |
| 1 | 165248182 | rs1833030 | -1.394 | -0.999 | -0.048 | -1.004 | -0.004 | -0.032 | -0.032 | 0 | 0.37 | 0.48 | 0.46 | 0.26 | 0.38 | 0.15 |
| 1 | 165250429 | rs1807798 | -1.4 | -0.793 | -0.329 | -0.944 | -0.001 | 0.001 | 0.001 | 0.063 | 0.37 | 0.38 | 0.48 | 0.26 | 0.48 | 0.22 |
| 1 | 165256014 | rs7540476 | -1.309 | -0.834 | -0.161 | -0.909 | 0.002 | -0.012 | -0.012 | 0 | 0.34 | 0.41 | 0.37 | 0.26 | 0.41 | 0.2 |
| 1 | 165256640 | rs11804125 | 1.809 | -0.647 | -0.503 | -0.893 | -0.053 | 0.057 | 0.057 | 0.181 | 0.1 | 0.02 | 0.09 | 0 | 0.21 | 0.02 |
| 1 | 165257255 | rs4361970 | 0.621 | -0.627 | -0.43 | -0.823 | -0.002 | -0.002 | -0.002 | 0 | 0.07 | 0.06 | 0.17 | 0.14 | 0.11 | 0.1 |
| 1 | 165263768 | rs10800080 | 0.475 | -0.855 | -0.452 | -0.498 | 0 | 0 | 0 | 0 | 0.64 | 0.41 | 0.59 | 0.68 | 0.61 | 0.71 |
| 1 | 165264856 | rs6662349 | 1.089 | -0.901 | -0.678 | -0.555 | -0.002 | -0.006 | -0.006 | 0 | 0.46 | 0.29 | 0.39 | 0.53 | 0.46 | 0.6 |
| 1 | 165265258 | rs10753670 | -0.194 | -0.359 | -0.802 | 0.355 | -0.004 | 0.005 | 0.005 | 0.003 | 0.86 | 0.83 | 0.87 | 0.8 | 0.89 | 0.85 |
| 1 | 165265537 | rs10800081 | 0.776 | -0.498 | -0.984 | 0.518 | 0.008 | -0.032 | -0.032 | 0.11 | 0.09 | 0 | 0.11 | 0.47 | 0.03 | 0.29 |
| 1 | 165267930 | rs10732283 | 0.034 | 0.078 | -0.45 | -0.131 | -0.003 | 0.005 | 0.005 | 0 | 0.86 | 0.82 | 0.87 | 0.8 | 0.89 | 0.85 |
| 1 | 165276634 | rs16842354 | 0.302 | 0.341 | -0.439 | -0.186 | 0 | -0.034 | -0.034 | 0.283 | 0.16 | 0.23 | 0.12 | 0 | 0.22 | 0.02 |
| 1 | 165284302 | rs1504698 | 0.478 | 0.057 | -0.397 | 0.541 | -0.009 | 0.013 | 0.013 | 0 | 0.85 | 0.79 | 0.86 | 0.79 | 0.88 | 0.84 |
| 1 | 165284723 | rs1546412 | 0.543 | 0.019 | -0.163 | 0.433 | -0.019 | 0.013 | 0.013 | 0.365 | 0.87 | 0.75 | 0.86 | 0.85 | 0.91 | 0.85 |
| 1 | 165286141 | rs906267 | 0.282 | -0.048 | -0.3 | 0.434 | -0.007 | -0.004 | -0.004 | 0 | 0.89 | 0.87 | 0.9 | 0.93 | 0.93 | 0.84 |
| 1 | 165286563 | rs875822 | -0.934 | 0.299 | 0.326 | 0.596 | 0.033 | 0.08 | 0.08 | 0.41 | 0.07 | 0.22 | 0.13 | 0.12 | 0.08 | 0.22 |
| 1 | 165287003 | rs7554002 | 0.824 | -0.187 | 0.628 | 0.375 | -0.042 | -0.031 | -0.031 | 0.118 | 0.84 | 0.72 | 0.88 | 0.93 | 0.92 | 0.84 |
| 1 | 165288357 | rs11809911 | 0.243 | -0.294 | 0.141 | -0.585 | 0 | -0.002 | -0.002 | 0 | 0.46 | 0.54 | 0.55 | 0.25 | 0.59 | 0.47 |

| | | | | | | | | | | | | | | | | |
|---|---|---|---|---|---|---|---|---|---|---|---|---|---|---|---|---|
| 1 | 165289385 | rs8179473 | 1.145 | 0.151 | -0.168 | -0.842 | -0.032 | -0.018 | -0.018 | 0.898 | 0.06 | 0 | 0.18 | 0.08 | 0.13 | 0.04 |
| 1 | 165290803 | rs10800084 | 0.202 | -0.289 | 0.076 | -0.744 | 0.004 | 0.02 | 0.02 | 0.069 | 0.44 | 0.38 | 0.54 | 0.23 | 0.59 | 0.47 |
| 1 | 165291536 | rs6688855 | 0.836 | -0.264 | 0.013 | -0.722 | 0.005 | 0.016 | 0.016 | 3.383 | 0.38 | 0.19 | 0.33 | 0.7 | 0.34 | 0.37 |
| 1 | 165293947 | rs4656444 | 0.367 | -0.223 | 0.085 | -0.703 | 0.021 | 0.023 | 0.023 | 4.48 | 0.44 | 0.35 | 0.35 | 0.68 | 0.34 | 0.37 |
| 1 | 165295689 | rs2055681 | 0.041 | -0.198 | 0.842 | -0.439 | -0.043 | -0.025 | -0.025 | 0 | 0.86 | 0.61 | 0.88 | 0.93 | 0.92 | 0.84 |
| 1 | 165297637 | rs2221246 | 0.26 | -0.432 | 0.358 | -0.855 | -0.012 | -0.129 | -0.129 | 0 | 0.25 | 0.36 | 0.14 | 0 | 0.22 | 0.02 |
| 1 | 165297703 | rs1948857 | -0.035 | -0.344 | 0.11 | -0.699 | -0.003 | 0.017 | 0.017 | 5.327 | 0.84 | 0.79 | 0.78 | 0.81 | 0.84 | 0.62 |
| 1 | 165313427 | rs1395548 | 0.715 | -0.319 | -0.142 | -0.543 | 0.02 | 0.039 | 0.039 | 0.089 | 0.58 | 0.32 | 0.64 | 0.78 | 0.62 | 0.54 |
| 1 | 165317732 | rs10918148 | 0.817 | -0.561 | 0.076 | -0.493 | 0.027 | 0.045 | 0.045 | 0.569 | 0.55 | 0.34 | 0.64 | 0.75 | 0.61 | 0.58 |
| 1 | 165317996 | rs1974080 | 0.609 | -0.553 | 0.539 | -0.418 | -0.019 | -0.098 | -0.098 | 0.519 | 0.2 | 0.31 | 0.11 | 0.04 | 0.18 | 0.12 |
| 1 | 165337200 | rs1506943 | 0.225 | -1.048 | 0.085 | 0.7 | -0.001 | 0.001 | 0.001 | 0.812 | 0.73 | 0.7 | 0.64 | 0.52 | 0.78 | 0.52 |
| 1 | 165337631 | rs12035029 | -0.137 | -0.718 | -0.185 | 0.591 | -0.008 | 0.042 | 0.042 | 0.486 | 0.19 | 0.3 | 0.23 | 0.4 | 0.19 | 0.16 |
| 1 | 165340884 | rs11590177 | 0.236 | -0.649 | -0.263 | 0.613 | -0.001 | 0.019 | 0.019 | 0.195 | 0.16 | 0.26 | 0.22 | 0.25 | 0.18 | 0.1 |
| 1 | 165341849 | rs12081765 | 0.093 | -0.621 | -0.361 | 0.462 | 0.026 | 0.022 | 0.022 | 0.055 | 0.52 | 0.62 | 0.45 | 0.44 | 0.57 | 0.37 |
| 1 | 165342963 | rs12057296 | 0.758 | -0.38 | -0.441 | 0.452 | 0.007 | -0.025 | -0.025 | 0.027 | 0.36 | 0.36 | 0.23 | 0.18 | 0.4 | 0.26 |
| 1 | 165349087 | rs965743 | -0.131 | -0.827 | -0.485 | 1.262 | -0.006 | 0.04 | 0.04 | 0.001 | 0.18 | 0.21 | 0.29 | 0.55 | 0.16 | 0.41 |
| 1 | 165352270 | rs6426914 | -0.512 | -1.019 | -0.593 | 1.056 | 0.001 | -0.001 | -0.001 | 0 | 0.9 | 0.92 | 0.91 | 0.89 | 0.93 | 0.88 |
| 1 | 165353548 | rs10489743 | -0.792 | -0.859 | -0.468 | 1.125 | -0.001 | 0 | 0 | 0 | 0.4 | 0.4 | 0.48 | 0.59 | 0.37 | 0.48 |
| 1 | 165355350 | rs1506951 | -0.443 | -0.841 | -0.876 | 1.376 | 0 | 0.005 | 0.005 | 0 | 0.19 | 0.18 | 0.27 | 0.5 | 0.15 | 0.33 |
| 1 | 165360114 | rs12739596 | 0.606 | -0.548 | -1.086 | 0.984 | -0.014 | 0.016 | 0.016 | 0 | 0.2 | 0.08 | 0.2 | 0.08 | 0.22 | 0.12 |
| 1 | 165366153 | rs16844019 | -1.082 | -0.813 | -0.23 | 1.4 | -0.01 | 0.147 | 0.147 | 0.347 | 0.06 | 0.24 | 0.01 | 0.21 | 0.02 | 0.24 |
| 1 | 165368861 | rs10800094 | -1.285 | -0.898 | -0.156 | 1.196 | -0.018 | 0.064 | 0.064 | 2.351 | 0.07 | 0.3 | 0.02 | 0.12 | 0.02 | 0.1 |
| 1 | 165370757 | rs283690 | -0.895 | -1.093 | -1.327 | 0.965 | -0.023 | 0.042 | 0.042 | 0.003 | 0.61 | 0.85 | 0.61 | 0.71 | 0.48 | 0.77 |
| 1 | 165372404 | rs3767333 | 0.374 | -1.635 | -0.681 | 1.518 | -0.041 | 0.018 | 0.018 | 0.117 | 0.27 | 0.37 | 0.26 | 0.31 | 0.22 | 0.3 |
| 1 | 165373146 | rs283700 | -0.04 | -1.39 | -1.808 | 0.977 | 0 | -0.042 | -0.042 | 0.001 | 0.81 | 0.91 | 0.8 | 0.74 | 0.85 | 0.87 |
| 1 | 165375835 | rs283698 | -1.777 | -1.603 | -1.675 | 0.827 | 0.02 | 0.03 | 0.03 | 0.057 | 0.81 | 0.88 | 0.83 | 0.93 | 0.8 | 0.91 |
| 1 | 165376499 | rs283695 | -1.579 | -1.102 | -1.751 | 1.044 | -0.003 | 0.125 | 0.125 | 0.044 | 0.52 | 0.82 | 0.57 | 0.79 | 0.43 | 0.62 |
| 1 | 165376685 | rs283694 | -0.989 | -1.353 | -1.322 | 1.253 | -0.004 | -0.017 | -0.017 | 0 | 0.31 | 0.25 | 0.33 | 0.48 | 0.3 | 0.46 |
| 1 | 165377552 | rs2134095 | 1.375 | -1.13 | -1.109 | 1.27 | 0.007 | -0.021 | -0.021 | 0 | 0.63 | 0.23 | 0.6 | 0.51 | 0.67 | 0.5 |
| 1 | 165379572 | rs746332 | 0.031 | -1.097 | -0.977 | 1.118 | -0.048 | 0.032 | 0.032 | 1.886 | 0.2 | 0.56 | 0.24 | 0.31 | 0.12 | 0.16 |
| 1 | 165381055 | rs2651860 | 0.036 | -0.696 | -0.972 | 1.086 | 0.003 | -0.004 | -0.004 | 0 | 0.15 | 0.12 | 0.15 | 0.1 | 0.21 | 0.19 |
| 1 | 165381666 | rs12069160 | 0.375 | -0.784 | -0.919 | 1.64 | -0.001 | 0.074 | 0.074 | 0.59 | 0.08 | 0.07 | 0.15 | 0.36 | 0.05 | 0.08 |
| 1 | 165383403 | rs157863 | 0.195 | -0.779 | -0.878 | 1.296 | 0.014 | 0.012 | 0.012 | 0 | 0.13 | 0.13 | 0.13 | 0.21 | 0.16 | 0.24 |
| 1 | 165384621 | rs188196 | 0.474 | -0.773 | -0.909 | 1.296 | -0.006 | -0.005 | -0.005 | 0 | 0.12 | 0.07 | 0.12 | 0.21 | 0.16 | 0.22 |
| 1 | 165386158 | rs2281985 | 1.007 | -1.023 | -1.31 | 0.998 | 0.022 | 0.065 | 0.065 | 0.23 | 0.47 | 0.23 | 0.38 | 0.32 | 0.53 | 0.55 |
| 1 | 165387533 | rs7533324 | 2.22 | -1.161 | -1.378 | 0.664 | 0.008 | 0.047 | 0.047 | 0 | 0.16 | 0.07 | 0.12 | 0 | 0.18 | 0.03 |
| 1 | 165388948 | rs157869 | -0.237 | -0.993 | -1.116 | 0.935 | 0.002 | -0.006 | -0.006 | 0.003 | 0.41 | 0.6 | 0.41 | 0.18 | 0.39 | 0.17 |
| 1 | 165389129 | rs1128977 | 1.939 | -0.906 | -1.179 | 0.668 | -0.11 | 0.118 | 0.118 | 0.03 | 0.35 | 0.07 | 0.26 | 0.12 | 0.4 | 0.34 |
| 1 | 165394882 | rs466639 | -0.073 | -0.304 | -0.501 | 1.058 | 0.003 | 0 | 0 | 3.627 | 0.88 | 0.86 | 0.95 | 0.88 | 0.88 | 0.77 |
| 1 | 165403903 | rs285480 | 0.83 | -0.284 | -0.261 | 1.056 | 0.043 | -0.001 | -0.001 | 0 | 0.83 | 0.62 | 0.82 | 0.86 | 0.79 | 0.68 |
| 1 | 165407290 | rs12043501 | -0.35 | -0.282 | -0.535 | 1.119 | -0.001 | -0.001 | -0.001 | 1.642 | 0.13 | 0.1 | 0.1 | 0.13 | 0.15 | 0.3 |
| 1 | 165409095 | rs10800098 | 0.529 | -0.249 | -0.79 | 1.622 | -0.019 | 0.006 | 0.006 | 1.709 | 0.06 | 0.01 | 0.15 | 0.35 | 0.05 | 0.07 |
| 1 | 165410468 | rs285481 | -1.617 | -0.507 | -0.622 | 1.445 | 0 | 0 | 0 | 0 | 0.37 | 0.58 | 0.49 | 0.71 | 0.34 | 0.4 |
| 1 | 165410723 | rs285482 | -0.985 | -0.504 | -0.609 | 1.365 | -0.003 | -0.001 | -0.001 | 0.061 | 0.31 | 0.6 | 0.35 | 0.36 | 0.28 | 0.34 |
| 1 | 165412959 | rs752739 | 0.868 | -0.659 | -0.961 | 0.974 | 0 | 0.004 | 0.004 | 0 | 0.19 | 0.09 | 0.18 | 0.15 | 0.23 | 0.19 |
| 1 | 165413644 | rs100537 | -1.255 | -0.596 | -0.891 | 0.941 | -0.022 | 0.046 | 0.046 | 0 | 0.56 | 0.73 | 0.66 | 0.72 | 0.53 | 0.49 |
| 1 | 165414933 | rs1467664 | 0.018 | -0.44 | 0.303 | 1.413 | 0.023 | 0.046 | 0.046 | 0 | 0.88 | 0.74 | 0.73 | 0.73 | 0.86 | 0.92 |
| 1 | 165422305 | rs16844995 | 0.037 | -0.518 | 0.199 | 1.493 | -0.001 | 0 | 0 | 0.024 | 0.21 | 0.21 | 0.19 | 0.28 | 0.19 | 0.24 |
| 1 | 165422460 | rs878180 | 1.422 | -0.715 | 0.137 | 1.367 | 0.018 | 0.03 | 0.03 | 0 | 0.54 | 0.36 | 0.45 | 0.29 | 0.62 | 0.55 |
| 1 | 165422559 | rs285428 | 0.312 | -0.726 | 0.137 | 1.368 | 0.002 | 0.007 | 0.007 | 0 | 0.11 | 0.12 | 0.25 | 0.27 | 0.14 | 0.09 |
| 1 | 165423876 | rs7536062 | 1.395 | -0.71 | 0.043 | 1.346 | 0.019 | 0.03 | 0.03 | 0 | 0.54 | 0.36 | 0.45 | 0.31 | 0.62 | 0.55 |
| 1 | 165425536 | rs7519141 | 1.53 | -0.577 | -0.711 | 1.133 | 0.024 | 0.183 | 0.183 | 0.007 | 0.49 | 0.21 | 0.37 | 0.14 | 0.56 | 0.36 |
| 1 | 165426193 | rs7542538 | 0.979 | -0.451 | -0.911 | 1.198 | -0.017 | 0.058 | 0.058 | 1.227 | 0.75 | 0.57 | 0.63 | 0.56 | 0.81 | 0.79 |
| 1 | 165427428 | rs10753678 | 0.174 | -0.076 | -1.146 | 0.46 | 0.017 | 0.038 | 0.038 | 0 | 0.88 | 0.8 | 0.75 | 0.71 | 0.86 | 0.91 |
| 1 | 165429012 | rs17430865 | -0.116 | 0.074 | -1.517 | -0.511 | 0.005 | 0.026 | 0.026 | 4.08 | 0.18 | 0.1 | 0.1 | 0.02 | 0.14 | 0.06 |
| 1 | 165430874 | rs1469267 | 0.49 | -0.134 | -1.373 | -0.013 | -0.013 | -0.03 | -0.03 | 0.026 | 0.25 | 0.14 | 0.44 | 0.67 | 0.28 | 0.35 |
| 1 | 165438912 | rs10800102 | 0.497 | -0.212 | -1.242 | 0.017 | -0.004 | -0.01 | -0.01 | 0.032 | 0.25 | 0.2 | 0.44 | 0.67 | 0.26 | 0.35 |
| 1 | 165440119 | rs185904 | -0.269 | -0.329 | -1.832 | -0.326 | -0.009 | 0.066 | 0.066 | 0 | 0.31 | 0.15 | 0.27 | 0.18 | 0.31 | 0.28 |
| 1 | 165440194 | rs285419 | -0.905 | 0.067 | -1.401 | 0.041 | -0.05 | 0.111 | 0.111 | 0 | 0.63 | 0.85 | 0.79 | 0.87 | 0.58 | 0.7 |
| 1 | 165448157 | rs10918196 | 0.375 | 0.016 | -1.507 | -0.384 | 0.017 | 0.084 | 0.084 | 0 | 0.4 | 0.23 | 0.51 | 0.81 | 0.34 | 0.54 |
| 1 | 165460351 | rs285488 | -1.119 | 0.015 | -0.903 | 0.986 | 0.004 | -0.011 | -0.011 | 0.035 | 0.21 | 0.16 | 0.19 | 0.27 | 0.19 | 0.36 |
| 1 | 165461566 | rs285492 | -1.3 | 0.07 | -0.877 | 1.001 | 0.003 | -0.003 | -0.003 | 0 | 0.25 | 0.23 | 0.24 | 0.3 | 0.2 | 0.45 |
| 1 | 165463932 | rs184042 | -0.979 | 0.326 | -1.079 | 0.87 | 0.003 | -0.012 | -0.012 | 0 | 0.16 | 0.14 | 0.2 | 0.22 | 0.13 | 0.29 |
| 1 | 165464051 | rs184043 | -0.977 | 0.354 | -1.077 | 0.873 | 0.002 | -0.012 | -0.012 | 0.832 | 0.16 | 0.14 | 0.2 | 0.22 | 0.14 | 0.27 |
| 1 | 165465281 | rs4657449 | -1.299 | -0.136 | -1.195 | 0.247 | 0.027 | 0.023 | 0.023 | 0 | 0.18 | 0.1 | 0.41 | 0.88 | 0.11 | 0.49 |
| 1 | 165465858 | rs373494 | -1.186 | 0.988 | -1.435 | 0.272 | -0.064 | -0.032 | -0.032 | 0 | 0.15 | 0.06 | 0.26 | 0.15 | 0.26 | 0.3 |
| 1 | 165466224 | rs12087952 | -0.655 | 0.852 | -1.343 | 0.125 | -0.001 | -0.002 | -0.002 | 0.176 | 0.07 | 0.09 | 0.03 | 0.07 | 0.05 | 0.07 |
| 1 | 165466540 | rs2433137 | -1.963 | 0.312 | -0.801 | -0.307 | -0.001 | 0.015 | 0.015 | 0 | 0.43 | 0.71 | 0.53 | 0.33 | 0.46 | 0.46 |
| 1 | 165467865 | rs17414421 | 0.588 | 0.334 | -0.018 | -0.601 | -0.001 | -0.026 | -0.026 | 0 | 0.09 | 0.1 | 0.06 | 0 | 0.09 | 0.01 |

| | | | | | | | | | | | | | | | | |
|---|---|---|---|---|---|---|---|---|---|---|---|---|---|---|---|---|
| 1 | 165470814 | rs185902 | -2.401 | 0.181 | 0.153 | -0.658 | 0 | 0.049 | 0.049 | 0.018 | 0.43 | 0.76 | 0.53 | 0.3 | 0.46 | 0.46 |
| 1 | 165471314 | rs12084656 | -0.33 | 0.311 | 0.279 | -0.634 | -0.001 | -0.003 | -0.003 | 0 | 0.08 | 0.09 | 0.03 | 0.07 | 0.05 | 0.07 |
| 1 | 165473592 | rs7549782 | 1.051 | 0.309 | 0.341 | -0.662 | 0 | -0.015 | -0.015 | 2.947 | 0.07 | 0.07 | 0.05 | 0 | 0.09 | 0.01 |
| 1 | 165475283 | rs1025159 | -0.92 | 0.221 | 0.519 | -0.684 | -0.01 | 0.003 | 0.003 | 0 | 0.17 | 0.1 | 0.19 | 0.15 | 0.18 | 0.23 |
| 1 | 165480347 | rs285473 | -0.843 | 0.253 | 1.256 | -0.678 | 0.009 | -0.002 | -0.002 | 0 | 0.15 | 0.2 | 0.26 | 0.16 | 0.23 | 0.27 |
| 1 | 165482762 | rs10753681 | 2.228 | 0.016 | 1.946 | -0.869 | -0.002 | -0.025 | -0.025 | 0.355 | 0.67 | 0.37 | 0.66 | 0.77 | 0.65 | 0.57 |
| 1 | 165486905 | rs285463 | 1.081 | 0.006 | 1.984 | -0.879 | -0.008 | -0.03 | -0.03 | 0.021 | 0.87 | 0.69 | 0.95 | 0.92 | 0.91 | 0.85 |
| 1 | 165489708 | rs10800108 | 2.557 | -0.078 | 1.931 | -0.903 | -0.017 | 0.083 | 0.083 | 0.185 | 0.56 | 0.1 | 0.47 | 0.7 | 0.55 | 0.52 |
| 1 | 165497692 | rs285461 | 1.69 | -0.086 | 1.963 | -0.908 | -0.028 | -0.061 | -0.061 | 0.31 | 0.83 | 0.61 | 0.92 | 0.92 | 0.87 | 0.84 |
| 1 | 165502862 | rs6669449 | -0.719 | 0.015 | 1.776 | -0.778 | 0 | 0.005 | 0.005 | 0 | 0.14 | 0.25 | 0.16 | 0.16 | 0.13 | 0.22 |
| 1 | 165513417 | rs7539622 | 0.056 | 0.05 | 1.629 | -0.764 | -0.036 | -0.039 | -0.039 | 0.099 | 0.11 | 0.01 | 0.13 | 0.15 | 0.13 | 0.23 |
| 1 | 165513439 | rs285446 | 0.719 | -0.074 | 1.795 | -0.777 | -0.028 | -0.041 | -0.041 | 0.301 | 0.93 | 0.81 | 0.96 | 1 | 0.96 | 0.94 |
| 1 | 165517729 | rs285439 | -1.59 | -0.017 | 1.567 | -0.765 | 0.011 | 0.003 | 0.003 | 0 | 0.06 | 0.07 | 0.24 | 0.07 | 0.15 | 0.16 |
| 1 | 165521770 | rs7528336 | 0.777 | -0.035 | 1.753 | -0.819 | 0.001 | -0.027 | -0.027 | 0.034 | 0.09 | 0.11 | 0.06 | 0 | 0.1 | 0.01 |
| 1 | 165523447 | rs4339837 | -0.758 | -0.045 | 1.95 | -0.839 | -0.005 | 0.009 | 0.009 | 0 | 0.14 | 0.31 | 0.15 | 0.15 | 0.12 | 0.23 |
| 1 | 165525069 | rs12750195 | 2.167 | -0.125 | 2.006 | -0.88 | 0.001 | -0.001 | -0.001 | 0 | 0.71 | 0.28 | 0.64 | 0.87 | 0.73 | 0.75 |
| 1 | 165525825 | rs11804904 | 0.631 | -0.081 | 1.75 | -0.855 | 0.002 | -0.03 | -0.03 | 0.005 | 0.09 | 0.14 | 0.06 | 0 | 0.1 | 0.02 |
| 1 | 165527669 | rs10800111 | 2.476 | -0.147 | 1.649 | -0.871 | -0.013 | 0.073 | 0.073 | 0.254 | 0.57 | 0.07 | 0.49 | 0.71 | 0.58 | 0.49 |
| 1 | 165530586 | rs10494444 | -0.866 | -0.097 | 1.812 | -0.822 | -0.02 | 0.008 | 0.008 | 0 | 0.06 | 0.14 | 0.16 | 0.06 | 0.04 | 0.07 |
| 1 | 165532746 | rs17407838 | 0.832 | -0.051 | 1.926 | -0.843 | 0.003 | -0.025 | -0.025 | 0.03 | 0.09 | 0.1 | 0.06 | 0 | 0.1 | 0.01 |
| 1 | 165533005 | rs453717 | -1.982 | -0.105 | 2.185 | -0.878 | 0.003 | -0.011 | -0.011 | 0 | 0.26 | 0.38 | 0.3 | 0.23 | 0.31 | 0.37 |
| 1 | 165533075 | rs6676948 | 0.836 | -0.105 | 2.244 | -0.879 | 0.005 | -0.031 | -0.031 | 0.158 | 0.09 | 0.11 | 0.06 | 0 | 0.1 | 0.01 |
| 1 | 165533188 | rs367199 | -1.992 | -0.104 | 2.21 | -0.861 | 0.003 | -0.011 | -0.011 | 1.511 | 0.26 | 0.38 | 0.3 | 0.23 | 0.31 | 0.37 |
| 1 | 165534130 | rs16847083 | -0.951 | -0.117 | 2.191 | -0.835 | -0.009 | -0.005 | -0.005 | 0 | 0.17 | 0.25 | 0.17 | 0.15 | 0.15 | 0.23 |
| 1 | 165534788 | rs2252583 | 1.484 | -0.148 | 2.261 | -0.841 | -0.002 | 0.076 | 0.076 | 3.142 | 0.92 | 0.6 | 0.85 | 0.94 | 0.99 | 0.88 |
| 1 | 165546671 | rs6657863 | -1.939 | -0.117 | 1.819 | -0.84 | -0.031 | -0.005 | -0.005 | 0 | 0.25 | 0.65 | 0.32 | 0.21 | 0.16 | 0.31 |
| 1 | 165550009 | rs1572542 | 2.229 | -0.52 | 1.211 | -1.072 | 0.006 | 0.101 | 0.101 | 0.225 | 0.54 | 0.12 | 0.48 | 0.71 | 0.57 | 0.52 |
| 1 | 165559225 | rs952375 | 0.947 | 0.075 | 1.589 | -1.443 | 0.09 | -0.029 | -0.029 | 0 | 0.89 | 0.63 | 0.74 | 0.98 | 0.86 | 0.87 |
| 1 | 165562701 | rs16847398 | 0.026 | -0.008 | 1.035 | -1.499 | -0.02 | -0.019 | -0.019 | 0 | 0.12 | 0.03 | 0.12 | 0.14 | 0.13 | 0.26 |
| 1 | 165563605 | rs6679871 | 2.153 | -0.045 | 1.139 | -1.607 | -0.015 | 0.15 | 0.15 | 0.239 | 0.54 | 0.2 | 0.5 | 0.8 | 0.53 | 0.51 |
| 1 | 165569064 | rs11806754 | -0.251 | 0.056 | 0.985 | -1.53 | 0.004 | 0.009 | 0.009 | 0 | 0.08 | 0.08 | 0.03 | 0.03 | 0.07 | 0.08 |
| 1 | 165569268 | rs11801981 | -0.027 | 0.036 | 1.043 | -1.53 | -0.002 | 0.002 | 0.002 | 0 | 0.16 | 0.14 | 0.09 | 0.03 | 0.17 | 0.09 |
| 1 | 165570840 | rs16847544 | -0.215 | 0.04 | 1.289 | -1.529 | 0.007 | -0.069 | -0.069 | 1.192 | 0.16 | 0.28 | 0.1 | 0.03 | 0.17 | 0.09 |
| 1 | 165576665 | rs12125420 | 0.176 | 0.141 | 0.854 | -1.46 | -0.019 | -0.033 | -0.033 | 0 | 0.12 | 0.03 | 0.08 | 0.15 | 0.12 | 0.26 |
| 1 | 165582470 | rs6426918 | 0.114 | 0.148 | 0.914 | -1.448 | -0.001 | -0.009 | -0.009 | 0 | 0.15 | 0.16 | 0.09 | 0.03 | 0.14 | 0.06 |
| 1 | 165589535 | rs1337444 | 2.412 | -0.242 | 0.373 | -1.711 | -0.092 | 0.177 | 0.177 | 0.622 | 0.48 | 0.02 | 0.54 | 0.79 | 0.53 | 0.49 |
| 1 | 165590676 | rs1913836 | 0.129 | -0.185 | 0.485 | -1.697 | -0.023 | -0.039 | -0.039 | 0 | 0.13 | 0.01 | 0.08 | 0.15 | 0.13 | 0.26 |
| 1 | 165592722 | rs11809495 | -1.472 | -0.162 | 1.218 | -1.687 | -0.034 | -0.089 | -0.089 | 0 | 0.28 | 0.61 | 0.26 | 0.06 | 0.18 | 0.15 |
| 1 | 165595883 | rs1415502 | -1.382 | -0.165 | 1.235 | -1.688 | -0.035 | -0.093 | -0.093 | 0.813 | 0.27 | 0.58 | 0.26 | 0.05 | 0.18 | 0.15 |
| 1 | 165597321 | rs6690679 | -1.527 | -0.172 | 1.277 | -1.692 | 0.003 | -0.014 | -0.014 | 0 | 0.29 | 0.7 | 0.26 | 0.05 | 0.18 | 0.15 |
| 1 | 165598854 | rs6426925 | 0.69 | -0.133 | 1.085 | -1.791 | -0.021 | 0.007 | 0.007 | 0.243 | 0.8 | 0.86 | 0.81 | 0.83 | 0.71 | 0.66 |
| 1 | 165599551 | rs6694924 | 0.734 | -0.14 | 1.086 | -1.791 | -0.023 | 0.001 | 0.001 | 0.338 | 0.8 | 0.86 | 0.72 | 0.81 | 0.71 | 0.6 |
| 1 | 165599804 | rs6657788 | -0.197 | -0.128 | 1.093 | -1.796 | 0.006 | -0.008 | -0.008 | 0 | 0.93 | 0.94 | 0.89 | 1 | 0.84 | 0.91 |
| 1 | 165599979 | rs4147591 | 0.842 | -0.033 | 1.198 | -1.797 | -0.011 | 0.001 | 0.001 | 0.026 | 0.79 | 0.81 | 0.71 | 0.81 | 0.71 | 0.6 |
| 1 | 165600146 | rs4147592 | 2.545 | 0.196 | 0.878 | -2.148 | -0.018 | 0.145 | 0.145 | 0 | 0.46 | 0.15 | 0.5 | 0.78 | 0.46 | 0.46 |
| 1 | 165600606 | rs4147594 | 2.545 | 0.196 | 0.878 | -2.148 | -0.018 | 0.145 | 0.145 | 0 | 0.46 | 0.15 | 0.5 | 0.78 | 0.45 | 0.46 |
| 1 | 165600766 | rs9333372 | -0.938 | 0.124 | 1.359 | -2.185 | -0.027 | -0.098 | -0.098 | 0 | 0.17 | 0.32 | 0.11 | 0.01 | 0.12 | 0.09 |
| 1 | 165601215 | rs4147595 | 0.636 | 0.157 | 1.029 | -2.166 | -0.012 | 0.002 | 0.002 | 0.26 | 0.79 | 0.82 | 0.71 | 0.81 | 0.71 | 0.6 |
| 1 | 165601365 | rs4147596 | 2.555 | 0.129 | 1.05 | -2.155 | -0.018 | 0.145 | 0.145 | 0 | 0.46 | 0.16 | 0.51 | 0.78 | 0.45 | 0.46 |
| 1 | 165601466 | rs9333378 | 1.801 | 0.161 | 1.026 | -2.155 | -0.007 | 0.021 | 0.021 | 0.265 | 0.61 | 0.33 | 0.68 | 0.96 | 0.61 | 0.79 |
| 1 | 165602524 | rs4147598 | 0.14 | 0.195 | 0.774 | -1.997 | -0.002 | -0.026 | -0.026 | 0 | 0.13 | 0.04 | 0.08 | 0.17 | 0.13 | 0.26 |
| 1 | 165603894 | rs1538114 | 0.153 | 0.258 | 0.923 | -2.004 | -0.012 | 0.006 | 0.006 | 0 | 0.91 | 0.95 | 0.87 | 1 | 0.85 | 0.93 |
| 1 | 165604991 | rs4147599 | 1.377 | 0.408 | 0.58 | -2.701 | 0.008 | 0.038 | 0.038 | 0 | 0.47 | 0.4 | 0.49 | 0.79 | 0.43 | 0.54 |
| 1 | 165605209 | rs4147600 | 0.393 | 0.449 | 0.348 | -2.694 | -0.036 | 0.085 | 0.085 | 0 | 0.64 | 0.85 | 0.7 | 0.84 | 0.56 | 0.66 |
| 1 | 165605310 | rs10800120 | 0.393 | 0.449 | 0.348 | -2.694 | -0.036 | 0.085 | 0.085 | 0 | 0.64 | 0.85 | 0.7 | 0.84 | 0.56 | 0.66 |
| 1 | 165609300 | rs9333413 | 2.328 | 0.614 | 0.064 | -2.712 | -0.009 | 0.063 | 0.063 | 0 | 0.36 | 0.14 | 0.43 | 0.8 | 0.34 | 0.47 |
| 1 | 165610464 | rs7554034 | -0.559 | 0.672 | 0.329 | -2.667 | -0.012 | -0.145 | -0.145 | 0 | 0.26 | 0.5 | 0.14 | 0.03 | 0.19 | 0.11 |
| 1 | 165616157 | rs957644 | -0.435 | 0.946 | 0.248 | -2.666 | 0.027 | -0.145 | -0.145 | 0.23 | 0.25 | 0.48 | 0.15 | 0.03 | 0.24 | 0.12 |
| 1 | 165616588 | rs6686643 | -0.518 | 0.956 | 0.172 | -2.655 | 0.019 | -0.148 | -0.148 | 3.074 | 0.26 | 0.48 | 0.15 | 0.03 | 0.23 | 0.12 |
| 1 | 165617320 | rs1878076 | 2.139 | 0.965 | -0.244 | -2.666 | 0.002 | 0.059 | 0.059 | 0 | 0.36 | 0.18 | 0.44 | 0.8 | 0.34 | 0.47 |
| 1 | 165619079 | rs6681 | -0.308 | 1.173 | -0.238 | -2.353 | 0.012 | 0.013 | 0.013 | 0 | 0.05 | 0.03 | 0.02 | 0 | 0.05 | 0.02 |
| 1 | 165619712 | rs9333471 | -1.22 | 1.346 | -0.098 | -2.383 | 0.008 | -0.106 | -0.106 | 0.116 | 0.29 | 0.48 | 0.18 | 0.03 | 0.3 | 0.15 |
| 1 | 165620792 | rs11799886 | 0.402 | 1.576 | -0.769 | -2.023 | -0.001 | -0.008 | -0.008 | 0 | 0.13 | 0.15 | 0.09 | 0.03 | 0.16 | 0.06 |
| 1 | 165621590 | rs7527279 | 0.667 | 1.373 | -1.346 | -2.22 | -0.029 | 0.119 | 0.119 | 1.149 | 0.71 | 0.9 | 0.69 | 0.97 | 0.68 | 0.78 |
| 1 | 165621603 | rs7549511 | 0.763 | 0.988 | -1.494 | -2.658 | 0.016 | 0.114 | 0.114 | 0.014 | 0.5 | 0.69 | 0.55 | 0.82 | 0.46 | 0.59 |
| 1 | 165621778 | rs4147602 | -0.279 | 0.476 | -1.232 | -2.243 | -0.005 | -0.01 | -0.01 | 0 | 0.7 | 0.57 | 0.55 | 0.26 | 0.66 | 0.52 |
| 1 | 165621807 | rs10918228 | 0.332 | 0.065 | -2.227 | -2.514 | -0.059 | 0.179 | 0.179 | 2.874 | 0.39 | 0.05 | 0.32 | 0.13 | 0.38 | 0.29 |
| 1 | 165622567 | rs3795540 | 0.738 | -0.367 | -1.234 | -2.13 | -0.027 | -0.016 | -0.016 | 0 | 0.21 | 0.33 | 0.39 | 0.81 | 0.13 | 0.39 |
| 1 | 165622662 | rs2297763 | 0.453 | -0.397 | -1.022 | -2.889 | -0.001 | -0.017 | -0.017 | 0 | 0.31 | 0.28 | 0.2 | 0.14 | 0.28 | 0.29 |

| | | | | | | | | | | | | | | | | |
|---|---|---|---|---|---|---|---|---|---|---|---|---|---|---|---|---|
| 1 | 165623994 | rs7533986 | -1.045 | -0.177 | -0.913 | -2.986 | -0.003 | -0.023 | -0.023 | 0.687 | 0.75 | 0.76 | 0.77 | 0.99 | 0.68 | 0.83 |
| 1 | 165624799 | rs8133 | 0.847 | -0.318 | -1.013 | -3.133 | -0.046 | 0.209 | 0.209 | 0.497 | 0.42 | 0.08 | 0.27 | 0.15 | 0.5 | 0.33 |
| 1 | 165625474 | rs7551594 | -1.417 | -0.274 | -0.997 | -3.102 | -0.007 | -0.001 | -0.001 | 0.101 | 0.76 | 0.81 | 0.78 | 0.99 | 0.69 | 0.84 |
| 1 | 165628921 | rs10800125 | 0.326 | -0.355 | -0.886 | -3.1 | -0.026 | -0.035 | -0.035 | 0 | 0.3 | 0.41 | 0.47 | 0.84 | 0.19 | 0.5 |
| 1 | 165631123 | rs6668102 | -1.425 | -0.293 | -0.929 | -3.116 | -0.006 | -0.002 | -0.002 | 2.086 | 0.76 | 0.81 | 0.77 | 0.99 | 0.69 | 0.84 |
| 1 | 165631506 | rs11506 | 1.54 | -0.398 | -0.955 | -3.042 | 0.018 | 0.02 | 0.02 | 0 | 0.09 | 0.06 | 0.04 | 0.02 | 0.06 | 0.06 |
| 1 | 165631670 | rs12670 | 1.326 | -0.231 | -0.682 | -2.906 | -0.01 | -0.006 | -0.006 | 0 | 0.2 | 0.26 | 0.38 | 0.82 | 0.12 | 0.36 |
| 1 | 165639264 | rs12065621 | 0.381 | 0.162 | -0.807 | -2.942 | -0.001 | -0.021 | -0.021 | 0 | 0.44 | 0.4 | 0.32 | 0.16 | 0.46 | 0.42 |
| 1 | 165644865 | rs10918239 | 1.184 | -0.339 | -0.814 | -2.924 | -0.006 | -0.004 | -0.004 | 0 | 0.22 | 0.27 | 0.39 | 0.82 | 0.13 | 0.38 |
| 1 | 165645370 | rs12081903 | 0.69 | -0.109 | -0.738 | -2.782 | -0.048 | 0.175 | 0.175 | 0.293 | 0.41 | 0.11 | 0.26 | 0.15 | 0.49 | 0.32 |
| 1 | 165646758 | rs7534437 | 0.263 | -0.083 | -0.736 | -2.777 | -0.001 | -0.018 | -0.018 | 0 | 0.44 | 0.37 | 0.33 | 0.15 | 0.5 | 0.41 |
| 1 | 165648886 | rs4646896 | 0.274 | -0.086 | -0.736 | -2.778 | -0.001 | -0.019 | -0.019 | 0 | 0.44 | 0.37 | 0.33 | 0.15 | 0.5 | 0.41 |
| 1 | 165649715 | rs16849374 | -0.645 | -0.105 | -0.126 | -2.425 | -0.035 | -0.055 | -0.055 | 0 | 0.11 | 0.39 | 0.1 | 0.01 | 0.05 | 0.16 |
| 1 | 165650616 | rs4578194 | 0.566 | -0.043 | -0.508 | -1.606 | 0.004 | -0.005 | -0.005 | 0 | 0.34 | 0.26 | 0.35 | 0.38 | 0.31 | 0.45 |
| 1 | 165651725 | rs1337442 | -0.553 | 0.058 | -0.253 | -1.849 | 0.001 | -0.063 | -0.063 | 0 | 0.59 | 0.61 | 0.77 | 0.99 | 0.64 | 0.78 |
| 1 | 165652273 | rs1143659 | -0.553 | 0.058 | -0.26 | -1.849 | 0.001 | -0.066 | -0.066 | 0 | 0.59 | 0.61 | 0.77 | 0.99 | 0.64 | 0.78 |
| 1 | 165652568 | rs4656457 | -0.553 | 0.058 | -0.26 | -1.849 | 0.001 | -0.066 | -0.066 | 0 | 0.59 | 0.61 | 0.77 | 0.99 | 0.64 | 0.78 |
| 1 | 165654291 | rs4646889 | -0.676 | 0.11 | -0.669 | -1.824 | 0.006 | 0.002 | 0.002 | 0 | 0.6 | 0.67 | 0.79 | 0.99 | 0.64 | 0.78 |
| 1 | 165654812 | rs4646887 | -0.805 | 0.196 | -0.58 | -1.752 | 0.003 | -0.018 | -0.018 | 0.276 | 0.62 | 0.68 | 0.8 | 0.99 | 0.65 | 0.83 |
| 1 | 165654869 | rs12033254 | 1.279 | 0.332 | -0.525 | -1.582 | -0.002 | 0.113 | 0.113 | 0 | 0.06 | 0.09 | 0.15 | 0.59 | 0.04 | 0.16 |
| 1 | 165658091 | rs4646886 | -0.771 | 0.299 | -0.644 | -1.703 | 0.002 | -0.037 | -0.037 | 0 | 0.62 | 0.65 | 0.8 | 0.99 | 0.65 | 0.82 |
| 1 | 165658598 | rs4646885 | 0.676 | 0.32 | -0.631 | -1.794 | 0.014 | 0 | 0 | 0 | 0.38 | 0.23 | 0.42 | 0.38 | 0.33 | 0.57 |
| 1 | 165658750 | rs1538115 | -0.869 | 0.32 | -0.632 | -1.802 | 0.002 | -0.037 | -0.037 | 0.308 | 0.62 | 0.65 | 0.8 | 0.99 | 0.65 | 0.82 |
| 1 | 165663798 | rs17482717 | 0.478 | 0.427 | -0.819 | -1.51 | 0.014 | -0.001 | -0.001 | 0 | 0.38 | 0.23 | 0.42 | 0.38 | 0.33 | 0.57 |
| 1 | 165666448 | rs10753688 | -0.418 | 0.382 | -1.06 | -1.343 | 0.003 | 0.006 | 0.006 | 0 | 0.58 | 0.62 | 0.65 | 0.38 | 0.61 | 0.66 |
| 1 | 165667781 | rs12408101 | 0.389 | 0.713 | -1.181 | -1.443 | -0.044 | 0.061 | 0.061 | 0 | 0.17 | 0.08 | 0.22 | 0 | 0.26 | 0.08 |
| 1 | 165670958 | rs1547725 | -0.926 | 0.856 | -1.21 | -1.084 | -0.017 | 0.015 | 0.015 | 0 | 0.92 | 0.95 | 0.88 | 0.99 | 0.87 | 0.96 |
| 1 | 165676606 | rs2090281 | 1.021 | 0.162 | -1.637 | -2.048 | -0.033 | 0.23 | 0.23 | 0.283 | 0.46 | 0.12 | 0.35 | 0.13 | 0.48 | 0.4 |
| 1 | 165677013 | rs10918256 | 0.203 | -0.321 | -0.832 | -2.045 | -0.009 | -0.008 | -0.008 | 0 | 0.29 | 0.31 | 0.3 | 0.22 | 0.25 | 0.43 |
| 1 | 165677216 | rs12043815 | 0.314 | -0.313 | -0.828 | -2.031 | -0.01 | -0.011 | -0.011 | 0 | 0.28 | 0.31 | 0.26 | 0.19 | 0.25 | 0.36 |
| 1 | 165677673 | rs12027066 | 1.144 | -0.099 | -0.435 | -2.004 | 0.001 | 0.043 | 0.043 | 0 | 0.06 | 0.04 | 0.13 | 0.64 | 0.05 | 0.12 |
| 1 | 165677933 | rs4341347 | 1.329 | -0.112 | -0.104 | -2.039 | -0.006 | -0.044 | -0.044 | 0 | 0.08 | 0.17 | 0.09 | 0 | 0.08 | 0.01 |
| 1 | 165683487 | rs12062372 | 0.087 | -0.105 | -0.153 | -1.967 | -0.016 | 0.16 | 0.16 | 0.644 | 0.16 | 0.45 | 0.23 | 0.63 | 0.13 | 0.13 |
| 1 | 165685067 | rs10800149 | -0.349 | -0.308 | -0.628 | -2.221 | -0.021 | 0.097 | 0.097 | 1.592 | 0.62 | 0.78 | 0.52 | 0.84 | 0.48 | 0.53 |
| 1 | 165685082 | rs10800150 | -0.17 | -0.034 | -0.299 | -1.556 | -0.021 | 0.157 | 0.157 | 1.262 | 0.17 | 0.46 | 0.23 | 0.63 | 0.13 | 0.13 |
| 1 | 165687205 | rs4656461 | -0.366 | 0.176 | -0.273 | -1.734 | 0.045 | -0.046 | -0.046 | 0 | 0.91 | 0.74 | 0.86 | 0.99 | 0.86 | 0.96 |
| 1 | 165695579 | rs1913845 | 0.963 | 0.306 | -0.266 | -2.225 | -0.011 | 0.017 | 0.017 | 1.606 | 0.24 | 0.16 | 0.23 | 0.14 | 0.27 | 0.34 |
| 1 | 165704732 | rs10918271 | 0.943 | 0.198 | -0.336 | -2.248 | 0.067 | 0.119 | 0.119 | 2.486 | 0.48 | 0.22 | 0.27 | 0.24 | 0.33 | 0.41 |
| 1 | 165734088 | rs10918281 | 0.454 | 0.367 | 0.078 | -1.936 | 0.016 | 0.096 | 0.096 | 0.288 | 0.17 | 0.28 | 0.32 | 0.58 | 0.26 | 0.14 |
| 1 | 165736880 | rs7518099 | -1.338 | 0.184 | -0.323 | -1.866 | 0.004 | -0.009 | -0.009 | 0.245 | 0.93 | 0.91 | 0.87 | 0.99 | 0.86 | 0.96 |
| 1 | 165739598 | rs2814471 | -1.037 | 0.181 | -0.313 | -1.868 | 0.016 | -0.023 | -0.023 | 0 | 0.93 | 0.86 | 0.87 | 0.99 | 0.86 | 0.96 |
| 1 | 165742416 | rs2790051 | 0.501 | -0.382 | -0.744 | -2.314 | 0.006 | 0.038 | 0.038 | 0 | 0.55 | 0.62 | 0.48 | 0.16 | 0.56 | 0.51 |
| 1 | 165743081 | rs12095969 | 0.78 | 0.289 | -1.012 | -1.82 | -0.003 | -0.021 | -0.021 | 0.028 | 0.05 | 0.19 | 0.01 | 0 | 0.01 | 0.03 |
| 1 | 165758106 | rs6674639 | 1.147 | 0.319 | -1.354 | -1.823 | -0.026 | 0.053 | 0.053 | 0 | 0.1 | 0 | 0.05 | 0.01 | 0.09 | 0.08 |
| 1 | 165765014 | rs6656194 | 1.154 | 0.731 | -1.887 | -1.222 | 0.004 | -0.036 | -0.036 | 0.727 | 0.1 | 0.05 | 0.21 | 0.58 | 0.12 | 0.14 |
| 1 | 165769653 | rs1934278 | 0.123 | 0.781 | -1.904 | -1.427 | -0.043 | 0.054 | 0.054 | 2.455 | 0.3 | 0.17 | 0.32 | 0.14 | 0.41 | 0.4 |
| 1 | 165787477 | rs2153264 | 0.254 | -0.878 | -1.442 | -1.026 | -0.015 | -0.003 | -0.003 | 0.27 | 0.65 | 0.67 | 0.46 | 0.38 | 0.54 | 0.59 |
| 1 | 165798011 | rs11585115 | -0.061 | -0.734 | -1.251 | -0.564 | 0.003 | 0 | 0 | 0 | 0.15 | 0.1 | 0.23 | 0.6 | 0.14 | 0.24 |
| 1 | 165799022 | rs10800160 | -0.271 | -0.373 | -1.199 | -0.433 | 0.003 | 0.004 | 0.004 | 0 | 0.31 | 0.26 | 0.3 | 0.31 | 0.31 | 0.5 |
| 1 | 165807812 | rs16851444 | -0.109 | -0.952 | -1.236 | -0.612 | 0.014 | 0.036 | 0.036 | 0 | 0.14 | 0.08 | 0.07 | 0.02 | 0.12 | 0.07 |
| 1 | 165808987 | rs10918291 | 0.005 | -1.242 | -1.217 | -0.626 | 0.005 | -0.003 | -0.003 | 0.014 | 0.45 | 0.38 | 0.51 | 0.53 | 0.42 | 0.56 |
| 1 | 165811729 | rs6677965 | 0.715 | -1.556 | -1.125 | -0.654 | -0.057 | -0.067 | -0.067 | 0.05 | 0.36 | 0.18 | 0.4 | 0.47 | 0.49 | 0.42 |
| 1 | 165812120 | rs6702462 | -0.463 | -1.287 | -0.446 | -0.094 | -0.001 | 0 | 0 | 0 | 0.48 | 0.49 | 0.52 | 0.52 | 0.41 | 0.53 |
| 1 | 165815939 | rs4535982 | -0.508 | -1.142 | -0.927 | 0.007 | 0.031 | -0.003 | -0.003 | 0 | 0.48 | 0.65 | 0.54 | 0.51 | 0.42 | 0.51 |
| 1 | 165819247 | rs10918293 | -0.246 | -0.779 | -1.181 | 0.6 | -0.033 | 0.002 | 0.002 | 0.087 | 0.22 | 0.03 | 0.16 | 0.22 | 0.23 | 0.37 |
| 1 | 165825166 | rs6426942 | 0.835 | -1.081 | -0.12 | 1.142 | -0.101 | 0.185 | 0.185 | 0 | 0.81 | 0.52 | 0.81 | 0.59 | 0.93 | 0.73 |
| 1 | 165826745 | rs946542 | 1.449 | -0.809 | 0.403 | 0.661 | 0.015 | 0.027 | 0.027 | 2.964 | 0.93 | 0.59 | 0.96 | 0.96 | 0.99 | 0.95 |
| 1 | 165827488 | rs6666086 | -0.67 | -0.797 | -0.299 | -0.012 | -0.007 | 0.057 | 0.057 | 0 | 0.22 | 0.1 | 0.13 | 0.13 | 0.24 | 0.47 |
| 1 | 165831093 | rs12075898 | -1.089 | -0.723 | -0.003 | -0.003 | 0.003 | 0.022 | 0.022 | 0 | 0.26 | 0.24 | 0.23 | 0.19 | 0.26 | 0.49 |
| 1 | 165831128 | rs10494448 | 0.749 | -0.557 | -0.387 | -0.038 | -0.01 | -0.006 | -0.006 | 0 | 0.37 | 0.27 | 0.34 | 0.34 | 0.4 | 0.24 |
| 1 | 165832085 | rs12066877 | -0.547 | -0.926 | -0.562 | -0.079 | 0.017 | 0.035 | 0.035 | 0 | 0.31 | 0.3 | 0.24 | 0.23 | 0.27 | 0.55 |
| 1 | 165835364 | rs4657487 | -1.006 | -1.007 | -0.525 | -0.101 | -0.001 | -0.043 | -0.043 | 0 | 0.66 | 0.8 | 0.6 | 0.56 | 0.66 | 0.79 |
| 1 | 165835455 | rs16851987 | 0.029 | -1.191 | -0.629 | -0.109 | -0.007 | 0.022 | 0.022 | 0.027 | 0.23 | 0.08 | 0.29 | 0.18 | 0.23 | 0.49 |
| 1 | 165840411 | rs6703280 | 0.857 | -1.128 | -0.616 | -0.003 | 0 | -0.025 | -0.025 | 0 | 0.34 | 0.24 | 0.41 | 0.45 | 0.35 | 0.2 |
| 1 | 165845042 | rs12080800 | 0.341 | -1.187 | -0.477 | 0.181 | -0.012 | 0.002 | 0.002 | 0 | 0.38 | 0.32 | 0.27 | 0.29 | 0.42 | 0.27 |
| 1 | 165856120 | rs4657489 | -0.035 | -1.16 | -0.663 | 0.13 | 0.009 | 0.034 | 0.034 | 0.009 | 0.23 | 0.11 | 0.28 | 0.18 | 0.22 | 0.49 |
| 1 | 165860813 | rs3762365 | -0.395 | -1.117 | 0.012 | 0.171 | -0.001 | 0 | 0 | 0 | 0.62 | 0.62 | 0.74 | 0.71 | 0.59 | 0.7 |
| 1 | 165871705 | rs4656467 | -0.328 | -1.022 | -0.296 | -0.248 | -0.002 | 0.056 | 0.056 | 0.2 | 0.29 | 0.19 | 0.26 | 0.16 | 0.33 | 0.52 |
| 1 | 165873392 | rs3790672 | 1.243 | -0.812 | -0.437 | -0.206 | -0.05 | -0.151 | -0.151 | 0.946 | 0.3 | 0.06 | 0.4 | 0.52 | 0.3 | 0.22 |

| | | | | | | | | | | | | | | | | |
|---|---|---|---|---|---|---|---|---|---|---|---|---|---|---|---|---|
| 1 | 165877751 | rs3748699 | 0.245 | -0.69 | -0.472 | -0.241 | -0.025 | 0.001 | 0.001 | 0 | 0.07 | 0.02 | 0.08 | 0.08 | 0.16 | 0.1 |
| 1 | 165878498 | rs506813 | 1.447 | -0.231 | -0.836 | -0.593 | -0.042 | -0.061 | -0.061 | 0.771 | 0.1 | 0 | 0.09 | 0.26 | 0.1 | 0.12 |
| 1 | 165889160 | rs475558 | 0.107 | -0.327 | -0.284 | -0.771 | 0 | -0.011 | -0.011 | 0.11 | 0.69 | 0.69 | 0.74 | 0.84 | 0.67 | 0.48 |
| 1 | 165895538 | rs10800167 | 1.215 | -0.273 | -0.173 | -0.465 | -0.002 | 0.017 | 0.017 | 0 | 0.32 | 0.37 | 0.43 | 0.48 | 0.28 | 0.2 |
| 1 | 165896757 | rs472888 | 0.29 | -0.59 | -0.805 | -1.467 | -0.001 | -0.02 | -0.02 | 0.326 | 0.54 | 0.46 | 0.54 | 0.82 | 0.57 | 0.48 |
| 1 | 165898768 | rs522854 | 0.273 | -0.557 | -0.759 | -1.512 | 0.026 | 0.055 | 0.055 | 2.906 | 0.94 | 0.63 | 0.88 | 0.98 | 1 | 0.98 |
| 1 | 165899981 | rs1663839 | -0.761 | -0.506 | -0.837 | -1.519 | 0.006 | -0.132 | -0.132 | 0 | 0.77 | 0.53 | 0.77 | 0.98 | 0.78 | 0.89 |
| 1 | 165903962 | rs2349106 | -0.112 | -0.095 | -1.14 | -1.5 | -0.002 | 0.015 | 0.015 | 0 | 0.42 | 0.37 | 0.44 | 0.18 | 0.48 | 0.51 |
| 1 | 165904700 | rs7525707 | -0.601 | 0.555 | -1.582 | -0.676 | 0.003 | 0.036 | 0.036 | 0 | 0.81 | 0.77 | 0.88 | 0.67 | 0.83 | 0.79 |
| 1 | 165906123 | rs12119052 | 0.227 | 0.736 | -1.924 | -0.844 | -0.065 | 0.078 | 0.078 | 0 | 0.22 | 0.04 | 0.19 | 0.08 | 0.31 | 0.16 |
| 1 | 165907256 | rs10800171 | 1.022 | 0.616 | -1.261 | -0.418 | -0.024 | 0.059 | 0.059 | 0 | 0.13 | 0.01 | 0.15 | 0.02 | 0.12 | 0.08 |
| 1 | 165908788 | rs12026896 | -0.653 | 1.046 | -0.343 | 0.425 | 0.011 | 0.149 | 0.149 | 0 | 0.21 | 0.44 | 0.15 | 0.37 | 0.18 | 0.29 |
| 1 | 165911600 | rs16853627 | -0.226 | 1.215 | -0.007 | -0.334 | 0.045 | -0.143 | -0.143 | 0 | 0.19 | 0.46 | 0.2 | 0.01 | 0.22 | 0.04 |
| 1 | 165913774 | rs6683842 | 0.197 | 1.401 | -0.763 | 0.241 | -0.022 | -0.052 | -0.052 | 0 | 0.71 | 0.79 | 0.73 | 0.52 | 0.68 | 0.58 |
| 1 | 165915221 | rs10918318 | 1.36 | 0.778 | -1.109 | -0.061 | 0.07 | 0.021 | 0.021 | 0 | 0.33 | 0.12 | 0.25 | 0.36 | 0.25 | 0.17 |
| 1 | 165917032 | rs6658459 | -0.374 | 0.991 | -1.114 | 0.135 | -0.022 | -0.054 | -0.054 | 0 | 0.72 | 0.79 | 0.68 | 0.51 | 0.69 | 0.58 |
| 1 | 165917109 | rs4657493 | 0.963 | 0.777 | -0.666 | 0.042 | 0.002 | -0.004 | -0.004 | 0 | 0.22 | 0.23 | 0.29 | 0.15 | 0.22 | 0.37 |
| 1 | 165928052 | rs10800173 | 0.534 | 0.878 | -0.714 | 0.156 | 0.01 | 0.053 | 0.053 | 0 | 0.53 | 0.23 | 0.49 | 0.46 | 0.46 | 0.49 |
| 1 | 165932506 | rs12083468 | -0.36 | 1.117 | 0.049 | 0.008 | -0.045 | -0.013 | -0.013 | 1.921 | 0.65 | 0.78 | 0.59 | 0.47 | 0.59 | 0.52 |
| 1 | 165953874 | rs7536863 | 0.696 | 1 | 0.508 | -0.083 | 0.016 | -0.035 | -0.035 | 0 | 0.07 | 0.1 | 0.05 | 0 | 0.11 | 0.02 |
| 1 | 165965211 | rs4657500 | 1.385 | 1.032 | 0.21 | 0.175 | -0.046 | 0.017 | 0.017 | 0 | 0.16 | 0.05 | 0.18 | 0.09 | 0.17 | 0.18 |
| 1 | 165967420 | rs732285 | 0.245 | 0.76 | 0.96 | 0.318 | 0.016 | -0.008 | -0.008 | 0 | 0.61 | 0.46 | 0.55 | 0.43 | 0.58 | 0.5 |
| 1 | 165967961 | rs9426841 | -0.18 | 0.753 | 0.809 | 0.682 | -0.074 | -0.071 | -0.071 | 1.885 | 0.64 | 0.79 | 0.57 | 0.51 | 0.58 | 0.5 |
| 1 | 165969492 | rs17422118 | 0.679 | 0.608 | 0.601 | 0.403 | -0.005 | 0.031 | 0.031 | 0 | 0.28 | 0.16 | 0.19 | 0.1 | 0.32 | 0.34 |
| 1 | 165988336 | rs6426957 | 0.108 | 0.62 | 0.368 | 0.442 | 0 | 0 | 0 | 0 | 0.6 | 0.36 | 0.53 | 0.51 | 0.57 | 0.49 |
| 1 | 165992161 | rs11585763 | 0.192 | 0.571 | 0.256 | 0.694 | -0.003 | -0.035 | -0.035 | 0.177 | 0.32 | 0.43 | 0.21 | 0.19 | 0.33 | 0.48 |
| 1 | 165992212 | rs6692476 | 1.362 | 0.53 | 0.291 | 0.727 | 0.002 | -0.001 | -0.001 | 0 | 0.34 | 0.27 | 0.34 | 0.4 | 0.26 | 0.14 |
| 1 | 165994034 | rs12040476 | 0.332 | 0.514 | 0.336 | 0.735 | -0.005 | 0.058 | 0.058 | 0.853 | 0.05 | 0.16 | 0.01 | 0.09 | 0.02 | 0.14 |
| 1 | 165994886 | rs6426960 | 1.214 | 0.542 | 0.016 | 0.649 | -0.037 | 0.005 | 0.005 | 0 | 0.34 | 0.45 | 0.36 | 0.41 | 0.25 | 0.15 |
| 1 | 165995470 | rs10918328 | 1.808 | 0.559 | -0.632 | 0.667 | 0.032 | -0.048 | -0.048 | 0 | 0.3 | 0.08 | 0.32 | 0.4 | 0.25 | 0.14 |
| 1 | 165996331 | rs6426962 | -0.213 | 0.791 | -1.173 | 0.611 | -0.139 | -0.122 | -0.122 | 0 | 0.67 | 0.88 | 0.57 | 0.6 | 0.58 | 0.63 |
| 1 | 166002131 | rs6672766 | -0.179 | 0.68 | -1.23 | 0.491 | -0.146 | -0.114 | -0.114 | 0 | 0.64 | 0.88 | 0.56 | 0.6 | 0.56 | 0.62 |
| 1 | 166002695 | rs867336 | 0.064 | 0.81 | -0.465 | 0.179 | 0.024 | -0.12 | -0.12 | 0.302 | 0.28 | 0.44 | 0.25 | 0.08 | 0.33 | 0.32 |
| 1 | 166003309 | rs1316805 | -0.779 | 0.862 | -1.004 | 0.982 | -0.06 | -0.09 | -0.09 | 0 | 0.85 | 0.97 | 0.85 | 0.61 | 0.81 | 0.76 |
| 1 | 166010221 | rs1867555 | -0.138 | 0.227 | -1.287 | -0.059 | -0.013 | 0.069 | 0.069 | 0.66 | 0.32 | 0.81 | 0.33 | 0.13 | 0.34 | 0.26 |
| 1 | 166010280 | rs12752269 | 1.028 | 0.036 | -1.51 | -0.07 | -0.044 | 0.013 | 0.013 | 0 | 0.19 | 0.04 | 0.15 | 0.19 | 0.23 | 0.27 |
| 1 | 166011666 | rs10753700 | 0.949 | 0.062 | -0.782 | -0.051 | -0.009 | 0.016 | 0.016 | 0 | 0.19 | 0.09 | 0.15 | 0.19 | 0.23 | 0.3 |
| 1 | 166018490 | rs4656477 | -0.927 | 0.006 | -0.897 | 0.5 | -0.049 | -0.096 | -0.096 | 0 | 0.85 | 0.97 | 0.87 | 0.59 | 0.84 | 0.76 |
| 1 | 166027277 | rs4657510 | -0.727 | -0.206 | -0.335 | 0.101 | -0.001 | -0.056 | -0.056 | 0 | 0.74 | 0.63 | 0.78 | 0.82 | 0.71 | 0.71 |
| 1 | 166028795 | rs11581600 | 1.419 | -0.38 | -0.323 | -0.033 | -0.028 | 0.013 | 0.013 | 0 | 0.2 | 0.13 | 0.23 | 0.12 | 0.26 | 0.15 |
| 1 | 166032197 | rs10918331 | 0.763 | -0.337 | -0.353 | -0.061 | -0.006 | 0.044 | 0.044 | 0 | 0.25 | 0.14 | 0.19 | 0.16 | 0.29 | 0.29 |
| 1 | 166032241 | rs3935592 | 0.672 | -0.8 | -0.477 | -0.217 | 0.01 | -0.059 | -0.059 | 0.086 | 0.34 | 0.23 | 0.23 | 0.41 | 0.29 | 0.35 |
| 1 | 166035856 | rs4262520 | 1.05 | -0.71 | -0.498 | -0.251 | -0.044 | 0.081 | 0.081 | 0 | 0.24 | 0.07 | 0.2 | 0.16 | 0.31 | 0.29 |
| 1 | 166036621 | rs4074017 | 2.334 | -0.791 | -0.504 | -0.275 | 0.019 | 0.004 | 0.004 | 1.576 | 0.2 | 0.21 | 0.12 | 0 | 0.13 | 0.11 |
| 1 | 166037909 | rs3856211 | 0.267 | -0.341 | 0.049 | 0.391 | -0.001 | -0.001 | -0.001 | 0.121 | 0.6 | 0.59 | 0.44 | 0.57 | 0.59 | 0.63 |
| 1 | 166041169 | rs6426964 | -0.319 | -1.088 | 0.812 | 1.548 | 0.026 | 0.155 | 0.155 | 0 | 0.17 | 0.29 | 0.15 | 0.43 | 0.17 | 0.25 |
| 1 | 166045424 | rs7524575 | 0.075 | -1.559 | 0.813 | 1.535 | 0.003 | -0.006 | -0.006 | 3.046 | 0.81 | 0.84 | 0.69 | 0.75 | 0.85 | 0.78 |
| 1 | 166054646 | rs10800183 | 0.314 | -1.276 | 1.109 | 1.553 | -0.009 | 0.024 | 0.024 | 0 | 0.78 | 0.69 | 0.8 | 0.67 | 0.8 | 0.64 |
| 1 | 166065145 | rs10918345 | -0.278 | -1.357 | 0.808 | 1.518 | 0.009 | -0.023 | -0.023 | 0.044 | 0.19 | 0.11 | 0.18 | 0.27 | 0.19 | 0.29 |
| 1 | 166086171 | rs4074897 | 1.46 | -1.76 | 1.054 | 1.494 | 0.042 | -0.009 | -0.009 | 0 | 0.61 | 0.3 | 0.54 | 0.55 | 0.71 | 0.48 |
| 1 | 166087578 | rs12756802 | 0.306 | -1.586 | 1.329 | 1.609 | -0.019 | 0.07 | 0.07 | 0 | 0.78 | 0.61 | 0.63 | 0.62 | 0.81 | 0.63 |
| 1 | 166098675 | rs4657520 | 0.103 | -1.587 | 1.404 | 1.609 | -0.012 | 0.055 | 0.055 | 0 | 0.79 | 0.72 | 0.63 | 0.63 | 0.81 | 0.66 |
| 1 | 166099990 | rs6676037 | 0.896 | -1.388 | 0.951 | 1.922 | -0.032 | -0.036 | -0.036 | 0 | 0.06 | 0 | 0.06 | 0.13 | 0.08 | 0.05 |
| 1 | 166104437 | rs10918373 | -0.288 | -1.626 | 1.23 | 1.642 | 0.004 | -0.007 | -0.007 | 0 | 0.19 | 0.17 | 0.18 | 0.26 | 0.18 | 0.35 |
| 1 | 166110181 | rs1470761 | 0.506 | -1.81 | 1.396 | 1.34 | -0.047 | -0.007 | -0.007 | 0 | 0.81 | 0.6 | 0.75 | 0.82 | 0.9 | 0.83 |
| 1 | 166114960 | rs17423848 | -0.193 | -1.813 | 1.301 | 1.302 | 0.021 | 0.042 | 0.042 | 0 | 0.15 | 0.02 | 0.06 | 0.07 | 0.09 | 0.14 |
| 1 | 166117868 | rs7548172 | -0.566 | -1.634 | 1.408 | 1.562 | 0.028 | -0.052 | -0.052 | 0 | 0.32 | 0.03 | 0.18 | 0.34 | 0.25 | 0.44 |
| 1 | 166130218 | rs2116519 | 0.856 | -1.641 | 1.569 | 1.566 | 0.039 | -0.024 | -0.024 | 0 | 0.62 | 0.32 | 0.57 | 0.51 | 0.73 | 0.49 |
| 1 | 166147182 | rs12402371 | 0.756 | -1.675 | 1.452 | 1.484 | 0.011 | 0 | 0 | 0.013 | 0.62 | 0.35 | 0.73 | 0.62 | 0.74 | 0.52 |
| 1 | 166166695 | rs6658513 | -0.571 | -1.658 | 1.567 | 1.529 | 0.037 | -0.023 | -0.023 | 0 | 0.32 | 0.23 | 0.23 | 0.41 | 0.22 | 0.36 |
| 1 | 166169203 | rs10800201 | 0.977 | -1.763 | 1.69 | 1.58 | 0.074 | -0.041 | -0.041 | 0 | 0.61 | 0.21 | 0.57 | 0.47 | 0.77 | 0.51 |
| 1 | 166175936 | rs896404 | 0.613 | -1.748 | 1.938 | 1.573 | -0.016 | 0.009 | 0.009 | 0.218 | 0.64 | 0.43 | 0.59 | 0.47 | 0.77 | 0.62 |
| 1 | 166177384 | rs4391646 | -0.544 | -1.746 | 1.914 | 1.573 | 0.024 | -0.015 | -0.015 | 0 | 0.32 | 0.25 | 0.23 | 0.41 | 0.22 | 0.34 |
| 1 | 166191677 | rs7543144 | 0.957 | -1.765 | 1.899 | 1.441 | 0.072 | -0.045 | -0.045 | 0 | 0.61 | 0.22 | 0.6 | 0.48 | 0.77 | 0.53 |
| 1 | 166206684 | rs10918405 | -0.552 | -1.618 | 1.788 | 1.467 | 0.029 | -0.019 | -0.019 | 0 | 0.32 | 0.25 | 0.21 | 0.41 | 0.22 | 0.34 |
| 1 | 166221811 | rs10918416 | -0.506 | -1.618 | 1.741 | 1.467 | 0.045 | -0.033 | -0.033 | 0 | 0.31 | 0.17 | 0.2 | 0.41 | 0.22 | 0.34 |
| 1 | 166226685 | rs6658303 | -0.739 | -1.615 | 1.781 | 1.469 | -0.048 | 0.021 | 0.021 | 0 | 0.33 | 0.43 | 0.22 | 0.41 | 0.22 | 0.34 |
| 1 | 166228386 | rs1432120 | 1.717 | -1.264 | 1.626 | 1.55 | 0.061 | 0.237 | 0.237 | 1.45 | 0.95 | 0.6 | 0.76 | 0.85 | 0.99 | 0.98 |
| 1 | 166235714 | rs2599513 | 0.073 | -0.816 | 0.805 | 1.642 | -0.082 | -0.104 | -0.104 | 0 | 0.13 | 0.01 | 0.13 | 0.41 | 0.19 | 0.3 |

| | | | | | | | | | | | | | | | | |
|---|---|---|---|---|---|---|---|---|---|---|---|---|---|---|---|---|
| 1 | 166250778 | rs1545007 | 0.801 | -1.882 | 1.486 | 1.665 | 0.043 | -0.023 | -0.023 | 0 | 0.64 | 0.26 | 0.61 | 0.52 | 0.77 | 0.63 |
| 1 | 166251059 | rs7524473 | -0.802 | -1.818 | 1.193 | 1.657 | 0.005 | -0.003 | -0.003 | 0.368 | 0.32 | 0.3 | 0.31 | 0.43 | 0.23 | 0.37 |
| 1 | 166262123 | rs10494457 | -0.645 | -1.725 | 0.999 | 1.659 | 0.031 | 0.012 | 0.012 | 0 | 0.34 | 0.21 | 0.2 | 0.33 | 0.3 | 0.38 |
| 1 | 166269296 | rs16856889 | -0.944 | -1.56 | 1.083 | 1.9 | -0.022 | 0.013 | 0.013 | 0 | 0.37 | 0.58 | 0.42 | 0.5 | 0.28 | 0.36 |
| 1 | 166270695 | rs4282794 | -1.241 | -1.525 | 1.084 | 2.199 | 0.034 | 0.009 | 0.009 | 0.161 | 0.39 | 0.75 | 0.46 | 0.68 | 0.31 | 0.42 |
| 1 | 166286486 | rs6426991 | -0.517 | -1.484 | 0.461 | 1.941 | 0.044 | 0.002 | 0.002 | 0 | 0.32 | 0.15 | 0.19 | 0.33 | 0.28 | 0.33 |
| 1 | 166299045 | rs9943075 | -1.255 | -1.51 | 0.549 | 1.958 | 0.077 | 0.03 | 0.03 | 0.902 | 0.42 | 0.85 | 0.47 | 0.68 | 0.32 | 0.42 |
| 1 | 166299138 | rs9943264 | -2.217 | -0.953 | 1.392 | 2.127 | 0.04 | 0.288 | 0.288 | 8 | 0.07 | 0.57 | 0.11 | 0.24 | 0.03 | 0.06 |
| 1 | 166304410 | rs10494458 | -2.221 | -0.85 | 1.258 | 2.028 | 0.047 | 0.266 | 0.266 | 0.205 | 0.05 | 0.27 | 0.21 | 0.29 | 0.04 | 0.09 |
| 1 | 166304467 | rs6426993 | -0.796 | -1.401 | 0.801 | 1.466 | -0.006 | 0.005 | 0.005 | 0 | 0.18 | 0.19 | 0.27 | 0.26 | 0.11 | 0.13 |
| 1 | 166305834 | rs9970244 | -1.56 | -1.32 | 0.768 | 1.492 | 0.04 | 0.046 | 0.046 | 0 | 0.4 | 0.76 | 0.46 | 0.71 | 0.33 | 0.42 |
| 1 | 166308210 | rs7543439 | -0.777 | -1.518 | 1.281 | 2.058 | -0.035 | 0.063 | 0.063 | 0 | 0.24 | 0.37 | 0.34 | 0.47 | 0.2 | 0.21 |
| 1 | 166308592 | rs12024710 | -0.441 | -1.528 | 1.485 | 2.115 | 0.004 | -0.005 | -0.005 | 0 | 0.21 | 0.18 | 0.12 | 0.37 | 0.19 | 0.19 |
| 1 | 166313850 | rs10800214 | -0.622 | -1.42 | 1.638 | 2.119 | -0.01 | -0.032 | -0.032 | 0 | 0.13 | 0.07 | 0.09 | 0.24 | 0.13 | 0.21 |
| 1 | 166318227 | rs10494459 | -1.818 | -1.426 | 1.65 | 2.103 | -0.006 | -0.008 | -0.008 | 0.002 | 0.36 | 0.4 | 0.43 | 0.71 | 0.33 | 0.42 |
| 1 | 166319377 | rs10494460 | -0.446 | -1.34 | 1.614 | 2.104 | 0.003 | -0.004 | -0.004 | 0 | 0.21 | 0.17 | 0.12 | 0.37 | 0.19 | 0.18 |
| 1 | 166325662 | rs6666649 | -1.972 | -1.398 | 1.565 | 1.921 | -0.017 | -0.03 | -0.03 | 0 | 0.38 | 0.51 | 0.44 | 0.74 | 0.33 | 0.42 |
| 1 | 166326750 | rs16856966 | -1.22 | -1.354 | 1.349 | 2.17 | -0.004 | 0.105 | 0.105 | 0 | 0.09 | 0.26 | 0.28 | 0.21 | 0.07 | 0.08 |
| 1 | 166334060 | rs7547839 | -2.132 | -1.281 | 1.344 | 2.144 | -0.012 | -0.072 | -0.072 | 0 | 0.34 | 0.47 | 0.45 | 0.83 | 0.3 | 0.5 |
| 1 | 166336254 | rs10800219 | -0.171 | -1.052 | 1.333 | 2.51 | -0.035 | -0.025 | -0.025 | 0 | 0.07 | 0.01 | 0.13 | 0.25 | 0.09 | 0.2 |
| 1 | 166336660 | rs3924947 | -0.907 | -1.02 | 1.41 | 2.483 | -0.017 | -0.031 | -0.031 | 0 | 0.13 | 0.07 | 0.19 | 0.38 | 0.16 | 0.3 |
| 1 | 166337053 | rs12095056 | -1.268 | -1.019 | 1.428 | 2.228 | 0.081 | 0.227 | 0.227 | 0.554 | 0.05 | 0.34 | 0.2 | 0.17 | 0.03 | 0.07 |
| 1 | 166352194 | rs4551573 | -2.108 | -1.175 | 1.11 | 2.047 | -0.011 | -0.054 | -0.054 | 0.72 | 0.34 | 0.54 | 0.45 | 0.83 | 0.29 | 0.49 |
| 1 | 166353381 | rs6426997 | -1.064 | -1.287 | 1.22 | 2.241 | 0 | 0.048 | 0.048 | 0 | 0.15 | 0.22 | 0.18 | 0.25 | 0.14 | 0.23 |
| 1 | 166355224 | rs12121870 | -0.622 | -1.24 | 0.732 | 2.263 | -0.035 | -0.078 | -0.078 | 0 | 0.13 | 0.02 | 0.16 | 0.25 | 0.14 | 0.23 |
| 1 | 166362651 | rs4626860 | -1.237 | -1.09 | 0.514 | 1.611 | -0.017 | 0.015 | 0.015 | 0.63 | 0.32 | 0.41 | 0.39 | 0.54 | 0.24 | 0.4 |
| 1 | 166364063 | rs4542197 | -1.342 | -1.195 | 0.116 | 1.168 | -0.005 | -0.004 | -0.004 | 0 | 0.37 | 0.63 | 0.43 | 0.69 | 0.25 | 0.39 |
| 1 | 166367325 | rs12562144 | -0.559 | 0.531 | -0.301 | 0.622 | -0.018 | 0 | 0 | 0 | 0.09 | 0.05 | 0.14 | 0.08 | 0.24 | 0.09 |
| 1 | 166369651 | rs4657543 | -0.871 | 0.757 | -0.336 | 0.115 | -0.013 | 0.012 | 0.012 | 0.743 | 0.34 | 0.72 | 0.36 | 0.3 | 0.5 | 0.35 |
| 1 | 166388391 | rs7545852 | 0.626 | -0.14 | -1.083 | -0.655 | 0.065 | 0.048 | 0.048 | 0.186 | 0.4 | 0.16 | 0.52 | 0.62 | 0.3 | 0.41 |
| 1 | 166389338 | rs1389779 | 1.176 | 0.057 | -0.431 | -0.709 | 0.032 | -0.064 | -0.064 | 0 | 0.21 | 0.4 | 0.2 | 0.15 | 0.24 | 0.24 |
| 1 | 166399575 | rs6427004 | -0.086 | 0.083 | -0.36 | -0.672 | 0.002 | 0.069 | 0.069 | 0 | 0.52 | 0.38 | 0.68 | 0.76 | 0.56 | 0.52 |
| 1 | 166405300 | rs2311322 | 1.089 | -0.281 | -0.909 | -0.299 | 0.046 | -0.147 | -0.147 | 0.371 | 0.24 | 0.03 | 0.35 | 0.39 | 0.14 | 0.2 |
| 1 | 166415259 | rs10918467 | -0.802 | 0.233 | -0.321 | -0.663 | 0.016 | 0.042 | 0.042 | 0.056 | 0.74 | 0.86 | 0.88 | 0.91 | 0.8 | 0.75 |
| 1 | 166418076 | rs11589054 | -0.046 | 0.388 | -0.34 | -0.676 | -0.037 | -0.017 | -0.017 | 0 | 0.11 | 0.05 | 0.15 | 0.14 | 0.27 | 0.13 |
| 1 | 166423222 | rs7544530 | -0.255 | 0.501 | -0.172 | -0.554 | 0.041 | -0.005 | -0.005 | 0 | 0.32 | 0.55 | 0.42 | 0.31 | 0.5 | 0.34 |
| 1 | 166431020 | rs7549089 | 0.306 | 0.806 | -0.185 | -0.668 | 0.001 | -0.009 | -0.009 | 0.005 | 0.61 | 0.6 | 0.71 | 0.78 | 0.5 | 0.63 |
| 1 | 166432265 | rs1908311 | -0.069 | 0.93 | -0.192 | -0.677 | -0.002 | -0.038 | -0.038 | 0.763 | 0.65 | 0.6 | 0.81 | 0.85 | 0.69 | 0.7 |
| 1 | 166435300 | rs10918474 | 0.863 | 0.513 | -0.458 | -0.614 | 0.073 | 0.022 | 0.022 | 0.193 | 0.41 | 0.12 | 0.45 | 0.63 | 0.3 | 0.4 |
| 1 | 166436973 | rs6427006 | -0.552 | 0.633 | -0.482 | -0.659 | -0.011 | 0.094 | 0.094 | 0 | 0.72 | 0.87 | 0.87 | 0.92 | 0.77 | 0.74 |
| 1 | 166440331 | rs16857149 | 1.054 | 0.869 | -0.139 | -0.481 | 0.009 | -0.025 | -0.025 | 0.772 | 0.19 | 0.37 | 0.25 | 0.14 | 0.2 | 0.22 |
| 1 | 166444011 | rs12566754 | -0.02 | 0.842 | 0.038 | -0.42 | 0.042 | -0.004 | -0.004 | 0.476 | 0.28 | 0.39 | 0.39 | 0.26 | 0.46 | 0.3 |
| 1 | 166444655 | rs10918477 | 0.904 | 0.619 | -0.085 | -0.527 | 0.057 | 0.016 | 0.016 | 1.024 | 0.43 | 0.26 | 0.46 | 0.59 | 0.3 | 0.41 |
| 1 | 166445013 | rs17428629 | 0.52 | 0.911 | -0.151 | -0.641 | -0.018 | -0.014 | -0.014 | 0 | 0.09 | 0.05 | 0.15 | 0.11 | 0.26 | 0.08 |
| 1 | 166445116 | rs1494430 | -0.22 | 0.751 | -0.113 | -0.826 | -0.001 | -0.034 | -0.034 | 0.818 | 0.72 | 0.7 | 0.86 | 0.91 | 0.77 | 0.74 |
| 1 | 166447321 | rs1027351 | 0.923 | 1.108 | -0.037 | -0.6 | 0.007 | -0.046 | -0.046 | 0.988 | 0.21 | 0.36 | 0.25 | 0.13 | 0.22 | 0.22 |
| 1 | 166447978 | rs1027353 | -0.227 | 1.262 | -0.163 | -0.63 | -0.013 | 0.057 | 0.057 | 0.363 | 0.78 | 0.85 | 0.9 | 0.92 | 0.8 | 0.77 |
| 1 | 166448963 | rs16857179 | 0.783 | 0.657 | -0.079 | -0.47 | -0.004 | -0.009 | -0.009 | 0 | 0.28 | 0.32 | 0.25 | 0.18 | 0.23 | 0.24 |
| 1 | 166450328 | rs12031195 | 1.042 | 0.775 | -0.445 | -0.416 | 0.052 | 0.031 | 0.031 | 1.981 | 0.38 | 0.17 | 0.45 | 0.65 | 0.29 | 0.44 |
| 1 | 166451443 | rs10918481 | 0.949 | 0.768 | -0.388 | -0.42 | 0.012 | 0.008 | 0.008 | 2.362 | 0.4 | 0.33 | 0.46 | 0.65 | 0.29 | 0.44 |
| 1 | 166452024 | rs2172714 | 1.094 | 0.847 | -1.038 | -0.314 | 0.047 | -0.022 | -0.022 | 0 | 0.37 | 0.17 | 0.45 | 0.54 | 0.29 | 0.4 |
| 1 | 166454802 | rs1494428 | 1.094 | 0.847 | -1.038 | -0.314 | 0.047 | -0.022 | -0.022 | 0 | 0.37 | 0.17 | 0.45 | 0.54 | 0.29 | 0.4 |
| 1 | 166456704 | rs7552777 | 0.206 | 0.892 | -1.028 | -0.37 | -0.016 | 0.014 | 0.014 | 1.963 | 0.68 | 0.68 | 0.71 | 0.82 | 0.52 | 0.67 |
| 1 | 166458160 | rs7544608 | 0.155 | 0.187 | -0.87 | 0.014 | 0.003 | 0.003 | 0.003 | 0 | 0.58 | 0.39 | 0.68 | 0.57 | 0.63 | 0.58 |
| 1 | 166458590 | rs10918487 | -0.34 | 0.3 | -0.819 | -0.768 | 0.001 | 0.001 | 0.001 | 0 | 0.41 | 0.61 | 0.32 | 0.24 | 0.37 | 0.37 |
| 1 | 166459849 | rs6680174 | 0.639 | 0.336 | -0.741 | -0.71 | -0.014 | -0.105 | -0.105 | 0.09 | 0.7 | 0.47 | 0.8 | 0.9 | 0.71 | 0.73 |
| 1 | 166460696 | rs6658665 | -0.255 | -0.077 | -1.901 | -0.692 | 0.044 | 0.095 | 0.095 | 0 | 0.4 | 0.04 | 0.23 | 0.21 | 0.35 | 0.37 |
| 1 | 166468429 | rs7514715 | 0.003 | 0.428 | -1.511 | -0.381 | -0.002 | -0.006 | -0.006 | 0 | 0.36 | 0.31 | 0.3 | 0.48 | 0.38 | 0.49 |
| 1 | 166479898 | rs10753736 | -0.553 | 0.564 | -1.131 | -0.403 | 0 | -0.012 | -0.012 | 0 | 0.43 | 0.42 | 0.66 | 0.8 | 0.51 | 0.58 |
| 1 | 166482753 | rs16857234 | 0.403 | 1.068 | -1.298 | 0.06 | -0.005 | -0.033 | -0.033 | 0 | 0.17 | 0.06 | 0.12 | 0.24 | 0.17 | 0.12 |
| 1 | 166483531 | rs12097144 | 0.504 | 1.17 | -0.925 | 0.152 | 0.031 | 0.218 | 0.218 | 0 | 0.06 | 0.19 | 0.3 | 0.28 | 0.04 | 0.05 |
| 1 | 166486813 | rs11579746 | 0.887 | 1.029 | -1.264 | -0.178 | -0.029 | 0.03 | 0.03 | 0 | 0.3 | 0.03 | 0.23 | 0.2 | 0.33 | 0.31 |
| 1 | 166489143 | rs6695015 | -0.05 | 0.897 | -1.246 | -0.684 | -0.027 | -0.01 | -0.01 | 0 | 0.56 | 0.26 | 0.6 | 0.48 | 0.54 | 0.44 |
| 1 | 166489372 | rs17429732 | 1.712 | 0.514 | -1.808 | -0.975 | 0.033 | 0.083 | 0.083 | 0.04 | 0.18 | 0.02 | 0.14 | 0 | 0.15 | 0.1 |
| 1 | 166489831 | rs17429774 | 0.701 | 1.091 | -1.29 | -0.308 | 0.002 | 0.077 | 0.077 | 0.315 | 0.21 | 0.11 | 0.08 | 0.01 | 0.16 | 0.09 |
| 1 | 166493260 | rs12118962 | -0.613 | 1.555 | -0.71 | 0.566 | -0.017 | -0.048 | -0.048 | 0.102 | 0.34 | 0.22 | 0.3 | 0.48 | 0.38 | 0.49 |
| 1 | 166493556 | rs12119773 | -0.598 | 1.576 | -0.7 | 0.576 | -0.017 | -0.048 | -0.048 | 0 | 0.34 | 0.22 | 0.3 | 0.48 | 0.38 | 0.49 |
| 1 | 166500023 | rs269676 | 1.602 | 1.608 | 0.071 | 0.634 | 0.093 | 0.078 | 0.078 | 0 | 0.91 | 0.83 | 0.85 | 0.81 | 0.8 | 0.84 |
| 1 | 166504879 | rs419212 | 0.855 | 0.628 | -0.155 | 0.175 | -0.009 | -0.034 | -0.034 | 0 | 0.94 | 0.85 | 0.95 | 1 | 0.96 | 0.99 |

| | | | | | | | | | | | | | | | | |
|---|---|---|---|---|---|---|---|---|---|---|---|---|---|---|---|---|
| 1 | 166511732 | rs1530321 | -0.979 | 0.143 | -0.084 | -0.021 | -0.028 | -0.041 | -0.041 | 0 | 0.23 | 0.36 | 0.1 | 0.14 | 0.13 | 0.07 |
| 1 | 166532565 | rs10918527 | -0.025 | 1.591 | 0.326 | 0.726 | 0.061 | 0.066 | 0.066 | 0 | 0.19 | 0.23 | 0.23 | 0.32 | 0.29 | 0.19 |
| 1 | 166536092 | rs2584523 | -1.276 | 0.519 | 0.023 | 0.194 | -0.047 | -0.055 | -0.055 | 0.34 | 0.77 | 0.87 | 0.49 | 0.59 | 0.66 | 0.41 |
| 1 | 166539001 | rs7516525 | 0.266 | 0.078 | 0.53 | -0.008 | 0 | 0 | 0 | 0.106 | 0.56 | 0.39 | 0.39 | 0.45 | 0.55 | 0.35 |
| 1 | 166545275 | rs10918532 | 0.749 | -0.2 | 0.603 | -0.278 | 0.028 | 0.08 | 0.08 | 2.65 | 0.43 | 0.3 | 0.27 | 0.15 | 0.41 | 0.21 |
| 1 | 166545748 | rs17431003 | 1.541 | -0.22 | 0.568 | -0.29 | 0.005 | 0.202 | 0.202 | 1.014 | 0.33 | 0.07 | 0.2 | 0 | 0.34 | 0.17 |
| 1 | 166546775 | rs12030083 | 1.162 | -0.202 | 0.602 | -0.245 | 0.004 | 0.006 | 0.006 | 0 | 0.16 | 0.13 | 0.15 | 0.19 | 0.18 | 0.12 |
| 1 | 166546851 | rs6670674 | -0.133 | -0.177 | 0.559 | -0.109 | -0.004 | 0 | 0 | 0.305 | 0.6 | 0.56 | 0.42 | 0.36 | 0.59 | 0.35 |
| 1 | 166549009 | rs2119902 | -0.734 | -0.102 | 0.177 | -0.073 | -0.037 | -0.033 | -0.033 | 0.822 | 0.67 | 0.78 | 0.47 | 0.59 | 0.62 | 0.39 |
| 1 | 166552723 | rs7539202 | -0.379 | -0.138 | 0.2 | -0.137 | 0.003 | 0.022 | 0.022 | 0 | 0.27 | 0.49 | 0.22 | 0.33 | 0.25 | 0.16 |
| 1 | 166553569 | rs1442500 | -1.515 | -0.042 | 0.175 | -0.181 | -0.011 | -0.053 | -0.053 | 0.011 | 0.88 | 0.83 | 0.94 | 1 | 0.92 | 0.97 |
| 1 | 166554578 | rs4657570 | 1.204 | -0.05 | 0.19 | -0.199 | -0.045 | -0.045 | -0.045 | 0 | 0.23 | 0.13 | 0.51 | 0.64 | 0.32 | 0.62 |
| 1 | 166555844 | rs565210 | -1.436 | -0.059 | 0.266 | -0.145 | -0.011 | -0.053 | -0.053 | 1.296 | 0.88 | 0.83 | 0.94 | 1 | 0.92 | 0.97 |
| 1 | 166556638 | rs509587 | -1.734 | -0.037 | 0.277 | -0.13 | -0.011 | -0.052 | -0.052 | 0 | 0.89 | 0.84 | 0.95 | 1 | 0.93 | 0.98 |
| 1 | 166560125 | rs497142 | -0.5 | -0.023 | 0.208 | -0.078 | 0.002 | 0.021 | 0.021 | 0 | 0.27 | 0.49 | 0.22 | 0.33 | 0.25 | 0.16 |
| 1 | 166565815 | rs528816 | -1.519 | -0.007 | 0.067 | 0.005 | 0.001 | -0.002 | -0.002 | 0.882 | 0.88 | 0.9 | 0.93 | 0.76 | 0.91 | 0.94 |
| 1 | 166568908 | rs510265 | 0.289 | -0.108 | 0.391 | -0.156 | -0.003 | 0.047 | 0.047 | 1.899 | 0.1 | 0.34 | 0.07 | 0.15 | 0.06 | 0.03 |
| 1 | 166571770 | rs6677448 | 0.462 | -0.814 | -0.125 | -0.783 | -0.033 | 0.04 | 0.04 | 0 | 0.34 | 0.15 | 0.25 | 0.19 | 0.38 | 0.23 |
| 1 | 166573130 | rs556269 | -1.115 | -0.91 | -0.392 | -0.868 | -0.053 | -0.013 | -0.013 | 1.606 | 0.66 | 0.84 | 0.46 | 0.35 | 0.61 | 0.34 |
| 1 | 166573473 | rs6427019 | -0.211 | -0.923 | -0.268 | -0.878 | 0 | -0.01 | -0.01 | 0 | 0.54 | 0.47 | 0.37 | 0.19 | 0.53 | 0.29 |
| 1 | 166573706 | rs12039131 | 0.249 | -0.949 | -0.268 | -0.917 | -0.003 | -0.005 | -0.005 | 0.886 | 0.36 | 0.31 | 0.26 | 0.19 | 0.32 | 0.17 |
| 1 | 166579080 | rs530690 | -1.003 | -0.933 | -0.268 | -0.906 | -0.026 | 0.002 | 0.002 | 1.788 | 0.64 | 0.7 | 0.45 | 0.34 | 0.61 | 0.32 |
| 1 | 166581706 | rs553928 | -1.045 | -0.933 | -0.28 | -0.906 | -0.045 | -0.002 | -0.002 | 0.318 | 0.64 | 0.8 | 0.45 | 0.34 | 0.61 | 0.32 |
| 1 | 166582292 | rs7546913 | -0.193 | -0.919 | -0.309 | -0.913 | -0.003 | 0.028 | 0.028 | 0.059 | 0.54 | 0.36 | 0.36 | 0.19 | 0.53 | 0.29 |
| 1 | 166591271 | rs17472444 | 1.205 | -0.931 | -0.308 | -0.892 | -0.002 | -0.002 | -0.002 | 0 | 0.16 | 0.1 | 0.14 | 0.19 | 0.18 | 0.1 |
| 1 | 166593754 | rs12741275 | 1.998 | -0.801 | -0.484 | -1.048 | -0.033 | 0.049 | 0.049 | 0 | 0.09 | 0 | 0.05 | 0 | 0.13 | 0.02 |
| 1 | 166595209 | rs714339 | 1.688 | -0.825 | -0.443 | -1.012 | 0.001 | -0.027 | -0.027 | 0.063 | 0.18 | 0.14 | 0.11 | 0 | 0.15 | 0.07 |
| 1 | 166602976 | rs12094885 | 1.539 | -0.714 | -0.452 | -0.904 | -0.027 | -0.028 | -0.028 | 0 | 0.27 | 0.2 | 0.51 | 0.65 | 0.31 | 0.64 |
| 1 | 166603023 | rs2065920 | 1.763 | -0.716 | -0.456 | -0.797 | -0.052 | -0.076 | -0.076 | 8.682 | 0.25 | 0.1 | 0.47 | 0.44 | 0.3 | 0.61 |
| 1 | 166603496 | rs481045 | -0.608 | -0.771 | -0.515 | -0.951 | 0.028 | 0.056 | 0.056 | 0.327 | 0.55 | 0.66 | 0.74 | 0.99 | 0.57 | 0.78 |
| 1 | 166604212 | rs509414 | -1.697 | -0.227 | -0.027 | -0.374 | 0.007 | 0.004 | 0.004 | 2.033 | 0.88 | 0.91 | 0.95 | 1 | 0.93 | 0.98 |
| 1 | 166606328 | rs1372682 | 1.113 | -0.231 | -0.013 | -0.364 | 0.018 | 0.014 | 0.014 | 1.589 | 0.59 | 0.3 | 0.71 | 0.66 | 0.66 | 0.83 |
| 1 | 166612213 | rs1582115 | 0.608 | -0.151 | -0.045 | -0.481 | -0.011 | -0.036 | -0.036 | 0 | 0.23 | 0.34 | 0.13 | 0.16 | 0.21 | 0.11 |
| 1 | 166613883 | rs529204 | 0.534 | -0.294 | 0.127 | -0.373 | 0.003 | 0.054 | 0.054 | 1.391 | 0.1 | 0.33 | 0.08 | 0.16 | 0.08 | 0.04 |
| 1 | 166616786 | rs3910715 | 1.197 | -0.282 | 0.029 | -0.391 | 0.019 | 0.013 | 0.013 | 1.243 | 0.58 | 0.3 | 0.71 | 0.65 | 0.66 | 0.81 |
| 1 | 166618118 | rs3845541 | 1.131 | -0.184 | -0.211 | -0.345 | -0.001 | -0.002 | -0.002 | 0.103 | 0.45 | 0.48 | 0.62 | 0.65 | 0.47 | 0.72 |
| 1 | 166623180 | rs530303 | -0.867 | -0.135 | -0.195 | -0.333 | 0.001 | -0.005 | -0.005 | 0 | 0.46 | 0.48 | 0.33 | 0.34 | 0.46 | 0.25 |
| 1 | 166627974 | rs3856213 | 1.423 | -0.112 | -0.273 | -0.362 | -0.022 | 0.072 | 0.072 | 0 | 0.19 | 0.08 | 0.11 | 0 | 0.21 | 0.11 |
| 1 | 166628260 | rs565064 | -1.528 | -0.118 | -0.236 | -0.334 | -0.001 | -0.025 | -0.025 | 0.022 | 0.88 | 0.87 | 0.95 | 1 | 0.92 | 0.98 |
| 1 | 166634216 | rs558843 | -1.318 | -0.245 | -0.16 | -0.585 | -0.021 | 0.078 | 0.078 | 0.795 | 0.81 | 0.94 | 0.9 | 1 | 0.8 | 0.96 |
| 1 | 166640279 | rs539482 | 0.314 | -0.226 | -0.003 | -0.541 | 0.002 | 0.044 | 0.044 | 1.6 | 0.1 | 0.34 | 0.08 | 0.15 | 0.08 | 0.03 |
| 1 | 166640635 | rs472293 | -0.644 | -0.23 | 0 | -0.52 | -0.002 | 0.156 | 0.156 | 0.003 | 0.18 | 0.41 | 0.17 | 0.34 | 0.13 | 0.12 |
| 1 | 166647034 | rs12033500 | 1.068 | -0.158 | -0.019 | -0.48 | 0.002 | 0.05 | 0.05 | 0 | 0.08 | 0.07 | 0.09 | 0.19 | 0.05 | 0.1 |
| 1 | 166652275 | rs4657583 | 0.882 | -0.121 | -0.331 | -0.58 | -0.003 | 0.008 | 0.008 | 0.501 | 0.43 | 0.39 | 0.62 | 0.66 | 0.46 | 0.72 |
| 1 | 166656798 | rs11807188 | 0.918 | 0.24 | -0.889 | -0.755 | -0.023 | 0.061 | 0.061 | 0 | 0.18 | 0.07 | 0.11 | 0 | 0.21 | 0.11 |
| 1 | 166671929 | rs562914 | -0.756 | 0.294 | -0.784 | -0.324 | -0.016 | 0.056 | 0.056 | 0.585 | 0.81 | 0.94 | 0.9 | 1 | 0.8 | 0.96 |
| 1 | 166674289 | rs579100 | -0.127 | 0.069 | 0.013 | 0.162 | -0.003 | -0.005 | -0.005 | 0.233 | 0.76 | 0.54 | 0.83 | 0.75 | 0.76 | 0.92 |
| 1 | 166674984 | rs12048648 | 0.577 | 0.055 | 0.322 | 0.542 | 0.008 | 0.072 | 0.072 | 0 | 0.08 | 0.14 | 0.08 | 0.18 | 0.05 | 0.1 |
| 1 | 166675316 | rs4656506 | 0.543 | -0.053 | 0.398 | 0.48 | -0.003 | 0.016 | 0.016 | 0.422 | 0.67 | 0.37 | 0.74 | 0.57 | 0.71 | 0.82 |
| 1 | 166693545 | rs1855834 | 0.756 | -0.152 | 0.016 | 0.325 | -0.016 | -0.01 | -0.01 | 0.026 | 0.42 | 0.17 | 0.57 | 0.4 | 0.44 | 0.66 |
| 1 | 166702558 | rs12029656 | 0.397 | -0.041 | 0.259 | 0.635 | 0.006 | 0.065 | 0.065 | 0 | 0.08 | 0.13 | 0.08 | 0.18 | 0.05 | 0.1 |
| 1 | 166707143 | rs487726 | -0.719 | -0.474 | 0.18 | 0.253 | -0.026 | 0.009 | 0.009 | 0.459 | 0.13 | 0.21 | 0.07 | 0.17 | 0.07 | 0.05 |
| 1 | 166709642 | rs477967 | -0.794 | -0.327 | 0.702 | 0.462 | -0.016 | 0.047 | 0.047 | 0 | 0.17 | 0.27 | 0.13 | 0.27 | 0.12 | 0.14 |
| 1 | 166711398 | rs12073480 | -0.262 | -0.238 | 0.224 | 0.509 | -0.002 | -0.01 | -0.01 | 0 | 0.15 | 0.13 | 0.1 | 0.27 | 0.13 | 0.25 |
| 1 | 166716641 | rs524612 | -0.692 | -0.299 | 0.309 | 0.015 | 0.015 | -0.002 | -0.002 | 0.196 | 0.47 | 0.69 | 0.34 | 0.54 | 0.42 | 0.49 |
| 1 | 166717206 | rs1815531 | -1.794 | -0.159 | -0.124 | -0.292 | -0.036 | 0.091 | 0.091 | 2.754 | 0.82 | 0.97 | 0.89 | 1 | 0.79 | 0.96 |
| 1 | 166718884 | rs10918573 | 0.246 | -0.152 | 0.103 | -0.203 | 0 | 0.007 | 0.007 | 0 | 0.14 | 0.16 | 0.1 | 0.27 | 0.13 | 0.25 |
| 1 | 166721207 | rs564138 | 0.45 | -0.219 | 0.315 | -0.252 | -0.018 | -0.013 | -0.013 | 0.292 | 0.33 | 0.53 | 0.24 | 0.27 | 0.29 | 0.24 |
| 1 | 166726772 | rs1548332 | 0.618 | -0.285 | 0.272 | -0.278 | -0.018 | -0.01 | -0.01 | 0 | 0.32 | 0.46 | 0.24 | 0.27 | 0.28 | 0.25 |
| 1 | 166729339 | rs7549229 | 1.388 | -0.573 | -0.106 | -0.751 | -0.016 | -0.017 | -0.017 | 0 | 0.18 | 0.1 | 0.43 | 0.47 | 0.23 | 0.4 |
| 1 | 166730775 | rs563244 | -0.082 | -0.594 | -0.072 | -0.724 | 0.015 | -0.004 | -0.004 | 0 | 0.46 | 0.67 | 0.33 | 0.53 | 0.41 | 0.49 |
| 1 | 166730812 | rs562445 | 0.786 | -0.584 | -0.092 | -0.422 | -0.015 | 0.002 | 0.002 | 0 | 0.33 | 0.55 | 0.24 | 0.31 | 0.28 | 0.28 |
| 1 | 166758542 | rs480820 | 0.452 | -0.539 | -0.036 | -0.462 | 0.003 | 0.02 | 0.02 | 0.13 | 0.5 | 0.44 | 0.35 | 0.27 | 0.49 | 0.29 |
| 1 | 166758737 | rs858792 | -0.812 | -0.44 | -0.408 | -0.331 | -0.005 | -0.021 | -0.021 | 0 | 0.87 | 0.91 | 0.92 | 0.75 | 0.87 | 0.75 |
| 1 | 166767050 | rs858800 | -0.092 | -0.477 | -0.415 | -0.2 | 0.026 | -0.003 | -0.003 | 9.489 | 0.71 | 0.82 | 0.8 | 0.7 | 0.72 | 0.7 |
| 1 | 166767772 | rs858802 | -0.031 | -0.492 | -0.38 | -0.167 | 0.029 | 0.005 | 0.005 | 0 | 0.69 | 0.79 | 0.79 | 0.7 | 0.72 | 0.7 |
| 1 | 166774303 | rs12131167 | 1.643 | -0.492 | -0.404 | -0.433 | 0.032 | 0.034 | 0.034 | 0 | 0.15 | 0.08 | 0.11 | 0 | 0.14 | 0.07 |
| 1 | 166774997 | rs858818 | 1.605 | -0.425 | -0.408 | -0.515 | -0.021 | 0.057 | 0.057 | 2.593 | 0.18 | 0.08 | 0.12 | 0 | 0.22 | 0.04 |
| 1 | 166778754 | rs2205647 | -0.449 | -0.46 | -0.409 | -0.315 | -0.021 | 0.061 | 0.061 | 0.775 | 0.69 | 0.82 | 0.81 | 0.85 | 0.66 | 0.75 |

| | | | | | | | | | | | | | | | | |
|---|---|---|---|---|---|---|---|---|---|---|---|---|---|---|---|---|
| 1 | 166782438 | rs2064067 | 0.246 | -0.34 | -0.233 | -0.291 | 0.015 | 0.069 | 0.069 | 0.013 | 0.51 | 0.6 | 0.68 | 0.88 | 0.51 | 0.68 |
| 1 | 166794217 | rs2312236 | -1.223 | -0.286 | -0.394 | -0.179 | -0.034 | 0.045 | 0.045 | 0 | 0.89 | 1 | 0.97 | 0.98 | 0.91 | 0.8 |
| 1 | 166798529 | rs12040139 | 0.626 | -0.529 | -0.038 | -0.076 | -0.002 | -0.001 | -0.001 | 0 | 0.33 | 0.32 | 0.23 | 0.28 | 0.26 | 0.25 |
| 1 | 166808253 | rs7548191 | 0.46 | -0.196 | 0.389 | 0.175 | 0.001 | 0 | 0 | 0 | 0.5 | 0.49 | 0.37 | 0.28 | 0.43 | 0.33 |
| 1 | 166824580 | rs12117020 | 0.431 | -0.245 | 0.321 | 0.834 | 0.039 | 0.028 | 0.028 | 0 | 0.51 | 0.37 | 0.37 | 0.69 | 0.42 | 0.52 |
| 1 | 166824846 | rs10800269 | 0.361 | -0.253 | 0.353 | 0.857 | 0.033 | 0.037 | 0.037 | 0.013 | 0.5 | 0.39 | 0.31 | 0.3 | 0.42 | 0.33 |
| 1 | 166826655 | rs6675033 | 0.386 | -0.26 | 0.342 | 0.853 | 0.043 | 0.047 | 0.047 | 0.009 | 0.49 | 0.37 | 0.31 | 0.3 | 0.42 | 0.33 |
| 1 | 166827224 | rs6675750 | 0.393 | -0.254 | 0.335 | 0.847 | 0.042 | 0.047 | 0.047 | 0.005 | 0.49 | 0.37 | 0.31 | 0.3 | 0.42 | 0.33 |
| 1 | 166828957 | rs12562794 | 0.483 | 0.721 | 0.386 | 0.856 | 0.012 | 0.081 | 0.081 | 1.45 | 0.13 | 0.2 | 0.17 | 0.58 | 0.16 | 0.34 |
| 1 | 166830976 | rs17331024 | 0.647 | 0.789 | 0.406 | 0.838 | 0.033 | 0.037 | 0.037 | 0 | 0.49 | 0.38 | 0.31 | 0.3 | 0.42 | 0.33 |
| 1 | 166832809 | rs12029367 | 0.669 | 0.967 | 0.328 | 0.764 | 0.03 | 0.034 | 0.034 | 0.006 | 0.49 | 0.39 | 0.31 | 0.3 | 0.42 | 0.33 |
| 1 | 166838281 | rs2294195 | 0.867 | 0.906 | -0.01 | 0.91 | -0.015 | 0 | 0 | 0 | 0.12 | 0.07 | 0.17 | 0.57 | 0.17 | 0.34 |
| 1 | 166850207 | rs11583927 | -0.128 | 1.14 | 0.212 | 0.78 | -0.017 | -0.034 | -0.034 | 0.43 | 0.33 | 0.2 | 0.3 | 0.57 | 0.37 | 0.38 |
| 1 | 166852434 | rs6690192 | 0.877 | 1.32 | 0.419 | 0.771 | 0.019 | 0.04 | 0.04 | 0.017 | 0.51 | 0.58 | 0.34 | 0.3 | 0.47 | 0.34 |
| 1 | 166856059 | rs4656517 | -1.042 | 1.324 | 0.557 | 0.754 | 0.002 | 0 | 0 | 0 | 0.86 | 0.86 | 0.65 | 0.87 | 0.84 | 0.72 |
| 1 | 166860471 | rs2223221 | -0.209 | 1.313 | 0.478 | 0.755 | -0.022 | -0.039 | -0.039 | 0 | 0.34 | 0.24 | 0.31 | 0.57 | 0.38 | 0.38 |
| 1 | 166882737 | rs1053407 | 0.781 | 1.328 | 0.805 | 0.444 | 0.023 | 0.023 | 0.023 | 0 | 0.52 | 0.37 | 0.38 | 0.69 | 0.46 | 0.51 |
| 1 | 166885604 | rs11108 | 1.469 | 1.409 | 0.706 | 0.397 | -0.004 | 0.059 | 0.059 | 5.814 | 0.2 | 0.07 | 0.12 | 0 | 0.21 | 0.04 |
| 1 | 166899261 | rs4657614 | 0.522 | 1.515 | 0.836 | 0.396 | 0.001 | 0.001 | 0.001 | 0.027 | 0.54 | 0.56 | 0.4 | 0.69 | 0.47 | 0.52 |
| 1 | 166910831 | rs10489574 | 1.555 | 1.518 | 0.513 | 0.217 | -0.007 | 0.052 | 0.052 | 4.421 | 0.19 | 0.07 | 0.12 | 0 | 0.21 | 0.04 |
| 1 | 166914258 | rs6427047 | 0.237 | 1.593 | 0.434 | 0.357 | 0 | 0.001 | 0.001 | 0.055 | 0.14 | 0.19 | 0.13 | 0.17 | 0.18 | 0.17 |
| 1 | 166925806 | rs2075981 | 0.99 | 1.753 | 0.421 | 0.51 | 0.036 | -0.005 | -0.005 | 0.08 | 0.51 | 0.32 | 0.48 | 0.59 | 0.44 | 0.46 |
| 1 | 166929527 | rs2075982 | 0.885 | 1.385 | 0.556 | 0.473 | 0.021 | -0.003 | -0.003 | 0 | 0.52 | 0.27 | 0.35 | 0.6 | 0.47 | 0.47 |
| 1 | 166936503 | rs10918603 | 1.065 | 1.403 | 0.618 | 0.323 | 0.006 | 0.012 | 0.012 | 1.457 | 0.11 | 0.16 | 0.11 | 0.17 | 0.16 | 0.17 |
| 1 | 166940095 | rs7520897 | -0.12 | 1.417 | 0.804 | 0.244 | -0.001 | 0.001 | 0.001 | 0 | 0.33 | 0.64 | 0.27 | 0.17 | 0.38 | 0.22 |
| 1 | 166947681 | rs7545911 | 0.001 | 1.713 | 0.79 | 0.301 | 0.099 | 0.099 | 0.099 | 0.133 | 0.15 | 0.56 | 0.17 | 0.25 | 0.22 | 0.18 |
| 1 | 166948628 | rs10918604 | 1.142 | 1.42 | 0.385 | 0.271 | 0.008 | -0.035 | -0.035 | 0 | 0.08 | 0.01 | 0.05 | 0.34 | 0.04 | 0.29 |
| 1 | 166950458 | rs402031 | 0.816 | 1.394 | 0.381 | 0.209 | 0.036 | 0.004 | 0.004 | 0.013 | 0.52 | 0.29 | 0.34 | 0.48 | 0.45 | 0.48 |
| 1 | 166951869 | rs6688613 | 0.023 | 1.459 | 0.814 | 0.309 | 0.09 | 0.09 | 0.09 | 0.118 | 0.14 | 0.37 | 0.15 | 0.25 | 0.22 | 0.18 |
| 1 | 166957803 | rs10918607 | 1.167 | 1.468 | 0.796 | 0.314 | 0.005 | 0.012 | 0.012 | 2.64 | 0.1 | 0.13 | 0.11 | 0.25 | 0.18 | 0.18 |
| 1 | 166958710 | rs11578336 | 0.019 | 1.452 | 0.831 | 0.307 | 0.09 | 0.09 | 0.09 | 0.118 | 0.14 | 0.37 | 0.15 | 0.25 | 0.23 | 0.18 |
| 1 | 166974420 | rs1041238 | 1.182 | 1.421 | 0.776 | 0.192 | 0.021 | 0.113 | 0.113 | 0.739 | 0.09 | 0.22 | 0.11 | 0.25 | 0.12 | 0.15 |
| 1 | 166978035 | rs6693185 | -0.611 | 1.178 | 0.191 | -0.268 | -0.001 | -0.044 | -0.044 | 0 | 0.71 | 0.79 | 0.78 | 0.52 | 0.72 | 0.64 |
| 1 | 166980583 | rs1952313 | 1.17 | 1.287 | -0.204 | -0.303 | -0.05 | -0.066 | -0.066 | 0 | 0.13 | 0.03 | 0.39 | 0.28 | 0.18 | 0.32 |
| 1 | 166983171 | rs10918609 | 1.258 | 1.26 | -0.049 | -0.25 | -0.03 | -0.034 | -0.034 | 0 | 0.08 | 0.01 | 0.08 | 0.35 | 0.08 | 0.27 |
| 1 | 166989908 | rs4657619 | 0.464 | 1.139 | 0.233 | -0.597 | -0.05 | 0.008 | 0.008 | 2.923 | 0.32 | 0.1 | 0.49 | 0.28 | 0.36 | 0.39 |
| 1 | 166997021 | rs10918610 | 0.278 | 0.896 | 0.283 | -0.312 | 0.002 | 0.004 | 0.004 | 0 | 0.38 | 0.64 | 0.28 | 0.24 | 0.36 | 0.26 |
| 1 | 167005881 | rs6661553 | 0.188 | 0.952 | 0.269 | -0.273 | 0.003 | -0.008 | -0.008 | 0.104 | 0.38 | 0.65 | 0.34 | 0.51 | 0.37 | 0.31 |
| 1 | 167011602 | rs12143525 | 1.309 | 1.013 | 0.506 | -0.161 | 0.011 | 0.086 | 0.086 | 0.78 | 0.09 | 0.19 | 0.1 | 0.25 | 0.12 | 0.14 |
| 1 | 167011971 | rs7541843 | 0.029 | 0.993 | 0.409 | -0.222 | 0.039 | 0.347 | 0.347 | 0.02 | 0.14 | 0.46 | 0.18 | 0.51 | 0.16 | 0.2 |
| 1 | 167015184 | rs6698931 | -0.261 | 0.883 | 0.113 | -0.313 | -0.014 | -0.01 | -0.01 | 0 | 0.58 | 0.66 | 0.42 | 0.51 | 0.52 | 0.37 |
| 1 | 167016853 | rs6427053 | 0.555 | 0.87 | -0.399 | 0.004 | 0.005 | 0.036 | 0.036 | 1.421 | 0.11 | 0.15 | 0.11 | 0.25 | 0.16 | 0.14 |
| 1 | 167016898 | rs17433906 | 0.813 | 0.737 | -0.61 | -0.529 | 0.012 | 0.044 | 0.044 | 0 | 0.24 | 0.12 | 0.15 | 0 | 0.21 | 0.11 |
| 1 | 167021041 | rs4233414 | 0.007 | 0.946 | -0.408 | -0.254 | -0.001 | 0 | 0 | 0 | 0.62 | 0.41 | 0.71 | 0.46 | 0.59 | 0.69 |
| 1 | 167022262 | rs11736 | 0.007 | 0.946 | -0.408 | -0.254 | -0.001 | 0 | 0 | 0 | 0.62 | 0.41 | 0.71 | 0.46 | 0.59 | 0.69 |
| 1 | 167022381 | rs12461 | 0.153 | 0.918 | -0.439 | -0.32 | -0.003 | -0.014 | -0.014 | 0 | 0.6 | 0.32 | 0.7 | 0.45 | 0.57 | 0.68 |
| 1 | 167022701 | rs3753914 | 0.505 | 0.917 | -0.722 | -0.461 | -0.017 | -0.043 | -0.043 | 2.91 | 0.36 | 0.23 | 0.57 | 0.46 | 0.38 | 0.58 |
| 1 | 167024631 | rs6666045 | 0.474 | 0.875 | -0.791 | -0.491 | -0.04 | -0.051 | -0.051 | 0.011 | 0.37 | 0.18 | 0.58 | 0.44 | 0.41 | 0.6 |
| 1 | 167026832 | rs4657625 | 0.196 | 0.821 | -0.249 | -0.424 | -0.009 | -0.006 | -0.006 | 1.146 | 0.42 | 0.35 | 0.6 | 0.42 | 0.43 | 0.61 |
| 1 | 167034382 | rs4657629 | -0.429 | 0.749 | -0.216 | -0.464 | 0.001 | 0.02 | 0.02 | 0 | 0.68 | 0.65 | 0.74 | 0.42 | 0.67 | 0.7 |
| 1 | 167035148 | rs1887537 | -0.544 | 0.946 | -0.251 | -1.075 | -0.002 | 0.002 | 0.002 | 0 | 0.72 | 0.71 | 0.76 | 0.26 | 0.75 | 0.75 |
| 1 | 167035702 | rs3767430 | 0.547 | 1.117 | -0.751 | -1.535 | -0.085 | 0.079 | 0.079 | 0.429 | 0.41 | 0.13 | 0.55 | 0.23 | 0.46 | 0.65 |
| 1 | 167036714 | rs12404508 | 0.975 | 1.124 | -0.783 | -1.658 | -0.058 | 0.231 | 0.231 | 1.071 | 0.37 | 0.09 | 0.49 | 0.02 | 0.36 | 0.58 |
| 1 | 167037025 | rs10918613 | 1.859 | 0.946 | -0.672 | -1.641 | 0.028 | 0.052 | 0.052 | 0.107 | 0.19 | 0.05 | 0.09 | 0 | 0.16 | 0.08 |
| 1 | 167040788 | rs2312489 | 0.325 | 0.862 | -0.399 | -1.491 | -0.004 | -0.006 | -0.006 | 0 | 0.27 | 0.31 | 0.24 | 0.77 | 0.24 | 0.25 |
| 1 | 167045333 | rs10918616 | -0.33 | 0.576 | -0.004 | -1.836 | -0.004 | -0.123 | -0.123 | 0.794 | 0.35 | 0.58 | 0.24 | 0 | 0.35 | 0.12 |
| 1 | 167050565 | rs12131781 | 0.124 | 0.759 | 0.061 | -1.721 | 0 | -0.097 | -0.097 | 0.087 | 0.3 | 0.34 | 0.2 | 0 | 0.33 | 0.12 |
| 1 | 167051092 | rs10918621 | 0.211 | 0.645 | 0.198 | -1.754 | -0.015 | -0.178 | -0.178 | 0.304 | 0.28 | 0.43 | 0.17 | 0 | 0.27 | 0.12 |
| 1 | 167054980 | rs1041236 | 0.265 | 0.743 | -0.221 | -1.583 | -0.004 | -0.004 | -0.004 | 0 | 0.28 | 0.2 | 0.27 | 0.73 | 0.29 | 0.34 |
| 1 | 167059760 | rs2281962 | 0.201 | 0.641 | -0.087 | -2.059 | 0.002 | 0.013 | 0.013 | 1.688 | 0.46 | 0.39 | 0.3 | 0.17 | 0.45 | 0.23 |
| 1 | 167059833 | rs2281960 | -0.27 | 0.546 | 0.097 | -2.097 | -0.001 | -0.002 | -0.002 | 0.116 | 0.52 | 0.55 | 0.34 | 0.17 | 0.53 | 0.23 |
| 1 | 167063122 | rs10918622 | -0.198 | 0.806 | 0.572 | -1.507 | 0.003 | 0.053 | 0.053 | 0 | 0.07 | 0.09 | 0.06 | 0.16 | 0.1 | 0.07 |
| 1 | 167069155 | rs11588401 | 0.818 | 0.67 | 0.573 | -1.693 | 0.009 | -0.067 | -0.067 | 0.02 | 0.17 | 0.21 | 0.16 | 0.01 | 0.2 | 0.07 |
| 1 | 167070658 | rs1590721 | -0.493 | 0.645 | 0.607 | -1.691 | 0.021 | -0.012 | -0.012 | 0 | 0.25 | 0.4 | 0.22 | 0.17 | 0.29 | 0.14 |
| 1 | 167071975 | rs9287077 | -0.42 | 0.699 | 0.555 | -1.786 | 0.012 | -0.012 | -0.012 | 0 | 0.24 | 0.32 | 0.21 | 0.16 | 0.3 | 0.14 |
| 1 | 167074750 | rs16858452 | 0.92 | 0.755 | 0.176 | -1.771 | -0.014 | 0.01 | 0.01 | 0 | 0.16 | 0.11 | 0.14 | 0.01 | 0.19 | 0.07 |
| 1 | 167077805 | rs6691731 | 0.43 | 0.747 | 0.244 | -1.968 | -0.004 | 0.007 | 0.007 | 1.867 | 0.27 | 0.56 | 0.28 | 0.74 | 0.25 | 0.34 |
| 1 | 167082869 | rs12122365 | -1.458 | 0.771 | -0.056 | -2.057 | 0.03 | 0.226 | 0.226 | 0 | 0.52 | 0.8 | 0.5 | 0.89 | 0.56 | 0.49 |
| 1 | 167084813 | rs4657630 | -0.902 | 0.8 | -0.034 | -1.949 | -0.007 | 0.025 | 0.025 | 1.018 | 0.43 | 0.65 | 0.41 | 0.75 | 0.44 | 0.41 |

| | | | | | | | | | | | | | | | | |
|---|---|---|---|---|---|---|---|---|---|---|---|---|---|---|---|---|
| 1 | 167088473 | rs1410244 | 0.467 | 0.714 | -0.64 | -1.673 | 0.002 | 0.013 | 0.013 | 1.139 | 0.25 | 0.39 | 0.25 | 0.74 | 0.24 | 0.36 |
| 1 | 167091859 | rs267741 | -1.033 | 0.568 | -1.581 | -1.829 | 0.013 | 0.226 | 0.226 | 0 | 0.57 | 0.87 | 0.52 | 0.89 | 0.61 | 0.51 |
| 1 | 167095163 | rs267745 | -1.117 | 0.24 | -1.721 | -1.781 | -0.038 | 0.216 | 0.216 | 3.453 | 0.55 | 0.89 | 0.46 | 0.87 | 0.48 | 0.37 |
| 1 | 167095881 | rs3795605 | 0.679 | 0.04 | -1.226 | -1.659 | -0.012 | 0.019 | 0.019 | 1.022 | 0.21 | 0.27 | 0.2 | 0.74 | 0.17 | 0.24 |
| 1 | 167096931 | rs267746 | -0.187 | 0.278 | -0.615 | -1.527 | 0.015 | 0.007 | 0.007 | 0.193 | 0.16 | 0.41 | 0.14 | 0.13 | 0.16 | 0.08 |
| 1 | 167097739 | rs2281959 | -0.298 | 0.206 | -1.288 | -1.644 | 0.018 | 0.017 | 0.017 | 0.657 | 0.37 | 0.28 | 0.29 | 0.74 | 0.31 | 0.29 |
| 1 | 167097945 | rs267747 | -0.995 | 0.209 | -0.455 | -1.323 | 0.004 | -0.003 | -0.003 | 0 | 0.17 | 0.58 | 0.17 | 0.13 | 0.16 | 0.08 |
| 1 | 167100361 | rs7519927 | -0.886 | 0.066 | -1.3 | -1.773 | -0.073 | 0.12 | 0.12 | 0.83 | 0.65 | 0.87 | 0.51 | 0.9 | 0.57 | 0.44 |
| 1 | 167102492 | rs11806848 | -0.375 | -0.384 | -1.731 | -2.805 | -0.02 | 0.238 | 0.238 | 0.101 | 0.33 | 0.04 | 0.16 | 0.02 | 0.26 | 0.25 |
| 1 | 167103023 | rs7540969 | 0.093 | -0.366 | -1.262 | -2.461 | -0.036 | -0.025 | -0.025 | 0 | 0.23 | 0.17 | 0.25 | 0.66 | 0.35 | 0.27 |
| 1 | 167103945 | rs988778 | -1.293 | -0.665 | -0.628 | -2.634 | -0.006 | 0.016 | 0.016 | 0.455 | 0.55 | 0.66 | 0.48 | 0.89 | 0.57 | 0.43 |
| 1 | 167104790 | rs12077809 | -0.207 | -0.456 | -0.363 | -2.302 | -0.012 | 0.068 | 0.068 | 0.897 | 0.18 | 0.29 | 0.1 | 0.53 | 0.13 | 0.21 |
| 1 | 167105038 | rs12723100 | 0.866 | -0.561 | -0.549 | -2.46 | -0.016 | 0.004 | 0.004 | 0.105 | 0.16 | 0.09 | 0.2 | 0.1 | 0.31 | 0.1 |
| 1 | 167110188 | rs12129059 | 0.661 | -0.74 | -0.373 | -2.519 | 0.004 | -0.005 | -0.005 | 0.221 | 0.16 | 0.11 | 0.08 | 0.01 | 0.12 | 0.08 |
| 1 | 167111293 | rs11808527 | 2.274 | -0.92 | -0.67 | -2.744 | -0.079 | 0.135 | 0.135 | 0.252 | 0.32 | 0.05 | 0.43 | 0.11 | 0.33 | 0.43 |
| 1 | 167111494 | rs10918638 | 0.685 | -0.825 | -0.399 | -2.773 | 0.011 | 0.024 | 0.024 | 0.752 | 0.6 | 0.31 | 0.64 | 0.22 | 0.64 | 0.57 |
| 1 | 167119523 | rs12140642 | -0.811 | -0.789 | -0.068 | -2.881 | -0.002 | 0.006 | 0.006 | 0 | 0.34 | 0.34 | 0.26 | 0.09 | 0.42 | 0.18 |
| 1 | 167121665 | rs4656525 | 0.699 | -0.837 | -0.355 | -2.986 | -0.039 | 0.093 | 0.093 | 0 | 0.15 | 0.01 | 0.09 | 0 | 0.13 | 0.14 |
| 1 | 167121814 | rs10753750 | 1.276 | -0.902 | 0.277 | -2.934 | -0.002 | 0.037 | 0.037 | 0 | 0.47 | 0.35 | 0.54 | 0.66 | 0.45 | 0.65 |
| 1 | 167122429 | rs12143783 | -0.303 | -0.936 | 0.37 | -3.052 | 0.008 | 0.062 | 0.062 | 0 | 0.27 | 0.12 | 0.14 | 0 | 0.22 | 0.19 |
| 1 | 167123154 | rs11590513 | 1.259 | -0.786 | 0.613 | -2.928 | -0.009 | -0.059 | -0.059 | 0.912 | 0.11 | 0.25 | 0.09 | 0.01 | 0.13 | 0.04 |
| 1 | 167124187 | rs11591049 | 0.904 | -0.728 | 0.466 | -2.927 | 0 | -0.001 | -0.001 | 1.049 | 0.11 | 0.12 | 0.12 | 0.06 | 0.18 | 0.07 |
| 1 | 167124834 | rs6427069 | 0.298 | -0.758 | 0.443 | -2.888 | -0.001 | -0.003 | -0.003 | 0.169 | 0.63 | 0.62 | 0.69 | 0.71 | 0.62 | 0.78 |
| 1 | 167126250 | rs869714 | 0.77 | -0.758 | 0.373 | -2.707 | 0 | -0.007 | -0.007 | 0 | 0.48 | 0.53 | 0.59 | 0.71 | 0.49 | 0.64 |
| 1 | 167129131 | rs6679415 | 1.189 | -0.771 | 0.603 | -2.648 | -0.005 | -0.033 | -0.033 | 0 | 0.11 | 0.17 | 0.09 | 0.01 | 0.13 | 0.04 |
| 1 | 167131321 | rs1032608 | -0.434 | -0.796 | 0.537 | -2.637 | -0.003 | 0.166 | 0.166 | 0 | 0.85 | 0.61 | 0.84 | 0.4 | 0.84 | 0.79 |
| 1 | 167132882 | rs1027493 | 1.525 | -0.505 | 0.023 | -2.594 | -0.022 | 0.11 | 0.11 | 0.354 | 0.33 | 0.14 | 0.43 | 0.11 | 0.32 | 0.42 |
| 1 | 167137645 | rs16858608 | -0.414 | 0.277 | -0.004 | -2.079 | 0 | 0.001 | 0.001 | 0.011 | 0.14 | 0.17 | 0.18 | 0.72 | 0.24 | 0.24 |
| 1 | 167137872 | rs12409740 | 1.597 | -0.061 | -0.183 | -2.485 | -0.007 | 0.027 | 0.027 | 0 | 0.32 | 0.25 | 0.48 | 0.05 | 0.34 | 0.45 |
| 1 | 167137949 | rs4657642 | -0.935 | -0.084 | -0.039 | -2.338 | -0.008 | 0.008 | 0.008 | 2.041 | 0.26 | 0.64 | 0.16 | 0.66 | 0.18 | 0.27 |
| 1 | 167138024 | rs4657643 | 0.131 | -0.27 | -0.375 | -2.514 | -0.018 | -0.074 | -0.074 | 0 | 0.12 | 0.34 | 0.06 | 0 | 0.05 | 0.04 |
| 1 | 167138099 | rs4657644 | -0.304 | -0.183 | -0.547 | -2.483 | -0.063 | -0.139 | -0.139 | 3.724 | 0.18 | 0.4 | 0.07 | 0 | 0.08 | 0.08 |
| 1 | 167140959 | rs6691080 | -0.983 | 0.413 | -1.316 | -2.233 | -0.019 | 0.066 | 0.066 | 0 | 0.61 | 0.73 | 0.42 | 0.88 | 0.47 | 0.48 |
| 1 | 167142822 | rs17348602 | 0.502 | 0.397 | -2.164 | -2.433 | -0.038 | 0.251 | 0.251 | 0.077 | 0.46 | 0.13 | 0.52 | 0.07 | 0.45 | 0.56 |
| 1 | 167143896 | rs10918653 | 1.242 | 0.584 | -2.096 | -2.247 | -0.071 | 0.147 | 0.147 | 0.038 | 0.3 | 0.08 | 0.46 | 0.06 | 0.33 | 0.45 |
| 1 | 167144171 | rs12048211 | 1.217 | 0.588 | -2.096 | -2.243 | -0.058 | 0.144 | 0.144 | 0.01 | 0.31 | 0.08 | 0.46 | 0.06 | 0.32 | 0.45 |
| 1 | 167146372 | rs12739886 | -0.237 | 0.651 | -1.258 | -1.602 | 0.029 | -0.004 | -0.004 | 0 | 0.21 | 0.3 | 0.28 | 0.24 | 0.33 | 0.16 |
| 1 | 167150538 | rs7548201 | 0.158 | 0.675 | -1.266 | -1.637 | 0.002 | -0.001 | -0.001 | 0 | 0.17 | 0.18 | 0.25 | 0.25 | 0.3 | 0.17 |
| 1 | 167150734 | rs7536570 | -0.551 | 0.351 | -1.447 | -1.909 | 0.02 | 0.039 | 0.039 | 0 | 0.86 | 0.9 | 0.93 | 1 | 0.9 | 0.95 |
| 1 | 167152574 | rs10494471 | 0.85 | 0.467 | -1.495 | -1.845 | -0.018 | -0.058 | -0.058 | 0 | 0.1 | 0.01 | 0.1 | 0.68 | 0.15 | 0.18 |
| 1 | 167154534 | rs12144916 | 1.601 | 0.281 | -1.42 | -1.921 | -0.033 | 0.211 | 0.211 | 0.267 | 0.42 | 0.08 | 0.49 | 0.07 | 0.39 | 0.53 |
| 1 | 167156500 | rs9803659 | -1.326 | 0.441 | -1.043 | -1.755 | 0.002 | 0.037 | 0.037 | 0 | 0.39 | 0.36 | 0.3 | 0.25 | 0.36 | 0.24 |
| 1 | 167156816 | rs9287078 | -0.035 | 0.459 | -1.042 | -1.745 | 0.005 | 0 | 0 | 0 | 0.17 | 0.21 | 0.22 | 0.25 | 0.23 | 0.15 |
| 1 | 167158183 | rs6658631 | 1.104 | 0.408 | -1.127 | -1.569 | 0 | 0.003 | 0.003 | 0 | 0.56 | 0.51 | 0.67 | 0.75 | 0.61 | 0.76 |
| 1 | 167158927 | rs16858688 | -0.748 | -0.235 | -1.249 | -0.669 | 0.019 | 0.008 | 0.008 | 0 | 0.22 | 0.21 | 0.16 | 0.22 | 0.13 | 0.11 |
| 1 | 167161095 | rs6699424 | -0.453 | 0.315 | -0.967 | -0.766 | -0.003 | -0.036 | -0.036 | 0.568 | 0.36 | 0.46 | 0.27 | 0.13 | 0.35 | 0.23 |
| 1 | 167168408 | rs7529557 | -0.479 | -0.129 | -1.471 | -1.262 | -0.048 | 0.034 | 0.034 | 0.129 | 0.62 | 0.85 | 0.36 | 0.29 | 0.49 | 0.36 |
| 1 | 167171571 | rs12127503 | -0.025 | -0.455 | -1.43 | -1.466 | -0.001 | -0.007 | -0.007 | 0.057 | 0.11 | 0.08 | 0.07 | 0.15 | 0.11 | 0.09 |
| 1 | 167178149 | rs2173298 | 0.629 | 0.169 | -2.011 | -1.975 | -0.129 | 0.006 | 0.006 | 0.662 | 0.36 | 0.06 | 0.61 | 0.7 | 0.48 | 0.65 |
| 1 | 167184952 | rs7549340 | -0.407 | 0.383 | -1.757 | -2.198 | 0.087 | 0.17 | 0.17 | 0.129 | 0.52 | 0.31 | 0.76 | 0.88 | 0.69 | 0.75 |
| 1 | 167198536 | rs1021621 | -0.199 | -0.133 | -1.171 | -2.415 | -0.001 | -0.007 | -0.007 | 0 | 0.52 | 0.48 | 0.32 | 0.27 | 0.49 | 0.33 |
| 1 | 167218385 | rs12068411 | 0.646 | -0.27 | -0.779 | -2.474 | 0.017 | 0.019 | 0.019 | 0.537 | 0.4 | 0.3 | 0.19 | 0.11 | 0.28 | 0.22 |
| 1 | 167226854 | rs607975 | -0.736 | -0.234 | -0.786 | -2.475 | 0.003 | -0.004 | -0.004 | 0 | 0.59 | 0.64 | 0.8 | 0.89 | 0.72 | 0.78 |
| 1 | 167233474 | rs4657652 | 1.086 | -0.199 | -0.925 | -2.475 | -0.149 | 0.004 | 0.004 | 0.649 | 0.35 | 0.05 | 0.61 | 0.72 | 0.49 | 0.66 |
| 1 | 167247473 | rs10800299 | -0.269 | -0.426 | -0.869 | -2.441 | 0.099 | 0.172 | 0.172 | 0.103 | 0.52 | 0.31 | 0.76 | 0.89 | 0.7 | 0.75 |
| 1 | 167248664 | rs11584337 | 0.242 | -0.366 | -0.865 | -2.425 | -0.028 | -0.004 | -0.004 | 1.641 | 0.11 | 0.06 | 0.11 | 0.14 | 0.21 | 0.1 |
| 1 | 167250080 | rs7532692 | 0.263 | -0.371 | -0.869 | -2.42 | -0.002 | -0.001 | -0.001 | 0 | 0.11 | 0.12 | 0.12 | 0.14 | 0.21 | 0.09 |
| 1 | 167256468 | rs266820 | 0.409 | -0.427 | -0.855 | -2.427 | 0.055 | 0.011 | 0.011 | 0.411 | 0.07 | 0.38 | 0.04 | 0.01 | 0.03 | 0.02 |
| 1 | 167259755 | rs4656538 | 1.032 | -0.39 | -1 | -2.423 | -0.117 | 0.013 | 0.013 | 0.476 | 0.36 | 0.08 | 0.62 | 0.7 | 0.49 | 0.66 |
| 1 | 167339632 | rs3736846 | -0.95 | -0.927 | -0.738 | -2.394 | 0.008 | 0.009 | 0.009 | 0 | 0.65 | 0.74 | 0.81 | 0.89 | 0.74 | 0.79 |
| 1 | 167355192 | rs2093658 | 0.309 | -0.821 | -1.13 | -2.234 | 0.096 | 0.169 | 0.169 | 0.442 | 0.46 | 0.2 | 0.73 | 0.86 | 0.69 | 0.73 |
| 1 | 167391904 | rs2949664 | 0.93 | -0.48 | -0.44 | -1.543 | -0.069 | -0.063 | -0.063 | 0 | 0.87 | 0.61 | 0.94 | 0.98 | 0.98 | 0.99 |
| 1 | 167397078 | rs6682013 | -0.636 | 0.575 | -1.524 | -1.899 | -0.106 | 0.174 | 0.174 | 0.782 | 0.27 | 0.03 | 0.25 | 0 | 0.34 | 0.14 |
| 1 | 167399503 | rs870873 | -0.924 | 0.238 | -0.317 | -1.01 | 0.017 | 0.054 | 0.054 | 0 | 0.09 | 0.16 | 0.1 | 0.26 | 0.12 | 0.09 |
| 1 | 167399734 | rs870875 | 0.141 | 0.487 | -0.126 | 0.787 | -0.004 | -0.005 | -0.005 | 0 | 0.28 | 0.3 | 0.53 | 0.42 | 0.37 | 0.51 |
| 1 | 167401254 | rs16859030 | -0.167 | 0.252 | 0.655 | 1.421 | -0.001 | 0.161 | 0.161 | 0.252 | 0.07 | 0.22 | 0.05 | 0.23 | 0.05 | 0.12 |
| 1 | 167403625 | rs953809 | 0.474 | 0.338 | 0.342 | 1.071 | 0.021 | 0.07 | 0.07 | 0 | 0.91 | 0.8 | 0.89 | 0.74 | 0.88 | 0.92 |
| 1 | 167408073 | rs2258497 | 0.8 | 0.271 | 0.238 | 1.02 | -0.001 | 0.02 | 0.02 | 0 | 0.88 | 0.82 | 0.89 | 0.75 | 0.87 | 0.91 |
| 1 | 167408565 | rs12141641 | -0.6 | 0.474 | 0.286 | 1.343 | 0.011 | 0.08 | 0.08 | 0 | 0.08 | 0.07 | 0.1 | 0.33 | 0.13 | 0.1 |

| | | | | | | | | | | | | | | | | |
|---|---|---|---|---|---|---|---|---|---|---|---|---|---|---|---|---|
| 1 | 167415791 | rs1554669 | -1.169 | -0.058 | 0.345 | 1.194 | -0.01 | 0.024 | 0.024 | 0 | 0.1 | 0.27 | 0.12 | 0.13 | 0.07 | 0.04 |
| 1 | 167427247 | rs7523907 | -0.835 | -0.33 | 0.085 | 0.819 | 0.001 | -0.035 | -0.035 | 0.005 | 0.52 | 0.58 | 0.71 | 0.94 | 0.58 | 0.75 |
| 1 | 167431857 | rs10918693 | -0.635 | 0.03 | 0.236 | 1.393 | -0.023 | 0.16 | 0.16 | 0 | 0.11 | 0.29 | 0.15 | 0.36 | 0.09 | 0.22 |
| 1 | 167432894 | rs1723015 | 0.434 | 0.105 | -0.599 | 1.01 | -0.012 | -0.012 | -0.012 | 1.358 | 0.55 | 0.38 | 0.43 | 0.52 | 0.5 | 0.39 |
| 1 | 167437953 | rs10918695 | -0.197 | 0.695 | -1.322 | 0.372 | -0.063 | -0.004 | -0.004 | 0 | 0.33 | 0.13 | 0.47 | 0.31 | 0.43 | 0.45 |
| 1 | 167443423 | rs2480678 | 0.446 | 0.124 | -1.643 | -0.117 | 0.085 | 0.001 | 0.001 | 0.492 | 0.45 | 0.13 | 0.29 | 0.52 | 0.36 | 0.34 |
| 1 | 167445001 | rs2995093 | -0.174 | 1.147 | -0.961 | -0.491 | 0.013 | -0.001 | -0.001 | 0.018 | 0.65 | 0.54 | 0.67 | 0.59 | 0.57 | 0.68 |
| 1 | 167448392 | rs7518703 | 0.287 | 1.443 | -1.132 | -0.396 | 0.02 | -0.007 | -0.007 | 0 | 0.05 | 0.05 | 0.31 | 0.02 | 0.12 | 0.15 |
| 1 | 167449608 | rs1214609 | -0.925 | 1.401 | -0.91 | -0.451 | 0.057 | -0.049 | -0.049 | 0 | 0.11 | 0.26 | 0.36 | 0.02 | 0.18 | 0.2 |
| 1 | 167451115 | rs12737372 | 1.16 | 0.606 | -1.983 | -0.484 | 0.039 | -0.091 | -0.091 | 0 | 0.33 | 0.06 | 0.25 | 0.49 | 0.27 | 0.29 |
| 1 | 167453334 | rs16859085 | -0.456 | 1.243 | -1.293 | -0.329 | 0.011 | -0.011 | -0.011 | 0 | 0.14 | 0.15 | 0.32 | 0.09 | 0.17 | 0.28 |
| 1 | 167455145 | rs858553 | 0.273 | 1.153 | -1.554 | -0.568 | -0.048 | 0.065 | 0.065 | 0.551 | 0.71 | 0.87 | 0.5 | 0.88 | 0.61 | 0.58 |
| 1 | 167457824 | rs863455 | 0.331 | -0.27 | -1.43 | -0.101 | 0.113 | 0.057 | 0.057 | 0.417 | 0.46 | 0.17 | 0.26 | 0.66 | 0.31 | 0.28 |
| 1 | 167463906 | rs6670426 | -0.728 | -0.047 | -0.593 | -0.096 | -0.033 | -0.031 | -0.031 | 0 | 0.21 | 0.33 | 0.08 | 0.12 | 0.11 | 0.38 |
| 1 | 167463916 | rs704852 | -1.18 | 0.945 | -0.51 | -0.327 | 0.012 | -0.024 | -0.024 | 0 | 0.17 | 0.25 | 0.31 | 0.08 | 0.2 | 0.26 |
| 1 | 167466121 | rs10918706 | 0.313 | 0.417 | -1.225 | -0.012 | 0.039 | 0.04 | 0.04 | 0.019 | 0.38 | 0.1 | 0.16 | 0.32 | 0.3 | 0.46 |
| 1 | 167467298 | rs858543 | 0.003 | 0.657 | -0.371 | 0.255 | 0.009 | -0.044 | -0.044 | 0 | 0.72 | 0.77 | 0.76 | 0.51 | 0.75 | 0.74 |
| 1 | 167467782 | rs858542 | -1.52 | 1.166 | 0.821 | -0.001 | 0.157 | 0.065 | 0.065 | 1.5 | 0.06 | 0.42 | 0.04 | 0 | 0.05 | 0.02 |
| 1 | 167468864 | rs704853 | -1.385 | 1.245 | 0.244 | 0.33 | 0.036 | 0.106 | 0.106 | 0 | 0.09 | 0.23 | 0.1 | 0.18 | 0.1 | 0.08 |
| 1 | 167475871 | rs1799704 | -0.999 | 1.965 | -0.022 | -0.215 | 0.034 | -0.013 | -0.013 | 0 | 0.15 | 0.25 | 0.46 | 0.08 | 0.32 | 0.22 |
| 1 | 167477937 | rs2982484 | 0.75 | 1.047 | -0.421 | -0.636 | 0.015 | 0.007 | 0.007 | 0 | 0.43 | 0.61 | 0.33 | 0.62 | 0.36 | 0.37 |
| 1 | 167479765 | rs858550 | -0.288 | 0.685 | -0.664 | -0.733 | 0.035 | 0.038 | 0.038 | 0 | 0.68 | 0.87 | 0.82 | 0.78 | 0.76 | 0.61 |
| 1 | 167479987 | rs12095738 | 0.286 | 0.584 | -0.556 | -0.603 | 0.021 | 0.003 | 0.003 | 0 | 0.23 | 0.1 | 0.11 | 0.22 | 0.12 | 0.34 |
| 1 | 167484888 | rs2982479 | -0.375 | 0.441 | 0.154 | -0.735 | 0 | 0.067 | 0.067 | 0 | 0.12 | 0.35 | 0.06 | 0.14 | 0.05 | 0.1 |
| 1 | 167485375 | rs1723012 | -0.384 | 0.263 | 0.665 | -0.794 | -0.054 | 0.067 | 0.067 | 0.23 | 0.11 | 0.56 | 0.05 | 0.14 | 0.02 | 0.09 |
| 1 | 167487880 | rs3820390 | 0.839 | 0.274 | 0.349 | -0.83 | -0.009 | -0.029 | -0.029 | 0 | 0.1 | 0.01 | 0.04 | 0.09 | 0.07 | 0.03 |
| 1 | 167490596 | rs12131537 | 0.725 | 0.211 | 0.303 | -1.05 | 0.027 | 0.095 | 0.095 | 0 | 0.22 | 0.06 | 0.11 | 0.03 | 0.14 | 0.24 |
| 1 | 167494139 | rs4657666 | -0.206 | 0.67 | 0.313 | -1.135 | -0.08 | 0.113 | 0.113 | 0 | 0.31 | 0.11 | 0.53 | 0.09 | 0.42 | 0.44 |
| 1 | 167494622 | rs2794984 | -1.164 | 0.574 | 0.457 | -1.148 | -0.037 | 0.118 | 0.118 | 0 | 0.45 | 0.77 | 0.6 | 0.23 | 0.45 | 0.54 |
| 1 | 167495240 | rs858484 | 0.198 | 0.627 | 0.95 | -1.055 | -0.029 | 0.075 | 0.075 | 0 | 0.1 | 0.3 | 0.05 | 0.14 | 0.03 | 0.09 |
| 1 | 167496303 | rs6664958 | -0.573 | 1.087 | 0.607 | -0.937 | 0.162 | -0.038 | -0.038 | 0 | 0.14 | 0.41 | 0.45 | 0.06 | 0.27 | 0.2 |
| 1 | 167497574 | rs2995062 | -0.891 | 1.074 | 0.326 | -0.788 | -0.032 | 0.056 | 0.056 | 0 | 0.45 | 0.75 | 0.62 | 0.33 | 0.45 | 0.53 |
| 1 | 167497812 | rs1618879 | -0.235 | 0.676 | 0.577 | -0.33 | -0.018 | -0.015 | -0.015 | 0.427 | 0.39 | 0.47 | 0.18 | 0.25 | 0.23 | 0.34 |
| 1 | 167499599 | rs1003815 | -0.158 | 1.767 | 0.074 | -0.052 | -0.002 | 0.031 | 0.031 | 0.069 | 0.89 | 0.91 | 0.92 | 1 | 0.87 | 0.93 |
| 1 | 167500815 | rs704859 | 0.74 | 2.077 | 0.188 | 0.004 | 0.053 | 0.001 | 0.001 | 0 | 0.83 | 0.66 | 0.74 | 0.83 | 0.69 | 0.77 |
| 1 | 167501611 | rs858493 | 0.646 | 1.137 | -0.05 | -0.35 | 0.001 | -0.082 | -0.082 | 0.043 | 0.76 | 0.45 | 0.83 | 0.92 | 0.78 | 0.91 |
| 1 | 167502112 | rs858492 | -0.123 | 1.279 | -0.991 | -0.47 | -0.044 | 0.036 | 0.036 | 1.182 | 0.35 | 0.19 | 0.64 | 0.8 | 0.5 | 0.58 |
| 1 | 167502598 | rs858490 | -0.204 | 1.007 | -1.121 | 0.005 | -0.001 | 0.002 | 0.002 | 0.321 | 0.5 | 0.53 | 0.53 | 0.56 | 0.56 | 0.5 |
| 1 | 167502914 | rs10918709 | -0.171 | 1.294 | -1.85 | -0.461 | -0.092 | -0.06 | -0.06 | 0 | 0.27 | 0.03 | 0.33 | 0.29 | 0.36 | 0.22 |
| 1 | 167503548 | rs7539688 | 0.606 | 0.639 | -1.551 | -1.077 | 0.001 | 0.027 | 0.027 | 0 | 0.55 | 0.32 | 0.56 | 0.67 | 0.57 | 0.71 |
| 1 | 167503778 | rs7527911 | 1.133 | 0.371 | -2.062 | -1.269 | -0.002 | 0 | 0 | 0 | 0.24 | 0.2 | 0.3 | 0.17 | 0.27 | 0.31 |
| 1 | 167507857 | rs16859159 | 0.748 | -0.175 | -1.918 | -1.407 | -0.011 | -0.025 | -0.025 | 0 | 0.15 | 0.1 | 0.28 | 0.28 | 0.21 | 0.15 |
| 1 | 167509563 | rs35591543 | -0.5 | -0.403 | -1.742 | -1.576 | 0.006 | -0.005 | -0.005 | 0 | 0.34 | 0.2 | 0.35 | 0.3 | 0.34 | 0.28 |
| 1 | 167510837 | rs2860662 | -1.013 | -0.417 | -1.176 | -1.595 | 0.012 | -0.038 | -0.038 | 0 | 0.92 | 0.85 | 0.96 | 1 | 0.95 | 0.99 |
| 1 | 167510996 | rs2143300 | -1.013 | -0.417 | -1.155 | -1.595 | 0.013 | -0.039 | -0.039 | 0 | 0.92 | 0.84 | 0.96 | 1 | 0.95 | 0.99 |
| 1 | 167511256 | rs2031131 | 1.613 | -0.381 | -1.091 | -1.594 | 0.01 | -0.032 | -0.032 | 0 | 0.12 | 0.07 | 0.09 | 0.24 | 0.11 | 0.08 |
| 1 | 167512558 | rs3820393 | 0.838 | -0.425 | -1.058 | -1.597 | -0.021 | -0.044 | -0.044 | 0 | 0.15 | 0.08 | 0.28 | 0.28 | 0.21 | 0.15 |
| 1 | 167512580 | rs4657667 | -0.983 | -0.397 | -0.898 | -1.565 | 0.012 | -0.038 | -0.038 | 0.007 | 0.92 | 0.86 | 0.96 | 1 | 0.95 | 0.99 |
| 1 | 167513159 | rs3753921 | 1.45 | -0.257 | -0.889 | -1.532 | -0.002 | 0.001 | 0.001 | 0 | 0.25 | 0.23 | 0.33 | 0.18 | 0.31 | 0.31 |
| 1 | 167513418 | rs1229430 | 0.782 | 0.406 | -0.921 | -1.249 | -0.001 | 0.015 | 0.015 | 0 | 0.59 | 0.65 | 0.6 | 0.7 | 0.6 | 0.7 |
| 1 | 167515272 | rs7513428 | -0.205 | 0.336 | -0.815 | -1.313 | -0.004 | -0.05 | -0.05 | 0 | 0.77 | 0.73 | 0.9 | 0.98 | 0.85 | 0.87 |
| 1 | 167520464 | rs3767443 | 1.351 | -0.162 | -1.189 | -1.254 | -0.068 | 0.042 | 0.042 | 0 | 0.29 | 0.05 | 0.31 | 0.19 | 0.3 | 0.34 |
| 1 | 167523387 | rs1737487 | 0.249 | -0.156 | -0.601 | -1.171 | -0.019 | -0.059 | -0.059 | 0.033 | 0.2 | 0.29 | 0.07 | 0.02 | 0.11 | 0.13 |
| 1 | 167525350 | rs1005298 | 0.893 | -0.073 | -0.57 | -1.074 | 0 | 0.018 | 0.018 | 0 | 0.47 | 0.41 | 0.53 | 0.7 | 0.52 | 0.7 |
| 1 | 167527216 | rs1737481 | -0.958 | -0.119 | -0.68 | -0.974 | -0.001 | -0.003 | -0.003 | 0 | 0.47 | 0.53 | 0.43 | 0.3 | 0.43 | 0.29 |
| 1 | 167529696 | rs10489192 | 0.369 | 0.477 | -1.185 | -1.268 | 0.001 | -0.017 | -0.017 | 0.003 | 0.11 | 0.09 | 0.06 | 0 | 0.11 | 0.01 |
| 1 | 167530546 | rs1737476 | -0.047 | 0.631 | -1.227 | -0.883 | 0 | 0.018 | 0.018 | 0 | 0.22 | 0.17 | 0.27 | 0.07 | 0.21 | 0.19 |
| 1 | 167532812 | rs12096911 | 0.519 | 0.919 | -1.13 | -0.577 | 0.007 | 0.04 | 0.04 | 0 | 0.38 | 0.3 | 0.33 | 0.21 | 0.34 | 0.32 |
| 1 | 167532944 | rs17477053 | 0.179 | 1.314 | -1.13 | 0.617 | -0.012 | -0.056 | -0.056 | 0.978 | 0.11 | 0.01 | 0.09 | 0.16 | 0.11 | 0.07 |
| 1 | 167533014 | rs909935 | 0.221 | 1.507 | -0.955 | -0.017 | -0.015 | 0.025 | 0.025 | 0 | 0.69 | 0.9 | 0.69 | 0.24 | 0.64 | 0.46 |
| 1 | 167537676 | rs16859250 | -1.097 | 1.996 | 0.787 | 0.88 | 0.021 | 0.043 | 0.043 | 0 | 0.16 | 0.23 | 0.17 | 0.27 | 0.21 | 0.2 |
| 1 | 167538798 | rs6697794 | -1.612 | 2.045 | 1.065 | 0.89 | 0.012 | 0.034 | 0.034 | 0 | 0.14 | 0.14 | 0.13 | 0.28 | 0.18 | 0.24 |
| 1 | 167542337 | rs7543683 | -1.696 | 2.333 | 0.787 | 0.583 | 0.008 | 0 | 0 | 0.498 | 0.45 | 0.52 | 0.53 | 0.46 | 0.63 | 0.39 |
| 1 | 167546460 | rs7513712 | -1.935 | 2.457 | 0.785 | 0.652 | -0.011 | -0.001 | -0.001 | 0 | 0.45 | 0.48 | 0.52 | 0.46 | 0.63 | 0.39 |
| 1 | 167547219 | rs6683618 | -0.797 | 2.593 | 0.666 | 0.631 | -0.045 | -0.034 | -0.034 | 0 | 0.3 | 0.2 | 0.45 | 0.43 | 0.53 | 0.27 |
| 1 | 167549747 | rs11584355 | -0.432 | 2.663 | 0.693 | 0.615 | -0.059 | 0.057 | 0.057 | 1.437 | 0.27 | 0.12 | 0.37 | 0.16 | 0.42 | 0.15 |
| 1 | 167553649 | rs10800319 | -0.911 | 2.542 | 0.739 | 0.63 | 0.009 | 0.006 | 0.006 | 0.685 | 0.31 | 0.34 | 0.46 | 0.43 | 0.54 | 0.28 |
| 1 | 167554230 | rs1935286 | -0.343 | 2.537 | 0.765 | 0.62 | 0.011 | -0.033 | -0.033 | 0 | 0.11 | 0.29 | 0.05 | 0.02 | 0.08 | 0.07 |
| 1 | 167554337 | rs7554445 | -1.978 | 2.507 | 0.747 | 0.608 | -0.01 | 0 | 0 | 0 | 0.44 | 0.49 | 0.52 | 0.45 | 0.64 | 0.35 |

| | | | | | | | | | | | | | | | | |
|---|---|---|---|---|---|---|---|---|---|---|---|---|---|---|---|---|
| 1 | 167560542 | rs767707 | -0.979 | 2.502 | 0.582 | 0.601 | -0.05 | -0.04 | -0.04 | 0.622 | 0.3 | 0.21 | 0.46 | 0.42 | 0.55 | 0.25 |
| 1 | 167562605 | rs12037105 | -1.711 | 2.943 | 0.624 | 0.495 | 0.023 | 0.01 | 0.01 | 0 | 0.35 | 0.48 | 0.33 | 0.39 | 0.41 | 0.3 |
| 1 | 167562816 | rs6701488 | -1.332 | 2.961 | 0.583 | 0.4 | -0.005 | -0.055 | -0.055 | 1.379 | 0.31 | 0.36 | 0.24 | 0.12 | 0.28 | 0.17 |
| 1 | 167569730 | rs7514302 | -2.156 | 2.789 | 0.528 | 0.764 | 0.007 | 0 | 0 | 0.096 | 0.46 | 0.53 | 0.67 | 0.52 | 0.64 | 0.35 |
| 1 | 167572161 | rs2143305 | -0.866 | 2.838 | 0.433 | 0.72 | -0.037 | 0.035 | 0.035 | 0 | 0.2 | 0.1 | 0.25 | 0.08 | 0.3 | 0.14 |
| 1 | 167573990 | rs6427095 | -1.12 | 2.854 | 0.368 | 0.721 | 0.017 | -0.029 | -0.029 | 0 | 0.22 | 0.27 | 0.26 | 0.08 | 0.3 | 0.14 |
| 1 | 167585884 | rs11809621 | 0.044 | 2.237 | 0.045 | 0.668 | -0.058 | -0.027 | -0.027 | 0.046 | 0.17 | 0.01 | 0.25 | 0.15 | 0.2 | 0.09 |
| 1 | 167589943 | rs1229403 | -1.904 | 2.124 | 0.121 | 0.649 | 0.005 | -0.001 | -0.001 | 2.661 | 0.46 | 0.55 | 0.66 | 0.52 | 0.63 | 0.41 |
| 1 | 167592532 | rs926521 | -1.652 | 2.475 | -0.042 | 0.704 | 0.014 | 0.007 | 0.007 | 0 | 0.24 | 0.27 | 0.41 | 0.36 | 0.43 | 0.31 |
| 1 | 167597473 | rs1890131 | -0.806 | 2.773 | -0.466 | 0.826 | -0.07 | -0.064 | -0.064 | 0.132 | 0.13 | 0.04 | 0.35 | 0.3 | 0.36 | 0.13 |
| 1 | 167603963 | rs1229393 | -0.224 | 1.452 | -0.17 | -0.443 | 0.005 | -0.059 | -0.059 | 0.128 | 0.65 | 0.49 | 0.67 | 0.82 | 0.68 | 0.77 |
| 1 | 167607340 | rs1229390 | -0.135 | 1.324 | 0.276 | -0.59 | -0.01 | -0.067 | -0.067 | 0 | 0.63 | 0.56 | 0.71 | 0.84 | 0.7 | 0.64 |
| 1 | 167610200 | rs1229387 | 0.466 | 1.282 | 0.107 | -0.645 | 0.001 | 0.002 | 0.002 | 0 | 0.07 | 0.06 | 0.04 | 0.02 | 0.06 | 0.09 |
| 1 | 167611221 | rs4657692 | 0.983 | 1.237 | -0.063 | -0.908 | 0.011 | 0.005 | 0.005 | 0 | 0.37 | 0.33 | 0.43 | 0.64 | 0.32 | 0.39 |
| 1 | 167611826 | rs3738226 | -0.443 | 1.247 | 0.222 | -1.129 | -0.001 | 0.044 | 0.044 | 0 | 0.62 | 0.72 | 0.7 | 0.78 | 0.67 | 0.57 |
| 1 | 167613419 | rs12062583 | -0.928 | 1.651 | 0.367 | -1.051 | 0.036 | -0.038 | -0.038 | 0 | 0.23 | 0.35 | 0.24 | 0.13 | 0.32 | 0.14 |
| 1 | 167614752 | rs742471 | 0.891 | 1.91 | 0.104 | -1.01 | -0.051 | -0.016 | -0.016 | 0.301 | 0.09 | 0.02 | 0.12 | 0.08 | 0.2 | 0.05 |
| 1 | 167615368 | rs761078 | -1.247 | 1.647 | 0.159 | -1.085 | -0.025 | 0.039 | 0.039 | 0 | 0.74 | 0.88 | 0.77 | 0.82 | 0.74 | 0.67 |
| 1 | 167617699 | rs10489193 | -1.094 | 1.504 | 0.547 | -0.77 | -0.018 | 0.008 | 0.008 | 1.348 | 0.09 | 0.34 | 0.1 | 0.06 | 0.01 | 0.07 |
| 1 | 167618170 | rs2008981 | -1.214 | 1.705 | 0.48 | -0.801 | 0.031 | -0.048 | -0.048 | 0 | 0.23 | 0.44 | 0.23 | 0.13 | 0.28 | 0.14 |
| 1 | 167621144 | rs6659046 | -1.249 | 1.849 | 0.345 | -0.861 | 0.039 | -0.052 | -0.052 | 0 | 0.23 | 0.42 | 0.26 | 0.14 | 0.34 | 0.18 |
| 1 | 167622119 | rs12067995 | -1.033 | 1.542 | 0.409 | -0.759 | 0.016 | -0.014 | -0.014 | 0.465 | 0.08 | 0.3 | 0.1 | 0.06 | 0.06 | 0.11 |
| 1 | 167623320 | rs16859384 | 0.65 | 1.418 | 0.112 | -0.754 | 0 | 0 | 0 | 0 | 0.38 | 0.38 | 0.44 | 0.61 | 0.32 | 0.35 |
| 1 | 167631806 | rs1229377 | 0.423 | 1.86 | -0.431 | -0.327 | -0.027 | -0.018 | -0.018 | 0 | 0.06 | 0.01 | 0.13 | 0.09 | 0.18 | 0.04 |
| 1 | 167632271 | rs1229373 | 0.328 | 0.976 | -0.614 | -0.694 | 0.003 | 0.088 | 0.088 | 0 | 0.4 | 0.24 | 0.29 | 0.19 | 0.38 | 0.21 |
| 1 | 167637763 | rs1229365 | -0.91 | 1.463 | -0.679 | -0.034 | -0.048 | 0.01 | 0.01 | 0 | 0.76 | 0.92 | 0.77 | 0.78 | 0.71 | 0.61 |
| 1 | 167638451 | rs3753928 | -0.927 | 1.149 | 0.642 | 0.776 | -0.006 | 0.106 | 0.106 | 0 | 0.18 | 0.39 | 0.36 | 0.56 | 0.18 | 0.29 |
| 1 | 167638461 | rs3753929 | -0.479 | 1.193 | 0.632 | 0.785 | -0.028 | 0.125 | 0.125 | 0 | 0.16 | 0.33 | 0.33 | 0.55 | 0.13 | 0.25 |
| 1 | 167638557 | rs6695906 | -0.914 | 1.185 | 0.61 | 0.792 | -0.005 | 0.097 | 0.097 | 0 | 0.18 | 0.35 | 0.35 | 0.56 | 0.18 | 0.29 |
| 1 | 167639855 | rs10918737 | -1.327 | 1.196 | 0.544 | 1.021 | -0.009 | 0.102 | 0.102 | 0 | 0.26 | 0.49 | 0.4 | 0.5 | 0.23 | 0.28 |
| 1 | 167641292 | rs1229355 | -1.409 | 1.401 | -0.089 | 0.849 | 0.013 | -0.004 | -0.004 | 0.091 | 0.37 | 0.63 | 0.59 | 0.63 | 0.46 | 0.46 |
| 1 | 167642494 | rs12081670 | -1.144 | 1.365 | 0.163 | 0.556 | 0.054 | -0.057 | -0.057 | 1.261 | 0.22 | 0.44 | 0.29 | 0.1 | 0.26 | 0.17 |
| 1 | 167652272 | rs7511792 | -1.056 | 0.671 | 0.241 | 1.338 | -0.019 | 0.166 | 0.166 | 0.251 | 0.18 | 0.61 | 0.29 | 0.46 | 0.17 | 0.22 |
| 1 | 167652660 | rs7514408 | -1.619 | 0.471 | -0.111 | 0.998 | -0.001 | 0.004 | 0.004 | 0 | 0.27 | 0.73 | 0.36 | 0.49 | 0.23 | 0.29 |
| 1 | 167666172 | rs12093287 | -0.722 | 0.43 | 0.042 | 0.591 | 0.002 | 0.002 | 0.002 | 0 | 0.1 | 0.1 | 0.07 | 0.03 | 0.07 | 0.07 |
| 1 | 167669487 | rs6427100 | -0.687 | 0.696 | -0.971 | -0.013 | -0.069 | 0.03 | 0.03 | 0.007 | 0.82 | 0.99 | 0.84 | 0.9 | 0.77 | 0.81 |
| 1 | 167672809 | rs1890129 | -0.712 | 1.096 | -1.275 | -0.238 | -0.03 | 0.016 | 0.016 | 0 | 0.29 | 0.14 | 0.25 | 0.23 | 0.3 | 0.31 |
| 1 | 167672969 | rs1890130 | 0.375 | 1.039 | -1.397 | -0.326 | -0.036 | -0.023 | -0.023 | 0 | 0.17 | 0.08 | 0.12 | 0.2 | 0.22 | 0.25 |
| 1 | 167673695 | rs2862866 | -0.967 | 1.067 | -1.376 | -0.299 | -0.042 | 0.014 | 0.014 | 0 | 0.29 | 0.12 | 0.25 | 0.23 | 0.3 | 0.31 |
| 1 | 167676364 | rs1933075 | -1.144 | 0.91 | -0.249 | -0.384 | 0.003 | -0.015 | -0.015 | 0.108 | 0.32 | 0.42 | 0.27 | 0.23 | 0.31 | 0.32 |
| 1 | 167679477 | rs4657699 | 0.617 | 0.894 | -0.302 | -0.288 | -0.013 | 0.03 | 0.03 | 0 | 0.09 | 0.01 | 0.07 | 0.02 | 0.07 | 0.06 |
| 1 | 167697059 | rs12566176 | 0.233 | 1.008 | 0.16 | -0.478 | 0.003 | 0.005 | 0.005 | 0 | 0.14 | 0.16 | 0.11 | 0.19 | 0.17 | 0.22 |
| 1 | 167700143 | rs3767445 | 1.275 | 0.767 | -0.378 | -0.975 | -0.021 | 0.027 | 0.027 | 0 | 0.18 | 0.07 | 0.22 | 0.11 | 0.22 | 0.18 |
| 1 | 167702255 | rs10918751 | -1.123 | 0.626 | 0.32 | -0.846 | 0.001 | -0.03 | -0.03 | 0.512 | 0.33 | 0.52 | 0.29 | 0.22 | 0.3 | 0.3 |
| 1 | 167706521 | rs10918754 | 0.52 | 0.727 | 0.101 | -0.747 | -0.002 | -0.002 | -0.002 | 0 | 0.14 | 0.15 | 0.11 | 0.18 | 0.16 | 0.18 |
| 1 | 167707655 | rs4657704 | -0.578 | 0.446 | 0.524 | -0.939 | -0.027 | -0.122 | -0.122 | 0 | 0.2 | 0.5 | 0.19 | 0.03 | 0.13 | 0.12 |
| 1 | 167709064 | rs7518652 | -0.512 | 1.165 | 0.324 | -0.975 | 0.002 | -0.041 | -0.041 | 0 | 0.19 | 0.26 | 0.18 | 0.02 | 0.18 | 0.13 |
| 1 | 167713516 | rs6427106 | -0.756 | 0.993 | 0.154 | -1.083 | 0 | -0.002 | -0.002 | 0.01 | 0.75 | 0.71 | 0.84 | 0.76 | 0.74 | 0.67 |
| 1 | 167715397 | rs2213883 | 1.103 | 0.626 | -0.385 | -1.111 | 0.011 | 0.02 | 0.02 | 0 | 0.21 | 0.05 | 0.23 | 0.15 | 0.18 | 0.19 |
| 1 | 167722003 | rs3753931 | 0.693 | 0.449 | 0.211 | -1.081 | -0.05 | 0.075 | 0.075 | 0.111 | 0.29 | 0.53 | 0.37 | 0.43 | 0.21 | 0.25 |
| 1 | 167724850 | rs12039424 | 2.012 | 0.414 | -0.084 | -0.919 | 0.003 | -0.007 | -0.007 | 0 | 0.17 | 0.15 | 0.3 | 0.35 | 0.16 | 0.15 |
| 1 | 167730783 | rs2097572 | -2.838 | 0.246 | -0.577 | -1.177 | -0.025 | 0.015 | 0.015 | 0.1 | 0.86 | 1 | 0.94 | 0.89 | 0.88 | 0.93 |
| 1 | 167732098 | rs10918760 | -1.155 | 0.246 | -0.466 | -1.159 | -0.029 | 0.019 | 0.019 | 0.011 | 0.73 | 0.87 | 0.83 | 0.77 | 0.68 | 0.66 |
| 1 | 167732984 | rs16859595 | 2.394 | 0.183 | -0.487 | -1.131 | 0.004 | -0.032 | -0.032 | 0 | 0.16 | 0.09 | 0.29 | 0.35 | 0.16 | 0.15 |
| 1 | 167739821 | rs4472738 | 0.232 | 0.262 | -0.33 | -1.079 | -0.017 | 0.009 | 0.009 | 0.128 | 0.36 | 0.48 | 0.49 | 0.55 | 0.3 | 0.44 |
| 1 | 167739861 | rs6680341 | 1.598 | 0.287 | -0.42 | -1.055 | -0.001 | 0.001 | 0.001 | 0 | 0.09 | 0.11 | 0.05 | 0.12 | 0.09 | 0.2 |
| 1 | 167744858 | rs16859650 | 1.623 | 0.375 | -0.124 | -0.901 | -0.017 | -0.016 | -0.016 | 0.074 | 0.1 | 0.18 | 0.06 | 0.07 | 0.04 | 0.09 |
| 1 | 167748385 | rs10918763 | 1.489 | 0.784 | -0.123 | -0.937 | 0.01 | 0.005 | 0.005 | 0 | 0.12 | 0.12 | 0.13 | 0.19 | 0.21 | 0.19 |
| 1 | 167749603 | rs6671664 | -0.811 | 0.796 | -0.623 | -1.014 | 0.001 | 0.033 | 0.033 | 0 | 0.63 | 0.71 | 0.77 | 0.7 | 0.63 | 0.56 |
| 1 | 167751269 | rs929137 | -0.491 | 0.883 | -0.61 | -0.982 | -0.01 | 0.015 | 0.015 | 0 | 0.44 | 0.3 | 0.38 | 0.33 | 0.47 | 0.4 |
| 1 | 167753387 | rs17481761 | 1.525 | 0.902 | -1.067 | -1.029 | -0.018 | 0.046 | 0.046 | 0 | 0.2 | 0.01 | 0.15 | 0.08 | 0.18 | 0.17 |
| 1 | 167758784 | rs3752606 | 1.064 | 0.918 | -0.744 | -0.847 | -0.005 | -0.011 | -0.011 | 0 | 0.12 | 0.22 | 0.1 | 0.08 | 0.11 | 0.15 |
| 1 | 167758901 | rs3088322 | -1.053 | 1.045 | -0.966 | -0.902 | -0.009 | 0.016 | 0.016 | 0 | 0.72 | 0.78 | 0.82 | 0.77 | 0.68 | 0.65 |
| 1 | 167759281 | rs7605 | 0.458 | 1.118 | -1.112 | -0.992 | 0.005 | -0.012 | -0.012 | 0 | 0.43 | 0.45 | 0.62 | 0.61 | 0.53 | 0.42 |
| 1 | 167760731 | rs10753764 | -1.149 | 0.606 | -1.769 | -0.781 | 0.006 | 0.019 | 0.019 | 0 | 0.73 | 0.82 | 0.85 | 0.77 | 0.75 | 0.67 |
| 1 | 167764925 | rs3830180 | -0.125 | 0.903 | -1.534 | -0.537 | 0.022 | 0.084 | 0.084 | 0 | 0.56 | 0.7 | 0.72 | 0.7 | 0.63 | 0.54 |
| 1 | 167777775 | rs10918774 | 1.03 | 1.511 | -1.639 | -0.028 | -0.059 | -0.049 | -0.049 | 0 | 0.1 | 0.02 | 0.25 | 0.23 | 0.24 | 0.21 |
| 1 | 167777989 | rs6427107 | -1.456 | 0.967 | -2.295 | -0.328 | 0.006 | 0.018 | 0.018 | 0 | 0.75 | 0.93 | 0.85 | 0.78 | 0.78 | 0.78 |
| 1 | 167778843 | rs3213588 | -0.225 | 0.229 | -2.548 | -0.415 | 0.035 | 0.237 | 0.237 | 1.874 | 0.51 | 0.84 | 0.38 | 0.18 | 0.43 | 0.38 |

| | | | | | | | | | | | | | | | | |
|---|---|---|---|---|---|---|---|---|---|---|---|---|---|---|---|---|
| 1 | 167782320 | rs12735367 | 0.788 | 0.191 | -1.703 | -0.366 | 0.024 | -0.026 | -0.026 | 0 | 0.18 | 0.38 | 0.18 | 0.11 | 0.19 | 0.08 |
| 1 | 167785060 | rs12021710 | 2.062 | -1.097 | -2.182 | -0.043 | 0.009 | -0.101 | -0.101 | 0 | 0.15 | 0.03 | 0.21 | 0.35 | 0.11 | 0.18 |
| 1 | 167785262 | rs1120604 | 1.501 | -0.981 | -1.925 | -0.414 | 0.029 | 0.076 | 0.076 | 0.972 | 0.15 | 0.02 | 0.1 | 0 | 0.09 | 0.02 |
| 1 | 167790618 | rs4656560 | -0.984 | -1.02 | -1.982 | -0.411 | 0.071 | 0.089 | 0.089 | 0.376 | 0.51 | 0.83 | 0.58 | 0.37 | 0.58 | 0.43 |
| 1 | 167793670 | rs1476074 | 0.296 | -0.998 | -1.4 | -0.11 | -0.005 | -0.005 | -0.005 | 0 | 0.25 | 0.3 | 0.47 | 0.42 | 0.42 | 0.3 |
| 1 | 167794406 | rs1476075 | -0.897 | -1.013 | -1.16 | -0.117 | 0.019 | 0.076 | 0.076 | 0.072 | 0.46 | 0.72 | 0.53 | 0.28 | 0.55 | 0.38 |
| 1 | 167796012 | rs12407873 | -0.256 | -0.857 | -0.73 | 0.123 | 0.003 | -0.009 | -0.009 | 0 | 0.38 | 0.48 | 0.49 | 0.28 | 0.49 | 0.26 |
| 1 | 167799520 | rs10800328 | 1.83 | -0.797 | -1.104 | 0.706 | 0.01 | -0.021 | -0.021 | 0 | 0.14 | 0.09 | 0.16 | 0.3 | 0.1 | 0.18 |
| 1 | 167801355 | rs17482924 | 2.032 | -0.064 | -0.892 | 0.664 | -0.007 | 0.014 | 0.014 | 0 | 0.09 | 0.06 | 0.09 | 0.04 | 0.1 | 0.02 |
| 1 | 167804738 | rs12409150 | 1.296 | 0.491 | -0.849 | 0.97 | -0.048 | -0.05 | -0.05 | 0 | 0.1 | 0.03 | 0.28 | 0.25 | 0.29 | 0.22 |
| 1 | 167810373 | rs204267 | -0.761 | -0.55 | -1.065 | 0.291 | 0.1 | 0.222 | 0.222 | 1.221 | 0.51 | 0.87 | 0.48 | 0.2 | 0.38 | 0.4 |
| 1 | 167812359 | rs204264 | 0.765 | -0.759 | -0.628 | 0.249 | 0.007 | 0.009 | 0.009 | 0 | 0.21 | 0.16 | 0.14 | 0.06 | 0.16 | 0.17 |
| 1 | 167814605 | rs16859798 | 0.366 | -1.04 | 0.12 | 0.22 | -0.075 | -0.1 | -0.1 | 2.932 | 0.22 | 0.43 | 0.16 | 0.07 | 0.13 | 0.11 |
| 1 | 167815167 | rs2269678 | 1.217 | -0.493 | -0.124 | 0.797 | -0.021 | -0.019 | -0.019 | 0 | 0.12 | 0.08 | 0.29 | 0.38 | 0.28 | 0.19 |
| 1 | 167815816 | rs2902467 | -0.043 | -0.638 | 0.172 | 0.649 | -0.045 | -0.088 | -0.088 | 0 | 0.28 | 0.58 | 0.2 | 0.08 | 0.19 | 0.25 |
| 1 | 167817639 | rs203795 | -1.949 | -0.627 | -0.542 | 0.785 | -0.004 | 0.126 | 0.126 | 0 | 0.67 | 0.93 | 0.84 | 0.86 | 0.71 | 0.74 |
| 1 | 167819017 | rs3767451 | -0.567 | 0.197 | 0.615 | 1.684 | -0.011 | -0.011 | -0.011 | 0 | 0.19 | 0.18 | 0.35 | 0.44 | 0.35 | 0.25 |
| 1 | 167822213 | rs3767457 | 0.263 | 0.217 | 0.672 | 1.833 | 0.043 | 0.027 | 0.027 | 0 | 0.14 | 0.18 | 0.29 | 0.31 | 0.29 | 0.2 |
| 1 | 167823505 | rs3738235 | 0.986 | 0.223 | -0.154 | 2.187 | -0.001 | -0.014 | -0.014 | 0 | 0.05 | 0.01 | 0.05 | 0.23 | 0.04 | 0.11 |
| 1 | 167823512 | rs3738236 | -1.612 | 0.14 | -0.138 | 1.702 | -0.004 | 0.053 | 0.053 | 0 | 0.46 | 0.76 | 0.63 | 0.67 | 0.52 | 0.55 |
| 1 | 167824112 | rs12032641 | 0.246 | 0.61 | -0.146 | 2.366 | 0.009 | 0.019 | 0.019 | 0.086 | 0.06 | 0.04 | 0.07 | 0.31 | 0.04 | 0.12 |
| 1 | 167825134 | rs7512378 | 0.27 | 1.153 | 0.236 | 1.87 | 0.161 | 0.044 | 0.044 | 0 | 0.12 | 0.29 | 0.27 | 0.24 | 0.27 | 0.16 |
| 1 | 167825485 | rs2071921 | -0.938 | 0.856 | 0.043 | 1.832 | 0.025 | -0.02 | -0.02 | 0 | 0.37 | 0.46 | 0.57 | 0.71 | 0.41 | 0.4 |
| 1 | 167825606 | rs2071922 | -0.182 | -0.603 | -0.666 | 1.813 | 0.061 | -0.106 | -0.106 | 0 | 0.23 | 0.08 | 0.28 | 0.42 | 0.13 | 0.21 |
| 1 | 167837285 | rs203812 | -1.089 | -0.196 | 0.403 | 1.828 | 0.014 | 0.342 | 0.342 | 0 | 0.06 | 0.35 | 0.07 | 0.27 | 0.01 | 0.1 |
| 1 | 167845787 | rs6427113 | 0.365 | 0.061 | -0.302 | 1.578 | -0.008 | 0.001 | 0.001 | 0 | 0.21 | 0.04 | 0.21 | 0.13 | 0.17 | 0.1 |
| 1 | 167847584 | rs203838 | -0.8 | 0.021 | 0.389 | 1.142 | -0.009 | -0.025 | -0.025 | 0 | 0.37 | 0.56 | 0.34 | 0.19 | 0.3 | 0.16 |
| 1 | 167847984 | rs203841 | -0.429 | -0.014 | 0.779 | 1.139 | 0.078 | 0.068 | 0.068 | 0.431 | 0.05 | 0.46 | 0.04 | 0 | 0.01 | 0.01 |
| 1 | 167848634 | rs11807288 | 0.808 | 0.239 | 0.371 | 1.029 | -0.017 | -0.025 | -0.025 | 0 | 0.07 | 0.01 | 0.1 | 0.23 | 0.08 | 0.08 |
| 1 | 167849327 | rs203848 | -0.171 | 0.232 | 0.492 | 0.559 | -0.039 | -0.114 | -0.114 | 0 | 0.15 | 0.52 | 0.11 | 0 | 0.09 | 0.02 |
| 1 | 167849414 | rs203849 | -0.637 | 0.065 | -0.053 | 0.164 | -0.008 | 0.01 | 0.01 | 0 | 0.59 | 0.67 | 0.46 | 0.32 | 0.5 | 0.44 |
| 1 | 167858881 | rs1829684 | 0.869 | -0.65 | -0.394 | 0.123 | 0.002 | 0.043 | 0.043 | 0 | 0.25 | 0.07 | 0.14 | 0.19 | 0.24 | 0.31 |
| 1 | 167863706 | rs437846 | 0.577 | -0.638 | 0.019 | 0.035 | 0.005 | 0.009 | 0.009 | 0 | 0.26 | 0.23 | 0.27 | 0.16 | 0.2 | 0.14 |
| 1 | 167865871 | rs16859886 | 1.105 | -0.668 | 0.287 | 0.048 | -0.008 | -0.049 | -0.049 | 0.222 | 0.09 | 0.17 | 0.07 | 0 | 0.09 | 0.02 |
| 1 | 167869334 | rs11590245 | 0.954 | -0.677 | 0.293 | 0.037 | -0.003 | -0.052 | -0.052 | 0 | 0.1 | 0.18 | 0.09 | 0 | 0.1 | 0.08 |
| 1 | 167873622 | rs203775 | -1.325 | -0.746 | 0.873 | 0.113 | -0.051 | -0.066 | -0.066 | 0 | 0.74 | 0.9 | 0.61 | 0.5 | 0.62 | 0.59 |
| 1 | 167876616 | rs203777 | -0.365 | -0.844 | 1.67 | -0.011 | 0.035 | 0.068 | 0.068 | 0 | 0.42 | 0.77 | 0.38 | 0.18 | 0.34 | 0.24 |
| 1 | 167879655 | rs11578481 | 1.175 | -0.865 | 1.865 | -0.027 | -0.001 | -0.055 | -0.055 | 0 | 0.1 | 0.29 | 0.08 | 0 | 0.11 | 0.05 |
| 1 | 167880085 | rs11579897 | 1.062 | -0.867 | 1.967 | -0.017 | 0.004 | -0.028 | -0.028 | 0.646 | 0.1 | 0.35 | 0.1 | 0.05 | 0.11 | 0.08 |
| 1 | 167881008 | rs11580404 | 1.683 | -0.868 | 1.989 | -0.018 | -0.002 | -0.024 | -0.024 | 0.013 | 0.08 | 0.07 | 0.07 | 0 | 0.11 | 0.02 |
| 1 | 167882386 | rs203782 | -0.424 | -0.88 | 2.075 | -0.02 | 0.03 | 0.053 | 0.053 | 0 | 0.42 | 0.74 | 0.52 | 0.18 | 0.34 | 0.24 |
| 1 | 167887416 | rs1050037 | 1.002 | -0.816 | 2.027 | 0.034 | 0 | -0.03 | -0.03 | 0 | 0.1 | 0.33 | 0.1 | 0.05 | 0.11 | 0.08 |
| 1 | 167887436 | rs1050036 | 1.122 | -0.815 | 1.992 | 0.024 | -0.004 | -0.056 | -0.056 | 0 | 0.1 | 0.27 | 0.08 | 0 | 0.11 | 0.05 |
| 1 | 167887642 | rs10918798 | 1.122 | -0.815 | 1.992 | 0.024 | -0.004 | -0.056 | -0.056 | 0 | 0.1 | 0.27 | 0.08 | 0 | 0.11 | 0.05 |
| 1 | 167903079 | rs10489202 | 1.072 | -0.875 | 1.772 | -0.012 | -0.035 | 0.013 | 0.013 | 0 | 0.25 | 0.04 | 0.39 | 0.13 | 0.23 | 0.16 |
| 1 | 167929037 | rs11578006 | 1.211 | -0.84 | 1.899 | 0.003 | -0.009 | -0.051 | -0.051 | 0 | 0.1 | 0.2 | 0.08 | 0 | 0.11 | 0.05 |
| 1 | 167961235 | rs12404574 | 1.007 | -0.87 | 1.91 | -0.005 | 0.005 | 0.024 | 0.024 | 0 | 0.26 | 0.19 | 0.39 | 0.13 | 0.23 | 0.16 |
| 1 | 167972023 | rs202248 | -1.338 | -0.93 | 1.811 | -0.009 | -0.064 | -0.069 | -0.069 | 0 | 0.73 | 0.9 | 0.75 | 0.53 | 0.62 | 0.56 |
| 1 | 167980218 | rs202252 | 0.03 | -0.977 | 1.926 | -0.054 | -0.029 | -0.077 | -0.077 | 0 | 0.16 | 0.48 | 0.12 | 0.06 | 0.12 | 0.08 |
| 1 | 168013850 | rs11558511 | 1.665 | -0.757 | 1.674 | 0.155 | -0.056 | 0.012 | 0.012 | 0 | 0.22 | 0.05 | 0.14 | 0.2 | 0.28 | 0.27 |
| 1 | 168018840 | rs17485889 | 0.905 | -1.541 | 1.9 | -0.191 | -0.02 | 0.021 | 0.021 | 0 | 0.26 | 0.06 | 0.4 | 0.14 | 0.23 | 0.16 |
| 1 | 168051266 | rs275137 | -0.709 | -1.53 | 1.961 | -0.245 | 0.033 | 0.058 | 0.058 | 0 | 0.43 | 0.77 | 0.54 | 0.22 | 0.36 | 0.25 |
| 1 | 168051609 | rs968020 | 1.463 | -1.555 | 1.863 | -0.314 | 0.003 | 0.006 | 0.006 | 0 | 0.08 | 0.05 | 0.07 | 0 | 0.11 | 0.02 |
| 1 | 168073601 | rs2294817 | -0.019 | -1.411 | 1.122 | -0.143 | 0.012 | 0.04 | 0.04 | 0 | 0.36 | 0.33 | 0.48 | 0.15 | 0.34 | 0.17 |
| 1 | 168074677 | rs3753934 | -0.224 | -1.451 | 1.076 | -0.101 | 0 | -0.001 | -0.001 | 0 | 0.38 | 0.44 | 0.5 | 0.22 | 0.34 | 0.24 |
| 1 | 168074746 | rs3820400 | -1.461 | -1.55 | 1.028 | -0.108 | -0.072 | -0.07 | -0.07 | 0.03 | 0.71 | 0.92 | 0.74 | 0.55 | 0.59 | 0.55 |
| 1 | 168075316 | rs10918826 | -0.422 | -1.395 | 1.001 | -0.1 | -0.001 | -0.003 | -0.003 | 0 | 0.39 | 0.46 | 0.5 | 0.22 | 0.34 | 0.24 |
| 1 | 168077636 | rs11582396 | -0.206 | -1.363 | 0.842 | -0.085 | 0.012 | 0.032 | 0.032 | 0 | 0.37 | 0.32 | 0.49 | 0.15 | 0.34 | 0.17 |
| 1 | 168078384 | rs16828093 | -0.321 | -1.35 | 0.72 | -0.083 | 0.005 | 0.009 | 0.009 | 0 | 0.38 | 0.42 | 0.49 | 0.22 | 0.34 | 0.24 |
| 1 | 168090899 | rs16828104 | -0.795 | -2.565 | 0.939 | 0.228 | -0.022 | -0.002 | -0.002 | 0 | 0.26 | 0.47 | 0.19 | 0.26 | 0.18 | 0.17 |
| 1 | 168095261 | rs11805261 | 0.449 | -2.312 | 0.459 | 0.306 | 0.003 | -0.023 | -0.023 | 0 | 0.12 | 0.03 | 0.08 | 0.19 | 0.08 | 0.09 |
| 1 | 168103545 | rs4540637 | -1.157 | -2.678 | 0.488 | 0.243 | -0.013 | -0.017 | -0.017 | 0.006 | 0.67 | 0.81 | 0.7 | 0.6 | 0.49 | 0.48 |
| 1 | 168112908 | rs2064329 | 0.627 | -2.62 | 0.926 | 0.079 | 0.044 | -0.013 | -0.013 | 0 | 0.16 | 0.1 | 0.35 | 0.15 | 0.11 | 0.07 |
| 1 | 168118292 | rs2902570 | -0.022 | -2.307 | 1.261 | 0.01 | 0.011 | -0.004 | -0.004 | 0.086 | 0.31 | 0.28 | 0.44 | 0.34 | 0.18 | 0.16 |
| 1 | 168126289 | rs12742613 | 0.091 | -2.339 | 1.216 | 0.003 | 0.031 | -0.01 | -0.01 | 2.218 | 0.3 | 0.25 | 0.45 | 0.36 | 0.18 | 0.16 |
| 1 | 168131592 | rs1543358 | -1.07 | -2.364 | 1.108 | -0.052 | 0.047 | 0.023 | 0.023 | 3.528 | 0.47 | 0.71 | 0.57 | 0.39 | 0.32 | 0.22 |
| 1 | 168138084 | rs12097383 | 0.409 | -2.147 | 0.99 | -0.287 | 0 | -0.051 | -0.051 | 0 | 0.09 | 0.17 | 0.08 | 0 | 0.1 | 0.02 |
| 1 | 168142489 | rs4657739 | -0.056 | -2.268 | 1.093 | -0.207 | -0.053 | 0.029 | 0.029 | 2.576 | 0.32 | 0.43 | 0.52 | 0.42 | 0.19 | 0.2 |
| 1 | 168147892 | rs2281016 | -0.377 | -2.286 | 0.816 | -0.215 | 0.012 | -0.008 | -0.008 | 0 | 0.35 | 0.69 | 0.53 | 0.42 | 0.19 | 0.2 |

| | | | | | | | | | | | | | | | | |
|---|---|---|---|---|---|---|---|---|---|---|---|---|---|---|---|---|
| 1 | 168148829 | rs10918838 | 0.127 | -2.226 | 0.302 | -0.186 | -0.006 | 0.001 | 0.001 | 2.944 | 0.31 | 0.31 | 0.51 | 0.42 | 0.19 | 0.16 |
| 1 | 168149142 | rs4657741 | -1.103 | -2.193 | 0.164 | -0.184 | 0.131 | 0.009 | 0.009 | 3.138 | 0.5 | 0.86 | 0.64 | 0.44 | 0.33 | 0.22 |
| 1 | 168159262 | rs2235207 | -0.925 | -2.172 | -0.291 | -0.212 | 0.148 | 0.004 | 0.004 | 1.246 | 0.48 | 0.86 | 0.63 | 0.42 | 0.32 | 0.22 |
| 1 | 168166616 | rs10753766 | -1.173 | -1.929 | -0.852 | -0.054 | 0.075 | -0.042 | -0.042 | 0.009 | 0.54 | 0.91 | 0.66 | 0.44 | 0.36 | 0.24 |
| 1 | 168167139 | rs4534357 | -0.683 | -1.363 | -0.201 | 0.824 | -0.029 | 0.017 | 0.017 | 2.412 | 0.31 | 0.38 | 0.52 | 0.42 | 0.2 | 0.16 |
| 1 | 168181725 | rs7551039 | -0.973 | -1.09 | -0.06 | 0.534 | -0.031 | -0.016 | -0.016 | 0 | 0.14 | 0.26 | 0.09 | 0.09 | 0.07 | 0.09 |
| 1 | 168182443 | rs9659234 | 0.023 | -1.58 | -0.113 | 0.657 | -0.007 | 0.007 | 0.007 | 0 | 0.66 | 0.67 | 0.58 | 0.61 | 0.72 | 0.62 |
| 1 | 168183278 | rs6702778 | -0.013 | -1.838 | -0.173 | 0.573 | 0.008 | -0.035 | -0.035 | 0 | 0.14 | 0.07 | 0.28 | 0.35 | 0.13 | 0.16 |
| 1 | 168186174 | rs12049577 | 1.439 | -2.132 | -0.374 | 0.197 | -0.018 | -0.001 | -0.001 | 0.619 | 0.39 | 0.29 | 0.42 | 0.4 | 0.55 | 0.29 |
| 1 | 168194927 | rs4656572 | 0.678 | -2.572 | -0.038 | 0.252 | -0.008 | 0.001 | 0.001 | 0 | 0.57 | 0.59 | 0.57 | 0.53 | 0.7 | 0.57 |
| 1 | 168202963 | rs2235178 | 0.908 | -3.044 | 0.311 | 0.05 | -0.011 | -0.003 | -0.003 | 0.104 | 0.55 | 0.57 | 0.54 | 0.44 | 0.67 | 0.42 |
| 1 | 168211256 | rs2280991 | 1.323 | -3.276 | 0.574 | -0.061 | 0.091 | 0.038 | 0.038 | 0 | 0.5 | 0.31 | 0.52 | 0.44 | 0.67 | 0.42 |
| 1 | 168211282 | rs2280990 | 0.159 | -3.265 | 0.809 | -0.048 | 0.011 | -0.014 | -0.014 | 0 | 0.64 | 0.72 | 0.6 | 0.53 | 0.72 | 0.61 |
| 1 | 168212234 | rs2280989 | 0.799 | -3.239 | 0.84 | -0.045 | -0.004 | -0.001 | -0.001 | 0 | 0.54 | 0.57 | 0.55 | 0.44 | 0.67 | 0.42 |
| 1 | 168214098 | rs10737541 | 0.627 | -3.382 | 0.729 | -0.137 | -0.027 | -0.002 | -0.002 | 0 | 0.59 | 0.53 | 0.58 | 0.44 | 0.71 | 0.5 |
| 1 | 168221225 | rs12608 | 0.938 | -3.233 | 0.832 | -0.044 | 0.022 | 0.009 | 0.009 | 0 | 0.53 | 0.46 | 0.54 | 0.44 | 0.67 | 0.42 |
| 1 | 168230181 | rs10489209 | 0.742 | -3.274 | 0.77 | -0.046 | -0.005 | -0.001 | -0.001 | 0.097 | 0.54 | 0.58 | 0.55 | 0.44 | 0.67 | 0.43 |
| 1 | 168240499 | rs1543311 | 0.336 | -3.41 | 0.624 | -0.109 | 0.017 | -0.011 | -0.011 | 0.28 | 0.6 | 0.7 | 0.57 | 0.49 | 0.67 | 0.52 |
| 1 | 168251535 | rs3767481 | -0.424 | -3.364 | 0.698 | -0.109 | -0.025 | 0.026 | 0.026 | 0 | 0.09 | 0.26 | 0.05 | 0.09 | 0.03 | 0.12 |
| 1 | 168254692 | rs1003503 | 0.308 | -3.553 | 0.561 | -0.106 | 0.028 | -0.005 | -0.005 | 0 | 0.58 | 0.67 | 0.57 | 0.46 | 0.67 | 0.51 |
| 1 | 168255020 | rs1003502 | 0.353 | -3.551 | 0.565 | -0.105 | 0.008 | -0.002 | -0.002 | 0 | 0.57 | 0.6 | 0.56 | 0.47 | 0.67 | 0.51 |
| 1 | 168257331 | rs3819818 | 1.164 | -3.603 | 0.423 | -0.168 | 0.063 | 0.04 | 0.04 | 0 | 0.45 | 0.3 | 0.51 | 0.38 | 0.62 | 0.36 |
| 1 | 168262289 | rs7543224 | 0.046 | -3.349 | 0.201 | -0.049 | 0.01 | -0.009 | -0.009 | 0.24 | 0.62 | 0.66 | 0.44 | 0.5 | 0.66 | 0.58 |
| 1 | 168262299 | rs7520743 | 0.899 | -3.011 | -0.214 | -0.12 | 0.058 | 0.041 | 0.041 | 0.334 | 0.46 | 0.2 | 0.35 | 0.41 | 0.6 | 0.34 |
| 1 | 168270110 | rs2072739 | -1.535 | -2.909 | 0.521 | 0.429 | -0.092 | 0.094 | 0.094 | 1.542 | 0.18 | 0.5 | 0.12 | 0.34 | 0.06 | 0.31 |
| 1 | 168274201 | rs4656579 | 1.216 | -3.299 | 0.5 | 0.637 | -0.014 | -0.001 | -0.001 | 0 | 0.58 | 0.44 | 0.56 | 0.38 | 0.71 | 0.39 |
| 1 | 168274691 | rs6427122 | 0.748 | -3.274 | -0.08 | 0.706 | 0.015 | -0.007 | -0.007 | 0.198 | 0.64 | 0.71 | 0.61 | 0.58 | 0.74 | 0.46 |
| 1 | 168275262 | rs2284766 | 0.05 | -3.194 | -0.075 | 0.681 | 0.001 | -0.001 | -0.001 | 0 | 0.76 | 0.78 | 0.67 | 0.72 | 0.79 | 0.68 |
| 1 | 168278264 | rs1883148 | 0.095 | -3.067 | -0.045 | 0.669 | 0 | 0 | 0 | 0 | 0.75 | 0.77 | 0.64 | 0.72 | 0.78 | 0.68 |
| 1 | 168282491 | rs1000533 | 0.766 | -0.706 | -0.731 | 0.351 | -0.117 | -0.043 | -0.043 | 0 | 0.2 | 0.04 | 0.18 | 0.28 | 0.35 | 0.14 |
| 1 | 168285045 | rs1008680 | 0.383 | -0.884 | -0.362 | 0.247 | 0 | 0 | 0 | 0.239 | 0.54 | 0.58 | 0.46 | 0.45 | 0.44 | 0.53 |
| 1 | 168285844 | rs2268568 | 2.152 | -1.232 | -0.758 | 0.086 | -0.063 | -0.079 | -0.079 | 0.663 | 0.33 | 0.03 | 0.36 | 0.31 | 0.38 | 0.3 |
| 1 | 168288835 | rs2300575 | 0.736 | -0.857 | -0.332 | 0.385 | 0.02 | 0.002 | 0.002 | 0 | 0.21 | 0.2 | 0.19 | 0.27 | 0.34 | 0.13 |
| 1 | 168294250 | rs2300573 | 0.416 | -0.534 | -0.348 | 0.515 | 0.008 | -0.018 | -0.018 | 0.446 | 0.23 | 0.14 | 0.33 | 0.28 | 0.21 | 0.35 |
| 1 | 168300075 | rs2179184 | -0.781 | -0.386 | -0.008 | 0.352 | -0.003 | -0.003 | -0.003 | 0 | 0.6 | 0.55 | 0.58 | 0.69 | 0.61 | 0.68 |
| 1 | 168309147 | rs7535315 | -1.791 | -0.015 | 0.978 | 0.481 | 0.077 | 0.031 | 0.031 | 0.065 | 0.06 | 0.43 | 0.06 | 0 | 0.03 | 0.02 |
| 1 | 168309388 | rs7512951 | -1.805 | -0.206 | 0.681 | 0.4 | -0.047 | -0.061 | -0.061 | 0.007 | 0.28 | 0.48 | 0.16 | 0.15 | 0.15 | 0.32 |
| 1 | 168317903 | rs10918873 | -1.9 | 0.268 | 1.173 | 0.822 | -0.007 | 0.032 | 0.032 | 0 | 0.26 | 0.42 | 0.3 | 0.27 | 0.24 | 0.31 |
| 1 | 168327555 | rs10800356 | -0.269 | 0.415 | 1.044 | 0.916 | -0.025 | -0.025 | -0.025 | 0 | 0.1 | 0.04 | 0.23 | 0.26 | 0.18 | 0.24 |
| 1 | 168329363 | rs16860434 | 0.04 | 0.047 | 1.083 | 0.936 | 0.01 | -0.012 | -0.012 | 0 | 0.12 | 0.04 | 0.08 | 0.15 | 0.08 | 0.25 |
| 1 | 168332753 | rs1886004 | -0.984 | 0.036 | 1.292 | 0.917 | -0.02 | 0 | 0 | 0.241 | 0.17 | 0.31 | 0.12 | 0.15 | 0.11 | 0.27 |
| 1 | 168336393 | rs760685 | -0.036 | 0.177 | 0.949 | 1.032 | -0.005 | -0.009 | -0.009 | 0.191 | 0.09 | 0.05 | 0.21 | 0.27 | 0.12 | 0.23 |
| 1 | 168336623 | rs2268558 | -1.6 | 0.133 | 1.204 | 1.009 | -0.02 | 0.086 | 0.086 | 0 | 0.24 | 0.39 | 0.33 | 0.43 | 0.2 | 0.49 |
| 1 | 168338962 | rs2078197 | -2.049 | -0.092 | 1.228 | 0.969 | -0.029 | 0.039 | 0.039 | 0 | 0.33 | 0.52 | 0.36 | 0.44 | 0.23 | 0.5 |
| 1 | 168341267 | rs2284759 | -1.005 | -0.014 | 1.134 | 0.842 | -0.022 | 0.02 | 0.02 | 0 | 0.15 | 0.28 | 0.11 | 0.16 | 0.09 | 0.26 |
| 1 | 168341835 | rs2268555 | -0.756 | 0.089 | 0.579 | 1.205 | -0.001 | -0.002 | -0.002 | 0 | 0.07 | 0.07 | 0.04 | 0.11 | 0.08 | 0.05 |
| 1 | 168341866 | rs2268554 | -1.953 | 0.024 | 0.567 | 1.171 | -0.006 | -0.019 | -0.019 | 0 | 0.14 | 0.67 | 0.11 | 0.11 | 0.11 | 0.07 |
| 1 | 168351648 | rs2179182 | -1.014 | 0.412 | 0.077 | 1.708 | 0.045 | 0.228 | 0.228 | 0.015 | 0.12 | 0.35 | 0.29 | 0.45 | 0.14 | 0.21 |
| 1 | 168352289 | rs7516573 | -0.486 | 0.407 | -0.808 | 1.165 | 0.01 | 0.019 | 0.019 | 0.11 | 0.06 | 0.12 | 0.06 | 0.19 | 0.1 | 0.05 |
| 1 | 168361867 | rs2255438 | -1.261 | 0.245 | -1.465 | 0.832 | -0.046 | -0.047 | -0.047 | 0 | 0.39 | 0.9 | 0.38 | 0.59 | 0.39 | 0.36 |
| 1 | 168369018 | rs2268545 | 0.467 | -0.802 | -0.84 | 0.268 | 0.015 | 0.015 | 0.015 | 0 | 0.65 | 0.28 | 0.55 | 0.73 | 0.66 | 0.81 |
| 1 | 168375973 | rs2300568 | -0.612 | -1.35 | -0.48 | -0.27 | 0.025 | 0.036 | 0.036 | 0.362 | 0.47 | 0.57 | 0.36 | 0.22 | 0.37 | 0.45 |
| 1 | 168378160 | rs12406610 | -0.841 | -1.461 | -0.721 | -0.088 | 0.061 | 0.015 | 0.015 | 0.144 | 0.53 | 0.83 | 0.54 | 0.45 | 0.42 | 0.52 |
| 1 | 168381150 | rs12740293 | -0.017 | -1.333 | -0.51 | -0.412 | 0.004 | -0.038 | -0.038 | 0.651 | 0.16 | 0.03 | 0.13 | 0.2 | 0.12 | 0.18 |
| 1 | 168386785 | rs2300565 | 0.19 | -1.461 | -0.216 | -0.615 | -0.031 | 0.04 | 0.04 | 0 | 0.17 | 0.01 | 0.15 | 0.05 | 0.19 | 0.22 |
| 1 | 168389917 | rs12691501 | -0.377 | -1.288 | 0.309 | -0.481 | 0.019 | 0.067 | 0.067 | 0 | 0.4 | 0.13 | 0.32 | 0.22 | 0.36 | 0.39 |
| 1 | 168392286 | rs9633330 | -1.205 | -0.909 | 1.674 | -0.247 | -0.038 | -0.008 | -0.008 | 0 | 0.08 | 0.25 | 0.22 | 0.12 | 0.05 | 0.09 |
| 1 | 168397414 | rs2268541 | -1.414 | -0.92 | 1.444 | -0.275 | -0.053 | -0.014 | -0.014 | 0.241 | 0.11 | 0.49 | 0.25 | 0.11 | 0.06 | 0.09 |
| 1 | 168405109 | rs4656588 | -1.558 | -0.902 | 0.756 | -0.299 | -0.067 | -0.044 | -0.044 | 0.201 | 0.13 | 0.63 | 0.27 | 0.09 | 0.07 | 0.08 |
| 1 | 168406461 | rs7549700 | -1.399 | -0.841 | 0.709 | -0.27 | -0.059 | -0.031 | -0.031 | 2.738 | 0.12 | 0.53 | 0.2 | 0.09 | 0.05 | 0.08 |
| 1 | 168407955 | rs742264 | -0.214 | -0.323 | -0.023 | -1.271 | -0.008 | 0.053 | 0.053 | 0 | 0.47 | 0.35 | 0.67 | 0.88 | 0.51 | 0.54 |
| 1 | 168412113 | rs2268539 | -0.469 | -1.173 | 0.139 | -1.768 | 0.023 | 0.015 | 0.015 | 0 | 0.18 | 0.14 | 0.06 | 0.04 | 0.08 | 0.21 |
| 1 | 168413221 | rs1810467 | -0.597 | -0.485 | 0.34 | -1.866 | -0.013 | -0.01 | -0.01 | 0 | 0.65 | 0.69 | 0.75 | 0.91 | 0.56 | 0.73 |
| 1 | 168418373 | rs7535780 | -0.441 | -0.228 | 0.268 | -0.158 | 0.003 | 0.12 | 0.12 | 0 | 0.09 | 0.12 | 0.27 | 0.44 | 0.11 | 0.21 |
| 1 | 168426153 | rs858099 | -0.106 | -0.134 | 0.162 | -0.334 | 0.013 | 0.046 | 0.046 | 0 | 0.66 | 0.76 | 0.76 | 0.81 | 0.67 | 0.71 |
| 1 | 168431914 | rs12564969 | -0.494 | -0.037 | 0.26 | -0.204 | -0.011 | 0.101 | 0.101 | 0 | 0.08 | 0.16 | 0.2 | 0.32 | 0.08 | 0.17 |
| 1 | 168436436 | rs7543621 | 0.835 | 0.439 | 0.337 | -0.661 | 0.008 | 0.106 | 0.106 | 0 | 0.5 | 0.26 | 0.43 | 0.22 | 0.49 | 0.44 |
| 1 | 168438268 | rs10918897 | 0.282 | 0.382 | 0.37 | -0.533 | 0 | 0.01 | 0.01 | 0 | 0.66 | 0.7 | 0.58 | 0.31 | 0.63 | 0.57 |
| 1 | 168443178 | rs1737168 | -1.079 | 0.558 | 0.485 | 0.877 | 0.024 | 0.079 | 0.079 | 0 | 0.12 | 0.22 | 0.28 | 0.37 | 0.17 | 0.25 |

| | | | | | | | | | | | | | | | |
|---|---|---|---|---|---|---|---|---|---|---|---|---|---|---|---|
| 1 | 168445500 | rs858114 | -0.805 | 0.639 | 0.647 | 0.996 | 0.036 | 0.093 | 0.093 | 0 | 0.11 | 0.22 | 0.27 | 0.34 | 0.17 | 0.24 |
| 1 | 168450775 | rs10918901 | -0.546 | -0.902 | 0.237 | 0.15 | -0.072 | 0.069 | 0.069 | 0 | 0.22 | 0.43 | 0.15 | 0.42 | 0.13 | 0.29 |
| 1 | 168451333 | rs6656805 | -0.534 | -0.9 | 0.244 | 0.151 | -0.073 | 0.073 | 0.073 | 0 | 0.22 | 0.43 | 0.15 | 0.43 | 0.13 | 0.29 |
| 1 | 168456819 | rs1014978 | -0.529 | -1.097 | -0.123 | -0.468 | -0.021 | 0.009 | 0.009 | 0 | 0.15 | 0.22 | 0.09 | 0.23 | 0.11 | 0.18 |
| 1 | 168460487 | rs10918911 | -0.497 | -1.097 | -0.111 | -0.401 | -0.021 | 0.021 | 0.021 | 0 | 0.14 | 0.22 | 0.09 | 0.25 | 0.11 | 0.18 |
| 1 | 168461703 | rs12094807 | -0.552 | -1.162 | -0.096 | -0.359 | -0.024 | 0.041 | 0.041 | 0 | 0.15 | 0.25 | 0.1 | 0.35 | 0.11 | 0.23 |
| 1 | 168463587 | rs7417366 | 0.373 | -0.738 | -1.151 | -1.418 | -0.049 | -0.027 | -0.027 | 0.416 | 0.35 | 0.07 | 0.47 | 0.56 | 0.36 | 0.48 |
| 1 | 168465557 | rs858120 | -1.695 | -0.541 | -1.349 | -2.138 | -0.044 | 0.104 | 0.104 | 0 | 0.74 | 0.95 | 0.81 | 0.95 | 0.68 | 0.83 |
| 1 | 168474310 | rs4656597 | -1.912 | -0.503 | -1.426 | -2.15 | -0.035 | 0.074 | 0.074 | 0.029 | 0.82 | 0.96 | 0.92 | 1 | 0.83 | 0.95 |
| 1 | 168475194 | rs10918912 | 1.068 | -0.564 | -1.293 | -1.945 | -0.02 | 0.037 | 0.037 | 0 | 0.07 | 0 | 0.13 | 0.04 | 0.08 | 0.13 |
| 1 | 168484505 | rs12126216 | 0.351 | -0.463 | -0.737 | -1.859 | 0.001 | -0.042 | -0.042 | 0.005 | 0.39 | 0.57 | 0.37 | 0.19 | 0.4 | 0.33 |
| 1 | 168487659 | rs6667542 | 0.553 | -0.459 | -0.404 | -1.785 | 0 | -0.081 | -0.081 | 2.649 | 0.38 | 0.44 | 0.35 | 0.1 | 0.4 | 0.3 |
| 1 | 168489847 | rs1323530 | -0.399 | -0.479 | -0.522 | -1.706 | 0.014 | 0.079 | 0.079 | 0 | 0.48 | 0.8 | 0.46 | 0.44 | 0.48 | 0.5 |
| 1 | 168492603 | rs16860761 | 0.293 | -1.104 | -0.356 | -1.365 | -0.001 | 0.141 | 0.141 | 0 | 0.09 | 0.32 | 0.1 | 0.25 | 0.07 | 0.14 |
| 1 | 168494042 | rs4656153 | 0.584 | -0.944 | -1.062 | -1.466 | -0.053 | -0.052 | -0.052 | 1.284 | 0.32 | 0.06 | 0.44 | 0.55 | 0.34 | 0.44 |
| 1 | 168496272 | rs1323532 | 0.99 | -0.658 | -0.819 | -1.813 | -0.027 | -0.034 | -0.034 | 0.03 | 0.17 | 0.07 | 0.38 | 0.63 | 0.21 | 0.32 |
| 1 | 168500219 | rs1323533 | 0.854 | -0.958 | -0.535 | -2.12 | 0.013 | 0.027 | 0.027 | 0 | 0.23 | 0.14 | 0.15 | 0.1 | 0.18 | 0.17 |
| 1 | 168500458 | rs7529824 | -0.444 | -1.039 | -0.066 | -2.165 | 0 | 0.002 | 0.002 | 0.171 | 0.41 | 0.4 | 0.56 | 0.75 | 0.41 | 0.5 |
| 1 | 168500750 | rs1040508 | -0.369 | -1.037 | -0.113 | -2.164 | 0.002 | 0.075 | 0.075 | 0.139 | 0.39 | 0.23 | 0.53 | 0.75 | 0.39 | 0.48 |
| 1 | 168501453 | rs10429892 | -1.222 | -1.069 | -0.192 | -2.443 | 0.007 | -0.032 | -0.032 | 0.509 | 0.67 | 0.66 | 0.66 | 0.87 | 0.54 | 0.65 |
| 1 | 168501594 | rs17572134 | 1.517 | -1.08 | -0.487 | -2.396 | 0.014 | 0.069 | 0.069 | 0.953 | 0.22 | 0.04 | 0.09 | 0.12 | 0.14 | 0.15 |
| 1 | 168503162 | rs1998510 | 1.34 | -0.895 | -0.32 | -2.172 | -0.02 | -0.025 | -0.025 | 0 | 0.16 | 0.09 | 0.38 | 0.64 | 0.21 | 0.32 |
| 1 | 168521417 | rs1933112 | -0.522 | -1.088 | 0.042 | -2.341 | 0.006 | 0.018 | 0.018 | 0 | 0.45 | 0.66 | 0.55 | 0.69 | 0.41 | 0.48 |
| 1 | 168521615 | rs12049040 | 1.376 | -1.068 | 0.179 | -2.35 | 0.001 | 0.002 | 0.002 | 0 | 0.17 | 0.19 | 0.39 | 0.66 | 0.21 | 0.31 |
| 1 | 168521747 | rs1933111 | 0.688 | -0.977 | 0.234 | -2.35 | -0.014 | -0.088 | -0.088 | 0 | 0.23 | 0.33 | 0.15 | 0.03 | 0.2 | 0.17 |
| 1 | 168524895 | rs1933107 | -1.027 | -0.996 | -0.18 | -2.481 | -0.019 | 0.02 | 0.02 | 0.015 | 0.67 | 0.78 | 0.65 | 0.8 | 0.53 | 0.63 |
| 1 | 168531765 | rs12058546 | 0.282 | -0.916 | 0.259 | -1.839 | -0.023 | -0.13 | -0.13 | 0 | 0.23 | 0.39 | 0.16 | 0 | 0.19 | 0.1 |
| 1 | 168537645 | rs2419114 | -0.844 | -0.71 | -0.227 | -1.505 | 0.008 | 0.027 | 0.027 | 0 | 0.47 | 0.65 | 0.55 | 0.67 | 0.38 | 0.51 |
| 1 | 168542041 | rs16860928 | 0.027 | -0.779 | -0.372 | -1.479 | -0.066 | -0.137 | -0.137 | 0 | 0.24 | 0.44 | 0.11 | 0 | 0.12 | 0.09 |
| 1 | 168544231 | rs3753938 | -0.084 | 0.371 | -0.777 | -0.755 | -0.044 | -0.025 | -0.025 | 0 | 0.21 | 0.12 | 0.42 | 0.67 | 0.26 | 0.42 |
| 1 | 168549720 | rs6687507 | 0.747 | 0.242 | -0.451 | -1.007 | 0.004 | -0.021 | -0.021 | 0 | 0.6 | 0.42 | 0.39 | 0.19 | 0.63 | 0.4 |
| 1 | 168549885 | rs6700487 | -0.591 | 0.277 | -0.395 | -0.511 | -0.037 | -0.043 | -0.043 | 0.543 | 0.78 | 0.84 | 0.56 | 0.39 | 0.74 | 0.53 |
| 1 | 168551236 | rs982143 | -0.107 | 0.047 | -0.317 | -0.578 | 0.005 | -0.019 | -0.019 | 0 | 0.14 | 0.06 | 0.15 | 0.19 | 0.1 | 0.13 |
| 1 | 168553057 | rs6662401 | -0.685 | 0.044 | -0.409 | -0.58 | -0.023 | 0.027 | 0.027 | 0.002 | 0.17 | 0.43 | 0.18 | 0.2 | 0.11 | 0.13 |
| 1 | 168557141 | rs2419115 | 0.184 | 0.283 | -0.656 | -0.845 | -0.055 | 0.07 | 0.07 | 0 | 0.66 | 0.81 | 0.43 | 0.16 | 0.65 | 0.43 |
| 1 | 168560132 | rs7556321 | 0.792 | 0.475 | -1.305 | -1.168 | -0.014 | -0.004 | -0.004 | 0 | 0.2 | 0.14 | 0.19 | 0.02 | 0.33 | 0.19 |
| 1 | 168563895 | rs6693141 | 0.594 | 0.281 | -0.871 | -0.572 | -0.004 | 0.019 | 0.019 | 0.023 | 0.58 | 0.34 | 0.37 | 0.21 | 0.63 | 0.43 |
| 1 | 168566850 | rs10918934 | -0.319 | 0.28 | -0.997 | -0.196 | -0.026 | -0.056 | -0.056 | 0 | 0.12 | 0.01 | 0.19 | 0.23 | 0.09 | 0.12 |
| 1 | 168568911 | rs1933090 | -0.029 | -0.227 | -0.665 | -0.884 | -0.002 | -0.005 | -0.005 | 0.491 | 0.71 | 0.72 | 0.72 | 0.8 | 0.78 | 0.85 |
| 1 | 168569112 | rs12137493 | 0.4 | -0.279 | -0.788 | -0.927 | -0.033 | -0.042 | -0.042 | 0 | 0.1 | 0 | 0.19 | 0.17 | 0.11 | 0.1 |
| 1 | 168569568 | rs10800360 | 0.727 | -0.139 | -0.505 | -1.179 | 0 | -0.008 | -0.008 | 1.852 | 0.43 | 0.44 | 0.35 | 0.26 | 0.49 | 0.46 |
| 1 | 168573041 | rs988724 | 0.174 | -0.2 | -0.446 | -1.105 | 0.013 | 0.029 | 0.029 | 0 | 0.3 | 0.45 | 0.39 | 0.55 | 0.3 | 0.38 |
| 1 | 168574698 | rs1933089 | 0.14 | -0.153 | -0.22 | -0.97 | 0.025 | 0.063 | 0.063 | 0 | 0.3 | 0.47 | 0.33 | 0.51 | 0.3 | 0.39 |
| 1 | 168575163 | rs10800363 | 0.05 | -0.153 | -0.217 | -0.953 | 0 | 0.002 | 0.002 | 5.158 | 0.53 | 0.48 | 0.6 | 0.46 | 0.6 | 0.55 |
| 1 | 168581220 | rs6427135 | 0.048 | -0.199 | 0.181 | -1.032 | 0 | 0.001 | 0.001 | 7.283 | 0.55 | 0.62 | 0.6 | 0.46 | 0.62 | 0.57 |
| 1 | 168582119 | rs10800364 | 0.725 | -0.286 | -0.102 | -1.12 | -0.035 | 0.063 | 0.063 | 3.97 | 0.4 | 0.22 | 0.32 | 0.26 | 0.52 | 0.45 |
| 1 | 168585289 | rs11589732 | 0.928 | -0.153 | -0.239 | -1.114 | -0.035 | -0.052 | -0.052 | 0.017 | 0.11 | 0.01 | 0.26 | 0.2 | 0.13 | 0.1 |
| 1 | 168587489 | rs12093734 | 0.711 | -0.276 | -0.015 | -1.184 | -0.005 | 0.016 | 0.016 | 3.858 | 0.4 | 0.35 | 0.33 | 0.26 | 0.5 | 0.44 |
| 1 | 168589619 | rs12401991 | 0.756 | -0.413 | 0.064 | -1.31 | -0.01 | -0.035 | -0.035 | 0 | 0.25 | 0.16 | 0.3 | 0.5 | 0.25 | 0.4 |
| 1 | 168589639 | rs2419118 | -2.27 | -0.378 | -0.059 | -1.334 | -0.005 | 0.072 | 0.072 | 5.097 | 0.84 | 0.98 | 0.92 | 0.97 | 0.88 | 0.95 |
| 1 | 168589988 | rs7355223 | 0.632 | -0.176 | 0.141 | -1.109 | -0.04 | 0.056 | 0.056 | 6.073 | 0.38 | 0.19 | 0.31 | 0.26 | 0.5 | 0.44 |
| 1 | 168594940 | rs2064298 | 0.141 | 0.161 | -0.876 | -1.104 | -0.006 | 0.003 | 0.003 | 0.061 | 0.45 | 0.17 | 0.47 | 0.51 | 0.58 | 0.6 |
| 1 | 168599216 | rs11580561 | -0.112 | 0.283 | -0.926 | -1.54 | -0.017 | -0.008 | -0.008 | 1.638 | 0.31 | 0.2 | 0.49 | 0.62 | 0.35 | 0.49 |
| 1 | 168599786 | rs4656609 | -2.606 | -0.009 | -1.026 | -1.674 | -0.003 | 0.057 | 0.057 | 2.814 | 0.87 | 0.98 | 0.93 | 0.97 | 0.9 | 0.95 |
| 1 | 168601255 | rs728354 | 0.064 | -0.378 | -0.39 | -1.844 | -0.01 | 0.059 | 0.059 | 0 | 0.27 | 0.52 | 0.42 | 0.54 | 0.23 | 0.38 |
| 1 | 168602862 | rs10489216 | 0.833 | -0.47 | -0.756 | -2.003 | -0.039 | -0.034 | -0.034 | 0 | 0.26 | 0.04 | 0.16 | 0.28 | 0.24 | 0.25 |
| 1 | 168606715 | rs10800374 | 0.782 | -0.444 | -0.455 | -2.012 | -0.014 | 0.043 | 0.043 | 1.703 | 0.32 | 0.23 | 0.33 | 0.14 | 0.44 | 0.32 |
| 1 | 168607373 | rs742479 | -0.029 | -0.388 | -0.294 | -1.871 | 0.001 | 0.001 | 0.001 | 2.358 | 0.59 | 0.4 | 0.5 | 0.43 | 0.68 | 0.57 |
| 1 | 168607663 | rs1076697 | -0.024 | -0.37 | -0.195 | -1.842 | -0.009 | 0.045 | 0.045 | 0 | 0.27 | 0.56 | 0.42 | 0.54 | 0.22 | 0.4 |
| 1 | 168609204 | rs6678237 | 0.721 | 0.108 | -0.716 | -1.712 | 0.031 | -0.004 | -0.004 | 0 | 0.09 | 0.27 | 0.06 | 0.06 | 0.07 | 0.07 |
| 1 | 168610065 | rs742483 | 1.07 | 0.083 | -1.198 | -1.489 | -0.004 | -0.045 | -0.045 | 0 | 0.15 | 0.06 | 0.35 | 0.48 | 0.15 | 0.33 |
| 1 | 168610817 | rs926523 | -2.033 | 0.111 | -1.039 | -1.458 | 0.005 | 0.036 | 0.036 | 3.463 | 0.86 | 0.95 | 0.92 | 0.97 | 0.9 | 0.95 |
| 1 | 168611846 | rs16861196 | 0.354 | 0.294 | -0.942 | -1.311 | -0.013 | -0.027 | -0.027 | 0.397 | 0.22 | 0.06 | 0.16 | 0.28 | 0.21 | 0.26 |
| 1 | 168612925 | rs6427139 | -1.236 | 0.51 | -1.145 | -1.794 | -0.017 | 0.015 | 0.015 | 0.271 | 0.59 | 0.76 | 0.44 | 0.41 | 0.53 | 0.53 |
| 1 | 168613176 | rs7550529 | 0.136 | 0.548 | -0.832 | -1.895 | 0.006 | -0.036 | -0.036 | 1.767 | 0.11 | 0.18 | 0.07 | 0.04 | 0.09 | 0.06 |
| 1 | 168616678 | rs10489218 | -1.376 | 0.466 | -0.966 | -1.948 | -0.016 | 0.022 | 0.022 | 1.112 | 0.58 | 0.77 | 0.44 | 0.41 | 0.52 | 0.52 |
| 1 | 168616738 | rs10918948 | 0.418 | 0.746 | -1.408 | -1.774 | -0.065 | 0.046 | 0.046 | 3.636 | 0.19 | 0.01 | 0.2 | 0.1 | 0.23 | 0.21 |
| 1 | 168618641 | rs1412337 | 0.17 | 0.58 | -0.799 | -1.501 | 0.006 | -0.007 | -0.007 | 0 | 0.23 | 0.13 | 0.14 | 0.27 | 0.2 | 0.25 |
| 1 | 168625843 | rs2064300 | 0.083 | 0.839 | -1.019 | -1.63 | -0.017 | 0.05 | 0.05 | 0.003 | 0.32 | 0.06 | 0.22 | 0.25 | 0.34 | 0.3 |

| | | | | | | | | | | | | | | | | |
|---|---|---|---|---|---|---|---|---|---|---|---|---|---|---|---|---|
| 1 | 168626707 | rs1015001 | 0.034 | 1.243 | -0.816 | -1.891 | 0.02 | -0.045 | -0.045 | 0.362 | 0.23 | 0.34 | 0.22 | 0.09 | 0.28 | 0.25 |
| 1 | 168633890 | rs1933115 | -0.954 | 1.209 | -0.773 | -1.841 | -0.004 | -0.01 | -0.01 | 0 | 0.56 | 0.52 | 0.45 | 0.35 | 0.62 | 0.55 |
| 1 | 168634922 | rs1407953 | 0.481 | 1.611 | -0.565 | -2.052 | -0.025 | 0.046 | 0.046 | 3.251 | 0.12 | 0.02 | 0.11 | 0.05 | 0.13 | 0.09 |
| 1 | 168635004 | rs1009434 | -0.471 | 1.407 | -0.149 | -1.7 | 0.019 | -0.043 | -0.043 | 0 | 0.22 | 0.32 | 0.21 | 0.09 | 0.28 | 0.25 |
| 1 | 168647094 | rs11808391 | -0.593 | 1.355 | -0.081 | -1.629 | 0.012 | -0.072 | -0.072 | 0 | 0.28 | 0.35 | 0.28 | 0.08 | 0.34 | 0.28 |
| 1 | 168653857 | rs7416358 | -0.944 | 1.032 | 0.132 | -1.709 | -0.001 | -0.029 | -0.029 | 0.013 | 0.31 | 0.68 | 0.27 | 0.03 | 0.33 | 0.23 |
| 1 | 168654331 | rs1415678 | -0.452 | 1.02 | -0.482 | -1.592 | 0.007 | -0.098 | -0.098 | 0.017 | 0.27 | 0.35 | 0.23 | 0.02 | 0.29 | 0.2 |
| 1 | 168655627 | rs17502304 | -0.452 | 1.02 | -0.482 | -1.592 | 0.007 | -0.098 | -0.098 | 2.165 | 0.27 | 0.35 | 0.23 | 0.02 | 0.29 | 0.2 |
| 1 | 168662302 | rs538060 | 0.61 | 0.95 | -1.485 | -1.738 | -0.068 | 0.255 | 0.255 | 0.323 | 0.55 | 0.07 | 0.66 | 0.88 | 0.53 | 0.66 |
| 1 | 168662352 | rs646441 | 0.153 | 0.159 | -0.467 | -1.555 | -0.05 | -0.087 | -0.087 | 0 | 0.77 | 0.54 | 0.87 | 0.96 | 0.85 | 0.91 |
| 1 | 168664450 | rs682367 | 0.324 | 0.25 | -0.535 | -0.95 | 0 | -0.001 | -0.001 | 0 | 0.44 | 0.44 | 0.49 | 0.51 | 0.41 | 0.46 |
| 1 | 168665974 | rs524705 | -0.287 | 0.49 | -0.662 | -1.252 | -0.05 | -0.042 | -0.042 | 0.009 | 0.59 | 0.83 | 0.58 | 0.58 | 0.54 | 0.62 |
| 1 | 168666850 | rs12116794 | 0.441 | 0.5 | -0.109 | -1.278 | 0.008 | -0.042 | -0.042 | 0.179 | 0.12 | 0.23 | 0.07 | 0.07 | 0.1 | 0.14 |
| 1 | 168667989 | rs10489360 | 1.019 | 0.427 | 0.044 | -1.64 | 0.004 | -0.063 | -0.063 | 0.599 | 0.17 | 0.24 | 0.13 | 0 | 0.19 | 0.13 |
| 1 | 168671797 | rs12122756 | 0.554 | 0.423 | 0.018 | -1.557 | 0.007 | -0.042 | -0.042 | 0 | 0.12 | 0.22 | 0.06 | 0.06 | 0.1 | 0.14 |
| 1 | 168672858 | rs12126497 | 0.261 | 0.414 | 0.082 | -1.562 | 0.009 | -0.052 | -0.052 | 0.593 | 0.13 | 0.32 | 0.07 | 0.06 | 0.1 | 0.14 |
| 1 | 168675971 | rs530022 | 0.313 | 0.413 | 0.005 | -1.553 | -0.002 | 0.001 | 0.001 | 0 | 0.46 | 0.51 | 0.51 | 0.51 | 0.44 | 0.48 |
| 1 | 168678324 | rs12133988 | 0.421 | 0.358 | 0.036 | -1.676 | 0.02 | -0.053 | -0.053 | 0.51 | 0.12 | 0.3 | 0.06 | 0.05 | 0.09 | 0.14 |
| 1 | 168679692 | rs581753 | -0.799 | 0.613 | -0.563 | -1.75 | -0.045 | -0.155 | -0.155 | 2.43 | 0.74 | 0.95 | 0.64 | 0.56 | 0.69 | 0.79 |
| 1 | 168687822 | rs7542028 | 0.979 | 0.611 | -0.464 | -1.704 | 0.014 | -0.044 | -0.044 | 0 | 0.11 | 0.21 | 0.05 | 0.05 | 0.09 | 0.13 |
| 1 | 168689940 | rs545833 | 1.049 | 0.677 | -0.587 | -1.693 | 0 | 0.001 | 0.001 | 0 | 0.34 | 0.28 | 0.4 | 0.54 | 0.28 | 0.44 |
| 1 | 168691293 | rs4656614 | 0.779 | 1.122 | -1.135 | -1.987 | -0.036 | 0.056 | 0.056 | 0.538 | 0.19 | 0.06 | 0.11 | 0.01 | 0.24 | 0.21 |
| 1 | 168692615 | rs11590043 | 0.299 | 1.069 | -0.906 | -1.722 | -0.006 | -0.041 | -0.041 | 2.26 | 0.35 | 0.3 | 0.2 | 0.01 | 0.39 | 0.33 |
| 1 | 168694212 | rs2207216 | 0.42 | 1.287 | -1.211 | -0.631 | -0.083 | -0.092 | -0.092 | 0.134 | 0.72 | 0.92 | 0.62 | 0.66 | 0.6 | 0.74 |
| 1 | 168695609 | rs736846 | -1.252 | 0.508 | -0.337 | -1.16 | -0.013 | -0.074 | -0.074 | 0 | 0.18 | 0.35 | 0.14 | 0.06 | 0.15 | 0.17 |
| 1 | 168697942 | rs628718 | -0.735 | 0.292 | -0.889 | -1.22 | 0.024 | 0.06 | 0.06 | 1.042 | 0.68 | 0.89 | 0.53 | 0.2 | 0.73 | 0.58 |
| 1 | 168698173 | rs1052591 | 0.234 | 0.429 | -0.632 | -1.559 | -0.016 | 0.029 | 0.029 | 0 | 0.37 | 0.25 | 0.31 | 0.16 | 0.51 | 0.37 |
| 1 | 168701088 | rs1555206 | 0.234 | 0.429 | -0.632 | -1.559 | -0.016 | 0.029 | 0.029 | 0 | 0.37 | 0.25 | 0.31 | 0.16 | 0.51 | 0.37 |
| 1 | 168707974 | rs610403 | 0.201 | 0.469 | -0.614 | -1.541 | -0.008 | 0.013 | 0.013 | 0 | 0.38 | 0.29 | 0.31 | 0.16 | 0.51 | 0.38 |
| 1 | 168711459 | rs655128 | 0.509 | 0.474 | -0.568 | -1.542 | -0.031 | 0.038 | 0.038 | 0 | 0.36 | 0.24 | 0.29 | 0.16 | 0.49 | 0.38 |
| 1 | 168720250 | rs3950134 | 0.102 | 0.291 | -0.64 | -1.687 | -0.02 | -0.129 | -0.129 | 0 | 0.31 | 0.62 | 0.2 | 0 | 0.22 | 0.18 |
| 1 | 168725876 | rs6427150 | -1.047 | 0.495 | -0.699 | -1.458 | 0.018 | 0.046 | 0.046 | 1.037 | 0.69 | 0.93 | 0.54 | 0.2 | 0.75 | 0.6 |
| 1 | 168732201 | rs10918977 | 0.616 | 0.232 | -0.4 | -1.563 | -0.022 | -0.156 | -0.156 | 0.128 | 0.24 | 0.43 | 0.17 | 0 | 0.2 | 0.18 |
| 1 | 168734457 | rs12065661 | 0.663 | 0.269 | -0.884 | -1.358 | -0.022 | -0.094 | -0.094 | 1.022 | 0.24 | 0.53 | 0.23 | 0.03 | 0.19 | 0.17 |
| 1 | 168737147 | rs2744666 | 0.644 | 0.465 | -1.084 | -1.251 | -0.034 | 0.005 | 0.005 | 0.12 | 0.34 | 0.04 | 0.46 | 0.75 | 0.3 | 0.41 |
| 1 | 168741270 | rs2786765 | -0.506 | 0.699 | -0.846 | -1.321 | -0.011 | 0.024 | 0.024 | 0 | 0.63 | 0.69 | 0.76 | 0.86 | 0.57 | 0.63 |
| 1 | 168741818 | rs10800380 | -0.035 | 0.652 | -0.032 | -0.801 | -0.007 | -0.102 | -0.102 | 0 | 0.26 | 0.49 | 0.28 | 0.06 | 0.23 | 0.21 |
| 1 | 168745136 | rs2206300 | -0.831 | 0.725 | 0.339 | 0.103 | 0.028 | 0.186 | 0.186 | 0 | 0.21 | 0.47 | 0.28 | 0.43 | 0.2 | 0.31 |
| 1 | 168748586 | rs17570453 | -1.002 | 0.795 | 0.845 | -0.438 | 0.107 | 0 | 0 | 0 | 0.13 | 0.49 | 0.17 | 0.06 | 0.17 | 0.14 |
| 1 | 168758306 | rs3738241 | -0.707 | 0.768 | 0.802 | -0.5 | 0.03 | -0.009 | -0.009 | 2.837 | 0.12 | 0.18 | 0.14 | 0.07 | 0.19 | 0.19 |
| 1 | 168762175 | rs2066895 | -0.789 | 0.767 | 0.804 | -0.501 | 0.064 | -0.012 | -0.012 | 0 | 0.12 | 0.27 | 0.15 | 0.07 | 0.19 | 0.18 |
| 1 | 168772393 | rs10918992 | -0.097 | -0.017 | -0.335 | -0.954 | 0.03 | 0.052 | 0.052 | 0 | 0.26 | 0.04 | 0.36 | 0.88 | 0.18 | 0.47 |
| 1 | 168812739 | rs7524938 | -0.679 | 0.036 | 0.072 | -1.716 | 0.013 | -0.117 | -0.117 | 0.263 | 0.69 | 0.58 | 0.81 | 0.97 | 0.68 | 0.87 |
| 1 | 168813241 | rs6695006 | 0.231 | -0.217 | -0.665 | -1.851 | 0.052 | 0.066 | 0.066 | 0 | 0.27 | 0.15 | 0.42 | 0.89 | 0.19 | 0.48 |
| 1 | 168816245 | rs2143091 | 0.837 | -0.328 | -0.831 | -2.035 | -0.057 | 0.097 | 0.097 | 0.005 | 0.34 | 0.13 | 0.27 | 0.06 | 0.41 | 0.35 |
| 1 | 168820779 | rs200558 | 0.542 | -0.401 | -0.851 | -2.042 | 0.054 | 0.058 | 0.058 | 0 | 0.25 | 0.08 | 0.41 | 0.88 | 0.19 | 0.47 |
| 1 | 168822447 | rs200561 | 0.023 | -0.358 | -0.762 | -2.045 | 0.068 | 0.147 | 0.147 | 0 | 0.32 | 0.15 | 0.5 | 0.92 | 0.26 | 0.52 |
| 1 | 168824111 | rs169266 | -0.649 | -0.326 | -0.796 | -2.059 | -0.002 | 0.035 | 0.035 | 0.175 | 0.66 | 0.29 | 0.78 | 0.97 | 0.68 | 0.87 |
| 1 | 168825618 | rs12124937 | 0.489 | 0.292 | -1.081 | -1.689 | -0.117 | 0.174 | 0.174 | 0.09 | 0.33 | 0.06 | 0.27 | 0.06 | 0.41 | 0.35 |
| 1 | 168837734 | rs1322494 | 0.101 | 0.736 | -0.242 | -1.169 | 0.047 | 0.003 | 0.003 | 0 | 0.1 | 0.25 | 0.14 | 0.11 | 0.15 | 0.16 |
| 1 | 168837845 | rs6702913 | 0.612 | 0.435 | -0.299 | -0.651 | -0.001 | 0.016 | 0.016 | 0 | 0.16 | 0.13 | 0.16 | 0.43 | 0.16 | 0.22 |
| 1 | 168849973 | rs7537225 | 0.728 | 0.561 | -0.32 | -1.197 | -0.017 | 0.062 | 0.062 | 0 | 0.31 | 0.44 | 0.42 | 0.43 | 0.25 | 0.3 |
| 1 | 168851940 | rs1885555 | 0.743 | 0.544 | 0.026 | -1.213 | -0.009 | 0.036 | 0.036 | 0 | 0.3 | 0.35 | 0.42 | 0.43 | 0.25 | 0.3 |
| 1 | 168852043 | rs1570334 | -1.591 | 0.559 | -0.483 | -1.18 | -0.016 | 0.015 | 0.015 | 0 | 0.93 | 0.99 | 0.93 | 0.97 | 0.9 | 0.93 |
| 1 | 168856283 | rs10919006 | 0.887 | 0.67 | -0.483 | -1.199 | 0 | 0.011 | 0.011 | 0 | 0.17 | 0.13 | 0.17 | 0.44 | 0.18 | 0.22 |
| 1 | 168862571 | rs10800393 | 0.887 | 0.869 | -0.023 | -0.712 | 0.01 | 0.04 | 0.04 | 0.373 | 0.49 | 0.19 | 0.36 | 0.44 | 0.5 | 0.47 |
| 1 | 168863309 | rs1407477 | -0.57 | 0.894 | 0.125 | -0.664 | -0.007 | -0.008 | -0.008 | 0 | 0.41 | 0.62 | 0.51 | 0.5 | 0.39 | 0.42 |
| 1 | 168869293 | rs6657987 | 0.972 | 1.156 | 0.323 | -0.496 | -0.025 | 0.004 | 0.004 | 0.022 | 0.56 | 0.26 | 0.37 | 0.47 | 0.53 | 0.55 |
| 1 | 168871028 | rs4142003 | 0.897 | 1.17 | 0.284 | -0.49 | -0.016 | 0.002 | 0.002 | 0 | 0.57 | 0.32 | 0.38 | 0.47 | 0.53 | 0.55 |
| 1 | 168874099 | rs2146204 | 0.736 | 1.242 | 0.507 | -0.159 | 0.032 | 0 | 0 | 0 | 0.08 | 0.09 | 0.08 | 0.08 | 0.15 | 0.11 |
| 1 | 168882428 | rs10919013 | -1.109 | 1.015 | 0.435 | -0.259 | 0.011 | 0.125 | 0.125 | 0.11 | 0.24 | 0.53 | 0.38 | 0.48 | 0.2 | 0.3 |
| 1 | 168888946 | rs973608 | 0.293 | 1.328 | -0.316 | -0.34 | -0.012 | 0 | 0 | 0.151 | 0.1 | 0.06 | 0.1 | 0.09 | 0.16 | 0.11 |
| 1 | 168889557 | rs17514245 | 0.704 | 1.65 | -0.224 | -0.605 | 0.046 | -0.032 | -0.032 | 0 | 0.06 | 0.14 | 0.05 | 0 | 0.09 | 0.04 |
| 1 | 168891735 | rs10737544 | -0.034 | 1.049 | -0.439 | -0.199 | -0.011 | -0.013 | -0.013 | 0 | 0.6 | 0.67 | 0.45 | 0.49 | 0.57 | 0.6 |
| 1 | 168892181 | rs6427160 | 0.171 | 0.454 | -1.21 | 0.002 | 0.05 | 0.031 | 0.031 | 0 | 0.49 | 0.23 | 0.3 | 0.46 | 0.45 | 0.5 |
| 1 | 168907072 | rs882126 | -0.431 | 0.122 | -0.897 | 0.106 | 0.004 | -0.004 | -0.004 | 1.197 | 0.21 | 0.19 | 0.2 | 0.49 | 0.12 | 0.27 |
| 1 | 168907616 | rs12743077 | 0.876 | 0.373 | -1.067 | -0.752 | -0.006 | 0.052 | 0.052 | 0 | 0.37 | 0.26 | 0.23 | 0.11 | 0.41 | 0.35 |
| 1 | 168907899 | rs3795611 | 0.744 | 0.544 | -0.94 | -0.713 | 0 | -0.001 | -0.001 | 0.705 | 0.07 | 0.07 | 0.14 | 0.04 | 0.1 | 0.09 |
| 1 | 168918766 | rs981257 | -1.047 | 0.407 | -1.146 | -1.413 | 0.006 | 0.01 | 0.01 | 0.268 | 0.42 | 0.64 | 0.47 | 0.63 | 0.34 | 0.52 |

| | | | | | | | | | | | | | | | | |
|---|---|---|---|---|---|---|---|---|---|---|---|---|---|---|---|---|
| 1 | 168922582 | rs12048750 | 1.318 | 0.45 | -1.242 | -1.517 | -0.001 | 0.024 | 0.024 | 0 | 0.39 | 0.32 | 0.24 | 0.11 | 0.42 | 0.33 |
| 1 | 168927614 | rs10489370 | 1.365 | 0.625 | -1.266 | -1.408 | -0.003 | 0.002 | 0.002 | 0.008 | 0.06 | 0.03 | 0.13 | 0.03 | 0.09 | 0.08 |
| 1 | 168930460 | rs12729624 | 1.121 | 0.869 | -1.331 | -1.622 | 0.002 | -0.001 | -0.001 | 0 | 0.12 | 0.12 | 0.12 | 0.11 | 0.22 | 0.2 |
| 1 | 168932500 | rs10489371 | 0.279 | 0.573 | -0.608 | -1.704 | -0.002 | -0.009 | -0.009 | 0 | 0.24 | 0.2 | 0.27 | 0.59 | 0.22 | 0.35 |
| 1 | 168933787 | rs16861817 | 0.674 | 0.304 | -0.501 | -1.688 | 0.005 | -0.006 | -0.006 | 0.961 | 0.14 | 0.2 | 0.12 | 0.1 | 0.23 | 0.21 |
| 1 | 168942829 | rs10800398 | 0.75 | 0.179 | -0.046 | -1.898 | 0.001 | -0.003 | -0.003 | 0.325 | 0.54 | 0.52 | 0.43 | 0.61 | 0.47 | 0.5 |
| 1 | 168943770 | rs2181210 | 0.991 | 0.118 | 0.223 | -1.927 | -0.005 | 0.03 | 0.03 | 0 | 0.53 | 0.3 | 0.41 | 0.61 | 0.46 | 0.5 |
| 1 | 168945426 | rs10919028 | 0.879 | 0.028 | 0.338 | -1.899 | -0.001 | -0.004 | -0.004 | 0 | 0.1 | 0.1 | 0.17 | 0.04 | 0.1 | 0.13 |
| 1 | 168950508 | rs3007381 | -1.073 | -0.025 | 0.217 | -1.964 | -0.001 | 0.008 | 0.008 | 0 | 0.35 | 0.48 | 0.48 | 0.38 | 0.4 | 0.5 |
| 1 | 168951141 | rs12030460 | 0.65 | 0.005 | -0.061 | -1.649 | 0 | -0.061 | -0.061 | 0 | 0.19 | 0.08 | 0.22 | 0.58 | 0.16 | 0.23 |
| 1 | 168956295 | rs4656634 | -1.493 | 0.113 | 0.259 | -1.76 | 0.004 | 0.004 | 0.004 | 0 | 0.49 | 0.62 | 0.55 | 0.95 | 0.48 | 0.66 |
| 1 | 168956784 | rs10800399 | 0.082 | 0.288 | 0.382 | -1.587 | 0.004 | -0.015 | -0.015 | 0 | 0.24 | 0.21 | 0.24 | 0.6 | 0.18 | 0.32 |
| 1 | 168960286 | rs1322487 | -0.607 | 0.111 | 0.194 | -1.436 | 0.001 | -0.012 | -0.012 | 0 | 0.39 | 0.48 | 0.36 | 0.7 | 0.4 | 0.53 |
| 1 | 168976442 | rs2984600 | 0.506 | 0.304 | 0.041 | -1.415 | -0.001 | -0.03 | -0.03 | 0 | 0.45 | 0.37 | 0.37 | 0.04 | 0.43 | 0.36 |
| 1 | 168982215 | rs6689418 | -0.713 | 0.296 | 0.601 | -0.65 | 0.021 | -0.023 | -0.023 | 0.18 | 0.2 | 0.51 | 0.18 | 0.15 | 0.24 | 0.2 |
| 1 | 168994513 | rs7544086 | 0.74 | 0.006 | 0.211 | -0.582 | -0.043 | 0.012 | 0.012 | 0.007 | 0.65 | 0.83 | 0.61 | 0.28 | 0.57 | 0.45 |
| 1 | 168996958 | rs4656640 | -0.222 | -1.035 | 0.331 | 0.03 | 0.009 | -0.099 | -0.099 | 0 | 0.86 | 0.98 | 0.83 | 0.41 | 0.9 | 0.74 |
| 1 | 169000292 | rs4486430 | -0.305 | -0.726 | 0.805 | -0.125 | -0.001 | 0.024 | 0.024 | 0 | 0.06 | 0.06 | 0.13 | 0.2 | 0.07 | 0.16 |
| 1 | 169000494 | rs10494476 | -0.4 | -0.816 | 0.93 | -0.303 | -0.001 | -0.003 | -0.003 | 0 | 0.09 | 0.21 | 0.21 | 0.06 | 0.08 | 0.09 |
| 1 | 169002599 | rs10919037 | -0.994 | -0.661 | 0.78 | -0.142 | 0.032 | 0.013 | 0.013 | 0.837 | 0.06 | 0.33 | 0.02 | 0 | 0.01 | 0 |
| 1 | 169005304 | rs11580269 | -0.357 | -0.347 | -0.343 | 0.076 | 0 | 0.022 | 0.022 | 0 | 0.06 | 0.04 | 0.13 | 0.2 | 0.07 | 0.16 |
| 1 | 169012985 | rs12403132 | 0.107 | -0.731 | -0.418 | 0.219 | 0.008 | -0.018 | -0.018 | 0 | 0.18 | 0.14 | 0.26 | 0.53 | 0.14 | 0.38 |
| 1 | 169017419 | rs2982467 | -0.962 | -1.221 | 0.01 | -0.478 | -0.024 | 0.006 | 0.006 | 0 | 0.37 | 0.47 | 0.43 | 0.61 | 0.24 | 0.49 |
| 1 | 169020222 | rs2982456 | 0.292 | -1.262 | -0.424 | -0.873 | 0.002 | 0.042 | 0.042 | 0 | 0.13 | 0.04 | 0.14 | 0.04 | 0.11 | 0.12 |
| 1 | 169026930 | rs12069869 | -1.053 | -1.429 | -0.12 | -1.099 | 0.002 | 0.002 | 0.002 | 0 | 0.25 | 0.26 | 0.27 | 0.11 | 0.21 | 0.2 |
| 1 | 169027073 | rs909928 | 0.461 | -1.44 | -0.136 | -1.103 | 0.004 | 0.044 | 0.044 | 0 | 0.14 | 0.04 | 0.13 | 0.03 | 0.11 | 0.12 |
| 1 | 169028438 | rs4656646 | 0.658 | -1.557 | -0.075 | -1.11 | 0.02 | -0.046 | -0.046 | 0 | 0.21 | 0.14 | 0.3 | 0.61 | 0.13 | 0.41 |
| 1 | 169029904 | rs12074062 | -0.519 | -1.441 | -0.523 | -1.061 | -0.006 | -0.008 | -0.008 | 0 | 0.12 | 0.22 | 0.14 | 0.07 | 0.1 | 0.08 |
| 1 | 169030431 | rs12078922 | 0.162 | -1.4 | -0.519 | -1.039 | 0.002 | 0.002 | 0.002 | 0.359 | 0.09 | 0.16 | 0.13 | 0.07 | 0.09 | 0.08 |
| 1 | 169030878 | rs1358713 | -1.411 | -1.422 | -0.261 | -1.043 | 0.077 | 0.088 | 0.088 | 1.057 | 0.5 | 0.77 | 0.6 | 0.72 | 0.34 | 0.61 |
| 1 | 169035485 | rs12032929 | 0.769 | -1.372 | -0.671 | -1.468 | -0.005 | -0.035 | -0.035 | 0 | 0.15 | 0.09 | 0.13 | 0.26 | 0.17 | 0.21 |
| 1 | 169041144 | rs10919051 | 0.408 | -1.421 | -0.529 | -1.335 | -0.055 | 0.123 | 0.123 | 1.068 | 0.24 | 0.52 | 0.32 | 0.61 | 0.13 | 0.41 |
| 1 | 169049881 | rs1200100 | 0.496 | -0.846 | -1.131 | -1.412 | 0.087 | 0.1 | 0.1 | 0.921 | 0.5 | 0.22 | 0.4 | 0.28 | 0.66 | 0.39 |
| 1 | 169056077 | rs909926 | -0.504 | -0.056 | -0.943 | -2.131 | -0.006 | 0.142 | 0.142 | 0 | 0.57 | 0.81 | 0.66 | 0.98 | 0.43 | 0.79 |
| 1 | 169061558 | rs1200115 | -0.878 | 0.471 | -1.021 | -1.665 | -0.131 | 0.124 | 0.124 | 0 | 0.8 | 0.99 | 0.81 | 0.97 | 0.69 | 0.9 |
| 1 | 169066351 | rs1200120 | -0.502 | -0.204 | -0.414 | -1.89 | -0.029 | -0.06 | -0.06 | 0 | 0.27 | 0.38 | 0.15 | 0.12 | 0.2 | 0.31 |
| 1 | 169068156 | rs1200121 | 0.544 | -0.024 | -0.537 | -1.898 | -0.002 | -0.024 | -0.024 | 0.03 | 0.09 | 0.11 | 0.03 | 0.02 | 0.07 | 0.17 |
| 1 | 169070332 | rs2051145 | 0.186 | -0.111 | -0.492 | -1.784 | -0.006 | 0.009 | 0.009 | 0.179 | 0.6 | 0.38 | 0.65 | 0.83 | 0.57 | 0.57 |
| 1 | 169070523 | rs2051146 | 0.158 | -0.12 | -0.473 | -1.791 | -0.002 | -0.01 | -0.01 | 0.046 | 0.61 | 0.42 | 0.66 | 0.86 | 0.58 | 0.67 |
| 1 | 169072731 | rs926517 | 0.231 | -0.019 | -0.677 | -1.74 | -0.007 | -0.018 | -0.018 | 0 | 0.6 | 0.38 | 0.71 | 0.92 | 0.57 | 0.63 |
| 1 | 169073050 | rs926516 | 1.034 | -0.177 | -0.712 | -1.821 | -0.023 | 0.112 | 0.112 | 0.483 | 0.42 | 0.09 | 0.55 | 0.76 | 0.44 | 0.52 |
| 1 | 169073346 | rs1320976 | -0.492 | -0.296 | -0.004 | -0.905 | -0.019 | -0.001 | -0.001 | 0 | 0.18 | 0.3 | 0.16 | 0.16 | 0.13 | 0.11 |
| 1 | 169078572 | rs1200130 | -0.811 | -0.122 | -0.451 | -0.909 | -0.001 | -0.004 | -0.004 | 0 | 0.31 | 0.33 | 0.21 | 0.18 | 0.24 | 0.27 |
| 1 | 169079664 | rs1200136 | 0.441 | 1.414 | -0.407 | -0.757 | 0.034 | -0.04 | -0.04 | 0 | 0.77 | 0.74 | 0.72 | 0.9 | 0.67 | 0.78 |
| 1 | 169080837 | rs1200138 | 0.762 | 1.488 | -0.361 | -0.536 | 0.09 | -0.028 | -0.028 | 0 | 0.83 | 0.54 | 0.77 | 0.89 | 0.8 | 0.81 |
| 1 | 169085119 | rs3766031 | -0.408 | 0.907 | -1.304 | -0.055 | -0.017 | -0.038 | -0.038 | 0.358 | 0.14 | 0.05 | 0.14 | 0.16 | 0.13 | 0.09 |
| 1 | 169086221 | rs1040503 | -0.208 | 0.492 | -1.754 | -0.985 | 0.039 | 0.223 | 0.223 | 0 | 0.54 | 0.86 | 0.56 | 0.88 | 0.39 | 0.76 |
| 1 | 169088132 | rs1040502 | -0.418 | 0.12 | -1.166 | -0.805 | 0.01 | -0.002 | -0.002 | 0 | 0.42 | 0.63 | 0.43 | 0.91 | 0.34 | 0.5 |
| 1 | 169088337 | rs1200150 | -0.375 | 0.115 | -1.168 | -0.787 | -0.005 | -0.007 | -0.007 | 2.82 | 0.39 | 0.55 | 0.33 | 0.34 | 0.31 | 0.35 |
| 1 | 169088679 | rs2143290 | 0.701 | 0.116 | -1.71 | -1.294 | -0.02 | 0.081 | 0.081 | 0 | 0.13 | 0 | 0.22 | 0.03 | 0.13 | 0.09 |
| 1 | 169090660 | rs12731646 | 1.033 | 0.306 | -1.912 | -1.354 | -0.06 | 0.106 | 0.106 | 0 | 0.3 | 0.13 | 0.25 | 0.08 | 0.4 | 0.25 |
| 1 | 169091466 | rs3753292 | -0.25 | 0.345 | -1.691 | -1.494 | 0.001 | 0.039 | 0.039 | 0 | 0.48 | 0.38 | 0.57 | 0.67 | 0.56 | 0.49 |
| 1 | 169093676 | rs7540067 | 0.151 | 0.361 | -1.586 | -1.619 | -0.043 | 0.154 | 0.154 | 0 | 0.43 | 0.22 | 0.47 | 0.09 | 0.52 | 0.31 |
| 1 | 169097165 | rs12128543 | 0.151 | 0.351 | -1.512 | -1.592 | -0.068 | 0.226 | 0.226 | 0 | 0.43 | 0.16 | 0.47 | 0.09 | 0.52 | 0.31 |
| 1 | 169100095 | rs11809180 | 0.437 | -0.1 | -1.687 | -1.662 | -0.017 | 0.016 | 0.016 | 0.071 | 0.12 | 0.16 | 0.14 | 0.73 | 0.07 | 0.23 |
| 1 | 169103195 | rs1200160 | 0.429 | -0.392 | -0.969 | -1.793 | 0.001 | -0.078 | -0.078 | 0 | 0.46 | 0.46 | 0.47 | 0.09 | 0.53 | 0.32 |
| 1 | 169105702 | rs4656651 | 0.329 | -0.354 | -0.979 | -1.66 | -0.002 | 0.032 | 0.032 | 0 | 0.42 | 0.37 | 0.38 | 0.79 | 0.47 | 0.44 |
| 1 | 169112504 | rs12724494 | 0.321 | -0.358 | -0.981 | -1.662 | -0.002 | 0.032 | 0.032 | 0 | 0.42 | 0.37 | 0.39 | 0.79 | 0.47 | 0.44 |
| 1 | 169112703 | rs12728466 | 0.321 | -0.358 | -0.966 | -1.662 | -0.002 | 0.03 | 0.03 | 0 | 0.42 | 0.37 | 0.39 | 0.79 | 0.47 | 0.44 |
| 1 | 169113282 | rs12084964 | 0.41 | -0.311 | -0.81 | -1.487 | 0.017 | -0.019 | -0.019 | 1.73 | 0.12 | 0.11 | 0.14 | 0.73 | 0.07 | 0.22 |
| 1 | 169115833 | rs10919075 | 1.875 | -0.395 | -1.056 | -1.776 | 0.023 | -0.085 | -0.085 | 0 | 0.21 | 0.29 | 0.18 | 0 | 0.35 | 0.09 |
| 1 | 169116017 | rs12082618 | 0.604 | -0.357 | -1.054 | -1.738 | 0 | -0.004 | -0.004 | 0 | 0.32 | 0.36 | 0.32 | 0.73 | 0.41 | 0.32 |
| 1 | 169121811 | rs1320967 | 0.596 | -0.359 | -1.045 | -1.738 | -0.006 | 0.017 | 0.017 | 0 | 0.32 | 0.31 | 0.32 | 0.73 | 0.41 | 0.32 |
| 1 | 169123592 | rs12758208 | 0.565 | -0.232 | -1.068 | -1.739 | 0.001 | -0.005 | -0.005 | 0.007 | 0.33 | 0.37 | 0.32 | 0.73 | 0.42 | 0.32 |
| 1 | 169129151 | rs12744184 | 0.62 | -0.292 | -1.072 | -1.747 | -0.004 | 0.013 | 0.013 | 0 | 0.32 | 0.32 | 0.32 | 0.72 | 0.41 | 0.32 |
| 1 | 169129271 | rs12744341 | 0.621 | -0.296 | -1.082 | -1.749 | -0.005 | 0.015 | 0.015 | 0 | 0.32 | 0.31 | 0.32 | 0.72 | 0.41 | 0.32 |
| 1 | 169133621 | rs12074013 | 1.435 | -0.338 | -1.138 | -1.642 | 0.019 | -0.025 | -0.025 | 0.089 | 0.08 | 0.03 | 0.04 | 0.15 | 0.04 | 0.08 |
| 1 | 169134027 | rs12023291 | 0.924 | -0.053 | -1.112 | -1.885 | 0.036 | 0.04 | 0.04 | 0 | 0.06 | 0.13 | 0.14 | 0.18 | 0.11 | 0.06 |
| 1 | 169136833 | rs10800408 | -0.28 | -0.269 | -0.794 | -2.124 | 0.002 | -0.042 | -0.042 | 0.866 | 0.51 | 0.58 | 0.68 | 0.93 | 0.61 | 0.46 |

| | | | | | | | | | | | | | | | | |
|---|---|---|---|---|---|---|---|---|---|---|---|---|---|---|---|---|
| 1 | 169138307 | rs4656656 | 1.025 | -0.286 | -0.82 | -2.09 | -0.024 | 0.035 | 0.035 | 0 | 0.31 | 0.2 | 0.31 | 0.72 | 0.41 | 0.32 |
| 1 | 169176590 | rs1080266 | 0.16 | -0.369 | -0.689 | -2.099 | -0.001 | 0.129 | 0.129 | 0.103 | 0.44 | 0.28 | 0.53 | 0.79 | 0.51 | 0.41 |
| 1 | 169255020 | rs12733626 | 1.893 | -0.338 | -0.691 | -2.069 | -0.039 | -0.028 | -0.028 | 0 | 0.23 | 0.15 | 0.28 | 0.63 | 0.37 | 0.24 |
| 1 | 169257750 | rs10442644 | 2.5 | -0.35 | -0.705 | -2.09 | -0.06 | 0.071 | 0.071 | 0 | 0.19 | 0.07 | 0.17 | 0 | 0.34 | 0.09 |
| 1 | 169258201 | rs12065030 | 1.55 | -0.387 | -0.637 | -2.058 | -0.014 | 0.008 | 0.008 | 3.702 | 0.09 | 0.15 | 0.05 | 0.13 | 0.04 | 0.08 |
| 1 | 169265794 | rs12561922 | 1.631 | -0.387 | -0.646 | -2.058 | -0.008 | 0.004 | 0.004 | 1.28 | 0.09 | 0.1 | 0.04 | 0.13 | 0.05 | 0.08 |
| 1 | 169265836 | rs1591733 | -0.371 | -0.391 | -0.633 | -2.13 | 0.009 | 0.023 | 0.023 | 0.268 | 0.52 | 0.66 | 0.68 | 0.92 | 0.62 | 0.46 |
| 1 | 169276816 | rs10732287 | -0.536 | -0.784 | -0.412 | -2.578 | -0.001 | -0.002 | -0.002 | 0 | 0.7 | 0.8 | 0.78 | 0.94 | 0.75 | 0.66 |
| 1 | 169280253 | rs10919132 | 0.167 | -0.565 | -0.448 | -2.344 | 0.006 | 0.025 | 0.025 | 1.085 | 0.48 | 0.58 | 0.58 | 0.29 | 0.59 | 0.31 |
| 1 | 169288770 | rs10753786 | 0.167 | -0.565 | -0.448 | -2.344 | 0.006 | 0.025 | 0.025 | 1.085 | 0.48 | 0.58 | 0.58 | 0.29 | 0.59 | 0.31 |
| 1 | 169291205 | rs10800431 | 1.405 | -0.494 | -0.525 | -2.37 | -0.029 | 0.065 | 0.065 | 0 | 0.32 | 0.17 | 0.39 | 0.02 | 0.44 | 0.18 |
| 1 | 169301175 | rs12043489 | 0.171 | -0.725 | -0.438 | -2.173 | 0.02 | 0.004 | 0.004 | 0 | 0.07 | 0.13 | 0.14 | 0.13 | 0.1 | 0.05 |
| 1 | 169315527 | rs10919147 | 1.601 | -0.762 | -0.497 | -2.195 | 0.004 | 0.068 | 0.068 | 0 | 0.12 | 0 | 0.21 | 0.02 | 0.1 | 0.08 |
| 1 | 169330585 | rs16862093 | 0.55 | -0.904 | -0.389 | -2.17 | -0.026 | 0.01 | 0.01 | 0 | 0.12 | 0.2 | 0.15 | 0.76 | 0.07 | 0.24 |
| 1 | 169332102 | rs12097151 | 0.55 | -0.904 | -0.389 | -2.17 | -0.026 | 0.01 | 0.01 | 0 | 0.12 | 0.2 | 0.15 | 0.77 | 0.08 | 0.24 |
| 1 | 169335034 | rs1400838 | 0.507 | -0.857 | -0.416 | -2.238 | 0.029 | 0.005 | 0.005 | 0 | 0.08 | 0.14 | 0.14 | 0.13 | 0.13 | 0.06 |
| 1 | 169336617 | rs4656671 | 2.674 | -0.935 | -0.548 | -2.488 | -0.076 | 0.099 | 0.099 | 0 | 0.22 | 0.07 | 0.19 | 0 | 0.37 | 0.13 |
| 1 | 169337376 | rs2275299 | 0.78 | -1.007 | -0.489 | -2.548 | 0.004 | 0.063 | 0.063 | 0 | 0.43 | 0.32 | 0.55 | 0.16 | 0.6 | 0.28 |
| 1 | 169354051 | rs4656672 | -0.005 | -1.051 | -0.576 | -2.54 | -0.005 | -0.076 | -0.076 | 0 | 0.55 | 0.57 | 0.72 | 0.92 | 0.67 | 0.51 |
| 1 | 169357144 | rs10919158 | 1.833 | -1.04 | -0.608 | -2.514 | 0.002 | 0.066 | 0.066 | 0 | 0.12 | 0 | 0.21 | 0.02 | 0.1 | 0.08 |
| 1 | 169363203 | rs17516734 | 1.913 | -1.282 | -0.69 | -2.63 | -0.004 | 0.043 | 0.043 | 0 | 0.08 | 0 | 0.05 | 0.01 | 0.06 | 0.12 |
| 1 | 169368301 | rs9783117 | 0.771 | -1.218 | -0.583 | -2.55 | -0.027 | 0.01 | 0.01 | 0 | 0.12 | 0.21 | 0.15 | 0.77 | 0.08 | 0.24 |
| 1 | 169384437 | rs10919164 | 1.041 | -1.211 | -0.592 | -2.57 | 0 | -0.021 | -0.021 | 0 | 0.44 | 0.44 | 0.56 | 0.16 | 0.59 | 0.28 |
| 1 | 169389288 | rs6427191 | 0.691 | -1.159 | -0.793 | -2.613 | 0.03 | 0.005 | 0.005 | 0 | 0.08 | 0.14 | 0.14 | 0.13 | 0.13 | 0.06 |
| 1 | 169390327 | rs10489188 | 1.403 | -1.153 | -1.145 | -2.609 | -0.037 | -0.025 | -0.025 | 0 | 0.06 | 0.01 | 0.14 | 0.13 | 0.12 | 0.06 |
| 1 | 169391154 | rs3820059 | 0.17 | -1.144 | -1.109 | -2.659 | 0.007 | -0.008 | -0.008 | 1.251 | 0.56 | 0.64 | 0.72 | 0.92 | 0.67 | 0.51 |
| 1 | 169403703 | rs12021580 | 1.517 | -0.968 | -1.297 | -2.437 | -0.075 | 0.178 | 0.178 | 0 | 0.35 | 0.11 | 0.41 | 0.02 | 0.47 | 0.21 |
| 1 | 169428944 | rs1208134 | -1.457 | -1.002 | -0.843 | -2.304 | -0.002 | -0.004 | -0.004 | 0 | 0.89 | 0.93 | 0.93 | 0.94 | 0.94 | 0.89 |
| 1 | 169431218 | rs12128350 | 2.508 | -0.752 | -0.932 | -2.688 | -0.059 | 0.07 | 0.07 | 0 | 0.23 | 0.1 | 0.19 | 0 | 0.37 | 0.13 |
| 1 | 169431363 | rs10800447 | -0.087 | -0.874 | -0.699 | -2.703 | 0.002 | 0.002 | 0.002 | 0.707 | 0.61 | 0.63 | 0.67 | 0.31 | 0.71 | 0.48 |
| 1 | 169434120 | rs16862199 | 0.117 | -0.91 | -0.63 | -2.743 | 0.039 | -0.044 | -0.044 | 0.27 | 0.18 | 0.41 | 0.23 | 0.15 | 0.19 | 0.17 |
| 1 | 169436546 | rs6656822 | -0.252 | -0.896 | -0.71 | -2.57 | 0.002 | -0.021 | -0.021 | 0 | 0.65 | 0.69 | 0.77 | 0.94 | 0.74 | 0.62 |
| 1 | 169446183 | rs1983546 | -0.795 | -0.744 | -0.516 | -2.717 | 0.004 | -0.032 | -0.032 | 0 | 0.43 | 0.65 | 0.59 | 0.94 | 0.37 | 0.5 |
| 1 | 169448377 | rs6427193 | 2.146 | -0.589 | -0.793 | -2.699 | -0.062 | 0.072 | 0.072 | 0 | 0.23 | 0.08 | 0.19 | 0 | 0.37 | 0.13 |
| 1 | 169450022 | rs17577986 | 0.793 | -0.77 | -0.418 | -2.368 | 0.003 | -0.058 | -0.058 | 0 | 0.11 | 0.28 | 0.09 | 0.01 | 0.09 | 0.12 |
| 1 | 169452090 | rs17518769 | 0.776 | -0.77 | -0.418 | -2.368 | 0.002 | -0.059 | -0.059 | 0.098 | 0.11 | 0.28 | 0.09 | 0.01 | 0.09 | 0.12 |
| 1 | 169455982 | rs2038024 | -0.161 | -1.386 | -0.953 | -2.542 | 0.019 | 0.034 | 0.034 | 0.012 | 0.76 | 0.95 | 0.87 | 0.86 | 0.85 | 0.81 |
| 1 | 169456046 | rs12408451 | 0.852 | -1.553 | -0.816 | -2.717 | 0.034 | 0.029 | 0.029 | 0 | 0.5 | 0.51 | 0.57 | 0.18 | 0.67 | 0.42 |
| 1 | 169460726 | rs6696217 | -1.042 | -1.326 | -0.81 | -2.594 | 0.008 | 0.005 | 0.005 | 0 | 0.88 | 0.91 | 0.94 | 1 | 0.91 | 0.94 |
| 1 | 169461406 | rs2285211 | 1.02 | -1.349 | -0.601 | -2.83 | -0.004 | 0.033 | 0.033 | 0 | 0.41 | 0.28 | 0.43 | 0.17 | 0.53 | 0.31 |
| 1 | 169465997 | rs723751 | -0.331 | -1.08 | -0.409 | -2.524 | 0 | -0.009 | -0.009 | 0 | 0.29 | 0.3 | 0.38 | 0.21 | 0.25 | 0.38 |
| 1 | 169473761 | rs2420134 | 1.729 | -1.23 | -0.407 | -2.347 | 0.018 | 0.031 | 0.031 | 0 | 0.08 | 0.01 | 0.09 | 0.03 | 0.07 | 0.06 |
| 1 | 169477574 | rs6687813 | -1.116 | -1.408 | -0.333 | -2.514 | -0.001 | -0.02 | -0.02 | 1.151 | 0.9 | 0.88 | 0.95 | 1 | 0.93 | 0.94 |
| 1 | 169478042 | rs12120904 | 1.046 | -1.157 | -0.411 | -2.485 | -0.009 | 0 | 0 | 0 | 0.07 | 0.04 | 0.07 | 0.05 | 0.1 | 0.11 |
| 1 | 169481950 | rs2187952 | 0.471 | -1.321 | -0.336 | -2.597 | 0.004 | 0.019 | 0.019 | 0 | 0.31 | 0.21 | 0.4 | 0.25 | 0.25 | 0.33 |
| 1 | 169482436 | rs2040444 | 1.353 | -1.424 | -0.123 | -2.609 | 0.002 | -0.008 | -0.008 | 0.644 | 0.41 | 0.37 | 0.38 | 0.65 | 0.54 | 0.45 |
| 1 | 169482678 | rs9332673 | 2.356 | -1.424 | -0.127 | -2.629 | 0.016 | 0.031 | 0.031 | 0 | 0.08 | 0.01 | 0.09 | 0.03 | 0.07 | 0.06 |
| 1 | 169483561 | rs6027 | 2.544 | -1.425 | -0.128 | -2.629 | 0.011 | 0.026 | 0.026 | 0 | 0.08 | 0 | 0.09 | 0.03 | 0.07 | 0.06 |
| 1 | 169484552 | rs3820060 | -0.196 | -1.422 | -0.097 | -2.627 | 0.02 | 0.052 | 0.052 | 0 | 0.39 | 0.25 | 0.49 | 0.28 | 0.32 | 0.39 |
| 1 | 169485400 | rs6670407 | 1.303 | -1.537 | -0.012 | -2.585 | 0.007 | -0.024 | -0.024 | 0.031 | 0.41 | 0.39 | 0.39 | 0.65 | 0.54 | 0.44 |
| 1 | 169488927 | rs3766103 | 1.299 | -1.538 | 0.016 | -2.586 | 0.009 | -0.031 | -0.031 | 0 | 0.41 | 0.42 | 0.39 | 0.65 | 0.54 | 0.44 |
| 1 | 169489585 | rs2213867 | 0.577 | -1.535 | -0.046 | -2.589 | 0.005 | 0.043 | 0.043 | 0 | 0.3 | 0.16 | 0.4 | 0.25 | 0.25 | 0.32 |
| 1 | 169490592 | rs9332655 | 0.555 | -1.535 | -0.046 | -2.589 | 0.007 | 0.04 | 0.04 | 0 | 0.31 | 0.17 | 0.4 | 0.25 | 0.25 | 0.32 |
| 1 | 169490772 | rs9332653 | -0.579 | -1.514 | 0.238 | -2.564 | 0.001 | -0.004 | -0.004 | 0.723 | 0.2 | 0.34 | 0.25 | 0.72 | 0.21 | 0.39 |
| 1 | 169493953 | rs12131397 | 1.713 | -1.961 | 0.331 | -2.118 | -0.011 | 0.002 | 0.002 | 0 | 0.38 | 0.26 | 0.28 | 0.04 | 0.51 | 0.27 |
| 1 | 169496253 | rs9332630 | 1.713 | -2.016 | 0.331 | -2.118 | -0.01 | 0.002 | 0.002 | 0 | 0.38 | 0.26 | 0.28 | 0.04 | 0.51 | 0.27 |
| 1 | 169498975 | rs6030 | -0.907 | -1.595 | 0.554 | -1.352 | 0.025 | 0.074 | 0.074 | 0 | 0.38 | 0.17 | 0.49 | 0.28 | 0.31 | 0.39 |
| 1 | 169500210 | rs6010 | 0.589 | -1.529 | 0.478 | -1.312 | 0.011 | -0.016 | -0.016 | 0.966 | 0.08 | 0.04 | 0.16 | 0.17 | 0.04 | 0.12 |
| 1 | 169505158 | rs4656687 | -1.481 | -1.56 | 0.469 | -1.311 | -0.001 | -0.021 | -0.021 | 0 | 0.43 | 0.39 | 0.6 | 0.89 | 0.34 | 0.56 |
| 1 | 169510118 | rs9332608 | 0.627 | -1.486 | 0.077 | -1.238 | 0.01 | -0.024 | -0.024 | 0 | 0.07 | 0.03 | 0.16 | 0.17 | 0.04 | 0.12 |
| 1 | 169511755 | rs4524 | -0.182 | -1.514 | 0.147 | -1.231 | 0.006 | 0.041 | 0.041 | 0 | 0.3 | 0.17 | 0.39 | 0.25 | 0.25 | 0.34 |
| 1 | 169511878 | rs6018 | 1.294 | -1.491 | 0.156 | -1.219 | 0.024 | 0.035 | 0.035 | 0.295 | 0.09 | 0 | 0.09 | 0.03 | 0.06 | 0.06 |
| 1 | 169512120 | rs6016 | -0.251 | -1.418 | -0.275 | -1.186 | 0.005 | 0.039 | 0.039 | 0 | 0.3 | 0.17 | 0.39 | 0.25 | 0.25 | 0.32 |
| 1 | 169512497 | rs2239851 | -0.227 | -1.416 | -0.274 | -1.184 | 0.004 | 0.038 | 0.038 | 0 | 0.3 | 0.17 | 0.39 | 0.25 | 0.25 | 0.32 |
| 1 | 169512524 | rs2239852 | -1.291 | -1.367 | -0.232 | -1.335 | 0.016 | 0.025 | 0.025 | 0 | 0.43 | 0.35 | 0.6 | 0.88 | 0.34 | 0.56 |
| 1 | 169513583 | rs6037 | 0.747 | -0.806 | -0.608 | -0.915 | 0.02 | 0.04 | 0.04 | 0 | 0.1 | 0.01 | 0.1 | 0.03 | 0.07 | 0.07 |
| 1 | 169515183 | rs3766110 | 0.093 | -0.844 | -0.554 | -0.782 | 0 | 0.002 | 0.002 | 0 | 0.24 | 0.2 | 0.37 | 0.22 | 0.23 | 0.31 |
| 1 | 169515726 | rs6036 | 0.778 | -0.867 | -0.569 | -0.806 | 0.02 | 0.041 | 0.041 | 0 | 0.1 | 0.01 | 0.1 | 0.03 | 0.07 | 0.07 |
| 1 | 169519894 | rs6015 | 0.793 | -0.867 | -0.569 | -0.806 | 0.02 | 0.04 | 0.04 | 0 | 0.1 | 0.01 | 0.1 | 0.03 | 0.07 | 0.07 |

| | | | | | | | | | | | | | | | | |
|---|---|---|---|---|---|---|---|---|---|---|---|---|---|---|---|---|
| 1 | 169520098 | rs10800456 | -1.347 | -0.919 | -0.402 | -0.786 | -0.006 | -0.025 | -0.025 | 1.55 | 0.4 | 0.58 | 0.58 | 0.88 | 0.34 | 0.56 |
| 1 | 169520364 | rs10158595 | -0.179 | -0.132 | -1.28 | -0.684 | -0.001 | 0.002 | 0.002 | 0 | 0.24 | 0.2 | 0.38 | 0.22 | 0.24 | 0.32 |
| 1 | 169521849 | rs6035 | -0.477 | -0.163 | -1.401 | -0.573 | -0.005 | 0.004 | 0.004 | 0 | 0.09 | 0.08 | 0.07 | 0.07 | 0.09 | 0.1 |
| 1 | 169521853 | rs6033 | -0.192 | -0.265 | -1.394 | -0.662 | 0.024 | 0.04 | 0.04 | 0 | 0.1 | 0.01 | 0.1 | 0.03 | 0.07 | 0.07 |
| 1 | 169523537 | rs6427199 | -0.318 | -0.154 | -1.187 | -0.597 | 0.008 | 0.017 | 0.017 | 0.089 | 0.41 | 0.61 | 0.36 | 0.69 | 0.39 | 0.44 |
| 1 | 169525312 | rs9332579 | -0.587 | 0.099 | -0.333 | -0.717 | 0.052 | -0.076 | -0.076 | 0.102 | 0.13 | 0.46 | 0.11 | 0 | 0.12 | 0.02 |
| 1 | 169528580 | rs6012 | -1.386 | -0.654 | -0.703 | -0.536 | -0.018 | 0.042 | 0.042 | 1.72 | 0.23 | 0.66 | 0.29 | 0.66 | 0.18 | 0.18 |
| 1 | 169528830 | rs6427202 | -1.852 | -0.555 | -1.286 | -1.199 | -0.073 | 0.189 | 0.189 | 1.993 | 0.6 | 0.89 | 0.7 | 0.88 | 0.58 | 0.57 |
| 1 | 169529826 | rs6022 | -0.916 | -0.531 | -1.267 | -1.177 | -0.018 | 0.042 | 0.042 | 1.72 | 0.23 | 0.65 | 0.29 | 0.66 | 0.18 | 0.18 |
| 1 | 169529973 | rs6029 | -0.826 | -0.544 | -1.233 | -1.183 | -0.018 | 0.061 | 0.061 | 1.036 | 0.22 | 0.58 | 0.29 | 0.66 | 0.18 | 0.18 |
| 1 | 169530837 | rs1894702 | 0.302 | -0.591 | -1.171 | -1.218 | -0.01 | 0.006 | 0.006 | 0 | 0.27 | 0.17 | 0.37 | 0.2 | 0.31 | 0.36 |
| 1 | 169531035 | rs3766120 | 0.436 | -0.657 | -1.184 | -1.25 | -0.078 | -0.028 | -0.028 | 2.703 | 0.25 | 0.03 | 0.36 | 0.2 | 0.31 | 0.36 |
| 1 | 169533266 | rs6678795 | -1.4 | -0.652 | -1.179 | -1.247 | -0.008 | 0.23 | 0.23 | 0 | 0.49 | 0.82 | 0.66 | 0.87 | 0.49 | 0.54 |
| 1 | 169535918 | rs12024736 | 0.28 | -0.703 | -1.186 | -1.182 | 0.008 | -0.031 | -0.031 | 0.039 | 0.08 | 0 | 0.17 | 0.66 | 0.06 | 0.15 |
| 1 | 169539348 | rs6663533 | 0.649 | -0.593 | -0.896 | -1.415 | 0.018 | -0.044 | -0.044 | 0 | 0.1 | 0.4 | 0.06 | 0 | 0.05 | 0.01 |
| 1 | 169540080 | rs2227245 | 0.018 | -0.407 | -1.53 | -1.157 | -0.009 | -0.009 | -0.009 | 1.203 | 0.27 | 0.18 | 0.37 | 0.28 | 0.31 | 0.36 |
| 1 | 169542135 | rs6691048 | -0.091 | -0.394 | -1.525 | -1.49 | -0.005 | 0.005 | 0.005 | 0 | 0.28 | 0.24 | 0.38 | 0.2 | 0.31 | 0.36 |
| 1 | 169557734 | rs1018828 | 0.083 | -0.357 | -1.226 | -1.756 | -0.024 | -0.06 | -0.06 | 1.018 | 0.11 | 0.21 | 0.06 | 0.01 | 0.09 | 0.04 |
| 1 | 169559791 | rs3917848 | 0.06 | -0.363 | -1.214 | -1.759 | -0.025 | -0.061 | -0.061 | 0 | 0.11 | 0.21 | 0.06 | 0.01 | 0.09 | 0.04 |
| 1 | 169563951 | rs6136 | 0.86 | 0.003 | -2.109 | -1.73 | -0.029 | 0.027 | 0.027 | 1.089 | 0.06 | 0 | 0.08 | 0 | 0.09 | 0.03 |
| 1 | 169565346 | rs6133 | 0.034 | -0.108 | -1.788 | -1.832 | -0.003 | -0.078 | -0.078 | 0 | 0.14 | 0.66 | 0.13 | 0 | 0.12 | 0.02 |
| 1 | 169565393 | rs3917811 | 0.034 | -0.108 | -1.788 | -1.832 | -0.003 | -0.078 | -0.078 | 0 | 0.14 | 0.66 | 0.13 | 0 | 0.12 | 0.02 |
| 1 | 169566313 | rs6127 | -1.403 | -0.218 | -2.091 | -2.152 | -0.061 | 0.207 | 0.207 | 1.623 | 0.58 | 0.87 | 0.7 | 0.96 | 0.59 | 0.57 |
| 1 | 169566326 | rs6135 | 0.18 | -0.193 | -2.002 | -2.056 | -0.003 | -0.078 | -0.078 | 0.01 | 0.14 | 0.66 | 0.13 | 0 | 0.12 | 0.02 |
| 1 | 169569285 | rs3917873 | 0.657 | -0.218 | -2.195 | -2.078 | 0.006 | 0.036 | 0.036 | 0.665 | 0.1 | 0.05 | 0.04 | 0.01 | 0.1 | 0.05 |
| 1 | 169570848 | rs3917779 | 0.544 | -0.212 | -2.137 | -2.09 | 0.013 | -0.072 | -0.072 | 1.338 | 0.13 | 0.59 | 0.12 | 0 | 0.12 | 0.02 |
| 1 | 169571076 | rs3917775 | -1.043 | -0.271 | -2.293 | -2.014 | 0.012 | 0.201 | 0.201 | 0 | 0.45 | 0.19 | 0.57 | 0.96 | 0.47 | 0.56 |
| 1 | 169572645 | rs2205895 | -0.823 | -0.022 | -2.123 | -1.772 | 0.006 | 0.07 | 0.07 | 0 | 0.39 | 0.27 | 0.55 | 0.95 | 0.39 | 0.52 |
| 1 | 169574887 | rs2223303 | -0.515 | 0.243 | -2.712 | -1.597 | -0.024 | 0.173 | 0.173 | 0 | 0.34 | 0.08 | 0.52 | 0.95 | 0.37 | 0.51 |
| 1 | 169577990 | rs3917744 | -0.568 | 0.747 | -2.475 | -1.138 | -0.041 | 0.203 | 0.203 | 1.63 | 0.63 | 0.93 | 0.8 | 0.94 | 0.7 | 0.64 |
| 1 | 169578586 | rs2076074 | -0.663 | 0.933 | -1.92 | 0.273 | -0.024 | -0.058 | -0.058 | 1.002 | 0.2 | 0.07 | 0.36 | 0.28 | 0.27 | 0.31 |
| 1 | 169579266 | rs3917740 | -1.003 | 1.666 | -0.728 | 0.551 | 0.086 | 0.058 | 0.058 | 0 | 0.16 | 0.44 | 0.14 | 0.21 | 0.19 | 0.2 |
| 1 | 169579378 | rs3917739 | 0.256 | 1.922 | -1.397 | 0.455 | -0.04 | -0.005 | -0.005 | 0 | 0.76 | 0.86 | 0.53 | 0.78 | 0.61 | 0.62 |
| 1 | 169579645 | rs2235305 | -0.093 | 1.227 | -0.914 | -0.376 | 0 | -0.017 | -0.017 | 0 | 0.46 | 0.47 | 0.27 | 0.19 | 0.4 | 0.35 |
| 1 | 169580885 | rs6131 | -0.348 | 0.874 | -0.267 | -0.65 | -0.003 | -0.011 | -0.011 | 0 | 0.23 | 0.34 | 0.14 | 0.2 | 0.19 | 0.27 |
| 1 | 169583197 | rs3917717 | 0.847 | 0.815 | 0.226 | -0.684 | 0.022 | -0.021 | -0.021 | 0 | 0.05 | 0.14 | 0.06 | 0.01 | 0.05 | 0.13 |
| 1 | 169583996 | rs2236868 | -0.312 | 0.536 | -0.061 | -1.204 | -0.016 | 0.094 | 0.094 | 0 | 0.45 | 0.22 | 0.29 | 0.21 | 0.44 | 0.37 |
| 1 | 169584545 | rs3917709 | 1.548 | 0.523 | -0.149 | -1.151 | 0.006 | -0.05 | -0.05 | 0.562 | 0.11 | 0.01 | 0.04 | 0.19 | 0.07 | 0.1 |
| 1 | 169586946 | rs2244526 | 0.203 | -0.09 | 0.332 | -1.053 | -0.015 | 0.007 | 0.007 | 0 | 0.12 | 0.19 | 0.15 | 0.13 | 0.1 | 0.12 |
| 1 | 169587032 | rs2244529 | -1.243 | -0.21 | 0.285 | -1.221 | 0.015 | 0.011 | 0.011 | 0.564 | 0.4 | 0.64 | 0.3 | 0.34 | 0.31 | 0.39 |
| 1 | 169590794 | rs3917691 | -1.716 | -0.229 | 0.142 | -0.498 | -0.005 | 0.005 | 0.005 | 0 | 0.87 | 0.88 | 0.62 | 0.71 | 0.75 | 0.74 |
| 1 | 169591164 | rs3917686 | -0.764 | -0.406 | 0.513 | -0.91 | -0.051 | -0.023 | -0.023 | 0 | 0.19 | 0.52 | 0.17 | 0.13 | 0.12 | 0.13 |
| 1 | 169599254 | rs732314 | -0.833 | -0.374 | 0.428 | -0.938 | -0.007 | 0.012 | 0.012 | 0 | 0.6 | 0.66 | 0.39 | 0.33 | 0.51 | 0.36 |
| 1 | 169601281 | rs1800805 | 0.52 | -0.31 | 0.05 | -1.164 | -0.051 | 0.108 | 0.108 | 0 | 0.4 | 0.12 | 0.23 | 0.2 | 0.39 | 0.24 |
| 1 | 169602109 | rs6666554 | -1.242 | -0.392 | 0.092 | -1.14 | -0.016 | 0.022 | 0.022 | 0 | 0.64 | 0.77 | 0.45 | 0.29 | 0.56 | 0.48 |
| 1 | 169602383 | rs6691334 | -0.965 | -0.371 | -0.177 | -1.129 | -0.031 | -0.071 | -0.071 | 0 | 0.23 | 0.59 | 0.22 | 0.09 | 0.18 | 0.24 |
| 1 | 169606008 | rs10800460 | -1.194 | -0.35 | -0.365 | -1.113 | -0.006 | 0.012 | 0.012 | 0 | 0.64 | 0.71 | 0.45 | 0.29 | 0.56 | 0.48 |
| 1 | 169613794 | rs10919202 | -0.813 | -0.297 | -0.367 | -1.085 | -0.026 | -0.065 | -0.065 | 0.637 | 0.22 | 0.39 | 0.21 | 0.09 | 0.18 | 0.26 |
| 1 | 169614949 | rs17589700 | 1.162 | 0.199 | -1.071 | -0.735 | -0.042 | 0.093 | 0.093 | 0 | 0.24 | 0.08 | 0.15 | 0 | 0.25 | 0.11 |
| 1 | 169623104 | rs10800463 | -0.396 | 0.211 | -0.837 | -0.491 | -0.003 | 0.021 | 0.021 | 0 | 0.52 | 0.42 | 0.38 | 0.27 | 0.48 | 0.36 |
| 1 | 169623680 | rs1569475 | -0.391 | 0.452 | -0.623 | 0.121 | 0.018 | -0.02 | -0.02 | 0.053 | 0.06 | 0.19 | 0.07 | 0.01 | 0.07 | 0.1 |
| 1 | 169647046 | rs1011266 | 0.336 | 0.542 | -1.46 | 0.288 | -0.027 | -0.049 | -0.049 | 0.094 | 0.15 | 0.01 | 0.08 | 0.19 | 0.13 | 0.11 |
| 1 | 169648469 | rs6663862 | -0.603 | 0.093 | -1.097 | 0.618 | 0.003 | 0.002 | 0.002 | 1.857 | 0.32 | 0.63 | 0.26 | 0.28 | 0.25 | 0.28 |
| 1 | 169650804 | rs17524608 | 0.889 | 0.282 | -1.809 | 0.215 | -0.044 | 0.058 | 0.058 | 0 | 0.3 | 0.11 | 0.19 | 0.16 | 0.31 | 0.24 |
| 1 | 169651044 | rs7531806 | -0.297 | 0.443 | -1.45 | 0.516 | -0.023 | -0.032 | -0.032 | 0 | 0.63 | 0.78 | 0.45 | 0.43 | 0.57 | 0.51 |
| 1 | 169658586 | rs4987388 | -0.763 | 1.249 | -1.409 | 1.247 | -0.055 | -0.049 | -0.049 | 0.551 | 0.91 | 0.99 | 0.79 | 0.63 | 0.72 | 0.82 |
| 1 | 169660002 | rs7553 | 0.407 | 0.189 | -0.876 | 0.81 | -0.002 | 0 | 0 | 0 | 0.53 | 0.49 | 0.37 | 0.39 | 0.38 | 0.47 |
| 1 | 169660781 | rs12938 | 1.146 | -0.215 | -0.961 | 0.143 | 0 | -0.006 | -0.006 | 0.025 | 0.3 | 0.3 | 0.18 | 0.05 | 0.24 | 0.22 |
| 1 | 169661936 | rs4987373 | 0.911 | -0.826 | -0.323 | -0.085 | -0.044 | -0.042 | -0.042 | 0 | 0.11 | 0.3 | 0.05 | 0.04 | 0.05 | 0.1 |
| 1 | 169663079 | rs2298902 | 1.244 | -0.886 | -0.498 | 0.14 | 0.041 | 0.007 | 0.007 | 0 | 0.2 | 0.08 | 0.13 | 0.11 | 0.13 | 0.2 |
| 1 | 169665632 | rs2223286 | 0.185 | -0.427 | -0.209 | 0.03 | 0.013 | 0.012 | 0.012 | 0.047 | 0.41 | 0.62 | 0.24 | 0.1 | 0.28 | 0.32 |
| 1 | 169667011 | rs4987352 | 0.529 | 0.033 | -0.165 | 0.671 | -0.041 | -0.04 | -0.04 | 0 | 0.1 | 0.21 | 0.04 | 0.04 | 0.05 | 0.1 |
| 1 | 169669020 | rs2298900 | -0.951 | 0.819 | -0.661 | 1.187 | -0.06 | -0.057 | -0.057 | 0.324 | 0.9 | 0.98 | 0.77 | 0.64 | 0.68 | 0.8 |
| 1 | 169669732 | rs2298899 | 1.083 | 0.147 | -0.678 | 1.19 | 0.009 | -0.016 | -0.016 | 0 | 0.12 | 0.08 | 0.24 | 0.2 | 0.13 | 0.15 |
| 1 | 169671088 | rs964555 | 0.495 | -0.015 | -0.16 | 1.109 | -0.018 | 0.021 | 0.021 | 0 | 0.14 | 0.22 | 0.25 | 0.2 | 0.13 | 0.15 |
| 1 | 169675210 | rs4987301 | 1.268 | -0.121 | 0.256 | 1.179 | 0.009 | -0.016 | -0.016 | 0 | 0.12 | 0.08 | 0.24 | 0.2 | 0.13 | 0.15 |
| 1 | 169675349 | rs4987299 | 1.854 | -0.281 | 0 | 0.865 | -0.016 | 0.05 | 0.05 | 0 | 0.11 | 0.03 | 0.06 | 0.01 | 0.1 | 0.09 |
| 1 | 169677709 | rs1051091 | -0.346 | -0.629 | 0.065 | 0.718 | 0.022 | 0.013 | 0.013 | 0 | 0.35 | 0.3 | 0.34 | 0.27 | 0.27 | 0.34 |
| 1 | 169678024 | rs4987285 | -0.689 | -0.617 | 0.046 | 0.71 | 0.004 | 0.001 | 0.001 | 0 | 0.38 | 0.37 | 0.36 | 0.33 | 0.29 | 0.45 |

| | | | | | | | | | | | | | | | | |
|---|---|---|---|---|---|---|---|---|---|---|---|---|---|---|---|---|
| 1 | 169688135 | rs4656703 | -0.688 | -0.622 | 0.041 | 0.707 | -0.009 | -0.002 | -0.002 | 1.009 | 0.38 | 0.43 | 0.36 | 0.33 | 0.29 | 0.45 |
| 1 | 169692132 | rs4786 | -0.883 | -0.821 | 0.784 | 0.966 | -0.02 | 0.002 | 0.002 | 0.005 | 0.4 | 0.57 | 0.42 | 0.56 | 0.3 | 0.49 |
| 1 | 169692522 | rs5359 | 0.025 | -0.85 | 0.769 | 0.96 | -0.014 | 0 | 0 | 0 | 0.24 | 0.27 | 0.28 | 0.26 | 0.17 | 0.25 |
| 1 | 169694695 | rs3917434 | -1.601 | -0.688 | 0.947 | 0.954 | -0.001 | -0.005 | -0.005 | 0 | 0.89 | 0.92 | 0.78 | 0.69 | 0.69 | 0.8 |
| 1 | 169694918 | rs5356 | 1.865 | -0.787 | 0.909 | 0.948 | 0.011 | -0.017 | -0.017 | 0.429 | 0.12 | 0.08 | 0.24 | 0.2 | 0.13 | 0.15 |
| 1 | 169695870 | rs5355 | 1.473 | -0.804 | 0.907 | 0.905 | 0.039 | 0.031 | 0.031 | 1.1 | 0.09 | 0.01 | 0.03 | 0.05 | 0.05 | 0.1 |
| 1 | 169696946 | rs5368 | 1.851 | -0.76 | 0.921 | 0.934 | 0.011 | -0.019 | -0.019 | 0 | 0.12 | 0.08 | 0.23 | 0.21 | 0.13 | 0.15 |
| 1 | 169698789 | rs5363 | 2.254 | -0.74 | 0.864 | 0.903 | -0.016 | 0.05 | 0.05 | 0 | 0.11 | 0.03 | 0.06 | 0.01 | 0.1 | 0.09 |
| 1 | 169698825 | rs1534904 | -1.931 | -0.688 | 0.848 | 0.935 | -0.059 | -0.058 | -0.058 | 0 | 0.9 | 0.98 | 0.78 | 0.69 | 0.69 | 0.8 |
| 1 | 169699819 | rs3917419 | 1.279 | -0.79 | 1.41 | 0.854 | 0.008 | 0.145 | 0.145 | 0 | 0.47 | 0.25 | 0.35 | 0.12 | 0.39 | 0.31 |
| 1 | 169701060 | rs5361 | 2.106 | -0.666 | 1.258 | 0.778 | -0.016 | 0.05 | 0.05 | 0 | 0.11 | 0.03 | 0.06 | 0.01 | 0.1 | 0.09 |
| 1 | 169701901 | rs1800016 | 2.106 | -0.666 | 1.258 | 0.778 | -0.016 | 0.05 | 0.05 | 0 | 0.11 | 0.03 | 0.06 | 0.01 | 0.1 | 0.09 |
| 1 | 169701904 | rs1800015 | 2.106 | -0.666 | 1.258 | 0.778 | -0.016 | 0.05 | 0.05 | 0 | 0.11 | 0.03 | 0.06 | 0.01 | 0.1 | 0.09 |
| 1 | 169710669 | rs10489182 | 0.153 | -0.696 | 1.204 | 0.853 | -0.019 | 0.041 | 0.041 | 0 | 0.28 | 0.41 | 0.36 | 0.46 | 0.24 | 0.28 |
| 1 | 169713146 | rs6427213 | 1.373 | -1.046 | 1.061 | 0.445 | 0.005 | 0.011 | 0.011 | 0 | 0.18 | 0.11 | 0.17 | 0.12 | 0.15 | 0.18 |
| 1 | 169719338 | rs969310 | -1.028 | -0.773 | 1.158 | 0.77 | 0.079 | 0.064 | 0.064 | 0.043 | 0.88 | 0.8 | 0.77 | 0.69 | 0.69 | 0.79 |
| 1 | 169733345 | rs2272818 | 0.447 | -1.03 | 1.012 | 0.728 | -0.048 | 0.157 | 0.157 | 0 | 0.18 | 0.41 | 0.31 | 0.44 | 0.14 | 0.2 |
| 1 | 169738202 | rs12027035 | -0.242 | -0.927 | 0.796 | 0.682 | -0.039 | 0.042 | 0.042 | 0 | 0.28 | 0.4 | 0.33 | 0.5 | 0.2 | 0.31 |
| 1 | 169739650 | rs2205851 | -0.955 | -0.599 | 0.887 | 0.72 | 0.107 | 0.1 | 0.1 | 0.309 | 0.88 | 0.76 | 0.77 | 0.69 | 0.7 | 0.8 |
| 1 | 169744647 | rs6673493 | -0.118 | -0.504 | 0.63 | 0.706 | -0.005 | 0.008 | 0.008 | 0 | 0.27 | 0.31 | 0.34 | 0.5 | 0.23 | 0.3 |
| 1 | 169749154 | rs12035144 | 0.556 | -0.592 | 0.709 | 0.676 | -0.001 | -0.002 | -0.002 | 0 | 0.62 | 0.6 | 0.45 | 0.19 | 0.5 | 0.5 |
| 1 | 169752700 | rs10919239 | -1.379 | -0.535 | 0.636 | 0.717 | -0.001 | -0.001 | -0.001 | 0 | 0.89 | 0.91 | 0.78 | 0.69 | 0.7 | 0.8 |
| 1 | 169755988 | rs10919242 | 0.522 | -0.592 | 0.757 | 0.52 | 0.004 | -0.03 | -0.03 | 0 | 0.61 | 0.52 | 0.45 | 0.19 | 0.5 | 0.5 |
| 1 | 169756389 | rs10157266 | 0.199 | -0.609 | 0.684 | 0.701 | -0.012 | -0.013 | -0.013 | 0.537 | 0.64 | 0.7 | 0.5 | 0.42 | 0.51 | 0.54 |
| 1 | 169756981 | rs6696884 | 0.612 | -0.923 | 0.378 | 0.249 | 0 | -0.01 | -0.01 | 0 | 0.54 | 0.51 | 0.42 | 0.16 | 0.47 | 0.43 |
| 1 | 169757316 | rs12130372 | 2.261 | -0.773 | -0.001 | 0.155 | -0.024 | 0.046 | 0.046 | 0.23 | 0.09 | 0 | 0.05 | 0.01 | 0.09 | 0.07 |
| 1 | 169757487 | rs10753796 | 0.389 | -0.81 | 0.337 | 0.139 | 0.006 | -0.035 | -0.035 | 3.508 | 0.6 | 0.45 | 0.43 | 0.18 | 0.49 | 0.48 |
| 1 | 169758843 | rs12135656 | 0.266 | -1.103 | 0.357 | -0.206 | -0.001 | -0.008 | -0.008 | 0 | 0.29 | 0.27 | 0.24 | 0.18 | 0.26 | 0.35 |
| 1 | 169761107 | rs10800477 | 0.88 | -1.09 | 0.361 | -0.208 | -0.006 | 0.001 | 0.001 | 0 | 0.2 | 0.27 | 0.2 | 0.17 | 0.16 | 0.28 |
| 1 | 169761591 | rs12135361 | 2.442 | -1.088 | 0.357 | -0.21 | -0.025 | 0.048 | 0.048 | 0.409 | 0.1 | 0.01 | 0.04 | 0.01 | 0.1 | 0.07 |
| 1 | 169762807 | rs10489177 | 0.841 | -1.092 | 0.365 | -0.205 | -0.007 | -0.001 | -0.001 | 0 | 0.2 | 0.27 | 0.2 | 0.17 | 0.16 | 0.28 |
| 1 | 169765340 | rs2205845 | -0.449 | -1.237 | 0.446 | 0.045 | 0.012 | 0.025 | 0.025 | 0.94 | 0.71 | 0.64 | 0.51 | 0.5 | 0.57 | 0.63 |
| 1 | 169768080 | rs6668114 | -0.602 | -1.309 | 0.424 | 0.215 | 0.026 | 0.011 | 0.011 | 0 | 0.77 | 0.72 | 0.7 | 0.66 | 0.58 | 0.73 |
| 1 | 169768372 | rs3766134 | -1.847 | -1.306 | 0.356 | 0.223 | -0.063 | -0.06 | -0.06 | 4.174 | 0.89 | 0.96 | 0.76 | 0.67 | 0.68 | 0.8 |
| 1 | 169773317 | rs11590604 | 1.781 | -1.451 | 0.33 | 0.146 | 0.028 | -0.011 | -0.011 | 0 | 0.16 | 0.09 | 0.24 | 0.17 | 0.11 | 0.17 |
| 1 | 169794359 | rs9803860 | 0.247 | -1.421 | 0.438 | 0.178 | 0.017 | -0.027 | -0.027 | 0 | 0.59 | 0.51 | 0.42 | 0.26 | 0.47 | 0.52 |
| 1 | 169802760 | rs7521700 | -0.205 | -1.363 | 0.47 | 0.186 | 0.037 | 0.036 | 0.036 | 0.001 | 0.74 | 0.67 | 0.66 | 0.43 | 0.57 | 0.69 |
| 1 | 169808651 | rs12133074 | 2.503 | -1.384 | 0.438 | 0.18 | -0.025 | 0.049 | 0.049 | 0.481 | 0.1 | 0.01 | 0.04 | 0.01 | 0.1 | 0.08 |
| 1 | 169818880 | rs2272918 | 1.251 | -1.567 | 0.422 | 0.066 | 0.018 | 0.187 | 0.187 | 0.14 | 0.39 | 0.14 | 0.22 | 0.1 | 0.31 | 0.24 |
| 1 | 169819065 | rs3817859 | 1.895 | -1.513 | 0.472 | 0.121 | -0.01 | 0.002 | 0.002 | 0 | 0.15 | 0.18 | 0.25 | 0.17 | 0.11 | 0.17 |
| 1 | 169823660 | rs1062976 | 1.995 | -1.524 | 0.404 | 0.103 | -0.009 | 0.121 | 0.121 | 0 | 0.3 | 0.13 | 0.19 | 0.01 | 0.26 | 0.12 |
| 1 | 169823718 | rs4656197 | -0.463 | -1.316 | 0.493 | 0.263 | 0.026 | 0.011 | 0.011 | 0.005 | 0.77 | 0.71 | 0.7 | 0.66 | 0.59 | 0.73 |
| 1 | 169824068 | rs12132384 | 2.542 | -1.325 | 0.432 | 0.255 | -0.025 | 0.049 | 0.049 | 0.481 | 0.1 | 0.01 | 0.04 | 0.01 | 0.1 | 0.08 |
| 1 | 169824104 | rs12135726 | 2.542 | -1.325 | 0.428 | 0.255 | -0.033 | 0.05 | 0.05 | 0.481 | 0.1 | 0 | 0.04 | 0.01 | 0.1 | 0.08 |
| 1 | 169825529 | rs12128754 | 0.959 | -1.348 | 0.518 | 0.2 | -0.007 | -0.002 | -0.002 | 0.106 | 0.2 | 0.26 | 0.2 | 0.17 | 0.16 | 0.28 |
| 1 | 169828327 | rs17602701 | 2.535 | -1.294 | 0.372 | 0.195 | -0.025 | 0.049 | 0.049 | 0.481 | 0.1 | 0.01 | 0.05 | 0.01 | 0.1 | 0.08 |
| 1 | 169834216 | rs4508080 | -0.979 | -1.273 | 0.442 | 0.245 | 0.084 | 0.085 | 0.085 | 0.264 | 0.86 | 0.74 | 0.75 | 0.67 | 0.68 | 0.8 |
| 1 | 169840185 | rs10489170 | 1.357 | -1.341 | 0.526 | 0.173 | 0 | 0.005 | 0.005 | 0 | 0.18 | 0.11 | 0.18 | 0.13 | 0.16 | 0.18 |
| 1 | 169844093 | rs12143057 | 2.442 | -1.197 | 0.486 | 0.226 | -0.025 | 0.049 | 0.049 | 0.481 | 0.1 | 0.01 | 0.04 | 0.01 | 0.1 | 0.07 |
| 1 | 169846883 | rs6692451 | 0.001 | -1.478 | 0.413 | -0.059 | -0.005 | -0.082 | -0.082 | 0 | 0.2 | 0.35 | 0.1 | 0.05 | 0.18 | 0.17 |
| 1 | 169847146 | rs10800485 | -1.293 | -1.187 | 0.157 | 0.378 | -0.02 | -0.012 | -0.012 | 5.693 | 0.87 | 0.9 | 0.73 | 0.43 | 0.68 | 0.76 |
| 1 | 169855576 | rs3766151 | -1.558 | -1.146 | 0.083 | 0.386 | -0.014 | -0.011 | -0.011 | 0 | 0.88 | 0.9 | 0.77 | 0.66 | 0.69 | 0.8 |
| 1 | 169858192 | rs11577641 | 2.078 | -1.455 | -0.023 | -0.102 | 0.017 | -0.002 | -0.002 | 0 | 0.15 | 0.11 | 0.24 | 0.16 | 0.11 | 0.16 |
| 1 | 169860199 | rs16862745 | 1.621 | -1.579 | -0.09 | -0.313 | -0.001 | 0.082 | 0.082 | 0 | 0.3 | 0.16 | 0.19 | 0.01 | 0.26 | 0.12 |
| 1 | 169860528 | rs3753312 | 1.88 | -1.352 | -0.007 | -0.032 | -0.002 | 0 | 0 | 2.076 | 0.16 | 0.15 | 0.24 | 0.18 | 0.11 | 0.16 |
| 1 | 169869344 | rs6670818 | -1.904 | -1.028 | -0.115 | 0.419 | -0.051 | -0.041 | -0.041 | 0 | 0.9 | 0.96 | 0.73 | 0.54 | 0.69 | 0.8 |
| 1 | 169875679 | rs12124561 | 2.279 | -1.274 | -0.183 | -0.205 | -0.032 | 0.048 | 0.048 | 0.904 | 0.1 | 0 | 0.04 | 0.01 | 0.1 | 0.07 |
| 1 | 169884386 | rs16862751 | 1.187 | -1.331 | 0.202 | -0.235 | -0.002 | -0.059 | -0.059 | 0.276 | 0.13 | 0.21 | 0.06 | 0.01 | 0.11 | 0.07 |
| 1 | 169886587 | rs10800488 | 1.624 | -1.686 | 0.146 | -0.202 | 0.028 | -0.007 | -0.007 | 0.67 | 0.17 | 0.12 | 0.23 | 0.16 | 0.11 | 0.27 |
| 1 | 169893204 | rs603246 | 0.127 | -1.725 | 0.152 | -0.304 | 0.029 | 0.004 | 0.004 | 0.035 | 0.68 | 0.61 | 0.63 | 0.29 | 0.54 | 0.58 |
| 1 | 169899427 | rs648576 | 1.6 | -1.857 | 0.011 | -0.516 | -0.011 | 0.083 | 0.083 | 0 | 0.3 | 0.16 | 0.19 | 0 | 0.26 | 0.15 |
| 1 | 169905053 | rs10919271 | 1.662 | -1.946 | 0.095 | -0.541 | 0 | 0.007 | 0.007 | 0 | 0.19 | 0.13 | 0.19 | 0.12 | 0.17 | 0.18 |
| 1 | 169907215 | rs516793 | 1.57 | -1.868 | -0.036 | -0.52 | -0.002 | 0.007 | 0.007 | 0 | 0.3 | 0.23 | 0.2 | 0 | 0.26 | 0.12 |
| 1 | 169916580 | rs1572551 | 2.044 | -1.683 | -0.195 | -0.297 | 0.027 | 0.005 | 0.005 | 0 | 0.16 | 0.09 | 0.23 | 0.13 | 0.11 | 0.16 |
| 1 | 169923676 | rs508063 | -0.129 | -1.355 | -0.075 | -0.103 | -0.019 | -0.008 | -0.008 | 1.793 | 0.72 | 0.76 | 0.65 | 0.29 | 0.55 | 0.57 |
| 1 | 169930761 | rs618692 | -0.888 | -1.239 | -0.048 | -0.014 | -0.001 | 0.001 | 0.001 | 0.566 | 0.84 | 0.85 | 0.86 | 0.85 | 0.88 | 0.77 |
| 1 | 169956450 | rs546894 | 0.948 | -1.019 | -0.746 | -0.953 | -0.022 | 0.042 | 0.042 | 0 | 0.24 | 0.16 | 0.19 | 0.01 | 0.25 | 0.13 |
| 1 | 169968152 | rs688100 | -0.954 | -1.371 | -0.72 | -1.099 | 0.001 | 0 | 0 | 0 | 0.66 | 0.7 | 0.75 | 0.85 | 0.69 | 0.66 |
| 1 | 169971492 | rs4656729 | -0.332 | -1.261 | -0.675 | -0.892 | -0.004 | -0.041 | -0.041 | 0 | 0.41 | 0.52 | 0.55 | 0.85 | 0.44 | 0.53 |

| | | | | | | | | | | | | | | | | |
|---|---|---|---|---|---|---|---|---|---|---|---|---|---|---|---|---|
| 1 | 169984446 | rs2152831 | 1.221 | -1.11 | -0.991 | -0.529 | -0.083 | -0.117 | -0.117 | 0 | 0.17 | 0.05 | 0.29 | 0.5 | 0.33 | 0.23 |
| 1 | 169985796 | rs16862933 | 1.586 | -0.974 | -1.046 | -0.501 | -0.087 | -0.126 | -0.126 | 0 | 0.16 | 0.01 | 0.27 | 0.48 | 0.3 | 0.18 |
| 1 | 169998200 | rs10800495 | 1.4 | -1.375 | -0.679 | -0.576 | 0.007 | 0.027 | 0.027 | 0.025 | 0.22 | 0.11 | 0.19 | 0.11 | 0.2 | 0.19 |
| 1 | 170004722 | rs4656732 | 1.874 | -1.477 | -0.681 | -0.578 | 0.032 | 0.004 | 0.004 | 0 | 0.16 | 0.05 | 0.24 | 0.13 | 0.11 | 0.16 |
| 1 | 170017067 | rs1330369 | 0.211 | -1.414 | -0.254 | -0.511 | -0.005 | 0.168 | 0.168 | 0.153 | 0.07 | 0.3 | 0.02 | 0.16 | 0.01 | 0.14 |
| 1 | 170037624 | rs12146137 | 1.477 | -1.503 | -0.25 | -0.625 | 0.001 | 0.004 | 0.004 | 0.039 | 0.22 | 0.17 | 0.2 | 0.11 | 0.2 | 0.19 |
| 1 | 170043851 | rs1360294 | 0.227 | -1.202 | 0.131 | -0.283 | -0.073 | 0.066 | 0.066 | 0 | 0.22 | 0.37 | 0.26 | 0.34 | 0.11 | 0.31 |
| 1 | 170048831 | rs11588736 | 1.404 | -0.932 | -0.512 | -0.234 | 0.023 | -0.014 | -0.014 | 1.517 | 0.15 | 0.07 | 0.24 | 0.19 | 0.11 | 0.16 |
| 1 | 170052537 | rs12146094 | 1.588 | -0.467 | -0.457 | -0.337 | -0.003 | -0.003 | -0.003 | 1.229 | 0.1 | 0.08 | 0.04 | 0 | 0.1 | 0.03 |
| 1 | 170053607 | rs4656204 | 1.033 | -0.437 | -0.482 | -0.288 | 0.028 | -0.002 | -0.002 | 0.176 | 0.18 | 0.07 | 0.25 | 0.19 | 0.12 | 0.2 |
| 1 | 170053959 | rs1330378 | -0.714 | 0.079 | -0.005 | -0.179 | 0.005 | 0.102 | 0.102 | 0 | 0.08 | 0.25 | 0.04 | 0.19 | 0.04 | 0.16 |
| 1 | 170057917 | rs12126443 | 1.485 | 0.235 | -0.427 | -0.378 | 0 | -0.018 | -0.018 | 0 | 0.07 | 0.09 | 0.03 | 0 | 0.06 | 0.03 |
| 1 | 170067296 | rs10919299 | 0.831 | 0.236 | -0.714 | -0.632 | -0.002 | -0.02 | -0.02 | 0 | 0.38 | 0.12 | 0.39 | 0.63 | 0.42 | 0.49 |
| 1 | 170084155 | rs12135070 | 1.115 | 0.403 | -0.8 | -0.621 | -0.002 | -0.008 | -0.008 | 0.03 | 0.09 | 0.11 | 0.05 | 0.01 | 0.09 | 0.07 |
| 1 | 170084238 | rs12131740 | 1.115 | 0.403 | -0.8 | -0.621 | -0.002 | -0.008 | -0.008 | 0 | 0.09 | 0.11 | 0.05 | 0.01 | 0.09 | 0.07 |
| 1 | 170085453 | rs12042109 | 0.056 | 1.003 | -0.348 | -0.603 | -0.002 | 0 | 0 | 0 | 0.38 | 0.34 | 0.47 | 0.35 | 0.36 | 0.39 |
| 1 | 170093425 | rs516935 | -1.162 | 1.137 | -0.762 | -0.984 | 0 | -0.025 | -0.025 | 0 | 0.9 | 0.86 | 0.93 | 1 | 0.89 | 0.96 |
| 1 | 170103379 | rs471569 | 0.806 | 1.152 | -1.438 | -0.866 | -0.031 | 0.068 | 0.068 | 0 | 0.44 | 0.09 | 0.43 | 0.69 | 0.48 | 0.47 |
| 1 | 170103732 | rs10800498 | 0.102 | 1.388 | -1.102 | -1.085 | 0.07 | -0.01 | -0.01 | 0 | 0.18 | 0.47 | 0.23 | 0.13 | 0.22 | 0.2 |
| 1 | 170112399 | rs575119 | 0.909 | 1.238 | -1.108 | -1.129 | 0.023 | 0.129 | 0.129 | 0 | 0.48 | 0.15 | 0.44 | 0.71 | 0.48 | 0.55 |
| 1 | 170118691 | rs10919307 | -0.189 | 1.37 | -1.054 | -1.127 | 0.082 | -0.028 | -0.028 | 3.261 | 0.21 | 0.5 | 0.25 | 0.13 | 0.27 | 0.22 |
| 1 | 170126618 | rs7354854 | 0.923 | 1.38 | -1.07 | -1.085 | 0 | -0.017 | -0.017 | 1.286 | 0.09 | 0.11 | 0.06 | 0 | 0.09 | 0.02 |
| 1 | 170131293 | rs504944 | -1.352 | 1.317 | -1.149 | -1.019 | 0.007 | 0.008 | 0.008 | 0 | 0.78 | 0.77 | 0.75 | 0.84 | 0.84 | 0.8 |
| 1 | 170139508 | rs532141 | 0.737 | 1.548 | -1.374 | -1 | -0.061 | 0.123 | 0.123 | 0 | 0.49 | 0.1 | 0.46 | 0.7 | 0.54 | 0.49 |
| 1 | 170141722 | rs11585754 | 1.077 | 1.946 | -1.608 | -1.008 | 0.068 | 0.16 | 0.16 | 0.226 | 0.41 | 0.15 | 0.23 | 0.12 | 0.33 | 0.31 |
| 1 | 170141746 | rs495127 | -0.656 | 1.963 | -1.4 | -1.007 | 0.055 | -0.006 | -0.006 | 5.659 | 0.83 | 0.5 | 0.79 | 0.88 | 0.81 | 0.82 |
| 1 | 170143426 | rs519459 | -0.621 | 1.961 | -1.412 | -1.008 | 0.052 | -0.009 | -0.009 | 0.436 | 0.83 | 0.5 | 0.79 | 0.88 | 0.81 | 0.82 |
| 1 | 170145099 | rs486427 | 0.05 | 2.111 | -1.468 | -1.025 | 0.003 | 0.021 | 0.021 | 0 | 0.38 | 0.32 | 0.52 | 0.75 | 0.39 | 0.44 |
| 1 | 170147431 | rs533550 | 1.487 | 1.84 | -1.275 | -0.705 | -0.052 | -0.036 | -0.036 | 0 | 0.09 | 0.01 | 0.24 | 0.57 | 0.2 | 0.13 |
| 1 | 170151517 | rs488150 | 1.329 | 1.883 | -1.255 | -0.613 | -0.056 | -0.073 | -0.073 | 0 | 0.11 | 0.01 | 0.24 | 0.56 | 0.2 | 0.13 |
| 1 | 170152927 | rs10919319 | -0.437 | 1.361 | -0.937 | -0.802 | 0.057 | -0.033 | -0.033 | 2.896 | 0.2 | 0.53 | 0.23 | 0.12 | 0.24 | 0.21 |
| 1 | 170168182 | rs7518785 | 0.269 | 1.271 | -0.937 | -0.458 | 0.01 | 0.045 | 0.045 | 0 | 0.53 | 0.6 | 0.42 | 0.25 | 0.46 | 0.5 |
| 1 | 170171083 | rs10919325 | 0.347 | 1.601 | -0.698 | -0.352 | 0.012 | 0.048 | 0.048 | 0 | 0.43 | 0.31 | 0.38 | 0.24 | 0.41 | 0.48 |
| 1 | 170171258 | rs10919326 | 0.511 | 1.633 | -0.75 | -0.487 | 0.033 | 0.08 | 0.08 | 0.364 | 0.41 | 0.22 | 0.35 | 0.24 | 0.38 | 0.37 |
| 1 | 170171702 | rs12043515 | 0.588 | 2.119 | -0.176 | -0.381 | -0.002 | -0.007 | -0.007 | 0 | 0.32 | 0.3 | 0.29 | 0.17 | 0.27 | 0.23 |
| 1 | 170172417 | rs10158534 | 0.535 | 2.435 | -0.127 | -0.385 | 0.004 | 0.04 | 0.04 | 0 | 0.47 | 0.41 | 0.4 | 0.26 | 0.44 | 0.63 |
| 1 | 170174298 | rs12144655 | -1.069 | 2.63 | -0.113 | -0.406 | 0.049 | -0.011 | -0.011 | 0 | 0.05 | 0.15 | 0.05 | 0.02 | 0.1 | 0.13 |
| 1 | 170174665 | rs1330385 | 0.716 | 2.573 | -0.184 | -0.417 | 0.012 | 0.053 | 0.053 | 0 | 0.43 | 0.33 | 0.37 | 0.25 | 0.4 | 0.51 |
| 1 | 170179064 | rs4656206 | 0.339 | 2.666 | -0.265 | -0.429 | -0.009 | 0.094 | 0.094 | 0.05 | 0.55 | 0.26 | 0.42 | 0.25 | 0.5 | 0.55 |
| 1 | 170186610 | rs7545084 | 0.956 | 2.326 | 0.017 | -0.394 | 0.027 | 0.052 | 0.052 | 0 | 0.42 | 0.26 | 0.32 | 0.26 | 0.36 | 0.51 |
| 1 | 170186747 | rs16863119 | 0.746 | 2.489 | 0.07 | -0.192 | -0.001 | -0.005 | -0.005 | 0 | 0.32 | 0.3 | 0.32 | 0.18 | 0.26 | 0.22 |
| 1 | 170187309 | rs6657002 | 0.999 | 2.437 | 0.045 | -0.253 | 0.028 | 0.054 | 0.054 | 0 | 0.42 | 0.26 | 0.32 | 0.26 | 0.36 | 0.51 |
| 1 | 170188648 | rs4656207 | 0.13 | 2.425 | 0.026 | -0.036 | 0.077 | 0.311 | 0.311 | 0.043 | 0.95 | 0.82 | 0.8 | 0.45 | 0.9 | 0.91 |
| 1 | 170192022 | rs4509636 | 0.352 | 1.752 | -0.308 | -0.693 | 0 | -0.012 | -0.012 | 1.358 | 0.34 | 0.31 | 0.32 | 0.11 | 0.26 | 0.22 |
| 1 | 170196342 | rs4437927 | -0.867 | 2.146 | -0.179 | -0.234 | 0.011 | -0.096 | -0.096 | 0 | 0.23 | 0.36 | 0.16 | 0.03 | 0.23 | 0.18 |
| 1 | 170202563 | rs2225588 | -0.667 | 2.254 | -0.674 | 0.374 | -0.028 | -0.025 | -0.025 | 0 | 0.33 | 0.24 | 0.36 | 0.61 | 0.41 | 0.38 |
| 1 | 170206902 | rs983118 | 0.64 | 2.118 | -0.085 | 0.48 | 0.012 | 0.016 | 0.016 | 0 | 0.64 | 0.54 | 0.62 | 0.41 | 0.58 | 0.62 |
| 1 | 170208425 | rs1412788 | -0.394 | 2.315 | -0.444 | 1.16 | -0.059 | -0.08 | -0.08 | 2.836 | 0.29 | 0.07 | 0.32 | 0.42 | 0.35 | 0.36 |
| 1 | 170208435 | rs12028052 | 1.047 | 1.828 | -0.925 | 0.589 | -0.047 | -0.04 | -0.04 | 0.098 | 0.13 | 0.02 | 0.13 | 0.17 | 0.22 | 0.1 |
| 1 | 170208724 | rs1412787 | 0.23 | 2.268 | -0.942 | 0.753 | -0.046 | -0.087 | -0.087 | 0.002 | 0.26 | 0.07 | 0.22 | 0.44 | 0.33 | 0.35 |
| 1 | 170210475 | rs12122745 | 0.151 | 2.277 | -0.959 | 0.683 | -0.052 | -0.107 | -0.107 | 0 | 0.26 | 0.05 | 0.22 | 0.45 | 0.33 | 0.36 |
| 1 | 170216618 | rs1332950 | 0.926 | 1.61 | -0.98 | 0.211 | -0.012 | -0.024 | -0.024 | 2.144 | 0.32 | 0.37 | 0.34 | 0.17 | 0.25 | 0.21 |
| 1 | 170216805 | rs10919336 | 1.494 | 1.778 | -1.239 | 0.175 | -0.09 | -0.057 | -0.057 | 0.278 | 0.15 | 0.02 | 0.24 | 0.19 | 0.26 | 0.1 |
| 1 | 170217002 | rs10919337 | 0.93 | 1.017 | -0.729 | 0.028 | 0.012 | 0.012 | 0.012 | 0.014 | 0.25 | 0.18 | 0.18 | 0.18 | 0.17 | 0.36 |
| 1 | 170220252 | rs2019438 | 0.428 | 1.107 | -0.551 | 0.068 | -0.016 | 0.002 | 0.002 | 0 | 0.41 | 0.27 | 0.45 | 0.64 | 0.57 | 0.43 |
| 1 | 170220451 | rs1018710 | 0.517 | 1.13 | -0.498 | 0.079 | -0.031 | -0.004 | -0.004 | 2.386 | 0.4 | 0.2 | 0.45 | 0.64 | 0.57 | 0.43 |
| 1 | 170223350 | rs1547282 | 0.015 | 0.776 | -0.343 | -0.277 | -0.025 | -0.047 | -0.047 | 0 | 0.33 | 0.49 | 0.35 | 0.17 | 0.26 | 0.21 |
| 1 | 170231780 | rs1556498 | -1.175 | 0.745 | 0.045 | -0.212 | 0.003 | -0.101 | -0.101 | 0 | 0.75 | 0.67 | 0.82 | 0.99 | 0.77 | 0.92 |
| 1 | 170233764 | rs2251095 | 0.305 | 0.735 | -0.062 | 0.02 | 0.003 | 0.004 | 0.004 | 0 | 0.56 | 0.6 | 0.58 | 0.53 | 0.42 | 0.64 |
| 1 | 170237860 | rs7517657 | -0.305 | 0.758 | -0.01 | 0.043 | -0.015 | -0.017 | -0.017 | 0 | 0.2 | 0.19 | 0.26 | 0.46 | 0.35 | 0.28 |
| 1 | 170240085 | rs2254605 | 0.334 | 0.696 | 0.072 | -0.012 | 0.003 | 0.004 | 0.004 | 0 | 0.54 | 0.43 | 0.56 | 0.53 | 0.42 | 0.64 |
| 1 | 170243444 | rs2990634 | -1.203 | 0.754 | -0.125 | -0.153 | 0.005 | -0.136 | -0.136 | 2.279 | 0.74 | 0.63 | 0.82 | 0.99 | 0.77 | 0.92 |
| 1 | 170254945 | rs2095134 | 0.014 | 0.844 | -0.686 | -0.054 | 0.003 | 0.018 | 0.018 | 0 | 0.58 | 0.68 | 0.59 | 0.57 | 0.42 | 0.64 |
| 1 | 170256887 | rs2184042 | -0.071 | 0.913 | -1.099 | -0.02 | -0.027 | 0.019 | 0.019 | 0.175 | 0.4 | 0.17 | 0.41 | 0.43 | 0.58 | 0.36 |
| 1 | 170265547 | rs12745248 | 1.694 | 0.899 | -1.105 | -0.064 | -0.081 | 0.135 | 0.135 | 0 | 0.21 | 0.02 | 0.16 | 0.01 | 0.23 | 0.08 |
| 1 | 170266921 | rs11582930 | 0.096 | 0.168 | -0.695 | 0.164 | -0.01 | 0.028 | 0.028 | 0 | 0.41 | 0.18 | 0.4 | 0.42 | 0.57 | 0.36 |
| 1 | 170280083 | rs4575137 | 0.907 | -0.072 | -0.07 | -0.404 | 0.013 | -0.133 | -0.133 | 0.141 | 0.3 | 0.47 | 0.27 | 0.01 | 0.3 | 0.11 |
| 1 | 170283044 | rs986032 | -0.969 | -0.835 | -0.591 | -0.318 | 0.001 | -0.032 | -0.032 | 0 | 0.81 | 0.94 | 0.75 | 0.8 | 0.8 | 0.88 |
| 1 | 170287952 | rs12732119 | 1.549 | -0.53 | -0.582 | -0.274 | -0.128 | 0.171 | 0.171 | 0.058 | 0.27 | 0.04 | 0.26 | 0.01 | 0.33 | 0.12 |

| | | | | | | | | | | | | | | | | |
|---|---|---|---|---|---|---|---|---|---|---|---|---|---|---|---|---|
| 1 | 170290129 | rs928462 | 0.499 | -0.849 | -0.651 | -0.131 | 0.006 | -0.081 | -0.081 | 0 | 0.24 | 0.1 | 0.24 | 0.45 | 0.18 | 0.42 |
| 1 | 170297878 | rs1332969 | 1.108 | -0.779 | -0.371 | -0.09 | 0.006 | -0.081 | -0.081 | 0.178 | 0.31 | 0.41 | 0.29 | 0.01 | 0.34 | 0.12 |
| 1 | 170299009 | rs4491109 | 0.401 | -0.696 | -0.203 | 0.078 | -0.026 | -0.027 | -0.027 | 0 | 0.15 | 0.06 | 0.15 | 0.32 | 0.23 | 0.24 |
| 1 | 170326065 | rs16863314 | 1.362 | -0.621 | -0.609 | 0.219 | 0.01 | 0.023 | 0.023 | 0 | 0.07 | 0.06 | 0.12 | 0.28 | 0.16 | 0.14 |
| 1 | 170330500 | rs1930537 | -0.63 | -0.875 | -0.57 | 0.118 | -0.002 | 0.018 | 0.018 | 0 | 0.37 | 0.26 | 0.39 | 0.76 | 0.36 | 0.63 |
| 1 | 170336440 | rs10919364 | 0.565 | -0.897 | -0.621 | 0.057 | 0.004 | -0.004 | -0.004 | 0.385 | 0.23 | 0.17 | 0.24 | 0.44 | 0.14 | 0.4 |
| 1 | 170339511 | rs6698352 | 1.746 | -0.864 | -0.84 | -0.18 | -0.13 | 0.173 | 0.173 | 0.297 | 0.27 | 0.03 | 0.27 | 0.01 | 0.34 | 0.12 |
| 1 | 170346164 | rs10919369 | 0.152 | -0.883 | -0.545 | -0.091 | 0.003 | 0.003 | 0.003 | 0 | 0.16 | 0.14 | 0.15 | 0.3 | 0.21 | 0.24 |
| 1 | 170356917 | rs1071521 | 0.217 | -0.877 | -0.538 | -0.088 | 0.021 | 0.022 | 0.022 | 0 | 0.8 | 0.43 | 0.82 | 0.69 | 0.79 | 0.76 |
| 1 | 170363183 | rs1112882 | 0.428 | -0.76 | -0.389 | 0.077 | 0.003 | 0.011 | 0.011 | 0 | 0.46 | 0.59 | 0.44 | 0.32 | 0.56 | 0.35 |
| 1 | 170365623 | rs7554931 | 0.435 | -0.757 | -0.387 | 0.078 | 0.003 | 0.01 | 0.01 | 0 | 0.46 | 0.59 | 0.44 | 0.32 | 0.56 | 0.35 |
| 1 | 170372503 | rs10919388 | 0.003 | -0.267 | -0.244 | -0.33 | -0.002 | -0.002 | -0.002 | 0 | 0.67 | 0.7 | 0.68 | 0.76 | 0.71 | 0.74 |
| 1 | 170377323 | rs12024348 | -0.352 | 0.473 | -0.581 | 0.124 | 0.005 | 0.038 | 0.038 | 0 | 0.15 | 0.14 | 0.24 | 0.58 | 0.21 | 0.27 |
| 1 | 170379727 | rs4656767 | 0.283 | 0.213 | -0.947 | -0.342 | 0.008 | -0.075 | -0.075 | 0.115 | 0.3 | 0.04 | 0.28 | 0.29 | 0.3 | 0.25 |
| 1 | 170379773 | rs4656768 | -0.363 | 0.365 | -0.514 | -0.405 | -0.018 | 0.092 | 0.092 | 0 | 0.39 | 0.25 | 0.61 | 0.87 | 0.5 | 0.49 |
| 1 | 170380480 | rs6427247 | 0.3 | 0.302 | -0.551 | -0.387 | 0.029 | -0.029 | -0.029 | 0 | 0.32 | 0.18 | 0.3 | 0.31 | 0.31 | 0.29 |
| 1 | 170387134 | rs2027020 | -0.721 | 0.135 | -0.517 | -0.788 | -0.003 | -0.028 | -0.028 | 0 | 0.58 | 0.42 | 0.67 | 0.97 | 0.59 | 0.81 |
| 1 | 170391041 | rs16863343 | 0.073 | 0.131 | -0.08 | -0.867 | -0.002 | -0.034 | -0.034 | 0 | 0.09 | 0.23 | 0.03 | 0.02 | 0.04 | 0.07 |
| 1 | 170402976 | rs12119356 | 0.002 | 0.045 | -0.151 | -0.933 | 0.003 | 0.101 | 0.101 | 0.033 | 0.56 | 0.29 | 0.36 | 0.16 | 0.51 | 0.51 |
| 1 | 170406261 | rs12032644 | 0.094 | 0.039 | -0.098 | -0.937 | -0.002 | 0.017 | 0.017 | 0 | 0.41 | 0.38 | 0.62 | 0.85 | 0.49 | 0.49 |
| 1 | 170407926 | rs1333141 | 0.094 | 0.039 | -0.098 | -0.937 | -0.002 | 0.017 | 0.017 | 0 | 0.41 | 0.38 | 0.62 | 0.85 | 0.49 | 0.49 |
| 1 | 170408985 | rs7512094 | 1.486 | 0.032 | -0.127 | -0.952 | -0.126 | 0.197 | 0.197 | 0.045 | 0.29 | 0.03 | 0.27 | 0.02 | 0.38 | 0.12 |
| 1 | 170413292 | rs1333147 | -0.055 | 0.178 | -0.203 | -0.765 | -0.002 | 0.012 | 0.012 | 0 | 0.42 | 0.41 | 0.62 | 0.85 | 0.48 | 0.49 |
| 1 | 170416126 | rs16863354 | 0.34 | -0.034 | -0.296 | -0.699 | 0.02 | -0.004 | -0.004 | 0 | 0.34 | 0.26 | 0.31 | 0.3 | 0.31 | 0.3 |
| 1 | 170433781 | rs4294468 | -1.18 | 0.177 | -0.258 | -0.923 | 0.007 | -0.098 | -0.098 | 0 | 0.68 | 0.64 | 0.7 | 0.98 | 0.62 | 0.88 |
| 1 | 170440359 | rs12747498 | 0.808 | 0.077 | -0.466 | -1.01 | -0.014 | 0.027 | 0.027 | 0 | 0.17 | 0.01 | 0.06 | 0.13 | 0.1 | 0.32 |
| 1 | 170443579 | rs12044570 | 0.047 | 0.046 | -0.179 | -1.067 | 0 | -0.03 | -0.03 | 0 | 0.43 | 0.46 | 0.63 | 0.85 | 0.48 | 0.49 |
| 1 | 170452449 | rs16863369 | -0.263 | 0.036 | -0.183 | -1.041 | 0.005 | 0.006 | 0.006 | 0 | 0.26 | 0.23 | 0.09 | 0.14 | 0.14 | 0.39 |
| 1 | 170480531 | rs10919423 | 1.378 | -0.028 | -0.34 | -1.014 | -0.126 | 0.197 | 0.197 | 0.045 | 0.29 | 0.03 | 0.27 | 0.02 | 0.38 | 0.12 |
| 1 | 170499706 | rs12408432 | 1.317 | 0.049 | -0.258 | -0.808 | 0.029 | 0.136 | 0.136 | 0 | 0.07 | 0.13 | 0.31 | 0.55 | 0.18 | 0.19 |
| 1 | 170501167 | rs16863397 | 1.236 | 0.044 | -0.256 | -0.869 | 0.009 | 0.092 | 0.092 | 0 | 0.07 | 0.11 | 0.31 | 0.57 | 0.18 | 0.21 |
| 1 | 170501456 | rs11578119 | 1.424 | 0.039 | -0.375 | -0.919 | -0.126 | 0.197 | 0.197 | 0.048 | 0.29 | 0.03 | 0.27 | 0.02 | 0.36 | 0.12 |
| 1 | 170514229 | rs12120764 | 1.448 | 0.035 | -0.373 | -0.917 | -0.124 | 0.195 | 0.195 | 0.053 | 0.29 | 0.03 | 0.26 | 0.02 | 0.36 | 0.12 |
| 1 | 170519884 | rs16863425 | 0.554 | 0.087 | -0.08 | -0.923 | 0.01 | -0.004 | -0.004 | 0 | 0.36 | 0.28 | 0.32 | 0.31 | 0.35 | 0.35 |
| 1 | 170521376 | rs913257 | -0.012 | 0.012 | -0.088 | -1.092 | 0.002 | -0.088 | -0.088 | 0.355 | 0.42 | 0.53 | 0.59 | 0.88 | 0.47 | 0.52 |
| 1 | 170521650 | rs7552922 | -0.02 | -0.917 | 0.034 | -0.862 | 0.003 | -0.002 | -0.002 | 0 | 0.23 | 0.23 | 0.12 | 0.12 | 0.14 | 0.35 |
| 1 | 170528547 | rs12564005 | 0.506 | -0.81 | -0.131 | -0.672 | 0.013 | -0.012 | -0.012 | 0 | 0.32 | 0.27 | 0.4 | 0.33 | 0.29 | 0.33 |
| 1 | 170528913 | rs6676293 | -0.184 | -0.585 | -0.398 | -0.608 | 0.011 | 0.036 | 0.036 | 0 | 0.22 | 0.15 | 0.11 | 0.12 | 0.14 | 0.35 |
| 1 | 170549087 | rs16863439 | -0.294 | -0.149 | -0.143 | -0.111 | -0.022 | -0.037 | -0.037 | 0 | 0.07 | 0.14 | 0.01 | 0 | 0.03 | 0.01 |
| 1 | 170553442 | rs6427265 | -0.507 | 0.195 | -0.313 | 0.014 | -0.016 | 0.005 | 0.005 | 0 | 0.37 | 0.46 | 0.45 | 0.31 | 0.35 | 0.4 |
| 1 | 170554924 | rs7535322 | -0.453 | 0.125 | -0.656 | -0.088 | 0.01 | 0 | 0 | 0.373 | 0.37 | 0.28 | 0.29 | 0.28 | 0.34 | 0.4 |
| 1 | 170560332 | rs6680785 | -0.395 | 0.562 | -1.016 | -0.334 | -0.02 | 0.031 | 0.031 | 0 | 0.44 | 0.33 | 0.61 | 0.7 | 0.49 | 0.44 |
| 1 | 170564545 | rs6697185 | -0.052 | -0.202 | -0.21 | -0.638 | 0.002 | -0.011 | -0.011 | 0 | 0.07 | 0.33 | 0.04 | 0 | 0.01 | 0.04 |
| 1 | 170565191 | rs2050000 | 0.167 | -0.455 | -0.761 | -0.943 | 0.073 | 0.104 | 0.104 | 0 | 0.3 | 0.1 | 0.32 | 0.08 | 0.15 | 0.23 |
| 1 | 170582464 | rs7540157 | 0.134 | -0.556 | -0.665 | -0.974 | 0.052 | 0.062 | 0.062 | 0 | 0.31 | 0.18 | 0.33 | 0.08 | 0.15 | 0.23 |
| 1 | 170585832 | rs619456 | -1.136 | -0.336 | -0.59 | -1.124 | -0.006 | 0.04 | 0.04 | 0 | 0.57 | 0.75 | 0.7 | 0.7 | 0.59 | 0.49 |
| 1 | 170586315 | rs736791 | 0.729 | -0.367 | -0.535 | -1.142 | -0.057 | -0.054 | -0.054 | 0 | 0.16 | 0.11 | 0.33 | 0.62 | 0.42 | 0.22 |
| 1 | 170587340 | rs577676 | -0.266 | -0.022 | -0.713 | -1.544 | -0.042 | -0.034 | -0.034 | 0 | 0.76 | 0.84 | 0.6 | 0.38 | 0.46 | 0.73 |
| 1 | 170588594 | rs2022373 | 1.144 | -0.027 | -0.721 | -1.532 | 0.016 | 0.004 | 0.004 | 0 | 0.36 | 0.21 | 0.22 | 0.3 | 0.3 | 0.46 |
| 1 | 170612873 | rs639652 | 0.056 | 0.058 | -0.54 | -1.494 | 0.092 | 0.075 | 0.075 | 0.022 | 0.75 | 0.51 | 0.63 | 0.38 | 0.56 | 0.75 |
| 1 | 170615833 | rs12129225 | 0.226 | 0.006 | -0.639 | -1.506 | 0.031 | 0.039 | 0.039 | 0 | 0.39 | 0.3 | 0.41 | 0.08 | 0.27 | 0.28 |
| 1 | 170617306 | rs608930 | -0.989 | 0.195 | -0.723 | -1.433 | 0.022 | 0.018 | 0.018 | 0 | 0.82 | 0.81 | 0.67 | 0.38 | 0.57 | 0.78 |
| 1 | 170618461 | rs12038255 | 1.697 | -0.215 | -1.589 | -1.85 | -0.073 | 0.017 | 0.017 | 0 | 0.2 | 0.01 | 0.11 | 0.08 | 0.22 | 0.22 |
| 1 | 170624572 | rs12132205 | 0.95 | -0.039 | -1.154 | -1.803 | 0.045 | 0.105 | 0.105 | 0.009 | 0.35 | 0.17 | 0.41 | 0.08 | 0.27 | 0.27 |
| 1 | 170625867 | rs501680 | 0.552 | -0.087 | -0.995 | -1.749 | 0.017 | 0.009 | 0.009 | 0 | 0.36 | 0.23 | 0.21 | 0.3 | 0.29 | 0.47 |
| 1 | 170632808 | rs12121907 | 1.224 | -0.274 | -1.116 | -1.743 | 0.088 | 0.126 | 0.126 | 0.066 | 0.31 | 0.07 | 0.33 | 0.09 | 0.16 | 0.24 |
| 1 | 170638658 | rs10919449 | -0.539 | 0.708 | -0.849 | -1.441 | 0.008 | 0.027 | 0.027 | 0 | 0.49 | 0.6 | 0.32 | 0.3 | 0.42 | 0.53 |
| 1 | 170642509 | rs591715 | 0.512 | 0.849 | -1.342 | -1.139 | 0.055 | -0.018 | -0.018 | 0 | 0.33 | 0.17 | 0.17 | 0.32 | 0.21 | 0.42 |
| 1 | 170642899 | rs593479 | -0.561 | 1.975 | -0.946 | -0.499 | -0.043 | -0.037 | -0.037 | 0 | 0.8 | 0.85 | 0.64 | 0.42 | 0.52 | 0.76 |
| 1 | 170649279 | rs4656794 | 0.182 | 0.776 | -1.6 | -0.682 | -0.008 | 0.026 | 0.026 | 0.045 | 0.45 | 0.14 | 0.29 | 0.59 | 0.36 | 0.53 |
| 1 | 170649992 | rs11576746 | 0.058 | 0.62 | -0.531 | -0.983 | 0.003 | -0.053 | -0.053 | 0 | 0.09 | 0.26 | 0.11 | 0 | 0.07 | 0.02 |
| 1 | 170652434 | rs12066968 | 1.353 | 0.556 | -0.249 | -1.223 | 0.001 | -0.042 | -0.042 | 0 | 0.07 | 0.17 | 0.05 | 0 | 0.06 | 0.03 |
| 1 | 170654118 | rs520131 | 0.423 | 1.026 | -0.063 | -1.027 | -0.005 | -0.084 | -0.084 | 1.294 | 0.41 | 0.42 | 0.46 | 0.12 | 0.42 | 0.28 |
| 1 | 170657335 | rs17550940 | 1.082 | 0.996 | -0.621 | -1.078 | -0.026 | 0.176 | 0.176 | 0.632 | 0.32 | 0.05 | 0.34 | 0.11 | 0.35 | 0.26 |
| 1 | 170657994 | rs531650 | -0.904 | 1.041 | -0.123 | -0.838 | 0.03 | 0.054 | 0.054 | 0 | 0.86 | 0.79 | 0.75 | 0.71 | 0.76 | 0.79 |
| 1 | 170658355 | rs12744534 | 1.338 | 0.498 | -0.632 | -0.668 | 0.032 | -0.082 | -0.082 | 0 | 0.21 | 0.07 | 0.18 | 0.42 | 0.1 | 0.32 |
| 1 | 170664237 | rs513287 | 0.005 | 0.293 | -0.138 | -0.6 | 0.008 | 0.008 | 0.008 | 1.482 | 0.58 | 0.46 | 0.61 | 0.51 | 0.5 | 0.6 |
| 1 | 170700758 | rs1476040 | -1.089 | 0.972 | 0.457 | -0.302 | 0.034 | 0.038 | 0.038 | 0 | 0.87 | 0.78 | 0.78 | 0.77 | 0.76 | 0.84 |
| 1 | 170709195 | rs4656799 | 0.296 | 0.787 | 0.137 | -0.702 | 0.019 | 0.014 | 0.014 | 0 | 0.44 | 0.32 | 0.43 | 0.12 | 0.38 | 0.29 |

| | | | | | | | | | | | | | | | | |
|---|---|---|---|---|---|---|---|---|---|---|---|---|---|---|---|---|
| 1 | 170715110 | rs6679313 | -0.323 | 0.244 | 0.272 | -0.754 | 0.005 | 0.028 | 0.028 | 0 | 0.36 | 0.43 | 0.38 | 0.5 | 0.32 | 0.48 |
| 1 | 170715531 | rs2421494 | 1.464 | 0.187 | -0.006 | -0.848 | 0.032 | 0.175 | 0.175 | 1.833 | 0.31 | 0.03 | 0.3 | 0.12 | 0.24 | 0.23 |
| 1 | 170719187 | rs991230 | 1.464 | 0.187 | -0.006 | -0.848 | 0.032 | 0.175 | 0.175 | 1.833 | 0.31 | 0.03 | 0.29 | 0.12 | 0.25 | 0.22 |
| 1 | 170719666 | rs6702078 | 0.069 | 0.221 | 0.039 | -0.869 | 0.124 | -0.062 | -0.062 | 0.048 | 0.1 | 0.25 | 0.13 | 0 | 0.19 | 0.07 |
| 1 | 170727442 | rs4656224 | -0.58 | 0.2 | 0.047 | -0.854 | 0 | 0.001 | 0.001 | 0 | 0.65 | 0.31 | 0.66 | 0.62 | 0.56 | 0.7 |
| 1 | 170728265 | rs12028481 | 0.858 | 0.062 | -0.011 | -0.59 | 0.046 | -0.133 | -0.133 | 0 | 0.24 | 0.05 | 0.23 | 0.49 | 0.13 | 0.4 |
| 1 | 170742228 | rs1029320 | -0.511 | 0.229 | 0.105 | -0.553 | 0 | 0.018 | 0.018 | 0 | 0.38 | 0.47 | 0.38 | 0.5 | 0.33 | 0.48 |
| 1 | 170762228 | rs17552579 | 1.723 | 0.346 | -0.003 | -0.483 | -0.011 | -0.01 | -0.01 | 0 | 0.09 | 0.07 | 0.09 | 0.12 | 0.15 | 0.1 |
| 1 | 170762680 | rs10919470 | 0.331 | 0.2 | 0.036 | -0.645 | -0.008 | -0.053 | -0.053 | 0.077 | 0.29 | 0.15 | 0.28 | 0.49 | 0.26 | 0.46 |
| 1 | 170764915 | rs10800542 | -0.381 | 0.07 | 0.18 | -0.638 | 0 | 0.008 | 0.008 | 0 | 0.37 | 0.4 | 0.38 | 0.5 | 0.32 | 0.49 |
| 1 | 170767276 | rs6677593 | 1.922 | 0.051 | -0.139 | -0.646 | -0.001 | -0.001 | -0.001 | 0 | 0.19 | 0.21 | 0.1 | 0.18 | 0.2 | 0.14 |
| 1 | 170773031 | rs2143109 | -0.523 | -0.07 | -0.138 | -0.549 | -0.001 | 0.012 | 0.012 | 0 | 0.37 | 0.4 | 0.39 | 0.5 | 0.34 | 0.49 |
| 1 | 170805627 | rs6664881 | 0.251 | -0.035 | -0.475 | -0.463 | 0.102 | 0.004 | 0.004 | 0 | 0.49 | 0.14 | 0.46 | 0.5 | 0.31 | 0.55 |
| 1 | 170807586 | rs10800546 | -1.019 | 0.082 | -0.447 | -0.492 | 0.022 | 0.021 | 0.021 | 0 | 0.66 | 0.46 | 0.62 | 0.62 | 0.52 | 0.69 |
| 1 | 170814360 | rs7551763 | 1.181 | 0.175 | -0.91 | -0.352 | 0.015 | 0.003 | 0.003 | 0 | 0.1 | 0.14 | 0.07 | 0.12 | 0.16 | 0.11 |
| 1 | 170818339 | rs6664433 | -0.411 | 0.085 | -0.856 | -0.365 | -0.01 | -0.003 | -0.003 | 0 | 0.6 | 0.38 | 0.54 | 0.63 | 0.47 | 0.66 |
| 1 | 170837041 | rs3765377 | 0.695 | -0.273 | -1.358 | -0.674 | 0.037 | 0.155 | 0.155 | 1.818 | 0.3 | 0.02 | 0.28 | 0.11 | 0.2 | 0.2 |
| 1 | 170847097 | rs16863701 | 1.963 | -0.244 | -1.751 | -0.682 | -0.035 | -0.058 | -0.058 | 0.709 | 0.19 | 0.01 | 0.11 | 0.18 | 0.23 | 0.15 |
| 1 | 170856852 | rs10489239 | -0.096 | -1.135 | -1.012 | -0.968 | -0.008 | 0.004 | 0.004 | 0 | 0.31 | 0.33 | 0.3 | 0.4 | 0.2 | 0.4 |
| 1 | 170858095 | rs10489241 | 0.614 | -1.177 | -1.023 | -0.979 | -0.022 | 0.04 | 0.04 | 0 | 0.23 | 0.32 | 0.22 | 0.39 | 0.13 | 0.35 |
| 1 | 170859317 | rs6667186 | -0.991 | -0.961 | -0.58 | -1.006 | -0.001 | -0.002 | -0.002 | 0.024 | 0.43 | 0.58 | 0.39 | 0.52 | 0.39 | 0.53 |
| 1 | 170859820 | rs16863739 | 1.543 | -1.098 | -0.702 | -0.923 | 0.034 | 0.118 | 0.118 | 3.225 | 0.25 | 0.02 | 0.25 | 0.1 | 0.16 | 0.16 |
| 1 | 170860531 | rs12038181 | 0.672 | -0.499 | 0 | -0.334 | 0.014 | -0.062 | -0.062 | 0 | 0.19 | 0.09 | 0.2 | 0.39 | 0.12 | 0.35 |
| 1 | 170897273 | rs979070 | 0.225 | 0.338 | 0.171 | -0.432 | 0.107 | 0.042 | 0.042 | 0.684 | 0.13 | 0.49 | 0.1 | 0.13 | 0.16 | 0.11 |
| 1 | 170898256 | rs6679669 | -0.149 | 0.363 | 0.055 | -0.44 | 0.09 | 0.008 | 0.008 | 0.56 | 0.16 | 0.44 | 0.12 | 0.13 | 0.19 | 0.13 |
| 1 | 170901445 | rs4656815 | -0.993 | -0.028 | -0.037 | -0.346 | -0.002 | -0.003 | -0.003 | 0 | 0.41 | 0.58 | 0.4 | 0.53 | 0.39 | 0.52 |
| 1 | 170903771 | rs6681553 | 0.304 | -0.059 | 0.004 | -0.03 | 0.015 | 0.054 | 0.054 | 0.484 | 0.49 | 0.39 | 0.37 | 0.25 | 0.38 | 0.39 |
| 1 | 170907494 | rs2143383 | -0.375 | 0.787 | -0.491 | 0.655 | 0.004 | 0.081 | 0.081 | 0.014 | 0.17 | 0.27 | 0.23 | 0.39 | 0.14 | 0.35 |
| 1 | 170908174 | rs6695615 | -0.384 | 0.626 | -0.012 | 0.369 | 0.031 | -0.014 | -0.014 | 1.281 | 0.06 | 0.25 | 0.08 | 0 | 0.06 | 0.02 |
| 1 | 170914071 | rs9426910 | -0.362 | 0.665 | -0.796 | 0.441 | -0.006 | -0.01 | -0.01 | 0 | 0.63 | 0.75 | 0.65 | 0.62 | 0.56 | 0.71 |
| 1 | 170915133 | rs4656233 | 0.003 | 0.648 | -0.715 | 0.468 | 0 | 0.002 | 0.002 | 0 | 0.57 | 0.61 | 0.61 | 0.62 | 0.5 | 0.69 |
| 1 | 170916201 | rs4656816 | -0.437 | 0.676 | -0.799 | 0.509 | -0.002 | -0.009 | -0.009 | 0.052 | 0.65 | 0.71 | 0.65 | 0.63 | 0.57 | 0.75 |
| 1 | 170916646 | rs4656817 | -0.602 | 0.61 | -0.606 | 1.154 | 0.008 | -0.014 | -0.014 | 1.996 | 0.26 | 0.14 | 0.27 | 0.32 | 0.2 | 0.37 |
| 1 | 170917459 | rs17620916 | -0.374 | 0.606 | -0.167 | 0.767 | 0.003 | -0.034 | -0.034 | 0 | 0.11 | 0.11 | 0.1 | 0 | 0.11 | 0.07 |
| 1 | 170917638 | rs1033481 | -0.117 | 0.69 | -0.398 | 1.7 | -0.017 | -0.094 | -0.094 | 0 | 0.15 | 0.03 | 0.17 | 0.43 | 0.13 | 0.41 |
| 1 | 170927615 | rs17563089 | 0.142 | 0.615 | -0.605 | 0.97 | -0.038 | 0.06 | 0.06 | 0 | 0.1 | 0.02 | 0.09 | 0 | 0.12 | 0.07 |
| 1 | 170929213 | rs6695676 | -0.586 | 1.057 | -0.69 | 1.088 | -0.015 | -0.015 | -0.015 | 0 | 0.94 | 0.97 | 0.73 | 0.85 | 0.85 | 0.9 |
| 1 | 170937288 | rs9427206 | -1.623 | 0.836 | 0.281 | 0.512 | 0 | -0.023 | -0.023 | 0.003 | 0.34 | 0.42 | 0.22 | 0.14 | 0.32 | 0.21 |
| 1 | 170939524 | rs28651462 | -0.206 | 0.698 | 0.307 | 0.567 | 0.008 | -0.009 | -0.009 | 0 | 0.16 | 0.12 | 0.07 | 0 | 0.08 | 0.06 |
| 1 | 170953552 | rs17621857 | -0.813 | 0.702 | 0.357 | 0.634 | -0.01 | -0.006 | -0.006 | 0 | 0.16 | 0.09 | 0.14 | 0.12 | 0.14 | 0.12 |
| 1 | 170954182 | rs16863898 | 1.766 | 0.94 | 0.839 | 0.636 | -0.007 | -0.005 | -0.005 | 0 | 0.41 | 0.46 | 0.28 | 0.44 | 0.39 | 0.25 |
| 1 | 170954492 | rs16863901 | -0.863 | 0.882 | 0.858 | 0.405 | 0.016 | 0.08 | 0.08 | 0 | 0.23 | 0.13 | 0.13 | 0 | 0.15 | 0.1 |
| 1 | 170958136 | rs2223610 | 1.219 | 0.82 | 1.225 | 0.644 | 0.068 | 0.075 | 0.075 | 0 | 0.87 | 0.74 | 0.65 | 0.81 | 0.78 | 0.86 |
| 1 | 170958464 | rs2050151 | 1.591 | 0.763 | 1.155 | 0.574 | 0.004 | -0.022 | -0.022 | 0 | 0.65 | 0.6 | 0.53 | 0.82 | 0.64 | 0.77 |
| 1 | 170959029 | rs16863922 | -1.705 | 0.53 | 0.751 | -0.275 | -0.015 | -0.129 | -0.129 | 0 | 0.31 | 0.4 | 0.19 | 0 | 0.22 | 0.15 |
| 1 | 170966813 | rs9427214 | -0.858 | 0.391 | -0.117 | -0.238 | 0.004 | -0.033 | -0.033 | 0 | 0.22 | 0.24 | 0.14 | 0 | 0.14 | 0.08 |
| 1 | 170974911 | rs3853181 | -1.773 | 0.36 | -0.146 | -0.262 | 0.008 | -0.039 | -0.039 | 2.288 | 0.33 | 0.66 | 0.2 | 0 | 0.21 | 0.14 |
| 1 | 170982001 | rs9426925 | -0.709 | 0.505 | -1.04 | 0.197 | -0.004 | 0.116 | 0.116 | 0.677 | 0.19 | 0.03 | 0.12 | 0 | 0.13 | 0.06 |
| 1 | 171007053 | rs1920142 | 0.873 | 0.828 | -1.418 | 1.048 | 0.005 | 0.012 | 0.012 | 0.499 | 0.58 | 0.17 | 0.49 | 0.48 | 0.63 | 0.79 |
| 1 | 171019772 | rs6423173 | 0.343 | 1.862 | -1.157 | 1.955 | -0.023 | 0.06 | 0.06 | 0 | 0.95 | 0.99 | 0.75 | 0.61 | 0.83 | 0.93 |
| 1 | 171034797 | rs16863955 | -0.614 | 1.397 | -0.772 | 0.927 | -0.014 | -0.066 | -0.066 | 0 | 0.18 | 0.27 | 0.13 | 0 | 0.11 | 0.06 |
| 1 | 171040083 | rs6608458 | 1.088 | 1.76 | -1.166 | 1.658 | -0.087 | -0.094 | -0.094 | 0.002 | 0.86 | 0.98 | 0.64 | 0.31 | 0.75 | 0.63 |
| 1 | 171045317 | rs2091657 | -1.279 | 1.982 | -0.343 | 1.663 | 0.143 | 0.097 | 0.097 | 0 | 0.05 | 0.18 | 0.09 | 0.16 | 0.15 | 0.03 |
| 1 | 171046434 | rs7878682 | -2.211 | 1.378 | 0.446 | 2.144 | 0.073 | 0.132 | 0.132 | 0 | 0.16 | 0.37 | 0.35 | 0.37 | 0.22 | 0.25 |
| 1 | 171050589 | rs7474070 | -1.769 | 1.317 | 0.549 | 2.126 | -0.035 | 0.067 | 0.067 | 0 | 0.13 | 0.31 | 0.27 | 0.21 | 0.07 | 0.23 |
| 1 | 171054550 | rs12118296 | -0.278 | 0.781 | 0.076 | 2.64 | 0.018 | -0.045 | -0.045 | 0 | 0.1 | 0.02 | 0.09 | 0.2 | 0.05 | 0.08 |
| 1 | 171055570 | rs16863990 | -0.699 | 0.741 | 0.097 | 2.615 | 0.008 | -0.019 | -0.019 | 0 | 0.06 | 0.03 | 0.22 | 0.21 | 0.04 | 0.22 |
| 1 | 171057312 | rs12402693 | -0.601 | 0.741 | 0.097 | 2.615 | 0.007 | -0.017 | -0.017 | 0 | 0.06 | 0.03 | 0.22 | 0.21 | 0.04 | 0.22 |
| 1 | 171058194 | rs1736554 | -0.252 | 0.768 | 0.095 | 2.621 | 0.015 | -0.044 | -0.044 | 0 | 0.1 | 0.02 | 0.09 | 0.2 | 0.06 | 0.08 |
| 1 | 171058236 | rs17564210 | -0.547 | 0.931 | -0.291 | 2.186 | -0.018 | 0.037 | 0.037 | 0 | 0.08 | 0.01 | 0.04 | 0.01 | 0.08 | 0.1 |
| 1 | 171058946 | rs12404183 | -1.468 | 0.892 | 0.104 | 2.198 | 0.001 | 0.001 | 0.001 | 0 | 0.1 | 0.12 | 0.25 | 0.21 | 0.06 | 0.22 |
| 1 | 171069959 | rs12076145 | -1.72 | 0.066 | 0.912 | 1.118 | 0.003 | -0.007 | -0.007 | 0 | 0.43 | 0.55 | 0.55 | 0.38 | 0.43 | 0.52 |
| 1 | 171071883 | rs10911192 | -1.806 | -0.026 | 1.09 | 1.073 | -0.01 | 0.032 | 0.032 | 0.27 | 0.46 | 0.75 | 0.56 | 0.38 | 0.43 | 0.52 |
| 1 | 171076935 | rs1800822 | -0.882 | 0.045 | 1.115 | 1.114 | 0.016 | -0.032 | -0.032 | 0.455 | 0.1 | 0.07 | 0.23 | 0.19 | 0.06 | 0.21 |
| 1 | 171076966 | rs2266782 | -1.017 | 0.025 | 1.195 | 1.102 | 0.021 | -0.042 | -0.042 | 0 | 0.34 | 0.5 | 0.32 | 0.19 | 0.37 | 0.31 |
| 1 | 171077198 | rs1920149 | -1.839 | 0.024 | 1.212 | 1.101 | -0.008 | 0.029 | 0.029 | 0.004 | 0.46 | 0.74 | 0.56 | 0.38 | 0.43 | 0.52 |
| 1 | 171077372 | rs2066534 | -0.466 | 0.011 | 1.245 | 1.089 | 0.009 | -0.009 | -0.009 | 0 | 0.28 | 0.27 | 0.21 | 0.02 | 0.19 | 0.28 |
| 1 | 171077782 | rs10797894 | -0.478 | 0.011 | 1.245 | 1.089 | 0.01 | -0.009 | -0.009 | 0 | 0.28 | 0.27 | 0.21 | 0.02 | 0.19 | 0.28 |
| 1 | 171079549 | rs2235192 | -1.113 | -0.017 | 1.445 | 1.082 | -0.034 | -0.042 | -0.042 | 0 | 0.36 | 0.52 | 0.45 | 0.22 | 0.24 | 0.48 |

| | | | | | | | | | | | | | | | | |
|---|---|---|---|---|---|---|---|---|---|---|---|---|---|---|---|---|
| 1 | 171080080 | rs1736557 | 0.718 | -0.069 | 1.394 | 1.118 | 0.01 | -0.042 | -0.042 | 0 | 0.1 | 0.03 | 0.1 | 0.21 | 0.06 | 0.12 |
| 1 | 171080485 | rs2075992 | -2.412 | -0.089 | 1.448 | 1.095 | -0.023 | 0.035 | 0.035 | 0.165 | 0.57 | 0.81 | 0.67 | 0.59 | 0.49 | 0.64 |
| 1 | 171080629 | rs2281002 | 0.039 | 0.099 | 1.548 | 1.194 | 0.006 | -0.015 | -0.015 | 0 | 0.06 | 0.03 | 0.21 | 0.19 | 0.04 | 0.21 |
| 1 | 171082896 | rs909529 | -1.052 | 0.087 | 1.714 | 1.188 | 0.001 | 0 | 0 | 0 | 0.09 | 0.07 | 0.23 | 0.19 | 0.06 | 0.21 |
| 1 | 171083174 | rs909530 | -2.294 | 0.111 | 1.837 | 1.284 | 0.12 | 0.167 | 0.167 | 0 | 0.17 | 0.46 | 0.35 | 0.35 | 0.24 | 0.24 |
| 1 | 171083714 | rs909532 | -1.047 | 0.112 | 1.834 | 1.284 | 0.001 | -0.001 | -0.001 | 0 | 0.09 | 0.07 | 0.23 | 0.18 | 0.06 | 0.21 |
| 1 | 171084242 | rs12406572 | -0.962 | 0.121 | 1.831 | 1.234 | 0.263 | 0.199 | 0.199 | 1.301 | 0.07 | 0.36 | 0.12 | 0.17 | 0.17 | 0.03 |
| 1 | 171085646 | rs1736559 | 0.66 | 0.105 | 1.778 | 1.241 | 0.009 | -0.041 | -0.041 | 0 | 0.1 | 0.02 | 0.09 | 0.2 | 0.06 | 0.12 |
| 1 | 171087660 | rs12086786 | -1.445 | 0.034 | 1.949 | 1.268 | -0.018 | 0.087 | 0.087 | 0 | 0.1 | 0.27 | 0.24 | 0.17 | 0.04 | 0.21 |
| 1 | 171088560 | rs12089338 | -1.445 | 0.034 | 1.949 | 1.268 | -0.018 | 0.087 | 0.087 | 0 | 0.1 | 0.27 | 0.24 | 0.17 | 0.04 | 0.21 |
| 1 | 171088641 | rs9970495 | -1.445 | 0.034 | 1.949 | 1.268 | -0.018 | 0.087 | 0.087 | 0 | 0.1 | 0.27 | 0.24 | 0.17 | 0.04 | 0.21 |
| 1 | 171088658 | rs9970392 | -2.018 | 0.027 | 1.955 | 1.263 | -0.025 | 0.083 | 0.083 | 0 | 0.13 | 0.37 | 0.27 | 0.17 | 0.06 | 0.22 |
| 1 | 171088854 | rs10911641 | -0.328 | -0.098 | 1.962 | 1.18 | -0.002 | -0.042 | -0.042 | 3.16 | 0.27 | 0.32 | 0.22 | 0.02 | 0.19 | 0.27 |
| 1 | 171090984 | rs742057 | -1.905 | -0.092 | 1.964 | 1.195 | -0.024 | 0.079 | 0.079 | 0 | 0.12 | 0.36 | 0.26 | 0.17 | 0.06 | 0.21 |
| 1 | 171091420 | rs742058 | -0.564 | -0.092 | 1.971 | 1.195 | -0.007 | 0.02 | 0.02 | 0 | 0.08 | 0.09 | 0.21 | 0.17 | 0.04 | 0.21 |
| 1 | 171091855 | rs10911664 | -2.017 | -0.093 | 1.97 | 1.194 | -0.027 | 0.076 | 0.076 | 0 | 0.13 | 0.37 | 0.26 | 0.17 | 0.06 | 0.21 |
| 1 | 171097561 | rs4916390 | -1.82 | -0.033 | 1.845 | 1.246 | -0.009 | 0.007 | 0.007 | 0.132 | 0.42 | 0.65 | 0.56 | 0.37 | 0.43 | 0.52 |
| 1 | 171103241 | rs2064080 | -1.977 | -0.026 | 1.911 | 1.251 | 0.039 | 0.063 | 0.063 | 0 | 0.15 | 0.25 | 0.34 | 0.34 | 0.24 | 0.24 |
| 1 | 171103607 | rs7882773 | -0.379 | -0.027 | 1.914 | 1.25 | -0.015 | -0.084 | -0.084 | 0.153 | 0.27 | 0.36 | 0.22 | 0.02 | 0.19 | 0.28 |
| 1 | 171103644 | rs7879505 | -0.405 | -0.027 | 1.879 | 1.25 | -0.024 | -0.111 | -0.111 | 0.35 | 0.27 | 0.39 | 0.22 | 0.02 | 0.19 | 0.28 |
| 1 | 171104052 | rs10911839 | -1.994 | 0.011 | 1.775 | 1.272 | 0.039 | 0.063 | 0.063 | 0 | 0.15 | 0.25 | 0.34 | 0.34 | 0.24 | 0.24 |
| 1 | 171104140 | rs10911841 | -1.994 | 0.011 | 1.775 | 1.272 | 0.039 | 0.063 | 0.063 | 0 | 0.15 | 0.25 | 0.34 | 0.34 | 0.24 | 0.24 |
| 1 | 171105390 | rs16864094 | -0.416 | 0.016 | 1.775 | 1.265 | 0.011 | -0.02 | -0.02 | 2.005 | 0.07 | 0.05 | 0.22 | 0.17 | 0.04 | 0.21 |
| 1 | 171110307 | rs1920144 | -1.982 | 0.197 | 1.895 | 1.172 | 0.042 | 0.069 | 0.069 | 0 | 0.16 | 0.25 | 0.34 | 0.34 | 0.23 | 0.24 |
| 1 | 171111365 | rs1795241 | -2.522 | 0.084 | 1.875 | 1.127 | -0.103 | 0.014 | 0.014 | 0.173 | 0.55 | 0.93 | 0.66 | 0.58 | 0.48 | 0.64 |
| 1 | 171112490 | rs1736565 | -2.341 | 0.096 | 1.957 | 1.133 | -0.014 | 0.057 | 0.057 | 0.06 | 0.54 | 0.79 | 0.65 | 0.58 | 0.48 | 0.64 |
| 1 | 171116130 | rs17565766 | -0.426 | 0.112 | 1.911 | 1.149 | 0.083 | 0.06 | 0.06 | 0 | 0.06 | 0.15 | 0.1 | 0.17 | 0.17 | 0.04 |
| 1 | 171116404 | rs1795243 | -2.346 | 0.102 | 1.945 | 1.165 | -0.013 | 0.054 | 0.054 | 0.022 | 0.54 | 0.79 | 0.65 | 0.58 | 0.48 | 0.64 |
| 1 | 171117113 | rs6608452 | -0.592 | 0.172 | 1.868 | 1.13 | 0.093 | 0.068 | 0.068 | 0 | 0.06 | 0.15 | 0.1 | 0.17 | 0.17 | 0.04 |
| 1 | 171118840 | rs2272797 | -0.196 | -0.162 | 2.026 | 1.13 | 0.116 | 0.114 | 0.114 | 0 | 0.05 | 0.17 | 0.09 | 0.17 | 0.14 | 0.03 |
| 1 | 171119021 | rs1795244 | -2.383 | -0.291 | 1.962 | 1.285 | -0.043 | 0.086 | 0.086 | 0.034 | 0.54 | 0.84 | 0.65 | 0.64 | 0.45 | 0.64 |
| 1 | 171119825 | rs10489229 | -0.238 | -0.137 | 2.062 | 1.204 | 0.134 | 0.132 | 0.132 | 0 | 0.05 | 0.19 | 0.08 | 0.17 | 0.14 | 0.03 |
| 1 | 171121173 | rs7889839 | -0.238 | -0.137 | 2.062 | 1.204 | 0.134 | 0.132 | 0.132 | 0 | 0.05 | 0.19 | 0.09 | 0.17 | 0.14 | 0.03 |
| 1 | 171121268 | rs7886938 | -0.238 | -0.137 | 2.062 | 1.204 | 0.134 | 0.132 | 0.132 | 0 | 0.05 | 0.19 | 0.09 | 0.17 | 0.14 | 0.03 |
| 1 | 171122296 | rs12754628 | -0.609 | -0.217 | 2.203 | 1.078 | -0.017 | -0.081 | -0.081 | 0.407 | 0.27 | 0.35 | 0.22 | 0.03 | 0.19 | 0.26 |
| 1 | 171125990 | rs7883894 | -0.643 | -0.197 | 2.051 | 1.088 | -0.016 | -0.076 | -0.076 | 0 | 0.27 | 0.34 | 0.22 | 0.03 | 0.19 | 0.26 |
| 1 | 171127032 | rs7881379 | -1.641 | -0.037 | 1.724 | 1.337 | 0.039 | 0.089 | 0.089 | 0 | 0.13 | 0.24 | 0.33 | 0.37 | 0.2 | 0.25 |
| 1 | 171127396 | rs7885012 | -1.051 | -0.014 | 1.668 | 1.358 | 0.043 | 0.103 | 0.103 | 0 | 0.11 | 0.2 | 0.31 | 0.37 | 0.18 | 0.25 |
| 1 | 171130346 | rs16864125 | -0.996 | -1.385 | 1.763 | 0.636 | -0.025 | -0.08 | -0.08 | 0 | 0.16 | 0.3 | 0.09 | 0.02 | 0.11 | 0.11 |
| 1 | 171131591 | rs12065048 | -1.037 | -1.362 | 1.786 | 0.649 | -0.014 | -0.049 | -0.049 | 0 | 0.14 | 0.23 | 0.07 | 0.01 | 0.1 | 0.1 |
| 1 | 171133251 | rs2421707 | -1.576 | -1.783 | 1.4 | 0.735 | -0.002 | 0.023 | 0.023 | 0 | 0.29 | 0.38 | 0.5 | 0.46 | 0.26 | 0.42 |
| 1 | 171133716 | rs11803918 | -0.924 | -1.828 | 1.483 | 0.807 | -0.001 | 0.017 | 0.017 | 0 | 0.18 | 0.18 | 0.35 | 0.4 | 0.15 | 0.33 |
| 1 | 171133889 | rs16864136 | 0.164 | -1.89 | 1.491 | 0.765 | -0.016 | -0.038 | -0.038 | 1.316 | 0.08 | 0.3 | 0.05 | 0 | 0.05 | 0.04 |
| 1 | 171135035 | rs12076944 | -0.281 | -1.838 | 1.476 | 0.75 | -0.022 | -0.061 | -0.061 | 0.137 | 0.1 | 0.3 | 0.07 | 0 | 0.07 | 0.05 |
| 1 | 171136125 | rs10912478 | -1.78 | -1.843 | 1.475 | 0.753 | -0.002 | 0.023 | 0.023 | 0 | 0.29 | 0.38 | 0.5 | 0.46 | 0.26 | 0.42 |
| 1 | 171137048 | rs12072579 | -0.285 | -1.836 | 1.471 | 0.745 | -0.022 | -0.061 | -0.061 | 0.172 | 0.1 | 0.3 | 0.07 | 0 | 0.07 | 0.05 |
| 1 | 171139823 | rs7887295 | -2.604 | -1.838 | 1.437 | 0.757 | 0.012 | -0.014 | -0.014 | 0.507 | 0.4 | 0.68 | 0.57 | 0.46 | 0.33 | 0.46 |
| 1 | 171145242 | rs11807206 | -0.796 | -1.84 | 1.457 | 0.762 | -0.001 | -0.025 | -0.025 | 0 | 0.17 | 0.13 | 0.35 | 0.4 | 0.15 | 0.33 |
| 1 | 171148521 | rs7545752 | -1.301 | -1.836 | 1.483 | 0.736 | -0.034 | -0.102 | -0.102 | 0 | 0.22 | 0.49 | 0.22 | 0.05 | 0.18 | 0.14 |
| 1 | 171151638 | rs12752688 | 0.031 | -1.825 | 1.467 | 0.745 | -0.001 | -0.016 | -0.016 | 2.681 | 0.12 | 0.2 | 0.15 | 0.05 | 0.11 | 0.09 |
| 1 | 171151991 | rs926268 | -2.856 | -2.435 | 1.052 | 0.498 | -0.054 | -0.021 | -0.021 | 0 | 0.59 | 0.94 | 0.63 | 0.6 | 0.39 | 0.62 |
| 1 | 171153448 | rs3754495 | -2.116 | -2.479 | 1.198 | 0.478 | -0.019 | -0.002 | -0.002 | 0 | 0.43 | 0.5 | 0.47 | 0.55 | 0.28 | 0.51 |
| 1 | 171154959 | rs2020870 | 0.456 | -2.483 | 1.22 | 0.48 | 0.032 | 0.003 | 0.003 | 0 | 0.15 | 0.06 | 0.04 | 0.15 | 0.06 | 0.14 |
| 1 | 171155103 | rs6657314 | -2.852 | -2.486 | 1.223 | 0.479 | -0.055 | -0.027 | -0.027 | 0 | 0.59 | 0.94 | 0.63 | 0.6 | 0.39 | 0.62 |
| 1 | 171156997 | rs11806072 | -2.657 | -2.462 | 1.243 | 0.49 | 0.019 | 0.028 | 0.028 | 0 | 0.55 | 0.69 | 0.62 | 0.61 | 0.39 | 0.58 |
| 1 | 171158284 | rs12566823 | -1.85 | -2.455 | 1.237 | 0.487 | -0.025 | 0.001 | 0.001 | 0.014 | 0.41 | 0.49 | 0.44 | 0.54 | 0.25 | 0.49 |
| 1 | 171158499 | rs12087971 | 0.368 | -2.431 | 1.262 | 0.489 | -0.016 | -0.038 | -0.038 | 0.502 | 0.08 | 0.3 | 0.05 | 0 | 0.04 | 0.04 |
| 1 | 171160698 | rs12091865 | 0.321 | -2.458 | 1.252 | 0.476 | -0.019 | -0.041 | -0.041 | 0.814 | 0.09 | 0.3 | 0.05 | 0 | 0.04 | 0.04 |
| 1 | 171162583 | rs2020860 | 0.345 | -2.458 | 1.252 | 0.476 | -0.017 | -0.039 | -0.039 | 0 | 0.09 | 0.3 | 0.05 | 0 | 0.04 | 0.04 |
| 1 | 171164111 | rs7556029 | 0.155 | -2.376 | 1.226 | 0.477 | 0.01 | 0.014 | 0.014 | 0 | 0.18 | 0.1 | 0.07 | 0.14 | 0.13 | 0.15 |
| 1 | 171164138 | rs7518693 | 0.155 | -2.428 | 1.226 | 0.477 | 0.011 | 0.014 | 0.014 | 0 | 0.18 | 0.1 | 0.07 | 0.14 | 0.13 | 0.15 |
| 1 | 171166532 | rs10912537 | -2.106 | -2.435 | 1.213 | 0.483 | -0.003 | -0.001 | -0.001 | 0.038 | 0.44 | 0.53 | 0.48 | 0.55 | 0.3 | 0.5 |
| 1 | 171168545 | rs2307492 | 0.35 | -2.435 | 1.223 | 0.477 | -0.003 | -0.012 | -0.012 | 0.696 | 0.11 | 0.19 | 0.14 | 0.06 | 0.09 | 0.08 |
| 1 | 171168584 | rs2020862 | -2.106 | -2.458 | 1.221 | 0.469 | -0.003 | -0.001 | -0.001 | 0 | 0.44 | 0.53 | 0.48 | 0.55 | 0.3 | 0.5 |
| 1 | 171168693 | rs7542361 | -0.01 | -2.47 | 1.241 | 0.466 | 0.01 | 0.012 | 0.012 | 0 | 0.19 | 0.13 | 0.08 | 0.14 | 0.13 | 0.15 |
| 1 | 171169436 | rs13376335 | 0.358 | -2.459 | 1.233 | 0.466 | -0.003 | -0.012 | -0.012 | 0 | 0.11 | 0.2 | 0.14 | 0.06 | 0.09 | 0.08 |
| 1 | 171169547 | rs6692347 | -2.116 | -2.462 | 1.236 | 0.468 | -0.003 | -0.001 | -0.001 | 0 | 0.44 | 0.53 | 0.48 | 0.55 | 0.3 | 0.5 |
| 1 | 171170209 | rs10798263 | -2.116 | -2.464 | 1.279 | 0.487 | -0.003 | -0.001 | -0.001 | 0 | 0.44 | 0.53 | 0.48 | 0.55 | 0.3 | 0.5 |
| 1 | 171170551 | rs10912552 | -2.116 | -2.464 | 1.279 | 0.487 | -0.003 | -0.001 | -0.001 | 0.086 | 0.44 | 0.53 | 0.48 | 0.55 | 0.3 | 0.5 |

| | | | | | | | | | | | | | | | | |
|---|---|---|---|---|---|---|---|---|---|---|---|---|---|---|---|---|
| 1 | 171170971 | rs12758408 | 0.344 | -2.424 | 1.238 | 0.474 | -0.003 | -0.012 | -0.012 | 0 | 0.11 | 0.19 | 0.14 | 0.06 | 0.09 | 0.08 |
| 1 | 171172147 | rs7527478 | -0.812 | -2.389 | 1.332 | 0.418 | -0.02 | -0.029 | -0.029 | 0 | 0.25 | 0.3 | 0.12 | 0.14 | 0.16 | 0.18 |
| 1 | 171173411 | rs2075988 | -0.067 | -2.424 | 1.307 | 0.52 | 0 | -0.063 | -0.063 | 0 | 0.13 | 0.04 | 0.23 | 0.24 | 0.09 | 0.26 |
| 1 | 171174152 | rs16864174 | -0.243 | -2.426 | 1.334 | 0.52 | 0.006 | -0.02 | -0.02 | 0 | 0.14 | 0.1 | 0.23 | 0.24 | 0.09 | 0.26 |
| 1 | 171174531 | rs2020863 | -0.067 | -2.426 | 1.334 | 0.52 | 0 | -0.063 | -0.063 | 0 | 0.13 | 0.04 | 0.23 | 0.24 | 0.09 | 0.26 |
| 1 | 171175586 | rs2049920 | -0.233 | -2.425 | 1.332 | 0.52 | 0.006 | -0.02 | -0.02 | 0.036 | 0.14 | 0.1 | 0.23 | 0.24 | 0.09 | 0.26 |
| 1 | 171176056 | rs2179105 | -3.404 | -2.674 | 1.07 | 0.248 | -0.1 | -0.054 | -0.054 | 0.127 | 0.64 | 0.93 | 0.65 | 0.6 | 0.41 | 0.63 |
| 1 | 171176235 | rs7527067 | 0.1 | -2.65 | 1.09 | 0.261 | 0.009 | 0.01 | 0.01 | 0 | 0.2 | 0.14 | 0.09 | 0.14 | 0.13 | 0.16 |
| 1 | 171176912 | rs2020865 | 0.53 | -2.654 | 1.141 | 0.259 | -0.012 | -0.035 | -0.035 | 0 | 0.09 | 0.33 | 0.05 | 0 | 0.05 | 0.04 |
| 1 | 171177119 | rs2075989 | 0.371 | -2.639 | 1.104 | 0.27 | 0.004 | 0.012 | 0.012 | 0 | 0.18 | 0.02 | 0.07 | 0.14 | 0.13 | 0.16 |
| 1 | 171177273 | rs2075991 | -2.434 | -2.637 | 1.102 | 0.271 | 0.003 | 0.006 | 0.006 | 0 | 0.55 | 0.58 | 0.6 | 0.6 | 0.37 | 0.59 |
| 1 | 171177525 | rs2072756 | 0.088 | -2.632 | 1.107 | 0.276 | 0.009 | 0.01 | 0.01 | 0 | 0.2 | 0.14 | 0.09 | 0.14 | 0.13 | 0.16 |
| 1 | 171178152 | rs2020869 | 0.14 | -2.59 | 1.181 | 0.312 | -0.001 | 0.002 | 0.002 | 1.154 | 0.15 | 0.18 | 0.25 | 0.22 | 0.12 | 0.15 |
| 1 | 171178490 | rs28369914 | -0.249 | -2.638 | 1.146 | 0.286 | 0.009 | -0.024 | -0.024 | 0.343 | 0.15 | 0.1 | 0.23 | 0.24 | 0.09 | 0.26 |
| 1 | 171179287 | rs7512785 | -1.062 | -2.565 | 1.103 | 0.31 | 0.007 | 0 | 0 | 0 | 0.36 | 0.33 | 0.34 | 0.36 | 0.25 | 0.3 |
| 1 | 171179477 | rs7515157 | -1.541 | -2.528 | 1.106 | 0.329 | -0.025 | -0.009 | -0.009 | 0 | 0.39 | 0.48 | 0.35 | 0.36 | 0.25 | 0.32 |
| 1 | 171187987 | rs10798271 | -2.269 | -2.466 | 0.788 | 0.333 | 0.107 | 0.025 | 0.025 | 0.273 | 0.48 | 0.74 | 0.41 | 0.38 | 0.3 | 0.48 |
| 1 | 171188057 | rs10912593 | -2.254 | -2.47 | 0.787 | 0.331 | 0.105 | 0.024 | 0.024 | 0.189 | 0.48 | 0.74 | 0.41 | 0.38 | 0.3 | 0.48 |
| 1 | 171190112 | rs10912598 | -1.309 | -2.526 | 0.883 | 0.342 | -0.024 | -0.011 | -0.011 | 0 | 0.28 | 0.62 | 0.34 | 0.23 | 0.19 | 0.2 |
| 1 | 171191583 | rs981460 | -1.209 | -2.508 | 0.749 | 0.35 | 0.034 | -0.04 | -0.04 | 0 | 0.34 | 0.15 | 0.33 | 0.38 | 0.24 | 0.42 |
| 1 | 171202164 | rs12072321 | 0.319 | -2.475 | 0.809 | 0.342 | -0.021 | -0.044 | -0.044 | 0 | 0.09 | 0.34 | 0.06 | 0 | 0.05 | 0.04 |
| 1 | 171205078 | rs7529080 | -3.137 | -2.53 | 0.712 | 0.344 | -0.112 | -0.063 | -0.063 | 1.004 | 0.63 | 0.94 | 0.66 | 0.61 | 0.41 | 0.63 |
| 1 | 171212746 | rs7545153 | -1.146 | -2.514 | 0.806 | 0.389 | 0.005 | 0 | 0 | 0 | 0.36 | 0.35 | 0.34 | 0.37 | 0.25 | 0.28 |
| 1 | 171214880 | rs7551531 | -1.273 | -2.48 | 0.47 | 0.397 | -0.008 | -0.002 | -0.002 | 0.056 | 0.37 | 0.42 | 0.34 | 0.37 | 0.26 | 0.28 |
| 1 | 171215242 | rs7551934 | 0.277 | -2.479 | 0.419 | 0.418 | 0.007 | -0.016 | -0.016 | 0 | 0.12 | 0.07 | 0.23 | 0.24 | 0.07 | 0.26 |
| 1 | 171217181 | rs2064078 | -2.969 | -2.487 | 0.409 | 0.392 | -0.079 | -0.031 | -0.031 | 0 | 0.61 | 0.91 | 0.66 | 0.61 | 0.41 | 0.61 |
| 1 | 171217445 | rs17350523 | -2.357 | -2.484 | 0.448 | 0.393 | 0.011 | 0.014 | 0.014 | 0 | 0.52 | 0.57 | 0.6 | 0.61 | 0.36 | 0.57 |
| 1 | 171217691 | rs12720462 | 0.002 | -2.464 | 0.331 | 0.407 | -0.001 | 0.012 | 0.012 | 0 | 0.13 | 0.12 | 0.24 | 0.24 | 0.08 | 0.26 |
| 1 | 171218287 | rs4916191 | -2.823 | -2.416 | 0.3 | 0.402 | -0.078 | -0.03 | -0.03 | 0 | 0.61 | 0.91 | 0.66 | 0.61 | 0.41 | 0.61 |
| 1 | 171218366 | rs4916192 | -2.894 | -2.415 | 0.3 | 0.402 | -0.079 | -0.032 | -0.032 | 0 | 0.62 | 0.91 | 0.66 | 0.61 | 0.41 | 0.62 |
| 1 | 171218925 | rs28384819 | 0.044 | -2.087 | 0.481 | 0.421 | 0.003 | 0.007 | 0.007 | 0 | 0.18 | 0.02 | 0.07 | 0.14 | 0.14 | 0.13 |
| 1 | 171219803 | rs6689616 | -0.302 | -1.855 | 1.072 | 0.45 | -0.022 | -0.046 | -0.046 | 0 | 0.09 | 0.34 | 0.05 | 0 | 0.05 | 0.04 |
| 1 | 171220236 | rs4631734 | -0.554 | -1.87 | 0.867 | 0.487 | -0.002 | 0.014 | 0.014 | 0 | 0.13 | 0.12 | 0.24 | 0.24 | 0.07 | 0.28 |
| 1 | 171220811 | rs13353034 | -0.204 | -1.794 | 0.944 | 0.375 | -0.011 | -0.043 | -0.043 | 0.617 | 0.09 | 0.34 | 0.05 | 0 | 0.05 | 0.04 |
| 1 | 171221917 | rs4073692 | -1.749 | -1.836 | 0.904 | 0.473 | -0.053 | 0.024 | 0.024 | 0 | 0.25 | 0.48 | 0.32 | 0.23 | 0.15 | 0.29 |
| 1 | 171222024 | rs4073693 | -0.858 | -1.22 | 0.645 | 0.89 | -0.002 | 0.005 | 0.005 | 0 | 0.16 | 0.13 | 0.25 | 0.23 | 0.08 | 0.29 |
| 1 | 171223015 | rs10912669 | -0.822 | -1.401 | -0.044 | 0.84 | -0.04 | 0.002 | 0.002 | 0 | 0.32 | 0.5 | 0.36 | 0.34 | 0.22 | 0.31 |
| 1 | 171223134 | rs4916273 | -1.021 | -1.018 | -0.31 | 1.047 | 0.011 | -0.018 | -0.018 | 0 | 0.13 | 0.07 | 0.23 | 0.24 | 0.07 | 0.24 |
| 1 | 171226034 | rs16864258 | -1.054 | -0.234 | 0.61 | 0.483 | 0.002 | 0 | 0 | 0 | 0.11 | 0.09 | 0.22 | 0.13 | 0.14 | 0.16 |
| 1 | 171226409 | rs6670639 | -1.081 | -0.573 | 1.231 | -0.275 | -0.022 | -0.115 | -0.115 | 0 | 0.22 | 0.54 | 0.12 | 0.04 | 0.15 | 0.23 |
| 1 | 171227487 | rs12141555 | -1.104 | -0.644 | 1.971 | -0.309 | -0.021 | -0.112 | -0.112 | 0.481 | 0.22 | 0.56 | 0.11 | 0.02 | 0.15 | 0.23 |
| 1 | 171228895 | rs10798297 | -1.149 | -0.692 | 2.262 | -0.332 | -0.017 | -0.084 | -0.084 | 0.145 | 0.22 | 0.58 | 0.12 | 0.02 | 0.15 | 0.23 |
| 1 | 171232213 | rs12040753 | -0.985 | -0.671 | 2.498 | -0.333 | -0.006 | -0.001 | -0.001 | 0.421 | 0.13 | 0.09 | 0.22 | 0.13 | 0.14 | 0.14 |
| 1 | 171232446 | rs17581251 | -0.266 | -0.735 | 2.889 | -0.383 | -0.002 | 0.001 | 0.001 | 0 | 0.11 | 0.08 | 0.06 | 0.02 | 0.12 | 0.17 |
| 1 | 171235742 | rs13376631 | -1.47 | -0.713 | 2.852 | -0.372 | -0.017 | -0.109 | -0.109 | 0.145 | 0.19 | 0.48 | 0.11 | 0.02 | 0.14 | 0.21 |
| 1 | 171236223 | rs10912694 | -3.06 | -0.676 | 2.771 | -0.264 | -0.004 | 0.051 | 0.051 | 0 | 0.35 | 0.76 | 0.34 | 0.18 | 0.28 | 0.36 |
| 1 | 171238786 | rs10912698 | -0.911 | -0.594 | 2.74 | -0.256 | -0.006 | -0.001 | -0.001 | 0 | 0.13 | 0.09 | 0.22 | 0.13 | 0.14 | 0.15 |
| 1 | 171238986 | rs12022278 | 0.07 | -0.56 | 2.732 | -0.284 | -0.024 | -0.024 | -0.024 | 0 | 0.06 | 0.02 | 0.2 | 0.13 | 0.1 | 0.12 |
| 1 | 171244759 | rs2076320 | -1.846 | -0.404 | 2.736 | -0.122 | -0.006 | -0.004 | -0.004 | 2.053 | 0.13 | 0.09 | 0.27 | 0.18 | 0.14 | 0.15 |
| 1 | 171244911 | rs16864302 | -1.064 | -0.412 | 2.894 | -0.129 | -0.005 | -0.055 | -0.055 | 0 | 0.17 | 0.25 | 0.1 | 0.02 | 0.14 | 0.21 |
| 1 | 171245579 | rs12062692 | -1.765 | -0.425 | 2.899 | -0.133 | -0.02 | -0.085 | -0.085 | 0.517 | 0.2 | 0.58 | 0.12 | 0.04 | 0.14 | 0.21 |
| 1 | 171248614 | rs7539057 | -1.804 | -0.42 | 2.867 | -0.131 | -0.019 | -0.069 | -0.069 | 0 | 0.21 | 0.66 | 0.12 | 0.04 | 0.14 | 0.21 |
| 1 | 171249094 | rs3766654 | -1.836 | -0.149 | 2.762 | -0.234 | -0.006 | -0.004 | -0.004 | 0 | 0.13 | 0.09 | 0.27 | 0.18 | 0.14 | 0.15 |
| 1 | 171250044 | rs742350 | -1.567 | -0.149 | 2.772 | -0.233 | -0.014 | -0.109 | -0.109 | 0.589 | 0.19 | 0.51 | 0.11 | 0.02 | 0.14 | 0.21 |
| 1 | 171251663 | rs10399952 | -1.806 | -0.163 | 2.797 | -0.24 | -0.019 | -0.069 | -0.069 | 3.721 | 0.21 | 0.66 | 0.12 | 0.04 | 0.14 | 0.21 |
| 1 | 171252013 | rs12049309 | 0.047 | -0.16 | 2.794 | -0.239 | -0.025 | -0.024 | -0.024 | 0 | 0.06 | 0.02 | 0.26 | 0.18 | 0.1 | 0.12 |
| 1 | 171252687 | rs12057955 | -1.447 | -0.159 | 2.796 | -0.239 | -0.008 | -0.107 | -0.107 | 1.57 | 0.18 | 0.46 | 0.11 | 0.02 | 0.14 | 0.21 |
| 1 | 171252868 | rs12354390 | -1.811 | -0.16 | 2.795 | -0.239 | -0.019 | -0.069 | -0.069 | 3.165 | 0.21 | 0.66 | 0.12 | 0.04 | 0.14 | 0.21 |
| 1 | 171253037 | rs10912714 | -1.811 | -0.16 | 2.795 | -0.239 | -0.019 | -0.069 | -0.069 | 0.062 | 0.21 | 0.66 | 0.12 | 0.04 | 0.14 | 0.21 |
| 1 | 171254209 | rs16864339 | 0.172 | -0.164 | 2.783 | -0.24 | -0.033 | -0.031 | -0.031 | 0 | 0.06 | 0 | 0.26 | 0.18 | 0.1 | 0.12 |
| 1 | 171254794 | rs12954 | -1.646 | -0.188 | 2.768 | -0.23 | -0.007 | -0.004 | -0.004 | 0 | 0.14 | 0.09 | 0.27 | 0.18 | 0.14 | 0.15 |
| 1 | 171254890 | rs7877 | -3.265 | -0.194 | 2.736 | -0.233 | -0.008 | 0.046 | 0.046 | 0 | 0.35 | 0.76 | 0.4 | 0.22 | 0.28 | 0.36 |
| 1 | 171256038 | rs12118836 | -1.531 | -0.188 | 2.683 | -0.229 | -0.006 | -0.004 | -0.004 | 1.004 | 0.13 | 0.09 | 0.27 | 0.18 | 0.14 | 0.15 |
| 1 | 171258131 | rs10912723 | -1.474 | -0.192 | 2.708 | -0.241 | -0.01 | -0.109 | -0.109 | 0 | 0.19 | 0.46 | 0.11 | 0.02 | 0.14 | 0.21 |
| 1 | 171258642 | rs10798304 | -3.243 | -0.204 | 2.688 | -0.245 | -0.009 | 0.048 | 0.048 | 0 | 0.35 | 0.77 | 0.4 | 0.21 | 0.29 | 0.36 |
| 1 | 171261193 | rs6660565 | -3.25 | -0.22 | 2.643 | -0.253 | -0.008 | 0.046 | 0.046 | 0 | 0.35 | 0.76 | 0.4 | 0.21 | 0.28 | 0.36 |
| 1 | 171270545 | rs12077755 | -1.859 | -0.214 | 2.635 | -0.25 | -0.021 | -0.073 | -0.073 | 0.347 | 0.21 | 0.64 | 0.13 | 0.04 | 0.14 | 0.21 |
| 1 | 171272500 | rs12070766 | -1.874 | -0.214 | 2.635 | -0.25 | -0.021 | -0.073 | -0.073 | 0 | 0.22 | 0.65 | 0.13 | 0.04 | 0.14 | 0.21 |
| 1 | 171273374 | rs10912745 | -1.472 | -0.212 | 2.633 | -0.249 | -0.005 | -0.004 | -0.004 | 0 | 0.13 | 0.09 | 0.27 | 0.18 | 0.14 | 0.15 |

| | | | | | | | | | | | | | | | | |
|---|---|---|---|---|---|---|---|---|---|---|---|---|---|---|---|---|
| 1 | 171275804 | rs12747080 | -2.052 | -0.432 | 2.596 | -0.313 | -0.022 | -0.073 | -0.073 | 0.199 | 0.22 | 0.64 | 0.13 | 0.04 | 0.14 | 0.21 |
| 1 | 171282853 | rs4607925 | -1.509 | -0.429 | 2.607 | -0.304 | -0.009 | -0.102 | -0.102 | 0 | 0.18 | 0.44 | 0.11 | 0.02 | 0.14 | 0.21 |
| 1 | 171289803 | rs12729885 | -1.002 | -0.425 | 2.61 | -0.306 | 0.001 | -0.054 | -0.054 | 0 | 0.16 | 0.26 | 0.1 | 0.02 | 0.14 | 0.21 |
| 1 | 171292017 | rs2076322 | -1.164 | -0.601 | 2.535 | -0.377 | -0.006 | -0.055 | -0.055 | 0 | 0.17 | 0.26 | 0.1 | 0.02 | 0.14 | 0.25 |
| 1 | 171295285 | rs2284131 | -1.848 | -0.584 | 1.996 | -0.369 | -0.023 | -0.073 | -0.073 | 0.426 | 0.23 | 0.64 | 0.13 | 0.04 | 0.14 | 0.25 |
| 1 | 171298940 | rs714839 | -1.972 | -0.423 | 0.771 | -0.178 | -0.003 | 0.006 | 0.006 | 0 | 0.46 | 0.56 | 0.46 | 0.34 | 0.42 | 0.6 |
| 1 | 171300348 | rs7541160 | -0.51 | -0.196 | 0.664 | 0.243 | -0.029 | -0.041 | -0.041 | 0 | 0.19 | 0.14 | 0.4 | 0.44 | 0.24 | 0.29 |
| 1 | 171301303 | rs12044691 | -0.51 | -0.196 | 0.665 | 0.253 | -0.031 | -0.043 | -0.043 | 0.345 | 0.19 | 0.14 | 0.4 | 0.44 | 0.24 | 0.29 |
| 1 | 171302256 | rs7534165 | -0.392 | -0.883 | 0.713 | -0.683 | -0.001 | -0.035 | -0.035 | 0 | 0.24 | 0.22 | 0.13 | 0.01 | 0.19 | 0.24 |
| 1 | 171308832 | rs2223477 | -1.499 | -1.488 | 0.731 | -0.073 | 0.003 | 0.007 | 0.007 | 0 | 0.43 | 0.66 | 0.31 | 0.41 | 0.35 | 0.46 |
| 1 | 171314142 | rs2235510 | -1.658 | -1.563 | 1.613 | -0.17 | -0.002 | -0.006 | -0.006 | 0 | 0.41 | 0.46 | 0.3 | 0.4 | 0.35 | 0.46 |
| 1 | 171322446 | rs1963273 | -1.241 | -1.562 | 1.602 | -0.169 | -0.002 | 0.001 | 0.001 | 0 | 0.38 | 0.36 | 0.29 | 0.4 | 0.34 | 0.44 |
| 1 | 171324017 | rs6670432 | -2.021 | -1.671 | 1.573 | -0.491 | -0.021 | 0.006 | 0.006 | 0 | 0.59 | 0.75 | 0.42 | 0.42 | 0.48 | 0.68 |
| 1 | 171324181 | rs6670627 | -0.754 | -1.833 | 1.631 | -0.582 | -0.064 | -0.056 | -0.056 | 0 | 0.29 | 0.1 | 0.26 | 0.4 | 0.32 | 0.4 |
| 1 | 171326508 | rs980120 | -2.06 | -1.742 | 1.561 | -0.526 | -0.021 | 0.006 | 0.006 | 0 | 0.59 | 0.75 | 0.42 | 0.42 | 0.47 | 0.68 |
| 1 | 171328083 | rs6425194 | -1.921 | -1.761 | 1.578 | -0.516 | 0.003 | 0.008 | 0.008 | 0 | 0.43 | 0.66 | 0.31 | 0.41 | 0.35 | 0.46 |
| 1 | 171340430 | rs1018391 | -2.053 | -1.74 | 1.397 | -0.517 | -0.023 | 0.005 | 0.005 | 0 | 0.59 | 0.75 | 0.42 | 0.42 | 0.47 | 0.68 |
| 1 | 171340692 | rs12407579 | -1.438 | -1.734 | 1.112 | -0.52 | -0.001 | 0.001 | 0.001 | 0 | 0.39 | 0.33 | 0.29 | 0.4 | 0.34 | 0.44 |
| 1 | 171348866 | rs7547112 | -2.041 | -1.678 | 1.095 | -0.518 | -0.063 | -0.01 | -0.01 | 2.794 | 0.59 | 0.91 | 0.43 | 0.42 | 0.47 | 0.68 |
| 1 | 171353864 | rs1018392 | -1.635 | -1.266 | 0.815 | -0.165 | -0.004 | 0.001 | 0.001 | 0 | 0.38 | 0.31 | 0.29 | 0.39 | 0.35 | 0.45 |
| 1 | 171355569 | rs12739998 | -0.822 | -1.064 | 0.387 | 0.007 | -0.048 | -0.062 | -0.062 | 0 | 0.27 | 0.1 | 0.26 | 0.4 | 0.29 | 0.38 |
| 1 | 171356755 | rs7515001 | -1.296 | -1.081 | 0.587 | 0.064 | -0.003 | -0.01 | -0.01 | 0 | 0.33 | 0.28 | 0.27 | 0.4 | 0.31 | 0.42 |
| 1 | 171357250 | rs12738952 | -0.104 | -1.092 | 0.401 | 0.062 | -0.016 | 0.02 | 0.02 | 0 | 0.12 | 0.21 | 0.07 | 0.1 | 0.06 | 0.13 |
| 1 | 171361138 | rs4916374 | -0.369 | -1.417 | -0.808 | -0.31 | 0.065 | 0.108 | 0.108 | 0 | 0.58 | 0.83 | 0.77 | 0.82 | 0.69 | 0.43 |
| 1 | 171362841 | rs4916375 | -0.377 | -1.417 | -0.808 | -0.32 | 0.065 | 0.111 | 0.111 | 0 | 0.58 | 0.83 | 0.77 | 0.82 | 0.69 | 0.43 |
| 1 | 171364141 | rs10912887 | -0.551 | -1.452 | -0.993 | -0.338 | 0.055 | 0.123 | 0.123 | 0 | 0.61 | 0.87 | 0.78 | 0.82 | 0.7 | 0.44 |
| 1 | 171371998 | rs2421809 | -0.412 | -0.312 | -0.749 | 0.035 | 0.002 | 0.001 | 0.001 | 0 | 0.38 | 0.55 | 0.37 | 0.65 | 0.39 | 0.33 |
| 1 | 171379002 | rs2207188 | 0.436 | -0.01 | -0.603 | 0.205 | -0.022 | -0.004 | -0.004 | 0 | 0.25 | 0.33 | 0.14 | 0.23 | 0.15 | 0.23 |
| 1 | 171388916 | rs2207189 | 1.003 | -0.291 | -0.958 | -0.226 | -0.022 | 0.03 | 0.03 | 0 | 0.41 | 0.19 | 0.47 | 0.28 | 0.41 | 0.5 |
| 1 | 171389475 | rs7519763 | 0.126 | -0.063 | -0.865 | -0.097 | 0 | 0 | 0 | 0 | 0.52 | 0.57 | 0.54 | 0.37 | 0.51 | 0.55 |
| 1 | 171391273 | rs7523796 | 1.182 | -0.067 | -0.875 | -0.043 | 0.062 | 0.063 | 0.063 | 0 | 0.06 | 0.2 | 0.05 | 0.07 | 0.09 | 0.07 |
| 1 | 171393060 | rs2143934 | 1.535 | -0.025 | -1.228 | -0.022 | -0.033 | -0.028 | -0.028 | 0 | 0.07 | 0.02 | 0.18 | 0.33 | 0.14 | 0.06 |
| 1 | 171393787 | rs7540116 | 0.088 | -0.078 | -1.125 | -0.298 | 0.019 | 0.012 | 0.012 | 0 | 0.54 | 0.42 | 0.67 | 0.61 | 0.57 | 0.54 |
| 1 | 171394149 | rs12122751 | 2.358 | -0.555 | -1.621 | -0.783 | -0.073 | 0.071 | 0.071 | 0 | 0.13 | 0.01 | 0.16 | 0 | 0.26 | 0.03 |
| 1 | 171394212 | rs4579793 | -0.163 | -0.602 | -1.216 | -0.429 | 0.007 | 0.006 | 0.006 | 0 | 0.6 | 0.69 | 0.73 | 0.71 | 0.67 | 0.6 |
| 1 | 171397553 | rs6682065 | -1.764 | 0.207 | -0.788 | 0.099 | 0.005 | -0.001 | -0.001 | 0 | 0.84 | 0.92 | 0.86 | 0.88 | 0.82 | 0.83 |
| 1 | 171398925 | rs10753107 | -1.764 | 0.207 | -0.788 | 0.099 | 0.005 | -0.001 | -0.001 | 0 | 0.84 | 0.92 | 0.86 | 0.88 | 0.82 | 0.83 |
| 1 | 171401212 | rs4916227 | -0.45 | 0.303 | -0.761 | 0.057 | -0.004 | -0.004 | -0.004 | 0.433 | 0.69 | 0.71 | 0.76 | 0.77 | 0.69 | 0.68 |
| 1 | 171402667 | rs12127476 | 0.642 | 1.316 | -1.143 | 0.895 | -0.105 | -0.092 | -0.092 | 0 | 0.2 | 0.02 | 0.37 | 0.25 | 0.3 | 0.13 |
| 1 | 171402862 | rs6691655 | -0.365 | 1.17 | -0.124 | 1.22 | -0.012 | -0.005 | -0.005 | 0 | 0.8 | 0.75 | 0.87 | 0.88 | 0.83 | 0.84 |
| 1 | 171404123 | rs12032308 | -1.048 | 2.171 | -0.166 | 1.474 | 0.04 | 0.064 | 0.064 | 0 | 0.14 | 0.18 | 0.23 | 0.44 | 0.23 | 0.12 |
| 1 | 171404591 | rs10912993 | -1.212 | 2.168 | -0.173 | 1.462 | 0.038 | 0.063 | 0.063 | 0 | 0.14 | 0.18 | 0.23 | 0.44 | 0.23 | 0.12 |
| 1 | 171405186 | rs12031933 | -1.201 | 2.228 | -0.291 | 1.491 | 0.074 | 0.115 | 0.115 | 0 | 0.14 | 0.31 | 0.25 | 0.44 | 0.24 | 0.12 |
| 1 | 171410532 | rs10913015 | 0.437 | 1.776 | -0.725 | 0.813 | -0.009 | 0.001 | 0.001 | 0.166 | 0.66 | 0.34 | 0.48 | 0.27 | 0.61 | 0.73 |
| 1 | 171413912 | rs7522366 | 0.538 | 1.934 | -0.631 | 0.739 | -0.014 | 0.004 | 0.004 | 0.047 | 0.65 | 0.32 | 0.46 | 0.27 | 0.6 | 0.73 |
| 1 | 171415856 | rs2207190 | 0.538 | 1.934 | -0.631 | 0.739 | -0.014 | 0.004 | 0.004 | 0.047 | 0.65 | 0.32 | 0.46 | 0.27 | 0.6 | 0.73 |
| 1 | 171417202 | rs10913039 | 0.206 | 2.132 | -0.607 | 1.088 | -0.001 | -0.001 | -0.001 | 0 | 0.7 | 0.37 | 0.65 | 0.38 | 0.68 | 0.78 |
| 1 | 171426448 | rs6670505 | -0.916 | 2.21 | -1.023 | 0.752 | -0.011 | -0.028 | -0.028 | 0.238 | 0.14 | 0.11 | 0.32 | 0.3 | 0.22 | 0.13 |
| 1 | 171433109 | rs12741782 | 0.924 | 1.775 | -0.554 | 0.448 | -0.039 | 0.128 | 0.128 | 0 | 0.61 | 0.78 | 0.37 | 0.08 | 0.46 | 0.68 |
| 1 | 171434331 | rs10489247 | 0.62 | 1.084 | -0.661 | -0.623 | 0.006 | -0.023 | -0.023 | 0 | 0.23 | 0.18 | 0.16 | 0 | 0.17 | 0.38 |
| 1 | 171435477 | rs7544906 | 1.329 | 1.039 | -0.132 | -0.631 | 0.008 | 0.003 | 0.003 | 0 | 0.37 | 0.32 | 0.19 | 0.07 | 0.29 | 0.31 |
| 1 | 171437669 | rs12145164 | 1.371 | 0.959 | 0.021 | -0.669 | 0.02 | 0.063 | 0.063 | 0 | 0.36 | 0.25 | 0.18 | 0.07 | 0.29 | 0.31 |
| 1 | 171439743 | rs10913090 | 0.233 | 1.054 | -0.029 | -0.588 | 0.008 | -0.09 | -0.09 | 0.558 | 0.6 | 0.49 | 0.35 | 0.08 | 0.46 | 0.68 |
| 1 | 171448397 | rs11576706 | -1.239 | 1.153 | -0.215 | -0.775 | 0.018 | 0.032 | 0.032 | 0 | 0.62 | 0.7 | 0.8 | 0.92 | 0.71 | 0.73 |
| 1 | 171449207 | rs4916386 | -0.761 | 0.935 | 0.116 | -1.161 | 0.029 | -0.029 | -0.029 | 0 | 0.89 | 0.66 | 0.91 | 0.97 | 0.89 | 0.9 |
| 1 | 171450816 | rs12239382 | 1.042 | 1.838 | 0.133 | -0.609 | 0.02 | 0.029 | 0.029 | 0 | 0.07 | 0.11 | 0.14 | 0.16 | 0.12 | 0.06 |
| 1 | 171458250 | rs10913129 | 1.379 | 1.821 | -0.173 | -0.812 | -0.038 | 0.126 | 0.126 | 0 | 0.19 | 0.02 | 0.15 | 0 | 0.18 | 0.39 |
| 1 | 171459507 | rs12759907 | 2.319 | 1.735 | -0.234 | -0.846 | 0.018 | 0.14 | 0.14 | 0 | 0.27 | 0.02 | 0.11 | 0.06 | 0.18 | 0.17 |
| 1 | 171463300 | rs4916388 | 2.061 | 2.105 | -0.305 | -0.406 | -0.045 | -0.105 | -0.105 | 0 | 0.16 | 0.01 | 0.18 | 0.43 | 0.19 | 0.07 |
| 1 | 171464091 | rs7536723 | -0.38 | 2.02 | -0.118 | -0.464 | -0.001 | -0.001 | -0.001 | 0.46 | 0.47 | 0.5 | 0.5 | 0.6 | 0.5 | 0.56 |
| 1 | 171472188 | rs10913149 | 1.09 | 2.069 | -0.081 | -0.174 | -0.005 | 0.003 | 0.003 | 0 | 0.1 | 0.06 | 0.11 | 0.24 | 0.13 | 0.12 |
| 1 | 171478523 | rs4384265 | -0.373 | 1.947 | -0.046 | -0.286 | 0.027 | -0.006 | -0.006 | 0 | 0.87 | 0.74 | 0.73 | 0.9 | 0.82 | 0.85 |
| 1 | 171481592 | rs2038042 | -0.353 | 1.947 | -0.05 | -0.286 | 0.024 | -0.006 | -0.006 | 0 | 0.86 | 0.75 | 0.73 | 0.9 | 0.82 | 0.85 |
| 1 | 171483717 | rs1055417 | 2.014 | 1.743 | -0.236 | -0.375 | 0.036 | 0.074 | 0.074 | 0 | 0.28 | 0.11 | 0.12 | 0.06 | 0.19 | 0.17 |
| 1 | 171486912 | rs10913157 | 1.091 | 1.901 | -0.215 | -0.214 | -0.005 | 0.003 | 0.003 | 0 | 0.1 | 0.06 | 0.11 | 0.24 | 0.13 | 0.12 |
| 1 | 171499108 | rs12142594 | 2.088 | 1.886 | -0.182 | -0.247 | 0.032 | 0.058 | 0.058 | 0 | 0.28 | 0.13 | 0.12 | 0.06 | 0.18 | 0.17 |
| 1 | 171500154 | rs7534557 | 2.101 | 1.886 | -0.182 | -0.247 | 0.032 | 0.057 | 0.057 | 0 | 0.28 | 0.12 | 0.12 | 0.06 | 0.18 | 0.17 |
| 1 | 171509014 | rs12734261 | 2.088 | 1.86 | -0.202 | -0.259 | 0.032 | 0.058 | 0.058 | 0 | 0.28 | 0.13 | 0.12 | 0.06 | 0.18 | 0.17 |
| 1 | 171509327 | rs760644 | -0.329 | 1.985 | -0.05 | -0.296 | 0.026 | -0.008 | -0.008 | 1.598 | 0.86 | 0.72 | 0.72 | 0.9 | 0.82 | 0.85 |

| | | | | | | | | | | | | | | | | |
|---|---|---|---|---|---|---|---|---|---|---|---|---|---|---|---|---|
| 1 | 171511442 | rs2272810 | 0.014 | 1.886 | 0.05 | -0.338 | -0.067 | -0.059 | -0.059 | 0.175 | 0.43 | 0.17 | 0.47 | 0.6 | 0.49 | 0.53 |
| 1 | 171519487 | rs235470 | -0.304 | 1.899 | 0.076 | -0.294 | 0.026 | -0.008 | -0.008 | 0 | 0.86 | 0.72 | 0.73 | 0.9 | 0.82 | 0.85 |
| 1 | 171526570 | rs1687057 | -0.482 | 1.898 | 0.083 | -0.295 | 0.025 | -0.006 | -0.006 | 0 | 0.87 | 0.75 | 0.73 | 0.9 | 0.82 | 0.85 |
| 1 | 171526910 | rs12025905 | 1.204 | 2.076 | 0.021 | -0.262 | 0.021 | 0.03 | 0.03 | 0 | 0.07 | 0.11 | 0.14 | 0.16 | 0.12 | 0.06 |
| 1 | 171528416 | rs235485 | -0.444 | 2.357 | 0.096 | -0.341 | 0.051 | -0.005 | -0.005 | 0 | 0.86 | 0.76 | 0.7 | 0.9 | 0.74 | 0.84 |
| 1 | 171531159 | rs235487 | 0.206 | 2.114 | -0.144 | -0.606 | 0.018 | -0.025 | -0.025 | 0 | 0.78 | 0.69 | 0.63 | 0.9 | 0.69 | 0.75 |
| 1 | 171543476 | rs12079162 | 0.785 | 1.934 | -0.181 | -0.637 | -0.032 | -0.07 | -0.07 | 0.172 | 0.24 | 0.16 | 0.33 | 0.59 | 0.31 | 0.14 |
| 1 | 171543863 | rs235509 | -0.15 | 1.721 | -0.067 | -0.709 | 0.02 | -0.012 | -0.012 | 0.048 | 0.85 | 0.7 | 0.73 | 0.9 | 0.82 | 0.82 |
| 1 | 171545051 | rs764206 | -0.75 | 1.71 | -0.08 | -0.718 | -0.004 | -0.024 | -0.024 | 0 | 0.56 | 0.56 | 0.61 | 0.83 | 0.63 | 0.65 |
| 1 | 171545469 | rs235511 | -1.691 | 1.865 | -0.217 | -0.663 | -0.032 | 0.007 | 0.007 | 0 | 0.93 | 0.99 | 0.93 | 0.97 | 0.9 | 0.9 |
| 1 | 171561225 | rs235907 | -0.14 | 1.462 | -0.145 | -0.612 | 0.019 | -0.015 | -0.015 | 1.1 | 0.85 | 0.68 | 0.72 | 0.9 | 0.82 | 0.82 |
| 1 | 171576573 | rs171005 | 1.323 | 0.667 | -0.096 | -0.631 | -0.046 | -0.114 | -0.114 | 0.268 | 0.32 | 0.44 | 0.14 | 0.06 | 0.19 | 0.17 |
| 1 | 171588583 | rs235864 | 0.3 | 0.911 | 0.191 | -0.129 | 0.013 | 0.029 | 0.029 | 0 | 0.72 | 0.65 | 0.54 | 0.59 | 0.63 | 0.67 |
| 1 | 171591189 | rs929115 | 0.605 | 0.783 | 0.035 | -0.182 | 0.003 | -0.006 | -0.006 | 0.297 | 0.25 | 0.13 | 0.18 | 0.32 | 0.22 | 0.09 |
| 1 | 171591818 | rs235852 | 1.868 | 0.827 | 0.297 | -0.231 | 0.011 | 0.003 | 0.003 | 0 | 0.3 | 0.26 | 0.11 | 0.1 | 0.16 | 0.19 |
| 1 | 171592553 | rs235853 | 1.235 | 0.653 | 0.207 | -0.167 | -0.068 | -0.067 | -0.067 | 0 | 0.19 | 0.03 | 0.18 | 0.32 | 0.24 | 0.1 |
| 1 | 171592869 | rs235854 | -0.626 | 0.492 | 0.488 | -0.17 | -0.01 | -0.002 | -0.002 | 0 | 0.43 | 0.38 | 0.44 | 0.48 | 0.48 | 0.48 |
| 1 | 171597996 | rs6425356 | -0.519 | 0.461 | 0.326 | -0.214 | -0.021 | -0.026 | -0.026 | 0 | 0.28 | 0.49 | 0.27 | 0.23 | 0.24 | 0.44 |
| 1 | 171599843 | rs235863 | -0.926 | 0.478 | 0.285 | -0.196 | -0.003 | -0.002 | -0.002 | 0 | 0.48 | 0.59 | 0.46 | 0.55 | 0.49 | 0.54 |
| 1 | 171599953 | rs2421853 | -0.558 | 0.441 | 0.331 | -0.19 | -0.019 | -0.023 | -0.023 | 0.315 | 0.29 | 0.53 | 0.27 | 0.23 | 0.24 | 0.44 |
| 1 | 171602202 | rs7547017 | 0.114 | 0.578 | -0.179 | -0.114 | -0.026 | -0.052 | -0.052 | 0 | 0.6 | 0.78 | 0.44 | 0.46 | 0.46 | 0.66 |
| 1 | 171603547 | rs235881 | -0.71 | 0.634 | -0.039 | -0.128 | 0.043 | 0.015 | 0.015 | 0 | 0.83 | 0.69 | 0.64 | 0.78 | 0.71 | 0.8 |
| 1 | 171606054 | rs12076134 | 0.651 | 0.601 | -0.494 | -0.161 | -0.004 | -0.004 | -0.004 | 0.941 | 0.18 | 0.16 | 0.21 | 0.22 | 0.21 | 0.41 |
| 1 | 171606739 | rs7545646 | 2.284 | 0.557 | -1.014 | -0.227 | 0.078 | 0.086 | 0.086 | 0 | 0.29 | 0.13 | 0.09 | 0.02 | 0.14 | 0.15 |
| 1 | 171607702 | rs2032555 | -1.139 | 0.639 | -1.006 | -0.182 | -0.022 | -0.013 | -0.013 | 0 | 0.85 | 0.89 | 0.65 | 0.79 | 0.71 | 0.8 |
| 1 | 171608466 | rs604864 | -1.032 | 0.676 | -0.884 | -0.148 | 0.006 | 0.023 | 0.023 | 0 | 0.55 | 0.62 | 0.56 | 0.77 | 0.58 | 0.65 |
| 1 | 171613756 | rs235875 | 1.414 | 1.12 | -0.913 | 0.178 | -0.077 | -0.088 | -0.088 | 3.106 | 0.17 | 0.01 | 0.16 | 0.31 | 0.21 | 0.09 |
| 1 | 171614683 | rs2236875 | 0.369 | 1.102 | -0.473 | 0.128 | 0.032 | 0.141 | 0.141 | 0 | 0.08 | 0.22 | 0.12 | 0.22 | 0.11 | 0.1 |
| 1 | 171615206 | rs12035960 | 1.033 | 1.15 | -0.815 | 0.152 | -0.005 | 0.028 | 0.028 | 0 | 0.07 | 0.09 | 0.11 | 0.22 | 0.1 | 0.1 |
| 1 | 171618656 | rs235913 | -0.209 | 1.155 | -0.854 | 0.084 | -0.021 | -0.021 | -0.021 | 0 | 0.79 | 0.79 | 0.56 | 0.63 | 0.62 | 0.71 |
| 1 | 171620579 | rs235917 | -0.896 | 1.068 | -0.614 | 0.039 | 0.024 | -0.011 | -0.011 | 0 | 0.84 | 0.82 | 0.67 | 0.88 | 0.72 | 0.83 |
| 1 | 171621834 | rs2075648 | 0.65 | 0.615 | -0.856 | -0.005 | -0.005 | 0.061 | 0.061 | 0 | 0.16 | 0.01 | 0.1 | 0.08 | 0.14 | 0.39 |
| 1 | 171621975 | rs235920 | 0.085 | 0.774 | -0.577 | 0.002 | 0.015 | 0.059 | 0.059 | 1.089 | 0.27 | 0.56 | 0.3 | 0.46 | 0.3 | 0.14 |
| 1 | 171623833 | rs2075537 | 0.471 | -0.576 | -0.874 | -0.187 | 0.075 | 0.021 | 0.021 | 0.044 | 0.5 | 0.32 | 0.27 | 0.41 | 0.34 | 0.65 |
| 1 | 171624145 | rs235922 | -0.006 | -0.217 | -0.55 | -0.064 | 0.039 | 0.021 | 0.021 | 0 | 0.73 | 0.65 | 0.47 | 0.63 | 0.57 | 0.61 |
| 1 | 171626941 | rs10157798 | 0.616 | -1.481 | -0.22 | -0.641 | -0.102 | -0.049 | -0.049 | 0.838 | 0.23 | 0.53 | 0.1 | 0.12 | 0.07 | 0.15 |
| 1 | 171633958 | rs12142786 | 0.7 | -1.381 | -0.697 | -0.709 | -0.021 | 0.085 | 0.085 | 0 | 0.15 | 0.01 | 0.11 | 0.01 | 0.14 | 0.38 |
| 1 | 171640888 | rs11799938 | 0.668 | -1.154 | 0.306 | -0.153 | -0.031 | -0.018 | -0.018 | 0 | 0.21 | 0.31 | 0.1 | 0.12 | 0.09 | 0.14 |
| 1 | 171645589 | rs10489248 | -0.282 | -1.167 | -0.179 | -0.195 | -0.04 | -0.034 | -0.034 | 0.114 | 0.3 | 0.56 | 0.15 | 0.12 | 0.12 | 0.17 |
| 1 | 171647475 | rs11800358 | 0.661 | -0.829 | -0.605 | 0.019 | 0.018 | 0.013 | 0.013 | 0 | 0.21 | 0.16 | 0.09 | 0.12 | 0.09 | 0.15 |
| 1 | 171648923 | rs2636177 | -0.222 | -0.243 | -0.537 | 0.259 | -0.003 | 0.003 | 0.003 | 0 | 0.57 | 0.56 | 0.41 | 0.64 | 0.48 | 0.35 |
| 1 | 171652583 | rs10913482 | 0.017 | -0.218 | -0.262 | 0.784 | -0.001 | 0.059 | 0.059 | 0 | 0.12 | 0.18 | 0.12 | 0.25 | 0.11 | 0.06 |
| 1 | 171652664 | rs9661596 | 1.136 | -0.513 | 0.414 | 0.519 | -0.012 | -0.02 | -0.02 | 0 | 0.25 | 0.26 | 0.14 | 0.12 | 0.18 | 0.16 |
| 1 | 171652978 | rs11799420 | 1.136 | -0.513 | 0.414 | 0.519 | -0.012 | -0.02 | -0.02 | 0 | 0.25 | 0.26 | 0.14 | 0.12 | 0.18 | 0.16 |
| 1 | 171654616 | rs6662720 | 1.136 | -0.513 | 0.414 | 0.519 | -0.012 | -0.02 | -0.02 | 0 | 0.25 | 0.26 | 0.14 | 0.12 | 0.18 | 0.15 |
| 1 | 171654968 | rs9943293 | -0.117 | -0.556 | 0.463 | 0.476 | -0.009 | 0.002 | 0.002 | 0 | 0.38 | 0.52 | 0.27 | 0.37 | 0.29 | 0.23 |
| 1 | 171656124 | rs9659578 | 0.01 | -0.552 | 0.379 | 0.49 | -0.023 | 0.011 | 0.011 | 0 | 0.37 | 0.44 | 0.26 | 0.38 | 0.29 | 0.21 |
| 1 | 171669823 | rs236894 | 1.045 | -0.64 | 0.078 | 0.553 | -0.036 | 0.018 | 0.018 | 0 | 0.18 | 0.02 | 0.11 | 0.16 | 0.2 | 0.09 |
| 1 | 171674774 | rs6685742 | 0.024 | -0.786 | 0.352 | 0.712 | -0.024 | 0.016 | 0.016 | 0 | 0.38 | 0.46 | 0.27 | 0.4 | 0.29 | 0.23 |
| 1 | 171684506 | rs1001373 | 1.17 | -0.83 | 0.512 | 0.634 | 0.018 | 0.018 | 0.018 | 0 | 0.25 | 0.13 | 0.13 | 0.15 | 0.18 | 0.15 |
| 1 | 171710405 | rs2298914 | 1.282 | -0.938 | 0.567 | 0.636 | 0.02 | 0.017 | 0.017 | 0 | 0.24 | 0.09 | 0.13 | 0.15 | 0.18 | 0.15 |
| 1 | 171712150 | rs10913576 | 0.262 | -0.978 | 0.776 | 0.559 | -0.019 | -0.048 | -0.048 | 0 | 0.09 | 0.3 | 0.06 | 0 | 0.03 | 0.02 |
| 1 | 171712868 | rs7556644 | -1.926 | -0.573 | 0.105 | 0.05 | -0.06 | 0.048 | 0.048 | 0 | 0.83 | 0.93 | 0.67 | 0.93 | 0.7 | 0.84 |
| 1 | 171713808 | rs17649050 | 1.611 | -0.699 | 0.212 | 0.214 | 0.006 | 0.005 | 0.005 | 0 | 0.25 | 0.18 | 0.13 | 0.16 | 0.18 | 0.16 |
| 1 | 171715585 | rs10913582 | 1.604 | -0.696 | 0.214 | 0.215 | 0.006 | 0.005 | 0.005 | 0 | 0.25 | 0.18 | 0.13 | 0.16 | 0.18 | 0.16 |
| 1 | 171721146 | rs6665581 | 0.146 | -1.158 | 0.008 | 0.187 | -0.008 | 0 | 0 | 0 | 0.42 | 0.52 | 0.27 | 0.39 | 0.29 | 0.25 |
| 1 | 171722990 | rs10157510 | 1.005 | -1.086 | 0.047 | 0.039 | -0.04 | -0.068 | -0.068 | 0 | 0.12 | 0.29 | 0.06 | 0 | 0.03 | 0.02 |
| 1 | 171731932 | rs7545663 | -0.721 | -1.286 | 0.136 | 0.14 | 0.011 | -0.022 | -0.022 | 0.122 | 0.65 | 0.55 | 0.5 | 0.81 | 0.53 | 0.4 |
| 1 | 171732363 | rs6675708 | 1.552 | -1.279 | -0.201 | 0.214 | -0.022 | 0.013 | 0.013 | 0 | 0.16 | 0.06 | 0.12 | 0.18 | 0.2 | 0.09 |
| 1 | 171732378 | rs10913625 | 0.859 | -1.3 | -0.17 | 0.246 | 0.021 | 0.014 | 0.014 | 0 | 0.37 | 0.22 | 0.32 | 0.63 | 0.29 | 0.3 |
| 1 | 171735198 | rs12032340 | -0.127 | -0.998 | 0.174 | 0.246 | 0.006 | 0.068 | 0.068 | 0 | 0.19 | 0.2 | 0.24 | 0.48 | 0.22 | 0.16 |
| 1 | 171735666 | rs7542116 | -0.093 | -0.998 | 0.175 | 0.246 | 0.006 | 0.064 | 0.064 | 0 | 0.19 | 0.17 | 0.24 | 0.48 | 0.22 | 0.16 |
| 1 | 171737431 | rs12754084 | -1.259 | -0.949 | 0.479 | 0.226 | 0 | 0 | 0 | 0 | 0.47 | 0.49 | 0.42 | 0.67 | 0.45 | 0.3 |
| 1 | 171738453 | rs17649609 | 1.42 | -0.881 | 0.097 | 0.233 | -0.06 | -0.025 | -0.025 | 1.123 | 0.15 | 0.01 | 0.11 | 0.19 | 0.2 | 0.09 |
| 1 | 171740946 | rs6656814 | -1.344 | -1.077 | 0.688 | 0.15 | 0.007 | 0.047 | 0.047 | 0 | 0.48 | 0.6 | 0.42 | 0.67 | 0.44 | 0.3 |
| 1 | 171743575 | rs11803889 | -0.225 | -1.137 | 1.01 | 0.268 | -0.027 | 0.003 | 0.003 | 0 | 0.25 | 0.29 | 0.21 | 0.24 | 0.17 | 0.13 |
| 1 | 171743865 | rs10798636 | -0.626 | -1.129 | 1.031 | 0.27 | -0.014 | 0.034 | 0.034 | 0 | 0.31 | 0.51 | 0.31 | 0.49 | 0.25 | 0.2 |
| 1 | 171744655 | rs2181216 | 1.102 | -1.11 | 0.969 | 0.271 | -0.013 | 0.003 | 0.003 | 0 | 0.17 | 0.09 | 0.12 | 0.18 | 0.2 | 0.09 |
| 1 | 171749665 | rs1555101 | -1.193 | -1.084 | 1.179 | -0.332 | 0.004 | 0.029 | 0.029 | 0 | 0.47 | 0.57 | 0.41 | 0.66 | 0.44 | 0.3 |

| | | | | | | | | | | | | | | | | |
|---|---|---|---|---|---|---|---|---|---|---|---|---|---|---|---|---|
| 1 | 171750181 | rs2232809 | -1.934 | -1.043 | 1.012 | -0.535 | -0.015 | 0.011 | 0.011 | 0 | 0.62 | 0.76 | 0.54 | 0.68 | 0.61 | 0.68 |
| 1 | 171750471 | rs2232811 | -1.194 | -1.045 | 1.03 | -0.525 | 0.007 | 0.038 | 0.038 | 0 | 0.48 | 0.6 | 0.42 | 0.66 | 0.44 | 0.3 |
| 1 | 171750557 | rs2232812 | -0.2 | -1.075 | 1.099 | -0.479 | -0.01 | 0.042 | 0.042 | 0 | 0.28 | 0.29 | 0.29 | 0.52 | 0.24 | 0.19 |
| 1 | 171751051 | rs2294719 | -1.087 | -1.075 | 1.108 | -0.479 | 0.003 | 0.027 | 0.027 | 0 | 0.47 | 0.57 | 0.41 | 0.66 | 0.44 | 0.3 |
| 1 | 171751236 | rs2294720 | -1.433 | -1.078 | 1.08 | -0.48 | 0.011 | 0.08 | 0.08 | 0 | 0.5 | 0.73 | 0.44 | 0.66 | 0.44 | 0.3 |
| 1 | 171753039 | rs2232816 | -0.205 | -1.06 | 1.034 | -0.472 | -0.009 | 0.041 | 0.041 | 0 | 0.28 | 0.28 | 0.29 | 0.52 | 0.24 | 0.19 |
| 1 | 171755170 | rs2232818 | -0.154 | -0.841 | 0.895 | -0.563 | -0.011 | 0.046 | 0.046 | 0 | 0.29 | 0.32 | 0.29 | 0.52 | 0.24 | 0.19 |
| 1 | 171755182 | rs2232819 | 1.026 | -0.845 | 0.807 | -0.556 | -0.035 | 0.018 | 0.018 | 0 | 0.16 | 0.02 | 0.11 | 0.13 | 0.2 | 0.09 |
| 1 | 171763549 | rs2232825 | 0.872 | -0.773 | 0.802 | -0.561 | -0.038 | 0.022 | 0.022 | 0 | 0.16 | 0.02 | 0.16 | 0.13 | 0.22 | 0.09 |
| 1 | 171764771 | rs909959 | -1.57 | -0.817 | 0.846 | -0.571 | 0.015 | 0.071 | 0.071 | 0 | 0.49 | 0.69 | 0.43 | 0.66 | 0.44 | 0.3 |
| 1 | 171765964 | rs1801898 | 0.279 | -0.793 | 0.854 | -0.56 | 0.005 | -0.018 | -0.018 | 0 | 0.19 | 0.26 | 0.12 | 0.13 | 0.2 | 0.11 |
| 1 | 171766580 | rs12140193 | 1.759 | -0.759 | 0.696 | -0.535 | -0.032 | 0.055 | 0.055 | 0.137 | 0.12 | 0.02 | 0.1 | 0.02 | 0.16 | 0.38 |
| 1 | 171766696 | rs7860 | -0.202 | -0.747 | 0.824 | -0.443 | -0.01 | 0.034 | 0.034 | 0 | 0.29 | 0.27 | 0.28 | 0.52 | 0.24 | 0.19 |
| 1 | 171781952 | rs6700184 | -1.131 | -0.839 | 0.759 | -0.476 | 0 | 0.002 | 0.002 | 0 | 0.45 | 0.32 | 0.4 | 0.67 | 0.43 | 0.28 |
| 1 | 171786475 | rs6689108 | -1.156 | -0.751 | 0.475 | -0.431 | 0.01 | 0.029 | 0.029 | 0 | 0.47 | 0.56 | 0.4 | 0.67 | 0.42 | 0.3 |
| 1 | 171786582 | rs6689213 | -1.221 | -0.753 | 0.466 | -0.432 | 0.016 | 0.042 | 0.042 | 0 | 0.47 | 0.65 | 0.41 | 0.67 | 0.42 | 0.3 |
| 1 | 171788782 | rs2206542 | -1.169 | -0.746 | 0.473 | -0.434 | 0.009 | 0.023 | 0.023 | 0 | 0.47 | 0.54 | 0.4 | 0.67 | 0.42 | 0.3 |
| 1 | 171791211 | rs6425438 | 0.039 | -0.82 | 0.184 | -0.242 | 0.004 | -0.006 | -0.006 | 0.039 | 0.26 | 0.18 | 0.25 | 0.51 | 0.15 | 0.18 |
| 1 | 171797731 | rs10753195 | -1.48 | -0.427 | -0.495 | -0.164 | -0.023 | 0.012 | 0.012 | 0 | 0.61 | 0.75 | 0.53 | 0.69 | 0.58 | 0.67 |
| 1 | 171808405 | rs2294723 | -0.438 | -0.422 | 0.247 | 0.279 | -0.008 | 0.063 | 0.063 | 0 | 0.23 | 0.32 | 0.26 | 0.48 | 0.21 | 0.15 |
| 1 | 171809976 | rs6701033 | -1.652 | 0.005 | -0.16 | -0.094 | -0.002 | 0.067 | 0.067 | 0.002 | 0.56 | 0.68 | 0.59 | 0.78 | 0.56 | 0.53 |
| 1 | 171815855 | rs4916401 | -2.484 | 0.153 | -0.26 | -0.231 | -0.015 | 0.047 | 0.047 | 0 | 0.69 | 0.88 | 0.64 | 0.83 | 0.65 | 0.79 |
| 1 | 171826306 | rs1040517 | -2.502 | 0.162 | -0.221 | -0.231 | -0.042 | 0.06 | 0.06 | 0 | 0.69 | 0.93 | 0.64 | 0.83 | 0.65 | 0.79 |
| 1 | 171829164 | rs10798722 | -2.462 | 0.157 | -0.054 | -0.234 | -0.008 | 0.034 | 0.034 | 0 | 0.69 | 0.84 | 0.64 | 0.83 | 0.65 | 0.79 |
| 1 | 171833688 | rs12136630 | 0.803 | 0.084 | 0.226 | -0.315 | -0.005 | -0.039 | -0.039 | 0 | 0.15 | 0.23 | 0.06 | 0.03 | 0.1 | 0.26 |
| 1 | 171835654 | rs2901600 | -1.497 | 0.077 | 0.24 | -0.277 | 0 | 0.024 | 0.024 | 0 | 0.54 | 0.56 | 0.57 | 0.8 | 0.54 | 0.53 |
| 1 | 171842607 | rs2206544 | -0.724 | -0.05 | 0.424 | -0.254 | 0.011 | -0.021 | -0.021 | 0 | 0.38 | 0.44 | 0.42 | 0.31 | 0.41 | 0.37 |
| 1 | 171847295 | rs12025368 | 0.393 | -0.031 | 0.453 | -0.199 | 0.004 | -0.003 | -0.003 | 0.181 | 0.21 | 0.19 | 0.16 | 0.5 | 0.14 | 0.16 |
| 1 | 171863472 | rs12121064 | 1.182 | -0.027 | 0.226 | -0.246 | 0.001 | 0.003 | 0.003 | 0 | 0.14 | 0.16 | 0.06 | 0.03 | 0.11 | 0.26 |
| 1 | 171869242 | rs10436867 | 0.268 | 0.04 | 0.31 | -0.172 | -0.034 | 0.081 | 0.081 | 0 | 0.22 | 0.3 | 0.17 | 0.5 | 0.14 | 0.16 |
| 1 | 171870596 | rs4916239 | 0.88 | 0.054 | 0.3 | -0.187 | 0.003 | 0.018 | 0.018 | 0 | 0.53 | 0.41 | 0.53 | 0.67 | 0.49 | 0.37 |
| 1 | 171871134 | rs7540873 | -0.644 | 0.136 | 0.028 | -0.214 | -0.054 | -0.004 | -0.004 | 0.274 | 0.84 | 0.91 | 0.66 | 0.86 | 0.69 | 0.72 |
| 1 | 171875628 | rs1535301 | 2.518 | 0.155 | -0.005 | -0.213 | -0.031 | 0.063 | 0.063 | 0 | 0.31 | 0.12 | 0.36 | 0.17 | 0.35 | 0.21 |
| 1 | 171881168 | rs6425479 | 0.272 | 0.395 | 0.092 | -0.15 | 0.036 | 0.017 | 0.017 | 0 | 0.75 | 0.63 | 0.65 | 0.69 | 0.67 | 0.76 |
| 1 | 171885747 | rs2901603 | -0.551 | -0.048 | 0.382 | 0.039 | 0.024 | -0.011 | -0.011 | 0 | 0.42 | 0.37 | 0.25 | 0.51 | 0.3 | 0.44 |
| 1 | 171897198 | rs6673646 | 1.582 | -0.02 | -0.052 | -0.217 | -0.018 | 0.093 | 0.093 | 0 | 0.55 | 0.22 | 0.47 | 0.22 | 0.55 | 0.49 |
| 1 | 171901805 | rs6673009 | 1.134 | 0.111 | -0.253 | -0.189 | -0.036 | 0.136 | 0.136 | 0 | 0.23 | 0.02 | 0.1 | 0.02 | 0.21 | 0.28 |
| 1 | 171903833 | rs12084926 | 0.615 | 0.158 | 0.236 | 0.484 | 0.029 | 0.033 | 0.033 | 0 | 0.76 | 0.7 | 0.6 | 0.56 | 0.64 | 0.66 |
| 1 | 171919618 | rs2208369 | 0.611 | 0.165 | 0.006 | 0.43 | 0.026 | 0.036 | 0.036 | 0 | 0.75 | 0.68 | 0.6 | 0.54 | 0.64 | 0.66 |
| 1 | 171927696 | rs12028426 | -0.072 | -0.528 | 0.06 | 0.522 | -0.074 | 0.091 | 0.091 | 0 | 0.19 | 0.42 | 0.12 | 0.34 | 0.08 | 0.16 |
| 1 | 171947018 | rs2180748 | 0.569 | 0.319 | -0.367 | 0.248 | 0.002 | 0.04 | 0.04 | 1.287 | 0.59 | 0.24 | 0.68 | 0.72 | 0.63 | 0.53 |
| 1 | 171955331 | rs2208370 | -0.842 | -0.171 | 0.092 | 0.189 | 0 | 0 | 0 | 0.037 | 0.37 | 0.62 | 0.33 | 0.22 | 0.35 | 0.42 |
| 1 | 171956803 | rs3736790 | -1.493 | 0.019 | 0.317 | 0.431 | 0.005 | 0.001 | 0.001 | 3.365 | 0.17 | 0.74 | 0.22 | 0.2 | 0.16 | 0.2 |
| 1 | 171960993 | rs10910863 | -1.405 | 0.04 | 0.482 | 0.453 | 0.008 | 0.003 | 0.003 | 2.646 | 0.16 | 0.7 | 0.22 | 0.2 | 0.16 | 0.2 |
| 1 | 171961138 | rs12077603 | -1.527 | 0.011 | 0.458 | 0.438 | 0.004 | 0 | 0 | 3.116 | 0.17 | 0.72 | 0.22 | 0.2 | 0.17 | 0.2 |
| 1 | 171972317 | rs12406353 | -0.136 | 0.052 | 0.85 | 0.429 | 0.082 | 0.015 | 0.015 | 0 | 0.06 | 0.23 | 0.11 | 0.05 | 0.09 | 0.14 |
| 1 | 171977284 | rs1073940 | -1.792 | 0.054 | 1.24 | 1.148 | 0.061 | 0.226 | 0.226 | 1.754 | 0.17 | 0.53 | 0.22 | 0.45 | 0.19 | 0.23 |
| 1 | 172004114 | rs9286842 | -1.745 | -0.146 | 0.984 | 0.747 | 0.031 | 0.038 | 0.038 | 0 | 0.28 | 0.48 | 0.42 | 0.62 | 0.32 | 0.37 |
| 1 | 172004280 | rs10910945 | -0.766 | -0.105 | 0.954 | 0.858 | 0.021 | 0.084 | 0.084 | 0.201 | 0.2 | 0.3 | 0.33 | 0.47 | 0.25 | 0.28 |
| 1 | 172019718 | rs12091015 | 1.196 | -0.003 | 0.631 | 0.436 | -0.005 | 0.084 | 0.084 | 0 | 0.46 | 0.26 | 0.4 | 0.16 | 0.47 | 0.37 |
| 1 | 172034439 | rs4916418 | -0.111 | -0.132 | 0.452 | 0.448 | 0.031 | 0.076 | 0.076 | 0 | 0.67 | 0.83 | 0.81 | 0.84 | 0.73 | 0.71 |
| 1 | 172039455 | rs4641331 | -0.788 | -0.153 | 0.294 | 0.587 | 0.021 | -0.01 | -0.01 | 0.244 | 0.89 | 0.95 | 0.96 | 0.86 | 0.95 | 0.9 |
| 1 | 172060763 | rs12119190 | -0.456 | 0.142 | 0.807 | 0.863 | 0.076 | 0.203 | 0.203 | 0 | 0.14 | 0.34 | 0.34 | 0.45 | 0.22 | 0.28 |
| 1 | 172063516 | rs4075021 | 1.753 | -0.183 | 0.423 | 0.614 | -0.04 | 0.11 | 0.11 | 0 | 0.4 | 0.1 | 0.37 | 0.19 | 0.42 | 0.34 |
| 1 | 172066592 | rs17350579 | 0.714 | -0.305 | 0.478 | 0.408 | 0.002 | -0.019 | -0.019 | 2.038 | 0.24 | 0.19 | 0.16 | 0.02 | 0.25 | 0.2 |
| 1 | 172069641 | rs6425638 | -0.848 | -0.324 | 0.186 | 0.474 | 0.011 | -0.01 | -0.01 | 0 | 0.88 | 0.92 | 0.96 | 0.79 | 0.95 | 0.9 |
| 1 | 172083881 | rs6694089 | 0.929 | -0.28 | 0.087 | 0.407 | -0.047 | 0.131 | 0.131 | 0 | 0.22 | 0.01 | 0.15 | 0.02 | 0.25 | 0.19 |
| 1 | 172087026 | rs10158753 | -1.182 | -0.243 | 0.494 | 0.484 | 0 | 0.031 | 0.031 | 0 | 0.47 | 0.76 | 0.59 | 0.6 | 0.53 | 0.56 |
| 1 | 172095226 | rs733190 | -1.131 | -1.193 | 0.485 | 0.463 | 0.017 | 0.064 | 0.064 | 0 | 0.48 | 0.76 | 0.59 | 0.67 | 0.48 | 0.55 |
| 1 | 172100831 | rs2421987 | -0.528 | -1.84 | 1.293 | 0.971 | 0.008 | 0.094 | 0.094 | 0 | 0.17 | 0.28 | 0.33 | 0.47 | 0.2 | 0.3 |
| 1 | 172101666 | rs10489299 | -2.028 | -1.869 | 1.311 | 0.916 | 0.012 | 0.193 | 0.193 | 3.287 | 0.07 | 0.4 | 0.07 | 0.19 | 0.01 | 0.1 |
| 1 | 172106806 | rs2761184 | 0.374 | -1.667 | 0.644 | 0.611 | 0.023 | -0.012 | -0.012 | 0 | 0.11 | 0.05 | 0.03 | 0.14 | 0.05 | 0.09 |
| 1 | 172119559 | rs2146901 | -1.033 | -1.338 | 1.042 | 0.813 | 0.007 | 0.137 | 0.137 | 0 | 0.2 | 0.42 | 0.34 | 0.43 | 0.2 | 0.34 |
| 1 | 172128536 | rs1475230 | -1.063 | -1.429 | 0.563 | 0.736 | -0.005 | 0.129 | 0.129 | 0 | 0.23 | 0.49 | 0.36 | 0.42 | 0.22 | 0.37 |
| 1 | 172134682 | rs11800549 | 0.183 | -1.679 | 0.372 | 0.166 | 0.006 | -0.004 | -0.004 | 0 | 0.11 | 0.08 | 0.05 | 0.14 | 0.06 | 0.08 |
| 1 | 172137019 | rs17361886 | 0.239 | -2.138 | 0.146 | -0.412 | 0.017 | 0.064 | 0.064 | 0 | 0.3 | 0.18 | 0.23 | 0.01 | 0.3 | 0.21 |
| 1 | 172137094 | rs16843944 | -0.251 | -2.216 | 0.454 | -0.079 | 0 | 0.074 | 0.074 | 0.078 | 0.16 | 0.48 | 0.1 | 0.22 | 0.12 | 0.22 |
| 1 | 172137119 | rs672740 | -1.682 | -2.288 | 0.361 | -0.118 | -0.037 | -0.007 | -0.007 | 0 | 0.59 | 0.88 | 0.38 | 0.38 | 0.47 | 0.51 |
| 1 | 172146648 | rs12129806 | 0.343 | -2.209 | -0.052 | -0.175 | 0.014 | 0.114 | 0.114 | 0 | 0.3 | 0.14 | 0.22 | 0.01 | 0.3 | 0.21 |

| | | | | | | | | | | | | | | | | |
|---|---|---|---|---|---|---|---|---|---|---|---|---|---|---|---|---|
| 1 | 172153224 | rs515060 | -0.799 | -2.251 | 0.067 | -0.128 | -0.021 | 0.017 | 0.017 | 1.384 | 0.19 | 0.64 | 0.11 | 0.22 | 0.12 | 0.22 |
| 1 | 172156886 | rs583578 | -1.09 | -2.246 | -0.099 | -0.137 | 0.054 | 0.127 | 0.127 | 0 | 0.48 | 0.78 | 0.33 | 0.23 | 0.42 | 0.43 |
| 1 | 172164499 | rs598562 | -1.109 | -2.247 | -0.121 | -0.138 | 0.06 | 0.151 | 0.151 | 0 | 0.49 | 0.81 | 0.34 | 0.23 | 0.42 | 0.43 |
| 1 | 172171850 | rs11805471 | 0.746 | -2.288 | -0.202 | -0.097 | 0.016 | -0.001 | -0.001 | 0 | 0.11 | 0.05 | 0.04 | 0.08 | 0.06 | 0.08 |
| 1 | 172172467 | rs11589142 | 0.525 | -2.321 | -0.244 | -0.153 | -0.007 | 0.164 | 0.164 | 0 | 0.28 | 0.06 | 0.22 | 0.01 | 0.3 | 0.21 |
| 1 | 172175115 | rs508742 | -1.657 | -2.269 | -0.217 | -0.085 | -0.073 | -0.036 | -0.036 | 0 | 0.6 | 0.95 | 0.39 | 0.38 | 0.48 | 0.51 |
| 1 | 172180396 | rs610587 | -1.033 | -2.268 | -0.016 | -0.01 | -0.018 | 0.006 | 0.006 | 1.716 | 0.2 | 0.7 | 0.11 | 0.22 | 0.12 | 0.22 |
| 1 | 172181377 | rs17277295 | 0.35 | -2.129 | -0.115 | -0.147 | -0.007 | 0.001 | 0.001 | 0 | 0.12 | 0.13 | 0.05 | 0.15 | 0.06 | 0.09 |
| 1 | 172185029 | rs6689455 | -1.25 | -2.185 | 0.225 | -0.103 | 0.001 | 0.002 | 0.002 | 0 | 0.54 | 0.46 | 0.36 | 0.36 | 0.47 | 0.51 |
| 1 | 172186321 | rs10489294 | -0.545 | -2.221 | 0.412 | -0.339 | -0.008 | 0.004 | 0.004 | 0 | 0.16 | 0.29 | 0.09 | 0.17 | 0.12 | 0.2 |
| 1 | 172186729 | rs633995 | -1.091 | -2.355 | 0.112 | -0.3 | 0.027 | 0.052 | 0.052 | 0 | 0.47 | 0.64 | 0.34 | 0.2 | 0.4 | 0.44 |
| 1 | 172186787 | rs10489293 | 0.122 | -2.256 | -0.502 | -0.675 | -0.004 | 0.192 | 0.192 | 0 | 0.27 | 0.01 | 0.18 | 0 | 0.26 | 0.21 |
| 1 | 172189889 | rs678962 | -0.129 | -2.574 | 0.181 | -0.078 | -0.028 | 0.056 | 0.056 | 0 | 0.19 | 0.32 | 0.26 | 0.27 | 0.19 | 0.22 |
| 1 | 172191394 | rs477289 | -1.177 | -2.98 | 0.704 | -0.216 | -0.034 | 0.027 | 0.027 | 0 | 0.29 | 0.61 | 0.32 | 0.34 | 0.23 | 0.31 |
| 1 | 172206026 | rs4916245 | 1.782 | -3.242 | -0.012 | -0.687 | -0.129 | -0.131 | -0.131 | 0 | 0.35 | 0.02 | 0.47 | 0.46 | 0.48 | 0.28 |
| 1 | 172214642 | rs12403844 | 0.546 | -3.21 | 0.136 | -0.589 | -0.01 | 0.011 | 0.011 | 0 | 0.19 | 0.22 | 0.25 | 0.23 | 0.19 | 0.24 |
| 1 | 172223403 | rs10489290 | 1.262 | -3.228 | 0.152 | -0.598 | 0.011 | -0.043 | -0.043 | 0.199 | 0.15 | 0.06 | 0.23 | 0.23 | 0.18 | 0.21 |
| 1 | 172234627 | rs12082431 | 0.418 | -3.218 | 0.229 | -0.593 | -0.027 | 0.035 | 0.035 | 0 | 0.19 | 0.3 | 0.26 | 0.23 | 0.19 | 0.24 |
| 1 | 172241251 | rs2421992 | 0.649 | -3.272 | 0.345 | -0.631 | 0.007 | 0.007 | 0.007 | 0 | 0.56 | 0.44 | 0.74 | 0.68 | 0.67 | 0.51 |
| 1 | 172247127 | rs2254810 | 0.913 | -3.133 | 0.139 | -0.554 | -0.01 | -0.001 | -0.001 | 0 | 0.08 | 0.1 | 0.04 | 0.07 | 0.04 | 0.1 |
| 1 | 172253585 | rs2757506 | 1.018 | -3.105 | 0.151 | -0.54 | -0.011 | -0.001 | -0.001 | 0 | 0.08 | 0.1 | 0.04 | 0.07 | 0.04 | 0.09 |
| 1 | 172254193 | rs2586392 | -0.41 | -3.181 | 0.196 | -0.549 | -0.064 | 0.05 | 0.05 | 0 | 0.27 | 0.48 | 0.31 | 0.31 | 0.23 | 0.34 |
| 1 | 172256116 | rs2586393 | -1.814 | -2.985 | -0.346 | -0.49 | -0.125 | -0.123 | -0.123 | 3.803 | 0.65 | 0.98 | 0.53 | 0.55 | 0.52 | 0.73 |
| 1 | 172264193 | rs2757489 | 0.401 | -2.713 | 0.057 | -0.352 | -0.024 | -0.004 | -0.004 | 0.093 | 0.1 | 0.19 | 0.05 | 0.07 | 0.06 | 0.11 |
| 1 | 172264425 | rs10911509 | 0.539 | -2.921 | 0.317 | -0.456 | 0.011 | 0.017 | 0.017 | 0 | 0.11 | 0.18 | 0.08 | 0.13 | 0.1 | 0.1 |
| 1 | 172267419 | rs2757493 | -0.145 | -2.852 | -0.108 | -0.369 | -0.01 | 0.009 | 0.009 | 0 | 0.25 | 0.3 | 0.28 | 0.31 | 0.25 | 0.32 |
| 1 | 172267554 | rs10752932 | 0.094 | -3.037 | -0.104 | -0.513 | -0.007 | -0.007 | -0.007 | 0.282 | 0.24 | 0.34 | 0.17 | 0.24 | 0.2 | 0.25 |
| 1 | 172269642 | rs2422077 | -0.552 | -2.288 | -0.293 | -0.109 | -0.046 | -0.015 | -0.015 | 0 | 0.13 | 0.37 | 0.07 | 0.07 | 0.07 | 0.13 |
| 1 | 172279021 | rs2757500 | 0.351 | -2.084 | -1.21 | -0.011 | -0.004 | -0.001 | -0.001 | 0.063 | 0.09 | 0.1 | 0.04 | 0.07 | 0.06 | 0.11 |
| 1 | 172279909 | rs2421994 | 0.88 | -2.083 | -1.261 | 0.022 | 0.007 | -0.016 | -0.016 | 0 | 0.16 | 0.13 | 0.23 | 0.24 | 0.18 | 0.22 |
| 1 | 172281341 | rs2757502 | -0.038 | -1.081 | -1.211 | 0.101 | -0.018 | -0.005 | -0.005 | 0.09 | 0.1 | 0.16 | 0.05 | 0.07 | 0.06 | 0.11 |
| 1 | 172292205 | rs6694387 | -0.596 | -0.471 | -1.721 | 0.223 | -0.136 | -0.13 | -0.13 | 0 | 0.65 | 0.98 | 0.56 | 0.54 | 0.52 | 0.73 |
| 1 | 172296601 | rs9425287 | -0.274 | -0.627 | -1.378 | 0 | -0.002 | -0.038 | -0.038 | 0 | 0.64 | 0.76 | 0.48 | 0.42 | 0.47 | 0.67 |
| 1 | 172300345 | rs10797969 | -0.369 | -0.504 | -1.763 | 0.045 | -0.036 | -0.093 | -0.093 | 0 | 0.66 | 0.88 | 0.5 | 0.42 | 0.47 | 0.67 |
| 1 | 172302544 | rs911713 | 0.386 | 0.295 | -1.327 | -0.832 | -0.003 | 0.038 | 0.038 | 0 | 0.6 | 0.3 | 0.6 | 0.83 | 0.58 | 0.63 |
| 1 | 172303509 | rs10911603 | -0.9 | 0.325 | -0.788 | -0.852 | 0.003 | -0.069 | -0.069 | 0 | 0.12 | 0.26 | 0.07 | 0.01 | 0.09 | 0.21 |
| 1 | 172305739 | rs12134640 | -1.11 | 0.294 | -0.698 | -0.867 | -0.001 | -0.073 | -0.073 | 0 | 0.13 | 0.28 | 0.07 | 0.01 | 0.09 | 0.19 |
| 1 | 172312769 | rs9425291 | -1.566 | 0.184 | -1.341 | -1.265 | -0.045 | 0.192 | 0.192 | 0.222 | 0.41 | 0.91 | 0.36 | 0.11 | 0.42 | 0.43 |
| 1 | 172324930 | rs12143497 | -0.214 | 0.276 | -1.116 | -1.081 | 0.001 | -0.007 | -0.007 | 0.948 | 0.09 | 0.07 | 0.04 | 0 | 0.07 | 0.15 |
| 1 | 172331059 | rs1894633 | -0.627 | 0.233 | -0.958 | -1.162 | 0.003 | -0.035 | -0.035 | 0 | 0.28 | 0.6 | 0.28 | 0.1 | 0.33 | 0.24 |
| 1 | 172334818 | rs4504922 | 0.428 | 0.112 | -0.918 | -1.305 | -0.014 | -0.015 | -0.015 | 0 | 0.8 | 0.32 | 0.8 | 0.87 | 0.85 | 0.82 |
| 1 | 172346548 | rs1011731 | 1.739 | 0.066 | -0.984 | -1.431 | -0.035 | 0.183 | 0.183 | 0.1 | 0.6 | 0.1 | 0.64 | 0.87 | 0.58 | 0.57 |
| 1 | 172349921 | rs3213564 | -0.5 | 0.073 | -0.88 | -1.397 | -0.03 | -0.03 | -0.03 | 0 | 0.2 | 0.58 | 0.18 | 0.12 | 0.15 | 0.19 |
| 1 | 172355132 | rs1023479 | 0.59 | 0.088 | -0.9 | -1.388 | -0.007 | -0.012 | -0.012 | 0.548 | 0.77 | 0.31 | 0.8 | 0.87 | 0.82 | 0.78 |
| 1 | 172355841 | rs17369123 | 0.11 | 0.195 | -0.931 | -1.199 | -0.015 | 0.013 | 0.013 | 0 | 0.08 | 0.06 | 0.11 | 0.02 | 0.16 | 0.06 |
| 1 | 172365330 | rs6424894 | -0.593 | -0.013 | -0.285 | -1.226 | 0.02 | 0.099 | 0.099 | 0.107 | 0.09 | 0.53 | 0.13 | 0.12 | 0.06 | 0.13 |
| 1 | 172369347 | rs9425596 | 0.6 | -0.357 | -1.697 | -1.18 | -0.135 | -0.091 | -0.091 | 0 | 0.43 | 0.04 | 0.41 | 0.55 | 0.47 | 0.41 |
| 1 | 172370049 | rs17369331 | 0.91 | -0.165 | -1.99 | -1.219 | -0.031 | -0.02 | -0.02 | 0 | 0.13 | 0.01 | 0.13 | 0.1 | 0.15 | 0.1 |
| 1 | 172371151 | rs9425304 | 1.028 | -0.492 | -1.485 | -1.263 | -0.109 | -0.129 | -0.129 | 3.693 | 0.38 | 0.06 | 0.4 | 0.51 | 0.41 | 0.26 |
| 1 | 172371922 | rs9425598 | 0.103 | -0.67 | -1.077 | -1.41 | -0.035 | -0.007 | -0.007 | 0 | 0.57 | 0.07 | 0.55 | 0.64 | 0.62 | 0.57 |
| 1 | 172374555 | rs978875 | -0.438 | -0.58 | -0.998 | -1.473 | 0.022 | -0.027 | -0.027 | 0 | 0.95 | 0.89 | 0.92 | 1 | 0.92 | 0.95 |
| 1 | 172377745 | rs16844417 | 0.783 | -0.599 | -1.152 | -1.53 | -0.002 | -0.011 | -0.011 | 0 | 0.07 | 0.05 | 0.06 | 0 | 0.11 | 0.05 |
| 1 | 172381284 | rs17278379 | 0.516 | -0.609 | -1.177 | -1.53 | -0.005 | -0.007 | -0.007 | 0 | 0.08 | 0.06 | 0.06 | 0 | 0.11 | 0.05 |
| 1 | 172381383 | rs7528296 | 1.416 | -0.599 | -1.213 | -1.522 | -0.147 | -0.166 | -0.166 | 0 | 0.37 | 0.02 | 0.39 | 0.51 | 0.4 | 0.26 |
| 1 | 172389198 | rs9425606 | 0.343 | -0.436 | -0.83 | -1.296 | 0.027 | -0.023 | -0.023 | 0 | 0.3 | 0.18 | 0.31 | 0.36 | 0.24 | 0.36 |
| 1 | 172390896 | rs2422139 | 0.533 | -0.355 | -0.859 | -1.278 | 0.095 | -0.038 | -0.038 | 1.033 | 0.93 | 0.8 | 0.9 | 1 | 0.84 | 0.93 |
| 1 | 172400484 | rs6702667 | -0.503 | -0.328 | -0.805 | -1.234 | 0.003 | 0.002 | 0.002 | 0.008 | 0.69 | 0.68 | 0.64 | 0.64 | 0.64 | 0.59 |
| 1 | 172410967 | rs1063412 | 0.576 | -0.295 | -1.287 | -1.247 | -0.043 | 0.011 | 0.011 | 1.745 | 0.55 | 0.06 | 0.54 | 0.66 | 0.6 | 0.57 |
| 1 | 172411768 | rs13932 | 0.685 | -0.31 | -1.057 | -1.03 | 0.028 | -0.038 | -0.038 | 0 | 0.26 | 0.13 | 0.27 | 0.34 | 0.2 | 0.37 |
| 1 | 172419651 | rs2269613 | 0.462 | -0.274 | -0.517 | -1.007 | -0.017 | 0.008 | 0.008 | 0 | 0.28 | 0.35 | 0.28 | 0.34 | 0.21 | 0.35 |
| 1 | 172426998 | rs6691993 | 0.66 | -0.336 | -0.327 | -1.401 | -0.008 | 0.032 | 0.032 | 0 | 0.45 | 0.36 | 0.51 | 0.7 | 0.5 | 0.42 |
| 1 | 172431333 | rs16844498 | 1.424 | -0.377 | -0.451 | -1.448 | 0.004 | 0.055 | 0.055 | 0 | 0.1 | 0.03 | 0.05 | 0 | 0.09 | 0.06 |
| 1 | 172434812 | rs2901656 | 0.747 | -0.339 | -0.356 | -1.441 | -0.015 | 0.042 | 0.042 | 0 | 0.45 | 0.32 | 0.5 | 0.69 | 0.49 | 0.42 |
| 1 | 172435451 | rs4244193 | -0.274 | -0.456 | -0.293 | -1.515 | 0.042 | -0.075 | -0.075 | 0 | 0.87 | 0.7 | 0.85 | 1 | 0.81 | 0.93 |
| 1 | 172435851 | rs9425329 | 0.499 | -0.454 | -0.225 | -1.4 | -0.002 | -0.001 | -0.001 | 1.798 | 0.31 | 0.29 | 0.29 | 0.3 | 0.22 | 0.45 |
| 1 | 172445883 | rs7551242 | -0.192 | -0.379 | -0.28 | -1.396 | 0.001 | -0.054 | -0.054 | 0 | 0.78 | 0.74 | 0.81 | 1 | 0.72 | 0.87 |
| 1 | 172447498 | rs6695139 | 0.079 | -0.328 | -0.429 | -1.226 | -0.016 | 0.016 | 0.016 | 0 | 0.34 | 0.41 | 0.38 | 0.43 | 0.27 | 0.53 |
| 1 | 172450689 | rs4916263 | -0.053 | -0.206 | -0.447 | -1.187 | 0.021 | -0.037 | -0.037 | 0 | 0.76 | 0.66 | 0.72 | 0.88 | 0.67 | 0.79 |
| 1 | 172450795 | rs4916266 | 0.902 | -0.059 | -0.741 | -0.883 | -0.108 | -0.095 | -0.095 | 0 | 0.34 | 0.06 | 0.37 | 0.57 | 0.4 | 0.22 |

| | | | | | | | | | | | | | | | | |
|---|---|---|---|---|---|---|---|---|---|---|---|---|---|---|---|---|
| 1 | 172453801 | rs16844588 | -0.933 | -0.27 | 0.143 | -0.837 | -0.043 | -0.086 | -0.086 | 0.06 | 0.14 | 0.34 | 0.06 | 0 | 0.06 | 0.13 |
| 1 | 172458743 | rs2157449 | -0.847 | 0.196 | -0.636 | -0.561 | 0.035 | 0.093 | 0.093 | 0.143 | 0.46 | 0.82 | 0.38 | 0.32 | 0.41 | 0.43 |
| 1 | 172462962 | rs3919435 | -0.841 | 0.171 | -0.379 | -0.515 | 0.034 | 0.07 | 0.07 | 0.107 | 0.46 | 0.82 | 0.38 | 0.35 | 0.41 | 0.42 |
| 1 | 172463344 | rs2213737 | -0.877 | -0.201 | -0.561 | -1.201 | -0.015 | -0.059 | -0.059 | 0.328 | 0.4 | 0.55 | 0.26 | 0.11 | 0.32 | 0.33 |
| 1 | 172463933 | rs16844620 | -0.313 | 0.182 | -0.792 | -0.773 | 0.001 | 0.008 | 0.008 | 0 | 0.17 | 0.15 | 0.09 | 0.1 | 0.18 | 0.17 |
| 1 | 172464497 | rs2213736 | -0.921 | 0.614 | -1.037 | -1.044 | -0.015 | 0.068 | 0.068 | 0 | 0.42 | 0.77 | 0.41 | 0.18 | 0.46 | 0.38 |
| 1 | 172464790 | rs1984404 | -0.633 | 0.258 | -0.882 | -1.419 | 0.001 | -0.017 | -0.017 | 0.325 | 0.22 | 0.26 | 0.21 | 0.14 | 0.28 | 0.18 |
| 1 | 172464899 | rs12068145 | -0.181 | -0.065 | -0.337 | -1.513 | -0.024 | -0.077 | -0.077 | 0 | 0.17 | 0.29 | 0.13 | 0 | 0.12 | 0.1 |
| 1 | 172467044 | rs10911830 | 0.305 | -0.068 | -0.232 | -1.459 | -0.011 | -0.013 | -0.013 | 0.947 | 0.27 | 0.39 | 0.17 | 0.19 | 0.16 | 0.34 |
| 1 | 172469707 | rs6424956 | -0.401 | -0.083 | -0.619 | -1.843 | 0.012 | 0.026 | 0.026 | 0 | 0.39 | 0.4 | 0.24 | 0.19 | 0.26 | 0.36 |
| 1 | 172485007 | rs10157519 | -0.272 | 0.613 | -0.163 | -1.729 | -0.001 | 0.005 | 0.005 | 0.255 | 0.46 | 0.64 | 0.39 | 0.38 | 0.48 | 0.71 |
| 1 | 172498216 | rs10489279 | -0.536 | 0.92 | 0.207 | -1.779 | 0.078 | 0.028 | 0.028 | 0.021 | 0.15 | 0.35 | 0.17 | 0.19 | 0.23 | 0.32 |
| 1 | 172501643 | rs2239816 | 0.841 | 0.794 | 0.29 | -1.897 | 0.011 | 0.041 | 0.041 | 0.153 | 0.31 | 0.21 | 0.21 | 0.19 | 0.25 | 0.39 |
| 1 | 172517567 | rs2472549 | 0.314 | 0.631 | 0.399 | -1.832 | -0.033 | -0.051 | -0.051 | 0 | 0.36 | 0.59 | 0.25 | 0.2 | 0.23 | 0.41 |
| 1 | 172518896 | rs12758548 | 0.104 | 0.635 | 0.46 | -1.808 | -0.009 | -0.009 | -0.009 | 0.258 | 0.11 | 0.09 | 0.14 | 0.18 | 0.23 | 0.29 |
| 1 | 172530687 | rs10911920 | 0.911 | 0.608 | 0.438 | -1.812 | 0.022 | 0.042 | 0.042 | 0.127 | 0.31 | 0.23 | 0.21 | 0.2 | 0.23 | 0.39 |
| 1 | 172559323 | rs3738736 | 0.668 | 0.635 | 0.318 | -1.798 | 0.001 | 0 | 0 | 7.502 | 0.33 | 0.38 | 0.23 | 0.2 | 0.23 | 0.39 |
| 1 | 172575959 | rs10489278 | 0.337 | 0.762 | 0.101 | -1.766 | -0.039 | -0.027 | -0.027 | 0 | 0.1 | 0.04 | 0.14 | 0.18 | 0.23 | 0.29 |
| 1 | 172595624 | rs9970543 | -0.254 | 0.669 | 0.067 | -1.829 | -0.003 | -0.016 | -0.016 | 0 | 0.47 | 0.56 | 0.31 | 0.2 | 0.33 | 0.44 |
| 1 | 172600841 | rs12065670 | 0.67 | 0.637 | 0.324 | -1.839 | 0 | -0.018 | -0.018 | 0.064 | 0.12 | 0.08 | 0.07 | 0 | 0.1 | 0.02 |
| 1 | 172603411 | rs10912080 | -0.992 | 0.596 | 0.194 | -1.8 | 0.001 | 0.002 | 0.002 | 0 | 0.59 | 0.71 | 0.46 | 0.37 | 0.57 | 0.73 |
| 1 | 172603940 | rs730278 | -0.887 | 0.84 | -0.189 | -1.795 | -0.041 | 0.018 | 0.018 | 0 | 0.24 | 0.15 | 0.21 | 0.17 | 0.33 | 0.35 |
| 1 | 172609760 | rs2639621 | -0.789 | 0.608 | 0.029 | -1.861 | -0.001 | -0.003 | -0.003 | 0 | 0.6 | 0.71 | 0.44 | 0.29 | 0.57 | 0.68 |
| 1 | 172613241 | rs2859228 | -0.085 | 0.628 | -0.309 | -1.948 | -0.011 | 0.039 | 0.039 | 0.026 | 0.45 | 0.45 | 0.36 | 0.29 | 0.47 | 0.63 |
| 1 | 172621538 | rs2021838 | -0.282 | 0.655 | -0.421 | -1.766 | -0.002 | 0.009 | 0.009 | 0 | 0.46 | 0.49 | 0.38 | 0.36 | 0.43 | 0.6 |
| 1 | 172627498 | rs763110 | -0.697 | 0.048 | -0.934 | -2.244 | 0.013 | 0.161 | 0.161 | 0 | 0.54 | 0.89 | 0.35 | 0.26 | 0.35 | 0.63 |
| 1 | 172630811 | rs6700734 | 0.643 | -0.203 | -0.542 | -2.523 | 0.009 | 0.031 | 0.031 | 0.442 | 0.34 | 0.28 | 0.21 | 0.11 | 0.26 | 0.41 |
| 1 | 172632603 | rs2639654 | -1.314 | -0.195 | -0.521 | -2.537 | 0.001 | 0.051 | 0.051 | 0.086 | 0.88 | 0.98 | 0.94 | 1 | 0.89 | 0.97 |
| 1 | 172638500 | rs12135884 | -0.09 | -0.315 | 0.065 | -2.468 | 0.001 | 0 | 0 | 0 | 0.42 | 0.65 | 0.29 | 0.26 | 0.3 | 0.61 |
| 1 | 172642118 | rs1492907 | -0.005 | -0.199 | -0.137 | -2.41 | -0.012 | -0.052 | -0.052 | 0 | 0.42 | 0.55 | 0.28 | 0.22 | 0.3 | 0.6 |
| 1 | 172643220 | rs2021840 | 0.595 | -0.175 | -0.121 | -2.454 | 0.01 | 0.032 | 0.032 | 0.102 | 0.35 | 0.29 | 0.21 | 0.11 | 0.26 | 0.41 |
| 1 | 172645898 | rs6673027 | 1.09 | -0.416 | -0.947 | -2.552 | 0.044 | 0.065 | 0.065 | 0 | 0.22 | 0.08 | 0.15 | 0.03 | 0.14 | 0.12 |
| 1 | 172647523 | rs2422207 | 0.169 | 0.014 | -0.571 | -2.257 | -0.011 | -0.042 | -0.042 | 0 | 0.41 | 0.55 | 0.27 | 0.25 | 0.3 | 0.6 |
| 1 | 172650388 | rs859656 | 0.037 | -0.266 | -0.23 | -2.135 | -0.019 | -0.081 | -0.081 | 0 | 0.83 | 0.78 | 0.91 | 1 | 0.88 | 0.95 |
| 1 | 172650685 | rs704842 | -0.818 | -0.308 | 0.225 | -2.151 | -0.014 | -0.048 | -0.048 | 0 | 0.22 | 0.48 | 0.19 | 0.15 | 0.16 | 0.22 |
| 1 | 172654377 | rs859652 | -0.493 | -0.594 | -0.141 | -4.071 | 0.007 | -0.066 | -0.066 | 0 | 0.47 | 0.57 | 0.32 | 0.01 | 0.37 | 0.17 |
| 1 | 172667967 | rs2071929 | 0.461 | -0.829 | -0.152 | -3.592 | -0.029 | -0.034 | -0.034 | 0 | 0.21 | 0.25 | 0.07 | 0.13 | 0.09 | 0.36 |
| 1 | 172670720 | rs859668 | -0.182 | -1.202 | -0.456 | -3.881 | 0.011 | -0.009 | -0.009 | 0 | 0.23 | 0.2 | 0.1 | 0.03 | 0.11 | 0.17 |
| 1 | 172673322 | rs859670 | 0.04 | -1.116 | -0.535 | -3.909 | 0.031 | 0.038 | 0.038 | 0 | 0.24 | 0.08 | 0.13 | 0.02 | 0.16 | 0.15 |
| 1 | 172674541 | rs7553067 | 0.156 | -1.072 | -0.07 | -4.116 | -0.022 | -0.153 | -0.153 | 0 | 0.22 | 0.48 | 0.14 | 0 | 0.2 | 0.1 |
| 1 | 172675525 | rs10798176 | 0.514 | -1.06 | -0.235 | -4.107 | 0.007 | 0.024 | 0.024 | 0 | 0.2 | 0.11 | 0.11 | 0 | 0.19 | 0.1 |
| 1 | 172699579 | rs1102706 | -0.583 | -0.749 | -0.316 | -4.189 | -0.008 | 0 | 0 | 0.013 | 0.65 | 0.69 | 0.37 | 0.09 | 0.54 | 0.58 |
| 1 | 172716461 | rs11587368 | 0.601 | -0.695 | -0.818 | -4.147 | -0.09 | 0.204 | 0.204 | 0 | 0.36 | 0.07 | 0.22 | 0.07 | 0.39 | 0.39 |
| 1 | 172724394 | rs2022168 | 0.628 | -0.964 | -0.846 | -4.108 | -0.135 | 0.2 | 0.2 | 0.047 | 0.36 | 0.08 | 0.24 | 0.09 | 0.43 | 0.37 |
| 1 | 172728946 | rs859627 | -0.038 | -1.05 | -0.381 | -4.16 | -0.013 | 0.081 | 0.081 | 0 | 0.45 | 0.32 | 0.28 | 0.09 | 0.45 | 0.37 |
| 1 | 172730829 | rs859624 | -0.714 | -0.571 | -0.662 | -4.282 | -0.011 | 0.013 | 0.013 | 0 | 0.78 | 0.89 | 0.84 | 0.98 | 0.75 | 0.82 |
| 1 | 172737172 | rs733684 | 0.255 | -0.659 | -0.298 | -4.364 | 0 | -0.064 | -0.064 | 0 | 0.46 | 0.45 | 0.27 | 0.08 | 0.43 | 0.35 |
| 1 | 172743727 | rs2077690 | 0.99 | -0.567 | -0.419 | -4.34 | 0.008 | 0.061 | 0.061 | 0 | 0.29 | 0.18 | 0.54 | 0.9 | 0.31 | 0.43 |
| 1 | 172745284 | rs859722 | 0.844 | -0.59 | -0.529 | -4.352 | 0.001 | -0.013 | -0.013 | 0 | 0.3 | 0.25 | 0.54 | 0.9 | 0.31 | 0.43 |
| 1 | 172777616 | rs1102737 | -1.201 | -0.066 | -0.587 | -4.142 | -0.037 | 0.021 | 0.021 | 0 | 0.94 | 0.99 | 0.92 | 0.98 | 0.88 | 0.91 |
| 1 | 172789765 | rs11810332 | 0.292 | -1.165 | -0.408 | -4.144 | 0.002 | -0.098 | -0.098 | 0 | 0.41 | 0.44 | 0.27 | 0.08 | 0.49 | 0.31 |
| 1 | 172800805 | rs10489277 | 0.653 | -1.578 | -0.59 | -3.67 | 0.005 | 0.112 | 0.112 | 0 | 0.2 | 0.03 | 0.09 | 0.02 | 0.16 | 0.15 |
| 1 | 172807254 | rs7519046 | -0.582 | -1.416 | -0.145 | -3.488 | 0.023 | 0.067 | 0.067 | 0.028 | 0.66 | 0.74 | 0.42 | 0.15 | 0.72 | 0.54 |
| 1 | 172814855 | rs12066143 | -0.607 | -1.421 | -0.32 | -3.49 | 0.024 | 0.073 | 0.073 | 0.03 | 0.66 | 0.76 | 0.42 | 0.15 | 0.73 | 0.55 |
| 1 | 172852118 | rs12037781 | -0.477 | -1.849 | -0.436 | -3.505 | -0.021 | 0.05 | 0.05 | 0 | 0.66 | 0.76 | 0.71 | 0.89 | 0.46 | 0.71 |
| 1 | 172852403 | rs12037853 | 0.891 | -2.111 | -0.296 | -3.646 | 0.008 | 0.01 | 0.01 | 0.239 | 0.31 | 0.34 | 0.53 | 0.86 | 0.23 | 0.41 |
| 1 | 172853460 | rs7517810 | 0.991 | -2.112 | -0.263 | -3.646 | 0.02 | 0.04 | 0.04 | 0.461 | 0.3 | 0.27 | 0.52 | 0.85 | 0.23 | 0.41 |
| 1 | 172854334 | rs4916283 | 0.962 | -2.102 | -0.258 | -3.745 | 0.021 | 0.043 | 0.043 | 0.361 | 0.31 | 0.27 | 0.52 | 0.87 | 0.23 | 0.41 |
| 1 | 172858580 | rs2422257 | -0.782 | -2.067 | -0.394 | -3.767 | -0.081 | 0.154 | 0.154 | 0 | 0.67 | 0.89 | 0.72 | 0.92 | 0.46 | 0.71 |
| 1 | 172862234 | rs9286879 | 0.806 | -2.02 | -0.281 | -3.622 | 0.004 | -0.003 | -0.003 | 0 | 0.31 | 0.36 | 0.53 | 0.88 | 0.23 | 0.42 |
| 1 | 172862939 | rs10489276 | 0.848 | -2.006 | -0.321 | -3.615 | 0.011 | 0.014 | 0.014 | 0.287 | 0.31 | 0.33 | 0.53 | 0.88 | 0.23 | 0.42 |
| 1 | 172871498 | rs12060919 | 0.379 | -1.115 | -0.071 | -3.085 | 0.031 | 0.031 | 0.031 | 0 | 0.13 | 0.07 | 0.03 | 0.03 | 0.04 | 0.08 |
| 1 | 172875119 | rs16845072 | 0.419 | -1.085 | 0.233 | -3.073 | -0.039 | -0.04 | -0.04 | 0 | 0.14 | 0.16 | 0.05 | 0.03 | 0.04 | 0.08 |
| 1 | 172879023 | rs2227203 | -0.856 | -0.854 | -0.247 | -3.23 | -0.148 | 0.066 | 0.066 | 0 | 0.81 | 0.97 | 0.76 | 0.92 | 0.56 | 0.77 |
| 1 | 172881827 | rs2213746 | -0.271 | -1.242 | 0.379 | -3.009 | 0.001 | -0.014 | -0.014 | 0 | 0.59 | 0.6 | 0.66 | 0.92 | 0.41 | 0.67 |
| 1 | 172884872 | rs4916288 | 0.632 | -1.241 | 0.191 | -3.008 | 0.021 | 0.077 | 0.077 | 0.014 | 0.44 | 0.36 | 0.6 | 0.89 | 0.37 | 0.58 |
| 1 | 172886500 | rs1883475 | 0.618 | -1.235 | 0.109 | -3.006 | 0.018 | 0.054 | 0.054 | 0.006 | 0.44 | 0.34 | 0.6 | 0.89 | 0.37 | 0.58 |
| 1 | 172898402 | rs12037606 | -0.293 | -1.144 | -0.478 | -2.964 | 0.002 | 0.022 | 0.022 | 0 | 0.6 | 0.65 | 0.67 | 0.92 | 0.41 | 0.67 |
| 1 | 172921727 | rs10912501 | 0.426 | -0.826 | -1.422 | -2.783 | 0.02 | 0.068 | 0.068 | 0.007 | 0.44 | 0.32 | 0.6 | 0.91 | 0.37 | 0.59 |

| | | | | | | | | | | | | | | | | |
|---|---|---|---|---|---|---|---|---|---|---|---|---|---|---|---|---|
| 1 | 172930064 | rs6425189 | 0.171 | 0.182 | -2.102 | -2.254 | -0.035 | 0.184 | 0.184 | 0 | 0.61 | 0.89 | 0.67 | 0.9 | 0.4 | 0.68 |
| 1 | 172936636 | rs972100 | -0.513 | 1.2 | -1.303 | -2.042 | 0.01 | -0.027 | -0.027 | 0 | 0.11 | 0.13 | 0.14 | 0 | 0.18 | 0.1 |
| 1 | 172936702 | rs972099 | -0.694 | 0.465 | -2.277 | -2.099 | 0.025 | 0.052 | 0.052 | 0.542 | 0.67 | 0.98 | 0.58 | 0.22 | 0.75 | 0.44 |
| 1 | 172940607 | rs10489273 | 0.068 | 0.683 | -2.28 | -2.459 | -0.104 | 0.089 | 0.089 | 0 | 0.23 | 0.06 | 0.21 | 0.03 | 0.41 | 0.17 |
| 1 | 172941713 | rs10489272 | 0.032 | -0.003 | -0.918 | -2.669 | 0.001 | -0.027 | -0.027 | 0.015 | 0.37 | 0.36 | 0.26 | 0.1 | 0.32 | 0.21 |
| 1 | 172944751 | rs4916291 | -0.284 | 0.612 | -1.249 | -2.472 | -0.108 | 0.111 | 0.111 | 0.039 | 0.27 | 0.05 | 0.22 | 0.07 | 0.47 | 0.22 |
| 1 | 172962496 | rs6697964 | -0.221 | 1.72 | -0.403 | -2.463 | -0.068 | 0.054 | 0.054 | 0.404 | 0.18 | 0.05 | 0.18 | 0.05 | 0.36 | 0.18 |
| 1 | 172985231 | rs1883478 | 0.03 | 1.2 | 0.006 | -2.656 | 0.004 | -0.015 | -0.015 | 0 | 0.42 | 0.41 | 0.3 | 0.12 | 0.37 | 0.2 |
| 1 | 173001583 | rs12067426 | -0.171 | 1.184 | 0.079 | -2.708 | 0.049 | 0.144 | 0.144 | 0.834 | 0.46 | 0.73 | 0.32 | 0.12 | 0.37 | 0.2 |
| 1 | 173009832 | rs16845308 | 0.663 | 0.617 | -0.073 | -2.745 | 0.057 | 0.081 | 0.081 | 0.526 | 0.42 | 0.24 | 0.42 | 0.77 | 0.27 | 0.59 |
| 1 | 173012804 | rs6425201 | -0.709 | 0.582 | -0.033 | -2.735 | -0.006 | -0.013 | -0.013 | 0 | 0.08 | 0.11 | 0.05 | 0.03 | 0.04 | 0.07 |
| 1 | 173013767 | rs975074 | -0.577 | 1.375 | 0.393 | -2.942 | 0.004 | -0.049 | -0.049 | 4.623 | 0.62 | 0.58 | 0.67 | 0.88 | 0.56 | 0.77 |
| 1 | 173017486 | rs2236876 | 0.473 | 1.305 | 0.471 | -2.948 | 0.019 | 0.021 | 0.021 | 0.341 | 0.44 | 0.37 | 0.44 | 0.77 | 0.27 | 0.6 |
| 1 | 173018680 | rs9286880 | 0.056 | 1.403 | 0.367 | -2.906 | 0.005 | -0.015 | -0.015 | 0 | 0.18 | 0.14 | 0.13 | 0.06 | 0.28 | 0.16 |
| 1 | 173021349 | rs7537126 | -0.631 | 1.329 | 0.292 | -2.826 | -0.006 | -0.013 | -0.013 | 0.387 | 0.08 | 0.11 | 0.05 | 0.03 | 0.04 | 0.07 |
| 1 | 173026851 | rs7546619 | 0.653 | 1.326 | 0.323 | -2.843 | 0.062 | 0.089 | 0.089 | 0.777 | 0.42 | 0.22 | 0.41 | 0.77 | 0.27 | 0.6 |
| 1 | 173049073 | rs2422272 | -0.002 | 1.393 | 0.526 | -2.753 | 0.045 | -0.081 | -0.081 | 0 | 0.82 | 0.59 | 0.85 | 0.94 | 0.75 | 0.86 |
| 1 | 173055352 | rs2207061 | -0.207 | 1.349 | 0.367 | -2.847 | -0.002 | 0.024 | 0.024 | 0.932 | 0.45 | 0.4 | 0.51 | 0.2 | 0.5 | 0.3 |
| 1 | 173062049 | rs16845401 | 1.39 | 1.329 | -0.333 | -2.809 | 0.024 | 0.025 | 0.025 | 2.909 | 0.34 | 0.02 | 0.33 | 0.74 | 0.25 | 0.56 |
| 1 | 173064279 | rs6425203 | 0.148 | 1.77 | -0.091 | -2.588 | 0.023 | -0.022 | -0.022 | 0 | 0.81 | 0.67 | 0.77 | 0.88 | 0.74 | 0.84 |
| 1 | 173064777 | rs12060864 | -0.258 | 1.984 | -0.843 | -2.665 | -0.063 | 0.146 | 0.146 | 0.065 | 0.4 | 0.18 | 0.36 | 0.12 | 0.47 | 0.24 |
| 1 | 173069244 | rs6425204 | 0.292 | 2.042 | -0.618 | -2.557 | 0.004 | -0.041 | -0.041 | 2.053 | 0.14 | 0.16 | 0.14 | 0 | 0.14 | 0.13 |
| 1 | 173079943 | rs10912533 | 0.144 | 1.562 | -0.36 | -2.19 | 0.036 | -0.056 | -0.056 | 0 | 0.83 | 0.75 | 0.87 | 0.94 | 0.76 | 0.86 |
| 1 | 173094569 | rs17372135 | -0.334 | 1.549 | -1.193 | -1.728 | -0.042 | 0.078 | 0.078 | 0.208 | 0.34 | 0.17 | 0.33 | 0.19 | 0.42 | 0.19 |
| 1 | 173096963 | rs11484630 | -0.896 | 1.273 | -0.746 | -1.677 | 0.004 | -0.048 | -0.048 | 0 | 0.13 | 0.22 | 0.08 | 0.01 | 0.1 | 0.08 |
| 1 | 173103273 | rs3861948 | -0.256 | 0.373 | -0.737 | -1.567 | -0.023 | -0.029 | -0.029 | 0 | 0.4 | 0.51 | 0.46 | 0.74 | 0.29 | 0.46 |
| 1 | 173104771 | rs10798262 | -0.191 | 0.415 | -0.405 | -1.598 | 0.012 | 0.018 | 0.018 | 0 | 0.41 | 0.58 | 0.47 | 0.8 | 0.31 | 0.46 |
| 1 | 173105414 | rs10912540 | -0.162 | 0.418 | -0.394 | -1.6 | 0.008 | 0.01 | 0.01 | 0 | 0.4 | 0.57 | 0.47 | 0.74 | 0.31 | 0.46 |
| 1 | 173105511 | rs6663669 | -0.481 | 0.898 | -0.405 | -1.571 | 0.027 | 0.093 | 0.093 | 0.145 | 0.44 | 0.71 | 0.64 | 0.78 | 0.35 | 0.51 |
| 1 | 173110953 | rs6663179 | 0.039 | 0.364 | -0.165 | -1.757 | -0.043 | -0.093 | -0.093 | 0 | 0.14 | 0.47 | 0.11 | 0.01 | 0.08 | 0.07 |
| 1 | 173117771 | rs1234305 | -0.707 | 0.952 | -1.283 | -2.381 | -0.039 | 0.025 | 0.025 | 0 | 0.73 | 0.8 | 0.62 | 0.84 | 0.59 | 0.74 |
| 1 | 173119967 | rs10912551 | 1.128 | 0.598 | -1.4 | -2.134 | -0.042 | -0.05 | -0.05 | 0.018 | 0.24 | 0.02 | 0.32 | 0.72 | 0.23 | 0.4 |
| 1 | 173131908 | rs7553711 | -0.926 | 0.066 | -1.177 | -2.643 | 0.068 | 0.171 | 0.171 | 0 | 0.43 | 0.8 | 0.34 | 0.11 | 0.29 | 0.26 |
| 1 | 173151156 | rs10127728 | -0.311 | -0.063 | -0.249 | -2.687 | -0.013 | -0.068 | -0.068 | 0 | 0.18 | 0.57 | 0.23 | 0.08 | 0.12 | 0.11 |
| 1 | 173153442 | rs3900307 | 0.406 | -0.167 | 0.212 | -2.732 | -0.038 | -0.076 | -0.076 | 0 | 0.12 | 0.54 | 0.09 | 0.01 | 0.05 | 0.06 |
| 1 | 173153468 | rs16845543 | 0.028 | -0.094 | 0.067 | -2.659 | 0.021 | 0.005 | 0.005 | 0 | 0.06 | 0.16 | 0.14 | 0.08 | 0.07 | 0.05 |
| 1 | 173156292 | rs3861950 | -1.447 | -0.154 | -0.081 | -2.803 | 0.071 | 0.228 | 0.228 | 0 | 0.44 | 0.87 | 0.38 | 0.12 | 0.32 | 0.26 |
| 1 | 173157201 | rs10158707 | 1.502 | -0.225 | 0.265 | -2.825 | -0.021 | -0.047 | -0.047 | 0.006 | 0.09 | 0.17 | 0.05 | 0 | 0.05 | 0.04 |
| 1 | 173159516 | rs4081545 | -1.483 | -0.195 | -0.096 | -2.817 | 0.072 | 0.229 | 0.229 | 0 | 0.44 | 0.87 | 0.38 | 0.12 | 0.32 | 0.27 |
| 1 | 173163512 | rs7518045 | 0.093 | -0.107 | -0.001 | -2.767 | 0.023 | 0.006 | 0.006 | 0 | 0.06 | 0.16 | 0.14 | 0.08 | 0.07 | 0.05 |
| 1 | 173164565 | rs10912561 | -0.617 | -0.157 | 0.297 | -2.796 | -0.02 | -0.077 | -0.077 | 0 | 0.35 | 0.57 | 0.21 | 0.03 | 0.24 | 0.19 |
| 1 | 173164811 | rs16845585 | 0.029 | 0.086 | -0.139 | -2.658 | 0.02 | 0.004 | 0.004 | 0 | 0.06 | 0.16 | 0.14 | 0.08 | 0.07 | 0.05 |
| 1 | 173166247 | rs1234313 | -2.097 | 0.627 | -0.761 | -2.397 | -0.057 | -0.053 | -0.053 | 0 | 0.75 | 0.89 | 0.57 | 0.35 | 0.69 | 0.56 |
| 1 | 173168397 | rs7525284 | -0.482 | 0.058 | -0.527 | -2.19 | -0.001 | -0.013 | -0.013 | 0 | 0.31 | 0.33 | 0.25 | 0.11 | 0.26 | 0.2 |
| 1 | 173175832 | rs3850641 | -0.872 | 0.543 | -0.411 | -1.302 | -0.045 | -0.032 | -0.032 | 0 | 0.15 | 0.05 | 0.15 | 0.18 | 0.17 | 0.2 |
| 1 | 173178453 | rs10489266 | -0.447 | 0.454 | -0.503 | -1.356 | 0.001 | 0.044 | 0.044 | 0 | 0.13 | 0.04 | 0.05 | 0.03 | 0.1 | 0.11 |
| 1 | 173178463 | rs1234315 | -1.602 | 0.331 | -0.666 | -1.478 | -0.056 | -0.011 | -0.011 | 0.056 | 0.54 | 0.84 | 0.57 | 0.42 | 0.48 | 0.44 |
| 1 | 173184897 | rs13374515 | -0.297 | 0.369 | 0.369 | -1.172 | 0.035 | 0 | 0 | 1.213 | 0.07 | 0.35 | 0.05 | 0 | 0.04 | 0.02 |
| 1 | 173189389 | rs2422334 | -1.271 | 0.443 | 0.134 | -1.065 | 0 | 0.001 | 0.001 | 0 | 0.2 | 0.26 | 0.17 | 0.2 | 0.18 | 0.2 |
| 1 | 173209495 | rs844644 | -1.708 | 0.054 | -0.19 | -1.144 | -0.079 | -0.064 | -0.064 | 0 | 0.57 | 0.87 | 0.63 | 0.43 | 0.53 | 0.48 |
| 1 | 173223863 | rs844648 | -1.659 | 0.868 | 0.874 | -0.509 | 0.004 | 0.008 | 0.008 | 0.796 | 0.39 | 0.5 | 0.52 | 0.45 | 0.41 | 0.4 |
| 1 | 173226195 | rs704840 | -0.5 | 0.914 | 0.625 | -0.496 | 0.005 | 0.016 | 0.016 | 0 | 0.24 | 0.24 | 0.38 | 0.36 | 0.28 | 0.24 |
| 1 | 173236065 | rs10489265 | 0.199 | 0.913 | 0.297 | -0.507 | -0.011 | -0.016 | -0.016 | 0 | 0.19 | 0.08 | 0.33 | 0.23 | 0.23 | 0.19 |
| 1 | 173246262 | rs12144635 | -0.274 | 0.852 | 0.401 | -0.296 | -0.014 | 0.021 | 0.021 | 0 | 0.1 | 0.06 | 0.06 | 0.03 | 0.11 | 0.11 |
| 1 | 173249439 | rs844666 | -1.499 | 0.906 | 0.528 | -0.244 | 0.001 | 0.015 | 0.015 | 0 | 0.08 | 0.13 | 0.1 | 0.18 | 0.07 | 0.1 |
| 1 | 173265908 | rs4916318 | -0.112 | 0.627 | 0.514 | -0.566 | 0.005 | -0.041 | -0.041 | 0 | 0.88 | 0.83 | 0.93 | 0.99 | 0.88 | 0.9 |
| 1 | 173266444 | rs10798267 | -0.112 | 0.627 | 0.514 | -0.566 | 0.005 | -0.041 | -0.041 | 0 | 0.88 | 0.82 | 0.93 | 0.99 | 0.88 | 0.9 |
| 1 | 173266578 | rs4916319 | 1.411 | 0.39 | 0.73 | -0.412 | -0.007 | 0.04 | 0.04 | 3.168 | 0.55 | 0.25 | 0.51 | 0.64 | 0.56 | 0.66 |
| 1 | 173270649 | rs1342032 | 1.096 | 0.785 | 0.354 | 0.015 | 0.007 | -0.002 | -0.002 | 1.144 | 0.64 | 0.45 | 0.57 | 0.64 | 0.62 | 0.67 |
| 1 | 173277164 | rs12691473 | 0.103 | 0.663 | 0.134 | -0.384 | -0.005 | 0.005 | 0.005 | 0.064 | 0.27 | 0.21 | 0.17 | 0.17 | 0.34 | 0.23 |
| 1 | 173278169 | rs6685520 | -0.221 | 0.865 | -0.418 | -0.263 | 0.007 | -0.037 | -0.037 | 0.024 | 0.33 | 0.39 | 0.2 | 0.18 | 0.35 | 0.23 |
| 1 | 173280999 | rs10912583 | -0.203 | 1.366 | -1.753 | 0.04 | -0.018 | 0.026 | 0.026 | 0 | 0.37 | 0.71 | 0.35 | 0.27 | 0.4 | 0.29 |
| 1 | 173286808 | rs10494489 | -0.492 | 1.28 | -1.557 | 0.145 | -0.015 | 0.029 | 0.029 | 0 | 0.1 | 0.06 | 0.07 | 0.01 | 0.11 | 0.1 |
| 1 | 173297704 | rs7546568 | 0.332 | 1.01 | -1.769 | 0.297 | -0.023 | 0.096 | 0.096 | 0 | 0.52 | 0.21 | 0.58 | 0.72 | 0.53 | 0.61 |
| 1 | 173300840 | rs17346536 | -0.307 | 0.62 | -1.79 | 0.716 | 0.031 | 0.029 | 0.029 | 0 | 0.49 | 0.36 | 0.46 | 0.57 | 0.39 | 0.45 |
| 1 | 173301349 | rs12120199 | 0.146 | 0.228 | -2.058 | 0.354 | 0.053 | 0.063 | 0.063 | 0 | 0.33 | 0.24 | 0.14 | 0.13 | 0.2 | 0.26 |
| 1 | 173301516 | rs1342038 | -0.739 | 0.196 | -1.326 | 0.327 | 0.004 | 0.004 | 0.004 | 0 | 0.51 | 0.48 | 0.47 | 0.57 | 0.39 | 0.48 |
| 1 | 173302097 | rs947505 | 0.188 | 0.167 | -0.131 | 0.705 | 0.055 | 0.072 | 0.072 | 0.308 | 0.87 | 0.68 | 0.89 | 0.82 | 0.85 | 0.85 |
| 1 | 173302136 | rs2096148 | -0.334 | -0.093 | -1.235 | -0.161 | -0.001 | 0.032 | 0.032 | 0 | 0.72 | 0.73 | 0.7 | 0.86 | 0.7 | 0.74 |

| | | | | | | | | | | | | | | | | |
|---|---|---|---|---|---|---|---|---|---|---|---|---|---|---|---|---|
| 1 | 173302306 | rs2096147 | -0.112 | -0.107 | -1.08 | -0.676 | 0.01 | -0.013 | -0.013 | 0 | 0.41 | 0.21 | 0.28 | 0.49 | 0.41 | 0.47 |
| 1 | 173302838 | rs1342036 | 0.358 | 0.293 | 0.06 | -1.097 | -0.012 | 0.007 | 0.007 | 0 | 0.9 | 0.69 | 0.95 | 0.9 | 0.93 | 0.97 |
| 1 | 173302872 | rs1342035 | 0.389 | 0.734 | -0.489 | -1.407 | -0.01 | 0.065 | 0.065 | 0 | 0.46 | 0.33 | 0.58 | 0.71 | 0.49 | 0.53 |
| 1 | 173305632 | rs4298771 | -0.251 | 1.506 | 0.031 | -0.558 | -0.003 | 0.008 | 0.008 | 0 | 0.1 | 0.06 | 0.09 | 0.1 | 0.1 | 0.05 |
| 1 | 173309713 | rs10798269 | 1.358 | 1.604 | 0.383 | -0.369 | 0.033 | 0.01 | 0.01 | 0 | 0.7 | 0.54 | 0.68 | 0.6 | 0.66 | 0.67 |
| 1 | 173312501 | rs1539259 | 0.553 | 1.905 | 0.118 | -0.489 | -0.02 | -0.047 | -0.047 | 0 | 0.17 | 0.1 | 0.4 | 0.41 | 0.22 | 0.26 |
| 1 | 173313834 | rs6425217 | 1.263 | 1.66 | -0.013 | -0.235 | 0.026 | 0.008 | 0.008 | 0 | 0.7 | 0.62 | 0.69 | 0.6 | 0.66 | 0.67 |
| 1 | 173315625 | rs17346550 | -0.703 | 1.708 | -0.056 | -0.277 | -0.024 | 0.04 | 0.04 | 0 | 0.22 | 0.11 | 0.24 | 0.11 | 0.25 | 0.17 |
| 1 | 173333829 | rs4916334 | 0.563 | 1.748 | -0.293 | -0.193 | -0.012 | -0.007 | -0.007 | 0.437 | 0.74 | 0.82 | 0.7 | 0.6 | 0.66 | 0.67 |
| 1 | 173362457 | rs1418191 | 0.092 | 0.981 | -0.369 | 0.19 | -0.017 | 0.137 | 0.137 | 0 | 0.17 | 0.31 | 0.35 | 0.4 | 0.12 | 0.23 |
| 1 | 173368454 | rs7517497 | 0.485 | 1.463 | -1.108 | -0.001 | -0.005 | 0.006 | 0.006 | 0 | 0.08 | 0.11 | 0.05 | 0.02 | 0.13 | 0.26 |
| 1 | 173368671 | rs1578624 | 0.662 | 1.404 | -1.098 | 0.142 | -0.038 | -0.01 | -0.01 | 0.717 | 0.68 | 0.78 | 0.64 | 0.57 | 0.57 | 0.7 |
| 1 | 173373183 | rs4916219 | 0.561 | 1.415 | -0.76 | 0.428 | -0.006 | -0.004 | -0.004 | 0 | 0.73 | 0.74 | 0.67 | 0.61 | 0.66 | 0.7 |
| 1 | 173379730 | rs4328106 | 0.809 | 0.828 | -0.432 | 0.696 | -0.005 | 0.047 | 0.047 | 0.988 | 0.7 | 0.44 | 0.49 | 0.51 | 0.75 | 0.72 |
| 1 | 173381535 | rs10753077 | -0.276 | 0.783 | -0.003 | 0.673 | 0.013 | 0.193 | 0.193 | 0 | 0.19 | 0.39 | 0.4 | 0.43 | 0.18 | 0.25 |
| 1 | 173381640 | rs10798280 | 0.992 | 0.791 | -0.174 | 0.671 | -0.003 | 0.015 | 0.015 | 1.116 | 0.7 | 0.38 | 0.49 | 0.51 | 0.75 | 0.72 |
| 1 | 173384422 | rs6681720 | 0.216 | 0.744 | -0.314 | 0.587 | 0 | 0.015 | 0.015 | 3.696 | 0.91 | 0.93 | 0.89 | 0.93 | 0.92 | 0.97 |
| 1 | 173385473 | rs4554754 | 1.079 | 1.051 | -0.125 | 0.808 | 0.004 | 0.034 | 0.034 | 2.584 | 0.85 | 0.76 | 0.84 | 0.67 | 0.83 | 0.85 |
| 1 | 173389640 | rs4294450 | 0.361 | 0.294 | -0.881 | 0.917 | 0.023 | -0.078 | -0.078 | 0 | 0.76 | 0.94 | 0.8 | 0.7 | 0.83 | 0.91 |
| 1 | 173391208 | rs6702339 | -1.169 | -0.048 | 0.17 | 0.822 | 0.034 | 0.207 | 0.207 | 0.46 | 0.08 | 0.34 | 0.08 | 0.28 | 0.09 | 0.12 |
| 1 | 173394054 | rs6425233 | 1.235 | -0.173 | 0.39 | 0.761 | -0.024 | 0.011 | 0.011 | 0.303 | 0.66 | 0.45 | 0.71 | 0.41 | 0.74 | 0.79 |
| 1 | 173396682 | rs10737301 | 0.402 | -0.171 | 0.354 | 0.743 | 0.005 | -0.005 | -0.005 | 0 | 0.74 | 0.78 | 0.8 | 0.7 | 0.83 | 0.91 |
| 1 | 173398436 | rs3934576 | 0.228 | -0.17 | 0.315 | 0.743 | 0.024 | -0.078 | -0.078 | 0.839 | 0.76 | 0.94 | 0.8 | 0.7 | 0.83 | 0.91 |
| 1 | 173399734 | rs10912618 | 1.271 | -0.253 | 0.619 | 0.749 | 0.007 | -0.004 | -0.004 | 0.625 | 0.65 | 0.32 | 0.71 | 0.41 | 0.74 | 0.79 |
| 1 | 173408212 | rs7537938 | -1.53 | -0.258 | 0.664 | 0.798 | -0.029 | 0.015 | 0.015 | 0.18 | 0.34 | 0.53 | 0.28 | 0.59 | 0.26 | 0.21 |
| 1 | 173410444 | rs4539167 | -0.604 | -0.275 | 0.602 | 0.791 | 0.011 | -0.012 | -0.012 | 0 | 0.24 | 0.17 | 0.2 | 0.3 | 0.17 | 0.08 |
| 1 | 173414992 | rs4405161 | 0.552 | 0.241 | 0.147 | -0.32 | -0.062 | 0.091 | 0.091 | 0.032 | 0.23 | 0.06 | 0.25 | 0.06 | 0.34 | 0.27 |
| 1 | 173423799 | rs4916358 | 0.639 | 0.216 | 0.364 | -0.287 | 0.029 | -0.008 | -0.008 | 0 | 0.64 | 0.26 | 0.71 | 0.41 | 0.75 | 0.78 |
| 1 | 173429995 | rs12140760 | 1.331 | 0.233 | 0.383 | -0.333 | -0.007 | 0.014 | 0.014 | 0 | 0.42 | 0.23 | 0.46 | 0.38 | 0.41 | 0.53 |
| 1 | 173437024 | rs10798282 | 0.674 | 0.225 | 0.4 | -0.234 | 0.029 | -0.008 | -0.008 | 0 | 0.64 | 0.26 | 0.71 | 0.42 | 0.75 | 0.78 |
| 1 | 173445678 | rs11576174 | 1.806 | -0.199 | -0.319 | -0.874 | -0.048 | 0.086 | 0.086 | 0 | 0.14 | 0.01 | 0.09 | 0.01 | 0.16 | 0.18 |
| 1 | 173457599 | rs7314 | -0.502 | -0.255 | -0.06 | -0.711 | 0.048 | -0.022 | -0.022 | 0.813 | 0.34 | 0.73 | 0.26 | 0.59 | 0.21 | 0.2 |
| 1 | 173465265 | rs7521536 | 0.115 | -0.279 | 0.332 | -0.726 | -0.007 | 0.002 | 0.002 | 0.308 | 0.28 | 0.35 | 0.23 | 0.3 | 0.17 | 0.09 |
| 1 | 173470105 | rs12065426 | 0.115 | -0.279 | 0.332 | -0.726 | -0.007 | 0.002 | 0.002 | 0.308 | 0.28 | 0.35 | 0.23 | 0.3 | 0.17 | 0.09 |
| 1 | 173474127 | rs1359460 | -0.717 | -0.497 | 0.129 | -1.06 | 0.041 | 0.127 | 0.127 | 0.033 | 0.48 | 0.71 | 0.57 | 0.94 | 0.38 | 0.52 |
| 1 | 173478877 | rs16846052 | -0.548 | -0.462 | 0.45 | -0.896 | 0.058 | 0.054 | 0.054 | 5.418 | 0.06 | 0.5 | 0.05 | 0 | 0 | 0 |
| 1 | 173507194 | rs12043475 | 0.793 | -0.404 | 0.175 | -0.888 | -0.023 | -0.054 | -0.054 | 0 | 0.17 | 0.06 | 0.32 | 0.64 | 0.21 | 0.43 |
| 1 | 173508785 | rs301696 | -1.161 | -0.656 | -0.153 | -1.23 | -0.058 | 0.2 | 0.2 | 0 | 0.71 | 0.98 | 0.82 | 0.99 | 0.72 | 0.76 |
| 1 | 173509365 | rs1925340 | 1.009 | -0.7 | 0.158 | -0.813 | -0.05 | -0.102 | -0.102 | 0.595 | 0.16 | 0.01 | 0.31 | 0.36 | 0.18 | 0.33 |
| 1 | 173516871 | rs16846206 | 0.567 | -0.649 | 0.235 | -0.979 | -0.007 | -0.017 | -0.017 | 0.426 | 0.18 | 0.13 | 0.33 | 0.64 | 0.21 | 0.43 |
| 1 | 173517540 | rs6425247 | -0.415 | -0.613 | 0.228 | -0.96 | 0.078 | 0.068 | 0.068 | 4.389 | 0.05 | 0.47 | 0.05 | 0 | 0 | 0 |
| 1 | 173532805 | rs12027862 | 0.36 | -0.419 | -0.726 | -0.899 | -0.045 | -0.075 | -0.075 | 0 | 0.2 | 0.04 | 0.36 | 0.64 | 0.26 | 0.42 |
| 1 | 173538503 | rs12040255 | 0.949 | -0.705 | -0.95 | -1.084 | 0.002 | -0.104 | -0.104 | 0 | 0.21 | 0.03 | 0.18 | 0.3 | 0.15 | 0.09 |
| 1 | 173538526 | rs12033786 | 0.935 | -0.71 | -0.952 | -1.087 | 0.002 | -0.104 | -0.104 | 0 | 0.21 | 0.02 | 0.18 | 0.3 | 0.15 | 0.09 |
| 1 | 173539032 | rs10912650 | -0.737 | -0.854 | -0.638 | -1.405 | 0.029 | 0.274 | 0.274 | 0.136 | 0.51 | 0.79 | 0.61 | 0.94 | 0.41 | 0.52 |
| 1 | 173546835 | rs2013387 | -0.824 | -0.761 | -0.23 | -1.014 | -0.033 | -0.036 | -0.036 | 3.205 | 0.07 | 0.68 | 0.05 | 0 | 0.01 | 0 |
| 1 | 173574871 | rs6678784 | -1.06 | -0.757 | -0.057 | -1.117 | -0.034 | -0.036 | -0.036 | 1.79 | 0.07 | 0.7 | 0.05 | 0 | 0.01 | 0 |
| 1 | 173579034 | rs12065033 | -1.355 | -0.772 | -0.017 | -1.124 | -0.043 | -0.045 | -0.045 | 0.65 | 0.08 | 0.73 | 0.06 | 0 | 0.01 | 0 |
| 1 | 173596273 | rs12059068 | -1.057 | -0.464 | -0.073 | -1.157 | -0.034 | -0.036 | -0.036 | 1.158 | 0.07 | 0.7 | 0.05 | 0 | 0.01 | 0 |
| 1 | 173603912 | rs10912656 | 0.99 | -0.709 | -0.334 | -1.056 | -0.047 | -0.104 | -0.104 | 0.192 | 0.17 | 0.01 | 0.32 | 0.37 | 0.18 | 0.33 |
| 1 | 173604527 | rs10158034 | -1.01 | -0.549 | -0.177 | -0.901 | -0.034 | -0.036 | -0.036 | 0 | 0.07 | 0.7 | 0.05 | 0 | 0.01 | 0 |
| 1 | 173615965 | rs7539322 | 0.43 | -0.489 | -0.245 | -0.802 | -0.002 | 0.056 | 0.056 | 0 | 0.05 | 0.03 | 0.04 | 0.32 | 0.08 | 0.1 |
| 1 | 173616525 | rs12075071 | -0.507 | -0.746 | -0.62 | -1.145 | 0.065 | 0.002 | 0.002 | 0 | 0.4 | 0.83 | 0.31 | 0.59 | 0.27 | 0.2 |
| 1 | 173672799 | rs6681766 | 0.573 | -0.776 | -0.82 | -1.094 | 0.03 | -0.044 | -0.044 | 0.356 | 0.28 | 0.12 | 0.21 | 0.27 | 0.19 | 0.1 |
| 1 | 173678039 | rs12564699 | 0.26 | 0.018 | -1.073 | -1.194 | 0.011 | -0.037 | -0.037 | 0.096 | 0.33 | 0.13 | 0.27 | 0.28 | 0.27 | 0.24 |
| 1 | 173683954 | rs10798295 | -0.702 | 0.023 | -0.738 | -1.22 | 0.027 | 0.053 | 0.053 | 0 | 0.46 | 0.82 | 0.37 | 0.59 | 0.36 | 0.34 |
| 1 | 173686353 | rs6413827 | -0.229 | 0.005 | -0.36 | -1.238 | -0.01 | -0.001 | -0.001 | 1.714 | 0.39 | 0.56 | 0.3 | 0.28 | 0.28 | 0.24 |
| 1 | 173690603 | rs12080389 | -0.253 | 0.02 | -0.304 | -1.223 | 0.027 | 0.023 | 0.023 | 4.607 | 0.06 | 0.4 | 0.03 | 0 | 0.01 | 0 |
| 1 | 173713130 | rs11578058 | 1.391 | -0.136 | -0.551 | -1.43 | -0.046 | 0.05 | 0.05 | 0 | 0.09 | 0 | 0.09 | 0 | 0.15 | 0.04 |
| 1 | 173746659 | rs7553539 | 1.799 | 0.187 | -0.483 | -0.918 | -0.021 | 0.012 | 0.012 | 0 | 0.11 | 0.03 | 0.13 | 0.05 | 0.16 | 0.18 |
| 1 | 173774952 | rs12561931 | 0.59 | 0.114 | -0.465 | -0.849 | -0.006 | -0.063 | -0.063 | 0 | 0.32 | 0.07 | 0.26 | 0.28 | 0.27 | 0.25 |
| 1 | 173789507 | rs12096832 | -0.143 | -0.339 | -0.35 | -0.799 | 0.028 | 0.025 | 0.025 | 4.607 | 0.06 | 0.4 | 0.03 | 0 | 0.01 | 0 |
| 1 | 173796169 | rs2068871 | 0.877 | -0.282 | -0.409 | -0.744 | -0.002 | 0.048 | 0.048 | 0 | 0.05 | 0.03 | 0.05 | 0.32 | 0.09 | 0.09 |
| 1 | 173813197 | rs3791020 | 0.576 | -0.178 | -0.475 | -0.843 | 0.037 | -0.051 | -0.051 | 0.685 | 0.32 | 0.09 | 0.26 | 0.28 | 0.24 | 0.25 |
| 1 | 173832806 | rs1322771 | -0.045 | -0.141 | -0.385 | -0.814 | 0.034 | 0.03 | 0.03 | 3.872 | 0.06 | 0.4 | 0.03 | 0 | 0.01 | 0 |
| 1 | 173835211 | rs2067079 | 0.523 | -0.365 | -0.684 | -0.752 | 0.014 | -0.093 | -0.093 | 0 | 0.29 | 0.04 | 0.24 | 0.28 | 0.23 | 0.23 |
| 1 | 173840799 | rs941987 | 0.765 | -0.318 | -0.524 | -0.349 | 0 | 0.006 | 0.006 | 0 | 0.09 | 0.05 | 0.21 | 0.34 | 0.09 | 0.1 |
| 1 | 173858524 | rs4652298 | 0.698 | -0.659 | -0.546 | -0.344 | -0.063 | 0.082 | 0.082 | 0.043 | 0.21 | 0.46 | 0.12 | 0.22 | 0.12 | 0.18 |
| 1 | 173858915 | rs6698314 | 1.072 | -0.743 | -0.596 | -0.405 | 0.005 | 0.012 | 0.012 | 4.752 | 0.15 | 0.09 | 0.17 | 0.06 | 0.13 | 0.08 |

| | | | | | | | | | | | | | | | | |
|---|---|---|---|---|---|---|---|---|---|---|---|---|---|---|---|---|
| 1 | 173860182 | rs16846546 | 0.466 | -0.775 | -0.556 | -0.385 | 0.013 | 0.135 | 0.135 | 0 | 0.11 | 0.29 | 0.08 | 0.31 | 0.1 | 0.12 |
| 1 | 173862626 | rs9425431 | 0.979 | -0.747 | -0.519 | -0.382 | -0.063 | 0.082 | 0.082 | 0.043 | 0.21 | 0.46 | 0.12 | 0.22 | 0.12 | 0.18 |
| 1 | 173863499 | rs4652328 | 0.963 | -0.747 | -0.53 | -0.382 | -0.065 | 0.085 | 0.085 | 0.058 | 0.22 | 0.46 | 0.12 | 0.22 | 0.12 | 0.18 |
| 1 | 173864944 | rs7525115 | 0.976 | -0.517 | -1.067 | -0.387 | 0.005 | 0.012 | 0.012 | 4.752 | 0.15 | 0.09 | 0.17 | 0.06 | 0.13 | 0.08 |
| 1 | 173875401 | rs2227612 | 1.884 | -0.606 | -1.287 | -0.285 | 0.016 | -0.028 | -0.028 | 0 | 0.16 | 0.07 | 0.09 | 0.22 | 0.12 | 0.18 |
| 1 | 173878832 | rs5878 | -1.099 | -0.54 | -1.402 | -0.312 | 0.089 | 0.07 | 0.07 | 0.119 | 0.48 | 0.84 | 0.37 | 0.59 | 0.35 | 0.37 |
| 1 | 173878862 | rs5877 | -1.057 | -0.579 | -1.391 | -0.287 | 0.09 | 0.084 | 0.084 | 0.064 | 0.47 | 0.84 | 0.43 | 0.62 | 0.34 | 0.37 |
| 1 | 173880752 | rs2227607 | 1.099 | -0.461 | -1.313 | -0.382 | 0.004 | 0.044 | 0.044 | 5.037 | 0.14 | 0.04 | 0.16 | 0.06 | 0.12 | 0.08 |
| 1 | 173885712 | rs2227592 | 0.853 | -0.46 | -1.355 | -0.239 | -0.001 | 0.001 | 0.001 | 0.087 | 0.1 | 0.06 | 0.06 | 0.31 | 0.1 | 0.12 |
| 1 | 173886216 | rs2227589 | 0.853 | -0.46 | -1.355 | -0.239 | -0.001 | 0.001 | 0.001 | 0.087 | 0.1 | 0.06 | 0.06 | 0.31 | 0.1 | 0.12 |
| 1 | 173892101 | rs1951626 | -0.179 | -0.56 | -1.509 | -0.493 | -0.015 | -0.015 | -0.015 | 0 | 0.28 | 0.18 | 0.31 | 0.4 | 0.33 | 0.33 |
| 1 | 173893018 | rs9425775 | -0.171 | -0.904 | -0.892 | -0.28 | 0.013 | -0.001 | -0.001 | 0 | 0.31 | 0.73 | 0.18 | 0.21 | 0.16 | 0.21 |
| 1 | 173896153 | rs7528380 | 0.312 | -0.649 | -0.838 | -0.112 | -0.016 | 0.002 | 0.002 | 1.112 | 0.24 | 0.74 | 0.15 | 0.2 | 0.12 | 0.18 |
| 1 | 173903737 | rs767053 | 0.085 | -0.678 | -1.212 | -0.119 | 0.032 | -0.013 | -0.013 | 0 | 0.25 | 0.81 | 0.16 | 0.2 | 0.12 | 0.18 |
| 1 | 173907812 | rs6703547 | 0.085 | -0.692 | -1.229 | -0.184 | 0.032 | -0.013 | -0.013 | 0 | 0.25 | 0.81 | 0.15 | 0.2 | 0.12 | 0.18 |
| 1 | 173929735 | rs7413166 | -0.842 | -0.839 | -1.713 | -0.158 | 0.114 | -0.056 | -0.056 | 0 | 0.4 | 0.9 | 0.24 | 0.52 | 0.24 | 0.33 |
| 1 | 173949332 | rs10798301 | 1.454 | -0.056 | -2.122 | -0.603 | -0.08 | -0.112 | -0.112 | 0 | 0.19 | 0.01 | 0.37 | 0.4 | 0.24 | 0.23 |
| 1 | 173987798 | rs10798302 | -0.004 | -0.406 | -1.332 | -0.324 | 0.073 | -0.061 | -0.061 | 0 | 0.32 | 0.86 | 0.21 | 0.53 | 0.18 | 0.26 |
| 1 | 174040594 | rs16846667 | 1.918 | -0.386 | -1.265 | -0.561 | 0.032 | -0.049 | -0.049 | 0 | 0.17 | 0.07 | 0.09 | 0.2 | 0.11 | 0.17 |
| 1 | 174061844 | rs6695327 | 0.55 | -0.351 | -0.626 | -0.604 | -0.032 | 0.042 | 0.042 | 0 | 0.27 | 0.52 | 0.17 | 0.51 | 0.18 | 0.26 |
| 1 | 174113342 | rs6670425 | 0.105 | -0.154 | -0.81 | -1 | -0.008 | -0.057 | -0.057 | 1.593 | 0.28 | 0.4 | 0.31 | 0.06 | 0.27 | 0.26 |
| 1 | 174126742 | rs6681627 | 2.154 | 0.138 | -0.992 | -1.205 | -0.041 | 0.04 | 0.04 | 0 | 0.07 | 0 | 0.1 | 0 | 0.16 | 0.04 |
| 1 | 174128140 | rs10912735 | -0.037 | -0.812 | -0.67 | -0.968 | -0.008 | -0.06 | -0.06 | 0 | 0.28 | 0.4 | 0.31 | 0.06 | 0.27 | 0.26 |
| 1 | 174197801 | rs10489254 | 1.538 | -0.809 | -0.615 | -0.842 | 0 | 0.04 | 0.04 | 0.036 | 0.07 | 0.05 | 0.05 | 0.31 | 0.06 | 0.09 |
| 1 | 174209978 | rs7535987 | -0.064 | -0.809 | -0.61 | -1.033 | -0.008 | -0.059 | -0.059 | 0.707 | 0.28 | 0.4 | 0.31 | 0.06 | 0.28 | 0.26 |
| 1 | 174218110 | rs6666588 | 0.449 | -0.82 | -0.63 | -1.038 | -0.002 | 0.073 | 0.073 | 0.204 | 0.24 | 0.12 | 0.28 | 0.06 | 0.27 | 0.26 |
| 1 | 174238786 | rs12072050 | 0.661 | -0.87 | -0.596 | -1.076 | 0.001 | -0.017 | -0.017 | 0 | 0.18 | 0.18 | 0.26 | 0.06 | 0.23 | 0.22 |
| 1 | 174246712 | rs6670617 | -1.832 | -1.067 | -0.68 | -1.106 | -0.018 | 0.027 | 0.027 | 0 | 0.55 | 0.92 | 0.48 | 0.58 | 0.45 | 0.52 |
| 1 | 174253651 | rs12060460 | 0.711 | -1.066 | -0.677 | -1.095 | 0.001 | -0.014 | -0.014 | 2.707 | 0.18 | 0.18 | 0.26 | 0.06 | 0.24 | 0.22 |
| 1 | 174254588 | rs6697959 | 0.15 | -1.069 | -0.613 | -1.097 | -0.001 | -0.077 | -0.077 | 4.358 | 0.21 | 0.43 | 0.29 | 0.06 | 0.24 | 0.22 |
| 1 | 174258877 | rs4650989 | -1.824 | -1.062 | -0.687 | -1.091 | -0.018 | 0.027 | 0.027 | 0 | 0.55 | 0.92 | 0.48 | 0.58 | 0.45 | 0.52 |
| 1 | 174265291 | rs6425277 | -0.439 | -1.099 | -0.343 | -1.112 | 0.009 | 0.006 | 0.006 | 0.123 | 0.31 | 0.77 | 0.33 | 0.06 | 0.28 | 0.26 |
| 1 | 174266642 | rs7555067 | -1.726 | -1.023 | -0.441 | -1.07 | -0.014 | 0.033 | 0.033 | 0 | 0.54 | 0.92 | 0.48 | 0.58 | 0.45 | 0.52 |
| 1 | 174270720 | rs2142863 | 0.845 | -1.005 | -0.411 | -1.063 | -0.01 | 0.027 | 0.027 | 0 | 0.17 | 0.12 | 0.26 | 0.06 | 0.24 | 0.22 |
| 1 | 174275837 | rs6700482 | 0.074 | -0.963 | -0.396 | -1.043 | -0.004 | -0.089 | -0.089 | 0.289 | 0.22 | 0.46 | 0.29 | 0.06 | 0.24 | 0.22 |
| 1 | 174312813 | rs17301013 | -1.623 | -0.793 | -0.787 | -0.948 | -0.017 | 0.029 | 0.029 | 0 | 0.55 | 0.92 | 0.48 | 0.58 | 0.45 | 0.52 |
| 1 | 174326318 | rs6684688 | -0.208 | -0.771 | -0.395 | -0.976 | -0.006 | -0.054 | -0.054 | 0 | 0.23 | 0.71 | 0.31 | 0.06 | 0.24 | 0.22 |
| 1 | 174343705 | rs10912773 | -0.865 | -0.997 | -0.552 | -0.893 | 0.097 | 0.064 | 0.064 | 0 | 0.48 | 0.85 | 0.43 | 0.27 | 0.38 | 0.44 |
| 1 | 174369767 | rs17373189 | 1.926 | -0.852 | -0.475 | -0.8 | -0.026 | 0.035 | 0.035 | 0.002 | 0.09 | 0.01 | 0.12 | 0.03 | 0.14 | 0.18 |
| 1 | 174400537 | rs12565430 | 1.186 | -0.924 | -0.392 | -0.732 | 0.013 | -0.015 | -0.015 | 0.114 | 0.17 | 0.09 | 0.09 | 0.2 | 0.1 | 0.17 |
| 1 | 174419672 | rs16847018 | 0.478 | -0.672 | -0.105 | -0.674 | 0 | 0.045 | 0.045 | 0 | 0.07 | 0.06 | 0.06 | 0.32 | 0.08 | 0.09 |
| 1 | 174438115 | rs12059051 | -0.272 | -0.693 | 0.267 | -0.792 | 0.069 | 0.071 | 0.071 | 4.555 | 0.05 | 0.51 | 0.04 | 0 | 0 | 0 |
| 1 | 174450715 | rs10912792 | -0.036 | -0.685 | 0.277 | -0.799 | 0.085 | 0.081 | 0.081 | 3.361 | 0.05 | 0.5 | 0.05 | 0 | 0 | 0 |
| 1 | 174468208 | rs1013769 | 0.215 | -0.672 | 0.046 | -0.939 | -0.024 | 0.038 | 0.038 | 0.029 | 0.15 | 0.09 | 0.26 | 0.06 | 0.24 | 0.22 |
| 1 | 174469794 | rs4652500 | -1.262 | -0.916 | 0.085 | -1.086 | 0.007 | -0.001 | -0.001 | 0.538 | 0.3 | 0.76 | 0.33 | 0.06 | 0.28 | 0.26 |
| 1 | 174469905 | rs12071794 | -1.262 | -0.916 | 0.085 | -1.086 | 0.007 | -0.001 | -0.001 | 0 | 0.3 | 0.76 | 0.33 | 0.06 | 0.28 | 0.26 |
| 1 | 174482433 | rs16847121 | 0.472 | -0.945 | 0.044 | -0.976 | -0.001 | 0.025 | 0.025 | 0.047 | 0.08 | 0.05 | 0.06 | 0.32 | 0.08 | 0.09 |
| 1 | 174484657 | rs4652512 | -0.048 | -0.947 | 0.063 | -0.993 | -0.01 | 0.026 | 0.026 | 0 | 0.17 | 0.12 | 0.26 | 0.06 | 0.24 | 0.22 |
| 1 | 174496267 | rs11487399 | -1.223 | -0.961 | 0.072 | -1 | 0.009 | 0.005 | 0.005 | 0.137 | 0.3 | 0.76 | 0.33 | 0.06 | 0.28 | 0.26 |
| 1 | 174497642 | rs12088073 | -1.901 | -1.104 | -0.123 | -1.046 | -0.033 | 0.021 | 0.021 | 0 | 0.56 | 0.92 | 0.49 | 0.59 | 0.47 | 0.52 |
| 1 | 174499977 | rs6425284 | -1.907 | -1.221 | -0.151 | -1.071 | -0.03 | 0.015 | 0.015 | 0 | 0.56 | 0.9 | 0.48 | 0.58 | 0.46 | 0.48 |
| 1 | 174500566 | rs909533 | -0.933 | -1.129 | -0.055 | -1.053 | -0.006 | -0.054 | -0.054 | 0 | 0.23 | 0.71 | 0.31 | 0.06 | 0.24 | 0.22 |
| 1 | 174501836 | rs6681970 | -1.123 | -1.128 | -0.084 | -1.052 | 0.007 | -0.002 | -0.002 | 0 | 0.3 | 0.76 | 0.33 | 0.06 | 0.28 | 0.26 |
| 1 | 174507529 | rs7547843 | 0.744 | -1.113 | -0.125 | -1.018 | -0.001 | 0.034 | 0.034 | 0.061 | 0.07 | 0.05 | 0.06 | 0.32 | 0.08 | 0.09 |
| 1 | 174533089 | rs11802744 | 1.832 | -1.088 | 0.081 | -1.037 | -0.003 | 0.003 | 0.003 | 0 | 0.06 | 0.08 | 0.13 | 0.03 | 0.09 | 0.04 |
| 1 | 174543850 | rs2038025 | -1.354 | -1.09 | 0.067 | -1.033 | 0.091 | 0.059 | 0.059 | 0 | 0.48 | 0.85 | 0.42 | 0.27 | 0.39 | 0.43 |
| 1 | 174546936 | rs6691118 | -1.108 | -1.08 | 0.072 | -1.031 | 0.009 | 0.005 | 0.005 | 0.137 | 0.3 | 0.77 | 0.33 | 0.06 | 0.28 | 0.26 |
| 1 | 174552811 | rs17301853 | 1.327 | -1.068 | 0.072 | -1.026 | -0.026 | 0.035 | 0.035 | 0.002 | 0.09 | 0.01 | 0.12 | 0.03 | 0.14 | 0.18 |
| 1 | 174592050 | rs6664864 | -1.638 | -1.126 | 0.118 | -0.951 | -0.058 | 0.001 | 0.001 | 0 | 0.56 | 0.97 | 0.5 | 0.58 | 0.47 | 0.51 |
| 1 | 174605170 | rs16847271 | 1.344 | -1.134 | 0.146 | -0.959 | 0.018 | -0.084 | -0.084 | 0 | 0.16 | 0.01 | 0.09 | 0.18 | 0.11 | 0.16 |
| 1 | 174611708 | rs3123641 | -0.474 | -1.169 | 0.207 | -1.017 | 0.025 | 0.021 | 0.021 | 5.467 | 0.06 | 0.59 | 0.05 | 0 | 0 | 0 |
| 1 | 174623077 | rs333454 | -0.559 | -1.169 | 0.198 | -1.017 | -0.002 | -0.028 | -0.028 | 0.766 | 0.24 | 0.77 | 0.31 | 0.06 | 0.24 | 0.22 |
| 1 | 174623922 | rs11806996 | 0.673 | -1.16 | 0.187 | -1.013 | -0.009 | 0.024 | 0.024 | 0 | 0.17 | 0.12 | 0.26 | 0.06 | 0.24 | 0.22 |
| 1 | 174681384 | rs332785 | -0.225 | -1.158 | 0.267 | -0.963 | 0.044 | 0.04 | 0.04 | 4.648 | 0.06 | 0.57 | 0.05 | 0 | 0 | 0 |
| 1 | 174715051 | rs16847402 | 1.19 | -1.179 | -0.006 | -0.96 | 0.021 | -0.088 | -0.088 | 3.006 | 0.16 | 0.01 | 0.09 | 0.21 | 0.11 | 0.19 |
| 1 | 174794144 | rs1793316 | 0.156 | -1.082 | 0.177 | -0.978 | -0.007 | -0.065 | -0.065 | 0 | 0.77 | 0.28 | 0.67 | 0.94 | 0.76 | 0.78 |
| 1 | 174818547 | rs3018410 | 1.458 | -1.14 | 0.071 | -0.965 | -0.064 | -0.004 | -0.004 | 0.03 | 0.44 | 0.03 | 0.49 | 0.42 | 0.53 | 0.49 |
| 1 | 174829296 | rs2987878 | -0.052 | -1.008 | 0.247 | -0.932 | 0.082 | 0.078 | 0.078 | 2.99 | 0.05 | 0.51 | 0.05 | 0 | 0 | 0 |
| 1 | 174835745 | rs913686 | -0.33 | -0.958 | -0.17 | -0.907 | -0.007 | -0.069 | -0.069 | 0 | 0.23 | 0.71 | 0.33 | 0.06 | 0.24 | 0.22 |

| | | | | | | | | | | | | | | | | |
|---|---|---|---|---|---|---|---|---|---|---|---|---|---|---|---|---|
| 1 | 174868328 | rs6700434 | 0.718 | -0.908 | -0.355 | -0.774 | -0.001 | 0.037 | 0.037 | 0.005 | 0.07 | 0.05 | 0.06 | 0.31 | 0.07 | 0.08 |
| 1 | 174884161 | rs6688577 | -1.37 | -1.062 | -0.442 | -0.95 | 0.095 | 0.063 | 0.063 | 0.1 | 0.5 | 0.92 | 0.45 | 0.27 | 0.39 | 0.43 |
| 1 | 174896299 | rs6675528 | -0.857 | -0.873 | -0.126 | -1.001 | 0.001 | -0.019 | -0.019 | 0 | 0.31 | 0.77 | 0.35 | 0.08 | 0.28 | 0.26 |
| 1 | 174903882 | rs6684515 | -0.633 | -0.701 | 0.048 | -0.806 | -0.001 | -0.02 | -0.02 | 2.279 | 0.24 | 0.79 | 0.33 | 0.07 | 0.25 | 0.22 |
| 1 | 174937352 | rs12072529 | -0.314 | -0.797 | 0.258 | -0.585 | 0.077 | 0.073 | 0.073 | 5.816 | 0.05 | 0.54 | 0.04 | 0 | 0.01 | 0 |
| 1 | 174969271 | rs1046439 | -2.014 | -0.59 | -0.594 | -0.859 | -0.106 | -0.069 | -0.069 | 0 | 0.69 | 0.98 | 0.62 | 0.62 | 0.69 | 0.67 |
| 1 | 174970505 | rs7537508 | 0.674 | -0.567 | -0.702 | -1.125 | -0.028 | 0.059 | 0.059 | 0 | 0.18 | 0.08 | 0.12 | 0.01 | 0.23 | 0.18 |
| 1 | 174971682 | rs6425310 | -1.995 | -0.549 | -0.68 | -1.052 | -0.106 | -0.069 | -0.069 | 0 | 0.69 | 0.98 | 0.62 | 0.63 | 0.69 | 0.68 |
| 1 | 174980323 | rs1034462 | 0.765 | -0.618 | -0.818 | -1.143 | -0.007 | 0.021 | 0.021 | 0.019 | 0.18 | 0.1 | 0.26 | 0.06 | 0.23 | 0.21 |
| 1 | 174983207 | rs16847706 | 1.072 | -0.788 | -0.828 | -1.008 | 0.015 | -0.025 | -0.025 | 0.78 | 0.16 | 0.07 | 0.1 | 0.23 | 0.11 | 0.17 |
| 1 | 174983633 | rs1058741 | 0.389 | -0.636 | -1.052 | -0.983 | -0.003 | 0.017 | 0.017 | 0 | 0.2 | 0.13 | 0.27 | 0.06 | 0.24 | 0.22 |
| 1 | 174984826 | rs10912867 | 0.729 | -0.554 | -1.045 | -0.949 | -0.037 | 0.085 | 0.085 | 0 | 0.19 | 0.06 | 0.12 | 0.01 | 0.23 | 0.18 |
| 1 | 174984885 | rs10489330 | 0.871 | -0.201 | -0.811 | -0.622 | 0.006 | 0.08 | 0.08 | 0.412 | 0.08 | 0.08 | 0.09 | 0.33 | 0.11 | 0.12 |
| 1 | 174988016 | rs10912870 | -0.325 | -0.372 | -0.839 | -0.742 | 0.026 | 0.029 | 0.029 | 0.482 | 0.42 | 0.78 | 0.26 | 0.24 | 0.34 | 0.35 |
| 1 | 174988736 | rs17302367 | 0.652 | -0.299 | -0.935 | -0.76 | -0.007 | 0.021 | 0.021 | 0.019 | 0.18 | 0.1 | 0.26 | 0.06 | 0.24 | 0.21 |
| 1 | 174988963 | rs10465483 | -0.678 | -0.107 | -0.921 | -0.418 | -0.055 | 0 | 0 | 0 | 0.6 | 0.9 | 0.52 | 0.3 | 0.58 | 0.54 |
| 1 | 174994790 | rs16847730 | 0.303 | -0.051 | -0.654 | -0.073 | -0.001 | 0.03 | 0.03 | 0.257 | 0.08 | 0.06 | 0.09 | 0.33 | 0.11 | 0.11 |
| 1 | 175010010 | rs997985 | -0.847 | -0.299 | -0.274 | -0.481 | 0.001 | 0.013 | 0.013 | 0 | 0.57 | 0.62 | 0.5 | 0.63 | 0.51 | 0.61 |
| 1 | 175022406 | rs10489329 | -0.034 | 0.17 | -0.127 | -0.358 | 0 | 0.001 | 0.001 | 0.118 | 0.22 | 0.24 | 0.34 | 0.32 | 0.25 | 0.31 |
| 1 | 175029932 | rs6672099 | -1.187 | -0.227 | -0.268 | -0.508 | -0.008 | -0.005 | -0.005 | 0.315 | 0.4 | 0.32 | 0.44 | 0.57 | 0.42 | 0.53 |
| 1 | 175033141 | rs6667611 | 0.318 | -0.022 | -0.623 | -0.35 | -0.005 | -0.049 | -0.049 | 0 | 0.18 | 0.06 | 0.1 | 0.25 | 0.16 | 0.22 |
| 1 | 175034437 | rs10798331 | -1.456 | 0.209 | -0.548 | -0.314 | -0.014 | -0.006 | -0.006 | 0.787 | 0.64 | 0.92 | 0.6 | 0.63 | 0.63 | 0.7 |
| 1 | 175041977 | rs6703740 | -0.699 | 0.232 | -0.548 | -0.296 | -0.024 | 0.038 | 0.038 | 0 | 0.53 | 0.87 | 0.52 | 0.38 | 0.51 | 0.49 |
| 1 | 175047382 | rs2269647 | 0.88 | 0.277 | -0.235 | -0.328 | 0.067 | -0.036 | -0.036 | 0 | 0.14 | 0.41 | 0.13 | 0.06 | 0.19 | 0.13 |
| 1 | 175049379 | rs16847812 | 0.771 | 0.023 | -0.874 | -0.161 | -0.04 | -0.061 | -0.061 | 0 | 0.09 | 0.01 | 0.07 | 0.25 | 0.13 | 0.2 |
| 1 | 175054548 | rs1573758 | -0.505 | -0.028 | -0.639 | -0.279 | -0.003 | 0.036 | 0.036 | 0 | 0.52 | 0.75 | 0.51 | 0.38 | 0.51 | 0.48 |
| 1 | 175057762 | rs6696640 | -0.53 | -0.022 | -0.689 | -0.313 | -0.005 | 0.041 | 0.041 | 0 | 0.52 | 0.76 | 0.51 | 0.38 | 0.51 | 0.48 |
| 1 | 175057782 | rs6694071 | -0.53 | -0.022 | -0.71 | -0.313 | -0.005 | 0.042 | 0.042 | 0 | 0.52 | 0.76 | 0.51 | 0.38 | 0.51 | 0.48 |
| 1 | 175083411 | rs7515773 | 0.79 | 0.311 | -0.751 | -0.702 | 0.046 | -0.034 | -0.034 | 0.363 | 0.15 | 0.25 | 0.14 | 0.05 | 0.23 | 0.13 |
| 1 | 175083952 | rs12083818 | 0.688 | 0.094 | -1.084 | -1.039 | -0.008 | -0.135 | -0.135 | 0 | 0.25 | 0.36 | 0.2 | 0.01 | 0.19 | 0.22 |
| 1 | 175086461 | rs6690229 | -1.147 | 0.355 | -1.615 | -0.632 | -0.082 | -0.129 | -0.129 | 0 | 0.62 | 0.96 | 0.55 | 0.42 | 0.58 | 0.6 |
| 1 | 175086965 | rs7546260 | 0.657 | 0.227 | -1.129 | -1.377 | 0.072 | -0.05 | -0.05 | 0 | 0.16 | 0.35 | 0.15 | 0.05 | 0.23 | 0.14 |
| 1 | 175087677 | rs6656284 | -0.176 | -0.493 | -1.87 | -0.82 | -0.079 | 0.327 | 0.327 | 0 | 0.51 | 0.91 | 0.38 | 0.08 | 0.45 | 0.39 |
| 1 | 175087729 | rs6696455 | -0.176 | -0.493 | -1.87 | -0.841 | -0.079 | 0.331 | 0.331 | 0 | 0.51 | 0.91 | 0.38 | 0.07 | 0.45 | 0.39 |
| 1 | 175090094 | rs6701127 | -0.258 | -0.529 | -0.8 | 0.017 | -0.006 | -0.009 | -0.009 | 0 | 0.45 | 0.44 | 0.62 | 0.64 | 0.42 | 0.55 |
| 1 | 175090414 | rs6703913 | -1.159 | -0.774 | -1.297 | -0.095 | 0.023 | -0.049 | -0.049 | 0.016 | 0.79 | 0.97 | 0.91 | 0.74 | 0.86 | 0.78 |
| 1 | 175091151 | rs7521981 | 0.858 | -0.501 | -0.822 | -0.279 | -0.022 | 0.018 | 0.018 | 0.108 | 0.3 | 0.27 | 0.27 | 0.1 | 0.43 | 0.19 |
| 1 | 175092840 | rs2072041 | 0.183 | -1.001 | -0.479 | 0.211 | -0.002 | 0 | 0 | 0.618 | 0.58 | 0.62 | 0.67 | 0.49 | 0.64 | 0.58 |
| 1 | 175102161 | rs12146041 | 0.879 | -1.321 | -0.433 | 0.326 | 0.009 | -0.017 | -0.017 | 0 | 0.26 | 0.23 | 0.39 | 0.39 | 0.2 | 0.38 |
| 1 | 175105553 | rs2072039 | 0.5 | -1.401 | -0.39 | 0.433 | 0.011 | -0.018 | -0.018 | 0 | 0.16 | 0.13 | 0.14 | 0.23 | 0.17 | 0.15 |
| 1 | 175105996 | rs2072036 | 1.037 | -1.518 | -0.164 | 0.565 | -0.01 | 0.031 | 0.031 | 0 | 0.09 | 0.1 | 0.11 | 0.21 | 0.09 | 0.11 |
| 1 | 175107826 | rs2157588 | 1.079 | -1.566 | -0.148 | 0.342 | -0.033 | 0.035 | 0.035 | 0.012 | 0.3 | 0.26 | 0.3 | 0.09 | 0.46 | 0.25 |
| 1 | 175109517 | rs12563833 | 0.742 | -1.681 | -0.325 | 0.415 | 0.021 | -0.038 | -0.038 | 0 | 0.32 | 0.15 | 0.43 | 0.4 | 0.28 | 0.38 |
| 1 | 175109722 | rs1018829 | -0.08 | -1.668 | -0.034 | 0.638 | 0.035 | -0.033 | -0.033 | 0.212 | 0.67 | 0.77 | 0.8 | 0.51 | 0.79 | 0.66 |
| 1 | 175110955 | rs2187958 | 0.349 | -1.333 | 0.185 | 0.414 | -0.011 | 0 | 0 | 0 | 0.25 | 0.23 | 0.13 | 0.28 | 0.15 | 0.25 |
| 1 | 175113118 | rs10489328 | -0.402 | -1.238 | 0.174 | 0.411 | -0.008 | -0.001 | -0.001 | 0 | 0.73 | 0.74 | 0.84 | 0.71 | 0.81 | 0.74 |
| 1 | 175118923 | rs4650704 | -0.825 | -1.196 | -0.317 | 0.454 | 0.032 | -0.029 | -0.029 | 1.851 | 0.77 | 0.9 | 0.89 | 0.65 | 0.84 | 0.75 |
| 1 | 175120715 | rs10912899 | -0.081 | -1.189 | -0.31 | 0.337 | 0.019 | 0.011 | 0.011 | 0.017 | 0.48 | 0.58 | 0.58 | 0.55 | 0.39 | 0.49 |
| 1 | 175125036 | rs6425323 | 0.662 | -0.896 | -0.488 | -0.024 | 0.015 | -0.026 | -0.026 | 0 | 0.28 | 0.33 | 0.31 | 0.1 | 0.45 | 0.24 |
| 1 | 175126644 | rs3737933 | -0.628 | -0.935 | -1.138 | 0.343 | 0.013 | -0.038 | -0.038 | 0.77 | 0.76 | 0.99 | 0.9 | 0.69 | 0.83 | 0.72 |
| 1 | 175129691 | rs2269650 | 0.105 | -1.061 | -0.603 | 0.19 | 0.008 | 0.004 | 0.004 | 0 | 0.41 | 0.34 | 0.55 | 0.52 | 0.36 | 0.42 |
| 1 | 175130204 | rs1057239 | 0.829 | -1.002 | -0.747 | 0.066 | -0.004 | -0.003 | -0.003 | 0.952 | 0.3 | 0.38 | 0.32 | 0.14 | 0.44 | 0.28 |
| 1 | 175135829 | rs2861158 | 0.883 | -1.211 | -1.354 | 0.506 | -0.037 | -0.107 | -0.107 | 2.371 | 0.28 | 0.02 | 0.44 | 0.29 | 0.26 | 0.31 |
| 1 | 175138289 | rs16847872 | 0.781 | -1.241 | -1.326 | 0.48 | 0.029 | -0.057 | -0.057 | 0.178 | 0.23 | 0 | 0.09 | 0.33 | 0.17 | 0.26 |
| 1 | 175142462 | rs3766696 | 0.701 | -0.686 | -0.434 | 0.91 | -0.02 | 0.066 | 0.066 | 0 | 0.12 | 0.18 | 0.11 | 0.22 | 0.1 | 0.11 |
| 1 | 175142638 | rs3766695 | 0.287 | -0.623 | -0.465 | 0.783 | -0.007 | -0.062 | -0.062 | 0 | 0.29 | 0.07 | 0.45 | 0.29 | 0.26 | 0.32 |
| 1 | 175151183 | rs1008459 | 0.481 | -0.265 | -0.404 | 0.82 | -0.014 | -0.052 | -0.052 | 0 | 0.25 | 0.11 | 0.51 | 0.33 | 0.25 | 0.32 |
| 1 | 175157838 | rs3766682 | 0.49 | -0.049 | -0.718 | 1.045 | -0.003 | 0.048 | 0.048 | 0 | 0.12 | 0.19 | 0.05 | 0.21 | 0.13 | 0.12 |
| 1 | 175161849 | rs2272784 | -0.01 | 0.354 | -1.203 | 0.693 | 0.001 | 0.024 | 0.024 | 0.162 | 0.55 | 0.34 | 0.37 | 0.45 | 0.56 | 0.55 |
| 1 | 175162805 | rs3753555 | -0.003 | 0.354 | -1.203 | 0.693 | 0.002 | 0.025 | 0.025 | 0.176 | 0.55 | 0.34 | 0.37 | 0.45 | 0.56 | 0.55 |
| 1 | 175173206 | rs1012876 | -0.056 | 0.631 | -1.569 | 0.895 | 0.021 | -0.04 | -0.04 | 0 | 0.73 | 0.87 | 0.6 | 0.68 | 0.79 | 0.77 |
| 1 | 175183939 | rs6659761 | -0.444 | 1.619 | -1.184 | 0.95 | -0.036 | -0.029 | -0.029 | 0 | 0.32 | 0.22 | 0.29 | 0.48 | 0.5 | 0.44 |
| 1 | 175187551 | rs10912915 | -0.201 | 2.322 | -1.073 | 1.04 | 0.075 | 0.02 | 0.02 | 0 | 0.13 | 0.17 | 0.15 | 0.16 | 0.3 | 0.19 |
| 1 | 175194390 | rs10798348 | -0.041 | 1.163 | -1.524 | 0.807 | 0.02 | -0.029 | -0.029 | 0 | 0.71 | 0.94 | 0.81 | 0.67 | 0.75 | 0.72 |
| 1 | 175198598 | rs6425326 | 0.037 | 1.209 | -1.494 | 0.787 | 0.031 | 0.001 | 0.001 | 0 | 0.7 | 0.85 | 0.79 | 0.7 | 0.73 | 0.72 |
| 1 | 175198786 | rs909467 | 0.339 | 2.064 | -1.375 | 0.836 | 0.012 | 0.004 | 0.004 | 0 | 0.11 | 0.1 | 0.1 | 0.17 | 0.21 | 0.15 |
| 1 | 175201727 | rs2012862 | 1.244 | 1.778 | -1.354 | 0.759 | 0 | -0.029 | -0.029 | 0.003 | 0.41 | 0.5 | 0.34 | 0.23 | 0.37 | 0.37 |
| 1 | 175201902 | rs998470 | -0.35 | 1.915 | -1.736 | 0.853 | -0.032 | -0.064 | -0.064 | 0 | 0.28 | 0.12 | 0.42 | 0.47 | 0.36 | 0.32 |
| 1 | 175202015 | rs3946168 | 0.448 | 1.942 | -1.268 | 0.764 | -0.028 | -0.056 | -0.056 | 1.443 | 0.2 | 0.02 | 0.16 | 0.29 | 0.21 | 0.25 |

| | | | | | | | | | | | | | | | | |
|---|---|---|---|---|---|---|---|---|---|---|---|---|---|---|---|---|
| 1 | 175218819 | rs4651418 | 0.904 | 1.897 | -1.17 | 0.496 | -0.002 | 0.018 | 0.018 | 0.307 | 0.58 | 0.72 | 0.71 | 0.62 | 0.54 | 0.49 |
| 1 | 175222943 | rs7540956 | 0.205 | 1.895 | -1.228 | 0.482 | 0.029 | 0.052 | 0.052 | 0 | 0.71 | 0.93 | 0.81 | 0.79 | 0.77 | 0.68 |
| 1 | 175233237 | rs6663906 | 1.11 | 1.925 | -1.177 | 0.469 | -0.001 | 0.009 | 0.009 | 0.043 | 0.58 | 0.62 | 0.7 | 0.62 | 0.54 | 0.49 |
| 1 | 175239758 | rs10157949 | -1.26 | 1.973 | -1.558 | 0.493 | 0.039 | 0.046 | 0.046 | 0.228 | 0.9 | 0.96 | 0.95 | 0.99 | 0.95 | 0.97 |
| 1 | 175241835 | rs12058559 | 0.355 | 1.988 | -1.529 | 0.5 | 0.005 | 0.011 | 0.011 | 0 | 0.2 | 0.08 | 0.14 | 0.2 | 0.18 | 0.29 |
| 1 | 175254593 | rs6674359 | 0.161 | 1.877 | -1.46 | 0.509 | -0.004 | -0.004 | -0.004 | 0 | 0.11 | 0.13 | 0.11 | 0.17 | 0.23 | 0.19 |
| 1 | 175257650 | rs1546844 | 0.974 | 1.919 | -1.464 | 0.502 | -0.002 | 0.025 | 0.025 | 0.178 | 0.59 | 0.75 | 0.71 | 0.62 | 0.55 | 0.49 |
| 1 | 175258634 | rs12744325 | 0.295 | 1.666 | -1.45 | 0.516 | -0.008 | -0.007 | -0.007 | 0 | 0.11 | 0.09 | 0.1 | 0.17 | 0.22 | 0.19 |
| 1 | 175268094 | rs11582340 | 0.342 | 2.146 | -1.397 | 0.582 | -0.049 | 0.008 | 0.008 | 0 | 0.2 | 0.04 | 0.16 | 0.2 | 0.24 | 0.3 |
| 1 | 175269065 | rs10912950 | 0.329 | 1.88 | -1.162 | 0.72 | -0.026 | 0.031 | 0.031 | 0.768 | 0.58 | 0.74 | 0.69 | 0.62 | 0.48 | 0.48 |
| 1 | 175283119 | rs12732088 | -0.758 | 2.135 | -0.218 | 0.801 | 0.092 | 0.081 | 0.081 | 0 | 0.15 | 0.25 | 0.24 | 0.28 | 0.29 | 0.33 |
| 1 | 175283481 | rs12133267 | -0.758 | 2.135 | -0.218 | 0.801 | 0.092 | 0.081 | 0.081 | 0 | 0.15 | 0.25 | 0.24 | 0.28 | 0.29 | 0.33 |
| 1 | 175286396 | rs10798380 | -0.251 | 1.949 | 0.157 | 0.707 | 0.043 | 0.143 | 0.143 | 1.498 | 0.15 | 0.17 | 0.37 | 0.49 | 0.25 | 0.35 |
| 1 | 175289399 | rs875326 | -0.384 | 1.447 | 0.491 | -0.107 | 0.06 | -0.1 | -0.1 | 0 | 0.16 | 0.53 | 0.14 | 0.03 | 0.16 | 0.07 |
| 1 | 175290566 | rs10489323 | -0.069 | 1.384 | 1.19 | -0.139 | 0.024 | -0.044 | -0.044 | 0.953 | 0.12 | 0.2 | 0.11 | 0.03 | 0.15 | 0.06 |
| 1 | 175290855 | rs11802001 | -0.078 | 1.385 | 1.203 | -0.139 | 0.025 | -0.045 | -0.045 | 0.027 | 0.12 | 0.21 | 0.11 | 0.03 | 0.15 | 0.06 |
| 1 | 175293424 | rs2213635 | -1.274 | 0.545 | 0.097 | -0.92 | 0.026 | 0.05 | 0.05 | 0 | 0.67 | 0.84 | 0.76 | 0.79 | 0.72 | 0.69 |
| 1 | 175294275 | rs7551235 | 0.764 | 0.576 | 0.182 | -0.919 | 0.026 | -0.037 | -0.037 | 0 | 0.11 | 0.16 | 0.11 | 0.03 | 0.15 | 0.06 |
| 1 | 175295662 | rs12753536 | 1.671 | 0.147 | -0.233 | -1.09 | 0.051 | -0.017 | -0.017 | 0 | 0.17 | 0.01 | 0.04 | 0.09 | 0.08 | 0.03 |
| 1 | 175298552 | rs6688599 | -1.443 | 0.098 | -0.001 | -1.122 | -0.008 | -0.016 | -0.016 | 0 | 0.64 | 0.6 | 0.74 | 0.79 | 0.71 | 0.67 |
| 1 | 175298757 | rs10489321 | -0.498 | 0.091 | 0.096 | -1.133 | 0.014 | 0.188 | 0.188 | 0.963 | 0.5 | 0.14 | 0.61 | 0.75 | 0.56 | 0.62 |
| 1 | 175308725 | rs2298918 | -0.021 | 0.082 | 0.175 | -1.109 | -0.097 | -0.024 | -0.024 | 0 | 0.32 | 0.09 | 0.56 | 0.66 | 0.47 | 0.57 |
| 1 | 175315600 | rs4650920 | -1.18 | 0.05 | 0.215 | -1.113 | -0.022 | -0.034 | -0.034 | 0 | 0.62 | 0.47 | 0.73 | 0.76 | 0.71 | 0.66 |
| 1 | 175321003 | rs2040425 | -0.079 | 0.149 | 0.027 | -1.023 | -0.061 | -0.009 | -0.009 | 0.888 | 0.33 | 0.15 | 0.57 | 0.66 | 0.47 | 0.57 |
| 1 | 175322695 | rs7547487 | -0.133 | -0.173 | 0.038 | -1.315 | 0.037 | 0.086 | 0.086 | 0 | 0.27 | 0.42 | 0.46 | 0.6 | 0.35 | 0.49 |
| 1 | 175322842 | rs16848281 | 1.591 | -0.053 | 0.058 | -1.378 | -0.003 | -0.004 | -0.004 | 0 | 0.08 | 0.07 | 0.07 | 0 | 0.09 | 0.05 |
| 1 | 175324651 | rs2228359 | -1.011 | -0.227 | -0.046 | -1.539 | -0.006 | -0.03 | -0.03 | 0.098 | 0.62 | 0.52 | 0.71 | 0.79 | 0.68 | 0.68 |
| 1 | 175325205 | rs723640 | 1.312 | -0.171 | -0.186 | -1.476 | 0.007 | 0.025 | 0.025 | 0 | 0.1 | 0.04 | 0.06 | 0.09 | 0.1 | 0.17 |
| 1 | 175326303 | rs10798390 | -0.878 | -0.239 | -0.118 | -1.536 | -0.006 | 0.033 | 0.033 | 0.101 | 0.54 | 0.51 | 0.58 | 0.69 | 0.52 | 0.58 |
| 1 | 175327147 | rs10489318 | -0.182 | -0.141 | -0.347 | -1.476 | -0.034 | 0.057 | 0.057 | 0 | 0.29 | 0.19 | 0.26 | 0.2 | 0.34 | 0.32 |
| 1 | 175338419 | rs2236883 | -0.144 | -0.576 | -0.425 | -1.548 | 0.018 | 0.002 | 0.002 | 1.297 | 0.34 | 0.41 | 0.57 | 0.61 | 0.49 | 0.48 |
| 1 | 175339021 | rs2236881 | -1.046 | -0.607 | -0.424 | -1.592 | -0.013 | -0.047 | -0.047 | 0.117 | 0.62 | 0.53 | 0.71 | 0.8 | 0.7 | 0.68 |
| 1 | 175341548 | rs16848353 | 1.87 | -0.59 | -0.679 | -1.527 | 0.032 | -0.02 | -0.02 | 0 | 0.17 | 0 | 0.05 | 0.09 | 0.09 | 0.04 |
| 1 | 175343297 | rs1351963 | -0.52 | -0.427 | -0.423 | -1.482 | 0.005 | 0.014 | 0.014 | 1.037 | 0.5 | 0.5 | 0.36 | 0.28 | 0.44 | 0.35 |
| 1 | 175348723 | rs859427 | -0.036 | -0.365 | -0.425 | -1.459 | 0.003 | -0.018 | -0.018 | 0 | 0.7 | 0.65 | 0.71 | 0.81 | 0.65 | 0.69 |
| 1 | 175352089 | rs4652084 | 1.992 | -0.629 | -0.686 | -1.704 | -0.01 | -0.034 | -0.034 | 4.946 | 0.09 | 0.02 | 0.1 | 0.2 | 0.08 | 0.2 |
| 1 | 175352218 | rs859433 | -1.381 | -0.179 | -0.388 | -1.419 | 0.014 | -0.016 | -0.016 | 0 | 0.83 | 0.75 | 0.79 | 0.88 | 0.76 | 0.84 |
| 1 | 175355391 | rs859437 | -1.581 | -0.209 | -0.329 | -1.417 | 0.009 | -0.015 | -0.015 | 0 | 0.84 | 0.8 | 0.86 | 0.89 | 0.77 | 0.85 |
| 1 | 175355618 | rs859438 | 0.027 | -0.165 | -0.328 | -1.408 | 0.003 | -0.025 | -0.025 | 0.571 | 0.69 | 0.64 | 0.77 | 0.81 | 0.65 | 0.69 |
| 1 | 175361190 | rs2021832 | 0.675 | -0.035 | -0.469 | -1.327 | -0.002 | 0.048 | 0.048 | 0 | 0.14 | 0.04 | 0.07 | 0.07 | 0.14 | 0.18 |
| 1 | 175362638 | rs10912968 | 0.53 | 0.087 | -1.02 | -1.27 | 0.069 | 0.107 | 0.107 | 1.386 | 0.22 | 0.34 | 0.44 | 0.52 | 0.34 | 0.38 |
| 1 | 175362736 | rs3752517 | -0.048 | 0.519 | -1.573 | -1.017 | -0.007 | 0.012 | 0.012 | 0 | 0.69 | 0.7 | 0.78 | 0.81 | 0.64 | 0.68 |
| 1 | 175363204 | rs12023965 | 0.219 | 0.48 | -1.589 | -1.037 | 0 | 0.016 | 0.016 | 0 | 0.43 | 0.55 | 0.51 | 0.6 | 0.43 | 0.41 |
| 1 | 175363509 | rs9425442 | 1.288 | 0.573 | -1.895 | -0.965 | 0.024 | 0.032 | 0.032 | 0 | 0.27 | 0.15 | 0.26 | 0.21 | 0.21 | 0.27 |
| 1 | 175364423 | rs6691883 | 1.288 | 0.557 | -1.895 | -0.965 | 0.019 | 0.032 | 0.032 | 0 | 0.27 | 0.15 | 0.26 | 0.21 | 0.22 | 0.27 |
| 1 | 175374390 | rs859401 | 0.517 | 1.678 | -1.589 | -0.201 | -0.015 | 0.03 | 0.03 | 0 | 0.7 | 0.76 | 0.78 | 0.82 | 0.63 | 0.65 |
| 1 | 175375469 | rs2239819 | 0.839 | 1.736 | -1.583 | -0.135 | -0.008 | 0.011 | 0.011 | 1.014 | 0.12 | 0.09 | 0.36 | 0.52 | 0.25 | 0.3 |
| 1 | 175375683 | rs859399 | 0.969 | 1.002 | -1.477 | -0.652 | -0.014 | 0.028 | 0.028 | 0 | 0.7 | 0.76 | 0.78 | 0.82 | 0.63 | 0.65 |
| 1 | 175375802 | rs859398 | 0.967 | 1.023 | -1.467 | -0.642 | -0.008 | 0.011 | 0.011 | 0 | 0.7 | 0.74 | 0.78 | 0.82 | 0.63 | 0.65 |
| 1 | 175378694 | rs12119177 | 0.004 | 1.228 | -1.349 | -0.227 | 0.008 | -0.014 | -0.014 | 0 | 0.83 | 0.8 | 0.86 | 0.89 | 0.78 | 0.87 |
| 1 | 175386594 | rs1485382 | -0.855 | 0.496 | -1.72 | -0.002 | -0.002 | 0.027 | 0.027 | 0 | 0.2 | 0.1 | 0.14 | 0.15 | 0.19 | 0.28 |
| 1 | 175387030 | rs12756192 | -0.176 | -0.041 | -2.2 | 0.281 | 0.001 | -0.037 | -0.037 | 1.319 | 0.21 | 0 | 0.09 | 0.15 | 0.14 | 0.05 |
| 1 | 175391741 | rs6666777 | -0.515 | -0.331 | -1.748 | 0.842 | -0.007 | 0.02 | 0.02 | 0 | 0.24 | 0.22 | 0.13 | 0.5 | 0.21 | 0.24 |
| 1 | 175394522 | rs984872 | 0.001 | 0.106 | -1.329 | 0.173 | 0.043 | -0.09 | -0.09 | 0 | 0.16 | 0.44 | 0.12 | 0.05 | 0.18 | 0.18 |
| 1 | 175403509 | rs6655978 | 0.401 | -0.102 | -1.404 | 0.819 | 0.005 | -0.018 | -0.018 | 0 | 0.1 | 0.04 | 0.03 | 0.27 | 0.09 | 0.06 |
| 1 | 175404089 | rs10912978 | 0.613 | 0.626 | -1.628 | 0.104 | -0.026 | 0.04 | 0.04 | 0 | 0.07 | 0.01 | 0.04 | 0 | 0.1 | 0.03 |
| 1 | 175411978 | rs12565878 | 0.401 | 0.292 | -1.878 | -0.238 | 0.017 | -0.016 | -0.016 | 0 | 0.37 | 0.24 | 0.2 | 0.51 | 0.27 | 0.25 |
| 1 | 175413886 | rs12735331 | 0.125 | 0.268 | -1.765 | 0.025 | 0.035 | -0.014 | -0.014 | 0 | 0.16 | 0.48 | 0.25 | 0.12 | 0.18 | 0.25 |
| 1 | 175416603 | rs859460 | -0.831 | 1.074 | -2.432 | -0.296 | -0.062 | -0.008 | -0.008 | 0 | 0.76 | 0.93 | 0.5 | 0.83 | 0.69 | 0.75 |
| 1 | 175417063 | rs4652090 | 0.158 | 0.563 | -2.119 | -0.245 | 0.062 | -0.015 | -0.015 | 0.041 | 0.43 | 0.67 | 0.26 | 0.55 | 0.32 | 0.3 |
| 1 | 175421100 | rs10912983 | 0.049 | 0.658 | -1.819 | -0.289 | 0.009 | 0.011 | 0.011 | 0 | 0.45 | 0.32 | 0.4 | 0.58 | 0.38 | 0.39 |
| 1 | 175422209 | rs10753106 | 0.3 | 0.78 | -1.214 | -0.402 | -0.013 | -0.036 | -0.036 | 0 | 0.69 | 0.42 | 0.78 | 0.81 | 0.69 | 0.61 |
| 1 | 175423386 | rs860906 | -0.738 | 0.989 | -1.48 | -0.024 | -0.015 | 0.022 | 0.022 | 2.941 | 0.53 | 0.9 | 0.64 | 0.37 | 0.58 | 0.63 |
| 1 | 175423905 | rs859455 | -0.744 | 0.99 | -1.48 | -0.024 | -0.016 | 0.021 | 0.021 | 2.935 | 0.53 | 0.9 | 0.64 | 0.37 | 0.58 | 0.63 |
| 1 | 175425083 | rs863519 | -1.021 | 0.761 | 0.203 | -0.249 | 0.071 | 0.05 | 0.05 | 1.094 | 0.05 | 0.55 | 0.06 | 0 | 0.01 | 0 |
| 1 | 175428957 | rs1016815 | -0.682 | 1.012 | -0.816 | -0.287 | 0.002 | 0.028 | 0.028 | 2.892 | 0.52 | 0.63 | 0.6 | 0.7 | 0.59 | 0.58 |
| 1 | 175430855 | rs7547439 | 0.287 | 1.262 | -1.69 | 0.067 | -0.031 | -0.047 | -0.047 | 0 | 0.11 | 0.04 | 0.32 | 0.55 | 0.19 | 0.21 |
| 1 | 175431064 | rs859374 | -1.058 | 0.896 | -1.416 | -0.289 | -0.066 | 0.046 | 0.046 | 3.48 | 0.61 | 0.96 | 0.65 | 0.71 | 0.65 | 0.63 |
| 1 | 175436833 | rs10912985 | -0.439 | 0.858 | -0.791 | -0.325 | -0.004 | 0.01 | 0.01 | 2.279 | 0.44 | 0.4 | 0.56 | 0.65 | 0.56 | 0.56 |

| | | | | | | | | | | | | | | | | |
|---|---|---|---|---|---|---|---|---|---|---|---|---|---|---|---|---|
| 1 | 175439176 | rs859369 | -0.92 | 0.652 | -0.986 | -0.504 | 0.039 | 0.082 | 0.082 | 3.468 | 0.55 | 0.81 | 0.7 | 0.69 | 0.64 | 0.53 |
| 1 | 175440469 | rs7548041 | 0.441 | 1.248 | -0.657 | -0.544 | -0.002 | -0.003 | -0.003 | 0 | 0.1 | 0.1 | 0.16 | 0.02 | 0.17 | 0.19 |
| 1 | 175442871 | rs12074467 | -0.344 | 1.324 | -0.017 | -0.415 | 0.023 | -0.021 | -0.021 | 0 | 0.18 | 0.31 | 0.21 | 0.11 | 0.21 | 0.23 |
| 1 | 175444933 | rs859413 | -1.745 | 1.086 | -0.188 | -0.586 | -0.007 | 0.072 | 0.072 | 0.997 | 0.56 | 0.84 | 0.67 | 0.68 | 0.57 | 0.53 |
| 1 | 175461844 | rs859395 | -1.272 | 1.323 | 0.019 | -0.605 | -0.01 | 0.036 | 0.036 | 0 | 0.36 | 0.73 | 0.33 | 0.13 | 0.4 | 0.34 |
| 1 | 175469011 | rs2861311 | -1.332 | 1.229 | -0.022 | -0.475 | -0.008 | 0.006 | 0.006 | 0.505 | 0.41 | 0.35 | 0.61 | 0.62 | 0.53 | 0.46 |
| 1 | 175476069 | rs859382 | -1.417 | 1.276 | 0.143 | -0.438 | -0.001 | 0.018 | 0.018 | 0.883 | 0.4 | 0.71 | 0.58 | 0.64 | 0.41 | 0.45 |
| 1 | 175476197 | rs12081663 | -0.214 | 1.488 | 0.368 | -0.287 | 0.014 | -0.024 | -0.024 | 0 | 0.2 | 0.31 | 0.22 | 0.1 | 0.24 | 0.22 |
| 1 | 175484720 | rs859353 | -0.892 | 0.914 | 0.597 | -0.063 | 0.018 | -0.05 | -0.05 | 0 | 0.15 | 0.32 | 0.14 | 0.05 | 0.18 | 0.14 |
| 1 | 175488409 | rs10912994 | -0.16 | 1.15 | 0.417 | 0.05 | -0.003 | -0.003 | -0.003 | 0 | 0.32 | 0.3 | 0.54 | 0.59 | 0.38 | 0.37 |
| 1 | 175492359 | rs864163 | -1.394 | 1.133 | 0.108 | 0.045 | 0.017 | 0.058 | 0.058 | 1.642 | 0.48 | 0.79 | 0.66 | 0.62 | 0.56 | 0.5 |
| 1 | 175493914 | rs859358 | -0.764 | 1.093 | 0.173 | -0.015 | 0.011 | -0.054 | -0.054 | 0.406 | 0.14 | 0.26 | 0.11 | 0.01 | 0.18 | 0.14 |
| 1 | 175494230 | rs859359 | -1.048 | 1.091 | 0.218 | -0.014 | 0.009 | -0.052 | -0.052 | 0.133 | 0.16 | 0.35 | 0.11 | 0.03 | 0.18 | 0.14 |
| 1 | 175495090 | rs859362 | -1.064 | 1.09 | 0.217 | -0.012 | 0.008 | -0.035 | -0.035 | 0 | 0.16 | 0.34 | 0.13 | 0.07 | 0.18 | 0.14 |
| 1 | 175495225 | rs859363 | -0.653 | 1.101 | 0.21 | -0.04 | 0.013 | -0.037 | -0.037 | 0.628 | 0.14 | 0.26 | 0.13 | 0.05 | 0.18 | 0.14 |
| 1 | 175505966 | rs7515041 | -1.122 | 1.101 | -0.451 | -0.208 | 0.019 | 0.032 | 0.032 | 0.152 | 0.53 | 0.43 | 0.67 | 0.7 | 0.63 | 0.57 |
| 1 | 175518442 | rs6667463 | 0.602 | 1.171 | -0.845 | -0.096 | -0.007 | -0.037 | -0.037 | 0 | 0.13 | 0.03 | 0.08 | 0.18 | 0.13 | 0.18 |
| 1 | 175521376 | rs1003964 | -1.015 | 1.048 | -0.594 | -0.554 | -0.002 | -0.005 | -0.005 | 0 | 0.6 | 0.62 | 0.75 | 0.72 | 0.68 | 0.65 |
| 1 | 175526526 | rs10489312 | 0.133 | 1.19 | -0.536 | -0.331 | 0.005 | -0.025 | -0.025 | 3.857 | 0.11 | 0.13 | 0.09 | 0 | 0.17 | 0.07 |
| 1 | 175526544 | rs6676780 | -0.062 | 1.096 | -0.647 | -0.122 | 0.004 | -0.014 | -0.014 | 0 | 0.41 | 0.45 | 0.64 | 0.69 | 0.44 | 0.48 |
| 1 | 175528029 | rs4652097 | -0.367 | 0.703 | -0.881 | 0.048 | 0.011 | 0.028 | 0.028 | 0.046 | 0.57 | 0.68 | 0.74 | 0.71 | 0.65 | 0.64 |
| 1 | 175528957 | rs7540211 | -0.503 | 0.266 | -1.104 | 0.818 | 0.002 | -0.052 | -0.052 | 0.526 | 0.23 | 0.14 | 0.14 | 0.43 | 0.21 | 0.33 |
| 1 | 175531634 | rs10913008 | 0.426 | 0.045 | -0.987 | 0.508 | 0.007 | -0.014 | -0.014 | 0 | 0.12 | 0.08 | 0.08 | 0.13 | 0.09 | 0.13 |
| 1 | 175534602 | rs17305076 | -0.281 | -0.055 | 0.228 | -0.147 | -0.001 | -0.02 | -0.02 | 0 | 0.29 | 0.29 | 0.29 | 0.03 | 0.34 | 0.3 |
| 1 | 175536944 | rs6693370 | 0.275 | -0.241 | 0.706 | 0.471 | 0.003 | 0 | 0 | 1.183 | 0.12 | 0.14 | 0.1 | 0.08 | 0.09 | 0.13 |
| 1 | 175539955 | rs6657672 | 0.031 | -0.341 | 0.759 | 0.45 | -0.025 | 0.049 | 0.049 | 0.206 | 0.13 | 0.34 | 0.11 | 0.13 | 0.07 | 0.13 |
| 1 | 175542257 | rs7354864 | 0.031 | -0.341 | 0.754 | 0.461 | -0.025 | 0.05 | 0.05 | 0.206 | 0.13 | 0.34 | 0.11 | 0.14 | 0.07 | 0.13 |
| 1 | 175552365 | rs10913019 | 0.136 | -0.231 | 0.94 | 0.453 | -0.013 | 0.024 | 0.024 | 0.061 | 0.13 | 0.27 | 0.1 | 0.15 | 0.09 | 0.13 |
| 1 | 175561555 | rs6688098 | -0.097 | -0.484 | 0.739 | 0.105 | -0.039 | -0.077 | -0.077 | 0 | 0.16 | 0.4 | 0.1 | 0.04 | 0.09 | 0.14 |
| 1 | 175561612 | rs10913025 | 0.128 | -0.484 | 0.733 | 0.105 | -0.031 | -0.067 | -0.067 | 0.884 | 0.15 | 0.35 | 0.1 | 0.04 | 0.09 | 0.14 |
| 1 | 175564860 | rs6690188 | -0.199 | -0.497 | 0.54 | 0.115 | -0.047 | -0.084 | -0.084 | 0 | 0.17 | 0.47 | 0.1 | 0.04 | 0.09 | 0.14 |
| 1 | 175567218 | rs7541077 | -0.612 | -1.26 | 0.14 | 0.465 | 0.01 | 0.001 | 0.001 | 3.57 | 0.37 | 0.63 | 0.19 | 0.43 | 0.19 | 0.33 |
| 1 | 175576828 | rs6692370 | -1.626 | -0.851 | -0.259 | 0.324 | -0.201 | -0.157 | -0.157 | 0.819 | 0.69 | 0.98 | 0.51 | 0.57 | 0.58 | 0.66 |
| 1 | 175578177 | rs10913029 | -0.245 | -0.733 | 0.069 | 0.406 | 0 | -0.001 | -0.001 | 0 | 0.26 | 0.73 | 0.16 | 0.43 | 0.19 | 0.2 |
| 1 | 175581072 | rs7546740 | -1.28 | -0.763 | 0.956 | 0.326 | 0.154 | 0.064 | 0.064 | 0.932 | 0.06 | 0.45 | 0.07 | 0.01 | 0.08 | 0.04 |
| 1 | 175581146 | rs7515728 | -1.306 | -0.983 | 0.996 | 0.174 | 0.025 | -0.047 | -0.047 | 0 | 0.42 | 0.83 | 0.38 | 0.51 | 0.39 | 0.36 |
| 1 | 175582788 | rs1359047 | -1.217 | -0.898 | 1.016 | 0.207 | 0.151 | 0.061 | 0.061 | 1.08 | 0.06 | 0.45 | 0.07 | 0.01 | 0.08 | 0.04 |
| 1 | 175585312 | rs2012430 | -1.176 | -0.962 | 0.694 | 0.182 | 0.01 | -0.033 | -0.033 | 0 | 0.4 | 0.75 | 0.37 | 0.51 | 0.39 | 0.36 |
| 1 | 175585503 | rs2179158 | -0.377 | -1.029 | 0.661 | 0.122 | 0.009 | -0.063 | -0.063 | 0.432 | 0.32 | 0.16 | 0.27 | 0.47 | 0.31 | 0.31 |
| 1 | 175590376 | rs10798409 | -1.161 | -1.064 | 0.35 | 0.148 | 0.008 | -0.015 | -0.015 | 0 | 0.39 | 0.71 | 0.37 | 0.48 | 0.39 | 0.36 |
| 1 | 175593305 | rs12037583 | 0.891 | -1.176 | -0.059 | 0.356 | 0.022 | -0.104 | -0.104 | 0 | 0.16 | 0 | 0.05 | 0.39 | 0.11 | 0.16 |
| 1 | 175594438 | rs10798411 | -0.65 | -1.297 | 0.512 | 0.262 | -0.068 | 0.023 | 0.023 | 0.171 | 0.29 | 0.57 | 0.14 | 0.42 | 0.16 | 0.3 |
| 1 | 175600179 | rs1323120 | -0.824 | -1.255 | 0.725 | 0.375 | -0.038 | 0.025 | 0.025 | 0 | 0.35 | 0.51 | 0.18 | 0.49 | 0.21 | 0.31 |
| 1 | 175604381 | rs17311923 | 0.515 | -1.262 | 0.756 | 0.275 | -0.002 | -0.009 | -0.009 | 0 | 0.08 | 0.07 | 0.06 | 0.07 | 0.09 | 0.03 |
| 1 | 175606606 | rs12082376 | -1.193 | -1.312 | 1.217 | 0.227 | -0.022 | 0.011 | 0.011 | 1.044 | 0.14 | 0.37 | 0.09 | 0.1 | 0.1 | 0.08 |
| 1 | 175607170 | rs6425354 | 0.421 | -1.334 | 1.223 | 0.252 | 0.006 | -0.008 | -0.008 | 0.042 | 0.23 | 0.23 | 0.1 | 0.42 | 0.11 | 0.22 |
| 1 | 175612924 | rs10913043 | 0.535 | -1.314 | 1.147 | 0.263 | 0.034 | -0.039 | -0.039 | 0.598 | 0.22 | 0.17 | 0.09 | 0.42 | 0.11 | 0.22 |
| 1 | 175615257 | rs10913045 | -1.006 | -1.292 | 1.275 | 0.208 | -0.03 | 0.018 | 0.018 | 0 | 0.37 | 0.61 | 0.19 | 0.51 | 0.21 | 0.3 |
| 1 | 175619177 | rs12088761 | -0.931 | -0.673 | 1.229 | 0.49 | -0.007 | 0.05 | 0.05 | 0.032 | 0.12 | 0.31 | 0.07 | 0.15 | 0.09 | 0.04 |
| 1 | 175623669 | rs942107 | 0.855 | 0.349 | 0.456 | 0.673 | -0.075 | -0.092 | -0.092 | 0.41 | 0.13 | 0.01 | 0.36 | 0.41 | 0.21 | 0.26 |
| 1 | 175624557 | rs12046065 | -1.68 | 0.36 | 0.387 | 0.394 | 0.012 | 0.043 | 0.043 | 0.005 | 0.59 | 0.68 | 0.75 | 0.73 | 0.66 | 0.71 |
| 1 | 175627395 | rs2057453 | -0.293 | 0.368 | 0.459 | 0.385 | -0.027 | -0.02 | -0.02 | 0.663 | 0.4 | 0.23 | 0.62 | 0.53 | 0.51 | 0.56 |
| 1 | 175629207 | rs4652099 | -0.418 | 0.35 | 0.451 | 0.374 | -0.033 | -0.027 | -0.027 | 1.271 | 0.4 | 0.21 | 0.62 | 0.53 | 0.51 | 0.56 |
| 1 | 175630097 | rs10913052 | -1.802 | 0.32 | 0.462 | 0.341 | 0.009 | 0.044 | 0.044 | 0 | 0.6 | 0.68 | 0.75 | 0.74 | 0.66 | 0.71 |
| 1 | 175636696 | rs4652101 | -0.698 | 0.715 | 0.04 | 0.543 | 0.006 | 0.08 | 0.08 | 0 | 0.11 | 0.31 | 0.08 | 0.18 | 0.09 | 0.03 |
| 1 | 175638336 | rs12025758 | 0.217 | 0.829 | -0.366 | 0.427 | 0.002 | 0.018 | 0.018 | 0.447 | 0.28 | 0.18 | 0.26 | 0.12 | 0.29 | 0.28 |
| 1 | 175639705 | rs12403473 | -0.134 | 0.508 | -0.142 | 0.092 | -0.018 | -0.016 | -0.016 | 0 | 0.4 | 0.27 | 0.62 | 0.53 | 0.49 | 0.55 |
| 1 | 175642887 | rs998677 | 0.71 | 0.616 | -0.532 | 0.337 | -0.025 | -0.031 | -0.031 | 1.481 | 0.05 | 0 | 0.05 | 0.11 | 0.09 | 0.03 |
| 1 | 175647041 | rs7544659 | -0.676 | 0.41 | -0.319 | -0.357 | 0.004 | 0.018 | 0.018 | 0 | 0.5 | 0.54 | 0.69 | 0.65 | 0.56 | 0.59 |
| 1 | 175650376 | rs6692146 | -1.286 | 0.374 | -0.085 | -0.932 | 0 | -0.041 | -0.041 | 0 | 0.43 | 0.56 | 0.57 | 0.94 | 0.46 | 0.55 |
| 1 | 175650399 | rs11582103 | -0.709 | 0.136 | -0.198 | -0.725 | 0.017 | -0.012 | -0.012 | 4.473 | 0.2 | 0.09 | 0.12 | 0.24 | 0.18 | 0.24 |
| 1 | 175651786 | rs4233164 | 0.163 | 0.33 | 0.126 | -0.571 | -0.012 | -0.031 | -0.031 | 0 | 0.21 | 0.12 | 0.44 | 0.68 | 0.24 | 0.31 |
| 1 | 175652529 | rs12401659 | -1.394 | 0.251 | 0.274 | -0.741 | -0.007 | -0.072 | -0.072 | 0 | 0.37 | 0.53 | 0.54 | 0.92 | 0.35 | 0.45 |
| 1 | 175652920 | rs17312104 | 0.077 | 0.055 | 1.067 | -0.061 | 0.014 | -0.041 | -0.041 | 0 | 0.13 | 0.19 | 0.17 | 0.03 | 0.16 | 0.25 |
| 1 | 175658531 | rs12404663 | -1.183 | -0.022 | 0.798 | 0.422 | -0.028 | 0.02 | 0.02 | 0 | 0.12 | 0.24 | 0.06 | 0.16 | 0.06 | 0.12 |
| 1 | 175664892 | rs1155910 | -0.882 | 0.001 | 0.758 | 0.448 | -0.006 | 0.034 | 0.034 | 0 | 0.11 | 0.14 | 0.07 | 0.25 | 0.1 | 0.13 |
| 1 | 175669902 | rs7539097 | -0.57 | -0.157 | 0.496 | -0.023 | 0.015 | -0.013 | -0.013 | 0 | 0.1 | 0.11 | 0.05 | 0.16 | 0.06 | 0.12 |
| 1 | 175675596 | rs10913063 | -2.029 | -0.197 | 0.659 | -0.075 | -0.038 | -0.002 | -0.002 | 0 | 0.18 | 0.43 | 0.1 | 0.16 | 0.12 | 0.13 |
| 1 | 175676896 | rs1323125 | -1.608 | 0.558 | 0.446 | -0.071 | 0.001 | -0.029 | -0.029 | 0 | 0.05 | 0.17 | 0.02 | 0 | 0.05 | 0.02 |

| | | | | | | | | | | | | | | | | |
|---|---|---|---|---|---|---|---|---|---|---|---|---|---|---|---|---|
| 1 | 175684583 | rs12144778 | -0.927 | 0.563 | 0.637 | -0.395 | -0.003 | -0.114 | -0.114 | 0.033 | 0.25 | 0.46 | 0.17 | 0.06 | 0.21 | 0.12 |
| 1 | 175687520 | rs10913064 | 1.274 | 0.503 | 1.013 | 0.038 | -0.005 | -0.011 | -0.011 | 0 | 0.67 | 0.46 | 0.79 | 0.71 | 0.7 | 0.81 |
| 1 | 175697716 | rs7534898 | -1.098 | 0.398 | 1.185 | 0.01 | -0.001 | 0.002 | 0.002 | 0.463 | 0.28 | 0.37 | 0.19 | 0.3 | 0.27 | 0.18 |
| 1 | 175700507 | rs6675876 | -1.05 | 0.428 | 1.261 | 0.025 | 0 | 0.002 | 0.002 | 0.646 | 0.27 | 0.37 | 0.19 | 0.3 | 0.27 | 0.17 |
| 1 | 175709171 | rs4565663 | -1.221 | 0.453 | 1.258 | 0.032 | 0 | 0.006 | 0.006 | 0.533 | 0.28 | 0.44 | 0.2 | 0.3 | 0.28 | 0.18 |
| 1 | 175709327 | rs909741 | -1.88 | 0.414 | 1.4 | 0.014 | 0.02 | 0.196 | 0.196 | 0 | 0.06 | 0.25 | 0.06 | 0.24 | 0.06 | 0.05 |
| 1 | 175711214 | rs12028534 | -0.293 | 0.419 | 1.067 | -0.023 | -0.012 | 0.051 | 0.051 | 2.433 | 0.21 | 0.12 | 0.13 | 0.06 | 0.21 | 0.12 |
| 1 | 175712239 | rs10913071 | -0.288 | 0.404 | 0.975 | -0.035 | -0.025 | 0.064 | 0.064 | 2.534 | 0.2 | 0.08 | 0.12 | 0.06 | 0.21 | 0.13 |
| 1 | 175717468 | rs546785 | 1.237 | 0.369 | 1.183 | -0.008 | -0.009 | -0.017 | -0.017 | 0 | 0.67 | 0.48 | 0.8 | 0.73 | 0.7 | 0.82 |
| 1 | 175722881 | rs645873 | 1.617 | 0.402 | 1.1 | 0.404 | -0.008 | 0.151 | 0.151 | 0 | 0.92 | 0.79 | 0.96 | 0.79 | 0.97 | 0.94 |
| 1 | 175724078 | rs10798415 | -2.115 | 0.84 | 0.703 | -0.108 | 0 | -0.016 | -0.016 | 0 | 0.14 | 0.79 | 0.21 | 0.04 | 0.13 | 0.09 |
| 1 | 175731846 | rs671000 | 0.441 | 0.79 | 0.139 | -0.726 | 0.002 | 0.023 | 0.023 | 0 | 0.73 | 0.85 | 0.74 | 0.89 | 0.74 | 0.69 |
| 1 | 175732289 | rs501915 | 0.745 | 0.723 | 0.752 | -0.759 | -0.003 | -0.028 | -0.028 | 0.093 | 0.71 | 0.64 | 0.73 | 0.89 | 0.73 | 0.69 |
| 1 | 175735380 | rs6659350 | -1.674 | 1.843 | 0.541 | -0.787 | 0.068 | -0.031 | -0.031 | 0 | 0.15 | 0.6 | 0.14 | 0.05 | 0.18 | 0.12 |
| 1 | 175739003 | rs640692 | 1.624 | 1.814 | 0.304 | -0.784 | -0.123 | 0.156 | 0.156 | 0 | 0.57 | 0.06 | 0.59 | 0.84 | 0.55 | 0.56 |
| 1 | 175744041 | rs1325914 | -0.849 | 1.652 | 1.003 | -0.775 | -0.007 | -0.035 | -0.035 | 0.206 | 0.29 | 0.35 | 0.27 | 0.13 | 0.27 | 0.35 |
| 1 | 175756706 | rs6425361 | 0.644 | 1.691 | 1.617 | -0.759 | -0.016 | -0.066 | -0.066 | 0 | 0.69 | 0.61 | 0.72 | 0.87 | 0.72 | 0.65 |
| 1 | 175757242 | rs12561921 | -0.497 | 1.738 | 1.509 | -0.898 | -0.009 | -0.023 | -0.023 | 0 | 0.12 | 0.18 | 0.04 | 0.05 | 0.09 | 0.13 |
| 1 | 175759323 | rs6667913 | -0.221 | 1.707 | 1.499 | -0.896 | -0.014 | 0.041 | 0.041 | 0 | 0.85 | 0.94 | 0.8 | 0.95 | 0.83 | 0.83 |
| 1 | 175769259 | rs7537684 | 0.745 | 1.658 | 1.562 | -0.917 | -0.011 | -0.053 | -0.053 | 0 | 0.79 | 0.65 | 0.75 | 0.92 | 0.81 | 0.8 |
| 1 | 175778102 | rs17353481 | -0.997 | 2.183 | 1.762 | -1.025 | 0.024 | -0.011 | -0.011 | 0 | 0.12 | 0.12 | 0.11 | 0.02 | 0.23 | 0.11 |
| 1 | 175779491 | rs1340998 | 0.381 | 1.624 | 1.539 | -0.898 | 0 | 0.041 | 0.041 | 0 | 0.82 | 0.89 | 0.79 | 0.95 | 0.83 | 0.83 |
| 1 | 175784378 | rs6656072 | 0.396 | 1.665 | 1.119 | -0.878 | 0.002 | 0.023 | 0.023 | 0 | 0.82 | 0.87 | 0.79 | 0.95 | 0.83 | 0.83 |
| 1 | 175791913 | rs7531762 | 1.04 | 1.676 | 1.133 | -0.839 | 0.004 | 0.012 | 0.012 | 0 | 0.72 | 0.77 | 0.75 | 0.89 | 0.74 | 0.7 |
| 1 | 175824579 | rs543445 | 0.078 | 1.907 | 0.792 | -1.108 | -0.022 | 0.049 | 0.049 | 0.429 | 0.75 | 0.87 | 0.7 | 0.86 | 0.75 | 0.67 |
| 1 | 175824630 | rs7548697 | -0.429 | 2.587 | 0.784 | 0.065 | -0.016 | -0.011 | -0.011 | 0.423 | 0.1 | 0.07 | 0.1 | 0.29 | 0.19 | 0.1 |
| 1 | 175824794 | rs571898 | -1.205 | 2.615 | 0.996 | 0.044 | 0.069 | 0.096 | 0.096 | 0 | 0.13 | 0.27 | 0.13 | 0.29 | 0.23 | 0.1 |
| 1 | 175827683 | rs11810084 | -0.981 | 2.717 | 0.808 | 0.21 | 0.028 | 0.053 | 0.053 | 0 | 0.13 | 0.2 | 0.18 | 0.31 | 0.25 | 0.16 |
| 1 | 175830514 | rs537298 | 0.192 | 2.73 | 0.076 | -0.406 | 0 | 0.093 | 0.093 | 1.049 | 0.66 | 0.87 | 0.69 | 0.86 | 0.73 | 0.69 |
| 1 | 175832217 | rs732292 | 1.507 | 1.838 | 0.164 | -0.087 | 0.016 | 0.011 | 0.011 | 0.293 | 0.38 | 0.19 | 0.37 | 0.38 | 0.37 | 0.38 |
| 1 | 175837332 | rs500894 | -0.349 | 1.566 | 0.499 | -0.138 | 0.003 | 0.072 | 0.072 | 0.012 | 0.26 | 0.59 | 0.33 | 0.39 | 0.3 | 0.21 |
| 1 | 175846158 | rs16849285 | 0.434 | 1.562 | 0.911 | -0.143 | 0.006 | 0.028 | 0.028 | 1.106 | 0.22 | 0.22 | 0.25 | 0.33 | 0.28 | 0.17 |
| 1 | 175848088 | rs578140 | -1.089 | 1.598 | 0.287 | -0.345 | -0.058 | 0.03 | 0.03 | 0 | 0.72 | 0.97 | 0.74 | 0.81 | 0.75 | 0.65 |
| 1 | 175848239 | rs4652137 | 0.832 | 1.653 | 0.403 | -0.316 | 0.004 | 0.013 | 0.013 | 0.603 | 0.46 | 0.37 | 0.4 | 0.42 | 0.45 | 0.44 |
| 1 | 175849782 | rs10913091 | 0.763 | 1.531 | 0.123 | -0.183 | -0.013 | -0.026 | -0.026 | 1.604 | 0.2 | 0.14 | 0.25 | 0.33 | 0.28 | 0.17 |
| 1 | 175852209 | rs532799 | 0.985 | 0.651 | 0.16 | -0.179 | -0.023 | 0.006 | 0.006 | 2.189 | 0.37 | 0.54 | 0.38 | 0.43 | 0.32 | 0.38 |
| 1 | 175859765 | rs12759426 | -0.287 | 0.128 | 0.462 | -0.432 | -0.015 | -0.037 | -0.037 | 0 | 0.35 | 0.51 | 0.3 | 0.2 | 0.25 | 0.43 |
| 1 | 175869554 | rs10753110 | 1.473 | 0.476 | -0.097 | -0.518 | -0.058 | 0.095 | 0.095 | 0 | 0.47 | 0.1 | 0.57 | 0.69 | 0.51 | 0.45 |
| 1 | 175874139 | rs10494494 | 0.64 | 0.448 | -0.046 | -0.627 | -0.017 | 0.029 | 0.029 | 0 | 0.14 | 0.05 | 0.07 | 0.06 | 0.15 | 0.07 |
| 1 | 175879489 | rs6425365 | 1.446 | 0.439 | 0.116 | -0.625 | -0.031 | 0.129 | 0.129 | 0 | 0.48 | 0.15 | 0.58 | 0.72 | 0.53 | 0.46 |
| 1 | 175881366 | rs12239742 | 0.673 | 0.883 | 0.113 | -0.536 | -0.002 | 0.001 | 0.001 | 0 | 0.25 | 0.25 | 0.27 | 0.17 | 0.23 | 0.28 |
| 1 | 175885411 | rs7525623 | 0.495 | 0.926 | 0.081 | -0.515 | 0.011 | -0.036 | -0.036 | 0 | 0.26 | 0.39 | 0.27 | 0.17 | 0.23 | 0.28 |
| 1 | 175886511 | rs6656281 | -0.329 | 0.611 | 0.153 | -0.341 | 0.008 | -0.031 | -0.031 | 4.277 | 0.41 | 0.47 | 0.35 | 0.22 | 0.39 | 0.36 |
| 1 | 175889627 | rs2030558 | -0.407 | 0.638 | -0.509 | -0.349 | -0.015 | 0.041 | 0.041 | 0 | 0.4 | 0.27 | 0.34 | 0.22 | 0.38 | 0.36 |
| 1 | 175891622 | rs7516327 | -0.661 | 0.568 | -0.39 | -0.56 | -0.018 | -0.028 | -0.028 | 0 | 0.39 | 0.27 | 0.4 | 0.49 | 0.41 | 0.42 |
| 1 | 175894857 | rs6695241 | -1.04 | 0.517 | -0.109 | -0.586 | 0.005 | 0.008 | 0.008 | 0 | 0.5 | 0.64 | 0.5 | 0.5 | 0.53 | 0.54 |
| 1 | 175902596 | rs6680839 | 0.896 | 0.071 | 0.648 | -0.548 | 0 | 0.005 | 0.005 | 0.044 | 0.53 | 0.42 | 0.53 | 0.44 | 0.53 | 0.37 |
| 1 | 175904809 | rs6687849 | 0.53 | -0.068 | 1.279 | -0.653 | 0.04 | 0.032 | 0.032 | 0.08 | 0.95 | 0.7 | 0.98 | 1 | 1 | 0.96 |
| 1 | 175911554 | rs10436856 | 1.363 | -0.041 | 1.298 | -0.724 | 0.003 | 0.018 | 0.018 | 0.011 | 0.47 | 0.31 | 0.44 | 0.44 | 0.45 | 0.31 |
| 1 | 175913828 | rs10913112 | 1.972 | -0.076 | 1.229 | -0.825 | -0.067 | -0.026 | -0.026 | 0.318 | 0.41 | 0.04 | 0.39 | 0.38 | 0.4 | 0.31 |
| 1 | 175936012 | rs4304533 | 0.624 | -0.298 | 1.393 | -0.805 | -0.056 | -0.013 | -0.013 | 0 | 0.88 | 0.54 | 0.94 | 0.92 | 0.98 | 0.79 |
| 1 | 175952542 | rs10798432 | 1.5 | -0.263 | 1.361 | -0.951 | -0.021 | -0.063 | -0.063 | 0.355 | 0.67 | 0.44 | 0.64 | 0.9 | 0.78 | 0.63 |
| 1 | 175983994 | rs4652143 | -0.635 | -0.261 | 1.502 | -0.997 | -0.026 | -0.017 | -0.017 | 0 | 0.12 | 0.25 | 0.07 | 0.08 | 0.06 | 0.21 |
| 1 | 175986057 | rs12041226 | 1.473 | -0.313 | 1.333 | -1.023 | -0.016 | -0.027 | -0.027 | 0 | 0.06 | 0 | 0.18 | 0.27 | 0.06 | 0.13 |
| 1 | 176000113 | rs12040241 | 2.872 | -0.217 | 1.767 | -0.892 | -0.066 | -0.025 | -0.025 | 0.338 | 0.41 | 0.04 | 0.39 | 0.38 | 0.4 | 0.31 |
| 1 | 176004673 | rs10913128 | 2.582 | -0.231 | 2.241 | -0.898 | -0.004 | 0.028 | 0.028 | 0.06 | 0.42 | 0.17 | 0.4 | 0.38 | 0.4 | 0.31 |
| 1 | 176008112 | rs7554913 | 0.998 | -0.208 | 2.206 | -0.879 | -0.025 | -0.088 | -0.088 | 0 | 0.79 | 0.67 | 0.77 | 0.99 | 0.88 | 0.87 |
| 1 | 176014659 | rs4129061 | 1.636 | -0.215 | 2.071 | -0.88 | -0.062 | -0.105 | -0.105 | 0.012 | 0.71 | 0.49 | 0.73 | 0.91 | 0.87 | 0.7 |
| 1 | 176018764 | rs12729524 | 0.238 | -0.198 | 2.059 | -0.874 | -0.051 | -0.081 | -0.081 | 0 | 0.19 | 0.11 | 0.19 | 0.48 | 0.33 | 0.32 |
| 1 | 176036351 | rs2502822 | 1.621 | -0.224 | 2.091 | -1.096 | -0.045 | -0.079 | -0.079 | 0 | 0.71 | 0.45 | 0.73 | 0.91 | 0.87 | 0.7 |
| 1 | 176051131 | rs4652148 | 0.461 | -0.213 | 2.061 | -1.077 | -0.029 | 0 | 0 | 0.005 | 0.07 | 0.16 | 0.04 | 0.07 | 0.02 | 0.17 |
| 1 | 176060829 | rs2481640 | 0.564 | -0.188 | 2.055 | -1.077 | 0.016 | 0.004 | 0.004 | 0.022 | 0.94 | 0.74 | 0.98 | 1 | 0.99 | 0.96 |
| 1 | 176088186 | rs2502841 | 1.67 | -0.293 | 2.123 | -1.097 | -0.033 | -0.06 | -0.06 | 0 | 0.71 | 0.42 | 0.72 | 0.91 | 0.87 | 0.71 |
| 1 | 176092668 | rs7517503 | 2.219 | -0.262 | 2.102 | -1.073 | 0.002 | 0.031 | 0.031 | 0.022 | 0.47 | 0.31 | 0.44 | 0.41 | 0.45 | 0.31 |
| 1 | 176111527 | rs2039232 | -1.091 | -0.27 | 2.111 | -1.074 | -0.059 | -0.016 | -0.016 | 1.831 | 0.13 | 0.46 | 0.06 | 0.07 | 0.02 | 0.2 |
| 1 | 176114487 | rs6665385 | -0.475 | -0.267 | 2.083 | -1.074 | -0.06 | -0.088 | -0.088 | 1.863 | 0.8 | 0.9 | 0.81 | 0.5 | 0.67 | 0.67 |
| 1 | 176118219 | rs3748618 | 2.759 | -0.273 | 2.065 | -1.082 | -0.067 | 0.014 | 0.014 | 0.188 | 0.41 | 0.04 | 0.39 | 0.35 | 0.4 | 0.31 |
| 1 | 176122170 | rs12239888 | 0.439 | -0.243 | 2.102 | -1.064 | -0.025 | 0.012 | 0.012 | 5.168 | 0.07 | 0.2 | 0.04 | 0.07 | 0.02 | 0.17 |
| 1 | 176129276 | rs520661 | 0.199 | -0.231 | 1.896 | -1.06 | -0.081 | -0.114 | -0.114 | 0.549 | 0.19 | 0.07 | 0.19 | 0.5 | 0.33 | 0.33 |

| | | | | | | | | | | | | | | | | |
|---|---|---|---|---|---|---|---|---|---|---|---|---|---|---|---|---|
| 1 | 176151494 | rs10494499 | -0.489 | -0.225 | 1.949 | -1.074 | 0.009 | 0.031 | 0.031 | 0.761 | 0.16 | 0.08 | 0.21 | 0.01 | 0.11 | 0.09 |
| 1 | 176172795 | rs10913150 | 1.654 | -0.213 | 2.012 | -1.069 | -0.041 | -0.075 | -0.075 | 0.017 | 0.71 | 0.44 | 0.73 | 0.91 | 0.87 | 0.7 |
| 1 | 176182722 | rs4526571 | 2.493 | -0.23 | 1.923 | -1 | -0.068 | 0.008 | 0.008 | 0.172 | 0.41 | 0.04 | 0.39 | 0.36 | 0.4 | 0.31 |
| 1 | 176183201 | rs704831 | 0.841 | -0.328 | 2.106 | -1.222 | -0.059 | -0.01 | -0.01 | 0 | 0.87 | 0.54 | 0.94 | 0.93 | 0.98 | 0.79 |
| 1 | 176186059 | rs704833 | 1.204 | -0.28 | 1.954 | -1.248 | 0 | -0.066 | -0.066 | 0.157 | 0.89 | 0.64 | 0.94 | 1 | 0.94 | 0.93 |
| 1 | 176186170 | rs12126441 | -0.127 | -0.143 | 1.803 | -1.191 | 0.003 | -0.01 | -0.01 | 0 | 0.17 | 0.13 | 0.28 | 0 | 0.16 | 0.12 |
| 1 | 176189141 | rs704834 | 0.004 | -0.064 | 1.83 | -1.049 | -0.034 | -0.062 | -0.062 | 0 | 0.19 | 0.15 | 0.18 | 0.46 | 0.3 | 0.32 |
| 1 | 176233932 | rs352314 | -1.711 | -0.092 | 1.905 | -1.087 | -0.014 | -0.005 | -0.005 | 0.019 | 0.42 | 0.33 | 0.49 | 0.55 | 0.47 | 0.62 |
| 1 | 176249709 | rs1503122 | -0.106 | -0.091 | 1.9 | -1.085 | 0.004 | -0.003 | -0.003 | 0 | 0.17 | 0.11 | 0.29 | 0 | 0.16 | 0.12 |
| 1 | 176261782 | rs2089015 | 0.133 | -0.087 | 1.9 | -1.083 | 0 | 0.045 | 0.045 | 0 | 0.16 | 0.03 | 0.27 | 0 | 0.16 | 0.12 |
| 1 | 176271944 | rs11589107 | 0.108 | -0.065 | 2.003 | -1.077 | 0.005 | -0.027 | -0.027 | 10.488 | 0.05 | 0.13 | 0.04 | 0 | 0.04 | 0.03 |
| 1 | 176273901 | rs12144020 | -0.972 | -0.073 | 2.011 | -1.076 | 0.007 | 0.006 | 0.006 | 0 | 0.24 | 0.21 | 0.31 | 0.09 | 0.17 | 0.3 |
| 1 | 176275821 | rs16849735 | -0.972 | -0.073 | 1.994 | -1.076 | 0.008 | 0.008 | 0.008 | 0 | 0.24 | 0.21 | 0.31 | 0.09 | 0.17 | 0.3 |
| 1 | 176277052 | rs2504473 | -1.937 | -0.02 | 2.03 | -1.031 | -0.001 | 0.006 | 0.006 | 0 | 0.48 | 0.51 | 0.53 | 0.55 | 0.52 | 0.65 |
| 1 | 176291703 | rs2455738 | -1.907 | -0.175 | 1.561 | -1.025 | -0.009 | 0.017 | 0.017 | 0 | 0.48 | 0.4 | 0.51 | 0.55 | 0.51 | 0.65 |
| 1 | 176295837 | rs2455730 | 0.2 | -0.163 | 1.091 | -1.022 | -0.053 | -0.091 | -0.091 | 0 | 0.18 | 0.1 | 0.18 | 0.46 | 0.3 | 0.32 |
| 1 | 176298497 | rs2455729 | -1.977 | -0.162 | 1.105 | -1.05 | -0.004 | 0.014 | 0.014 | 0 | 0.49 | 0.45 | 0.51 | 0.55 | 0.51 | 0.65 |
| 1 | 176311597 | rs10913158 | -1.2 | -0.147 | 0.749 | -1.133 | 0.025 | 0.081 | 0.081 | 0 | 0.28 | 0.14 | 0.31 | 0.08 | 0.21 | 0.33 |
| 1 | 176345142 | rs2504491 | 0.071 | -0.198 | 0.741 | -1.024 | -0.053 | -0.092 | -0.092 | 0 | 0.19 | 0.09 | 0.17 | 0.45 | 0.3 | 0.33 |
| 1 | 176345256 | rs4652171 | -0.209 | -0.281 | 0.775 | -1.043 | -0.019 | -0.001 | -0.001 | 0.199 | 0.08 | 0.13 | 0.02 | 0.08 | 0.01 | 0.16 |
| 1 | 176350944 | rs4147204 | 2.212 | -0.275 | 0.537 | -1.08 | -0.063 | -0.036 | -0.036 | 0.291 | 0.43 | 0.07 | 0.42 | 0.41 | 0.44 | 0.36 |
| 1 | 176354126 | rs1503124 | -2.003 | -0.225 | 0.504 | -0.992 | -0.001 | 0.012 | 0.012 | 0 | 0.49 | 0.41 | 0.49 | 0.54 | 0.51 | 0.63 |
| 1 | 176402381 | rs1393762 | -0.453 | 0.22 | 0.532 | -0.34 | -0.069 | -0.112 | -0.112 | 0 | 0.19 | 0.06 | 0.17 | 0.45 | 0.31 | 0.33 |
| 1 | 176405993 | rs16849826 | 0.21 | 0.347 | 0.603 | -0.238 | -0.043 | -0.049 | -0.049 | 0.068 | 0.13 | 0.08 | 0.09 | 0.41 | 0.25 | 0.23 |
| 1 | 176411602 | rs4652174 | -0.354 | -0.169 | 0.023 | -0.087 | 0.015 | -0.004 | -0.004 | 0 | 0.07 | 0.07 | 0.02 | 0.09 | 0.01 | 0.12 |
| 1 | 176414781 | rs4650943 | -0.762 | 0.091 | 0.163 | -0.339 | -0.002 | -0.004 | -0.004 | 0.106 | 0.4 | 0.42 | 0.46 | 0.54 | 0.45 | 0.6 |
| 1 | 176422273 | rs4233167 | 1.533 | -0.486 | 0.485 | -0.867 | 0.001 | -0.013 | -0.013 | 0 | 0.75 | 0.23 | 0.84 | 0.92 | 0.87 | 0.73 |
| 1 | 176422595 | rs13374369 | -1.305 | -0.481 | 1.303 | -0.882 | 0.024 | 0 | 0 | 0 | 0.1 | 0.3 | 0.1 | 0.08 | 0.07 | 0.22 |
| 1 | 176422849 | rs12096483 | -1.252 | -0.53 | 1.285 | -0.907 | 0.01 | 0.021 | 0.021 | 0.01 | 0.09 | 0.09 | 0.04 | 0 | 0.06 | 0.06 |
| 1 | 176423140 | rs4471226 | 1.367 | -0.432 | 1.687 | -0.747 | 0.034 | 0.002 | 0.002 | 0.01 | 0.89 | 0.54 | 0.9 | 0.92 | 0.93 | 0.78 |
| 1 | 176429364 | rs7552857 | -1.191 | -0.249 | 1.163 | -0.745 | 0.034 | -0.023 | -0.023 | 0 | 0.08 | 0.23 | 0.1 | 0.05 | 0.11 | 0.04 |
| 1 | 176435410 | rs7527035 | -2.384 | -0.697 | 1.004 | -0.994 | 0.021 | 0.024 | 0.024 | 0.237 | 0.27 | 0.79 | 0.18 | 0.07 | 0.17 | 0.28 |
| 1 | 176435567 | rs7515586 | -1.298 | -0.291 | 0.93 | -0.7 | 0.003 | -0.033 | -0.033 | 2.14 | 0.1 | 0.23 | 0.1 | 0.04 | 0.1 | 0.02 |
| 1 | 176440645 | rs7536647 | -1.61 | -0.722 | 0.916 | -0.996 | -0.048 | -0.096 | -0.096 | 0 | 0.17 | 0.63 | 0.13 | 0.01 | 0.09 | 0.19 |
| 1 | 176445702 | rs6425386 | -0.972 | -0.807 | -0.305 | -1.442 | 0.068 | 0.182 | 0.182 | 0 | 0.5 | 0.81 | 0.42 | 0.2 | 0.4 | 0.42 |
| 1 | 176453688 | rs2223579 | 0.486 | -1.41 | 0.264 | -2.072 | -0.038 | -0.101 | -0.101 | 0 | 0.7 | 0.5 | 0.8 | 0.97 | 0.81 | 0.67 |
| 1 | 176457223 | rs6676641 | -1.493 | -1.589 | 0.245 | -2.272 | 0.052 | 0.073 | 0.073 | 0.037 | 0.46 | 0.66 | 0.27 | 0.04 | 0.31 | 0.41 |
| 1 | 176462695 | rs7535501 | 0.081 | -1.768 | 0.584 | -2.408 | -0.001 | -0.015 | -0.015 | 0 | 0.91 | 0.9 | 0.92 | 1 | 0.93 | 0.95 |
| 1 | 176465969 | rs6667264 | -1.312 | -1.857 | 0.982 | -2.439 | -0.031 | -0.118 | -0.118 | 0 | 0.33 | 0.52 | 0.21 | 0.01 | 0.23 | 0.22 |
| 1 | 176470980 | rs7521130 | -1.857 | -1.675 | 0.949 | -2.458 | 0.058 | 0.092 | 0.092 | 0.039 | 0.45 | 0.69 | 0.27 | 0.05 | 0.31 | 0.39 |
| 1 | 176475180 | rs11584882 | -0.393 | -1.737 | 0.816 | -2.465 | 0.003 | -0.022 | -0.022 | 0 | 0.24 | 0.22 | 0.14 | 0.01 | 0.18 | 0.14 |
| 1 | 176487700 | rs10489473 | 0.159 | -1.703 | 0.723 | -2.44 | 0.002 | 0.001 | 0.001 | 0 | 0.13 | 0.14 | 0.06 | 0.04 | 0.08 | 0.17 |
| 1 | 176507187 | rs2206508 | 1.365 | -1.701 | 1.058 | -2.442 | -0.002 | -0.045 | -0.045 | 0 | 0.57 | 0.57 | 0.77 | 0.95 | 0.7 | 0.69 |
| 1 | 176508293 | rs992922 | -0.612 | -1.623 | 0.6 | -2.419 | -0.014 | -0.032 | -0.032 | 0 | 0.19 | 0.23 | 0.1 | 0.04 | 0.12 | 0.18 |
| 1 | 176513537 | rs2206510 | 0.923 | -0.759 | -0.612 | -2.048 | -0.1 | -0.08 | -0.08 | 0.113 | 0.2 | 0.01 | 0.24 | 0.25 | 0.26 | 0.22 |
| 1 | 176520317 | rs6671694 | 0.389 | -1.037 | 0.081 | -2.202 | -0.042 | -0.154 | -0.154 | 0.347 | 0.73 | 0.54 | 0.83 | 0.99 | 0.82 | 0.81 |
| 1 | 176535567 | rs6659102 | -0.718 | -0.804 | -0.892 | -2.261 | 0.011 | 0.1 | 0.1 | 0 | 0.23 | 0.06 | 0.17 | 0.03 | 0.19 | 0.27 |
| 1 | 176547669 | rs240099 | 0.723 | -0.976 | -0.8 | -2.247 | 0.023 | 0.076 | 0.076 | 0 | 0.73 | 0.9 | 0.81 | 0.94 | 0.79 | 0.54 |
| 1 | 176548900 | rs12094283 | 0.853 | -1.324 | -0.713 | -3.051 | 0.005 | -0.035 | -0.035 | 0.045 | 0.26 | 0.22 | 0.1 | 0 | 0.18 | 0.15 |
| 1 | 176551626 | rs11587239 | -0.628 | -0.901 | -0.497 | -2.822 | 0.028 | 0.053 | 0.053 | 0 | 0.52 | 0.32 | 0.28 | 0.06 | 0.39 | 0.61 |
| 1 | 176565307 | rs909941 | -1.029 | -0.818 | -0.589 | -2.171 | -0.03 | 0.092 | 0.092 | 0 | 0.61 | 0.84 | 0.41 | 0.22 | 0.46 | 0.6 |
| 1 | 176567941 | rs10913214 | 1.107 | -1.269 | -0.223 | -2.43 | 0.023 | 0.034 | 0.034 | 0 | 0.28 | 0.15 | 0.11 | 0 | 0.2 | 0.17 |
| 1 | 176571396 | rs12066567 | -1.313 | -1.263 | -0.229 | -2.444 | -0.035 | 0.094 | 0.094 | 0 | 0.61 | 0.85 | 0.41 | 0.22 | 0.46 | 0.6 |
| 1 | 176571431 | rs240104 | 1.48 | 0.036 | -0.708 | -2.225 | -0.096 | -0.007 | -0.007 | 1.024 | 0.16 | 0.01 | 0.24 | 0.14 | 0.24 | 0.13 |
| 1 | 176571578 | rs10798463 | 0.417 | -0.65 | 0.068 | -2.284 | 0.011 | 0.023 | 0.023 | 0 | 0.33 | 0.23 | 0.19 | 0.2 | 0.29 | 0.36 |
| 1 | 176603840 | rs7529392 | 0.359 | -0.446 | 0.05 | -2.576 | -0.001 | -0.029 | -0.029 | 0.004 | 0.34 | 0.31 | 0.17 | 0.08 | 0.28 | 0.35 |
| 1 | 176611275 | rs10913227 | 0.359 | -0.444 | 0.047 | -2.579 | -0.002 | -0.029 | -0.029 | 0 | 0.34 | 0.31 | 0.17 | 0.08 | 0.28 | 0.35 |
| 1 | 176630817 | rs1015901 | -1.216 | -0.494 | -0.134 | -2.619 | -0.012 | 0.131 | 0.131 | 0 | 0.6 | 0.8 | 0.36 | 0.11 | 0.45 | 0.59 |
| 1 | 176632533 | rs12071948 | 0.583 | -0.091 | -0.203 | -2.174 | 0.007 | -0.035 | -0.035 | 0 | 0.29 | 0.22 | 0.12 | 0 | 0.2 | 0.17 |
| 1 | 176649181 | rs10454444 | 0.583 | -0.042 | -0.171 | -2.19 | 0.007 | -0.035 | -0.035 | 0 | 0.29 | 0.22 | 0.12 | 0 | 0.2 | 0.17 |
| 1 | 176656255 | rs7540152 | 0.084 | -0.256 | 0.108 | -2.541 | -0.011 | -0.079 | -0.079 | 0 | 0.19 | 0.3 | 0.16 | 0.02 | 0.16 | 0.23 |
| 1 | 176659790 | rs4492574 | 0.394 | -0.254 | 0.008 | -2.533 | 0.004 | -0.009 | -0.009 | 0 | 0.34 | 0.27 | 0.17 | 0.08 | 0.28 | 0.35 |
| 1 | 176667810 | rs791031 | -0.283 | -0.163 | -0.014 | -2.369 | -0.005 | -0.049 | -0.049 | 0 | 0.82 | 0.74 | 0.84 | 0.98 | 0.84 | 0.77 |
| 1 | 176684432 | rs4507975 | -0.072 | 0.032 | 0.287 | -2.484 | -0.013 | -0.092 | -0.092 | 0.26 | 0.39 | 0.46 | 0.2 | 0.07 | 0.34 | 0.35 |
| 1 | 176684803 | rs1086520 | -0.049 | 0.206 | -0.095 | -2.588 | 0.002 | 0.069 | 0.069 | 0 | 0.81 | 0.91 | 0.86 | 0.98 | 0.84 | 0.76 |
| 1 | 176684851 | rs1086519 | 0.936 | 0.175 | 0.059 | -2.638 | 0 | 0.013 | 0.013 | 0 | 0.75 | 0.79 | 0.81 | 0.94 | 0.76 | 0.74 |
| 1 | 176686086 | rs791033 | 0.499 | 0.05 | 0.083 | -2.669 | -0.002 | -0.018 | -0.018 | 0 | 0.86 | 0.84 | 0.92 | 0.98 | 0.9 | 0.81 |
| 1 | 176690586 | rs7526821 | -0.931 | 0.078 | 0.302 | -2.976 | 0.002 | 0.022 | 0.022 | 0 | 0.56 | 0.65 | 0.32 | 0.06 | 0.42 | 0.39 |
| 1 | 176695533 | rs10442600 | -0.101 | 0.109 | 0.237 | -2.963 | -0.029 | 0.019 | 0.019 | 0 | 0.09 | 0.02 | 0.07 | 0.04 | 0.12 | 0.03 |

| | | | | | | | | | | | | | | | | |
|---|---|---|---|---|---|---|---|---|---|---|---|---|---|---|---|---|
| 1 | 176703743 | rs7528287 | -0.907 | 0.037 | 0.371 | -2.992 | 0.003 | 0.012 | 0.012 | 0 | 0.55 | 0.64 | 0.3 | 0.06 | 0.42 | 0.39 |
| 1 | 176703822 | rs12117539 | -1.353 | -0.004 | 0.387 | -3.022 | -0.001 | 0.016 | 0.016 | 0 | 0.23 | 0.18 | 0.15 | 0.06 | 0.21 | 0.22 |
| 1 | 176705147 | rs10913245 | 1.047 | -0.096 | 0.364 | -3.029 | -0.032 | -0.141 | -0.141 | 0.046 | 0.32 | 0.41 | 0.15 | 0 | 0.21 | 0.17 |
| 1 | 176716118 | rs10798479 | -0.848 | -0.054 | 0.388 | -2.93 | 0.005 | 0.02 | 0.02 | 0 | 0.56 | 0.67 | 0.3 | 0.06 | 0.42 | 0.39 |
| 1 | 176727102 | rs7529487 | -0.746 | -0.043 | 0.419 | -2.949 | -0.002 | -0.065 | -0.065 | 0.153 | 0.54 | 0.51 | 0.29 | 0.06 | 0.42 | 0.4 |
| 1 | 176738253 | rs1325601 | -0.649 | 0.229 | 0.546 | -2.953 | 0.014 | 0.044 | 0.044 | 0 | 0.49 | 0.64 | 0.27 | 0.05 | 0.4 | 0.37 |
| 1 | 176743400 | rs1325600 | -1.489 | 0.682 | 0.276 | -2.587 | -0.052 | 0.015 | 0.015 | 0 | 0.64 | 0.91 | 0.55 | 0.28 | 0.56 | 0.53 |
| 1 | 176747993 | rs16850193 | 0.023 | 0.57 | 0.442 | -2.634 | -0.01 | -0.013 | -0.013 | 0 | 0.08 | 0.13 | 0.05 | 0.03 | 0.05 | 0.17 |
| 1 | 176759336 | rs2076336 | -0.495 | 0.663 | 0.361 | -2.646 | 0.011 | 0.043 | 0.043 | 0 | 0.5 | 0.64 | 0.3 | 0.05 | 0.41 | 0.38 |
| 1 | 176769358 | rs10913256 | 0.76 | 0.59 | 0.407 | -2.715 | -0.035 | -0.16 | -0.16 | 0 | 0.34 | 0.47 | 0.17 | 0 | 0.22 | 0.17 |
| 1 | 176772029 | rs17378160 | -0.485 | 0.648 | 0.232 | -2.641 | -0.002 | 0.034 | 0.034 | 0.001 | 0.13 | 0.05 | 0.08 | 0.02 | 0.1 | 0.18 |
| 1 | 176772360 | rs1325597 | 0.592 | 0.777 | 0.147 | -2.582 | -0.042 | -0.021 | -0.021 | 0 | 0.79 | 0.93 | 0.66 | 0.77 | 0.76 | 0.66 |
| 1 | 176791571 | rs7543049 | 1.529 | 0.551 | -0.099 | -2.683 | 0.023 | 0.061 | 0.061 | 0 | 0.27 | 0.14 | 0.08 | 0 | 0.2 | 0.17 |
| 1 | 176795135 | rs12035200 | 0.535 | 0.702 | -0.101 | -2.701 | 0.015 | 0.106 | 0.106 | 0 | 0.46 | 0.72 | 0.51 | 0.74 | 0.45 | 0.46 |
| 1 | 176795627 | rs12131804 | -0.182 | 0.705 | -0.115 | -2.667 | 0.037 | -0.011 | -0.011 | 1.154 | 0.05 | 0.11 | 0.06 | 0.03 | 0.11 | 0.02 |
| 1 | 176811873 | rs1044299 | 0.011 | 0.675 | -0.354 | -2.633 | -0.045 | 0.142 | 0.142 | 0.053 | 0.52 | 0.89 | 0.57 | 0.74 | 0.55 | 0.47 |
| 1 | 176811918 | rs12137517 | -0.177 | 0.917 | -0.234 | -2.635 | 0.009 | -0.004 | -0.004 | 0.46 | 0.05 | 0.09 | 0.06 | 0.03 | 0.11 | 0.02 |
| 1 | 176816410 | rs2038114 | 0.708 | 0.749 | 0.119 | -2.719 | 0.004 | 0.034 | 0.034 | 0 | 0.42 | 0.32 | 0.45 | 0.75 | 0.43 | 0.46 |
| 1 | 176832613 | rs6425398 | -1.422 | 0.697 | 0.334 | -2.545 | -0.044 | -0.048 | -0.048 | 0 | 0.92 | 0.99 | 0.74 | 0.8 | 0.85 | 0.86 |
| 1 | 176842230 | rs16850282 | -0.89 | 0.519 | 0.506 | -2.754 | -0.015 | -0.084 | -0.084 | 0 | 0.17 | 0.38 | 0.13 | 0.02 | 0.11 | 0.18 |
| 1 | 176842745 | rs10913267 | 1.756 | 0.64 | 0.211 | -2.946 | -0.012 | 0.174 | 0.174 | 0 | 0.28 | 0.03 | 0.11 | 0 | 0.25 | 0.21 |
| 1 | 176844170 | rs7532411 | -0.61 | 0.804 | 0.729 | -2.838 | 0.121 | 0.016 | 0.016 | 0 | 0.06 | 0.25 | 0.07 | 0.03 | 0.11 | 0.02 |
| 1 | 176845386 | rs2076339 | 1.357 | 0.48 | 0.19 | -3.016 | -0.051 | 0.016 | 0.016 | 0.016 | 0.31 | 0.08 | 0.36 | 0.75 | 0.36 | 0.44 |
| 1 | 176859116 | rs6679170 | -1.053 | 0.023 | 0.596 | -2.951 | 0.01 | 0.021 | 0.021 | 1.535 | 0.91 | 0.86 | 0.72 | 0.8 | 0.86 | 0.86 |
| 1 | 176863867 | rs2228956 | -1.024 | 0.033 | 0.499 | -2.946 | 0.009 | 0.02 | 0.02 | 1.432 | 0.91 | 0.86 | 0.72 | 0.8 | 0.86 | 0.86 |
| 1 | 176867860 | rs726517 | 0.746 | -0.137 | 0.575 | -3.134 | -0.022 | -0.152 | -0.152 | 0.577 | 0.37 | 0.38 | 0.17 | 0 | 0.29 | 0.22 |
| 1 | 176878679 | rs1016612 | -0.406 | 0.04 | -0.124 | -2.995 | 0.031 | 0.169 | 0.169 | 1.156 | 0.54 | 0.75 | 0.3 | 0.02 | 0.39 | 0.39 |
| 1 | 176885306 | rs2038047 | -0.789 | -0.034 | -0.299 | -2.884 | -0.096 | 0.142 | 0.142 | 1.02 | 0.6 | 0.9 | 0.51 | 0.18 | 0.51 | 0.49 |
| 1 | 176887445 | rs11587780 | -1.23 | 0.212 | 0.243 | -2.538 | 0.003 | -0.018 | -0.018 | 2.091 | 0.18 | 0.21 | 0.1 | 0.02 | 0.12 | 0.17 |
| 1 | 176890692 | rs16850406 | 1.121 | 0.264 | 0.515 | -2.545 | 0.029 | 0.039 | 0.039 | 0.364 | 0.08 | 0.15 | 0.24 | 0.16 | 0.12 | 0.11 |
| 1 | 176891234 | rs6670777 | -1.689 | 0.235 | 0.577 | -2.556 | 0.001 | -0.02 | -0.02 | 0 | 0.26 | 0.37 | 0.34 | 0.18 | 0.24 | 0.28 |
| 1 | 176898702 | rs11804030 | 0.553 | 0.114 | 0.465 | -2.253 | -0.007 | -0.02 | -0.02 | 0 | 0.06 | 0.1 | 0.04 | 0 | 0.03 | 0.01 |
| 1 | 176901432 | rs4652207 | -1.853 | -0.042 | 1.019 | -2.353 | -0.018 | -0.03 | -0.03 | 0.026 | 0.22 | 0.41 | 0.3 | 0.15 | 0.16 | 0.12 |
| 1 | 176906304 | rs17313493 | 0.521 | -0.03 | 0.867 | -2.34 | -0.037 | -0.039 | -0.039 | 0 | 0.09 | 0.02 | 0.22 | 0.15 | 0.12 | 0.11 |
| 1 | 176918675 | rs2067038 | -1.713 | -0.053 | 1.084 | -2.32 | -0.015 | -0.024 | -0.024 | 0 | 0.22 | 0.45 | 0.31 | 0.19 | 0.16 | 0.12 |
| 1 | 176919799 | rs4652209 | -1.527 | -0.012 | 0.747 | -2.302 | -0.008 | -0.014 | -0.014 | 0.65 | 0.21 | 0.36 | 0.29 | 0.19 | 0.16 | 0.12 |
| 1 | 176925539 | rs4652214 | -0.058 | 0.035 | 0.748 | -2.317 | -0.004 | -0.04 | -0.04 | 0 | 0.08 | 0.26 | 0.06 | 0 | 0.04 | 0.01 |
| 1 | 176930714 | rs4652216 | -1.363 | 0.035 | 0.496 | -2.304 | -0.007 | -0.016 | -0.016 | 0 | 0.21 | 0.34 | 0.29 | 0.2 | 0.17 | 0.11 |
| 1 | 176945623 | rs227992 | -2.042 | -0.015 | 0.998 | -2.019 | -0.009 | -0.015 | -0.015 | 0 | 0.25 | 0.66 | 0.31 | 0.22 | 0.21 | 0.16 |
| 1 | 176952890 | rs2142756 | 0.159 | 0.08 | 1.557 | -1.919 | 0.017 | 0.023 | 0.023 | 0.895 | 0.1 | 0.16 | 0.23 | 0.2 | 0.13 | 0.14 |
| 1 | 176954713 | rs16850555 | -0.344 | 0.059 | 1.544 | -1.91 | 0.02 | 0.03 | 0.03 | 0.245 | 0.11 | 0.25 | 0.23 | 0.2 | 0.13 | 0.14 |
| 1 | 176963207 | rs228008 | 0.01 | 0.007 | 0.795 | -2.172 | -0.02 | 0.09 | 0.09 | 0.876 | 0.6 | 0.77 | 0.49 | 0.2 | 0.57 | 0.39 |
| 1 | 176976782 | rs735607 | 1.27 | 0.009 | 0.698 | -2.18 | -0.002 | 0.099 | 0.099 | 0.846 | 0.43 | 0.27 | 0.22 | 0 | 0.43 | 0.25 |
| 1 | 176978465 | rs6661023 | 0.532 | 0.099 | 0.333 | -1.967 | 0 | 0.003 | 0.003 | 1.227 | 0.56 | 0.61 | 0.47 | 0.2 | 0.56 | 0.39 |
| 1 | 176992553 | rs172917 | 0.019 | 0.515 | -0.089 | -1.67 | -0.031 | 0.087 | 0.087 | 0 | 0.63 | 0.79 | 0.5 | 0.21 | 0.59 | 0.39 |
| 1 | 177000503 | rs2223301 | 0.288 | 0.745 | -0.416 | -1.346 | -0.002 | 0.015 | 0.015 | 0 | 0.59 | 0.42 | 0.46 | 0.26 | 0.6 | 0.41 |
| 1 | 177015606 | rs16850689 | -0.116 | 1.001 | -0.792 | -0.748 | -0.036 | -0.051 | -0.051 | 0 | 0.11 | 0.03 | 0.2 | 0.25 | 0.15 | 0.09 |
| 1 | 177016368 | rs6667588 | 0.162 | 0.926 | -0.208 | -1.001 | 0.011 | -0.038 | -0.038 | 0 | 0.65 | 0.61 | 0.48 | 0.22 | 0.56 | 0.32 |
| 1 | 177016829 | rs16850697 | -1.279 | 0.748 | 0.552 | -0.177 | 0.062 | 0.198 | 0.198 | 1.945 | 0.07 | 0.31 | 0.14 | 0.18 | 0.06 | 0.02 |
| 1 | 177021726 | rs2861885 | -1.036 | 0.401 | 0.265 | -0.953 | -0.002 | -0.049 | -0.049 | 0.208 | 0.15 | 0.26 | 0.11 | 0.05 | 0.11 | 0.14 |
| 1 | 177026983 | rs7548151 | -1.02 | 0.369 | 0.266 | -0.953 | -0.006 | -0.048 | -0.048 | 0 | 0.14 | 0.26 | 0.11 | 0.05 | 0.1 | 0.14 |
| 1 | 177054771 | rs6678480 | 0.374 | 0.352 | -0.836 | -1.856 | -0.056 | 0.128 | 0.128 | 0.041 | 0.38 | 0.15 | 0.26 | 0.01 | 0.48 | 0.17 |
| 1 | 177058201 | rs227530 | -0.672 | 0.341 | -0.435 | -1.712 | -0.001 | 0.08 | 0.08 | 0 | 0.58 | 0.73 | 0.36 | 0.14 | 0.57 | 0.32 |
| 1 | 177071733 | rs12722726 | 0.455 | 0.617 | -1.09 | -1.668 | -0.057 | 0.168 | 0.168 | 0.253 | 0.34 | 0.12 | 0.19 | 0.12 | 0.42 | 0.17 |
| 1 | 177074669 | rs7519595 | -0.471 | 0.569 | -1.014 | -1.643 | 0.02 | 0.103 | 0.103 | 0 | 0.51 | 0.34 | 0.34 | 0.14 | 0.59 | 0.32 |
| 1 | 177075950 | rs17314157 | -0.227 | 0.447 | -0.955 | -0.864 | 0.001 | 0.05 | 0.05 | 0 | 0.13 | 0.04 | 0.06 | 0.01 | 0.09 | 0.13 |
| 1 | 177081676 | rs12141039 | -0.415 | 0.297 | -1.436 | -0.263 | 0 | -0.065 | -0.065 | 0 | 0.39 | 0.1 | 0.46 | 0.42 | 0.35 | 0.43 |
| 1 | 177093387 | rs227511 | -0.775 | 0.023 | -0.248 | -0.088 | -0.013 | 0.049 | 0.049 | 4.686 | 0.16 | 0.58 | 0.2 | 0.23 | 0.12 | 0.12 |
| 1 | 177096557 | rs10798497 | 0.149 | -0.045 | -0.571 | -0.383 | 0.002 | 0 | 0 | 0 | 0.12 | 0.13 | 0.08 | 0.04 | 0.09 | 0.13 |
| 1 | 177097442 | rs10798499 | -1.05 | -0.148 | 0.3 | -0.029 | -0.061 | 0.098 | 0.098 | 3.283 | 0.17 | 0.45 | 0.13 | 0.25 | 0.07 | 0.16 |
| 1 | 177103690 | rs10047146 | -0.513 | -0.171 | 0.311 | -0.159 | -0.049 | 0.088 | 0.088 | 6.087 | 0.13 | 0.47 | 0.09 | 0.19 | 0.05 | 0.11 |
| 1 | 177104130 | rs10047083 | -0.49 | -0.166 | 0.315 | -0.157 | -0.048 | 0.085 | 0.085 | 5.895 | 0.13 | 0.47 | 0.09 | 0.18 | 0.05 | 0.11 |
| 1 | 177106260 | rs972484 | -1.598 | -0.185 | 0.413 | -0.198 | 0 | 0 | 0 | 0 | 0.2 | 0.73 | 0.2 | 0.19 | 0.14 | 0.16 |
| 1 | 177129230 | rs11577465 | -1.159 | -0.154 | 0.677 | -0.093 | 0.051 | -0.034 | -0.034 | 0.013 | 0.07 | 0.23 | 0.09 | 0.01 | 0.09 | 0.05 |
| 1 | 177132067 | rs12021769 | -1.505 | -0.149 | 0.177 | -0.082 | -0.002 | -0.001 | -0.001 | 0 | 0.2 | 0.72 | 0.19 | 0.19 | 0.14 | 0.16 |
| 1 | 177148159 | rs725416 | -1.494 | -0.123 | 0.145 | -0.051 | -0.007 | -0.003 | -0.003 | 0.064 | 0.2 | 0.7 | 0.17 | 0.19 | 0.14 | 0.16 |
| 1 | 177152360 | rs12090000 | -0.238 | -0.168 | 0.143 | 0.263 | -0.028 | 0.049 | 0.049 | 3.633 | 0.09 | 0.17 | 0.05 | 0.19 | 0.03 | 0.1 |
| 1 | 177156815 | rs556482 | 0.131 | 0.36 | -0.454 | -0.697 | -0.01 | -0.007 | -0.007 | 0.186 | 0.25 | 0.17 | 0.16 | 0 | 0.28 | 0.13 |
| 1 | 177158553 | rs12117541 | 0.731 | 0.238 | -0.268 | -0.03 | 0.015 | 0.027 | 0.027 | 0.579 | 0.48 | 0.3 | 0.43 | 0.47 | 0.44 | 0.51 |

| | | | | | | | | | | | | | | | | |
|---|---|---|---|---|---|---|---|---|---|---|---|---|---|---|---|---|
| 1 | 177158631 | rs12128612 | 0.649 | 0.17 | -0.254 | -0.108 | 0.008 | 0.027 | 0.027 | 0.199 | 0.51 | 0.36 | 0.44 | 0.47 | 0.46 | 0.55 |
| 1 | 177162809 | rs12031932 | 0.211 | 0.164 | 0.109 | 0.22 | -0.034 | 0.097 | 0.097 | 5.666 | 0.12 | 0.31 | 0.07 | 0.23 | 0.05 | 0.1 |
| 1 | 177165301 | rs10753144 | -0.393 | 0.206 | -0.443 | 0.227 | -0.081 | -0.011 | -0.011 | 0.423 | 0.66 | 0.87 | 0.57 | 0.7 | 0.51 | 0.67 |
| 1 | 177173363 | rs4652232 | 0.699 | 0.216 | -0.531 | 0.113 | 0.016 | 0.028 | 0.028 | 0.445 | 0.48 | 0.3 | 0.43 | 0.46 | 0.44 | 0.52 |
| 1 | 177178708 | rs2294254 | 0.686 | 0.6 | -0.709 | -0.274 | -0.037 | 0.08 | 0.08 | 0 | 0.21 | 0.07 | 0.16 | 0.01 | 0.24 | 0.1 |
| 1 | 177181347 | rs556287 | 0.003 | -0.086 | -0.77 | -0.384 | -0.004 | 0 | 0 | 0 | 0.36 | 0.39 | 0.29 | 0.37 | 0.29 | 0.38 |
| 1 | 177188408 | rs510635 | -0.415 | -0.025 | -0.543 | -0.215 | 0.007 | -0.003 | -0.003 | 0.206 | 0.57 | 0.52 | 0.45 | 0.38 | 0.53 | 0.48 |
| 1 | 177193697 | rs932311 | -0.021 | -0.083 | -0.356 | -0.2 | 0 | -0.001 | -0.001 | 0 | 0.45 | 0.52 | 0.49 | 0.58 | 0.46 | 0.58 |
| 1 | 177195050 | rs760688 | -0.504 | -0.389 | -0.474 | -0.343 | 0.008 | -0.007 | -0.007 | 0.075 | 0.65 | 0.66 | 0.66 | 0.58 | 0.7 | 0.66 |
| 1 | 177197448 | rs989536 | 1.279 | -0.147 | -0.586 | -0.344 | -0.018 | 0.024 | 0.024 | 0 | 0.19 | 0.12 | 0.16 | 0 | 0.23 | 0.08 |
| 1 | 177198998 | rs3176441 | 0.998 | -0.193 | -0.268 | -0.368 | 0.005 | -0.069 | -0.069 | 0 | 0.21 | 0.26 | 0.17 | 0 | 0.23 | 0.09 |
| 1 | 177204902 | rs1879563 | 0.043 | -0.405 | -0.356 | -0.56 | -0.005 | -0.001 | -0.001 | 0 | 0.52 | 0.47 | 0.59 | 0.53 | 0.61 | 0.61 |
| 1 | 177208980 | rs982875 | 0.63 | -0.305 | -0.523 | -0.503 | -0.023 | -0.027 | -0.027 | 0.1 | 0.32 | 0.22 | 0.42 | 0.53 | 0.38 | 0.52 |
| 1 | 177217942 | rs11577869 | 0.692 | -0.338 | -0.553 | -0.517 | -0.065 | -0.074 | -0.074 | 0 | 0.31 | 0.14 | 0.41 | 0.53 | 0.38 | 0.52 |
| 1 | 177230782 | rs12083865 | 1.419 | -0.337 | -0.455 | -0.686 | -0.03 | 0.054 | 0.054 | 0 | 0.2 | 0.12 | 0.16 | 0 | 0.24 | 0.09 |
| 1 | 177231743 | rs4650961 | -1.111 | -0.785 | -0.166 | -0.809 | -0.007 | -0.025 | -0.025 | 0 | 0.84 | 0.68 | 0.88 | 0.88 | 0.83 | 0.8 |
| 1 | 177237533 | rs17380778 | 0.9 | -0.421 | -0.181 | -0.416 | 0.006 | 0.027 | 0.027 | 0.68 | 0.23 | 0.13 | 0.14 | 0.16 | 0.15 | 0.21 |
| 1 | 177250670 | rs1045736 | 1.501 | -0.201 | -0.235 | -0.412 | -0.039 | 0.069 | 0.069 | 0 | 0.19 | 0.06 | 0.15 | 0 | 0.24 | 0.11 |
| 1 | 177258204 | rs16851031 | 0.663 | -0.555 | -0.152 | -0.069 | -0.122 | -0.081 | -0.081 | 0.187 | 0.32 | 0.1 | 0.43 | 0.39 | 0.47 | 0.43 |
| 1 | 177261108 | rs11587000 | 0.834 | -0.648 | 0.062 | -0.134 | 0.004 | 0.009 | 0.009 | 0.572 | 0.23 | 0.17 | 0.14 | 0.16 | 0.16 | 0.2 |
| 1 | 177269474 | rs766021 | -0.89 | -0.761 | 0.55 | -0.167 | -0.014 | 0.007 | 0.007 | 0 | 0.65 | 0.54 | 0.64 | 0.6 | 0.69 | 0.65 |
| 1 | 177271313 | rs16851037 | -0.305 | -0.736 | 0.648 | -0.438 | -0.03 | -0.051 | -0.051 | 0 | 0.34 | 0.43 | 0.21 | 0.21 | 0.22 | 0.22 |
| 1 | 177279412 | rs12569209 | 0.1 | -0.636 | 0.501 | -0.272 | 0.002 | 0 | 0 | 0.841 | 0.37 | 0.34 | 0.44 | 0.52 | 0.48 | 0.43 |
| 1 | 177280491 | rs760687 | 0.428 | -0.633 | 0.482 | -0.279 | -0.082 | -0.048 | -0.048 | 0 | 0.35 | 0.15 | 0.42 | 0.52 | 0.48 | 0.43 |
| 1 | 177281746 | rs16851050 | 0.581 | -0.643 | 0.587 | -0.313 | -0.013 | -0.026 | -0.026 | 0.021 | 0.24 | 0.27 | 0.18 | 0.14 | 0.14 | 0.21 |
| 1 | 177286000 | rs12401643 | -1.023 | -0.64 | 0.584 | -0.342 | -0.017 | -0.02 | -0.02 | 0 | 0.66 | 0.54 | 0.64 | 0.71 | 0.68 | 0.66 |
| 1 | 177300379 | rs1123317 | 0.136 | -0.191 | 0.669 | -0.112 | -0.022 | -0.045 | -0.045 | 0 | 0.24 | 0.36 | 0.19 | 0.13 | 0.15 | 0.21 |
| 1 | 177300809 | rs4453002 | 0.239 | -0.139 | -0.006 | -0.087 | 0.018 | 0.034 | 0.034 | 1.051 | 0.24 | 0.14 | 0.16 | 0.14 | 0.15 | 0.21 |
| 1 | 177303628 | rs10798510 | -1.2 | 0.039 | 0.392 | -0.038 | -0.016 | -0.023 | -0.023 | 0.014 | 0.61 | 0.61 | 0.63 | 0.7 | 0.65 | 0.61 |
| 1 | 177304312 | rs6667899 | 1.778 | 0.063 | 0.125 | -0.069 | -0.009 | 0.016 | 0.016 | 0 | 0.06 | 0 | 0.04 | 0.03 | 0.06 | 0.04 |
| 1 | 177305805 | rs7528533 | 0.211 | -0.007 | 0.169 | -0.041 | -0.103 | -0.076 | -0.076 | 2.08 | 0.31 | 0.09 | 0.36 | 0.5 | 0.45 | 0.37 |
| 1 | 177306387 | rs17659437 | 0.488 | 0.51 | 0.752 | 0.438 | 0.005 | 0.012 | 0.012 | 0.26 | 0.21 | 0.15 | 0.19 | 0.17 | 0.14 | 0.17 |
| 1 | 177309473 | rs6425419 | 0.146 | 0.611 | 0.746 | 0.556 | -0.043 | -0.042 | -0.042 | 0.502 | 0.2 | 0.11 | 0.34 | 0.33 | 0.39 | 0.31 |
| 1 | 177309490 | rs4650963 | 0.611 | 0.705 | 0.704 | -0.371 | 0.014 | 0.026 | 0.026 | 0.358 | 0.2 | 0.13 | 0.19 | 0.13 | 0.14 | 0.17 |
| 1 | 177317366 | rs6678710 | -0.989 | 1.237 | 0.941 | -0.458 | 0.004 | -0.003 | -0.003 | 0 | 0.35 | 0.65 | 0.47 | 0.65 | 0.37 | 0.47 |
| 1 | 177320470 | rs6664016 | -1.617 | 1.174 | 0.938 | -0.472 | 0.008 | 0.032 | 0.032 | 0 | 0.45 | 0.66 | 0.57 | 0.68 | 0.54 | 0.57 |
| 1 | 177321917 | rs9425805 | 0.036 | 1.358 | 0.831 | -0.38 | -0.033 | -0.029 | -0.029 | 0.09 | 0.19 | 0.11 | 0.33 | 0.53 | 0.39 | 0.33 |
| 1 | 177326069 | rs10489219 | -0.393 | 0.45 | 1.224 | -0.232 | -0.026 | -0.049 | -0.049 | 0.398 | 0.18 | 0.59 | 0.2 | 0.12 | 0.09 | 0.15 |
| 1 | 177328296 | rs4652247 | -1.326 | 0.429 | 1.142 | -0.285 | -0.028 | -0.063 | -0.063 | 0 | 0.29 | 0.61 | 0.26 | 0.15 | 0.19 | 0.25 |
| 1 | 177337444 | rs10913310 | 0.325 | 0.738 | 0.868 | -0.164 | 0.011 | 0.015 | 0.015 | 0 | 0.17 | 0.15 | 0.32 | 0.53 | 0.28 | 0.29 |
| 1 | 177341802 | rs2902103 | -0.305 | 0.733 | 0.813 | -0.253 | -0.012 | -0.038 | -0.038 | 0.646 | 0.13 | 0.2 | 0.08 | 0.03 | 0.1 | 0.09 |
| 1 | 177355133 | rs10429832 | -0.904 | 0.744 | 0.563 | -0.191 | -0.049 | -0.037 | -0.037 | 0.066 | 0.28 | 0.13 | 0.37 | 0.53 | 0.37 | 0.39 |
| 1 | 177363013 | rs736840 | 0.763 | 0.663 | 0.621 | -0.229 | -0.001 | -0.001 | -0.001 | 0 | 0.12 | 0.18 | 0.17 | 0.14 | 0.09 | 0.16 |
| 1 | 177376746 | rs12135765 | 0.386 | 0.662 | 0.575 | -0.266 | -0.008 | 0.047 | 0.047 | 0.893 | 0.11 | 0 | 0.06 | 0.03 | 0.1 | 0.09 |
| 1 | 177381322 | rs2142754 | -0.133 | 0.735 | 0.764 | -0.278 | 0.034 | 0.049 | 0.049 | 0.345 | 0.2 | 0.26 | 0.32 | 0.51 | 0.27 | 0.3 |
| 1 | 177383443 | rs11581304 | 0.831 | 0.649 | 0.84 | -0.224 | -0.001 | 0.001 | 0.001 | 0 | 0.12 | 0.08 | 0.16 | 0.14 | 0.09 | 0.16 |
| 1 | 177384036 | rs9662701 | -1.732 | 0.663 | 0.828 | -0.211 | -0.003 | 0.021 | 0.021 | 0 | 0.45 | 0.63 | 0.56 | 0.68 | 0.46 | 0.55 |
| 1 | 177386423 | rs2207029 | -1.904 | 0.491 | 0.832 | -0.255 | 0.012 | 0.054 | 0.054 | 0 | 0.53 | 0.68 | 0.6 | 0.68 | 0.51 | 0.56 |
| 1 | 177395665 | rs16851250 | -0.089 | 0.121 | 0.699 | 0.28 | -0.004 | -0.005 | -0.005 | 0 | 0.14 | 0.2 | 0.17 | 0.14 | 0.09 | 0.16 |
| 1 | 177403769 | rs7543021 | -1.386 | 0.274 | 0.447 | 0.177 | 0.011 | -0.001 | -0.001 | 0.051 | 0.34 | 0.56 | 0.33 | 0.63 | 0.32 | 0.43 |
| 1 | 177413184 | rs1883903 | -0.862 | 0.266 | 0.464 | 0.119 | 0.01 | 0.042 | 0.042 | 0.05 | 0.22 | 0.14 | 0.17 | 0.09 | 0.15 | 0.24 |
| 1 | 177414054 | rs1883902 | -0.587 | 0.71 | 0.713 | 0.491 | 0.026 | 0.089 | 0.089 | 0 | 0.21 | 0.39 | 0.21 | 0.55 | 0.23 | 0.27 |
| 1 | 177418397 | rs4650965 | -0.135 | 0.73 | 0.141 | 0.503 | -0.027 | -0.048 | -0.048 | 0 | 0.17 | 0.1 | 0.18 | 0.56 | 0.23 | 0.29 |
| 1 | 177427162 | rs11578222 | 0.193 | 0.781 | -0.218 | 0.508 | -0.001 | 0.006 | 0.006 | 0 | 0.12 | 0.11 | 0.12 | 0.07 | 0.08 | 0.17 |
| 1 | 177431168 | rs715359 | -1.013 | 0.769 | 0.044 | 1.066 | 0.033 | 0.12 | 0.12 | 0 | 0.22 | 0.51 | 0.22 | 0.54 | 0.23 | 0.29 |
| 1 | 177433277 | rs6675650 | -1.065 | 0.555 | -0.889 | 1.073 | -0.007 | -0.059 | -0.059 | 0 | 0.27 | 0.13 | 0.22 | 0.55 | 0.28 | 0.35 |
| 1 | 177434054 | rs7521346 | -1.351 | -0.127 | 0.103 | 1.46 | -0.05 | 0.156 | 0.156 | 0 | 0.21 | 0.5 | 0.16 | 0.44 | 0.16 | 0.31 |
| 1 | 177436018 | rs6425426 | -1.613 | -0.396 | -0.034 | 0.802 | 0.002 | -0.004 | -0.004 | 3.434 | 0.37 | 0.37 | 0.3 | 0.48 | 0.3 | 0.42 |
| 1 | 177443057 | rs10798522 | -1.788 | -0.584 | 0.041 | 0.665 | 0.009 | -0.012 | -0.012 | 0 | 0.38 | 0.35 | 0.28 | 0.48 | 0.31 | 0.43 |
| 1 | 177450874 | rs1885021 | -1.436 | -0.631 | 0.074 | 0.656 | 0.012 | -0.109 | -0.109 | 0.245 | 0.34 | 0.13 | 0.23 | 0.48 | 0.29 | 0.42 |
| 1 | 177458170 | rs12038173 | -1.783 | -1.148 | 0.838 | 0.647 | -0.022 | 0.018 | 0.018 | 0.378 | 0.36 | 0.52 | 0.22 | 0.53 | 0.28 | 0.4 |
| 1 | 177463978 | rs2902113 | -1.937 | -1.485 | 0.134 | 0.049 | -0.095 | 0.052 | 0.052 | 0 | 0.62 | 0.89 | 0.59 | 0.69 | 0.51 | 0.63 |
| 1 | 177466529 | rs10737324 | -1.375 | -1.497 | 0.407 | 0.048 | 0.024 | -0.018 | -0.018 | 0 | 0.38 | 0.64 | 0.23 | 0.53 | 0.28 | 0.4 |
| 1 | 177468716 | rs12089461 | 0.291 | -1.72 | 0.779 | 0.367 | -0.002 | 0.015 | 0.015 | 0 | 0.17 | 0.16 | 0.12 | 0.38 | 0.13 | 0.3 |
| 1 | 177470662 | rs12567599 | 0.305 | -1.72 | 0.779 | 0.367 | -0.003 | 0.015 | 0.015 | 0 | 0.17 | 0.16 | 0.12 | 0.38 | 0.14 | 0.31 |
| 1 | 177480934 | rs12047881 | -1.154 | -1.459 | 0.703 | 0.073 | -0.02 | 0.019 | 0.019 | 0.219 | 0.35 | 0.42 | 0.23 | 0.53 | 0.28 | 0.4 |
| 1 | 177482911 | rs12047065 | -1.071 | -1.432 | 0.393 | 0.084 | -0.026 | 0.023 | 0.023 | 0.281 | 0.36 | 0.5 | 0.23 | 0.53 | 0.28 | 0.4 |
| 1 | 177486382 | rs1321774 | 0.118 | -1.249 | 0.486 | 0.168 | -0.001 | 0.006 | 0.006 | 0.063 | 0.23 | 0.17 | 0.32 | 0.16 | 0.23 | 0.23 |
| 1 | 177493799 | rs841789 | -0.179 | -1.256 | 0.187 | 0.12 | 0.001 | 0.001 | 0.001 | 0.003 | 0.15 | 0.26 | 0.26 | 0.13 | 0.14 | 0.16 |

| | | | | | | | | | | | | | | | | |
|---|---|---|---|---|---|---|---|---|---|---|---|---|---|---|---|---|
| 1 | 177508044 | rs1098777 | -1.839 | -0.739 | -0.707 | -0.195 | -0.087 | 0.043 | 0.043 | 0 | 0.64 | 0.89 | 0.7 | 0.7 | 0.56 | 0.67 |
| 1 | 177512260 | rs10737325 | -0.697 | -0.825 | -0.112 | -0.2 | 0 | 0 | 0 | 2.482 | 0.48 | 0.49 | 0.42 | 0.56 | 0.42 | 0.51 |
| 1 | 177525531 | rs2093203 | -0.979 | -0.83 | -0.078 | -0.206 | 0.018 | 0.017 | 0.017 | 0 | 0.53 | 0.65 | 0.44 | 0.56 | 0.44 | 0.51 |
| 1 | 177537041 | rs841775 | 0.452 | -0.787 | -0.024 | -0.201 | 0.003 | 0.003 | 0.003 | 0 | 0.13 | 0.15 | 0.26 | 0.13 | 0.14 | 0.17 |
| 1 | 177560435 | rs1415339 | -0.175 | -1.164 | -0.213 | -0.359 | -0.006 | 0.007 | 0.007 | 0 | 0.36 | 0.59 | 0.33 | 0.49 | 0.26 | 0.45 |
| 1 | 177566659 | rs1890035 | -1.587 | -1.185 | -0.337 | -0.388 | 0.012 | 0.011 | 0.011 | 0.037 | 0.55 | 0.72 | 0.45 | 0.56 | 0.44 | 0.51 |
| 1 | 177586110 | rs868715 | -0.276 | -0.93 | -0.089 | -0.193 | -0.023 | 0.039 | 0.039 | 0 | 0.34 | 0.47 | 0.32 | 0.49 | 0.26 | 0.45 |
| 1 | 177586974 | rs867541 | -0.706 | -0.817 | -0.856 | -0.135 | 0.009 | 0.008 | 0.008 | 0 | 0.87 | 0.8 | 0.75 | 0.87 | 0.87 | 0.84 |
| 1 | 177587339 | rs10913387 | -0.565 | -0.757 | -0.654 | -0.043 | 0.027 | -0.025 | -0.025 | 0.005 | 0.37 | 0.72 | 0.34 | 0.49 | 0.26 | 0.45 |
| 1 | 177591270 | rs1415334 | 1.814 | -0.037 | -0.372 | -0.365 | 0.002 | -0.012 | -0.012 | 0 | 0.34 | 0.35 | 0.33 | 0.2 | 0.41 | 0.3 |
| 1 | 177613863 | rs12140710 | 0.775 | -0.092 | -0.519 | -0.518 | 0.012 | 0.152 | 0.152 | 0.184 | 0.51 | 0.24 | 0.37 | 0.12 | 0.57 | 0.34 |
| 1 | 177621201 | rs1572511 | 0.885 | -0.121 | -0.361 | -0.529 | -0.001 | 0.031 | 0.031 | 0 | 0.49 | 0.44 | 0.4 | 0.2 | 0.52 | 0.36 |
| 1 | 177625201 | rs4652252 | 1.802 | -0.133 | 0.083 | -0.544 | 0.004 | -0.021 | -0.021 | 0 | 0.35 | 0.41 | 0.33 | 0.2 | 0.41 | 0.31 |
| 1 | 177626285 | rs1252066 | -0.128 | -0.203 | 0.289 | -0.403 | -0.009 | -0.007 | -0.007 | 0 | 0.32 | 0.44 | 0.31 | 0.69 | 0.29 | 0.44 |
| 1 | 177629723 | rs17625634 | 1.195 | -0.223 | 0.31 | -0.16 | -0.005 | -0.003 | -0.003 | 1.796 | 0.13 | 0.14 | 0.11 | 0.14 | 0.18 | 0.16 |
| 1 | 177631002 | rs10798559 | 1.213 | -0.095 | 0.418 | -0.292 | 0 | 0.057 | 0.057 | 0.07 | 0.53 | 0.38 | 0.37 | 0.12 | 0.55 | 0.33 |
| 1 | 177639015 | rs7529907 | -0.747 | -0.269 | 0.005 | -0.218 | -0.019 | 0.075 | 0.075 | 0 | 0.29 | 0.12 | 0.37 | 0.11 | 0.29 | 0.24 |
| 1 | 177648733 | rs12035381 | 0.317 | -0.017 | 0.563 | 0.303 | 0.006 | 0.011 | 0.011 | 0.297 | 0.19 | 0.33 | 0.16 | 0.22 | 0.18 | 0.19 |
| 1 | 177648870 | rs16851688 | 0.037 | -0.053 | -0.006 | 0.149 | 0.003 | 0.009 | 0.009 | 0 | 0.09 | 0.16 | 0.28 | 0.11 | 0.11 | 0.13 |
| 1 | 177649849 | rs1923630 | 1.163 | -0.588 | -0.141 | -0.099 | -0.043 | 0.218 | 0.218 | 0 | 0.46 | 0.18 | 0.34 | 0.12 | 0.51 | 0.32 |
| 1 | 177651210 | rs2104212 | 0.281 | -1.122 | 0.722 | 0.695 | -0.02 | 0.12 | 0.12 | 0 | 0.8 | 0.71 | 0.81 | 0.49 | 0.84 | 0.7 |
| 1 | 177652462 | rs1885646 | 0.719 | -1.51 | 0.089 | -0.137 | 0.053 | 0.056 | 0.056 | 0.379 | 0.73 | 0.83 | 0.89 | 0.86 | 0.84 | 0.81 |
| 1 | 177656039 | rs2224639 | 0.262 | -1.356 | 0.001 | -0.025 | 0.024 | -0.014 | -0.014 | 4.999 | 0.08 | 0.03 | 0.02 | 0.14 | 0.05 | 0.12 |
| 1 | 177669302 | rs12139234 | 1.749 | -1.253 | -0.217 | -0.355 | -0.045 | 0.123 | 0.123 | 0 | 0.32 | 0.1 | 0.34 | 0.09 | 0.35 | 0.25 |
| 1 | 177670577 | rs2181263 | 0.416 | -1.124 | -0.116 | 0.093 | -0.001 | 0.029 | 0.029 | 0 | 0.11 | 0.08 | 0.16 | 0.39 | 0.13 | 0.23 |
| 1 | 177671478 | rs4509537 | -0.321 | -1.193 | 0.337 | 0.03 | -0.024 | 0.017 | 0.017 | 0 | 0.41 | 0.28 | 0.41 | 0.34 | 0.44 | 0.36 |
| 1 | 177680031 | rs7542551 | 0.354 | -0.704 | 0.712 | 0.305 | -0.008 | -0.006 | -0.006 | 0.056 | 0.26 | 0.23 | 0.34 | 0.31 | 0.34 | 0.29 |
| 1 | 177700306 | rs3131308 | -0.632 | -0.637 | 0.553 | 0.15 | 0.007 | -0.023 | -0.023 | 0 | 0.39 | 0.43 | 0.52 | 0.72 | 0.49 | 0.51 |
| 1 | 177702841 | rs1923643 | -0.281 | -1.093 | 0.546 | 0.401 | -0.023 | 0.146 | 0.146 | 0.644 | 0.14 | 0.28 | 0.18 | 0.39 | 0.11 | 0.25 |
| 1 | 177710595 | rs2065140 | -0.105 | -1.331 | 0.149 | 0.165 | 0.044 | 0.094 | 0.094 | 0 | 0.25 | 0.04 | 0.08 | 0.14 | 0.18 | 0.19 |
| 1 | 177710610 | rs4652262 | 0.019 | -1.192 | 0.377 | 0.272 | -0.007 | 0.06 | 0.06 | 0.003 | 0.13 | 0.16 | 0.19 | 0.4 | 0.1 | 0.25 |
| 1 | 177714468 | rs2902197 | 0.146 | -1.123 | 0.196 | 0.233 | -0.021 | 0.131 | 0.131 | 0 | 0.13 | 0.21 | 0.2 | 0.39 | 0.11 | 0.21 |
| 1 | 177719196 | rs3131323 | 1.031 | -1.204 | -1.116 | 0.181 | 0.136 | -0.035 | -0.035 | 0.007 | 0.55 | 0.19 | 0.66 | 0.46 | 0.71 | 0.57 |
| 1 | 177723089 | rs3131325 | 0.893 | -0.837 | -1.308 | 0.069 | -0.047 | -0.039 | -0.039 | 1.112 | 0.23 | 0.14 | 0.32 | 0.36 | 0.34 | 0.28 |
| 1 | 177737303 | rs1885644 | -0.395 | -0.541 | -1.582 | -0.43 | 0.042 | 0.134 | 0.134 | 0 | 0.28 | 0.04 | 0.1 | 0.13 | 0.21 | 0.21 |
| 1 | 177739231 | rs932676 | 0.011 | -0.682 | -1.156 | 0.094 | 0.011 | 0.028 | 0.028 | 0 | 0.48 | 0.56 | 0.58 | 0.71 | 0.56 | 0.54 |
| 1 | 177742007 | rs3131333 | -0.423 | 0.625 | -1.079 | 0.668 | -0.004 | 0.016 | 0.016 | 0 | 0.68 | 0.73 | 0.65 | 0.83 | 0.66 | 0.68 |
| 1 | 177748866 | rs878833 | -0.749 | 0.472 | -0.96 | 1.647 | -0.008 | 0.07 | 0.07 | 0 | 0.08 | 0.1 | 0.13 | 0.28 | 0.09 | 0.16 |
| 1 | 177749363 | rs16851861 | -0.604 | 1.052 | -1.534 | 1.479 | -0.039 | -0.043 | -0.043 | 0 | 0.21 | 0.07 | 0.18 | 0.29 | 0.27 | 0.22 |
| 1 | 177750489 | rs2902199 | -0.25 | 0.626 | -2.108 | 0.959 | 0.014 | 0.05 | 0.05 | 1.969 | 0.39 | 0.81 | 0.44 | 0.28 | 0.35 | 0.32 |
| 1 | 177751488 | rs746584 | 0.022 | 0.23 | -1.911 | 1.025 | -0.014 | -0.017 | -0.017 | 0.164 | 0.23 | 0.11 | 0.3 | 0.31 | 0.27 | 0.24 |
| 1 | 177752701 | rs2093519 | -0.308 | 0.059 | -0.672 | 0.922 | 0.011 | 0.003 | 0.003 | 0 | 0.26 | 0.35 | 0.34 | 0.31 | 0.31 | 0.24 |
| 1 | 177755464 | rs10913419 | -0.574 | -0.221 | -1.371 | 1.355 | -0.012 | -0.001 | -0.001 | 0 | 0.39 | 0.2 | 0.43 | 0.67 | 0.41 | 0.48 |
| 1 | 177758118 | rs10737331 | -0.752 | -0.686 | -0.75 | 0.962 | 0.029 | -0.077 | -0.077 | 1.407 | 0.3 | 0.11 | 0.22 | 0.41 | 0.23 | 0.32 |
| 1 | 177760918 | rs9970077 | 0.514 | -0.58 | -0.656 | 1.113 | 0 | -0.02 | -0.02 | 0 | 0.17 | 0.12 | 0.14 | 0.41 | 0.15 | 0.25 |
| 1 | 177762061 | rs6425445 | -0.007 | -0.838 | -0.423 | 0.765 | 0.007 | 0.006 | 0.006 | 0.911 | 0.15 | 0.12 | 0.09 | 0.05 | 0.09 | 0.08 |
| 1 | 177762940 | rs7411090 | -0.145 | -1.09 | -0.747 | 1.163 | -0.015 | -0.048 | -0.048 | 0 | 0.39 | 0.16 | 0.35 | 0.55 | 0.42 | 0.35 |
| 1 | 177766616 | rs3922649 | -0.612 | -1.546 | -0.328 | 0.835 | 0.013 | 0.096 | 0.096 | 0.253 | 0.48 | 0.69 | 0.55 | 0.76 | 0.45 | 0.62 |
| 1 | 177769687 | rs4072161 | -0.09 | -1.483 | 0.078 | 0.814 | 0 | -0.02 | -0.02 | 1.009 | 0.42 | 0.45 | 0.45 | 0.76 | 0.41 | 0.5 |
| 1 | 177770655 | rs10913425 | 0.468 | -1.447 | -0.068 | 0.838 | -0.01 | -0.021 | -0.021 | 0 | 0.06 | 0.2 | 0.09 | 0 | 0.03 | 0.12 |
| 1 | 177770978 | rs4652267 | 1.339 | -1.35 | -0.31 | 1.023 | -0.012 | 0.051 | 0.051 | 0 | 0.07 | 0.12 | 0.12 | 0.22 | 0.03 | 0.17 |
| 1 | 177772815 | rs4442331 | -1.029 | -0.446 | -1.036 | 0.547 | -0.055 | 0.068 | 0.068 | 2.726 | 0.67 | 0.95 | 0.73 | 0.81 | 0.65 | 0.75 |
| 1 | 177778091 | rs12135579 | 1.06 | -0.158 | -0.662 | 0.618 | -0.062 | 0.029 | 0.029 | 0 | 0.17 | 0.03 | 0.16 | 0.05 | 0.2 | 0.12 |
| 1 | 177778970 | rs10913431 | 0.909 | -0.365 | -0.269 | 0.775 | -0.015 | 0.115 | 0.115 | 0 | 0.08 | 0.24 | 0.13 | 0.2 | 0.03 | 0.17 |
| 1 | 177782904 | rs6674700 | -0.971 | -0.501 | -1.022 | 0.351 | -0.013 | -0.007 | -0.007 | 0 | 0.9 | 0.99 | 0.87 | 0.91 | 0.89 | 0.97 |
| 1 | 177783644 | rs4652270 | 0.784 | -1.127 | -1.121 | 0.127 | 0.029 | -0.029 | -0.029 | 0.554 | 0.42 | 0.19 | 0.37 | 0.55 | 0.38 | 0.38 |
| 1 | 177783670 | rs4652271 | 2.298 | -0.97 | -1.238 | 0.407 | -0.03 | -0.053 | -0.053 | 0 | 0.09 | 0.01 | 0.1 | 0.17 | 0.11 | 0.05 |
| 1 | 177784137 | rs4081366 | -0.45 | -0.906 | -1.031 | 0.138 | 0 | 0.029 | 0.029 | 0.028 | 0.38 | 0.78 | 0.39 | 0.3 | 0.33 | 0.43 |
| 1 | 177788008 | rs10798570 | 0.468 | -1.098 | -0.339 | 0.356 | -0.04 | 0.051 | 0.051 | 0 | 0.17 | 0.4 | 0.19 | 0.2 | 0.09 | 0.29 |
| 1 | 177790062 | rs4081367 | 0.452 | -1.1 | -0.331 | 0.355 | -0.041 | 0.061 | 0.061 | 0 | 0.17 | 0.43 | 0.19 | 0.2 | 0.09 | 0.29 |
| 1 | 177791110 | rs10798572 | 0.055 | -1.068 | -1.182 | 0.237 | 0.103 | 0.034 | 0.034 | 0 | 0.48 | 0.82 | 0.48 | 0.65 | 0.34 | 0.6 |
| 1 | 177791775 | rs12748219 | 1.022 | -0.7 | -0.847 | -0.026 | 0.024 | 0.04 | 0.04 | 0.52 | 0.19 | 0.08 | 0.15 | 0.07 | 0.14 | 0.08 |
| 1 | 177792715 | rs2068973 | -0.173 | -0.249 | -0.327 | -0.292 | 0.009 | -0.086 | -0.086 | 1.993 | 0.34 | 0.51 | 0.31 | 0.1 | 0.33 | 0.2 |
| 1 | 177793990 | rs12735595 | 2.066 | -0.144 | -0.361 | -0.243 | -0.046 | 0.041 | 0.041 | 0 | 0.11 | 0.03 | 0.12 | 0.03 | 0.18 | 0.12 |
| 1 | 177797447 | rs7542718 | 2.125 | -0.453 | -0.046 | 0.055 | -0.01 | 0.04 | 0.04 | 0 | 0.07 | 0.1 | 0.11 | 0.19 | 0.03 | 0.17 |
| 1 | 177803087 | rs1854288 | 0.219 | -0.507 | 0.072 | -0.003 | 0.01 | -0.046 | -0.046 | 0 | 0.31 | 0.53 | 0.3 | 0.18 | 0.31 | 0.2 |
| 1 | 177806838 | rs2902200 | 2.146 | -0.48 | 0.146 | -0.02 | 0.003 | -0.005 | -0.005 | 0 | 0.11 | 0.08 | 0.13 | 0.03 | 0.18 | 0.12 |
| 1 | 177807335 | rs12043001 | 1.395 | -0.552 | 0.152 | -0.029 | 0.02 | 0.03 | 0.03 | 0.56 | 0.19 | 0.11 | 0.15 | 0.07 | 0.14 | 0.08 |
| 1 | 177809627 | rs10798574 | 0.347 | -0.521 | 0.004 | -0.02 | -0.007 | 0.013 | 0.013 | 0 | 0.29 | 0.23 | 0.27 | 0.18 | 0.31 | 0.2 |

| | | | | | | | | | | | | | | | | |
|---|---|---|---|---|---|---|---|---|---|---|---|---|---|---|---|---|
| 1 | 177811499 | rs34612022 | 1.371 | -0.541 | -0.008 | -0.035 | 0.02 | 0.031 | 0.031 | 0.505 | 0.19 | 0.11 | 0.15 | 0.07 | 0.14 | 0.08 |
| 1 | 177811939 | rs12737338 | 1.739 | -0.484 | 0.011 | -0.027 | 0.013 | -0.019 | -0.019 | 0 | 0.13 | 0.13 | 0.14 | 0.03 | 0.18 | 0.12 |
| 1 | 177814914 | rs3131310 | 1.402 | -0.427 | -0.467 | -0.055 | -0.092 | -0.012 | -0.012 | 0 | 0.19 | 0.01 | 0.23 | 0.16 | 0.29 | 0.2 |
| 1 | 177815433 | rs12091553 | 0.253 | -0.711 | 0.056 | 0.012 | -0.008 | 0.013 | 0.013 | 0 | 0.29 | 0.23 | 0.28 | 0.18 | 0.32 | 0.2 |
| 1 | 177817796 | rs12022841 | 1.404 | -1.556 | 0.222 | 0.418 | 0.013 | 0.005 | 0.005 | 0 | 0.16 | 0.1 | 0.14 | 0.15 | 0.13 | 0.08 |
| 1 | 177820480 | rs3131312 | -0.657 | -1.692 | 0.097 | 0.414 | -0.02 | 0.018 | 0.018 | 0 | 0.86 | 0.76 | 0.84 | 0.81 | 0.91 | 0.71 |
| 1 | 177821366 | rs3131313 | 0.598 | -1.726 | -0.12 | 0.368 | 0.027 | -0.001 | -0.001 | 0.324 | 0.66 | 0.28 | 0.66 | 0.66 | 0.79 | 0.63 |
| 1 | 177822050 | rs3131314 | -1.59 | -1.573 | -0.134 | 0.101 | -0.01 | -0.031 | -0.031 | 0 | 0.92 | 0.87 | 0.95 | 1 | 0.95 | 0.89 |
| 1 | 177828915 | rs12747989 | 0.949 | -1.609 | -0.467 | 0.146 | 0.032 | 0.045 | 0.045 | 0.347 | 0.19 | 0.07 | 0.15 | 0.08 | 0.14 | 0.08 |
| 1 | 177830530 | rs3131322 | 0.441 | -1.587 | -0.338 | 0.238 | 0.024 | 0.007 | 0.007 | 0 | 0.66 | 0.28 | 0.66 | 0.73 | 0.78 | 0.65 |
| 1 | 177836264 | rs943763 | -0.047 | -1.268 | -0.093 | 0.338 | -0.043 | 0.015 | 0.015 | 0 | 0.82 | 0.38 | 0.79 | 0.81 | 0.91 | 0.71 |
| 1 | 177839612 | rs10489881 | 0.391 | -1.413 | -0.511 | 0.475 | 0.001 | -0.001 | -0.001 | 0.565 | 0.56 | 0.55 | 0.63 | 0.53 | 0.73 | 0.53 |
| 1 | 177841895 | rs2902202 | -0.455 | -0.783 | -0.566 | 0.37 | 0.007 | -0.003 | -0.003 | 0 | 0.77 | 0.75 | 0.75 | 0.77 | 0.72 | 0.7 |
| 1 | 177842162 | rs943764 | 0.543 | -0.878 | -0.799 | 0.129 | -0.022 | 0.067 | 0.067 | 0.03 | 0.39 | 0.17 | 0.39 | 0.26 | 0.44 | 0.38 |
| 1 | 177843620 | rs2862278 | 1.314 | -0.598 | -0.478 | -0.013 | 0.005 | 0.051 | 0.051 | 0.568 | 0.19 | 0.08 | 0.12 | 0.05 | 0.17 | 0.07 |
| 1 | 177844043 | rs12040283 | -0.063 | -0.535 | 0.015 | 0.295 | -0.029 | 0.038 | 0.038 | 0 | 0.86 | 0.65 | 0.92 | 0.79 | 0.93 | 0.83 |
| 1 | 177845657 | rs16852055 | 0.932 | -0.498 | -0.379 | 0.066 | -0.004 | -0.019 | -0.019 | 0 | 0.16 | 0.18 | 0.18 | 0.04 | 0.17 | 0.18 |
| 1 | 177852580 | rs633715 | 1.432 | 0.111 | -0.43 | 0.557 | -0.035 | -0.03 | -0.03 | 0 | 0.13 | 0.08 | 0.18 | 0.15 | 0.2 | 0.15 |
| 1 | 177856925 | rs10798578 | 0.053 | 0.053 | -0.108 | 0.811 | 0.003 | 0.005 | 0.005 | 0.529 | 0.72 | 0.72 | 0.59 | 0.51 | 0.65 | 0.64 |
| 1 | 177864554 | rs10798580 | 1.055 | -0.224 | -0.289 | 0.514 | 0.004 | 0.096 | 0.096 | 0 | 0.4 | 0.23 | 0.4 | 0.14 | 0.42 | 0.42 |
| 1 | 177867608 | rs6663622 | 0.536 | -0.096 | 0.095 | 0.736 | 0.009 | 0.001 | 0.001 | 0.18 | 0.67 | 0.41 | 0.54 | 0.43 | 0.61 | 0.62 |
| 1 | 177868882 | rs527065 | -0.133 | -0.1 | 0.169 | 0.724 | -0.019 | -0.016 | -0.016 | 0 | 0.24 | 0.21 | 0.4 | 0.57 | 0.33 | 0.28 |
| 1 | 177873333 | rs10157480 | 0.462 | -0.107 | 0.27 | 0.729 | 0.011 | 0.001 | 0.001 | 0.037 | 0.68 | 0.64 | 0.56 | 0.42 | 0.61 | 0.62 |
| 1 | 177885762 | rs630372 | 0.589 | -0.051 | 0.048 | 0.754 | 0.045 | 0.051 | 0.051 | 0 | 0.19 | 0.38 | 0.22 | 0.23 | 0.22 | 0.16 |
| 1 | 177887214 | rs480603 | -1.632 | -0.116 | -0.024 | 0.668 | -0.007 | -0.031 | -0.031 | 0.065 | 0.92 | 0.84 | 0.96 | 0.99 | 0.95 | 0.9 |
| 1 | 177888756 | rs17275640 | 2.181 | -0.128 | -0.374 | 0.628 | -0.014 | 0.011 | 0.011 | 0 | 0.1 | 0.01 | 0.13 | 0.03 | 0.12 | 0.14 |
| 1 | 177888809 | rs16852100 | 1.219 | 0.158 | -0.22 | 0.988 | -0.009 | 0.044 | 0.044 | 0 | 0.06 | 0.03 | 0.19 | 0.36 | 0.11 | 0.12 |
| 1 | 177889480 | rs543874 | 1.329 | 0.239 | -0.101 | 0.909 | 0.033 | 0.025 | 0.025 | 0 | 0.15 | 0.28 | 0.19 | 0.15 | 0.19 | 0.15 |
| 1 | 177890153 | rs570762 | 0.16 | -0.05 | -0.179 | 0.682 | 0 | 0 | 0 | 0.375 | 0.63 | 0.66 | 0.63 | 0.33 | 0.63 | 0.62 |
| 1 | 177898799 | rs623479 | 0.132 | 0.111 | -0.222 | 0.672 | 0.024 | 0.041 | 0.041 | 0 | 0.5 | 0.46 | 0.64 | 0.71 | 0.59 | 0.53 |
| 1 | 177900098 | rs575908 | 0.459 | 0.23 | -0.188 | 0.733 | 0.018 | 0.029 | 0.029 | 0 | 0.2 | 0.29 | 0.37 | 0.54 | 0.29 | 0.27 |
| 1 | 177901619 | rs7531399 | 1.663 | 0.151 | -0.265 | 0.392 | 0 | 0.052 | 0.052 | 0 | 0.27 | 0.16 | 0.25 | 0.11 | 0.26 | 0.24 |
| 1 | 177901713 | rs522367 | 0.571 | 0.176 | -0.115 | 0.54 | 0.006 | -0.011 | -0.011 | 4.384 | 0.42 | 0.49 | 0.44 | 0.28 | 0.44 | 0.4 |
| 1 | 177902725 | rs3813649 | 1.49 | 0.13 | -0.251 | 0.306 | 0.002 | 0.019 | 0.019 | 0 | 0.28 | 0.24 | 0.26 | 0.11 | 0.26 | 0.24 |
| 1 | 177904495 | rs10489882 | 0.842 | 0.283 | -0.614 | 0.336 | 0.005 | 0.138 | 0.138 | 0 | 0.46 | 0.2 | 0.37 | 0.23 | 0.44 | 0.3 |
| 1 | 177905174 | rs2902210 | 0.548 | 0.383 | -0.237 | 0.569 | 0.016 | -0.017 | -0.017 | 0.257 | 0.65 | 0.42 | 0.55 | 0.27 | 0.61 | 0.53 |
| 1 | 177905469 | rs7522194 | 1.487 | 0.383 | -0.683 | 0.722 | -0.005 | 0.046 | 0.046 | 0 | 0.27 | 0.1 | 0.23 | 0.17 | 0.26 | 0.25 |
| 1 | 177908852 | rs943762 | -0.113 | 0.934 | -0.216 | 1.42 | -0.013 | 0.001 | 0.001 | 0 | 0.08 | 0.05 | 0.18 | 0.33 | 0.12 | 0.16 |
| 1 | 177915474 | rs842192 | 0.646 | 0.87 | -0.027 | 1.272 | -0.005 | 0.024 | 0.024 | 0 | 0.83 | 0.63 | 0.88 | 0.82 | 0.86 | 0.84 |
| 1 | 177918165 | rs1984159 | -0.325 | 1.025 | -0.895 | 1.363 | -0.012 | -0.038 | -0.038 | 0 | 0.87 | 0.96 | 0.76 | 0.7 | 0.84 | 0.75 |
| 1 | 177918499 | rs10798583 | -0.031 | 1.201 | -0.935 | 1.038 | -0.039 | -0.058 | -0.058 | 0 | 0.75 | 0.86 | 0.63 | 0.51 | 0.67 | 0.57 |
| 1 | 177920754 | rs3828137 | -0.335 | 0.857 | -0.893 | 0.219 | 0.01 | 0.035 | 0.035 | 0.128 | 0.42 | 0.7 | 0.39 | 0.13 | 0.41 | 0.28 |
| 1 | 177923440 | rs12129817 | 0.418 | 1.018 | -1.032 | 0.313 | -0.001 | 0.009 | 0.009 | 0 | 0.19 | 0.18 | 0.2 | 0.03 | 0.23 | 0.21 |
| 1 | 177929554 | rs10913472 | -0.416 | 1.136 | -1.024 | 0.758 | -0.029 | -0.01 | -0.01 | 1.237 | 0.39 | 0.26 | 0.53 | 0.57 | 0.46 | 0.6 |
| 1 | 177929940 | rs7413442 | 0.198 | 0.591 | -0.844 | 0.713 | -0.023 | -0.027 | -0.027 | 0.255 | 0.25 | 0.57 | 0.2 | 0.19 | 0.18 | 0.12 |
| 1 | 177930096 | rs867275 | 0.813 | 0.876 | -1.114 | 1.135 | -0.018 | -0.038 | -0.038 | 1.385 | 0.23 | 0.11 | 0.34 | 0.31 | 0.27 | 0.37 |
| 1 | 177930319 | rs945639 | 0.941 | 0.844 | -1.112 | 1.274 | -0.019 | -0.024 | -0.024 | 0 | 0.23 | 0.11 | 0.34 | 0.27 | 0.28 | 0.37 |
| 1 | 177930894 | rs945641 | -1.806 | 1.102 | -1.247 | 0.962 | 0.013 | 0.03 | 0.03 | 0 | 0.7 | 0.89 | 0.78 | 0.77 | 0.74 | 0.77 |
| 1 | 177931122 | rs10798585 | 0.687 | 1.131 | -1.097 | 1.018 | -0.009 | -0.013 | -0.013 | 0.13 | 0.18 | 0.12 | 0.21 | 0.33 | 0.22 | 0.26 |
| 1 | 177931508 | rs12747656 | -0.703 | 1.067 | -0.773 | 1.02 | 0.007 | 0.005 | 0.005 | 0 | 0.42 | 0.48 | 0.38 | 0.42 | 0.44 | 0.35 |
| 1 | 177935443 | rs954503 | -0.635 | 1.085 | -0.84 | 1.029 | 0.005 | 0.003 | 0.003 | 0 | 0.41 | 0.42 | 0.38 | 0.42 | 0.44 | 0.35 |
| 1 | 177936525 | rs12137761 | 0.92 | 0.843 | -0.746 | 0.508 | -0.001 | 0.033 | 0.033 | 0.401 | 0.17 | 0.09 | 0.13 | 0.06 | 0.16 | 0.07 |
| 1 | 177936536 | rs6667659 | -0.291 | 0.921 | -0.512 | 0.941 | -0.001 | -0.013 | -0.013 | 0 | 0.31 | 0.42 | 0.48 | 0.28 | 0.34 | 0.4 |
| 1 | 177937441 | rs6425457 | -1.46 | 1.064 | -0.38 | 1.492 | -0.012 | -0.037 | -0.037 | 0 | 0.37 | 0.79 | 0.47 | 0.52 | 0.37 | 0.39 |
| 1 | 177937755 | rs7523508 | -1.99 | 0.644 | -0.106 | 1.721 | 0.033 | -0.072 | -0.072 | 0 | 0.33 | 0.85 | 0.41 | 0.52 | 0.25 | 0.37 |
| 1 | 177938437 | rs6682862 | -1.527 | 0.716 | 0.99 | 1.997 | 0.01 | 0.168 | 0.168 | 0 | 0.18 | 0.42 | 0.2 | 0.42 | 0.16 | 0.27 |
| 1 | 177938712 | rs10158522 | -0.886 | 0.694 | 1.225 | 2.018 | -0.001 | 0.038 | 0.038 | 0.001 | 0.09 | 0.3 | 0.08 | 0.06 | 0.05 | 0.03 |
| 1 | 177940209 | rs10798588 | -0.407 | 0.655 | 1.216 | 2.053 | -0.026 | -0.041 | -0.041 | 0.891 | 0.11 | 0.05 | 0.24 | 0.42 | 0.12 | 0.22 |
| 1 | 177940514 | rs10913477 | -0.651 | 0.653 | 1.378 | 2.061 | -0.001 | 0.035 | 0.035 | 0 | 0.11 | 0.13 | 0.25 | 0.42 | 0.12 | 0.22 |
| 1 | 177940612 | rs10158853 | -2.368 | 0.632 | 1.563 | 1.89 | -0.01 | 0.052 | 0.052 | 0 | 0.26 | 0.56 | 0.36 | 0.51 | 0.21 | 0.31 |
| 1 | 177943812 | rs869286 | -1.585 | 0.656 | 1.383 | 1.905 | 0.003 | 0.167 | 0.167 | 0 | 0.21 | 0.43 | 0.33 | 0.48 | 0.17 | 0.26 |
| 1 | 177965223 | rs10913485 | -2.358 | 0.639 | 0.719 | 1.882 | 0.011 | -0.028 | -0.028 | 0.024 | 0.29 | 0.71 | 0.37 | 0.51 | 0.21 | 0.31 |
| 1 | 177968616 | rs16852317 | -1.931 | 0.723 | 0.873 | 2 | 0 | 0.044 | 0.044 | 0 | 0.26 | 0.34 | 0.42 | 0.56 | 0.26 | 0.37 |
| 1 | 177969239 | rs16852325 | 0.16 | 0.733 | 0.827 | 2.124 | 0.001 | 0.043 | 0.043 | 0 | 0.05 | 0.07 | 0.12 | 0.26 | 0.09 | 0.11 |
| 1 | 177970998 | rs12409266 | -2.559 | 0.583 | 1.538 | 2.23 | -0.038 | 0.073 | 0.073 | 0.212 | 0.22 | 0.55 | 0.24 | 0.35 | 0.2 | 0.2 |
| 1 | 177971147 | rs12409286 | -2.813 | 0.987 | 1.292 | 1.974 | -0.008 | 0.012 | 0.012 | 0.918 | 0.29 | 0.63 | 0.34 | 0.35 | 0.28 | 0.3 |
| 1 | 177971866 | rs7340104 | -3.179 | 0.908 | 1.33 | 2.042 | 0.003 | -0.001 | -0.001 | 0 | 0.4 | 0.79 | 0.57 | 0.6 | 0.38 | 0.52 |
| 1 | 177974569 | rs12074053 | -3 | 1.492 | 1.129 | 1.745 | -0.005 | 0 | 0 | 0 | 0.38 | 0.65 | 0.47 | 0.35 | 0.47 | 0.38 |
| 1 | 177976440 | rs11588190 | -0.014 | 1.538 | 1.122 | 1.771 | -0.015 | -0.015 | -0.015 | 0.07 | 0.1 | 0.07 | 0.19 | 0.26 | 0.17 | 0.16 |

| | | | | | | | | | | | | | | | | |
|---|---|---|---|---|---|---|---|---|---|---|---|---|---|---|---|---|
| 1 | 177981232 | rs4147158 | -1.925 | 1.51 | 1.026 | 1.645 | 0.037 | 0.021 | 0.021 | 0.974 | 0.25 | 0.55 | 0.35 | 0.31 | 0.29 | 0.32 |
| 1 | 177989198 | rs17360705 | -1.029 | 1.168 | 0.829 | 1.172 | -0.007 | 0.007 | 0.007 | 0 | 0.13 | 0.08 | 0.12 | 0.04 | 0.18 | 0.07 |
| 1 | 177989327 | rs7529702 | -0.228 | 1.002 | 0.953 | 1.062 | 0.004 | -0.009 | -0.009 | 0 | 0.06 | 0.09 | 0.1 | 0.01 | 0.08 | 0.1 |
| 1 | 177991255 | rs16852358 | -1.1 | 0.923 | 1.009 | 1.321 | -0.016 | 0.139 | 0.139 | 0 | 0.06 | 0.16 | 0.17 | 0.24 | 0.03 | 0.18 |
| 1 | 177997709 | rs946817 | -1.681 | 1.492 | 1.432 | 0.705 | 0.075 | 0.082 | 0.082 | 0 | 0.16 | 0.39 | 0.21 | 0.23 | 0.25 | 0.11 |
| 1 | 177999767 | rs12410468 | -2.444 | 1.12 | 0.703 | 0.224 | -0.078 | 0.052 | 0.052 | 0.449 | 0.43 | 0.89 | 0.47 | 0.27 | 0.43 | 0.29 |
| 1 | 178001395 | rs7530170 | -0.386 | 1.123 | 1.067 | 0.217 | 0.009 | -0.079 | -0.079 | 0 | 0.15 | 0.41 | 0.14 | 0.02 | 0.13 | 0.11 |
| 1 | 178018617 | rs7540551 | -1.325 | 1.218 | 0.849 | 0.312 | 0.012 | -0.037 | -0.037 | 0.016 | 0.25 | 0.36 | 0.27 | 0.06 | 0.31 | 0.19 |
| 1 | 178019772 | rs7543930 | -0.86 | 0.852 | 0.515 | 0.551 | 0.001 | 0.026 | 0.026 | 0 | 0.11 | 0.05 | 0.13 | 0.02 | 0.1 | 0.07 |
| 1 | 178021124 | rs11803039 | -1.385 | 0.858 | 0.554 | 0.652 | 0.007 | 0.032 | 0.032 | 0 | 0.14 | 0.25 | 0.17 | 0.21 | 0.13 | 0.1 |
| 1 | 178025632 | rs16852393 | -1.105 | 1.039 | 0.103 | 0.519 | -0.005 | 0.006 | 0.006 | 0 | 0.09 | 0.07 | 0.12 | 0.04 | 0.14 | 0.06 |
| 1 | 178027683 | rs4652289 | -2.229 | 0.663 | 1.002 | 0.603 | 0.002 | 0.048 | 0.048 | 1.511 | 0.17 | 0.47 | 0.3 | 0.21 | 0.16 | 0.23 |
| 1 | 178029944 | rs17360740 | -2.539 | 0.316 | 0.116 | 0.211 | -0.011 | 0.012 | 0.012 | 3.252 | 0.34 | 0.93 | 0.47 | 0.25 | 0.28 | 0.28 |
| 1 | 178037791 | rs6703721 | -1.768 | 0.243 | 0.878 | 0.09 | -0.014 | -0.066 | -0.066 | 0.017 | 0.26 | 0.39 | 0.24 | 0.07 | 0.2 | 0.11 |
| 1 | 178039670 | rs12081217 | -1.044 | 0.327 | 1.661 | 0.126 | 0.051 | 0.072 | 0.072 | 0 | 0.06 | 0.24 | 0.2 | 0.09 | 0.07 | 0.18 |
| 1 | 178041772 | rs16852409 | -1.201 | 0.196 | 1.601 | 0.025 | 0.004 | 0.01 | 0.01 | 0.214 | 0.16 | 0.13 | 0.15 | 0.04 | 0.12 | 0.05 |
| 1 | 178042145 | rs11583320 | -2.221 | 0.016 | 1.828 | -0.069 | -0.011 | -0.05 | -0.05 | 0 | 0.27 | 0.4 | 0.26 | 0.07 | 0.22 | 0.11 |
| 1 | 178045388 | rs2862303 | -2.346 | 0.006 | 1.981 | -0.02 | -0.012 | -0.034 | -0.034 | 0.843 | 0.25 | 0.45 | 0.36 | 0.16 | 0.23 | 0.22 |
| 1 | 178062359 | rs964993 | -2.467 | -0.067 | 1.997 | 0.062 | -0.015 | -0.037 | -0.037 | 0 | 0.23 | 0.41 | 0.24 | 0.1 | 0.19 | 0.11 |
| 1 | 178092233 | rs10913506 | -2.505 | -0.072 | 2.147 | 0.118 | -0.019 | -0.041 | -0.041 | 0.621 | 0.23 | 0.45 | 0.25 | 0.11 | 0.19 | 0.11 |
| 1 | 178095048 | rs728569 | -1.102 | -0.084 | 2.152 | 0.086 | -0.001 | -0.023 | -0.023 | 0.001 | 0.1 | 0.23 | 0.09 | 0.04 | 0.09 | 0.06 |
| 1 | 178156816 | rs6701848 | -1.052 | -0.085 | 2.134 | 0.121 | -0.001 | 0 | 0 | 4.84 | 0.12 | 0.11 | 0.13 | 0.1 | 0.1 | 0.05 |
| 1 | 178159362 | rs6665850 | -0.831 | -0.14 | 1.004 | 0.144 | -0.005 | 0.019 | 0.019 | 0.166 | 0.37 | 0.48 | 0.44 | 0.51 | 0.38 | 0.46 |
| 1 | 178160268 | rs12037925 | -0.094 | -0.541 | 1.204 | -0.257 | -0.001 | 0.004 | 0.004 | 0 | 0.11 | 0.06 | 0.13 | 0.08 | 0.1 | 0.05 |
| 1 | 178163144 | rs389563 | 2.434 | -0.579 | 1.418 | -0.305 | 0.013 | 0.028 | 0.028 | 2.136 | 0.7 | 0.2 | 0.64 | 0.81 | 0.73 | 0.69 |
| 1 | 178163651 | rs12036491 | -0.077 | -0.554 | 1.257 | -0.271 | -0.001 | 0.004 | 0.004 | 0 | 0.11 | 0.06 | 0.13 | 0.08 | 0.1 | 0.05 |
| 1 | 178181554 | rs178397 | 1.27 | -0.81 | 0.472 | -0.597 | -0.104 | -0.019 | -0.019 | 0 | 0.32 | 0.01 | 0.2 | 0.26 | 0.35 | 0.23 |
| 1 | 178206664 | rs1412784 | -2.015 | -1.145 | 0.988 | -0.515 | -0.021 | -0.029 | -0.029 | 0 | 0.25 | 0.49 | 0.27 | 0.19 | 0.21 | 0.21 |
| 1 | 178211322 | rs623225 | 1.752 | -1.151 | 1.022 | -0.518 | -0.014 | -0.017 | -0.017 | 0 | 0.74 | 0.33 | 0.7 | 0.81 | 0.78 | 0.79 |
| 1 | 178232469 | rs7531227 | -1.932 | -1.132 | 0.959 | -0.516 | -0.019 | -0.031 | -0.031 | 0 | 0.25 | 0.49 | 0.27 | 0.19 | 0.22 | 0.21 |
| 1 | 178254568 | rs10798604 | -1.684 | -1.052 | 0.955 | -0.506 | -0.036 | -0.034 | -0.034 | 0 | 0.22 | 0.64 | 0.22 | 0.11 | 0.13 | 0.1 |
| 1 | 178261750 | rs12037817 | 0.097 | -1.008 | 0.869 | -0.491 | 0 | -0.001 | -0.001 | 0 | 0.11 | 0.11 | 0.13 | 0.08 | 0.1 | 0.05 |
| 1 | 178319105 | rs2175254 | 0.483 | -0.989 | 0.917 | -0.482 | -0.021 | -0.007 | -0.007 | 0 | 0.1 | 0.02 | 0.13 | 0.08 | 0.1 | 0.05 |
| 1 | 178322757 | rs11590951 | 0.222 | -0.991 | 0.972 | -0.483 | 0 | -0.003 | -0.003 | 0 | 0.11 | 0.12 | 0.13 | 0.08 | 0.1 | 0.05 |
| 1 | 178350980 | rs10913544 | 0.238 | -0.992 | 0.976 | -0.483 | 0 | -0.003 | -0.003 | 0 | 0.11 | 0.12 | 0.13 | 0.08 | 0.1 | 0.05 |
| 1 | 178353532 | rs977034 | -0.441 | -1.027 | 0.994 | -0.489 | -0.027 | -0.009 | -0.009 | 0.054 | 0.08 | 0.22 | 0.04 | 0.04 | 0.02 | 0.05 |
| 1 | 178374117 | rs4342816 | 1.26 | -0.977 | 0.608 | -0.586 | -0.147 | -0.122 | -0.122 | 0.306 | 0.36 | 0.02 | 0.43 | 0.58 | 0.39 | 0.47 |
| 1 | 178408795 | rs10913550 | -2.142 | -0.91 | 1.398 | -0.692 | -0.038 | -0.037 | -0.037 | 0 | 0.23 | 0.63 | 0.22 | 0.11 | 0.13 | 0.1 |
| 1 | 178410425 | rs9803679 | -2.037 | -0.9 | 1.403 | -0.687 | -0.037 | -0.035 | -0.035 | 0 | 0.22 | 0.63 | 0.22 | 0.11 | 0.13 | 0.1 |
| 1 | 178419417 | rs2017349 | -2.927 | -0.92 | 1.403 | -0.724 | 0.017 | 0.026 | 0.026 | 2.832 | 0.29 | 0.79 | 0.3 | 0.16 | 0.21 | 0.29 |
| 1 | 178427933 | rs12073428 | -1.643 | -0.927 | 1.43 | -0.666 | -0.028 | -0.024 | -0.024 | 0.119 | 0.2 | 0.35 | 0.17 | 0.11 | 0.12 | 0.1 |
| 1 | 178444907 | rs12038910 | -0.224 | -0.963 | 1.253 | -0.635 | 0 | -0.001 | -0.001 | 0 | 0.12 | 0.11 | 0.13 | 0.08 | 0.1 | 0.05 |
| 1 | 178454768 | rs6672334 | -2.09 | -0.844 | 1.048 | -0.582 | -0.031 | -0.039 | -0.039 | 0 | 0.24 | 0.55 | 0.25 | 0.12 | 0.17 | 0.18 |
| 1 | 178458708 | rs1008495 | -0.728 | -1.032 | 0.529 | -0.75 | -0.002 | -0.005 | -0.005 | 0.013 | 0.28 | 0.34 | 0.24 | 0.11 | 0.19 | 0.11 |
| 1 | 178463591 | rs2746325 | -1.316 | -0.311 | 0.687 | -0.369 | 0.035 | -0.051 | -0.051 | 0 | 0.21 | 0.43 | 0.28 | 0.12 | 0.26 | 0.26 |
| 1 | 178466769 | rs6666248 | -0.233 | -0.483 | 0.183 | 0.141 | -0.005 | -0.015 | -0.015 | 0.005 | 0.59 | 0.72 | 0.49 | 0.44 | 0.48 | 0.33 |
| 1 | 178476554 | rs4304537 | -0.112 | -0.511 | 0.323 | 0.261 | -0.047 | -0.029 | -0.029 | 0 | 0.6 | 0.75 | 0.58 | 0.44 | 0.54 | 0.4 |
| 1 | 178477432 | rs4076972 | 0.004 | -0.6 | 1.107 | 0.218 | -0.002 | -0.001 | -0.001 | 0 | 0.58 | 0.41 | 0.54 | 0.44 | 0.54 | 0.4 |
| 1 | 178481750 | rs10913562 | -0.135 | -0.769 | 0.672 | -0.026 | 0.005 | 0.017 | 0.017 | 0 | 0.13 | 0.11 | 0.05 | 0.06 | 0.09 | 0.06 |
| 1 | 178491966 | rs12033565 | -0.904 | -0.846 | 0.786 | -0.017 | -0.006 | -0.004 | -0.004 | 0.275 | 0.19 | 0.23 | 0.08 | 0.1 | 0.09 | 0.1 |
| 1 | 178492635 | rs7527363 | -0.577 | -0.699 | 0.738 | 0.155 | -0.039 | -0.031 | -0.031 | 0 | 0.64 | 0.73 | 0.59 | 0.5 | 0.55 | 0.45 |
| 1 | 178494428 | rs6664493 | 0.753 | -0.498 | 0.523 | 0.272 | -0.053 | -0.048 | -0.048 | 0.451 | 0.31 | 0.16 | 0.27 | 0.4 | 0.36 | 0.39 |
| 1 | 178504880 | rs12734345 | 1.073 | -0.75 | 0.715 | -0.068 | -0.009 | -0.035 | -0.035 | 0 | 0.69 | 0.57 | 0.73 | 0.8 | 0.75 | 0.69 |
| 1 | 178506594 | rs4075631 | 0.816 | -0.654 | 0.905 | 0.022 | 0.005 | 0.005 | 0.005 | 0.033 | 0.31 | 0.3 | 0.3 | 0.42 | 0.36 | 0.37 |
| 1 | 178510497 | rs4233174 | 0.003 | -0.678 | 0.53 | -0.08 | 0.003 | -0.002 | -0.002 | 0 | 0.92 | 0.88 | 0.85 | 0.9 | 0.88 | 0.81 |
| 1 | 178511941 | rs10913567 | 0.909 | -0.633 | 0.495 | -0.017 | -0.067 | -0.068 | -0.068 | 0 | 0.3 | 0.15 | 0.29 | 0.43 | 0.35 | 0.36 |
| 1 | 178514622 | rs12568310 | 1.168 | -0.789 | 0.599 | -0.087 | 0.019 | -0.024 | -0.024 | 1.584 | 0.37 | 0.17 | 0.34 | 0.37 | 0.4 | 0.32 |
| 1 | 178515187 | rs4650993 | 0.221 | -0.957 | 0.581 | -0.186 | -0.054 | -0.081 | -0.081 | 0 | 0.42 | 0.12 | 0.36 | 0.54 | 0.46 | 0.52 |
| 1 | 178515312 | rs4650994 | 0.221 | -0.957 | 0.581 | -0.186 | -0.054 | -0.081 | -0.081 | 0 | 0.42 | 0.12 | 0.36 | 0.54 | 0.46 | 0.52 |
| 1 | 178517316 | rs4650995 | 0.221 | -0.957 | 0.581 | -0.186 | -0.054 | -0.081 | -0.081 | 0 | 0.42 | 0.12 | 0.36 | 0.54 | 0.46 | 0.52 |
| 1 | 178517365 | rs4650996 | 1.155 | -0.94 | 0.569 | -0.185 | -0.107 | -0.106 | -0.106 | 0 | 0.29 | 0.06 | 0.27 | 0.43 | 0.35 | 0.37 |
| 1 | 178517591 | rs4233175 | 1.425 | -0.955 | 0.955 | -0.191 | -0.032 | -0.074 | -0.074 | 0.046 | 0.8 | 0.43 | 0.72 | 0.91 | 0.86 | 0.84 |
| 1 | 178518008 | rs10913569 | 1.5 | -0.952 | 0.92 | -0.19 | -0.017 | -0.038 | -0.038 | 0 | 0.79 | 0.29 | 0.7 | 0.91 | 0.86 | 0.84 |
| 1 | 178529757 | rs7540745 | 1.058 | -0.907 | 0.85 | -0.148 | 0.009 | -0.004 | -0.004 | 0 | 0.39 | 0.31 | 0.35 | 0.37 | 0.4 | 0.33 |
| 1 | 178531117 | rs12046110 | 0.589 | -0.856 | 0.716 | -0.136 | -0.031 | -0.007 | -0.007 | 0 | 0.13 | 0.04 | 0.09 | 0.1 | 0.11 | 0.16 |
| 1 | 178532188 | rs4650999 | -0.161 | -0.932 | 1.16 | -0.176 | -0.002 | -0.019 | -0.019 | 0 | 0.45 | 0.31 | 0.38 | 0.54 | 0.46 | 0.51 |
| 1 | 178532223 | rs4651000 | 0.972 | -0.927 | 1.119 | -0.169 | -0.036 | -0.036 | -0.036 | 0.02 | 0.31 | 0.2 | 0.29 | 0.44 | 0.35 | 0.36 |
| 1 | 178534221 | rs4072643 | -0.216 | -0.967 | 1.322 | -0.184 | -0.024 | -0.027 | -0.027 | 1.583 | 0.08 | 0.29 | 0.04 | 0 | 0.01 | 0.01 |
| 1 | 178551472 | rs2811297 | 1.268 | -0.95 | 1.288 | -0.174 | 0.019 | -0.026 | -0.026 | 0 | 0.37 | 0.17 | 0.34 | 0.37 | 0.4 | 0.32 |

| | | | | | | | | | | | | | | | | |
|---|---|---|---|---|---|---|---|---|---|---|---|---|---|---|---|---|
| 1 | 178553480 | rs2492279 | -1.758 | -0.866 | 1.304 | -0.176 | -0.005 | -0.048 | -0.048 | 0 | 0.3 | 0.52 | 0.37 | 0.2 | 0.25 | 0.31 |
| 1 | 178563987 | rs2493856 | -0.146 | -0.825 | 1.177 | -0.158 | 0.005 | -0.004 | -0.004 | 0.113 | 0.09 | 0.13 | 0.23 | 0.09 | 0.12 | 0.15 |
| 1 | 178565860 | rs2492276 | -1.949 | -0.857 | 1.306 | -0.175 | -0.005 | -0.05 | -0.05 | 0 | 0.3 | 0.49 | 0.37 | 0.2 | 0.25 | 0.31 |
| 1 | 178567934 | rs12032055 | -0.872 | -0.776 | 1.102 | -0.164 | -0.017 | -0.04 | -0.04 | 0 | 0.21 | 0.36 | 0.13 | 0.1 | 0.13 | 0.16 |
| 1 | 178572265 | rs2147030 | -1.051 | -0.784 | 0.51 | -0.126 | -0.042 | -0.05 | -0.05 | 0 | 0.68 | 0.85 | 0.72 | 0.55 | 0.65 | 0.64 |
| 1 | 178572978 | rs2811290 | -0.267 | -0.928 | 0.628 | -0.129 | 0.004 | -0.002 | -0.002 | 0 | 0.55 | 0.78 | 0.63 | 0.43 | 0.53 | 0.48 |
| 1 | 178575265 | rs2476557 | -1.477 | -0.934 | 0.937 | -0.151 | -0.017 | -0.063 | -0.063 | 0 | 0.17 | 0.57 | 0.29 | 0.09 | 0.13 | 0.17 |
| 1 | 178590394 | rs2811300 | -0.105 | -1.009 | 0.799 | -0.204 | 0.032 | 0.017 | 0.017 | 0 | 0.5 | 0.66 | 0.6 | 0.43 | 0.45 | 0.46 |
| 1 | 178594196 | rs2811301 | 0.055 | -1.013 | 0.819 | -0.188 | 0.038 | 0.018 | 0.018 | 0 | 0.5 | 0.69 | 0.59 | 0.43 | 0.45 | 0.46 |
| 1 | 178597882 | rs10798623 | -0.103 | -0.996 | 1.068 | -0.194 | 0.021 | 0.008 | 0.008 | 0.088 | 0.08 | 0.26 | 0.06 | 0.08 | 0.09 | 0.07 |
| 1 | 178598873 | rs6659392 | 0.173 | -0.953 | 0.697 | -0.168 | -0.017 | 0.003 | 0.003 | 0 | 0.17 | 0.1 | 0.11 | 0.13 | 0.19 | 0.18 |
| 1 | 178599348 | rs2761471 | 0 | -0.994 | 0.783 | -0.155 | 0.048 | 0.022 | 0.022 | 0 | 0.5 | 0.73 | 0.6 | 0.43 | 0.45 | 0.46 |
| 1 | 178605501 | rs10913584 | 1.242 | -0.811 | -0.034 | -0.071 | 0.024 | -0.016 | -0.016 | 0 | 0.33 | 0.19 | 0.32 | 0.34 | 0.32 | 0.29 |
| 1 | 178617359 | rs7552657 | -0.823 | -0.748 | 0.013 | -0.032 | -0.001 | 0.002 | 0.002 | 0 | 0.23 | 0.2 | 0.38 | 0.16 | 0.21 | 0.26 |
| 1 | 178618316 | rs10913588 | -0.564 | -0.713 | -0.231 | -0.015 | -0.012 | 0.002 | 0.002 | 0 | 0.21 | 0.1 | 0.37 | 0.16 | 0.21 | 0.26 |
| 1 | 178618726 | rs6704082 | -0.384 | -0.71 | -0.271 | -0.014 | -0.036 | -0.016 | -0.016 | 0 | 0.21 | 0.05 | 0.37 | 0.16 | 0.2 | 0.26 |
| 1 | 178619183 | rs2811314 | -0.995 | -0.782 | -0.401 | 0.135 | -0.121 | -0.121 | -0.121 | 0 | 0.71 | 0.94 | 0.8 | 0.5 | 0.64 | 0.62 |
| 1 | 178625499 | rs2761476 | 0.766 | -0.625 | 0.625 | 0.177 | -0.034 | -0.05 | -0.05 | 0 | 0.74 | 0.55 | 0.77 | 0.87 | 0.82 | 0.79 |
| 1 | 178626526 | rs2761477 | 0.746 | -0.581 | 0.629 | 0.221 | -0.032 | -0.042 | -0.042 | 0.434 | 0.75 | 0.54 | 0.77 | 0.86 | 0.82 | 0.79 |
| 1 | 178633373 | rs10798625 | 0.059 | -0.035 | -0.093 | 0.415 | -0.041 | -0.016 | -0.016 | 0 | 0.3 | 0.17 | 0.23 | 0.38 | 0.37 | 0.34 |
| 1 | 178641945 | rs4652309 | -0.79 | -0.259 | -0.229 | 0.306 | -0.062 | -0.034 | -0.034 | 0 | 0.43 | 0.16 | 0.29 | 0.5 | 0.49 | 0.43 |
| 1 | 178645560 | rs10913603 | -0.181 | -0.356 | 0.146 | 0.228 | -0.003 | -0.002 | -0.002 | 0 | 0.11 | 0.15 | 0.21 | 0.1 | 0.08 | 0.08 |
| 1 | 178657038 | rs12407833 | -0.338 | -0.401 | 0.733 | 0.204 | -0.005 | -0.003 | -0.003 | 0 | 0.12 | 0.15 | 0.21 | 0.1 | 0.08 | 0.08 |
| 1 | 178657616 | rs2773091 | 1.195 | -0.504 | 0.977 | 0.164 | -0.033 | -0.022 | -0.022 | 2.829 | 0.72 | 0.56 | 0.7 | 0.76 | 0.81 | 0.81 |
| 1 | 178657678 | rs2811250 | 1.195 | -0.504 | 0.968 | 0.164 | -0.033 | -0.022 | -0.022 | 2.829 | 0.72 | 0.56 | 0.7 | 0.76 | 0.81 | 0.81 |
| 1 | 178659410 | rs4532794 | 1.674 | -0.552 | 1.322 | 0.168 | 0.005 | -0.004 | -0.004 | 0 | 0.38 | 0.22 | 0.45 | 0.34 | 0.39 | 0.4 |
| 1 | 178659933 | rs3922812 | 1.748 | -0.573 | 1.345 | 0.158 | 0.003 | -0.007 | -0.007 | 0 | 0.38 | 0.2 | 0.45 | 0.34 | 0.39 | 0.4 |
| 1 | 178661341 | rs16853243 | 0.404 | -0.533 | 1.491 | 0.151 | 0.004 | 0.008 | 0.008 | 0 | 0.05 | 0.11 | 0.02 | 0.06 | 0.05 | 0.1 |
| 1 | 178661629 | rs6425496 | 0.422 | -0.542 | 1.381 | 0.145 | 0.007 | 0.001 | 0.001 | 0 | 0.58 | 0.49 | 0.55 | 0.54 | 0.54 | 0.6 |
| 1 | 178666488 | rs2811267 | 0.986 | -0.553 | 1.374 | 0.136 | -0.027 | -0.018 | -0.018 | 0.012 | 0.73 | 0.58 | 0.7 | 0.76 | 0.81 | 0.81 |
| 1 | 178668341 | rs12045804 | 1.11 | -0.487 | 1.394 | 0.166 | -0.003 | 0.001 | 0.001 | 0 | 0.44 | 0.42 | 0.48 | 0.4 | 0.43 | 0.49 |
| 1 | 178669915 | rs12141498 | -0.425 | -0.43 | 1.312 | 0.133 | -0.017 | -0.016 | -0.016 | 0.859 | 0.13 | 0.25 | 0.22 | 0.1 | 0.08 | 0.08 |
| 1 | 178671750 | rs16853281 | 0.024 | 0.369 | 0.956 | 0.298 | 0.037 | 0.042 | 0.042 | 0 | 0.13 | 0.21 | 0.08 | 0.26 | 0.15 | 0.24 |
| 1 | 178674074 | rs9286919 | 1.856 | 0.224 | 0.47 | 0.299 | 0 | -0.01 | -0.01 | 0 | 0.38 | 0.21 | 0.45 | 0.35 | 0.39 | 0.41 |
| 1 | 178676113 | rs867556 | 0.582 | 0.471 | 0.152 | 0.341 | -0.049 | 0.007 | 0.007 | 0 | 0.23 | 0.04 | 0.15 | 0.15 | 0.27 | 0.17 |
| 1 | 178679194 | rs7543277 | -1.244 | 0.449 | 0.573 | 0.445 | 0.014 | 0.026 | 0.026 | 0 | 0.25 | 0.41 | 0.29 | 0.36 | 0.23 | 0.33 |
| 1 | 178686247 | rs2811280 | -0.397 | 0.64 | 0.533 | 0.292 | 0.005 | -0.005 | -0.005 | 0 | 0.31 | 0.4 | 0.35 | 0.25 | 0.35 | 0.24 |
| 1 | 178686299 | rs10913618 | -0.05 | 0.228 | 0.329 | 0.11 | 0.023 | 0.01 | 0.01 | 0 | 0.16 | 0.07 | 0.09 | 0.14 | 0.11 | 0.08 |
| 1 | 178686331 | rs16853302 | 0.156 | 0.284 | 0.451 | 0.144 | 0.002 | 0.005 | 0.005 | 0 | 0.06 | 0.11 | 0.02 | 0.06 | 0.04 | 0.1 |
| 1 | 178686809 | rs2988268 | 0.948 | 0.562 | 0.496 | 0.069 | -0.039 | -0.008 | -0.008 | 0 | 0.19 | 0.07 | 0.14 | 0.15 | 0.25 | 0.15 |
| 1 | 178687459 | rs1018669 | 1.123 | 0.389 | 0.927 | 0.031 | 0.015 | 0.055 | 0.055 | 0 | 0.54 | 0.28 | 0.7 | 0.71 | 0.57 | 0.74 |
| 1 | 178696750 | rs2773080 | 1.122 | 0.422 | 1.022 | -0.471 | 0.013 | -0.035 | -0.035 | 0.796 | 0.63 | 0.48 | 0.72 | 0.78 | 0.57 | 0.78 |
| 1 | 178702875 | rs2862517 | -1.487 | 0.418 | 1.116 | -0.473 | -0.005 | 0.009 | 0.009 | 0 | 0.38 | 0.7 | 0.3 | 0.22 | 0.43 | 0.22 |
| 1 | 178709966 | rs4652319 | 0.761 | 0.414 | 1.097 | -0.522 | -0.05 | 0.014 | 0.014 | 0 | 0.16 | 0.1 | 0.17 | 0.11 | 0.32 | 0.13 |
| 1 | 178719306 | rs4651005 | 0.854 | 0.398 | 0.948 | -0.526 | -0.063 | 0.012 | 0.012 | 0 | 0.16 | 0.07 | 0.17 | 0.11 | 0.32 | 0.13 |
| 1 | 178721805 | rs7519811 | -0.123 | 0.353 | 1.069 | -0.549 | 0 | 0 | 0 | 0 | 0.11 | 0.11 | 0.06 | 0.1 | 0.07 | 0.07 |
| 1 | 178724419 | rs4540610 | -0.166 | 0.353 | 1.069 | -0.548 | 0.001 | 0 | 0 | 0 | 0.11 | 0.11 | 0.06 | 0.1 | 0.08 | 0.07 |
| 1 | 178740427 | rs6693283 | -2.413 | 0.346 | 0.918 | -0.548 | 0.049 | -0.021 | -0.021 | 0 | 0.54 | 0.81 | 0.58 | 0.54 | 0.58 | 0.43 |
| 1 | 178752762 | rs883561 | 0.942 | 0.411 | 0.904 | -0.518 | -0.056 | 0.008 | 0.008 | 0 | 0.16 | 0.07 | 0.16 | 0.12 | 0.32 | 0.13 |
| 1 | 178765148 | rs16853350 | 0.797 | 0.415 | 0.721 | -0.502 | -0.064 | -0.083 | -0.083 | 0.901 | 0.14 | 0.01 | 0.27 | 0.32 | 0.16 | 0.22 |
| 1 | 178782625 | rs965558 | 0.442 | 0.432 | 0.781 | -0.494 | -0.009 | -0.018 | -0.018 | 0 | 0.16 | 0.1 | 0.29 | 0.32 | 0.16 | 0.21 |
| 1 | 178800481 | rs12064728 | -2.088 | 0.418 | 0.83 | -0.507 | 0.051 | 0 | 0 | 0 | 0.52 | 0.74 | 0.56 | 0.54 | 0.58 | 0.43 |
| 1 | 178813531 | rs3766642 | -1.512 | 0.97 | 1.204 | -0.156 | -0.051 | -0.074 | -0.074 | 0 | 0.21 | 0.47 | 0.12 | 0.1 | 0.1 | 0.09 |
| 1 | 178817723 | rs16853405 | 0.459 | 0.984 | 1.121 | -0.149 | 0.001 | -0.031 | -0.031 | 0 | 0.08 | 0.32 | 0.05 | 0 | 0.03 | 0.02 |
| 1 | 178819553 | rs10913632 | -0.609 | 1.016 | 0.561 | -0.085 | -0.003 | -0.004 | -0.004 | 0 | 0.12 | 0.14 | 0.06 | 0.1 | 0.08 | 0.07 |
| 1 | 178820159 | rs1544124 | -2.342 | 1.011 | 0.515 | -0.081 | 0.054 | -0.01 | -0.01 | 0 | 0.53 | 0.81 | 0.57 | 0.54 | 0.58 | 0.43 |
| 1 | 178822365 | rs12117953 | 0.216 | 0.965 | 0.872 | -0.258 | -0.009 | 0.033 | 0.033 | 0 | 0.07 | 0.02 | 0.03 | 0 | 0.06 | 0.03 |
| 1 | 178826282 | rs10913634 | -1.945 | 1.014 | 0.876 | -0.234 | -0.009 | 0.017 | 0.017 | 0 | 0.38 | 0.71 | 0.29 | 0.22 | 0.43 | 0.22 |
| 1 | 178841980 | rs6696204 | -0.572 | 0.753 | -0.062 | -0.118 | -0.002 | -0.003 | -0.003 | 0 | 0.12 | 0.14 | 0.07 | 0.1 | 0.08 | 0.07 |
| 1 | 178848223 | rs3766637 | -0.572 | 0.753 | -0.062 | -0.118 | -0.002 | -0.003 | -0.003 | 2.981 | 0.12 | 0.14 | 0.07 | 0.1 | 0.08 | 0.07 |
| 1 | 178871889 | rs12139850 | -0.463 | 0.766 | -0.103 | -0.112 | 0.001 | 0.001 | 0.001 | 0 | 0.11 | 0.11 | 0.06 | 0.1 | 0.08 | 0.07 |
| 1 | 178903637 | rs871631 | -0.982 | 1.303 | -0.018 | -0.074 | 0.055 | -0.062 | -0.062 | 0 | 0.25 | 0.55 | 0.2 | 0.11 | 0.32 | 0.15 |
| 1 | 178907461 | rs10913645 | -0.964 | 1.228 | -0.654 | 0.246 | 0.004 | -0.002 | -0.002 | 0 | 0.31 | 0.69 | 0.24 | 0.31 | 0.36 | 0.22 |
| 1 | 178922451 | rs6425507 | -0.162 | 1.392 | -1.125 | 0.425 | 0.003 | 0.012 | 0.012 | 0 | 0.18 | 0.19 | 0.25 | 0.52 | 0.26 | 0.29 |
| 1 | 178924427 | rs2039536 | -0.643 | -0.102 | -1.599 | -0.458 | 0.042 | 0.081 | 0.081 | 0 | 0.73 | 0.89 | 0.86 | 0.91 | 0.8 | 0.83 |
| 1 | 178927427 | rs7522133 | -0.464 | 0.292 | -1.742 | -0.305 | -0.016 | 0.083 | 0.083 | 0 | 0.43 | 0.18 | 0.59 | 0.7 | 0.54 | 0.49 |
| 1 | 178933590 | rs7539025 | -0.062 | -0.294 | -0.934 | 0.02 | -0.056 | 0.007 | 0.007 | 0.123 | 0.17 | 0.55 | 0.15 | 0.18 | 0.1 | 0.15 |
| 1 | 178934776 | rs10798640 | -0.164 | 0.455 | -1.394 | 0.206 | -0.001 | 0 | 0 | 0 | 0.15 | 0.14 | 0.2 | 0.09 | 0.19 | 0.11 |
| 1 | 178943012 | rs2031470 | -0.497 | 0.126 | -1.933 | 0.358 | 0.019 | 0.041 | 0.041 | 0 | 0.32 | 0.2 | 0.19 | 0.23 | 0.28 | 0.2 |

| | | | | | | | | | | | | | | | | |
|---|---|---|---|---|---|---|---|---|---|---|---|---|---|---|---|---|
| 1 | 178943576 | rs10913662 | -0.343 | -0.153 | -2.19 | 0.46 | 0.036 | 0.019 | 0.019 | 0.012 | 0.2 | 0.02 | 0.07 | 0.11 | 0.11 | 0.08 |
| 1 | 178956553 | rs4652341 | -0.738 | 0.714 | -1.966 | 0.928 | -0.039 | -0.05 | -0.05 | 0.081 | 0.09 | 0.01 | 0.11 | 0.15 | 0.12 | 0.13 |
| 1 | 178964604 | rs1012066 | -0.183 | 1.022 | -2.472 | 1.075 | -0.038 | -0.044 | -0.044 | 0 | 0.71 | 0.9 | 0.64 | 0.62 | 0.67 | 0.75 |
| 1 | 178967229 | rs6697180 | -0.654 | 1.07 | -1.465 | 0.903 | 0.024 | 0.032 | 0.032 | 0.269 | 0.19 | 0.37 | 0.23 | 0.25 | 0.22 | 0.2 |
| 1 | 178967996 | rs997419 | 0.412 | 0.416 | -2.024 | 0.312 | -0.002 | 0.049 | 0.049 | 0 | 0.21 | 0.06 | 0.16 | 0.1 | 0.21 | 0.23 |
| 1 | 178976556 | rs10798648 | -0.285 | 0.569 | -1.376 | 0.379 | 0.017 | 0.036 | 0.036 | 0.665 | 0.53 | 0.41 | 0.39 | 0.31 | 0.41 | 0.48 |
| 1 | 178976731 | rs12137038 | 0.344 | -0.182 | -1.873 | -0.101 | 0.043 | 0.063 | 0.063 | 0.688 | 0.25 | 0.07 | 0.11 | 0.1 | 0.16 | 0.18 |
| 1 | 178984086 | rs12725164 | -0.687 | 0.014 | -0.85 | -0.117 | 0.028 | 0.052 | 0.052 | 0 | 0.47 | 0.65 | 0.37 | 0.32 | 0.37 | 0.39 |
| 1 | 178991490 | rs7516491 | -0.07 | 0.458 | 0.016 | -0.359 | 0.022 | 0 | 0 | 1.472 | 0.12 | 0.31 | 0.09 | 0.09 | 0.1 | 0.13 |
| 1 | 178994644 | rs3813638 | -2.363 | 0.786 | -0.743 | 0.096 | -0.069 | -0.093 | -0.093 | 0.278 | 0.81 | 0.97 | 0.63 | 0.66 | 0.7 | 0.72 |
| 1 | 179006462 | rs6425513 | -1.386 | 0.747 | -0.911 | 0.26 | -0.037 | -0.077 | -0.077 | 0.117 | 0.69 | 0.93 | 0.53 | 0.6 | 0.65 | 0.62 |
| 1 | 179013204 | rs11802142 | 2.797 | 0.723 | -0.924 | 0.224 | 0.016 | 0.027 | 0.027 | 0 | 0.05 | 0 | 0.03 | 0 | 0.03 | 0.02 |
| 1 | 179016398 | rs3766636 | 0.126 | 0.662 | -0.876 | 0.174 | 0 | -0.008 | -0.008 | 0 | 0.49 | 0.58 | 0.4 | 0.24 | 0.44 | 0.5 |
| 1 | 179016483 | rs3766635 | 1.053 | 0.633 | -0.787 | 0.334 | -0.026 | -0.039 | -0.039 | 1.481 | 0.12 | 0.06 | 0.08 | 0.31 | 0.17 | 0.08 |
| 1 | 179019186 | rs12733622 | 1.324 | 0.042 | -0.674 | -0.158 | 0.003 | 0.007 | 0.007 | 0 | 0.27 | 0.23 | 0.25 | 0.2 | 0.24 | 0.23 |
| 1 | 179022236 | rs7527432 | 0.053 | 0.111 | -0.499 | 0.028 | 0.009 | 0.118 | 0.118 | 0 | 0.2 | 0.33 | 0.13 | 0.38 | 0.2 | 0.11 |
| 1 | 179038155 | rs12083856 | 1.363 | 0.017 | -0.288 | 0.005 | -0.01 | 0.052 | 0.052 | 0.147 | 0.09 | 0.31 | 0.05 | 0.08 | 0.03 | 0.02 |
| 1 | 179039044 | rs12022201 | -0.254 | 0.54 | -0.16 | 0.465 | -0.003 | 0.03 | 0.03 | 0 | 0.53 | 0.38 | 0.42 | 0.33 | 0.54 | 0.54 |
| 1 | 179042092 | rs16853612 | 1.336 | 0.785 | -0.448 | 0.344 | -0.022 | -0.033 | -0.033 | 0 | 0.11 | 0.06 | 0.07 | 0.3 | 0.16 | 0.08 |
| 1 | 179042227 | rs9725887 | -0.647 | 0.759 | -0.441 | 0.294 | -0.003 | 0.008 | 0.008 | 0 | 0.61 | 0.67 | 0.48 | 0.54 | 0.61 | 0.59 |
| 1 | 179042300 | rs4652352 | 0.23 | 0.766 | -0.446 | 0.297 | 0.001 | 0.025 | 0.025 | 0 | 0.48 | 0.51 | 0.4 | 0.24 | 0.44 | 0.5 |
| 1 | 179043553 | rs3766625 | 1.007 | 0.736 | -0.538 | 0.288 | -0.013 | -0.024 | -0.024 | 0 | 0.12 | 0.07 | 0.07 | 0.3 | 0.16 | 0.08 |
| 1 | 179043939 | rs16853619 | 1.878 | 0.755 | -0.52 | 0.293 | -0.013 | 0.044 | 0.044 | 0.431 | 0.09 | 0.25 | 0.05 | 0.08 | 0.03 | 0.02 |
| 1 | 179044723 | rs16853621 | 2.001 | 1.009 | -1.116 | 0.345 | 0.008 | 0.01 | 0.01 | 0 | 0.12 | 0.01 | 0.1 | 0.05 | 0.07 | 0.11 |
| 1 | 179045500 | rs1220 | -1.096 | 1.122 | -1.093 | 0.433 | -0.057 | -0.037 | -0.037 | 1.665 | 0.68 | 0.93 | 0.5 | 0.66 | 0.6 | 0.61 |
| 1 | 179050478 | rs3813639 | 1.505 | 1.187 | -0.948 | 0.518 | -0.03 | -0.042 | -0.042 | 0 | 0.1 | 0.02 | 0.06 | 0.26 | 0.14 | 0.07 |
| 1 | 179050593 | rs3813640 | 1.258 | 1.177 | -0.86 | 0.509 | -0.001 | 0.005 | 0.005 | 0 | 0.11 | 0.11 | 0.07 | 0.25 | 0.14 | 0.07 |
| 1 | 179051300 | rs2296377 | -2.754 | 1.259 | -1.016 | 0.492 | -0.08 | -0.102 | -0.102 | 0 | 0.82 | 0.99 | 0.64 | 0.67 | 0.71 | 0.72 |
| 1 | 179055020 | rs2296376 | 1.361 | 1.246 | -0.728 | 0.523 | -0.001 | 0.005 | 0.005 | 0 | 0.11 | 0.11 | 0.07 | 0.26 | 0.14 | 0.07 |
| 1 | 179058010 | rs16853640 | 2.312 | 1 | -0.489 | 0.099 | -0.015 | -0.071 | -0.071 | 0.084 | 0.14 | 0.17 | 0.07 | 0.01 | 0.1 | 0.07 |
| 1 | 179061108 | rs1570807 | 0.404 | 0.962 | -0.412 | 0.129 | 0 | 0 | 0 | 0 | 0.33 | 0.37 | 0.3 | 0.23 | 0.27 | 0.33 |
| 1 | 179061427 | rs10798652 | -1.347 | 0.926 | -0.552 | 0.154 | -0.009 | -0.006 | -0.006 | 0 | 0.64 | 0.71 | 0.54 | 0.58 | 0.54 | 0.53 |
| 1 | 179062497 | rs16853647 | 1.037 | 0.917 | -0.576 | 0.013 | -0.002 | -0.003 | -0.003 | 0 | 0.23 | 0.21 | 0.15 | 0.27 | 0.24 | 0.15 |
| 1 | 179064875 | rs1046385 | -2.785 | 1.046 | -0.775 | 0.074 | -0.064 | -0.089 | -0.089 | 0.011 | 0.81 | 0.97 | 0.64 | 0.68 | 0.71 | 0.71 |
| 1 | 179065125 | rs1128952 | 0.17 | 1.105 | -0.609 | -0.051 | -0.019 | -0.024 | -0.024 | 0 | 0.41 | 0.51 | 0.32 | 0.29 | 0.28 | 0.35 |
| 1 | 179065563 | rs12143004 | 1.501 | 0.931 | -0.444 | 0.037 | -0.028 | 0.023 | 0.023 | 0 | 0.14 | 0.11 | 0.09 | 0.08 | 0.18 | 0.18 |
| 1 | 179067071 | rs10913690 | -1.439 | 1.459 | 0.171 | 0.329 | 0.03 | 0.098 | 0.098 | 0 | 0.77 | 0.71 | 0.53 | 0.54 | 0.62 | 0.68 |
| 1 | 179069429 | rs3210839 | -1.901 | 1.45 | 0.199 | 0.335 | -0.008 | -0.009 | -0.009 | 0 | 0.8 | 0.88 | 0.6 | 0.6 | 0.62 | 0.7 |
| 1 | 179070334 | rs3813643 | -1.329 | 1.435 | 0.154 | 0.333 | 0.03 | 0.103 | 0.103 | 0 | 0.77 | 0.7 | 0.53 | 0.54 | 0.62 | 0.68 |
| 1 | 179071918 | rs1325194 | -2.141 | 1.498 | 0.114 | 0.345 | -0.056 | -0.107 | -0.107 | 0.031 | 0.81 | 0.99 | 0.65 | 0.68 | 0.72 | 0.71 |
| 1 | 179072728 | rs3829794 | -0.098 | 1.343 | 0.173 | 0.165 | 0 | -0.007 | -0.007 | 1.974 | 0.58 | 0.48 | 0.42 | 0.53 | 0.5 | 0.59 |
| 1 | 179074869 | rs10913692 | -0.044 | 1.354 | 0.166 | 0.159 | 0.001 | -0.008 | -0.008 | 0 | 0.58 | 0.48 | 0.42 | 0.54 | 0.5 | 0.59 |
| 1 | 179075755 | rs1490311 | -2.117 | 1.801 | 0.169 | 0.275 | -0.008 | -0.001 | -0.001 | 0.635 | 0.87 | 0.9 | 0.91 | 0.99 | 0.85 | 0.91 |
| 1 | 179089491 | rs9728480 | -1.358 | 1.376 | 0.307 | 0.477 | 0.035 | 0.103 | 0.103 | 0 | 0.77 | 0.72 | 0.53 | 0.54 | 0.62 | 0.68 |
| 1 | 179103701 | rs2636287 | 1.341 | 1.362 | 0.235 | 0.459 | 0.016 | 0.031 | 0.031 | 0 | 0.21 | 0.18 | 0.41 | 0.4 | 0.38 | 0.31 |
| 1 | 179120795 | rs2636278 | 0.805 | 1.241 | -0.059 | 0.286 | -0.004 | -0.025 | -0.025 | 0 | 0.3 | 0.14 | 0.42 | 0.33 | 0.37 | 0.37 |
| 1 | 179130359 | rs6672395 | 1.372 | 1.303 | -0.092 | 0.137 | -0.025 | -0.029 | -0.029 | 0 | 0.13 | 0.06 | 0.18 | 0.25 | 0.15 | 0.16 |
| 1 | 179148739 | rs12126360 | 2.118 | 0.936 | 0.265 | 0.12 | 0 | -0.019 | -0.019 | 0 | 0.11 | 0.13 | 0.08 | 0.01 | 0.09 | 0.07 |
| 1 | 179153697 | rs2636293 | 0.237 | 0.977 | 0.385 | 0.148 | 0.016 | 0.039 | 0.039 | 0 | 0.33 | 0.4 | 0.55 | 0.46 | 0.48 | 0.39 |
| 1 | 179199652 | rs2636266 | 1.651 | 1.047 | 0.132 | 0.259 | -0.057 | -0.105 | -0.105 | 0 | 0.19 | 0.01 | 0.35 | 0.32 | 0.29 | 0.29 |
| 1 | 179222840 | rs451088 | 0.874 | 0.838 | 0.395 | 0.021 | 0.048 | 0.14 | 0.14 | 0 | 0.23 | 0.38 | 0.48 | 0.46 | 0.39 | 0.32 |
| 1 | 179226551 | rs2816173 | -0.887 | 0.516 | 0.521 | -0.216 | -0.041 | -0.114 | -0.114 | 0.32 | 0.8 | 0.66 | 0.87 | 0.99 | 0.88 | 0.91 |
| 1 | 179233083 | rs12128920 | 1.878 | 0.522 | 0.395 | -0.156 | -0.01 | -0.043 | -0.043 | 0.026 | 0.12 | 0.22 | 0.08 | 0.01 | 0.08 | 0.07 |
| 1 | 179236814 | rs2783387 | 0.292 | 0.645 | 0.198 | -0.164 | 0.003 | 0.043 | 0.043 | 0 | 0.51 | 0.42 | 0.26 | 0.29 | 0.39 | 0.49 |
| 1 | 179239244 | rs2636270 | -1.231 | 0.876 | 0.144 | 0.24 | 0.002 | 0.015 | 0.015 | 0 | 0.91 | 0.89 | 0.84 | 0.76 | 0.88 | 0.9 |
| 1 | 179248952 | rs2816177 | 0.315 | 0.736 | 0.176 | 0.104 | 0.003 | 0.017 | 0.017 | 0 | 0.29 | 0.23 | 0.5 | 0.46 | 0.42 | 0.34 |
| 1 | 179252540 | rs34415902 | 0.537 | 0.784 | 0.559 | 0.172 | 0.04 | 0.046 | 0.046 | 1.028 | 0.11 | 0.28 | 0.15 | 0.13 | 0.13 | 0.05 |
| 1 | 179255957 | rs2493117 | 0.823 | 0.243 | 0.247 | -0.617 | 0 | -0.017 | -0.017 | 0 | 0.27 | 0.3 | 0.16 | 0.13 | 0.24 | 0.22 |
| 1 | 179262334 | rs12408349 | 2.046 | 0.269 | -0.005 | -0.492 | -0.051 | -0.103 | -0.103 | 0 | 0.19 | 0.01 | 0.35 | 0.31 | 0.28 | 0.22 |
| 1 | 179264397 | rs10913708 | 0.482 | 0.499 | -0.002 | -0.857 | -0.005 | -0.039 | -0.039 | 0 | 0.3 | 0.22 | 0.28 | 0.48 | 0.27 | 0.37 |
| 1 | 179267783 | rs2265932 | -1.54 | 0.427 | 0.014 | -0.938 | 0 | -0.072 | -0.072 | 0 | 0.79 | 0.74 | 0.87 | 0.96 | 0.79 | 0.83 |
| 1 | 179268323 | rs4652362 | 1.63 | 0.443 | -0.402 | -0.932 | -0.009 | 0.037 | 0.037 | 0.622 | 0.19 | 0.12 | 0.11 | 0.04 | 0.21 | 0.17 |
| 1 | 179269788 | rs7522641 | -0.419 | 0.506 | -0.274 | -0.705 | 0 | 0.071 | 0.071 | 0.321 | 0.51 | 0.35 | 0.7 | 0.84 | 0.55 | 0.6 |
| 1 | 179275259 | rs10913713 | -0.365 | 0.51 | -0.27 | -0.703 | 0.002 | 0.08 | 0.08 | 0.17 | 0.51 | 0.34 | 0.7 | 0.84 | 0.55 | 0.6 |
| 1 | 179275579 | rs12022539 | 1.898 | 0.715 | -0.192 | -0.537 | -0.018 | -0.089 | -0.089 | 0.038 | 0.19 | 0.05 | 0.42 | 0.36 | 0.28 | 0.23 |
| 1 | 179302003 | rs7531307 | 0.65 | 0.722 | -0.14 | -0.533 | 0.004 | 0.019 | 0.019 | 0 | 0.23 | 0.25 | 0.25 | 0.48 | 0.24 | 0.35 |
| 1 | 179308803 | rs3753524 | 1.531 | 0.472 | -0.543 | -0.399 | -0.002 | -0.055 | -0.055 | 0 | 0.2 | 0.09 | 0.42 | 0.36 | 0.28 | 0.23 |
| 1 | 179311602 | rs3753525 | -0.699 | -0.071 | -0.478 | -0.512 | 0.023 | 0.045 | 0.045 | 0.277 | 0.68 | 0.75 | 0.85 | 0.88 | 0.78 | 0.75 |
| 1 | 179312102 | rs4651024 | 0.282 | -0.162 | 0.013 | -0.376 | 0.001 | 0.001 | 0.001 | 0 | 0.39 | 0.38 | 0.43 | 0.49 | 0.55 | 0.45 |

| | | | | | | | | | | | | | | | | |
|---|---|---|---|---|---|---|---|---|---|---|---|---|---|---|---|---|
| 1 | 179312235 | rs10913728 | 0.173 | -0.31 | -0.03 | -0.49 | -0.015 | 0.016 | 0.016 | 0 | 0.37 | 0.49 | 0.48 | 0.42 | 0.34 | 0.39 |
| 1 | 179312752 | rs10753191 | 0.346 | -0.293 | -0.039 | -0.397 | -0.038 | 0.127 | 0.127 | 0.973 | 0.17 | 0.35 | 0.29 | 0.35 | 0.13 | 0.17 |
| 1 | 179315357 | rs2152319 | -0.16 | -0.27 | 0.061 | -0.484 | 0 | -0.027 | -0.027 | 0 | 0.47 | 0.46 | 0.66 | 0.87 | 0.62 | 0.65 |
| 1 | 179317069 | rs10913733 | 1.228 | -0.22 | -0.383 | -0.233 | -0.01 | 0.093 | 0.093 | 0 | 0.07 | 0.07 | 0.17 | 0.33 | 0.07 | 0.18 |
| 1 | 179319580 | rs3753527 | -0.739 | -0.209 | -0.472 | -0.284 | 0.031 | 0.116 | 0.116 | 0.099 | 0.57 | 0.73 | 0.72 | 0.87 | 0.67 | 0.66 |
| 1 | 179320578 | rs13306731 | 1.893 | -0.111 | -0.494 | -0.072 | -0.002 | -0.006 | -0.006 | 1.033 | 0.06 | 0.01 | 0.15 | 0.3 | 0.07 | 0.14 |
| 1 | 179334934 | rs10798670 | 1.453 | -0.107 | -0.405 | -0.201 | -0.006 | 0.077 | 0.077 | 0 | 0.07 | 0.06 | 0.17 | 0.33 | 0.07 | 0.18 |
| 1 | 179334984 | rs10753193 | -1.37 | -0.315 | -0.49 | -0.358 | 0.046 | 0.048 | 0.048 | 0 | 0.77 | 0.88 | 0.91 | 0.91 | 0.89 | 0.84 |
| 1 | 179335138 | rs17369102 | 2.015 | 0.011 | -0.74 | -0.385 | -0.084 | -0.003 | -0.003 | 0 | 0.18 | 0.01 | 0.26 | 0.16 | 0.3 | 0.2 |
| 1 | 179335366 | rs10913736 | 0.182 | -0.351 | -0.029 | -0.271 | 0.038 | 0.013 | 0.013 | 0.058 | 0.5 | 0.67 | 0.55 | 0.54 | 0.6 | 0.48 |
| 1 | 179339334 | rs6666358 | -0.152 | -0.396 | -0.032 | -0.195 | 0.022 | -0.018 | -0.018 | 0 | 0.67 | 0.74 | 0.66 | 0.52 | 0.8 | 0.62 |
| 1 | 179347832 | rs10798674 | 0.429 | -0.291 | 0.05 | -0.234 | -0.014 | 0.03 | 0.03 | 0 | 0.44 | 0.28 | 0.53 | 0.7 | 0.52 | 0.62 |
| 1 | 179352736 | rs4077551 | 1.009 | -0.217 | 0.06 | -0.219 | -0.012 | 0.089 | 0.089 | 0 | 0.09 | 0.1 | 0.18 | 0.33 | 0.09 | 0.2 |
| 1 | 179353439 | rs4424451 | 1.766 | -0.11 | -0.011 | -0.404 | -0.013 | 0.026 | 0.026 | 0.457 | 0.18 | 0.12 | 0.19 | 0.05 | 0.22 | 0.16 |
| 1 | 179356378 | rs12749461 | 0.726 | -0.148 | 0.195 | -0.452 | -0.001 | -0.017 | -0.017 | 1.646 | 0.27 | 0.27 | 0.24 | 0.05 | 0.27 | 0.18 |
| 1 | 179361178 | rs4297245 | -0.742 | -0.226 | 0.443 | -0.466 | -0.008 | -0.016 | -0.016 | 0 | 0.73 | 0.68 | 0.84 | 0.8 | 0.79 | 0.82 |
| 1 | 179363954 | rs4606280 | -1.221 | -0.273 | 0.258 | -0.552 | 0.037 | 0.038 | 0.038 | 0 | 0.77 | 0.85 | 0.9 | 0.9 | 0.89 | 0.84 |
| 1 | 179367301 | rs12760534 | 0.699 | -0.182 | 0.171 | -0.519 | 0 | -0.019 | -0.019 | 0 | 0.28 | 0.29 | 0.24 | 0.05 | 0.27 | 0.18 |
| 1 | 179371973 | rs4651025 | 0.001 | -0.24 | 0.195 | -0.399 | -0.015 | 0.041 | 0.041 | 0.035 | 0.66 | 0.55 | 0.65 | 0.48 | 0.69 | 0.63 |
| 1 | 179396444 | rs4652380 | 1.063 | -0.249 | 0.176 | -0.405 | -0.023 | -0.029 | -0.029 | 0 | 0.37 | 0.26 | 0.4 | 0.43 | 0.42 | 0.44 |
| 1 | 179399253 | rs7555722 | 1.42 | -0.51 | 0.003 | -0.704 | -0.003 | -0.005 | -0.005 | 0 | 0.22 | 0.2 | 0.2 | 0.05 | 0.22 | 0.17 |
| 1 | 179404013 | rs6659294 | -1.164 | -0.499 | 0.092 | -0.656 | 0.01 | 0.01 | 0.01 | 0 | 0.78 | 0.83 | 0.9 | 0.9 | 0.9 | 0.86 |
| 1 | 179414105 | rs11577579 | 0.278 | -0.034 | -0.119 | -0.474 | 0.001 | -0.006 | -0.006 | 0 | 0.3 | 0.29 | 0.24 | 0.05 | 0.28 | 0.19 |
| 1 | 179415532 | rs4130266 | 0.189 | -0.044 | -0.098 | -0.479 | -0.001 | -0.054 | -0.054 | 0 | 0.31 | 0.39 | 0.25 | 0.05 | 0.28 | 0.19 |
| 1 | 179416956 | rs10913766 | 0.264 | 0.008 | -0.205 | -0.454 | 0.001 | -0.01 | -0.01 | 0 | 0.3 | 0.29 | 0.25 | 0.05 | 0.28 | 0.19 |
| 1 | 179423620 | rs12403933 | 1.025 | 0.074 | -0.24 | -0.219 | -0.001 | 0.008 | 0.008 | 0 | 0.22 | 0.21 | 0.26 | 0.11 | 0.21 | 0.19 |
| 1 | 179426390 | rs12740455 | 0.05 | -0.129 | 0.032 | 0.111 | -0.016 | 0.112 | 0.112 | 0.198 | 0.12 | 0.27 | 0.2 | 0.33 | 0.12 | 0.19 |
| 1 | 179427632 | rs12725126 | -0.461 | -0.217 | 0.027 | 0.009 | -0.025 | 0.04 | 0.04 | 0.701 | 0.2 | 0.33 | 0.25 | 0.33 | 0.17 | 0.21 |
| 1 | 179452275 | rs10494520 | 0.567 | -0.201 | 0.06 | 0.026 | -0.006 | 0.046 | 0.046 | 0 | 0.11 | 0.14 | 0.2 | 0.33 | 0.12 | 0.21 |
| 1 | 179452381 | rs10494519 | 1.038 | -0.222 | 0.088 | -0.002 | -0.016 | -0.049 | -0.049 | 0.196 | 0.09 | 0.19 | 0.05 | 0 | 0.05 | 0.02 |
| 1 | 179452695 | rs962429 | 0.305 | -0.25 | -0.098 | -0.226 | -0.017 | 0.099 | 0.099 | 0 | 0.46 | 0.29 | 0.58 | 0.75 | 0.53 | 0.66 |
| 1 | 179458821 | rs2225213 | -0.298 | -0.262 | -0.102 | -0.231 | 0 | 0.004 | 0.004 | 0 | 0.55 | 0.49 | 0.64 | 0.75 | 0.59 | 0.68 |
| 1 | 179459554 | rs6674086 | 1.087 | -0.253 | -0.128 | -0.184 | -0.068 | -0.022 | -0.022 | 0 | 0.35 | 0.15 | 0.32 | 0.37 | 0.41 | 0.45 |
| 1 | 179460125 | rs6425567 | 0.246 | -0.246 | -0.12 | -0.313 | -0.016 | 0.102 | 0.102 | 0 | 0.46 | 0.29 | 0.58 | 0.75 | 0.53 | 0.66 |
| 1 | 179460524 | rs16828724 | 1.812 | -0.186 | -0.035 | -0.231 | 0.002 | -0.008 | -0.008 | 1.003 | 0.08 | 0.07 | 0.05 | 0 | 0.05 | 0.02 |
| 1 | 179460806 | rs17369441 | 2.068 | -0.19 | -0.054 | -0.231 | -0.003 | 0.04 | 0.04 | 0 | 0.07 | 0 | 0.04 | 0 | 0.05 | 0.02 |
| 1 | 179469314 | rs12047808 | -0.5 | -0.197 | 0.43 | -0.314 | 0.142 | 0.229 | 0.229 | 0 | 0.06 | 0.32 | 0.08 | 0.16 | 0.11 | 0.03 |
| 1 | 179471750 | rs10913801 | 0.276 | -0.424 | 0.363 | -0.458 | -0.005 | -0.012 | -0.012 | 0 | 0.68 | 0.43 | 0.78 | 0.8 | 0.74 | 0.82 |
| 1 | 179473513 | rs1329276 | 0.639 | -0.43 | 0.352 | -0.453 | 0.005 | -0.002 | -0.002 | 0 | 0.32 | 0.28 | 0.39 | 0.38 | 0.33 | 0.36 |
| 1 | 179474053 | rs1928004 | -1.353 | -0.422 | 0.331 | -0.459 | -0.014 | -0.049 | -0.049 | 0 | 0.82 | 0.8 | 0.91 | 0.96 | 0.91 | 0.87 |
| 1 | 179479012 | rs10913805 | -0.356 | -0.414 | 0.339 | -0.451 | -0.006 | -0.012 | -0.012 | 0 | 0.76 | 0.48 | 0.83 | 0.8 | 0.8 | 0.84 |
| 1 | 179479758 | rs12131400 | -1.029 | -0.543 | 0.238 | -0.574 | -0.038 | -0.097 | -0.097 | 0 | 0.8 | 0.66 | 0.9 | 0.96 | 0.91 | 0.87 |
| 1 | 179479803 | rs10913807 | -0.428 | -0.516 | 0.062 | -0.57 | -0.009 | -0.055 | -0.055 | 0 | 0.74 | 0.74 | 0.86 | 0.96 | 0.85 | 0.85 |
| 1 | 179492085 | rs7551440 | -0.292 | -0.358 | -0.302 | -0.441 | -0.001 | -0.002 | -0.002 | 2.056 | 0.43 | 0.39 | 0.49 | 0.54 | 0.49 | 0.4 |
| 1 | 179514777 | rs2296960 | 0.906 | -0.16 | -0.842 | -0.239 | -0.085 | -0.101 | -0.101 | 0 | 0.35 | 0.11 | 0.39 | 0.44 | 0.41 | 0.47 |
| 1 | 179516738 | rs10913815 | -0.366 | -0.011 | -0.735 | -0.381 | 0.038 | -0.02 | -0.02 | 0 | 0.55 | 0.73 | 0.59 | 0.53 | 0.61 | 0.47 |
| 1 | 179517140 | rs10494515 | 0.482 | 0.203 | -0.198 | -0.549 | 0.044 | 0.092 | 0.092 | 0 | 0.21 | 0.23 | 0.3 | 0.48 | 0.31 | 0.27 |
| 1 | 179518858 | rs10913816 | -0.319 | 0.049 | 0.173 | -0.538 | 0.029 | 0.054 | 0.054 | 0.171 | 0.3 | 0.54 | 0.36 | 0.48 | 0.37 | 0.29 |
| 1 | 179520050 | rs2274622 | 1.561 | -0.073 | -0.078 | -0.63 | -0.016 | -0.062 | -0.062 | 0 | 0.28 | 0.11 | 0.29 | 0.37 | 0.31 | 0.4 |
| 1 | 179520108 | rs2274623 | 1.185 | 0.156 | -0.088 | -0.4 | -0.066 | -0.09 | -0.09 | 0 | 0.17 | 0.02 | 0.22 | 0.32 | 0.21 | 0.25 |
| 1 | 179520151 | rs1410590 | -1.592 | 0.107 | -0.061 | -0.449 | 0.001 | -0.016 | -0.016 | 2.286 | 0.83 | 0.84 | 0.94 | 0.97 | 0.93 | 0.88 |
| 1 | 179520506 | rs1410592 | 0.083 | 0.132 | 0.309 | -0.352 | 0.011 | 0.024 | 0.024 | 1.032 | 0.63 | 0.59 | 0.63 | 0.49 | 0.62 | 0.62 |
| 1 | 179521932 | rs2274626 | 0.457 | 0.052 | 0.304 | -0.389 | -0.012 | 0.011 | 0.011 | 0.145 | 0.4 | 0.57 | 0.43 | 0.45 | 0.37 | 0.44 |
| 1 | 179529150 | rs10913817 | -0.285 | 0.083 | 0.282 | -0.337 | 0 | -0.001 | -0.001 | 0.006 | 0.67 | 0.75 | 0.7 | 0.66 | 0.71 | 0.64 |
| 1 | 179545669 | rs3829795 | 0.146 | -0.001 | 0.166 | -0.588 | -0.015 | -0.022 | -0.022 | 0 | 0.28 | 0.2 | 0.32 | 0.48 | 0.37 | 0.32 |
| 1 | 179551381 | rs1887815 | -1.134 | -0.142 | 0.133 | -0.727 | 0.008 | 0.066 | 0.066 | 0 | 0.72 | 0.84 | 0.77 | 0.93 | 0.75 | 0.76 |
| 1 | 179551652 | rs1887814 | 0.06 | -0.134 | 0.159 | -0.655 | -0.013 | -0.013 | -0.013 | 0 | 0.3 | 0.22 | 0.4 | 0.55 | 0.39 | 0.34 |
| 1 | 179553724 | rs3930537 | -1.091 | -0.071 | 0.084 | -0.661 | 0.007 | 0.061 | 0.061 | 0 | 0.72 | 0.83 | 0.77 | 0.93 | 0.75 | 0.76 |
| 1 | 179558801 | rs913514 | 0.993 | -0.486 | 0.128 | -0.78 | 0.007 | -0.022 | -0.022 | 0 | 0.31 | 0.3 | 0.45 | 0.45 | 0.31 | 0.39 |
| 1 | 179559632 | rs10753199 | 0.263 | -0.468 | 0.26 | -0.845 | -0.001 | -0.001 | -0.001 | 0 | 0.48 | 0.44 | 0.68 | 0.83 | 0.52 | 0.65 |
| 1 | 179560154 | rs6691184 | 0.449 | -0.454 | -0.147 | -0.829 | -0.017 | 0.112 | 0.112 | 0 | 0.47 | 0.25 | 0.61 | 0.76 | 0.51 | 0.63 |
| 1 | 179562707 | rs12070589 | 1.808 | -0.545 | 0.016 | -0.541 | -0.005 | -0.044 | -0.044 | 0.756 | 0.08 | 0.13 | 0.04 | 0 | 0.06 | 0.02 |
| 1 | 179563717 | rs4652421 | -0.87 | -0.64 | 0.03 | -0.555 | -0.029 | -0.018 | -0.018 | 0 | 0.92 | 0.75 | 0.96 | 0.95 | 0.99 | 0.94 |
| 1 | 179574943 | rs10913831 | 1.289 | -0.505 | 0.003 | -0.46 | -0.01 | -0.001 | -0.001 | 3.221 | 0.11 | 0.14 | 0.04 | 0.11 | 0.08 | 0.05 |
| 1 | 179583165 | rs1410594 | -1.002 | -0.483 | -0.25 | -0.586 | 0.001 | -0.024 | -0.024 | 0 | 0.73 | 0.73 | 0.84 | 0.9 | 0.76 | 0.76 |
| 1 | 179598179 | rs4610966 | 0.633 | -0.36 | -0.438 | -0.233 | -0.026 | -0.062 | -0.062 | 0.242 | 0.28 | 0.1 | 0.3 | 0.36 | 0.29 | 0.31 |
| 1 | 179600001 | rs6704505 | 1.374 | -0.382 | -0.559 | -0.594 | 0.001 | 0.001 | 0.001 | 0.301 | 0.11 | 0.09 | 0.04 | 0.11 | 0.09 | 0.05 |
| 1 | 179604514 | rs4474210 | 0.794 | -0.449 | -0.275 | -0.7 | -0.01 | -0.008 | -0.008 | 0.449 | 0.32 | 0.28 | 0.44 | 0.37 | 0.4 | 0.39 |
| 1 | 179604864 | rs4628468 | -1.326 | -0.483 | -0.085 | -0.729 | 0 | -0.032 | -0.032 | 0 | 0.73 | 0.72 | 0.86 | 0.9 | 0.78 | 0.77 |

| | | | | | | | | | | | | | | | | |
|---|---|---|---|---|---|---|---|---|---|---|---|---|---|---|---|---|
| 1 | 179606157 | rs10494522 | -0.088 | -0.432 | -0.172 | -0.667 | -0.01 | -0.01 | -0.01 | 0 | 0.43 | 0.35 | 0.48 | 0.48 | 0.49 | 0.44 |
| 1 | 179608461 | rs4468120 | 1.898 | -0.441 | -0.665 | -0.631 | -0.049 | -0.007 | -0.007 | 0 | 0.17 | 0.01 | 0.22 | 0.13 | 0.24 | 0.24 |
| 1 | 179611021 | rs11806102 | -0.186 | -0.668 | -0.388 | -0.556 | -0.014 | -0.013 | -0.013 | 0.679 | 0.43 | 0.32 | 0.47 | 0.48 | 0.49 | 0.44 |
| 1 | 179633040 | rs12035247 | 0.351 | -0.515 | -0.341 | -0.544 | -0.007 | -0.006 | -0.006 | 1.586 | 0.32 | 0.29 | 0.43 | 0.37 | 0.4 | 0.39 |
| 1 | 179636209 | rs4652433 | 0.593 | -0.503 | -0.424 | -0.549 | -0.091 | -0.079 | -0.079 | 0 | 0.29 | 0.08 | 0.41 | 0.37 | 0.4 | 0.39 |
| 1 | 179638838 | rs4651045 | 0.459 | -0.463 | -0.441 | -0.519 | -0.023 | -0.019 | -0.019 | 0.503 | 0.31 | 0.22 | 0.43 | 0.37 | 0.4 | 0.39 |
| 1 | 179647658 | rs10798708 | -0.516 | -0.537 | -0.268 | -0.715 | -0.016 | -0.094 | -0.094 | 2.863 | 0.73 | 0.65 | 0.85 | 0.9 | 0.78 | 0.77 |
| 1 | 179669906 | rs10733002 | -0.066 | -0.522 | -0.214 | -0.034 | -0.029 | 0.051 | 0.051 | 0.345 | 0.92 | 0.75 | 0.96 | 0.86 | 0.99 | 0.93 |
| 1 | 179678643 | rs4652445 | -0.011 | -0.022 | -1.274 | -0.251 | -0.017 | -0.024 | -0.024 | 0 | 0.56 | 0.83 | 0.6 | 0.41 | 0.53 | 0.51 |
| 1 | 179680642 | rs10798713 | -0.188 | 0.005 | -1.07 | -0.884 | -0.009 | 0.024 | 0.024 | 0 | 0.6 | 0.75 | 0.63 | 0.65 | 0.57 | 0.6 |
| 1 | 179687226 | rs10913869 | 0.661 | 0.117 | -1.01 | -0.898 | 0.006 | -0.001 | -0.001 | 0 | 0.11 | 0.09 | 0.05 | 0 | 0.08 | 0.05 |
| 1 | 179687259 | rs2504057 | -0.479 | -0.068 | -0.877 | -1.262 | 0.007 | 0.041 | 0.041 | 0.08 | 0.53 | 0.76 | 0.42 | 0.4 | 0.42 | 0.56 |
| 1 | 179687646 | rs10913870 | 0.613 | 0.379 | -0.696 | -0.888 | 0.001 | 0.005 | 0.005 | 0 | 0.21 | 0.18 | 0.41 | 0.58 | 0.22 | 0.27 |
| 1 | 179695998 | rs12128834 | 0.783 | 0.151 | -0.816 | -0.943 | 0.01 | 0.02 | 0.02 | 0 | 0.11 | 0.05 | 0.05 | 0 | 0.08 | 0.05 |
| 1 | 179700133 | rs2504031 | 0.429 | 0.003 | -1.071 | -1.198 | -0.007 | -0.031 | -0.031 | 0 | 0.3 | 0.19 | 0.3 | 0.4 | 0.29 | 0.45 |
| 1 | 179701692 | rs10798715 | 1.345 | -0.203 | -1.002 | -1.369 | -0.004 | -0.021 | -0.021 | 0 | 0.08 | 0.08 | 0.03 | 0 | 0.04 | 0.07 |
| 1 | 179703019 | rs2504033 | 0.492 | -0.2 | -0.706 | -1.566 | 0.002 | 0.007 | 0.007 | 0 | 0.3 | 0.36 | 0.3 | 0.38 | 0.3 | 0.47 |
| 1 | 179705637 | rs2454190 | -0.927 | -0.093 | -1.221 | -1.99 | -0.044 | 0.214 | 0.214 | 0 | 0.63 | 0.93 | 0.78 | 0.97 | 0.55 | 0.78 |
| 1 | 179711649 | rs1052447 | 1.245 | -0.241 | -0.833 | -2.022 | -0.002 | -0.003 | -0.003 | 0 | 0.21 | 0.17 | 0.4 | 0.6 | 0.2 | 0.24 |
| 1 | 179716507 | rs2245195 | -0.292 | -0.309 | -0.927 | -2.248 | -0.01 | 0.005 | 0.005 | 0 | 0.62 | 0.79 | 0.49 | 0.4 | 0.59 | 0.71 |
| 1 | 179723197 | rs17370084 | 1.917 | -0.236 | -0.942 | -2.186 | -0.021 | 0.069 | 0.069 | 0 | 0.2 | 0.05 | 0.12 | 0.03 | 0.24 | 0.17 |
| 1 | 179726432 | rs10913879 | 0.898 | -0.34 | -0.396 | -1.687 | 0.008 | 0.034 | 0.034 | 0 | 0.21 | 0.23 | 0.4 | 0.61 | 0.2 | 0.24 |
| 1 | 179727752 | rs2487742 | -0.887 | -0.591 | -0.759 | -1.794 | 0.004 | 0.049 | 0.049 | 0 | 0.82 | 0.93 | 0.89 | 1 | 0.8 | 0.94 |
| 1 | 179729126 | rs2504053 | 0.752 | -0.59 | -0.701 | -1.96 | 0.043 | 0.035 | 0.035 | 0 | 0.52 | 0.3 | 0.45 | 0.41 | 0.6 | 0.67 |
| 1 | 179729554 | rs2454199 | 0.106 | -0.198 | -0.682 | -2.287 | 0 | -0.052 | -0.052 | 0 | 0.73 | 0.69 | 0.83 | 1 | 0.75 | 0.87 |
| 1 | 179733193 | rs17279065 | -0.163 | 0.091 | -0.817 | -2.455 | -0.011 | 0.022 | 0.022 | 0 | 0.31 | 0.26 | 0.2 | 0.08 | 0.34 | 0.25 |
| 1 | 179735387 | rs2318393 | -0.1 | -0.043 | -0.7 | -2.241 | 0.001 | 0.001 | 0.001 | 0 | 0.13 | 0.09 | 0.08 | 0.06 | 0.11 | 0.07 |
| 1 | 179743876 | rs4652453 | -0.616 | -0.07 | -0.095 | -2.349 | -0.02 | -0.067 | -0.067 | 2.226 | 0.16 | 0.34 | 0.13 | 0.06 | 0.12 | 0.12 |
| 1 | 179745671 | rs7514717 | -0.616 | -0.07 | -0.105 | -2.349 | -0.02 | -0.066 | -0.066 | 0.059 | 0.16 | 0.34 | 0.13 | 0.06 | 0.12 | 0.12 |
| 1 | 179748224 | rs4448483 | -0.741 | -0.085 | -0.245 | -2.388 | -0.001 | -0.001 | -0.001 | 0 | 0.41 | 0.64 | 0.28 | 0.25 | 0.39 | 0.31 |
| 1 | 179748514 | rs4448484 | -0.333 | -0.2 | -0.687 | -3.018 | 0 | -0.05 | -0.05 | 0 | 0.58 | 0.44 | 0.71 | 0.96 | 0.6 | 0.63 |
| 1 | 179750647 | rs10913887 | 0.703 | 0.092 | -0.636 | -2.7 | -0.01 | 0.013 | 0.013 | 0 | 0.59 | 0.36 | 0.65 | 0.87 | 0.51 | 0.59 |
| 1 | 179753147 | rs6680679 | -1.483 | -0.623 | 0.04 | -2.332 | -0.004 | -0.04 | -0.04 | 0 | 0.3 | 0.64 | 0.22 | 0.1 | 0.27 | 0.28 |
| 1 | 179755583 | rs11583659 | 0.324 | -0.618 | 0.082 | -2.311 | -0.033 | -0.081 | -0.081 | 0 | 0.13 | 0.28 | 0.09 | 0 | 0.06 | 0.12 |
| 1 | 179756639 | rs6678978 | -0.702 | -0.495 | -0.51 | -2.251 | -0.003 | -0.079 | -0.079 | 0 | 0.27 | 0.58 | 0.21 | 0.09 | 0.27 | 0.26 |
| 1 | 179757536 | rs10913888 | -1.561 | -0.447 | -1.647 | -2.267 | -0.022 | 0.192 | 0.192 | 0 | 0.53 | 0.84 | 0.37 | 0.19 | 0.52 | 0.45 |
| 1 | 179757855 | rs12756748 | 0.388 | -0.066 | -1.318 | -1.694 | -0.088 | 0.04 | 0.04 | 0 | 0.21 | 0.01 | 0.13 | 0.1 | 0.25 | 0.15 |
| 1 | 179761601 | rs12120743 | -1.019 | -0.46 | -0.488 | -1.269 | -0.042 | 0.032 | 0.032 | 0.492 | 0.2 | 0.1 | 0.17 | 0.14 | 0.26 | 0.21 |
| 1 | 179763611 | rs16854720 | -0.243 | -1.319 | 0.146 | -1.541 | -0.043 | -0.044 | -0.044 | 0 | 0.25 | 0.37 | 0.16 | 0.14 | 0.17 | 0.21 |
| 1 | 179766086 | rs10798724 | -0.275 | -1.323 | 0.165 | -1.546 | -0.054 | -0.045 | -0.045 | 1.566 | 0.26 | 0.37 | 0.15 | 0.14 | 0.15 | 0.21 |
| 1 | 179766172 | rs17370290 | -0.674 | -1.34 | 0.262 | -1.67 | -0.015 | 0.038 | 0.038 | 0.153 | 0.19 | 0.1 | 0.14 | 0.06 | 0.18 | 0.17 |
| 1 | 179768464 | rs6425594 | -1.37 | -1.23 | 0.61 | -1.288 | 0.009 | 0.022 | 0.022 | 0 | 0.45 | 0.37 | 0.28 | 0.27 | 0.36 | 0.39 |
| 1 | 179769066 | rs6701652 | -1.38 | -1.247 | 0.644 | -1.525 | 0.011 | 0.021 | 0.021 | 0 | 0.45 | 0.38 | 0.28 | 0.19 | 0.34 | 0.38 |
| 1 | 179770256 | rs2647225 | -1.201 | -1.237 | 0.523 | -1.52 | 0.017 | 0.045 | 0.045 | 0 | 0.43 | 0.32 | 0.26 | 0.19 | 0.33 | 0.38 |
| 1 | 179776841 | rs2777920 | -1.533 | -1.467 | 0.011 | -1.554 | 0.01 | 0.143 | 0.143 | 0.673 | 0.56 | 0.87 | 0.34 | 0.27 | 0.4 | 0.45 |
| 1 | 179785891 | rs2252384 | -1.46 | -1.387 | 0.923 | -1.57 | -0.004 | -0.048 | -0.048 | 0 | 0.38 | 0.47 | 0.28 | 0.2 | 0.35 | 0.34 |
| 1 | 179787027 | rs2794579 | -1.529 | -1.48 | 1.033 | -1.588 | 0.016 | 0.119 | 0.119 | 1.682 | 0.4 | 0.84 | 0.3 | 0.21 | 0.35 | 0.34 |
| 1 | 179788333 | rs12118565 | 0.042 | -1.468 | 0.832 | -1.674 | -0.002 | -0.018 | -0.018 | 0 | 0.17 | 0.16 | 0.14 | 0.01 | 0.19 | 0.17 |
| 1 | 179790518 | rs10913893 | 0.042 | -1.468 | 0.782 | -1.674 | -0.009 | 0.002 | 0.002 | 0 | 0.17 | 0.13 | 0.13 | 0.01 | 0.19 | 0.17 |
| 1 | 179794602 | rs872376 | -0.218 | -1.492 | 0.831 | -1.644 | -0.001 | 0.003 | 0.003 | 0 | 0.11 | 0.12 | 0.08 | 0.05 | 0.11 | 0.08 |
| 1 | 179795505 | rs1148821 | -1.918 | -1.547 | 1.092 | -1.63 | 0.008 | 0.034 | 0.034 | 1.404 | 0.4 | 0.73 | 0.31 | 0.21 | 0.35 | 0.33 |
| 1 | 179802378 | rs1148819 | -0.89 | -1.575 | 1.032 | -1.384 | -0.007 | 0.005 | 0.005 | 0 | 0.14 | 0.22 | 0.1 | 0.15 | 0.09 | 0.07 |
| 1 | 179813613 | rs1281410 | -2.036 | -1.171 | 0.476 | -1.543 | -0.128 | 0.055 | 0.055 | 0.107 | 0.56 | 0.97 | 0.38 | 0.31 | 0.44 | 0.44 |
| 1 | 179814457 | rs1281411 | -1.761 | -1.088 | 0.985 | -1.601 | -0.01 | 0.06 | 0.06 | 0.455 | 0.4 | 0.78 | 0.34 | 0.22 | 0.39 | 0.32 |
| 1 | 179817654 | rs1281432 | -1.78 | -1.079 | 0.848 | -1.597 | -0.003 | 0.004 | 0.004 | 0.215 | 0.39 | 0.67 | 0.33 | 0.22 | 0.39 | 0.32 |
| 1 | 179826051 | rs1281414 | -1.266 | -1.043 | 1.213 | -1.643 | -0.009 | -0.039 | -0.039 | 0 | 0.3 | 0.38 | 0.15 | 0.09 | 0.19 | 0.19 |
| 1 | 179827381 | rs1281415 | -0.576 | -1.066 | 1.297 | -1.658 | -0.008 | -0.002 | -0.002 | 0 | 0.1 | 0.26 | 0.06 | 0.08 | 0.05 | 0.04 |
| 1 | 179830012 | rs3737813 | -0.82 | -1.043 | 1.266 | -1.656 | -0.007 | -0.012 | -0.012 | 0.077 | 0.06 | 0.04 | 0.15 | 0.13 | 0.19 | 0.12 |
| 1 | 179831248 | rs1048418 | 0.196 | -1.085 | 1.268 | -1.673 | 0.003 | 0.024 | 0.024 | 0 | 0.18 | 0.1 | 0.09 | 0.01 | 0.14 | 0.13 |
| 1 | 179841753 | rs7411457 | -1.978 | -1.095 | 1.367 | -1.632 | 0.082 | 0.017 | 0.017 | 0 | 0.11 | 0.24 | 0.17 | 0.13 | 0.22 | 0.16 |
| 1 | 179843291 | rs668813 | -0.763 | -1.126 | 1.299 | -1.636 | -0.01 | -0.003 | -0.003 | 0 | 0.1 | 0.3 | 0.07 | 0.09 | 0.05 | 0.04 |
| 1 | 179847402 | rs1144575 | -2.165 | -1.109 | 1.23 | -1.632 | 0.03 | 0.006 | 0.006 | 0 | 0.19 | 0.34 | 0.21 | 0.21 | 0.25 | 0.19 |
| 1 | 179847635 | rs2249346 | -1.961 | -1.088 | 1.125 | -1.612 | -0.005 | 0.015 | 0.015 | 0.205 | 0.39 | 0.68 | 0.33 | 0.22 | 0.39 | 0.32 |
| 1 | 179852074 | rs1281378 | 1.556 | -1.071 | 1 | -1.604 | 0.003 | -0.018 | -0.018 | 0.461 | 0.62 | 0.38 | 0.68 | 0.78 | 0.61 | 0.69 |
| 1 | 179861042 | rs7530362 | -1.203 | -1.034 | 0.982 | -1.591 | 0.044 | -0.071 | -0.071 | 0 | 0.29 | 0.46 | 0.27 | 0.14 | 0.35 | 0.28 |
| 1 | 179887788 | rs531814 | 1.86 | -1.128 | 0.163 | -1.548 | -0.124 | 0.075 | 0.075 | 0.105 | 0.44 | 0.03 | 0.62 | 0.72 | 0.56 | 0.56 |
| 1 | 179900199 | rs539181 | 1.567 | -1.285 | 0.895 | -0.981 | 0.015 | 0.054 | 0.054 | 0 | 0.83 | 0.62 | 0.83 | 0.71 | 0.83 | 0.87 |
| 1 | 179900978 | rs2777910 | 0.575 | -1.231 | 0.961 | -1.254 | 0.012 | 0.014 | 0.014 | 0 | 0.89 | 0.8 | 0.91 | 0.83 | 0.86 | 0.94 |
| 1 | 179906900 | rs561233 | 1.391 | -1.499 | 1.163 | -1.477 | -0.001 | -0.033 | -0.033 | 0 | 0.65 | 0.51 | 0.8 | 0.82 | 0.71 | 0.78 |

| | | | | | | | | | | | | | | | | |
|---|---|---|---|---|---|---|---|---|---|---|---|---|---|---|---|---|
| 1 | 179909076 | rs2281805 | 0.089 | -1.481 | 0.947 | -1.461 | 0.011 | -0.046 | -0.046 | 0 | 0.08 | 0 | 0.02 | 0.17 | 0.03 | 0.05 |
| 1 | 179910422 | rs10913912 | 0.997 | -1.539 | 0.971 | -1.518 | -0.007 | -0.02 | -0.02 | 0 | 0.08 | 0.09 | 0.04 | 0 | 0.04 | 0.07 |
| 1 | 179911891 | rs528350 | 2.032 | -1.864 | 1.114 | -1.679 | 0.042 | 0.099 | 0.099 | 0 | 0.49 | 0.25 | 0.74 | 0.76 | 0.65 | 0.64 |
| 1 | 179918897 | rs12141134 | 0.719 | -1.799 | 0.975 | -1.632 | -0.039 | 0.065 | 0.065 | 0.308 | 0.11 | 0 | 0.06 | 0.01 | 0.11 | 0.1 |
| 1 | 179929388 | rs12124175 | 0.66 | -1.894 | 1.345 | -1.659 | -0.029 | -0.052 | -0.052 | 0 | 0.1 | 0.2 | 0.05 | 0 | 0.04 | 0.07 |
| 1 | 179942388 | rs10913918 | 0.665 | -1.841 | 1.423 | -1.644 | 0.017 | 0.026 | 0.026 | 0 | 0.89 | 0.73 | 0.92 | 0.84 | 0.87 | 0.95 |
| 1 | 179960787 | rs12132694 | -0.218 | -2.002 | 1.474 | -1.559 | -0.007 | 0.021 | 0.021 | 0.479 | 0.14 | 0.09 | 0.09 | 0.08 | 0.13 | 0.14 |
| 1 | 179964879 | rs2501615 | 2.323 | -2.127 | 1.512 | -1.585 | 0.032 | 0.072 | 0.072 | 0 | 0.48 | 0.25 | 0.73 | 0.7 | 0.65 | 0.65 |
| 1 | 179985156 | rs16855107 | -0.214 | -2.084 | 1.483 | -1.58 | -0.004 | 0.013 | 0.013 | 0.55 | 0.14 | 0.11 | 0.09 | 0.08 | 0.13 | 0.14 |
| 1 | 179994628 | rs16855130 | -0.349 | -2.085 | 1.464 | -1.58 | 0.001 | -0.017 | -0.017 | 0 | 0.15 | 0.18 | 0.1 | 0.08 | 0.13 | 0.15 |
| 1 | 179995975 | rs7521948 | 0.745 | -2.1 | 1.445 | -1.585 | -0.003 | -0.002 | -0.002 | 0 | 0.75 | 0.68 | 0.89 | 0.78 | 0.82 | 0.86 |
| 1 | 180000542 | rs12125245 | 0.718 | -2.082 | 1.325 | -1.574 | -0.042 | 0.065 | 0.065 | 0.108 | 0.11 | 0 | 0.06 | 0.01 | 0.11 | 0.08 |
| 1 | 180010911 | rs16855164 | -0.357 | -2.093 | 1.392 | -1.563 | 0.04 | -0.006 | -0.006 | 0 | 0.2 | 0.09 | 0.04 | 0.22 | 0.07 | 0.12 |
| 1 | 180037987 | rs6692394 | 1.485 | -2.127 | 1.414 | -1.564 | -0.033 | -0.037 | -0.037 | 0 | 0.63 | 0.43 | 0.83 | 0.78 | 0.78 | 0.79 |
| 1 | 180055985 | rs7540856 | 1.064 | -2.135 | 1.36 | -1.583 | 0.034 | 0.032 | 0.032 | 0 | 0.12 | 0.05 | 0.02 | 0.05 | 0.03 | 0.07 |
| 1 | 180059811 | rs10798729 | 2.15 | -2.061 | 1.347 | -1.551 | 0.033 | 0.076 | 0.076 | 0 | 0.49 | 0.25 | 0.73 | 0.71 | 0.65 | 0.65 |
| 1 | 180067988 | rs12121845 | -0.331 | -1.878 | 1.182 | -1.452 | 0 | -0.013 | -0.013 | 0.033 | 0.15 | 0.17 | 0.1 | 0.08 | 0.13 | 0.15 |
| 1 | 180075388 | rs6425607 | 0.228 | -1.742 | 0.974 | -1.341 | 0.014 | 0.018 | 0.018 | 0 | 0.89 | 0.76 | 0.93 | 0.84 | 0.87 | 0.95 |
| 1 | 180080115 | rs3767202 | 1.817 | -1.816 | 0.886 | -1.407 | 0.039 | 0.102 | 0.102 | 0 | 0.48 | 0.2 | 0.72 | 0.71 | 0.65 | 0.65 |
| 1 | 180085620 | rs12075450 | -0.691 | -2.031 | 0.941 | -1.473 | 0.008 | 0.005 | 0.005 | 0 | 0.26 | 0.28 | 0.1 | 0.21 | 0.17 | 0.13 |
| 1 | 180089858 | rs7544467 | 1.23 | -2.032 | 1.256 | -1.484 | -0.037 | -0.044 | -0.044 | 0.195 | 0.63 | 0.45 | 0.83 | 0.79 | 0.77 | 0.79 |
| 1 | 180095348 | rs6425609 | 2.085 | -2.066 | 1.202 | -1.637 | 0.016 | 0.051 | 0.051 | 0 | 0.47 | 0.29 | 0.75 | 0.76 | 0.64 | 0.65 |
| 1 | 180096149 | rs9425483 | 1.412 | -2.059 | 1.239 | -1.599 | 0.01 | 0.01 | 0.01 | 0 | 0.6 | 0.31 | 0.81 | 0.78 | 0.75 | 0.74 |
| 1 | 180096531 | rs9425484 | -0.784 | -1.836 | 1.414 | -1.456 | -0.032 | -0.057 | -0.057 | 0.022 | 0.89 | 0.77 | 0.94 | 1 | 0.96 | 0.93 |
| 1 | 180097729 | rs4652469 | -0.065 | -1.833 | 1.39 | -1.453 | -0.035 | -0.027 | -0.027 | 0 | 0.87 | 0.71 | 0.93 | 0.93 | 0.94 | 0.87 |
| 1 | 180099463 | rs7551181 | 1.661 | -1.745 | 1.206 | -1.457 | 0.026 | 0.084 | 0.084 | 0 | 0.49 | 0.28 | 0.75 | 0.77 | 0.64 | 0.67 |
| 1 | 180103851 | rs4336810 | 2.002 | -2.173 | 1.095 | -1.593 | 0.006 | 0.08 | 0.08 | 0 | 0.43 | 0.2 | 0.72 | 0.75 | 0.64 | 0.57 |
| 1 | 180105578 | rs2874053 | 0.684 | -2.084 | 1.011 | -1.552 | -0.063 | -0.063 | -0.063 | 0.431 | 0.81 | 0.59 | 0.9 | 0.91 | 0.92 | 0.77 |
| 1 | 180107859 | rs6691435 | 1.625 | -1.863 | 0.837 | -1.422 | 0.007 | 0.025 | 0.025 | 0 | 0.46 | 0.33 | 0.75 | 0.75 | 0.64 | 0.57 |
| 1 | 180114023 | rs7551902 | -1.294 | -1.715 | 0.535 | -1.339 | -0.001 | -0.005 | -0.005 | 0 | 0.46 | 0.51 | 0.21 | 0.27 | 0.32 | 0.38 |
| 1 | 180119349 | rs17371566 | -0.305 | -1.425 | -0.025 | -1.187 | 0.008 | 0.005 | 0.005 | 0 | 0.1 | 0.18 | 0.08 | 0.14 | 0.14 | 0.06 |
| 1 | 180119544 | rs3845391 | 0.829 | -1.671 | -0.001 | -1.266 | 0.013 | -0.001 | -0.001 | 0 | 0.54 | 0.46 | 0.8 | 0.86 | 0.77 | 0.62 |
| 1 | 180121480 | rs3753814 | 0.981 | -1.361 | -0.185 | -1.205 | 0.037 | 0.117 | 0.117 | 0 | 0.54 | 0.25 | 0.75 | 0.85 | 0.68 | 0.73 |
| 1 | 180122812 | rs7516257 | -0.265 | -1.458 | 0.683 | -1.051 | -0.033 | -0.058 | -0.058 | 0.033 | 0.1 | 0.27 | 0.04 | 0 | 0.05 | 0.07 |
| 1 | 180124667 | rs3767199 | -1.713 | -1.273 | 0.311 | -1.428 | 0 | -0.024 | -0.024 | 0 | 0.39 | 0.58 | 0.29 | 0.11 | 0.35 | 0.38 |
| 1 | 180124872 | rs3767196 | -1.954 | -1.365 | 0.312 | -0.715 | 0.004 | 0.062 | 0.062 | 0.032 | 0.43 | 0.81 | 0.36 | 0.34 | 0.37 | 0.41 |
| 1 | 180126141 | rs3753812 | 1.709 | -1.402 | 0.3 | -0.744 | 0 | 0.075 | 0.075 | 0.081 | 0.56 | 0.14 | 0.64 | 0.66 | 0.63 | 0.59 |
| 1 | 180127107 | rs6425611 | -1.608 | -1.415 | 0.492 | -0.854 | -0.018 | -0.114 | -0.114 | 0.692 | 0.29 | 0.58 | 0.14 | 0.01 | 0.2 | 0.2 |
| 1 | 180129205 | rs3842888 | 1.619 | -1.756 | 0.088 | -0.638 | 0.073 | 0.042 | 0.042 | 0.298 | 0.63 | 0.26 | 0.81 | 0.75 | 0.84 | 0.69 |
| 1 | 180138318 | rs3820308 | 1.715 | -1.714 | 0.253 | -0.666 | 0.034 | 0.05 | 0.05 | 0 | 0.62 | 0.26 | 0.78 | 0.77 | 0.74 | 0.67 |
| 1 | 180138800 | rs6703180 | 2.306 | -1.977 | 0.215 | -0.641 | 0.103 | 0.151 | 0.151 | 0 | 0.51 | 0.17 | 0.7 | 0.76 | 0.7 | 0.53 |
| 1 | 180141367 | rs3767184 | 1.571 | -1.984 | 0.298 | -0.534 | -0.002 | -0.002 | -0.002 | 0 | 0.63 | 0.4 | 0.85 | 0.59 | 0.84 | 0.68 |
| 1 | 180144485 | rs10913939 | 0.446 | -1.847 | 0.179 | -0.493 | -0.009 | 0.003 | 0.003 | 0 | 0.11 | 0.09 | 0.11 | 0 | 0.14 | 0.15 |
| 1 | 180145088 | rs2298206 | -0.413 | -1.905 | 0.326 | -0.488 | -0.011 | 0.038 | 0.038 | 1.156 | 0.9 | 0.81 | 0.97 | 0.85 | 0.96 | 0.9 |
| 1 | 180145769 | rs12075418 | 0.007 | -1.906 | 0.324 | -0.479 | -0.007 | 0.093 | 0.093 | 0.767 | 0.11 | 0.15 | 0.07 | 0.39 | 0.06 | 0.13 |
| 1 | 180154776 | rs3842890 | 0.224 | -1.9 | 0.314 | -0.51 | -0.011 | 0.012 | 0.012 | 0 | 0.12 | 0.08 | 0.15 | 0.07 | 0.16 | 0.18 |
| 1 | 180156251 | rs3767175 | 1.01 | -1.924 | 0.293 | -0.505 | -0.012 | -0.046 | -0.046 | 0 | 0.08 | 0 | 0.03 | 0.16 | 0.04 | 0.11 |
| 1 | 180157567 | rs3767170 | 2.373 | -2.09 | 0.263 | -0.603 | 0.079 | -0.051 | -0.051 | 0 | 0.49 | 0.12 | 0.66 | 0.51 | 0.68 | 0.5 |
| 1 | 180157593 | rs3767169 | 1.112 | -1.821 | 0.422 | -0.449 | 0.008 | 0.079 | 0.079 | 0 | 0.77 | 0.65 | 0.76 | 0.51 | 0.78 | 0.69 |
| 1 | 180159654 | rs7521513 | 0.104 | -1.713 | 0.257 | -0.477 | -0.011 | 0.013 | 0.013 | 0 | 0.12 | 0.08 | 0.16 | 0.07 | 0.16 | 0.18 |
| 1 | 180162364 | rs10913943 | -1.717 | -2.015 | 0.781 | -0.616 | -0.013 | -0.02 | -0.02 | 0 | 0.31 | 0.35 | 0.15 | 0.17 | 0.22 | 0.31 |
| 1 | 180162555 | rs7542624 | -0.954 | -1.914 | -0.114 | 0.238 | 0.006 | -0.003 | -0.003 | 0.353 | 0.41 | 0.39 | 0.32 | 0.45 | 0.26 | 0.38 |
| 1 | 180165091 | rs3767165 | 2.255 | -1.148 | 0.144 | 0.647 | 0.049 | 0.004 | 0.004 | 0 | 0.54 | 0.3 | 0.68 | 0.48 | 0.69 | 0.53 |
| 1 | 180167090 | rs7597 | -0.146 | -0.964 | 0.179 | 0.784 | 0.005 | 0.01 | 0.01 | 1.645 | 0.15 | 0.11 | 0.07 | 0.45 | 0.06 | 0.18 |
| 1 | 180167440 | rs3843273 | -1.293 | -0.989 | 0.203 | 0.736 | -0.078 | -0.086 | -0.086 | 0 | 0.21 | 0.49 | 0.14 | 0.07 | 0.1 | 0.14 |
| 1 | 180168238 | rs6425612 | -0.27 | -0.84 | 0.025 | 0.71 | 0.036 | 0.011 | 0.011 | 0.186 | 0.07 | 0.34 | 0.05 | 0 | 0.05 | 0.05 |
| 1 | 180169108 | rs10913945 | 0.173 | -0.762 | -0.862 | 0.78 | 0.015 | -0.026 | -0.026 | 0 | 0.13 | 0.07 | 0.03 | 0.2 | 0.05 | 0.16 |
| 1 | 180169330 | rs11126 | 1.296 | -0.783 | -0.886 | 0.81 | -0.002 | 0.005 | 0.005 | 0.01 | 0.66 | 0.38 | 0.77 | 0.53 | 0.74 | 0.58 |
| 1 | 180169405 | rs7513063 | -0.286 | -0.746 | -0.902 | 0.823 | 0.015 | -0.012 | -0.012 | 0 | 0.08 | 0.42 | 0.06 | 0.01 | 0.05 | 0.08 |
| 1 | 180171254 | rs6704314 | 0.759 | -0.094 | -1.65 | 1.211 | -0.004 | -0.036 | -0.036 | 0 | 0.75 | 0.82 | 0.84 | 0.55 | 0.79 | 0.66 |
| 1 | 180173484 | rs3845393 | 0.542 | -0.028 | -1.614 | 0.773 | -0.004 | -0.006 | -0.006 | 0 | 0.76 | 0.83 | 0.88 | 0.79 | 0.81 | 0.69 |
| 1 | 180177266 | rs7521602 | -0.763 | 0.23 | -2.237 | 0.599 | -0.084 | 0.075 | 0.075 | 0.384 | 0.33 | 0.11 | 0.18 | 0.17 | 0.35 | 0.46 |
| 1 | 180178695 | rs10913949 | -0.042 | 1.198 | -1.837 | 1.021 | -0.028 | -0.058 | -0.058 | 0.016 | 0.22 | 0.1 | 0.32 | 0.3 | 0.27 | 0.17 |
| 1 | 180186713 | rs6697992 | -1.033 | 1.031 | -1.239 | 0.838 | 0 | -0.033 | -0.033 | 0 | 0.21 | 0.18 | 0.24 | 0.34 | 0.22 | 0.25 |
| 1 | 180194443 | rs355623 | 0.713 | 1.849 | -0.905 | 0.446 | 0.007 | -0.001 | -0.001 | 0.316 | 0.91 | 0.89 | 0.93 | 0.86 | 0.83 | 0.92 |
| 1 | 180194498 | rs355622 | 1.235 | 1.001 | -0.595 | 0.01 | -0.01 | 0.013 | 0.013 | 0 | 0.57 | 0.33 | 0.7 | 0.68 | 0.57 | 0.43 |
| 1 | 180201104 | rs355627 | -1.619 | 0.685 | -0.371 | -0.186 | -0.004 | 0.052 | 0.052 | 0 | 0.5 | 0.86 | 0.35 | 0.37 | 0.48 | 0.62 |
| 1 | 180201184 | rs6425616 | -1.553 | 0.621 | 0.28 | -0.296 | 0.036 | -0.039 | -0.039 | 0 | 0.34 | 0.45 | 0.29 | 0.2 | 0.44 | 0.41 |
| 1 | 180206814 | rs7525475 | -0.211 | 0.437 | 0.48 | -0.157 | -0.053 | 0.028 | 0.028 | 0 | 0.17 | 0.34 | 0.05 | 0.18 | 0.06 | 0.21 |

| | | | | | | | | | | | | | | | | |
|---|---|---|---|---|---|---|---|---|---|---|---|---|---|---|---|---|
| 1 | 180214469 | rs3911335 | -1.448 | 0.848 | -0.124 | -0.343 | 0.016 | -0.021 | -0.021 | 0.344 | 0.32 | 0.38 | 0.45 | 0.19 | 0.48 | 0.41 |
| 1 | 180217250 | rs4652490 | 0.108 | 1.101 | -0.042 | -0.298 | 0.068 | 0.017 | 0.017 | 0 | 0.11 | 0.2 | 0.1 | 0.18 | 0.21 | 0.09 |
| 1 | 180218442 | rs4652492 | -1.321 | 1.011 | -0.068 | -0.355 | -0.003 | -0.006 | -0.006 | 0.234 | 0.47 | 0.48 | 0.48 | 0.37 | 0.52 | 0.62 |
| 1 | 180219900 | rs7539244 | -1.007 | 1.109 | -0.535 | -0.293 | -0.006 | 0.001 | 0.001 | 0 | 0.39 | 0.35 | 0.4 | 0.36 | 0.41 | 0.54 |
| 1 | 180221084 | rs357069 | -0.682 | 1.076 | -1.301 | 0.098 | -0.003 | 0.003 | 0.003 | 0 | 0.38 | 0.72 | 0.24 | 0.37 | 0.33 | 0.39 |
| 1 | 180225223 | rs357065 | -0.684 | 0.834 | -2.052 | 0.25 | -0.059 | -0.116 | -0.116 | 0.046 | 0.63 | 0.97 | 0.58 | 0.42 | 0.6 | 0.75 |
| 1 | 180225664 | rs99174 | 0.01 | 0.739 | -1.597 | 0.467 | -0.011 | -0.008 | -0.008 | 0.315 | 0.57 | 0.73 | 0.4 | 0.44 | 0.54 | 0.58 |
| 1 | 180225837 | rs357064 | -0.68 | 0.45 | -1.361 | 0.742 | 0.018 | -0.036 | -0.036 | 0 | 0.3 | 0.2 | 0.28 | 0.55 | 0.26 | 0.43 |
| 1 | 180226390 | rs357062 | -1.435 | 1.116 | -0.257 | 0.247 | 0.059 | -0.005 | -0.005 | 0 | 0.2 | 0.48 | 0.32 | 0.21 | 0.22 | 0.24 |
| 1 | 180226900 | rs6661125 | -1.467 | 0.567 | -0.135 | -0.362 | -0.006 | -0.146 | -0.146 | 1.812 | 0.22 | 0.56 | 0.13 | 0.01 | 0.18 | 0.17 |
| 1 | 180227658 | rs357058 | -1.397 | 0.4 | -0.001 | 0.341 | -0.022 | 0.036 | 0.036 | 0.425 | 0.17 | 0.67 | 0.11 | 0.25 | 0.11 | 0.1 |
| 1 | 180230081 | rs704336 | -1.106 | 0.618 | 0.288 | -0.159 | 0.025 | -0.043 | -0.043 | 0 | 0.09 | 0.13 | 0.07 | 0.02 | 0.12 | 0.07 |
| 1 | 180232750 | rs7520941 | -0.631 | 1.238 | 0.914 | -0.524 | 0.035 | -0.02 | -0.02 | 0 | 0.11 | 0.23 | 0.18 | 0.05 | 0.17 | 0.06 |
| 1 | 180232890 | rs7528949 | 1.252 | 0.813 | 0.465 | -1.483 | -0.006 | 0.015 | 0.015 | 0 | 0.56 | 0.63 | 0.67 | 0.89 | 0.55 | 0.57 |
| 1 | 180233602 | rs357052 | 1.041 | 0.536 | 0.348 | -1.613 | 0.021 | 0.021 | 0.021 | 0.368 | 0.14 | 0.08 | 0.05 | 0 | 0.07 | 0.03 |
| 1 | 180233647 | rs357051 | -0.491 | 0.524 | 0.34 | -1.623 | -0.004 | -0.06 | -0.06 | 0 | 0.24 | 0.23 | 0.12 | 0.02 | 0.2 | 0.17 |
| 1 | 180234458 | rs3911332 | 0.008 | 0.547 | 0.397 | -1.616 | 0.025 | -0.018 | -0.018 | 1.104 | 0.11 | 0.23 | 0.17 | 0.06 | 0.17 | 0.06 |
| 1 | 180234642 | rs2764448 | 0.332 | 0.529 | 0.352 | -1.511 | -0.033 | 0.026 | 0.026 | 0 | 0.07 | 0.01 | 0.04 | 0.02 | 0.11 | 0.12 |
| 1 | 180234729 | rs357050 | -0.979 | 0.782 | 0.345 | -1.554 | -0.002 | 0.014 | 0.014 | 0 | 0.32 | 0.22 | 0.2 | 0.04 | 0.3 | 0.35 |
| 1 | 180236235 | rs2794961 | 0.437 | 0.4 | 0.847 | -1.562 | 0.006 | 0.022 | 0.022 | 0 | 0.69 | 0.78 | 0.82 | 0.97 | 0.77 | 0.67 |
| 1 | 180238442 | rs17372142 | 0.396 | 0.562 | 0.998 | -1.429 | 0.009 | -0.003 | -0.003 | 0 | 0.07 | 0.12 | 0.1 | 0.06 | 0.09 | 0.02 |
| 1 | 180238495 | rs2290478 | 1.756 | 0.036 | 0.14 | -1.538 | -0.126 | -0.093 | -0.093 | 0 | 0.32 | 0.03 | 0.25 | 0.55 | 0.43 | 0.34 |
| 1 | 180239473 | rs357044 | 0.87 | 0.266 | 1.217 | -1.035 | 0.001 | 0.122 | 0.122 | 0 | 0.9 | 0.68 | 0.82 | 0.67 | 0.92 | 0.82 |
| 1 | 180239670 | rs119465 | -0.13 | 0.251 | 1.13 | -1.067 | 0.017 | -0.031 | -0.031 | 0 | 0.07 | 0.15 | 0.05 | 0.01 | 0.09 | 0.07 |
| 1 | 180240752 | rs3845397 | 1.665 | 0.005 | 1.141 | -1.121 | -0.046 | 0.066 | 0.066 | 0 | 0.51 | 0.1 | 0.31 | 0.56 | 0.53 | 0.6 |
| 1 | 180243524 | rs7536561 | 1.671 | -0.331 | 1.213 | -1.276 | 0 | 0.108 | 0.108 | 0 | 0.51 | 0.12 | 0.33 | 0.59 | 0.55 | 0.62 |
| 1 | 180246261 | rs6680361 | 1.253 | -0.524 | 1.567 | -1.094 | 0.005 | -0.065 | -0.065 | 0 | 0.86 | 0.66 | 0.88 | 0.97 | 0.83 | 0.8 |
| 1 | 180249395 | rs2764454 | -0.429 | -0.527 | 1.595 | -1.029 | -0.007 | -0.033 | -0.033 | 0 | 0.07 | 0.21 | 0.05 | 0.02 | 0.07 | 0.08 |
| 1 | 180254865 | rs2794968 | -0.817 | -0.54 | 1.829 | -1.035 | 0.005 | -0.03 | -0.03 | 0 | 0.08 | 0.32 | 0.06 | 0.02 | 0.07 | 0.08 |
| 1 | 180268476 | rs2296378 | 1.413 | -0.562 | 1.795 | -0.971 | 0.017 | 0.037 | 0.037 | 0 | 0.56 | 0.16 | 0.48 | 0.56 | 0.64 | 0.66 |
| 1 | 180310005 | rs6658310 | 0.284 | -0.541 | 1.747 | -0.931 | -0.003 | -0.007 | -0.007 | 0 | 0.89 | 0.86 | 0.88 | 0.93 | 0.88 | 0.91 |
| 1 | 180317247 | rs4259605 | -0.8 | -0.554 | 1.686 | -0.929 | 0 | -0.002 | -0.002 | 0 | 0.95 | 0.96 | 0.97 | 0.99 | 0.93 | 0.93 |
| 1 | 180321192 | rs6425622 | -0.8 | -0.554 | 1.686 | -0.929 | 0 | -0.002 | -0.002 | 0 | 0.95 | 0.96 | 0.97 | 0.99 | 0.93 | 0.93 |
| 1 | 180323689 | rs17372362 | 0.889 | -0.577 | 1.66 | -0.982 | -0.021 | 0.062 | 0.062 | 0 | 0.19 | 0.07 | 0.12 | 0 | 0.17 | 0.09 |
| 1 | 180332045 | rs6425625 | 1.757 | -0.576 | 1.802 | -0.94 | -0.038 | 0.069 | 0.069 | 0 | 0.81 | 0.4 | 0.73 | 0.61 | 0.87 | 0.85 |
| 1 | 180335264 | rs4333819 | 0.462 | -0.559 | 1.822 | -0.949 | -0.001 | -0.033 | -0.033 | 0.018 | 0.92 | 0.73 | 0.94 | 0.98 | 0.93 | 0.92 |
| 1 | 180340706 | rs1418102 | 0.849 | -0.542 | 1.778 | -0.939 | -0.008 | 0.097 | 0.097 | 0 | 0.86 | 0.73 | 0.76 | 0.61 | 0.87 | 0.85 |
| 1 | 180356025 | rs4642852 | 0.535 | -0.543 | 1.747 | -0.96 | -0.002 | -0.001 | -0.001 | 0 | 0.67 | 0.62 | 0.64 | 0.61 | 0.7 | 0.76 |
| 1 | 180369966 | rs1361628 | -0.845 | -0.522 | 1.641 | -0.946 | -0.009 | 0.021 | 0.021 | 0 | 0.95 | 1 | 0.97 | 0.99 | 0.93 | 0.93 |
| 1 | 180381923 | rs7532806 | 1.432 | -0.564 | 1.75 | -0.972 | 0.011 | 0.042 | 0.042 | 0.046 | 0.55 | 0.1 | 0.47 | 0.56 | 0.64 | 0.66 |
| 1 | 180393514 | rs6425635 | -1.084 | -0.522 | 1.775 | -0.943 | -0.001 | -0.03 | -0.03 | 0 | 0.09 | 0.27 | 0.14 | 0.05 | 0.06 | 0.1 |
| 1 | 180437525 | rs6670950 | -1.169 | -0.467 | 1.508 | -0.934 | 0 | 0.082 | 0.082 | 0 | 0.09 | 0.19 | 0.24 | 0.37 | 0.09 | 0.12 |
| 1 | 180445616 | rs10914005 | 1.244 | -0.433 | 1.449 | -0.889 | 0.004 | 0.006 | 0.006 | 0 | 0.58 | 0.35 | 0.5 | 0.56 | 0.64 | 0.67 |
| 1 | 180448546 | rs10914007 | -0.528 | -0.35 | 1.498 | -0.637 | -0.004 | 0.164 | 0.164 | 0 | 0.09 | 0.23 | 0.2 | 0.38 | 0.06 | 0.08 |
| 1 | 180462683 | rs12093073 | -1.101 | -0.264 | 1.137 | -0.68 | 0.001 | -0.03 | -0.03 | 1.529 | 0.09 | 0.27 | 0.14 | 0.05 | 0.06 | 0.1 |
| 1 | 180472298 | rs3806301 | -1.428 | -0.279 | 1.163 | -0.608 | -0.01 | 0.092 | 0.092 | 0.05 | 0.17 | 0.37 | 0.33 | 0.43 | 0.12 | 0.18 |
| 1 | 180472370 | rs3806300 | -0.904 | -0.261 | 1.13 | -0.601 | 0 | 0.036 | 0.036 | 0 | 0.13 | 0.17 | 0.23 | 0.39 | 0.12 | 0.17 |
| 1 | 180472511 | rs17300712 | 0.839 | -0.301 | 1.112 | -0.879 | -0.009 | 0.016 | 0.016 | 0 | 0.19 | 0.11 | 0.13 | 0 | 0.17 | 0.09 |
| 1 | 180491338 | rs7515052 | -0.4 | -0.365 | 1.243 | -0.935 | -0.005 | -0.022 | -0.022 | 0 | 0.94 | 0.91 | 0.96 | 0.99 | 0.93 | 0.93 |
| 1 | 180493944 | rs16856086 | -0.705 | -0.365 | 1.28 | -0.879 | 0.001 | 0.044 | 0.044 | 0 | 0.13 | 0.17 | 0.23 | 0.4 | 0.12 | 0.18 |
| 1 | 180494062 | rs4651067 | -0.258 | -0.368 | 1.187 | -0.911 | -0.002 | -0.018 | -0.018 | 0 | 0.75 | 0.8 | 0.83 | 0.99 | 0.76 | 0.84 |
| 1 | 180495670 | rs2331739 | -0.302 | -0.336 | 1.203 | -0.865 | -0.006 | -0.025 | -0.025 | 0 | 0.94 | 0.89 | 0.96 | 0.99 | 0.93 | 0.93 |
| 1 | 180496261 | rs11806607 | 0.974 | -0.564 | 0.487 | -1.38 | -0.007 | 0.081 | 0.081 | 0 | 0.15 | 0.01 | 0.06 | 0 | 0.13 | 0.1 |
| 1 | 180496582 | rs1416823 | -0.895 | -0.51 | 1.458 | -0.782 | 0.003 | 0.001 | 0.001 | 0 | 0.33 | 0.46 | 0.37 | 0.4 | 0.29 | 0.27 |
| 1 | 180501196 | rs12039331 | -0.602 | -0.509 | 1.322 | -0.731 | -0.003 | 0.005 | 0.005 | 0 | 0.3 | 0.41 | 0.35 | 0.37 | 0.24 | 0.18 |
| 1 | 180506038 | rs7543366 | 1.552 | -0.549 | 1.209 | -0.752 | 0.001 | 0.025 | 0.025 | 0.155 | 0.55 | 0.1 | 0.46 | 0.56 | 0.63 | 0.66 |
| 1 | 180510172 | rs6425646 | 0.129 | -0.491 | 1.441 | -0.719 | 0.001 | 0.013 | 0.013 | 0 | 0.23 | 0.42 | 0.27 | 0.3 | 0.18 | 0.15 |
| 1 | 180512764 | rs1538622 | 1.287 | -0.407 | 1.311 | -0.623 | 0.003 | 0.015 | 0.015 | 0 | 0.58 | 0.26 | 0.55 | 0.58 | 0.63 | 0.66 |
| 1 | 180513655 | rs4652507 | 1.206 | -0.389 | 1.187 | -0.617 | -0.001 | -0.035 | -0.035 | 0 | 0.81 | 0.72 | 0.82 | 0.92 | 0.82 | 0.81 |
| 1 | 180516881 | rs10798774 | -0.712 | -0.517 | 0.924 | -0.369 | -0.007 | 0.017 | 0.017 | 0 | 0.31 | 0.51 | 0.35 | 0.4 | 0.24 | 0.17 |
| 1 | 180521281 | rs7552084 | -0.476 | -0.425 | 0.324 | -0.325 | -0.003 | -0.019 | -0.019 | 0.217 | 0.28 | 0.2 | 0.32 | 0.4 | 0.24 | 0.16 |
| 1 | 180523777 | rs7513944 | -0.518 | -0.415 | 0.343 | -0.328 | -0.001 | -0.012 | -0.012 | 0 | 0.28 | 0.23 | 0.33 | 0.4 | 0.24 | 0.16 |
| 1 | 180530439 | rs3845398 | 0.562 | -0.443 | 0.507 | -0.135 | 0.002 | 0.002 | 0.002 | 0 | 0.87 | 0.85 | 0.89 | 0.94 | 0.91 | 0.97 |
| 1 | 180534407 | rs12562357 | -1.715 | -0.336 | 0.996 | 0.041 | -0.014 | 0.167 | 0.167 | 0.936 | 0.14 | 0.46 | 0.21 | 0.35 | 0.1 | 0.15 |
| 1 | 180535972 | rs2331545 | 1.476 | -0.399 | 0.008 | 0.356 | -0.178 | -0.004 | -0.004 | 0 | 0.38 | 0.02 | 0.26 | 0.26 | 0.46 | 0.37 |
| 1 | 180536954 | rs10798778 | 1.035 | -0.489 | 0.345 | 0.314 | -0.011 | 0.062 | 0.062 | 0 | 0.44 | 0.17 | 0.3 | 0.28 | 0.53 | 0.48 |
| 1 | 180540896 | rs10733006 | -0.517 | -0.241 | 0.225 | 0 | -0.015 | 0.09 | 0.09 | 0 | 0.89 | 0.78 | 0.92 | 0.72 | 0.93 | 0.97 |
| 1 | 180541363 | rs12030731 | -0.517 | -0.241 | 0.225 | 0 | -0.015 | 0.09 | 0.09 | 0 | 0.89 | 0.78 | 0.92 | 0.72 | 0.93 | 0.97 |
| 1 | 180546296 | rs10798780 | 0.452 | -0.309 | 0.362 | -0.051 | 0.01 | 0.007 | 0.007 | 0 | 0.55 | 0.59 | 0.48 | 0.64 | 0.58 | 0.6 |

| | | | | | | | | | | | | | | | |
|---|---|---|---|---|---|---|---|---|---|---|---|---|---|---|---|
| 1 | 180546563 | rs10798781 | 0.443 | -0.284 | -0.013 | -0.04 | 0.013 | 0.007 | 0.007 | 0 | 0.56 | 0.7 | 0.49 | 0.64 | 0.58 | 0.6 |
| 1 | 180549164 | rs4652519 | 0.515 | -0.328 | -0.046 | -0.071 | -0.011 | 0.118 | 0.118 | 0 | 0.3 | 0.1 | 0.39 | 0.08 | 0.29 | 0.31 |
| 1 | 180551151 | rs12735662 | 0.027 | -0.144 | -0.004 | 0.018 | -0.018 | 0.025 | 0.025 | 0 | 0.07 | 0.04 | 0.08 | 0.01 | 0.13 | 0.1 |
| 1 | 180552260 | rs10737346 | -0.511 | -0.637 | 0.088 | 0.196 | -0.013 | 0.054 | 0.054 | 0 | 0.89 | 0.82 | 0.92 | 0.72 | 0.94 | 0.97 |
| 1 | 180560692 | rs7519155 | 0.943 | -0.191 | 0.377 | 0.143 | -0.004 | -0.003 | -0.003 | 0 | 0.43 | 0.3 | 0.41 | 0.61 | 0.52 | 0.51 |
| 1 | 180561644 | rs10737348 | -0.425 | -0.256 | 0.255 | -0.032 | 0.042 | -0.004 | -0.004 | 0 | 0.62 | 0.7 | 0.69 | 0.66 | 0.76 | 0.71 |
| 1 | 180562366 | rs10494531 | -0.286 | -0.021 | -0.06 | 0.034 | -0.019 | 0.029 | 0.029 | 0 | 0.09 | 0.04 | 0.2 | 0.01 | 0.15 | 0.1 |
| 1 | 180564513 | rs12078544 | -0.85 | -0.464 | 0.179 | 0.334 | -0.001 | -0.003 | -0.003 | 0 | 0.08 | 0.24 | 0.13 | 0.05 | 0.08 | 0.1 |
| 1 | 180567428 | rs10798783 | 0.679 | -0.502 | -0.102 | 0.373 | 0.051 | 0.013 | 0.013 | 0 | 0.5 | 0.36 | 0.53 | 0.61 | 0.66 | 0.61 |
| 1 | 180567596 | rs16856269 | -1.126 | 0.153 | 0.245 | 0.214 | -0.006 | -0.015 | -0.015 | 0 | 0.11 | 0.27 | 0.15 | 0.05 | 0.1 | 0.1 |
| 1 | 180570408 | rs3843276 | -0.118 | 0.743 | -0.237 | -0.064 | 0.008 | -0.01 | -0.01 | 0.051 | 0.32 | 0.34 | 0.33 | 0.19 | 0.43 | 0.39 |
| 1 | 180575168 | rs9970363 | -0.552 | 0.531 | -0.209 | 0.581 | 0.002 | 0.002 | 0.002 | 0 | 0.41 | 0.44 | 0.4 | 0.45 | 0.5 | 0.46 |
| 1 | 180576443 | rs12038863 | 0.231 | 0.036 | -0.293 | 0.353 | -0.004 | 0.004 | 0.004 | 0 | 0.15 | 0.18 | 0.15 | 0.18 | 0.1 | 0.19 |
| 1 | 180587744 | rs16856306 | 0.098 | -0.01 | -0.316 | 0.343 | -0.004 | 0.003 | 0.003 | 0 | 0.16 | 0.18 | 0.16 | 0.19 | 0.1 | 0.2 |
| 1 | 180591237 | rs3908502 | -1.923 | -0.515 | -0.893 | -0.309 | 0.019 | 0.095 | 0.095 | 0 | 0.8 | 0.95 | 0.88 | 1 | 0.85 | 0.91 |
| 1 | 180596196 | rs3842895 | 0.028 | -0.017 | -0.426 | -0.036 | 0.021 | -0.045 | -0.045 | 0 | 0.37 | 0.53 | 0.39 | 0.17 | 0.45 | 0.41 |
| 1 | 180600599 | rs4651073 | -0.299 | -0.153 | -0.21 | -0.105 | 0.03 | -0.005 | -0.005 | 0.012 | 0.52 | 0.31 | 0.54 | 0.44 | 0.62 | 0.56 |
| 1 | 180657965 | rs7556184 | -0.552 | -0.136 | 0.437 | -0.094 | 0.016 | -0.07 | -0.07 | 0 | 0.13 | 0.35 | 0.1 | 0.01 | 0.11 | 0.08 |
| 1 | 180672792 | rs10914074 | 1.214 | -0.02 | 0.457 | -0.213 | -0.015 | 0.011 | 0.011 | 0 | 0.26 | 0.19 | 0.31 | 0.16 | 0.39 | 0.35 |
| 1 | 180685921 | rs10914077 | 0.936 | -0.14 | 0.666 | -0.295 | 0.002 | -0.002 | -0.002 | 0 | 0.27 | 0.24 | 0.33 | 0.16 | 0.4 | 0.35 |
| 1 | 180690420 | rs10494532 | 1.242 | -0.059 | 0.695 | -0.259 | -0.026 | 0.018 | 0.018 | 0 | 0.25 | 0.16 | 0.31 | 0.15 | 0.39 | 0.35 |
| 1 | 180692815 | rs12408176 | -0.029 | -0.247 | 0.903 | -0.156 | -0.005 | 0.006 | 0.006 | 0 | 0.27 | 0.31 | 0.25 | 0.28 | 0.24 | 0.22 |
| 1 | 180700133 | rs10914083 | -0.162 | -0.26 | 0.929 | -0.185 | 0.021 | -0.046 | -0.046 | 0 | 0.38 | 0.51 | 0.4 | 0.16 | 0.5 | 0.42 |
| 1 | 180706924 | rs12401315 | 1.795 | -0.308 | 0.777 | -0.249 | -0.002 | 0.033 | 0.033 | 0 | 0.13 | 0.04 | 0.17 | 0 | 0.15 | 0.11 |
| 1 | 180721279 | rs7550031 | -2.28 | -0.426 | 0.824 | -0.265 | 0.01 | 0.119 | 0.119 | 0 | 0.72 | 0.93 | 0.85 | 0.95 | 0.77 | 0.79 |
| 1 | 180722353 | rs6700926 | -0.152 | -0.414 | 0.922 | -0.255 | -0.016 | -0.012 | -0.012 | 0.746 | 0.4 | 0.66 | 0.49 | 0.53 | 0.5 | 0.48 |
| 1 | 180736428 | rs16856420 | 1.397 | -0.417 | 0.747 | -0.262 | -0.007 | -0.013 | -0.013 | 0.381 | 0.07 | 0.09 | 0.03 | 0 | 0.03 | 0.05 |
| 1 | 180742308 | rs1339768 | -1.403 | -0.301 | 0.707 | -0.182 | 0.03 | -0.07 | -0.07 | 0 | 0.65 | 0.84 | 0.65 | 0.44 | 0.74 | 0.64 |
| 1 | 180757056 | rs3761904 | 1.297 | -0.172 | 0.847 | -0.378 | -0.026 | 0.015 | 0.015 | 0 | 0.26 | 0.16 | 0.31 | 0.17 | 0.39 | 0.35 |
| 1 | 180760180 | rs6661417 | 0.768 | -0.188 | 0.916 | -0.401 | 0.036 | -0.047 | -0.047 | 0.474 | 0.29 | 0.43 | 0.35 | 0.15 | 0.39 | 0.35 |
| 1 | 180761873 | rs1554185 | 1.038 | -0.18 | 0.876 | -0.397 | -0.001 | -0.001 | -0.001 | 0 | 0.27 | 0.24 | 0.32 | 0.15 | 0.39 | 0.35 |
| 1 | 180770320 | rs12047845 | 1.261 | -0.198 | 0.868 | -0.358 | -0.002 | 0.023 | 0.023 | 0 | 0.05 | 0.09 | 0.11 | 0.14 | 0.02 | 0.09 |
| 1 | 180781318 | rs12120511 | 1.302 | -0.151 | 0.589 | -0.403 | -0.026 | 0.018 | 0.018 | 0 | 0.26 | 0.16 | 0.31 | 0.15 | 0.39 | 0.35 |
| 1 | 180784878 | rs2944259 | 2.013 | -0.206 | 0.528 | -0.439 | 0.01 | 0.117 | 0.117 | 0 | 0.28 | 0.07 | 0.16 | 0.05 | 0.24 | 0.21 |
| 1 | 180820794 | rs7536258 | -0.204 | 0.431 | 0.995 | 0.095 | 0.029 | -0.047 | -0.047 | 0 | 0.34 | 0.43 | 0.38 | 0.16 | 0.47 | 0.41 |
| 1 | 180833665 | rs3856060 | 0.454 | 0.421 | 0.81 | 0.111 | -0.002 | 0 | 0 | 0 | 0.28 | 0.25 | 0.32 | 0.15 | 0.39 | 0.33 |
| 1 | 180849081 | rs2271668 | 0.58 | 0.745 | 1.04 | 0.438 | 0.036 | 0.166 | 0.166 | 0 | 0.07 | 0.17 | 0.1 | 0.32 | 0.11 | 0.07 |
| 1 | 180849494 | rs10914123 | 0.203 | 0.74 | 1.051 | -0.403 | 0.014 | -0.027 | -0.027 | 0 | 0.63 | 0.57 | 0.63 | 0.84 | 0.51 | 0.58 |
| 1 | 180852364 | rs4550006 | 0.374 | 0.658 | 0.636 | -0.363 | -0.001 | -0.001 | -0.001 | 0 | 0.28 | 0.25 | 0.32 | 0.16 | 0.38 | 0.35 |
| 1 | 180853719 | rs1061015 | 0.726 | 0.708 | -0.166 | -0.36 | -0.022 | 0.02 | 0.02 | 0 | 0.26 | 0.16 | 0.31 | 0.16 | 0.38 | 0.35 |
| 1 | 180868330 | rs12562489 | 0.495 | 0.626 | -0.574 | -0.277 | -0.022 | 0.066 | 0.066 | 0 | 0.55 | 0.23 | 0.48 | 0.7 | 0.48 | 0.49 |
| 1 | 180873153 | rs17374151 | 0.847 | 0.79 | -0.658 | -0.299 | 0.003 | -0.024 | -0.024 | 0.062 | 0.06 | 0.07 | 0.03 | 0 | 0.08 | 0.07 |
| 1 | 180875730 | rs12118635 | 0.604 | 1.063 | -0.882 | -0.341 | -0.007 | -0.023 | -0.023 | 0 | 0.13 | 0.04 | 0.13 | 0.15 | 0.16 | 0.31 |
| 1 | 180877409 | rs7536392 | -0.235 | 0.959 | -0.488 | -0.737 | 0 | -0.047 | -0.047 | 0.134 | 0.47 | 0.52 | 0.62 | 0.94 | 0.48 | 0.56 |
| 1 | 180881148 | rs6672415 | 0.297 | 0.87 | -0.702 | -0.707 | 0.006 | 0.016 | 0.016 | 0 | 0.44 | 0.34 | 0.34 | 0.05 | 0.42 | 0.37 |
| 1 | 180887289 | rs7551076 | 0.178 | 0.954 | -1.362 | -0.048 | -0.031 | -0.002 | -0.002 | 0 | 0.17 | 0.08 | 0.21 | 0.15 | 0.21 | 0.2 |
| 1 | 180888910 | rs12034554 | 0.259 | 0.897 | -0.876 | 0.308 | 0.007 | 0.058 | 0.058 | 0 | 0.08 | 0.23 | 0.16 | 0.14 | 0.08 | 0.07 |
| 1 | 180890173 | rs7515553 | -0.404 | 0.942 | -1.716 | 0.518 | -0.005 | -0.027 | -0.027 | 0 | 0.53 | 0.19 | 0.51 | 0.53 | 0.51 | 0.65 |
| 1 | 180893135 | rs12735084 | 0.284 | 0.739 | -1.723 | 0.291 | 0.001 | 0.009 | 0.009 | 0 | 0.35 | 0.14 | 0.29 | 0.33 | 0.29 | 0.46 |
| 1 | 180894304 | rs2331901 | 0.228 | 0.76 | -2.023 | 0.167 | -0.009 | -0.082 | -0.082 | 0 | 0.23 | 0.11 | 0.26 | 0.46 | 0.26 | 0.24 |
| 1 | 180895678 | rs12128456 | 0.779 | 0.758 | -1.718 | -0.438 | -0.043 | 0.13 | 0.13 | 0.171 | 0.3 | 0.12 | 0.23 | 0.06 | 0.35 | 0.24 |
| 1 | 180897569 | rs3747958 | -0.296 | 0.693 | -1.635 | -0.001 | 0.007 | -0.007 | -0.007 | 1.503 | 0.4 | 0.27 | 0.41 | 0.55 | 0.41 | 0.35 |
| 1 | 180905352 | rs3795505 | 0.151 | 0.567 | -1.719 | -0.534 | -0.046 | 0.126 | 0.126 | 0 | 0.39 | 0.15 | 0.4 | 0.26 | 0.42 | 0.51 |
| 1 | 180905448 | rs3795504 | -0.004 | 0.606 | -1.377 | -0.431 | -0.005 | 0.063 | 0.063 | 0 | 0.46 | 0.35 | 0.43 | 0.27 | 0.45 | 0.53 |
| 1 | 180915596 | rs4652539 | 0.418 | 0.966 | -0.906 | 0.095 | 0.009 | 0.022 | 0.022 | 0.005 | 0.52 | 0.32 | 0.52 | 0.6 | 0.56 | 0.48 |
| 1 | 180916130 | rs2331996 | -0.395 | 1.032 | -0.732 | 0.024 | -0.005 | 0.014 | 0.014 | 0.344 | 0.38 | 0.66 | 0.42 | 0.25 | 0.37 | 0.19 |
| 1 | 180919151 | rs2297184 | 0.748 | 0.74 | -0.635 | 0.004 | -0.031 | 0.004 | 0.004 | 0 | 0.34 | 0.43 | 0.31 | 0.35 | 0.32 | 0.28 |
| 1 | 180919255 | rs2297185 | 0.484 | 0.825 | -0.85 | -0.144 | -0.046 | 0.084 | 0.084 | 1.706 | 0.28 | 0.1 | 0.32 | 0.09 | 0.29 | 0.17 |
| 1 | 180920226 | rs10753229 | 1.866 | 0.664 | -0.958 | -0.12 | 0.001 | 0.02 | 0.02 | 0 | 0.24 | 0.01 | 0.22 | 0.17 | 0.25 | 0.24 |
| 1 | 180931508 | rs3737062 | 0.298 | 0.77 | -0.43 | -0.005 | 0 | 0.007 | 0.007 | 0.035 | 0.51 | 0.42 | 0.52 | 0.49 | 0.54 | 0.68 |
| 1 | 180932175 | rs6691233 | -1.102 | 0.921 | -0.973 | -0.022 | -0.009 | 0.057 | 0.057 | 3.812 | 0.77 | 0.9 | 0.81 | 0.91 | 0.8 | 0.88 |
| 1 | 180934047 | rs12022368 | -0.836 | 0.894 | -0.914 | 0.012 | -0.001 | 0.05 | 0.05 | 0 | 0.75 | 0.66 | 0.73 | 0.58 | 0.75 | 0.8 |
| 1 | 180934163 | rs4651080 | 1.42 | 0.821 | -1.469 | -0.449 | -0.019 | 0.057 | 0.057 | 0 | 0.3 | 0.16 | 0.36 | 0.18 | 0.34 | 0.29 |
| 1 | 180936062 | rs10429833 | -1.194 | 0.974 | -1.425 | 0.098 | 0.003 | -0.003 | -0.003 | 0 | 0.86 | 0.91 | 0.85 | 0.86 | 0.86 | 0.87 |
| 1 | 180938117 | rs2002041 | -0.072 | 1.021 | -1.226 | 0.098 | -0.034 | 0.084 | 0.084 | 3.876 | 0.23 | 0.04 | 0.19 | 0.09 | 0.21 | 0.13 |
| 1 | 180940912 | rs12740112 | 0.32 | 0.956 | -0.724 | 0.399 | -0.001 | 0.015 | 0.015 | 0 | 0.14 | 0.09 | 0.07 | 0.06 | 0.11 | 0.11 |
| 1 | 180953853 | rs3747957 | 0.262 | 0.886 | -0.283 | 0.324 | 0 | 0.012 | 0.012 | 0 | 0.51 | 0.45 | 0.63 | 0.63 | 0.57 | 0.4 |
| 1 | 180956195 | rs7556005 | -0.71 | 0.825 | 0.166 | 0.307 | -0.001 | 0.019 | 0.019 | 0 | 0.67 | 0.76 | 0.72 | 0.69 | 0.68 | 0.5 |
| 1 | 180957962 | rs12754041 | 0.256 | 0.849 | 0.094 | 0.333 | 0 | 0.012 | 0.012 | 0 | 0.51 | 0.45 | 0.63 | 0.63 | 0.57 | 0.4 |

| | | | | | | | | | | | | | | | | |
|---|---|---|---|---|---|---|---|---|---|---|---|---|---|---|---|---|
| 1 | 180972918 | rs17374409 | 0.563 | 0.769 | 0.342 | 0.435 | 0.003 | 0.012 | 0.012 | 0 | 0.08 | 0.16 | 0.06 | 0.11 | 0.08 | 0.02 |
| 1 | 180975865 | rs10494536 | -0.835 | 0.825 | 0.036 | 0.569 | -0.007 | 0.015 | 0.015 | 0 | 0.67 | 0.82 | 0.73 | 0.7 | 0.68 | 0.52 |
| 1 | 180976091 | rs2331992 | 1.219 | 0.911 | -0.158 | 0.748 | -0.004 | 0.047 | 0.047 | 0.103 | 0.32 | 0.18 | 0.37 | 0.2 | 0.34 | 0.28 |
| 1 | 180994590 | rs2331902 | 1.354 | 0.773 | -0.237 | 0.779 | -0.025 | 0.027 | 0.027 | 0 | 0.13 | 0.01 | 0.05 | 0.04 | 0.11 | 0.11 |
| 1 | 180999351 | rs10159067 | -0.913 | 0.75 | -0.156 | 0.798 | 0.001 | 0.023 | 0.023 | 0.025 | 0.34 | 0.53 | 0.34 | 0.5 | 0.33 | 0.23 |
| 1 | 181002479 | rs3806284 | -0.564 | 0.668 | 0.211 | 1.235 | 0.008 | 0.183 | 0.183 | 0.862 | 0.21 | 0.48 | 0.28 | 0.43 | 0.21 | 0.13 |
| 1 | 181005644 | rs6692913 | 0.7 | 0.797 | 0.078 | 1.096 | 0.009 | 0.043 | 0.043 | 0.336 | 0.32 | 0.2 | 0.36 | 0.2 | 0.33 | 0.29 |
| 1 | 181012229 | rs12134601 | -1.048 | 0.78 | 0.679 | 1.707 | 0.01 | 0.189 | 0.189 | 0.311 | 0.2 | 0.45 | 0.27 | 0.43 | 0.21 | 0.12 |
| 1 | 181020317 | rs3789357 | -0.932 | 0.352 | 0.501 | 1.606 | -0.02 | 0.097 | 0.097 | 0 | 0.09 | 0.17 | 0.2 | 0.37 | 0.06 | 0.3 |
| 1 | 181026704 | rs4652553 | -0.836 | 0.681 | 0.928 | 0.702 | -0.005 | 0.016 | 0.016 | 0.403 | 0.59 | 0.64 | 0.72 | 0.89 | 0.57 | 0.79 |
| 1 | 181027559 | rs2282347 | -0.14 | 0.77 | 1.17 | 0.942 | 0.008 | 0.074 | 0.074 | 0 | 0.12 | 0.16 | 0.22 | 0.34 | 0.14 | 0.12 |
| 1 | 181027841 | rs6686208 | -1.254 | 0.887 | 1.234 | 0.808 | -0.03 | 0.135 | 0.135 | 0.305 | 0.62 | 0.84 | 0.76 | 0.91 | 0.59 | 0.81 |
| 1 | 181027925 | rs2282348 | 0.72 | 1.023 | 1.31 | 0.798 | -0.003 | -0.001 | -0.001 | 0 | 0.05 | 0.03 | 0.06 | 0.05 | 0.08 | 0.06 |
| 1 | 181031783 | rs3936939 | -0.788 | 0.918 | 1.322 | 0.767 | -0.039 | 0.139 | 0.139 | 0.027 | 0.56 | 0.77 | 0.69 | 0.86 | 0.51 | 0.74 |
| 1 | 181032155 | rs4652554 | 0.105 | 0.789 | 1.691 | 0.775 | 0.011 | -0.002 | -0.002 | 0.118 | 0.42 | 0.34 | 0.45 | 0.51 | 0.37 | 0.62 |
| 1 | 181033591 | rs12240122 | 0.49 | 0.832 | 1.648 | 0.262 | -0.003 | -0.001 | -0.001 | 0 | 0.05 | 0.11 | 0.04 | 0.05 | 0.05 | 0.06 |
| 1 | 181035754 | rs10797668 | -1.421 | 0.942 | 1.908 | 0.513 | 0.046 | 0.097 | 0.097 | 0 | 0.19 | 0.44 | 0.3 | 0.3 | 0.22 | 0.2 |
| 1 | 181039039 | rs12144382 | -1.51 | 0.921 | 1.979 | 0.516 | 0.053 | 0.113 | 0.113 | 0.008 | 0.19 | 0.47 | 0.31 | 0.3 | 0.22 | 0.2 |
| 1 | 181039628 | rs2332059 | -1.475 | 0.909 | 1.952 | 0.402 | -0.038 | 0.154 | 0.154 | 0.774 | 0.63 | 0.88 | 0.76 | 0.91 | 0.61 | 0.81 |
| 1 | 181045421 | rs10910862 | -0.6 | 0.939 | 2.114 | 0.395 | 0.058 | 0.034 | 0.034 | 0 | 0.07 | 0.33 | 0.09 | 0.05 | 0.07 | 0.06 |
| 1 | 181045443 | rs3856067 | -0.358 | 0.948 | 2.004 | 0.42 | 0.004 | 0.019 | 0.019 | 0 | 0.12 | 0.15 | 0.22 | 0.24 | 0.15 | 0.14 |
| 1 | 181050713 | rs4651082 | 0.249 | 0.977 | 1.994 | 0.355 | 0.018 | -0.014 | -0.014 | 0 | 0.4 | 0.25 | 0.45 | 0.6 | 0.38 | 0.61 |
| 1 | 181056160 | rs6662507 | 0.605 | 0.862 | 1.807 | 0.302 | -0.022 | -0.091 | -0.091 | 0 | 0.36 | 0.06 | 0.42 | 0.6 | 0.36 | 0.6 |
| 1 | 181060618 | rs4287133 | -0.954 | 0.827 | 2.121 | 0.073 | -0.002 | 0.027 | 0.027 | 0.346 | 0.58 | 0.65 | 0.71 | 0.9 | 0.59 | 0.82 |
| 1 | 181060791 | rs6658698 | -1.174 | 0.821 | 1.916 | 0.083 | -0.006 | 0.09 | 0.09 | 0.305 | 0.61 | 0.77 | 0.75 | 0.91 | 0.61 | 0.82 |
| 1 | 181079712 | rs2877547 | -0.918 | 1.123 | 1.999 | 0.266 | 0.002 | 0.032 | 0.032 | 0 | 0.12 | 0.25 | 0.21 | 0.21 | 0.1 | 0.12 |
| 1 | 181086687 | rs7541438 | 0.384 | 0.93 | 1.737 | 0.152 | 0.012 | -0.008 | -0.008 | 0 | 0.36 | 0.16 | 0.42 | 0.65 | 0.38 | 0.62 |
| 1 | 181095779 | rs12025764 | 2.416 | 1.088 | 2.369 | 0.454 | -0.005 | -0.002 | -0.002 | 0 | 0.76 | 0.2 | 0.67 | 0.77 | 0.78 | 0.83 |
| 1 | 181106211 | rs10910872 | 1.197 | 1.021 | 2.363 | 0.612 | 0.016 | 0.079 | 0.079 | 0 | 0.88 | 0.7 | 0.8 | 0.8 | 0.89 | 0.88 |
| 1 | 181108029 | rs10910875 | 1.697 | 1.019 | 1.915 | 0.59 | 0.017 | 0.069 | 0.069 | 0.255 | 0.83 | 0.55 | 0.76 | 0.79 | 0.87 | 0.86 |
| 1 | 181109478 | rs881964 | 1.441 | 1.102 | 1.605 | 0.635 | 0.022 | 0.071 | 0.071 | 0.059 | 0.85 | 0.63 | 0.78 | 0.79 | 0.87 | 0.87 |
| 1 | 181116023 | rs10752838 | -1.354 | 1.454 | 0.131 | 0.165 | -0.018 | 0.004 | 0.004 | 0.461 | 0.39 | 0.69 | 0.46 | 0.42 | 0.45 | 0.32 |
| 1 | 181119835 | rs6425668 | 0.733 | 1.431 | -0.111 | 0.156 | 0.002 | -0.001 | -0.001 | 0.161 | 0.65 | 0.3 | 0.65 | 0.59 | 0.66 | 0.77 |
| 1 | 181126503 | rs11582767 | 0.684 | 1.107 | 0.094 | -0.123 | -0.008 | -0.011 | -0.011 | 0 | 0.72 | 0.32 | 0.72 | 0.77 | 0.78 | 0.85 |
| 1 | 181128206 | rs4652572 | 0.827 | 1.291 | 0.023 | -0.031 | -0.001 | 0 | 0 | 0 | 0.94 | 0.93 | 0.9 | 0.99 | 0.9 | 0.9 |
| 1 | 181140881 | rs4651091 | 0.877 | 0.847 | 0.017 | 0.061 | 0.006 | -0.008 | -0.008 | 0 | 0.93 | 0.88 | 0.89 | 0.96 | 0.94 | 0.91 |
| 1 | 181142796 | rs2013836 | -1.494 | 0.536 | 0.119 | -0.01 | -0.003 | -0.029 | -0.029 | 0 | 0.07 | 0.24 | 0.05 | 0 | 0.04 | 0.01 |
| 1 | 181145529 | rs12566430 | 0.132 | 0.085 | -0.803 | -0.263 | 0.015 | 0.02 | 0.02 | 0 | 0.62 | 0.32 | 0.67 | 0.7 | 0.67 | 0.71 |
| 1 | 181147045 | rs10910888 | 0.169 | -0.02 | -0.775 | -0.48 | 0.007 | 0.012 | 0.012 | 0 | 0.62 | 0.33 | 0.66 | 0.7 | 0.67 | 0.63 |
| 1 | 181147914 | rs7523662 | 0.152 | 0.422 | -0.753 | -0.438 | 0 | -0.002 | -0.002 | 0 | 0.6 | 0.37 | 0.66 | 0.69 | 0.61 | 0.61 |
| 1 | 181150358 | rs6681270 | -0.312 | 0.643 | -0.632 | -0.18 | 0.002 | -0.005 | -0.005 | 0 | 0.31 | 0.36 | 0.28 | 0.27 | 0.34 | 0.28 |
| 1 | 181154835 | rs946646 | 0.453 | -0.157 | -0.123 | -1.306 | -0.003 | -0.015 | -0.015 | 0 | 0.67 | 0.33 | 0.75 | 0.78 | 0.73 | 0.78 |
| 1 | 181159459 | rs3827707 | 1.036 | 0.309 | -0.779 | -1.323 | -0.03 | -0.004 | -0.004 | 0 | 0.36 | 0.14 | 0.47 | 0.57 | 0.41 | 0.24 |
| 1 | 181160271 | rs3827706 | 0.059 | 0.265 | -0.492 | -1.415 | -0.002 | 0.003 | 0.003 | 0 | 0.3 | 0.29 | 0.29 | 0.22 | 0.31 | 0.5 |
| 1 | 181160639 | rs10910892 | -0.336 | 0.093 | 0.209 | -1.558 | 0.004 | -0.036 | -0.036 | 0.138 | 0.36 | 0.54 | 0.36 | 0.22 | 0.37 | 0.51 |
| 1 | 181160693 | rs7417573 | -0.051 | -0.003 | 0.931 | -1.817 | -0.005 | -0.014 | -0.014 | 0 | 0.75 | 0.72 | 0.82 | 0.76 | 0.78 | 0.79 |
| 1 | 181171768 | rs10157919 | -0.035 | 0.215 | 1.127 | -1.926 | -0.001 | 0.026 | 0.026 | 0 | 0.17 | 0.07 | 0.14 | 0.08 | 0.11 | 0.2 |
| 1 | 181173577 | rs6675445 | 0.188 | 0.281 | 1.815 | -1.823 | 0.056 | 0.008 | 0.008 | 0.179 | 0.94 | 0.71 | 0.94 | 1 | 0.96 | 0.97 |
| 1 | 181177854 | rs4652591 | 1.273 | 0.26 | 1.698 | -1.862 | 0.053 | -0.025 | -0.025 | 0 | 0.91 | 0.49 | 0.91 | 1 | 0.94 | 0.97 |
| 1 | 181179048 | rs1544174 | 1.34 | 0.214 | 1.734 | -1.852 | 0.025 | -0.028 | -0.028 | 0.1 | 0.91 | 0.44 | 0.92 | 1 | 0.95 | 0.97 |
| 1 | 181180236 | rs4651093 | 0.275 | 0.263 | 1.666 | -1.829 | 0.055 | 0.006 | 0.006 | 0.549 | 0.94 | 0.7 | 0.94 | 1 | 0.96 | 0.97 |
| 1 | 181181995 | rs10797688 | 0.857 | 0.518 | 1.527 | -2.111 | 0.001 | -0.066 | -0.066 | 0 | 0.69 | 0.58 | 0.72 | 0.9 | 0.71 | 0.7 |
| 1 | 181185861 | rs6425678 | 0.412 | 0.562 | 1.562 | -2.16 | 0.003 | -0.076 | -0.076 | 0 | 0.81 | 0.73 | 0.84 | 0.98 | 0.8 | 0.88 |
| 1 | 181199735 | rs10797698 | 0.162 | 0.561 | 1.617 | -2.161 | 0.002 | -0.05 | -0.05 | 0 | 0.82 | 0.79 | 0.85 | 0.98 | 0.8 | 0.88 |
| 1 | 181204194 | rs10797699 | 0.71 | 0.511 | 1.742 | -2.203 | -0.003 | -0.106 | -0.106 | 0 | 0.79 | 0.63 | 0.83 | 0.98 | 0.79 | 0.87 |
| 1 | 181205284 | rs4652618 | 0.521 | 0.513 | 1.666 | -2.202 | -0.001 | -0.082 | -0.082 | 0 | 0.8 | 0.69 | 0.84 | 0.98 | 0.79 | 0.87 |
| 1 | 181209210 | rs10910900 | -2.123 | 0.359 | 2.043 | -2.249 | 0.045 | 0.021 | 0.021 | 0.804 | 0.05 | 0.35 | 0.05 | 0 | 0.02 | 0.01 |
| 1 | 181210746 | rs3911582 | 0.485 | 0.396 | 2.034 | -2.233 | -0.003 | -0.082 | -0.082 | 0 | 0.81 | 0.69 | 0.85 | 0.98 | 0.81 | 0.88 |
| 1 | 181214224 | rs11582429 | 1.731 | 0.34 | 1.979 | -2.287 | 0.008 | 0.031 | 0.031 | 0.447 | 0.65 | 0.25 | 0.69 | 0.91 | 0.72 | 0.71 |
| 1 | 181225293 | rs10797708 | 0.244 | 0.441 | 1.809 | -2.286 | -0.001 | -0.06 | -0.06 | 0.008 | 0.81 | 0.75 | 0.86 | 0.98 | 0.81 | 0.88 |
| 1 | 181227982 | rs4652630 | 0.263 | 0.446 | 1.766 | -2.284 | -0.001 | -0.073 | -0.073 | 0 | 0.81 | 0.74 | 0.85 | 0.98 | 0.81 | 0.88 |
| 1 | 181240106 | rs12027236 | 2.118 | 0.621 | 1.084 | -2.751 | -0.011 | 0.021 | 0.021 | 0 | 0.35 | 0.24 | 0.52 | 0.71 | 0.43 | 0.48 |
| 1 | 181240950 | rs7544832 | 0.205 | 0.559 | 1.358 | -2.825 | -0.002 | -0.018 | -0.018 | 0.035 | 0.12 | 0.24 | 0.14 | 0.06 | 0.09 | 0.17 |
| 1 | 181242019 | rs1458844 | 0.243 | 0.433 | 1.384 | -2.91 | -0.014 | -0.12 | -0.12 | 1.851 | 0.77 | 0.6 | 0.83 | 0.98 | 0.8 | 0.87 |
| 1 | 181243896 | rs2501540 | -0.03 | 0.458 | 1.363 | -2.879 | -0.001 | 0.055 | 0.055 | 0 | 0.3 | 0.18 | 0.19 | 0.22 | 0.28 | 0.22 |
| 1 | 181260215 | rs6674511 | 0.636 | 0.476 | 1.311 | -2.832 | -0.002 | 0.008 | 0.008 | 0 | 0.11 | 0.07 | 0.12 | 0.06 | 0.09 | 0.17 |
| 1 | 181263259 | rs2804682 | 1.201 | 0.334 | 1.453 | -2.89 | 0.005 | 0.157 | 0.157 | 0 | 0.51 | 0.73 | 0.68 | 0.77 | 0.52 | 0.65 |
| 1 | 181265245 | rs10910917 | 1.405 | 0.332 | 1.477 | -2.892 | 0 | -0.013 | -0.013 | 0 | 0.47 | 0.43 | 0.64 | 0.77 | 0.51 | 0.64 |
| 1 | 181266420 | rs2332203 | 1.55 | 0.335 | 1.336 | -2.891 | -0.009 | 0.089 | 0.089 | 0.106 | 0.45 | 0.25 | 0.62 | 0.77 | 0.51 | 0.64 |

| | | | | | | | | | | | | | | | | |
|---|---|---|---|---|---|---|---|---|---|---|---|---|---|---|---|---|
| 1 | 181267758 | rs3911589 | -0.119 | 0.4 | 0.935 | -2.857 | -0.001 | -0.043 | -0.043 | 0 | 0.8 | 0.75 | 0.85 | 0.99 | 0.81 | 0.88 |
| 1 | 181270232 | rs2609500 | -0.28 | 0.416 | 0.922 | -2.85 | 0.001 | -0.002 | -0.002 | 0 | 0.81 | 0.85 | 0.86 | 0.99 | 0.81 | 0.88 |
| 1 | 181274415 | rs7523745 | 0.843 | 0.643 | 1.024 | -2.671 | 0.006 | 0.113 | 0.113 | 0 | 0.49 | 0.64 | 0.67 | 0.77 | 0.52 | 0.63 |
| 1 | 181291866 | rs2609483 | -0.187 | 0.999 | 0.832 | -2.562 | -0.018 | -0.158 | -0.158 | 0.183 | 0.29 | 0.54 | 0.23 | 0.02 | 0.26 | 0.2 |
| 1 | 181296112 | rs7525299 | -1.503 | 0.968 | 0.429 | -2.447 | -0.003 | 0.113 | 0.113 | 0.348 | 0.52 | 0.74 | 0.42 | 0.21 | 0.55 | 0.35 |
| 1 | 181296409 | rs7548242 | -1.493 | 1 | 0.446 | -2.44 | -0.002 | 0.115 | 0.115 | 0.457 | 0.52 | 0.74 | 0.4 | 0.22 | 0.55 | 0.34 |
| 1 | 181299951 | rs12094454 | 0.582 | 1.006 | -0.02 | -2.413 | -0.001 | -0.007 | -0.007 | 0 | 0.07 | 0.07 | 0.05 | 0.17 | 0.09 | 0.05 |
| 1 | 181300447 | rs12139587 | 0.402 | 1.076 | 0.154 | -2.499 | 0.008 | -0.007 | -0.007 | 0 | 0.24 | 0.2 | 0.15 | 0.02 | 0.19 | 0.19 |
| 1 | 181301619 | rs6670140 | -1.255 | 1.075 | 0.176 | -2.591 | -0.005 | -0.026 | -0.026 | 0 | 0.42 | 0.52 | 0.28 | 0.22 | 0.39 | 0.34 |
| 1 | 181307883 | rs12119916 | 0.118 | 1.31 | 0.197 | -2.216 | -0.008 | -0.058 | -0.058 | 0 | 0.25 | 0.25 | 0.16 | 0.02 | 0.2 | 0.21 |
| 1 | 181309400 | rs9803794 | -0.898 | 1.149 | 0.285 | -2.042 | -0.009 | 0.051 | 0.051 | 0 | 0.44 | 0.36 | 0.31 | 0.19 | 0.47 | 0.29 |
| 1 | 181310418 | rs12129769 | -0.869 | 1.155 | 0.265 | -2.038 | -0.011 | 0.059 | 0.059 | 0 | 0.43 | 0.33 | 0.31 | 0.19 | 0.47 | 0.28 |
| 1 | 181310522 | rs12756919 | -1.033 | 1.116 | 0.096 | -1.72 | -0.005 | -0.038 | -0.038 | 0 | 0.56 | 0.51 | 0.43 | 0.26 | 0.58 | 0.48 |
| 1 | 181311555 | rs3845428 | -0.786 | 1.27 | -0.255 | -1.956 | 0.008 | 0.022 | 0.022 | 0.03 | 0.51 | 0.61 | 0.43 | 0.11 | 0.53 | 0.5 |
| 1 | 181313404 | rs16857131 | -0.64 | 1.281 | 0.093 | -0.934 | -0.001 | -0.01 | -0.01 | 0.109 | 0.31 | 0.31 | 0.24 | 0.21 | 0.31 | 0.25 |
| 1 | 181317690 | rs16857133 | -0.837 | 1.476 | 0.516 | -0.511 | 0.034 | -0.022 | -0.022 | 6.579 | 0.05 | 0.18 | 0.05 | 0 | 0.05 | 0.04 |
| 1 | 181329186 | rs12408625 | -1.529 | 1.769 | 0.249 | -0.312 | 0.029 | -0.028 | -0.028 | 0.503 | 0.23 | 0.35 | 0.24 | 0.17 | 0.28 | 0.43 |
| 1 | 181335507 | rs2609473 | -1.206 | 1.359 | 0.07 | -0.027 | -0.016 | 0.008 | 0.008 | 0 | 0.37 | 0.7 | 0.56 | 0.4 | 0.55 | 0.48 |
| 1 | 181336389 | rs2609475 | -1.28 | 1.326 | 0.967 | 0.172 | 0.082 | 0.083 | 0.083 | 0 | 0.11 | 0.28 | 0.21 | 0.23 | 0.16 | 0.26 |
| 1 | 181336528 | rs6658713 | -0.752 | 1.333 | 1.1 | 0.247 | 0.01 | -0.038 | -0.038 | 0.027 | 0.23 | 0.5 | 0.32 | 0.2 | 0.3 | 0.18 |
| 1 | 181338339 | rs12039057 | -0.756 | 1.317 | 1.084 | 0.258 | 0.015 | -0.037 | -0.037 | 0.598 | 0.23 | 0.5 | 0.32 | 0.2 | 0.3 | 0.17 |
| 1 | 181341867 | rs10910926 | -0.391 | 1.307 | 1.53 | 0.248 | 0.023 | -0.024 | -0.024 | 0.356 | 0.21 | 0.31 | 0.31 | 0.19 | 0.3 | 0.17 |
| 1 | 181343029 | rs2609497 | -2.158 | 0.478 | 1 | 0.326 | -0.037 | -0.033 | -0.033 | 0 | 0.49 | 0.92 | 0.61 | 0.57 | 0.5 | 0.51 |
| 1 | 181358407 | rs617322 | -2.004 | 1.217 | 1.428 | 0.269 | -0.041 | -0.011 | -0.011 | 0.049 | 0.35 | 0.79 | 0.53 | 0.43 | 0.44 | 0.41 |
| 1 | 181359880 | rs634225 | -1.599 | 1.192 | 1.63 | 0.242 | 0.025 | 0.008 | 0.008 | 0 | 0.31 | 0.55 | 0.51 | 0.43 | 0.44 | 0.4 |
| 1 | 181364473 | rs10797715 | -1.416 | 1.294 | 1.63 | 0.318 | 0.064 | 0.036 | 0.036 | 0 | 0.27 | 0.43 | 0.49 | 0.42 | 0.43 | 0.38 |
| 1 | 181370365 | rs10910931 | -1.008 | 1.354 | 1.605 | 0.121 | 0.027 | 0.008 | 0.008 | 0 | 0.24 | 0.25 | 0.47 | 0.36 | 0.43 | 0.4 |
| 1 | 181372624 | rs667586 | -2.056 | 1.35 | 1.037 | 0.137 | -0.045 | 0.019 | 0.019 | 0.057 | 0.36 | 0.8 | 0.55 | 0.36 | 0.46 | 0.42 |
| 1 | 181374969 | rs673671 | -2.059 | 1.251 | 1.022 | 0.112 | -0.016 | 0.011 | 0.011 | 0 | 0.37 | 0.67 | 0.54 | 0.37 | 0.46 | 0.5 |
| 1 | 181381242 | rs596424 | -1.006 | 1.184 | 1.196 | 0.219 | 0.025 | -0.013 | -0.013 | 0 | 0.05 | 0.24 | 0.02 | 0 | 0.03 | 0.02 |
| 1 | 181384078 | rs591650 | -2.114 | 1.153 | 0.754 | 0.205 | -0.035 | -0.07 | -0.07 | 0 | 0.13 | 0.64 | 0.09 | 0.02 | 0.05 | 0.02 |
| 1 | 181386651 | rs650635 | -1.354 | 1.728 | 0.074 | 0.556 | 0.014 | 0.008 | 0.008 | 0 | 0.11 | 0.18 | 0.18 | 0.2 | 0.14 | 0.3 |
| 1 | 181389092 | rs10752856 | -0.761 | 1.607 | 0.004 | 0.448 | 0.004 | -0.004 | -0.004 | 0.802 | 0.2 | 0.22 | 0.28 | 0.21 | 0.29 | 0.2 |
| 1 | 181390870 | rs602992 | -1.61 | 1.409 | 0.607 | 0.573 | 0.035 | 0.048 | 0.048 | 0.627 | 0.13 | 0.35 | 0.24 | 0.18 | 0.14 | 0.26 |
| 1 | 181393238 | rs6696547 | -0.511 | 1.632 | 0.544 | -0.226 | 0.008 | -0.053 | -0.053 | 0 | 0.36 | 0.48 | 0.36 | 0.14 | 0.39 | 0.27 |
| 1 | 181394765 | rs3898528 | -0.996 | 1.659 | 0.484 | 0.458 | 0.001 | -0.003 | -0.003 | 0 | 0.46 | 0.47 | 0.66 | 0.41 | 0.55 | 0.42 |
| 1 | 181397134 | rs601354 | -1.59 | 1.321 | 0.038 | -0.03 | -0.02 | 0.125 | 0.125 | 0 | 0.54 | 0.86 | 0.61 | 0.25 | 0.56 | 0.59 |
| 1 | 181397224 | rs10910937 | -0.908 | 1.119 | 0.118 | 0.052 | -0.033 | -0.016 | -0.016 | 0 | 0.09 | 0.01 | 0.19 | 0.12 | 0.14 | 0.31 |
| 1 | 181401738 | rs673236 | -1.944 | 1.103 | 0.254 | 0.048 | 0 | 0 | 0 | 0.02 | 0.17 | 0.21 | 0.23 | 0.12 | 0.18 | 0.33 |
| 1 | 181408925 | rs681590 | -0.012 | 1.121 | 0.358 | 0.185 | -0.014 | -0.031 | -0.031 | 0 | 0.06 | 0.15 | 0.04 | 0 | 0.03 | 0.02 |
| 1 | 181412487 | rs3845431 | -0.79 | 1.336 | 0.408 | 0.088 | 0.003 | -0.033 | -0.033 | 0 | 0.39 | 0.48 | 0.49 | 0.25 | 0.44 | 0.44 |
| 1 | 181415083 | rs581582 | 0.077 | 1.527 | 0.374 | 0.069 | 0 | -0.012 | -0.012 | 0 | 0.05 | 0.08 | 0.03 | 0 | 0.04 | 0.02 |
| 1 | 181417470 | rs12565562 | -0.076 | 1.574 | 0.513 | 0.073 | 0.001 | -0.042 | -0.042 | 1.363 | 0.32 | 0.42 | 0.32 | 0.13 | 0.32 | 0.24 |
| 1 | 181419678 | rs588789 | -1.304 | 1.552 | -0.31 | 0.062 | -0.036 | 0.158 | 0.158 | 0 | 0.48 | 0.86 | 0.55 | 0.25 | 0.48 | 0.48 |
| 1 | 181428298 | rs6689920 | -1.837 | 1.537 | -0.343 | 0.234 | -0.126 | -0.097 | -0.097 | 0 | 0.57 | 0.97 | 0.61 | 0.45 | 0.55 | 0.6 |
| 1 | 181433468 | rs6424813 | -0.48 | 1.735 | -0.337 | -0.108 | 0.009 | 0.036 | 0.036 | 0.115 | 0.33 | 0.72 | 0.34 | 0.16 | 0.31 | 0.26 |
| 1 | 181437147 | rs629760 | -1.776 | 1.81 | -0.431 | -0.052 | -0.003 | 0.009 | 0.009 | 0 | 0.13 | 0.14 | 0.2 | 0.09 | 0.17 | 0.22 |
| 1 | 181441744 | rs16857368 | -0.83 | 1.673 | -0.641 | 0.026 | -0.012 | -0.002 | -0.002 | 0 | 0.06 | 0.02 | 0.17 | 0.09 | 0.11 | 0.2 |
| 1 | 181443014 | rs633125 | -0.982 | 1.613 | -0.51 | 0.131 | 0.002 | 0.077 | 0.077 | 0 | 0.46 | 0.73 | 0.53 | 0.27 | 0.47 | 0.51 |
| 1 | 181449485 | rs633143 | -0.505 | 1.789 | -0.386 | 0.213 | -0.008 | 0.009 | 0.009 | 0 | 0.14 | 0.11 | 0.11 | 0.06 | 0.14 | 0.06 |
| 1 | 181456700 | rs553042 | -1.496 | 1.489 | -0.177 | 0.607 | -0.004 | -0.001 | -0.001 | 0 | 0.29 | 0.3 | 0.36 | 0.31 | 0.26 | 0.32 |
| 1 | 181462926 | rs943795 | -1.55 | 1.385 | 0.281 | 0.537 | 0.007 | -0.004 | -0.004 | 0.317 | 0.32 | 0.51 | 0.39 | 0.31 | 0.27 | 0.32 |
| 1 | 181466550 | rs650707 | -1.285 | 1.678 | -0.131 | 0.75 | -0.001 | -0.001 | -0.001 | 0.036 | 0.26 | 0.74 | 0.32 | 0.26 | 0.3 | 0.3 |
| 1 | 181473020 | rs576006 | -1.349 | 1.109 | -0.513 | 0.423 | -0.001 | 0.027 | 0.027 | 0 | 0.52 | 0.82 | 0.37 | 0.37 | 0.41 | 0.35 |
| 1 | 181476430 | rs4652661 | 0.056 | 0.98 | -0.527 | 0.308 | -0.018 | 0.047 | 0.047 | 0 | 0.18 | 0.08 | 0.12 | 0.07 | 0.17 | 0.09 |
| 1 | 181477525 | rs558994 | -0.91 | 0.926 | 0.085 | 0.216 | -0.011 | 0.008 | 0.008 | 0 | 0.36 | 0.45 | 0.26 | 0.35 | 0.29 | 0.15 |
| 1 | 181480183 | rs17494681 | 0.33 | 0.901 | -0.092 | 0.056 | -0.03 | 0.054 | 0.054 | 0 | 0.18 | 0.05 | 0.13 | 0.07 | 0.19 | 0.12 |
| 1 | 181483786 | rs681271 | -2.478 | 0.738 | -0.405 | 0.203 | -0.139 | -0.097 | -0.097 | 0 | 0.64 | 0.95 | 0.52 | 0.5 | 0.5 | 0.63 |
| 1 | 181489802 | rs683927 | -1.901 | 0.972 | 0.605 | 0.365 | 0 | 0.005 | 0.005 | 0.008 | 0.46 | 0.43 | 0.54 | 0.49 | 0.5 | 0.59 |
| 1 | 181490018 | rs12402123 | 0.145 | 0.943 | 0.563 | 0.382 | 0.013 | -0.02 | -0.02 | 1.742 | 0.12 | 0.07 | 0.14 | 0.13 | 0.08 | 0.25 |
| 1 | 181494775 | rs16857509 | -0.145 | 0.886 | 0.412 | 0.231 | 0 | -0.014 | -0.014 | 0 | 0.15 | 0.12 | 0.08 | 0.02 | 0.12 | 0.19 |
| 1 | 181494858 | rs593080 | -2.412 | 0.883 | 0.464 | 0.32 | 0.015 | 0.047 | 0.047 | 0 | 0.5 | 0.77 | 0.53 | 0.33 | 0.5 | 0.58 |
| 1 | 181499238 | rs6685263 | 0.109 | 0.89 | 0.54 | 0.329 | -0.013 | -0.007 | -0.007 | 0 | 0.19 | 0.16 | 0.27 | 0.19 | 0.3 | 0.12 |
| 1 | 181500324 | rs625226 | -0.709 | 0.884 | 0.524 | 0.34 | -0.004 | 0.003 | 0.003 | 0 | 0.17 | 0.21 | 0.13 | 0.17 | 0.13 | 0.21 |
| 1 | 181503903 | rs4146634 | 2.175 | 0.894 | 0.331 | 0.336 | -0.036 | -0.002 | -0.002 | 0.208 | 0.48 | 0.09 | 0.4 | 0.51 | 0.49 | 0.41 |
| 1 | 181506017 | rs6424814 | -0.354 | 1.012 | 0.392 | 0.388 | 0.079 | 0.037 | 0.037 | 0 | 0.22 | 0.5 | 0.31 | 0.19 | 0.3 | 0.12 |
| 1 | 181508326 | rs6681017 | -0.78 | 1.206 | 0.245 | 0.555 | 0.239 | 0.079 | 0.079 | 0 | 0.07 | 0.36 | 0.24 | 0.06 | 0.19 | 0.08 |
| 1 | 181513867 | rs12024842 | -0.291 | 0.336 | 0.292 | 0.495 | 0.013 | -0.033 | -0.033 | 0 | 0.12 | 0.08 | 0.19 | 0.28 | 0.09 | 0.27 |
| 1 | 181516101 | rs4652663 | -0.987 | 0.406 | 0.259 | 0.507 | -0.011 | 0.088 | 0.088 | 0 | 0.11 | 0.37 | 0.13 | 0.15 | 0.08 | 0.04 |

| | | | | | | | | | | | | | | | | |
|---|---|---|---|---|---|---|---|---|---|---|---|---|---|---|---|---|
| 1 | 181517638 | rs7524309 | -0.034 | 0.6 | -0.068 | 0.7 | -0.008 | -0.001 | -0.001 | 2.593 | 0.26 | 0.22 | 0.31 | 0.21 | 0.35 | 0.28 |
| 1 | 181519521 | rs10910955 | 0.99 | 0.724 | -0.687 | 0.431 | 0.005 | 0.039 | 0.039 | 0 | 0.5 | 0.17 | 0.41 | 0.51 | 0.52 | 0.43 |
| 1 | 181525755 | rs2332429 | 0.9 | 0.363 | 0.164 | 0.406 | -0.023 | 0.014 | 0.014 | 0 | 0.82 | 0.64 | 0.78 | 0.84 | 0.89 | 0.77 |
| 1 | 181526369 | rs16857648 | -0.702 | 0.532 | -0.654 | 0.488 | 0.006 | -0.017 | -0.017 | 1.129 | 0.15 | 0.07 | 0.17 | 0.15 | 0.11 | 0.24 |
| 1 | 181529199 | rs10910958 | -0.291 | 0.92 | -1.109 | 0.013 | -0.037 | -0.012 | -0.012 | 0 | 0.23 | 0.08 | 0.32 | 0.27 | 0.29 | 0.26 |
| 1 | 181532332 | rs10910960 | 1.09 | 0.62 | -2.156 | -0.989 | -0.101 | -0.001 | -0.001 | 0 | 0.37 | 0.05 | 0.35 | 0.66 | 0.47 | 0.29 |
| 1 | 181534435 | rs1953690 | -0.211 | 0.099 | -1.514 | -1.35 | 0.004 | 0.012 | 0.012 | 0 | 0.32 | 0.28 | 0.37 | 0.27 | 0.31 | 0.26 |
| 1 | 181534471 | rs1574781 | -0.068 | 0.216 | -0.202 | -1.424 | -0.017 | -0.066 | -0.066 | 0 | 0.27 | 0.32 | 0.21 | 0.07 | 0.16 | 0.44 |
| 1 | 181539747 | rs10797729 | 0.488 | 0.355 | -0.226 | -1.357 | -0.033 | -0.026 | -0.026 | 0 | 0.25 | 0.07 | 0.31 | 0.29 | 0.3 | 0.24 |
| 1 | 181541159 | rs10797730 | -0.472 | 0.106 | 0.657 | -1.576 | 0.006 | 0.017 | 0.017 | 0 | 0.4 | 0.33 | 0.48 | 0.34 | 0.38 | 0.49 |
| 1 | 181541353 | rs1999838 | -0.464 | 0.113 | 0.7 | -1.572 | 0.005 | 0.013 | 0.013 | 0 | 0.4 | 0.34 | 0.48 | 0.34 | 0.38 | 0.49 |
| 1 | 181545037 | rs7511748 | 0.334 | 0.089 | 0.813 | -1.614 | 0.009 | -0.023 | -0.023 | 0 | 0.59 | 0.51 | 0.58 | 0.69 | 0.57 | 0.57 |
| 1 | 181545124 | rs12135959 | 1.17 | 0.096 | 0.804 | -1.608 | -0.002 | 0.047 | 0.047 | 0 | 0.44 | 0.24 | 0.41 | 0.64 | 0.49 | 0.32 |
| 1 | 181549547 | rs10494540 | 1.151 | 0.1 | 0.95 | -1.606 | 0.003 | 0.023 | 0.023 | 0 | 0.45 | 0.39 | 0.43 | 0.64 | 0.49 | 0.32 |
| 1 | 181550465 | rs10494541 | -0.556 | 0.058 | 0.978 | -1.573 | 0.045 | -0.083 | -0.083 | 0.238 | 0.85 | 0.61 | 0.9 | 0.98 | 0.87 | 0.82 |
| 1 | 181553070 | rs12239392 | 0.667 | 0.009 | 0.953 | -1.536 | -0.015 | -0.041 | -0.041 | 0 | 0.14 | 0.23 | 0.16 | 0.05 | 0.08 | 0.25 |
| 1 | 181561783 | rs3856094 | -0.163 | -0.153 | 0.852 | -1.559 | 0.008 | 0.014 | 0.014 | 0 | 0.33 | 0.24 | 0.37 | 0.29 | 0.32 | 0.26 |
| 1 | 181562198 | rs4652667 | 0.521 | 0.006 | 1.416 | -1.964 | -0.008 | -0.045 | -0.045 | 0.541 | 0.13 | 0.23 | 0.07 | 0.02 | 0.08 | 0.17 |
| 1 | 181562269 | rs16857762 | 0.506 | -0.032 | 1.341 | -1.989 | -0.032 | -0.058 | -0.058 | 0.18 | 0.14 | 0.28 | 0.14 | 0.05 | 0.07 | 0.24 |
| 1 | 181569891 | rs4651109 | 1.234 | 0.017 | 1.144 | -2.002 | 0.015 | 0.045 | 0.045 | 0.009 | 0.65 | 0.25 | 0.72 | 0.93 | 0.78 | 0.56 |
| 1 | 181572088 | rs12742236 | 0.798 | 0.086 | 1.169 | -1.934 | 0.005 | 0.002 | 0.002 | 0 | 0.69 | 0.26 | 0.74 | 0.93 | 0.8 | 0.57 |
| 1 | 181579231 | rs199960 | 0.274 | -0.047 | 1.085 | -1.924 | 0.001 | 0.045 | 0.045 | 0 | 0.3 | 0.18 | 0.28 | 0.1 | 0.29 | 0.22 |
| 1 | 181586946 | rs16857808 | 0.549 | 0.115 | 1.486 | -1.452 | -0.006 | -0.015 | -0.015 | 0 | 0.13 | 0.11 | 0.12 | 0.05 | 0.07 | 0.25 |
| 1 | 181590858 | rs199950 | 1.041 | 0.088 | 1.7 | -1.482 | -0.013 | -0.109 | -0.109 | 1.453 | 0.71 | 0.45 | 0.75 | 0.93 | 0.8 | 0.59 |
| 1 | 181595979 | rs199939 | 0.844 | 0.239 | 1.851 | -1.37 | -0.001 | -0.097 | -0.097 | 2.278 | 0.76 | 0.43 | 0.76 | 0.93 | 0.83 | 0.61 |
| 1 | 181596223 | rs6691980 | -0.56 | 0.293 | 1.694 | -1.345 | 0.005 | -0.047 | -0.047 | 0 | 0.21 | 0.28 | 0.2 | 0.07 | 0.17 | 0.4 |
| 1 | 181605165 | rs175338 | 0.853 | 0.433 | 1.871 | -1.473 | 0.009 | -0.099 | -0.099 | 0.697 | 0.73 | 0.42 | 0.74 | 0.93 | 0.76 | 0.58 |
| 1 | 181609567 | rs3845439 | -0.365 | 0.444 | 2.098 | -1.473 | -0.002 | -0.047 | -0.047 | 0 | 0.22 | 0.27 | 0.22 | 0.07 | 0.18 | 0.28 |
| 1 | 181611287 | rs175337 | 0.483 | 0.436 | 2.195 | -1.465 | -0.009 | -0.085 | -0.085 | 0 | 0.76 | 0.6 | 0.77 | 0.92 | 0.81 | 0.59 |
| 1 | 181612041 | rs199955 | 0.538 | 0.435 | 2.192 | -1.482 | -0.011 | -0.096 | -0.096 | 0 | 0.76 | 0.59 | 0.84 | 0.94 | 0.81 | 0.59 |
| 1 | 181612753 | rs10910977 | -0.711 | 0.505 | 1.832 | -1.46 | 0.026 | -0.101 | -0.101 | 0 | 0.24 | 0.41 | 0.18 | 0.06 | 0.23 | 0.28 |
| 1 | 181613691 | rs6697349 | -0.781 | 0.506 | 1.824 | -1.46 | 0.024 | -0.105 | -0.105 | 0 | 0.24 | 0.43 | 0.18 | 0.06 | 0.23 | 0.29 |
| 1 | 181614143 | rs6675785 | -0.781 | 0.506 | 1.824 | -1.46 | 0.024 | -0.105 | -0.105 | 0.006 | 0.24 | 0.43 | 0.18 | 0.06 | 0.23 | 0.29 |
| 1 | 181629618 | rs6680280 | -1.778 | 0.425 | 1.949 | -1.136 | 0.113 | 0.009 | 0.009 | 2.129 | 0.07 | 0.37 | 0.06 | 0 | 0.07 | 0.02 |
| 1 | 181635954 | rs199916 | -0.866 | 0.174 | 1.313 | -1.297 | -0.027 | -0.006 | -0.006 | 0.502 | 0.86 | 0.98 | 0.85 | 0.82 | 0.83 | 0.91 |
| 1 | 181638511 | rs3820260 | 0.333 | -0.289 | 1.315 | -1.695 | 0.011 | 0.008 | 0.008 | 0 | 0.2 | 0.14 | 0.11 | 0.05 | 0.13 | 0.26 |
| 1 | 181643711 | rs3766980 | 1 | -0.725 | 1.132 | -2.074 | 0.006 | 0.007 | 0.007 | 0 | 0.49 | 0.44 | 0.52 | 0.73 | 0.58 | 0.36 |
| 1 | 181653111 | rs10910979 | 1.363 | -0.518 | 1.06 | -2.037 | -0.015 | 0.152 | 0.152 | 0 | 0.45 | 0.13 | 0.47 | 0.72 | 0.53 | 0.36 |
| 1 | 181667283 | rs3766988 | 0.7 | -0.606 | 1.178 | -2.083 | -0.001 | 0.005 | 0.005 | 0 | 0.12 | 0.09 | 0.15 | 0.02 | 0.13 | 0.1 |
| 1 | 181672682 | rs7513540 | 0.842 | -0.698 | 1.559 | -2.148 | -0.008 | -0.035 | -0.035 | 0 | 0.66 | 0.39 | 0.6 | 0.77 | 0.68 | 0.63 |
| 1 | 181674891 | rs3845444 | 0.323 | -0.685 | 1.703 | -2.142 | -0.001 | -0.023 | -0.023 | 3.348 | 0.7 | 0.64 | 0.63 | 0.77 | 0.68 | 0.63 |
| 1 | 181686216 | rs7540850 | 0.792 | -0.844 | 1.733 | -2.092 | 0.03 | 0.087 | 0.087 | 0 | 0.6 | 0.2 | 0.63 | 0.76 | 0.68 | 0.64 |
| 1 | 181688483 | rs4651112 | -1.188 | -0.775 | 1.664 | -2.001 | 0.005 | -0.001 | -0.001 | 0.555 | 0.85 | 0.78 | 0.8 | 0.81 | 0.83 | 0.89 |
| 1 | 181704788 | rs7418539 | 0.242 | -0.842 | 1.64 | -2.085 | -0.012 | -0.072 | -0.072 | 0 | 0.22 | 0.28 | 0.15 | 0.05 | 0.16 | 0.27 |
| 1 | 181712290 | rs3767000 | 1.697 | -1.094 | 0.59 | -1.386 | -0.024 | -0.018 | -0.018 | 0 | 0.34 | 0.23 | 0.29 | 0.3 | 0.4 | 0.23 |
| 1 | 181712376 | rs3767001 | -1.715 | -1.112 | 0.688 | -1.348 | 0.001 | 0 | 0 | 0 | 0.85 | 0.82 | 0.83 | 0.8 | 0.83 | 0.9 |
| 1 | 181714184 | rs16858051 | 1.072 | -1.155 | 0.636 | -1.363 | -0.001 | -0.004 | -0.004 | 0 | 0.11 | 0.08 | 0.16 | 0.03 | 0.12 | 0.1 |
| 1 | 181715146 | rs3753748 | -0.029 | -1.176 | 0.726 | -1.352 | 0 | -0.028 | -0.028 | 0 | 0.72 | 0.63 | 0.66 | 0.77 | 0.71 | 0.8 |
| 1 | 181716658 | rs3767003 | 0.463 | -1.112 | 0.658 | -1.254 | 0.006 | 0.025 | 0.025 | 0 | 0.52 | 0.46 | 0.51 | 0.72 | 0.55 | 0.53 |
| 1 | 181717047 | rs3767004 | 0.259 | -1.131 | 0.269 | -0.721 | -0.001 | 0.069 | 0.069 | 0.236 | 0.07 | 0.13 | 0.14 | 0.33 | 0.07 | 0.06 |
| 1 | 181717867 | rs12131032 | 1.912 | -1.097 | 0.117 | -0.734 | -0.065 | -0.037 | -0.037 | 0 | 0.32 | 0.13 | 0.27 | 0.29 | 0.39 | 0.23 |
| 1 | 181725110 | rs4652678 | -0.306 | -1.051 | 0.258 | -0.821 | -0.023 | -0.135 | -0.135 | 2.189 | 0.34 | 0.49 | 0.33 | 0.08 | 0.29 | 0.37 |
| 1 | 181727273 | rs704329 | 1.583 | -1.025 | -0.484 | -0.709 | -0.055 | -0.066 | -0.066 | 0 | 0.36 | 0.16 | 0.32 | 0.38 | 0.41 | 0.44 |
| 1 | 181734600 | rs199920 | 0.112 | -0.93 | -0.253 | -0.715 | 0.006 | 0.004 | 0.004 | 0 | 0.3 | 0.27 | 0.29 | 0.07 | 0.26 | 0.35 |
| 1 | 181736189 | rs697259 | 1.049 | -1.007 | -0.388 | -0.543 | -0.02 | 0.017 | 0.017 | 0 | 0.46 | 0.25 | 0.51 | 0.51 | 0.47 | 0.5 |
| 1 | 181741301 | rs199930 | 0.034 | -0.904 | -0.33 | -0.639 | 0.006 | 0.001 | 0.001 | 0 | 0.3 | 0.28 | 0.28 | 0.08 | 0.26 | 0.34 |
| 1 | 181753030 | rs546191 | -0.086 | -1.037 | -0.475 | -0.554 | -0.012 | -0.063 | -0.063 | 0 | 0.72 | 0.51 | 0.7 | 0.83 | 0.74 | 0.79 |
| 1 | 181754803 | rs473200 | 0.381 | -0.781 | -0.859 | 0.113 | 0.007 | 0 | 0 | 0.939 | 0.26 | 0.25 | 0.19 | 0.27 | 0.21 | 0.29 |
| 1 | 181759657 | rs704326 | 1.225 | -0.266 | -1.169 | 0.506 | -0.103 | -0.078 | -0.078 | 0.331 | 0.34 | 0.12 | 0.32 | 0.39 | 0.43 | 0.18 |
| 1 | 181766437 | rs3845446 | -0.089 | -0.052 | -0.892 | 1.136 | -0.002 | 0.045 | 0.045 | 0.025 | 0.06 | 0.06 | 0.05 | 0.31 | 0.05 | 0.12 |
| 1 | 181768186 | rs2280869 | -0.134 | -0.056 | -0.856 | 1.131 | -0.001 | 0.057 | 0.057 | 0.032 | 0.06 | 0.08 | 0.16 | 0.31 | 0.05 | 0.12 |
| 1 | 181768985 | rs638132 | -0.031 | -0.256 | -0.7 | 0.537 | 0.002 | -0.005 | -0.005 | 0 | 0.75 | 0.74 | 0.64 | 0.74 | 0.73 | 0.82 |
| 1 | 181771992 | rs12045458 | 1.22 | -0.27 | -1.057 | 0.538 | -0.062 | 0.008 | 0.008 | 0 | 0.15 | 0.02 | 0.27 | 0.15 | 0.22 | 0.09 |
| 1 | 181773948 | rs3681 | 1.649 | 0.101 | -1.766 | 0.659 | -0.104 | -0.065 | -0.065 | 0 | 0.2 | 0.01 | 0.23 | 0.25 | 0.29 | 0.19 |
| 1 | 181779469 | rs3902279 | -0.127 | -0.119 | -1.448 | 0.795 | -0.008 | 0.004 | 0.004 | 0 | 0.07 | 0.12 | 0.09 | 0.08 | 0.03 | 0.06 |
| 1 | 181782017 | rs509391 | -0.072 | -0.366 | -0.92 | 0.874 | 0 | 0.005 | 0.005 | 0.689 | 0.68 | 0.43 | 0.6 | 0.45 | 0.69 | 0.61 |
| 1 | 181787784 | rs480752 | 1.835 | -0.759 | -1.26 | 0.468 | -0.004 | -0.038 | -0.038 | 1.321 | 0.25 | 0.29 | 0.13 | 0.05 | 0.18 | 0.23 |
| 1 | 181788646 | rs605863 | 0.401 | -0.097 | -0.738 | 1.022 | 0.033 | 0.063 | 0.063 | 0.066 | 0.49 | 0.42 | 0.38 | 0.17 | 0.41 | 0.42 |
| 1 | 181804790 | rs511136 | -0.001 | -0.046 | -0.471 | 0.937 | 0.011 | 0.057 | 0.057 | 0.524 | 0.7 | 0.5 | 0.61 | 0.44 | 0.7 | 0.74 |

| | | | | | | | | | | | | | | | | |
|---|---|---|---|---|---|---|---|---|---|---|---|---|---|---|---|---|
| 1 | 181808610 | rs659234 | -1.439 | -0.035 | -0.37 | 0.887 | -0.007 | 0.007 | 0.007 | 0 | 0.8 | 0.87 | 0.72 | 0.84 | 0.75 | 0.87 |
| 1 | 181813549 | rs12071191 | 0.664 | -0.041 | -0.285 | 0.929 | 0.038 | 0.28 | 0.28 | 1.612 | 0.07 | 0.29 | 0.08 | 0.28 | 0.03 | 0.06 |
| 1 | 181813769 | rs486003 | -0.61 | -0.019 | -0.585 | 0.564 | 0.026 | 0.078 | 0.078 | 0.26 | 0.74 | 0.57 | 0.63 | 0.56 | 0.71 | 0.81 |
| 1 | 181816509 | rs695072 | 0.6 | -0.173 | -0.617 | 0.494 | 0.034 | 0.096 | 0.096 | 0.062 | 0.44 | 0.32 | 0.31 | 0.09 | 0.39 | 0.37 |
| 1 | 181824882 | rs7533297 | 0.17 | 0.303 | -0.533 | 0.634 | 0.017 | 0.029 | 0.029 | 0.044 | 0.29 | 0.39 | 0.43 | 0.46 | 0.32 | 0.2 |
| 1 | 181826928 | rs598714 | -0.689 | 0.495 | -0.602 | 0.798 | -0.003 | -0.006 | -0.006 | 0.003 | 0.75 | 0.78 | 0.72 | 0.57 | 0.7 | 0.69 |
| 1 | 181827111 | rs677618 | -1.216 | 0.666 | -0.992 | 0.759 | -0.031 | -0.051 | -0.051 | 0 | 0.79 | 0.93 | 0.76 | 0.68 | 0.72 | 0.77 |
| 1 | 181827563 | rs9286844 | -0.737 | 0.677 | -0.653 | 0.361 | 0.008 | 0.018 | 0.018 | 0.943 | 0.35 | 0.55 | 0.45 | 0.49 | 0.35 | 0.43 |
| 1 | 181827989 | rs630341 | -0.53 | 0.909 | -1.24 | 0.088 | -0.042 | 0.097 | 0.097 | 0.044 | 0.36 | 0.14 | 0.33 | 0.15 | 0.44 | 0.33 |
| 1 | 181828178 | rs10910993 | 0.238 | -0.101 | -1.074 | 0.472 | 0.017 | -0.059 | -0.059 | 0 | 0.21 | 0.11 | 0.19 | 0.44 | 0.18 | 0.24 |
| 1 | 181841744 | rs12408910 | 0.328 | -0.04 | -0.894 | 0.719 | -0.006 | -0.02 | -0.02 | 1.473 | 0.09 | 0.04 | 0.1 | 0.33 | 0.13 | 0.21 |
| 1 | 181843583 | rs12410953 | -0.415 | -0.143 | -0.82 | 0.67 | 0 | 0.012 | 0.012 | 0 | 0.11 | 0.1 | 0.11 | 0.33 | 0.13 | 0.21 |
| 1 | 181850780 | rs6701187 | -1.598 | -0.533 | -0.174 | 0.489 | -0.039 | 0.104 | 0.104 | 0.019 | 0.2 | 0.48 | 0.19 | 0.38 | 0.15 | 0.28 |
| 1 | 181869716 | rs12086274 | -0.92 | -0.543 | 0.166 | 0.482 | -0.012 | 0.092 | 0.092 | 0 | 0.13 | 0.3 | 0.13 | 0.33 | 0.13 | 0.21 |
| 1 | 181870217 | rs498212 | 0.77 | -0.273 | -0.115 | 0.213 | 0.011 | 0.002 | 0.002 | 0 | 0.55 | 0.38 | 0.59 | 0.54 | 0.64 | 0.43 |
| 1 | 181876870 | rs16858326 | 0.596 | 0 | -0.185 | 0.15 | -0.014 | 0.002 | 0.002 | 0 | 0.06 | 0.13 | 0.05 | 0.04 | 0.02 | 0.07 |
| 1 | 181883902 | rs12565283 | 0.408 | -0.002 | -0.132 | 0.149 | -0.001 | 0.022 | 0.022 | 0 | 0.06 | 0.24 | 0.06 | 0.04 | 0.02 | 0.07 |
| 1 | 181888775 | rs4651117 | 1.033 | 0.114 | -0.692 | 0.078 | -0.013 | 0.089 | 0.089 | 0 | 0.23 | 0.06 | 0.22 | 0.09 | 0.21 | 0.13 |
| 1 | 181895558 | rs615014 | 0.607 | 0.101 | -0.982 | -0.04 | -0.038 | -0.005 | -0.005 | 0.967 | 0.14 | 0.01 | 0.08 | 0.1 | 0.17 | 0.2 |
| 1 | 181897939 | rs1281294 | 0.779 | 0.01 | -0.7 | -0.008 | 0.007 | 0.052 | 0.052 | 0 | 0.25 | 0.11 | 0.23 | 0.09 | 0.21 | 0.29 |
| 1 | 181904093 | rs10494543 | -0.42 | -0.174 | -0.454 | 0.533 | 0 | 0.002 | 0.002 | 0 | 0.15 | 0.18 | 0.14 | 0.34 | 0.18 | 0.25 |
| 1 | 181906592 | rs6424828 | -1.434 | -0.202 | -0.416 | 0.512 | -0.018 | 0.037 | 0.037 | 0 | 0.23 | 0.43 | 0.2 | 0.38 | 0.2 | 0.33 |
| 1 | 181914287 | rs2262513 | -1.438 | -0.165 | -0.36 | 0.439 | -0.027 | 0.048 | 0.048 | 0 | 0.24 | 0.49 | 0.21 | 0.33 | 0.19 | 0.49 |
| 1 | 181938516 | rs16858396 | 0.413 | -0.162 | -0.45 | 0.397 | -0.018 | -0.003 | -0.003 | 0 | 0.07 | 0.12 | 0.05 | 0.05 | 0.02 | 0.24 |
| 1 | 181951768 | rs10494544 | 1.612 | -0.13 | -0.675 | 0.503 | -0.001 | 0.003 | 0.003 | 0 | 0.05 | 0.04 | 0.05 | 0.1 | 0.07 | 0.12 |
| 1 | 181952711 | rs12039940 | 1.318 | 0.181 | -1.234 | 0.287 | -0.093 | 0.03 | 0.03 | 0.101 | 0.26 | 0.03 | 0.25 | 0.18 | 0.3 | 0.14 |
| 1 | 181954718 | rs10911006 | 1.117 | 0.307 | -1.12 | 0.443 | -0.017 | -0.032 | -0.032 | 0.048 | 0.07 | 0.03 | 0.08 | 0.19 | 0.09 | 0.12 |
| 1 | 181965454 | rs1281317 | -0.868 | 0.327 | -1.2 | 0.33 | -0.014 | 0.013 | 0.013 | 0 | 0.84 | 0.96 | 0.89 | 0.86 | 0.81 | 0.85 |
| 1 | 181972879 | rs3904616 | -0.71 | 0.177 | -0.766 | 0.753 | -0.009 | 0.034 | 0.034 | 0.223 | 0.15 | 0.25 | 0.14 | 0.25 | 0.12 | 0.42 |
| 1 | 182005722 | rs6700658 | 1.039 | 0.108 | -0.651 | 0.536 | -0.009 | 0.001 | 0.001 | 0 | 0.4 | 0.31 | 0.49 | 0.33 | 0.45 | 0.22 |
| 1 | 182010689 | rs6695585 | -0.314 | 0.138 | -0.845 | 0.587 | -0.028 | 0.002 | 0.002 | 0 | 0.09 | 0.31 | 0.07 | 0.05 | 0.02 | 0.29 |
| 1 | 182022938 | rs3856101 | -0.08 | 0.32 | -1.002 | 0.563 | -0.006 | -0.018 | -0.018 | 0.184 | 0.14 | 0.1 | 0.12 | 0.29 | 0.16 | 0.25 |
| 1 | 182023046 | rs3908801 | -1.591 | 0.284 | -0.965 | 0.549 | -0.02 | 0.03 | 0.03 | 0 | 0.24 | 0.45 | 0.21 | 0.34 | 0.19 | 0.54 |
| 1 | 182024515 | rs11802562 | -0.12 | 0.309 | -1.198 | 0.623 | -0.027 | -0.009 | -0.009 | 0.779 | 0.08 | 0.29 | 0.07 | 0.04 | 0.02 | 0.29 |
| 1 | 182027022 | rs12564283 | -0.281 | 0.966 | -1.971 | 0.832 | -0.004 | -0.019 | -0.019 | 0 | 0.13 | 0.09 | 0.12 | 0.28 | 0.15 | 0.25 |
| 1 | 182039442 | rs11803436 | 0.066 | 0.753 | -2.214 | 0.629 | -0.075 | 0.027 | 0.027 | 0 | 0.29 | 0.03 | 0.27 | 0.19 | 0.32 | 0.16 |
| 1 | 182042119 | rs12569254 | -0.747 | 0.531 | -1.443 | 0.999 | 0.002 | -0.018 | -0.018 | 0 | 0.14 | 0.1 | 0.12 | 0.28 | 0.13 | 0.23 |
| 1 | 182053214 | rs10797752 | 0.254 | 0.68 | -1.269 | 0.837 | -0.002 | 0 | 0 | 0 | 0.36 | 0.35 | 0.41 | 0.31 | 0.42 | 0.16 |
| 1 | 182054142 | rs7540941 | 0.009 | -0.112 | -1.215 | 1.109 | -0.001 | 0.001 | 0.001 | 0 | 0.8 | 0.82 | 0.81 | 0.69 | 0.84 | 0.78 |
| 1 | 182054977 | rs3856106 | 0.422 | -0.365 | -0.965 | 1.121 | -0.001 | -0.005 | -0.005 | 0 | 0.39 | 0.39 | 0.27 | 0.27 | 0.38 | 0.45 |
| 1 | 182057506 | rs7413116 | -1.325 | -0.096 | -0.666 | 1.051 | -0.035 | 0.002 | 0.002 | 1.803 | 0.65 | 0.79 | 0.59 | 0.69 | 0.59 | 0.84 |
| 1 | 182058556 | rs7537211 | 0.435 | -0.084 | -0.75 | 1.045 | -0.001 | -0.004 | -0.004 | 0 | 0.4 | 0.39 | 0.28 | 0.27 | 0.39 | 0.45 |
| 1 | 182058955 | rs2236537 | -0.576 | -0.067 | -0.76 | 1.047 | 0.013 | -0.025 | -0.025 | 4.496 | 0.23 | 0.17 | 0.29 | 0.42 | 0.18 | 0.38 |
| 1 | 182062812 | rs10911018 | -1.154 | -0.044 | -0.778 | 0.963 | 0.001 | -0.001 | -0.001 | 0 | 0.65 | 0.63 | 0.59 | 0.69 | 0.6 | 0.84 |
| 1 | 182069992 | rs3856109 | -1.19 | -0.108 | -0.809 | 0.918 | 0.002 | -0.001 | -0.001 | 0 | 0.66 | 0.63 | 0.59 | 0.69 | 0.6 | 0.84 |
| 1 | 182077680 | rs17498867 | 1.088 | -0.652 | -1.573 | 0.511 | -0.008 | 0.025 | 0.025 | 3.542 | 0.25 | 0.01 | 0.09 | 0.15 | 0.18 | 0.23 |
| 1 | 182079178 | rs17498888 | 1.286 | -0.692 | -1.503 | 0.789 | -0.014 | -0.031 | -0.031 | 0.105 | 0.08 | 0 | 0.12 | 0.09 | 0.08 | 0.14 |
| 1 | 182079986 | rs7517310 | -0.587 | -0.9 | -1.453 | 0.659 | 0.004 | -0.001 | -0.001 | 0.169 | 0.26 | 0.2 | 0.28 | 0.28 | 0.27 | 0.32 |
| 1 | 182081960 | rs10911021 | -0.893 | -0.832 | -0.972 | 0.606 | 0.003 | -0.009 | -0.009 | 0 | 0.28 | 0.52 | 0.32 | 0.28 | 0.29 | 0.31 |
| 1 | 182094598 | rs1281331 | 1.323 | -0.641 | -1.091 | 0.582 | 0.001 | 0.002 | 0.002 | 0.042 | 0.38 | 0.38 | 0.49 | 0.42 | 0.39 | 0.34 |
| 1 | 182096856 | rs2236539 | -1.447 | -0.621 | -1.063 | 0.578 | -0.003 | -0.003 | -0.003 | 0 | 0.62 | 0.63 | 0.5 | 0.58 | 0.58 | 0.63 |
| 1 | 182098096 | rs16858559 | 0.861 | -0.723 | -1.173 | 0.529 | 0.018 | 0.001 | 0.001 | 1.871 | 0.32 | 0.11 | 0.17 | 0.3 | 0.27 | 0.31 |
| 1 | 182101577 | rs1281322 | 0.792 | -0.742 | -1.121 | 0.502 | -0.003 | 0.008 | 0.008 | 0.558 | 0.46 | 0.47 | 0.56 | 0.62 | 0.48 | 0.47 |
| 1 | 182101740 | rs1281323 | -0.011 | -0.735 | -1.111 | 0.453 | -0.013 | 0.05 | 0.05 | 0.738 | 0.77 | 0.87 | 0.76 | 0.9 | 0.73 | 0.76 |
| 1 | 182104189 | rs3845452 | -0.989 | -0.466 | -1.205 | 0.791 | -0.003 | -0.001 | -0.001 | 0.281 | 0.58 | 0.58 | 0.48 | 0.58 | 0.55 | 0.61 |
| 1 | 182106702 | rs3795490 | 1.044 | -0.543 | -1.443 | 0.812 | 0.003 | -0.004 | -0.004 | 0 | 0.1 | 0.07 | 0.13 | 0.09 | 0.11 | 0.14 |
| 1 | 182108102 | rs6674234 | -1.333 | -0.271 | -1.218 | 0.777 | -0.042 | -0.016 | -0.016 | 0 | 0.67 | 0.81 | 0.63 | 0.69 | 0.61 | 0.82 |
| 1 | 182110058 | rs4652690 | 0.464 | -0.334 | -1.104 | 0.741 | 0.001 | 0.011 | 0.011 | 1.191 | 0.43 | 0.38 | 0.54 | 0.6 | 0.42 | 0.46 |
| 1 | 182111188 | rs4112312 | 0.726 | -0.364 | -1.029 | 0.801 | -0.001 | -0.001 | -0.001 | 0 | 0.4 | 0.42 | 0.5 | 0.41 | 0.4 | 0.41 |
| 1 | 182111988 | rs1281337 | 0.383 | -0.544 | -1.158 | 0.946 | -0.014 | 0.027 | 0.027 | 1.422 | 0.48 | 0.75 | 0.57 | 0.38 | 0.44 | 0.45 |
| 1 | 182112106 | rs1281338 | 0.255 | -0.77 | -1.507 | 0.634 | -0.006 | -0.028 | -0.028 | 0 | 0.7 | 0.86 | 0.71 | 0.67 | 0.68 | 0.66 |
| 1 | 182115278 | rs3845453 | 0.892 | -0.994 | -1.6 | 0.391 | 0.021 | 0.008 | 0.008 | 0 | 0.18 | 0.06 | 0.14 | 0.14 | 0.14 | 0.3 |
| 1 | 182115787 | rs1281340 | 0.069 | -0.882 | -1.041 | 0.779 | 0.001 | -0.005 | -0.005 | 0 | 0.43 | 0.35 | 0.4 | 0.49 | 0.38 | 0.32 |
| 1 | 182119516 | rs6694132 | 0.66 | -0.896 | -1.071 | 0.559 | 0.013 | 0.031 | 0.031 | 0.157 | 0.36 | 0.14 | 0.33 | 0.25 | 0.32 | 0.23 |
| 1 | 182121598 | rs1689804 | -0.372 | -0.75 | -0.748 | 0.491 | -0.011 | 0.023 | 0.023 | 0 | 0.53 | 0.73 | 0.52 | 0.62 | 0.5 | 0.5 |
| 1 | 182131294 | rs10911029 | 1.975 | -0.517 | -0.754 | 0.575 | -0.017 | 0.007 | 0.007 | 0.043 | 0.23 | 0.18 | 0.33 | 0.18 | 0.34 | 0.2 |
| 1 | 182133412 | rs7414231 | 0.872 | -0.316 | -1.02 | 0.77 | -0.001 | 0.009 | 0.009 | 0 | 0.14 | 0.15 | 0.13 | 0.37 | 0.15 | 0.26 |
| 1 | 182151909 | rs1689791 | 0.451 | -0.469 | -1.11 | 0.624 | 0.002 | 0.006 | 0.006 | 0 | 0.36 | 0.25 | 0.34 | 0.26 | 0.32 | 0.22 |
| 1 | 182157235 | rs2243976 | 0.362 | -0.586 | -1.132 | 0.634 | -0.001 | 0.002 | 0.002 | 0 | 0.36 | 0.3 | 0.35 | 0.26 | 0.35 | 0.22 |

| | | | | | | | | | | | | | | | | |
|---|---|---|---|---|---|---|---|---|---|---|---|---|---|---|---|---|
| 1 | 182173711 | rs10911031 | -0.021 | -0.372 | -1.235 | 1.017 | -0.001 | -0.003 | -0.003 | 0 | 0.15 | 0.16 | 0.14 | 0.37 | 0.15 | 0.26 |
| 1 | 182176656 | rs1779821 | -0.596 | -0.528 | -1.413 | -0.388 | -0.009 | -0.051 | -0.051 | 0 | 0.84 | 0.98 | 0.87 | 0.84 | 0.86 | 0.73 |
| 1 | 182177802 | rs4147173 | -0.312 | -0.252 | -0.669 | -1.023 | 0.035 | 0.061 | 0.061 | 0 | 0.65 | 0.82 | 0.77 | 0.87 | 0.72 | 0.64 |
| 1 | 182178482 | rs1779819 | -0.343 | -0.283 | -0.696 | -1.026 | 0.022 | 0.114 | 0.114 | 0 | 0.66 | 0.93 | 0.78 | 0.87 | 0.71 | 0.64 |
| 1 | 182178847 | rs4113814 | -0.056 | -0.16 | -0.482 | -1.051 | 0.009 | 0.024 | 0.024 | 0 | 0.63 | 0.71 | 0.75 | 0.87 | 0.68 | 0.64 |
| 1 | 182182651 | rs1419326 | -0.413 | -0.055 | -0.576 | -1.103 | 0.034 | 0.14 | 0.14 | 0 | 0.66 | 0.9 | 0.78 | 0.92 | 0.71 | 0.63 |
| 1 | 182184243 | rs6662373 | 0.288 | -0.024 | -0.628 | -1.18 | -0.004 | -0.009 | -0.009 | 0.223 | 0.39 | 0.68 | 0.47 | 0.54 | 0.44 | 0.39 |
| 1 | 182185360 | rs10911032 | 0.255 | -0.141 | -0.658 | -1.167 | -0.02 | -0.018 | -0.018 | 0 | 0.41 | 0.73 | 0.48 | 0.54 | 0.46 | 0.39 |
| 1 | 182200659 | rs10494545 | 0.805 | -0.063 | -1.059 | -1.12 | -0.034 | -0.1 | -0.1 | 0 | 0.24 | 0.06 | 0.28 | 0.38 | 0.25 | 0.24 |
| 1 | 182204828 | rs10911041 | 0.379 | -0.126 | -0.565 | -1.097 | -0.012 | -0.01 | -0.01 | 0 | 0.36 | 0.37 | 0.44 | 0.54 | 0.46 | 0.38 |
| 1 | 182218476 | rs2489285 | 0.097 | 0.065 | -1.027 | -1.006 | -0.079 | -0.069 | -0.069 | 0 | 0.41 | 0.87 | 0.5 | 0.54 | 0.45 | 0.37 |
| 1 | 182224790 | rs1018611 | 0.896 | 0.135 | -0.671 | -0.528 | -0.004 | -0.071 | -0.071 | 0.536 | 0.17 | 0.07 | 0.21 | 0.36 | 0.15 | 0.23 |
| 1 | 182226964 | rs2251093 | 0.405 | 0.321 | -0.168 | -0.554 | 0.001 | 0.001 | 0.001 | 0 | 0.49 | 0.52 | 0.65 | 0.61 | 0.58 | 0.45 |
| 1 | 182237719 | rs7546440 | 0.569 | 0.381 | -0.08 | -0.705 | -0.021 | 0.061 | 0.061 | 0 | 0.13 | 0.23 | 0.08 | 0.31 | 0.07 | 0.15 |
| 1 | 182244786 | rs7524031 | 1.02 | 0.705 | -0.546 | -0.643 | -0.142 | -0.01 | -0.01 | 0.746 | 0.29 | 0.04 | 0.4 | 0.25 | 0.45 | 0.22 |
| 1 | 182256110 | rs10911044 | 1.279 | 0.338 | -0.851 | -0.709 | -0.092 | -0.044 | -0.044 | 0 | 0.28 | 0.06 | 0.32 | 0.26 | 0.39 | 0.26 |
| 1 | 182259826 | rs6659875 | 0.773 | -0.232 | -0.629 | -0.52 | -0.003 | -0.036 | -0.036 | 0 | 0.18 | 0.12 | 0.23 | 0.41 | 0.16 | 0.26 |
| 1 | 182262763 | rs4600010 | 1.061 | 0.15 | -0.777 | -0.425 | -0.124 | -0.012 | -0.012 | 0.477 | 0.33 | 0.06 | 0.39 | 0.27 | 0.46 | 0.32 |
| 1 | 182264659 | rs2494466 | -0.643 | 0.041 | -0.566 | -0.49 | -0.008 | -0.012 | -0.012 | 0 | 0.45 | 0.52 | 0.34 | 0.32 | 0.37 | 0.46 |
| 1 | 182266517 | rs2477185 | 0.484 | 0.063 | -0.093 | 0.191 | 0.001 | 0 | 0 | 0.812 | 0.63 | 0.32 | 0.63 | 0.29 | 0.77 | 0.59 |
| 1 | 182267259 | rs3118146 | 0.494 | 0.061 | -0.094 | 0.19 | 0.001 | 0 | 0 | 1.318 | 0.63 | 0.32 | 0.63 | 0.29 | 0.77 | 0.59 |
| 1 | 182270803 | rs4652696 | 0.548 | 0.107 | -0.369 | 0.157 | 0.003 | -0.009 | -0.009 | 0.244 | 0.14 | 0.09 | 0.08 | 0 | 0.12 | 0.06 |
| 1 | 182282817 | rs10911051 | -0.294 | 0.497 | 0.104 | 1.056 | -0.025 | 0.065 | 0.065 | 0 | 0.15 | 0.35 | 0.12 | 0.29 | 0.08 | 0.19 |
| 1 | 182283352 | rs2494467 | -0.158 | 0.534 | 0.05 | 1.08 | -0.003 | 0.046 | 0.046 | 0.319 | 0.77 | 0.66 | 0.75 | 0.58 | 0.82 | 0.77 |
| 1 | 182291435 | rs6703548 | -1.155 | 0.728 | 0.007 | 0.949 | -0.024 | 0.065 | 0.065 | 0 | 0.15 | 0.35 | 0.13 | 0.29 | 0.08 | 0.19 |
| 1 | 182301508 | rs4075837 | -1.373 | 1.873 | -0.276 | 0.679 | 0.042 | 0.005 | 0.005 | 0.121 | 0.14 | 0.25 | 0.21 | 0.15 | 0.25 | 0.21 |
| 1 | 182305147 | rs2985441 | -1.744 | 2.083 | -0.417 | 0.434 | 0.007 | -0.04 | -0.04 | 0.003 | 0.1 | 0.16 | 0.07 | 0 | 0.13 | 0.08 |
| 1 | 182305938 | rs2985440 | -0.789 | 1.748 | -0.55 | -0.069 | 0.003 | 0.018 | 0.018 | 0.008 | 0.43 | 0.64 | 0.37 | 0.21 | 0.38 | 0.54 |
| 1 | 182305990 | rs2985439 | -1.235 | 1.742 | -0.346 | -0.05 | 0.033 | 0.061 | 0.061 | 0.08 | 0.51 | 0.83 | 0.55 | 0.38 | 0.56 | 0.68 |
| 1 | 182308962 | rs12022482 | 0.894 | 1.793 | -1.018 | 0.07 | -0.018 | -0.128 | -0.128 | 0 | 0.23 | 0.01 | 0.27 | 0.36 | 0.15 | 0.17 |
| 1 | 182309477 | rs16858860 | 0.612 | 1.841 | -0.918 | 0.064 | 0.018 | -0.032 | -0.032 | 0 | 0.06 | 0.14 | 0.05 | 0 | 0.1 | 0.08 |
| 1 | 182309601 | rs2477179 | -0.121 | 1.77 | -0.897 | -0.021 | -0.005 | -0.009 | -0.009 | 0 | 0.68 | 0.73 | 0.54 | 0.47 | 0.68 | 0.7 |
| 1 | 182312476 | rs10797768 | 0.403 | 1.555 | -1.185 | -0.596 | 0 | -0.003 | -0.003 | 0 | 0.47 | 0.49 | 0.46 | 0.63 | 0.53 | 0.51 |
| 1 | 182315534 | rs2246119 | -0.188 | 1.558 | -1.286 | -0.563 | 0.008 | 0.001 | 0.001 | 0 | 0.42 | 0.23 | 0.45 | 0.37 | 0.35 | 0.38 |
| 1 | 182318783 | rs10911063 | 1.061 | 1.521 | -1.314 | -0.569 | 0.01 | -0.052 | -0.052 | 1.478 | 0.23 | 0.09 | 0.27 | 0.36 | 0.16 | 0.15 |
| 1 | 182319534 | rs11587233 | 1.07 | 1.515 | -1.453 | -0.59 | -0.045 | 0.105 | 0.105 | 0 | 0.17 | 0.02 | 0.17 | 0.01 | 0.16 | 0.23 |
| 1 | 182319935 | rs2256956 | -0.518 | 1.645 | -1.421 | -0.555 | -0.022 | -0.032 | -0.032 | 0 | 0.7 | 0.79 | 0.55 | 0.47 | 0.66 | 0.71 |
| 1 | 182327842 | rs2494461 | -0.176 | 1.737 | -1.48 | -0.547 | -0.082 | 0.031 | 0.031 | 0 | 0.76 | 0.9 | 0.66 | 0.83 | 0.66 | 0.61 |
| 1 | 182331836 | rs2252633 | -0.609 | 1.74 | -1.405 | -0.448 | -0.024 | -0.02 | -0.02 | 0 | 0.92 | 0.94 | 0.83 | 0.84 | 0.82 | 0.83 |
| 1 | 182333131 | rs2252483 | 0.532 | 1.419 | -1.312 | -0.542 | 0.029 | 0.014 | 0.014 | 0 | 0.47 | 0.67 | 0.34 | 0.45 | 0.44 | 0.41 |
| 1 | 182358944 | rs12136955 | 0.99 | 1.431 | 0.136 | -0.646 | -0.007 | 0 | 0 | 0 | 0.39 | 0.39 | 0.28 | 0.44 | 0.33 | 0.35 |
| 1 | 182360315 | rs12403634 | 1.028 | 1.607 | -0.41 | -0.703 | -0.031 | -0.131 | -0.131 | 0 | 0.23 | 0.01 | 0.29 | 0.39 | 0.19 | 0.19 |
| 1 | 182367548 | rs2038891 | 0.376 | 1.704 | -0.187 | -0.714 | -0.048 | 0.043 | 0.043 | 0 | 0.68 | 0.76 | 0.61 | 0.83 | 0.54 | 0.6 |
| 1 | 182367893 | rs6424853 | 0.376 | 1.704 | -0.185 | -0.714 | -0.05 | 0.045 | 0.045 | 0 | 0.68 | 0.76 | 0.61 | 0.83 | 0.54 | 0.6 |
| 1 | 182369078 | rs6698722 | -0.534 | 1.847 | -0.22 | -0.759 | -0.03 | -0.028 | -0.028 | 0 | 0.91 | 0.96 | 0.82 | 0.84 | 0.82 | 0.8 |
| 1 | 182369233 | rs6674281 | 0.527 | 1.84 | -0.223 | -0.764 | -0.048 | 0.043 | 0.043 | 0 | 0.68 | 0.76 | 0.61 | 0.83 | 0.54 | 0.6 |
| 1 | 182369738 | rs7551829 | -0.656 | 1.844 | -0.29 | -0.756 | -0.029 | -0.027 | -0.027 | 3.134 | 0.91 | 0.96 | 0.82 | 0.84 | 0.82 | 0.79 |
| 1 | 182370774 | rs1925829 | -0.227 | 1.841 | -0.297 | -0.764 | -0.042 | -0.03 | -0.03 | 0.262 | 0.88 | 0.96 | 0.81 | 0.84 | 0.79 | 0.79 |
| 1 | 182372452 | rs2148026 | 0.498 | 1.751 | -0.304 | -0.77 | -0.074 | 0.103 | 0.103 | 0 | 0.19 | 0.07 | 0.18 | 0.01 | 0.25 | 0.19 |
| 1 | 182376260 | rs3818458 | 0.923 | 1.879 | -0.35 | -0.792 | -0.019 | -0.133 | -0.133 | 0.872 | 0.25 | 0.01 | 0.3 | 0.38 | 0.2 | 0.21 |
| 1 | 182383133 | rs6704116 | 0.447 | 1.986 | -0.167 | -0.768 | -0.057 | 0.076 | 0.076 | 0 | 0.19 | 0.1 | 0.19 | 0.01 | 0.25 | 0.19 |
| 1 | 182386012 | rs267856 | -1.134 | 2.396 | -0.159 | -0.841 | -0.024 | 0.025 | 0.025 | 0 | 0.3 | 0.24 | 0.32 | 0.17 | 0.39 | 0.38 |
| 1 | 182388872 | rs267854 | -1.246 | 2.01 | -0.05 | -1.021 | -0.009 | -0.052 | -0.052 | 0 | 0.38 | 0.5 | 0.39 | 0.18 | 0.33 | 0.37 |
| 1 | 182392412 | rs267873 | -1.073 | 1.719 | -0.139 | -0.664 | -0.011 | 0.05 | 0.05 | 0.083 | 0.77 | 0.89 | 0.66 | 0.9 | 0.77 | 0.84 |
| 1 | 182397765 | rs6692352 | 1.688 | 1.584 | 0.056 | -0.549 | 0.009 | -0.027 | -0.027 | 0.188 | 0.33 | 0.2 | 0.21 | 0.46 | 0.34 | 0.34 |
| 1 | 182398201 | rs6659412 | 1.208 | 1.582 | 0.049 | -0.623 | 0.003 | 0.004 | 0.004 | 0.573 | 0.39 | 0.54 | 0.27 | 0.72 | 0.35 | 0.43 |
| 1 | 182400213 | rs11585935 | 0.313 | 1.545 | -0.017 | -0.487 | -0.051 | 0.076 | 0.076 | 0 | 0.19 | 0.02 | 0.2 | 0.05 | 0.17 | 0.17 |
| 1 | 182400696 | rs169154 | -1.103 | 1.497 | 0.098 | -0.459 | -0.012 | 0.024 | 0.024 | 0 | 0.38 | 0.37 | 0.4 | 0.18 | 0.43 | 0.4 |
| 1 | 182404101 | rs6667166 | 1.332 | 1.486 | -0.732 | -0.436 | 0.002 | -0.027 | -0.027 | 0 | 0.38 | 0.23 | 0.25 | 0.47 | 0.39 | 0.4 |
| 1 | 182406654 | rs497915 | -1.185 | 1.498 | -1.36 | -0.512 | -0.003 | 0.03 | 0.03 | 0.331 | 0.8 | 0.92 | 0.69 | 0.89 | 0.82 | 0.83 |
| 1 | 182410409 | rs6677117 | 1.052 | 1.599 | -1.318 | -0.181 | 0.01 | -0.076 | -0.076 | 0 | 0.37 | 0.15 | 0.26 | 0.53 | 0.38 | 0.49 |
| 1 | 182420448 | rs267866 | 0.305 | 2.106 | -0.518 | 0.339 | 0.055 | 0.108 | 0.108 | 0 | 0.92 | 0.79 | 0.92 | 0.76 | 0.88 | 0.88 |
| 1 | 182422667 | rs12744324 | -0.754 | 1.495 | -1.004 | -0.019 | -0.026 | 0.018 | 0.018 | 0 | 0.12 | 0.03 | 0.11 | 0.07 | 0.17 | 0.1 |
| 1 | 182424985 | rs155552 | 0.501 | 1.336 | -0.77 | 0.718 | 0.001 | -0.007 | -0.007 | 0 | 0.46 | 0.34 | 0.35 | 0.53 | 0.48 | 0.49 |
| 1 | 182429728 | rs266508 | -0.098 | 1.078 | -0.639 | 0.424 | -0.021 | 0.06 | 0.06 | 0.281 | 0.72 | 0.84 | 0.63 | 0.94 | 0.68 | 0.82 |
| 1 | 182433987 | rs2477875 | -0.239 | 1.125 | -0.517 | 0.831 | 0.002 | 0.093 | 0.093 | 0 | 0.06 | 0.11 | 0.15 | 0.25 | 0.06 | 0.07 |
| 1 | 182443462 | rs649437 | -1.124 | 0.553 | -0.13 | 0.575 | 0.001 | 0 | 0 | 0 | 0.45 | 0.55 | 0.53 | 0.46 | 0.35 | 0.48 |
| 1 | 182446188 | rs4271175 | 1.135 | 0.467 | -0.266 | 0.492 | -0.008 | 0.097 | 0.097 | 0 | 0.23 | 0.02 | 0.27 | 0.08 | 0.2 | 0.21 |
| 1 | 182457029 | rs12024567 | 0.217 | 0.407 | 0.382 | 0.532 | -0.023 | 0.007 | 0.007 | 0 | 0.11 | 0.17 | 0.09 | 0.14 | 0.08 | 0.19 |

| | | | | | | | | | | | | | | | | |
|---|---|---|---|---|---|---|---|---|---|---|---|---|---|---|---|---|
| 1 | 182474029 | rs10797777 | 1.472 | 0.668 | 0.345 | 0.448 | 0.002 | 0.011 | 0.011 | 0.168 | 0.5 | 0.42 | 0.39 | 0.54 | 0.52 | 0.49 |
| 1 | 182474160 | rs563988 | 0.55 | 0.589 | 0.49 | 0.564 | -0.012 | 0.057 | 0.057 | 0 | 0.84 | 0.81 | 0.77 | 0.62 | 0.87 | 0.74 |
| 1 | 182474704 | rs471910 | 0.55 | 0.589 | 0.49 | 0.564 | -0.012 | 0.057 | 0.057 | 0 | 0.84 | 0.81 | 0.77 | 0.62 | 0.87 | 0.74 |
| 1 | 182494063 | rs513553 | 0.498 | 0.615 | 0.406 | 0.581 | 0.005 | -0.007 | -0.007 | 0 | 0.85 | 0.88 | 0.79 | 0.61 | 0.87 | 0.74 |
| 1 | 182496829 | rs7535533 | 1.804 | 0.855 | 0.344 | 0.522 | 0.003 | -0.039 | -0.039 | 0 | 0.36 | 0.19 | 0.24 | 0.43 | 0.39 | 0.32 |
| 1 | 182497021 | rs266511 | 0.788 | 0.746 | 0.345 | 0.583 | -0.004 | -0.001 | -0.001 | 0.262 | 0.55 | 0.45 | 0.47 | 0.54 | 0.66 | 0.56 |
| 1 | 182511398 | rs4119585 | 0.684 | 0.824 | 0.435 | 0.73 | -0.001 | -0.001 | -0.001 | 3.476 | 0.56 | 0.43 | 0.4 | 0.44 | 0.54 | 0.54 |
| 1 | 182519861 | rs185580 | -0.184 | 0.877 | -0.013 | 0.934 | 0 | 0.074 | 0.074 | 0 | 0.93 | 0.92 | 0.85 | 0.65 | 0.92 | 0.9 |
| 1 | 182520612 | rs266539 | -1.527 | 0.852 | 0.441 | 0.745 | 0.02 | -0.052 | -0.052 | 0.195 | 0.2 | 0.4 | 0.17 | 0.12 | 0.22 | 0.16 |
| 1 | 182523591 | rs266547 | -0.892 | 0.949 | 0.387 | 0.689 | -0.002 | -0.029 | -0.029 | 0 | 0.44 | 0.47 | 0.44 | 0.21 | 0.42 | 0.4 |
| 1 | 182526839 | rs501085 | 1.076 | 0.92 | 0.408 | 0.699 | -0.002 | 0.005 | 0.005 | 0.025 | 0.48 | 0.39 | 0.4 | 0.45 | 0.5 | 0.5 |
| 1 | 182527409 | rs10911097 | 1.216 | 1.015 | -0.559 | 0.75 | 0.005 | 0.107 | 0.107 | 0.232 | 0.26 | 0.02 | 0.27 | 0.08 | 0.21 | 0.24 |
| 1 | 182528269 | rs266549 | -0.209 | 1.036 | -0.523 | 0.788 | 0 | 0.072 | 0.072 | 0 | 0.93 | 0.92 | 0.85 | 0.65 | 0.92 | 0.9 |
| 1 | 182536498 | rs10911098 | -0.479 | 1.029 | -0.537 | 0.249 | 0.002 | 0.091 | 0.091 | 0 | 0.42 | 0.26 | 0.37 | 0.2 | 0.42 | 0.39 |
| 1 | 182537494 | rs17568993 | 0.486 | 1.19 | -0.581 | 0.363 | 0.042 | -0.028 | -0.028 | 0 | 0.06 | 0.12 | 0.07 | 0 | 0.13 | 0.02 |
| 1 | 182538063 | rs12034992 | -0.182 | 0.673 | -0.972 | 0.518 | 0.009 | -0.001 | -0.001 | 2.936 | 0.1 | 0.04 | 0.08 | 0.12 | 0.09 | 0.14 |
| 1 | 182538615 | rs474939 | 0.042 | 0.744 | -0.933 | 1.053 | 0.001 | 0.107 | 0.107 | 0 | 0.92 | 0.9 | 0.84 | 0.51 | 0.93 | 0.9 |
| 1 | 182540149 | rs10911099 | -0.18 | 0.475 | -1.172 | 0.249 | 0.004 | 0.004 | 0.004 | 0 | 0.11 | 0.04 | 0.1 | 0.12 | 0.11 | 0.17 |
| 1 | 182543331 | rs11072 | 0.148 | 0.721 | -1.042 | 0.264 | 0.036 | 0.091 | 0.091 | 0 | 0.38 | 0.16 | 0.37 | 0.21 | 0.31 | 0.38 |
| 1 | 182545221 | rs672527 | 0.862 | 0.798 | -1.299 | 0.224 | -0.049 | 0.071 | 0.071 | 0 | 0.26 | 0.04 | 0.19 | 0.08 | 0.29 | 0.18 |
| 1 | 182549019 | rs533259 | 0.613 | 0.624 | -0.06 | 0.311 | 0.083 | 0.068 | 0.068 | 0 | 0.93 | 0.68 | 0.95 | 0.95 | 0.94 | 0.93 |
| 1 | 182551337 | rs627928 | -0.49 | 0.616 | -0.843 | 0.137 | 0.007 | 0.07 | 0.07 | 0 | 0.44 | 0.26 | 0.44 | 0.71 | 0.57 | 0.47 |
| 1 | 182554557 | rs486907 | 0.688 | 1.205 | -1.458 | 0.524 | -0.075 | -0.018 | -0.018 | 0 | 0.28 | 0.07 | 0.22 | 0.24 | 0.37 | 0.3 |
| 1 | 182556036 | rs3738579 | 0.727 | 1.207 | -1.48 | 0.536 | -0.076 | -0.026 | -0.026 | 0 | 0.26 | 0.07 | 0.21 | 0.25 | 0.36 | 0.3 |
| 1 | 182559510 | rs682585 | 1.254 | 0.687 | -1.546 | 0.335 | 0.022 | 0.166 | 0.166 | 0.391 | 0.45 | 0.08 | 0.47 | 0.24 | 0.38 | 0.46 |
| 1 | 182565274 | rs942436 | 0.02 | 0.836 | -0.536 | 0.378 | 0.037 | -0.001 | -0.001 | 0 | 0.69 | 0.55 | 0.73 | 0.71 | 0.6 | 0.67 |
| 1 | 182567440 | rs4652733 | -0.097 | 0.997 | -1.349 | 0.348 | -0.075 | -0.012 | -0.012 | 0 | 0.28 | 0.07 | 0.21 | 0.24 | 0.37 | 0.3 |
| 1 | 182567739 | rs3795485 | -0.097 | 0.997 | -1.349 | 0.348 | -0.075 | -0.012 | -0.012 | 0 | 0.28 | 0.07 | 0.21 | 0.24 | 0.37 | 0.3 |
| 1 | 182575376 | rs10489966 | 0.093 | 1.01 | -1.259 | 0.437 | -0.064 | -0.012 | -0.012 | 1.401 | 0.26 | 0.08 | 0.21 | 0.25 | 0.36 | 0.3 |
| 1 | 182579035 | rs516567 | -0.017 | 0.128 | -0.836 | 0.437 | -0.001 | -0.001 | -0.001 | 0 | 0.42 | 0.41 | 0.5 | 0.3 | 0.55 | 0.57 |
| 1 | 182589991 | rs1287816 | 1.581 | 0.129 | -0.305 | 0.795 | 0.019 | 0.069 | 0.069 | 0.782 | 0.8 | 0.39 | 0.74 | 0.63 | 0.78 | 0.82 |
| 1 | 182592336 | rs528846 | -1.353 | 0.06 | -0.129 | 0.834 | 0.044 | -0.015 | -0.015 | 0.062 | 0.1 | 0.39 | 0.09 | 0.04 | 0.11 | 0.05 |
| 1 | 182596728 | rs672684 | -1.275 | -0.537 | -0.39 | 0.941 | -0.013 | 0.051 | 0.051 | 0 | 0.11 | 0.18 | 0.16 | 0.33 | 0.11 | 0.11 |
| 1 | 182600851 | rs661506 | -1.403 | -0.648 | -0.3 | 0.616 | -0.004 | 0.006 | 0.006 | 0 | 0.11 | 0.13 | 0.16 | 0.15 | 0.12 | 0.07 |
| 1 | 182623509 | rs10797788 | -0.373 | -1.147 | -0.836 | 0.708 | 0.03 | 0.014 | 0.014 | 0 | 0.65 | 0.73 | 0.73 | 0.72 | 0.76 | 0.78 |
| 1 | 182626738 | rs4652741 | -0.148 | -1.036 | -0.886 | 0.519 | -0.002 | -0.001 | -0.001 | 0 | 0.61 | 0.55 | 0.63 | 0.61 | 0.74 | 0.75 |
| 1 | 182642857 | rs6670735 | -0.259 | 0.363 | -1.311 | 0.093 | -0.164 | 0.071 | 0.071 | 0 | 0.28 | 0.02 | 0.33 | 0.13 | 0.36 | 0.34 |
| 1 | 182652564 | rs6424866 | -1.338 | -0.41 | -0.167 | 0.617 | -0.052 | 0.038 | 0.038 | 0 | 0.18 | 0.48 | 0.12 | 0.18 | 0.09 | 0.1 |
| 1 | 182664258 | rs6672020 | -0.746 | -0.129 | -0.373 | 0.853 | -0.022 | 0.081 | 0.081 | 0 | 0.05 | 0.16 | 0.03 | 0.15 | 0.02 | 0.04 |
| 1 | 182675034 | rs6680242 | -0.664 | -0.061 | -0.581 | 0.41 | 0 | -0.009 | -0.009 | 0 | 0.07 | 0.09 | 0.02 | 0 | 0.05 | 0.03 |
| 1 | 182679351 | rs508607 | -0.183 | -0.279 | -0.816 | 0.44 | 0.003 | -0.012 | -0.012 | 2.227 | 0.53 | 0.76 | 0.44 | 0.48 | 0.49 | 0.49 |
| 1 | 182684087 | rs2482806 | 0.294 | -0.371 | -0.356 | 0.458 | 0.02 | -0.01 | -0.01 | 0 | 0.46 | 0.67 | 0.42 | 0.5 | 0.44 | 0.46 |
| 1 | 182700570 | rs2482808 | -0.564 | 0.044 | -0.676 | 0.353 | -0.058 | -0.036 | -0.036 | 0.414 | 0.68 | 0.86 | 0.69 | 0.61 | 0.64 | 0.7 |
| 1 | 182705001 | rs490285 | 0.942 | -0.24 | -0.458 | 0.473 | -0.004 | 0.002 | 0.002 | 0 | 0.4 | 0.48 | 0.48 | 0.43 | 0.43 | 0.51 |
| 1 | 182727061 | rs11805728 | 0.39 | -0.23 | -0.673 | 0.47 | -0.028 | -0.014 | -0.014 | 0 | 0.34 | 0.19 | 0.3 | 0.38 | 0.35 | 0.31 |
| 1 | 182727253 | rs10911129 | 0.381 | -0.229 | -0.673 | 0.47 | -0.028 | -0.013 | -0.013 | 0.483 | 0.34 | 0.19 | 0.3 | 0.38 | 0.35 | 0.31 |
| 1 | 182728887 | rs16859619 | 0.515 | -0.247 | -0.743 | 0.538 | -0.003 | 0 | 0 | 0 | 0.08 | 0.06 | 0.04 | 0.05 | 0.08 | 0.07 |
| 1 | 182731008 | rs6660584 | -0.062 | -0.134 | -0.444 | 0.456 | -0.028 | 0.006 | 0.006 | 0 | 0.4 | 0.27 | 0.32 | 0.38 | 0.4 | 0.34 |
| 1 | 182740586 | rs12118059 | -1.097 | -0.147 | -0.13 | 0.539 | -0.019 | 0.012 | 0.012 | 0 | 0.33 | 0.58 | 0.25 | 0.34 | 0.23 | 0.24 |
| 1 | 182740955 | rs3904695 | 1.343 | -0.399 | 0.135 | 0.519 | 0 | -0.002 | -0.002 | 0 | 0.3 | 0.19 | 0.26 | 0.32 | 0.36 | 0.33 |
| 1 | 182744205 | rs7540658 | -1.857 | 0.246 | 0.638 | 0.567 | -0.001 | 0.007 | 0.007 | 0 | 0.35 | 0.58 | 0.28 | 0.35 | 0.3 | 0.25 |
| 1 | 182746231 | rs10797793 | -0.96 | -0.098 | -0.792 | 0.391 | 0.005 | -0.019 | -0.019 | 0 | 0.2 | 0.06 | 0.19 | 0.18 | 0.16 | 0.17 |
| 1 | 182759592 | rs6678876 | -1.221 | -0.257 | -0.013 | 0.177 | -0.035 | -0.028 | -0.028 | 0.545 | 0.22 | 0.64 | 0.17 | 0.13 | 0.11 | 0.24 |
| 1 | 182760730 | rs12066158 | -0.913 | -0.328 | 0.657 | 0.128 | -0.068 | -0.004 | -0.004 | 0.776 | 0.17 | 0.5 | 0.11 | 0.13 | 0.06 | 0.21 |
| 1 | 182760859 | rs3904696 | -0.93 | -0.31 | 0.58 | 0.133 | 0.016 | -0.026 | -0.026 | 0 | 0.05 | 0.14 | 0.06 | 0 | 0.04 | 0.03 |
| 1 | 182766498 | rs7554927 | -1.566 | 0.068 | 0.765 | 0.136 | -0.029 | -0.03 | -0.03 | 0 | 0.21 | 0.62 | 0.18 | 0.13 | 0.13 | 0.25 |
| 1 | 182784022 | rs2275171 | -0.913 | -0.17 | 0.016 | -0.048 | -0.002 | 0.008 | 0.008 | 0 | 0.56 | 0.76 | 0.51 | 0.38 | 0.57 | 0.63 |
| 1 | 182786953 | rs12080837 | 0.049 | -0.27 | 0.823 | -0.11 | 0.001 | -0.005 | -0.005 | 0.054 | 0.06 | 0.27 | 0.04 | 0.01 | 0.01 | 0.06 |
| 1 | 182798924 | rs1045971 | -1.43 | -0.285 | 0.811 | -0.118 | -0.056 | -0.029 | -0.029 | 0 | 0.2 | 0.61 | 0.16 | 0.13 | 0.08 | 0.23 |
| 1 | 182834817 | rs13373977 | 0.101 | -0.295 | 1.24 | -0.121 | -0.013 | 0.001 | 0.001 | 0 | 0.11 | 0.14 | 0.06 | 0.12 | 0.05 | 0.16 |
| 1 | 182839174 | rs13375165 | 0.081 | -0.294 | 1.271 | -0.121 | -0.018 | 0.002 | 0.002 | 0 | 0.11 | 0.16 | 0.06 | 0.12 | 0.05 | 0.16 |
| 1 | 182854017 | rs3130492 | -0.799 | -0.108 | 0.986 | -0.145 | -0.121 | 0.134 | 0.134 | 0 | 0.68 | 0.94 | 0.78 | 0.9 | 0.56 | 0.68 |
| 1 | 182865717 | rs6424873 | -1.355 | -0.422 | 1.901 | 0.551 | -0.041 | -0.003 | -0.003 | 0 | 0.15 | 0.33 | 0.12 | 0.12 | 0.07 | 0.17 |
| 1 | 182867528 | rs12073553 | -0.462 | -0.398 | 1.841 | 0.578 | 0.007 | 0 | 0 | 0.002 | 0.06 | 0.28 | 0.04 | 0.01 | 0.01 | 0.06 |
| 1 | 182878158 | rs6681606 | -0.608 | -0.012 | 0.871 | 0.796 | -0.028 | 0.005 | 0.005 | 0 | 0.12 | 0.2 | 0.07 | 0.12 | 0.05 | 0.16 |
| 1 | 182922975 | rs3814325 | -1.166 | -0.047 | 0.629 | 0.608 | -0.04 | -0.002 | -0.002 | 0.038 | 0.15 | 0.33 | 0.13 | 0.12 | 0.08 | 0.16 |
| 1 | 182931205 | rs10797795 | -1.615 | 0.299 | 0.46 | 1.036 | 0.004 | 0.036 | 0.036 | 0 | 0.14 | 0.17 | 0.28 | 0.32 | 0.15 | 0.29 |
| 1 | 182938102 | rs6664353 | -2.041 | -0.079 | 2.34 | 0.929 | -0.004 | 0.115 | 0.115 | 0 | 0.16 | 0.59 | 0.32 | 0.32 | 0.14 | 0.26 |
| 1 | 182971816 | rs10737235 | -0.66 | -0.143 | 0.871 | 0.364 | 0.019 | 0.016 | 0.016 | 3.967 | 0.5 | 0.32 | 0.61 | 0.63 | 0.53 | 0.6 |

| | | | | | | | | | | | | | | | | |
|---|---|---|---|---|---|---|---|---|---|---|---|---|---|---|---|---|
| 1 | 182973024 | rs10911187 | 0.144 | -0.046 | 0.863 | 0.443 | -0.004 | -0.002 | -0.002 | 0.22 | 0.07 | 0.03 | 0.22 | 0.11 | 0.11 | 0.2 |
| 1 | 182979560 | rs12138053 | 0.084 | -0.332 | 0.751 | 0.296 | -0.028 | -0.059 | -0.059 | 0.047 | 0.38 | 0.14 | 0.36 | 0.52 | 0.41 | 0.33 |
| 1 | 183018062 | rs16860009 | -0.383 | -0.316 | 0.815 | 0.309 | -0.007 | -0.019 | -0.019 | 0 | 0.06 | 0.13 | 0.03 | 0 | 0.02 | 0.07 |
| 1 | 183025785 | rs10911214 | 0.15 | -0.315 | 0.796 | 0.301 | -0.017 | -0.008 | -0.008 | 3.347 | 0.07 | 0.02 | 0.22 | 0.1 | 0.11 | 0.2 |
| 1 | 183028226 | rs5015713 | 0.02 | -0.317 | 0.813 | 0.299 | 0.012 | 0.003 | 0.003 | 0 | 0.07 | 0.07 | 0.22 | 0.11 | 0.11 | 0.2 |
| 1 | 183037695 | rs6424883 | 0.127 | -0.293 | 0.792 | 0.32 | -0.047 | -0.076 | -0.076 | 0.188 | 0.37 | 0.13 | 0.35 | 0.52 | 0.41 | 0.33 |
| 1 | 183046065 | rs6666744 | 0.019 | -0.301 | 0.818 | 0.348 | 0.012 | 0.003 | 0.003 | 0 | 0.07 | 0.07 | 0.22 | 0.11 | 0.11 | 0.2 |
| 1 | 183061890 | rs12073936 | -0.013 | -0.155 | 0.629 | 0.531 | -0.018 | 0.034 | 0.034 | 0 | 0.06 | 0 | 0.13 | 0.43 | 0.08 | 0.09 |
| 1 | 183071024 | rs10752897 | -0.757 | -0.243 | 0.464 | 0.299 | 0.014 | 0.005 | 0.005 | 3.513 | 0.48 | 0.2 | 0.6 | 0.63 | 0.54 | 0.53 |
| 1 | 183072590 | rs2296288 | -1.18 | -0.219 | 0.489 | 0.335 | 0.014 | -0.001 | -0.001 | 0.003 | 0.54 | 0.33 | 0.63 | 0.63 | 0.56 | 0.6 |
| 1 | 183072701 | rs2296289 | 0.272 | -0.165 | 0.485 | 0.353 | -0.017 | -0.008 | -0.008 | 0.918 | 0.07 | 0.02 | 0.22 | 0.11 | 0.11 | 0.2 |
| 1 | 183086757 | rs2296292 | -1.067 | -0.141 | 0.52 | 0.358 | 0.016 | 0.002 | 0.002 | 2.668 | 0.54 | 0.33 | 0.63 | 0.63 | 0.56 | 0.6 |
| 1 | 183090011 | rs3768622 | -1.172 | -0.165 | 0.771 | 0.346 | 0 | -0.001 | -0.001 | 0.021 | 0.55 | 0.46 | 0.64 | 0.63 | 0.56 | 0.6 |
| 1 | 183094547 | rs20558 | -1.061 | -0.158 | 0.666 | 0.349 | 0.016 | 0.002 | 0.002 | 2.668 | 0.54 | 0.33 | 0.63 | 0.63 | 0.56 | 0.6 |
| 1 | 183106739 | rs10797854 | -0.947 | -0.317 | 0.714 | 0.377 | 0.018 | -0.005 | -0.005 | 0 | 0.53 | 0.3 | 0.63 | 0.62 | 0.56 | 0.59 |
| 1 | 183112412 | rs1062044 | -0.963 | -0.333 | 0.726 | 0.389 | 0.007 | -0.006 | -0.006 | 0 | 0.55 | 0.4 | 0.64 | 0.62 | 0.57 | 0.59 |
| 1 | 183112505 | rs944971 | -0.963 | -0.333 | 0.726 | 0.389 | 0.007 | -0.006 | -0.006 | 0 | 0.55 | 0.4 | 0.64 | 0.62 | 0.57 | 0.59 |
| 1 | 183113862 | rs3738829 | -0.97 | -0.301 | 0.708 | 0.38 | 0.007 | -0.007 | -0.007 | 0 | 0.55 | 0.4 | 0.64 | 0.62 | 0.57 | 0.59 |
| 1 | 183113952 | rs1547715 | -0.961 | -0.3 | 0.708 | 0.381 | 0.007 | -0.006 | -0.006 | 0 | 0.55 | 0.4 | 0.64 | 0.62 | 0.57 | 0.59 |
| 1 | 183114119 | rs1547714 | -0.961 | -0.3 | 0.708 | 0.381 | 0.007 | -0.006 | -0.006 | 0 | 0.55 | 0.4 | 0.64 | 0.62 | 0.57 | 0.59 |
| 1 | 183114634 | rs7473 | -0.961 | -0.3 | 0.708 | 0.381 | 0.007 | -0.006 | -0.006 | 0 | 0.55 | 0.4 | 0.64 | 0.62 | 0.57 | 0.59 |
| 1 | 183121028 | rs2333618 | -0.013 | -0.436 | 0.899 | 0.284 | -0.004 | 0.007 | 0.007 | 0.809 | 0.31 | 0.36 | 0.3 | 0.48 | 0.33 | 0.25 |
| 1 | 183134719 | rs10911269 | -1.242 | -0.579 | 0.293 | 0.687 | 0.041 | 0.057 | 0.057 | 5.115 | 0.5 | 0.79 | 0.6 | 0.68 | 0.45 | 0.54 |
| 1 | 183136196 | rs4652781 | -1.691 | -0.488 | 0.065 | 0.874 | 0.001 | -0.007 | -0.007 | 0 | 0.2 | 0.2 | 0.29 | 0.26 | 0.16 | 0.32 |
| 1 | 183137568 | rs16860401 | -0.287 | -0.434 | -0.003 | 0.885 | -0.006 | -0.018 | -0.018 | 0 | 0.05 | 0.06 | 0.1 | 0.01 | 0.01 | 0.09 |
| 1 | 183143422 | rs17479287 | -0.037 | -0.328 | -0.17 | 0.992 | -0.012 | -0.008 | -0.008 | 0 | 0.07 | 0.03 | 0.05 | 0.08 | 0.05 | 0.16 |
| 1 | 183148228 | rs16860429 | 0.432 | -0.312 | -0.068 | 1.062 | -0.011 | -0.015 | -0.015 | 0 | 0.05 | 0.02 | 0.13 | 0.17 | 0.08 | 0.07 |
| 1 | 183155305 | rs2276543 | -0.201 | -0.589 | -0.12 | 1.306 | 0.006 | -0.006 | -0.006 | 0 | 0.29 | 0.18 | 0.39 | 0.38 | 0.29 | 0.32 |
| 1 | 183157924 | rs547905 | -1.264 | -0.56 | 0.484 | 1.304 | 0.004 | 0.045 | 0.045 | 1.7 | 0.49 | 0.71 | 0.61 | 0.7 | 0.49 | 0.54 |
| 1 | 183159377 | rs1005716 | -1.491 | -0.393 | 0.831 | 1.379 | 0.019 | 0.116 | 0.116 | 2.05 | 0.15 | 0.5 | 0.21 | 0.26 | 0.15 | 0.21 |
| 1 | 183169690 | rs661451 | -0.275 | -0.864 | 0.485 | 1.218 | 0.001 | 0.001 | 0.001 | 0 | 0.41 | 0.39 | 0.41 | 0.42 | 0.37 | 0.34 |
| 1 | 183172548 | rs10494560 | 0.539 | -0.654 | 0.479 | 1.475 | 0.006 | 0.061 | 0.061 | 0 | 0.06 | 0.11 | 0.15 | 0.2 | 0.1 | 0.08 |
| 1 | 183176430 | rs525410 | -0.332 | -0.855 | 0.365 | 1.228 | -0.013 | 0.002 | 0.002 | 0 | 0.42 | 0.33 | 0.47 | 0.61 | 0.51 | 0.35 |
| 1 | 183177218 | rs647347 | -0.712 | -0.978 | 0.344 | 1.203 | 0.001 | -0.001 | -0.001 | 1.129 | 0.48 | 0.47 | 0.5 | 0.57 | 0.46 | 0.49 |
| 1 | 183177454 | rs635796 | -1.184 | -1.118 | 0.656 | 1.044 | -0.026 | 0.044 | 0.044 | 0 | 0.56 | 0.78 | 0.65 | 0.73 | 0.55 | 0.56 |
| 1 | 183184616 | rs2274980 | -1.796 | -1.068 | 1.156 | 1.079 | -0.01 | 0.093 | 0.093 | 0.46 | 0.17 | 0.4 | 0.32 | 0.33 | 0.16 | 0.33 |
| 1 | 183189331 | rs6673920 | -1.54 | -0.65 | 1.407 | 1.307 | -0.002 | 0.047 | 0.047 | 0.183 | 0.12 | 0.26 | 0.26 | 0.18 | 0.11 | 0.19 |
| 1 | 183192304 | rs1047980 | -2.583 | -0.687 | 1.513 | 1.3 | -0.028 | 0.086 | 0.086 | 1.057 | 0.2 | 0.55 | 0.33 | 0.34 | 0.16 | 0.33 |
| 1 | 183192496 | rs3738824 | 0.314 | -0.666 | 1.534 | 1.304 | 0.007 | 0.024 | 0.024 | 0 | 0.06 | 0.11 | 0.14 | 0.14 | 0.1 | 0.08 |
| 1 | 183197305 | rs6658628 | -2.283 | -0.669 | 1.416 | 1.279 | -0.026 | 0.089 | 0.089 | 0.848 | 0.2 | 0.54 | 0.33 | 0.34 | 0.16 | 0.34 |
| 1 | 183198013 | rs3768597 | -1.051 | -0.626 | 1.148 | 1.221 | -0.011 | 0.02 | 0.02 | 0.007 | 0.07 | 0.25 | 0.06 | 0.08 | 0.04 | 0.14 |
| 1 | 183212548 | rs3768593 | -1.593 | -0.653 | 0.962 | 1.414 | -0.023 | 0.089 | 0.089 | 1.713 | 0.19 | 0.37 | 0.31 | 0.37 | 0.15 | 0.33 |
| 1 | 183220937 | rs549191 | -1.825 | -0.656 | 1.002 | 1.343 | -0.035 | 0.133 | 0.133 | 2.506 | 0.2 | 0.51 | 0.32 | 0.37 | 0.16 | 0.33 |
| 1 | 183253213 | rs607332 | 0.68 | 0.208 | 0.146 | 0.759 | 0.01 | -0.011 | -0.011 | 0.023 | 0.35 | 0.4 | 0.3 | 0.25 | 0.36 | 0.39 |
| 1 | 183257218 | rs2480767 | 0.608 | 0.203 | 0.042 | 0.757 | -0.012 | 0.01 | 0.01 | 0.232 | 0.35 | 0.28 | 0.29 | 0.25 | 0.36 | 0.39 |
| 1 | 183259125 | rs2301877 | 0.973 | -0.277 | -0.714 | 0.244 | -0.045 | -0.021 | -0.021 | 0 | 0.33 | 0.08 | 0.29 | 0.3 | 0.38 | 0.29 |
| 1 | 183260659 | rs946173 | 1.278 | -0.439 | -0.76 | -0.047 | 0.003 | 0.049 | 0.049 | 0 | 0.13 | 0.05 | 0.08 | 0 | 0.12 | 0.03 |
| 1 | 183260894 | rs11589702 | 1.442 | 0.17 | -0.385 | -0.042 | 0.008 | 0.015 | 0.015 | 0.065 | 0.29 | 0.23 | 0.2 | 0.06 | 0.19 | 0.29 |
| 1 | 183263356 | rs10752907 | 0.227 | 0.086 | -0.463 | 0.247 | -0.017 | 0.051 | 0.051 | 0.375 | 0.48 | 0.19 | 0.39 | 0.38 | 0.52 | 0.31 |
| 1 | 183266182 | rs4652795 | -0.468 | 0.215 | -0.122 | 0.397 | -0.008 | -0.016 | -0.016 | 0 | 0.58 | 0.31 | 0.61 | 0.52 | 0.63 | 0.46 |
| 1 | 183272295 | rs6687056 | 1.133 | -0.003 | 0.239 | 0.303 | -0.003 | -0.024 | -0.024 | 1.136 | 0.32 | 0.36 | 0.26 | 0.1 | 0.24 | 0.37 |
| 1 | 183272486 | rs12748895 | 0.547 | -0.067 | 0.299 | 0.284 | -0.041 | 0.028 | 0.028 | 0.042 | 0.4 | 0.15 | 0.36 | 0.33 | 0.46 | 0.33 |
| 1 | 183273751 | rs2276879 | 0.201 | -0.212 | 0.483 | 0.418 | -0.002 | -0.058 | -0.058 | 0 | 0.18 | 0.1 | 0.22 | 0.36 | 0.2 | 0.23 |
| 1 | 183279029 | rs602182 | 0.078 | -0.083 | 0.553 | 0.1 | 0.002 | -0.01 | -0.01 | 0 | 0.43 | 0.48 | 0.38 | 0.35 | 0.45 | 0.31 |
| 1 | 183281919 | rs2050701 | -1.729 | -0.538 | 0.777 | -0.751 | -0.004 | 0.011 | 0.011 | 0 | 0.69 | 0.75 | 0.73 | 0.92 | 0.7 | 0.71 |
| 1 | 183286777 | rs10911305 | -0.03 | -0.521 | 0.879 | -0.719 | -0.018 | 0.154 | 0.154 | 0 | 0.47 | 0.23 | 0.45 | 0.83 | 0.51 | 0.63 |
| 1 | 183298150 | rs616545 | -1.75 | -0.526 | 0.866 | -0.783 | -0.002 | 0.001 | 0.001 | 0 | 0.68 | 0.73 | 0.72 | 0.89 | 0.7 | 0.72 |
| 1 | 183298502 | rs10797875 | -0.004 | -0.539 | 1.026 | -0.796 | -0.015 | 0.128 | 0.128 | 0 | 0.46 | 0.27 | 0.45 | 0.81 | 0.51 | 0.63 |
| 1 | 183302111 | rs675220 | -0.575 | -0.574 | 1.005 | -0.728 | -0.015 | -0.082 | -0.082 | 0.134 | 0.22 | 0.46 | 0.26 | 0.1 | 0.19 | 0.09 |
| 1 | 183307942 | rs2788053 | -1.637 | -0.501 | 0.904 | -0.722 | -0.003 | 0.001 | 0.001 | 0 | 0.68 | 0.74 | 0.72 | 0.9 | 0.7 | 0.71 |
| 1 | 183317819 | rs3924298 | -0.007 | -0.592 | 0.45 | -0.671 | -0.012 | 0.14 | 0.14 | 0 | 0.44 | 0.14 | 0.47 | 0.77 | 0.58 | 0.49 |
| 1 | 183325722 | rs664422 | -1.157 | -0.731 | 0.69 | -0.66 | 0.005 | -0.055 | -0.055 | 2.7 | 0.63 | 0.54 | 0.58 | 0.82 | 0.62 | 0.65 |
| 1 | 183330413 | rs2811558 | 0.217 | -0.751 | 0.696 | -0.614 | -0.009 | 0.028 | 0.028 | 0 | 0.43 | 0.31 | 0.44 | 0.69 | 0.5 | 0.42 |
| 1 | 183337933 | rs542349 | -1.624 | -0.629 | 0.757 | -0.663 | -0.002 | -0.021 | -0.021 | 0.121 | 0.68 | 0.69 | 0.72 | 0.91 | 0.7 | 0.71 |
| 1 | 183339847 | rs17556785 | 1.142 | -0.737 | 0.451 | -0.614 | -0.001 | 0.073 | 0.073 | 0 | 0.14 | 0.01 | 0.11 | 0.01 | 0.12 | 0.04 |
| 1 | 183346669 | rs2811563 | -0.382 | -0.825 | 0.627 | -0.532 | 0.006 | 0.058 | 0.058 | 0.867 | 0.55 | 0.28 | 0.52 | 0.66 | 0.61 | 0.45 |
| 1 | 183353853 | rs2022013 | -0.409 | -0.829 | 0.616 | -0.561 | 0.005 | 0.061 | 0.061 | 0.873 | 0.55 | 0.26 | 0.52 | 0.66 | 0.61 | 0.45 |
| 1 | 183358405 | rs2078087 | -1.283 | -0.833 | 0.639 | -0.579 | -0.012 | 0.026 | 0.026 | 0.011 | 0.13 | 0.26 | 0.18 | 0.25 | 0.1 | 0.26 |
| 1 | 183358440 | rs1819628 | -1.552 | -0.722 | 0.511 | -0.606 | 0 | -0.045 | -0.045 | 0 | 0.7 | 0.7 | 0.72 | 0.92 | 0.7 | 0.72 |

| | | | | | | | | | | | | | | | | |
|---|---|---|---|---|---|---|---|---|---|---|---|---|---|---|---|---|
| 1 | 183373665 | rs3120798 | -1.536 | -0.704 | 0.377 | -0.586 | 0.009 | -0.102 | -0.102 | 0 | 0.69 | 0.58 | 0.71 | 0.9 | 0.7 | 0.71 |
| 1 | 183383643 | rs4652800 | 0.579 | -0.837 | 0.304 | -0.586 | -0.069 | 0.037 | 0.037 | 0.112 | 0.37 | 0.1 | 0.39 | 0.64 | 0.47 | 0.41 |
| 1 | 183384796 | rs2788045 | -1.017 | -0.743 | 0.407 | -0.434 | -0.001 | -0.057 | -0.057 | 0 | 0.64 | 0.57 | 0.69 | 0.86 | 0.69 | 0.63 |
| 1 | 183397899 | rs12144629 | -0.326 | -0.952 | 0.263 | -0.534 | -0.027 | 0.002 | 0.002 | 0.022 | 0.15 | 0.21 | 0.06 | 0.2 | 0.03 | 0.23 |
| 1 | 183400890 | rs2788056 | -1.337 | -0.507 | 0.283 | -0.649 | 0.006 | -0.086 | -0.086 | 0.558 | 0.68 | 0.6 | 0.71 | 0.9 | 0.7 | 0.72 |
| 1 | 183404466 | rs10752913 | -0.865 | -0.757 | 0.418 | -0.259 | -0.038 | -0.019 | -0.019 | 0.898 | 0.34 | 0.47 | 0.21 | 0.25 | 0.21 | 0.21 |
| 1 | 183415318 | rs4047801 | 0.135 | -0.486 | 0.189 | -0.121 | -0.004 | 0.009 | 0.009 | 0.556 | 0.54 | 0.69 | 0.51 | 0.47 | 0.49 | 0.52 |
| 1 | 183426758 | rs4652802 | 1.198 | -0.521 | 0.451 | -0.134 | -0.006 | 0.003 | 0.003 | 0.055 | 0.38 | 0.33 | 0.41 | 0.2 | 0.43 | 0.37 |
| 1 | 183439483 | rs2275675 | -0.989 | -0.511 | 0.385 | -0.121 | -0.021 | -0.031 | -0.031 | 1.92 | 0.73 | 0.76 | 0.61 | 0.47 | 0.63 | 0.54 |
| 1 | 183440720 | rs4047761 | 0.535 | -0.544 | 0.442 | -0.161 | -0.001 | -0.034 | -0.034 | 1.281 | 0.18 | 0.18 | 0.1 | 0 | 0.13 | 0.04 |
| 1 | 183449357 | rs2226099 | 1.875 | -0.546 | 0.233 | -0.319 | -0.084 | 0.182 | 0.182 | 0 | 0.32 | 0.06 | 0.19 | 0.09 | 0.34 | 0.28 |
| 1 | 183457396 | rs2702204 | 1.069 | -0.548 | 0.252 | -0.282 | -0.046 | -0.067 | -0.067 | 0 | 0.26 | 0.12 | 0.38 | 0.53 | 0.37 | 0.46 |
| 1 | 183486859 | rs789169 | -1.237 | -0.517 | 0.045 | -0.304 | 0.024 | 0.083 | 0.083 | 0 | 0.84 | 0.96 | 0.91 | 1 | 0.87 | 0.96 |
| 1 | 183496658 | rs2877978 | 1.884 | -0.492 | 0.002 | -0.305 | -0.101 | 0.193 | 0.193 | 0 | 0.32 | 0.04 | 0.19 | 0.09 | 0.34 | 0.28 |
| 1 | 183497627 | rs12144253 | 0.45 | -0.365 | 0.117 | -0.173 | 0.008 | -0.01 | -0.01 | 0 | 0.13 | 0.12 | 0.09 | 0.25 | 0.06 | 0.14 |
| 1 | 183515428 | rs2298083 | -0.614 | -0.363 | 0.212 | -0.462 | 0.077 | 0.102 | 0.102 | 0.49 | 0.07 | 0.29 | 0.22 | 0.11 | 0.1 | 0.08 |
| 1 | 183515777 | rs789177 | -0.796 | -0.364 | 0.246 | -0.462 | 0.103 | 0.131 | 0.131 | 0.878 | 0.08 | 0.35 | 0.23 | 0.11 | 0.1 | 0.08 |
| 1 | 183524855 | rs796860 | -0.921 | -0.389 | 0.089 | -0.543 | 0.029 | 0.057 | 0.057 | 0 | 0.83 | 0.95 | 0.91 | 1 | 0.87 | 0.96 |
| 1 | 183533910 | rs11811630 | -0.358 | 0.348 | 0.782 | 0.052 | 0.009 | -0.001 | -0.001 | 0.871 | 0.33 | 0.32 | 0.6 | 0.62 | 0.47 | 0.47 |
| 1 | 183542387 | rs2274064 | -0.409 | 0.336 | 0.791 | -0.068 | 0.008 | -0.001 | -0.001 | 0.552 | 0.33 | 0.33 | 0.6 | 0.62 | 0.47 | 0.53 |
| 1 | 183546061 | rs12753665 | -0.067 | 0.004 | 0.812 | 0.044 | 0.004 | 0.011 | 0.011 | 0 | 0.28 | 0.32 | 0.4 | 0.54 | 0.34 | 0.43 |
| 1 | 183548992 | rs10911362 | -1.067 | -0.228 | 0.418 | 0.43 | -0.048 | 0.046 | 0.046 | 1.173 | 0.15 | 0.27 | 0.1 | 0.27 | 0.06 | 0.15 |
| 1 | 183551760 | rs3845466 | -1.082 | -0.107 | 0.428 | 0.457 | -0.032 | 0.053 | 0.053 | 2.15 | 0.16 | 0.29 | 0.16 | 0.27 | 0.09 | 0.16 |
| 1 | 183554281 | rs789185 | 0.296 | 0.255 | 0.421 | -0.419 | 0.001 | -0.012 | -0.012 | 0 | 0.42 | 0.57 | 0.37 | 0.16 | 0.4 | 0.26 |
| 1 | 183555779 | rs11578964 | 0.27 | 0.405 | 0.399 | -0.713 | -0.009 | 0.166 | 0.166 | 3.622 | 0.43 | 0.2 | 0.47 | 0.16 | 0.43 | 0.31 |
| 1 | 183559704 | rs2274065 | -1.677 | 0.176 | 1.289 | -0.597 | 0.047 | 0.031 | 0.031 | 2.203 | 0.1 | 0.49 | 0.23 | 0.07 | 0.09 | 0.06 |
| 1 | 183561560 | rs12066019 | -0.39 | 0.263 | 1.194 | -0.555 | 0.013 | 0.007 | 0.007 | 0.161 | 0.06 | 0.14 | 0.2 | 0.07 | 0.08 | 0.07 |
| 1 | 183562782 | rs3754515 | 0.108 | 0.232 | 1.028 | -0.716 | -0.001 | -0.037 | -0.037 | 1.646 | 0.46 | 0.53 | 0.51 | 0.16 | 0.45 | 0.32 |
| 1 | 183565989 | rs2333685 | 0.239 | 0.033 | 0.985 | -0.674 | 0 | -0.022 | -0.022 | 0 | 0.47 | 0.44 | 0.35 | 0.19 | 0.4 | 0.34 |
| 1 | 183581914 | rs12069669 | -0.574 | 0.224 | 1.21 | -0.562 | 0.03 | -0.03 | -0.03 | 0.09 | 0.08 | 0.27 | 0.21 | 0.03 | 0.07 | 0.08 |
| 1 | 183596136 | rs11755 | -0.37 | 0.076 | 1.226 | -0.744 | -0.032 | 0.154 | 0.154 | 5.201 | 0.55 | 0.8 | 0.57 | 0.27 | 0.47 | 0.4 |
| 1 | 183596703 | rs2231241 | -0.48 | -0.048 | 1.697 | -0.809 | -0.014 | 0.009 | 0.009 | 0.153 | 0.08 | 0.2 | 0.04 | 0.11 | 0.03 | 0.07 |
| 1 | 183597537 | rs10911382 | -0.345 | -0.042 | 1.703 | -0.807 | 0.031 | -0.029 | -0.029 | 0 | 0.08 | 0.28 | 0.22 | 0.03 | 0.07 | 0.08 |
| 1 | 183602308 | rs12032504 | -0.755 | -0.129 | 1.706 | -0.844 | -0.013 | -0.003 | -0.003 | 0 | 0.09 | 0.14 | 0.03 | 0.11 | 0.03 | 0.07 |
| 1 | 183602795 | rs2254972 | -0.422 | -0.103 | 1.748 | -0.849 | -0.027 | 0.146 | 0.146 | 5.158 | 0.54 | 0.79 | 0.57 | 0.27 | 0.47 | 0.4 |
| 1 | 183604258 | rs11811536 | -0.701 | -0.11 | 1.8 | -0.808 | -0.018 | 0.003 | 0.003 | 0.227 | 0.08 | 0.18 | 0.04 | 0.11 | 0.03 | 0.07 |
| 1 | 183614855 | rs12025732 | -0.756 | -0.106 | 1.816 | -0.806 | -0.009 | 0.02 | 0.02 | 0.175 | 0.08 | 0.24 | 0.04 | 0.11 | 0.03 | 0.07 |
| 1 | 183616263 | rs16861371 | -0.675 | -0.107 | 1.816 | -0.807 | -0.016 | 0.005 | 0.005 | 0.1 | 0.08 | 0.18 | 0.04 | 0.11 | 0.03 | 0.07 |
| 1 | 183616687 | rs1174656 | 1.111 | -0.132 | 1.871 | -0.821 | -0.004 | 0.087 | 0.087 | 1.26 | 0.36 | 0.2 | 0.3 | 0.08 | 0.37 | 0.25 |
| 1 | 183616781 | rs10911388 | -0.439 | -0.123 | 1.88 | -0.818 | -0.015 | 0.004 | 0.004 | 0.028 | 0.08 | 0.16 | 0.03 | 0.11 | 0.03 | 0.07 |
| 1 | 183616805 | rs10911389 | -0.672 | -0.122 | 1.86 | -0.818 | -0.016 | 0.005 | 0.005 | 0.122 | 0.08 | 0.18 | 0.04 | 0.11 | 0.03 | 0.07 |
| 1 | 183616926 | rs1174657 | 0.308 | -0.123 | 1.852 | -0.817 | -0.004 | -0.033 | -0.033 | 0 | 0.46 | 0.49 | 0.35 | 0.24 | 0.4 | 0.32 |
| 1 | 183617693 | rs16861394 | -0.737 | -0.084 | 1.811 | -0.63 | -0.017 | 0.004 | 0.004 | 0.122 | 0.08 | 0.18 | 0.04 | 0.11 | 0.03 | 0.07 |
| 1 | 183622314 | rs6661715 | -0.808 | -0.084 | 1.811 | -0.63 | -0.009 | 0.02 | 0.02 | 0.175 | 0.08 | 0.24 | 0.04 | 0.11 | 0.03 | 0.07 |
| 1 | 183622379 | rs6699011 | -0.738 | -0.078 | 1.661 | -0.627 | -0.017 | 0.004 | 0.004 | 0.122 | 0.08 | 0.18 | 0.04 | 0.11 | 0.03 | 0.07 |
| 1 | 183667375 | rs10911417 | -0.756 | -0.024 | 1.031 | -0.754 | 0.023 | -0.063 | -0.063 | 0 | 0.12 | 0.4 | 0.25 | 0.02 | 0.1 | 0.12 |
| 1 | 183667908 | rs10494565 | -0.756 | -0.024 | 1.012 | -0.754 | 0.024 | -0.063 | -0.063 | 7.453 | 0.12 | 0.4 | 0.25 | 0.02 | 0.1 | 0.12 |
| 1 | 183668132 | rs6424904 | -0.705 | -0.083 | 0.919 | -0.916 | 0.014 | 0.089 | 0.089 | 2.815 | 0.56 | 0.72 | 0.68 | 0.76 | 0.6 | 0.66 |
| 1 | 183684569 | rs17559937 | -0.43 | -0.116 | 0.85 | -1.01 | 0.007 | 0.046 | 0.046 | 0.712 | 0.51 | 0.39 | 0.46 | 0.89 | 0.54 | 0.61 |
| 1 | 183687038 | rs12045314 | -0.051 | -0.055 | 0.17 | -0.838 | 0.01 | -0.014 | -0.014 | 0 | 0.07 | 0.02 | 0.02 | 0.11 | 0.03 | 0.07 |
| 1 | 183696656 | rs6424909 | -1.259 | -0.052 | 0.188 | -0.84 | -0.005 | 0.117 | 0.117 | 0.753 | 0.64 | 0.84 | 0.71 | 0.92 | 0.64 | 0.74 |
| 1 | 183707595 | rs2491430 | 1.114 | -0.067 | 0.181 | -0.848 | -0.02 | 0.165 | 0.165 | 0.962 | 0.36 | 0.12 | 0.28 | 0.08 | 0.36 | 0.27 |
| 1 | 183725820 | rs12130615 | 0.698 | 0.342 | -0.299 | -0.082 | -0.05 | -0.118 | -0.118 | 0.167 | 0.16 | 0.01 | 0.19 | 0.38 | 0.18 | 0.24 |
| 1 | 183743599 | rs12024301 | -0.311 | 0.508 | 0.08 | -0.636 | 0.002 | -0.001 | -0.001 | 0 | 0.08 | 0.09 | 0.04 | 0.11 | 0.06 | 0.08 |
| 1 | 183754672 | rs1952235 | -0.63 | 0.497 | 0.111 | -0.672 | 0.011 | 0.106 | 0.106 | 2.785 | 0.55 | 0.7 | 0.66 | 0.81 | 0.58 | 0.65 |
| 1 | 183754706 | rs1952234 | -1.193 | 0.522 | 0.056 | -0.667 | 0.001 | 0.09 | 0.09 | 0.688 | 0.63 | 0.78 | 0.7 | 0.92 | 0.63 | 0.73 |
| 1 | 183759410 | rs4609388 | -0.724 | 0.661 | 0.075 | -0.528 | 0.008 | 0.04 | 0.04 | 2.485 | 0.54 | 0.63 | 0.66 | 0.81 | 0.58 | 0.65 |
| 1 | 183765059 | rs2500097 | -0.304 | 1.01 | 0.066 | 0.087 | -0.01 | -0.024 | -0.024 | 0 | 0.37 | 0.25 | 0.37 | 0.49 | 0.4 | 0.59 |
| 1 | 183769285 | rs16861532 | 0.036 | 1.263 | 0.025 | -0.307 | 0.001 | -0.039 | -0.039 | 0 | 0.16 | 0.17 | 0.2 | 0.02 | 0.19 | 0.12 |
| 1 | 183772418 | rs10082229 | 0.077 | 0.459 | 0.185 | -0.131 | -0.032 | 0.161 | 0.161 | 2.128 | 0.1 | 0.24 | 0.14 | 0.4 | 0.05 | 0.11 |
| 1 | 183781391 | rs11580938 | 1.227 | 0.519 | -0.208 | -0.344 | -0.058 | 0.174 | 0.174 | 0.308 | 0.32 | 0.05 | 0.25 | 0.05 | 0.34 | 0.19 |
| 1 | 183785174 | rs10797912 | -0.627 | 0.31 | 0.248 | -0.079 | -0.052 | 0.191 | 0.191 | 1.424 | 0.13 | 0.3 | 0.13 | 0.41 | 0.05 | 0.12 |
| 1 | 183785842 | rs2500125 | -0.781 | 1.092 | -0.236 | -0.036 | 0 | -0.001 | -0.001 | 0 | 0.59 | 0.44 | 0.6 | 0.56 | 0.6 | 0.7 |
| 1 | 183787675 | rs2500123 | -0.209 | 1.602 | -1.102 | -0.059 | -0.001 | -0.066 | -0.066 | 0.037 | 0.4 | 0.08 | 0.27 | 0.45 | 0.34 | 0.57 |
| 1 | 183788570 | rs10494568 | -0.772 | 2.149 | 0.043 | -0.239 | 0.057 | -0.126 | -0.126 | 0.417 | 0.21 | 0.35 | 0.32 | 0.02 | 0.34 | 0.17 |
| 1 | 183792288 | rs10911443 | -0.426 | 2.045 | -0.649 | 0.668 | -0.045 | -0.088 | -0.088 | 0.313 | 0.23 | 0.09 | 0.37 | 0.44 | 0.31 | 0.31 |
| 1 | 183793323 | rs4652825 | 0.313 | 1.331 | -0.464 | 0.793 | 0.007 | 0.044 | 0.044 | 0 | 0.8 | 0.66 | 0.7 | 0.68 | 0.79 | 0.65 |
| 1 | 183798877 | rs6424913 | -0.398 | 1.733 | -1.338 | 0.798 | -0.073 | -0.041 | -0.041 | 0 | 0.28 | 0.13 | 0.43 | 0.44 | 0.4 | 0.39 |
| 1 | 183801920 | rs2500116 | -0.507 | 1.061 | -1.583 | 0.658 | 0.006 | 0.001 | 0.001 | 0 | 0.79 | 0.81 | 0.83 | 0.84 | 0.84 | 0.67 |

| | | | | | | | | | | | | | | | | |
|---|---|---|---|---|---|---|---|---|---|---|---|---|---|---|---|---|
| 1 | 183804680 | rs4388664 | 0.961 | -0.096 | -1.379 | 0.02 | -0.067 | -0.1 | -0.1 | 0.207 | 0.21 | 0.47 | 0.1 | 0.07 | 0.12 | 0.07 |
| 1 | 183811400 | rs4262497 | -0.125 | 0.146 | -1.418 | 0.01 | 0.005 | 0.018 | 0.018 | 0 | 0.49 | 0.59 | 0.56 | 0.51 | 0.56 | 0.49 |
| 1 | 183811777 | rs2500103 | -1.083 | 0.075 | -1.377 | -0.01 | 0.001 | 0 | 0 | 0 | 0.78 | 0.78 | 0.83 | 0.84 | 0.84 | 0.67 |
| 1 | 183811931 | rs2500102 | 1.608 | -0.169 | -1.57 | -0.086 | -0.028 | -0.064 | -0.064 | 0 | 0.18 | 0.05 | 0.15 | 0.28 | 0.19 | 0.08 |
| 1 | 183813858 | rs6424914 | 1.093 | -0.107 | -1.548 | -0.059 | -0.001 | -0.002 | -0.002 | 0.152 | 0.1 | 0.12 | 0.12 | 0.06 | 0.08 | 0.1 |
| 1 | 183817289 | rs3010055 | -0.329 | -0.136 | -1.565 | -0.073 | 0.041 | 0.014 | 0.014 | 0 | 0.5 | 0.65 | 0.36 | 0.41 | 0.4 | 0.25 |
| 1 | 183820769 | rs3010058 | -0.193 | -0.181 | -1.428 | -0.097 | 0.001 | 0.001 | 0.001 | 0 | 0.68 | 0.66 | 0.71 | 0.78 | 0.75 | 0.6 |
| 1 | 183826941 | rs4650644 | 0.131 | -0.293 | -1.489 | -0.527 | 0.027 | 0.026 | 0.026 | 1.845 | 0.62 | 0.36 | 0.76 | 0.87 | 0.79 | 0.82 |
| 1 | 183830469 | rs3002652 | 0.752 | -0.112 | -1.645 | -0.347 | -0.011 | -0.034 | -0.034 | 1.893 | 0.36 | 0.25 | 0.31 | 0.43 | 0.36 | 0.38 |
| 1 | 183831184 | rs12116846 | 0.899 | -0.47 | -1.64 | -0.824 | 0.047 | 0.108 | 0.108 | 3.172 | 0.22 | 0.01 | 0.07 | 0.05 | 0.12 | 0.1 |
| 1 | 183831786 | rs11591044 | 0.923 | -0.412 | -1.442 | -0.755 | 0.002 | 0.016 | 0.016 | 0 | 0.09 | 0.04 | 0.11 | 0.06 | 0.08 | 0.09 |
| 1 | 183834061 | rs9425322 | 0.713 | -0.186 | -1.462 | -0.778 | -0.029 | 0.024 | 0.024 | 0.02 | 0.11 | 0.05 | 0.14 | 0.05 | 0.15 | 0.09 |
| 1 | 183835393 | rs9425324 | -0.317 | 0.521 | -1.745 | -0.99 | -0.082 | 0.078 | 0.078 | 0.039 | 0.27 | 0.1 | 0.42 | 0.13 | 0.41 | 0.26 |
| 1 | 183835838 | rs7540012 | 0.419 | -0.136 | -0.59 | -0.779 | -0.035 | -0.052 | -0.052 | 0 | 0.24 | 0.32 | 0.08 | 0.05 | 0.12 | 0.1 |
| 1 | 183837032 | rs4651156 | 0.351 | 0.084 | -1.001 | -0.625 | 0.034 | 0.058 | 0.058 | 1.482 | 0.6 | 0.32 | 0.73 | 0.88 | 0.71 | 0.8 |
| 1 | 183841796 | rs10797917 | 0.295 | 0.086 | -1.072 | -0.627 | 0.041 | 0.075 | 0.075 | 1.71 | 0.61 | 0.29 | 0.73 | 0.88 | 0.72 | 0.83 |
| 1 | 183842979 | rs12070135 | -0.648 | 0.087 | -1.158 | -0.094 | 0.007 | 0.018 | 0.018 | 0 | 0.5 | 0.7 | 0.52 | 0.6 | 0.52 | 0.53 |
| 1 | 183843104 | rs10911457 | -0.61 | 0.164 | -1.006 | -0.068 | 0 | 0.014 | 0.014 | 0 | 0.49 | 0.65 | 0.51 | 0.6 | 0.5 | 0.52 |
| 1 | 183844367 | rs12563514 | 0.832 | 0.308 | -1.335 | 0.102 | -0.071 | -0.116 | -0.116 | 0 | 0.16 | 0.01 | 0.19 | 0.42 | 0.2 | 0.33 |
| 1 | 183850901 | rs11585126 | -0.657 | 0.341 | -0.858 | -0.032 | 0.051 | 0.059 | 0.059 | 0 | 0.31 | 0.49 | 0.43 | 0.49 | 0.42 | 0.42 |
| 1 | 183861007 | rs3814329 | 0.44 | 0.271 | -0.759 | -0.106 | -0.014 | 0.012 | 0.012 | 0 | 0.12 | 0.11 | 0.2 | 0.07 | 0.22 | 0.11 |
| 1 | 183866849 | rs12031722 | -0.46 | 0.263 | -0.871 | -0.238 | -0.011 | 0.015 | 0.015 | 0 | 0.18 | 0.16 | 0.22 | 0.07 | 0.26 | 0.15 |
| 1 | 183876287 | rs16861715 | 0.317 | 0.065 | -0.591 | -0.196 | -0.004 | -0.041 | -0.041 | 0 | 0.3 | 0.21 | 0.25 | 0.5 | 0.27 | 0.35 |
| 1 | 183886016 | rs10797921 | -1.209 | -0.154 | -0.735 | -0.439 | 0.007 | 0.001 | 0.001 | 0 | 0.53 | 0.61 | 0.56 | 0.57 | 0.56 | 0.54 |
| 1 | 183887602 | rs6658545 | 0.675 | -0.009 | -0.622 | -0.359 | 0.003 | 0.034 | 0.034 | 0 | 0.27 | 0.31 | 0.27 | 0.5 | 0.25 | 0.32 |
| 1 | 183896961 | rs4650 | -0.984 | -0.001 | -0.599 | -0.322 | 0.002 | 0.001 | 0.001 | 0 | 0.55 | 0.63 | 0.56 | 0.57 | 0.56 | 0.58 |
| 1 | 183898790 | rs7522001 | -0.641 | 0.085 | -0.705 | -0.361 | -0.011 | 0.029 | 0.029 | 0.039 | 0.24 | 0.2 | 0.28 | 0.08 | 0.31 | 0.22 |
| 1 | 183905073 | rs1053093 | 0.681 | 0.025 | -0.766 | -0.308 | -0.001 | 0.021 | 0.021 | 0 | 0.3 | 0.31 | 0.26 | 0.5 | 0.26 | 0.35 |
| 1 | 183907252 | rs3828027 | -1.129 | 0.045 | -0.73 | -0.335 | 0.001 | 0.002 | 0.002 | 0 | 0.55 | 0.59 | 0.55 | 0.57 | 0.56 | 0.58 |
| 1 | 183907291 | rs3814332 | -1.105 | 0.074 | -0.727 | -0.32 | 0.002 | 0.004 | 0.004 | 0 | 0.55 | 0.59 | 0.55 | 0.57 | 0.56 | 0.57 |
| 1 | 183908222 | rs3010040 | 1.815 | 0.3 | -1.217 | -0.49 | -0.001 | -0.013 | -0.013 | 0 | 0.2 | 0.08 | 0.25 | 0.18 | 0.2 | 0.23 |
| 1 | 183908813 | rs10911472 | 0.611 | 0.294 | -1.204 | -0.526 | -0.009 | 0.013 | 0.013 | 0.062 | 0.18 | 0.16 | 0.25 | 0.07 | 0.26 | 0.18 |
| 1 | 183909717 | rs2296713 | 1.635 | 0.375 | -1.193 | -0.364 | -0.001 | -0.013 | -0.013 | 0 | 0.2 | 0.07 | 0.25 | 0.18 | 0.2 | 0.22 |
| 1 | 183912784 | rs11803131 | 0.237 | 0.515 | -0.568 | 0.304 | -0.006 | 0.005 | 0.005 | 0.031 | 0.35 | 0.36 | 0.35 | 0.6 | 0.29 | 0.38 |
| 1 | 183913540 | rs17504460 | -0.34 | 0.93 | -0.403 | 0.6 | 0.017 | -0.019 | -0.019 | 0.004 | 0.13 | 0.18 | 0.17 | 0.06 | 0.2 | 0.15 |
| 1 | 183918161 | rs2378788 | 1.128 | 0.724 | -0.4 | 0.623 | 0.026 | -0.029 | -0.029 | 0 | 0.81 | 0.71 | 0.72 | 0.93 | 0.74 | 0.85 |
| 1 | 183920191 | rs16861794 | -0.257 | 0.754 | -0.594 | 0.657 | -0.002 | 0.003 | 0.003 | 0 | 0.05 | 0.04 | 0.02 | 0.06 | 0.03 | 0.05 |
| 1 | 183922435 | rs2986550 | -0.098 | 0.904 | -0.796 | 0.489 | 0.032 | -0.049 | -0.049 | 1.431 | 0.88 | 0.81 | 0.75 | 1 | 0.78 | 0.88 |
| 1 | 183925087 | rs1926870 | -0.641 | 0.681 | -1.415 | 0.955 | 0.001 | -0.008 | -0.008 | 0.004 | 0.37 | 0.67 | 0.44 | 0.44 | 0.37 | 0.51 |
| 1 | 183929771 | rs12096645 | -0.65 | 0.597 | -1.559 | 0.693 | 0.003 | -0.024 | -0.024 | 0 | 0.37 | 0.69 | 0.45 | 0.49 | 0.37 | 0.52 |
| 1 | 183932632 | rs17504880 | -0.057 | 0.485 | -1.764 | 0.325 | -0.044 | 0.05 | 0.05 | 0 | 0.28 | 0.05 | 0.19 | 0.15 | 0.25 | 0.24 |
| 1 | 183933498 | rs12725649 | -0.215 | 0.491 | -1.717 | 0.757 | -0.014 | -0.018 | -0.018 | 0.121 | 0.14 | 0.09 | 0.3 | 0.18 | 0.19 | 0.17 |
| 1 | 183937066 | rs10911474 | 0.066 | 0.304 | -1.832 | 0.579 | -0.002 | 0.082 | 0.082 | 0 | 0.37 | 0.13 | 0.27 | 0.22 | 0.34 | 0.3 |
| 1 | 183938304 | rs4651158 | -1.558 | -0.24 | -1.632 | 0.457 | -0.028 | 0.024 | 0.024 | 0 | 0.74 | 0.85 | 0.76 | 0.84 | 0.66 | 0.71 |
| 1 | 183939080 | rs12736977 | 0.016 | -0.314 | -1.011 | 0.729 | -0.017 | -0.023 | -0.023 | 0 | 0.14 | 0.09 | 0.29 | 0.18 | 0.19 | 0.16 |
| 1 | 183941554 | rs7530464 | -1.23 | -0.427 | -1.126 | 0.487 | 0.054 | 0.057 | 0.057 | 0 | 0.42 | 0.88 | 0.51 | 0.7 | 0.34 | 0.4 |
| 1 | 183942271 | rs7521638 | -1.134 | -0.403 | 0.073 | 0.847 | -0.011 | 0.019 | 0.019 | 0.003 | 0.31 | 0.49 | 0.45 | 0.62 | 0.29 | 0.38 |
| 1 | 183942950 | rs2296714 | 1.055 | -1.17 | -0.268 | 0.809 | 0.065 | 0.013 | 0.013 | 0 | 0.64 | 0.18 | 0.54 | 0.31 | 0.77 | 0.61 |
| 1 | 183945447 | rs2182626 | 0.993 | -1.109 | -0.212 | 0.728 | -0.006 | 0 | 0 | 0 | 0.65 | 0.37 | 0.55 | 0.36 | 0.77 | 0.62 |
| 1 | 183949169 | rs12021841 | 0.517 | -1.077 | -0.953 | 0.97 | 0.032 | -0.039 | -0.039 | 0 | 0.13 | 0.06 | 0.1 | 0.45 | 0.06 | 0.19 |
| 1 | 183954003 | rs3930248 | -0.731 | -1.135 | -0.966 | 0.895 | 0.023 | -0.043 | -0.043 | 0 | 0.22 | 0.14 | 0.33 | 0.45 | 0.17 | 0.28 |
| 1 | 183955574 | rs1926868 | 0.548 | -1.108 | -1.158 | 0.914 | 0.033 | -0.04 | -0.04 | 0 | 0.13 | 0.06 | 0.1 | 0.44 | 0.06 | 0.19 |
| 1 | 183958442 | rs6698088 | -1.737 | -1.303 | -1.304 | 0.784 | 0.105 | -0.024 | -0.024 | 0 | 0.39 | 0.93 | 0.48 | 0.65 | 0.24 | 0.41 |
| 1 | 183962308 | rs12748062 | 0.474 | -0.925 | -1.117 | 0.7 | -0.029 | 0.045 | 0.045 | 0 | 0.08 | 0.01 | 0.22 | 0 | 0.11 | 0.09 |
| 1 | 183966166 | rs7519097 | -0.952 | -0.995 | -0.773 | 0.781 | -0.003 | 0 | 0 | 0.237 | 0.25 | 0.72 | 0.36 | 0.2 | 0.18 | 0.23 |
| 1 | 183977285 | rs1887282 | 0.363 | -0.964 | -1.155 | 1.053 | 0.027 | -0.03 | -0.03 | 0 | 0.13 | 0.06 | 0.1 | 0.47 | 0.06 | 0.19 |
| 1 | 183979729 | rs10737249 | -1.075 | -0.229 | -1.343 | 0.606 | -0.135 | 0.016 | 0.016 | 0 | 0.75 | 0.98 | 0.73 | 0.84 | 0.62 | 0.7 |
| 1 | 183979966 | rs7529873 | -0.834 | -0.588 | -0.326 | 0.78 | -0.053 | 0.066 | 0.066 | 2.342 | 0.16 | 0.54 | 0.13 | 0.17 | 0.07 | 0.14 |
| 1 | 183982487 | rs1952251 | -0.339 | -0.216 | -0.983 | 0.617 | 0.001 | -0.003 | -0.003 | 0.628 | 0.35 | 0.33 | 0.29 | 0.3 | 0.35 | 0.36 |
| 1 | 183982631 | rs12405373 | -0.797 | -0.703 | -1.543 | 0.433 | 0.037 | -0.003 | -0.003 | 0 | 0.26 | 0.07 | 0.15 | 0.21 | 0.16 | 0.19 |
| 1 | 183983087 | rs12089250 | -0.248 | -0.656 | -0.866 | 0.811 | -0.013 | 0.116 | 0.116 | 1.49 | 0.17 | 0.23 | 0.22 | 0.49 | 0.13 | 0.2 |
| 1 | 183983580 | rs10494572 | -0.653 | -1.007 | -0.217 | 0.424 | -0.013 | 0.144 | 0.144 | 1.317 | 0.22 | 0.39 | 0.22 | 0.45 | 0.18 | 0.29 |
| 1 | 183985203 | rs10911492 | -0.209 | -1.188 | -0.316 | 0.339 | -0.006 | 0.05 | 0.05 | 0.095 | 0.2 | 0.23 | 0.19 | 0.44 | 0.16 | 0.26 |
| 1 | 183986897 | rs1926880 | -1.312 | -1.024 | -1.189 | -0.404 | -0.107 | 0.128 | 0.128 | 0 | 0.68 | 0.94 | 0.77 | 0.87 | 0.61 | 0.68 |
| 1 | 183990746 | rs10911494 | -0.466 | -1.316 | -1.052 | -0.391 | -0.047 | 0.055 | 0.055 | 0.446 | 0.54 | 0.83 | 0.47 | 0.58 | 0.46 | 0.43 |
| 1 | 183995354 | rs10158437 | -1.251 | -1.147 | -0.256 | -0.549 | -0.009 | 0.042 | 0.042 | 0 | 0.65 | 0.75 | 0.75 | 0.86 | 0.62 | 0.65 |
| 1 | 183997429 | rs10494573 | 1.584 | -1.013 | -0.797 | -0.52 | -0.067 | -0.013 | -0.013 | 0 | 0.2 | 0.02 | 0.15 | 0.12 | 0.22 | 0.08 |
| 1 | 184001499 | rs10911498 | -1.233 | -0.767 | 0.05 | -0.154 | -0.023 | 0.019 | 0.019 | 0.268 | 0.22 | 0.54 | 0.34 | 0.29 | 0.18 | 0.26 |
| 1 | 184002332 | rs10797934 | 0.25 | -0.505 | -0.402 | 0.015 | -0.005 | -0.014 | -0.014 | 0.362 | 0.12 | 0.1 | 0.3 | 0.29 | 0.15 | 0.18 |

| | | | | | | | | | | | | | | | | |
|---|---|---|---|---|---|---|---|---|---|---|---|---|---|---|---|---|
| 1 | 184002874 | rs756199 | 0.128 | -0.378 | -0.279 | -0.014 | 0 | -0.016 | -0.016 | 3.502 | 0.23 | 0.19 | 0.25 | 0.44 | 0.21 | 0.31 |
| 1 | 184010727 | rs10494574 | -0.219 | 0.068 | -0.325 | -0.016 | -0.018 | -0.045 | -0.045 | 0.674 | 0.3 | 0.2 | 0.29 | 0.43 | 0.33 | 0.36 |
| 1 | 184018475 | rs1926872 | 0.113 | 0.3 | -0.298 | 0.071 | -0.039 | -0.062 | -0.062 | 0 | 0.26 | 0.12 | 0.3 | 0.45 | 0.33 | 0.38 |
| 1 | 184020945 | rs2274432 | 0.122 | 0.301 | -0.297 | 0.071 | -0.039 | -0.061 | -0.061 | 0.333 | 0.26 | 0.12 | 0.3 | 0.45 | 0.33 | 0.37 |
| 1 | 184023059 | rs11582961 | 0.057 | 0.29 | -0.175 | -0.026 | 0 | -0.007 | -0.007 | 0 | 0.08 | 0.1 | 0.23 | 0.05 | 0.09 | 0.09 |
| 1 | 184023529 | rs1046934 | 0.196 | 0.282 | -0.253 | -0.019 | -0.039 | -0.062 | -0.062 | 0 | 0.26 | 0.12 | 0.3 | 0.45 | 0.33 | 0.37 |
| 1 | 184027676 | rs16822379 | -0.776 | 0.43 | 0.429 | -0.268 | 0.039 | -0.044 | -0.044 | 2.223 | 0.11 | 0.34 | 0.27 | 0.05 | 0.12 | 0.1 |
| 1 | 184031561 | rs1409812 | -0.776 | 0.43 | 0.429 | -0.268 | 0.039 | -0.044 | -0.044 | 0 | 0.11 | 0.34 | 0.27 | 0.05 | 0.12 | 0.1 |
| 1 | 184053769 | rs10911506 | -1.123 | -0.199 | 0.023 | -0.421 | 0.019 | 0.011 | 0.011 | 0 | 0.69 | 0.77 | 0.78 | 0.74 | 0.72 | 0.6 |
| 1 | 184058359 | rs1409813 | -1.15 | -0.208 | 0.067 | -0.42 | 0.034 | 0.014 | 0.014 | 0 | 0.69 | 0.82 | 0.79 | 0.74 | 0.72 | 0.6 |
| 1 | 184067280 | rs10752931 | -0.371 | -0.365 | 0.522 | -0.64 | 0.004 | 0.004 | 0.004 | 0 | 0.59 | 0.42 | 0.72 | 0.73 | 0.67 | 0.58 |
| 1 | 184068508 | rs971572 | 0.473 | 0.4 | 0.522 | -0.329 | -0.053 | -0.076 | -0.076 | 0.101 | 0.27 | 0.13 | 0.31 | 0.57 | 0.34 | 0.39 |
| 1 | 184069215 | rs16822450 | 1.13 | 0.393 | 0.835 | -0.643 | -0.002 | -0.041 | -0.041 | 0.029 | 0.07 | 0.19 | 0.05 | 0.01 | 0.04 | 0.02 |
| 1 | 184082630 | rs12725276 | 0.383 | 0.442 | 0.623 | -0.637 | 0.005 | -0.021 | -0.021 | 0 | 0.08 | 0.12 | 0.23 | 0.04 | 0.09 | 0.08 |
| 1 | 184099405 | rs6703693 | 0.327 | 0.282 | 0.878 | -0.735 | -0.015 | -0.062 | -0.062 | 5.728 | 0.1 | 0.28 | 0.06 | 0.01 | 0.05 | 0.02 |
| 1 | 184110720 | rs11585756 | 0.039 | 0.301 | 0.671 | -0.728 | 0.004 | -0.033 | -0.033 | 0.039 | 0.09 | 0.21 | 0.24 | 0.04 | 0.09 | 0.12 |
| 1 | 184111365 | rs10911518 | 0.633 | 0.456 | -0.104 | -0.359 | -0.071 | -0.078 | -0.078 | 0 | 0.26 | 0.12 | 0.3 | 0.44 | 0.34 | 0.4 |
| 1 | 184113245 | rs732812 | -1.209 | 0.315 | 0.037 | -0.418 | -0.012 | 0.018 | 0.018 | 0.127 | 0.47 | 0.75 | 0.63 | 0.49 | 0.48 | 0.54 |
| 1 | 184113568 | rs890063 | 0.972 | 0.18 | -0.266 | -0.474 | -0.019 | -0.04 | -0.04 | 0 | 0.22 | 0.12 | 0.19 | 0.4 | 0.27 | 0.33 |
| 1 | 184115890 | rs1409814 | 0.279 | 0.688 | 0.078 | -0.366 | -0.078 | -0.086 | -0.086 | 0 | 0.27 | 0.08 | 0.32 | 0.45 | 0.34 | 0.38 |
| 1 | 184116980 | rs10797941 | -0.356 | 0.679 | 0.234 | -0.359 | -0.008 | -0.084 | -0.084 | 0 | 0.17 | 0.34 | 0.29 | 0.04 | 0.14 | 0.07 |
| 1 | 184128264 | rs6424925 | -0.987 | 0.661 | -0.507 | -0.16 | 0.001 | 0.014 | 0.014 | 0 | 0.48 | 0.68 | 0.64 | 0.49 | 0.49 | 0.51 |
| 1 | 184144995 | rs11810038 | -1.26 | 1.229 | -0.294 | -0.632 | 0.004 | -0.034 | -0.034 | 0.007 | 0.23 | 0.68 | 0.45 | 0.07 | 0.25 | 0.16 |
| 1 | 184145642 | rs12119930 | -0.935 | 1.28 | 0.478 | -0.678 | 0.068 | -0.083 | -0.083 | 1.217 | 0.16 | 0.5 | 0.38 | 0.06 | 0.18 | 0.14 |
| 1 | 184148645 | rs4650656 | -1.573 | 1.387 | -0.129 | -0.558 | 0.034 | -0.023 | -0.023 | 2.399 | 0.47 | 0.91 | 0.7 | 0.56 | 0.58 | 0.49 |
| 1 | 184158796 | rs6424930 | -0.462 | 1.412 | 0.592 | -0.35 | 0.048 | -0.062 | -0.062 | 0 | 0.12 | 0.37 | 0.29 | 0.04 | 0.12 | 0.07 |
| 1 | 184161757 | rs2025808 | 0.534 | 1.797 | 0.178 | -0.331 | 0.006 | 0.043 | 0.043 | 0 | 0.17 | 0.22 | 0.21 | 0.49 | 0.24 | 0.29 |
| 1 | 184169492 | rs10911524 | -1.104 | 1.237 | 0.262 | -0.592 | -0.001 | 0 | 0 | 0 | 0.43 | 0.56 | 0.67 | 0.57 | 0.55 | 0.55 |
| 1 | 184174866 | rs12136030 | 0.315 | 1.289 | 0.074 | -0.591 | -0.037 | -0.054 | -0.054 | 0 | 0.27 | 0.13 | 0.3 | 0.5 | 0.37 | 0.4 |
| 1 | 184177245 | rs1863155 | -1.099 | 1.204 | 0.038 | -0.568 | -0.015 | 0.025 | 0.025 | 0 | 0.46 | 0.87 | 0.71 | 0.57 | 0.55 | 0.55 |
| 1 | 184189782 | rs1409808 | -0.699 | 0.879 | 0.688 | -0.466 | 0.071 | -0.067 | -0.067 | 0.077 | 0.15 | 0.46 | 0.39 | 0.06 | 0.18 | 0.14 |
| 1 | 184192465 | rs6696916 | 0.326 | 1.015 | 0.618 | -0.505 | -0.014 | -0.019 | -0.019 | 0.168 | 0.3 | 0.27 | 0.34 | 0.5 | 0.4 | 0.38 |
| 1 | 184192838 | rs1854085 | -0.663 | 0.849 | 0.971 | -0.402 | 0.046 | -0.054 | -0.054 | 0.02 | 0.14 | 0.36 | 0.36 | 0.06 | 0.17 | 0.13 |
| 1 | 184195075 | rs10911530 | -0.93 | 0.708 | 0.662 | -0.584 | 0.052 | 0.029 | 0.029 | 0.273 | 0.49 | 0.7 | 0.73 | 0.56 | 0.61 | 0.53 |
| 1 | 184198012 | rs7517993 | 0.465 | 0.699 | 0.604 | -0.596 | -0.093 | -0.092 | -0.092 | 0.094 | 0.33 | 0.09 | 0.34 | 0.5 | 0.43 | 0.4 |
| 1 | 184208586 | rs714625 | -0.918 | 0.69 | 0.657 | -0.592 | 0.065 | 0.034 | 0.034 | 0.29 | 0.49 | 0.75 | 0.73 | 0.56 | 0.61 | 0.54 |
| 1 | 184210805 | rs2840274 | -1.409 | 0.66 | 0.587 | -0.626 | 0.057 | -0.028 | -0.028 | 0 | 0.54 | 0.96 | 0.78 | 0.56 | 0.65 | 0.56 |
| 1 | 184215054 | rs12754974 | -0.897 | 0.541 | 1.064 | -0.643 | 0.049 | -0.041 | -0.041 | 1.699 | 0.13 | 0.26 | 0.37 | 0.04 | 0.21 | 0.14 |
| 1 | 184217979 | rs1930285 | 0.411 | 0.558 | 0.796 | -0.64 | -0.064 | -0.056 | -0.056 | 0.023 | 0.34 | 0.16 | 0.35 | 0.5 | 0.44 | 0.4 |
| 1 | 184223494 | rs1889026 | -0.884 | 0.591 | 0.698 | -0.603 | 0.064 | 0.031 | 0.031 | 0.483 | 0.49 | 0.73 | 0.71 | 0.55 | 0.61 | 0.56 |
| 1 | 184235480 | rs4650664 | -1.16 | 0.581 | 0.67 | -0.609 | 0.077 | 0.016 | 0.016 | 0.376 | 0.53 | 0.81 | 0.74 | 0.57 | 0.65 | 0.58 |
| 1 | 184241399 | rs10911551 | 0.299 | 0.642 | 1.017 | -0.639 | 0.004 | 0.004 | 0.004 | 0.462 | 0.35 | 0.31 | 0.37 | 0.5 | 0.45 | 0.43 |
| 1 | 184245631 | rs7549909 | -1.091 | 0.689 | 0.936 | -0.613 | 0.078 | 0.027 | 0.027 | 0.326 | 0.53 | 0.8 | 0.74 | 0.58 | 0.65 | 0.58 |
| 1 | 184254001 | rs12079081 | -1.431 | 0.684 | 1.018 | -0.64 | 0.004 | -0.097 | -0.097 | 0 | 0.18 | 0.68 | 0.33 | 0.03 | 0.17 | 0.15 |
| 1 | 184269036 | rs7530243 | 0.474 | 0.664 | 1.021 | -0.628 | -0.109 | -0.084 | -0.084 | 0 | 0.35 | 0.07 | 0.36 | 0.51 | 0.46 | 0.46 |
| 1 | 184282979 | rs10911555 | 0.187 | 0.442 | 0.95 | -0.668 | -0.022 | -0.005 | -0.005 | 0 | 0.39 | 0.28 | 0.51 | 0.53 | 0.47 | 0.48 |
| 1 | 184283376 | rs10911556 | -0.391 | 0.958 | 1.357 | -0.623 | 0.055 | -0.065 | -0.065 | 0 | 0.11 | 0.28 | 0.17 | 0 | 0.18 | 0.11 |
| 1 | 184287575 | rs12128914 | -1.722 | 0.95 | 1.324 | -0.631 | 0.061 | -0.115 | -0.115 | 2.177 | 0.17 | 0.55 | 0.19 | 0.04 | 0.2 | 0.18 |
| 1 | 184290625 | rs4618934 | -0.436 | 1.024 | 1.124 | -0.532 | 0.083 | -0.068 | -0.068 | 0.069 | 0.11 | 0.34 | 0.18 | 0 | 0.18 | 0.11 |
| 1 | 184294167 | rs1930293 | -0.665 | 1.077 | 0.821 | -0.517 | 0.092 | -0.08 | -0.08 | 0.404 | 0.12 | 0.4 | 0.19 | 0 | 0.18 | 0.12 |
| 1 | 184297478 | rs2779286 | 0.264 | 0.223 | 0.167 | -0.687 | 0.011 | -0.035 | -0.035 | 0 | 0.63 | 0.48 | 0.66 | 0.78 | 0.56 | 0.68 |
| 1 | 184298164 | rs2151171 | -0.992 | 0.027 | -0.543 | -0.473 | -0.01 | -0.01 | -0.01 | 0.584 | 0.75 | 0.95 | 0.85 | 0.79 | 0.74 | 0.8 |
| 1 | 184304073 | rs12119892 | -0.154 | -0.037 | 0.174 | -0.459 | -0.057 | -0.049 | -0.049 | 0 | 0.25 | 0.5 | 0.22 | 0.23 | 0.16 | 0.22 |
| 1 | 184314411 | rs4568789 | -1.302 | 0.199 | 0.193 | -0.142 | 0.01 | 0.011 | 0.011 | 1.64 | 0.35 | 0.76 | 0.33 | 0.35 | 0.29 | 0.33 |
| 1 | 184318553 | rs10797957 | -0.651 | 0.232 | 0.03 | -0.096 | 0.002 | 0.013 | 0.013 | 0 | 0.8 | 0.8 | 0.83 | 0.73 | 0.81 | 0.81 |
| 1 | 184319293 | rs12407200 | 1.286 | 0.152 | -0.31 | -0.312 | -0.001 | 0.035 | 0.035 | 1.184 | 0.49 | 0.23 | 0.56 | 0.54 | 0.53 | 0.54 |
| 1 | 184321542 | rs4338323 | 1.138 | -0.029 | -0.407 | -0.345 | 0.015 | 0.002 | 0.002 | 0.212 | 0.17 | 0.1 | 0.14 | 0.21 | 0.08 | 0.14 |
| 1 | 184322625 | rs7554617 | -0.208 | -0.053 | -0.247 | -0.33 | -0.06 | -0.006 | -0.006 | 0.103 | 0.27 | 0.5 | 0.21 | 0.29 | 0.15 | 0.27 |
| 1 | 184323468 | rs9970847 | -1.489 | -0.034 | -0.406 | -0.336 | -0.043 | -0.033 | -0.033 | 0.573 | 0.85 | 0.98 | 0.86 | 0.85 | 0.82 | 0.87 |
| 1 | 184324403 | rs4390140 | 0.972 | -0.016 | -0.387 | -0.341 | 0.001 | -0.006 | -0.006 | 0 | 0.05 | 0.07 | 0.07 | 0 | 0.12 | 0.04 |
| 1 | 184324486 | rs4288563 | -0.856 | -0.021 | -0.493 | -0.31 | -0.011 | -0.014 | -0.014 | 0 | 0.33 | 0.6 | 0.29 | 0.3 | 0.27 | 0.31 |
| 1 | 184324652 | rs4356011 | -1.158 | -0.106 | -1.17 | -0.127 | -0.043 | -0.033 | -0.033 | 0 | 0.84 | 0.98 | 0.86 | 0.85 | 0.82 | 0.87 |
| 1 | 184326198 | rs12406462 | 0.601 | -0.374 | -0.728 | 0.006 | 0.027 | 0.003 | 0.003 | 0.212 | 0.17 | 0.09 | 0.13 | 0.21 | 0.08 | 0.14 |
| 1 | 184332991 | rs10911583 | 0.347 | 0.022 | -0.701 | 0.022 | 0.006 | 0.002 | 0.002 | 0 | 0.42 | 0.38 | 0.43 | 0.57 | 0.35 | 0.48 |
| 1 | 184333114 | rs4539094 | 0.876 | -0.628 | -1.388 | -0.214 | -0.04 | -0.019 | -0.019 | 1.174 | 0.56 | 0.05 | 0.63 | 0.63 | 0.59 | 0.57 |
| 1 | 184335001 | rs6692317 | 0.776 | -0.951 | -0.209 | -0.06 | 0.011 | 0.001 | 0.001 | 0.695 | 0.72 | 0.14 | 0.74 | 0.74 | 0.74 | 0.74 |
| 1 | 184335888 | rs6655949 | -0.379 | -1.063 | 0.282 | -0.096 | -0.001 | -0.001 | -0.001 | 0 | 0.83 | 0.82 | 0.84 | 0.84 | 0.81 | 0.87 |
| 1 | 184336331 | rs6683308 | -0.529 | -1.211 | 0.865 | -0.079 | 0.018 | 0.136 | 0.136 | 16.288 | 0.07 | 0.42 | 0.05 | 0.09 | 0.01 | 0.06 |
| 1 | 184336748 | rs6683726 | -0.201 | -1.156 | 0.808 | -0.093 | 0 | -0.003 | -0.003 | 0 | 0.82 | 0.76 | 0.83 | 0.84 | 0.81 | 0.87 |

| | | | | | | | | | | | | | | | | |
|---|---|---|---|---|---|---|---|---|---|---|---|---|---|---|---|---|
| 1 | 184338037 | rs10797961 | 0.218 | -1.164 | 0.927 | -0.095 | 0.013 | -0.013 | -0.013 | 0 | 0.8 | 0.65 | 0.81 | 0.83 | 0.75 | 0.81 |
| 1 | 184341562 | rs7536226 | 1.58 | -1.294 | 0.472 | -0.187 | -0.013 | -0.057 | -0.057 | 0 | 0.52 | 0.05 | 0.56 | 0.5 | 0.62 | 0.59 |
| 1 | 184341849 | rs6424949 | -0.557 | -0.8 | 1.035 | -0.123 | 0.008 | -0.008 | -0.008 | 0 | 0.86 | 0.93 | 0.9 | 0.84 | 0.88 | 0.89 |
| 1 | 184352288 | rs10911592 | 1.345 | -0.812 | 1.076 | -0.128 | 0.008 | 0.002 | 0.002 | 1.143 | 0.72 | 0.18 | 0.75 | 0.74 | 0.75 | 0.75 |
| 1 | 184364513 | rs16823097 | 0.307 | -0.812 | 1.365 | -0.087 | 0.006 | -0.007 | -0.007 | 3.067 | 0.14 | 0.09 | 0.11 | 0.16 | 0.12 | 0.1 |
| 1 | 184369713 | rs2207524 | -0.122 | -0.812 | 1.572 | -0.086 | 0.053 | 0.184 | 0.184 | 6.668 | 0.05 | 0.37 | 0.04 | 0.1 | 0 | 0.04 |
| 1 | 184374910 | rs732959 | -1.35 | -0.887 | 1.539 | -0.113 | 0.029 | -0.012 | -0.012 | 0.16 | 0.24 | 0.73 | 0.2 | 0.26 | 0.14 | 0.15 |
| 1 | 184375503 | rs6696161 | 0.063 | -0.83 | 1.587 | -0.142 | -0.008 | 0.009 | 0.009 | 0 | 0.15 | 0.17 | 0.13 | 0.17 | 0.13 | 0.11 |
| 1 | 184386071 | rs6666696 | -1.18 | -0.842 | 1.598 | -0.155 | -0.021 | -0.012 | -0.012 | 0 | 0.22 | 0.54 | 0.19 | 0.18 | 0.19 | 0.18 |
| 1 | 184391279 | rs11799574 | -1.572 | -0.819 | 1.872 | -0.141 | 0.093 | 0.03 | 0.03 | 0.12 | 0.05 | 0.38 | 0.05 | 0 | 0.05 | 0.07 |
| 1 | 184398346 | rs10158710 | -1.5 | -0.673 | 1.605 | -0.153 | 0.007 | -0.014 | -0.014 | 0 | 0.24 | 0.64 | 0.25 | 0.19 | 0.29 | 0.19 |
| 1 | 184409965 | rs7531736 | -1.45 | -0.295 | 1.328 | -0.138 | 0.139 | 0.003 | 0.003 | 0 | 0.05 | 0.3 | 0.07 | 0 | 0.12 | 0.08 |
| 1 | 184422553 | rs10489722 | 0.006 | -0.467 | 0.976 | -0.128 | 0 | -0.002 | -0.002 | 0.153 | 0.14 | 0.12 | 0.15 | 0.17 | 0.15 | 0.11 |
| 1 | 184430842 | rs16823195 | 0.025 | -0.736 | 1.043 | -0.221 | 0.01 | -0.049 | -0.049 | 0.332 | 0.14 | 0.04 | 0.11 | 0.2 | 0.12 | 0.15 |
| 1 | 184432880 | rs2207526 | -1.622 | -0.906 | 1.151 | -0.364 | -0.001 | -0.005 | -0.005 | 0 | 0.28 | 0.64 | 0.24 | 0.21 | 0.28 | 0.2 |
| 1 | 184442054 | rs4651208 | -1.72 | -1.245 | 0.83 | -0.438 | 0.023 | 0.008 | 0.008 | 0 | 0.45 | 0.63 | 0.51 | 0.46 | 0.36 | 0.39 |
| 1 | 184443829 | rs12043012 | 0.326 | -1.27 | 0.819 | -0.432 | 0.007 | 0.004 | 0.004 | 1.109 | 0.2 | 0.16 | 0.31 | 0.26 | 0.14 | 0.21 |
| 1 | 184447624 | rs6683932 | -0.086 | -1.153 | 1.029 | -0.242 | -0.029 | 0.039 | 0.039 | 0.112 | 0.17 | 0.32 | 0.13 | 0.2 | 0.12 | 0.15 |
| 1 | 184452580 | rs2013889 | 0.65 | -0.953 | 0.553 | -0.35 | 0.001 | 0.011 | 0.011 | 0 | 0.2 | 0.09 | 0.35 | 0.25 | 0.2 | 0.18 |
| 1 | 184466179 | rs4651213 | -1.755 | -0.957 | 0.421 | -0.158 | -0.042 | -0.052 | -0.052 | 0 | 0.53 | 0.84 | 0.63 | 0.52 | 0.52 | 0.43 |
| 1 | 184476621 | rs9633308 | -1.552 | -0.945 | 0.428 | -0.169 | 0.022 | -0.038 | -0.038 | 0 | 0.51 | 0.84 | 0.57 | 0.52 | 0.46 | 0.41 |
| 1 | 184488963 | rs7513091 | -0.25 | -0.967 | 0.65 | -0.18 | -0.002 | 0.007 | 0.007 | 0.083 | 0.22 | 0.4 | 0.14 | 0.19 | 0.21 | 0.17 |
| 1 | 184498780 | rs1416751 | -1.442 | -0.779 | 0.433 | -0.105 | -0.039 | -0.113 | -0.113 | 0 | 0.52 | 0.97 | 0.58 | 0.52 | 0.46 | 0.41 |
| 1 | 184503803 | rs1339563 | -1.286 | -0.833 | 0.562 | -0.112 | 0.007 | 0.038 | 0.038 | 0 | 0.49 | 0.67 | 0.54 | 0.45 | 0.48 | 0.39 |
| 1 | 184533939 | rs4651221 | -1.457 | -0.914 | -0.706 | 0.053 | -0.061 | -0.042 | -0.042 | 0 | 0.54 | 0.95 | 0.59 | 0.63 | 0.47 | 0.44 |
| 1 | 184536171 | rs749214 | -1.33 | -0.791 | -0.647 | 0.019 | -0.047 | -0.07 | -0.07 | 2.162 | 0.52 | 0.95 | 0.59 | 0.59 | 0.47 | 0.43 |
| 1 | 184539342 | rs10911612 | 0.76 | -0.829 | -0.608 | 0.005 | -0.003 | -0.005 | -0.005 | 0 | 0.19 | 0.1 | 0.35 | 0.3 | 0.19 | 0.19 |
| 1 | 184552327 | rs1856562 | -1.16 | -0.613 | -0.548 | 0.089 | -0.009 | -0.023 | -0.023 | 0.226 | 0.52 | 0.91 | 0.58 | 0.6 | 0.47 | 0.44 |
| 1 | 184558656 | rs6424958 | -1.163 | -0.615 | -0.548 | 0.088 | -0.01 | -0.023 | -0.023 | 1.098 | 0.52 | 0.91 | 0.59 | 0.6 | 0.48 | 0.44 |
| 1 | 184573082 | rs10911619 | 0.392 | -0.458 | -0.709 | 0.099 | 0.004 | 0.003 | 0.003 | 0 | 0.23 | 0.24 | 0.37 | 0.3 | 0.21 | 0.2 |
| 1 | 184576095 | rs6696816 | -1.695 | -1.103 | -1.336 | -0.067 | -0.161 | -0.147 | -0.147 | 0.362 | 0.7 | 0.98 | 0.69 | 0.61 | 0.63 | 0.55 |
| 1 | 184577317 | rs6695173 | -0.231 | -0.879 | -0.249 | 0.689 | -0.025 | 0.006 | 0.006 | 0 | 0.22 | 0.26 | 0.1 | 0.23 | 0.14 | 0.18 |
| 1 | 184580250 | rs4651223 | -0.669 | -0.117 | -0.592 | 0.373 | 0.004 | 0.005 | 0.005 | 0.224 | 0.29 | 0.4 | 0.46 | 0.38 | 0.33 | 0.26 |
| 1 | 184595020 | rs6701227 | -0.858 | -0.725 | -0.619 | 0.13 | -0.053 | -0.027 | -0.027 | 0.495 | 0.18 | 0.33 | 0.11 | 0.09 | 0.11 | 0.11 |
| 1 | 184597999 | rs8972 | -0.31 | -0.467 | -1.512 | -0.254 | -0.011 | 0.069 | 0.069 | 0.623 | 0.17 | 0.04 | 0.12 | 0.02 | 0.16 | 0.12 |
| 1 | 184602236 | rs1572766 | 0.031 | 0.466 | -1.757 | -0.232 | -0.01 | 0 | 0 | 0 | 0.84 | 0.89 | 0.75 | 0.78 | 0.79 | 0.73 |
| 1 | 184603098 | rs7540093 | 0.44 | 0.266 | -2.794 | -0.491 | -0.007 | -0.062 | -0.062 | 0 | 0.31 | 0.06 | 0.32 | 0.66 | 0.3 | 0.36 |
| 1 | 184621930 | rs6671374 | -0.394 | 0.448 | -2.973 | 0.231 | -0.105 | -0.127 | -0.127 | 0.688 | 0.68 | 0.97 | 0.66 | 0.52 | 0.63 | 0.51 |
| 1 | 184622136 | rs10797982 | 0.307 | 0.144 | -2.191 | -0.343 | -0.024 | 0.043 | 0.043 | 0 | 0.28 | 0.21 | 0.24 | 0.08 | 0.32 | 0.2 |
| 1 | 184639886 | rs9286853 | -0.47 | -0.336 | -0.727 | 0.04 | 0.022 | 0.103 | 0.103 | 0.027 | 0.08 | 0.42 | 0.07 | 0.07 | 0.04 | 0.06 |
| 1 | 184644321 | rs12131285 | -0.046 | -0.706 | -1.943 | 0.143 | -0.132 | -0.185 | -0.185 | 0 | 0.35 | 0.03 | 0.35 | 0.58 | 0.37 | 0.51 |
| 1 | 184653633 | rs2024766 | -0.413 | -1.121 | -1.531 | -0.015 | -0.022 | -0.006 | -0.006 | 0 | 0.46 | 0.2 | 0.41 | 0.63 | 0.44 | 0.59 |
| 1 | 184655649 | rs6673341 | -1.565 | -1.272 | -1.903 | -0.044 | -0.15 | 0.012 | 0.012 | 0 | 0.64 | 0.97 | 0.71 | 0.8 | 0.53 | 0.69 |
| 1 | 184656801 | rs6677073 | -0.882 | -0.936 | -1.219 | 0.052 | -0.003 | 0.029 | 0.029 | 0 | 0.52 | 0.6 | 0.47 | 0.7 | 0.47 | 0.61 |
| 1 | 184661642 | rs6663178 | -1.186 | -1.007 | -0.95 | 0.024 | -0.049 | 0.029 | 0.029 | 0 | 0.56 | 0.82 | 0.51 | 0.7 | 0.48 | 0.62 |
| 1 | 184662181 | rs1061508 | -0.46 | -0.997 | -0.949 | 0.039 | -0.036 | -0.026 | -0.026 | 1.093 | 0.44 | 0.19 | 0.4 | 0.63 | 0.43 | 0.59 |
| 1 | 184663537 | rs9425343 | 0.379 | -1.01 | -0.972 | 0.02 | -0.117 | -0.168 | -0.168 | 0 | 0.33 | 0.04 | 0.34 | 0.58 | 0.37 | 0.54 |
| 1 | 184667731 | rs10489697 | 0.685 | -0.761 | -0.811 | -0.068 | -0.008 | -0.033 | -0.033 | 0 | 0.09 | 0.13 | 0.06 | 0.01 | 0.06 | 0.03 |
| 1 | 184677464 | rs3736757 | -0.954 | -0.669 | -0.935 | 0.01 | -0.071 | 0.025 | 0.025 | 0 | 0.56 | 0.85 | 0.51 | 0.7 | 0.48 | 0.61 |
| 1 | 184705437 | rs2024767 | -0.683 | -0.551 | -0.632 | 0.067 | -0.002 | 0.014 | 0.014 | 0 | 0.51 | 0.53 | 0.46 | 0.7 | 0.47 | 0.61 |
| 1 | 184725099 | rs861575 | -0.316 | -0.623 | -0.556 | 0.017 | 0 | -0.002 | -0.002 | 0 | 0.42 | 0.41 | 0.4 | 0.69 | 0.41 | 0.59 |
| 1 | 184727100 | rs35779894 | 0.59 | -0.113 | -0.641 | -0.135 | 0.001 | -0.019 | -0.019 | 0 | 0.13 | 0.12 | 0.08 | 0.01 | 0.13 | 0.05 |
| 1 | 184728874 | rs3895196 | 0.573 | -0.178 | -0.679 | -0.045 | 0.003 | 0.004 | 0.004 | 0 | 0.45 | 0.47 | 0.52 | 0.3 | 0.45 | 0.37 |
| 1 | 184738494 | rs861588 | -0.051 | 0.03 | -0.542 | 0.38 | -0.01 | -0.013 | -0.013 | 0 | 0.25 | 0.27 | 0.35 | 0.47 | 0.29 | 0.37 |
| 1 | 184747087 | rs10752942 | -0.137 | -0.314 | -0.185 | 0.328 | -0.041 | -0.023 | -0.023 | 0 | 0.22 | 0.39 | 0.11 | 0.13 | 0.14 | 0.13 |
| 1 | 184757199 | rs861620 | -0.509 | -0.056 | -0.021 | 0.279 | -0.012 | -0.054 | -0.054 | 0.252 | 0.36 | 0.16 | 0.37 | 0.56 | 0.35 | 0.46 |
| 1 | 184760001 | rs570441 | -1.651 | -0.154 | 0.07 | 0.316 | 0.007 | 0.026 | 0.026 | 0 | 0.5 | 0.73 | 0.46 | 0.65 | 0.42 | 0.51 |
| 1 | 184760433 | rs7360 | -1.402 | -0.118 | 0.078 | 0.314 | 0.003 | 0.002 | 0.002 | 0 | 0.48 | 0.71 | 0.45 | 0.6 | 0.43 | 0.5 |
| 1 | 184761365 | rs682331 | -1.5 | -0.082 | 0.075 | 0.298 | 0.007 | 0.023 | 0.023 | 0 | 0.5 | 0.73 | 0.46 | 0.65 | 0.43 | 0.51 |
| 1 | 184795779 | rs520605 | 0.2 | 0.873 | -0.448 | 0.868 | -0.087 | -0.079 | -0.079 | 0 | 0.83 | 0.94 | 0.71 | 0.59 | 0.76 | 0.63 |
| 1 | 184801065 | rs487675 | -0.763 | 0.644 | -0.116 | 0.453 | 0 | -0.017 | -0.017 | 0 | 0.36 | 0.38 | 0.24 | 0.16 | 0.34 | 0.26 |
| 1 | 184809353 | rs626576 | 0.328 | 0.433 | 0.167 | 0.489 | -0.043 | -0.043 | -0.043 | 0 | 0.62 | 0.77 | 0.6 | 0.44 | 0.58 | 0.44 |
| 1 | 184810494 | rs642027 | -0.326 | 0.379 | 0.192 | 0.473 | 0.022 | 0.006 | 0.006 | 0 | 0.09 | 0.2 | 0.09 | 0.03 | 0.08 | 0.05 |
| 1 | 184810895 | rs491291 | 0.435 | 0.389 | 0.214 | 0.515 | -0.032 | -0.033 | -0.033 | 1.427 | 0.61 | 0.71 | 0.59 | 0.39 | 0.58 | 0.44 |
| 1 | 184812165 | rs10911651 | 1.432 | 0.084 | 0.529 | 0.435 | 0.007 | 0.047 | 0.047 | 0.086 | 0.46 | 0.17 | 0.47 | 0.3 | 0.49 | 0.37 |
| 1 | 184824457 | rs591890 | -1.551 | 0.124 | 0.605 | 0.176 | 0.017 | -0.001 | -0.001 | 0 | 0.52 | 0.8 | 0.49 | 0.6 | 0.5 | 0.62 |
| 1 | 184830048 | rs234669 | -0.452 | 0.256 | 0.986 | 0.328 | -0.007 | 0.022 | 0.022 | 0.23 | 0.06 | 0.28 | 0.04 | 0.05 | 0.02 | 0.03 |
| 1 | 184833344 | rs234644 | -1.489 | 0.13 | 0.861 | 0.191 | 0.016 | 0.004 | 0.004 | 0.02 | 0.49 | 0.77 | 0.46 | 0.6 | 0.49 | 0.6 |
| 1 | 184838335 | rs170915 | -1.524 | 0.183 | 0.596 | 0.191 | 0.016 | -0.002 | -0.002 | 0.085 | 0.51 | 0.82 | 0.46 | 0.6 | 0.49 | 0.6 |

| | | | | | | | | | | | | | | | | |
|---|---|---|---|---|---|---|---|---|---|---|---|---|---|---|---|---|
| 1 | 184842453 | rs555206 | -0.753 | 0.322 | 0.779 | 0.253 | -0.034 | -0.034 | -0.034 | 0 | 0.35 | 0.21 | 0.38 | 0.49 | 0.4 | 0.53 |
| 1 | 184848690 | rs10797991 | 0.151 | 0.65 | 0.528 | 0.228 | -0.097 | 0.009 | 0.009 | 0 | 0.23 | 0.01 | 0.1 | 0.1 | 0.25 | 0.3 |
| 1 | 184851655 | rs532092 | -0.446 | 0.368 | 0.829 | 0.279 | -0.06 | -0.06 | -0.06 | 0 | 0.35 | 0.14 | 0.38 | 0.49 | 0.4 | 0.53 |
| 1 | 184852414 | rs234660 | -0.492 | 0.363 | 1.038 | 0.293 | -0.001 | -0.001 | -0.001 | 0 | 0.36 | 0.33 | 0.4 | 0.49 | 0.4 | 0.53 |
| 1 | 184859611 | rs476539 | -0.394 | 0.412 | 0.536 | 0.362 | -0.052 | -0.052 | -0.052 | 0 | 0.36 | 0.15 | 0.38 | 0.49 | 0.4 | 0.53 |
| 1 | 184878307 | rs234670 | -0.636 | 0.407 | 0.549 | 0.31 | 0.005 | 0.014 | 0.014 | 0 | 0.38 | 0.43 | 0.4 | 0.56 | 0.4 | 0.53 |
| 1 | 184888653 | rs234680 | -0.44 | 0.227 | 0.757 | 0.597 | 0.081 | 0.142 | 0.142 | 1.771 | 0.08 | 0.51 | 0.07 | 0.11 | 0.06 | 0.03 |
| 1 | 184890756 | rs234683 | -0.989 | 0.148 | 0.327 | 0.592 | -0.009 | 0.059 | 0.059 | 4.516 | 0.12 | 0.66 | 0.09 | 0.17 | 0.06 | 0.05 |
| 1 | 184898266 | rs234686 | -1.194 | 0.271 | -0.988 | 0.288 | -0.045 | -0.149 | -0.149 | 0 | 0.67 | 0.94 | 0.68 | 0.59 | 0.68 | 0.68 |
| 1 | 184905261 | rs10911659 | -0.035 | 0.51 | -0.884 | -0.004 | -0.062 | 0.032 | 0.032 | 0 | 0.36 | 0.12 | 0.4 | 0.33 | 0.5 | 0.56 |
| 1 | 184910915 | rs234089 | 0.35 | 0.447 | -0.164 | 0.264 | -0.012 | 0.011 | 0.011 | 0 | 0.39 | 0.51 | 0.31 | 0.38 | 0.33 | 0.35 |
| 1 | 184914948 | rs6686749 | 0.378 | 0.599 | -0.077 | -0.022 | -0.016 | 0.041 | 0.041 | 0 | 0.36 | 0.28 | 0.37 | 0.2 | 0.4 | 0.34 |
| 1 | 184916682 | rs118188 | 0.449 | 0.657 | 0.636 | -0.03 | -0.001 | 0.002 | 0.002 | 0 | 0.44 | 0.47 | 0.34 | 0.39 | 0.39 | 0.39 |
| 1 | 184921581 | rs7512071 | 0.664 | 0.204 | 0.508 | -0.008 | 0.025 | 0.127 | 0.127 | 1.067 | 0.48 | 0.28 | 0.49 | 0.21 | 0.57 | 0.47 |
| 1 | 184922863 | rs4651243 | 0.33 | 0.196 | 0.588 | -0.019 | -0.004 | -0.012 | -0.012 | 0 | 0.55 | 0.53 | 0.53 | 0.21 | 0.64 | 0.51 |
| 1 | 184928165 | rs234123 | 0.11 | 0.193 | 0.199 | 0.153 | -0.015 | 0.025 | 0.025 | 0.037 | 0.31 | 0.42 | 0.23 | 0.33 | 0.23 | 0.2 |
| 1 | 184937385 | rs234098 | -0.02 | 0.097 | 0.275 | 0.23 | -0.02 | 0.095 | 0.095 | 0 | 0.78 | 0.57 | 0.7 | 0.53 | 0.8 | 0.73 |
| 1 | 184938222 | rs234100 | 0.03 | 0.426 | 0.482 | -0.531 | -0.005 | 0.027 | 0.027 | 0.242 | 0.48 | 0.64 | 0.47 | 0.09 | 0.48 | 0.49 |
| 1 | 184941749 | rs170885 | -0.11 | 0.472 | 0.277 | -0.465 | -0.008 | 0.143 | 0.143 | 0.153 | 0.51 | 0.75 | 0.43 | 0.15 | 0.49 | 0.52 |
| 1 | 184951676 | rs234117 | 0.046 | 0.616 | 0.176 | -0.532 | -0.001 | -0.046 | -0.046 | 0.321 | 0.47 | 0.53 | 0.39 | 0.09 | 0.48 | 0.48 |
| 1 | 184961804 | rs12029679 | -0.718 | 0.578 | 0.901 | 0.163 | 0.052 | 0.08 | 0.08 | 0.27 | 0.06 | 0.33 | 0.11 | 0.04 | 0.04 | 0.04 |
| 1 | 184962749 | rs10911661 | -1.103 | 0.327 | 0.948 | 0.137 | -0.01 | -0.002 | -0.002 | 0 | 0.25 | 0.7 | 0.25 | 0.24 | 0.18 | 0.22 |
| 1 | 184967014 | rs7516808 | 0.712 | 0.309 | 0.731 | 0.101 | -0.02 | -0.057 | -0.057 | 0 | 0.11 | 0.24 | 0.06 | 0 | 0.07 | 0.11 |
| 1 | 184986725 | rs12040650 | -0.214 | 0.487 | 0.919 | 0.562 | 0.027 | 0.104 | 0.104 | 0.013 | 0.05 | 0.18 | 0.09 | 0.12 | 0.06 | 0.09 |
| 1 | 184999431 | rs6668714 | 0.883 | 0.536 | 1.066 | 0.488 | -0.001 | -0.026 | -0.026 | 0 | 0.1 | 0.13 | 0.05 | 0 | 0.08 | 0.11 |
| 1 | 185030547 | rs12127009 | 0.408 | 0.616 | 0.577 | 1.045 | -0.032 | -0.034 | -0.034 | 2.418 | 0.1 | 0 | 0.1 | 0.1 | 0.13 | 0.14 |
| 1 | 185041630 | rs12038919 | -0.464 | 0.828 | 0.539 | 0.764 | 0.019 | 0.087 | 0.087 | 0 | 0.09 | 0.16 | 0.16 | 0.27 | 0.14 | 0.11 |
| 1 | 185046574 | rs10911674 | -0.437 | 0.833 | 0.54 | 0.766 | 0.02 | 0.088 | 0.088 | 0 | 0.08 | 0.16 | 0.16 | 0.27 | 0.14 | 0.11 |
| 1 | 185067805 | rs2057222 | 0.346 | 0.591 | 0.161 | 0.268 | 0.014 | -0.016 | -0.016 | 0.687 | 0.41 | 0.78 | 0.47 | 0.41 | 0.36 | 0.45 |
| 1 | 185071831 | rs10911676 | -0.808 | 0.19 | 0.499 | 0.325 | 0.024 | 0.307 | 0.307 | 0.346 | 0.08 | 0.28 | 0.15 | 0.35 | 0.06 | 0.12 |
| 1 | 185110367 | rs6424971 | -0.171 | 0.509 | 0.341 | 0.174 | -0.062 | 0.008 | 0.008 | 0 | 0.52 | 0.9 | 0.54 | 0.39 | 0.49 | 0.55 |
| 1 | 185114837 | rs6684586 | 0.302 | 0.603 | 0.632 | 0.445 | -0.007 | 0.012 | 0.012 | 1.134 | 0.39 | 0.54 | 0.45 | 0.3 | 0.41 | 0.35 |
| 1 | 185121752 | rs10489581 | 0.13 | 0.558 | 0.525 | 0.433 | 0.014 | 0.168 | 0.168 | 0 | 0.06 | 0.16 | 0.14 | 0.25 | 0.06 | 0.08 |
| 1 | 185123630 | rs10737256 | 0.099 | 0.585 | 0.542 | 0.467 | -0.021 | -0.009 | -0.009 | 0.672 | 0.42 | 0.78 | 0.49 | 0.39 | 0.42 | 0.45 |
| 1 | 185125043 | rs1544052 | -0.131 | 0.593 | 0.534 | 0.477 | 0.006 | 0.203 | 0.203 | 0 | 0.06 | 0.16 | 0.14 | 0.34 | 0.06 | 0.11 |
| 1 | 185128250 | rs7548237 | 0.702 | 0.639 | 0.545 | 0.326 | 0.003 | -0.032 | -0.032 | 0.047 | 0.35 | 0.53 | 0.4 | 0.2 | 0.35 | 0.35 |
| 1 | 185135745 | rs950327 | 0.779 | 0.614 | 0.581 | 0.313 | 0.009 | -0.062 | -0.062 | 0 | 0.34 | 0.42 | 0.39 | 0.19 | 0.35 | 0.36 |
| 1 | 185136458 | rs6674490 | 0.382 | 0.277 | 0.154 | 0.38 | -0.027 | 0.04 | 0.04 | 1.531 | 0.41 | 0.78 | 0.49 | 0.3 | 0.42 | 0.43 |
| 1 | 185143721 | rs10489579 | 0.748 | 0.356 | 0.385 | 0.318 | -0.001 | 0 | 0 | 0 | 0.28 | 0.7 | 0.35 | 0.27 | 0.34 | 0.4 |
| 1 | 185151378 | rs1200610 | 1.744 | -0.079 | -0.189 | 0.664 | -0.004 | -0.078 | -0.078 | 0.084 | 0.19 | 0.04 | 0.18 | 0.28 | 0.19 | 0.15 |
| 1 | 185157131 | rs10798000 | 0.747 | 0.038 | 0.149 | 1.006 | -0.008 | -0.012 | -0.012 | 0.016 | 0.3 | 0.75 | 0.35 | 0.37 | 0.35 | 0.43 |
| 1 | 185171869 | rs6698109 | 1.643 | -0.034 | 0.674 | 1.021 | 0.004 | -0.001 | -0.001 | 4.773 | 0.23 | 0.22 | 0.25 | 0.2 | 0.34 | 0.39 |
| 1 | 185176395 | rs7543962 | 1.519 | -0.024 | 0.768 | 0.998 | 0.026 | -0.009 | -0.009 | 0 | 0.24 | 0.3 | 0.26 | 0.2 | 0.34 | 0.39 |
| 1 | 185201061 | rs12096987 | 0.803 | -0.074 | 0.37 | 1.18 | -0.013 | -0.028 | -0.028 | 0 | 0.3 | 0.78 | 0.35 | 0.41 | 0.34 | 0.42 |
| 1 | 185211545 | rs16823859 | 1.294 | -0.226 | 0.067 | 1.05 | -0.021 | -0.068 | -0.068 | 0 | 0.13 | 0.01 | 0.18 | 0.29 | 0.14 | 0.12 |
| 1 | 185212686 | rs2383269 | 0.74 | -0.1 | 0.438 | 0.893 | -0.006 | -0.012 | -0.012 | 0 | 0.3 | 0.73 | 0.34 | 0.4 | 0.34 | 0.41 |
| 1 | 185219680 | rs12135172 | 1.185 | -0.115 | 0.649 | 0.885 | 0.035 | 0.065 | 0.065 | 1.613 | 0.27 | 0.55 | 0.33 | 0.4 | 0.34 | 0.42 |
| 1 | 185227886 | rs6666742 | 0.734 | -0.109 | 0.413 | 0.888 | -0.006 | -0.011 | -0.011 | 0 | 0.3 | 0.73 | 0.34 | 0.4 | 0.34 | 0.42 |
| 1 | 185230850 | rs883753 | 0.568 | -0.062 | 0.658 | 1.091 | -0.006 | 0.022 | 0.022 | 3.271 | 0.41 | 0.65 | 0.44 | 0.24 | 0.43 | 0.46 |
| 1 | 185240474 | rs12041704 | 1.178 | -0.036 | 0.825 | 1.083 | -0.016 | 0.012 | 0.012 | 4.264 | 0.36 | 0.24 | 0.4 | 0.25 | 0.44 | 0.46 |
| 1 | 185253790 | rs7537341 | -0.115 | -0.084 | 0.244 | 1.018 | -0.009 | -0.012 | -0.012 | 0 | 0.49 | 0.83 | 0.52 | 0.43 | 0.48 | 0.54 |
| 1 | 185265149 | rs10798004 | 0.216 | 0.033 | 0.317 | 1.056 | -0.004 | -0.019 | -0.019 | 0 | 0.46 | 0.78 | 0.41 | 0.42 | 0.45 | 0.54 |
| 1 | 185265711 | rs7123 | 0.745 | 0.033 | 0.418 | 0.978 | -0.004 | 0.007 | 0.007 | 1.918 | 0.39 | 0.62 | 0.34 | 0.25 | 0.43 | 0.48 |
| 1 | 185267114 | rs10174 | 0.745 | -0.008 | 0.412 | 0.978 | -0.003 | 0.006 | 0.006 | 1.918 | 0.39 | 0.62 | 0.33 | 0.25 | 0.42 | 0.48 |
| 1 | 185286189 | rs1064902 | 1.525 | 0.052 | 0.227 | 0.947 | 0.006 | 0.027 | 0.027 | 0 | 0.05 | 0.01 | 0.01 | 0 | 0.02 | 0.03 |
| 1 | 185291426 | rs10911704 | 1.107 | 0.044 | 0.577 | 1.02 | 0.005 | -0.007 | -0.007 | 0 | 0.35 | 0.35 | 0.31 | 0.25 | 0.42 | 0.49 |
| 1 | 185295754 | rs726567 | 1.111 | 0.037 | 0.574 | 1.017 | 0.005 | -0.007 | -0.007 | 0 | 0.35 | 0.35 | 0.31 | 0.25 | 0.42 | 0.49 |
| 1 | 185298185 | rs11803823 | -0.703 | -0.002 | 0.707 | 1.166 | 0.06 | 0.326 | 0.326 | 1.567 | 0.06 | 0.4 | 0.09 | 0.19 | 0.01 | 0.03 |
| 1 | 185304859 | rs10911707 | 0.145 | 0.036 | -0.061 | 0.893 | -0.02 | 0.067 | 0.067 | 0.872 | 0.47 | 0.86 | 0.36 | 0.32 | 0.45 | 0.55 |
| 1 | 185317357 | rs6424976 | -0.357 | 0.021 | -0.465 | 0.606 | -0.088 | 0.04 | 0.04 | 0.735 | 0.59 | 0.96 | 0.55 | 0.31 | 0.58 | 0.73 |
| 1 | 185325547 | rs6689206 | -0.261 | 0.032 | -0.457 | 0.711 | -0.087 | 0.089 | 0.089 | 1.153 | 0.58 | 0.94 | 0.55 | 0.24 | 0.56 | 0.7 |
| 1 | 185328658 | rs2224027 | 1.404 | -0.08 | 0.076 | 0.66 | 0.012 | -0.022 | -0.022 | 0.059 | 0.35 | 0.41 | 0.35 | 0.24 | 0.42 | 0.51 |
| 1 | 185331197 | rs1074229 | 0.331 | 0.009 | -0.297 | 0.602 | 0.002 | -0.019 | -0.019 | 0 | 0.15 | 0.15 | 0.15 | 0 | 0.1 | 0.17 |
| 1 | 185338550 | rs10489486 | 1.239 | 0.065 | -0.848 | 1.058 | -0.054 | -0.098 | -0.098 | 0.056 | 0.2 | 0.02 | 0.16 | 0.26 | 0.23 | 0.11 |
| 1 | 185341597 | rs4651254 | -0.381 | 0.138 | -0.864 | 0.942 | -0.025 | -0.022 | -0.022 | 0 | 0.67 | 0.86 | 0.62 | 0.36 | 0.65 | 0.75 |
| 1 | 185344764 | rs977004 | -0.208 | 0.143 | -0.858 | 0.816 | -0.021 | 0.073 | 0.073 | 0.817 | 0.66 | 0.84 | 0.57 | 0.17 | 0.63 | 0.73 |
| 1 | 185347488 | rs10911713 | 1.587 | 0.191 | -0.776 | 0.712 | -0.013 | 0.04 | 0.04 | 3.153 | 0.36 | 0.21 | 0.36 | 0.25 | 0.42 | 0.55 |
| 1 | 185359467 | rs10489483 | -0.103 | 0.307 | -0.606 | 0.808 | -0.02 | -0.133 | -0.133 | 0 | 0.18 | 0.41 | 0.18 | 0 | 0.12 | 0.16 |
| 1 | 185371940 | rs2144291 | 0.012 | 0.637 | -1.322 | 1.449 | -0.046 | -0.044 | -0.044 | 0 | 0.67 | 0.89 | 0.63 | 0.36 | 0.65 | 0.76 |

| | | | | | | | | | | | | | | | | |
|---|---|---|---|---|---|---|---|---|---|---|---|---|---|---|---|---|
| 1 | 185374848 | rs4651256 | -0.582 | 0.575 | -1.028 | 1.779 | -0.024 | -0.087 | -0.087 | 0.538 | 0.87 | 0.99 | 0.79 | 0.63 | 0.88 | 0.89 |
| 1 | 185375966 | rs9803977 | 0.286 | 0.623 | -0.83 | 1.587 | 0.016 | -0.035 | -0.035 | 0 | 0.6 | 0.76 | 0.69 | 0.46 | 0.67 | 0.79 |
| 1 | 185381662 | rs1543409 | -0.022 | 0.969 | -0.243 | 1.241 | 0.002 | 0.008 | 0.008 | 0.696 | 0.32 | 0.35 | 0.26 | 0.39 | 0.3 | 0.16 |
| 1 | 185386615 | rs10489482 | 0.061 | 0.411 | -0.414 | 0.902 | 0.036 | 0.003 | 0.003 | 0 | 0.41 | 0.69 | 0.48 | 0.4 | 0.29 | 0.36 |
| 1 | 185389564 | rs6424982 | -0.391 | 0.693 | -0.434 | 0.625 | -0.004 | -0.013 | -0.013 | 0 | 0.6 | 0.72 | 0.65 | 0.56 | 0.63 | 0.81 |
| 1 | 185394176 | rs10737261 | -0.455 | 0.672 | -0.426 | 0.603 | -0.004 | 0.005 | 0.005 | 0 | 0.6 | 0.74 | 0.69 | 0.61 | 0.63 | 0.81 |
| 1 | 185394834 | rs10798014 | 0.741 | 0.746 | -0.022 | 0.465 | 0.001 | -0.002 | -0.002 | 1.281 | 0.34 | 0.3 | 0.45 | 0.42 | 0.3 | 0.44 |
| 1 | 185395182 | rs10911725 | 0.111 | 0.8 | -0.102 | 0.414 | 0.017 | -0.05 | -0.05 | 0.007 | 0.25 | 0.38 | 0.2 | 0.15 | 0.33 | 0.37 |
| 1 | 185403172 | rs6701364 | 0.11 | 0.93 | -0.279 | 0.529 | -0.008 | 0.003 | 0.003 | 0 | 0.4 | 0.25 | 0.31 | 0.39 | 0.37 | 0.2 |
| 1 | 185410975 | rs4651259 | -0.428 | 0.912 | -0.158 | 0.443 | -0.004 | 0.002 | 0.002 | 0 | 0.61 | 0.56 | 0.52 | 0.57 | 0.66 | 0.6 |
| 1 | 185416257 | rs4651260 | 0.123 | 1.365 | -0.037 | 0.619 | 0.001 | 0.001 | 0.001 | 0.025 | 0.42 | 0.34 | 0.32 | 0.41 | 0.34 | 0.3 |
| 1 | 185422346 | rs7519702 | 0.131 | 1.402 | -0.406 | 0.622 | 0.007 | 0.004 | 0.004 | 0 | 0.42 | 0.29 | 0.31 | 0.41 | 0.33 | 0.3 |
| 1 | 185422424 | rs10911734 | -0.043 | 1.544 | -0.63 | 0.662 | 0.006 | -0.006 | -0.006 | 0 | 0.18 | 0.22 | 0.17 | 0.11 | 0.31 | 0.28 |
| 1 | 185422604 | rs10911736 | -0.006 | 1.568 | -0.839 | 0.674 | -0.003 | 0.003 | 0.003 | 0 | 0.18 | 0.18 | 0.17 | 0.11 | 0.31 | 0.28 |
| 1 | 185426382 | rs232295 | -0.035 | 0.798 | -1.087 | 0.907 | 0.029 | 0.008 | 0.008 | 0 | 0.44 | 0.25 | 0.51 | 0.44 | 0.39 | 0.46 |
| 1 | 185430494 | rs4323662 | -0.341 | 1.125 | -0.325 | 1.383 | 0.011 | 0.077 | 0.077 | 0 | 0.26 | 0.54 | 0.32 | 0.57 | 0.25 | 0.22 |
| 1 | 185432942 | rs1010863 | 0.009 | 0.964 | -0.203 | 1.139 | 0 | 0.002 | 0.002 | 0 | 0.44 | 0.58 | 0.43 | 0.31 | 0.54 | 0.38 |
| 1 | 185438020 | rs1504323 | 0.147 | 0.768 | -0.049 | 1.281 | 0.019 | 0.102 | 0.102 | 0.013 | 0.52 | 0.75 | 0.54 | 0.67 | 0.55 | 0.53 |
| 1 | 185439258 | rs7416136 | 0.793 | 0.554 | -0.048 | 1.09 | -0.042 | 0.046 | 0.046 | 0 | 0.41 | 0.14 | 0.35 | 0.3 | 0.51 | 0.38 |
| 1 | 185444202 | rs10911739 | 0.155 | 0.903 | 0.111 | 0.139 | -0.011 | 0.138 | 0.138 | 0.114 | 0.56 | 0.77 | 0.58 | 0.79 | 0.57 | 0.62 |
| 1 | 185451824 | rs6685999 | -0.579 | 0.627 | 0.903 | -0.119 | 0.001 | -0.106 | -0.106 | 0.106 | 0.38 | 0.55 | 0.38 | 0.04 | 0.4 | 0.19 |
| 1 | 185453884 | rs12128031 | -0.333 | 0.599 | 1.108 | -0.149 | 0.001 | -0.077 | -0.077 | 0 | 0.36 | 0.36 | 0.34 | 0.02 | 0.4 | 0.18 |
| 1 | 185455790 | rs232298 | 1.037 | 0.741 | 0.85 | 0.393 | -0.061 | -0.064 | -0.064 | 0.863 | 0.46 | 0.11 | 0.43 | 0.51 | 0.47 | 0.57 |
| 1 | 185463490 | rs6424984 | -0.725 | 0.233 | 1.378 | 0.602 | -0.014 | 0.108 | 0.108 | 0 | 0.07 | 0.17 | 0.08 | 0.21 | 0.02 | 0.12 |
| 1 | 185472032 | rs6696736 | -0.484 | 0.281 | 1.674 | 0.504 | -0.02 | 0.059 | 0.059 | 0 | 0.42 | 0.23 | 0.4 | 0.26 | 0.46 | 0.31 |
| 1 | 185484309 | rs12063048 | -2.398 | 0.167 | 2.383 | 0.968 | -0.038 | 0.127 | 0.127 | 3.669 | 0.1 | 0.6 | 0.13 | 0.23 | 0.03 | 0.14 |
| 1 | 185489589 | rs10911745 | -0.897 | 0.238 | 1.919 | 0.986 | -0.016 | 0.088 | 0.088 | 0 | 0.06 | 0.12 | 0.06 | 0.21 | 0.02 | 0.11 |
| 1 | 185489752 | rs6670762 | -1.473 | 0.238 | 1.942 | 0.986 | 0.008 | 0.187 | 0.187 | 0.282 | 0.07 | 0.3 | 0.07 | 0.21 | 0.02 | 0.11 |
| 1 | 185495843 | rs7544587 | -1.237 | 0.529 | 1.391 | 0.608 | 0.076 | -0.022 | -0.022 | 0 | 0.09 | 0.53 | 0.11 | 0.01 | 0.1 | 0.03 |
| 1 | 185498284 | rs7536974 | -1.268 | 0.502 | -0.087 | 0.617 | 0.01 | -0.012 | -0.012 | 1.339 | 0.44 | 0.82 | 0.5 | 0.48 | 0.42 | 0.41 |
| 1 | 185500084 | rs1407277 | -1.272 | 0.54 | -0.149 | 0.643 | 0.01 | -0.007 | -0.007 | 0 | 0.45 | 0.79 | 0.5 | 0.47 | 0.43 | 0.41 |
| 1 | 185510752 | rs10489702 | -1.173 | 0.341 | -0.024 | 0.714 | 0.013 | 0.088 | 0.088 | 1.187 | 0.13 | 0.51 | 0.18 | 0.2 | 0.11 | 0.14 |
| 1 | 185515461 | rs1321996 | 0.589 | 0.464 | -1.571 | 0.2 | -0.017 | 0.141 | 0.141 | 0 | 0.52 | 0.09 | 0.52 | 0.75 | 0.54 | 0.66 |
| 1 | 185518666 | rs1321997 | -0.764 | -0.041 | -0.583 | 0.565 | -0.025 | 0.063 | 0.063 | 0 | 0.09 | 0.2 | 0.11 | 0.19 | 0.04 | 0.11 |
| 1 | 185520600 | rs534817 | -1.263 | 0.117 | -1.693 | 0.249 | -0.049 | 0.101 | 0.101 | 0.022 | 0.47 | 0.95 | 0.38 | 0.27 | 0.45 | 0.48 |
| 1 | 185522687 | rs10798019 | -0.208 | -0.39 | -1.92 | -0.353 | -0.008 | 0.082 | 0.082 | 0 | 0.25 | 0.09 | 0.15 | 0.09 | 0.23 | 0.26 |
| 1 | 185524918 | rs1321999 | -0.447 | -0.4 | -1.008 | -0.329 | -0.014 | 0.018 | 0.018 | 0 | 0.08 | 0.16 | 0.05 | 0.11 | 0.06 | 0.07 |
| 1 | 185527738 | rs11605 | -1.345 | -0.492 | -0.069 | -0.408 | 0 | -0.004 | -0.004 | 0 | 0.43 | 0.58 | 0.32 | 0.16 | 0.41 | 0.43 |
| 1 | 185533785 | rs556035 | -1.405 | -0.508 | 0.132 | -0.411 | 0.007 | 0.034 | 0.034 | 0 | 0.43 | 0.64 | 0.33 | 0.16 | 0.41 | 0.43 |
| 1 | 185540618 | rs11583193 | -0.376 | -0.576 | -0.144 | -0.34 | -0.008 | 0.003 | 0.003 | 0 | 0.08 | 0.08 | 0.04 | 0.08 | 0.06 | 0.06 |
| 1 | 185543071 | rs4651272 | -0.245 | -0.712 | 0.062 | -0.258 | 0.012 | 0.071 | 0.071 | 0 | 0.28 | 0.12 | 0.16 | 0.12 | 0.22 | 0.26 |
| 1 | 185543344 | rs333722 | -1.105 | -0.605 | 0.375 | -0.166 | 0.005 | 0.024 | 0.024 | 0 | 0.41 | 0.64 | 0.32 | 0.15 | 0.39 | 0.4 |
| 1 | 185545130 | rs333724 | -2.003 | -0.561 | 0.371 | -0.168 | -0.001 | 0.097 | 0.097 | 0 | 0.53 | 0.83 | 0.39 | 0.28 | 0.47 | 0.5 |
| 1 | 185548517 | rs2208600 | -1.48 | -0.57 | 0.391 | -0.138 | 0.018 | 0.084 | 0.084 | 0 | 0.45 | 0.73 | 0.34 | 0.19 | 0.4 | 0.43 |
| 1 | 185554904 | rs567021 | -1.644 | -0.436 | 0.493 | -0.099 | 0.019 | 0.079 | 0.079 | 0 | 0.51 | 0.72 | 0.38 | 0.27 | 0.46 | 0.5 |
| 1 | 185565704 | rs4344284 | -1.508 | -0.93 | 0.403 | -0.05 | 0.061 | 0.116 | 0.116 | 0 | 0.48 | 0.74 | 0.32 | 0.25 | 0.37 | 0.45 |
| 1 | 185571933 | rs971224 | -1.463 | -0.912 | 0.425 | -0.049 | 0.048 | 0.096 | 0.096 | 0 | 0.48 | 0.69 | 0.31 | 0.25 | 0.37 | 0.45 |
| 1 | 185597494 | rs12089482 | -2.299 | -0.971 | 1.011 | 0.839 | -0.035 | 0.195 | 0.195 | 0 | 0.1 | 0.33 | 0.12 | 0.28 | 0.04 | 0.19 |
| 1 | 185625971 | rs1321991 | -2.525 | -0.887 | 1.136 | 0.878 | -0.01 | 0.235 | 0.235 | 0 | 0.1 | 0.34 | 0.14 | 0.29 | 0.06 | 0.17 |
| 1 | 185675592 | rs4651286 | -0.931 | -1.041 | 1.176 | -0.155 | -0.026 | -0.148 | -0.148 | 0 | 0.32 | 0.49 | 0.19 | 0.05 | 0.27 | 0.3 |
| 1 | 185685544 | rs10127540 | -1.694 | -0.845 | 0.906 | -0.239 | -0.049 | 0.207 | 0.207 | 0.021 | 0.42 | 0.96 | 0.27 | 0.12 | 0.35 | 0.41 |
| 1 | 185689139 | rs900775 | -0.718 | -0.852 | 1.125 | -0.291 | -0.017 | -0.125 | -0.125 | 0 | 0.31 | 0.42 | 0.19 | 0.05 | 0.28 | 0.3 |
| 1 | 185707213 | rs35425624 | -0.194 | -0.614 | 1.074 | -0.327 | -0.003 | 0.011 | 0.011 | 0 | 0.26 | 0.21 | 0.17 | 0.04 | 0.27 | 0.22 |
| 1 | 185716964 | rs4651287 | -0.913 | -0.699 | 1.824 | -0.366 | -0.01 | -0.086 | -0.086 | 0.131 | 0.32 | 0.58 | 0.21 | 0.05 | 0.28 | 0.26 |
| 1 | 185717714 | rs11583867 | -0.143 | -0.707 | 1.802 | -0.369 | -0.001 | 0.003 | 0.003 | 0 | 0.25 | 0.23 | 0.17 | 0.04 | 0.26 | 0.22 |
| 1 | 185742967 | rs6696804 | -1.925 | -0.663 | 2.045 | -0.323 | 0.014 | 0.011 | 0.011 | 0.918 | 0.06 | 0.31 | 0.04 | 0.01 | 0.01 | 0.04 |
| 1 | 185781037 | rs6657554 | -2.092 | -0.667 | 2.124 | -0.325 | 0.039 | 0.035 | 0.035 | 1.232 | 0.06 | 0.41 | 0.04 | 0.01 | 0.01 | 0.05 |
| 1 | 185794315 | rs4651288 | -0.543 | -0.923 | 1.525 | -0.447 | 0.007 | 0.07 | 0.07 | 0 | 0.29 | 0.16 | 0.17 | 0.05 | 0.28 | 0.27 |
| 1 | 185827128 | rs17448230 | -0.696 | -0.875 | 1.565 | -0.453 | -0.001 | -0.045 | -0.045 | 1.116 | 0.31 | 0.34 | 0.19 | 0.05 | 0.29 | 0.27 |
| 1 | 185827238 | rs1407431 | -0.575 | -0.882 | 1.555 | -0.457 | -0.004 | -0.087 | -0.087 | 0 | 0.29 | 0.38 | 0.19 | 0.04 | 0.27 | 0.23 |
| 1 | 185833565 | rs7556537 | -2.204 | -0.936 | 1.253 | 0.309 | -0.101 | -0.143 | -0.143 | 0.116 | 0.54 | 0.97 | 0.44 | 0.48 | 0.45 | 0.54 |
| 1 | 185849364 | rs6698721 | -1.982 | -1.026 | 2.122 | -0.066 | -0.006 | -0.01 | -0.01 | 0.23 | 0.08 | 0.58 | 0.06 | 0.01 | 0.01 | 0.05 |
| 1 | 185855289 | rs12078444 | -1.205 | -1.004 | 1.858 | -0.139 | 0.034 | 0.04 | 0.04 | 0.42 | 0.35 | 0.72 | 0.21 | 0.05 | 0.24 | 0.31 |
| 1 | 185886151 | rs1555494 | -2.439 | -0.651 | 1.554 | 0.37 | -0.046 | -0.052 | -0.052 | 1.119 | 0.1 | 0.68 | 0.07 | 0.01 | 0.02 | 0.06 |
| 1 | 185902978 | rs12068599 | -0.943 | -0.587 | 1.767 | 0.397 | -0.008 | -0.019 | -0.019 | 0 | 0.05 | 0.23 | 0.04 | 0 | 0.02 | 0 |
| 1 | 185906206 | rs16824765 | -0.449 | -0.87 | 0.701 | 0.266 | -0.074 | -0.128 | -0.128 | 0 | 0.24 | 0.04 | 0.2 | 0.39 | 0.26 | 0.22 |
| 1 | 185940266 | rs10911789 | -1.81 | -0.878 | 1.509 | -0.006 | -0.004 | -0.015 | -0.015 | 0.887 | 0.09 | 0.48 | 0.06 | 0.01 | 0.02 | 0.06 |
| 1 | 185958737 | rs7539719 | -2.143 | -0.955 | 1.805 | -0.026 | -0.033 | -0.036 | -0.036 | 0 | 0.1 | 0.6 | 0.07 | 0.01 | 0.03 | 0.06 |
| 1 | 185984517 | rs6665753 | 1.471 | -1.018 | 0.251 | -0.077 | -0.165 | -0.066 | -0.066 | 0 | 0.37 | 0.03 | 0.43 | 0.42 | 0.48 | 0.48 |

| | | | | | | | | | | | | | | | | |
|---|---|---|---|---|---|---|---|---|---|---|---|---|---|---|---|---|
| 1 | 185987152 | rs10911798 | 0.428 | -1.462 | -0.294 | -0.014 | -0.018 | 0.011 | 0.011 | 0 | 0.23 | 0.04 | 0.13 | 0.18 | 0.22 | 0.14 |
| 1 | 185995616 | rs4141459 | -1.778 | -1.307 | 0.143 | -0.28 | -0.061 | 0 | 0 | 3.549 | 0.62 | 0.82 | 0.65 | 0.56 | 0.52 | 0.5 |
| 1 | 186026474 | rs12129650 | -1.169 | -1.344 | 0.421 | -0.269 | 0.001 | -0.003 | -0.003 | 0 | 0.57 | 0.54 | 0.61 | 0.57 | 0.5 | 0.46 |
| 1 | 186050417 | rs10798035 | -1.484 | -1.49 | 0.041 | -0.563 | -0.02 | 0.017 | 0.017 | 0 | 0.59 | 0.7 | 0.63 | 0.57 | 0.5 | 0.5 |
| 1 | 186055371 | rs6425017 | -1.828 | -1.452 | -0.222 | -0.539 | -0.12 | -0.016 | -0.016 | 7.722 | 0.63 | 0.92 | 0.66 | 0.58 | 0.52 | 0.54 |
| 1 | 186073181 | rs10911820 | -1.558 | -1.467 | -0.228 | -0.548 | -0.087 | 0.013 | 0.013 | 0 | 0.61 | 0.89 | 0.65 | 0.58 | 0.5 | 0.5 |
| 1 | 186090370 | rs2146098 | -1.386 | -1.61 | -0.087 | -0.681 | 0.05 | 0.057 | 0.057 | 0 | 0.52 | 0.77 | 0.55 | 0.57 | 0.35 | 0.51 |
| 1 | 186097908 | rs6673390 | -0.916 | -1.659 | 0.36 | -0.674 | 0.002 | 0.002 | 0.002 | 0 | 0.45 | 0.56 | 0.52 | 0.54 | 0.32 | 0.44 |
| 1 | 186101539 | rs10911825 | -0.992 | -1.816 | 0.245 | -0.848 | 0.008 | 0.001 | 0.001 | 0.018 | 0.38 | 0.65 | 0.48 | 0.57 | 0.26 | 0.4 |
| 1 | 186104488 | rs10489755 | -0.563 | -1.816 | 0.213 | -0.834 | -0.02 | 0.008 | 0.008 | 0 | 0.34 | 0.53 | 0.46 | 0.55 | 0.24 | 0.37 |
| 1 | 186114325 | rs1322374 | -1.466 | -2.127 | 0.079 | -1.026 | -0.02 | 0.011 | 0.011 | 0.017 | 0.34 | 0.53 | 0.46 | 0.54 | 0.25 | 0.38 |
| 1 | 186120139 | rs1322376 | -0.576 | -2.351 | -0.335 | -1.25 | -0.026 | -0.116 | -0.116 | 0 | 0.28 | 0.41 | 0.16 | 0.05 | 0.22 | 0.14 |
| 1 | 186126099 | rs7545010 | -2.284 | -2.201 | 0.156 | -1.141 | -0.013 | 0.011 | 0.011 | 0 | 0.61 | 0.7 | 0.6 | 0.59 | 0.47 | 0.57 |
| 1 | 186133290 | rs2208692 | -1.419 | -2.287 | -0.088 | -1.164 | -0.007 | -0.066 | -0.066 | 0 | 0.47 | 0.51 | 0.23 | 0.1 | 0.36 | 0.28 |
| 1 | 186136572 | rs2383457 | -1.745 | -2.215 | -0.252 | -1.159 | 0.006 | -0.031 | -0.031 | 0 | 0.5 | 0.47 | 0.24 | 0.1 | 0.37 | 0.32 |
| 1 | 186139951 | rs16824998 | 0.368 | -2.086 | -0.724 | -1.047 | 0.009 | 0.006 | 0.006 | 0 | 0.1 | 0.06 | 0.04 | 0.01 | 0.05 | 0.08 |
| 1 | 186149032 | rs1407434 | 0.25 | -1.793 | -0.452 | -0.65 | 0.002 | 0.166 | 0.166 | 0 | 0.12 | 0.24 | 0.36 | 0.48 | 0.1 | 0.23 |
| 1 | 186149235 | rs10911836 | -1.406 | -1.745 | -1.164 | -1.211 | 0.028 | 0.06 | 0.06 | 0 | 0.38 | 0.27 | 0.16 | 0.09 | 0.31 | 0.24 |
| 1 | 186149704 | rs17466882 | -0.395 | -0.792 | -1.392 | -0.555 | -0.042 | 0.055 | 0.055 | 0 | 0.14 | 0.01 | 0.08 | 0.02 | 0.17 | 0.07 |
| 1 | 186154816 | rs10489750 | -0.686 | -0.625 | -1.174 | 0.366 | 0 | 0.007 | 0.007 | 0 | 0.2 | 0.22 | 0.34 | 0.5 | 0.19 | 0.28 |
| 1 | 186156052 | rs682568 | 0.155 | 0.013 | -1.738 | -0.561 | -0.015 | 0.035 | 0.035 | 0 | 0.24 | 0.11 | 0.24 | 0.01 | 0.28 | 0.21 |
| 1 | 186156514 | rs680638 | -1.243 | 0.201 | -2.285 | -0.369 | -0.059 | -0.103 | -0.103 | 1.09 | 0.83 | 0.99 | 0.79 | 0.63 | 0.77 | 0.72 |
| 1 | 186163076 | rs1322375 | 0.63 | 0.245 | -1.935 | -0.311 | -0.001 | 0 | 0 | 0 | 0.13 | 0.15 | 0.15 | 0.1 | 0.1 | 0.09 |
| 1 | 186164899 | rs651967 | 1.016 | 0.413 | -2.156 | -0.454 | -0.03 | 0.068 | 0.068 | 0 | 0.2 | 0.07 | 0.21 | 0.01 | 0.23 | 0.17 |
| 1 | 186167933 | rs10752963 | 0.591 | 0.307 | -1.597 | -0.366 | 0.002 | 0.01 | 0.01 | 0 | 0.39 | 0.47 | 0.36 | 0.52 | 0.37 | 0.45 |
| 1 | 186168281 | rs857753 | -0.607 | 0.441 | -1.692 | 0.777 | -0.066 | -0.098 | -0.098 | 0.48 | 0.84 | 0.98 | 0.77 | 0.63 | 0.76 | 0.73 |
| 1 | 186172443 | rs1885522 | 0.074 | 0.669 | -1.439 | 0.424 | 0.006 | -0.003 | -0.003 | 0 | 0.12 | 0.1 | 0.13 | 0.11 | 0.1 | 0.08 |
| 1 | 186173178 | rs676236 | -0.656 | 1.143 | -1.252 | 1.01 | -0.063 | -0.106 | -0.106 | 0 | 0.87 | 0.99 | 0.68 | 0.47 | 0.78 | 0.76 |
| 1 | 186175300 | rs6696879 | -0.996 | 0.686 | -0.45 | 0.113 | -0.003 | -0.006 | -0.006 | 0 | 0.09 | 0.26 | 0.15 | 0.06 | 0.06 | 0.08 |
| 1 | 186182156 | rs6666859 | -0.599 | 1.136 | -0.869 | -0.088 | -0.001 | -0.018 | -0.018 | 0 | 0.11 | 0.13 | 0.18 | 0 | 0.16 | 0.05 |
| 1 | 186189394 | rs643128 | -0.334 | 2.009 | -1.308 | -0.28 | -0.028 | 0.021 | 0.021 | 0 | 0.14 | 0.1 | 0.09 | 0.13 | 0.21 | 0.19 |
| 1 | 186190163 | rs6676055 | 0.945 | 1.611 | -1.095 | -1.191 | 0.024 | 0.032 | 0.032 | 0 | 0.49 | 0.36 | 0.47 | 0.81 | 0.36 | 0.63 |
| 1 | 186190474 | rs628066 | 0.535 | 1.701 | -1.116 | -1.053 | -0.053 | 0.027 | 0.027 | 0 | 0.14 | 0.03 | 0.08 | 0.13 | 0.21 | 0.19 |
| 1 | 186190564 | rs650797 | 0.535 | 1.748 | -1.116 | -1.053 | -0.054 | 0.027 | 0.027 | 0 | 0.14 | 0.03 | 0.08 | 0.13 | 0.21 | 0.19 |
| 1 | 186199641 | rs585738 | 0.541 | 1.741 | -1.125 | -1.052 | -0.061 | 0.026 | 0.026 | 0 | 0.14 | 0.03 | 0.08 | 0.13 | 0.21 | 0.2 |
| 1 | 186205867 | rs1743691 | 0.415 | 1.724 | -1.079 | -1.061 | -0.044 | 0.028 | 0.028 | 0 | 0.14 | 0.06 | 0.09 | 0.13 | 0.21 | 0.2 |
| 1 | 186225257 | rs639524 | -0.975 | 1.594 | -1.115 | -1.293 | -0.04 | 0.104 | 0.104 | 0 | 0.77 | 0.95 | 0.83 | 0.97 | 0.73 | 0.91 |
| 1 | 186237510 | rs1285115 | 0.753 | 1.561 | -0.782 | -1.34 | -0.009 | 0.019 | 0.019 | 0 | 0.58 | 0.33 | 0.7 | 0.83 | 0.52 | 0.67 |
| 1 | 186243582 | rs2146060 | 0.765 | 1.525 | -0.752 | -1.309 | -0.03 | 0.019 | 0.019 | 0 | 0.13 | 0.06 | 0.08 | 0.13 | 0.2 | 0.2 |
| 1 | 186247924 | rs1795022 | 0.169 | 1.456 | -1.114 | -0.103 | 0.002 | 0.006 | 0.006 | 0 | 0.64 | 0.37 | 0.59 | 0.54 | 0.59 | 0.68 |
| 1 | 186255294 | rs857590 | 0.523 | 1.413 | -0.704 | -0.13 | -0.018 | -0.012 | -0.012 | 2.612 | 0.57 | 0.69 | 0.43 | 0.51 | 0.47 | 0.58 |
| 1 | 186257518 | rs857591 | 0.461 | 1.051 | -0.666 | 0.226 | 0.008 | 0.044 | 0.044 | 0.35 | 0.52 | 0.7 | 0.38 | 0.37 | 0.42 | 0.56 |
| 1 | 186258229 | rs1293970 | -0.052 | 1.035 | -0.732 | 0.088 | -0.018 | 0.097 | 0.097 | 0.81 | 0.61 | 0.8 | 0.56 | 0.93 | 0.56 | 0.75 |
| 1 | 186261061 | rs857587 | 1.481 | 0.85 | -0.658 | 0.258 | 0.02 | 0 | 0 | 0 | 0.37 | 0.31 | 0.27 | 0.32 | 0.22 | 0.37 |
| 1 | 186264721 | rs35877985 | 0.839 | 0.905 | -0.877 | 0.449 | -0.027 | 0.014 | 0.014 | 0.093 | 0.08 | 0.03 | 0.18 | 0.49 | 0.13 | 0.19 |
| 1 | 186266293 | rs1293988 | 0.17 | 0.842 | -0.802 | 0.007 | -0.02 | 0.11 | 0.11 | 0.836 | 0.61 | 0.81 | 0.56 | 0.93 | 0.56 | 0.75 |
| 1 | 186267100 | rs1293989 | 0.176 | 0.833 | -0.809 | 0.002 | -0.017 | 0.095 | 0.095 | 0.856 | 0.61 | 0.8 | 0.56 | 0.93 | 0.56 | 0.75 |
| 1 | 186270894 | rs1547084 | -0.36 | 1.016 | -0.896 | 0.051 | -0.02 | 0.048 | 0.048 | 0 | 0.29 | 0.15 | 0.41 | 0.13 | 0.33 | 0.22 |
| 1 | 186273994 | rs2273779 | -0.447 | 1.014 | -0.899 | 0.048 | -0.019 | 0.052 | 0.052 | 0.065 | 0.3 | 0.15 | 0.41 | 0.13 | 0.33 | 0.22 |
| 1 | 186275760 | rs16825150 | -0.086 | 1.213 | -0.439 | -0.1 | 0.129 | 0.04 | 0.04 | 0 | 0.09 | 0.31 | 0.09 | 0.13 | 0.15 | 0.14 |
| 1 | 186277989 | rs3737940 | -0.147 | 1.201 | -0.404 | -0.105 | -0.034 | 0.068 | 0.068 | 0 | 0.28 | 0.11 | 0.4 | 0.13 | 0.33 | 0.22 |
| 1 | 186281696 | rs7756 | -1.604 | 1.243 | -0.346 | 0.29 | 0.047 | 0.018 | 0.018 | 0 | 0.49 | 0.62 | 0.6 | 0.66 | 0.62 | 0.53 |
| 1 | 186290521 | rs10494584 | 0.315 | 1.28 | -0.287 | 0.368 | 0.021 | 0.002 | 0.002 | 0 | 0.08 | 0.12 | 0.07 | 0.13 | 0.15 | 0.14 |
| 1 | 186307064 | rs16825185 | -0.129 | 1.329 | -0.342 | 0.437 | 0.129 | 0.044 | 0.044 | 0 | 0.09 | 0.31 | 0.09 | 0.13 | 0.15 | 0.14 |
| 1 | 186315707 | rs3766703 | -0.129 | 1.322 | -0.332 | 0.353 | 0.129 | 0.04 | 0.04 | 0 | 0.09 | 0.31 | 0.09 | 0.13 | 0.15 | 0.14 |
| 1 | 186316488 | rs3753565 | 0.617 | 1.318 | -0.322 | 0.351 | -0.011 | -0.008 | -0.008 | 0 | 0.07 | 0.02 | 0.06 | 0.13 | 0.15 | 0.14 |
| 1 | 186330643 | rs10494586 | -1.353 | 1.297 | -0.356 | 0.33 | 0.001 | 0 | 0 | 0.472 | 0.43 | 0.53 | 0.58 | 0.65 | 0.58 | 0.48 |
| 1 | 186343759 | rs2297226 | -0.112 | 1.319 | -0.354 | 0.339 | 0.129 | 0.04 | 0.04 | 0 | 0.09 | 0.31 | 0.09 | 0.13 | 0.15 | 0.14 |
| 1 | 186345281 | rs6425028 | 0.369 | 1.33 | -0.422 | 0.344 | 0.022 | 0.002 | 0.002 | 0 | 0.08 | 0.12 | 0.07 | 0.13 | 0.15 | 0.14 |
| 1 | 186351104 | rs2383477 | 0.87 | 1.309 | -0.597 | 0.224 | -0.047 | 0.03 | 0.03 | 0.054 | 0.15 | 0.01 | 0.27 | 0.05 | 0.14 | 0.14 |
| 1 | 186358588 | rs1003127 | -0.139 | 1.357 | -0.47 | 0.32 | -0.02 | 0.048 | 0.048 | 0 | 0.3 | 0.15 | 0.41 | 0.13 | 0.33 | 0.22 |
| 1 | 186359750 | rs3115752 | 1.003 | 0.864 | -0.734 | -0.041 | 0.023 | 0.05 | 0.05 | 0.056 | 0.12 | 0.02 | 0.03 | 0 | 0.07 | 0.05 |
| 1 | 186379265 | rs6665157 | -0.991 | 1.72 | -0.173 | 0.409 | 0.114 | 0.258 | 0.258 | 0 | 0.14 | 0.38 | 0.18 | 0.53 | 0.25 | 0.25 |
| 1 | 186382460 | rs10911858 | -0.32 | 1.577 | -0.37 | 0.329 | -0.02 | 0.048 | 0.048 | 0 | 0.3 | 0.15 | 0.41 | 0.13 | 0.33 | 0.22 |
| 1 | 186390303 | rs8824 | -1.397 | 1.321 | -0.326 | 0.472 | 0.006 | 0 | 0 | 0 | 0.44 | 0.54 | 0.58 | 0.63 | 0.58 | 0.48 |
| 1 | 186403648 | rs4567244 | -0.431 | 1.225 | -0.247 | 0.515 | 0.01 | 0.123 | 0.123 | 0.015 | 0.23 | 0.49 | 0.4 | 0.43 | 0.23 | 0.27 |
| 1 | 186411579 | rs1882926 | 0.766 | 1.265 | -0.187 | 0.499 | -0.001 | -0.012 | -0.012 | 0 | 0.15 | 0.14 | 0.17 | 0.04 | 0.11 | 0.14 |
| 1 | 186412913 | rs35066734 | 0.766 | 1.265 | -0.187 | 0.499 | -0.001 | -0.012 | -0.012 | 0 | 0.15 | 0.14 | 0.17 | 0.04 | 0.11 | 0.14 |
| 1 | 186418600 | rs12407957 | 0.22 | 1.261 | -0.185 | 0.498 | -0.001 | -0.08 | -0.08 | 0 | 0.18 | 0.44 | 0.2 | 0.04 | 0.12 | 0.14 |

| | | | | | | | | | | | | | | | | |
|---|---|---|---|---|---|---|---|---|---|---|---|---|---|---|---|---|
| 1 | 186418798 | rs11801527 | 0.227 | 1.272 | -0.229 | 0.553 | -0.004 | 0.076 | 0.076 | 0.527 | 0.06 | 0.04 | 0.2 | 0.39 | 0.11 | 0.13 |
| 1 | 186425526 | rs10798041 | 0.214 | 1.246 | -0.182 | 0.43 | -0.001 | -0.08 | -0.08 | 0 | 0.18 | 0.44 | 0.2 | 0.04 | 0.12 | 0.14 |
| 1 | 186441585 | rs3131557 | -1.504 | 1.447 | -0.212 | 0.387 | 0.002 | 0.002 | 0.002 | 0.036 | 0.36 | 0.58 | 0.48 | 0.56 | 0.43 | 0.45 |
| 1 | 186447591 | rs12062743 | 0.508 | 1.367 | -0.16 | 0.421 | -0.004 | -0.047 | -0.047 | 0 | 0.16 | 0.28 | 0.18 | 0.04 | 0.12 | 0.14 |
| 1 | 186467274 | rs3901950 | -0.549 | 1.379 | -0.156 | 0.43 | 0.009 | 0.122 | 0.122 | 0.028 | 0.23 | 0.5 | 0.4 | 0.43 | 0.23 | 0.27 |
| 1 | 186474889 | rs12072014 | 0.673 | 1.389 | -0.177 | 0.282 | -0.001 | -0.041 | -0.041 | 0.577 | 0.16 | 0.26 | 0.18 | 0.04 | 0.12 | 0.15 |
| 1 | 186483596 | rs3863719 | 0.722 | 1.389 | -0.179 | 0.281 | 0 | -0.027 | -0.027 | 0 | 0.15 | 0.17 | 0.17 | 0.04 | 0.12 | 0.15 |
| 1 | 186504772 | rs16825519 | 0.388 | 1.442 | -0.311 | 0.41 | -0.026 | 0.036 | 0.036 | 0.428 | 0.05 | 0.01 | 0.19 | 0.38 | 0.12 | 0.13 |
| 1 | 186506889 | rs10911878 | -0.612 | 1.407 | -0.214 | 0.388 | 0.006 | 0.113 | 0.113 | 0.031 | 0.23 | 0.53 | 0.4 | 0.43 | 0.24 | 0.28 |
| 1 | 186509790 | rs7550953 | 0.45 | 1.401 | -0.207 | 0.198 | 0.007 | -0.068 | -0.068 | 3.592 | 0.16 | 0.42 | 0.2 | 0.04 | 0.12 | 0.15 |
| 1 | 186532231 | rs7542810 | 0.825 | 1.417 | -0.281 | 0.206 | 0.001 | -0.023 | -0.023 | 0 | 0.14 | 0.15 | 0.17 | 0.04 | 0.12 | 0.15 |
| 1 | 186547952 | rs1569879 | 0.908 | 1.419 | -0.339 | 0.207 | -0.002 | 0.014 | 0.014 | 0 | 0.14 | 0.07 | 0.16 | 0.04 | 0.12 | 0.15 |
| 1 | 186557453 | rs4651315 | 0.161 | 1.495 | -0.597 | 0.219 | -0.039 | -0.092 | -0.092 | 0 | 0.2 | 0.04 | 0.24 | 0.28 | 0.26 | 0.27 |
| 1 | 186576930 | rs11811335 | -0.319 | 1.541 | -0.314 | 0.52 | 0.006 | 0.134 | 0.134 | 0 | 0.06 | 0.12 | 0.21 | 0.45 | 0.12 | 0.09 |
| 1 | 186581005 | rs12038031 | 1.284 | 1.593 | -0.439 | 0.539 | -0.008 | -0.024 | -0.024 | 1.19 | 0.07 | 0 | 0.12 | 0.23 | 0.08 | 0.2 |
| 1 | 186598526 | rs6658482 | -0.288 | 1.688 | -0.182 | 0.399 | -0.058 | -0.007 | -0.007 | 0 | 0.12 | 0.02 | 0.08 | 0.14 | 0.19 | 0.18 |
| 1 | 186599524 | rs2383513 | -0.411 | 1.219 | 0.009 | 0.447 | 0.058 | 0.353 | 0.353 | 0 | 0.06 | 0.26 | 0.22 | 0.45 | 0.12 | 0.07 |
| 1 | 186612593 | rs10911898 | -0.91 | 1.146 | -0.021 | 0.331 | -0.002 | 0.061 | 0.061 | 0.035 | 0.24 | 0.63 | 0.41 | 0.49 | 0.24 | 0.27 |
| 1 | 186623470 | rs12405141 | 0.54 | 0.733 | 0.135 | -0.879 | 0 | -0.002 | -0.002 | 0.221 | 0.15 | 0.11 | 0.17 | 0.04 | 0.12 | 0.16 |
| 1 | 186641058 | rs689470 | -1.767 | 0.722 | 0.182 | -0.885 | 0.035 | 0.019 | 0.019 | 0.893 | 0.07 | 0.5 | 0.05 | 0.01 | 0.03 | 0.07 |
| 1 | 186643058 | rs5275 | -1.721 | 1.118 | 0.468 | -0.61 | -0.001 | -0.004 | -0.004 | 0.062 | 0.34 | 0.64 | 0.37 | 0.2 | 0.31 | 0.4 |
| 1 | 186650846 | rs689465 | 0.371 | 0.954 | 0.707 | -0.74 | 0 | -0.008 | -0.008 | 0 | 0.16 | 0.16 | 0.18 | 0.05 | 0.13 | 0.16 |
| 1 | 186654076 | rs2179555 | -0.264 | 0.945 | 0.745 | -0.745 | -0.006 | -0.078 | -0.078 | 0 | 0.19 | 0.39 | 0.19 | 0.04 | 0.13 | 0.2 |
| 1 | 186659125 | rs12064238 | -0.352 | 0.932 | 0.751 | -0.754 | -0.008 | -0.089 | -0.089 | 0.058 | 0.2 | 0.43 | 0.22 | 0.05 | 0.15 | 0.19 |
| 1 | 186669092 | rs2383529 | -1.193 | 0.987 | 0.698 | -0.77 | -0.003 | -0.067 | -0.067 | 0 | 0.27 | 0.62 | 0.24 | 0.05 | 0.2 | 0.28 |
| 1 | 186684977 | rs964570 | 1.16 | 1.111 | 0.632 | -0.696 | 0.027 | -0.05 | -0.05 | 0 | 0.91 | 0.71 | 0.97 | 1 | 0.93 | 0.88 |
| 1 | 186685818 | rs6685280 | 0.706 | 1.144 | 0.559 | -0.699 | 0.007 | -0.092 | -0.092 | 0.315 | 0.76 | 0.61 | 0.79 | 0.95 | 0.81 | 0.72 |
| 1 | 186687347 | rs766400 | 0.293 | 1.003 | 0.183 | -0.442 | 0.003 | -0.02 | -0.02 | 0 | 0.94 | 0.91 | 0.98 | 1 | 0.93 | 0.92 |
| 1 | 186704555 | rs4650701 | -0.004 | 1.053 | 0.001 | -0.418 | 0.033 | -0.033 | -0.033 | 0 | 0.94 | 0.82 | 0.97 | 1 | 0.93 | 0.92 |
| 1 | 186722430 | rs760707 | -0.119 | 1.09 | -0.067 | -0.389 | 0.011 | -0.028 | -0.028 | 0.798 | 0.95 | 0.91 | 0.98 | 1 | 0.95 | 0.93 |
| 1 | 186725003 | rs4140564 | -0.42 | 1.1 | -0.084 | -0.385 | 0.013 | -0.025 | -0.025 | 0 | 0.95 | 0.91 | 0.98 | 1 | 0.95 | 0.93 |
| 1 | 186742947 | rs10489401 | 0.6 | 1.111 | -0.554 | -0.669 | -0.062 | 0.104 | 0.104 | 0 | 0.3 | 0.03 | 0.22 | 0.11 | 0.36 | 0.3 |
| 1 | 186744874 | rs10752974 | 0.993 | 1.127 | 0.409 | -0.347 | 0.031 | -0.069 | -0.069 | 0 | 0.88 | 0.53 | 0.94 | 0.99 | 0.92 | 0.9 |
| 1 | 186746556 | rs10489402 | 0.113 | 1.134 | 0.202 | -0.327 | 0.03 | 0.012 | 0.012 | 0 | 0.06 | 0.33 | 0.04 | 0.01 | 0.03 | 0.03 |
| 1 | 186763179 | rs7521061 | -0.297 | 1.209 | -0.042 | -0.301 | 0.013 | -0.026 | -0.026 | 0 | 0.95 | 0.91 | 0.98 | 1 | 0.95 | 0.93 |
| 1 | 186767992 | rs7554410 | 0.957 | 1.139 | 0.043 | -0.338 | 0.02 | -0.078 | -0.078 | 0 | 0.88 | 0.56 | 0.94 | 0.99 | 0.92 | 0.89 |
| 1 | 186776884 | rs4651329 | -0.003 | 1.56 | -0.541 | -0.054 | 0.009 | -0.025 | -0.025 | 0 | 0.95 | 0.91 | 0.98 | 1 | 0.95 | 0.93 |
| 1 | 186784505 | rs926324 | -0.485 | 1.81 | -1.032 | 0.176 | -0.001 | -0.001 | -0.001 | 0 | 0.24 | 0.2 | 0.31 | 0.25 | 0.25 | 0.33 |
| 1 | 186797479 | rs979924 | 0.442 | 1.52 | -1.25 | 0.713 | -0.044 | -0.034 | -0.034 | 0.508 | 0.91 | 0.99 | 0.82 | 0.83 | 0.89 | 0.84 |
| 1 | 186808343 | rs3820185 | -0.165 | 0.73 | -0.966 | 0.846 | -0.001 | -0.001 | -0.001 | 0.171 | 0.37 | 0.39 | 0.39 | 0.58 | 0.36 | 0.32 |
| 1 | 186823284 | rs2076075 | -1.032 | 1.277 | -1.031 | 1.767 | -0.011 | 0.044 | 0.044 | 0 | 0.08 | 0.07 | 0.28 | 0.48 | 0.1 | 0.09 |
| 1 | 186830092 | rs12404877 | -0.146 | 0.461 | -1.744 | 0.731 | 0.016 | 0.077 | 0.077 | 0 | 0.27 | 0.08 | 0.16 | 0.09 | 0.2 | 0.23 |
| 1 | 186831073 | rs10737270 | -0.099 | 0.28 | -1.637 | 0.666 | -0.005 | -0.015 | -0.015 | 0.977 | 0.84 | 0.92 | 0.86 | 0.78 | 0.84 | 0.83 |
| 1 | 186831728 | rs6685652 | 0.307 | 0.101 | -0.545 | 0.574 | 0.005 | 0.032 | 0.032 | 0 | 0.82 | 0.56 | 0.84 | 0.78 | 0.83 | 0.83 |
| 1 | 186844926 | rs2223307 | 0.89 | -0.61 | -0.432 | 0.65 | 0.001 | 0 | 0 | 0 | 0.68 | 0.37 | 0.5 | 0.34 | 0.78 | 0.58 |
| 1 | 186849186 | rs12746200 | -0.158 | -0.718 | -0.606 | 0.628 | -0.011 | 0.018 | 0.018 | 0 | 0.09 | 0.01 | 0.15 | 0.05 | 0.08 | 0.07 |
| 1 | 186850960 | rs1980444 | 1.04 | -1.739 | 0.255 | 0.617 | -0.017 | 0.037 | 0.037 | 0.73 | 0.75 | 0.32 | 0.64 | 0.43 | 0.83 | 0.72 |
| 1 | 186851157 | rs17526478 | 1.966 | -1.386 | -0.022 | 0.568 | 0.004 | 0.115 | 0.115 | 1.113 | 0.55 | 0.08 | 0.44 | 0.3 | 0.68 | 0.46 |
| 1 | 186851216 | rs2049963 | -1.373 | -1.65 | 0.416 | 0.866 | -0.068 | 0.13 | 0.13 | 0.231 | 0.22 | 0.54 | 0.3 | 0.57 | 0.1 | 0.25 |
| 1 | 186852597 | rs10911945 | 0.28 | -1.673 | 0.577 | 0.823 | -0.033 | -0.019 | -0.019 | 0.396 | 0.11 | 0.17 | 0.04 | 0.04 | 0.05 | 0.11 |
| 1 | 186855555 | rs16826060 | -0.746 | -0.979 | 0.295 | 1.212 | 0.012 | 0.159 | 0.159 | 0 | 0.1 | 0.22 | 0.27 | 0.46 | 0.08 | 0.14 |
| 1 | 186858135 | rs12042344 | -0.673 | -0.975 | 0.297 | 1.433 | 0.007 | 0.089 | 0.089 | 0 | 0.09 | 0.15 | 0.25 | 0.35 | 0.08 | 0.13 |
| 1 | 186868740 | rs10911952 | -0.678 | -1.046 | -0.246 | 0.523 | -0.015 | 0.024 | 0.024 | 0 | 0.25 | 0.21 | 0.5 | 0.14 | 0.26 | 0.39 |
| 1 | 186870263 | rs6695515 | -0.747 | -1.897 | -0.678 | 0.405 | -0.005 | -0.002 | -0.002 | 0 | 0.32 | 0.6 | 0.2 | 0.37 | 0.18 | 0.28 |
| 1 | 186873704 | rs1569479 | -1.733 | -1.882 | -0.712 | 0.409 | -0.009 | -0.05 | -0.05 | 0 | 0.57 | 0.81 | 0.7 | 0.51 | 0.44 | 0.67 |
| 1 | 186875459 | rs10798069 | 1.788 | -1.768 | -0.782 | 0.385 | -0.091 | -0.096 | -0.096 | 0.225 | 0.41 | 0.1 | 0.28 | 0.48 | 0.51 | 0.31 |
| 1 | 186879043 | rs6683515 | 0.065 | -1.754 | -0.764 | 0.375 | -0.059 | 0.035 | 0.035 | 0 | 0.23 | 0.08 | 0.38 | 0.13 | 0.24 | 0.37 |
| 1 | 186887107 | rs6686848 | -0.098 | -1.701 | -0.476 | 0.403 | -0.05 | 0.057 | 0.057 | 0 | 0.14 | 0.41 | 0.09 | 0.2 | 0.07 | 0.09 |
| 1 | 186894402 | rs4336803 | -1.08 | -1.736 | -0.518 | 0.375 | -0.026 | 0.004 | 0.004 | 0.186 | 0.32 | 0.59 | 0.2 | 0.38 | 0.18 | 0.29 |
| 1 | 186898065 | rs12726519 | -1.144 | -1.736 | -0.555 | 0.375 | -0.003 | -0.002 | -0.002 | 0 | 0.32 | 0.64 | 0.21 | 0.37 | 0.18 | 0.29 |
| 1 | 186910591 | rs6697145 | -2.069 | -1.712 | -0.594 | 0.356 | -0.002 | -0.038 | -0.038 | 0 | 0.57 | 0.8 | 0.7 | 0.51 | 0.44 | 0.67 |
| 1 | 186913263 | rs2207209 | 2.185 | -1.692 | -0.819 | 0.353 | -0.103 | -0.105 | -0.105 | 0 | 0.41 | 0.07 | 0.27 | 0.48 | 0.51 | 0.31 |
| 1 | 186917520 | rs7555140 | -0.008 | -1.592 | -0.798 | 0.365 | -0.041 | 0.045 | 0.045 | 0 | 0.73 | 0.84 | 0.49 | 0.86 | 0.69 | 0.6 |
| 1 | 186920114 | rs10489409 | 0.141 | -1.592 | -0.725 | 0.421 | 0.035 | -0.015 | -0.015 | 0 | 0.17 | 0.08 | 0.1 | 0.18 | 0.1 | 0.19 |
| 1 | 186933555 | rs932476 | -1.546 | -0.788 | -0.18 | 0.647 | 0.016 | 0.019 | 0.019 | 0 | 0.4 | 0.71 | 0.59 | 0.33 | 0.33 | 0.49 |
| 1 | 186934187 | rs12144159 | -0.409 | -0.641 | -0.603 | 0.184 | -0.015 | 0.018 | 0.018 | 0 | 0.25 | 0.17 | 0.49 | 0.17 | 0.26 | 0.39 |
| 1 | 186941897 | rs12353944 | -0.134 | -0.72 | -0.364 | 0.253 | 0 | -0.002 | -0.002 | 0 | 0.2 | 0.25 | 0.3 | 0.1 | 0.2 | 0.31 |
| 1 | 186946386 | rs2076675 | 1.699 | -0.829 | -1.226 | 0.287 | -0.118 | -0.097 | -0.097 | 0.063 | 0.45 | 0.03 | 0.36 | 0.62 | 0.54 | 0.42 |
| 1 | 186951429 | rs7545121 | 1.766 | -0.807 | -1.147 | 0.31 | -0.057 | -0.041 | -0.041 | 0 | 0.45 | 0.07 | 0.36 | 0.62 | 0.54 | 0.42 |

| | | | | | | | | | | | | | | | | |
|---|---|---|---|---|---|---|---|---|---|---|---|---|---|---|---|---|
| 1 | 186952409 | rs4402086 | -0.694 | -0.846 | -1.128 | 0.326 | -0.02 | -0.034 | -0.034 | 1.115 | 0.27 | 0.16 | 0.34 | 0.26 | 0.25 | 0.44 |
| 1 | 186953521 | rs12564966 | 0.035 | -0.756 | -0.876 | 0.2 | -0.035 | -0.022 | -0.022 | 0 | 0.18 | 0.58 | 0.2 | 0.12 | 0.11 | 0.1 |
| 1 | 186961387 | rs761517 | -0.138 | -0.415 | -1.216 | 0.415 | -0.022 | -0.042 | -0.042 | 0 | 0.75 | 0.45 | 0.72 | 0.88 | 0.79 | 0.86 |
| 1 | 186967242 | rs2144000 | 0.093 | -0.06 | -1.371 | 0.78 | 0.01 | 0.063 | 0.063 | 0 | 0.55 | 0.28 | 0.43 | 0.77 | 0.59 | 0.53 |
| 1 | 186967702 | rs10798078 | 0.348 | 0.256 | -1.248 | 0.709 | -0.002 | -0.005 | -0.005 | 0 | 0.47 | 0.25 | 0.3 | 0.6 | 0.54 | 0.4 |
| 1 | 186981339 | rs10752993 | -0.408 | 0.678 | -0.917 | 0.628 | 0.006 | 0.002 | 0.002 | 0 | 0.37 | 0.43 | 0.36 | 0.65 | 0.36 | 0.34 |
| 1 | 186984283 | rs12735309 | 0.388 | 0.007 | -0.852 | 0.184 | -0.007 | 0.002 | 0.002 | 0 | 0.18 | 0.21 | 0.22 | 0.24 | 0.14 | 0.19 |
| 1 | 186985141 | rs2207215 | -0.127 | -0.271 | -1.066 | -0.074 | 0.002 | -0.004 | -0.004 | 0 | 0.42 | 0.5 | 0.37 | 0.65 | 0.37 | 0.37 |
| 1 | 186996815 | rs6665890 | -1.142 | -0.148 | -1.092 | -0.112 | -0.072 | 0.185 | 0.185 | 0 | 0.63 | 0.91 | 0.67 | 0.95 | 0.54 | 0.61 |
| 1 | 187002419 | rs1884022 | 0.53 | -0.299 | -0.805 | -0.122 | -0.011 | -0.006 | -0.006 | 0 | 0.35 | 0.27 | 0.35 | 0.65 | 0.36 | 0.34 |
| 1 | 187005310 | rs7540422 | 1.009 | 0.149 | -1.663 | 0.652 | -0.037 | -0.044 | -0.044 | 0.991 | 0.1 | 0.02 | 0.23 | 0.35 | 0.17 | 0.14 |
| 1 | 187007181 | rs12566615 | 0.841 | 0.093 | -1.616 | 0.381 | -0.045 | -0.07 | -0.07 | 0 | 0.23 | 0.03 | 0.1 | 0.29 | 0.19 | 0.18 |
| 1 | 187013915 | rs1160832 | -1.176 | 0.258 | -1.342 | 0.261 | -0.046 | 0.161 | 0.161 | 0 | 0.63 | 0.85 | 0.67 | 0.95 | 0.54 | 0.61 |
| 1 | 187032137 | rs12569046 | 0.97 | 0.464 | -1.473 | 0.728 | -0.041 | -0.065 | -0.065 | 0 | 0.15 | 0.06 | 0.1 | 0.3 | 0.17 | 0.17 |
| 1 | 187033639 | rs1297671 | -1.451 | 0.763 | -1.958 | 0.029 | -0.028 | 0.082 | 0.082 | 0 | 0.77 | 0.92 | 0.79 | 0.95 | 0.71 | 0.68 |
| 1 | 187039464 | rs10911992 | 0.259 | 0.76 | -1.203 | 0.384 | 0.018 | 0.04 | 0.04 | 0 | 0.13 | 0.2 | 0.29 | 0.32 | 0.19 | 0.15 |
| 1 | 187039770 | rs12048590 | 0.405 | 0.786 | -1.554 | 0.388 | -0.02 | -0.028 | -0.028 | 0 | 0.13 | 0.07 | 0.28 | 0.32 | 0.19 | 0.15 |
| 1 | 187041011 | rs1297676 | -0.035 | 0.112 | -1.207 | -0.101 | 0.013 | 0.027 | 0.027 | 0 | 0.36 | 0.24 | 0.29 | 0.24 | 0.3 | 0.27 |
| 1 | 187051532 | rs971145 | -0.15 | 0.07 | -1.291 | -0.296 | 0.013 | 0.065 | 0.065 | 0 | 0.65 | 0.77 | 0.74 | 0.86 | 0.72 | 0.76 |
| 1 | 187054897 | rs2144438 | 0.284 | 0.373 | -0.766 | -0.167 | 0.047 | 0.021 | 0.021 | 0 | 0.75 | 0.57 | 0.63 | 0.64 | 0.7 | 0.69 |
| 1 | 187055323 | rs17545182 | 1.236 | 0.398 | -2.004 | -0.78 | -0.075 | 0.153 | 0.153 | 0 | 0.25 | 0.03 | 0.18 | 0.06 | 0.3 | 0.25 |
| 1 | 187055561 | rs7511950 | 1.096 | 0.19 | -1.869 | -0.205 | -0.001 | -0.056 | -0.056 | 0 | 0.21 | 0.13 | 0.15 | 0.4 | 0.2 | 0.26 |
| 1 | 187057636 | rs10911997 | -0.009 | 0.302 | -1.399 | 0.016 | 0.004 | -0.004 | -0.004 | 0.53 | 0.15 | 0.08 | 0.2 | 0.19 | 0.12 | 0.12 |
| 1 | 187070355 | rs118654 | -1.02 | 0.913 | -1.43 | -0.167 | -0.05 | -0.01 | -0.01 | 0 | 0.85 | 0.91 | 0.79 | 0.82 | 0.8 | 0.83 |
| 1 | 187073950 | rs230629 | -0.013 | 1.003 | -0.676 | -0.007 | 0.036 | 0.021 | 0.021 | 0 | 0.78 | 0.74 | 0.64 | 0.58 | 0.69 | 0.69 |
| 1 | 187074490 | rs10798086 | -0.027 | 0.468 | -0.516 | -0.553 | -0.004 | -0.002 | -0.002 | 0 | 0.38 | 0.42 | 0.41 | 0.55 | 0.33 | 0.35 |
| 1 | 187078276 | rs230624 | 0.662 | 0.334 | -0.492 | -0.358 | 0.02 | 0.025 | 0.025 | 0 | 0.63 | 0.57 | 0.5 | 0.48 | 0.57 | 0.59 |
| 1 | 187078671 | rs10912005 | 1.823 | 0.323 | -0.743 | -0.237 | 0.001 | -0.122 | -0.122 | 0.093 | 0.16 | 0.04 | 0.18 | 0.44 | 0.13 | 0.2 |
| 1 | 187081199 | rs2383574 | 1.404 | 0.355 | -0.636 | -0.241 | -0.003 | 0.041 | 0.041 | 0 | 0.18 | 0.22 | 0.21 | 0.44 | 0.16 | 0.21 |
| 1 | 187082330 | rs230619 | 1.086 | 0.379 | -0.734 | -0.743 | -0.003 | 0.195 | 0.195 | 0 | 0.4 | 0.14 | 0.21 | 0.02 | 0.37 | 0.34 |
| 1 | 187092482 | rs1924736 | -0.312 | 0.374 | -0.197 | -0.364 | -0.015 | 0.003 | 0.003 | 0 | 0.28 | 0.37 | 0.31 | 0.3 | 0.23 | 0.27 |
| 1 | 187093377 | rs16826435 | 0.674 | 0.397 | -0.233 | -0.362 | 0.02 | 0.08 | 0.08 | 0 | 0.11 | 0.18 | 0.22 | 0.28 | 0.14 | 0.17 |
| 1 | 187095959 | rs12047450 | -0.269 | 0.376 | -0.204 | -0.379 | -0.002 | 0 | 0 | 0 | 0.27 | 0.21 | 0.3 | 0.3 | 0.23 | 0.27 |
| 1 | 187097401 | rs391391 | -1.129 | 0.629 | 0.068 | -0.267 | 0.022 | -0.002 | -0.002 | 0.245 | 0.84 | 0.74 | 0.75 | 0.8 | 0.79 | 0.83 |
| 1 | 187100011 | rs10798089 | 0.377 | 0.53 | -0.112 | -0.139 | 0.023 | 0.076 | 0.076 | 0 | 0.12 | 0.18 | 0.24 | 0.3 | 0.15 | 0.18 |
| 1 | 187114753 | rs12065268 | -0.404 | 0.525 | -0.119 | -0.272 | 0.028 | 0.016 | 0.016 | 0.003 | 0.28 | 0.47 | 0.33 | 0.33 | 0.29 | 0.29 |
| 1 | 187131218 | rs7545246 | 1.223 | 0.208 | -0.211 | -0.348 | -0.025 | 0.072 | 0.072 | 0.991 | 0.2 | 0.33 | 0.15 | 0.37 | 0.14 | 0.2 |
| 1 | 187135612 | rs12133085 | 1.028 | 0.19 | -0.726 | -0.409 | -0.001 | -0.051 | -0.051 | 0 | 0.11 | 0.01 | 0.13 | 0.13 | 0.13 | 0.1 |
| 1 | 187154470 | rs1924724 | -1.258 | 0.209 | -0.676 | -0.491 | -0.056 | 0.175 | 0.175 | 0 | 0.59 | 0.82 | 0.62 | 0.83 | 0.56 | 0.59 |
| 1 | 187158604 | rs10494592 | 0.731 | 0.457 | 0.118 | 0.017 | 0.042 | 0.057 | 0.057 | 0 | 0.1 | 0.11 | 0.23 | 0.31 | 0.18 | 0.17 |
| 1 | 187158671 | rs10494593 | 0.805 | 0.103 | 0.118 | -0.357 | -0.006 | 0.001 | 0.001 | 0 | 0.14 | 0.15 | 0.22 | 0.14 | 0.16 | 0.17 |
| 1 | 187163851 | rs2209363 | 0.191 | 0.08 | 0.136 | -0.381 | -0.018 | -0.022 | -0.022 | 0 | 0.77 | 0.48 | 0.64 | 0.81 | 0.81 | 0.76 |
| 1 | 187173215 | rs2025513 | -0.768 | 0.09 | 0.12 | -0.394 | 0.013 | -0.004 | -0.004 | 0.671 | 0.84 | 0.51 | 0.77 | 0.86 | 0.88 | 0.87 |
| 1 | 187177513 | rs12134977 | 0.491 | 0.05 | 0.16 | -0.377 | -0.013 | -0.011 | -0.011 | 0.185 | 0.08 | 0.2 | 0.14 | 0.03 | 0.07 | 0.1 |
| 1 | 187181021 | rs17524475 | 1.359 | -0.079 | -0.19 | -0.423 | 0.019 | -0.005 | -0.005 | 0 | 0.14 | 0.01 | 0.09 | 0.06 | 0.1 | 0.09 |
| 1 | 187183135 | rs330708 | -1.075 | -0.082 | -0.174 | -0.37 | 0.006 | 0.01 | 0.01 | 0 | 0.31 | 0.69 | 0.41 | 0.21 | 0.29 | 0.27 |
| 1 | 187196272 | rs330712 | 0.021 | 0.171 | -0.497 | -0.213 | 0.029 | 0.026 | 0.026 | 0 | 0.31 | 0.15 | 0.3 | 0.2 | 0.25 | 0.22 |
| 1 | 187208014 | rs330725 | -1.044 | 0.227 | -0.379 | -0.14 | 0.009 | 0.012 | 0.012 | 0 | 0.46 | 0.48 | 0.49 | 0.29 | 0.41 | 0.38 |
| 1 | 187212988 | rs10912035 | 0.019 | 0.847 | -0.307 | 0.298 | 0.002 | 0.003 | 0.003 | 0 | 0.08 | 0.08 | 0.16 | 0.05 | 0.09 | 0.12 |
| 1 | 187223137 | rs330723 | -0.132 | 1.391 | -0.022 | 0.826 | 0.125 | 0.248 | 0.248 | 0 | 0.9 | 0.71 | 0.76 | 0.66 | 0.85 | 0.84 |
| 1 | 187229039 | rs7547443 | 0.557 | 1.232 | -0.732 | 0.479 | -0.027 | 0.003 | 0.003 | 0 | 0.16 | 0.1 | 0.2 | 0.18 | 0.18 | 0.17 |
| 1 | 187232021 | rs330744 | -0.497 | 1.19 | -0.846 | 0.376 | -0.078 | 0.039 | 0.039 | 0 | 0.81 | 0.93 | 0.77 | 0.88 | 0.73 | 0.79 |
| 1 | 187236958 | rs17590592 | 0.711 | 1.036 | -0.525 | 0.183 | -0.003 | -0.013 | -0.013 | 0 | 0.06 | 0.07 | 0.05 | 0.03 | 0.07 | 0.04 |
| 1 | 187239230 | rs10494595 | -0.13 | 0.917 | 0.183 | 0.445 | 0.097 | 0.424 | 0.424 | 0 | 0.08 | 0.42 | 0.22 | 0.34 | 0.07 | 0.15 |
| 1 | 187239318 | rs4650724 | 0.379 | 0.719 | -0.372 | 0.138 | 0.01 | 0.042 | 0.042 | 0 | 0.24 | 0.46 | 0.34 | 0.69 | 0.21 | 0.31 |
| 1 | 187239405 | rs330736 | 1.791 | 0.296 | -0.69 | -0.074 | -0.038 | 0.237 | 0.237 | 0.024 | 0.4 | 0.03 | 0.28 | 0.08 | 0.38 | 0.35 |
| 1 | 187239429 | rs330737 | 1.744 | 0.296 | -0.688 | -0.094 | 0.001 | 0.22 | 0.22 | 0.001 | 0.41 | 0.11 | 0.28 | 0.08 | 0.38 | 0.35 |
| 1 | 187239805 | rs170708 | 1.791 | 0.299 | -0.735 | -0.093 | -0.038 | 0.243 | 0.243 | 0.024 | 0.4 | 0.03 | 0.28 | 0.08 | 0.38 | 0.35 |
| 1 | 187243378 | rs12139370 | 0.788 | 0.306 | -0.628 | 0.023 | 0.02 | -0.005 | -0.005 | 0 | 0.08 | 0.02 | 0.07 | 0.07 | 0.07 | 0.1 |
| 1 | 187247150 | rs2453727 | 0.407 | 0.628 | -0.667 | 0.182 | -0.001 | -0.032 | -0.032 | 0.078 | 0.57 | 0.48 | 0.42 | 0.18 | 0.52 | 0.48 |
| 1 | 187258353 | rs6658044 | -0.304 | 0.635 | -0.987 | 0.367 | -0.027 | 0.01 | 0.01 | 0 | 0.76 | 0.84 | 0.73 | 0.82 | 0.69 | 0.72 |
| 1 | 187265734 | rs17591285 | 1.428 | 0.266 | -1.305 | 0.029 | 0.004 | 0.033 | 0.033 | 0 | 0.15 | 0.01 | 0.12 | 0.05 | 0.12 | 0.1 |
| 1 | 187276553 | rs1390496 | -0.792 | 0.749 | -0.854 | 0.398 | -0.017 | 0.011 | 0.011 | 0 | 0.8 | 0.85 | 0.76 | 0.87 | 0.74 | 0.8 |
| 1 | 187279324 | rs1874414 | 0.489 | 0.638 | -0.977 | 0.541 | -0.055 | -0.022 | -0.022 | 2.048 | 0.6 | 0.8 | 0.61 | 0.46 | 0.57 | 0.55 |
| 1 | 187290395 | rs12131731 | 0.596 | 0.638 | -0.922 | 0.669 | -0.006 | -0.097 | -0.097 | 0 | 0.2 | 0.09 | 0.16 | 0.43 | 0.18 | 0.25 |
| 1 | 187295508 | rs1032538 | -0.693 | 0.679 | -0.765 | 0.58 | -0.005 | 0.003 | 0.003 | 0 | 0.78 | 0.76 | 0.77 | 0.86 | 0.73 | 0.77 |
| 1 | 187295740 | rs1285496 | -0.679 | 0.673 | -0.768 | 0.577 | -0.005 | 0.003 | 0.003 | 0 | 0.78 | 0.76 | 0.77 | 0.86 | 0.73 | 0.77 |
| 1 | 187309604 | rs10798106 | 0.619 | 0.661 | -0.811 | 0.617 | -0.018 | -0.121 | -0.121 | 0.082 | 0.2 | 0.08 | 0.16 | 0.43 | 0.18 | 0.24 |
| 1 | 187312447 | rs1148759 | 1.007 | 0.672 | -0.293 | 0.439 | 0.019 | 0.093 | 0.093 | 0 | 0.51 | 0.61 | 0.43 | 0.12 | 0.48 | 0.39 |

| | | | | | | | | | | | | | | | | |
|---|---|---|---|---|---|---|---|---|---|---|---|---|---|---|---|---|
| 1 | 187326714 | rs2800913 | -0.687 | 0.772 | -0.384 | 0.688 | -0.007 | 0.004 | 0.004 | 0 | 0.78 | 0.76 | 0.76 | 0.84 | 0.73 | 0.77 |
| 1 | 187327396 | rs2802475 | 0.676 | 0.614 | -0.584 | 0.676 | -0.006 | -0.005 | -0.005 | 0 | 0.63 | 0.33 | 0.47 | 0.43 | 0.56 | 0.5 |
| 1 | 187328545 | rs2802474 | 0.388 | 0.564 | -0.521 | 0.725 | 0.005 | -0.009 | -0.009 | 0 | 0.67 | 0.37 | 0.65 | 0.74 | 0.63 | 0.63 |
| 1 | 187329185 | rs10912054 | 0.897 | 1.051 | -1.036 | 0.739 | -0.057 | -0.107 | -0.107 | 0 | 0.17 | 0.04 | 0.14 | 0.34 | 0.17 | 0.17 |
| 1 | 187333447 | rs1285513 | -0.105 | 0.967 | -0.533 | 0.945 | 0.011 | 0.023 | 0.023 | 0 | 0.75 | 0.66 | 0.58 | 0.63 | 0.67 | 0.66 |
| 1 | 187352093 | rs12027637 | -0.171 | 0.926 | -0.448 | 1.087 | -0.005 | -0.006 | -0.006 | 0 | 0.25 | 0.24 | 0.4 | 0.35 | 0.33 | 0.33 |
| 1 | 187353527 | rs4409608 | -0.151 | 0.915 | -0.415 | 1.084 | 0.006 | 0.011 | 0.011 | 0 | 0.25 | 0.31 | 0.4 | 0.34 | 0.33 | 0.35 |
| 1 | 187354887 | rs10912059 | -1.342 | 0.618 | -0.234 | 1.039 | 0.002 | 0.101 | 0.101 | 0 | 0.49 | 0.75 | 0.54 | 0.72 | 0.5 | 0.54 |
| 1 | 187355994 | rs11583351 | 0.523 | 0.701 | -0.371 | 0.688 | -0.014 | 0.004 | 0.004 | 0 | 0.21 | 0.16 | 0.25 | 0.18 | 0.29 | 0.27 |
| 1 | 187362785 | rs10798107 | 0.694 | 0.339 | -1.138 | -0.006 | 0.052 | 0.204 | 0.204 | 0.941 | 0.49 | 0.17 | 0.42 | 0.21 | 0.59 | 0.46 |
| 1 | 187362820 | rs10912060 | 1.63 | 0.399 | -1.156 | 0.435 | -0.014 | -0.035 | -0.035 | 0.622 | 0.15 | 0.03 | 0.07 | 0.21 | 0.13 | 0.1 |
| 1 | 187363393 | rs12728510 | 1.916 | 0.83 | -1.266 | 0.302 | 0.024 | 0.014 | 0.014 | 0 | 0.1 | 0.15 | 0.18 | 0.14 | 0.17 | 0.15 |
| 1 | 187363621 | rs4651374 | -0.137 | 0.197 | -0.718 | 0.331 | -0.022 | 0.07 | 0.07 | 0.36 | 0.73 | 0.48 | 0.57 | 0.59 | 0.76 | 0.67 |
| 1 | 187374362 | rs7534916 | 0.856 | 0.043 | -0.651 | 0.393 | -0.005 | 0.035 | 0.035 | 0 | 0.13 | 0.21 | 0.09 | 0.3 | 0.11 | 0.11 |
| 1 | 187379326 | rs4650729 | -0.51 | 0.003 | -0.624 | 0.293 | -0.017 | 0.093 | 0.093 | 0.123 | 0.77 | 0.65 | 0.61 | 0.57 | 0.8 | 0.7 |
| 1 | 187384477 | rs4651378 | 0.525 | 0.101 | -0.868 | 0.242 | 0.004 | 0.024 | 0.024 | 0 | 0.11 | 0.18 | 0.08 | 0.19 | 0.1 | 0.15 |
| 1 | 187386201 | rs12145400 | 1.568 | -0.256 | -0.809 | 0.458 | 0.061 | 0.179 | 0.179 | 0.047 | 0.52 | 0.19 | 0.45 | 0.2 | 0.61 | 0.49 |
| 1 | 187390668 | rs12142409 | 0.19 | 0.318 | -0.812 | 0.424 | 0.001 | 0.006 | 0.006 | 0 | 0.17 | 0.13 | 0.09 | 0.19 | 0.16 | 0.11 |
| 1 | 187397003 | rs12121531 | 0.743 | 0.704 | -0.597 | 0.736 | -0.045 | -0.044 | -0.044 | 0 | 0.09 | 0.01 | 0.06 | 0.19 | 0.15 | 0.09 |
| 1 | 187397277 | rs6696686 | -0.115 | 0.714 | -0.46 | 0.795 | 0.021 | 0.094 | 0.094 | 0 | 0.13 | 0.31 | 0.14 | 0.36 | 0.17 | 0.19 |
| 1 | 187406532 | rs2490407 | -0.406 | 0.451 | -0.589 | 0.729 | -0.01 | 0.054 | 0.054 | 0.213 | 0.23 | 0.28 | 0.37 | 0.51 | 0.19 | 0.31 |
| 1 | 187416536 | rs2490327 | -0.983 | 1.034 | -0.773 | -0.261 | 0.01 | -0.014 | -0.014 | 0 | 0.81 | 0.8 | 0.78 | 0.93 | 0.76 | 0.74 |
| 1 | 187418404 | rs4651384 | 0.578 | 0.639 | -1.391 | -0.343 | -0.021 | -0.105 | -0.105 | 0.514 | 0.28 | 0.07 | 0.28 | 0.48 | 0.28 | 0.23 |
| 1 | 187422872 | rs1938512 | -0.202 | 0.258 | -1.308 | -0.74 | -0.039 | 0.048 | 0.048 | 0 | 0.22 | 0.41 | 0.22 | 0.3 | 0.13 | 0.28 |
| 1 | 187428231 | rs12137534 | 0.683 | 0.78 | -0.988 | -1.616 | -0.008 | 0.065 | 0.065 | 0 | 0.3 | 0.12 | 0.24 | 0.14 | 0.33 | 0.21 |
| 1 | 187432059 | rs10798113 | 0.538 | 0.835 | -1.178 | -1.561 | -0.003 | -0.015 | -0.015 | 3.977 | 0.39 | 0.07 | 0.34 | 0.63 | 0.31 | 0.4 |
| 1 | 187435132 | rs2249521 | 0.168 | 1.297 | 0.2 | -1.326 | 0.05 | -0.035 | -0.035 | 0.253 | 0.74 | 0.48 | 0.64 | 0.81 | 0.66 | 0.6 |
| 1 | 187435580 | rs1938522 | 0.528 | 1.283 | 0.22 | -1.334 | 0.001 | -0.006 | -0.006 | 0 | 0.69 | 0.35 | 0.62 | 0.81 | 0.65 | 0.59 |
| 1 | 187436592 | rs2249371 | -0.352 | 0.952 | -0.073 | -1.386 | 0.028 | -0.054 | -0.054 | 1.896 | 0.8 | 0.7 | 0.78 | 0.94 | 0.77 | 0.74 |
| 1 | 187443751 | rs10753005 | -0.527 | 0.58 | -0.26 | -0.999 | 0.002 | -0.078 | -0.078 | 1.849 | 0.82 | 0.75 | 0.89 | 0.98 | 0.79 | 0.92 |
| 1 | 187444151 | rs7349120 | -0.25 | 0.243 | -0.512 | -1.467 | -0.013 | -0.023 | -0.023 | 0 | 0.38 | 0.56 | 0.4 | 0.27 | 0.29 | 0.39 |
| 1 | 187444628 | rs4607826 | 0.711 | 0.316 | 0.142 | -1.295 | 0.001 | -0.007 | -0.007 | 0.045 | 0.07 | 0.15 | 0.14 | 0.04 | 0.06 | 0.12 |
| 1 | 187447280 | rs1892537 | -0.684 | 0.468 | -0.146 | -1.153 | 0 | -0.045 | -0.045 | 0 | 0.82 | 0.83 | 0.89 | 0.98 | 0.79 | 0.92 |
| 1 | 187461719 | rs7554573 | -0.491 | 0.388 | -0.139 | -1.202 | 0.002 | -0.023 | -0.023 | 0 | 0.77 | 0.78 | 0.87 | 0.98 | 0.76 | 0.94 |
| 1 | 187479735 | rs7555613 | -0.202 | -0.133 | 0.697 | -1.184 | -0.024 | -0.015 | -0.015 | 1.232 | 0.1 | 0.16 | 0.06 | 0.05 | 0.04 | 0.06 |
| 1 | 187483206 | rs4593773 | -1.068 | -0.016 | 0.489 | -1.202 | -0.013 | 0.008 | 0.008 | 0 | 0.61 | 0.7 | 0.5 | 0.37 | 0.52 | 0.48 |
| 1 | 187492543 | rs7535036 | -0.018 | -0.166 | 0.618 | -1.173 | -0.032 | -0.019 | -0.019 | 0 | 0.11 | 0.27 | 0.07 | 0.05 | 0.04 | 0.06 |
| 1 | 187498538 | rs1361298 | 0.443 | -0.338 | 0.16 | -1.341 | -0.05 | -0.054 | -0.054 | 0 | 0.25 | 0.05 | 0.29 | 0.3 | 0.27 | 0.37 |
| 1 | 187500314 | rs10494596 | 0.051 | -0.584 | 0.84 | -1.521 | 0.003 | 0.001 | 0.001 | 0 | 0.07 | 0.06 | 0.03 | 0.05 | 0.03 | 0.06 |
| 1 | 187500338 | rs1416476 | -0.603 | -0.599 | 0.806 | -1.554 | 0.018 | 0.028 | 0.028 | 0 | 0.76 | 0.83 | 0.89 | 0.98 | 0.8 | 0.94 |
| 1 | 187501925 | rs10798116 | -0.629 | -0.457 | 0.771 | -1.349 | 0.023 | 0.024 | 0.024 | 0 | 0.36 | 0.7 | 0.2 | 0.09 | 0.25 | 0.12 |
| 1 | 187522674 | rs2891285 | -0.708 | 0.208 | 0.812 | -0.428 | 0.007 | 0.06 | 0.06 | 0.881 | 0.06 | 0.24 | 0.04 | 0.06 | 0.02 | 0.1 |
| 1 | 187524263 | rs10912094 | -0.689 | 0.1 | 0.742 | -0.695 | -0.005 | -0.033 | -0.033 | 0.775 | 0.15 | 0.37 | 0.35 | 0.11 | 0.12 | 0.28 |
| 1 | 187531473 | rs2802752 | -0.006 | 0.573 | 0.829 | -0.637 | 0 | 0 | 0 | 0 | 0.2 | 0.17 | 0.32 | 0.7 | 0.34 | 0.38 |
| 1 | 187533898 | rs2789391 | 0.398 | 0.671 | 0.758 | -0.637 | 0.003 | 0.041 | 0.041 | 0 | 0.09 | 0.08 | 0.18 | 0.64 | 0.16 | 0.33 |
| 1 | 187543154 | rs2789394 | -0.607 | 0.54 | 0.974 | -0.748 | 0.015 | 0.001 | 0.001 | 0.279 | 0.23 | 0.29 | 0.34 | 0.76 | 0.36 | 0.4 |
| 1 | 187557096 | rs12127474 | 0.382 | 0.227 | 0.722 | -1.075 | -0.033 | -0.008 | -0.008 | 0 | 0.17 | 0.02 | 0.28 | 0.13 | 0.17 | 0.15 |
| 1 | 187573295 | rs12075120 | 0.595 | 0.761 | 0.635 | -1.328 | 0.014 | -0.083 | -0.083 | 0.163 | 0.22 | 0.51 | 0.21 | 0.1 | 0.2 | 0.31 |
| 1 | 187575386 | rs2789405 | 1.02 | 0.875 | 0.435 | -1.288 | 0.004 | 0.041 | 0.041 | 0 | 0.09 | 0.08 | 0.18 | 0.63 | 0.16 | 0.34 |
| 1 | 187581303 | rs2383658 | 0.602 | 0.123 | 0.155 | -1.492 | -0.006 | 0.075 | 0.075 | 0 | 0.46 | 0.33 | 0.49 | 0.76 | 0.59 | 0.52 |
| 1 | 187583469 | rs17604852 | 0.052 | 0.528 | 0.518 | -0.677 | -0.032 | 0.048 | 0.048 | 0 | 0.18 | 0.07 | 0.09 | 0.1 | 0.2 | 0.31 |
| 1 | 187584447 | rs11809135 | -0.277 | 0.528 | 0.531 | -0.735 | 0.006 | -0.041 | -0.041 | 0 | 0.2 | 0.27 | 0.11 | 0.08 | 0.2 | 0.31 |
| 1 | 187588239 | rs12746529 | 0.004 | 0.701 | -0.576 | -0.648 | -0.034 | 0.061 | 0.061 | 0 | 0.18 | 0.07 | 0.09 | 0.08 | 0.2 | 0.31 |
| 1 | 187596559 | rs12741706 | 0.002 | 0.847 | -1.403 | -0.618 | -0.026 | 0.053 | 0.053 | 0 | 0.18 | 0.08 | 0.09 | 0.08 | 0.2 | 0.31 |
| 1 | 187599437 | rs16827381 | 0.341 | 0.666 | -1.335 | 0.019 | -0.012 | -0.027 | -0.027 | 0.026 | 0.15 | 0.05 | 0.18 | 0.2 | 0.22 | 0.1 |
| 1 | 187601186 | rs10798126 | -0.014 | 0.659 | -1.714 | -0.338 | -0.014 | -0.042 | -0.042 | 1.433 | 0.22 | 0.11 | 0.26 | 0.63 | 0.24 | 0.4 |
| 1 | 187601249 | rs6678774 | 0.787 | 0.176 | -1.304 | -1.026 | 0.003 | 0.057 | 0.057 | 0.601 | 0.41 | 0.27 | 0.36 | 0.18 | 0.35 | 0.21 |
| 1 | 187602253 | rs12058310 | 0.849 | 0.095 | -1.364 | -1.07 | 0 | 0.044 | 0.044 | 0.874 | 0.38 | 0.27 | 0.36 | 0.16 | 0.34 | 0.17 |
| 1 | 187604765 | rs6703330 | -0.087 | -0.354 | -0.112 | -0.882 | -0.008 | 0.025 | 0.025 | 0.442 | 0.25 | 0.41 | 0.27 | 0.67 | 0.22 | 0.41 |
| 1 | 187607223 | rs6686651 | 0.612 | -0.119 | -0.242 | -0.994 | 0.001 | 0.068 | 0.068 | 0.012 | 0.37 | 0.22 | 0.36 | 0.16 | 0.34 | 0.2 |
| 1 | 187609850 | rs955835 | 0.03 | -0.133 | 0.109 | -1.102 | -0.001 | 0.024 | 0.024 | 1.4 | 0.42 | 0.31 | 0.42 | 0.69 | 0.46 | 0.6 |
| 1 | 187611003 | rs12076164 | 1.759 | -0.72 | -0.105 | -1.403 | -0.005 | 0.015 | 0.015 | 0 | 0.17 | 0.11 | 0.11 | 0 | 0.16 | 0.09 |
| 1 | 187611232 | rs10753015 | -0.917 | -0.718 | 0.101 | -1.367 | 0.007 | -0.026 | -0.026 | 0 | 0.78 | 0.56 | 0.78 | 0.84 | 0.81 | 0.8 |
| 1 | 187615141 | rs2066081 | 0.226 | -0.8 | 0.021 | -1.442 | 0.024 | 0.061 | 0.061 | 0 | 0.55 | 0.31 | 0.57 | 0.16 | 0.65 | 0.38 |
| 1 | 187623942 | rs10912115 | 0.604 | -0.868 | 0.179 | -1.41 | -0.025 | 0.02 | 0.02 | 0.069 | 0.23 | 0.31 | 0.22 | 0.66 | 0.16 | 0.35 |
| 1 | 187623971 | rs12749748 | 1.5 | -0.862 | 0.214 | -1.42 | 0.01 | -0.002 | -0.002 | 0 | 0.17 | 0.25 | 0.2 | 0.17 | 0.17 | 0.21 |
| 1 | 187626149 | rs10737282 | -0.064 | -0.864 | 0.044 | -1.405 | -0.003 | -0.021 | -0.021 | 0 | 0.4 | 0.57 | 0.42 | 0.83 | 0.34 | 0.56 |
| 1 | 187630186 | rs10494603 | 0.649 | -0.888 | -0.003 | -1.38 | -0.032 | 0.077 | 0.077 | 0 | 0.32 | 0.17 | 0.2 | 0.14 | 0.32 | 0.3 |
| 1 | 187642686 | rs10912138 | 0.671 | -0.893 | -0.011 | -1.383 | -0.047 | 0.096 | 0.096 | 0 | 0.31 | 0.13 | 0.2 | 0.14 | 0.32 | 0.3 |

| | | | | | | | | | | | | | | | | |
|---|---|---|---|---|---|---|---|---|---|---|---|---|---|---|---|---|
| 1 | 187644756 | rs17547862 | 1.219 | -0.907 | -0.046 | -1.426 | -0.008 | 0.004 | 0.004 | 0.184 | 0.18 | 0.06 | 0.26 | 0.16 | 0.19 | 0.11 |
| 1 | 187647751 | rs10912140 | 0.882 | -1.061 | 0.083 | -1.456 | -0.013 | 0.014 | 0.014 | 0.205 | 0.21 | 0.24 | 0.21 | 0.66 | 0.16 | 0.34 |
| 1 | 187651919 | rs17613702 | 1.985 | -1.048 | -0.012 | -1.479 | -0.072 | 0.102 | 0.102 | 0.25 | 0.2 | 0.04 | 0.2 | 0.01 | 0.31 | 0.19 |
| 1 | 187652296 | rs2891292 | -1.174 | -1.042 | 0.1 | -1.475 | 0.017 | -0.09 | -0.09 | 0 | 0.74 | 0.6 | 0.68 | 0.97 | 0.67 | 0.74 |
| 1 | 187661530 | rs953554 | 1.621 | -1.49 | 0.543 | -0.852 | -0.073 | 0.14 | 0.14 | 0 | 0.37 | 0.14 | 0.31 | 0.01 | 0.47 | 0.29 |
| 1 | 187662721 | rs3127505 | 1.077 | -1.588 | 0.558 | -0.832 | 0.093 | 0.214 | 0.214 | 0 | 0.52 | 0.2 | 0.4 | 0.14 | 0.64 | 0.49 |
| 1 | 187663473 | rs3127507 | 1.077 | -1.588 | 0.558 | -0.832 | 0.093 | 0.214 | 0.214 | 0 | 0.52 | 0.2 | 0.4 | 0.14 | 0.64 | 0.49 |
| 1 | 187668728 | rs16827441 | 0.749 | -1.597 | 0.541 | -0.825 | -0.001 | 0.008 | 0.008 | 0.07 | 0.19 | 0.09 | 0.27 | 0.16 | 0.19 | 0.11 |
| 1 | 187669092 | rs2050741 | 1.078 | -1.602 | 0.552 | -0.828 | 0.09 | 0.205 | 0.205 | 0 | 0.52 | 0.21 | 0.4 | 0.14 | 0.64 | 0.49 |
| 1 | 187672247 | rs3120966 | 0.278 | -1.6 | 0.763 | -0.821 | -0.016 | -0.001 | -0.001 | 0 | 0.72 | 0.33 | 0.67 | 0.3 | 0.83 | 0.6 |
| 1 | 187676040 | rs1538478 | 0.875 | -1.54 | 0.659 | -0.853 | 0.014 | 0.007 | 0.007 | 0 | 0.53 | 0.41 | 0.43 | 0.14 | 0.64 | 0.49 |
| 1 | 187677162 | rs6685655 | 1.519 | -1.436 | 0.538 | -0.922 | -0.071 | 0.131 | 0.131 | 0.005 | 0.37 | 0.15 | 0.31 | 0.01 | 0.47 | 0.29 |
| 1 | 187681623 | rs4521954 | 0.214 | -1.461 | 0.766 | -0.853 | -0.014 | -0.041 | -0.041 | 1.69 | 0.63 | 0.49 | 0.67 | 0.21 | 0.68 | 0.46 |
| 1 | 187684427 | rs11800915 | 0.075 | -1.471 | 0.739 | -0.879 | -0.003 | -0.005 | -0.005 | 0 | 0.23 | 0.27 | 0.28 | 0.18 | 0.2 | 0.17 |
| 1 | 187684475 | rs4074556 | 0.075 | -1.475 | 0.554 | -0.884 | 0.007 | 0.028 | 0.028 | 0 | 0.63 | 0.7 | 0.7 | 0.2 | 0.68 | 0.45 |
| 1 | 187687844 | rs10912148 | -0.235 | -1.375 | 0.723 | -0.797 | -0.025 | -0.039 | -0.039 | 0 | 0.25 | 0.47 | 0.31 | 0.18 | 0.21 | 0.17 |
| 1 | 187689619 | rs4498762 | -0.35 | -1.524 | 0.627 | -0.754 | -0.029 | -0.007 | -0.007 | 0 | 0.28 | 0.4 | 0.31 | 0.72 | 0.22 | 0.4 |
| 1 | 187693731 | rs4233131 | 0.042 | -1.457 | 0.629 | -0.618 | -0.013 | 0.007 | 0.007 | 0 | 0.25 | 0.28 | 0.31 | 0.69 | 0.21 | 0.34 |
| 1 | 187694044 | rs4319280 | -1.027 | -1.268 | 0.527 | -0.569 | -0.003 | 0.007 | 0.007 | 0 | 0.8 | 0.82 | 0.86 | 0.91 | 0.8 | 0.76 |
| 1 | 187699152 | rs10798137 | 0.313 | -1.278 | 0.448 | -0.586 | -0.004 | -0.023 | -0.023 | 0 | 0.62 | 0.54 | 0.59 | 0.73 | 0.67 | 0.67 |
| 1 | 187700076 | rs6661294 | 0.313 | -1.278 | 0.448 | -0.586 | -0.004 | -0.023 | -0.023 | 0.019 | 0.62 | 0.54 | 0.59 | 0.73 | 0.67 | 0.67 |
| 1 | 187703817 | rs4622024 | -1.096 | -1.277 | 0.448 | -0.586 | -0.009 | 0.025 | 0.025 | 0 | 0.8 | 0.85 | 0.87 | 0.92 | 0.8 | 0.76 |
| 1 | 187704353 | rs10912155 | 0.309 | -1.322 | 0.534 | -0.371 | -0.021 | 0.005 | 0.005 | 0.023 | 0.19 | 0.31 | 0.28 | 0.2 | 0.13 | 0.09 |
| 1 | 187711717 | rs10912160 | 0.879 | -1.142 | -0.203 | -0.285 | -0.005 | -0.039 | -0.039 | 0 | 0.16 | 0.02 | 0.24 | 0.18 | 0.12 | 0.09 |
| 1 | 187715194 | rs4418543 | -0.733 | -1.472 | 0.026 | -0.08 | 0.024 | 0.024 | 0.024 | 1.06 | 0.4 | 0.28 | 0.51 | 0.72 | 0.32 | 0.43 |
| 1 | 187725317 | rs7546941 | 0.388 | -1.313 | -0.024 | 0.498 | -0.018 | 0.069 | 0.069 | 0 | 0.12 | 0.25 | 0.12 | 0.18 | 0.1 | 0.09 |
| 1 | 187731782 | rs12125012 | 0.89 | -1.263 | -0.564 | 0.619 | 0.002 | -0.008 | -0.008 | 0 | 0.11 | 0.05 | 0.16 | 0.39 | 0.1 | 0.26 |
| 1 | 187732601 | rs2495445 | 0.661 | -1.387 | -0.731 | 0.474 | 0.048 | -0.022 | -0.022 | 0 | 0.21 | 0.08 | 0.23 | 0.21 | 0.13 | 0.13 |
| 1 | 187741568 | rs3123042 | -0.326 | -0.664 | -0.414 | 0.028 | -0.005 | -0.023 | -0.023 | 0.004 | 0.31 | 0.22 | 0.33 | 0.59 | 0.31 | 0.51 |
| 1 | 187744623 | rs1158512 | -0.546 | -0.843 | -0.258 | -0.034 | -0.034 | 0.034 | 0.034 | 0 | 0.69 | 0.56 | 0.63 | 0.6 | 0.78 | 0.78 |
| 1 | 187748399 | rs967778 | 0.056 | -0.644 | -0.602 | -0.016 | 0.034 | 0.083 | 0.083 | 0.644 | 0.51 | 0.73 | 0.5 | 0.4 | 0.57 | 0.54 |
| 1 | 187748759 | rs967776 | 0.373 | -0.652 | -0.606 | 0.098 | -0.001 | 0.107 | 0.107 | 0 | 0.13 | 0.39 | 0.2 | 0.31 | 0.1 | 0.27 |
| 1 | 187749447 | rs10912174 | 0.381 | -0.382 | -0.825 | -0.08 | -0.01 | -0.002 | -0.002 | 2.234 | 0.38 | 0.32 | 0.3 | 0.09 | 0.47 | 0.28 |
| 1 | 187757027 | rs6664686 | -0.563 | 0.062 | -0.961 | 0.795 | -0.001 | -0.017 | -0.017 | 0 | 0.34 | 0.72 | 0.37 | 0.51 | 0.31 | 0.52 |
| 1 | 187760345 | rs1476901 | -0.357 | 0.162 | -1.182 | 0.807 | -0.008 | 0.026 | 0.026 | 0 | 0.42 | 0.23 | 0.53 | 0.58 | 0.47 | 0.58 |
| 1 | 187767589 | rs2495421 | -0.704 | -0.201 | -0.968 | 0.732 | -0.009 | 0.005 | 0.005 | 0 | 0.91 | 0.89 | 0.88 | 0.88 | 0.96 | 0.94 |
| 1 | 187768381 | rs2495419 | -0.671 | -0.206 | -0.97 | 0.729 | -0.009 | 0.005 | 0.005 | 0 | 0.91 | 0.89 | 0.88 | 0.88 | 0.96 | 0.94 |
| 1 | 187779509 | rs10798148 | -0.062 | -0.238 | -1.143 | 1.598 | -0.012 | -0.04 | -0.04 | 0 | 0.38 | 0.18 | 0.46 | 0.42 | 0.44 | 0.49 |
| 1 | 187783367 | rs2430081 | 0.354 | -0.891 | -0.32 | 1.066 | -0.078 | -0.032 | -0.032 | 0 | 0.81 | 0.52 | 0.84 | 0.88 | 0.93 | 0.91 |
| 1 | 187812024 | rs1860919 | 0.33 | -0.732 | 0.029 | 1.268 | 0 | -0.003 | -0.003 | 0.178 | 0.48 | 0.46 | 0.38 | 0.3 | 0.51 | 0.43 |
| 1 | 187813938 | rs2495474 | 0.479 | -0.695 | 0.152 | 1.246 | -0.008 | 0.086 | 0.086 | 0 | 0.46 | 0.3 | 0.37 | 0.29 | 0.5 | 0.45 |
| 1 | 187817247 | rs2495469 | 0.433 | -0.787 | 0.165 | 1.244 | -0.001 | 0.05 | 0.05 | 0 | 0.46 | 0.37 | 0.37 | 0.29 | 0.51 | 0.43 |
| 1 | 187823537 | rs2430144 | 0.147 | -0.791 | -0.252 | 1.477 | -0.003 | 0.039 | 0.039 | 0 | 0.67 | 0.51 | 0.67 | 0.56 | 0.71 | 0.79 |
| 1 | 187832346 | rs10489379 | 0.231 | -0.934 | -0.235 | 1.706 | -0.001 | -0.01 | -0.01 | 0 | 0.36 | 0.23 | 0.42 | 0.35 | 0.37 | 0.43 |
| 1 | 187834475 | rs2430132 | 0.063 | -0.657 | -0.14 | 1.846 | -0.044 | 0.103 | 0.103 | 0 | 0.81 | 0.48 | 0.78 | 0.65 | 0.87 | 0.86 |
| 1 | 187835370 | rs2192114 | -0.056 | -0.614 | -0.264 | 1.867 | -0.031 | 0.065 | 0.065 | 0 | 0.81 | 0.61 | 0.8 | 0.65 | 0.87 | 0.86 |
| 1 | 187840501 | rs2430130 | -0.047 | 0.058 | -0.749 | 2.034 | 0.002 | -0.008 | -0.008 | 0 | 0.46 | 0.27 | 0.39 | 0.47 | 0.54 | 0.5 |
| 1 | 187846237 | rs800821 | 0.092 | 0.265 | -0.582 | 0.803 | -0.031 | 0.039 | 0.039 | 0 | 0.41 | 0.25 | 0.36 | 0.32 | 0.49 | 0.41 |
| 1 | 187852709 | rs16827680 | -0.733 | -0.037 | 0.334 | 1.149 | 0.02 | 0.11 | 0.11 | 0.167 | 0.11 | 0.46 | 0.09 | 0.12 | 0.06 | 0.17 |
| 1 | 187852822 | rs10798153 | -0.785 | 0.178 | 0.11 | 1.148 | 0.004 | -0.002 | -0.002 | 0.614 | 0.2 | 0.24 | 0.19 | 0.2 | 0.21 | 0.22 |
| 1 | 187859825 | rs800805 | -0.376 | 0.189 | -0.633 | 1.018 | -0.034 | -0.096 | -0.096 | 0 | 0.26 | 0.08 | 0.32 | 0.4 | 0.29 | 0.37 |
| 1 | 187862656 | rs800801 | 0.173 | 0.175 | -0.565 | 0.907 | 0 | 0.021 | 0.021 | 0.022 | 0.17 | 0.15 | 0.12 | 0.06 | 0.18 | 0.32 |
| 1 | 187866906 | rs12024761 | 1.408 | 0.186 | -0.421 | 0.92 | 0.007 | 0.041 | 0.041 | 2.523 | 0.61 | 0.23 | 0.66 | 0.77 | 0.62 | 0.44 |
| 1 | 187870452 | rs2311063 | 0.677 | 0.183 | -0.413 | 0.872 | -0.013 | 0.067 | 0.067 | 0.374 | 0.71 | 0.82 | 0.76 | 0.88 | 0.68 | 0.63 |
| 1 | 187872751 | rs11591211 | 2.162 | 0.135 | -0.483 | 0.978 | -0.014 | 0.074 | 0.074 | 0.83 | 0.43 | 0.17 | 0.37 | 0.3 | 0.41 | 0.22 |
| 1 | 187873230 | rs800788 | -0.453 | 0.286 | -0.539 | 0.983 | 0.006 | 0.012 | 0.012 | 0.546 | 0.93 | 0.97 | 0.94 | 1 | 0.93 | 0.98 |
| 1 | 187882589 | rs887738 | -0.096 | 0.314 | -0.532 | 1.01 | -0.016 | 0.036 | 0.036 | 0 | 0.22 | 0.15 | 0.18 | 0.12 | 0.25 | 0.35 |
| 1 | 187884372 | rs1572507 | 0.032 | 0.437 | -0.483 | 1.144 | 0.016 | 0.116 | 0.116 | 1.59 | 0.1 | 0.52 | 0.09 | 0.11 | 0.06 | 0.15 |
| 1 | 187886683 | rs585964 | -0.035 | 0.442 | -0.518 | 1.221 | -0.017 | -0.064 | -0.064 | 0.734 | 0.83 | 0.92 | 0.71 | 0.52 | 0.79 | 0.79 |
| 1 | 187887030 | rs12075837 | -0.272 | 0.407 | -0.436 | 1.182 | 0.044 | 0.061 | 0.061 | 1.889 | 0.13 | 0.5 | 0.11 | 0.11 | 0.12 | 0.19 |
| 1 | 187888876 | rs776798 | 0.21 | 0.503 | -0.508 | 1.11 | -0.001 | 0.006 | 0.006 | 0 | 0.82 | 0.85 | 0.7 | 0.51 | 0.79 | 0.75 |
| 1 | 187888914 | rs776797 | -0.477 | -0.045 | -0.157 | 0.959 | -0.001 | -0.017 | -0.017 | 0 | 0.31 | 0.53 | 0.25 | 0.22 | 0.31 | 0.5 |
| 1 | 187891919 | rs16827734 | -0.457 | 0.149 | 0.184 | 1.092 | -0.011 | 0.067 | 0.067 | 0 | 0.06 | 0.07 | 0.15 | 0.28 | 0.04 | 0.03 |
| 1 | 187902981 | rs655167 | 1.347 | 0.451 | 0.564 | 0.073 | 0.011 | -0.111 | -0.111 | 0 | 0.86 | 0.53 | 0.87 | 1 | 0.89 | 0.9 |
| 1 | 187903176 | rs627163 | 0.537 | 0.725 | 0.399 | -0.137 | 0.019 | -0.035 | -0.035 | 0.387 | 0.63 | 0.52 | 0.68 | 0.78 | 0.56 | 0.62 |
| 1 | 187903798 | rs1368175 | 1.177 | -0.071 | 0.884 | 0.209 | -0.026 | 0.024 | 0.024 | 0 | 0.9 | 0.73 | 0.9 | 0.89 | 0.94 | 0.9 |
| 1 | 187904692 | rs612355 | -0.593 | -0.333 | 0.657 | 0.003 | 0.003 | 0.017 | 0.017 | 0 | 0.13 | 0.04 | 0.11 | 0.07 | 0.14 | 0.13 |
| 1 | 187912763 | rs1572508 | 1.907 | -0.59 | 1.171 | 0.961 | -0.021 | 0.04 | 0.04 | 0 | 0.72 | 0.63 | 0.6 | 0.54 | 0.74 | 0.69 |
| 1 | 187920237 | rs4650748 | 2.321 | -0.933 | 1.614 | 0.82 | 0.059 | -0.054 | -0.054 | 0 | 0.67 | 0.17 | 0.55 | 0.53 | 0.74 | 0.65 |

| | | | | | | | | | | | | | | | | |
|---|---|---|---|---|---|---|---|---|---|---|---|---|---|---|---|---|
| 1 | 187933982 | rs635926 | 0.7 | -0.901 | 1.846 | 0.847 | -0.001 | 0.029 | 0.029 | 0 | 0.84 | 0.82 | 0.71 | 0.61 | 0.85 | 0.82 |
| 1 | 187934869 | rs12733186 | 2.217 | -0.941 | 2.12 | 0.823 | -0.01 | 0.01 | 0.01 | 0 | 0.68 | 0.37 | 0.57 | 0.54 | 0.75 | 0.66 |
| 1 | 187937724 | rs6674919 | 2.033 | -0.931 | 2.314 | 0.827 | -0.036 | 0.065 | 0.065 | 0 | 0.71 | 0.57 | 0.59 | 0.54 | 0.75 | 0.69 |
| 1 | 187938431 | rs776795 | -0.205 | -0.905 | 2.239 | 0.761 | 0.009 | 0.015 | 0.015 | 0 | 0.12 | 0.04 | 0.09 | 0.07 | 0.11 | 0.13 |
| 1 | 187946548 | rs10912216 | 1.433 | -0.854 | 2.5 | 0.766 | -0.042 | -0.024 | -0.024 | 0 | 0.78 | 0.65 | 0.83 | 0.82 | 0.82 | 0.77 |
| 1 | 187947140 | rs1432432 | 2.355 | -0.882 | 2.444 | 0.983 | 0.051 | -0.051 | -0.051 | 0 | 0.67 | 0.21 | 0.53 | 0.55 | 0.74 | 0.66 |
| 1 | 187950418 | rs12564137 | -0.571 | -0.848 | 2.427 | 0.921 | 0.006 | 0.263 | 0.263 | 0 | 0.1 | 0.37 | 0.3 | 0.28 | 0.08 | 0.08 |
| 1 | 187967161 | rs7543364 | 2.516 | -0.859 | 2.088 | 0.856 | 0.064 | -0.062 | -0.062 | 0 | 0.63 | 0.16 | 0.47 | 0.57 | 0.69 | 0.57 |
| 1 | 187967988 | rs603811 | -1.809 | -0.882 | 2.258 | 0.843 | -0.051 | -0.101 | -0.101 | 0.242 | 0.23 | 0.43 | 0.17 | 0.12 | 0.18 | 0.21 |
| 1 | 187968134 | rs657814 | 1.026 | -0.729 | 2.231 | 0.909 | -0.002 | 0.209 | 0.209 | 0 | 0.86 | 0.6 | 0.64 | 0.68 | 0.87 | 0.75 |
| 1 | 187976838 | rs10912219 | 0.844 | -0.593 | 0.799 | 0.259 | -0.023 | -0.014 | -0.014 | 1.788 | 0.35 | 0.29 | 0.55 | 0.51 | 0.44 | 0.23 |
| 1 | 187978644 | rs10798156 | 0.952 | -1.285 | 0.589 | -0.211 | -0.026 | -0.026 | -0.026 | 0 | 0.18 | 0.31 | 0.09 | 0.12 | 0.1 | 0.12 |
| 1 | 187983850 | rs683527 | -0.446 | -1.126 | 0.513 | -0.247 | 0.018 | -0.016 | -0.016 | 0.083 | 0.92 | 0.86 | 0.88 | 0.96 | 0.88 | 0.84 |
| 1 | 187988077 | rs613118 | 0.691 | -1.284 | 0.493 | -0.345 | -0.021 | -0.055 | -0.055 | 0.003 | 0.64 | 0.5 | 0.72 | 0.79 | 0.7 | 0.58 |
| 1 | 187992993 | rs11801907 | 0.786 | -1.254 | 0.455 | -0.362 | 0 | 0.014 | 0.014 | 0.01 | 0.29 | 0.16 | 0.17 | 0.29 | 0.27 | 0.35 |
| 1 | 188003233 | rs671564 | -1.52 | -1.264 | 0.481 | -0.368 | 0.002 | 0.006 | 0.006 | 0 | 0.91 | 0.96 | 0.93 | 0.95 | 0.92 | 0.88 |
| 1 | 188017479 | rs10442612 | 0.774 | -1.203 | 0.398 | -0.337 | 0.024 | 0.047 | 0.047 | 0.095 | 0.62 | 0.31 | 0.7 | 0.79 | 0.7 | 0.57 |
| 1 | 188018355 | rs12137769 | 0.802 | -1.24 | 0.358 | -0.337 | -0.008 | 0.016 | 0.016 | 0 | 0.28 | 0.11 | 0.16 | 0.26 | 0.27 | 0.35 |
| 1 | 188023515 | rs1432427 | 2.523 | -1.199 | 0.265 | -0.151 | -0.137 | -0.024 | -0.024 | 0 | 0.24 | 0.02 | 0.23 | 0.2 | 0.31 | 0.15 |
| 1 | 188023780 | rs17477137 | 0.358 | -1.111 | 0.4 | -0.069 | -0.001 | 0.001 | 0.001 | 0 | 0.08 | 0.08 | 0.11 | 0.05 | 0.12 | 0.16 |
| 1 | 188024831 | rs1122919 | 1.175 | -1.132 | 0.408 | -0.051 | -0.005 | 0.08 | 0.08 | 0 | 0.09 | 0.14 | 0.3 | 0.3 | 0.12 | 0.07 |
| 1 | 188039590 | rs12023936 | -0.002 | -1.089 | 0.406 | -0.125 | -0.038 | -0.055 | -0.055 | 0 | 0.72 | 0.58 | 0.83 | 0.83 | 0.82 | 0.76 |
| 1 | 188071966 | rs4650754 | 1.243 | -1.483 | -0.448 | -0.352 | -0.018 | 0.016 | 0.016 | 0 | 0.41 | 0.37 | 0.6 | 0.63 | 0.48 | 0.39 |
| 1 | 188073361 | rs12024781 | 1.395 | -1.458 | -0.398 | -0.234 | 0 | 0.036 | 0.036 | 0 | 0.1 | 0.1 | 0.32 | 0.29 | 0.15 | 0.1 |
| 1 | 188081057 | rs12410224 | 0.866 | -1.373 | -0.727 | -0.29 | -0.059 | -0.062 | -0.062 | 0 | 0.14 | 0.02 | 0.11 | 0.3 | 0.2 | 0.27 |
| 1 | 188085833 | rs10489764 | 3.101 | -1.382 | -0.686 | -0.417 | -0.043 | 0.027 | 0.027 | 0 | 0.28 | 0.15 | 0.24 | 0.19 | 0.3 | 0.16 |
| 1 | 188086156 | rs7546596 | 0.777 | -1.441 | -0.564 | -0.451 | 0.013 | 0.018 | 0.018 | 0 | 0.55 | 0.46 | 0.69 | 0.79 | 0.66 | 0.54 |
| 1 | 188099917 | rs12137000 | -0.028 | -1.389 | -0.593 | -0.424 | 0.002 | 0.048 | 0.048 | 0 | 0.17 | 0.09 | 0.14 | 0.04 | 0.19 | 0.22 |
| 1 | 188103345 | rs12042467 | -1.536 | -1.42 | -0.477 | -0.425 | -0.001 | 0.001 | 0.001 | 1.08 | 0.45 | 0.46 | 0.6 | 0.64 | 0.54 | 0.6 |
| 1 | 188114531 | rs12048287 | -0.299 | -1.394 | -0.609 | -0.411 | 0 | 0.004 | 0.004 | 0.081 | 0.27 | 0.31 | 0.46 | 0.58 | 0.35 | 0.38 |
| 1 | 188116416 | rs10489766 | -0.019 | -1.381 | -0.652 | -0.422 | -0.005 | 0.061 | 0.061 | 0 | 0.17 | 0.08 | 0.14 | 0.04 | 0.19 | 0.22 |
| 1 | 188124227 | rs354246 | -0.355 | -1.37 | -0.471 | -0.34 | -0.048 | -0.047 | -0.047 | 0.249 | 0.72 | 0.6 | 0.84 | 0.82 | 0.84 | 0.79 |
| 1 | 188124446 | rs1032215 | 0.615 | -0.985 | -0.405 | -0.048 | -0.007 | -0.007 | -0.007 | 0 | 0.16 | 0.14 | 0.14 | 0.28 | 0.2 | 0.28 |
| 1 | 188124666 | rs7542386 | 0.768 | -0.916 | -0.303 | -0.114 | -0.01 | 0.102 | 0.102 | 0 | 0.12 | 0.24 | 0.33 | 0.29 | 0.16 | 0.1 |
| 1 | 188145787 | rs354255 | -1.525 | -1.068 | -0.406 | -0.308 | -0.002 | 0 | 0 | 0 | 0.93 | 0.93 | 0.92 | 0.93 | 0.94 | 0.91 |
| 1 | 188154257 | rs12140392 | 0.193 | -1.17 | -0.592 | -0.303 | -0.02 | -0.04 | -0.04 | 0 | 0.2 | 0.11 | 0.15 | 0.37 | 0.22 | 0.35 |
| 1 | 188154357 | rs10912259 | 1.68 | -1.361 | -0.104 | -0.426 | -0.016 | -0.027 | -0.027 | 0.021 | 0.14 | 0.3 | 0.07 | 0.07 | 0.07 | 0.05 |
| 1 | 188162030 | rs1487492 | 0.428 | -1.381 | -0.188 | -0.435 | -0.028 | -0.027 | -0.027 | 1.494 | 0.22 | 0.4 | 0.16 | 0.14 | 0.13 | 0.17 |
| 1 | 188168004 | rs17352839 | 0.331 | -1.294 | -0.101 | -0.456 | 0 | -0.005 | -0.005 | 0.075 | 0.1 | 0.11 | 0.11 | 0.05 | 0.12 | 0.16 |
| 1 | 188169106 | rs7414437 | 0.208 | -1.303 | -0.012 | -0.46 | 0.008 | -0.02 | -0.02 | 0.487 | 0.1 | 0.2 | 0.12 | 0.04 | 0.12 | 0.16 |
| 1 | 188172047 | rs1906255 | 2.828 | -1.337 | -0.411 | -0.551 | -0.114 | 0.083 | 0.083 | 0 | 0.28 | 0.03 | 0.25 | 0.13 | 0.33 | 0.16 |
| 1 | 188172757 | rs12044617 | 1.091 | -1.324 | -0.22 | -0.411 | -0.016 | 0.063 | 0.063 | 0 | 0.13 | 0.18 | 0.29 | 0.32 | 0.12 | 0.11 |
| 1 | 188187947 | rs12734405 | 0.399 | -1.138 | -0.241 | -0.486 | 0 | -0.003 | -0.003 | 0 | 0.1 | 0.09 | 0.11 | 0.04 | 0.15 | 0.16 |
| 1 | 188190768 | rs1825798 | -0.317 | -1.313 | -0.359 | -0.409 | -0.004 | -0.036 | -0.036 | 0 | 0.27 | 0.17 | 0.21 | 0.45 | 0.25 | 0.5 |
| 1 | 188193927 | rs1843860 | 0.597 | -1.314 | -0.337 | -0.414 | -0.001 | -0.009 | -0.009 | 0.067 | 0.67 | 0.76 | 0.74 | 0.55 | 0.67 | 0.45 |
| 1 | 188202082 | rs16828303 | 0.949 | -1.231 | -0.567 | -0.328 | -0.056 | -0.093 | -0.093 | 0 | 0.18 | 0.02 | 0.13 | 0.38 | 0.19 | 0.41 |
| 1 | 188202548 | rs10494610 | 0.953 | -1.248 | -0.35 | -0.28 | -0.015 | 0.06 | 0.06 | 0 | 0.13 | 0.17 | 0.29 | 0.31 | 0.12 | 0.1 |
| 1 | 188215107 | rs1994423 | 0.436 | -1.143 | -0.493 | -0.347 | 0.006 | -0.01 | -0.01 | 0 | 0.65 | 0.77 | 0.73 | 0.55 | 0.67 | 0.47 |
| 1 | 188215137 | rs1386565 | 2.424 | -1.178 | -0.809 | -0.45 | -0.052 | 0.07 | 0.07 | 0 | 0.29 | 0.11 | 0.26 | 0.12 | 0.34 | 0.16 |
| 1 | 188216544 | rs10912271 | 1.324 | -1.552 | -0.506 | -0.241 | -0.011 | 0.08 | 0.08 | 0 | 0.11 | 0.17 | 0.27 | 0.32 | 0.12 | 0.12 |
| 1 | 188216715 | rs10912272 | 1.375 | -1.531 | -0.497 | -0.232 | -0.011 | 0.083 | 0.083 | 0 | 0.11 | 0.17 | 0.27 | 0.32 | 0.12 | 0.12 |
| 1 | 188248649 | rs3010654 | -0.783 | -1.577 | -0.861 | -0.323 | 0.038 | -0.064 | -0.064 | 0.327 | 0.37 | 0.81 | 0.48 | 0.46 | 0.33 | 0.31 |
| 1 | 188249558 | rs10798187 | -0.498 | -1.154 | -0.694 | -0.127 | -0.003 | 0.048 | 0.048 | 0 | 0.84 | 0.76 | 0.83 | 0.75 | 0.83 | 0.84 |
| 1 | 188250062 | rs1386560 | 0.099 | -1.184 | -0.719 | -0.196 | -0.006 | 0.021 | 0.021 | 0 | 0.79 | 0.74 | 0.8 | 0.75 | 0.79 | 0.81 |
| 1 | 188250201 | rs2871869 | 0.358 | -1.41 | -0.592 | -0.346 | -0.017 | -0.019 | -0.019 | 0 | 0.32 | 0.36 | 0.16 | 0.28 | 0.2 | 0.35 |
| 1 | 188250810 | rs1487489 | -0.666 | -1.047 | -0.029 | 0.159 | -0.008 | 0.05 | 0.05 | 0 | 0.22 | 0.36 | 0.4 | 0.37 | 0.26 | 0.28 |
| 1 | 188267144 | rs12030453 | 1.114 | -1.046 | -0.061 | 0.202 | -0.011 | 0.077 | 0.077 | 0 | 0.11 | 0.19 | 0.28 | 0.33 | 0.12 | 0.12 |
| 1 | 188273214 | rs2067988 | -0.604 | -1.056 | -0.007 | 0.178 | 0.003 | 0.065 | 0.065 | 0 | 0.87 | 0.79 | 0.89 | 0.76 | 0.87 | 0.86 |
| 1 | 188274620 | rs6694771 | -0.072 | -1.053 | -0.015 | 0.172 | -0.005 | 0.034 | 0.034 | 0 | 0.83 | 0.76 | 0.86 | 0.76 | 0.83 | 0.83 |
| 1 | 188283839 | rs12564813 | -0.708 | -0.998 | -0.076 | 0.218 | -0.006 | 0.038 | 0.038 | 0 | 0.22 | 0.34 | 0.39 | 0.36 | 0.26 | 0.28 |
| 1 | 188300868 | rs4463623 | 0.146 | -0.898 | -0.044 | 0.107 | -0.014 | 0.048 | 0.048 | 0 | 0.25 | 0.38 | 0.38 | 0.53 | 0.24 | 0.44 |
| 1 | 188301971 | rs7538400 | -1.058 | -0.904 | -0.03 | 0.092 | -0.006 | 0.011 | 0.011 | 0 | 0.35 | 0.45 | 0.47 | 0.57 | 0.37 | 0.6 |
| 1 | 188308289 | rs4434808 | 0.116 | -0.892 | -0.047 | 0.08 | -0.014 | 0.05 | 0.05 | 0 | 0.25 | 0.38 | 0.38 | 0.53 | 0.24 | 0.44 |
| 1 | 188324516 | rs1489256 | 1.375 | -0.952 | 0.085 | 0.04 | 0.003 | 0.093 | 0.093 | 0 | 0.48 | 0.32 | 0.36 | 0.2 | 0.46 | 0.24 |
| 1 | 188328650 | rs1387343 | -0.696 | -0.938 | 0.044 | 0.146 | 0 | 0 | 0 | 0 | 0.34 | 0.39 | 0.48 | 0.56 | 0.37 | 0.6 |
| 1 | 188338756 | rs12036328 | -0.91 | -0.891 | 0.106 | 0.126 | -0.004 | 0.005 | 0.005 | 0 | 0.38 | 0.45 | 0.5 | 0.56 | 0.42 | 0.63 |
| 1 | 188376810 | rs4399114 | 0.902 | -0.912 | 0.05 | 0.261 | -0.015 | 0.08 | 0.08 | 0 | 0.12 | 0.21 | 0.28 | 0.32 | 0.12 | 0.12 |
| 1 | 188378902 | rs6425181 | 0.108 | -0.889 | 0.118 | 0.179 | -0.008 | 0.048 | 0.048 | 0 | 0.82 | 0.75 | 0.85 | 0.75 | 0.84 | 0.84 |
| 1 | 188379428 | rs12125250 | 0.669 | -0.908 | 0.076 | 0.134 | -0.027 | 0.006 | 0.006 | 0 | 0.21 | 0.1 | 0.18 | 0.24 | 0.25 | 0.45 |

| | | | | | | | | | | | | | | | | |
|---|---|---|---|---|---|---|---|---|---|---|---|---|---|---|---|---|
| 1 | 188383316 | rs12142669 | 0.261 | -0.743 | 0.348 | 0.357 | -0.008 | 0.058 | 0.058 | 0 | 0.82 | 0.72 | 0.84 | 0.75 | 0.84 | 0.84 |
| 1 | 188384464 | rs10798210 | 0.261 | -0.743 | 0.348 | 0.357 | -0.008 | 0.058 | 0.058 | 0 | 0.82 | 0.72 | 0.84 | 0.75 | 0.84 | 0.84 |
| 1 | 188391229 | rs7541884 | -0.457 | -0.737 | 0.244 | 0.48 | -0.01 | 0.045 | 0.045 | 0 | 0.87 | 0.84 | 0.88 | 0.76 | 0.9 | 0.87 |
| 1 | 188396320 | rs12029171 | 0.934 | -0.702 | 0.2 | 0.562 | -0.012 | 0.096 | 0.096 | 0 | 0.11 | 0.2 | 0.28 | 0.36 | 0.12 | 0.12 |
| 1 | 188413159 | rs10912385 | -1.359 | -0.719 | 0.422 | 0.092 | -0.006 | 0.032 | 0.032 | 0 | 0.56 | 0.67 | 0.63 | 0.85 | 0.54 | 0.78 |
| 1 | 188423382 | rs9988491 | 0.584 | -0.779 | 0.486 | 0.161 | -0.022 | 0.095 | 0.095 | 0 | 0.13 | 0.26 | 0.3 | 0.32 | 0.13 | 0.16 |
| 1 | 188424374 | rs12086010 | 1.521 | -0.879 | 0.242 | -0.234 | 0.015 | 0.024 | 0.024 | 1.111 | 0.19 | 0.15 | 0.1 | 0.04 | 0.13 | 0.05 |
| 1 | 188428780 | rs12138047 | 0.82 | -0.909 | 0.377 | -0.107 | -0.01 | 0.02 | 0.02 | 0 | 0.24 | 0.13 | 0.18 | 0.24 | 0.26 | 0.44 |
| 1 | 188436062 | rs6680786 | 1.287 | -0.45 | -0.032 | -0.119 | -0.024 | 0.108 | 0.108 | 0 | 0.42 | 0.25 | 0.34 | 0.15 | 0.41 | 0.24 |
| 1 | 188440351 | rs12091277 | 1.287 | -0.45 | -0.032 | -0.133 | -0.024 | 0.109 | 0.109 | 0 | 0.42 | 0.25 | 0.34 | 0.15 | 0.41 | 0.24 |
| 1 | 188455409 | rs10912414 | -0.695 | -0.484 | 0.146 | -0.065 | -0.016 | 0.016 | 0.016 | 0 | 0.26 | 0.46 | 0.44 | 0.61 | 0.27 | 0.29 |
| 1 | 188456430 | rs1489267 | 0.488 | -0.496 | 0.168 | 0.037 | -0.01 | 0.075 | 0.075 | 0 | 0.12 | 0.18 | 0.12 | 0.28 | 0.1 | 0.12 |
| 1 | 188458988 | rs6656158 | 0.468 | -0.438 | 0.192 | 0.024 | -0.001 | 0.091 | 0.091 | 0 | 0.15 | 0.28 | 0.32 | 0.33 | 0.17 | 0.17 |
| 1 | 188465568 | rs284076 | -0.499 | -0.441 | 0.161 | 0.103 | -0.007 | 0.033 | 0.033 | 0 | 0.22 | 0.34 | 0.35 | 0.29 | 0.22 | 0.2 |
| 1 | 188465943 | rs284077 | 0.334 | -0.431 | 0.16 | 0.044 | 0.001 | -0.002 | -0.002 | 0 | 0.3 | 0.3 | 0.21 | 0.32 | 0.28 | 0.52 |
| 1 | 188467273 | rs2220034 | 0.282 | -0.557 | 0.194 | 0.051 | 0 | 0.012 | 0.012 | 0.024 | 0.13 | 0.14 | 0.14 | 0.25 | 0.15 | 0.08 |
| 1 | 188469106 | rs12038869 | 1.26 | -0.529 | 0.197 | 0.064 | 0.001 | 0.043 | 0.043 | 0 | 0.12 | 0.18 | 0.31 | 0.28 | 0.15 | 0.12 |
| 1 | 188477749 | rs284113 | -0.874 | -0.374 | 0.192 | 0.036 | 0.003 | 0.009 | 0.009 | 0 | 0.44 | 0.62 | 0.53 | 0.6 | 0.45 | 0.67 |
| 1 | 188484287 | rs284131 | -1.073 | -0.403 | 0.159 | 0.038 | -0.001 | -0.005 | -0.005 | 1.03 | 0.32 | 0.31 | 0.46 | 0.52 | 0.36 | 0.23 |
| 1 | 188485563 | rs12732082 | -0.173 | -0.399 | 0.185 | 0.023 | -0.001 | -0.005 | -0.005 | 0 | 0.36 | 0.36 | 0.3 | 0.57 | 0.37 | 0.57 |
| 1 | 188486435 | rs1489241 | -0.928 | -0.385 | 0.181 | -0.015 | 0 | -0.054 | -0.054 | 0 | 0.48 | 0.55 | 0.6 | 0.85 | 0.52 | 0.69 |
| 1 | 188490154 | rs1906767 | -1.372 | -0.111 | -0.127 | 0.143 | -0.002 | -0.025 | -0.025 | 0 | 0.92 | 0.92 | 0.97 | 1 | 0.93 | 0.97 |
| 1 | 188502148 | rs12727900 | 0.359 | -0.119 | -0.209 | 0.156 | -0.007 | -0.026 | -0.026 | 0.002 | 0.13 | 0.08 | 0.13 | 0.24 | 0.14 | 0.08 |
| 1 | 188512805 | rs17356003 | -1.337 | -0.189 | -0.162 | 0.156 | 0.002 | -0.003 | -0.003 | 0 | 0.92 | 0.96 | 0.97 | 1 | 0.94 | 0.97 |
| 1 | 188515145 | rs1586059 | 0.744 | -0.217 | -0.154 | 0.156 | 0 | 0.016 | 0.016 | 0 | 0.53 | 0.35 | 0.62 | 0.44 | 0.54 | 0.37 |
| 1 | 188520376 | rs6662623 | -0.268 | -0.371 | -0.468 | 0.125 | -0.005 | -0.042 | -0.042 | 0.831 | 0.89 | 0.87 | 0.96 | 1 | 0.92 | 0.95 |
| 1 | 188524167 | rs1384351 | -0.767 | -0.351 | -0.458 | 0.145 | -0.006 | -0.04 | -0.04 | 0.158 | 0.91 | 0.89 | 0.97 | 1 | 0.93 | 0.97 |
| 1 | 188532630 | rs10494614 | -0.625 | -0.34 | -0.428 | 0.15 | -0.008 | -0.047 | -0.047 | 0 | 0.91 | 0.86 | 0.96 | 1 | 0.93 | 0.97 |
| 1 | 188536146 | rs10737293 | -1.07 | -0.323 | -0.477 | 0.159 | -0.004 | -0.038 | -0.038 | 0.233 | 0.92 | 0.9 | 0.97 | 1 | 0.93 | 0.97 |
| 1 | 188536685 | rs2128368 | 2.09 | -0.32 | -0.877 | 0.03 | -0.118 | 0.079 | 0.079 | 0 | 0.22 | 0.02 | 0.23 | 0.11 | 0.29 | 0.2 |
| 1 | 188553599 | rs1609772 | 0.75 | -0.411 | -0.305 | 0.439 | -0.001 | -0.002 | -0.002 | 0 | 0.23 | 0.24 | 0.17 | 0.28 | 0.26 | 0.44 |
| 1 | 188561149 | rs2312286 | 1.096 | -0.491 | -0.255 | 0.248 | 0.016 | 0.003 | 0.003 | 0 | 0.52 | 0.36 | 0.64 | 0.46 | 0.54 | 0.36 |
| 1 | 188562024 | rs10798231 | 0.546 | -0.508 | -0.188 | 0.278 | -0.023 | 0.103 | 0.103 | 0 | 0.82 | 0.62 | 0.85 | 0.72 | 0.86 | 0.83 |
| 1 | 188572755 | rs16829161 | 0.545 | -0.431 | -0.216 | 0.268 | 0.004 | -0.007 | -0.007 | 1.826 | 0.59 | 0.39 | 0.68 | 0.46 | 0.6 | 0.39 |
| 1 | 188573091 | rs16829150 | 0.163 | -0.329 | -0.981 | 0.376 | -0.013 | -0.055 | -0.055 | 0.149 | 0.13 | 0.02 | 0.1 | 0.23 | 0.13 | 0.08 |
| 1 | 188597187 | rs12079114 | 0.757 | -0.264 | -0.944 | 0.356 | -0.001 | 0 | 0 | 0.306 | 0.23 | 0.24 | 0.17 | 0.25 | 0.26 | 0.44 |
| 1 | 188601792 | rs10912458 | 0.562 | 0.152 | -1.246 | 0.689 | -0.007 | 0.036 | 0.036 | 0 | 0.84 | 0.8 | 0.86 | 0.72 | 0.86 | 0.83 |
| 1 | 188609111 | rs1902875 | -0.24 | 0.401 | -1.32 | 0.162 | 0.002 | 0.009 | 0.009 | 0 | 0.09 | 0.05 | 0.04 | 0 | 0.07 | 0.05 |
| 1 | 188613133 | rs7415825 | 0.411 | 0.527 | -1.15 | 0.466 | 0.004 | 0 | 0 | 0 | 0.92 | 0.87 | 0.95 | 0.95 | 0.94 | 0.88 |
| 1 | 188617210 | rs7418999 | -0.332 | 0.358 | -1.55 | 0.507 | 0.007 | 0.017 | 0.017 | 0 | 0.57 | 0.34 | 0.7 | 0.67 | 0.62 | 0.51 |
| 1 | 188620700 | rs12082345 | -0.332 | 0.358 | -1.55 | 0.51 | 0.007 | 0.012 | 0.012 | 0 | 0.57 | 0.34 | 0.7 | 0.66 | 0.63 | 0.51 |
| 1 | 188620903 | rs10912463 | 0.323 | 0.853 | -1.912 | 0.203 | -0.081 | 0 | 0 | 0 | 0.22 | 0.04 | 0.24 | 0.15 | 0.29 | 0.21 |
| 1 | 188622706 | rs12038742 | 0.673 | -0.132 | -1.305 | 0.068 | -0.009 | -0.003 | -0.003 | 0 | 0.81 | 0.23 | 0.87 | 0.83 | 0.88 | 0.87 |
| 1 | 188624613 | rs12033740 | -0.019 | 0.436 | -2.158 | -0.581 | -0.069 | -0.011 | -0.011 | 0 | 0.21 | 0.01 | 0.13 | 0.16 | 0.25 | 0.24 |
| 1 | 188629757 | rs2312283 | 0.432 | 0.828 | -2.087 | -1.04 | -0.007 | 0.005 | 0.005 | 0.031 | 0.11 | 0.09 | 0.09 | 0 | 0.16 | 0.17 |
| 1 | 188635050 | rs12024989 | 0.013 | 0.839 | -2.55 | -1.178 | -0.082 | 0.094 | 0.094 | 0.021 | 0.43 | 0.09 | 0.46 | 0.68 | 0.5 | 0.44 |
| 1 | 188635456 | rs10912467 | -0.657 | 0.634 | -1.434 | -1.655 | 0 | -0.093 | -0.093 | 0 | 0.79 | 0.67 | 0.82 | 0.95 | 0.82 | 0.7 |
| 1 | 188640013 | rs6692148 | -1.453 | 0.766 | -1.234 | -1.625 | 0.013 | -0.025 | -0.025 | 0 | 0.89 | 0.85 | 0.88 | 0.95 | 0.87 | 0.81 |
| 1 | 188650160 | rs7539920 | -0.115 | 0.591 | -1.01 | -1.532 | -0.031 | 0.09 | 0.09 | 0 | 0.44 | 0.19 | 0.45 | 0.32 | 0.45 | 0.44 |
| 1 | 188659420 | rs7416924 | -0.02 | 0.601 | -0.911 | -1.533 | -0.012 | 0.05 | 0.05 | 0 | 0.45 | 0.31 | 0.47 | 0.32 | 0.47 | 0.45 |
| 1 | 188664093 | rs1902895 | -1.369 | 0.575 | -0.929 | -1.464 | 0.024 | 0.136 | 0.136 | 0 | 0.83 | 0.66 | 0.68 | 0.44 | 0.8 | 0.82 |
| 1 | 188677169 | rs4474247 | -1.387 | 0.565 | -0.935 | -1.469 | 0.024 | 0.136 | 0.136 | 0 | 0.83 | 0.66 | 0.68 | 0.44 | 0.8 | 0.82 |
| 1 | 188683442 | rs4589114 | 0.462 | 0.621 | -1.67 | -1.58 | -0.05 | 0.018 | 0.018 | 0.036 | 0.33 | 0.06 | 0.37 | 0.32 | 0.32 | 0.32 |
| 1 | 188686273 | rs12028712 | 0.772 | 0.523 | -1.857 | -1.584 | -0.006 | -0.094 | -0.094 | 0 | 0.25 | 0.06 | 0.33 | 0.56 | 0.24 | 0.29 |
| 1 | 188688134 | rs11587440 | 0.896 | 0.601 | -1.744 | -1.262 | -0.054 | 0.004 | 0.004 | 0.239 | 0.21 | 0.01 | 0.13 | 0.12 | 0.24 | 0.2 |
| 1 | 188694491 | rs6665576 | 0.2 | 0.769 | -0.616 | -0.968 | -0.002 | 0.007 | 0.007 | 0.073 | 0.65 | 0.63 | 0.61 | 0.69 | 0.68 | 0.68 |
| 1 | 188700069 | rs12758150 | 0.509 | 0.967 | -1.052 | -0.938 | 0.004 | -0.003 | -0.003 | 0.504 | 0.22 | 0.13 | 0.25 | 0.26 | 0.15 | 0.12 |
| 1 | 188705079 | rs12134565 | -0.047 | 1.152 | -0.727 | -0.892 | -0.056 | 0.035 | 0.035 | 0.004 | 0.11 | 0.01 | 0.12 | 0.06 | 0.16 | 0.19 |
| 1 | 188735336 | rs10754227 | 0.03 | 0.973 | -0.376 | -0.45 | -0.02 | 0.007 | 0.007 | 0 | 0.38 | 0.24 | 0.5 | 0.62 | 0.42 | 0.4 |
| 1 | 188751380 | rs4439360 | 0.097 | 0.609 | -0.194 | -0.236 | 0.012 | 0.027 | 0.027 | 0 | 0.51 | 0.4 | 0.42 | 0.38 | 0.45 | 0.53 |
| 1 | 188760628 | rs10737570 | 1.262 | 0.183 | 0.696 | -0.334 | -0.011 | -0.065 | -0.065 | 0 | 0.89 | 0.47 | 0.91 | 0.99 | 0.94 | 0.98 |
| 1 | 188761359 | rs4417067 | -0.581 | 0.504 | 0.139 | 0.033 | 0 | 0 | 0 | 0 | 0.43 | 0.41 | 0.41 | 0.58 | 0.45 | 0.44 |
| 1 | 188764115 | rs12048906 | -0.24 | 0.417 | -0.322 | 0.185 | -0.003 | -0.111 | -0.111 | 0 | 0.34 | 0.07 | 0.32 | 0.47 | 0.32 | 0.34 |
| 1 | 188764543 | rs4551597 | -0.429 | 0.396 | 0.17 | 0.187 | -0.016 | 0.016 | 0.016 | 0 | 0.4 | 0.48 | 0.37 | 0.47 | 0.35 | 0.35 |
| 1 | 188770589 | rs4255388 | -0.626 | 0.539 | -0.113 | 0.131 | -0.008 | 0.02 | 0.02 | 0 | 0.46 | 0.7 | 0.45 | 0.57 | 0.47 | 0.44 |
| 1 | 188773111 | rs7537197 | -1.897 | -0.331 | 1.1 | -0.2 | -0.036 | -0.025 | -0.025 | 2.528 | 0.14 | 0.58 | 0.08 | 0.09 | 0.06 | 0.08 |
| 1 | 188788576 | rs12029028 | 0.297 | -0.438 | 0.404 | -0.334 | -0.057 | -0.039 | -0.039 | 0 | 0.32 | 0.08 | 0.44 | 0.39 | 0.38 | 0.41 |
| 1 | 188825570 | rs1445660 | -1.881 | -1.101 | 2.22 | -0.46 | -0.027 | 0.015 | 0.015 | 1.112 | 0.12 | 0.46 | 0.05 | 0.11 | 0.04 | 0.07 |
| 1 | 188835473 | rs1031191 | -2.46 | -1.109 | 2.765 | -0.464 | -0.048 | -0.02 | -0.02 | 0.617 | 0.13 | 0.64 | 0.07 | 0.11 | 0.04 | 0.07 |

| | | | | | | | | | | | | | | | | |
|---|---|---|---|---|---|---|---|---|---|---|---|---|---|---|---|---|
| 1 | 188836078 | rs16829453 | -0.613 | -1.074 | 2.685 | -0.448 | -0.012 | -0.03 | -0.03 | 0 | 0.07 | 0.14 | 0.02 | 0 | 0.02 | 0.01 |
| 1 | 188836200 | rs16829456 | -1.966 | -1.071 | 2.541 | -0.446 | -0.024 | 0.017 | 0.017 | 1.208 | 0.12 | 0.51 | 0.06 | 0.11 | 0.04 | 0.07 |
| 1 | 188857624 | rs7414565 | -0.463 | -1.402 | 2.022 | -0.723 | -0.003 | -0.017 | -0.017 | 0 | 0.89 | 0.85 | 0.94 | 0.98 | 0.91 | 0.86 |
| 1 | 188870483 | rs16829527 | -2.164 | -1.191 | 2.706 | -0.543 | -0.057 | -0.014 | -0.014 | 0 | 0.12 | 0.53 | 0.04 | 0.07 | 0.01 | 0.08 |
| 1 | 188884571 | rs6428035 | -2.206 | -1.151 | 2.43 | -0.524 | 0.053 | 0.102 | 0.102 | 2.768 | 0.06 | 0.42 | 0.03 | 0.07 | 0.01 | 0.07 |
| 1 | 188895696 | rs1445661 | -2.002 | -1.109 | 2.179 | -0.527 | -0.036 | -0.004 | -0.004 | 0 | 0.11 | 0.46 | 0.04 | 0.07 | 0.02 | 0.08 |
| 1 | 188946679 | rs10920995 | 0.306 | -0.856 | 1.232 | -0.473 | -0.001 | -0.001 | -0.001 | 0 | 0.58 | 0.44 | 0.62 | 0.51 | 0.58 | 0.52 |
| 1 | 188983626 | rs7550535 | -0.066 | -0.924 | 1.409 | -0.53 | -0.002 | 0.001 | 0.001 | 0 | 0.64 | 0.69 | 0.65 | 0.53 | 0.58 | 0.53 |
| 1 | 188984321 | rs2861427 | 0.518 | -0.991 | 1.813 | -0.564 | 0.005 | 0.007 | 0.007 | 0 | 0.57 | 0.39 | 0.62 | 0.52 | 0.57 | 0.53 |
| 1 | 188993946 | rs12738395 | 0.361 | -1.082 | 2.103 | -0.575 | -0.003 | -0.004 | -0.004 | 0 | 0.58 | 0.53 | 0.62 | 0.53 | 0.57 | 0.53 |
| 1 | 189002771 | rs10921121 | -0.189 | -1.05 | 1.939 | -0.66 | -0.037 | 0.02 | 0.02 | 0.107 | 0.63 | 0.84 | 0.64 | 0.6 | 0.54 | 0.62 |
| 1 | 189012899 | rs1912920 | 0.534 | -1.089 | 2.346 | -0.645 | 0.002 | 0.012 | 0.012 | 0 | 0.56 | 0.66 | 0.6 | 0.5 | 0.52 | 0.49 |
| 1 | 189022610 | rs7546105 | 0.1 | -1.067 | 2.3 | -0.623 | -0.023 | 0.001 | 0.001 | 0 | 0.07 | 0.12 | 0.03 | 0.08 | 0.01 | 0.1 |
| 1 | 189034234 | rs4388707 | 0.595 | -1.138 | 2.595 | -0.678 | -0.001 | -0.003 | -0.003 | 0 | 0.55 | 0.49 | 0.59 | 0.58 | 0.52 | 0.59 |
| 1 | 189045079 | rs1912915 | -0.114 | -1.164 | 2.2 | -0.759 | -0.021 | 0.019 | 0.019 | 0.046 | 0.62 | 0.79 | 0.64 | 0.58 | 0.53 | 0.62 |
| 1 | 189069537 | rs7518946 | 0.455 | -1.189 | 2.353 | -0.768 | 0.002 | 0.015 | 0.015 | 0 | 0.56 | 0.64 | 0.61 | 0.51 | 0.52 | 0.49 |
| 1 | 189100707 | rs1912937 | 0.497 | -1.456 | 2.52 | -0.869 | -0.003 | -0.005 | -0.005 | 0 | 0.52 | 0.53 | 0.59 | 0.51 | 0.52 | 0.48 |
| 1 | 189108008 | rs10754056 | -0.015 | -1.467 | 2.526 | -0.875 | -0.002 | 0.009 | 0.009 | 0 | 0.58 | 0.68 | 0.62 | 0.51 | 0.53 | 0.49 |
| 1 | 189112184 | rs10921341 | 0.348 | -1.525 | 3.001 | -0.967 | 0.008 | 0.018 | 0.018 | 0 | 0.57 | 0.23 | 0.61 | 0.51 | 0.56 | 0.5 |
| 1 | 189123194 | rs1912933 | 0.694 | -1.548 | 2.924 | -0.98 | 0.022 | 0.053 | 0.053 | 0 | 0.53 | 0.2 | 0.59 | 0.54 | 0.54 | 0.5 |
| 1 | 189159261 | rs1891457 | -0.215 | -1.602 | 2.897 | -1.065 | -0.026 | -0.013 | -0.013 | 0 | 0.65 | 0.83 | 0.66 | 0.54 | 0.57 | 0.54 |
| 1 | 189172684 | rs7527682 | 0.643 | -1.611 | 3.078 | -1.071 | -0.003 | -0.006 | -0.006 | 0 | 0.55 | 0.51 | 0.61 | 0.54 | 0.55 | 0.53 |
| 1 | 189181097 | rs1342555 | -0.752 | -1.637 | 2.737 | -1.2 | -0.016 | 0.019 | 0.019 | 0 | 0.95 | 0.98 | 0.95 | 1 | 0.92 | 0.95 |
| 1 | 189188538 | rs10458404 | 0.957 | -1.672 | 2.751 | -1.149 | -0.071 | -0.05 | -0.05 | 0 | 0.26 | 0.03 | 0.26 | 0.33 | 0.35 | 0.28 |
| 1 | 189238665 | rs10921627 | -0.345 | -1.656 | 2.927 | -0.953 | -0.008 | -0.013 | -0.013 | 0.086 | 0.65 | 0.73 | 0.65 | 0.51 | 0.56 | 0.53 |
| 1 | 189240528 | rs1418547 | -0.052 | -1.657 | 2.953 | -0.954 | 0.003 | 0.007 | 0.007 | 0 | 0.63 | 0.49 | 0.63 | 0.51 | 0.56 | 0.52 |
| 1 | 189240602 | rs1418548 | 0.614 | -1.672 | 2.85 | -0.947 | 0.008 | 0.006 | 0.006 | 0 | 0.55 | 0.36 | 0.6 | 0.5 | 0.54 | 0.5 |
| 1 | 189252528 | rs7541670 | -0.304 | -1.639 | 2.594 | -0.949 | -0.021 | -0.029 | -0.029 | 0 | 0.65 | 0.8 | 0.65 | 0.51 | 0.56 | 0.53 |
| 1 | 189255779 | rs1113560 | -0.387 | -1.624 | 2.511 | -0.942 | -0.015 | -0.022 | -0.022 | 0.433 | 0.65 | 0.78 | 0.65 | 0.51 | 0.56 | 0.52 |
| 1 | 189258909 | rs10921664 | 0.582 | -1.639 | 2.623 | -0.944 | 0.016 | 0.013 | 0.013 | 0 | 0.54 | 0.25 | 0.59 | 0.51 | 0.54 | 0.49 |
| 1 | 189266248 | rs12130800 | 0.527 | -1.639 | 2.741 | -0.945 | -0.004 | -0.003 | -0.003 | 0 | 0.56 | 0.49 | 0.61 | 0.51 | 0.54 | 0.49 |
| 1 | 189270440 | rs1336885 | 0.481 | -1.639 | 2.569 | -0.944 | 0.01 | 0.008 | 0.008 | 0 | 0.55 | 0.32 | 0.6 | 0.51 | 0.54 | 0.49 |
| 1 | 189276962 | rs12135892 | 0.064 | -1.639 | 2.566 | -0.945 | 0 | 0.004 | 0.004 | 0 | 0.62 | 0.47 | 0.62 | 0.51 | 0.56 | 0.51 |
| 1 | 189282802 | rs1361011 | 0.733 | -1.587 | 2.266 | -0.92 | 0.025 | 0.021 | 0.021 | 0 | 0.52 | 0.25 | 0.59 | 0.51 | 0.54 | 0.49 |
| 1 | 189284520 | rs10921762 | -0.502 | -1.621 | 2.12 | -0.967 | -0.005 | -0.02 | -0.02 | 0 | 0.66 | 0.78 | 0.67 | 0.51 | 0.59 | 0.59 |
| 1 | 189286113 | rs6678596 | 0.071 | -1.557 | 1.786 | -1.1 | 0 | 0.001 | 0.001 | 0 | 0.6 | 0.29 | 0.61 | 0.55 | 0.56 | 0.51 |
| 1 | 189286816 | rs3909054 | 0.616 | -1.505 | 1.656 | -1.071 | 0.022 | 0.035 | 0.035 | 0 | 0.52 | 0.22 | 0.59 | 0.55 | 0.54 | 0.48 |
| 1 | 189293928 | rs7517688 | 0.627 | -1.661 | 1.649 | -1.058 | 0.022 | 0.032 | 0.032 | 0 | 0.52 | 0.22 | 0.6 | 0.55 | 0.54 | 0.48 |
| 1 | 189301911 | rs12747712 | 0.632 | -1.671 | 1.566 | -1.063 | 0.023 | 0.035 | 0.035 | 0 | 0.52 | 0.21 | 0.59 | 0.55 | 0.54 | 0.48 |
| 1 | 189310658 | rs2000219 | 0.073 | -1.725 | 1.718 | -1.093 | 0.001 | 0.001 | 0.001 | 0 | 0.59 | 0.3 | 0.62 | 0.55 | 0.56 | 0.5 |
| 1 | 189317485 | rs1336893 | 0.672 | -1.731 | 1.717 | -1.097 | 0.024 | 0.036 | 0.036 | 0 | 0.52 | 0.2 | 0.59 | 0.55 | 0.54 | 0.48 |
| 1 | 189317621 | rs1336891 | -0.227 | -1.758 | 2.064 | -1.108 | -0.004 | -0.003 | -0.003 | 0 | 0.63 | 0.7 | 0.65 | 0.55 | 0.57 | 0.51 |
| 1 | 189319775 | rs10921850 | 0.15 | -1.744 | 1.855 | -1.12 | -0.001 | 0.004 | 0.004 | 0 | 0.58 | 0.25 | 0.62 | 0.55 | 0.56 | 0.5 |
| 1 | 189319892 | rs10801417 | 0.795 | -1.738 | 1.857 | -1.116 | 0.019 | 0.043 | 0.043 | 0 | 0.51 | 0.16 | 0.59 | 0.55 | 0.54 | 0.47 |
| 1 | 189337997 | rs10921875 | 1.215 | -2.387 | 1.327 | -1.414 | -0.131 | -0.071 | -0.071 | 5.908 | 0.41 | 0.04 | 0.57 | 0.56 | 0.51 | 0.45 |
| 1 | 189343623 | rs12408416 | 1.189 | -2.383 | 1.329 | -1.412 | -0.13 | -0.071 | -0.071 | 0 | 0.41 | 0.05 | 0.57 | 0.56 | 0.51 | 0.45 |
| 1 | 189348462 | rs2152731 | -0.717 | -2.218 | 1.848 | -1.308 | 0 | 0 | 0 | 0 | 0.64 | 0.63 | 0.66 | 0.56 | 0.59 | 0.55 |
| 1 | 189372882 | rs10921990 | -0.335 | -1.936 | 2.058 | -1.28 | 0.004 | -0.028 | -0.028 | 0 | 0.05 | 0.16 | 0.03 | 0.01 | 0.03 | 0.05 |
| 1 | 189378725 | rs1148604 | -0.742 | -2.008 | 2.069 | -1.302 | -0.002 | -0.003 | -0.003 | 0 | 0.64 | 0.68 | 0.67 | 0.57 | 0.59 | 0.55 |
| 1 | 189383411 | rs7543574 | -0.752 | -2.008 | 2.053 | -1.302 | -0.003 | -0.005 | -0.005 | 0 | 0.64 | 0.73 | 0.67 | 0.57 | 0.59 | 0.55 |
| 1 | 189389416 | rs1148606 | -0.752 | -1.988 | 1.841 | -1.297 | -0.005 | -0.006 | -0.006 | 0.45 | 0.64 | 0.74 | 0.67 | 0.57 | 0.59 | 0.55 |
| 1 | 189430002 | rs6695079 | 1.007 | -2.02 | 1.839 | -1.263 | -0.03 | -0.038 | -0.038 | 0.283 | 0.08 | 0.12 | 0.02 | 0 | 0.02 | 0.03 |
| 1 | 189435354 | rs816812 | -0.518 | -2.142 | 1.91 | -1.239 | -0.007 | -0.002 | -0.002 | 0 | 0.62 | 0.68 | 0.65 | 0.56 | 0.56 | 0.51 |
| 1 | 189471770 | rs269737 | -0.383 | -2.192 | 1.876 | -1.246 | -0.018 | -0.004 | -0.004 | 0 | 0.61 | 0.77 | 0.65 | 0.56 | 0.54 | 0.49 |
| 1 | 189478530 | rs547918 | -0.379 | -2.185 | 1.896 | -1.242 | -0.013 | -0.002 | -0.002 | 0.088 | 0.61 | 0.73 | 0.65 | 0.56 | 0.54 | 0.49 |
| 1 | 189478630 | rs12137197 | 0.948 | -1.999 | 2.077 | -1.159 | -0.039 | -0.049 | -0.049 | 0 | 0.09 | 0.17 | 0.03 | 0 | 0.02 | 0.03 |
| 1 | 189491402 | rs269720 | -0.493 | -2.044 | 2.03 | -1.178 | -0.026 | -0.007 | -0.007 | 0 | 0.62 | 0.8 | 0.69 | 0.57 | 0.54 | 0.51 |
| 1 | 189496903 | rs269741 | -0.486 | -2.083 | 2.058 | -1.195 | -0.026 | -0.007 | -0.007 | 0 | 0.62 | 0.8 | 0.7 | 0.57 | 0.54 | 0.51 |
| 1 | 189503853 | rs269743 | -0.49 | -2.08 | 2.06 | -1.192 | -0.026 | -0.005 | -0.005 | 1.055 | 0.62 | 0.8 | 0.7 | 0.57 | 0.54 | 0.51 |
| 1 | 189512884 | rs12071479 | 0.543 | -2.058 | 1.84 | -1.184 | -0.046 | -0.056 | -0.056 | 0 | 0.1 | 0.26 | 0.04 | 0 | 0.02 | 0.02 |
| 1 | 189518210 | rs12126281 | 0.514 | -2.023 | 1.391 | -1.159 | -0.016 | 0.036 | 0.036 | 0 | 0.48 | 0.13 | 0.63 | 0.57 | 0.52 | 0.48 |
| 1 | 189523135 | rs1393495 | -0.658 | -1.676 | 2.08 | -0.921 | -0.067 | -0.075 | -0.075 | 0.072 | 0.12 | 0.56 | 0.06 | 0 | 0.02 | 0.03 |
| 1 | 189526779 | rs12127810 | -0.887 | -1.626 | 2.022 | -0.955 | -0.038 | -0.096 | -0.096 | 0.199 | 0.14 | 0.34 | 0.07 | 0.01 | 0.05 | 0.08 |
| 1 | 189527261 | rs12128689 | -0.623 | -1.639 | 2.046 | -0.966 | -0.02 | -0.055 | -0.055 | 0 | 0.13 | 0.18 | 0.05 | 0.01 | 0.05 | 0.09 |
| 1 | 189527897 | rs367510 | -0.024 | -1.525 | 2.009 | -1.008 | -0.001 | 0 | 0 | 0.096 | 0.49 | 0.52 | 0.48 | 0.59 | 0.49 | 0.41 |
| 1 | 189546726 | rs483402 | -0.561 | -1.812 | 1.671 | -0.936 | -0.014 | 0.012 | 0.012 | 0 | 0.57 | 0.82 | 0.52 | 0.58 | 0.52 | 0.48 |
| 1 | 189548411 | rs421604 | -0.467 | -1.805 | 1.666 | -0.932 | -0.004 | 0.016 | 0.016 | 0 | 0.56 | 0.76 | 0.52 | 0.58 | 0.52 | 0.48 |
| 1 | 189548619 | rs12131655 | 0.248 | -1.81 | 0.947 | -0.937 | -0.019 | 0.016 | 0.016 | 0 | 0.45 | 0.18 | 0.41 | 0.58 | 0.5 | 0.46 |
| 1 | 189550729 | rs1502595 | 0.432 | -1.809 | 0.626 | -0.848 | -0.092 | -0.098 | -0.098 | 0 | 0.43 | 0.05 | 0.4 | 0.44 | 0.5 | 0.42 |

| | | | | | | | | | | | | | | | | |
|---|---|---|---|---|---|---|---|---|---|---|---|---|---|---|---|---|
| 1 | 189552579 | rs445857 | 0.09 | -1.822 | 1.24 | -0.852 | 0.003 | 0.001 | 0.001 | 0 | 0.48 | 0.54 | 0.4 | 0.44 | 0.51 | 0.43 |
| 1 | 189558666 | rs366839 | -0.781 | -1.713 | 1.14 | -0.761 | -0.01 | -0.016 | -0.016 | 0.07 | 0.5 | 0.76 | 0.46 | 0.44 | 0.47 | 0.39 |
| 1 | 189566655 | rs155916 | -0.781 | -1.713 | 1.14 | -0.761 | -0.01 | -0.016 | -0.016 | 0 | 0.5 | 0.76 | 0.46 | 0.44 | 0.47 | 0.39 |
| 1 | 189568547 | rs10922329 | 0.45 | -1.766 | 1.263 | -0.772 | -0.088 | -0.084 | -0.084 | 2.78 | 0.37 | 0.12 | 0.33 | 0.44 | 0.45 | 0.35 |
| 1 | 189580639 | rs10489760 | -0.731 | -1.748 | 1.226 | -0.781 | -0.017 | -0.023 | -0.023 | 0.369 | 0.51 | 0.79 | 0.46 | 0.44 | 0.47 | 0.39 |
| 1 | 189586353 | rs13374026 | -1.998 | -1.753 | 1.643 | -0.783 | 0.073 | 0.075 | 0.075 | 0.417 | 0.05 | 0.49 | 0.03 | 0 | 0 | 0.01 |
| 1 | 189587034 | rs7532675 | -0.623 | -1.636 | 1.002 | -0.735 | -0.006 | -0.011 | -0.011 | 0 | 0.49 | 0.71 | 0.46 | 0.44 | 0.47 | 0.39 |
| 1 | 189595647 | rs2863819 | -0.798 | -1.487 | 0.724 | -0.687 | -0.017 | -0.016 | -0.016 | 0 | 0.5 | 0.79 | 0.45 | 0.44 | 0.47 | 0.39 |
| 1 | 189602917 | rs6672054 | 0.058 | -1.485 | 0.968 | -0.683 | -0.004 | -0.004 | -0.004 | 0 | 0.39 | 0.4 | 0.35 | 0.44 | 0.45 | 0.37 |
| 1 | 189608672 | rs540093 | -0.759 | -1.492 | 0.743 | -0.681 | -0.008 | -0.011 | -0.011 | 0 | 0.49 | 0.75 | 0.46 | 0.44 | 0.47 | 0.39 |
| 1 | 189612335 | rs534695 | -0.759 | -1.492 | 0.743 | -0.681 | -0.008 | -0.011 | -0.011 | 0 | 0.49 | 0.75 | 0.46 | 0.44 | 0.47 | 0.39 |
| 1 | 189620278 | rs532905 | 0.014 | -1.512 | 1.04 | -0.693 | -0.001 | -0.001 | -0.001 | 0 | 0.41 | 0.54 | 0.37 | 0.44 | 0.45 | 0.37 |
| 1 | 189622286 | rs508382 | 0.004 | -1.512 | 1.04 | -0.693 | -0.001 | -0.001 | -0.001 | 2.386 | 0.41 | 0.54 | 0.37 | 0.44 | 0.45 | 0.37 |
| 1 | 189623964 | rs495240 | 0.012 | -1.507 | 1.042 | -0.691 | -0.001 | -0.001 | -0.001 | 0 | 0.41 | 0.54 | 0.37 | 0.44 | 0.45 | 0.37 |
| 1 | 189627183 | rs473773 | 0.012 | -1.507 | 1.042 | -0.691 | -0.001 | -0.001 | -0.001 | 0 | 0.41 | 0.54 | 0.37 | 0.44 | 0.45 | 0.37 |
| 1 | 189653244 | rs479137 | -0.017 | -1.546 | 1.233 | -0.693 | 0.008 | 0.008 | 0.008 | 0 | 0.41 | 0.52 | 0.36 | 0.44 | 0.45 | 0.37 |
| 1 | 189658034 | rs518342 | 0.075 | -1.55 | 1.241 | -0.695 | 0.006 | 0.005 | 0.005 | 0 | 0.41 | 0.46 | 0.36 | 0.44 | 0.45 | 0.39 |
| 1 | 189667459 | rs485028 | -0.7 | -1.511 | 1.179 | -0.72 | -0.002 | -0.001 | -0.001 | 0 | 0.49 | 0.58 | 0.39 | 0.44 | 0.47 | 0.41 |
| 1 | 189678231 | rs2902583 | 0.105 | -1.481 | 1.262 | -0.705 | -0.003 | -0.003 | -0.003 | 0.056 | 0.39 | 0.38 | 0.35 | 0.44 | 0.45 | 0.39 |
| 1 | 189688790 | rs10465587 | 1.038 | -0.878 | 0.905 | -0.908 | 0 | -0.028 | -0.028 | 0 | 0.14 | 0.13 | 0.14 | 0 | 0.21 | 0.03 |
| 1 | 189695297 | rs528001 | -0.233 | -1.149 | 0.718 | -0.38 | -0.011 | 0.021 | 0.021 | 0 | 0.49 | 0.73 | 0.46 | 0.39 | 0.47 | 0.42 |
| 1 | 189721806 | rs10800627 | -0.209 | -1.004 | -0.132 | -0.31 | -0.016 | 0.02 | 0.02 | 0 | 0.5 | 0.79 | 0.47 | 0.39 | 0.47 | 0.42 |
| 1 | 189736833 | rs12566065 | 0.539 | -0.885 | -0.864 | -0.231 | -0.146 | -0.18 | -0.18 | 0 | 0.34 | 0.04 | 0.48 | 0.52 | 0.47 | 0.32 |
| 1 | 189752408 | rs6666578 | -0.209 | -0.261 | 0.265 | -0.007 | 0 | 0 | 0 | 0 | 0.45 | 0.43 | 0.37 | 0.37 | 0.48 | 0.45 |
| 1 | 189755774 | rs11586173 | -0.217 | -0.118 | 0.191 | 0.435 | -0.001 | -0.001 | -0.001 | 0.808 | 0.63 | 0.67 | 0.69 | 0.67 | 0.56 | 0.54 |
| 1 | 189761051 | rs2418508 | 1.099 | -0.996 | 0.84 | 0.568 | -0.062 | -0.01 | -0.01 | 2.184 | 0.81 | 0.34 | 0.72 | 0.85 | 0.92 | 0.83 |
| 1 | 189763755 | rs7551425 | 0.986 | -0.64 | 0.772 | 0.832 | 0.003 | -0.008 | -0.008 | 2.014 | 0.9 | 0.92 | 0.89 | 0.78 | 0.93 | 0.8 |
| 1 | 189772005 | rs4845149 | 1.334 | -0.478 | 1.777 | 0.739 | 0.034 | 0.051 | 0.051 | 0.401 | 0.93 | 0.67 | 0.94 | 0.96 | 0.96 | 0.92 |
| 1 | 189775505 | rs10919787 | -0.71 | -0.451 | 1.671 | 0.752 | 0.01 | 0.024 | 0.024 | 0 | 0.05 | 0.17 | 0.04 | 0.04 | 0.04 | 0.08 |
| 1 | 189777241 | rs11800026 | -0.71 | -0.451 | 1.671 | 0.752 | 0.01 | 0.024 | 0.024 | 0 | 0.05 | 0.17 | 0.04 | 0.04 | 0.04 | 0.08 |
| 1 | 189779629 | rs2418511 | 1.93 | -0.454 | 1.881 | 1.197 | 0.035 | 0.181 | 0.181 | 0 | 0.83 | 0.57 | 0.67 | 0.64 | 0.84 | 0.76 |
| 1 | 189780142 | rs12036655 | -2.038 | -0.454 | 1.919 | 1.197 | 0.023 | 0.126 | 0.126 | 0 | 0.16 | 0.32 | 0.33 | 0.36 | 0.16 | 0.24 |
| 1 | 189781569 | rs9427700 | 1.166 | -0.455 | 1.937 | 1.203 | 0.051 | 0.189 | 0.189 | 0 | 0.89 | 0.67 | 0.7 | 0.68 | 0.88 | 0.84 |
| 1 | 189785489 | rs2039776 | 2.291 | -0.543 | 2.193 | 1.131 | -0.007 | 0.066 | 0.066 | 0 | 0.76 | 0.36 | 0.6 | 0.6 | 0.79 | 0.6 |
| 1 | 189786558 | rs9427447 | 1.175 | -0.533 | 2.136 | 1.134 | 0.049 | 0.184 | 0.184 | 0 | 0.88 | 0.67 | 0.7 | 0.68 | 0.88 | 0.84 |
| 1 | 189790182 | rs12137220 | 2.252 | -0.541 | 2.054 | 1.163 | -0.007 | 0.066 | 0.066 | 0 | 0.76 | 0.36 | 0.6 | 0.6 | 0.79 | 0.6 |
| 1 | 189795996 | rs10919831 | -0.813 | -0.509 | 1.769 | 0.992 | -0.006 | -0.004 | -0.004 | 0.067 | 0.11 | 0.15 | 0.08 | 0.08 | 0.08 | 0.25 |
| 1 | 189805990 | rs10919878 | -1.209 | -0.44 | 1.723 | 1.025 | 0.008 | 0.053 | 0.053 | 0 | 0.1 | 0.15 | 0.29 | 0.31 | 0.12 | 0.16 |
| 1 | 189807634 | rs12038154 | -1.041 | -0.413 | 1.771 | 1.036 | 0.021 | 0.094 | 0.094 | 0 | 0.1 | 0.2 | 0.29 | 0.31 | 0.12 | 0.16 |
| 1 | 189812397 | rs12141679 | -1.088 | -0.402 | 1.714 | 1.041 | 0.015 | 0.074 | 0.074 | 0 | 0.1 | 0.16 | 0.28 | 0.3 | 0.12 | 0.16 |
| 1 | 189816957 | rs2176360 | -2.046 | -0.418 | 1.388 | 1.029 | 0.01 | 0.119 | 0.119 | 0 | 0.21 | 0.57 | 0.37 | 0.38 | 0.2 | 0.41 |
| 1 | 189825109 | rs11801703 | 0.285 | -0.423 | -0.129 | -0.053 | 0.01 | 0.089 | 0.089 | 0 | 0.39 | 0.11 | 0.15 | 0.28 | 0.3 | 0.24 |
| 1 | 189836689 | rs12062955 | -1.037 | -0.396 | 0.515 | 0.191 | -0.009 | -0.006 | -0.006 | 0 | 0.1 | 0.16 | 0.09 | 0.08 | 0.08 | 0.24 |
| 1 | 189841077 | rs950569 | -1.028 | -0.241 | 0.632 | 0.341 | 0.006 | 0.003 | 0.003 | 0 | 0.06 | 0.17 | 0.09 | 0.04 | 0.05 | 0.05 |
| 1 | 189841516 | rs1517085 | -1.293 | -0.586 | 0.845 | -0.145 | -0.005 | -0.019 | -0.019 | 0 | 0.33 | 0.37 | 0.21 | 0.14 | 0.27 | 0.32 |
| 1 | 189843320 | rs12743001 | -0.653 | -0.441 | 1.371 | -0.034 | 0.01 | -0.042 | -0.042 | 0.061 | 0.2 | 0.32 | 0.17 | 0.09 | 0.2 | 0.12 |
| 1 | 189846139 | rs6693137 | -1.01 | -0.522 | 1.221 | -0.084 | 0.008 | -0.061 | -0.061 | 0.067 | 0.22 | 0.39 | 0.18 | 0.1 | 0.21 | 0.18 |
| 1 | 189855850 | rs12402777 | 0.22 | -0.664 | 0.795 | 0.108 | -0.005 | -0.052 | -0.052 | 0 | 0.11 | 0.03 | 0.19 | 0.23 | 0.09 | 0.07 |
| 1 | 189865412 | rs7539739 | -1.444 | -0.725 | 0.975 | 0.126 | -0.013 | -0.044 | -0.044 | 0 | 0.31 | 0.47 | 0.2 | 0.15 | 0.24 | 0.32 |
| 1 | 189868708 | rs10920065 | -1.725 | -0.816 | 1.128 | 0.082 | 0.011 | 0.024 | 0.024 | 0 | 0.36 | 0.77 | 0.39 | 0.32 | 0.29 | 0.24 |
| 1 | 189878303 | rs10920093 | -0.315 | -0.621 | 0.822 | 0.176 | 0.005 | -0.021 | -0.021 | 0 | 0.19 | 0.22 | 0.16 | 0.08 | 0.2 | 0.12 |
| 1 | 189878594 | rs6413913 | -1.61 | -0.407 | 0.395 | 0.288 | 0.011 | 0.006 | 0.006 | 0 | 0.42 | 0.66 | 0.4 | 0.38 | 0.33 | 0.38 |
| 1 | 189879087 | rs12138080 | -1.003 | -0.039 | 0.612 | 0.016 | -0.008 | -0.038 | -0.038 | 0 | 0.29 | 0.43 | 0.19 | 0.14 | 0.24 | 0.27 |
| 1 | 189883637 | rs1852906 | -1.079 | -0.048 | 0.592 | 0.036 | -0.009 | -0.035 | -0.035 | 0 | 0.3 | 0.56 | 0.2 | 0.15 | 0.24 | 0.32 |
| 1 | 189887715 | rs10920116 | -1.017 | -0.046 | 0.594 | 0.034 | -0.008 | -0.034 | -0.034 | 0 | 0.29 | 0.43 | 0.19 | 0.14 | 0.24 | 0.27 |
| 1 | 189916279 | rs2699385 | 0.315 | 0.72 | 0.001 | 0.373 | 0.011 | -0.008 | -0.008 | 0 | 0.81 | 0.75 | 0.83 | 0.88 | 0.78 | 0.82 |
| 1 | 189918985 | rs815742 | -0.171 | 0.728 | -0.433 | 0.565 | 0.004 | -0.051 | -0.051 | 0.5 | 0.3 | 0.22 | 0.43 | 0.5 | 0.24 | 0.41 |
| 1 | 189922408 | rs860107 | -0.588 | 0.712 | -0.255 | 0.559 | 0.004 | -0.004 | -0.004 | 0 | 0.82 | 0.82 | 0.83 | 0.88 | 0.78 | 0.82 |
| 1 | 189929364 | rs13375157 | 0.188 | 0.509 | 0.008 | 0.087 | -0.005 | -0.032 | -0.032 | 0 | 0.08 | 0.09 | 0.05 | 0 | 0.07 | 0.01 |
| 1 | 189931733 | rs1341976 | 0.642 | 0.518 | 0.024 | 0.11 | -0.001 | 0.005 | 0.005 | 0 | 0.51 | 0.53 | 0.39 | 0.38 | 0.54 | 0.41 |
| 1 | 189951042 | rs815724 | 0.299 | 0.643 | 0.48 | 0.385 | 0.002 | 0.001 | 0.001 | 0 | 0.21 | 0.19 | 0.19 | 0.18 | 0.27 | 0.24 |
| 1 | 189952609 | rs10920249 | 0.816 | 0.763 | 0.411 | 0.462 | 0.002 | 0 | 0 | 0 | 0.06 | 0.06 | 0.05 | 0.06 | 0.07 | 0.12 |
| 1 | 189953701 | rs815730 | -1.405 | 0.317 | 0.743 | 0.557 | -0.008 | 0.002 | 0.002 | 0 | 0.54 | 0.77 | 0.67 | 0.54 | 0.49 | 0.51 |
| 1 | 189954040 | rs815731 | -1.709 | 0.303 | 0.732 | 0.551 | -0.049 | 0.001 | 0.001 | 0 | 0.59 | 0.84 | 0.71 | 0.61 | 0.56 | 0.62 |
| 1 | 189958783 | rs16829304 | 0.698 | -0.027 | 0.78 | 0.351 | -0.002 | -0.047 | -0.047 | 0 | 0.16 | 0.09 | 0.4 | 0.36 | 0.16 | 0.25 |
| 1 | 189958874 | rs1730734 | 0.791 | -0.017 | 0.828 | 0.355 | 0.003 | 0.002 | 0.002 | 0 | 0.21 | 0.21 | 0.19 | 0.18 | 0.27 | 0.24 |
| 1 | 189960204 | rs1320748 | -1.944 | 0.003 | 0.816 | 0.353 | -0.051 | -0.001 | -0.001 | 0.566 | 0.59 | 0.83 | 0.71 | 0.61 | 0.56 | 0.62 |
| 1 | 189962502 | rs845629 | -1.612 | -0.02 | 0.858 | 0.483 | -0.006 | 0.014 | 0.014 | 0 | 0.53 | 0.71 | 0.68 | 0.56 | 0.51 | 0.51 |
| 1 | 189963444 | rs6684090 | -0.394 | -0.055 | 1.203 | 0.496 | -0.026 | -0.055 | -0.055 | 0.658 | 0.1 | 0.23 | 0.05 | 0.01 | 0.04 | 0.01 |

| | | | | | | | | | | | | | | | | |
|---|---|---|---|---|---|---|---|---|---|---|---|---|---|---|---|---|
| 1 | 189968691 | rs4845208 | 0.258 | -0.006 | 0.869 | 0.621 | 0.006 | -0.016 | -0.016 | 0 | 0.15 | 0.11 | 0.38 | 0.36 | 0.12 | 0.24 |
| 1 | 189970400 | rs10920278 | -0.707 | -0.048 | 1.082 | 0.492 | 0.007 | -0.017 | -0.017 | 0.094 | 0.36 | 0.47 | 0.28 | 0.26 | 0.38 | 0.4 |
| 1 | 189971441 | rs12117595 | 0.07 | -0.029 | 0.895 | 0.473 | 0.002 | -0.002 | -0.002 | 0.366 | 0.16 | 0.13 | 0.4 | 0.36 | 0.13 | 0.25 |
| 1 | 189973857 | rs1529315 | -0.69 | 0.261 | -0.117 | 0.065 | -0.104 | -0.009 | -0.009 | 0 | 0.51 | 0.95 | 0.58 | 0.41 | 0.52 | 0.6 |
| 1 | 189983720 | rs10737578 | -1.686 | 0.069 | -0.255 | 0.446 | -0.062 | -0.179 | -0.179 | 6.897 | 0.65 | 0.95 | 0.65 | 0.51 | 0.66 | 0.66 |
| 1 | 189984428 | rs10800809 | -0.884 | -0.151 | 0.326 | 0.277 | 0.005 | 0.001 | 0.001 | 0 | 0.54 | 0.51 | 0.58 | 0.45 | 0.6 | 0.49 |
| 1 | 190005340 | rs1534745 | -1.514 | -0.4 | 0.434 | 0.476 | 0.003 | 0.003 | 0.003 | 0 | 0.61 | 0.63 | 0.58 | 0.5 | 0.53 | 0.65 |
| 1 | 190010100 | rs6427944 | -1.616 | -0.421 | 0.713 | 0.474 | -0.041 | -0.046 | -0.046 | 0 | 0.61 | 0.79 | 0.59 | 0.5 | 0.53 | 0.65 |
| 1 | 190010245 | rs17374565 | 0.151 | -0.535 | 0.572 | 0.415 | -0.04 | 0.131 | 0.131 | 0 | 0.36 | 0.1 | 0.23 | 0.15 | 0.35 | 0.41 |
| 1 | 190020770 | rs2037871 | -0.932 | -0.518 | 0.778 | 0.532 | -0.02 | 0.032 | 0.032 | 0 | 0.24 | 0.39 | 0.35 | 0.34 | 0.17 | 0.24 |
| 1 | 190034469 | rs6427956 | -1.806 | -0.409 | 0.63 | 0.423 | -0.123 | -0.131 | -0.131 | 1.288 | 0.63 | 0.93 | 0.63 | 0.5 | 0.54 | 0.65 |
| 1 | 190036140 | rs10920424 | 0.796 | -0.405 | 0.947 | 0.374 | -0.012 | 0.061 | 0.061 | 0 | 0.28 | 0.18 | 0.21 | 0.1 | 0.31 | 0.24 |
| 1 | 190037310 | rs7516991 | 0.796 | -0.405 | 0.947 | 0.374 | -0.012 | 0.061 | 0.061 | 0 | 0.28 | 0.18 | 0.21 | 0.1 | 0.31 | 0.24 |
| 1 | 190042706 | rs10920441 | -0.796 | -0.387 | 0.958 | 0.411 | 0.002 | -0.001 | -0.001 | 0 | 0.23 | 0.27 | 0.34 | 0.22 | 0.17 | 0.23 |
| 1 | 190050376 | rs4585977 | -1.626 | -0.269 | 0.602 | 0.436 | -0.035 | -0.039 | -0.039 | 1.143 | 0.61 | 0.77 | 0.62 | 0.5 | 0.53 | 0.65 |
| 1 | 190057680 | rs2419374 | -1.368 | -0.29 | 0.886 | 0.393 | -0.022 | -0.003 | -0.003 | 0 | 0.28 | 0.39 | 0.14 | 0.27 | 0.21 | 0.29 |
| 1 | 190064165 | rs1418811 | -1.076 | -0.194 | 0.861 | 0.404 | 0.001 | -0.001 | -0.001 | 0 | 0.24 | 0.22 | 0.11 | 0.26 | 0.2 | 0.26 |
| 1 | 190066386 | rs1935881 | 0.551 | -0.196 | 0.986 | 0.226 | -0.016 | 0.097 | 0.097 | 0 | 0.31 | 0.14 | 0.21 | 0.1 | 0.33 | 0.27 |
| 1 | 190070919 | rs17375108 | 0.1 | -0.189 | 0.935 | 0.253 | -0.008 | 0.003 | 0.003 | 0 | 0.14 | 0.07 | 0.06 | 0.09 | 0.14 | 0.08 |
| 1 | 190073382 | rs6427982 | -1.031 | -0.105 | 1.218 | 0.417 | -0.011 | 0.02 | 0.02 | 0 | 0.56 | 0.62 | 0.34 | 0.38 | 0.52 | 0.53 |
| 1 | 190086179 | rs2226167 | -0.817 | -0.409 | 0.29 | 0.113 | -0.019 | 0.01 | 0.01 | 0 | 0.6 | 0.72 | 0.61 | 0.39 | 0.55 | 0.55 |
| 1 | 190099810 | rs2211430 | -0.399 | -0.876 | 0.263 | -0.018 | -0.04 | 0.03 | 0.03 | 0.236 | 0.12 | 0.34 | 0.3 | 0.19 | 0.05 | 0.2 |
| 1 | 190102031 | rs10753936 | 0.12 | -0.738 | -0.718 | -0.14 | -0.006 | 0.141 | 0.141 | 0 | 0.47 | 0.23 | 0.29 | 0.2 | 0.46 | 0.33 |
| 1 | 190104227 | rs1418808 | -0.584 | -0.294 | -1.197 | 0.038 | -0.059 | 0.007 | 0.007 | 0 | 0.61 | 0.77 | 0.61 | 0.4 | 0.51 | 0.53 |
| 1 | 190108881 | rs2419370 | -0.67 | 0.346 | -1.531 | 0.482 | -0.114 | -0.095 | -0.095 | 2.049 | 0.63 | 0.89 | 0.6 | 0.53 | 0.49 | 0.54 |
| 1 | 190112653 | rs815342 | 0.466 | 0.296 | -1.05 | 0.07 | 0.014 | 0.038 | 0.038 | 0 | 0.4 | 0.31 | 0.24 | 0.24 | 0.35 | 0.28 |
| 1 | 190117274 | rs1557188 | 0.569 | 0.032 | -0.397 | 0.709 | -0.058 | 0.042 | 0.042 | 0 | 0.18 | 0.31 | 0.15 | 0.25 | 0.05 | 0.28 |
| 1 | 190118911 | rs2990998 | 1.094 | 0.029 | -0.184 | 0.687 | -0.006 | -0.016 | -0.016 | 0 | 0.4 | 0.46 | 0.26 | 0.24 | 0.35 | 0.3 |
| 1 | 190131750 | rs10920654 | -0.251 | 0.325 | -0.23 | 0.663 | -0.04 | 0.078 | 0.078 | 0 | 0.22 | 0.38 | 0.19 | 0.37 | 0.15 | 0.33 |
| 1 | 190135811 | rs1093086 | -2.42 | 0.937 | -0.552 | 0.736 | -0.05 | -0.097 | -0.097 | 0 | 0.78 | 0.97 | 0.73 | 0.65 | 0.79 | 0.79 |
| 1 | 190149198 | rs12041138 | 2.255 | 0.421 | -0.573 | 0.436 | 0.014 | 0.081 | 0.081 | 0 | 0.28 | 0.08 | 0.15 | 0.12 | 0.25 | 0.22 |
| 1 | 190150037 | rs2253618 | -0.743 | 0.49 | -0.469 | 0.442 | -0.004 | -0.044 | -0.044 | 0 | 0.6 | 0.79 | 0.43 | 0.49 | 0.42 | 0.59 |
| 1 | 190152217 | rs815344 | -0.722 | 0.464 | -0.409 | 0.446 | 0.003 | -0.029 | -0.029 | 0 | 0.59 | 0.74 | 0.43 | 0.49 | 0.42 | 0.59 |
| 1 | 190155633 | rs815343 | 1.118 | 0.447 | -0.341 | 0.442 | 0.008 | 0.02 | 0.02 | 0 | 0.4 | 0.37 | 0.27 | 0.24 | 0.36 | 0.28 |
| 1 | 190158234 | rs1592245 | 0.929 | 0.464 | -0.324 | 0.485 | -0.047 | 0.03 | 0.03 | 0 | 0.18 | 0.26 | 0.15 | 0.25 | 0.05 | 0.29 |
| 1 | 190164775 | rs815330 | 0.355 | 0.558 | -0.263 | 0.409 | 0.01 | 0.01 | 0.01 | 0 | 0.1 | 0.21 | 0.12 | 0.11 | 0.12 | 0.06 |
| 1 | 190170694 | rs1592251 | 2.113 | 0.563 | -0.278 | 0.388 | 0.012 | 0.084 | 0.084 | 0 | 0.28 | 0.07 | 0.16 | 0.13 | 0.25 | 0.23 |
| 1 | 190176008 | rs10800936 | -1.116 | 0.552 | -0.098 | 0.402 | -0.006 | -0.009 | -0.009 | 0 | 0.61 | 0.52 | 0.72 | 0.65 | 0.57 | 0.69 |
| 1 | 190194909 | rs4633293 | -1.112 | 0.549 | -0.093 | 0.401 | -0.003 | -0.004 | -0.004 | 0 | 0.61 | 0.55 | 0.72 | 0.65 | 0.57 | 0.69 |
| 1 | 190201724 | rs4399162 | 0.257 | 0.467 | -0.093 | 0.463 | -0.025 | 0.006 | 0.006 | 0 | 0.23 | 0.26 | 0.19 | 0.25 | 0.09 | 0.29 |
| 1 | 190213532 | rs17368045 | 1.017 | 0.673 | -0.326 | 0.425 | -0.083 | 0.021 | 0.021 | 0.526 | 0.13 | 0.02 | 0.24 | 0.05 | 0.25 | 0.18 |
| 1 | 190234436 | rs10920675 | 1.082 | 0.212 | -0.07 | 0.541 | -0.057 | -0.014 | -0.014 | 0 | 0.36 | 0.15 | 0.52 | 0.39 | 0.48 | 0.37 |
| 1 | 190239919 | rs10920679 | -1.146 | 0.164 | 0.003 | 0.533 | 0.002 | 0.005 | 0.005 | 0 | 0.61 | 0.65 | 0.73 | 0.65 | 0.57 | 0.67 |
| 1 | 190259661 | rs480692 | 1.061 | 0.224 | -0.031 | 0.557 | -0.043 | -0.01 | -0.01 | 1.368 | 0.36 | 0.19 | 0.52 | 0.39 | 0.48 | 0.39 |
| 1 | 190263214 | rs1093040 | -0.13 | 0.29 | -0.048 | 0.512 | 0.039 | 0.091 | 0.091 | 0 | 0.91 | 0.77 | 0.89 | 0.77 | 0.82 | 0.9 |
| 1 | 190287713 | rs655598 | -0.939 | 0.22 | 0.154 | 0.661 | -0.003 | -0.002 | -0.002 | 0 | 0.6 | 0.52 | 0.72 | 0.63 | 0.57 | 0.69 |
| 1 | 190288297 | rs652953 | 0.992 | 0.242 | -0.297 | 0.619 | -0.036 | 0.009 | 0.009 | 0 | 0.37 | 0.2 | 0.52 | 0.37 | 0.48 | 0.38 |
| 1 | 190291298 | rs1759762 | 1.077 | 0.248 | -0.445 | 0.622 | -0.065 | 0.002 | 0.002 | 0 | 0.36 | 0.14 | 0.51 | 0.37 | 0.48 | 0.38 |
| 1 | 190307072 | rs718609 | 1.891 | 0.107 | -0.418 | 0.289 | 0.002 | 0.054 | 0.054 | 0 | 0.28 | 0.05 | 0.16 | 0.14 | 0.25 | 0.22 |
| 1 | 190318694 | rs12030570 | 1.746 | -0.284 | -0.229 | 0.319 | 0.017 | 0.012 | 0.012 | 0.588 | 0.23 | 0.16 | 0.1 | 0.14 | 0.15 | 0.19 |
| 1 | 190337198 | rs1148612 | -0.133 | 0.116 | -0.078 | 0.405 | -0.007 | -0.018 | -0.018 | 0.309 | 0.73 | 0.63 | 0.78 | 0.8 | 0.79 | 0.7 |
| 1 | 190362935 | rs1031076 | 0.944 | 0.26 | -0.469 | 0.56 | -0.06 | -0.03 | -0.03 | 0 | 0.4 | 0.15 | 0.57 | 0.41 | 0.45 | 0.4 |
| 1 | 190377988 | rs1171148 | 0.967 | 0.25 | -0.525 | 0.562 | -0.074 | -0.035 | -0.035 | 0 | 0.4 | 0.13 | 0.55 | 0.4 | 0.45 | 0.4 |
| 1 | 190383667 | rs1171045 | -0.086 | -0.175 | -0.082 | 0.516 | -0.009 | -0.02 | -0.02 | 0 | 0.74 | 0.67 | 0.78 | 0.78 | 0.79 | 0.7 |
| 1 | 190385136 | rs12065497 | 1.947 | -0.22 | -0.179 | 0.464 | 0.032 | 0.01 | 0.01 | 0.427 | 0.22 | 0.09 | 0.09 | 0.15 | 0.16 | 0.2 |
| 1 | 190385304 | rs1171044 | 0.181 | -0.123 | 0.117 | 0.544 | -0.014 | 0.012 | 0.012 | 0.015 | 0.72 | 0.62 | 0.78 | 0.64 | 0.78 | 0.68 |
| 1 | 190385714 | rs1171042 | 0.212 | -0.077 | 0.155 | 0.577 | -0.008 | 0.02 | 0.02 | 0.314 | 0.72 | 0.61 | 0.78 | 0.62 | 0.79 | 0.73 |
| 1 | 190388053 | rs13374312 | 0.881 | -0.183 | -0.077 | 0.469 | 0.001 | -0.001 | -0.001 | 0 | 0.14 | 0.12 | 0.07 | 0.14 | 0.16 | 0.08 |
| 1 | 190394746 | rs17377527 | 0.587 | -0.221 | 0.025 | 0.331 | -0.001 | -0.01 | -0.01 | 0 | 0.15 | 0.17 | 0.09 | 0.07 | 0.16 | 0.07 |
| 1 | 190399946 | rs9970018 | 0.408 | -0.529 | 0.255 | 0.416 | -0.011 | -0.01 | -0.01 | 0.885 | 0.38 | 0.42 | 0.22 | 0.26 | 0.31 | 0.31 |
| 1 | 190406331 | rs12091129 | 1.836 | -0.575 | 0.414 | 0.356 | -0.01 | -0.001 | -0.001 | 0.461 | 0.24 | 0.25 | 0.14 | 0.18 | 0.15 | 0.24 |
| 1 | 190412944 | rs6660355 | 2.205 | -0.651 | 0.388 | 0.319 | 0.031 | -0.007 | -0.007 | 0.722 | 0.21 | 0.09 | 0.13 | 0.18 | 0.15 | 0.21 |
| 1 | 190416298 | rs12087474 | 0.306 | -0.658 | 0.411 | 0.274 | -0.003 | -0.005 | -0.005 | 0 | 0.4 | 0.44 | 0.26 | 0.26 | 0.34 | 0.32 |
| 1 | 190418653 | rs1171023 | 1.19 | -0.218 | 0.314 | 0.45 | -0.063 | -0.007 | -0.007 | 0 | 0.38 | 0.14 | 0.38 | 0.38 | 0.46 | 0.34 |
| 1 | 190424966 | rs12058205 | 1.747 | -0.225 | 0.336 | 0.52 | 0.018 | 0.001 | 0.001 | 1.374 | 0.26 | 0.17 | 0.16 | 0.2 | 0.18 | 0.21 |
| 1 | 190441481 | rs12121097 | 0.69 | -0.211 | 0.729 | 0.622 | 0.02 | -0.009 | -0.009 | 0.725 | 0.12 | 0.18 | 0.26 | 0.09 | 0.18 | 0.17 |
| 1 | 190442971 | rs2490271 | -0.396 | -0.245 | 0.846 | 0.632 | -0.003 | 0.033 | 0.033 | 0 | 0.79 | 0.73 | 0.82 | 0.67 | 0.83 | 0.75 |
| 1 | 190446098 | rs1171049 | 0.776 | -0.377 | 0.46 | 0.612 | 0.015 | 0.018 | 0.018 | 0 | 0.66 | 0.5 | 0.56 | 0.59 | 0.65 | 0.58 |
| 1 | 190455544 | rs12034237 | 1.884 | -0.279 | 0.248 | 0.66 | 0.026 | -0.007 | -0.007 | 0.363 | 0.22 | 0.13 | 0.13 | 0.23 | 0.15 | 0.21 |

| | | | | | | | | | | | | | | | | |
|---|---|---|---|---|---|---|---|---|---|---|---|---|---|---|---|---|
| 1 | 190466822 | rs814941 | 0.161 | -0.349 | 0.557 | 0.703 | 0.103 | 0.032 | 0.032 | 0.486 | 0.79 | 0.58 | 0.58 | 0.76 | 0.67 | 0.73 |
| 1 | 190467826 | rs703931 | 0.149 | -0.418 | 0.466 | 0.699 | -0.008 | -0.009 | -0.009 | 0.456 | 0.55 | 0.4 | 0.46 | 0.53 | 0.52 | 0.48 |
| 1 | 190472727 | rs10920731 | 0.859 | -0.406 | 0.288 | 0.706 | -0.035 | 0.037 | 0.037 | 0 | 0.43 | 0.16 | 0.41 | 0.36 | 0.49 | 0.35 |
| 1 | 190476185 | rs814977 | 1.151 | -0.145 | 0.523 | 0.763 | 0.045 | 0.057 | 0.057 | 0 | 0.72 | 0.54 | 0.57 | 0.59 | 0.66 | 0.61 |
| 1 | 190476831 | rs12405769 | 1.261 | -0.037 | 0.167 | 0.876 | -0.003 | 0.003 | 0.003 | 0 | 0.1 | 0.1 | 0.03 | 0.18 | 0.03 | 0.13 |
| 1 | 190483784 | rs814972 | 0.516 | -0.025 | 0.222 | 0.789 | 0.099 | 0.062 | 0.062 | 0 | 0.81 | 0.58 | 0.62 | 0.73 | 0.71 | 0.66 |
| 1 | 190485500 | rs814923 | 0.516 | -0.025 | 0.222 | 0.789 | 0.099 | 0.062 | 0.062 | 0 | 0.81 | 0.58 | 0.62 | 0.73 | 0.71 | 0.66 |
| 1 | 190492280 | rs814914 | 0.864 | -0.367 | 0.018 | 0.719 | -0.042 | 0.053 | 0.053 | 0 | 0.47 | 0.16 | 0.43 | 0.33 | 0.5 | 0.33 |
| 1 | 190492543 | rs814913 | 1.349 | -0.284 | 0.157 | 0.771 | 0.004 | 0.012 | 0.012 | 0.039 | 0.69 | 0.41 | 0.57 | 0.55 | 0.67 | 0.53 |
| 1 | 190494536 | rs17378706 | 0.225 | -0.004 | 0.252 | 0.874 | 0.015 | -0.004 | -0.004 | 0 | 0.12 | 0.21 | 0.13 | 0.1 | 0.15 | 0.22 |
| 1 | 190501742 | rs7541423 | 1.435 | 0.104 | -0.193 | 0.945 | 0.013 | -0.013 | -0.013 | 0.262 | 0.1 | 0.09 | 0.03 | 0.18 | 0.04 | 0.11 |
| 1 | 190503966 | rs755805 | 1.479 | 0.517 | -0.096 | 0.844 | -0.025 | -0.026 | -0.026 | 9.14 | 0.05 | 0.01 | 0.24 | 0.16 | 0.13 | 0.07 |
| 1 | 190506603 | rs814968 | -0.083 | 0.382 | -0.083 | 0.788 | -0.013 | -0.019 | -0.019 | 0 | 0.61 | 0.36 | 0.49 | 0.51 | 0.55 | 0.49 |
| 1 | 190514183 | rs814958 | -0.098 | 0.839 | -0.047 | 0.713 | 0.109 | 0.099 | 0.099 | 0.016 | 0.93 | 0.81 | 0.75 | 0.82 | 0.86 | 0.89 |
| 1 | 190522315 | rs10920743 | 0.776 | 0.839 | -0.636 | 0.669 | -0.018 | -0.014 | -0.014 | 0.026 | 0.08 | 0.03 | 0.21 | 0.16 | 0.15 | 0.1 |
| 1 | 190533202 | rs12036325 | 0.977 | 1.007 | -1.551 | 0.514 | -0.069 | -0.066 | -0.066 | 0.062 | 0.24 | 0.03 | 0.23 | 0.27 | 0.27 | 0.23 |
| 1 | 190533700 | rs1327872 | 0.54 | 0.771 | -1.368 | 0.618 | -0.027 | -0.011 | -0.011 | 0.404 | 0.12 | 0.04 | 0.21 | 0.16 | 0.16 | 0.1 |
| 1 | 190543473 | rs12136925 | 0.805 | 0.795 | -0.531 | 0.662 | 0.049 | 0.034 | 0.034 | 0.423 | 0.08 | 0.19 | 0.12 | 0.11 | 0.15 | 0.21 |
| 1 | 190543495 | rs17370725 | 0.765 | 0.68 | -1.103 | 0.368 | -0.098 | 0.003 | 0.003 | 0 | 0.36 | 0.03 | 0.38 | 0.27 | 0.39 | 0.27 |
| 1 | 190546931 | rs11582878 | -0.185 | 0.406 | -0.78 | 0.329 | 0.011 | 0.029 | 0.029 | 0.042 | 0.51 | 0.62 | 0.57 | 0.39 | 0.57 | 0.51 |
| 1 | 190548529 | rs16832476 | 1.118 | 0.355 | -0.915 | 0.26 | 0.065 | -0.088 | -0.088 | 0 | 0.32 | 0.15 | 0.14 | 0.43 | 0.19 | 0.33 |
| 1 | 190549233 | rs16832477 | 0.917 | 0.523 | -0.556 | 0.223 | 0.061 | 0.053 | 0.053 | 0 | 0.08 | 0.29 | 0.12 | 0.1 | 0.14 | 0.2 |
| 1 | 190551418 | rs642748 | -0.029 | 0.681 | -0.795 | 0.238 | -0.005 | 0.037 | 0.037 | 0 | 0.46 | 0.62 | 0.53 | 0.29 | 0.52 | 0.5 |
| 1 | 190558106 | rs49314 | 1.248 | 0.698 | -0.57 | 0.24 | 0.027 | -0.036 | -0.036 | 0 | 0.09 | 0.19 | 0.16 | 0.01 | 0.09 | 0.07 |
| 1 | 190560021 | rs749390 | 1.354 | 0.575 | -0.689 | 0.436 | 0.052 | -0.128 | -0.128 | 0 | 0.3 | 0.09 | 0.13 | 0.43 | 0.18 | 0.28 |
| 1 | 190566014 | rs694168 | -0.161 | 0.67 | -0.616 | 0.442 | -0.001 | -0.001 | -0.001 | 0 | 0.79 | 0.8 | 0.68 | 0.72 | 0.71 | 0.83 |
| 1 | 190567035 | rs17379742 | 1.14 | 0.706 | -0.714 | 0.444 | -0.064 | -0.035 | -0.035 | 0.438 | 0.11 | 0.01 | 0.2 | 0.17 | 0.17 | 0.1 |
| 1 | 190570848 | rs2149028 | 2.463 | 0.409 | -0.879 | 0.69 | -0.015 | -0.087 | -0.087 | 0 | 0.2 | 0.01 | 0.12 | 0.23 | 0.16 | 0.18 |
| 1 | 190570878 | rs12144398 | 1.287 | 0.698 | -0.642 | 0.438 | 0.031 | 0.01 | 0.01 | 0.144 | 0.09 | 0.18 | 0.12 | 0.09 | 0.17 | 0.21 |
| 1 | 190573073 | rs6668224 | 1.778 | 0.648 | -0.764 | 0.439 | 0.037 | -0.128 | -0.128 | 5.79 | 0.27 | 0.09 | 0.13 | 0.43 | 0.18 | 0.28 |
| 1 | 190577336 | rs584161 | -1.659 | 0.757 | -0.523 | 0.461 | 0.001 | 0.003 | 0.003 | 0 | 0.9 | 0.85 | 0.89 | 0.89 | 0.89 | 0.93 |
| 1 | 190581372 | rs7531899 | 0.465 | 0.776 | -0.377 | 0.365 | 0.037 | 0.002 | 0.002 | 0.015 | 0.12 | 0.32 | 0.15 | 0.09 | 0.18 | 0.23 |
| 1 | 190583153 | rs694698 | 0.602 | 0.647 | -0.453 | 0.297 | -0.002 | 0.014 | 0.014 | 0 | 0.36 | 0.31 | 0.39 | 0.19 | 0.36 | 0.3 |
| 1 | 190596095 | rs604813 | -0.312 | 0.936 | 0.098 | 0.907 | 0.011 | 0.028 | 0.028 | 0 | 0.88 | 0.85 | 0.78 | 0.72 | 0.84 | 0.87 |
| 1 | 190604337 | rs12079787 | 0.633 | 0.962 | 0.106 | 0.746 | 0.022 | -0.043 | -0.043 | 0 | 0.11 | 0.25 | 0.14 | 0.04 | 0.16 | 0.05 |
| 1 | 190607833 | rs16832542 | 1.077 | 0.997 | 0.038 | 0.762 | 0.017 | -0.028 | -0.028 | 0.005 | 0.09 | 0.16 | 0.12 | 0.04 | 0.16 | 0.05 |
| 1 | 190608808 | rs2149025 | 0.712 | 1.035 | -0.136 | 0.87 | -0.049 | -0.054 | -0.054 | 0 | 0.1 | 0.01 | 0.2 | 0.24 | 0.16 | 0.08 |
| 1 | 190632794 | rs10800956 | -1.253 | 0.858 | 0.29 | 0.76 | 0.035 | 0.017 | 0.017 | 0 | 0.26 | 0.49 | 0.37 | 0.32 | 0.32 | 0.2 |
| 1 | 190633980 | rs12022777 | -0.747 | 0.839 | 0.434 | 0.821 | 0.059 | 0.053 | 0.053 | 0 | 0.19 | 0.39 | 0.34 | 0.31 | 0.31 | 0.19 |
| 1 | 190634131 | rs16832590 | 1.784 | 0.577 | -0.093 | 0.879 | 0.033 | -0.069 | -0.069 | 0.063 | 0.22 | 0.02 | 0.12 | 0.22 | 0.13 | 0.13 |
| 1 | 190645553 | rs1325222 | -0.38 | 0.63 | 0.055 | 0.842 | 0 | -0.002 | -0.002 | 0.002 | 0.44 | 0.57 | 0.48 | 0.53 | 0.44 | 0.32 |
| 1 | 190647315 | rs6678095 | -0.391 | 0.63 | 0.055 | 0.842 | 0 | -0.002 | -0.002 | 0 | 0.44 | 0.57 | 0.48 | 0.53 | 0.44 | 0.32 |
| 1 | 190650449 | rs6428024 | 1.207 | 0.71 | 0.19 | 0.869 | 0.016 | -0.018 | -0.018 | 0 | 0.09 | 0.18 | 0.12 | 0.05 | 0.16 | 0.06 |
| 1 | 190651533 | rs10753952 | 0.187 | 0.642 | 0.143 | 0.92 | -0.012 | 0.023 | 0.023 | 0 | 0.35 | 0.4 | 0.36 | 0.48 | 0.28 | 0.26 |
| 1 | 190653512 | rs2025585 | -1.03 | 0.726 | 0.178 | 0.577 | 0.015 | -0.025 | -0.025 | 1.584 | 0.91 | 0.8 | 0.85 | 0.98 | 0.92 | 0.9 |
| 1 | 190671240 | rs6675319 | -0.504 | 0.558 | -0.05 | 1.072 | -0.005 | 0 | 0 | 0.01 | 0.46 | 0.54 | 0.31 | 0.45 | 0.36 | 0.41 |
| 1 | 190673080 | rs1325227 | 0.261 | 0.395 | -0.38 | 1.105 | 0.076 | -0.034 | -0.034 | 0 | 0.33 | 0.11 | 0.17 | 0.37 | 0.2 | 0.31 |
| 1 | 190681824 | rs1408840 | -0.006 | 0.754 | -0.379 | 1.19 | 0.005 | -0.069 | -0.069 | 0 | 0.37 | 0.1 | 0.35 | 0.61 | 0.34 | 0.38 |
| 1 | 190687205 | rs10800961 | -0.771 | 0.756 | 0.056 | 1.126 | 0.024 | 0.01 | 0.01 | 0 | 0.2 | 0.3 | 0.34 | 0.27 | 0.32 | 0.16 |
| 1 | 190689478 | rs10920786 | -0.616 | 0.764 | -0.073 | 1.13 | -0.003 | -0.002 | -0.002 | 0 | 0.19 | 0.2 | 0.33 | 0.28 | 0.32 | 0.16 |
| 1 | 190694813 | rs12024738 | -0.117 | 0.65 | -0.051 | 1.123 | -0.019 | 0.001 | 0.001 | 0 | 0.37 | 0.49 | 0.37 | 0.64 | 0.29 | 0.37 |
| 1 | 190697042 | rs1359454 | 1.551 | 0.719 | -0.178 | 1.213 | -0.026 | -0.014 | -0.014 | 0.014 | 0.05 | 0.01 | 0.19 | 0.24 | 0.13 | 0.06 |
| 1 | 190713300 | rs573068 | -0.196 | 0.617 | -0.161 | 1.099 | -0.01 | -0.012 | -0.012 | 0 | 0.42 | 0.47 | 0.27 | 0.25 | 0.34 | 0.3 |
| 1 | 190713567 | rs12036438 | 1.217 | 0.658 | -0.162 | 1.093 | 0.001 | 0.019 | 0.019 | 0 | 0.06 | 0.06 | 0.2 | 0.24 | 0.12 | 0.06 |
| 1 | 190713980 | rs490184 | -0.996 | 0.421 | -0.178 | 1.097 | 0 | 0.009 | 0.009 | 0 | 0.53 | 0.6 | 0.5 | 0.68 | 0.49 | 0.47 |
| 1 | 190718173 | rs10494640 | 1.137 | 0.656 | -0.403 | 1.145 | -0.032 | -0.026 | -0.026 | 0 | 0.07 | 0.01 | 0.1 | 0.09 | 0.11 | 0.16 |
| 1 | 190735457 | rs12087744 | -0.244 | 0.352 | -0.202 | 1.172 | -0.007 | -0.003 | -0.003 | 0 | 0.2 | 0.18 | 0.31 | 0.29 | 0.3 | 0.15 |
| 1 | 190750491 | rs16832764 | 1.29 | 0.193 | -0.339 | 1.155 | 0.047 | -0.027 | -0.027 | 0 | 0.22 | 0.05 | 0.13 | 0.19 | 0.13 | 0.13 |
| 1 | 190756262 | rs513810 | -0.379 | 0.223 | 0.203 | 1.261 | 0.006 | -0.004 | -0.004 | 0 | 0.47 | 0.61 | 0.48 | 0.5 | 0.44 | 0.34 |
| 1 | 190756370 | rs511362 | -0.307 | 0.18 | 0.491 | 1.234 | -0.008 | 0.007 | 0.007 | 0.18 | 0.11 | 0.43 | 0.08 | 0.06 | 0.07 | 0.13 |
| 1 | 190760788 | rs16832771 | 0.247 | 0.255 | 0.143 | 1.265 | -0.017 | 0.034 | 0.034 | 0 | 0.12 | 0.21 | 0.07 | 0.23 | 0.1 | 0.17 |
| 1 | 190777205 | rs660849 | -0.452 | 0.341 | 0.261 | 1.155 | 0.005 | 0.021 | 0.021 | 0 | 0.51 | 0.59 | 0.49 | 0.68 | 0.47 | 0.45 |
| 1 | 190781850 | rs625802 | -0.513 | 0.381 | 0.103 | 1.158 | 0.006 | 0.028 | 0.028 | 0 | 0.51 | 0.63 | 0.5 | 0.68 | 0.47 | 0.45 |
| 1 | 190785818 | rs12029687 | 0.959 | 0.591 | -0.117 | 1.365 | -0.013 | -0.007 | -0.007 | 0 | 0.06 | 0.03 | 0.19 | 0.24 | 0.1 | 0.07 |
| 1 | 190786827 | rs12564225 | -0.226 | 0.459 | -0.019 | 1.276 | 0.006 | 0.03 | 0.03 | 0 | 0.46 | 0.28 | 0.46 | 0.66 | 0.46 | 0.39 |
| 1 | 190799545 | rs7522517 | 0.613 | 0.397 | -0.045 | 1.356 | -0.008 | 0.016 | 0.016 | 0 | 0.11 | 0.17 | 0.07 | 0.23 | 0.1 | 0.17 |
| 1 | 190804630 | rs2489676 | -1.106 | 0.358 | 0.023 | 1.172 | -0.011 | 0.024 | 0.024 | 0 | 0.6 | 0.68 | 0.62 | 0.75 | 0.54 | 0.54 |
| 1 | 190813289 | rs10494649 | 0.652 | 0.362 | -0.179 | 1.262 | -0.043 | -0.072 | -0.072 | 0 | 0.35 | 0.11 | 0.39 | 0.43 | 0.36 | 0.22 |
| 1 | 190818192 | rs12143564 | 1.133 | 0.743 | 0.011 | 1.025 | -0.006 | 0.038 | 0.038 | 0 | 0.36 | 0.21 | 0.33 | 0.24 | 0.38 | 0.42 |

| | | | | | | | | | | | | | | | | |
|---|---|---|---|---|---|---|---|---|---|---|---|---|---|---|---|---|
| 1 | 190818216 | rs10920810 | -0.167 | 0.759 | 0.247 | 0.975 | 0.043 | 0 | 0 | 0.374 | 0.83 | 0.53 | 0.79 | 0.9 | 0.84 | 0.82 |
| 1 | 190818256 | rs10800972 | -0.167 | 0.759 | 0.247 | 0.975 | 0.043 | 0 | 0 | 0.374 | 0.83 | 0.53 | 0.79 | 0.9 | 0.84 | 0.82 |
| 1 | 190834267 | rs1438532 | -0.156 | 0.766 | -0.254 | 0.934 | 0.03 | -0.019 | -0.019 | 0 | 0.82 | 0.49 | 0.8 | 0.92 | 0.84 | 0.83 |
| 1 | 190840017 | rs12136008 | 0.687 | 0.683 | -0.45 | 0.967 | 0.013 | -0.022 | -0.022 | 0.546 | 0.12 | 0.07 | 0.07 | 0.23 | 0.11 | 0.18 |
| 1 | 190849399 | rs10800988 | 0.541 | 0.704 | -0.5 | 0.934 | -0.02 | -0.011 | -0.011 | 0 | 0.41 | 0.28 | 0.51 | 0.5 | 0.48 | 0.49 |
| 1 | 190860691 | rs6699971 | 0.164 | 0.632 | -0.354 | 0.845 | 0.006 | -0.025 | -0.025 | 0 | 0.81 | 0.4 | 0.79 | 0.92 | 0.83 | 0.81 |
| 1 | 190863011 | rs12135498 | 0.718 | 0.534 | -0.501 | 0.929 | 0.009 | -0.014 | -0.014 | 0 | 0.12 | 0.09 | 0.07 | 0.23 | 0.11 | 0.18 |
| 1 | 190872645 | rs2419932 | 0.841 | 0.643 | -0.462 | 0.981 | -0.001 | -0.001 | -0.001 | 0 | 0.37 | 0.38 | 0.34 | 0.25 | 0.38 | 0.44 |
| 1 | 190874331 | rs12120220 | 0.869 | 0.59 | -0.47 | 1.024 | 0.011 | -0.018 | -0.018 | 3.235 | 0.12 | 0.09 | 0.07 | 0.23 | 0.11 | 0.18 |
| 1 | 190875644 | rs12411096 | 1.605 | 0.668 | -0.595 | 1.089 | -0.045 | -0.031 | -0.031 | 0.085 | 0.28 | 0.02 | 0.2 | 0.2 | 0.24 | 0.14 |
| 1 | 190876071 | rs6669436 | 0.968 | 0.747 | -0.468 | 1.041 | -0.003 | 0.017 | 0.017 | 0 | 0.37 | 0.31 | 0.33 | 0.25 | 0.38 | 0.44 |
| 1 | 190876141 | rs1936980 | -0.549 | 0.791 | 0.104 | 0.997 | 0.008 | -0.019 | -0.019 | 0 | 0.85 | 0.77 | 0.82 | 0.95 | 0.84 | 0.87 |
| 1 | 190885399 | rs1936988 | 0.508 | 0.749 | 0.051 | 0.977 | 0.046 | 0.064 | 0.064 | 0 | 0.77 | 0.51 | 0.62 | 0.68 | 0.73 | 0.75 |
| 1 | 190894392 | rs4372256 | 1.089 | 0.877 | -0.132 | 0.749 | -0.001 | -0.005 | -0.005 | 0 | 0.29 | 0.32 | 0.25 | 0.15 | 0.29 | 0.26 |
| 1 | 190895591 | rs11811680 | 1.262 | 0.847 | -0.085 | 0.735 | -0.002 | 0.028 | 0.028 | 0 | 0.28 | 0.2 | 0.24 | 0.15 | 0.3 | 0.25 |
| 1 | 190903602 | rs12134172 | 0.288 | 0.797 | -0.024 | 0.896 | 0.004 | -0.03 | -0.03 | 0.049 | 0.19 | 0.11 | 0.16 | 0.32 | 0.19 | 0.35 |
| 1 | 190913478 | rs12567544 | -0.06 | 0.632 | -0.063 | 0.652 | 0.007 | -0.046 | -0.046 | 1.41 | 0.78 | 0.49 | 0.77 | 0.92 | 0.81 | 0.82 |
| 1 | 190920467 | rs1011726 | 0.212 | 0.649 | -0.194 | 0.747 | 0.015 | -0.018 | -0.018 | 0 | 0.45 | 0.24 | 0.36 | 0.53 | 0.4 | 0.49 |
| 1 | 190923837 | rs1431139 | -0.099 | 0.668 | -0.192 | 0.728 | -0.008 | 0.104 | 0.104 | 0 | 0.51 | 0.3 | 0.54 | 0.77 | 0.51 | 0.58 |
| 1 | 190934017 | rs1431141 | -0.741 | 0.759 | -0.199 | 0.706 | -0.022 | 0.04 | 0.04 | 0 | 0.83 | 0.94 | 0.81 | 0.95 | 0.82 | 0.87 |
| 1 | 190934383 | rs1431142 | -0.027 | 0.687 | 0.117 | 0.802 | -0.001 | 0 | 0 | 0 | 0.48 | 0.5 | 0.37 | 0.52 | 0.41 | 0.5 |
| 1 | 190940126 | rs7546929 | 0.12 | 0.677 | 0.043 | 0.792 | 0.016 | -0.002 | -0.002 | 0.249 | 0.47 | 0.37 | 0.36 | 0.52 | 0.41 | 0.5 |
| 1 | 190946081 | rs1469568 | 0.363 | 0.679 | 0.005 | 0.818 | 0.003 | -0.014 | -0.014 | 0 | 0.19 | 0.15 | 0.16 | 0.32 | 0.19 | 0.35 |
| 1 | 190952999 | rs11802112 | 1.341 | 0.551 | -0.397 | 0.794 | -0.026 | -0.03 | -0.03 | 0 | 0.06 | 0.01 | 0.19 | 0.25 | 0.12 | 0.08 |
| 1 | 190953471 | rs2025509 | 1.784 | 0.542 | -0.349 | 0.621 | 0.002 | 0.004 | 0.004 | 0.02 | 0.26 | 0.18 | 0.2 | 0.2 | 0.22 | 0.14 |
| 1 | 190965934 | rs12145021 | 0.15 | 0.445 | -0.104 | 0.673 | -0.01 | 0.046 | 0.046 | 0.695 | 0.21 | 0.39 | 0.17 | 0.32 | 0.2 | 0.36 |
| 1 | 190976940 | rs7534290 | -0.615 | 0.536 | -0.398 | 0.483 | -0.025 | 0.094 | 0.094 | 0 | 0.58 | 0.73 | 0.58 | 0.82 | 0.54 | 0.63 |
| 1 | 190977609 | rs6683411 | -0.011 | 0.464 | -0.385 | 0.526 | 0.032 | 0.008 | 0.008 | 0 | 0.5 | 0.66 | 0.38 | 0.52 | 0.42 | 0.51 |
| 1 | 190988857 | rs10737592 | -0.509 | 0.579 | -0.448 | 0.4 | -0.011 | 0.031 | 0.031 | 0 | 0.82 | 0.89 | 0.81 | 0.99 | 0.81 | 0.86 |
| 1 | 190989756 | rs1431144 | -0.621 | 0.582 | -0.464 | 0.402 | -0.03 | 0.074 | 0.074 | 0 | 0.83 | 0.95 | 0.82 | 0.99 | 0.81 | 0.86 |
| 1 | 190990146 | rs10801004 | 0.193 | 0.538 | -0.391 | 0.479 | 0.018 | 0.019 | 0.019 | 0 | 0.5 | 0.58 | 0.41 | 0.55 | 0.42 | 0.54 |
| 1 | 191016101 | rs1886414 | 1.459 | 0.471 | -0.439 | 0.509 | 0.009 | -0.004 | -0.004 | 0.088 | 0.29 | 0.2 | 0.23 | 0.23 | 0.22 | 0.18 |
| 1 | 191016342 | rs10920877 | 0.114 | 0.45 | -0.341 | 0.533 | -0.019 | 0.041 | 0.041 | 1.127 | 0.22 | 0.52 | 0.18 | 0.33 | 0.19 | 0.36 |
| 1 | 191025899 | rs12047686 | 0.011 | 0.512 | -0.48 | 0.557 | -0.011 | -0.018 | -0.018 | 0.47 | 0.1 | 0.06 | 0.19 | 0.29 | 0.14 | 0.14 |
| 1 | 191040861 | rs6685319 | 1.768 | 0.503 | -0.495 | 0.432 | -0.005 | 0.019 | 0.019 | 0 | 0.25 | 0.2 | 0.23 | 0.16 | 0.28 | 0.23 |
| 1 | 191062388 | rs7521746 | -0.589 | 0.45 | -0.599 | 0.403 | -0.014 | 0.082 | 0.082 | 0.316 | 0.57 | 0.82 | 0.6 | 0.79 | 0.59 | 0.74 |
| 1 | 191071885 | rs9427751 | 0.261 | 0.295 | -0.473 | 0.489 | -0.048 | 0.086 | 0.086 | 0 | 0.16 | 0.02 | 0.18 | 0.01 | 0.19 | 0.13 |
| 1 | 191076486 | rs1080901 | 0.742 | 0.231 | -0.317 | 0.548 | -0.001 | 0.053 | 0.053 | 0.038 | 0.7 | 0.52 | 0.64 | 0.37 | 0.68 | 0.49 |
| 1 | 191093816 | rs1454353 | 0.58 | -0.015 | -0.146 | 0.594 | 0 | -0.001 | -0.001 | 0.735 | 0.64 | 0.65 | 0.64 | 0.65 | 0.63 | 0.51 |
| 1 | 191106183 | rs6428044 | 1.616 | -0.286 | -0.128 | 0.524 | 0.005 | 0.045 | 0.045 | 0 | 0.5 | 0.34 | 0.5 | 0.35 | 0.5 | 0.36 |
| 1 | 191109973 | rs12067987 | 1.52 | -0.252 | -0.187 | 0.542 | 0.002 | 0.028 | 0.028 | 0 | 0.51 | 0.41 | 0.51 | 0.35 | 0.5 | 0.36 |
| 1 | 191145367 | rs1580087 | 0.972 | -0.227 | -0.389 | 0.61 | -0.006 | -0.002 | -0.002 | 2.566 | 0.56 | 0.63 | 0.54 | 0.4 | 0.49 | 0.34 |
| 1 | 191146188 | rs10737595 | 0.048 | -0.224 | -0.458 | 0.621 | 0.013 | 0.01 | 0.01 | 0 | 0.75 | 0.69 | 0.71 | 0.66 | 0.7 | 0.47 |
| 1 | 191150709 | rs10920908 | -0.271 | -0.148 | -0.595 | 0.786 | -0.03 | -0.071 | -0.071 | 0 | 0.19 | 0.04 | 0.17 | 0.26 | 0.21 | 0.14 |
| 1 | 191151059 | rs11801275 | 0.015 | 0.038 | 0.038 | 0.623 | -0.006 | 0.016 | 0.016 | 1.002 | 0.05 | 0.17 | 0.02 | 0.02 | 0.01 | 0.05 |
| 1 | 191154092 | rs12143616 | -0.026 | -0.191 | -0.382 | 0.522 | -0.016 | 0.002 | 0.002 | 0 | 0.42 | 0.25 | 0.51 | 0.38 | 0.48 | 0.39 |
| 1 | 191158206 | rs2045121 | -0.4 | -0.311 | 0.046 | 0.407 | 0 | 0.002 | 0.002 | 0 | 0.48 | 0.51 | 0.54 | 0.38 | 0.48 | 0.4 |
| 1 | 191161284 | rs16833288 | 0.791 | -0.339 | -0.13 | 0.603 | -0.024 | -0.057 | -0.057 | 0 | 0.12 | 0.04 | 0.16 | 0.36 | 0.19 | 0.22 |
| 1 | 191164972 | rs12121657 | 1.213 | -0.227 | -0.273 | 0.602 | -0.041 | 0.031 | 0.031 | 0 | 0.29 | 0.06 | 0.33 | 0.18 | 0.31 | 0.22 |
| 1 | 191167624 | rs7554791 | 0.675 | -0.193 | -0.143 | 0.619 | 0.003 | 0.008 | 0.008 | 0.018 | 0.33 | 0.31 | 0.36 | 0.19 | 0.31 | 0.22 |
| 1 | 191173607 | rs7517109 | -0.192 | -0.182 | -0.153 | 0.579 | -0.003 | -0.003 | -0.003 | 0 | 0.13 | 0.11 | 0.16 | 0.19 | 0.16 | 0.16 |
| 1 | 191180003 | rs2045119 | -1.229 | -0.193 | -0.034 | 0.561 | -0.001 | -0.009 | -0.009 | 1.398 | 0.6 | 0.56 | 0.69 | 0.73 | 0.67 | 0.61 |
| 1 | 191195800 | rs7418163 | -0.992 | -0.046 | -0.097 | 0.792 | -0.003 | -0.005 | -0.005 | 0.096 | 0.26 | 0.23 | 0.33 | 0.52 | 0.36 | 0.38 |
| 1 | 191199033 | rs7515438 | -0.356 | -0.023 | -0.096 | 0.822 | 0.013 | 0.018 | 0.018 | 0 | 0.13 | 0.17 | 0.18 | 0.19 | 0.17 | 0.16 |
| 1 | 191223734 | rs12120479 | 0.685 | -0.085 | -0.141 | 0.759 | -0.009 | -0.018 | -0.018 | 0.051 | 0.36 | 0.39 | 0.38 | 0.21 | 0.32 | 0.23 |
| 1 | 191231120 | rs1629378 | 0.816 | -0.054 | -0.161 | 0.681 | -0.001 | -0.012 | -0.012 | 1.674 | 0.39 | 0.44 | 0.3 | 0.27 | 0.32 | 0.38 |
| 1 | 191243984 | rs1246839 | 0.864 | -0.04 | -0.176 | 0.688 | -0.001 | -0.009 | -0.009 | 2.657 | 0.38 | 0.44 | 0.3 | 0.27 | 0.32 | 0.38 |
| 1 | 191271830 | rs3128539 | -0.517 | 0.089 | -0.353 | 0.714 | 0 | 0.002 | 0.002 | 0 | 0.9 | 0.91 | 0.86 | 0.81 | 0.86 | 0.84 |
| 1 | 191277082 | rs1931204 | -0.517 | 0.089 | -0.355 | 0.714 | -0.003 | -0.002 | -0.002 | 0 | 0.9 | 0.92 | 0.86 | 0.81 | 0.86 | 0.84 |
| 1 | 191316583 | rs12029728 | 1.184 | 0.01 | -0.492 | 0.533 | -0.036 | 0.015 | 0.015 | 0 | 0.31 | 0.06 | 0.34 | 0.2 | 0.31 | 0.23 |
| 1 | 191321186 | rs10159191 | 0.428 | 0.082 | -0.104 | 0.57 | -0.005 | 0.051 | 0.051 | 0 | 0.1 | 0.15 | 0.05 | 0.26 | 0.09 | 0.26 |
| 1 | 191340943 | rs6690512 | 1.102 | 0.178 | -0.075 | 0.404 | 0.025 | 0.107 | 0.107 | 0.042 | 0.05 | 0.23 | 0.04 | 0.06 | 0.01 | 0.07 |
| 1 | 191349614 | rs1246726 | 0.178 | 0.127 | -0.042 | 0.337 | -0.001 | 0.002 | 0.002 | 0.157 | 0.49 | 0.46 | 0.43 | 0.47 | 0.45 | 0.57 |
| 1 | 191358099 | rs1923812 | 1.766 | 0.365 | -0.217 | 0.259 | -0.009 | 0.054 | 0.054 | 0 | 0.23 | 0.02 | 0.29 | 0.13 | 0.21 | 0.12 |
| 1 | 191362050 | rs10920932 | -0.153 | 0.199 | 0.082 | 0.549 | 0.005 | 0.086 | 0.086 | 0.034 | 0.19 | 0.28 | 0.21 | 0.39 | 0.22 | 0.23 |
| 1 | 191362545 | rs12145952 | 1.347 | 0.218 | -0.096 | 0.551 | -0.061 | 0.107 | 0.107 | 0 | 0.3 | 0.04 | 0.34 | 0.14 | 0.32 | 0.2 |
| 1 | 191377664 | rs10920936 | 0.563 | 0.288 | 0.009 | 0.614 | 0.004 | -0.002 | -0.002 | 0 | 0.42 | 0.35 | 0.28 | 0.45 | 0.34 | 0.48 |
| 1 | 191388293 | rs598834 | -0.472 | 0.479 | -0.859 | 0.362 | -0.006 | 0.12 | 0.12 | 0 | 0.59 | 0.76 | 0.52 | 0.84 | 0.58 | 0.71 |
| 1 | 191391663 | rs9651063 | 0.37 | 0.461 | -0.8 | 0.419 | 0.005 | -0.004 | -0.004 | 0 | 0.45 | 0.59 | 0.32 | 0.5 | 0.35 | 0.55 |

| | | | | | | | | | | | | | | | | |
|---|---|---|---|---|---|---|---|---|---|---|---|---|---|---|---|---|
| 1 | 191395497 | rs535553 | -0.583 | 0.584 | -0.956 | 0.453 | 0.002 | -0.016 | -0.016 | 1.642 | 0.91 | 0.9 | 0.88 | 0.99 | 0.89 | 0.91 |
| 1 | 191403411 | rs2135943 | 0.492 | 0.586 | -0.961 | 0.49 | 0.007 | 0.015 | 0.015 | 0 | 0.76 | 0.73 | 0.69 | 0.64 | 0.67 | 0.75 |
| 1 | 191406367 | rs10801043 | 0.161 | 0.521 | -0.997 | 0.706 | -0.016 | -0.002 | -0.002 | 0 | 0.45 | 0.26 | 0.56 | 0.48 | 0.53 | 0.35 |
| 1 | 191424963 | rs7548615 | -0.241 | 0.247 | -0.912 | 0.605 | -0.003 | -0.034 | -0.034 | 0 | 0.1 | 0.03 | 0.05 | 0.12 | 0.08 | 0.1 |
| 1 | 191428481 | rs535509 | 0.019 | 0.008 | -0.074 | 0.295 | 0.008 | -0.013 | -0.013 | 0 | 0.11 | 0.46 | 0.06 | 0.04 | 0.06 | 0.16 |
| 1 | 191429227 | rs650957 | -0.217 | 0.149 | -0.642 | 0.318 | 0.022 | 0.058 | 0.058 | 0 | 0.51 | 0.67 | 0.55 | 0.31 | 0.44 | 0.48 |
| 1 | 191430223 | rs11800005 | -0.319 | 0.185 | -0.538 | 0.705 | -0.006 | 0.01 | 0.01 | 0 | 0.1 | 0.17 | 0.04 | 0.09 | 0.06 | 0.05 |
| 1 | 191431169 | rs10920948 | 0.096 | 0.307 | -0.909 | 0.482 | -0.014 | 0.031 | 0.031 | 0 | 0.32 | 0.11 | 0.32 | 0.24 | 0.33 | 0.21 |
| 1 | 191433463 | rs613836 | -0.937 | 0.5 | 0.181 | 0.916 | 0.128 | 0.277 | 0.277 | 0.007 | 0.06 | 0.47 | 0.22 | 0.16 | 0.05 | 0.23 |
| 1 | 191437899 | rs583439 | -0.331 | 0.153 | -0.787 | 0.642 | 0.012 | -0.011 | -0.011 | 0 | 0.73 | 0.89 | 0.82 | 0.75 | 0.74 | 0.84 |
| 1 | 191438591 | rs17398140 | 0.684 | 0.836 | -1.305 | 0.073 | -0.062 | 0.096 | 0.096 | 0 | 0.28 | 0.11 | 0.31 | 0.09 | 0.35 | 0.17 |
| 1 | 191446589 | rs623950 | -0.02 | 1.016 | -0.618 | -0.043 | 0.055 | -0.025 | -0.025 | 0 | 0.05 | 0.19 | 0.07 | 0 | 0.09 | 0.07 |
| 1 | 191449983 | rs669885 | -0.965 | 0.384 | -0.748 | -0.32 | 0.062 | 0.05 | 0.05 | 0 | 0.6 | 0.82 | 0.79 | 0.72 | 0.67 | 0.69 |
| 1 | 191453259 | rs12071696 | 0.009 | 0.269 | -0.539 | -0.163 | -0.003 | 0.016 | 0.016 | 0 | 0.4 | 0.35 | 0.38 | 0.18 | 0.4 | 0.35 |
| 1 | 191459828 | rs10920956 | 0.344 | 0.372 | -0.507 | -0.132 | -0.005 | 0.012 | 0.012 | 0 | 0.36 | 0.3 | 0.37 | 0.16 | 0.39 | 0.3 |
| 1 | 191467283 | rs10920959 | 0.849 | 0.591 | -0.952 | 0.291 | -0.022 | 0.013 | 0.013 | 0 | 0.08 | 0.02 | 0.12 | 0.46 | 0.14 | 0.19 |
| 1 | 191472500 | rs639721 | -0.484 | 0.358 | -0.967 | -0.076 | 0.011 | 0.115 | 0.115 | 0.001 | 0.47 | 0.91 | 0.67 | 0.27 | 0.52 | 0.48 |
| 1 | 191481432 | rs648348 | -0.395 | 0.291 | -1.018 | 0.039 | -0.043 | 0.044 | 0.044 | 0 | 0.45 | 0.91 | 0.66 | 0.31 | 0.5 | 0.46 |
| 1 | 191491194 | rs1338029 | 0.999 | 0.323 | -0.956 | -0.055 | -0.062 | 0.107 | 0.107 | 0 | 0.28 | 0.08 | 0.29 | 0.09 | 0.34 | 0.16 |
| 1 | 191491491 | rs591536 | 0.188 | 0.304 | -0.017 | -0.034 | 0.007 | -0.013 | -0.013 | 0.525 | 0.36 | 0.38 | 0.37 | 0.22 | 0.38 | 0.3 |
| 1 | 191502099 | rs1892294 | 0.855 | 0.164 | -0.081 | -0.112 | -0.057 | 0.072 | 0.072 | 0 | 0.29 | 0.07 | 0.3 | 0.13 | 0.35 | 0.17 |
| 1 | 191511510 | rs661322 | -0.676 | 0.152 | -0.1 | -0.101 | -0.019 | 0.125 | 0.125 | 0 | 0.47 | 0.85 | 0.61 | 0.23 | 0.49 | 0.37 |
| 1 | 191517119 | rs682333 | -1.65 | -0.042 | 0.246 | -0.086 | 0.048 | 0.017 | 0.017 | 1.15 | 0.12 | 0.59 | 0.29 | 0.1 | 0.12 | 0.19 |
| 1 | 191525674 | rs652772 | -1.308 | -0.049 | 0.461 | -0.094 | 0.097 | 0.046 | 0.046 | 0.655 | 0.1 | 0.53 | 0.27 | 0.09 | 0.12 | 0.16 |
| 1 | 191542315 | rs2065987 | 0.567 | 0.246 | 0.169 | -0.12 | -0.016 | 0.08 | 0.08 | 0 | 0.32 | 0.13 | 0.3 | 0.13 | 0.33 | 0.17 |
| 1 | 191546478 | rs2095726 | 1.105 | 0.368 | 0.032 | 0.369 | 0.021 | 0.091 | 0.091 | 0 | 0.42 | 0.1 | 0.26 | 0.24 | 0.38 | 0.36 |
| 1 | 191555290 | rs10920970 | 0.871 | 0.412 | 0.046 | 0.563 | -0.023 | -0.004 | -0.004 | 0.012 | 0.08 | 0.02 | 0.12 | 0.43 | 0.13 | 0.17 |
| 1 | 191561185 | rs12144096 | 1.107 | 0.374 | 0.039 | 0.407 | 0.02 | 0.094 | 0.094 | 0 | 0.42 | 0.1 | 0.26 | 0.24 | 0.38 | 0.37 |
| 1 | 191564468 | rs6657223 | 0.583 | 0.351 | 0.359 | 0.382 | 0 | 0.004 | 0.004 | 0 | 0.48 | 0.55 | 0.49 | 0.34 | 0.45 | 0.52 |
| 1 | 191571287 | rs12403168 | -0.923 | 0.347 | 0.365 | 0.4 | 0.003 | 0.043 | 0.043 | 0 | 0.17 | 0.21 | 0.19 | 0.53 | 0.22 | 0.31 |
| 1 | 191577392 | rs2050664 | 1.059 | 0.356 | 0.312 | 0.308 | 0.02 | 0.082 | 0.082 | 0 | 0.42 | 0.12 | 0.26 | 0.24 | 0.38 | 0.37 |
| 1 | 191590362 | rs7547370 | 0.453 | 0.355 | 0.353 | 0.21 | 0.006 | 0.049 | 0.049 | 0 | 0.48 | 0.68 | 0.49 | 0.34 | 0.45 | 0.52 |
| 1 | 191599645 | rs1856294 | 1.031 | 0.483 | 0.015 | 0.031 | 0.025 | 0.039 | 0.039 | 0 | 0.4 | 0.15 | 0.36 | 0.28 | 0.34 | 0.35 |
| 1 | 191606863 | rs2814611 | -1.994 | 0.995 | -0.136 | -0.382 | -0.009 | 0.006 | 0.006 | 0 | 0.85 | 0.88 | 0.83 | 1 | 0.82 | 0.92 |
| 1 | 191610760 | rs2814580 | -0.372 | 0.938 | -0.106 | -0.45 | 0.009 | -0.07 | -0.07 | 0 | 0.76 | 0.37 | 0.67 | 0.93 | 0.75 | 0.77 |
| 1 | 191615655 | rs945672 | -2.012 | 0.946 | -0.096 | -0.36 | -0.011 | 0.013 | 0.013 | 0 | 0.85 | 0.9 | 0.84 | 1 | 0.82 | 0.92 |
| 1 | 191618129 | rs12047092 | -0.313 | 0.967 | 0.067 | -0.177 | 0.193 | 0.158 | 0.158 | 0 | 0.05 | 0.5 | 0.17 | 0.07 | 0.07 | 0.15 |
| 1 | 191619165 | rs7551667 | 0.468 | 0.806 | -0.263 | -0.17 | 0.006 | 0.011 | 0.011 | 0 | 0.17 | 0.12 | 0.11 | 0.11 | 0.18 | 0.14 |
| 1 | 191619550 | rs10753990 | -0.468 | 0.826 | -0.307 | 0.025 | -0.012 | -0.039 | -0.039 | 0 | 0.31 | 0.18 | 0.27 | 0.55 | 0.37 | 0.33 |
| 1 | 191624744 | rs11582040 | -0.188 | 0.766 | -0.113 | 0.202 | 0.002 | 0.09 | 0.09 | 0 | 0.06 | 0.15 | 0.04 | 0.11 | 0.03 | 0.07 |
| 1 | 191627004 | rs1338035 | -0.034 | 0.926 | -0.061 | 0.021 | 0.085 | 0.016 | 0.016 | 0.105 | 0.1 | 0.38 | 0.21 | 0.06 | 0.15 | 0.16 |
| 1 | 191627282 | rs1338034 | -0.459 | 0.887 | -0.486 | -0.077 | -0.044 | -0.036 | -0.036 | 0.521 | 0.72 | 0.85 | 0.69 | 0.56 | 0.63 | 0.71 |
| 1 | 191632198 | rs1336864 | 0.599 | 0.676 | -0.383 | -0.204 | 0.006 | -0.013 | -0.013 | 0 | 0.57 | 0.46 | 0.54 | 0.72 | 0.5 | 0.61 |
| 1 | 191633346 | rs12036028 | 1.165 | 0.73 | -0.839 | -0.012 | -0.03 | -0.041 | -0.041 | 0 | 0.12 | 0.04 | 0.14 | 0.48 | 0.16 | 0.19 |
| 1 | 191640891 | rs12564198 | 0.61 | 0.618 | -0.861 | 0.071 | 0.009 | 0.001 | 0.001 | 0 | 0.13 | 0.05 | 0.06 | 0.13 | 0.13 | 0.14 |
| 1 | 191641336 | rs2785776 | 2.195 | 0.582 | -1.035 | -0.178 | 0.039 | 0.079 | 0.079 | 0 | 0.35 | 0.08 | 0.34 | 0.21 | 0.28 | 0.34 |
| 1 | 191641732 | rs4142414 | 1.328 | 0.698 | -1.046 | -0.053 | -0.039 | -0.058 | -0.058 | 0.118 | 0.12 | 0.03 | 0.14 | 0.48 | 0.16 | 0.2 |
| 1 | 191642616 | rs10801053 | 1.311 | 0.7 | -1.045 | -0.052 | -0.039 | -0.059 | -0.059 | 0 | 0.12 | 0.03 | 0.14 | 0.48 | 0.16 | 0.2 |
| 1 | 191658150 | rs12080045 | 0.813 | 0.673 | -0.828 | -0.133 | 0.017 | -0.02 | -0.02 | 0 | 0.06 | 0.17 | 0.05 | 0.02 | 0.06 | 0.04 |
| 1 | 191670376 | rs1889994 | -1.11 | 0.942 | -0.555 | -0.23 | 0.017 | 0.001 | 0.001 | 0 | 0.31 | 0.37 | 0.42 | 0.66 | 0.4 | 0.48 |
| 1 | 191673544 | rs1509703 | 2.392 | 0.768 | -0.648 | -0.337 | 0.027 | 0.074 | 0.074 | 0 | 0.35 | 0.06 | 0.33 | 0.21 | 0.28 | 0.33 |
| 1 | 191677851 | rs4575057 | 1.252 | 0.96 | -0.567 | -0.198 | -0.026 | -0.041 | -0.041 | 0 | 0.13 | 0.04 | 0.14 | 0.48 | 0.16 | 0.2 |
| 1 | 191689375 | rs10494658 | 0.467 | 1.096 | -0.516 | -0.232 | -0.003 | -0.001 | -0.001 | 0 | 0.11 | 0.11 | 0.23 | 0.15 | 0.18 | 0.15 |
| 1 | 191689401 | rs12059896 | 0.728 | 1.056 | -0.482 | -0.265 | 0.023 | -0.018 | -0.018 | 0 | 0.06 | 0.2 | 0.05 | 0.02 | 0.06 | 0.04 |
| 1 | 191693460 | rs833959 | -0.601 | 1.028 | -0.367 | -0.328 | 0.006 | -0.098 | -0.098 | 0 | 0.76 | 0.64 | 0.79 | 0.98 | 0.75 | 0.89 |
| 1 | 191703596 | rs833976 | 1.268 | 0.912 | -0.459 | -0.288 | -0.004 | -0.006 | -0.006 | 1.046 | 0.51 | 0.49 | 0.42 | 0.34 | 0.41 | 0.47 |
| 1 | 191719427 | rs2785761 | 0.413 | 0.934 | -0.412 | -0.337 | 0.018 | -0.031 | -0.031 | 0 | 0.64 | 0.52 | 0.56 | 0.83 | 0.57 | 0.73 |
| 1 | 191729205 | rs2814584 | -2.011 | 0.994 | -0.711 | -0.4 | -0.005 | -0.015 | -0.015 | 0.016 | 0.86 | 0.88 | 0.85 | 1 | 0.81 | 0.93 |
| 1 | 191755912 | rs833986 | 0.896 | 0.778 | -0.726 | -0.294 | 0 | -0.002 | -0.002 | 0 | 0.55 | 0.5 | 0.51 | 0.7 | 0.45 | 0.55 |
| 1 | 191758108 | rs7546487 | 3.006 | -0.241 | -1.39 | -0.746 | 0.041 | 0.036 | 0.036 | 0 | 0.14 | 0.02 | 0.03 | 0.08 | 0.06 | 0.25 |
| 1 | 191759743 | rs833982 | -2.898 | -0.209 | -1.419 | -0.829 | -0.014 | 0.003 | 0.003 | 0 | 0.86 | 0.9 | 0.86 | 1 | 0.81 | 0.9 |
| 1 | 191776078 | rs10494663 | 0.412 | -0.316 | -1.384 | -0.81 | 0.026 | 0.169 | 0.169 | 1.447 | 0.19 | 0.42 | 0.17 | 0.42 | 0.17 | 0.21 |
| 1 | 191778671 | rs2000054 | -2.188 | -0.282 | -1.71 | -0.8 | 0.044 | 0.056 | 0.056 | 0 | 0.64 | 0.93 | 0.66 | 0.73 | 0.73 | 0.66 |
| 1 | 191784175 | rs16833826 | 1.78 | -0.29 | -1.696 | -0.791 | -0.039 | -0.059 | -0.059 | 0.27 | 0.12 | 0.03 | 0.14 | 0.41 | 0.17 | 0.16 |
| 1 | 191788524 | rs17402518 | 3.05 | -0.258 | -1.656 | -0.793 | 0.041 | 0.036 | 0.036 | 0 | 0.14 | 0.02 | 0.03 | 0.09 | 0.06 | 0.25 |
| 1 | 191792216 | rs7528316 | -2.071 | -0.234 | -1.638 | -0.78 | 0.046 | 0.058 | 0.058 | 0 | 0.63 | 0.93 | 0.66 | 0.73 | 0.73 | 0.66 |
| 1 | 191798964 | rs12023209 | 1.922 | -0.119 | -1.562 | -0.67 | -0.002 | -0.02 | -0.02 | 0 | 0.07 | 0.01 | 0.13 | 0.07 | 0.07 | 0.11 |
| 1 | 191804265 | rs6696942 | 2.647 | -0.042 | -1.532 | -0.73 | -0.011 | -0.003 | -0.003 | 0.005 | 0.14 | 0.11 | 0.14 | 0 | 0.19 | 0.1 |
| 1 | 191809160 | rs7556435 | 2.142 | -0.233 | -1.664 | -0.816 | 0.049 | 0.063 | 0.063 | 0 | 0.18 | 0.03 | 0.04 | 0.09 | 0.1 | 0.32 |

| | | | | | | | | | | | | | | | | |
|---|---|---|---|---|---|---|---|---|---|---|---|---|---|---|---|---|
| 1 | 191822724 | rs9427533 | 2.26 | 0.681 | -1.012 | -0.337 | -0.011 | -0.003 | -0.003 | 0 | 0.14 | 0.11 | 0.14 | 0 | 0.19 | 0.1 |
| 1 | 191833819 | rs3908964 | -0.285 | 0.153 | -0.762 | -0.049 | 0.003 | -0.001 | -0.001 | 0 | 0.13 | 0.22 | 0.19 | 0.11 | 0.15 | 0.16 |
| 1 | 191836128 | rs2369435 | -2.226 | 0.078 | -0.852 | -0.065 | 0.058 | 0.054 | 0.054 | 0 | 0.64 | 0.92 | 0.66 | 0.73 | 0.74 | 0.65 |
| 1 | 191836316 | rs2211195 | -0.615 | 0.113 | -0.771 | 0.014 | 0.029 | 0.096 | 0.096 | 1.358 | 0.23 | 0.61 | 0.22 | 0.47 | 0.25 | 0.24 |
| 1 | 191848287 | rs4845294 | 1.531 | 0.077 | -0.882 | 0.074 | -0.004 | -0.02 | -0.02 | 0 | 0.07 | 0.01 | 0.13 | 0.07 | 0.07 | 0.12 |
| 1 | 191855740 | rs9427802 | -1.383 | 0.204 | -0.777 | 0 | 0.092 | 0.069 | 0.069 | 0 | 0.54 | 0.85 | 0.52 | 0.65 | 0.66 | 0.55 |
| 1 | 191859736 | rs9427545 | -0.482 | 0.458 | -0.864 | -0.022 | -0.012 | 0.012 | 0.012 | 0.209 | 0.39 | 0.41 | 0.35 | 0.25 | 0.49 | 0.4 |
| 1 | 191865100 | rs12081603 | 0.863 | 0.622 | -1.061 | 0.262 | 0.016 | -0.003 | -0.003 | 0 | 0.07 | 0.18 | 0.06 | 0.05 | 0.08 | 0.12 |
| 1 | 191865391 | rs1320604 | -1.241 | 0.366 | -1.653 | 0.25 | 0.034 | 0.04 | 0.04 | 0 | 0.54 | 0.95 | 0.49 | 0.63 | 0.65 | 0.55 |
| 1 | 191868116 | rs1340202 | -1.285 | 0.329 | -1.264 | 0.288 | 0.053 | 0.029 | 0.029 | 0 | 0.6 | 0.88 | 0.62 | 0.64 | 0.72 | 0.56 |
| 1 | 191868313 | rs9427808 | -0.326 | 1.093 | -0.554 | 0.467 | 0.042 | -0.033 | -0.033 | 0 | 0.29 | 0.53 | 0.28 | 0.2 | 0.41 | 0.28 |
| 1 | 191870504 | rs9427809 | -0.496 | 0.912 | -0.82 | 1.299 | 0.004 | 0.053 | 0.053 | 0 | 0.09 | 0.14 | 0.12 | 0.3 | 0.08 | 0.15 |
| 1 | 191872334 | rs2369527 | -0.288 | 0.815 | -0.372 | 1.288 | 0.052 | 0.094 | 0.094 | 0 | 0.11 | 0.41 | 0.12 | 0.14 | 0.08 | 0.19 |
| 1 | 191875752 | rs9427811 | -0.455 | 0.882 | -0.334 | 1.3 | 0.013 | 0.101 | 0.101 | 0 | 0.09 | 0.2 | 0.1 | 0.3 | 0.07 | 0.16 |
| 1 | 191887735 | rs10921043 | -0.417 | 0.658 | -1.464 | 0.67 | -0.036 | 0.017 | 0.017 | 0 | 0.6 | 0.75 | 0.41 | 0.64 | 0.47 | 0.75 |
| 1 | 191913535 | rs10801082 | -0.573 | 0.598 | -1.677 | 0.374 | -0.026 | 0.012 | 0.012 | 2.689 | 0.67 | 0.74 | 0.45 | 0.73 | 0.51 | 0.73 |
| 1 | 191916122 | rs12405594 | -0.586 | 0.591 | -1.715 | 0.371 | -0.032 | 0.015 | 0.015 | 2.619 | 0.67 | 0.75 | 0.45 | 0.73 | 0.51 | 0.73 |
| 1 | 191919402 | rs10801083 | -0.591 | 0.593 | -1.714 | 0.372 | -0.032 | 0.014 | 0.014 | 2.647 | 0.68 | 0.75 | 0.45 | 0.73 | 0.51 | 0.73 |
| 1 | 191923076 | rs7530235 | 1.26 | 0.352 | -1.878 | 0.338 | 0.054 | 0.097 | 0.097 | 0.168 | 0.42 | 0.15 | 0.2 | 0.22 | 0.32 | 0.41 |
| 1 | 191932047 | rs12064769 | -0.432 | 0.428 | -1.888 | 0.367 | -0.031 | 0.017 | 0.017 | 1.668 | 0.67 | 0.76 | 0.47 | 0.73 | 0.5 | 0.74 |
| 1 | 191932822 | rs12144141 | 0.525 | 0.541 | -2.07 | 0.399 | -0.002 | 0.02 | 0.02 | 0.426 | 0.5 | 0.18 | 0.26 | 0.56 | 0.41 | 0.54 |
| 1 | 191934144 | rs946962 | -0.246 | 0.621 | -1.843 | 0.3 | -0.024 | 0.04 | 0.04 | 0.017 | 0.38 | 0.25 | 0.59 | 0.27 | 0.51 | 0.27 |
| 1 | 191941669 | rs9427554 | 0.463 | 0.625 | -1.93 | 0.297 | -0.084 | 0.01 | 0.01 | 0.67 | 0.31 | 0.07 | 0.55 | 0.27 | 0.48 | 0.26 |
| 1 | 191961497 | rs10429898 | 1.415 | 0.657 | -1.986 | 0.409 | -0.038 | -0.067 | -0.067 | 1.893 | 0.1 | 0.01 | 0.06 | 0.35 | 0.09 | 0.13 |
| 1 | 191962527 | rs16833949 | 0.784 | 0.66 | -1.968 | 0.409 | -0.021 | -0.053 | -0.053 | 0.011 | 0.12 | 0.05 | 0.08 | 0.35 | 0.11 | 0.13 |
| 1 | 191979285 | rs7416935 | 0.784 | 0.66 | -1.968 | 0.409 | -0.021 | -0.053 | -0.053 | 0 | 0.12 | 0.04 | 0.08 | 0.35 | 0.11 | 0.13 |
| 1 | 191983048 | rs12134799 | 1.22 | 0.553 | -2.068 | 0.387 | -0.005 | 0.101 | 0.101 | 0 | 0.41 | 0.07 | 0.2 | 0.17 | 0.33 | 0.43 |
| 1 | 191983858 | rs6428079 | 1.104 | 1.249 | -2.105 | 0.298 | -0.053 | -0.057 | -0.057 | 2.183 | 0.23 | 0.03 | 0.46 | 0.26 | 0.33 | 0.24 |
| 1 | 192005194 | rs6688496 | -0.61 | 1.041 | -1.627 | 0.49 | 0.011 | 0.078 | 0.078 | 0 | 0.89 | 0.4 | 0.85 | 0.79 | 0.93 | 0.81 |
| 1 | 192011987 | rs7553698 | -1.11 | 1.131 | -1.553 | 0.557 | 0.004 | 0.062 | 0.062 | 0 | 0.88 | 0.81 | 0.94 | 0.65 | 0.91 | 0.87 |
| 1 | 192012070 | rs10801086 | 0.507 | 1.061 | -1.665 | 0.226 | 0.012 | 0.019 | 0.019 | 0 | 0.51 | 0.6 | 0.36 | 0.37 | 0.41 | 0.62 |
| 1 | 192033140 | rs12567153 | 1.061 | 1.061 | -2.322 | 0.171 | -0.004 | 0.112 | 0.112 | 0 | 0.41 | 0.07 | 0.2 | 0.17 | 0.34 | 0.43 |
| 1 | 192047263 | rs3012194 | -0.559 | 1.185 | -1.966 | -0.139 | 0.007 | 0.057 | 0.057 | 0 | 0.88 | 0.39 | 0.83 | 0.8 | 0.9 | 0.82 |
| 1 | 192085298 | rs12569272 | 0.607 | 1.273 | -2.382 | -0.19 | 0.001 | -0.013 | -0.013 | 0 | 0.72 | 0.3 | 0.74 | 0.42 | 0.79 | 0.67 |
| 1 | 192088638 | rs12119436 | 0.051 | 0.916 | -2.815 | -0.545 | -0.034 | 0.059 | 0.059 | 0 | 0.4 | 0.23 | 0.6 | 0.25 | 0.54 | 0.28 |
| 1 | 192089215 | rs17521643 | 1.416 | 1.014 | -3.096 | -0.415 | -0.007 | 0.043 | 0.043 | 0 | 0.32 | 0.07 | 0.14 | 0.17 | 0.25 | 0.4 |
| 1 | 192091291 | rs12736318 | 1.938 | 1.337 | -3.332 | -0.575 | -0.055 | -0.07 | -0.07 | 1.134 | 0.15 | 0.01 | 0.4 | 0.21 | 0.24 | 0.12 |
| 1 | 192096142 | rs12024380 | 0.262 | 1.417 | -2.62 | -0.234 | -0.015 | -0.086 | -0.086 | 0.221 | 0.16 | 0.07 | 0.15 | 0.38 | 0.12 | 0.13 |
| 1 | 192096277 | rs12026956 | 0.172 | 1.417 | -2.338 | -0.496 | 0.004 | -0.019 | -0.019 | 2.496 | 0.48 | 0.14 | 0.3 | 0.55 | 0.36 | 0.53 |
| 1 | 192100668 | rs10801098 | 0.528 | 1.595 | -1.323 | -0.337 | 0.017 | -0.002 | -0.002 | 0 | 0.77 | 0.33 | 0.7 | 0.8 | 0.71 | 0.73 |
| 1 | 192108478 | rs11581394 | 0.528 | 1.165 | -1.595 | -0.892 | -0.006 | 0.066 | 0.066 | 0 | 0.43 | 0.17 | 0.55 | 0.72 | 0.48 | 0.38 |
| 1 | 192121369 | rs17587884 | 1.037 | 1.076 | -1.658 | -0.961 | 0.012 | 0.147 | 0.147 | 0 | 0.31 | 0.04 | 0.12 | 0.08 | 0.21 | 0.33 |
| 1 | 192123711 | rs7526348 | 0.442 | 1.096 | -1.638 | -0.938 | 0.005 | 0.069 | 0.069 | 0 | 0.46 | 0.26 | 0.58 | 0.72 | 0.49 | 0.4 |
| 1 | 192128183 | rs4454527 | -0.099 | 1.049 | -1.583 | -0.995 | 0.067 | 0.007 | 0.007 | 0.096 | 0.8 | 0.39 | 0.71 | 0.8 | 0.71 | 0.73 |
| 1 | 192136366 | rs16829396 | 1.328 | 1.203 | -2.024 | -0.759 | -0.047 | -0.076 | -0.076 | 0 | 0.12 | 0.01 | 0.13 | 0.47 | 0.14 | 0.1 |
| 1 | 192151174 | rs1937235 | 0.798 | 0.969 | -1.507 | -1.349 | 0.004 | 0.084 | 0.084 | 0 | 0.45 | 0.25 | 0.57 | 0.72 | 0.49 | 0.43 |
| 1 | 192152345 | rs10921107 | 1.122 | 0.986 | -1.547 | -1.335 | -0.013 | 0.073 | 0.073 | 0 | 0.43 | 0.15 | 0.54 | 0.72 | 0.47 | 0.38 |
| 1 | 192170402 | rs1856492 | 0.79 | 1.011 | -1.557 | -1.336 | 0.008 | 0.149 | 0.149 | 0 | 0.31 | 0.06 | 0.12 | 0.08 | 0.22 | 0.31 |
| 1 | 192170415 | rs1175174 | 0.835 | 1.027 | -1.353 | -1.179 | -0.018 | 0.01 | 0.01 | 0.132 | 0.74 | 0.21 | 0.67 | 0.8 | 0.69 | 0.73 |
| 1 | 192176651 | rs1768306 | -0.502 | 1.031 | -0.594 | -1.258 | 0.088 | -0.001 | -0.001 | 0 | 0.17 | 0.55 | 0.21 | 0.13 | 0.24 | 0.16 |
| 1 | 192182559 | rs2801415 | -1.331 | 0.902 | -0.749 | -1.419 | 0.003 | 0.076 | 0.076 | 0 | 0.54 | 0.7 | 0.38 | 0.21 | 0.5 | 0.5 |
| 1 | 192206032 | rs1175111 | -1.014 | 0.8 | -0.711 | -1.414 | 0.02 | 0.124 | 0.124 | 0 | 0.5 | 0.76 | 0.36 | 0.21 | 0.48 | 0.49 |
| 1 | 192211842 | rs946421 | -1.35 | 0.934 | -0.207 | -1.365 | -0.017 | 0.04 | 0.04 | 0 | 0.41 | 0.67 | 0.35 | 0.16 | 0.43 | 0.46 |
| 1 | 192219415 | rs1175142 | -0.466 | 1.12 | 0.621 | -1.411 | 0.158 | 0.017 | 0.017 | 0 | 0.16 | 0.48 | 0.2 | 0.13 | 0.24 | 0.16 |
| 1 | 192225979 | rs7517414 | 0.315 | 1.118 | 0.556 | -1.417 | 0.015 | 0.039 | 0.039 | 0.008 | 0.25 | 0.17 | 0.15 | 0.03 | 0.19 | 0.31 |
| 1 | 192230089 | rs1175154 | -1.604 | 1.102 | 0.549 | -1.422 | -0.017 | 0.086 | 0.086 | 0 | 0.45 | 0.75 | 0.38 | 0.16 | 0.45 | 0.47 |
| 1 | 192245030 | rs17523810 | 0.161 | 1.158 | 0.351 | -1.393 | 0.018 | 0.094 | 0.094 | 0 | 0.25 | 0.13 | 0.14 | 0.03 | 0.19 | 0.31 |
| 1 | 192249116 | rs1175119 | -0.883 | 1.173 | 0.382 | -1.389 | 0.105 | -0.009 | -0.009 | 0.032 | 0.18 | 0.55 | 0.2 | 0.13 | 0.24 | 0.16 |
| 1 | 192253353 | rs6704532 | -1.601 | 1.151 | 0.365 | -1.381 | -0.01 | 0.048 | 0.048 | 0 | 0.45 | 0.7 | 0.37 | 0.16 | 0.45 | 0.49 |
| 1 | 192275796 | rs10921130 | 0.291 | 1.277 | 0.333 | -1.278 | 0.016 | 0.056 | 0.056 | 0 | 0.24 | 0.16 | 0.14 | 0.03 | 0.19 | 0.28 |
| 1 | 192281152 | rs6697334 | 0.335 | 1.276 | 0.322 | -1.278 | 0.011 | 0.111 | 0.111 | 0 | 0.23 | 0.07 | 0.13 | 0.03 | 0.19 | 0.28 |
| 1 | 192284703 | rs10921131 | 0.515 | 1.286 | 0.327 | -1.293 | 0.018 | 0.046 | 0.046 | 0 | 0.22 | 0.15 | 0.14 | 0.03 | 0.17 | 0.28 |
| 1 | 192294011 | rs17413507 | 0.286 | 1.29 | 0.319 | -1.292 | 0.016 | 0.067 | 0.067 | 0 | 0.23 | 0.14 | 0.14 | 0.03 | 0.19 | 0.28 |
| 1 | 192296038 | rs6663865 | -1.769 | 1.317 | 0.42 | -1.264 | -0.012 | 0.046 | 0.046 | 0 | 0.44 | 0.7 | 0.36 | 0.17 | 0.45 | 0.49 |
| 1 | 192313535 | rs11805712 | 0.009 | 1.77 | 0.106 | -1.057 | 0.007 | -0.013 | -0.013 | 0.018 | 0.07 | 0.06 | 0.09 | 0.01 | 0.13 | 0.03 |
| 1 | 192314681 | rs7514892 | -0.3 | 1.363 | -0.094 | -0.981 | 0.163 | 0.013 | 0.013 | 0 | 0.15 | 0.48 | 0.19 | 0.13 | 0.23 | 0.18 |
| 1 | 192323753 | rs1936671 | 0.304 | 1.227 | -0.428 | -1.087 | 0.025 | 0.105 | 0.105 | 0 | 0.27 | 0.13 | 0.17 | 0.03 | 0.25 | 0.29 |
| 1 | 192324175 | rs10494671 | -0.596 | 1.361 | -0.424 | -0.978 | 0.084 | -0.033 | -0.033 | 0.257 | 0.2 | 0.53 | 0.22 | 0.12 | 0.25 | 0.19 |
| 1 | 192335792 | rs7528947 | -0.531 | 1.036 | -0.585 | -0.468 | -0.001 | 0.049 | 0.049 | 0 | 0.47 | 0.66 | 0.39 | 0.16 | 0.52 | 0.49 |

| | | | | | | | | | | | | | | | | |
|---|---|---|---|---|---|---|---|---|---|---|---|---|---|---|---|---|
| 1 | 192352776 | rs1936712 | -0.539 | 1.352 | -0.189 | 0.586 | 0.029 | 0.009 | 0.009 | 0 | 0.21 | 0.29 | 0.29 | 0.21 | 0.24 | 0.13 |
| 1 | 192356989 | rs1936709 | -0.902 | 1.455 | 0.181 | 1.332 | 0.088 | 0.101 | 0.101 | 0 | 0.09 | 0.26 | 0.15 | 0.22 | 0.16 | 0.06 |
| 1 | 192358256 | rs12753442 | 0.103 | 1.07 | -0.276 | 1.176 | -0.002 | 0.027 | 0.027 | 0 | 0.31 | 0.12 | 0.27 | 0.24 | 0.29 | 0.47 |
| 1 | 192366691 | rs1936706 | 0.698 | 1.32 | 0.799 | 1.309 | 0.085 | 0.2 | 0.2 | 1.121 | 0.92 | 0.56 | 0.83 | 0.87 | 0.93 | 0.94 |
| 1 | 192370110 | rs2369810 | 0.013 | 1.32 | 0.822 | 1.327 | 0.018 | 0.066 | 0.066 | 0.044 | 0.94 | 0.83 | 0.85 | 0.88 | 0.93 | 0.95 |
| 1 | 192372674 | rs7545533 | 1.553 | 1.316 | 0.863 | 1.321 | 0.069 | 0.144 | 0.144 | 0 | 0.84 | 0.49 | 0.69 | 0.64 | 0.77 | 0.88 |
| 1 | 192390633 | rs10754006 | 0.634 | 0.981 | 0.395 | 0.715 | 0.078 | 0.193 | 0.193 | 0.877 | 0.93 | 0.61 | 0.82 | 0.87 | 0.93 | 0.94 |
| 1 | 192405661 | rs6676076 | -0.597 | 0.783 | -0.091 | 0.297 | -0.002 | -0.004 | -0.004 | 0 | 0.45 | 0.51 | 0.45 | 0.36 | 0.51 | 0.61 |
| 1 | 192410674 | rs12048806 | 0.39 | 1.226 | -0.198 | 0.424 | 0.038 | -0.001 | -0.001 | 0 | 0.05 | 0.12 | 0.05 | 0.04 | 0.11 | 0.05 |
| 1 | 192411280 | rs12568515 | 0.798 | 1.172 | -0.271 | 0.429 | 0 | 0.002 | 0.002 | 0 | 0.2 | 0.15 | 0.22 | 0.15 | 0.18 | 0.13 |
| 1 | 192414282 | rs12725829 | 0.044 | 0.672 | -0.337 | 0.513 | -0.003 | 0.001 | 0.001 | 0 | 0.19 | 0.19 | 0.16 | 0.16 | 0.19 | 0.43 |
| 1 | 192414988 | rs1936688 | -0.605 | 0.487 | -0.337 | 0.6 | 0.029 | 0.034 | 0.034 | 0 | 0.44 | 0.22 | 0.52 | 0.35 | 0.54 | 0.61 |
| 1 | 192417416 | rs2887558 | -0.28 | 0.337 | 0.323 | 0.548 | 0.013 | 0.054 | 0.054 | 0 | 0.29 | 0.46 | 0.44 | 0.36 | 0.29 | 0.17 |
| 1 | 192417494 | rs12568724 | 0.184 | 0.332 | 0.333 | 0.567 | -0.044 | 0.041 | 0.041 | 0 | 0.25 | 0.07 | 0.28 | 0.13 | 0.28 | 0.14 |
| 1 | 192418922 | rs1892149 | 0.087 | 0.323 | 0.141 | 0.395 | -0.019 | 0.026 | 0.026 | 0 | 0.26 | 0.14 | 0.29 | 0.15 | 0.28 | 0.14 |
| 1 | 192427901 | rs10754009 | 0.066 | 0.347 | -0.385 | 0.373 | -0.024 | 0.042 | 0.042 | 0 | 0.26 | 0.12 | 0.29 | 0.13 | 0.28 | 0.14 |
| 1 | 192430085 | rs2186024 | -0.205 | 0.081 | -0.163 | 0.197 | 0.001 | 0.001 | 0.001 | 0.043 | 0.45 | 0.58 | 0.52 | 0.41 | 0.43 | 0.29 |
| 1 | 192461456 | rs7515864 | -0.705 | -0.393 | 0.069 | -0.285 | -0.042 | -0.089 | -0.089 | 0.225 | 0.28 | 0.47 | 0.18 | 0.08 | 0.19 | 0.2 |
| 1 | 192462312 | rs6699851 | -1.558 | -0.107 | 0.923 | 0.262 | 0.049 | 0.087 | 0.087 | 3.593 | 0.06 | 0.54 | 0.05 | 0.02 | 0.01 | 0 |
| 1 | 192476716 | rs2476312 | 0.622 | -0.362 | 0.382 | 0.152 | -0.023 | -0.013 | -0.013 | 0 | 0.76 | 0.63 | 0.84 | 0.81 | 0.79 | 0.89 |
| 1 | 192481285 | rs4657830 | 0.177 | 0.296 | 0.706 | -0.235 | 0.026 | -0.012 | -0.012 | 0.055 | 0.74 | 0.62 | 0.7 | 0.69 | 0.71 | 0.81 |
| 1 | 192502783 | rs10754013 | 0.489 | 0.233 | 1.206 | -0.377 | 0.017 | -0.022 | -0.022 | 1.58 | 0.7 | 0.49 | 0.7 | 0.69 | 0.67 | 0.79 |
| 1 | 192510011 | rs12033611 | 0.334 | 0.431 | 1.066 | -0.447 | -0.029 | -0.007 | -0.007 | 2.011 | 0.06 | 0 | 0.03 | 0.11 | 0.07 | 0.09 |
| 1 | 192510030 | rs1323299 | 0.148 | 0.404 | 1.448 | -0.46 | 0.005 | -0.008 | -0.008 | 0 | 0.77 | 0.77 | 0.74 | 0.8 | 0.75 | 0.88 |
| 1 | 192511183 | rs10489868 | 1.041 | 0.692 | 0.901 | -0.351 | -0.026 | -0.015 | -0.015 | 2.072 | 0.19 | 0.09 | 0.26 | 0.19 | 0.24 | 0.38 |
| 1 | 192514029 | rs10921191 | 0.964 | 0.511 | 1.149 | 0.116 | 0 | -0.001 | -0.001 | 1.806 | 0.68 | 0.36 | 0.59 | 0.6 | 0.66 | 0.77 |
| 1 | 192519865 | rs1323298 | -0.839 | 0.273 | 1.236 | 0.168 | -0.042 | -0.019 | -0.019 | 0 | 0.83 | 0.98 | 0.8 | 0.81 | 0.82 | 0.93 |
| 1 | 192521591 | rs10801129 | 0.7 | 0.264 | 1.309 | 0.162 | 0.011 | -0.023 | -0.023 | 0 | 0.73 | 0.49 | 0.74 | 0.7 | 0.74 | 0.84 |
| 1 | 192522392 | rs10921192 | 0.126 | 0.294 | 1.388 | 0.141 | -0.004 | 0.001 | 0.001 | 0 | 0.07 | 0.06 | 0.03 | 0.11 | 0.07 | 0.09 |
| 1 | 192523807 | rs2760519 | -1.154 | 0.281 | 1.367 | 0.101 | 0.062 | 0.008 | 0.008 | 0.063 | 0.1 | 0.43 | 0.05 | 0.11 | 0.07 | 0.09 |
| 1 | 192524049 | rs2760520 | 0.442 | 0.136 | 0.932 | -0.061 | -0.001 | 0.05 | 0.05 | 0 | 0.7 | 0.88 | 0.71 | 0.8 | 0.75 | 0.9 |
| 1 | 192536813 | rs2816316 | -0.402 | 0.119 | 0.395 | 0.019 | 0.002 | -0.001 | -0.001 | 4.271 | 0.81 | 0.71 | 0.75 | 0.8 | 0.82 | 0.93 |
| 1 | 192538496 | rs2760530 | 0.255 | 0.069 | 0.363 | -0.026 | 0.003 | 0.026 | 0.026 | 0 | 0.7 | 0.8 | 0.71 | 0.8 | 0.75 | 0.9 |
| 1 | 192545099 | rs7535818 | 0.405 | 0.11 | -0.756 | 0.546 | -0.002 | 0.003 | 0.003 | 0.151 | 0.76 | 0.29 | 0.65 | 0.64 | 0.81 | 0.88 |
| 1 | 192547052 | rs10921202 | -0.698 | 0.123 | -1.032 | 0.208 | -0.016 | 0.022 | 0.022 | 0 | 0.07 | 0.02 | 0.03 | 0.04 | 0.06 | 0.1 |
| 1 | 192547710 | rs1923949 | -0.704 | -0.386 | -1.641 | 0.362 | 0.071 | -0.112 | -0.112 | 0.365 | 0.38 | 0.92 | 0.44 | 0.42 | 0.27 | 0.55 |
| 1 | 192548543 | rs1323291 | -0.748 | -0.058 | -0.193 | 0.399 | 0.107 | 0.006 | 0.006 | 0 | 0.07 | 0.4 | 0.08 | 0 | 0.1 | 0.07 |
| 1 | 192549912 | rs2760535 | -0.694 | -0.053 | -0.174 | 0.402 | 0.086 | -0.006 | -0.006 | 0 | 0.07 | 0.37 | 0.08 | 0 | 0.09 | 0.07 |
| 1 | 192551624 | rs2984921 | -0.329 | -0.096 | -0.729 | 0.627 | -0.006 | 0.057 | 0.057 | 0 | 0.32 | 0.47 | 0.31 | 0.46 | 0.28 | 0.24 |
| 1 | 192551676 | rs10489871 | -0.359 | -0.151 | -0.521 | 0.567 | -0.021 | 0.029 | 0.029 | 0 | 0.31 | 0.49 | 0.38 | 0.38 | 0.25 | 0.44 |
| 1 | 192556309 | rs6701700 | -1.005 | -0.39 | 0.129 | 0.39 | 0.001 | -0.018 | -0.018 | 0 | 0.12 | 0.21 | 0.09 | 0.07 | 0.1 | 0.27 |
| 1 | 192559646 | rs10489873 | -1.206 | -0.514 | 0.771 | 0.817 | -0.01 | 0.059 | 0.059 | 0 | 0.12 | 0.21 | 0.16 | 0.24 | 0.07 | 0.27 |
| 1 | 192561472 | rs2760511 | -0.604 | -0.816 | 0.413 | -0.013 | -0.024 | -0.019 | -0.019 | 0 | 0.36 | 0.58 | 0.25 | 0.27 | 0.22 | 0.3 |
| 1 | 192561545 | rs2760512 | -1.104 | -0.581 | 0.346 | 0.048 | 0.022 | 0.025 | 0.025 | 0 | 0.46 | 0.63 | 0.33 | 0.35 | 0.32 | 0.57 |
| 1 | 192561712 | rs1407818 | -0.463 | 0.484 | 0.282 | 0.169 | -0.001 | 0.004 | 0.004 | 0 | 0.21 | 0.16 | 0.15 | 0.19 | 0.2 | 0.09 |
| 1 | 192567674 | rs10921206 | -0.852 | 0.214 | -0.043 | 0.301 | 0.004 | 0.006 | 0.006 | 0 | 0.18 | 0.15 | 0.08 | 0.12 | 0.17 | 0.38 |
| 1 | 192568943 | rs2984918 | -0.821 | 0.307 | -0.73 | 0.437 | 0.055 | 0.034 | 0.034 | 0 | 0.55 | 0.88 | 0.53 | 0.66 | 0.64 | 0.66 |
| 1 | 192570150 | rs10489877 | 0.186 | 1.259 | -0.291 | 0.937 | -0.039 | -0.07 | -0.07 | 2.166 | 0.2 | 0.12 | 0.19 | 0.4 | 0.27 | 0.2 |
| 1 | 192598635 | rs7553821 | -0.216 | 0.53 | 0.362 | 0.572 | -0.001 | 0.009 | 0.009 | 0 | 0.25 | 0.25 | 0.28 | 0.2 | 0.22 | 0.26 |
| 1 | 192605325 | rs6658382 | 0.262 | 0.164 | -0.833 | 0.298 | -0.175 | -0.148 | -0.148 | 0.604 | 0.37 | 0.04 | 0.35 | 0.61 | 0.49 | 0.42 |
| 1 | 192609268 | rs12044559 | 0.073 | -0.374 | -0.182 | -0.022 | 0 | 0 | 0 | 0 | 0.51 | 0.43 | 0.53 | 0.33 | 0.45 | 0.51 |
| 1 | 192616680 | rs4147217 | -0.162 | -0.911 | 0.323 | -0.245 | 0.002 | 0.063 | 0.063 | 0 | 0.53 | 0.78 | 0.49 | 0.72 | 0.54 | 0.57 |
| 1 | 192621580 | rs4568808 | 0.149 | -0.973 | 0.106 | -0.28 | 0.011 | -0.004 | -0.004 | 0 | 0.39 | 0.78 | 0.58 | 0.33 | 0.35 | 0.31 |
| 1 | 192625530 | rs9287126 | 0.039 | -1.138 | 0.164 | -0.196 | -0.033 | 0.03 | 0.03 | 0 | 0.52 | 0.13 | 0.35 | 0.66 | 0.52 | 0.55 |
| 1 | 192626826 | rs2296020 | -0.388 | -1.177 | 0.825 | -0.23 | -0.004 | 0.014 | 0.014 | 0 | 0.57 | 0.61 | 0.5 | 0.73 | 0.54 | 0.59 |
| 1 | 192627952 | rs12118386 | 0.101 | -1.091 | 0.218 | -0.235 | -0.051 | 0.049 | 0.049 | 0 | 0.5 | 0.08 | 0.34 | 0.67 | 0.49 | 0.55 |
| 1 | 192638215 | rs1933709 | 0.388 | -1.381 | 0.044 | -0.146 | -0.091 | -0.052 | -0.052 | 0.194 | 0.41 | 0.07 | 0.32 | 0.63 | 0.45 | 0.35 |
| 1 | 192660401 | rs1933705 | -0.396 | -1.359 | 0.18 | -0.147 | 0 | 0.023 | 0.023 | 0 | 0.53 | 0.35 | 0.45 | 0.7 | 0.49 | 0.56 |
| 1 | 192682671 | rs2146609 | -0.456 | -1.207 | 0.494 | -0.264 | -0.044 | -0.016 | -0.016 | 0 | 0.11 | 0.33 | 0.14 | 0.07 | 0.03 | 0.21 |
| 1 | 192696582 | rs16834698 | -0.299 | -1.208 | 0.482 | -0.262 | -0.04 | -0.011 | -0.011 | 0 | 0.11 | 0.33 | 0.14 | 0.07 | 0.03 | 0.2 |
| 1 | 192739041 | rs10921257 | -0.112 | -1.115 | -0.217 | 0.059 | -0.002 | -0.006 | -0.006 | 0.015 | 0.49 | 0.32 | 0.35 | 0.53 | 0.5 | 0.4 |
| 1 | 192755994 | rs17646713 | -0.169 | -0.906 | -0.235 | 0.136 | 0.013 | 0.016 | 0.016 | 0 | 0.13 | 0.09 | 0.04 | 0.05 | 0.07 | 0.23 |
| 1 | 192757432 | rs6703040 | 0.508 | -1.511 | -0.269 | -0.136 | -0.002 | -0.003 | -0.003 | 0 | 0.33 | 0.34 | 0.39 | 0.53 | 0.41 | 0.32 |
| 1 | 192758615 | rs6670601 | -0.595 | -1.851 | -0.082 | -0.263 | -0.001 | -0.002 | -0.002 | 2.322 | 0.49 | 0.56 | 0.45 | 0.61 | 0.48 | 0.41 |
| 1 | 192758841 | rs6670801 | 0.707 | -1.839 | -0.013 | -0.232 | -0.036 | -0.046 | -0.046 | 0 | 0.33 | 0.19 | 0.3 | 0.49 | 0.41 | 0.32 |
| 1 | 192759186 | rs16834831 | 1.005 | -1.862 | 0.002 | -0.214 | -0.015 | -0.003 | -0.003 | 0.081 | 0.12 | 0.17 | 0.15 | 0.09 | 0.06 | 0.05 |
| 1 | 192763948 | rs10801153 | 0.969 | -0.923 | -0.43 | -0.315 | 0.001 | -0.023 | -0.023 | 0.022 | 0.28 | 0.22 | 0.22 | 0.48 | 0.29 | 0.3 |
| 1 | 192764820 | rs1933695 | 0.59 | -0.878 | -0.68 | -0.338 | -0.052 | 0.051 | 0.051 | 4.049 | 0.16 | 0.02 | 0.12 | 0.06 | 0.18 | 0.07 |
| 1 | 192764888 | rs6428136 | 1.043 | -0.842 | -0.581 | -0.303 | -0.011 | -0.087 | -0.087 | 0 | 0.27 | 0.08 | 0.2 | 0.48 | 0.29 | 0.3 |

| | | | | | | | | | | | | | | | | |
|---|---|---|---|---|---|---|---|---|---|---|---|---|---|---|---|---|
| 1 | 192765943 | rs7531013 | -1.446 | -0.781 | -0.684 | -0.288 | 0.004 | -0.001 | -0.001 | 0 | 0.61 | 0.64 | 0.49 | 0.66 | 0.54 | 0.64 |
| 1 | 192770933 | rs1890397 | -1.347 | -0.673 | -0.818 | -0.212 | 0.006 | -0.003 | -0.003 | 1.592 | 0.59 | 0.57 | 0.48 | 0.64 | 0.54 | 0.6 |
| 1 | 192777564 | rs2746071 | 0.372 | -0.757 | -0.276 | 0.124 | -0.008 | 0.066 | 0.066 | 0 | 0.3 | 0.48 | 0.32 | 0.53 | 0.29 | 0.3 |
| 1 | 192787743 | rs17587185 | -0.275 | -0.486 | -0.401 | -0.282 | 0.012 | 0.02 | 0.02 | 0 | 0.13 | 0.08 | 0.04 | 0.05 | 0.08 | 0.23 |
| 1 | 192795419 | rs710146 | 0.495 | 0.533 | 0.177 | -0.311 | 0.006 | -0.016 | -0.016 | 0 | 0.75 | 0.69 | 0.73 | 0.86 | 0.71 | 0.79 |
| 1 | 192796240 | rs710145 | 0.535 | 0.522 | 0.309 | -0.313 | -0.013 | 0.021 | 0.021 | 0 | 0.75 | 0.81 | 0.74 | 0.86 | 0.71 | 0.79 |
| 1 | 192796532 | rs710143 | 0.193 | 0.316 | 0.452 | -0.455 | -0.011 | -0.025 | -0.025 | 0 | 0.08 | 0.11 | 0.03 | 0 | 0.03 | 0.05 |
| 1 | 192800241 | rs10489514 | -1.222 | 0.2 | 0.52 | -0.532 | -0.056 | -0.089 | -0.089 | 0.155 | 0.14 | 0.38 | 0.05 | 0.01 | 0.05 | 0.08 |
| 1 | 192802381 | rs2153084 | -1.086 | 0.099 | 0.643 | -0.577 | -0.047 | -0.076 | -0.076 | 0 | 0.12 | 0.59 | 0.07 | 0 | 0.03 | 0.05 |
| 1 | 192810167 | rs1775686 | -0.627 | 0.533 | -0.128 | -0.164 | 0.01 | 0.024 | 0.024 | 0 | 0.43 | 0.75 | 0.46 | 0.43 | 0.39 | 0.51 |
| 1 | 192818018 | rs1231728 | 0.792 | 0.361 | -0.19 | -0.118 | 0.01 | -0.019 | -0.019 | 0.012 | 0.74 | 0.36 | 0.66 | 0.81 | 0.71 | 0.79 |
| 1 | 192822069 | rs1231741 | -0.528 | 0.706 | -0.904 | 0.073 | -0.04 | -0.013 | -0.013 | 3.018 | 0.84 | 0.93 | 0.71 | 0.82 | 0.73 | 0.82 |
| 1 | 192845807 | rs4140450 | 0.046 | 0.312 | -0.978 | 0.005 | -0.048 | -0.041 | -0.041 | 0 | 0.32 | 0.09 | 0.35 | 0.39 | 0.34 | 0.46 |
| 1 | 192880701 | rs7517778 | -0.389 | 0.638 | -0.082 | 0.075 | 0.045 | -0.016 | -0.016 | 0 | 0.18 | 0.42 | 0.26 | 0.13 | 0.25 | 0.19 |
| 1 | 192883411 | rs7527185 | -0.823 | 0.212 | -0.288 | 0.068 | -0.009 | 0.013 | 0.013 | 0 | 0.4 | 0.53 | 0.39 | 0.51 | 0.37 | 0.6 |
| 1 | 192886814 | rs10801161 | -0.173 | 0.15 | -0.121 | 0.028 | -0.006 | -0.057 | -0.057 | 0 | 0.32 | 0.18 | 0.36 | 0.51 | 0.35 | 0.56 |
| 1 | 192888345 | rs6671680 | -1.013 | 0.002 | 0.257 | -0.041 | 0.032 | -0.031 | -0.031 | 0 | 0.42 | 0.8 | 0.42 | 0.52 | 0.37 | 0.59 |
| 1 | 192891048 | rs2370026 | -0.365 | 0.326 | 0.239 | -0.009 | 0.004 | -0.019 | -0.019 | 0.096 | 0.08 | 0.35 | 0.03 | 0 | 0.02 | 0.03 |
| 1 | 192894881 | rs2887613 | -0.319 | 0.266 | -0.459 | 0.412 | 0.012 | -0.002 | -0.002 | 0 | 0.31 | 0.25 | 0.38 | 0.68 | 0.26 | 0.43 |
| 1 | 192896434 | rs17525308 | 0.122 | 0.508 | -0.438 | 0.076 | 0.001 | -0.021 | -0.021 | 0.008 | 0.08 | 0.12 | 0.09 | 0 | 0.1 | 0.02 |
| 1 | 192900182 | rs1417416 | 0.151 | 0.562 | -1.226 | 0.194 | -0.093 | -0.039 | -0.039 | 0 | 0.42 | 0.08 | 0.43 | 0.42 | 0.42 | 0.56 |
| 1 | 192902415 | rs842796 | -0.753 | 0.377 | -0.536 | 0.312 | -0.005 | -0.003 | -0.003 | 0 | 0.6 | 0.69 | 0.59 | 0.45 | 0.58 | 0.63 |
| 1 | 192910766 | rs10801165 | -0.289 | 0.266 | -0.084 | 0.09 | 0.02 | -0.069 | -0.069 | 0.293 | 0.11 | 0.33 | 0.11 | 0 | 0.12 | 0.04 |
| 1 | 192911972 | rs842814 | -1.525 | 0.313 | -0.485 | 0.326 | -0.048 | -0.051 | -0.051 | 0.329 | 0.76 | 0.95 | 0.77 | 0.74 | 0.73 | 0.81 |
| 1 | 192912108 | rs710140 | -0.525 | 0.302 | -0.202 | -0.271 | 0.02 | -0.085 | -0.085 | 1.693 | 0.13 | 0.43 | 0.12 | 0 | 0.12 | 0.04 |
| 1 | 192940642 | rs12025826 | 0.003 | 0.324 | -0.083 | -0.16 | -0.007 | 0.01 | 0.01 | 0.039 | 0.57 | 0.31 | 0.55 | 0.43 | 0.57 | 0.62 |
| 1 | 192946832 | rs7512241 | -0.871 | 0.401 | 0.451 | -0.18 | 0.017 | -0.08 | -0.08 | 1.465 | 0.12 | 0.49 | 0.12 | 0 | 0.11 | 0.04 |
| 1 | 193027834 | rs2887640 | 1.09 | 0.916 | -0.478 | -0.401 | 0 | 0.02 | 0.02 | 2.69 | 0.06 | 0.03 | 0.05 | 0 | 0.05 | 0.01 |
| 1 | 193030600 | rs7554496 | 0.92 | 1.043 | -0.857 | -1.156 | -0.004 | 0.037 | 0.037 | 0 | 0.23 | 0.15 | 0.21 | 0.07 | 0.21 | 0.14 |
| 1 | 193035013 | rs12131585 | 2.053 | 1.431 | -1.049 | -1.264 | -0.013 | 0.027 | 0.027 | 0 | 0.05 | 0 | 0.02 | 0 | 0.04 | 0.01 |
| 1 | 193051685 | rs2275444 | 0.463 | 1.195 | -0.809 | -1.061 | 0.034 | 0.065 | 0.065 | 0 | 0.74 | 0.52 | 0.66 | 0.44 | 0.68 | 0.7 |
| 1 | 193052205 | rs10921308 | 0.104 | 1.154 | -0.817 | -1.085 | 0.042 | 0.072 | 0.072 | 0 | 0.77 | 0.58 | 0.68 | 0.44 | 0.69 | 0.74 |
| 1 | 193054154 | rs10921309 | 1.74 | 0.309 | -1.475 | -1.618 | -0.122 | -0.041 | -0.041 | 0.68 | 0.31 | 0.02 | 0.36 | 0.34 | 0.38 | 0.46 |
| 1 | 193054485 | rs10801173 | 0.871 | 0.775 | -0.895 | -1.018 | 0.015 | 0.028 | 0.028 | 0 | 0.69 | 0.48 | 0.63 | 0.44 | 0.65 | 0.69 |
| 1 | 193075108 | rs912071 | 0.222 | 0.646 | -0.72 | -0.96 | 0.007 | 0.013 | 0.013 | 0 | 0.77 | 0.73 | 0.69 | 0.44 | 0.68 | 0.78 |
| 1 | 193077301 | rs10921312 | -0.289 | 0.654 | -0.717 | -0.957 | 0.014 | 0.025 | 0.025 | 0 | 0.79 | 0.75 | 0.7 | 0.45 | 0.69 | 0.82 |
| 1 | 193078985 | rs2181408 | 0.142 | 0.657 | -0.716 | -0.954 | 0.007 | 0.014 | 0.014 | 0 | 0.77 | 0.73 | 0.69 | 0.45 | 0.68 | 0.78 |
| 1 | 193083501 | rs12123511 | 2.198 | 0.646 | -0.737 | -0.964 | -0.001 | -0.002 | -0.002 | 0.178 | 0.05 | 0.05 | 0.05 | 0 | 0.05 | 0.01 |
| 1 | 193084045 | rs10733078 | 0.895 | 0.637 | -0.702 | -0.953 | 0.027 | 0.031 | 0.031 | 0 | 0.7 | 0.46 | 0.62 | 0.45 | 0.63 | 0.77 |
| 1 | 193140246 | rs10754052 | -0.295 | 0.662 | -0.679 | -0.918 | 0.015 | 0.028 | 0.028 | 0 | 0.79 | 0.74 | 0.69 | 0.45 | 0.7 | 0.82 |
| 1 | 193150873 | rs10494675 | 2.1 | 0.656 | -0.665 | -0.922 | -0.001 | -0.001 | -0.001 | 0 | 0.05 | 0.06 | 0.05 | 0 | 0.05 | 0.01 |
| 1 | 193161959 | rs6676878 | -0.298 | 0.659 | -0.661 | -0.918 | 0.015 | 0.028 | 0.028 | 0 | 0.79 | 0.74 | 0.69 | 0.45 | 0.7 | 0.82 |
| 1 | 193193597 | rs12410213 | 0.973 | 0.67 | -0.601 | -0.976 | 0.029 | 0.034 | 0.034 | 0 | 0.7 | 0.51 | 0.63 | 0.45 | 0.63 | 0.77 |
| 1 | 193210415 | rs6667670 | 1.658 | 0.45 | -0.832 | -1.292 | -0.016 | 0.063 | 0.063 | 0.062 | 0.18 | 0.05 | 0.16 | 0 | 0.18 | 0.02 |
| 1 | 193236656 | rs1578242 | 0.713 | 0.678 | -0.495 | -0.815 | 0.032 | 0.038 | 0.038 | 0 | 0.7 | 0.47 | 0.62 | 0.45 | 0.63 | 0.77 |
| 1 | 193282883 | rs560900 | -0.19 | 0.772 | -0.441 | -0.596 | 0.038 | 0.05 | 0.05 | 0 | 0.21 | 0.37 | 0.31 | 0.59 | 0.3 | 0.18 |
| 1 | 193288856 | rs1770000 | -1.027 | 0.826 | -0.28 | -0.333 | 0.051 | 0.053 | 0.053 | 0 | 0.27 | 0.52 | 0.35 | 0.54 | 0.35 | 0.21 |
| 1 | 193290358 | rs1749429 | -0.493 | 0.863 | -0.241 | -0.231 | 0.054 | 0.104 | 0.104 | 0 | 0.22 | 0.45 | 0.31 | 0.52 | 0.3 | 0.21 |
| 1 | 193291213 | rs1769999 | -0.284 | 0.911 | -0.162 | -0.21 | 0.056 | 0.088 | 0.088 | 0 | 0.2 | 0.38 | 0.3 | 0.54 | 0.29 | 0.18 |
| 1 | 193334309 | rs16835319 | -0.119 | 0.924 | -0.226 | -0.247 | 0.015 | 0.034 | 0.034 | 0 | 0.19 | 0.23 | 0.29 | 0.49 | 0.29 | 0.19 |
| 1 | 193356684 | rs725209 | -1.411 | 0.468 | -0.591 | -0.411 | -0.002 | -0.001 | -0.001 | 0 | 0.61 | 0.63 | 0.61 | 0.56 | 0.63 | 0.39 |
| 1 | 193383564 | rs6667545 | -0.971 | -0.016 | 0.18 | 0.159 | -0.052 | 0.09 | 0.09 | 0 | 0.25 | 0.44 | 0.33 | 0.48 | 0.16 | 0.21 |
| 1 | 193391298 | rs16829482 | 0.321 | 0.295 | 0.323 | 0.49 | 0.117 | 0.014 | 0.014 | 0 | 0.09 | 0.22 | 0.13 | 0.07 | 0.22 | 0.05 |
| 1 | 193391538 | rs12043960 | 1.569 | 0.045 | -0.092 | 0.642 | 0.001 | 0.079 | 0.079 | 0 | 0.06 | 0.04 | 0.24 | 0.48 | 0.05 | 0.09 |
| 1 | 193418907 | rs16835376 | 0.168 | -0.008 | 0.014 | 0.93 | -0.014 | 0.134 | 0.134 | 0 | 0.13 | 0.2 | 0.25 | 0.37 | 0.09 | 0.13 |
| 1 | 193421360 | rs7528241 | -0.097 | 0.005 | -0.044 | 0.98 | -0.02 | 0.136 | 0.136 | 0 | 0.14 | 0.26 | 0.27 | 0.35 | 0.1 | 0.15 |
| 1 | 193424164 | rs2038750 | -1.538 | 0.126 | -0.426 | 0.812 | -0.002 | 0.075 | 0.075 | 0 | 0.3 | 0.45 | 0.39 | 0.43 | 0.28 | 0.26 |
| 1 | 193435334 | rs10921350 | -0.798 | -0.269 | -0.834 | -0.123 | -0.003 | 0 | 0 | 0.005 | 0.4 | 0.57 | 0.37 | 0.38 | 0.25 | 0.25 |
| 1 | 193438444 | rs12033145 | 0.722 | -0.366 | -0.869 | -0.659 | 0.003 | -0.012 | -0.012 | 0.025 | 0.11 | 0.08 | 0.29 | 0.35 | 0.1 | 0.14 |
| 1 | 193442375 | rs7518647 | -0.978 | -0.385 | -0.915 | -0.933 | -0.001 | -0.001 | -0.001 | 0 | 0.4 | 0.59 | 0.44 | 0.4 | 0.25 | 0.25 |
| 1 | 193442605 | rs942439 | 0.474 | -0.381 | -1.439 | -1.366 | -0.068 | 0.044 | 0.044 | 0 | 0.16 | 0.01 | 0.18 | 0.09 | 0.23 | 0.07 |
| 1 | 193442639 | rs7549962 | 0.937 | -0.524 | -1.476 | -1.423 | -0.018 | 0.14 | 0.14 | 0.99 | 0.22 | 0.04 | 0.1 | 0 | 0.15 | 0.09 |
| 1 | 193443157 | rs752276 | 1.018 | -0.357 | -1.081 | -1.139 | -0.013 | -0.017 | -0.017 | 0 | 0.43 | 0.32 | 0.37 | 0.51 | 0.51 | 0.68 |
| 1 | 193443701 | rs11808338 | -0.634 | -0.052 | -0.274 | -0.513 | -0.032 | 0.2 | 0.2 | 0 | 0.17 | 0.54 | 0.33 | 0.39 | 0.1 | 0.15 |
| 1 | 193446704 | rs2224874 | 0.535 | 0.039 | -0.496 | -2.097 | 0.001 | -0.024 | -0.024 | 0 | 0.44 | 0.45 | 0.57 | 0.85 | 0.46 | 0.69 |
| 1 | 193449584 | rs2147305 | 0.889 | 0.287 | -0.383 | -2.081 | -0.001 | -0.013 | -0.013 | 0 | 0.42 | 0.42 | 0.54 | 0.8 | 0.41 | 0.64 |
| 1 | 193450009 | rs10494682 | 0.913 | 0.273 | -0.645 | -2.443 | -0.019 | 0.101 | 0.101 | 1.67 | 0.24 | 0.08 | 0.12 | 0.03 | 0.24 | 0.08 |
| 1 | 193451373 | rs10921353 | 1.026 | 0.31 | -0.514 | -2.265 | 0 | -0.012 | -0.012 | 0.184 | 0.41 | 0.44 | 0.52 | 0.75 | 0.41 | 0.64 |
| 1 | 193457511 | rs12406893 | 0.16 | 0.512 | -0.664 | -2.532 | 0.029 | -0.007 | -0.007 | 0 | 0.15 | 0.19 | 0.26 | 0.14 | 0.27 | 0.17 |

| | | | | | | | | | | | | | | | | |
|---|---|---|---|---|---|---|---|---|---|---|---|---|---|---|---|---|
| 1 | 193459219 | rs12757080 | 0.864 | 0.335 | -0.725 | -2.709 | 0 | 0.072 | 0.072 | 1.681 | 0.24 | 0.11 | 0.12 | 0.03 | 0.23 | 0.08 |
| 1 | 193461282 | rs12729450 | -0.759 | 0.228 | -0.757 | -2.762 | -0.01 | 0.047 | 0.047 | 0 | 0.44 | 0.33 | 0.4 | 0.17 | 0.54 | 0.3 |
| 1 | 193463330 | rs12139329 | 0.783 | 0.357 | -0.755 | -2.63 | -0.007 | 0.073 | 0.073 | 1.628 | 0.24 | 0.11 | 0.13 | 0.03 | 0.25 | 0.08 |
| 1 | 193464485 | rs6683479 | -1.222 | 0.445 | -0.49 | -2.581 | 0.003 | -0.049 | -0.049 | 0 | 0.6 | 0.46 | 0.47 | 0.19 | 0.6 | 0.37 |
| 1 | 193469347 | rs7538476 | -0.112 | 0.281 | -0.296 | -2.473 | 0.003 | -0.01 | -0.01 | 0 | 0.3 | 0.24 | 0.16 | 0.04 | 0.29 | 0.13 |
| 1 | 193477045 | rs1555649 | -0.684 | 0.783 | -0.208 | -2.017 | -0.003 | 0.088 | 0.088 | 0 | 0.56 | 0.28 | 0.48 | 0.18 | 0.6 | 0.39 |
| 1 | 193493591 | rs10218732 | 0.087 | 0.845 | -0.42 | -2.193 | -0.028 | 0.006 | 0.006 | 0.08 | 0.16 | 0.08 | 0.28 | 0.17 | 0.28 | 0.21 |
| 1 | 193493617 | rs1408408 | -1.231 | 0.647 | -0.366 | -2.203 | -0.004 | 0.017 | 0.017 | 0 | 0.6 | 0.64 | 0.51 | 0.18 | 0.62 | 0.43 |
| 1 | 193494656 | rs17591577 | 0.748 | 0.659 | -0.441 | -2.216 | -0.01 | 0.026 | 0.026 | 0 | 0.05 | 0.01 | 0.04 | 0 | 0.05 | 0.05 |
| 1 | 193499345 | rs12125262 | -0.169 | 0.877 | -0.219 | -2.015 | -0.023 | 0.008 | 0.008 | 0.033 | 0.17 | 0.11 | 0.3 | 0.17 | 0.3 | 0.21 |
| 1 | 193502609 | rs6692705 | -0.199 | 0.667 | 0.242 | -1.377 | -0.003 | 0.043 | 0.043 | 0 | 0.57 | 0.34 | 0.49 | 0.17 | 0.61 | 0.4 |
| 1 | 193510827 | rs1007460 | 0.504 | 1.294 | -0.099 | -0.361 | -0.018 | 0.013 | 0.013 | 0 | 0.65 | 0.73 | 0.54 | 0.75 | 0.54 | 0.66 |
| 1 | 193521335 | rs9660530 | 0.922 | 0.998 | 0.445 | -0.667 | 0.015 | 0.028 | 0.028 | 0 | 0.5 | 0.41 | 0.45 | 0.7 | 0.43 | 0.54 |
| 1 | 193523636 | rs4657871 | 0.411 | 0.85 | 0.618 | -0.771 | 0.006 | -0.013 | -0.013 | 0 | 0.59 | 0.57 | 0.52 | 0.74 | 0.52 | 0.64 |
| 1 | 193528905 | rs17546014 | 0.032 | 0.701 | 0.957 | -0.721 | 0.009 | -0.037 | -0.037 | 0 | 0.08 | 0.12 | 0.06 | 0 | 0.08 | 0.07 |
| 1 | 193536949 | rs7544296 | -1.344 | 0.649 | 1.204 | -0.815 | -0.004 | -0.1 | -0.1 | 0.691 | 0.16 | 0.37 | 0.12 | 0.01 | 0.13 | 0.11 |
| 1 | 193547909 | rs16835564 | -1.132 | 0.67 | 0.898 | -0.78 | -0.007 | -0.032 | -0.032 | 0 | 0.15 | 0.22 | 0.12 | 0.04 | 0.13 | 0.12 |
| 1 | 193559230 | rs731361 | -0.575 | 0.577 | 0.779 | -1.015 | -0.012 | -0.082 | -0.082 | 0 | 0.16 | 0.24 | 0.12 | 0 | 0.13 | 0.11 |
| 1 | 193569748 | rs979497 | 0.773 | 1.557 | 0.661 | -0.623 | 0.002 | -0.009 | -0.009 | 0.054 | 0.62 | 0.63 | 0.56 | 0.76 | 0.57 | 0.63 |
| 1 | 193576625 | rs7529129 | -1.755 | 2.182 | 0.858 | -0.688 | 0.004 | -0.023 | -0.023 | 0.194 | 0.29 | 0.62 | 0.33 | 0.12 | 0.37 | 0.3 |
| 1 | 193581076 | rs17376294 | -0.394 | 3.08 | 0.631 | -0.539 | -0.036 | -0.024 | -0.024 | 0 | 0.06 | 0.01 | 0.06 | 0.05 | 0.12 | 0.09 |
| 1 | 193581678 | rs12131305 | -0.927 | 3.319 | 1.294 | -0.652 | 0.006 | -0.041 | -0.041 | 0.309 | 0.19 | 0.24 | 0.11 | 0.03 | 0.22 | 0.19 |
| 1 | 193584062 | rs6668266 | -1.773 | 3.198 | 1.634 | -0.676 | 0.008 | -0.012 | -0.012 | 0 | 0.25 | 0.29 | 0.17 | 0.08 | 0.34 | 0.28 |
| 1 | 193592394 | rs6428182 | -0.825 | 3.259 | 1.798 | -0.683 | 0.001 | -0.046 | -0.046 | 0 | 0.18 | 0.24 | 0.09 | 0.03 | 0.18 | 0.18 |
| 1 | 193600999 | rs2370438 | 2.321 | 3.237 | 2.1 | -0.669 | 0.115 | 0.028 | 0.028 | 0 | 0.82 | 0.57 | 0.68 | 0.84 | 0.69 | 0.8 |
| 1 | 193609764 | rs10801214 | 1.419 | 3.193 | 2.059 | -0.665 | 0.052 | -0.054 | -0.054 | 0 | 0.75 | 0.59 | 0.82 | 0.92 | 0.66 | 0.72 |
| 1 | 193622690 | rs1416601 | -1.039 | 3.133 | 1.98 | -0.702 | -0.009 | 0.02 | 0.02 | 0 | 0.07 | 0.02 | 0.11 | 0 | 0.14 | 0.04 |
| 1 | 193625278 | rs10921400 | 1.976 | 3.208 | 2.088 | -0.71 | -0.001 | 0.004 | 0.004 | 0 | 0.69 | 0.31 | 0.66 | 0.87 | 0.65 | 0.7 |
| 1 | 193627682 | rs1410397 | 2.49 | 3.223 | 2.046 | -0.708 | -0.051 | 0.054 | 0.054 | 0.341 | 0.63 | 0.24 | 0.58 | 0.82 | 0.51 | 0.62 |
| 1 | 193630577 | rs6695669 | -1.091 | 3.217 | 2.079 | -0.708 | 0.004 | -0.069 | -0.069 | 0 | 0.2 | 0.31 | 0.1 | 0.03 | 0.19 | 0.19 |
| 1 | 193634651 | rs7550917 | 2.442 | 3.168 | 2.003 | -0.699 | -0.05 | 0.053 | 0.053 | 0.363 | 0.63 | 0.23 | 0.59 | 0.82 | 0.51 | 0.62 |
| 1 | 193649198 | rs1410399 | 2.371 | 3.194 | 2.244 | -0.686 | -0.04 | 0.041 | 0.041 | 0.28 | 0.63 | 0.26 | 0.59 | 0.82 | 0.51 | 0.62 |
| 1 | 193655428 | rs17376588 | -0.294 | 3.275 | 2.292 | -0.685 | 0.032 | 0.004 | 0.004 | 0 | 0.05 | 0.04 | 0.07 | 0.05 | 0.15 | 0.08 |
| 1 | 193661626 | rs10921412 | 1.936 | 3.268 | 2.373 | -0.646 | -0.001 | -0.002 | -0.002 | 1.015 | 0.69 | 0.35 | 0.67 | 0.87 | 0.65 | 0.7 |
| 1 | 193667563 | rs17376699 | -0.055 | 3.094 | 2.358 | -0.643 | -0.026 | -0.024 | -0.024 | 0.286 | 0.05 | 0.01 | 0.07 | 0.05 | 0.15 | 0.08 |
| 1 | 193678599 | rs10801229 | 2.408 | 3.113 | 2.376 | -0.652 | -0.039 | 0.043 | 0.043 | 0.288 | 0.63 | 0.28 | 0.61 | 0.82 | 0.54 | 0.63 |
| 1 | 193684171 | rs12075525 | -1.052 | 3.141 | 2.562 | -0.655 | 0 | -0.03 | -0.03 | 0.386 | 0.2 | 0.24 | 0.1 | 0.03 | 0.19 | 0.19 |
| 1 | 193695077 | rs2031832 | 0.7 | 3.146 | 2.833 | -0.678 | 0.108 | 0.029 | 0.029 | 0 | 0.93 | 0.81 | 0.91 | 0.95 | 0.85 | 0.92 |
| 1 | 193700250 | rs10494690 | 2.25 | 3.047 | 2.622 | -0.687 | -0.014 | 0.013 | 0.013 | 0.187 | 0.64 | 0.34 | 0.62 | 0.82 | 0.54 | 0.63 |
| 1 | 193700387 | rs10801243 | 2.321 | 3.073 | 2.593 | -0.674 | -0.039 | 0.04 | 0.04 | 0.39 | 0.64 | 0.28 | 0.61 | 0.82 | 0.54 | 0.63 |
| 1 | 193702204 | rs1999034 | 2.193 | 3.084 | 2.326 | -0.658 | 0.034 | -0.023 | -0.023 | 0 | 0.67 | 0.5 | 0.63 | 0.78 | 0.54 | 0.63 |
| 1 | 193709733 | rs10921423 | 2.171 | 3.326 | 2.236 | -0.88 | 0.042 | -0.055 | -0.055 | 0 | 0.7 | 0.59 | 0.78 | 0.89 | 0.61 | 0.72 |
| 1 | 193711172 | rs4344313 | 2.631 | 2.925 | 2.068 | -0.801 | -0.001 | 0.001 | 0.001 | 0 | 0.59 | 0.57 | 0.6 | 0.75 | 0.48 | 0.62 |
| 1 | 193715323 | rs10921426 | -2.483 | 2.961 | 2.069 | -0.798 | 0.009 | 0.001 | 0.001 | 0 | 0.24 | 0.26 | 0.35 | 0.23 | 0.4 | 0.21 |
| 1 | 193720321 | rs1887628 | -1.669 | 2.977 | 2.127 | -0.799 | 0.013 | -0.003 | -0.003 | 0 | 0.16 | 0.17 | 0.17 | 0.08 | 0.29 | 0.12 |
| 1 | 193721143 | rs1359937 | 0.016 | 2.96 | 2.112 | -0.81 | -0.002 | -0.023 | -0.023 | 0 | 0.86 | 0.83 | 0.95 | 0.98 | 0.89 | 0.84 |
| 1 | 193722542 | rs6697603 | -3.029 | 2.968 | 2.165 | -0.803 | 0.066 | 0.002 | 0.002 | 0 | 0.29 | 0.44 | 0.37 | 0.23 | 0.42 | 0.22 |
| 1 | 193733125 | rs10921430 | 0.267 | 2.963 | 2.136 | -0.821 | -0.003 | -0.074 | -0.074 | 0 | 0.86 | 0.71 | 0.94 | 0.98 | 0.89 | 0.84 |
| 1 | 193740390 | rs6661798 | 2.056 | 2.985 | 2.102 | -0.827 | -0.015 | 0.012 | 0.012 | 0 | 0.71 | 0.7 | 0.78 | 0.89 | 0.6 | 0.72 |
| 1 | 193741207 | rs1328177 | -0.913 | 2.991 | 2.126 | -0.819 | 0.008 | 0.025 | 0.025 | 0 | 0.09 | 0.11 | 0.19 | 0.14 | 0.13 | 0.1 |
| 1 | 193742890 | rs6669264 | 0.094 | 3.02 | 2.064 | -0.803 | 0 | -0.02 | -0.02 | 0 | 0.86 | 0.83 | 0.94 | 0.97 | 0.88 | 0.84 |
| 1 | 193749358 | rs1854037 | 2.962 | 2.996 | 2.106 | -0.784 | -0.001 | -0.001 | -0.001 | 0 | 0.57 | 0.47 | 0.57 | 0.75 | 0.48 | 0.62 |
| 1 | 193749837 | rs3753949 | 0.951 | 3.009 | 1.867 | -0.811 | -0.006 | 0.01 | 0.01 | 0 | 0.79 | 0.83 | 0.86 | 0.97 | 0.77 | 0.82 |
| 1 | 193756440 | rs6687160 | 2.676 | 2.77 | 1.798 | -0.774 | -0.001 | 0.001 | 0.001 | 0 | 0.58 | 0.45 | 0.57 | 0.75 | 0.47 | 0.62 |
| 1 | 193757853 | rs1328180 | 1.822 | 2.81 | 1.623 | -0.744 | 0.044 | -0.052 | -0.052 | 0 | 0.7 | 0.6 | 0.77 | 0.89 | 0.6 | 0.72 |
| 1 | 193763292 | rs990812 | 0.701 | 2.724 | 1.155 | -0.68 | 0.008 | -0.006 | -0.006 | 0 | 0.8 | 0.74 | 0.88 | 0.89 | 0.74 | 0.73 |
| 1 | 193764013 | rs2370633 | 2.028 | 2.237 | 0.5 | -0.54 | 0.036 | 0.074 | 0.074 | 0 | 0.5 | 0.31 | 0.53 | 0.74 | 0.44 | 0.53 |
| 1 | 193772147 | rs12085891 | -1.387 | 2.289 | 0.411 | 0.219 | -0.033 | -0.035 | -0.035 | 2.668 | 0.09 | 0.02 | 0.17 | 0.29 | 0.13 | 0.18 |
| 1 | 193774083 | rs10801256 | 1.567 | 2.307 | 0.994 | 0.024 | 0.021 | -0.076 | -0.076 | 0.141 | 0.69 | 0.41 | 0.7 | 0.85 | 0.61 | 0.82 |
| 1 | 193777572 | rs1326853 | 1.781 | 2.279 | 0.624 | 0.036 | -0.055 | -0.001 | -0.001 | 1.432 | 0.67 | 0.12 | 0.56 | 0.67 | 0.62 | 0.78 |
| 1 | 193783317 | rs10921443 | 2.832 | 2.183 | 0.31 | -0.273 | 0.031 | -0.067 | -0.067 | 1.262 | 0.47 | 0.03 | 0.38 | 0.55 | 0.33 | 0.54 |
| 1 | 193788485 | rs6693857 | -1.008 | 2.362 | 0.348 | -0.23 | -0.023 | -0.002 | -0.002 | 0 | 0.14 | 0.08 | 0.1 | 0.12 | 0.2 | 0.23 |
| 1 | 193792484 | rs1575938 | 1.513 | 2.346 | 0.785 | -0.338 | -0.001 | -0.001 | -0.001 | 0.342 | 0.76 | 0.33 | 0.73 | 0.8 | 0.84 | 0.81 |
| 1 | 193808170 | rs10801260 | 2.636 | 2.329 | 0.405 | -0.387 | -0.067 | 0.041 | 0.041 | 0 | 0.6 | 0.11 | 0.56 | 0.67 | 0.54 | 0.76 |
| 1 | 193808664 | rs10737633 | 1.189 | 1.78 | 0.907 | -0.236 | -0.005 | -0.06 | -0.06 | 0 | 0.8 | 0.76 | 0.88 | 0.98 | 0.84 | 0.88 |
| 1 | 193824708 | rs12036144 | 1.647 | 1.817 | 0.385 | -0.117 | 0.058 | -0.023 | -0.023 | 3.136 | 0.72 | 0.13 | 0.76 | 0.67 | 0.84 | 0.79 |
| 1 | 193832544 | rs650465 | 1.338 | 2.424 | 1.415 | -0.046 | 0.058 | 0.019 | 0.019 | 0.078 | 0.84 | 0.61 | 0.77 | 0.85 | 0.8 | 0.73 |
| 1 | 193833027 | rs1887401 | -1.384 | 2.37 | 1.603 | -0.052 | 0.031 | 0.008 | 0.008 | 0.167 | 0.15 | 0.22 | 0.2 | 0.15 | 0.21 | 0.28 |
| 1 | 193834315 | rs592371 | 2.117 | 2.789 | 2.238 | -0.299 | 0.046 | -0.032 | -0.032 | 1.877 | 0.7 | 0.41 | 0.56 | 0.84 | 0.52 | 0.69 |

| | | | | | | | | | | | | | | | |
|---|---|---|---|---|---|---|---|---|---|---|---|---|---|---|---|
| 1 | 193836086 | rs600023 | 2.101 | 2.789 | 2.245 | -0.299 | 0.05 | -0.033 | -0.033 | 0 | 0.7 | 0.42 | 0.56 | 0.84 | 0.52 | 0.69 |
| 1 | 193849656 | rs517206 | 1.183 | 2.7 | 2.289 | -0.284 | 0.087 | 0.04 | 0.04 | 0.033 | 0.84 | 0.54 | 0.77 | 0.85 | 0.8 | 0.74 |
| 1 | 193859025 | rs670747 | 1.547 | 2.698 | 2.425 | -0.29 | 0.042 | 0.013 | 0.013 | 0.255 | 0.83 | 0.39 | 0.76 | 0.85 | 0.8 | 0.73 |
| 1 | 193864579 | rs12057376 | -1.452 | 2.702 | 2.556 | -0.288 | 0.062 | 0.027 | 0.027 | 0.037 | 0.16 | 0.35 | 0.22 | 0.15 | 0.2 | 0.27 |
| 1 | 193866782 | rs12744347 | -1.123 | 2.695 | 2.525 | -0.291 | -0.013 | -0.007 | -0.007 | 0 | 0.14 | 0.11 | 0.2 | 0.15 | 0.2 | 0.27 |
| 1 | 193871478 | rs544511 | 1.439 | 2.699 | 2.442 | -0.29 | 0.06 | 0.022 | 0.022 | 0.106 | 0.84 | 0.48 | 0.76 | 0.85 | 0.81 | 0.74 |
| 1 | 193878858 | rs4326606 | -0.876 | 2.818 | 1.973 | -0.282 | -0.078 | 0.084 | 0.084 | 0 | 0.14 | 0.01 | 0.19 | 0.01 | 0.28 | 0.05 |
| 1 | 193883110 | rs550899 | 3.129 | 2.636 | 2.171 | -0.087 | 0.064 | 0.008 | 0.008 | 0 | 0.47 | 0.16 | 0.38 | 0.57 | 0.35 | 0.45 |
| 1 | 193886619 | rs6662524 | -1.495 | 2.66 | 2.559 | -0.08 | -0.005 | -0.01 | -0.01 | 0 | 0.19 | 0.28 | 0.14 | 0.15 | 0.16 | 0.2 |
| 1 | 193889470 | rs12747764 | -1.762 | 2.637 | 2.775 | -0.074 | 0 | 0 | 0 | 0 | 0.18 | 0.18 | 0.24 | 0.26 | 0.21 | 0.3 |
| 1 | 193890360 | rs7541852 | -3.647 | 2.574 | 2.742 | -0.111 | 0.004 | 0.001 | 0.001 | 0 | 0.52 | 0.47 | 0.58 | 0.43 | 0.64 | 0.55 |
| 1 | 193890660 | rs11805075 | -1.741 | 2.574 | 2.741 | -0.111 | 0 | 0 | 0 | 0 | 0.18 | 0.18 | 0.24 | 0.26 | 0.21 | 0.3 |
| 1 | 193895333 | rs4330903 | -2.426 | 2.575 | 2.711 | -0.11 | -0.01 | 0.023 | 0.023 | 0 | 0.35 | 0.29 | 0.34 | 0.16 | 0.43 | 0.29 |
| 1 | 193896826 | rs2771108 | 3.341 | 2.56 | 2.495 | -0.098 | 0.045 | 0.014 | 0.014 | 0 | 0.46 | 0.25 | 0.4 | 0.58 | 0.36 | 0.46 |
| 1 | 193901506 | rs574986 | 2.603 | 2.712 | 2.538 | 0.012 | -0.006 | 0.055 | 0.055 | 0 | 0.58 | 0.25 | 0.59 | 0.71 | 0.54 | 0.69 |
| 1 | 193901655 | rs4657892 | -1.773 | 2.872 | 2.479 | 0.003 | 0.008 | -0.014 | -0.014 | 0 | 0.21 | 0.33 | 0.19 | 0.17 | 0.25 | 0.27 |
| 1 | 193917413 | rs6680840 | -1.699 | 2.849 | 2.484 | 0.029 | 0 | -0.006 | -0.006 | 3.36 | 0.17 | 0.32 | 0.19 | 0.15 | 0.18 | 0.13 |
| 1 | 193921548 | rs606149 | 3.162 | 2.846 | 2.371 | 0.025 | -0.038 | 0.145 | 0.145 | 0 | 0.51 | 0.05 | 0.52 | 0.72 | 0.44 | 0.62 |
| 1 | 193923694 | rs1570442 | -1.79 | 2.639 | 2.583 | 0.16 | 0.001 | -0.012 | -0.012 | 2.064 | 0.18 | 0.3 | 0.2 | 0.14 | 0.21 | 0.13 |
| 1 | 193929977 | rs642668 | 1.53 | 2.607 | 2.824 | 0.152 | 0.02 | -0.013 | -0.013 | 0.621 | 0.83 | 0.54 | 0.75 | 0.87 | 0.79 | 0.95 |
| 1 | 193932788 | rs17613894 | -1.803 | 2.591 | 2.366 | 0.05 | 0.007 | -0.047 | -0.047 | 0.711 | 0.19 | 0.27 | 0.15 | 0.03 | 0.22 | 0.11 |
| 1 | 193937947 | rs1110217 | -2.268 | 2.143 | 1.844 | 0.456 | -0.001 | 0 | 0 | 0 | 0.29 | 0.31 | 0.4 | 0.19 | 0.4 | 0.21 |
| 1 | 193942375 | rs1323086 | -1.082 | 1.649 | 1.878 | 0.571 | 0.004 | -0.014 | -0.014 | 0 | 0.06 | 0.09 | 0.05 | 0.01 | 0.09 | 0.06 |
| 1 | 193946970 | rs16835994 | -1.225 | 1.614 | 1.972 | 0.649 | 0.001 | -0.002 | -0.002 | 0 | 0.09 | 0.08 | 0.14 | 0.13 | 0.06 | 0.12 |
| 1 | 193952069 | rs12403843 | -1.081 | 1.641 | 2.081 | 0.706 | 0.006 | 0.012 | 0.012 | 0 | 0.05 | 0.1 | 0.11 | 0.1 | 0.06 | 0.07 |
| 1 | 193956727 | rs669015 | 3.111 | 1.515 | 2.544 | 0.669 | 0 | 0.001 | 0.001 | 0 | 0.75 | 0.25 | 0.66 | 0.77 | 0.79 | 0.73 |
| 1 | 193957678 | rs654447 | 2.863 | 1.51 | 2.786 | 0.696 | 0.005 | 0.007 | 0.007 | 0 | 0.77 | 0.37 | 0.67 | 0.75 | 0.79 | 0.73 |
| 1 | 193958320 | rs651691 | 3.233 | 1.678 | 2.676 | 0.573 | -0.1 | 0.054 | 0.054 | 0 | 0.62 | 0.06 | 0.52 | 0.74 | 0.58 | 0.62 |
| 1 | 193960753 | rs1590324 | -2.619 | 0.903 | 3.119 | 1.073 | 0.01 | 0.001 | 0.001 | 0 | 0.17 | 0.33 | 0.2 | 0.15 | 0.15 | 0.19 |
| 1 | 193965213 | rs12749556 | -3.303 | 0.901 | 3.113 | 1.072 | 0.019 | 0.016 | 0.016 | 0 | 0.23 | 0.47 | 0.32 | 0.24 | 0.21 | 0.26 |
| 1 | 193969210 | rs16836062 | -1.394 | 0.906 | 2.685 | 1.112 | 0.002 | -0.002 | -0.002 | 0 | 0.1 | 0.08 | 0.14 | 0.16 | 0.06 | 0.13 |
| 1 | 193973521 | rs617952 | 2.781 | 0.918 | 2.693 | 1.111 | 0 | 0.004 | 0.004 | 0 | 0.76 | 0.32 | 0.66 | 0.75 | 0.79 | 0.77 |
| 1 | 193982142 | rs598015 | 2.396 | 1.1 | 2.076 | 1.005 | 0.009 | -0.028 | -0.028 | 0.963 | 0.78 | 0.42 | 0.73 | 0.86 | 0.79 | 0.78 |
| 1 | 193984170 | rs1951744 | -1.476 | 1.134 | 1.627 | 0.889 | -0.002 | 0.003 | 0.003 | 0 | 0.15 | 0.19 | 0.25 | 0.21 | 0.12 | 0.2 |
| 1 | 193994024 | rs10737634 | 1.557 | 1.074 | 1.092 | 0.939 | 0.018 | -0.005 | -0.005 | 0 | 0.77 | 0.48 | 0.69 | 0.79 | 0.79 | 0.75 |
| 1 | 194011689 | rs9943194 | 1.474 | 0.361 | 0.38 | 1.577 | 0.008 | -0.066 | -0.066 | 0 | 0.6 | 0.16 | 0.49 | 0.5 | 0.63 | 0.54 |
| 1 | 194011835 | rs12731751 | -1.308 | 0.282 | 1.611 | 1.607 | -0.022 | -0.056 | -0.056 | 0 | 0.24 | 0.33 | 0.29 | 0.08 | 0.19 | 0.23 |
| 1 | 194014041 | rs11803098 | -0.621 | 0.167 | 1.682 | 1.518 | -0.005 | 0.02 | 0.02 | 0 | 0.13 | 0.05 | 0.08 | 0 | 0.13 | 0.14 |
| 1 | 194025452 | rs1442421 | 2.501 | 0.035 | 2.535 | 1.478 | -0.005 | -0.037 | -0.037 | 0 | 0.66 | 0.16 | 0.65 | 0.58 | 0.67 | 0.65 |
| 1 | 194032486 | rs10429908 | -2.194 | 0.035 | 2.656 | 1.545 | 0.011 | 0.008 | 0.008 | 0 | 0.26 | 0.35 | 0.25 | 0.32 | 0.3 | 0.34 |
| 1 | 194033705 | rs2584257 | 3.043 | 0.132 | 2.536 | 1.449 | -0.01 | -0.086 | -0.086 | 0 | 0.6 | 0.11 | 0.48 | 0.5 | 0.63 | 0.54 |
| 1 | 194036654 | rs648371 | 1.796 | 0.16 | 2.65 | 1.434 | -0.02 | -0.062 | -0.062 | 0 | 0.77 | 0.52 | 0.73 | 0.89 | 0.83 | 0.77 |
| 1 | 194041058 | rs488612 | 2.402 | 0.182 | 2.579 | 1.508 | 0 | 0.039 | 0.039 | 0 | 0.81 | 0.27 | 0.73 | 0.61 | 0.81 | 0.79 |
| 1 | 194053792 | rs12402555 | -0.308 | 0.195 | 2.486 | 1.497 | 0 | -0.023 | -0.023 | 0 | 0.07 | 0.01 | 0.16 | 0.08 | 0.04 | 0.09 |
| 1 | 194061306 | rs7543008 | -1.624 | 0.364 | 2.772 | 1.96 | -0.013 | -0.028 | -0.028 | 0 | 0.05 | 0.12 | 0.04 | 0 | 0.03 | 0.01 |
| 1 | 194062066 | rs16836246 | -2.55 | 0.36 | 2.862 | 1.958 | 0.014 | -0.01 | -0.01 | 0 | 0.07 | 0.33 | 0.05 | 0 | 0.03 | 0.01 |
| 1 | 194064837 | rs685089 | 1.597 | 0.405 | 2.787 | 2.031 | 0.076 | 0.233 | 0.233 | 0 | 0.87 | 0.53 | 0.78 | 0.63 | 0.84 | 0.81 |
| 1 | 194065895 | rs520472 | 1.973 | 0.389 | 2.756 | 2.033 | 0.019 | 0.035 | 0.035 | 0 | 0.75 | 0.36 | 0.69 | 0.63 | 0.71 | 0.67 |
| 1 | 194077406 | rs12022618 | -1.26 | 0.307 | 2.764 | 2.198 | 0.002 | 0.06 | 0.06 | 0 | 0.08 | 0.07 | 0.27 | 0.38 | 0.06 | 0.12 |
| 1 | 194080827 | rs719813 | -3.385 | 0.233 | 2.937 | 2.182 | 0.014 | 0.146 | 0.146 | 0 | 0.18 | 0.62 | 0.37 | 0.39 | 0.2 | 0.21 |
| 1 | 194083056 | rs592817 | -2.898 | 0.227 | 3.004 | 2.173 | -0.04 | 0.173 | 0.173 | 0 | 0.14 | 0.37 | 0.31 | 0.35 | 0.09 | 0.13 |
| 1 | 194084026 | rs490363 | -3.407 | -0.002 | 2.812 | 2.484 | -0.008 | -0.064 | -0.064 | 0 | 0.42 | 0.94 | 0.59 | 0.56 | 0.36 | 0.44 |
| 1 | 194084297 | rs2584256 | -2.294 | 0.396 | 3.184 | 2.567 | 0.045 | 0.195 | 0.195 | 0 | 0.12 | 0.26 | 0.32 | 0.4 | 0.17 | 0.15 |
| 1 | 194089817 | rs693833 | -2.106 | 0.005 | 3.538 | 1.607 | 0.057 | 0.042 | 0.042 | 0 | 0.06 | 0.35 | 0.05 | 0 | 0.02 | 0.01 |
| 1 | 194094713 | rs10494696 | -0.96 | 0.009 | 3.123 | 1.539 | -0.005 | 0.034 | 0.034 | 0 | 0.13 | 0.05 | 0.08 | 0.01 | 0.12 | 0.15 |
| 1 | 194100972 | rs2506081 | -2.356 | 0.01 | 2.112 | 1.603 | 0.025 | 0.023 | 0.023 | 0.049 | 0.23 | 0.82 | 0.19 | 0.12 | 0.15 | 0.23 |
| 1 | 194106280 | rs2506058 | -2.358 | -0.028 | 2.245 | 1.557 | -0.005 | -0.005 | -0.005 | 1.625 | 0.23 | 0.73 | 0.17 | 0.12 | 0.15 | 0.23 |
| 1 | 194113102 | rs2485045 | -3.078 | -0.129 | 2.85 | 1.412 | -0.03 | 0.025 | 0.025 | 0.451 | 0.1 | 0.67 | 0.09 | 0.07 | 0.04 | 0.04 |
| 1 | 194118455 | rs2485041 | -3.125 | 0.365 | 2.603 | 1.586 | -0.023 | -0.023 | -0.023 | 0.17 | 0.28 | 0.84 | 0.41 | 0.3 | 0.29 | 0.3 |
| 1 | 194120062 | rs10754088 | 2.222 | 0.445 | 3.253 | 1.547 | 0.029 | -0.12 | -0.12 | 0 | 0.82 | 0.49 | 0.83 | 0.97 | 0.83 | 0.83 |
| 1 | 194121249 | rs16836431 | -1.794 | 0.343 | 3.646 | 1.667 | 0.033 | 0.028 | 0.028 | 0 | 0.06 | 0.13 | 0.19 | 0.14 | 0.09 | 0.1 |
| 1 | 194130204 | rs1487528 | -3.739 | 0.334 | 3.88 | 1.709 | 0.07 | 0.228 | 0.228 | 0 | 0.13 | 0.43 | 0.26 | 0.27 | 0.13 | 0.15 |
| 1 | 194145754 | rs16836463 | -0.843 | 0.11 | 3.755 | 1.888 | -0.017 | 0.056 | 0.056 | 0 | 0.06 | 0.09 | 0.09 | 0.18 | 0.02 | 0.08 |
| 1 | 194147683 | rs6670222 | -3.29 | 0.356 | 3.984 | 1.207 | 0.045 | 0.04 | 0.04 | 0 | 0.11 | 0.33 | 0.21 | 0.15 | 0.11 | 0.11 |
| 1 | 194157220 | rs660889 | -1.606 | 0.297 | 4.054 | 1.175 | 0.013 | -0.078 | -0.078 | 0.098 | 0.12 | 0.3 | 0.12 | 0 | 0.12 | 0.08 |
| 1 | 194158003 | rs6678341 | -2.504 | 0.255 | 3.978 | 1.24 | -0.008 | -0.006 | -0.006 | 0 | 0.05 | 0.18 | 0.02 | 0.01 | 0.01 | 0.02 |
| 1 | 194201170 | rs4657769 | -1.661 | 0.287 | 3.72 | 1.257 | -0.006 | -0.005 | -0.005 | 0.179 | 0.06 | 0.04 | 0.18 | 0.14 | 0.09 | 0.1 |
| 1 | 194213693 | rs12067383 | -1.675 | 0.35 | 3.78 | 1.277 | 0.042 | 0.04 | 0.04 | 0 | 0.06 | 0.13 | 0.19 | 0.14 | 0.09 | 0.1 |
| 1 | 194251044 | rs302224 | 3.882 | 0.444 | 3.945 | 1.277 | 0.115 | 0.246 | 0.246 | 0.04 | 0.91 | 0.57 | 0.9 | 0.84 | 0.91 | 0.86 |

| | | | | | | | | | | | | | | | | |
|---|---|---|---|---|---|---|---|---|---|---|---|---|---|---|---|---|
| 1 | 194258816 | rs2506048 | 3.324 | 0.528 | 3.832 | 1.113 | 0.082 | -0.012 | -0.012 | 0 | 0.92 | 0.68 | 0.93 | 0.96 | 0.91 | 0.9 |
| 1 | 194266914 | rs7550778 | 3.295 | 0.512 | 3.84 | 1.081 | 0.01 | -0.044 | -0.044 | 4 | 0.75 | 0.42 | 0.68 | 0.83 | 0.75 | 0.75 |
| 1 | 194278945 | rs1325426 | 3.357 | 0.496 | 3.867 | 1.095 | -0.007 | -0.012 | -0.012 | 0 | 0.73 | 0.21 | 0.62 | 0.72 | 0.73 | 0.71 |
| 1 | 194288364 | rs16836678 | -2.209 | 0.507 | 3.836 | 1.044 | -0.002 | -0.013 | -0.013 | 0 | 0.17 | 0.25 | 0.25 | 0.13 | 0.16 | 0.15 |
| 1 | 194299382 | rs2400329 | 3.445 | 0.509 | 3.835 | 1.057 | 0.096 | -0.001 | -0.001 | 1.557 | 0.92 | 0.62 | 0.92 | 0.96 | 0.91 | 0.89 |
| 1 | 194301753 | rs17562861 | -1.265 | 0.5 | 3.722 | 1.031 | -0.022 | 0.064 | 0.064 | 0 | 0.12 | 0.02 | 0.08 | 0 | 0.13 | 0.1 |
| 1 | 194307301 | rs12083179 | -1.539 | 0.708 | 3.672 | 0.907 | 0.057 | -0.039 | -0.039 | 0 | 0.08 | 0.22 | 0.07 | 0.03 | 0.11 | 0.19 |
| 1 | 194308202 | rs6671783 | 2.497 | 0.671 | 3.682 | 0.906 | 0.052 | -0.08 | -0.08 | 0.111 | 0.86 | 0.59 | 0.89 | 0.96 | 0.85 | 0.79 |
| 1 | 194309053 | rs6700148 | -1.753 | 0.66 | 3.345 | 0.92 | -0.002 | -0.013 | -0.013 | 0.251 | 0.17 | 0.25 | 0.25 | 0.13 | 0.16 | 0.15 |
| 1 | 194322167 | rs4390154 | 2.995 | 0.646 | 3.389 | 0.888 | 0.01 | -0.056 | -0.056 | 0.008 | 0.71 | 0.45 | 0.65 | 0.83 | 0.7 | 0.61 |
| 1 | 194324826 | rs7515996 | 2.151 | 0.671 | 2.857 | 0.91 | 0.039 | -0.063 | -0.063 | 3.492 | 0.88 | 0.69 | 0.9 | 0.96 | 0.86 | 0.76 |
| 1 | 194329561 | rs10921544 | -1.096 | 0.662 | 2.795 | 0.906 | 0.006 | -0.013 | -0.013 | 0 | 0.07 | 0.07 | 0.05 | 0.03 | 0.11 | 0.19 |
| 1 | 194329737 | rs6428220 | 2.151 | 0.666 | 2.831 | 0.914 | 0.047 | -0.065 | -0.065 | 0.689 | 0.88 | 0.69 | 0.9 | 0.96 | 0.86 | 0.76 |
| 1 | 194334907 | rs4417046 | 1.564 | 0.239 | 2.296 | 1.108 | 0.037 | 0.105 | 0.105 | 0 | 0.89 | 0.63 | 0.81 | 0.84 | 0.88 | 0.76 |
| 1 | 194348173 | rs7548680 | -2.903 | 0.175 | 1.904 | 1.101 | -0.005 | 0.008 | 0.008 | 0 | 0.27 | 0.65 | 0.36 | 0.29 | 0.24 | 0.37 |
| 1 | 194351596 | rs10921548 | -1.702 | 0.18 | 0.885 | 0.812 | 0 | 0 | 0 | 0 | 0.16 | 0.17 | 0.16 | 0.14 | 0.14 | 0.15 |
| 1 | 194365120 | rs10754101 | 1.635 | -0.253 | 1.695 | 0.959 | -0.027 | 0.018 | 0.018 | 0 | 0.74 | 0.39 | 0.68 | 0.69 | 0.84 | 0.71 |
| 1 | 194368842 | rs12088170 | 1.964 | 0.314 | 1.689 | 0.601 | 0 | 0 | 0 | 0 | 0.65 | 0.39 | 0.61 | 0.67 | 0.69 | 0.5 |
| 1 | 194377156 | rs4460614 | -1.542 | 0.174 | 2.238 | 0.662 | 0.019 | 0.277 | 0.277 | 0 | 0.07 | 0.27 | 0.21 | 0.31 | 0.06 | 0.19 |
| 1 | 194403061 | rs12401747 | -2.092 | -0.278 | 2.471 | 0.423 | -0.033 | 0.006 | 0.006 | 0 | 0.24 | 0.33 | 0.3 | 0.28 | 0.16 | 0.4 |
| 1 | 194405611 | rs6428233 | 1.44 | 0.274 | 2.501 | 0.231 | -0.004 | -0.017 | -0.017 | 0.05 | 0.86 | 0.86 | 0.85 | 0.99 | 0.8 | 0.77 |
| 1 | 194407083 | rs4291483 | 2.804 | 0.272 | 2.575 | 0.233 | -0.011 | -0.018 | -0.018 | 0 | 0.61 | 0.47 | 0.55 | 0.71 | 0.65 | 0.37 |
| 1 | 194412218 | rs12121745 | -1.191 | 0.327 | 2.548 | 0.218 | -0.016 | 0.031 | 0.031 | 0 | 0.09 | 0.05 | 0.1 | 0.01 | 0.1 | 0.19 |
| 1 | 194416810 | rs4381164 | 2.936 | 0.343 | 2.655 | 0.379 | 0.013 | 0.016 | 0.016 | 0 | 0.61 | 0.3 | 0.53 | 0.69 | 0.65 | 0.37 |
| 1 | 194422151 | rs12130142 | 2.033 | -0.978 | 0.992 | -1.021 | 0.054 | 0.155 | 0.155 | 1.613 | 0.24 | 0.01 | 0.1 | 0 | 0.13 | 0.03 |
| 1 | 194445466 | rs6699952 | 2.269 | -0.538 | 1.278 | -0.512 | -0.015 | -0.018 | -0.018 | 0 | 0.62 | 0.48 | 0.55 | 0.69 | 0.65 | 0.36 |
| 1 | 194448945 | rs7514959 | -1.627 | -0.5 | 1.114 | -0.409 | -0.035 | 0.013 | 0.013 | 0 | 0.25 | 0.36 | 0.3 | 0.31 | 0.16 | 0.4 |
| 1 | 194486186 | rs16837077 | -1.33 | -0.5 | 0.972 | -0.381 | -0.001 | -0.002 | -0.002 | 0 | 0.22 | 0.25 | 0.29 | 0.3 | 0.15 | 0.4 |
| 1 | 194492493 | rs2494316 | 1.642 | -0.48 | 0.998 | -0.376 | -0.027 | 0.004 | 0.004 | 0 | 0.7 | 0.54 | 0.65 | 0.69 | 0.75 | 0.55 |
| 1 | 194506982 | rs2494164 | 1.918 | -0.465 | 0.596 | -0.433 | 0.027 | 0.046 | 0.046 | 0.042 | 0.58 | 0.18 | 0.5 | 0.69 | 0.62 | 0.37 |
| 1 | 194530914 | rs3001167 | -2.249 | -0.824 | 1.848 | -0.264 | -0.05 | 0.019 | 0.019 | 0 | 0.2 | 0.66 | 0.28 | 0.28 | 0.13 | 0.27 |
| 1 | 194537085 | rs3009382 | 1.006 | -0.737 | 1.647 | -0.483 | 0.01 | -0.033 | -0.033 | 0 | 0.9 | 0.87 | 0.82 | 0.99 | 0.85 | 0.81 |
| 1 | 194546767 | rs16837141 | -0.929 | -0.801 | 1.954 | -0.441 | -0.012 | 0.011 | 0.011 | 0 | 0.09 | 0.14 | 0.15 | 0.14 | 0.09 | 0.17 |
| 1 | 194548819 | rs10921612 | -1.609 | -1.2 | 2.259 | -0.675 | -0.034 | -0.018 | -0.018 | 0 | 0.14 | 0.29 | 0.16 | 0.1 | 0.12 | 0.18 |
| 1 | 194555331 | rs10801339 | 2.708 | -1.259 | 2.366 | -0.711 | 0.029 | 0.036 | 0.036 | 0 | 0.69 | 0.18 | 0.54 | 0.76 | 0.73 | 0.54 |
| 1 | 194558050 | rs12406807 | 2.782 | -1.227 | 2.628 | -0.695 | -0.018 | -0.022 | -0.022 | 0 | 0.71 | 0.41 | 0.55 | 0.75 | 0.73 | 0.53 |
| 1 | 194560576 | rs7523452 | 1.74 | -1.203 | 2.542 | -0.683 | 0.007 | -0.017 | -0.017 | 0.697 | 0.85 | 0.69 | 0.72 | 0.86 | 0.85 | 0.71 |
| 1 | 194564447 | rs12128318 | -0.754 | -1.273 | 2.379 | -0.563 | -0.027 | -0.004 | -0.004 | 4.182 | 0.11 | 0.17 | 0.15 | 0.1 | 0.09 | 0.16 |
| 1 | 194565350 | rs12124558 | -2.132 | -1.283 | 2.338 | -0.568 | -0.055 | 0.017 | 0.017 | 0.237 | 0.17 | 0.39 | 0.26 | 0.23 | 0.12 | 0.21 |
| 1 | 194569085 | rs3001223 | -2.575 | -0.267 | 1.743 | -0.965 | -0.001 | -0.004 | -0.004 | 1.04 | 0.32 | 0.64 | 0.41 | 0.24 | 0.33 | 0.37 |
| 1 | 194572285 | rs6691362 | -1.889 | -0.151 | 1.907 | -0.619 | 0 | 0.015 | 0.015 | 0.518 | 0.16 | 0.36 | 0.2 | 0.19 | 0.14 | 0.18 |
| 1 | 194580669 | rs3009322 | -2.757 | -0.274 | 1.871 | -0.579 | -0.005 | -0.005 | -0.005 | 0 | 0.32 | 0.62 | 0.41 | 0.31 | 0.31 | 0.42 |
| 1 | 194583153 | rs3009329 | -2.942 | -0.49 | 1.685 | -0.685 | 0.024 | 0.033 | 0.033 | 0.096 | 0.36 | 0.9 | 0.43 | 0.31 | 0.31 | 0.43 |
| 1 | 194584977 | rs10494709 | -0.928 | -0.47 | 1.922 | -0.682 | -0.003 | 0.01 | 0.01 | 0.004 | 0.1 | 0.19 | 0.14 | 0.14 | 0.08 | 0.06 |
| 1 | 194590401 | rs3001199 | -2.82 | -0.468 | 2.085 | -0.679 | -0.006 | -0.007 | -0.007 | 0 | 0.33 | 0.61 | 0.41 | 0.31 | 0.31 | 0.43 |
| 1 | 194591753 | rs13376197 | -0.279 | -0.388 | 1.872 | -0.616 | -0.008 | 0.002 | 0.002 | 0 | 0.07 | 0.18 | 0.07 | 0.07 | 0.04 | 0.13 |
| 1 | 194617533 | rs3009358 | -0.931 | -0.799 | 1.545 | -0.846 | -0.007 | -0.02 | -0.02 | 0.614 | 0.19 | 0.23 | 0.21 | 0.1 | 0.19 | 0.26 |
| 1 | 194621839 | rs3009361 | -0.931 | -0.799 | 1.545 | -0.846 | -0.007 | -0.02 | -0.02 | 0 | 0.19 | 0.23 | 0.21 | 0.1 | 0.19 | 0.26 |
| 1 | 194621938 | rs16837245 | -0.494 | -0.843 | 1.85 | -0.849 | -0.001 | 0.013 | 0.013 | 0 | 0.09 | 0.31 | 0.08 | 0.07 | 0.04 | 0.13 |
| 1 | 194623450 | rs3009362 | -2.969 | -0.885 | 1.798 | -0.869 | 0.038 | 0.044 | 0.044 | 0.021 | 0.39 | 0.92 | 0.43 | 0.31 | 0.31 | 0.44 |
| 1 | 194625139 | rs3009364 | -2.296 | -0.881 | 1.838 | -0.866 | -0.019 | -0.037 | -0.037 | 0 | 0.31 | 0.56 | 0.36 | 0.2 | 0.28 | 0.4 |
| 1 | 194635480 | rs12074264 | 0.12 | -0.185 | 1.054 | -0.115 | -0.022 | -0.03 | -0.03 | 1.703 | 0.14 | 0.06 | 0.22 | 0.39 | 0.21 | 0.25 |
| 1 | 194637756 | rs12737828 | -0.43 | -0.199 | 1.092 | -0.136 | 0.007 | 0.018 | 0.018 | 0 | 0.18 | 0.11 | 0.2 | 0.1 | 0.19 | 0.26 |
| 1 | 194638931 | rs16837273 | -0.806 | -0.206 | 1.154 | -0.14 | -0.002 | -0.005 | -0.005 | 0.213 | 0.12 | 0.32 | 0.15 | 0.1 | 0.08 | 0.15 |
| 1 | 194639330 | rs10801343 | -3.074 | -0.212 | 0.99 | -0.149 | -0.086 | 0.088 | 0.088 | 0.312 | 0.52 | 0.91 | 0.65 | 0.7 | 0.52 | 0.69 |
| 1 | 194639414 | rs10157433 | -2.384 | -0.202 | 1.014 | -0.14 | -0.004 | -0.013 | -0.013 | 0 | 0.34 | 0.73 | 0.45 | 0.6 | 0.33 | 0.43 |
| 1 | 194650446 | rs12141663 | -0.305 | -0.217 | 0.984 | -0.145 | 0.005 | 0.019 | 0.019 | 0.001 | 0.17 | 0.09 | 0.2 | 0.1 | 0.19 | 0.26 |
| 1 | 194660227 | rs7555373 | 0.091 | -0.193 | 1.109 | -0.099 | -0.002 | 0.035 | 0.035 | 0 | 0.07 | 0.21 | 0.07 | 0.11 | 0.04 | 0.03 |
| 1 | 194674393 | rs2400382 | -0.212 | -0.218 | 1.14 | -0.122 | -0.005 | 0.009 | 0.009 | 0 | 0.09 | 0.3 | 0.08 | 0.07 | 0.04 | 0.13 |
| 1 | 194676825 | rs1929237 | -3.247 | -0.018 | 0.128 | -0.867 | -0.033 | 0.122 | 0.122 | 0.126 | 0.81 | 0.98 | 0.91 | 1 | 0.83 | 0.97 |
| 1 | 194682062 | rs10801344 | -0.547 | 0.17 | 0.642 | -0.423 | 0.029 | 0.143 | 0.143 | 0 | 0.22 | 0.41 | 0.3 | 0.5 | 0.25 | 0.28 |
| 1 | 194683889 | rs6428262 | 0.688 | 0.144 | 0.745 | -0.428 | -0.003 | 0.011 | 0.011 | 0 | 0.09 | 0.29 | 0.08 | 0.07 | 0.04 | 0.13 |
| 1 | 194683991 | rs12140285 | -1.316 | 0.13 | 0.673 | -0.47 | -0.024 | -0.048 | -0.048 | 0 | 0.3 | 0.5 | 0.36 | 0.2 | 0.28 | 0.41 |
| 1 | 194691302 | rs4657928 | 0.493 | 0.144 | 0.598 | -0.424 | -0.008 | 0.005 | 0.005 | 0 | 0.09 | 0.29 | 0.07 | 0.07 | 0.04 | 0.13 |
| 1 | 194702451 | rs6665228 | 0.776 | 0.255 | 0.612 | -0.394 | 0.007 | 0.05 | 0.05 | 0 | 0.15 | 0.2 | 0.23 | 0.38 | 0.21 | 0.25 |
| 1 | 194702869 | rs12138052 | -1.178 | 0.28 | 0.67 | -0.453 | -0.011 | -0.031 | -0.031 | 0 | 0.28 | 0.44 | 0.35 | 0.24 | 0.27 | 0.31 |
| 1 | 194703530 | rs10801345 | 0.653 | 0.36 | 1.155 | -0.111 | 0.008 | 0.033 | 0.033 | 0 | 0.07 | 0.18 | 0.07 | 0.11 | 0.04 | 0.04 |
| 1 | 194706180 | rs12040494 | 0.653 | 0.361 | 1.152 | -0.111 | 0.012 | 0.039 | 0.039 | 0 | 0.07 | 0.21 | 0.07 | 0.11 | 0.04 | 0.04 |
| 1 | 194707591 | rs995805 | 0.393 | 0.368 | 1.125 | -0.105 | 0.016 | 0.086 | 0.086 | 0 | 0.16 | 0.26 | 0.23 | 0.38 | 0.21 | 0.25 |

| | | | | | | | | | | | | | | | |
|---|---|---|---|---|---|---|---|---|---|---|---|---|---|---|---|
| 1 | 194709891 | rs914273 | 0.178 | 0.015 | 0.927 | -0.268 | -0.013 | -0.002 | -0.002 | 5.211 | 0.09 | 0.25 | 0.07 | 0.07 | 0.04 | 0.13 |
| 1 | 194729943 | rs1929219 | -0.991 | 0.239 | 0.712 | -0.202 | 0.024 | 0.041 | 0.041 | 0.059 | 0.11 | 0.35 | 0.15 | 0.14 | 0.08 | 0.05 |
| 1 | 194736589 | rs2131384 | -0.202 | 0.269 | -0.07 | -0.385 | 0.017 | 0.019 | 0.019 | 0 | 0.48 | 0.39 | 0.49 | 0.57 | 0.56 | 0.43 |
| 1 | 194749075 | rs6669812 | -1.63 | -0.038 | -0.545 | -0.539 | -0.069 | -0.005 | -0.005 | 0 | 0.7 | 0.97 | 0.72 | 0.79 | 0.69 | 0.72 |
| 1 | 194760481 | rs6702232 | 0.817 | -0.271 | 0.158 | -0.547 | 0.017 | 0.008 | 0.008 | 0 | 0.05 | 0.15 | 0.05 | 0.04 | 0.04 | 0.04 |
| 1 | 194765326 | rs764552 | -0.142 | -0.591 | 0.781 | -0.651 | -0.081 | -0.001 | -0.001 | 0 | 0.16 | 0.37 | 0.08 | 0.16 | 0.07 | 0.15 |
| 1 | 194772457 | rs10754115 | -0.551 | -0.341 | 0.764 | -0.559 | 0.048 | -0.026 | -0.026 | 0 | 0.54 | 0.82 | 0.5 | 0.49 | 0.39 | 0.46 |
| 1 | 194775338 | rs6656153 | -1.185 | -0.371 | 0.967 | -0.505 | -0.012 | -0.001 | -0.001 | 0 | 0.23 | 0.7 | 0.17 | 0.23 | 0.12 | 0.24 |
| 1 | 194783067 | rs7550967 | -0.177 | -0.343 | 0.905 | -0.493 | 0.019 | 0.033 | 0.033 | 0.018 | 0.07 | 0.28 | 0.08 | 0.07 | 0.05 | 0.08 |
| 1 | 194792340 | rs12033267 | -0.041 | -0.348 | 1.194 | -0.515 | -0.046 | -0.003 | -0.003 | 0 | 0.15 | 0.19 | 0.07 | 0.16 | 0.07 | 0.15 |
| 1 | 194802115 | rs10494714 | -1.009 | -0.106 | 0.822 | -0.694 | -0.013 | -0.01 | -0.01 | 0 | 0.22 | 0.31 | 0.21 | 0.22 | 0.19 | 0.32 |
| 1 | 194804211 | rs10921671 | -1.009 | -0.068 | 0.818 | -0.694 | -0.016 | -0.014 | -0.014 | 0 | 0.22 | 0.35 | 0.21 | 0.22 | 0.19 | 0.32 |
| 1 | 194810589 | rs6686114 | 1.789 | 0.032 | 0.181 | -0.724 | -0.052 | -0.034 | -0.034 | 0 | 0.3 | 0.03 | 0.28 | 0.26 | 0.29 | 0.24 |
| 1 | 194811972 | rs1410470 | -0.707 | 0.028 | 0.218 | -0.654 | -0.035 | -0.031 | -0.031 | 0 | 0.58 | 0.81 | 0.54 | 0.48 | 0.48 | 0.57 |
| 1 | 194819196 | rs163401 | 0.234 | 0.011 | 0.18 | -0.49 | 0.029 | -0.007 | -0.007 | 2.092 | 0.12 | 0.06 | 0.06 | 0.16 | 0.06 | 0.16 |
| 1 | 194839289 | rs339620 | -0.616 | 0.565 | 0.291 | -0.658 | -0.004 | -0.005 | -0.005 | 0 | 0.72 | 0.71 | 0.6 | 0.57 | 0.65 | 0.72 |
| 1 | 194842734 | rs339567 | -0.118 | 0.202 | 0.306 | -0.82 | 0.016 | 0.02 | 0.02 | 0 | 0.31 | 0.18 | 0.24 | 0.3 | 0.24 | 0.34 |
| 1 | 194843625 | rs339565 | -0.725 | 0.299 | 0.352 | -1.008 | 0.017 | 0.028 | 0.028 | 0 | 0.42 | 0.33 | 0.31 | 0.3 | 0.33 | 0.36 |
| 1 | 194846016 | rs4657943 | 0.149 | 0.416 | 0.72 | -1.047 | 0.117 | 0.072 | 0.072 | 0 | 0.07 | 0.36 | 0.1 | 0.07 | 0.13 | 0.14 |
| 1 | 194846662 | rs339560 | -1.292 | 0.417 | 0.586 | -1.094 | 0.018 | 0.076 | 0.076 | 0 | 0.49 | 0.65 | 0.38 | 0.31 | 0.45 | 0.5 |
| 1 | 194863270 | rs1927707 | 0.756 | 0.253 | 0.615 | -1.037 | 0.081 | -0.009 | -0.009 | 0 | 0.06 | 0.21 | 0.07 | 0.01 | 0.12 | 0.13 |
| 1 | 194868558 | rs4657944 | 0.932 | 0.108 | 0.875 | -1.114 | -0.018 | -0.069 | -0.069 | 0 | 0.13 | 0.2 | 0.08 | 0 | 0.08 | 0.02 |
| 1 | 194888936 | rs339653 | 0.161 | 0.466 | 0.67 | -1.138 | -0.011 | 0.022 | 0.022 | 0 | 0.3 | 0.14 | 0.26 | 0.3 | 0.3 | 0.33 |
| 1 | 194889402 | rs7515328 | 1.384 | 0.576 | 0.578 | -0.931 | 0.001 | -0.009 | -0.009 | 0 | 0.38 | 0.4 | 0.35 | 0.27 | 0.4 | 0.39 |
| 1 | 194899193 | rs339649 | 0.721 | 0.306 | -0.067 | -0.751 | -0.011 | 0.027 | 0.027 | 0 | 0.08 | 0.12 | 0.06 | 0.22 | 0.06 | 0.16 |
| 1 | 194900910 | rs6656502 | 2.089 | 0.369 | -0.347 | -0.785 | -0.033 | 0.023 | 0.023 | 0 | 0.31 | 0.04 | 0.27 | 0.22 | 0.27 | 0.25 |
| 1 | 194902278 | rs560517 | -1.216 | 0.315 | -0.096 | -0.958 | -0.001 | 0.088 | 0.088 | 0 | 0.49 | 0.7 | 0.41 | 0.35 | 0.5 | 0.49 |
| 1 | 194908303 | rs339639 | -1.161 | 0.442 | -0.148 | -0.865 | -0.004 | 0.081 | 0.081 | 0 | 0.49 | 0.72 | 0.42 | 0.38 | 0.52 | 0.56 |
| 1 | 194909372 | rs4292939 | 1.737 | 0.341 | -0.259 | -0.65 | 0.021 | -0.003 | -0.003 | 0.073 | 0.3 | 0.13 | 0.3 | 0.3 | 0.24 | 0.19 |
| 1 | 194914012 | rs339597 | -0.629 | 0.641 | 0.505 | -0.67 | 0.011 | 0.024 | 0.024 | 0 | 0.16 | 0.34 | 0.14 | 0.22 | 0.2 | 0.29 |
| 1 | 194919249 | rs339601 | -0.541 | 0.598 | 0.484 | -0.767 | 0.008 | -0.01 | -0.01 | 0 | 0.32 | 0.52 | 0.32 | 0.3 | 0.37 | 0.49 |
| 1 | 194924039 | rs541688 | -0.293 | 0.605 | 0.57 | -0.742 | 0.025 | 0.092 | 0.092 | 0 | 0.13 | 0.4 | 0.13 | 0.23 | 0.15 | 0.23 |
| 1 | 194938031 | rs9659629 | 1.93 | 0.112 | 0.018 | -1.011 | -0.046 | -0.048 | -0.048 | 0 | 0.19 | 0.03 | 0.25 | 0.25 | 0.22 | 0.18 |
| 1 | 194940905 | rs339570 | 0.02 | 0.235 | 0.21 | -0.864 | 0.028 | 0.095 | 0.095 | 0 | 0.13 | 0.39 | 0.13 | 0.23 | 0.15 | 0.23 |
| 1 | 194941213 | rs339571 | 0.02 | 0.235 | 0.21 | -0.864 | 0.028 | 0.095 | 0.095 | 0 | 0.13 | 0.39 | 0.13 | 0.23 | 0.15 | 0.23 |
| 1 | 194945275 | rs572607 | -1.512 | -0.039 | 0.151 | -1.127 | -0.016 | -0.053 | -0.053 | 0 | 0.75 | 0.81 | 0.68 | 0.56 | 0.73 | 0.72 |
| 1 | 194950013 | rs576187 | 1.259 | -0.058 | 0.05 | -1.083 | -0.003 | 0.015 | 0.015 | 0 | 0.07 | 0.09 | 0.06 | 0.22 | 0.06 | 0.12 |
| 1 | 194955350 | rs6693848 | 0.389 | -0.054 | -0.02 | -1.222 | 0.019 | 0.054 | 0.054 | 0 | 0.43 | 0.27 | 0.36 | 0.25 | 0.36 | 0.23 |
| 1 | 194965856 | rs11583374 | 1.451 | 0.136 | -0.327 | -1.085 | -0.013 | 0.029 | 0.029 | 0.11 | 0.19 | 0.12 | 0.19 | 0.08 | 0.22 | 0.26 |
| 1 | 194975253 | rs4525024 | 1.488 | 0.104 | -0.465 | -1.123 | -0.021 | 0.055 | 0.055 | 0.156 | 0.19 | 0.06 | 0.17 | 0.06 | 0.21 | 0.25 |
| 1 | 194980508 | rs2210331 | -0.051 | 0.166 | 0.098 | -1.165 | -0.002 | 0.01 | 0.01 | 0 | 0.6 | 0.32 | 0.53 | 0.32 | 0.58 | 0.49 |
| 1 | 194993511 | rs10754124 | -1.051 | 0.142 | 0.502 | -1.095 | 0.01 | 0.084 | 0.084 | 0 | 0.72 | 0.52 | 0.65 | 0.53 | 0.73 | 0.7 |
| 1 | 195001536 | rs12121014 | 1.314 | -0.006 | 0.344 | -1.293 | 0.004 | -0.024 | -0.024 | 0 | 0.19 | 0.15 | 0.1 | 0 | 0.13 | 0.04 |
| 1 | 195015798 | rs12032648 | 0.683 | 0.052 | 0.365 | -1.199 | 0.001 | 0.021 | 0.021 | 0.05 | 0.12 | 0.16 | 0.1 | 0.21 | 0.15 | 0.21 |
| 1 | 195019104 | rs10801376 | 0.461 | 0.025 | 0.512 | -1.322 | 0.004 | 0.019 | 0.019 | 0 | 0.4 | 0.3 | 0.36 | 0.26 | 0.36 | 0.23 |
| 1 | 195031417 | rs12040659 | 0.848 | 0.051 | 0.591 | -0.839 | 0 | 0.012 | 0.012 | 0.071 | 0.12 | 0.11 | 0.12 | 0.25 | 0.15 | 0.22 |
| 1 | 195032861 | rs1934206 | -0.987 | 0.061 | 0.523 | -0.865 | 0.003 | 0.045 | 0.045 | 0 | 0.71 | 0.52 | 0.65 | 0.57 | 0.72 | 0.7 |
| 1 | 195041705 | rs17649943 | 1.566 | 0.141 | 0.421 | -0.974 | -0.021 | 0.063 | 0.063 | 0 | 0.19 | 0.05 | 0.17 | 0.06 | 0.21 | 0.25 |
| 1 | 195044251 | rs16837755 | 0.504 | 0.262 | 0.417 | -0.925 | 0 | 0 | 0 | 0 | 0.18 | 0.16 | 0.19 | 0.23 | 0.25 | 0.22 |
| 1 | 195050507 | rs696393 | -1.782 | 0.161 | 0.301 | -0.81 | -0.051 | -0.15 | -0.15 | 0 | 0.8 | 0.97 | 0.75 | 0.55 | 0.83 | 0.71 |
| 1 | 195052656 | rs980422 | -1.348 | 0.184 | 0.54 | -0.799 | -0.001 | 0.076 | 0.076 | 0 | 0.77 | 0.6 | 0.72 | 0.55 | 0.83 | 0.71 |
| 1 | 195053221 | rs696392 | -0.327 | 0.165 | 0.424 | -0.883 | 0.004 | -0.007 | -0.007 | 0.186 | 0.6 | 0.64 | 0.64 | 0.55 | 0.7 | 0.67 |
| 1 | 195053488 | rs696390 | -0.863 | 0.449 | 0.486 | -0.802 | -0.001 | 0.001 | 0.001 | 0 | 0.59 | 0.59 | 0.48 | 0.3 | 0.6 | 0.53 |
| 1 | 195058871 | rs12067603 | 1.487 | 0.255 | 0.081 | -0.83 | 0.009 | -0.048 | -0.048 | 0 | 0.23 | 0.15 | 0.26 | 0.45 | 0.17 | 0.31 |
| 1 | 195072336 | rs2785795 | -1.696 | 0.16 | -0.152 | -1.115 | -0.035 | 0.078 | 0.078 | 0.065 | 0.8 | 0.94 | 0.79 | 0.95 | 0.78 | 0.75 |
| 1 | 195082738 | rs1542731 | 1.64 | 0.196 | -0.243 | -1.137 | 0.001 | -0.026 | -0.026 | 0.52 | 0.17 | 0.14 | 0.09 | 0 | 0.12 | 0.06 |
| 1 | 195085692 | rs6684597 | 0.34 | 0.26 | -0.277 | -1.187 | 0 | 0.007 | 0.007 | 0 | 0.2 | 0.19 | 0.2 | 0.28 | 0.26 | 0.2 |
| 1 | 195095179 | rs822456 | 0.298 | 0.286 | -0.404 | -1.147 | 0.003 | -0.004 | -0.004 | 0 | 0.4 | 0.38 | 0.35 | 0.46 | 0.3 | 0.38 |
| 1 | 195111306 | rs635815 | 0.075 | 0.313 | -0.449 | -1.145 | 0 | -0.001 | -0.001 | 0.234 | 0.57 | 0.46 | 0.54 | 0.67 | 0.52 | 0.53 |
| 1 | 195113428 | rs12067771 | 1.782 | 0.238 | -0.432 | -1.187 | 0.001 | -0.003 | -0.003 | 0.227 | 0.23 | 0.21 | 0.27 | 0.45 | 0.17 | 0.28 |
| 1 | 195116480 | rs10921803 | 2.019 | 0.226 | -0.444 | -1.213 | -0.027 | -0.005 | -0.005 | 0 | 0.18 | 0.09 | 0.21 | 0.22 | 0.23 | 0.17 |
| 1 | 195122910 | rs581571 | 0.497 | 0.258 | -0.443 | -1.178 | -0.007 | 0.045 | 0.045 | 0 | 0.34 | 0.25 | 0.28 | 0.22 | 0.35 | 0.25 |
| 1 | 195132593 | rs598224 | 0.115 | 0.325 | -0.457 | -1.128 | 0 | 0 | 0 | 0.447 | 0.57 | 0.46 | 0.55 | 0.67 | 0.52 | 0.53 |
| 1 | 195134340 | rs643532 | -1.386 | 0.296 | -0.398 | -1.219 | 0.004 | -0.087 | -0.087 | 0.348 | 0.77 | 0.62 | 0.75 | 0.95 | 0.78 | 0.75 |
| 1 | 195139365 | rs603495 | -1.283 | 0.351 | -0.715 | -1.193 | 0.001 | -0.109 | -0.109 | 1.16 | 0.76 | 0.58 | 0.75 | 0.95 | 0.78 | 0.75 |
| 1 | 195141706 | rs17607167 | 1.28 | 0.656 | -1.21 | -0.878 | 0.008 | 0.006 | 0.006 | 0.233 | 0.17 | 0.11 | 0.07 | 0 | 0.12 | 0.06 |
| 1 | 195142993 | rs11581486 | -0.102 | 1.177 | -0.997 | -0.146 | -0.006 | -0.02 | -0.02 | 0 | 0.19 | 0.12 | 0.2 | 0.27 | 0.26 | 0.22 |
| 1 | 195144412 | rs592350 | 1.289 | 1.188 | -1.005 | -0.136 | -0.019 | -0.005 | -0.005 | 0.635 | 0.17 | 0.13 | 0.21 | 0.22 | 0.23 | 0.17 |
| 1 | 195145500 | rs3845553 | -0.599 | 1.094 | -0.605 | 0.077 | -0.006 | -0.02 | -0.02 | 0 | 0.19 | 0.11 | 0.2 | 0.27 | 0.26 | 0.22 |

| | | | | | | | | | | | | | | | | |
|---|---|---|---|---|---|---|---|---|---|---|---|---|---|---|---|---|
| 1 | 195155161 | rs11581720 | -0.613 | 1.063 | -0.59 | 0.063 | -0.003 | -0.018 | -0.018 | 0 | 0.19 | 0.11 | 0.2 | 0.27 | 0.27 | 0.21 |
| 1 | 195155293 | rs656518 | -0.211 | 1.371 | 0.054 | -0.36 | 0.027 | -0.091 | -0.091 | 0.407 | 0.8 | 0.59 | 0.67 | 0.94 | 0.8 | 0.73 |
| 1 | 195164476 | rs613938 | -0.196 | 1.476 | -0.348 | 0.066 | -0.002 | -0.009 | -0.009 | 0 | 0.64 | 0.73 | 0.72 | 0.59 | 0.64 | 0.71 |
| 1 | 195166911 | rs12755016 | 0.882 | 2.025 | -0.725 | -0.121 | -0.044 | 0.045 | 0.045 | 0.147 | 0.1 | 0.02 | 0.1 | 0 | 0.2 | 0.03 |
| 1 | 195170004 | rs687523 | 0.152 | 2.122 | -0.543 | -0.338 | 0.014 | -0.029 | -0.029 | 2.434 | 0.13 | 0.15 | 0.16 | 0.01 | 0.15 | 0.13 |
| 1 | 195191693 | rs6688980 | 0.927 | 2.043 | -0.256 | -0.149 | 0.005 | 0.007 | 0.007 | 0 | 0.47 | 0.57 | 0.49 | 0.39 | 0.44 | 0.46 |
| 1 | 195195201 | rs4657974 | -0.279 | 2.051 | -0.479 | -0.052 | -0.027 | -0.031 | -0.031 | 0 | 0.65 | 0.82 | 0.72 | 0.59 | 0.62 | 0.7 |
| 1 | 195202201 | rs2031953 | -0.29 | 2.012 | -0.585 | -0.047 | -0.037 | -0.047 | -0.047 | 0 | 0.65 | 0.85 | 0.73 | 0.59 | 0.62 | 0.7 |
| 1 | 195219984 | rs2183227 | -0.956 | 1.934 | -0.699 | -0.019 | 0.009 | -0.105 | -0.105 | 0 | 0.74 | 0.93 | 0.82 | 0.56 | 0.81 | 0.73 |
| 1 | 195224639 | rs4657975 | -0.143 | 2.045 | -0.77 | -0.1 | -0.036 | -0.071 | -0.071 | 0 | 0.63 | 0.86 | 0.73 | 0.56 | 0.62 | 0.69 |
| 1 | 195239349 | rs12024167 | -0.208 | 2.154 | -0.644 | -0.162 | -0.012 | -0.011 | -0.011 | 0 | 0.16 | 0.11 | 0.23 | 0.2 | 0.21 | 0.24 |
| 1 | 195241106 | rs16838214 | 1.463 | 1.895 | -0.191 | -0.475 | 0.014 | 0.018 | 0.018 | 0 | 0.38 | 0.28 | 0.43 | 0.36 | 0.37 | 0.44 |
| 1 | 195241681 | rs17609129 | -0.046 | 2.173 | -0.495 | -0.322 | -0.012 | -0.011 | -0.011 | 0 | 0.16 | 0.11 | 0.23 | 0.2 | 0.21 | 0.24 |
| 1 | 195242066 | rs7551364 | -1.585 | 2.122 | -0.814 | -0.36 | 0.001 | -0.093 | -0.093 | 0.907 | 0.77 | 0.93 | 0.82 | 0.61 | 0.83 | 0.75 |
| 1 | 195247438 | rs2026082 | 1.427 | 2.12 | -1.261 | -0.615 | -0.03 | 0.03 | 0.03 | 0.04 | 0.11 | 0.06 | 0.1 | 0 | 0.2 | 0.04 |
| 1 | 195262063 | rs17661321 | 0.666 | 1.741 | -1.164 | -0.367 | -0.054 | 0.034 | 0.034 | 0 | 0.1 | 0.01 | 0.15 | 0.01 | 0.15 | 0.12 |
| 1 | 195267072 | rs6428282 | 0.14 | 1.352 | -0.81 | -0.257 | -0.003 | 0.01 | 0.01 | 0 | 0.55 | 0.44 | 0.55 | 0.36 | 0.57 | 0.48 |
| 1 | 195277736 | rs1324707 | -1.202 | 1.725 | -1.454 | -0.615 | -0.083 | 0.045 | 0.045 | 0 | 0.39 | 0.85 | 0.43 | 0.22 | 0.45 | 0.4 |
| 1 | 195279977 | rs16838261 | 0.551 | 2.002 | -1.285 | -0.736 | 0 | -0.008 | -0.008 | 0 | 0.08 | 0.07 | 0.1 | 0 | 0.1 | 0.11 |
| 1 | 195280424 | rs12068912 | -0.812 | 2.389 | -1.143 | -0.781 | 0.03 | -0.008 | -0.008 | 1.583 | 0.14 | 0.68 | 0.19 | 0.08 | 0.16 | 0.1 |
| 1 | 195288287 | rs11587011 | 1.282 | 2.106 | -1.36 | -0.803 | -0.021 | 0.026 | 0.026 | 0 | 0.12 | 0.06 | 0.09 | 0 | 0.16 | 0.05 |
| 1 | 195291956 | rs12409559 | -1.495 | 1.724 | -1.4 | -0.624 | -0.085 | 0.033 | 0.033 | 0 | 0.39 | 0.85 | 0.43 | 0.22 | 0.44 | 0.4 |
| 1 | 195303475 | rs10921835 | -1.114 | 1.811 | -1.312 | -0.73 | -0.089 | 0.113 | 0.113 | 0.047 | 0.34 | 0.84 | 0.37 | 0.08 | 0.41 | 0.25 |
| 1 | 195316969 | rs12731553 | 0.981 | 1.034 | -1.503 | -0.744 | -0.008 | 0.008 | 0.008 | 0.713 | 0.25 | 0.19 | 0.3 | 0.28 | 0.32 | 0.24 |
| 1 | 195318164 | rs4657806 | -1.1 | 0.655 | -1.743 | -0.886 | 0.041 | -0.115 | -0.115 | 0.485 | 0.68 | 0.9 | 0.71 | 0.5 | 0.77 | 0.68 |
| 1 | 195341727 | rs12125032 | -0.218 | 0.927 | -1.558 | -1.122 | -0.015 | -0.038 | -0.038 | 0.446 | 0.22 | 0.08 | 0.26 | 0.22 | 0.21 | 0.38 |
| 1 | 195343788 | rs17668176 | 1.981 | 0.79 | -1.921 | -1.303 | -0.048 | 0.048 | 0.048 | 0.054 | 0.2 | 0.04 | 0.16 | 0.12 | 0.28 | 0.19 |
| 1 | 195346285 | rs10754139 | -1.329 | 0.776 | -0.896 | -1.275 | 0.006 | -0.009 | -0.009 | 0.343 | 0.68 | 0.7 | 0.7 | 0.48 | 0.77 | 0.69 |
| 1 | 195348938 | rs12729977 | -0.198 | 0.75 | -1.145 | -1.668 | 0.006 | 0.013 | 0.013 | 0 | 0.52 | 0.58 | 0.47 | 0.27 | 0.61 | 0.37 |
| 1 | 195350153 | rs12563798 | 0.218 | 1.007 | -0.652 | -1.425 | -0.011 | 0.051 | 0.051 | 0 | 0.42 | 0.32 | 0.4 | 0.27 | 0.55 | 0.31 |
| 1 | 195363004 | rs6690599 | 1.717 | 1.016 | -0.674 | -1.435 | -0.076 | 0.09 | 0.09 | 0 | 0.15 | 0.02 | 0.09 | 0 | 0.23 | 0.05 |
| 1 | 195363412 | rs6428283 | 0.285 | 0.797 | -0.466 | -1.342 | -0.007 | 0.034 | 0.034 | 0 | 0.42 | 0.37 | 0.4 | 0.27 | 0.55 | 0.3 |
| 1 | 195376972 | rs6702266 | -0.487 | 0.734 | -0.449 | -1.307 | -0.014 | -0.011 | -0.011 | 0 | 0.54 | 0.53 | 0.6 | 0.36 | 0.69 | 0.52 |
| 1 | 195384254 | rs6692973 | 0.234 | 0.771 | -0.47 | -1.318 | -0.01 | 0.057 | 0.057 | 0 | 0.43 | 0.33 | 0.4 | 0.27 | 0.55 | 0.3 |
| 1 | 195385150 | rs1998647 | 0.082 | 1.005 | 0.046 | -1.38 | 0.05 | 0.002 | 0.002 | 0 | 0.87 | 0.41 | 0.79 | 0.9 | 0.88 | 0.83 |
| 1 | 195386897 | rs7538519 | 0.414 | 0.994 | -0.031 | -1.424 | 0.039 | 0.016 | 0.016 | 0 | 0.1 | 0.22 | 0.18 | 0.1 | 0.12 | 0.18 |
| 1 | 195388740 | rs12754009 | 1.502 | 1.175 | -0.278 | -1.566 | -0.036 | 0.039 | 0.039 | 0 | 0.15 | 0.07 | 0.1 | 0 | 0.22 | 0.05 |
| 1 | 195390646 | rs16838352 | 0.835 | 1.178 | -0.318 | -1.583 | 0.05 | -0.119 | -0.119 | 0 | 0.19 | 0.38 | 0.13 | 0 | 0.24 | 0.07 |
| 1 | 195392852 | rs2119441 | -1.522 | 0.923 | -0.638 | -1.612 | 0.035 | -0.084 | -0.084 | 0 | 0.71 | 0.92 | 0.68 | 0.39 | 0.79 | 0.68 |
| 1 | 195393613 | rs7523327 | 0.223 | 0.929 | -0.644 | -1.66 | -0.043 | 0.062 | 0.062 | 1.434 | 0.39 | 0.25 | 0.48 | 0.27 | 0.49 | 0.44 |
| 1 | 195393642 | rs7515446 | -1.48 | 0.976 | -0.862 | -1.556 | 0.032 | -0.094 | -0.094 | 0 | 0.71 | 0.94 | 0.68 | 0.39 | 0.79 | 0.68 |
| 1 | 195397391 | rs11803658 | 1.466 | 1.029 | -0.747 | -1.509 | 0.047 | -0.1 | -0.1 | 0.156 | 0.16 | 0.33 | 0.12 | 0 | 0.22 | 0.05 |
| 1 | 195399786 | rs16838372 | 0.829 | 0.313 | -1.417 | -1.145 | -0.025 | -0.016 | -0.016 | 0 | 0.09 | 0.05 | 0.16 | 0.1 | 0.12 | 0.18 |
| 1 | 195416104 | rs12403940 | 0.842 | 0.321 | -1.407 | -1.173 | -0.019 | -0.012 | -0.012 | 0 | 0.09 | 0.05 | 0.16 | 0.1 | 0.12 | 0.18 |
| 1 | 195420070 | rs12128390 | 0.919 | -0.16 | -1.374 | -1.051 | 0.011 | -0.049 | -0.049 | 0 | 0.11 | 0 | 0.05 | 0.13 | 0.07 | 0.17 |
| 1 | 195424142 | rs7515652 | -0.398 | -0.084 | -1.322 | -1.166 | 0.077 | 0.087 | 0.087 | 0.094 | 0.6 | 0.88 | 0.64 | 0.26 | 0.72 | 0.51 |
| 1 | 195442756 | rs12732243 | 0.282 | 0.495 | -0.724 | -1.111 | 0.008 | 0.014 | 0.014 | 0.154 | 0.49 | 0.59 | 0.44 | 0.6 | 0.61 | 0.42 |
| 1 | 195443569 | rs1361350 | -0.854 | 0.373 | -0.972 | -1.289 | 0.026 | 0.053 | 0.053 | 1.04 | 0.69 | 0.81 | 0.81 | 0.88 | 0.78 | 0.74 |
| 1 | 195444261 | rs6695362 | -1.662 | 0.279 | -1.034 | -1.372 | 0.028 | 0.044 | 0.044 | 3.93 | 0.78 | 0.89 | 0.87 | 0.93 | 0.85 | 0.89 |
| 1 | 195448403 | rs10921854 | 0.045 | 0.525 | -0.795 | -1.089 | -0.032 | -0.07 | -0.07 | 0.206 | 0.18 | 0.04 | 0.36 | 0.28 | 0.17 | 0.33 |
| 1 | 195448770 | rs10754140 | -0.727 | 0.84 | -0.373 | -0.948 | 0.012 | 0.007 | 0.007 | 1.558 | 0.68 | 0.75 | 0.81 | 0.88 | 0.78 | 0.74 |
| 1 | 195454557 | rs4657983 | -0.359 | 1.245 | -0.328 | -0.99 | -0.001 | -0.048 | -0.048 | 0 | 0.65 | 0.63 | 0.74 | 0.88 | 0.64 | 0.71 |
| 1 | 195456425 | rs2888961 | -0.823 | 1.075 | -0.83 | -0.964 | 0.017 | 0.028 | 0.028 | 3.482 | 0.71 | 0.79 | 0.82 | 0.88 | 0.8 | 0.75 |
| 1 | 195467306 | rs7541185 | -0.844 | 1.084 | -0.826 | -0.96 | 0.016 | 0.027 | 0.027 | 2.579 | 0.71 | 0.79 | 0.82 | 0.88 | 0.8 | 0.75 |
| 1 | 195473977 | rs10921859 | -0.99 | 1.082 | -0.882 | -0.961 | 0.025 | 0.078 | 0.078 | 1.833 | 0.72 | 0.86 | 0.82 | 0.88 | 0.8 | 0.75 |
| 1 | 195477551 | rs10754143 | 0.564 | 1.232 | -0.755 | -0.901 | 0.001 | 0.014 | 0.014 | 0.631 | 0.52 | 0.43 | 0.38 | 0.6 | 0.48 | 0.42 |
| 1 | 195487796 | rs1339556 | 0.749 | 1.129 | -0.859 | -0.936 | -0.047 | 0.049 | 0.049 | 0.679 | 0.5 | 0.13 | 0.35 | 0.6 | 0.48 | 0.42 |
| 1 | 195506117 | rs12038043 | 0.382 | 1.014 | -0.842 | -0.902 | -0.051 | -0.075 | -0.075 | 0 | 0.12 | 0.02 | 0.34 | 0.28 | 0.15 | 0.28 |
| 1 | 195507334 | rs1416750 | 0.708 | 0.994 | -0.689 | -0.931 | 0.01 | 0.045 | 0.045 | 0.424 | 0.51 | 0.36 | 0.37 | 0.62 | 0.47 | 0.43 |
| 1 | 195515431 | rs2773454 | 0.702 | 1.019 | -0.671 | -1.013 | 0.009 | 0.048 | 0.048 | 0.477 | 0.51 | 0.37 | 0.37 | 0.6 | 0.47 | 0.43 |
| 1 | 195527078 | rs2790068 | 0.643 | 0.974 | -0.653 | -1.009 | 0.007 | 0.029 | 0.029 | 0.477 | 0.51 | 0.42 | 0.38 | 0.6 | 0.47 | 0.42 |
| 1 | 195531423 | rs1890645 | -0.685 | 1.13 | -0.604 | -0.941 | 0.069 | 0.047 | 0.047 | 0 | 0.18 | 0.32 | 0.44 | 0.27 | 0.31 | 0.32 |
| 1 | 195537767 | rs2480756 | -0.8 | 0.844 | -1.188 | -0.705 | 0.024 | 0.084 | 0.084 | 1.504 | 0.71 | 0.85 | 0.82 | 0.88 | 0.78 | 0.74 |
| 1 | 195546330 | rs12024378 | 0.117 | 0.862 | -1.184 | -0.701 | -0.016 | -0.031 | -0.031 | 0 | 0.13 | 0.09 | 0.36 | 0.27 | 0.15 | 0.27 |
| 1 | 195551814 | rs2480761 | -0.396 | 0.8 | -2.28 | -0.943 | 0.02 | 0.08 | 0.08 | 1.348 | 0.71 | 0.86 | 0.81 | 0.86 | 0.78 | 0.74 |
| 1 | 195552999 | rs1538595 | -0.401 | 0.722 | -2.265 | -1.031 | 0.025 | 0.086 | 0.086 | 1.339 | 0.71 | 0.85 | 0.82 | 0.88 | 0.78 | 0.74 |
| 1 | 195567234 | rs2993831 | -0.258 | 0.505 | -2.309 | -0.944 | -0.023 | 0.26 | 0.26 | 0 | 0.52 | 0.81 | 0.68 | 0.88 | 0.49 | 0.68 |
| 1 | 195568383 | rs12078156 | -0.425 | -0.37 | -2.373 | 0.446 | 0.048 | -0.123 | -0.123 | 0.656 | 0.26 | 0.09 | 0.22 | 0.46 | 0.2 | 0.35 |
| 1 | 195569614 | rs10494727 | -0.299 | -0.318 | -2.783 | 0.63 | -0.062 | -0.11 | -0.11 | 0 | 0.32 | 0.05 | 0.19 | 0.44 | 0.33 | 0.41 |

| | | | | | | | | | | | | | | | | |
|---|---|---|---|---|---|---|---|---|---|---|---|---|---|---|---|---|
| 1 | 195576106 | rs6699138 | -0.206 | -0.83 | -1.555 | 0.655 | 0.006 | -0.003 | -0.003 | 0 | 0.58 | 0.45 | 0.55 | 0.55 | 0.69 | 0.66 |
| 1 | 195606741 | rs10921881 | -0.026 | -0.92 | -1.4 | 0.646 | 0.11 | -0.047 | -0.047 | 0.475 | 0.56 | 0.17 | 0.55 | 0.55 | 0.7 | 0.66 |
| 1 | 195629083 | rs1556455 | -0.582 | -1.138 | -1.11 | 0.587 | -0.07 | 0.028 | 0.028 | 2.012 | 0.64 | 0.58 | 0.63 | 0.45 | 0.8 | 0.69 |
| 1 | 195632382 | rs2150899 | 0.268 | -1.149 | -1.171 | 0.564 | 0.124 | -0.062 | -0.062 | 0 | 0.55 | 0.13 | 0.56 | 0.45 | 0.7 | 0.66 |
| 1 | 195632577 | rs2183858 | -0.467 | -1.128 | -0.922 | 0.576 | 0.021 | -0.012 | -0.012 | 0 | 0.63 | 0.37 | 0.63 | 0.45 | 0.8 | 0.69 |
| 1 | 195646909 | rs10921887 | 0.227 | -1.124 | -0.965 | 0.577 | 0.122 | -0.051 | -0.051 | 0 | 0.56 | 0.15 | 0.56 | 0.45 | 0.7 | 0.66 |
| 1 | 195658978 | rs7526474 | -0.137 | -0.541 | -0.774 | 0.769 | 0.028 | -0.036 | -0.036 | 0 | 0.07 | 0.2 | 0.07 | 0 | 0.1 | 0.03 |
| 1 | 195659664 | rs12122249 | 0.39 | -0.917 | -0.902 | 1.363 | -0.001 | 0.071 | 0.071 | 0 | 0.06 | 0.05 | 0.08 | 0.32 | 0.05 | 0.06 |
| 1 | 195672584 | rs10754147 | -0.29 | -1.186 | -0.741 | 0.873 | -0.05 | 0.022 | 0.022 | 3.039 | 0.63 | 0.56 | 0.65 | 0.45 | 0.79 | 0.68 |
| 1 | 195685800 | rs10921897 | -0.147 | -0.709 | -0.329 | 0.829 | -0.001 | 0.014 | 0.014 | 0 | 0.07 | 0.07 | 0.13 | 0.24 | 0.05 | 0.06 |
| 1 | 195686601 | rs17672688 | 0.31 | -0.137 | -0.364 | 0.167 | -0.025 | 0.034 | 0.034 | 0.589 | 0.29 | 0.23 | 0.35 | 0.11 | 0.37 | 0.26 |
| 1 | 195687201 | rs12035852 | -0.714 | -0.201 | 0.228 | 0.259 | 0.007 | -0.013 | -0.013 | 0 | 0.4 | 0.59 | 0.46 | 0.3 | 0.5 | 0.54 |
| 1 | 195702820 | rs2895786 | -0.392 | -0.22 | 0.338 | 0.259 | -0.016 | 0.014 | 0.014 | 0.054 | 0.37 | 0.34 | 0.44 | 0.3 | 0.5 | 0.54 |
| 1 | 195707490 | rs12048401 | 0.064 | -0.194 | 0.315 | 0.269 | -0.008 | -0.014 | -0.014 | 0 | 0.08 | 0.07 | 0.08 | 0.19 | 0.13 | 0.28 |
| 1 | 195714307 | rs12034483 | -0.286 | -0.24 | 0.4 | 0.252 | -0.057 | 0.07 | 0.07 | 0 | 0.36 | 0.17 | 0.43 | 0.24 | 0.49 | 0.5 |
| 1 | 195724306 | rs9943101 | -1.362 | 0.041 | 0.842 | 0.902 | 0.047 | 0.181 | 0.181 | 0 | 0.11 | 0.45 | 0.11 | 0.26 | 0.13 | 0.33 |
| 1 | 195726127 | rs12028579 | -0.128 | 0.144 | 0.35 | 0.892 | -0.03 | -0.041 | -0.041 | 0.394 | 0.08 | 0.01 | 0.08 | 0.18 | 0.12 | 0.28 |
| 1 | 195736070 | rs10494730 | -0.128 | -0.02 | 0.486 | 0.569 | -0.006 | 0.023 | 0.023 | 0.907 | 0.43 | 0.37 | 0.34 | 0.3 | 0.5 | 0.56 |
| 1 | 195738953 | rs12035482 | -0.313 | -0.141 | 0.947 | 0.528 | -0.001 | 0.055 | 0.055 | 0.113 | 0.46 | 0.68 | 0.38 | 0.3 | 0.5 | 0.56 |
| 1 | 195745830 | rs2400664 | 0.017 | -0.111 | 0.839 | 0.545 | -0.002 | 0.036 | 0.036 | 0.102 | 0.55 | 0.36 | 0.63 | 0.7 | 0.5 | 0.45 |
| 1 | 195745968 | rs2989965 | 0.094 | -0.112 | 0.705 | 0.545 | -0.003 | 0.073 | 0.073 | 0.09 | 0.54 | 0.29 | 0.62 | 0.7 | 0.5 | 0.43 |
| 1 | 195747936 | rs2942921 | -0.758 | 0.119 | -0.185 | 0.546 | -0.023 | 0.064 | 0.064 | 0.342 | 0.64 | 0.74 | 0.73 | 0.88 | 0.61 | 0.69 |
| 1 | 195749012 | rs12042375 | 0.596 | 0.286 | -0.437 | 0.688 | -0.027 | -0.028 | -0.028 | 0 | 0.06 | 0.01 | 0.08 | 0.19 | 0.1 | 0.26 |
| 1 | 195757610 | rs2225518 | 0.737 | 0.069 | -0.59 | 0.512 | -0.075 | -0.141 | -0.141 | 0.98 | 0.43 | 0.04 | 0.45 | 0.43 | 0.45 | 0.37 |
| 1 | 195763875 | rs2942926 | -0.143 | 0.287 | 0.342 | -0.141 | -0.017 | 0.07 | 0.07 | 0 | 0.62 | 0.75 | 0.72 | 0.88 | 0.61 | 0.7 |
| 1 | 195765740 | rs2989984 | 0.041 | 0.353 | 0.323 | -0.109 | 0 | -0.035 | -0.035 | 0 | 0.59 | 0.41 | 0.69 | 0.88 | 0.61 | 0.69 |
| 1 | 195773441 | rs3010167 | -0.028 | 0.383 | 0.084 | -0.114 | -0.006 | 0.022 | 0.022 | 0 | 0.6 | 0.66 | 0.7 | 0.89 | 0.61 | 0.7 |
| 1 | 195774752 | rs3010171 | -0.034 | 0.388 | -0.071 | -0.112 | -0.012 | 0.062 | 0.062 | 0 | 0.6 | 0.68 | 0.71 | 0.89 | 0.61 | 0.7 |
| 1 | 195797849 | rs4604678 | -0.3 | 0.378 | -0.045 | -0.103 | -0.054 | 0.151 | 0.151 | 0.04 | 0.63 | 0.84 | 0.72 | 0.88 | 0.61 | 0.71 |
| 1 | 195804144 | rs10754156 | -0.006 | 0.419 | -0.018 | -0.097 | -0.006 | 0.01 | 0.01 | 0.017 | 0.6 | 0.59 | 0.71 | 0.88 | 0.6 | 0.69 |
| 1 | 195811203 | rs7519009 | -0.23 | 0.449 | -0.368 | -0.083 | -0.07 | 0.164 | 0.164 | 0.088 | 0.62 | 0.85 | 0.72 | 0.88 | 0.6 | 0.69 |
| 1 | 195825304 | rs7530086 | 0.117 | 0.43 | 0.047 | -0.113 | 0.002 | -0.062 | -0.062 | 0 | 0.58 | 0.42 | 0.69 | 0.88 | 0.6 | 0.69 |
| 1 | 195826632 | rs6679783 | -0.085 | 0.421 | 0.071 | -0.118 | -0.012 | 0.058 | 0.058 | 0 | 0.6 | 0.68 | 0.7 | 0.88 | 0.6 | 0.69 |
| 1 | 195826738 | rs6679891 | -0.085 | 0.421 | 0.071 | -0.118 | -0.012 | 0.058 | 0.058 | 0 | 0.6 | 0.68 | 0.7 | 0.88 | 0.6 | 0.69 |
| 1 | 195832997 | rs7542834 | -0.115 | 0.649 | -1.303 | 0.591 | 0.082 | 0.067 | 0.067 | 0.218 | 0.5 | 0.8 | 0.57 | 0.64 | 0.36 | 0.55 |
| 1 | 195849028 | rs7546529 | 0.038 | 0.831 | -1.619 | 0.326 | 0.011 | 0.013 | 0.013 | 0 | 0.36 | 0.18 | 0.24 | 0.31 | 0.34 | 0.35 |
| 1 | 195852964 | rs2923891 | 0.356 | 0.927 | -1.978 | 0.55 | -0.104 | -0.054 | -0.054 | 0 | 0.19 | 0.01 | 0.32 | 0.24 | 0.3 | 0.11 |
| 1 | 195853219 | rs2923889 | 0.27 | 0.752 | -1.863 | 0.598 | -0.018 | 0.027 | 0.027 | 0 | 0.5 | 0.12 | 0.59 | 0.46 | 0.5 | 0.31 |
| 1 | 195854637 | rs921516 | 0.317 | 0.197 | -1.292 | 0.024 | 0.011 | 0.018 | 0.018 | 0 | 0.54 | 0.27 | 0.48 | 0.53 | 0.52 | 0.54 |
| 1 | 195859022 | rs12121336 | -0.413 | 0.086 | -0.553 | -0.204 | 0.01 | 0.009 | 0.009 | 0 | 0.1 | 0.2 | 0.12 | 0.12 | 0.13 | 0.16 |
| 1 | 195859179 | rs2923884 | 1.521 | 0.686 | -0.697 | -0.409 | -0.054 | 0.049 | 0.049 | 0 | 0.1 | 0.03 | 0.15 | 0 | 0.22 | 0.02 |
| 1 | 195859677 | rs3010143 | -0.647 | 0.381 | 0.117 | -0.607 | 0.012 | -0.118 | -0.118 | 0.782 | 0.77 | 0.58 | 0.84 | 0.94 | 0.81 | 0.8 |
| 1 | 195862859 | rs10494737 | 1.387 | -0.504 | -0.326 | -0.827 | -0.005 | -0.031 | -0.031 | 0 | 0.27 | 0.18 | 0.25 | 0.36 | 0.3 | 0.26 |
| 1 | 195864347 | rs1869042 | 0.437 | -0.523 | -0.324 | -0.839 | 0.012 | 0.045 | 0.045 | 0 | 0.57 | 0.34 | 0.57 | 0.82 | 0.45 | 0.61 |
| 1 | 195866450 | rs1376803 | 0.936 | -0.776 | -0.355 | -0.774 | 0.092 | -0.053 | -0.053 | 1.075 | 0.31 | 0.14 | 0.32 | 0.46 | 0.16 | 0.35 |
| 1 | 195869123 | rs3008576 | -0.845 | -0.111 | 0.415 | -0.665 | 0.007 | -0.089 | -0.089 | 0 | 0.77 | 0.65 | 0.84 | 0.94 | 0.81 | 0.8 |
| 1 | 195871839 | rs3010147 | 0.254 | -0.105 | 0.337 | -0.656 | 0.01 | -0.06 | -0.06 | 0.434 | 0.67 | 0.46 | 0.73 | 0.82 | 0.68 | 0.63 |
| 1 | 195881459 | rs3008552 | -0.589 | 0.134 | 0.4 | -0.865 | 0.011 | -0.114 | -0.114 | 0 | 0.77 | 0.57 | 0.84 | 0.94 | 0.81 | 0.8 |
| 1 | 195883861 | rs2923911 | -0.957 | 0.154 | 0.493 | -0.853 | -0.008 | 0.019 | 0.019 | 0 | 0.79 | 0.83 | 0.88 | 0.94 | 0.81 | 0.8 |
| 1 | 195891127 | rs10921984 | 0.222 | 0.276 | 0.723 | -0.76 | 0.004 | 0.004 | 0.004 | 0 | 0.63 | 0.43 | 0.47 | 0.51 | 0.55 | 0.67 |
| 1 | 195892617 | rs2923875 | 1.458 | 0.515 | 0.803 | -0.811 | 0.101 | -0.062 | -0.062 | 0 | 0.1 | 0.24 | 0.16 | 0 | 0.21 | 0.02 |
| 1 | 195901676 | rs984081 | 0.176 | 0.497 | 0.873 | -0.83 | 0.189 | -0.114 | -0.114 | 0 | 0.16 | 0.51 | 0.22 | 0 | 0.33 | 0.09 |
| 1 | 195903057 | rs3010159 | 1.466 | 0.463 | 0.626 | -0.765 | 0.103 | -0.061 | -0.061 | 3.175 | 0.1 | 0.24 | 0.16 | 0 | 0.21 | 0.02 |
| 1 | 195908262 | rs1901425 | -1.208 | -0.11 | 0.142 | -0.533 | 0.04 | -0.127 | -0.127 | 0.039 | 0.8 | 0.99 | 0.7 | 0.58 | 0.88 | 0.85 |
| 1 | 195909031 | rs1451924 | -0.182 | 0.005 | 0.462 | -0.592 | -0.031 | 0.081 | 0.081 | 0 | 0.73 | 0.56 | 0.62 | 0.53 | 0.77 | 0.7 |
| 1 | 195909700 | rs16839132 | 0.465 | -0.352 | -0.051 | -1.048 | 0.022 | 0.044 | 0.044 | 0 | 0.31 | 0.24 | 0.13 | 0.07 | 0.24 | 0.36 |
| 1 | 195914273 | rs7517337 | 0.246 | -0.218 | -0.086 | -1.042 | 0.024 | 0.041 | 0.041 | 0 | 0.5 | 0.56 | 0.56 | 0.86 | 0.42 | 0.48 |
| 1 | 195915483 | rs5024503 | 0.364 | -0.154 | -0.187 | -0.877 | 0 | 0 | 0 | 0 | 0.42 | 0.54 | 0.47 | 0.54 | 0.42 | 0.57 |
| 1 | 195915580 | rs1376811 | -0.21 | 0.282 | -0.543 | -0.823 | 0.035 | -0.001 | -0.001 | 0 | 0.46 | 0.37 | 0.44 | 0.52 | 0.37 | 0.47 |
| 1 | 195915909 | rs7537893 | -0.266 | 0.055 | -0.736 | -0.893 | -0.036 | -0.065 | -0.065 | 0 | 0.7 | 0.9 | 0.61 | 0.56 | 0.65 | 0.82 |
| 1 | 195916112 | rs7516958 | 0.355 | 0.194 | -0.662 | -0.808 | 0.001 | -0.005 | -0.005 | 0 | 0.45 | 0.55 | 0.4 | 0.47 | 0.42 | 0.49 |
| 1 | 195916500 | rs984557 | -1.312 | 0.101 | -0.478 | -0.307 | -0.005 | 0.001 | 0.001 | 0 | 0.91 | 0.95 | 0.84 | 1 | 0.78 | 0.98 |
| 1 | 195916887 | rs4295871 | 0.003 | -0.415 | -0.655 | -0.231 | 0.014 | -0.018 | -0.018 | 0 | 0.25 | 0.25 | 0.25 | 0.44 | 0.14 | 0.16 |
| 1 | 195919285 | rs2878283 | 0.295 | 1.099 | -0.083 | -0.617 | 0.13 | -0.139 | -0.139 | 0.142 | 0.8 | 0.61 | 0.68 | 1 | 0.68 | 0.87 |
| 1 | 195927611 | rs10921993 | -0.221 | -0.184 | -0.858 | 0.019 | 0.066 | -0.031 | -0.031 | 0.488 | 0.39 | 0.28 | 0.37 | 0.54 | 0.23 | 0.41 |
| 1 | 195930996 | rs11580534 | 0.31 | -0.665 | -1.122 | 0.155 | 0.023 | -0.037 | -0.037 | 0 | 0.15 | 0.09 | 0.26 | 0.33 | 0.11 | 0.34 |
| 1 | 195932083 | rs4658011 | -0.78 | -0.03 | -1.18 | 0.079 | 0.033 | 0.102 | 0.102 | 0 | 0.5 | 0.38 | 0.49 | 0.85 | 0.39 | 0.64 |
| 1 | 195934150 | rs10801499 | -0.84 | 0.057 | -0.762 | -0.595 | 0.014 | -0.086 | -0.086 | 0.766 | 0.72 | 0.7 | 0.71 | 1 | 0.58 | 0.88 |
| 1 | 195934627 | rs10921998 | 1.802 | -0.071 | -0.917 | -0.624 | -0.043 | -0.006 | -0.006 | 0 | 0.19 | 0.01 | 0.2 | 0.14 | 0.19 | 0.24 |

| | | | | | | | | | | | | | | | | |
|---|---|---|---|---|---|---|---|---|---|---|---|---|---|---|---|---|
| 1 | 195935240 | rs2923905 | 1.159 | -0.027 | -1.044 | -0.777 | -0.114 | 0.134 | 0.134 | 0 | 0.21 | 0.02 | 0.21 | 0 | 0.36 | 0.05 |
| 1 | 195936137 | rs6683264 | 1.545 | -0.08 | -0.995 | -0.771 | 0.041 | 0.008 | 0.008 | 0 | 0.23 | 0.12 | 0.1 | 0.21 | 0.12 | 0.07 |
| 1 | 195939912 | rs896516 | 0.614 | -0.071 | -0.724 | -0.604 | -0.06 | 0.081 | 0.081 | 0 | 0.27 | 0.13 | 0.27 | 0.02 | 0.42 | 0.12 |
| 1 | 195941818 | rs4657818 | -0.001 | 0.137 | -1.069 | -0.442 | -0.003 | -0.012 | -0.012 | 0 | 0.27 | 0.26 | 0.39 | 0.65 | 0.28 | 0.57 |
| 1 | 195945395 | rs1597730 | 1.778 | 0.117 | -1.088 | -0.455 | -0.043 | -0.007 | -0.007 | 0 | 0.19 | 0.01 | 0.2 | 0.14 | 0.19 | 0.24 |
| 1 | 195953642 | rs10494739 | 0.038 | 0.116 | -1.087 | -0.455 | -0.003 | -0.012 | -0.012 | 0 | 0.27 | 0.26 | 0.39 | 0.65 | 0.28 | 0.57 |
| 1 | 195954813 | rs1597732 | -0.213 | 0.073 | -1.03 | -0.563 | -0.017 | 0.132 | 0.132 | 0 | 0.46 | 0.28 | 0.59 | 0.79 | 0.47 | 0.8 |
| 1 | 195977493 | rs3008584 | 0.79 | 0.254 | -1.232 | -0.237 | 0.013 | -0.043 | -0.043 | 0 | 0.15 | 0.1 | 0.26 | 0.4 | 0.14 | 0.34 |
| 1 | 195977640 | rs10922008 | 1.262 | 0.487 | -1.157 | -0.851 | -0.044 | -0.046 | -0.046 | 0 | 0.2 | 0.02 | 0.2 | 0.2 | 0.2 | 0.23 |
| 1 | 195985616 | rs12029379 | 1.176 | 0.839 | -0.769 | -0.496 | 0.016 | 0.005 | 0.005 | 0 | 0.17 | 0.12 | 0.09 | 0.14 | 0.09 | 0.06 |
| 1 | 195987908 | rs3008588 | -0.45 | 2.002 | -0.384 | 0.019 | 0.009 | -0.058 | -0.058 | 0 | 0.79 | 0.8 | 0.79 | 1 | 0.66 | 0.96 |
| 1 | 195989351 | rs6428320 | -0.809 | 1.666 | -0.345 | 1.031 | 0.005 | 0.011 | 0.011 | 0 | 0.29 | 0.38 | 0.42 | 0.65 | 0.29 | 0.57 |
| 1 | 196007974 | rs12136731 | -0.5 | 3.098 | -1.279 | 0.159 | -0.051 | 0.049 | 0.049 | 0 | 0.1 | 0.02 | 0.11 | 0 | 0.21 | 0.04 |
| 1 | 196018808 | rs10801509 | -1.97 | 2.528 | -0.532 | 0.269 | 0.008 | -0.011 | -0.011 | 0.036 | 0.28 | 0.28 | 0.28 | 0.1 | 0.38 | 0.19 |
| 1 | 196020060 | rs17688413 | -1.168 | 2.339 | -0.373 | 0.19 | -0.078 | 0.075 | 0.075 | 0 | 0.14 | 0.03 | 0.13 | 0 | 0.27 | 0.05 |
| 1 | 196027872 | rs10922013 | -0.804 | 1.913 | -0.013 | 0.338 | 0.019 | -0.029 | -0.029 | 0 | 0.11 | 0.16 | 0.12 | 0 | 0.23 | 0.03 |
| 1 | 196030268 | rs12740444 | 0.436 | 0.56 | -0.224 | 0.312 | -0.03 | 0.019 | 0.019 | 0 | 0.43 | 0.21 | 0.42 | 0.36 | 0.5 | 0.44 |
| 1 | 196050876 | rs7536964 | 0.712 | -0.345 | 0.277 | 0.203 | 0.042 | 0.06 | 0.06 | 0 | 0.22 | 0.06 | 0.19 | 0.11 | 0.12 | 0.17 |
| 1 | 196053435 | rs4658015 | -0.266 | -0.478 | 0.668 | 0.224 | -0.071 | 0.236 | 0.236 | 0.022 | 0.83 | 0.5 | 0.79 | 0.47 | 0.93 | 0.68 |
| 1 | 196057313 | rs12145838 | -0.114 | -0.854 | 0.529 | 0.106 | -0.013 | -0.003 | -0.003 | 0 | 0.18 | 0.25 | 0.14 | 0.16 | 0.13 | 0.19 |
| 1 | 196059593 | rs6428323 | 0.262 | -0.816 | 0.879 | 0.198 | 0.011 | 0.044 | 0.044 | 0 | 0.76 | 0.7 | 0.7 | 0.44 | 0.7 | 0.74 |
| 1 | 196061711 | rs11579497 | -0.577 | -0.7 | 0.772 | -0.099 | 0.001 | 0.002 | 0.002 | 0 | 0.24 | 0.22 | 0.17 | 0.16 | 0.18 | 0.3 |
| 1 | 196070782 | rs4453019 | -0.238 | -0.667 | 0.977 | -0.095 | -0.008 | -0.002 | -0.002 | 0 | 0.83 | 0.77 | 0.88 | 0.84 | 0.89 | 0.73 |
| 1 | 196080988 | rs12405380 | 1.779 | -0.843 | 1.004 | -0.172 | -0.028 | 0.039 | 0.039 | 0 | 0.3 | 0.16 | 0.33 | 0.17 | 0.4 | 0.26 |
| 1 | 196081615 | rs4498784 | 0.624 | -0.945 | 0.976 | 0.5 | 0.038 | 0.011 | 0.011 | 0 | 0.51 | 0.62 | 0.58 | 0.49 | 0.69 | 0.43 |
| 1 | 196089818 | rs12036795 | -0.66 | -0.888 | 0.902 | 0.311 | 0.06 | 0.028 | 0.028 | 0 | 0.47 | 0.28 | 0.41 | 0.37 | 0.3 | 0.57 |
| 1 | 196096194 | rs12567147 | 0.939 | -1.105 | 0.81 | 0.187 | 0.033 | -0.007 | -0.007 | 0 | 0.21 | 0.14 | 0.11 | 0.22 | 0.09 | 0.22 |
| 1 | 196099023 | rs10922038 | -1.873 | -1.032 | 0.976 | 0.147 | 0.006 | -0.022 | -0.022 | 0 | 0.67 | 0.44 | 0.65 | 0.82 | 0.6 | 0.72 |
| 1 | 196099926 | rs6676015 | 0.048 | -0.414 | 1 | -0.079 | -0.034 | 0.023 | 0.023 | 0 | 0.13 | 0.07 | 0.24 | 0 | 0.28 | 0.09 |
| 1 | 196103959 | rs7543048 | 0.71 | -0.519 | 0.802 | 0.144 | 0.026 | -0.008 | -0.008 | 0 | 0.2 | 0.14 | 0.11 | 0.21 | 0.09 | 0.18 |
| 1 | 196110486 | rs7513153 | 0.654 | -0.474 | 0.798 | 0.152 | -0.044 | 0.017 | 0.017 | 0 | 0.2 | 0.29 | 0.09 | 0.22 | 0.08 | 0.18 |
| 1 | 196111734 | rs33964123 | -0.104 | -0.18 | 0.422 | 0.2 | 0.001 | -0.013 | -0.013 | 0 | 0.14 | 0.06 | 0.1 | 0.17 | 0.11 | 0.27 |
| 1 | 196119310 | rs12727479 | 0.546 | -0.156 | 0.677 | 0.11 | 0.002 | -0.036 | -0.036 | 1.895 | 0.07 | 0.17 | 0.04 | 0 | 0.06 | 0.1 |
| 1 | 196120294 | rs10754178 | -1.38 | -0.399 | 0.39 | 0.055 | 0.037 | 0.011 | 0.011 | 0 | 0.83 | 0.92 | 0.93 | 0.88 | 0.91 | 0.87 |
| 1 | 196120613 | rs16839556 | 0.968 | -0.415 | 0.42 | 0.141 | 0.004 | -0.009 | -0.009 | 1.142 | 0.09 | 0.05 | 0.07 | 0.2 | 0.05 | 0.16 |
| 1 | 196125008 | rs12122832 | 0.45 | -0.216 | 0.325 | 0.153 | 0.065 | 0.013 | 0.013 | 0.01 | 0.1 | 0.21 | 0.15 | 0.09 | 0.25 | 0.11 |
| 1 | 196136924 | rs10754180 | 0.63 | -0.235 | 0.107 | 0.155 | 0.011 | -0.001 | -0.001 | 0 | 0.1 | 0.14 | 0.15 | 0.09 | 0.25 | 0.11 |
| 1 | 196142986 | rs4658018 | -1.382 | -0.238 | -0.157 | 0.161 | 0.028 | 0.013 | 0.013 | 0 | 0.82 | 0.89 | 0.93 | 0.88 | 0.91 | 0.87 |
| 1 | 196144843 | rs10801520 | 1.033 | -0.244 | -0.396 | 0.021 | 0.002 | -0.003 | -0.003 | 0.249 | 0.48 | 0.34 | 0.5 | 0.56 | 0.49 | 0.52 |
| 1 | 196163897 | rs1033928 | 0.585 | -0.43 | -1.199 | 0.074 | -0.015 | -0.005 | -0.005 | 0 | 0.58 | 0.55 | 0.66 | 0.63 | 0.74 | 0.63 |
| 1 | 196166278 | rs2094029 | 0.328 | -0.182 | -2.417 | -0.042 | 0.004 | 0.006 | 0.006 | 0 | 0.08 | 0.05 | 0.15 | 0 | 0.05 | 0.06 |
| 1 | 196167779 | rs1325300 | 0.997 | -0.045 | -2.378 | 0.267 | -0.02 | 0.035 | 0.035 | 0 | 0.07 | 0 | 0.04 | 0 | 0.07 | 0.03 |
| 1 | 196170439 | rs10754182 | 0.398 | -0.136 | -2.136 | 0.574 | -0.054 | -0.104 | -0.104 | 0 | 0.39 | 0.08 | 0.43 | 0.5 | 0.43 | 0.42 |
| 1 | 196174790 | rs927724 | -0.181 | -0.158 | -1.48 | 0.374 | 0.03 | 0.016 | 0.016 | 0 | 0.82 | 0.92 | 0.93 | 0.87 | 0.9 | 0.87 |
| 1 | 196177193 | rs6683628 | -0.416 | 0.278 | -1.306 | 0.037 | -0.001 | 0.002 | 0.002 | 0.434 | 0.48 | 0.51 | 0.33 | 0.26 | 0.53 | 0.25 |
| 1 | 196186750 | rs10801528 | 0.923 | 0.436 | -0.783 | -0.312 | -0.002 | -0.02 | -0.02 | 0 | 0.86 | 0.23 | 0.83 | 0.95 | 0.83 | 0.84 |
| 1 | 196187665 | rs1998712 | 0.943 | 0.418 | -0.293 | -0.321 | 0.005 | -0.028 | -0.028 | 0 | 0.87 | 0.31 | 0.84 | 0.95 | 0.83 | 0.84 |
| 1 | 196236425 | rs1998652 | 0.948 | 0.369 | 1.233 | -0.393 | 0.065 | 0.013 | 0.013 | 0 | 0.91 | 0.56 | 0.87 | 0.97 | 0.88 | 0.85 |
| 1 | 196240120 | rs912760 | 1.832 | 0.357 | 1.165 | -0.398 | -0.002 | -0.021 | -0.021 | 0 | 0.86 | 0.23 | 0.83 | 0.95 | 0.82 | 0.83 |
| 1 | 196243012 | rs2025599 | 1.244 | 0.37 | 1.391 | -0.392 | 0.052 | -0.007 | -0.007 | 0 | 0.88 | 0.43 | 0.83 | 0.95 | 0.82 | 0.84 |
| 1 | 196256498 | rs12058062 | -1.737 | 0.369 | 1.393 | -0.392 | 0.049 | -0.01 | -0.01 | 0 | 0.12 | 0.59 | 0.17 | 0.05 | 0.18 | 0.16 |
| 1 | 196304863 | rs7554267 | 0.263 | 0.446 | 0.229 | -0.271 | -0.007 | -0.101 | -0.101 | 0 | 0.39 | 0.03 | 0.22 | 0.6 | 0.28 | 0.26 |
| 1 | 196329362 | rs11801881 | -1.164 | 0.703 | 0.663 | -0.416 | 0.051 | -0.009 | -0.009 | 3.056 | 0.12 | 0.59 | 0.17 | 0.05 | 0.18 | 0.16 |
| 1 | 196331833 | rs7544651 | -1.602 | 0.704 | 0.664 | -0.411 | 0.004 | -0.033 | -0.033 | 0 | 0.13 | 0.7 | 0.17 | 0.05 | 0.17 | 0.13 |
| 1 | 196351150 | rs2027368 | 0.152 | 0.845 | 0.08 | -0.489 | -0.006 | -0.014 | -0.014 | 0 | 0.27 | 0.13 | 0.35 | 0.32 | 0.31 | 0.29 |
| 1 | 196359665 | rs946750 | -0.052 | 0.926 | 0.808 | -0.524 | -0.006 | 0.006 | 0.006 | 0 | 0.71 | 0.54 | 0.61 | 0.68 | 0.69 | 0.71 |
| 1 | 196369947 | rs2878551 | 0.961 | 0.515 | 0.306 | -0.679 | 0.009 | -0.125 | -0.125 | 0.039 | 0.34 | 0.03 | 0.25 | 0.56 | 0.22 | 0.24 |
| 1 | 196385649 | rs3927686 | -0.239 | 0.539 | 0.79 | -0.31 | 0 | 0 | 0 | 0 | 0.73 | 0.76 | 0.64 | 0.68 | 0.69 | 0.71 |
| 1 | 196387632 | rs12729649 | 0.017 | 0.528 | 0.807 | -0.315 | -0.008 | -0.003 | -0.003 | 0 | 0.69 | 0.43 | 0.6 | 0.68 | 0.68 | 0.71 |
| 1 | 196397819 | rs1538686 | 0.616 | 0.557 | 0.894 | -0.33 | -0.001 | -0.001 | -0.001 | 0 | 0.65 | 0.22 | 0.52 | 0.64 | 0.59 | 0.6 |
| 1 | 196402773 | rs2878557 | -0.003 | 0.534 | 0.922 | -0.324 | -0.009 | -0.003 | -0.003 | 0 | 0.69 | 0.45 | 0.61 | 0.68 | 0.68 | 0.71 |
| 1 | 196418527 | rs9659527 | -0.306 | 0.525 | 0.781 | -0.291 | 0.003 | 0.002 | 0.002 | 0 | 0.29 | 0.6 | 0.42 | 0.33 | 0.38 | 0.3 |
| 1 | 196425133 | rs10494743 | 0.538 | 0.451 | 0.348 | -0.373 | -0.01 | -0.023 | -0.023 | 0 | 0.64 | 0.14 | 0.51 | 0.63 | 0.61 | 0.61 |
| 1 | 196432158 | rs6682225 | 0.44 | 0.423 | 0.701 | -0.386 | 0.001 | -0.004 | -0.004 | 0 | 0.66 | 0.28 | 0.52 | 0.63 | 0.61 | 0.61 |
| 1 | 196433870 | rs10922071 | -0.178 | 0.463 | 0.643 | -0.38 | -0.004 | 0.003 | 0.003 | 0 | 0.29 | 0.33 | 0.4 | 0.34 | 0.31 | 0.28 |
| 1 | 196455824 | rs10737671 | 1.128 | 0.464 | 0.85 | -0.377 | 0.087 | 0.054 | 0.054 | 0.289 | 0.93 | 0.47 | 0.91 | 0.98 | 0.92 | 0.9 |
| 1 | 196494547 | rs4388642 | 1.232 | 0.641 | -0.299 | -0.609 | -0.118 | 0.057 | 0.057 | 0 | 0.26 | 0.02 | 0.21 | 0.08 | 0.31 | 0.28 |
| 1 | 196519023 | rs10754192 | 1.053 | 0.607 | 0.369 | -0.511 | 0.037 | 0.011 | 0.011 | 0.173 | 0.93 | 0.37 | 0.9 | 0.97 | 0.91 | 0.9 |
| 1 | 196525314 | rs10922084 | 0.04 | 0.518 | 0.2 | -0.487 | 0.006 | -0.004 | -0.004 | 0 | 0.29 | 0.22 | 0.39 | 0.36 | 0.3 | 0.28 |

| | | | | | | | | | | | | | | | | |
|---|---|---|---|---|---|---|---|---|---|---|---|---|---|---|---|---|
| 1 | 196542567 | rs10494744 | 1.831 | 0.974 | 0.009 | -0.751 | -0.057 | 0.053 | 0.053 | 0 | 0.1 | 0.01 | 0.08 | 0 | 0.16 | 0.07 |
| 1 | 196557329 | rs12128544 | -0.885 | 0.426 | 0.593 | -0.513 | 0.058 | 0.06 | 0.06 | 0.436 | 0.05 | 0.55 | 0.06 | 0.02 | 0.01 | 0.08 |
| 1 | 196567597 | rs4462080 | 0.353 | 0.484 | -0.199 | -0.494 | 0.014 | 0.007 | 0.007 | 0 | 0.64 | 0.15 | 0.5 | 0.64 | 0.61 | 0.62 |
| 1 | 196620917 | rs3753394 | 0.59 | 0.392 | 0.864 | -0.532 | -0.021 | -0.118 | -0.118 | 0.115 | 0.32 | 0.04 | 0.27 | 0.55 | 0.27 | 0.25 |
| 1 | 196623337 | rs7524776 | 1.076 | 0.605 | 0.828 | -0.462 | 0.008 | -0.024 | -0.024 | 0 | 0.12 | 0.16 | 0.1 | 0.03 | 0.17 | 0.09 |
| 1 | 196635470 | rs34388368 | 0.512 | 0.523 | 0.071 | -0.519 | -0.008 | -0.019 | -0.019 | 0 | 0.17 | 0.12 | 0.34 | 0.33 | 0.25 | 0.23 |
| 1 | 196646176 | rs1329424 | -0.881 | 0.688 | -0.219 | -1.159 | -0.015 | 0.055 | 0.055 | 0 | 0.65 | 0.71 | 0.77 | 0.95 | 0.64 | 0.71 |
| 1 | 196646261 | rs572515 | -1.082 | 0.635 | -0.179 | -1.185 | -0.035 | 0.1 | 0.1 | 0 | 0.67 | 0.82 | 0.78 | 0.95 | 0.64 | 0.71 |
| 1 | 196651832 | rs3766404 | 0.033 | 0.277 | -0.088 | -1.394 | -0.026 | -0.047 | -0.047 | 0.081 | 0.19 | 0.46 | 0.15 | 0.06 | 0.14 | 0.3 |
| 1 | 196654324 | rs1061147 | -1.341 | 0.288 | 0.118 | -1.389 | 0.001 | -0.039 | -0.039 | 0.014 | 0.65 | 0.57 | 0.76 | 0.95 | 0.64 | 0.71 |
| 1 | 196655743 | rs10801553 | -1.246 | 0.289 | 0.067 | -1.401 | -0.009 | 0.022 | 0.022 | 0 | 0.65 | 0.65 | 0.77 | 0.95 | 0.64 | 0.71 |
| 1 | 196661613 | rs12405238 | 1.023 | 0.216 | -0.177 | -1.152 | -0.018 | -0.106 | -0.106 | 0 | 0.31 | 0.08 | 0.25 | 0.54 | 0.26 | 0.2 |
| 1 | 196662459 | rs10922096 | 0.18 | 0.243 | -0.104 | -1.188 | -0.042 | 0.195 | 0.195 | 0.434 | 0.46 | 0.16 | 0.62 | 0.89 | 0.5 | 0.41 |
| 1 | 196670839 | rs10754199 | -1.239 | 0.402 | 0.032 | -1.076 | -0.009 | 0.021 | 0.021 | 0 | 0.65 | 0.64 | 0.77 | 0.95 | 0.64 | 0.71 |
| 1 | 196677163 | rs1410997 | -1.323 | 0.263 | 0.047 | -1.061 | -0.016 | 0.034 | 0.034 | 0 | 0.64 | 0.64 | 0.75 | 0.94 | 0.6 | 0.7 |
| 1 | 196677945 | rs203685 | -1.429 | 0.253 | 0.034 | -1.069 | -0.05 | 0.107 | 0.107 | 0 | 0.66 | 0.81 | 0.75 | 0.95 | 0.6 | 0.7 |
| 1 | 196686918 | rs6677604 | 0.14 | -0.119 | -0.375 | -1.468 | -0.037 | -0.107 | -0.107 | 0 | 0.3 | 0.39 | 0.17 | 0.05 | 0.19 | 0.36 |
| 1 | 196692148 | rs11801630 | 1.014 | -0.069 | -0.497 | -1.36 | 0 | -0.086 | -0.086 | 1.899 | 0.18 | 0.08 | 0.19 | 0.51 | 0.18 | 0.15 |
| 1 | 196695742 | rs3753396 | 1.208 | -0.057 | -0.498 | -1.355 | -0.017 | -0.112 | -0.112 | 0.097 | 0.17 | 0.04 | 0.19 | 0.51 | 0.18 | 0.15 |
| 1 | 196696933 | rs1410996 | -0.594 | -0.096 | -0.556 | -1.389 | 0.012 | 0.025 | 0.025 | 0.022 | 0.46 | 0.56 | 0.55 | 0.42 | 0.43 | 0.56 |
| 1 | 196698272 | rs395544 | -1.988 | -0.072 | -0.568 | -1.395 | -0.035 | 0.081 | 0.081 | 0 | 0.65 | 0.76 | 0.75 | 0.95 | 0.6 | 0.71 |
| 1 | 196701284 | rs7540032 | -0.465 | -0.09 | -0.525 | -1.364 | 0 | 0.001 | 0.001 | 0.133 | 0.45 | 0.47 | 0.54 | 0.44 | 0.42 | 0.56 |
| 1 | 196702525 | rs2284664 | 1.66 | -0.092 | -0.538 | -1.377 | -0.025 | -0.046 | -0.046 | 0 | 0.15 | 0.06 | 0.36 | 0.38 | 0.24 | 0.2 |
| 1 | 196702810 | rs1329428 | -0.604 | -0.033 | -0.512 | -1.295 | 0 | 0.001 | 0.001 | 0.129 | 0.45 | 0.47 | 0.54 | 0.44 | 0.43 | 0.56 |
| 1 | 196704997 | rs70620 | 0.382 | -0.172 | -0.164 | -0.905 | -0.014 | 0.072 | 0.072 | 0.054 | 0.2 | 0.36 | 0.22 | 0.51 | 0.18 | 0.15 |
| 1 | 196705520 | rs742855 | 0.81 | -0.17 | -0.252 | -0.909 | 0 | -0.085 | -0.085 | 0.013 | 0.18 | 0.08 | 0.19 | 0.51 | 0.18 | 0.15 |
| 1 | 196709774 | rs1065489 | 0.961 | -0.135 | -0.237 | -0.858 | -0.016 | -0.116 | -0.116 | 0 | 0.17 | 0.04 | 0.2 | 0.51 | 0.18 | 0.15 |
| 1 | 196823613 | rs7542235 | -0.026 | -0.122 | -0.056 | -1.58 | -0.034 | -0.103 | -0.103 | 0 | 0.29 | 0.38 | 0.17 | 0.05 | 0.19 | 0.36 |
| 1 | 196824773 | rs16840639 | -0.026 | -0.122 | -0.049 | -1.505 | -0.033 | -0.085 | -0.085 | 0.352 | 0.29 | 0.37 | 0.17 | 0.06 | 0.19 | 0.36 |
| 1 | 196825628 | rs16840658 | 1.154 | -0.273 | -0.342 | -1.172 | 0.005 | -0.006 | -0.006 | 0 | 0.13 | 0.11 | 0.08 | 0.13 | 0.14 | 0.05 |
| 1 | 196840173 | rs7519501 | 0.424 | -0.305 | -0.33 | -1.784 | -0.013 | -0.115 | -0.115 | 3.007 | 0.16 | 0.5 | 0.13 | 0 | 0.13 | 0.16 |
| 1 | 196843711 | rs6428363 | -0.857 | -0.326 | -0.443 | -1.784 | 0.048 | 0.071 | 0.071 | 0 | 0.44 | 0.69 | 0.26 | 0.19 | 0.33 | 0.4 |
| 1 | 196852781 | rs10801575 | 0.778 | 0.178 | -0.501 | -1.784 | 0.035 | 0.086 | 0.086 | 0 | 0.6 | 0.2 | 0.73 | 0.83 | 0.68 | 0.65 |
| 1 | 196862569 | rs12404243 | 0.219 | 0.188 | -0.533 | -1.77 | 0.001 | 0 | 0 | 0 | 0.24 | 0.27 | 0.13 | 0.17 | 0.18 | 0.2 |
| 1 | 196881420 | rs4044578 | -1.423 | 0.612 | -1.164 | -2.03 | -0.105 | 0.182 | 0.182 | 0.265 | 0.66 | 0.96 | 0.79 | 0.95 | 0.62 | 0.75 |
| 1 | 196882344 | rs7522952 | 1.247 | 0.512 | -0.771 | -1.815 | -0.001 | 0.054 | 0.054 | 0.204 | 0.11 | 0.03 | 0.08 | 0 | 0.11 | 0.14 |
| 1 | 196892322 | rs7548070 | 0.248 | 0.677 | 0.316 | -1.597 | -0.006 | -0.093 | -0.093 | 0.151 | 0.15 | 0.33 | 0.12 | 0 | 0.13 | 0.15 |
| 1 | 196901753 | rs7538501 | 0.227 | 0.676 | 0.327 | -1.598 | -0.006 | -0.093 | -0.093 | 0 | 0.15 | 0.33 | 0.12 | 0 | 0.13 | 0.14 |
| 1 | 196922860 | rs9427934 | -0.738 | 0.626 | 0.519 | -1.627 | -0.015 | -0.042 | -0.042 | 0 | 0.37 | 0.47 | 0.24 | 0.17 | 0.31 | 0.35 |
| 1 | 196929310 | rs7531555 | 0.984 | 0.756 | 0.345 | -1.634 | 0 | -0.009 | -0.009 | 0 | 0.22 | 0.19 | 0.48 | 0.7 | 0.25 | 0.37 |
| 1 | 196933245 | rs9427938 | 0.676 | 0.81 | 0.568 | -1.59 | -0.017 | -0.083 | -0.083 | 2.071 | 0.13 | 0.27 | 0.1 | 0 | 0.11 | 0.14 |
| 1 | 196937536 | rs6428379 | -1.265 | 0.824 | 0.45 | -1.599 | -0.069 | 0.163 | 0.163 | 0 | 0.61 | 0.86 | 0.73 | 0.87 | 0.56 | 0.71 |
| 1 | 196941493 | rs6667243 | -1.099 | 0.927 | 0.45 | -1.469 | -0.044 | 0.21 | 0.21 | 0.095 | 0.56 | 0.86 | 0.71 | 0.87 | 0.52 | 0.71 |
| 1 | 196943321 | rs9427657 | 0.438 | 0.664 | 0.86 | -1.311 | -0.02 | -0.085 | -0.085 | 0 | 0.13 | 0.37 | 0.1 | 0 | 0.1 | 0.14 |
| 1 | 196946546 | rs9427661 | 0.562 | 0.919 | 0.507 | -1.155 | 0.003 | -0.073 | -0.073 | 0 | 0.12 | 0.48 | 0.11 | 0 | 0.08 | 0.1 |
| 1 | 196946775 | rs9427662 | 0.562 | 0.919 | 0.507 | -1.155 | 0.003 | -0.073 | -0.073 | 0 | 0.12 | 0.48 | 0.11 | 0 | 0.08 | 0.1 |
| 1 | 196954225 | rs1750311 | -0.818 | 1.126 | 0.229 | -1.159 | 0.017 | 0.031 | 0.031 | 0 | 0.37 | 0.75 | 0.26 | 0.18 | 0.33 | 0.31 |
| 1 | 196978615 | rs10922153 | 0.969 | 1.149 | -0.057 | -1.132 | -0.015 | 0.231 | 0.231 | 0.212 | 0.48 | 0.14 | 0.3 | 0.15 | 0.49 | 0.35 |
| 1 | 196993973 | rs1360558 | 0.162 | 1.213 | 0.013 | -1.013 | -0.117 | 0.041 | 0.041 | 0 | 0.34 | 0.04 | 0.22 | 0.17 | 0.42 | 0.28 |
| 1 | 196995848 | rs1332667 | 0.418 | 1.182 | 0.148 | -1.032 | -0.014 | 0.113 | 0.113 | 0 | 0.54 | 0.27 | 0.65 | 0.82 | 0.5 | 0.62 |
| 1 | 197008365 | rs698859 | -0.007 | 1.267 | 0.459 | -1.572 | -0.057 | 0.066 | 0.066 | 0 | 0.34 | 0.09 | 0.23 | 0.17 | 0.41 | 0.29 |
| 1 | 197009798 | rs5998 | 0.538 | 1.27 | 0.491 | -1.577 | -0.023 | 0.141 | 0.141 | 0 | 0.53 | 0.19 | 0.65 | 0.82 | 0.5 | 0.61 |
| 1 | 197012111 | rs10754210 | 1.15 | 1.3 | 0.468 | -1.592 | -0.021 | 0.132 | 0.132 | 0 | 0.38 | 0.13 | 0.22 | 0.15 | 0.33 | 0.29 |
| 1 | 197012194 | rs857026 | -0.535 | 1.211 | 0.561 | -1.553 | -0.021 | -0.041 | -0.041 | 0.266 | 0.87 | 0.31 | 0.87 | 0.98 | 0.91 | 0.9 |
| 1 | 197027606 | rs1759007 | -1.247 | 1.1 | 0.713 | -1.537 | -0.002 | -0.063 | -0.063 | 0 | 0.9 | 0.69 | 0.91 | 1 | 0.91 | 0.9 |
| 1 | 197031021 | rs6003 | -0.561 | 1.041 | 0.115 | -1.534 | -0.02 | -0.045 | -0.045 | 0 | 0.87 | 0.32 | 0.88 | 0.98 | 0.91 | 0.9 |
| 1 | 197053373 | rs12677 | -0.656 | 1.041 | 0.141 | -1.537 | -0.019 | -0.074 | -0.074 | 0.306 | 0.88 | 0.38 | 0.88 | 1 | 0.91 | 0.9 |
| 1 | 197055925 | rs10754213 | -0.656 | 1.059 | 0.117 | -1.531 | -0.022 | -0.074 | -0.074 | 1.186 | 0.88 | 0.38 | 0.88 | 1 | 0.91 | 0.9 |
| 1 | 197070442 | rs3762271 | 0.158 | 1.069 | 0.016 | -1.571 | -0.115 | 0.041 | 0.041 | 0 | 0.33 | 0.04 | 0.23 | 0.16 | 0.41 | 0.29 |
| 1 | 197070776 | rs10922162 | 1.601 | 1.08 | -0.006 | -1.566 | -0.03 | -0.105 | -0.105 | 0.065 | 0.15 | 0.02 | 0.43 | 0.67 | 0.17 | 0.32 |
| 1 | 197070815 | rs1412640 | -1.026 | 1.083 | 0.088 | -1.602 | 0.008 | -0.069 | -0.069 | 1.574 | 0.89 | 0.5 | 0.89 | 1 | 0.91 | 0.9 |
| 1 | 197073932 | rs2878749 | 0.165 | 1.069 | -0.049 | -1.61 | -0.115 | 0.038 | 0.038 | 0 | 0.33 | 0.04 | 0.23 | 0.16 | 0.41 | 0.29 |
| 1 | 197079671 | rs2026429 | -0.957 | 1.052 | 0.004 | -1.612 | -0.014 | -0.07 | -0.07 | 0 | 0.89 | 0.38 | 0.88 | 1 | 0.91 | 0.9 |
| 1 | 197123043 | rs1127661 | 0.121 | 1.1 | -0.12 | -1.608 | -0.113 | 0.046 | 0.046 | 0 | 0.34 | 0.04 | 0.23 | 0.16 | 0.41 | 0.29 |
| 1 | 197126649 | rs4639796 | 1.379 | 0.669 | 0.164 | -1.34 | -0.03 | -0.105 | -0.105 | 0.32 | 0.16 | 0.03 | 0.43 | 0.67 | 0.17 | 0.32 |
| 1 | 197127491 | rs12028827 | 1.379 | 0.669 | 0.164 | -1.338 | -0.03 | -0.104 | -0.104 | 0.166 | 0.16 | 0.03 | 0.43 | 0.67 | 0.17 | 0.32 |
| 1 | 197128726 | rs12128631 | 1.388 | 0.676 | 0.168 | -1.34 | -0.03 | -0.104 | -0.104 | 0 | 0.15 | 0.03 | 0.43 | 0.67 | 0.17 | 0.32 |
| 1 | 197146144 | rs10429911 | -0.095 | 0.432 | 0.852 | -1.543 | -0.018 | -0.044 | -0.044 | 1.227 | 0.87 | 0.34 | 0.87 | 0.98 | 0.91 | 0.9 |
| 1 | 197151246 | rs1415217 | -0.095 | 0.432 | 0.878 | -1.543 | -0.019 | -0.048 | -0.048 | 0 | 0.87 | 0.35 | 0.87 | 0.98 | 0.91 | 0.9 |

| | | | | | | | | | | | | | | | |
|---|---|---|---|---|---|---|---|---|---|---|---|---|---|---|---|
| 1 | 197152149 | rs1556763 | -0.157 | 0.429 | 0.89 | -1.544 | -0.017 | -0.048 | -0.048 | 0 | 0.88 | 0.38 | 0.88 | 0.98 | 0.91 | 0.9 |
| 1 | 197161212 | rs4915378 | -0.319 | 0.438 | 0.894 | -1.54 | -0.014 | -0.047 | -0.047 | 0 | 0.88 | 0.36 | 0.88 | 0.98 | 0.91 | 0.9 |
| 1 | 197161428 | rs4915379 | -0.464 | 0.44 | 0.942 | -1.54 | 0.012 | -0.071 | -0.071 | 0.182 | 0.89 | 0.5 | 0.89 | 1 | 0.91 | 0.91 |
| 1 | 197165014 | rs10754218 | 0.422 | 0.51 | 0.395 | -1.597 | 0.02 | 0.094 | 0.094 | 0.201 | 0.67 | 0.19 | 0.75 | 0.92 | 0.74 | 0.69 |
| 1 | 197187605 | rs10922187 | 0.454 | 0.395 | 0.542 | -1.573 | 0.027 | 0.094 | 0.094 | 0.034 | 0.65 | 0.18 | 0.74 | 0.92 | 0.71 | 0.66 |
| 1 | 197192918 | rs12125062 | 0.43 | 0.392 | 0.576 | -1.604 | 0.002 | 0.053 | 0.053 | 0 | 0.22 | 0.1 | 0.14 | 0.06 | 0.19 | 0.22 |
| 1 | 197221248 | rs12119142 | 0.532 | 0.386 | 0.591 | -1.607 | -0.006 | 0.058 | 0.058 | 0 | 0.21 | 0.08 | 0.13 | 0.06 | 0.19 | 0.22 |
| 1 | 197227033 | rs517222 | -0.014 | 0.388 | 0.747 | -1.605 | -0.02 | -0.058 | -0.058 | 0 | 0.25 | 0.53 | 0.17 | 0.08 | 0.19 | 0.24 |
| 1 | 197235926 | rs11802085 | 0.434 | 0.404 | 0.383 | -1.603 | 0.002 | 0.052 | 0.052 | 0 | 0.22 | 0.1 | 0.14 | 0.06 | 0.19 | 0.22 |
| 1 | 197257090 | rs1009188 | 0.684 | 0.541 | -0.044 | -1.745 | -0.042 | 0.11 | 0.11 | 0 | 0.34 | 0.09 | 0.19 | 0.16 | 0.29 | 0.25 |
| 1 | 197266535 | rs10801601 | 0.728 | 0.687 | 0.098 | -1.413 | -0.108 | -0.028 | -0.028 | 0.235 | 0.31 | 0.04 | 0.54 | 0.76 | 0.42 | 0.4 |
| 1 | 197302402 | rs2821131 | -0.529 | 0.358 | 0.247 | -1.391 | -0.004 | -0.082 | -0.082 | 0 | 0.62 | 0.6 | 0.77 | 0.92 | 0.67 | 0.72 |
| 1 | 197306099 | rs2786111 | -0.543 | 0.433 | -1.056 | -1.432 | 0 | -0.026 | -0.026 | 0 | 0.63 | 0.64 | 0.77 | 0.93 | 0.67 | 0.72 |
| 1 | 197308507 | rs17552989 | -0.492 | 0.283 | -1.309 | -0.507 | 0.019 | 0.048 | 0.048 | 0 | 0.2 | 0.06 | 0.12 | 0.06 | 0.16 | 0.21 |
| 1 | 197311228 | rs2786106 | 0.533 | 0.591 | -0.525 | -0.733 | -0.002 | -0.088 | -0.088 | 0 | 0.77 | 0.4 | 0.79 | 0.95 | 0.78 | 0.8 |
| 1 | 197313189 | rs949571 | 0.544 | 1.191 | -1.22 | -0.571 | -0.041 | -0.002 | -0.002 | 0 | 0.72 | 0.11 | 0.67 | 0.75 | 0.67 | 0.81 |
| 1 | 197316408 | rs1572514 | -1.2 | 1.436 | -0.628 | -0.611 | 0.042 | -0.047 | -0.047 | 0 | 0.09 | 0.68 | 0.12 | 0.01 | 0.1 | 0.09 |
| 1 | 197321440 | rs10922200 | -0.983 | 1.422 | 0.704 | -0.618 | 0.126 | 0.004 | 0.004 | 0 | 0.07 | 0.35 | 0.1 | 0.01 | 0.1 | 0.09 |
| 1 | 197329275 | rs12126335 | 0.75 | 1.414 | 0.631 | -0.627 | -0.037 | -0.004 | -0.004 | 0 | 0.73 | 0.13 | 0.68 | 0.75 | 0.68 | 0.81 |
| 1 | 197329535 | rs2786119 | -0.035 | 1.37 | 1.245 | -0.613 | 0.012 | 0.016 | 0.016 | 0 | 0.79 | 0.68 | 0.77 | 0.75 | 0.77 | 0.84 |
| 1 | 197342686 | rs2821108 | -0.961 | 0.654 | 0.468 | -1.319 | 0.001 | -0.004 | -0.004 | 0 | 0.35 | 0.71 | 0.27 | 0.11 | 0.3 | 0.24 |
| 1 | 197342725 | rs10801604 | -0.215 | 0.623 | 0.535 | -1.334 | -0.003 | 0.053 | 0.053 | 0 | 0.31 | 0.2 | 0.24 | 0.1 | 0.3 | 0.24 |
| 1 | 197347920 | rs2494271 | -0.898 | 0.626 | 0.437 | -1.341 | -0.004 | -0.045 | -0.045 | 0 | 0.34 | 0.64 | 0.26 | 0.11 | 0.29 | 0.26 |
| 1 | 197359999 | rs4418639 | 0.597 | 0.657 | -0.126 | -1.45 | -0.104 | -0.047 | -0.047 | 1.27 | 0.61 | 0.04 | 0.56 | 0.65 | 0.59 | 0.68 |
| 1 | 197361571 | rs2759662 | -1.2 | 0.656 | 0.239 | -1.368 | -0.001 | -0.004 | -0.004 | 0 | 0.34 | 0.68 | 0.28 | 0.23 | 0.29 | 0.25 |
| 1 | 197365105 | rs4604725 | 1.47 | 0.659 | 0.058 | -1.367 | -0.083 | 0.014 | 0.014 | 0 | 0.48 | 0.03 | 0.5 | 0.65 | 0.48 | 0.62 |
| 1 | 197368281 | rs2476020 | -0.576 | 0.701 | 0.634 | -1.349 | -0.001 | -0.026 | -0.026 | 0 | 0.32 | 0.46 | 0.27 | 0.23 | 0.29 | 0.27 |
| 1 | 197372250 | rs11802794 | -0.344 | 0.697 | 0.703 | -1.351 | -0.011 | 0.041 | 0.041 | 0 | 0.3 | 0.17 | 0.25 | 0.22 | 0.29 | 0.24 |
| 1 | 197392067 | rs12134409 | 1.299 | 0.878 | 0.61 | -1.307 | -0.054 | -0.018 | -0.018 | 0.364 | 0.16 | 0.04 | 0.2 | 0.12 | 0.23 | 0.14 |
| 1 | 197396108 | rs17554990 | 1.343 | 0.885 | 0.613 | -1.303 | -0.054 | -0.019 | -0.019 | 0 | 0.15 | 0.04 | 0.2 | 0.12 | 0.22 | 0.14 |
| 1 | 197414097 | rs1923759 | -1.097 | 0.821 | 0.646 | -1.35 | -0.025 | 0.045 | 0.045 | 0 | 0.44 | 0.15 | 0.4 | 0.33 | 0.43 | 0.37 |
| 1 | 197438606 | rs9659183 | 1.764 | 0.247 | -0.242 | -1.017 | -0.107 | -0.011 | -0.011 | 0.878 | 0.3 | 0.02 | 0.26 | 0.17 | 0.35 | 0.29 |
| 1 | 197446702 | rs1135810 | -1.105 | 0.359 | 0.496 | -1.021 | 0.006 | 0.046 | 0.046 | 0 | 0.45 | 0.19 | 0.4 | 0.34 | 0.42 | 0.38 |
| 1 | 197463936 | rs10494754 | -1.156 | 0.355 | 0.439 | -1.215 | 0.011 | 0.071 | 0.071 | 0.008 | 0.51 | 0.26 | 0.47 | 0.34 | 0.52 | 0.4 |
| 1 | 197472070 | rs9658810 | 1.187 | 0.351 | 0.581 | -1.135 | -0.001 | 0.009 | 0.009 | 0 | 0.45 | 0.29 | 0.5 | 0.63 | 0.48 | 0.6 |
| 1 | 197482265 | rs2287537 | 1.048 | 0.28 | 0.584 | -1.291 | 0.006 | 0.021 | 0.021 | 0 | 0.47 | 0.34 | 0.56 | 0.65 | 0.57 | 0.62 |
| 1 | 197485094 | rs10494756 | 1.036 | 0.276 | 0.582 | -1.293 | 0.006 | 0.021 | 0.021 | 0 | 0.48 | 0.34 | 0.56 | 0.65 | 0.57 | 0.62 |
| 1 | 197508901 | rs4915551 | -0.526 | 0.667 | 0.531 | -1.325 | 0.005 | 0.022 | 0.022 | 0 | 0.76 | 0.19 | 0.72 | 0.87 | 0.77 | 0.83 |
| 1 | 197520273 | rs10801614 | -1.706 | 0.433 | 0.721 | -1.254 | 0 | -0.001 | -0.001 | 0 | 0.83 | 0.75 | 0.78 | 0.87 | 0.8 | 0.87 |
| 1 | 197521877 | rs6677361 | -1.45 | 0.429 | 0.826 | -1.256 | -0.001 | -0.003 | -0.003 | 0 | 0.82 | 0.67 | 0.77 | 0.87 | 0.8 | 0.87 |
| 1 | 197529218 | rs2111931 | -1.671 | 0.414 | 0.742 | -1.249 | 0 | 0 | 0 | 0.473 | 0.83 | 0.77 | 0.78 | 0.87 | 0.79 | 0.86 |
| 1 | 197546426 | rs6676073 | 1.679 | 0.415 | 0.672 | -1.249 | -0.031 | -0.009 | -0.009 | 0 | 0.16 | 0.05 | 0.21 | 0.12 | 0.21 | 0.14 |
| 1 | 197546891 | rs6428410 | -1.675 | 0.417 | 0.69 | -1.248 | 0 | 0 | 0 | 0 | 0.83 | 0.77 | 0.78 | 0.87 | 0.79 | 0.86 |
| 1 | 197559324 | rs6702421 | -0.33 | 0.277 | 0.532 | -1.298 | 0.039 | 0.062 | 0.062 | 0 | 0.32 | 0.13 | 0.2 | 0.22 | 0.21 | 0.24 |
| 1 | 197582780 | rs11799915 | -0.531 | 0.409 | 0.784 | -1.188 | 0.001 | 0.009 | 0.009 | 0 | 0.32 | 0.29 | 0.26 | 0.22 | 0.29 | 0.25 |
| 1 | 197586232 | rs2111930 | 1.525 | 0.446 | 0.576 | -1.224 | -0.117 | 0 | 0 | 0 | 0.46 | 0.03 | 0.48 | 0.65 | 0.49 | 0.61 |
| 1 | 197590190 | rs4282851 | 1.235 | 0.46 | 0.964 | -1.199 | 0.021 | 0.034 | 0.034 | 0 | 0.51 | 0.31 | 0.56 | 0.65 | 0.59 | 0.62 |
| 1 | 197601092 | rs12127378 | 1.553 | 0.534 | 0.809 | -1.146 | -0.039 | -0.008 | -0.008 | 0.068 | 0.16 | 0.06 | 0.21 | 0.12 | 0.21 | 0.14 |
| 1 | 197606708 | rs10442656 | 1.273 | 0.524 | 0.93 | -1.149 | 0.001 | -0.001 | -0.001 | 0 | 0.18 | 0.22 | 0.23 | 0.13 | 0.21 | 0.14 |
| 1 | 197644773 | rs9633354 | 1.404 | 0.514 | 1.09 | -1.186 | 0 | 0.004 | 0.004 | 0 | 0.5 | 0.44 | 0.51 | 0.66 | 0.49 | 0.61 |
| 1 | 197646544 | rs12751508 | 1.336 | 0.457 | 0.917 | -1.21 | -0.028 | 0.067 | 0.067 | 0 | 0.11 | 0.01 | 0.07 | 0 | 0.1 | 0.05 |
| 1 | 197674344 | rs12354257 | -0.192 | 0.586 | 0.954 | -1.057 | -0.01 | 0.058 | 0.058 | 0 | 0.32 | 0.14 | 0.25 | 0.22 | 0.29 | 0.25 |
| 1 | 197698103 | rs1747815 | -1.912 | 0.636 | 0.714 | -0.973 | -0.056 | -0.025 | -0.025 | 0.579 | 0.85 | 0.98 | 0.8 | 0.88 | 0.8 | 0.85 |
| 1 | 197702401 | rs2488401 | 0.09 | 0.499 | 0.82 | -1.032 | -0.002 | -0.002 | -0.002 | 0 | 0.28 | 0.31 | 0.2 | 0.22 | 0.18 | 0.23 |
| 1 | 197704717 | rs17641524 | 0.404 | 0.566 | 0.391 | -1.027 | 0.002 | 0.098 | 0.098 | 0 | 0.27 | 0.01 | 0.17 | 0.09 | 0.18 | 0.2 |
| 1 | 197727298 | rs1775454 | -1.248 | 0.78 | 0.704 | -0.918 | 0 | 0 | 0 | 0 | 0.84 | 0.85 | 0.79 | 0.87 | 0.81 | 0.86 |
| 1 | 197735497 | rs2488394 | -1.025 | 0.739 | 0.753 | -0.918 | 0.004 | 0 | 0 | 0 | 0.83 | 0.79 | 0.79 | 0.87 | 0.8 | 0.86 |
| 1 | 197781198 | rs12134279 | 0.136 | 0.516 | 0.552 | -0.93 | 0.043 | 0.042 | 0.042 | 0 | 0.28 | 0.09 | 0.19 | 0.22 | 0.18 | 0.22 |
| 1 | 197794699 | rs6667560 | 0.598 | 0.568 | 0.631 | -1.343 | -0.003 | 0.039 | 0.039 | 0.077 | 0.49 | 0.29 | 0.48 | 0.65 | 0.53 | 0.61 |
| 1 | 197808778 | rs17642632 | 0.22 | 0.408 | 0.324 | -1.445 | 0.045 | 0.137 | 0.137 | 0 | 0.3 | 0.05 | 0.21 | 0.1 | 0.18 | 0.21 |
| 1 | 197811282 | rs6678151 | -0.348 | 0.428 | 0.754 | -1.442 | 0.005 | -0.008 | -0.008 | 0 | 0.68 | 0.68 | 0.77 | 0.91 | 0.82 | 0.79 |
| 1 | 197811566 | rs10922298 | 0.517 | 0.527 | 0.778 | -1.353 | -0.001 | 0.015 | 0.015 | 0 | 0.49 | 0.4 | 0.59 | 0.75 | 0.55 | 0.63 |
| 1 | 197813032 | rs12026183 | 0.997 | 0.56 | 0.677 | -1.273 | -0.048 | -0.043 | -0.043 | 0 | 0.15 | 0.03 | 0.11 | 0.14 | 0.18 | 0.14 |
| 1 | 197823360 | rs10922306 | -0.303 | 0.667 | 0.923 | -1.283 | 0.075 | 0.015 | 0.015 | 0 | 0.79 | 0.54 | 0.79 | 0.84 | 0.7 | 0.85 |
| 1 | 197830485 | rs2047540 | -2.951 | 0.488 | 1.147 | -1.168 | 0.095 | 0.105 | 0.105 | 0.468 | 0.08 | 0.55 | 0.12 | 0.13 | 0.09 | 0.03 |
| 1 | 197830838 | rs2047541 | -1.108 | 0.502 | 1.269 | -1.171 | 0.187 | 0.063 | 0.063 | 0.094 | 0.06 | 0.35 | 0.09 | 0.01 | 0.09 | 0.02 |
| 1 | 197836347 | rs10922313 | -0.75 | 0.463 | 0.411 | -1.385 | -0.004 | 0.007 | 0.007 | 0 | 0.55 | 0.59 | 0.41 | 0.38 | 0.51 | 0.46 |
| 1 | 197843204 | rs1499602 | -0.19 | 0.129 | 0.231 | -1.408 | -0.011 | 0.105 | 0.105 | 0 | 0.46 | 0.07 | 0.29 | 0.26 | 0.4 | 0.4 |
| 1 | 197844367 | rs10801629 | -0.685 | 0.12 | 0.42 | -1.397 | 0.007 | 0.011 | 0.011 | 0 | 0.52 | 0.53 | 0.34 | 0.35 | 0.42 | 0.44 |

| | | | | | | | | | | | | | | | | |
|---|---|---|---|---|---|---|---|---|---|---|---|---|---|---|---|---|
| 1 | 197851893 | rs6677863 | -0.672 | 0.134 | 0.474 | -1.39 | 0.004 | 0.006 | 0.006 | 0 | 0.52 | 0.52 | 0.34 | 0.35 | 0.42 | 0.44 |
| 1 | 197862321 | rs10922323 | -0.672 | 0.134 | 0.413 | -1.422 | 0.012 | 0.02 | 0.02 | 0 | 0.52 | 0.56 | 0.34 | 0.35 | 0.42 | 0.44 |
| 1 | 197866357 | rs12138550 | -0.259 | 0.331 | -0.321 | -1.395 | 0.065 | 0.076 | 0.076 | 0 | 0.42 | 0.07 | 0.25 | 0.26 | 0.3 | 0.39 |
| 1 | 197867662 | rs10922326 | 0.512 | 0.574 | 0.246 | -0.658 | -0.037 | -0.049 | -0.049 | 0.276 | 0.14 | 0.03 | 0.1 | 0.17 | 0.18 | 0.1 |
| 1 | 197874949 | rs2270763 | 0.024 | 0.002 | 0.551 | -0.437 | -0.004 | -0.02 | -0.02 | 0.49 | 0.36 | 0.41 | 0.39 | 0.26 | 0.37 | 0.36 |
| 1 | 197887980 | rs12024850 | 0.016 | -0.35 | 0.786 | 0.345 | -0.003 | -0.003 | -0.003 | 0.004 | 0.16 | 0.15 | 0.35 | 0.39 | 0.19 | 0.24 |
| 1 | 197890802 | rs13376365 | 0.64 | -0.139 | 1.259 | 0.306 | 0.011 | 0.006 | 0.006 | 0 | 0.29 | 0.25 | 0.26 | 0.29 | 0.21 | 0.34 |
| 1 | 197897332 | rs12568860 | 0.654 | -0.083 | 1.224 | 0.339 | 0.03 | 0.014 | 0.014 | 0 | 0.28 | 0.2 | 0.25 | 0.29 | 0.21 | 0.34 |
| 1 | 197906399 | rs10754237 | 0.426 | -0.067 | 0.855 | 0.24 | 0.06 | 0.011 | 0.011 | 0.113 | 0.94 | 0.67 | 0.96 | 1 | 0.96 | 0.99 |
| 1 | 197912389 | rs17644383 | 0.914 | -0.087 | 0.355 | 0.28 | -0.111 | 0.086 | 0.086 | 0 | 0.44 | 0.03 | 0.31 | 0.2 | 0.53 | 0.35 |
| 1 | 197913101 | rs6671069 | -0.723 | -0.273 | 0.697 | 0.5 | -0.008 | -0.009 | -0.009 | 0 | 0.44 | 0.55 | 0.62 | 0.7 | 0.41 | 0.55 |
| 1 | 197919693 | rs1909507 | 0.184 | -0.063 | 0.103 | 0.588 | 0.001 | 0.03 | 0.03 | 0.353 | 0.13 | 0.15 | 0.35 | 0.41 | 0.18 | 0.23 |
| 1 | 197922395 | rs2047539 | -0.748 | 0.098 | -0.598 | 0.621 | 0.018 | 0.016 | 0.016 | 0 | 0.45 | 0.64 | 0.62 | 0.69 | 0.39 | 0.54 |
| 1 | 197927281 | rs4319370 | -0.88 | 0.599 | -0.759 | 0.126 | -0.021 | -0.002 | -0.002 | 0 | 0.35 | 0.42 | 0.3 | 0.39 | 0.26 | 0.42 |
| 1 | 197929497 | rs549890 | 0.19 | 0.667 | -1.195 | 0.078 | 0.002 | 0.014 | 0.014 | 0 | 0.67 | 0.74 | 0.7 | 0.72 | 0.69 | 0.66 |
| 1 | 197949082 | rs435062 | -0.339 | 0.673 | -1.221 | -0.199 | 0.001 | 0.033 | 0.033 | 0.063 | 0.37 | 0.26 | 0.46 | 0.14 | 0.37 | 0.29 |
| 1 | 197959936 | rs1475145 | -0.363 | 0.783 | -1.935 | -0.107 | -0.016 | 0.001 | 0.001 | 0.022 | 0.75 | 0.92 | 0.84 | 0.78 | 0.76 | 0.57 |
| 1 | 197960889 | rs17304978 | 0.525 | 0.088 | -1.608 | -0.511 | -0.011 | 0.072 | 0.072 | 0 | 0.33 | 0.22 | 0.43 | 0.1 | 0.36 | 0.15 |
| 1 | 197966051 | rs10801637 | 0.451 | -0.365 | -1.071 | 0.059 | -0.01 | 0.002 | 0.002 | 0.057 | 0.56 | 0.52 | 0.65 | 0.52 | 0.62 | 0.38 |
| 1 | 197966427 | rs12045836 | 0.244 | -0.416 | -0.929 | 0.045 | 0.005 | -0.013 | -0.013 | 0.042 | 0.12 | 0.1 | 0.13 | 0.22 | 0.11 | 0.05 |
| 1 | 197968019 | rs1543511 | -0.119 | -0.398 | -0.806 | 0.046 | 0.011 | -0.024 | -0.024 | 1.371 | 0.63 | 0.75 | 0.72 | 0.56 | 0.65 | 0.53 |
| 1 | 197968386 | rs1125953 | 1.099 | -0.437 | -0.389 | 0.015 | -0.022 | -0.048 | -0.048 | 0 | 0.25 | 0.1 | 0.22 | 0.42 | 0.27 | 0.24 |
| 1 | 197970270 | rs2359374 | 1.193 | -0.412 | -0.427 | 0.018 | -0.023 | 0.058 | 0.058 | 0.564 | 0.28 | 0.19 | 0.4 | 0.1 | 0.33 | 0.14 |
| 1 | 197971220 | rs10801638 | 0.681 | -0.488 | -0.414 | 0.004 | -0.04 | 0.006 | 0.006 | 0.083 | 0.35 | 0.2 | 0.5 | 0.32 | 0.43 | 0.18 |
| 1 | 197971280 | rs10801639 | 0.089 | -0.85 | -0.414 | -0.02 | -0.028 | -0.027 | -0.027 | 0 | 0.64 | 0.52 | 0.75 | 0.74 | 0.72 | 0.42 |
| 1 | 197972312 | rs2181288 | -0.939 | -0.742 | -1.098 | -0.007 | -0.028 | -0.029 | -0.029 | 0 | 0.85 | 0.99 | 0.72 | 0.84 | 0.83 | 0.63 |
| 1 | 197973573 | rs2224648 | 0.505 | -0.878 | -1.635 | 0.072 | -0.006 | -0.118 | -0.118 | 0 | 0.18 | 0.02 | 0.14 | 0.37 | 0.13 | 0.16 |
| 1 | 197976444 | rs12561995 | -0.573 | -1.111 | -0.831 | -0.242 | -0.02 | 0.022 | 0.022 | 0 | 0.29 | 0.33 | 0.18 | 0.38 | 0.18 | 0.28 |
| 1 | 197977954 | rs2065150 | 1.294 | -1.285 | -1.558 | -0.805 | -0.133 | 0.179 | 0.179 | 0.216 | 0.38 | 0.03 | 0.32 | 0.11 | 0.51 | 0.21 |
| 1 | 197982554 | rs1114740 | -1.064 | -1.424 | -1.58 | -0.61 | -0.038 | -0.054 | -0.054 | 0 | 0.84 | 0.97 | 0.66 | 0.75 | 0.78 | 0.56 |
| 1 | 197987125 | rs2359376 | 1.851 | -1.525 | -2.147 | -1.138 | -0.135 | 0.113 | 0.113 | 0.175 | 0.23 | 0.03 | 0.22 | 0.04 | 0.35 | 0.13 |
| 1 | 197988207 | rs1120608 | 0.427 | -1.618 | -1.78 | -0.754 | -0.002 | 0.001 | 0.001 | 0 | 0.07 | 0.07 | 0.06 | 0.08 | 0.05 | 0.16 |
| 1 | 197995095 | rs12085890 | 1.247 | -1.691 | -2.022 | -0.485 | -0.025 | -0.117 | -0.117 | 2.632 | 0.17 | 0.02 | 0.15 | 0.44 | 0.15 | 0.15 |
| 1 | 197995961 | rs7551861 | 1.277 | -1.823 | -2.157 | -0.688 | -0.127 | 0.227 | 0.227 | 0.002 | 0.42 | 0.04 | 0.31 | 0.11 | 0.51 | 0.24 |
| 1 | 198000253 | rs6667378 | 0.499 | -1.679 | -1.428 | -0.535 | 0.012 | -0.009 | -0.009 | 0.051 | 0.64 | 0.33 | 0.56 | 0.62 | 0.73 | 0.5 |
| 1 | 198003122 | rs2359377 | 0.514 | -0.805 | -1.83 | 0.661 | -0.043 | 0.115 | 0.115 | 0.254 | 0.58 | 0.93 | 0.48 | 0.25 | 0.6 | 0.42 |
| 1 | 198011397 | rs2093531 | -0.436 | -0.135 | -1.435 | 0.783 | -0.049 | -0.068 | -0.068 | 0 | 0.81 | 0.99 | 0.69 | 0.77 | 0.81 | 0.56 |
| 1 | 198026675 | rs2146334 | 0.944 | -0.077 | -0.654 | 0.984 | -0.036 | -0.077 | -0.077 | 0 | 0.13 | 0.01 | 0.13 | 0.35 | 0.15 | 0.15 |
| 1 | 198036134 | rs1233833 | -0.915 | -0.132 | -0.03 | 0.199 | -0.01 | -0.036 | -0.036 | 0 | 0.07 | 0.23 | 0.05 | 0.02 | 0.07 | 0.07 |
| 1 | 198043697 | rs1233827 | 0.03 | -0.355 | 0.133 | 0.603 | 0.004 | -0.031 | -0.031 | 1.599 | 0.19 | 0.08 | 0.19 | 0.52 | 0.14 | 0.25 |
| 1 | 198044170 | rs12125644 | -0.092 | -0.092 | 0.219 | 0.27 | -0.001 | -0.019 | -0.019 | 0.753 | 0.14 | 0.16 | 0.09 | 0.05 | 0.14 | 0.09 |
| 1 | 198051161 | rs7529161 | -0.548 | -0.203 | 0.237 | 0.168 | 0 | 0.032 | 0.032 | 0.463 | 0.16 | 0.09 | 0.1 | 0.03 | 0.19 | 0.11 |
| 1 | 198055493 | rs1233817 | 1.448 | 0.034 | 0.592 | -0.138 | -0.001 | 0.001 | 0.001 | 0.024 | 0.6 | 0.4 | 0.44 | 0.58 | 0.57 | 0.44 |
| 1 | 198066365 | rs4409685 | 0.989 | 0.059 | 0.333 | -0.136 | 0.005 | 0.006 | 0.006 | 0.138 | 0.09 | 0.07 | 0.05 | 0.03 | 0.07 | 0.02 |
| 1 | 198068036 | rs1233812 | 1.295 | 0.034 | 0.368 | 0.05 | 0.006 | -0.022 | -0.022 | 0 | 0.8 | 0.86 | 0.82 | 0.71 | 0.75 | 0.77 |
| 1 | 198068767 | rs1233811 | 1.188 | 0.029 | 0.468 | 0.047 | -0.002 | 0.005 | 0.005 | 0 | 0.8 | 0.77 | 0.82 | 0.71 | 0.75 | 0.78 |
| 1 | 198069021 | rs1233810 | -0.388 | 0.032 | 0.351 | 0.093 | -0.002 | 0.029 | 0.029 | 0.071 | 0.94 | 0.95 | 0.91 | 0.75 | 0.89 | 0.85 |
| 1 | 198071724 | rs12046946 | 0.448 | -0.018 | 0.332 | 0.021 | -0.019 | -0.006 | -0.006 | 0.762 | 0.18 | 0.06 | 0.36 | 0.15 | 0.18 | 0.33 |
| 1 | 198077025 | rs12136149 | -0.27 | -0.028 | 0.199 | 0.002 | 0.002 | -0.008 | -0.008 | 0 | 0.15 | 0.14 | 0.15 | 0.26 | 0.18 | 0.16 |
| 1 | 198081181 | rs16842401 | -0.393 | 0.216 | 0.507 | 0.307 | 0.003 | 0.058 | 0.058 | 0 | 0.07 | 0.11 | 0.11 | 0.22 | 0.12 | 0.15 |
| 1 | 198083440 | rs10158950 | 0.172 | 0.098 | 0.51 | 0.429 | -0.006 | 0.003 | 0.003 | 0 | 0.18 | 0.13 | 0.37 | 0.14 | 0.18 | 0.33 |
| 1 | 198083850 | rs10922367 | 1.236 | 0.372 | 0.394 | 0.315 | 0 | 0.023 | 0.023 | 0.152 | 0.6 | 0.35 | 0.4 | 0.31 | 0.61 | 0.34 |
| 1 | 198085033 | rs4288631 | -0.16 | 0.833 | 0.173 | 0.9 | -0.003 | 0.006 | 0.006 | 0 | 0.85 | 0.92 | 0.83 | 0.45 | 0.86 | 0.73 |
| 1 | 198097178 | rs16842420 | -1.203 | 0.742 | 0.938 | 0.795 | 0.034 | 0.15 | 0.15 | 0 | 0.08 | 0.32 | 0.09 | 0.2 | 0.08 | 0.1 |
| 1 | 198105828 | rs6690826 | -1.359 | 0.261 | 0.657 | 0.298 | 0.007 | 0.003 | 0.003 | 0.126 | 0.3 | 0.28 | 0.45 | 0.33 | 0.26 | 0.42 |
| 1 | 198117538 | rs11585160 | 1.599 | 1.146 | 0.675 | 0.004 | -0.028 | 0.028 | 0.028 | 0 | 0.05 | 0.01 | 0.07 | 0 | 0.11 | 0.01 |
| 1 | 198128859 | rs3762382 | -1.219 | 0.202 | 0.826 | 0.168 | -0.004 | 0.009 | 0.009 | 0.027 | 0.13 | 0.18 | 0.14 | 0.25 | 0.14 | 0.12 |
| 1 | 198140938 | rs6671879 | -2.05 | -0.151 | 1.054 | -0.074 | 0.01 | 0.015 | 0.015 | 0.839 | 0.48 | 0.53 | 0.66 | 0.9 | 0.41 | 0.76 |
| 1 | 198150496 | rs10158410 | -0.543 | -0.138 | 0.957 | -0.054 | -0.008 | 0.056 | 0.056 | 0 | 0.08 | 0.17 | 0.1 | 0.25 | 0.06 | 0.1 |
| 1 | 198155858 | rs6670073 | 0.017 | -0.145 | 1.198 | -0.058 | 0.001 | -0.005 | -0.005 | 0 | 0.18 | 0.12 | 0.19 | 0.56 | 0.14 | 0.33 |
| 1 | 198161614 | rs2359665 | -1.494 | -0.361 | 1.217 | 0.084 | -0.012 | -0.006 | -0.006 | 0 | 0.31 | 0.39 | 0.46 | 0.34 | 0.26 | 0.45 |
| 1 | 198224252 | rs12060034 | 0.156 | -0.53 | 1.504 | 0.006 | -0.026 | -0.046 | -0.046 | 1.73 | 0.17 | 0.34 | 0.32 | 0.09 | 0.1 | 0.32 |
| 1 | 198233907 | rs1927252 | -1.529 | -0.569 | 1.452 | -0.066 | -0.031 | -0.051 | -0.051 | 0 | 0.31 | 0.46 | 0.33 | 0.8 | 0.27 | 0.42 |
| 1 | 198249195 | rs6668469 | -1.519 | -0.512 | 1.519 | 0.071 | -0.009 | 0.03 | 0.03 | 0.251 | 0.15 | 0.32 | 0.16 | 0.24 | 0.16 | 0.14 |
| 1 | 198269416 | rs12090377 | -0.277 | -0.529 | 1.476 | 0.002 | -0.038 | -0.065 | -0.065 | 1.373 | 0.18 | 0.43 | 0.32 | 0.09 | 0.1 | 0.31 |
| 1 | 198299144 | rs4525073 | -1.924 | -0.535 | 1.366 | 0.653 | 0.029 | 0.031 | 0.031 | 0.748 | 0.44 | 0.67 | 0.64 | 0.69 | 0.36 | 0.68 |
| 1 | 198300755 | rs6428452 | 0.001 | -0.683 | 1.399 | 0.799 | -0.002 | 0.036 | 0.036 | 0 | 0.16 | 0.15 | 0.14 | 0.42 | 0.14 | 0.26 |
| 1 | 198315934 | rs1556017 | -1.246 | -0.751 | 1.542 | 0.751 | -0.049 | 0.087 | 0.087 | 0.137 | 0.24 | 0.61 | 0.46 | 0.47 | 0.14 | 0.46 |
| 1 | 198324701 | rs10922405 | -0.149 | -0.744 | 1.333 | 0.878 | -0.002 | 0.033 | 0.033 | 0 | 0.17 | 0.15 | 0.14 | 0.43 | 0.14 | 0.26 |

| | | | | | | | | | | | | | | | |
|---|---|---|---|---|---|---|---|---|---|---|---|---|---|---|---|
| 1 | 198327168 | rs6702048 | -0.877 | -0.741 | 1.347 | 0.853 | -0.003 | 0.023 | 0.023 | 0.072 | 0.22 | 0.2 | 0.18 | 0.43 | 0.22 | 0.27 |
| 1 | 198338612 | rs12040677 | 0.094 | -0.736 | 1.316 | 0.928 | -0.004 | 0.042 | 0.042 | 0 | 0.15 | 0.13 | 0.13 | 0.43 | 0.11 | 0.23 |
| 1 | 198358514 | rs12087101 | -0.703 | -0.734 | 1.455 | 0.826 | -0.07 | 0.101 | 0.101 | 0 | 0.17 | 0.46 | 0.32 | 0.3 | 0.06 | 0.36 |
| 1 | 198389958 | rs7545645 | 0.211 | -0.602 | 1.28 | 1.166 | -0.008 | 0.055 | 0.055 | 0.1 | 0.12 | 0.17 | 0.12 | 0.33 | 0.09 | 0.19 |
| 1 | 198393815 | rs12078566 | -0.45 | -0.616 | 1.235 | 1.098 | 0.003 | -0.011 | -0.011 | 10.945 | 0.17 | 0.13 | 0.16 | 0.33 | 0.17 | 0.22 |
| 1 | 198408321 | rs16843078 | -1.226 | -0.931 | 1.638 | 1.18 | -0.009 | 0.198 | 0.198 | 0 | 0.13 | 0.32 | 0.18 | 0.48 | 0.13 | 0.2 |
| 1 | 198410156 | rs12407744 | -0.822 | -0.884 | 1.574 | 1.16 | -0.012 | 0.083 | 0.083 | 0.914 | 0.22 | 0.37 | 0.3 | 0.51 | 0.21 | 0.32 |
| 1 | 198417152 | rs10922421 | 0.204 | -0.993 | 1.426 | 0.958 | -0.031 | -0.025 | -0.025 | 0 | 0.13 | 0.25 | 0.28 | 0.09 | 0.04 | 0.31 |
| 1 | 198421267 | rs2640175 | -1.387 | -0.975 | 1.249 | 1.055 | -0.026 | -0.051 | -0.051 | 0 | 0.3 | 0.53 | 0.34 | 0.81 | 0.29 | 0.42 |
| 1 | 198433606 | rs12732681 | 1.19 | -0.74 | 0.042 | 0.384 | -0.183 | 0.22 | 0.22 | 0.088 | 0.3 | 0.03 | 0.23 | 0 | 0.41 | 0.15 |
| 1 | 198436809 | rs4477310 | -1.487 | -0.861 | 1.278 | 1.475 | -0.002 | 0.285 | 0.285 | 0.022 | 0.14 | 0.43 | 0.2 | 0.47 | 0.14 | 0.19 |
| 1 | 198450612 | rs10801671 | -1.41 | -0.968 | 1.286 | 1.489 | -0.015 | 0.294 | 0.294 | 0 | 0.14 | 0.43 | 0.2 | 0.47 | 0.13 | 0.19 |
| 1 | 198453920 | rs4915287 | -1.864 | -0.907 | 1.125 | 1.148 | -0.043 | 0.121 | 0.121 | 1.326 | 0.25 | 0.5 | 0.34 | 0.44 | 0.16 | 0.41 |
| 1 | 198454280 | rs4915149 | -1.502 | -0.937 | 1.133 | 1.13 | -0.038 | 0.185 | 0.185 | 1.142 | 0.2 | 0.41 | 0.29 | 0.44 | 0.1 | 0.4 |
| 1 | 198454949 | rs4582831 | -1.134 | -0.93 | 1.146 | 1.142 | -0.038 | 0.223 | 0.223 | 0.387 | 0.18 | 0.41 | 0.28 | 0.44 | 0.08 | 0.39 |
| 1 | 198457891 | rs10922436 | -0.175 | -0.908 | 0.948 | 1.002 | 0.002 | 0.003 | 0.003 | 0.041 | 0.11 | 0.09 | 0.23 | 0.06 | 0.03 | 0.28 |
| 1 | 198461064 | rs4915292 | 0.08 | -0.905 | 0.919 | 1.004 | 0.009 | 0.013 | 0.013 | 0.71 | 0.1 | 0.02 | 0.23 | 0.06 | 0.03 | 0.28 |
| 1 | 198469162 | rs10922438 | -0.293 | -0.903 | 0.957 | 1.013 | 0.007 | 0.009 | 0.009 | 0.631 | 0.13 | 0.09 | 0.25 | 0.08 | 0.05 | 0.27 |
| 1 | 198474448 | rs12048578 | -1.606 | -1.107 | 1.063 | 0.951 | -0.025 | 0.166 | 0.166 | 0 | 0.21 | 0.44 | 0.14 | 0.38 | 0.21 | 0.18 |
| 1 | 198480661 | rs11587248 | 0.179 | -1.097 | 0.981 | 0.913 | 0.009 | 0.021 | 0.021 | 1.866 | 0.12 | 0.03 | 0.24 | 0.08 | 0.05 | 0.27 |
| 1 | 198481371 | rs12058080 | -2.003 | -1.169 | 1.482 | 0.87 | -0.038 | -0.119 | -0.119 | 1.965 | 0.28 | 0.47 | 0.36 | 0.08 | 0.22 | 0.34 |
| 1 | 198484209 | rs2359847 | -2.003 | -1.169 | 1.482 | 0.87 | -0.038 | -0.119 | -0.119 | 2.955 | 0.28 | 0.47 | 0.36 | 0.08 | 0.22 | 0.34 |
| 1 | 198484427 | rs2249320 | -0.099 | -1.177 | 1.512 | 0.915 | -0.005 | 0.26 | 0.26 | 0 | 0.94 | 0.85 | 0.97 | 0.62 | 0.96 | 0.89 |
| 1 | 198484784 | rs16843226 | -2.005 | -1.161 | 1.429 | 0.861 | -0.038 | -0.119 | -0.119 | 1.42 | 0.28 | 0.47 | 0.36 | 0.08 | 0.22 | 0.34 |
| 1 | 198485706 | rs12083381 | -0.088 | -1.132 | 1.443 | 0.885 | -0.011 | -0.017 | -0.017 | 0 | 0.13 | 0.19 | 0.25 | 0.08 | 0.05 | 0.27 |
| 1 | 198489457 | rs10494777 | 0.122 | -1.124 | 1.424 | 0.889 | 0.003 | 0.005 | 0.005 | 1.524 | 0.12 | 0.09 | 0.24 | 0.08 | 0.05 | 0.27 |
| 1 | 198492454 | rs1055160 | -0.128 | -1.147 | 1.435 | 0.869 | -0.008 | -0.051 | -0.051 | 0 | 0.09 | 0.2 | 0.05 | 0 | 0.08 | 0.05 |
| 1 | 198494473 | rs10922446 | -1.081 | -1.144 | 1.287 | 0.872 | -0.021 | -0.055 | -0.055 | 0.579 | 0.21 | 0.3 | 0.3 | 0.08 | 0.14 | 0.31 |
| 1 | 198505546 | rs3736929 | -2.278 | -1.166 | 1.099 | 0.91 | 0.022 | -0.024 | -0.024 | 0 | 0.35 | 0.68 | 0.39 | 0.46 | 0.26 | 0.45 |
| 1 | 198511122 | rs6428467 | -2.105 | -0.87 | 1.226 | 0.998 | -0.011 | 0.024 | 0.024 | 0.301 | 0.27 | 0.64 | 0.35 | 0.45 | 0.18 | 0.43 |
| 1 | 198513826 | rs16843286 | 0.038 | -0.787 | 1.206 | 1.056 | -0.005 | -0.047 | -0.047 | 0 | 0.08 | 0.19 | 0.05 | 0 | 0.08 | 0.05 |
| 1 | 198517308 | rs6675366 | -1.051 | -0.628 | 0.725 | 1.201 | 0.014 | 0.417 | 0.417 | 0 | 0.07 | 0.32 | 0.05 | 0.37 | 0.04 | 0.11 |
| 1 | 198520125 | rs2359848 | -0.326 | -0.493 | 0.026 | 1.264 | -0.005 | 0.264 | 0.264 | 0 | 0.06 | 0.14 | 0.03 | 0.38 | 0.04 | 0.11 |
| 1 | 198521135 | rs17659788 | 0.685 | 0.005 | -0.945 | 0.677 | -0.092 | 0.126 | 0.126 | 0 | 0.19 | 0.01 | 0.14 | 0 | 0.25 | 0.06 |
| 1 | 198540028 | rs4915303 | -1.111 | -0.681 | 0.194 | 1.401 | -0.029 | 0.079 | 0.079 | 0.34 | 0.27 | 0.5 | 0.34 | 0.45 | 0.19 | 0.46 |
| 1 | 198542032 | rs2359852 | 0.496 | -0.487 | 0.174 | 1.449 | 0.002 | -0.007 | -0.007 | 0 | 0.91 | 0.91 | 0.81 | 0.86 | 0.94 | 0.8 |
| 1 | 198544730 | rs12742531 | -0.382 | -0.97 | 0.068 | 1.504 | 0.004 | -0.004 | -0.004 | 0 | 0.1 | 0.08 | 0.1 | 0.14 | 0.07 | 0.1 |
| 1 | 198561100 | rs2359854 | 0.931 | -1.631 | -0.384 | 1.115 | 0.024 | 0.031 | 0.031 | 0 | 0.48 | 0.45 | 0.46 | 0.26 | 0.64 | 0.33 |
| 1 | 198574902 | rs16843430 | -0.107 | -1.137 | -0.876 | 1.306 | 0.024 | -0.067 | -0.067 | 0.227 | 0.26 | 0.08 | 0.21 | 0.38 | 0.23 | 0.31 |
| 1 | 198601197 | rs2038926 | 0.044 | -0.279 | -0.653 | 0.309 | -0.004 | 0.001 | 0.001 | 0.043 | 0.33 | 0.41 | 0.4 | 0.69 | 0.36 | 0.48 |
| 1 | 198607328 | rs3754094 | 1.028 | -0.562 | -0.869 | 0.277 | 0.015 | -0.034 | -0.034 | 0 | 0.06 | 0 | 0.08 | 0.15 | 0.03 | 0.09 |
| 1 | 198609305 | rs6673408 | 0.514 | -0.774 | -0.31 | 0.109 | 0.004 | 0.021 | 0.021 | 0 | 0.1 | 0.26 | 0.21 | 0.05 | 0.05 | 0.14 |
| 1 | 198618707 | rs6686725 | 0.598 | -0.828 | -0.339 | 0.055 | -0.003 | 0.003 | 0.003 | 0 | 0.32 | 0.39 | 0.39 | 0.69 | 0.35 | 0.47 |
| 1 | 198619686 | rs1326270 | -1.451 | -0.989 | -0.437 | -0.286 | -0.098 | 0.146 | 0.146 | 0 | 0.64 | 0.9 | 0.76 | 0.89 | 0.56 | 0.82 |
| 1 | 198619759 | rs1326271 | -1.919 | -1.03 | -0.443 | -0.293 | -0.066 | 0.096 | 0.096 | 0 | 0.71 | 0.9 | 0.82 | 0.9 | 0.65 | 0.86 |
| 1 | 198628483 | rs10801677 | 0.822 | -1.042 | -0.335 | -0.245 | -0.024 | 0.036 | 0.036 | 1.959 | 0.07 | 0.18 | 0.09 | 0.15 | 0.03 | 0.09 |
| 1 | 198632356 | rs1998843 | 0.884 | -0.932 | -0.037 | -0.238 | 0.001 | -0.01 | -0.01 | 0 | 0.31 | 0.24 | 0.39 | 0.69 | 0.35 | 0.48 |
| 1 | 198634625 | rs1052238 | 0.698 | -0.83 | -0.284 | -0.366 | -0.002 | 0 | 0 | 0.103 | 0.39 | 0.4 | 0.49 | 0.69 | 0.54 | 0.51 |
| 1 | 198634997 | rs3767742 | 0.358 | -0.925 | 0.449 | -0.346 | -0.001 | 0.004 | 0.004 | 0 | 0.83 | 0.88 | 0.74 | 0.95 | 0.85 | 0.81 |
| 1 | 198644124 | rs9803750 | -1.823 | -1.027 | 0.553 | -0.354 | 0.023 | 0.048 | 0.048 | 0 | 0.38 | 0.7 | 0.42 | 0.18 | 0.3 | 0.36 |
| 1 | 198645649 | rs9803978 | 0.402 | -1.05 | 0.717 | -0.407 | 0.016 | -0.045 | -0.045 | 0.007 | 0.08 | 0.13 | 0.07 | 0 | 0.1 | 0.05 |
| 1 | 198656407 | rs2359952 | -1.937 | -1.021 | 0.781 | -0.023 | -0.012 | 0.001 | 0.001 | 0 | 0.38 | 0.51 | 0.47 | 0.35 | 0.3 | 0.42 |
| 1 | 198658856 | rs3754098 | -0.636 | -0.993 | 0.626 | 0.006 | -0.005 | 0.004 | 0.004 | 0.21 | 0.15 | 0.19 | 0.27 | 0.17 | 0.08 | 0.21 |
| 1 | 198660724 | rs16843712 | 0.332 | -0.959 | 0.641 | 0.009 | -0.005 | -0.01 | -0.01 | 0 | 0.1 | 0.17 | 0.19 | 0.04 | 0.06 | 0.14 |
| 1 | 198663661 | rs10494783 | -0.635 | -0.683 | 0.315 | 0.091 | -0.014 | 0.099 | 0.099 | 1.247 | 0.07 | 0.12 | 0.16 | 0.3 | 0.04 | 0.14 |
| 1 | 198664018 | rs12124872 | 0.368 | -0.564 | 0.324 | -0.034 | 0.011 | -0.037 | -0.037 | 0 | 0.08 | 0.09 | 0.06 | 0 | 0.11 | 0.05 |
| 1 | 198670555 | rs17623914 | 0.568 | -0.528 | -0.062 | -0.01 | 0.01 | 0.018 | 0.018 | 0 | 0.12 | 0.06 | 0.06 | 0 | 0.1 | 0.09 |
| 1 | 198672299 | rs16843742 | -0.646 | -0.49 | 0.021 | 0.06 | -0.002 | -0.003 | -0.003 | 4.868 | 0.22 | 0.21 | 0.2 | 0.17 | 0.22 | 0.2 |
| 1 | 198699793 | rs12120762 | -0.235 | -0.586 | -0.394 | -0.323 | 0.022 | 0.009 | 0.009 | 1.045 | 0.58 | 0.69 | 0.57 | 0.33 | 0.63 | 0.41 |
| 1 | 198704294 | rs7540378 | -0.117 | -0.789 | -0.172 | -0.596 | -0.015 | 0.022 | 0.022 | 0 | 0.19 | 0.27 | 0.3 | 0.24 | 0.12 | 0.21 |
| 1 | 198705081 | rs11589894 | 0.798 | -0.792 | -0.133 | -0.726 | -0.028 | 0.089 | 0.089 | 1.555 | 0.37 | 0.22 | 0.27 | 0.09 | 0.5 | 0.19 |
| 1 | 198719973 | rs874909 | 1.727 | -0.632 | -0.345 | -0.886 | -0.077 | 0.083 | 0.083 | 0.474 | 0.27 | 0.12 | 0.18 | 0.09 | 0.35 | 0.14 |
| 1 | 198739870 | rs1932436 | -0.562 | -1.614 | -0.605 | -1.259 | 0.089 | 0.133 | 0.133 | 3.304 | 0.43 | 0.77 | 0.44 | 0.86 | 0.25 | 0.64 |
| 1 | 198745866 | rs7538393 | 0.151 | -1.453 | -0.141 | -0.936 | -0.043 | 0.087 | 0.087 | 0.03 | 0.16 | 0.43 | 0.14 | 0.2 | 0.1 | 0.16 |
| 1 | 198751401 | rs2784115 | -0.007 | -1.261 | -0.397 | -1.144 | 0.005 | 0.104 | 0.104 | 0 | 0.26 | 0.09 | 0.31 | 0.06 | 0.27 | 0.24 |
| 1 | 198758216 | rs16843838 | 0.82 | -1.481 | -0.398 | -1.064 | -0.029 | 0.044 | 0.044 | 0 | 0.1 | 0.31 | 0.07 | 0.14 | 0.03 | 0.12 |
| 1 | 198760554 | rs2784103 | -2.151 | -1.318 | -0.866 | -1.097 | -0.072 | 0.073 | 0.073 | 0.322 | 0.72 | 0.85 | 0.8 | 0.91 | 0.62 | 0.87 |
| 1 | 198769899 | rs10919581 | -0.289 | -1.363 | -0.647 | -0.799 | 0.014 | 0.019 | 0.019 | 0.002 | 0.34 | 0.19 | 0.4 | 0.72 | 0.31 | 0.49 |
| 1 | 198780386 | rs2784110 | -0.804 | -1.365 | -0.368 | -0.671 | -0.001 | 0 | 0 | 2.067 | 0.53 | 0.44 | 0.62 | 0.59 | 0.54 | 0.7 |

| | | | | | | | | | | | | | | | | |
|---|---|---|---|---|---|---|---|---|---|---|---|---|---|---|---|---|
| 1 | 198785494 | rs1575394 | -0.164 | -1.523 | -0.519 | -0.789 | -0.007 | -0.012 | -0.012 | 0 | 0.4 | 0.28 | 0.55 | 0.59 | 0.44 | 0.61 |
| 1 | 198786953 | rs16843867 | 1.629 | -1.579 | -0.522 | -0.736 | 0.005 | -0.001 | -0.001 | 0 | 0.07 | 0.05 | 0.07 | 0.16 | 0.03 | 0.06 |
| 1 | 198788077 | rs12144017 | -0.278 | -1.38 | -0.414 | -0.689 | -0.004 | -0.007 | -0.007 | 0.099 | 0.4 | 0.33 | 0.55 | 0.59 | 0.44 | 0.61 |
| 1 | 198790120 | rs1359020 | -1.65 | -1.329 | -0.433 | -0.814 | -0.011 | 0.004 | 0.004 | 3.429 | 0.63 | 0.67 | 0.7 | 0.71 | 0.61 | 0.83 |
| 1 | 198791103 | rs7537080 | 0.985 | -1.168 | -0.609 | -0.522 | 0.009 | 0.005 | 0.005 | 0 | 0.11 | 0.09 | 0.07 | 0 | 0.11 | 0.08 |
| 1 | 198807259 | rs4141743 | 0.169 | -0.959 | -0.332 | -0.167 | -0.017 | 0.024 | 0.024 | 0 | 0.1 | 0.26 | 0.07 | 0.12 | 0.06 | 0.13 |
| 1 | 198807862 | rs12042054 | 0.067 | -1.037 | -0.885 | -0.129 | -0.003 | -0.089 | -0.089 | 1.694 | 0.26 | 0.08 | 0.3 | 0.51 | 0.24 | 0.4 |
| 1 | 198811372 | rs10919593 | 0.949 | -1.408 | -0.918 | -0.008 | 0.015 | -0.082 | -0.082 | 0 | 0.15 | 0.03 | 0.22 | 0.5 | 0.11 | 0.36 |
| 1 | 198821238 | rs7537170 | -0.087 | -1.684 | -0.672 | -0.668 | 0.014 | 0.004 | 0.004 | 0 | 0.31 | 0.25 | 0.29 | 0.72 | 0.19 | 0.5 |
| 1 | 198825960 | rs12406238 | 0.068 | -1.69 | -0.111 | -0.897 | -0.002 | -0.065 | -0.065 | 0 | 0.11 | 0.26 | 0.07 | 0 | 0.1 | 0.04 |
| 1 | 198837151 | rs6701884 | 0.441 | -1.749 | -0.163 | -1.418 | -0.003 | 0.002 | 0.002 | 0 | 0.21 | 0.2 | 0.31 | 0.1 | 0.24 | 0.29 |
| 1 | 198844315 | rs1007939 | 0.568 | -2.001 | -0.137 | -1.217 | -0.038 | 0 | 0 | 0.744 | 0.3 | 0.38 | 0.29 | 0.71 | 0.2 | 0.49 |
| 1 | 198845784 | rs10800596 | -1.955 | -1.536 | -0.434 | -1.192 | -0.104 | 0.016 | 0.016 | 3.337 | 0.72 | 0.94 | 0.76 | 0.81 | 0.64 | 0.88 |
| 1 | 198876258 | rs1322916 | 0.396 | -1.514 | -0.21 | -1.177 | -0.006 | 0.016 | 0.016 | 0 | 0.16 | 0.1 | 0.24 | 0.1 | 0.19 | 0.21 |
| 1 | 198880267 | rs6662894 | -2.454 | -1.908 | -0.093 | -1.46 | -0.044 | 0.017 | 0.017 | 0.487 | 0.79 | 0.89 | 0.76 | 0.86 | 0.64 | 0.89 |
| 1 | 198886404 | rs1275161 | -2.454 | -1.909 | -0.112 | -1.458 | -0.074 | 0.021 | 0.021 | 0.541 | 0.79 | 0.91 | 0.76 | 0.86 | 0.64 | 0.89 |
| 1 | 198894457 | rs1275164 | 0.428 | -2.439 | 0.085 | -1.653 | 0.001 | -0.024 | -0.024 | 0 | 0.26 | 0.27 | 0.18 | 0.07 | 0.24 | 0.16 |
| 1 | 198898427 | rs10919605 | -0.408 | -2.405 | 0.246 | -1.611 | 0.03 | 0.078 | 0.078 | 0.473 | 0.51 | 0.38 | 0.55 | 0.79 | 0.41 | 0.7 |
| 1 | 198914510 | rs10800602 | -0.629 | -2.397 | 0.301 | -1.592 | -0.023 | -0.1 | -0.1 | 0 | 0.74 | 0.61 | 0.81 | 0.93 | 0.77 | 0.85 |
| 1 | 198915057 | rs12047947 | 0.547 | -2.566 | 0.132 | -1.651 | 0.037 | 0.01 | 0.01 | 0.305 | 0.34 | 0.2 | 0.31 | 0.69 | 0.22 | 0.5 |
| 1 | 198920036 | rs6703089 | -0.928 | -2.525 | 0.289 | -1.651 | -0.002 | -0.029 | -0.029 | 9.021 | 0.76 | 0.74 | 0.82 | 0.93 | 0.77 | 0.82 |
| 1 | 198925918 | rs12758891 | 2.366 | -2.141 | 0.337 | -1.434 | -0.073 | 0.018 | 0.018 | 0.459 | 0.21 | 0.09 | 0.25 | 0.14 | 0.36 | 0.11 |
| 1 | 198929441 | rs6680666 | 0.326 | -2.215 | 0.381 | -1.454 | 0 | 0 | 0 | 0.013 | 0.36 | 0.33 | 0.36 | 0.69 | 0.22 | 0.53 |
| 1 | 198934601 | rs7550308 | 2.26 | -2.072 | 0.315 | -1.342 | 0.014 | -0.003 | -0.003 | 0.122 | 0.21 | 0.19 | 0.25 | 0.14 | 0.36 | 0.11 |
| 1 | 198939403 | rs1938042 | 0.887 | -2.125 | 0.049 | -1.354 | 0.024 | -0.001 | -0.001 | 0.012 | 0.31 | 0.21 | 0.29 | 0.69 | 0.19 | 0.52 |
| 1 | 198942452 | rs2154327 | 2.191 | -1.634 | -0.013 | -1.231 | -0.107 | 0.011 | 0.011 | 1.533 | 0.2 | 0.06 | 0.25 | 0.14 | 0.35 | 0.11 |
| 1 | 198961568 | rs1892433 | -0.51 | -1.843 | 0.307 | -1.304 | 0.037 | 0.067 | 0.067 | 1.541 | 0.5 | 0.39 | 0.56 | 0.74 | 0.4 | 0.72 |
| 1 | 198970728 | rs4915165 | -0.537 | -1.952 | 0.171 | -1.306 | 0.004 | -0.013 | -0.013 | 0 | 0.71 | 0.75 | 0.82 | 0.93 | 0.75 | 0.81 |
| 1 | 198976492 | rs12239179 | -0.168 | -1.72 | 0.069 | -1.202 | -0.027 | -0.053 | -0.053 | 0 | 0.65 | 0.57 | 0.76 | 0.86 | 0.75 | 0.71 |
| 1 | 198976520 | rs12240178 | 2.243 | -1.526 | -0.753 | -1.122 | -0.09 | 0.005 | 0.005 | 2.022 | 0.2 | 0.07 | 0.19 | 0.16 | 0.34 | 0.09 |
| 1 | 198981241 | rs12757682 | 2.034 | -1.558 | -0.724 | -1.157 | -0.069 | 0.045 | 0.045 | 3.237 | 0.22 | 0.11 | 0.21 | 0.12 | 0.34 | 0.09 |
| 1 | 198981329 | rs12737296 | 2.281 | -1.547 | -0.766 | -1.151 | -0.102 | 0.045 | 0.045 | 2.193 | 0.2 | 0.06 | 0.19 | 0.12 | 0.34 | 0.09 |
| 1 | 198991093 | rs2186188 | 0.84 | -1.823 | -0.793 | -1.196 | 0.016 | -0.014 | -0.014 | 0 | 0.27 | 0.21 | 0.24 | 0.66 | 0.21 | 0.46 |
| 1 | 198992043 | rs12125200 | -0.482 | -1.593 | -1.146 | -1.179 | 0.046 | 0.091 | 0.091 | 0 | 0.68 | 0.88 | 0.74 | 0.85 | 0.74 | 0.73 |
| 1 | 198995597 | rs12756679 | 1.661 | -0.687 | -0.918 | -0.765 | -0.037 | 0.026 | 0.026 | 3.199 | 0.22 | 0.17 | 0.23 | 0.12 | 0.34 | 0.09 |
| 1 | 198998535 | rs11581099 | -0.365 | -1.13 | -0.597 | 0.277 | -0.004 | -0.043 | -0.043 | 3.722 | 0.07 | 0 | 0.06 | 0.17 | 0.07 | 0.1 |
| 1 | 199015253 | rs7550394 | 0.347 | -0.793 | -0.282 | -0.12 | -0.003 | -0.002 | -0.002 | 0.052 | 0.84 | 0.87 | 0.77 | 0.83 | 0.81 | 0.76 |
| 1 | 199018143 | rs10919618 | 0.282 | -1.371 | -0.116 | -0.251 | 0.005 | -0.011 | -0.011 | 0 | 0.2 | 0.21 | 0.23 | 0.63 | 0.15 | 0.44 |
| 1 | 199019855 | rs322931 | -0.527 | -1.183 | 0.603 | -0.222 | 0 | -0.001 | -0.001 | 0 | 0.16 | 0.15 | 0.23 | 0.17 | 0.18 | 0.23 |
| 1 | 199020987 | rs322929 | -0.608 | -1.187 | 0.538 | -0.223 | -0.002 | 0.002 | 0.002 | 0.092 | 0.16 | 0.23 | 0.24 | 0.18 | 0.19 | 0.24 |
| 1 | 199033440 | rs322883 | 0.006 | -1.432 | 0.521 | -0.625 | -0.054 | -0.163 | -0.163 | 0 | 0.3 | 0.48 | 0.15 | 0 | 0.21 | 0.07 |
| 1 | 199034906 | rs962793 | -0.46 | -1.535 | 0.406 | -0.644 | 0.013 | 0.051 | 0.051 | 2.023 | 0.32 | 0.17 | 0.4 | 0.8 | 0.3 | 0.65 |
| 1 | 199038043 | rs10919619 | 0.688 | -1.5 | 0.304 | -0.498 | -0.032 | -0.053 | -0.053 | 0 | 0.1 | 0.01 | 0.19 | 0.19 | 0.14 | 0.2 |
| 1 | 199038249 | rs322879 | -1.406 | -1.291 | 0.683 | -0.752 | -0.01 | 0.036 | 0.036 | 0 | 0.63 | 0.75 | 0.54 | 0.81 | 0.49 | 0.73 |
| 1 | 199054666 | rs16844162 | -0.399 | -1.095 | 0.876 | -0.656 | 0.003 | 0.012 | 0.012 | 1.736 | 0.33 | 0.31 | 0.42 | 0.83 | 0.32 | 0.62 |
| 1 | 199075025 | rs4915342 | -0.653 | -1.171 | 0.925 | -0.638 | -0.001 | -0.017 | -0.017 | 0 | 0.36 | 0.36 | 0.42 | 0.84 | 0.34 | 0.62 |
| 1 | 199080141 | rs7538550 | -0.159 | -1.147 | 0.994 | -0.504 | 0.002 | 0.006 | 0.006 | 0.924 | 0.31 | 0.25 | 0.41 | 0.77 | 0.3 | 0.61 |
| 1 | 199086009 | rs322904 | 0.444 | -1.28 | 1.126 | -0.633 | -0.025 | -0.091 | -0.091 | 0 | 0.26 | 0.34 | 0.1 | 0 | 0.17 | 0.09 |
| 1 | 199097277 | rs427790 | -1.266 | -1.141 | 0.463 | -0.558 | -0.012 | 0.059 | 0.059 | 0 | 0.61 | 0.76 | 0.53 | 0.79 | 0.48 | 0.7 |
| 1 | 199102610 | rs322875 | -1.402 | -1.154 | 0.461 | -0.564 | -0.021 | 0.069 | 0.069 | 0 | 0.62 | 0.81 | 0.53 | 0.79 | 0.48 | 0.71 |
| 1 | 199109802 | rs13374545 | -0.009 | -1.106 | 0.539 | -0.569 | -0.061 | -0.153 | -0.153 | 0 | 0.29 | 0.5 | 0.14 | 0.02 | 0.21 | 0.1 |
| 1 | 199114986 | rs10753846 | -0.461 | -1.016 | 0.507 | -0.496 | 0.011 | 0.02 | 0.02 | 0 | 0.34 | 0.29 | 0.39 | 0.77 | 0.28 | 0.61 |
| 1 | 199115592 | rs10800611 | -0.347 | -0.955 | 0.517 | -0.469 | 0.006 | 0.018 | 0.018 | 0 | 0.33 | 0.28 | 0.4 | 0.77 | 0.29 | 0.61 |
| 1 | 199116609 | rs3861923 | -0.278 | -0.944 | 0.467 | -0.507 | 0.008 | 0.049 | 0.049 | 0 | 0.33 | 0.22 | 0.4 | 0.82 | 0.29 | 0.61 |
| 1 | 199118713 | rs16844287 | 0.215 | -1.103 | 0.497 | -0.458 | -0.056 | -0.153 | -0.153 | 2.558 | 0.25 | 0.42 | 0.12 | 0 | 0.17 | 0.09 |
| 1 | 199122765 | rs4915347 | -0.78 | 0.096 | -0.049 | 0.218 | -0.011 | 0.068 | 0.068 | 0 | 0.6 | 0.75 | 0.52 | 0.8 | 0.44 | 0.69 |
| 1 | 199134034 | rs16844329 | -0.802 | 0.455 | 0.366 | 1.279 | -0.018 | -0.027 | -0.027 | 0 | 0.07 | 0.01 | 0.29 | 0.23 | 0.08 | 0.19 |
| 1 | 199134239 | rs1471646 | 0.508 | 0.746 | 0.652 | 1.193 | 0.016 | 0.016 | 0.016 | 0.066 | 0.51 | 0.33 | 0.59 | 0.4 | 0.58 | 0.47 |
| 1 | 199134987 | rs12068990 | -1.382 | 1.365 | 0.935 | 1.982 | -0.028 | -0.038 | -0.038 | 0 | 0.07 | 0.01 | 0.3 | 0.22 | 0.1 | 0.18 |
| 1 | 199145557 | rs6667273 | -2.275 | 1.084 | 1.534 | 1.79 | 0.075 | 0.214 | 0.214 | 0.128 | 0.05 | 0.3 | 0.25 | 0.19 | 0.04 | 0.13 |
| 1 | 199181905 | rs6427760 | 0.749 | 1.002 | 0.993 | 1.936 | -0.013 | -0.044 | -0.044 | 0 | 0.69 | 0.81 | 0.68 | 0.45 | 0.67 | 0.59 |
| 1 | 199188907 | rs6427761 | -1.104 | 0.85 | 1.088 | 2.195 | -0.039 | 0.029 | 0.029 | 0 | 0.1 | 0.2 | 0.1 | 0.16 | 0.02 | 0.19 |
| 1 | 199193889 | rs10800619 | -2.215 | 0.78 | 1.271 | 2.208 | -0.06 | 0.101 | 0.101 | 0 | 0.16 | 0.57 | 0.34 | 0.3 | 0.08 | 0.39 |
| 1 | 199204697 | rs1482659 | 0.764 | 1.088 | 0.841 | 2.242 | -0.091 | -0.088 | -0.088 | 0 | 0.68 | 0.92 | 0.51 | 0.44 | 0.57 | 0.54 |
| 1 | 199204879 | rs1482658 | 0.526 | 1.092 | 0.839 | 2.172 | -0.112 | -0.142 | -0.142 | 0.566 | 0.72 | 0.95 | 0.73 | 0.59 | 0.63 | 0.75 |
| 1 | 199220833 | rs12076262 | -0.624 | 1.002 | 0.729 | 1.882 | -0.028 | 0.03 | 0.03 | 0 | 0.08 | 0.14 | 0.09 | 0.16 | 0.02 | 0.18 |
| 1 | 199221398 | rs6667933 | 0.302 | 1.127 | 0.35 | 2.021 | -0.127 | -0.099 | -0.099 | 1.249 | 0.74 | 0.98 | 0.76 | 0.67 | 0.64 | 0.75 |
| 1 | 199231736 | rs7538098 | -1.33 | 0.323 | 1.603 | 1.874 | -0.018 | 0.083 | 0.083 | 0 | 0.08 | 0.29 | 0.1 | 0.15 | 0.04 | 0.16 |
| 1 | 199240161 | rs7537542 | -1.519 | 0.895 | 0.434 | 1.381 | 0.005 | 0.091 | 0.091 | 2.2 | 0.14 | 0.56 | 0.41 | 0.29 | 0.12 | 0.28 |

| | | | | | | | | | | | | | | | | |
|---|---|---|---|---|---|---|---|---|---|---|---|---|---|---|---|---|
| 1 | 199242481 | rs12145889 | -0.5 | 2.073 | -0.408 | 0.632 | -0.068 | 0.071 | 0.071 | 0 | 0.12 | 0.01 | 0.11 | 0 | 0.21 | 0.03 |
| 1 | 199242683 | rs9663039 | -1.808 | 1.622 | 0.264 | 0.615 | 0.017 | -0.053 | -0.053 | 0 | 0.27 | 0.55 | 0.24 | 0.15 | 0.3 | 0.1 |
| 1 | 199244301 | rs1325193 | -2.122 | 2.407 | -0.161 | 1.104 | 0.019 | 0.004 | 0.004 | 0 | 0.38 | 0.55 | 0.52 | 0.34 | 0.48 | 0.31 |
| 1 | 199248661 | rs10919655 | -1.088 | 2.219 | -0.54 | 1.023 | -0.09 | -0.095 | -0.095 | 2.12 | 0.26 | 0.09 | 0.4 | 0.29 | 0.34 | 0.42 |
| 1 | 199254953 | rs6697569 | -0.194 | 2.495 | -0.234 | 0.863 | -0.003 | -0.011 | -0.011 | 0.19 | 0.11 | 0.12 | 0.11 | 0 | 0.2 | 0.03 |
| 1 | 199257785 | rs881576 | -1.783 | 2.255 | 0.105 | 1.123 | 0.004 | 0.002 | 0.002 | 0 | 0.4 | 0.51 | 0.53 | 0.35 | 0.54 | 0.45 |
| 1 | 199260653 | rs17682977 | -0.265 | 2.678 | -0.326 | 0.714 | -0.029 | 0.028 | 0.028 | 0 | 0.05 | 0 | 0.09 | 0.01 | 0.15 | 0 |
| 1 | 199263161 | rs7538555 | -0.845 | 2.045 | -1.805 | -0.787 | -0.048 | 0.252 | 0.252 | 0 | 0.67 | 0.98 | 0.78 | 1 | 0.67 | 0.7 |
| 1 | 199265270 | rs12075138 | -0.762 | 1.922 | -0.766 | 0.124 | 0.011 | 0.051 | 0.051 | 0 | 0.25 | 0.36 | 0.44 | 0.36 | 0.27 | 0.18 |
| 1 | 199266874 | rs11579522 | -0.704 | 1.869 | -0.63 | 0.246 | 0.035 | 0.019 | 0.019 | 0 | 0.17 | 0.28 | 0.23 | 0.19 | 0.21 | 0.14 |
| 1 | 199270943 | rs723692 | -2.026 | 1.324 | -0.211 | 0.864 | 0.013 | -0.018 | -0.018 | 0.084 | 0.34 | 0.75 | 0.52 | 0.57 | 0.31 | 0.28 |
| 1 | 199271533 | rs953051 | -0.518 | 1.413 | 0.117 | 0.9 | 0.017 | 0.007 | 0.007 | 0 | 0.17 | 0.23 | 0.22 | 0.19 | 0.21 | 0.14 |
| 1 | 199274467 | rs7546413 | -1.488 | 1.346 | 0.466 | 0.856 | -0.003 | 0.014 | 0.014 | 0 | 0.31 | 0.59 | 0.5 | 0.57 | 0.31 | 0.28 |
| 1 | 199281617 | rs2164524 | 0.491 | 1.075 | 0.565 | 0.748 | 0 | 0.067 | 0.067 | 0 | 0.1 | 0.21 | 0.12 | 0.18 | 0.06 | 0.13 |
| 1 | 199291722 | rs10800624 | 0.763 | 1.229 | -0.3 | 0.526 | 0 | -0.02 | -0.02 | 0 | 0.23 | 0.18 | 0.15 | 0.35 | 0.19 | 0.14 |
| 1 | 199298683 | rs1561144 | -1.463 | 1.267 | -0.458 | 0.138 | -0.069 | 0.251 | 0.251 | 0 | 0.54 | 0.89 | 0.66 | 0.89 | 0.49 | 0.44 |
| 1 | 199301087 | rs6413909 | -1.295 | 1.293 | -0.272 | 0.269 | 0.004 | -0.027 | -0.027 | 1.163 | 0.31 | 0.72 | 0.51 | 0.54 | 0.29 | 0.3 |
| 1 | 199304892 | rs6656521 | -0.581 | 1.152 | 0.068 | -0.088 | 0.026 | 0.209 | 0.209 | 0.023 | 0.1 | 0.45 | 0.22 | 0.23 | 0.07 | 0.13 |
| 1 | 199312280 | rs1325184 | -0.3 | 0.52 | -0.096 | -0.351 | -0.022 | 0.02 | 0.02 | 0.457 | 0.25 | 0.32 | 0.3 | 0.3 | 0.2 | 0.19 |
| 1 | 199315719 | rs12726043 | 0.783 | 0.955 | 0.629 | 0.088 | -0.001 | -0.015 | -0.015 | 0 | 0.23 | 0.18 | 0.15 | 0.35 | 0.2 | 0.14 |
| 1 | 199316227 | rs11580692 | -0.237 | 0.868 | 0.953 | 0.084 | 0.017 | 0.167 | 0.167 | 0 | 0.08 | 0.27 | 0.2 | 0.23 | 0.07 | 0.13 |
| 1 | 199316461 | rs645943 | 1.774 | 0.89 | 0.774 | -0.103 | 0.019 | 0.059 | 0.059 | 0 | 0.41 | 0.25 | 0.33 | 0.17 | 0.34 | 0.36 |
| 1 | 199316935 | rs17633331 | 1.924 | 1.102 | 0.794 | -0.133 | -0.034 | -0.003 | -0.003 | 0.14 | 0.08 | 0.01 | 0.1 | 0.06 | 0.13 | 0.25 |
| 1 | 199320785 | rs590258 | 1.336 | 1.099 | 0.855 | -0.119 | 0.044 | 0.051 | 0.051 | 0 | 0.48 | 0.29 | 0.39 | 0.3 | 0.35 | 0.37 |
| 1 | 199328001 | rs6669877 | -0.812 | 1.162 | 0.837 | -0.103 | 0.098 | 0.154 | 0.154 | 0.038 | 0.13 | 0.42 | 0.3 | 0.25 | 0.22 | 0.1 |
| 1 | 199329274 | rs576141 | 0.201 | 1.157 | 0.755 | -0.117 | -0.025 | -0.011 | -0.011 | 0 | 0.61 | 0.74 | 0.7 | 0.55 | 0.58 | 0.47 |
| 1 | 199342144 | rs473023 | 1.923 | 1.119 | 0.782 | -0.147 | 0.022 | 0.091 | 0.091 | 0 | 0.4 | 0.18 | 0.32 | 0.17 | 0.34 | 0.36 |
| 1 | 199342535 | rs9887912 | -1.093 | 1.209 | 0.86 | -0.133 | 0.099 | 0.098 | 0.098 | 0 | 0.17 | 0.45 | 0.33 | 0.25 | 0.27 | 0.13 |
| 1 | 199346031 | rs670197 | 0.133 | 0.917 | 1.004 | -0.034 | 0.013 | 0.045 | 0.045 | 0 | 0.78 | 0.64 | 0.8 | 0.67 | 0.75 | 0.54 |
| 1 | 199347080 | rs1898240 | 0.889 | 0.907 | 0.99 | -0.035 | -0.001 | -0.012 | -0.012 | 5.461 | 0.23 | 0.18 | 0.15 | 0.35 | 0.2 | 0.14 |
| 1 | 199348223 | rs477569 | -0.823 | 0.915 | 0.987 | -0.048 | -0.03 | 0.027 | 0.027 | 0 | 0.84 | 0.92 | 0.85 | 0.9 | 0.78 | 0.61 |
| 1 | 199348842 | rs12065723 | 0.869 | 0.962 | 1.075 | -0.021 | 0.044 | -0.006 | -0.006 | 0 | 0.05 | 0.16 | 0.11 | 0.02 | 0.15 | 0 |
| 1 | 199354034 | rs589437 | 0.372 | 0.921 | 1.083 | -0.017 | 0.001 | -0.004 | -0.004 | 0.489 | 0.57 | 0.56 | 0.66 | 0.32 | 0.56 | 0.41 |
| 1 | 199357512 | rs1819293 | 0.306 | 1.147 | 1.525 | 0.465 | 0.035 | 0.282 | 0.282 | 0 | 0.05 | 0.28 | 0.05 | 0.23 | 0.02 | 0.07 |
| 1 | 199360836 | rs611739 | 1.738 | 1.077 | 1.457 | 0.38 | 0.027 | 0.135 | 0.135 | 0 | 0.46 | 0.18 | 0.34 | 0.17 | 0.39 | 0.43 |
| 1 | 199362216 | rs571609 | 1.309 | 1.147 | 1.415 | 0.41 | 0.011 | 0.022 | 0.022 | 0 | 0.53 | 0.36 | 0.41 | 0.3 | 0.4 | 0.44 |
| 1 | 199362820 | rs571754 | 0.048 | 1.166 | 1.359 | 0.445 | -0.019 | -0.02 | -0.02 | 0 | 0.66 | 0.73 | 0.72 | 0.55 | 0.63 | 0.55 |
| 1 | 199363883 | rs7515853 | -0.27 | 0.864 | 1.438 | 0.7 | 0.009 | 0.06 | 0.06 | 0 | 0.83 | 0.74 | 0.83 | 0.62 | 0.8 | 0.61 |
| 1 | 199374488 | rs661480 | 0.91 | 0.954 | 1.011 | 0.626 | 0 | -0.012 | -0.012 | 0 | 0.57 | 0.48 | 0.44 | 0.3 | 0.43 | 0.44 |
| 1 | 199377000 | rs585476 | 1.228 | 0.907 | 1.052 | 0.563 | -0.004 | -0.01 | -0.01 | 0.124 | 0.54 | 0.47 | 0.42 | 0.3 | 0.41 | 0.42 |
| 1 | 199386880 | rs528227 | 1.511 | 0.999 | 1.003 | 0.589 | 0.013 | 0.043 | 0.043 | 0 | 0.48 | 0.36 | 0.35 | 0.17 | 0.4 | 0.42 |
| 1 | 199394582 | rs1069784 | 1.599 | 0.991 | 1.011 | 0.585 | 0.02 | 0.086 | 0.086 | 0 | 0.47 | 0.29 | 0.35 | 0.17 | 0.4 | 0.42 |
| 1 | 199405454 | rs11579495 | 0.015 | 1.028 | 1.006 | 0.6 | 0.013 | -0.031 | -0.031 | 0 | 0.06 | 0.1 | 0.17 | 0 | 0.08 | 0.05 |
| 1 | 199418899 | rs2152872 | -0.491 | 0.938 | 1.047 | 0.765 | 0.002 | 0.012 | 0.012 | 0 | 0.81 | 0.76 | 0.81 | 0.62 | 0.78 | 0.58 |
| 1 | 199419004 | rs16844836 | 0.98 | 0.958 | 0.989 | 0.762 | 0 | -0.006 | -0.006 | 2.131 | 0.19 | 0.18 | 0.15 | 0.31 | 0.17 | 0.12 |
| 1 | 199424523 | rs2184965 | 0.556 | 0.983 | 0.952 | 0.804 | -0.008 | -0.006 | -0.006 | 0.725 | 0.62 | 0.66 | 0.55 | 0.55 | 0.59 | 0.5 |
| 1 | 199425806 | rs12037316 | -0.179 | 0.993 | 0.916 | 0.829 | -0.011 | -0.047 | -0.047 | 0 | 0.29 | 0.22 | 0.29 | 0.5 | 0.33 | 0.14 |
| 1 | 199426535 | rs6676168 | -0.404 | 1.002 | 0.918 | 0.856 | 0.018 | 0.03 | 0.03 | 0.056 | 0.1 | 0.11 | 0.27 | 0.25 | 0.23 | 0.1 |
| 1 | 199427140 | rs10800628 | -0.761 | 0.983 | 1.015 | 0.863 | 0.014 | 0.082 | 0.082 | 0 | 0.17 | 0.25 | 0.35 | 0.38 | 0.24 | 0.11 |
| 1 | 199428249 | rs6658450 | 1.868 | 0.936 | 0.942 | 0.787 | -0.037 | 0.039 | 0.039 | 1.395 | 0.07 | 0.14 | 0.07 | 0.13 | 0.01 | 0.01 |
| 1 | 199433291 | rs1590636 | -0.605 | 0.983 | 0.896 | 0.789 | 0.012 | 0.037 | 0.037 | 0 | 0.82 | 0.77 | 0.81 | 0.67 | 0.78 | 0.58 |
| 1 | 199434760 | rs1327068 | 1.28 | 0.989 | 0.799 | 0.755 | 0 | 0.019 | 0.019 | 0 | 0.53 | 0.44 | 0.45 | 0.19 | 0.55 | 0.42 |
| 1 | 199437760 | rs6427769 | 0.716 | 0.972 | 0.819 | 0.732 | 0.023 | -0.007 | -0.007 | 0 | 0.7 | 0.63 | 0.53 | 0.76 | 0.58 | 0.61 |
| 1 | 199438213 | rs12144397 | 1.525 | 0.96 | 0.815 | 0.738 | 0.011 | 0.009 | 0.009 | 0 | 0.51 | 0.42 | 0.38 | 0.45 | 0.42 | 0.49 |
| 1 | 199441521 | rs1571147 | 1.243 | 1.006 | 0.82 | 0.757 | 0.03 | 0.032 | 0.032 | 0 | 0.65 | 0.5 | 0.47 | 0.49 | 0.55 | 0.53 |
| 1 | 199442312 | rs10494793 | 0.929 | 0.93 | 0.668 | 0.535 | 0 | -0.006 | -0.006 | 0 | 0.19 | 0.19 | 0.14 | 0.35 | 0.17 | 0.12 |
| 1 | 199449355 | rs1415665 | 0.295 | 1.039 | 0.779 | -0.597 | 0.003 | -0.005 | -0.005 | 0.683 | 0.59 | 0.55 | 0.63 | 0.32 | 0.56 | 0.46 |
| 1 | 199450063 | rs12057228 | -0.032 | 1.086 | 0.801 | -0.796 | 0.001 | -0.001 | -0.001 | 0 | 0.13 | 0.17 | 0.13 | 0.11 | 0.17 | 0.01 |
| 1 | 199454878 | rs1890133 | -0.666 | 1.108 | 0.857 | -0.818 | 0.011 | -0.014 | -0.014 | 0.899 | 0.16 | 0.27 | 0.33 | 0.11 | 0.22 | 0.05 |
| 1 | 199457527 | rs7540552 | 0.167 | 1.055 | 0.881 | -0.844 | -0.004 | 0.009 | 0.009 | 0 | 0.59 | 0.62 | 0.68 | 0.3 | 0.58 | 0.45 |
| 1 | 199464611 | rs2095683 | 0.946 | 0.905 | 0.779 | -0.963 | 0 | -0.05 | -0.05 | 0 | 0.48 | 0.4 | 0.5 | 0.02 | 0.55 | 0.4 |
| 1 | 199471616 | rs12124166 | 1.662 | 0.992 | 0.78 | -0.96 | 0.011 | -0.006 | -0.006 | 0.138 | 0.38 | 0.27 | 0.21 | 0 | 0.33 | 0.35 |
| 1 | 199473534 | rs12065500 | 1.229 | 0.825 | 0.595 | -0.819 | -0.046 | 0.002 | 0.002 | 1.332 | 0.09 | 0.22 | 0.04 | 0.09 | 0.01 | 0.01 |
| 1 | 199474027 | rs1415667 | 0.775 | 0.968 | 0.366 | -0.798 | -0.004 | -0.03 | -0.03 | 0 | 0.23 | 0.17 | 0.18 | 0.61 | 0.21 | 0.22 |
| 1 | 199478315 | rs2050643 | 2 | 0.931 | 0.06 | -1.009 | -0.031 | 0.248 | 0.248 | 0 | 0.33 | 0.02 | 0.19 | 0 | 0.32 | 0.26 |
| 1 | 199480321 | rs1361106 | -1.388 | 0.887 | 0.397 | -0.676 | -0.022 | 0.025 | 0.025 | 0 | 0.83 | 0.9 | 0.87 | 0.9 | 0.79 | 0.67 |
| 1 | 199486414 | rs6683501 | 0.64 | 0.85 | 0.631 | -0.716 | -0.002 | -0.001 | -0.001 | 0 | 0.55 | 0.57 | 0.52 | 0.12 | 0.57 | 0.41 |
| 1 | 199496362 | rs10494797 | 1.002 | 1.317 | 0.657 | -0.492 | 0 | -0.023 | -0.023 | 0.904 | 0.45 | 0.54 | 0.43 | 0.14 | 0.41 | 0.43 |
| 1 | 199519488 | rs2225880 | 0.418 | 1.144 | 0.772 | -0.363 | -0.005 | 0.014 | 0.014 | 0 | 0.23 | 0.31 | 0.21 | 0.69 | 0.19 | 0.22 |

| | | | | | | | | | | | | | | | | |
|---|---|---|---|---|---|---|---|---|---|---|---|---|---|---|---|---|
| 1 | 199521371 | rs2786723 | -0.36 | 1.155 | 0.711 | -0.409 | 0.008 | -0.001 | -0.001 | 0 | 0.73 | 0.84 | 0.72 | 0.75 | 0.76 | 0.65 |
| 1 | 199531025 | rs6691743 | 0.314 | 1.129 | 0.79 | -0.356 | -0.005 | 0.01 | 0.01 | 0 | 0.24 | 0.31 | 0.21 | 0.69 | 0.19 | 0.22 |
| 1 | 199539063 | rs1415661 | 0.091 | 1.25 | 0.76 | -0.298 | -0.007 | 0.005 | 0.005 | 0 | 0.67 | 0.74 | 0.63 | 0.73 | 0.62 | 0.65 |
| 1 | 199544080 | rs945878 | 0.385 | 1.182 | 0.686 | -0.279 | 0.002 | -0.006 | -0.006 | 0.438 | 0.64 | 0.64 | 0.46 | 0.73 | 0.58 | 0.62 |
| 1 | 199544559 | rs2812261 | 1.373 | 1.209 | 0.27 | -0.298 | 0.009 | 0.067 | 0.067 | 0 | 0.4 | 0.26 | 0.24 | 0.04 | 0.37 | 0.4 |
| 1 | 199552831 | rs2812265 | 1.38 | 1.217 | 0.019 | -0.285 | 0.005 | 0.01 | 0.01 | 0 | 0.4 | 0.3 | 0.24 | 0.04 | 0.37 | 0.39 |
| 1 | 199555143 | rs4367802 | 0.98 | 0.942 | -0.569 | 0.069 | 0.017 | -0.01 | -0.01 | 0 | 0.05 | 0.01 | 0.14 | 0.19 | 0.01 | 0.03 |
| 1 | 199555572 | rs1337728 | -0.542 | 1.209 | -0.437 | 0.11 | 0.025 | 0.005 | 0.005 | 0.408 | 0.72 | 0.89 | 0.73 | 0.75 | 0.77 | 0.64 |
| 1 | 199557306 | rs16845170 | 0.316 | 1.259 | -0.506 | 0.273 | -0.004 | -0.016 | -0.016 | 1.856 | 0.21 | 0.19 | 0.19 | 0.59 | 0.21 | 0.22 |
| 1 | 199557772 | rs2786738 | 0.88 | 1.288 | -0.53 | 0.198 | -0.002 | -0.045 | -0.045 | 0 | 0.45 | 0.45 | 0.43 | 0.15 | 0.43 | 0.44 |
| 1 | 199571190 | rs7521829 | -0.579 | 1.068 | -0.692 | 0.538 | 0 | 0.001 | 0.001 | 0.136 | 0.34 | 0.32 | 0.42 | 0.29 | 0.37 | 0.34 |
| 1 | 199582376 | rs6662178 | -0.777 | 1.295 | -0.502 | 0.99 | 0.023 | 0.055 | 0.055 | 0.292 | 0.29 | 0.54 | 0.23 | 0.49 | 0.32 | 0.26 |
| 1 | 199582872 | rs9628679 | -0.27 | 1.296 | -1.412 | 0.837 | 0.006 | 0.015 | 0.015 | 0.861 | 0.47 | 0.88 | 0.54 | 0.44 | 0.54 | 0.31 |
| 1 | 199584037 | rs4915364 | -0.672 | 0.728 | -1.993 | 1 | 0.057 | -0.015 | -0.015 | 0 | 0.65 | 0.92 | 0.76 | 0.67 | 0.7 | 0.6 |
| 1 | 199587072 | rs10800646 | -0.152 | 1.373 | -0.769 | 0.935 | -0.004 | -0.003 | -0.003 | 0 | 0.57 | 0.5 | 0.55 | 0.55 | 0.56 | 0.57 |
| 1 | 199588458 | rs4612653 | 0.735 | 1.453 | -0.495 | 0.903 | 0.001 | 0.033 | 0.033 | 1.159 | 0.4 | 0.29 | 0.44 | 0.26 | 0.41 | 0.27 |
| 1 | 199589986 | rs6693206 | 0.425 | 1.668 | 0.366 | 1.048 | 0.11 | 0.117 | 0.117 | 0.271 | 0.05 | 0.28 | 0.15 | 0.12 | 0.05 | 0.02 |
| 1 | 199590921 | rs12730081 | 0.206 | 1.632 | 0.234 | 1.049 | -0.054 | -0.09 | -0.09 | 0.11 | 0.16 | 0.02 | 0.1 | 0.28 | 0.16 | 0.3 |
| 1 | 199593842 | rs12725952 | -1.311 | 1.641 | 0.321 | 1.053 | 0.036 | 0.042 | 0.042 | 0 | 0.26 | 0.6 | 0.33 | 0.4 | 0.3 | 0.33 |
| 1 | 199598783 | rs4290087 | -1.169 | 1.491 | 0.373 | 0.78 | 0.169 | 0.073 | 0.073 | 0.26 | 0.09 | 0.43 | 0.22 | 0.13 | 0.14 | 0.03 |
| 1 | 199608286 | rs4301659 | -0.598 | 1.548 | 0.278 | 0.808 | 0.172 | 0.081 | 0.081 | 0.215 | 0.07 | 0.36 | 0.21 | 0.13 | 0.13 | 0.03 |
| 1 | 199610237 | rs4369252 | -1.017 | 1.353 | 0.555 | 0.764 | 0.09 | 0.057 | 0.057 | 0.271 | 0.08 | 0.36 | 0.19 | 0.12 | 0.1 | 0.04 |
| 1 | 199611059 | rs4431873 | -0.876 | 1.232 | -0.005 | 0.773 | 0.054 | -0.012 | -0.012 | 0.051 | 0.64 | 0.92 | 0.75 | 0.68 | 0.69 | 0.6 |
| 1 | 199611305 | rs4604723 | -0.876 | 1.232 | -0.005 | 0.773 | 0.054 | -0.012 | -0.012 | 0.019 | 0.64 | 0.92 | 0.75 | 0.68 | 0.69 | 0.6 |
| 1 | 199629438 | rs4623735 | -1.239 | 1.304 | 1.207 | 0.746 | 0.1 | 0.065 | 0.065 | 0.23 | 0.08 | 0.4 | 0.18 | 0.12 | 0.1 | 0.02 |
| 1 | 199634402 | rs16845384 | 0.066 | 1.486 | 0.996 | 0.945 | -0.022 | -0.05 | -0.05 | 0 | 0.17 | 0.08 | 0.12 | 0.28 | 0.17 | 0.27 |
| 1 | 199634598 | rs10919736 | -1.311 | 1.461 | 1.044 | 0.931 | 0.031 | 0.058 | 0.058 | 0 | 0.27 | 0.6 | 0.31 | 0.4 | 0.27 | 0.29 |
| 1 | 199636944 | rs16845393 | -0.994 | 1.448 | 0.576 | 0.788 | 0.071 | 0.044 | 0.044 | 0 | 0.07 | 0.3 | 0.18 | 0.12 | 0.1 | 0.02 |
| 1 | 199688536 | rs1429700 | 0.917 | 0.751 | -0.489 | 0.08 | -0.005 | 0.072 | 0.072 | 0 | 0.17 | 0 | 0.12 | 0.03 | 0.19 | 0.13 |
| 1 | 199690702 | rs2435240 | -0.845 | 0.919 | -0.077 | -0.019 | 0.01 | -0.013 | -0.013 | 0 | 0.58 | 0.5 | 0.41 | 0.62 | 0.5 | 0.68 |
| 1 | 199691414 | rs16845432 | 0.291 | 0.381 | -0.779 | -0.151 | 0.056 | -0.043 | -0.043 | 0 | 0.38 | 0.12 | 0.23 | 0.38 | 0.31 | 0.38 |
| 1 | 199691458 | rs2435242 | -1.014 | 0.692 | -0.379 | -0.348 | 0.022 | -0.016 | -0.016 | 0 | 0.67 | 0.53 | 0.51 | 0.71 | 0.56 | 0.74 |
| 1 | 199707262 | rs17655162 | 1.101 | 1.2 | -0.69 | -0.525 | -0.003 | 0.003 | 0.003 | 0 | 0.3 | 0.29 | 0.3 | 0.24 | 0.37 | 0.21 |
| 1 | 199764832 | rs4465240 | -1.042 | 1.086 | -0.796 | -0.608 | 0.002 | -0.011 | -0.011 | 0 | 0.6 | 0.57 | 0.64 | 0.72 | 0.57 | 0.71 |
| 1 | 199772234 | rs1346680 | 1.559 | 1.126 | -0.79 | -0.61 | -0.011 | 0.009 | 0.009 | 0 | 0.09 | 0.03 | 0.07 | 0.04 | 0.09 | 0.08 |
| 1 | 199778956 | rs1366978 | -0.007 | 0.926 | -0.999 | -0.498 | -0.003 | -0.015 | -0.015 | 0 | 0.43 | 0.5 | 0.45 | 0.35 | 0.43 | 0.41 |
| 1 | 199779779 | rs265450 | 0.953 | 0.909 | -1.009 | -0.5 | -0.006 | -0.066 | -0.066 | 1.246 | 0.17 | 0.07 | 0.2 | 0.38 | 0.14 | 0.29 |
| 1 | 199782570 | rs10800652 | -0.276 | 0.772 | -0.795 | -0.419 | -0.004 | -0.017 | -0.017 | 0 | 0.43 | 0.49 | 0.44 | 0.35 | 0.42 | 0.41 |
| 1 | 199783024 | rs987648 | 0.845 | 0.39 | -0.397 | -0.465 | 0.01 | 0.006 | 0.006 | 0.015 | 0.09 | 0.02 | 0.04 | 0.04 | 0.05 | 0.08 |
| 1 | 199785642 | rs17719305 | 0.41 | 0.444 | -0.416 | -0.483 | -0.003 | 0.012 | 0.012 | 0.163 | 0.1 | 0.02 | 0.07 | 0.04 | 0.09 | 0.08 |
| 1 | 199787801 | rs4915387 | -0.456 | 0.115 | -0.246 | -0.63 | 0.007 | 0.033 | 0.033 | 0 | 0.51 | 0.41 | 0.61 | 0.66 | 0.5 | 0.63 |
| 1 | 199792844 | rs12027362 | 0.244 | -0.009 | -0.264 | -0.8 | -0.057 | 0.021 | 0.021 | 0.195 | 0.39 | 0.17 | 0.53 | 0.64 | 0.43 | 0.57 |
| 1 | 199797615 | rs7528943 | -0.942 | 0.567 | 0.086 | -0.564 | -0.003 | -0.015 | -0.015 | 0 | 0.63 | 0.62 | 0.69 | 0.76 | 0.59 | 0.72 |
| 1 | 199801373 | rs6700960 | -1.008 | 0.342 | 0.002 | -0.542 | 0.002 | 0.004 | 0.004 | 0 | 0.63 | 0.66 | 0.7 | 0.76 | 0.6 | 0.73 |
| 1 | 199804080 | rs12751057 | 0.8 | 0.368 | 0.23 | -0.311 | 0.007 | -0.002 | -0.002 | 0.295 | 0.08 | 0.02 | 0.06 | 0.05 | 0.05 | 0.07 |
| 1 | 199809292 | rs2553731 | 0.03 | 0.35 | 0.257 | -0.298 | 0.001 | 0.014 | 0.014 | 0 | 0.5 | 0.42 | 0.41 | 0.37 | 0.54 | 0.38 |
| 1 | 199814415 | rs10919745 | 0.439 | 0.605 | 0.388 | -0.134 | 0.002 | -0.003 | -0.003 | 0.078 | 0.09 | 0.08 | 0.06 | 0.13 | 0.05 | 0.09 |
| 1 | 199822974 | rs12096427 | -0.358 | 0.57 | 0.389 | -0.182 | 0.007 | 0.025 | 0.025 | 0 | 0.12 | 0.27 | 0.1 | 0.12 | 0.09 | 0.12 |
| 1 | 199831111 | rs1346098 | -0.186 | 0.543 | 0.45 | -0.258 | 0.004 | 0.029 | 0.029 | 0 | 0.5 | 0.37 | 0.57 | 0.64 | 0.48 | 0.63 |
| 1 | 199832023 | rs1425791 | -0.437 | 0.567 | 0.461 | -0.243 | 0.008 | 0.014 | 0.014 | 0 | 0.13 | 0.34 | 0.1 | 0.09 | 0.09 | 0.12 |
| 1 | 199839780 | rs7550378 | 0.267 | 0.363 | 0.399 | -0.226 | -0.015 | 0.011 | 0.011 | 0 | 0.1 | 0.2 | 0.07 | 0.09 | 0.05 | 0.12 |
| 1 | 199840991 | rs16845628 | 1.147 | 0.441 | 0.247 | -0.137 | -0.039 | -0.086 | -0.086 | 0 | 0.13 | 0.01 | 0.17 | 0.31 | 0.13 | 0.18 |
| 1 | 199844057 | rs10919757 | -0.091 | 0.636 | 0.931 | -0.803 | -0.016 | 0.012 | 0.012 | 0 | 0.1 | 0.22 | 0.07 | 0.09 | 0.05 | 0.12 |
| 1 | 199881715 | rs12759783 | 0.265 | 0.647 | -0.131 | -0.593 | 0 | -0.003 | -0.003 | 0 | 0.52 | 0.5 | 0.59 | 0.64 | 0.48 | 0.62 |
| 1 | 199889671 | rs6677214 | 0.149 | 0.682 | -0.764 | -0.68 | -0.001 | 0.032 | 0.032 | 0 | 0.53 | 0.61 | 0.6 | 0.68 | 0.49 | 0.65 |
| 1 | 199891278 | rs12745887 | 0.015 | 0.557 | -0.708 | -0.503 | 0.008 | -0.01 | -0.01 | 0.138 | 0.08 | 0.02 | 0.06 | 0.06 | 0.05 | 0.11 |
| 1 | 199902929 | rs755556 | -0.362 | 0.806 | -0.497 | -1.038 | 0 | -0.004 | -0.004 | 0 | 0.7 | 0.68 | 0.73 | 0.8 | 0.65 | 0.75 |
| 1 | 199925306 | rs1991245 | 0.349 | 0.974 | -1.149 | -1.5 | -0.032 | 0.098 | 0.098 | 0 | 0.45 | 0.07 | 0.52 | 0.72 | 0.39 | 0.65 |
| 1 | 199926184 | rs4915398 | -0.624 | 0.987 | -0.253 | -1.57 | -0.006 | -0.035 | -0.035 | 0.087 | 0.63 | 0.48 | 0.7 | 0.78 | 0.63 | 0.74 |
| 1 | 199931323 | rs6662512 | 0.159 | 0.871 | -0.243 | -1.33 | 0.014 | -0.022 | -0.022 | 0.027 | 0.06 | 0.1 | 0.06 | 0.01 | 0.09 | 0.06 |
| 1 | 199940193 | rs9662614 | 0.227 | 0.895 | -0.235 | -1.329 | 0.006 | -0.013 | -0.013 | 0 | 0.06 | 0.06 | 0.06 | 0.01 | 0.1 | 0.06 |
| 1 | 199963622 | rs4465241 | 0.786 | 0.854 | -0.14 | -1.415 | 0.015 | -0.037 | -0.037 | 0 | 0.88 | 0.81 | 0.88 | 0.99 | 0.83 | 0.94 |
| 1 | 199964344 | rs4147452 | -0.395 | 0.501 | -0.139 | -1.404 | 0.021 | -0.026 | -0.026 | 0 | 0.94 | 0.86 | 0.94 | 0.99 | 0.91 | 0.94 |
| 1 | 199965168 | rs10919791 | 1.173 | 0.136 | -0.666 | -1.698 | -0.02 | -0.068 | -0.068 | 0 | 0.25 | 0.09 | 0.36 | 0.67 | 0.22 | 0.48 |
| 1 | 199971261 | rs4244145 | 0.294 | 0.253 | -0.319 | -1.636 | 0.021 | -0.061 | -0.061 | 0 | 0.88 | 0.75 | 0.9 | 0.99 | 0.85 | 0.94 |
| 1 | 199985700 | rs2816939 | -0.683 | 0.245 | -0.083 | -1.624 | 0.053 | -0.069 | -0.069 | 0 | 0.13 | 0.36 | 0.11 | 0.02 | 0.16 | 0.07 |
| 1 | 199995418 | rs9427440 | -0.796 | 0.008 | -0.587 | -1.774 | -0.063 | 0.146 | 0.146 | 0 | 0.52 | 0.93 | 0.62 | 0.76 | 0.48 | 0.61 |
| 1 | 199997778 | rs2816949 | -0.98 | 0.018 | -0.52 | -1.778 | 0.003 | 0.067 | 0.067 | 2.985 | 0.23 | 0.82 | 0.19 | 0.06 | 0.22 | 0.09 |
| 1 | 200000125 | rs3790854 | 0.78 | 0.048 | -0.504 | -1.628 | -0.027 | -0.038 | -0.038 | 0 | 0.28 | 0.11 | 0.43 | 0.7 | 0.26 | 0.52 |

| | | | | | | | | | | | | | | | | |
|---|---|---|---|---|---|---|---|---|---|---|---|---|---|---|---|---|
| 1 | 200002126 | rs3762399 | 0.683 | -0.213 | 0.126 | -1.633 | 0.019 | -0.03 | -0.03 | 1.663 | 0.1 | 0.29 | 0.08 | 0.04 | 0.09 | 0.02 |
| 1 | 200007432 | rs3790844 | 1.07 | -0.079 | -0.861 | -1.596 | -0.008 | -0.064 | -0.064 | 0.084 | 0.26 | 0.11 | 0.4 | 0.65 | 0.24 | 0.47 |
| 1 | 200008271 | rs2821361 | -0.834 | -0.029 | -0.83 | -1.781 | -0.038 | 0.123 | 0.123 | 0.317 | 0.55 | 0.87 | 0.64 | 0.75 | 0.49 | 0.62 |
| 1 | 200014762 | rs2817008 | -1.908 | -0.126 | 0.013 | -1.72 | 0.022 | 0.18 | 0.18 | 5.724 | 0.39 | 0.77 | 0.28 | 0.06 | 0.33 | 0.21 |
| 1 | 200014790 | rs2817009 | -0.964 | -0.154 | 0.428 | -1.74 | -0.038 | -0.132 | -0.132 | 0.001 | 0.25 | 0.51 | 0.17 | 0.04 | 0.18 | 0.13 |
| 1 | 200015228 | rs2248673 | 0.007 | 0.034 | 0.26 | -1.477 | -0.001 | -0.029 | -0.029 | 0 | 0.15 | 0.11 | 0.08 | 0 | 0.09 | 0.11 |
| 1 | 200016460 | rs3790840 | -0.896 | 0.177 | 0.239 | -1.258 | 0.02 | -0.077 | -0.077 | 0 | 0.15 | 0.27 | 0.11 | 0.02 | 0.15 | 0.08 |
| 1 | 200020261 | rs2737640 | -0.254 | -0.177 | -0.123 | -1.09 | -0.002 | -0.029 | -0.029 | 0 | 0.15 | 0.11 | 0.08 | 0 | 0.08 | 0.11 |
| 1 | 200021261 | rs2821372 | -1.312 | -0.342 | 0.349 | -1.38 | -0.043 | -0.142 | -0.142 | 0 | 0.29 | 0.44 | 0.18 | 0.05 | 0.2 | 0.15 |
| 1 | 200022667 | rs2737641 | -1.149 | -0.784 | 0.477 | -1.091 | -0.049 | -0.05 | -0.05 | 0.021 | 0.3 | 0.54 | 0.19 | 0.18 | 0.21 | 0.18 |
| 1 | 200031107 | rs2816913 | -1.214 | -0.499 | 0.188 | -1.296 | 0.019 | 0.063 | 0.063 | 0 | 0.31 | 0.83 | 0.27 | 0.2 | 0.27 | 0.3 |
| 1 | 200036139 | rs12069549 | -1.037 | -0.283 | 0.939 | -1.332 | 0.018 | 0.013 | 0.013 | 0 | 0.16 | 0.46 | 0.14 | 0.18 | 0.15 | 0.16 |
| 1 | 200037007 | rs2363570 | 0.74 | -0.276 | 0.406 | 0.032 | -0.009 | 0.012 | 0.012 | 0 | 0.06 | 0 | 0.1 | 0.35 | 0.05 | 0.21 |
| 1 | 200037727 | rs10919806 | 1.111 | -0.064 | -0.104 | 0.022 | 0.002 | -0.008 | -0.008 | 0 | 0.2 | 0.16 | 0.27 | 0.26 | 0.18 | 0.26 |
| 1 | 200042031 | rs3828112 | -0.228 | -0.112 | -0.428 | -0.278 | 0 | 0.003 | 0.003 | 0 | 0.34 | 0.28 | 0.24 | 0.18 | 0.33 | 0.26 |
| 1 | 200046444 | rs2737654 | -0.525 | -0.023 | -0.176 | -0.129 | -0.002 | 0.002 | 0.002 | 0 | 0.55 | 0.53 | 0.52 | 0.44 | 0.51 | 0.52 |
| 1 | 200051274 | rs2821313 | -0.485 | -0.018 | -0.202 | -0.127 | 0.001 | -0.004 | -0.004 | 0 | 0.54 | 0.49 | 0.51 | 0.44 | 0.51 | 0.52 |
| 1 | 200061873 | rs3790826 | -0.051 | 0.097 | -0.059 | 0.651 | -0.001 | 0.029 | 0.029 | 0 | 0.07 | 0.1 | 0.15 | 0.28 | 0.06 | 0.22 |
| 1 | 200062389 | rs2737665 | -0.385 | -0.08 | -0.133 | 0.19 | -0.03 | 0.066 | 0.066 | 0.014 | 0.32 | 0.48 | 0.43 | 0.54 | 0.23 | 0.42 |
| 1 | 200066752 | rs2816999 | -0.734 | 0.474 | -0.523 | -0.286 | -0.028 | 0.011 | 0.011 | 0.31 | 0.39 | 0.69 | 0.53 | 0.68 | 0.44 | 0.48 |
| 1 | 200068658 | rs2253771 | 1.042 | -0.097 | -0.981 | -0.716 | -0.002 | -0.01 | -0.01 | 0.625 | 0.29 | 0.22 | 0.31 | 0.29 | 0.34 | 0.35 |
| 1 | 200069216 | rs2816992 | 0.702 | 0.003 | -0.658 | -0.656 | 0.027 | 0.083 | 0.083 | 0 | 0.44 | 0.28 | 0.4 | 0.67 | 0.4 | 0.53 |
| 1 | 200070765 | rs2816989 | 0.704 | 0.01 | -0.643 | -0.652 | 0.027 | 0.086 | 0.086 | 0 | 0.44 | 0.28 | 0.4 | 0.67 | 0.4 | 0.53 |
| 1 | 200078006 | rs2821324 | 0.817 | -0.148 | -0.339 | -0.622 | 0.013 | 0.083 | 0.083 | 0.206 | 0.46 | 0.32 | 0.41 | 0.67 | 0.44 | 0.55 |
| 1 | 200080463 | rs3828110 | -0.85 | -0.36 | 0.097 | -0.838 | -0.024 | 0.018 | 0.018 | 0.062 | 0.34 | 0.65 | 0.52 | 0.29 | 0.43 | 0.32 |
| 1 | 200083654 | rs2249150 | 1.017 | -0.337 | -0.001 | -0.82 | 0.016 | 0.086 | 0.086 | 0.183 | 0.47 | 0.36 | 0.42 | 0.68 | 0.44 | 0.52 |
| 1 | 200085004 | rs2816972 | -1.251 | -0.34 | 0.068 | -0.826 | 0.023 | 0.009 | 0.009 | 0 | 0.79 | 0.88 | 0.92 | 0.97 | 0.88 | 0.87 |
| 1 | 200086424 | rs12041297 | 0.538 | 0.333 | 0.189 | -0.882 | -0.024 | 0.022 | 0.022 | 0.002 | 0.12 | 0.06 | 0.09 | 0.06 | 0.16 | 0.07 |
| 1 | 200090120 | rs3762398 | 0.535 | 0.031 | 0.504 | -0.75 | -0.011 | -0.001 | -0.001 | 0.741 | 0.17 | 0.19 | 0.39 | 0.23 | 0.27 | 0.27 |
| 1 | 200098537 | rs2821338 | 0.918 | 0.084 | 0.457 | -0.822 | 0.007 | 0.039 | 0.039 | 0 | 0.55 | 0.42 | 0.5 | 0.69 | 0.56 | 0.55 |
| 1 | 200100159 | rs3790800 | -0.012 | 0.073 | 0.429 | -0.807 | -0.012 | -0.003 | -0.003 | 0.158 | 0.21 | 0.18 | 0.39 | 0.28 | 0.29 | 0.31 |
| 1 | 200101481 | rs2816959 | -1.142 | 0.235 | 0.079 | -0.825 | 0.016 | 0.015 | 0.015 | 0 | 0.78 | 0.88 | 0.91 | 0.97 | 0.83 | 0.84 |
| 1 | 200101920 | rs2816958 | -1.085 | -0.209 | 0 | -0.79 | -0.049 | -0.114 | -0.114 | 0.457 | 0.78 | 0.71 | 0.9 | 0.97 | 0.88 | 0.87 |
| 1 | 200104972 | rs10919814 | 0.263 | 0.446 | -0.169 | 0.031 | -0.055 | -0.024 | -0.024 | 0.149 | 0.16 | 0.08 | 0.38 | 0.24 | 0.27 | 0.27 |
| 1 | 200107578 | rs2737619 | 0.623 | 0.524 | -0.155 | 0.413 | 0 | 0.008 | 0.008 | 0 | 0.67 | 0.43 | 0.49 | 0.45 | 0.65 | 0.5 |
| 1 | 200108094 | rs12049278 | 1.397 | 0.372 | -0.485 | 0.346 | 0.026 | -0.109 | -0.109 | 1.496 | 0.15 | 0.01 | 0.09 | 0.38 | 0.07 | 0.17 |
| 1 | 200113077 | rs7514543 | 1.036 | 1.419 | 0.353 | 0.504 | -0.002 | 0.017 | 0.017 | 0 | 0.67 | 0.35 | 0.55 | 0.69 | 0.66 | 0.59 |
| 1 | 200118439 | rs871446 | -0.54 | 1.158 | 0.627 | 0.557 | 0.036 | -0.041 | -0.041 | 0.197 | 0.08 | 0.13 | 0.08 | 0.01 | 0.13 | 0.03 |
| 1 | 200128676 | rs7551367 | -1.918 | 1.275 | 1.224 | 1.111 | 0.059 | -0.017 | -0.017 | 0 | 0.15 | 0.3 | 0.11 | 0.11 | 0.18 | 0.11 |
| 1 | 200129258 | rs3790791 | -2.182 | 1.055 | 0.698 | 0.893 | -0.004 | -0.025 | -0.025 | 0 | 0.32 | 0.76 | 0.28 | 0.36 | 0.27 | 0.34 |
| 1 | 200132084 | rs10494809 | -1.991 | 0.897 | 0.76 | 0.682 | -0.004 | -0.001 | -0.001 | 0.061 | 0.28 | 0.67 | 0.26 | 0.25 | 0.23 | 0.32 |
| 1 | 200135564 | rs12138728 | -0.743 | 1.033 | 0.604 | 0.857 | 0.001 | 0.053 | 0.053 | 0 | 0.13 | 0.22 | 0.16 | 0.24 | 0.09 | 0.22 |
| 1 | 200136582 | rs10494810 | -1.624 | 1.015 | 0.104 | 0.795 | -0.003 | -0.006 | -0.006 | 0 | 0.29 | 0.74 | 0.27 | 0.32 | 0.27 | 0.33 |
| 1 | 200137649 | rs2690039 | -1.002 | 0.315 | 0.201 | 0.752 | -0.035 | 0.06 | 0.06 | 0.792 | 0.3 | 0.46 | 0.5 | 0.44 | 0.26 | 0.27 |
| 1 | 200138774 | rs7556049 | -0.506 | 0.336 | 0.791 | -0.078 | 0.021 | -0.085 | -0.085 | 0 | 0.13 | 0.43 | 0.1 | 0.02 | 0.12 | 0.01 |
| 1 | 200139350 | rs2246923 | -1.461 | 0.457 | 0.727 | 0 | -0.005 | 0.011 | 0.011 | 0 | 0.27 | 0.43 | 0.32 | 0.44 | 0.3 | 0.29 |
| 1 | 200139792 | rs1524163 | -1.753 | 0.398 | 0.436 | -0.282 | -0.01 | -0.011 | -0.011 | 0.504 | 0.37 | 0.54 | 0.39 | 0.46 | 0.38 | 0.45 |
| 1 | 200142317 | rs2690035 | -1.637 | 0.192 | 0.702 | -0.403 | -0.004 | -0.005 | -0.005 | 0 | 0.36 | 0.43 | 0.37 | 0.46 | 0.37 | 0.45 |
| 1 | 200143281 | rs1060060 | -0.101 | 0.269 | -0.122 | -0.671 | 0.002 | 0.006 | 0.006 | 0 | 0.3 | 0.37 | 0.32 | 0.52 | 0.29 | 0.26 |
| 1 | 200143442 | rs1060061 | 0.816 | 0.372 | 0.031 | -1.245 | -0.004 | 0.04 | 0.04 | 0 | 0.57 | 0.69 | 0.51 | 0.09 | 0.57 | 0.4 |
| 1 | 200145533 | rs2246209 | 0.155 | 0.935 | -0.4 | -0.511 | 0.008 | -0.018 | -0.018 | 0 | 0.69 | 0.64 | 0.65 | 0.34 | 0.65 | 0.74 |
| 1 | 200148319 | rs6665426 | 0.329 | 0.329 | -0.496 | -0.324 | -0.001 | -0.002 | -0.002 | 0.035 | 0.69 | 0.68 | 0.72 | 0.34 | 0.75 | 0.67 |
| 1 | 200150738 | rs4915195 | -0.674 | -0.167 | -0.186 | -0.162 | -0.007 | -0.02 | -0.02 | 1.698 | 0.41 | 0.48 | 0.64 | 0.77 | 0.35 | 0.5 |
| 1 | 200152167 | rs2885820 | -0.975 | -0.345 | -0.302 | -0.251 | 0.043 | 0.103 | 0.103 | 0.012 | 0.45 | 0.74 | 0.66 | 0.77 | 0.36 | 0.5 |
| 1 | 200154474 | rs10919830 | -0.752 | -0.717 | 0.07 | 0.342 | -0.01 | 0.144 | 0.144 | 0.07 | 0.14 | 0.23 | 0.28 | 0.47 | 0.12 | 0.23 |
| 1 | 200161148 | rs12088130 | -1.34 | -0.673 | -0.023 | 0.245 | 0.047 | 0.109 | 0.109 | 0.037 | 0.48 | 0.74 | 0.67 | 0.74 | 0.39 | 0.53 |
| 1 | 200162670 | rs12562944 | -0.084 | -0.832 | 0.052 | 0.324 | -0.01 | -0.012 | -0.012 | 0 | 0.28 | 0.33 | 0.34 | 0.27 | 0.23 | 0.29 |
| 1 | 200162783 | rs2885821 | -0.968 | -0.764 | -0.193 | 0.871 | -0.002 | 0.052 | 0.052 | 0 | 0.13 | 0.13 | 0.27 | 0.46 | 0.12 | 0.23 |
| 1 | 200167465 | rs10753870 | -0.798 | -1.369 | -0.602 | 0.568 | 0.045 | -0.039 | -0.039 | 0.022 | 0.36 | 0.72 | 0.48 | 0.44 | 0.22 | 0.43 |
| 1 | 200167658 | rs6694617 | -0.806 | -1.435 | 0.186 | 0.543 | -0.02 | 0.022 | 0.022 | 0 | 0.34 | 0.51 | 0.47 | 0.45 | 0.22 | 0.44 |
| 1 | 200174695 | rs16846260 | -0.731 | -1.56 | 0.358 | 0.531 | -0.014 | 0.01 | 0.01 | 0 | 0.35 | 0.52 | 0.46 | 0.45 | 0.22 | 0.45 |
| 1 | 200175042 | rs4915415 | -0.979 | -1.869 | 0.063 | 0.351 | 0.02 | 0.016 | 0.016 | 0 | 0.46 | 0.52 | 0.55 | 0.67 | 0.31 | 0.53 |
| 1 | 200176068 | rs10919843 | -1.225 | -1.931 | -0.104 | 0.316 | 0.105 | 0.073 | 0.073 | 0 | 0.48 | 0.81 | 0.57 | 0.66 | 0.31 | 0.53 |
| 1 | 200178964 | rs10919845 | -0.977 | -1.939 | -0.037 | 0.312 | 0.036 | 0.026 | 0.026 | 0 | 0.46 | 0.56 | 0.55 | 0.66 | 0.31 | 0.53 |
| 1 | 200182664 | rs6427806 | -1.185 | -2.329 | -0.46 | 0.215 | 0.062 | 0.193 | 0.193 | 0 | 0.56 | 0.85 | 0.6 | 0.85 | 0.35 | 0.6 |
| 1 | 200184939 | rs7529495 | -1.181 | -2.336 | -0.464 | 0.192 | 0.057 | 0.15 | 0.15 | 0 | 0.56 | 0.75 | 0.59 | 0.85 | 0.35 | 0.6 |
| 1 | 200187504 | rs10919851 | -1.316 | -2.237 | -0.41 | 0.095 | 0.019 | 0.029 | 0.029 | 0 | 0.35 | 0.66 | 0.42 | 0.78 | 0.26 | 0.46 |
| 1 | 200187645 | rs7521914 | -0.779 | -2.251 | -0.127 | 0.185 | -0.014 | 0.008 | 0.008 | 0 | 0.3 | 0.58 | 0.38 | 0.61 | 0.22 | 0.36 |
| 1 | 200189305 | rs6427807 | 0.978 | -2.136 | -0.701 | -0.113 | 0.037 | 0.13 | 0.13 | 0.048 | 0.54 | 0.29 | 0.49 | 0.16 | 0.63 | 0.42 |

| | | | | | | | | | | | | | | | | |
|---|---|---|---|---|---|---|---|---|---|---|---|---|---|---|---|---|
| 1 | 200192024 | rs7547703 | 0.347 | -2.302 | -0.657 | 0.326 | 0.028 | -0.013 | -0.013 | 0 | 0.58 | 0.37 | 0.62 | 0.52 | 0.68 | 0.66 |
| 1 | 200193823 | rs1111642 | 0.271 | -2.307 | -0.23 | 0.321 | -0.014 | 0.01 | 0.01 | 0 | 0.61 | 0.53 | 0.63 | 0.52 | 0.68 | 0.67 |
| 1 | 200197538 | rs12128213 | 1.721 | -2.508 | -0.61 | 0.017 | -0.187 | 0.244 | 0.244 | 0.007 | 0.33 | 0.02 | 0.38 | 0.01 | 0.51 | 0.3 |
| 1 | 200197618 | rs12128246 | 1.202 | -2.074 | 0.111 | 0.731 | 0.062 | 0.145 | 0.145 | 0 | 0.56 | 0.28 | 0.53 | 0.06 | 0.67 | 0.44 |
| 1 | 200199414 | rs3861927 | 1.084 | -1.586 | -0.127 | 0.785 | 0.034 | 0.062 | 0.062 | 0 | 0.53 | 0.28 | 0.48 | 0.35 | 0.58 | 0.55 |
| 1 | 200199971 | rs2204309 | 0.333 | -1.586 | 0.182 | 0.932 | -0.001 | 0.025 | 0.025 | 0 | 0.71 | 0.6 | 0.68 | 0.71 | 0.7 | 0.74 |
| 1 | 200204364 | rs10919858 | 0.648 | -1.56 | -0.31 | 0.912 | 0.008 | -0.033 | -0.033 | 2.49 | 0.68 | 0.29 | 0.58 | 0.55 | 0.7 | 0.63 |
| 1 | 200220794 | rs12045034 | -0.206 | -1.172 | -0.553 | 1.34 | -0.025 | -0.138 | -0.138 | 0 | 0.19 | 0.02 | 0.14 | 0.5 | 0.17 | 0.21 |
| 1 | 200221769 | rs10919862 | -0.734 | -1.206 | -0.283 | 1.389 | -0.01 | 0.041 | 0.041 | 0 | 0.22 | 0.3 | 0.25 | 0.65 | 0.17 | 0.33 |
| 1 | 200222777 | rs12047019 | -0.422 | -1.195 | -0.266 | 1.394 | 0.004 | -0.012 | -0.012 | 0 | 0.21 | 0.17 | 0.16 | 0.49 | 0.17 | 0.21 |
| 1 | 200234381 | rs10919870 | 0.33 | -1.093 | -0.315 | 1.08 | -0.014 | 0.016 | 0.016 | 0 | 0.14 | 0.16 | 0.1 | 0.25 | 0.1 | 0.07 |
| 1 | 200247025 | rs10919876 | -1.004 | -0.982 | -0.202 | 1.169 | 0 | 0.027 | 0.027 | 0 | 0.25 | 0.25 | 0.28 | 0.63 | 0.28 | 0.36 |
| 1 | 200247910 | rs16846402 | 0.565 | -1.144 | -0.275 | 1.096 | 0.007 | -0.004 | -0.004 | 0 | 0.14 | 0.1 | 0.1 | 0.18 | 0.11 | 0.08 |
| 1 | 200251050 | rs4915419 | 1.337 | -1.378 | -0.052 | 0.954 | -0.003 | -0.002 | -0.002 | 0.034 | 0.57 | 0.49 | 0.59 | 0.32 | 0.58 | 0.46 |
| 1 | 200255788 | rs6427814 | 1.395 | -1.371 | -0.046 | 0.959 | 0.001 | 0 | 0 | 0 | 0.56 | 0.41 | 0.59 | 0.34 | 0.58 | 0.45 |
| 1 | 200259607 | rs7515411 | 1.344 | -1.561 | 0.173 | 1.194 | 0 | -0.001 | -0.001 | 0 | 0.61 | 0.41 | 0.65 | 0.35 | 0.66 | 0.45 |
| 1 | 200259938 | rs34959467 | 0.939 | -1.565 | 0.267 | 1.137 | 0 | 0.018 | 0.018 | 0 | 0.66 | 0.68 | 0.74 | 0.8 | 0.67 | 0.69 |
| 1 | 200261547 | rs10429903 | 0.664 | -1.549 | 0.223 | 1.154 | 0.012 | -0.019 | -0.019 | 0.235 | 0.13 | 0.04 | 0.09 | 0.17 | 0.11 | 0.08 |
| 1 | 200263697 | rs2153279 | 0.393 | -1.271 | 0.13 | 0.553 | 0.009 | -0.074 | -0.074 | 0 | 0.78 | 0.68 | 0.83 | 0.96 | 0.78 | 0.76 |
| 1 | 200271692 | rs819635 | 0.648 | -1.098 | -0.65 | 0.886 | 0.003 | 0.006 | 0.006 | 0 | 0.6 | 0.63 | 0.74 | 0.66 | 0.66 | 0.64 |
| 1 | 200285880 | rs2790125 | 0.637 | -0.919 | -0.65 | 1.334 | 0.001 | 0.003 | 0.003 | 0 | 0.63 | 0.67 | 0.6 | 0.26 | 0.67 | 0.46 |
| 1 | 200293248 | rs10733062 | 0.312 | -0.983 | -0.886 | 1.152 | 0.01 | 0.025 | 0.025 | 0 | 0.66 | 0.81 | 0.72 | 0.71 | 0.73 | 0.73 |
| 1 | 200300299 | rs2808477 | 0.798 | -1.113 | -0.87 | 1.274 | 0.012 | 0.033 | 0.033 | 0 | 0.61 | 0.75 | 0.62 | 0.24 | 0.67 | 0.4 |
| 1 | 200308381 | rs7553682 | 0.217 | -1.119 | -0.893 | 1.262 | 0.012 | 0.021 | 0.021 | 0 | 0.67 | 0.81 | 0.72 | 0.7 | 0.74 | 0.71 |
| 1 | 200314618 | rs12117979 | 0.93 | -0.622 | -1.129 | 0.639 | -0.003 | -0.037 | -0.037 | 0.51 | 0.29 | 0.51 | 0.31 | 0.16 | 0.28 | 0.1 |
| 1 | 200324163 | rs12086509 | 1.096 | -0.666 | -1.188 | 0.631 | -0.115 | 0.146 | 0.146 | 0 | 0.31 | 0.05 | 0.29 | 0.08 | 0.39 | 0.29 |
| 1 | 200324744 | rs2790129 | -0.131 | -0.627 | -1.108 | 0.633 | -0.008 | -0.011 | -0.011 | 2.482 | 0.66 | 0.65 | 0.71 | 0.7 | 0.74 | 0.71 |
| 1 | 200330479 | rs2808500 | 0.693 | -0.598 | -1.116 | 0.659 | -0.106 | -0.146 | -0.146 | 0 | 0.34 | 0.05 | 0.36 | 0.53 | 0.4 | 0.57 |
| 1 | 200342825 | rs2153280 | -0.01 | -0.314 | -0.896 | 0.778 | -0.003 | -0.004 | -0.004 | 0.022 | 0.66 | 0.67 | 0.71 | 0.71 | 0.74 | 0.71 |
| 1 | 200365655 | rs16846614 | 1.095 | -0.332 | -1.077 | 0.643 | -0.001 | -0.01 | -0.01 | 0 | 0.13 | 0.11 | 0.1 | 0.03 | 0.12 | 0.16 |
| 1 | 200365963 | rs4915434 | -0.101 | -0.341 | -1.097 | 0.678 | 0.027 | 0.01 | 0.01 | 0 | 0.71 | 0.85 | 0.76 | 0.71 | 0.79 | 0.75 |
| 1 | 200366091 | rs12141206 | 1.141 | -0.511 | -1.111 | 0.583 | -0.011 | -0.033 | -0.033 | 0 | 0.32 | 0.48 | 0.31 | 0.16 | 0.29 | 0.12 |
| 1 | 200369696 | rs2790108 | 0.993 | -0.372 | -1.146 | 0.77 | -0.104 | 0.136 | 0.136 | 0 | 0.29 | 0.05 | 0.28 | 0.07 | 0.38 | 0.3 |
| 1 | 200375209 | rs1128817 | -0.25 | -0.517 | -1.102 | 0.755 | 0.012 | 0.008 | 0.008 | 0 | 0.3 | 0.19 | 0.24 | 0.29 | 0.22 | 0.25 |
| 1 | 200380314 | rs4244146 | -0.222 | -0.492 | -1.098 | 0.809 | 0.011 | 0.003 | 0.003 | 0 | 0.3 | 0.19 | 0.25 | 0.32 | 0.22 | 0.25 |
| 1 | 200391561 | rs4414069 | 0.733 | -0.399 | -1.088 | 0.857 | 0 | -0.011 | -0.011 | 0.581 | 0.38 | 0.62 | 0.38 | 0.17 | 0.39 | 0.23 |
| 1 | 200403095 | rs10494814 | 0.808 | -0.33 | -1.168 | 0.89 | 0.003 | -0.036 | -0.036 | 0 | 0.38 | 0.53 | 0.38 | 0.17 | 0.38 | 0.23 |
| 1 | 200404343 | rs7544904 | -0.36 | -0.262 | -1.454 | 0.957 | -0.005 | -0.003 | -0.003 | 0 | 0.31 | 0.3 | 0.26 | 0.32 | 0.22 | 0.25 |
| 1 | 200405735 | rs16846692 | -0.286 | -0.257 | -1.472 | 0.959 | 0.018 | 0.007 | 0.007 | 0 | 0.3 | 0.17 | 0.25 | 0.32 | 0.22 | 0.25 |
| 1 | 200421466 | rs4427435 | 0.527 | 0.176 | -1.551 | 1.601 | 0 | 0.096 | 0.096 | 0 | 0.54 | 0.74 | 0.5 | 0.2 | 0.52 | 0.39 |
| 1 | 200423841 | rs12095036 | 0.138 | -0.06 | -1.268 | 1.756 | -0.018 | 0.015 | 0.015 | 0.027 | 0.16 | 0.18 | 0.15 | 0.29 | 0.09 | 0.09 |
| 1 | 200437506 | rs12058730 | 0.171 | -0.064 | -1.303 | 1.773 | -0.021 | 0.053 | 0.053 | 0.015 | 0.16 | 0.19 | 0.16 | 0.46 | 0.08 | 0.1 |
| 1 | 200450197 | rs7546093 | 0.462 | 0.088 | -1.897 | 1.655 | 0.03 | -0.048 | -0.048 | 0 | 0.14 | 0.03 | 0.13 | 0.37 | 0.08 | 0.11 |
| 1 | 200453320 | rs4506486 | 0.046 | 1.206 | -1.689 | 1.321 | -0.101 | -0.111 | -0.111 | 0 | 0.71 | 0.96 | 0.63 | 0.59 | 0.6 | 0.49 |
| 1 | 200456424 | rs10919946 | -1.021 | 1.163 | -1.21 | 1.639 | -0.006 | 0.175 | 0.175 | 0.155 | 0.1 | 0.24 | 0.15 | 0.34 | 0.06 | 0.2 |
| 1 | 200464803 | rs6427830 | -0.125 | 1.795 | -2.065 | 0.933 | -0.124 | 0.088 | 0.088 | 0 | 0.77 | 0.98 | 0.74 | 0.93 | 0.65 | 0.68 |
| 1 | 200466004 | rs6427832 | 0.601 | 1.821 | -1.468 | 1.941 | -0.036 | -0.057 | -0.057 | 0 | 0.86 | 0.92 | 0.72 | 0.58 | 0.78 | 0.8 |
| 1 | 200469505 | rs6427833 | -1.011 | 1.362 | -1.003 | 2.142 | 0.008 | 0.085 | 0.085 | 0.051 | 0.12 | 0.17 | 0.38 | 0.4 | 0.19 | 0.16 |
| 1 | 200471464 | rs6680807 | 0.453 | 0.719 | -0.514 | 1.806 | -0.028 | 0.073 | 0.073 | 0 | 0.83 | 0.7 | 0.78 | 0.62 | 0.88 | 0.66 |
| 1 | 200473718 | rs4417099 | 0.569 | 1.103 | -0.552 | 2.123 | -0.012 | 0.017 | 0.017 | 0 | 0.89 | 0.94 | 0.67 | 0.53 | 0.79 | 0.81 |
| 1 | 200475235 | rs6675963 | 0.457 | 0.687 | -0.839 | 0.704 | -0.003 | 0.08 | 0.08 | 0 | 0.49 | 0.68 | 0.33 | 0.22 | 0.47 | 0.39 |
| 1 | 200479973 | rs10919960 | 1.009 | -0.211 | -1.061 | 0.114 | -0.042 | 0.171 | 0.171 | 0.11 | 0.45 | 0.17 | 0.26 | 0.2 | 0.48 | 0.34 |
| 1 | 200481555 | rs7549548 | 0.07 | -0.542 | -0.27 | 0.193 | -0.107 | 0.003 | 0.003 | 0 | 0.28 | 0.5 | 0.31 | 0.31 | 0.11 | 0.37 |
| 1 | 200481756 | rs4335405 | 0.256 | -0.33 | -0.372 | 0.425 | 0.002 | 0.011 | 0.011 | 0 | 0.81 | 0.83 | 0.64 | 0.61 | 0.68 | 0.79 |
| 1 | 200481934 | rs4418638 | -0.611 | -0.29 | -0.504 | 0.501 | -0.002 | 0.033 | 0.033 | 0 | 0.88 | 0.91 | 0.72 | 0.61 | 0.8 | 0.81 |
| 1 | 200483591 | rs9659917 | 1.569 | -0.392 | -0.639 | 0.192 | -0.106 | 0.089 | 0.089 | 0.075 | 0.37 | 0.11 | 0.22 | 0.24 | 0.4 | 0.31 |
| 1 | 200494416 | rs12567052 | 1.604 | -0.425 | -0.797 | 0.313 | -0.144 | 0.024 | 0.024 | 0.072 | 0.36 | 0.04 | 0.22 | 0.25 | 0.4 | 0.31 |
| 1 | 200496095 | rs10800706 | -0.088 | -0.608 | -0.621 | 0.943 | -0.008 | -0.019 | -0.019 | 0 | 0.85 | 0.91 | 0.7 | 0.68 | 0.79 | 0.81 |
| 1 | 200502030 | rs11811022 | 0.497 | -0.951 | -0.312 | 0.713 | 0 | -0.007 | -0.007 | 0 | 0.22 | 0.2 | 0.28 | 0.45 | 0.1 | 0.34 |
| 1 | 200502671 | rs10919967 | -0.05 | -0.57 | -0.444 | 0.804 | -0.008 | -0.02 | -0.02 | 0 | 0.85 | 0.91 | 0.69 | 0.74 | 0.79 | 0.81 |
| 1 | 200511466 | rs7516247 | -0.843 | -0.929 | -0.094 | 0.513 | 0.026 | -0.002 | -0.002 | 1.358 | 0.38 | 0.74 | 0.42 | 0.45 | 0.3 | 0.4 |
| 1 | 200521131 | rs10737572 | -0.187 | -0.713 | -0.326 | 0.449 | -0.011 | -0.034 | -0.034 | 5.794 | 0.82 | 0.9 | 0.67 | 0.69 | 0.75 | 0.75 |
| 1 | 200521412 | rs12060793 | 1.382 | -0.896 | 0.241 | 0.24 | 0.015 | 0.002 | 0.002 | 0 | 0.6 | 0.48 | 0.5 | 0.63 | 0.49 | 0.62 |
| 1 | 200529259 | rs10919976 | 1.872 | -0.748 | -0.439 | 0.073 | -0.155 | 0.02 | 0.02 | 0.226 | 0.38 | 0.02 | 0.2 | 0.23 | 0.39 | 0.28 |
| 1 | 200531693 | rs4332382 | 0.763 | -0.906 | -0.15 | -0.122 | 0.063 | -0.076 | -0.076 | 0.002 | 0.12 | 0.34 | 0.1 | 0 | 0.13 | 0.08 |
| 1 | 200534248 | rs7543730 | 1.012 | -0.965 | -0.091 | -0.115 | -0.005 | -0.001 | -0.001 | 0 | 0.64 | 0.41 | 0.55 | 0.69 | 0.58 | 0.64 |
| 1 | 200535312 | rs6688734 | 0.552 | -0.918 | -0.047 | -0.112 | 0.079 | -0.087 | -0.087 | 0 | 0.13 | 0.47 | 0.1 | 0 | 0.13 | 0.08 |
| 1 | 200545187 | rs10800712 | -0.484 | -0.931 | -0.061 | -0.109 | -0.011 | -0.034 | -0.034 | 0 | 0.82 | 0.9 | 0.67 | 0.69 | 0.75 | 0.75 |
| 1 | 200557767 | rs4504918 | -0.974 | -1.011 | -0.095 | 0.19 | -0.013 | -0.008 | -0.008 | 0.77 | 0.85 | 0.91 | 0.74 | 0.82 | 0.75 | 0.83 |

| | | | | | | | | | | | | | | | | |
|---|---|---|---|---|---|---|---|---|---|---|---|---|---|---|---|---|
| 1 | 200558363 | rs6665951 | -0.298 | -0.999 | -0.088 | 0.192 | -0.012 | -0.037 | -0.037 | 0 | 0.82 | 0.9 | 0.67 | 0.69 | 0.75 | 0.75 |
| 1 | 200561456 | rs16846948 | -0.144 | -1.004 | -0.06 | 0.179 | 0.032 | -0.135 | -0.135 | 0 | 0.18 | 0.47 | 0.12 | 0 | 0.17 | 0.11 |
| 1 | 200563971 | rs10494824 | 0.683 | -1.055 | 0.155 | 0.455 | -0.039 | 0.038 | 0.038 | 0 | 0.18 | 0.25 | 0.27 | 0.4 | 0.09 | 0.28 |
| 1 | 200569392 | rs2809357 | -1.761 | -0.574 | 0.164 | 0.493 | -0.01 | 0.062 | 0.062 | 0.292 | 0.95 | 0.91 | 0.9 | 0.87 | 0.97 | 0.91 |
| 1 | 200571961 | rs10494823 | 0.991 | -0.444 | 0.08 | 0.181 | -0.002 | 0.002 | 0.002 | 0 | 0.07 | 0.07 | 0.07 | 0.06 | 0.09 | 0.08 |
| 1 | 200575794 | rs2794410 | -0.662 | -0.306 | 0.173 | 0.193 | 0.033 | -0.017 | -0.017 | 0.377 | 0.25 | 0.34 | 0.32 | 0.18 | 0.35 | 0.23 |
| 1 | 200592275 | rs2808248 | -1.628 | -0.417 | -0.724 | 0.019 | -0.039 | 0.044 | 0.044 | 0.963 | 0.6 | 0.83 | 0.76 | 0.72 | 0.58 | 0.7 |
| 1 | 200592295 | rs11807125 | -0.349 | -0.771 | -0.329 | 0.045 | 0.029 | -0.011 | -0.011 | 1.049 | 0.29 | 0.42 | 0.34 | 0.22 | 0.38 | 0.25 |
| 1 | 200593325 | rs12744809 | -0.802 | -0.925 | -0.39 | -0.051 | 0.014 | -0.017 | -0.017 | 0 | 0.35 | 0.43 | 0.37 | 0.22 | 0.42 | 0.26 |
| 1 | 200596263 | rs2809345 | -1.647 | -0.899 | -0.519 | -0.033 | -0.037 | 0.079 | 0.079 | 0.048 | 0.55 | 0.82 | 0.74 | 0.72 | 0.53 | 0.69 |
| 1 | 200596345 | rs10919991 | 1.988 | -0.947 | -0.541 | -0.047 | -0.094 | 0.031 | 0.031 | 0.786 | 0.39 | 0.07 | 0.23 | 0.28 | 0.42 | 0.3 |
| 1 | 200597650 | rs2809342 | 0.5 | -0.906 | -0.329 | -0.017 | 0.015 | -0.02 | -0.02 | 0 | 0.64 | 0.45 | 0.63 | 0.78 | 0.58 | 0.73 |
| 1 | 200607466 | rs10919992 | 1.892 | -0.707 | -0.291 | 0.029 | -0.049 | 0.027 | 0.027 | 0.975 | 0.37 | 0.11 | 0.24 | 0.28 | 0.41 | 0.28 |
| 1 | 200611710 | rs6695836 | 0.415 | -0.936 | -0.146 | -0.016 | -0.051 | 0.077 | 0.077 | 0.068 | 0.74 | 0.6 | 0.6 | 0.5 | 0.83 | 0.62 |
| 1 | 200618238 | rs6665604 | -1.408 | -0.835 | -0.214 | -0.062 | -0.005 | -0.029 | -0.029 | 0 | 0.94 | 0.91 | 0.97 | 1 | 0.95 | 0.98 |
| 1 | 200622353 | rs2809361 | 0.162 | -0.951 | -0.249 | 0.067 | 0.003 | -0.001 | -0.001 | 0 | 0.08 | 0.1 | 0.13 | 0.07 | 0.05 | 0.17 |
| 1 | 200625693 | rs16847097 | 1.656 | -0.944 | -0.015 | 0.048 | -0.003 | -0.031 | -0.031 | 0 | 0.06 | 0.11 | 0.03 | 0 | 0.04 | 0.01 |
| 1 | 200631885 | rs16847099 | 0.977 | -1.006 | -0.056 | 0.118 | -0.023 | 0.07 | 0.07 | 0 | 0.12 | 0.17 | 0.24 | 0.43 | 0.07 | 0.2 |
| 1 | 200639683 | rs3806358 | 1.295 | -0.901 | -0.104 | -0.07 | 0.011 | 0.068 | 0.068 | 0.156 | 0.49 | 0.28 | 0.48 | 0.72 | 0.48 | 0.48 |
| 1 | 200648439 | rs4437908 | -0.552 | -0.903 | -0.003 | -0.078 | 0.008 | -0.014 | -0.014 | 0 | 0.37 | 0.41 | 0.36 | 0.21 | 0.42 | 0.34 |
| 1 | 200650063 | rs6665979 | 0.118 | -0.902 | 0.009 | -0.028 | -0.02 | -0.023 | -0.023 | 0 | 0.86 | 0.69 | 0.83 | 0.93 | 0.9 | 0.83 |
| 1 | 200650490 | rs12082129 | -0.138 | -0.643 | 0.1 | -0.188 | 0.001 | -0.002 | -0.002 | 0.069 | 0.29 | 0.31 | 0.31 | 0.21 | 0.33 | 0.32 |
| 1 | 200652600 | rs4378238 | 0.325 | -0.68 | 0.017 | -0.047 | -0.021 | -0.036 | -0.036 | 1.358 | 0.85 | 0.68 | 0.82 | 0.93 | 0.89 | 0.81 |
| 1 | 200655233 | rs11581339 | -0.524 | -0.812 | -0.111 | -0.165 | -0.001 | -0.002 | -0.002 | 3.077 | 0.39 | 0.39 | 0.37 | 0.21 | 0.44 | 0.33 |
| 1 | 200655272 | rs7519923 | 0.033 | -0.386 | -0.456 | -0.346 | -0.009 | -0.003 | -0.003 | 0 | 0.91 | 0.82 | 0.85 | 0.93 | 0.93 | 0.82 |
| 1 | 200659854 | rs4915451 | -0.64 | -0.428 | -0.621 | -0.228 | -0.001 | 0.007 | 0.007 | 0 | 0.51 | 0.53 | 0.59 | 0.64 | 0.51 | 0.54 |
| 1 | 200662215 | rs7542210 | 1.306 | 0.059 | -1.273 | 0.035 | -0.039 | -0.002 | -0.002 | 0 | 0.33 | 0.15 | 0.22 | 0.29 | 0.38 | 0.23 |
| 1 | 200669180 | rs6427847 | 0.379 | 0.11 | -0.939 | 0.232 | 0.001 | 0.002 | 0.002 | 0 | 0.65 | 0.65 | 0.67 | 0.57 | 0.63 | 0.6 |
| 1 | 200672699 | rs7535122 | 0.665 | -0.069 | -0.83 | 0.159 | 0.002 | 0.007 | 0.007 | 0 | 0.62 | 0.56 | 0.66 | 0.57 | 0.63 | 0.6 |
| 1 | 200679613 | rs12137256 | 1.716 | 0.168 | -1.168 | -0.072 | -0.028 | 0.056 | 0.056 | 0 | 0.33 | 0.13 | 0.28 | 0.21 | 0.38 | 0.19 |
| 1 | 200682721 | rs1389644 | 1.564 | 0.018 | -1.068 | -0.209 | -0.016 | 0.023 | 0.023 | 0 | 0.12 | 0.06 | 0.08 | 0.01 | 0.14 | 0.07 |
| 1 | 200692168 | rs6664413 | 0.848 | 0.11 | -1.229 | -0.091 | 0 | -0.015 | -0.015 | 0 | 0.57 | 0.55 | 0.68 | 0.74 | 0.6 | 0.63 |
| 1 | 200703541 | rs6660197 | 1.397 | 0.17 | -0.863 | 0.054 | 0.016 | 0.063 | 0.063 | 0 | 0.1 | 0.17 | 0.12 | 0.2 | 0.13 | 0.11 |
| 1 | 200704754 | rs10494819 | 0.953 | 0.027 | -0.939 | -0.015 | -0.007 | 0.007 | 0.007 | 0 | 0.48 | 0.37 | 0.42 | 0.4 | 0.52 | 0.31 |
| 1 | 200707090 | rs10920017 | 1.063 | -0.034 | -0.815 | 0.027 | -0.028 | 0.089 | 0.089 | 0 | 0.14 | 0.24 | 0.28 | 0.36 | 0.1 | 0.33 |
| 1 | 200709796 | rs7517998 | -1.175 | 0.007 | -0.895 | -0.015 | 0 | 0.005 | 0.005 | 0 | 0.38 | 0.52 | 0.5 | 0.55 | 0.37 | 0.52 |
| 1 | 200718422 | rs7529518 | 1.082 | -0.058 | -0.805 | 0.061 | -0.021 | 0.068 | 0.068 | 0 | 0.13 | 0.22 | 0.28 | 0.36 | 0.1 | 0.32 |
| 1 | 200727873 | rs7519005 | 1.792 | 0.077 | -0.938 | -0.056 | -0.015 | 0.026 | 0.026 | 0 | 0.12 | 0.04 | 0.08 | 0.01 | 0.13 | 0.07 |
| 1 | 200728818 | rs4382754 | 2.289 | 0.044 | -1.031 | -0.087 | -0.049 | 0.068 | 0.068 | 0 | 0.33 | 0.09 | 0.28 | 0.2 | 0.39 | 0.19 |
| 1 | 200750576 | rs7520287 | -0.518 | 0.061 | -0.874 | -0.028 | -0.001 | -0.007 | -0.007 | 10.101 | 0.74 | 0.71 | 0.79 | 0.75 | 0.76 | 0.7 |
| 1 | 200776641 | rs7552353 | 1.109 | -0.007 | -0.858 | 0.069 | -0.021 | 0.067 | 0.067 | 0 | 0.14 | 0.22 | 0.27 | 0.36 | 0.1 | 0.33 |
| 1 | 200780153 | rs6427859 | -0.432 | 0.012 | -0.929 | 0.055 | -0.018 | 0.076 | 0.076 | 0 | 0.26 | 0.47 | 0.42 | 0.53 | 0.24 | 0.44 |
| 1 | 200791954 | rs6697899 | 1.582 | 0.03 | -1.205 | 0.014 | 0.006 | -0.012 | -0.012 | 0.215 | 0.1 | 0.06 | 0.05 | 0.15 | 0.08 | 0.03 |
| 1 | 200827806 | rs12733378 | 1.534 | 0.054 | -1.026 | 0.192 | -0.015 | -0.022 | -0.022 | 0.009 | 0.09 | 0.06 | 0.13 | 0.18 | 0.13 | 0.11 |
| 1 | 200829992 | rs7514015 | 1.781 | 0.069 | -1.06 | -0.102 | -0.022 | 0.085 | 0.085 | 0 | 0.36 | 0.17 | 0.28 | 0.2 | 0.39 | 0.19 |
| 1 | 200841534 | rs12120209 | -0.27 | 0.239 | -0.563 | -0.061 | -0.03 | 0.04 | 0.04 | 0 | 0.28 | 0.13 | 0.18 | 0.18 | 0.29 | 0.11 |
| 1 | 200843768 | rs2292097 | 0.582 | -0.117 | -0.309 | 0.185 | -0.021 | 0.075 | 0.075 | 0 | 0.13 | 0.21 | 0.26 | 0.37 | 0.1 | 0.32 |
| 1 | 200849360 | rs2062847 | -1.76 | 0.052 | -0.704 | 0.028 | -0.004 | -0.005 | -0.005 | 0.964 | 0.55 | 0.62 | 0.55 | 0.56 | 0.51 | 0.46 |
| 1 | 200849676 | rs10800745 | 0.437 | 0.441 | -0.94 | 0.291 | -0.034 | 0.145 | 0.145 | 0 | 0.14 | 0.31 | 0.27 | 0.38 | 0.1 | 0.33 |
| 1 | 200852820 | rs12564878 | -0.719 | 0.832 | -1.646 | 0.05 | -0.067 | 0.081 | 0.081 | 0 | 0.35 | 0.12 | 0.23 | 0.19 | 0.39 | 0.12 |
| 1 | 200857774 | rs3753955 | 0.167 | 0.933 | -1.482 | 0.306 | -0.014 | 0.003 | 0.003 | 0 | 0.16 | 0.07 | 0.09 | 0.14 | 0.16 | 0.05 |
| 1 | 200859413 | rs6427864 | -0.743 | 0.643 | -1.035 | 0.412 | 0 | 0.002 | 0.002 | 0 | 0.52 | 0.58 | 0.53 | 0.56 | 0.51 | 0.46 |
| 1 | 200861003 | rs2271019 | -0.734 | 0.64 | -1.051 | 0.413 | 0 | 0.003 | 0.003 | 0 | 0.52 | 0.6 | 0.53 | 0.57 | 0.51 | 0.46 |
| 1 | 200862557 | rs885197 | 0.461 | 0.349 | -1.03 | 0.48 | 0.009 | 0.004 | 0.004 | 0 | 0.43 | 0.44 | 0.47 | 0.34 | 0.35 | 0.47 |
| 1 | 200865768 | rs296533 | 0.308 | 0.544 | -1.647 | -0.392 | -0.061 | 0.102 | 0.102 | 0 | 0.27 | 0.08 | 0.21 | 0.02 | 0.34 | 0.17 |
| 1 | 200874375 | rs2134757 | 1.083 | 0.236 | -1.926 | -0.361 | -0.054 | 0.069 | 0.069 | 0.089 | 0.3 | 0.06 | 0.42 | 0.15 | 0.35 | 0.24 |
| 1 | 200877562 | rs7554511 | 0.739 | 0.043 | -1.494 | -0.224 | -0.008 | -0.019 | -0.019 | 0 | 0.21 | 0.14 | 0.16 | 0 | 0.27 | 0.12 |
| 1 | 200881595 | rs7522462 | 0.795 | 0.054 | -1.553 | -0.222 | -0.025 | 0.022 | 0.022 | 0 | 0.21 | 0.11 | 0.16 | 0 | 0.27 | 0.13 |
| 1 | 200889833 | rs296543 | -0.482 | -0.189 | -1.504 | 0.516 | -0.013 | 0.002 | 0.002 | 0 | 0.73 | 0.82 | 0.8 | 0.8 | 0.71 | 0.84 |
| 1 | 200892137 | rs296547 | 0.667 | -0.735 | -0.99 | 0.694 | 0.027 | 0.024 | 0.024 | 0 | 0.55 | 0.29 | 0.62 | 0.38 | 0.63 | 0.42 |
| 1 | 200896708 | rs7536488 | 1.044 | -0.498 | -0.662 | 0.889 | 0.034 | 0.029 | 0.029 | 0 | 0.05 | 0.12 | 0.05 | 0.1 | 0.1 | 0.04 |
| 1 | 200901416 | rs425079 | -0.242 | -0.877 | -0.239 | 0.893 | -0.04 | 0.098 | 0.098 | 0.087 | 0.87 | 0.77 | 0.85 | 0.58 | 0.94 | 0.58 |
| 1 | 200906473 | rs296513 | -0.308 | -0.787 | -0.673 | 0.512 | 0.023 | -0.049 | -0.049 | 0 | 0.27 | 0.09 | 0.25 | 0.44 | 0.22 | 0.31 |
| 1 | 200921385 | rs396029 | 0.36 | -0.699 | -0.634 | 0.534 | 0.016 | -0.001 | -0.001 | 0.045 | 0.21 | 0.07 | 0.13 | 0.26 | 0.17 | 0.25 |
| 1 | 200929144 | rs169850 | -0.246 | -0.665 | -0.665 | 0.437 | 0.003 | -0.01 | -0.01 | 0 | 0.65 | 0.28 | 0.71 | 0.66 | 0.68 | 0.72 |
| 1 | 200930005 | rs502658 | 0.485 | -0.57 | -0.656 | 0.446 | 0.002 | 0.012 | 0.012 | 0 | 0.2 | 0.14 | 0.14 | 0.13 | 0.18 | 0.14 |
| 1 | 200934432 | rs4915471 | 1.608 | -0.477 | -0.851 | 0.48 | -0.03 | 0.025 | 0.025 | 0.16 | 0.33 | 0.1 | 0.42 | 0.24 | 0.35 | 0.45 |
| 1 | 200935866 | rs11584383 | 1.353 | -0.481 | -0.576 | 0.268 | -0.011 | -0.014 | -0.014 | 0 | 0.22 | 0.18 | 0.16 | 0 | 0.3 | 0.12 |
| 1 | 200938029 | rs10494829 | 0.112 | -0.519 | -0.295 | 0.422 | 0.036 | 0.119 | 0.119 | 0.498 | 0.15 | 0.44 | 0.2 | 0.29 | 0.16 | 0.13 |

| | | | | | | | | | | | | | | | | |
|---|---|---|---|---|---|---|---|---|---|---|---|---|---|---|---|---|
| 1 | 200941755 | rs296569 | 0.668 | -0.529 | -0.589 | 0.333 | -0.006 | 0.053 | 0.053 | 0 | 0.2 | 0.03 | 0.12 | 0.13 | 0.18 | 0.14 |
| 1 | 200956288 | rs2275485 | -0.258 | -0.332 | 0.467 | 0.779 | 0.032 | 0.113 | 0.113 | 0.665 | 0.16 | 0.45 | 0.2 | 0.29 | 0.15 | 0.13 |
| 1 | 200957032 | rs10494828 | 1.404 | -0.262 | 0.1 | 0.563 | -0.014 | -0.006 | -0.006 | 0 | 0.2 | 0.16 | 0.16 | 0 | 0.28 | 0.09 |
| 1 | 200964578 | rs10800750 | -1.196 | -0.389 | -0.311 | 0.731 | -0.065 | 0.052 | 0.052 | 0.092 | 0.47 | 0.87 | 0.47 | 0.63 | 0.47 | 0.39 |
| 1 | 200974888 | rs11584527 | 0.664 | -0.348 | -0.09 | 0.405 | 0.017 | -0.003 | -0.003 | 0.093 | 0.07 | 0.05 | 0.04 | 0.22 | 0.01 | 0.06 |
| 1 | 200975064 | rs10920091 | -0.3 | -0.328 | 0.097 | 0.445 | -0.032 | 0.016 | 0.016 | 0 | 0.34 | 0.14 | 0.21 | 0.22 | 0.35 | 0.17 |
| 1 | 200984480 | rs12122721 | 0.92 | -0.231 | 0.097 | 0.076 | -0.008 | 0.009 | 0.009 | 0 | 0.22 | 0.17 | 0.19 | 0.13 | 0.29 | 0.17 |
| 1 | 200996189 | rs6427870 | -1.485 | -0.326 | -0.146 | 0.176 | -0.082 | 0.009 | 0.009 | 0.366 | 0.49 | 0.9 | 0.37 | 0.37 | 0.46 | 0.3 |
| 1 | 200997930 | rs12140231 | 0.322 | -0.318 | 0.222 | 0.273 | 0.018 | 0.003 | 0.003 | 0.122 | 0.1 | 0.22 | 0.07 | 0.1 | 0.1 | 0.06 |
| 1 | 200998358 | rs12081096 | -0.1 | -0.331 | -0.335 | 0.326 | 0.015 | -0.02 | -0.02 | 1.399 | 0.13 | 0.07 | 0.06 | 0.14 | 0.06 | 0.07 |
| 1 | 201003606 | rs10920099 | -1.068 | -0.142 | -0.226 | 0.232 | -0.031 | 0.009 | 0.009 | 0.482 | 0.49 | 0.85 | 0.38 | 0.4 | 0.46 | 0.29 |
| 1 | 201004512 | rs12132696 | -0.511 | -0.038 | 0.313 | 0.25 | 0.022 | -0.002 | -0.002 | 0.134 | 0.13 | 0.59 | 0.1 | 0.09 | 0.1 | 0.06 |
| 1 | 201009182 | rs12139527 | -0.545 | -0.052 | 0.309 | 0.275 | 0.019 | 0.011 | 0.011 | 1.822 | 0.13 | 0.61 | 0.1 | 0.11 | 0.11 | 0.06 |
| 1 | 201009556 | rs1546416 | -0.374 | -0.198 | 0.488 | 0.189 | 0.002 | -0.004 | -0.004 | 0 | 0.38 | 0.42 | 0.29 | 0.3 | 0.36 | 0.25 |
| 1 | 201016296 | rs3850625 | 1.368 | -0.442 | 0.204 | 0.056 | -0.018 | -0.005 | -0.005 | 0 | 0.09 | 0.01 | 0.09 | 0.04 | 0.12 | 0.2 |
| 1 | 201016646 | rs2297905 | -0.009 | -0.223 | 0.81 | 0.217 | 0.048 | 0.043 | 0.043 | 1.446 | 0.11 | 0.4 | 0.1 | 0.11 | 0.1 | 0.05 |
| 1 | 201020727 | rs3767498 | -1.112 | -0.372 | 0.25 | -0.034 | -0.092 | 0.162 | 0.162 | 0 | 0.73 | 0.96 | 0.82 | 0.99 | 0.67 | 0.89 |
| 1 | 201020851 | rs3767499 | 1.037 | -0.999 | 0.424 | 0.15 | -0.005 | -0.018 | -0.018 | 0 | 0.46 | 0.17 | 0.59 | 0.58 | 0.48 | 0.68 |
| 1 | 201023436 | rs3818873 | -0.027 | -0.924 | 0.546 | 0.125 | -0.002 | -0.012 | -0.012 | 0 | 0.6 | 0.37 | 0.72 | 0.87 | 0.56 | 0.84 |
| 1 | 201025682 | rs3767503 | 0.744 | -0.911 | 0.387 | 0.095 | 0.017 | 0.02 | 0.02 | 0.061 | 0.49 | 0.25 | 0.58 | 0.59 | 0.47 | 0.68 |
| 1 | 201029431 | rs4915475 | 0.627 | -0.768 | 0.474 | 0.104 | 0.015 | 0.009 | 0.009 | 0.098 | 0.5 | 0.29 | 0.59 | 0.57 | 0.47 | 0.67 |
| 1 | 201030668 | rs2297904 | -1.239 | -0.858 | 0.638 | 0.05 | -0.048 | 0.086 | 0.086 | 0 | 0.21 | 0.35 | 0.18 | 0.43 | 0.11 | 0.21 |
| 1 | 201031307 | rs2297903 | 0.513 | -0.823 | 0.635 | 0.097 | 0.001 | -0.033 | -0.033 | 0 | 0.55 | 0.49 | 0.66 | 0.76 | 0.5 | 0.71 |
| 1 | 201032737 | rs10920103 | -0.838 | -1.186 | 0.911 | 0.927 | -0.03 | -0.03 | -0.03 | 0 | 0.14 | 0.25 | 0.11 | 0.07 | 0.07 | 0.06 |
| 1 | 201033720 | rs16847613 | -1.464 | -1.175 | 0.768 | 0.981 | -0.039 | 0.041 | 0.041 | 0 | 0.2 | 0.31 | 0.18 | 0.26 | 0.1 | 0.1 |
| 1 | 201035607 | rs2297901 | -1.334 | -0.703 | 0.476 | 0.814 | -0.032 | 0.139 | 0.139 | 0.674 | 0.2 | 0.54 | 0.14 | 0.35 | 0.13 | 0.2 |
| 1 | 201040180 | rs9427711 | -0.407 | -0.828 | 0.455 | -0.656 | -0.002 | -0.043 | -0.043 | 0 | 0.81 | 0.77 | 0.78 | 0.97 | 0.77 | 0.91 |
| 1 | 201041198 | rs9427712 | 1.246 | -0.547 | -0.468 | -1.584 | -0.107 | 0.118 | 0.118 | 0 | 0.22 | 0.04 | 0.21 | 0 | 0.34 | 0.12 |
| 1 | 201042268 | rs3767507 | 1.032 | -1.149 | -0.848 | -1.498 | 0.023 | -0.032 | -0.032 | 0.117 | 0.12 | 0.06 | 0.12 | 0.19 | 0.07 | 0.04 |
| 1 | 201042983 | rs2296386 | 1.018 | -1.531 | -1.218 | -1.761 | -0.04 | -0.068 | -0.068 | 0 | 0.25 | 0.03 | 0.32 | 0.77 | 0.24 | 0.69 |
| 1 | 201044097 | rs3767508 | -0.795 | -1.612 | -1.04 | -1.884 | -0.025 | -0.11 | -0.11 | 0 | 0.76 | 0.5 | 0.75 | 0.96 | 0.76 | 0.91 |
| 1 | 201047075 | rs4915477 | -0.731 | -1.48 | -1.244 | -1.799 | -0.051 | -0.026 | -0.026 | 1.23 | 0.69 | 0.94 | 0.65 | 0.23 | 0.66 | 0.28 |
| 1 | 201047888 | rs11811602 | 0.528 | -1.317 | -0.815 | -1.717 | -0.006 | -0.001 | -0.001 | 0 | 0.3 | 0.32 | 0.33 | 0.77 | 0.24 | 0.69 |
| 1 | 201049320 | rs942707 | -0.897 | -1.422 | -0.791 | -1.775 | -0.009 | -0.084 | -0.084 | 0.515 | 0.86 | 0.7 | 0.92 | 1 | 0.88 | 0.96 |
| 1 | 201050046 | rs942705 | -0.147 | -1.439 | -0.916 | -1.866 | 0.01 | 0.059 | 0.059 | 0 | 0.52 | 0.63 | 0.6 | 0.81 | 0.48 | 0.8 |
| 1 | 201050233 | rs942704 | -1.014 | -1.434 | -0.916 | -1.865 | -0.009 | -0.08 | -0.08 | 3.506 | 0.87 | 0.71 | 0.92 | 1 | 0.88 | 0.96 |
| 1 | 201050284 | rs734881 | 0.559 | -1.164 | -1.396 | -1.642 | -0.133 | 0.03 | 0.03 | 0.044 | 0.34 | 0.07 | 0.32 | 0.19 | 0.4 | 0.16 |
| 1 | 201050568 | rs7539618 | 0.683 | -1.305 | -0.58 | -0.914 | 0.03 | -0.044 | -0.044 | 3.781 | 0.13 | 0.04 | 0.12 | 0.19 | 0.07 | 0.04 |
| 1 | 201052199 | rs3767511 | -0.064 | -0.706 | -0.297 | -0.626 | -0.002 | -0.05 | -0.05 | 0 | 0.23 | 0.3 | 0.23 | 0.03 | 0.24 | 0.11 |
| 1 | 201059084 | rs3753960 | -0.366 | -0.365 | -0.364 | -0.023 | -0.005 | 0.092 | 0.092 | 0 | 0.87 | 0.82 | 0.8 | 0.44 | 0.89 | 0.58 |
| 1 | 201059787 | rs10159219 | 0.151 | -0.644 | -0.599 | -0.605 | -0.005 | -0.049 | -0.049 | 0 | 0.29 | 0.28 | 0.29 | 0.03 | 0.28 | 0.11 |
| 1 | 201060865 | rs2296383 | 0.912 | -0.218 | -1.308 | -0.686 | -0.104 | -0.046 | -0.046 | 0.099 | 0.41 | 0.05 | 0.4 | 0.39 | 0.41 | 0.38 |
| 1 | 201067011 | rs17454870 | 0.165 | -0.021 | -0.966 | -0.228 | -0.003 | -0.014 | -0.014 | 3.271 | 0.16 | 0.12 | 0.13 | 0.13 | 0.14 | 0.11 |
| 1 | 201067784 | rs12409138 | -0.221 | 0.108 | -0.301 | -0.532 | -0.005 | 0.022 | 0.022 | 0 | 0.33 | 0.23 | 0.31 | 0.04 | 0.38 | 0.15 |
| 1 | 201068298 | rs6699862 | -0.417 | 0.204 | -0.742 | -0.356 | -0.018 | 0.061 | 0.061 | 0 | 0.82 | 0.73 | 0.8 | 0.56 | 0.85 | 0.58 |
| 1 | 201068787 | rs1574408 | 0.141 | 0.745 | -1.222 | -0.275 | -0.008 | -0.007 | -0.007 | 1.115 | 0.44 | 0.3 | 0.44 | 0.52 | 0.49 | 0.44 |
| 1 | 201069761 | rs17454947 | -1.085 | 1.417 | -1.087 | -0.505 | 0.001 | -0.027 | -0.027 | 0.945 | 0.13 | 0.13 | 0.11 | 0 | 0.23 | 0.1 |
| 1 | 201070740 | rs12405259 | -0.744 | 0.996 | -0.982 | -0.148 | 0.004 | -0.024 | -0.024 | 2.961 | 0.07 | 0.12 | 0.08 | 0 | 0.12 | 0.05 |
| 1 | 201071790 | rs10449267 | -0.318 | 0.399 | -1.252 | -0.105 | 0.013 | 0.029 | 0.029 | 0 | 0.46 | 0.69 | 0.58 | 0.66 | 0.45 | 0.54 |
| 1 | 201071884 | rs10920115 | -0.384 | -0.01 | -1.457 | -0.209 | 0.012 | 0.027 | 0.027 | 0 | 0.47 | 0.69 | 0.52 | 0.63 | 0.47 | 0.51 |
| 1 | 201073555 | rs12135240 | -0.774 | -0.427 | -1.05 | 0.027 | -0.003 | -0.002 | -0.002 | 0 | 0.14 | 0.16 | 0.23 | 0.11 | 0.11 | 0.19 |
| 1 | 201073691 | rs12561765 | -0.601 | -1.108 | -0.711 | 0.117 | -0.046 | 0.085 | 0.085 | 0.036 | 0.23 | 0.35 | 0.15 | 0.5 | 0.16 | 0.35 |
| 1 | 201074293 | rs12022389 | -0.609 | -1.098 | -0.709 | 0.119 | -0.044 | 0.087 | 0.087 | 0.012 | 0.23 | 0.35 | 0.16 | 0.5 | 0.17 | 0.35 |
| 1 | 201075231 | rs12239772 | 0.141 | -0.953 | -0.799 | -0.108 | 0.004 | 0.013 | 0.013 | 0 | 0.1 | 0.07 | 0.07 | 0.01 | 0.09 | 0.03 |
| 1 | 201075245 | rs2365293 | -0.688 | -1.12 | -0.485 | -0.229 | 0.004 | 0.003 | 0.003 | 0 | 0.29 | 0.37 | 0.47 | 0.66 | 0.35 | 0.48 |
| 1 | 201075796 | rs998135 | -0.882 | -1.737 | -0.795 | -0.431 | 0.064 | 0.13 | 0.13 | 1.53 | 0.46 | 0.83 | 0.56 | 0.77 | 0.37 | 0.6 |
| 1 | 201077212 | rs4915215 | -0.821 | -1.952 | 0.178 | -0.554 | -0.002 | -0.015 | -0.015 | 0 | 0.45 | 0.47 | 0.56 | 0.82 | 0.42 | 0.55 |
| 1 | 201079088 | rs1325311 | -0.837 | -1.961 | 0.364 | -0.55 | -0.009 | -0.032 | -0.032 | 0 | 0.46 | 0.52 | 0.55 | 0.82 | 0.39 | 0.54 |
| 1 | 201081943 | rs2281845 | -0.931 | -1.973 | 0.527 | -0.06 | 0.054 | 0.072 | 0.072 | 0.241 | 0.47 | 0.73 | 0.54 | 0.74 | 0.41 | 0.47 |
| 1 | 201083343 | rs3738262 | -1.172 | -2.038 | 0.436 | -0.105 | 0.074 | 0.137 | 0.137 | 1.005 | 0.49 | 0.83 | 0.54 | 0.77 | 0.41 | 0.55 |
| 1 | 201085420 | rs12079819 | 0.039 | -2.292 | 0.839 | 0.657 | 0.006 | 0.059 | 0.059 | 0 | 0.22 | 0.31 | 0.38 | 0.51 | 0.27 | 0.36 |
| 1 | 201085996 | rs10494826 | 0.515 | -2.67 | 0.46 | 0.078 | -0.001 | -0.063 | -0.063 | 2.57 | 0.14 | 0.31 | 0.16 | 0.03 | 0.11 | 0.1 |
| 1 | 201088889 | rs942702 | -2.042 | -1.995 | -0.497 | -0.055 | -0.14 | 0.105 | 0.105 | 0 | 0.66 | 0.96 | 0.76 | 0.86 | 0.59 | 0.64 |
| 1 | 201092550 | rs10753889 | -0.141 | -2.008 | 0.172 | 0.174 | -0.071 | -0.01 | -0.01 | 0 | 0.25 | 0.44 | 0.17 | 0.23 | 0.17 | 0.11 |
| 1 | 201095714 | rs10920125 | -0.126 | -2.024 | 0.155 | 0.173 | -0.081 | -0.01 | -0.01 | 0 | 0.25 | 0.46 | 0.16 | 0.23 | 0.15 | 0.11 |
| 1 | 201097881 | rs12029703 | 0.827 | -1.99 | -0.401 | 0.233 | 0.006 | -0.029 | -0.029 | 1.794 | 0.08 | 0.03 | 0.09 | 0.11 | 0.07 | 0.12 |
| 1 | 201099275 | rs2057579 | -0.906 | -2.284 | -0.416 | 0.129 | 0.082 | 0.117 | 0.117 | 0.696 | 0.48 | 0.81 | 0.54 | 0.74 | 0.41 | 0.46 |
| 1 | 201100439 | rs10800760 | 0.432 | -2.203 | -0.081 | 0.306 | -0.002 | -0.036 | -0.036 | 0 | 0.22 | 0.17 | 0.35 | 0.51 | 0.25 | 0.35 |
| 1 | 201100482 | rs10753891 | -0.32 | -2.157 | -0.122 | 0.141 | -0.022 | 0.015 | 0.015 | 0 | 0.32 | 0.54 | 0.43 | 0.62 | 0.28 | 0.39 |

| | | | | | | | | | | | | | | | | |
|---|---|---|---|---|---|---|---|---|---|---|---|---|---|---|---|---|
| 1 | 201101413 | rs6697303 | -1.267 | -2.131 | -0.512 | -0.12 | 0.015 | 0.112 | 0.112 | 0.138 | 0.54 | 0.8 | 0.68 | 0.76 | 0.47 | 0.64 |
| 1 | 201103404 | rs17455455 | 1.358 | -0.694 | -1.031 | -0.25 | -0.035 | -0.017 | -0.017 | 0.37 | 0.16 | 0.08 | 0.17 | 0.17 | 0.25 | 0.28 |
| 1 | 201104589 | rs1059625 | 0.256 | -1.105 | -0.995 | -0.493 | -0.023 | 0.02 | 0.02 | 1.924 | 0.37 | 0.25 | 0.38 | 0.25 | 0.48 | 0.37 |
| 1 | 201105488 | rs6693143 | 0.192 | -1.684 | -0.625 | -0.326 | -0.02 | 0.016 | 0.016 | 0 | 0.3 | 0.35 | 0.31 | 0.49 | 0.18 | 0.4 |
| 1 | 201105708 | rs754924 | 0.119 | -1.711 | -0.594 | -0.34 | -0.047 | 0.033 | 0.033 | 0 | 0.31 | 0.46 | 0.31 | 0.49 | 0.18 | 0.4 |
| 1 | 201108096 | rs11590996 | 0.261 | -1.823 | -0.588 | -0.022 | -0.025 | 0.025 | 0.025 | 0 | 0.25 | 0.32 | 0.28 | 0.44 | 0.13 | 0.32 |
| 1 | 201108503 | rs17421734 | 0.557 | -1.119 | -0.899 | -0.267 | 0.002 | 0.005 | 0.005 | 0 | 0.13 | 0.1 | 0.13 | 0.03 | 0.11 | 0.13 |
| 1 | 201109903 | rs10920129 | 0.638 | -1.107 | -1.043 | -0.314 | -0.011 | 0.027 | 0.027 | 2.566 | 0.21 | 0.14 | 0.21 | 0.12 | 0.24 | 0.25 |
| 1 | 201109984 | rs1408869 | 0.087 | -1.89 | -0.956 | -0.291 | -0.005 | 0.004 | 0.004 | 0 | 0.67 | 0.61 | 0.68 | 0.62 | 0.71 | 0.66 |
| 1 | 201112204 | rs7545363 | -0.035 | -1.857 | -0.918 | -0.279 | -0.009 | 0.003 | 0.003 | 2.78 | 0.67 | 0.64 | 0.71 | 0.63 | 0.76 | 0.75 |
| 1 | 201112981 | rs8158 | -0.035 | -1.857 | -0.918 | -0.279 | -0.009 | 0.003 | 0.003 | 2.78 | 0.67 | 0.64 | 0.71 | 0.63 | 0.76 | 0.75 |
| 1 | 201113118 | rs3738266 | -2.158 | -1.963 | -0.879 | -0.186 | 0.013 | -0.013 | -0.013 | 0 | 0.92 | 0.94 | 0.96 | 0.87 | 0.95 | 0.97 |
| 1 | 201113318 | rs10920130 | 1.845 | -2.081 | -0.961 | -0.388 | 0.001 | -0.007 | -0.007 | 0.948 | 0.27 | 0.25 | 0.26 | 0.39 | 0.32 | 0.32 |
| 1 | 201115288 | rs3752975 | 2.312 | -1.939 | -1.11 | -0.402 | -0.05 | -0.01 | -0.01 | 0.707 | 0.16 | 0.08 | 0.2 | 0.14 | 0.29 | 0.23 |
| 1 | 201115822 | rs3752978 | 1.194 | -2.241 | -0.792 | -0.2 | -0.017 | 0.011 | 0.011 | 0 | 0.33 | 0.42 | 0.29 | 0.39 | 0.32 | 0.32 |
| 1 | 201116502 | rs4915489 | -0.016 | -1.838 | -0.659 | 0.04 | 0.004 | -0.003 | -0.003 | 0 | 0.68 | 0.68 | 0.72 | 0.62 | 0.76 | 0.75 |
| 1 | 201117133 | rs3767514 | 1.544 | -1.784 | -0.921 | 0.047 | -0.009 | 0.014 | 0.014 | 5.076 | 0.16 | 0.11 | 0.18 | 0.09 | 0.18 | 0.08 |
| 1 | 201121523 | rs6694122 | -0.226 | -1.43 | -1.128 | 0.113 | -0.011 | 0.004 | 0.004 | 2.58 | 0.67 | 0.63 | 0.71 | 0.62 | 0.76 | 0.75 |
| 1 | 201122816 | rs16847895 | 1.383 | -0.419 | -1.422 | -0.212 | -0.028 | 0.012 | 0.012 | 0 | 0.16 | 0.04 | 0.22 | 0.13 | 0.17 | 0.11 |
| 1 | 201124410 | rs3753967 | -0.254 | -0.042 | -0.999 | -0.188 | -0.049 | 0.035 | 0.035 | 0.154 | 0.36 | 0.2 | 0.43 | 0.22 | 0.44 | 0.44 |
| 1 | 201125326 | rs7542768 | -0.011 | 0.225 | -1.481 | 0.05 | -0.053 | 0.02 | 0.02 | 0.004 | 0.31 | 0.17 | 0.4 | 0.22 | 0.38 | 0.36 |
| 1 | 201133774 | rs863826 | 0.035 | 0.048 | -1.442 | 0.105 | -0.019 | 0.042 | 0.042 | 0 | 0.54 | 0.27 | 0.61 | 0.7 | 0.5 | 0.58 |
| 1 | 201133955 | rs831748 | 0.701 | -0.414 | -1.97 | 0.345 | -0.077 | -0.078 | -0.078 | 0.253 | 0.38 | 0.04 | 0.5 | 0.44 | 0.37 | 0.39 |
| 1 | 201137500 | rs875495 | 0.326 | -0.461 | -1.514 | 0.014 | -0.015 | -0.009 | -0.009 | 0 | 0.61 | 0.23 | 0.65 | 0.47 | 0.57 | 0.52 |
| 1 | 201137527 | rs6699983 | 0.618 | -0.085 | -0.871 | 0.108 | 0.118 | 0.114 | 0.114 | 0 | 0.1 | 0.27 | 0.15 | 0.23 | 0.23 | 0.25 |
| 1 | 201138428 | rs1122396 | -0.476 | -0.403 | -0.678 | -0.022 | -0.026 | -0.037 | -0.037 | 0.14 | 0.35 | 0.74 | 0.36 | 0.46 | 0.45 | 0.41 |
| 1 | 201138712 | rs1722755 | -1.03 | -0.743 | -0.622 | 0.109 | 0.023 | 0.026 | 0.026 | 0 | 0.83 | 0.91 | 0.9 | 0.92 | 0.88 | 0.9 |
| 1 | 201139616 | rs6413916 | -0.895 | -0.748 | -0.557 | 0.106 | 0.021 | 0.024 | 0.024 | 0 | 0.82 | 0.89 | 0.9 | 0.92 | 0.88 | 0.9 |
| 1 | 201142554 | rs10920138 | 0.833 | -0.743 | -0.442 | 0.154 | -0.026 | 0.161 | 0.161 | 0 | 0.11 | 0.41 | 0.1 | 0.24 | 0.08 | 0.12 |
| 1 | 201143774 | rs2066284 | 0.649 | -0.611 | -0.883 | 0.223 | -0.065 | -0.03 | -0.03 | 0 | 0.13 | 0.04 | 0.16 | 0.13 | 0.28 | 0.31 |
| 1 | 201149033 | rs831768 | 0.572 | -1.309 | -0.579 | 0.314 | 0.015 | 0.019 | 0.019 | 0 | 0.18 | 0.13 | 0.1 | 0.07 | 0.12 | 0.1 |
| 1 | 201150096 | rs16847947 | 1.024 | -0.995 | -0.717 | 0.498 | -0.032 | 0.07 | 0.07 | 0 | 0.1 | 0.2 | 0.08 | 0.24 | 0.05 | 0.12 |
| 1 | 201155654 | rs10920139 | 1.041 | -1.137 | -1.229 | 0.407 | 0.046 | -0.016 | -0.016 | 0.274 | 0.27 | 0.14 | 0.29 | 0.33 | 0.18 | 0.32 |
| 1 | 201155702 | rs831756 | -0.362 | -0.601 | -0.871 | 0.334 | -0.012 | 0.005 | 0.005 | 0 | 0.69 | 0.53 | 0.68 | 0.65 | 0.73 | 0.78 |
| 1 | 201157146 | rs831752 | 0.393 | -0.27 | -1.055 | 0.433 | 0.007 | 0.005 | 0.005 | 0 | 0.12 | 0.09 | 0.09 | 0.07 | 0.09 | 0.09 |
| 1 | 201158976 | rs831751 | -0.681 | -0.12 | -1.113 | 0.511 | 0.001 | -0.003 | -0.003 | 0 | 0.88 | 0.87 | 0.8 | 0.85 | 0.94 | 0.86 |
| 1 | 201159461 | rs831750 | 0.718 | 0.617 | -1.14 | 0.748 | -0.01 | -0.009 | -0.009 | 3.011 | 0.17 | 0.14 | 0.2 | 0.35 | 0.32 | 0.36 |
| 1 | 201163584 | rs1568809 | -0.394 | 0.055 | -1.506 | -0.149 | -0.057 | 0.137 | 0.137 | 0 | 0.61 | 0.81 | 0.57 | 0.86 | 0.55 | 0.79 |
| 1 | 201164099 | rs1007227 | 0.713 | -0.309 | -1.153 | -0.029 | 0.018 | 0.06 | 0.06 | 0 | 0.26 | 0.4 | 0.24 | 0.46 | 0.33 | 0.41 |
| 1 | 201164930 | rs4915219 | 2.095 | -0.652 | -1.67 | -0.066 | 0.014 | 0.075 | 0.075 | 0 | 0.13 | 0.01 | 0.1 | 0 | 0.07 | 0.09 |
| 1 | 201166281 | rs7533096 | 0.005 | -0.455 | -1.151 | 0.199 | 0.004 | 0.006 | 0.006 | 0 | 0.36 | 0.62 | 0.32 | 0.53 | 0.41 | 0.54 |
| 1 | 201166383 | rs4915221 | -0.734 | -0.39 | -1.435 | -0.078 | -0.047 | 0.138 | 0.138 | 0 | 0.6 | 0.81 | 0.57 | 0.86 | 0.55 | 0.8 |
| 1 | 201169554 | rs2365299 | 0.312 | -0.249 | -1.072 | 0.518 | -0.002 | -0.001 | -0.001 | 0.11 | 0.24 | 0.19 | 0.2 | 0.26 | 0.14 | 0.25 |
| 1 | 201170571 | rs10157719 | 0.272 | 0.563 | -0.935 | 1.046 | 0.002 | 0.001 | 0.001 | 0.084 | 0.19 | 0.23 | 0.18 | 0.23 | 0.31 | 0.29 |
| 1 | 201171283 | rs947379 | -0.242 | 0.536 | -0.864 | 0.92 | 0.005 | 0.008 | 0.008 | 0 | 0.64 | 0.58 | 0.54 | 0.53 | 0.57 | 0.59 |
| 1 | 201175274 | rs9651057 | -0.315 | -0.248 | -1.494 | 0.777 | -0.041 | -0.024 | -0.024 | 0 | 0.4 | 0.15 | 0.41 | 0.59 | 0.48 | 0.33 |
| 1 | 201176711 | rs12568690 | 0.733 | -0.2 | -1.582 | 0.909 | -0.027 | -0.018 | -0.018 | 0 | 0.07 | 0.01 | 0.09 | 0.08 | 0.11 | 0.06 |
| 1 | 201177278 | rs7551098 | 1.524 | -0.723 | -1.504 | 0.424 | -0.018 | -0.008 | -0.008 | 0.069 | 0.29 | 0.21 | 0.32 | 0.31 | 0.35 | 0.41 |
| 1 | 201181217 | rs12070918 | 1.491 | -0.732 | -1.281 | 0.419 | -0.007 | -0.003 | -0.003 | 0 | 0.29 | 0.25 | 0.32 | 0.31 | 0.35 | 0.42 |
| 1 | 201181994 | rs6688782 | 0.024 | -1.043 | -0.849 | 0.243 | -0.045 | 0.023 | 0.023 | 0 | 0.1 | 0.36 | 0.07 | 0.1 | 0.01 | 0.14 |
| 1 | 201183363 | rs2282415 | 1.46 | -0.771 | -1.417 | 0.248 | -0.007 | -0.003 | -0.003 | 0 | 0.29 | 0.25 | 0.32 | 0.31 | 0.35 | 0.42 |
| 1 | 201183402 | rs1568810 | 1.527 | -0.78 | -1.419 | 0.372 | -0.022 | -0.022 | -0.022 | 0.048 | 0.28 | 0.19 | 0.33 | 0.34 | 0.35 | 0.42 |
| 1 | 201183773 | rs4915495 | 0.733 | -0.753 | -1.595 | 0.434 | -0.003 | -0.006 | -0.006 | 0 | 0.14 | 0.1 | 0.15 | 0.22 | 0.16 | 0.19 |
| 1 | 201184275 | rs2275673 | 1.54 | -0.694 | -1.554 | 0.258 | -0.011 | -0.011 | -0.011 | 0 | 0.29 | 0.24 | 0.33 | 0.35 | 0.35 | 0.42 |
| 1 | 201187828 | rs3738268 | -1.948 | -0.581 | -1.332 | -0.377 | -0.017 | 0.079 | 0.079 | 0 | 0.84 | 0.96 | 0.85 | 1 | 0.89 | 0.91 |
| 1 | 201187954 | rs6667020 | -1.928 | -0.581 | -1.326 | -0.377 | -0.01 | 0.067 | 0.067 | 0 | 0.84 | 0.95 | 0.85 | 1 | 0.89 | 0.91 |
| 1 | 201190732 | rs3738269 | -0.547 | -0.732 | -1.395 | -0.099 | -0.018 | -0.02 | -0.02 | 0 | 0.2 | 0.11 | 0.25 | 0.26 | 0.27 | 0.26 |
| 1 | 201193499 | rs6687300 | 0.669 | -0.93 | -1.466 | -0.438 | 0.038 | 0.087 | 0.087 | 1.82 | 0.61 | 0.23 | 0.6 | 0.83 | 0.71 | 0.66 |
| 1 | 201195119 | rs3738270 | 0.759 | -1.042 | -1.563 | -0.558 | 0.057 | 0.123 | 0.123 | 1.045 | 0.58 | 0.2 | 0.59 | 0.83 | 0.7 | 0.64 |
| 1 | 201197331 | rs3753971 | 0.199 | -1.516 | -1.847 | -0.354 | -0.092 | 0.091 | 0.091 | 0 | 0.52 | 0.93 | 0.6 | 0.67 | 0.53 | 0.73 |
| 1 | 201197471 | rs3753973 | 0.026 | -1.582 | -1.889 | -0.393 | -0.09 | -0.037 | -0.037 | 0 | 0.33 | 0.04 | 0.26 | 0.33 | 0.37 | 0.18 |
| 1 | 201197781 | rs14061 | 0.015 | -1.619 | -1.387 | -0.409 | -0.09 | -0.036 | -0.036 | 1.785 | 0.33 | 0.04 | 0.26 | 0.33 | 0.37 | 0.18 |
| 1 | 201198425 | rs1578531 | 1.696 | -1.613 | -1.435 | -0.411 | -0.047 | -0.063 | -0.063 | 0 | 0.2 | 0.01 | 0.12 | 0.26 | 0.19 | 0.08 |
| 1 | 201202490 | rs1917348 | 0.23 | -1.566 | -1.362 | -0.361 | -0.092 | 0.091 | 0.091 | 0 | 0.52 | 0.93 | 0.6 | 0.66 | 0.52 | 0.73 |
| 1 | 201216597 | rs10800765 | 0.229 | -1.557 | -1.357 | -0.349 | -0.093 | 0.069 | 0.069 | 0 | 0.52 | 0.93 | 0.57 | 0.66 | 0.52 | 0.73 |
| 1 | 201217148 | rs10920155 | 1.8 | -1.554 | -1.282 | -0.235 | -0.053 | -0.075 | -0.075 | 0.13 | 0.33 | 0.16 | 0.36 | 0.49 | 0.4 | 0.48 |
| 1 | 201218342 | rs6427893 | 0.391 | -1.491 | -0.959 | -0.217 | 0.051 | 0.056 | 0.056 | 0.842 | 0.66 | 0.22 | 0.65 | 0.83 | 0.77 | 0.67 |
| 1 | 201219233 | rs10157947 | 0.573 | -1.258 | -0.25 | 0.027 | 0.002 | 0.057 | 0.057 | 0 | 0.11 | 0.2 | 0.14 | 0.15 | 0.11 | 0.18 |
| 1 | 201233751 | rs2078570 | 0.299 | -1.225 | -1.068 | -0.023 | -0.04 | 0.015 | 0.015 | 0.019 | 0.11 | 0.02 | 0.14 | 0.08 | 0.17 | 0.11 |

| | | | | | | | | | | | | | | | | |
|---|---|---|---|---|---|---|---|---|---|---|---|---|---|---|---|---|
| 1 | 201235047 | rs4244148 | -1.274 | -1.268 | -0.776 | -0.017 | -0.009 | -0.006 | -0.006 | 0.073 | 0.28 | 0.49 | 0.31 | 0.22 | 0.29 | 0.32 |
| 1 | 201239651 | rs4915501 | -0.847 | -1.814 | -1.433 | -0.014 | 0.06 | -0.054 | -0.054 | 0 | 0.41 | 0.83 | 0.33 | 0.42 | 0.3 | 0.3 |
| 1 | 201240182 | rs10800771 | -0.911 | -1.837 | -1.425 | -0.042 | 0.059 | -0.011 | -0.011 | 0 | 0.42 | 0.75 | 0.34 | 0.59 | 0.3 | 0.38 |
| 1 | 201240763 | rs4915503 | -0.057 | -1.686 | -1.48 | -0.75 | -0.023 | 0.001 | 0.001 | 0 | 0.67 | 0.93 | 0.75 | 0.74 | 0.72 | 0.83 |
| 1 | 201241279 | rs6687886 | -1.277 | -1.584 | -1.429 | -0.733 | -0.006 | 0.072 | 0.072 | 0 | 0.85 | 0.96 | 0.87 | 1 | 0.89 | 0.9 |
| 1 | 201244429 | rs16848196 | 0.493 | -1.489 | -0.946 | 0.045 | -0.04 | -0.085 | -0.085 | 0 | 0.18 | 0.01 | 0.12 | 0.26 | 0.17 | 0.07 |
| 1 | 201246636 | rs832179 | 0.963 | -1.662 | -0.553 | -0.273 | 0.011 | 0.021 | 0.021 | 0 | 0.62 | 0.62 | 0.72 | 0.75 | 0.67 | 0.86 |
| 1 | 201246862 | rs12143238 | -0.131 | -1.614 | -0.632 | -0.564 | -0.013 | 0.054 | 0.054 | 0 | 0.09 | 0.01 | 0.07 | 0 | 0.09 | 0.02 |
| 1 | 201247121 | rs16848203 | -1.001 | -1.849 | -0.098 | -0.736 | -0.016 | -0.05 | -0.05 | 5.262 | 0.12 | 0.12 | 0.07 | 0 | 0.08 | 0.07 |
| 1 | 201248438 | rs854493 | -0.947 | -1.885 | 0.073 | -0.617 | -0.013 | -0.065 | -0.065 | 0.984 | 0.16 | 0.19 | 0.11 | 0 | 0.12 | 0.06 |
| 1 | 201251316 | rs854488 | 1.9 | -2.125 | -0.245 | -0.67 | 0.042 | 0.211 | 0.211 | 0.295 | 0.47 | 0.08 | 0.6 | 0.75 | 0.62 | 0.69 |
| 1 | 201252866 | rs2268147 | -0.23 | -2.135 | 0.741 | -0.573 | -0.006 | -0.024 | -0.024 | 0 | 0.1 | 0.38 | 0.11 | 0.09 | 0.07 | 0.17 |
| 1 | 201254116 | rs1257394 | 0.72 | -2.142 | 0.55 | -0.586 | -0.062 | -0.119 | -0.119 | 0.423 | 0.8 | 0.56 | 0.84 | 0.91 | 0.87 | 0.79 |
| 1 | 201255199 | rs832164 | 1.863 | -2.263 | 0.118 | -0.438 | 0.085 | 0.115 | 0.115 | 3.832 | 0.68 | 0.1 | 0.73 | 0.85 | 0.81 | 0.73 |
| 1 | 201256121 | rs832168 | 1.853 | -2.235 | 0.246 | -0.62 | 0.072 | 0.109 | 0.109 | 3.085 | 0.67 | 0.15 | 0.75 | 0.9 | 0.8 | 0.74 |
| 1 | 201257016 | rs832170 | 0.611 | -1.949 | 0.126 | -0.583 | -0.071 | -0.017 | -0.017 | 0 | 0.2 | 0.01 | 0.14 | 0.15 | 0.21 | 0.08 |
| 1 | 201257373 | rs16848229 | -1.184 | -1.983 | 0.696 | -0.612 | -0.013 | -0.095 | -0.095 | 0 | 0.13 | 0.48 | 0.1 | 0 | 0.09 | 0.02 |
| 1 | 201258519 | rs832174 | 1.774 | -1.996 | 0.371 | -0.615 | 0.02 | 0.019 | 0.019 | 1.96 | 0.7 | 0.24 | 0.76 | 0.9 | 0.81 | 0.74 |
| 1 | 201262432 | rs860554 | -0.436 | -1.797 | 0.106 | -0.499 | -0.062 | 0.063 | 0.063 | 0.41 | 0.29 | 0.05 | 0.21 | 0.15 | 0.3 | 0.1 |
| 1 | 201264902 | rs2300293 | 1.367 | -1.829 | 0.41 | -0.463 | 0.002 | -0.001 | -0.001 | 0.434 | 0.52 | 0.57 | 0.63 | 0.75 | 0.6 | 0.65 |
| 1 | 201265770 | rs713292 | 1.532 | -1.875 | -0.217 | -0.408 | 0.015 | 0.195 | 0.195 | 0.304 | 0.52 | 0.04 | 0.64 | 0.77 | 0.63 | 0.67 |
| 1 | 201271222 | rs706494 | 0.581 | -1.762 | 0.198 | 0.077 | -0.014 | -0.02 | -0.02 | 0.356 | 0.17 | 0.09 | 0.19 | 0.24 | 0.2 | 0.11 |
| 1 | 201278862 | rs1772847 | 0.708 | -1.806 | 0.354 | 0.069 | -0.022 | -0.017 | -0.017 | 0.293 | 0.17 | 0.09 | 0.15 | 0.18 | 0.18 | 0.08 |
| 1 | 201279398 | rs1779304 | 0.007 | -2.035 | 1.08 | -0.079 | -0.033 | -0.077 | -0.077 | 0.091 | 0.88 | 0.71 | 0.93 | 1 | 0.94 | 0.96 |
| 1 | 201280649 | rs832152 | 0.526 | -1.954 | 1.039 | 0.012 | -0.024 | -0.036 | -0.036 | 0.354 | 0.18 | 0.09 | 0.19 | 0.23 | 0.2 | 0.11 |
| 1 | 201280956 | rs832151 | 0.225 | -1.973 | 1.332 | 0.014 | 0.007 | 0.021 | 0.021 | 0.808 | 0.19 | 0.25 | 0.2 | 0.25 | 0.2 | 0.11 |
| 1 | 201285712 | rs7514146 | -1.056 | -2.097 | 1.322 | -0.108 | -0.036 | -0.081 | -0.081 | 0 | 0.15 | 0.31 | 0.08 | 0.01 | 0.1 | 0.19 |
| 1 | 201288955 | rs1626370 | 0.533 | -2.06 | 1.234 | -0.073 | -0.023 | -0.036 | -0.036 | 0 | 0.18 | 0.09 | 0.19 | 0.23 | 0.2 | 0.11 |
| 1 | 201292473 | rs10920173 | -1.073 | -2.145 | 1.395 | -0.136 | -0.037 | -0.082 | -0.082 | 0 | 0.15 | 0.33 | 0.09 | 0.01 | 0.1 | 0.19 |
| 1 | 201292794 | rs1543805 | 1.181 | -2.153 | 1.342 | -0.103 | -0.012 | -0.015 | -0.015 | 0.148 | 0.65 | 0.38 | 0.71 | 0.76 | 0.7 | 0.7 |
| 1 | 201296112 | rs3820427 | -0.022 | -2.19 | 1.63 | -0.144 | -0.028 | -0.045 | -0.045 | 0 | 0.11 | 0.23 | 0.07 | 0.02 | 0.06 | 0.04 |
| 1 | 201297059 | rs1341944 | -0.949 | -2.189 | 1.573 | -0.147 | -0.027 | -0.066 | -0.066 | 0.779 | 0.14 | 0.26 | 0.08 | 0.01 | 0.1 | 0.19 |
| 1 | 201298352 | rs1572968 | 1.871 | -2.128 | 1.428 | 0.023 | 0.044 | 0.05 | 0.05 | 2.36 | 0.69 | 0.21 | 0.78 | 0.88 | 0.8 | 0.7 |
| 1 | 201298866 | rs1361902 | 2.227 | -2.168 | 1.327 | 0.03 | 0.017 | 0.041 | 0.041 | 0.107 | 0.51 | 0.03 | 0.64 | 0.7 | 0.62 | 0.61 |
| 1 | 201299470 | rs947376 | 1.865 | -2.3 | 1.406 | -0.119 | 0.043 | 0.044 | 0.044 | 3.375 | 0.71 | 0.2 | 0.83 | 0.94 | 0.84 | 0.76 |
| 1 | 201299785 | rs2365648 | 0.938 | -2.31 | 1.698 | -0.107 | -0.031 | -0.023 | -0.023 | 0 | 0.84 | 0.54 | 0.87 | 0.91 | 0.9 | 0.89 |
| 1 | 201299852 | rs1105331 | 2.079 | -2.342 | 1.642 | -0.148 | 0.054 | 0.06 | 0.06 | 2.731 | 0.68 | 0.19 | 0.78 | 0.87 | 0.79 | 0.69 |
| 1 | 201300025 | rs1104755 | -1.036 | -2.276 | 1.623 | -0.112 | -0.032 | -0.05 | -0.05 | 0 | 0.15 | 0.31 | 0.09 | 0.04 | 0.11 | 0.19 |
| 1 | 201300842 | rs10920175 | 1.101 | -2.253 | 1.727 | -0.101 | 0.026 | 0.049 | 0.049 | 0 | 0.94 | 0.8 | 0.94 | 0.93 | 0.96 | 0.93 |
| 1 | 201301053 | rs17425876 | 0.966 | -2.274 | 1.797 | -0.081 | -0.012 | 0.006 | 0.006 | 0.733 | 0.16 | 0.09 | 0.14 | 0.14 | 0.18 | 0.08 |
| 1 | 201301111 | rs12120834 | -1.125 | -2.276 | 1.91 | -0.089 | -0.031 | -0.036 | -0.036 | 0.928 | 0.14 | 0.37 | 0.09 | 0.06 | 0.1 | 0.2 |
| 1 | 201301696 | rs12143423 | -2.065 | -2.227 | 1.597 | -0.121 | -0.034 | -0.071 | -0.071 | 0 | 0.3 | 0.64 | 0.2 | 0.06 | 0.21 | 0.25 |
| 1 | 201303231 | rs10800773 | 1.067 | -2.183 | 1.487 | -0.074 | 0 | 0 | 0 | 2.922 | 0.64 | 0.37 | 0.71 | 0.73 | 0.69 | 0.66 |
| 1 | 201310733 | rs4915517 | -1.26 | -2.018 | 1.546 | 0.128 | -0.002 | 0.001 | 0.001 | 2.504 | 0.84 | 0.89 | 0.89 | 0.83 | 0.83 | 0.92 |
| 1 | 201311165 | rs2365650 | -0.39 | -2.069 | 1.642 | -0.026 | -0.011 | -0.057 | -0.057 | 0 | 0.13 | 0.16 | 0.1 | 0 | 0.11 | 0.05 |
| 1 | 201312033 | rs12126907 | 0.832 | -2.197 | 1.624 | 0.622 | -0.016 | -0.008 | -0.008 | 0.021 | 0.65 | 0.52 | 0.78 | 0.65 | 0.71 | 0.85 |
| 1 | 201312346 | rs4915518 | 0.749 | -2.152 | 1.597 | 0.476 | 0 | -0.055 | -0.055 | 0 | 0.66 | 0.57 | 0.74 | 0.8 | 0.67 | 0.8 |
| 1 | 201316482 | rs1539694 | 0.389 | -2.101 | 1.281 | 0.558 | -0.002 | -0.043 | -0.043 | 0 | 0.71 | 0.69 | 0.79 | 0.84 | 0.71 | 0.87 |
| 1 | 201322209 | rs12073852 | -1.542 | -1.842 | 1.225 | 0.739 | 0.041 | 0.041 | 0.041 | 0.135 | 0.07 | 0.24 | 0.08 | 0.08 | 0.09 | 0.06 |
| 1 | 201325462 | rs7540834 | -2.258 | -2.267 | 1.072 | 0.696 | -0.015 | 0.079 | 0.079 | 0 | 0.14 | 0.3 | 0.1 | 0.29 | 0.1 | 0.11 |
| 1 | 201327894 | rs10920180 | -0.865 | -2.383 | 1.251 | 0.563 | -0.042 | -0.044 | -0.044 | 0 | 0.16 | 0.32 | 0.09 | 0.07 | 0.09 | 0.2 |
| 1 | 201328824 | rs2275863 | -2.1 | -2.38 | 1.29 | 0.616 | -0.102 | 0.039 | 0.039 | 0.194 | 0.22 | 0.54 | 0.1 | 0.28 | 0.09 | 0.22 |
| 1 | 201332020 | rs1892027 | -0.36 | -2.253 | 1.394 | 0.699 | 0.004 | 0.017 | 0.017 | 1.385 | 0.78 | 0.8 | 0.8 | 0.53 | 0.74 | 0.84 |
| 1 | 201334382 | rs3729547 | 0.115 | -2.248 | 1.405 | 0.698 | 0.019 | 0.069 | 0.069 | 1.35 | 0.75 | 0.63 | 0.8 | 0.53 | 0.74 | 0.84 |
| 1 | 201334795 | rs3729845 | 0.491 | -2.305 | 1.439 | 0.516 | -0.022 | -0.054 | -0.054 | 0.46 | 0.09 | 0.19 | 0.06 | 0 | 0.07 | 0.05 |
| 1 | 201336984 | rs3729843 | 2.943 | -2.294 | 1.042 | 0.859 | -0.192 | 0.058 | 0.058 | 0 | 0.37 | 0.02 | 0.48 | 0.18 | 0.47 | 0.42 |
| 1 | 201337768 | rs3753997 | -0.923 | -1.503 | 1.745 | 0.576 | -0.032 | -0.034 | -0.034 | 0 | 0.15 | 0.26 | 0.07 | 0.07 | 0.09 | 0.19 |
| 1 | 201338107 | rs16848462 | -0.923 | -1.503 | 1.728 | 0.576 | -0.033 | -0.035 | -0.035 | 0 | 0.15 | 0.26 | 0.07 | 0.07 | 0.09 | 0.19 |
| 1 | 201340813 | rs6427901 | -0.99 | -1.504 | 1.735 | 0.576 | -0.042 | -0.044 | -0.044 | 0.354 | 0.16 | 0.32 | 0.08 | 0.07 | 0.09 | 0.19 |
| 1 | 201341341 | rs868407 | -0.476 | -1.46 | 1.678 | 0.619 | 0.021 | 0.011 | 0.011 | 0.165 | 0.8 | 0.74 | 0.81 | 0.74 | 0.75 | 0.84 |
| 1 | 201343638 | rs2996494 | 0.026 | -1.46 | 1.524 | 0.556 | 0.006 | 0.041 | 0.041 | 0 | 0.05 | 0.23 | 0.03 | 0.05 | 0.02 | 0.17 |
| 1 | 201345352 | rs947485 | 1.512 | -1.498 | 1.166 | 0.57 | 0.007 | -0.022 | -0.022 | 0.348 | 0.6 | 0.31 | 0.72 | 0.49 | 0.65 | 0.63 |
| 1 | 201345487 | rs12564445 | -2.134 | -1.367 | 1.315 | 0.737 | -0.028 | 0.074 | 0.074 | 0.124 | 0.22 | 0.43 | 0.16 | 0.35 | 0.18 | 0.27 |
| 1 | 201347315 | rs2799686 | 0.481 | -1.25 | 0.716 | 0.427 | -0.014 | -0.032 | -0.032 | 0.039 | 0.88 | 0.83 | 0.94 | 0.96 | 0.91 | 0.97 |
| 1 | 201350082 | rs16848493 | 0.142 | -0.988 | 0.466 | 0.66 | -0.027 | -0.046 | -0.046 | 0.021 | 0.09 | 0.13 | 0.05 | 0 | 0.05 | 0.04 |
| 1 | 201355522 | rs4128458 | -1.953 | -0.851 | 0.04 | 0.705 | -0.036 | -0.04 | -0.04 | 0 | 0.54 | 0.88 | 0.38 | 0.52 | 0.46 | 0.36 |
| 1 | 201355653 | rs11805972 | -0.103 | -0.985 | 0.432 | 0.664 | -0.024 | -0.014 | -0.014 | 0 | 0.08 | 0.25 | 0.04 | 0.04 | 0.03 | 0.02 |
| 1 | 201355943 | rs2799677 | -1.764 | -1.005 | 0.467 | 0.651 | 0.006 | 0.005 | 0.005 | 0.195 | 0.51 | 0.75 | 0.38 | 0.52 | 0.46 | 0.36 |
| 1 | 201358304 | rs3738281 | -0.489 | -1.014 | 0.359 | 0.553 | 0.025 | 0.052 | 0.052 | 0.652 | 0.11 | 0.36 | 0.07 | 0.11 | 0.09 | 0.17 |

| | | | | | | | | | | | | | | | | |
|---|---|---|---|---|---|---|---|---|---|---|---|---|---|---|---|---|
| 1 | 201359697 | rs6671391 | -1.279 | -0.973 | -0.114 | 0.567 | -0.029 | -0.006 | -0.006 | 0 | 0.19 | 0.64 | 0.11 | 0.14 | 0.12 | 0.19 |
| 1 | 201360281 | rs2719184 | 0.291 | -0.334 | -0.507 | 0.494 | -0.038 | 0.042 | 0.042 | 4.67 | 0.21 | 0.04 | 0.14 | 0.14 | 0.22 | 0.11 |
| 1 | 201364434 | rs3923236 | -0.119 | -0.468 | -0.159 | 0.436 | -0.029 | -0.044 | -0.044 | 0 | 0.09 | 0.13 | 0.04 | 0 | 0.05 | 0.04 |
| 1 | 201364570 | rs4606345 | -0.82 | -0.219 | -0.307 | 0.796 | 0.003 | -0.006 | -0.006 | 0 | 0.44 | 0.6 | 0.31 | 0.48 | 0.39 | 0.34 |
| 1 | 201369279 | rs2799685 | 0.034 | -0.158 | -0.665 | 0.666 | 0.023 | 0.008 | 0.008 | 0 | 0.52 | 0.31 | 0.63 | 0.52 | 0.59 | 0.67 |
| 1 | 201373844 | rs1107256 | 0.921 | -0.357 | -0.684 | 0.582 | -0.032 | 0.048 | 0.048 | 0 | 0.26 | 0.04 | 0.11 | 0.13 | 0.25 | 0.15 |
| 1 | 201374677 | rs2799680 | -0.598 | -0.112 | -0.675 | 0.688 | 0.027 | 0.008 | 0.008 | 0 | 0.49 | 0.7 | 0.36 | 0.47 | 0.41 | 0.33 |
| 1 | 201375658 | rs1256945 | -0.598 | -0.155 | -0.675 | 0.685 | 0.029 | 0.006 | 0.006 | 0 | 0.49 | 0.69 | 0.36 | 0.48 | 0.41 | 0.33 |
| 1 | 201376261 | rs1256948 | -0.678 | -0.258 | -0.713 | 0.588 | 0.026 | 0 | 0 | 0 | 0.52 | 0.76 | 0.37 | 0.48 | 0.43 | 0.34 |
| 1 | 201377885 | rs4915521 | -0.085 | 0.022 | -0.768 | 0.893 | 0.003 | 0.004 | 0.004 | 0 | 0.53 | 0.47 | 0.55 | 0.65 | 0.62 | 0.61 |
| 1 | 201378488 | rs12048410 | 0.151 | -0.984 | -0.522 | 1.222 | 0.001 | -0.003 | -0.003 | 0 | 0.16 | 0.18 | 0.17 | 0.27 | 0.14 | 0.16 |
| 1 | 201383643 | rs2026594 | 0.209 | -1.158 | -0.296 | 1.249 | -0.019 | 0.037 | 0.037 | 0 | 0.84 | 0.73 | 0.85 | 0.72 | 0.86 | 0.76 |
| 1 | 201385061 | rs1819291 | 0.118 | -1.183 | -0.422 | 0.969 | -0.017 | 0.005 | 0.005 | 0 | 0.84 | 0.75 | 0.87 | 0.8 | 0.88 | 0.78 |
| 1 | 201385868 | rs7513481 | -0.115 | -0.765 | -0.834 | 1.085 | -0.008 | 0.001 | 0.001 | 0 | 0.3 | 0.31 | 0.26 | 0.27 | 0.22 | 0.41 |
| 1 | 201386916 | rs3738286 | -0.293 | -1.033 | -1.389 | 0.907 | 0.021 | -0.044 | -0.044 | 0 | 0.1 | 0 | 0.03 | 0.12 | 0.05 | 0.23 |
| 1 | 201388148 | rs3767548 | 0.503 | -1.09 | -1.393 | 0.238 | 0.004 | 0.049 | 0.049 | 1.358 | 0.4 | 0.33 | 0.43 | 0.15 | 0.57 | 0.21 |
| 1 | 201389299 | rs2799670 | 0.107 | -1.774 | -1.794 | 1.037 | 0.02 | -0.057 | -0.057 | 0 | 0.72 | 0.87 | 0.63 | 0.54 | 0.76 | 0.86 |
| 1 | 201390540 | rs3767551 | -0.598 | -1.998 | -2.014 | 1.346 | -0.001 | -0.091 | -0.091 | 0 | 0.19 | 0.02 | 0.15 | 0.28 | 0.15 | 0.28 |
| 1 | 201391030 | rs2799671 | 0.091 | -1.767 | -1.115 | 1.235 | -0.004 | 0.042 | 0.042 | 0 | 0.06 | 0.12 | 0.02 | 0.1 | 0.04 | 0.25 |
| 1 | 201392764 | rs1256941 | 0.466 | -1.818 | -1.487 | 1.705 | -0.032 | -0.044 | -0.044 | 0.606 | 0.91 | 0.98 | 0.89 | 0.7 | 0.86 | 0.77 |
| 1 | 201400088 | rs10800777 | 1.539 | -2.966 | -2.126 | 0.928 | -0.002 | 0.149 | 0.149 | 0 | 0.42 | 0.11 | 0.47 | 0.22 | 0.6 | 0.26 |
| 1 | 201400200 | rs10920194 | 1.501 | -2.983 | -2.136 | 0.923 | 0 | 0.15 | 0.15 | 0 | 0.43 | 0.11 | 0.48 | 0.22 | 0.6 | 0.26 |
| 1 | 201401025 | rs10800779 | -1.318 | -3.086 | -1.417 | 0.949 | 0.083 | 0.023 | 0.023 | 0 | 0.44 | 0.71 | 0.41 | 0.63 | 0.27 | 0.64 |
| 1 | 201407782 | rs6697016 | -0.54 | -3.265 | -0.662 | 1.216 | -0.026 | 0.142 | 0.142 | 0 | 0.22 | 0.47 | 0.36 | 0.48 | 0.18 | 0.48 |
| 1 | 201415592 | rs10494830 | -0.382 | -3.455 | -0.426 | 0.582 | -0.035 | -0.042 | -0.042 | 0 | 0.14 | 0.19 | 0.04 | 0.05 | 0.05 | 0.06 |
| 1 | 201419375 | rs7525711 | 0.744 | -3.654 | -0.558 | 0.626 | -0.013 | -0.019 | -0.019 | 0 | 0.67 | 0.52 | 0.55 | 0.27 | 0.69 | 0.46 |
| 1 | 201422201 | rs10920205 | 1.478 | -3.516 | -0.98 | 0.377 | 0.029 | 0.099 | 0.099 | 0 | 0.54 | 0.34 | 0.5 | 0.23 | 0.62 | 0.29 |
| 1 | 201422464 | rs10920206 | -0.03 | -3.05 | -0.992 | 0.485 | 0.022 | 0.055 | 0.055 | 1.167 | 0.12 | 0.01 | 0.02 | 0.05 | 0.06 | 0.06 |
| 1 | 201423658 | rs16848610 | -0.398 | -3.244 | -0.958 | 0.509 | 0.003 | 0.018 | 0.018 | 0 | 0.1 | 0.04 | 0.03 | 0 | 0.07 | 0.12 |
| 1 | 201424899 | rs10920207 | -0.29 | -2.989 | -0.835 | 0.801 | -0.002 | 0 | 0 | 0.345 | 0.12 | 0.12 | 0.03 | 0.1 | 0.05 | 0.12 |
| 1 | 201428511 | rs4025021 | 0.586 | -2.572 | -1.064 | 0.649 | 0.001 | 0.007 | 0.007 | 0 | 0.67 | 0.86 | 0.78 | 0.74 | 0.71 | 0.71 |
| 1 | 201429381 | rs12090950 | 1.643 | -2.38 | -1.323 | 0.026 | -0.045 | 0.105 | 0.105 | 0 | 0.38 | 0.17 | 0.33 | 0.13 | 0.53 | 0.18 |
| 1 | 201430285 | rs11591030 | 1.69 | -2.289 | -1.823 | -0.057 | -0.119 | 0.151 | 0.151 | 0 | 0.37 | 0.07 | 0.33 | 0.11 | 0.53 | 0.21 |
| 1 | 201430572 | rs6690874 | -0.529 | -1.875 | -0.61 | 0.519 | -0.024 | -0.026 | -0.026 | 0 | 0.1 | 0.26 | 0.14 | 0.04 | 0.06 | 0.13 |
| 1 | 201439473 | rs654072 | 1.059 | -1.75 | -1.336 | 0.752 | 0.032 | 0.051 | 0.051 | 0 | 0.55 | 0.71 | 0.62 | 0.29 | 0.64 | 0.44 |
| 1 | 201440012 | rs3753993 | -0.795 | -1.661 | -1.618 | 1.241 | 0.007 | -0.059 | -0.059 | 0 | 0.25 | 0.09 | 0.27 | 0.51 | 0.21 | 0.36 |
| 1 | 201441035 | rs3753990 | 0.529 | -1.719 | -1.692 | 1.661 | -0.029 | -0.046 | -0.046 | 0.132 | 0.08 | 0.01 | 0.21 | 0.35 | 0.08 | 0.17 |
| 1 | 201441635 | rs12030255 | -0.723 | -1.64 | -1.611 | 1.212 | 0.008 | -0.06 | -0.06 | 0 | 0.25 | 0.09 | 0.27 | 0.51 | 0.21 | 0.37 |
| 1 | 201441734 | rs1256929 | 0.056 | -1.367 | -1.425 | 0.814 | 0.007 | -0.018 | -0.018 | 0 | 0.85 | 0.89 | 0.92 | 0.8 | 0.88 | 0.81 |
| 1 | 201451852 | rs10800783 | -0.978 | -1.293 | -1.486 | 0.853 | 0.019 | -0.013 | -0.013 | 0 | 0.31 | 0.15 | 0.39 | 0.71 | 0.24 | 0.61 |
| 1 | 201454923 | rs645390 | 0.423 | -0.871 | -1.726 | 0.628 | -0.046 | -0.014 | -0.014 | 0 | 0.86 | 0.99 | 0.91 | 0.88 | 0.84 | 0.85 |
| 1 | 201455815 | rs3767542 | -0.404 | -1.529 | -1.944 | 0.768 | 0.02 | -0.032 | -0.032 | 0.111 | 0.29 | 0.22 | 0.34 | 0.45 | 0.21 | 0.33 |
| 1 | 201455910 | rs3767541 | 1.409 | -1.791 | -2.154 | 0.107 | 0.033 | 0.252 | 0.252 | 0.002 | 0.48 | 0.08 | 0.37 | 0.16 | 0.61 | 0.36 |
| 1 | 201457054 | rs675508 | -0.02 | -1.517 | -2.035 | 0.081 | -0.048 | -0.024 | -0.024 | 0 | 0.84 | 0.99 | 0.91 | 0.86 | 0.84 | 0.82 |
| 1 | 201461809 | rs495886 | 0.862 | -1.062 | -1.06 | -0.228 | 0 | -0.015 | -0.015 | 0 | 0.55 | 0.5 | 0.4 | 0.18 | 0.63 | 0.44 |
| 1 | 201465944 | rs4915529 | 1.03 | 0.125 | -0.895 | -0.115 | -0.01 | 0.009 | 0.009 | 0 | 0.54 | 0.35 | 0.48 | 0.4 | 0.5 | 0.45 |
| 1 | 201466057 | rs10800784 | -0.23 | -0.437 | -0.154 | 0.268 | -0.04 | 0.096 | 0.096 | 0 | 0.2 | 0.32 | 0.28 | 0.5 | 0.13 | 0.32 |
| 1 | 201471723 | rs685023 | 0.534 | -0.087 | -0.696 | 0.347 | -0.006 | -0.014 | -0.014 | 0 | 0.06 | 0.02 | 0.04 | 0.1 | 0.08 | 0.09 |
| 1 | 201475727 | rs2038972 | -0.117 | -0.302 | -0.736 | 0.264 | -0.007 | 0.017 | 0.017 | 0 | 0.77 | 0.72 | 0.78 | 0.69 | 0.81 | 0.67 |
| 1 | 201480039 | rs12021779 | 0.925 | -0.422 | -1.331 | 0.366 | 0.025 | -0.016 | -0.016 | 0 | 0.1 | 0.05 | 0.31 | 0.46 | 0.04 | 0.24 |
| 1 | 201486928 | rs11586401 | 1.612 | 0.307 | -1.612 | -0.812 | -0.097 | 0.164 | 0.164 | 0.095 | 0.35 | 0.03 | 0.19 | 0.09 | 0.34 | 0.14 |
| 1 | 201500825 | rs4915537 | 0.092 | -0.373 | -0.669 | 0.053 | 0.026 | -0.054 | -0.054 | 0.735 | 0.17 | 0.11 | 0.36 | 0.56 | 0.13 | 0.43 |
| 1 | 201504189 | rs1932023 | -0.392 | 0.069 | -0.007 | 0.312 | -0.031 | 0.016 | 0.016 | 0 | 0.2 | 0.3 | 0.21 | 0.25 | 0.16 | 0.23 |
| 1 | 201517349 | rs11577209 | 0.917 | 0.249 | -0.467 | 0.283 | -0.011 | 0.017 | 0.017 | 0 | 0.71 | 0.48 | 0.7 | 0.64 | 0.72 | 0.61 |
| 1 | 201523953 | rs662755 | -0.563 | -0.063 | -0.963 | 0.399 | -0.123 | 0.061 | 0.061 | 0 | 0.8 | 0.97 | 0.82 | 0.94 | 0.67 | 0.79 |
| 1 | 201524412 | rs12030585 | -0.076 | -0.397 | -0.061 | 0.672 | -0.043 | 0.067 | 0.067 | 0.033 | 0.23 | 0.31 | 0.36 | 0.56 | 0.15 | 0.43 |
| 1 | 201524471 | rs556903 | 0.258 | 0.63 | 0.46 | 0.953 | 0.021 | -0.009 | -0.009 | 0 | 0.65 | 0.53 | 0.59 | 0.68 | 0.53 | 0.6 |
| 1 | 201528243 | rs7536246 | -1.839 | 0.255 | 1.379 | 1.647 | 0.038 | 0.2 | 0.2 | 0.984 | 0.1 | 0.37 | 0.3 | 0.3 | 0.1 | 0.3 |
| 1 | 201530119 | rs654897 | -1.967 | -0.291 | 1.782 | 1.356 | -0.046 | 0.042 | 0.042 | 4.391 | 0.24 | 0.48 | 0.35 | 0.31 | 0.16 | 0.32 |
| 1 | 201531663 | rs1751342 | -0.946 | -0.251 | 1.713 | 1.396 | -0.03 | -0.071 | -0.071 | 0 | 0.14 | 0.22 | 0.06 | 0.01 | 0.07 | 0.13 |
| 1 | 201532404 | rs2148619 | -1.837 | -0.192 | 1.94 | 1.471 | 0.071 | 0.365 | 0.365 | 1.897 | 0.09 | 0.46 | 0.3 | 0.3 | 0.07 | 0.19 |
| 1 | 201537485 | rs4915236 | -1.601 | -0.018 | 1.258 | 1.68 | -0.024 | 0.117 | 0.117 | 1.392 | 0.26 | 0.49 | 0.43 | 0.48 | 0.25 | 0.48 |
| 1 | 201540509 | rs2205595 | -1.693 | 0.556 | 1.171 | 1.698 | 0.081 | 0.081 | 0.081 | 0.003 | 0.09 | 0.3 | 0.17 | 0.13 | 0.13 | 0.13 |
| 1 | 201550615 | rs544127 | -0.887 | 0.413 | 1.133 | 2.343 | -0.026 | 0.036 | 0.036 | 0 | 0.06 | 0.17 | 0.02 | 0.15 | 0.01 | 0.13 |
| 1 | 201553563 | rs614452 | -1.676 | 0.389 | 1.047 | 2.227 | -0.017 | 0 | 0 | 0 | 0.12 | 0.2 | 0.06 | 0.15 | 0.06 | 0.14 |
| 1 | 201554323 | rs16865416 | -0.262 | 0.393 | 1.024 | 2.218 | 0.009 | 0.01 | 0.01 | 0 | 0.07 | 0.04 | 0.05 | 0 | 0.05 | 0.02 |
| 1 | 201560163 | rs608356 | -2.99 | 0.345 | 1.109 | 2.227 | -0.004 | 0.063 | 0.063 | 0 | 0.26 | 0.62 | 0.42 | 0.5 | 0.24 | 0.42 |
| 1 | 201561620 | rs545372 | -1.665 | 0.319 | -0.06 | 1.498 | -0.015 | 0 | 0 | 0 | 0.12 | 0.19 | 0.06 | 0.15 | 0.06 | 0.14 |
| 1 | 201563914 | rs873260 | -0.914 | 0.33 | -0.017 | 1.433 | -0.016 | -0.05 | -0.05 | 0 | 0.09 | 0.21 | 0.07 | 0 | 0.06 | 0.02 |

| | | | | | | | | | | | | | | | | |
|---|---|---|---|---|---|---|---|---|---|---|---|---|---|---|---|---|
| 1 | 201564909 | rs609391 | -2.381 | 0.333 | -0.028 | 1.467 | -0.009 | 0.021 | 0.021 | 0.113 | 0.21 | 0.38 | 0.22 | 0.28 | 0.19 | 0.26 |
| 1 | 201567030 | rs620078 | -0.819 | 0.638 | -0.463 | 1.644 | -0.023 | 0.027 | 0.027 | 0 | 0.06 | 0.15 | 0.02 | 0.14 | 0.01 | 0.12 |
| 1 | 201572071 | rs483489 | 0.125 | 1.294 | -1.211 | 0.941 | -0.06 | 0.096 | 0.096 | 0 | 0.76 | 1 | 0.91 | 0.94 | 0.79 | 0.82 |
| 1 | 201575999 | rs681068 | -1.322 | 1.404 | -0.219 | 1.709 | 0.014 | 0.018 | 0.018 | 0 | 0.08 | 0.13 | 0.17 | 0.15 | 0.13 | 0.13 |
| 1 | 201603081 | rs679352 | 0.148 | 0.996 | -0.334 | 1.832 | 0.024 | 0 | 0 | 0.004 | 0.5 | 0.36 | 0.54 | 0.46 | 0.56 | 0.5 |
| 1 | 201605716 | rs10920222 | -0.031 | 0.56 | 0.064 | 1.926 | 0.005 | 0.006 | 0.006 | 0 | 0.16 | 0.2 | 0.4 | 0.18 | 0.18 | 0.06 |
| 1 | 201607272 | rs631225 | -0.157 | 0.086 | 0.424 | 1.798 | 0.027 | -0.006 | -0.006 | 0 | 0.74 | 0.82 | 0.89 | 0.69 | 0.82 | 0.66 |
| 1 | 201616627 | rs7364578 | -0.048 | 0.056 | -0.193 | 1.136 | -0.034 | -0.011 | -0.011 | 0.222 | 0.23 | 0.05 | 0.43 | 0.2 | 0.27 | 0.09 |
| 1 | 201616835 | rs1736450 | 0.778 | 0.359 | -0.093 | 1.3 | -0.019 | 0.007 | 0.007 | 0 | 0.45 | 0.71 | 0.38 | 0.38 | 0.44 | 0.48 |
| 1 | 201623215 | rs967392 | -0.057 | 0.911 | -0.18 | 1.036 | 0.001 | 0 | 0 | 0.197 | 0.67 | 0.79 | 0.81 | 0.63 | 0.72 | 0.55 |
| 1 | 201625823 | rs478472 | -0.346 | 0.99 | -0.095 | 0.856 | 0.022 | -0.042 | -0.042 | 0.133 | 0.08 | 0.13 | 0.05 | 0 | 0.07 | 0.04 |
| 1 | 201626598 | rs2819395 | 0.87 | 1.017 | 0.04 | 1.069 | -0.005 | 0.001 | 0.001 | 0 | 0.37 | 0.49 | 0.33 | 0.34 | 0.37 | 0.43 |
| 1 | 201628577 | rs652930 | -0.741 | 0.733 | -0.726 | 1.023 | -0.016 | -0.004 | -0.004 | 0.015 | 0.78 | 0.99 | 0.88 | 0.85 | 0.82 | 0.76 |
| 1 | 201628990 | rs664896 | 0.575 | 0.781 | 0.028 | 1.082 | -0.007 | 0.006 | 0.006 | 0 | 0.44 | 0.36 | 0.36 | 0.65 | 0.46 | 0.64 |
| 1 | 201630461 | rs724220 | 0.041 | 1.04 | -0.207 | 1.572 | -0.048 | -0.072 | -0.072 | 0 | 0.24 | 0.11 | 0.46 | 0.41 | 0.34 | 0.2 |
| 1 | 201630517 | rs724221 | -1.175 | 0.7 | 0.468 | 1.227 | -0.008 | -0.008 | -0.008 | 0 | 0.4 | 0.64 | 0.58 | 0.45 | 0.47 | 0.26 |
| 1 | 201630955 | rs586688 | -1.018 | 0.781 | 0.999 | 1.169 | 0.014 | -0.094 | -0.094 | 0 | 0.16 | 0.51 | 0.11 | 0.04 | 0.15 | 0.07 |
| 1 | 201634364 | rs1569403 | 0.384 | 1.095 | 0.654 | 1.22 | -0.029 | -0.059 | -0.059 | 0 | 0.22 | 0.11 | 0.44 | 0.4 | 0.28 | 0.19 |
| 1 | 201636724 | rs12045814 | 0.311 | 1.067 | 0.839 | 1.268 | -0.003 | -0.001 | -0.001 | 0 | 0.23 | 0.17 | 0.45 | 0.41 | 0.28 | 0.19 |
| 1 | 201637442 | rs7367640 | 0.145 | 1.046 | 0.872 | 1.256 | 0.002 | 0.009 | 0.009 | 0 | 0.24 | 0.22 | 0.46 | 0.41 | 0.29 | 0.19 |
| 1 | 201638858 | rs525492 | -1.009 | 0.635 | 1.564 | 1.109 | 0 | -0.094 | -0.094 | 0 | 0.16 | 0.3 | 0.09 | 0.02 | 0.14 | 0.06 |
| 1 | 201640821 | rs6427916 | -0.063 | 0.853 | 0.39 | 1.059 | -0.012 | -0.017 | -0.017 | 0 | 0.29 | 0.25 | 0.49 | 0.46 | 0.37 | 0.19 |
| 1 | 201644752 | rs10800790 | -0.095 | 0.719 | 0.401 | 1.018 | -0.007 | -0.011 | -0.011 | 0 | 0.3 | 0.28 | 0.48 | 0.49 | 0.37 | 0.21 |
| 1 | 201645660 | rs531606 | 0.447 | -0.975 | 0.099 | 0.669 | 0.002 | -0.006 | -0.006 | 0 | 0.78 | 0.79 | 0.9 | 0.7 | 0.92 | 0.81 |
| 1 | 201651484 | rs3000763 | -0.23 | -0.643 | 0.368 | 0.628 | -0.012 | 0.023 | 0.023 | 0 | 0.37 | 0.31 | 0.5 | 0.23 | 0.4 | 0.16 |
| 1 | 201657709 | rs602668 | 0.056 | -0.411 | 0.612 | 0.439 | 0.026 | -0.023 | -0.023 | 0 | 0.85 | 0.71 | 0.91 | 0.88 | 0.83 | 0.91 |
| 1 | 201661705 | rs1317704 | 1.029 | -0.433 | 0.604 | 0.606 | 0.019 | 0.01 | 0.01 | 0 | 0.48 | 0.38 | 0.4 | 0.56 | 0.43 | 0.68 |
| 1 | 201664101 | rs606114 | -0.343 | -0.371 | 0.355 | 0.569 | 0.017 | -0.026 | -0.026 | 0 | 0.87 | 0.81 | 0.92 | 0.92 | 0.84 | 0.93 |
| 1 | 201680451 | rs675141 | -0.457 | -0.991 | 0.037 | 0.678 | -0.014 | 0.003 | 0.003 | 0 | 0.39 | 0.27 | 0.37 | 0.35 | 0.4 | 0.24 |
| 1 | 201684221 | rs10920236 | -0.754 | -1.47 | 0.181 | 0.546 | -0.001 | -0.002 | -0.002 | 1.021 | 0.47 | 0.5 | 0.51 | 0.35 | 0.47 | 0.29 |
| 1 | 201684691 | rs722807 | -0.357 | -1.669 | 0.272 | 0.404 | -0.011 | -0.007 | -0.007 | 0 | 0.24 | 0.23 | 0.15 | 0.11 | 0.12 | 0.13 |
| 1 | 201689201 | rs732813 | -0.731 | -1.628 | -0.103 | 0.224 | 0.002 | -0.003 | -0.003 | 0.361 | 0.58 | 0.46 | 0.71 | 0.59 | 0.56 | 0.56 |
| 1 | 201692871 | rs239967 | 0.474 | -1.858 | -0.127 | 0.084 | -0.025 | 0.066 | 0.066 | 0 | 0.45 | 0.26 | 0.26 | 0.32 | 0.49 | 0.4 |
| 1 | 201699395 | rs239995 | -0.326 | -1.506 | 0.178 | 0.496 | -0.013 | -0.022 | -0.022 | 0 | 0.88 | 0.75 | 0.92 | 0.91 | 0.94 | 0.9 |
| 1 | 201705650 | rs4950838 | 1.015 | -1.598 | 0.063 | 0.537 | -0.006 | 0.009 | 0.009 | 0 | 0.18 | 0.16 | 0.3 | 0.35 | 0.11 | 0.32 |
| 1 | 201708422 | rs12026046 | 0.452 | -1.588 | 0.017 | 0.443 | -0.001 | 0.011 | 0.011 | 2.072 | 0.41 | 0.43 | 0.42 | 0.24 | 0.44 | 0.21 |
| 1 | 201709880 | rs239983 | -1.437 | -1.574 | 0.091 | 0.444 | -0.004 | -0.033 | -0.033 | 0 | 0.93 | 0.9 | 0.94 | 1 | 0.96 | 0.93 |
| 1 | 201711704 | rs4950846 | -0.445 | -1.552 | -0.089 | 0.493 | 0.002 | -0.002 | -0.002 | 0 | 0.59 | 0.6 | 0.72 | 0.59 | 0.55 | 0.53 |
| 1 | 201712748 | rs10920247 | 0.525 | -1.083 | 0.36 | 0.87 | -0.013 | 0.019 | 0.019 | 0 | 0.18 | 0.16 | 0.28 | 0.35 | 0.1 | 0.32 |
| 1 | 201717721 | rs239987 | 0.561 | -0.028 | 0.373 | 0.458 | -0.011 | 0.003 | 0.003 | 0 | 0.88 | 0.75 | 0.91 | 0.86 | 0.93 | 0.89 |
| 1 | 201722102 | rs239980 | 1.16 | 0.37 | 0.112 | 0.443 | 0.019 | -0.019 | -0.019 | 0 | 0.83 | 0.64 | 0.88 | 0.84 | 0.81 | 0.91 |
| 1 | 201732694 | rs7535989 | -0.129 | 0.602 | -0.885 | 0.302 | -0.109 | 0.033 | 0.033 | 0.153 | 0.2 | 0.03 | 0.31 | 0.08 | 0.34 | 0.1 |
| 1 | 201733944 | rs7527583 | -1.233 | 0.33 | -0.45 | 0.35 | 0.004 | -0.007 | -0.007 | 0 | 0.32 | 0.4 | 0.23 | 0.2 | 0.39 | 0.13 |
| 1 | 201735730 | rs507068 | -1.705 | 0.148 | -0.553 | 0.534 | 0.004 | -0.005 | -0.005 | 0 | 0.41 | 0.51 | 0.49 | 0.32 | 0.44 | 0.31 |
| 1 | 201735913 | rs882624 | 0.291 | 0.743 | -1.028 | 0.947 | -0.081 | 0.021 | 0.021 | 0 | 0.16 | 0.05 | 0.17 | 0.07 | 0.35 | 0.08 |
| 1 | 201749156 | rs650720 | -0.744 | 0.699 | -1.289 | 0.679 | -0.071 | -0.031 | -0.031 | 0.301 | 0.28 | 0.13 | 0.43 | 0.31 | 0.4 | 0.25 |
| 1 | 201754528 | rs16849345 | 0.03 | -0.024 | -0.507 | 0.197 | -0.009 | -0.038 | -0.038 | 0 | 0.09 | 0.13 | 0.06 | 0 | 0.05 | 0.01 |
| 1 | 201754720 | rs903676 | -0.084 | 0.05 | -0.005 | 0.011 | 0.015 | -0.02 | -0.02 | 0 | 0.18 | 0.55 | 0.23 | 0.15 | 0.22 | 0.12 |
| 1 | 201761301 | rs504988 | 0.743 | -0.619 | -0.975 | -0.044 | 0.049 | 0.103 | 0.103 | 0 | 0.52 | 0.11 | 0.63 | 0.67 | 0.63 | 0.66 |
| 1 | 201763912 | rs481530 | 0.354 | -0.733 | -0.876 | -0.079 | 0.021 | 0.097 | 0.097 | 0 | 0.59 | 0.16 | 0.65 | 0.75 | 0.64 | 0.74 |
| 1 | 201770373 | rs12401827 | 0.454 | -0.851 | -0.995 | 0.742 | 0.001 | 0.016 | 0.016 | 0 | 0.13 | 0.12 | 0.25 | 0.36 | 0.07 | 0.08 |
| 1 | 201770929 | rs2678209 | 1.044 | -0.161 | -1.739 | 0.249 | -0.076 | -0.038 | -0.038 | 0 | 0.21 | 0.05 | 0.35 | 0.19 | 0.27 | 0.31 |
| 1 | 201775746 | rs4379719 | 0.092 | -0.496 | -1.014 | 0.296 | 0.009 | 0.006 | 0.006 | 0 | 0.5 | 0.58 | 0.45 | 0.58 | 0.43 | 0.38 |
| 1 | 201777250 | rs2820289 | 1.55 | -0.021 | -1.197 | 0.346 | -0.03 | -0.026 | -0.026 | 0 | 0.06 | 0 | 0.14 | 0.18 | 0.06 | 0.09 |
| 1 | 201790270 | rs1022361 | 0.172 | -0.59 | -0.863 | -0.128 | -0.016 | 0.003 | 0.003 | 0 | 0.35 | 0.5 | 0.2 | 0.36 | 0.31 | 0.22 |
| 1 | 201790826 | rs1032524 | 0.443 | -0.814 | -0.931 | -0.214 | -0.043 | 0.056 | 0.056 | 0.071 | 0.38 | 0.21 | 0.37 | 0.24 | 0.52 | 0.55 |
| 1 | 201792306 | rs10464 | 1.386 | -0.935 | -0.612 | -0.245 | -0.008 | 0.043 | 0.043 | 0 | 0.25 | 0.2 | 0.17 | 0.09 | 0.29 | 0.15 |
| 1 | 201797696 | rs10920259 | 0.152 | -0.747 | -0.649 | -0.339 | 0.001 | 0.001 | 0.001 | 0 | 0.5 | 0.49 | 0.35 | 0.49 | 0.52 | 0.5 |
| 1 | 201800582 | rs10920260 | 1.45 | -0.551 | -0.15 | -0.274 | -0.002 | -0.031 | -0.031 | 1.779 | 0.18 | 0.25 | 0.16 | 0.14 | 0.21 | 0.27 |
| 1 | 201800868 | rs2820295 | 0.777 | -0.492 | -0.118 | -0.246 | 0.035 | -0.02 | -0.02 | 0.314 | 0.22 | 0.28 | 0.23 | 0.1 | 0.3 | 0.28 |
| 1 | 201814994 | rs12562614 | 0.848 | -0.644 | -0.355 | -0.191 | -0.03 | -0.073 | -0.073 | 1.105 | 0.35 | 0.13 | 0.39 | 0.42 | 0.36 | 0.25 |
| 1 | 201816572 | rs12042456 | 1.086 | -0.687 | -0.318 | -0.157 | -0.016 | 0.055 | 0.055 | 0 | 0.08 | 0.12 | 0.03 | 0.27 | 0.02 | 0.07 |
| 1 | 201840210 | rs950114 | 1.265 | -0.686 | -0.326 | -0.166 | 0.028 | -0.028 | -0.028 | 0 | 0.08 | 0.02 | 0.02 | 0.27 | 0.02 | 0.07 |
| 1 | 201845432 | rs1043823 | 1.693 | -0.648 | -0.131 | -0.223 | 0.004 | 0.045 | 0.045 | 1.387 | 0.15 | 0.03 | 0.17 | 0.07 | 0.1 | 0.12 |
| 1 | 201845575 | rs8024 | 0.855 | -0.277 | 0.241 | -0.053 | -0.005 | 0.003 | 0.003 | 0.019 | 0.21 | 0.2 | 0.22 | 0.1 | 0.3 | 0.27 |
| 1 | 201846689 | rs10920269 | 1.606 | -0.297 | 0.225 | -0.065 | -0.041 | 0.015 | 0.015 | 0 | 0.16 | 0.01 | 0.15 | 0.14 | 0.2 | 0.28 |
| 1 | 201850057 | rs1517810 | 0.693 | -0.243 | 0.26 | -0.039 | -0.005 | 0.003 | 0.003 | 0.231 | 0.22 | 0.21 | 0.23 | 0.1 | 0.3 | 0.27 |
| 1 | 201855412 | rs6427922 | 1.153 | -0.201 | 0.266 | -0.016 | -0.002 | -0.046 | -0.046 | 0.285 | 0.19 | 0.34 | 0.17 | 0.14 | 0.21 | 0.29 |
| 1 | 201857560 | rs13374557 | 1.262 | -0.193 | 0.12 | -0.012 | -0.003 | -0.031 | -0.031 | 0.541 | 0.18 | 0.25 | 0.16 | 0.14 | 0.21 | 0.29 |

| | | | | | | | | | | | | | | | | |
|---|---|---|---|---|---|---|---|---|---|---|---|---|---|---|---|---|
| 1 | 201860626 | rs2250377 | -1.114 | -0.15 | 0.129 | -0.107 | -0.005 | 0.003 | 0.003 | 0.018 | 0.79 | 0.8 | 0.78 | 0.9 | 0.7 | 0.73 |
| 1 | 201865763 | rs8028 | -1.108 | -0.145 | 0.134 | -0.111 | -0.005 | 0.003 | 0.003 | 0.019 | 0.79 | 0.8 | 0.77 | 0.9 | 0.7 | 0.74 |
| 1 | 201868333 | rs3738290 | 1.258 | -0.161 | 0.048 | -0.173 | -0.002 | -0.059 | -0.059 | 0 | 0.18 | 0.24 | 0.16 | 0.08 | 0.2 | 0.28 |
| 1 | 201869257 | rs2820312 | 0.816 | -0.166 | -0.006 | -0.146 | -0.001 | 0.001 | 0.001 | 0.019 | 0.21 | 0.21 | 0.22 | 0.1 | 0.3 | 0.26 |
| 1 | 201883199 | rs11580908 | 1.171 | -0.32 | -0.12 | -0.126 | -0.006 | -0.055 | -0.055 | 0.182 | 0.18 | 0.24 | 0.16 | 0.08 | 0.19 | 0.28 |
| 1 | 201884668 | rs4494199 | -0.01 | -0.41 | -0.354 | 0.141 | 0.002 | -0.007 | -0.007 | 0 | 0.54 | 0.31 | 0.59 | 0.56 | 0.48 | 0.39 |
| 1 | 201886402 | rs2047264 | -0.243 | -0.157 | -0.528 | -0.039 | -0.039 | 0.1 | 0.1 | 0 | 0.59 | 0.25 | 0.61 | 0.81 | 0.5 | 0.46 |
| 1 | 201887721 | rs17432675 | 0.389 | -0.429 | -0.606 | 0.107 | 0 | -0.024 | -0.024 | 3.14 | 0.35 | 0.25 | 0.39 | 0.42 | 0.35 | 0.23 |
| 1 | 201888672 | rs2790903 | 0.1 | -0.647 | -0.264 | 0.412 | -0.014 | -0.008 | -0.008 | 0 | 0.64 | 0.62 | 0.62 | 0.33 | 0.78 | 0.68 |
| 1 | 201888969 | rs2360546 | 0.23 | -0.665 | -0.271 | 0.451 | 0.03 | -0.06 | -0.06 | 0 | 0.29 | 0.1 | 0.3 | 0.39 | 0.19 | 0.22 |
| 1 | 201891254 | rs1390398 | 0.248 | -0.64 | -0.071 | 0.317 | -0.025 | -0.029 | -0.029 | 0.001 | 0.62 | 0.54 | 0.59 | 0.29 | 0.75 | 0.65 |
| 1 | 201892257 | rs16849442 | 0.155 | -0.571 | -0.587 | 0.325 | 0.019 | -0.083 | -0.083 | 1.028 | 0.29 | 0.1 | 0.32 | 0.43 | 0.22 | 0.25 |
| 1 | 201897670 | rs9787358 | -0.336 | -0.57 | -0.416 | 0.242 | 0.017 | 0.02 | 0.02 | 0.027 | 0.36 | 0.27 | 0.4 | 0.7 | 0.25 | 0.36 |
| 1 | 201904589 | rs2820327 | 0.41 | -0.268 | -0.462 | 0.15 | 0.007 | -0.002 | -0.002 | 0 | 0.46 | 0.39 | 0.48 | 0.45 | 0.58 | 0.5 |
| 1 | 201908597 | rs10800799 | 0.64 | -0.473 | -0.672 | 0.551 | -0.002 | -0.026 | -0.026 | 0.548 | 0.15 | 0.1 | 0.16 | 0.26 | 0.12 | 0.12 |
| 1 | 201910609 | rs2819369 | -0.083 | -0.174 | -0.236 | 0.34 | -0.002 | 0.003 | 0.003 | 0 | 0.37 | 0.39 | 0.37 | 0.45 | 0.29 | 0.3 |
| 1 | 201912501 | rs16849455 | 0.526 | 0.025 | -0.194 | 0.939 | 0.009 | -0.013 | -0.013 | 0 | 0.11 | 0.07 | 0.14 | 0.26 | 0.1 | 0.11 |
| 1 | 201918527 | rs2820304 | -0.005 | 0.913 | -1.432 | 0.06 | -0.102 | 0.08 | 0.08 | 0.055 | 0.33 | 0.08 | 0.23 | 0.2 | 0.42 | 0.56 |
| 1 | 201922759 | rs17473434 | -0.032 | 1.529 | -0.993 | 0.343 | 0.001 | 0 | 0 | 0 | 0.17 | 0.16 | 0.28 | 0.19 | 0.3 | 0.08 |
| 1 | 201923383 | rs4950744 | -0.021 | 0.593 | -1.123 | 0.948 | 0.014 | -0.027 | -0.027 | 0.225 | 0.22 | 0.11 | 0.19 | 0.27 | 0.15 | 0.2 |
| 1 | 201931219 | rs2819376 | -0.007 | 0.234 | -0.437 | 0.937 | -0.012 | 0.003 | 0.003 | 0 | 0.76 | 0.64 | 0.8 | 0.78 | 0.83 | 0.78 |
| 1 | 201936165 | rs4950802 | -0.483 | 0.206 | -0.276 | 0.923 | 0.003 | -0.002 | -0.002 | 0 | 0.77 | 0.7 | 0.81 | 0.78 | 0.83 | 0.78 |
| 1 | 201936324 | rs12090635 | 0.526 | 0.94 | 0.167 | 1.165 | 0.031 | 0.025 | 0.025 | 0 | 0.14 | 0.26 | 0.26 | 0.29 | 0.2 | 0.06 |
| 1 | 201937414 | rs10920285 | 0.991 | 0.644 | -0.234 | 0.845 | -0.012 | -0.015 | -0.015 | 0 | 0.34 | 0.21 | 0.2 | 0.34 | 0.38 | 0.49 |
| 1 | 201938995 | rs7513 | 0.66 | 0.408 | -0.232 | 0.783 | -0.015 | -0.004 | -0.004 | 0 | 0.38 | 0.25 | 0.27 | 0.33 | 0.47 | 0.51 |
| 1 | 201939752 | rs1045893 | 0.811 | 0.47 | -0.34 | 0.75 | -0.011 | -0.01 | -0.01 | 0 | 0.34 | 0.23 | 0.24 | 0.33 | 0.4 | 0.48 |
| 1 | 201940216 | rs12754196 | 0.283 | 0.237 | 0.086 | 0.813 | 0.008 | 0.012 | 0.012 | 0 | 0.13 | 0.25 | 0.22 | 0.2 | 0.15 | 0.05 |
| 1 | 201950038 | rs7529387 | 0.352 | 0.062 | 0.083 | 0.841 | 0 | 0 | 0 | 0 | 0.52 | 0.52 | 0.47 | 0.53 | 0.61 | 0.6 |
| 1 | 201954697 | rs6689324 | 0.464 | 0.087 | 0.014 | 0.881 | 0.011 | 0.013 | 0.013 | 0 | 0.51 | 0.41 | 0.45 | 0.51 | 0.61 | 0.6 |
| 1 | 201954987 | rs6689634 | 0.851 | 0.314 | -0.353 | 1.105 | -0.053 | -0.059 | -0.059 | 0 | 0.21 | 0.12 | 0.24 | 0.3 | 0.3 | 0.16 |
| 1 | 201955614 | rs11586632 | -0.042 | 0.297 | -0.273 | 1.083 | 0.008 | 0.009 | 0.009 | 0 | 0.52 | 0.57 | 0.46 | 0.51 | 0.61 | 0.6 |
| 1 | 201959455 | rs16849483 | 0.666 | 0.12 | -0.339 | 1.005 | -0.006 | -0.008 | -0.008 | 0 | 0.29 | 0.29 | 0.21 | 0.22 | 0.3 | 0.44 |
| 1 | 201959490 | rs12734984 | -0.022 | 0.499 | -0.577 | 1.086 | 0.008 | 0.018 | 0.018 | 0 | 0.09 | 0.12 | 0.21 | 0.21 | 0.15 | 0.05 |
| 1 | 201966774 | rs10920300 | -0.294 | 0.406 | -0.699 | 0.874 | -0.004 | -0.006 | -0.006 | 0 | 0.11 | 0.08 | 0.29 | 0.18 | 0.16 | 0.06 |
| 1 | 201969082 | rs1130790 | 0.24 | 0.096 | -1.201 | 0.713 | -0.085 | -0.082 | -0.082 | 0 | 0.33 | 0.16 | 0.44 | 0.4 | 0.43 | 0.21 |
| 1 | 201970748 | rs4131469 | 0.208 | 0.054 | -0.849 | 0.693 | -0.079 | -0.077 | -0.077 | 0 | 0.33 | 0.16 | 0.44 | 0.4 | 0.43 | 0.21 |
| 1 | 201973073 | rs6427928 | 0.151 | -0.473 | -0.359 | 0.286 | -0.025 | -0.075 | -0.075 | 0 | 0.12 | 0.37 | 0.05 | 0.02 | 0.06 | 0.12 |
| 1 | 201973565 | rs3820439 | 1.331 | -0.734 | -0.913 | 0.118 | 0.013 | -0.01 | -0.01 | 0 | 0.33 | 0.22 | 0.24 | 0.4 | 0.29 | 0.51 |
| 1 | 201973961 | rs3795621 | -0.221 | -0.192 | -1.113 | 0.334 | -0.001 | 0.013 | 0.013 | 0 | 0.21 | 0.12 | 0.25 | 0.18 | 0.22 | 0.16 |
| 1 | 201974442 | rs10920303 | -0.221 | -0.192 | -1.113 | 0.334 | -0.001 | 0.013 | 0.013 | 0 | 0.21 | 0.12 | 0.25 | 0.18 | 0.22 | 0.16 |
| 1 | 201977254 | rs2819360 | -0.399 | 0.248 | -1.187 | 0.655 | -0.071 | -0.037 | -0.037 | 0 | 0.67 | 0.84 | 0.57 | 0.65 | 0.58 | 0.73 |
| 1 | 201977827 | rs9427715 | 0.175 | 0.369 | -0.698 | 0.616 | 0 | -0.013 | -0.013 | 0 | 0.58 | 0.65 | 0.53 | 0.52 | 0.57 | 0.66 |
| 1 | 201978323 | rs10920304 | 0.518 | 0.21 | -0.868 | -0.046 | 0 | -0.012 | -0.012 | 0.297 | 0.56 | 0.44 | 0.52 | 0.79 | 0.47 | 0.58 |
| 1 | 201979455 | rs3806357 | 1.439 | 0.395 | -1.525 | 0.658 | -0.002 | -0.057 | -0.057 | 0 | 0.1 | 0.02 | 0.06 | 0.23 | 0.09 | 0.06 |
| 1 | 201981862 | rs2819362 | -0.083 | 0.987 | -1.396 | 0.017 | -0.07 | 0.062 | 0.062 | 0 | 0.67 | 0.85 | 0.6 | 0.78 | 0.55 | 0.67 |
| 1 | 201983242 | rs2819365 | 0.594 | 1.351 | -1.556 | 0.135 | -0.13 | 0.016 | 0.016 | 0 | 0.19 | 0.01 | 0.32 | 0.13 | 0.32 | 0.23 |
| 1 | 201986418 | rs12756809 | 0.535 | 0.021 | -0.985 | 0.24 | 0.031 | -0.028 | -0.028 | 0.278 | 0.38 | 0.2 | 0.26 | 0.54 | 0.32 | 0.49 |
| 1 | 201995426 | rs4950760 | -0.039 | 0.35 | -0.959 | 0.194 | 0.003 | 0.009 | 0.009 | 0 | 0.48 | 0.17 | 0.4 | 0.54 | 0.48 | 0.54 |
| 1 | 201996215 | rs10920308 | -0.325 | 0.545 | -0.65 | 0.318 | -0.002 | 0 | 0 | 0 | 0.55 | 0.57 | 0.43 | 0.54 | 0.49 | 0.54 |
| 1 | 202003269 | rs12139192 | 0.569 | 0.792 | -1.315 | 0.206 | 0 | -0.009 | -0.009 | 0 | 0.1 | 0.08 | 0.05 | 0 | 0.09 | 0.03 |
| 1 | 202013757 | rs12731187 | 0.08 | 0.83 | -1.233 | 0.75 | 0.012 | -0.022 | -0.022 | 0.03 | 0.39 | 0.28 | 0.36 | 0.48 | 0.36 | 0.53 |
| 1 | 202017400 | rs6656470 | 0.325 | 1.542 | -1.243 | 0.818 | -0.071 | -0.023 | -0.023 | 0 | 0.76 | 0.91 | 0.62 | 0.68 | 0.61 | 0.75 |
| 1 | 202027170 | rs10920315 | -0.411 | 0.553 | -1.257 | 0.091 | -0.053 | 0.087 | 0.087 | 0.904 | 0.25 | 0.06 | 0.2 | 0.15 | 0.3 | 0.16 |
| 1 | 202032235 | rs9427477 | 0.721 | -0.345 | -0.895 | 0.062 | -0.041 | 0.022 | 0.022 | 0 | 0.22 | 0.35 | 0.28 | 0.16 | 0.15 | 0.35 |
| 1 | 202036251 | rs12037174 | 0.925 | -1.084 | -1.242 | -0.407 | 0.041 | 0.051 | 0.051 | 0.454 | 0.23 | 0.12 | 0.2 | 0.09 | 0.15 | 0.16 |
| 1 | 202037092 | rs12071151 | 0.555 | -1.09 | -1.127 | 0.113 | -0.05 | 0.055 | 0.055 | 0 | 0.29 | 0.43 | 0.37 | 0.33 | 0.2 | 0.47 |
| 1 | 202038945 | rs9427727 | -0.759 | -0.422 | -1.169 | 0.154 | -0.037 | -0.027 | -0.027 | 0 | 0.77 | 0.86 | 0.8 | 0.64 | 0.63 | 0.78 |
| 1 | 202045541 | rs12724599 | 0.969 | -0.504 | -1.115 | 0.19 | -0.041 | 0.073 | 0.073 | 0.124 | 0.27 | 0.09 | 0.17 | 0.2 | 0.29 | 0.11 |
| 1 | 202045985 | rs9427478 | -0.189 | -0.545 | -0.922 | 0.053 | 0.038 | -0.01 | -0.01 | 0 | 0.56 | 0.7 | 0.61 | 0.45 | 0.39 | 0.68 |
| 1 | 202052416 | rs7535074 | 0.516 | -0.695 | -1.165 | 0.107 | 0 | 0.001 | 0.001 | 0 | 0.31 | 0.32 | 0.29 | 0.26 | 0.2 | 0.29 |
| 1 | 202055938 | rs10800812 | -1.461 | -0.028 | -1.753 | 0.296 | -0.054 | -0.05 | -0.05 | 0 | 0.85 | 0.94 | 0.81 | 0.65 | 0.72 | 0.79 |
| 1 | 202055955 | rs10800813 | -0.115 | -0.198 | -1.756 | 0.206 | 0.062 | -0.008 | -0.008 | 0 | 0.55 | 0.81 | 0.63 | 0.5 | 0.4 | 0.68 |
| 1 | 202056808 | rs12122896 | 0.391 | -0.336 | -1.667 | -0.127 | -0.022 | 0.111 | 0.111 | 0.025 | 0.3 | 0.1 | 0.18 | 0.15 | 0.32 | 0.11 |
| 1 | 202066564 | rs4576706 | 0.245 | -0.488 | -0.654 | 0.542 | -0.009 | 0.008 | 0.008 | 0 | 0.22 | 0.26 | 0.3 | 0.17 | 0.2 | 0.34 |
| 1 | 202075669 | rs4290117 | 0.328 | -1.521 | -0.769 | 0.223 | -0.006 | -0.003 | -0.003 | 0 | 0.6 | 0.57 | 0.73 | 0.7 | 0.76 | 0.72 |
| 1 | 202077927 | rs4950826 | 0.666 | -1.197 | -0.875 | 0.305 | -0.001 | -0.01 | -0.01 | 0 | 0.34 | 0.32 | 0.47 | 0.37 | 0.38 | 0.49 |
| 1 | 202078075 | rs4245703 | -0.421 | -1.542 | -1.192 | 0.164 | 0.041 | 0.004 | 0.004 | 0 | 0.24 | 0.29 | 0.37 | 0.72 | 0.39 | 0.33 |
| 1 | 202079202 | rs12745308 | 0.596 | -1.437 | -1.522 | 0.13 | -0.097 | -0.072 | -0.072 | 0 | 0.18 | 0.03 | 0.2 | 0.38 | 0.35 | 0.22 |
| 1 | 202081212 | rs10920333 | 0.611 | -1.617 | -1.326 | 0.176 | 0.112 | 0.014 | 0.014 | 1.106 | 0.45 | 0.08 | 0.46 | 0.41 | 0.69 | 0.61 |

| | | | | | | | | | | | | | | | | |
|---|---|---|---|---|---|---|---|---|---|---|---|---|---|---|---|---|
| 1 | 202087936 | rs4436441 | 0.404 | -2.211 | -0.565 | 0.109 | 0.114 | -0.014 | -0.014 | 0 | 0.57 | 0.25 | 0.57 | 0.49 | 0.79 | 0.75 |
| 1 | 202091660 | rs3922999 | 0.001 | -2.523 | -0.301 | -0.077 | -0.025 | 0.008 | 0.008 | 0.781 | 0.67 | 0.39 | 0.57 | 0.49 | 0.85 | 0.8 |
| 1 | 202092046 | rs3795593 | -0.012 | -2.456 | -0.308 | -0.098 | -0.02 | 0.008 | 0.008 | 0.475 | 0.67 | 0.39 | 0.56 | 0.49 | 0.84 | 0.77 |
| 1 | 202092415 | rs3795596 | -0.796 | -2.761 | 0.052 | -0.165 | -0.01 | -0.004 | -0.004 | 0 | 0.78 | 0.78 | 0.83 | 0.84 | 0.91 | 0.92 |
| 1 | 202093628 | rs6676433 | 0.183 | -2.766 | -0.007 | -0.189 | -0.01 | -0.036 | -0.036 | 0 | 0.12 | 0.14 | 0.03 | 0 | 0.07 | 0.05 |
| 1 | 202093735 | rs6701064 | 0.739 | -2.622 | -0.117 | -0.272 | 0.007 | 0.117 | 0.117 | 0.211 | 0.49 | 0.34 | 0.37 | 0.07 | 0.5 | 0.57 |
| 1 | 202095590 | rs4376792 | -0.783 | -2.573 | -0.129 | -0.196 | -0.013 | -0.003 | -0.003 | 0.006 | 0.78 | 0.78 | 0.83 | 0.82 | 0.91 | 0.92 |
| 1 | 202096815 | rs3820380 | 0.745 | -2.344 | -0.165 | -0.184 | 0.15 | 0.002 | 0.002 | 1.495 | 0.55 | 0.18 | 0.54 | 0.48 | 0.78 | 0.74 |
| 1 | 202099045 | rs16849688 | 0.336 | -1.327 | 0.334 | 0.344 | -0.034 | -0.031 | -0.031 | 0 | 0.18 | 0.18 | 0.2 | 0.43 | 0.35 | 0.22 |
| 1 | 202100239 | rs4950829 | -0.767 | -1.459 | 0.792 | 0.557 | -0.017 | 0.18 | 0.18 | 0 | 0.09 | 0.23 | 0.23 | 0.32 | 0.05 | 0.13 |
| 1 | 202106797 | rs16849710 | 0.392 | -1.191 | 0.519 | 0.084 | 0.012 | 0.052 | 0.052 | 0 | 0.52 | 0.41 | 0.51 | 0.27 | 0.55 | 0.53 |
| 1 | 202115945 | rs10920336 | 0.201 | -1.185 | 0.715 | 0.264 | 0.001 | -0.01 | -0.01 | 0 | 0.55 | 0.71 | 0.65 | 0.44 | 0.54 | 0.6 |
| 1 | 202116238 | rs9077 | 1.795 | -1.27 | 0.372 | -0.845 | -0.111 | 0.157 | 0.157 | 0.071 | 0.3 | 0.02 | 0.27 | 0.08 | 0.35 | 0.37 |
| 1 | 202117002 | rs2070257 | -0.615 | -1.237 | 0.714 | -0.665 | -0.003 | 0.004 | 0.004 | 0 | 0.22 | 0.44 | 0.23 | 0.19 | 0.19 | 0.16 |
| 1 | 202119308 | rs7551879 | 0.424 | -1.002 | -0.189 | -0.765 | 0.001 | -0.001 | -0.001 | 0.381 | 0.22 | 0.22 | 0.24 | 0.35 | 0.16 | 0.26 |
| 1 | 202121643 | rs10920337 | 0.037 | -0.458 | -0.355 | -0.458 | 0.011 | -0.003 | -0.003 | 0 | 0.35 | 0.72 | 0.36 | 0.37 | 0.26 | 0.37 |
| 1 | 202122613 | rs10920338 | -0.141 | 0.319 | -0.419 | -0.386 | 0.04 | 0.076 | 0.076 | 0 | 0.73 | 0.64 | 0.64 | 0.51 | 0.63 | 0.81 |
| 1 | 202122783 | rs11590076 | 0.115 | 0.561 | -0.996 | -1.085 | -0.03 | 0.089 | 0.089 | 0.13 | 0.29 | 0.12 | 0.27 | 0.07 | 0.32 | 0.35 |
| 1 | 202125994 | rs12141644 | -0.842 | 0.419 | -0.267 | -0.186 | -0.005 | -0.052 | -0.052 | 0 | 0.11 | 0.18 | 0.05 | 0 | 0.06 | 0.01 |
| 1 | 202128179 | rs10920345 | -1.294 | 0.331 | -0.262 | -0.378 | -0.008 | -0.067 | -0.067 | 0 | 0.2 | 0.29 | 0.1 | 0 | 0.13 | 0.04 |
| 1 | 202129112 | rs4359077 | -1.103 | 0.947 | 0.4 | -0.113 | 0.08 | 0.017 | 0.017 | 0 | 0.07 | 0.26 | 0.12 | 0.06 | 0.12 | 0.04 |
| 1 | 202129277 | rs4631763 | 0.248 | 0.908 | -0.25 | 0.399 | -0.007 | -0.027 | -0.027 | 0 | 0.93 | 0.98 | 0.96 | 0.84 | 0.94 | 0.91 |
| 1 | 202130754 | rs3934648 | -0.016 | 1.34 | -1.178 | 0.746 | -0.029 | 0.03 | 0.03 | 0 | 0.73 | 0.87 | 0.8 | 0.77 | 0.77 | 0.74 |
| 1 | 202131108 | rs6660999 | -0.018 | 1.38 | -0.531 | 0.512 | 0.005 | -0.016 | -0.016 | 0.013 | 0.15 | 0.29 | 0.13 | 0.09 | 0.16 | 0.06 |
| 1 | 202132777 | rs12737056 | -0.361 | 2.031 | 0.083 | 0.376 | 0.076 | 0.088 | 0.088 | 0 | 0.1 | 0.34 | 0.16 | 0.21 | 0.18 | 0.06 |
| 1 | 202138239 | rs3935008 | 0.151 | 2.126 | 0.538 | 0.328 | 0.027 | -0.014 | -0.014 | 0 | 0.15 | 0.34 | 0.14 | 0.08 | 0.16 | 0.07 |
| 1 | 202140400 | rs7550551 | -0.772 | 2.298 | 0.362 | 0.348 | -0.017 | 0.035 | 0.035 | 0 | 0.73 | 0.87 | 0.79 | 0.8 | 0.77 | 0.74 |
| 1 | 202148042 | rs10920352 | 0.776 | 2.109 | 0.421 | 0.361 | 0.007 | 0.018 | 0.018 | 0 | 0.51 | 0.42 | 0.55 | 0.67 | 0.44 | 0.63 |
| 1 | 202151566 | rs10920357 | -0.168 | 2.111 | -0.155 | 0.356 | -0.034 | 0.055 | 0.055 | 0 | 0.66 | 0.8 | 0.69 | 0.77 | 0.6 | 0.69 |
| 1 | 202152129 | rs10920358 | 1.082 | 1.848 | -0.28 | 0.282 | -0.011 | -0.13 | -0.13 | 0 | 0.42 | 0.04 | 0.33 | 0.43 | 0.37 | 0.58 |
| 1 | 202159922 | rs4950833 | 0.618 | 1.841 | -0.021 | 0.426 | -0.001 | -0.006 | -0.006 | 0 | 0.76 | 0.25 | 0.64 | 0.67 | 0.77 | 0.9 |
| 1 | 202161644 | rs2993446 | 0.328 | 1.565 | 0.07 | 0.392 | -0.005 | 0.003 | 0.003 | 1.108 | 0.79 | 0.29 | 0.81 | 0.72 | 0.83 | 0.91 |
| 1 | 202167176 | rs1823696 | -0.017 | 1.228 | -0.487 | 0.385 | -0.002 | -0.001 | -0.001 | 0 | 0.55 | 0.47 | 0.67 | 0.58 | 0.59 | 0.68 |
| 1 | 202171339 | rs3010096 | 0.2 | 1.249 | -1.052 | 0.479 | 0.016 | 0.017 | 0.017 | 0.148 | 0.48 | 0.21 | 0.35 | 0.42 | 0.37 | 0.62 |
| 1 | 202172594 | rs10920360 | 0.712 | 0.888 | -1.32 | 0.539 | 0.006 | -0.031 | -0.031 | 0 | 0.2 | 0.07 | 0.2 | 0.26 | 0.14 | 0.36 |
| 1 | 202172769 | rs12122827 | 0.227 | 0.488 | -1.842 | 0.193 | -0.04 | -0.092 | -0.092 | 0 | 0.25 | 0.03 | 0.25 | 0.35 | 0.29 | 0.29 |
| 1 | 202180401 | rs12033508 | -0.078 | 0.662 | -1.91 | 0.09 | -0.066 | 0.018 | 0.018 | 0 | 0.3 | 0.1 | 0.26 | 0.28 | 0.43 | 0.28 |
| 1 | 202183308 | rs10920362 | -0.588 | 0.681 | -1.055 | -0.034 | 0.024 | -0.024 | -0.024 | 1.552 | 0.37 | 0.47 | 0.44 | 0.31 | 0.48 | 0.29 |
| 1 | 202186178 | rs10920364 | -0.388 | 0.662 | -0.974 | -0.083 | 0.021 | -0.014 | -0.014 | 0 | 0.36 | 0.41 | 0.44 | 0.31 | 0.49 | 0.3 |
| 1 | 202188616 | rs2860229 | -0.322 | 0.089 | -0.995 | -0.514 | -0.016 | -0.09 | -0.09 | 1.652 | 0.79 | 0.47 | 0.75 | 0.95 | 0.86 | 0.92 |
| 1 | 202189711 | rs10920365 | 0.306 | 0.066 | -0.92 | -0.535 | -0.006 | 0.004 | 0.004 | 0 | 0.31 | 0.26 | 0.29 | 0.27 | 0.43 | 0.28 |
| 1 | 202190546 | rs4950774 | 0.448 | 0.054 | -1.031 | -0.55 | -0.072 | 0.026 | 0.026 | 0 | 0.29 | 0.11 | 0.27 | 0.26 | 0.43 | 0.28 |
| 1 | 202190742 | rs2993433 | 0.795 | 0.073 | -1.026 | -0.532 | 0.009 | 0.111 | 0.111 | 0.298 | 0.48 | 0.19 | 0.46 | 0.68 | 0.43 | 0.64 |
| 1 | 202193434 | rs3010069 | -0.651 | -0.073 | -1.04 | -0.686 | -0.043 | -0.102 | -0.102 | 0 | 0.79 | 0.65 | 0.89 | 0.99 | 0.88 | 0.94 |
| 1 | 202193981 | rs2993437 | 0.898 | 0.007 | -1.298 | -0.654 | 0.024 | 0.108 | 0.108 | 0.332 | 0.47 | 0.22 | 0.46 | 0.69 | 0.4 | 0.65 |
| 1 | 202197161 | rs3795598 | 0.54 | -0.075 | -1.442 | -0.525 | -0.091 | 0.017 | 0.017 | 0 | 0.27 | 0.1 | 0.27 | 0.25 | 0.42 | 0.26 |
| 1 | 202197551 | rs3820383 | 1.001 | -1.332 | -1.416 | -0.23 | 0.045 | -0.07 | -0.07 | 1.872 | 0.2 | 0.09 | 0.21 | 0.32 | 0.09 | 0.26 |
| 1 | 202201905 | rs2361450 | -0.068 | -0.767 | -0.829 | -0.982 | -0.03 | -0.106 | -0.106 | 0.013 | 0.75 | 0.65 | 0.85 | 0.99 | 0.84 | 0.94 |
| 1 | 202203119 | rs2123138 | 0.402 | -0.415 | -0.999 | -0.739 | 0.031 | 0.1 | 0.1 | 0 | 0.53 | 0.24 | 0.64 | 0.71 | 0.56 | 0.59 |
| 1 | 202203337 | rs10920369 | -0.051 | -0.374 | -0.819 | -0.83 | 0 | 0.008 | 0.008 | 0 | 0.61 | 0.63 | 0.72 | 0.81 | 0.57 | 0.61 |
| 1 | 202205323 | rs896549 | 0.281 | -0.319 | -1.012 | -0.752 | 0.007 | 0.025 | 0.025 | 0 | 0.56 | 0.41 | 0.69 | 0.75 | 0.56 | 0.58 |
| 1 | 202209967 | rs7524808 | -0.367 | -0.575 | -0.824 | -0.862 | -0.004 | -0.019 | -0.019 | 0 | 0.47 | 0.51 | 0.3 | 0.17 | 0.37 | 0.37 |
| 1 | 202219472 | rs880766 | 0.049 | -1.054 | -1.368 | -1.223 | 0.006 | 0.012 | 0.012 | 1.214 | 0.66 | 0.36 | 0.76 | 0.84 | 0.69 | 0.77 |
| 1 | 202224520 | rs12123327 | 0.414 | -1.104 | -1.407 | -1.146 | 0.029 | 0.044 | 0.044 | 0 | 0.2 | 0.05 | 0.12 | 0.12 | 0.13 | 0.12 |
| 1 | 202228042 | rs3010077 | 0.011 | -1.292 | -1.674 | -1.375 | 0.025 | 0.048 | 0.048 | 2.749 | 0.59 | 0.36 | 0.75 | 0.83 | 0.66 | 0.73 |
| 1 | 202233116 | rs12739853 | 0.419 | -1.182 | -1.793 | -1.358 | -0.038 | 0.009 | 0.009 | 0 | 0.16 | 0.01 | 0.07 | 0.07 | 0.18 | 0.05 |
| 1 | 202239771 | rs2993444 | 0.839 | -0.938 | -1.461 | -1.153 | -0.022 | 0.031 | 0.031 | 4.954 | 0.38 | 0.23 | 0.63 | 0.71 | 0.53 | 0.61 |
| 1 | 202248554 | rs2993447 | 0.836 | -0.808 | -1.493 | -1.138 | -0.055 | 0.036 | 0.036 | 5.357 | 0.38 | 0.16 | 0.63 | 0.72 | 0.52 | 0.61 |
| 1 | 202266547 | rs788827 | -0.093 | -0.861 | -1.175 | -1.218 | -0.004 | -0.012 | -0.012 | 2.255 | 0.6 | 0.46 | 0.73 | 0.77 | 0.64 | 0.7 |
| 1 | 202266607 | rs788826 | 0.573 | -0.82 | -1.617 | -1.192 | -0.005 | 0.043 | 0.043 | 4.227 | 0.43 | 0.3 | 0.65 | 0.72 | 0.55 | 0.62 |
| 1 | 202273023 | rs7532633 | 1.235 | -0.828 | -2.063 | -0.628 | -0.109 | -0.046 | -0.046 | 0.355 | 0.28 | 0.09 | 0.42 | 0.33 | 0.49 | 0.37 |
| 1 | 202279709 | rs705766 | -0.109 | -0.645 | -1.712 | -1.091 | 0.014 | 0.037 | 0.037 | 2.273 | 0.59 | 0.36 | 0.72 | 0.77 | 0.64 | 0.7 |
| 1 | 202281902 | rs788640 | 0.681 | -0.455 | -1.807 | -1.099 | -0.034 | 0.059 | 0.059 | 4.065 | 0.42 | 0.21 | 0.62 | 0.72 | 0.53 | 0.61 |
| 1 | 202286109 | rs788796 | -0.102 | -0.402 | -1.844 | -1.097 | 0.018 | 0.044 | 0.044 | 2.48 | 0.59 | 0.34 | 0.72 | 0.77 | 0.64 | 0.71 |
| 1 | 202288462 | rs788792 | -0.089 | -0.406 | -1.846 | -1.098 | 0.019 | 0.045 | 0.045 | 2.378 | 0.59 | 0.34 | 0.72 | 0.77 | 0.64 | 0.71 |
| 1 | 202290303 | rs12126278 | 0.547 | 0.054 | -1.623 | -0.777 | 0.016 | 0.07 | 0.07 | 0 | 0.16 | 0.06 | 0.1 | 0.05 | 0.11 | 0.08 |
| 1 | 202299085 | rs4079028 | -0.084 | 0.324 | -1.585 | -0.678 | -0.022 | 0.022 | 0.022 | 0 | 0.17 | 0.06 | 0.1 | 0.1 | 0.19 | 0.08 |
| 1 | 202300127 | rs35873832 | 0.498 | -0.024 | -2.046 | -0.934 | 0.017 | 0.063 | 0.063 | 0 | 0.2 | 0.01 | 0.13 | 0.07 | 0.15 | 0.09 |
| 1 | 202308344 | rs10465591 | 1.043 | 0.458 | -2.212 | -1.367 | -0.037 | 0.031 | 0.031 | 3.186 | 0.36 | 0.2 | 0.61 | 0.72 | 0.49 | 0.62 |

| | | | | | | | | | | | | | | | | |
|---|---|---|---|---|---|---|---|---|---|---|---|---|---|---|---|---|
| 1 | 202322113 | rs6658708 | 1.066 | 0.074 | -2.292 | -1.399 | -0.098 | -0.001 | -0.001 | 0 | 0.35 | 0.09 | 0.6 | 0.72 | 0.49 | 0.61 |
| 1 | 202356935 | rs17438379 | 1.002 | 0.072 | -2.315 | -1.416 | 0.017 | 0.063 | 0.063 | 0.001 | 0.2 | 0.01 | 0.13 | 0.07 | 0.15 | 0.1 |
| 1 | 202364505 | rs2843410 | -0.278 | -0.109 | -1.182 | -1.448 | 0.007 | 0.012 | 0.012 | 2.017 | 0.4 | 0.37 | 0.22 | 0.21 | 0.35 | 0.28 |
| 1 | 202370997 | rs11583656 | 0.981 | -0.109 | -0.92 | -1.418 | 0.035 | 0.058 | 0.058 | 0 | 0.2 | 0.05 | 0.13 | 0.07 | 0.15 | 0.09 |
| 1 | 202373394 | rs2741849 | 0.73 | 0.184 | -0.138 | -0.961 | -0.011 | -0.014 | -0.014 | 0 | 0.19 | 0.2 | 0.11 | 0.11 | 0.15 | 0.19 |
| 1 | 202381531 | rs2696961 | 0.746 | 0.188 | -0.071 | -0.957 | -0.009 | -0.01 | -0.01 | 0 | 0.19 | 0.19 | 0.11 | 0.12 | 0.15 | 0.19 |
| 1 | 202405671 | rs2741853 | 0.381 | 0.211 | 0.297 | -0.892 | 0.007 | 0.011 | 0.011 | 2.115 | 0.6 | 0.63 | 0.78 | 0.8 | 0.65 | 0.72 |
| 1 | 202547101 | rs3767392 | 1.302 | 0.229 | -0.193 | -0.016 | -0.114 | -0.019 | -0.019 | 0 | 0.43 | 0.06 | 0.43 | 0.38 | 0.51 | 0.36 |
| 1 | 202550272 | rs2008455 | 0.351 | 0.032 | 0.323 | 0.053 | -0.015 | -0.031 | -0.031 | 0 | 0.61 | 0.52 | 0.76 | 0.79 | 0.66 | 0.72 |
| 1 | 202550547 | rs2362941 | 0.568 | -0.042 | 0.276 | 0.108 | 0.048 | 0.088 | 0.088 | 1.108 | 0.55 | 0.14 | 0.7 | 0.71 | 0.62 | 0.66 |
| 1 | 202551071 | rs747640 | 0.626 | -0.103 | 0.259 | 0.05 | 0.037 | 0.082 | 0.082 | 0.756 | 0.56 | 0.12 | 0.7 | 0.71 | 0.64 | 0.67 |
| 1 | 202552100 | rs4950858 | 0.574 | -0.079 | 0.312 | 0.038 | -0.005 | -0.01 | -0.01 | 0 | 0.57 | 0.46 | 0.72 | 0.71 | 0.62 | 0.67 |
| 1 | 202552839 | rs4950859 | 0.605 | 0.026 | 0.367 | 0.196 | 0.037 | 0.078 | 0.078 | 0.567 | 0.58 | 0.14 | 0.7 | 0.72 | 0.64 | 0.69 |
| 1 | 202559753 | rs1046269 | 0.662 | 0.08 | 0.02 | 0.156 | 0.002 | 0.002 | 0.002 | 0 | 0.14 | 0.03 | 0.08 | 0.1 | 0.11 | 0.06 |
| 1 | 202560412 | rs1040010 | 0.043 | 0.095 | 0.357 | 0.264 | -0.011 | -0.02 | -0.02 | 0 | 0.6 | 0.49 | 0.73 | 0.72 | 0.65 | 0.69 |
| 1 | 202562427 | rs6427957 | 0.301 | 0.505 | 0.607 | 0.647 | -0.001 | 0.022 | 0.022 | 0 | 0.59 | 0.3 | 0.67 | 0.68 | 0.55 | 0.61 |
| 1 | 202563966 | rs12033091 | 0.379 | 0.212 | 0.349 | 0.895 | 0.002 | 0.232 | 0.232 | 0.22 | 0.12 | 0.39 | 0.27 | 0.34 | 0.06 | 0.2 |
| 1 | 202564040 | rs12033827 | 0.379 | 0.216 | 0.349 | 0.857 | -0.002 | 0.236 | 0.236 | 0.22 | 0.12 | 0.39 | 0.27 | 0.34 | 0.06 | 0.22 |
| 1 | 202564221 | rs4520471 | 0.922 | 0.235 | 0.201 | 0.899 | -0.017 | 0.066 | 0.066 | 0.048 | 0.5 | 0.08 | 0.46 | 0.36 | 0.57 | 0.44 |
| 1 | 202566200 | rs9633344 | 0.289 | 0.415 | 0.769 | 0.454 | -0.009 | -0.013 | -0.013 | 0 | 0.65 | 0.54 | 0.73 | 0.71 | 0.65 | 0.68 |
| 1 | 202566430 | rs10494834 | 0.302 | 0.435 | 0.778 | 0.442 | -0.009 | -0.005 | -0.005 | 1.199 | 0.65 | 0.54 | 0.73 | 0.67 | 0.65 | 0.68 |
| 1 | 202595153 | rs16830648 | -1.035 | 1.471 | 0.851 | 1.061 | 0.063 | 0.069 | 0.069 | 0 | 0.14 | 0.34 | 0.19 | 0.25 | 0.25 | 0.12 |
| 1 | 202604386 | rs12354333 | -0.922 | 0.695 | 0.34 | 0.69 | 0 | -0.003 | -0.003 | 0.48 | 0.27 | 0.69 | 0.25 | 0.22 | 0.31 | 0.1 |
| 1 | 202615197 | rs11585565 | 0.52 | 0.358 | -1.297 | 0.256 | -0.126 | 0.131 | 0.131 | 0.148 | 0.43 | 0.05 | 0.51 | 0.21 | 0.48 | 0.5 |
| 1 | 202616107 | rs12409072 | -0.268 | -0.007 | -1.207 | 0.702 | 0.039 | 0.037 | 0.037 | 0 | 0.67 | 0.79 | 0.85 | 0.75 | 0.79 | 0.6 |
| 1 | 202616789 | rs12121078 | -0.059 | 0.34 | -2.232 | 0.416 | -0.079 | 0.001 | 0.001 | 0 | 0.37 | 0.09 | 0.39 | 0.4 | 0.51 | 0.47 |
| 1 | 202617807 | rs867313 | -0.352 | 0.968 | -2.231 | 0.215 | -0.028 | 0.034 | 0.034 | 0 | 0.38 | 0.17 | 0.43 | 0.76 | 0.42 | 0.46 |
| 1 | 202618162 | rs867315 | -0.253 | 0.946 | -2.22 | 0.251 | -0.074 | 0.003 | 0.003 | 0 | 0.37 | 0.07 | 0.43 | 0.74 | 0.42 | 0.46 |
| 1 | 202618202 | rs867316 | -0.602 | 1.175 | -2.252 | 0.042 | -0.033 | 0.119 | 0.119 | 0 | 0.45 | 0.21 | 0.47 | 0.81 | 0.5 | 0.56 |
| 1 | 202618673 | rs946857 | -0.287 | 1.112 | -2.217 | 0.093 | -0.033 | 0.035 | 0.035 | 0 | 0.38 | 0.16 | 0.43 | 0.76 | 0.42 | 0.46 |
| 1 | 202622302 | rs12404945 | 0.985 | 1.386 | -2.563 | 0.078 | -0.047 | -0.021 | -0.021 | 0 | 0.19 | 0.02 | 0.2 | 0.21 | 0.17 | 0.13 |
| 1 | 202623455 | rs10737582 | -0.582 | 1.356 | -1.896 | 0.104 | -0.008 | 0.022 | 0.022 | 0 | 0.39 | 0.3 | 0.44 | 0.75 | 0.43 | 0.46 |
| 1 | 202623622 | rs10920443 | 0.846 | 1.27 | -1.884 | 0.222 | -0.039 | -0.088 | -0.088 | 0 | 0.13 | 0.02 | 0.16 | 0.53 | 0.19 | 0.32 |
| 1 | 202638960 | rs6662116 | -0.235 | 0.696 | -2.074 | -0.112 | 0.003 | 0.043 | 0.043 | 1.385 | 0.62 | 0.25 | 0.54 | 0.8 | 0.62 | 0.66 |
| 1 | 202643007 | rs4950866 | -0.195 | 0.731 | -2.104 | -0.095 | 0 | 0.063 | 0.063 | 1.234 | 0.61 | 0.19 | 0.54 | 0.8 | 0.62 | 0.66 |
| 1 | 202652939 | rs6698441 | 0.181 | 0.973 | -2.11 | 0.365 | -0.006 | -0.021 | -0.021 | 0 | 0.32 | 0.15 | 0.31 | 0.41 | 0.29 | 0.26 |
| 1 | 202658132 | rs10920457 | 0.79 | 0.966 | -1.403 | 0.419 | -0.035 | -0.074 | -0.074 | 0 | 0.31 | 0.07 | 0.25 | 0.38 | 0.34 | 0.41 |
| 1 | 202662356 | rs4362037 | 0.662 | 1.009 | -1.243 | 0.497 | 0 | -0.023 | -0.023 | 0 | 0.33 | 0.19 | 0.25 | 0.38 | 0.35 | 0.43 |
| 1 | 202675394 | rs4400672 | 0.562 | 1.082 | -1.195 | 0.491 | -0.001 | -0.021 | -0.021 | 0.432 | 0.35 | 0.19 | 0.27 | 0.4 | 0.36 | 0.42 |
| 1 | 202675543 | rs4364933 | -0.666 | 0.965 | -1.292 | 0.388 | 0.006 | -0.01 | -0.01 | 0 | 0.69 | 0.38 | 0.72 | 0.79 | 0.65 | 0.72 |
| 1 | 202675629 | rs4418670 | 0.202 | 0.948 | -1.476 | 0.343 | -0.004 | -0.016 | -0.016 | 0.629 | 0.33 | 0.13 | 0.44 | 0.39 | 0.29 | 0.3 |
| 1 | 202676881 | rs12141884 | 0.514 | 1.22 | -0.684 | 0.075 | 0.001 | -0.014 | -0.014 | 4.921 | 0.24 | 0.26 | 0.22 | 0.1 | 0.3 | 0.17 |
| 1 | 202678105 | rs4325195 | 0.746 | 1.02 | -0.926 | 0.459 | -0.026 | -0.003 | -0.003 | 0 | 0.34 | 0.18 | 0.31 | 0.32 | 0.39 | 0.41 |
| 1 | 202685803 | rs4561090 | -0.177 | 0.409 | -0.476 | 0.339 | 0.003 | 0.007 | 0.007 | 0.847 | 0.67 | 0.3 | 0.75 | 0.82 | 0.69 | 0.72 |
| 1 | 202688077 | rs7529811 | -0.053 | 0.351 | -0.444 | 0.858 | -0.097 | 0.079 | 0.079 | 0.37 | 0.43 | 0.05 | 0.44 | 0.33 | 0.52 | 0.42 |
| 1 | 202689028 | rs4570460 | 2.33 | 0.287 | -0.487 | 0.818 | 0.002 | -0.027 | -0.027 | 0 | 0.05 | 0 | 0.03 | 0.08 | 0.03 | 0.13 |
| 1 | 202695165 | rs4950740 | -0.546 | 0.303 | -0.089 | 0.771 | -0.032 | -0.002 | -0.002 | 0 | 0.77 | 0.56 | 0.87 | 0.81 | 0.83 | 0.72 |
| 1 | 202696247 | rs4348778 | 0.196 | 0.3 | -0.074 | 0.861 | -0.004 | -0.055 | -0.055 | 0 | 0.7 | 0.2 | 0.66 | 0.52 | 0.74 | 0.57 |
| 1 | 202702353 | rs4950791 | -0.378 | 0.352 | -0.616 | 0.881 | -0.028 | -0.005 | -0.005 | 0 | 0.76 | 0.46 | 0.86 | 0.81 | 0.83 | 0.72 |
| 1 | 202715284 | rs1141108 | 0.006 | 0.358 | -0.678 | 0.885 | 0 | 0 | 0 | 0.219 | 0.73 | 0.32 | 0.83 | 0.78 | 0.77 | 0.64 |
| 1 | 202718202 | rs1892164 | 0.334 | 0.37 | -0.763 | 0.903 | -0.003 | -0.04 | -0.04 | 0 | 0.7 | 0.24 | 0.67 | 0.52 | 0.74 | 0.57 |
| 1 | 202721148 | rs10920473 | 1.529 | 0.237 | -0.925 | 0.771 | 0.001 | -0.012 | -0.012 | 0 | 0.18 | 0.01 | 0.18 | 0.16 | 0.14 | 0.16 |
| 1 | 202726145 | rs12132801 | 1.813 | 0.547 | -0.511 | 1.095 | 0.023 | -0.022 | -0.022 | 0.141 | 0.25 | 0.04 | 0.22 | 0.18 | 0.22 | 0.14 |
| 1 | 202733238 | rs3196669 | 0.396 | 0.721 | -0.216 | 1.223 | 0 | 0 | 0 | 0 | 0.72 | 0.32 | 0.83 | 0.77 | 0.77 | 0.64 |
| 1 | 202760658 | rs4351714 | 0.628 | 0.747 | -0.244 | 1.173 | -0.002 | -0.04 | -0.04 | 0.51 | 0.7 | 0.24 | 0.67 | 0.52 | 0.75 | 0.58 |
| 1 | 202765279 | rs6427967 | 0.205 | 0.496 | -0.26 | 1.173 | -0.019 | 0.058 | 0.058 | 0.016 | 0.74 | 0.4 | 0.7 | 0.55 | 0.8 | 0.64 |
| 1 | 202778261 | rs4245713 | -0.738 | 0.701 | -0.113 | 1.256 | -0.004 | 0.037 | 0.037 | 0 | 0.83 | 0.59 | 0.88 | 0.85 | 0.85 | 0.73 |
| 1 | 202780530 | rs2275895 | 1.311 | 0.83 | -0.62 | 1.3 | 0.013 | -0.02 | -0.02 | 0 | 0.26 | 0.04 | 0.21 | 0.19 | 0.22 | 0.14 |
| 1 | 202780663 | rs2275896 | -0.222 | 1.085 | -0.626 | 1.239 | 0.005 | 0.021 | 0.021 | 0 | 0.49 | 0.33 | 0.48 | 0.61 | 0.55 | 0.46 |
| 1 | 202781164 | rs2246083 | 0.724 | 1.385 | -0.717 | 1.196 | -0.058 | 0.021 | 0.021 | 0.062 | 0.18 | 0.05 | 0.21 | 0.16 | 0.31 | 0.24 |
| 1 | 202788043 | rs12095873 | -0.573 | 0.852 | 0.004 | 1.722 | 0.053 | 0.169 | 0.169 | 0 | 0.06 | 0.25 | 0.07 | 0.3 | 0.07 | 0.12 |
| 1 | 202794787 | rs12139944 | 0.775 | 0.62 | -0.163 | 1.64 | 0.019 | -0.02 | -0.02 | 0 | 0.27 | 0.06 | 0.23 | 0.24 | 0.25 | 0.18 |
| 1 | 202810562 | rs4950949 | -0.318 | 0.642 | -0.155 | 1.47 | -0.004 | 0.007 | 0.007 | 0 | 0.63 | 0.42 | 0.68 | 0.47 | 0.66 | 0.69 |
| 1 | 202812654 | rs11576909 | 0.563 | 0.346 | -0.911 | 1.476 | -0.104 | 0.095 | 0.095 | 0 | 0.45 | 0.05 | 0.47 | 0.32 | 0.5 | 0.42 |
| 1 | 202815986 | rs10920503 | -0.184 | -0.287 | -0.008 | 1.813 | -0.034 | 0.034 | 0.034 | 0 | 0.67 | 0.56 | 0.71 | 0.48 | 0.73 | 0.75 |
| 1 | 202827191 | rs6663182 | -0.599 | 1.292 | 0.369 | 1.541 | 0.033 | -0.001 | -0.001 | 0 | 0.25 | 0.3 | 0.27 | 0.22 | 0.4 | 0.24 |
| 1 | 202842333 | rs12094889 | -0.772 | 1.215 | 0.892 | 1.588 | 0.001 | -0.001 | -0.001 | 0 | 0.27 | 0.26 | 0.4 | 0.3 | 0.41 | 0.3 |
| 1 | 202843127 | rs11590469 | -1.151 | 1.084 | 1.512 | 1.638 | 0.037 | 0.028 | 0.028 | 0 | 0.31 | 0.58 | 0.43 | 0.4 | 0.44 | 0.43 |
| 1 | 202847719 | rs6703138 | 0.384 | 0.749 | 0.99 | 1.538 | -0.03 | -0.087 | -0.087 | 1.536 | 0.16 | 0.02 | 0.31 | 0.43 | 0.2 | 0.29 |

| | | | | | | | | | | | | | | | | |
|---|---|---|---|---|---|---|---|---|---|---|---|---|---|---|---|---|
| 1 | 202849723 | rs1131906 | 0.017 | -0.292 | 0.33 | 0.66 | 0.005 | 0.002 | 0.002 | 0 | 0.54 | 0.47 | 0.53 | 0.41 | 0.65 | 0.51 |
| 1 | 202849770 | rs1131905 | 0.017 | -0.292 | 0.33 | 0.66 | 0.005 | 0.002 | 0.002 | 0 | 0.54 | 0.47 | 0.53 | 0.41 | 0.65 | 0.51 |
| 1 | 202853101 | rs6698793 | -0.793 | -0.437 | 0.558 | 0.525 | -0.032 | -0.065 | -0.065 | 0 | 0.72 | 0.62 | 0.87 | 0.88 | 0.85 | 0.81 |
| 1 | 202859600 | rs3806346 | 0.303 | -0.34 | 0.531 | 0.638 | -0.014 | -0.057 | -0.057 | 0 | 0.2 | 0.07 | 0.33 | 0.52 | 0.23 | 0.4 |
| 1 | 202870330 | rs4950887 | -0.708 | -0.41 | 0.674 | 0.608 | -0.044 | -0.088 | -0.088 | 0 | 0.71 | 0.55 | 0.86 | 0.88 | 0.85 | 0.81 |
| 1 | 202874730 | rs6686801 | 0.407 | -0.416 | 0.693 | 0.606 | 0.026 | 0.048 | 0.048 | 0 | 0.5 | 0.4 | 0.51 | 0.31 | 0.61 | 0.4 |
| 1 | 202877397 | rs9919219 | -0.966 | -0.412 | 0.745 | 0.608 | -0.007 | -0.022 | -0.022 | 0.79 | 0.72 | 0.72 | 0.87 | 0.88 | 0.85 | 0.81 |
| 1 | 202881302 | rs6692075 | 0.55 | -0.409 | 0.462 | 0.608 | 0.022 | 0.04 | 0.04 | 0 | 0.5 | 0.42 | 0.53 | 0.32 | 0.62 | 0.4 |
| 1 | 202884063 | rs10800877 | 0.311 | -0.375 | 0.38 | 0.628 | -0.014 | -0.057 | -0.057 | 0 | 0.2 | 0.07 | 0.33 | 0.52 | 0.23 | 0.39 |
| 1 | 202887295 | rs2275734 | 0.382 | -0.383 | 0.392 | 0.618 | 0.026 | 0.048 | 0.048 | 0 | 0.5 | 0.39 | 0.51 | 0.31 | 0.62 | 0.4 |
| 1 | 202892917 | rs10800880 | 0.145 | -0.133 | 0.311 | 0.794 | -0.014 | -0.058 | -0.058 | 0 | 0.2 | 0.07 | 0.33 | 0.52 | 0.23 | 0.39 |
| 1 | 202899361 | rs2363772 | 0.243 | -0.121 | 0.268 | 1.339 | -0.007 | -0.027 | -0.027 | 0 | 0.18 | 0.09 | 0.14 | 0.34 | 0.2 | 0.34 |
| 1 | 202908836 | rs10920531 | 0.239 | -0.396 | 0.122 | 1.075 | 0.007 | 0.01 | 0.01 | 0.059 | 0.54 | 0.46 | 0.53 | 0.35 | 0.64 | 0.43 |
| 1 | 202914356 | rs1342387 | -1.092 | 0.387 | 0.364 | 1.271 | 0.006 | -0.007 | -0.007 | 0 | 0.42 | 0.5 | 0.48 | 0.63 | 0.53 | 0.54 |
| 1 | 202916806 | rs12045862 | -0.397 | 0.491 | 0.134 | 1.408 | -0.011 | -0.08 | -0.08 | 0.313 | 0.24 | 0.06 | 0.46 | 0.59 | 0.26 | 0.43 |
| 1 | 202919851 | rs16850799 | 0.145 | 0.51 | 0.152 | 1.413 | -0.023 | -0.109 | -0.109 | 0.829 | 0.19 | 0.05 | 0.44 | 0.46 | 0.22 | 0.3 |
| 1 | 202922194 | rs10494839 | 0.103 | 0.679 | 0.412 | 1.546 | -0.048 | 0.042 | 0.042 | 0 | 0.15 | 0.13 | 0.18 | 0.05 | 0.29 | 0.06 |
| 1 | 202923178 | rs10753929 | -0.301 | 0.712 | 0.614 | 1.453 | 0.041 | 0.017 | 0.017 | 0 | 0.92 | 0.81 | 0.87 | 0.91 | 0.86 | 0.85 |
| 1 | 202928703 | rs6666089 | 0.089 | 0.583 | -0.095 | 1.452 | -0.047 | 0.042 | 0.042 | 0 | 0.15 | 0.13 | 0.18 | 0.05 | 0.29 | 0.06 |
| 1 | 202935911 | rs2232842 | -1.119 | 0.208 | 0.087 | 1.876 | -0.022 | 0.248 | 0.248 | 0 | 0.06 | 0.14 | 0.22 | 0.41 | 0.03 | 0.19 |
| 1 | 202937634 | rs12758184 | -0.337 | 0.385 | -0.393 | 1.491 | -0.065 | -0.046 | -0.046 | 0 | 0.25 | 0.09 | 0.35 | 0.28 | 0.34 | 0.37 |
| 1 | 202948992 | rs2363770 | 1.469 | -0.016 | 1.297 | 2.087 | -0.001 | 0.366 | 0.366 | 0 | 0.91 | 0.53 | 0.92 | 0.65 | 0.97 | 0.78 |
| 1 | 202957262 | rs16850859 | -0.434 | -0.269 | 1.274 | 1.224 | -0.014 | -0.02 | -0.02 | 0 | 0.05 | 0.22 | 0.02 | 0 | 0 | 0 |
| 1 | 202967491 | rs12121955 | -0.765 | 0.052 | 1.129 | 1.128 | -0.001 | 0.01 | 0.01 | 0 | 0.12 | 0.16 | 0.21 | 0.18 | 0.12 | 0.15 |
| 1 | 202969332 | rs10800895 | -1.461 | -0.13 | 1.131 | 1.322 | -0.043 | 0.363 | 0.363 | 0.36 | 0.11 | 0.4 | 0.24 | 0.49 | 0.03 | 0.18 |
| 1 | 202974481 | rs1361734 | 0.758 | 0.125 | 1.016 | 1.015 | 0.017 | 0.056 | 0.056 | 0 | 0.86 | 0.64 | 0.76 | 0.82 | 0.88 | 0.86 |
| 1 | 202983294 | rs11582753 | 0.995 | 0.83 | -0.061 | 0.464 | -0.002 | -0.03 | -0.03 | 0 | 0.49 | 0.49 | 0.29 | 0.13 | 0.41 | 0.35 |
| 1 | 202994268 | rs1774847 | 0.51 | 1.408 | -0.004 | 0.828 | 0.005 | -0.004 | -0.004 | 0 | 0.63 | 0.62 | 0.5 | 0.33 | 0.51 | 0.48 |
| 1 | 202998819 | rs782791 | 0.147 | 0.907 | -0.267 | 1.02 | -0.005 | 0.047 | 0.047 | 0 | 0.71 | 0.61 | 0.6 | 0.43 | 0.69 | 0.7 |
| 1 | 203003641 | rs10920550 | -0.332 | 1.259 | -0.316 | 0.972 | 0.018 | 0.026 | 0.026 | 0 | 0.09 | 0.2 | 0.1 | 0.09 | 0.11 | 0.17 |
| 1 | 203003870 | rs782800 | 0.567 | 0.83 | -0.71 | 1.098 | 0.014 | 0.002 | 0.002 | 0 | 0.54 | 0.35 | 0.47 | 0.53 | 0.59 | 0.55 |
| 1 | 203004367 | rs10920551 | -0.162 | 0.532 | -0.703 | 1.2 | -0.024 | -0.002 | -0.002 | 2.243 | 0.16 | 0.08 | 0.29 | 0.22 | 0.21 | 0.2 |
| 1 | 203005571 | rs4950907 | 0.763 | 0.042 | -0.668 | 1.522 | -0.001 | 0.025 | 0.025 | 0 | 0.06 | 0.04 | 0.07 | 0.23 | 0.07 | 0.07 |
| 1 | 203005821 | rs782802 | -0.33 | -0.227 | -1.134 | 1.022 | -0.033 | 0.01 | 0.01 | 0.384 | 0.37 | 0.17 | 0.44 | 0.42 | 0.38 | 0.44 |
| 1 | 203006098 | rs782803 | -0.467 | -0.206 | -1.135 | 1.081 | -0.04 | 0.009 | 0.009 | 0.711 | 0.38 | 0.16 | 0.46 | 0.44 | 0.41 | 0.44 |
| 1 | 203012906 | rs4950908 | -0.486 | -0.289 | -0.296 | 0.934 | -0.002 | -0.018 | -0.018 | 2.915 | 0.73 | 0.48 | 0.7 | 0.8 | 0.78 | 0.67 |
| 1 | 203014196 | rs4472812 | 0.727 | -0.277 | -0.641 | 0.884 | -0.008 | 0.03 | 0.03 | 0.151 | 0.51 | 0.21 | 0.36 | 0.33 | 0.56 | 0.45 |
| 1 | 203017087 | rs754894 | 0.548 | -0.662 | -0.028 | 0.967 | -0.029 | 0.02 | 0.02 | 0 | 0.73 | 0.59 | 0.58 | 0.59 | 0.78 | 0.64 |
| 1 | 203019387 | rs12122472 | 0.716 | -0.459 | -0.426 | 0.955 | 0.013 | -0.007 | -0.007 | 0 | 0.13 | 0.05 | 0.09 | 0.15 | 0.11 | 0.15 |
| 1 | 203019650 | rs12130133 | 0.723 | -0.466 | -0.423 | 1.044 | 0.012 | -0.021 | -0.021 | 0.19 | 0.13 | 0.05 | 0.08 | 0.2 | 0.11 | 0.15 |
| 1 | 203019747 | rs12755165 | -0.027 | -0.386 | -0.645 | 0.543 | 0.002 | -0.005 | -0.005 | 0 | 0.21 | 0.17 | 0.16 | 0.18 | 0.25 | 0.16 |
| 1 | 203021807 | rs4950876 | -0.488 | -0.319 | -0.484 | 0.418 | 0 | 0.002 | 0.002 | 0 | 0.6 | 0.66 | 0.53 | 0.46 | 0.62 | 0.63 |
| 1 | 203023122 | rs2291691 | 0.967 | -0.488 | -0.458 | 0.218 | 0 | -0.004 | -0.004 | 0.037 | 0.29 | 0.29 | 0.18 | 0.11 | 0.25 | 0.25 |
| 1 | 203024767 | rs3736314 | 0.904 | -0.677 | -0.415 | 0.285 | 0.002 | 0.004 | 0.004 | 0 | 0.12 | 0.22 | 0.11 | 0.13 | 0.15 | 0.12 |
| 1 | 203025428 | rs12097602 | 0.932 | -0.672 | -0.49 | 0.286 | 0 | 0 | 0 | 0.008 | 0.12 | 0.19 | 0.11 | 0.13 | 0.16 | 0.12 |
| 1 | 203026591 | rs17461925 | 0.344 | -0.408 | -0.419 | 0.396 | -0.005 | -0.019 | -0.019 | 0 | 0.41 | 0.53 | 0.29 | 0.23 | 0.38 | 0.36 |
| 1 | 203034906 | rs3737883 | 0.019 | -0.499 | -0.758 | 0.338 | 0.015 | 0.062 | 0.062 | 0 | 0.5 | 0.26 | 0.39 | 0.39 | 0.56 | 0.54 |
| 1 | 203035551 | rs6670637 | 0.419 | -0.326 | -0.907 | 0.396 | 0.004 | 0.024 | 0.024 | 0 | 0.39 | 0.4 | 0.28 | 0.21 | 0.38 | 0.35 |
| 1 | 203041457 | rs4950914 | 0.982 | -0.03 | -1.223 | 0.548 | -0.034 | -0.008 | -0.008 | 0 | 0.13 | 0.03 | 0.12 | 0.19 | 0.18 | 0.18 |
| 1 | 203041492 | rs4950915 | -0.39 | -0.092 | -0.653 | 0.73 | 0.002 | 0.006 | 0.006 | 0 | 0.63 | 0.67 | 0.54 | 0.46 | 0.62 | 0.62 |
| 1 | 203045583 | rs6682652 | 0.506 | 0.125 | -1.605 | 0.387 | 0.001 | -0.015 | -0.015 | 0 | 0.32 | 0.45 | 0.34 | 0.29 | 0.37 | 0.34 |
| 1 | 203046534 | rs3820151 | 0.35 | 0.816 | -2.247 | 0.231 | -0.021 | 0.049 | 0.049 | 0 | 0.24 | 0.02 | 0.15 | 0.09 | 0.17 | 0.23 |
| 1 | 203047621 | rs3087949 | 0.111 | 1.369 | -1.066 | 0.905 | -0.028 | 0.002 | 0.002 | 0 | 0.78 | 0.89 | 0.77 | 0.75 | 0.72 | 0.77 |
| 1 | 203055101 | rs4950877 | -0.238 | 0.316 | -0.38 | 1.413 | 0.002 | 0.003 | 0.003 | 0 | 0.48 | 0.52 | 0.62 | 0.53 | 0.54 | 0.41 |
| 1 | 203058842 | rs1038984 | -0.138 | -0.313 | -1.092 | 0.724 | -0.025 | 0.078 | 0.078 | 0 | 0.32 | 0.13 | 0.27 | 0.14 | 0.33 | 0.36 |
| 1 | 203059728 | rs6664108 | 0.413 | -1.007 | -0.668 | 0.658 | 0.022 | 0.04 | 0.04 | 0.315 | 0.59 | 0.69 | 0.71 | 0.78 | 0.67 | 0.5 |
| 1 | 203064086 | rs10920564 | -0.2 | -1.132 | -0.637 | 0.464 | 0.002 | 0.037 | 0.037 | 0 | 0.34 | 0.24 | 0.28 | 0.15 | 0.34 | 0.38 |
| 1 | 203080264 | rs11590405 | -0.172 | -1.211 | -0.287 | 0.318 | -0.003 | -0.023 | -0.023 | 0 | 0.36 | 0.36 | 0.3 | 0.15 | 0.34 | 0.37 |
| 1 | 203082811 | rs11582098 | -0.118 | -1.206 | -0.293 | 0.32 | -0.002 | -0.018 | -0.018 | 1.707 | 0.35 | 0.35 | 0.3 | 0.15 | 0.34 | 0.37 |
| 1 | 203090976 | rs7555556 | 0.229 | -1.223 | -0.498 | 0.342 | -0.045 | 0.094 | 0.094 | 0 | 0.3 | 0.08 | 0.25 | 0.12 | 0.34 | 0.37 |
| 1 | 203091274 | rs903361 | 0.462 | -1.326 | -0.424 | 0.763 | 0.011 | 0 | 0 | 0.259 | 0.6 | 0.65 | 0.71 | 0.62 | 0.66 | 0.48 |
| 1 | 203098818 | rs10920570 | 0.119 | -1.154 | -0.722 | 0.657 | -0.026 | 0.073 | 0.073 | 0.02 | 0.31 | 0.1 | 0.26 | 0.15 | 0.34 | 0.38 |
| 1 | 203101029 | rs3766568 | 0.166 | -1.193 | -0.559 | 0.622 | -0.026 | 0.088 | 0.088 | 0 | 0.31 | 0.12 | 0.25 | 0.12 | 0.34 | 0.37 |
| 1 | 203102632 | rs12123037 | 1.328 | -1.008 | -0.878 | 1.105 | -0.067 | -0.113 | -0.113 | 0.679 | 0.15 | 0.01 | 0.28 | 0.37 | 0.23 | 0.13 |
| 1 | 203102728 | rs6701725 | -0.757 | -1.437 | -0.538 | 0.903 | -0.001 | 0.006 | 0.006 | 0 | 0.58 | 0.61 | 0.73 | 0.77 | 0.61 | 0.6 |
| 1 | 203103428 | rs4950921 | -0.049 | -1.519 | -0.198 | 1.318 | 0 | 0.012 | 0.012 | 0 | 0.94 | 0.94 | 0.98 | 0.8 | 0.99 | 0.87 |
| 1 | 203104064 | rs6427993 | 0.199 | -1.314 | 0.409 | 1.003 | -0.015 | 0.089 | 0.089 | 1.644 | 0.1 | 0.37 | 0.19 | 0.2 | 0.04 | 0.07 |
| 1 | 203106390 | rs7549561 | -0.764 | -1.236 | 0.116 | 0.95 | 0.015 | 0.023 | 0.023 | 0.323 | 0.44 | 0.65 | 0.47 | 0.4 | 0.37 | 0.47 |
| 1 | 203108963 | rs7534251 | 0.163 | -0.828 | 0.117 | 1.02 | 0.001 | -0.002 | -0.002 | 0 | 0.08 | 0.12 | 0.05 | 0.05 | 0.08 | 0.18 |

| | | | | | | | | | | | | | | | | |
|---|---|---|---|---|---|---|---|---|---|---|---|---|---|---|---|---|
| 1 | 203111304 | rs10800901 | 0.116 | -1.055 | -0.518 | 0.952 | 0 | -0.001 | -0.001 | 0 | 0.67 | 0.66 | 0.63 | 0.37 | 0.69 | 0.64 |
| 1 | 203111695 | rs3766563 | -0.458 | -1.129 | -0.513 | 1.386 | -0.003 | 0.007 | 0.007 | 0.05 | 0.91 | 0.92 | 0.94 | 0.8 | 0.98 | 0.95 |
| 1 | 203115832 | rs6686206 | 0.662 | -1.154 | -0.523 | 1.11 | -0.024 | 0.022 | 0.022 | 0 | 0.72 | 0.39 | 0.67 | 0.6 | 0.84 | 0.8 |
| 1 | 203119491 | rs17511192 | 1.473 | -1.13 | -0.921 | 0.727 | -0.136 | 0.276 | 0.276 | 0.089 | 0.38 | 0.04 | 0.27 | 0.01 | 0.48 | 0.23 |
| 1 | 203123258 | rs6677137 | -0.342 | -1.264 | -0.71 | 0.782 | 0.001 | -0.032 | -0.032 | 0 | 0.4 | 0.51 | 0.49 | 0.2 | 0.4 | 0.31 |
| 1 | 203129179 | rs10920576 | 0.375 | -1.255 | -0.667 | 0.777 | -0.014 | -0.031 | -0.031 | 0 | 0.18 | 0.47 | 0.24 | 0.11 | 0.13 | 0.1 |
| 1 | 203130671 | rs17465037 | 1.79 | -0.905 | -1.26 | 0.279 | -0.091 | 0.227 | 0.227 | 1.259 | 0.33 | 0.04 | 0.25 | 0.03 | 0.41 | 0.17 |
| 1 | 203131533 | rs3753472 | 0.495 | -0.832 | -1.203 | 0.389 | -0.093 | 0.091 | 0.091 | 0 | 0.26 | 0.04 | 0.19 | 0.1 | 0.33 | 0.27 |
| 1 | 203132307 | rs3766555 | 1.122 | -0.831 | -1.202 | 0.389 | -0.064 | 0.084 | 0.084 | 0 | 0.21 | 0.04 | 0.18 | 0.06 | 0.27 | 0.2 |
| 1 | 203133493 | rs11587584 | 1.073 | -0.628 | -0.93 | 0.798 | -0.021 | 0.002 | 0.002 | 0.168 | 0.05 | 0.01 | 0.02 | 0.04 | 0.06 | 0.1 |
| 1 | 203133652 | rs16851020 | 0.574 | -0.807 | -0.772 | 0.792 | -0.009 | -0.01 | -0.01 | 0 | 0.17 | 0.42 | 0.24 | 0.14 | 0.13 | 0.1 |
| 1 | 203135674 | rs12744240 | 0.571 | -0.558 | -0.898 | 1.095 | -0.003 | 0.006 | 0.006 | 0.244 | 0.09 | 0.08 | 0.05 | 0.08 | 0.06 | 0.23 |
| 1 | 203138970 | rs2250509 | 0.594 | -0.543 | -0.995 | 1.117 | -0.012 | 0.06 | 0.06 | 0 | 0.83 | 0.77 | 0.74 | 0.41 | 0.89 | 0.81 |
| 1 | 203141689 | rs3766550 | 0.391 | -0.399 | -1.081 | 1.199 | 0.006 | 0.136 | 0.136 | 0.054 | 0.06 | 0.1 | 0.12 | 0.4 | 0.09 | 0.14 |
| 1 | 203143395 | rs3737872 | 1.533 | -0.61 | -1.279 | 0.851 | -0.094 | 0.149 | 0.149 | 0.015 | 0.32 | 0.08 | 0.26 | 0.11 | 0.42 | 0.22 |
| 1 | 203145301 | rs762625 | 1.148 | -0.793 | -0.516 | 1.246 | -0.018 | 0.008 | 0.008 | 0.52 | 0.75 | 0.65 | 0.68 | 0.32 | 0.82 | 0.53 |
| 1 | 203145665 | rs12139072 | -0.183 | -0.759 | -0.784 | 1.151 | -0.006 | 0.008 | 0.008 | 0 | 0.91 | 0.93 | 0.85 | 0.81 | 0.98 | 0.95 |
| 1 | 203147139 | rs903358 | 0.191 | -1.01 | -0.494 | 1.051 | -0.043 | 0.013 | 0.013 | 0 | 0.84 | 0.76 | 0.81 | 0.83 | 0.95 | 0.91 |
| 1 | 203151176 | rs7542294 | -0.242 | -0.828 | -0.459 | 1.469 | -0.001 | 0.086 | 0.086 | 0 | 0.22 | 0.54 | 0.34 | 0.44 | 0.16 | 0.24 |
| 1 | 203152801 | rs880633 | 1.358 | -0.562 | -1.163 | 1.015 | -0.041 | 0.015 | 0.015 | 0.075 | 0.43 | 0.17 | 0.33 | 0.38 | 0.54 | 0.48 |
| 1 | 203154532 | rs1538372 | 0.544 | -0.555 | -1.2 | 0.935 | 0.021 | 0 | 0 | 0 | 0.63 | 0.71 | 0.65 | 0.66 | 0.68 | 0.65 |
| 1 | 203155998 | rs10399805 | 0.181 | -0.554 | -0.431 | 1.214 | 0.001 | 0.09 | 0.09 | 0.065 | 0.19 | 0.44 | 0.3 | 0.29 | 0.13 | 0.16 |
| 1 | 203156080 | rs10399931 | 0.294 | -0.836 | -0.904 | 1.039 | -0.003 | 0.015 | 0.015 | 0 | 0.76 | 0.71 | 0.71 | 0.68 | 0.79 | 0.73 |
| 1 | 203157873 | rs946261 | 1.229 | -0.887 | -1.465 | 1.002 | 0.026 | 0.049 | 0.049 | 0.83 | 0.55 | 0.13 | 0.38 | 0.37 | 0.65 | 0.56 |
| 1 | 203158972 | rs10920579 | -0.521 | -0.618 | -1.166 | 1.267 | -0.029 | -0.002 | -0.002 | 0.713 | 0.9 | 0.94 | 0.88 | 0.92 | 0.82 | 0.85 |
| 1 | 203167682 | rs12562024 | 0.377 | -0.88 | -1.079 | 1.329 | 0.03 | -0.033 | -0.033 | 0 | 0.13 | 0.07 | 0.18 | 0.24 | 0.03 | 0.13 |
| 1 | 203168764 | rs2486064 | -0.193 | -0.632 | -1.76 | 1.061 | -0.117 | -0.116 | -0.116 | 0 | 0.68 | 0.98 | 0.76 | 0.62 | 0.62 | 0.63 |
| 1 | 203169391 | rs872129 | -0.952 | -1.277 | -0.775 | 1.341 | -0.03 | 0.038 | 0.038 | 0 | 0.23 | 0.3 | 0.33 | 0.51 | 0.09 | 0.26 |
| 1 | 203173356 | rs2486949 | -1.369 | -0.672 | -0.613 | 0.406 | -0.017 | 0.089 | 0.089 | 0 | 0.55 | 0.7 | 0.7 | 0.96 | 0.48 | 0.75 |
| 1 | 203173656 | rs12042412 | -0.029 | -0.787 | -0.484 | 0.746 | -0.022 | 0.128 | 0.128 | 0 | 0.11 | 0.33 | 0.13 | 0.25 | 0.04 | 0.06 |
| 1 | 203174186 | rs883389 | 0.345 | -0.611 | -0.418 | 0.936 | -0.029 | 0.014 | 0.014 | 0 | 0.19 | 0.12 | 0.26 | 0.14 | 0.28 | 0.24 |
| 1 | 203174864 | rs4950932 | -0.053 | -0.628 | -0.484 | 0.996 | -0.01 | -0.179 | -0.179 | 0.009 | 0.27 | 0.02 | 0.3 | 0.53 | 0.19 | 0.46 |
| 1 | 203175878 | rs2486062 | -1.429 | -0.586 | -0.378 | 0.67 | 0 | -0.063 | -0.063 | 0 | 0.53 | 0.49 | 0.68 | 0.96 | 0.48 | 0.75 |
| 1 | 203179943 | rs2494281 | -1.31 | -0.47 | -0.657 | 0.739 | -0.001 | 0.046 | 0.046 | 0 | 0.51 | 0.35 | 0.67 | 0.96 | 0.48 | 0.75 |
| 1 | 203181362 | rs12141375 | -0.139 | -0.457 | -0.783 | 0.923 | 0.002 | -0.142 | -0.142 | 0 | 0.27 | 0.03 | 0.3 | 0.57 | 0.19 | 0.47 |
| 1 | 203183093 | rs17466945 | 0.789 | 0.045 | -0.674 | 0.869 | -0.033 | 0.018 | 0.018 | 0 | 0.06 | 0.01 | 0.16 | 0.02 | 0.12 | 0.07 |
| 1 | 203188692 | rs2486958 | -1.648 | 0.121 | -0.502 | 0.824 | 0 | -0.056 | -0.056 | 0 | 0.54 | 0.54 | 0.71 | 0.93 | 0.5 | 0.75 |
| 1 | 203190240 | rs946258 | -0.529 | 0.115 | -0.478 | 0.992 | 0.011 | 0.009 | 0.009 | 0 | 0.25 | 0.28 | 0.38 | 0.36 | 0.33 | 0.27 |
| 1 | 203191866 | rs2486067 | -0.025 | 0.082 | -0.474 | 1.018 | 0.033 | -0.115 | -0.115 | 0.094 | 0.27 | 0.08 | 0.3 | 0.53 | 0.16 | 0.45 |
| 1 | 203191904 | rs2486961 | -0.341 | 0.08 | -0.461 | 1.017 | 0.015 | -0.015 | -0.015 | 0 | 0.29 | 0.25 | 0.32 | 0.58 | 0.16 | 0.48 |
| 1 | 203193261 | rs3766539 | 0.755 | 0.102 | -0.815 | 0.964 | 0.031 | -0.077 | -0.077 | 0.954 | 0.2 | 0.01 | 0.12 | 0.27 | 0.1 | 0.31 |
| 1 | 203193736 | rs2486069 | -1.45 | 0.343 | -0.601 | -0.047 | 0 | -0.02 | -0.02 | 1.294 | 0.53 | 0.5 | 0.69 | 0.89 | 0.49 | 0.72 |
| 1 | 203194186 | rs2297950 | -1.007 | 1.059 | -0.925 | 0.343 | 0.006 | 0.001 | 0.001 | 0 | 0.25 | 0.27 | 0.33 | 0.28 | 0.32 | 0.26 |
| 1 | 203195877 | rs3766537 | -0.184 | 0.953 | -0.944 | 0.326 | 0.009 | 0.013 | 0.013 | 0 | 0.19 | 0.24 | 0.17 | 0.26 | 0.2 | 0.19 |
| 1 | 203196281 | rs1340237 | -0.716 | 0.653 | -0.886 | 0.329 | 0.004 | 0.002 | 0.002 | 0 | 0.23 | 0.27 | 0.29 | 0.27 | 0.27 | 0.25 |
| 1 | 203197257 | rs2486072 | -0.46 | 0.472 | -0.846 | -0.321 | -0.003 | -0.027 | -0.027 | 0 | 0.49 | 0.47 | 0.55 | 0.88 | 0.36 | 0.66 |
| 1 | 203200196 | rs17533071 | -0.446 | 0.878 | -1.242 | -0.551 | -0.049 | 0.092 | 0.092 | 0.101 | 0.31 | 0.17 | 0.27 | 0.06 | 0.41 | 0.26 |
| 1 | 203207520 | rs2494287 | -0.097 | 0.547 | -1.323 | 0.298 | -0.008 | -0.011 | -0.011 | 0 | 0.91 | 0.96 | 0.7 | 0.7 | 0.91 | 0.82 |
| 1 | 203208696 | rs10920589 | -0.909 | -0.162 | -0.877 | 0.384 | 0.014 | 0.001 | 0.001 | 0 | 0.45 | 0.66 | 0.37 | 0.47 | 0.38 | 0.46 |
| 1 | 203208848 | rs10920590 | 0.376 | -0.211 | -0.911 | 0.387 | -0.043 | 0.106 | 0.106 | 0.228 | 0.27 | 0.09 | 0.22 | 0.04 | 0.33 | 0.24 |
| 1 | 203209481 | rs7547716 | 0.37 | -0.34 | -0.057 | 0.818 | -0.037 | 0.2 | 0.2 | 0.108 | 0.1 | 0.29 | 0.1 | 0.27 | 0.03 | 0.12 |
| 1 | 203218176 | rs2494295 | -0.858 | -0.084 | -0.335 | 0.391 | -0.009 | 0.043 | 0.043 | 0 | 0.42 | 0.66 | 0.34 | 0.21 | 0.44 | 0.36 |
| 1 | 203238568 | rs2486928 | 1.442 | -0.318 | 0.012 | 0.558 | -0.021 | 0.106 | 0.106 | 0 | 0.45 | 0.24 | 0.33 | 0.22 | 0.52 | 0.36 |
| 1 | 203238707 | rs6427997 | -0.072 | -0.265 | 0.054 | 0.664 | 0.001 | -0.017 | -0.017 | 0 | 0.32 | 0.39 | 0.26 | 0.24 | 0.33 | 0.31 |
| 1 | 203244113 | rs7551146 | 0.096 | -0.01 | 0.356 | 1.093 | -0.034 | 0.202 | 0.202 | 0.032 | 0.1 | 0.29 | 0.1 | 0.27 | 0.03 | 0.12 |
| 1 | 203245214 | rs2494302 | -1.313 | -0.109 | 0.476 | 1.074 | -0.074 | 0.173 | 0.173 | 3.672 | 0.19 | 0.58 | 0.15 | 0.45 | 0.05 | 0.22 |
| 1 | 203250860 | rs1934657 | -0.598 | -0.119 | 0.348 | 1.085 | -0.065 | 0.235 | 0.235 | 1.95 | 0.14 | 0.61 | 0.13 | 0.38 | 0.04 | 0.18 |
| 1 | 203253226 | rs871850 | 1.651 | -0.169 | 0.026 | 0.944 | -0.032 | 0.151 | 0.151 | 0 | 0.45 | 0.18 | 0.33 | 0.17 | 0.53 | 0.34 |
| 1 | 203254197 | rs2486939 | -0.468 | -0.192 | -0.025 | 0.945 | -0.01 | -0.018 | -0.018 | 0.012 | 0.91 | 0.97 | 0.7 | 0.65 | 0.91 | 0.78 |
| 1 | 203255818 | rs2494279 | -0.558 | -0.377 | 0.129 | 0.929 | -0.066 | 0.178 | 0.178 | 3.005 | 0.15 | 0.67 | 0.14 | 0.38 | 0.04 | 0.18 |
| 1 | 203256800 | rs4971226 | 0.021 | -0.334 | -0.044 | 0.928 | -0.074 | -0.027 | -0.027 | 0 | 0.64 | 0.87 | 0.49 | 0.6 | 0.58 | 0.56 |
| 1 | 203261445 | rs4483473 | 1.293 | -0.22 | -0.008 | 0.911 | -0.047 | 0.175 | 0.175 | 0 | 0.45 | 0.13 | 0.33 | 0.17 | 0.53 | 0.35 |
| 1 | 203268658 | rs12080252 | -1.139 | -0.397 | 0.139 | 1.321 | -0.067 | 0.201 | 0.201 | 2.607 | 0.15 | 0.62 | 0.14 | 0.38 | 0.04 | 0.16 |
| 1 | 203271930 | rs12136280 | 1.279 | -0.256 | -0.774 | 0.37 | -0.009 | 0.127 | 0.127 | 0 | 0.4 | 0.12 | 0.29 | 0.19 | 0.43 | 0.37 |
| 1 | 203273366 | rs4971234 | 0.337 | -0.501 | -0.364 | 0.633 | -0.015 | -0.008 | -0.008 | 0 | 0.81 | 0.75 | 0.9 | 0.88 | 0.86 | 0.86 |
| 1 | 203277053 | rs12085417 | 0.158 | -0.893 | 0.205 | 1.28 | -0.004 | 0.005 | 0.005 | 0 | 0.11 | 0.13 | 0.08 | 0.13 | 0.06 | 0.04 |
| 1 | 203288743 | rs7513835 | 0.026 | -0.942 | 0.104 | 1.227 | -0.029 | 0.007 | 0.007 | 0 | 0.88 | 0.85 | 0.97 | 0.87 | 0.95 | 0.89 |
| 1 | 203291579 | rs12406229 | -0.865 | -0.426 | 0.275 | 1.192 | 0.044 | -0.058 | -0.058 | 0 | 0.14 | 0.44 | 0.14 | 0.05 | 0.14 | 0.18 |
| 1 | 203291735 | rs10920610 | 0.08 | -0.908 | -0.186 | 1.361 | 0 | -0.002 | -0.002 | 0 | 0.11 | 0.11 | 0.08 | 0.13 | 0.06 | 0.04 |

| | | | | | | | | | | | | | | | | |
|---|---|---|---|---|---|---|---|---|---|---|---|---|---|---|---|---|
| 1 | 203295794 | rs884926 | -0.109 | -0.633 | -0.476 | 1.641 | -0.069 | -0.079 | -0.079 | 0.644 | 0.11 | 0.01 | 0.37 | 0.46 | 0.17 | 0.22 |
| 1 | 203300280 | rs16851314 | -1.023 | -0.802 | 0.04 | 1.421 | -0.008 | -0.002 | -0.002 | 0.447 | 0.23 | 0.33 | 0.19 | 0.17 | 0.18 | 0.21 |
| 1 | 203304679 | rs2050931 | -0.004 | -0.808 | -0.268 | 1.512 | 0.03 | -0.028 | -0.028 | 0 | 0.1 | 0.04 | 0.04 | 0.13 | 0.05 | 0.11 |
| 1 | 203308968 | rs12757745 | -0.548 | -0.6 | -0.204 | 1.524 | -0.038 | -0.07 | -0.07 | 0 | 0.16 | 0.06 | 0.39 | 0.5 | 0.18 | 0.27 |
| 1 | 203309752 | rs2886220 | -0.574 | -0.627 | -0.216 | 1.496 | -0.038 | -0.071 | -0.071 | 0 | 0.16 | 0.07 | 0.39 | 0.48 | 0.18 | 0.28 |
| 1 | 203309880 | rs7208 | -0.488 | -0.615 | -0.247 | 1.501 | -0.056 | -0.097 | -0.097 | 0 | 0.16 | 0.03 | 0.38 | 0.48 | 0.18 | 0.28 |
| 1 | 203312556 | rs3820224 | 0.724 | -1.065 | 0.258 | 1.27 | -0.018 | 0.01 | 0.01 | 0 | 0.85 | 0.77 | 0.92 | 0.84 | 0.89 | 0.84 |
| 1 | 203314594 | rs2105309 | 0.459 | -1.004 | 0.538 | 1.263 | -0.002 | 0.079 | 0.079 | 0 | 0.82 | 0.62 | 0.61 | 0.63 | 0.86 | 0.74 |
| 1 | 203315893 | rs10800913 | 0.431 | -1.31 | 0.567 | 1.269 | -0.01 | 0.03 | 0.03 | 0 | 0.84 | 0.77 | 0.85 | 0.72 | 0.9 | 0.82 |
| 1 | 203317162 | rs7543148 | 1.726 | -1.476 | 0.765 | 1.004 | 0.01 | -0.009 | -0.009 | 0 | 0.7 | 0.27 | 0.79 | 0.65 | 0.86 | 0.68 |
| 1 | 203317708 | rs12077300 | -0.674 | -1.104 | 0.901 | 0.599 | -0.007 | -0.02 | -0.02 | 0 | 0.08 | 0.22 | 0.04 | 0.03 | 0.04 | 0.08 |
| 1 | 203318375 | rs12093676 | -0.711 | -1.108 | 0.758 | 0.548 | -0.005 | -0.017 | -0.017 | 0 | 0.1 | 0.17 | 0.05 | 0.03 | 0.06 | 0.1 |
| 1 | 203319739 | rs10128007 | 0.049 | -1.087 | 0.834 | 0.515 | -0.001 | 0 | 0 | 0 | 0.09 | 0.13 | 0.08 | 0.1 | 0.1 | 0.12 |
| 1 | 203319816 | rs6697630 | 1.274 | -1.043 | 0.627 | 0.598 | -0.031 | -0.072 | -0.072 | 0 | 0.86 | 0.56 | 0.94 | 0.97 | 0.94 | 0.89 |
| 1 | 203328740 | rs10494841 | 0.398 | -0.812 | -0.277 | 1.209 | -0.034 | 0.116 | 0.116 | 0.805 | 0.85 | 0.73 | 0.87 | 0.58 | 0.94 | 0.86 |
| 1 | 203332622 | rs1977812 | 0.349 | -0.245 | -1.024 | 0.155 | -0.016 | -0.033 | -0.033 | 0.235 | 0.81 | 0.89 | 0.86 | 0.77 | 0.81 | 0.84 |
| 1 | 203333064 | rs6690853 | -0.6 | -0.961 | -0.447 | -0.164 | -0.055 | 0.019 | 0.019 | 0 | 0.29 | 0.47 | 0.45 | 0.66 | 0.18 | 0.38 |
| 1 | 203334114 | rs10800916 | 0.519 | -0.273 | -0.783 | -0.989 | 0.007 | 0.109 | 0.109 | 0 | 0.42 | 0.21 | 0.35 | 0.07 | 0.56 | 0.47 |
| 1 | 203334320 | rs10800917 | 0.155 | -0.118 | -0.816 | -0.914 | 0.002 | 0.06 | 0.06 | 0.263 | 0.44 | 0.6 | 0.35 | 0.08 | 0.55 | 0.38 |
| 1 | 203335702 | rs1891175 | 0.198 | -0.434 | -0.736 | -0.33 | 0.025 | 0.009 | 0.009 | 0 | 0.51 | 0.57 | 0.57 | 0.47 | 0.68 | 0.57 |
| 1 | 203336808 | rs10800919 | -0.838 | 0.492 | -0.714 | -0.29 | 0.001 | 0 | 0 | 0.007 | 0.38 | 0.39 | 0.46 | 0.44 | 0.49 | 0.45 |
| 1 | 203339985 | rs12139058 | -0.092 | -0.33 | -0.686 | -0.04 | 0.011 | 0.019 | 0.019 | 0 | 0.16 | 0.1 | 0.09 | 0.08 | 0.11 | 0.14 |
| 1 | 203340424 | rs1417926 | -0.533 | -0.345 | -0.207 | 0.347 | -0.013 | 0.035 | 0.035 | 0 | 0.24 | 0.26 | 0.4 | 0.42 | 0.18 | 0.26 |
| 1 | 203349524 | rs6428005 | -0.621 | -0.039 | -0.134 | -0.047 | -0.002 | -0.021 | -0.021 | 0 | 0.12 | 0.1 | 0.14 | 0.02 | 0.09 | 0.14 |
| 1 | 203351887 | rs7540920 | -1.5 | 0.493 | 0.564 | 0.513 | 0.014 | 0.04 | 0.04 | 0 | 0.23 | 0.53 | 0.33 | 0.29 | 0.29 | 0.26 |
| 1 | 203352061 | rs10920623 | -0.606 | 0.776 | 0.245 | 0.654 | 0.017 | 0.031 | 0.031 | 0 | 0.13 | 0.15 | 0.18 | 0.25 | 0.22 | 0.14 |
| 1 | 203365232 | rs10800925 | 0.897 | 0.585 | -0.109 | 0.745 | 0.033 | 0.083 | 0.083 | 0 | 0.5 | 0.24 | 0.51 | 0.31 | 0.54 | 0.39 |
| 1 | 203366863 | rs2185644 | 0.89 | 0.544 | 0.133 | 1.015 | 0.01 | 0.01 | 0.01 | 0 | 0.57 | 0.4 | 0.56 | 0.33 | 0.64 | 0.4 |
| 1 | 203367047 | rs10920631 | -0.494 | 1.316 | 0.097 | 1.436 | 0.01 | 0.045 | 0.045 | 0 | 0.08 | 0.1 | 0.14 | 0.24 | 0.13 | 0.11 |
| 1 | 203370193 | rs7521975 | -0.696 | 1.739 | 0.025 | 1.368 | -0.003 | -0.002 | -0.002 | 0 | 0.18 | 0.13 | 0.35 | 0.28 | 0.23 | 0.13 |
| 1 | 203371570 | rs879446 | 1.434 | 1.372 | -0.574 | 0.626 | -0.023 | 0.227 | 0.227 | 0.028 | 0.41 | 0.11 | 0.32 | 0.07 | 0.44 | 0.34 |
| 1 | 203372186 | rs1539144 | -0.906 | 1.435 | -0.554 | 1.031 | 0 | 0.004 | 0.004 | 0.035 | 0.26 | 0.26 | 0.48 | 0.5 | 0.32 | 0.26 |
| 1 | 203372485 | rs1539143 | -0.099 | 1.231 | 0.376 | 1.101 | -0.019 | 0.033 | 0.033 | 0 | 0.69 | 0.52 | 0.81 | 0.58 | 0.76 | 0.61 |
| 1 | 203374751 | rs16851527 | -0.538 | 1.395 | 0.601 | 1.317 | 0.036 | 0.064 | 0.064 | 0.298 | 0.09 | 0.19 | 0.13 | 0.23 | 0.1 | 0.14 |
| 1 | 203375246 | rs10920638 | -1.03 | 0.867 | 1.033 | 1.286 | 0.003 | 0.027 | 0.027 | 0 | 0.31 | 0.51 | 0.41 | 0.37 | 0.32 | 0.33 |
| 1 | 203378754 | rs1891181 | 1.434 | 0.77 | 1.339 | 1.22 | -0.023 | 0.028 | 0.028 | 0 | 0.68 | 0.51 | 0.6 | 0.37 | 0.76 | 0.56 |
| 1 | 203379696 | rs4971222 | -0.173 | 0.751 | 1.36 | 1.789 | 0.007 | 0.047 | 0.047 | 0 | 0.1 | 0.13 | 0.09 | 0.25 | 0.1 | 0.19 |
| 1 | 203388521 | rs12127756 | -1.779 | 0.732 | 1.484 | 1.661 | -0.001 | 0.001 | 0.001 | 0 | 0.31 | 0.36 | 0.39 | 0.61 | 0.24 | 0.44 |
| 1 | 203393566 | rs10920650 | 1.688 | 0.632 | 1.624 | 1.967 | -0.006 | 0.006 | 0.006 | 0 | 0.56 | 0.26 | 0.64 | 0.39 | 0.58 | 0.5 |
| 1 | 203395863 | rs10920651 | 2.075 | 0.828 | 1.39 | 1.934 | -0.001 | 0.156 | 0.156 | 0.316 | 0.49 | 0.13 | 0.41 | 0.19 | 0.49 | 0.41 |
| 1 | 203402652 | rs10920652 | 1.371 | 0.924 | 1.708 | 2.086 | 0.04 | 0.022 | 0.022 | 0.04 | 0.66 | 0.52 | 0.56 | 0.37 | 0.59 | 0.57 |
| 1 | 203402716 | rs4971229 | -0.665 | 0.943 | 1.752 | 2.14 | -0.037 | -0.05 | -0.05 | 0.737 | 0.23 | 0.14 | 0.18 | 0.42 | 0.3 | 0.29 |
| 1 | 203403117 | rs1417925 | -1.376 | 0.909 | 1.698 | 2.128 | 0.018 | 0.046 | 0.046 | 0.111 | 0.12 | 0.26 | 0.29 | 0.23 | 0.12 | 0.18 |
| 1 | 203403981 | rs12757606 | 2.272 | 0.734 | 1.421 | 1.919 | 0.021 | 0.106 | 0.106 | 0.325 | 0.51 | 0.21 | 0.42 | 0.2 | 0.48 | 0.41 |
| 1 | 203404635 | rs6681328 | -0.674 | 0.69 | 1.163 | 2.02 | -0.013 | 0.014 | 0.014 | 0 | 0.12 | 0.26 | 0.1 | 0.16 | 0.09 | 0.11 |
| 1 | 203411859 | rs999051 | -1.401 | 0.929 | 1.041 | 2.13 | 0.013 | 0.035 | 0.035 | 0 | 0.13 | 0.17 | 0.13 | 0.24 | 0.15 | 0.21 |
| 1 | 203430997 | rs2802791 | -0.495 | 0.796 | 1.36 | 2.329 | -0.016 | 0.064 | 0.064 | 0 | 0.07 | 0.17 | 0.21 | 0.21 | 0.04 | 0.1 |
| 1 | 203435718 | rs880878 | -2.387 | 0.718 | 1.61 | 2.322 | 0.007 | 0.117 | 0.117 | 0 | 0.22 | 0.39 | 0.34 | 0.45 | 0.2 | 0.31 |
| 1 | 203440733 | rs2802796 | -1.776 | 0.819 | 1.553 | 2.238 | -0.009 | 0.019 | 0.019 | 0 | 0.39 | 0.51 | 0.54 | 0.58 | 0.38 | 0.42 |
| 1 | 203444384 | rs2794448 | -1.239 | 0.996 | 1.518 | 2.268 | 0.006 | 0.045 | 0.045 | 0 | 0.33 | 0.49 | 0.5 | 0.56 | 0.37 | 0.38 |
| 1 | 203449615 | rs734591 | -1.765 | 0.959 | 1.415 | 2.154 | 0.071 | 0.244 | 0.244 | 0.74 | 0.16 | 0.47 | 0.31 | 0.44 | 0.19 | 0.28 |
| 1 | 203456221 | rs1166 | -0.289 | 1.24 | 0.891 | 2.218 | 0.067 | 0.119 | 0.119 | 0 | 0.06 | 0.15 | 0.09 | 0.24 | 0.14 | 0.17 |
| 1 | 203458407 | rs9475 | -0.384 | 1.252 | 0.889 | 2.393 | 0.006 | 0.001 | 0.001 | 0 | 0.25 | 0.31 | 0.29 | 0.31 | 0.32 | 0.27 |
| 1 | 203459718 | rs16851710 | -0.384 | 1.252 | 0.893 | 2.393 | 0.006 | 0.001 | 0.001 | 0 | 0.25 | 0.31 | 0.29 | 0.31 | 0.32 | 0.27 |
| 1 | 203460673 | rs16851713 | -1.878 | 1.014 | 0.327 | 2.403 | 0.017 | 0.042 | 0.042 | 0.008 | 0.11 | 0.16 | 0.13 | 0.26 | 0.15 | 0.22 |
| 1 | 203462917 | rs1078839 | -0.779 | 1.047 | 0.269 | 2.448 | 0.07 | 0.159 | 0.159 | 0 | 0.06 | 0.16 | 0.09 | 0.3 | 0.15 | 0.18 |
| 1 | 203474617 | rs4971243 | -1.685 | 0.988 | 0.261 | 2.354 | 0.016 | 0.046 | 0.046 | 0.006 | 0.12 | 0.16 | 0.13 | 0.31 | 0.15 | 0.22 |
| 1 | 203476659 | rs4246546 | -1.685 | 0.988 | 0.261 | 2.354 | 0.016 | 0.046 | 0.046 | 0.006 | 0.12 | 0.17 | 0.13 | 0.31 | 0.15 | 0.22 |
| 1 | 203478370 | rs3766902 | -2.298 | 0.941 | 0.154 | 2.559 | -0.002 | 0.015 | 0.015 | 0 | 0.13 | 0.13 | 0.15 | 0.42 | 0.14 | 0.28 |
| 1 | 203481101 | rs4971215 | -1.736 | 0.971 | 0.281 | 2.645 | 0.021 | 0.086 | 0.086 | 0 | 0.09 | 0.15 | 0.12 | 0.39 | 0.13 | 0.24 |
| 1 | 203485142 | rs13303252 | -0.366 | -0.706 | -0.33 | 2.315 | 0.037 | -0.028 | -0.028 | 0.509 | 0.29 | 0.2 | 0.3 | 0.36 | 0.15 | 0.36 |
| 1 | 203485284 | rs16851750 | -1.558 | -0.317 | -0.109 | 2.176 | -0.002 | 0.072 | 0.072 | 0 | 0.09 | 0.23 | 0.07 | 0.23 | 0.07 | 0.21 |
| 1 | 203488951 | rs4971240 | -1.553 | -0.37 | -0.096 | 2.18 | -0.001 | 0.094 | 0.094 | 0 | 0.09 | 0.24 | 0.06 | 0.24 | 0.06 | 0.22 |
| 1 | 203491392 | rs4971212 | -0.355 | -0.342 | -0.391 | 1.918 | -0.007 | 0.013 | 0.013 | 0.417 | 0.27 | 0.32 | 0.44 | 0.42 | 0.18 | 0.52 |
| 1 | 203492121 | rs6688116 | 0.23 | -0.236 | -0.865 | 1.317 | -0.001 | 0 | 0 | 0 | 0.61 | 0.57 | 0.72 | 0.63 | 0.59 | 0.66 |
| 1 | 203508174 | rs6670438 | -0.463 | -0.12 | -1.361 | 0.978 | -0.105 | -0.132 | -0.132 | 0 | 0.67 | 0.92 | 0.35 | 0.53 | 0.48 | 0.54 |
| 1 | 203514662 | rs2821225 | -0.686 | -0.351 | -0.263 | 0.476 | -0.003 | 0.012 | 0.012 | 0 | 0.1 | 0.31 | 0.07 | 0.08 | 0.03 | 0.05 |
| 1 | 203527111 | rs2821233 | 0.149 | -0.55 | -1.022 | 0.092 | -0.023 | -0.032 | -0.032 | 0.91 | 0.37 | 0.54 | 0.22 | 0.17 | 0.28 | 0.22 |
| 1 | 203527812 | rs2802774 | -0.595 | 0.118 | -1.883 | 0.554 | -0.149 | -0.136 | -0.136 | 0 | 0.65 | 0.92 | 0.36 | 0.49 | 0.51 | 0.52 |

| | | | | | | | | | | | | | | | | |
|---|---|---|---|---|---|---|---|---|---|---|---|---|---|---|---|---|
| 1 | 203538470 | rs2821218 | -0.08 | -0.538 | -0.811 | 0.073 | -0.039 | -0.068 | -0.068 | 0.226 | 0.36 | 0.44 | 0.23 | 0.16 | 0.28 | 0.23 |
| 1 | 203544404 | rs1541251 | -0.528 | -0.563 | -0.625 | 0.09 | 0.009 | 0.022 | 0.022 | 0.234 | 0.41 | 0.68 | 0.36 | 0.81 | 0.36 | 0.56 |
| 1 | 203554848 | rs2798625 | -1.003 | -0.783 | 0.215 | 1.007 | -0.026 | 0.152 | 0.152 | 0.011 | 0.14 | 0.39 | 0.07 | 0.33 | 0.09 | 0.26 |
| 1 | 203566485 | rs2798634 | -0.887 | -0.757 | 0.162 | 0.512 | 0.032 | 0.073 | 0.073 | 0 | 0.49 | 0.63 | 0.53 | 0.88 | 0.43 | 0.66 |
| 1 | 203571766 | rs2802782 | -1.08 | -0.677 | -0.339 | 0.977 | -0.132 | 0.249 | 0.249 | 0.258 | 0.56 | 0.97 | 0.58 | 0.91 | 0.48 | 0.71 |
| 1 | 203575667 | rs4951272 | -0.028 | -0.977 | -0.263 | 1.552 | -0.01 | -0.02 | -0.02 | 0 | 0.34 | 0.06 | 0.27 | 0.27 | 0.33 | 0.39 |
| 1 | 203583705 | rs3900967 | 0.147 | -0.968 | 0.713 | 1.448 | -0.017 | 0.307 | 0.307 | 0 | 0.9 | 0.62 | 0.8 | 0.59 | 0.96 | 0.77 |
| 1 | 203584850 | rs6593903 | 0.772 | -0.945 | 0.775 | 1.435 | 0.017 | 0.273 | 0.273 | 0.712 | 0.87 | 0.49 | 0.77 | 0.59 | 0.9 | 0.77 |
| 1 | 203587538 | rs12120868 | 0.874 | -0.918 | 0.799 | 1.312 | 0.021 | 0.102 | 0.102 | 0 | 0.51 | 0.29 | 0.47 | 0.17 | 0.56 | 0.37 |
| 1 | 203595146 | rs4245719 | 0.059 | -0.803 | 0.457 | 1.604 | -0.012 | -0.068 | -0.068 | 0 | 0.28 | 0.06 | 0.24 | 0.37 | 0.27 | 0.35 |
| 1 | 203596318 | rs4600103 | -0.087 | -0.779 | 0.777 | 1.379 | -0.045 | 0.133 | 0.133 | 0 | 0.8 | 0.61 | 0.74 | 0.57 | 0.89 | 0.75 |
| 1 | 203600654 | rs12749310 | 0.717 | -0.735 | 0.876 | 1.309 | 0.024 | 0.032 | 0.032 | 0 | 0.51 | 0.38 | 0.47 | 0.25 | 0.64 | 0.4 |
| 1 | 203605690 | rs10793746 | 0.014 | -0.36 | 0.566 | 1.163 | -0.001 | 0.006 | 0.006 | 0 | 0.66 | 0.6 | 0.69 | 0.58 | 0.66 | 0.62 |
| 1 | 203608872 | rs10793748 | 0.171 | -0.41 | 0.577 | 1.165 | -0.003 | 0.007 | 0.007 | 0 | 0.65 | 0.54 | 0.67 | 0.59 | 0.66 | 0.62 |
| 1 | 203634403 | rs7520864 | 1.25 | -0.165 | 0.848 | 1.082 | 0.05 | 0.135 | 0.135 | 1.116 | 0.95 | 0.67 | 0.96 | 0.9 | 0.98 | 0.95 |
| 1 | 203635699 | rs10900561 | 1.813 | -0.193 | 0.984 | 1.007 | 0.005 | 0.005 | 0.005 | 2.117 | 0.88 | 0.46 | 0.77 | 0.88 | 0.89 | 0.89 |
| 1 | 203636551 | rs12568960 | 1.879 | -0.268 | 0.799 | 1.416 | -0.017 | 0.066 | 0.066 | 3.254 | 0.85 | 0.35 | 0.69 | 0.72 | 0.88 | 0.79 |
| 1 | 203636660 | rs4297354 | 0.461 | -0.136 | -0.143 | 0.374 | -0.158 | 0.103 | 0.103 | 0 | 0.39 | 0.03 | 0.39 | 0.17 | 0.51 | 0.31 |
| 1 | 203636917 | rs12066685 | -1.435 | 0.102 | 1.217 | 0.847 | 0.031 | -0.009 | -0.009 | 0 | 0.07 | 0.22 | 0.18 | 0.02 | 0.08 | 0.06 |
| 1 | 203642721 | rs4073264 | 0.192 | 0.185 | -0.413 | 0.799 | 0.051 | -0.009 | -0.009 | 0 | 0.46 | 0.35 | 0.38 | 0.46 | 0.28 | 0.59 |
| 1 | 203643053 | rs11576343 | 0.148 | 1.033 | -0.116 | 0.996 | 0.084 | 0.149 | 0.149 | 0 | 0.85 | 0.54 | 0.83 | 0.71 | 0.82 | 0.83 |
| 1 | 203644549 | rs11580986 | 0.204 | 1.324 | -0.364 | 0.826 | -0.01 | 0.045 | 0.045 | 0 | 0.4 | 0.33 | 0.45 | 0.26 | 0.5 | 0.26 |
| 1 | 203644628 | rs4951357 | 0.357 | 1.38 | 0.519 | 1.302 | 0.157 | 0.262 | 0.262 | 0 | 0.88 | 0.59 | 0.83 | 0.68 | 0.79 | 0.83 |
| 1 | 203651824 | rs11240734 | 0.037 | 0.374 | 0.195 | 0.466 | -0.027 | -0.116 | -0.116 | 0.001 | 0.84 | 0.63 | 0.94 | 1 | 0.9 | 0.88 |
| 1 | 203651927 | rs1541252 | 0.037 | 0.374 | 0.195 | 0.466 | -0.027 | -0.116 | -0.116 | 0.287 | 0.84 | 0.63 | 0.94 | 1 | 0.9 | 0.88 |
| 1 | 203652444 | rs1419114 | 0.037 | 0.374 | 0.195 | 0.466 | -0.027 | -0.116 | -0.116 | 3.531 | 0.84 | 0.64 | 0.94 | 1 | 0.9 | 0.88 |
| 1 | 203656814 | rs10900588 | 0.037 | 0.374 | 0.195 | 0.466 | -0.027 | -0.116 | -0.116 | 0 | 0.84 | 0.63 | 0.94 | 1 | 0.9 | 0.87 |
| 1 | 203658486 | rs4951073 | 0.922 | 0.238 | -0.151 | 0.799 | -0.007 | -0.006 | -0.006 | 0.19 | 0.65 | 0.39 | 0.79 | 0.81 | 0.71 | 0.59 |
| 1 | 203659355 | rs7529482 | 0.17 | -0.306 | -0.619 | 0.396 | 0.033 | -0.004 | -0.004 | 0 | 0.58 | 0.33 | 0.49 | 0.36 | 0.71 | 0.52 |
| 1 | 203670179 | rs3766752 | 0.21 | -0.181 | -0.12 | 1.157 | -0.008 | 0.026 | 0.026 | 0 | 0.26 | 0.31 | 0.45 | 0.6 | 0.19 | 0.34 |
| 1 | 203671877 | rs4951081 | -1.127 | -0.152 | -0.156 | 1.106 | 0.005 | 0.004 | 0.004 | 0 | 0.45 | 0.63 | 0.6 | 0.82 | 0.37 | 0.63 |
| 1 | 203675105 | rs16852191 | 0.439 | -0.559 | 0.062 | 1.86 | -0.043 | -0.094 | -0.094 | 0.441 | 0.16 | 0.01 | 0.13 | 0.21 | 0.18 | 0.29 |
| 1 | 203679934 | rs2032018 | -1.403 | -0.462 | -0.083 | 1.725 | 0.004 | 0.048 | 0.048 | 0.282 | 0.55 | 0.69 | 0.67 | 0.8 | 0.46 | 0.65 |
| 1 | 203680492 | rs6655965 | -1.19 | -0.424 | -0.007 | 1.716 | 0.019 | 0.066 | 0.066 | 0.398 | 0.52 | 0.67 | 0.65 | 0.83 | 0.43 | 0.65 |
| 1 | 203689314 | rs11808688 | -0.173 | -0.282 | -0.154 | 1.855 | -0.007 | 0.058 | 0.058 | 0 | 0.2 | 0.33 | 0.19 | 0.53 | 0.18 | 0.39 |
| 1 | 203692179 | rs10494845 | -0.057 | -0.09 | -0.194 | 1.085 | 0.008 | 0.009 | 0.009 | 0 | 0.29 | 0.23 | 0.44 | 0.22 | 0.24 | 0.23 |
| 1 | 203695836 | rs4951130 | -0.381 | 0.414 | -0.076 | 1.497 | -0.001 | 0.018 | 0.018 | 0 | 0.18 | 0.28 | 0.18 | 0.28 | 0.18 | 0.35 |
| 1 | 203699716 | rs12410036 | -0.033 | 0.54 | -0.618 | 1.249 | -0.004 | -0.026 | -0.026 | 0 | 0.17 | 0.14 | 0.13 | 0.22 | 0.17 | 0.3 |
| 1 | 203709034 | rs955865 | -1.25 | 0.489 | -0.719 | 0.799 | 0.004 | -0.003 | -0.003 | 0 | 0.53 | 0.67 | 0.73 | 0.5 | 0.58 | 0.54 |
| 1 | 203713133 | rs568 | -0.819 | -0.204 | -0.925 | 0.565 | 0.009 | 0.013 | 0.013 | 1.802 | 0.26 | 0.23 | 0.43 | 0.18 | 0.22 | 0.2 |
| 1 | 203719848 | rs11240347 | -0.566 | -0.212 | -0.927 | 0.774 | -0.003 | 0 | 0 | 1.744 | 0.2 | 0.23 | 0.42 | 0.24 | 0.14 | 0.16 |
| 1 | 203733579 | rs11579602 | -0.605 | -0.184 | -1.093 | 0.792 | 0.015 | -0.002 | -0.002 | 0 | 0.23 | 0.14 | 0.13 | 0.2 | 0.2 | 0.28 |
| 1 | 203734985 | rs7540760 | -0.79 | -0.518 | -0.8 | 0.735 | 0.013 | 0.002 | 0.002 | 0 | 0.25 | 0.21 | 0.16 | 0.21 | 0.21 | 0.3 |
| 1 | 203738765 | rs4951180 | 0.794 | -0.691 | -0.221 | 0.258 | -0.006 | -0.008 | -0.008 | 0 | 0.65 | 0.58 | 0.83 | 0.76 | 0.65 | 0.64 |
| 1 | 203738781 | rs4951181 | 1.356 | -1.194 | 0.285 | 0.656 | 0.048 | 0.11 | 0.11 | 0 | 0.53 | 0.3 | 0.45 | 0.14 | 0.59 | 0.41 |
| 1 | 203740615 | rs3737976 | 1.41 | -1.319 | 0.375 | 0.733 | 0.045 | 0.097 | 0.097 | 0.064 | 0.52 | 0.32 | 0.46 | 0.1 | 0.6 | 0.41 |
| 1 | 203743122 | rs3820194 | -0.691 | -1.381 | 0.758 | 0.849 | 0 | 0 | 0 | 0 | 0.32 | 0.32 | 0.17 | 0.23 | 0.33 | 0.37 |
| 1 | 203743653 | rs3766748 | -0.691 | -1.381 | 0.726 | 0.849 | 0.002 | 0.004 | 0.004 | 0 | 0.32 | 0.29 | 0.16 | 0.23 | 0.33 | 0.37 |
| 1 | 203747752 | rs2185080 | -0.675 | -1.457 | 0.782 | 0.752 | 0.001 | 0.002 | 0.002 | 0 | 0.33 | 0.3 | 0.17 | 0.23 | 0.34 | 0.37 |
| 1 | 203749270 | rs9787160 | -1.476 | -1.655 | 1.045 | 0.741 | -0.001 | -0.012 | -0.012 | 0 | 0.47 | 0.49 | 0.54 | 0.86 | 0.41 | 0.6 |
| 1 | 203753454 | rs6593939 | 0.06 | -1.447 | 0.927 | 1.093 | 0.002 | 0.029 | 0.029 | 0.497 | 0.12 | 0.11 | 0.36 | 0.63 | 0.07 | 0.22 |
| 1 | 203762306 | rs6593945 | 0.835 | -0.716 | 1.122 | 0.888 | -0.003 | 0.091 | 0.091 | 0.505 | 0.81 | 0.63 | 0.65 | 0.33 | 0.84 | 0.71 |
| 1 | 203766331 | rs4951011 | -0.215 | -0.783 | 1.051 | 1.679 | -0.022 | -0.063 | -0.063 | 0 | 0.14 | 0.05 | 0.33 | 0.31 | 0.15 | 0.23 |
| 1 | 203766395 | rs6693481 | 1.497 | -0.633 | 1.027 | 1.555 | -0.001 | 0.001 | 0.001 | 0 | 0.71 | 0.59 | 0.6 | 0.29 | 0.72 | 0.64 |
| 1 | 203766827 | rs872032 | 1.621 | -1.068 | 1.036 | 1.273 | 0.001 | -0.002 | -0.002 | 0 | 0.46 | 0.4 | 0.41 | 0.06 | 0.5 | 0.38 |
| 1 | 203768328 | rs7552670 | 0.504 | -0.964 | 1.214 | 1.344 | -0.003 | -0.047 | -0.047 | 0 | 0.12 | 0.05 | 0.32 | 0.29 | 0.12 | 0.23 |
| 1 | 203769532 | rs7540041 | 0.504 | -0.964 | 1.221 | 1.344 | -0.001 | -0.039 | -0.039 | 0 | 0.12 | 0.05 | 0.32 | 0.29 | 0.12 | 0.23 |
| 1 | 203769891 | rs10494844 | 0.504 | -0.964 | 1.204 | 1.344 | -0.003 | -0.047 | -0.047 | 0.22 | 0.12 | 0.05 | 0.32 | 0.29 | 0.12 | 0.23 |
| 1 | 203777741 | rs12145049 | 0.854 | -0.305 | 1.429 | 0.991 | 0.053 | -0.038 | -0.038 | 0 | 0.08 | 0.16 | 0.16 | 0.01 | 0.16 | 0.04 |
| 1 | 203784993 | rs4951022 | -1.144 | -0.489 | 1.354 | 1.205 | -0.002 | -0.006 | -0.006 | 0 | 0.24 | 0.32 | 0.39 | 0.78 | 0.26 | 0.38 |
| 1 | 203800426 | rs12069734 | 0.53 | -0.471 | 1.23 | 1.192 | -0.011 | 0.011 | 0.011 | 0 | 0.1 | 0.05 | 0.05 | 0.04 | 0.12 | 0.09 |
| 1 | 203819440 | rs10494847 | 0.634 | -0.504 | 1.253 | 1.276 | -0.003 | -0.039 | -0.039 | 0 | 0.12 | 0.05 | 0.32 | 0.29 | 0.12 | 0.23 |
| 1 | 203829556 | rs4951031 | -0.128 | -0.5 | 1.257 | 1.026 | 0.003 | 0.003 | 0.003 | 0 | 0.7 | 0.7 | 0.79 | 0.83 | 0.76 | 0.77 |
| 1 | 203831261 | rs11240609 | 0.489 | -0.573 | 1.36 | 0.998 | 0.001 | -0.008 | -0.008 | 0.26 | 0.1 | 0.13 | 0.05 | 0.04 | 0.11 | 0.08 |
| 1 | 203837577 | rs6696091 | 0.68 | -0.52 | 1.352 | 0.994 | 0.009 | 0.013 | 0.013 | 0 | 0.6 | 0.63 | 0.74 | 0.8 | 0.65 | 0.69 |
| 1 | 203840037 | rs4598553 | -0.138 | -0.507 | 1.346 | 0.994 | 0.003 | 0.003 | 0.003 | 0 | 0.7 | 0.7 | 0.79 | 0.83 | 0.76 | 0.77 |
| 1 | 203850090 | rs6702219 | 1.234 | -0.544 | 1.341 | 1.061 | 0.002 | -0.001 | -0.001 | 1.703 | 0.48 | 0.52 | 0.41 | 0.55 | 0.53 | 0.45 |
| 1 | 203871055 | rs2796375 | -1.12 | -0.51 | 1.152 | 0.88 | 0.004 | -0.021 | -0.021 | 0 | 0.05 | 0.11 | 0.03 | 0 | 0.07 | 0.02 |
| 1 | 203877662 | rs1891432 | -0.745 | -0.473 | 1.055 | 0.99 | 0.03 | -0.032 | -0.032 | 0.205 | 0.39 | 0.12 | 0.52 | 0.41 | 0.36 | 0.47 |

| | | | | | | | | | | | | | | | | |
|---|---|---|---|---|---|---|---|---|---|---|---|---|---|---|---|---|
| 1 | 203884915 | rs11584733 | -0.881 | -0.489 | 1.103 | 0.982 | 0.006 | -0.036 | -0.036 | 0.067 | 0.42 | 0.18 | 0.54 | 0.44 | 0.43 | 0.5 |
| 1 | 203905103 | rs16852507 | 0.043 | -0.607 | 1.172 | 1.258 | 0.003 | -0.004 | -0.004 | 0.507 | 0.14 | 0.11 | 0.32 | 0.34 | 0.13 | 0.21 |
| 1 | 203907135 | rs7545327 | -1.467 | -0.65 | 1.323 | 1.228 | -0.018 | 0.093 | 0.093 | 3.272 | 0.23 | 0.47 | 0.38 | 0.34 | 0.2 | 0.24 |
| 1 | 203910220 | rs1935588 | 0.183 | -0.744 | 1.294 | 1.168 | 0 | 0 | 0 | 0.154 | 0.57 | 0.58 | 0.62 | 0.54 | 0.56 | 0.61 |
| 1 | 203911557 | rs2366257 | 0.559 | -0.787 | 1.135 | 1.16 | 0.012 | 0.017 | 0.017 | 0.138 | 0.5 | 0.41 | 0.58 | 0.54 | 0.48 | 0.58 |
| 1 | 203913823 | rs12410049 | -0.394 | -0.604 | 1.385 | 1.433 | -0.013 | 0.067 | 0.067 | 0.021 | 0.16 | 0.29 | 0.36 | 0.34 | 0.13 | 0.21 |
| 1 | 203915958 | rs11240629 | 1.53 | -0.831 | 1.18 | 1.168 | -0.04 | 0.176 | 0.176 | 0.239 | 0.47 | 0.15 | 0.26 | 0.2 | 0.46 | 0.4 |
| 1 | 203918185 | rs9438454 | -0.025 | -0.8 | 1.384 | 1.248 | 0.001 | 0.018 | 0.018 | 0 | 0.69 | 0.59 | 0.66 | 0.57 | 0.66 | 0.64 |
| 1 | 203945059 | rs13374988 | 1.274 | -0.636 | 0.524 | 1.056 | -0.027 | 0.173 | 0.173 | 0.074 | 0.48 | 0.2 | 0.26 | 0.2 | 0.46 | 0.4 |
| 1 | 203948660 | rs12408223 | -0.63 | -0.477 | 1.039 | 1.798 | 0.002 | 0.011 | 0.011 | 0 | 0.14 | 0.12 | 0.34 | 0.39 | 0.13 | 0.23 |
| 1 | 203955957 | rs2796421 | -1.931 | -0.237 | 1.458 | 1.433 | 0.024 | 0.068 | 0.068 | 5.057 | 0.06 | 0.35 | 0.03 | 0.05 | 0.01 | 0.05 |
| 1 | 203963678 | rs11588335 | 0.499 | 0.059 | 0.893 | 1.317 | -0.024 | -0.004 | -0.004 | 0.068 | 0.43 | 0.29 | 0.24 | 0.44 | 0.46 | 0.5 |
| 1 | 203967091 | rs6674546 | -0.439 | -0.015 | 0.164 | 0.487 | -0.026 | 0.172 | 0.172 | 1.646 | 0.6 | 0.89 | 0.63 | 0.84 | 0.58 | 0.73 |
| 1 | 203972021 | rs2796424 | 0.297 | 0.178 | 0.024 | 0.633 | 0 | 0.001 | 0.001 | 2.251 | 0.45 | 0.55 | 0.26 | 0.64 | 0.4 | 0.55 |
| 1 | 203972175 | rs12120627 | -0.318 | 0.574 | -0.319 | 0.893 | 0.018 | 0.063 | 0.063 | 0 | 0.15 | 0.28 | 0.37 | 0.23 | 0.19 | 0.18 |
| 1 | 203972649 | rs7556505 | 0.235 | 0.348 | -0.962 | 1.113 | -0.015 | 0.009 | 0.009 | 3.094 | 0.4 | 0.23 | 0.22 | 0.64 | 0.39 | 0.55 |
| 1 | 203979519 | rs11240647 | -0.067 | 0.504 | -0.945 | 0.938 | -0.006 | 0.006 | 0.006 | 0 | 0.54 | 0.38 | 0.54 | 0.44 | 0.49 | 0.63 |
| 1 | 203982102 | rs2810616 | 0.355 | 0.252 | -1.401 | 1.329 | 0.005 | -0.112 | -0.112 | 0.044 | 0.83 | 0.97 | 0.78 | 0.54 | 0.88 | 0.78 |
| 1 | 203982333 | rs2796433 | 1.268 | 0.465 | -0.672 | 0.596 | -0.003 | 0.022 | 0.022 | 0 | 0.66 | 0.46 | 0.63 | 0.52 | 0.67 | 0.73 |
| 1 | 203982534 | rs2810615 | 0.11 | -0.018 | -0.853 | -0.45 | -0.02 | -0.065 | -0.065 | 0 | 0.17 | 0.25 | 0.08 | 0 | 0.1 | 0.11 |
| 1 | 203985706 | rs2796435 | -1.137 | -0.069 | 0.111 | -0.48 | 0.002 | -0.122 | -0.122 | 0 | 0.2 | 0.47 | 0.15 | 0.02 | 0.2 | 0.03 |
| 1 | 203997926 | rs10900541 | 1.644 | 0.53 | 0.072 | -0.364 | -0.008 | 0.014 | 0.014 | 0.149 | 0.72 | 0.24 | 0.69 | 0.82 | 0.69 | 0.85 |
| 1 | 204000843 | rs7525696 | 1.35 | 0.446 | 0.598 | -0.273 | 0 | -0.03 | -0.03 | 0 | 0.77 | 0.38 | 0.73 | 0.87 | 0.77 | 0.86 |
| 1 | 204005018 | rs4301679 | -1.176 | 0.763 | 0.571 | -0.17 | -0.005 | -0.033 | -0.033 | 0 | 0.33 | 0.41 | 0.34 | 0.23 | 0.31 | 0.28 |
| 1 | 204006597 | rs4951039 | -0.099 | 1.099 | 0.192 | 0.457 | -0.016 | -0.013 | -0.013 | 0 | 0.92 | 0.94 | 0.94 | 0.76 | 0.87 | 0.82 |
| 1 | 204011313 | rs2366009 | -0.417 | 0.969 | -0.262 | 0.366 | -0.045 | -0.054 | -0.054 | 0.008 | 0.91 | 0.99 | 0.88 | 0.8 | 0.86 | 0.81 |
| 1 | 204014607 | rs12133634 | 0.106 | 0.748 | -0.484 | 0.058 | -0.019 | 0.021 | 0.021 | 0.626 | 0.17 | 0.05 | 0.12 | 0.12 | 0.18 | 0.1 |
| 1 | 204020481 | rs11240655 | 0.772 | 0.958 | 0.22 | 0.207 | 0.06 | 0.085 | 0.085 | 0.026 | 0.8 | 0.47 | 0.77 | 0.75 | 0.74 | 0.79 |
| 1 | 204028177 | rs6660981 | 0.146 | 0.57 | -0.38 | -0.057 | -0.001 | 0.037 | 0.037 | 0 | 0.54 | 0.28 | 0.46 | 0.32 | 0.45 | 0.57 |
| 1 | 204030721 | rs3911890 | -0.23 | 0.479 | 0.089 | 0.155 | 0.068 | 0.103 | 0.103 | 0 | 0.84 | 0.65 | 0.79 | 0.73 | 0.74 | 0.8 |
| 1 | 204031363 | rs6673836 | 1.109 | 0.323 | -0.787 | -0.056 | 0.021 | 0.046 | 0.046 | 0 | 0.33 | 0.1 | 0.32 | 0.22 | 0.26 | 0.48 |
| 1 | 204041346 | rs11240657 | 0.107 | 0.221 | -0.765 | 0.089 | -0.002 | -0.032 | -0.032 | 0 | 0.83 | 0.91 | 0.91 | 0.72 | 0.85 | 0.79 |
| 1 | 204044566 | rs4951300 | 0.091 | 0.224 | -0.764 | 0.099 | -0.002 | -0.032 | -0.032 | 0.02 | 0.83 | 0.91 | 0.91 | 0.72 | 0.85 | 0.79 |
| 1 | 204045170 | rs6683103 | 1.454 | 0.191 | -0.692 | 0.008 | -0.011 | -0.014 | -0.014 | 0.648 | 0.05 | 0.06 | 0.02 | 0 | 0.02 | 0.01 |
| 1 | 204045948 | rs12023823 | 1.67 | 0.392 | -0.79 | 0.097 | 0.005 | -0.011 | -0.011 | 0 | 0.19 | 0.14 | 0.24 | 0.36 | 0.17 | 0.14 |
| 1 | 204049354 | rs4951302 | 1.342 | 0.532 | -1.08 | -0.122 | 0.001 | 0.058 | 0.058 | 0.029 | 0.31 | 0.11 | 0.39 | 0.2 | 0.28 | 0.52 |
| 1 | 204050926 | rs12122693 | -0.329 | 0.792 | -0.381 | 0.057 | 0.022 | -0.012 | -0.012 | 0.001 | 0.14 | 0.32 | 0.1 | 0.09 | 0.16 | 0.08 |
| 1 | 204052408 | rs1554678 | -1.475 | 0.963 | -0.776 | 0.41 | -0.041 | -0.033 | -0.033 | 0 | 0.93 | 0.99 | 0.94 | 0.79 | 0.87 | 0.84 |
| 1 | 204053090 | rs940657 | 0.839 | 0.763 | -0.805 | 0.181 | -0.003 | -0.002 | -0.002 | 0 | 0.54 | 0.3 | 0.71 | 0.61 | 0.57 | 0.67 |
| 1 | 204054323 | rs7550796 | 0.222 | 0.664 | -0.577 | 0.056 | -0.001 | -0.001 | -0.001 | 0 | 0.25 | 0.2 | 0.11 | 0.14 | 0.17 | 0.13 |
| 1 | 204054521 | rs11577055 | 1.177 | 0.834 | -0.801 | 0.566 | -0.017 | 0.012 | 0.012 | 0 | 0.29 | 0.15 | 0.41 | 0.24 | 0.31 | 0.52 |
| 1 | 204057993 | rs12239182 | 0.73 | 0.53 | -0.298 | 0.723 | 0.001 | 0.002 | 0.002 | 0 | 0.31 | 0.26 | 0.17 | 0.14 | 0.27 | 0.14 |
| 1 | 204058458 | rs11240662 | 0.045 | 0.787 | -0.319 | 0.867 | -0.001 | 0.003 | 0.003 | 0.085 | 0.56 | 0.58 | 0.77 | 0.65 | 0.59 | 0.62 |
| 1 | 204062992 | rs6593993 | -0.596 | 0.699 | -0.37 | 1.158 | -0.01 | -0.006 | -0.006 | 1.045 | 0.58 | 0.67 | 0.47 | 0.59 | 0.53 | 0.35 |
| 1 | 204068110 | rs16852826 | 0.511 | 0.705 | -0.495 | 1.021 | 0 | -0.019 | -0.019 | 0.208 | 0.39 | 0.38 | 0.24 | 0.12 | 0.36 | 0.18 |
| 1 | 204069838 | rs3795579 | 1.069 | 0.609 | -0.51 | 1.058 | 0 | 0 | 0 | 0 | 0.35 | 0.33 | 0.47 | 0.26 | 0.33 | 0.47 |
| 1 | 204070007 | rs1534098 | 0.319 | 0.798 | -0.228 | 2.119 | -0.051 | -0.057 | -0.057 | 0 | 0.91 | 0.99 | 0.94 | 0.81 | 0.86 | 0.79 |
| 1 | 204070224 | rs11240666 | 1.236 | 0.831 | -0.077 | 1.829 | 0.003 | 0.075 | 0.075 | 0 | 0.73 | 0.53 | 0.72 | 0.43 | 0.74 | 0.67 |
| 1 | 204074283 | rs11240668 | -1.054 | 0.802 | -0.317 | 1.956 | -0.003 | 0.012 | 0.012 | 0 | 0.12 | 0.1 | 0.21 | 0.44 | 0.16 | 0.13 |
| 1 | 204074532 | rs12563446 | -1.919 | 0.317 | 0.493 | 1.801 | -0.047 | 0.057 | 0.057 | 1.257 | 0.15 | 0.65 | 0.16 | 0.25 | 0.08 | 0.13 |
| 1 | 204075303 | rs4951304 | 0.955 | 0.68 | 0.077 | 1.23 | -0.007 | 0.007 | 0.007 | 0 | 0.67 | 0.77 | 0.67 | 0.27 | 0.59 | 0.67 |
| 1 | 204076846 | rs6689899 | -0.248 | 0.78 | -0.241 | 1.537 | -0.064 | -0.056 | -0.056 | 0 | 0.86 | 0.98 | 0.89 | 0.79 | 0.79 | 0.8 |
| 1 | 204079268 | rs7549807 | 1.894 | 0.355 | -0.672 | 0.227 | -0.076 | 0.083 | 0.083 | 1.105 | 0.42 | 0.04 | 0.52 | 0.25 | 0.41 | 0.52 |
| 1 | 204081196 | rs940654 | 0.89 | 0.376 | 0.503 | 0.643 | 0.012 | 0.03 | 0.03 | 0.994 | 0.92 | 0.53 | 0.95 | 0.97 | 0.98 | 0.93 |
| 1 | 204081892 | rs2250538 | -0.862 | 0.518 | 0.47 | 0.762 | -0.008 | -0.011 | -0.011 | 1.873 | 0.27 | 0.22 | 0.3 | 0.58 | 0.37 | 0.28 |
| 1 | 204082938 | rs2037679 | 1.016 | 0.077 | 0.591 | 0.67 | -0.02 | 0.02 | 0.02 | 0 | 0.87 | 0.58 | 0.92 | 0.86 | 0.93 | 0.91 |
| 1 | 204094781 | rs16852929 | -0.4 | 0.254 | 0.269 | 0.527 | -0.007 | -0.034 | -0.034 | 0.325 | 0.08 | 0.14 | 0.04 | 0 | 0.06 | 0.03 |
| 1 | 204095478 | rs16852933 | -0.571 | 0.235 | 0.467 | 0.518 | -0.01 | -0.048 | -0.048 | 0.017 | 0.09 | 0.19 | 0.04 | 0 | 0.06 | 0.03 |
| 1 | 204095573 | rs3737658 | 0.291 | 0.245 | -0.079 | 0.355 | -0.015 | 0.015 | 0.015 | 0.602 | 0.09 | 0.04 | 0.1 | 0.06 | 0.1 | 0.04 |
| 1 | 204095687 | rs3737659 | 0.376 | 0.232 | -0.241 | 0.834 | -0.005 | -0.063 | -0.063 | 0.026 | 0.13 | 0.03 | 0.09 | 0.35 | 0.1 | 0.23 |
| 1 | 204096630 | rs1138617 | 0.035 | -0.007 | 0.248 | 0.491 | -0.032 | -0.089 | -0.089 | 0 | 0.87 | 0.6 | 0.91 | 1 | 0.92 | 0.96 |
| 1 | 204097746 | rs10900550 | 2.051 | 0.669 | -0.793 | 0.424 | -0.056 | 0.069 | 0.069 | 0.1 | 0.39 | 0.03 | 0.47 | 0.22 | 0.39 | 0.45 |
| 1 | 204097859 | rs10900551 | 1.229 | -0.079 | -0.084 | 0.474 | 0.019 | -0.045 | -0.045 | 0.124 | 0.62 | 0.21 | 0.74 | 0.47 | 0.68 | 0.6 |
| 1 | 204103618 | rs2293335 | 1.84 | -0.836 | -0.396 | 0.394 | 0.089 | -0.04 | -0.04 | 0 | 0.53 | 0.15 | 0.67 | 0.53 | 0.69 | 0.59 |
| 1 | 204108163 | rs10900553 | -0.339 | -0.015 | -0.128 | 0.953 | -0.007 | 0.045 | 0.045 | 0 | 0.16 | 0.23 | 0.11 | 0.37 | 0.12 | 0.27 |
| 1 | 204108493 | rs4951307 | -1.003 | 0.35 | -0.258 | 0.263 | -0.003 | 0.028 | 0.028 | 0.638 | 0.13 | 0.05 | 0.12 | 0.08 | 0.11 | 0.05 |
| 1 | 204110660 | rs3737656 | -1.237 | 0.195 | 0.229 | 0.259 | -0.026 | -0.056 | -0.056 | 0 | 0.17 | 0.34 | 0.13 | 0.07 | 0.09 | 0.08 |
| 1 | 204112406 | rs4951309 | 0.386 | 0.335 | -0.212 | 0.418 | 0.015 | -0.008 | -0.008 | 0 | 0.47 | 0.38 | 0.55 | 0.48 | 0.43 | 0.41 |
| 1 | 204112984 | rs3795578 | -0.62 | 0.635 | 0.173 | 0.262 | -0.014 | 0.008 | 0.008 | 0 | 0.4 | 0.34 | 0.31 | 0.41 | 0.48 | 0.48 |

| | | | | | | | | | | | | | | | | |
|---|---|---|---|---|---|---|---|---|---|---|---|---|---|---|---|---|
| 1 | 204114846 | rs1917542 | 0.87 | 0.635 | 0.365 | 0.444 | -0.003 | 0 | 0 | 0 | 0.48 | 0.53 | 0.56 | 0.45 | 0.41 | 0.45 |
| 1 | 204115477 | rs940652 | -0.438 | 0.267 | 0.372 | 0.45 | 0.013 | 0.001 | 0.001 | 0.133 | 0.89 | 0.84 | 0.87 | 0.92 | 0.91 | 0.94 |
| 1 | 204117810 | rs4951310 | -0.507 | 0.282 | 0.414 | 0.388 | 0.001 | -0.014 | -0.014 | 0 | 0.9 | 0.86 | 0.89 | 0.95 | 0.93 | 0.95 |
| 1 | 204118360 | rs1997034 | 0.792 | 0.451 | 0.213 | 0.382 | 0 | 0 | 0 | 0.122 | 0.68 | 0.72 | 0.79 | 0.67 | 0.75 | 0.65 |
| 1 | 204125089 | rs3795575 | -0.608 | 0.203 | 0.178 | 0.12 | -0.001 | -0.005 | -0.005 | 0 | 0.17 | 0.18 | 0.18 | 0.07 | 0.14 | 0.08 |
| 1 | 204129671 | rs5707 | -0.708 | 0.074 | 0.22 | 0.559 | 0 | 0 | 0 | 0 | 0.26 | 0.28 | 0.22 | 0.4 | 0.25 | 0.23 |
| 1 | 204131186 | rs5705 | -0.631 | -0.224 | 0.824 | 0.371 | -0.018 | -0.04 | -0.04 | 0 | 0.19 | 0.32 | 0.18 | 0.08 | 0.14 | 0.09 |
| 1 | 204131631 | rs11571080 | -1.442 | -0.348 | 0.944 | 0.265 | -0.014 | -0.001 | -0.001 | 0 | 0.41 | 0.58 | 0.4 | 0.48 | 0.33 | 0.28 |
| 1 | 204136628 | rs6681776 | 1.197 | -0.952 | 0.386 | 0.077 | -0.008 | -0.001 | -0.001 | 0 | 0.19 | 0.21 | 0.09 | 0.13 | 0.12 | 0.13 |
| 1 | 204140906 | rs7539596 | -1.222 | -1.099 | 0.536 | 0.054 | -0.03 | -0.027 | -0.027 | 0 | 0.6 | 0.8 | 0.48 | 0.54 | 0.46 | 0.38 |
| 1 | 204143007 | rs6702587 | 0.887 | -1.076 | 0.941 | 0.068 | -0.028 | -0.006 | -0.006 | 0 | 0.21 | 0.3 | 0.1 | 0.13 | 0.12 | 0.15 |
| 1 | 204158132 | rs11240695 | -0.825 | -1.244 | 0.961 | 0.057 | -0.015 | 0.003 | 0.003 | 0 | 0.33 | 0.39 | 0.3 | 0.43 | 0.25 | 0.25 |
| 1 | 204161490 | rs4951316 | -0.84 | -1.581 | 0.661 | -0.094 | -0.01 | -0.001 | -0.001 | 0 | 0.36 | 0.38 | 0.32 | 0.42 | 0.26 | 0.24 |
| 1 | 204162142 | rs10158616 | 0.786 | -1.319 | 0.672 | 0.078 | -0.003 | -0.028 | -0.028 | 0 | 0.11 | 0.14 | 0.06 | 0.02 | 0.11 | 0.05 |
| 1 | 204164691 | rs12568353 | -0.719 | -1.474 | 0.591 | 0.099 | -0.007 | 0.075 | 0.075 | 0 | 0.13 | 0.21 | 0.21 | 0.41 | 0.11 | 0.13 |
| 1 | 204166043 | rs3924586 | 0.209 | -1.015 | 0.348 | -0.473 | 0.003 | 0.003 | 0.003 | 0 | 0.58 | 0.61 | 0.42 | 0.29 | 0.62 | 0.5 |
| 1 | 204166082 | rs3924585 | 0.603 | -0.636 | 0.826 | -0.629 | 0.012 | -0.051 | -0.051 | 0.106 | 0.12 | 0.27 | 0.07 | 0.02 | 0.11 | 0.06 |
| 1 | 204170993 | rs6700426 | -1.392 | -0.427 | 0.792 | -0.547 | 0.026 | -0.014 | -0.014 | 0 | 0.09 | 0.39 | 0.1 | 0.03 | 0.05 | 0.06 |
| 1 | 204175579 | rs4373805 | -0.597 | -0.448 | 0.165 | -0.61 | -0.017 | -0.092 | -0.092 | 0 | 0.22 | 0.41 | 0.15 | 0.06 | 0.17 | 0.13 |
| 1 | 204176954 | rs11240698 | -1.145 | -0.712 | -0.752 | -0.317 | 0.066 | 0.081 | 0.081 | 0 | 0.51 | 0.18 | 0.34 | 0.28 | 0.58 | 0.56 |
| 1 | 204180540 | rs10900559 | -0.166 | -0.72 | -0.152 | -0.512 | 0.01 | -0.01 | -0.01 | 0.474 | 0.2 | 0.15 | 0.08 | 0 | 0.13 | 0.08 |
| 1 | 204188517 | rs16853105 | 0.313 | -0.053 | 0.489 | -0.257 | -0.001 | -0.021 | -0.021 | 0 | 0.12 | 0.15 | 0.06 | 0 | 0.1 | 0.05 |
| 1 | 204190508 | rs4631764 | 1.014 | 0.275 | 0.754 | -0.068 | 0.004 | -0.002 | -0.002 | 0 | 0.42 | 0.36 | 0.46 | 0.48 | 0.35 | 0.31 |
| 1 | 204190576 | rs4489623 | 1.014 | 0.275 | 0.754 | -0.068 | 0.004 | -0.002 | -0.002 | 0 | 0.42 | 0.36 | 0.46 | 0.48 | 0.35 | 0.31 |
| 1 | 204190873 | rs10793758 | 0.138 | 0.276 | 0.825 | -0.078 | -0.004 | -0.001 | -0.001 | 0 | 0.55 | 0.62 | 0.54 | 0.49 | 0.46 | 0.36 |
| 1 | 204196327 | rs4581328 | 0.114 | 0.324 | 0.758 | -0.067 | -0.005 | -0.002 | -0.002 | 0.915 | 0.56 | 0.69 | 0.54 | 0.48 | 0.44 | 0.36 |
| 1 | 204198209 | rs3795571 | 0.093 | 0.267 | 0.772 | -0.07 | -0.005 | -0.002 | -0.002 | 0.484 | 0.56 | 0.7 | 0.54 | 0.48 | 0.43 | 0.36 |
| 1 | 204210750 | rs4412655 | 1.303 | 0.167 | 1.158 | -0.126 | 0.019 | -0.026 | -0.026 | 1.004 | 0.39 | 0.18 | 0.45 | 0.48 | 0.33 | 0.28 |
| 1 | 204211925 | rs12744138 | 0.734 | 0.307 | 1.039 | 0.045 | -0.007 | 0 | 0 | 0 | 0.11 | 0.07 | 0.24 | 0.33 | 0.19 | 0.19 |
| 1 | 204216410 | rs4951329 | -0.609 | 0.332 | 1.034 | 0.018 | -0.014 | 0.024 | 0.024 | 0 | 0.66 | 0.77 | 0.78 | 0.81 | 0.62 | 0.55 |
| 1 | 204216468 | rs4512713 | -0.343 | 0.323 | 1.103 | 0.013 | -0.002 | 0.009 | 0.009 | 0.001 | 0.64 | 0.66 | 0.76 | 0.81 | 0.62 | 0.53 |
| 1 | 204218852 | rs12040324 | -0.383 | 0.321 | 1.044 | 0.018 | -0.006 | 0.026 | 0.026 | 0 | 0.64 | 0.75 | 0.77 | 0.81 | 0.62 | 0.53 |
| 1 | 204221842 | rs7520626 | -0.473 | 0.43 | 0.939 | 0.074 | -0.009 | 0.018 | 0.018 | 0 | 0.66 | 0.72 | 0.77 | 0.81 | 0.62 | 0.55 |
| 1 | 204230052 | rs4951338 | 0.269 | 0.299 | 0.882 | 0.141 | -0.005 | 0.04 | 0.04 | 0 | 0.54 | 0.61 | 0.71 | 0.81 | 0.54 | 0.51 |
| 1 | 204231164 | rs4338423 | 0.46 | 0.386 | 0.71 | 0.102 | -0.005 | 0.022 | 0.022 | 0.047 | 0.14 | 0.07 | 0.07 | 0 | 0.13 | 0.04 |
| 1 | 204232972 | rs4951056 | 0.3 | 0.49 | 0.809 | 0.266 | -0.003 | -0.001 | -0.001 | 0 | 0.57 | 0.54 | 0.53 | 0.49 | 0.48 | 0.35 |
| 1 | 204237416 | rs10900571 | -0.467 | 0.54 | 0.527 | 0.243 | 0.001 | -0.022 | -0.022 | 0.826 | 0.67 | 0.48 | 0.76 | 0.8 | 0.66 | 0.55 |
| 1 | 204244473 | rs12145347 | 0.396 | 0.578 | 0.186 | 0.505 | 0.003 | 0.017 | 0.017 | 0 | 0.11 | 0.13 | 0.24 | 0.29 | 0.18 | 0.2 |
| 1 | 204245347 | rs4951347 | 1.276 | 0.438 | -0.072 | 0.453 | 0.008 | -0.021 | -0.021 | 0 | 0.39 | 0.27 | 0.46 | 0.51 | 0.35 | 0.31 |
| 1 | 204251754 | rs12124529 | 0.57 | 0.582 | -0.155 | 0.573 | -0.005 | 0.003 | 0.003 | 0 | 0.11 | 0.08 | 0.24 | 0.32 | 0.18 | 0.2 |
| 1 | 204254288 | rs4245726 | 1.007 | 0.353 | -0.138 | 0.508 | 0.008 | -0.015 | -0.015 | 0.039 | 0.38 | 0.29 | 0.45 | 0.5 | 0.34 | 0.3 |
| 1 | 204255848 | rs4951353 | 0.328 | 0.554 | -0.447 | 0.641 | -0.026 | -0.03 | -0.03 | 0 | 0.56 | 0.76 | 0.56 | 0.57 | 0.5 | 0.4 |
| 1 | 204258019 | rs4951059 | -0.056 | 0.174 | -0.033 | 0.397 | -0.017 | 0.079 | 0.079 | 0.593 | 0.89 | 0.57 | 0.96 | 0.93 | 0.97 | 0.96 |
| 1 | 204259763 | rs4132008 | 0.433 | 0.596 | -1.349 | 0.159 | 0.017 | 0.11 | 0.11 | 0.204 | 0.55 | 0.22 | 0.71 | 0.82 | 0.63 | 0.53 |
| 1 | 204260710 | rs4951354 | 0.995 | 0.782 | -1.718 | 0.267 | -0.01 | -0.104 | -0.104 | 0.245 | 0.33 | 0.07 | 0.41 | 0.49 | 0.31 | 0.32 |
| 1 | 204262260 | rs16853237 | -0.118 | 1.045 | -1.132 | 0.12 | -0.016 | -0.051 | -0.051 | 0 | 0.15 | 0.08 | 0.24 | 0.58 | 0.18 | 0.19 |
| 1 | 204263809 | rs11240716 | -0.214 | 1.048 | -1.094 | 0.075 | -0.016 | -0.045 | -0.045 | 1.317 | 0.15 | 0.08 | 0.23 | 0.59 | 0.18 | 0.19 |
| 1 | 204263864 | rs7525113 | 1.276 | 1.183 | -0.577 | -0.385 | 0 | 0.102 | 0.102 | 0 | 0.45 | 0.28 | 0.49 | 0.22 | 0.44 | 0.38 |
| 1 | 204264047 | rs17405517 | 0.339 | 1.134 | -0.55 | -0.571 | -0.03 | 0.039 | 0.039 | 0.464 | 0.22 | 0.07 | 0.19 | 0.14 | 0.28 | 0.35 |
| 1 | 204266980 | rs6594001 | 0.245 | 1.149 | -0.028 | -0.553 | 0.001 | 0.014 | 0.014 | 0.264 | 0.15 | 0.17 | 0.24 | 0.59 | 0.18 | 0.19 |
| 1 | 204269242 | rs4619033 | 0.249 | 1.136 | 0.189 | -0.588 | -0.021 | 0.038 | 0.038 | 0.61 | 0.23 | 0.11 | 0.19 | 0.14 | 0.28 | 0.36 |
| 1 | 204274256 | rs6660437 | -0.911 | 1.121 | 0.203 | -0.597 | -0.017 | 0.044 | 0.044 | 0 | 0.38 | 0.29 | 0.44 | 0.73 | 0.46 | 0.54 |
| 1 | 204281004 | rs4951064 | 1.703 | 1.014 | 0.293 | -0.63 | -0.033 | 0.147 | 0.147 | 0 | 0.44 | 0.16 | 0.48 | 0.22 | 0.44 | 0.39 |
| 1 | 204282936 | rs17333794 | 0.279 | 1.008 | 0.53 | -0.646 | -0.004 | 0.021 | 0.021 | 0.315 | 0.22 | 0.16 | 0.19 | 0.14 | 0.27 | 0.35 |
| 1 | 204283820 | rs4457633 | 0.034 | 0.909 | 1.337 | -0.709 | -0.019 | -0.053 | -0.053 | 0 | 0.09 | 0.24 | 0.04 | 0 | 0.02 | 0.02 |
| 1 | 204285075 | rs4245728 | 1.822 | 0.911 | 1.498 | -0.706 | -0.005 | 0.095 | 0.095 | 0 | 0.46 | 0.28 | 0.5 | 0.22 | 0.44 | 0.39 |
| 1 | 204289051 | rs10900579 | 0.485 | 0.93 | 1.563 | -0.693 | 0.006 | 0.015 | 0.015 | 0 | 0.78 | 0.73 | 0.73 | 0.36 | 0.74 | 0.76 |
| 1 | 204289393 | rs4245729 | 1.747 | 0.912 | 1.578 | -0.718 | 0 | 0.063 | 0.063 | 0 | 0.46 | 0.33 | 0.5 | 0.22 | 0.44 | 0.39 |
| 1 | 204289780 | rs17333933 | -0.731 | 0.897 | 1.487 | -1.043 | -0.007 | -0.044 | -0.044 | 0 | 0.31 | 0.4 | 0.22 | 0.14 | 0.3 | 0.37 |
| 1 | 204290800 | rs4951359 | -0.682 | 0.958 | 1.072 | -0.992 | -0.001 | -0.009 | -0.009 | 1.639 | 0.93 | 0.93 | 0.97 | 0.95 | 0.92 | 0.95 |
| 1 | 204298123 | rs12044614 | 0.358 | 0.899 | 1.029 | -0.98 | -0.01 | 0.028 | 0.028 | 0.205 | 0.21 | 0.13 | 0.18 | 0.14 | 0.27 | 0.35 |
| 1 | 204301822 | rs17405885 | 0.362 | 0.885 | 0.914 | -0.987 | -0.025 | 0.04 | 0.04 | 0.468 | 0.22 | 0.07 | 0.18 | 0.14 | 0.27 | 0.27 |
| 1 | 204306109 | rs17334002 | 0.747 | 0.792 | 0.77 | -1.033 | -0.039 | 0.02 | 0.02 | 0 | 0.18 | 0.06 | 0.17 | 0.13 | 0.27 | 0.27 |
| 1 | 204312033 | rs6594005 | 2.082 | 0.77 | 0.734 | -1.051 | -0.089 | 0.205 | 0.205 | 0 | 0.42 | 0.08 | 0.48 | 0.14 | 0.44 | 0.39 |
| 1 | 204312363 | rs6656057 | 0.258 | 0.403 | 1.429 | -0.892 | 0.011 | -0.055 | -0.055 | 0 | 0.9 | 0.7 | 0.94 | 0.95 | 0.92 | 0.95 |
| 1 | 204314219 | rs4951362 | -0.483 | 0.39 | 1.355 | -0.893 | -0.007 | -0.026 | -0.026 | 1.97 | 0.93 | 0.9 | 0.96 | 1 | 0.97 | 0.97 |
| 1 | 204317155 | rs4951365 | 0.652 | 0.376 | 1.289 | -0.911 | 0.002 | -0.057 | -0.057 | 0.002 | 0.7 | 0.62 | 0.78 | 0.82 | 0.66 | 0.74 |
| 1 | 204317650 | rs1398150 | 1.519 | 0.383 | 0.813 | -0.91 | 0.01 | 0.193 | 0.193 | 2.761 | 0.58 | 0.15 | 0.73 | 0.82 | 0.63 | 0.64 |
| 1 | 204322935 | rs2942135 | -1.732 | -0.538 | 1.032 | -0.799 | -0.003 | -0.047 | -0.047 | 2.032 | 0.22 | 0.71 | 0.09 | 0.06 | 0.12 | 0.14 |

| | | | | | | | | | | | | | | | |
|---|---|---|---|---|---|---|---|---|---|---|---|---|---|---|---|
| 1 | 204323363 | rs10494855 | -0.799 | -0.677 | 1.537 | -0.785 | 0.006 | -0.047 | -0.047 | 0 | 0.1 | 0.4 | 0.04 | 0 | 0.02 | 0.09 |
| 1 | 204323874 | rs10494854 | 1.387 | -0.624 | 0.95 | -0.802 | 0.07 | 0.12 | 0.12 | 5.174 | 0.59 | 0.23 | 0.79 | 0.82 | 0.72 | 0.6 |
| 1 | 204324483 | rs2942132 | -1.394 | -0.374 | 1.457 | -0.37 | -0.013 | -0.08 | -0.08 | 0.142 | 0.13 | 0.43 | 0.05 | 0.05 | 0.08 | 0.06 |
| 1 | 204327412 | rs3125168 | -1.827 | -0.431 | 1.101 | 0.113 | -0.077 | 0.162 | 0.162 | 0.615 | 0.2 | 0.5 | 0.26 | 0.62 | 0.11 | 0.31 |
| 1 | 204334765 | rs955323 | -1.296 | 0.281 | 0.725 | -0.481 | -0.003 | -0.109 | -0.109 | 2.214 | 0.24 | 0.34 | 0.1 | 0.01 | 0.2 | 0.16 |
| 1 | 204344375 | rs955514 | -1.649 | 0.404 | 1.333 | -0.085 | 0.041 | -0.06 | -0.06 | 0 | 0.13 | 0.25 | 0.09 | 0.06 | 0.16 | 0.1 |
| 1 | 204344685 | rs2942147 | -2.073 | 0.132 | 1.629 | 0.403 | -0.013 | 0.156 | 0.156 | 0 | 0.23 | 0.47 | 0.31 | 0.54 | 0.17 | 0.23 |
| 1 | 204345392 | rs11240732 | -1.064 | -0.03 | 1.26 | 0.288 | 0.009 | -0.043 | -0.043 | 0 | 0.29 | 0.43 | 0.22 | 0.22 | 0.35 | 0.38 |
| 1 | 204345923 | rs1980051 | 1.539 | -0.006 | 0.909 | 0.139 | 0.019 | 0.03 | 0.03 | 0 | 0.55 | 0.31 | 0.55 | 0.26 | 0.59 | 0.42 |
| 1 | 204353050 | rs10736846 | 1.526 | 0.116 | 1.075 | 0.488 | -0.013 | 0.086 | 0.086 | 0 | 0.71 | 0.49 | 0.69 | 0.43 | 0.75 | 0.67 |
| 1 | 204354548 | rs2942121 | 0.512 | 0.162 | 0.969 | 0.143 | -0.009 | -0.028 | -0.028 | 0.884 | 0.93 | 0.9 | 0.99 | 1 | 0.97 | 0.99 |
| 1 | 204355244 | rs3106364 | -1.89 | 0.252 | 1.056 | 0.13 | 0.033 | -0.022 | -0.022 | 0.029 | 0.08 | 0.25 | 0.07 | 0.07 | 0.11 | 0.15 |
| 1 | 204355646 | rs11808492 | -0.092 | 0.136 | 0.297 | 0.28 | -0.002 | -0.014 | -0.014 | 0.233 | 0.05 | 0.06 | 0.01 | 0 | 0.03 | 0.01 |
| 1 | 204359333 | rs11240735 | 0.418 | 0.805 | 0.552 | 0.331 | 0.093 | -0.017 | -0.017 | 0 | 0.71 | 0.56 | 0.67 | 0.8 | 0.58 | 0.78 |
| 1 | 204359597 | rs1512113 | 1.212 | -0.789 | 1.132 | 1.039 | -0.071 | 0.073 | 0.073 | 0.662 | 0.74 | 0.38 | 0.8 | 0.47 | 0.9 | 0.67 |
| 1 | 204365749 | rs7530308 | -1.351 | -0.443 | 1.419 | 0.556 | 0.02 | -0.064 | -0.064 | 0 | 0.11 | 0.38 | 0.07 | 0 | 0.13 | 0.03 |
| 1 | 204368977 | rs6694389 | -1.172 | -0.449 | 1.464 | 0.563 | 0.01 | -0.057 | -0.057 | 0 | 0.11 | 0.31 | 0.07 | 0.01 | 0.13 | 0.07 |
| 1 | 204369875 | rs2942133 | 1.387 | -0.573 | 1.248 | 0.844 | 0.026 | -0.011 | -0.011 | 0.336 | 0.65 | 0.22 | 0.71 | 0.39 | 0.73 | 0.57 |
| 1 | 204389314 | rs12061474 | -1.862 | 0.083 | 1.6 | 0.967 | 0.07 | 0.057 | 0.057 | 0.031 | 0.06 | 0.28 | 0.1 | 0.08 | 0.13 | 0.08 |
| 1 | 204392257 | rs4951373 | -1.035 | -0.758 | 1.633 | 1.329 | 0.001 | -0.002 | -0.002 | 0.111 | 0.21 | 0.22 | 0.15 | 0.58 | 0.13 | 0.31 |
| 1 | 204396728 | rs17406271 | -1.857 | -0.632 | 1.431 | 1.143 | 0.05 | 0.046 | 0.046 | 0.191 | 0.07 | 0.3 | 0.11 | 0.08 | 0.11 | 0.06 |
| 1 | 204396830 | rs17334387 | -1.818 | -0.628 | 1.422 | 1.145 | 0.052 | 0.048 | 0.048 | 0.236 | 0.07 | 0.28 | 0.1 | 0.08 | 0.11 | 0.06 |
| 1 | 204400783 | rs2271421 | -0.88 | -1.153 | 0.698 | 1.882 | 0.048 | -0.111 | -0.111 | 0.126 | 0.19 | 0.02 | 0.13 | 0.46 | 0.11 | 0.28 |
| 1 | 204403659 | rs3747636 | -2.171 | -1.111 | 1.738 | 2.02 | -0.036 | 0.063 | 0.063 | 0 | 0.32 | 0.53 | 0.27 | 0.51 | 0.26 | 0.38 |
| 1 | 204404363 | rs12031854 | 1.072 | -0.479 | 0.626 | 1.485 | -0.079 | -0.001 | -0.001 | 0 | 0.14 | 0.02 | 0.1 | 0.07 | 0.22 | 0.17 |
| 1 | 204408386 | rs9658849 | -0.744 | -0.756 | 1.193 | 1.663 | 0.007 | 0.005 | 0.005 | 0 | 0.08 | 0.06 | 0.02 | 0.03 | 0.04 | 0.05 |
| 1 | 204413297 | rs2292459 | 1.515 | -0.97 | 2.208 | 1.617 | -0.021 | 0.031 | 0.031 | 0 | 0.71 | 0.42 | 0.77 | 0.49 | 0.78 | 0.7 |
| 1 | 204413845 | rs2999495 | 1.657 | -0.999 | 2.732 | 1.768 | 0.055 | 0.237 | 0.237 | 6.879 | 0.95 | 0.59 | 0.97 | 0.89 | 1 | 0.95 |
| 1 | 204416501 | rs3827534 | -1.206 | -1.009 | 2.677 | 1.787 | 0.007 | -0.014 | -0.014 | 0 | 0.24 | 0.14 | 0.15 | 0.43 | 0.22 | 0.29 |
| 1 | 204417972 | rs3106366 | 1.55 | -1.018 | 2.71 | 1.789 | 0.006 | -0.004 | -0.004 | 0 | 0.69 | 0.35 | 0.76 | 0.39 | 0.77 | 0.65 |
| 1 | 204418198 | rs2271427 | 1.385 | -1.012 | 2.721 | 1.79 | -0.048 | 0.061 | 0.061 | 0 | 0.72 | 0.57 | 0.84 | 0.57 | 0.78 | 0.7 |
| 1 | 204420629 | rs3014638 | 2.478 | -1.022 | 2.387 | 1.702 | -0.01 | 0.22 | 0.22 | 7.11 | 0.92 | 0.45 | 0.92 | 0.82 | 0.99 | 0.94 |
| 1 | 204426295 | rs1008833 | 0.283 | -1.061 | 2.004 | 1.912 | -0.029 | -0.088 | -0.088 | 0.235 | 0.14 | 0.01 | 0.07 | 0.37 | 0.17 | 0.24 |
| 1 | 204427980 | rs2999487 | 1.241 | -1.033 | 2.294 | 1.805 | 0.013 | 0.005 | 0.005 | 4.445 | 0.94 | 0.7 | 0.98 | 1 | 1 | 0.99 |
| 1 | 204430462 | rs2999484 | 1.804 | -1.072 | 1.672 | 1.775 | 0.056 | -0.03 | -0.03 | 0.177 | 0.67 | 0.2 | 0.81 | 0.43 | 0.77 | 0.66 |
| 1 | 204432236 | rs16853770 | -0.894 | -1.098 | 1.721 | 1.837 | 0.017 | -0.062 | -0.062 | 0 | 0.23 | 0.11 | 0.13 | 0.46 | 0.21 | 0.29 |
| 1 | 204433093 | rs2271415 | 1.606 | -1.086 | 1.511 | 1.768 | 0.036 | -0.02 | -0.02 | 0 | 0.68 | 0.25 | 0.81 | 0.42 | 0.78 | 0.66 |
| 1 | 204438334 | rs1124777 | 1.972 | -0.884 | 1.67 | 1.624 | -0.052 | 0.047 | 0.047 | 11.165 | 0.88 | 0.24 | 0.89 | 0.82 | 0.98 | 0.91 |
| 1 | 204438643 | rs1553921 | 1.982 | -0.887 | 1.655 | 1.623 | -0.051 | 0.044 | 0.044 | 11.515 | 0.88 | 0.24 | 0.9 | 0.82 | 0.98 | 0.91 |
| 1 | 204446313 | rs2999479 | 2.01 | -0.909 | 1.633 | 1.639 | -0.065 | 0.089 | 0.089 | 10.608 | 0.88 | 0.3 | 0.9 | 0.82 | 0.98 | 0.91 |
| 1 | 204452779 | rs12402641 | 1.189 | -0.536 | 0.012 | 1.274 | -0.111 | 0.014 | 0.014 | 1.636 | 0.4 | 0.07 | 0.52 | 0.32 | 0.47 | 0.38 |
| 1 | 204453320 | rs11240748 | -0.4 | -0.634 | 0.52 | 0.958 | -0.01 | 0.017 | 0.017 | 0 | 0.27 | 0.29 | 0.13 | 0.41 | 0.26 | 0.28 |
| 1 | 204457336 | rs7556371 | 0.059 | -0.65 | 0.562 | 0.909 | -0.001 | -0.003 | -0.003 | 0 | 0.67 | 0.36 | 0.65 | 0.73 | 0.73 | 0.64 |
| 1 | 204462050 | rs11240751 | 0.893 | -0.752 | 0.56 | 0.849 | 0.036 | 0.053 | 0.053 | 1.859 | 0.58 | 0.16 | 0.61 | 0.72 | 0.7 | 0.61 |
| 1 | 204494576 | rs4252675 | 0.578 | -0.753 | 1.264 | 0.633 | 0.002 | -0.021 | -0.021 | 0.049 | 0.05 | 0.1 | 0.04 | 0 | 0.05 | 0.12 |
| 1 | 204498364 | rs4252690 | 0.466 | -0.85 | 1.362 | 0.642 | -0.025 | -0.033 | -0.033 | 0.498 | 0.08 | 0.08 | 0.02 | 0.01 | 0.03 | 0.04 |
| 1 | 204505487 | rs10458588 | 1.843 | -0.822 | 1.174 | 0.718 | -0.099 | 0.023 | 0.023 | 0.609 | 0.4 | 0.07 | 0.51 | 0.31 | 0.46 | 0.37 |
| 1 | 204511704 | rs2290853 | -0.804 | -0.837 | 1.3 | 0.82 | 0.024 | 0.239 | 0.239 | 1.77 | 0.06 | 0.35 | 0.02 | 0.16 | 0.01 | 0.07 |
| 1 | 204516025 | rs2290854 | 0.986 | -0.852 | 1.298 | 0.81 | 0.024 | 0.028 | 0.028 | 0.539 | 0.59 | 0.22 | 0.61 | 0.7 | 0.69 | 0.6 |
| 1 | 204516485 | rs3789050 | -0.416 | -0.843 | 1.268 | 0.815 | 0.055 | 0.298 | 0.298 | 2.965 | 0.05 | 0.4 | 0.03 | 0.16 | 0 | 0.07 |
| 1 | 204518842 | rs4245739 | -0.561 | -0.839 | 0.827 | 0.816 | -0.01 | 0.013 | 0.013 | 0.23 | 0.75 | 0.77 | 0.71 | 0.95 | 0.74 | 0.74 |
| 1 | 204522489 | rs10900597 | 1.165 | -0.743 | 0.531 | 0.857 | 0.031 | 0.045 | 0.045 | 0.596 | 0.57 | 0.12 | 0.6 | 0.7 | 0.68 | 0.59 |
| 1 | 204551830 | rs16854023 | 0.95 | -0.646 | 0.535 | 1.056 | -0.017 | -0.059 | -0.059 | 0 | 0.17 | 0.06 | 0.09 | 0.37 | 0.22 | 0.22 |
| 1 | 204566183 | rs6682208 | 1.006 | -0.767 | 0.337 | 0.869 | 0.038 | 0.037 | 0.037 | 0.26 | 0.56 | 0.22 | 0.61 | 0.69 | 0.67 | 0.57 |
| 1 | 204566999 | rs11240762 | 1.774 | -0.67 | 0.284 | 0.794 | -0.059 | 0.024 | 0.024 | 0.055 | 0.37 | 0.13 | 0.51 | 0.33 | 0.44 | 0.34 |
| 1 | 204572050 | rs10900601 | -0.688 | -0.732 | 0.457 | 0.959 | -0.008 | 0.013 | 0.013 | 0 | 0.76 | 0.74 | 0.73 | 0.88 | 0.74 | 0.75 |
| 1 | 204572071 | rs12143943 | 1.487 | -0.787 | 0.152 | 0.989 | -0.062 | 0.031 | 0.031 | 0 | 0.37 | 0.16 | 0.52 | 0.33 | 0.45 | 0.31 |
| 1 | 204586184 | rs3789047 | 1.634 | -0.658 | -0.69 | 0.98 | -0.139 | -0.022 | -0.022 | 0.148 | 0.35 | 0.05 | 0.5 | 0.34 | 0.43 | 0.32 |
| 1 | 204586805 | rs3789046 | 1.634 | -0.658 | -0.69 | 0.98 | -0.139 | -0.022 | -0.022 | 0.948 | 0.35 | 0.05 | 0.5 | 0.34 | 0.43 | 0.32 |
| 1 | 204587047 | rs11588857 | 0.794 | -0.698 | -0.793 | 1.032 | -0.013 | -0.048 | -0.048 | 0 | 0.2 | 0.06 | 0.09 | 0.31 | 0.22 | 0.25 |
| 1 | 204587240 | rs3747632 | 1.771 | -0.654 | -0.846 | 1.037 | -0.162 | -0.042 | -0.042 | 0 | 0.34 | 0.03 | 0.5 | 0.34 | 0.43 | 0.32 |
| 1 | 204589101 | rs3789044 | 0.522 | -0.501 | -0.433 | 1.071 | -0.02 | -0.067 | -0.067 | 0 | 0.21 | 0.04 | 0.09 | 0.31 | 0.22 | 0.24 |
| 1 | 204609194 | rs4951087 | 0.769 | -0.372 | -0.045 | 0.706 | -0.008 | 0.003 | 0.003 | 1.078 | 0.62 | 0.27 | 0.63 | 0.71 | 0.59 | 0.63 |
| 1 | 204612679 | rs7533494 | -0.065 | -0.315 | 0.455 | 0.567 | 0.005 | -0.041 | -0.041 | 0.988 | 0.1 | 0.27 | 0.05 | 0.01 | 0.05 | 0.1 |
| 1 | 204614475 | rs11801229 | -0.097 | -0.83 | 0.206 | 0.426 | -0.012 | 0.008 | 0.008 | 0 | 0.28 | 0.31 | 0.14 | 0.42 | 0.15 | 0.34 |
| 1 | 204616390 | rs1329631 | -0.613 | -0.414 | 0.448 | 0.497 | 0.024 | -0.06 | -0.06 | 0.453 | 0.31 | 0.46 | 0.32 | 0.13 | 0.39 | 0.3 |
| 1 | 204617240 | rs2815832 | 0.853 | -0.451 | 0.3 | 0.485 | -0.051 | 0.024 | 0.024 | 0.174 | 0.18 | 0.07 | 0.26 | 0.11 | 0.34 | 0.16 |
| 1 | 204618533 | rs4951090 | 2.138 | -0.455 | 0.032 | 0.345 | -0.052 | -0.035 | -0.035 | 1.398 | 0.43 | 0.14 | 0.56 | 0.46 | 0.46 | 0.42 |
| 1 | 204618573 | rs4951092 | 0.746 | -0.361 | 0.166 | 0.19 | 0.018 | -0.062 | -0.062 | 0 | 0.64 | 0.48 | 0.66 | 0.87 | 0.57 | 0.7 |

| | | | | | | | | | | | | | | | | |
|---|---|---|---|---|---|---|---|---|---|---|---|---|---|---|---|---|
| 1 | 204626549 | rs6667578 | 1.691 | -0.15 | -0.123 | 0.424 | 0.008 | -0.024 | -0.024 | 0 | 0.45 | 0.15 | 0.57 | 0.42 | 0.43 | 0.46 |
| 1 | 204627052 | rs2772229 | 0.65 | 0.15 | 0.753 | 0.379 | 0.01 | -0.03 | -0.03 | 0 | 0.93 | 0.74 | 0.96 | 1 | 0.95 | 1 |
| 1 | 204628101 | rs12079489 | -0.543 | 0.168 | 0.27 | 0.362 | -0.007 | -0.061 | -0.061 | 0.116 | 0.1 | 0.26 | 0.07 | 0 | 0.05 | 0.09 |
| 1 | 204628601 | rs1411109 | 1.421 | 0.16 | -0.182 | 0.591 | -0.016 | 0 | 0 | 0 | 0.77 | 0.3 | 0.87 | 0.82 | 0.9 | 0.82 |
| 1 | 204632065 | rs2772233 | 0.279 | 0.973 | -0.525 | 0.535 | -0.003 | 0.001 | 0.001 | 0 | 0.16 | 0.15 | 0.26 | 0.12 | 0.31 | 0.19 |
| 1 | 204633340 | rs17413714 | -0.382 | -0.024 | -1.295 | 0.239 | 0.005 | 0.015 | 0.015 | 0.026 | 0.3 | 0.08 | 0.11 | 0.32 | 0.24 | 0.29 |
| 1 | 204633526 | rs1610400 | 0.815 | 0.05 | -0.794 | -0.01 | 0.013 | 0.047 | 0.047 | 0.436 | 0.53 | 0.34 | 0.56 | 0.18 | 0.63 | 0.53 |
| 1 | 204633858 | rs1610399 | 1.305 | -0.358 | -0.281 | 0.438 | 0.029 | 0.021 | 0.021 | 0 | 0.47 | 0.26 | 0.6 | 0.42 | 0.43 | 0.55 |
| 1 | 204646295 | rs10900421 | -0.959 | -1.487 | 0.216 | 0.996 | 0.016 | -0.075 | -0.075 | 0 | 0.25 | 0.1 | 0.15 | 0.37 | 0.2 | 0.17 |
| 1 | 204656803 | rs2782524 | 1.669 | -1.98 | 0.51 | 0.608 | -0.009 | -0.001 | -0.001 | 0.581 | 0.58 | 0.42 | 0.79 | 0.68 | 0.79 | 0.78 |
| 1 | 204668280 | rs10751426 | 1.552 | -1.791 | 0.746 | 0.791 | -0.038 | 0.021 | 0.021 | 0.892 | 0.69 | 0.36 | 0.71 | 0.51 | 0.88 | 0.77 |
| 1 | 204671283 | rs1014818 | 0.746 | -1.675 | 0.883 | 0.613 | -0.037 | -0.057 | -0.057 | 0.596 | 0.9 | 0.73 | 0.97 | 1 | 0.98 | 0.95 |
| 1 | 204672660 | rs1610402 | 1.354 | -1.701 | 0.798 | 0.599 | -0.064 | -0.094 | -0.094 | 0 | 0.86 | 0.65 | 0.95 | 1 | 0.98 | 0.95 |
| 1 | 204674668 | rs16854206 | 0.903 | -1.321 | 0.961 | 0.817 | -0.045 | -0.068 | -0.068 | 0 | 0.89 | 0.62 | 0.95 | 1 | 0.98 | 0.95 |
| 1 | 204679248 | rs1411112 | -0.946 | -0.694 | 0.262 | 1.374 | -0.036 | 0.123 | 0.123 | 0 | 0.17 | 0.32 | 0.25 | 0.48 | 0.1 | 0.18 |
| 1 | 204681324 | rs4951095 | -0.859 | 0.267 | 0.317 | 0.653 | 0.071 | -0.016 | -0.016 | 0.399 | 0.27 | 0.41 | 0.29 | 0.25 | 0.35 | 0.17 |
| 1 | 204682011 | rs4075338 | -0.571 | -1.739 | 0.65 | -0.049 | 0.017 | 0.013 | 0.013 | 5.473 | 0.4 | 0.67 | 0.19 | 0.14 | 0.19 | 0.19 |
| 1 | 204682267 | rs868507 | -0.047 | -1.917 | 1.163 | -0.069 | 0.012 | 0.012 | 0.012 | 4.91 | 0.32 | 0.33 | 0.13 | 0.15 | 0.15 | 0.17 |
| 1 | 204682411 | rs868506 | -0.241 | -1.992 | 1.09 | -0.105 | 0.059 | 0.063 | 0.063 | 3.01 | 0.34 | 0.21 | 0.14 | 0.18 | 0.18 | 0.22 |
| 1 | 204684237 | rs3738158 | -0.743 | -2.355 | 1.626 | 0.326 | 0.005 | 0.007 | 0.007 | 3.793 | 0.4 | 0.69 | 0.28 | 0.3 | 0.17 | 0.24 |
| 1 | 204685624 | rs6695611 | -1.491 | -2.327 | 1.747 | 0.301 | -0.013 | 0.111 | 0.111 | 0.377 | 0.08 | 0.35 | 0.15 | 0.15 | 0.02 | 0.07 |
| 1 | 204688365 | rs11803257 | -0.806 | -2.373 | 1.782 | 0.284 | 0.078 | 0.033 | 0.033 | 2.806 | 0.41 | 0.77 | 0.28 | 0.33 | 0.17 | 0.24 |
| 1 | 204691325 | rs11240252 | -0.464 | -2.3 | 1.791 | 0.152 | -0.011 | 0.086 | 0.086 | 0.444 | 0.06 | 0.2 | 0.13 | 0.15 | 0.01 | 0.07 |
| 1 | 204693036 | rs12075982 | -1.024 | -2.294 | 1.836 | 0.151 | -0.014 | 0.092 | 0.092 | 0.703 | 0.07 | 0.28 | 0.14 | 0.15 | 0.01 | 0.07 |
| 1 | 204696179 | rs16854270 | 0.471 | -2.428 | 1.832 | 0.084 | 0.044 | 0.05 | 0.05 | 2.512 | 0.27 | 0.14 | 0.11 | 0.15 | 0.15 | 0.17 |
| 1 | 204699500 | rs6689270 | -1.592 | -2.375 | 1.885 | 0.114 | -0.022 | 0.113 | 0.113 | 0.31 | 0.1 | 0.44 | 0.14 | 0.15 | 0.02 | 0.05 |
| 1 | 204703066 | rs12408321 | 0.131 | -1.884 | 1.036 | 0.439 | -0.06 | -0.101 | -0.101 | 0 | 0.13 | 0.01 | 0.4 | 0.51 | 0.23 | 0.13 |
| 1 | 204703480 | rs12239515 | -0.07 | -1.884 | 1.106 | 0.436 | -0.038 | -0.013 | -0.013 | 0.036 | 0.36 | 0.58 | 0.26 | 0.3 | 0.17 | 0.25 |
| 1 | 204717211 | rs2012130 | -0.819 | -1.606 | 0.91 | 0.3 | 0.012 | 0.023 | 0.023 | 1.653 | 0.53 | 0.64 | 0.68 | 0.81 | 0.4 | 0.38 |
| 1 | 204733391 | rs6672027 | -1.293 | -1.918 | 1.819 | 0.631 | -0.037 | 0.023 | 0.023 | 0.261 | 0.23 | 0.67 | 0.49 | 0.63 | 0.11 | 0.11 |
| 1 | 204734495 | rs7551603 | -0.256 | -1.895 | 1.73 | 0.645 | 0.016 | -0.047 | -0.047 | 0.098 | 0.14 | 0.06 | 0.34 | 0.53 | 0.1 | 0.08 |
| 1 | 204735453 | rs9438425 | 0.701 | -2.138 | 1.571 | 0.448 | 0.01 | 0.016 | 0.016 | 2.188 | 0.07 | 0.02 | 0.03 | 0 | 0.04 | 0.04 |
| 1 | 204740078 | rs11240265 | -0.371 | -2.014 | 1.726 | 0.533 | -0.007 | 0.049 | 0.049 | 0 | 0.15 | 0.17 | 0.35 | 0.53 | 0.11 | 0.08 |
| 1 | 204747491 | rs11240271 | -0.981 | -2.054 | 1.39 | 0.363 | -0.063 | 0.074 | 0.074 | 0 | 0.21 | 0.52 | 0.5 | 0.65 | 0.11 | 0.12 |
| 1 | 204753539 | rs11240277 | 1.217 | -1.902 | 0.287 | -0.061 | -0.136 | 0.037 | 0.037 | 8.909 | 0.33 | 0.03 | 0.26 | 0.18 | 0.55 | 0.54 |
| 1 | 204777269 | rs12047135 | -0.474 | -2.007 | 1.259 | 0.633 | -0.02 | 0.094 | 0.094 | 0 | 0.15 | 0.28 | 0.35 | 0.68 | 0.1 | 0.11 |
| 1 | 204779073 | rs4314941 | 1.744 | -2.27 | 0.816 | 0.453 | 0.135 | 0.066 | 0.066 | 1.54 | 0.6 | 0.04 | 0.44 | 0.2 | 0.83 | 0.77 |
| 1 | 204788314 | rs16854428 | -1.585 | -1.408 | 1.574 | 0.894 | -0.047 | 0.025 | 0.025 | 0 | 0.2 | 0.45 | 0.44 | 0.78 | 0.1 | 0.12 |
| 1 | 204788772 | rs7413575 | -1.073 | -1.409 | 1.604 | 1.245 | -0.025 | 0.103 | 0.103 | 0 | 0.17 | 0.33 | 0.34 | 0.61 | 0.1 | 0.09 |
| 1 | 204789624 | rs2208843 | -0.73 | -1.409 | 1.62 | 1.247 | -0.003 | 0.035 | 0.035 | 0 | 0.15 | 0.19 | 0.33 | 0.52 | 0.1 | 0.09 |
| 1 | 204807500 | rs11801589 | -1.578 | -1.385 | 1.545 | 1.217 | -0.046 | 0.043 | 0.043 | 0 | 0.2 | 0.45 | 0.37 | 0.74 | 0.1 | 0.12 |
| 1 | 204810636 | rs911990 | -1.35 | -1.384 | 1.542 | 1.186 | -0.041 | 0.183 | 0.183 | 0 | 0.18 | 0.45 | 0.29 | 0.56 | 0.09 | 0.09 |
| 1 | 204815729 | rs4951136 | -1.283 | -1.366 | 1.346 | 1.195 | -0.035 | 0.148 | 0.148 | 0 | 0.18 | 0.41 | 0.29 | 0.56 | 0.09 | 0.09 |
| 1 | 204823161 | rs4951140 | -1.321 | -1.365 | 1.251 | 1.178 | -0.04 | 0.043 | 0.043 | 0 | 0.18 | 0.4 | 0.37 | 0.74 | 0.08 | 0.12 |
| 1 | 204829234 | rs1548272 | -0.847 | -1.283 | 0.921 | 1.055 | -0.018 | 0.076 | 0.076 | 0 | 0.16 | 0.26 | 0.26 | 0.55 | 0.08 | 0.1 |
| 1 | 204833446 | rs11806216 | -0.921 | -1.258 | 1.063 | 1.069 | -0.035 | 0.155 | 0.155 | 0 | 0.17 | 0.34 | 0.28 | 0.48 | 0.07 | 0.09 |
| 1 | 204842180 | rs4951142 | -1.288 | -1.437 | 1.675 | 0.956 | -0.058 | 0.08 | 0.08 | 0 | 0.19 | 0.49 | 0.38 | 0.68 | 0.08 | 0.1 |
| 1 | 204846772 | rs2759282 | -0.856 | -1.756 | 1.554 | 0.511 | -0.03 | -0.079 | -0.079 | 1.97 | 0.14 | 0.33 | 0.08 | 0.02 | 0.06 | 0.07 |
| 1 | 204849035 | rs11240302 | 1.028 | -1.734 | 0.297 | -0.229 | -0.162 | 0.012 | 0.012 | 9.705 | 0.34 | 0.03 | 0.36 | 0.22 | 0.57 | 0.55 |
| 1 | 204856039 | rs11240305 | -0.986 | -1.732 | 1.013 | 0.594 | -0.055 | 0.055 | 0.055 | 0.287 | 0.2 | 0.52 | 0.38 | 0.69 | 0.08 | 0.11 |
| 1 | 204861371 | rs12405378 | -1.227 | -1.805 | 0.277 | 0.576 | -0.068 | 0.031 | 0.031 | 1.409 | 0.21 | 0.61 | 0.38 | 0.72 | 0.07 | 0.1 |
| 1 | 204862644 | rs7550921 | -0.116 | -1.719 | -0.975 | 0.575 | 0.008 | 0.01 | 0.01 | 0 | 0.13 | 0.12 | 0.32 | 0.51 | 0.06 | 0.08 |
| 1 | 204866365 | rs2759281 | -1.962 | -1.665 | -1.379 | 0.412 | 0.118 | 0.043 | 0.043 | 2.735 | 0.34 | 0.95 | 0.45 | 0.82 | 0.13 | 0.18 |
| 1 | 204869559 | rs12025272 | -0.078 | -0.862 | -0.936 | 0.694 | 0.018 | -0.046 | -0.046 | 0 | 0.11 | 0.03 | 0.35 | 0.52 | 0.07 | 0.08 |
| 1 | 204870988 | rs2997505 | -0.606 | -0.324 | -1.301 | -0.78 | -0.006 | 0.055 | 0.055 | 0.091 | 0.28 | 0.17 | 0.15 | 0 | 0.3 | 0.16 |
| 1 | 204871146 | rs2759280 | 0.153 | 0.469 | -1.456 | -0.561 | 0.01 | -0.001 | -0.001 | 0 | 0.68 | 0.62 | 0.5 | 0.31 | 0.56 | 0.67 |
| 1 | 204871981 | rs11240308 | 0.579 | 0.279 | -2.655 | -0.623 | -0.037 | -0.045 | -0.045 | 0 | 0.36 | 0.02 | 0.33 | 0.3 | 0.34 | 0.48 |
| 1 | 204876966 | rs17336419 | 0.035 | 0.985 | -2.023 | -0.707 | -0.004 | -0.005 | -0.005 | 0 | 0.15 | 0.13 | 0.15 | 0.02 | 0.26 | 0.09 |
| 1 | 204877652 | rs2247208 | -0.312 | 1.058 | -2.68 | -0.864 | -0.043 | -0.032 | -0.032 | 0 | 0.85 | 0.98 | 0.93 | 0.81 | 0.85 | 0.75 |
| 1 | 204877670 | rs2369423 | -0.499 | 0.271 | -2.928 | -0.773 | -0.003 | -0.011 | -0.011 | 0 | 0.45 | 0.32 | 0.59 | 0.49 | 0.45 | 0.39 |
| 1 | 204878704 | rs11240309 | 0.37 | 0.072 | -2.881 | -0.681 | 0.041 | -0.053 | -0.053 | 0 | 0.3 | 0.18 | 0.45 | 0.46 | 0.19 | 0.29 |
| 1 | 204888536 | rs4950972 | 0.93 | 0.203 | -2.788 | -0.746 | 0.001 | 0.08 | 0.08 | 0 | 0.45 | 0.21 | 0.41 | 0.19 | 0.63 | 0.41 |
| 1 | 204889532 | rs11240310 | 0.929 | 0.205 | -2.788 | -0.745 | 0.001 | 0.08 | 0.08 | 0 | 0.45 | 0.21 | 0.41 | 0.19 | 0.63 | 0.42 |
| 1 | 204891547 | rs4951146 | 0.135 | 0.141 | -1.351 | -0.63 | -0.012 | 0.011 | 0.011 | 0 | 0.91 | 0.81 | 0.96 | 0.87 | 0.96 | 0.9 |
| 1 | 204892329 | rs10900434 | 0.4 | -0.094 | -1.17 | -0.827 | -0.007 | -0.012 | -0.012 | 0 | 0.69 | 0.45 | 0.84 | 0.67 | 0.79 | 0.69 |
| 1 | 204895216 | rs2802851 | 1.283 | 0.161 | -1.313 | -1.03 | -0.048 | -0.005 | -0.005 | 0 | 0.32 | 0.08 | 0.26 | 0.18 | 0.37 | 0.35 |
| 1 | 204904835 | rs2625221 | 0.72 | -0.502 | -0.595 | -1.256 | -0.006 | 0.035 | 0.035 | 0 | 0.41 | 0.35 | 0.3 | 0.17 | 0.43 | 0.39 |
| 1 | 204905311 | rs2625220 | -1.032 | -0.8 | -0.153 | -1.261 | 0 | -0.025 | -0.025 | 3.437 | 0.86 | 0.84 | 0.92 | 1 | 0.91 | 0.92 |
| 1 | 204905920 | rs2595955 | -0.259 | -0.986 | -0.122 | -1.342 | -0.011 | -0.055 | -0.055 | 1.171 | 0.22 | 0.28 | 0.09 | 0.01 | 0.11 | 0.09 |

| | | | | | | | | | | | | | | | | |
|---|---|---|---|---|---|---|---|---|---|---|---|---|---|---|---|---|
| 1 | 204906922 | rs2802814 | -0.702 | -1.119 | 0.092 | -1.414 | -0.035 | -0.098 | -0.098 | 1.052 | 0.25 | 0.4 | 0.1 | 0.01 | 0.11 | 0.12 |
| 1 | 204906939 | rs2802815 | 0.752 | -1.093 | -0.173 | -1.401 | 0.001 | -0.023 | -0.023 | 0 | 0.15 | 0.16 | 0.08 | 0.01 | 0.1 | 0.09 |
| 1 | 204910576 | rs2802829 | -0.581 | -0.945 | 0.203 | -1.329 | -0.056 | -0.133 | -0.133 | 0.143 | 0.27 | 0.58 | 0.12 | 0.01 | 0.11 | 0.12 |
| 1 | 204912843 | rs12737855 | 0.664 | -0.753 | 0.241 | -1.086 | 0 | 0 | 0 | 0 | 0.18 | 0.17 | 0.18 | 0.24 | 0.3 | 0.19 |
| 1 | 204913652 | rs2595961 | -1.397 | -0.83 | 0.013 | -1.061 | -0.003 | 0.014 | 0.014 | 0 | 0.81 | 0.8 | 0.6 | 0.42 | 0.86 | 0.68 |
| 1 | 204917680 | rs2068667 | 0.717 | -0.644 | -0.418 | -1.576 | 0 | 0 | 0 | 0 | 0.18 | 0.16 | 0.18 | 0.23 | 0.3 | 0.17 |
| 1 | 204919232 | rs4951147 | 0.413 | -0.933 | -0.325 | -1.565 | 0.069 | 0.072 | 0.072 | 0 | 0.55 | 0.32 | 0.49 | 0.3 | 0.75 | 0.56 |
| 1 | 204919494 | rs2066306 | 0.398 | -0.344 | -0.02 | -1.703 | 0.022 | 0.022 | 0.022 | 0.813 | 0.19 | 0.12 | 0.08 | 0 | 0.1 | 0.09 |
| 1 | 204924685 | rs7534993 | 0.888 | 0.386 | 0.268 | -1.338 | -0.003 | -0.005 | -0.005 | 0 | 0.31 | 0.31 | 0.25 | 0.1 | 0.35 | 0.32 |
| 1 | 204927136 | rs2275696 | 0.367 | 0.453 | 0.248 | -1.342 | 0.035 | -0.063 | -0.063 | 0 | 0.16 | 0.29 | 0.16 | 0.05 | 0.25 | 0.06 |
| 1 | 204928459 | rs4950974 | 0.248 | 0.05 | -0.107 | -1.005 | 0.003 | 0.033 | 0.033 | 0 | 0.45 | 0.34 | 0.39 | 0.13 | 0.58 | 0.35 |
| 1 | 204930143 | rs7543481 | 0.061 | -0.019 | -0.208 | -0.78 | 0.008 | 0.007 | 0.007 | 0 | 0.52 | 0.43 | 0.52 | 0.24 | 0.67 | 0.46 |
| 1 | 204931137 | rs16854813 | -0.405 | 0.223 | 0.215 | -0.392 | 0.015 | 0.001 | 0.001 | 0.125 | 0.09 | 0.28 | 0.07 | 0.09 | 0.09 | 0.11 |
| 1 | 204932805 | rs6685118 | -0.327 | 0.15 | 0.394 | -0.596 | 0.012 | -0.044 | -0.044 | 0.022 | 0.09 | 0.23 | 0.07 | 0 | 0.08 | 0.11 |
| 1 | 204936835 | rs12744678 | -0.338 | -0.093 | 0.201 | -1.265 | 0 | -0.006 | -0.006 | 0 | 0.24 | 0.17 | 0.2 | 0.03 | 0.27 | 0.1 |
| 1 | 204940972 | rs12036722 | 0.949 | -0.356 | -0.111 | -1.086 | -0.013 | 0.002 | 0.002 | 0 | 0.32 | 0.06 | 0.22 | 0.25 | 0.3 | 0.28 |
| 1 | 204946764 | rs6663324 | -0.018 | -0.591 | 0.346 | -1.342 | -0.034 | -0.075 | -0.075 | 0.269 | 0.18 | 0.23 | 0.07 | 0.01 | 0.08 | 0.08 |
| 1 | 204947702 | rs3892248 | 1.104 | 0.047 | 0.252 | -1.27 | 0.008 | -0.024 | -0.024 | 0.136 | 0.12 | 0.11 | 0.13 | 0.03 | 0.2 | 0.04 |
| 1 | 204954993 | rs2278795 | 0.239 | -0.095 | 0.107 | -0.993 | 0.007 | 0.009 | 0.009 | 1.396 | 0.51 | 0.46 | 0.3 | 0.3 | 0.38 | 0.37 |
| 1 | 204955329 | rs11240325 | 0.682 | 0.008 | -0.656 | -0.863 | 0.042 | 0.047 | 0.047 | 0.022 | 0.42 | 0.09 | 0.25 | 0.29 | 0.32 | 0.34 |
| 1 | 204960239 | rs3820336 | 0.637 | 0.975 | -0.571 | 0.101 | -0.019 | -0.024 | -0.024 | 0 | 0.05 | 0.01 | 0.11 | 0.1 | 0.07 | 0.18 |
| 1 | 204960930 | rs867450 | -0.106 | 0.896 | -0.375 | 0.14 | 0.005 | 0.038 | 0.038 | 0.376 | 0.22 | 0.35 | 0.38 | 0.59 | 0.23 | 0.31 |
| 1 | 204966170 | rs16854920 | 0.31 | 0.779 | -1.146 | -0.134 | -0.037 | -0.01 | -0.01 | 0 | 0.32 | 0.13 | 0.35 | 0.31 | 0.34 | 0.36 |
| 1 | 204966428 | rs2802808 | -1.136 | 1.503 | -0.974 | 0.802 | 0.014 | 0.048 | 0.048 | 0.685 | 0.18 | 0.21 | 0.29 | 0.43 | 0.25 | 0.4 |
| 1 | 204968945 | rs6700069 | 0.68 | 1.024 | -1.245 | 0.056 | -0.024 | -0.013 | -0.013 | 2.211 | 0.27 | 0.13 | 0.18 | 0.24 | 0.27 | 0.29 |
| 1 | 204969586 | rs2802809 | 0.055 | 1.07 | -0.942 | 0.027 | 0.054 | 0.104 | 0.104 | 0.522 | 0.27 | 0.42 | 0.39 | 0.55 | 0.35 | 0.34 |
| 1 | 204971485 | rs6697344 | 0.32 | 0.585 | -1.467 | 0.06 | -0.014 | -0.003 | -0.003 | 0.322 | 0.28 | 0.14 | 0.18 | 0.24 | 0.27 | 0.33 |
| 1 | 204974508 | rs16854930 | -0.431 | -0.378 | -1.139 | -0.079 | 0.018 | 0.016 | 0.016 | 0.01 | 0.44 | 0.56 | 0.27 | 0.26 | 0.26 | 0.47 |
| 1 | 204975368 | rs10900436 | 0.071 | -0.717 | -1.01 | -0.397 | -0.061 | -0.133 | -0.133 | 0 | 0.33 | 0.51 | 0.15 | 0.01 | 0.12 | 0.17 |
| 1 | 204975735 | rs2802841 | 1.185 | 0.091 | -1.714 | -0.684 | -0.151 | 0.036 | 0.036 | 0 | 0.32 | 0.02 | 0.27 | 0.18 | 0.47 | 0.27 |
| 1 | 204976138 | rs2802842 | 0.334 | 0.054 | -1.677 | -0.652 | -0.032 | -0.084 | -0.084 | 0 | 0.43 | 0.1 | 0.39 | 0.44 | 0.61 | 0.57 |
| 1 | 204977799 | rs2595968 | -0.226 | -0.54 | -1.756 | -0.911 | 0.013 | -0.028 | -0.028 | 0 | 0.65 | 0.67 | 0.52 | 0.45 | 0.73 | 0.72 |
| 1 | 204980360 | rs2595967 | 0.914 | -0.365 | -1.237 | -0.855 | -0.033 | 0.058 | 0.058 | 0 | 0.38 | 0.2 | 0.3 | 0.21 | 0.48 | 0.32 |
| 1 | 204980907 | rs2802845 | -0.065 | -0.078 | -1.165 | -0.787 | 0.02 | 0.096 | 0.096 | 0.03 | 0.75 | 0.54 | 0.53 | 0.45 | 0.74 | 0.75 |
| 1 | 204982713 | rs2802846 | -0.051 | -0.238 | -1.563 | -0.879 | -0.003 | -0.006 | -0.006 | 0.955 | 0.73 | 0.75 | 0.48 | 0.42 | 0.67 | 0.67 |
| 1 | 204984572 | rs2802847 | 0.362 | -0.286 | -1.985 | -1.788 | -0.006 | 0.301 | 0.301 | 0.043 | 0.47 | 0.15 | 0.4 | 0.06 | 0.62 | 0.33 |
| 1 | 204987629 | rs16854941 | -0.404 | -0.812 | -0.75 | -1.132 | 0.028 | 0.025 | 0.025 | 0.575 | 0.06 | 0.29 | 0.04 | 0.01 | 0.01 | 0.01 |
| 1 | 204988535 | rs2794866 | 0.396 | -1.272 | -2.197 | -1.338 | 0.147 | 0.126 | 0.126 | 0.544 | 0.51 | 0.16 | 0.68 | 0.75 | 0.71 | 0.53 |
| 1 | 204989163 | rs2841633 | 0.402 | -1.27 | -2.196 | -1.337 | 0.148 | 0.127 | 0.127 | 0.541 | 0.51 | 0.16 | 0.68 | 0.75 | 0.71 | 0.53 |
| 1 | 204998375 | rs4951155 | 0.437 | -0.897 | -2.248 | -1.111 | 0.066 | 0.104 | 0.104 | 1.27 | 0.56 | 0.08 | 0.65 | 0.78 | 0.68 | 0.6 |
| 1 | 205007950 | rs11240339 | 0.007 | -0.117 | -1.648 | -1.115 | -0.001 | 0.091 | 0.091 | 1.227 | 0.52 | 0.34 | 0.71 | 0.85 | 0.43 | 0.67 |
| 1 | 205011659 | rs4951158 | -0.351 | -0.516 | -1.321 | -0.758 | 0 | -0.001 | -0.001 | 0 | 0.14 | 0.1 | 0.14 | 0.18 | 0.13 | 0.15 |
| 1 | 205013871 | rs6660593 | 0.387 | -1.336 | -1.071 | -0.39 | 0.001 | 0 | 0 | 0.006 | 0.24 | 0.24 | 0.28 | 0.47 | 0.11 | 0.28 |
| 1 | 205014616 | rs3753848 | 0.383 | -1.505 | -0.561 | -0.428 | -0.034 | 0.07 | 0.07 | 2.139 | 0.24 | 0.31 | 0.29 | 0.49 | 0.11 | 0.27 |
| 1 | 205015284 | rs11240341 | -0.213 | -1.385 | -1.554 | -1.222 | 0.014 | 0.005 | 0.005 | 0 | 0.65 | 0.85 | 0.66 | 0.69 | 0.66 | 0.63 |
| 1 | 205016018 | rs1006054 | 0.164 | -1.442 | -1.405 | -1.057 | -0.009 | -0.001 | -0.001 | 0 | 0.07 | 0.06 | 0.03 | 0.05 | 0.02 | 0.02 |
| 1 | 205019488 | rs10900438 | 0.355 | -1.893 | -1.524 | -1.284 | 0.05 | -0.016 | -0.016 | 0 | 0.3 | 0.21 | 0.34 | 0.65 | 0.15 | 0.47 |
| 1 | 205019571 | rs10793724 | 0.549 | -1.895 | -1.531 | -1.273 | 0.061 | -0.067 | -0.067 | 0.475 | 0.28 | 0.15 | 0.31 | 0.58 | 0.15 | 0.46 |
| 1 | 205020566 | rs4951161 | 0.364 | -1.882 | -1.52 | -1.282 | 0.05 | -0.018 | -0.018 | 0 | 0.3 | 0.21 | 0.34 | 0.65 | 0.15 | 0.47 |
| 1 | 205023316 | rs9787334 | 0.19 | -1.938 | -1.687 | -1.437 | 0.037 | 0.097 | 0.097 | 0.249 | 0.41 | 0.19 | 0.51 | 0.81 | 0.3 | 0.51 |
| 1 | 205027737 | rs2275697 | 0.63 | -1.92 | -1.839 | -1.625 | 0.07 | -0.126 | -0.126 | 0.077 | 0.3 | 0.14 | 0.28 | 0.48 | 0.15 | 0.47 |
| 1 | 205028915 | rs10793725 | 0.943 | -2.157 | -1.978 | -1.94 | -0.094 | 0.12 | 0.12 | 0.217 | 0.3 | 0.12 | 0.3 | 0.05 | 0.51 | 0.17 |
| 1 | 205031769 | rs2242000 | 1.51 | -2.118 | -1.817 | -1.805 | -0.002 | -0.012 | -0.012 | 0 | 0.16 | 0.09 | 0.27 | 0.4 | 0.18 | 0.28 |
| 1 | 205033283 | rs1470637 | -0.533 | -2.174 | -1.088 | -1.897 | 0.01 | -0.021 | -0.021 | 0.174 | 0.55 | 0.44 | 0.59 | 0.95 | 0.35 | 0.76 |
| 1 | 205033374 | rs2305277 | -0.177 | -2.161 | -1.322 | -1.862 | 0.066 | 0.138 | 0.138 | 0 | 0.51 | 0.35 | 0.56 | 0.88 | 0.34 | 0.74 |
| 1 | 205034434 | rs3901740 | 1.253 | -1.908 | -1.473 | -1.562 | 0.031 | -0.158 | -0.158 | 0 | 0.24 | 0.01 | 0.25 | 0.46 | 0.11 | 0.3 |
| 1 | 205041158 | rs2229868 | 0.38 | -1.824 | -1.179 | -1.639 | 0.096 | -0.041 | -0.041 | 0 | 0.39 | 0.24 | 0.32 | 0.55 | 0.2 | 0.5 |
| 1 | 205041465 | rs6662930 | 0.833 | -1.677 | -1.175 | -1.88 | -0.102 | 0.096 | 0.096 | 0.117 | 0.26 | 0.11 | 0.28 | 0.05 | 0.46 | 0.15 |
| 1 | 205041542 | rs3903399 | 0.871 | -1.832 | -0.995 | -1.966 | -0.001 | 0 | 0 | 0 | 0.19 | 0.15 | 0.31 | 0.43 | 0.21 | 0.31 |
| 1 | 205041952 | rs6696846 | 0.398 | -1.793 | -0.915 | -2.013 | -0.069 | 0.064 | 0.064 | 0 | 0.29 | 0.18 | 0.31 | 0.13 | 0.47 | 0.17 |
| 1 | 205044339 | rs11240351 | 0.467 | -1.592 | -0.744 | -2.021 | 0.008 | 0.027 | 0.027 | 0 | 0.56 | 0.42 | 0.6 | 0.87 | 0.41 | 0.8 |
| 1 | 205044502 | rs16855036 | 0.207 | -1.767 | -0.809 | -2.124 | -0.03 | -0.046 | -0.046 | 0.174 | 0.1 | 0.28 | 0.04 | 0.02 | 0.05 | 0.16 |
| 1 | 205044757 | rs16855040 | 0.207 | -1.767 | -0.809 | -2.125 | -0.03 | -0.054 | -0.054 | 0 | 0.1 | 0.28 | 0.04 | 0.01 | 0.05 | 0.16 |
| 1 | 205045087 | rs1572993 | 0.084 | -1.768 | -0.812 | -2.134 | 0 | -0.033 | -0.033 | 0.486 | 0.59 | 0.45 | 0.63 | 0.93 | 0.42 | 0.83 |
| 1 | 205046894 | rs16855045 | 1.595 | -1.775 | -0.809 | -2.037 | -0.002 | 0.003 | 0.003 | 0.408 | 0.16 | 0.12 | 0.28 | 0.39 | 0.19 | 0.28 |
| 1 | 205048383 | rs7523477 | -0.9 | -1.704 | -0.732 | -2.109 | -0.019 | -0.111 | -0.111 | 0.479 | 0.81 | 0.56 | 0.89 | 0.98 | 0.85 | 0.93 |
| 1 | 205052723 | rs10494858 | 0.17 | -1.711 | -0.757 | -2.106 | -0.03 | -0.054 | -0.054 | 0 | 0.1 | 0.28 | 0.04 | 0.01 | 0.05 | 0.15 |
| 1 | 205052939 | rs4950979 | -0.549 | -1.838 | -0.786 | -2.198 | -0.027 | -0.1 | -0.1 | 2.548 | 0.78 | 0.5 | 0.87 | 0.98 | 0.84 | 0.92 |
| 1 | 205053150 | rs16855059 | 0.137 | -1.678 | -0.706 | -2.103 | -0.03 | -0.054 | -0.054 | 0 | 0.1 | 0.28 | 0.04 | 0.01 | 0.05 | 0.16 |

| | | | | | | | | | | | | | | | | |
|---|---|---|---|---|---|---|---|---|---|---|---|---|---|---|---|---|
| 1 | 205053219 | rs4951168 | -0.887 | -1.676 | -0.701 | -2.102 | -0.019 | -0.11 | -0.11 | 0.652 | 0.81 | 0.55 | 0.89 | 0.98 | 0.85 | 0.93 |
| 1 | 205054114 | rs7520752 | 0.951 | -1.671 | -0.748 | -2.138 | -0.099 | 0.087 | 0.087 | 0.022 | 0.22 | 0.11 | 0.26 | 0.06 | 0.43 | 0.1 |
| 1 | 205054406 | rs2369526 | -0.165 | -1.667 | -0.743 | -2.134 | -0.014 | -0.011 | -0.011 | 0 | 0.32 | 0.39 | 0.3 | 0.07 | 0.48 | 0.26 |
| 1 | 205055162 | rs2197909 | -1.02 | -1.668 | -0.658 | -2.094 | -0.013 | -0.11 | -0.11 | 0 | 0.82 | 0.6 | 0.89 | 0.98 | 0.85 | 0.93 |
| 1 | 205063949 | rs3738154 | -0.532 | -1.224 | -0.771 | -2.352 | 0.001 | -0.028 | -0.028 | 0 | 0.31 | 0.35 | 0.28 | 0.09 | 0.45 | 0.23 |
| 1 | 205065733 | rs11577406 | 0.805 | -1.738 | -0.625 | -2.039 | -0.094 | 0.083 | 0.083 | 0.023 | 0.22 | 0.11 | 0.26 | 0.06 | 0.43 | 0.1 |
| 1 | 205070573 | rs11240358 | 0.824 | -1.783 | -0.63 | -2.052 | 0.058 | 0.251 | 0.251 | 0.062 | 0.5 | 0.16 | 0.59 | 0.91 | 0.38 | 0.67 |
| 1 | 205072625 | rs17416419 | 0.846 | -1.667 | -0.59 | -1.97 | -0.003 | -0.015 | -0.015 | 0 | 0.08 | 0.16 | 0.03 | 0.01 | 0.04 | 0.15 |
| 1 | 205084120 | rs7515178 | -0.344 | -1.674 | -0.598 | -1.886 | -0.014 | -0.011 | -0.011 | 0 | 0.33 | 0.4 | 0.3 | 0.07 | 0.47 | 0.25 |
| 1 | 205099959 | rs3862948 | 0.246 | -1.661 | -0.539 | -1.715 | 0.001 | -0.005 | -0.005 | 0.657 | 0.24 | 0.29 | 0.3 | 0.4 | 0.24 | 0.43 |
| 1 | 205110929 | rs11240368 | -1.073 | -1.776 | -0.578 | -1.792 | -0.018 | -0.111 | -0.111 | 0 | 0.82 | 0.62 | 0.89 | 0.98 | 0.86 | 0.93 |
| 1 | 205111761 | rs2836 | 0.871 | -1.799 | -0.726 | -1.858 | -0.126 | 0.111 | 0.111 | 0.016 | 0.22 | 0.07 | 0.26 | 0.04 | 0.43 | 0.1 |
| 1 | 205112911 | rs3795556 | 0.204 | -1.801 | -0.811 | -1.835 | -0.006 | 0.037 | 0.037 | 0.145 | 0.26 | 0.42 | 0.32 | 0.4 | 0.26 | 0.44 |
| 1 | 205113595 | rs12037322 | 1.452 | -1.814 | -0.815 | -1.838 | 0.065 | -0.134 | -0.134 | 3.16 | 0.34 | 0.05 | 0.31 | 0.52 | 0.18 | 0.39 |
| 1 | 205113833 | rs3183297 | 0.193 | -1.814 | -0.814 | -1.838 | -0.007 | 0.037 | 0.037 | 0.15 | 0.26 | 0.42 | 0.32 | 0.4 | 0.26 | 0.44 |
| 1 | 205115428 | rs10900452 | 1.375 | -1.779 | -0.993 | -1.75 | 0.065 | -0.133 | -0.133 | 0 | 0.34 | 0.05 | 0.31 | 0.52 | 0.18 | 0.39 |
| 1 | 205128059 | rs10751431 | 0.707 | -1.71 | -0.997 | -1.714 | 0.023 | -0.031 | -0.031 | 0 | 0.56 | 0.15 | 0.57 | 0.58 | 0.61 | 0.49 |
| 1 | 205139804 | rs2887527 | -0.913 | -1.605 | -0.804 | -1.776 | -0.026 | -0.106 | -0.106 | 0 | 0.81 | 0.53 | 0.88 | 0.98 | 0.87 | 0.93 |
| 1 | 205145005 | rs10900456 | 0.894 | -1.697 | -1.122 | -1.711 | -0.009 | -0.077 | -0.077 | 0.891 | 0.52 | 0.04 | 0.53 | 0.49 | 0.59 | 0.45 |
| 1 | 205152596 | rs3851287 | 1.551 | -1.732 | -1.066 | -1.72 | -0.007 | -0.01 | -0.01 | 0.569 | 0.16 | 0.11 | 0.28 | 0.4 | 0.21 | 0.28 |
| 1 | 205153208 | rs10900461 | 0.567 | -1.54 | -1.14 | -1.609 | 0.017 | -0.059 | -0.059 | 0 | 0.55 | 0.11 | 0.56 | 0.57 | 0.61 | 0.48 |
| 1 | 205157903 | rs2369634 | -1.609 | -1.422 | -0.653 | -1.433 | 0.006 | 0.014 | 0.014 | 0 | 0.85 | 0.93 | 0.92 | 1 | 0.87 | 0.93 |
| 1 | 205195831 | rs7513642 | 1.446 | -1.397 | -0.965 | -1.435 | 0.041 | -0.134 | -0.134 | 0 | 0.3 | 0.01 | 0.28 | 0.45 | 0.17 | 0.35 |
| 1 | 205209530 | rs1078362 | 0.362 | -1.229 | -1.018 | -1.491 | 0.025 | -0.039 | -0.039 | 0 | 0.51 | 0.11 | 0.54 | 0.57 | 0.58 | 0.47 |
| 1 | 205213883 | rs10900473 | 1.356 | -1.218 | -0.971 | -1.514 | 0.046 | -0.093 | -0.093 | 0.195 | 0.32 | 0.12 | 0.33 | 0.51 | 0.2 | 0.39 |
| 1 | 205216110 | rs17417252 | 0.341 | -0.976 | -0.649 | -1.405 | 0.005 | -0.032 | -0.032 | 0 | 0.28 | 0.24 | 0.33 | 0.41 | 0.28 | 0.45 |
| 1 | 205220241 | rs1172155 | -1.213 | -0.878 | -0.603 | -1.367 | -0.017 | -0.111 | -0.111 | 0 | 0.82 | 0.6 | 0.89 | 0.98 | 0.87 | 0.93 |
| 1 | 205224758 | rs1172149 | -0.016 | -0.512 | -1.223 | -1.122 | 0.037 | -0.005 | -0.005 | 0 | 0.54 | 0.24 | 0.56 | 0.57 | 0.59 | 0.47 |
| 1 | 205225457 | rs1172147 | 0.296 | 0.335 | -1.077 | -1.119 | -0.06 | 0.05 | 0.05 | 0 | 0.18 | 0.11 | 0.22 | 0.06 | 0.38 | 0.08 |
| 1 | 205236233 | rs4951182 | 0.385 | -0.066 | -1.084 | -1.327 | 0.002 | 0.132 | 0.132 | 0 | 0.54 | 0.26 | 0.63 | 0.86 | 0.44 | 0.65 |
| 1 | 205238275 | rs1768586 | 0.242 | 1.005 | -1.063 | -0.792 | -0.047 | 0.033 | 0.033 | 0 | 0.14 | 0.1 | 0.19 | 0.05 | 0.35 | 0.07 |
| 1 | 205244953 | rs1172130 | -0.325 | 0.99 | -1.007 | -0.789 | -0.031 | 0.012 | 0.012 | 0 | 0.17 | 0.15 | 0.21 | 0.11 | 0.37 | 0.11 |
| 1 | 205257338 | rs1562866 | 0.325 | 0.336 | -1.232 | -0.527 | 0.049 | -0.134 | -0.134 | 0.016 | 0.34 | 0.05 | 0.33 | 0.49 | 0.21 | 0.35 |
| 1 | 205263122 | rs17345837 | -0.128 | 1.827 | -0.764 | -0.55 | -0.1 | 0.033 | 0.033 | 0 | 0.13 | 0.02 | 0.18 | 0.04 | 0.35 | 0.07 |
| 1 | 205272292 | rs7550061 | -1.002 | 0.292 | 0.366 | -0.601 | -0.005 | -0.042 | -0.042 | 0 | 0.19 | 0.41 | 0.13 | 0.12 | 0.13 | 0.19 |
| 1 | 205272926 | rs1106201 | 0.72 | 0.16 | -0.427 | -0.503 | 0 | -0.063 | -0.063 | 0.58 | 0.31 | 0.14 | 0.42 | 0.46 | 0.3 | 0.39 |
| 1 | 205274606 | rs2275869 | -0.155 | 0.067 | -0.171 | -0.542 | -0.029 | 0 | 0 | 1.103 | 0.07 | 0.19 | 0.06 | 0.08 | 0.01 | 0.12 |
| 1 | 205274675 | rs1539695 | 0.702 | 0.265 | -0.335 | -0.488 | -0.005 | -0.068 | -0.068 | 0.406 | 0.31 | 0.13 | 0.42 | 0.46 | 0.3 | 0.39 |
| 1 | 205276025 | rs1110304 | -0.274 | 0.447 | -0.002 | -1.309 | 0 | -0.116 | -0.116 | 0 | 0.63 | 0.52 | 0.64 | 0.93 | 0.63 | 0.77 |
| 1 | 205277781 | rs2275868 | 0.668 | 0.077 | -0.535 | -1.191 | 0.019 | 0.013 | 0.013 | 0.044 | 0.08 | 0.03 | 0.05 | 0.06 | 0.02 | 0.11 |
| 1 | 205282979 | rs4950989 | 0.979 | 0.14 | -0.694 | -1.14 | -0.013 | -0.071 | -0.071 | 0 | 0.29 | 0.14 | 0.41 | 0.49 | 0.3 | 0.35 |
| 1 | 205284082 | rs12074332 | 0.639 | 0.625 | -0.16 | -1.464 | 0.013 | -0.094 | -0.094 | 0 | 0.18 | 0.26 | 0.22 | 0 | 0.2 | 0.07 |
| 1 | 205292420 | rs11240414 | -0.471 | 0.702 | -0.161 | -1.411 | 0.017 | -0.067 | -0.067 | 0 | 0.21 | 0.25 | 0.13 | 0.03 | 0.28 | 0.17 |
| 1 | 205292964 | rs13374813 | -1.505 | -0.085 | 0.081 | -1.736 | -0.003 | -0.116 | -0.116 | 0.195 | 0.34 | 0.49 | 0.2 | 0.09 | 0.3 | 0.29 |
| 1 | 205305048 | rs10900479 | -0.044 | -0.068 | 0.013 | -1.761 | 0.003 | -0.032 | -0.032 | 0 | 0.19 | 0.2 | 0.11 | 0.03 | 0.26 | 0.14 |
| 1 | 205305684 | rs13375059 | -0.459 | -0.28 | 0.01 | -1.883 | -0.031 | -0.045 | -0.045 | 0.025 | 0.15 | 0.19 | 0.05 | 0.01 | 0.04 | 0.16 |
| 1 | 205306102 | rs10900482 | -0.155 | 0.191 | -0.114 | -1.648 | 0.003 | -0.032 | -0.032 | 0.585 | 0.19 | 0.2 | 0.11 | 0.03 | 0.26 | 0.14 |
| 1 | 205307789 | rs11240441 | -0.184 | 0.227 | -0.204 | -1.647 | 0.002 | -0.031 | -0.031 | 0.946 | 0.19 | 0.2 | 0.11 | 0.03 | 0.27 | 0.14 |
| 1 | 205313647 | rs4951199 | 0.059 | -0.273 | 0.111 | -1.523 | -0.034 | -0.047 | -0.047 | 0 | 0.91 | 0.83 | 0.97 | 1 | 0.98 | 0.98 |
| 1 | 205317503 | rs12059162 | -0.072 | -0.771 | -0.403 | -1.885 | 0.012 | 0.035 | 0.035 | 0 | 0.23 | 0.14 | 0.09 | 0.21 | 0.17 | 0.35 |
| 1 | 205318321 | rs10751435 | 0.646 | -0.872 | -0.189 | -1.69 | 0 | -0.001 | -0.001 | 0 | 0.58 | 0.37 | 0.7 | 0.48 | 0.6 | 0.48 |
| 1 | 205318983 | rs913722 | -0.124 | -0.797 | -0.117 | -1.975 | -0.002 | -0.01 | -0.01 | 0 | 0.59 | 0.45 | 0.57 | 0.69 | 0.56 | 0.73 |
| 1 | 205322091 | rs7516786 | 0.722 | 0.026 | -0.304 | -1.838 | -0.003 | 0.007 | 0.007 | 0 | 0.37 | 0.33 | 0.2 | 0.21 | 0.35 | 0.42 |
| 1 | 205322809 | rs2864859 | -0.566 | -0.14 | -0.333 | -1.55 | 0.004 | -0.022 | -0.022 | 0 | 0.77 | 0.88 | 0.8 | 0.7 | 0.75 | 0.72 |
| 1 | 205322934 | rs2864860 | 1.326 | -0.146 | -0.044 | -1.576 | 0.002 | 0.041 | 0.041 | 0 | 0.15 | 0.16 | 0.36 | 0.49 | 0.18 | 0.2 |
| 1 | 205324725 | rs2183230 | 0.04 | -0.414 | 0.045 | -1.95 | -0.002 | 0.011 | 0.011 | 0 | 0.62 | 0.68 | 0.44 | 0.21 | 0.56 | 0.52 |
| 1 | 205327237 | rs12140669 | 0.5 | -0.448 | 0.084 | -1.91 | -0.005 | 0.023 | 0.023 | 0 | 0.38 | 0.32 | 0.21 | 0.18 | 0.36 | 0.45 |
| 1 | 205327281 | rs4951209 | 0.45 | -0.43 | -0.079 | -2.211 | -0.051 | 0.075 | 0.075 | 0 | 0.33 | 0.17 | 0.54 | 0.8 | 0.48 | 0.38 |
| 1 | 205330243 | rs963550 | -0.856 | -0.313 | 0.646 | -1.767 | -0.003 | -0.022 | -0.022 | 0 | 0.64 | 0.63 | 0.81 | 0.79 | 0.73 | 0.62 |
| 1 | 205336875 | rs6676552 | -0.193 | -0.061 | 0.536 | -1.437 | 0.008 | -0.002 | -0.002 | 0 | 0.51 | 0.36 | 0.62 | 0.47 | 0.56 | 0.47 |
| 1 | 205339119 | rs6661074 | 0.502 | -0.153 | 0.325 | -1.251 | -0.027 | -0.025 | -0.025 | 0 | 0.44 | 0.17 | 0.55 | 0.43 | 0.47 | 0.33 |
| 1 | 205340269 | rs6664679 | 1.324 | -0.109 | 0.128 | -1.098 | 0 | 0.026 | 0.026 | 0 | 0.13 | 0.16 | 0.19 | 0.32 | 0.18 | 0.15 |
| 1 | 205343648 | rs12402771 | 0.652 | -0.326 | 0.175 | -1.196 | 0.018 | 0.083 | 0.083 | 0.046 | 0.34 | 0.16 | 0.31 | 0.21 | 0.26 | 0.37 |
| 1 | 205348215 | rs7551359 | -0.443 | -0.196 | 0.345 | -1.145 | -0.002 | -0.018 | -0.018 | 0.004 | 0.57 | 0.58 | 0.64 | 0.77 | 0.72 | 0.61 |
| 1 | 205354025 | rs2149733 | 0.552 | -0.183 | -0.173 | -1.032 | -0.052 | -0.046 | -0.046 | 0 | 0.39 | 0.16 | 0.42 | 0.42 | 0.48 | 0.32 |
| 1 | 205354073 | rs11581020 | 0.944 | 0.146 | 0.07 | -0.827 | 0.001 | -0.002 | -0.002 | 0 | 0.13 | 0.12 | 0.17 | 0.23 | 0.14 | 0.17 |
| 1 | 205358594 | rs11240465 | -1.134 | 0.02 | -0.327 | -1.001 | -0.002 | -0.045 | -0.045 | 0 | 0.74 | 0.83 | 0.73 | 0.6 | 0.75 | 0.77 |
| 1 | 205358622 | rs11240466 | -0.233 | -0.192 | -0.182 | -1.268 | -0.002 | -0.01 | -0.01 | 0 | 0.43 | 0.47 | 0.39 | 0.22 | 0.43 | 0.25 |
| 1 | 205358741 | rs17346510 | 0.594 | -0.737 | -0.204 | -1.758 | 0.002 | 0.01 | 0.01 | 1.968 | 0.28 | 0.2 | 0.29 | 0.25 | 0.23 | 0.4 |

| | | | | | | | | | | | | | | | | |
|---|---|---|---|---|---|---|---|---|---|---|---|---|---|---|---|---|
| 1 | 205366268 | rs1928444 | -0.649 | -0.73 | -0.232 | -1.855 | 0 | 0.003 | 0.003 | 1.649 | 0.54 | 0.54 | 0.51 | 0.38 | 0.53 | 0.63 |
| 1 | 205369075 | rs1928442 | 0.017 | -0.567 | -0.241 | -1.833 | 0.006 | -0.069 | -0.069 | 0 | 0.31 | 0.52 | 0.24 | 0.13 | 0.3 | 0.21 |
| 1 | 205377069 | rs6421773 | 0.077 | -0.533 | -0.306 | -1.817 | 0.005 | -0.051 | -0.051 | 2.234 | 0.3 | 0.42 | 0.22 | 0.13 | 0.3 | 0.21 |
| 1 | 205377965 | rs7533222 | 0.954 | -0.768 | -0.842 | -1.717 | -0.076 | -0.006 | -0.006 | 0.207 | 0.33 | 0.08 | 0.43 | 0.62 | 0.45 | 0.35 |
| 1 | 205381940 | rs1338624 | 1.414 | -1.303 | -0.201 | -2.001 | 0.085 | 0.191 | 0.191 | 0 | 0.54 | 0.12 | 0.77 | 0.88 | 0.69 | 0.64 |
| 1 | 205388023 | rs913728 | 0.199 | -1.236 | 0.057 | -1.998 | -0.037 | -0.17 | -0.17 | 0.342 | 0.73 | 0.48 | 0.87 | 1 | 0.83 | 0.93 |
| 1 | 205389460 | rs16855560 | -0.889 | -0.989 | -0.261 | -1.58 | -0.018 | -0.01 | -0.01 | 0 | 0.19 | 0.3 | 0.1 | 0.13 | 0.14 | 0.29 |
| 1 | 205404878 | rs10900490 | 0.877 | -1.098 | -0.554 | -1.643 | -0.034 | -0.075 | -0.075 | 0 | 0.67 | 0.54 | 0.83 | 0.92 | 0.77 | 0.87 |
| 1 | 205405358 | rs1999132 | 0.578 | -1.239 | -0.501 | -1.714 | -0.059 | -0.102 | -0.102 | 0 | 0.82 | 0.69 | 0.95 | 0.99 | 0.93 | 0.97 |
| 1 | 205405964 | rs3738744 | 0.404 | -0.683 | -0.376 | -1.281 | -0.008 | -0.041 | -0.041 | 0 | 0.8 | 0.57 | 0.86 | 0.93 | 0.82 | 0.9 |
| 1 | 205415348 | rs10793730 | 0.219 | -0.764 | -1.053 | -0.878 | -0.057 | -0.021 | -0.021 | 0 | 0.79 | 0.36 | 0.92 | 0.88 | 0.93 | 0.82 |
| 1 | 205418134 | rs1999131 | -0.452 | 0.764 | -1.28 | -0.777 | -0.012 | 0.008 | 0.008 | 0.521 | 0.34 | 0.3 | 0.29 | 0.23 | 0.41 | 0.26 |
| 1 | 205423893 | rs10900493 | 0.13 | 0.713 | -1.616 | -0.913 | -0.066 | 0.107 | 0.107 | 0 | 0.45 | 0.13 | 0.64 | 0.76 | 0.55 | 0.6 |
| 1 | 205428066 | rs12086423 | -0.223 | 0.127 | -0.837 | -1.757 | -0.008 | -0.075 | -0.075 | 0 | 0.43 | 0.51 | 0.2 | 0.08 | 0.28 | 0.22 |
| 1 | 205434743 | rs6692041 | -0.442 | 0.212 | -1.453 | -1.735 | 0.063 | 0.29 | 0.29 | 0 | 0.49 | 0.86 | 0.27 | 0.08 | 0.35 | 0.23 |
| 1 | 205438914 | rs10900494 | -0.604 | -0.296 | -0.516 | -1.395 | -0.046 | -0.073 | -0.073 | 0 | 0.29 | 0.38 | 0.14 | 0.11 | 0.21 | 0.16 |
| 1 | 205439769 | rs4279903 | 0.18 | 0.361 | -0.224 | -1.35 | -0.001 | -0.015 | -0.015 | 0.712 | 0.78 | 0.77 | 0.83 | 0.93 | 0.81 | 0.91 |
| 1 | 205442194 | rs1109953 | 1.068 | 0.416 | -0.282 | -1.15 | 0.003 | 0.043 | 0.043 | 0 | 0.6 | 0.26 | 0.69 | 0.73 | 0.65 | 0.8 |
| 1 | 205444187 | rs2027288 | 0.599 | 0.486 | -0.035 | -1.067 | 0.01 | -0.025 | -0.025 | 1.171 | 0.66 | 0.52 | 0.72 | 0.73 | 0.67 | 0.83 |
| 1 | 205447508 | rs1338610 | 0.612 | 0.526 | -0.175 | -1.321 | 0.009 | -0.073 | -0.073 | 0.438 | 0.64 | 0.45 | 0.6 | 0.86 | 0.64 | 0.8 |
| 1 | 205451659 | rs4951226 | -0.919 | 0.204 | -0.029 | -0.856 | 0.001 | 0.006 | 0.006 | 0.084 | 0.23 | 0.4 | 0.23 | 0.19 | 0.21 | 0.17 |
| 1 | 205452302 | rs4951002 | -1.54 | -0.108 | -0.227 | -1.025 | -0.02 | -0.018 | -0.018 | 0.154 | 0.29 | 0.56 | 0.26 | 0.19 | 0.24 | 0.17 |
| 1 | 205455368 | rs7555999 | -1.104 | -0.156 | -0.63 | -1.036 | -0.01 | -0.007 | -0.007 | 0 | 0.27 | 0.68 | 0.25 | 0.19 | 0.22 | 0.19 |
| 1 | 205457566 | rs949710 | -0.622 | 0.031 | -1.955 | -1.282 | 0.051 | 0.179 | 0.179 | 0 | 0.49 | 0.91 | 0.34 | 0.19 | 0.37 | 0.35 |
| 1 | 205459889 | rs2027285 | 0.339 | 0.684 | -1.685 | -0.916 | -0.023 | 0.01 | 0.01 | 0.839 | 0.6 | 0.21 | 0.6 | 0.33 | 0.61 | 0.64 |
| 1 | 205461250 | rs11240493 | -0.221 | 1.207 | -1.736 | -0.978 | -0.101 | -0.072 | -0.072 | 0 | 0.27 | 0.12 | 0.28 | 0.61 | 0.48 | 0.37 |
| 1 | 205461563 | rs1041309 | 0.324 | 0.427 | -1.167 | -1.259 | -0.022 | 0.033 | 0.033 | 0 | 0.77 | 0.44 | 0.54 | 0.65 | 0.83 | 0.84 |
| 1 | 205465032 | rs746110 | 0.077 | 0.461 | -0.437 | -1.141 | -0.025 | 0.001 | 0.001 | 0 | 0.83 | 0.77 | 0.61 | 0.8 | 0.92 | 0.9 |
| 1 | 205467785 | rs10900497 | 0.705 | 0.376 | -0.05 | -1.187 | -0.071 | -0.02 | -0.02 | 0 | 0.79 | 0.54 | 0.58 | 0.79 | 0.89 | 0.9 |
| 1 | 205467891 | rs12119834 | -0.249 | -0.215 | -0.486 | -1.74 | 0.046 | 0.047 | 0.047 | 0 | 0.5 | 0.41 | 0.48 | 0.78 | 0.75 | 0.66 |
| 1 | 205468289 | rs6681714 | 0.483 | -0.153 | -0.421 | -1.78 | -0.032 | -0.013 | -0.013 | 0 | 0.76 | 0.62 | 0.55 | 0.78 | 0.82 | 0.86 |
| 1 | 205468612 | rs4951231 | 0.108 | -0.141 | -0.406 | -1.783 | 0.04 | 0.057 | 0.057 | 0 | 0.43 | 0.31 | 0.42 | 0.76 | 0.66 | 0.62 |
| 1 | 205470633 | rs12738728 | 1.326 | 0.097 | -0.665 | -1.434 | 0.044 | 0.045 | 0.045 | 0 | 0.28 | 0.15 | 0.1 | 0.02 | 0.14 | 0.24 |
| 1 | 205472184 | rs1538365 | 0.129 | 0.613 | -0.828 | -1.143 | -0.002 | -0.009 | -0.009 | 0 | 0.59 | 0.51 | 0.39 | 0.63 | 0.62 | 0.61 |
| 1 | 205473207 | rs10900498 | -0.222 | 0.655 | -0.778 | -1.018 | -0.004 | -0.003 | -0.003 | 0 | 0.66 | 0.57 | 0.45 | 0.65 | 0.71 | 0.65 |
| 1 | 205482189 | rs4074998 | -0.074 | 1.456 | -1.174 | -0.694 | 0 | 0 | 0 | 0 | 0.24 | 0.27 | 0.22 | 0.61 | 0.34 | 0.35 |
| 1 | 205482946 | rs1105822 | 0.561 | 0.502 | -1.587 | -0.835 | -0.021 | 0.027 | 0.027 | 0 | 0.06 | 0.01 | 0.05 | 0.01 | 0.09 | 0.05 |
| 1 | 205483124 | rs16855698 | 0.126 | -0.023 | -1.396 | -0.794 | -0.003 | 0 | 0 | 0 | 0.12 | 0.12 | 0.36 | 0.2 | 0.06 | 0.09 |
| 1 | 205483795 | rs12561888 | -0.315 | 0.413 | -1.443 | -0.751 | 0.034 | 0.082 | 0.082 | 0 | 0.49 | 0.28 | 0.3 | 0.7 | 0.43 | 0.5 |
| 1 | 205484073 | rs11240503 | -0.122 | 1.527 | -1.303 | 0.165 | -0.069 | -0.082 | -0.082 | 0 | 0.18 | 0.05 | 0.19 | 0.67 | 0.28 | 0.26 |
| 1 | 205499495 | rs12035825 | -0.124 | -0.609 | -1.168 | 0.09 | 0.038 | -0.022 | -0.022 | 0.874 | 0.27 | 0.17 | 0.25 | 0.31 | 0.12 | 0.35 |
| 1 | 205500543 | rs11240507 | 0.088 | -0.212 | -1.534 | -0.6 | -0.024 | 0.047 | 0.047 | 0 | 0.2 | 0.09 | 0.16 | 0.01 | 0.27 | 0.1 |
| 1 | 205500829 | rs3795547 | 0.189 | -0.845 | -1.265 | -0.677 | -0.022 | -0.009 | -0.009 | 0 | 0.07 | 0.14 | 0.13 | 0.03 | 0.04 | 0.06 |
| 1 | 205501521 | rs14028 | 0.191 | -0.543 | -0.879 | -1.222 | 0.027 | 0.005 | 0.005 | 0 | 0.38 | 0.23 | 0.35 | 0.36 | 0.31 | 0.51 |
| 1 | 205501843 | rs8027 | 0.134 | -0.546 | -0.88 | -1.224 | 0.033 | -0.004 | -0.004 | 0 | 0.4 | 0.23 | 0.44 | 0.39 | 0.33 | 0.56 |
| 1 | 205503250 | rs12727646 | 0.172 | -0.604 | -0.893 | -1.242 | 0.031 | 0.007 | 0.007 | 0 | 0.39 | 0.23 | 0.35 | 0.37 | 0.31 | 0.51 |
| 1 | 205504159 | rs4951236 | 1.421 | -0.547 | -1.016 | -1.229 | 0.047 | -0.057 | -0.057 | 0 | 0.22 | 0.05 | 0.11 | 0.29 | 0.11 | 0.28 |
| 1 | 205510242 | rs12751472 | 0.057 | -0.644 | -0.125 | -1.349 | -0.006 | -0.095 | -0.095 | 0 | 0.69 | 0.52 | 0.68 | 0.97 | 0.66 | 0.84 |
| 1 | 205510376 | rs12041365 | 1.009 | -0.709 | 0.109 | -1.379 | -0.001 | -0.001 | -0.001 | 0 | 0.31 | 0.27 | 0.42 | 0.63 | 0.37 | 0.39 |
| 1 | 205515921 | rs10900509 | 0.081 | -0.659 | 0.181 | -1.371 | -0.007 | -0.11 | -0.11 | 0 | 0.68 | 0.43 | 0.67 | 0.97 | 0.66 | 0.84 |
| 1 | 205525798 | rs1980 | -0.641 | -0.517 | 0.101 | -1.317 | 0.002 | -0.063 | -0.063 | 0.016 | 0.76 | 0.73 | 0.82 | 0.99 | 0.7 | 0.9 |
| 1 | 205533082 | rs11240521 | 0.113 | -0.872 | -0.134 | -1.037 | 0.02 | 0.009 | 0.009 | 0 | 0.41 | 0.33 | 0.36 | 0.36 | 0.31 | 0.51 |
| 1 | 205541515 | rs12035608 | 0.7 | -0.307 | -0.368 | -0.887 | -0.034 | 0.006 | 0.006 | 0 | 0.29 | 0.14 | 0.29 | 0.75 | 0.37 | 0.28 |
| 1 | 205543767 | rs7534154 | 0.52 | -0.855 | 0.074 | -1.511 | -0.001 | -0.079 | -0.079 | 0.12 | 0.41 | 0.44 | 0.28 | 0.02 | 0.42 | 0.23 |
| 1 | 205543862 | rs1891506 | 0.271 | -1.084 | 0.153 | -1.296 | -0.015 | -0.03 | -0.03 | 0 | 0.7 | 0.59 | 0.65 | 0.78 | 0.79 | 0.55 |
| 1 | 205550898 | rs7523059 | 0.405 | -1.121 | 0.224 | -1.313 | 0 | -0.018 | -0.018 | 0 | 0.17 | 0.16 | 0.24 | 0.06 | 0.16 | 0.2 |
| 1 | 205552864 | rs10494871 | 1.018 | -1.461 | -0.001 | -1.547 | 0.029 | 0.011 | 0.011 | 0 | 0.23 | 0.1 | 0.22 | 0.19 | 0.13 | 0.32 |
| 1 | 205554095 | rs7515820 | -0.472 | -1.472 | 0.226 | -1.723 | 0.003 | -0.089 | -0.089 | 0.017 | 0.38 | 0.55 | 0.4 | 0.06 | 0.45 | 0.29 |
| 1 | 205555145 | rs7531623 | -0.342 | -1.524 | 0.371 | -1.488 | -0.004 | -0.016 | -0.016 | 0 | 0.43 | 0.52 | 0.47 | 0.21 | 0.42 | 0.45 |
| 1 | 205556426 | rs6593954 | 0.022 | -0.958 | 0.434 | -1.338 | 0.001 | -0.169 | -0.169 | 1.322 | 0.24 | 0.54 | 0.19 | 0 | 0.31 | 0.09 |
| 1 | 205558200 | rs7530988 | -0.298 | -0.715 | -0.066 | -1.34 | -0.003 | -0.022 | -0.022 | 0 | 0.24 | 0.25 | 0.33 | 0.04 | 0.35 | 0.17 |
| 1 | 205558675 | rs11240528 | 0.774 | -0.95 | -0.251 | -0.51 | 0.005 | 0 | 0 | 0.602 | 0.4 | 0.49 | 0.28 | 0.34 | 0.42 | 0.26 |
| 1 | 205558976 | rs6677703 | 1.025 | -1.025 | 0.091 | -0.361 | -0.018 | -0.001 | -0.001 | 0.166 | 0.4 | 0.26 | 0.28 | 0.35 | 0.48 | 0.27 |
| 1 | 205560119 | rs7539751 | -1.625 | -1.237 | -0.629 | -0.604 | 0.019 | 0.027 | 0.027 | 0 | 0.79 | 0.89 | 0.88 | 0.95 | 0.89 | 0.83 |
| 1 | 205561372 | rs7529077 | 1.131 | -1.187 | -0.624 | -0.548 | 0.002 | -0.03 | -0.03 | 0 | 0.15 | 0.22 | 0.13 | 0.01 | 0.22 | 0.06 |
| 1 | 205562202 | rs7553871 | -0.237 | -1.061 | -0.616 | 0.437 | -0.009 | 0 | 0 | 0 | 0.67 | 0.45 | 0.67 | 0.72 | 0.78 | 0.69 |
| 1 | 205563162 | rs3795550 | 0.356 | -1.088 | -0.921 | 0.521 | 0.04 | 0.022 | 0.022 | 0 | 0.57 | 0.71 | 0.48 | 0.32 | 0.7 | 0.37 |
| 1 | 205564044 | rs11240530 | 0.37 | -0.773 | -1.638 | 0.738 | 0.011 | -0.033 | -0.033 | 0 | 0.16 | 0.12 | 0.22 | 0.41 | 0.11 | 0.37 |
| 1 | 205566236 | rs12728206 | 1.239 | -0.746 | -1.637 | 0.582 | -0.101 | -0.066 | -0.066 | 0 | 0.37 | 0.11 | 0.36 | 0.46 | 0.46 | 0.31 |

| | | | | | | | | | | | | | | | | |
|---|---|---|---|---|---|---|---|---|---|---|---|---|---|---|---|---|
| 1 | 205568213 | rs2282449 | -0.459 | -0.476 | -1.247 | 0.724 | 0.008 | -0.056 | -0.056 | 0 | 0.21 | 0.14 | 0.39 | 0.64 | 0.18 | 0.49 |
| 1 | 205569170 | rs12080053 | 0.745 | -0.448 | -1.208 | 0.683 | -0.001 | -0.005 | -0.005 | 0 | 0.15 | 0.17 | 0.13 | 0.01 | 0.22 | 0.06 |
| 1 | 205569973 | rs3818970 | -0.827 | -0.189 | -1.119 | 1.301 | 0.008 | -0.053 | -0.053 | 0.181 | 0.22 | 0.15 | 0.39 | 0.64 | 0.18 | 0.46 |
| 1 | 205588637 | rs3789461 | -0.444 | 0.095 | -0.814 | 1.445 | -0.002 | 0.099 | 0.099 | 0 | 0.11 | 0.23 | 0.13 | 0.38 | 0.1 | 0.31 |
| 1 | 205596870 | rs2185811 | -0.267 | 0.03 | -1.059 | 1.428 | -0.005 | 0.042 | 0.042 | 0 | 0.1 | 0.15 | 0.12 | 0.37 | 0.07 | 0.27 |
| 1 | 205602553 | rs6593958 | 0.406 | 0.343 | -1.561 | 0.817 | -0.084 | 0.011 | 0.011 | 0 | 0.37 | 0.09 | 0.3 | 0.32 | 0.41 | 0.26 |
| 1 | 205606172 | rs1114126 | -0.546 | -0.266 | -0.826 | 1.012 | -0.046 | 0.08 | 0.08 | 0 | 0.29 | 0.51 | 0.28 | 0.5 | 0.15 | 0.32 |
| 1 | 205607227 | rs11579421 | 0.546 | 0.094 | -0.718 | 0.675 | -0.007 | -0.005 | -0.005 | 0.312 | 0.26 | 0.21 | 0.38 | 0.25 | 0.34 | 0.25 |
| 1 | 205607514 | rs10900518 | -0.537 | 0.436 | -0.182 | 0.311 | 0.014 | -0.072 | -0.072 | 0 | 0.87 | 0.76 | 0.92 | 1 | 0.89 | 0.91 |
| 1 | 205609010 | rs12039255 | 0.241 | 0.243 | 0.147 | 0.671 | 0 | 0.202 | 0.202 | 0 | 0.08 | 0.26 | 0.11 | 0.38 | 0.06 | 0.26 |
| 1 | 205609410 | rs17347710 | 0.614 | 0.157 | -0.189 | 0.063 | 0.005 | 0.008 | 0.008 | 0 | 0.15 | 0.11 | 0.11 | 0.05 | 0.09 | 0.1 |
| 1 | 205610774 | rs6661827 | 0.041 | 0.142 | -0.008 | 0.111 | 0.007 | 0.004 | 0.004 | 0 | 0.35 | 0.44 | 0.49 | 0.63 | 0.39 | 0.51 |
| 1 | 205616849 | rs2153903 | 1.278 | 0.051 | -0.527 | 0.063 | 0.019 | 0.044 | 0.044 | 0 | 0.4 | 0.2 | 0.32 | 0.31 | 0.4 | 0.3 |
| 1 | 205622253 | rs7514718 | 0.645 | 0.05 | -0.275 | 0.111 | 0.002 | -0.05 | -0.05 | 0.286 | 0.76 | 0.73 | 0.83 | 0.95 | 0.82 | 0.81 |
| 1 | 205622275 | rs11589772 | -0.091 | 0.243 | -0.421 | 0.242 | 0.019 | -0.062 | -0.062 | 0.001 | 0.88 | 0.8 | 0.91 | 1 | 0.88 | 0.91 |
| 1 | 205627886 | rs12565987 | -0.126 | -0.773 | -1.23 | 0.211 | 0.032 | 0.062 | 0.062 | 0.012 | 0.17 | 0.03 | 0.09 | 0.05 | 0.07 | 0.1 |
| 1 | 205630893 | rs3761916 | -1.027 | -0.865 | -0.986 | 0.967 | -0.004 | -0.04 | -0.04 | 0 | 0.27 | 0.19 | 0.44 | 0.45 | 0.26 | 0.48 |
| 1 | 205631767 | rs16856110 | -0.26 | -0.78 | -1.202 | 1.22 | -0.064 | -0.083 | -0.083 | 0 | 0.14 | 0.04 | 0.38 | 0.35 | 0.21 | 0.28 |
| 1 | 205632932 | rs11240547 | 1.68 | -1.529 | -1.566 | 0.884 | 0.014 | -0.063 | -0.063 | 1.044 | 0.51 | 0.02 | 0.35 | 0.45 | 0.66 | 0.33 |
| 1 | 205639045 | rs3902968 | -1.731 | -1.436 | -1.032 | 0.701 | 0.035 | 0.067 | 0.067 | 3.196 | 0.44 | 0.85 | 0.63 | 0.65 | 0.33 | 0.66 |
| 1 | 205642790 | rs2153904 | -0.194 | -2.27 | -0.529 | -0.161 | -0.102 | 0.034 | 0.034 | 0 | 0.65 | 0.5 | 0.79 | 0.52 | 0.84 | 0.57 |
| 1 | 205648707 | rs1775147 | 0.975 | -2.283 | -0.414 | -0.271 | 0.164 | 0.053 | 0.053 | 0.066 | 0.45 | 0.21 | 0.63 | 0.36 | 0.71 | 0.38 |
| 1 | 205652472 | rs4245718 | -0.487 | -1.635 | -0.29 | -0.26 | 0.033 | 0.044 | 0.044 | 0.402 | 0.15 | 0.23 | 0.38 | 0.35 | 0.27 | 0.24 |
| 1 | 205655725 | rs16856148 | -0.321 | -1.925 | -0.191 | -0.394 | -0.003 | -0.028 | -0.028 | 0 | 0.14 | 0.12 | 0.05 | 0.01 | 0.11 | 0.09 |
| 1 | 205658939 | rs1172199 | 0.772 | -1.949 | -0.213 | -0.318 | 0.078 | 0.04 | 0.04 | 0.049 | 0.46 | 0.33 | 0.63 | 0.35 | 0.7 | 0.37 |
| 1 | 205661614 | rs12752037 | -0.681 | -1.997 | -0.101 | -0.37 | 0.009 | 0.011 | 0.011 | 0.015 | 0.49 | 0.37 | 0.32 | 0.65 | 0.3 | 0.63 |
| 1 | 205662977 | rs6676110 | -0.205 | -2.058 | 0.025 | -0.21 | -0.021 | 0.001 | 0.001 | 0.51 | 0.42 | 0.34 | 0.2 | 0.5 | 0.25 | 0.51 |
| 1 | 205684067 | rs10900522 | 0.015 | -1.684 | -0.197 | -0.06 | -0.029 | -0.03 | -0.03 | 0 | 0.13 | 0.09 | 0.35 | 0.34 | 0.24 | 0.22 |
| 1 | 205695695 | rs1772145 | -0.449 | -2.256 | 0.157 | -0.381 | -0.087 | -0.003 | -0.003 | 0 | 0.65 | 0.56 | 0.71 | 0.36 | 0.84 | 0.45 |
| 1 | 205698026 | rs7513645 | -0.228 | -1.935 | 0.313 | -0.159 | -0.034 | -0.036 | -0.036 | 0 | 0.14 | 0.1 | 0.37 | 0.34 | 0.25 | 0.22 |
| 1 | 205713378 | rs823128 | -0.547 | -1.964 | 0.432 | -0.023 | -0.024 | 0.141 | 0.141 | 0.015 | 0.91 | 0.64 | 0.84 | 0.85 | 0.97 | 0.88 |
| 1 | 205719532 | rs823114 | 2.16 | -1.693 | 0.283 | -0.152 | -0.038 | -0.038 | -0.038 | 0 | 0.56 | 0.15 | 0.37 | 0.5 | 0.54 | 0.56 |
| 1 | 205731824 | rs2027527 | 0.31 | -1.602 | 0.289 | -0.175 | 0 | -0.004 | -0.004 | 0 | 0.09 | 0.06 | 0.04 | 0.01 | 0.09 | 0.05 |
| 1 | 205735229 | rs823149 | 0.3 | -1.724 | 0.507 | -0.158 | -0.049 | 0.029 | 0.029 | 0 | 0.84 | 0.62 | 0.82 | 0.85 | 0.9 | 0.83 |
| 1 | 205735891 | rs7522056 | -0.91 | -1.675 | 0.516 | -0.123 | -0.019 | -0.016 | -0.016 | 0 | 0.21 | 0.17 | 0.39 | 0.34 | 0.33 | 0.25 |
| 1 | 205738166 | rs823135 | -0.444 | -1.707 | 0.548 | -0.044 | -0.03 | 0.126 | 0.126 | 0.021 | 0.9 | 0.63 | 0.84 | 0.86 | 0.97 | 0.88 |
| 1 | 205738251 | rs823136 | -0.444 | -1.707 | 0.548 | -0.044 | -0.03 | 0.126 | 0.126 | 0.021 | 0.9 | 0.63 | 0.84 | 0.86 | 0.97 | 0.88 |
| 1 | 205739266 | rs708723 | 2.281 | -1.714 | 0.515 | -0.095 | -0.034 | -0.037 | -0.037 | 0.808 | 0.56 | 0.17 | 0.38 | 0.51 | 0.53 | 0.56 |
| 1 | 205739396 | rs823139 | 1.326 | -1.988 | 0.669 | -0.027 | -0.051 | -0.012 | -0.012 | 0 | 0.78 | 0.44 | 0.78 | 0.85 | 0.86 | 0.79 |
| 1 | 205744218 | rs1572931 | 0.183 | -1.729 | 0.273 | 0.058 | -0.009 | -0.026 | -0.026 | 0.118 | 0.12 | 0.09 | 0.33 | 0.34 | 0.15 | 0.21 |
| 1 | 205744546 | rs823144 | 1.403 | -1.68 | 0.203 | -0.169 | -0.01 | 0.016 | 0.016 | 0 | 0.67 | 0.36 | 0.48 | 0.51 | 0.75 | 0.63 |
| 1 | 205748309 | rs10900524 | 1.152 | -1.657 | 0.196 | -0.149 | -0.023 | 0.08 | 0.08 | 0 | 0.71 | 0.44 | 0.5 | 0.51 | 0.76 | 0.65 |
| 1 | 205755046 | rs12565968 | 1.087 | -1.621 | -0.332 | -0.12 | 0.003 | -0.007 | -0.007 | 0 | 0.69 | 0.25 | 0.48 | 0.51 | 0.76 | 0.65 |
| 1 | 205756484 | rs6679073 | 1.245 | -1.805 | -0.601 | -0.343 | 0.013 | -0.142 | -0.142 | 0 | 0.27 | 0.03 | 0.17 | 0.5 | 0.25 | 0.45 |
| 1 | 205759876 | rs1341249 | 0.344 | -1.735 | -0.303 | -0.225 | -0.032 | 0.163 | 0.163 | 1.169 | 0.78 | 0.42 | 0.51 | 0.51 | 0.82 | 0.65 |
| 1 | 205762139 | rs708726 | 1.087 | -1.715 | -0.426 | -0.256 | 0.001 | -0.002 | -0.002 | 0.401 | 0.69 | 0.29 | 0.46 | 0.52 | 0.75 | 0.63 |
| 1 | 205764640 | rs823156 | 1.029 | -1.991 | -0.516 | -0.408 | -0.007 | -0.001 | -0.001 | 0 | 0.7 | 0.34 | 0.75 | 0.76 | 0.81 | 0.73 |
| 1 | 205771173 | rs823066 | -0.465 | -2.09 | -0.782 | -0.303 | -0.04 | 0.106 | 0.106 | 0.013 | 0.86 | 0.61 | 0.82 | 0.76 | 0.93 | 0.81 |
| 1 | 205777780 | rs708730 | 1 | -2.008 | -1.116 | -0.276 | 0.063 | 0.006 | 0.006 | 1.276 | 0.68 | 0.19 | 0.74 | 0.75 | 0.81 | 0.74 |
| 1 | 205785932 | rs1891094 | 0.61 | -1.761 | -1.022 | 0.044 | -0.004 | 0.009 | 0.009 | 0.101 | 0.08 | 0.08 | 0.31 | 0.32 | 0.11 | 0.16 |
| 1 | 205786935 | rs823082 | 0.596 | -1.813 | -0.986 | -0.122 | -0.001 | -0.048 | -0.048 | 0 | 0.43 | 0.34 | 0.31 | 0.01 | 0.52 | 0.2 |
| 1 | 205794175 | rs823088 | -0.041 | -1.819 | -0.975 | -0.098 | -0.003 | -0.006 | -0.006 | 0 | 0.52 | 0.52 | 0.63 | 0.33 | 0.63 | 0.36 |
| 1 | 205795960 | rs7529407 | 0.636 | -1.749 | -1.205 | -0.05 | -0.003 | 0.008 | 0.008 | 0.101 | 0.08 | 0.08 | 0.31 | 0.32 | 0.11 | 0.16 |
| 1 | 205801872 | rs1361754 | 0.449 | -1.595 | -1.221 | -0.418 | -0.001 | -0.043 | -0.043 | 0 | 0.43 | 0.35 | 0.31 | 0.01 | 0.52 | 0.2 |
| 1 | 205806179 | rs6694490 | -0.564 | -1.39 | -0.558 | 0.118 | -0.014 | -0.099 | -0.099 | 0 | 0.16 | 0.32 | 0.08 | 0.01 | 0.12 | 0.08 |
| 1 | 205811828 | rs4951025 | -0.689 | -0.798 | -0.481 | 0.489 | -0.047 | 0.06 | 0.06 | 0 | 0.23 | 0.42 | 0.24 | 0.29 | 0.14 | 0.28 |
| 1 | 205816798 | rs12756994 | -0.491 | -2.061 | -0.212 | -0.436 | -0.013 | 0.065 | 0.065 | 0 | 0.9 | 0.87 | 0.84 | 0.74 | 0.93 | 0.83 |
| 1 | 205817574 | rs1539394 | -0.274 | -2.126 | -0.074 | -0.474 | -0.024 | 0.181 | 0.181 | 0.386 | 0.88 | 0.53 | 0.8 | 0.75 | 0.93 | 0.84 |
| 1 | 205857654 | rs7364618 | -0.774 | -2.312 | -0.203 | -0.56 | -0.056 | 0.063 | 0.063 | 0.064 | 0.21 | 0.54 | 0.23 | 0.25 | 0.12 | 0.21 |
| 1 | 206720207 | rs11119007 | -0.02 | 0.342 | 0.357 | 0.273 | 0.007 | -0.051 | -0.051 | 0 | 0.88 | 0.81 | 0.89 | 1 | 0.85 | 0.98 |
| 1 | 206734044 | rs6669685 | 0.825 | -0.076 | 0.137 | 0.253 | -0.072 | -0.005 | -0.005 | 0.202 | 0.24 | 0.08 | 0.3 | 0.2 | 0.33 | 0.21 |
| 1 | 206736573 | rs4844540 | -0.692 | -0.114 | 0.506 | 0.329 | -0.001 | 0.009 | 0.009 | 0 | 0.25 | 0.23 | 0.35 | 0.57 | 0.26 | 0.48 |
| 1 | 206737276 | rs17434403 | 0.024 | -0.056 | 0.543 | 0.466 | 0.001 | -0.002 | -0.002 | 0 | 0.08 | 0.06 | 0.07 | 0.06 | 0.12 | 0.06 |
| 1 | 206741917 | rs944780 | 0.767 | -0.282 | 0.595 | 0.454 | 0 | -0.008 | -0.008 | 0 | 0.64 | 0.68 | 0.55 | 0.43 | 0.59 | 0.49 |
| 1 | 206744191 | rs10779478 | 0.721 | -0.219 | 0.485 | 0.568 | 0.002 | 0.031 | 0.031 | 0 | 0.65 | 0.51 | 0.56 | 0.46 | 0.6 | 0.57 |
| 1 | 206748424 | rs7512541 | 0.916 | -0.276 | 0.112 | 0.597 | -0.004 | -0.018 | -0.018 | 0 | 0.61 | 0.34 | 0.5 | 0.47 | 0.55 | 0.56 |
| 1 | 206757857 | rs4845112 | -0.248 | 0.25 | 0.469 | 1.072 | -0.019 | 0.046 | 0.046 | 0 | 0.07 | 0.14 | 0.1 | 0.15 | 0.06 | 0.07 |
| 1 | 206759106 | rs4501842 | -1.014 | 0.268 | 0.547 | 1.008 | 0.012 | 0.003 | 0.003 | 0.031 | 0.5 | 0.59 | 0.57 | 0.45 | 0.58 | 0.46 |
| 1 | 206759622 | rs4240840 | -0.074 | 0.311 | 0.682 | 0.987 | -0.006 | -0.036 | -0.036 | 0 | 0.37 | 0.23 | 0.43 | 0.4 | 0.39 | 0.32 |

| | | | | | | | | | | | | | | | | |
|---|---|---|---|---|---|---|---|---|---|---|---|---|---|---|---|---|
| 1 | 206759783 | rs7535580 | -0.917 | 0.22 | 0.634 | 0.977 | 0.001 | 0.004 | 0.004 | 0 | 0.86 | 0.84 | 0.77 | 0.7 | 0.83 | 0.71 |
| 1 | 206759864 | rs7531197 | 0.732 | 0.203 | 0.632 | 1.017 | -0.022 | -0.014 | -0.014 | 0.28 | 0.3 | 0.14 | 0.37 | 0.28 | 0.33 | 0.24 |
| 1 | 206760474 | rs7541244 | -0.858 | 0.317 | 0.238 | 0.331 | 0.001 | 0.005 | 0.005 | 0 | 0.86 | 0.83 | 0.8 | 0.7 | 0.83 | 0.71 |
| 1 | 206765672 | rs12075916 | 0.021 | 0.284 | 0.298 | 0.288 | -0.018 | 0.043 | 0.043 | 0 | 0.07 | 0.17 | 0.06 | 0.12 | 0.06 | 0.07 |
| 1 | 206767938 | rs11119107 | 0.621 | 0.326 | 0.301 | 0.44 | -0.026 | -0.063 | -0.063 | 1.553 | 0.3 | 0.12 | 0.36 | 0.32 | 0.32 | 0.24 |
| 1 | 206772753 | rs4845116 | -0.966 | 0.338 | 0.344 | 0.368 | 0.001 | 0.008 | 0.008 | 0 | 0.85 | 0.82 | 0.78 | 0.66 | 0.83 | 0.71 |
| 1 | 206775390 | rs3813973 | -0.744 | 0.367 | 0.356 | 0.61 | 0.006 | -0.004 | -0.004 | 0 | 0.17 | 0.25 | 0.19 | 0.12 | 0.25 | 0.12 |
| 1 | 206775427 | rs3813972 | 0.937 | -0.012 | 0.316 | 0.193 | 0.006 | 0.012 | 0.012 | 0.015 | 0.52 | 0.58 | 0.45 | 0.52 | 0.43 | 0.55 |
| 1 | 206776097 | rs6658181 | -1.125 | 0.004 | 0.084 | 0.168 | 0.006 | 0.015 | 0.015 | 0.014 | 0.48 | 0.41 | 0.56 | 0.48 | 0.57 | 0.45 |
| 1 | 206783307 | rs11119136 | -0.123 | -0.463 | 0.178 | 0.31 | -0.002 | 0.017 | 0.017 | 0 | 0.24 | 0.27 | 0.29 | 0.35 | 0.21 | 0.23 |
| 1 | 206784809 | rs6685813 | -0.625 | -0.084 | 0.806 | -0.232 | 0.019 | -0.035 | -0.035 | 0 | 0.06 | 0.14 | 0.06 | 0 | 0.09 | 0.03 |
| 1 | 206788458 | rs6677934 | -1.424 | -0.461 | 0.681 | 0.001 | -0.044 | -0.054 | -0.054 | 0.096 | 0.14 | 0.27 | 0.1 | 0.06 | 0.07 | 0.17 |
| 1 | 206788556 | rs6681092 | -1.054 | -0.28 | 0.032 | 0.088 | -0.014 | -0.019 | -0.019 | 0.02 | 0.12 | 0.14 | 0.09 | 0.06 | 0.07 | 0.17 |
| 1 | 206795267 | rs12096152 | -0.67 | 0.701 | -1.17 | -0.065 | -0.033 | 0.079 | 0.079 | 0 | 0.23 | 0.09 | 0.22 | 0.09 | 0.24 | 0.36 |
| 1 | 206796165 | rs6659397 | 0.816 | 0.736 | -2.467 | -0.624 | -0.096 | 0.062 | 0.062 | 0.207 | 0.35 | 0.03 | 0.35 | 0.32 | 0.37 | 0.34 |
| 1 | 206796179 | rs4845119 | -0.547 | 0.58 | -1.408 | -0.395 | 0.011 | 0.021 | 0.021 | 0 | 0.49 | 0.45 | 0.44 | 0.54 | 0.44 | 0.54 |
| 1 | 206803101 | rs7549114 | 0.936 | 0.504 | -1.28 | -0.602 | -0.025 | 0.045 | 0.045 | 0 | 0.14 | 0.08 | 0.1 | 0 | 0.17 | 0.07 |
| 1 | 206807914 | rs3813965 | -0.251 | 1.162 | -1.571 | -0.95 | -0.08 | 0.101 | 0.101 | 0.058 | 0.27 | 0.12 | 0.24 | 0.09 | 0.39 | 0.46 |
| 1 | 206810946 | rs17013271 | -0.513 | 1.03 | -0.668 | -1.009 | -0.022 | 0.031 | 0.031 | 1.375 | 0.3 | 0.26 | 0.32 | 0.09 | 0.4 | 0.47 |
| 1 | 206813737 | rs6540512 | -0.985 | 0.968 | 0.092 | -1.039 | -0.015 | 0.018 | 0.018 | 0 | 0.74 | 0.74 | 0.81 | 0.53 | 0.82 | 0.84 |
| 1 | 206825605 | rs10863766 | -0.569 | 0.95 | 0.299 | -1.034 | 0.003 | -0.003 | -0.003 | 0 | 0.33 | 0.36 | 0.38 | 0.24 | 0.4 | 0.5 |
| 1 | 206832961 | rs7512726 | 1.069 | 0.874 | 0.309 | -1.093 | -0.014 | 0.074 | 0.074 | 0.805 | 0.38 | 0.2 | 0.4 | 0.29 | 0.41 | 0.36 |
| 1 | 206835985 | rs12140926 | 0.624 | 1.02 | 0.524 | -1.07 | 0.004 | -0.009 | -0.009 | 0 | 0.63 | 0.37 | 0.66 | 0.77 | 0.58 | 0.52 |
| 1 | 206840403 | rs17041922 | 0.775 | 0.717 | 0.393 | -1.114 | 0.019 | -0.024 | -0.024 | 0.556 | 0.24 | 0.16 | 0.24 | 0.47 | 0.14 | 0.17 |
| 1 | 206840713 | rs11119265 | 0.884 | 1.035 | 0.351 | -1.248 | 0.002 | 0.004 | 0.004 | 2.35 | 0.58 | 0.4 | 0.53 | 0.76 | 0.4 | 0.45 |
| 1 | 206841857 | rs17014035 | 0.626 | 1.108 | 0.535 | -1.187 | 0.032 | 0.028 | 0.028 | 0 | 0.08 | 0.11 | 0.13 | 0.14 | 0.17 | 0.1 |
| 1 | 206857407 | rs4844550 | 1.767 | 0.874 | 0.523 | -1.372 | 0.033 | 0.069 | 0.069 | 1.979 | 0.31 | 0.09 | 0.25 | 0.3 | 0.24 | 0.28 |
| 1 | 206866203 | rs12756861 | 1.784 | 0.774 | 0.458 | -1.429 | 0.027 | 0.06 | 0.06 | 1.882 | 0.3 | 0.06 | 0.24 | 0.3 | 0.24 | 0.29 |
| 1 | 206872487 | rs4845130 | 0.266 | 0.766 | 0.647 | -1.486 | 0.073 | -0.007 | -0.007 | 0 | 0.85 | 0.66 | 0.83 | 0.88 | 0.79 | 0.86 |
| 1 | 206877337 | rs12060808 | 0.024 | 0.578 | 0.109 | -1.549 | -0.053 | 0.065 | 0.065 | 0 | 0.29 | 0.22 | 0.29 | 0.1 | 0.41 | 0.44 |
| 1 | 206890435 | rs6669284 | 0.279 | 1.106 | 0.178 | -1.308 | -0.048 | -0.043 | -0.043 | 0 | 0.27 | 0.08 | 0.34 | 0.54 | 0.32 | 0.21 |
| 1 | 206890449 | rs4256810 | 0.185 | 1.114 | 0.213 | -1.3 | -0.015 | -0.011 | -0.011 | 0 | 0.27 | 0.16 | 0.35 | 0.54 | 0.32 | 0.21 |
| 1 | 206896622 | rs4240847 | 0.195 | 1.198 | 0.22 | -1.305 | 0.108 | 0.001 | 0.001 | 0 | 0.86 | 0.63 | 0.82 | 0.9 | 0.8 | 0.86 |
| 1 | 206904464 | rs10863805 | 0.053 | 1.042 | -0.522 | -1.118 | -0.015 | -0.012 | -0.012 | 0 | 0.27 | 0.17 | 0.35 | 0.54 | 0.32 | 0.21 |
| 1 | 206911566 | rs4311892 | 0.993 | 0.337 | -0.905 | -1.104 | 0.045 | 0.092 | 0.092 | 0 | 0.37 | 0.2 | 0.31 | 0.32 | 0.29 | 0.55 |
| 1 | 206912501 | rs4147011 | 0.216 | 0.272 | -0.756 | 0.132 | 0.066 | -0.116 | -0.116 | 0 | 0.21 | 0.07 | 0.18 | 0.49 | 0.12 | 0.14 |
| 1 | 206915809 | rs10494878 | -0.083 | 0.88 | -0.422 | -0.225 | 0.017 | -0.033 | -0.033 | 0 | 0.11 | 0.1 | 0.15 | 0.01 | 0.2 | 0.09 |
| 1 | 206923478 | rs11119447 | 0.619 | 0.569 | -0.269 | 0.551 | 0 | 0.004 | 0.004 | 0 | 0.72 | 0.76 | 0.69 | 0.68 | 0.71 | 0.8 |
| 1 | 206927516 | rs6692511 | 0.618 | 0.532 | -0.326 | 0.533 | 0 | -0.004 | -0.004 | 0 | 0.72 | 0.79 | 0.7 | 0.67 | 0.71 | 0.79 |
| 1 | 206932451 | rs4390174 | -0.471 | 0.344 | -0.845 | 0.194 | -0.004 | 0.004 | 0.004 | 0 | 0.32 | 0.34 | 0.27 | 0.25 | 0.36 | 0.28 |
| 1 | 206933490 | rs7548373 | 0.339 | -0.162 | -0.316 | -0.203 | -0.011 | -0.053 | -0.053 | 0 | 0.09 | 0.24 | 0.05 | 0 | 0.06 | 0.03 |
| 1 | 206939904 | rs3024505 | 0.914 | -0.07 | -0.956 | -0.752 | -0.042 | 0.078 | 0.078 | 0 | 0.15 | 0.03 | 0.08 | 0.03 | 0.17 | 0.14 |
| 1 | 206940310 | rs3024502 | -0.704 | 0.294 | -0.679 | -1.183 | -0.029 | 0.016 | 0.016 | 0 | 0.36 | 0.26 | 0.29 | 0.03 | 0.45 | 0.23 |
| 1 | 206941529 | rs3024498 | 1.064 | 0.279 | -0.676 | -1.188 | -0.019 | 0.004 | 0.004 | 0 | 0.14 | 0.1 | 0.17 | 0 | 0.23 | 0.08 |
| 1 | 206941864 | rs3024496 | -0.883 | 0.222 | -0.649 | -1.223 | 0 | -0.07 | -0.07 | 0 | 0.38 | 0.39 | 0.3 | 0.03 | 0.46 | 0.23 |
| 1 | 206944952 | rs3021094 | 0.939 | -0.335 | -0.765 | 0.016 | -0.01 | 0.081 | 0.081 | 0 | 0.05 | 0.01 | 0.11 | 0.47 | 0.09 | 0.08 |
| 1 | 206946634 | rs1800871 | -0.721 | -0.759 | -0.541 | -0.392 | -0.036 | -0.019 | -0.019 | 0.045 | 0.68 | 0.56 | 0.67 | 0.32 | 0.76 | 0.54 |
| 1 | 206946897 | rs1800896 | -0.032 | -0.472 | -0.724 | -1.025 | -0.02 | 0.012 | 0.012 | 0 | 0.36 | 0.31 | 0.3 | 0.05 | 0.45 | 0.24 |
| 1 | 206947167 | rs1800893 | -0.349 | -0.455 | -0.751 | -1.013 | -0.015 | -0.004 | -0.004 | 0.154 | 0.37 | 0.35 | 0.3 | 0.05 | 0.45 | 0.25 |
| 1 | 206955651 | rs6686931 | -0.135 | -0.625 | -0.365 | -0.419 | -0.023 | 0.012 | 0.012 | 0 | 0.67 | 0.6 | 0.68 | 0.57 | 0.78 | 0.58 |
| 1 | 206957092 | rs6667202 | -0.167 | -0.037 | -0.332 | -1.107 | 0.077 | -0.156 | -0.156 | 0 | 0.72 | 0.48 | 0.76 | 0.97 | 0.61 | 0.78 |
| 1 | 206957558 | rs4072227 | -0.659 | -0.919 | -0.3 | -0.501 | -0.008 | 0.06 | 0.06 | 3.148 | 0.9 | 0.82 | 0.84 | 0.73 | 0.94 | 0.91 |
| 1 | 206960216 | rs880790 | -0.31 | -1.044 | -0.337 | -1.006 | -0.03 | -0.021 | -0.021 | 0 | 0.66 | 0.48 | 0.62 | 0.69 | 0.78 | 0.53 |
| 1 | 206962416 | rs885334 | 0.159 | -1.016 | -0.379 | -1.206 | 0.005 | 0.005 | 0.005 | 0 | 0.58 | 0.42 | 0.58 | 0.7 | 0.69 | 0.48 |
| 1 | 206967152 | rs6683473 | 1.383 | -0.605 | -0.441 | -1.18 | 0.024 | -0.035 | -0.035 | 2.952 | 0.11 | 0.17 | 0.11 | 0.01 | 0.2 | 0.1 |
| 1 | 206968893 | rs11583394 | -0.409 | -0.937 | -0.074 | -1.102 | -0.027 | -0.021 | -0.021 | 2.203 | 0.66 | 0.59 | 0.62 | 0.71 | 0.78 | 0.52 |
| 1 | 206973239 | rs11119584 | -0.426 | -0.655 | 0.169 | -0.455 | 0.012 | 0.019 | 0.019 | 0 | 0.39 | 0.24 | 0.42 | 0.68 | 0.39 | 0.31 |
| 1 | 206974779 | rs12042283 | -0.45 | -0.608 | 0.175 | -0.444 | -0.002 | -0.003 | -0.003 | 0 | 0.41 | 0.38 | 0.43 | 0.68 | 0.39 | 0.3 |
| 1 | 206977937 | rs11119597 | -0.925 | -0.274 | 0.564 | -0.063 | -0.012 | -0.038 | -0.038 | 0.106 | 0.07 | 0.2 | 0.04 | 0 | 0.06 | 0.04 |
| 1 | 206978340 | rs4347211 | -0.037 | -0.3 | 0.818 | -0.233 | 0.02 | 0.005 | 0.005 | 4.444 | 0.2 | 0.31 | 0.28 | 0.23 | 0.27 | 0.17 |
| 1 | 206980977 | rs3950619 | -1.264 | -0.354 | 0.393 | -0.595 | -0.024 | 0.127 | 0.127 | 0.892 | 0.45 | 0.74 | 0.59 | 0.9 | 0.48 | 0.34 |
| 1 | 206986545 | rs7540516 | 0.045 | -0.19 | 0.214 | 0.025 | 0.047 | 0.098 | 0.098 | 5.158 | 0.2 | 0.36 | 0.27 | 0.42 | 0.27 | 0.16 |
| 1 | 206986926 | rs7521798 | -0.528 | -0.561 | 0.116 | -0.614 | 0.023 | -0.129 | -0.129 | 0 | 0.39 | 0.5 | 0.53 | 0.9 | 0.44 | 0.3 |
| 1 | 206988533 | rs12044804 | -0.682 | -0.478 | 0.179 | -0.55 | 0.01 | -0.117 | -0.117 | 0.112 | 0.42 | 0.51 | 0.55 | 0.88 | 0.46 | 0.31 |
| 1 | 207005321 | rs11119670 | 0.82 | -1.459 | 0.554 | 0.539 | -0.034 | -0.004 | -0.004 | 0 | 0.66 | 0.57 | 0.65 | 0.41 | 0.76 | 0.71 |
| 1 | 207007641 | rs2243158 | -0.225 | -1.561 | 0.788 | 0.986 | -0.028 | 0.042 | 0.042 | 0.641 | 0.85 | 0.84 | 0.87 | 0.8 | 0.92 | 0.92 |
| 1 | 207009910 | rs2243170 | -0.281 | -1.564 | 0.859 | 0.955 | -0.043 | 0.109 | 0.109 | 0.53 | 0.85 | 0.68 | 0.86 | 0.8 | 0.92 | 0.93 |
| 1 | 207012444 | rs2243176 | -0.332 | -1.541 | 0.789 | 0.72 | 0 | -0.047 | -0.047 | 0 | 0.18 | 0.11 | 0.22 | 0.39 | 0.16 | 0.2 |
| 1 | 207015957 | rs2243191 | 0.802 | -1.446 | 0.76 | 0.543 | 0.008 | 0.003 | 0.003 | 0.243 | 0.71 | 0.82 | 0.67 | 0.34 | 0.77 | 0.72 |

| | | | | | | | | | | | | | | | | |
|---|---|---|---|---|---|---|---|---|---|---|---|---|---|---|---|---|
| 1 | 207016125 | rs1798 | -0.379 | -1.45 | 0.718 | 0.669 | -0.025 | -0.109 | -0.109 | 0.024 | 0.19 | 0.03 | 0.24 | 0.35 | 0.15 | 0.22 |
| 1 | 207016225 | rs2243193 | 1.25 | -1.444 | 0.785 | 0.622 | -0.031 | -0.029 | -0.029 | 0 | 0.65 | 0.54 | 0.64 | 0.33 | 0.76 | 0.7 |
| 1 | 207024762 | rs2883036 | 1.415 | -1.758 | 0.256 | 0.265 | 0.022 | 0.129 | 0.129 | 0 | 0.44 | 0.27 | 0.38 | 0.16 | 0.58 | 0.29 |
| 1 | 207032751 | rs574773 | 0.243 | -1.202 | 0.109 | 0.593 | -0.02 | 0.071 | 0.071 | 0.108 | 0.14 | 0.15 | 0.1 | 0.3 | 0.08 | 0.06 |
| 1 | 207037475 | rs1713239 | -0.206 | -0.48 | -0.033 | 0.79 | -0.028 | -0.097 | -0.097 | 0 | 0.84 | 0.97 | 0.77 | 0.64 | 0.85 | 0.8 |
| 1 | 207038686 | rs1400986 | -0.593 | -0.464 | 0.796 | 0.022 | -0.033 | -0.036 | -0.036 | 0 | 0.22 | 0.41 | 0.26 | 0.18 | 0.17 | 0.44 |
| 1 | 207064400 | rs12022348 | -0.473 | 0.258 | -0.116 | -0.437 | 0.018 | 0.116 | 0.116 | 0.17 | 0.51 | 0.62 | 0.46 | 0.2 | 0.45 | 0.23 |
| 1 | 207068672 | rs3762344 | -0.422 | 0.224 | -0.033 | -0.454 | 0.019 | 0.108 | 0.108 | 0.225 | 0.51 | 0.61 | 0.46 | 0.2 | 0.45 | 0.22 |
| 1 | 207072959 | rs1150255 | -0.402 | 0.227 | 0.848 | -0.571 | -0.001 | 0.125 | 0.125 | 0.64 | 0.48 | 0.27 | 0.44 | 0.21 | 0.45 | 0.23 |
| 1 | 207074905 | rs1150258 | -0.402 | 0.227 | 0.848 | -0.571 | -0.001 | 0.125 | 0.125 | 0.64 | 0.48 | 0.29 | 0.44 | 0.21 | 0.45 | 0.23 |
| 1 | 207075171 | rs291107 | -0.867 | 0.114 | 0.904 | -0.603 | -0.041 | 0.155 | 0.155 | 0.251 | 0.54 | 0.84 | 0.5 | 0.24 | 0.48 | 0.24 |
| 1 | 207077210 | rs3093438 | -0.928 | -0.049 | 0.82 | -0.646 | 0.007 | 0.032 | 0.032 | 1.029 | 0.05 | 0.19 | 0.15 | 0.08 | 0.02 | 0.04 |
| 1 | 207080565 | rs12121499 | -0.278 | -0.073 | 0.752 | -0.681 | -0.025 | 0.175 | 0.175 | 0.782 | 0.44 | 0.18 | 0.3 | 0.13 | 0.45 | 0.19 |
| 1 | 207086194 | rs749860 | -1.453 | -0.31 | 0.795 | -0.055 | 0.018 | 0.217 | 0.217 | 2.741 | 0.07 | 0.52 | 0.05 | 0.19 | 0.02 | 0.02 |
| 1 | 207086845 | rs94872 | -1.002 | -0.215 | 0.479 | -0.125 | -0.114 | -0.009 | -0.009 | 0 | 0.57 | 0.93 | 0.52 | 0.42 | 0.49 | 0.26 |
| 1 | 207101264 | rs12129555 | 0.374 | -0.576 | 0.068 | -0.701 | 0.025 | 0.171 | 0.171 | 0.112 | 0.25 | 0.01 | 0.11 | 0 | 0.18 | 0.1 |
| 1 | 207105641 | rs908702 | -0.648 | -0.602 | 0.226 | -0.61 | -0.024 | 0.081 | 0.081 | 1.786 | 0.46 | 0.17 | 0.32 | 0.33 | 0.46 | 0.22 |
| 1 | 207106444 | rs172361 | -1.471 | -0.618 | 0.38 | -0.477 | -0.03 | -0.003 | -0.003 | 1.055 | 0.1 | 0.74 | 0.19 | 0.12 | 0.03 | 0.05 |
| 1 | 207106478 | rs291102 | -1.588 | -0.619 | 0.38 | -0.478 | -0.023 | -0.004 | -0.004 | 0 | 0.1 | 0.76 | 0.19 | 0.12 | 0.03 | 0.05 |
| 1 | 207107696 | rs908701 | -1.455 | -0.602 | 0.396 | -0.469 | -0.025 | 0.027 | 0.027 | 7.979 | 0.09 | 0.63 | 0.18 | 0.12 | 0.02 | 0.05 |
| 1 | 207109116 | rs2275531 | -0.67 | -0.632 | 0.087 | -0.506 | -0.012 | 0.084 | 0.084 | 1.495 | 0.46 | 0.19 | 0.33 | 0.33 | 0.46 | 0.21 |
| 1 | 207110888 | rs291097 | -1.659 | -0.613 | 0.324 | -0.486 | -0.03 | -0.028 | -0.028 | 0 | 0.1 | 0.69 | 0.18 | 0.05 | 0.02 | 0.04 |
| 1 | 207119853 | rs2275529 | -0.811 | -0.527 | 0.27 | -0.748 | 0.025 | 0.027 | 0.027 | 0.878 | 0.42 | 0.6 | 0.35 | 0.12 | 0.3 | 0.17 |
| 1 | 207130317 | rs6697345 | -0.961 | -0.753 | 0.147 | -0.819 | 0.054 | 0.077 | 0.077 | 1.77 | 0.44 | 0.63 | 0.35 | 0.07 | 0.3 | 0.17 |
| 1 | 207135991 | rs11119925 | -0.828 | -0.775 | 0.147 | -0.831 | 0.079 | 0.137 | 0.137 | 2.145 | 0.45 | 0.69 | 0.35 | 0.13 | 0.3 | 0.17 |
| 1 | 207139046 | rs11576146 | -0.622 | -0.448 | 0.057 | -0.714 | 0.018 | 0.055 | 0.055 | 0 | 0.12 | 0.21 | 0.15 | 0.32 | 0.16 | 0.08 |
| 1 | 207143422 | rs1856746 | -1.501 | -0.505 | 0.718 | -0.717 | -0.071 | -0.007 | -0.007 | 0 | 0.55 | 0.93 | 0.51 | 0.44 | 0.47 | 0.26 |
| 1 | 207146837 | rs2353550 | -0.744 | -0.343 | 0.812 | -0.722 | -0.005 | 0.123 | 0.123 | 0 | 0.45 | 0.21 | 0.27 | 0.24 | 0.45 | 0.2 |
| 1 | 207156555 | rs11119981 | -0.976 | -0.2 | 0.522 | -0.873 | 0.018 | 0.131 | 0.131 | 0 | 0.49 | 0.3 | 0.28 | 0.25 | 0.45 | 0.2 |
| 1 | 207159378 | rs6672060 | -0.81 | -0.538 | 0.463 | -1.026 | 0.044 | 0.106 | 0.106 | 4.222 | 0.45 | 0.26 | 0.25 | 0.25 | 0.38 | 0.18 |
| 1 | 207162433 | rs2791708 | -1.268 | -0.385 | 0.418 | -0.967 | 0.047 | 0.045 | 0.045 | 3.689 | 0.52 | 0.8 | 0.41 | 0.44 | 0.41 | 0.22 |
| 1 | 207178900 | rs2629646 | -1.022 | -0.252 | 1.415 | -1.048 | 0.025 | 0.034 | 0.034 | 4.11 | 0.47 | 0.7 | 0.39 | 0.39 | 0.41 | 0.22 |
| 1 | 207179782 | rs11586424 | 0.662 | 0.365 | 0.883 | -0.896 | -0.046 | -0.036 | -0.036 | 0 | 0.1 | 0.01 | 0.09 | 0.14 | 0.13 | 0.15 |
| 1 | 207179822 | rs4844564 | -0.32 | -0.202 | 0.985 | -1.241 | 0.005 | -0.013 | -0.013 | 0.193 | 0.21 | 0.24 | 0.13 | 0.12 | 0.21 | 0.09 |
| 1 | 207187406 | rs764317 | 0.057 | -0.528 | 0.887 | -1.179 | 0.045 | 0.061 | 0.061 | 0.804 | 0.25 | 0.15 | 0.21 | 0.07 | 0.19 | 0.12 |
| 1 | 207188207 | rs2842721 | -1.16 | -0.547 | 0.819 | -1.182 | 0.038 | 0.009 | 0.009 | 3.271 | 0.37 | 0.76 | 0.32 | 0.37 | 0.26 | 0.17 |
| 1 | 207189574 | rs12409177 | 0.068 | -0.446 | 0.378 | -1.436 | 0.063 | 0.111 | 0.111 | 0.813 | 0.24 | 0.07 | 0.21 | 0.07 | 0.19 | 0.12 |
| 1 | 207191164 | rs12138969 | 0.103 | -0.472 | 0.187 | -1.448 | 0.061 | 0.11 | 0.11 | 0.781 | 0.24 | 0.07 | 0.21 | 0.07 | 0.19 | 0.12 |
| 1 | 207191936 | rs3020 | -1.591 | -0.356 | 0.586 | -1.361 | 0.013 | 0.053 | 0.053 | 0 | 0.14 | 0.3 | 0.08 | 0.19 | 0.11 | 0.07 |
| 1 | 207192180 | rs10877 | -0.845 | -0.77 | 0.682 | -1.467 | -0.009 | -0.004 | -0.004 | 0.211 | 0.3 | 0.34 | 0.17 | 0.25 | 0.21 | 0.17 |
| 1 | 207193177 | rs901107 | -0.698 | -0.979 | 0.486 | -2.275 | -0.017 | 0.085 | 0.085 | 0.173 | 0.38 | 0.25 | 0.24 | 0.08 | 0.38 | 0.24 |
| 1 | 207200549 | rs6696573 | 0.386 | -0.986 | 0.673 | -2.347 | -0.01 | 0.035 | 0.035 | 0.126 | 0.14 | 0.04 | 0.19 | 0.07 | 0.16 | 0.1 |
| 1 | 207209364 | rs11799614 | 0.244 | -0.994 | 0.878 | -2.423 | 0 | -0.002 | -0.002 | 0 | 0.1 | 0.13 | 0.06 | 0.07 | 0.1 | 0.1 |
| 1 | 207211161 | rs11120092 | -0.707 | -1.143 | 0.865 | -2.404 | -0.007 | 0.138 | 0.138 | 0 | 0.45 | 0.24 | 0.26 | 0.17 | 0.42 | 0.27 |
| 1 | 207217359 | rs1044145 | -1.881 | -1.225 | 1.062 | -2.424 | -0.021 | 0.049 | 0.049 | 3.591 | 0.63 | 0.85 | 0.5 | 0.24 | 0.58 | 0.37 |
| 1 | 207220800 | rs2629665 | -0.686 | -1.18 | 0.886 | -2.404 | 0.001 | 0.028 | 0.028 | 0.004 | 0.46 | 0.66 | 0.3 | 0.17 | 0.42 | 0.27 |
| 1 | 207221612 | rs2054780 | 0.412 | -1.163 | 0.851 | -2.391 | -0.003 | 0.028 | 0.028 | 0.075 | 0.14 | 0.06 | 0.19 | 0.07 | 0.16 | 0.1 |
| 1 | 207224992 | rs2306714 | -0.676 | -1.091 | 0.891 | -2.352 | -0.001 | 0.065 | 0.065 | 0 | 0.44 | 0.37 | 0.26 | 0.17 | 0.41 | 0.27 |
| 1 | 207235954 | rs2075863 | -0.524 | -1.07 | 0.571 | -2.36 | -0.015 | 0.131 | 0.131 | 0 | 0.43 | 0.2 | 0.25 | 0.17 | 0.42 | 0.26 |
| 1 | 207241281 | rs2243816 | 0.144 | -0.971 | 0.038 | -2.226 | -0.001 | 0.05 | 0.05 | 0.118 | 0.17 | 0.05 | 0.2 | 0.07 | 0.16 | 0.11 |
| 1 | 207249855 | rs11120148 | 1.724 | -0.983 | 0.042 | -2.396 | -0.099 | 0.048 | 0.048 | 1.419 | 0.34 | 0.07 | 0.53 | 0.76 | 0.44 | 0.6 |
| 1 | 207250300 | rs1060286 | 0.296 | -1.089 | 0.245 | -2.479 | 0.012 | 0.092 | 0.092 | 0.006 | 0.54 | 0.7 | 0.75 | 0.83 | 0.6 | 0.71 |
| 1 | 207250456 | rs1060287 | 1.472 | -0.703 | -0.405 | -2.202 | -0.094 | 0.051 | 0.051 | 1.334 | 0.34 | 0.07 | 0.53 | 0.76 | 0.44 | 0.6 |
| 1 | 207250570 | rs1565 | 1.376 | -0.719 | -0.459 | -2.19 | -0.022 | 0.031 | 0.031 | 0.514 | 0.35 | 0.29 | 0.56 | 0.76 | 0.44 | 0.6 |
| 1 | 207253031 | rs3748671 | 0.607 | -0.282 | -0.62 | -2.161 | 0.009 | 0.193 | 0.193 | 0 | 0.52 | 0.17 | 0.68 | 0.88 | 0.55 | 0.78 |
| 1 | 207253165 | rs3748672 | 0.24 | -0.167 | 0.07 | -2.101 | 0.001 | -0.034 | -0.034 | 0.019 | 0.6 | 0.56 | 0.72 | 0.88 | 0.57 | 0.79 |
| 1 | 207253487 | rs6673422 | -0.319 | -0.861 | 0.082 | -1.74 | -0.007 | -0.024 | -0.024 | 0 | 0.17 | 0.26 | 0.16 | 0.12 | 0.16 | 0.1 |
| 1 | 207257303 | rs2842754 | -0.866 | -0.911 | -0.152 | -1.868 | 0 | 0.001 | 0.001 | 2.196 | 0.29 | 0.25 | 0.18 | 0.07 | 0.25 | 0.13 |
| 1 | 207262980 | rs12711513 | 0.589 | -0.711 | -0.124 | -1.581 | 0.004 | -0.048 | -0.048 | 0.552 | 0.43 | 0.52 | 0.64 | 0.84 | 0.48 | 0.62 |
| 1 | 207263829 | rs6690037 | 1.094 | -0.371 | -1.197 | -1.455 | -0.127 | 0.046 | 0.046 | 1.803 | 0.36 | 0.07 | 0.54 | 0.75 | 0.47 | 0.55 |
| 1 | 207269858 | rs3813948 | -0.721 | -0.901 | -0.151 | -0.934 | 0.007 | 0.001 | 0.001 | 0.146 | 0.17 | 0.15 | 0.07 | 0.16 | 0.07 | 0.14 |
| 1 | 207276379 | rs11120211 | -0.464 | -0.828 | -0.001 | -0.712 | -0.013 | 0.005 | 0.005 | 0 | 0.15 | 0.17 | 0.08 | 0.2 | 0.07 | 0.14 |
| 1 | 207278451 | rs11120218 | -1.686 | -1.066 | 0.601 | -0.923 | -0.056 | -0.025 | -0.025 | 0 | 0.25 | 0.47 | 0.15 | 0.2 | 0.14 | 0.17 |
| 1 | 207287187 | rs2808470 | -1.939 | -1.13 | 0.253 | -1.034 | 0.056 | 0.046 | 0.046 | 0 | 0.31 | 0.84 | 0.28 | 0.2 | 0.18 | 0.25 |
| 1 | 207288297 | rs17020983 | -1.194 | -0.993 | 0.417 | -0.679 | -0.056 | 0.001 | 0.001 | 0 | 0.2 | 0.56 | 0.12 | 0.2 | 0.08 | 0.13 |
| 1 | 207289436 | rs2842704 | 0.475 | -0.868 | -0.007 | -0.552 | -0.023 | -0.004 | -0.004 | 0.32 | 0.73 | 0.74 | 0.86 | 0.76 | 0.84 | 0.81 |
| 1 | 207293496 | rs4259650 | 0.325 | -0.361 | -0.524 | -0.598 | 0.044 | 0.065 | 0.065 | 1.884 | 0.54 | 0.34 | 0.63 | 0.74 | 0.63 | 0.75 |
| 1 | 207293780 | rs4425986 | 0.548 | 0.046 | -1.292 | -0.295 | -0.064 | 0.019 | 0.019 | 2.729 | 0.44 | 0.09 | 0.46 | 0.66 | 0.51 | 0.57 |
| 1 | 207300259 | rs2491393 | -0.649 | 0.267 | -0.334 | -0.055 | -0.002 | -0.005 | -0.005 | 0 | 0.37 | 0.6 | 0.43 | 0.27 | 0.37 | 0.38 |

| | | | | | | | | | | | | | | | | |
|---|---|---|---|---|---|---|---|---|---|---|---|---|---|---|---|---|
| 1 | 207304900 | rs4844573 | -0.757 | 0.208 | -0.19 | -0.141 | 0.005 | 0.026 | 0.026 | 0 | 0.4 | 0.84 | 0.46 | 0.27 | 0.38 | 0.38 |
| 1 | 207305588 | rs4571969 | -0.402 | 0.194 | 0.546 | -0.156 | -0.016 | 0.003 | 0.003 | 0 | 0.32 | 0.5 | 0.31 | 0.27 | 0.29 | 0.3 |
| 1 | 207311481 | rs7524207 | -1.916 | 0.243 | 1.616 | 0.039 | -0.027 | -0.05 | -0.05 | 0 | 0.1 | 0.62 | 0.07 | 0 | 0.03 | 0.04 |
| 1 | 207314356 | rs12091603 | -0.608 | 0.277 | 2.018 | 0.056 | -0.007 | -0.022 | -0.022 | 0 | 0.05 | 0.21 | 0.03 | 0 | 0.03 | 0 |
| 1 | 207345819 | rs6666604 | 0.261 | 0.481 | 1.625 | -0.314 | -0.003 | 0 | 0 | 0 | 0.18 | 0.13 | 0.12 | 0.06 | 0.18 | 0.14 |
| 1 | 207353449 | rs7529677 | -1.595 | 0.46 | 1.922 | -0.275 | 0.005 | -0.021 | -0.021 | 0 | 0.08 | 0.42 | 0.05 | 0 | 0.03 | 0.01 |
| 1 | 207360067 | rs6672985 | 0.019 | 0.437 | 1.297 | -0.536 | 0.001 | 0.002 | 0.002 | 0 | 0.73 | 0.69 | 0.65 | 0.32 | 0.66 | 0.61 |
| 1 | 207369066 | rs1429909 | 1.002 | 0.297 | 1.523 | -0.108 | 0.017 | 0.015 | 0.015 | 0.381 | 0.51 | 0.31 | 0.51 | 0.36 | 0.47 | 0.39 |
| 1 | 207402853 | rs10864144 | -0.017 | 0.59 | 1.58 | -0.152 | 0.008 | 0.029 | 0.029 | 0 | 0.73 | 0.62 | 0.65 | 0.42 | 0.68 | 0.55 |
| 1 | 207414732 | rs17258843 | 0.211 | 0.599 | 1.494 | 0.369 | 0.01 | -0.054 | -0.054 | 0 | 0.22 | 0.28 | 0.17 | 0.06 | 0.25 | 0.16 |
| 1 | 207418996 | rs1864369 | 1.095 | 0.559 | 1.518 | 0.393 | 0.032 | 0.037 | 0.037 | 0.766 | 0.49 | 0.14 | 0.48 | 0.37 | 0.46 | 0.39 |
| 1 | 207440851 | rs6540896 | -0.238 | 0.639 | 1.411 | 0.423 | 0.01 | 0.046 | 0.046 | 0.054 | 0.76 | 0.63 | 0.69 | 0.43 | 0.72 | 0.56 |
| 1 | 207441975 | rs11584921 | 0.25 | 0.684 | 1.434 | 0.402 | -0.006 | -0.003 | -0.003 | 1.908 | 0.22 | 0.2 | 0.16 | 0.06 | 0.25 | 0.17 |
| 1 | 207457845 | rs12134133 | -0.294 | 0.717 | 1.324 | 0.454 | 0.019 | -0.133 | -0.133 | 0 | 0.28 | 0.5 | 0.2 | 0.06 | 0.27 | 0.17 |
| 1 | 207470460 | rs1652333 | -0.801 | 0.704 | 1.265 | 0.558 | -0.005 | -0.005 | -0.005 | 0.052 | 0.8 | 0.82 | 0.71 | 0.42 | 0.74 | 0.56 |
| 1 | 207471799 | rs2802235 | -0.078 | 0.702 | 1.275 | 0.524 | 0.013 | -0.088 | -0.088 | 0.046 | 0.26 | 0.42 | 0.19 | 0.06 | 0.27 | 0.17 |
| 1 | 207491489 | rs891376 | -0.719 | 0.771 | 1.238 | 0.651 | 0.013 | 0.049 | 0.049 | 0.896 | 0.8 | 0.74 | 0.71 | 0.41 | 0.74 | 0.56 |
| 1 | 207500293 | rs4844591 | -0.554 | 0.764 | 1.248 | 0.633 | 0.017 | 0.066 | 0.066 | 1.05 | 0.79 | 0.7 | 0.7 | 0.41 | 0.74 | 0.55 |
| 1 | 207509022 | rs7544288 | 0.93 | 0.685 | 1.105 | 0.541 | 0.037 | 0.038 | 0.038 | 0.535 | 0.5 | 0.19 | 0.49 | 0.35 | 0.47 | 0.38 |
| 1 | 207509799 | rs1507761 | 0.452 | 0.778 | 1.001 | 0.63 | -0.002 | 0.006 | 0.006 | 0.679 | 0.19 | 0.16 | 0.29 | 0.59 | 0.25 | 0.44 |
| 1 | 207536971 | rs11117564 | -0.485 | 0.547 | 0.203 | 0.487 | -0.004 | 0.021 | 0.021 | 0.305 | 0.44 | 0.38 | 0.46 | 0.42 | 0.51 | 0.53 |
| 1 | 207556770 | rs12116783 | 0.439 | 0.126 | -0.463 | 0.125 | 0.042 | 0.076 | 0.076 | 0 | 0.34 | 0.11 | 0.19 | 0.18 | 0.26 | 0.17 |
| 1 | 207569884 | rs10779277 | 0.373 | 0.67 | 0.191 | 0.425 | -0.029 | 0.008 | 0.008 | 0.095 | 0.41 | 0.2 | 0.6 | 0.44 | 0.45 | 0.61 |
| 1 | 207571043 | rs2490255 | 0.133 | 0.631 | 0.323 | 0.2 | -0.024 | 0.052 | 0.052 | 0.203 | 0.44 | 0.25 | 0.62 | 0.71 | 0.47 | 0.7 |
| 1 | 207572294 | rs17186722 | 1.256 | 0.432 | 0.169 | -0.005 | 0.028 | 0.119 | 0.119 | 0.216 | 0.31 | 0.08 | 0.17 | 0.1 | 0.24 | 0.16 |
| 1 | 207573387 | rs2940252 | -0.124 | 0.598 | 0.358 | -0.103 | 0.037 | 0.076 | 0.076 | 0 | 0.52 | 0.31 | 0.7 | 0.8 | 0.61 | 0.73 |
| 1 | 207578194 | rs2661361 | -0.926 | 0.604 | 0.956 | -0.042 | -0.005 | -0.03 | -0.03 | 0 | 0.63 | 0.61 | 0.77 | 0.89 | 0.68 | 0.8 |
| 1 | 207578284 | rs6664001 | 1.379 | 0.371 | 0.86 | -0.18 | 0.003 | 0.009 | 0.009 | 0 | 0.23 | 0.21 | 0.15 | 0.11 | 0.2 | 0.15 |
| 1 | 207594943 | rs2135924 | 1.379 | 0.365 | 0.966 | -0.185 | 0.003 | 0.009 | 0.009 | 0 | 0.23 | 0.21 | 0.15 | 0.11 | 0.2 | 0.15 |
| 1 | 207595822 | rs6656123 | 0.669 | 0.357 | 1.015 | -0.141 | -0.015 | 0.092 | 0.092 | 0 | 0.52 | 0.22 | 0.66 | 0.72 | 0.52 | 0.71 |
| 1 | 207596484 | rs311299 | -0.588 | 0.257 | 1.156 | -0.16 | -0.006 | -0.067 | -0.067 | 0.077 | 0.7 | 0.56 | 0.8 | 0.89 | 0.71 | 0.8 |
| 1 | 207634670 | rs17044576 | 0.55 | 0.277 | 1.004 | -0.152 | -0.01 | 0.013 | 0.013 | 10.958 | 0.07 | 0.15 | 0.05 | 0.09 | 0.06 | 0.07 |
| 1 | 207640345 | rs1032980 | -0.606 | 0.43 | 0.924 | -0.252 | -0.008 | -0.084 | -0.084 | 0.033 | 0.7 | 0.54 | 0.8 | 0.89 | 0.71 | 0.8 |
| 1 | 207646462 | rs17615 | 0.402 | 0.429 | 0.909 | -0.252 | -0.005 | -0.074 | -0.074 | 0 | 0.29 | 0.43 | 0.2 | 0.11 | 0.29 | 0.2 |
| 1 | 207646898 | rs4308977 | 0.271 | 0.444 | 0.925 | -0.256 | -0.01 | -0.072 | -0.072 | 1.498 | 0.31 | 0.54 | 0.21 | 0.11 | 0.29 | 0.2 |
| 1 | 207646923 | rs17616 | 0.376 | 0.485 | 0.872 | -0.218 | -0.008 | -0.074 | -0.074 | 0 | 0.29 | 0.42 | 0.2 | 0.11 | 0.29 | 0.2 |
| 1 | 207650938 | rs7549152 | -0.259 | 0.694 | -0.006 | -0.098 | 0.023 | -0.022 | -0.022 | 0 | 0.05 | 0.12 | 0.04 | 0 | 0.09 | 0.04 |
| 1 | 207653106 | rs2182909 | 0.684 | 0.555 | -0.072 | -0.109 | -0.001 | 0.006 | 0.006 | 0 | 0.27 | 0.21 | 0.18 | 0.11 | 0.29 | 0.19 |
| 1 | 207656050 | rs6690215 | 0.968 | 0.197 | -0.267 | 0.072 | 0.006 | 0.01 | 0.01 | 0 | 0.53 | 0.4 | 0.7 | 0.48 | 0.58 | 0.57 |
| 1 | 207658145 | rs12021671 | 0.702 | 0.186 | -0.378 | -0.164 | 0.003 | 0.001 | 0.001 | 0 | 0.57 | 0.41 | 0.75 | 0.48 | 0.64 | 0.61 |
| 1 | 207660071 | rs2182911 | 0.323 | 0.306 | -0.361 | -0.873 | -0.002 | -0.012 | -0.012 | 0 | 0.82 | 0.84 | 0.88 | 0.96 | 0.8 | 0.94 |
| 1 | 207661640 | rs4618970 | -0.312 | 0.214 | -0.254 | -0.77 | 0.007 | -0.027 | -0.027 | 0 | 0.87 | 0.85 | 0.92 | 0.96 | 0.86 | 0.98 |
| 1 | 207662912 | rs9429940 | -0.312 | 0.214 | -0.277 | -0.77 | 0.006 | -0.026 | -0.026 | 2.048 | 0.87 | 0.85 | 0.92 | 0.96 | 0.86 | 0.98 |
| 1 | 207670928 | rs1571344 | 0.159 | -0.222 | -0.14 | -1.004 | -0.057 | -0.131 | -0.131 | 0.155 | 0.2 | 0.4 | 0.1 | 0 | 0.11 | 0.06 |
| 1 | 207673631 | rs7533408 | 0.436 | -0.091 | -0.429 | -0.918 | -0.011 | -0.092 | -0.092 | 0 | 0.66 | 0.45 | 0.82 | 0.96 | 0.75 | 0.92 |
| 1 | 207675655 | rs2025935 | 0.998 | 0.104 | -0.629 | -0.708 | -0.012 | 0.033 | 0.033 | 0 | 0.27 | 0.22 | 0.17 | 0.1 | 0.29 | 0.13 |
| 1 | 207684359 | rs11117956 | 0.062 | -0.434 | -0.46 | -0.623 | -0.017 | -0.02 | -0.02 | 0 | 0.29 | 0.33 | 0.32 | 0.32 | 0.22 | 0.5 |
| 1 | 207684519 | rs11117959 | 0.444 | -0.467 | -0.369 | -0.65 | 0.018 | 0.014 | 0.014 | 0.03 | 0.26 | 0.15 | 0.27 | 0.32 | 0.18 | 0.5 |
| 1 | 207748107 | rs10127904 | 1.832 | -0.484 | -0.455 | -1.263 | -0.095 | 0.142 | 0.142 | 0.005 | 0.26 | 0.04 | 0.17 | 0.03 | 0.3 | 0.1 |
| 1 | 207753345 | rs2274566 | 1.071 | -0.029 | -0.134 | -0.474 | -0.023 | -0.052 | -0.052 | 0 | 0.51 | 0.12 | 0.56 | 0.35 | 0.6 | 0.42 |
| 1 | 207762358 | rs3738468 | 0.581 | -0.101 | 0.013 | -0.418 | 0.014 | 0.011 | 0.011 | 0.073 | 0.27 | 0.16 | 0.27 | 0.33 | 0.18 | 0.49 |
| 1 | 207782707 | rs17259045 | 1.842 | -0.129 | -0.313 | -0.5 | -0.024 | 0.027 | 0.027 | 0 | 0.05 | 0.01 | 0.16 | 0 | 0.1 | 0.07 |
| 1 | 207782889 | rs17047661 | -0.92 | -0.269 | -0.101 | -0.266 | -0.012 | -0.015 | -0.015 | 0 | 0.07 | 0.71 | 0.05 | 0 | 0.01 | 0 |
| 1 | 207782931 | rs6691117 | -0.409 | -0.27 | -0.155 | -0.204 | 0.087 | 0.074 | 0.074 | 0.45 | 0.34 | 0.87 | 0.38 | 0.34 | 0.23 | 0.49 |
| 1 | 207783507 | rs12032275 | 0.665 | -0.152 | -0.141 | -0.135 | 0.033 | 0.017 | 0.017 | 0.087 | 0.26 | 0.12 | 0.28 | 0.34 | 0.18 | 0.49 |
| 1 | 207784968 | rs3818361 | 0.182 | -0.073 | -0.339 | -0.217 | 0.073 | 0.11 | 0.11 | 0 | 0.81 | 0.55 | 0.89 | 0.69 | 0.82 | 0.92 |
| 1 | 207793964 | rs12734030 | 0.621 | 0.097 | -1.202 | -0.044 | 0.032 | 0.023 | 0.023 | 0.047 | 0.25 | 0.11 | 0.26 | 0.32 | 0.17 | 0.49 |
| 1 | 207803595 | rs12034383 | 0.371 | 0.349 | -1.443 | -0.105 | -0.024 | -0.053 | -0.053 | 0.11 | 0.55 | 0.17 | 0.57 | 0.37 | 0.6 | 0.44 |
| 1 | 207814451 | rs10779339 | 0.025 | 0.319 | -1.052 | -0.087 | -0.012 | -0.002 | -0.002 | 0 | 0.58 | 0.66 | 0.4 | 0.68 | 0.5 | 0.59 |
| 1 | 207819729 | rs6696840 | 0.433 | 0.173 | -0.373 | -0.328 | 0.003 | 0.016 | 0.016 | 0 | 0.52 | 0.35 | 0.37 | 0.68 | 0.45 | 0.47 |
| 1 | 207820612 | rs12080578 | 0.188 | 0.312 | 0.006 | -0.288 | 0.048 | -0.009 | -0.009 | 0 | 0.05 | 0.18 | 0.03 | 0.01 | 0.05 | 0.13 |
| 1 | 207823240 | rs1323721 | 0.276 | -0.069 | -0.709 | -0.203 | -0.004 | 0.02 | 0.02 | 0 | 0.41 | 0.64 | 0.34 | 0.35 | 0.43 | 0.45 |
| 1 | 207840776 | rs10863461 | 0.757 | -0.112 | -0.69 | -0.226 | 0.001 | -0.023 | -0.023 | 1.056 | 0.37 | 0.51 | 0.32 | 0.33 | 0.38 | 0.32 |
| 1 | 207841934 | rs12567945 | 1.038 | -0.125 | -1.008 | -0.162 | 0.021 | -0.092 | -0.092 | 0 | 0.15 | 0.02 | 0.05 | 0.35 | 0.07 | 0.14 |
| 1 | 207847876 | rs10494884 | 0.454 | -0.102 | -0.566 | -0.25 | 0.005 | -0.018 | -0.018 | 0 | 0.39 | 0.42 | 0.33 | 0.35 | 0.43 | 0.45 |
| 1 | 207872290 | rs7527798 | 1.783 | 0.014 | -1.232 | -0.402 | -0.064 | 0.114 | 0.114 | 0 | 0.24 | 0.05 | 0.14 | 0.03 | 0.25 | 0.1 |
| 1 | 207874272 | rs1830762 | 0.707 | 0.255 | -0.783 | -0.175 | 0.007 | -0.004 | -0.004 | 0.073 | 0.4 | 0.24 | 0.22 | 0.38 | 0.32 | 0.25 |
| 1 | 207875175 | rs4844614 | 0.648 | 0.186 | -0.765 | -0.272 | 0.001 | 0.043 | 0.043 | 0.073 | 0.23 | 0.26 | 0.5 | 0.32 | 0.27 | 0.34 |
| 1 | 207875431 | rs12073783 | 0.467 | 0.249 | -1.352 | -0.273 | -0.015 | 0.006 | 0.006 | 0 | 0.43 | 0.22 | 0.23 | 0.4 | 0.37 | 0.38 |

| | | | | | | | | | | | | | | | | |
|---|---|---|---|---|---|---|---|---|---|---|---|---|---|---|---|---|
| 1 | 207880683 | rs1977265 | -0.024 | 0.808 | -1.322 | -0.658 | -0.099 | 0.103 | 0.103 | 0 | 0.69 | 0.91 | 0.49 | 0.88 | 0.57 | 0.74 |
| 1 | 207886487 | rs1998538 | 0.151 | 0.69 | -0.605 | -0.713 | -0.015 | 0.014 | 0.014 | 0 | 0.68 | 0.74 | 0.47 | 0.88 | 0.56 | 0.74 |
| 1 | 207893782 | rs2761424 | 0.448 | 0.955 | -0.057 | -0.891 | -0.029 | 0.028 | 0.028 | 0 | 0.2 | 0.11 | 0.31 | 0.01 | 0.27 | 0.15 |
| 1 | 207894416 | rs11118475 | 1.132 | 0.574 | 0.168 | -0.943 | -0.001 | -0.015 | -0.015 | 0 | 0.6 | 0.43 | 0.41 | 0.88 | 0.51 | 0.67 |
| 1 | 207923081 | rs2761437 | -0.544 | 1.067 | 0.545 | -0.832 | 0.012 | -0.038 | -0.038 | 1.673 | 0.86 | 0.79 | 0.72 | 0.99 | 0.8 | 0.86 |
| 1 | 207924066 | rs1970530 | 1.137 | 1.254 | 0.702 | -0.829 | 0.036 | -0.046 | -0.046 | 0 | 0.73 | 0.6 | 0.48 | 0.92 | 0.61 | 0.72 |
| 1 | 207930203 | rs2724384 | -0.587 | 1.228 | 0.703 | -0.841 | 0.004 | -0.027 | -0.027 | 0.996 | 0.86 | 0.83 | 0.72 | 0.99 | 0.8 | 0.86 |
| 1 | 207932617 | rs2724387 | 0.991 | 1.146 | 0.716 | -0.816 | 0.031 | -0.04 | -0.04 | 0.512 | 0.73 | 0.61 | 0.48 | 0.92 | 0.6 | 0.72 |
| 1 | 207936103 | rs17006738 | 1.006 | 0.664 | 0.275 | -1.314 | -0.007 | -0.031 | -0.031 | 0 | 0.12 | 0.17 | 0.08 | 0.04 | 0.07 | 0.06 |
| 1 | 207938236 | rs7545126 | 1.141 | 0.011 | -0.154 | -0.578 | 0.014 | -0.023 | -0.023 | 0.011 | 0.15 | 0.08 | 0.1 | 0.29 | 0.1 | 0.2 |
| 1 | 207940046 | rs2466571 | 1.543 | -0.237 | 0.286 | -0.692 | -0.022 | 0.104 | 0.104 | 0 | 0.55 | 0.24 | 0.38 | 0.85 | 0.52 | 0.65 |
| 1 | 207955823 | rs2796278 | 1.131 | -0.219 | 0.366 | -0.475 | 0.008 | 0.005 | 0.005 | 0 | 0.58 | 0.52 | 0.37 | 0.6 | 0.5 | 0.53 |
| 1 | 207957555 | rs7541230 | -0.144 | 0.33 | 0.582 | -0.661 | -0.012 | 0.008 | 0.008 | 0 | 0.21 | 0.2 | 0.46 | 0.09 | 0.33 | 0.22 |
| 1 | 207961058 | rs10449303 | -0.585 | 0.126 | 0.673 | -0.677 | 0.013 | -0.008 | -0.008 | 0.09 | 0.23 | 0.29 | 0.5 | 0.09 | 0.39 | 0.28 |
| 1 | 207962270 | rs859706 | -0.663 | 0.133 | 0.737 | -0.672 | 0.023 | -0.011 | -0.011 | 0 | 0.24 | 0.34 | 0.54 | 0.12 | 0.39 | 0.28 |
| 1 | 207981419 | rs761276 | -1.845 | 0.02 | 0.528 | -0.723 | -0.027 | 0.085 | 0.085 | 0 | 0.43 | 0.76 | 0.64 | 0.16 | 0.48 | 0.33 |
| 1 | 207997973 | rs2724375 | -0.889 | 0.269 | 0.819 | -0.45 | 0.076 | -0.04 | -0.04 | 0 | 0.25 | 0.45 | 0.51 | 0.11 | 0.38 | 0.27 |
| 1 | 208018367 | rs2745980 | -0.845 | 0.272 | 0.576 | -0.449 | 0.071 | -0.044 | -0.044 | 0 | 0.26 | 0.53 | 0.51 | 0.11 | 0.38 | 0.27 |
| 1 | 208024820 | rs12731740 | 1.4 | 0.281 | 0.016 | -0.311 | -0.032 | -0.033 | -0.033 | 0 | 0.06 | 0.01 | 0.17 | 0.08 | 0.12 | 0.08 |
| 1 | 208026977 | rs926631 | 0.938 | 0.033 | 0.097 | -0.064 | 0.005 | -0.015 | -0.015 | 0 | 0.59 | 0.45 | 0.49 | 0.65 | 0.58 | 0.6 |
| 1 | 208027459 | rs2796241 | 1.84 | 0.052 | 0.066 | 0.224 | 0.022 | 0.008 | 0.008 | 0 | 0.51 | 0.25 | 0.3 | 0.48 | 0.44 | 0.52 |
| 1 | 208029947 | rs7550821 | 0.896 | 0.053 | 0.179 | -0.017 | -0.012 | 0.004 | 0.004 | 0 | 0.15 | 0.12 | 0.28 | 0.01 | 0.22 | 0.14 |
| 1 | 208030852 | rs2796245 | -0.061 | -0.087 | 0.212 | 0.054 | -0.03 | 0.055 | 0.055 | 0 | 0.77 | 0.54 | 0.81 | 0.66 | 0.84 | 0.81 |
| 1 | 208033037 | rs859709 | 1.397 | 0.015 | -0.271 | -0.025 | 0 | -0.042 | -0.042 | 0 | 0.11 | 0.17 | 0.08 | 0 | 0.1 | 0.04 |
| 1 | 208035088 | rs984984 | 0.677 | -0.032 | -0.504 | 0.253 | -0.016 | 0.002 | 0.002 | 0 | 0.57 | 0.26 | 0.46 | 0.59 | 0.54 | 0.6 |
| 1 | 208036976 | rs2796247 | 0.677 | -0.032 | -0.504 | 0.245 | -0.016 | 0.002 | 0.002 | 0 | 0.57 | 0.26 | 0.46 | 0.59 | 0.54 | 0.6 |
| 1 | 208038684 | rs2796249 | 1.947 | -0.701 | -1.33 | -0.47 | -0.056 | 0.176 | 0.176 | 0 | 0.3 | 0.05 | 0.18 | 0.04 | 0.32 | 0.13 |
| 1 | 208044432 | rs1204672 | 0.167 | -0.633 | -0.83 | -0.125 | -0.014 | 0.019 | 0.019 | 0 | 0.66 | 0.63 | 0.44 | 0.83 | 0.56 | 0.69 |
| 1 | 208049077 | rs2952 | -0.043 | -0.535 | -0.85 | -0.14 | -0.021 | 0.025 | 0.025 | 0 | 0.71 | 0.69 | 0.49 | 0.83 | 0.63 | 0.72 |
| 1 | 208050360 | rs17009223 | 0.928 | -0.833 | -0.565 | -0.468 | -0.023 | -0.088 | -0.088 | 0 | 0.16 | 0.25 | 0.09 | 0 | 0.1 | 0.05 |
| 1 | 208051059 | rs2745948 | 2.513 | -1.152 | -0.995 | -0.622 | -0.075 | 0.147 | 0.147 | 0 | 0.25 | 0.04 | 0.18 | 0.01 | 0.31 | 0.13 |
| 1 | 208053785 | rs607952 | 0.05 | -1.314 | -0.756 | -0.146 | 0.005 | -0.003 | -0.003 | 0 | 0.62 | 0.43 | 0.39 | 0.69 | 0.55 | 0.59 |
| 1 | 208055666 | rs694748 | -1.07 | -1.186 | -0.652 | 0.128 | -0.009 | 0 | 0 | 0 | 0.36 | 0.37 | 0.21 | 0.67 | 0.24 | 0.45 |
| 1 | 208060009 | rs7572 | -0.744 | -1.445 | -0.323 | 1.022 | -0.002 | 0.005 | 0.005 | 0 | 0.25 | 0.23 | 0.29 | 0.51 | 0.2 | 0.28 |
| 1 | 208060359 | rs2556 | 0.904 | -1.148 | -0.547 | 1.146 | 0.003 | 0.041 | 0.041 | 0 | 0.07 | 0.05 | 0.05 | 0.37 | 0.07 | 0.12 |
| 1 | 208060464 | rs3820521 | -0.498 | -1.129 | -0.369 | 0.765 | -0.006 | 0.059 | 0.059 | 0 | 0.07 | 0.13 | 0.04 | 0.25 | 0.04 | 0.22 |
| 1 | 208065610 | rs2250321 | 0.133 | -1.217 | -1.006 | 0.583 | -0.033 | -0.114 | -0.114 | 0 | 0.31 | 0.08 | 0.4 | 0.4 | 0.35 | 0.37 |
| 1 | 208068579 | rs2259397 | 1.16 | -1.304 | -0.717 | 0.506 | 0 | 0.016 | 0.016 | 0 | 0.51 | 0.58 | 0.5 | 0.44 | 0.49 | 0.41 |
| 1 | 208069137 | rs2796254 | 2.73 | -1.455 | -0.782 | 0.353 | -0.088 | 0.143 | 0.143 | 0 | 0.28 | 0.09 | 0.21 | 0.02 | 0.35 | 0.11 |
| 1 | 208069877 | rs2488250 | 1.263 | -1.483 | -0.794 | 0.405 | 0 | 0.034 | 0.034 | 0.309 | 0.47 | 0.58 | 0.32 | 0.24 | 0.43 | 0.24 |
| 1 | 208071105 | rs3738460 | 0.986 | -1.562 | -1.008 | 0.428 | -0.02 | -0.021 | -0.021 | 0 | 0.09 | 0.03 | 0.05 | 0.13 | 0.12 | 0.09 |
| 1 | 208072626 | rs12029239 | -0.196 | -1.533 | -0.795 | 0.458 | -0.058 | 0.071 | 0.071 | 0 | 0.21 | 0.39 | 0.25 | 0.39 | 0.12 | 0.28 |
| 1 | 208077006 | rs1204682 | 0.01 | -1.371 | -0.716 | -0.191 | -0.056 | -0.073 | -0.073 | 0.316 | 0.35 | 0.03 | 0.57 | 0.61 | 0.4 | 0.53 |
| 1 | 208077643 | rs656801 | 0.578 | -1.167 | -0.196 | 0.245 | 0.028 | 0.007 | 0.007 | 0 | 0.82 | 0.55 | 0.89 | 0.85 | 0.84 | 0.78 |
| 1 | 208078238 | rs1932819 | -0.15 | -1.189 | 0.131 | 0.142 | -0.072 | -0.005 | -0.005 | 0 | 0.18 | 0.39 | 0.08 | 0.19 | 0.06 | 0.11 |
| 1 | 208085735 | rs2267894 | -0.323 | -1.106 | 0.001 | 0.086 | -0.06 | 0.002 | 0.002 | 0 | 0.19 | 0.38 | 0.11 | 0.21 | 0.08 | 0.13 |
| 1 | 208086939 | rs3754171 | -0.258 | -1.069 | -0.238 | 0.104 | -0.056 | 0.005 | 0.005 | 0 | 0.18 | 0.37 | 0.11 | 0.21 | 0.08 | 0.13 |
| 1 | 208097796 | rs12041505 | 0.34 | -0.984 | -1.202 | -0.011 | -0.001 | -0.098 | -0.098 | 0 | 0.14 | 0.02 | 0.24 | 0.38 | 0.13 | 0.25 |
| 1 | 208101656 | rs2473653 | -0.183 | -1.041 | -1.073 | -0.45 | 0.013 | 0.03 | 0.03 | 0.073 | 0.64 | 0.57 | 0.54 | 0.54 | 0.56 | 0.5 |
| 1 | 208112125 | rs614732 | -0.055 | -1.027 | -0.98 | -0.721 | 0.064 | 0.024 | 0.024 | 2.155 | 0.92 | 0.51 | 0.93 | 0.98 | 0.97 | 0.96 |
| 1 | 208113731 | rs1028400 | -0.734 | -1.081 | -1.12 | -0.786 | 0.005 | 0.143 | 0.143 | 0.93 | 0.49 | 0.28 | 0.66 | 0.93 | 0.55 | 0.81 |
| 1 | 208127490 | rs1074327 | 1.729 | -0.912 | -1.384 | -0.728 | -0.04 | 0.06 | 0.06 | 0 | 0.31 | 0.22 | 0.23 | 0.05 | 0.36 | 0.14 |
| 1 | 208128855 | rs11579848 | 1.772 | -0.91 | -1.442 | -0.727 | -0.051 | 0.079 | 0.079 | 0 | 0.31 | 0.19 | 0.23 | 0.05 | 0.36 | 0.14 |
| 1 | 208130627 | rs11579568 | 1.952 | -0.859 | -1.901 | -0.725 | -0.122 | 0.179 | 0.179 | 0 | 0.29 | 0.05 | 0.19 | 0.02 | 0.35 | 0.12 |
| 1 | 208140468 | rs916508 | 0.122 | -0.504 | -1.686 | 0.046 | -0.1 | -0.157 | -0.157 | 0 | 0.3 | 0.03 | 0.4 | 0.45 | 0.41 | 0.45 |
| 1 | 208142232 | rs2184820 | -0.615 | -0.383 | -0.841 | -0.891 | -0.02 | 0.146 | 0.146 | 0 | 0.54 | 0.27 | 0.67 | 0.81 | 0.58 | 0.72 |
| 1 | 208145073 | rs560311 | -0.265 | 0.028 | -1.058 | -0.426 | -0.13 | 0.201 | 0.201 | 1.401 | 0.4 | 0.08 | 0.52 | 0.83 | 0.54 | 0.64 |
| 1 | 208151575 | rs606714 | 1.295 | -0.089 | 0.529 | 0.802 | -0.002 | -0.033 | -0.033 | 1.309 | 0.88 | 0.36 | 0.78 | 0.95 | 0.91 | 0.88 |
| 1 | 208163349 | rs637746 | -1.686 | -0.173 | 0.689 | 1.445 | -0.004 | 0.158 | 0.158 | 0.148 | 0.11 | 0.36 | 0.24 | 0.22 | 0.11 | 0.24 |
| 1 | 208164369 | rs652727 | 0.961 | -0.052 | 0.464 | 1.694 | -0.001 | 0.007 | 0.007 | 0 | 0.64 | 0.52 | 0.62 | 0.36 | 0.62 | 0.44 |
| 1 | 208164620 | rs545947 | 0.594 | -0.135 | 0.656 | 1.69 | 0 | 0.006 | 0.006 | 0 | 0.68 | 0.65 | 0.82 | 0.41 | 0.7 | 0.57 |
| 1 | 208168613 | rs586683 | 1.655 | -0.207 | 0.673 | 1.567 | 0.013 | 0.178 | 0.178 | 0 | 0.9 | 0.59 | 0.72 | 0.79 | 0.89 | 0.69 |
| 1 | 208170487 | rs586030 | 0.915 | -0.23 | 0.741 | 1.429 | -0.001 | 0.03 | 0.03 | 0 | 0.65 | 0.49 | 0.58 | 0.54 | 0.62 | 0.47 |
| 1 | 208171368 | rs651480 | -0.938 | -0.941 | 0.706 | 0.594 | -0.058 | -0.006 | -0.006 | 0 | 0.21 | 0.46 | 0.27 | 0.19 | 0.13 | 0.2 |
| 1 | 208173117 | rs7535719 | -0.617 | -0.335 | 0.662 | 0.739 | -0.002 | 0.024 | 0.024 | 0 | 0.13 | 0.26 | 0.11 | 0.17 | 0.09 | 0.21 |
| 1 | 208175991 | rs567125 | -0.898 | -0.224 | 0.795 | 0.643 | 0.023 | -0.021 | -0.021 | 0 | 0.07 | 0.32 | 0.18 | 0.01 | 0.08 | 0.08 |
| 1 | 208177525 | rs6666839 | -0.826 | -0.078 | 0.067 | 0.926 | -0.003 | 0.021 | 0.021 | 0 | 0.14 | 0.27 | 0.11 | 0.18 | 0.1 | 0.27 |
| 1 | 208178218 | rs3901316 | -0.753 | 0.048 | -0.837 | 1.146 | -0.003 | -0.084 | -0.084 | 0 | 0.26 | 0.09 | 0.25 | 0.48 | 0.28 | 0.24 |
| 1 | 208183804 | rs11811665 | -0.91 | -0.248 | -0.398 | 0.272 | -0.018 | -0.057 | -0.057 | 0.778 | 0.23 | 0.41 | 0.16 | 0.1 | 0.26 | 0.23 |

| | | | | | | | | | | | | | | | | |
|---|---|---|---|---|---|---|---|---|---|---|---|---|---|---|---|---|
| 1 | 208184956 | rs481764 | 0.504 | -0.277 | -0.64 | 0.427 | 0.003 | 0.036 | 0.036 | 0 | 0.7 | 0.8 | 0.77 | 0.81 | 0.77 | 0.78 |
| 1 | 208186905 | rs6684965 | -0.469 | -0.219 | -0.377 | 0.421 | 0.024 | -0.017 | -0.017 | 2.978 | 0.2 | 0.35 | 0.21 | 0.15 | 0.22 | 0.29 |
| 1 | 208188196 | rs2745945 | -0.234 | -0.313 | -0.492 | 0.416 | -0.006 | 0 | 0 | 0.066 | 0.87 | 0.83 | 0.96 | 0.84 | 0.96 | 0.95 |
| 1 | 208190623 | rs16808 | 0.511 | -0.074 | -0.868 | 0.513 | -0.009 | 0.002 | 0.002 | 0 | 0.63 | 0.42 | 0.69 | 0.64 | 0.73 | 0.63 |
| 1 | 208193732 | rs1152832 | 0.519 | -0.07 | -1.36 | 0.526 | -0.001 | 0.005 | 0.005 | 0 | 0.45 | 0.22 | 0.45 | 0.48 | 0.59 | 0.48 |
| 1 | 208195516 | rs1193096 | -0.22 | -0.556 | -0.813 | 0.845 | -0.01 | 0.038 | 0.038 | 0 | 0.94 | 0.81 | 0.9 | 0.92 | 0.98 | 0.97 |
| 1 | 208195981 | rs556625 | 0.35 | -1.245 | -0.989 | -0.056 | 0.004 | 0.009 | 0.009 | 0 | 0.41 | 0.37 | 0.34 | 0.13 | 0.61 | 0.28 |
| 1 | 208196081 | rs2767567 | -0.881 | -1.245 | -0.887 | 0.232 | -0.016 | 0.057 | 0.057 | 0 | 0.93 | 0.8 | 0.85 | 0.91 | 0.98 | 0.97 |
| 1 | 208197785 | rs591752 | 0.823 | -2.029 | -1.445 | 0.603 | -0.002 | -0.087 | -0.087 | 0 | 0.17 | 0.03 | 0.11 | 0.38 | 0.14 | 0.38 |
| 1 | 208204399 | rs3811386 | 0.961 | -2.424 | -1.295 | 0.366 | 0.003 | 0.008 | 0.008 | 0 | 0.41 | 0.36 | 0.33 | 0.15 | 0.6 | 0.28 |
| 1 | 208207839 | rs3736963 | 0.469 | -2.642 | -0.802 | 0.413 | 0.007 | 0.007 | 0.007 | 0.007 | 0.48 | 0.54 | 0.45 | 0.26 | 0.62 | 0.31 |
| 1 | 208209385 | rs4844633 | 0.853 | -3.166 | -0.873 | 0.488 | 0.021 | -0.029 | -0.029 | 0 | 0.21 | 0.14 | 0.31 | 0.32 | 0.14 | 0.23 |
| 1 | 208210864 | rs12087306 | 0.654 | -3.119 | -0.768 | 0.498 | -0.015 | 0.046 | 0.046 | 0 | 0.06 | 0.15 | 0.12 | 0.11 | 0.02 | 0.04 |
| 1 | 208212397 | rs2767566 | -0.846 | -3.498 | -0.899 | 0.363 | 0.048 | 0.047 | 0.047 | 0.952 | 0.44 | 0.22 | 0.53 | 0.74 | 0.37 | 0.66 |
| 1 | 208213692 | rs1536472 | 0.923 | -3.45 | -0.84 | 0.426 | -0.026 | 0.045 | 0.045 | 0.005 | 0.71 | 0.63 | 0.69 | 0.46 | 0.82 | 0.55 |
| 1 | 208214504 | rs2785622 | 1.429 | -3.444 | -0.946 | 0.448 | -0.034 | -0.077 | -0.077 | 0 | 0.12 | 0.02 | 0.14 | 0.37 | 0.15 | 0.36 |
| 1 | 208216304 | rs2274448 | -0.678 | -3.428 | -0.853 | 0.399 | -0.044 | 0.022 | 0.022 | 0.391 | 0.14 | 0.22 | 0.16 | 0.16 | 0.03 | 0.09 |
| 1 | 208218841 | rs11588627 | 1.341 | -3.31 | -1.145 | 0.608 | -0.021 | -0.052 | -0.052 | 0 | 0.08 | 0.01 | 0.27 | 0.25 | 0.09 | 0.16 |
| 1 | 208225357 | rs6540451 | 2.072 | -3.55 | -1.194 | 0.301 | -0.002 | -0.02 | -0.02 | 0 | 0.4 | 0.39 | 0.35 | 0.15 | 0.59 | 0.28 |
| 1 | 208225909 | rs841865 | 1.421 | -3.622 | -1.348 | 0.915 | -0.041 | -0.076 | -0.076 | 1.786 | 0.12 | 0.01 | 0.12 | 0.31 | 0.16 | 0.34 |
| 1 | 208233998 | rs2251465 | 1.669 | -3.561 | -1.292 | 0.997 | -0.003 | 0.02 | 0.02 | 0 | 0.06 | 0.08 | 0.13 | 0.11 | 0.05 | 0.04 |
| 1 | 208234388 | rs4844634 | -0.548 | -3.713 | -1.2 | 0.938 | 0.037 | -0.017 | -0.017 | 0 | 0.34 | 0.25 | 0.39 | 0.44 | 0.22 | 0.31 |
| 1 | 208236164 | rs12057548 | -0.005 | -3.606 | -1.201 | 1.061 | 0.022 | -0.01 | -0.01 | 0 | 0.3 | 0.23 | 0.35 | 0.34 | 0.22 | 0.26 |
| 1 | 208237677 | rs752016 | -0.174 | -3.665 | -1.271 | 1.041 | 0.031 | -0.101 | -0.101 | 0 | 0.21 | 0.05 | 0.22 | 0.45 | 0.14 | 0.36 |
| 1 | 208239176 | rs6656034 | 0.525 | -3.886 | -1.301 | 0.924 | 0 | -0.002 | -0.002 | 0 | 0.61 | 0.61 | 0.5 | 0.55 | 0.68 | 0.63 |
| 1 | 208239519 | rs12057794 | -1.598 | -3.878 | -1.17 | 0.897 | 0.048 | 0.053 | 0.053 | 0.509 | 0.51 | 0.39 | 0.5 | 0.73 | 0.36 | 0.67 |
| 1 | 208243860 | rs10863692 | 0.172 | -3.592 | -1.095 | 0.93 | -0.001 | -0.001 | -0.001 | 2.236 | 0.22 | 0.24 | 0.35 | 0.2 | 0.24 | 0.27 |
| 1 | 208244039 | rs6673084 | 1.719 | -3.445 | -1.027 | 0.89 | 0.013 | 0.016 | 0.016 | 0 | 0.5 | 0.4 | 0.39 | 0.35 | 0.6 | 0.33 |
| 1 | 208244853 | rs2478816 | -0.089 | -3.325 | -1.042 | 0.958 | -0.023 | 0.024 | 0.024 | 0 | 0.81 | 0.74 | 0.89 | 0.66 | 0.91 | 0.68 |
| 1 | 208253883 | rs7540179 | 2.17 | -3.042 | -1.306 | 0.66 | -0.001 | -0.005 | -0.005 | 0 | 0.44 | 0.4 | 0.34 | 0.31 | 0.55 | 0.31 |
| 1 | 208255877 | rs6540453 | 1.031 | -3.042 | -1.28 | 0.711 | -0.012 | -0.004 | -0.004 | 0.171 | 0.61 | 0.43 | 0.46 | 0.66 | 0.68 | 0.6 |
| 1 | 208262786 | rs10863694 | 0.273 | -2.181 | -1.559 | 0.082 | -0.012 | -0.003 | -0.003 | 0 | 0.23 | 0.14 | 0.39 | 0.23 | 0.26 | 0.27 |
| 1 | 208263150 | rs2498024 | 0.775 | -2.223 | -0.945 | 0.106 | 0.006 | -0.013 | -0.013 | 1.418 | 0.16 | 0.07 | 0.12 | 0.34 | 0.13 | 0.27 |
| 1 | 208263222 | rs2498023 | -0.147 | -2.331 | -0.768 | 0.001 | -0.015 | -0.033 | -0.033 | 2.627 | 0.8 | 0.63 | 0.81 | 0.9 | 0.85 | 0.84 |
| 1 | 208265612 | rs2478806 | 0.951 | -2.189 | -0.653 | 0.213 | -0.001 | -0.006 | -0.006 | 0.019 | 0.57 | 0.5 | 0.42 | 0.66 | 0.59 | 0.56 |
| 1 | 208267578 | rs2148699 | 2.021 | -2.293 | -0.735 | 0.078 | 0.001 | -0.003 | -0.003 | 0 | 0.4 | 0.43 | 0.29 | 0.32 | 0.43 | 0.28 |
| 1 | 208270244 | rs3818517 | -0.53 | -2.197 | -0.683 | -0.012 | -0.011 | 0.062 | 0.062 | 1.495 | 0.38 | 0.23 | 0.43 | 0.18 | 0.38 | 0.35 |
| 1 | 208274114 | rs2478819 | 0.986 | -2.15 | -0.447 | 0.082 | -0.001 | -0.002 | -0.002 | 0.315 | 0.56 | 0.51 | 0.43 | 0.68 | 0.59 | 0.57 |
| 1 | 208281097 | rs12032925 | 0.369 | -1.025 | -0.622 | 0.281 | -0.001 | 0.016 | 0.016 | 0 | 0.14 | 0.11 | 0.16 | 0.38 | 0.12 | 0.3 |
| 1 | 208281939 | rs4844406 | -0.349 | -1.136 | -0.794 | -0.061 | -0.015 | 0.058 | 0.058 | 2.43 | 0.34 | 0.16 | 0.42 | 0.18 | 0.35 | 0.36 |
| 1 | 208293513 | rs4844645 | -0.765 | -1.414 | -0.329 | -0.256 | -0.016 | -0.059 | -0.059 | 0.361 | 0.89 | 0.73 | 0.93 | 1 | 0.93 | 0.96 |
| 1 | 208296083 | rs11118988 | 0.787 | -1.502 | -0.555 | -0.234 | -0.012 | -0.018 | -0.018 | 0 | 0.18 | 0.1 | 0.37 | 0.23 | 0.21 | 0.23 |
| 1 | 208299199 | rs10863698 | 1.292 | -1.752 | -0.441 | -0.267 | 0.008 | 0.03 | 0.03 | 0.549 | 0.49 | 0.57 | 0.37 | 0.33 | 0.55 | 0.29 |
| 1 | 208300491 | rs681241 | 1.724 | -1.805 | -0.469 | -0.266 | 0.005 | -0.033 | -0.033 | 0 | 0.12 | 0.07 | 0.13 | 0.35 | 0.11 | 0.3 |
| 1 | 208302633 | rs895241 | 1.186 | -1.709 | -0.395 | -0.23 | 0.008 | 0.012 | 0.012 | 0.551 | 0.49 | 0.57 | 0.43 | 0.4 | 0.55 | 0.29 |
| 1 | 208303998 | rs3811389 | 1.222 | -1.728 | -0.175 | -0.239 | 0.002 | 0.003 | 0.003 | 0.424 | 0.49 | 0.49 | 0.42 | 0.4 | 0.55 | 0.29 |
| 1 | 208304053 | rs3811390 | 2.506 | -2.118 | -0.482 | -0.447 | 0.026 | -0.004 | -0.004 | 1.632 | 0.15 | 0.08 | 0.05 | 0.16 | 0.08 | 0.05 |
| 1 | 208306480 | rs17187133 | 2.458 | -1.321 | -0.453 | -1.096 | 0.002 | -0.005 | -0.005 | 0.011 | 0.11 | 0.09 | 0.07 | 0.03 | 0.1 | 0.13 |
| 1 | 208309240 | rs859499 | 0.055 | -0.89 | -0.236 | -0.845 | 0.001 | 0.032 | 0.032 | 1.354 | 0.36 | 0.24 | 0.45 | 0.25 | 0.35 | 0.38 |
| 1 | 208310909 | rs473500 | 1.352 | -1.186 | -0.51 | -0.973 | -0.064 | 0.127 | 0.127 | 0.408 | 0.19 | 0.02 | 0.08 | 0 | 0.21 | 0.17 |
| 1 | 208313310 | rs668878 | -1.964 | -1.318 | -0.054 | -0.773 | -0.006 | 0 | 0 | 0.884 | 0.9 | 0.79 | 0.88 | 0.92 | 0.93 | 0.89 |
| 1 | 208314807 | rs686065 | 0.641 | -1.347 | 0.043 | -0.86 | -0.006 | 0.022 | 0.022 | 0 | 0.29 | 0.12 | 0.14 | 0 | 0.3 | 0.21 |
| 1 | 208321118 | rs1875431 | 0.031 | -1.08 | -0.051 | -0.502 | 0 | -0.002 | -0.002 | 0 | 0.56 | 0.49 | 0.27 | 0.33 | 0.46 | 0.42 |
| 1 | 208323140 | rs11578181 | 3.17 | -0.818 | -0.207 | -0.543 | -0.066 | 0.054 | 0.054 | 0 | 0.13 | 0.06 | 0.17 | 0 | 0.3 | 0.04 |
| 1 | 208325244 | rs600031 | -1.014 | -1.025 | -0.017 | -0.325 | -0.013 | 0.075 | 0.075 | 0 | 0.87 | 0.75 | 0.85 | 0.67 | 0.91 | 0.7 |
| 1 | 208325802 | rs6671923 | 0.801 | -1.105 | -0.123 | -0.174 | -0.072 | 0.015 | 0.015 | 0 | 0.31 | 0.16 | 0.52 | 0.24 | 0.45 | 0.27 |
| 1 | 208326212 | rs4844411 | 0.264 | -0.991 | 0.462 | -0.384 | -0.01 | 0.121 | 0.121 | 0.093 | 0.15 | 0.21 | 0.43 | 0.58 | 0.13 | 0.43 |
| 1 | 208327039 | rs6697350 | 1.04 | -1.081 | -0.15 | -0.517 | 0.001 | -0.004 | -0.004 | 0 | 0.46 | 0.49 | 0.33 | 0.36 | 0.5 | 0.24 |
| 1 | 208327712 | rs550948 | -0.001 | -1.122 | -0.182 | -0.561 | -0.034 | 0.133 | 0.133 | 0 | 0.35 | 0.1 | 0.23 | 0.08 | 0.36 | 0.28 |
| 1 | 208328224 | rs2590688 | -0.11 | -0.958 | -0.212 | -0.467 | -0.004 | 0.051 | 0.051 | 0 | 0.38 | 0.25 | 0.25 | 0.08 | 0.37 | 0.33 |
| 1 | 208328749 | rs486753 | -0.218 | -0.242 | -0.075 | 0.036 | -0.015 | 0.099 | 0.099 | 0.022 | 0.83 | 0.7 | 0.58 | 0.44 | 0.87 | 0.57 |
| 1 | 208329768 | rs1604364 | 0.684 | 0.147 | -0.459 | 0.146 | -0.01 | 0.013 | 0.013 | 0.01 | 0.42 | 0.35 | 0.3 | 0.36 | 0.48 | 0.24 |
| 1 | 208339232 | rs2590685 | -0.654 | 0.342 | -0.286 | -0.483 | -0.004 | -0.052 | -0.052 | 0.039 | 0.69 | 0.69 | 0.79 | 1 | 0.65 | 0.93 |
| 1 | 208339601 | rs11118997 | -0.237 | 0.208 | -0.07 | 0.275 | -0.044 | -0.014 | -0.014 | 0 | 0.41 | 0.23 | 0.61 | 0.57 | 0.47 | 0.48 |
| 1 | 208341500 | rs895237 | 0.074 | 0.092 | 0.067 | -0.192 | -0.001 | -0.088 | -0.088 | 0 | 0.56 | 0.51 | 0.73 | 1 | 0.54 | 0.73 |
| 1 | 208349259 | rs12033110 | 0.199 | 0.184 | 0.097 | 0.561 | 0.001 | 0.095 | 0.095 | 0.051 | 0.11 | 0.16 | 0.41 | 0.5 | 0.13 | 0.39 |
| 1 | 208353855 | rs6673048 | 0.702 | 0.425 | 0.316 | 0.327 | 0.003 | 0.014 | 0.014 | 0 | 0.42 | 0.3 | 0.27 | 0.3 | 0.42 | 0.27 |
| 1 | 208366353 | rs2218481 | -0.839 | 0.671 | -0.128 | 0.393 | -0.082 | 0.092 | 0.092 | 0 | 0.68 | 0.85 | 0.74 | 0.87 | 0.59 | 0.67 |
| 1 | 208367614 | rs1609841 | 0.379 | 0.39 | -0.071 | 0.819 | -0.006 | -0.003 | -0.003 | 3.4 | 0.2 | 0.21 | 0.35 | 0.18 | 0.16 | 0.21 |

| | | | | | | | | | | | | | | | | |
|---|---|---|---|---|---|---|---|---|---|---|---|---|---|---|---|---|
| 1 | 208368201 | rs11590686 | 0.616 | 0.52 | 0.219 | 1.122 | 0.001 | 0.013 | 0.013 | 0.758 | 0.43 | 0.36 | 0.28 | 0.29 | 0.43 | 0.28 |
| 1 | 208369825 | rs17012158 | 0.46 | 0.593 | 0.189 | 0.648 | -0.002 | -0.013 | -0.013 | 0 | 0.45 | 0.49 | 0.37 | 0.7 | 0.44 | 0.46 |
| 1 | 208375738 | rs10863703 | -0.916 | 0.735 | 0.065 | 0.674 | -0.009 | 0.015 | 0.015 | 0 | 0.73 | 0.76 | 0.77 | 0.86 | 0.68 | 0.74 |
| 1 | 208376078 | rs1249865 | -1.352 | 0.732 | -0.199 | 0.444 | -0.03 | 0.05 | 0.05 | 0 | 0.77 | 0.85 | 0.84 | 0.99 | 0.7 | 0.86 |
| 1 | 208378625 | rs756098 | 0.617 | 0.451 | -0.146 | 1.186 | 0.031 | 0.342 | 0.342 | 0.288 | 0.93 | 0.62 | 0.88 | 0.66 | 0.98 | 0.72 |
| 1 | 208382040 | rs1664232 | 0.568 | 0.603 | -1.075 | 0.926 | -0.038 | 0.014 | 0.014 | 0.034 | 0.42 | 0.23 | 0.31 | 0.3 | 0.48 | 0.39 |
| 1 | 208382156 | rs10863704 | 1.082 | 0.981 | -1.624 | 0.142 | -0.036 | 0.048 | 0.048 | 0 | 0.24 | 0.07 | 0.22 | 0.15 | 0.3 | 0.15 |
| 1 | 208383009 | rs1318276 | -0.23 | 0.699 | -1.4 | 0.367 | 0.015 | 0.018 | 0.018 | 0 | 0.25 | 0.16 | 0.33 | 0.2 | 0.21 | 0.24 |
| 1 | 208390155 | rs4844657 | 0.21 | 1.042 | -1.634 | 0.243 | -0.026 | 0.006 | 0.006 | 0.086 | 0.2 | 0.05 | 0.2 | 0.14 | 0.21 | 0.08 |
| 1 | 208390162 | rs4844658 | 0.303 | 1.055 | -1.661 | 0.239 | -0.058 | -0.019 | -0.019 | 0 | 0.19 | 0.01 | 0.2 | 0.14 | 0.21 | 0.08 |
| 1 | 208390469 | rs3748735 | 1.303 | 1.003 | -1.963 | 0.003 | 0.023 | 0.042 | 0.042 | 0 | 0.23 | 0.07 | 0.17 | 0.15 | 0.2 | 0.15 |
| 1 | 208391085 | rs12120681 | 0.718 | 1.444 | -2.205 | -0.165 | -0.046 | 0.007 | 0.007 | 0 | 0.07 | 0.01 | 0.14 | 0.02 | 0.16 | 0.12 |
| 1 | 208393548 | rs881758 | 1.455 | 1.294 | -2.128 | -0.142 | -0.003 | 0.042 | 0.042 | 0 | 0.23 | 0.06 | 0.19 | 0.14 | 0.23 | 0.16 |
| 1 | 208397575 | rs17259450 | 1.87 | 0.874 | -2.368 | -0.511 | 0.021 | 0.043 | 0.043 | 0.025 | 0.14 | 0.03 | 0.03 | 0.02 | 0.07 | 0.02 |
| 1 | 208403258 | rs875044 | 0.981 | 0.97 | -2.408 | -0.46 | -0.039 | -0.013 | -0.013 | 0 | 0.19 | 0.03 | 0.21 | 0.18 | 0.21 | 0.09 |
| 1 | 208404011 | rs11586367 | 1.501 | 1.301 | -2.236 | -0.306 | 0.004 | 0.054 | 0.054 | 0 | 0.23 | 0.09 | 0.19 | 0.11 | 0.23 | 0.16 |
| 1 | 208404823 | rs1537693 | -0.848 | 1.297 | -1.18 | 0.31 | 0.028 | 0.012 | 0.012 | 0 | 0.86 | 0.77 | 0.86 | 0.86 | 0.82 | 0.83 |
| 1 | 208408503 | rs2782934 | -0.293 | 1.374 | -0.754 | 0.519 | 0.031 | 0.099 | 0.099 | 0.057 | 0.22 | 0.37 | 0.32 | 0.4 | 0.27 | 0.41 |
| 1 | 208408890 | rs11119017 | 0.149 | 0.953 | -1.067 | -0.017 | 0.008 | 0.037 | 0.037 | 0.332 | 0.42 | 0.32 | 0.4 | 0.3 | 0.42 | 0.24 |
| 1 | 208410160 | rs6685511 | -1.205 | 0.929 | -0.47 | 0.095 | 0 | -0.007 | -0.007 | 0 | 0.13 | 0.25 | 0.21 | 0.12 | 0.12 | 0.19 |
| 1 | 208410510 | rs12739500 | -0.526 | 1.156 | -0.768 | -0.712 | -0.016 | 0.012 | 0.012 | 0.217 | 0.29 | 0.23 | 0.19 | 0.01 | 0.36 | 0.18 |
| 1 | 208414033 | rs1335045 | -0.536 | 0.305 | -0.323 | -0.093 | 0.001 | 0.005 | 0.005 | 0 | 0.52 | 0.66 | 0.53 | 0.56 | 0.5 | 0.4 |
| 1 | 208415871 | rs7533778 | -0.21 | 0.157 | -0.37 | -0.066 | 0.002 | -0.003 | -0.003 | 0.42 | 0.14 | 0.12 | 0.07 | 0.25 | 0.08 | 0.2 |
| 1 | 208419288 | rs1620298 | 0.119 | 0.128 | 0.157 | -0.085 | -0.002 | 0.001 | 0.001 | 0 | 0.37 | 0.42 | 0.44 | 0.27 | 0.42 | 0.19 |
| 1 | 208419966 | rs1664248 | -0.971 | 0.103 | 0.22 | -0.01 | 0.008 | 0.175 | 0.175 | 0 | 0.09 | 0.4 | 0.26 | 0.27 | 0.08 | 0.06 |
| 1 | 208420588 | rs1781038 | -1.318 | 0.104 | 0.004 | -0.001 | -0.031 | 0.082 | 0.082 | 0 | 0.24 | 0.62 | 0.35 | 0.52 | 0.16 | 0.26 |
| 1 | 208424205 | rs1781030 | -1.033 | 0.11 | -0.274 | -0.227 | -0.003 | -0.054 | -0.054 | 0 | 0.6 | 0.82 | 0.56 | 0.54 | 0.57 | 0.42 |
| 1 | 208424905 | rs12058868 | -0.62 | -0.125 | 0.265 | -0.111 | -0.033 | 0.027 | 0.027 | 0 | 0.2 | 0.5 | 0.12 | 0.21 | 0.11 | 0.21 |
| 1 | 208425671 | rs2801193 | -1.152 | -0.096 | 0.2 | -0.188 | -0.001 | 0.005 | 0.005 | 0 | 0.25 | 0.76 | 0.34 | 0.54 | 0.17 | 0.26 |
| 1 | 208432764 | rs1445355 | -1.012 | -0.109 | 0.315 | -0.172 | -0.021 | 0.1 | 0.1 | 0 | 0.23 | 0.64 | 0.34 | 0.54 | 0.16 | 0.27 |
| 1 | 208433883 | rs11119027 | 0.522 | -0.006 | 0.186 | -0.234 | -0.063 | 0.136 | 0.136 | 0 | 0.36 | 0.11 | 0.21 | 0 | 0.4 | 0.16 |
| 1 | 208443170 | rs6703732 | 0.592 | -0.12 | 0.207 | -0.196 | -0.029 | 0.132 | 0.132 | 0 | 0.33 | 0.09 | 0.17 | 0 | 0.33 | 0.16 |
| 1 | 208443456 | rs2154229 | -0.702 | -0.072 | 0.385 | -0.014 | 0.005 | 0.206 | 0.206 | 0 | 0.21 | 0.5 | 0.28 | 0.54 | 0.15 | 0.24 |
| 1 | 208445800 | rs2590691 | -1.229 | -0.164 | 0.208 | -0.055 | 0.013 | -0.039 | -0.039 | 0 | 0.27 | 0.81 | 0.3 | 0.55 | 0.15 | 0.24 |
| 1 | 208446188 | rs6689034 | -0.833 | -0.165 | 0.315 | 0.016 | -0.008 | 0 | 0 | 0 | 0.23 | 0.55 | 0.15 | 0.21 | 0.15 | 0.22 |
| 1 | 208447524 | rs1617731 | -1.048 | -0.161 | -0.329 | -0.096 | -0.039 | -0.066 | -0.066 | 3.104 | 0.58 | 0.86 | 0.47 | 0.55 | 0.48 | 0.4 |
| 1 | 208449483 | rs12121098 | 0.513 | 0.116 | -0.402 | -0.038 | -0.056 | 0.17 | 0.17 | 0.217 | 0.31 | 0.04 | 0.17 | 0 | 0.33 | 0.15 |
| 1 | 208461343 | rs6704329 | -1.11 | -0.44 | 0.022 | 0.522 | -0.002 | 0 | 0 | 0.326 | 0.25 | 0.73 | 0.25 | 0.46 | 0.11 | 0.27 |
| 1 | 208464118 | rs3121647 | -1.682 | -0.84 | 0.074 | 0.198 | -0.063 | -0.028 | -0.028 | 0 | 0.18 | 0.68 | 0.08 | 0.13 | 0.03 | 0.2 |
| 1 | 208469340 | rs3121651 | -1.641 | -0.599 | 0.31 | 0.202 | -0.014 | -0.098 | -0.098 | 0 | 0.24 | 0.65 | 0.15 | 0 | 0.13 | 0.06 |
| 1 | 208477423 | rs1762463 | -0.413 | -0.595 | 0.903 | 0.226 | -0.014 | -0.047 | -0.047 | 0 | 0.08 | 0.26 | 0.04 | 0 | 0.03 | 0.01 |
| 1 | 208485874 | rs6702300 | 0.881 | -0.567 | 0.321 | 0.546 | -0.038 | -0.07 | -0.07 | 0 | 0.11 | 0.01 | 0.28 | 0.44 | 0.17 | 0.29 |
| 1 | 208490895 | rs2897064 | -2.131 | -0.706 | 0.506 | 0.205 | 0.024 | -0.024 | -0.024 | 0 | 0.3 | 0.82 | 0.28 | 0.4 | 0.19 | 0.2 |
| 1 | 208493971 | rs1499587 | -1.108 | -0.628 | 0.569 | 0.23 | -0.014 | -0.071 | -0.071 | 0 | 0.2 | 0.45 | 0.15 | 0.04 | 0.15 | 0.13 |
| 1 | 208505326 | rs4844673 | -0.537 | -0.849 | 0.372 | 0.297 | 0.008 | -0.009 | -0.009 | 0 | 0.16 | 0.15 | 0.19 | 0.4 | 0.11 | 0.19 |
| 1 | 208507471 | rs6699314 | 1.208 | -0.872 | -0.233 | -0.143 | -0.099 | 0.152 | 0.152 | 0.083 | 0.3 | 0.05 | 0.2 | 0 | 0.38 | 0.11 |
| 1 | 208510805 | rs4844419 | -2.187 | -0.954 | 0.857 | 1.095 | -0.025 | 0.017 | 0.017 | 0.873 | 0.26 | 0.64 | 0.25 | 0.36 | 0.14 | 0.16 |
| 1 | 208511351 | rs1499577 | -2.601 | -0.964 | 0.849 | 1.199 | 0.01 | -0.013 | -0.013 | 0 | 0.29 | 0.75 | 0.26 | 0.44 | 0.16 | 0.21 |
| 1 | 208515347 | rs7543864 | -1.856 | -1.038 | 0.535 | 1.227 | -0.059 | 0.125 | 0.125 | 0.666 | 0.23 | 0.47 | 0.2 | 0.44 | 0.11 | 0.13 |
| 1 | 208520153 | rs6689464 | -0.215 | -0.92 | 0.547 | 1.034 | 0.008 | 0.012 | 0.012 | 0.045 | 0.12 | 0.09 | 0.06 | 0 | 0.07 | 0.05 |
| 1 | 208520568 | rs2217357 | -0.849 | -0.906 | 0.297 | 1.083 | -0.051 | 0.246 | 0.246 | 0.215 | 0.17 | 0.44 | 0.19 | 0.47 | 0.07 | 0.12 |
| 1 | 208526097 | rs6661433 | -1.41 | -0.752 | -0.535 | 0.814 | 0.064 | -0.087 | -0.087 | 0.015 | 0.3 | 0.85 | 0.35 | 0.53 | 0.17 | 0.27 |
| 1 | 208535774 | rs6677524 | -1.125 | 0.145 | -0.636 | -0.002 | -0.001 | -0.005 | -0.005 | 0.015 | 0.13 | 0.19 | 0.15 | 0.11 | 0.12 | 0.1 |
| 1 | 208547582 | rs696985 | 0.507 | 0.247 | -1.352 | -1.294 | -0.017 | 0.039 | 0.039 | 0 | 0.38 | 0.3 | 0.41 | 0.86 | 0.42 | 0.6 |
| 1 | 208549846 | rs696981 | -0.257 | -0.143 | -0.899 | -1.578 | -0.001 | -0.006 | -0.006 | 0 | 0.52 | 0.41 | 0.53 | 0.14 | 0.51 | 0.38 |
| 1 | 208552368 | rs12138270 | 0.926 | 0.043 | -1.18 | -0.679 | -0.082 | -0.117 | -0.117 | 1.448 | 0.31 | 0.08 | 0.2 | 0.41 | 0.36 | 0.44 |
| 1 | 208554185 | rs701641 | -0.201 | -0.246 | -0.614 | -0.878 | 0.001 | -0.024 | -0.024 | 0 | 0.24 | 0.27 | 0.16 | 0.13 | 0.24 | 0.12 |
| 1 | 208557123 | rs1499588 | 0.104 | -0.582 | -0.539 | -0.907 | 0.013 | 0.002 | 0.002 | 0 | 0.25 | 0.15 | 0.33 | 0.01 | 0.23 | 0.35 |
| 1 | 208558547 | rs696979 | 1.023 | -0.181 | -1.073 | -1.001 | -0.043 | 0.091 | 0.091 | 0 | 0.35 | 0.16 | 0.24 | 0.12 | 0.37 | 0.17 |
| 1 | 208561353 | rs2163039 | -0.166 | 0.474 | -0.437 | 0.15 | -0.006 | -0.02 | -0.02 | 0 | 0.3 | 0.22 | 0.28 | 0.41 | 0.34 | 0.38 |
| 1 | 208562158 | rs696977 | 0.525 | 0.445 | -0.275 | 0.043 | 0.003 | -0.047 | -0.047 | 0 | 0.63 | 0.56 | 0.58 | 0.13 | 0.61 | 0.53 |
| 1 | 208567416 | rs11119064 | -0.628 | 0.491 | -0.594 | 0.066 | -0.007 | 0.034 | 0.034 | 0 | 0.36 | 0.25 | 0.4 | 0.87 | 0.39 | 0.47 |
| 1 | 208568131 | rs17012470 | 0.117 | 0.413 | -0.497 | 0.71 | -0.001 | -0.014 | -0.014 | 0 | 0.07 | 0.06 | 0.06 | 0.01 | 0.09 | 0.17 |
| 1 | 208569243 | rs17187394 | -0.247 | 0.362 | -0.515 | 1.135 | 0.001 | -0.025 | -0.025 | 0 | 0.05 | 0.01 | 0.07 | 0.11 | 0.05 | 0.03 |
| 1 | 208578200 | rs710417 | 0.015 | 0.483 | -0.263 | 1.001 | -0.001 | 0.208 | 0.208 | 0 | 0.93 | 0.84 | 0.86 | 0.53 | 0.95 | 0.9 |
| 1 | 208583530 | rs696944 | 0.002 | 0.857 | -0.548 | 0.66 | 0.011 | 0.042 | 0.042 | 0 | 0.69 | 0.48 | 0.71 | 0.58 | 0.69 | 0.84 |
| 1 | 208589905 | rs696948 | -0.079 | 0.86 | -0.947 | 0.83 | -0.013 | -0.012 | -0.012 | 0.455 | 0.34 | 0.24 | 0.33 | 0.41 | 0.46 | 0.33 |
| 1 | 208590432 | rs696172 | 0.223 | 1 | -1.354 | 0.568 | -0.125 | -0.035 | -0.035 | 0 | 0.27 | 0.03 | 0.22 | 0.25 | 0.38 | 0.29 |
| 1 | 208593803 | rs710420 | 0.274 | 0.15 | -0.296 | 0.433 | -0.031 | 0.099 | 0.099 | 0.014 | 0.78 | 0.61 | 0.78 | 0.54 | 0.83 | 0.66 |

| | | | | | | | | | | | | | | | | |
|---|---|---|---|---|---|---|---|---|---|---|---|---|---|---|---|---|
| 1 | 208598450 | rs696964 | 0.723 | -0.016 | 0.231 | 0.346 | -0.007 | 0.017 | 0.017 | 0.312 | 0.75 | 0.27 | 0.73 | 0.51 | 0.8 | 0.66 |
| 1 | 208605807 | rs12047825 | 1.009 | -0.011 | 0.031 | 0.093 | -0.005 | 0.06 | 0.06 | 0 | 0.05 | 0.01 | 0.13 | 0.49 | 0.04 | 0.1 |
| 1 | 208606109 | rs6662504 | -0.079 | -0.03 | 0.327 | 0.04 | -0.018 | -0.078 | -0.078 | 0.146 | 0.15 | 0.24 | 0.09 | 0 | 0.13 | 0.25 |
| 1 | 208611123 | rs701618 | 0.789 | -0.027 | 0.07 | 0.063 | -0.027 | 0.076 | 0.076 | 0.12 | 0.75 | 0.4 | 0.74 | 0.51 | 0.8 | 0.65 |
| 1 | 208619842 | rs701610 | 0.19 | -0.14 | -0.795 | -0.823 | 0.007 | 0.058 | 0.058 | 0 | 0.3 | 0.14 | 0.15 | 0.13 | 0.24 | 0.46 |
| 1 | 208619858 | rs4844678 | 0.73 | 0.296 | -0.892 | -0.967 | -0.055 | 0.024 | 0.024 | 0 | 0.35 | 0.14 | 0.46 | 0.25 | 0.39 | 0.36 |
| 1 | 208622869 | rs12089705 | 0.473 | 0.138 | 0.326 | -0.288 | -0.003 | 0.373 | 0.373 | 0 | 0.06 | 0.22 | 0.14 | 0.44 | 0.03 | 0.08 |
| 1 | 208623242 | rs12122106 | 0.547 | 0.173 | -0.399 | -0.573 | -0.026 | 0.096 | 0.096 | 0 | 0.37 | 0.15 | 0.21 | 0.17 | 0.35 | 0.5 |
| 1 | 208623475 | rs701608 | 0.575 | 0.348 | -0.79 | -0.32 | -0.026 | -0.037 | -0.037 | 0.297 | 0.37 | 0.14 | 0.51 | 0.39 | 0.41 | 0.38 |
| 1 | 208625731 | rs1499585 | -0.4 | 0.269 | -0.696 | -0.211 | -0.007 | 0.089 | 0.089 | 0 | 0.1 | 0.16 | 0.1 | 0.43 | 0.09 | 0.08 |
| 1 | 208626075 | rs1499586 | 0.19 | 0.752 | -0.425 | -1.652 | 0.004 | 0.108 | 0.108 | 0 | 0.51 | 0.38 | 0.7 | 0.85 | 0.52 | 0.47 |
| 1 | 208629207 | rs17012564 | -1.158 | 0.624 | -0.482 | -0.609 | 0.029 | -0.085 | -0.085 | 0 | 0.13 | 0.41 | 0.1 | 0 | 0.16 | 0.02 |
| 1 | 208634002 | rs696958 | -0.368 | 0.548 | -1.901 | -0.684 | 0.005 | 0.15 | 0.15 | 0.242 | 0.36 | 0.08 | 0.3 | 0.14 | 0.34 | 0.48 |
| 1 | 208636607 | rs17259644 | -0.506 | 1.015 | -1.774 | -1.113 | -0.02 | 0.054 | 0.054 | 0 | 0.22 | 0.11 | 0.12 | 0 | 0.24 | 0.06 |
| 1 | 208637976 | rs7536351 | -0.589 | 0.954 | -1.43 | -0.849 | 0.001 | -0.001 | -0.001 | 0 | 0.14 | 0.19 | 0.2 | 0.12 | 0.13 | 0.26 |
| 1 | 208640150 | rs10863712 | 0.779 | 0.528 | -1.975 | -0.996 | -0.086 | 0.224 | 0.224 | 0 | 0.45 | 0.09 | 0.47 | 0.14 | 0.51 | 0.32 |
| 1 | 208644820 | rs696951 | 0.591 | 0.305 | 0.029 | 0.968 | 0.021 | 0.288 | 0.288 | 0 | 0.87 | 0.61 | 0.81 | 0.52 | 0.88 | 0.79 |
| 1 | 208646090 | rs701627 | 0.876 | 0.715 | -0.558 | 0.211 | 0.019 | 0.133 | 0.133 | 0.049 | 0.48 | 0.16 | 0.53 | 0.27 | 0.41 | 0.4 |
| 1 | 208658283 | rs701615 | 0.218 | 0.785 | 0.057 | 0.321 | 0.044 | 0.342 | 0.342 | 0 | 0.87 | 0.54 | 0.82 | 0.5 | 0.87 | 0.79 |
| 1 | 208676383 | rs7551719 | 0.821 | 0.694 | 0.09 | 0.33 | 0.023 | 0.107 | 0.107 | 0.007 | 0.47 | 0.21 | 0.54 | 0.27 | 0.4 | 0.4 |
| 1 | 208676718 | rs11119075 | -0.975 | 0.779 | 0.404 | 0.67 | -0.001 | -0.008 | -0.008 | 0 | 0.94 | 0.93 | 0.97 | 1 | 0.93 | 0.98 |
| 1 | 208700733 | rs2405104 | -0.487 | 0.815 | 0.336 | -0.218 | 0.004 | -0.066 | -0.066 | 0 | 0.36 | 0.57 | 0.26 | 0.13 | 0.39 | 0.28 |
| 1 | 208701114 | rs4844686 | 0.564 | 0.518 | 0.478 | 0.172 | -0.002 | 0.003 | 0.003 | 1.342 | 0.6 | 0.58 | 0.69 | 0.72 | 0.55 | 0.58 |
| 1 | 208701134 | rs4844687 | 1.346 | 0.804 | -0.103 | 0.296 | 0.029 | 0.12 | 0.12 | 1.676 | 0.43 | 0.24 | 0.44 | 0.17 | 0.35 | 0.24 |
| 1 | 208701907 | rs1417953 | 0.697 | 0.507 | -0.256 | 0.151 | -0.002 | 0.023 | 0.023 | 0 | 0.61 | 0.66 | 0.73 | 0.71 | 0.61 | 0.57 |
| 1 | 208706936 | rs11576725 | -0.209 | 0.882 | -0.776 | -0.127 | -0.054 | 0.101 | 0.101 | 0 | 0.24 | 0.08 | 0.15 | 0.05 | 0.34 | 0.2 |
| 1 | 208710349 | rs7512748 | -0.601 | 0.814 | -0.818 | -0.574 | -0.06 | 0.114 | 0.114 | 0 | 0.37 | 0.16 | 0.3 | 0.01 | 0.46 | 0.22 |
| 1 | 208710593 | rs10863714 | 0.699 | 0.56 | 0.049 | -0.334 | -0.003 | -0.004 | -0.004 | 3.252 | 0.58 | 0.59 | 0.61 | 0.79 | 0.53 | 0.6 |
| 1 | 208711001 | rs12752047 | -0.388 | 0.569 | -0.16 | -0.404 | -0.098 | 0.177 | 0.177 | 0 | 0.36 | 0.1 | 0.29 | 0.01 | 0.46 | 0.22 |
| 1 | 208713165 | rs10863715 | 1.851 | 0.039 | -1.134 | -0.26 | -0.028 | -0.052 | -0.052 | 0 | 0.29 | 0.01 | 0.25 | 0.3 | 0.24 | 0.2 |
| 1 | 208717605 | rs10863718 | -1.033 | 0.088 | -0.481 | 0.457 | 0.02 | -0.007 | -0.007 | 15.008 | 0.61 | 0.33 | 0.57 | 0.39 | 0.68 | 0.66 |
| 1 | 208719848 | rs7548249 | -0.073 | 0.284 | -0.584 | 0.422 | -0.029 | 0.089 | 0.089 | 0 | 0.43 | 0.26 | 0.36 | 0.23 | 0.47 | 0.43 |
| 1 | 208720185 | rs7527585 | 0.641 | 0.35 | -0.845 | 0.484 | -0.055 | -0.059 | -0.059 | 0 | 0.16 | 0.01 | 0.18 | 0.15 | 0.2 | 0.21 |
| 1 | 208722372 | rs6656598 | -0.275 | 0.45 | -0.575 | 0.504 | -0.006 | 0.016 | 0.016 | 0 | 0.45 | 0.37 | 0.37 | 0.23 | 0.48 | 0.43 |
| 1 | 208732621 | rs17012827 | -0.118 | 0.598 | -0.132 | 1.299 | -0.003 | -0.012 | -0.012 | 0.052 | 0.2 | 0.23 | 0.23 | 0.35 | 0.25 | 0.22 |
| 1 | 208739034 | rs10779485 | -0.3 | 0.65 | 0.546 | 1.446 | 0.027 | 0.153 | 0.153 | 2.552 | 0.06 | 0.25 | 0.12 | 0.15 | 0.03 | 0.11 |
| 1 | 208739309 | rs10779486 | -0.424 | 0.639 | 0.561 | 1.44 | 0.025 | 0.15 | 0.15 | 2.752 | 0.06 | 0.25 | 0.13 | 0.15 | 0.03 | 0.11 |
| 1 | 208750816 | rs17187568 | 1.255 | 1.115 | 0.072 | 1.144 | -0.039 | 0.044 | 0.044 | 0 | 0.11 | 0.04 | 0.14 | 0.02 | 0.14 | 0.06 |
| 1 | 208755605 | rs6540478 | -0.858 | 0.816 | -0.227 | 0.296 | 0 | -0.002 | -0.002 | 0 | 0.44 | 0.51 | 0.38 | 0.56 | 0.49 | 0.55 |
| 1 | 208755804 | rs17187582 | 1.245 | 0.82 | -0.493 | 0.049 | -0.046 | 0.086 | 0.086 | 0 | 0.16 | 0.03 | 0.08 | 0 | 0.19 | 0.15 |
| 1 | 208761593 | rs17259791 | 2.049 | 0.604 | -0.642 | -0.001 | -0.042 | 0.016 | 0.016 | 0 | 0.34 | 0.02 | 0.22 | 0.27 | 0.29 | 0.25 |
| 1 | 208766610 | rs12029495 | 0.518 | 0.403 | -0.091 | 0.39 | 0.001 | 0.002 | 0.002 | 0.033 | 0.19 | 0.21 | 0.2 | 0.24 | 0.26 | 0.23 |
| 1 | 208766895 | rs951279 | 1.103 | 0.379 | -0.346 | 0.214 | -0.031 | 0.166 | 0.166 | 0 | 0.49 | 0.13 | 0.45 | 0.28 | 0.47 | 0.34 |
| 1 | 208772958 | rs10779489 | 0.903 | 0.752 | -0.176 | 0.23 | -0.003 | 0.004 | 0.004 | 0 | 0.55 | 0.41 | 0.57 | 0.4 | 0.48 | 0.49 |
| 1 | 208777606 | rs6694783 | 0.561 | 0.863 | -0.225 | 0.571 | -0.007 | -0.015 | -0.015 | 0 | 0.16 | 0.11 | 0.19 | 0.19 | 0.21 | 0.21 |
| 1 | 208777749 | rs946801 | -1.276 | 1.021 | 0.023 | 0.347 | 0.002 | -0.049 | -0.049 | 0 | 0.87 | 0.85 | 0.95 | 1 | 0.9 | 0.98 |
| 1 | 208777895 | rs869521 | 1.508 | 1.24 | 0.176 | 0.283 | 0.02 | 0.009 | 0.009 | 0 | 0.44 | 0.34 | 0.36 | 0.46 | 0.42 | 0.52 |
| 1 | 208779683 | rs2185383 | 0.58 | 1.215 | 0.117 | 0.222 | -0.06 | 0.064 | 0.064 | 0 | 0.17 | 0.06 | 0.17 | 0.03 | 0.24 | 0.09 |
| 1 | 208781486 | rs17259861 | 1.653 | 1.001 | -0.104 | 0.476 | 0.042 | 0.014 | 0.014 | 0.019 | 0.42 | 0.25 | 0.34 | 0.39 | 0.34 | 0.49 |
| 1 | 208787740 | rs1340113 | -0.149 | 1.114 | -0.095 | 0.299 | -0.002 | -0.007 | -0.007 | 0 | 0.24 | 0.37 | 0.34 | 0.21 | 0.27 | 0.27 |
| 1 | 208789949 | rs12122043 | -0.017 | 0.873 | -0.493 | 0.019 | -0.052 | 0.064 | 0.064 | 0 | 0.24 | 0.09 | 0.18 | 0.03 | 0.28 | 0.1 |
| 1 | 208804121 | rs7546488 | 1.04 | 0.645 | -0.291 | 0.017 | 0 | -0.007 | -0.007 | 0 | 0.45 | 0.35 | 0.39 | 0.52 | 0.49 | 0.53 |
| 1 | 208804925 | rs4391668 | -0.51 | 0.438 | -0.98 | 0.107 | -0.036 | 0.002 | 0.002 | 0.156 | 0.43 | 0.16 | 0.37 | 0.34 | 0.4 | 0.26 |
| 1 | 208805287 | rs2404678 | 0.567 | 0.049 | -0.106 | -0.108 | -0.008 | -0.013 | -0.013 | 0.13 | 0.77 | 0.7 | 0.71 | 0.84 | 0.82 | 0.73 |
| 1 | 208806940 | rs6685613 | -0.894 | 0.435 | -0.118 | 0.039 | -0.007 | -0.076 | -0.076 | 0 | 0.2 | 0.26 | 0.11 | 0 | 0.15 | 0.07 |
| 1 | 208809622 | rs1166861 | 0.958 | 0.6 | 0.278 | 1.052 | 0.011 | 0.117 | 0.117 | 0.747 | 0.83 | 0.6 | 0.74 | 0.67 | 0.84 | 0.71 |
| 1 | 208810027 | rs10779491 | 0.058 | 0.367 | -0.139 | 0.919 | 0.011 | 0.007 | 0.007 | 0.027 | 0.49 | 0.3 | 0.35 | 0.44 | 0.45 | 0.33 |
| 1 | 208810098 | rs1166864 | -0.294 | 0.281 | -0.004 | 0.918 | 0 | 0.003 | 0.003 | 0.057 | 0.48 | 0.6 | 0.63 | 0.51 | 0.53 | 0.59 |
| 1 | 208813119 | rs1166868 | -0.824 | 1.072 | 0.033 | 0.761 | 0.001 | 0 | 0 | 0 | 0.23 | 0.7 | 0.29 | 0.26 | 0.26 | 0.25 |
| 1 | 208830611 | rs1340117 | -1.44 | -0.688 | 0.554 | 0.589 | -0.068 | 0.011 | 0.011 | 0.001 | 0.2 | 0.72 | 0.15 | 0.26 | 0.05 | 0.22 |
| 1 | 208830888 | rs11589583 | -0.91 | -0.569 | 0.828 | 0.312 | -0.02 | -0.041 | -0.041 | 0 | 0.14 | 0.21 | 0.05 | 0 | 0.04 | 0.08 |
| 1 | 208840386 | rs17013019 | -1.253 | -0.456 | 1.073 | 0.256 | -0.077 | -0.026 | -0.026 | 1.074 | 0.22 | 0.55 | 0.17 | 0.2 | 0.07 | 0.15 |
| 1 | 208849634 | rs17259875 | 0.096 | -0.373 | 0.669 | -0.012 | -0.001 | -0.016 | -0.016 | 0 | 0.09 | 0.07 | 0.07 | 0 | 0.12 | 0.02 |
| 1 | 208855395 | rs6664062 | -1.345 | -0.705 | 0.766 | 0.351 | -0.077 | -0.033 | -0.033 | 0 | 0.25 | 0.57 | 0.19 | 0.2 | 0.08 | 0.24 |
| 1 | 208860293 | rs10863723 | -1.065 | 0.308 | 0.897 | 0.543 | 0.001 | 0.012 | 0.012 | 0.164 | 0.19 | 0.38 | 0.17 | 0.16 | 0.17 | 0.2 |
| 1 | 208863701 | rs17013082 | -0.391 | 0.14 | 0.387 | 0.381 | 0.009 | 0.015 | 0.015 | 0 | 0.07 | 0 | 0.03 | 0.03 | 0.04 | 0.03 |
| 1 | 208864258 | rs11119111 | -1.516 | 0.066 | 0.777 | 0.791 | -0.014 | 0.064 | 0.064 | 0.84 | 0.25 | 0.59 | 0.27 | 0.33 | 0.23 | 0.34 |
| 1 | 208866218 | rs6668364 | -1.567 | -0.078 | 0.864 | 0.585 | -0.018 | -0.036 | -0.036 | 0 | 0.12 | 0.15 | 0.06 | 0.03 | 0.06 | 0.05 |
| 1 | 208872435 | rs1166872 | 0.624 | 0.668 | 0.428 | 0.288 | -0.027 | 0.033 | 0.033 | 0.168 | 0.83 | 0.9 | 0.9 | 1 | 0.76 | 0.91 |

| | | | | | | | | | | | | | | | | |
|---|---|---|---|---|---|---|---|---|---|---|---|---|---|---|---|---|
| 1 | 208872494 | rs1166873 | 1.951 | 0.505 | 1.004 | 0.614 | -0.013 | -0.007 | -0.007 | 0.104 | 0.69 | 0.44 | 0.76 | 0.74 | 0.7 | 0.75 |
| 1 | 208893836 | rs17259881 | 0.088 | 0.659 | 0.55 | 0.708 | -0.03 | -0.04 | -0.04 | 0.465 | 0.08 | 0.01 | 0.12 | 0.11 | 0.09 | 0.13 |
| 1 | 208897416 | rs1166879 | 2.052 | 0.422 | 1.075 | 0.479 | -0.011 | -0.005 | -0.005 | 0 | 0.69 | 0.42 | 0.77 | 0.74 | 0.71 | 0.76 |
| 1 | 208897975 | rs12063248 | -2.077 | 0.003 | 1.24 | 0.91 | -0.075 | 0.161 | 0.161 | 0 | 0.14 | 0.43 | 0.13 | 0.26 | 0.05 | 0.15 |
| 1 | 208909872 | rs11119123 | -1.567 | 0.021 | 1.156 | 0.882 | -0.048 | 0.015 | 0.015 | 0 | 0.12 | 0.43 | 0.07 | 0.07 | 0.04 | 0.04 |
| 1 | 208917791 | rs4844721 | -1.852 | 0.034 | 1.119 | 0.861 | -0.076 | 0.17 | 0.17 | 0 | 0.14 | 0.45 | 0.13 | 0.26 | 0.04 | 0.14 |
| 1 | 208925929 | rs17013209 | -0.185 | 0.651 | 0.566 | 0.634 | 0.008 | -0.023 | -0.023 | 0 | 0.05 | 0.08 | 0.05 | 0 | 0.07 | 0.03 |
| 1 | 208926042 | rs17013211 | -1.834 | 0.582 | 1.106 | 0.519 | -0.007 | -0.041 | -0.041 | 0 | 0.13 | 0.29 | 0.16 | 0.04 | 0.1 | 0.11 |
| 1 | 208929732 | rs12404464 | -0.89 | 1.224 | 0.645 | 0.55 | -0.054 | -0.06 | -0.06 | 0.411 | 0.1 | 0 | 0.18 | 0.17 | 0.14 | 0.11 |
| 1 | 208930265 | rs12076253 | -1.837 | 1.239 | 0.487 | 0.482 | -0.052 | -0.05 | -0.05 | 0 | 0.2 | 0.05 | 0.26 | 0.23 | 0.24 | 0.2 |
| 1 | 208935416 | rs12565978 | -1.714 | 1.306 | 0.502 | 0.507 | -0.039 | -0.067 | -0.067 | 0.188 | 0.17 | 0.04 | 0.23 | 0.21 | 0.16 | 0.11 |
| 1 | 208935974 | rs17013220 | -2.562 | 1.201 | 1.179 | 0.464 | 0.01 | 0.01 | 0.01 | 0 | 0.23 | 0.37 | 0.34 | 0.22 | 0.25 | 0.2 |
| 1 | 208937403 | rs17259916 | -2.991 | 1.01 | 1.025 | 0.368 | -0.001 | 0.008 | 0.008 | 0 | 0.36 | 0.29 | 0.42 | 0.27 | 0.35 | 0.29 |
| 1 | 208938670 | rs1933566 | -2.498 | 0.918 | 1.155 | 0.372 | -0.019 | -0.052 | -0.052 | 0 | 0.27 | 0.37 | 0.24 | 0.09 | 0.21 | 0.18 |
| 1 | 208941395 | rs6690048 | -1.967 | 0.947 | 1.008 | 0.414 | -0.011 | -0.043 | -0.043 | 0 | 0.19 | 0.27 | 0.18 | 0.06 | 0.16 | 0.17 |
| 1 | 208946799 | rs17013249 | -2.255 | 0.857 | 1.067 | 0.62 | 0.013 | 0.032 | 0.032 | 0 | 0.22 | 0.42 | 0.34 | 0.23 | 0.2 | 0.21 |
| 1 | 208947435 | rs6681543 | -1.835 | 0.795 | 1.043 | 0.489 | 0.001 | 0.002 | 0.002 | 0 | 0.18 | 0.17 | 0.17 | 0.07 | 0.16 | 0.15 |
| 1 | 208948015 | rs12079674 | -1.977 | 0.638 | 1.081 | 0.413 | -0.042 | -0.068 | -0.068 | 3.071 | 0.18 | 0.37 | 0.1 | 0.05 | 0.11 | 0.1 |
| 1 | 208948034 | rs11119127 | -0.6 | 0.779 | 0.929 | 0.509 | -0.044 | -0.051 | -0.051 | 0.501 | 0.1 | 0.01 | 0.13 | 0.15 | 0.11 | 0.14 |
| 1 | 208948194 | rs11119129 | -0.6 | 0.779 | 0.954 | 0.51 | -0.039 | -0.048 | -0.048 | 1.135 | 0.1 | 0.03 | 0.13 | 0.15 | 0.11 | 0.14 |
| 1 | 208949367 | rs12076644 | -3.108 | 0.74 | 1.014 | 0.515 | -0.007 | -0.011 | -0.011 | 0.487 | 0.36 | 0.52 | 0.37 | 0.26 | 0.31 | 0.29 |
| 1 | 208950832 | rs10863725 | -2.879 | 0.873 | 0.954 | 0.531 | -0.002 | -0.012 | -0.012 | 0 | 0.3 | 0.49 | 0.32 | 0.22 | 0.27 | 0.29 |
| 1 | 208951613 | rs7536675 | -1.437 | 0.765 | 0.672 | 0.543 | 0.008 | 0.015 | 0.015 | 0 | 0.14 | 0.1 | 0.08 | 0.05 | 0.12 | 0.1 |
| 1 | 208954872 | rs12085219 | -0.752 | 1.029 | 0.656 | 1.03 | -0.004 | 0.004 | 0.004 | 0 | 0.07 | 0.05 | 0.07 | 0.05 | 0.09 | 0.08 |
| 1 | 208956621 | rs1166928 | 1.895 | 1.124 | 0.963 | 1.004 | 0.012 | 0.014 | 0.014 | 0 | 0.74 | 0.48 | 0.75 | 0.76 | 0.77 | 0.85 |
| 1 | 208960876 | rs1166923 | 2.028 | 1.146 | 0.967 | 0.978 | 0.011 | 0.008 | 0.008 | 0.034 | 0.73 | 0.47 | 0.75 | 0.76 | 0.76 | 0.85 |
| 1 | 208968409 | rs17013392 | -1.732 | 1.159 | 1.111 | 0.968 | 0 | -0.002 | -0.002 | 0.044 | 0.2 | 0.21 | 0.2 | 0.2 | 0.13 | 0.12 |
| 1 | 208969750 | rs12403109 | -1.675 | 1.147 | 1.12 | 0.964 | -0.01 | -0.025 | -0.025 | 0 | 0.19 | 0.11 | 0.19 | 0.2 | 0.13 | 0.12 |
| 1 | 208970709 | rs17013396 | -0.762 | 1.151 | 1.096 | 0.966 | -0.006 | -0.009 | -0.009 | 0.258 | 0.1 | 0.14 | 0.07 | 0.05 | 0.04 | 0.01 |
| 1 | 208971775 | rs1316354 | -0.717 | 1.156 | 1.094 | 0.97 | -0.01 | -0.017 | -0.017 | 0 | 0.1 | 0.07 | 0.13 | 0.15 | 0.1 | 0.12 |
| 1 | 208971963 | rs1182870 | 1.345 | 1.173 | 1.097 | 1.057 | 0.093 | 0.062 | 0.062 | 0.16 | 0.93 | 0.65 | 0.94 | 0.95 | 0.89 | 0.97 |
| 1 | 208971979 | rs1317167 | -1.054 | 1.371 | 0.623 | 1.249 | -0.004 | -0.004 | -0.004 | 0 | 0.09 | 0.08 | 0.08 | 0.16 | 0.13 | 0.09 |
| 1 | 208972144 | rs1166900 | 2.004 | 1.408 | 0.582 | 1.376 | 0.008 | 0.018 | 0.018 | 0.017 | 0.69 | 0.36 | 0.71 | 0.59 | 0.69 | 0.78 |
| 1 | 208973877 | rs12145483 | -0.726 | 1.58 | 0.118 | 1.354 | -0.04 | -0.029 | -0.029 | 0 | 0.06 | 0 | 0.14 | 0.2 | 0.13 | 0.13 |
| 1 | 208974905 | rs1166906 | 1.628 | 1.46 | 0.354 | 1.199 | 0 | 0 | 0 | 1.091 | 0.69 | 0.33 | 0.64 | 0.72 | 0.65 | 0.72 |
| 1 | 208974935 | rs1166907 | 1.662 | 1.442 | 0.42 | 0.935 | 0.007 | -0.007 | -0.007 | 0 | 0.72 | 0.34 | 0.66 | 0.85 | 0.68 | 0.74 |
| 1 | 208976796 | rs1166917 | 1.932 | 1.427 | 0.451 | 0.864 | 0.011 | -0.011 | -0.011 | 0 | 0.67 | 0.4 | 0.63 | 0.79 | 0.64 | 0.74 |
| 1 | 208986593 | rs10863726 | -1.915 | 1.489 | 0.479 | 1.09 | 0.043 | 0.046 | 0.046 | 0 | 0.14 | 0.22 | 0.2 | 0.2 | 0.21 | 0.16 |
| 1 | 208994195 | rs1320540 | 0.481 | 1.062 | 0.251 | 0.794 | -0.01 | -0.033 | -0.033 | 1.303 | 0.89 | 0.84 | 0.96 | 1 | 0.95 | 0.94 |
| 1 | 208994644 | rs17187764 | -0.394 | 1.567 | -0.016 | 1.145 | -0.033 | -0.03 | -0.03 | 0 | 0.06 | 0 | 0.14 | 0.13 | 0.13 | 0.14 |
| 1 | 208995966 | rs4844443 | -0.063 | 1.106 | -0.404 | 0.231 | 0.007 | -0.01 | -0.01 | 0.121 | 0.41 | 0.54 | 0.53 | 0.29 | 0.43 | 0.46 |
| 1 | 208996828 | rs11119134 | 0.366 | 0.665 | -0.849 | 0.54 | -0.023 | 0.029 | 0.029 | 1.276 | 0.69 | 0.45 | 0.8 | 0.59 | 0.76 | 0.82 |
| 1 | 208998375 | rs1933544 | 0.396 | 0.478 | -0.899 | 0.649 | -0.022 | 0.01 | 0.01 | 0.782 | 0.71 | 0.43 | 0.77 | 0.68 | 0.78 | 0.69 |
| 1 | 209004061 | rs12035745 | 0.977 | 0.894 | -1.89 | -0.042 | -0.063 | 0.094 | 0.094 | 0.384 | 0.44 | 0.11 | 0.44 | 0.31 | 0.48 | 0.38 |
| 1 | 209007778 | rs12408989 | 1.038 | 1.334 | -2.258 | -0.2 | -0.022 | 0.024 | 0.024 | 0 | 0.28 | 0.12 | 0.34 | 0.16 | 0.29 | 0.21 |
| 1 | 209012975 | rs1890340 | 1.898 | 1.367 | -1.743 | -0.092 | 0.022 | 0.028 | 0.028 | 0 | 0.06 | 0.11 | 0.08 | 0.09 | 0.09 | 0.03 |
| 1 | 209025861 | rs17187785 | 1.423 | 1.325 | -1.879 | -0.014 | -0.029 | -0.026 | -0.026 | 0 | 0.06 | 0.01 | 0.14 | 0.16 | 0.09 | 0.14 |
| 1 | 209031988 | rs1933564 | -0.384 | 0.757 | -1.525 | -0.228 | -0.004 | 0.006 | 0.006 | 0 | 0.68 | 0.62 | 0.74 | 0.6 | 0.7 | 0.61 |
| 1 | 209034761 | rs12569218 | -0.485 | 0.557 | -1.324 | -0.357 | -0.002 | 0.022 | 0.022 | 0 | 0.49 | 0.56 | 0.5 | 0.29 | 0.45 | 0.3 |
| 1 | 209049429 | rs11119145 | 0.344 | 0.573 | -1.334 | -0.244 | -0.03 | 0.018 | 0.018 | 0 | 0.43 | 0.16 | 0.43 | 0.44 | 0.48 | 0.59 |
| 1 | 209052333 | rs7515208 | 0.356 | 0.599 | -1.289 | -0.214 | -0.022 | 0.019 | 0.019 | 0.047 | 0.42 | 0.2 | 0.43 | 0.44 | 0.48 | 0.58 |
| 1 | 209054775 | rs2211045 | 0.84 | 0.953 | -1.39 | -0.607 | -0.006 | -0.004 | -0.004 | 0 | 0.25 | 0.14 | 0.23 | 0 | 0.28 | 0.07 |
| 1 | 209055813 | rs11119146 | 1.275 | 0.725 | -1.582 | -0.165 | 0.008 | -0.011 | -0.011 | 0.123 | 0.08 | 0.01 | 0.04 | 0.13 | 0.05 | 0.11 |
| 1 | 209055852 | rs4844730 | 0.496 | 0.57 | -1.646 | 0.355 | -0.002 | 0.004 | 0.004 | 0.094 | 0.1 | 0.09 | 0.2 | 0.24 | 0.09 | 0.18 |
| 1 | 209059239 | rs11119148 | 1.516 | 0.554 | -1.662 | 0.27 | 0.01 | -0.009 | -0.009 | 0.399 | 0.08 | 0.01 | 0.03 | 0.12 | 0.05 | 0.1 |
| 1 | 209062504 | rs1933572 | 0.552 | 1.24 | -2.013 | -0.196 | -0.03 | 0.055 | 0.055 | 0 | 0.25 | 0.17 | 0.16 | 0.1 | 0.29 | 0.17 |
| 1 | 209063680 | rs6668834 | 0.223 | 1.009 | -1.617 | 0.079 | -0.021 | 0.048 | 0.048 | 0 | 0.32 | 0.49 | 0.42 | 0.5 | 0.31 | 0.57 |
| 1 | 209064416 | rs17260007 | 1.064 | 1.217 | -2.203 | -0.071 | -0.005 | 0.152 | 0.152 | 0 | 0.26 | 0.03 | 0.15 | 0 | 0.25 | 0.07 |
| 1 | 209067110 | rs1933575 | 0.653 | 1.173 | -1.992 | 0.381 | -0.015 | -0.04 | -0.04 | 0.001 | 0.09 | 0.02 | 0.19 | 0.25 | 0.08 | 0.16 |
| 1 | 209073907 | rs12130305 | 0.491 | 1.252 | -2.105 | 0.024 | -0.058 | 0.089 | 0.089 | 0.401 | 0.42 | 0.15 | 0.44 | 0.28 | 0.48 | 0.42 |
| 1 | 209079535 | rs17187806 | 1.111 | 1.18 | -2.394 | -0.154 | 0.012 | 0.123 | 0.123 | 0 | 0.24 | 0.04 | 0.13 | 0 | 0.21 | 0.06 |
| 1 | 209081012 | rs1933554 | 0.173 | 1.587 | -2.16 | 0.435 | -0.107 | 0.095 | 0.095 | 0.503 | 0.41 | 0.1 | 0.44 | 0.26 | 0.47 | 0.42 |
| 1 | 209083609 | rs4844732 | -0.116 | 1.453 | -0.914 | 1.272 | 0.001 | 0 | 0 | 0.151 | 0.08 | 0.05 | 0.19 | 0.24 | 0.06 | 0.15 |
| 1 | 209084106 | rs12039361 | -0.042 | 1.491 | -0.893 | 1.27 | 0.002 | 0.01 | 0.01 | 0.073 | 0.08 | 0.05 | 0.22 | 0.23 | 0.05 | 0.15 |
| 1 | 209088514 | rs7368359 | -0.935 | 0.922 | -0.929 | 0.1 | -0.003 | 0.007 | 0.007 | 0.471 | 0.39 | 0.33 | 0.37 | 0.24 | 0.38 | 0.26 |
| 1 | 209092756 | rs17260014 | -0.272 | 1.187 | -1.44 | -0.419 | -0.049 | 0.119 | 0.119 | 0 | 0.27 | 0.1 | 0.17 | 0 | 0.31 | 0.12 |
| 1 | 209097788 | rs10779495 | 0.497 | 0.816 | -0.49 | 0.357 | 0.015 | 0.04 | 0.04 | 2.287 | 0.45 | 0.36 | 0.26 | 0.24 | 0.39 | 0.28 |
| 1 | 209100322 | rs7516842 | 0.135 | 0.593 | 0.118 | 0.521 | 0.019 | 0.054 | 0.054 | 0.634 | 0.53 | 0.67 | 0.37 | 0.32 | 0.45 | 0.39 |
| 1 | 209101899 | rs11119168 | 0.734 | 0.556 | 0.199 | 0.512 | 0.015 | 0.039 | 0.039 | 2.48 | 0.45 | 0.34 | 0.26 | 0.24 | 0.39 | 0.28 |

| | | | | | | | | | | | | | | | | |
|---|---|---|---|---|---|---|---|---|---|---|---|---|---|---|---|---|
| 1 | 209101942 | rs1933569 | 0 | 0.6 | 0.429 | 0.531 | -0.008 | 0.03 | 0.03 | 0 | 0.07 | 0.24 | 0.1 | 0.08 | 0.06 | 0.1 |
| 1 | 209102552 | rs12410555 | -0.799 | 1.002 | 0.452 | 0.914 | 0.031 | 0.071 | 0.071 | 0.094 | 0.1 | 0.27 | 0.26 | 0.19 | 0.14 | 0.18 |
| 1 | 209105367 | rs11577857 | 0.029 | 0.936 | 0.499 | 0.819 | -0.013 | 0.021 | 0.021 | 0.051 | 0.08 | 0.23 | 0.1 | 0.08 | 0.06 | 0.11 |
| 1 | 209108493 | rs1320539 | 0.316 | 0.94 | 0.283 | 1.004 | 0.004 | 0.008 | 0.008 | 0.119 | 0.14 | 0.07 | 0.08 | 0.14 | 0.11 | 0.12 |
| 1 | 209113459 | rs12738015 | -0.575 | 0.83 | 0.369 | 1.014 | -0.009 | 0.006 | 0.006 | 0.282 | 0.43 | 0.33 | 0.49 | 0.58 | 0.48 | 0.54 |
| 1 | 209114372 | rs4129434 | 0.894 | 0.975 | 0.188 | 0.932 | 0.028 | 0.094 | 0.094 | 3.831 | 0.44 | 0.22 | 0.25 | 0.24 | 0.38 | 0.28 |
| 1 | 209117500 | rs10735503 | -0.644 | 1.08 | -0.108 | 1.297 | -0.023 | -0.026 | -0.026 | 0 | 0.75 | 0.8 | 0.6 | 0.64 | 0.64 | 0.74 |
| 1 | 209118300 | rs10863733 | -0.763 | 1.075 | -0.104 | 1.298 | -0.035 | -0.038 | -0.038 | 0 | 0.76 | 0.85 | 0.61 | 0.65 | 0.64 | 0.75 |
| 1 | 209118786 | rs7540998 | -0.338 | 1.107 | -0.109 | 1.324 | -0.022 | -0.022 | -0.022 | 0 | 0.72 | 0.76 | 0.44 | 0.57 | 0.56 | 0.68 |
| 1 | 209118888 | rs7517671 | -1.112 | 0.571 | 0.356 | 1.482 | -0.014 | 0.027 | 0.027 | 0.246 | 0.29 | 0.44 | 0.35 | 0.39 | 0.26 | 0.46 |
| 1 | 209121677 | rs11119182 | -0.588 | 0.219 | 0.462 | 1.596 | -0.006 | 0.037 | 0.037 | 0 | 0.13 | 0.21 | 0.12 | 0.17 | 0.14 | 0.12 |
| 1 | 209121854 | rs6697933 | -0.694 | 0.182 | 0.923 | 1.578 | 0 | 0.064 | 0.064 | 0 | 0.13 | 0.3 | 0.13 | 0.17 | 0.15 | 0.12 |
| 1 | 209126301 | rs12073738 | -0.379 | -0.28 | 0.975 | 2.168 | 0.006 | -0.006 | -0.006 | 0.205 | 0.11 | 0.1 | 0.05 | 0.2 | 0.04 | 0.07 |
| 1 | 209129752 | rs1924250 | -2.017 | 0.206 | 1.049 | 2.328 | 0 | 0.033 | 0.033 | 0 | 0.43 | 0.68 | 0.56 | 0.68 | 0.42 | 0.49 |
| 1 | 209133898 | rs2807391 | -2.041 | 0.193 | 1.14 | 2.331 | -0.001 | 0.043 | 0.043 | 0 | 0.43 | 0.7 | 0.6 | 0.71 | 0.42 | 0.49 |
| 1 | 209139480 | rs10746417 | -2.058 | 0.166 | 1.377 | 2.328 | 0 | 0.008 | 0.008 | 0 | 0.43 | 0.59 | 0.55 | 0.71 | 0.41 | 0.49 |
| 1 | 209141386 | rs10863734 | -0.054 | 0.175 | 1.438 | 2.432 | -0.01 | -0.01 | -0.01 | 0.935 | 0.16 | 0.12 | 0.33 | 0.34 | 0.21 | 0.25 |
| 1 | 209141842 | rs11119189 | -1.512 | 0.174 | 1.469 | 2.421 | -0.043 | 0.054 | 0.054 | 0 | 0.27 | 0.44 | 0.19 | 0.34 | 0.2 | 0.24 |
| 1 | 209146127 | rs12037214 | -0.493 | 0.35 | 1.486 | 2.381 | 0.011 | 0.055 | 0.055 | 0.573 | 0.14 | 0.34 | 0.13 | 0.16 | 0.15 | 0.16 |
| 1 | 209148991 | rs6696639 | -1.122 | 0.335 | 1.468 | 2.44 | -0.015 | -0.049 | -0.049 | 0.391 | 0.26 | 0.19 | 0.37 | 0.49 | 0.26 | 0.31 |
| 1 | 209149149 | rs6685924 | 0.161 | 0.334 | 1.447 | 2.442 | 0.025 | -0.037 | -0.037 | 0 | 0.1 | 0.06 | 0.04 | 0.18 | 0.04 | 0.08 |
| 1 | 209152380 | rs2807394 | -2.35 | 0.336 | 1.491 | 2.446 | -0.005 | 0.035 | 0.035 | 0 | 0.43 | 0.65 | 0.59 | 0.71 | 0.42 | 0.48 |
| 1 | 209162412 | rs7549301 | -0.503 | 0.352 | 1.475 | 2.458 | 0.012 | 0.06 | 0.06 | 0.535 | 0.14 | 0.36 | 0.17 | 0.17 | 0.15 | 0.15 |
| 1 | 209162928 | rs10863739 | -1.211 | 0.353 | 1.472 | 2.459 | -0.011 | -0.041 | -0.041 | 0 | 0.27 | 0.2 | 0.34 | 0.51 | 0.26 | 0.32 |
| 1 | 209177088 | rs10863744 | -1.469 | 0.447 | 1.414 | 2.381 | 0.034 | 0.067 | 0.067 | 0 | 0.3 | 0.49 | 0.47 | 0.5 | 0.36 | 0.39 |
| 1 | 209181761 | rs1078550 | -2.145 | 0.369 | 1.317 | 2.325 | -0.006 | 0.051 | 0.051 | 0 | 0.43 | 0.7 | 0.61 | 0.72 | 0.42 | 0.46 |
| 1 | 209182220 | rs4554722 | -1.353 | 0.387 | 1.36 | 2.371 | 0.018 | 0.04 | 0.04 | 0 | 0.29 | 0.34 | 0.46 | 0.51 | 0.36 | 0.36 |
| 1 | 209182651 | rs2807403 | -2.087 | 0.403 | 1.368 | 2.395 | 0.001 | -0.007 | -0.007 | 0.767 | 0.42 | 0.52 | 0.52 | 0.66 | 0.42 | 0.44 |
| 1 | 209188309 | rs4240828 | -1.356 | 0.412 | 1.341 | 2.47 | 0.042 | 0.034 | 0.034 | 0 | 0.29 | 0.42 | 0.39 | 0.38 | 0.36 | 0.31 |
| 1 | 209192139 | rs4844743 | -0.574 | -0.094 | 1.187 | 2.066 | 0.001 | 0.025 | 0.025 | 0.453 | 0.14 | 0.23 | 0.2 | 0.15 | 0.15 | 0.14 |
| 1 | 209195248 | rs6670957 | -0.544 | 0.005 | 0.768 | 1.872 | 0.016 | -0.013 | -0.013 | 0 | 0.12 | 0.09 | 0.05 | 0.16 | 0.06 | 0.08 |
| 1 | 209211154 | rs17013951 | -0.718 | 0.316 | 1.447 | 1.913 | 0.022 | 0.116 | 0.116 | 0 | 0.1 | 0.27 | 0.18 | 0.2 | 0.11 | 0.14 |
| 1 | 209211775 | rs4844747 | 1.007 | 0.756 | 1.385 | 1.69 | -0.013 | 0.014 | 0.014 | 0 | 0.6 | 0.33 | 0.34 | 0.32 | 0.5 | 0.56 |
| 1 | 209211968 | rs11119208 | 0.219 | 0.735 | 1.554 | 1.596 | -0.069 | -0.072 | -0.072 | 0 | 0.71 | 0.85 | 0.54 | 0.53 | 0.61 | 0.7 |
| 1 | 209214390 | rs17013958 | 1.037 | 0.699 | 1.614 | 1.603 | -0.017 | -0.001 | -0.001 | 0 | 0.6 | 0.3 | 0.34 | 0.38 | 0.5 | 0.56 |
| 1 | 209214739 | rs10218724 | -0.13 | 0.867 | 1.491 | 1.689 | 0.012 | 0.045 | 0.045 | 0 | 0.09 | 0.21 | 0.16 | 0.15 | 0.11 | 0.12 |
| 1 | 209220739 | rs1010692 | 1.167 | 0.493 | 1.003 | 1.412 | -0.008 | 0.014 | 0.014 | 0 | 0.55 | 0.32 | 0.33 | 0.38 | 0.48 | 0.54 |
| 1 | 209222499 | rs6685963 | -0.3 | 0.581 | 0.782 | 1.163 | 0.039 | 0.071 | 0.071 | 0 | 0.1 | 0.29 | 0.17 | 0.16 | 0.12 | 0.13 |
| 1 | 209233483 | rs10735505 | 0.299 | 0.816 | -0.202 | 1.161 | -0.058 | -0.104 | -0.104 | 1.295 | 0.67 | 0.86 | 0.55 | 0.5 | 0.63 | 0.7 |
| 1 | 209234757 | rs1360279 | 0.777 | 0.498 | 0.23 | 0.839 | 0.002 | -0.004 | -0.004 | 0.068 | 0.56 | 0.53 | 0.41 | 0.38 | 0.52 | 0.51 |
| 1 | 209237451 | rs17014015 | -0.318 | 0.807 | 0.247 | 0.416 | 0.001 | -0.001 | -0.001 | 0 | 0.12 | 0.13 | 0.12 | 0.11 | 0.1 | 0.08 |
| 1 | 209237470 | rs12410473 | -0.919 | 0.131 | 0.632 | 0.149 | -0.016 | -0.011 | -0.011 | 0 | 0.15 | 0.2 | 0.14 | 0.11 | 0.11 | 0.25 |
| 1 | 209239011 | rs17260119 | -0.727 | 0.079 | 0.799 | -0.114 | 0.011 | 0.021 | 0.021 | 0 | 0.14 | 0.06 | 0.07 | 0.03 | 0.11 | 0.25 |
| 1 | 209239811 | rs6540503 | -0.34 | 0.283 | 0.4 | -0.528 | -0.072 | 0.08 | 0.08 | 0.016 | 0.78 | 0.94 | 0.75 | 0.9 | 0.69 | 0.75 |
| 1 | 209245859 | rs10494916 | 1.603 | -0.61 | 0.273 | -1.546 | 0.003 | -0.002 | -0.002 | 0 | 0.28 | 0.31 | 0.47 | 0.73 | 0.36 | 0.29 |
| 1 | 209247368 | rs7415860 | -0.949 | -0.38 | 0.065 | -1.611 | -0.054 | 0.072 | 0.072 | 0 | 0.78 | 0.93 | 0.76 | 0.9 | 0.73 | 0.77 |
| 1 | 209256181 | rs6692331 | 0.344 | -0.117 | 0.458 | -1.344 | 0.023 | 0.002 | 0.002 | 0.013 | 0.15 | 0.29 | 0.15 | 0.13 | 0.14 | 0.11 |
| 1 | 209262232 | rs12121667 | -1.595 | -0.115 | -0.021 | -1.265 | -0.03 | 0.01 | 0.01 | 0 | 0.73 | 0.83 | 0.54 | 0.26 | 0.64 | 0.71 |
| 1 | 209268696 | rs17014039 | -0.08 | -0.226 | 0.165 | -1.24 | 0.053 | -0.013 | -0.013 | 0 | 0.17 | 0.41 | 0.17 | 0.12 | 0.15 | 0.11 |
| 1 | 209269627 | rs6540507 | -1.459 | -0.161 | -0.285 | -1.339 | -0.091 | 0.009 | 0.009 | 0 | 0.73 | 0.89 | 0.47 | 0.24 | 0.55 | 0.64 |
| 1 | 209271004 | rs12138003 | 1.29 | -0.794 | -0.258 | -1.725 | 0.032 | 0.041 | 0.041 | 0 | 0.22 | 0.11 | 0.11 | 0.07 | 0.16 | 0.1 |
| 1 | 209273939 | rs10494917 | 0.181 | -0.712 | 0.153 | -1.636 | -0.031 | -0.066 | -0.066 | 0 | 0.14 | 0.33 | 0.08 | 0.03 | 0.11 | 0.25 |
| 1 | 209276472 | rs1174556 | 1.515 | -0.227 | -0.749 | -1.617 | -0.066 | 0.019 | 0.019 | 0 | 0.26 | 0.09 | 0.48 | 0.78 | 0.35 | 0.34 |
| 1 | 209282540 | rs4258224 | 0.939 | -0.53 | -0.898 | -1.761 | -0.07 | 0.071 | 0.071 | 0 | 0.21 | 0.02 | 0.24 | 0.08 | 0.26 | 0.21 |
| 1 | 209285053 | rs1330233 | 1.409 | -0.173 | -0.488 | -1.374 | -0.062 | 0.018 | 0.018 | 0 | 0.26 | 0.09 | 0.47 | 0.78 | 0.34 | 0.33 |
| 1 | 209291638 | rs6694118 | -0.052 | -0.241 | -0.206 | -1.471 | -0.01 | -0.082 | -0.082 | 0 | 0.16 | 0.3 | 0.09 | 0.02 | 0.14 | 0.29 |
| 1 | 209295363 | rs4844751 | 1.211 | 0.015 | -0.502 | -1.099 | -0.081 | 0.003 | 0.003 | 0 | 0.26 | 0.08 | 0.47 | 0.76 | 0.35 | 0.34 |
| 1 | 209317220 | rs6660522 | 0.969 | -1.069 | 0.26 | -1.378 | -0.017 | -0.015 | -0.015 | 0 | 0.16 | 0.15 | 0.08 | 0.07 | 0.1 | 0.07 |
| 1 | 209317968 | rs10779502 | 0.558 | -1.189 | -0.605 | -1.855 | -0.012 | 0.025 | 0.025 | 0.458 | 0.38 | 0.34 | 0.56 | 0.83 | 0.5 | 0.56 |
| 1 | 209318792 | rs34104149 | 0.73 | -1.287 | -0.612 | -1.9 | -0.038 | 0.221 | 0.221 | 0 | 0.4 | 0.12 | 0.34 | 0.09 | 0.41 | 0.35 |
| 1 | 209323995 | rs4844769 | 0.035 | -0.371 | -0.393 | -1.331 | -0.006 | 0.009 | 0.009 | 0.52 | 0.39 | 0.4 | 0.55 | 0.82 | 0.49 | 0.56 |
| 1 | 209325038 | rs7514220 | 0.264 | -0.305 | -0.681 | -1.211 | -0.017 | 0.005 | 0.005 | 0.015 | 0.11 | 0.29 | 0.13 | 0.1 | 0.08 | 0.19 |
| 1 | 209327341 | rs12067269 | 1.043 | 0.109 | -1.439 | -0.372 | -0.176 | -0.122 | -0.122 | 0 | 0.27 | 0.03 | 0.42 | 0.64 | 0.41 | 0.35 |
| 1 | 209331789 | rs6672714 | -0.417 | -2.04 | -0.734 | -1.311 | -0.01 | 0 | 0 | 0 | 0.15 | 0.2 | 0.2 | 0.15 | 0.12 | 0.25 |
| 1 | 209335123 | rs6540521 | -1.338 | -2.498 | 0.16 | -1.52 | -0.059 | -0.019 | -0.019 | 0 | 0.22 | 0.45 | 0.23 | 0.16 | 0.12 | 0.27 |
| 1 | 209336246 | rs11119231 | 1.435 | -3.431 | -0.139 | -2.233 | 0.061 | 0.089 | 0.089 | 0 | 0.48 | 0.32 | 0.54 | 0.78 | 0.64 | 0.53 |
| 1 | 209342622 | rs1933612 | 0.572 | -3.5 | -0.159 | -2.266 | -0.003 | 0.136 | 0.136 | 0 | 0.28 | 0.02 | 0.22 | 0.06 | 0.24 | 0.2 |
| 1 | 209345380 | rs4844779 | 1.356 | -3.437 | -0.033 | -2.237 | 0.032 | 0.043 | 0.043 | 0 | 0.49 | 0.4 | 0.54 | 0.78 | 0.64 | 0.53 |
| 1 | 209363016 | rs10494918 | 0.166 | -2.997 | 0.097 | -2.167 | -0.007 | -0.012 | -0.012 | 0 | 0.11 | 0.17 | 0.05 | 0.06 | 0.05 | 0.1 |

| | | | | | | | | | | | | | | | | |
|---|---|---|---|---|---|---|---|---|---|---|---|---|---|---|---|---|
| 1 | 209371538 | rs2788170 | 0.074 | -3.478 | -0.011 | -2.431 | 0.041 | 0.064 | 0.064 | 0.271 | 0.31 | 0.21 | 0.23 | 0.06 | 0.23 | 0.19 |
| 1 | 209380481 | rs2970558 | 0.405 | -3.492 | -0.094 | -2.437 | 0.035 | 0.147 | 0.147 | 0.864 | 0.29 | 0.08 | 0.22 | 0.06 | 0.23 | 0.19 |
| 1 | 209383412 | rs1933616 | 0.945 | -3.641 | 0.3 | -2.436 | 0.068 | 0.1 | 0.1 | 0.477 | 0.54 | 0.3 | 0.7 | 0.88 | 0.72 | 0.7 |
| 1 | 209397138 | rs2250701 | 0.103 | -3.62 | 0.19 | -2.421 | 0.047 | 0.072 | 0.072 | 0.414 | 0.31 | 0.19 | 0.23 | 0.06 | 0.22 | 0.19 |
| 1 | 209400118 | rs10863764 | 0.918 | -3.584 | -0.014 | -2.399 | 0.047 | 0.063 | 0.063 | 0.559 | 0.54 | 0.35 | 0.7 | 0.88 | 0.72 | 0.7 |
| 1 | 209405753 | rs2800919 | 0.446 | -3.185 | -0.615 | -2.238 | 0.009 | 0.101 | 0.101 | 2.518 | 0.24 | 0.02 | 0.18 | 0.06 | 0.2 | 0.17 |
| 1 | 209420098 | rs2788154 | -0.153 | -3.134 | -0.35 | -2.119 | 0.05 | 0.076 | 0.076 | 1.367 | 0.32 | 0.16 | 0.17 | 0.07 | 0.22 | 0.18 |
| 1 | 209431182 | rs4844798 | 0.231 | -2.216 | -0.158 | -1.623 | 0 | 0 | 0 | 0 | 0.14 | 0.14 | 0.21 | 0.12 | 0.13 | 0.24 |
| 1 | 209436177 | rs2800930 | -0.647 | -2.291 | -0.205 | -1.501 | -0.001 | 0.018 | 0.018 | 0.017 | 0.9 | 0.79 | 0.95 | 0.87 | 0.95 | 0.89 |
| 1 | 209456475 | rs2843171 | 0.665 | -2.047 | -0.246 | -1.507 | -0.004 | 0.014 | 0.014 | 2.384 | 0.73 | 0.61 | 0.72 | 0.68 | 0.75 | 0.58 |
| 1 | 209456703 | rs2843170 | 0.899 | -2.086 | -0.058 | -1.509 | -0.008 | 0.016 | 0.016 | 2.284 | 0.72 | 0.4 | 0.7 | 0.67 | 0.75 | 0.57 |
| 1 | 209460346 | rs1553623 | -0.711 | -2.07 | -0.24 | -1.476 | -0.002 | 0.051 | 0.051 | 0 | 0.9 | 0.72 | 0.95 | 0.83 | 0.95 | 0.88 |
| 1 | 209472850 | rs1507341 | 0.321 | -2.088 | -0.331 | -1.607 | 0.001 | -0.002 | -0.002 | 0 | 0.73 | 0.69 | 0.71 | 0.7 | 0.79 | 0.63 |
| 1 | 209482201 | rs2660635 | 1.268 | -2.349 | -0.716 | -1.831 | -0.034 | 0.077 | 0.077 | 3.707 | 0.24 | 0.02 | 0.16 | 0.08 | 0.25 | 0.1 |
| 1 | 209482984 | rs11119253 | 1.453 | -1.326 | -0.138 | -1.292 | -0.14 | -0.143 | -0.143 | 0 | 0.27 | 0.05 | 0.44 | 0.54 | 0.42 | 0.33 |
| 1 | 209489688 | rs2495295 | 1.248 | -0.812 | 0.419 | -1.017 | -0.008 | 0.007 | 0.007 | 0.569 | 0.67 | 0.44 | 0.67 | 0.67 | 0.71 | 0.55 |
| 1 | 209501407 | rs1542981 | 0.215 | -0.6 | 0.787 | -0.586 | -0.012 | 0.002 | 0.002 | 0.021 | 0.86 | 0.72 | 0.93 | 0.86 | 0.93 | 0.87 |
| 1 | 209512173 | rs17014320 | 0.27 | -0.599 | 0.783 | -0.585 | -0.024 | -0.032 | -0.032 | 0 | 0.13 | 0.02 | 0.22 | 0.13 | 0.14 | 0.24 |
| 1 | 209512373 | rs1021993 | 0.93 | -0.582 | 0.802 | -0.577 | 0 | -0.001 | -0.001 | 0 | 0.73 | 0.69 | 0.72 | 0.73 | 0.78 | 0.64 |
| 1 | 209527706 | rs10494924 | -1.598 | -0.284 | 0.707 | -0.477 | 0.005 | 0.008 | 0.008 | 1.729 | 0.29 | 0.42 | 0.32 | 0.3 | 0.29 | 0.47 |
| 1 | 209540091 | rs17014370 | -0.138 | -0.241 | 0.358 | -0.46 | -0.005 | -0.002 | -0.002 | 0 | 0.14 | 0.21 | 0.22 | 0.13 | 0.14 | 0.21 |
| 1 | 209548546 | rs2055698 | -1.64 | -0.226 | 0.42 | -0.454 | 0 | 0 | 0 | 0.866 | 0.32 | 0.65 | 0.33 | 0.29 | 0.29 | 0.45 |
| 1 | 209552636 | rs1006317 | -0.309 | -0.021 | -0.018 | -0.207 | -0.003 | -0.002 | -0.002 | 0 | 0.15 | 0.16 | 0.22 | 0.13 | 0.15 | 0.23 |
| 1 | 209560036 | rs12142846 | -0.704 | 0.063 | -0.115 | -0.155 | -0.006 | -0.009 | -0.009 | 0 | 0.22 | 0.31 | 0.28 | 0.17 | 0.23 | 0.32 |
| 1 | 209575391 | rs7548177 | -0.189 | 1.064 | 0.428 | 0.441 | -0.021 | -0.03 | -0.03 | 0 | 0.68 | 0.76 | 0.53 | 0.46 | 0.62 | 0.73 |
| 1 | 209577743 | rs6665154 | -0.268 | 1.087 | 0.077 | 0.452 | -0.045 | -0.06 | -0.06 | 0.834 | 0.68 | 0.84 | 0.53 | 0.46 | 0.62 | 0.73 |
| 1 | 209581483 | rs1988163 | -0.05 | 0.74 | -0.115 | 0.322 | 0.014 | 0.002 | 0.002 | 1.88 | 0.63 | 0.56 | 0.47 | 0.4 | 0.54 | 0.65 |
| 1 | 209581629 | rs1354855 | 0.271 | 0.263 | -0.462 | 0.056 | 0.02 | 0.048 | 0.048 | 3.834 | 0.52 | 0.43 | 0.4 | 0.26 | 0.42 | 0.5 |
| 1 | 209588509 | rs742678 | -1.068 | 0.378 | 0.132 | 0.465 | -0.004 | -0.013 | -0.013 | 0.206 | 0.23 | 0.58 | 0.18 | 0.19 | 0.21 | 0.3 |
| 1 | 209590546 | rs6691406 | -1.273 | 0.218 | 0.364 | 0.953 | -0.009 | 0.03 | 0.03 | 0.924 | 0.24 | 0.59 | 0.26 | 0.27 | 0.22 | 0.3 |
| 1 | 209592106 | rs11119272 | -0.983 | 0.226 | 0.366 | 1.103 | 0.001 | 0.01 | 0.01 | 0.255 | 0.22 | 0.56 | 0.17 | 0.22 | 0.21 | 0.3 |
| 1 | 209603996 | rs2294938 | 1.073 | -0.196 | -0.076 | 0.385 | 0.049 | 0.073 | 0.073 | 0.481 | 0.59 | 0.13 | 0.71 | 0.76 | 0.68 | 0.67 |
| 1 | 209608020 | rs6690362 | 1.073 | -0.146 | -0.01 | 0.412 | 0.032 | 0.047 | 0.047 | 0.258 | 0.61 | 0.22 | 0.72 | 0.76 | 0.68 | 0.67 |
| 1 | 209611172 | rs10863770 | 1.066 | -0.153 | -0.26 | 0.442 | 0.046 | 0.063 | 0.063 | 0.263 | 0.6 | 0.13 | 0.71 | 0.76 | 0.68 | 0.67 |
| 1 | 209617663 | rs11119275 | -1.481 | -0.217 | 0.75 | 0.736 | 0.026 | 0.09 | 0.09 | 0.014 | 0.08 | 0.27 | 0.03 | 0.12 | 0.06 | 0.14 |
| 1 | 209619557 | rs6679530 | -1.629 | -0.131 | 0.51 | 0.873 | -0.006 | -0.015 | -0.015 | 0 | 0.4 | 0.74 | 0.44 | 0.42 | 0.39 | 0.37 |
| 1 | 209619604 | rs6688014 | -0.583 | -0.207 | 1.087 | 0.826 | -0.011 | -0.002 | -0.002 | 0 | 0.25 | 0.26 | 0.38 | 0.26 | 0.33 | 0.2 |
| 1 | 209621776 | rs2179655 | -1.089 | -0.518 | 1.561 | 0.493 | 0.026 | 0.003 | 0.003 | 0.012 | 0.29 | 0.47 | 0.31 | 0.29 | 0.33 | 0.2 |
| 1 | 209628156 | rs11577136 | -0.912 | -0.625 | 1.072 | 0.487 | -0.019 | 0.068 | 0.068 | 1.229 | 0.09 | 0.22 | 0.16 | 0.21 | 0.05 | 0.09 |
| 1 | 209632204 | rs7526722 | 0.521 | -0.946 | -0.167 | 0.323 | 0.001 | 0.072 | 0.072 | 1.039 | 0.47 | 0.19 | 0.4 | 0.33 | 0.52 | 0.4 |
| 1 | 209637764 | rs9919172 | -0.723 | -1.057 | -0.112 | 0.352 | 0 | 0.003 | 0.003 | 0.425 | 0.13 | 0.11 | 0.08 | 0.09 | 0.12 | 0.16 |
| 1 | 209641025 | rs12031184 | 0.175 | -1.034 | -0.162 | 0.35 | 0.015 | -0.013 | -0.013 | 0 | 0.07 | 0 | 0.02 | 0.06 | 0.03 | 0.11 |
| 1 | 209643416 | rs10863774 | -1.635 | -1.109 | -0.057 | 0.361 | -0.029 | 0.025 | 0.025 | 0.106 | 0.22 | 0.53 | 0.24 | 0.28 | 0.16 | 0.2 |
| 1 | 209651535 | rs6684558 | 0.23 | -1.221 | -0.794 | 0.037 | -0.039 | 0.071 | 0.071 | 1.807 | 0.21 | 0.07 | 0.15 | 0.01 | 0.28 | 0.07 |
| 1 | 209657070 | rs7521084 | -0.749 | -1.686 | -0.299 | 0.562 | -0.021 | 0.143 | 0.143 | 4.48 | 0.11 | 0.47 | 0.22 | 0.22 | 0.05 | 0.09 |
| 1 | 209660134 | rs17014581 | -1.231 | -1.737 | -0.336 | 0.59 | -0.024 | -0.02 | -0.02 | 0.124 | 0.19 | 0.29 | 0.09 | 0.14 | 0.12 | 0.15 |
| 1 | 209661527 | rs4844840 | -2.598 | -1.756 | -0.584 | 0.647 | 0.052 | -0.035 | -0.035 | 0 | 0.33 | 0.79 | 0.32 | 0.41 | 0.18 | 0.29 |
| 1 | 209661908 | rs12737310 | -0.485 | -1.744 | -0.596 | 0.658 | 0 | -0.001 | -0.001 | 0 | 0.12 | 0.1 | 0.07 | 0.14 | 0.11 | 0.14 |
| 1 | 209662378 | rs17014586 | -0.866 | -1.901 | -0.756 | 0.691 | -0.03 | 0.194 | 0.194 | 4.767 | 0.12 | 0.47 | 0.22 | 0.27 | 0.05 | 0.13 |
| 1 | 209682162 | rs17014649 | -1.211 | -1.747 | -0.732 | 0.676 | -0.056 | 0.014 | 0.014 | 0 | 0.19 | 0.33 | 0.09 | 0.2 | 0.09 | 0.2 |
| 1 | 209694672 | rs1883356 | -1.366 | -1.582 | -2.002 | 0.673 | 0.083 | -0.027 | -0.027 | 0 | 0.36 | 0.76 | 0.31 | 0.41 | 0.17 | 0.3 |
| 1 | 209698543 | rs11119302 | 0.408 | -0.997 | -2.427 | 0.28 | 0.016 | -0.001 | -0.001 | 0 | 0.14 | 0.02 | 0.08 | 0.1 | 0.08 | 0.08 |
| 1 | 209705895 | rs1011398 | -1.592 | -0.489 | -2.964 | 0.054 | -0.049 | 0.086 | 0.086 | 0.051 | 0.86 | 0.98 | 0.93 | 1 | 0.83 | 0.86 |
| 1 | 209706685 | rs2356932 | -1.076 | -0.698 | -2.553 | 0.618 | 0.1 | 0.008 | 0.008 | 1.607 | 0.39 | 0.83 | 0.35 | 0.63 | 0.23 | 0.34 |
| 1 | 209712889 | rs6703183 | -0.896 | -0.417 | -2.365 | 0.846 | 0.072 | 0.083 | 0.083 | 0 | 0.48 | 0.83 | 0.4 | 0.63 | 0.35 | 0.37 |
| 1 | 209713138 | rs6703522 | -0.255 | 0.358 | -2.397 | 0.646 | 0.005 | 0.004 | 0.004 | 0 | 0.33 | 0.21 | 0.19 | 0.28 | 0.28 | 0.24 |
| 1 | 209717109 | rs4844851 | -0.077 | 0.738 | -2.28 | 0.52 | -0.095 | 0 | 0 | 0 | 0.72 | 0.98 | 0.77 | 0.79 | 0.69 | 0.66 |
| 1 | 209720842 | rs996146 | -0.867 | 0.171 | -1.243 | 0.763 | -0.01 | 0 | 0 | 0 | 0.19 | 0.2 | 0.15 | 0.21 | 0.11 | 0.12 |
| 1 | 209722772 | rs11119306 | 0.529 | 0.905 | -1.542 | 0.822 | -0.02 | -0.001 | -0.001 | 0 | 0.62 | 0.76 | 0.6 | 0.58 | 0.56 | 0.47 |
| 1 | 209723420 | rs6673693 | 0.624 | 1.461 | -1.356 | 0.761 | -0.003 | 0 | 0 | 0.879 | 0.7 | 0.73 | 0.59 | 0.64 | 0.7 | 0.55 |
| 1 | 209728590 | rs10746420 | -0.052 | 1.044 | -1.726 | 0.219 | 0.039 | 0.083 | 0.083 | 0 | 0.52 | 0.7 | 0.57 | 0.66 | 0.42 | 0.55 |
| 1 | 209739903 | rs7525534 | -0.65 | 1.535 | -1.37 | 0.098 | 0.028 | 0.052 | 0.052 | 0 | 0.29 | 0.48 | 0.39 | 0.42 | 0.4 | 0.29 |
| 1 | 209741946 | rs7536585 | 0.699 | 1.628 | -1.308 | 0.105 | -0.004 | 0 | 0 | 0 | 0.16 | 0.12 | 0.15 | 0.04 | 0.21 | 0.2 |
| 1 | 209745395 | rs6695896 | -0.711 | 1.72 | -1.235 | 0.141 | 0.026 | 0.05 | 0.05 | 0 | 0.29 | 0.46 | 0.39 | 0.42 | 0.38 | 0.31 |
| 1 | 209756419 | rs17014820 | 0.529 | 1.661 | -1.287 | 0.124 | 0.002 | -0.034 | -0.034 | 0 | 0.17 | 0.2 | 0.15 | 0.04 | 0.21 | 0.2 |
| 1 | 209758758 | rs926386 | -1.076 | 1.872 | -0.747 | -0.179 | 0.026 | -0.021 | -0.021 | 0.019 | 0.59 | 0.75 | 0.61 | 0.46 | 0.68 | 0.55 |
| 1 | 209758914 | rs4556360 | -0.86 | 1.87 | -0.573 | -0.18 | -0.001 | 0 | 0 | 0.505 | 0.58 | 0.65 | 0.59 | 0.46 | 0.68 | 0.55 |
| 1 | 209765087 | rs7528698 | 0.9 | 1.803 | -0.822 | -0.363 | -0.038 | 0.094 | 0.094 | 0 | 0.23 | 0.07 | 0.17 | 0.05 | 0.25 | 0.25 |
| 1 | 209768979 | rs6690557 | 0.617 | 1.785 | -0.983 | -0.327 | -0.079 | -0.053 | -0.053 | 0.047 | 0.25 | 0.04 | 0.23 | 0.31 | 0.28 | 0.29 |

| | | | | | | | | | | | | | | | | |
|---|---|---|---|---|---|---|---|---|---|---|---|---|---|---|---|---|
| 1 | 209771347 | rs9429821 | 0.576 | 1.663 | -0.972 | -0.267 | -0.104 | -0.055 | -0.055 | 0.106 | 0.27 | 0.05 | 0.2 | 0.31 | 0.33 | 0.31 |
| 1 | 209774889 | rs2281139 | -1.593 | 1.453 | -1.372 | -0.376 | -0.058 | 0.014 | 0.014 | 0 | 0.89 | 0.99 | 0.88 | 0.94 | 0.82 | 0.89 |
| 1 | 209775800 | rs9430004 | -0.529 | 1.529 | -0.836 | -0.173 | -0.001 | -0.002 | -0.002 | 0 | 0.42 | 0.63 | 0.43 | 0.45 | 0.51 | 0.4 |
| 1 | 209779355 | rs12402938 | 0.397 | 1.501 | -0.753 | 0.524 | 0.022 | 0.021 | 0.021 | 0.004 | 0.12 | 0.26 | 0.2 | 0.12 | 0.18 | 0.06 |
| 1 | 209781125 | rs760833 | -0.073 | 1.437 | -0.787 | 0.376 | 0.03 | 0.106 | 0.106 | 0.013 | 0.14 | 0.33 | 0.22 | 0.37 | 0.21 | 0.1 |
| 1 | 209782121 | rs6658827 | 0.214 | 0.722 | -1.201 | -0.139 | -0.003 | 0.06 | 0.06 | 2.065 | 0.17 | 0.04 | 0.14 | 0.05 | 0.16 | 0.17 |
| 1 | 209784927 | rs2076230 | -0.333 | 0.402 | -0.614 | -0.091 | 0.017 | -0.069 | -0.069 | 0.549 | 0.29 | 0.47 | 0.29 | 0.11 | 0.34 | 0.33 |
| 1 | 209785206 | rs11119315 | 0.104 | 0.46 | -0.233 | -0.212 | 0.035 | 0.014 | 0.014 | 0 | 0.14 | 0.31 | 0.19 | 0.11 | 0.18 | 0.1 |
| 1 | 209786078 | rs2272880 | -1.382 | 0.088 | -1.078 | 0.158 | -0.021 | 0.069 | 0.069 | 0 | 0.59 | 0.91 | 0.78 | 0.74 | 0.64 | 0.68 |
| 1 | 209786883 | rs2206107 | -0.243 | 0.048 | -1.002 | 0.131 | 0.021 | -0.051 | -0.051 | 0 | 0.27 | 0.36 | 0.27 | 0.11 | 0.34 | 0.33 |
| 1 | 209788213 | rs926387 | -1.596 | -0.045 | -1.146 | 0.345 | -0.064 | -0.031 | -0.031 | 0 | 0.61 | 0.96 | 0.79 | 0.7 | 0.66 | 0.71 |
| 1 | 209788514 | rs2566 | 0.551 | -0.296 | -0.561 | 1.19 | -0.004 | -0.013 | -0.013 | 0 | 0.24 | 0.24 | 0.41 | 0.36 | 0.27 | 0.24 |
| 1 | 209795302 | rs742257 | -0.081 | -0.345 | -0.508 | 0.992 | 0.022 | -0.052 | -0.052 | 0 | 0.27 | 0.37 | 0.27 | 0.11 | 0.34 | 0.32 |
| 1 | 209795909 | rs3179860 | 1.245 | -0.434 | -0.425 | 1.139 | -0.019 | 0.041 | 0.041 | 0 | 0.14 | 0.04 | 0.09 | 0 | 0.16 | 0.24 |
| 1 | 209799253 | rs2179402 | -0.471 | -0.072 | -0.028 | 2.021 | -0.005 | 0.005 | 0.005 | 0 | 0.78 | 0.78 | 0.71 | 0.83 | 0.73 | 0.75 |
| 1 | 209800230 | rs2076349 | -0.994 | -0.173 | 1.042 | 2.911 | -0.008 | 0.461 | 0.461 | 0 | 0.09 | 0.32 | 0.21 | 0.58 | 0.03 | 0.11 |
| 1 | 209804095 | rs4844863 | 0.39 | 0.076 | 0.542 | 2.173 | 0.017 | 0.026 | 0.026 | 0 | 0.91 | 0.83 | 0.78 | 0.82 | 0.87 | 0.81 |
| 1 | 209807438 | rs12145126 | 0.389 | 0.2 | 0.241 | 2.023 | -0.011 | -0.006 | -0.006 | 0 | 0.09 | 0.06 | 0.09 | 0.06 | 0.14 | 0.1 |
| 1 | 209807506 | rs12137442 | 0.245 | 0.19 | 0.322 | 2.009 | 0.009 | 0.002 | 0.002 | 0.009 | 0.1 | 0.18 | 0.11 | 0.06 | 0.14 | 0.1 |
| 1 | 209812063 | rs2076357 | 1.324 | -0.051 | 0.052 | 1.557 | -0.001 | 0.003 | 0.003 | 0.605 | 0.71 | 0.34 | 0.6 | 0.65 | 0.68 | 0.71 |
| 1 | 209817981 | rs2236896 | -0.071 | 0.139 | -0.255 | 1.496 | 0.013 | 0.025 | 0.025 | 0 | 0.11 | 0.22 | 0.15 | 0.11 | 0.14 | 0.09 |
| 1 | 209822859 | rs2298928 | 0.835 | 0.187 | -0.803 | 1.462 | -0.004 | -0.006 | -0.006 | 0 | 0.57 | 0.52 | 0.43 | 0.35 | 0.61 | 0.44 |
| 1 | 209823936 | rs2076358 | -0.499 | 0.661 | -1.225 | 1.337 | 0.001 | -0.003 | -0.003 | 0 | 0.28 | 0.21 | 0.41 | 0.34 | 0.27 | 0.33 |
| 1 | 209830347 | rs12059601 | -0.12 | 1.506 | -1.715 | 0.691 | -0.08 | 0.03 | 0.03 | 0.023 | 0.3 | 0.03 | 0.32 | 0.23 | 0.3 | 0.24 |
| 1 | 209831544 | rs742375 | -0.362 | 1.178 | -1.391 | 0.818 | -0.058 | 0.045 | 0.045 | 0 | 0.41 | 0.14 | 0.43 | 0.31 | 0.46 | 0.35 |
| 1 | 209837683 | rs4844484 | 0.722 | 1.954 | -0.307 | 1.452 | 0.101 | 0.104 | 0.104 | 0 | 0.78 | 0.51 | 0.8 | 0.59 | 0.71 | 0.74 |
| 1 | 209839374 | rs11580294 | -1.073 | 1.739 | 0.176 | 1.3 | 0.017 | -0.01 | -0.01 | 0.115 | 0.15 | 0.2 | 0.24 | 0.09 | 0.21 | 0.15 |
| 1 | 209844517 | rs4844486 | 1.693 | 1.731 | 0.725 | 2.069 | 0.038 | 0.1 | 0.1 | 0.611 | 0.78 | 0.42 | 0.71 | 0.55 | 0.73 | 0.75 |
| 1 | 209845370 | rs6540546 | 1.972 | 1.495 | 0.868 | 2.31 | 0.012 | 0.058 | 0.058 | 0.893 | 0.75 | 0.34 | 0.61 | 0.5 | 0.72 | 0.71 |
| 1 | 209848432 | rs932374 | 1.6 | 1.502 | 1.025 | 2.331 | 0.047 | 0.132 | 0.132 | 0.366 | 0.78 | 0.45 | 0.62 | 0.54 | 0.74 | 0.72 |
| 1 | 209856626 | rs17317033 | 0.226 | 0.219 | 0.382 | 1.335 | 0.02 | 0.023 | 0.023 | 0 | 0.16 | 0.06 | 0.07 | 0.02 | 0.1 | 0.1 |
| 1 | 209861525 | rs760951 | 1.488 | 0.717 | 0.991 | 1.991 | -0.014 | 0.146 | 0.146 | 0 | 0.81 | 0.51 | 0.79 | 0.59 | 0.88 | 0.68 |
| 1 | 209872590 | rs10863782 | 1.451 | 0.832 | 0.989 | 1.917 | -0.01 | 0.11 | 0.11 | 0 | 0.8 | 0.44 | 0.77 | 0.57 | 0.84 | 0.67 |
| 1 | 209876251 | rs846911 | -0.898 | 0.518 | 1.33 | 1.356 | -0.03 | -0.004 | -0.004 | 0 | 0.08 | 0.31 | 0.02 | 0.03 | 0 | 0.09 |
| 1 | 209879599 | rs11799643 | -0.587 | 0.525 | 1.634 | 1.357 | 0.018 | 0.048 | 0.048 | 0.198 | 0.07 | 0.35 | 0.03 | 0.04 | 0 | 0.08 |
| 1 | 209885318 | rs11119328 | -0.542 | 0.449 | 1.117 | 2.454 | -0.001 | 0.012 | 0.012 | 0.018 | 0.16 | 0.2 | 0.21 | 0.23 | 0.17 | 0.18 |
| 1 | 209888435 | rs12094117 | -0.591 | 0.998 | 0.992 | 2.28 | 0.002 | 0.006 | 0.006 | 0 | 0.15 | 0.18 | 0.2 | 0.23 | 0.21 | 0.17 |
| 1 | 209902110 | rs1415542 | -0.649 | 0.992 | 1.147 | 2.375 | 0.009 | 0.017 | 0.017 | 0 | 0.16 | 0.23 | 0.21 | 0.22 | 0.21 | 0.17 |
| 1 | 209906675 | rs12059226 | -0.608 | 0.992 | 1.146 | 2.375 | 0.003 | 0.006 | 0.006 | 0 | 0.16 | 0.19 | 0.21 | 0.22 | 0.21 | 0.17 |
| 1 | 209908876 | rs10863785 | -0.662 | 0.992 | 1.147 | 2.37 | 0.009 | 0.018 | 0.018 | 0 | 0.16 | 0.23 | 0.21 | 0.23 | 0.21 | 0.18 |
| 1 | 209916884 | rs9430016 | -1.801 | 0.826 | 1.669 | 2.296 | -0.012 | 0.008 | 0.008 | 0 | 0.24 | 0.59 | 0.25 | 0.26 | 0.22 | 0.26 |
| 1 | 209918090 | rs7532324 | -0.819 | 0.826 | 1.69 | 2.466 | 0.011 | 0.073 | 0.073 | 0.025 | 0.16 | 0.24 | 0.4 | 0.49 | 0.23 | 0.25 |
| 1 | 209918151 | rs3765240 | -1.71 | 0.825 | 1.636 | 2.386 | -0.01 | -0.002 | -0.002 | 0.132 | 0.23 | 0.59 | 0.25 | 0.22 | 0.22 | 0.22 |
| 1 | 209922058 | rs17389198 | -0.572 | 0.918 | 1.447 | 2.364 | 0.004 | 0.003 | 0.003 | 0 | 0.16 | 0.19 | 0.21 | 0.18 | 0.23 | 0.15 |
| 1 | 209929579 | rs2272867 | -0.534 | 0.869 | 1.405 | 2.073 | -0.005 | -0.005 | -0.005 | 0 | 0.15 | 0.13 | 0.2 | 0.18 | 0.22 | 0.15 |
| 1 | 209932121 | rs10489341 | -0.195 | 0.98 | 1.502 | 1.971 | 0.005 | 0.017 | 0.017 | 0.973 | 0.08 | 0.32 | 0.04 | 0.03 | 0.02 | 0.07 |
| 1 | 209936914 | rs17015169 | -1.625 | 1.052 | 1.066 | 2.291 | -0.003 | 0.005 | 0.005 | 0.127 | 0.26 | 0.69 | 0.28 | 0.31 | 0.23 | 0.27 |
| 1 | 209937382 | rs654984 | 1.4 | 1.024 | 1.029 | 2.301 | -0.001 | 0 | 0 | 0.103 | 0.74 | 0.3 | 0.72 | 0.69 | 0.75 | 0.73 |
| 1 | 209943489 | rs11119344 | 0.201 | 0.997 | -0.132 | 1.321 | -0.045 | 0.034 | 0.034 | 1.66 | 0.2 | 0.06 | 0.16 | 0.11 | 0.25 | 0.15 |
| 1 | 209950681 | rs649275 | 0.875 | 0.854 | -0.023 | 1.025 | -0.005 | -0.006 | -0.006 | 2.946 | 0.61 | 0.59 | 0.67 | 0.63 | 0.62 | 0.57 |
| 1 | 209953571 | rs2076153 | 1.437 | 0.531 | -0.056 | 0.934 | 0 | 0.001 | 0.001 | 0 | 0.47 | 0.42 | 0.45 | 0.55 | 0.42 | 0.44 |
| 1 | 209955848 | rs651141 | 1.814 | 0.552 | 0.374 | 1.122 | -0.015 | 0.132 | 0.132 | 0 | 0.46 | 0.2 | 0.26 | 0.23 | 0.42 | 0.36 |
| 1 | 209956542 | rs7550857 | 0.179 | 0.485 | 0.73 | 1.103 | -0.008 | -0.022 | -0.022 | 0.041 | 0.17 | 0.22 | 0.1 | 0.07 | 0.14 | 0.2 |
| 1 | 209959614 | rs1044516 | -0.118 | 0.485 | 0.87 | 1.276 | -0.007 | 0.082 | 0.082 | 0 | 0.18 | 0.25 | 0.29 | 0.56 | 0.16 | 0.31 |
| 1 | 209968684 | rs2013162 | -0.922 | 0.45 | 0.675 | 1.089 | -0.005 | -0.023 | -0.023 | 0.417 | 0.32 | 0.24 | 0.48 | 0.58 | 0.36 | 0.43 |
| 1 | 209975386 | rs861019 | 1.871 | 0.402 | 0.805 | 1.061 | 0.002 | 0.008 | 0.008 | 0 | 0.48 | 0.55 | 0.3 | 0.23 | 0.42 | 0.35 |
| 1 | 209977111 | rs861020 | -0.466 | 0.448 | 0.399 | 1.087 | -0.007 | -0.003 | -0.003 | 0 | 0.81 | 0.89 | 0.79 | 0.81 | 0.78 | 0.78 |
| 1 | 209982923 | rs17015259 | -0.364 | 0.481 | 0.504 | 1.104 | -0.018 | 0.153 | 0.153 | 0.194 | 0.2 | 0.49 | 0.31 | 0.5 | 0.15 | 0.3 |
| 1 | 209987172 | rs17389541 | 0.526 | 0.617 | 0.17 | 1.173 | -0.005 | 0.007 | 0.007 | 0.142 | 0.15 | 0.1 | 0.21 | 0.08 | 0.22 | 0.13 |
| 1 | 209989270 | rs642961 | -0.401 | 0.604 | -0.071 | 1.159 | -0.011 | -0.007 | -0.007 | 0.423 | 0.82 | 0.91 | 0.79 | 0.81 | 0.79 | 0.78 |
| 1 | 209996542 | rs9430018 | 0.588 | 0.524 | -0.195 | 1.002 | -0.041 | 0.012 | 0.012 | 0 | 0.64 | 0.78 | 0.36 | 0.3 | 0.53 | 0.55 |
| 1 | 209997448 | rs11584081 | 0.738 | 0.652 | -0.286 | 1.033 | -0.045 | 0.001 | 0.001 | 0 | 0.13 | 0.01 | 0.2 | 0.08 | 0.2 | 0.12 |
| 1 | 209998662 | rs630065 | 0.37 | 0.47 | 0.244 | 0.984 | 0.007 | 0.034 | 0.034 | 0.288 | 0.73 | 0.61 | 0.54 | 0.39 | 0.73 | 0.67 |
| 1 | 209999790 | rs3753513 | 0.845 | 0.474 | 0.189 | 0.971 | -0.045 | 0.001 | 0.001 | 0 | 0.13 | 0.02 | 0.2 | 0.08 | 0.21 | 0.12 |
| 1 | 210001048 | rs660975 | 1.015 | 0.457 | 0.296 | 0.964 | -0.001 | 0 | 0 | 0 | 0.59 | 0.55 | 0.34 | 0.3 | 0.52 | 0.55 |
| 1 | 210005821 | rs10127937 | 0.656 | 0.392 | 0.281 | 0.925 | -0.029 | 0.013 | 0.013 | 0 | 0.14 | 0.05 | 0.2 | 0.08 | 0.21 | 0.12 |
| 1 | 210006603 | rs656566 | 0.4 | 0.419 | 0.305 | 0.952 | 0.006 | 0.032 | 0.032 | 0.27 | 0.73 | 0.61 | 0.54 | 0.39 | 0.73 | 0.67 |
| 1 | 210007218 | rs17015309 | 0.687 | 0.32 | 0.189 | 0.798 | 0.002 | 0.017 | 0.017 | 0 | 0.13 | 0.08 | 0.09 | 0.08 | 0.13 | 0.21 |
| 1 | 210019701 | rs3766614 | -1.08 | 0.27 | 0.514 | 0.778 | -0.011 | -0.051 | -0.051 | 0.472 | 0.33 | 0.5 | 0.32 | 0.16 | 0.34 | 0.33 |

| | | | | | | | | | | | | | | | | |
|---|---|---|---|---|---|---|---|---|---|---|---|---|---|---|---|---|
| 1 | 210019824 | rs126280 | -0.449 | 0.27 | 0.482 | 0.941 | -0.007 | 0.002 | 0.002 | 0.987 | 0.79 | 0.82 | 0.74 | 0.8 | 0.75 | 0.76 |
| 1 | 210020971 | rs583814 | 2.002 | 0.269 | 0.47 | 0.919 | 0.003 | 0.024 | 0.024 | 0 | 0.45 | 0.31 | 0.24 | 0.22 | 0.39 | 0.34 |
| 1 | 210023765 | rs17015379 | 0.196 | 0.37 | 0.476 | 0.852 | -0.001 | -0.017 | -0.017 | 0 | 0.15 | 0.19 | 0.1 | 0.08 | 0.15 | 0.21 |
| 1 | 210025751 | rs7539638 | 0.028 | 0.341 | 0.601 | 0.837 | 0.001 | -0.045 | -0.045 | 0 | 0.16 | 0.4 | 0.12 | 0.08 | 0.15 | 0.21 |
| 1 | 210027026 | rs6689839 | 0.584 | 0.353 | 0.427 | 0.947 | -0.009 | 0.021 | 0.021 | 0.434 | 0.64 | 0.7 | 0.65 | 0.75 | 0.61 | 0.56 |
| 1 | 210028454 | rs2357231 | -1.557 | 0.315 | 0.011 | 1.033 | 0 | 0.004 | 0.004 | 0.284 | 0.4 | 0.68 | 0.56 | 0.67 | 0.38 | 0.51 |
| 1 | 210040672 | rs7548533 | -2.096 | -0.031 | -0.347 | 0.715 | -0.005 | 0.026 | 0.026 | 0 | 0.59 | 0.72 | 0.75 | 0.75 | 0.59 | 0.66 |
| 1 | 210043565 | rs7537070 | -1.252 | -0.096 | -0.403 | 0.635 | 0.001 | 0.003 | 0.003 | 0 | 0.41 | 0.64 | 0.56 | 0.64 | 0.38 | 0.47 |
| 1 | 210050794 | rs9430019 | -2.267 | -0.065 | -0.494 | 0.538 | -0.005 | 0.026 | 0.026 | 0 | 0.59 | 0.72 | 0.75 | 0.75 | 0.59 | 0.66 |
| 1 | 210055630 | rs932347 | 0.051 | -0.119 | -0.846 | 0.546 | -0.015 | 0.006 | 0.006 | 0.001 | 0.24 | 0.03 | 0.23 | 0.19 | 0.21 | 0.21 |
| 1 | 210057927 | rs1883725 | 0.133 | 0.046 | -0.844 | 0.486 | -0.074 | 0.034 | 0.034 | 0 | 0.68 | 0.92 | 0.68 | 0.75 | 0.57 | 0.61 |
| 1 | 210066145 | rs6674507 | 1.049 | -0.103 | -0.618 | 0.44 | 0.001 | 0.01 | 0.01 | 0 | 0.6 | 0.39 | 0.65 | 0.74 | 0.58 | 0.61 |
| 1 | 210068117 | rs4844913 | 0.586 | -0.065 | -0.916 | 0.227 | 0.002 | 0.003 | 0.003 | 0 | 0.45 | 0.38 | 0.53 | 0.56 | 0.43 | 0.47 |
| 1 | 210086539 | rs12140373 | -0.794 | -0.079 | -1.106 | 0.26 | 0.001 | 0.002 | 0.002 | 0 | 0.56 | 0.72 | 0.47 | 0.45 | 0.57 | 0.53 |
| 1 | 210108473 | rs6540570 | -0.363 | -0.183 | -0.995 | 0.203 | 0.019 | 0.023 | 0.023 | 0 | 0.49 | 0.24 | 0.44 | 0.45 | 0.57 | 0.53 |
| 1 | 210110529 | rs9429829 | -0.259 | -0.197 | -1.237 | 0.196 | -0.008 | -0.001 | -0.001 | 0 | 0.48 | 0.12 | 0.43 | 0.45 | 0.57 | 0.53 |
| 1 | 210119135 | rs1016479 | -2.085 | -0.101 | -1.487 | 0.047 | -0.025 | -0.002 | -0.002 | 0 | 0.89 | 0.99 | 0.93 | 0.93 | 0.86 | 0.86 |
| 1 | 210122322 | rs7528647 | -1.011 | -0.121 | -1.391 | 0.152 | -0.001 | 0.119 | 0.119 | 0 | 0.58 | 0.78 | 0.76 | 0.88 | 0.59 | 0.64 |
| 1 | 210145298 | rs12034991 | 1.153 | -0.077 | -1.684 | 0.094 | -0.12 | -0.012 | -0.012 | 0 | 0.27 | 0.03 | 0.21 | 0.22 | 0.32 | 0.36 |
| 1 | 210156770 | rs7525980 | -1.076 | -0.031 | -1.3 | 0.139 | -0.007 | 0.154 | 0.154 | 0 | 0.58 | 0.79 | 0.76 | 0.88 | 0.58 | 0.64 |
| 1 | 210216946 | rs227178 | 0.042 | -0.04 | -1.38 | 0.409 | 0.001 | 0.034 | 0.034 | 0 | 0.52 | 0.87 | 0.58 | 0.58 | 0.43 | 0.47 |
| 1 | 210252888 | rs227185 | 1.789 | 0.085 | -1.427 | 0.298 | -0.031 | -0.003 | -0.003 | 0.99 | 0.11 | 0.01 | 0.07 | 0.07 | 0.15 | 0.14 |
| 1 | 210272790 | rs1334569 | -0.911 | -0.026 | -0.874 | 0.331 | 0.003 | 0.01 | 0.01 | 0 | 0.56 | 0.67 | 0.48 | 0.44 | 0.58 | 0.55 |
| 1 | 210291476 | rs2130628 | 0.942 | 0.046 | -0.65 | 0.368 | 0.013 | 0.004 | 0.004 | 1.31 | 0.07 | 0.53 | 0.04 | 0 | 0.01 | 0 |
| 1 | 210300491 | rs13376651 | 1.512 | 0.054 | -0.434 | 0.372 | 0.038 | 0.029 | 0.029 | 0.231 | 0.06 | 0.39 | 0.03 | 0 | 0.01 | 0 |
| 1 | 210300915 | rs6689269 | 1.132 | 0.059 | -0.715 | 0.374 | 0.027 | 0.017 | 0.017 | 1.203 | 0.07 | 0.51 | 0.04 | 0 | 0.01 | 0 |
| 1 | 210304319 | rs2046850 | 1.62 | 0.184 | -1.41 | 0.373 | -0.031 | -0.032 | -0.032 | 0.22 | 0.13 | 0.04 | 0.21 | 0.24 | 0.23 | 0.15 |
| 1 | 210306846 | rs4844949 | 0.804 | -0.118 | -1.463 | 0.371 | -0.117 | -0.003 | -0.003 | 0 | 0.28 | 0.04 | 0.21 | 0.22 | 0.32 | 0.37 |
| 1 | 210311800 | rs12117075 | -0.098 | 0 | -1.614 | 0.211 | -0.107 | -0.032 | -0.032 | 0 | 0.4 | 0.08 | 0.35 | 0.44 | 0.52 | 0.37 |
| 1 | 210312760 | rs1002382 | -0.861 | 0.004 | -1.341 | 0.129 | -0.004 | 0.172 | 0.172 | 0 | 0.49 | 0.76 | 0.68 | 0.87 | 0.54 | 0.47 |
| 1 | 210331377 | rs10494925 | 1.77 | 0.067 | -1.283 | 0.135 | -0.04 | 0.057 | 0.057 | 0 | 0.13 | 0.02 | 0.1 | 0.02 | 0.15 | 0.23 |
| 1 | 210331585 | rs4537554 | 0.661 | 0.073 | -1.36 | 0.645 | -0.069 | -0.11 | -0.11 | 0 | 0.23 | 0.03 | 0.24 | 0.44 | 0.36 | 0.22 |
| 1 | 210337043 | rs4844954 | -2.186 | -0.262 | -1.094 | 0.15 | 0.015 | 0.053 | 0.053 | 0 | 0.73 | 0.84 | 0.85 | 0.94 | 0.83 | 0.83 |
| 1 | 210337691 | rs17015701 | 0.68 | -0.188 | -0.8 | 0.329 | -0.004 | 0.072 | 0.072 | 0 | 0.25 | 0.55 | 0.43 | 0.62 | 0.18 | 0.25 |
| 1 | 210338196 | rs4421592 | -1.198 | -0.194 | -0.85 | 0.311 | 0.019 | 0.074 | 0.074 | 0 | 0.61 | 0.76 | 0.78 | 0.89 | 0.69 | 0.71 |
| 1 | 210338397 | rs12133399 | 0.167 | -0.177 | -0.955 | 0.213 | -0.108 | 0.044 | 0.044 | 0 | 0.35 | 0.06 | 0.34 | 0.27 | 0.51 | 0.46 |
| 1 | 210340969 | rs11119426 | 0.857 | -0.047 | -0.717 | 0.319 | -0.004 | -0.021 | -0.021 | 0 | 0.23 | 0.49 | 0.15 | 0.2 | 0.17 | 0.14 |
| 1 | 210348155 | rs2485893 | -1.045 | -0.113 | -1.008 | 0.483 | 0.013 | -0.023 | -0.023 | 0 | 0.61 | 0.77 | 0.5 | 0.47 | 0.68 | 0.59 |
| 1 | 210354442 | rs12567108 | -0.688 | -0.5 | -1.427 | -0.007 | 0.028 | 0.166 | 0.166 | 0.029 | 0.53 | 0.13 | 0.71 | 0.76 | 0.68 | 0.67 |
| 1 | 210354586 | rs11119431 | -1.309 | -0.252 | -1.27 | -0.019 | -0.002 | -0.011 | -0.011 | 0 | 0.67 | 0.71 | 0.5 | 0.47 | 0.67 | 0.62 |
| 1 | 210356238 | rs1014370 | 1.7 | -0.165 | -0.708 | 0.149 | -0.001 | 0.001 | 0.001 | 0 | 0.17 | 0.14 | 0.11 | 0.19 | 0.14 | 0.13 |
| 1 | 210369136 | rs12094311 | 1.25 | -0.128 | -0.721 | 0.226 | 0 | 0.052 | 0.052 | 0 | 0.12 | 0.09 | 0.36 | 0.47 | 0.15 | 0.2 |
| 1 | 210370700 | rs11812030 | 1.278 | -0.222 | -0.095 | 0.16 | -0.005 | -0.026 | -0.026 | 0 | 0.19 | 0.33 | 0.13 | 0.15 | 0.14 | 0.12 |
| 1 | 210370761 | rs2485901 | -0.074 | -0.256 | -0.242 | -0.07 | -0.022 | 0.037 | 0.037 | 0 | 0.39 | 0.22 | 0.35 | 0.28 | 0.51 | 0.45 |
| 1 | 210370851 | rs2485902 | -2.03 | -0.252 | -0.317 | -0.069 | 0.003 | 0.089 | 0.089 | 0 | 0.72 | 0.87 | 0.85 | 0.9 | 0.79 | 0.78 |
| 1 | 210371949 | rs2485903 | -0.127 | -0.183 | -0.229 | -0.09 | -0.006 | 0.013 | 0.013 | 0 | 0.41 | 0.32 | 0.36 | 0.29 | 0.51 | 0.45 |
| 1 | 210374337 | rs2485888 | -0.917 | -0.155 | -0.57 | 0.003 | 0.005 | 0.009 | 0.009 | 0 | 0.6 | 0.77 | 0.49 | 0.41 | 0.64 | 0.57 |
| 1 | 210381257 | rs11119439 | 1.003 | 0.042 | -0.377 | -0.005 | -0.06 | 0.03 | 0.03 | 0 | 0.2 | 0.03 | 0.12 | 0.13 | 0.24 | 0.16 |
| 1 | 210386561 | rs2185059 | 0.242 | 0.043 | -0.145 | -0.15 | -0.005 | -0.09 | -0.09 | 0 | 0.21 | 0.36 | 0.13 | 0.04 | 0.18 | 0.29 |
| 1 | 210386892 | rs2494184 | -1.421 | 0.104 | -0.319 | 0.072 | -0.042 | 0.035 | 0.035 | 0.097 | 0.58 | 0.85 | 0.64 | 0.65 | 0.56 | 0.65 |
| 1 | 210389417 | rs1342135 | -0.83 | 0.112 | -0.16 | 0.09 | 0.01 | 0.094 | 0.094 | 0 | 0.5 | 0.28 | 0.31 | 0.23 | 0.54 | 0.53 |
| 1 | 210394270 | rs591594 | -1.127 | 0.158 | 0.17 | 0.397 | -0.021 | 0.032 | 0.032 | 0 | 0.8 | 0.92 | 0.87 | 0.87 | 0.76 | 0.85 |
| 1 | 210397552 | rs1473696 | 0.123 | -0.046 | 0.789 | 0.694 | -0.007 | -0.003 | -0.003 | 0 | 0.35 | 0.51 | 0.31 | 0.35 | 0.32 | 0.31 |
| 1 | 210398242 | rs10863816 | 0.504 | -0.058 | 0.635 | 0.689 | -0.001 | -0.009 | -0.009 | 0.1 | 0.32 | 0.22 | 0.3 | 0.35 | 0.31 | 0.31 |
| 1 | 210399652 | rs677520 | -0.192 | -0.095 | -0.067 | 0.582 | 0.01 | 0.023 | 0.023 | 0 | 0.38 | 0.72 | 0.62 | 0.77 | 0.33 | 0.43 |
| 1 | 210413739 | rs654523 | -0.004 | 0.351 | -0.984 | 0.938 | -0.05 | 0.052 | 0.052 | 0 | 0.38 | 0.14 | 0.64 | 0.78 | 0.43 | 0.54 |
| 1 | 210417098 | rs701944 | 2.052 | 0.535 | -0.675 | 1.283 | -0.025 | -0.037 | -0.037 | 0.59 | 0.06 | 0.01 | 0.33 | 0.17 | 0.1 | 0.06 |
| 1 | 210421594 | rs634601 | 1.693 | 0.518 | -0.668 | 0.754 | -0.032 | -0.009 | -0.009 | 0 | 0.07 | 0.01 | 0.33 | 0.43 | 0.11 | 0.2 |
| 1 | 210429102 | rs845451 | 0.063 | 0.353 | -0.557 | 0.247 | -0.015 | 0.035 | 0.035 | 0 | 0.38 | 0.29 | 0.65 | 0.78 | 0.43 | 0.53 |
| 1 | 210443474 | rs685532 | -0.457 | 0.269 | -0.474 | 0.389 | -0.014 | 0.085 | 0.085 | 0.602 | 0.55 | 0.76 | 0.75 | 0.79 | 0.51 | 0.6 |
| 1 | 210443546 | rs6666540 | 0.53 | 0.388 | -0.997 | 0.458 | -0.025 | 0.021 | 0.021 | 0 | 0.4 | 0.19 | 0.38 | 0.33 | 0.39 | 0.36 |
| 1 | 210458861 | rs10489391 | 1.366 | 0.802 | -0.744 | 0.992 | -0.032 | 0.002 | 0.002 | 0 | 0.07 | 0.01 | 0.33 | 0.5 | 0.11 | 0.21 |
| 1 | 210459159 | rs649009 | -1.083 | 0.802 | -1.011 | 0.261 | -0.011 | 0.018 | 0.018 | 1.7 | 0.75 | 0.81 | 0.85 | 0.95 | 0.7 | 0.81 |
| 1 | 210460602 | rs7527633 | 0.473 | 0.788 | -1.29 | 0.331 | -0.008 | -0.02 | -0.02 | 0 | 0.11 | 0.03 | 0.07 | 0.14 | 0.14 | 0.11 |
| 1 | 210461837 | rs616413 | -0.551 | 0.549 | -0.878 | 0.016 | -0.004 | 0.052 | 0.052 | 0 | 0.64 | 0.75 | 0.82 | 0.93 | 0.65 | 0.68 |
| 1 | 210463122 | rs689248 | 0.806 | 0.651 | -0.628 | 0.656 | 0.013 | 0.153 | 0.153 | 0 | 0.09 | 0.14 | 0.35 | 0.49 | 0.13 | 0.21 |
| 1 | 210467047 | rs633352 | -0.765 | 0.415 | -0.43 | 0.376 | 0.01 | 0.054 | 0.054 | 0.962 | 0.29 | 0.51 | 0.45 | 0.61 | 0.32 | 0.44 |
| 1 | 210475254 | rs1338285 | 0.585 | 0.745 | -1.527 | -0.102 | -0.068 | 0.078 | 0.078 | 0 | 0.33 | 0.11 | 0.28 | 0.19 | 0.38 | 0.34 |
| 1 | 210482140 | rs742690 | 0.07 | 1.027 | -1.216 | 0.215 | -0.008 | -0.022 | -0.022 | 0 | 0.31 | 0.27 | 0.5 | 0.49 | 0.34 | 0.3 |

| | | | | | | | | | | | | | | | | |
|---|---|---|---|---|---|---|---|---|---|---|---|---|---|---|---|---|
| 1 | 210486746 | rs12117063 | 0.409 | 0.937 | -1.678 | -0.515 | -0.01 | 0.169 | 0.169 | 0 | 0.39 | 0.21 | 0.22 | 0.01 | 0.34 | 0.22 |
| 1 | 210486895 | rs4844997 | 1.798 | 1.001 | -1.476 | 0.555 | -0.036 | -0.029 | -0.029 | 0 | 0.07 | 0.01 | 0.3 | 0.25 | 0.12 | 0.09 |
| 1 | 210494973 | rs6666552 | -0.007 | 0.654 | -1.235 | 0.18 | -0.004 | -0.009 | -0.009 | 0 | 0.56 | 0.54 | 0.35 | 0.34 | 0.55 | 0.48 |
| 1 | 210499394 | rs4430327 | 0.584 | 0.827 | -0.897 | 0.202 | 0.002 | -0.005 | -0.005 | 0 | 0.07 | 0.13 | 0.03 | 0.04 | 0.07 | 0.09 |
| 1 | 210500594 | rs6697682 | -0.799 | 0.661 | -0.537 | 0.044 | 0.025 | -0.038 | -0.038 | 0 | 0.22 | 0.47 | 0.15 | 0.16 | 0.23 | 0.27 |
| 1 | 210505726 | rs589027 | -1.336 | 0.538 | -1.572 | 0.24 | -0.08 | -0.13 | -0.13 | 0 | 0.67 | 0.98 | 0.81 | 0.59 | 0.67 | 0.81 |
| 1 | 210510894 | rs1999890 | -0.633 | 0.387 | -1.409 | 0.241 | -0.016 | 0.018 | 0.018 | 0.57 | 0.56 | 0.83 | 0.5 | 0.36 | 0.55 | 0.64 |
| 1 | 210518082 | rs6696657 | -0.365 | 0.447 | -0.599 | 0.15 | 0.001 | 0.003 | 0.003 | 0 | 0.47 | 0.5 | 0.4 | 0.22 | 0.42 | 0.48 |
| 1 | 210519658 | rs1338289 | 1.932 | 0.872 | -1.116 | 0.54 | -0.034 | -0.035 | -0.035 | 0.176 | 0.07 | 0.01 | 0.3 | 0.25 | 0.11 | 0.09 |
| 1 | 210528907 | rs4845004 | -0.6 | 0.61 | -1.054 | 0.208 | 0.022 | -0.01 | -0.01 | 0 | 0.5 | 0.24 | 0.64 | 0.47 | 0.53 | 0.62 |
| 1 | 210529582 | rs4845006 | 0.172 | 0.689 | -1.134 | 0.03 | -0.008 | 0.077 | 0.077 | 0 | 0.35 | 0.14 | 0.34 | 0.16 | 0.33 | 0.37 |
| 1 | 210529833 | rs17708168 | 1.342 | 0.702 | -1.065 | 0.159 | -0.014 | 0.01 | 0.01 | 0 | 0.07 | 0.02 | 0.02 | 0.04 | 0.08 | 0.1 |
| 1 | 210531572 | rs17759116 | 1.259 | 0.691 | -1.057 | 0.154 | -0.012 | 0.009 | 0.009 | 0 | 0.07 | 0.02 | 0.02 | 0.04 | 0.08 | 0.1 |
| 1 | 210536025 | rs7527939 | 0.316 | 0.601 | -0.917 | -0.086 | 0 | -0.034 | -0.034 | 0 | 0.2 | 0.68 | 0.22 | 0.12 | 0.2 | 0.15 |
| 1 | 210536554 | rs12141729 | 1.792 | 0.768 | -1.312 | -0.401 | -0.039 | 0.08 | 0.08 | 0 | 0.14 | 0.03 | 0.09 | 0 | 0.14 | 0.1 |
| 1 | 210537248 | rs17016036 | 1.316 | 0.786 | -1.397 | -0.383 | -0.015 | -0.018 | -0.018 | 0 | 0.14 | 0.08 | 0.31 | 0.14 | 0.2 | 0.2 |
| 1 | 210543246 | rs3765874 | -0.678 | 0.564 | -1.279 | -0.469 | -0.03 | 0.078 | 0.078 | 0 | 0.43 | 0.23 | 0.45 | 0.2 | 0.43 | 0.46 |
| 1 | 210547213 | rs3765871 | -0.501 | 0.424 | -1.243 | -0.405 | 0.005 | 0.082 | 0.082 | 0.152 | 0.48 | 0.79 | 0.63 | 0.27 | 0.53 | 0.46 |
| 1 | 210550034 | rs3819987 | -2.109 | 0.402 | -1.54 | -0.13 | -0.018 | -0.127 | -0.127 | 0 | 0.72 | 0.99 | 0.86 | 0.59 | 0.75 | 0.78 |
| 1 | 210557308 | rs717659 | -0.297 | 0.661 | -1.206 | -0.079 | -0.009 | 0.012 | 0.012 | 1.005 | 0.34 | 0.72 | 0.53 | 0.24 | 0.38 | 0.35 |
| 1 | 210559076 | rs3765867 | -1.533 | 0.518 | -1.388 | -0.005 | 0.026 | -0.05 | -0.05 | 0.124 | 0.64 | 0.93 | 0.84 | 0.57 | 0.72 | 0.71 |
| 1 | 210559381 | rs3765865 | 1.978 | 0.587 | -1.403 | 0.137 | -0.029 | -0.047 | -0.047 | 0.434 | 0.08 | 0.01 | 0.17 | 0.28 | 0.13 | 0.18 |
| 1 | 210561713 | rs3765863 | -0.941 | 0.629 | -1.121 | -0.195 | 0.052 | 0.056 | 0.056 | 0.442 | 0.54 | 0.78 | 0.65 | 0.28 | 0.61 | 0.53 |
| 1 | 210567430 | rs1076708 | -0.334 | 0.891 | -0.486 | -0.15 | 0.023 | -0.03 | -0.03 | 0 | 0.34 | 0.41 | 0.45 | 0.16 | 0.41 | 0.37 |
| 1 | 210572044 | rs2073038 | 0.306 | 0.88 | -0.796 | 0.023 | -0.024 | -0.011 | -0.011 | 0 | 0.21 | 0.09 | 0.34 | 0.17 | 0.26 | 0.28 |
| 1 | 210572449 | rs6540592 | -0.838 | 0.519 | -0.663 | 0.101 | 0.041 | 0.016 | 0.016 | 1.241 | 0.55 | 0.71 | 0.65 | 0.39 | 0.61 | 0.58 |
| 1 | 210575777 | rs2143440 | -0.446 | 0.597 | -0.515 | -0.088 | -0.001 | 0 | 0 | 0 | 0.44 | 0.36 | 0.47 | 0.21 | 0.45 | 0.53 |
| 1 | 210577134 | rs10863825 | 1.426 | 0.482 | -0.777 | 0.792 | -0.029 | -0.048 | -0.048 | 0.927 | 0.08 | 0.01 | 0.17 | 0.25 | 0.13 | 0.11 |
| 1 | 210580381 | rs1033491 | -0.209 | 0.434 | -0.338 | 0.77 | 0.017 | 0.013 | 0.013 | 1.508 | 0.33 | 0.51 | 0.49 | 0.35 | 0.36 | 0.36 |
| 1 | 210581223 | rs7527094 | 0.29 | 0.263 | -0.807 | 0.484 | -0.007 | -0.01 | -0.01 | 0.079 | 0.15 | 0.16 | 0.05 | 0.1 | 0.09 | 0.16 |
| 1 | 210582538 | rs3765857 | -0.858 | 0.386 | -1.193 | 0.609 | -0.056 | 0.031 | 0.031 | 2.651 | 0.56 | 0.9 | 0.73 | 0.66 | 0.56 | 0.65 |
| 1 | 210585575 | rs2092449 | -0.091 | 0.518 | -0.475 | 0.844 | 0 | 0.041 | 0.041 | 0 | 0.91 | 0.73 | 0.83 | 0.83 | 0.88 | 0.91 |
| 1 | 210587818 | rs4845015 | 0.855 | 0.893 | -1.145 | 0.762 | -0.06 | -0.031 | -0.031 | 0 | 0.14 | 0.03 | 0.35 | 0.11 | 0.2 | 0.19 |
| 1 | 210589297 | rs4601592 | 0.235 | 0.273 | -0.925 | 0.61 | -0.004 | 0.013 | 0.013 | 0.182 | 0.6 | 0.45 | 0.57 | 0.5 | 0.55 | 0.59 |
| 1 | 210590153 | rs926579 | 0.237 | 0.411 | -0.824 | 0.589 | -0.004 | 0.004 | 0.004 | 0.243 | 0.6 | 0.42 | 0.58 | 0.51 | 0.59 | 0.59 |
| 1 | 210592577 | rs2357587 | 0.098 | 0.278 | -1.051 | 0.275 | 0.002 | -0.001 | -0.001 | 0 | 0.51 | 0.44 | 0.52 | 0.5 | 0.48 | 0.56 |
| 1 | 210598837 | rs4845021 | 0.353 | 0.257 | -0.971 | 0.481 | -0.006 | 0.016 | 0.016 | 1.58 | 0.45 | 0.37 | 0.5 | 0.33 | 0.46 | 0.48 |
| 1 | 210605563 | rs4845025 | -0.077 | 0.108 | -1.133 | 0.449 | 0.014 | 0.023 | 0.023 | 2.887 | 0.54 | 0.4 | 0.64 | 0.72 | 0.6 | 0.7 |
| 1 | 210607991 | rs926578 | 1.278 | 0.453 | -0.969 | 0.016 | 0.022 | 0.007 | 0.007 | 0 | 0.16 | 0.24 | 0.37 | 0.11 | 0.2 | 0.19 |
| 1 | 210611882 | rs910032 | 1.752 | 0.475 | -1.092 | 0.027 | -0.062 | -0.041 | -0.041 | 0 | 0.14 | 0.03 | 0.36 | 0.11 | 0.2 | 0.2 |
| 1 | 210612528 | rs910029 | -1.419 | 0.321 | -0.994 | 0.023 | 0.013 | -0.054 | -0.054 | 0 | 0.78 | 0.86 | 0.77 | 0.56 | 0.82 | 0.81 |
| 1 | 210613616 | rs3765851 | 0.005 | 0.219 | -1.136 | -0.35 | -0.001 | 0.012 | 0.012 | 0 | 0.47 | 0.43 | 0.5 | 0.32 | 0.47 | 0.47 |
| 1 | 210615064 | rs12121798 | -0.41 | 0.219 | -1.147 | -0.39 | 0.002 | -0.003 | -0.003 | 0 | 0.56 | 0.42 | 0.63 | 0.54 | 0.6 | 0.68 |
| 1 | 210620492 | rs12025108 | 0.642 | 0.485 | -1.033 | 0.118 | -0.002 | -0.002 | -0.002 | 0 | 0.25 | 0.26 | 0.5 | 0.31 | 0.34 | 0.33 |
| 1 | 210622527 | rs742563 | -0.668 | 0.189 | -1.141 | 0.329 | -0.001 | 0.005 | 0.005 | 0.436 | 0.73 | 0.69 | 0.73 | 0.55 | 0.75 | 0.78 |
| 1 | 210622790 | rs3819985 | 1.43 | 0.568 | -1.299 | 0.289 | -0.035 | -0.047 | -0.047 | 0 | 0.08 | 0.01 | 0.13 | 0.2 | 0.13 | 0.13 |
| 1 | 210632175 | rs12128618 | 1.45 | 0.273 | -1.123 | -0.216 | -0.004 | -0.027 | -0.027 | 0.093 | 0.1 | 0.1 | 0.04 | 0.01 | 0.08 | 0.13 |
| 1 | 210633518 | rs7555818 | 1.403 | 0.212 | -1.203 | -0.627 | 0 | 0.001 | 0.001 | 0.046 | 0.14 | 0.1 | 0.08 | 0.03 | 0.15 | 0.08 |
| 1 | 210637550 | rs7554508 | -0.188 | 0.229 | -1.546 | -0.599 | -0.011 | 0.013 | 0.013 | 0 | 0.56 | 0.7 | 0.47 | 0.39 | 0.54 | 0.48 |
| 1 | 210641438 | rs981754 | 1.181 | 0.33 | -1.266 | -0.361 | -0.011 | -0.007 | -0.007 | 0.291 | 0.14 | 0.05 | 0.15 | 0.16 | 0.15 | 0.21 |
| 1 | 210644714 | rs2223630 | -0.467 | 0.507 | -1.367 | -0.086 | -0.012 | -0.052 | -0.052 | 0.157 | 0.68 | 0.78 | 0.59 | 0.4 | 0.67 | 0.64 |
| 1 | 210647296 | rs7530369 | -1.608 | 0.34 | -1.287 | 0.047 | 0.002 | -0.02 | -0.02 | 0 | 0.82 | 0.85 | 0.74 | 0.56 | 0.83 | 0.84 |
| 1 | 210650464 | rs17016283 | 2.336 | 0.448 | -1.388 | -0.086 | 0.013 | -0.029 | -0.029 | 3.569 | 0.06 | 0.1 | 0.04 | 0 | 0.09 | 0.03 |
| 1 | 210651567 | rs761174 | -1.968 | 0.175 | -1.769 | -0.547 | -0.017 | 0.107 | 0.107 | 2.85 | 0.72 | 0.92 | 0.85 | 0.99 | 0.73 | 0.74 |
| 1 | 210652685 | rs12031180 | -0.296 | 0.146 | -0.597 | -0.045 | 0.009 | 0.03 | 0.03 | 0 | 0.31 | 0.41 | 0.39 | 0.39 | 0.36 | 0.34 |
| 1 | 210653374 | rs2050187 | -0.993 | 0.057 | -1.206 | -0.288 | -0.007 | 0.196 | 0.196 | 0.458 | 0.51 | 0.83 | 0.67 | 0.83 | 0.53 | 0.49 |
| 1 | 210654164 | rs2206610 | -0.447 | -0.367 | -0.227 | 0.533 | -0.017 | 0.098 | 0.098 | 0 | 0.12 | 0.29 | 0.13 | 0.19 | 0.08 | 0.08 |
| 1 | 210659966 | rs2473623 | -1.59 | -0.199 | -1.439 | 0.027 | -0.114 | 0.066 | 0.066 | 0.298 | 0.55 | 0.94 | 0.55 | 0.72 | 0.5 | 0.62 |
| 1 | 210660195 | rs2473621 | -0.529 | -0.38 | -0.751 | 0.284 | 0 | 0 | 0 | 0 | 0.32 | 0.39 | 0.23 | 0.24 | 0.29 | 0.47 |
| 1 | 210662359 | rs6682994 | -0.374 | -0.438 | -0.703 | 0.244 | -0.002 | 0.009 | 0.009 | 0 | 0.92 | 0.88 | 0.92 | 0.84 | 0.94 | 0.89 |
| 1 | 210664959 | rs6675248 | 2.548 | -0.363 | -0.425 | 0.116 | -0.008 | 0.001 | 0.001 | 0.572 | 0.33 | 0.38 | 0.31 | 0.34 | 0.31 | 0.19 |
| 1 | 210665583 | rs7546331 | 0.432 | -0.399 | -0.526 | 0.022 | -0.006 | -0.009 | -0.009 | 0 | 0.71 | 0.66 | 0.58 | 0.58 | 0.66 | 0.57 |
| 1 | 210666439 | rs4478810 | 0.946 | -0.427 | -0.457 | 0.004 | -0.008 | -0.001 | -0.001 | 1.369 | 0.84 | 0.63 | 0.86 | 0.84 | 0.86 | 0.77 |
| 1 | 210670422 | rs7514691 | 0.739 | -0.424 | -0.317 | -0.084 | -0.017 | -0.005 | -0.005 | 0 | 0.55 | 0.66 | 0.5 | 0.55 | 0.52 | 0.35 |
| 1 | 210671640 | rs7534803 | 2.867 | -0.338 | -0.264 | -0.29 | -0.003 | 0.117 | 0.117 | 1.445 | 0.31 | 0.11 | 0.18 | 0 | 0.3 | 0.17 |
| 1 | 210672483 | rs12748886 | 0.444 | -0.363 | -0.291 | -0.272 | -0.006 | 0.002 | 0.002 | 0.434 | 0.59 | 0.55 | 0.55 | 0.6 | 0.55 | 0.36 |
| 1 | 210672629 | rs17016349 | -0.858 | -0.334 | -0.125 | -0.12 | -0.007 | 0.025 | 0.025 | 0.004 | 0.28 | 0.51 | 0.38 | 0.59 | 0.25 | 0.19 |
| 1 | 210674209 | rs12411254 | 0.231 | 0.337 | 0.585 | 0.517 | -0.018 | -0.018 | -0.018 | 0.263 | 0.2 | 0.13 | 0.31 | 0.26 | 0.26 | 0.32 |
| 1 | 210674600 | rs12759944 | 1.325 | 0.343 | 0.249 | 0.62 | -0.075 | -0.091 | -0.091 | 0 | 0.18 | 0.02 | 0.28 | 0.26 | 0.22 | 0.21 |

| | | | | | | | | | | | | | | | | |
|---|---|---|---|---|---|---|---|---|---|---|---|---|---|---|---|---|
| 1 | 210677551 | rs4537555 | 0.913 | 0.513 | 0.248 | 0.631 | -0.029 | 0.017 | 0.017 | 0 | 0.08 | 0.03 | 0.07 | 0.05 | 0.12 | 0.07 |
| 1 | 210679612 | rs6672183 | -0.203 | 0.472 | 0.532 | 0.621 | 0.008 | 0.009 | 0.009 | 0 | 0.69 | 0.4 | 0.41 | 0.37 | 0.66 | 0.69 |
| 1 | 210683732 | rs10863837 | 0.434 | 0.541 | 0.621 | 0.612 | -0.001 | -0.001 | -0.001 | 1.801 | 0.58 | 0.42 | 0.55 | 0.56 | 0.59 | 0.39 |
| 1 | 210687396 | rs10746425 | 0.572 | -0.063 | 0.698 | 0.715 | 0 | 0 | 0 | 0 | 0.53 | 0.45 | 0.5 | 0.56 | 0.49 | 0.38 |
| 1 | 210688180 | rs10735507 | -0.563 | 0.176 | 0.598 | 0.88 | 0.025 | 0.012 | 0.012 | 0 | 0.72 | 0.6 | 0.61 | 0.68 | 0.67 | 0.72 |
| 1 | 210688784 | rs12145163 | 0.866 | -0.362 | 0.03 | 0.656 | -0.038 | 0.037 | 0.037 | 0 | 0.14 | 0.01 | 0.07 | 0.05 | 0.14 | 0.17 |
| 1 | 210696806 | rs6540602 | 0.767 | -0.555 | -0.087 | -0.087 | 0.012 | 0.026 | 0.026 | 1.009 | 0.36 | 0.23 | 0.21 | 0.01 | 0.37 | 0.34 |
| 1 | 210699389 | rs6691759 | -0.057 | -0.455 | -0.127 | 0.332 | 0.012 | 0.091 | 0.091 | 0 | 0.51 | 0.29 | 0.39 | 0.24 | 0.49 | 0.44 |
| 1 | 210707065 | rs4845045 | 0.128 | -0.998 | -0.143 | 0.486 | 0.058 | 0.05 | 0.05 | 0 | 0.39 | 0.17 | 0.31 | 0.32 | 0.3 | 0.41 |
| 1 | 210710747 | rs1316508 | 0.61 | -0.956 | 0.196 | 0.648 | -0.019 | 0.01 | 0.01 | 0 | 0.4 | 0.24 | 0.44 | 0.32 | 0.51 | 0.44 |
| 1 | 210717147 | rs2069151 | 0.308 | -0.855 | 0.254 | 0.648 | 0.03 | 0.024 | 0.024 | 0 | 0.4 | 0.31 | 0.32 | 0.32 | 0.3 | 0.29 |
| 1 | 210721874 | rs12130516 | 0.234 | -0.712 | -0.37 | 0.642 | -0.054 | -0.031 | -0.031 | 0.002 | 0.15 | 0.02 | 0.14 | 0.11 | 0.17 | 0.11 |
| 1 | 210727515 | rs2490215 | 0.138 | -0.859 | -0.09 | 0.568 | -0.012 | 0.006 | 0.006 | 0.11 | 0.44 | 0.6 | 0.6 | 0.6 | 0.48 | 0.48 |
| 1 | 210729242 | rs4844531 | -0.149 | -0.767 | -0.528 | 0.776 | -0.001 | -0.013 | -0.013 | 0.028 | 0.56 | 0.34 | 0.46 | 0.46 | 0.46 | 0.47 |
| 1 | 210729897 | rs12040304 | -0.224 | -1.475 | 0.145 | 0.495 | 0.027 | 0.014 | 0.014 | 0 | 0.4 | 0.34 | 0.31 | 0.34 | 0.28 | 0.28 |
| 1 | 210730635 | rs17016427 | -0.869 | -0.635 | 1.032 | 1.5 | 0 | -0.001 | -0.001 | 0 | 0.06 | 0.06 | 0.05 | 0.02 | 0.07 | 0.08 |
| 1 | 210731236 | rs11807403 | -1.269 | -0.494 | 1.382 | 1.785 | -0.027 | 0.25 | 0.25 | 0.312 | 0.12 | 0.3 | 0.25 | 0.6 | 0.06 | 0.17 |
| 1 | 210747424 | rs2501907 | 0.967 | 0.457 | 1.214 | 0.621 | 0.072 | -0.026 | -0.026 | 0 | 0.89 | 0.62 | 0.91 | 0.99 | 0.87 | 0.94 |
| 1 | 210749312 | rs12567982 | -0.133 | -0.047 | 0.602 | 0.261 | -0.002 | 0.013 | 0.013 | 0 | 0.24 | 0.14 | 0.12 | 0.16 | 0.26 | 0.16 |
| 1 | 210750063 | rs12217125 | 0.22 | 0.1 | 0.757 | -0.016 | -0.012 | 0.005 | 0.005 | 0 | 0.37 | 0.25 | 0.47 | 0.67 | 0.44 | 0.38 |
| 1 | 210772300 | rs11119547 | 0.414 | -0.092 | 0.784 | -0.211 | -0.016 | 0.086 | 0.086 | 0.433 | 0.46 | 0.27 | 0.33 | 0.26 | 0.47 | 0.47 |
| 1 | 210775500 | rs12745924 | 0.799 | 0.026 | 0.041 | -0.476 | -0.113 | -0.126 | -0.126 | 0 | 0.26 | 0.02 | 0.39 | 0.68 | 0.33 | 0.34 |
| 1 | 210775735 | rs6693576 | 0.283 | -0.022 | 0.405 | -0.583 | -0.006 | 0.045 | 0.045 | 0 | 0.57 | 0.74 | 0.41 | 0.27 | 0.56 | 0.52 |
| 1 | 210780516 | rs4245753 | 0.58 | -0.183 | 0.573 | -0.543 | 0.041 | -0.034 | -0.034 | 0 | 0.9 | 0.52 | 0.93 | 0.99 | 0.92 | 0.96 |
| 1 | 210781750 | rs6540610 | 0.934 | -0.1 | 0.359 | -0.476 | -0.03 | -0.07 | -0.07 | 0 | 0.73 | 0.42 | 0.72 | 0.94 | 0.81 | 0.82 |
| 1 | 210789792 | rs6540614 | -1.256 | 0.281 | 0.361 | -0.56 | -0.009 | -0.106 | -0.106 | 0 | 0.22 | 0.49 | 0.21 | 0.06 | 0.14 | 0.22 |
| 1 | 210789920 | rs7541034 | 0.261 | 0.22 | -0.482 | -0.601 | -0.002 | -0.004 | -0.004 | 0 | 0.57 | 0.55 | 0.54 | 0.63 | 0.62 | 0.48 |
| 1 | 210790214 | rs4488079 | -0.657 | 0.835 | -1.097 | 0.724 | -0.018 | -0.037 | -0.037 | 0 | 0.14 | 0.07 | 0.15 | 0.37 | 0.18 | 0.19 |
| 1 | 210793972 | rs11119552 | -0.438 | 0.689 | -1.485 | 1.13 | -0.027 | -0.071 | -0.071 | 0 | 0.23 | 0.12 | 0.22 | 0.36 | 0.26 | 0.24 |
| 1 | 210794866 | rs12133319 | 0.957 | 0.495 | -2.412 | 0.851 | -0.113 | -0.035 | -0.035 | 0 | 0.38 | 0.06 | 0.27 | 0.38 | 0.39 | 0.42 |
| 1 | 210794975 | rs12409512 | 0.589 | 0.339 | -2.334 | 0.676 | 0.007 | 0.035 | 0.035 | 0 | 0.11 | 0.03 | 0.13 | 0.05 | 0.08 | 0.05 |
| 1 | 210798142 | rs10465688 | -0.449 | 0.86 | -2.136 | 0.977 | -0.027 | -0.031 | -0.031 | 0 | 0.15 | 0.09 | 0.21 | 0.28 | 0.26 | 0.15 |
| 1 | 210799183 | rs11590767 | 0.867 | 0.453 | -1.97 | 0.899 | 0.002 | -0.003 | -0.003 | 1.103 | 0.13 | 0.15 | 0.15 | 0.19 | 0.1 | 0.17 |
| 1 | 210800547 | rs4951675 | -0.224 | 0.579 | -1.5 | 1.156 | 0.009 | 0.004 | 0.004 | 0 | 0.15 | 0.14 | 0.22 | 0.25 | 0.29 | 0.15 |
| 1 | 210801879 | rs4504959 | -1.611 | 0.371 | -0.746 | 1.23 | -0.003 | 0.035 | 0.035 | 0 | 0.89 | 0.75 | 0.92 | 0.85 | 0.9 | 0.86 |
| 1 | 210803032 | rs12410510 | 0.115 | -0.041 | -0.695 | 0.855 | 0.044 | 0.089 | 0.089 | 0 | 0.27 | 0.13 | 0.3 | 0.1 | 0.19 | 0.22 |
| 1 | 210804430 | rs4951682 | 0.106 | 0.134 | -0.686 | 0.974 | 0.01 | 0.017 | 0.017 | 0 | 0.14 | 0.2 | 0.2 | 0.26 | 0.21 | 0.14 |
| 1 | 210807437 | rs3935665 | 1.894 | -0.357 | -0.74 | 0.834 | -0.074 | -0.031 | -0.031 | 0.535 | 0.36 | 0.08 | 0.25 | 0.38 | 0.34 | 0.41 |
| 1 | 210808270 | rs4951488 | -1.399 | -0.383 | -0.507 | 0.805 | -0.004 | 0.032 | 0.032 | 0.014 | 0.52 | 0.63 | 0.59 | 0.4 | 0.56 | 0.44 |
| 1 | 210810054 | rs12090004 | 1.172 | -0.642 | -0.474 | 0.606 | -0.002 | -0.031 | -0.031 | 0 | 0.14 | 0.12 | 0.05 | 0.02 | 0.09 | 0.11 |
| 1 | 210810170 | rs6658987 | 0.331 | -0.51 | -0.498 | 0.772 | 0.012 | 0.01 | 0.01 | 0 | 0.16 | 0.17 | 0.22 | 0.31 | 0.3 | 0.16 |
| 1 | 210815017 | rs12745898 | -0.075 | -1.128 | -0.542 | 0.285 | -0.006 | -0.026 | -0.026 | 0.001 | 0.14 | 0.14 | 0.07 | 0.02 | 0.07 | 0.06 |
| 1 | 210816382 | rs6540616 | -1.905 | -1.115 | -0.319 | 0.483 | -0.009 | 0.044 | 0.044 | 0 | 0.88 | 0.8 | 0.83 | 0.78 | 0.9 | 0.86 |
| 1 | 210818347 | rs6683572 | 0.298 | -1.291 | -0.587 | 0.567 | 0.011 | -0.007 | -0.007 | 0.043 | 0.61 | 0.35 | 0.62 | 0.51 | 0.69 | 0.62 |
| 1 | 210819646 | rs12096661 | -0.847 | -0.972 | -0.556 | 0.697 | -0.005 | -0.003 | -0.003 | 2.203 | 0.36 | 0.31 | 0.46 | 0.44 | 0.44 | 0.31 |
| 1 | 210819791 | rs12097792 | 0.214 | -0.589 | 0.064 | 1.343 | 0.014 | 0.021 | 0.021 | 0 | 0.17 | 0.18 | 0.38 | 0.43 | 0.33 | 0.18 |
| 1 | 210822016 | rs10863847 | -0.893 | -0.696 | 0.024 | 1.196 | 0.01 | 0.006 | 0.006 | 0 | 0.26 | 0.27 | 0.4 | 0.37 | 0.37 | 0.23 |
| 1 | 210829477 | rs4951508 | -0.096 | -0.927 | -0.743 | 0.831 | -0.003 | 0 | 0 | 0.253 | 0.7 | 0.33 | 0.66 | 0.47 | 0.79 | 0.66 |
| 1 | 210830320 | rs4472816 | -0.09 | -0.637 | -0.846 | 0.868 | -0.014 | -0.013 | -0.013 | 0 | 0.18 | 0.15 | 0.36 | 0.37 | 0.32 | 0.22 |
| 1 | 210830431 | rs12121954 | 2.134 | -0.756 | -1.058 | 0.557 | -0.101 | 0.197 | 0.197 | 2.372 | 0.38 | 0.05 | 0.21 | 0.1 | 0.37 | 0.38 |
| 1 | 210831820 | rs12116836 | 2.134 | -0.756 | -1.058 | 0.557 | -0.101 | 0.197 | 0.197 | 2.372 | 0.38 | 0.05 | 0.21 | 0.1 | 0.37 | 0.38 |
| 1 | 210836847 | rs1890843 | 1.352 | -0.231 | -0.044 | 1.366 | -0.011 | 0.04 | 0.04 | 0 | 0.09 | 0.16 | 0.07 | 0.2 | 0.08 | 0.13 |
| 1 | 210837691 | rs10746432 | 1.408 | -0.291 | -0.106 | 1.281 | -0.001 | 0 | 0 | 0 | 0.67 | 0.67 | 0.55 | 0.63 | 0.57 | 0.72 |
| 1 | 210839705 | rs7413508 | -0.128 | 0.051 | 0.006 | 1.465 | 0.029 | 0.024 | 0.024 | 0 | 0.86 | 0.78 | 0.66 | 0.65 | 0.69 | 0.83 |
| 1 | 210840755 | rs1970425 | -0.019 | -0.183 | 0.123 | 1.541 | -0.034 | 0.303 | 0.303 | 0.023 | 0.14 | 0.55 | 0.23 | 0.5 | 0.08 | 0.17 |
| 1 | 210848224 | rs10863850 | -0.445 | 0.132 | -0.084 | 1.588 | 0.028 | 0.029 | 0.029 | 0 | 0.89 | 0.85 | 0.7 | 0.67 | 0.73 | 0.84 |
| 1 | 210849262 | rs17016653 | 1.216 | 0.068 | -0.195 | 1.505 | -0.013 | -0.041 | -0.041 | 2.735 | 0.09 | 0.03 | 0.07 | 0.2 | 0.1 | 0.12 |
| 1 | 210857457 | rs1135317 | 0.594 | -0.018 | -0.022 | 1.552 | 0.025 | 0.024 | 0.024 | 0.018 | 0.84 | 0.81 | 0.69 | 0.65 | 0.71 | 0.8 |
| 1 | 210862656 | rs6540619 | 0.631 | -0.366 | -0.247 | 0.941 | -0.04 | -0.055 | -0.055 | 8.991 | 0.73 | 0.82 | 0.61 | 0.46 | 0.56 | 0.65 |
| 1 | 210871341 | rs17260690 | -0.003 | -0.678 | -0.021 | 0.885 | -0.01 | -0.044 | -0.044 | 0.925 | 0.15 | 0.15 | 0.09 | 0 | 0.08 | 0.06 |
| 1 | 210873142 | rs1890842 | -0.615 | -0.534 | 0.093 | 1.142 | 0.087 | 0.684 | 0.684 | 4.188 | 0.05 | 0.53 | 0.18 | 0.31 | 0 | 0.05 |
| 1 | 210884699 | rs9430044 | -1.316 | -0.65 | -0.336 | 1.135 | -0.051 | 0.051 | 0.051 | 1.449 | 0.16 | 0.69 | 0.16 | 0.3 | 0.06 | 0.12 |
| 1 | 210885735 | rs4951542 | 0.168 | 0.282 | -0.313 | 1.248 | -0.01 | -0.008 | -0.008 | 0.725 | 0.8 | 0.81 | 0.63 | 0.47 | 0.6 | 0.71 |
| 1 | 210885868 | rs4951543 | -0.776 | 0.204 | -0.18 | 1.327 | -0.065 | 0.118 | 0.118 | 0.642 | 0.24 | 0.58 | 0.28 | 0.36 | 0.14 | 0.14 |
| 1 | 210886979 | rs1934618 | 0.405 | 1.29 | -0.461 | 1.659 | 0.004 | 0.016 | 0.016 | 0.056 | 0.85 | 0.84 | 0.59 | 0.44 | 0.61 | 0.73 |
| 1 | 210887739 | rs17016764 | -1.061 | 0.56 | 0.042 | 1.446 | 0 | 0.012 | 0.012 | 1.166 | 0.09 | 0.22 | 0.08 | 0.14 | 0.08 | 0.05 |
| 1 | 210893635 | rs6681860 | -1.103 | 0.209 | 0.672 | 1.548 | -0.029 | -0.021 | -0.021 | 0 | 0.18 | 0.43 | 0.27 | 0.09 | 0.13 | 0.14 |
| 1 | 210896693 | rs7555360 | -1.074 | 0.269 | 0.82 | 1.848 | -0.027 | -0.018 | -0.018 | 0 | 0.18 | 0.43 | 0.27 | 0.09 | 0.13 | 0.14 |
| 1 | 210898263 | rs1934620 | -0.996 | 0.251 | 1.047 | 1.838 | -0.015 | -0.011 | -0.011 | 0 | 0.17 | 0.3 | 0.26 | 0.09 | 0.13 | 0.14 |

| | | | | | | | | | | | | | | | | |
|---|---|---|---|---|---|---|---|---|---|---|---|---|---|---|---|---|
| 1 | 210900481 | rs1934623 | -0.729 | 0.386 | 1.017 | 1.81 | 0.006 | 0.045 | 0.045 | 0 | 0.15 | 0.15 | 0.19 | 0.45 | 0.18 | 0.29 |
| 1 | 210901459 | rs1953951 | -0.849 | 0.562 | 0.47 | 0.918 | 0 | -0.009 | -0.009 | 0 | 0.39 | 0.5 | 0.45 | 0.23 | 0.43 | 0.39 |
| 1 | 210903194 | rs11119583 | 0.845 | 0.514 | 0.473 | 0.918 | -0.009 | 0.064 | 0.064 | 1.146 | 0.76 | 0.48 | 0.65 | 0.71 | 0.76 | 0.76 |
| 1 | 210907239 | rs1890844 | 0.827 | 0.442 | 0.602 | 0.72 | -0.001 | 0.009 | 0.009 | 0 | 0.56 | 0.38 | 0.53 | 0.86 | 0.5 | 0.65 |
| 1 | 210907463 | rs1890845 | 0.205 | 0.413 | 0.38 | 0.694 | -0.012 | 0.046 | 0.046 | 0 | 0.64 | 0.75 | 0.65 | 0.89 | 0.55 | 0.71 |
| 1 | 210912467 | rs1934628 | -0.251 | 0.382 | 0.242 | 0.858 | -0.001 | -0.014 | -0.014 | 0 | 0.86 | 0.89 | 0.8 | 0.9 | 0.83 | 0.88 |
| 1 | 210923061 | rs4951563 | -0.788 | 0.453 | 0.7 | 0.948 | 0.034 | 0.13 | 0.13 | 0.072 | 0.09 | 0.41 | 0.16 | 0.17 | 0.07 | 0.1 |
| 1 | 210929893 | rs1934636 | -0.298 | 0.325 | 0.425 | 0.846 | 0.001 | -0.004 | -0.004 | 0 | 0.85 | 0.85 | 0.79 | 0.89 | 0.82 | 0.89 |
| 1 | 210939931 | rs1777264 | -0.453 | 0.331 | 0.572 | 0.859 | 0.009 | 0.017 | 0.017 | 0 | 0.17 | 0.4 | 0.23 | 0.11 | 0.19 | 0.11 |
| 1 | 210941930 | rs1777245 | 0.021 | 0.354 | 0.302 | 0.87 | -0.006 | -0.009 | -0.009 | 0 | 0.14 | 0.11 | 0.21 | 0.11 | 0.19 | 0.11 |
| 1 | 210955200 | rs1777255 | -0.992 | 0.414 | 0.31 | 0.867 | 0.005 | 0.02 | 0.02 | 0 | 0.46 | 0.68 | 0.56 | 0.29 | 0.6 | 0.43 |
| 1 | 210966290 | rs10863856 | -0.244 | 0.347 | 0.502 | 1.156 | -0.011 | 0.073 | 0.073 | 0 | 0.13 | 0.15 | 0.11 | 0.37 | 0.09 | 0.25 |
| 1 | 210967608 | rs6694502 | -1.205 | 0.469 | 0.479 | 0.66 | 0.078 | 0.075 | 0.075 | 0.608 | 0.14 | 0.53 | 0.2 | 0.18 | 0.16 | 0.13 |
| 1 | 210978485 | rs7553708 | 0.487 | 0.263 | -0.038 | 0.416 | -0.001 | 0.004 | 0.004 | 0 | 0.23 | 0.14 | 0.17 | 0.5 | 0.23 | 0.36 |
| 1 | 210982084 | rs1573159 | -1.418 | -0.009 | 0.088 | 0.118 | -0.022 | -0.026 | -0.026 | 0 | 0.7 | 0.65 | 0.72 | 0.83 | 0.82 | 0.79 |
| 1 | 210984565 | rs7532949 | 0.767 | 0.122 | 0.066 | -0.023 | -0.056 | 0.078 | 0.078 | 0 | 0.26 | 0.12 | 0.28 | 0.03 | 0.39 | 0.24 |
| 1 | 210985460 | rs6658580 | 0.277 | 0.097 | 0.135 | 0.118 | 0 | 0 | 0 | 0 | 0.16 | 0.18 | 0.23 | 0.13 | 0.2 | 0.11 |
| 1 | 210990713 | rs4282878 | -0.014 | 0.135 | 0.121 | 0.128 | 0 | -0.001 | -0.001 | 1.584 | 0.36 | 0.44 | 0.4 | 0.29 | 0.46 | 0.35 |
| 1 | 210994339 | rs1418973 | -1.028 | 0.017 | -0.018 | 0.047 | -0.005 | -0.026 | -0.026 | 0 | 0.65 | 0.59 | 0.6 | 0.81 | 0.7 | 0.74 |
| 1 | 210994395 | rs1112269 | 0.48 | 0.014 | -0.018 | 0.088 | -0.005 | -0.061 | -0.061 | 0 | 0.28 | 0.11 | 0.2 | 0.51 | 0.24 | 0.38 |
| 1 | 210994910 | rs12034491 | -0.763 | 0.036 | 0.005 | 0.077 | 0 | -0.01 | -0.01 | 2.435 | 0.45 | 0.44 | 0.44 | 0.63 | 0.44 | 0.49 |
| 1 | 210996703 | rs4951641 | -1.691 | 0.048 | -0.271 | 0.186 | 0.043 | 0.023 | 0.023 | 0.178 | 0.73 | 0.84 | 0.73 | 0.8 | 0.84 | 0.8 |
| 1 | 210997462 | rs11119611 | -0.339 | 0.231 | -0.074 | 0.272 | -0.006 | -0.009 | -0.009 | 0 | 0.34 | 0.28 | 0.31 | 0.5 | 0.37 | 0.43 |
| 1 | 211000020 | rs4098807 | -0.881 | 0.397 | -0.409 | 0.1 | 0.006 | 0.147 | 0.147 | 0.077 | 0.49 | 0.71 | 0.47 | 0.84 | 0.46 | 0.56 |
| 1 | 211000868 | rs11119613 | -0.871 | 0.645 | -0.509 | 0.316 | -0.001 | 0 | 0 | 0.022 | 0.45 | 0.43 | 0.44 | 0.64 | 0.47 | 0.51 |
| 1 | 211002077 | rs7545932 | 1.041 | 0.776 | -0.709 | 0.207 | -0.016 | 0.031 | 0.031 | 0 | 0.18 | 0.31 | 0.2 | 0.18 | 0.11 | 0.09 |
| 1 | 211002232 | rs6695591 | 1.43 | 1.001 | -0.774 | 0.204 | -0.086 | 0.109 | 0.109 | 0 | 0.2 | 0.04 | 0.18 | 0.01 | 0.34 | 0.19 |
| 1 | 211006126 | rs7556358 | -0.153 | 0.464 | -0.793 | 0.22 | 0.018 | 0.009 | 0.009 | 0.104 | 0.52 | 0.85 | 0.58 | 0.38 | 0.6 | 0.38 |
| 1 | 211010936 | rs4509646 | -0.166 | 0.471 | -0.756 | 0.222 | 0.021 | 0.003 | 0.003 | 0.078 | 0.52 | 0.83 | 0.58 | 0.38 | 0.6 | 0.38 |
| 1 | 211014027 | rs17016986 | 0.808 | 0.419 | -0.638 | 0.287 | 0.003 | -0.001 | -0.001 | 0.756 | 0.23 | 0.35 | 0.26 | 0.18 | 0.23 | 0.11 |
| 1 | 211016418 | rs10863862 | 1.531 | 0.477 | -0.917 | 0.538 | -0.034 | 0.008 | 0.008 | 0 | 0.06 | 0.01 | 0.05 | 0.36 | 0.12 | 0.16 |
| 1 | 211021787 | rs4532912 | -1.545 | 0.029 | -0.833 | 0.043 | 0.039 | 0.046 | 0.046 | 0 | 0.73 | 0.87 | 0.71 | 0.83 | 0.85 | 0.77 |
| 1 | 211041590 | rs9701252 | 0.404 | 0.672 | -0.778 | 0.635 | -0.064 | -0.102 | -0.102 | 0 | 0.15 | 0.04 | 0.11 | 0.45 | 0.19 | 0.32 |
| 1 | 211047777 | rs12122965 | 0.547 | 0.599 | -0.72 | -0.311 | -0.042 | 0.048 | 0.048 | 0 | 0.09 | 0.01 | 0.12 | 0.02 | 0.16 | 0.09 |
| 1 | 211047846 | rs11119624 | -0.738 | -0.047 | -0.298 | -0.132 | 0.022 | 0.034 | 0.034 | 0 | 0.48 | 0.39 | 0.49 | 0.66 | 0.59 | 0.52 |
| 1 | 211073277 | rs1697618 | -0.083 | 0.366 | -0.155 | 0.182 | -0.001 | -0.012 | -0.012 | 0 | 0.22 | 0.21 | 0.17 | 0.12 | 0.22 | 0.12 |
| 1 | 211079618 | rs924569 | -0.426 | 0.423 | -0.453 | 0.325 | -0.012 | -0.024 | -0.024 | 0 | 0.31 | 0.45 | 0.34 | 0.22 | 0.27 | 0.31 |
| 1 | 211082285 | rs1777250 | 0.059 | 0.564 | -0.678 | 0.423 | 0.003 | -0.002 | -0.002 | 6.867 | 0.57 | 0.5 | 0.56 | 0.53 | 0.56 | 0.46 |
| 1 | 211085683 | rs1160575 | 0.463 | 0.505 | -0.984 | 0.615 | -0.001 | -0.02 | -0.02 | 0.196 | 0.06 | 0.01 | 0.04 | 0.26 | 0.05 | 0.12 |
| 1 | 211088400 | rs4951487 | -0.414 | -0.232 | -0.907 | 0.114 | -0.013 | 0.104 | 0.104 | 0.513 | 0.52 | 0.72 | 0.54 | 0.29 | 0.49 | 0.56 |
| 1 | 211089156 | rs1875435 | 0.644 | -0.562 | -0.465 | -0.048 | -0.007 | 0.01 | 0.01 | 0.028 | 0.33 | 0.3 | 0.38 | 0.21 | 0.38 | 0.37 |
| 1 | 211093590 | rs1501569 | 0.625 | -0.569 | -0.484 | -0.018 | -0.005 | 0.005 | 0.005 | 0.034 | 0.34 | 0.31 | 0.38 | 0.21 | 0.38 | 0.37 |
| 1 | 211094848 | rs10494930 | 0.289 | -0.635 | -0.563 | -0.075 | 0.006 | 0.035 | 0.035 | 1.495 | 0.14 | 0.01 | 0.13 | 0.08 | 0.11 | 0.19 |
| 1 | 211097125 | rs1501555 | -1.096 | -0.622 | -0.49 | -0.029 | -0.022 | -0.031 | -0.031 | 0.12 | 0.59 | 0.75 | 0.58 | 0.54 | 0.54 | 0.68 |
| 1 | 211098350 | rs1354112 | -1.458 | -0.871 | -0.998 | -0.43 | -0.064 | 0.11 | 0.11 | 0 | 0.75 | 0.92 | 0.84 | 0.93 | 0.68 | 0.85 |
| 1 | 211108733 | rs7529770 | 0.068 | -0.923 | -0.941 | -0.454 | -0.005 | -0.004 | -0.004 | 0 | 0.56 | 0.54 | 0.68 | 0.59 | 0.57 | 0.58 |
| 1 | 211112257 | rs12036999 | 1.036 | -0.983 | -0.972 | -0.401 | 0 | 0.028 | 0.028 | 0 | 0.08 | 0.05 | 0.19 | 0.38 | 0.05 | 0.15 |
| 1 | 211120598 | rs6540636 | 0.043 | -0.976 | -0.786 | -0.357 | -0.002 | 0.053 | 0.053 | 0 | 0.15 | 0.17 | 0.26 | 0.38 | 0.12 | 0.16 |
| 1 | 211135613 | rs12566456 | 1.314 | -0.908 | -0.828 | -0.166 | -0.004 | -0.035 | -0.035 | 3.701 | 0.06 | 0.01 | 0.04 | 0.19 | 0.05 | 0.12 |
| 1 | 211153359 | rs1393026 | 1.238 | -1.077 | -0.894 | -0.357 | -0.012 | 0.013 | 0.013 | 0.243 | 0.33 | 0.29 | 0.33 | 0.21 | 0.42 | 0.37 |
| 1 | 211163369 | rs6680997 | 0.081 | -1.158 | -0.889 | -0.352 | 0 | 0.01 | 0.01 | 0 | 0.18 | 0.15 | 0.26 | 0.38 | 0.11 | 0.17 |
| 1 | 211176922 | rs1501551 | 0.02 | -1.191 | -1.31 | -0.77 | -0.012 | 0.036 | 0.036 | 0 | 0.44 | 0.31 | 0.45 | 0.36 | 0.48 | 0.58 |
| 1 | 211178828 | rs11806852 | 0.421 | -1.212 | -1.308 | -0.761 | 0.004 | -0.007 | -0.007 | 0 | 0.18 | 0.12 | 0.25 | 0.38 | 0.11 | 0.17 |
| 1 | 211185917 | rs7526579 | -0.669 | -1.182 | -1.288 | -0.769 | 0.032 | -0.058 | -0.058 | 0 | 0.29 | 0.17 | 0.38 | 0.46 | 0.19 | 0.35 |
| 1 | 211187663 | rs924568 | -1.465 | -1.191 | -1.242 | -0.908 | -0.073 | 0.028 | 0.028 | 0 | 0.68 | 0.91 | 0.74 | 0.74 | 0.61 | 0.76 |
| 1 | 211187730 | rs10494931 | 0.738 | -1.084 | -0.905 | -0.872 | -0.002 | -0.015 | -0.015 | 0.675 | 0.32 | 0.65 | 0.32 | 0.09 | 0.37 | 0.29 |
| 1 | 211187783 | rs17188797 | 1.764 | -0.747 | -1.444 | -1.274 | -0.067 | 0.027 | 0.027 | 0 | 0.14 | 0.01 | 0.19 | 0.08 | 0.19 | 0.11 |
| 1 | 211188937 | rs12760791 | 0.704 | -0.812 | -0.954 | -1.14 | 0.024 | -0.068 | -0.068 | 0 | 0.28 | 0.39 | 0.3 | 0.09 | 0.35 | 0.28 |
| 1 | 211191894 | rs11582507 | 0.819 | -0.788 | -1.084 | -1.128 | 0.004 | -0.025 | -0.025 | 0 | 0.27 | 0.28 | 0.29 | 0.09 | 0.35 | 0.28 |
| 1 | 211201592 | rs11119666 | -0.622 | -0.903 | -1.348 | -1.007 | 0.033 | 0.002 | 0.002 | 0 | 0.32 | 0.22 | 0.39 | 0.65 | 0.2 | 0.42 |
| 1 | 211202705 | rs12134819 | 0.301 | -0.882 | -1.55 | -0.833 | 0.034 | -0.042 | -0.042 | 4.14 | 0.14 | 0.07 | 0.13 | 0.28 | 0.09 | 0.25 |
| 1 | 211203217 | rs4951491 | -1.984 | -0.854 | -1.586 | -1.005 | -0.036 | 0.073 | 0.073 | 0 | 0.74 | 0.88 | 0.84 | 0.9 | 0.68 | 0.85 |
| 1 | 211208401 | rs6540647 | -0.66 | -0.709 | -1.48 | -0.927 | -0.013 | -0.036 | -0.036 | 0 | 0.54 | 0.79 | 0.6 | 0.47 | 0.52 | 0.5 |
| 1 | 211218816 | rs12124026 | -0.162 | -0.083 | -0.889 | -0.417 | 0.032 | 0.154 | 0.154 | 0 | 0.79 | 0.63 | 0.67 | 0.52 | 0.79 | 0.83 |
| 1 | 211222788 | rs12122529 | 0.116 | -0.157 | -1.177 | -0.725 | 0.016 | -0.01 | -0.01 | 0 | 0.46 | 0.24 | 0.49 | 0.42 | 0.45 | 0.59 |
| 1 | 211230021 | rs6540652 | 0.327 | -0.075 | -1.221 | -0.692 | -0.003 | 0.022 | 0.022 | 0 | 0.39 | 0.24 | 0.4 | 0.3 | 0.4 | 0.47 |
| 1 | 211255524 | rs17188839 | 1.646 | 0.06 | -1.224 | -0.827 | -0.016 | 0.066 | 0.066 | 0 | 0.23 | 0.13 | 0.12 | 0.09 | 0.25 | 0.15 |
| 1 | 211255936 | rs4620600 | -0.452 | -0.002 | -1.117 | -0.81 | 0.012 | 0.133 | 0.133 | 0 | 0.77 | 0.61 | 0.69 | 0.52 | 0.8 | 0.78 |
| 1 | 211258161 | rs946143 | 0.23 | -0.076 | -1.178 | -0.853 | 0.002 | 0.033 | 0.033 | 0.162 | 0.44 | 0.26 | 0.43 | 0.32 | 0.46 | 0.46 |

| | | | | | | | | | | | | | | | | |
|---|---|---|---|---|---|---|---|---|---|---|---|---|---|---|---|---|
| 1 | 211263253 | rs9970961 | 0.492 | -0.003 | -1.218 | -0.846 | -0.001 | 0.012 | 0.012 | 0 | 0.34 | 0.32 | 0.27 | 0.21 | 0.36 | 0.32 |
| 1 | 211263614 | rs3828159 | 2.2 | 0.266 | -1.636 | -1.204 | -0.018 | 0.008 | 0.008 | 0 | 0.1 | 0.03 | 0.05 | 0.09 | 0.11 | 0.06 |
| 1 | 211264402 | rs4951502 | -0.79 | 0.084 | -1.228 | -0.378 | 0.01 | 0.077 | 0.077 | 0 | 0.77 | 0.65 | 0.69 | 0.52 | 0.79 | 0.8 |
| 1 | 211265284 | rs1338348 | 1.223 | 0.103 | -1.404 | -0.401 | -0.027 | -0.039 | -0.039 | 0 | 0.09 | 0 | 0.13 | 0.11 | 0.11 | 0.17 |
| 1 | 211267168 | rs12735839 | 1.046 | 0.065 | -1.174 | -0.58 | -0.001 | 0.008 | 0.008 | 0 | 0.24 | 0.21 | 0.13 | 0.09 | 0.24 | 0.16 |
| 1 | 211276828 | rs3762452 | 1.131 | 0.038 | -1.232 | -0.561 | -0.008 | 0.05 | 0.05 | 0 | 0.24 | 0.13 | 0.12 | 0.09 | 0.24 | 0.14 |
| 1 | 211279739 | rs4578265 | 0.251 | 0.042 | -0.861 | -0.263 | 0.008 | 0.03 | 0.03 | 0 | 0.67 | 0.56 | 0.54 | 0.41 | 0.67 | 0.61 |
| 1 | 211282681 | rs6660415 | 0.346 | 0.149 | -0.884 | -0.575 | -0.029 | -0.072 | -0.072 | 0.37 | 0.25 | 0.42 | 0.25 | 0.1 | 0.17 | 0.23 |
| 1 | 211288097 | rs4512717 | 1.253 | 0.389 | -0.744 | -0.525 | 0 | 0.011 | 0.011 | 0 | 0.4 | 0.3 | 0.22 | 0.24 | 0.42 | 0.28 |
| 1 | 211296202 | rs6540658 | 1.411 | 0.315 | -0.753 | -0.611 | -0.013 | -0.049 | -0.049 | 0 | 0.21 | 0.28 | 0.17 | 0.06 | 0.17 | 0.18 |
| 1 | 211308116 | rs3811395 | 1.375 | 0.331 | -0.959 | -0.517 | -0.001 | 0.032 | 0.032 | 0 | 0.39 | 0.22 | 0.22 | 0.24 | 0.42 | 0.27 |
| 1 | 211311215 | rs2185069 | 1.603 | 0.467 | -1.16 | -0.063 | -0.031 | -0.059 | -0.059 | 6.188 | 0.1 | 0.01 | 0.14 | 0.16 | 0.12 | 0.21 |
| 1 | 211321345 | rs17188895 | 2.156 | 0.327 | -1.191 | -0.252 | 0.006 | 0.054 | 0.054 | 0 | 0.16 | 0 | 0.12 | 0.06 | 0.09 | 0.16 |
| 1 | 211331893 | rs6540664 | 0.821 | 0.481 | -1.337 | -0.168 | -0.001 | 0.037 | 0.037 | 0 | 0.47 | 0.37 | 0.33 | 0.28 | 0.5 | 0.37 |
| 1 | 211339584 | rs9430058 | 1.002 | 0.425 | -1.25 | -0.207 | -0.033 | -0.105 | -0.105 | 0.086 | 0.23 | 0.43 | 0.19 | 0.07 | 0.16 | 0.19 |
| 1 | 211344081 | rs12036302 | 1.654 | 0.448 | -1.274 | -0.269 | -0.047 | 0.034 | 0.034 | 0 | 0.37 | 0.07 | 0.2 | 0.25 | 0.41 | 0.27 |
| 1 | 211351513 | rs12134651 | 1.903 | 0.571 | -1.328 | -0.211 | -0.005 | 0.015 | 0.015 | 0 | 0.08 | 0.05 | 0.11 | 0.02 | 0.08 | 0.1 |
| 1 | 211351594 | rs1572650 | -1.164 | 0.451 | -1.175 | 0.037 | -0.024 | -0.047 | -0.047 | 0 | 0.77 | 0.85 | 0.63 | 0.51 | 0.73 | 0.73 |
| 1 | 211352625 | rs1338356 | 0.163 | 0.106 | -1.024 | 0.019 | -0.03 | -0.003 | -0.003 | 0 | 0.29 | 0.45 | 0.3 | 0.23 | 0.23 | 0.36 |
| 1 | 211357154 | rs1572648 | -1.186 | 0.63 | -1.671 | 0.086 | -0.031 | 0.036 | 0.036 | 0.529 | 0.82 | 0.93 | 0.78 | 0.9 | 0.8 | 0.79 |
| 1 | 211358608 | rs4347267 | 1.958 | 0.539 | -1.624 | 0.231 | -0.013 | -0.027 | -0.027 | 0 | 0.06 | 0.01 | 0.1 | 0.15 | 0.06 | 0.16 |
| 1 | 211358755 | rs1538291 | 1.811 | 0.562 | -2.169 | -0.178 | -0.085 | 0.121 | 0.121 | 0 | 0.25 | 0.05 | 0.16 | 0.03 | 0.34 | 0.2 |
| 1 | 211363628 | rs3111229 | -0.118 | -0.159 | -0.8 | 0.23 | -0.033 | 0.027 | 0.027 | 0 | 0.3 | 0.46 | 0.3 | 0.45 | 0.19 | 0.23 |
| 1 | 211372063 | rs1538286 | -0.612 | -0.029 | -1.436 | -0.16 | 0.043 | -0.018 | -0.018 | 0 | 0.39 | 0.74 | 0.42 | 0.6 | 0.26 | 0.4 |
| 1 | 211383129 | rs1361193 | -0.876 | 0.107 | -1.181 | -0.686 | 0.059 | 0.074 | 0.074 | 0.289 | 0.43 | 0.84 | 0.39 | 0.36 | 0.34 | 0.42 |
| 1 | 211408176 | rs11586573 | 0.557 | 0.199 | -0.955 | -0.763 | -0.015 | 0.018 | 0.018 | 0 | 0.12 | 0 | 0.11 | 0.07 | 0.07 | 0.14 |
| 1 | 211425658 | rs11811405 | 1.072 | 0.471 | -0.82 | -0.843 | 0.022 | 0.087 | 0.087 | 0 | 0.5 | 0.22 | 0.56 | 0.62 | 0.55 | 0.45 |
| 1 | 211432029 | rs12118279 | 0.939 | 0.453 | -0.851 | -0.852 | 0.006 | 0.098 | 0.098 | 0 | 0.5 | 0.13 | 0.56 | 0.62 | 0.55 | 0.45 |
| 1 | 211433672 | rs6679980 | 0.759 | 0.245 | -0.231 | -0.93 | -0.001 | -0.012 | -0.012 | 0.016 | 0.2 | 0.23 | 0.09 | 0.04 | 0.16 | 0.22 |
| 1 | 211447625 | rs11580084 | 1.369 | 0.27 | -0.268 | -0.919 | 0.031 | 0.087 | 0.087 | 0 | 0.52 | 0.15 | 0.58 | 0.62 | 0.6 | 0.47 |
| 1 | 211451467 | rs11577985 | -0.066 | 0.352 | 0.241 | -0.863 | -0.006 | -0.057 | -0.057 | 0.022 | 0.18 | 0.3 | 0.16 | 0.07 | 0.14 | 0.15 |
| 1 | 211491130 | rs17267540 | 1.365 | 0.386 | 0.184 | -0.817 | -0.016 | 0.006 | 0.006 | 0 | 0.07 | 0.03 | 0.03 | 0.03 | 0.06 | 0.1 |
| 1 | 211491152 | rs6540679 | -0.4 | 0.349 | 0.193 | -0.806 | 0.002 | 0.048 | 0.048 | 0 | 0.9 | 0.81 | 0.84 | 0.73 | 0.9 | 0.84 |
| 1 | 211491284 | rs17017394 | -0.077 | 0.386 | 0.338 | -0.885 | -0.006 | -0.057 | -0.057 | 0.139 | 0.18 | 0.3 | 0.16 | 0.07 | 0.14 | 0.15 |
| 1 | 211493573 | rs6540681 | -0.945 | 0.414 | 0.096 | -0.856 | 0.004 | 0.011 | 0.011 | 0.415 | 0.8 | 0.81 | 0.9 | 0.95 | 0.81 | 0.77 |
| 1 | 211529569 | rs7550702 | 1.324 | 0.392 | 0.234 | -0.882 | 0.024 | 0.061 | 0.061 | 0 | 0.53 | 0.2 | 0.58 | 0.62 | 0.6 | 0.49 |
| 1 | 211530480 | rs12752670 | 1.002 | 0.364 | 0.232 | -0.914 | 0.007 | 0.028 | 0.028 | 0.235 | 0.18 | 0.11 | 0.07 | 0.05 | 0.15 | 0.21 |
| 1 | 211531329 | rs11119729 | 1.5 | 0.842 | -0.221 | -1.423 | -0.023 | 0.031 | 0.031 | 0 | 0.18 | 0.09 | 0.18 | 0.01 | 0.21 | 0.17 |
| 1 | 211533027 | rs2292483 | 1.215 | 0.537 | 0.391 | -0.776 | 0.002 | 0 | 0 | 0 | 0.71 | 0.28 | 0.65 | 0.66 | 0.75 | 0.7 |
| 1 | 211548387 | rs3738200 | 0.894 | 0.016 | 0.679 | -0.609 | -0.01 | 0.068 | 0.068 | 0.214 | 0.18 | 0.07 | 0.06 | 0.05 | 0.17 | 0.22 |
| 1 | 211557908 | rs17017451 | 0.08 | 0.106 | 1.088 | -0.391 | 0 | -0.02 | -0.02 | 0 | 0.1 | 0.24 | 0.14 | 0.06 | 0.1 | 0.07 |
| 1 | 211570581 | rs34418448 | 0.779 | 0.113 | 0.755 | -0.401 | -0.023 | 0.106 | 0.106 | 0.151 | 0.24 | 0.07 | 0.08 | 0.05 | 0.21 | 0.27 |
| 1 | 211591687 | rs12739630 | 0.823 | 0.105 | 0.596 | -0.405 | -0.043 | 0.108 | 0.108 | 0.108 | 0.23 | 0.05 | 0.08 | 0.05 | 0.21 | 0.27 |
| 1 | 211603026 | rs12733930 | 0.835 | 0.104 | 0.302 | -0.406 | -0.052 | 0.102 | 0.102 | 0.104 | 0.23 | 0.03 | 0.08 | 0.05 | 0.21 | 0.27 |
| 1 | 211605697 | rs1055565 | -0.897 | 0.083 | 0.146 | -0.409 | 0.007 | 0.018 | 0.018 | 0 | 0.42 | 0.37 | 0.35 | 0.38 | 0.37 | 0.47 |
| 1 | 211614280 | rs11580728 | 0.646 | 0.231 | -0.304 | -0.353 | 0.005 | 0.024 | 0.024 | 0.284 | 0.26 | 0.21 | 0.1 | 0.05 | 0.21 | 0.27 |
| 1 | 211621170 | rs1663517 | 0.657 | 0.471 | -0.875 | -0.08 | 0.022 | 0.076 | 0.076 | 0 | 0.52 | 0.14 | 0.59 | 0.62 | 0.59 | 0.51 |
| 1 | 211641950 | rs1419090 | -0.683 | 0.462 | -0.071 | 0.411 | 0 | -0.021 | -0.021 | 0 | 0.1 | 0.24 | 0.14 | 0.06 | 0.1 | 0.07 |
| 1 | 211645082 | rs1361031 | -0.826 | 0.563 | 0.139 | 0.874 | 0.053 | 0.22 | 0.22 | 0 | 0.11 | 0.36 | 0.24 | 0.32 | 0.12 | 0.14 |
| 1 | 211649327 | rs12130155 | 0.16 | 0.275 | -0.722 | 1.083 | -0.009 | -0.11 | -0.11 | 0 | 0.14 | 0.02 | 0.23 | 0.48 | 0.14 | 0.16 |
| 1 | 211657371 | rs17189035 | 1.003 | 0.139 | -0.779 | 0.692 | -0.003 | 0.01 | 0.01 | 0 | 0.06 | 0 | 0.03 | 0.04 | 0.05 | 0.07 |
| 1 | 211662836 | rs10863897 | 0.535 | 0.137 | -0.824 | 1.489 | -0.02 | -0.076 | -0.076 | 0 | 0.13 | 0.02 | 0.15 | 0.27 | 0.13 | 0.16 |
| 1 | 211663221 | rs11119751 | -0.175 | 0.04 | -0.488 | 1.006 | 0.055 | 0.251 | 0.251 | 0 | 0.08 | 0.35 | 0.16 | 0.27 | 0.08 | 0.1 |
| 1 | 211663632 | rs1530436 | 0.527 | 0.028 | -0.535 | 1.132 | -0.018 | -0.093 | -0.093 | 0 | 0.13 | 0.02 | 0.23 | 0.44 | 0.13 | 0.16 |
| 1 | 211663679 | rs17017564 | -0.094 | -0.124 | -0.308 | 1.041 | -0.003 | -0.001 | -0.001 | 0.062 | 0.11 | 0.13 | 0.11 | 0.11 | 0.09 | 0.11 |
| 1 | 211666218 | rs10863899 | -0.36 | 0.121 | -0.897 | 0.106 | 0 | -0.002 | -0.002 | 0.836 | 0.43 | 0.67 | 0.37 | 0.44 | 0.43 | 0.4 |
| 1 | 211668437 | rs11587632 | 1.022 | 0.09 | -0.876 | 0.084 | -0.002 | 0.001 | 0.001 | 0.209 | 0.28 | 0.27 | 0.18 | 0.11 | 0.3 | 0.23 |
| 1 | 211671118 | rs1578838 | -1.384 | 0.116 | -0.88 | 0.081 | -0.014 | 0.114 | 0.114 | 0 | 0.61 | 0.82 | 0.68 | 0.93 | 0.61 | 0.6 |
| 1 | 211672506 | rs755448 | -0.283 | -0.029 | -0.79 | 0.316 | -0.002 | 0.138 | 0.138 | 0 | 0.14 | 0.4 | 0.18 | 0.33 | 0.13 | 0.15 |
| 1 | 211675125 | rs894984 | -2.153 | 0.039 | -1.449 | -0.121 | -0.021 | 0.077 | 0.077 | 0 | 0.82 | 0.99 | 0.88 | 0.96 | 0.84 | 0.85 |
| 1 | 211676378 | rs12403299 | -0.299 | 0.082 | -0.868 | 0.466 | 0 | -0.012 | -0.012 | 0.262 | 0.17 | 0.15 | 0.31 | 0.5 | 0.18 | 0.2 |
| 1 | 211686020 | rs753647 | -0.322 | 0.096 | -0.876 | 0.437 | 0 | -0.013 | -0.013 | 0 | 0.17 | 0.14 | 0.32 | 0.51 | 0.18 | 0.2 |
| 1 | 211686121 | rs753646 | -0.738 | 0.167 | -1.003 | -0.013 | 0.004 | 0.019 | 0.019 | 0 | 0.52 | 0.44 | 0.6 | 0.88 | 0.55 | 0.57 |
| 1 | 211686860 | rs10863911 | -0.008 | 0.206 | -0.991 | 0.014 | -0.013 | -0.019 | -0.019 | 0.579 | 0.37 | 0.32 | 0.33 | 0.44 | 0.39 | 0.42 |
| 1 | 211690631 | rs4951758 | -1.321 | 0.208 | -0.951 | -0.161 | -0.015 | 0.082 | 0.082 | 0 | 0.63 | 0.79 | 0.68 | 0.94 | 0.62 | 0.66 |
| 1 | 211693071 | rs1020334 | 0.409 | 0.014 | -0.566 | 0.688 | -0.031 | -0.078 | -0.078 | 0.024 | 0.11 | 0.01 | 0.23 | 0.41 | 0.13 | 0.16 |
| 1 | 211694012 | rs4951761 | -0.992 | -0.115 | -0.455 | -0.296 | -0.001 | -0.004 | -0.004 | 0 | 0.27 | 0.33 | 0.46 | 0.86 | 0.27 | 0.36 |
| 1 | 211694508 | rs17017668 | -0.435 | -0.057 | -0.321 | 0.251 | -0.006 | -0.014 | -0.014 | 0.307 | 0.1 | 0.11 | 0.07 | 0.06 | 0.1 | 0.1 |
| 1 | 211705317 | rs7525277 | -0.992 | -0.235 | 0.006 | 0.735 | -0.042 | 0.207 | 0.207 | 1.918 | 0.16 | 0.51 | 0.31 | 0.59 | 0.11 | 0.2 |

| | | | | | | | | | | | | | | | | |
|---|---|---|---|---|---|---|---|---|---|---|---|---|---|---|---|---|
| 1 | 211710139 | rs17695731 | 0.332 | -0.251 | -0.381 | 0.57 | -0.01 | 0.029 | 0.029 | 0 | 0.07 | 0 | 0.06 | 0.03 | 0.05 | 0.1 |
| 1 | 211711946 | rs12033819 | -0.184 | 0.13 | -0.611 | 0.278 | -0.016 | -0.03 | -0.03 | 0 | 0.13 | 0.07 | 0.17 | 0.28 | 0.16 | 0.17 |
| 1 | 211712690 | rs6679498 | -1.709 | -0.36 | -0.202 | 0.523 | -0.043 | 0.045 | 0.045 | 0.399 | 0.28 | 0.63 | 0.39 | 0.6 | 0.18 | 0.39 |
| 1 | 211712719 | rs4951768 | -1.141 | -0.104 | -0.095 | 0.797 | -0.032 | 0.104 | 0.104 | 0 | 0.21 | 0.42 | 0.37 | 0.46 | 0.16 | 0.28 |
| 1 | 211722446 | rs11119770 | 0.264 | 0.151 | -0.598 | 0.898 | 0.002 | 0.038 | 0.038 | 0 | 0.07 | 0.03 | 0.29 | 0.41 | 0.07 | 0.12 |
| 1 | 211726975 | rs9430069 | 0.809 | 0.155 | -0.756 | -0.105 | -0.005 | -0.04 | -0.04 | 0 | 0.34 | 0.38 | 0.17 | 0.09 | 0.25 | 0.11 |
| 1 | 211730036 | rs2165993 | 0.022 | 1.021 | -1.143 | 0.135 | -0.033 | -0.04 | -0.04 | 0 | 0.68 | 0.83 | 0.51 | 0.54 | 0.53 | 0.57 |
| 1 | 211736031 | rs12026206 | 0.013 | 0.706 | -0.586 | 1.159 | -0.023 | -0.047 | -0.047 | 0 | 0.08 | 0 | 0.08 | 0.26 | 0.08 | 0.14 |
| 1 | 211736470 | rs7530862 | -0.904 | 0.486 | -0.159 | 1.014 | -0.014 | 0.011 | 0.011 | 0 | 0.17 | 0.23 | 0.11 | 0.19 | 0.12 | 0.22 |
| 1 | 211737647 | rs9430071 | -0.131 | 0.633 | 0.078 | 1.126 | -0.005 | 0.174 | 0.174 | 0 | 0.08 | 0.09 | 0.31 | 0.42 | 0.07 | 0.16 |
| 1 | 211744536 | rs17017761 | -0.406 | 0.728 | 0.156 | 0.36 | -0.004 | 0.065 | 0.065 | 0 | 0.08 | 0.22 | 0.06 | 0.14 | 0.06 | 0.14 |
| 1 | 211750362 | rs12562288 | -0.377 | 0.732 | 0.158 | 0.362 | -0.003 | 0.066 | 0.066 | 0 | 0.08 | 0.22 | 0.06 | 0.14 | 0.06 | 0.14 |
| 1 | 211754866 | rs7546852 | 0.419 | 1.348 | -0.06 | 0.257 | 0.026 | -0.019 | -0.019 | 0 | 0.93 | 0.91 | 0.93 | 0.97 | 0.84 | 0.9 |
| 1 | 211755204 | rs12030478 | -0.83 | 1.014 | 0.172 | 0.475 | 0.016 | 0.046 | 0.046 | 0 | 0.12 | 0.3 | 0.12 | 0.21 | 0.1 | 0.14 |
| 1 | 211763053 | rs17017782 | -0.455 | 1.01 | -1.058 | 0.229 | -0.017 | -0.045 | -0.045 | 0 | 0.09 | 0.01 | 0.05 | 0.14 | 0.07 | 0.09 |
| 1 | 211772031 | rs6692675 | 0.809 | 1.165 | -1.695 | -0.938 | -0.026 | 0.211 | 0.211 | 0.014 | 0.52 | 0.12 | 0.64 | 0.86 | 0.46 | 0.55 |
| 1 | 211774350 | rs6702177 | -0.175 | 1.555 | -0.686 | -1.326 | 0.03 | -0.119 | -0.119 | 0.356 | 0.18 | 0.42 | 0.13 | 0.03 | 0.16 | 0.18 |
| 1 | 211775742 | rs7527059 | -0.303 | 1.827 | -0.554 | -1.422 | -0.005 | 0.001 | 0.001 | 0 | 0.11 | 0.12 | 0.13 | 0.07 | 0.19 | 0.12 |
| 1 | 211776439 | rs4951536 | 1.062 | 1.634 | -0.246 | -1.463 | -0.002 | 0.081 | 0.081 | 0 | 0.64 | 0.24 | 0.76 | 0.93 | 0.65 | 0.66 |
| 1 | 211785039 | rs7546959 | 1.656 | 1.443 | -0.503 | -1.273 | -0.029 | 0.113 | 0.113 | 0.065 | 0.55 | 0.15 | 0.67 | 0.76 | 0.48 | 0.53 |
| 1 | 211785065 | rs7525247 | 1.236 | 1.32 | -0.468 | -1.512 | -0.047 | 0.079 | 0.079 | 0.646 | 0.64 | 0.24 | 0.75 | 0.92 | 0.54 | 0.57 |
| 1 | 211797879 | rs4951537 | 1.875 | 0.961 | -0.289 | -1.75 | 0.019 | 0.151 | 0.151 | 0.033 | 0.52 | 0.22 | 0.68 | 0.77 | 0.46 | 0.49 |
| 1 | 211797917 | rs12751701 | -0.988 | 0.949 | -0.303 | -1.855 | -0.085 | 0.087 | 0.087 | 0 | 0.28 | 0.13 | 0.19 | 0.04 | 0.45 | 0.26 |
| 1 | 211798430 | rs12736496 | -1.297 | 0.932 | 0.084 | -1.864 | -0.019 | -0.013 | -0.013 | 0.547 | 0.32 | 0.31 | 0.22 | 0.04 | 0.46 | 0.3 |
| 1 | 211803723 | rs11119778 | -1.9 | 0.92 | 0.201 | -1.868 | 0.004 | -0.034 | -0.034 | 0.073 | 0.41 | 0.49 | 0.29 | 0.19 | 0.48 | 0.4 |
| 1 | 211804676 | rs7523748 | 1.938 | 0.634 | -0.026 | -2.12 | 0.029 | 0.177 | 0.177 | 0.003 | 0.47 | 0.15 | 0.62 | 0.78 | 0.42 | 0.48 |
| 1 | 211805625 | rs4951420 | 0.719 | 0.595 | 0.164 | -2.121 | -0.02 | -0.041 | -0.041 | 0.903 | 0.87 | 0.54 | 0.93 | 0.96 | 0.93 | 0.79 |
| 1 | 211806784 | rs6660850 | 1.111 | 0.594 | 0.005 | -2.195 | -0.012 | 0.024 | 0.024 | 0 | 0.61 | 0.32 | 0.67 | 0.81 | 0.51 | 0.68 |
| 1 | 211809647 | rs6670986 | 1.876 | 0.589 | 0.011 | -2.129 | 0.024 | 0.171 | 0.171 | 0 | 0.47 | 0.16 | 0.62 | 0.78 | 0.41 | 0.47 |
| 1 | 211813010 | rs4951541 | 0.374 | 0.62 | 0.093 | -2.165 | -0.007 | 0.093 | 0.093 | 0 | 0.89 | 0.55 | 0.9 | 0.86 | 0.96 | 0.91 |
| 1 | 211815287 | rs1373293 | -1.505 | 0.813 | 0.012 | -2.119 | -0.002 | -0.014 | -0.014 | 0 | 0.42 | 0.38 | 0.27 | 0.08 | 0.54 | 0.44 |
| 1 | 211817646 | rs11576789 | -1.403 | 0.806 | 0.146 | -2.108 | -0.009 | 0.034 | 0.034 | 0 | 0.41 | 0.31 | 0.27 | 0.08 | 0.54 | 0.44 |
| 1 | 211824797 | rs7553718 | -0.919 | 0.331 | -0.361 | -2.301 | -0.003 | 0.057 | 0.057 | 0 | 0.43 | 0.35 | 0.33 | 0.19 | 0.51 | 0.29 |
| 1 | 211826546 | rs1530400 | -1.362 | 0.227 | -0.283 | -2.384 | 0.045 | 0.171 | 0.171 | 0 | 0.51 | 0.8 | 0.39 | 0.19 | 0.58 | 0.35 |
| 1 | 211828134 | rs7516189 | -0.537 | 0.386 | -0.161 | -2.344 | -0.038 | 0.055 | 0.055 | 0 | 0.36 | 0.25 | 0.27 | 0.03 | 0.48 | 0.25 |
| 1 | 211831380 | rs2028706 | 0.18 | -0.234 | 0.068 | -1.889 | -0.011 | 0.013 | 0.013 | 0 | 0.24 | 0.39 | 0.35 | 0.26 | 0.23 | 0.32 |
| 1 | 211832827 | rs6695691 | 0.454 | 0.286 | -0.021 | -2.121 | -0.046 | 0.05 | 0.05 | 0 | 0.26 | 0.15 | 0.19 | 0 | 0.34 | 0.14 |
| 1 | 211835786 | rs894852 | -2.342 | 0.053 | -0.144 | -2.001 | 0.032 | -0.056 | -0.056 | 0 | 0.7 | 0.85 | 0.69 | 0.54 | 0.76 | 0.68 |
| 1 | 211837911 | rs10158205 | 1.452 | 0.044 | -0.216 | -2.044 | 0.001 | 0.041 | 0.041 | 0 | 0.12 | 0.17 | 0.21 | 0.22 | 0.13 | 0.13 |
| 1 | 211842808 | rs697004 | -0.823 | 0.072 | -0.499 | -2.071 | -0.062 | 0.082 | 0.082 | 0 | 0.74 | 0.87 | 0.81 | 1 | 0.67 | 0.87 |
| 1 | 211845859 | rs11119790 | 0.919 | 0.065 | -0.068 | -1.398 | 0.007 | -0.047 | -0.047 | 0 | 0.08 | 0 | 0.05 | 0.24 | 0.04 | 0.08 |
| 1 | 211856601 | rs10494940 | 0.605 | 0.21 | -0.242 | -1.409 | -0.009 | -0.026 | -0.026 | 0.008 | 0.12 | 0.22 | 0.15 | 0.04 | 0.1 | 0.19 |
| 1 | 211859913 | rs7516751 | 0.83 | 0.487 | -0.288 | -1.298 | -0.001 | 0.024 | 0.024 | 0 | 0.12 | 0.14 | 0.2 | 0.23 | 0.13 | 0.13 |
| 1 | 211859925 | rs6661565 | 0.224 | 0.62 | -0.433 | -1.38 | 0.01 | -0.029 | -0.029 | 0.02 | 0.1 | 0.17 | 0.08 | 0.03 | 0.15 | 0.14 |
| 1 | 211870825 | rs7513065 | 1.037 | 0.2 | -0.716 | -1.018 | -0.016 | 0.027 | 0.027 | 0.089 | 0.1 | 0.05 | 0.05 | 0 | 0.11 | 0.05 |
| 1 | 211873190 | rs10494941 | -0.895 | -0.229 | -0.244 | -0.569 | -0.03 | -0.003 | -0.003 | 0 | 0.22 | 0.38 | 0.21 | 0.27 | 0.14 | 0.23 |
| 1 | 211877481 | rs12096414 | -0.792 | -0.342 | -0.314 | -0.243 | -0.036 | 0.007 | 0.007 | 0 | 0.15 | 0.24 | 0.08 | 0.24 | 0.05 | 0.09 |
| 1 | 211886246 | rs12569332 | -0.084 | 0.291 | -0.538 | 0.054 | -0.002 | 0.014 | 0.014 | 0 | 0.13 | 0.17 | 0.2 | 0.22 | 0.13 | 0.13 |
| 1 | 211887918 | rs7533652 | -0.878 | 0.477 | -0.953 | 0.074 | -0.003 | -0.004 | -0.004 | 0.001 | 0.32 | 0.26 | 0.35 | 0.45 | 0.33 | 0.23 |
| 1 | 211892568 | rs17494100 | 0.482 | 0.033 | -1.394 | 0.689 | 0.019 | -0.103 | -0.103 | 0 | 0.14 | 0.01 | 0.16 | 0.37 | 0.09 | 0.16 |
| 1 | 211893655 | rs7512315 | -0.072 | 0.525 | -0.793 | 0.398 | -0.002 | -0.013 | -0.013 | 0 | 0.23 | 0.28 | 0.18 | 0.22 | 0.2 | 0.14 |
| 1 | 211898313 | rs1773353 | 0.316 | 1.192 | -0.844 | 0.293 | 0.003 | 0 | 0 | 0 | 0.76 | 0.71 | 0.72 | 0.75 | 0.68 | 0.75 |
| 1 | 211905584 | rs701911 | -0.604 | 0.794 | -0.831 | -0.179 | -0.002 | 0 | 0 | 0.1 | 0.35 | 0.36 | 0.47 | 0.67 | 0.31 | 0.28 |
| 1 | 211909230 | rs1157168 | 0.781 | 0.037 | -1.683 | -1.224 | -0.11 | 0.248 | 0.248 | 0 | 0.4 | 0.09 | 0.26 | 0.07 | 0.53 | 0.42 |
| 1 | 211914587 | rs3862952 | 1.12 | 0.158 | -1.625 | -1.509 | -0.037 | 0.17 | 0.17 | 0 | 0.36 | 0.12 | 0.22 | 0.06 | 0.41 | 0.33 |
| 1 | 211916142 | rs10494942 | 0.314 | 0.133 | -1.869 | -1.678 | -0.022 | 0.002 | 0.002 | 0 | 0.1 | 0.02 | 0.13 | 0.04 | 0.14 | 0.11 |
| 1 | 211919937 | rs3795834 | -0.18 | -0.091 | -1.145 | -1.641 | -0.008 | -0.021 | -0.021 | 0.121 | 0.37 | 0.54 | 0.45 | 0.75 | 0.27 | 0.4 |
| 1 | 211920427 | rs12566620 | 1.09 | -0.157 | -1.139 | -1.677 | 0.007 | -0.046 | -0.046 | 0 | 0.1 | 0.01 | 0.16 | 0.14 | 0.1 | 0.11 |
| 1 | 211920624 | rs10465691 | 0.32 | -0.231 | -1.075 | -1.666 | 0.001 | 0.003 | 0.003 | 0 | 0.32 | 0.5 | 0.43 | 0.71 | 0.24 | 0.29 |
| 1 | 211921229 | rs11119806 | 0.225 | -0.234 | -1.092 | -1.668 | -0.001 | 0.002 | 0.002 | 0 | 0.32 | 0.54 | 0.44 | 0.71 | 0.24 | 0.29 |
| 1 | 211946375 | rs4951426 | 0.381 | -0.253 | -1.168 | -1.634 | 0.014 | -0.036 | -0.036 | 0.175 | 0.81 | 0.96 | 0.82 | 0.82 | 0.83 | 0.74 |
| 1 | 211955125 | rs11119810 | 0.05 | -0.263 | -1.153 | -1.615 | 0.023 | -0.05 | -0.05 | 0.046 | 0.15 | 0.02 | 0.17 | 0.18 | 0.13 | 0.22 |
| 1 | 211972775 | rs1065607 | 1.166 | -0.264 | -0.728 | -1.614 | 0.007 | -0.046 | -0.046 | 0 | 0.1 | 0.01 | 0.16 | 0.14 | 0.1 | 0.11 |
| 1 | 211986052 | rs4951553 | -0.344 | -0.276 | -0.61 | -1.701 | -0.006 | -0.013 | -0.013 | 1.026 | 0.37 | 0.58 | 0.45 | 0.75 | 0.27 | 0.4 |
| 1 | 211994671 | rs12038038 | -0.174 | -0.197 | -0.583 | -1.692 | 0.003 | -0.013 | -0.013 | 0 | 0.37 | 0.55 | 0.52 | 0.71 | 0.31 | 0.35 |
| 1 | 212010527 | rs4364936 | 1.076 | -0.199 | -0.722 | -1.665 | 0.009 | -0.046 | -0.046 | 0.23 | 0.11 | 0.01 | 0.16 | 0.14 | 0.1 | 0.11 |
| 1 | 212027436 | rs28668719 | 0.557 | -0.239 | -0.714 | -1.879 | -0.006 | 0.008 | 0.008 | 0 | 0.91 | 0.94 | 0.95 | 0.97 | 0.89 | 0.92 |
| 1 | 212042686 | rs28688444 | -0.847 | -0.178 | -0.56 | -2.086 | 0 | 0.099 | 0.099 | 0.094 | 0.45 | 0.8 | 0.57 | 0.77 | 0.37 | 0.47 |
| 1 | 212046710 | rs11119820 | -1.517 | -0.145 | -0.579 | -2.096 | -0.033 | 0.166 | 0.166 | 0 | 0.6 | 0.84 | 0.77 | 0.94 | 0.54 | 0.62 |

| | | | | | | | | | | | | | | | | |
|---|---|---|---|---|---|---|---|---|---|---|---|---|---|---|---|---|
| 1 | 212079778 | rs7540019 | 1.342 | -0.071 | -0.658 | -2.024 | -0.066 | 0.217 | 0.217 | 0 | 0.4 | 0.09 | 0.23 | 0.09 | 0.46 | 0.38 |
| 1 | 212080831 | rs2970582 | -0.081 | -0.111 | -0.317 | -1.981 | 0.004 | -0.01 | -0.01 | 0 | 0.63 | 0.53 | 0.47 | 0.27 | 0.66 | 0.66 |
| 1 | 212087682 | rs7550801 | 0.718 | -0.127 | -0.16 | -1.941 | 0.058 | -0.021 | -0.021 | 0.049 | 0.06 | 0.19 | 0.12 | 0.01 | 0.11 | 0.04 |
| 1 | 212090822 | rs12135924 | 0.86 | -0.121 | -0.782 | -1.899 | -0.001 | -0.045 | -0.045 | 0 | 0.11 | 0.01 | 0.17 | 0.13 | 0.1 | 0.11 |
| 1 | 212092918 | rs2788124 | -0.047 | -0.142 | -0.591 | -1.781 | -0.002 | -0.007 | -0.007 | 0.4 | 0.64 | 0.54 | 0.56 | 0.22 | 0.72 | 0.57 |
| 1 | 212097521 | rs2970580 | -0.072 | -0.181 | -0.699 | -1.497 | 0.002 | -0.002 | -0.002 | 0 | 0.16 | 0.1 | 0.19 | 0.14 | 0.14 | 0.14 |
| 1 | 212098460 | rs2970578 | 1.077 | -0.153 | -0.584 | -2.071 | -0.013 | 0.05 | 0.05 | 0 | 0.36 | 0.22 | 0.19 | 0.06 | 0.4 | 0.32 |
| 1 | 212102871 | rs7534669 | -0.221 | 0.036 | 0.502 | -1.722 | 0.033 | 0.017 | 0.017 | 0 | 0.07 | 0.27 | 0.07 | 0.02 | 0.07 | 0.08 |
| 1 | 212104854 | rs2788140 | -0.038 | -0.48 | 0.467 | -1.404 | -0.003 | 0.017 | 0.017 | 0 | 0.64 | 0.54 | 0.54 | 0.42 | 0.67 | 0.62 |
| 1 | 212113995 | rs2788139 | -0.775 | -0.672 | 1.019 | -1.458 | -0.001 | -0.01 | -0.01 | 0 | 0.75 | 0.76 | 0.65 | 0.46 | 0.78 | 0.81 |
| 1 | 212117205 | rs12022699 | 0.92 | -0.637 | 1.13 | -1.472 | 0 | 0.009 | 0.009 | 0 | 0.08 | 0.05 | 0.08 | 0.04 | 0.09 | 0.17 |
| 1 | 212123221 | rs2808388 | 0.18 | -0.614 | 1.062 | -1.45 | 0.044 | -0.014 | -0.014 | 0 | 0.05 | 0.22 | 0.02 | 0 | 0.06 | 0.05 |
| 1 | 212149699 | rs3795836 | 1.478 | -0.666 | 0.82 | -1.462 | 0.003 | -0.047 | -0.047 | 0 | 0.1 | 0.01 | 0.14 | 0.13 | 0.07 | 0.11 |
| 1 | 212152173 | rs3010004 | -0.844 | -0.68 | 0.851 | -1.475 | -0.002 | -0.007 | -0.007 | 0 | 0.75 | 0.76 | 0.66 | 0.46 | 0.78 | 0.81 |
| 1 | 212190116 | rs2993531 | 0.354 | -0.557 | 0.574 | -1.323 | -0.003 | -0.002 | -0.002 | 0.133 | 0.58 | 0.47 | 0.49 | 0.41 | 0.61 | 0.54 |
| 1 | 212211284 | rs1578993 | -0.135 | -0.444 | 0.291 | -1.359 | -0.001 | 0.019 | 0.019 | 0 | 0.64 | 0.54 | 0.54 | 0.42 | 0.67 | 0.62 |
| 1 | 212237798 | rs10863936 | -0.719 | -0.112 | 0.241 | -1.529 | -0.017 | 0.07 | 0.07 | 0 | 0.55 | 0.72 | 0.71 | 0.73 | 0.51 | 0.58 |
| 1 | 212277107 | rs1387815 | 0.819 | -0.077 | -0.578 | -1.477 | -0.095 | 0.076 | 0.076 | 0.304 | 0.39 | 0.05 | 0.27 | 0.26 | 0.43 | 0.37 |
| 1 | 212295571 | rs6690208 | 1.007 | -0.554 | -0.377 | -0.902 | 0.055 | 0.153 | 0.153 | 0.005 | 0.5 | 0.2 | 0.32 | 0.23 | 0.57 | 0.49 |
| 1 | 212298804 | rs12087287 | 0.736 | -0.688 | -0.479 | -0.209 | -0.005 | 0.017 | 0.017 | 0.716 | 0.07 | 0.01 | 0.27 | 0.55 | 0.05 | 0.1 |
| 1 | 212300741 | rs9804001 | 0.215 | -0.786 | -0.378 | -0.357 | 0.015 | -0.005 | -0.005 | 0 | 0.16 | 0.09 | 0.16 | 0.13 | 0.1 | 0.21 |
| 1 | 212301643 | rs11809969 | -0.86 | -0.64 | 0.272 | -0.315 | -0.011 | 0.026 | 0.026 | 0 | 0.27 | 0.34 | 0.38 | 0.63 | 0.23 | 0.25 |
| 1 | 212320694 | rs923851 | 0.043 | -0.552 | 0.735 | -0.451 | -0.017 | -0.054 | -0.054 | 0 | 0.86 | 0.56 | 0.92 | 0.94 | 0.91 | 0.96 |
| 1 | 212322447 | rs12125796 | 0.433 | -0.579 | 0.071 | -0.427 | -0.004 | -0.05 | -0.05 | 1.4 | 0.13 | 0.01 | 0.15 | 0.14 | 0.1 | 0.19 |
| 1 | 212324488 | rs4951438 | -0.168 | -0.657 | 0.162 | -0.458 | -0.003 | 0.041 | 0.041 | 0.082 | 0.18 | 0.2 | 0.34 | 0.59 | 0.18 | 0.21 |
| 1 | 212324535 | rs7540143 | 0.001 | -0.762 | 0.036 | -0.663 | -0.033 | 0.011 | 0.011 | 0 | 0.22 | 0.4 | 0.15 | 0.21 | 0.15 | 0.18 |
| 1 | 212332294 | rs11119860 | 1.381 | -0.508 | -0.67 | -0.578 | 0.006 | 0.063 | 0.063 | 0 | 0.46 | 0.31 | 0.35 | 0.06 | 0.52 | 0.39 |
| 1 | 212338360 | rs11119861 | -0.533 | -0.065 | -1.046 | 0.489 | -0.021 | -0.118 | -0.118 | 0.92 | 0.17 | 0.03 | 0.32 | 0.58 | 0.18 | 0.22 |
| 1 | 212344323 | rs11119865 | -0.924 | 0.105 | -1.723 | -0.063 | -0.011 | -0.027 | -0.027 | 0 | 0.3 | 0.05 | 0.48 | 0.72 | 0.28 | 0.41 |
| 1 | 212345669 | rs7550051 | 0.261 | 1.025 | -1.242 | -0.349 | -0.001 | -0.003 | -0.003 | 0 | 0.68 | 0.69 | 0.62 | 0.86 | 0.56 | 0.7 |
| 1 | 212347352 | rs6680653 | -0.02 | 0.367 | -1.365 | -1.417 | 0.018 | 0.032 | 0.032 | 0 | 0.47 | 0.34 | 0.36 | 0.04 | 0.57 | 0.45 |
| 1 | 212348372 | rs903124 | -0.3 | -0.279 | -1.173 | -0.632 | -0.022 | 0.06 | 0.06 | 0 | 0.26 | 0.43 | 0.44 | 0.61 | 0.25 | 0.21 |
| 1 | 212350402 | rs10863949 | -1.221 | -0.385 | -1.493 | -0.42 | 0.005 | -0.003 | -0.003 | 0.283 | 0.76 | 0.78 | 0.82 | 0.75 | 0.84 | 0.66 |
| 1 | 212350975 | rs6703980 | 0.323 | -0.619 | -1.212 | -0.314 | -0.007 | -0.004 | -0.004 | 0 | 0.15 | 0.17 | 0.13 | 0.1 | 0.06 | 0.13 |
| 1 | 212351181 | rs4325202 | 0.696 | -0.577 | -1.264 | -0.346 | 0.023 | 0.063 | 0.063 | 0 | 0.51 | 0.59 | 0.38 | 0.13 | 0.58 | 0.47 |
| 1 | 212353818 | rs17719838 | 1.483 | -0.185 | -1.768 | -0.981 | -0.012 | 0.064 | 0.064 | 0 | 0.2 | 0.06 | 0.1 | 0 | 0.18 | 0.18 |
| 1 | 212355971 | rs12039815 | -0.261 | -0.096 | -1.604 | -0.787 | -0.002 | 0.002 | 0.002 | 0.597 | 0.42 | 0.56 | 0.59 | 0.64 | 0.59 | 0.35 |
| 1 | 212357017 | rs11119874 | 0.437 | 0.08 | -1.673 | -0.507 | -0.043 | -0.021 | -0.021 | 0 | 0.08 | 0.01 | 0.28 | 0.51 | 0.18 | 0.06 |
| 1 | 212370745 | rs1962127 | -0.905 | -0.215 | -1.899 | -0.793 | -0.016 | 0.111 | 0.111 | 0.966 | 0.58 | 0.9 | 0.72 | 0.78 | 0.65 | 0.49 |
| 1 | 212378485 | rs11119880 | 0.605 | -0.054 | -1.211 | -1.04 | -0.007 | 0.006 | 0.006 | 0.894 | 0.3 | 0.26 | 0.28 | 0.17 | 0.41 | 0.3 |
| 1 | 212379399 | rs7546833 | 1.416 | -0.044 | -1.305 | -1.06 | -0.035 | 0.109 | 0.109 | 0 | 0.19 | 0.03 | 0.1 | 0 | 0.18 | 0.18 |
| 1 | 212400146 | rs11119884 | -0.005 | 0.049 | -0.606 | -0.668 | 0.014 | -0.017 | -0.017 | 0 | 0.33 | 0.38 | 0.31 | 0.2 | 0.44 | 0.3 |
| 1 | 212402580 | rs1472226 | -0.261 | 0.113 | -0.902 | -0.681 | -0.082 | 0.077 | 0.077 | 2.444 | 0.36 | 0.88 | 0.41 | 0.2 | 0.43 | 0.31 |
| 1 | 212409008 | rs7544225 | 0.203 | 0.657 | -0.805 | -0.682 | 0.003 | -0.001 | -0.001 | 0.059 | 0.25 | 0.23 | 0.35 | 0.2 | 0.37 | 0.29 |
| 1 | 212412332 | rs1009622 | 0.444 | 0.984 | -0.62 | -0.824 | 0.005 | 0.013 | 0.013 | 0.414 | 0.1 | 0.14 | 0.2 | 0.19 | 0.12 | 0.22 |
| 1 | 212416078 | rs4951583 | -1.578 | 0.552 | -0.884 | -0.718 | -0.055 | 0.05 | 0.05 | 0.419 | 0.44 | 0.86 | 0.62 | 0.66 | 0.58 | 0.33 |
| 1 | 212418044 | rs10863957 | -0.304 | 0.579 | -0.555 | -0.722 | 0.003 | 0.064 | 0.064 | 0 | 0.16 | 0.22 | 0.24 | 0.47 | 0.22 | 0.05 |
| 1 | 212421771 | rs351386 | -0.002 | 0.638 | -0.814 | -0.746 | 0.006 | -0.003 | -0.003 | 0 | 0.25 | 0.24 | 0.35 | 0.2 | 0.36 | 0.28 |
| 1 | 212429259 | rs10494951 | 1.709 | 0.529 | -1.245 | -1.024 | -0.027 | 0.138 | 0.138 | 0 | 0.23 | 0.03 | 0.09 | 0 | 0.18 | 0.24 |
| 1 | 212431398 | rs351405 | -1.855 | 0.36 | -1.327 | -0.967 | -0.046 | -0.05 | -0.05 | 0.033 | 0.77 | 0.96 | 0.81 | 0.76 | 0.78 | 0.69 |
| 1 | 212434815 | rs7531219 | 0.006 | 0.161 | -1.099 | -1.142 | 0.074 | 0.137 | 0.137 | 0 | 0.42 | 0.79 | 0.26 | 0.1 | 0.24 | 0.35 |
| 1 | 212447142 | rs726558 | 0.281 | 0.325 | -1.427 | -1.148 | -0.01 | 0.203 | 0.203 | 0 | 0.39 | 0.06 | 0.16 | 0.12 | 0.29 | 0.44 |
| 1 | 212457414 | rs351413 | -0.519 | 0.201 | -0.417 | -0.622 | 0.003 | -0.013 | -0.013 | 0 | 0.6 | 0.43 | 0.57 | 0.39 | 0.57 | 0.64 |
| 1 | 212468974 | rs411278 | -0.217 | 0.216 | -0.053 | -0.615 | 0.107 | 0.067 | 0.067 | 0 | 0.05 | 0.4 | 0.05 | 0 | 0.04 | 0 |
| 1 | 212474288 | rs351392 | -1.139 | 0.152 | -0.192 | -0.656 | -0.1 | -0.06 | -0.06 | 0 | 0.67 | 0.95 | 0.62 | 0.39 | 0.6 | 0.64 |
| 1 | 212500433 | rs4951588 | -0.162 | 0.127 | -0.175 | -0.77 | 0.032 | -0.02 | -0.02 | 0.024 | 0.31 | 0.41 | 0.39 | 0.29 | 0.4 | 0.32 |
| 1 | 212515613 | rs1774249 | 0.441 | 0.201 | -0.273 | -0.776 | -0.007 | 0 | 0 | 0 | 0.25 | 0.19 | 0.33 | 0.28 | 0.34 | 0.31 |
| 1 | 212516875 | rs12736502 | 0.92 | 0.111 | -0.374 | -0.816 | 0.033 | 0.14 | 0.14 | 0 | 0.31 | 0.07 | 0.2 | 0.1 | 0.2 | 0.32 |
| 1 | 212526288 | rs792442 | -1.971 | 0.243 | -0.459 | -0.636 | -0.042 | -0.017 | -0.017 | 0.039 | 0.74 | 0.95 | 0.8 | 0.78 | 0.75 | 0.68 |
| 1 | 212534448 | rs14123 | -0.692 | 0.321 | -0.071 | -0.525 | 0.034 | 0.233 | 0.233 | 1.224 | 0.15 | 0.45 | 0.23 | 0.4 | 0.19 | 0.04 |
| 1 | 212537553 | rs3767861 | 0.318 | 0.279 | -0.306 | -0.578 | -0.025 | 0.002 | 0.002 | 0 | 0.26 | 0.14 | 0.35 | 0.28 | 0.37 | 0.32 |
| 1 | 212537702 | rs3767862 | 0.338 | 0.299 | -0.297 | -0.568 | -0.025 | 0.001 | 0.001 | 0 | 0.26 | 0.14 | 0.35 | 0.28 | 0.37 | 0.32 |
| 1 | 212538456 | rs12073615 | 1.309 | -0.061 | -0.247 | -0.764 | 0.028 | 0.007 | 0.007 | 0.087 | 0.1 | 0 | 0.03 | 0.07 | 0.04 | 0.09 |
| 1 | 212556678 | rs792446 | -0.531 | 0.268 | 0.321 | -0.624 | 0.001 | -0.025 | -0.025 | 0.198 | 0.57 | 0.46 | 0.49 | 0.26 | 0.57 | 0.46 |
| 1 | 212556744 | rs792447 | 1.199 | 0.473 | 0.07 | -0.88 | -0.001 | -0.01 | -0.01 | 0 | 0.17 | 0.14 | 0.14 | 0.06 | 0.19 | 0.11 |
| 1 | 212559906 | rs12119779 | 0.586 | 0.288 | 0.183 | -0.585 | -0.052 | -0.037 | -0.037 | 0 | 0.09 | 0.01 | 0.28 | 0.41 | 0.18 | 0.04 |
| 1 | 212562737 | rs1086892 | -1.904 | 0.265 | 0.134 | -0.633 | -0.062 | -0.107 | -0.107 | 0 | 0.72 | 0.97 | 0.8 | 0.63 | 0.74 | 0.47 |
| 1 | 212562975 | rs6676208 | 1.272 | 0.237 | 0.226 | -0.674 | -0.013 | -0.101 | -0.101 | 0.542 | 0.25 | 0.34 | 0.11 | 0 | 0.16 | 0.2 |
| 1 | 212571575 | rs12060029 | 0.296 | 0.264 | -0.681 | -0.618 | 0.02 | 0.001 | 0.001 | 0.555 | 0.11 | 0.03 | 0.13 | 0.1 | 0.06 | 0.07 |

| | | | | | | | | | | | | | | | | |
|---|---|---|---|---|---|---|---|---|---|---|---|---|---|---|---|---|
| 1 | 212580922 | rs1539206 | -0.888 | 0.292 | -0.66 | -0.618 | 0.026 | 0.007 | 0.007 | 0.013 | 0.48 | 0.81 | 0.56 | 0.51 | 0.4 | 0.32 |
| 1 | 212588893 | rs512458 | -0.944 | 0.427 | -0.554 | -0.555 | 0.005 | -0.067 | -0.067 | 1.167 | 0.59 | 0.9 | 0.67 | 0.53 | 0.68 | 0.4 |
| 1 | 212590678 | rs4336916 | 0.708 | 0.497 | -0.069 | -0.44 | -0.029 | -0.154 | -0.154 | 1.976 | 0.26 | 0.53 | 0.12 | 0 | 0.16 | 0.2 |
| 1 | 212601099 | rs10863965 | -0.859 | 0.74 | -0.32 | -0.069 | 0.044 | 0.247 | 0.247 | 1.591 | 0.14 | 0.61 | 0.33 | 0.41 | 0.18 | 0.05 |
| 1 | 212602176 | rs612414 | 0.244 | 0.607 | -0.5 | -0.179 | -0.005 | 0.012 | 0.012 | 0 | 0.29 | 0.23 | 0.27 | 0.12 | 0.38 | 0.16 |
| 1 | 212603619 | rs10863966 | -0.436 | 0.648 | -0.431 | -0.063 | 0.002 | -0.007 | -0.007 | 0 | 0.4 | 0.66 | 0.5 | 0.5 | 0.35 | 0.33 |
| 1 | 212607404 | rs3767868 | -0.236 | 0.756 | 0.027 | 0.853 | -0.03 | -0.008 | -0.008 | 2.412 | 0.08 | 0 | 0.24 | 0.41 | 0.1 | 0.04 |
| 1 | 212610755 | rs583058 | -1.282 | 0.612 | 0.419 | 0.303 | -0.004 | -0.016 | -0.016 | 0 | 0.43 | 0.49 | 0.56 | 0.68 | 0.38 | 0.27 |
| 1 | 212614788 | rs791720 | -0.707 | 0.604 | 0.558 | 0.47 | 0.014 | 0.238 | 0.238 | 0 | 0.09 | 0.15 | 0.27 | 0.42 | 0.09 | 0.03 |
| 1 | 212617423 | rs12567517 | -0.529 | 0.589 | 0.615 | 0.43 | 0.009 | 0.026 | 0.026 | 0.811 | 0.34 | 0.3 | 0.3 | 0.25 | 0.29 | 0.24 |
| 1 | 212619339 | rs4804 | -1.371 | 0.649 | 0.57 | 0.473 | -0.004 | -0.016 | -0.016 | 0 | 0.43 | 0.49 | 0.56 | 0.67 | 0.38 | 0.3 |
| 1 | 212620214 | rs483954 | -0.882 | 0.623 | 0.557 | 0.471 | 0 | -0.002 | -0.002 | 0 | 0.37 | 0.38 | 0.45 | 0.56 | 0.36 | 0.18 |
| 1 | 212620938 | rs4951446 | 0.186 | 0.504 | 0.427 | 0.68 | -0.024 | 0.045 | 0.045 | 0 | 0.06 | 0 | 0.23 | 0.42 | 0.07 | 0.03 |
| 1 | 212621251 | rs7520772 | -0.367 | 0.512 | 0.599 | 0.557 | -0.017 | 0.025 | 0.025 | 0 | 0.06 | 0.14 | 0.11 | 0.11 | 0.02 | 0.12 |
| 1 | 212628393 | rs731075 | -1.111 | 0.536 | 0.44 | 0.59 | -0.004 | 0.088 | 0.088 | 0 | 0.11 | 0.17 | 0.34 | 0.53 | 0.09 | 0.16 |
| 1 | 212629935 | rs479564 | -1.454 | 0.694 | 0.34 | 0.419 | 0.006 | 0.046 | 0.046 | 0.094 | 0.45 | 0.65 | 0.61 | 0.68 | 0.44 | 0.34 |
| 1 | 212634583 | rs675065 | -0.748 | 0.982 | 0.285 | 0.714 | 0.071 | 0.462 | 0.462 | 0 | 0.08 | 0.28 | 0.27 | 0.43 | 0.1 | 0.03 |
| 1 | 212636824 | rs12746015 | -0.41 | 0.836 | -0.615 | -0.071 | -0.015 | 0.072 | 0.072 | 0.224 | 0.29 | 0.13 | 0.22 | 0.14 | 0.32 | 0.18 |
| 1 | 212639003 | rs902438 | -0.859 | 0.651 | -0.085 | -0.333 | -0.006 | -0.073 | -0.073 | 0.466 | 0.35 | 0.44 | 0.21 | 0.14 | 0.31 | 0.19 |
| 1 | 212640367 | rs2292480 | 0.243 | 0.638 | -0.485 | -0.234 | -0.01 | -0.022 | -0.022 | 2.027 | 0.09 | 0.02 | 0.06 | 0.14 | 0.1 | 0.17 |
| 1 | 212640982 | rs734976 | -1.166 | 1.115 | -0.139 | -0.002 | -0.001 | -0.003 | -0.003 | 1.546 | 0.47 | 0.55 | 0.5 | 0.69 | 0.5 | 0.44 |
| 1 | 212646225 | rs6697091 | 0.294 | 0.747 | -0.121 | -0.732 | -0.021 | -0.09 | -0.09 | 0 | 0.19 | 0.31 | 0.07 | 0 | 0.11 | 0.13 |
| 1 | 212648135 | rs748517 | 0.279 | 0.429 | 0.008 | -0.919 | -0.002 | -0.026 | -0.026 | 2.282 | 0.29 | 0.36 | 0.3 | 0.17 | 0.28 | 0.2 |
| 1 | 212652970 | rs12129531 | 1.452 | 0.401 | -0.675 | -1.041 | 0.007 | 0.037 | 0.037 | 0 | 0.16 | 0 | 0.06 | 0.05 | 0.1 | 0.09 |
| 1 | 212656471 | rs11119947 | 1.074 | 0.608 | -0.305 | -0.826 | -0.01 | -0.049 | -0.049 | 0 | 0.16 | 0.2 | 0.06 | 0 | 0.1 | 0.13 |
| 1 | 212660681 | rs7544479 | -0.913 | 0.679 | -0.497 | -0.439 | -0.085 | -0.039 | -0.039 | 0.011 | 0.56 | 0.85 | 0.63 | 0.5 | 0.5 | 0.39 |
| 1 | 212668552 | rs12729438 | 0.521 | 0.641 | -0.243 | -0.544 | 0 | -0.008 | -0.008 | 0 | 0.3 | 0.27 | 0.19 | 0.13 | 0.29 | 0.2 |
| 1 | 212670865 | rs6540742 | -0.636 | 0.646 | -0.427 | -0.549 | 0.025 | 0.274 | 0.274 | 0.05 | 0.5 | 0.83 | 0.39 | 0.17 | 0.39 | 0.33 |
| 1 | 212675482 | rs597459 | -1.08 | 1.423 | -0.338 | -0.416 | 0.042 | 0.098 | 0.098 | 0 | 0.57 | 0.72 | 0.73 | 0.84 | 0.66 | 0.6 |
| 1 | 212677319 | rs626178 | 0.379 | 1.547 | -0.139 | -0.317 | -0.001 | -0.058 | -0.058 | 0 | 0.33 | 0.51 | 0.33 | 0.19 | 0.3 | 0.2 |
| 1 | 212678727 | rs653006 | 0.519 | 1.545 | -0.161 | -0.341 | 0.001 | -0.034 | -0.034 | 1.523 | 0.32 | 0.36 | 0.21 | 0.13 | 0.3 | 0.2 |
| 1 | 212683565 | rs878947 | 0.111 | 1.609 | -0.852 | -0.272 | -0.002 | -0.001 | -0.001 | 0.01 | 0.16 | 0.15 | 0.17 | 0.32 | 0.26 | 0.34 |
| 1 | 212683888 | rs562683 | -0.555 | 1.139 | -1.169 | -0.328 | 0.063 | 0.163 | 0.163 | 0 | 0.55 | 0.79 | 0.73 | 0.83 | 0.66 | 0.6 |
| 1 | 212693423 | rs10863978 | 0.503 | 1.411 | -1.069 | 0.253 | -0.047 | -0.017 | -0.017 | 0 | 0.08 | 0.01 | 0.26 | 0.32 | 0.14 | 0.05 |
| 1 | 212700618 | rs1021378 | -0.221 | 1.387 | -1.197 | -0.311 | -0.036 | -0.013 | -0.013 | 3.331 | 0.61 | 0.72 | 0.49 | 0.44 | 0.56 | 0.67 |
| 1 | 212702656 | rs1331060 | 0.646 | 1.791 | -1.432 | -0.28 | -0.041 | -0.042 | -0.042 | 0.145 | 0.07 | 0 | 0.07 | 0.15 | 0.13 | 0.14 |
| 1 | 212711012 | rs650229 | -0.5 | 1.676 | -1.207 | -0.39 | -0.044 | 0.015 | 0.015 | 0 | 0.63 | 0.81 | 0.65 | 0.63 | 0.56 | 0.57 |
| 1 | 212712720 | rs1512847 | 0.153 | 1.677 | -1.207 | -0.391 | 0.007 | 0.014 | 0.014 | 0 | 0.09 | 0.11 | 0.11 | 0.15 | 0.15 | 0.14 |
| 1 | 212713653 | rs11802726 | 1.45 | 1.567 | -1.036 | -0.526 | -0.003 | 0.023 | 0.023 | 0 | 0.08 | 0.14 | 0.15 | 0.24 | 0.08 | 0.23 |
| 1 | 212714455 | rs599028 | -1.486 | 1.558 | -1.098 | -0.549 | -0.041 | 0.073 | 0.073 | 0 | 0.72 | 0.92 | 0.77 | 0.78 | 0.71 | 0.71 |
| 1 | 212716033 | rs616408 | -0.027 | 1.517 | -1.007 | -0.534 | 0.002 | -0.004 | -0.004 | 0.523 | 0.53 | 0.59 | 0.49 | 0.39 | 0.46 | 0.32 |
| 1 | 212716491 | rs17666978 | 1.533 | 1.146 | -0.999 | -0.812 | 0.004 | -0.006 | -0.006 | 0 | 0.15 | 0.14 | 0.09 | 0 | 0.14 | 0.12 |
| 1 | 212716579 | rs11119967 | 0.354 | 1.238 | -0.9 | -0.743 | -0.019 | -0.089 | -0.089 | 0 | 0.44 | 0.45 | 0.24 | 0.05 | 0.33 | 0.27 |
| 1 | 212723176 | rs791851 | -0.67 | 1.51 | -1.256 | -0.519 | -0.069 | -0.036 | -0.036 | 0.252 | 0.63 | 0.83 | 0.51 | 0.44 | 0.56 | 0.65 |
| 1 | 212727262 | rs7522039 | 0.364 | 1.309 | -1.031 | -0.46 | -0.012 | -0.011 | -0.011 | 0 | 0.37 | 0.5 | 0.28 | 0.29 | 0.27 | 0.38 |
| 1 | 212730334 | rs2884515 | -0.89 | 1.966 | -0.898 | 0.094 | 0.01 | 0.015 | 0.015 | 0 | 0.17 | 0.18 | 0.37 | 0.49 | 0.29 | 0.19 |
| 1 | 212733925 | rs11119972 | 0.034 | 1.828 | -0.736 | 0.224 | 0.042 | 0.177 | 0.177 | 0 | 0.1 | 0.22 | 0.26 | 0.36 | 0.14 | 0.07 |
| 1 | 212734994 | rs493894 | 0.744 | 1.722 | -1.399 | 0.011 | -0.031 | 0.016 | 0.016 | 0 | 0.19 | 0.08 | 0.19 | 0.23 | 0.25 | 0.22 |
| 1 | 212747117 | rs2137424 | 0.571 | 1.357 | -1.216 | -0.017 | -0.001 | 0.099 | 0.099 | 0.043 | 0.35 | 0.2 | 0.27 | 0.24 | 0.33 | 0.38 |
| 1 | 212752129 | rs3122720 | -1.127 | -0.042 | -0.813 | -0.044 | 0 | -0.001 | -0.001 | 0 | 0.81 | 0.82 | 0.78 | 0.76 | 0.84 | 0.7 |
| 1 | 212757658 | rs4543871 | 0.205 | 0.091 | -0.206 | 0.484 | -0.037 | -0.075 | -0.075 | 0.27 | 0.28 | 0.38 | 0.16 | 0.08 | 0.22 | 0.17 |
| 1 | 212757891 | rs1877474 | -0.742 | 0.292 | -0.686 | 0.633 | 0.001 | -0.003 | -0.003 | 0 | 0.39 | 0.55 | 0.42 | 0.39 | 0.37 | 0.21 |
| 1 | 212759493 | rs6674072 | -0.485 | 0.746 | -0.873 | 1.163 | -0.049 | -0.064 | -0.064 | 1.409 | 0.1 | 0.01 | 0.27 | 0.3 | 0.15 | 0.06 |
| 1 | 212761057 | rs12739624 | -0.982 | 0.614 | -0.531 | 1.118 | 0.004 | 0.03 | 0.03 | 0 | 0.12 | 0.13 | 0.28 | 0.25 | 0.15 | 0.05 |
| 1 | 212764085 | rs10494953 | -0.13 | -0.143 | -0.128 | 0.312 | -0.016 | -0.04 | -0.04 | 1.391 | 0.23 | 0.33 | 0.18 | 0.17 | 0.21 | 0.24 |
| 1 | 212764844 | rs11119982 | 0.826 | -0.783 | -0.821 | -0.433 | -0.046 | 0.134 | 0.134 | 0 | 0.38 | 0.18 | 0.27 | 0.19 | 0.5 | 0.46 |
| 1 | 212767840 | rs6682064 | 0.631 | -0.798 | -0.224 | -0.228 | -0.025 | -0.035 | -0.035 | 0 | 0.22 | 0.44 | 0.18 | 0.21 | 0.2 | 0.26 |
| 1 | 212768428 | rs1567710 | 0.181 | -0.959 | -0.516 | 0.17 | 0.016 | -0.026 | -0.026 | 3.22 | 0.33 | 0.25 | 0.28 | 0.42 | 0.26 | 0.22 |
| 1 | 212769411 | rs11119985 | 0.977 | -0.667 | -0.913 | -0.129 | -0.148 | 0.155 | 0.155 | 0 | 0.33 | 0.07 | 0.23 | 0.18 | 0.47 | 0.42 |
| 1 | 212771881 | rs9430097 | -0.114 | -1.491 | -0.313 | -0.045 | -0.012 | 0.01 | 0.01 | 0 | 0.33 | 0.35 | 0.37 | 0.59 | 0.3 | 0.38 |
| 1 | 212772937 | rs17019438 | -0.552 | -1.969 | -0.134 | 0.004 | -0.032 | 0.028 | 0.028 | 0 | 0.37 | 0.46 | 0.43 | 0.49 | 0.25 | 0.4 |
| 1 | 212773328 | rs3795837 | 1.449 | -1.991 | 0.089 | 0.005 | -0.008 | 0.039 | 0.039 | 0 | 0.12 | 0.21 | 0.1 | 0.27 | 0.11 | 0.22 |
| 1 | 212774340 | rs17019442 | 0.738 | -2.132 | 0.07 | -0.108 | 0.027 | 0.053 | 0.053 | 0.007 | 0.19 | 0.08 | 0.06 | 0 | 0.1 | 0.15 |
| 1 | 212776328 | rs10746435 | 0.685 | -2.289 | 0.262 | -0.114 | -0.009 | -0.036 | -0.036 | 0.059 | 0.57 | 0.49 | 0.32 | 0.25 | 0.6 | 0.57 |
| 1 | 212778286 | rs3125296 | -0.29 | -2.298 | 0.155 | -0.162 | 0.001 | 0.001 | 0.001 | 0 | 0.74 | 0.72 | 0.78 | 0.72 | 0.74 | 0.7 |
| 1 | 212780471 | rs4406706 | 0.242 | -1.986 | 0.234 | -0.206 | 0.017 | 0.235 | 0.235 | 0 | 0.06 | 0.17 | 0.27 | 0.23 | 0.04 | 0.07 |
| 1 | 212782500 | rs6666492 | 0.765 | -2.049 | -0.242 | -0.637 | -0.012 | -0.037 | -0.037 | 0 | 0.17 | 0.17 | 0.04 | 0 | 0.07 | 0.09 |
| 1 | 212788206 | rs11119988 | 1.468 | -1.815 | -0.57 | -0.549 | -0.014 | 0.038 | 0.038 | 0 | 0.1 | 0.04 | 0.1 | 0 | 0.12 | 0.07 |
| 1 | 212790335 | rs12070345 | 1.475 | -1.912 | -0.583 | -0.689 | -0.046 | 0.066 | 0.066 | 0 | 0.35 | 0.23 | 0.25 | 0.21 | 0.48 | 0.47 |

| | | | | | | | | | | | | | | | | |
|---|---|---|---|---|---|---|---|---|---|---|---|---|---|---|---|---|
| 1 | 212792191 | rs11571552 | 1.791 | -2.032 | -0.691 | -0.731 | -0.007 | 0.038 | 0.038 | 0 | 0.08 | 0 | 0.03 | 0 | 0.06 | 0.06 |
| 1 | 212792580 | rs11571553 | 2.251 | -2 | -0.554 | -0.667 | 0.004 | -0.003 | -0.003 | 0 | 0.06 | 0.03 | 0.01 | 0 | 0.01 | 0.01 |
| 1 | 212793505 | rs1126526 | 1.072 | -1.994 | -0.757 | -0.706 | 0.002 | 0.029 | 0.029 | 0 | 0.15 | 0.11 | 0.12 | 0 | 0.15 | 0.08 |
| 1 | 212793613 | rs10475 | -0.609 | -1.871 | -0.538 | -0.286 | -0.003 | 0.014 | 0.014 | 0 | 0.75 | 0.73 | 0.7 | 0.43 | 0.76 | 0.69 |
| 1 | 212797890 | rs7533306 | -0.52 | -1.548 | -0.915 | -0.593 | 0 | -0.001 | -0.001 | 0 | 0.72 | 0.72 | 0.68 | 0.44 | 0.73 | 0.65 |
| 1 | 212798977 | rs3122712 | -0.554 | -1.814 | -0.574 | -0.633 | -0.025 | -0.063 | -0.063 | 0.021 | 0.38 | 0.46 | 0.2 | 0.08 | 0.26 | 0.19 |
| 1 | 212799870 | rs3122713 | -0.833 | -1.632 | -0.784 | -0.077 | -0.005 | 0.028 | 0.028 | 0 | 0.81 | 0.77 | 0.76 | 0.68 | 0.84 | 0.87 |
| 1 | 212799949 | rs3795842 | 0.37 | -1.43 | -1.181 | -0.538 | 0.044 | 0.062 | 0.062 | 0 | 0.54 | 0.34 | 0.32 | 0.45 | 0.65 | 0.68 |
| 1 | 212804825 | rs3123534 | 0.676 | -0.977 | -1.179 | -0.413 | 0.035 | -0.013 | -0.013 | 0 | 0.46 | 0.23 | 0.37 | 0.57 | 0.63 | 0.65 |
| 1 | 212809606 | rs3123536 | -0.108 | -1.362 | -0.621 | -1.154 | -0.071 | -0.131 | -0.131 | 2.009 | 0.73 | 0.56 | 0.92 | 0.92 | 0.88 | 0.84 |
| 1 | 212811875 | rs3923950 | 1.001 | -1.39 | -1.27 | -1.023 | -0.077 | -0.08 | -0.08 | 0.804 | 0.22 | 0.01 | 0.26 | 0.24 | 0.25 | 0.18 |
| 1 | 212811947 | rs3123537 | 1.363 | -1.405 | -1.25 | -1.05 | 0.024 | 0.07 | 0.07 | 0 | 0.46 | 0.23 | 0.61 | 0.67 | 0.62 | 0.65 |
| 1 | 212813503 | rs3890801 | 1.074 | -0.941 | -1.408 | -0.827 | 0 | -0.019 | -0.019 | 0.621 | 0.14 | 0.12 | 0.03 | 0 | 0.08 | 0.12 |
| 1 | 212817909 | rs6674102 | 0.646 | -0.712 | -1.311 | -0.584 | -0.033 | -0.034 | -0.034 | 0 | 0.22 | 0.05 | 0.26 | 0.24 | 0.25 | 0.18 |
| 1 | 212827841 | rs2501841 | -0.341 | -0.418 | -0.955 | -0.599 | -0.004 | -0.038 | -0.038 | 0 | 0.82 | 0.75 | 0.93 | 0.91 | 0.85 | 0.83 |
| 1 | 212828748 | rs2501842 | 0.728 | -0.659 | -1.391 | -0.837 | 0.006 | 0.052 | 0.052 | 0 | 0.45 | 0.23 | 0.6 | 0.67 | 0.59 | 0.61 |
| 1 | 212829330 | rs6540751 | 0.563 | -0.193 | -1.417 | -0.926 | 0 | -0.006 | -0.006 | 3.595 | 0.14 | 0.1 | 0.05 | 0 | 0.12 | 0.12 |
| 1 | 212838716 | rs12119575 | -0.231 | 0.189 | -1.531 | -0.944 | -0.006 | 0.085 | 0.085 | 0 | 0.48 | 0.32 | 0.28 | 0.2 | 0.45 | 0.35 |
| 1 | 212842330 | rs4951633 | 0.28 | 0.51 | -1.61 | -0.143 | -0.011 | -0.006 | -0.006 | 0 | 0.17 | 0.09 | 0.25 | 0.2 | 0.15 | 0.25 |
| 1 | 212844187 | rs2492788 | 0.505 | 0.782 | -1.278 | -0.001 | -0.002 | -0.022 | -0.022 | 0 | 0.48 | 0.2 | 0.61 | 0.51 | 0.46 | 0.47 |
| 1 | 212845034 | rs2492789 | -0.633 | 0.905 | -0.762 | -0.015 | 0.006 | 0.048 | 0.048 | 0 | 0.79 | 0.67 | 0.84 | 0.7 | 0.79 | 0.59 |
| 1 | 212846165 | rs17019556 | 0.189 | 0.647 | -1.241 | -0.179 | -0.002 | 0.049 | 0.049 | 0 | 0.23 | 0.15 | 0.14 | 0.04 | 0.21 | 0.06 |
| 1 | 212846338 | rs2501847 | 0.061 | 0.665 | -0.863 | 0.048 | 0.014 | 0.063 | 0.063 | 0 | 0.72 | 0.52 | 0.78 | 0.55 | 0.67 | 0.53 |
| 1 | 212858797 | rs906363 | 0.142 | 0.095 | -1.509 | -0.17 | 0.008 | 0.077 | 0.077 | 0 | 0.19 | 0.09 | 0.12 | 0.04 | 0.16 | 0.06 |
| 1 | 212863052 | rs2246053 | 0.644 | 0.172 | -1.703 | 0 | 0.001 | 0.022 | 0.022 | 0 | 0.32 | 0.1 | 0.52 | 0.24 | 0.29 | 0.26 |
| 1 | 212868263 | rs2884573 | 0.279 | 0.376 | -1.269 | 0.152 | -0.021 | 0.002 | 0.002 | 0 | 0.16 | 0.01 | 0.11 | 0.17 | 0.15 | 0.07 |
| 1 | 212873431 | rs2221593 | -0.229 | 0.279 | -0.851 | 0.582 | -0.007 | -0.026 | -0.026 | 0 | 0.13 | 0.08 | 0.09 | 0.19 | 0.12 | 0.34 |
| 1 | 212885805 | rs7555918 | -0.428 | 0.523 | -0.182 | 0.318 | 0.057 | -0.02 | -0.02 | 0.253 | 0.06 | 0.19 | 0.03 | 0 | 0.06 | 0.01 |
| 1 | 212887790 | rs12753524 | -0.386 | 0.154 | 0.287 | 0.304 | -0.024 | 0.03 | 0.03 | 0.075 | 0.64 | 0.54 | 0.44 | 0.56 | 0.68 | 0.66 |
| 1 | 212889867 | rs11120002 | 0.87 | 0.152 | 0.36 | 0.316 | -0.045 | 0.055 | 0.055 | 0.151 | 0.47 | 0.16 | 0.31 | 0.41 | 0.5 | 0.42 |
| 1 | 212892113 | rs6681171 | -0.843 | 0.32 | 0.559 | 0.114 | 0.082 | 0.046 | 0.046 | 0.291 | 0.15 | 0.48 | 0.11 | 0.14 | 0.15 | 0.21 |
| 1 | 212894578 | rs11800991 | -0.975 | 0.341 | 0.509 | 0.1 | 0.011 | -0.022 | -0.022 | 0 | 0.09 | 0.13 | 0.05 | 0.02 | 0.1 | 0.04 |
| 1 | 212900487 | rs2244935 | -1.174 | 0.236 | 0.637 | 0.234 | -0.004 | 0.016 | 0.016 | 0 | 0.93 | 0.91 | 0.96 | 0.89 | 0.92 | 0.89 |
| 1 | 212903089 | rs12747251 | 0.936 | 0.186 | 0.534 | 0.219 | -0.04 | 0.093 | 0.093 | 0.037 | 0.47 | 0.16 | 0.31 | 0.38 | 0.51 | 0.43 |
| 1 | 212909744 | rs11800642 | -0.179 | 0.127 | 0.853 | 0.232 | 0.064 | 0.036 | 0.036 | 0 | 0.12 | 0.3 | 0.11 | 0.14 | 0.14 | 0.19 |
| 1 | 212911836 | rs15702 | -0.449 | 0.157 | 0.926 | 0.221 | 0.004 | -0.008 | -0.008 | 0 | 0.66 | 0.65 | 0.44 | 0.53 | 0.69 | 0.66 |
| 1 | 212913168 | rs2271148 | 0.878 | 0.176 | 0.971 | 0.221 | 0.023 | 0.052 | 0.052 | 0 | 0.08 | 0.16 | 0.07 | 0.14 | 0.07 | 0.18 |
| 1 | 212913590 | rs6702101 | -0.765 | 0.076 | 0.92 | 0.242 | 0.102 | 0.032 | 0.032 | 0 | 0.17 | 0.4 | 0.13 | 0.15 | 0.18 | 0.23 |
| 1 | 212925857 | rs6540762 | -0.746 | -0.004 | 0.92 | 0.242 | 0.099 | 0.034 | 0.034 | 0 | 0.17 | 0.4 | 0.13 | 0.15 | 0.18 | 0.23 |
| 1 | 212928157 | rs1032965 | 1.156 | -0.051 | 0.899 | 0.219 | -0.024 | -0.117 | -0.117 | 0.806 | 0.25 | 0.03 | 0.49 | 0.36 | 0.23 | 0.23 |
| 1 | 212931215 | rs4951638 | 1.733 | -0.272 | 0.701 | -0.016 | -0.13 | -0.062 | -0.062 | 0 | 0.31 | 0.02 | 0.19 | 0.37 | 0.4 | 0.37 |
| 1 | 212943581 | rs11808846 | -0.587 | -0.278 | 0.704 | -0.011 | 0.099 | 0.034 | 0.034 | 0 | 0.17 | 0.4 | 0.13 | 0.15 | 0.18 | 0.23 |
| 1 | 212947520 | rs10863997 | 0.273 | -0.266 | 0.679 | 0.004 | 0.064 | 0.036 | 0.036 | 0 | 0.12 | 0.3 | 0.11 | 0.14 | 0.15 | 0.19 |
| 1 | 212955231 | rs11120025 | 0.592 | -0.26 | 0.485 | -0.218 | -0.016 | 0.098 | 0.098 | 0.339 | 0.51 | 0.15 | 0.32 | 0.39 | 0.54 | 0.47 |
| 1 | 212962207 | rs11120027 | 1.601 | -0.292 | 0.477 | -0.18 | -0.062 | -0.009 | -0.009 | 1.701 | 0.32 | 0.1 | 0.2 | 0.37 | 0.4 | 0.37 |
| 1 | 212972599 | rs12128410 | 0.238 | -0.038 | 0.428 | 0.147 | 0.041 | 0.079 | 0.079 | 0 | 0.12 | 0.23 | 0.09 | 0.25 | 0.15 | 0.2 |
| 1 | 212975778 | rs4951639 | -0.32 | -0.017 | 0.489 | 0.1 | 0.003 | -0.002 | -0.002 | 0.939 | 0.66 | 0.65 | 0.44 | 0.64 | 0.69 | 0.67 |
| 1 | 212978197 | rs1532951 | 0.159 | 0.024 | 0.514 | 0.152 | 0.021 | 0.057 | 0.057 | 0 | 0.12 | 0.22 | 0.09 | 0.28 | 0.15 | 0.2 |
| 1 | 212986554 | rs7515115 | -0.017 | 0.033 | 0.513 | 0.091 | 0.065 | 0.119 | 0.119 | 0 | 0.13 | 0.29 | 0.1 | 0.25 | 0.15 | 0.23 |
| 1 | 212997385 | rs10864002 | -0.238 | 0.067 | 0.532 | 0.121 | 0.003 | -0.002 | -0.002 | 0 | 0.66 | 0.64 | 0.44 | 0.64 | 0.69 | 0.67 |
| 1 | 212999701 | rs10746436 | -0.32 | 0.064 | 0.534 | 0.112 | 0.016 | -0.058 | -0.058 | 0 | 0.9 | 0.68 | 0.92 | 1 | 0.86 | 0.88 |
| 1 | 213002088 | rs2884675 | -0.178 | 0.059 | 0.517 | 0.115 | 0.003 | -0.002 | -0.002 | 0 | 0.66 | 0.64 | 0.44 | 0.64 | 0.69 | 0.67 |
| 1 | 213003428 | rs12038225 | 0.909 | 0.08 | 0.24 | 0.116 | 0.006 | -0.122 | -0.122 | 0 | 0.25 | 0.02 | 0.48 | 0.36 | 0.18 | 0.22 |
| 1 | 213005012 | rs6694190 | 0.124 | 0.266 | 0.369 | 0.255 | 0.024 | 0.045 | 0.045 | 0 | 0.12 | 0.21 | 0.1 | 0.25 | 0.15 | 0.21 |
| 1 | 213006397 | rs6700410 | 0.527 | 0.177 | 0.204 | 0.259 | -0.002 | 0.035 | 0.035 | 0 | 0.52 | 0.35 | 0.33 | 0.39 | 0.52 | 0.44 |
| 1 | 213009469 | rs10864004 | -0.082 | -0.635 | -0.086 | -0.042 | 0.034 | -0.024 | -0.024 | 0 | 0.66 | 0.79 | 0.46 | 0.54 | 0.74 | 0.66 |
| 1 | 213031158 | rs1047881 | 0.588 | -0.448 | -0.368 | -0.118 | 0.051 | 0.003 | 0.003 | 0.136 | 0.46 | 0.23 | 0.31 | 0.55 | 0.56 | 0.61 |
| 1 | 213037624 | rs963327 | 0.374 | -0.794 | 0.128 | -0.398 | -0.025 | -0.138 | -0.138 | 2.791 | 0.73 | 0.44 | 0.81 | 0.99 | 0.76 | 0.83 |
| 1 | 213038375 | rs2792561 | -0.101 | -0.843 | 0.68 | -0.451 | -0.013 | -0.064 | -0.064 | 0 | 0.78 | 0.75 | 0.86 | 0.99 | 0.82 | 0.85 |
| 1 | 213046236 | rs1890932 | 0.714 | -0.769 | 0.719 | -0.362 | 0.009 | -0.014 | -0.014 | 0 | 0.26 | 0.17 | 0.49 | 0.44 | 0.19 | 0.23 |
| 1 | 213064209 | rs7553536 | -0.532 | -0.553 | 0.861 | -0.272 | 0.02 | 0.03 | 0.03 | 0 | 0.05 | 0.18 | 0.04 | 0.09 | 0.06 | 0.02 |
| 1 | 213068595 | rs3207090 | 1.096 | -0.5 | 0.889 | -0.271 | -0.023 | -0.047 | -0.047 | 0 | 0.43 | 0.13 | 0.28 | 0.46 | 0.5 | 0.58 |
| 1 | 213069783 | rs6682947 | 0.838 | -0.568 | 0.771 | -0.305 | -0.004 | -0.038 | -0.038 | 3.301 | 0.69 | 0.3 | 0.77 | 0.9 | 0.69 | 0.81 |
| 1 | 213071341 | rs1284855 | -0.386 | -0.55 | 0.814 | -0.227 | -0.003 | 0.009 | 0.009 | 0 | 0.35 | 0.64 | 0.58 | 0.54 | 0.32 | 0.27 |
| 1 | 213073013 | rs1284852 | -0.591 | -0.398 | 0.765 | -0.261 | 0 | -0.002 | -0.002 | 0 | 0.39 | 0.65 | 0.6 | 0.55 | 0.36 | 0.31 |
| 1 | 213073366 | rs1284851 | -1.363 | 0.164 | 1.192 | 0.05 | 0.105 | 0.05 | 0.05 | 0.141 | 0.1 | 0.48 | 0.09 | 0.09 | 0.13 | 0.04 |
| 1 | 213080601 | rs4951651 | 1.286 | 0.107 | 0.853 | 0.013 | 0.014 | -0.033 | -0.033 | 0.004 | 0.82 | 0.32 | 0.86 | 0.91 | 0.78 | 0.85 |
| 1 | 213088494 | rs12042181 | 0.225 | -0.21 | 0.603 | 0.083 | 0.064 | -0.015 | -0.015 | 0.007 | 0.39 | 0.17 | 0.58 | 0.45 | 0.28 | 0.27 |
| 1 | 213089037 | rs1770778 | -0.06 | -0.119 | 0.649 | 0.095 | 0.005 | -0.038 | -0.038 | 0 | 0.92 | 0.76 | 0.94 | 1 | 0.91 | 0.89 |

| | | | | | | | | | | | | | | | | |
|---|---|---|---|---|---|---|---|---|---|---|---|---|---|---|---|---|
| 1 | 213091573 | rs12140722 | 0.62 | -0.067 | 0.709 | 0.208 | -0.01 | -0.007 | -0.007 | 0.002 | 0.42 | 0.34 | 0.27 | 0.42 | 0.47 | 0.54 |
| 1 | 213093069 | rs6679951 | 0.711 | -0.03 | 0.541 | 0.215 | -0.022 | -0.013 | -0.013 | 2.054 | 0.41 | 0.25 | 0.27 | 0.41 | 0.47 | 0.54 |
| 1 | 213094034 | rs1284864 | 0.079 | -0.054 | 0.675 | 0.198 | -0.01 | -0.048 | -0.048 | 0 | 0.91 | 0.77 | 0.94 | 1 | 0.91 | 0.89 |
| 1 | 213094557 | rs11120067 | 0.56 | 0.069 | 0.126 | 0.337 | -0.022 | -0.013 | -0.013 | 0 | 0.41 | 0.25 | 0.27 | 0.41 | 0.47 | 0.54 |
| 1 | 213097217 | rs1890933 | 0.75 | 0.335 | 0.722 | 0.416 | 0.009 | 0.037 | 0.037 | 0 | 0.91 | 0.79 | 0.94 | 0.9 | 0.9 | 0.96 |
| 1 | 213097929 | rs937348 | 1.257 | 0.413 | 0.841 | 0.19 | 0.047 | -0.035 | -0.035 | 0 | 0.84 | 0.56 | 0.89 | 0.9 | 0.81 | 0.9 |
| 1 | 213100605 | rs2792564 | 0.197 | -0.135 | 0.296 | 0.304 | 0.002 | -0.009 | -0.009 | 0 | 0.62 | 0.65 | 0.51 | 0.55 | 0.63 | 0.63 |
| 1 | 213101844 | rs970081 | 0.319 | -0.205 | 0.713 | 0.274 | -0.005 | 0.01 | 0.01 | 0 | 0.61 | 0.53 | 0.49 | 0.55 | 0.63 | 0.63 |
| 1 | 213109096 | rs3124665 | 0.119 | -0.343 | 0.226 | 0.088 | -0.038 | 0.011 | 0.011 | 0 | 0.58 | 0.84 | 0.61 | 0.63 | 0.55 | 0.62 |
| 1 | 213111959 | rs3104212 | 0.29 | -0.376 | 0.242 | -0.029 | 0.005 | -0.037 | -0.037 | 0 | 0.58 | 0.77 | 0.53 | 0.53 | 0.6 | 0.66 |
| 1 | 213120103 | rs6656706 | -0.684 | -0.879 | 0.101 | -0.14 | -0.032 | -0.024 | -0.024 | 0 | 0.19 | 0.7 | 0.24 | 0.14 | 0.11 | 0.17 |
| 1 | 213120185 | rs6656738 | -0.295 | -0.907 | 0.487 | -0.13 | -0.035 | -0.052 | -0.052 | 0 | 0.17 | 0.52 | 0.21 | 0.08 | 0.1 | 0.17 |
| 1 | 213123172 | rs6667192 | -0.665 | -0.915 | 0.998 | -0.194 | -0.033 | -0.078 | -0.078 | 1.467 | 0.17 | 0.48 | 0.12 | 0.06 | 0.1 | 0.08 |
| 1 | 213126541 | rs2935214 | 1.592 | -0.907 | 0.524 | -0.232 | -0.088 | -0.058 | -0.058 | 0 | 0.35 | 0.06 | 0.24 | 0.42 | 0.44 | 0.47 |
| 1 | 213127101 | rs17020046 | -0.253 | -0.663 | 0.937 | -0.347 | -0.018 | -0.085 | -0.085 | 0 | 0.14 | 0.3 | 0.09 | 0 | 0.09 | 0.07 |
| 1 | 213129663 | rs3010785 | 0.86 | -0.844 | 0.379 | -0.343 | -0.015 | -0.02 | -0.02 | 0.127 | 0.39 | 0.23 | 0.28 | 0.45 | 0.45 | 0.51 |
| 1 | 213132001 | rs903803 | -1.964 | -1.229 | 1.08 | -0.071 | -0.091 | -0.011 | -0.011 | 0 | 0.2 | 0.59 | 0.33 | 0.14 | 0.07 | 0.15 |
| 1 | 213138075 | rs6679892 | 0.18 | -0.715 | 0.254 | -0.284 | -0.028 | 0.033 | 0.033 | 0 | 0.57 | 0.76 | 0.59 | 0.66 | 0.54 | 0.59 |
| 1 | 213140398 | rs10779598 | 0.726 | -0.744 | 0.292 | -0.325 | 0.019 | 0.004 | 0.004 | 0 | 0.49 | 0.21 | 0.35 | 0.49 | 0.56 | 0.55 |
| 1 | 213144851 | rs11586518 | 1.692 | -0.773 | 0.107 | -0.347 | -0.093 | -0.085 | -0.085 | 0.204 | 0.37 | 0.07 | 0.27 | 0.49 | 0.46 | 0.44 |
| 1 | 213146944 | rs7511930 | 0.082 | -0.788 | 0.194 | -0.382 | -0.011 | 0.021 | 0.021 | 0 | 0.6 | 0.79 | 0.66 | 0.67 | 0.61 | 0.61 |
| 1 | 213146952 | rs11809007 | -0.929 | -0.92 | 0.497 | -0.106 | 0.024 | 0.05 | 0.05 | 0 | 0.05 | 0.22 | 0.03 | 0.05 | 0.02 | 0.01 |
| 1 | 213161171 | rs10864032 | 1.237 | -0.876 | -0.588 | -0.288 | 0.005 | -0.117 | -0.117 | 0 | 0.48 | 0.04 | 0.37 | 0.53 | 0.64 | 0.59 |
| 1 | 213164918 | rs3010780 | -0.259 | -1.192 | -0.816 | -0.545 | -0.046 | -0.071 | -0.071 | 0 | 0.32 | 0.08 | 0.48 | 0.37 | 0.34 | 0.34 |
| 1 | 213174934 | rs6692011 | -1.171 | -1.66 | 0.089 | -0.537 | -0.064 | -0.092 | -0.092 | 0 | 0.15 | 0.58 | 0.05 | 0 | 0.04 | 0.05 |
| 1 | 213181916 | rs3002284 | -1.562 | -1.675 | 0.068 | -0.536 | 0.071 | -0.027 | -0.027 | 0 | 0.44 | 0.78 | 0.56 | 0.44 | 0.3 | 0.4 |
| 1 | 213184233 | rs3124668 | 0.167 | -1.592 | 0.033 | -0.518 | -0.015 | -0.075 | -0.075 | 0 | 0.27 | 0.08 | 0.42 | 0.37 | 0.27 | 0.29 |
| 1 | 213247854 | rs12065249 | -0.016 | -1.612 | 0.47 | -0.542 | 0 | 0 | 0 | 0 | 0.06 | 0.08 | 0.05 | 0.03 | 0.05 | 0.01 |
| 1 | 213248758 | rs6697921 | 0.7 | -1.565 | 0.535 | -0.549 | 0.003 | -0.01 | -0.01 | 0 | 0.62 | 0.62 | 0.43 | 0.52 | 0.67 | 0.6 |
| 1 | 213273769 | rs6540777 | 1.772 | -1.58 | 0.261 | -0.561 | -0.015 | -0.112 | -0.112 | 0.776 | 0.48 | 0.04 | 0.37 | 0.52 | 0.63 | 0.56 |
| 1 | 213311427 | rs6657234 | -0.289 | -1.604 | 0.281 | -0.568 | -0.013 | -0.059 | -0.059 | 0 | 0.72 | 0.87 | 0.57 | 0.62 | 0.72 | 0.7 |
| 1 | 213316546 | rs10864037 | 1.713 | -1.624 | 0.201 | -0.577 | -0.008 | -0.107 | -0.107 | 1.59 | 0.49 | 0.04 | 0.35 | 0.52 | 0.63 | 0.56 |
| 1 | 213318765 | rs6671872 | -0.271 | -1.567 | 0.256 | -0.546 | -0.023 | -0.079 | -0.079 | 0 | 0.72 | 0.91 | 0.55 | 0.62 | 0.72 | 0.7 |
| 1 | 213348514 | rs10864038 | -0.284 | -1.565 | 0.257 | -0.545 | -0.024 | -0.079 | -0.079 | 0 | 0.72 | 0.91 | 0.57 | 0.62 | 0.72 | 0.7 |
| 1 | 213378760 | rs2647387 | -0.274 | -1.563 | 0.253 | -0.544 | -0.019 | -0.071 | -0.071 | 4.121 | 0.72 | 0.9 | 0.57 | 0.62 | 0.72 | 0.7 |
| 1 | 213416302 | rs10494957 | -0.238 | -1.429 | 0.17 | -0.501 | -0.014 | -0.033 | -0.033 | 0 | 0.07 | 0.1 | 0.03 | 0 | 0.03 | 0.05 |
| 1 | 213437680 | rs10779599 | 1.583 | -1.357 | -0.194 | -0.182 | -0.015 | -0.111 | -0.111 | 0 | 0.47 | 0.04 | 0.37 | 0.52 | 0.63 | 0.56 |
| 1 | 213449418 | rs10864039 | 1.498 | -1.342 | -0.186 | -0.174 | -0.008 | -0.106 | -0.106 | 0.122 | 0.48 | 0.04 | 0.37 | 0.52 | 0.63 | 0.56 |
| 1 | 213451958 | rs12040029 | -0.184 | -1.533 | -0.347 | -0.971 | -0.01 | -0.013 | -0.013 | 0 | 0.17 | 0.02 | 0.24 | 0.14 | 0.12 | 0.1 |
| 1 | 213452267 | rs1628854 | 1.165 | -1.32 | 0.691 | -0.83 | 0.013 | 0.111 | 0.111 | 0.15 | 0.9 | 0.54 | 0.84 | 0.86 | 0.95 | 0.92 |
| 1 | 213456220 | rs1187802 | 1.199 | -1.35 | 0.392 | -0.849 | -0.017 | 0.032 | 0.032 | 0 | 0.91 | 0.75 | 0.85 | 0.87 | 0.95 | 0.92 |
| 1 | 213464445 | rs12026921 | -0.57 | -1.365 | -0.005 | -1.149 | 0.018 | 0.052 | 0.052 | 0 | 0.38 | 0.25 | 0.34 | 0.2 | 0.33 | 0.36 |
| 1 | 213468908 | rs10864040 | -1.802 | -1.363 | -0.056 | -1.154 | 0.03 | 0.121 | 0.121 | 0 | 0.53 | 0.8 | 0.45 | 0.24 | 0.42 | 0.45 |
| 1 | 213484976 | rs6696434 | -0.548 | -1.777 | -0.16 | -1.128 | 0.02 | 0.053 | 0.053 | 0 | 0.38 | 0.26 | 0.35 | 0.2 | 0.34 | 0.36 |
| 1 | 213491399 | rs1655996 | 1.6 | -1.581 | -0.642 | -1.11 | -0.064 | -0.024 | -0.024 | 2.08 | 0.42 | 0.04 | 0.45 | 0.67 | 0.58 | 0.5 |
| 1 | 213493727 | rs4359010 | 1.005 | -1.214 | 0.072 | -0.905 | 0.007 | 0.008 | 0.008 | 0 | 0.56 | 0.41 | 0.5 | 0.68 | 0.63 | 0.58 |
| 1 | 213499232 | rs1699185 | 0.836 | -1.319 | 0.23 | -0.53 | -0.031 | 0.012 | 0.012 | 0.833 | 0.88 | 0.59 | 0.87 | 0.9 | 0.96 | 0.83 |
| 1 | 213511231 | rs1434889 | 1.237 | -1.375 | -0.979 | -0.736 | 0.108 | 0.079 | 0.079 | 0.063 | 0.53 | 0.2 | 0.54 | 0.63 | 0.71 | 0.56 |
| 1 | 213513059 | rs11580058 | 0.366 | -1.162 | -1.522 | -0.746 | -0.012 | 0.001 | 0.001 | 0 | 0.06 | 0.04 | 0.08 | 0.02 | 0.1 | 0.05 |
| 1 | 213524560 | rs1187767 | -0.142 | -1.293 | -1.548 | -0.799 | 0.026 | 0.004 | 0.004 | 0.144 | 0.23 | 0.06 | 0.27 | 0.21 | 0.2 | 0.27 |
| 1 | 213527242 | rs1656022 | 1.076 | -0.965 | -1.519 | -0.622 | 0.069 | 0.022 | 0.022 | 0.026 | 0.48 | 0.07 | 0.46 | 0.61 | 0.66 | 0.53 |
| 1 | 213536227 | rs1656015 | -1.65 | -0.361 | -0.856 | -0.954 | 0.035 | 0.093 | 0.093 | 0 | 0.45 | 0.82 | 0.35 | 0.27 | 0.37 | 0.33 |
| 1 | 213536488 | rs1656014 | -1.572 | -0.401 | -0.752 | -0.952 | 0.026 | 0.134 | 0.134 | 0 | 0.43 | 0.77 | 0.26 | 0.06 | 0.37 | 0.23 |
| 1 | 213538617 | rs1656011 | -1.556 | -0.398 | -0.767 | -0.951 | 0.024 | 0.135 | 0.135 | 0 | 0.43 | 0.77 | 0.26 | 0.06 | 0.37 | 0.23 |
| 1 | 213542560 | rs753369 | -0.923 | -0.429 | -0.197 | -0.287 | 0.001 | 0.022 | 0.022 | 0 | 0.17 | 0.16 | 0.09 | 0.01 | 0.15 | 0.1 |
| 1 | 213558937 | rs11809986 | -1.127 | -0.036 | -0.153 | -0.404 | -0.02 | -0.008 | -0.008 | 0 | 0.1 | 0.17 | 0.05 | 0.03 | 0.03 | 0.03 |
| 1 | 213563818 | rs1187770 | 0.591 | 0.221 | 0.038 | -0.736 | 0.023 | -0.058 | -0.058 | 1.482 | 0.88 | 0.78 | 0.92 | 0.99 | 0.86 | 0.91 |
| 1 | 213565567 | rs320427 | 1.174 | -0.084 | 0.209 | -0.905 | -0.019 | -0.12 | -0.12 | 0 | 0.78 | 0.47 | 0.87 | 0.99 | 0.84 | 0.89 |
| 1 | 213568239 | rs7551878 | 1.218 | -0.185 | -0.739 | -0.939 | 0.024 | 0.086 | 0.086 | 0.251 | 0.75 | 0.2 | 0.85 | 0.98 | 0.83 | 0.9 |
| 1 | 213578006 | rs4655383 | 1.322 | 0.071 | -1.085 | -1.131 | -0.006 | 0.183 | 0.183 | 0 | 0.63 | 0.15 | 0.76 | 0.96 | 0.68 | 0.77 |
| 1 | 213580800 | rs11807941 | -0.802 | -0.316 | -1.053 | -1.457 | 0.015 | 0.011 | 0.011 | 0 | 0.34 | 0.25 | 0.15 | 0.07 | 0.22 | 0.31 |
| 1 | 213583547 | rs458411 | -0.977 | 0.458 | -2.012 | -0.675 | -0.074 | -0.077 | -0.077 | 0 | 0.64 | 0.89 | 0.54 | 0.61 | 0.58 | 0.64 |
| 1 | 213589359 | rs320464 | -0.196 | 0.662 | -1.617 | -0.549 | 0.001 | -0.006 | -0.006 | 0.213 | 0.67 | 0.68 | 0.71 | 0.64 | 0.67 | 0.44 |
| 1 | 213594118 | rs748795 | 1.2 | 0.532 | -2.127 | -0.32 | -0.113 | -0.157 | -0.157 | 0 | 0.36 | 0.06 | 0.52 | 0.46 | 0.45 | 0.38 |
| 1 | 213595546 | rs12035134 | 0.313 | 0.901 | -2.031 | -0.273 | -0.055 | 0.005 | 0.005 | 0 | 0.15 | 0.01 | 0.09 | 0.09 | 0.16 | 0.07 |
| 1 | 213597841 | rs320461 | -0.381 | 0.658 | -1.325 | -0.14 | 0 | 0 | 0 | 0.534 | 0.73 | 0.71 | 0.75 | 0.64 | 0.74 | 0.58 |
| 1 | 213599316 | rs3910657 | 0.57 | 0.82 | -1.759 | -0.035 | -0.016 | -0.031 | -0.031 | 2.599 | 0.38 | 0.24 | 0.55 | 0.56 | 0.46 | 0.48 |
| 1 | 213602132 | rs12023455 | 0.29 | 0.827 | -1.518 | -0.221 | -0.006 | -0.006 | -0.006 | 0 | 0.21 | 0.16 | 0.14 | 0.23 | 0.2 | 0.1 |
| 1 | 213603077 | rs11120129 | 1.102 | 0.501 | -1.741 | -0.191 | -0.064 | -0.061 | -0.061 | 1.301 | 0.21 | 0.01 | 0.14 | 0.24 | 0.22 | 0.09 |

| | | | | | | | | | | | | | | | | |
|---|---|---|---|---|---|---|---|---|---|---|---|---|---|---|---|---|
| 1 | 213603105 | rs12045355 | 1.783 | 0.644 | -1.947 | 0.356 | -0.042 | -0.056 | -0.056 | 0.396 | 0.1 | 0.01 | 0.29 | 0.4 | 0.12 | 0.27 |
| 1 | 213611635 | rs6540789 | 0.559 | -0.288 | -1.537 | -0.801 | -0.025 | -0.018 | -0.018 | 0 | 0.35 | 0.26 | 0.6 | 0.57 | 0.44 | 0.45 |
| 1 | 213612779 | rs12049265 | 1.995 | -0.198 | -1.572 | -0.786 | -0.048 | -0.053 | -0.053 | 0.036 | 0.2 | 0.01 | 0.12 | 0.24 | 0.17 | 0.08 |
| 1 | 213617738 | rs320451 | -0.457 | -0.23 | -0.75 | -0.844 | -0.002 | -0.04 | -0.04 | 0 | 0.65 | 0.66 | 0.82 | 0.87 | 0.67 | 0.57 |
| 1 | 213625292 | rs320442 | -1.143 | -0.354 | -0.691 | -1.062 | 0.005 | -0.071 | -0.071 | 0 | 0.77 | 0.73 | 0.86 | 0.98 | 0.74 | 0.73 |
| 1 | 213626376 | rs4655282 | -0.153 | -0.269 | -1.281 | -1.003 | -0.002 | 0.027 | 0.027 | 0 | 0.48 | 0.4 | 0.69 | 0.68 | 0.52 | 0.62 |
| 1 | 213629078 | rs320439 | 0.903 | 0.215 | -1.48 | -1.578 | 0.007 | -0.01 | -0.01 | 0 | 0.27 | 0.2 | 0.16 | 0.29 | 0.22 | 0.1 |
| 1 | 213636801 | rs12036568 | 0.02 | 0.742 | -1.4 | -1.03 | -0.033 | -0.02 | -0.02 | 0.16 | 0.34 | 0.21 | 0.59 | 0.58 | 0.44 | 0.45 |
| 1 | 213641943 | rs320480 | -0.924 | 0.684 | -0.61 | -1.051 | 0.004 | -0.022 | -0.022 | 0.495 | 0.89 | 0.82 | 0.91 | 0.98 | 0.83 | 0.83 |
| 1 | 213657633 | rs10864046 | 1.103 | 0.524 | -1.264 | -1.045 | -0.084 | -0.094 | -0.094 | 0.015 | 0.21 | 0.01 | 0.23 | 0.28 | 0.21 | 0.08 |
| 1 | 213657863 | rs320471 | 0.053 | 0.726 | -0.791 | -0.955 | -0.02 | -0.031 | -0.031 | 0.495 | 0.28 | 0.22 | 0.44 | 0.59 | 0.37 | 0.47 |
| 1 | 213659931 | rs4655421 | 0.732 | -0.016 | -0.407 | -1.233 | -0.02 | 0.012 | 0.012 | 1.769 | 0.34 | 0.19 | 0.4 | 0.26 | 0.29 | 0.14 |
| 1 | 213660162 | rs320476 | -0.718 | -0.277 | 0.036 | -1.526 | -0.029 | -0.133 | -0.133 | 0.057 | 0.72 | 0.59 | 0.82 | 0.99 | 0.8 | 0.69 |
| 1 | 213661444 | rs11120139 | 0.899 | -0.094 | -0.156 | -1.361 | -0.02 | 0.012 | 0.012 | 1.811 | 0.33 | 0.19 | 0.4 | 0.26 | 0.29 | 0.14 |
| 1 | 213664461 | rs2165129 | 0.943 | -0.169 | -0.258 | -1.362 | -0.001 | 0 | 0 | 2.089 | 0.25 | 0.29 | 0.43 | 0.62 | 0.38 | 0.46 |
| 1 | 213666937 | rs320374 | 0.465 | -0.133 | -0.11 | -1.423 | 0.013 | 0.011 | 0.011 | 1.584 | 0.41 | 0.3 | 0.42 | 0.37 | 0.29 | 0.25 |
| 1 | 213670585 | rs11120142 | 0.662 | -0.165 | -0.188 | -1.443 | -0.021 | -0.027 | -0.027 | 0 | 0.38 | 0.1 | 0.41 | 0.37 | 0.29 | 0.22 |
| 1 | 213674732 | rs965121 | 1.53 | -0.706 | 0.876 | -1.374 | 0.058 | 0.183 | 0.183 | 0 | 0.55 | 0.26 | 0.62 | 0.87 | 0.65 | 0.57 |
| 1 | 213676893 | rs10494959 | -0.376 | 0.52 | 1.145 | -1.26 | 0.04 | -0.034 | -0.034 | 0.183 | 0.06 | 0.12 | 0.07 | 0 | 0.13 | 0.04 |
| 1 | 213677091 | rs320379 | 0.538 | 0.381 | 0.776 | -1.129 | 0 | -0.092 | -0.092 | 0.398 | 0.68 | 0.65 | 0.76 | 0.99 | 0.67 | 0.72 |
| 1 | 213685950 | rs430997 | 0.845 | 0.376 | 0.167 | -0.095 | 0.001 | -0.013 | -0.013 | 2.51 | 0.83 | 0.87 | 0.89 | 0.99 | 0.86 | 0.81 |
| 1 | 213692263 | rs893665 | 1.87 | 0.525 | 0.966 | -0.554 | 0 | -0.061 | -0.061 | 0 | 0.71 | 0.31 | 0.79 | 0.99 | 0.72 | 0.76 |
| 1 | 213702665 | rs320425 | -0.788 | 0.529 | 1.692 | -0.54 | 0.075 | -0.03 | -0.03 | 0 | 0.06 | 0.15 | 0.06 | 0 | 0.13 | 0.05 |
| 1 | 213703223 | rs320424 | 1.314 | 0.444 | 2.39 | -0.577 | -0.03 | -0.143 | -0.143 | 0 | 0.78 | 0.54 | 0.85 | 0.97 | 0.85 | 0.8 |
| 1 | 213707153 | rs10864051 | -2.337 | 0.309 | 2.426 | 0.637 | -0.031 | 0.129 | 0.129 | 0.328 | 0.11 | 0.25 | 0.24 | 0.27 | 0.05 | 0.06 |
| 1 | 213711548 | rs320414 | 1.386 | 0.142 | 3.106 | 0.5 | -0.012 | -0.073 | -0.073 | 0.104 | 0.78 | 0.65 | 0.86 | 0.97 | 0.85 | 0.81 |
| 1 | 213714381 | rs11120157 | 2.444 | 0.096 | 3.197 | 0.518 | 0.003 | -0.014 | -0.014 | 0 | 0.65 | 0.24 | 0.59 | 0.7 | 0.69 | 0.7 |
| 1 | 213715274 | rs6680294 | 2.044 | 0.177 | 3.175 | 0.438 | 0.003 | -0.111 | -0.111 | 0 | 0.71 | 0.56 | 0.8 | 0.97 | 0.72 | 0.75 |
| 1 | 213720321 | rs320395 | -1.71 | 0.557 | 2.683 | 0.795 | -0.019 | -0.002 | -0.002 | 0.531 | 0.1 | 0.06 | 0.11 | 0.1 | 0.2 | 0.15 |
| 1 | 213726328 | rs7530543 | -2.441 | 0.56 | 2.642 | 0.937 | 0.095 | 0.236 | 0.236 | 0 | 0.14 | 0.27 | 0.21 | 0.38 | 0.21 | 0.16 |
| 1 | 213752902 | rs4475718 | -1.612 | 0.429 | 1.942 | 0.876 | -0.007 | -0.001 | -0.001 | 0 | 0.1 | 0.08 | 0.1 | 0.1 | 0.18 | 0.12 |
| 1 | 213754770 | rs6540792 | -2.539 | 0.281 | 2.005 | 0.857 | -0.004 | -0.017 | -0.017 | 0 | 0.37 | 0.69 | 0.36 | 0.4 | 0.37 | 0.36 |
| 1 | 213759712 | rs11120162 | -1.851 | 0.249 | 1.891 | 1.527 | 0.002 | 0.065 | 0.065 | 0 | 0.08 | 0.06 | 0.06 | 0.44 | 0.06 | 0.1 |
| 1 | 213760845 | rs12062324 | -2.162 | 0.163 | 2.233 | 1.309 | -0.001 | 0.028 | 0.028 | 0 | 0.1 | 0.27 | 0.05 | 0.1 | 0.06 | 0.08 |
| 1 | 213762107 | rs17021041 | -1.541 | 0.189 | 2.119 | 1.325 | 0.006 | 0.036 | 0.036 | 0 | 0.06 | 0.16 | 0.04 | 0.1 | 0.04 | 0.07 |
| 1 | 213772821 | rs11120168 | 1.675 | -0.259 | 1.667 | 0.997 | -0.012 | 0.006 | 0.006 | 0 | 0.67 | 0.35 | 0.69 | 0.69 | 0.72 | 0.76 |
| 1 | 213776679 | rs4561011 | 1.82 | -0.269 | 1.4 | 0.896 | 0.011 | -0.015 | -0.015 | 0.07 | 0.64 | 0.29 | 0.66 | 0.61 | 0.69 | 0.7 |
| 1 | 213777711 | rs10779604 | 0.857 | -0.116 | 0.974 | 0.541 | -0.021 | -0.081 | -0.081 | 0 | 0.84 | 0.7 | 0.88 | 0.97 | 0.89 | 0.88 |
| 1 | 213778619 | rs10864054 | 0.992 | -0.068 | 0.582 | 0.516 | -0.015 | -0.071 | -0.071 | 0 | 0.82 | 0.77 | 0.87 | 0.97 | 0.88 | 0.88 |
| 1 | 213779196 | rs11120170 | -0.668 | 0.275 | -0.303 | 1.141 | -0.003 | 0.003 | 0.003 | 0 | 0.42 | 0.47 | 0.64 | 0.39 | 0.37 | 0.4 |
| 1 | 213786241 | rs885179 | -0.779 | 0.293 | -1.076 | 2.13 | -0.01 | -0.026 | -0.026 | 0.538 | 0.13 | 0.04 | 0.2 | 0.17 | 0.09 | 0.2 |
| 1 | 213796830 | rs11805334 | 0.351 | 0.508 | -1.547 | 1.913 | -0.027 | -0.014 | -0.014 | 0 | 0.71 | 0.84 | 0.61 | 0.66 | 0.62 | 0.69 |
| 1 | 213797431 | rs12740651 | 0.485 | 0.367 | -1.344 | 1.909 | -0.012 | -0.008 | -0.008 | 0.737 | 0.7 | 0.77 | 0.61 | 0.64 | 0.62 | 0.67 |
| 1 | 213805502 | rs4655454 | 0.604 | -0.311 | -1.638 | 0.826 | -0.054 | 0.047 | 0.047 | 0.145 | 0.39 | 0.15 | 0.29 | 0.33 | 0.45 | 0.44 |
| 1 | 213811721 | rs7529857 | 0.401 | -0.621 | -1.178 | 0.544 | -0.006 | -0.019 | -0.019 | 0.004 | 0.1 | 0.11 | 0.03 | 0.01 | 0.05 | 0.16 |
| 1 | 213812489 | rs12045301 | 0.399 | 0.076 | -1.404 | 0.759 | -0.049 | -0.034 | -0.034 | 0 | 0.09 | 0 | 0.15 | 0.13 | 0.18 | 0.19 |
| 1 | 213817311 | rs12144639 | -0.359 | -0.43 | -0.783 | 0.738 | -0.042 | -0.054 | -0.054 | 1.869 | 0.33 | 0.54 | 0.23 | 0.2 | 0.23 | 0.22 |
| 1 | 213821667 | rs6540796 | 0.201 | -0.055 | -1.034 | 0.891 | -0.023 | -0.027 | -0.027 | 0 | 0.79 | 0.85 | 0.56 | 0.47 | 0.72 | 0.64 |
| 1 | 213823471 | rs12128947 | 1.636 | -0.254 | -1.11 | 0.917 | -0.034 | 0.077 | 0.077 | 0.013 | 0.41 | 0.13 | 0.3 | 0.31 | 0.46 | 0.42 |
| 1 | 213825023 | rs11120179 | 1.317 | -0.242 | -1.013 | 0.902 | -0.004 | 0.083 | 0.083 | 0 | 0.46 | 0.28 | 0.33 | 0.27 | 0.48 | 0.42 |
| 1 | 213826374 | rs12023700 | 0.62 | -0.33 | -1.527 | 0.796 | 0.031 | 0.093 | 0.093 | 0 | 0.21 | 0.09 | 0.11 | 0.04 | 0.16 | 0.15 |
| 1 | 213827269 | rs6693655 | 0.203 | -0.112 | -1.553 | 1.308 | -0.049 | -0.068 | -0.068 | 0.194 | 0.13 | 0.02 | 0.28 | 0.26 | 0.2 | 0.24 |
| 1 | 213827795 | rs1509863 | -0.448 | -0.081 | -1.303 | 1.305 | 0.009 | 0.036 | 0.036 | 0 | 0.87 | 0.83 | 0.83 | 0.72 | 0.88 | 0.87 |
| 1 | 213828800 | rs10779605 | -0.062 | -0.111 | -1.512 | 1.27 | -0.024 | -0.024 | -0.024 | 0.082 | 0.33 | 0.17 | 0.4 | 0.25 | 0.35 | 0.37 |
| 1 | 213834509 | rs2884830 | 1.075 | 0.211 | -1.652 | 1.121 | -0.03 | 0.085 | 0.085 | 0 | 0.42 | 0.11 | 0.31 | 0.3 | 0.46 | 0.43 |
| 1 | 213834902 | rs1509861 | -0.142 | 0.464 | -1.549 | 1.098 | 0.007 | 0.014 | 0.014 | 0.067 | 0.87 | 0.85 | 0.83 | 0.78 | 0.87 | 0.89 |
| 1 | 213835207 | rs6662813 | 0.388 | 0.228 | -1.885 | 0.753 | -0.004 | 0.049 | 0.049 | 0.09 | 0.49 | 0.17 | 0.38 | 0.42 | 0.5 | 0.55 |
| 1 | 213843800 | rs12033150 | 0.346 | 0.482 | -2.607 | 0.454 | -0.09 | -0.049 | -0.049 | 3.508 | 0.53 | 0.07 | 0.43 | 0.43 | 0.53 | 0.74 |
| 1 | 213849457 | rs17021216 | 0.363 | 0.509 | -2.61 | 0.424 | -0.094 | -0.044 | -0.044 | 0 | 0.53 | 0.09 | 0.43 | 0.43 | 0.52 | 0.74 |
| 1 | 213853968 | rs4518871 | -0.274 | 0.627 | -1.52 | 0.378 | -0.009 | -0.024 | -0.024 | 0 | 0.19 | 0.27 | 0.09 | 0.13 | 0.15 | 0.1 |
| 1 | 213854151 | rs12035846 | 0.451 | 0.562 | -1.359 | 0.324 | -0.02 | -0.017 | -0.017 | 0 | 0.15 | 0.11 | 0.45 | 0.43 | 0.28 | 0.11 |
| 1 | 213859249 | rs11585232 | 0.07 | 0.248 | -1.237 | 0.129 | 0 | -0.004 | -0.004 | 0 | 0.12 | 0.13 | 0.06 | 0.05 | 0.1 | 0.07 |
| 1 | 213861309 | rs1509867 | -0.398 | -0.114 | -2.116 | -0.609 | 0.009 | 0.098 | 0.098 | 0.129 | 0.59 | 0.8 | 0.75 | 0.82 | 0.61 | 0.43 |
| 1 | 213861648 | rs1509868 | 0.277 | -0.57 | -1.499 | -0.915 | 0 | 0 | 0 | 0 | 0.63 | 0.47 | 0.47 | 0.52 | 0.54 | 0.78 |
| 1 | 213862164 | rs12136457 | 0.999 | -0.733 | -1.353 | -0.835 | -0.045 | -0.076 | -0.076 | 0 | 0.26 | 0.04 | 0.21 | 0.32 | 0.27 | 0.31 |
| 1 | 213863315 | rs11120194 | -0.786 | -0.69 | -0.109 | -1.012 | -0.039 | -0.068 | -0.068 | 0 | 0.17 | 0.57 | 0.09 | 0.06 | 0.11 | 0.06 |
| 1 | 213866694 | rs12759209 | 0.67 | -0.714 | -0.348 | -1.229 | -0.018 | -0.052 | -0.052 | 0 | 0.19 | 0.21 | 0.09 | 0.03 | 0.11 | 0.14 |
| 1 | 213867407 | rs937973 | 0.584 | -0.684 | -1.076 | -1.405 | -0.132 | -0.059 | -0.059 | 0 | 0.31 | 0.09 | 0.62 | 0.68 | 0.53 | 0.5 |
| 1 | 213868299 | rs11120195 | 0.613 | -0.668 | -1.052 | -1.348 | -0.118 | -0.071 | -0.071 | 0 | 0.31 | 0.1 | 0.59 | 0.64 | 0.52 | 0.49 |

| | | | | | | | | | | | | | | | | |
|---|---|---|---|---|---|---|---|---|---|---|---|---|---|---|---|---|
| 1 | 213873500 | rs1588374 | -1.265 | -1.023 | -0.906 | -1.552 | 0.007 | 0.015 | 0.015 | 0.23 | 0.73 | 0.83 | 0.8 | 0.81 | 0.78 | 0.81 |
| 1 | 213877776 | rs10864064 | -1.232 | -0.985 | -0.908 | -1.553 | 0.006 | 0.018 | 0.018 | 0.158 | 0.72 | 0.83 | 0.8 | 0.81 | 0.78 | 0.81 |
| 1 | 213886423 | rs11588107 | 0.136 | -1.076 | -0.635 | -1.59 | 0.01 | 0.013 | 0.013 | 0 | 0.18 | 0.17 | 0.07 | 0.02 | 0.12 | 0.12 |
| 1 | 213887109 | rs12044591 | -0.056 | -0.703 | -0.835 | -1.634 | -0.033 | -0.043 | -0.043 | 0 | 0.35 | 0.22 | 0.46 | 0.6 | 0.41 | 0.3 |
| 1 | 213888861 | rs11120203 | -0.17 | -0.825 | -0.569 | -1.743 | -0.019 | -0.078 | -0.078 | 0 | 0.38 | 0.43 | 0.14 | 0.06 | 0.23 | 0.27 |
| 1 | 213889563 | rs1553794 | -2.071 | -0.956 | -0.283 | -1.833 | 0.001 | -0.002 | -0.002 | 0 | 0.77 | 0.78 | 0.8 | 0.81 | 0.78 | 0.81 |
| 1 | 213889697 | rs6692320 | 1.559 | -0.917 | -0.225 | -1.817 | -0.019 | -0.051 | -0.051 | 0 | 0.2 | 0.21 | 0.07 | 0.03 | 0.11 | 0.14 |
| 1 | 213890404 | rs1509869 | -0.303 | -0.49 | -0.218 | -1.529 | -0.001 | 0.002 | 0.002 | 0 | 0.52 | 0.48 | 0.38 | 0.24 | 0.49 | 0.63 |
| 1 | 213898329 | rs861229 | 0.529 | -0.442 | -0.334 | -1.499 | -0.021 | 0.029 | 0.029 | 0 | 0.19 | 0.15 | 0.26 | 0.11 | 0.25 | 0.34 |
| 1 | 213898344 | rs3008203 | -0.679 | -0.333 | -0.31 | -1.345 | -0.023 | 0.045 | 0.045 | 0 | 0.59 | 0.75 | 0.41 | 0.24 | 0.49 | 0.64 |
| 1 | 213905747 | rs1771901 | -1.177 | -0.236 | -0.337 | -1.254 | -0.007 | -0.001 | -0.001 | 0 | 0.84 | 0.84 | 0.75 | 0.82 | 0.75 | 0.64 |
| 1 | 213907945 | rs1704199 | -1.445 | -0.21 | -0.533 | -1.271 | -0.056 | 0.018 | 0.018 | 0 | 0.86 | 0.95 | 0.76 | 0.9 | 0.75 | 0.67 |
| 1 | 213913001 | rs4655466 | 1.193 | -0.108 | -0.355 | -1.073 | -0.006 | -0.007 | -0.007 | 0 | 0.18 | 0.13 | 0.39 | 0.57 | 0.28 | 0.18 |
| 1 | 213915710 | rs2793276 | -0.028 | -0.059 | -0.566 | -1.479 | 0 | -0.031 | -0.031 | 0 | 0.48 | 0.49 | 0.62 | 0.83 | 0.52 | 0.4 |
| 1 | 213918920 | rs11120205 | 0.162 | 0.182 | -0.536 | -0.998 | 0.017 | -0.066 | -0.066 | 0.037 | 0.13 | 0.23 | 0.1 | 0 | 0.14 | 0.06 |
| 1 | 213923233 | rs701903 | -0.127 | -0.158 | -1.325 | -1.072 | 0.019 | 0.014 | 0.014 | 0.005 | 0.38 | 0.68 | 0.16 | 0.06 | 0.22 | 0.24 |
| 1 | 213924960 | rs3008204 | -0.974 | 0.27 | -1.282 | -0.814 | -0.107 | 0.009 | 0.009 | 0.123 | 0.67 | 0.9 | 0.54 | 0.71 | 0.56 | 0.52 |
| 1 | 213925444 | rs5011573 | -0.002 | 0.919 | -0.508 | 0.022 | -0.003 | -0.008 | -0.008 | 0 | 0.25 | 0.2 | 0.37 | 0.6 | 0.33 | 0.25 |
| 1 | 213926607 | rs1704203 | -0.767 | 0.775 | -0.067 | -0.02 | 0.028 | -0.076 | -0.076 | 0 | 0.12 | 0.38 | 0.07 | 0 | 0.09 | 0.05 |
| 1 | 213932821 | rs12041173 | -0.373 | 1.41 | -1.076 | 0.606 | -0.039 | -0.038 | -0.038 | 0 | 0.21 | 0.12 | 0.37 | 0.7 | 0.29 | 0.25 |
| 1 | 213937037 | rs2359934 | 0.371 | 1.278 | -2.065 | 0.397 | -0.125 | -0.089 | -0.089 | 0.063 | 0.28 | 0.07 | 0.38 | 0.35 | 0.47 | 0.42 |
| 1 | 213937382 | rs1891058 | 0.138 | 0.345 | -1.459 | 0.029 | 0.023 | 0.089 | 0.089 | 0.255 | 0.47 | 0.26 | 0.62 | 0.79 | 0.62 | 0.63 |
| 1 | 213942168 | rs12141140 | -0.508 | -0.568 | -0.931 | -0.703 | -0.004 | -0.057 | -0.057 | 0.239 | 0.79 | 0.7 | 0.88 | 0.98 | 0.85 | 0.85 |
| 1 | 213945761 | rs3008206 | -1.086 | -0.606 | -0.889 | -0.773 | -0.001 | -0.03 | -0.03 | 0.181 | 0.86 | 0.82 | 0.92 | 0.98 | 0.9 | 0.86 |
| 1 | 213947269 | rs10864069 | 0.634 | -0.561 | -0.976 | -0.648 | 0.039 | 0.127 | 0.127 | 0.187 | 0.48 | 0.27 | 0.65 | 0.84 | 0.63 | 0.64 |
| 1 | 213947964 | rs4233314 | -0.419 | -0.605 | -0.759 | -0.977 | -0.01 | -0.086 | -0.086 | 0.499 | 0.78 | 0.63 | 0.88 | 0.98 | 0.85 | 0.84 |
| 1 | 213952574 | rs2359936 | 0.076 | -0.136 | -1.173 | -0.539 | 0.05 | 0.134 | 0.134 | 0.133 | 0.49 | 0.3 | 0.75 | 0.91 | 0.64 | 0.69 |
| 1 | 213961900 | rs10864070 | -0.685 | -0.669 | -0.734 | 0.228 | -0.018 | 0.004 | 0.004 | 0.06 | 0.18 | 0.21 | 0.28 | 0.31 | 0.1 | 0.28 |
| 1 | 213965246 | rs11120214 | -1.395 | -1.289 | -0.301 | -0.076 | -0.006 | 0 | 0 | 0.189 | 0.27 | 0.71 | 0.34 | 0.27 | 0.15 | 0.33 |
| 1 | 213966887 | rs10494964 | -1.696 | -1.937 | -0.98 | 0.052 | 0.135 | 0.004 | 0.004 | 0 | 0.49 | 0.81 | 0.48 | 0.53 | 0.28 | 0.48 |
| 1 | 213967677 | rs2282386 | -0.52 | -1.844 | -0.255 | -0.006 | -0.069 | -0.015 | -0.015 | 0 | 0.22 | 0.49 | 0.3 | 0.23 | 0.1 | 0.27 |
| 1 | 213969354 | rs1891060 | -1.477 | -1.614 | -0.861 | -0.209 | 0.05 | 0.039 | 0.039 | 0.793 | 0.44 | 0.76 | 0.46 | 0.46 | 0.31 | 0.45 |
| 1 | 213970960 | rs6702664 | -0.65 | -1.554 | 0.008 | 0.17 | -0.031 | 0.044 | 0.044 | 0 | 0.16 | 0.37 | 0.08 | 0.24 | 0.06 | 0.16 |
| 1 | 213974734 | rs17709142 | 1.007 | -1.417 | -0.114 | 0.107 | 0.002 | 0.011 | 0.011 | 0 | 0.11 | 0.06 | 0.1 | 0.04 | 0.09 | 0.04 |
| 1 | 213977989 | rs12403122 | -0.592 | -1.199 | 0.253 | 0.29 | -0.034 | 0.038 | 0.038 | 0 | 0.15 | 0.32 | 0.08 | 0.23 | 0.06 | 0.15 |
| 1 | 213978359 | rs11120217 | -0.437 | -1.147 | 0.24 | 0.323 | -0.002 | 0.084 | 0.084 | 0.022 | 0.05 | 0.16 | 0.17 | 0.23 | 0.03 | 0.2 |
| 1 | 213985718 | rs17021481 | -1.793 | -1.232 | 0.081 | 0.239 | -0.034 | 0.032 | 0.032 | 0.45 | 0.23 | 0.69 | 0.25 | 0.46 | 0.1 | 0.37 |
| 1 | 213991857 | rs1578435 | -1.828 | -1.237 | -0.107 | 0.266 | -0.018 | 0.016 | 0.016 | 0.687 | 0.23 | 0.73 | 0.27 | 0.46 | 0.1 | 0.38 |
| 1 | 213999434 | rs1341152 | -0.665 | -0.886 | -0.214 | 0.299 | -0.005 | -0.008 | -0.008 | 0 | 0.16 | 0.14 | 0.21 | 0.26 | 0.1 | 0.3 |
| 1 | 214001594 | rs10494967 | -2.114 | -1.411 | -0.112 | -0.006 | -0.014 | -0.001 | -0.001 | 0 | 0.28 | 0.68 | 0.28 | 0.36 | 0.15 | 0.35 |
| 1 | 214002260 | rs6693625 | -0.364 | -1.344 | 0.158 | 0.019 | -0.013 | -0.048 | -0.048 | 0 | 0.1 | 0.36 | 0.07 | 0 | 0.04 | 0.01 |
| 1 | 214008627 | rs7515689 | -1.892 | -1.419 | -0.962 | -0.527 | -0.045 | 0.057 | 0.057 | 0.109 | 0.78 | 0.99 | 0.9 | 0.93 | 0.81 | 0.8 |
| 1 | 214009019 | rs6540803 | -1.901 | -1.311 | -0.638 | -0.282 | -0.004 | 0 | 0 | 0 | 0.24 | 0.78 | 0.27 | 0.36 | 0.12 | 0.28 |
| 1 | 214021214 | rs10864074 | -0.75 | -1.37 | -0.081 | -0.64 | 0.003 | 0.006 | 0.006 | 0 | 0.21 | 0.21 | 0.31 | 0.17 | 0.14 | 0.29 |
| 1 | 214023872 | rs12118140 | 0.641 | -1.378 | -0.035 | -0.65 | -0.002 | -0.03 | -0.03 | 0 | 0.11 | 0.12 | 0.15 | 0 | 0.11 | 0.07 |
| 1 | 214026324 | rs2236580 | 1.244 | -0.932 | -0.415 | -0.016 | -0.031 | -0.05 | -0.05 | 0.021 | 0.22 | 0.11 | 0.32 | 0.46 | 0.33 | 0.3 |
| 1 | 214043216 | rs853752 | 1.216 | -1.209 | 0.048 | -0.019 | 0.031 | -0.009 | -0.009 | 0.964 | 0.8 | 0.13 | 0.75 | 0.72 | 0.92 | 0.7 |
| 1 | 214047014 | rs860505 | 0.626 | -1.12 | 0.68 | 0.099 | 0.038 | 0.097 | 0.097 | 0 | 0.92 | 0.67 | 0.82 | 0.83 | 0.95 | 0.81 |
| 1 | 214051497 | rs701899 | 0.492 | -1.11 | 0.99 | 0.116 | 0.023 | 0.079 | 0.079 | 0 | 0.92 | 0.72 | 0.82 | 0.83 | 0.95 | 0.81 |
| 1 | 214051639 | rs853757 | 1.961 | -1.183 | 0.912 | 0.094 | -0.031 | 0.078 | 0.078 | 0.972 | 0.86 | 0.41 | 0.78 | 0.72 | 0.94 | 0.71 |
| 1 | 214057330 | rs1341146 | -0.429 | -1.344 | 0.235 | -0.13 | 0.053 | 0.029 | 0.029 | 0.985 | 0.6 | 0.11 | 0.65 | 0.64 | 0.73 | 0.49 |
| 1 | 214063635 | rs17710111 | 0.522 | -0.993 | 0.609 | 0.124 | 0.004 | 0.025 | 0.025 | 0 | 0.05 | 0.11 | 0.16 | 0.18 | 0.04 | 0.2 |
| 1 | 214067353 | rs853733 | 0.272 | -1.004 | 0.753 | 0.255 | 0.033 | 0.111 | 0.111 | 0 | 0.93 | 0.71 | 0.82 | 0.8 | 0.95 | 0.81 |
| 1 | 214077644 | rs10864079 | -0.843 | -1.058 | 0.437 | 0.098 | -0.013 | 0.003 | 0.003 | 0.834 | 0.1 | 0.25 | 0.15 | 0.16 | 0.06 | 0.23 |
| 1 | 214081498 | rs12089197 | -0.601 | -0.999 | 0.451 | 0.138 | 0.005 | 0.031 | 0.031 | 0 | 0.08 | 0.26 | 0.13 | 0.14 | 0.04 | 0.17 |
| 1 | 214083466 | rs1891062 | 0.868 | -0.808 | 0.331 | 0.64 | 0.025 | 0.048 | 0.048 | 0 | 0.24 | 0.33 | 0.3 | 0.38 | 0.33 | 0.32 |
| 1 | 214084384 | rs12087028 | -0.737 | -0.945 | 0.707 | 0.81 | 0.006 | 0.041 | 0.041 | 0 | 0.08 | 0.27 | 0.13 | 0.14 | 0.04 | 0.17 |
| 1 | 214089860 | rs6681892 | -1.63 | -0.962 | 0.59 | 0.971 | 0 | 0.023 | 0.023 | 2.249 | 0.11 | 0.62 | 0.14 | 0.11 | 0.04 | 0.18 |
| 1 | 214094291 | rs17761554 | -0.305 | -0.984 | 0.191 | 0.815 | -0.003 | 0.086 | 0.086 | 0 | 0.39 | 0.2 | 0.34 | 0.16 | 0.4 | 0.17 |
| 1 | 214095868 | rs1073569 | -0.359 | -0.772 | 0.487 | 1.121 | 0.02 | 0.183 | 0.183 | 0 | 0.07 | 0.21 | 0.22 | 0.32 | 0.05 | 0.22 |
| 1 | 214099716 | rs7550337 | -1.58 | -0.89 | 0.062 | 0.991 | -0.015 | -0.012 | -0.012 | 0 | 0.55 | 0.87 | 0.62 | 0.46 | 0.47 | 0.47 |
| 1 | 214102041 | rs1991879 | -0.587 | -0.133 | 0.192 | 0.776 | 0.019 | 0.081 | 0.081 | 0.241 | 0.48 | 0.35 | 0.38 | 0.16 | 0.46 | 0.21 |
| 1 | 214108641 | rs1159179 | 0.659 | -0.262 | 0.321 | 1.064 | -0.025 | 0.083 | 0.083 | 2.141 | 0.85 | 0.36 | 0.71 | 0.68 | 0.93 | 0.72 |
| 1 | 214110152 | rs11120231 | 1.39 | -0.161 | 0.22 | 1.153 | -0.104 | -0.089 | -0.089 | 0 | 0.4 | 0.12 | 0.36 | 0.48 | 0.47 | 0.5 |
| 1 | 214118343 | rs6540809 | -0.424 | -0.08 | 0.189 | 0.998 | 0.012 | 0.107 | 0.107 | 0 | 0.46 | 0.24 | 0.35 | 0.18 | 0.46 | 0.22 |
| 1 | 214121995 | rs12130674 | -0.427 | -0.103 | 0.105 | 0.973 | 0.01 | 0.086 | 0.086 | 0 | 0.46 | 0.26 | 0.35 | 0.19 | 0.46 | 0.22 |
| 1 | 214130413 | rs6698217 | 1.577 | 0.11 | -0.136 | 0.952 | -0.032 | 0.023 | 0.023 | 0 | 0.1 | 0.01 | 0.08 | 0.04 | 0.13 | 0.03 |
| 1 | 214135660 | rs6700065 | 0.009 | 0.333 | 0.541 | 1.146 | 0.001 | 0.005 | 0.005 | 0 | 0.39 | 0.37 | 0.26 | 0.17 | 0.38 | 0.19 |
| 1 | 214140993 | rs12567631 | -0.86 | -0.128 | 0.582 | 1.55 | 0.003 | 0.129 | 0.129 | 1.4 | 0.12 | 0.34 | 0.27 | 0.34 | 0.08 | 0.27 |

| | | | | | | | | | | | | | | | | |
|---|---|---|---|---|---|---|---|---|---|---|---|---|---|---|---|---|
| 1 | 214145389 | rs340884 | 0.601 | -0.01 | 0.031 | 0.897 | 0.065 | 0.012 | 0.012 | 0.122 | 0.49 | 0.29 | 0.6 | 0.59 | 0.6 | 0.64 |
| 1 | 214145706 | rs340883 | 1.195 | 0.358 | -0.136 | 1.047 | -0.079 | 0.017 | 0.017 | 0.406 | 0.41 | 0.1 | 0.34 | 0.42 | 0.47 | 0.52 |
| 1 | 214147889 | rs7529073 | -1.184 | 0.299 | 0.024 | 1.005 | -0.032 | -0.016 | -0.016 | 0 | 0.57 | 0.82 | 0.61 | 0.53 | 0.49 | 0.44 |
| 1 | 214152475 | rs1431979 | -0.848 | 0.44 | 0.258 | 1.165 | -0.049 | 0.108 | 0.108 | 0.018 | 0.18 | 0.48 | 0.2 | 0.28 | 0.1 | 0.19 |
| 1 | 214152590 | rs1431978 | -0.848 | 0.44 | 0.258 | 1.165 | -0.049 | 0.108 | 0.108 | 0.018 | 0.18 | 0.48 | 0.2 | 0.28 | 0.1 | 0.19 |
| 1 | 214156514 | rs340879 | 0.346 | 1.09 | -0.457 | 1.123 | 0.059 | 0.014 | 0.014 | 0.06 | 0.48 | 0.22 | 0.6 | 0.61 | 0.6 | 0.62 |
| 1 | 214160529 | rs340873 | 0.382 | 1.008 | -0.379 | 1.19 | 0.015 | 0.001 | 0.001 | 0 | 0.47 | 0.23 | 0.53 | 0.6 | 0.54 | 0.61 |
| 1 | 214161820 | rs340839 | 0.529 | 0.851 | -0.364 | 0.966 | -0.016 | -0.001 | -0.001 | 0 | 0.46 | 0.22 | 0.48 | 0.57 | 0.48 | 0.58 |
| 1 | 214163675 | rs340835 | 0.496 | 0.841 | -0.338 | 0.928 | -0.023 | 0.054 | 0.054 | 0.182 | 0.45 | 0.17 | 0.35 | 0.4 | 0.47 | 0.51 |
| 1 | 214167508 | rs4655314 | -1.43 | 1.281 | 0.636 | 0.7 | -0.001 | -0.004 | -0.004 | 0 | 0.17 | 0.18 | 0.3 | 0.17 | 0.18 | 0.13 |
| 1 | 214173840 | rs3767848 | -2.037 | 0.981 | 0.865 | 1.128 | 0.019 | 0.089 | 0.089 | 0 | 0.24 | 0.45 | 0.44 | 0.3 | 0.27 | 0.17 |
| 1 | 214179972 | rs3754138 | -1.86 | 0.94 | 0.81 | 1.127 | 0.001 | 0.009 | 0.009 | 0 | 0.24 | 0.27 | 0.46 | 0.29 | 0.27 | 0.17 |
| 1 | 214181251 | rs4655480 | -1.847 | 0.941 | 0.807 | 1.128 | 0.002 | 0.011 | 0.011 | 0 | 0.24 | 0.27 | 0.45 | 0.29 | 0.27 | 0.17 |
| 1 | 214187713 | rs399887 | 1.413 | 0.494 | 1.339 | 0.6 | -0.015 | 0.059 | 0.059 | 0 | 0.94 | 0.83 | 0.79 | 0.87 | 0.96 | 0.89 |
| 1 | 214196634 | rs6686424 | -1.501 | 0.447 | 1.64 | 0.406 | 0.044 | 0.021 | 0.021 | 0.317 | 0.05 | 0.32 | 0.03 | 0.02 | 0.03 | 0 |
| 1 | 214197679 | rs7543057 | -2.866 | 0.388 | 1.653 | 0.428 | -0.038 | 0.046 | 0.046 | 0 | 0.13 | 0.48 | 0.23 | 0.13 | 0.06 | 0.12 |
| 1 | 214230381 | rs4655316 | -2.411 | 0.536 | 1.325 | 0.433 | 0.003 | 0.05 | 0.05 | 0 | 0.11 | 0.38 | 0.23 | 0.13 | 0.07 | 0.1 |
| 1 | 214240736 | rs4655317 | -2.434 | 0.616 | 0.777 | 0.507 | -0.008 | 0.046 | 0.046 | 0 | 0.11 | 0.35 | 0.22 | 0.14 | 0.06 | 0.11 |
| 1 | 214268998 | rs1223760 | -2.681 | 0.774 | 0.618 | 0.435 | 0.019 | 0.018 | 0.018 | 0 | 0.13 | 0.42 | 0.14 | 0.14 | 0.11 | 0.09 |
| 1 | 214274759 | rs1223751 | -1.732 | 0.816 | 0.529 | 0.44 | 0.002 | 0.004 | 0.004 | 0 | 0.08 | 0.13 | 0.11 | 0.11 | 0.08 | 0.08 |
| 1 | 214276204 | rs1023763 | -2.211 | 0.815 | 0.344 | 0.458 | 0.017 | 0.017 | 0.017 | 0 | 0.13 | 0.39 | 0.14 | 0.13 | 0.11 | 0.09 |
| 1 | 214280532 | rs1223742 | -0.069 | 0.896 | -0.936 | 0.264 | -0.002 | 0.04 | 0.04 | 0 | 0.72 | 0.91 | 0.72 | 0.82 | 0.74 | 0.75 |
| 1 | 214287420 | rs920489 | -0.651 | 1.525 | -0.258 | 0.702 | 0.056 | 0.032 | 0.032 | 0.049 | 0.23 | 0.39 | 0.39 | 0.32 | 0.28 | 0.17 |
| 1 | 214288429 | rs1795040 | -0.037 | 0.829 | 0.056 | 0.053 | 0.001 | -0.021 | -0.021 | 0.692 | 0.5 | 0.51 | 0.28 | 0.1 | 0.38 | 0.17 |
| 1 | 214292309 | rs1795036 | -1.642 | 0.766 | 0.981 | 0.954 | 0.059 | 0.144 | 0.144 | 2.672 | 0.05 | 0.29 | 0.04 | 0.08 | 0.03 | 0.1 |
| 1 | 214301559 | rs7540655 | -0.704 | 0.444 | 0.142 | 1.635 | -0.003 | -0.004 | -0.004 | 0.126 | 0.14 | 0.11 | 0.19 | 0.36 | 0.18 | 0.31 |
| 1 | 214302012 | rs1795052 | -1.173 | -0.277 | 0.119 | 1.307 | 0 | -0.001 | -0.001 | 0 | 0.33 | 0.65 | 0.48 | 0.55 | 0.31 | 0.45 |
| 1 | 214304491 | rs723098 | 0.48 | -0.549 | 0.049 | 1.458 | -0.026 | -0.013 | -0.013 | 0 | 0.87 | 0.94 | 0.71 | 0.84 | 0.82 | 0.9 |
| 1 | 214304616 | rs723099 | -1.296 | -0.949 | 0.543 | 2.04 | -0.028 | 0.081 | 0.081 | 0.027 | 0.12 | 0.3 | 0.08 | 0.33 | 0.07 | 0.28 |
| 1 | 214306040 | rs1223786 | -1.282 | -0.728 | 0.662 | 2.025 | -0.03 | 0.28 | 0.28 | 0.044 | 0.14 | 0.52 | 0.24 | 0.44 | 0.11 | 0.23 |
| 1 | 214312495 | rs7535649 | -1.311 | -0.697 | 1.059 | 1.935 | -0.035 | 0.204 | 0.204 | 0 | 0.15 | 0.31 | 0.23 | 0.46 | 0.11 | 0.21 |
| 1 | 214318748 | rs10864086 | 1.118 | -0.755 | 1.285 | 1.524 | 0.001 | 0.027 | 0.027 | 0 | 0.75 | 0.7 | 0.63 | 0.47 | 0.75 | 0.59 |
| 1 | 214322247 | rs1228534 | 0.696 | -0.769 | 1.472 | 1.91 | 0.007 | 0.027 | 0.027 | 0 | 0.89 | 0.85 | 0.74 | 0.73 | 0.82 | 0.86 |
| 1 | 214332987 | rs1223761 | 0.533 | -0.779 | 1.387 | 1.895 | -0.047 | -0.052 | -0.052 | 0.003 | 0.9 | 0.97 | 0.77 | 0.76 | 0.84 | 0.87 |
| 1 | 214344393 | rs1223817 | -0.584 | -0.795 | 1.661 | 1.882 | -0.019 | 0.031 | 0.031 | 0 | 0.15 | 0.26 | 0.15 | 0.29 | 0.1 | 0.28 |
| 1 | 214351618 | rs17022092 | -2.856 | -0.854 | 2.171 | 1.817 | 0.034 | 0.096 | 0.096 | 0.653 | 0.05 | 0.37 | 0.04 | 0.04 | 0.02 | 0.02 |
| 1 | 214359179 | rs11120266 | -0.42 | -0.971 | 2.115 | 1.791 | -0.015 | 0.087 | 0.087 | 0 | 0.13 | 0.16 | 0.16 | 0.41 | 0.09 | 0.21 |
| 1 | 214361661 | rs2919490 | 2.014 | -1.103 | 2.329 | 1.445 | -0.018 | -0.016 | -0.016 | 0.131 | 0.7 | 0.47 | 0.56 | 0.31 | 0.72 | 0.56 |
| 1 | 214363164 | rs12026597 | -0.332 | -1.096 | 2.391 | 1.411 | -0.01 | 0.042 | 0.042 | 0 | 0.13 | 0.23 | 0.15 | 0.3 | 0.1 | 0.27 |
| 1 | 214364640 | rs10864087 | -0.881 | -1.101 | 2.388 | 1.472 | -0.039 | 0.174 | 0.174 | 0 | 0.15 | 0.29 | 0.24 | 0.48 | 0.1 | 0.23 |
| 1 | 214368330 | rs6699206 | 1.686 | -1.035 | 2.427 | 1.357 | -0.033 | -0.002 | -0.002 | 0 | 0.73 | 0.41 | 0.79 | 0.69 | 0.83 | 0.65 |
| 1 | 214375999 | rs2033923 | -0.357 | -0.739 | 2.438 | 1.45 | -0.005 | 0.023 | 0.023 | 0 | 0.14 | 0.19 | 0.16 | 0.27 | 0.11 | 0.27 |
| 1 | 214384723 | rs11120275 | -0.404 | -0.638 | 2.365 | 1.514 | -0.015 | 0.187 | 0.187 | 0.454 | 0.12 | 0.27 | 0.2 | 0.44 | 0.09 | 0.23 |
| 1 | 214392240 | rs9787247 | -0.121 | -0.631 | 2.269 | 1.464 | -0.011 | 0.163 | 0.163 | 0.991 | 0.11 | 0.18 | 0.2 | 0.45 | 0.09 | 0.19 |
| 1 | 214395861 | rs1223777 | 1.772 | -0.681 | 2.435 | 1.447 | -0.058 | 0.153 | 0.153 | 0 | 0.77 | 0.52 | 0.73 | 0.5 | 0.84 | 0.71 |
| 1 | 214398058 | rs7539006 | 1.949 | -0.786 | 1.825 | 1.319 | -0.021 | -0.018 | -0.018 | 0 | 0.62 | 0.4 | 0.6 | 0.26 | 0.73 | 0.45 |
| 1 | 214402245 | rs10864089 | -0.893 | -0.383 | 1.47 | 1.505 | 0.003 | 0.053 | 0.053 | 0 | 0.18 | 0.3 | 0.23 | 0.34 | 0.18 | 0.32 |
| 1 | 214402742 | rs1223672 | -1.776 | -0.485 | 1.273 | 1.427 | -0.013 | -0.006 | -0.006 | 0 | 0.3 | 0.6 | 0.38 | 0.73 | 0.22 | 0.47 |
| 1 | 214403709 | rs4655320 | -0.658 | -0.007 | 0.696 | 1.831 | -0.002 | 0.006 | 0.006 | 0 | 0.16 | 0.21 | 0.15 | 0.28 | 0.12 | 0.28 |
| 1 | 214405194 | rs1795065 | 0.949 | 0.004 | 0.149 | 1.385 | 0.009 | 0.06 | 0.06 | 0 | 0.69 | 0.48 | 0.6 | 0.47 | 0.67 | 0.5 |
| 1 | 214418139 | rs6682783 | 0.114 | -0.844 | -0.47 | 1.06 | 0.01 | -0.006 | -0.006 | 0 | 0.83 | 0.84 | 0.8 | 0.81 | 0.93 | 0.88 |
| 1 | 214420058 | rs4655323 | -0.661 | 0.238 | -0.08 | 1.213 | 0.066 | 0.148 | 0.148 | 0 | 0.24 | 0.46 | 0.43 | 0.53 | 0.31 | 0.23 |
| 1 | 214425225 | rs7531751 | 0.879 | 0.034 | -0.656 | 0.003 | -0.004 | 0.074 | 0.074 | 0 | 0.43 | 0.23 | 0.24 | 0.19 | 0.4 | 0.35 |
| 1 | 214430221 | rs4265402 | 0.917 | 0.076 | -0.623 | -0.012 | 0 | 0.066 | 0.066 | 0 | 0.42 | 0.22 | 0.25 | 0.2 | 0.39 | 0.35 |
| 1 | 214432191 | rs1391551 | 0.199 | 0.478 | 0.093 | -0.294 | 0.001 | -0.005 | -0.005 | 0 | 0.65 | 0.6 | 0.52 | 0.26 | 0.65 | 0.72 |
| 1 | 214436619 | rs6540814 | -0.208 | 0.63 | 0.182 | -0.281 | -0.008 | -0.072 | -0.072 | 0 | 0.23 | 0.36 | 0.28 | 0.06 | 0.25 | 0.38 |
| 1 | 214437315 | rs12755490 | 0.518 | 0.998 | -0.534 | -0.317 | -0.003 | -0.002 | -0.002 | 0.279 | 0.06 | 0.04 | 0.06 | 0.1 | 0.1 | 0.08 |
| 1 | 214457009 | rs9308433 | -0.221 | 1.184 | -0.27 | -0.901 | 0.007 | -0.07 | -0.07 | 0 | 0.31 | 0.61 | 0.32 | 0.07 | 0.35 | 0.38 |
| 1 | 214457286 | rs9970083 | -1.629 | 0.432 | -1.18 | -0.528 | 0.05 | -0.138 | -0.138 | 0.219 | 0.78 | 0.99 | 0.76 | 0.63 | 0.85 | 0.88 |
| 1 | 214461354 | rs6660726 | -0.27 | 0.885 | -1.366 | -0.66 | 0.001 | 0.024 | 0.024 | 0.323 | 0.63 | 0.75 | 0.53 | 0.25 | 0.63 | 0.56 |
| 1 | 214462988 | rs11120288 | 0.559 | 0.517 | -1.325 | -0.693 | -0.036 | -0.035 | -0.035 | 0 | 0.39 | 0.24 | 0.39 | 0.48 | 0.45 | 0.38 |
| 1 | 214465791 | rs7519743 | 0.781 | 0.463 | -1.025 | -0.917 | -0.001 | -0.008 | -0.008 | 0 | 0.1 | 0.12 | 0.04 | 0.07 | 0.08 | 0.24 |
| 1 | 214466068 | rs4655330 | 1.055 | 0.473 | -0.579 | -0.955 | 0.017 | -0.098 | -0.098 | 0 | 0.22 | 0.35 | 0.27 | 0 | 0.28 | 0.15 |
| 1 | 214466932 | rs7544984 | -0.643 | 0.456 | -0.802 | -0.913 | -0.004 | 0.082 | 0.082 | 0 | 0.38 | 0.76 | 0.37 | 0.16 | 0.4 | 0.43 |
| 1 | 214469508 | rs6701764 | -0.997 | 0.206 | -1.002 | -1.049 | 0.06 | -0.083 | -0.083 | 2.211 | 0.68 | 0.95 | 0.69 | 0.62 | 0.77 | 0.72 |
| 1 | 214473363 | rs12057464 | -1.099 | 0.235 | -1.031 | -1.049 | 0.029 | -0.124 | -0.124 | 0.243 | 0.69 | 0.98 | 0.7 | 0.62 | 0.77 | 0.77 |
| 1 | 214476561 | rs10779613 | 0.077 | 0.3 | -0.712 | -1.235 | -0.007 | 0.012 | 0.012 | 0.142 | 0.45 | 0.71 | 0.42 | 0.62 | 0.47 | 0.57 |
| 1 | 214482888 | rs2133175 | 2.461 | 0.606 | -0.48 | -1.247 | -0.001 | -0.002 | -0.002 | 0 | 0.17 | 0.19 | 0.14 | 0.09 | 0.15 | 0.1 |
| 1 | 214489193 | rs6676709 | 0.615 | 0.567 | -0.658 | -1.375 | -0.001 | -0.001 | -0.001 | 0.869 | 0.34 | 0.33 | 0.36 | 0.57 | 0.39 | 0.35 |

| | | | | | | | | | | | | | | | | |
|---|---|---|---|---|---|---|---|---|---|---|---|---|---|---|---|---|
| 1 | 214492286 | rs1134647 | -2.412 | 0.446 | -0.698 | -1.413 | 0.082 | -0.138 | -0.138 | 0.184 | 0.79 | 1 | 0.76 | 0.62 | 0.91 | 0.8 |
| 1 | 214492596 | rs12059328 | 1.867 | 1.067 | -0.586 | -1.034 | -0.001 | 0.013 | 0.013 | 0 | 0.05 | 0.04 | 0.15 | 0.26 | 0.15 | 0.09 |
| 1 | 214494006 | rs12737638 | 0.523 | 1.01 | -0.584 | -1.067 | 0.048 | 0.104 | 0.104 | 0.135 | 0.19 | 0.32 | 0.25 | 0.48 | 0.24 | 0.25 |
| 1 | 214495139 | rs6687629 | 0.574 | 1.006 | -0.641 | -1.091 | 0.044 | -0.162 | -0.162 | 0 | 0.29 | 0.47 | 0.34 | 0 | 0.42 | 0.17 |
| 1 | 214498226 | rs3767843 | 0.506 | 0.954 | -0.54 | -0.989 | 0.03 | 0.068 | 0.068 | 0.114 | 0.18 | 0.29 | 0.25 | 0.48 | 0.24 | 0.25 |
| 1 | 214503489 | rs2291830 | -0.289 | 0.521 | -0.707 | -1 | 0 | 0 | 0 | 0.106 | 0.48 | 0.52 | 0.46 | 0.62 | 0.59 | 0.61 |
| 1 | 214504604 | rs2270704 | -1.441 | 0.482 | -1.69 | -0.904 | 0.127 | -0.118 | -0.118 | 0 | 0.72 | 0.96 | 0.74 | 0.62 | 0.89 | 0.78 |
| 1 | 214511519 | rs1048791 | 1.203 | 1.281 | -1.345 | -0.626 | -0.059 | 0.106 | 0.106 | 0 | 0.21 | 0.05 | 0.26 | 0 | 0.31 | 0.15 |
| 1 | 214514361 | rs7512594 | 1.092 | 1.545 | -0.886 | 0.021 | -0.036 | -0.052 | -0.052 | 0.869 | 0.22 | 0.14 | 0.28 | 0.36 | 0.29 | 0.19 |
| 1 | 214527252 | rs1472441 | -0.502 | 0.987 | -1.394 | -0.228 | -0.074 | -0.046 | -0.046 | 1.469 | 0.64 | 0.9 | 0.5 | 0.4 | 0.58 | 0.69 |
| 1 | 214529786 | rs7543972 | 0.863 | 0.322 | -1.113 | -0.373 | 0.005 | 0.004 | 0.004 | 0.729 | 0.2 | 0.19 | 0.1 | 0.18 | 0.13 | 0.19 |
| 1 | 214531239 | rs2291831 | 1.928 | 0.405 | -0.949 | -0.438 | 0 | -0.001 | -0.001 | 0 | 0.19 | 0.19 | 0.13 | 0.16 | 0.14 | 0.11 |
| 1 | 214536061 | rs3002297 | -1.149 | 0.618 | -1.132 | -0.325 | 0.044 | -0.035 | -0.035 | 1.077 | 0.68 | 0.85 | 0.67 | 0.65 | 0.76 | 0.71 |
| 1 | 214541336 | rs11120305 | 0.197 | 0.655 | -0.815 | -0.371 | 0.069 | -0.002 | -0.002 | 0 | 0.29 | 0.43 | 0.42 | 0.32 | 0.5 | 0.42 |
| 1 | 214542819 | rs1135352 | -1.136 | 0.546 | -1.083 | -0.187 | 0.041 | -0.102 | -0.102 | 2.161 | 0.7 | 0.96 | 0.67 | 0.65 | 0.78 | 0.78 |
| 1 | 214543098 | rs3738425 | 1.1 | 0.878 | -1.055 | -0.196 | 0.015 | 0.015 | 0.015 | 0 | 0.07 | 0.06 | 0.18 | 0.26 | 0.2 | 0.08 |
| 1 | 214544632 | rs3002309 | -0.968 | -0.452 | -0.936 | -0.221 | 0.042 | -0.101 | -0.101 | 0 | 0.7 | 0.96 | 0.67 | 0.65 | 0.78 | 0.78 |
| 1 | 214544854 | rs6668557 | 2.33 | -0.476 | -1.014 | -0.238 | -0.016 | -0.058 | -0.058 | 0 | 0.16 | 0 | 0.12 | 0.15 | 0.13 | 0.1 |
| 1 | 214547938 | rs7532953 | 0.112 | -0.561 | -0.817 | -0.309 | 0.006 | 0.028 | 0.028 | 0 | 0.39 | 0.72 | 0.49 | 0.3 | 0.56 | 0.44 |
| 1 | 214549501 | rs1125452 | 1.119 | -0.547 | -0.842 | -0.27 | -0.016 | 0 | 0 | 0.259 | 0.15 | 0.2 | 0.07 | 0.19 | 0.09 | 0.17 |
| 1 | 214550568 | rs2936014 | -0.945 | -0.464 | -0.961 | -0.196 | 0.057 | -0.077 | -0.077 | 0 | 0.7 | 0.96 | 0.68 | 0.65 | 0.79 | 0.77 |
| 1 | 214554966 | rs6676318 | 1.145 | -0.187 | -0.74 | -0.164 | -0.003 | -0.001 | -0.001 | 0 | 0.21 | 0.3 | 0.15 | 0.17 | 0.18 | 0.11 |
| 1 | 214554980 | rs1472444 | 1.259 | -0.19 | -1.135 | 0.032 | -0.008 | 0.001 | 0.001 | 0.001 | 0.13 | 0.18 | 0.06 | 0.17 | 0.09 | 0.17 |
| 1 | 214556082 | rs3002296 | 0.445 | -0.189 | -1.146 | 0.027 | 0.013 | -0.007 | -0.007 | 0.974 | 0.3 | 0.26 | 0.2 | 0.35 | 0.26 | 0.27 |
| 1 | 214558228 | rs3013446 | -0.537 | -0.664 | -1.358 | 0.143 | 0.022 | -0.116 | -0.116 | 0 | 0.71 | 0.99 | 0.68 | 0.64 | 0.79 | 0.8 |
| 1 | 214558658 | rs3013445 | 0.205 | 0.541 | -0.547 | 0.603 | 0.001 | 0 | 0 | 0 | 0.14 | 0.18 | 0.07 | 0.17 | 0.11 | 0.17 |
| 1 | 214569107 | rs4129186 | 0.11 | 1.646 | -1.376 | 0.896 | -0.037 | 0 | 0 | 0 | 0.06 | 0 | 0.23 | 0.36 | 0.12 | 0.1 |
| 1 | 214572600 | rs12753105 | -0.178 | 1.478 | -1.302 | -0.007 | -0.022 | 0.034 | 0.034 | 0.336 | 0.31 | 0.16 | 0.32 | 0.17 | 0.34 | 0.22 |
| 1 | 214576482 | rs4636427 | -0.849 | 1.309 | -0.991 | -0.145 | -0.009 | 0.017 | 0.017 | 0 | 0.27 | 0.19 | 0.19 | 0.13 | 0.27 | 0.2 |
| 1 | 214576513 | rs6657749 | -0.894 | 0.694 | -0.413 | 0.26 | -0.001 | 0.007 | 0.007 | 0 | 0.19 | 0.22 | 0.25 | 0.22 | 0.18 | 0.11 |
| 1 | 214576614 | rs6673035 | -0.119 | -0.143 | -0.512 | -0.397 | -0.034 | -0.007 | -0.007 | 0 | 0.27 | 0.31 | 0.17 | 0.22 | 0.19 | 0.15 |
| 1 | 214577101 | rs12129366 | -1.02 | 1.005 | -0.577 | -0.489 | 0 | -0.025 | -0.025 | 0 | 0.18 | 0.18 | 0.09 | 0.06 | 0.18 | 0.18 |
| 1 | 214581206 | rs11120313 | -0.767 | 0.66 | -0.186 | -0.329 | -0.008 | -0.049 | -0.049 | 0 | 0.12 | 0.18 | 0.12 | 0.01 | 0.1 | 0.09 |
| 1 | 214582984 | rs4655249 | -0.002 | 0.943 | -0.489 | -0.022 | -0.012 | -0.017 | -0.017 | 0 | 0.95 | 0.98 | 0.87 | 0.86 | 0.9 | 0.83 |
| 1 | 214588645 | rs12078874 | -0.572 | 0.866 | -0.314 | -0.075 | -0.003 | -0.022 | -0.022 | 0 | 0.12 | 0.1 | 0.11 | 0.01 | 0.1 | 0.09 |
| 1 | 214589914 | rs10779614 | -0.383 | 0.244 | -0.515 | -0.42 | 0.017 | 0.002 | 0.002 | 0 | 0.67 | 0.74 | 0.74 | 0.64 | 0.73 | 0.72 |
| 1 | 214593855 | rs4540625 | 0.256 | 0.686 | -0.288 | -0.367 | 0.002 | -0.005 | -0.005 | 0 | 0.44 | 0.44 | 0.53 | 0.57 | 0.49 | 0.47 |
| 1 | 214601209 | rs4655344 | 0.125 | 0.294 | -0.789 | 0.022 | 0.022 | -0.038 | -0.038 | 0.185 | 0.34 | 0.75 | 0.37 | 0.46 | 0.26 | 0.34 |
| 1 | 214601540 | rs12047119 | 0.887 | 0.041 | -1.257 | -0.156 | 0.051 | 0.069 | 0.069 | 0.117 | 0.31 | 0.07 | 0.16 | 0.17 | 0.21 | 0.11 |
| 1 | 214601710 | rs6682627 | 0.228 | 0.518 | -0.57 | 0.11 | 0.001 | -0.002 | -0.002 | 0 | 0.84 | 0.86 | 0.84 | 0.82 | 0.79 | 0.73 |
| 1 | 214604236 | rs4615815 | 0.874 | 0.37 | -0.042 | 0.317 | 0.008 | -0.002 | -0.002 | 0 | 0.54 | 0.63 | 0.49 | 0.51 | 0.42 | 0.35 |
| 1 | 214608879 | rs4465172 | -0.793 | 0.403 | -0.071 | 0.372 | -0.02 | -0.033 | -0.033 | 0 | 0.94 | 0.99 | 0.87 | 0.82 | 0.9 | 0.89 |
| 1 | 214609822 | rs4253890 | 0.932 | 0.398 | 0.037 | 0.344 | -0.001 | 0 | 0 | 0.747 | 0.78 | 0.74 | 0.76 | 0.67 | 0.74 | 0.58 |
| 1 | 214611723 | rs6698662 | -0.067 | 0.265 | 0.733 | 0.277 | 0.012 | 0.062 | 0.062 | 1.796 | 0.09 | 0.38 | 0.05 | 0.1 | 0.04 | 0.19 |
| 1 | 214612807 | rs10864096 | 0.61 | 0.29 | 0.664 | 0.265 | -0.047 | 0.018 | 0.018 | 2.17 | 0.24 | 0.11 | 0.27 | 0.16 | 0.33 | 0.22 |
| 1 | 214622818 | rs4394613 | 0.446 | 0.248 | 0.477 | 0.27 | -0.066 | -0.017 | -0.017 | 0 | 0.62 | 0.86 | 0.54 | 0.61 | 0.46 | 0.54 |
| 1 | 214629616 | rs12757404 | 1.241 | 0.286 | 0.421 | 0.282 | 0.018 | 0.003 | 0.003 | 0.325 | 0.51 | 0.29 | 0.47 | 0.51 | 0.42 | 0.37 |
| 1 | 214630783 | rs4375234 | -1.189 | 0.388 | 0.667 | 0.376 | -0.017 | -0.024 | -0.024 | 0 | 0.19 | 0.55 | 0.12 | 0.15 | 0.16 | 0.3 |
| 1 | 214632398 | rs17733405 | 1.187 | 0.488 | 0.212 | 0.421 | 0.004 | -0.015 | -0.015 | 0 | 0.06 | 0.03 | 0.05 | 0.11 | 0.04 | 0.19 |
| 1 | 214636442 | rs4284238 | 0.257 | 0.507 | 0.381 | 0.388 | -0.009 | -0.012 | -0.012 | 0 | 0.86 | 0.92 | 0.82 | 0.78 | 0.8 | 0.78 |
| 1 | 214641548 | rs6540838 | 0.438 | 0.499 | 0.513 | 0.384 | 0 | 0 | 0 | 0.541 | 0.84 | 0.83 | 0.8 | 0.78 | 0.8 | 0.77 |
| 1 | 214644771 | rs10864104 | 0.539 | 0.311 | 0.255 | 0.209 | -0.059 | 0.039 | 0.039 | 5.146 | 0.28 | 0.11 | 0.29 | 0.16 | 0.35 | 0.24 |
| 1 | 214645263 | rs10158490 | 0.605 | 0.322 | 0.35 | 0.209 | -0.001 | -0.001 | -0.001 | 0 | 0.09 | 0.07 | 0.06 | 0.05 | 0.1 | 0.11 |
| 1 | 214645307 | rs4655351 | 1.06 | 0.28 | 0.625 | 0.205 | 0.006 | 0.014 | 0.014 | 0.294 | 0.81 | 0.48 | 0.79 | 0.78 | 0.8 | 0.77 |
| 1 | 214647628 | rs17790562 | 0.58 | 0.412 | 0.184 | 0.237 | -0.001 | -0.001 | -0.001 | 0 | 0.09 | 0.07 | 0.05 | 0.05 | 0.1 | 0.11 |
| 1 | 214658211 | rs10494979 | 0.208 | 0.337 | 0.298 | 0.346 | -0.031 | 0.028 | 0.028 | 2.556 | 0.34 | 0.21 | 0.33 | 0.26 | 0.39 | 0.42 |
| 1 | 214662822 | rs10864105 | 1.306 | 0.705 | 0.643 | 1.009 | 0.028 | 0.067 | 0.067 | 1.341 | 0.73 | 0.53 | 0.51 | 0.4 | 0.66 | 0.58 |
| 1 | 214664556 | rs11120342 | -0.786 | 0.47 | 0.598 | 1.181 | 0.038 | 0.211 | 0.211 | 0 | 0.11 | 0.29 | 0.29 | 0.4 | 0.13 | 0.19 |
| 1 | 214674925 | rs17042140 | 0.433 | -0.579 | 0.273 | 1.002 | 0.014 | -0.049 | -0.049 | 0.294 | 0.75 | 0.88 | 0.58 | 0.59 | 0.8 | 0.79 |
| 1 | 214676121 | rs4129059 | -0.259 | -0.988 | 0.875 | 1.079 | -0.03 | 0.058 | 0.058 | 0 | 0.18 | 0.52 | 0.17 | 0.22 | 0.14 | 0.23 |
| 1 | 214677026 | rs6680964 | -0.909 | -1.753 | 0.378 | 0.452 | -0.038 | -0.023 | -0.023 | 0 | 0.34 | 0.57 | 0.23 | 0.23 | 0.2 | 0.35 |
| 1 | 214679264 | rs4077549 | 0.585 | -1.813 | 0.716 | 0.296 | 0.002 | -0.014 | -0.014 | 0 | 0.16 | 0.13 | 0.06 | 0.02 | 0.07 | 0.13 |
| 1 | 214679285 | rs6690349 | -0.363 | -1.855 | 1.026 | 0.281 | 0.06 | 0.045 | 0.045 | 0.279 | 0.06 | 0.44 | 0.03 | 0 | 0.01 | 0.02 |
| 1 | 214682931 | rs6701315 | 0.766 | -1.91 | 0.814 | 0.247 | -0.024 | -0.039 | -0.039 | 0 | 0.14 | 0.2 | 0.05 | 0.02 | 0.06 | 0.12 |
| 1 | 214683833 | rs6540847 | -0.614 | -1.886 | 0.792 | 0.257 | -0.072 | -0.122 | -0.122 | 0 | 0.22 | 0.59 | 0.1 | 0.02 | 0.08 | 0.15 |
| 1 | 214685978 | rs6693492 | -1.43 | -1.752 | 0.685 | 0.44 | 0.06 | -0.046 | -0.046 | 0 | 0.37 | 0.81 | 0.25 | 0.52 | 0.23 | 0.43 |
| 1 | 214710841 | rs4326598 | 0.303 | -1.494 | 0.957 | 0.327 | -0.002 | 0.012 | 0.012 | 0 | 0.15 | 0.25 | 0.15 | 0.23 | 0.14 | 0.22 |
| 1 | 214712251 | rs17023155 | -0.564 | -1.941 | 1.238 | 0.025 | -0.029 | -0.064 | -0.064 | 0 | 0.22 | 0.3 | 0.1 | 0.02 | 0.11 | 0.16 |
| 1 | 214716479 | rs7541263 | -0.642 | -1.916 | 1.515 | 0.127 | -0.017 | 0.007 | 0.007 | 0 | 0.2 | 0.62 | 0.17 | 0.22 | 0.16 | 0.27 |

| | | | | | | | | | | | | | | | | |
|---|---|---|---|---|---|---|---|---|---|---|---|---|---|---|---|---|
| 1 | 214719346 | rs12747035 | 0.573 | -1.886 | 1.19 | -0.361 | -0.003 | -0.063 | -0.063 | 0 | 0.57 | 0.46 | 0.39 | 0.07 | 0.63 | 0.44 |
| 1 | 214727383 | rs11120350 | 0.668 | -1.832 | 1.092 | -0.25 | 0.018 | 0.015 | 0.015 | 0 | 0.62 | 0.73 | 0.42 | 0.32 | 0.7 | 0.6 |
| 1 | 214729875 | rs4311856 | 1.845 | -1.842 | 0.961 | -0.374 | -0.032 | 0.185 | 0.185 | 0 | 0.41 | 0.14 | 0.32 | 0.09 | 0.55 | 0.31 |
| 1 | 214730401 | rs35329321 | 0.679 | -1.854 | 1.131 | -0.231 | -0.05 | -0.062 | -0.062 | 4.334 | 0.14 | 0.25 | 0.07 | 0.02 | 0.05 | 0.11 |
| 1 | 214742612 | rs10864111 | 0.789 | -1.935 | 0.99 | 0.205 | -0.018 | 0.071 | 0.071 | 0.005 | 0.06 | 0.06 | 0.11 | 0.24 | 0.04 | 0.08 |
| 1 | 214743309 | rs9699681 | -0.578 | -1.811 | 1.099 | 0.204 | -0.017 | -0.032 | -0.032 | 0.128 | 0.83 | 0.89 | 0.6 | 0.58 | 0.8 | 0.8 |
| 1 | 214746812 | rs2666838 | -0.417 | -1.383 | 1.466 | -0.017 | 0.011 | 0.155 | 0.155 | 0 | 0.14 | 0.48 | 0.16 | 0.25 | 0.1 | 0.12 |
| 1 | 214747137 | rs6688339 | 0.811 | -1.385 | 1.236 | -0.044 | -0.001 | 0.005 | 0.005 | 0 | 0.11 | 0.06 | 0.14 | 0.25 | 0.11 | 0.12 |
| 1 | 214747257 | rs2807644 | -1.499 | -1.524 | 1.088 | -0.194 | 0.083 | -0.015 | -0.015 | 0 | 0.47 | 0.78 | 0.31 | 0.48 | 0.33 | 0.55 |
| 1 | 214747718 | rs17735873 | 1.565 | -1.486 | 0.897 | -0.21 | 0.035 | 0.037 | 0.037 | 0 | 0.12 | 0.07 | 0.05 | 0.02 | 0.05 | 0.1 |
| 1 | 214751786 | rs6656590 | 0.989 | -1.502 | 1.192 | -0.07 | -0.003 | -0.012 | -0.012 | 0 | 0.61 | 0.55 | 0.4 | 0.32 | 0.65 | 0.61 |
| 1 | 214756410 | rs12563432 | -0.291 | -1.631 | 0.846 | 0.008 | 0.039 | 0.026 | 0.026 | 0 | 0.29 | 0.11 | 0.12 | 0.23 | 0.2 | 0.38 |
| 1 | 214762003 | rs2807650 | -1.462 | -1.643 | 0.632 | 0.073 | 0.028 | -0.006 | -0.006 | 0.15 | 0.45 | 0.63 | 0.29 | 0.48 | 0.31 | 0.54 |
| 1 | 214763986 | rs2807653 | -0.535 | -1.395 | 0.954 | 0.101 | -0.012 | 0.123 | 0.123 | 0.036 | 0.16 | 0.48 | 0.16 | 0.25 | 0.11 | 0.17 |
| 1 | 214770895 | rs11487873 | 0.462 | -1.222 | 0.377 | -0.131 | -0.023 | -0.027 | -0.027 | 0 | 0.14 | 0.16 | 0.07 | 0.04 | 0.06 | 0.09 |
| 1 | 214786635 | rs2807663 | 0.193 | -1.261 | 0.408 | -0.092 | -0.013 | 0.007 | 0.007 | 0 | 0.63 | 0.7 | 0.53 | 0.28 | 0.61 | 0.49 |
| 1 | 214789375 | rs2666834 | -0.827 | -1.695 | 0.248 | -0.263 | -0.046 | -0.007 | -0.007 | 0 | 0.25 | 0.62 | 0.21 | 0.16 | 0.1 | 0.21 |
| 1 | 214804409 | rs12408153 | -0.247 | -1.715 | 0.492 | -0.242 | -0.008 | 0.008 | 0.008 | 0 | 0.09 | 0.07 | 0.11 | 0.13 | 0.05 | 0.15 |
| 1 | 214814125 | rs3795522 | -0.537 | -1.737 | 0.51 | -0.254 | -0.012 | 0.008 | 0.008 | 0 | 0.1 | 0.1 | 0.11 | 0.13 | 0.05 | 0.15 |
| 1 | 214814582 | rs3795520 | -0.537 | -1.737 | 0.51 | -0.254 | -0.012 | 0.008 | 0.008 | 0 | 0.1 | 0.1 | 0.11 | 0.13 | 0.05 | 0.15 |
| 1 | 214814733 | rs3795519 | -0.537 | -1.708 | 0.51 | -0.254 | -0.011 | 0.008 | 0.008 | 0 | 0.1 | 0.1 | 0.11 | 0.13 | 0.05 | 0.15 |
| 1 | 214814995 | rs12067133 | -0.537 | -1.742 | 0.51 | -0.254 | -0.012 | 0.008 | 0.008 | 0 | 0.1 | 0.1 | 0.11 | 0.13 | 0.05 | 0.15 |
| 1 | 214816142 | rs3795516 | -0.537 | -1.742 | 0.53 | -0.254 | -0.014 | 0.01 | 0.01 | 0 | 0.1 | 0.1 | 0.11 | 0.13 | 0.05 | 0.15 |
| 1 | 214816224 | rs2666839 | -1.157 | -1.811 | 0.644 | -0.288 | -0.027 | -0.003 | -0.003 | 0.046 | 0.26 | 0.68 | 0.21 | 0.16 | 0.1 | 0.2 |
| 1 | 214816297 | rs3795514 | 0.431 | -1.662 | 0.243 | -0.233 | 0.025 | -0.041 | -0.041 | 0 | 0.07 | 0 | 0.1 | 0.13 | 0.04 | 0.12 |
| 1 | 214819328 | rs3748695 | -0.698 | -1.683 | 0.279 | -0.357 | -0.015 | 0.007 | 0.007 | 0 | 0.1 | 0.11 | 0.11 | 0.12 | 0.05 | 0.15 |
| 1 | 214819587 | rs12058704 | -0.698 | -1.683 | 0.279 | -0.357 | -0.015 | 0.007 | 0.007 | 0 | 0.1 | 0.11 | 0.11 | 0.12 | 0.05 | 0.15 |
| 1 | 214820099 | rs3748697 | 1.739 | -1.635 | 0.157 | -0.458 | -0.165 | 0.159 | 0.159 | 0 | 0.37 | 0.05 | 0.31 | 0.12 | 0.48 | 0.29 |
| 1 | 214820299 | rs3748698 | -0.784 | -1.439 | 0.403 | -0.361 | -0.015 | 0.007 | 0.007 | 0 | 0.1 | 0.11 | 0.11 | 0.12 | 0.05 | 0.15 |
| 1 | 214820494 | rs3790648 | -0.784 | -1.439 | 0.403 | -0.361 | -0.015 | 0.007 | 0.007 | 0 | 0.1 | 0.11 | 0.11 | 0.12 | 0.05 | 0.15 |
| 1 | 214820524 | rs3790649 | -0.784 | -1.439 | 0.403 | -0.361 | -0.015 | 0.007 | 0.007 | 0 | 0.1 | 0.11 | 0.11 | 0.12 | 0.05 | 0.15 |
| 1 | 214826196 | rs335524 | -1.044 | -1.871 | 0.276 | -0.926 | 0.019 | 0.096 | 0.096 | 0 | 0.52 | 0.29 | 0.59 | 0.88 | 0.44 | 0.7 |
| 1 | 214828812 | rs435043 | -1.964 | -1.772 | 0.512 | -1.053 | -0.045 | 0.075 | 0.075 | 0 | 0.66 | 0.79 | 0.67 | 0.89 | 0.56 | 0.74 |
| 1 | 214830617 | rs438034 | -1.544 | -1.774 | 0.519 | -1.054 | 0.002 | -0.037 | -0.037 | 0 | 0.62 | 0.35 | 0.64 | 0.89 | 0.56 | 0.74 |
| 1 | 214831405 | rs12744637 | 1.205 | -2.124 | -0.269 | -0.744 | -0.014 | -0.073 | -0.073 | 0.002 | 0.13 | 0.01 | 0.08 | 0.23 | 0.11 | 0.23 |
| 1 | 214833425 | rs417774 | -1.111 | -2.126 | -0.228 | -0.894 | 0.005 | 0.098 | 0.098 | 0 | 0.47 | 0.27 | 0.55 | 0.84 | 0.45 | 0.64 |
| 1 | 214839175 | rs335546 | -1.936 | -2.122 | -0.238 | -0.813 | 0.006 | -0.041 | -0.041 | 0 | 0.62 | 0.36 | 0.67 | 0.87 | 0.56 | 0.76 |
| 1 | 214842590 | rs335544 | -0.408 | -2.168 | -0.198 | -0.762 | -0.001 | -0.004 | -0.004 | 0 | 0.32 | 0.24 | 0.39 | 0.57 | 0.34 | 0.31 |
| 1 | 214847804 | rs6687966 | 0.1 | -2.084 | -0.043 | -1.989 | 0.014 | 0.014 | 0.014 | 0.009 | 0.16 | 0.12 | 0.09 | 0.05 | 0.1 | 0.11 |
| 1 | 214849923 | rs11120375 | 0.039 | -2.165 | -0.108 | -2.027 | 0.031 | 0.04 | 0.04 | 0 | 0.17 | 0.08 | 0.08 | 0.05 | 0.1 | 0.11 |
| 1 | 214855222 | rs430311 | 0.711 | -2.065 | -0.183 | -2.286 | 0 | 0.005 | 0.005 | 1.481 | 0.08 | 0 | 0.1 | 0.06 | 0.09 | 0.08 |
| 1 | 214859231 | rs374482 | 0.711 | -2.082 | 0.063 | -2.301 | -0.004 | 0.001 | 0.001 | 1.481 | 0.08 | 0 | 0.1 | 0.06 | 0.09 | 0.08 |
| 1 | 214859676 | rs759 | -0.706 | -2.186 | -0.048 | -2.507 | -0.001 | 0.053 | 0.053 | 0 | 0.43 | 0.26 | 0.48 | 0.83 | 0.39 | 0.62 |
| 1 | 214863137 | rs1511047 | -0.747 | -1.974 | -0.032 | -2.634 | 0 | -0.032 | -0.032 | 0 | 0.45 | 0.35 | 0.49 | 0.82 | 0.42 | 0.64 |
| 1 | 214863736 | rs2597661 | -0.698 | -1.892 | 0.108 | -2.476 | -0.035 | -0.096 | -0.096 | 0.014 | 0.87 | 0.48 | 0.93 | 1 | 0.93 | 0.96 |
| 1 | 214868909 | rs335565 | 0.285 | -1.497 | -0.497 | -2.284 | -0.021 | 0.022 | 0.022 | 0.267 | 0.1 | 0 | 0.12 | 0.06 | 0.12 | 0.1 |
| 1 | 214883583 | rs11120384 | -0.538 | -1.637 | -0.173 | -2.517 | -0.002 | 0.187 | 0.187 | 0 | 0.44 | 0.13 | 0.47 | 0.81 | 0.39 | 0.68 |
| 1 | 214888381 | rs335554 | -1.393 | -1.878 | 0.57 | -2.551 | -0.027 | -0.114 | -0.114 | 0 | 0.22 | 0.44 | 0.16 | 0.07 | 0.2 | 0.08 |
| 1 | 214889859 | rs335555 | -2.367 | -1.328 | 0.621 | -2.635 | -0.025 | 0.05 | 0.05 | 0 | 0.64 | 0.69 | 0.65 | 0.88 | 0.56 | 0.74 |
| 1 | 214892608 | rs1450597 | -0.276 | -1.359 | 0.502 | -2.484 | -0.01 | 0.173 | 0.173 | 0.008 | 0.41 | 0.1 | 0.47 | 0.81 | 0.36 | 0.67 |
| 1 | 214912720 | rs12385729 | -1.126 | -0.869 | 1.045 | -2.181 | -0.033 | -0.097 | -0.097 | 0 | 0.14 | 0.35 | 0.06 | 0 | 0.08 | 0.04 |
| 1 | 214920883 | rs6659655 | -2.226 | -0.811 | 0.212 | -2.091 | -0.058 | 0.168 | 0.168 | 0 | 0.55 | 0.81 | 0.55 | 0.79 | 0.42 | 0.71 |
| 1 | 214921207 | rs6699978 | -1.279 | -0.71 | 0.388 | -2 | 0.002 | 0.006 | 0.006 | 0 | 0.46 | 0.38 | 0.52 | 0.79 | 0.37 | 0.7 |
| 1 | 214939196 | rs4655371 | -0.194 | -0.611 | -0.029 | -1.954 | -0.005 | 0.013 | 0.013 | 0 | 0.3 | 0.21 | 0.43 | 0.73 | 0.28 | 0.64 |
| 1 | 214941018 | rs11120394 | -0.23 | -0.863 | -0.044 | -1.461 | 0 | 0.002 | 0.002 | 0.064 | 0.08 | 0.04 | 0.07 | 0.06 | 0.08 | 0.04 |
| 1 | 214950361 | rs11800020 | -0.921 | -0.843 | -0.034 | -1.483 | 0.002 | -0.005 | -0.005 | 0 | 0.14 | 0.13 | 0.08 | 0.06 | 0.09 | 0.05 |
| 1 | 214959995 | rs11120400 | -0.451 | -0.757 | -0.483 | -1.837 | 0.001 | -0.022 | -0.022 | 0.105 | 0.39 | 0.42 | 0.49 | 0.73 | 0.38 | 0.65 |
| 1 | 214962771 | rs12734277 | -0.896 | -0.739 | -0.462 | -1.818 | -0.001 | -0.015 | -0.015 | 1.222 | 0.48 | 0.46 | 0.56 | 0.79 | 0.47 | 0.69 |
| 1 | 214963300 | rs992228 | -0.903 | -0.739 | -0.467 | -1.821 | -0.001 | -0.015 | -0.015 | 0.866 | 0.48 | 0.46 | 0.56 | 0.79 | 0.47 | 0.69 |
| 1 | 214965101 | rs2154119 | -1.303 | -0.772 | -0.452 | -1.843 | -0.103 | 0.157 | 0.157 | 1.3 | 0.55 | 0.86 | 0.58 | 0.79 | 0.47 | 0.7 |
| 1 | 214980895 | rs11120417 | -0.985 | -0.502 | -0.431 | -1.722 | -0.001 | -0.015 | -0.015 | 0.032 | 0.48 | 0.47 | 0.56 | 0.79 | 0.47 | 0.69 |
| 1 | 214991610 | rs12407130 | -0.14 | -0.359 | -0.367 | -1.33 | -0.022 | 0.03 | 0.03 | 0 | 0.33 | 0.19 | 0.46 | 0.73 | 0.36 | 0.64 |
| 1 | 214999519 | rs1272551 | 0.516 | -0.372 | -0.206 | -1.007 | 0.016 | 0.076 | 0.076 | 0 | 0.57 | 0.23 | 0.5 | 0.3 | 0.63 | 0.34 |
| 1 | 215005621 | rs1856050 | 1.247 | -0.509 | -0.338 | -1.141 | -0.015 | 0.142 | 0.142 | 1.781 | 0.45 | 0.14 | 0.43 | 0.23 | 0.55 | 0.28 |
| 1 | 215006939 | rs11587867 | 0.493 | -0.374 | -0.191 | -1.014 | 0.016 | 0.065 | 0.065 | 0 | 0.57 | 0.26 | 0.5 | 0.3 | 0.63 | 0.34 |
| 1 | 215014275 | rs1779665 | -0.104 | -0.233 | -0.196 | -0.881 | 0.001 | 0.008 | 0.008 | 0 | 0.67 | 0.68 | 0.55 | 0.3 | 0.68 | 0.38 |
| 1 | 215024396 | rs7414883 | -0.503 | -0.626 | -0.194 | -0.675 | 0.018 | 0.032 | 0.032 | 3.551 | 0.12 | 0.05 | 0.07 | 0.07 | 0.08 | 0.07 |
| 1 | 215025255 | rs12128520 | -0.088 | -0.578 | -0.014 | -0.761 | -0.001 | -0.014 | -0.014 | 0 | 0.07 | 0.07 | 0.02 | 0 | 0.06 | 0.03 |
| 1 | 215030041 | rs6699503 | -0.423 | -0.832 | 0.502 | -0.843 | -0.022 | -0.029 | -0.029 | 0 | 0.34 | 0.41 | 0.49 | 0.72 | 0.29 | 0.63 |

| | | | | | | | | | | | | | | | | |
|---|---|---|---|---|---|---|---|---|---|---|---|---|---|---|---|---|
| 1 | 215031151 | rs1415181 | -0.993 | -0.374 | 0.591 | -0.996 | 0 | -0.007 | -0.007 | 0.231 | 0.46 | 0.51 | 0.59 | 0.81 | 0.47 | 0.73 |
| 1 | 215044315 | rs869380 | 2.166 | -0.636 | 1.997 | -0.273 | -0.01 | -0.018 | -0.018 | 0 | 0.7 | 0.36 | 0.78 | 0.76 | 0.78 | 0.64 |
| 1 | 215048949 | rs7547694 | -1.024 | -0.458 | 2.329 | -0.224 | 0.047 | -0.03 | -0.03 | 1.671 | 0.07 | 0.19 | 0.13 | 0.01 | 0.12 | 0.12 |
| 1 | 215058414 | rs1268910 | 2.711 | -0.512 | 2.219 | -0.22 | 0.015 | 0.022 | 0.022 | 0.07 | 0.63 | 0.24 | 0.74 | 0.66 | 0.72 | 0.56 |
| 1 | 215064388 | rs11807086 | -1.038 | -0.804 | 2.304 | -0.163 | -0.001 | 0.002 | 0.002 | 0.102 | 0.1 | 0.11 | 0.04 | 0.22 | 0.05 | 0.21 |
| 1 | 215068688 | rs1336948 | -0.911 | -0.837 | 2.52 | -0.343 | -0.007 | -0.009 | -0.009 | 0 | 0.06 | 0 | 0.05 | 0.11 | 0.05 | 0.06 |
| 1 | 215073104 | rs12750128 | -2.25 | -0.661 | 2.842 | -0.219 | -0.004 | -0.054 | -0.054 | 0 | 0.22 | 0.36 | 0.2 | 0.12 | 0.2 | 0.23 |
| 1 | 215080890 | rs7524233 | -1.747 | -0.657 | 2.745 | -0.22 | -0.003 | -0.073 | -0.073 | 0 | 0.18 | 0.3 | 0.15 | 0.01 | 0.15 | 0.15 |
| 1 | 215081934 | rs10779629 | -2.381 | -0.696 | 2.785 | -0.124 | -0.02 | -0.011 | -0.011 | 0 | 0.26 | 0.4 | 0.22 | 0.23 | 0.17 | 0.35 |
| 1 | 215093013 | rs17601260 | -0.57 | -0.818 | 2.725 | -0.239 | 0.001 | -0.004 | -0.004 | 0.031 | 0.06 | 0.07 | 0.11 | 0.01 | 0.09 | 0.11 |
| 1 | 215093394 | rs12025511 | -0.957 | -0.817 | 2.713 | -0.221 | -0.007 | 0.003 | 0.003 | 0.182 | 0.09 | 0.11 | 0.07 | 0.19 | 0.03 | 0.2 |
| 1 | 215094201 | rs554343 | -2.875 | -0.823 | 2.806 | -0.206 | -0.02 | -0.012 | -0.012 | 0 | 0.32 | 0.66 | 0.26 | 0.32 | 0.21 | 0.4 |
| 1 | 215102445 | rs573277 | -2.123 | -0.877 | 2.81 | -0.185 | -0.089 | -0.009 | -0.009 | 0 | 0.22 | 0.58 | 0.18 | 0.2 | 0.08 | 0.25 |
| 1 | 215107372 | rs520982 | -2.662 | -0.8 | 2.716 | -0.193 | 0.052 | 0.031 | 0.031 | 0.052 | 0.34 | 0.88 | 0.29 | 0.32 | 0.23 | 0.35 |
| 1 | 215110998 | rs1963938 | -1.38 | -0.789 | 2.809 | -0.188 | -0.015 | -0.129 | -0.129 | 0 | 0.2 | 0.51 | 0.15 | 0.01 | 0.14 | 0.15 |
| 1 | 215112145 | rs497870 | -2.688 | -0.791 | 2.803 | -0.185 | 0.049 | 0.028 | 0.028 | 0 | 0.34 | 0.87 | 0.29 | 0.32 | 0.23 | 0.35 |
| 1 | 215113661 | rs6684902 | -2.28 | -0.616 | 2.895 | 0.493 | -0.032 | 0.07 | 0.07 | 0 | 0.14 | 0.34 | 0.14 | 0.31 | 0.09 | 0.2 |
| 1 | 215124414 | rs566170 | -2.025 | -0.408 | 1.274 | 0.607 | -0.052 | 0.085 | 0.085 | 0 | 0.17 | 0.62 | 0.15 | 0.33 | 0.1 | 0.28 |
| 1 | 215126369 | rs11120451 | -1.262 | 0.62 | -0.224 | 0.317 | -0.012 | -0.024 | -0.024 | 0.481 | 0.06 | 0 | 0.06 | 0.13 | 0.07 | 0.07 |
| 1 | 215130169 | rs476847 | -1.407 | 0.683 | 0.273 | 0.02 | 0.015 | -0.07 | -0.07 | 0 | 0.11 | 0.29 | 0.06 | 0 | 0.09 | 0.05 |
| 1 | 215130661 | rs11120453 | -1.233 | 0.765 | 0.035 | 0.038 | 0.004 | -0.047 | -0.047 | 2.358 | 0.1 | 0.15 | 0.06 | 0 | 0.09 | 0.05 |
| 1 | 215134041 | rs1077323 | -0.76 | 1.606 | -0.255 | 0.527 | -0.015 | -0.013 | -0.013 | 0 | 0.25 | 0.17 | 0.44 | 0.37 | 0.31 | 0.53 |
| 1 | 215136657 | rs10494987 | -0.876 | 1.632 | 0.789 | 0.425 | 0.043 | 0.031 | 0.031 | 2.251 | 0.24 | 0.35 | 0.39 | 0.31 | 0.31 | 0.51 |
| 1 | 215140251 | rs476557 | -1.247 | 0.612 | 0.377 | -0.346 | 0.003 | 0.016 | 0.016 | 0 | 0.57 | 0.73 | 0.56 | 0.44 | 0.56 | 0.76 |
| 1 | 215141570 | rs10494988 | -0.693 | 0.292 | 0.874 | -0.606 | 0.009 | 0.003 | 0.003 | 0.463 | 0.36 | 0.41 | 0.44 | 0.43 | 0.39 | 0.69 |
| 1 | 215150317 | rs2077982 | 0.607 | 0.303 | 0.778 | -0.552 | -0.004 | -0.003 | -0.003 | 1.257 | 0.21 | 0.16 | 0.37 | 0.29 | 0.29 | 0.48 |
| 1 | 215153348 | rs495296 | -1.642 | -0.531 | 0.406 | -1.02 | -0.051 | -0.05 | -0.05 | 0 | 0.65 | 0.91 | 0.61 | 0.44 | 0.59 | 0.79 |
| 1 | 215154120 | rs1452632 | -0.786 | -0.637 | 0.734 | -1.042 | 0.014 | 0.04 | 0.04 | 0 | 0.52 | 0.56 | 0.54 | 0.33 | 0.55 | 0.66 |
| 1 | 215164399 | rs2841623 | -1.439 | -0.834 | 1.226 | -0.649 | 0.073 | 0.278 | 0.278 | 2.278 | 0.06 | 0.45 | 0.03 | 0.14 | 0 | 0.11 |
| 1 | 215165307 | rs1452623 | 0.521 | -0.51 | 0.791 | -0.555 | -0.054 | -0.052 | -0.052 | 0 | 0.22 | 0.05 | 0.38 | 0.34 | 0.26 | 0.48 |
| 1 | 215165487 | rs2841622 | -1.83 | -1.181 | 0.849 | -1.027 | 0.017 | 0.019 | 0.019 | 0 | 0.66 | 0.6 | 0.59 | 0.52 | 0.53 | 0.79 |
| 1 | 215165561 | rs10864142 | -0.472 | -1.299 | 0.836 | -1.101 | 0.06 | 0.111 | 0.111 | 0.139 | 0.34 | 0.15 | 0.16 | 0.01 | 0.24 | 0.14 |
| 1 | 215166503 | rs6659170 | -0.988 | -1.213 | 0.895 | -0.98 | 0.05 | 0.239 | 0.239 | 2.494 | 0.05 | 0.39 | 0.04 | 0.14 | 0 | 0.09 |
| 1 | 215171545 | rs12031300 | 0.794 | -1.191 | 0.762 | -0.954 | -0.055 | -0.054 | -0.054 | 0 | 0.22 | 0.05 | 0.38 | 0.33 | 0.26 | 0.48 |
| 1 | 215174180 | rs1452634 | -1.225 | -1.192 | 1.077 | -0.905 | 0.07 | 0.259 | 0.259 | 1.412 | 0.05 | 0.41 | 0.03 | 0.14 | 0 | 0.11 |
| 1 | 215178483 | rs10779632 | -1.094 | -1.547 | 0.869 | -1.131 | -0.036 | -0.054 | -0.054 | 0 | 0.39 | 0.5 | 0.17 | 0.16 | 0.24 | 0.2 |
| 1 | 215178824 | rs7538655 | -1.184 | -1.605 | 0.66 | -1.16 | 0.031 | 0.036 | 0.036 | 3.843 | 0.4 | 0.35 | 0.19 | 0.18 | 0.25 | 0.3 |
| 1 | 215186044 | rs11120462 | -0.764 | -1.892 | 0.128 | -1.45 | -0.019 | 0.042 | 0.042 | 0 | 0.55 | 0.14 | 0.5 | 0.36 | 0.4 | 0.61 |
| 1 | 215186972 | rs12131478 | -0.311 | -2.275 | -0.061 | -1.664 | 0.014 | 0.014 | 0.014 | 0 | 0.41 | 0.15 | 0.45 | 0.35 | 0.33 | 0.57 |
| 1 | 215187250 | rs2841612 | -2.359 | -2.327 | 0.357 | -1.714 | -0.016 | -0.018 | -0.018 | 0 | 0.7 | 0.76 | 0.59 | 0.53 | 0.5 | 0.82 |
| 1 | 215193196 | rs11120466 | -0.279 | -2.341 | 0.297 | -1.719 | 0.069 | 0.102 | 0.102 | 5.86 | 0.27 | 0.1 | 0.1 | 0.01 | 0.13 | 0.09 |
| 1 | 215206914 | rs12141327 | -0.303 | -2.133 | 0.565 | -1.561 | 0.001 | -0.003 | -0.003 | 0 | 0.4 | 0.38 | 0.46 | 0.51 | 0.33 | 0.72 |
| 1 | 215219587 | rs12097900 | 0.309 | -2.101 | 0.314 | -1.492 | 0.011 | 0.023 | 0.023 | 0 | 0.21 | 0.11 | 0.1 | 0.01 | 0.16 | 0.07 |
| 1 | 215220345 | rs2123331 | -0.038 | -2.349 | -0.496 | -1.881 | 0.012 | -0.075 | -0.075 | 0 | 0.44 | 0.09 | 0.49 | 0.49 | 0.37 | 0.72 |
| 1 | 215222115 | rs2841615 | 2.002 | -2.218 | -0.378 | -1.76 | -0.044 | -0.048 | -0.048 | 0 | 0.24 | 0.16 | 0.37 | 0.47 | 0.45 | 0.12 |
| 1 | 215228234 | rs12131054 | -0.085 | -2.115 | -0.436 | -1.751 | 0.044 | 0.022 | 0.022 | 0 | 0.43 | 0.07 | 0.49 | 0.33 | 0.34 | 0.62 |
| 1 | 215231468 | rs6665177 | -0.242 | -2.329 | 0.248 | -2.028 | -0.037 | -0.139 | -0.139 | 0 | 0.74 | 0.57 | 0.87 | 0.96 | 0.82 | 0.83 |
| 1 | 215236196 | rs12038094 | 0.012 | -2.328 | 0.178 | -2.036 | 0.073 | 0.063 | 0.063 | 0 | 0.43 | 0.12 | 0.36 | 0.31 | 0.33 | 0.62 |
| 1 | 215238904 | rs1157492 | -1.592 | -2.223 | 0.628 | -1.954 | 0.057 | 0.035 | 0.035 | 0.576 | 0.72 | 0.64 | 0.48 | 0.45 | 0.51 | 0.87 |
| 1 | 215244510 | rs10779637 | 0.034 | -2.288 | 0.614 | -2.08 | -0.044 | -0.11 | -0.11 | 0 | 0.26 | 0.55 | 0.12 | 0.11 | 0.16 | 0.16 |
| 1 | 215244538 | rs1947364 | -0.147 | -2.367 | 0.333 | -2.037 | -0.02 | -0.052 | -0.052 | 0 | 0.82 | 0.77 | 0.91 | 0.97 | 0.88 | 0.81 |
| 1 | 215251227 | rs12136349 | -1.11 | -2.454 | 0.285 | -2.092 | 0.004 | -0.026 | -0.026 | 0 | 0.41 | 0.62 | 0.18 | 0.11 | 0.26 | 0.26 |
| 1 | 215255307 | rs2363561 | 0.524 | -2.431 | -0.216 | -1.484 | -0.002 | -0.001 | -0.001 | 0 | 0.44 | 0.53 | 0.58 | 0.53 | 0.58 | 0.25 |
| 1 | 215264486 | rs12133857 | 0.817 | -2.284 | -0.206 | -1.478 | 0.01 | 0.024 | 0.024 | 0 | 0.21 | 0.17 | 0.1 | 0.08 | 0.15 | 0.16 |
| 1 | 215265676 | rs7547655 | -0.779 | -2.361 | -0.149 | -1.411 | 0.01 | -0.011 | -0.011 | 0 | 0.71 | 0.79 | 0.7 | 0.61 | 0.75 | 0.51 |
| 1 | 215269410 | rs10864152 | -0.196 | -2.372 | -0.24 | -1.489 | 0.042 | -0.007 | -0.007 | 0.346 | 0.64 | 0.24 | 0.65 | 0.61 | 0.73 | 0.42 |
| 1 | 215271773 | rs6703032 | 0.649 | -2.378 | -0.267 | -1.575 | 0.042 | 0.07 | 0.07 | 0 | 0.16 | 0.05 | 0.07 | 0.01 | 0.1 | 0.11 |
| 1 | 215288919 | rs4411107 | 0.686 | -2.301 | -0.251 | -1.505 | 0.082 | 0.064 | 0.064 | 0 | 0.49 | 0.3 | 0.59 | 0.63 | 0.63 | 0.34 |
| 1 | 215307777 | rs11120504 | 2.322 | -2.173 | -0.397 | -1.485 | -0.187 | -0.141 | -0.141 | 0.812 | 0.27 | 0.02 | 0.48 | 0.56 | 0.47 | 0.16 |
| 1 | 215309924 | rs12031308 | 0.764 | -2.216 | -0.115 | -1.428 | 0.082 | 0.067 | 0.067 | 0 | 0.49 | 0.3 | 0.59 | 0.63 | 0.63 | 0.32 |
| 1 | 215317651 | rs4509553 | -0.394 | -2.302 | -0.052 | -1.477 | -0.005 | 0 | 0 | 0 | 0.67 | 0.41 | 0.66 | 0.64 | 0.73 | 0.45 |
| 1 | 215322806 | rs10779645 | 0.969 | -2.312 | -0.063 | -1.483 | 0.088 | 0.067 | 0.067 | 0 | 0.48 | 0.24 | 0.59 | 0.63 | 0.63 | 0.32 |
| 1 | 215324031 | rs12138417 | 0.124 | -2.177 | -0.528 | -1.402 | 0.024 | 0.017 | 0.017 | 0 | 0.17 | 0.1 | 0.07 | 0.01 | 0.1 | 0.11 |
| 1 | 215345938 | rs10779651 | 0.212 | -2.134 | -0.613 | -1.336 | 0.051 | 0.027 | 0.027 | 0 | 0.59 | 0.77 | 0.65 | 0.66 | 0.69 | 0.47 |
| 1 | 215359972 | rs1339407 | 0.234 | -1.613 | -0.254 | -0.966 | 0.012 | 0.002 | 0.002 | 0.007 | 0.13 | 0.07 | 0.05 | 0 | 0.06 | 0.05 |
| 1 | 215361592 | rs1556905 | -0.701 | -1.358 | -0.131 | -0.902 | 0.013 | 0.031 | 0.031 | 0 | 0.57 | 0.26 | 0.37 | 0.35 | 0.42 | 0.62 |
| 1 | 215362207 | rs10494994 | 0.509 | -1.557 | -0.341 | -0.914 | 0.014 | -0.019 | -0.019 | 0.463 | 0.3 | 0.02 | 0.2 | 0.27 | 0.24 | 0.48 |
| 1 | 215370762 | rs1538546 | 0.635 | -1.402 | 1.228 | -0.619 | 0.063 | 0.005 | 0.005 | 0 | 0.92 | 0.58 | 0.92 | 0.98 | 0.95 | 0.86 |
| 1 | 215377026 | rs7530450 | 1.126 | -1.403 | 1.3 | -0.625 | 0.018 | -0.038 | -0.038 | 0 | 0.9 | 0.39 | 0.9 | 0.97 | 0.94 | 0.88 |

| | | | | | | | | | | | | | | | | |
|---|---|---|---|---|---|---|---|---|---|---|---|---|---|---|---|---|
| 1 | 215380352 | rs12753507 | 0.787 | -1.39 | 0.822 | -0.698 | 0.002 | 0.007 | 0.007 | 0 | 0.4 | 0.38 | 0.65 | 0.66 | 0.57 | 0.39 |
| 1 | 215394162 | rs17024500 | -0.834 | -1.389 | 1.719 | -0.55 | -0.001 | -0.001 | -0.001 | 0 | 0.57 | 0.59 | 0.37 | 0.4 | 0.43 | 0.67 |
| 1 | 215396543 | rs12080135 | 0.229 | -1.785 | 1.567 | -0.732 | -0.028 | -0.059 | -0.059 | 0 | 0.28 | 0.37 | 0.14 | 0.08 | 0.19 | 0.12 |
| 1 | 215412906 | rs7549184 | -0.865 | -1.816 | 1.544 | -0.662 | -0.028 | 0.006 | 0.006 | 1.452 | 0.72 | 0.56 | 0.77 | 0.72 | 0.76 | 0.52 |
| 1 | 215414522 | rs12094438 | 0.88 | -1.817 | 1.575 | -0.612 | 0.002 | -0.002 | -0.002 | 0 | 0.27 | 0.27 | 0.22 | 0.29 | 0.23 | 0.47 |
| 1 | 215415487 | rs946488 | 0.707 | -1.876 | 1.654 | -0.675 | 0.015 | 0.015 | 0.015 | 0 | 0.45 | 0.39 | 0.64 | 0.64 | 0.57 | 0.4 |
| 1 | 215418210 | rs3933352 | -1.04 | -1.877 | 1.606 | -0.675 | -0.008 | -0.008 | -0.008 | 0 | 0.55 | 0.51 | 0.35 | 0.36 | 0.43 | 0.59 |
| 1 | 215426570 | rs6540890 | -0.011 | -0.994 | 1.314 | -0.714 | -0.016 | -0.037 | -0.037 | 0.082 | 0.3 | 0.33 | 0.14 | 0.08 | 0.19 | 0.12 |
| 1 | 215443789 | rs9662592 | 1.04 | -0.914 | 1.171 | -0.677 | 0.01 | 0.007 | 0.007 | 0.234 | 0.42 | 0.3 | 0.63 | 0.6 | 0.57 | 0.34 |
| 1 | 215453286 | rs1339400 | 1.819 | -1.159 | 0.796 | -0.835 | -0.01 | 0.013 | 0.013 | 0 | 0.13 | 0.01 | 0.05 | 0.08 | 0.09 | 0.05 |
| 1 | 215457085 | rs4442346 | 0.999 | -1.081 | 0.839 | -0.782 | -0.012 | -0.073 | -0.073 | 0 | 0.27 | 0.05 | 0.21 | 0.32 | 0.24 | 0.55 |
| 1 | 215460047 | rs10494997 | 0.174 | -1.056 | 0.976 | -0.801 | 0.025 | 0.036 | 0.036 | 0 | 0.27 | 0.17 | 0.13 | 0.08 | 0.19 | 0.12 |
| 1 | 215474906 | rs11120546 | -1.217 | -1.063 | 1.09 | -0.782 | 0.006 | 0.006 | 0.006 | 0.041 | 0.58 | 0.62 | 0.38 | 0.39 | 0.44 | 0.64 |
| 1 | 215480395 | rs6540891 | 0.232 | -1.038 | 1.134 | -0.786 | 0.036 | 0.061 | 0.061 | 0 | 0.27 | 0.11 | 0.12 | 0.08 | 0.19 | 0.12 |
| 1 | 215511379 | rs952844 | -1.347 | -0.983 | 1.16 | -0.733 | -0.007 | -0.004 | -0.004 | 0 | 0.6 | 0.81 | 0.39 | 0.4 | 0.44 | 0.64 |
| 1 | 215561548 | rs2135811 | -0.332 | -1.081 | -1.054 | -0.846 | 0.042 | 0.054 | 0.054 | 0.146 | 0.29 | 0.79 | 0.16 | 0.12 | 0.15 | 0.21 |
| 1 | 215570581 | rs2089427 | -1.384 | -0.759 | -1.109 | -0.538 | -0.128 | -0.115 | -0.115 | 0.02 | 0.64 | 0.98 | 0.43 | 0.43 | 0.46 | 0.78 |
| 1 | 215592251 | rs973197 | 0.051 | -0.932 | -1.033 | -0.53 | 0.022 | 0.015 | 0.015 | 0 | 0.37 | 0.7 | 0.32 | 0.31 | 0.28 | 0.55 |
| 1 | 215694431 | rs1557174 | 0.027 | -0.841 | -1.508 | -0.387 | 0.031 | 0.014 | 0.014 | 0.07 | 0.4 | 0.69 | 0.3 | 0.37 | 0.29 | 0.66 |
| 1 | 215721124 | rs11120592 | 0.781 | -2.012 | -1.586 | -0.312 | 0.029 | -0.093 | -0.093 | 0.757 | 0.33 | 0.05 | 0.23 | 0.41 | 0.29 | 0.49 |
| 1 | 215747924 | rs10495002 | 1.45 | -1.919 | -1.771 | -0.297 | 0.002 | -0.025 | -0.025 | 0 | 0.06 | 0 | 0.05 | 0.09 | 0.05 | 0.08 |
| 1 | 215753355 | rs11803705 | 0.702 | -1.914 | -1.832 | -0.321 | 0.025 | -0.127 | -0.127 | 0 | 0.35 | 0.04 | 0.24 | 0.44 | 0.3 | 0.52 |
| 1 | 215765548 | rs12080379 | 0.919 | -1.872 | -1.801 | -0.355 | -0.019 | -0.06 | -0.06 | 0.44 | 0.18 | 0.22 | 0.07 | 0.03 | 0.11 | 0.06 |
| 1 | 215773825 | rs2797393 | 0.021 | -1.896 | -1.896 | -0.345 | 0.049 | -0.001 | -0.001 | 0 | 0.45 | 0.71 | 0.35 | 0.55 | 0.35 | 0.68 |
| 1 | 215793834 | rs14137 | -0.128 | -1.756 | -1.844 | -0.338 | 0.062 | 0.041 | 0.041 | 0.574 | 0.53 | 0.29 | 0.61 | 0.35 | 0.64 | 0.28 |
| 1 | 215796350 | rs6667509 | 0.857 | -1.815 | -1.818 | -0.369 | 0.009 | 0.005 | 0.005 | 0 | 0.18 | 0.13 | 0.06 | 0.07 | 0.11 | 0.12 |
| 1 | 215797021 | rs17024993 | 1.035 | -1.69 | -2.109 | -0.244 | 0.01 | -0.017 | -0.017 | 0 | 0.07 | 0.02 | 0.1 | 0.18 | 0.05 | 0.09 |
| 1 | 215805813 | rs2797386 | 1.015 | -1.375 | -2.627 | -0.335 | -0.158 | -0.08 | -0.08 | 0.148 | 0.32 | 0.03 | 0.47 | 0.29 | 0.51 | 0.21 |
| 1 | 215816688 | rs2677111 | 0.461 | -1.227 | -2.09 | 0.146 | 0.084 | 0.066 | 0.066 | 0.492 | 0.51 | 0.26 | 0.55 | 0.32 | 0.64 | 0.28 |
| 1 | 215828499 | rs4655413 | -0.122 | -0.934 | -2.688 | 0.295 | 0.031 | -0.121 | -0.121 | 0 | 0.35 | 0.03 | 0.28 | 0.45 | 0.29 | 0.52 |
| 1 | 215845147 | rs7548730 | -0.234 | -0.646 | -2.337 | 0.206 | 0.05 | -0.009 | -0.009 | 0 | 0.4 | 0.26 | 0.3 | 0.42 | 0.27 | 0.57 |
| 1 | 215845310 | rs12132749 | -0.055 | -0.656 | -2.321 | 0.208 | 0.046 | -0.065 | -0.065 | 0 | 0.37 | 0.06 | 0.28 | 0.37 | 0.27 | 0.51 |
| 1 | 215845984 | rs2066405 | -0.259 | 0.093 | -2.166 | 0.382 | 0.002 | 0.009 | 0.009 | 0.761 | 0.69 | 0.29 | 0.73 | 0.75 | 0.69 | 0.74 |
| 1 | 215848062 | rs2009923 | -0.101 | 0.094 | -2.248 | 0.719 | -0.019 | -0.113 | -0.113 | 0 | 0.21 | 0.03 | 0.18 | 0.35 | 0.19 | 0.31 |
| 1 | 215848587 | rs2797234 | 0.514 | 0.291 | -2.097 | 0.472 | -0.041 | 0.012 | 0.012 | 0 | 0.31 | 0.18 | 0.45 | 0.32 | 0.37 | 0.28 |
| 1 | 215848641 | rs2797235 | 0.435 | -0.05 | -0.75 | 0.696 | -0.019 | 0.051 | 0.051 | 0 | 0.77 | 0.61 | 0.8 | 0.64 | 0.81 | 0.64 |
| 1 | 215849589 | rs10864182 | -0.58 | 0.204 | -1.306 | 0.79 | 0 | -0.049 | -0.049 | 0 | 0.23 | 0.11 | 0.19 | 0.38 | 0.22 | 0.39 |
| 1 | 215849648 | rs10495004 | -0.743 | 0.07 | -1.178 | 0.758 | 0.007 | -0.027 | -0.027 | 1.879 | 0.24 | 0.16 | 0.21 | 0.39 | 0.22 | 0.45 |
| 1 | 215851464 | rs2797236 | -0.189 | 0.337 | -0.917 | 0.678 | -0.003 | -0.007 | -0.007 | 1.362 | 0.64 | 0.39 | 0.57 | 0.52 | 0.63 | 0.71 |
| 1 | 215853116 | rs11578710 | -0.144 | -0.334 | -0.785 | 0.229 | -0.002 | 0.007 | 0.007 | 3.318 | 0.26 | 0.3 | 0.29 | 0.42 | 0.22 | 0.55 |
| 1 | 215854582 | rs2797238 | 1.013 | -0.652 | -0.842 | -0.425 | 0.007 | 0.042 | 0.042 | 5.097 | 0.21 | 0.12 | 0.09 | 0 | 0.17 | 0.04 |
| 1 | 215857876 | rs2797239 | -0.87 | -1.005 | -0.317 | -0.641 | -0.006 | -0.047 | -0.047 | 0.022 | 0.8 | 0.61 | 0.8 | 0.92 | 0.76 | 0.88 |
| 1 | 215859895 | rs2797246 | -0.879 | -1.011 | -0.32 | -0.644 | -0.006 | -0.047 | -0.047 | 0.007 | 0.8 | 0.61 | 0.8 | 0.92 | 0.76 | 0.88 |
| 1 | 215861411 | rs2820691 | -0.347 | -1.028 | -0.271 | -0.288 | 0.015 | 0.047 | 0.047 | 0 | 0.45 | 0.39 | 0.27 | 0.3 | 0.38 | 0.56 |
| 1 | 215861649 | rs2820693 | -0.374 | -1.029 | -0.271 | -0.28 | 0.012 | 0.018 | 0.018 | 0 | 0.46 | 0.39 | 0.39 | 0.42 | 0.39 | 0.59 |
| 1 | 215863578 | rs2820699 | 0.825 | -0.872 | -0.271 | -0.267 | 0.01 | 0.035 | 0.035 | 4.815 | 0.22 | 0.13 | 0.1 | 0 | 0.17 | 0.04 |
| 1 | 215868952 | rs4628479 | 0.514 | -0.826 | -0.221 | -0.163 | 0.002 | 0.034 | 0.034 | 0.004 | 0.13 | 0.23 | 0.14 | 0.31 | 0.13 | 0.35 |
| 1 | 215878243 | rs10864190 | 0.02 | -1.082 | 0.064 | -0.426 | -0.006 | -0.026 | -0.026 | 1.621 | 0.58 | 0.45 | 0.59 | 0.79 | 0.59 | 0.77 |
| 1 | 215880177 | rs7546433 | 0.309 | -1.092 | 0.228 | -0.491 | -0.004 | -0.002 | -0.002 | 0 | 0.61 | 0.47 | 0.64 | 0.57 | 0.66 | 0.52 |
| 1 | 215883350 | rs2364860 | 0.76 | -1.083 | 0.231 | -0.487 | -0.003 | -0.002 | -0.002 | 0 | 0.47 | 0.35 | 0.46 | 0.48 | 0.46 | 0.43 |
| 1 | 215883434 | rs2246472 | -0.565 | -1.064 | 0.333 | -0.504 | 0.006 | -0.016 | -0.016 | 0.04 | 0.95 | 0.83 | 0.97 | 1 | 0.97 | 0.93 |
| 1 | 215885296 | rs6692669 | -0.765 | -1.02 | 0.327 | -0.462 | -0.004 | -0.017 | -0.017 | 0 | 0.74 | 0.68 | 0.78 | 0.87 | 0.8 | 0.84 |
| 1 | 215885838 | rs2886199 | 0.977 | -1.007 | 0.309 | -0.44 | -0.002 | 0.001 | 0.001 | 0 | 0.11 | 0.1 | 0.13 | 0.31 | 0.13 | 0.34 |
| 1 | 215886975 | rs4465171 | 0.702 | -1.026 | 0.293 | -0.455 | -0.002 | -0.001 | -0.001 | 0 | 0.47 | 0.36 | 0.46 | 0.48 | 0.46 | 0.44 |
| 1 | 215887929 | rs6656554 | -0.403 | -1.092 | 0.53 | -0.469 | -0.001 | 0.022 | 0.022 | 0 | 0.29 | 0.56 | 0.33 | 0.39 | 0.34 | 0.38 |
| 1 | 215889357 | rs4655423 | 0.519 | -0.791 | 0.294 | -0.397 | -0.003 | -0.003 | -0.003 | 0.041 | 0.13 | 0.16 | 0.15 | 0.31 | 0.17 | 0.32 |
| 1 | 215889410 | rs12029094 | 0.645 | -0.657 | 0.248 | -0.448 | -0.004 | -0.023 | -0.023 | 0 | 0.42 | 0.17 | 0.42 | 0.48 | 0.4 | 0.43 |
| 1 | 215890030 | rs2257059 | -1.046 | -0.57 | 0.549 | -0.513 | -0.021 | -0.061 | -0.061 | 0 | 0.89 | 0.8 | 0.96 | 1 | 0.94 | 0.88 |
| 1 | 215891174 | rs17620843 | 0.892 | -0.526 | 0.016 | 0.221 | -0.095 | 0.043 | 0.043 | 0.922 | 0.43 | 0.07 | 0.31 | 0.28 | 0.4 | 0.34 |
| 1 | 215893063 | rs4655426 | 0.713 | -0.534 | 0.149 | 0.396 | -0.002 | -0.002 | -0.002 | 0 | 0.13 | 0.15 | 0.16 | 0.21 | 0.18 | 0.36 |
| 1 | 215893303 | rs2797218 | 0.315 | -0.479 | 0.467 | 0.237 | -0.007 | -0.088 | -0.088 | 0 | 0.25 | 0.34 | 0.11 | 0 | 0.2 | 0.12 |
| 1 | 215893559 | rs17562778 | 1.604 | -0.542 | 0.379 | 0.235 | -0.075 | 0.002 | 0.002 | 0.585 | 0.38 | 0.07 | 0.28 | 0.28 | 0.36 | 0.28 |
| 1 | 215893818 | rs2820706 | -0.587 | -0.695 | 0.461 | 0.183 | -0.002 | -0.023 | -0.023 | 0 | 0.38 | 0.47 | 0.26 | 0.21 | 0.37 | 0.45 |
| 1 | 215894886 | rs4449992 | 1.1 | -0.867 | 0.686 | 0.316 | -0.003 | 0.008 | 0.008 | 0 | 0.16 | 0.12 | 0.19 | 0.08 | 0.23 | 0.06 |
| 1 | 215897099 | rs2820710 | 0.239 | -0.869 | 0.775 | 0.3 | 0.011 | 0.015 | 0.015 | 0 | 0.23 | 0.17 | 0.1 | 0 | 0.19 | 0.12 |
| 1 | 215900309 | rs2797213 | 0.552 | -0.852 | 0.888 | 0.48 | 0.028 | 0.414 | 0.414 | 0 | 0.92 | 0.62 | 0.84 | 0.65 | 0.97 | 0.88 |
| 1 | 215903837 | rs2797261 | 0.207 | -0.866 | 0.812 | 0.49 | -0.001 | 0.315 | 0.315 | 0 | 0.93 | 0.74 | 0.85 | 0.65 | 0.98 | 0.88 |
| 1 | 215903907 | rs17025267 | 1.024 | -0.884 | -0.024 | 0.535 | 0.001 | -0.037 | -0.037 | 1.276 | 0.49 | 0.09 | 0.43 | 0.47 | 0.54 | 0.64 |
| 1 | 215912320 | rs4233312 | 0.469 | -0.721 | 0.154 | 0.458 | 0.023 | -0.005 | -0.005 | 0 | 0.67 | 0.27 | 0.72 | 0.64 | 0.78 | 0.75 |

| | | | | | | | | | | | | | | | | |
|---|---|---|---|---|---|---|---|---|---|---|---|---|---|---|---|---|
| 1 | 215915378 | rs2797222 | -0.558 | -0.497 | 0.219 | 0.526 | 0 | 0 | 0 | 0 | 0.37 | 0.38 | 0.49 | 0.4 | 0.43 | 0.27 |
| 1 | 215916563 | rs11120616 | 0.817 | -0.497 | 0.287 | 0.511 | -0.001 | -0.001 | -0.001 | 0.01 | 0.16 | 0.15 | 0.19 | 0.08 | 0.23 | 0.07 |
| 1 | 215919848 | rs2364865 | -0.579 | -0.504 | 0.197 | 0.443 | 0 | 0 | 0 | 0 | 0.37 | 0.38 | 0.49 | 0.4 | 0.43 | 0.27 |
| 1 | 215921679 | rs2820724 | 0.768 | -0.506 | 0.071 | 0.371 | -0.001 | 0.027 | 0.027 | 0 | 0.19 | 0.12 | 0.08 | 0 | 0.18 | 0.09 |
| 1 | 215921956 | rs12141500 | 0.425 | -0.442 | 0.026 | 0.605 | 0.005 | 0.046 | 0.046 | 0 | 0.18 | 0.21 | 0.4 | 0.39 | 0.24 | 0.18 |
| 1 | 215922860 | rs4595347 | -0.908 | -0.553 | -0.194 | 0.575 | 0.002 | -0.018 | -0.018 | 0 | 0.29 | 0.68 | 0.48 | 0.48 | 0.3 | 0.34 |
| 1 | 215928321 | rs2820727 | 0.793 | -0.563 | -0.956 | 0.051 | 0.009 | 0.129 | 0.129 | 0.033 | 0.48 | 0.14 | 0.56 | 0.23 | 0.56 | 0.27 |
| 1 | 215928483 | rs11120620 | 0.036 | -0.597 | -0.193 | 0.209 | -0.012 | -0.045 | -0.045 | 0 | 0.23 | 0.32 | 0.12 | 0.11 | 0.2 | 0.17 |
| 1 | 215929795 | rs2488418 | 0.164 | -0.54 | -1.035 | 0.497 | 0.02 | -0.062 | -0.062 | 0.01 | 0.72 | 0.2 | 0.75 | 0.49 | 0.79 | 0.73 |
| 1 | 215933460 | rs2797227 | 0.124 | -0.373 | -1.018 | 0.451 | 0.014 | -0.036 | -0.036 | 0 | 0.72 | 0.26 | 0.73 | 0.49 | 0.78 | 0.78 |
| 1 | 215935660 | rs7521489 | 0.863 | 0.162 | -1.571 | 0.11 | -0.056 | 0.067 | 0.067 | 0.17 | 0.42 | 0.16 | 0.5 | 0.26 | 0.46 | 0.29 |
| 1 | 215936900 | rs12146110 | 0.679 | 0.265 | -1.178 | 0.347 | -0.001 | -0.035 | -0.035 | 0 | 0.19 | 0.11 | 0.22 | 0.39 | 0.21 | 0.15 |
| 1 | 215955937 | rs6665799 | 0.035 | 0.493 | -1.081 | 0.142 | -0.014 | -0.022 | -0.022 | 0 | 0.37 | 0.49 | 0.27 | 0.27 | 0.33 | 0.34 |
| 1 | 215956304 | rs7518466 | -0.518 | 0.53 | -1.703 | -0.366 | -0.02 | 0.031 | 0.031 | 0 | 0.57 | 0.75 | 0.5 | 0.65 | 0.54 | 0.49 |
| 1 | 215958222 | rs7527334 | -1.336 | 0.599 | -1.806 | -0.43 | -0.017 | -0.009 | -0.009 | 0 | 0.85 | 0.9 | 0.82 | 0.8 | 0.81 | 0.68 |
| 1 | 215960167 | rs10864198 | -0.483 | 0.688 | -1.833 | -0.331 | -0.014 | 0.025 | 0.025 | 0 | 0.57 | 0.71 | 0.49 | 0.65 | 0.54 | 0.49 |
| 1 | 215963035 | rs4462116 | 1.289 | 0.821 | -2.335 | -0.469 | -0.077 | 0.004 | 0.004 | 4.77 | 0.23 | 0.02 | 0.3 | 0.15 | 0.25 | 0.18 |
| 1 | 215963374 | rs11120629 | 0.64 | 0.586 | -1.34 | -0.2 | 0.002 | -0.014 | -0.014 | 0 | 0.15 | 0.13 | 0.07 | 0 | 0.14 | 0.04 |
| 1 | 215968551 | rs10864199 | -0.246 | 0.864 | -1.294 | 0.318 | -0.006 | 0.008 | 0.008 | 0 | 0.17 | 0.12 | 0.17 | 0.12 | 0.19 | 0.11 |
| 1 | 215968999 | rs10864200 | -0.466 | 0.794 | -1.5 | 0.443 | 0.009 | 0.074 | 0.074 | 0.297 | 0.5 | 0.22 | 0.48 | 0.74 | 0.54 | 0.58 |
| 1 | 215970985 | rs4314862 | -0.719 | 0.626 | -1.304 | 0.756 | -0.005 | -0.009 | -0.009 | 0 | 0.06 | 0.03 | 0.07 | 0.16 | 0.08 | 0.09 |
| 1 | 215972795 | rs4363405 | 0.484 | 0.058 | -0.882 | 0.628 | -0.001 | -0.005 | -0.005 | 0 | 0.42 | 0.4 | 0.65 | 0.31 | 0.52 | 0.29 |
| 1 | 215974257 | rs6540911 | -0.288 | -0.567 | -0.426 | 0.698 | 0.015 | 0.005 | 0.005 | 0 | 0.58 | 0.61 | 0.72 | 0.57 | 0.59 | 0.51 |
| 1 | 215976180 | rs10779660 | 0.405 | -0.937 | -0.292 | 0.737 | 0.008 | 0.003 | 0.003 | 0 | 0.54 | 0.54 | 0.75 | 0.64 | 0.61 | 0.48 |
| 1 | 215978775 | rs6688650 | -0.675 | -0.975 | -0.251 | 0.717 | 0.003 | 0.016 | 0.016 | 0 | 0.67 | 0.84 | 0.79 | 0.73 | 0.67 | 0.63 |
| 1 | 215979386 | rs4405115 | 1.426 | -0.89 | 0.036 | 0.726 | -0.01 | 0.009 | 0.009 | 0 | 0.39 | 0.28 | 0.62 | 0.31 | 0.48 | 0.28 |
| 1 | 215984094 | rs11120641 | -0.281 | -0.757 | 0.059 | 0.872 | -0.002 | -0.004 | -0.004 | 0 | 0.07 | 0.05 | 0.07 | 0.16 | 0.08 | 0.09 |
| 1 | 215985959 | rs11120643 | 0.867 | -0.758 | 0.016 | 0.918 | 0.001 | -0.001 | -0.001 | 0 | 0.09 | 0.12 | 0.06 | 0.16 | 0.08 | 0.12 |
| 1 | 216003557 | rs7536020 | 0.392 | -0.81 | 0.349 | 0.871 | -0.017 | 0.013 | 0.013 | 0 | 0.11 | 0.27 | 0.07 | 0.16 | 0.08 | 0.12 |
| 1 | 216005035 | rs10864208 | 1.258 | -0.65 | 0.116 | 0.643 | 0.006 | -0.011 | -0.011 | 0 | 0.42 | 0.44 | 0.62 | 0.31 | 0.46 | 0.26 |
| 1 | 216011803 | rs17025482 | 0.592 | -0.753 | -0.011 | 0.686 | 0.002 | -0.004 | -0.004 | 0.057 | 0.11 | 0.08 | 0.01 | 0.11 | 0.02 | 0.15 |
| 1 | 216016747 | rs17623850 | 1.217 | -0.608 | -0.292 | 0.584 | -0.028 | -0.071 | -0.071 | 0.416 | 0.12 | 0.01 | 0.07 | 0.22 | 0.13 | 0.26 |
| 1 | 216035149 | rs970464 | 0.151 | -0.749 | 0.104 | 0.514 | 0 | 0 | 0 | 0.017 | 0.62 | 0.62 | 0.69 | 0.58 | 0.57 | 0.52 |
| 1 | 216036382 | rs1028796 | 0.777 | -0.699 | 0.197 | 0.556 | -0.002 | -0.001 | -0.001 | 0 | 0.47 | 0.47 | 0.67 | 0.46 | 0.5 | 0.33 |
| 1 | 216041451 | rs12733601 | 0.415 | -0.293 | 0.069 | 0.552 | 0.004 | 0.003 | 0.003 | 0 | 0.11 | 0.19 | 0.12 | 0.17 | 0.11 | 0.15 |
| 1 | 216042210 | rs9804120 | -0.23 | -0.336 | 0.145 | 0.43 | -0.006 | 0.013 | 0.013 | 0.838 | 0.62 | 0.7 | 0.72 | 0.73 | 0.55 | 0.6 |
| 1 | 216047519 | rs11120682 | -0.378 | -0.301 | 0.227 | 0.516 | 0.009 | 0.026 | 0.026 | 0 | 0.16 | 0.26 | 0.17 | 0.31 | 0.16 | 0.23 |
| 1 | 216050785 | rs11120686 | 1.23 | -0.35 | 0.19 | 0.521 | -0.001 | -0.002 | -0.002 | 0 | 0.41 | 0.38 | 0.59 | 0.46 | 0.41 | 0.29 |
| 1 | 216051287 | rs7518358 | -1.588 | -0.173 | -0.018 | 0.609 | -0.014 | 0.019 | 0.019 | 0 | 0.28 | 0.37 | 0.19 | 0.42 | 0.19 | 0.39 |
| 1 | 216053787 | rs17025539 | 0.547 | -0.176 | -0.231 | 0.433 | 0.021 | -0.015 | -0.015 | 0.024 | 0.09 | 0.05 | 0.02 | 0.11 | 0.02 | 0.16 |
| 1 | 216054086 | rs7544464 | -0.502 | -0.059 | -0.149 | 0.397 | -0.01 | 0.004 | 0.004 | 0 | 0.22 | 0.27 | 0.14 | 0.27 | 0.14 | 0.31 |
| 1 | 216055286 | rs7516705 | 0.832 | -0.07 | -0.272 | -0.207 | -0.002 | 0.005 | 0.005 | 0.259 | 0.35 | 0.3 | 0.49 | 0.28 | 0.33 | 0.21 |
| 1 | 216057058 | rs5022147 | -0.516 | -0.066 | -0.155 | -1.028 | 0.007 | -0.066 | -0.066 | 0 | 0.34 | 0.43 | 0.27 | 0.1 | 0.41 | 0.33 |
| 1 | 216063246 | rs11120692 | 1.571 | 0.04 | -0.239 | -1.075 | 0.005 | 0.005 | 0.005 | 0.882 | 0.07 | 0.05 | 0.01 | 0.05 | 0.03 | 0.03 |
| 1 | 216064898 | rs17025561 | 0.831 | -0.066 | 0.129 | -1.012 | 0 | 0.028 | 0.028 | 0 | 0.11 | 0.18 | 0.12 | 0.17 | 0.11 | 0.36 |
| 1 | 216069531 | rs6540916 | -1.271 | -0.386 | 0.13 | -1.459 | 0.008 | 0.012 | 0.012 | 0 | 0.4 | 0.64 | 0.27 | 0.29 | 0.34 | 0.56 |
| 1 | 216074778 | rs7528148 | -1.367 | 0.11 | 0.766 | -2.19 | 0.127 | 0.058 | 0.058 | 0 | 0.07 | 0.31 | 0.12 | 0.06 | 0.09 | 0.09 |
| 1 | 216076167 | rs12046356 | 1.769 | -0.32 | 0.419 | -2.534 | -0.014 | 0.056 | 0.056 | 0 | 0.44 | 0.32 | 0.66 | 0.68 | 0.49 | 0.58 |
| 1 | 216076263 | rs4564111 | -0.281 | -0.685 | 0.433 | -2.509 | 0.028 | 0.043 | 0.043 | 0 | 0.76 | 0.86 | 0.89 | 0.8 | 0.8 | 0.83 |
| 1 | 216080155 | rs1934424 | 0.732 | -0.523 | 0.575 | -2.529 | -0.008 | -0.059 | -0.059 | 0.357 | 0.57 | 0.49 | 0.71 | 0.79 | 0.59 | 0.68 |
| 1 | 216082763 | rs12034100 | 1.83 | -0.461 | 0.311 | -2.524 | -0.005 | 0.054 | 0.054 | 0 | 0.44 | 0.32 | 0.67 | 0.68 | 0.52 | 0.57 |
| 1 | 216096735 | rs7519354 | 1.059 | -0.261 | -0.241 | -2.331 | -0.001 | -0.003 | -0.003 | 0.709 | 0.15 | 0.13 | 0.09 | 0 | 0.16 | 0.11 |
| 1 | 216096763 | rs927744 | -0.312 | -0.418 | -0.402 | -2.188 | 0.03 | 0.044 | 0.044 | 0 | 0.77 | 0.92 | 0.9 | 0.8 | 0.83 | 0.83 |
| 1 | 216104980 | rs498911 | 0.301 | -0.451 | -0.461 | -2.155 | 0.031 | 0.03 | 0.03 | 0 | 0.73 | 0.87 | 0.88 | 0.74 | 0.79 | 0.73 |
| 1 | 216112208 | rs492474 | -0.602 | -0.438 | -0.432 | -2.148 | 0.025 | 0.025 | 0.025 | 0 | 0.28 | 0.16 | 0.12 | 0.27 | 0.22 | 0.27 |
| 1 | 216115551 | rs509655 | 0.468 | -0.359 | 0.237 | -1.947 | -0.01 | -0.082 | -0.082 | 0 | 0.7 | 0.46 | 0.79 | 0.91 | 0.74 | 0.73 |
| 1 | 216115708 | rs683804 | -1.098 | -0.358 | 0.382 | -1.989 | -0.012 | -0.064 | -0.064 | 0 | 0.84 | 0.74 | 0.91 | 0.98 | 0.86 | 0.89 |
| 1 | 216117840 | rs531780 | 0.697 | 0.043 | -0.067 | -2.36 | -0.068 | 0.077 | 0.077 | 1.663 | 0.28 | 0.09 | 0.2 | 0.2 | 0.32 | 0.19 |
| 1 | 216129082 | rs17025707 | 0.404 | 0.158 | 0.066 | -2.245 | 0.019 | -0.057 | -0.057 | 0 | 0.1 | 0.22 | 0.11 | 0 | 0.09 | 0.06 |
| 1 | 216132878 | rs12410350 | 0.493 | 0.167 | 0.027 | -2.243 | 0.015 | -0.051 | -0.051 | 0 | 0.1 | 0.18 | 0.11 | 0 | 0.09 | 0.06 |
| 1 | 216136620 | rs3845529 | 1.151 | -0.218 | 0.139 | -2.233 | 0 | 0.002 | 0.002 | 0 | 0.35 | 0.29 | 0.56 | 0.67 | 0.37 | 0.52 |
| 1 | 216136873 | rs571005 | -1.303 | 0.095 | -0.142 | -2.124 | -0.006 | -0.045 | -0.045 | 0.084 | 0.84 | 0.78 | 0.91 | 0.98 | 0.85 | 0.89 |
| 1 | 216138953 | rs12069955 | 1.255 | 0.209 | -0.503 | -1.675 | 0.005 | 0.005 | 0.005 | 0.052 | 0.08 | 0.05 | 0.01 | 0.04 | 0.03 | 0.03 |
| 1 | 216139209 | rs2576002 | -0.526 | 0.411 | -0.319 | -1.524 | -0.022 | 0.136 | 0.136 | 1.733 | 0.46 | 0.27 | 0.32 | 0.24 | 0.45 | 0.29 |
| 1 | 216143270 | rs479826 | -0.58 | 0.603 | -0.452 | -1.424 | -0.007 | 0.053 | 0.053 | 0.703 | 0.46 | 0.38 | 0.32 | 0.25 | 0.45 | 0.29 |
| 1 | 216151789 | rs634691 | 0.328 | 0.565 | -0.133 | -1.055 | -0.009 | -0.05 | -0.05 | 1.962 | 0.89 | 0.84 | 0.94 | 1 | 0.93 | 0.93 |
| 1 | 216152301 | rs12403934 | -0.885 | 1.103 | -0.399 | -0.669 | -0.046 | -0.044 | -0.044 | 0.048 | 0.08 | 0 | 0.11 | 0.16 | 0.13 | 0.1 |
| 1 | 216154283 | rs2813727 | 0.829 | 0.885 | 0.406 | -1.07 | 0.019 | -0.033 | -0.033 | 0 | 0.86 | 0.79 | 0.87 | 0.93 | 0.85 | 0.85 |
| 1 | 216157829 | rs747808 | 1.159 | 1.34 | 0.684 | -1.406 | 0 | -0.009 | -0.009 | 0 | 0.72 | 0.73 | 0.73 | 0.88 | 0.65 | 0.76 |

| | | | | | | | | | | | | | | | | |
|---|---|---|---|---|---|---|---|---|---|---|---|---|---|---|---|---|
| 1 | 216159935 | rs9308440 | 0.423 | 0.778 | 0.403 | -1.678 | -0.029 | -0.061 | -0.061 | 0.716 | 0.24 | 0.33 | 0.09 | 0.03 | 0.16 | 0.12 |
| 1 | 216163612 | rs565454 | 1.118 | 0.837 | 0.254 | -1.684 | -0.04 | 0.033 | 0.033 | 0 | 0.13 | 0.07 | 0.14 | 0.06 | 0.2 | 0.07 |
| 1 | 216165271 | rs1821763 | 0.44 | 0.757 | 0.623 | -1.745 | 0.01 | 0.045 | 0.045 | 0 | 0.65 | 0.74 | 0.8 | 0.91 | 0.7 | 0.73 |
| 1 | 216166005 | rs3856186 | 0.274 | 0.782 | 0.566 | -1.757 | -0.029 | 0.046 | 0.046 | 0 | 0.77 | 0.83 | 0.82 | 0.94 | 0.75 | 0.86 |
| 1 | 216167644 | rs1992099 | 1.134 | 0.781 | 0.553 | -1.746 | 0.025 | 0.038 | 0.038 | 0 | 0.21 | 0.15 | 0.08 | 0.03 | 0.15 | 0.11 |
| 1 | 216172380 | rs10864219 | 1.322 | 0.781 | 0.765 | -1.686 | 0.002 | 0.003 | 0.003 | 0 | 0.49 | 0.42 | 0.65 | 0.83 | 0.5 | 0.64 |
| 1 | 216172585 | rs1436780 | -1.307 | 0.841 | 0.363 | -1.624 | 0.003 | 0.004 | 0.004 | 0 | 0.92 | 0.95 | 0.96 | 1 | 0.95 | 0.96 |
| 1 | 216172740 | rs12128250 | 0.852 | 0.849 | 0.098 | -1.587 | -0.04 | 0.037 | 0.037 | 0 | 0.14 | 0.07 | 0.14 | 0.06 | 0.2 | 0.09 |
| 1 | 216179808 | rs4471233 | 1.68 | 0.917 | -0.031 | -1.07 | -0.013 | 0.019 | 0.019 | 0 | 0.42 | 0.19 | 0.64 | 0.67 | 0.47 | 0.4 |
| 1 | 216187482 | rs4537521 | -0.479 | 1.026 | -0.506 | -1.055 | -0.057 | 0.004 | 0.004 | 0 | 0.88 | 0.99 | 0.89 | 0.93 | 0.86 | 0.87 |
| 1 | 216188600 | rs7519838 | 0.754 | 1.045 | -0.423 | -1.013 | 0.023 | 0.037 | 0.037 | 0 | 0.22 | 0.18 | 0.07 | 0.04 | 0.18 | 0.11 |
| 1 | 216188698 | rs10779662 | 1.525 | 1.122 | -0.368 | -0.979 | 0.001 | 0.011 | 0.011 | 0 | 0.43 | 0.35 | 0.64 | 0.68 | 0.46 | 0.41 |
| 1 | 216194948 | rs2317936 | 1.485 | 1.999 | 0.03 | -0.212 | 0.036 | -0.024 | -0.024 | 0 | 0.66 | 0.53 | 0.72 | 0.72 | 0.63 | 0.53 |
| 1 | 216205110 | rs12121718 | -0.148 | 1.234 | -0.531 | -0.092 | -0.041 | -0.023 | -0.023 | 0.266 | 0.17 | 0.08 | 0.15 | 0.19 | 0.21 | 0.32 |
| 1 | 216209672 | rs6692241 | 0.727 | 0.523 | 0.105 | -0.165 | -0.017 | -0.036 | -0.036 | 0 | 0.86 | 0.7 | 0.94 | 0.93 | 0.92 | 0.89 |
| 1 | 216215741 | rs11120726 | 0.207 | 0.729 | -0.077 | -0.072 | 0.023 | 0.01 | 0.01 | 1.301 | 0.37 | 0.52 | 0.44 | 0.41 | 0.48 | 0.3 |
| 1 | 216217169 | rs17025913 | -0.031 | 0.542 | -0.252 | -0.062 | -0.009 | -0.002 | -0.002 | 0.925 | 0.11 | 0.3 | 0.06 | 0.05 | 0.1 | 0.05 |
| 1 | 216219781 | rs6657250 | -0.19 | 0.573 | -1.165 | -0.242 | -0.01 | -0.021 | -0.021 | 0 | 0.67 | 0.8 | 0.47 | 0.45 | 0.66 | 0.5 |
| 1 | 216220364 | rs4655443 | -0.786 | -0.177 | -1.437 | -0.212 | 0.061 | -0.013 | -0.013 | 0 | 0.44 | 0.81 | 0.53 | 0.38 | 0.32 | 0.33 |
| 1 | 216229053 | rs12067972 | 0.011 | 0.638 | -0.597 | -0.209 | 0.051 | -0.064 | -0.064 | 0 | 0.11 | 0.34 | 0.1 | 0.02 | 0.1 | 0.08 |
| 1 | 216232533 | rs7539701 | -0.359 | 0.602 | -2.068 | -0.128 | -0.065 | -0.092 | -0.092 | 0 | 0.59 | 0.91 | 0.44 | 0.55 | 0.6 | 0.48 |
| 1 | 216235074 | rs1436776 | 0.427 | 0.519 | -1.982 | -0.179 | 0.001 | -0.03 | -0.03 | 0 | 0.44 | 0.25 | 0.31 | 0.52 | 0.51 | 0.4 |
| 1 | 216235233 | rs12126435 | 0.37 | 0.602 | -1.954 | -0.15 | 0 | -0.026 | -0.026 | 0 | 0.45 | 0.25 | 0.32 | 0.52 | 0.51 | 0.41 |
| 1 | 216236311 | rs6419476 | -0.02 | 0.735 | -2.512 | 0.32 | -0.056 | -0.088 | -0.088 | 0.2 | 0.59 | 0.88 | 0.44 | 0.54 | 0.61 | 0.48 |
| 1 | 216240301 | rs4655445 | -0.264 | 0.656 | -2.066 | 0.142 | -0.056 | 0.017 | 0.017 | 0.53 | 0.54 | 0.88 | 0.5 | 0.44 | 0.49 | 0.6 |
| 1 | 216246044 | rs17026017 | -0.65 | 0.039 | -0.719 | -0.422 | -0.038 | -0.108 | -0.108 | 0 | 0.16 | 0.55 | 0.07 | 0.02 | 0.09 | 0.07 |
| 1 | 216249570 | rs11587260 | -0.107 | 0.345 | -0.953 | -0.862 | -0.009 | 0.045 | 0.045 | 0 | 0.43 | 0.27 | 0.56 | 0.77 | 0.48 | 0.54 |
| 1 | 216252742 | rs2068721 | -0.086 | 0.775 | -0.703 | -1.492 | -0.012 | 0.062 | 0.062 | 0 | 0.34 | 0.24 | 0.57 | 0.85 | 0.37 | 0.51 |
| 1 | 216264294 | rs1544298 | -0.471 | 0.386 | -0.519 | -2.198 | -0.001 | -0.015 | -0.015 | 0 | 0.53 | 0.52 | 0.68 | 0.88 | 0.53 | 0.58 |
| 1 | 216264388 | rs1544299 | 0.438 | 0.408 | -0.398 | -2.172 | -0.01 | 0.047 | 0.047 | 0 | 0.35 | 0.27 | 0.58 | 0.86 | 0.37 | 0.5 |
| 1 | 216269176 | rs17026081 | 0.624 | 0.491 | -0.073 | -2.074 | 0.038 | -0.085 | -0.085 | 2.172 | 0.13 | 0.25 | 0.1 | 0 | 0.17 | 0.06 |
| 1 | 216277327 | rs12123638 | 0.972 | 0.279 | -0.304 | -2.13 | -0.004 | 0.051 | 0.051 | 0 | 0.32 | 0.22 | 0.21 | 0.14 | 0.3 | 0.33 |
| 1 | 216281673 | rs11120735 | -0.544 | 0.216 | -0.352 | -2.21 | -0.001 | 0.089 | 0.089 | 0.199 | 0.82 | 0.93 | 0.9 | 0.99 | 0.83 | 0.94 |
| 1 | 216282322 | rs4511139 | 1.015 | -0.363 | -0.647 | -2.536 | 0.025 | -0.007 | -0.007 | 0.025 | 0.09 | 0.04 | 0.04 | 0.08 | 0.03 | 0.13 |
| 1 | 216282495 | rs421821 | 0.17 | -0.322 | -0.252 | -2.572 | -0.002 | -0.122 | -0.122 | 0.217 | 0.68 | 0.5 | 0.79 | 0.99 | 0.67 | 0.88 |
| 1 | 216286427 | rs12562620 | 0.708 | -0.235 | -0.368 | -2.374 | -0.016 | 0.087 | 0.087 | 0 | 0.34 | 0.17 | 0.57 | 0.84 | 0.36 | 0.52 |
| 1 | 216292566 | rs301734 | -0.051 | -0.221 | -0.31 | -1.818 | -0.003 | 0.013 | 0.013 | 0 | 0.45 | 0.51 | 0.34 | 0.24 | 0.45 | 0.4 |
| 1 | 216295188 | rs10864233 | 0.754 | -0.217 | -0.414 | -1.757 | -0.01 | 0.019 | 0.019 | 0 | 0.35 | 0.19 | 0.55 | 0.75 | 0.37 | 0.5 |
| 1 | 216298635 | rs532342 | 0.36 | -0.272 | -0.305 | -1.558 | -0.017 | -0.099 | -0.099 | 0.084 | 0.75 | 0.57 | 0.87 | 0.99 | 0.79 | 0.9 |
| 1 | 216305479 | rs301736 | 0.634 | -0.191 | -0.548 | -1.353 | 0.023 | 0.016 | 0.016 | 0 | 0.72 | 0.77 | 0.79 | 0.82 | 0.77 | 0.75 |
| 1 | 216313208 | rs448588 | -0.151 | -0.094 | -0.514 | -1.346 | 0.005 | 0.015 | 0.015 | 0 | 0.8 | 0.81 | 0.9 | 0.98 | 0.82 | 0.91 |
| 1 | 216347556 | rs439940 | -0.185 | -0.286 | -0.701 | -1.334 | 0.019 | -0.018 | -0.018 | 0.011 | 0.22 | 0.16 | 0.19 | 0.21 | 0.21 | 0.22 |
| 1 | 216348764 | rs1805049 | 0.924 | -0.506 | 0.257 | -1.494 | -0.004 | -0.008 | -0.008 | 0 | 0.61 | 0.55 | 0.7 | 0.78 | 0.61 | 0.71 |
| 1 | 216362024 | rs301748 | -1.281 | -0.546 | 0.095 | -1.54 | -0.014 | 0.037 | 0.037 | 0 | 0.91 | 1 | 0.97 | 0.99 | 0.92 | 0.97 |
| 1 | 216368181 | rs400358 | -1.32 | -0.569 | 0.093 | -1.542 | -0.005 | 0.036 | 0.036 | 0 | 0.92 | 1 | 0.97 | 0.99 | 0.93 | 0.97 |
| 1 | 216384908 | rs3754064 | 1.005 | -0.525 | 0.33 | -1.617 | -0.023 | -0.02 | -0.02 | 0 | 0.62 | 0.53 | 0.72 | 0.77 | 0.68 | 0.73 |
| 1 | 216390694 | rs1324330 | 1.4 | -0.765 | 0.965 | -1.754 | 0.025 | 0.045 | 0.045 | 0.876 | 0.57 | 0.3 | 0.68 | 0.77 | 0.61 | 0.7 |
| 1 | 216405772 | rs378340 | -0.082 | -0.866 | 0.862 | -1.742 | 0.007 | -0.005 | -0.005 | 2.017 | 0.23 | 0.22 | 0.22 | 0.21 | 0.23 | 0.21 |
| 1 | 216408083 | rs619803 | -0.496 | -0.839 | 0.741 | -1.707 | 0.012 | 0.072 | 0.072 | 0 | 0.86 | 1 | 0.96 | 0.99 | 0.91 | 0.95 |
| 1 | 216410939 | rs3767693 | 1.162 | -1.025 | 0.237 | -2.152 | -0.116 | -0.047 | -0.047 | 2.015 | 0.34 | 0.05 | 0.51 | 0.68 | 0.4 | 0.49 |
| 1 | 216413725 | rs2926178 | -0.722 | -1.016 | 0.343 | -2.087 | 0.011 | 0.071 | 0.071 | 0 | 0.86 | 1 | 0.96 | 0.99 | 0.91 | 0.95 |
| 1 | 216417675 | rs682319 | -0.503 | -1.396 | 0.135 | -2.23 | 0.023 | 0.097 | 0.097 | 0.258 | 0.84 | 1 | 0.95 | 0.99 | 0.89 | 0.92 |
| 1 | 216417771 | rs696720 | 0.695 | -1.568 | 0.614 | -1.981 | -0.031 | -0.016 | -0.016 | 0.432 | 0.6 | 0.49 | 0.7 | 0.72 | 0.66 | 0.71 |
| 1 | 216431962 | rs386654 | 0.263 | -1.489 | 0.628 | -1.929 | -0.005 | 0.007 | 0.007 | 0 | 0.23 | 0.27 | 0.24 | 0.25 | 0.23 | 0.21 |
| 1 | 216440105 | rs2168924 | 1.5 | -1.554 | 0.046 | -0.732 | -0.042 | -0.05 | -0.05 | 0 | 0.24 | 0.11 | 0.33 | 0.43 | 0.28 | 0.39 |
| 1 | 216446653 | rs12134474 | 1.095 | -1.904 | 0.508 | -0.892 | 0.062 | 0.041 | 0.041 | 1.107 | 0.58 | 0.29 | 0.7 | 0.75 | 0.69 | 0.69 |
| 1 | 216454483 | rs1159143 | 0.368 | -2.119 | 0.73 | -0.952 | -0.013 | -0.001 | -0.001 | 0 | 0.71 | 0.33 | 0.71 | 0.78 | 0.78 | 0.75 |
| 1 | 216455107 | rs1512581 | -0.122 | -2.12 | 0.68 | -0.935 | -0.004 | 0 | 0 | 0 | 0.76 | 0.77 | 0.75 | 0.82 | 0.78 | 0.75 |
| 1 | 216473745 | rs17042212 | 0.13 | -2.081 | 0.011 | -0.921 | 0.024 | -0.063 | -0.063 | 0.044 | 0.25 | 0.02 | 0.21 | 0.18 | 0.21 | 0.24 |
| 1 | 216475249 | rs1398382 | 1.243 | -2.021 | 0.18 | -0.991 | 0.096 | 0.094 | 0.094 | 2.835 | 0.54 | 0.25 | 0.67 | 0.77 | 0.67 | 0.66 |
| 1 | 216488762 | rs10864238 | -1.488 | -2.082 | 0.516 | -1.009 | 0.008 | 0.006 | 0.006 | 2.021 | 0.44 | 0.56 | 0.32 | 0.17 | 0.33 | 0.32 |
| 1 | 216496947 | rs1805050 | -0.466 | -2.115 | 0.687 | -1.028 | -0.02 | -0.031 | -0.031 | 0.085 | 0.31 | 0.39 | 0.28 | 0.13 | 0.24 | 0.24 |
| 1 | 216505468 | rs12030122 | 0.442 | -2.068 | 0.567 | -1.145 | -0.016 | -0.033 | -0.033 | 0.066 | 0.14 | 0.17 | 0.04 | 0.04 | 0.09 | 0.08 |
| 1 | 216521091 | rs1606357 | -0.415 | -2.095 | 0.628 | -1.127 | -0.021 | -0.031 | -0.031 | 0.623 | 0.3 | 0.38 | 0.28 | 0.13 | 0.24 | 0.24 |
| 1 | 216537936 | rs1588657 | 0.9 | -1.85 | 0.454 | -0.985 | 0.031 | 0.041 | 0.041 | 2.949 | 0.56 | 0.36 | 0.67 | 0.83 | 0.68 | 0.68 |
| 1 | 216549082 | rs1588656 | 1.492 | -1.124 | -0.264 | -0.406 | -0.127 | -0.115 | -0.115 | 0 | 0.18 | 0.04 | 0.36 | 0.44 | 0.39 | 0.4 |
| 1 | 216552091 | rs2669053 | 1.059 | -1.395 | -0.064 | -0.802 | 0.061 | 0.087 | 0.087 | 1.186 | 0.51 | 0.33 | 0.64 | 0.78 | 0.66 | 0.63 |
| 1 | 216552367 | rs17026635 | -0.563 | -1.551 | 0.11 | -0.81 | 0.027 | 0.017 | 0.017 | 1.584 | 0.07 | 0.42 | 0.04 | 0 | 0.02 | 0 |
| 1 | 216571742 | rs4531347 | 1.152 | -1.531 | -0.234 | -0.792 | 0.049 | 0.06 | 0.06 | 0.985 | 0.51 | 0.36 | 0.65 | 0.78 | 0.66 | 0.64 |

| | | | | | | | | | | | | | | | | |
|---|---|---|---|---|---|---|---|---|---|---|---|---|---|---|---|---|
| 1 | 216576160 | rs4420153 | -0.254 | -1.616 | -0.293 | -0.765 | 0.033 | 0.05 | 0.05 | 0 | 0.68 | 0.82 | 0.71 | 0.82 | 0.73 | 0.78 |
| 1 | 216591419 | rs6682779 | -0.419 | -1.652 | -0.112 | -0.713 | 0.002 | 0.03 | 0.03 | 1.23 | 0.07 | 0.33 | 0.03 | 0.04 | 0.02 | 0 |
| 1 | 216595306 | rs10779261 | -0.007 | -1.551 | -0.648 | -0.737 | 0.029 | 0.037 | 0.037 | 0 | 0.67 | 0.76 | 0.71 | 0.81 | 0.73 | 0.77 |
| 1 | 216605071 | rs12723493 | 0.855 | -1.272 | -1.261 | -0.535 | 0.026 | 0.008 | 0.008 | 0.057 | 0.51 | 0.2 | 0.54 | 0.56 | 0.47 | 0.47 |
| 1 | 216608268 | rs11582099 | 1.327 | -1.3 | -1.471 | -0.507 | 0.019 | -0.019 | -0.019 | 0 | 0.22 | 0.02 | 0.18 | 0.18 | 0.15 | 0.17 |
| 1 | 216613761 | rs11117581 | 1.195 | -0.981 | -1.011 | -0.351 | -0.018 | -0.015 | -0.015 | 0 | 0.26 | 0.26 | 0.34 | 0.4 | 0.38 | 0.35 |
| 1 | 216622208 | rs11117582 | 0.152 | -0.715 | -1.251 | -0.3 | -0.036 | -0.027 | -0.027 | 0 | 0.39 | 0.76 | 0.44 | 0.45 | 0.44 | 0.45 |
| 1 | 216630596 | rs4240918 | 0.021 | 0.192 | -1.558 | -0.24 | 0.014 | 0.049 | 0.049 | 0 | 0.63 | 0.72 | 0.75 | 0.81 | 0.69 | 0.62 |
| 1 | 216630824 | rs4548509 | -0.928 | 0.738 | -1.713 | 0.175 | -0.025 | 0.069 | 0.069 | 0 | 0.88 | 1 | 0.94 | 0.99 | 0.89 | 0.92 |
| 1 | 216632415 | rs11117584 | 0.079 | 0.981 | -1.157 | 0.777 | 0 | -0.001 | -0.001 | 0 | 0.27 | 0.25 | 0.33 | 0.39 | 0.27 | 0.24 |
| 1 | 216632948 | rs10495024 | -0.733 | 1.843 | -1.507 | 0.925 | 0.006 | 0.012 | 0.012 | 0 | 0.21 | 0.23 | 0.32 | 0.51 | 0.34 | 0.25 |
| 1 | 216633391 | rs4846466 | 0.19 | 1.283 | -1.936 | 0.251 | -0.068 | 0.005 | 0.005 | 4.552 | 0.65 | 0.86 | 0.63 | 0.65 | 0.52 | 0.49 |
| 1 | 216638249 | rs12142867 | -0.852 | 0.589 | -1.12 | 0.209 | -0.019 | 0.05 | 0.05 | 0 | 0.09 | 0.11 | 0.1 | 0.19 | 0.05 | 0.2 |
| 1 | 216638273 | rs12725653 | 0.285 | 0.786 | -1.623 | -0.171 | -0.089 | -0.101 | -0.101 | 0 | 0.24 | 0.04 | 0.42 | 0.7 | 0.3 | 0.31 |
| 1 | 216641894 | rs7533687 | -0.203 | 0.312 | -1.083 | -0.331 | 0.034 | 0.069 | 0.069 | 0.547 | 0.07 | 0.54 | 0.04 | 0.04 | 0.02 | 0.09 |
| 1 | 216649667 | rs4338426 | -1.192 | 0.325 | -1.058 | -0.665 | 0.055 | 0.137 | 0.137 | 0 | 0.51 | 0.3 | 0.65 | 0.9 | 0.6 | 0.78 |
| 1 | 216650304 | rs4256876 | -1.227 | 0.326 | -1.061 | -0.668 | 0.06 | 0.156 | 0.156 | 0 | 0.52 | 0.26 | 0.64 | 0.9 | 0.6 | 0.78 |
| 1 | 216655122 | rs6691953 | 0.162 | 0.176 | -0.736 | -0.247 | 0.091 | 0.074 | 0.074 | 0 | 0.06 | 0.43 | 0.04 | 0.01 | 0.02 | 0.03 |
| 1 | 216655251 | rs6604611 | -1.534 | 0.118 | -1.155 | -0.255 | 0.014 | -0.015 | -0.015 | 0 | 0.18 | 0.81 | 0.16 | 0.22 | 0.08 | 0.29 |
| 1 | 216657848 | rs4348785 | -0.061 | 0.682 | -0.977 | 0.042 | -0.056 | -0.073 | -0.073 | 0.673 | 0.25 | 0.09 | 0.42 | 0.65 | 0.3 | 0.31 |
| 1 | 216665455 | rs12031665 | -0.453 | 0.565 | -1.063 | -0.02 | 0.011 | 0.008 | 0.008 | 0 | 0.31 | 0.57 | 0.46 | 0.68 | 0.32 | 0.4 |
| 1 | 216670203 | rs11582478 | -0.505 | 0.56 | -1.23 | 0.188 | -0.119 | 0.106 | 0.106 | 0 | 0.24 | 0.01 | 0.17 | 0.04 | 0.32 | 0.27 |
| 1 | 216670895 | rs11584377 | -0.428 | 0.578 | -1.223 | 0.197 | -0.115 | 0.102 | 0.102 | 0 | 0.23 | 0.01 | 0.16 | 0.04 | 0.3 | 0.27 |
| 1 | 216671992 | rs4360581 | -0.428 | 0.644 | -1.02 | 0.576 | -0.005 | 0.118 | 0.118 | 0 | 0.05 | 0.08 | 0.15 | 0.24 | 0.04 | 0.15 |
| 1 | 216672805 | rs6666194 | -0.745 | 0.907 | -1.032 | 0.411 | -0.018 | -0.01 | -0.01 | 0.561 | 0.35 | 0.7 | 0.37 | 0.28 | 0.37 | 0.43 |
| 1 | 216690332 | rs7529338 | -0.903 | 1.835 | 0.691 | -0.002 | -0.087 | -0.009 | -0.009 | 0 | 0.12 | 0.02 | 0.17 | 0.08 | 0.29 | 0.18 |
| 1 | 216691055 | rs17042719 | -0.461 | 1.101 | 0.418 | 0.077 | 0.017 | -0.012 | -0.012 | 0 | 0.13 | 0.07 | 0.19 | 0.16 | 0.08 | 0.09 |
| 1 | 216691305 | rs6604627 | -0.498 | 1.072 | 0.06 | -0.42 | -0.008 | 0.014 | 0.014 | 0 | 0.76 | 0.84 | 0.81 | 0.82 | 0.73 | 0.77 |
| 1 | 216692002 | rs10779265 | 0.296 | 0.343 | 0.223 | -0.345 | -0.001 | 0.021 | 0.021 | 0 | 0.63 | 0.73 | 0.64 | 0.74 | 0.43 | 0.59 |
| 1 | 216692669 | rs945453 | 0.291 | 0.342 | 0.223 | -0.345 | -0.001 | 0.021 | 0.021 | 0 | 0.63 | 0.73 | 0.64 | 0.74 | 0.43 | 0.59 |
| 1 | 216693791 | rs12027943 | 0.589 | 0.407 | -0.271 | -0.304 | 0.027 | 0.109 | 0.109 | 0 | 0.56 | 0.2 | 0.58 | 0.74 | 0.42 | 0.57 |
| 1 | 216700824 | rs12058696 | 0.218 | 0.311 | -0.025 | -0.43 | -0.007 | -0.023 | -0.023 | 0 | 0.1 | 0.16 | 0.07 | 0.01 | 0.08 | 0.01 |
| 1 | 216701242 | rs11117608 | -0.984 | 0.873 | -0.152 | -0.686 | -0.016 | 0.016 | 0.016 | 0 | 0.79 | 0.86 | 0.81 | 0.86 | 0.73 | 0.78 |
| 1 | 216702685 | rs4846514 | -0.154 | 0.48 | 0.502 | -0.27 | -0.002 | 0.009 | 0.009 | 0 | 0.09 | 0.08 | 0.19 | 0.19 | 0.07 | 0.09 |
| 1 | 216702964 | rs6683007 | -1.139 | 0.49 | 1.004 | -0.405 | -0.002 | -0.038 | -0.038 | 0 | 0.31 | 0.36 | 0.32 | 0.16 | 0.34 | 0.2 |
| 1 | 216705965 | rs6696225 | -2.276 | 0.165 | 0.724 | -0.51 | 0.004 | 0.081 | 0.081 | 0.098 | 0.49 | 0.86 | 0.58 | 0.35 | 0.49 | 0.33 |
| 1 | 216706093 | rs9651260 | -1.075 | 0.182 | 1.217 | -0.547 | 0.051 | -0.039 | -0.039 | 0 | 0.08 | 0.34 | 0.07 | 0.01 | 0.07 | 0.03 |
| 1 | 216707012 | rs1833036 | -1.972 | 0.079 | 1.014 | -0.532 | 0.008 | 0.009 | 0.009 | 0.239 | 0.43 | 0.56 | 0.52 | 0.34 | 0.42 | 0.3 |
| 1 | 216707519 | rs6604631 | -1.021 | 0.178 | 0.765 | -0.505 | -0.003 | 0.032 | 0.032 | 1.669 | 0.29 | 0.18 | 0.29 | 0.16 | 0.35 | 0.21 |
| 1 | 216712891 | rs6694260 | -0.059 | 0.252 | 0.619 | -0.452 | -0.028 | -0.045 | -0.045 | 2.509 | 0.13 | 0.24 | 0.08 | 0.01 | 0.08 | 0.01 |
| 1 | 216715514 | rs2377098 | -1.802 | 0.774 | 0.351 | -0.145 | -0.006 | 0.003 | 0.003 | 0 | 0.33 | 0.37 | 0.47 | 0.33 | 0.39 | 0.32 |
| 1 | 216716320 | rs12145418 | -1.449 | 0.804 | -0.012 | -0.103 | -0.028 | -0.044 | -0.044 | 0 | 0.28 | 0.06 | 0.41 | 0.32 | 0.33 | 0.29 |
| 1 | 216724215 | rs1846224 | -0.484 | 0.453 | -1.173 | 0.059 | -0.068 | -0.006 | -0.006 | 0.896 | 0.18 | 0.02 | 0.21 | 0.14 | 0.26 | 0.16 |
| 1 | 216726333 | rs10442688 | -1.302 | 0.536 | -0.975 | 0.078 | 0 | 0 | 0 | 0.686 | 0.29 | 0.27 | 0.44 | 0.31 | 0.34 | 0.27 |
| 1 | 216727323 | rs4846528 | -0.995 | 0.8 | -2.019 | 0.189 | -0.069 | -0.068 | -0.068 | 0 | 0.27 | 0.03 | 0.4 | 0.29 | 0.33 | 0.26 |
| 1 | 216729434 | rs10863252 | -0.6 | 0.893 | -1.95 | 0.289 | -0.085 | -0.005 | -0.005 | 0.03 | 0.25 | 0.01 | 0.17 | 0.16 | 0.29 | 0.16 |
| 1 | 216730036 | rs1566224 | 0.333 | 1.003 | -1.432 | 1.325 | 0.016 | 0.12 | 0.12 | 0.002 | 0.93 | 0.87 | 0.81 | 0.65 | 0.91 | 0.78 |
| 1 | 216731498 | rs17668665 | 0.726 | 0.943 | -2.474 | 0.385 | -0.039 | 0.111 | 0.111 | 0 | 0.18 | 0.02 | 0.09 | 0 | 0.17 | 0.12 |
| 1 | 216732718 | rs7520795 | 0.272 | 0.513 | -1.444 | 0.177 | -0.005 | -0.064 | -0.064 | 0 | 0.22 | 0.25 | 0.11 | 0 | 0.17 | 0.12 |
| 1 | 216741314 | rs11572766 | 0.839 | 0.69 | -1.681 | 0.159 | -0.077 | 0.13 | 0.13 | 0.071 | 0.21 | 0.01 | 0.1 | 0 | 0.2 | 0.14 |
| 1 | 216743264 | rs11117625 | 0.787 | 0.761 | -2.277 | -0.04 | -0.036 | -0.014 | -0.014 | 0 | 0.21 | 0.01 | 0.18 | 0.14 | 0.24 | 0.14 |
| 1 | 216745245 | rs1984137 | -0.193 | 0.706 | -2.258 | -0.029 | -0.084 | 0.167 | 0.167 | 0 | 0.43 | 0.11 | 0.28 | 0.15 | 0.44 | 0.28 |
| 1 | 216754098 | rs1335964 | 0.083 | 0.615 | -1.757 | -0.119 | -0.004 | -0.014 | -0.014 | 0.132 | 0.84 | 0.87 | 0.67 | 0.61 | 0.83 | 0.73 |
| 1 | 216755753 | rs6701062 | 0.538 | 0.634 | -1.624 | -0.082 | 0.027 | -0.026 | -0.026 | 1.329 | 0.44 | 0.74 | 0.39 | 0.46 | 0.39 | 0.44 |
| 1 | 216756821 | rs17669622 | 1.426 | 0.351 | -1.698 | 0.051 | -0.073 | -0.008 | -0.008 | 0.236 | 0.24 | 0.01 | 0.17 | 0.15 | 0.24 | 0.15 |
| 1 | 216761107 | rs10218694 | 0.393 | 0.551 | -1.334 | 0.004 | 0.025 | -0.02 | -0.02 | 0.694 | 0.41 | 0.6 | 0.37 | 0.46 | 0.34 | 0.44 |
| 1 | 216770716 | rs2201838 | -0.351 | 0.747 | -1.345 | 0.134 | -0.003 | 0.004 | 0.004 | 0 | 0.86 | 0.87 | 0.67 | 0.61 | 0.83 | 0.73 |
| 1 | 216771649 | rs1833039 | 1.457 | 0.873 | -1.375 | 0.014 | -0.071 | 0.138 | 0.138 | 0.069 | 0.21 | 0.03 | 0.11 | 0 | 0.21 | 0.15 |
| 1 | 216777687 | rs7544758 | 0.169 | 0.542 | -0.7 | 0.172 | -0.001 | 0 | 0 | 0 | 0.36 | 0.34 | 0.36 | 0.41 | 0.34 | 0.4 |
| 1 | 216783528 | rs1826977 | -0.267 | 0.794 | -0.465 | 0.317 | 0.011 | 0.041 | 0.041 | 0.076 | 0.86 | 0.82 | 0.7 | 0.64 | 0.82 | 0.67 |
| 1 | 216787169 | rs10863259 | -0.281 | 0.632 | -0.552 | 0.103 | -0.017 | 0 | 0 | 0.012 | 0.61 | 0.7 | 0.59 | 0.57 | 0.61 | 0.53 |
| 1 | 216787802 | rs11572727 | 1.351 | 0.496 | -0.718 | 0.072 | -0.063 | 0.075 | 0.075 | 2.308 | 0.26 | 0.02 | 0.19 | 0.08 | 0.29 | 0.14 |
| 1 | 216788184 | rs7553463 | 0.233 | 0.531 | -0.561 | 0.063 | 0.007 | -0.02 | -0.02 | 0.043 | 0.36 | 0.65 | 0.39 | 0.49 | 0.32 | 0.39 |
| 1 | 216789563 | rs6673703 | -0.564 | 0.533 | -0.576 | 0.067 | -0.003 | 0.004 | 0.004 | 0.092 | 0.86 | 0.87 | 0.7 | 0.65 | 0.82 | 0.68 |
| 1 | 216791605 | rs1857407 | 0.626 | 0.533 | -0.318 | 0.032 | -0.006 | 0.154 | 0.154 | 0.055 | 0.51 | 0.31 | 0.32 | 0.15 | 0.5 | 0.29 |
| 1 | 216795695 | rs11577585 | 1.45 | 0.503 | -0.36 | 0.014 | -0.018 | 0.122 | 0.122 | 0 | 0.24 | 0.11 | 0.11 | 0.01 | 0.21 | 0.14 |
| 1 | 216798806 | rs1414529 | 1.163 | 0.45 | -0.104 | 0.288 | 0 | -0.027 | -0.027 | 0.062 | 0.24 | 0.15 | 0.26 | 0.39 | 0.2 | 0.28 |
| 1 | 216798830 | rs1414530 | 0.093 | 0.455 | -0.25 | 0.295 | -0.006 | 0.015 | 0.015 | 0.039 | 0.35 | 0.52 | 0.38 | 0.49 | 0.32 | 0.4 |
| 1 | 216800699 | rs12089054 | 0.499 | 0.551 | 0.511 | 0.424 | -0.002 | 0.021 | 0.021 | 0 | 0.11 | 0.27 | 0.11 | 0.1 | 0.12 | 0.12 |

| | | | | | | | | | | | | | | | | |
|---|---|---|---|---|---|---|---|---|---|---|---|---|---|---|---|---|
| 1 | 216801205 | rs10779269 | -0.052 | 0.759 | 0.045 | 0.73 | 0.014 | 0.062 | 0.062 | 0.426 | 0.86 | 0.8 | 0.69 | 0.62 | 0.82 | 0.68 |
| 1 | 216807520 | rs1846228 | 0.517 | 0.77 | -0.215 | 0.714 | 0.003 | 0.001 | 0.001 | 2.259 | 0.5 | 0.3 | 0.46 | 0.51 | 0.49 | 0.42 |
| 1 | 216813244 | rs1817998 | -0.069 | 0.804 | -0.214 | 0.742 | 0.002 | 0.026 | 0.026 | 0.071 | 0.86 | 0.85 | 0.69 | 0.62 | 0.82 | 0.68 |
| 1 | 216814307 | rs1498282 | -0.211 | 0.841 | -0.261 | 0.765 | -0.003 | 0.012 | 0.012 | 0.138 | 0.87 | 0.87 | 0.69 | 0.62 | 0.82 | 0.68 |
| 1 | 216816881 | rs2047429 | 0.08 | 0.558 | -0.138 | 0.223 | -0.012 | -0.071 | -0.071 | 0 | 0.36 | 0.57 | 0.24 | 0.11 | 0.33 | 0.26 |
| 1 | 216822615 | rs4846617 | -1.247 | 1.057 | -0.003 | 1.452 | 0.02 | 0.186 | 0.186 | 0 | 0.09 | 0.22 | 0.34 | 0.43 | 0.13 | 0.22 |
| 1 | 216827026 | rs11117644 | 0.286 | 1.135 | 0.261 | 1.029 | -0.018 | 0.109 | 0.109 | 3.19 | 0.48 | 0.21 | 0.42 | 0.22 | 0.51 | 0.27 |
| 1 | 216829908 | rs7538424 | 0.858 | 1.226 | 0.027 | 0.741 | -0.001 | -0.004 | -0.004 | 0 | 0.18 | 0.2 | 0.14 | 0.11 | 0.18 | 0.08 |
| 1 | 216833926 | rs1339344 | -0.268 | 1.245 | 0.168 | 0.56 | 0.01 | 0.02 | 0.02 | 0 | 0.86 | 0.8 | 0.89 | 0.78 | 0.86 | 0.82 |
| 1 | 216836576 | rs946468 | 1.302 | 1.224 | 0.215 | 0.537 | -0.001 | -0.011 | -0.011 | 0 | 0.18 | 0.2 | 0.14 | 0.11 | 0.18 | 0.08 |
| 1 | 216839426 | rs7550534 | 1.409 | 1.292 | 0.083 | 0.489 | -0.001 | -0.002 | -0.002 | 0 | 0.17 | 0.16 | 0.14 | 0.11 | 0.18 | 0.08 |
| 1 | 216840371 | rs946465 | 0.697 | 1.17 | 0.19 | 0.795 | 0.005 | 0.066 | 0.066 | 0 | 0.81 | 0.37 | 0.79 | 0.63 | 0.8 | 0.79 |
| 1 | 216842535 | rs1416612 | -0.029 | 1.154 | 0.314 | 0.742 | 0.004 | 0.007 | 0.007 | 0 | 0.85 | 0.84 | 0.81 | 0.75 | 0.8 | 0.8 |
| 1 | 216844423 | rs10127480 | 0.196 | 1.173 | 0.132 | 0.707 | 0 | 0.009 | 0.009 | 0.255 | 0.53 | 0.36 | 0.59 | 0.61 | 0.53 | 0.64 |
| 1 | 216848771 | rs12121367 | 0.994 | 1.124 | -0.113 | 0.64 | -0.056 | 0.055 | 0.055 | 2.661 | 0.29 | 0.01 | 0.2 | 0.08 | 0.27 | 0.17 |
| 1 | 216852176 | rs4426040 | 0.282 | 1.063 | 0.236 | 0.739 | 0.031 | 0.051 | 0.051 | 0 | 0.83 | 0.6 | 0.8 | 0.75 | 0.8 | 0.79 |
| 1 | 216853179 | rs12754372 | 0.054 | 1.626 | 0.102 | 1.056 | 0.03 | 0.065 | 0.065 | 0 | 0.09 | 0.14 | 0.37 | 0.33 | 0.18 | 0.22 |
| 1 | 216854001 | rs6695824 | 1.207 | 1.479 | 0.057 | 0.941 | 0.025 | 0.051 | 0.051 | 1.036 | 0.28 | 0.14 | 0.19 | 0.08 | 0.23 | 0.15 |
| 1 | 216854148 | rs11580756 | 2.173 | 1.488 | -0.089 | 0.883 | -0.008 | 0.07 | 0.07 | 1.094 | 0.25 | 0.02 | 0.1 | 0.12 | 0.2 | 0.33 |
| 1 | 216856115 | rs4846641 | -0.046 | 1.548 | 0.204 | 1.002 | -0.001 | 0.002 | 0.002 | 0.097 | 0.4 | 0.62 | 0.57 | 0.46 | 0.41 | 0.37 |
| 1 | 216858013 | rs4085154 | -0.01 | 1.813 | -0.023 | 1.028 | 0.004 | 0.002 | 0.002 | 0.086 | 0.55 | 0.62 | 0.61 | 0.61 | 0.55 | 0.64 |
| 1 | 216859096 | rs17043350 | 0.078 | 1.838 | -0.017 | 1.112 | 0.033 | 0.095 | 0.095 | 0 | 0.09 | 0.14 | 0.37 | 0.38 | 0.18 | 0.23 |
| 1 | 216864218 | rs2211118 | 0.316 | 1.613 | -0.365 | 0.975 | 0.017 | 0.041 | 0.041 | 0 | 0.83 | 0.73 | 0.8 | 0.68 | 0.78 | 0.79 |
| 1 | 216872201 | rs11117663 | 1.143 | 1.783 | -0.627 | 0.832 | -0.049 | 0.058 | 0.058 | 0.826 | 0.28 | 0.01 | 0.18 | 0.07 | 0.26 | 0.15 |
| 1 | 216872461 | rs2820883 | -0.189 | 1.842 | -0.236 | 0.958 | 0.035 | 0.034 | 0.034 | 0 | 0.86 | 0.7 | 0.88 | 0.77 | 0.83 | 0.81 |
| 1 | 216873939 | rs17627954 | 1.553 | 1.676 | -0.47 | 0.839 | 0.013 | 0.041 | 0.041 | 0.736 | 0.28 | 0.18 | 0.11 | 0.12 | 0.22 | 0.35 |
| 1 | 216875154 | rs17679272 | 1.792 | 1.776 | -0.797 | 0.871 | -0.009 | 0.067 | 0.067 | 1.475 | 0.27 | 0.03 | 0.1 | 0.13 | 0.22 | 0.35 |
| 1 | 216877492 | rs2066108 | -1.229 | 1.632 | -0.664 | 0.993 | 0.01 | 0.014 | 0.014 | 0.163 | 0.8 | 0.84 | 0.86 | 0.91 | 0.84 | 0.93 |
| 1 | 216878458 | rs1416611 | 0.189 | 2.015 | -0.436 | 1.448 | 0.006 | 0.025 | 0.025 | 0 | 0.11 | 0.1 | 0.12 | 0.3 | 0.16 | 0.19 |
| 1 | 216880662 | rs11572672 | 0.167 | 2.031 | -0.444 | 1.344 | 0.007 | 0.009 | 0.009 | 0 | 0.11 | 0.09 | 0.12 | 0.23 | 0.16 | 0.18 |
| 1 | 216886001 | rs2789718 | -0.321 | 1.694 | -0.642 | 1.419 | -0.006 | 0.022 | 0.022 | 0 | 0.07 | 0.09 | 0.31 | 0.33 | 0.14 | 0.22 |
| 1 | 216897131 | rs1416620 | 1.478 | 1.251 | -0.756 | 0.81 | -0.008 | 0.058 | 0.058 | 0.84 | 0.25 | 0.03 | 0.1 | 0.11 | 0.2 | 0.35 |
| 1 | 216900811 | rs2820878 | -0.578 | 1.161 | -0.294 | 0.643 | -0.009 | -0.02 | -0.02 | 0 | 0.74 | 0.65 | 0.84 | 0.86 | 0.79 | 0.91 |
| 1 | 216901955 | rs12144183 | -1.319 | 1.33 | -0.41 | 0.715 | -0.014 | -0.029 | -0.029 | 0 | 0.75 | 0.85 | 0.57 | 0.54 | 0.68 | 0.83 |
| 1 | 216903455 | rs7521774 | 0.274 | 1.352 | -0.005 | 0.728 | 0.006 | 0.028 | 0.028 | 0 | 0.11 | 0.16 | 0.16 | 0.27 | 0.14 | 0.28 |
| 1 | 216904564 | rs17043567 | 0.274 | 1.352 | -0.005 | 0.715 | 0.006 | 0.051 | 0.051 | 0 | 0.11 | 0.16 | 0.17 | 0.32 | 0.14 | 0.28 |
| 1 | 216905611 | rs2999504 | -0.806 | 1.266 | -0.066 | 0.543 | -0.009 | -0.019 | -0.019 | 0 | 0.74 | 0.65 | 0.84 | 0.86 | 0.79 | 0.91 |
| 1 | 216908871 | rs1572740 | 0.84 | 1.506 | -0.151 | 0.913 | -0.102 | -0.133 | -0.133 | 0 | 0.27 | 0.04 | 0.44 | 0.56 | 0.37 | 0.24 |
| 1 | 216910585 | rs954276 | 0.545 | 1.476 | -0.095 | 0.912 | 0.01 | -0.007 | -0.007 | 1.649 | 0.09 | 0.16 | 0.12 | 0.09 | 0.1 | 0.2 |
| 1 | 216913261 | rs2813703 | -0.831 | 1.408 | 0.126 | 0.827 | 0.004 | 0.049 | 0.049 | 0 | 0.45 | 0.28 | 0.71 | 0.75 | 0.56 | 0.56 |
| 1 | 216915514 | rs1339352 | -1.442 | 1.399 | 0.183 | 0.754 | -0.019 | -0.059 | -0.059 | 0 | 0.68 | 0.46 | 0.85 | 0.85 | 0.73 | 0.63 |
| 1 | 216919617 | rs12070175 | 0.878 | 1.3 | 0.162 | 0.783 | -0.029 | -0.045 | -0.045 | 0 | 0.25 | 0.41 | 0.14 | 0.09 | 0.16 | 0.06 |
| 1 | 216920286 | rs2820875 | -0.993 | 1.302 | -0.088 | 0.808 | -0.047 | -0.048 | -0.048 | 3.304 | 0.74 | 0.52 | 0.85 | 0.85 | 0.84 | 0.93 |
| 1 | 216921580 | rs6671659 | 0.128 | 1.872 | 0.28 | 1.299 | 0.011 | 0.016 | 0.016 | 0 | 0.09 | 0.17 | 0.14 | 0.17 | 0.1 | 0.22 |
| 1 | 216922557 | rs11572654 | -0.342 | 1.931 | 0.272 | 1.424 | 0.018 | 0.064 | 0.064 | 0 | 0.11 | 0.18 | 0.18 | 0.32 | 0.16 | 0.29 |
| 1 | 216923473 | rs2789731 | 0.604 | 1.757 | 0.251 | 1.168 | -0.032 | -0.051 | -0.051 | 0.721 | 0.24 | 0.09 | 0.42 | 0.41 | 0.32 | 0.16 |
| 1 | 216925972 | rs10779279 | 0.057 | 1.415 | -0.06 | 0.69 | -0.027 | 0.008 | 0.008 | 0 | 0.08 | 0 | 0.11 | 0.02 | 0.09 | 0.11 |
| 1 | 216926414 | rs2185237 | -0.999 | 1.39 | 0.095 | 0.798 | 0 | -0.001 | -0.001 | 0 | 0.17 | 0.18 | 0.3 | 0.19 | 0.19 | 0.28 |
| 1 | 216927091 | rs1361332 | -1.31 | 1.487 | 0.695 | 0.463 | 0.035 | -0.073 | -0.073 | 0 | 0.2 | 0.48 | 0.24 | 0.05 | 0.23 | 0.16 |
| 1 | 216927468 | rs6604643 | -0.253 | 1.302 | 0.045 | 0.567 | 0.01 | 0.032 | 0.032 | 0 | 0.5 | 0.66 | 0.43 | 0.32 | 0.41 | 0.52 |
| 1 | 216936881 | rs7547186 | 0.628 | 1.448 | -0.532 | 0.725 | -0.009 | 0.017 | 0.017 | 0 | 0.36 | 0.18 | 0.17 | 0.29 | 0.32 | 0.4 |
| 1 | 216938541 | rs6693633 | -0.342 | 1.028 | -0.633 | 0.691 | 0.013 | 0.013 | 0.013 | 0 | 0.63 | 0.59 | 0.51 | 0.41 | 0.52 | 0.7 |
| 1 | 216941624 | rs17681575 | 0.404 | 0.667 | -1.434 | 0.677 | 0.013 | 0.028 | 0.028 | 0 | 0.34 | 0.12 | 0.14 | 0.26 | 0.27 | 0.34 |
| 1 | 216942781 | rs2813694 | -0.946 | 0.63 | -1.574 | 1.571 | -0.022 | -0.085 | -0.085 | 2.542 | 0.13 | 0.02 | 0.12 | 0.42 | 0.16 | 0.23 |
| 1 | 216942929 | rs11117671 | -0.157 | -0.246 | -1.147 | 0.737 | -0.04 | -0.039 | -0.039 | 0 | 0.32 | 0.5 | 0.25 | 0.28 | 0.25 | 0.22 |
| 1 | 216944098 | rs6683574 | 1.112 | 0.047 | -0.956 | 0.794 | 0.009 | 0.024 | 0.024 | 0.145 | 0.21 | 0.25 | 0.37 | 0.26 | 0.23 | 0.24 |
| 1 | 216944772 | rs17043767 | 1.509 | -0.012 | -1.024 | 0.765 | -0.081 | -0.105 | -0.105 | 2.206 | 0.19 | 0.01 | 0.35 | 0.26 | 0.23 | 0.24 |
| 1 | 216945100 | rs946470 | -0.38 | -0.349 | -1.038 | 0.57 | -0.042 | -0.026 | -0.026 | 0 | 0.6 | 0.8 | 0.62 | 0.54 | 0.5 | 0.53 |
| 1 | 216949851 | rs6687061 | 1.223 | -0.263 | -0.548 | 0.882 | -0.003 | -0.006 | -0.006 | 0.214 | 0.22 | 0.17 | 0.36 | 0.26 | 0.23 | 0.24 |
| 1 | 216949936 | rs6658214 | 0.15 | -0.25 | -0.192 | 0.716 | 0.019 | 0.011 | 0.011 | 1.962 | 0.74 | 0.65 | 0.74 | 0.65 | 0.64 | 0.79 |
| 1 | 216950504 | rs9970445 | 1.524 | -0.533 | -0.036 | 0.584 | 0.012 | 0.011 | 0.011 | 1.283 | 0.15 | 0.12 | 0.07 | 0.07 | 0.07 | 0.14 |
| 1 | 216956685 | rs6702277 | 1.645 | -0.445 | 0.09 | 0.609 | 0.003 | 0.012 | 0.012 | 0 | 0.22 | 0.23 | 0.37 | 0.27 | 0.23 | 0.24 |
| 1 | 216958033 | rs7522593 | 0.127 | -0.491 | 0.133 | 0.552 | -0.055 | -0.074 | -0.074 | 0.054 | 0.23 | 0.44 | 0.16 | 0.13 | 0.13 | 0.18 |
| 1 | 216964401 | rs10157225 | 0.797 | -0.375 | -0.541 | 0.575 | 0.007 | 0.021 | 0.021 | 0 | 0.29 | 0.44 | 0.44 | 0.33 | 0.28 | 0.28 |
| 1 | 216965808 | rs12023645 | -0.1 | -0.197 | 0.176 | 0.794 | 0.038 | 0.019 | 0.019 | 0 | 0.08 | 0.37 | 0.08 | 0.06 | 0.06 | 0.04 |
| 1 | 216965834 | rs10218498 | 0.664 | -0.024 | -0.559 | 0.995 | -0.019 | 0.002 | 0.002 | 0.972 | 0.16 | 0.18 | 0.1 | 0.21 | 0.07 | 0.16 |
| 1 | 216966601 | rs7355008 | 0.273 | 1.033 | -0.771 | 0.92 | -0.031 | -0.024 | -0.024 | 0.641 | 0.81 | 0.91 | 0.81 | 0.72 | 0.7 | 0.82 |
| 1 | 216973196 | rs2789726 | 0.664 | 0.593 | -0.247 | 1.148 | 0 | 0.1 | 0.1 | 0.674 | 0.92 | 0.83 | 0.9 | 0.73 | 0.92 | 0.89 |
| 1 | 216974886 | rs2813663 | 1.437 | 0.504 | 0.207 | 1.186 | 0.01 | 0.132 | 0.132 | 0 | 0.83 | 0.6 | 0.8 | 0.62 | 0.83 | 0.83 |

| | | | | | | | | | | | | | | | | |
|---|---|---|---|---|---|---|---|---|---|---|---|---|---|---|---|---|
| 1 | 216980277 | rs11117683 | -1.271 | 0.308 | 0.414 | 1.18 | 0.021 | 0.057 | 0.057 | 0.016 | 0.07 | 0.31 | 0.06 | 0.06 | 0.04 | 0.02 |
| 1 | 216981412 | rs17041521 | 0.448 | 0.018 | 0.298 | 0.318 | -0.002 | -0.04 | -0.04 | 0 | 0.19 | 0.19 | 0.1 | 0 | 0.15 | 0.11 |
| 1 | 216983034 | rs12760372 | -0.137 | -0.013 | -0.09 | 0.287 | 0 | -0.002 | -0.002 | 0 | 0.56 | 0.42 | 0.57 | 0.26 | 0.6 | 0.56 |
| 1 | 216986215 | rs2576236 | 0.976 | -0.039 | -0.164 | 0.46 | -0.01 | -0.02 | -0.02 | 0 | 0.33 | 0.13 | 0.27 | 0.36 | 0.32 | 0.27 |
| 1 | 216987005 | rs2576234 | 0.835 | -0.046 | -0.148 | 0.444 | -0.003 | -0.048 | -0.048 | 0 | 0.34 | 0.18 | 0.28 | 0.48 | 0.32 | 0.36 |
| 1 | 216991606 | rs12563692 | 1.231 | 0.07 | -0.65 | 0.22 | -0.065 | -0.02 | -0.02 | 0.698 | 0.15 | 0.01 | 0.17 | 0.14 | 0.21 | 0.32 |
| 1 | 216994412 | rs2032028 | -1.246 | -0.066 | -0.326 | 0.387 | -0.005 | -0.01 | -0.01 | 0 | 0.67 | 0.84 | 0.73 | 0.65 | 0.68 | 0.74 |
| 1 | 216995157 | rs12753723 | -0.26 | 0.112 | -0.151 | 0.234 | -0.001 | -0.015 | -0.015 | 0 | 0.55 | 0.47 | 0.57 | 0.26 | 0.58 | 0.55 |
| 1 | 216997113 | rs11117687 | -0.376 | 0.078 | -0.109 | 0.218 | -0.002 | 0.036 | 0.036 | 0 | 0.57 | 0.68 | 0.59 | 0.26 | 0.58 | 0.54 |
| 1 | 216998464 | rs3738490 | -0.248 | 0.029 | -0.217 | 0.25 | -0.001 | -0.016 | -0.016 | 0 | 0.12 | 0.11 | 0.14 | 0.27 | 0.13 | 0.13 |
| 1 | 217000074 | rs17044015 | 1.351 | -0.077 | -0.194 | 0.164 | -0.005 | -0.045 | -0.045 | 0 | 0.18 | 0.19 | 0.08 | 0 | 0.13 | 0.11 |
| 1 | 217001114 | rs17044026 | 0.224 | -0.056 | -0.525 | 0.189 | 0 | 0.011 | 0.011 | 0.554 | 0.1 | 0.12 | 0.12 | 0.26 | 0.1 | 0.11 |
| 1 | 217001587 | rs2164614 | 0.506 | -0.045 | -0.325 | -0.299 | -0.067 | 0.03 | 0.03 | 0 | 0.35 | 0.12 | 0.47 | 0.26 | 0.45 | 0.43 |
| 1 | 217003419 | rs2164613 | -0.332 | 0.172 | -0.136 | 0.011 | 0 | 0.008 | 0.008 | 0 | 0.56 | 0.58 | 0.58 | 0.26 | 0.58 | 0.53 |
| 1 | 217005202 | rs1436897 | -0.522 | 0.348 | -0.272 | 0.105 | -0.001 | -0.003 | -0.003 | 0 | 0.57 | 0.54 | 0.59 | 0.39 | 0.61 | 0.59 |
| 1 | 217006749 | rs17044062 | 0.477 | 0.349 | -0.018 | -0.294 | 0.026 | -0.106 | -0.106 | 0 | 0.16 | 0.35 | 0.11 | 0 | 0.18 | 0.1 |
| 1 | 217007348 | rs1561186 | 1.488 | 0.221 | -0.091 | -0.441 | 0 | 0.003 | 0.003 | 0 | 0.17 | 0.19 | 0.29 | 0.1 | 0.18 | 0.16 |
| 1 | 217007399 | rs1561185 | 0.849 | 0.043 | 0.158 | -0.573 | -0.003 | -0.021 | -0.021 | 0.351 | 0.21 | 0.36 | 0.33 | 0.1 | 0.21 | 0.17 |
| 1 | 217007912 | rs1561182 | -0.29 | 0.222 | 0.148 | -0.505 | -0.001 | -0.122 | -0.122 | 0 | 0.2 | 0.51 | 0.12 | 0 | 0.18 | 0.11 |
| 1 | 217008849 | rs7537081 | -0.306 | 0.106 | 0.013 | -0.471 | -0.001 | -0.004 | -0.004 | 0 | 0.35 | 0.68 | 0.41 | 0.14 | 0.34 | 0.26 |
| 1 | 217009130 | rs7548403 | 0.919 | 0.089 | 0.027 | -0.419 | -0.003 | -0.002 | -0.002 | 0 | 0.22 | 0.31 | 0.32 | 0.14 | 0.19 | 0.17 |
| 1 | 217016670 | rs2576258 | -1.445 | -0.002 | -0.428 | -0.307 | -0.006 | -0.045 | -0.045 | 0 | 0.67 | 0.83 | 0.74 | 0.49 | 0.68 | 0.67 |
| 1 | 217020295 | rs1156007 | 1.739 | -0.001 | -0.264 | -0.395 | 0.03 | -0.067 | -0.067 | 0 | 0.11 | 0.23 | 0.08 | 0 | 0.14 | 0.09 |
| 1 | 217020992 | rs12086903 | 0.824 | -0.065 | -0.388 | -0.324 | -0.019 | -0.031 | -0.031 | 0.008 | 0.26 | 0.47 | 0.31 | 0.14 | 0.23 | 0.18 |
| 1 | 217022982 | rs12410353 | 1.376 | 0.01 | -0.514 | -0.221 | 0.007 | 0.004 | 0.004 | 0 | 0.22 | 0.19 | 0.31 | 0.15 | 0.19 | 0.17 |
| 1 | 217026271 | rs4338425 | -1.454 | 0.119 | -0.747 | -0.124 | -0.005 | -0.043 | -0.043 | 0 | 0.67 | 0.83 | 0.74 | 0.49 | 0.69 | 0.67 |
| 1 | 217026858 | rs10779283 | 0.867 | 0.052 | -0.488 | -0.397 | -0.005 | -0.004 | -0.004 | 0 | 0.23 | 0.3 | 0.33 | 0.15 | 0.19 | 0.17 |
| 1 | 217034049 | rs7532370 | 0.493 | 0.773 | -0.523 | -0.251 | -0.031 | 0.001 | 0.001 | 0.022 | 0.13 | 0.05 | 0.15 | 0.11 | 0.18 | 0.21 |
| 1 | 217048338 | rs11572547 | 0.187 | 0.278 | -0.144 | -0.191 | 0.03 | -0.078 | -0.078 | 0 | 0.12 | 0.27 | 0.23 | 0 | 0.16 | 0.09 |
| 1 | 217053037 | rs17044239 | 0.69 | -0.303 | -0.196 | -0.125 | -0.011 | -0.061 | -0.061 | 0.337 | 0.1 | 0.22 | 0.2 | 0 | 0.09 | 0.08 |
| 1 | 217054551 | rs2576262 | -0.181 | -0.338 | -0.632 | -0.079 | -0.007 | -0.055 | -0.055 | 0 | 0.62 | 0.63 | 0.66 | 0.86 | 0.66 | 0.73 |
| 1 | 217058479 | rs1436900 | 0.03 | -0.461 | -0.988 | 0.733 | -0.017 | -0.003 | -0.003 | 0 | 0.66 | 0.7 | 0.57 | 0.26 | 0.62 | 0.48 |
| 1 | 217059640 | rs1370150 | -0.308 | -0.162 | -0.854 | 1.05 | -0.012 | -0.002 | -0.002 | 0.01 | 0.72 | 0.7 | 0.66 | 0.61 | 0.64 | 0.73 |
| 1 | 217059931 | rs2576267 | -0.462 | -0.905 | -0.109 | 0.862 | -0.062 | -0.019 | -0.019 | 0.148 | 0.27 | 0.44 | 0.26 | 0.18 | 0.13 | 0.17 |
| 1 | 217062140 | rs11117717 | -0.538 | 0.384 | 0.231 | 1.407 | 0.019 | 0.147 | 0.147 | 0 | 0.07 | 0.21 | 0.19 | 0.4 | 0.1 | 0.12 |
| 1 | 217069513 | rs3795704 | 0.205 | 0.188 | 0.353 | 0.416 | -0.007 | 0.021 | 0.021 | 0.043 | 0.07 | 0.12 | 0.12 | 0.08 | 0.06 | 0.07 |
| 1 | 217070040 | rs4436447 | -0.056 | 0.299 | -0.202 | -0.006 | -0.088 | 0.098 | 0.098 | 0 | 0.72 | 0.96 | 0.77 | 0.92 | 0.66 | 0.75 |
| 1 | 217073060 | rs2576203 | -1.267 | 0.12 | 0.527 | -0.112 | -0.014 | 0.008 | 0.008 | 0 | 0.16 | 0.56 | 0.19 | 0.12 | 0.13 | 0.08 |
| 1 | 217073889 | rs1436883 | -0.206 | 0.113 | 0.723 | -0.111 | 0.01 | -0.003 | -0.003 | 0.205 | 0.06 | 0.3 | 0.03 | 0 | 0.01 | 0.01 |
| 1 | 217074421 | rs1898312 | -1.314 | 0.048 | 0.705 | -0.143 | -0.029 | 0.004 | 0.004 | 0 | 0.15 | 0.63 | 0.18 | 0.11 | 0.12 | 0.08 |
| 1 | 217076563 | rs11117722 | 0.891 | 0.214 | 0.847 | -0.076 | 0.002 | -0.03 | -0.03 | 0 | 0.57 | 0.5 | 0.59 | 0.81 | 0.53 | 0.66 |
| 1 | 217081526 | rs1025653 | 1.098 | 0.267 | 0.588 | 0.038 | 0.004 | 0.035 | 0.035 | 0 | 0.54 | 0.33 | 0.52 | 0.72 | 0.53 | 0.48 |
| 1 | 217082283 | rs4348786 | 1.042 | 0.372 | 0.001 | 0.114 | -0.001 | -0.012 | -0.012 | 0 | 0.56 | 0.54 | 0.55 | 0.72 | 0.53 | 0.48 |
| 1 | 217082385 | rs1436879 | 0.077 | 0.408 | 0.058 | 0.266 | -0.012 | 0.042 | 0.042 | 0 | 0.08 | 0.13 | 0.18 | 0.17 | 0.07 | 0.23 |
| 1 | 217084292 | rs4147268 | 0.304 | 0.758 | -0.28 | -0.189 | -0.001 | -0.007 | -0.007 | 0 | 0.68 | 0.7 | 0.79 | 0.93 | 0.66 | 0.75 |
| 1 | 217088341 | rs2576269 | 0.165 | 1.184 | -0.079 | 0.047 | 0.055 | -0.04 | -0.04 | 0 | 0.72 | 0.59 | 0.77 | 0.9 | 0.63 | 0.75 |
| 1 | 217088584 | rs2818747 | -0.044 | 0.666 | -0.13 | 0.478 | -0.012 | 0.124 | 0.124 | 0 | 0.19 | 0.4 | 0.43 | 0.44 | 0.16 | 0.26 |
| 1 | 217088750 | rs2818748 | -0.498 | 1.138 | -0.045 | 0.44 | 0.019 | -0.093 | -0.093 | 0 | 0.75 | 0.65 | 0.87 | 0.97 | 0.72 | 0.84 |
| 1 | 217092655 | rs10779284 | 0.337 | 0.997 | -0.147 | 0.579 | -0.003 | -0.005 | -0.005 | 4.581 | 0.57 | 0.38 | 0.45 | 0.54 | 0.56 | 0.57 |
| 1 | 217093310 | rs10863278 | 1.176 | 0.633 | -0.953 | 0.273 | -0.03 | -0.031 | -0.031 | 0.786 | 0.33 | 0.02 | 0.18 | 0.3 | 0.29 | 0.28 |
| 1 | 217094044 | rs11572505 | 0.48 | 0.389 | -0.536 | 0.303 | 0.027 | 0.021 | 0.021 | 0 | 0.11 | 0.02 | 0.06 | 0.07 | 0.07 | 0.1 |
| 1 | 217100192 | rs2576241 | 0.125 | 0.878 | -0.102 | 0.493 | 0.009 | 0.011 | 0.011 | 0 | 0.59 | 0.51 | 0.45 | 0.54 | 0.56 | 0.57 |
| 1 | 217102061 | rs2099557 | 0.209 | 1.163 | -0.043 | 0.633 | 0.001 | 0 | 0 | 0 | 0.59 | 0.45 | 0.53 | 0.65 | 0.62 | 0.53 |
| 1 | 217102472 | rs12562202 | -0.569 | 0.924 | -0.166 | 0.134 | 0.002 | -0.01 | -0.01 | 0 | 0.23 | 0.22 | 0.38 | 0.33 | 0.22 | 0.37 |
| 1 | 217103658 | rs6682369 | 1.081 | 1.43 | 0.066 | 0.617 | 0.022 | 0.049 | 0.049 | 0.114 | 0.51 | 0.16 | 0.28 | 0.47 | 0.47 | 0.45 |
| 1 | 217104215 | rs6677596 | -0.246 | 1.623 | 0.346 | 0.306 | -0.007 | -0.063 | -0.063 | 0 | 0.73 | 0.74 | 0.86 | 0.97 | 0.76 | 0.84 |
| 1 | 217108951 | rs7551571 | -1.09 | 1.935 | 0.045 | 1.008 | 0.121 | 0.118 | 0.118 | 0.269 | 0.07 | 0.24 | 0.25 | 0.19 | 0.15 | 0.09 |
| 1 | 217110899 | rs11572495 | -0.071 | 1.396 | -0.559 | 0.882 | -0.047 | -0.042 | -0.042 | 0 | 0.08 | 0.01 | 0.21 | 0.14 | 0.15 | 0.04 |
| 1 | 217112871 | rs7417600 | 0.009 | 0.976 | -0.405 | 0.303 | 0.002 | 0.016 | 0.016 | 0 | 0.21 | 0.18 | 0.13 | 0.09 | 0.22 | 0.18 |
| 1 | 217116732 | rs1030845 | 0.253 | 0.238 | -0.49 | 0.589 | -0.006 | 0.006 | 0.006 | 0 | 0.74 | 0.34 | 0.62 | 0.65 | 0.82 | 0.58 |
| 1 | 217122207 | rs2576197 | -0.921 | 0.085 | -0.245 | 0.65 | -0.003 | -0.003 | -0.003 | 0 | 0.87 | 0.91 | 0.72 | 0.66 | 0.86 | 0.64 |
| 1 | 217122449 | rs11117734 | 0.4 | 0.079 | -0.162 | 0.778 | -0.023 | 0.062 | 0.062 | 0 | 0.77 | 0.4 | 0.63 | 0.63 | 0.84 | 0.62 |
| 1 | 217124015 | rs10492958 | 1.02 | 0.353 | -0.788 | 0.895 | -0.041 | -0.044 | -0.044 | 0 | 0.07 | 0.01 | 0.23 | 0.16 | 0.15 | 0.03 |
| 1 | 217124731 | rs10492957 | 1.073 | 0.241 | -0.793 | 0.677 | -0.058 | 0.095 | 0.095 | 0.067 | 0.48 | 0.09 | 0.27 | 0.35 | 0.49 | 0.41 |
| 1 | 217126354 | rs11117737 | 1.193 | 0.334 | -0.839 | 0.636 | -0.086 | 0.13 | 0.13 | 0.009 | 0.46 | 0.08 | 0.26 | 0.32 | 0.47 | 0.39 |
| 1 | 217131368 | rs2576226 | 0.1 | 0.114 | 0.869 | 1.159 | -0.02 | 0.099 | 0.099 | 0 | 0.79 | 0.43 | 0.69 | 0.57 | 0.84 | 0.61 |
| 1 | 217136990 | rs2576213 | 0.714 | 0.026 | 0.61 | 0.843 | 0.008 | 0.053 | 0.053 | 0 | 0.19 | 0.09 | 0.12 | 0 | 0.19 | 0.15 |
| 1 | 217154258 | rs2789550 | 0.375 | 0.038 | 1.022 | 1.059 | 0 | 0.12 | 0.12 | 0 | 0.15 | 0.28 | 0.19 | 0.41 | 0.13 | 0.39 |
| 1 | 217159448 | rs867410 | 0.219 | 0.278 | 0.986 | 1.134 | -0.004 | -0.001 | -0.001 | 0 | 0.11 | 0.11 | 0.34 | 0.2 | 0.2 | 0.07 |

| | | | | | | | | | | | | | | | | |
|---|---|---|---|---|---|---|---|---|---|---|---|---|---|---|---|---|
| 1 | 217162443 | rs17044788 | -0.572 | 0.233 | 1.149 | 1.107 | 0.056 | 0.098 | 0.098 | 0 | 0.15 | 0.36 | 0.35 | 0.2 | 0.21 | 0.07 |
| 1 | 217162879 | rs3000609 | 0.377 | 0.225 | 1.193 | 1.111 | 0.001 | 0.128 | 0.128 | 0 | 0.15 | 0.29 | 0.19 | 0.41 | 0.13 | 0.39 |
| 1 | 217167698 | rs5017053 | 1.494 | 0.144 | 0.914 | 1.026 | -0.043 | 0.137 | 0.137 | 0 | 0.48 | 0.15 | 0.28 | 0.29 | 0.47 | 0.39 |
| 1 | 217168377 | rs1556891 | 0.476 | 0.136 | 0.942 | 1.028 | 0.002 | -0.015 | -0.015 | 0 | 0.66 | 0.64 | 0.53 | 0.8 | 0.6 | 0.79 |
| 1 | 217168483 | rs2818765 | -0.446 | 0.131 | 0.875 | 1.012 | -0.012 | 0.211 | 0.211 | 0 | 0.19 | 0.5 | 0.26 | 0.51 | 0.13 | 0.4 |
| 1 | 217172606 | rs1339221 | 0.291 | 0.2 | 0.836 | 1.05 | -0.011 | 0.008 | 0.008 | 0.048 | 0.64 | 0.53 | 0.72 | 0.59 | 0.69 | 0.47 |
| 1 | 217174168 | rs10863285 | 0.55 | 0.22 | 0.654 | 0.945 | -0.013 | 0.015 | 0.015 | 0.054 | 0.61 | 0.42 | 0.66 | 0.5 | 0.68 | 0.46 |
| 1 | 217174535 | rs10863286 | 1.025 | -0.512 | -0.024 | 0.183 | -0.005 | -0.001 | -0.001 | 0.179 | 0.39 | 0.3 | 0.27 | 0.16 | 0.46 | 0.34 |
| 1 | 217175860 | rs2185226 | -0.93 | -0.955 | -0.397 | 0.309 | 0.039 | -0.03 | -0.03 | 0 | 0.36 | 0.72 | 0.29 | 0.51 | 0.23 | 0.43 |
| 1 | 217177525 | rs2789567 | -0.931 | -0.973 | -0.292 | 0.303 | 0.022 | -0.017 | -0.017 | 0 | 0.36 | 0.67 | 0.29 | 0.52 | 0.23 | 0.43 |
| 1 | 217180260 | rs1339222 | -1.391 | -0.816 | -0.416 | 0.266 | 0.035 | 0.096 | 0.096 | 0.212 | 0.47 | 0.73 | 0.53 | 0.74 | 0.4 | 0.48 |
| 1 | 217180432 | rs2789571 | -1.409 | -0.819 | -0.409 | 0.264 | 0.038 | 0.105 | 0.105 | 0.25 | 0.47 | 0.78 | 0.54 | 0.74 | 0.4 | 0.48 |
| 1 | 217181451 | rs2818760 | -0.873 | -0.83 | -0.303 | 0.435 | 0.026 | -0.016 | -0.016 | 0 | 0.34 | 0.7 | 0.23 | 0.45 | 0.18 | 0.36 |
| 1 | 217183535 | rs2789573 | 0.037 | -0.973 | 0.122 | 0.364 | -0.005 | 0.015 | 0.015 | 0 | 0.25 | 0.24 | 0.18 | 0.45 | 0.17 | 0.36 |
| 1 | 217184151 | rs2789574 | -1.811 | -0.777 | -0.631 | -0.233 | -0.048 | 0.007 | 0.007 | 0 | 0.75 | 0.92 | 0.86 | 0.76 | 0.69 | 0.74 |
| 1 | 217184643 | rs946438 | -0.083 | -0.907 | -0.365 | -0.241 | -0.001 | -0.007 | -0.007 | 0.04 | 0.48 | 0.48 | 0.61 | 0.26 | 0.52 | 0.33 |
| 1 | 217190815 | rs1890554 | -1.677 | -0.794 | -0.649 | -0.243 | -0.053 | -0.01 | -0.01 | 0.514 | 0.74 | 0.9 | 0.81 | 0.7 | 0.66 | 0.69 |
| 1 | 217192021 | rs11117750 | 1.524 | -0.862 | -0.699 | -0.424 | -0.001 | 0.085 | 0.085 | 0 | 0.29 | 0.13 | 0.32 | 0.05 | 0.29 | 0.28 |
| 1 | 217194760 | rs2789551 | -2.033 | -0.864 | -0.729 | -0.374 | -0.066 | 0.061 | 0.061 | 0.336 | 0.76 | 0.93 | 0.86 | 0.85 | 0.68 | 0.77 |
| 1 | 217195463 | rs6604651 | -0.17 | -0.884 | -0.529 | -0.316 | 0.002 | 0.014 | 0.014 | 0 | 0.48 | 0.55 | 0.56 | 0.26 | 0.51 | 0.34 |
| 1 | 217199141 | rs4846802 | -0.095 | -0.876 | -0.521 | -0.319 | 0 | 0.001 | 0.001 | 0 | 0.48 | 0.53 | 0.56 | 0.26 | 0.51 | 0.33 |
| 1 | 217199613 | rs11572427 | 1.549 | -0.756 | -0.6 | -0.112 | -0.057 | -0.066 | -0.066 | 0.093 | 0.1 | 0.01 | 0.19 | 0.2 | 0.2 | 0.06 |
| 1 | 217199626 | rs2818805 | -0.948 | -0.79 | -0.474 | -0.052 | -0.021 | 0.085 | 0.085 | 4.912 | 0.9 | 0.67 | 0.9 | 0.86 | 0.95 | 0.91 |
| 1 | 217200883 | rs2377360 | -0.577 | -1.028 | -0.682 | -0.331 | -0.016 | -0.027 | -0.027 | 0 | 0.66 | 0.4 | 0.62 | 0.82 | 0.73 | 0.68 |
| 1 | 217200953 | rs6684076 | 1.224 | -0.696 | -0.648 | -0.05 | -0.054 | -0.06 | -0.06 | 0 | 0.1 | 0.01 | 0.19 | 0.16 | 0.2 | 0.06 |
| 1 | 217203091 | rs7527281 | 0.362 | -0.798 | -0.663 | -0.071 | -0.019 | 0.036 | 0.036 | 0 | 0.26 | 0.32 | 0.25 | 0.42 | 0.15 | 0.32 |
| 1 | 217206190 | rs6688120 | -1.017 | -0.819 | -0.504 | -0.118 | 0 | -0.003 | -0.003 | 0 | 0.61 | 0.64 | 0.72 | 0.67 | 0.58 | 0.64 |
| 1 | 217206705 | rs11117754 | 1.296 | -0.808 | -0.572 | -0.146 | 0.003 | 0.021 | 0.021 | 0 | 0.24 | 0.21 | 0.28 | 0.05 | 0.23 | 0.23 |
| 1 | 217207089 | rs4846806 | -0.323 | -0.782 | -0.677 | -0.118 | 0.003 | 0 | 0 | 0 | 0.49 | 0.55 | 0.53 | 0.47 | 0.38 | 0.56 |
| 1 | 217215465 | rs4360582 | 1.267 | -0.699 | -0.707 | -0.259 | 0.003 | 0.063 | 0.063 | 0 | 0.24 | 0.13 | 0.27 | 0.05 | 0.23 | 0.23 |
| 1 | 217217013 | rs11807917 | 0.481 | -0.032 | -0.408 | 0.123 | 0.002 | -0.039 | -0.039 | 0 | 0.38 | 0.38 | 0.23 | 0.18 | 0.4 | 0.29 |
| 1 | 217222233 | rs7513573 | -0.253 | -0.263 | -0.58 | 0.015 | 0.004 | 0.005 | 0.005 | 0 | 0.49 | 0.52 | 0.43 | 0.41 | 0.6 | 0.36 |
| 1 | 217223816 | rs7520126 | 0.204 | 0.562 | -0.129 | -0.488 | -0.001 | -0.001 | -0.001 | 0 | 0.45 | 0.43 | 0.42 | 0.63 | 0.41 | 0.55 |
| 1 | 217224368 | rs17045075 | 0.907 | 0.238 | -0.4 | -0.72 | 0.003 | 0 | 0 | 0 | 0.09 | 0.08 | 0.03 | 0.08 | 0.05 | 0.07 |
| 1 | 217227540 | rs11117762 | 0.923 | 0.606 | -0.431 | -0.886 | -0.021 | 0.066 | 0.066 | 0 | 0.33 | 0.19 | 0.39 | 0.21 | 0.38 | 0.23 |
| 1 | 217228039 | rs6678584 | 0.726 | 0.987 | -0.511 | -0.923 | 0.015 | -0.002 | -0.002 | 0 | 0.07 | 0.09 | 0.15 | 0.04 | 0.14 | 0.07 |
| 1 | 217228116 | rs7532636 | 0.401 | 0.774 | -0.396 | -0.905 | 0.005 | 0.027 | 0.027 | 0 | 0.43 | 0.29 | 0.36 | 0.64 | 0.41 | 0.53 |
| 1 | 217230015 | rs6680670 | -0.085 | 0.891 | -0.113 | -0.468 | 0.008 | -0.005 | -0.005 | 0.038 | 0.17 | 0.22 | 0.2 | 0.12 | 0.2 | 0.15 |
| 1 | 217234774 | rs4846399 | -0.532 | 0.212 | -0.153 | -0.518 | -0.02 | -0.03 | -0.03 | 0 | 0.8 | 0.72 | 0.9 | 0.91 | 0.9 | 0.86 |
| 1 | 217235542 | rs6684903 | 0.564 | 0.842 | -0.669 | -0.51 | 0 | 0.014 | 0.014 | 0 | 0.25 | 0.18 | 0.35 | 0.21 | 0.28 | 0.21 |
| 1 | 217236801 | rs1416527 | -0.566 | 1.19 | -0.198 | -0.294 | -0.002 | -0.01 | -0.01 | 0 | 0.55 | 0.55 | 0.54 | 0.73 | 0.61 | 0.63 |
| 1 | 217236895 | rs1416526 | 0.687 | 1.3 | -0.37 | -0.186 | -0.003 | -0.004 | -0.004 | 0 | 0.21 | 0.21 | 0.14 | 0.12 | 0.17 | 0.27 |
| 1 | 217237052 | rs2225967 | -1.287 | 1.287 | -0.318 | -0.553 | 0.021 | 0.034 | 0.034 | 0.104 | 0.59 | 0.68 | 0.59 | 0.74 | 0.66 | 0.66 |
| 1 | 217237449 | rs11117763 | 0.59 | 1.713 | -0.739 | -0.127 | -0.005 | -0.027 | -0.027 | 0 | 0.2 | 0.23 | 0.14 | 0.08 | 0.18 | 0.25 |
| 1 | 217237998 | rs17045141 | -0.645 | 1.951 | -0.798 | -0.162 | -0.026 | 0.006 | 0.006 | 0 | 0.32 | 0.24 | 0.33 | 0.66 | 0.39 | 0.34 |
| 1 | 217238186 | rs4537652 | -0.645 | 1.951 | -0.798 | -0.162 | -0.026 | 0.006 | 0.006 | 0 | 0.32 | 0.23 | 0.33 | 0.66 | 0.39 | 0.34 |
| 1 | 217238322 | rs4539203 | -0.296 | 2.165 | -1.316 | -0.041 | -0.005 | -0.004 | -0.004 | 0 | 0.13 | 0.11 | 0.22 | 0.62 | 0.16 | 0.24 |
| 1 | 217240584 | rs4846809 | 0.193 | 2.373 | -1.21 | 0.285 | -0.011 | 0.023 | 0.023 | 0 | 0.08 | 0.07 | 0.19 | 0.56 | 0.12 | 0.2 |
| 1 | 217245008 | rs12402289 | 0.147 | 2.206 | -1.227 | 0.062 | -0.083 | 0.057 | 0.057 | 0 | 0.18 | 0.02 | 0.11 | 0.04 | 0.23 | 0.1 |
| 1 | 217245278 | rs4846400 | -0.502 | 1.532 | -0.513 | 0.668 | 0.003 | 0.007 | 0.007 | 0 | 0.28 | 0.32 | 0.34 | 0.63 | 0.3 | 0.45 |
| 1 | 217252549 | rs1553347 | -0.72 | 1.728 | -0.69 | 0.972 | -0.01 | -0.012 | -0.012 | 0 | 0.1 | 0.07 | 0.23 | 0.2 | 0.15 | 0.09 |
| 1 | 217255780 | rs7539648 | -0.244 | 1.54 | -0.36 | 0.64 | -0.002 | -0.008 | -0.008 | 0 | 0.55 | 0.5 | 0.6 | 0.76 | 0.55 | 0.49 |
| 1 | 217256257 | rs6604652 | 0.067 | 1.316 | -0.296 | 0.566 | 0.004 | -0.003 | -0.003 | 0 | 0.44 | 0.41 | 0.35 | 0.56 | 0.39 | 0.41 |
| 1 | 217264291 | rs11572381 | -0.167 | 0.796 | -0.278 | 0.446 | -0.002 | 0 | 0 | 0 | 0.41 | 0.47 | 0.29 | 0.46 | 0.3 | 0.35 |
| 1 | 217266496 | rs1502357 | 0.224 | -0.517 | -0.542 | 0.21 | -0.03 | -0.007 | -0.007 | 0.011 | 0.44 | 0.18 | 0.46 | 0.55 | 0.51 | 0.37 |
| 1 | 217270667 | rs830307 | -0.001 | -1.172 | -0.081 | -0.089 | -0.003 | -0.016 | -0.016 | 0 | 0.63 | 0.43 | 0.6 | 0.69 | 0.66 | 0.6 |
| 1 | 217272170 | rs7514298 | 0.137 | -1.352 | 0.024 | -0.229 | -0.001 | -0.001 | -0.001 | 0 | 0.25 | 0.24 | 0.31 | 0.29 | 0.25 | 0.23 |
| 1 | 217272629 | rs12033461 | 0.962 | -1.338 | 0.033 | -0.209 | -0.01 | 0.021 | 0.021 | 0 | 0.13 | 0.22 | 0.09 | 0.23 | 0.08 | 0.11 |
| 1 | 217275575 | rs11808388 | 0.447 | -1.233 | 0.176 | -0.295 | 0.001 | -0.043 | -0.043 | 0 | 0.1 | 0.24 | 0.08 | 0.02 | 0.09 | 0.17 |
| 1 | 217277611 | rs11587946 | 0.895 | -1.344 | 0.029 | -0.052 | -0.04 | -0.013 | -0.013 | 0 | 0.08 | 0 | 0.17 | 0.05 | 0.11 | 0.09 |
| 1 | 217279717 | rs6604654 | 0.106 | -1.609 | 0.349 | 0.032 | -0.04 | -0.015 | -0.015 | 0 | 0.2 | 0.33 | 0.15 | 0.14 | 0.15 | 0.22 |
| 1 | 217286487 | rs1166331 | -0.236 | -1.628 | -0.258 | 0.144 | 0.012 | -0.04 | -0.04 | 0 | 0.27 | 0.18 | 0.25 | 0.45 | 0.18 | 0.43 |
| 1 | 217287741 | rs1166332 | -0.885 | -1.779 | -0.049 | -0.246 | -0.02 | 0.028 | 0.028 | 0.048 | 0.57 | 0.7 | 0.49 | 0.42 | 0.54 | 0.47 |
| 1 | 217288723 | rs4846815 | 0.866 | -1.806 | -0.088 | -0.259 | -0.044 | 0.03 | 0.03 | 0.07 | 0.41 | 0.18 | 0.5 | 0.58 | 0.46 | 0.53 |
| 1 | 217293917 | rs12040162 | 0.877 | -1.794 | -0.012 | -0.254 | 0.001 | 0.01 | 0.01 | 0 | 0.06 | 0.08 | 0.06 | 0.21 | 0.06 | 0.09 |
| 1 | 217298122 | rs10495041 | 1.349 | -1.916 | -0.105 | -0.375 | -0.039 | 0.015 | 0.015 | 0 | 0.12 | 0 | 0.21 | 0.06 | 0.15 | 0.14 |
| 1 | 217302383 | rs830321 | -1.385 | -2.003 | -0.009 | -0.368 | -0.025 | 0.017 | 0.017 | 0.248 | 0.59 | 0.74 | 0.5 | 0.44 | 0.54 | 0.59 |
| 1 | 217302743 | rs830322 | -0.742 | -1.738 | 0.193 | -0.298 | 0 | -0.003 | -0.003 | 0 | 0.36 | 0.59 | 0.37 | 0.32 | 0.35 | 0.37 |
| 1 | 217308047 | rs6696627 | -1.429 | -1.289 | 0.069 | -0.16 | 0.019 | 0.063 | 0.063 | 0.009 | 0.46 | 0.74 | 0.46 | 0.39 | 0.44 | 0.47 |

| | | | | | | | | | | | | | | | | |
|---|---|---|---|---|---|---|---|---|---|---|---|---|---|---|---|---|
| 1 | 217333272 | rs12732971 | 1.031 | -1.584 | -0.152 | -0.466 | -0.029 | 0.025 | 0.025 | 0 | 0.12 | 0 | 0.21 | 0.04 | 0.13 | 0.14 |
| 1 | 217334278 | rs17696298 | 1.554 | -1.633 | -0.249 | -0.526 | -0.037 | -0.004 | -0.004 | 0 | 0.1 | 0 | 0.08 | 0.1 | 0.14 | 0.08 |
| 1 | 217334839 | rs11117781 | 0.586 | -1.89 | 0.181 | -0.435 | -0.004 | -0.004 | -0.004 | 0 | 0.58 | 0.59 | 0.55 | 0.77 | 0.67 | 0.62 |
| 1 | 217337735 | rs1502352 | 1.1 | -1.783 | 0.319 | -0.377 | 0.004 | 0.023 | 0.023 | 0 | 0.44 | 0.26 | 0.44 | 0.66 | 0.52 | 0.41 |
| 1 | 217340451 | rs1553346 | -1.162 | -1.456 | -0.43 | -0.333 | -0.003 | 0 | 0 | 0 | 0.8 | 0.81 | 0.82 | 0.81 | 0.88 | 0.8 |
| 1 | 217341609 | rs7514342 | -0.967 | -1.44 | -0.594 | -0.329 | -0.023 | 0.011 | 0.011 | 0 | 0.79 | 0.68 | 0.79 | 0.82 | 0.84 | 0.78 |
| 1 | 217342298 | rs1354220 | -1.155 | -1.423 | -0.853 | -0.304 | -0.019 | 0.008 | 0.008 | 0 | 0.8 | 0.76 | 0.82 | 0.82 | 0.88 | 0.8 |
| 1 | 217344002 | rs12407512 | 0.851 | -1.25 | -1.674 | -0.546 | -0.029 | 0.009 | 0.009 | 0 | 0.12 | 0 | 0.26 | 0.06 | 0.13 | 0.14 |
| 1 | 217349340 | rs2377477 | 0.866 | -0.802 | -1.512 | -0.266 | -0.053 | 0.013 | 0.013 | 0 | 0.41 | 0.11 | 0.4 | 0.64 | 0.52 | 0.53 |
| 1 | 217349392 | rs2377478 | -0.257 | -0.31 | -0.395 | 0.291 | -0.028 | -0.02 | -0.02 | 0.829 | 0.79 | 0.55 | 0.81 | 0.87 | 0.85 | 0.79 |
| 1 | 217350383 | rs7555255 | 0.038 | -0.148 | -1.011 | 0.475 | 0.013 | 0.005 | 0.005 | 0.006 | 0.16 | 0.12 | 0.1 | 0.2 | 0.17 | 0.12 |
| 1 | 217351406 | rs11117784 | -0.571 | 0.912 | -0.768 | 0.424 | 0.011 | 0.001 | 0.001 | 1.731 | 0.28 | 0.3 | 0.47 | 0.68 | 0.38 | 0.45 |
| 1 | 217352068 | rs1892259 | 0.028 | 1.116 | -0.928 | 1.193 | 0.023 | 0.054 | 0.054 | 0.011 | 0.17 | 0.19 | 0.37 | 0.38 | 0.28 | 0.35 |
| 1 | 217352414 | rs11117785 | -0.248 | 0.753 | -1.158 | 0.964 | -0.018 | -0.056 | -0.056 | 0 | 0.09 | 0.01 | 0.06 | 0.32 | 0.08 | 0.1 |
| 1 | 217355588 | rs11117791 | -0.289 | 0.905 | -1.096 | 0.37 | -0.015 | 0.011 | 0.011 | 0 | 0.34 | 0.2 | 0.23 | 0.3 | 0.37 | 0.24 |
| 1 | 217356401 | rs2221285 | -0.255 | 1.491 | -0.859 | 0.686 | -0.028 | -0.023 | -0.023 | 0.279 | 0.12 | 0.09 | 0.23 | 0.27 | 0.28 | 0.12 |
| 1 | 217356792 | rs17045418 | 0.542 | 1.077 | -0.723 | 0.385 | -0.008 | -0.029 | -0.029 | 0 | 0.06 | 0.11 | 0.01 | 0 | 0.03 | 0.08 |
| 1 | 217357160 | rs12038637 | -0.462 | 1.311 | -0.589 | 0.926 | 0.011 | 0.1 | 0.1 | 0 | 0.07 | 0.05 | 0.12 | 0.42 | 0.13 | 0.08 |
| 1 | 217359519 | rs11117794 | -0.917 | 1.272 | 0.143 | 0.886 | 0.09 | 0.273 | 0.273 | 0 | 0.08 | 0.22 | 0.14 | 0.42 | 0.14 | 0.08 |
| 1 | 217362297 | rs12143637 | 1.143 | 1.285 | -0.4 | 0.332 | -0.078 | 0.107 | 0.107 | 0 | 0.18 | 0.01 | 0.18 | 0.01 | 0.21 | 0.13 |
| 1 | 217362375 | rs1502349 | -0.236 | 1.299 | -0.331 | 0.415 | 0.038 | 0.16 | 0.16 | 0 | 0.07 | 0.11 | 0.13 | 0.42 | 0.14 | 0.08 |
| 1 | 217362648 | rs1502348 | -1.545 | 0.321 | -0.605 | -0.184 | 0.023 | 0.102 | 0.102 | 0.396 | 0.68 | 0.86 | 0.78 | 0.9 | 0.7 | 0.81 |
| 1 | 217364103 | rs17045459 | 0.185 | 0.454 | -0.489 | -0.003 | 0.013 | 0.1 | 0.1 | 0 | 0.07 | 0.06 | 0.13 | 0.42 | 0.13 | 0.08 |
| 1 | 217365587 | rs6669865 | 0.466 | -0.077 | 0.065 | -0.452 | -0.001 | -0.001 | -0.001 | 0 | 0.28 | 0.27 | 0.41 | 0.14 | 0.32 | 0.4 |
| 1 | 217366425 | rs17045466 | 1.081 | -0.103 | 0.44 | -0.427 | 0.001 | 0.01 | 0.01 | 0 | 0.11 | 0.17 | 0.23 | 0.13 | 0.1 | 0.28 |
| 1 | 217368088 | rs1393385 | -1.076 | -0.246 | 0.482 | -0.386 | 0.001 | -0.001 | -0.001 | 0 | 0.61 | 0.61 | 0.66 | 0.44 | 0.64 | 0.73 |
| 1 | 217370116 | rs2134314 | -0.508 | -0.13 | 0.488 | -0.135 | -0.011 | 0.117 | 0.117 | 0 | 0.14 | 0.2 | 0.13 | 0.45 | 0.1 | 0.15 |
| 1 | 217374423 | rs12728997 | 0.752 | 0.166 | 0.32 | 0.031 | 0.002 | 0.143 | 0.143 | 0 | 0.05 | 0.05 | 0.11 | 0.45 | 0.06 | 0.08 |
| 1 | 217375824 | rs1910147 | 1.045 | 0.125 | 0.202 | -0.043 | -0.032 | -0.034 | -0.034 | 0 | 0.11 | 0.06 | 0.15 | 0.24 | 0.18 | 0.13 |
| 1 | 217386144 | rs10429948 | -1.208 | -0.141 | 0.295 | -0.325 | 0.04 | 0.141 | 0.141 | 0.035 | 0.54 | 0.8 | 0.72 | 0.83 | 0.6 | 0.69 |
| 1 | 217392390 | rs17045519 | 1.33 | -0.046 | 0.85 | -0.285 | -0.013 | -0.03 | -0.03 | 0 | 0.06 | 0.14 | 0.01 | 0 | 0.03 | 0.07 |
| 1 | 217393675 | rs17045558 | 0.691 | -0.046 | 0.675 | -0.28 | -0.001 | 0 | 0 | 0 | 0.12 | 0.17 | 0.24 | 0.14 | 0.1 | 0.28 |
| 1 | 217402330 | rs1000796 | -1.841 | -0.212 | 0.428 | -0.419 | 0.028 | 0.084 | 0.084 | 0.217 | 0.67 | 0.85 | 0.78 | 0.89 | 0.7 | 0.81 |
| 1 | 217403176 | rs11117815 | -0.503 | 0.097 | 0.468 | -0.196 | 0.017 | 0.046 | 0.046 | 0.163 | 0.24 | 0.48 | 0.41 | 0.38 | 0.28 | 0.41 |
| 1 | 217406858 | rs4332417 | 0.915 | 0.083 | 0.366 | -0.135 | -0.01 | -0.016 | -0.016 | 0 | 0.23 | 0.14 | 0.29 | 0.46 | 0.28 | 0.18 |
| 1 | 217412147 | rs12085534 | 0.184 | 0.16 | 0.407 | 0.079 | -0.005 | 0.019 | 0.019 | 0.316 | 0.13 | 0.25 | 0.25 | 0.14 | 0.1 | 0.28 |
| 1 | 217414922 | rs4846824 | -0.658 | 0.087 | 0.347 | 0.083 | -0.004 | 0.017 | 0.017 | 0.308 | 0.31 | 0.49 | 0.35 | 0.41 | 0.3 | 0.32 |
| 1 | 217417838 | rs10495045 | -0.812 | -0.211 | 0.416 | -0.321 | -0.001 | 0.013 | 0.013 | 0.843 | 0.27 | 0.58 | 0.32 | 0.69 | 0.28 | 0.31 |
| 1 | 217418118 | rs4846826 | -1.344 | -0.474 | -0.053 | -0.927 | 0.007 | 0.077 | 0.077 | 1.151 | 0.64 | 0.83 | 0.68 | 0.88 | 0.65 | 0.61 |
| 1 | 217420660 | rs6701766 | 1.61 | -0.619 | -0.054 | -0.892 | -0.028 | 0.022 | 0.022 | 0 | 0.06 | 0.15 | 0.11 | 0.11 | 0.01 | 0.11 |
| 1 | 217421415 | rs11586825 | 0.914 | -0.652 | 0.223 | -1.01 | -0.004 | -0.007 | -0.007 | 0 | 0.13 | 0.18 | 0.09 | 0.09 | 0.11 | 0.1 |
| 1 | 217421644 | rs4388759 | -1.975 | -0.699 | 0.185 | -1.15 | 0.014 | 0.05 | 0.05 | 0.273 | 0.7 | 0.85 | 0.78 | 0.88 | 0.72 | 0.8 |
| 1 | 217423277 | rs4339926 | -1.86 | -0.709 | 0.384 | -1.155 | 0.001 | -0.005 | -0.005 | 0 | 0.69 | 0.73 | 0.77 | 0.88 | 0.72 | 0.8 |
| 1 | 217424810 | rs3923328 | -0.459 | -0.741 | 0.559 | -1.235 | 0.001 | -0.005 | -0.005 | 0.04 | 0.32 | 0.35 | 0.37 | 0.76 | 0.36 | 0.33 |
| 1 | 217427087 | rs10495047 | 1.591 | -0.616 | 0.284 | -1.168 | -0.027 | 0.053 | 0.053 | 0 | 0.21 | 0.11 | 0.15 | 0.01 | 0.22 | 0.1 |
| 1 | 217429135 | rs10495049 | -1.189 | -0.575 | 0.612 | -0.994 | 0.003 | -0.018 | -0.018 | 0 | 0.35 | 0.45 | 0.54 | 0.78 | 0.38 | 0.59 |
| 1 | 217435636 | rs4146799 | -0.256 | -0.704 | 0.694 | -0.974 | -0.016 | -0.013 | -0.013 | 0 | 0.16 | 0.29 | 0.25 | 0.11 | 0.11 | 0.29 |
| 1 | 217438561 | rs7541546 | -1.479 | -0.773 | 0.585 | -1.146 | -0.017 | 0.086 | 0.086 | 0.534 | 0.66 | 0.88 | 0.67 | 0.88 | 0.63 | 0.62 |
| 1 | 217443063 | rs4436446 | -0.812 | -0.765 | 0.792 | -1.179 | 0.001 | 0.012 | 0.012 | 0 | 0.5 | 0.59 | 0.49 | 0.22 | 0.45 | 0.5 |
| 1 | 217445254 | rs11589117 | 0.462 | -0.837 | 0.746 | -1.26 | -0.025 | 0.101 | 0.101 | 0 | 0.33 | 0.19 | 0.23 | 0.11 | 0.34 | 0.22 |
| 1 | 217450293 | rs7518401 | -0.937 | -0.811 | 0.819 | -1.22 | -0.03 | -0.036 | -0.036 | 5.537 | 0.29 | 0.48 | 0.34 | 0.19 | 0.22 | 0.37 |
| 1 | 217455903 | rs10158471 | -1.142 | -0.808 | 0.686 | -1.208 | -0.017 | 0.057 | 0.057 | 0.009 | 0.56 | 0.73 | 0.51 | 0.21 | 0.48 | 0.58 |
| 1 | 217456447 | rs7535028 | -1.151 | -0.807 | 0.517 | -1.208 | -0.031 | 0.093 | 0.093 | 0.027 | 0.56 | 0.8 | 0.52 | 0.21 | 0.48 | 0.58 |
| 1 | 217463023 | rs12074270 | -0.331 | -0.703 | 0.204 | -0.795 | -0.015 | -0.015 | -0.015 | 0 | 0.16 | 0.27 | 0.25 | 0.09 | 0.11 | 0.27 |
| 1 | 217466024 | rs2377567 | 0.873 | -0.461 | -0.16 | -0.636 | -0.004 | -0.004 | -0.004 | 0 | 0.12 | 0.06 | 0.08 | 0.1 | 0.11 | 0.1 |
| 1 | 217471874 | rs7518650 | 0.171 | -0.302 | 0.151 | -0.615 | 0.006 | 0.002 | 0.002 | 0 | 0.11 | 0.17 | 0.26 | 0.1 | 0.12 | 0.28 |
| 1 | 217471995 | rs12758898 | 1.429 | -0.381 | -0.402 | -0.937 | -0.072 | 0.116 | 0.116 | 0 | 0.22 | 0.03 | 0.15 | 0.02 | 0.24 | 0.11 |
| 1 | 217472501 | rs17045910 | -0.028 | -0.609 | 0.332 | -1 | -0.023 | -0.033 | -0.033 | 0 | 0.2 | 0.35 | 0.11 | 0.12 | 0.15 | 0.18 |
| 1 | 217473650 | rs6703333 | -0.124 | -0.502 | 0.66 | -0.933 | -0.001 | -0.05 | -0.05 | 0 | 0.42 | 0.52 | 0.27 | 0.14 | 0.38 | 0.29 |
| 1 | 217473967 | rs2815259 | 0.667 | -0.483 | 0.386 | -0.787 | -0.02 | -0.023 | -0.023 | 0.058 | 0.14 | 0.1 | 0.27 | 0.63 | 0.25 | 0.22 |
| 1 | 217474593 | rs17700529 | 1.501 | -0.668 | 0.394 | -0.831 | -0.013 | -0.016 | -0.016 | 0.228 | 0.1 | 0.06 | 0.08 | 0.11 | 0.11 | 0.1 |
| 1 | 217476282 | rs2646811 | -0.438 | -0.59 | 0.475 | -0.31 | 0 | 0.001 | 0.001 | 0 | 0.52 | 0.56 | 0.41 | 0.35 | 0.55 | 0.43 |
| 1 | 217486401 | rs12083375 | 0.674 | -0.994 | 0.66 | -0.594 | -0.069 | 0.012 | 0.012 | 1.298 | 0.12 | 0.28 | 0.23 | 0.11 | 0.02 | 0.1 |
| 1 | 217487940 | rs11117853 | 0.591 | -0.903 | 0.158 | -0.594 | -0.058 | -0.096 | -0.096 | 0 | 0.2 | 0.09 | 0.24 | 0.54 | 0.3 | 0.36 |
| 1 | 217488788 | rs2088457 | 0.171 | -1.309 | -0.133 | -0.869 | 0.021 | -0.034 | -0.034 | 0 | 0.47 | 0.18 | 0.4 | 0.56 | 0.55 | 0.48 |
| 1 | 217492666 | rs2646860 | -0.616 | -1.424 | 0.111 | -0.914 | 0.028 | 0.02 | 0.02 | 0 | 0.62 | 0.38 | 0.57 | 0.77 | 0.75 | 0.7 |
| 1 | 217494099 | rs2060 | -0.192 | -1.198 | 0.037 | -0.825 | -0.035 | 0.013 | 0.013 | 1.561 | 0.35 | 0.3 | 0.41 | 0.75 | 0.5 | 0.58 |
| 1 | 217497526 | rs2646800 | -1.479 | -0.67 | 0.096 | -0.723 | 0.007 | 0.024 | 0.024 | 0 | 0.76 | 0.89 | 0.81 | 0.88 | 0.77 | 0.8 |
| 1 | 217500820 | rs2815217 | -1.11 | -0.537 | 0.065 | -0.658 | 0.008 | 0.011 | 0.011 | 0 | 0.72 | 0.82 | 0.8 | 0.87 | 0.77 | 0.8 |

| | | | | | | | | | | | | | | | | |
|---|---|---|---|---|---|---|---|---|---|---|---|---|---|---|---|---|
| 1 | 217506787 | rs2815220 | -1.099 | -0.599 | 0.055 | -0.657 | 0.009 | 0.01 | 0.01 | 0 | 0.72 | 0.82 | 0.82 | 0.87 | 0.78 | 0.8 |
| 1 | 217507115 | rs1032601 | -0.246 | -0.481 | 0.342 | -0.615 | -0.004 | -0.003 | -0.003 | 0 | 0.6 | 0.47 | 0.64 | 0.67 | 0.62 | 0.65 |
| 1 | 217507258 | rs2646852 | -0.091 | -0.507 | 0.421 | -0.716 | -0.001 | -0.001 | -0.001 | 1.516 | 0.4 | 0.4 | 0.34 | 0.22 | 0.42 | 0.28 |
| 1 | 217509430 | rs11800230 | -1.047 | -0.468 | 1.028 | -0.607 | 0.083 | 0.073 | 0.073 | 0.057 | 0.05 | 0.42 | 0.03 | 0 | 0.01 | 0.01 |
| 1 | 217511930 | rs964347 | -1.516 | -0.517 | 1.212 | -0.611 | -0.005 | -0.008 | -0.008 | 0 | 0.06 | 0.58 | 0.04 | 0 | 0.01 | 0.01 |
| 1 | 217523112 | rs2646810 | -0.554 | -0.554 | -0.205 | -0.77 | 0.043 | 0.046 | 0.046 | 0 | 0.64 | 0.36 | 0.74 | 0.83 | 0.77 | 0.67 |
| 1 | 217538165 | rs2815231 | -0.413 | 0.026 | 0.071 | -0.692 | 0.012 | 0.014 | 0.014 | 0 | 0.66 | 0.43 | 0.74 | 0.85 | 0.77 | 0.77 |
| 1 | 217542361 | rs2646828 | 0.401 | 0.013 | 0.309 | -0.698 | 0.001 | 0 | 0 | 0 | 0.32 | 0.25 | 0.23 | 0.15 | 0.23 | 0.23 |
| 1 | 217548036 | rs2132994 | 0.432 | 0.011 | 0.28 | -0.708 | -0.001 | -0.003 | -0.003 | 0.078 | 0.32 | 0.25 | 0.23 | 0.15 | 0.23 | 0.23 |
| 1 | 217549909 | rs2815241 | 0.195 | 0.037 | 0.101 | -0.696 | 0.038 | 0.047 | 0.047 | 0 | 0.35 | 0.64 | 0.26 | 0.16 | 0.23 | 0.34 |
| 1 | 217559619 | rs2810777 | 0.214 | 0.05 | 0.021 | -0.69 | 0.027 | 0.033 | 0.033 | 0 | 0.35 | 0.61 | 0.26 | 0.16 | 0.23 | 0.34 |
| 1 | 217573623 | rs2815249 | 0.06 | 0.936 | -0.389 | 0.098 | -0.132 | -0.112 | -0.112 | 0 | 0.78 | 0.93 | 0.71 | 0.33 | 0.59 | 0.57 |
| 1 | 217577346 | rs12403652 | -0.173 | 0.322 | -0.751 | -0.445 | 0.002 | 0.046 | 0.046 | 0 | 0.64 | 0.83 | 0.71 | 0.78 | 0.66 | 0.69 |
| 1 | 217583807 | rs2815251 | -0.498 | 0.179 | -0.224 | -0.339 | 0.004 | 0.001 | 0.001 | 0 | 0.46 | 0.37 | 0.35 | 0.47 | 0.41 | 0.54 |
| 1 | 217583937 | rs2646850 | -0.216 | -0.088 | -0.272 | -0.499 | -0.008 | -0.018 | -0.018 | 0 | 0.38 | 0.23 | 0.33 | 0.47 | 0.37 | 0.45 |
| 1 | 217584510 | rs4543873 | -0.796 | -0.067 | 0.28 | -0.496 | 0.016 | 0.009 | 0.009 | 0 | 0.48 | 0.65 | 0.38 | 0.48 | 0.42 | 0.55 |
| 1 | 217589433 | rs557114 | 0.179 | -1.042 | 0.621 | -1.544 | -0.04 | -0.148 | -0.148 | 0 | 0.2 | 0.45 | 0.11 | 0 | 0.11 | 0.08 |
| 1 | 217589894 | rs561609 | 0.561 | -1.051 | 0.754 | -1.549 | -0.013 | -0.071 | -0.071 | 0 | 0.18 | 0.25 | 0.09 | 0 | 0.1 | 0.08 |
| 1 | 217592010 | rs629840 | -0.456 | -1.107 | 0.752 | -1.586 | -0.049 | -0.153 | -0.153 | 0 | 0.24 | 0.43 | 0.12 | 0 | 0.12 | 0.11 |
| 1 | 217595347 | rs12127692 | 0.735 | -0.981 | 0.454 | -1.861 | 0.012 | 0.011 | 0.011 | 0 | 0.4 | 0.34 | 0.22 | 0.09 | 0.31 | 0.27 |
| 1 | 217600228 | rs171946 | -0.204 | -0.961 | 0.589 | -1.857 | -0.001 | 0.022 | 0.022 | 0.4 | 0.58 | 0.66 | 0.32 | 0.09 | 0.41 | 0.35 |
| 1 | 217603847 | rs1048126 | 0.763 | -1.008 | 0.671 | -1.873 | 0.011 | 0.011 | 0.011 | 0 | 0.4 | 0.35 | 0.23 | 0.09 | 0.31 | 0.27 |
| 1 | 217603884 | rs265140 | 0.918 | -0.991 | 0.681 | -1.873 | -0.009 | -0.061 | -0.061 | 0.358 | 0.18 | 0.23 | 0.09 | 0 | 0.1 | 0.08 |
| 1 | 217604124 | rs166449 | -0.231 | -0.994 | 0.649 | -1.874 | -0.002 | 0.012 | 0.012 | 0.383 | 0.59 | 0.66 | 0.32 | 0.09 | 0.42 | 0.35 |
| 1 | 217604777 | rs7520483 | 0.656 | -0.842 | 0.235 | -1.754 | 0.012 | 0.013 | 0.013 | 0 | 0.4 | 0.34 | 0.23 | 0.09 | 0.31 | 0.27 |
| 1 | 217616744 | rs265129 | -0.736 | -1.039 | 0.89 | -1.138 | -0.051 | -0.136 | -0.136 | 0 | 0.27 | 0.5 | 0.14 | 0.04 | 0.16 | 0.14 |
| 1 | 217617702 | rs2911709 | -1.271 | -1.102 | 0.765 | -1.148 | -0.025 | -0.08 | -0.08 | 0 | 0.32 | 0.57 | 0.16 | 0.04 | 0.17 | 0.18 |
| 1 | 217620430 | rs265125 | -0.577 | -0.569 | 0.324 | -0.835 | -0.025 | -0.086 | -0.086 | 0.122 | 0.23 | 0.33 | 0.11 | 0.04 | 0.14 | 0.13 |
| 1 | 217625140 | rs12067356 | -1.091 | -0.573 | 0.648 | -0.522 | -0.027 | 0.001 | 0.001 | 0.082 | 0.11 | 0.38 | 0.06 | 0.09 | 0.04 | 0.07 |
| 1 | 217637075 | rs2818973 | -1.544 | -1.287 | 0.745 | -1.041 | -0.056 | -0.098 | -0.098 | 0 | 0.29 | 0.51 | 0.15 | 0.09 | 0.15 | 0.17 |
| 1 | 217658496 | rs13375527 | -1.28 | -1.132 | 0.992 | -0.807 | -0.046 | -0.021 | -0.021 | 0.293 | 0.12 | 0.37 | 0.06 | 0.09 | 0.04 | 0.08 |
| 1 | 217671223 | rs3790729 | -0.28 | 0.152 | 0.128 | -0.242 | 0.023 | 0.018 | 0.018 | 1.466 | 0.18 | 0.19 | 0.19 | 0.35 | 0.34 | 0.33 |
| 1 | 217677256 | rs12041730 | -0.411 | -0.743 | -0.436 | -1.141 | 0.006 | -0.014 | -0.014 | 0 | 0.56 | 0.41 | 0.71 | 0.92 | 0.67 | 0.72 |
| 1 | 217680035 | rs4846415 | 0.826 | -0.637 | -1.041 | -0.7 | -0.044 | -0.155 | -0.155 | 0 | 0.19 | 0.04 | 0.37 | 0.51 | 0.17 | 0.25 |
| 1 | 217704670 | rs1930301 | -0.9 | -0.881 | -0.929 | -0.967 | 0.021 | 0.021 | 0.021 | 0.091 | 0.52 | 0.39 | 0.68 | 0.92 | 0.64 | 0.69 |
| 1 | 217710863 | rs11806776 | -0.17 | -0.367 | -0.596 | -0.405 | -0.016 | 0 | 0 | 0 | 0.05 | 0.05 | 0.01 | 0.07 | 0.02 | 0.05 |
| 1 | 217715564 | rs4846419 | -0.499 | -0.406 | -0.882 | -0.852 | 0.005 | -0.007 | -0.007 | 0.019 | 0.56 | 0.4 | 0.67 | 0.94 | 0.63 | 0.63 |
| 1 | 217727217 | rs10495060 | 0.132 | 0.575 | -0.943 | 0.285 | 0.009 | 0.025 | 0.025 | 0.849 | 0.12 | 0.07 | 0.2 | 0.37 | 0.3 | 0.25 |
| 1 | 217728354 | rs17666128 | 1.095 | 0.176 | -0.76 | -0.199 | -0.003 | -0.023 | -0.023 | 0 | 0.05 | 0.09 | 0.03 | 0 | 0.04 | 0.03 |
| 1 | 217743949 | rs11117883 | -0.435 | 0.186 | -1.043 | -0.359 | -0.024 | 0.092 | 0.092 | 0 | 0.42 | 0.3 | 0.6 | 0.93 | 0.51 | 0.49 |
| 1 | 217746802 | rs17666266 | 0.591 | 0.631 | -1.118 | -0.146 | -0.033 | -0.031 | -0.031 | 0.708 | 0.08 | 0.02 | 0.11 | 0.16 | 0.23 | 0.13 |
| 1 | 217753698 | rs12097653 | 0.76 | 0.324 | -0.665 | 0.023 | 0.001 | -0.006 | -0.006 | 0 | 0.27 | 0.29 | 0.4 | 0.46 | 0.23 | 0.25 |
| 1 | 217754783 | rs11117887 | 0.975 | 0.638 | -0.52 | -0.584 | 0 | -0.036 | -0.036 | 0 | 0.19 | 0.18 | 0.13 | 0 | 0.17 | 0.25 |
| 1 | 217756427 | rs4846425 | 1.088 | 0.596 | -0.48 | -0.428 | 0.003 | -0.032 | -0.032 | 0 | 0.27 | 0.24 | 0.39 | 0.5 | 0.23 | 0.25 |
| 1 | 217756963 | rs4846244 | -0.365 | 0.838 | -0.361 | -0.414 | -0.006 | 0.006 | 0.006 | 0 | 0.44 | 0.44 | 0.6 | 0.93 | 0.51 | 0.49 |
| 1 | 217760264 | rs10863327 | -0.276 | 0.851 | -0.354 | -0.408 | -0.006 | 0.001 | 0.001 | 0 | 0.43 | 0.44 | 0.6 | 0.93 | 0.51 | 0.49 |
| 1 | 217762620 | rs11583496 | 0.31 | 1.036 | -0.017 | 0.372 | 0.019 | 0.02 | 0.02 | 0 | 0.1 | 0.1 | 0.12 | 0.26 | 0.23 | 0.19 |
| 1 | 217765449 | rs12045640 | -0.334 | -0.204 | 0.128 | -0.312 | -0.03 | -0.107 | -0.107 | 0 | 0.64 | 0.51 | 0.77 | 0.93 | 0.72 | 0.75 |
| 1 | 217767109 | rs11117891 | -0.305 | -0.207 | 0.057 | -0.244 | -0.02 | -0.021 | -0.021 | 0 | 0.64 | 0.48 | 0.7 | 0.77 | 0.7 | 0.72 |
| 1 | 217769684 | rs10779300 | -0.354 | -0.053 | -0.499 | -0.839 | -0.008 | -0.046 | -0.046 | 0 | 0.65 | 0.66 | 0.77 | 0.93 | 0.72 | 0.75 |
| 1 | 217780472 | rs11117896 | -0.5 | -0.116 | -0.317 | -1.158 | 0.002 | 0.037 | 0.037 | 0 | 0.58 | 0.68 | 0.38 | 0.19 | 0.63 | 0.43 |
| 1 | 217783810 | rs12749938 | 0.415 | -0.111 | -0.813 | -1.748 | -0.006 | 0.003 | 0.003 | 0.474 | 0.27 | 0.27 | 0.21 | 0.1 | 0.38 | 0.26 |
| 1 | 217784092 | rs17723306 | 0.175 | -1.08 | -0.865 | -1.861 | 0.003 | -0.006 | -0.006 | 0.369 | 0.12 | 0.08 | 0.04 | 0 | 0.1 | 0.07 |
| 1 | 217796299 | rs12124165 | -0.144 | -1.095 | -0.944 | -1.824 | -0.029 | 0.062 | 0.062 | 2.42 | 0.22 | 0.07 | 0.14 | 0.1 | 0.24 | 0.16 |
| 1 | 217830178 | rs10863333 | -0.493 | -1.097 | -0.795 | -1.75 | -0.015 | -0.053 | -0.053 | 0.202 | 0.27 | 0.38 | 0.16 | 0.1 | 0.24 | 0.17 |
| 1 | 217835210 | rs7555851 | -1.353 | -0.887 | -0.955 | -1.884 | 0.002 | 0.001 | 0.001 | 0.01 | 0.59 | 0.63 | 0.42 | 0.2 | 0.69 | 0.44 |
| 1 | 217842887 | rs10863335 | 1.599 | -1.048 | -1.708 | -1.247 | -0.025 | -0.162 | -0.162 | 0.015 | 0.23 | 0.02 | 0.29 | 0.56 | 0.18 | 0.33 |
| 1 | 217849797 | rs17724824 | 0.773 | -0.265 | -1.021 | -1.081 | -0.001 | 0.016 | 0.016 | 0 | 0.07 | 0.03 | 0.03 | 0 | 0.06 | 0.03 |
| 1 | 217889346 | rs17725494 | -0.529 | -0.049 | -0.855 | -0.771 | -0.046 | 0.022 | 0.022 | 0 | 0.33 | 0.26 | 0.33 | 0.26 | 0.46 | 0.3 |
| 1 | 217893504 | rs5017213 | 0.354 | 1.138 | -0.742 | -0.478 | -0.062 | 0.015 | 0.015 | 0 | 0.14 | 0.06 | 0.19 | 0.1 | 0.36 | 0.16 |
| 1 | 217911017 | rs4846431 | 0.45 | -0.253 | -0.574 | 0.213 | -0.005 | 0.041 | 0.041 | 0 | 0.22 | 0.23 | 0.31 | 0.52 | 0.17 | 0.39 |
| 1 | 217914116 | rs12120516 | 0.353 | -0.882 | -0.999 | -0.573 | -0.013 | 0.051 | 0.051 | 0.112 | 0.4 | 0.25 | 0.34 | 0.23 | 0.6 | 0.32 |
| 1 | 217916357 | rs6673733 | 0.477 | -0.888 | -1.002 | -0.575 | -0.012 | 0.05 | 0.05 | 0.098 | 0.4 | 0.25 | 0.34 | 0.23 | 0.6 | 0.32 |
| 1 | 217922972 | rs10863340 | 0.993 | -0.719 | -0.345 | -0.621 | 0.001 | 0.033 | 0.033 | 0 | 0.22 | 0.22 | 0.32 | 0.53 | 0.2 | 0.4 |
| 1 | 217924096 | rs4344379 | -0.543 | -0.995 | -0.063 | -0.682 | 0.009 | 0.007 | 0.007 | 0 | 0.66 | 0.69 | 0.68 | 0.73 | 0.81 | 0.73 |
| 1 | 217931977 | rs11117926 | 0.219 | -0.945 | -0.034 | -0.69 | 0.004 | 0.053 | 0.053 | 0.542 | 0.43 | 0.33 | 0.35 | 0.23 | 0.61 | 0.34 |
| 1 | 217933988 | rs11583816 | 1.691 | -0.701 | -0.418 | -0.956 | -0.101 | 0.074 | 0.074 | 0 | 0.24 | 0.07 | 0.21 | 0.1 | 0.4 | 0.17 |
| 1 | 217947738 | rs6604561 | -0.665 | -0.843 | -0.272 | -0.913 | 0.011 | 0.008 | 0.008 | 0 | 0.66 | 0.69 | 0.68 | 0.73 | 0.81 | 0.73 |
| 1 | 217951156 | rs17726462 | 0.48 | -0.844 | -0.309 | -0.927 | -0.033 | 0.038 | 0.038 | 0.191 | 0.38 | 0.25 | 0.32 | 0.23 | 0.57 | 0.3 |

| | | | | | | | | | | | | | | | | |
|---|---|---|---|---|---|---|---|---|---|---|---|---|---|---|---|---|
| 1 | 217952876 | rs10863348 | -1.766 | -1.055 | -0.736 | -0.974 | 0.001 | -0.013 | -0.013 | 0.176 | 0.87 | 0.98 | 0.81 | 0.89 | 0.9 | 0.9 |
| 1 | 217954378 | rs4846440 | 0.421 | -0.8 | -0.424 | -0.786 | 0.007 | 0.075 | 0.075 | 0 | 0.27 | 0.35 | 0.36 | 0.5 | 0.24 | 0.43 |
| 1 | 217967094 | rs6694862 | 1.057 | -0.807 | -0.438 | -0.648 | 0.005 | 0.091 | 0.091 | 0 | 0.22 | 0.31 | 0.21 | 0.47 | 0.2 | 0.41 |
| 1 | 217971980 | rs7532704 | -0.462 | -0.959 | -0.439 | -0.844 | 0.001 | -0.005 | -0.005 | 0 | 0.4 | 0.45 | 0.34 | 0.6 | 0.38 | 0.54 |
| 1 | 217975830 | rs6604563 | -0.318 | -0.956 | -0.342 | -0.981 | -0.021 | -0.015 | -0.015 | 0.234 | 0.67 | 0.61 | 0.68 | 0.72 | 0.79 | 0.78 |
| 1 | 217982896 | rs2377703 | 0.35 | -0.866 | -0.394 | -0.928 | 0 | 0.001 | 0.001 | 0 | 0.55 | 0.43 | 0.54 | 0.63 | 0.58 | 0.69 |
| 1 | 217988772 | rs4846441 | -0.397 | -0.648 | -0.313 | -0.96 | 0.009 | 0.015 | 0.015 | 0 | 0.39 | 0.58 | 0.37 | 0.24 | 0.27 | 0.22 |
| 1 | 217989437 | rs1510263 | -0.375 | -0.772 | -0.384 | -1.087 | -0.021 | -0.006 | -0.006 | 6.334 | 0.65 | 0.54 | 0.72 | 0.36 | 0.7 | 0.46 |
| 1 | 217991773 | rs8179429 | 0.487 | -1.015 | -0.396 | -1.207 | -0.013 | -0.042 | -0.042 | 0 | 0.28 | 0.33 | 0.26 | 0.08 | 0.2 | 0.13 |
| 1 | 217992744 | rs4332416 | 1.408 | -1.011 | -0.448 | -1.206 | -0.024 | -0.026 | -0.026 | 0.26 | 0.05 | 0.01 | 0.09 | 0.16 | 0.07 | 0.07 |
| 1 | 218003361 | rs11117946 | -0.343 | -1.025 | -0.439 | -1.209 | -0.022 | -0.009 | -0.009 | 0 | 0.64 | 0.54 | 0.71 | 0.36 | 0.7 | 0.46 |
| 1 | 218006850 | rs12040010 | -0.148 | -1.03 | -0.436 | -1.205 | -0.011 | -0.031 | -0.031 | 0.466 | 0.34 | 0.43 | 0.36 | 0.24 | 0.27 | 0.21 |
| 1 | 218008002 | rs4846443 | 1.838 | -0.938 | -0.456 | -1.145 | -0.09 | 0.08 | 0.08 | 0 | 0.26 | 0.07 | 0.32 | 0.12 | 0.39 | 0.23 |
| 1 | 218008788 | rs6674883 | -1.314 | -0.973 | -0.395 | -1.131 | -0.006 | 0.031 | 0.031 | 0 | 0.53 | 0.72 | 0.5 | 0.37 | 0.45 | 0.35 |
| 1 | 218010172 | rs10495081 | -0.647 | -0.964 | -0.311 | -1.097 | -0.021 | 0.015 | 0.015 | 0 | 0.69 | 0.54 | 0.7 | 0.33 | 0.77 | 0.47 |
| 1 | 218010441 | rs4846444 | -1.812 | -0.927 | -0.299 | -0.944 | -0.004 | 0.022 | 0.022 | 0 | 0.79 | 0.79 | 0.82 | 0.49 | 0.85 | 0.58 |
| 1 | 218018089 | rs17041601 | 1.016 | -0.915 | -0.295 | -0.973 | -0.002 | -0.005 | -0.005 | 0 | 0.11 | 0.15 | 0.09 | 0.13 | 0.15 | 0.09 |
| 1 | 218018956 | rs1567449 | -1.659 | -1.003 | -0.152 | -0.921 | -0.004 | 0.023 | 0.023 | 0 | 0.79 | 0.79 | 0.82 | 0.47 | 0.85 | 0.58 |
| 1 | 218019292 | rs1567448 | 1.369 | -0.674 | -0.002 | -0.965 | -0.004 | -0.032 | -0.032 | 0 | 0.11 | 0.15 | 0.04 | 0 | 0.07 | 0.06 |
| 1 | 218021137 | rs12759918 | -0.326 | -0.492 | 0.08 | -0.702 | -0.017 | -0.003 | -0.003 | 0.087 | 0.57 | 0.55 | 0.57 | 0.38 | 0.68 | 0.48 |
| 1 | 218024308 | rs12748013 | -0.876 | 0.216 | 0.344 | -0.63 | 0.115 | -0.005 | -0.005 | 0 | 0.07 | 0.29 | 0.06 | 0 | 0.11 | 0.04 |
| 1 | 218030584 | rs10863350 | -0.43 | -0.049 | -0.059 | -1.113 | 0.012 | 0.08 | 0.08 | 0 | 0.5 | 0.68 | 0.51 | 0.11 | 0.43 | 0.31 |
| 1 | 218036817 | rs11117953 | 0.661 | 0.019 | -0.288 | -0.858 | 0.007 | 0.012 | 0.012 | 0 | 0.13 | 0.07 | 0.05 | 0.06 | 0.07 | 0.08 |
| 1 | 218038479 | rs12756739 | 1.576 | -0.401 | -0.462 | -1.124 | -0.059 | 0.05 | 0.05 | 0 | 0.21 | 0.05 | 0.36 | 0.11 | 0.3 | 0.16 |
| 1 | 218044052 | rs12405333 | 0.612 | -0.506 | -0.282 | -1.174 | -0.016 | 0.08 | 0.08 | 0.086 | 0.29 | 0.14 | 0.39 | 0.11 | 0.32 | 0.2 |
| 1 | 218047636 | rs4846248 | -0.718 | -0.602 | -0.694 | -1.065 | 0.004 | 0.008 | 0.008 | 0 | 0.78 | 0.75 | 0.78 | 0.72 | 0.74 | 0.81 |
| 1 | 218048010 | rs17739437 | 1.385 | -0.569 | -0.738 | -1.024 | 0.003 | -0.008 | -0.008 | 0 | 0.11 | 0.08 | 0.04 | 0 | 0.06 | 0.07 |
| 1 | 218050263 | rs11117962 | 1.664 | -0.503 | -1.14 | -0.91 | -0.053 | -0.165 | -0.165 | 0 | 0.29 | 0.04 | 0.27 | 0.53 | 0.31 | 0.49 |
| 1 | 218054687 | rs17739539 | 1.218 | -0.151 | -0.929 | -1.336 | -0.012 | 0.007 | 0.007 | 0 | 0.12 | 0.06 | 0.09 | 0.08 | 0.17 | 0.08 |
| 1 | 218058598 | rs907057 | -0.451 | -0.458 | -0.596 | -1.134 | -0.006 | 0.013 | 0.013 | 0.9 | 0.57 | 0.68 | 0.42 | 0.65 | 0.44 | 0.65 |
| 1 | 218058946 | rs872171 | -0.966 | -0.222 | -0.854 | -1.022 | 0.009 | 0.033 | 0.033 | 0 | 0.92 | 0.84 | 0.89 | 0.84 | 0.92 | 0.9 |
| 1 | 218067292 | rs7546839 | -0.086 | -0.217 | -1.377 | -1.194 | 0.004 | 0.011 | 0.011 | 0 | 0.2 | 0.12 | 0.1 | 0.02 | 0.13 | 0.16 |
| 1 | 218068893 | rs7538903 | 1.18 | -0.059 | -1.601 | -0.747 | -0.026 | -0.155 | -0.155 | 0 | 0.31 | 0.04 | 0.26 | 0.52 | 0.3 | 0.49 |
| 1 | 218077622 | rs4573568 | 0.614 | 0.149 | -1.384 | -0.911 | -0.067 | 0.066 | 0.066 | 0.695 | 0.34 | 0.13 | 0.47 | 0.18 | 0.48 | 0.24 |
| 1 | 218084087 | rs12744967 | 0.62 | 0.105 | -0.708 | -0.929 | -0.005 | 0.005 | 0.005 | 0.111 | 0.36 | 0.36 | 0.5 | 0.18 | 0.48 | 0.24 |
| 1 | 218084926 | rs11591117 | 0.363 | 0.009 | -0.215 | -0.968 | 0.015 | -0.041 | -0.041 | 0.71 | 0.37 | 0.46 | 0.52 | 0.19 | 0.48 | 0.24 |
| 1 | 218084991 | rs871379 | -0.379 | -0.149 | 0.045 | -0.984 | 0.002 | 0.012 | 0.012 | 0 | 0.82 | 0.73 | 0.85 | 0.79 | 0.86 | 0.84 |
| 1 | 218085255 | rs871378 | 0.393 | -0.463 | -0.016 | -1.09 | -0.006 | -0.041 | -0.041 | 0 | 0.11 | 0.15 | 0.06 | 0.02 | 0.07 | 0.09 |
| 1 | 218087593 | rs10779305 | -0.458 | -0.281 | -0.11 | -0.986 | -0.011 | 0.013 | 0.013 | 0 | 0.82 | 0.76 | 0.86 | 0.78 | 0.88 | 0.84 |
| 1 | 218090930 | rs17046996 | 0.141 | -0.369 | -0.292 | -1.137 | 0.002 | 0.003 | 0.003 | 0 | 0.46 | 0.46 | 0.57 | 0.17 | 0.57 | 0.36 |
| 1 | 218094151 | rs6604568 | -0.851 | -0.161 | -0.581 | -0.928 | 0.014 | -0.027 | -0.027 | 0 | 0.65 | 0.75 | 0.74 | 0.47 | 0.71 | 0.53 |
| 1 | 218099577 | rs1397178 | 0.572 | 0.376 | -0.976 | -1.022 | -0.004 | 0.017 | 0.017 | 0.156 | 0.36 | 0.14 | 0.49 | 0.32 | 0.38 | 0.3 |
| 1 | 218105019 | rs10495085 | 0.363 | 0.012 | -0.805 | -0.699 | 0.01 | -0.002 | -0.002 | 0 | 0.09 | 0.01 | 0.08 | 0.07 | 0.07 | 0.06 |
| 1 | 218107388 | rs2136685 | 0.504 | 0.205 | -0.404 | -0.854 | 0.01 | 0 | 0 | 0 | 0.08 | 0.18 | 0.06 | 0.06 | 0.07 | 0.07 |
| 1 | 218108088 | rs966209 | 0.052 | 0.052 | -0.381 | -0.932 | 0.011 | 0.012 | 0.012 | 0 | 0.49 | 0.38 | 0.62 | 0.43 | 0.59 | 0.43 |
| 1 | 218108164 | rs966208 | 0.524 | 0.075 | -0.681 | -1.118 | -0.049 | 0.028 | 0.028 | 0.561 | 0.39 | 0.14 | 0.51 | 0.34 | 0.47 | 0.32 |
| 1 | 218111251 | rs17047029 | 0.109 | -0.284 | -0.176 | -1.002 | -0.02 | -0.039 | -0.039 | 0 | 0.1 | 0.09 | 0.04 | 0 | 0.04 | 0.05 |
| 1 | 218113797 | rs7550363 | 0.356 | -0.091 | 0.167 | -1.303 | -0.025 | 0.049 | 0.049 | 1.708 | 0.39 | 0.32 | 0.52 | 0.13 | 0.47 | 0.31 |
| 1 | 218124633 | rs1890608 | -1.547 | -0.035 | 0.184 | -1.298 | 0.009 | -0.026 | -0.026 | 0 | 0.94 | 0.94 | 0.97 | 1 | 0.95 | 0.96 |
| 1 | 218125630 | rs4846450 | -0.561 | 0.195 | 0.29 | -1.443 | 0.009 | 0.01 | 0.01 | 0.512 | 0.69 | 0.46 | 0.5 | 0.62 | 0.68 | 0.71 |
| 1 | 218126726 | rs1567445 | -0.216 | 0.178 | 0.422 | -1.424 | 0.063 | 0.175 | 0.175 | 0.06 | 0.91 | 0.64 | 0.89 | 0.76 | 0.93 | 0.94 |
| 1 | 218127794 | rs4846249 | -0.343 | 0.18 | 0.415 | -1.423 | 0.056 | 0.16 | 0.16 | 0.037 | 0.91 | 0.66 | 0.89 | 0.76 | 0.93 | 0.94 |
| 1 | 218129861 | rs12031059 | 0.507 | 0.057 | 0.202 | -1.598 | 0.029 | 0.01 | 0.01 | 0 | 0.41 | 0.3 | 0.33 | 0.62 | 0.34 | 0.52 |
| 1 | 218133537 | rs1832544 | 0.131 | 0.063 | 0.234 | -1.613 | 0.011 | -0.029 | -0.029 | 0 | 0.63 | 0.46 | 0.71 | 0.76 | 0.59 | 0.74 |
| 1 | 218136962 | rs2225975 | 0.053 | 0.058 | 0.121 | -1.601 | 0.027 | 0.097 | 0.097 | 1.145 | 0.5 | 0.33 | 0.55 | 0.14 | 0.59 | 0.42 |
| 1 | 218139006 | rs4428953 | 1.995 | -0.047 | 0.082 | -1.589 | 0.029 | -0.093 | -0.093 | 0.46 | 0.24 | 0.05 | 0.22 | 0.54 | 0.21 | 0.39 |
| 1 | 218146855 | rs4475815 | 1.017 | 0.127 | 0.675 | -1.437 | 0.034 | 0.008 | 0.008 | 0 | 0.41 | 0.16 | 0.33 | 0.62 | 0.35 | 0.53 |
| 1 | 218146954 | rs17742354 | 1.602 | 0.16 | 0.692 | -1.462 | -0.024 | 0.024 | 0.024 | 0 | 0.15 | 0.07 | 0.12 | 0 | 0.22 | 0.1 |
| 1 | 218147297 | rs7546902 | 1.161 | 0.132 | 0.682 | -1.454 | 0 | -0.021 | -0.021 | 0 | 0.11 | 0.13 | 0.04 | 0 | 0.1 | 0.09 |
| 1 | 218148677 | rs4846453 | 0.379 | 0.155 | 0.689 | -1.432 | 0.029 | 0.047 | 0.047 | 0.734 | 0.52 | 0.28 | 0.37 | 0.64 | 0.45 | 0.62 |
| 1 | 218149826 | rs7520319 | 0.186 | 0.17 | 0.649 | -1.438 | -0.042 | 0.069 | 0.069 | 2.453 | 0.37 | 0.2 | 0.49 | 0.14 | 0.47 | 0.32 |
| 1 | 218154463 | rs12043516 | 1.368 | 0.201 | 0.603 | -1.393 | -0.03 | 0.019 | 0.019 | 2.705 | 0.21 | 0.06 | 0.37 | 0.13 | 0.24 | 0.22 |
| 1 | 218155651 | rs1889607 | -0.704 | 0.202 | 0.615 | -1.399 | -0.005 | 0.015 | 0.015 | 0 | 0.43 | 0.65 | 0.54 | 0.36 | 0.42 | 0.38 |
| 1 | 218156176 | rs1335006 | -0.907 | 0.159 | 0.569 | -1.398 | 0 | 0.023 | 0.023 | 0 | 0.44 | 0.67 | 0.54 | 0.36 | 0.42 | 0.37 |
| 1 | 218158712 | rs17742891 | 0.824 | 0.221 | 0.121 | -2.018 | 0.001 | -0.017 | -0.017 | 0 | 0.12 | 0.13 | 0.05 | 0 | 0.11 | 0.1 |
| 1 | 218162375 | rs12401951 | 1.239 | 0.269 | -0.177 | -1.911 | -0.032 | 0.021 | 0.021 | 2.168 | 0.21 | 0.06 | 0.38 | 0.13 | 0.24 | 0.21 |
| 1 | 218162502 | rs10495090 | 0.867 | 0.566 | 0.133 | -1.54 | -0.076 | 0.09 | 0.09 | 0 | 0.15 | 0.02 | 0.11 | 0 | 0.22 | 0.1 |
| 1 | 218168100 | rs7548817 | 0.867 | 0.648 | 0.311 | -1.46 | -0.004 | -0.042 | -0.042 | 0 | 0.61 | 0.4 | 0.76 | 0.98 | 0.59 | 0.77 |
| 1 | 218188851 | rs4282881 | 0.125 | 0.619 | 0.334 | -0.751 | 0.016 | -0.15 | -0.15 | 0 | 0.78 | 0.51 | 0.83 | 1 | 0.75 | 0.9 |

| | | | | | | | | | | | | | | | | |
|---|---|---|---|---|---|---|---|---|---|---|---|---|---|---|---|---|
| 1 | 218198027 | rs1490331 | -0.09 | 0.662 | 0.157 | -0.731 | 0.003 | -0.067 | -0.067 | 0.627 | 0.79 | 0.7 | 0.84 | 1 | 0.76 | 0.9 |
| 1 | 218198668 | rs7512034 | 0.062 | 0.971 | 0.642 | -0.048 | 0.046 | 0.016 | 0.016 | 0 | 0.78 | 0.57 | 0.8 | 0.76 | 0.73 | 0.84 |
| 1 | 218201025 | rs12142065 | 0.626 | 0.515 | -0.236 | 0.071 | -0.005 | -0.037 | -0.037 | 0 | 0.3 | 0.13 | 0.25 | 0.42 | 0.3 | 0.35 |
| 1 | 218201594 | rs6604578 | -0.351 | 0.346 | 0.065 | 0.049 | 0.046 | 0.022 | 0.022 | 0.441 | 0.74 | 0.53 | 0.78 | 0.71 | 0.71 | 0.76 |
| 1 | 218204695 | rs11118016 | 1.008 | -0.162 | -0.583 | -0.335 | -0.023 | 0.029 | 0.029 | 1.055 | 0.24 | 0.08 | 0.4 | 0.13 | 0.27 | 0.18 |
| 1 | 218208472 | rs6661310 | 0.217 | -0.163 | -0.299 | -0.328 | -0.003 | 0.004 | 0.004 | 0 | 0.63 | 0.37 | 0.73 | 0.71 | 0.63 | 0.7 |
| 1 | 218209224 | rs1556913 | 0.043 | -0.269 | -0.304 | -0.41 | -0.028 | 0.063 | 0.063 | 0 | 0.82 | 0.43 | 0.84 | 0.71 | 0.88 | 0.81 |
| 1 | 218209667 | rs12138324 | 0.703 | -0.293 | -0.388 | -0.421 | -0.008 | 0.014 | 0.014 | 0 | 0.6 | 0.34 | 0.71 | 0.69 | 0.61 | 0.62 |
| 1 | 218214914 | rs11118017 | 0.37 | -0.235 | -0.393 | -0.426 | -0.003 | 0.004 | 0.004 | 0 | 0.63 | 0.37 | 0.73 | 0.71 | 0.63 | 0.7 |
| 1 | 218214924 | rs17693785 | 1.218 | 0.108 | -0.954 | -0.49 | -0.099 | 0.116 | 0.116 | 0 | 0.18 | 0.02 | 0.1 | 0 | 0.25 | 0.11 |
| 1 | 218223205 | rs12729245 | 0.444 | -0.456 | -0.879 | -0.393 | -0.094 | 0.16 | 0.16 | 0.879 | 0.41 | 0.08 | 0.5 | 0.15 | 0.53 | 0.29 |
| 1 | 218224279 | rs992010 | -0.147 | -0.809 | -0.294 | -0.641 | 0.008 | 0.029 | 0.029 | 0 | 0.49 | 0.58 | 0.45 | 0.84 | 0.43 | 0.58 |
| 1 | 218237411 | rs12405427 | -0.891 | -0.811 | -0.336 | -0.652 | -0.079 | 0.156 | 0.156 | 0.914 | 0.59 | 0.89 | 0.49 | 0.85 | 0.47 | 0.71 |
| 1 | 218238773 | rs6697579 | 0.883 | -0.783 | -0.096 | -0.335 | -0.001 | -0.047 | -0.047 | 0.258 | 0.31 | 0.16 | 0.3 | 0.57 | 0.32 | 0.39 |
| 1 | 218241796 | rs1981212 | 0.016 | -0.63 | 0.09 | -0.217 | 0.012 | -0.005 | -0.005 | 0.191 | 0.44 | 0.32 | 0.35 | 0.57 | 0.4 | 0.45 |
| 1 | 218251692 | rs10863373 | 1.114 | -0.622 | -0.301 | 0.274 | -0.013 | 0.053 | 0.053 | 0.011 | 0.76 | 0.4 | 0.78 | 0.53 | 0.84 | 0.67 |
| 1 | 218253106 | rs2067836 | 1.141 | -0.861 | -0.778 | 0.061 | 0.013 | 0.007 | 0.007 | 2.256 | 0.14 | 0.04 | 0.3 | 0.13 | 0.12 | 0.11 |
| 1 | 218253210 | rs2067837 | 0.369 | -0.888 | -0.569 | 0.207 | 0.005 | 0.236 | 0.236 | 0.069 | 0.87 | 0.44 | 0.83 | 0.57 | 0.92 | 0.84 |
| 1 | 218253312 | rs11118029 | 0.369 | -0.888 | -0.597 | 0.223 | 0.004 | 0.221 | 0.221 | 0.069 | 0.87 | 0.44 | 0.83 | 0.57 | 0.92 | 0.84 |
| 1 | 218253812 | rs2046842 | 0.907 | -1.138 | -0.503 | -0.034 | -0.033 | 0 | 0 | 0 | 0.79 | 0.54 | 0.88 | 0.79 | 0.9 | 0.72 |
| 1 | 218259988 | rs7539193 | -0.175 | -0.831 | -0.951 | 0.174 | -0.001 | -0.002 | -0.002 | 3.235 | 0.42 | 0.53 | 0.28 | 0.49 | 0.4 | 0.29 |
| 1 | 218261632 | rs877850 | -0.373 | -0.919 | -1.579 | 0.178 | -0.028 | 0.023 | 0.023 | 0 | 0.19 | 0.6 | 0.13 | 0.25 | 0.1 | 0.1 |
| 1 | 218265683 | rs6670238 | 1.387 | -1.02 | -1.737 | 0.389 | 0.023 | -0.022 | -0.022 | 0 | 0.14 | 0.05 | 0.25 | 0.24 | 0.1 | 0.18 |
| 1 | 218267766 | rs6604589 | -1.346 | -1.059 | -1.763 | 0.302 | 0.034 | -0.015 | -0.015 | 0 | 0.39 | 0.64 | 0.48 | 0.5 | 0.24 | 0.38 |
| 1 | 218268137 | rs11118031 | 0.945 | -0.906 | -1.543 | 0.409 | -0.007 | 0.031 | 0.031 | 1.875 | 0.16 | 0.19 | 0.3 | 0.45 | 0.12 | 0.16 |
| 1 | 218271784 | rs12083333 | -0.89 | -0.841 | -1.888 | -0.203 | -0.025 | 0.018 | 0.018 | 0 | 0.58 | 0.7 | 0.55 | 0.69 | 0.51 | 0.43 |
| 1 | 218272851 | rs6699736 | 1.328 | -0.821 | -1.818 | -0.174 | 0 | 0 | 0 | 0.02 | 0.14 | 0.12 | 0.26 | 0.19 | 0.1 | 0.15 |
| 1 | 218278650 | rs4639829 | -1.327 | -0.232 | -2.17 | -0.431 | -0.09 | -0.004 | -0.004 | 0 | 0.68 | 0.93 | 0.68 | 0.77 | 0.58 | 0.63 |
| 1 | 218281230 | rs6672959 | 0.976 | -1.185 | -2.063 | -0.361 | 0.007 | 0.006 | 0.006 | 0.509 | 0.1 | 0.06 | 0.06 | 0 | 0.06 | 0.07 |
| 1 | 218284056 | rs1873326 | -0.771 | -0.196 | -1.925 | -0.132 | -0.009 | 0.023 | 0.023 | 0 | 0.62 | 0.72 | 0.61 | 0.74 | 0.4 | 0.54 |
| 1 | 218284122 | rs2377781 | -0.259 | -0.294 | -1.896 | -0.102 | 0.035 | 0.03 | 0.03 | 0 | 0.56 | 0.71 | 0.5 | 0.69 | 0.36 | 0.41 |
| 1 | 218287335 | rs7527372 | 0.075 | -0.778 | -1.615 | -0.02 | 0.019 | -0.006 | -0.006 | 0.537 | 0.12 | 0.17 | 0.2 | 0.08 | 0.17 | 0.22 |
| 1 | 218292715 | rs580733 | -0.425 | -1.503 | -1.664 | -0.006 | 0.045 | 0.022 | 0.022 | 0 | 0.55 | 0.71 | 0.37 | 0.7 | 0.33 | 0.39 |
| 1 | 218294968 | rs570822 | 0.275 | -0.94 | -1.707 | -0.153 | 0.026 | 0.072 | 0.072 | 0 | 0.21 | 0.1 | 0.08 | 0.02 | 0.15 | 0.14 |
| 1 | 218295180 | rs568909 | 0.261 | -0.919 | -1.657 | -0.11 | 0.019 | 0.053 | 0.053 | 0 | 0.19 | 0.11 | 0.08 | 0.02 | 0.14 | 0.12 |
| 1 | 218299276 | rs12125587 | -0.162 | -1.671 | -2.002 | 0.939 | 0.02 | -0.059 | -0.059 | 0 | 0.22 | 0.04 | 0.18 | 0.28 | 0.18 | 0.14 |
| 1 | 218302651 | rs12033297 | 0.527 | -2.138 | -2.357 | 0.613 | 0.051 | -0.006 | -0.006 | 0 | 0.16 | 0.03 | 0.1 | 0.16 | 0.11 | 0.12 |
| 1 | 218304258 | rs17047439 | 0.16 | -1.662 | -1.525 | 0.288 | 0.049 | 0.019 | 0.019 | 0 | 0.13 | 0.32 | 0.18 | 0.11 | 0.12 | 0.15 |
| 1 | 218304346 | rs575962 | -1.149 | -1.601 | -2.279 | 0.002 | -0.103 | 0.072 | 0.072 | 0 | 0.67 | 0.88 | 0.58 | 0.83 | 0.49 | 0.55 |
| 1 | 218328002 | rs11805614 | 0.466 | -1.826 | -1.822 | -0.079 | -0.005 | -0.028 | -0.028 | 0 | 0.15 | 0.16 | 0.08 | 0.02 | 0.11 | 0.1 |
| 1 | 218331504 | rs12742498 | 0.561 | -1.839 | -2.351 | -0.211 | -0.078 | 0.044 | 0.044 | 0 | 0.23 | 0.02 | 0.18 | 0.1 | 0.26 | 0.27 |
| 1 | 218333601 | rs683968 | -0.723 | -2.03 | -2.469 | 0.31 | 0.121 | -0.046 | -0.046 | 0.144 | 0.42 | 0.9 | 0.42 | 0.59 | 0.25 | 0.29 |
| 1 | 218333622 | rs12133828 | 0.605 | -1.563 | -2.274 | 0.211 | 0.005 | 0.001 | 0.001 | 0 | 0.15 | 0.11 | 0.07 | 0.03 | 0.11 | 0.1 |
| 1 | 218333741 | rs511223 | -0.747 | -0.888 | -2.094 | 0.563 | -0.184 | -0.042 | -0.042 | 0 | 0.66 | 0.97 | 0.6 | 0.7 | 0.52 | 0.56 |
| 1 | 218337199 | rs621070 | -0.659 | -0.957 | -0.464 | 0.874 | -0.074 | 0.088 | 0.088 | 0.799 | 0.19 | 0.48 | 0.23 | 0.22 | 0.1 | 0.09 |
| 1 | 218347150 | rs6658076 | -0.584 | -0.264 | -1.428 | 0.231 | 0.002 | 0.041 | 0.041 | 0 | 0.34 | 0.16 | 0.3 | 0.29 | 0.32 | 0.4 |
| 1 | 218348909 | rs10495096 | -0.137 | -0.511 | -1.572 | 0.221 | 0.014 | -0.016 | -0.016 | 0 | 0.14 | 0.12 | 0.15 | 0.27 | 0.09 | 0.19 |
| 1 | 218352449 | rs1338451 | -0.53 | 1.592 | -1.166 | -0.322 | 0.043 | -0.032 | -0.032 | 0.011 | 0.89 | 0.89 | 0.88 | 1 | 0.69 | 0.92 |
| 1 | 218353211 | rs6694166 | -0.025 | 1.189 | -1.225 | -0.698 | 0.002 | -0.022 | -0.022 | 0.508 | 0.2 | 0.19 | 0.09 | 0.01 | 0.14 | 0.11 |
| 1 | 218361784 | rs7528076 | 1.43 | 0.751 | -0.937 | -0.521 | 0.062 | -0.004 | -0.004 | 0 | 0.5 | 0.31 | 0.46 | 0.51 | 0.38 | 0.52 |
| 1 | 218362082 | rs12116939 | 0.371 | 0.68 | -0.762 | -0.585 | 0.001 | 0.159 | 0.159 | 0 | 0.09 | 0.19 | 0.28 | 0.48 | 0.1 | 0.22 |
| 1 | 218367181 | rs12757063 | 0.204 | 0.536 | -0.789 | -1.455 | 0.005 | 0.028 | 0.028 | 0 | 0.08 | 0.01 | 0.04 | 0 | 0.07 | 0.07 |
| 1 | 218371223 | rs7556454 | 0.975 | 0.509 | -0.539 | -1.399 | 0.001 | -0.089 | -0.089 | 0 | 0.59 | 0.48 | 0.73 | 0.99 | 0.49 | 0.75 |
| 1 | 218372797 | rs11118050 | 0.04 | 0.367 | -1.169 | -0.792 | 0.013 | 0.037 | 0.037 | 0 | 0.19 | 0.08 | 0.08 | 0.01 | 0.14 | 0.11 |
| 1 | 218388769 | rs2039711 | -0.341 | 1.364 | -1.042 | -0.546 | -0.033 | 0.02 | 0.02 | 0 | 0.87 | 0.94 | 0.88 | 1 | 0.71 | 0.89 |
| 1 | 218391681 | rs1556836 | -0.329 | 1.221 | -1.02 | -0.64 | -0.026 | 0.008 | 0.008 | 0 | 0.85 | 0.9 | 0.87 | 1 | 0.69 | 0.88 |
| 1 | 218392447 | rs11118054 | 0.526 | 0.285 | -1.922 | -0.758 | -0.061 | -0.031 | -0.031 | 0 | 0.33 | 0.06 | 0.4 | 0.35 | 0.33 | 0.34 |
| 1 | 218397580 | rs1416156 | -0.014 | 0.056 | -1.6 | -0.83 | -0.018 | 0.016 | 0.016 | 0.059 | 0.72 | 0.74 | 0.78 | 0.92 | 0.57 | 0.74 |
| 1 | 218418085 | rs4445537 | 0.019 | -0.04 | -1.12 | -0.844 | 0.006 | -0.023 | -0.023 | 0 | 0.72 | 0.68 | 0.78 | 0.92 | 0.59 | 0.76 |
| 1 | 218418911 | rs12087406 | -0.084 | -0.729 | -0.73 | -0.296 | -0.004 | 0.009 | 0.009 | 0 | 0.08 | 0.17 | 0.06 | 0.08 | 0.07 | 0.11 |
| 1 | 218422950 | rs12563602 | -0.555 | -1.133 | -0.342 | -0.544 | -0.031 | -0.036 | -0.036 | 0 | 0.18 | 0.37 | 0.12 | 0.08 | 0.14 | 0.2 |
| 1 | 218424514 | rs12082932 | -0.252 | -0.676 | -0.223 | -0.115 | -0.004 | 0.008 | 0.008 | 0 | 0.08 | 0.18 | 0.06 | 0.08 | 0.07 | 0.11 |
| 1 | 218428101 | rs10863384 | 0.027 | -0.757 | -0.191 | 0.642 | -0.015 | 0.248 | 0.248 | 0 | 0.12 | 0.34 | 0.18 | 0.38 | 0.07 | 0.19 |
| 1 | 218432309 | rs12128367 | 0.147 | -0.659 | -1.166 | 0.489 | -0.014 | -0.054 | -0.054 | 0 | 0.14 | 0.01 | 0.16 | 0.2 | 0.11 | 0.12 |
| 1 | 218439928 | rs1338445 | -0.378 | -0.517 | -0.413 | 1.038 | -0.024 | 0.121 | 0.121 | 0.062 | 0.16 | 0.3 | 0.21 | 0.36 | 0.11 | 0.2 |
| 1 | 218448425 | rs12027949 | -0.239 | -0.448 | -1.165 | 0.421 | 0.024 | -0.027 | -0.027 | 0 | 0.32 | 0.08 | 0.23 | 0.38 | 0.23 | 0.33 |
| 1 | 218458058 | rs3753592 | -0.664 | 0.223 | -0.441 | 0.242 | -0.003 | 0.004 | 0.004 | 0 | 0.37 | 0.7 | 0.33 | 0.38 | 0.41 | 0.43 |
| 1 | 218458227 | rs2788193 | 1.183 | -0.233 | -0.982 | 0.127 | -0.026 | -0.054 | -0.054 | 0 | 0.29 | 0.16 | 0.39 | 0.41 | 0.32 | 0.28 |
| 1 | 218492121 | rs3121580 | -0.147 | -1.475 | 0.013 | 0.235 | 0.032 | 0.018 | 0.018 | 0 | 0.75 | 0.95 | 0.8 | 0.83 | 0.81 | 0.74 |

| | | | | | | | | | | | | | | | | |
|---|---|---|---|---|---|---|---|---|---|---|---|---|---|---|---|---|
| 1 | 218506743 | rs6604603 | -0.85 | -0.514 | 1.29 | 0.767 | 0.081 | 0.146 | 0.146 | 0.029 | 0.07 | 0.5 | 0.07 | 0.07 | 0.04 | 0.01 |
| 1 | 218517668 | rs6604604 | -0.549 | -0.49 | 1.304 | 0.778 | 0.067 | 0.129 | 0.129 | 0 | 0.06 | 0.31 | 0.05 | 0.07 | 0.04 | 0.01 |
| 1 | 218523025 | rs17047682 | 0.393 | -0.494 | 0.986 | 0.804 | -0.004 | -0.031 | -0.031 | 0 | 0.06 | 0 | 0.09 | 0.14 | 0.06 | 0.11 |
| 1 | 218523650 | rs1418555 | -0.562 | -0.297 | 1.229 | 0.707 | -0.01 | -0.006 | -0.006 | 0 | 0.29 | 0.36 | 0.23 | 0.31 | 0.22 | 0.35 |
| 1 | 218523730 | rs1417488 | -0.766 | -0.293 | 0.957 | 0.716 | -0.019 | 0.008 | 0.008 | 0 | 0.31 | 0.43 | 0.26 | 0.38 | 0.25 | 0.36 |
| 1 | 218524632 | rs2799097 | -0.245 | -0.607 | 0.542 | 0.75 | -0.009 | -0.021 | -0.021 | 0 | 0.87 | 0.95 | 0.82 | 0.81 | 0.84 | 0.78 |
| 1 | 218525588 | rs17047703 | -0.561 | -0.572 | 0.822 | 0.344 | 0.02 | 0.009 | 0.009 | 0 | 0.27 | 0.16 | 0.22 | 0.31 | 0.22 | 0.35 |
| 1 | 218542059 | rs6662137 | -1.107 | -0.326 | -0.104 | 0.404 | -0.004 | -0.003 | -0.003 | 0 | 0.37 | 0.62 | 0.29 | 0.39 | 0.31 | 0.42 |
| 1 | 218548521 | rs17558745 | -0.834 | 0.151 | -1.664 | 0.274 | -0.067 | -0.01 | -0.01 | 0.135 | 0.3 | 0.01 | 0.2 | 0.31 | 0.3 | 0.4 |
| 1 | 218553297 | rs3892225 | -0.942 | -0.154 | -0.988 | 0.236 | -0.001 | 0.004 | 0.004 | 0 | 0.18 | 0.14 | 0.16 | 0.16 | 0.18 | 0.22 |
| 1 | 218556036 | rs10482750 | -1.095 | -0.499 | -0.593 | 0.359 | -0.001 | -0.003 | -0.003 | 0 | 0.11 | 0.08 | 0.12 | 0.19 | 0.12 | 0.11 |
| 1 | 218556297 | rs10482751 | 0.586 | -1.462 | -1.555 | 0.292 | 0.11 | -0.046 | -0.046 | 0.326 | 0.51 | 0.09 | 0.57 | 0.49 | 0.66 | 0.52 |
| 1 | 218557609 | rs4335431 | -0.146 | -1.621 | -1.598 | 0.258 | 0 | 0.003 | 0.003 | 0 | 0.1 | 0.07 | 0.08 | 0.13 | 0.09 | 0.1 |
| 1 | 218558327 | rs2027566 | 0.75 | -1.486 | -1.845 | 0.295 | 0.003 | -0.114 | -0.114 | 0.012 | 0.47 | 0.02 | 0.55 | 0.48 | 0.63 | 0.49 |
| 1 | 218558531 | rs11466383 | 0.354 | -1.975 | -1.484 | 0.053 | -0.004 | -0.017 | -0.017 | 0 | 0.06 | 0.08 | 0.06 | 0 | 0.04 | 0.04 |
| 1 | 218558588 | rs2027567 | 0.469 | -1.885 | -0.99 | 0.312 | 0.048 | 0.004 | 0.004 | 0 | 0.59 | 0.34 | 0.68 | 0.57 | 0.74 | 0.58 |
| 1 | 218559280 | rs12058490 | -1.069 | -0.89 | 0.522 | 0.2 | -0.003 | -0.027 | -0.027 | 0 | 0.14 | 0.35 | 0.11 | 0.13 | 0.1 | 0.12 |
| 1 | 218564881 | rs947712 | -0.851 | -0.993 | 0.389 | 0.52 | -0.002 | 0.006 | 0.006 | 0 | 0.39 | 0.52 | 0.49 | 0.54 | 0.34 | 0.52 |
| 1 | 218565324 | rs1539399 | -0.83 | -1.001 | 0.371 | 0.669 | -0.001 | 0.009 | 0.009 | 0 | 0.39 | 0.54 | 0.49 | 0.48 | 0.34 | 0.52 |
| 1 | 218568829 | rs2799090 | -0.848 | -1.366 | 0.958 | 0.361 | 0.011 | 0.038 | 0.038 | 0 | 0.36 | 0.74 | 0.3 | 0.29 | 0.28 | 0.33 |
| 1 | 218574789 | rs6703224 | -0.53 | -1.326 | 1.242 | 0.324 | 0 | 0.063 | 0.063 | 0.187 | 0.11 | 0.44 | 0.15 | 0.13 | 0.06 | 0.06 |
| 1 | 218575202 | rs1317681 | -0.297 | -1.46 | 1.028 | 0.316 | 0.002 | -0.003 | -0.003 | 0.085 | 0.16 | 0.15 | 0.36 | 0.5 | 0.17 | 0.29 |
| 1 | 218579644 | rs11118103 | -0.589 | -1.497 | 1.004 | 0.24 | -0.018 | 0.016 | 0.016 | 0 | 0.12 | 0.29 | 0.14 | 0.14 | 0.08 | 0.09 |
| 1 | 218579985 | rs1891467 | -1.752 | -1.589 | 0.926 | 0.216 | -0.032 | 0.013 | 0.013 | 0 | 0.29 | 0.52 | 0.5 | 0.64 | 0.24 | 0.38 |
| 1 | 218581617 | rs2799083 | -0.474 | -1.475 | 0.996 | 0.068 | -0.001 | -0.048 | -0.048 | 0.115 | 0.32 | 0.39 | 0.31 | 0.21 | 0.26 | 0.32 |
| 1 | 218584419 | rs2799103 | 0.3 | -1.473 | 1.119 | 0.01 | 0.007 | -0.093 | -0.093 | 0 | 0.23 | 0.29 | 0.16 | 0.07 | 0.2 | 0.23 |
| 1 | 218587636 | rs2000220 | -1.727 | -1.458 | 0.646 | 0.056 | -0.055 | 0.132 | 0.132 | 0.237 | 0.49 | 0.96 | 0.64 | 0.73 | 0.4 | 0.6 |
| 1 | 218596461 | rs6657275 | -1.677 | -1.201 | 1.018 | -0.065 | -0.017 | -0.008 | -0.008 | 0 | 0.3 | 0.49 | 0.51 | 0.69 | 0.29 | 0.36 |
| 1 | 218597859 | rs1342586 | -0.215 | -1.228 | 1.059 | 0.003 | -0.01 | -0.039 | -0.039 | 0.622 | 0.16 | 0.14 | 0.32 | 0.56 | 0.22 | 0.26 |
| 1 | 218598410 | rs4846479 | -1.317 | -1.174 | 1.074 | 0.001 | -0.003 | 0 | 0 | 0 | 0.26 | 0.35 | 0.49 | 0.69 | 0.28 | 0.33 |
| 1 | 218604709 | rs1890994 | -0.154 | -1.284 | 1.027 | -0.034 | -0.052 | -0.116 | -0.116 | 2.104 | 0.15 | 0.01 | 0.31 | 0.56 | 0.21 | 0.27 |
| 1 | 218611878 | rs2798631 | -1.852 | -1.232 | 1.019 | -0.132 | -0.129 | 0.132 | 0.132 | 0.306 | 0.53 | 0.97 | 0.69 | 0.75 | 0.49 | 0.58 |
| 1 | 218617900 | rs6683598 | -1.534 | -1.2 | 1.096 | -0.084 | -0.008 | -0.003 | -0.003 | 0 | 0.29 | 0.41 | 0.49 | 0.69 | 0.29 | 0.37 |
| 1 | 218620747 | rs1473526 | -1.188 | -1.196 | 1.061 | -0.083 | -0.003 | 0 | 0 | 0 | 0.26 | 0.35 | 0.49 | 0.69 | 0.28 | 0.33 |
| 1 | 218622319 | rs1473527 | -1.703 | -1.363 | 1.179 | -0.175 | -0.013 | -0.013 | -0.013 | 0 | 0.33 | 0.62 | 0.52 | 0.72 | 0.29 | 0.37 |
| 1 | 218632368 | rs4846483 | -1.228 | -1.272 | 1.256 | -0.128 | -0.006 | -0.003 | -0.003 | 0 | 0.28 | 0.36 | 0.48 | 0.7 | 0.28 | 0.35 |
| 1 | 218632925 | rs11118109 | 0.579 | -1.217 | 0.978 | -0.184 | 0.009 | -0.1 | -0.1 | 1.089 | 0.23 | 0.35 | 0.17 | 0.05 | 0.21 | 0.23 |
| 1 | 218643940 | rs725033 | -0.929 | -1.218 | 0.741 | 0.236 | -0.039 | 0.038 | 0.038 | 0 | 0.21 | 0.43 | 0.27 | 0.31 | 0.14 | 0.26 |
| 1 | 218646954 | rs6604616 | -0.275 | -1.163 | 0.772 | 0.284 | -0.034 | 0.018 | 0.018 | 0.115 | 0.13 | 0.27 | 0.18 | 0.14 | 0.06 | 0.06 |
| 1 | 218663189 | rs3009935 | -0.385 | -1 | 0.422 | 0.318 | -0.004 | -0.011 | -0.011 | 0 | 0.67 | 0.71 | 0.6 | 0.55 | 0.63 | 0.63 |
| 1 | 218689155 | rs3009947 | 0.495 | -0.913 | 0.139 | -0.008 | -0.002 | 0.029 | 0.029 | 0 | 0.49 | 0.43 | 0.37 | 0.24 | 0.5 | 0.42 |
| 1 | 218690577 | rs6665024 | -0.373 | -1.076 | 0.678 | 0.042 | -0.014 | 0.011 | 0.011 | 0 | 0.18 | 0.27 | 0.23 | 0.25 | 0.13 | 0.22 |
| 1 | 218692641 | rs12034138 | -1.216 | -1.103 | 0.674 | 0.028 | -0.048 | 0.012 | 0.012 | 0 | 0.23 | 0.55 | 0.27 | 0.26 | 0.14 | 0.25 |
| 1 | 218693872 | rs11118119 | -1 | -1.102 | 0.55 | 0.028 | -0.041 | 0.016 | 0.016 | 0 | 0.22 | 0.56 | 0.27 | 0.26 | 0.14 | 0.24 |
| 1 | 218698641 | rs11805322 | -0.768 | -0.575 | 0.183 | 0.265 | 0.019 | 0.021 | 0.021 | 0.033 | 0.05 | 0.27 | 0.04 | 0.01 | 0.01 | 0.03 |
| 1 | 218702876 | rs528797 | -0.385 | -0.194 | -1.083 | -0.019 | 0.033 | 0.058 | 0.058 | 0 | 0.4 | 0.26 | 0.36 | 0.24 | 0.33 | 0.29 |
| 1 | 218705761 | rs11118120 | 0.089 | 0.11 | -1.001 | 0.774 | 0 | 0.014 | 0.014 | 0 | 0.15 | 0.18 | 0.27 | 0.3 | 0.14 | 0.23 |
| 1 | 218706772 | rs12563275 | -0.443 | 0.396 | -1.513 | 0.445 | -0.033 | -0.054 | -0.054 | 0 | 0.25 | 0.07 | 0.3 | 0.29 | 0.25 | 0.26 |
| 1 | 218708110 | rs512589 | -0.514 | -0.113 | -1.823 | 0.429 | 0.005 | -0.003 | -0.003 | 0 | 0.83 | 0.86 | 0.9 | 0.82 | 0.91 | 0.78 |
| 1 | 218708603 | rs2047163 | 0.035 | -0.02 | -1.832 | 0.522 | -0.022 | -0.039 | -0.039 | 0 | 0.25 | 0.1 | 0.3 | 0.29 | 0.25 | 0.28 |
| 1 | 218714208 | rs604045 | 1.385 | 0.287 | -2.611 | -0.081 | -0.111 | -0.131 | -0.131 | 0.01 | 0.26 | 0.04 | 0.37 | 0.44 | 0.39 | 0.33 |
| 1 | 218717742 | rs4846488 | -0.822 | 0.147 | -1.543 | -0.3 | -0.002 | 0.001 | 0.001 | 0 | 0.45 | 0.42 | 0.46 | 0.4 | 0.44 | 0.41 |
| 1 | 218719400 | rs12038116 | -0.025 | -0.023 | -1.067 | -0.552 | -0.008 | -0.046 | -0.046 | 0.14 | 0.19 | 0.31 | 0.15 | 0.06 | 0.17 | 0.12 |
| 1 | 218726797 | rs1764694 | -0.146 | -0.085 | -1.548 | -0.442 | 0.023 | 0.029 | 0.029 | 0 | 0.82 | 0.94 | 0.92 | 0.88 | 0.89 | 0.78 |
| 1 | 218727857 | rs1603059 | -0.347 | -0.491 | -0.791 | -1.109 | -0.018 | -0.109 | -0.109 | 0 | 0.3 | 0.44 | 0.16 | 0.05 | 0.25 | 0.19 |
| 1 | 218729548 | rs472905 | 0.139 | -0.649 | -0.616 | -0.834 | -0.014 | -0.015 | -0.015 | 6.208 | 0.65 | 0.42 | 0.69 | 0.8 | 0.71 | 0.69 |
| 1 | 218732057 | rs11585026 | 1.217 | -0.284 | -0.787 | -0.97 | -0.032 | 0.014 | 0.014 | 0 | 0.06 | 0.01 | 0.06 | 0.02 | 0.12 | 0.09 |
| 1 | 218737104 | rs10863401 | 0.701 | -0.684 | -0.16 | -1.16 | -0.017 | 0.012 | 0.012 | 0 | 0.2 | 0.38 | 0.24 | 0.14 | 0.17 | 0.21 |
| 1 | 218739482 | rs682483 | 1.167 | -0.771 | -0.651 | -1.242 | -0.087 | -0.034 | -0.034 | 2.675 | 0.37 | 0.12 | 0.52 | 0.66 | 0.48 | 0.39 |
| 1 | 218741263 | rs11118129 | 0.087 | -1.077 | -0.365 | -1.375 | -0.056 | -0.034 | -0.034 | 0.993 | 0.33 | 0.45 | 0.28 | 0.18 | 0.22 | 0.23 |
| 1 | 218741315 | rs10863402 | -0.525 | -0.975 | -0.748 | -1.303 | 0.054 | 0.076 | 0.076 | 0 | 0.42 | 0.76 | 0.38 | 0.2 | 0.35 | 0.32 |
| 1 | 218744152 | rs17048147 | -1.185 | -0.063 | -0.887 | -0.595 | -0.027 | -0.068 | -0.068 | 1.006 | 0.69 | 0.85 | 0.57 | 0.51 | 0.62 | 0.72 |
| 1 | 218748958 | rs1932971 | -0.322 | -0.137 | -0.627 | -0.703 | -0.004 | -0.001 | -0.001 | 0 | 0.39 | 0.43 | 0.32 | 0.47 | 0.33 | 0.35 |
| 1 | 218749231 | rs259464 | -2.056 | -0.175 | -0.789 | -0.767 | -0.019 | -0.035 | -0.035 | 0 | 0.94 | 0.88 | 0.97 | 1 | 0.97 | 0.98 |
| 1 | 218758944 | rs1337110 | 1.078 | -0.291 | -0.774 | -0.888 | -0.001 | 0.016 | 0.016 | 0 | 0.12 | 0.19 | 0.16 | 0.13 | 0.11 | 0.2 |
| 1 | 218761681 | rs17048238 | 0.374 | -0.484 | -0.838 | -1.136 | 0.03 | 0.076 | 0.076 | 0 | 0.21 | 0.38 | 0.27 | 0.44 | 0.23 | 0.28 |
| 1 | 218762925 | rs10495099 | 0.993 | -0.227 | -0.946 | -0.748 | -0.001 | 0.008 | 0.008 | 0 | 0.12 | 0.17 | 0.16 | 0.13 | 0.11 | 0.2 |
| 1 | 218763810 | rs28459747 | 1.343 | 0.195 | -1.195 | -0.843 | -0.037 | 0.028 | 0.028 | 15.905 | 0.06 | 0 | 0.13 | 0.02 | 0.12 | 0.11 |
| 1 | 218766058 | rs1481347 | -1.212 | 0.132 | -1.185 | -0.578 | -0.013 | -0.032 | -0.032 | 0.021 | 0.71 | 0.81 | 0.71 | 0.62 | 0.65 | 0.74 |

| | | | | | | | | | | | | | | | | |
|---|---|---|---|---|---|---|---|---|---|---|---|---|---|---|---|---|
| 1 | 218777698 | rs17048282 | 1.175 | 0.016 | -1.125 | -0.595 | -0.002 | -0.003 | -0.003 | 0 | 0.26 | 0.21 | 0.32 | 0.44 | 0.28 | 0.36 |
| 1 | 218786549 | rs2965012 | -0.826 | 0.425 | -1.085 | -0.51 | -0.002 | -0.061 | -0.061 | 0.052 | 0.84 | 0.79 | 0.84 | 0.98 | 0.86 | 0.86 |
| 1 | 218786572 | rs3009943 | 1.223 | 0.496 | -1.699 | -0.631 | -0.126 | -0.143 | -0.143 | 0 | 0.25 | 0.04 | 0.28 | 0.38 | 0.37 | 0.25 |
| 1 | 218794670 | rs729283 | 0.88 | 0.223 | -1.496 | -0.701 | 0.004 | 0.003 | 0.003 | 1.082 | 0.42 | 0.35 | 0.37 | 0.45 | 0.34 | 0.39 |
| 1 | 218810253 | rs1481358 | 1.716 | 0.387 | -1.999 | -0.373 | 0 | -0.026 | -0.026 | 0 | 0.11 | 0.11 | 0.05 | 0 | 0.11 | 0.06 |
| 1 | 218811824 | rs17048343 | 1.178 | 0.316 | -2.248 | 0.008 | -0.079 | -0.115 | -0.115 | 0.002 | 0.19 | 0.04 | 0.24 | 0.39 | 0.29 | 0.22 |
| 1 | 218813191 | rs1026841 | -0.044 | 0.094 | -2.15 | -0.215 | -0.016 | 0.017 | 0.017 | 0 | 0.39 | 0.12 | 0.42 | 0.57 | 0.4 | 0.41 |
| 1 | 218813701 | rs1318827 | -0.058 | -0.051 | -1.262 | 0.021 | 0.029 | -0.006 | -0.006 | 0 | 0.57 | 0.72 | 0.66 | 0.44 | 0.68 | 0.49 |
| 1 | 218827855 | rs1481345 | 0.521 | 0.217 | -1.046 | -0.102 | 0.02 | 0.013 | 0.013 | 0 | 0.5 | 0.32 | 0.44 | 0.49 | 0.53 | 0.46 |
| 1 | 218832257 | rs6690218 | 0.068 | -0.038 | -0.463 | -0.307 | 0 | 0.032 | 0.032 | 0 | 0.66 | 0.54 | 0.54 | 0.49 | 0.64 | 0.56 |
| 1 | 218833214 | rs11485142 | 1.082 | 0.174 | -1.388 | -0.639 | -0.01 | 0.09 | 0.09 | 0.054 | 0.23 | 0.13 | 0.16 | 0 | 0.22 | 0.18 |
| 1 | 218841163 | rs11583395 | 0.285 | 0.457 | -0.966 | -0.284 | -0.006 | -0.006 | -0.006 | 0.7 | 0.39 | 0.35 | 0.37 | 0.48 | 0.43 | 0.35 |
| 1 | 218856744 | rs4601642 | -0.287 | 0.405 | -0.888 | -0.21 | -0.004 | 0.019 | 0.019 | 0 | 0.54 | 0.4 | 0.41 | 0.07 | 0.49 | 0.45 |
| 1 | 218858100 | rs922715 | -0.692 | 0.566 | -0.872 | 0.434 | 0.001 | -0.001 | -0.001 | 0 | 0.46 | 0.48 | 0.56 | 0.56 | 0.5 | 0.52 |
| 1 | 218860068 | rs993925 | -0.128 | 0.716 | -0.711 | 0.888 | 0.006 | 0.021 | 0.021 | 0 | 0.28 | 0.34 | 0.29 | 0.46 | 0.31 | 0.28 |
| 1 | 218866367 | rs1481354 | -0.466 | -0.227 | -0.786 | 1.386 | 0.01 | -0.022 | -0.022 | 0 | 0.18 | 0.15 | 0.12 | 0.39 | 0.12 | 0.2 |
| 1 | 218870348 | rs4078315 | -0.49 | -0.286 | -0.537 | 1.511 | 0.002 | 0.092 | 0.092 | 0.276 | 0.13 | 0.18 | 0.38 | 0.44 | 0.13 | 0.4 |
| 1 | 218879235 | rs17512031 | 0.739 | -0.554 | -0.763 | 0.531 | -0.006 | 0.002 | 0.002 | 0.006 | 0.15 | 0.13 | 0.13 | 0 | 0.17 | 0.11 |
| 1 | 218891136 | rs17048442 | 0.255 | -1.692 | -0.671 | 0.159 | -0.004 | -0.01 | -0.01 | 0 | 0.15 | 0.16 | 0.04 | 0.04 | 0.06 | 0.13 |
| 1 | 218891564 | rs17048445 | 1.087 | -1.945 | -0.161 | 0.155 | 0.075 | 0.1 | 0.1 | 0 | 0.52 | 0.33 | 0.45 | 0.14 | 0.71 | 0.26 |
| 1 | 218902223 | rs6604859 | 0.447 | -1.904 | 0.416 | 0.863 | -0.009 | 0.07 | 0.07 | 0 | 0.82 | 0.71 | 0.63 | 0.63 | 0.86 | 0.57 |
| 1 | 218903928 | rs1031961 | -0.022 | -1.913 | 0.415 | 0.86 | -0.012 | 0.106 | 0.106 | 0.036 | 0.85 | 0.71 | 0.64 | 0.64 | 0.88 | 0.6 |
| 1 | 218906453 | rs4372320 | 1.092 | -2.026 | 0.252 | 1.082 | 0.038 | -0.079 | -0.079 | 0 | 0.13 | 0.02 | 0.08 | 0.42 | 0.06 | 0.17 |
| 1 | 218908668 | rs1481356 | -1.591 | -2.259 | 0.516 | 0.941 | 0.081 | 0.083 | 0.083 | 0 | 0.46 | 0.7 | 0.52 | 0.81 | 0.26 | 0.73 |
| 1 | 218912366 | rs1351472 | -1.024 | -2.307 | 0.701 | 1.375 | -0.009 | 0.084 | 0.084 | 0 | 0.1 | 0.24 | 0.09 | 0.24 | 0.06 | 0.2 |
| 1 | 218912421 | rs1552122 | -0.784 | -2.33 | 0.662 | 1.424 | -0.064 | 0.05 | 0.05 | 0 | 0.32 | 0.45 | 0.31 | 0.53 | 0.18 | 0.44 |
| 1 | 218920738 | rs2378003 | -1.003 | -2.256 | 0.714 | 1.351 | -0.067 | 0.025 | 0.025 | 0 | 0.23 | 0.43 | 0.15 | 0.27 | 0.11 | 0.33 |
| 1 | 218931976 | rs7542694 | 1.496 | -2.244 | 0.437 | 0.987 | -0.008 | -0.007 | -0.007 | 0 | 0.13 | 0.14 | 0.14 | 0 | 0.19 | 0.11 |
| 1 | 218932794 | rs6692647 | -1.46 | -2.44 | 0.492 | 1.103 | 0.052 | 0.027 | 0.027 | 0 | 0.41 | 0.68 | 0.38 | 0.75 | 0.21 | 0.65 |
| 1 | 218932807 | rs6665999 | -0.956 | -2.413 | 0.311 | 1.258 | -0.062 | 0.076 | 0.076 | 0.202 | 0.23 | 0.45 | 0.27 | 0.34 | 0.14 | 0.45 |
| 1 | 218934929 | rs17048517 | 1.177 | -2.209 | 0.319 | 1.83 | 0.039 | -0.03 | -0.03 | 0 | 0.13 | 0.04 | 0.07 | 0.23 | 0.04 | 0.16 |
| 1 | 218936752 | rs9431471 | -0.879 | -2.083 | 0.615 | 1.852 | 0 | 0.187 | 0.187 | 0 | 0.1 | 0.28 | 0.22 | 0.32 | 0.08 | 0.33 |
| 1 | 218938244 | rs9431472 | -1.493 | -1.925 | 0.088 | 1.038 | -0.012 | 0.006 | 0.006 | 0.139 | 0.64 | 0.71 | 0.59 | 0.76 | 0.55 | 0.79 |
| 1 | 218938412 | rs7512685 | -1.113 | -2.276 | -0.219 | 1.185 | 0.009 | 0.009 | 0.009 | 1.114 | 0.47 | 0.65 | 0.59 | 0.63 | 0.39 | 0.72 |
| 1 | 218938961 | rs6541199 | -0.303 | -2.197 | -0.154 | 1.297 | -0.023 | 0.03 | 0.03 | 0.03 | 0.35 | 0.57 | 0.31 | 0.5 | 0.26 | 0.44 |
| 1 | 218940037 | rs12746518 | -0.495 | -2.184 | -0.187 | 1.151 | -0.004 | -0.009 | -0.009 | 0 | 0.11 | 0.04 | 0.16 | 0.1 | 0.09 | 0.23 |
| 1 | 218940225 | rs6698925 | -0.413 | -2.404 | -0.116 | 1.261 | -0.088 | 0 | 0 | 0 | 0.24 | 0.57 | 0.12 | 0.3 | 0.08 | 0.22 |
| 1 | 218940944 | rs12026846 | -0.604 | -2.201 | -0.653 | 1.407 | 0.013 | -0.011 | -0.011 | 0 | 0.27 | 0.24 | 0.24 | 0.41 | 0.14 | 0.41 |
| 1 | 218958105 | rs6683441 | -0.535 | -2.27 | -0.845 | 1.449 | 0.028 | 0.054 | 0.054 | 0.9 | 0.5 | 0.72 | 0.5 | 0.38 | 0.44 | 0.61 |
| 1 | 218960310 | rs6655980 | -1.452 | -2.269 | -0.857 | 1.459 | -0.046 | -0.028 | -0.028 | 0.007 | 0.67 | 0.83 | 0.59 | 0.68 | 0.52 | 0.8 |
| 1 | 218970504 | rs10495107 | -0.473 | -2.211 | -0.811 | 1.459 | -0.002 | 0.013 | 0.013 | 0 | 0.06 | 0.06 | 0.1 | 0.16 | 0.04 | 0.22 |
| 1 | 218975899 | rs11118155 | 1.564 | -2.229 | -0.843 | 1.695 | -0.005 | -0.008 | -0.008 | 0 | 0.05 | 0 | 0.15 | 0.17 | 0.05 | 0.09 |
| 1 | 218982922 | rs7415744 | 0.196 | -2.3 | -0.751 | 1.461 | -0.006 | -0.071 | -0.071 | 0 | 0.39 | 0.57 | 0.3 | 0.15 | 0.37 | 0.25 |
| 1 | 218988754 | rs17514738 | -0.306 | -2.323 | -0.771 | 1.516 | 0.011 | 0.038 | 0.038 | 0.924 | 0.45 | 0.69 | 0.39 | 0.33 | 0.41 | 0.44 |
| 1 | 218992800 | rs12033283 | 1.117 | -2.213 | -0.689 | 1.477 | 0.013 | -0.01 | -0.01 | 0.534 | 0.15 | 0.08 | 0.07 | 0.25 | 0.06 | 0.15 |
| 1 | 218999505 | rs11118161 | 0.289 | -2.239 | -0.66 | 1.493 | 0.006 | -0.009 | -0.009 | 0 | 0.2 | 0.15 | 0.15 | 0.4 | 0.08 | 0.37 |
| 1 | 219001663 | rs6541234 | -0.845 | -2.33 | -0.695 | 1.451 | -0.006 | -0.006 | -0.006 | 0 | 0.59 | 0.7 | 0.45 | 0.57 | 0.47 | 0.6 |
| 1 | 219009835 | rs2647116 | 0.142 | -2.323 | -0.689 | 1.446 | -0.011 | -0.042 | -0.042 | 0 | 0.4 | 0.55 | 0.36 | 0.29 | 0.37 | 0.4 |
| 1 | 219012713 | rs2647115 | -1.453 | -2.323 | -0.701 | 1.397 | -0.048 | -0.034 | -0.034 | 0 | 0.66 | 0.83 | 0.59 | 0.66 | 0.52 | 0.78 |
| 1 | 219013166 | rs4846513 | -0.852 | -2.326 | -0.692 | 1.389 | -0.012 | -0.013 | -0.013 | 0 | 0.59 | 0.75 | 0.45 | 0.57 | 0.47 | 0.58 |
| 1 | 219013302 | rs4846281 | -0.408 | -2.348 | -0.676 | 1.389 | 0.001 | 0 | 0 | 0 | 0.55 | 0.61 | 0.42 | 0.54 | 0.43 | 0.57 |
| 1 | 219020707 | rs12045234 | 1.056 | -2.26 | -1.007 | 1.44 | 0.025 | -0.011 | -0.011 | 0 | 0.15 | 0.07 | 0.06 | 0.24 | 0.06 | 0.15 |
| 1 | 219024035 | rs17574573 | 0.62 | -2.131 | -1.079 | 1.418 | 0.004 | -0.015 | -0.015 | 1.283 | 0.07 | 0.05 | 0.13 | 0.1 | 0.06 | 0.21 |
| 1 | 219036330 | rs617385 | 0.614 | -2.023 | -1.064 | 1.386 | -0.01 | 0.006 | 0.006 | 0 | 0.69 | 0.69 | 0.71 | 0.53 | 0.82 | 0.45 |
| 1 | 219040605 | rs637991 | 0.208 | -2.035 | -1.225 | 1.225 | -0.021 | 0 | 0 | 0 | 0.73 | 0.76 | 0.81 | 0.65 | 0.89 | 0.63 |
| 1 | 219042817 | rs617696 | -0.526 | -1.985 | -1.199 | 1.131 | -0.016 | 0.001 | 0.001 | 0 | 0.79 | 0.8 | 0.9 | 0.74 | 0.88 | 0.83 |
| 1 | 219050679 | rs1909099 | 1.157 | -1.951 | -1.147 | 1.118 | 0.019 | -0.008 | -0.008 | 0 | 0.15 | 0.06 | 0.06 | 0.23 | 0.06 | 0.15 |
| 1 | 219051070 | rs924270 | 0.81 | -1.616 | -1.157 | 1.219 | 0.011 | -0.06 | -0.06 | 0.373 | 0.3 | 0.51 | 0.26 | 0.18 | 0.34 | 0.21 |
| 1 | 219053292 | rs650681 | -0.737 | -1.493 | -0.928 | 1.001 | -0.023 | -0.004 | -0.004 | 0.575 | 0.77 | 0.81 | 0.9 | 0.76 | 0.91 | 0.84 |
| 1 | 219053400 | rs17516059 | 0.229 | -1.409 | -1.005 | 1.035 | -0.014 | -0.015 | -0.015 | 0.194 | 0.11 | 0.06 | 0.16 | 0.12 | 0.13 | 0.21 |
| 1 | 219053465 | rs649752 | 1.324 | -0.89 | -0.955 | 0.976 | -0.013 | 0.004 | 0.004 | 0 | 0.26 | 0.16 | 0.23 | 0.19 | 0.35 | 0.11 |
| 1 | 219053976 | rs647485 | 0.795 | -1.288 | -1.011 | 0.957 | 0.008 | -0.002 | -0.002 | 0 | 0.63 | 0.69 | 0.69 | 0.54 | 0.76 | 0.42 |
| 1 | 219054054 | rs636764 | 0.795 | -1.288 | -1.011 | 0.957 | 0.008 | -0.002 | -0.002 | 0 | 0.63 | 0.69 | 0.69 | 0.54 | 0.76 | 0.42 |
| 1 | 219055281 | rs676416 | 0.56 | -1.084 | -1.028 | 0.76 | 0.002 | 0 | 0 | 0 | 0.66 | 0.71 | 0.73 | 0.63 | 0.78 | 0.63 |
| 1 | 219060101 | rs882373 | -0.437 | -0.78 | -1.879 | 0.009 | 0.018 | 0.114 | 0.114 | 0 | 0.28 | 0.07 | 0.2 | 0.07 | 0.25 | 0.24 |
| 1 | 219062526 | rs3897852 | -0.991 | -0.587 | -1.694 | 1.007 | 0.011 | -0.034 | -0.034 | 0 | 0.34 | 0.15 | 0.27 | 0.43 | 0.31 | 0.39 |
| 1 | 219071000 | rs11118172 | -1.243 | -0.911 | -0.91 | 1.053 | 0.006 | 0.002 | 0.002 | 0.023 | 0.45 | 0.59 | 0.44 | 0.55 | 0.38 | 0.56 |
| 1 | 219072116 | rs1338457 | 0.427 | -1.037 | -1.011 | 0.675 | -0.027 | 0.121 | 0.121 | 1.849 | 0.2 | 0.01 | 0.12 | 0.01 | 0.19 | 0.19 |
| 1 | 219072415 | rs1338458 | -0.221 | -0.968 | -0.855 | 0.732 | 0.024 | 0.096 | 0.096 | 0.016 | 0.28 | 0.09 | 0.2 | 0.07 | 0.25 | 0.25 |

| | | | | | | | | | | | | | | | | |
|---|---|---|---|---|---|---|---|---|---|---|---|---|---|---|---|---|
| 1 | 219073958 | rs878394 | -1.604 | -1.098 | -1.175 | 0.677 | -0.001 | 0.004 | 0.004 | 0 | 0.56 | 0.59 | 0.61 | 0.8 | 0.43 | 0.63 |
| 1 | 219083060 | rs9661984 | -0.836 | -1.223 | -1.46 | 0.774 | 0.033 | -0.041 | -0.041 | 0 | 0.39 | 0.16 | 0.27 | 0.51 | 0.28 | 0.41 |
| 1 | 219085469 | rs11118174 | -0.899 | -1.166 | -1.691 | 0.243 | 0.037 | -0.02 | -0.02 | 0.046 | 0.52 | 0.17 | 0.36 | 0.58 | 0.41 | 0.6 |
| 1 | 219088165 | rs7540766 | -1.962 | -1.199 | -1.18 | -0.114 | -0.009 | -0.019 | -0.019 | 0.202 | 0.69 | 0.69 | 0.7 | 0.96 | 0.58 | 0.83 |
| 1 | 219088578 | rs17517067 | 0.944 | -1.104 | -1.417 | -0.012 | 0.018 | -0.003 | -0.003 | 0 | 0.05 | 0.04 | 0.06 | 0.03 | 0.09 | 0.06 |
| 1 | 219088779 | rs10495112 | -0.618 | -1.21 | -1.492 | 0.152 | 0.056 | 0.15 | 0.15 | 0 | 0.48 | 0.16 | 0.3 | 0.27 | 0.39 | 0.49 |
| 1 | 219089006 | rs10495114 | -0.115 | -1.216 | -1.511 | 0.149 | 0.104 | 0.149 | 0.149 | 0 | 0.43 | 0.11 | 0.24 | 0.24 | 0.31 | 0.43 |
| 1 | 219089109 | rs10495115 | 0.872 | -1.254 | -1.571 | 0.124 | 0.026 | 0.038 | 0.038 | 0 | 0.21 | 0.07 | 0.16 | 0.15 | 0.2 | 0.25 |
| 1 | 219089114 | rs17576463 | 0.768 | -1.015 | -1.444 | 0.217 | -0.002 | 0.002 | 0.002 | 0 | 0.05 | 0.02 | 0.06 | 0.03 | 0.09 | 0.06 |
| 1 | 219089313 | rs11118175 | 0.4 | -1.497 | -0.903 | 0.466 | 0.048 | 0.083 | 0.083 | 0 | 0.06 | 0.22 | 0.16 | 0.12 | 0.09 | 0.15 |
| 1 | 219094943 | rs1933661 | -0.812 | -1.537 | -1.057 | 0.288 | 0.022 | 0.017 | 0.017 | 0 | 0.53 | 0.36 | 0.46 | 0.68 | 0.4 | 0.69 |
| 1 | 219095423 | rs7525547 | -0.621 | -1.558 | -1.02 | 0.613 | -0.034 | 0.027 | 0.027 | 0 | 0.33 | 0.56 | 0.37 | 0.41 | 0.29 | 0.35 |
| 1 | 219099712 | rs12122701 | 0.737 | -0.792 | -1.96 | 0.602 | -0.026 | -0.057 | -0.057 | 0.93 | 0.32 | 0.02 | 0.29 | 0.37 | 0.25 | 0.28 |
| 1 | 219101174 | rs639907 | -0.334 | -1.013 | -1.048 | 0.448 | 0.056 | 0.052 | 0.052 | 4.379 | 0.95 | 0.46 | 0.97 | 1 | 1 | 1 |
| 1 | 219101848 | rs626518 | 0.498 | -0.768 | -1.834 | 0.849 | -0.081 | -0.025 | -0.025 | 0.386 | 0.39 | 0.13 | 0.49 | 0.33 | 0.51 | 0.52 |
| 1 | 219104226 | rs11118178 | 0.632 | -0.826 | -1.621 | 0.741 | -0.001 | 0.005 | 0.005 | 0 | 0.17 | 0.21 | 0.1 | 0.27 | 0.18 | 0.18 |
| 1 | 219106144 | rs12124340 | 1.709 | -0.808 | -2.061 | -0.072 | 0.009 | 0.116 | 0.116 | 0 | 0.22 | 0.02 | 0.16 | 0.02 | 0.16 | 0.13 |
| 1 | 219106550 | rs10746381 | -0.991 | -0.815 | -1.888 | -0.095 | 0.006 | -0.069 | -0.069 | 1.205 | 0.89 | 0.36 | 0.92 | 1 | 0.93 | 0.98 |
| 1 | 219110621 | rs631060 | 0.274 | -0.98 | -2.1 | -0.079 | 0.015 | 0.015 | 0.015 | 1.91 | 0.8 | 0.16 | 0.88 | 0.92 | 0.91 | 0.84 |
| 1 | 219117743 | rs2788271 | 1.339 | -1.063 | -2.217 | 0.208 | 0.051 | -0.07 | -0.07 | 0 | 0.68 | 0.15 | 0.64 | 0.5 | 0.84 | 0.69 |
| 1 | 219117863 | rs11118180 | 0.079 | -0.954 | -2.216 | 0.251 | -0.006 | -0.069 | -0.069 | 0 | 0.12 | 0.02 | 0.24 | 0.42 | 0.08 | 0.15 |
| 1 | 219126448 | rs638699 | 1.388 | -1.355 | -1.929 | 0.446 | -0.027 | 0.178 | 0.178 | 1.99 | 0.4 | 0.1 | 0.36 | 0.14 | 0.59 | 0.42 |
| 1 | 219135058 | rs10495117 | 1.863 | -1.303 | -1.956 | 0.467 | 0.024 | 0.09 | 0.09 | 0 | 0.22 | 0.04 | 0.16 | 0.02 | 0.16 | 0.13 |
| 1 | 219135985 | rs11118183 | 1.506 | -1.36 | -1.955 | 0.614 | 0.007 | 0.043 | 0.043 | 0 | 0.06 | 0.02 | 0.22 | 0.42 | 0.02 | 0.13 |
| 1 | 219138400 | rs662150 | 1.361 | -1.376 | -1.976 | 0.559 | -0.021 | 0.177 | 0.177 | 1.353 | 0.41 | 0.12 | 0.36 | 0.14 | 0.59 | 0.42 |
| 1 | 219141960 | rs601175 | 0.874 | -1.413 | -1.94 | 0.601 | 0 | -0.001 | -0.001 | 0.168 | 0.75 | 0.22 | 0.66 | 0.51 | 0.89 | 0.71 |
| 1 | 219143598 | rs629222 | 0.329 | -1.372 | -1.968 | 0.616 | 0.087 | -0.007 | -0.007 | 0.078 | 0.53 | 0.16 | 0.6 | 0.57 | 0.67 | 0.57 |
| 1 | 219149298 | rs618171 | 0.284 | -1.391 | -1.961 | 0.612 | 0.109 | -0.027 | -0.027 | 0 | 0.53 | 0.17 | 0.5 | 0.5 | 0.73 | 0.58 |
| 1 | 219150241 | rs17048938 | 1.565 | -1.323 | -1.926 | 0.654 | 0.006 | 0.048 | 0.048 | 0 | 0.06 | 0.01 | 0.22 | 0.41 | 0.02 | 0.13 |
| 1 | 219150622 | rs11581158 | 1.59 | -1.278 | -1.898 | 0.464 | -0.011 | 0.018 | 0.018 | 0 | 0.06 | 0 | 0.02 | 0.02 | 0.05 | 0.02 |
| 1 | 219163360 | rs6703753 | 0.575 | -1.354 | -1.791 | 0.404 | -0.001 | -0.043 | -0.043 | 0 | 0.33 | 0.66 | 0.24 | 0.02 | 0.23 | 0.16 |
| 1 | 219176608 | rs7540160 | -0.393 | -1.237 | -1.754 | 0.703 | 0.044 | -0.027 | -0.027 | 0 | 0.42 | 0.23 | 0.34 | 0.5 | 0.31 | 0.43 |
| 1 | 219187143 | rs17048983 | -0.074 | -1.131 | -1.718 | 0.813 | -0.018 | -0.05 | -0.05 | 0 | 0.12 | 0.04 | 0.14 | 0.36 | 0.14 | 0.16 |
| 1 | 219189306 | rs17048992 | 0.591 | -1.246 | -1.665 | 0.803 | -0.033 | 0.003 | 0.003 | 0 | 0.08 | 0.18 | 0.04 | 0.12 | 0.02 | 0.14 |
| 1 | 219191704 | rs1256590 | 0.192 | -1.155 | -1.618 | 0.81 | -0.022 | 0.146 | 0.146 | 2.255 | 0.17 | 0.63 | 0.3 | 0.35 | 0.09 | 0.15 |
| 1 | 219221780 | rs6690339 | 0.201 | -1.232 | -1.759 | 0.684 | 0.024 | 0.034 | 0.034 | 0.207 | 0.36 | 0.25 | 0.22 | 0.16 | 0.23 | 0.29 |
| 1 | 219222948 | rs17041707 | 0.994 | -1.15 | -1.805 | 0.792 | -0.007 | 0.03 | 0.03 | 0.096 | 0.06 | 0.02 | 0.1 | 0.35 | 0.08 | 0.13 |
| 1 | 219223317 | rs871107 | -0.91 | -1.193 | -1.821 | 0.785 | 0.074 | -0.04 | -0.04 | 0 | 0.53 | 0.87 | 0.54 | 0.51 | 0.32 | 0.43 |
| 1 | 219240970 | rs1256603 | -0.497 | -0.712 | -1.266 | 0.758 | 0.036 | 0.062 | 0.062 | 0.321 | 0.45 | 0.69 | 0.58 | 0.71 | 0.33 | 0.41 |
| 1 | 219247211 | rs12041410 | -0.729 | -0.182 | -0.336 | 1.454 | -0.022 | 0.228 | 0.228 | 1.888 | 0.17 | 0.59 | 0.39 | 0.43 | 0.09 | 0.14 |
| 1 | 219273881 | rs12141207 | 0.449 | 0.577 | -0.766 | 0.392 | -0.007 | -0.006 | -0.006 | 0 | 0.6 | 0.36 | 0.45 | 0.59 | 0.49 | 0.58 |
| 1 | 219298539 | rs7545152 | -0.664 | 0.833 | -0.314 | 0.223 | 0 | 0.023 | 0.023 | 0 | 0.17 | 0.21 | 0.33 | 0.33 | 0.16 | 0.2 |
| 1 | 219306923 | rs12067803 | -0.782 | 0.67 | -0.373 | -0.011 | 0 | 0.016 | 0.016 | 0.696 | 0.19 | 0.2 | 0.4 | 0.34 | 0.18 | 0.15 |
| 1 | 219325260 | rs17521251 | 0.721 | 0.796 | -0.302 | -0.071 | -0.029 | 0.03 | 0.03 | 0 | 0.07 | 0.01 | 0.05 | 0 | 0.11 | 0.03 |
| 1 | 219337294 | rs9308387 | -0.152 | 0.763 | -0.011 | -0.146 | 0.058 | -0.099 | -0.099 | 0 | 0.75 | 0.5 | 0.76 | 0.91 | 0.63 | 0.74 |
| 1 | 219350309 | rs7536671 | 0.367 | 0.643 | 0.572 | -0.202 | 0.057 | -0.064 | -0.064 | 0 | 0.21 | 0.33 | 0.23 | 0.08 | 0.36 | 0.21 |
| 1 | 219350856 | rs7536586 | 1.778 | 0.277 | -0.198 | -0.493 | 0.044 | -0.073 | -0.073 | 0.693 | 0.44 | 0.02 | 0.45 | 0.59 | 0.34 | 0.41 |
| 1 | 219355789 | rs6699816 | -0.333 | 0.583 | 0.1 | -0.239 | 0.001 | 0.023 | 0.023 | 0 | 0.14 | 0.17 | 0.15 | 0.27 | 0.12 | 0.12 |
| 1 | 219374030 | rs12402039 | -0.511 | 1.653 | 0.071 | -0.29 | 0.007 | 0.007 | 0.007 | 0 | 0.16 | 0.17 | 0.22 | 0.27 | 0.26 | 0.14 |
| 1 | 219383192 | rs6692892 | 0.997 | 1.316 | 0.156 | -0.354 | -0.027 | 0.005 | 0.005 | 0 | 0.61 | 0.3 | 0.6 | 0.64 | 0.51 | 0.61 |
| 1 | 219392381 | rs4846539 | 0.882 | 1.284 | 0.41 | -0.37 | -0.007 | 0 | 0 | 0 | 0.62 | 0.35 | 0.61 | 0.64 | 0.51 | 0.62 |
| 1 | 219412715 | rs17588395 | 1.218 | 1.238 | -0.024 | -0.404 | -0.051 | 0.017 | 0.017 | 0 | 0.1 | 0.01 | 0.1 | 0.05 | 0.13 | 0.13 |
| 1 | 219446253 | rs11118250 | 0.972 | 1.18 | 0.221 | -0.488 | -0.023 | 0.01 | 0.01 | 0 | 0.6 | 0.26 | 0.59 | 0.64 | 0.5 | 0.59 |
| 1 | 219464144 | rs10863444 | 0.857 | 1.159 | 0.259 | -0.58 | -0.011 | 0.043 | 0.043 | 0 | 0.61 | 0.32 | 0.67 | 0.79 | 0.48 | 0.62 |
| 1 | 219466670 | rs12080306 | 0.562 | 1.022 | 0.501 | -0.581 | 0.021 | -0.04 | -0.04 | 0 | 0.08 | 0.13 | 0.06 | 0 | 0.11 | 0.03 |
| 1 | 219466796 | rs12030447 | 1.457 | 0.829 | 0.427 | -0.61 | 0.09 | 0.027 | 0.027 | 0 | 0.48 | 0.23 | 0.36 | 0.59 | 0.35 | 0.5 |
| 1 | 219470069 | rs6657322 | 0.799 | 0.774 | 0.576 | -0.578 | -0.031 | 0.015 | 0.015 | 1.646 | 0.09 | 0.15 | 0.02 | 0.14 | 0.01 | 0.11 |
| 1 | 219470417 | rs6665677 | -1.4 | 0.967 | 0.571 | -0.702 | 0.001 | -0.011 | -0.011 | 0 | 0.92 | 0.89 | 0.93 | 0.98 | 0.92 | 0.9 |
| 1 | 219477443 | rs6671444 | 0.199 | 0.979 | 0.629 | -0.729 | 0.022 | -0.063 | -0.063 | 0 | 0.84 | 0.73 | 0.88 | 0.98 | 0.81 | 0.87 |
| 1 | 219479238 | rs12086585 | 0.832 | 0.896 | 0.622 | -0.647 | -0.031 | 0.05 | 0.05 | 1.32 | 0.09 | 0.15 | 0.08 | 0.24 | 0.01 | 0.11 |
| 1 | 219480797 | rs7538186 | -1.373 | 1.09 | 0.582 | -0.714 | 0.002 | -0.013 | -0.013 | 0 | 0.92 | 0.88 | 0.93 | 0.98 | 0.92 | 0.9 |
| 1 | 219482935 | rs10158029 | 0.05 | 1.111 | 0.558 | -0.74 | 0.029 | -0.014 | -0.014 | 0.442 | 0.26 | 0.3 | 0.44 | 0.16 | 0.45 | 0.26 |
| 1 | 219484416 | rs4846547 | 0.41 | 1.136 | 0.671 | -0.81 | 0.002 | 0.001 | 0.001 | 0 | 0.21 | 0.2 | 0.39 | 0.16 | 0.4 | 0.23 |
| 1 | 219484671 | rs4846293 | -0.496 | 1.005 | 0.581 | -0.855 | 0.008 | -0.007 | -0.007 | 0 | 0.3 | 0.3 | 0.45 | 0.16 | 0.47 | 0.31 |
| 1 | 219490301 | rs11118272 | 1.407 | 0.94 | 0.773 | -0.895 | 0.046 | 0.069 | 0.069 | 0.152 | 0.54 | 0.31 | 0.42 | 0.81 | 0.33 | 0.56 |
| 1 | 219497096 | rs11118275 | 0.286 | 1.045 | 0.729 | -0.755 | 0.045 | -0.092 | -0.092 | 6.336 | 0.83 | 0.63 | 0.87 | 0.97 | 0.8 | 0.87 |
| 1 | 219507866 | rs11118277 | 1.07 | 0.894 | 0.708 | -0.656 | 0.026 | -0.031 | -0.031 | 0 | 0.08 | 0.21 | 0.07 | 0.03 | 0.09 | 0.1 |
| 1 | 219508319 | rs1531090 | 1.869 | 0.534 | 0.549 | -0.845 | 0.046 | -0.002 | -0.002 | 0 | 0.41 | 0.23 | 0.34 | 0.58 | 0.31 | 0.41 |
| 1 | 219516224 | rs4846295 | 0.385 | 0.572 | 0.54 | -0.953 | -0.012 | 0.007 | 0.007 | 0.05 | 0.24 | 0.22 | 0.44 | 0.13 | 0.41 | 0.27 |

| | | | | | | | | | | | | | | | | |
|---|---|---|---|---|---|---|---|---|---|---|---|---|---|---|---|---|
| 1 | 219520895 | rs13376087 | 0.871 | 0.802 | 0.594 | -0.966 | -0.025 | 0.01 | 0.01 | 0.527 | 0.2 | 0.07 | 0.36 | 0.13 | 0.25 | 0.25 |
| 1 | 219533817 | rs10779349 | 0.792 | 0.711 | 0.839 | -0.981 | -0.001 | -0.005 | -0.005 | 0 | 0.59 | 0.38 | 0.49 | 0.84 | 0.57 | 0.54 |
| 1 | 219539066 | rs7553882 | 0.598 | 0.833 | 0.616 | -0.693 | -0.012 | -0.013 | -0.013 | 0 | 0.11 | 0.07 | 0.13 | 0.27 | 0.17 | 0.11 |
| 1 | 219544174 | rs17005342 | 0.598 | 0.833 | 0.617 | -0.693 | -0.016 | -0.019 | -0.019 | 0 | 0.11 | 0.07 | 0.13 | 0.28 | 0.17 | 0.11 |
| 1 | 219547050 | rs6670103 | 1.86 | 0.727 | 0.563 | -0.669 | 0.056 | -0.019 | -0.019 | 0 | 0.42 | 0.21 | 0.3 | 0.51 | 0.31 | 0.42 |
| 1 | 219548373 | rs6666109 | -0.53 | 0.82 | 0.441 | -0.654 | -0.012 | 0.039 | 0.039 | 0 | 0.17 | 0.12 | 0.1 | 0.06 | 0.19 | 0.14 |
| 1 | 219557659 | rs10495123 | 0.647 | 0.833 | 0.483 | -0.639 | -0.001 | -0.007 | -0.007 | 1.404 | 0.1 | 0.12 | 0.06 | 0.06 | 0.11 | 0.06 |
| 1 | 219557811 | rs6671991 | -0.065 | 0.847 | 0.558 | -0.666 | -0.018 | 0.037 | 0.037 | 0.399 | 0.27 | 0.16 | 0.41 | 0.13 | 0.32 | 0.3 |
| 1 | 219564357 | rs1547502 | -1.556 | 0.755 | 0.811 | -0.67 | 0.003 | 0.001 | 0.001 | 0 | 0.5 | 0.5 | 0.62 | 0.46 | 0.6 | 0.47 |
| 1 | 219565545 | rs1340101 | -0.076 | 0.81 | 0.889 | -0.662 | 0.038 | -0.076 | -0.076 | 0 | 0.82 | 0.62 | 0.86 | 0.91 | 0.8 | 0.84 |
| 1 | 219570103 | rs10863457 | -0.392 | 0.872 | 0.746 | -0.602 | 0.015 | 0.014 | 0.014 | 0 | 0.33 | 0.37 | 0.51 | 0.4 | 0.42 | 0.33 |
| 1 | 219577106 | rs12759274 | 1.753 | 0.318 | 0.537 | -1.031 | 0.03 | 0.09 | 0.09 | 0 | 0.34 | 0.14 | 0.19 | 0.09 | 0.28 | 0.29 |
| 1 | 219577622 | rs12123734 | 1.764 | 0.318 | 0.532 | -1.032 | 0.032 | 0.126 | 0.126 | 0 | 0.34 | 0.11 | 0.19 | 0.09 | 0.28 | 0.29 |
| 1 | 219577777 | rs11118293 | 0.785 | 0.491 | 0.468 | -0.989 | -0.005 | -0.053 | -0.053 | 0 | 0.24 | 0.11 | 0.46 | 0.53 | 0.26 | 0.34 |
| 1 | 219578414 | rs2820461 | -0.915 | 0.503 | 0.57 | -1.008 | 0 | -0.035 | -0.035 | 0 | 0.48 | 0.47 | 0.66 | 0.81 | 0.52 | 0.54 |
| 1 | 219579930 | rs17005406 | 0.677 | 0.736 | 0.181 | -0.823 | 0.01 | -0.023 | -0.023 | 0.692 | 0.1 | 0.19 | 0.07 | 0.07 | 0.11 | 0.08 |
| 1 | 219580434 | rs12081796 | 0.035 | 0.65 | 0.066 | -0.794 | -0.003 | -0.003 | -0.003 | 0.126 | 0.17 | 0.19 | 0.14 | 0.1 | 0.2 | 0.17 |
| 1 | 219583791 | rs2791558 | -0.353 | 0.673 | -0.152 | -0.746 | -0.012 | -0.014 | -0.014 | 0 | 0.2 | 0.15 | 0.15 | 0.28 | 0.24 | 0.2 |
| 1 | 219586124 | rs1001329 | -0.36 | 1.197 | 0.445 | -0.294 | 0.026 | -0.055 | -0.055 | 0 | 0.18 | 0.37 | 0.16 | 0.11 | 0.2 | 0.17 |
| 1 | 219586415 | rs12032159 | 0.014 | 1.154 | -0.168 | 0.145 | 0.001 | -0.012 | -0.012 | 0 | 0.25 | 0.2 | 0.48 | 0.52 | 0.25 | 0.34 |
| 1 | 219588803 | rs2791559 | -0.66 | 1.09 | -0.74 | -0.103 | -0.016 | 0.024 | 0.024 | 0 | 0.33 | 0.25 | 0.59 | 0.8 | 0.42 | 0.45 |
| 1 | 219590047 | rs4846300 | -0.672 | 0.547 | -1.151 | 0.176 | -0.009 | -0.016 | -0.016 | 0 | 0.34 | 0.21 | 0.54 | 0.59 | 0.36 | 0.41 |
| 1 | 219591948 | rs4846301 | -0.305 | -0.223 | -1.188 | -0.089 | 0.025 | 0.031 | 0.031 | 0.77 | 0.18 | 0.08 | 0.15 | 0.07 | 0.12 | 0.16 |
| 1 | 219596070 | rs2820455 | -1.512 | -1.081 | -1.225 | 0.203 | 0.023 | -0.111 | -0.111 | 0 | 0.74 | 0.96 | 0.73 | 0.54 | 0.79 | 0.74 |
| 1 | 219598894 | rs12119837 | 0.32 | -0.313 | -1.351 | 0.045 | -0.094 | 0.072 | 0.072 | 3.028 | 0.3 | 0.11 | 0.39 | 0.15 | 0.4 | 0.35 |
| 1 | 219602093 | rs2605087 | 0.03 | -0.394 | -0.681 | 0.025 | -0.025 | 0.042 | 0.042 | 1.65 | 0.33 | 0.33 | 0.42 | 0.15 | 0.41 | 0.38 |
| 1 | 219604453 | rs17005517 | 0.369 | -0.454 | 0.001 | 0.004 | 0.04 | 0.032 | 0.032 | 0 | 0.17 | 0.26 | 0.14 | 0.26 | 0.26 | 0.15 |
| 1 | 219604950 | rs2085261 | -0.632 | -0.477 | -0.185 | 0.059 | 0.026 | 0.012 | 0.012 | 0.362 | 0.5 | 0.6 | 0.55 | 0.41 | 0.67 | 0.52 |
| 1 | 219619538 | rs2605082 | -0.947 | -0.012 | 0.196 | 0.54 | 0.018 | 0.027 | 0.027 | 0 | 0.28 | 0.5 | 0.26 | 0.37 | 0.31 | 0.2 |
| 1 | 219622596 | rs6669442 | 0.524 | 0.31 | 0.442 | 0.47 | 0.012 | 0.007 | 0.007 | 0.002 | 0.67 | 0.4 | 0.5 | 0.45 | 0.64 | 0.76 |
| 1 | 219625808 | rs6541226 | 0.062 | 0.293 | 0.425 | 0.322 | 0.002 | -0.007 | -0.007 | 0 | 0.11 | 0.16 | 0.08 | 0.11 | 0.11 | 0.13 |
| 1 | 219636951 | rs2820437 | 0.438 | 0.158 | 0.778 | 0.55 | -0.021 | 0.174 | 0.174 | 0.143 | 0.12 | 0.26 | 0.16 | 0.41 | 0.05 | 0.06 |
| 1 | 219641900 | rs2605096 | 0.219 | -0.141 | 0.504 | 0.316 | 0.009 | -0.014 | -0.014 | 0.094 | 0.7 | 0.48 | 0.57 | 0.82 | 0.66 | 0.71 |
| 1 | 219642109 | rs2605097 | -0.713 | -0.153 | 0.596 | 0.319 | 0.024 | -0.005 | -0.005 | 0.343 | 0.78 | 0.65 | 0.66 | 0.85 | 0.69 | 0.82 |
| 1 | 219644224 | rs2605100 | -1.02 | -0.149 | 0.649 | 0.302 | -0.049 | -0.007 | -0.007 | 2.022 | 0.8 | 0.91 | 0.68 | 0.85 | 0.7 | 0.84 |
| 1 | 219648119 | rs11804949 | 0.941 | -0.137 | 0.644 | 0.296 | -0.028 | 0 | 0 | 3.76 | 0.08 | 0.17 | 0.09 | 0.03 | 0.03 | 0.11 |
| 1 | 219651133 | rs12022722 | 1.816 | -0.013 | 0.288 | 0.28 | -0.132 | 0.161 | 0.161 | 1.367 | 0.46 | 0.02 | 0.33 | 0.3 | 0.5 | 0.52 |
| 1 | 219653101 | rs1563355 | 0.374 | -0.068 | 0.794 | 0.504 | 0 | -0.007 | -0.007 | 0 | 0.69 | 0.41 | 0.57 | 0.82 | 0.67 | 0.7 |
| 1 | 219660535 | rs2820426 | 1.016 | 0.014 | 0.504 | 0.314 | 0.007 | 0.034 | 0.034 | 0 | 0.59 | 0.3 | 0.43 | 0.4 | 0.62 | 0.67 |
| 1 | 219668252 | rs1538742 | 1.552 | -0.07 | 0.369 | 0.162 | -0.001 | 0.123 | 0.123 | 0 | 0.53 | 0.13 | 0.4 | 0.38 | 0.59 | 0.62 |
| 1 | 219669226 | rs765751 | 0.89 | 0.026 | 0.72 | 0.18 | -0.001 | -0.007 | -0.007 | 0 | 0.61 | 0.56 | 0.44 | 0.39 | 0.62 | 0.69 |
| 1 | 219675016 | rs2820466 | -0.299 | 0.134 | 0.845 | 0.155 | 0.052 | 0.007 | 0.007 | 0 | 0.06 | 0.27 | 0.03 | 0 | 0.03 | 0.08 |
| 1 | 219686440 | rs12031785 | 1.577 | 0.599 | 0.39 | 0.039 | -0.011 | 0.165 | 0.165 | 0 | 0.49 | 0.09 | 0.31 | 0.34 | 0.59 | 0.59 |
| 1 | 219697819 | rs6696581 | 0.339 | 0.428 | 0.874 | 0.336 | -0.048 | 0.21 | 0.21 | 2.528 | 0.15 | 0.36 | 0.16 | 0.42 | 0.06 | 0.09 |
| 1 | 219700063 | rs2820460 | -0.462 | 0.391 | 0.647 | 0.233 | -0.021 | -0.011 | -0.011 | 0 | 0.29 | 0.11 | 0.49 | 0.25 | 0.33 | 0.23 |
| 1 | 219700334 | rs6704014 | 2.431 | 0.473 | 0.383 | 0.032 | -0.013 | 0.133 | 0.133 | 0 | 0.41 | 0.13 | 0.19 | 0.26 | 0.35 | 0.47 |
| 1 | 219700471 | rs2785979 | -1.9 | 0.49 | 0.384 | 0.051 | 0.007 | 0.14 | 0.14 | 0 | 0.51 | 0.71 | 0.76 | 0.74 | 0.55 | 0.43 |
| 1 | 219705205 | rs4500327 | -0.063 | 0.564 | 0.204 | -0.608 | -0.066 | -0.023 | -0.023 | 3.348 | 0.27 | 0.05 | 0.46 | 0.2 | 0.32 | 0.22 |
| 1 | 219706428 | rs10746385 | 1.411 | 0.493 | 0.486 | -0.694 | 0.003 | 0.019 | 0.019 | 0.304 | 0.58 | 0.34 | 0.41 | 0.75 | 0.57 | 0.63 |
| 1 | 219707074 | rs2485301 | 1.029 | 0.85 | 0.117 | -0.755 | -0.047 | 0.035 | 0.035 | 2.626 | 0.12 | 0.03 | 0.18 | 0.05 | 0.25 | 0.15 |
| 1 | 219708773 | rs1572505 | -0.099 | 0.832 | 0.279 | -0.625 | -0.067 | -0.055 | -0.055 | 0 | 0.27 | 0.05 | 0.48 | 0.24 | 0.33 | 0.23 |
| 1 | 219714999 | rs2820449 | -0.162 | 0.832 | 0.347 | -0.625 | -0.025 | -0.017 | -0.017 | 0.646 | 0.27 | 0.12 | 0.49 | 0.24 | 0.33 | 0.22 |
| 1 | 219723599 | rs1932963 | 2.434 | 0.814 | 0.192 | -0.651 | -0.057 | -0.029 | -0.029 | 0 | 0.41 | 0.04 | 0.27 | 0.68 | 0.35 | 0.48 |
| 1 | 219724937 | rs1337100 | 0.572 | 0.705 | 0.564 | -0.631 | -0.002 | -0.041 | -0.041 | 0.092 | 0.09 | 0.15 | 0.04 | 0 | 0.09 | 0.1 |
| 1 | 219727779 | rs3001032 | -0.77 | 0.782 | 0.793 | -0.608 | -0.007 | -0.004 | -0.004 | 0 | 0.31 | 0.48 | 0.53 | 0.25 | 0.32 | 0.24 |
| 1 | 219736063 | rs9431182 | -0.349 | 0.866 | 0.442 | -0.59 | -0.126 | -0.048 | -0.048 | 0.092 | 0.27 | 0.05 | 0.54 | 0.24 | 0.46 | 0.22 |
| 1 | 219739907 | rs11118343 | 2.244 | 1.081 | 0.57 | -0.487 | -0.016 | -0.022 | -0.022 | 0 | 0.45 | 0.06 | 0.27 | 0.66 | 0.37 | 0.5 |
| 1 | 219739966 | rs4428898 | 1.522 | 1.075 | 0.62 | -0.49 | -0.054 | 0.004 | 0.004 | 0.588 | 0.54 | 0.13 | 0.3 | 0.67 | 0.46 | 0.58 |
| 1 | 219742768 | rs1415289 | 0.545 | 1.193 | 0.98 | -0.531 | 0.012 | -0.055 | -0.055 | 0 | 0.78 | 0.41 | 0.81 | 0.91 | 0.75 | 0.78 |
| 1 | 219748779 | rs2785991 | 1.258 | 1.193 | 1.118 | -0.53 | 0.062 | 0.014 | 0.014 | 0.137 | 0.9 | 0.75 | 0.88 | 0.91 | 0.81 | 0.82 |
| 1 | 219751388 | rs11118350 | 0.805 | 1.169 | 1.132 | -0.549 | 0.006 | -0.057 | -0.057 | 0 | 0.77 | 0.35 | 0.8 | 0.91 | 0.75 | 0.78 |
| 1 | 219759378 | rs17006168 | 0.929 | 1.134 | 1.124 | -0.536 | -0.028 | -0.076 | -0.076 | 0 | 0.13 | 0.29 | 0.08 | 0 | 0.06 | 0.04 |
| 1 | 219759682 | rs4373767 | 2.231 | 1.141 | 1.035 | -0.54 | 0.01 | 0.011 | 0.011 | 0.621 | 0.46 | 0.09 | 0.27 | 0.67 | 0.38 | 0.53 |
| 1 | 219760467 | rs3897377 | 0.865 | 0.815 | 1.018 | -0.521 | -0.026 | -0.086 | -0.086 | 0 | 0.13 | 0.41 | 0.08 | 0 | 0.06 | 0.04 |
| 1 | 219762581 | rs2494196 | 0.035 | 1.037 | 0.834 | -0.444 | -0.001 | -0.001 | -0.001 | 1.862 | 0.23 | 0.2 | 0.49 | 0.23 | 0.28 | 0.2 |
| 1 | 219770722 | rs6696190 | 1.174 | 1.026 | 0.582 | -0.377 | 0.044 | 0.02 | 0.02 | 0 | 0.91 | 0.79 | 0.82 | 0.88 | 0.82 | 0.84 |
| 1 | 219786890 | rs10779363 | 1.291 | 0.679 | 0.488 | -0.463 | -0.01 | 0.001 | 0.001 | 0.982 | 0.6 | 0.63 | 0.38 | 0.67 | 0.46 | 0.55 |
| 1 | 219791385 | rs1572506 | 0.436 | -0.581 | 0.296 | -1.001 | -0.03 | -0.091 | -0.091 | 0.015 | 0.75 | 0.6 | 0.8 | 0.96 | 0.84 | 0.84 |
| 1 | 219792920 | rs7553734 | 0.845 | -0.405 | 0.68 | -0.215 | 0.033 | 0.065 | 0.065 | 3.767 | 0.93 | 0.5 | 0.88 | 0.96 | 0.98 | 0.92 |

| | | | | | | | | | | | | | | | | |
|---|---|---|---|---|---|---|---|---|---|---|---|---|---|---|---|---|
| 1 | 219798632 | rs1415991 | -0.613 | -0.013 | -0.391 | -0.211 | -0.032 | 0.05 | 0.05 | 0 | 0.23 | 0.04 | 0.24 | 0.12 | 0.25 | 0.18 |
| 1 | 219812901 | rs11118371 | 1.145 | -0.709 | -0.67 | -0.048 | -0.102 | -0.044 | -0.044 | 0 | 0.32 | 0.02 | 0.38 | 0.3 | 0.36 | 0.42 |
| 1 | 219821067 | rs4472763 | 0.47 | -1.26 | -0.03 | 0.006 | -0.063 | -0.048 | -0.048 | 0.056 | 0.78 | 0.44 | 0.8 | 0.88 | 0.9 | 0.82 |
| 1 | 219822453 | rs17006225 | -0.183 | -1.328 | 0.131 | -0.03 | -0.02 | -0.027 | -0.027 | 1.11 | 0.07 | 0.29 | 0.04 | 0 | 0.01 | 0.11 |
| 1 | 219825554 | rs10495132 | 0.716 | -1.341 | 0.097 | -0.035 | -0.005 | -0.036 | -0.036 | 0 | 0.09 | 0.01 | 0.1 | 0.12 | 0.05 | 0.07 |
| 1 | 219825766 | rs2378282 | 0.46 | -1.332 | 0.165 | -0.035 | -0.046 | -0.085 | -0.085 | 0 | 0.88 | 0.54 | 0.91 | 1 | 0.95 | 0.89 |
| 1 | 219829351 | rs17006249 | 0.47 | -1.332 | 0.091 | -0.027 | 0.005 | -0.004 | -0.004 | 0 | 0.1 | 0.1 | 0.11 | 0.12 | 0.05 | 0.07 |
| 1 | 219829958 | rs6675807 | -0.488 | -1.293 | 0.475 | -0.023 | -0.01 | -0.02 | -0.02 | 0.09 | 0.08 | 0.39 | 0.06 | 0 | 0.01 | 0.11 |
| 1 | 219838104 | rs945717 | -0.553 | -1.464 | -0.036 | 0.125 | -0.017 | 0.003 | 0.003 | 0.029 | 0.34 | 0.59 | 0.25 | 0.38 | 0.17 | 0.31 |
| 1 | 219842739 | rs12405272 | 0.223 | -0.857 | -0.295 | 0.117 | -0.018 | -0.03 | -0.03 | 0.117 | 0.06 | 0.03 | 0.13 | 0.2 | 0.23 | 0.13 |
| 1 | 219848628 | rs1415998 | 1.219 | -3.088 | 0.124 | 0.278 | 0.146 | -0.001 | -0.001 | 0 | 0.51 | 0.19 | 0.61 | 0.5 | 0.74 | 0.61 |
| 1 | 219859158 | rs1538027 | 1.494 | -2.995 | 0.306 | 0.051 | 0.025 | 0.024 | 0.024 | 0 | 0.58 | 0.31 | 0.64 | 0.68 | 0.68 | 0.62 |
| 1 | 219865236 | rs12096993 | -0.69 | -3.306 | 0.117 | -0.137 | 0.014 | 0.066 | 0.066 | 0 | 0.32 | 0.11 | 0.19 | 0.2 | 0.27 | 0.22 |
| 1 | 219866299 | rs6657605 | 0.356 | -3.275 | 0.148 | -0.075 | 0.017 | 0.035 | 0.035 | 0.202 | 0.17 | 0.11 | 0.11 | 0.06 | 0.11 | 0.08 |
| 1 | 219881894 | rs10779377 | 1.16 | -3.267 | 0.36 | -0.064 | 0.02 | 0.03 | 0.03 | 0.27 | 0.54 | 0.63 | 0.65 | 0.68 | 0.64 | 0.72 |
| 1 | 219886534 | rs12048860 | 1.808 | -3.273 | 0.278 | -0.06 | 0.015 | 0.023 | 0.023 | 0.345 | 0.46 | 0.33 | 0.6 | 0.67 | 0.63 | 0.59 |
| 1 | 219890503 | rs4532841 | -0.481 | -3.207 | 0.242 | -0.09 | -0.024 | -0.039 | -0.039 | 0.295 | 0.23 | 0.38 | 0.13 | 0.15 | 0.17 | 0.25 |
| 1 | 219911743 | rs10157552 | 0.172 | -3.061 | 0.17 | -0.014 | 0 | -0.003 | -0.003 | 0.102 | 0.17 | 0.17 | 0.12 | 0.06 | 0.11 | 0.08 |
| 1 | 219923104 | rs1776037 | 0.439 | -3.199 | 0.158 | 0.081 | -0.049 | -0.026 | -0.026 | 1.291 | 0.79 | 0.57 | 0.8 | 0.86 | 0.89 | 0.8 |
| 1 | 219923989 | rs2797548 | 1.417 | -3.267 | 0.178 | 0.056 | -0.005 | -0.004 | -0.004 | 0.818 | 0.61 | 0.36 | 0.69 | 0.8 | 0.78 | 0.72 |
| 1 | 219926135 | rs12035718 | 0.606 | -3.277 | 0.195 | 0.383 | -0.045 | 0.075 | 0.075 | 0 | 0.69 | 0.51 | 0.68 | 0.52 | 0.8 | 0.73 |
| 1 | 219926717 | rs17006538 | -0.36 | -2.868 | 0.252 | 0.528 | 0.005 | 0.016 | 0.016 | 0 | 0.05 | 0.18 | 0.07 | 0.06 | 0.05 | 0.06 |
| 1 | 219931145 | rs12090457 | -0.455 | -2.872 | 0.246 | 0.526 | 0.008 | 0.019 | 0.019 | 0 | 0.06 | 0.2 | 0.07 | 0.06 | 0.05 | 0.06 |
| 1 | 219931448 | rs4846586 | 0.764 | -2.998 | 0.141 | 0.452 | -0.006 | 0 | 0 | 1.856 | 0.57 | 0.56 | 0.65 | 0.52 | 0.75 | 0.71 |
| 1 | 219934116 | rs1776018 | -1.862 | -2.975 | 0.301 | 0.407 | -0.005 | 0.007 | 0.007 | 0 | 0.62 | 0.66 | 0.46 | 0.7 | 0.44 | 0.53 |
| 1 | 219934247 | rs17006588 | -0.297 | -3.088 | 0.395 | 0.477 | -0.003 | 0.001 | 0.001 | 0 | 0.16 | 0.23 | 0.17 | 0.16 | 0.13 | 0.19 |
| 1 | 219934317 | rs946109 | -0.397 | -3.084 | 0.431 | 0.474 | -0.015 | 0.01 | 0.01 | 0 | 0.87 | 0.72 | 0.87 | 0.87 | 0.92 | 0.86 |
| 1 | 219936765 | rs17006599 | -0.829 | -3.158 | 0.409 | 0.462 | -0.004 | 0.001 | 0.001 | 0 | 0.3 | 0.35 | 0.21 | 0.34 | 0.22 | 0.27 |
| 1 | 219937300 | rs1776059 | -0.448 | -3.247 | 0.462 | 0.439 | -0.003 | 0 | 0 | 0 | 0.33 | 0.3 | 0.25 | 0.37 | 0.21 | 0.25 |
| 1 | 219937351 | rs1766406 | -2.053 | -3.252 | 0.464 | 0.437 | -0.003 | 0.003 | 0.003 | 0 | 0.62 | 0.65 | 0.46 | 0.71 | 0.44 | 0.52 |
| 1 | 219938676 | rs1338243 | -0.486 | -3.188 | 0.333 | 0.448 | -0.003 | 0 | 0 | 0 | 0.33 | 0.3 | 0.25 | 0.37 | 0.21 | 0.25 |
| 1 | 219943715 | rs17006614 | 0.523 | -2.954 | -0.044 | 0.527 | -0.015 | -0.061 | -0.061 | 0.681 | 0.16 | 0.03 | 0.12 | 0.25 | 0.15 | 0.14 |
| 1 | 219945686 | rs12022944 | 0.469 | -2.955 | 0.103 | 0.546 | 0.002 | -0.026 | -0.026 | 0.046 | 0.16 | 0.1 | 0.13 | 0.26 | 0.15 | 0.14 |
| 1 | 219947840 | rs7535102 | 0.079 | -2.987 | 0.187 | 0.364 | 0 | -0.009 | -0.009 | 0 | 0.93 | 0.9 | 0.94 | 0.98 | 0.92 | 0.94 |
| 1 | 219954267 | rs6664688 | 0.474 | -3.079 | 0.234 | 0.372 | -0.038 | 0.007 | 0.007 | 1.18 | 0.61 | 0.54 | 0.73 | 0.56 | 0.79 | 0.69 |
| 1 | 219963912 | rs4565719 | 1.768 | -3.043 | 0.216 | 0.4 | -0.008 | 0.001 | 0.001 | 0 | 0.37 | 0.32 | 0.54 | 0.39 | 0.57 | 0.49 |
| 1 | 219967632 | rs4846599 | 0.528 | -3.204 | 0.468 | 0.477 | -0.035 | 0.006 | 0.006 | 1.315 | 0.61 | 0.54 | 0.73 | 0.56 | 0.8 | 0.68 |
| 1 | 219967811 | rs10779385 | 0.528 | -3.204 | 0.468 | 0.477 | -0.035 | 0.006 | 0.006 | 1.315 | 0.61 | 0.54 | 0.73 | 0.56 | 0.8 | 0.68 |
| 1 | 219968010 | rs4846603 | 1.158 | -3.227 | 0.429 | 0.462 | 0.063 | 0.003 | 0.003 | 0.737 | 0.53 | 0.32 | 0.66 | 0.54 | 0.72 | 0.62 |
| 1 | 219968306 | rs13375416 | -0.672 | -3.019 | 0.275 | 0.539 | 0.008 | -0.025 | -0.025 | 0.161 | 0.08 | 0.2 | 0.06 | 0.02 | 0.08 | 0.06 |
| 1 | 219969405 | rs1857237 | -0.463 | -3.181 | 0.138 | 0.525 | 0.056 | -0.082 | -0.082 | 1.617 | 0.77 | 0.94 | 0.84 | 0.66 | 0.87 | 0.92 |
| 1 | 219978259 | rs6692539 | -1.203 | -2.686 | 0.646 | 0.45 | 0.032 | -0.019 | -0.019 | 0.063 | 0.1 | 0.37 | 0.08 | 0.02 | 0.08 | 0.06 |
| 1 | 219980552 | rs2066418 | -0.644 | -2.789 | 0.533 | 0.46 | 0 | -0.022 | -0.022 | 0 | 0.08 | 0.18 | 0.06 | 0.02 | 0.07 | 0.06 |
| 1 | 219983092 | rs12083182 | -0.64 | -3.019 | 0.327 | 0.476 | -0.008 | -0.003 | -0.003 | 0 | 0.23 | 0.28 | 0.19 | 0.17 | 0.2 | 0.19 |
| 1 | 219984995 | rs11118423 | 1.234 | -2.744 | 0.135 | 0.867 | -0.003 | -0.01 | -0.01 | 0.533 | 0.38 | 0.23 | 0.59 | 0.46 | 0.59 | 0.5 |
| 1 | 220006085 | rs6541105 | -0.113 | -2.969 | -0.211 | 1.05 | 0.002 | -0.002 | -0.002 | 0 | 0.84 | 0.78 | 0.82 | 0.74 | 0.87 | 0.9 |
| 1 | 220008308 | rs6664521 | 1.524 | -3.161 | 0.04 | 1.091 | 0.018 | 0.02 | 0.02 | 0.695 | 0.34 | 0.41 | 0.5 | 0.46 | 0.55 | 0.46 |
| 1 | 220009078 | rs6667735 | -0.195 | -3.018 | -0.178 | 1.327 | 0.013 | -0.034 | -0.034 | 0.016 | 0.85 | 0.88 | 0.83 | 0.74 | 0.87 | 0.9 |
| 1 | 220032318 | rs2008602 | 0.893 | -3.399 | -0.692 | 1.101 | 0.12 | 0.133 | 0.133 | 0.257 | 0.5 | 0.07 | 0.53 | 0.29 | 0.67 | 0.56 |
| 1 | 220052720 | rs11118448 | 1.298 | -2.684 | -0.798 | 0.726 | -0.033 | 0.14 | 0.14 | 1.026 | 0.4 | 0.03 | 0.42 | 0.18 | 0.62 | 0.5 |
| 1 | 220056396 | rs7514869 | -0.399 | -1.379 | 0.235 | 1.676 | 0.026 | 0.12 | 0.12 | 0 | 0.11 | 0.37 | 0.14 | 0.15 | 0.08 | 0.12 |
| 1 | 220068681 | rs4384244 | 0.195 | -1.162 | -0.764 | 1.906 | -0.006 | -0.064 | -0.064 | 0 | 0.9 | 0.98 | 0.95 | 0.78 | 0.91 | 0.94 |
| 1 | 220069860 | rs17006810 | -1.19 | -1.185 | -0.277 | 1.855 | -0.023 | 0.094 | 0.094 | 0.007 | 0.14 | 0.21 | 0.26 | 0.49 | 0.06 | 0.25 |
| 1 | 220070518 | rs9787289 | 1.205 | -0.821 | 0.044 | 1.295 | 0.001 | 0.01 | 0.01 | 0.076 | 0.83 | 0.6 | 0.75 | 0.86 | 0.85 | 0.91 |
| 1 | 220070855 | rs6674051 | 0.334 | -0.559 | -0.726 | 1.382 | -0.03 | -0.05 | -0.05 | 0 | 0.87 | 0.99 | 0.85 | 0.88 | 0.86 | 0.9 |
| 1 | 220076195 | rs6694002 | -0.14 | -0.505 | -0.566 | 1.306 | -0.012 | 0.016 | 0.016 | 0.784 | 0.33 | 0.49 | 0.39 | 0.39 | 0.29 | 0.31 |
| 1 | 220077320 | rs11118454 | 0.781 | -0.753 | -1.235 | 0.59 | -0.079 | -0.027 | -0.027 | 0 | 0.3 | 0.15 | 0.27 | 0.44 | 0.42 | 0.38 |
| 1 | 220078188 | rs17006829 | 1.116 | -0.934 | -1.259 | 0.458 | 0.013 | 0.013 | 0.013 | 0.215 | 0.19 | 0.18 | 0.14 | 0.07 | 0.09 | 0.15 |
| 1 | 220078622 | rs2378330 | 0.854 | -1.036 | -0.849 | 0.478 | -0.022 | 0.035 | 0.035 | 0.6 | 0.33 | 0.43 | 0.39 | 0.39 | 0.29 | 0.31 |
| 1 | 220082150 | rs884127 | 0.041 | -0.895 | -0.875 | 0.551 | 0.061 | -0.003 | -0.003 | 0 | 0.52 | 0.68 | 0.52 | 0.44 | 0.36 | 0.46 |
| 1 | 220088047 | rs2275707 | -0.844 | -0.809 | -1.032 | 0.488 | -0.061 | -0.021 | -0.021 | 0 | 0.83 | 0.99 | 0.81 | 0.88 | 0.79 | 0.82 |
| 1 | 220093188 | rs4846607 | 0.376 | -0.893 | -0.054 | 0.326 | -0.008 | -0.001 | -0.001 | 1.153 | 0.56 | 0.5 | 0.53 | 0.47 | 0.39 | 0.5 |
| 1 | 220098329 | rs6700061 | 1.014 | -1.318 | -0.116 | 0.131 | -0.015 | -0.009 | -0.009 | 0 | 0.24 | 0.27 | 0.16 | 0.09 | 0.09 | 0.15 |
| 1 | 220107917 | rs12129498 | 2.535 | -0.349 | -0.988 | 0.363 | -0.036 | -0.038 | -0.038 | 0 | 0.06 | 0 | 0.07 | 0.16 | 0.12 | 0.03 |
| 1 | 220108918 | rs950455 | 0.98 | -0.909 | -0.877 | 0.302 | 0.012 | -0.068 | -0.068 | 0.358 | 0.31 | 0.1 | 0.36 | 0.38 | 0.3 | 0.36 |
| 1 | 220110560 | rs1776046 | -1.079 | -0.868 | -1.09 | 0.281 | -0.061 | -0.018 | -0.018 | 1.291 | 0.83 | 0.99 | 0.82 | 0.88 | 0.79 | 0.83 |
| 1 | 220111321 | rs7530353 | 1.212 | -0.899 | -1.269 | 0.302 | -0.054 | -0.02 | -0.02 | 0 | 0.15 | 0.02 | 0.17 | 0.1 | 0.21 | 0.23 |
| 1 | 220113915 | rs1498384 | 0.872 | -1.012 | -0.561 | 0.228 | -0.074 | -0.049 | -0.049 | 0.005 | 0.23 | 0.37 | 0.15 | 0.09 | 0.06 | 0.12 |
| 1 | 220119384 | rs11118465 | 0.057 | -0.056 | -0.662 | 0.595 | 0.012 | 0 | 0 | 0 | 0.28 | 0.33 | 0.3 | 0.41 | 0.42 | 0.32 |

| | | | | | | | | | | | | | | | | |
|---|---|---|---|---|---|---|---|---|---|---|---|---|---|---|---|---|
| 1 | 220154768 | rs1061160 | 0.069 | -0.781 | -0.754 | 0.774 | 0.019 | 0.001 | 0.001 | 0.675 | 0.44 | 0.4 | 0.43 | 0.34 | 0.31 | 0.3 |
| 1 | 220156090 | rs2647462 | -0.736 | -0.472 | -0.33 | 0.785 | 0.004 | -0.01 | -0.01 | 1.613 | 0.83 | 0.8 | 0.8 | 0.92 | 0.81 | 0.85 |
| 1 | 220159585 | rs2789794 | -0.142 | -0.75 | -0.252 | 0.659 | 0.001 | 0.008 | 0.008 | 0.081 | 0.78 | 0.77 | 0.84 | 0.95 | 0.79 | 0.82 |
| 1 | 220160262 | rs2789796 | 0.878 | -0.775 | -0.513 | 0.902 | -0.002 | -0.001 | -0.001 | 0.493 | 0.63 | 0.4 | 0.64 | 0.58 | 0.67 | 0.59 |
| 1 | 220161360 | rs2647459 | 1.003 | -1.212 | -0.463 | 0.973 | 0.019 | 0.001 | 0.001 | 0 | 0.59 | 0.34 | 0.63 | 0.52 | 0.66 | 0.55 |
| 1 | 220167371 | rs10495143 | -0.079 | -1.246 | -0.022 | 1.32 | 0.065 | 0.371 | 0.371 | 0.328 | 0.06 | 0.37 | 0.16 | 0.35 | 0.04 | 0.21 |
| 1 | 220175777 | rs2647447 | -0.177 | -1.255 | -0.09 | 1.25 | 0 | -0.01 | -0.01 | 0 | 0.79 | 0.75 | 0.84 | 0.94 | 0.79 | 0.85 |
| 1 | 220185726 | rs2248717 | -0.344 | -1.268 | -0.223 | 1.243 | -0.02 | 0.058 | 0.058 | 0.035 | 0.8 | 0.83 | 0.85 | 0.94 | 0.79 | 0.85 |
| 1 | 220190663 | rs12048112 | 0.063 | -1.239 | 0.153 | 1.306 | 0.015 | 0.233 | 0.233 | 0.026 | 0.06 | 0.23 | 0.15 | 0.35 | 0.04 | 0.21 |
| 1 | 220197625 | rs2230301 | -0.927 | -1.239 | -0.085 | 1.243 | -0.062 | 0.03 | 0.03 | 0.009 | 0.83 | 0.99 | 0.87 | 0.94 | 0.79 | 0.86 |
| 1 | 220199852 | rs2647458 | 0.855 | -1.717 | 0.04 | 1.304 | 0.072 | -0.02 | -0.02 | 0 | 0.32 | 0.14 | 0.35 | 0.25 | 0.2 | 0.2 |
| 1 | 220214212 | rs12022181 | -0.548 | -1.088 | 0.619 | 1.632 | -0.028 | 0.259 | 0.259 | 0.05 | 0.11 | 0.42 | 0.17 | 0.41 | 0.05 | 0.25 |
| 1 | 220226732 | rs7515501 | -0.461 | -0.838 | 0.117 | 1.706 | -0.01 | -0.032 | -0.032 | 0 | 0.88 | 0.94 | 0.81 | 0.63 | 0.87 | 0.72 |
| 1 | 220228658 | rs10863529 | 1.938 | -1.254 | 0.191 | 1.774 | 0.025 | -0.01 | -0.01 | 0 | 0.63 | 0.32 | 0.62 | 0.39 | 0.71 | 0.52 |
| 1 | 220232797 | rs904328 | 1.595 | -1.221 | 0.283 | 1.804 | -0.003 | 0.003 | 0.003 | 0.137 | 0.72 | 0.34 | 0.69 | 0.43 | 0.82 | 0.63 |
| 1 | 220239448 | rs1146773 | 0.457 | -1.077 | 0.559 | 1.833 | -0.019 | 0.17 | 0.17 | 0 | 0.81 | 0.56 | 0.74 | 0.52 | 0.86 | 0.67 |
| 1 | 220240862 | rs17563262 | 1.571 | -0.848 | 0.101 | 1.66 | 0.016 | -0.019 | -0.019 | 0 | 0.18 | 0.03 | 0.25 | 0.12 | 0.15 | 0.12 |
| 1 | 220240997 | rs2577152 | -0.468 | -0.751 | 0.173 | 1.532 | -0.006 | 0.002 | 0.002 | 0 | 0.16 | 0.23 | 0.12 | 0.16 | 0.12 | 0.16 |
| 1 | 220264913 | rs34672561 | 2.348 | -0.462 | -0.184 | 1.549 | -0.178 | 0.068 | 0.068 | 1.232 | 0.36 | 0.02 | 0.29 | 0.2 | 0.46 | 0.31 |
| 1 | 220273564 | rs2289190 | 0.614 | -0.806 | 0.379 | 1.883 | -0.021 | 0.038 | 0.038 | 0 | 0.23 | 0.27 | 0.32 | 0.39 | 0.18 | 0.19 |
| 1 | 220276333 | rs2789789 | -1.274 | -0.765 | 0.344 | 1.835 | 0.062 | 0.02 | 0.02 | 0 | 0.44 | 0.69 | 0.46 | 0.61 | 0.31 | 0.43 |
| 1 | 220289803 | rs1465816 | -0.832 | -0.79 | 0.404 | 1.886 | -0.028 | 0 | 0 | 0 | 0.39 | 0.46 | 0.39 | 0.33 | 0.28 | 0.39 |
| 1 | 220300035 | rs2289191 | 1.706 | -0.679 | 0.13 | 1.647 | -0.018 | -0.011 | -0.011 | 0 | 0.06 | 0 | 0.05 | 0.03 | 0.07 | 0.05 |
| 1 | 220314945 | rs10442693 | 2.081 | -0.736 | 0.175 | 1.63 | -0.026 | 0.106 | 0.106 | 0.341 | 0.4 | 0.16 | 0.33 | 0.23 | 0.56 | 0.33 |
| 1 | 220319110 | rs3738336 | 1.367 | -0.925 | 0.434 | 1.827 | 0.009 | -0.034 | -0.034 | 0 | 0.18 | 0.01 | 0.25 | 0.12 | 0.16 | 0.15 |
| 1 | 220319301 | rs11806086 | 1.017 | -0.961 | 0.621 | 1.95 | -0.012 | 0.178 | 0.178 | 0 | 0.07 | 0.28 | 0.1 | 0.21 | 0.01 | 0.05 |
| 1 | 220321023 | rs11095 | 1.302 | -0.946 | 0.494 | 1.718 | 0.016 | -0.032 | -0.032 | 0 | 0.19 | 0.02 | 0.25 | 0.13 | 0.15 | 0.15 |
| 1 | 220330672 | rs11547779 | 0.55 | -0.862 | 0.513 | 1.74 | -0.018 | -0.009 | -0.009 | 0 | 0.09 | 0.01 | 0.05 | 0.05 | 0.09 | 0.12 |
| 1 | 220332876 | rs2808026 | -1.339 | -0.864 | 0.744 | 1.85 | 0.097 | 0.105 | 0.105 | 0.25 | 0.46 | 0.75 | 0.52 | 0.69 | 0.31 | 0.42 |
| 1 | 220339479 | rs2808027 | 0.843 | -0.668 | 0.687 | 1.952 | -0.014 | 0.212 | 0.212 | 0 | 0.07 | 0.29 | 0.11 | 0.22 | 0.01 | 0.05 |
| 1 | 220345860 | rs7527410 | 1.832 | -0.633 | 0.33 | 1.921 | -0.022 | 0.103 | 0.103 | 0 | 0.05 | 0.11 | 0.09 | 0.22 | 0.01 | 0.05 |
| 1 | 220351723 | rs11805285 | -0.718 | -0.566 | 0.225 | 1.681 | 0.005 | 0.005 | 0.005 | 0.019 | 0.15 | 0.12 | 0.09 | 0.09 | 0.12 | 0.14 |
| 1 | 220380819 | rs12035921 | 2.208 | -0.288 | -0.326 | 1.376 | -0.178 | 0.118 | 0.118 | 1.15 | 0.36 | 0.02 | 0.29 | 0.2 | 0.45 | 0.33 |
| 1 | 220385443 | rs4351641 | -0.179 | -0.571 | 0.732 | 1.562 | -0.014 | 0.011 | 0.011 | 0.491 | 0.26 | 0.3 | 0.36 | 0.34 | 0.16 | 0.2 |
| 1 | 220392412 | rs17565134 | 0.501 | -0.586 | 0.618 | 1.623 | 0.013 | 0.001 | 0.001 | 0.013 | 0.21 | 0.18 | 0.26 | 0.13 | 0.15 | 0.15 |
| 1 | 220401009 | rs6541127 | -1.802 | -0.303 | 0.535 | 1.365 | 0.094 | -0.022 | -0.022 | 0 | 0.44 | 0.8 | 0.49 | 0.44 | 0.3 | 0.39 |
| 1 | 220435999 | rs10779401 | -0.349 | 0.021 | 0.595 | 1.614 | 0.03 | 0.071 | 0.071 | 0 | 0.07 | 0.23 | 0.1 | 0.09 | 0.06 | 0.06 |
| 1 | 220448330 | rs1511695 | -0.524 | 0.208 | -0.099 | 1.6 | -0.021 | 0.022 | 0.022 | 0.445 | 0.25 | 0.31 | 0.33 | 0.33 | 0.15 | 0.21 |
| 1 | 220454174 | rs10779402 | 0.515 | 1.949 | 0.17 | 1.383 | 0.07 | 0.072 | 0.072 | 0.104 | 0.86 | 0.51 | 0.79 | 0.86 | 0.83 | 0.9 |
| 1 | 220457101 | rs17565835 | -0.224 | 1.383 | -0.097 | 1.093 | -0.014 | 0.015 | 0.015 | 0 | 0.12 | 0.07 | 0.16 | 0.09 | 0.13 | 0.14 |
| 1 | 220471610 | rs925445 | 0.087 | 1.354 | 0.061 | 1.035 | -0.009 | 0.009 | 0.009 | 0 | 0.1 | 0.07 | 0.12 | 0.07 | 0.12 | 0.14 |
| 1 | 220472295 | rs4846310 | -0.36 | 1.239 | 0.384 | 1.277 | -0.034 | 0.045 | 0.045 | 0 | 0.14 | 0.25 | 0.15 | 0.21 | 0.07 | 0.09 |
| 1 | 220473570 | rs17566172 | 0.161 | 1.437 | 0.295 | 1.231 | -0.031 | -0.027 | -0.027 | 0 | 0.05 | 0.01 | 0.05 | 0.05 | 0.12 | 0.1 |
| 1 | 220476572 | rs907416 | -0.8 | 1.288 | 0.498 | 1.306 | -0.024 | 0.106 | 0.106 | 0 | 0.17 | 0.39 | 0.18 | 0.27 | 0.12 | 0.16 |
| 1 | 220486349 | rs4272630 | -0.077 | 1.265 | 0.484 | 1.298 | -0.035 | 0.059 | 0.059 | 0 | 0.13 | 0.25 | 0.14 | 0.21 | 0.06 | 0.09 |
| 1 | 220489544 | rs6541128 | -0.808 | 1.002 | 0.027 | 1.141 | 0.002 | 0.001 | 0.001 | 0.399 | 0.79 | 0.8 | 0.74 | 0.83 | 0.85 | 0.84 |
| 1 | 220490497 | rs7540014 | -0.486 | 1.144 | 0.206 | 1.234 | -0.019 | 0.087 | 0.087 | 0 | 0.17 | 0.35 | 0.17 | 0.28 | 0.12 | 0.16 |
| 1 | 220495396 | rs2175480 | -1.773 | 1.166 | 0.142 | 1.114 | 0.007 | 0.011 | 0.011 | 0 | 0.36 | 0.44 | 0.35 | 0.37 | 0.37 | 0.44 |
| 1 | 220496232 | rs6541137 | -0.523 | 1.2 | 0.211 | 1.185 | -0.02 | 0.076 | 0.076 | 0 | 0.17 | 0.34 | 0.17 | 0.26 | 0.12 | 0.16 |
| 1 | 220497813 | rs1983248 | 1.858 | 1.111 | -0.038 | 1.092 | -0.001 | 0.014 | 0.014 | 0 | 0.45 | 0.32 | 0.4 | 0.46 | 0.5 | 0.43 |
| 1 | 220498513 | rs17007467 | -0.525 | 1.207 | 0.075 | 1.143 | -0.019 | 0.104 | 0.104 | 0 | 0.17 | 0.41 | 0.18 | 0.26 | 0.12 | 0.16 |
| 1 | 220499020 | rs6673738 | -0.342 | 1.215 | 0.056 | 1.137 | -0.033 | 0.013 | 0.013 | 0.014 | 0.17 | 0.1 | 0.17 | 0.11 | 0.24 | 0.27 |
| 1 | 220499314 | rs12058934 | -1.763 | 1.204 | 0.049 | 1.126 | 0.015 | 0.024 | 0.024 | 0 | 0.34 | 0.47 | 0.34 | 0.38 | 0.37 | 0.43 |
| 1 | 220504423 | rs11118527 | 0.357 | 1.289 | -0.117 | 1.062 | -0.006 | 0.011 | 0.011 | 0 | 0.13 | 0.1 | 0.13 | 0.07 | 0.15 | 0.15 |
| 1 | 220506871 | rs11118528 | -1.039 | 1.32 | -0.057 | 1.092 | -0.009 | 0.004 | 0.004 | 0.938 | 0.21 | 0.18 | 0.2 | 0.18 | 0.3 | 0.34 |
| 1 | 220507580 | rs17641611 | -0.64 | 1.42 | -0.275 | 1.114 | -0.003 | -0.004 | -0.004 | 0 | 0.09 | 0.08 | 0.07 | 0.11 | 0.16 | 0.19 |
| 1 | 220512519 | rs1567602 | 0.263 | 1.271 | -0.514 | 1.006 | 0.001 | -0.014 | -0.014 | 0 | 0.23 | 0.05 | 0.27 | 0.17 | 0.16 | 0.19 |
| 1 | 220523521 | rs9803782 | -0.947 | 1.497 | -0.194 | 1.294 | 0.026 | -0.004 | -0.004 | 0 | 0.21 | 0.25 | 0.21 | 0.18 | 0.3 | 0.34 |
| 1 | 220525993 | rs10863546 | 0.085 | 0.738 | -0.011 | 1.381 | -0.033 | 0.033 | 0.033 | 0 | 0.16 | 0.22 | 0.15 | 0.2 | 0.08 | 0.12 |
| 1 | 220527820 | rs7544009 | -0.547 | 1.304 | -0.3 | 1.05 | -0.003 | -0.002 | -0.002 | 0 | 0.45 | 0.51 | 0.49 | 0.48 | 0.39 | 0.45 |
| 1 | 220530560 | rs6541139 | -0.493 | 0.844 | -0.39 | 1.045 | -0.003 | 0.002 | 0.002 | 0 | 0.42 | 0.46 | 0.46 | 0.42 | 0.34 | 0.36 |
| 1 | 220539533 | rs12754802 | -0.369 | 1.5 | -0.759 | 0.778 | -0.068 | 0.029 | 0.029 | 0.061 | 0.13 | 0.01 | 0.11 | 0.06 | 0.24 | 0.24 |
| 1 | 220546421 | rs1338770 | 1.18 | 1.011 | 0.682 | 0.887 | -0.027 | 0.059 | 0.059 | 0.167 | 0.54 | 0.25 | 0.43 | 0.34 | 0.49 | 0.51 |
| 1 | 220556658 | rs6702655 | 0.108 | 0.982 | 1.333 | 1.671 | -0.037 | -0.088 | -0.088 | 0.559 | 0.15 | 0.04 | 0.17 | 0.36 | 0.18 | 0.07 |
| 1 | 220559400 | rs11118542 | 1.678 | 0.961 | 1.338 | 1.676 | -0.028 | 0.063 | 0.063 | 0.167 | 0.54 | 0.25 | 0.43 | 0.34 | 0.49 | 0.51 |
| 1 | 220563560 | rs12732389 | -0.28 | 0.991 | 1.535 | 1.631 | 0.005 | 0.074 | 0.074 | 0 | 0.17 | 0.26 | 0.19 | 0.36 | 0.18 | 0.07 |
| 1 | 220565331 | rs7531705 | 1.079 | 0.547 | 1.428 | 1.596 | 0.002 | 0.004 | 0.004 | 0.183 | 0.64 | 0.41 | 0.68 | 0.53 | 0.56 | 0.6 |
| 1 | 220567226 | rs1578046 | 1.2 | 0.553 | 1.308 | 1.589 | -0.02 | -0.036 | -0.036 | 0.377 | 0.63 | 0.24 | 0.66 | 0.52 | 0.56 | 0.6 |
| 1 | 220580172 | rs972980 | 0.694 | -0.296 | 1.278 | 2.055 | -0.05 | 0.093 | 0.093 | 0 | 0.82 | 0.65 | 0.85 | 0.65 | 0.89 | 0.86 |

| | | | | | | | | | | | | | | | | |
|---|---|---|---|---|---|---|---|---|---|---|---|---|---|---|---|---|
| 1 | 220583501 | rs4846323 | -1.002 | 0.187 | 0.957 | 1.236 | 0.02 | -0.025 | -0.025 | 0 | 0.05 | 0.16 | 0.07 | 0 | 0.04 | 0.03 |
| 1 | 220590437 | rs1578049 | -1.219 | -0.155 | 0.834 | 1.534 | -0.016 | 0.012 | 0.012 | 0 | 0.32 | 0.36 | 0.53 | 0.67 | 0.24 | 0.3 |
| 1 | 220594269 | rs1538419 | 0.771 | -0.408 | 1.15 | 1.249 | -0.022 | 0.013 | 0.013 | 0 | 0.93 | 0.75 | 0.97 | 0.95 | 0.99 | 0.92 |
| 1 | 220599845 | rs1338763 | 1.189 | -0.729 | 1.465 | 1.814 | -0.039 | 0.06 | 0.06 | 0.2 | 0.69 | 0.55 | 0.66 | 0.56 | 0.79 | 0.73 |
| 1 | 220601623 | rs7512221 | 1.234 | -0.848 | 2.16 | 1.979 | -0.037 | 0.033 | 0.033 | 0 | 0.88 | 0.53 | 0.93 | 0.89 | 0.95 | 0.87 |
| 1 | 220604048 | rs11576868 | 1.207 | -0.851 | 2.159 | 1.991 | -0.036 | 0.043 | 0.043 | 0 | 0.88 | 0.53 | 0.93 | 0.88 | 0.95 | 0.87 |
| 1 | 220608142 | rs12032166 | -0.354 | -0.905 | 1.917 | 2.075 | -0.026 | -0.129 | -0.129 | 0 | 0.19 | 0.02 | 0.27 | 0.34 | 0.16 | 0.13 |
| 1 | 220611332 | rs6670633 | 2.008 | -1.633 | 1.451 | 1.751 | 0.103 | 0.123 | 0.123 | 0.275 | 0.52 | 0.12 | 0.44 | 0.28 | 0.69 | 0.6 |
| 1 | 220613513 | rs1338775 | 1.067 | -1.649 | 1.598 | 1.748 | 0.101 | 0.226 | 0.226 | 0.589 | 0.95 | 0.61 | 0.97 | 0.91 | 0.97 | 0.95 |
| 1 | 220614557 | rs1953809 | 1.263 | -1.649 | 1.272 | 1.743 | 0.003 | -0.005 | -0.005 | 0.623 | 0.67 | 0.38 | 0.65 | 0.53 | 0.79 | 0.73 |
| 1 | 220621750 | rs6672068 | 0.122 | -1.475 | 1.191 | 1.756 | -0.011 | 0.004 | 0.004 | 0 | 0.16 | 0.25 | 0.22 | 0.25 | 0.11 | 0.14 |
| 1 | 220622474 | rs12408337 | -0.081 | -1.268 | 0.99 | 1.736 | -0.027 | -0.129 | -0.129 | 0 | 0.19 | 0.01 | 0.27 | 0.34 | 0.16 | 0.13 |
| 1 | 220624026 | rs6694188 | 1.08 | -1.293 | 1.292 | 1.624 | -0.031 | 0.047 | 0.047 | 0.373 | 0.69 | 0.51 | 0.66 | 0.53 | 0.79 | 0.74 |
| 1 | 220625992 | rs7555789 | -1.244 | -1.567 | 1.143 | 1.449 | -0.027 | -0.008 | -0.008 | 0.049 | 0.16 | 0.21 | 0.05 | 0.12 | 0.05 | 0.16 |
| 1 | 220626086 | rs7553380 | -1.244 | -1.567 | 1.143 | 1.437 | -0.027 | -0.009 | -0.009 | 0.071 | 0.16 | 0.21 | 0.05 | 0.11 | 0.05 | 0.16 |
| 1 | 220647006 | rs12561793 | -0.115 | -1.687 | 1.582 | 1.554 | -0.025 | -0.113 | -0.113 | 0.487 | 0.15 | 0.02 | 0.26 | 0.36 | 0.14 | 0.12 |
| 1 | 220658704 | rs12563560 | 1.195 | -1.713 | 1.592 | 1.532 | -0.009 | 0.005 | 0.005 | 0.219 | 0.06 | 0.06 | 0.01 | 0.09 | 0.02 | 0.06 |
| 1 | 220658889 | rs3842967 | 1.84 | -1.632 | 1.685 | 1.524 | -0.02 | 0.014 | 0.014 | 1.663 | 0.67 | 0.41 | 0.66 | 0.53 | 0.81 | 0.72 |
| 1 | 220664952 | rs7530302 | -1.364 | -1.522 | 1.606 | 1.414 | -0.037 | -0.017 | -0.017 | 0 | 0.16 | 0.25 | 0.05 | 0.11 | 0.05 | 0.15 |
| 1 | 220667154 | rs13376101 | -1.783 | -1.92 | 1.35 | 1.141 | -0.018 | -0.01 | -0.01 | 0.022 | 0.22 | 0.25 | 0.06 | 0.13 | 0.08 | 0.2 |
| 1 | 220671035 | rs3860322 | 1.919 | -1.989 | 0.768 | 0.74 | 0.034 | -0.009 | -0.009 | 0 | 0.54 | 0.39 | 0.61 | 0.55 | 0.72 | 0.67 |
| 1 | 220683696 | rs12753927 | -1.162 | -2.088 | 1.077 | 0.734 | -0.001 | 0 | 0 | 0 | 0.31 | 0.26 | 0.33 | 0.36 | 0.23 | 0.23 |
| 1 | 220684386 | rs10863557 | 1.967 | -2.234 | 1.204 | 0.673 | 0.046 | -0.01 | -0.01 | 0 | 0.55 | 0.36 | 0.61 | 0.53 | 0.72 | 0.61 |
| 1 | 220725132 | rs2589586 | -1.779 | -2.244 | 0.867 | 0.604 | 0.026 | -0.114 | -0.114 | 0 | 0.38 | 0.1 | 0.34 | 0.46 | 0.3 | 0.35 |
| 1 | 220748832 | rs1933002 | -0.233 | -1.982 | 0.213 | 0.51 | -0.059 | -0.156 | -0.156 | 0.87 | 0.23 | 0.01 | 0.3 | 0.36 | 0.23 | 0.22 |
| 1 | 220755159 | rs17007981 | -0.28 | -2.242 | 0.189 | 0.431 | 0.041 | 0.009 | 0.009 | 0 | 0.15 | 0 | 0.03 | 0.1 | 0.07 | 0.13 |
| 1 | 220801974 | rs10495150 | -0.941 | -2.356 | 0.168 | 0.3 | -0.058 | -0.061 | -0.061 | 1.506 | 0.11 | 0.61 | 0.06 | 0.01 | 0.01 | 0.07 |
| 1 | 220817803 | rs6656145 | 0.611 | -2.567 | -0.338 | 0.543 | -0.021 | -0.073 | -0.073 | 0 | 0.71 | 0.93 | 0.6 | 0.69 | 0.75 | 0.73 |
| 1 | 220825543 | rs3737297 | -0.206 | -2.394 | -0.236 | 0.46 | 0.046 | 0.022 | 0.022 | 0 | 0.17 | 0.05 | 0.05 | 0.11 | 0.08 | 0.13 |
| 1 | 220828130 | rs2889895 | 1.498 | -2.319 | -0.117 | 0.49 | 0.033 | -0.017 | -0.017 | 0 | 0.59 | 0.31 | 0.68 | 0.54 | 0.73 | 0.6 |
| 1 | 220849101 | rs366414 | -1.281 | -2.509 | 0.272 | 0.633 | -0.018 | -0.018 | -0.018 | 0 | 0.38 | 0.53 | 0.47 | 0.77 | 0.23 | 0.4 |
| 1 | 220849330 | rs2994525 | -1.686 | -2.628 | 0.25 | 0.572 | 0.065 | 0.066 | 0.066 | 0 | 0.45 | 0.64 | 0.49 | 0.78 | 0.24 | 0.46 |
| 1 | 220851520 | rs12410279 | 0.593 | -2.65 | -0.132 | 0.823 | -0.003 | -0.137 | -0.137 | 0.163 | 0.18 | 0.01 | 0.31 | 0.38 | 0.14 | 0.22 |
| 1 | 220863423 | rs3806329 | 0.509 | -2.796 | 0.07 | 1.03 | -0.002 | -0.14 | -0.14 | 0.319 | 0.18 | 0.02 | 0.31 | 0.37 | 0.15 | 0.22 |
| 1 | 220864976 | rs364360 | 0.346 | -2.744 | 0.551 | 1.108 | 0.024 | 0.231 | 0.231 | 1.135 | 0.07 | 0.26 | 0.11 | 0.31 | 0.04 | 0.09 |
| 1 | 220867903 | rs9431295 | -1.144 | -2.799 | 0.535 | 1.078 | -0.001 | 0 | 0 | 0 | 0.31 | 0.3 | 0.43 | 0.68 | 0.19 | 0.32 |
| 1 | 220868627 | rs437078 | -0.563 | -2.825 | 0.515 | 1.067 | -0.04 | 0.054 | 0.054 | 0 | 0.19 | 0.26 | 0.12 | 0.42 | 0.07 | 0.17 |
| 1 | 220871123 | rs337154 | -1.698 | -2.666 | 0.614 | 0.951 | -0.026 | -0.04 | -0.04 | 0.132 | 0.45 | 0.51 | 0.47 | 0.83 | 0.25 | 0.41 |
| 1 | 220871320 | rs337153 | -1.698 | -2.666 | 0.569 | 0.951 | -0.022 | -0.035 | -0.035 | 0.58 | 0.45 | 0.52 | 0.47 | 0.83 | 0.25 | 0.41 |
| 1 | 220874101 | rs426020 | -1.931 | -2.623 | 0.578 | 0.923 | -0.022 | -0.031 | -0.031 | 0.018 | 0.45 | 0.52 | 0.47 | 0.81 | 0.25 | 0.41 |
| 1 | 220879617 | rs6675203 | -0.565 | -2.507 | 0.345 | 1.018 | 0.052 | 0.044 | 0.044 | 0.542 | 0.2 | 0.05 | 0.1 | 0.11 | 0.08 | 0.15 |
| 1 | 220884498 | rs439513 | -1.371 | -2.63 | 0.861 | 1.234 | -0.072 | 0.045 | 0.045 | 0.067 | 0.27 | 0.41 | 0.16 | 0.44 | 0.11 | 0.23 |
| 1 | 220885612 | rs796486 | -0.389 | -2.834 | 0.715 | 1.445 | -0.018 | 0.022 | 0.022 | 0 | 0.27 | 0.31 | 0.37 | 0.61 | 0.16 | 0.27 |
| 1 | 220887035 | rs368664 | 0.922 | -2.708 | 0.595 | 1.485 | -0.007 | 0.172 | 0.172 | 0 | 0.06 | 0.13 | 0.05 | 0.32 | 0.02 | 0.08 |
| 1 | 220887172 | rs17008345 | -0.519 | -2.622 | 0.06 | 1.247 | 0.047 | -0.007 | -0.007 | 0 | 0.2 | 0.08 | 0.1 | 0.16 | 0.08 | 0.14 |
| 1 | 220900396 | rs337170 | -1.369 | -2.457 | 0.312 | 1.214 | 0.009 | -0.006 | -0.006 | 0 | 0.41 | 0.39 | 0.44 | 0.45 | 0.24 | 0.34 |
| 1 | 220906624 | rs337182 | 0.468 | -2.407 | 0.239 | 1.326 | 0.003 | -0.004 | -0.004 | 0 | 0.19 | 0.17 | 0.32 | 0.26 | 0.15 | 0.19 |
| 1 | 220907887 | rs35297838 | 0.23 | -2.367 | 0.064 | 1.233 | 0 | -0.031 | -0.031 | 0.464 | 0.08 | 0 | 0.07 | 0.07 | 0.05 | 0.06 |
| 1 | 220914920 | rs337161 | -0.828 | -2.372 | 0.301 | 1.387 | 0.015 | -0.017 | -0.017 | 0.003 | 0.33 | 0.3 | 0.39 | 0.47 | 0.2 | 0.32 |
| 1 | 220918953 | rs189231 | -1.379 | -2.183 | 0.078 | 1.308 | 0.065 | 0.041 | 0.041 | 0 | 0.42 | 0.73 | 0.48 | 0.68 | 0.25 | 0.4 |
| 1 | 220920857 | rs337190 | -1.141 | -2.364 | 0.063 | 1.294 | 0.045 | -0.001 | -0.001 | 0 | 0.41 | 0.68 | 0.43 | 0.62 | 0.22 | 0.36 |
| 1 | 220925376 | rs393585 | -1.153 | -2.324 | 0.01 | 1.207 | 0.051 | 0.014 | 0.014 | 0 | 0.39 | 0.71 | 0.43 | 0.71 | 0.21 | 0.35 |
| 1 | 220931227 | rs6659246 | -0.241 | -2.491 | 0.608 | 1.456 | -0.035 | 0.081 | 0.081 | 0 | 0.25 | 0.37 | 0.38 | 0.46 | 0.17 | 0.3 |
| 1 | 220934077 | rs337139 | -1.297 | -2.324 | -0.285 | 0.668 | -0.075 | 0.196 | 0.196 | 0 | 0.66 | 0.98 | 0.73 | 0.92 | 0.66 | 0.7 |
| 1 | 220935081 | rs2275575 | 0.901 | -2.661 | 0.364 | 1.363 | 0.011 | -0.011 | -0.011 | 0.122 | 0.19 | 0.16 | 0.33 | 0.23 | 0.14 | 0.22 |
| 1 | 220935808 | rs10779415 | -0.71 | -2.721 | 0.343 | 1.373 | -0.038 | 0.002 | 0.002 | 0 | 0.36 | 0.41 | 0.4 | 0.69 | 0.21 | 0.35 |
| 1 | 220939821 | rs17008587 | 1.176 | -2.643 | 0.113 | 1.333 | 0.005 | -0.058 | -0.058 | 0 | 0.18 | 0.04 | 0.32 | 0.22 | 0.14 | 0.19 |
| 1 | 220940411 | rs337147 | -1.09 | -2.789 | 0.247 | 1.31 | 0.027 | 0.019 | 0.019 | 0 | 0.4 | 0.63 | 0.47 | 0.74 | 0.21 | 0.36 |
| 1 | 220940834 | rs17008595 | 1.3 | -2.567 | -0.181 | 1.375 | 0.02 | -0.015 | -0.015 | 0.164 | 0.11 | 0.04 | 0.02 | 0.22 | 0.04 | 0.05 |
| 1 | 220941444 | rs11118598 | -0.094 | -2.556 | -0.191 | 1.366 | -0.071 | 0.155 | 0.155 | 0 | 0.19 | 0.4 | 0.14 | 0.46 | 0.06 | 0.14 |
| 1 | 220945483 | rs7519414 | -0.9 | -2.403 | -0.162 | 1.111 | -0.032 | -0.007 | -0.007 | 0 | 0.43 | 0.43 | 0.46 | 0.68 | 0.22 | 0.32 |
| 1 | 220951135 | rs6657944 | -1.017 | -2.419 | -0.11 | 1.095 | 0.013 | 0.012 | 0.012 | 0.021 | 0.46 | 0.52 | 0.47 | 0.73 | 0.23 | 0.37 |
| 1 | 220956012 | rs7537305 | 0.199 | -2.313 | -0.071 | 1.123 | -0.045 | 0.044 | 0.044 | 0.302 | 0.14 | 0.27 | 0.13 | 0.24 | 0.04 | 0.1 |
| 1 | 220959517 | rs868494 | 0.237 | -2.31 | -0.068 | 1.124 | -0.04 | 0.036 | 0.036 | 0.359 | 0.14 | 0.23 | 0.13 | 0.25 | 0.04 | 0.1 |
| 1 | 220960915 | rs7536276 | -0.932 | -2.394 | -0.109 | 1.109 | -0.014 | -0.009 | -0.009 | 0 | 0.45 | 0.41 | 0.38 | 0.73 | 0.23 | 0.38 |
| 1 | 220964537 | rs6682431 | 0.194 | -2.267 | -0.336 | 1.192 | -0.049 | 0.051 | 0.051 | 0.311 | 0.14 | 0.31 | 0.13 | 0.24 | 0.04 | 0.1 |
| 1 | 220965474 | rs4131748 | -0.112 | -1.537 | -1.04 | 1.395 | 0.034 | -0.034 | -0.034 | 0 | 0.29 | 0.14 | 0.25 | 0.44 | 0.18 | 0.22 |
| 1 | 220967683 | rs1389742 | -0.048 | -0.453 | -1.079 | 0.822 | 0.009 | -0.026 | -0.026 | 0.65 | 0.45 | 0.2 | 0.63 | 0.48 | 0.61 | 0.55 |
| 1 | 220970499 | rs2642439 | -0.547 | -0.794 | -1.338 | 0.33 | 0.003 | 0.111 | 0.111 | 0 | 0.64 | 0.89 | 0.77 | 0.78 | 0.71 | 0.74 |

| | | | | | | | | | | | | | | | | |
|---|---|---|---|---|---|---|---|---|---|---|---|---|---|---|---|---|
| 1 | 220972343 | rs867772 | -0.513 | -0.84 | -1.43 | 0.328 | -0.003 | 0.118 | 0.118 | 0.001 | 0.64 | 0.94 | 0.78 | 0.78 | 0.71 | 0.74 |
| 1 | 220973426 | rs2642441 | -0.027 | 0.584 | -0.523 | 0.637 | 0.062 | 0.008 | 0.008 | 0 | 0.17 | 0.34 | 0.23 | 0.16 | 0.3 | 0.23 |
| 1 | 220974174 | rs12046182 | -0.566 | 0.507 | -0.594 | 0.892 | 0 | 0.027 | 0.027 | 0 | 0.22 | 0.27 | 0.24 | 0.45 | 0.22 | 0.39 |
| 1 | 220974202 | rs2807837 | -1.149 | 0.495 | -0.839 | 0.732 | -0.002 | -0.012 | -0.012 | 0 | 0.92 | 0.97 | 0.93 | 0.86 | 0.9 | 0.95 |
| 1 | 220974218 | rs2642444 | -1.078 | 0.255 | -0.158 | 0.682 | -0.018 | 0.039 | 0.039 | 0.177 | 0.29 | 0.44 | 0.29 | 0.44 | 0.25 | 0.38 |
| 1 | 220976257 | rs17008806 | 1.205 | 0.294 | -0.117 | 0.293 | 0.021 | 0.013 | 0.013 | 0.002 | 0.26 | 0.39 | 0.34 | 0.29 | 0.29 | 0.26 |
| 1 | 220977615 | rs2642421 | 0.324 | -0.053 | -0.317 | -0.123 | 0 | -0.013 | -0.013 | 0 | 0.45 | 0.48 | 0.47 | 0.32 | 0.46 | 0.45 |
| 1 | 220977650 | rs2642420 | 0.885 | -0.171 | -0.417 | -0.207 | 0.01 | 0.026 | 0.026 | 0 | 0.2 | 0.13 | 0.13 | 0.03 | 0.16 | 0.18 |
| 1 | 220979429 | rs17008835 | 1.176 | 0.365 | -0.052 | -0.104 | -0.022 | 0.01 | 0.01 | 0 | 0.28 | 0.29 | 0.28 | 0.34 | 0.16 | 0.33 |
| 1 | 220979994 | rs1874113 | -0.875 | 0.312 | -0.438 | -0.186 | 0.065 | 0.02 | 0.02 | 0 | 0.57 | 0.77 | 0.64 | 0.37 | 0.7 | 0.5 |
| 1 | 220981358 | rs6541160 | 1.052 | 0.688 | -0.313 | -0.255 | -0.006 | -0.012 | -0.012 | 0 | 0.19 | 0.17 | 0.19 | 0.02 | 0.3 | 0.18 |
| 1 | 220983402 | rs17649913 | 1.463 | 0.195 | -0.498 | -0.223 | -0.025 | 0.053 | 0.053 | 0.402 | 0.17 | 0.08 | 0.12 | 0.03 | 0.2 | 0.19 |
| 1 | 220984364 | rs11118607 | 0.861 | -0.06 | -0.208 | -0.189 | -0.004 | -0.017 | -0.017 | 0 | 0.2 | 0.31 | 0.1 | 0.15 | 0.16 | 0.08 |
| 1 | 220985515 | rs12129004 | 1.881 | -0.14 | -0.525 | -0.019 | -0.025 | -0.054 | -0.054 | 0 | 0.25 | 0.15 | 0.32 | 0.34 | 0.29 | 0.32 |
| 1 | 220985677 | rs2642449 | -1.365 | -0.13 | -0.369 | 0.049 | 0.023 | -0.018 | -0.018 | 0 | 0.81 | 0.93 | 0.88 | 0.71 | 0.87 | 0.85 |
| 1 | 220987155 | rs2807840 | -1.163 | -0.134 | -0.169 | 0.045 | -0.007 | 0.002 | 0.002 | 0 | 0.81 | 0.79 | 0.87 | 0.71 | 0.86 | 0.85 |
| 1 | 220988346 | rs4143772 | 0.739 | -0.15 | -0.198 | 0.051 | -0.008 | 0.009 | 0.009 | 0 | 0.47 | 0.34 | 0.43 | 0.44 | 0.46 | 0.35 |
| 1 | 220992933 | rs2642431 | -1.368 | -0.063 | -0.322 | 0.257 | 0.02 | -0.023 | -0.023 | 0 | 0.82 | 0.94 | 0.89 | 0.72 | 0.87 | 0.85 |
| 1 | 221007855 | rs2131887 | 0.079 | 0.066 | 0.034 | 0.194 | 0 | 0.015 | 0.015 | 0 | 0.66 | 0.59 | 0.62 | 0.52 | 0.62 | 0.59 |
| 1 | 221010205 | rs12073837 | 1.933 | -0.142 | -0.35 | 0.073 | -0.02 | -0.021 | -0.021 | 0 | 0.18 | 0.09 | 0.31 | 0.28 | 0.27 | 0.26 |
| 1 | 221010513 | rs2807848 | 0.402 | -0.359 | -0.403 | -0.119 | 0.003 | 0.021 | 0.021 | 0 | 0.37 | 0.35 | 0.28 | 0.2 | 0.32 | 0.26 |
| 1 | 221011546 | rs11118618 | 0.999 | 0.15 | -0.405 | 0.219 | -0.002 | -0.002 | -0.002 | 0.013 | 0.15 | 0.14 | 0.24 | 0.18 | 0.24 | 0.27 |
| 1 | 221022916 | rs1603046 | 0.877 | -0.155 | -0.453 | -0.117 | -0.02 | -0.003 | -0.003 | 0.005 | 0.21 | 0.32 | 0.11 | 0.24 | 0.13 | 0.15 |
| 1 | 221029841 | rs2807854 | -0.449 | -0.01 | -0.812 | -0.356 | 0.017 | 0.082 | 0.082 | 0.159 | 0.59 | 0.71 | 0.77 | 0.91 | 0.65 | 0.72 |
| 1 | 221042175 | rs11118622 | -0.036 | 0.465 | -0.901 | 0.132 | 0.002 | 0.016 | 0.016 | 0 | 0.23 | 0.23 | 0.41 | 0.41 | 0.27 | 0.4 |
| 1 | 221043003 | rs2807880 | -1.248 | 0.268 | -1.197 | -0.458 | -0.032 | 0.083 | 0.083 | 0.7 | 0.84 | 0.98 | 0.89 | 0.98 | 0.83 | 0.81 |
| 1 | 221051792 | rs3806325 | 0.552 | 0.367 | -1.175 | -0.312 | -0.045 | 0.04 | 0.04 | 0 | 0.12 | 0.01 | 0.15 | 0.07 | 0.17 | 0.15 |
| 1 | 221052640 | rs2807857 | -0.527 | 0.231 | -1.216 | -0.303 | -0.006 | -0.014 | -0.014 | 0.009 | 0.75 | 0.79 | 0.64 | 0.6 | 0.67 | 0.58 |
| 1 | 221055463 | rs2247213 | -0.338 | -0.564 | -0.934 | -1.357 | -0.012 | -0.103 | -0.103 | 0.466 | 0.39 | 0.47 | 0.29 | 0.1 | 0.34 | 0.24 |
| 1 | 221057646 | rs2738755 | -0.338 | -0.564 | -0.934 | -1.357 | -0.012 | -0.103 | -0.103 | 0 | 0.39 | 0.47 | 0.29 | 0.1 | 0.34 | 0.24 |
| 1 | 221057662 | rs3738182 | 0.637 | -0.67 | -0.603 | -1.416 | 0.01 | -0.034 | -0.034 | 0 | 0.15 | 0.26 | 0.18 | 0.09 | 0.18 | 0.2 |
| 1 | 221057739 | rs11578466 | 0.999 | -0.863 | -0.355 | -1.209 | 0.031 | 0.058 | 0.058 | 0.094 | 0.23 | 0.07 | 0.11 | 0 | 0.16 | 0.04 |
| 1 | 221058059 | rs2738756 | 0.819 | -0.841 | -0.404 | -1.199 | -0.001 | -0.049 | -0.049 | 1.642 | 0.24 | 0.21 | 0.11 | 0 | 0.16 | 0.04 |
| 1 | 221062684 | rs2784252 | 0.78 | -0.846 | -0.345 | -1.21 | -0.009 | -0.07 | -0.07 | 0 | 0.26 | 0.26 | 0.13 | 0 | 0.18 | 0.04 |
| 1 | 221063887 | rs6678259 | 1.817 | -1.082 | -0.535 | -1.149 | 0.02 | -0.022 | -0.022 | 0 | 0.15 | 0.1 | 0.07 | 0.25 | 0.09 | 0.12 |
| 1 | 221064481 | rs2807862 | 0.531 | -1.111 | -0.411 | -1.178 | -0.059 | -0.165 | -0.165 | 0 | 0.28 | 0.5 | 0.16 | 0.02 | 0.18 | 0.09 |
| 1 | 221067103 | rs2807863 | -0.347 | -0.837 | -0.666 | -1.042 | -0.02 | -0.002 | -0.002 | 0 | 0.61 | 0.77 | 0.51 | 0.43 | 0.52 | 0.44 |
| 1 | 221069370 | rs6541166 | 0.234 | -0.912 | -0.658 | -1.111 | 0.009 | 0.009 | 0.009 | 2.326 | 0.41 | 0.36 | 0.19 | 0.25 | 0.26 | 0.16 |
| 1 | 221084222 | rs1537806 | 0.031 | -0.762 | -0.839 | -1.172 | -0.004 | 0.036 | 0.036 | 0 | 0.45 | 0.62 | 0.44 | 0.07 | 0.43 | 0.24 |
| 1 | 221094068 | rs2807869 | -0.499 | -0.701 | -1.111 | -1.023 | -0.018 | -0.013 | -0.013 | 0.165 | 0.65 | 0.79 | 0.53 | 0.38 | 0.55 | 0.49 |
| 1 | 221099886 | rs11587049 | 2.043 | -0.754 | -1.022 | -1.202 | -0.004 | 0.002 | 0.002 | 0.239 | 0.17 | 0.16 | 0.27 | 0.06 | 0.26 | 0.15 |
| 1 | 221103744 | rs1335915 | 0.653 | -0.815 | -1.204 | -1.235 | -0.002 | -0.06 | -0.06 | 0 | 0.29 | 0.24 | 0.15 | 0 | 0.2 | 0.1 |
| 1 | 221107246 | rs12567705 | 1.925 | -0.815 | -1.507 | -1.192 | 0.009 | -0.087 | -0.087 | 0.054 | 0.15 | 0 | 0.05 | 0.3 | 0.09 | 0.13 |
| 1 | 221107796 | rs10495157 | 0.462 | -0.549 | -1.002 | -1.007 | 0.026 | 0.025 | 0.025 | 0 | 0.28 | 0.12 | 0.15 | 0 | 0.2 | 0.1 |
| 1 | 221111780 | rs7542375 | 0.638 | -0.145 | -0.816 | -0.584 | 0.006 | 0.009 | 0.009 | 0 | 0.36 | 0.53 | 0.38 | 0.41 | 0.35 | 0.39 |
| 1 | 221113623 | rs2807873 | -0.614 | 0.244 | -1.045 | -0.824 | 0.003 | 0.009 | 0.009 | 0 | 0.29 | 0.18 | 0.15 | 0 | 0.25 | 0.13 |
| 1 | 221126914 | rs10779417 | 0.807 | 0.397 | -1.678 | -0.062 | 0.011 | 0.045 | 0.045 | 0.365 | 0.53 | 0.26 | 0.48 | 0.38 | 0.54 | 0.33 |
| 1 | 221136672 | rs10495161 | -0.452 | 0.965 | -1.332 | -0.045 | -0.058 | 0.033 | 0.033 | 0 | 0.17 | 0 | 0.16 | 0.11 | 0.2 | 0.13 |
| 1 | 221151806 | rs2808220 | -0.489 | 0.921 | -0.701 | 0.623 | -0.007 | -0.01 | -0.01 | 0 | 0.37 | 0.27 | 0.56 | 0.5 | 0.42 | 0.6 |
| 1 | 221152881 | rs17513397 | -0.266 | 0.291 | -0.189 | 0.277 | 0 | -0.005 | -0.005 | 0.009 | 0.06 | 0.08 | 0.05 | 0.02 | 0.06 | 0.08 |
| 1 | 221155819 | rs2784264 | 0.179 | 0.303 | -0.443 | -0.146 | 0.035 | 0.041 | 0.041 | 2.527 | 0.49 | 0.71 | 0.46 | 0.57 | 0.42 | 0.41 |
| 1 | 221171310 | rs2484696 | 1.022 | 0.152 | -0.093 | 0.326 | -0.003 | 0.005 | 0.005 | 0 | 0.33 | 0.34 | 0.25 | 0.37 | 0.31 | 0.18 |
| 1 | 221174133 | rs2808212 | -0.937 | 0.725 | -0.739 | 0.376 | -0.028 | 0.04 | 0.04 | 0.142 | 0.78 | 0.9 | 0.71 | 0.89 | 0.68 | 0.8 |
| 1 | 221194793 | rs4142350 | -0.084 | 0.068 | -0.437 | 0.592 | 0.04 | 0.036 | 0.036 | 2.113 | 0.49 | 0.72 | 0.45 | 0.56 | 0.43 | 0.41 |
| 1 | 221200621 | rs7536397 | -0.461 | 0.316 | -0.345 | 0.243 | -0.004 | -0.019 | -0.019 | 0 | 0.06 | 0.1 | 0.05 | 0.02 | 0.06 | 0.08 |
| 1 | 221209184 | rs7542237 | 0.192 | 0.171 | -0.482 | 0.12 | -0.02 | 0.053 | 0.053 | 0 | 0.54 | 0.81 | 0.53 | 0.59 | 0.55 | 0.39 |
| 1 | 221228069 | rs13375635 | -0.01 | 0.075 | -0.591 | 0.819 | 0.001 | -0.013 | -0.013 | 0 | 0.08 | 0.07 | 0.21 | 0.21 | 0.09 | 0.1 |
| 1 | 221244372 | rs11118642 | 0.506 | -0.613 | -0.129 | 0.165 | 0.001 | 0.001 | 0.001 | 0 | 0.54 | 0.38 | 0.35 | 0.39 | 0.55 | 0.32 |
| 1 | 221255989 | rs7523927 | -0.154 | -0.581 | -0.095 | 0.13 | -0.007 | -0.004 | -0.004 | 0 | 0.36 | 0.43 | 0.43 | 0.41 | 0.33 | 0.56 |
| 1 | 221261287 | rs1992480 | -0.171 | -0.214 | -0.223 | 0.366 | -0.009 | -0.036 | -0.036 | 0 | 0.91 | 0.96 | 0.78 | 0.8 | 0.89 | 0.9 |
| 1 | 221280094 | rs10218738 | -0.304 | -0.203 | -0.217 | 0.382 | -0.01 | -0.033 | -0.033 | 0 | 0.91 | 0.96 | 0.79 | 0.8 | 0.89 | 0.9 |
| 1 | 221280235 | rs10218741 | 0.411 | -0.438 | -0.043 | 0.237 | 0.003 | 0.007 | 0.007 | 0 | 0.62 | 0.47 | 0.4 | 0.39 | 0.58 | 0.33 |
| 1 | 221289044 | rs1965817 | -0.4 | -0.414 | -0.124 | 0.25 | -0.008 | -0.029 | -0.029 | 0 | 0.91 | 0.95 | 0.79 | 0.79 | 0.89 | 0.88 |
| 1 | 221298122 | rs4595370 | 0.109 | -0.41 | 0.022 | 0.186 | 0.004 | 0.011 | 0.011 | 0.069 | 0.28 | 0.41 | 0.38 | 0.4 | 0.31 | 0.49 |
| 1 | 221313677 | rs4484929 | 0.321 | -0.48 | 0.015 | 0.245 | -0.002 | 0 | 0 | 0 | 0.62 | 0.56 | 0.4 | 0.32 | 0.59 | 0.36 |
| 1 | 221325413 | rs6667902 | 0.213 | -0.448 | 0.024 | 0.356 | -0.002 | -0.005 | -0.005 | 0.016 | 0.28 | 0.26 | 0.35 | 0.4 | 0.29 | 0.52 |
| 1 | 221328346 | rs7515181 | 0.124 | -0.5 | 0.037 | 0.431 | -0.001 | -0.001 | -0.001 | 0.165 | 0.29 | 0.35 | 0.37 | 0.4 | 0.29 | 0.48 |
| 1 | 221339474 | rs17009536 | 0.253 | -0.496 | 0.005 | 0.421 | -0.003 | -0.005 | -0.005 | 0.165 | 0.28 | 0.26 | 0.35 | 0.4 | 0.29 | 0.48 |
| 1 | 221352843 | rs12032422 | 0.927 | -0.621 | -0.825 | -0.202 | -0.056 | -0.075 | -0.075 | 0 | 0.28 | 0.07 | 0.17 | 0.32 | 0.31 | 0.27 |

| | | | | | | | | | | | | | | | |
|---|---|---|---|---|---|---|---|---|---|---|---|---|---|---|---|
| 1 | 221365849 | rs6541181 | -0.42 | -0.662 | -0.309 | -0.039 | -0.001 | -0.001 | -0.001 | 0.09 | 0.65 | 0.59 | 0.51 | 0.58 | 0.64 | 0.49 |
| 1 | 221377435 | rs4846689 | 0.102 | -0.572 | -0.359 | 0.063 | 0.001 | -0.033 | -0.033 | 0 | 0.12 | 0.04 | 0.15 | 0.19 | 0.08 | 0.12 |
| 1 | 221388783 | rs10779428 | 0.162 | -0.967 | 0.012 | -0.13 | -0.001 | -0.006 | -0.006 | 0 | 0.63 | 0.31 | 0.43 | 0.4 | 0.64 | 0.5 |
| 1 | 221393875 | rs10863595 | 0.089 | -0.891 | -0.162 | -0.126 | -0.001 | 0.004 | 0.004 | 0 | 0.63 | 0.47 | 0.43 | 0.32 | 0.64 | 0.48 |
| 1 | 221397750 | rs17009664 | 1.231 | -0.81 | -0.549 | 0.373 | -0.007 | -0.017 | -0.017 | 0 | 0.06 | 0.02 | 0.13 | 0.24 | 0.05 | 0.09 |
| 1 | 221399536 | rs6687985 | -0.502 | -0.895 | -0.366 | 0.341 | -0.002 | -0.002 | -0.002 | 0.107 | 0.71 | 0.74 | 0.59 | 0.53 | 0.68 | 0.56 |
| 1 | 221403841 | rs2889909 | 1.209 | -0.38 | -0.854 | 0.015 | -0.041 | -0.095 | -0.095 | 0 | 0.22 | 0.01 | 0.14 | 0.26 | 0.2 | 0.26 |
| 1 | 221413423 | rs9431087 | 0.358 | -0.125 | 0.054 | 0.555 | 0 | 0 | 0 | 0 | 0.63 | 0.35 | 0.43 | 0.29 | 0.63 | 0.47 |
| 1 | 221425021 | rs9431457 | -0.449 | 0.12 | -0.623 | 0.586 | -0.015 | -0.009 | -0.009 | 0 | 0.73 | 0.81 | 0.59 | 0.61 | 0.68 | 0.58 |
| 1 | 221425311 | rs6664072 | -0.404 | 0.136 | -0.682 | 0.498 | -0.006 | -0.007 | -0.007 | 0.069 | 0.72 | 0.78 | 0.59 | 0.54 | 0.68 | 0.56 |
| 1 | 221430342 | rs11802047 | 0.741 | 0.025 | -0.836 | 0.904 | 0.003 | 0.006 | 0.006 | 0 | 0.07 | 0.07 | 0.11 | 0.19 | 0.05 | 0.09 |
| 1 | 221440464 | rs3924909 | -0.012 | -0.304 | -1.121 | 0.447 | 0.022 | 0.012 | 0.012 | 0 | 0.45 | 0.25 | 0.27 | 0.37 | 0.37 | 0.34 |
| 1 | 221442575 | rs3923935 | -0.086 | -0.45 | -1.036 | 0.472 | 0.021 | 0.003 | 0.003 | 0 | 0.46 | 0.3 | 0.3 | 0.42 | 0.37 | 0.37 |
| 1 | 221443609 | rs4609432 | 0.136 | -0.411 | -1.116 | 0.728 | 0 | 0 | 0 | 0.092 | 0.69 | 0.64 | 0.64 | 0.6 | 0.72 | 0.72 |
| 1 | 221443624 | rs4609433 | 0.496 | -0.337 | -1.379 | 0.727 | 0.012 | -0.031 | -0.031 | 2.549 | 0.63 | 0.7 | 0.6 | 0.53 | 0.67 | 0.64 |
| 1 | 221448322 | rs4846704 | 0.431 | -0.082 | -1.389 | 0.487 | -0.008 | -0.012 | -0.012 | 1.307 | 0.22 | 0.08 | 0.21 | 0.19 | 0.21 | 0.23 |
| 1 | 221451217 | rs10489952 | -1.437 | 0.614 | -1.295 | 0.019 | -0.018 | 0.009 | 0.009 | 0 | 0.87 | 0.86 | 0.88 | 1 | 0.82 | 0.91 |
| 1 | 221456218 | rs3924520 | 0.724 | 0.464 | -1.558 | 0.111 | -0.019 | -0.004 | -0.004 | 0 | 0.22 | 0.15 | 0.17 | 0.23 | 0.26 | 0.22 |
| 1 | 221472290 | rs3890716 | 1.432 | 0.269 | -0.979 | 0.107 | 0.006 | -0.047 | -0.047 | 0 | 0.17 | 0.27 | 0.15 | 0.07 | 0.16 | 0.06 |
| 1 | 221480857 | rs11118691 | -0.834 | -0.09 | -1.041 | 0.215 | -0.007 | 0 | 0 | 0.303 | 0.62 | 0.54 | 0.6 | 0.36 | 0.71 | 0.54 |
| 1 | 221484024 | rs6660101 | 2.17 | -0.342 | -1.235 | 0.525 | -0.003 | 0.009 | 0.009 | 0.052 | 0.2 | 0.22 | 0.27 | 0.36 | 0.19 | 0.29 |
| 1 | 221486559 | rs10863602 | 0.924 | -0.152 | -1.344 | 0.441 | -0.012 | -0.01 | -0.01 | 0.421 | 0.17 | 0.13 | 0.17 | 0.17 | 0.19 | 0.22 |
| 1 | 221493742 | rs2998397 | 0.166 | -0.178 | -1.274 | 0.544 | 0.005 | -0.028 | -0.028 | 0 | 0.68 | 0.7 | 0.75 | 0.62 | 0.71 | 0.65 |
| 1 | 221495334 | rs17521587 | 2.03 | -0.019 | -1.516 | 0.451 | 0.006 | -0.088 | -0.088 | 0 | 0.17 | 0.2 | 0.13 | 0 | 0.15 | 0.06 |
| 1 | 221496601 | rs2996005 | -0.006 | 0.129 | -1.871 | 0.434 | 0 | 0.009 | 0.009 | 1.166 | 0.48 | 0.48 | 0.47 | 0.26 | 0.51 | 0.37 |
| 1 | 221500943 | rs2996003 | 0.728 | 0.511 | -1.962 | 0.279 | -0.005 | -0.07 | -0.07 | 0 | 0.35 | 0.32 | 0.24 | 0.04 | 0.29 | 0.16 |
| 1 | 221511715 | rs4313406 | -0.966 | 0.488 | -1.977 | 0.337 | 0.001 | 0.006 | 0.006 | 0.886 | 0.62 | 0.5 | 0.54 | 0.37 | 0.62 | 0.48 |
| 1 | 221515711 | rs17587612 | 1.548 | 0.507 | -1.982 | 0.298 | -0.018 | -0.012 | -0.012 | 0.078 | 0.15 | 0.11 | 0.16 | 0.16 | 0.19 | 0.22 |
| 1 | 221523403 | rs10863605 | -1.488 | 0.519 | -1.71 | 0.047 | 0.001 | 0.013 | 0.013 | 0.65 | 0.62 | 0.53 | 0.54 | 0.48 | 0.62 | 0.49 |
| 1 | 221523673 | rs4517352 | 0.384 | 0.579 | -1.733 | -0.451 | -0.005 | -0.07 | -0.07 | 0 | 0.35 | 0.32 | 0.24 | 0.04 | 0.29 | 0.16 |
| 1 | 221526452 | rs12044275 | -0.878 | 0.784 | -2.184 | 0.427 | -0.015 | -0.034 | -0.034 | 0 | 0.27 | 0.18 | 0.3 | 0.34 | 0.32 | 0.33 |
| 1 | 221533350 | rs12410731 | 0.059 | 1.353 | -1.597 | 0.368 | 0.002 | 0.001 | 0.001 | 1.5 | 0.13 | 0.13 | 0.24 | 0.19 | 0.22 | 0.21 |
| 1 | 221535733 | rs12027086 | 0.602 | 1.34 | -1.608 | 0.362 | -0.067 | -0.062 | -0.062 | 0 | 0.11 | 0.01 | 0.23 | 0.19 | 0.22 | 0.21 |
| 1 | 221543543 | rs4846715 | 0.14 | 1.37 | -1.59 | 0.377 | 0.002 | 0.001 | 0.001 | 1.5 | 0.13 | 0.13 | 0.24 | 0.19 | 0.22 | 0.21 |
| 1 | 221550655 | rs731372 | 0.479 | 0.647 | -1.531 | 0.003 | -0.005 | 0.057 | 0.057 | 1.424 | 0.17 | 0.02 | 0.1 | 0.04 | 0.15 | 0.1 |
| 1 | 221556115 | rs17009942 | -0.827 | 1.287 | -0.839 | 0.275 | 0.004 | -0.017 | -0.017 | 0.485 | 0.14 | 0.19 | 0.14 | 0.05 | 0.17 | 0.14 |
| 1 | 221556157 | rs12029534 | -0.658 | 1.176 | -0.766 | 0.885 | 0.002 | -0.005 | -0.005 | 0 | 0.08 | 0.08 | 0.08 | 0.2 | 0.07 | 0.12 |
| 1 | 221556266 | rs17436982 | -0.309 | 1.639 | -0.511 | 0.614 | 0.023 | -0.019 | -0.019 | 0 | 0.1 | 0.15 | 0.11 | 0.03 | 0.15 | 0.1 |
| 1 | 221556290 | rs3123739 | 0.031 | 1.287 | -0.086 | 0.607 | -0.02 | 0 | 0 | 0.113 | 0.76 | 0.62 | 0.82 | 0.82 | 0.83 | 0.87 |
| 1 | 221563143 | rs12139598 | -1.028 | 1.186 | -0.362 | 0.821 | 0.004 | 0.012 | 0.012 | 0 | 0.28 | 0.23 | 0.28 | 0.23 | 0.24 | 0.22 |
| 1 | 221568175 | rs12736367 | 0.087 | 1.268 | -0.299 | 0.827 | 0.018 | -0.023 | -0.023 | 0 | 0.1 | 0.16 | 0.1 | 0.03 | 0.13 | 0.1 |
| 1 | 221569322 | rs9729478 | 0.357 | 1.636 | -0.431 | 1.017 | -0.011 | -0.011 | -0.011 | 0 | 0.13 | 0.07 | 0.21 | 0.22 | 0.22 | 0.24 |
| 1 | 221577849 | rs1903136 | 1.86 | 1.056 | -0.576 | 0.574 | 0.059 | 0.082 | 0.082 | 0.084 | 0.5 | 0.2 | 0.45 | 0.44 | 0.42 | 0.47 |
| 1 | 221579628 | rs3131533 | 0.169 | 1.152 | 0.144 | 0.608 | 0.083 | 0.027 | 0.027 | 0 | 0.82 | 0.62 | 0.83 | 0.85 | 0.77 | 0.83 |
| 1 | 221581041 | rs3131534 | -1.772 | 1.086 | 0.206 | 0.651 | 0.064 | 0.095 | 0.095 | 0 | 0.92 | 0.79 | 0.93 | 0.89 | 0.9 | 0.93 |
| 1 | 221582028 | rs1459223 | 1.403 | 1.06 | 0.416 | 0.616 | 0 | -0.007 | -0.007 | 1.23 | 0.58 | 0.47 | 0.56 | 0.6 | 0.46 | 0.55 |
| 1 | 221583559 | rs3131537 | 2.676 | 1.01 | 0.406 | 0.484 | 0.021 | 0.011 | 0.011 | 0 | 0.37 | 0.17 | 0.36 | 0.39 | 0.34 | 0.43 |
| 1 | 221585043 | rs10779435 | -1.644 | 1.271 | 0.587 | 0.442 | 0.064 | 0.088 | 0.088 | 0.007 | 0.92 | 0.79 | 0.93 | 0.89 | 0.9 | 0.93 |
| 1 | 221603164 | rs9727891 | -1.246 | 1.25 | -0.162 | 0.336 | -0.006 | -0.002 | -0.002 | 0 | 0.33 | 0.29 | 0.37 | 0.29 | 0.44 | 0.37 |
| 1 | 221604980 | rs9726422 | 0.524 | 0.96 | -0.264 | 0.483 | 0.008 | 0.008 | 0.008 | 0 | 0.16 | 0.17 | 0.12 | 0.07 | 0.08 | 0.05 |
| 1 | 221605304 | rs1031675 | -0.594 | 1.68 | -0.15 | 0.753 | -0.004 | -0.005 | -0.005 | 0 | 0.2 | 0.15 | 0.26 | 0.26 | 0.29 | 0.22 |
| 1 | 221615451 | rs10863614 | 0.674 | 1.273 | -0.442 | 0.377 | -0.023 | 0.024 | 0.024 | 0 | 0.53 | 0.78 | 0.51 | 0.45 | 0.49 | 0.53 |
| 1 | 221624817 | rs11118717 | 1.52 | 1.266 | 0.099 | 0.414 | 0.002 | 0.001 | 0.001 | 0 | 0.34 | 0.28 | 0.34 | 0.39 | 0.34 | 0.39 |
| 1 | 221627469 | rs6655923 | -0.763 | 1.305 | -0.016 | 0.374 | -0.02 | -0.035 | -0.035 | 0 | 0.21 | 0.17 | 0.27 | 0.45 | 0.33 | 0.31 |
| 1 | 221631938 | rs12757436 | -1.11 | 1.3 | -0.182 | 0.427 | -0.1 | -0.141 | -0.141 | 1.98 | 0.24 | 0.01 | 0.3 | 0.4 | 0.35 | 0.33 |
| 1 | 221632498 | rs1459229 | -0.468 | 1.242 | 0.309 | 0.265 | -0.002 | -0.027 | -0.027 | 0 | 0.66 | 0.68 | 0.7 | 0.91 | 0.74 | 0.82 |
| 1 | 221652503 | rs1903138 | -0.328 | 1.401 | 0.217 | 0.389 | -0.085 | -0.115 | -0.115 | 0.117 | 0.17 | 0.01 | 0.25 | 0.35 | 0.28 | 0.23 |
| 1 | 221654528 | rs6694450 | 0.733 | 1.251 | 0.38 | 0.413 | -0.002 | -0.001 | -0.001 | 0 | 0.45 | 0.51 | 0.44 | 0.55 | 0.43 | 0.54 |
| 1 | 221657634 | rs11118731 | 0.857 | 1.24 | 0.369 | 0.406 | 0.001 | -0.004 | -0.004 | 0 | 0.43 | 0.39 | 0.41 | 0.55 | 0.43 | 0.54 |
| 1 | 221659411 | rs997927 | 0.701 | 1.289 | -0.39 | 0.431 | -0.001 | -0.001 | -0.001 | 0 | 0.46 | 0.57 | 0.44 | 0.55 | 0.44 | 0.54 |
| 1 | 221674514 | rs966769 | 0.864 | 1.408 | -0.676 | 0.648 | -0.04 | -0.016 | -0.016 | 0.028 | 0.07 | 0.01 | 0.21 | 0.32 | 0.19 | 0.18 |
| 1 | 221677434 | rs12406123 | 0.332 | 1.014 | -0.672 | 0.406 | 0.012 | -0.005 | -0.005 | 0 | 0.09 | 0 | 0.08 | 0.08 | 0.06 | 0.1 |
| 1 | 221677672 | rs10779438 | 0.595 | 1.113 | -0.573 | 0.333 | 0.047 | 0.079 | 0.079 | 0 | 0.52 | 0.3 | 0.59 | 0.73 | 0.63 | 0.63 |
| 1 | 221682007 | rs10863628 | 0.595 | 1.113 | -0.573 | 0.333 | 0.047 | 0.079 | 0.079 | 0 | 0.52 | 0.3 | 0.59 | 0.73 | 0.63 | 0.63 |
| 1 | 221684604 | rs2005574 | -0.04 | 1.072 | -0.557 | 0.332 | 0.023 | 0.048 | 0.048 | 0.238 | 0.59 | 0.31 | 0.64 | 0.79 | 0.69 | 0.72 |
| 1 | 221685360 | rs9725646 | 1.65 | 1.089 | -0.562 | 0.294 | 0.007 | 0.003 | 0.003 | 0 | 0.35 | 0.17 | 0.32 | 0.38 | 0.32 | 0.38 |
| 1 | 221690507 | rs1858566 | -0.893 | 1.103 | -0.489 | 0.292 | 0.007 | 0.1 | 0.1 | 0.34 | 0.68 | 0.9 | 0.73 | 0.91 | 0.75 | 0.81 |
| 1 | 221690855 | rs9308395 | 0.584 | 1.404 | -0.207 | 0.523 | 0.017 | 0.02 | 0.02 | 0 | 0.44 | 0.37 | 0.38 | 0.44 | 0.39 | 0.48 |
| 1 | 221694627 | rs9730262 | -0.251 | 0.867 | 0.153 | 0.567 | 0.022 | 0.039 | 0.039 | 0.35 | 0.07 | 0.37 | 0.07 | 0.05 | 0.04 | 0.05 |

| | | | | | | | | | | | | | | | | |
|---|---|---|---|---|---|---|---|---|---|---|---|---|---|---|---|---|
| 1 | 221696159 | rs10863631 | 0.771 | 0.873 | 0.035 | 0.593 | 0.01 | -0.025 | -0.025 | 0 | 0.4 | 0.33 | 0.34 | 0.49 | 0.36 | 0.47 |
| 1 | 221696457 | rs10863632 | -0.818 | 0.897 | -0.147 | 0.519 | 0.008 | 0.111 | 0.111 | 0 | 0.65 | 0.89 | 0.71 | 0.91 | 0.71 | 0.81 |
| 1 | 221710721 | rs2889911 | -0.022 | 0.941 | -0.33 | 0.617 | -0.009 | -0.018 | -0.018 | 6.312 | 0.57 | 0.53 | 0.63 | 0.72 | 0.64 | 0.7 |
| 1 | 221712011 | rs12025155 | -0.239 | 1.091 | -0.541 | 0.665 | -0.104 | -0.123 | -0.123 | 0 | 0.18 | 0.02 | 0.29 | 0.35 | 0.31 | 0.28 |
| 1 | 221715719 | rs1026514 | -0.714 | 1.003 | -0.696 | 0.277 | 0.001 | 0.096 | 0.096 | 0.021 | 0.7 | 0.92 | 0.73 | 0.9 | 0.75 | 0.8 |
| 1 | 221723419 | rs12411238 | 0.098 | 0.99 | -0.414 | 0.526 | 0.005 | 0 | 0 | 0.941 | 0.05 | 0.04 | 0.03 | 0.06 | 0.03 | 0.05 |
| 1 | 221723524 | rs11118750 | 0.665 | 0.936 | -0.391 | 0.365 | 0.02 | 0.057 | 0.057 | 0.686 | 0.31 | 0.16 | 0.26 | 0.09 | 0.23 | 0.17 |
| 1 | 221734237 | rs9725124 | -0.583 | 1.158 | -0.369 | 0.638 | -0.013 | -0.027 | -0.027 | 0.674 | 0.23 | 0.22 | 0.3 | 0.42 | 0.32 | 0.42 |
| 1 | 221737354 | rs17041853 | 0.413 | 1.124 | -0.407 | 0.638 | 0.003 | -0.002 | -0.002 | 0.218 | 0.05 | 0.04 | 0.03 | 0.08 | 0.03 | 0.05 |
| 1 | 221743047 | rs12564947 | -0.222 | 1.113 | -0.44 | 0.851 | -0.108 | -0.095 | -0.095 | 0 | 0.19 | 0.02 | 0.28 | 0.3 | 0.32 | 0.4 |
| 1 | 221752376 | rs12133543 | 0.951 | 1.052 | -0.395 | 0.765 | 0.018 | 0.054 | 0.054 | 0.537 | 0.3 | 0.15 | 0.27 | 0.09 | 0.23 | 0.16 |
| 1 | 221753645 | rs9659823 | -1.257 | 0.791 | -0.339 | 0.898 | 0.001 | 0.005 | 0.005 | 1.383 | 0.29 | 0.3 | 0.34 | 0.49 | 0.36 | 0.54 |
| 1 | 221771162 | rs11118768 | -1.034 | 0.939 | -0.154 | 0.902 | -0.001 | 0 | 0 | 0 | 0.59 | 0.6 | 0.62 | 0.6 | 0.63 | 0.63 |
| 1 | 221787416 | rs4498809 | -1.033 | 0.957 | -0.091 | 0.831 | -0.001 | 0.001 | 0.001 | 0 | 0.59 | 0.6 | 0.62 | 0.6 | 0.63 | 0.63 |
| 1 | 221809552 | rs3860324 | -0.436 | 0.887 | -0.426 | 0.893 | 0.012 | 0.041 | 0.041 | 0.11 | 0.51 | 0.15 | 0.54 | 0.4 | 0.57 | 0.57 |
| 1 | 221811055 | rs6697462 | -0.804 | 1.006 | -0.348 | 0.76 | -0.001 | 0.011 | 0.011 | 0 | 0.57 | 0.33 | 0.59 | 0.6 | 0.6 | 0.62 |
| 1 | 221817308 | rs2176690 | 0.627 | 0.754 | -0.472 | 0.303 | 0.022 | 0.075 | 0.075 | 0.225 | 0.31 | 0.14 | 0.27 | 0.08 | 0.25 | 0.17 |
| 1 | 221817785 | rs12744368 | -0.492 | 0.832 | -0.235 | 0.418 | 0.02 | 0.018 | 0.018 | 0 | 0.47 | 0.59 | 0.38 | 0.29 | 0.37 | 0.28 |
| 1 | 221818681 | rs2176691 | -0.443 | 0.834 | -0.2 | 0.419 | 0.014 | 0.012 | 0.012 | 0 | 0.47 | 0.54 | 0.38 | 0.29 | 0.37 | 0.28 |
| 1 | 221821302 | rs7520263 | -0.443 | 0.827 | -0.156 | 0.441 | 0.014 | 0.012 | 0.012 | 0 | 0.47 | 0.54 | 0.38 | 0.29 | 0.37 | 0.28 |
| 1 | 221824935 | rs12027361 | -0.269 | 0.849 | -0.125 | 0.452 | 0.003 | 0.003 | 0.003 | 0 | 0.45 | 0.41 | 0.36 | 0.29 | 0.37 | 0.28 |
| 1 | 221830871 | rs12136173 | 0.292 | 1.033 | -0.25 | -0.311 | 0.017 | 0.09 | 0.09 | 0 | 0.36 | 0.19 | 0.3 | 0.09 | 0.31 | 0.24 |
| 1 | 221831403 | rs4504898 | -0.226 | 1.295 | -0.118 | -0.307 | -0.004 | -0.026 | -0.026 | 0.018 | 0.46 | 0.5 | 0.39 | 0.17 | 0.4 | 0.27 |
| 1 | 221837700 | rs6701049 | -0.37 | 1.332 | -0.281 | -0.3 | 0.027 | 0.072 | 0.072 | 0 | 0.47 | 0.62 | 0.4 | 0.17 | 0.4 | 0.28 |
| 1 | 221838876 | rs11807554 | 0.446 | 1.255 | -0.284 | -0.348 | 0.013 | 0.071 | 0.071 | 0 | 0.36 | 0.19 | 0.31 | 0.1 | 0.31 | 0.25 |
| 1 | 221847292 | rs7523649 | -0.671 | 1.286 | -0.533 | -0.333 | 0.01 | 0.074 | 0.074 | 0 | 0.52 | 0.62 | 0.44 | 0.18 | 0.47 | 0.35 |
| 1 | 221848970 | rs13376168 | -0.572 | 1.371 | -0.373 | -0.016 | 0.009 | -0.055 | -0.055 | 0.714 | 0.19 | 0.33 | 0.14 | 0.09 | 0.22 | 0.12 |
| 1 | 221856899 | rs10863647 | -1.063 | 1.332 | -0.929 | 0.663 | -0.025 | -0.019 | -0.019 | 0 | 0.16 | 0.1 | 0.2 | 0.21 | 0.28 | 0.38 |
| 1 | 221862428 | rs953111 | -0.071 | 0.043 | -1.767 | 0.224 | -0.037 | -0.056 | -0.056 | 0.406 | 0.45 | 0.1 | 0.45 | 0.54 | 0.46 | 0.56 |
| 1 | 221864710 | rs1815577 | -0.323 | 0.088 | -1.223 | -0.066 | 0.002 | -0.007 | -0.007 | 1.71 | 0.55 | 0.39 | 0.55 | 0.54 | 0.58 | 0.63 |
| 1 | 221867661 | rs11118827 | -0.302 | 0.073 | -1.291 | 0.092 | 0.001 | -0.008 | -0.008 | 0 | 0.55 | 0.39 | 0.54 | 0.59 | 0.57 | 0.67 |
| 1 | 221868406 | rs6687398 | 0.287 | -0.379 | -1.041 | -0.057 | 0 | 0.002 | 0.002 | 0.271 | 0.39 | 0.54 | 0.43 | 0.44 | 0.42 | 0.34 |
| 1 | 221868913 | rs6655992 | -0.488 | -0.876 | -1.241 | -0.799 | 0.013 | 0 | 0 | 0 | 0.31 | 0.35 | 0.19 | 0.03 | 0.23 | 0.14 |
| 1 | 221869434 | rs6604665 | -0.487 | -1.011 | -1.115 | -0.718 | -0.008 | -0.078 | -0.078 | 0 | 0.31 | 0.37 | 0.21 | 0.04 | 0.3 | 0.17 |
| 1 | 221874684 | rs908857 | -0.127 | -1.041 | -1.761 | -0.514 | -0.059 | -0.015 | -0.015 | 0.82 | 0.5 | 0.9 | 0.51 | 0.5 | 0.51 | 0.41 |
| 1 | 221876222 | rs2280339 | 0.991 | -0.802 | -1.289 | -0.288 | -0.033 | -0.058 | -0.058 | 0 | 0.31 | 0.16 | 0.36 | 0.43 | 0.37 | 0.3 |
| 1 | 221883225 | rs908858 | -0.873 | -0.596 | -0.889 | -0.281 | 0.002 | 0.006 | 0.006 | 0 | 0.61 | 0.59 | 0.54 | 0.51 | 0.61 | 0.68 |
| 1 | 221883496 | rs12749729 | -0.214 | 0.045 | -0.322 | 0.345 | 0.003 | 0.003 | 0.003 | 0 | 0.28 | 0.3 | 0.31 | 0.4 | 0.28 | 0.45 |
| 1 | 221884371 | rs11118832 | -0.574 | 0.218 | -0.267 | 0.192 | 0.006 | -0.042 | -0.042 | 3.056 | 0.09 | 0.4 | 0.1 | 0 | 0.08 | 0.04 |
| 1 | 221894739 | rs1474159 | 0.07 | 0.684 | -1.529 | 0.662 | -0.083 | -0.137 | -0.137 | 0.94 | 0.73 | 0.94 | 0.54 | 0.43 | 0.69 | 0.42 |
| 1 | 221895393 | rs17010629 | 0.794 | 0.175 | -0.767 | 0.474 | -0.007 | 0.013 | 0.013 | 0 | 0.58 | 0.37 | 0.37 | 0.31 | 0.54 | 0.33 |
| 1 | 221898783 | rs12091463 | -0.18 | -0.53 | -1.627 | -0.142 | -0.116 | 0.081 | 0.081 | 0 | 0.52 | 0.96 | 0.43 | 0.26 | 0.49 | 0.3 |
| 1 | 221899355 | rs17490808 | 0.792 | -0.541 | -1.401 | -0.262 | -0.007 | 0.032 | 0.032 | 0 | 0.36 | 0.29 | 0.25 | 0.14 | 0.34 | 0.2 |
| 1 | 221902183 | rs4275462 | 0.098 | -0.467 | -0.528 | -0.076 | -0.006 | -0.013 | -0.013 | 0 | 0.11 | 0.15 | 0.14 | 0.06 | 0.14 | 0.08 |
| 1 | 221906395 | rs6679673 | -0.522 | -0.619 | -0.638 | -0.46 | -0.012 | -0.096 | -0.096 | 0 | 0.16 | 0.28 | 0.12 | 0 | 0.16 | 0.14 |
| 1 | 221909193 | rs10863651 | -0.492 | 0.185 | -0.695 | -0.522 | 0.002 | -0.013 | -0.013 | 0 | 0.58 | 0.57 | 0.38 | 0.17 | 0.57 | 0.3 |
| 1 | 221912361 | rs1141714 | -0.34 | -0.357 | -1.499 | -0.612 | 0 | -0.019 | -0.019 | 2.39 | 0.69 | 0.83 | 0.68 | 0.67 | 0.64 | 0.73 |
| 1 | 221915967 | rs4589119 | 0.493 | -0.343 | -1.216 | -0.404 | 0.04 | 0.08 | 0.08 | 0.942 | 0.13 | 0.45 | 0.18 | 0.16 | 0.13 | 0.14 |
| 1 | 221923042 | rs10863652 | -0.538 | -0.657 | -1.381 | -0.547 | 0.004 | 0.003 | 0.003 | 0.152 | 0.26 | 0.71 | 0.26 | 0.23 | 0.24 | 0.24 |
| 1 | 221927143 | rs4603139 | 0.936 | -0.664 | -1.159 | -0.573 | -0.012 | -0.034 | -0.034 | 0 | 0.13 | 0.19 | 0.06 | 0.06 | 0.12 | 0.1 |
| 1 | 221928305 | rs17491176 | 1.934 | -0.557 | -1.718 | -0.992 | -0.009 | 0.032 | 0.032 | 2.361 | 0.15 | 0.08 | 0.1 | 0 | 0.12 | 0.02 |
| 1 | 221929177 | rs6690778 | -0.774 | -0.533 | -1.864 | -1.009 | 0.008 | 0.002 | 0.002 | 0 | 0.64 | 0.83 | 0.67 | 0.68 | 0.62 | 0.72 |
| 1 | 221931688 | rs7550075 | 1.071 | -0.107 | -1.629 | -0.896 | 0.001 | -0.001 | -0.001 | 0.209 | 0.2 | 0.18 | 0.16 | 0.2 | 0.21 | 0.1 |
| 1 | 221934718 | rs11581623 | 1.685 | 0.056 | -1.741 | -0.831 | -0.046 | -0.088 | -0.088 | 0 | 0.18 | 0.06 | 0.17 | 0.39 | 0.2 | 0.35 |
| 1 | 221935450 | rs7546398 | 0.046 | 0.13 | -1.631 | -1.116 | 0.001 | -0.001 | -0.001 | 0 | 0.51 | 0.45 | 0.31 | 0.26 | 0.46 | 0.29 |
| 1 | 221935951 | rs12058360 | 0.772 | -0.105 | -1.528 | -0.812 | -0.013 | -0.057 | -0.057 | 2.328 | 0.19 | 0.15 | 0.07 | 0 | 0.14 | 0.13 |
| 1 | 221939222 | rs7551561 | -1.766 | 0.214 | -1.795 | -0.574 | -0.041 | -0.021 | -0.021 | 0.058 | 0.87 | 0.96 | 0.82 | 0.87 | 0.84 | 0.88 |
| 1 | 221940271 | rs11118852 | -0.23 | -0.351 | -1.472 | -0.624 | 0.042 | 0.055 | 0.055 | 0 | 0.35 | 0.68 | 0.28 | 0.15 | 0.26 | 0.26 |
| 1 | 221943162 | rs7526577 | 1.233 | 0.04 | -1.678 | -0.325 | -0.05 | -0.065 | -0.065 | 0.859 | 0.3 | 0.13 | 0.28 | 0.37 | 0.38 | 0.42 |
| 1 | 221945714 | rs12026940 | 0.691 | -0.276 | -1.149 | 0.033 | 0.001 | 0.045 | 0.045 | 0 | 0.07 | 0.07 | 0.18 | 0.3 | 0.07 | 0.16 |
| 1 | 221946172 | rs4433402 | 0.595 | -0.443 | -0.918 | -0.075 | -0.018 | 0.015 | 0.015 | 0.897 | 0.18 | 0.48 | 0.21 | 0.2 | 0.13 | 0.13 |
| 1 | 221946207 | rs3010852 | -0.864 | -0.367 | -1.437 | -0.122 | 0.067 | -0.03 | -0.03 | 0 | 0.42 | 0.77 | 0.46 | 0.5 | 0.33 | 0.41 |
| 1 | 221948024 | rs3003902 | -0.877 | -0.361 | -1.413 | -0.125 | 0.067 | -0.029 | -0.029 | 0.215 | 0.43 | 0.77 | 0.46 | 0.5 | 0.34 | 0.41 |
| 1 | 221951794 | rs12036282 | 0.544 | -0.05 | -1.288 | 0.166 | -0.006 | 0.011 | 0.011 | 0 | 0.07 | 0.04 | 0.17 | 0.32 | 0.08 | 0.17 |
| 1 | 221952958 | rs11118858 | -0.247 | -0.536 | -0.923 | -0.151 | -0.017 | 0.008 | 0.008 | 0 | 0.26 | 0.3 | 0.26 | 0.32 | 0.21 | 0.29 |
| 1 | 221953558 | rs11118860 | 0.784 | -0.659 | -0.268 | 0.089 | -0.013 | -0.01 | -0.01 | 0 | 0.16 | 0.27 | 0.13 | 0.13 | 0.13 | 0.09 |
| 1 | 221953846 | rs4623704 | 0.799 | -0.659 | -0.158 | -0.031 | -0.012 | -0.029 | -0.029 | 0 | 0.16 | 0.24 | 0.13 | 0.1 | 0.13 | 0.09 |
| 1 | 221958189 | rs4240927 | -0.024 | -0.204 | -0.724 | -0.232 | 0.005 | 0.007 | 0.007 | 0 | 0.49 | 0.56 | 0.43 | 0.43 | 0.52 | 0.56 |
| 1 | 221958998 | rs10495177 | 1.961 | -0.174 | -1.059 | -0.248 | -0.033 | -0.047 | -0.047 | 0 | 0.26 | 0.15 | 0.25 | 0.38 | 0.37 | 0.39 |

| | | | | | | | | | | | | | | | | |
|---|---|---|---|---|---|---|---|---|---|---|---|---|---|---|---|---|
| 1 | 221966781 | rs4290069 | 0.625 | -0.156 | -1.01 | -0.884 | -0.018 | -0.063 | -0.063 | 0 | 0.22 | 0.21 | 0.09 | 0 | 0.15 | 0.16 |
| 1 | 221968224 | rs12089785 | 0.594 | -0.155 | -1.014 | -0.884 | -0.019 | -0.065 | -0.065 | 0.848 | 0.22 | 0.22 | 0.09 | 0 | 0.15 | 0.16 |
| 1 | 221977513 | rs9441867 | -0.965 | 0.31 | -1.366 | -0.576 | -0.075 | 0.092 | 0.092 | 0 | 0.69 | 0.92 | 0.73 | 0.87 | 0.7 | 0.8 |
| 1 | 221982925 | rs3003888 | -0.957 | 0.304 | -1.385 | -0.587 | -0.104 | 0.09 | 0.09 | 0 | 0.69 | 0.92 | 0.71 | 0.87 | 0.68 | 0.79 |
| 1 | 221985915 | rs4317815 | 1.147 | -0.122 | -1.244 | -0.519 | 0.004 | 0.03 | 0.03 | 0 | 0.26 | 0.08 | 0.28 | 0.23 | 0.2 | 0.2 |
| 1 | 221986751 | rs4551601 | 1.081 | -0.177 | -1.087 | -0.817 | -0.001 | -0.025 | -0.025 | 1.967 | 0.22 | 0.22 | 0.14 | 0.04 | 0.22 | 0.2 |
| 1 | 221991112 | rs6604673 | 2.239 | -0.568 | -0.85 | -0.726 | -0.053 | 0.011 | 0.011 | 1.211 | 0.13 | 0.26 | 0.1 | 0.13 | 0.02 | 0.05 |
| 1 | 221991428 | rs12042853 | 2.292 | -0.209 | -1.165 | -0.767 | -0.044 | -0.056 | -0.056 | 0 | 0.14 | 0.05 | 0.21 | 0.48 | 0.27 | 0.35 |
| 1 | 221994043 | rs3003892 | -1.828 | -0.225 | -1.137 | -0.878 | 0.02 | 0.048 | 0.048 | 0.937 | 0.7 | 0.89 | 0.72 | 0.77 | 0.79 | 0.8 |
| 1 | 221994525 | rs10863663 | 1.759 | -0.134 | -0.91 | -0.844 | -0.016 | -0.003 | -0.003 | 0 | 0.19 | 0.16 | 0.3 | 0.2 | 0.28 | 0.19 |
| 1 | 221997281 | rs11580024 | 2.349 | -0.226 | -1.012 | -0.897 | 0.018 | 0.043 | 0.043 | 0 | 0.15 | 0.05 | 0.05 | 0.01 | 0.14 | 0.14 |
| 1 | 222006376 | rs9441977 | -1.63 | -0.1 | -0.861 | -0.763 | 0.011 | 0.037 | 0.037 | 0.384 | 0.71 | 0.89 | 0.71 | 0.77 | 0.79 | 0.81 |
| 1 | 222007116 | rs4846354 | -0.492 | -0.038 | -0.637 | -0.692 | 0.001 | 0.014 | 0.014 | 0 | 0.51 | 0.58 | 0.46 | 0.65 | 0.51 | 0.62 |
| 1 | 222008026 | rs4418610 | 1.079 | -0.391 | -0.382 | -0.959 | -0.006 | -0.039 | -0.039 | 0 | 0.23 | 0.3 | 0.15 | 0.07 | 0.22 | 0.23 |
| 1 | 222008757 | rs17010904 | 2.084 | -0.765 | -0.467 | -0.778 | -0.053 | 0.002 | 0.002 | 0.774 | 0.13 | 0.24 | 0.1 | 0.1 | 0.02 | 0.04 |
| 1 | 222010866 | rs12125690 | 1.262 | -0.564 | -0.783 | -0.879 | 0.007 | -0.003 | -0.003 | 0 | 0.22 | 0.17 | 0.11 | 0 | 0.22 | 0.18 |
| 1 | 222012300 | rs7519761 | 1.035 | -0.51 | -0.664 | -0.717 | 0.01 | 0.053 | 0.053 | 0 | 0.31 | 0.22 | 0.23 | 0.05 | 0.31 | 0.26 |
| 1 | 222012950 | rs4579763 | -1.954 | -0.476 | -0.829 | -0.669 | 0.011 | 0.005 | 0.005 | 2.488 | 0.79 | 0.89 | 0.81 | 0.77 | 0.88 | 0.87 |
| 1 | 222014275 | rs12026361 | 0.614 | -0.542 | -0.712 | -0.596 | -0.049 | -0.022 | -0.022 | 0 | 0.34 | 0.2 | 0.46 | 0.6 | 0.54 | 0.56 |
| 1 | 222015508 | rs3003900 | -1.854 | -0.514 | -0.503 | -0.747 | 0.011 | 0.005 | 0.005 | 1.853 | 0.79 | 0.89 | 0.81 | 0.77 | 0.88 | 0.87 |
| 1 | 222019354 | rs6673674 | 0.23 | 0.19 | -0.623 | -0.362 | -0.048 | 0.024 | 0.024 | 1.482 | 0.33 | 0.21 | 0.49 | 0.68 | 0.49 | 0.56 |
| 1 | 222021079 | rs10495181 | 1.524 | 0.631 | -0.465 | 0.095 | -0.014 | -0.013 | -0.013 | 0 | 0.13 | 0.07 | 0.18 | 0.32 | 0.22 | 0.31 |
| 1 | 222032485 | rs12144971 | -0.268 | 0.495 | -0.569 | -0.083 | 0.001 | 0.002 | 0.002 | 0 | 0.47 | 0.49 | 0.41 | 0.38 | 0.59 | 0.55 |
| 1 | 222039055 | rs4240931 | 0.062 | 0.495 | -0.529 | -0.315 | -0.001 | -0.003 | -0.003 | 0 | 0.52 | 0.47 | 0.59 | 0.64 | 0.41 | 0.45 |
| 1 | 222039967 | rs11118875 | 0.414 | 0.495 | -0.8 | -0.491 | -0.001 | -0.051 | -0.051 | 0.114 | 0.34 | 0.35 | 0.23 | 0.03 | 0.37 | 0.26 |
| 1 | 222042179 | rs6604676 | 0.414 | 0.495 | -0.8 | -0.491 | -0.001 | -0.051 | -0.051 | 0.027 | 0.34 | 0.35 | 0.23 | 0.03 | 0.37 | 0.26 |
| 1 | 222053234 | rs11118882 | 1.202 | 0.899 | -0.918 | 0.401 | -0.006 | 0 | 0 | 0 | 0.12 | 0.1 | 0.17 | 0.32 | 0.19 | 0.29 |
| 1 | 222055403 | rs11811613 | 0.615 | 0.468 | -1.227 | 0.316 | -0.071 | -0.107 | -0.107 | 0.42 | 0.19 | 0.06 | 0.39 | 0.4 | 0.28 | 0.13 |
| 1 | 222061022 | rs11118883 | 0.585 | -0.32 | -0.893 | -1.06 | -0.004 | -0.018 | -0.018 | 0 | 0.29 | 0.22 | 0.21 | 0 | 0.35 | 0.21 |
| 1 | 222065709 | rs7547751 | -0.44 | -0.572 | -0.474 | -1.113 | -0.008 | 0.031 | 0.031 | 0 | 0.59 | 0.62 | 0.35 | 0.26 | 0.47 | 0.48 |
| 1 | 222070603 | rs6704182 | -0.066 | -0.479 | -0.689 | -1.094 | 0.004 | 0.009 | 0.009 | 0 | 0.51 | 0.43 | 0.32 | 0.16 | 0.46 | 0.44 |
| 1 | 222070893 | rs17493593 | 0.703 | -0.617 | -0.896 | -0.515 | 0.028 | 0.028 | 0.028 | 0.11 | 0.25 | 0.15 | 0.12 | 0.16 | 0.13 | 0.25 |
| 1 | 222072819 | rs11579212 | -0.031 | -0.064 | -0.786 | -0.793 | 0.012 | 0.071 | 0.071 | 0 | 0.36 | 0.23 | 0.2 | 0.1 | 0.36 | 0.26 |
| 1 | 222072998 | rs4846734 | 0.828 | 0.671 | -0.531 | -0.099 | -0.006 | -0.004 | -0.004 | 0 | 0.17 | 0.15 | 0.29 | 0.32 | 0.21 | 0.13 |
| 1 | 222076208 | rs2378489 | 1.276 | 0.434 | -0.542 | 0.059 | -0.043 | -0.057 | -0.057 | 0.218 | 0.15 | 0.07 | 0.35 | 0.37 | 0.27 | 0.31 |
| 1 | 222078040 | rs6604689 | -0.39 | -0.173 | -0.286 | -0.245 | -0.014 | -0.005 | -0.005 | 0 | 0.6 | 0.46 | 0.83 | 0.7 | 0.83 | 0.69 |
| 1 | 222078514 | rs17011057 | 2.282 | -0.164 | -0.3 | -0.255 | -0.017 | -0.02 | -0.02 | 0 | 0.06 | 0.08 | 0.01 | 0.01 | 0.02 | 0.05 |
| 1 | 222078560 | rs2378494 | 0.795 | -0.241 | 0.138 | -0.251 | 0.003 | 0.004 | 0.004 | 2.694 | 0.33 | 0.27 | 0.14 | 0.2 | 0.16 | 0.3 |
| 1 | 222078600 | rs4457591 | -0.189 | -0.169 | 0.057 | -0.144 | 0.038 | 0.079 | 0.079 | 0.552 | 0.49 | 0.33 | 0.44 | 0.28 | 0.58 | 0.43 |
| 1 | 222084135 | rs7520693 | -1.144 | -0.11 | 0.205 | -0.021 | 0.026 | 0.082 | 0.082 | 0 | 0.83 | 0.69 | 0.63 | 0.63 | 0.74 | 0.75 |
| 1 | 222086099 | rs17011091 | 1.86 | -0.089 | -0.035 | -0.06 | -0.023 | -0.024 | -0.024 | 0 | 0.06 | 0.09 | 0.02 | 0.01 | 0.02 | 0.05 |
| 1 | 222087406 | rs10863667 | 0.882 | -0.28 | -0.028 | -0.164 | -0.018 | -0.015 | -0.015 | 0 | 0.31 | 0.3 | 0.14 | 0.2 | 0.16 | 0.29 |
| 1 | 222087833 | rs3828088 | 0.16 | -0.061 | -0.136 | -0.19 | -0.009 | -0.014 | -0.014 | 0 | 0.24 | 0.24 | 0.27 | 0.32 | 0.25 | 0.24 |
| 1 | 222092700 | rs12132356 | 1.143 | -0.044 | -0.178 | -0.183 | -0.057 | -0.056 | -0.056 | 0 | 0.17 | 0.09 | 0.25 | 0.28 | 0.23 | 0.19 |
| 1 | 222093006 | rs11118895 | -0.507 | -0.116 | -0.117 | -0.217 | 0 | 0 | 0 | 0.195 | 0.55 | 0.49 | 0.41 | 0.58 | 0.41 | 0.55 |
| 1 | 222096695 | rs7527906 | 0.959 | -0.084 | -0.079 | -0.336 | -0.018 | -0.015 | -0.015 | 0 | 0.31 | 0.31 | 0.14 | 0.2 | 0.17 | 0.3 |
| 1 | 222097527 | rs10863669 | -0.391 | 0.442 | -0.02 | -0.229 | 0.004 | 0.005 | 0.005 | 0 | 0.62 | 0.55 | 0.43 | 0.6 | 0.51 | 0.59 |
| 1 | 222102362 | rs7544501 | 0.773 | 0.485 | -0.184 | -0.196 | -0.033 | -0.039 | -0.039 | 1.975 | 0.18 | 0.16 | 0.26 | 0.32 | 0.24 | 0.23 |
| 1 | 222102582 | rs7554115 | -0.868 | 0.405 | 0.012 | -0.223 | 0.023 | 0.034 | 0.034 | 0.008 | 0.58 | 0.72 | 0.84 | 0.85 | 0.73 | 0.67 |
| 1 | 222109111 | rs1416894 | -0.9 | 0.732 | 0.045 | 0.037 | 0.007 | 0.033 | 0.033 | 0 | 0.8 | 0.73 | 0.57 | 0.63 | 0.74 | 0.73 |
| 1 | 222111006 | rs2800853 | -0.934 | 0.734 | 0.038 | 0.038 | 0.003 | 0.019 | 0.019 | 0 | 0.8 | 0.8 | 0.58 | 0.63 | 0.74 | 0.73 |
| 1 | 222113648 | rs17011146 | 1.088 | 0.75 | -0.064 | -0.052 | 0 | 0.005 | 0.005 | 0 | 0.2 | 0.18 | 0.14 | 0.07 | 0.22 | 0.16 |
| 1 | 222113940 | rs949618 | 1.085 | 0.801 | -0.029 | -0.049 | 0 | -0.005 | -0.005 | 0.201 | 0.2 | 0.21 | 0.15 | 0.17 | 0.24 | 0.17 |
| 1 | 222120767 | rs12122784 | 0.596 | 1.065 | -0.128 | 0.081 | -0.029 | -0.03 | -0.03 | 0 | 0.13 | 0.09 | 0.24 | 0.27 | 0.23 | 0.23 |
| 1 | 222122590 | rs2800846 | 0.234 | 0.766 | -0.111 | -0.031 | 0.017 | -0.009 | -0.009 | 2.017 | 0.66 | 0.58 | 0.33 | 0.32 | 0.52 | 0.49 |
| 1 | 222152402 | rs12036536 | 0.061 | 0.719 | -0.485 | 0.083 | 0.015 | 0.007 | 0.007 | 0 | 0.67 | 0.63 | 0.45 | 0.38 | 0.54 | 0.46 |
| 1 | 222163217 | rs1416896 | -0.072 | 0.886 | -0.63 | -0.103 | -0.016 | 0 | 0 | 0 | 0.31 | 0.24 | 0.34 | 0.31 | 0.4 | 0.32 |
| 1 | 222164948 | rs6687758 | 1.124 | 0.882 | -0.648 | -0.115 | -0.001 | -0.003 | -0.003 | 0 | 0.19 | 0.18 | 0.23 | 0.2 | 0.22 | 0.13 |
| 1 | 222168110 | rs11118924 | 0.675 | 0.898 | -0.613 | -0.156 | 0.034 | 0.073 | 0.073 | 0.017 | 0.47 | 0.3 | 0.21 | 0.17 | 0.32 | 0.33 |
| 1 | 222168464 | rs7546586 | -0.774 | 0.996 | -0.635 | 0.018 | 0.024 | 0.067 | 0.067 | 0 | 0.79 | 0.71 | 0.57 | 0.61 | 0.71 | 0.67 |
| 1 | 222174101 | rs2800856 | -0.021 | 0.889 | -0.704 | -0.042 | -0.007 | -0.004 | -0.004 | 0.056 | 0.6 | 0.65 | 0.45 | 0.6 | 0.5 | 0.52 |
| 1 | 222183226 | rs2800860 | 0.729 | 0.889 | -0.644 | -0.087 | 0.009 | 0.008 | 0.008 | 0 | 0.12 | 0.18 | 0.12 | 0.23 | 0.17 | 0.2 |
| 1 | 222187794 | rs17442148 | 0.855 | 0.243 | -1.189 | -0.534 | 0.033 | -0.007 | -0.007 | 0.04 | 0.21 | 0.01 | 0.07 | 0.16 | 0.1 | 0.2 |
| 1 | 222190318 | rs2225999 | 0.55 | 0.685 | -0.656 | -0.007 | 0.005 | 0.004 | 0.004 | 0.081 | 0.48 | 0.44 | 0.33 | 0.38 | 0.32 | 0.33 |
| 1 | 222195002 | rs2378497 | 0.869 | 0.903 | -0.716 | 0.115 | -0.038 | -0.003 | -0.003 | 0 | 0.11 | 0.06 | 0.11 | 0.11 | 0.17 | 0.2 |
| 1 | 222195599 | rs1815631 | -1.468 | 0.709 | -0.556 | 0.239 | -0.065 | -0.116 | -0.116 | 0.084 | 0.85 | 0.99 | 0.64 | 0.66 | 0.8 | 0.76 |
| 1 | 222204361 | rs2584314 | -0.019 | 0.316 | -0.279 | -0.103 | -0.002 | 0.001 | 0.001 | 0 | 0.37 | 0.36 | 0.35 | 0.42 | 0.4 | 0.35 |
| 1 | 222205808 | rs1156668 | -0.828 | 0.369 | -0.279 | -0.107 | 0.012 | 0.056 | 0.056 | 0 | 0.78 | 0.73 | 0.58 | 0.58 | 0.72 | 0.66 |
| 1 | 222206887 | rs2584315 | -0.454 | 0.148 | -0.15 | 0.224 | -0.011 | 0.025 | 0.025 | 0 | 0.36 | 0.26 | 0.34 | 0.3 | 0.4 | 0.34 |

| | | | | | | | | | | | | | | | | |
|---|---|---|---|---|---|---|---|---|---|---|---|---|---|---|---|---|
| 1 | 222216000 | rs2790746 | -0.459 | 0.239 | -0.846 | 0.265 | -0.07 | 0.003 | 0.003 | 0 | 0.35 | 0.09 | 0.33 | 0.31 | 0.41 | 0.34 |
| 1 | 222222851 | rs7538392 | -0.425 | 0.702 | -0.729 | 0.342 | 0 | 0 | 0 | 0 | 0.35 | 0.39 | 0.35 | 0.42 | 0.4 | 0.34 |
| 1 | 222230715 | rs2066136 | -0.16 | 0.582 | -0.943 | 0.401 | 0.013 | 0.02 | 0.02 | 0 | 0.59 | 0.67 | 0.65 | 0.65 | 0.67 | 0.49 |
| 1 | 222241688 | rs7548749 | -0.271 | 1.023 | -0.182 | 0.655 | 0.023 | 0.031 | 0.031 | 0.638 | 0.33 | 0.55 | 0.37 | 0.43 | 0.41 | 0.25 |
| 1 | 222250020 | rs10495185 | -0.245 | 0.716 | 0.041 | 0.559 | 0.087 | 0.045 | 0.045 | 3.68 | 0.06 | 0.49 | 0.05 | 0 | 0.03 | 0.07 |
| 1 | 222252021 | rs12138480 | 0.076 | 1.066 | -0.221 | 0.785 | -0.146 | -0.124 | -0.124 | 0.201 | 0.29 | 0.02 | 0.33 | 0.33 | 0.42 | 0.23 |
| 1 | 222255923 | rs10863685 | -0.558 | 0.999 | -0.151 | 0.845 | 0.008 | 0.007 | 0.007 | 0.932 | 0.37 | 0.58 | 0.38 | 0.43 | 0.45 | 0.32 |
| 1 | 222266287 | rs11586620 | -0.155 | 0.461 | -0.957 | 0.412 | 0.005 | -0.02 | -0.02 | 1.026 | 0.28 | 0.21 | 0.35 | 0.55 | 0.28 | 0.32 |
| 1 | 222270724 | rs6702063 | 0.079 | 0.214 | -0.747 | 0.213 | 0.007 | -0.061 | -0.061 | 0 | 0.36 | 0.6 | 0.28 | 0.1 | 0.44 | 0.25 |
| 1 | 222271325 | rs7517086 | -1.47 | 0.232 | -1.205 | 0.226 | 0.015 | 0.005 | 0.005 | 3.352 | 0.64 | 0.82 | 0.63 | 0.65 | 0.72 | 0.57 |
| 1 | 222273249 | rs753961 | -0.147 | 0.277 | -1.408 | 0.847 | 0.007 | 0.023 | 0.023 | 0 | 0.22 | 0.12 | 0.12 | 0.13 | 0.21 | 0.18 |
| 1 | 222282882 | rs10779468 | -0.213 | 0.587 | -1.246 | 0.07 | -0.086 | 0.168 | 0.168 | 1.004 | 0.7 | 0.93 | 0.78 | 1 | 0.59 | 0.83 |
| 1 | 222289598 | rs17444339 | -0.137 | 0.588 | -1.12 | 0.519 | -0.028 | 0.101 | 0.101 | 1.199 | 0.69 | 0.88 | 0.82 | 1 | 0.62 | 0.84 |
| 1 | 222298970 | rs12036840 | 1.545 | 0.25 | -0.308 | 0.182 | 0.009 | -0.003 | -0.003 | 0.429 | 0.37 | 0.11 | 0.31 | 0.37 | 0.3 | 0.46 |
| 1 | 222301317 | rs2378522 | 1.166 | 0.237 | -0.293 | 0.175 | 0.002 | 0.002 | 0.002 | 0.567 | 0.41 | 0.37 | 0.34 | 0.37 | 0.31 | 0.47 |
| 1 | 222302968 | rs11118953 | -0.445 | 0.135 | 0.341 | 0.085 | -0.008 | -0.012 | -0.012 | 0 | 0.28 | 0.23 | 0.46 | 0.62 | 0.32 | 0.38 |
| 1 | 222304374 | rs12752144 | 1.654 | 0.123 | 0.3 | 0.079 | 0.005 | 0.149 | 0.149 | 0.244 | 0.36 | 0.08 | 0.22 | 0.12 | 0.3 | 0.45 |
| 1 | 222358676 | rs17162898 | 0.725 | -0.288 | 0.363 | -0.046 | 0.049 | 0.038 | 0.038 | 0.407 | 0.45 | 0.66 | 0.38 | 0.39 | 0.31 | 0.5 |
| 1 | 222360246 | rs1537852 | 1.838 | -0.391 | 0.446 | -0.064 | 0.029 | 0.071 | 0.071 | 0 | 0.34 | 0.12 | 0.27 | 0.23 | 0.28 | 0.49 |
| 1 | 222362840 | rs12750381 | 1.79 | -0.395 | 0.436 | -0.075 | 0.029 | 0.074 | 0.074 | 0 | 0.34 | 0.12 | 0.27 | 0.23 | 0.28 | 0.49 |
| 1 | 222365644 | rs17162930 | 1.935 | -0.386 | 0.441 | -0.07 | 0.025 | 0.067 | 0.067 | 0 | 0.33 | 0.12 | 0.26 | 0.23 | 0.26 | 0.48 |
| 1 | 222366608 | rs158735 | -0.694 | -0.336 | 0.39 | 0.32 | -0.066 | -0.125 | -0.125 | 0 | 0.71 | 0.9 | 0.64 | 0.53 | 0.61 | 0.8 |
| 1 | 222380752 | rs11487839 | -0.3 | -0.279 | 0.749 | 0.38 | 0.053 | 0.213 | 0.213 | 0.853 | 0.06 | 0.37 | 0.1 | 0.16 | 0.02 | 0.01 |
| 1 | 222396017 | rs158710 | -0.417 | -0.43 | 0.467 | 0.305 | -0.014 | -0.056 | -0.056 | 0 | 0.67 | 0.79 | 0.64 | 0.52 | 0.61 | 0.79 |
| 1 | 222426667 | rs1028861 | 1.527 | -0.499 | 0.415 | 0.312 | 0.029 | 0.056 | 0.056 | 0 | 0.33 | 0.11 | 0.21 | 0.24 | 0.26 | 0.5 |
| 1 | 222439668 | rs110046 | 0.089 | 0.236 | 0.928 | 1.202 | -0.152 | 0.032 | 0.032 | 0.055 | 0.72 | 0.97 | 0.72 | 0.9 | 0.53 | 0.76 |
| 1 | 222443578 | rs158731 | -0.085 | -0.099 | 1.031 | 0.873 | -0.168 | 0.072 | 0.072 | 0 | 0.72 | 0.96 | 0.7 | 0.94 | 0.52 | 0.76 |
| 1 | 222446350 | rs983816 | 1.821 | -0.061 | 1.258 | 0.743 | 0.005 | 0.025 | 0.025 | 0 | 0.4 | 0.3 | 0.28 | 0.21 | 0.34 | 0.44 |
| 1 | 222447753 | rs2172915 | 1.825 | -0.052 | 1.232 | 0.747 | 0.005 | 0.022 | 0.022 | 0 | 0.4 | 0.3 | 0.28 | 0.2 | 0.34 | 0.43 |
| 1 | 222455646 | rs158758 | -0.153 | -0.075 | 1.266 | 0.825 | -0.006 | -0.003 | -0.003 | 0 | 0.12 | 0.17 | 0.09 | 0.11 | 0.09 | 0.21 |
| 1 | 222456692 | rs35869320 | 1.806 | 0.035 | 1.052 | 0.679 | 0.009 | 0.071 | 0.071 | 0 | 0.39 | 0.21 | 0.27 | 0.2 | 0.34 | 0.44 |
| 1 | 222466162 | rs11487844 | 1.698 | 0.042 | 1.162 | 0.682 | -0.004 | -0.029 | -0.029 | 0 | 0.4 | 0.38 | 0.29 | 0.2 | 0.34 | 0.43 |
| 1 | 222484745 | rs11485173 | 1.825 | 0.118 | 1.161 | 0.708 | 0.012 | 0.075 | 0.075 | 0 | 0.39 | 0.2 | 0.26 | 0.2 | 0.34 | 0.45 |
| 1 | 222490196 | rs1034020 | 1.822 | 0.122 | 1.144 | 0.709 | -0.008 | -0.047 | -0.047 | 0 | 0.4 | 0.41 | 0.29 | 0.2 | 0.34 | 0.44 |
| 1 | 222497526 | rs35082873 | 1.172 | 0.091 | 1.097 | 0.672 | -0.004 | -0.031 | -0.031 | 0 | 0.53 | 0.46 | 0.33 | 0.21 | 0.4 | 0.49 |
| 1 | 222504752 | rs1155471 | 0.126 | 0.182 | 0.925 | 0.579 | -0.157 | 0.147 | 0.147 | 0 | 0.26 | 0.04 | 0.29 | 0.02 | 0.44 | 0.22 |
| 1 | 222506713 | rs17506627 | 1.166 | 0.171 | 1.003 | 0.63 | -0.003 | -0.036 | -0.036 | 1.024 | 0.55 | 0.44 | 0.35 | 0.21 | 0.44 | 0.51 |
| 1 | 222511433 | rs4384246 | 0.492 | 0.191 | 1.086 | 0.857 | -0.03 | 0.152 | 0.152 | 0 | 0.82 | 0.56 | 0.68 | 0.44 | 0.88 | 0.74 |
| 1 | 222513180 | rs4511115 | 1.506 | 0.579 | 0.83 | 0.583 | 0.005 | 0.029 | 0.029 | 0 | 0.42 | 0.3 | 0.29 | 0.2 | 0.39 | 0.46 |
| 1 | 222519003 | rs4364889 | 0.121 | -0.157 | 0.683 | 0.842 | -0.023 | -0.028 | -0.028 | 0 | 0.74 | 0.72 | 0.87 | 0.91 | 0.87 | 0.76 |
| 1 | 222538556 | rs34559460 | 0.997 | 0.214 | 0.074 | 0.818 | -0.001 | -0.002 | -0.002 | 0.066 | 0.47 | 0.33 | 0.43 | 0.44 | 0.47 | 0.55 |
| 1 | 222547295 | rs4459116 | -0.077 | 0.39 | -0.272 | 0.452 | -0.052 | 0.059 | 0.059 | 0 | 0.23 | 0.1 | 0.19 | 0.01 | 0.38 | 0.16 |
| 1 | 222550292 | rs4846364 | 0.696 | 0.282 | 0.291 | 0.96 | -0.012 | 0.086 | 0.086 | 0 | 0.46 | 0.26 | 0.37 | 0.22 | 0.46 | 0.53 |
| 1 | 222553902 | rs17507835 | 0.526 | 0.406 | 0.483 | 1.241 | -0.001 | -0.001 | -0.001 | 0 | 0.47 | 0.43 | 0.43 | 0.42 | 0.46 | 0.58 |
| 1 | 222560139 | rs4846748 | 1.075 | 0.141 | 1.331 | 1.746 | -0.039 | 0.107 | 0.107 | 0 | 0.86 | 0.77 | 0.9 | 0.71 | 0.93 | 0.87 |
| 1 | 222561006 | rs10495189 | -1.324 | 1.279 | 1.316 | 1.441 | 0.01 | 0.009 | 0.009 | 0 | 0.1 | 0.11 | 0.1 | 0.12 | 0.19 | 0.08 |
| 1 | 222562019 | rs17163084 | -0.514 | 1.018 | 1.205 | 0.876 | -0.023 | -0.025 | -0.025 | 0 | 0.06 | 0.28 | 0.03 | 0 | 0.01 | 0 |
| 1 | 222578271 | rs1108870 | -0.901 | 1.617 | 0.638 | 1.075 | 0.062 | 0.055 | 0.055 | 0.19 | 0.1 | 0.21 | 0.16 | 0.23 | 0.19 | 0.16 |
| 1 | 222584505 | rs17461806 | -1.045 | 1.508 | 0.563 | 0.833 | -0.001 | -0.008 | -0.008 | 1.638 | 0.16 | 0.14 | 0.13 | 0.01 | 0.25 | 0.14 |
| 1 | 222603792 | rs1843913 | -0.852 | 1.712 | 0.404 | 1.093 | 0.056 | 0.055 | 0.055 | 0.107 | 0.1 | 0.21 | 0.15 | 0.23 | 0.18 | 0.16 |
| 1 | 222625645 | rs17163136 | -1.614 | 1.337 | 0.165 | 1.072 | 0.004 | -0.003 | -0.003 | 0 | 0.31 | 0.62 | 0.26 | 0.25 | 0.39 | 0.31 |
| 1 | 222718314 | rs17463531 | -1.378 | 1.147 | -0.076 | 1.946 | -0.017 | 0.037 | 0.037 | 0 | 0.12 | 0.04 | 0.08 | 0 | 0.16 | 0.1 |
| 1 | 222718718 | rs17011628 | -0.98 | 1.079 | 0.994 | 2.061 | 0.046 | 0.125 | 0.125 | 0 | 0.1 | 0.35 | 0.1 | 0.15 | 0.1 | 0.04 |
| 1 | 222719037 | rs17463586 | 1.59 | 1.038 | 0.713 | 1.194 | -0.005 | 0.073 | 0.073 | 0.51 | 0.56 | 0.28 | 0.61 | 0.83 | 0.59 | 0.66 |
| 1 | 222721288 | rs4846382 | 1.365 | 0.468 | 0.933 | 1.262 | 0.012 | 0.021 | 0.021 | 0.08 | 0.54 | 0.31 | 0.72 | 0.64 | 0.6 | 0.77 |
| 1 | 222725850 | rs17011636 | -1.45 | 0.447 | 1.331 | 1.305 | 0 | 0.002 | 0.002 | 0 | 0.23 | 0.28 | 0.21 | 0.22 | 0.25 | 0.16 |
| 1 | 222726036 | rs17163203 | -1.348 | 0.539 | 1.403 | 0.906 | 0.033 | -0.049 | -0.049 | 0 | 0.27 | 0.47 | 0.23 | 0.13 | 0.33 | 0.33 |
| 1 | 222727556 | rs11485178 | -1.504 | 0.488 | 0.954 | 0.795 | 0.025 | 0.016 | 0.016 | 0 | 0.14 | 0.43 | 0.1 | 0.14 | 0.14 | 0.08 |
| 1 | 222729986 | rs4400619 | -0.509 | 0.526 | 0.604 | 0.733 | -0.033 | 0.057 | 0.057 | 0 | 0.1 | 0.01 | 0.07 | 0 | 0.14 | 0.08 |
| 1 | 222732656 | rs4481859 | 1.713 | 0.368 | 0.265 | 1.019 | -0.14 | -0.096 | -0.096 | 0 | 0.43 | 0.02 | 0.35 | 0.37 | 0.51 | 0.41 |
| 1 | 222738970 | rs2133185 | -0.938 | 0.317 | 0.901 | 0.946 | 0.034 | 0.029 | 0.029 | 0 | 0.12 | 0.42 | 0.1 | 0.14 | 0.14 | 0.08 |
| 1 | 222748085 | rs3002124 | -1.297 | 0.188 | 0.914 | 0.866 | -0.009 | -0.026 | -0.026 | 0.36 | 0.34 | 0.59 | 0.19 | 0.22 | 0.27 | 0.17 |
| 1 | 222751034 | rs2936033 | 0.849 | 0.155 | 1.015 | 1.07 | -0.002 | -0.001 | -0.001 | 0.198 | 0.64 | 0.37 | 0.51 | 0.57 | 0.76 | 0.56 |
| 1 | 222761771 | rs2270705 | 0.393 | 0.421 | 0.603 | 0.82 | -0.018 | 0.002 | 0.002 | 0 | 0.11 | 0.06 | 0.07 | 0.06 | 0.11 | 0.07 |
| 1 | 222762128 | rs17464836 | 0.023 | 0.434 | 0.918 | 0.783 | 0.001 | -0.002 | -0.002 | 1.827 | 0.07 | 0.06 | 0.08 | 0.04 | 0.09 | 0.2 |
| 1 | 222762709 | rs17464857 | -0.033 | 0.062 | 1.003 | 0.563 | 0.013 | 0.034 | 0.034 | 0 | 0.21 | 0.09 | 0.08 | 0 | 0.13 | 0.08 |
| 1 | 222762958 | rs17511318 | -0.033 | 0.062 | 1.003 | 0.563 | 0.013 | 0.034 | 0.034 | 0 | 0.21 | 0.09 | 0.08 | 0 | 0.13 | 0.08 |
| 1 | 222763026 | rs4846767 | 1.037 | 0.099 | 1.443 | 0.679 | -0.01 | -0.028 | -0.028 | 0.543 | 0.66 | 0.42 | 0.82 | 0.78 | 0.73 | 0.83 |
| 1 | 222769593 | rs1995152 | -0.36 | 0.109 | 1.573 | 0.669 | 0.02 | 0.02 | 0.02 | 0 | 0.11 | 0.24 | 0.08 | 0.14 | 0.14 | 0.08 |

| | | | | | | | | | | | | | | | | |
|---|---|---|---|---|---|---|---|---|---|---|---|---|---|---|---|---|
| 1 | 222772139 | rs3002130 | -0.204 | -0.086 | 1.447 | 0.653 | 0.01 | 0.009 | 0.009 | 0 | 0.79 | 0.87 | 0.92 | 1 | 0.86 | 0.92 |
| 1 | 222788062 | rs3002142 | 0.727 | -0.014 | 1.664 | 0.676 | -0.002 | 0.078 | 0.078 | 0.058 | 0.85 | 0.6 | 0.88 | 0.74 | 0.88 | 0.85 |
| 1 | 222790366 | rs904323 | -0.396 | 0.028 | 1.289 | 0.273 | 0.029 | 0.067 | 0.067 | 0 | 0.2 | 0.06 | 0.08 | 0 | 0.15 | 0.11 |
| 1 | 222794232 | rs3008611 | -0.454 | 0.208 | 1.358 | 0.496 | -0.014 | 0.007 | 0.007 | 0 | 0.95 | 0.95 | 0.98 | 0.89 | 0.98 | 0.89 |
| 1 | 222798965 | rs17011666 | 1.29 | 0.174 | 1.636 | 0.609 | -0.033 | 0.032 | 0.032 | 0.165 | 0.73 | 0.63 | 0.59 | 0.59 | 0.81 | 0.58 |
| 1 | 222823529 | rs17465637 | 1.679 | 0.197 | 1.598 | 0.616 | 0.04 | -0.01 | -0.01 | 0 | 0.65 | 0.19 | 0.51 | 0.61 | 0.74 | 0.58 |
| 1 | 222826481 | rs2291832 | 1.73 | 0.194 | 1.598 | 0.598 | 0.044 | -0.019 | -0.019 | 0 | 0.64 | 0.16 | 0.51 | 0.58 | 0.74 | 0.55 |
| 1 | 222831372 | rs2088514 | -0.838 | 0.188 | 1.716 | 0.618 | -0.016 | 0.014 | 0.014 | 0 | 0.21 | 0.32 | 0.39 | 0.28 | 0.16 | 0.27 |
| 1 | 222836894 | rs2936025 | -0.503 | 0.184 | 1.727 | 0.618 | -0.014 | 0.009 | 0.009 | 0 | 0.95 | 0.95 | 0.98 | 0.89 | 0.98 | 0.87 |
| 1 | 222837594 | rs2270707 | 1.239 | 0.18 | 1.732 | 0.617 | -0.033 | 0.029 | 0.029 | 0.144 | 0.73 | 0.62 | 0.59 | 0.61 | 0.82 | 0.61 |
| 1 | 222839838 | rs1053316 | 0.791 | 0.096 | 1.779 | 0.622 | -0.019 | -0.022 | -0.022 | 0.014 | 0.85 | 0.38 | 0.89 | 0.88 | 0.89 | 0.84 |
| 1 | 222845338 | rs17011686 | -0.483 | 0.012 | 1.734 | 0.585 | -0.02 | 0.089 | 0.089 | 0 | 0.08 | 0.15 | 0.32 | 0.27 | 0.03 | 0.18 |
| 1 | 222845411 | rs3008634 | 0.762 | 0.043 | 1.746 | 0.598 | -0.022 | -0.016 | -0.016 | 0 | 0.85 | 0.35 | 0.89 | 0.86 | 0.89 | 0.83 |
| 1 | 222848029 | rs1391558 | -0.723 | 0.144 | 1.727 | 0.589 | -0.002 | 0.001 | 0.001 | 0 | 0.2 | 0.19 | 0.37 | 0.28 | 0.15 | 0.28 |
| 1 | 222855196 | rs11487815 | 0.421 | 0.176 | 1.869 | 0.518 | 0.046 | -0.031 | -0.031 | 0 | 0.08 | 0.29 | 0.07 | 0.01 | 0.08 | 0.03 |
| 1 | 222855415 | rs17163429 | 0.242 | -0.145 | 1.405 | 0.486 | -0.006 | -0.01 | -0.01 | 0 | 0.23 | 0.08 | 0.13 | 0.16 | 0.2 | 0.07 |
| 1 | 222858398 | rs17163431 | 0.515 | -0.068 | 1.498 | 0.51 | 0.046 | -0.031 | -0.031 | 0.038 | 0.08 | 0.29 | 0.07 | 0.01 | 0.08 | 0.03 |
| 1 | 222878687 | rs17163437 | 0.029 | 0.111 | 1.288 | 0.447 | 0.074 | -0.052 | -0.052 | 0.164 | 0.1 | 0.46 | 0.11 | 0.01 | 0.12 | 0.07 |
| 1 | 222882892 | rs17163441 | 0.262 | 0.147 | 1.065 | 0.464 | 0.047 | -0.051 | -0.051 | 0 | 0.09 | 0.31 | 0.1 | 0.01 | 0.12 | 0.07 |
| 1 | 222886119 | rs3811465 | 0.3 | 0.099 | 0.912 | 0.487 | -0.002 | 0.001 | 0.001 | 0 | 0.8 | 0.81 | 0.63 | 0.73 | 0.85 | 0.72 |
| 1 | 222886588 | rs3748631 | 0.526 | 0.149 | 0.965 | -0.204 | 0.005 | -0.031 | -0.031 | 0 | 0.09 | 0.14 | 0.09 | 0.01 | 0.13 | 0.07 |
| 1 | 222897664 | rs17163453 | 0.17 | 0.113 | 0.703 | -0.125 | -0.014 | 0.013 | 0.013 | 0 | 0.06 | 0.13 | 0.07 | 0.08 | 0.02 | 0.03 |
| 1 | 222904298 | rs4240934 | -0.729 | 0.183 | 0.383 | -0.161 | 0.004 | 0.043 | 0.043 | 0 | 0.87 | 0.96 | 0.94 | 1 | 0.88 | 0.89 |
| 1 | 222913107 | rs12058171 | 0.173 | 0.41 | 0.613 | -0.043 | 0.051 | -0.051 | -0.051 | 0 | 0.09 | 0.31 | 0.1 | 0.01 | 0.13 | 0.07 |
| 1 | 222919682 | rs9286760 | 0.173 | 0.385 | 0.615 | -0.043 | 0.045 | -0.051 | -0.051 | 0 | 0.09 | 0.3 | 0.1 | 0.01 | 0.13 | 0.07 |
| 1 | 222922168 | rs17011714 | 0.262 | 0.324 | -0.139 | -0.092 | -0.017 | -0.02 | -0.02 | 0 | 0.22 | 0.04 | 0.13 | 0.16 | 0.19 | 0.07 |
| 1 | 222923001 | rs2068814 | 1.272 | -0.054 | 0.328 | -0.537 | 0.042 | 0.074 | 0.074 | 1.536 | 0.62 | 0.23 | 0.74 | 0.77 | 0.7 | 0.66 |
| 1 | 222923351 | rs6683071 | 0.575 | -0.08 | 0.114 | -0.41 | 0.006 | 0.005 | 0.005 | 0 | 0.73 | 0.75 | 0.85 | 0.8 | 0.83 | 0.76 |
| 1 | 222930485 | rs35806535 | 0.513 | -0.38 | -0.185 | -0.676 | -0.024 | -0.019 | -0.019 | 0 | 0.72 | 0.64 | 0.84 | 0.8 | 0.83 | 0.76 |
| 1 | 222939905 | rs2378603 | 0.934 | -0.324 | -0.671 | -0.712 | 0.046 | 0.071 | 0.071 | 1.546 | 0.62 | 0.21 | 0.73 | 0.75 | 0.7 | 0.66 |
| 1 | 222946204 | rs3922385 | 0.015 | -0.532 | -0.362 | -0.667 | -0.019 | -0.017 | -0.017 | 0 | 0.8 | 0.76 | 0.93 | 0.89 | 0.9 | 0.81 |
| 1 | 222951972 | rs3922772 | 0.381 | -0.201 | -0.363 | -1.011 | -0.003 | -0.047 | -0.047 | 0 | 0.53 | 0.5 | 0.72 | 0.86 | 0.57 | 0.67 |
| 1 | 222957080 | rs17011733 | -0.318 | -0.449 | -0.207 | -0.781 | -0.038 | -0.004 | -0.004 | 0 | 0.11 | 0.27 | 0.09 | 0.1 | 0.04 | 0.04 |
| 1 | 222962336 | rs17534083 | -0.027 | -0.338 | -0.363 | -1.021 | -0.02 | 0.062 | 0.062 | 0 | 0.32 | 0.21 | 0.24 | 0.13 | 0.37 | 0.22 |
| 1 | 222970177 | rs2378598 | 0.355 | -1.158 | -0.698 | -0.334 | -0.014 | -0.006 | -0.006 | 0 | 0.44 | 0.27 | 0.52 | 0.5 | 0.5 | 0.7 |
| 1 | 222971968 | rs4846778 | -1.055 | -1.455 | -0.86 | -0.168 | 0.025 | -0.011 | -0.011 | 0 | 0.86 | 0.96 | 0.93 | 0.89 | 0.9 | 0.95 |
| 1 | 222974926 | rs17162912 | 0.248 | -1.238 | -0.007 | -0.483 | 0.006 | -0.057 | -0.057 | 0 | 0.1 | 0.24 | 0.07 | 0 | 0.1 | 0.02 |
| 1 | 222978580 | rs34830798 | -0.875 | -0.522 | -0.794 | -1.083 | -0.054 | 0.025 | 0.025 | 0 | 0.88 | 0.99 | 0.83 | 0.95 | 0.83 | 0.81 |
| 1 | 222987443 | rs34419230 | 1.329 | -0.57 | -1.833 | -1.442 | -0.002 | 0.035 | 0.035 | 0.936 | 0.25 | 0.06 | 0.21 | 0.19 | 0.22 | 0.14 |
| 1 | 222991691 | rs4609435 | 0.715 | -0.228 | -1.894 | -1.7 | -0.064 | 0.051 | 0.051 | 0.257 | 0.14 | 0.04 | 0.22 | 0.05 | 0.25 | 0.25 |
| 1 | 222992740 | rs34229647 | 1.877 | -0.237 | -1.898 | -1.704 | -0.051 | 0.017 | 0.017 | 0 | 0.1 | 0.01 | 0.17 | 0.05 | 0.16 | 0.2 |
| 1 | 223003498 | rs17475774 | 1.917 | -0.182 | -1.901 | -1.707 | -0.049 | 0.035 | 0.035 | 0 | 0.1 | 0.01 | 0.06 | 0.04 | 0.15 | 0.18 |
| 1 | 223004049 | rs11487861 | 1.475 | -0.229 | -1.132 | -1.76 | 0.019 | -0.029 | -0.029 | 0.244 | 0.05 | 0.13 | 0.07 | 0 | 0.09 | 0.06 |
| 1 | 223022931 | rs34194482 | 1.991 | -0.103 | -1.173 | -1.78 | -0.056 | 0.042 | 0.042 | 0 | 0.09 | 0.01 | 0.06 | 0.02 | 0.17 | 0.18 |
| 1 | 223034487 | rs34764163 | 0.786 | 0.002 | -1.113 | -1.782 | -0.05 | 0.049 | 0.049 | 0.099 | 0.14 | 0.06 | 0.12 | 0.02 | 0.27 | 0.22 |
| 1 | 223035354 | rs17163530 | 1.956 | 0.573 | -1.658 | -0.841 | -0.025 | -0.064 | -0.064 | 1.184 | 0.1 | 0 | 0.1 | 0.26 | 0.08 | 0.04 |
| 1 | 223042940 | rs34892353 | -0.08 | 0.743 | -0.799 | 0.039 | 0.03 | 0.028 | 0.028 | 1.108 | 0.3 | 0.4 | 0.28 | 0.38 | 0.38 | 0.55 |
| 1 | 223057033 | rs35506390 | 3.359 | 0.671 | -0.876 | 0.167 | -0.05 | 0.015 | 0.015 | 0 | 0.08 | 0.01 | 0.06 | 0.06 | 0.16 | 0.17 |
| 1 | 223068480 | rs17163542 | -3.613 | 0.759 | -0.847 | 0.043 | 0.035 | 0.063 | 0.063 | 0.087 | 0.73 | 0.93 | 0.79 | 0.82 | 0.79 | 0.88 |
| 1 | 223078802 | rs34715037 | 1.583 | 0.79 | -0.373 | 0.029 | -0.004 | 0.023 | 0.023 | 0 | 0.15 | 0.15 | 0.1 | 0.31 | 0.09 | 0.28 |
| 1 | 223082306 | rs2068483 | 2.447 | 0.543 | -0.43 | 0.163 | -0.015 | -0.069 | -0.069 | 0 | 0.11 | 0.01 | 0.1 | 0.26 | 0.08 | 0.05 |
| 1 | 223085331 | rs35947998 | -1.537 | 0.591 | 0.608 | 0.085 | -0.009 | -0.001 | -0.001 | 0 | 0.58 | 0.61 | 0.65 | 0.58 | 0.68 | 0.78 |
| 1 | 223096578 | rs2609398 | 0.558 | 1.081 | 0.151 | 0.363 | 0.003 | -0.007 | -0.007 | 0 | 0.27 | 0.25 | 0.3 | 0.45 | 0.22 | 0.49 |
| 1 | 223100293 | rs2789966 | 1.967 | 0.773 | 0.047 | 0.399 | 0 | 0.002 | 0.002 | 0 | 0.17 | 0.16 | 0.18 | 0.3 | 0.14 | 0.32 |
| 1 | 223102867 | rs1538560 | 2.267 | 0.772 | 0.058 | 0.397 | -0.007 | -0.006 | -0.006 | 0 | 0.05 | 0.05 | 0.14 | 0.09 | 0.11 | 0.03 |
| 1 | 223104829 | rs2609406 | 1.812 | 0.534 | 0.295 | 0.254 | 0.001 | -0.003 | -0.003 | 0 | 0.17 | 0.15 | 0.18 | 0.3 | 0.14 | 0.32 |
| 1 | 223108966 | rs2254329 | -1.861 | 0.543 | 0.372 | 0.24 | 0.015 | 0.005 | 0.005 | 0.212 | 0.52 | 0.44 | 0.51 | 0.49 | 0.57 | 0.75 |
| 1 | 223110870 | rs34232799 | 3.687 | 0.567 | 0.258 | 0.304 | -0.049 | 0.014 | 0.014 | 0 | 0.08 | 0.01 | 0.06 | 0.03 | 0.16 | 0.16 |
| 1 | 223114549 | rs17536944 | 3.527 | 0.386 | 0.558 | 0.495 | 0.007 | 0.013 | 0.013 | 0 | 0.3 | 0.15 | 0.34 | 0.26 | 0.32 | 0.14 |
| 1 | 223116472 | rs2609383 | 1.166 | 0.728 | 1.618 | 0.317 | 0 | 0.001 | 0.001 | 0 | 0.17 | 0.16 | 0.17 | 0.29 | 0.14 | 0.31 |
| 1 | 223117960 | rs9651262 | -0.033 | 0.518 | 1.595 | 0.349 | -0.049 | -0.026 | -0.026 | 0 | 0.71 | 0.79 | 0.65 | 0.65 | 0.58 | 0.5 |
| 1 | 223118253 | rs2039780 | 2.11 | 0.358 | 1.772 | 0.49 | 0.002 | -0.034 | -0.034 | 0 | 0.45 | 0.15 | 0.44 | 0.5 | 0.42 | 0.26 |
| 1 | 223123348 | rs10495205 | 0.816 | 0.228 | 1.941 | 0.603 | 0.002 | -0.01 | -0.01 | 0 | 0.19 | 0.14 | 0.19 | 0.29 | 0.16 | 0.34 |
| 1 | 223124916 | rs2609379 | -0.445 | -0.158 | 1.697 | 0.493 | -0.006 | -0.032 | -0.032 | 0 | 0.39 | 0.73 | 0.45 | 0.51 | 0.37 | 0.61 |
| 1 | 223130319 | rs2789931 | 1.265 | 0.064 | 1.233 | 0.464 | 0.002 | -0.008 | -0.008 | 0.504 | 0.25 | 0.23 | 0.31 | 0.2 | 0.27 | 0.34 |
| 1 | 223141268 | rs728006 | 1.184 | 0.079 | 1.24 | 0.468 | 0.001 | -0.006 | -0.006 | 0.02 | 0.26 | 0.23 | 0.31 | 0.19 | 0.27 | 0.35 |
| 1 | 223147097 | rs36017274 | 0.426 | 0.06 | 0.845 | 0.467 | 0 | 0.019 | 0.019 | 0 | 0.1 | 0.11 | 0.13 | 0.27 | 0.09 | 0.14 |
| 1 | 223148727 | rs34684409 | 1.181 | 0.074 | 0.854 | 0.477 | 0.012 | -0.018 | -0.018 | 1.159 | 0.45 | 0.21 | 0.42 | 0.46 | 0.43 | 0.31 |
| 1 | 223159380 | rs2789954 | 0.279 | 0.167 | 1.559 | 1.265 | 0.006 | -0.009 | -0.009 | 1.066 | 0.28 | 0.36 | 0.33 | 0.27 | 0.29 | 0.46 |

| | | | | | | | | | | | | | | | | |
|---|---|---|---|---|---|---|---|---|---|---|---|---|---|---|---|---|
| 1 | 223172695 | rs17163660 | -0.595 | 0.41 | -0.59 | 1.496 | 0.001 | 0.011 | 0.011 | 0 | 0.1 | 0.11 | 0.13 | 0.27 | 0.09 | 0.15 |
| 1 | 223180844 | rs2789912 | 0.18 | 0.67 | -0.616 | 0.713 | 0 | -0.002 | -0.002 | 1.214 | 0.29 | 0.29 | 0.31 | 0.22 | 0.28 | 0.29 |
| 1 | 223184459 | rs2789948 | 0.174 | 0.584 | -0.623 | 0.733 | 0 | -0.001 | -0.001 | 0 | 0.29 | 0.3 | 0.34 | 0.26 | 0.28 | 0.3 |
| 1 | 223188116 | rs34558820 | 0.431 | 0.225 | -1.249 | 0.792 | 0.019 | 0.003 | 0.003 | 0 | 0.54 | 0.14 | 0.49 | 0.44 | 0.61 | 0.41 |
| 1 | 223192017 | rs17538288 | 0.003 | 0.186 | -1.086 | 0.314 | 0.01 | 0.037 | 0.037 | 0 | 0.5 | 0.42 | 0.52 | 0.7 | 0.47 | 0.58 |
| 1 | 223194033 | rs2609348 | -0.037 | 0.417 | -1.186 | 0.364 | 0.004 | 0.004 | 0.004 | 0 | 0.46 | 0.57 | 0.45 | 0.58 | 0.36 | 0.61 |
| 1 | 223194696 | rs17538400 | 0.833 | 0.682 | -1.948 | 0.364 | -0.066 | -0.033 | -0.033 | 0 | 0.27 | 0.03 | 0.35 | 0.24 | 0.31 | 0.15 |
| 1 | 223209450 | rs2789921 | -0.273 | 0.369 | -1.382 | 0.482 | -0.026 | 0.037 | 0.037 | 1.954 | 0.68 | 0.43 | 0.79 | 0.52 | 0.78 | 0.55 |
| 1 | 223213967 | rs946511 | -0.01 | 1.154 | -1.879 | 0.121 | -0.012 | -0.004 | -0.004 | 1.699 | 0.27 | 0.19 | 0.32 | 0.28 | 0.27 | 0.32 |
| 1 | 223214800 | rs755352 | 0.289 | 1.158 | -1.902 | -0.04 | 0.005 | -0.054 | -0.054 | 0 | 0.16 | 0.2 | 0.12 | 0 | 0.21 | 0.12 |
| 1 | 223282150 | rs3795743 | 0.127 | 1.512 | -2.649 | 0.346 | -0.045 | 0.077 | 0.077 | 0 | 0.34 | 0.1 | 0.35 | 0.2 | 0.41 | 0.46 |
| 1 | 223284528 | rs5744174 | 0.126 | 1.48 | -2.642 | 0.346 | -0.038 | 0.077 | 0.077 | 0 | 0.34 | 0.11 | 0.35 | 0.2 | 0.41 | 0.47 |
| 1 | 223284599 | rs2072493 | 0.839 | 1.689 | -2.784 | 0.893 | -0.051 | -0.066 | -0.066 | 0.008 | 0.1 | 0 | 0.21 | 0.27 | 0.16 | 0.12 |
| 1 | 223294851 | rs851193 | -0.447 | 1.509 | -2.708 | 0.416 | -0.038 | 0.077 | 0.077 | 0 | 0.66 | 0.89 | 0.65 | 0.8 | 0.59 | 0.53 |
| 1 | 223300713 | rs2241097 | 1.026 | 1.295 | -3 | 0.296 | 0.001 | 0.065 | 0.065 | 0 | 0.28 | 0.09 | 0.26 | 0.11 | 0.26 | 0.19 |
| 1 | 223306431 | rs851178 | -0.277 | 1.8 | -2.567 | 0.521 | -0.024 | 0.064 | 0.064 | 0 | 0.66 | 0.84 | 0.65 | 0.8 | 0.59 | 0.54 |
| 1 | 223310048 | rs2241096 | 0.261 | 1.333 | -2.763 | 1.016 | -0.029 | -0.081 | -0.081 | 0 | 0.13 | 0 | 0.06 | 0.27 | 0.1 | 0.19 |
| 1 | 223311607 | rs1640827 | -0.19 | 1.691 | -2.487 | 0.535 | -0.124 | 0.107 | 0.107 | 0.255 | 0.66 | 0.94 | 0.65 | 0.85 | 0.56 | 0.61 |
| 1 | 223311845 | rs2096142 | -0.071 | 1.654 | -2.268 | 0.533 | -0.056 | 0.06 | 0.06 | 0.002 | 0.64 | 0.79 | 0.63 | 0.76 | 0.53 | 0.55 |
| 1 | 223339689 | rs2247421 | 0.053 | 1.399 | -2.056 | 0.649 | -0.001 | 0.008 | 0.008 | 0 | 0.31 | 0.27 | 0.2 | 0.24 | 0.33 | 0.28 |
| 1 | 223349020 | rs851171 | 0.253 | 1.342 | -2.051 | 0.672 | 0.002 | 0.009 | 0.009 | 0 | 0.29 | 0.22 | 0.19 | 0.24 | 0.29 | 0.26 |
| 1 | 223353343 | rs851168 | -0.367 | 2.165 | -2.354 | 1.123 | -0.103 | -0.075 | -0.075 | 0 | 0.85 | 0.99 | 0.64 | 0.76 | 0.68 | 0.73 |
| 1 | 223369577 | rs17161975 | -0.651 | 1.531 | -2.075 | 0.977 | -0.061 | -0.079 | -0.079 | 0 | 0.18 | 0.01 | 0.17 | 0.26 | 0.2 | 0.27 |
| 1 | 223374948 | rs851165 | 0.394 | 1.032 | -2.05 | 0.584 | 0.002 | 0.003 | 0.003 | 0.081 | 0.31 | 0.29 | 0.18 | 0.23 | 0.28 | 0.24 |
| 1 | 223399328 | rs4661250 | -0.958 | 1.329 | -2.056 | 0.988 | -0.05 | -0.049 | -0.049 | 0 | 0.19 | 0.01 | 0.15 | 0.19 | 0.15 | 0.17 |
| 1 | 223404483 | rs1418151 | -0.115 | 1.443 | -1.856 | 1.192 | -0.034 | -0.046 | -0.046 | 0 | 0.08 | 0 | 0.13 | 0.16 | 0.08 | 0.08 |
| 1 | 223424645 | rs851206 | -0.158 | 1.199 | -1.875 | 0.564 | -0.006 | 0.016 | 0.016 | 0.018 | 0.32 | 0.23 | 0.22 | 0.23 | 0.39 | 0.28 |
| 1 | 223428569 | rs851201 | -0.199 | 1.315 | -0.966 | 0.922 | 0.029 | 0.01 | 0.01 | 0 | 0.31 | 0.44 | 0.3 | 0.37 | 0.42 | 0.31 |
| 1 | 223434633 | rs712857 | -0.385 | 1.307 | -0.939 | 0.842 | 0.017 | -0.027 | -0.027 | 0.091 | 0.34 | 0.44 | 0.32 | 0.25 | 0.44 | 0.31 |
| 1 | 223436336 | rs1640849 | -0.135 | 1.301 | -0.936 | 0.822 | -0.004 | -0.001 | -0.001 | 0 | 0.32 | 0.3 | 0.26 | 0.1 | 0.42 | 0.27 |
| 1 | 223438771 | rs1418150 | 0.102 | 1.311 | -1.04 | 0.826 | -0.139 | 0.086 | 0.086 | 0.002 | 0.29 | 0.04 | 0.24 | 0.1 | 0.42 | 0.26 |
| 1 | 223441672 | rs3820596 | -0.459 | 1.338 | -0.937 | 0.945 | 0.011 | 0.003 | 0.003 | 0 | 0.34 | 0.43 | 0.31 | 0.38 | 0.43 | 0.33 |
| 1 | 223442673 | rs885279 | -0.405 | 1.343 | -0.947 | 0.993 | 0.004 | 0.002 | 0.002 | 0 | 0.33 | 0.37 | 0.3 | 0.38 | 0.43 | 0.33 |
| 1 | 223444263 | rs17161988 | -0.076 | 1.38 | -1.028 | 0.958 | -0.04 | 0.028 | 0.028 | 0 | 0.31 | 0.18 | 0.29 | 0.25 | 0.42 | 0.3 |
| 1 | 223449236 | rs17517806 | 0.32 | 1.488 | -1.546 | 1.071 | -0.093 | -0.019 | -0.019 | 0.096 | 0.77 | 0.99 | 0.73 | 0.84 | 0.78 | 0.75 |
| 1 | 223458980 | rs9442041 | 0.335 | 1.489 | -1.545 | 1.202 | -0.093 | -0.102 | -0.102 | 3.394 | 0.77 | 0.99 | 0.73 | 0.71 | 0.78 | 0.72 |
| 1 | 223466400 | rs2795063 | 0.944 | 1.297 | -1.444 | 1.185 | 0.069 | -0.001 | -0.001 | 0.433 | 0.46 | 0.83 | 0.44 | 0.58 | 0.36 | 0.45 |
| 1 | 223467316 | rs17163774 | 1.013 | 1.304 | -1.351 | 1.132 | 0.013 | -0.005 | -0.005 | 0.389 | 0.44 | 0.6 | 0.44 | 0.46 | 0.37 | 0.44 |
| 1 | 223469630 | rs17518279 | 0.954 | 1.293 | -1.344 | 1.144 | 0.05 | 0.001 | 0.001 | 0 | 0.45 | 0.75 | 0.49 | 0.59 | 0.37 | 0.47 |
| 1 | 223479813 | rs1518044 | 0.575 | 1.236 | -1.28 | 1.353 | 0.005 | 0.014 | 0.014 | 0.022 | 0.78 | 0.69 | 0.7 | 0.7 | 0.81 | 0.72 |
| 1 | 223482641 | rs34468460 | 0.906 | 1.184 | -1.423 | 1.18 | -0.003 | 0 | 0 | 0 | 0.51 | 0.58 | 0.47 | 0.54 | 0.53 | 0.55 |
| 1 | 223486421 | rs17487717 | 0.786 | 1.246 | -1.374 | 1.236 | 0.012 | 0.051 | 0.051 | 0.276 | 0.76 | 0.62 | 0.65 | 0.64 | 0.79 | 0.66 |
| 1 | 223488310 | rs17518517 | 0.286 | 1.136 | -1.383 | 1.134 | -0.024 | 0.016 | 0.016 | 0.376 | 0.13 | 0.07 | 0.12 | 0.01 | 0.23 | 0.22 |
| 1 | 223497150 | rs1518040 | 1.14 | 1.21 | -1.034 | 1.819 | -0.002 | 0.012 | 0.012 | 0 | 0.52 | 0.57 | 0.45 | 0.42 | 0.54 | 0.5 |
| 1 | 223502811 | rs1400960 | -0.309 | 1.207 | -1.014 | 1.831 | -0.039 | -0.046 | -0.046 | 3.268 | 0.17 | 0.05 | 0.27 | 0.22 | 0.16 | 0.24 |
| 1 | 223503982 | rs17163816 | 1.115 | 1.263 | -0.957 | 1.879 | -0.002 | 0.009 | 0.009 | 0 | 0.52 | 0.57 | 0.45 | 0.43 | 0.54 | 0.51 |
| 1 | 223523854 | rs17163825 | -0.714 | 0.766 | -0.735 | 1.304 | -0.014 | 0.003 | 0.003 | 0.327 | 0.35 | 0.48 | 0.32 | 0.39 | 0.32 | 0.31 |
| 1 | 223523973 | rs17578203 | 0.274 | 0.938 | -1.244 | 0.996 | -0.051 | 0.066 | 0.066 | 2.559 | 0.25 | 0.02 | 0.18 | 0.11 | 0.28 | 0.12 |
| 1 | 223561802 | rs2487521 | 0.173 | 0.978 | -0.851 | 1.061 | -0.005 | -0.002 | -0.002 | 0 | 0.68 | 0.61 | 0.72 | 0.67 | 0.69 | 0.75 |
| 1 | 223567803 | rs10907376 | 0.048 | 1.049 | -1.271 | 1.038 | 0.006 | 0.039 | 0.039 | 1.835 | 0.27 | 0.18 | 0.19 | 0.11 | 0.27 | 0.12 |
| 1 | 223579329 | rs1772290 | -1.485 | 1.073 | -1.331 | 1.391 | -0.002 | -0.003 | -0.003 | 0.128 | 0.85 | 0.87 | 0.84 | 0.68 | 0.85 | 0.94 |
| 1 | 223584719 | rs825130 | 0.364 | 0.98 | -1.352 | 1.217 | 0.002 | 0.003 | 0.003 | 3.072 | 0.55 | 0.58 | 0.58 | 0.45 | 0.54 | 0.73 |
| 1 | 223587515 | rs825116 | -0.657 | 1.186 | -1.248 | 1.214 | 0 | 0.046 | 0.046 | 0 | 0.81 | 0.76 | 0.78 | 0.56 | 0.81 | 0.86 |
| 1 | 223587815 | rs866096 | -0.357 | 1.786 | -0.855 | 0.707 | -0.055 | 0.007 | 0.007 | 0 | 0.14 | 0.02 | 0.25 | 0.1 | 0.15 | 0.22 |
| 1 | 223590774 | rs12134824 | -0.192 | 1.59 | -0.514 | 0.685 | -0.002 | 0.02 | 0.02 | 0 | 0.23 | 0.19 | 0.19 | 0.1 | 0.26 | 0.11 |
| 1 | 223594234 | rs2176507 | 0.678 | 1.583 | -0.33 | 0.756 | -0.006 | 0 | 0 | 0 | 0.69 | 0.56 | 0.73 | 0.65 | 0.69 | 0.78 |
| 1 | 223597253 | rs866637 | 0.605 | 1.784 | -0.323 | 0.777 | 0.039 | 0.14 | 0.14 | 0 | 0.95 | 0.71 | 0.92 | 0.85 | 0.96 | 0.92 |
| 1 | 223603809 | rs12141995 | 0.137 | 1.813 | -0.444 | 0.589 | -0.047 | 0.102 | 0.102 | 0.048 | 0.22 | 0.04 | 0.18 | 0.05 | 0.26 | 0.1 |
| 1 | 223604499 | rs825133 | 0.579 | 1.74 | -0.283 | 0.707 | 0.001 | 0.181 | 0.181 | 0 | 0.91 | 0.65 | 0.9 | 0.71 | 0.94 | 0.88 |
| 1 | 223607389 | rs12040106 | 0.611 | 1.792 | -0.349 | 0.594 | -0.006 | 0.001 | 0.001 | 0.043 | 0.68 | 0.46 | 0.7 | 0.65 | 0.67 | 0.75 |
| 1 | 223607825 | rs7526539 | -0.229 | 1.908 | -0.759 | 0.327 | -0.006 | 0.023 | 0.023 | 0.128 | 0.25 | 0.23 | 0.2 | 0.07 | 0.3 | 0.13 |
| 1 | 223609583 | rs999017 | 0.073 | 2.028 | -0.789 | 0.553 | -0.066 | -0.045 | -0.045 | 0 | 0.12 | 0.01 | 0.25 | 0.16 | 0.17 | 0.24 |
| 1 | 223611033 | rs1417749 | -0.649 | 1.11 | -0.548 | 0.682 | 0.003 | 0 | 0 | 0 | 0.3 | 0.27 | 0.23 | 0.3 | 0.27 | 0.36 |
| 1 | 223613551 | rs1417748 | -0.617 | 1.111 | -0.595 | 0.683 | 0 | 0 | 0 | 0 | 0.31 | 0.3 | 0.24 | 0.3 | 0.27 | 0.35 |
| 1 | 223619066 | rs11261272 | 0.937 | 1.134 | -0.769 | 0.638 | -0.007 | 0.002 | 0.002 | 0 | 0.45 | 0.43 | 0.4 | 0.51 | 0.37 | 0.27 |
| 1 | 223629454 | rs12402934 | -1.216 | 0.794 | -0.437 | 0.668 | -0.005 | -0.004 | -0.004 | 0 | 0.28 | 0.39 | 0.25 | 0.28 | 0.23 | 0.34 |
| 1 | 223632045 | rs11261280 | -1.113 | 0.752 | -0.599 | 1.148 | 0.001 | 0.001 | 0.001 | 0 | 0.09 | 0.09 | 0.08 | 0.23 | 0.07 | 0.22 |
| 1 | 223632876 | rs4233516 | 0.972 | 1.088 | 0.262 | 0.943 | 0.079 | 0.059 | 0.059 | 0 | 0.89 | 0.63 | 0.88 | 0.88 | 0.87 | 0.87 |
| 1 | 223635192 | rs4481842 | 0.298 | 1.042 | -0.522 | 0.571 | -0.011 | 0.039 | 0.039 | 1.131 | 0.56 | 0.33 | 0.29 | 0.29 | 0.47 | 0.46 |

| | | | | | | | | | | | | | | | | |
|---|---|---|---|---|---|---|---|---|---|---|---|---|---|---|---|---|
| 1 | 223636166 | rs4233518 | 0.13 | 1.031 | 0.067 | 0.895 | 0.001 | 0 | 0 | 0 | 0.58 | 0.42 | 0.43 | 0.39 | 0.48 | 0.53 |
| 1 | 223638165 | rs6689716 | -0.32 | 1.004 | 0.132 | 0.882 | 0.005 | 0 | 0 | 0.909 | 0.41 | 0.56 | 0.57 | 0.61 | 0.51 | 0.48 |
| 1 | 223638214 | rs12748479 | 0.605 | 1.008 | -0.224 | 1.621 | -0.002 | -0.082 | -0.082 | 0 | 0.13 | 0.01 | 0.29 | 0.34 | 0.14 | 0.12 |
| 1 | 223639469 | rs11261281 | -0.992 | 0.536 | -0.698 | 0.659 | -0.002 | -0.037 | -0.037 | 0 | 0.88 | 0.87 | 0.95 | 1 | 0.9 | 0.95 |
| 1 | 223641440 | rs6685346 | -0.04 | 0.917 | -0.93 | 0.292 | 0.03 | 0.106 | 0.106 | 0 | 0.27 | 0.5 | 0.4 | 0.53 | 0.33 | 0.26 |
| 1 | 223642646 | rs6657657 | 1.277 | 0.976 | -0.946 | 0.306 | 0.018 | 0.002 | 0.002 | 1.973 | 0.43 | 0.31 | 0.37 | 0.39 | 0.37 | 0.46 |
| 1 | 223645256 | rs10430272 | -0.055 | 0.024 | -0.709 | 0.198 | 0.005 | -0.005 | -0.005 | 0 | 0.24 | 0.12 | 0.21 | 0.21 | 0.22 | 0.3 |
| 1 | 223646781 | rs12757009 | -0.51 | 0.063 | -0.396 | 0.16 | -0.005 | 0.001 | 0.001 | 2.195 | 0.43 | 0.33 | 0.57 | 0.62 | 0.54 | 0.5 |
| 1 | 223647537 | rs6687878 | 0.579 | -0.006 | -0.433 | 0.128 | 0.011 | 0.006 | 0.006 | 0 | 0.12 | 0.28 | 0.2 | 0.11 | 0.1 | 0.12 |
| 1 | 223647907 | rs6604717 | 0.975 | -0.45 | -1.16 | -0.618 | 0.009 | -0.018 | -0.018 | 0 | 0.23 | 0.16 | 0.13 | 0.28 | 0.2 | 0.33 |
| 1 | 223648361 | rs17162164 | 0.959 | -0.705 | -0.818 | -0.588 | -0.003 | 0.006 | 0.006 | 13.718 | 0.07 | 0.06 | 0.03 | 0.13 | 0.03 | 0.07 |
| 1 | 223648847 | rs6691566 | 0.276 | -0.795 | -0.827 | -0.672 | -0.017 | -0.017 | -0.017 | 0 | 0.27 | 0.32 | 0.13 | 0.13 | 0.18 | 0.13 |
| 1 | 223649032 | rs4323709 | 1.157 | -0.571 | -1.029 | -0.602 | 0.003 | -0.044 | -0.044 | 0.212 | 0.1 | 0.15 | 0.05 | 0 | 0.1 | 0.05 |
| 1 | 223649473 | rs10796476 | 0.647 | -0.777 | -0.92 | -0.724 | 0.01 | 0.017 | 0.017 | 2.664 | 0.55 | 0.37 | 0.65 | 0.79 | 0.65 | 0.68 |
| 1 | 223649811 | rs12745781 | 0.788 | -0.615 | -1.139 | -0.813 | -0.031 | 0.15 | 0.15 | 1.122 | 0.23 | 0.02 | 0.11 | 0.01 | 0.21 | 0.14 |
| 1 | 223649878 | rs17161952 | 0.456 | -0.877 | -0.896 | -0.598 | 0 | -0.017 | -0.017 | 0 | 0.12 | 0.24 | 0.23 | 0.08 | 0.11 | 0.15 |
| 1 | 223651266 | rs12134930 | 0.214 | -0.831 | -1.165 | -0.615 | 0.007 | -0.008 | -0.008 | 0.072 | 0.28 | 0.24 | 0.12 | 0.01 | 0.24 | 0.14 |
| 1 | 223651783 | rs7517790 | 0.121 | -0.812 | -1.341 | -0.097 | -0.006 | 0.027 | 0.027 | 0 | 0.25 | 0.31 | 0.22 | 0.37 | 0.22 | 0.29 |
| 1 | 223653181 | rs11261288 | 1.041 | -0.706 | -1.377 | -0.665 | 0.01 | 0.128 | 0.128 | 1.666 | 0.21 | 0.03 | 0.09 | 0.01 | 0.16 | 0.13 |
| 1 | 223653326 | rs4478802 | -0.262 | -0.581 | -1.092 | -0.317 | -0.006 | 0.001 | 0.001 | 0 | 0.75 | 0.69 | 0.79 | 0.8 | 0.8 | 0.72 |
| 1 | 223653362 | rs4512640 | 0.006 | 0.112 | -0.657 | 0.166 | 0 | 0.002 | 0.002 | 0 | 0.46 | 0.53 | 0.55 | 0.75 | 0.56 | 0.49 |
| 1 | 223655061 | rs12138067 | -1.285 | 0.504 | 0.328 | 0.632 | 0.046 | 0.172 | 0.172 | 0.135 | 0.13 | 0.41 | 0.18 | 0.2 | 0.11 | 0.25 |
| 1 | 223656288 | rs10907383 | -0.272 | 0.604 | -0.215 | 0.362 | -0.019 | -0.049 | -0.049 | 0.391 | 0.62 | 0.82 | 0.58 | 0.53 | 0.62 | 0.6 |
| 1 | 223656537 | rs10495209 | 0.761 | 0.555 | 0.157 | 0.344 | 0.006 | 0.056 | 0.056 | 0.089 | 0.51 | 0.33 | 0.46 | 0.39 | 0.5 | 0.45 |
| 1 | 223657162 | rs3935462 | 0.867 | 0.516 | 0.168 | 0.254 | 0.003 | 0.1 | 0.1 | 0.355 | 0.48 | 0.22 | 0.4 | 0.33 | 0.48 | 0.37 |
| 1 | 223657340 | rs3935463 | -0.051 | 0.554 | 0.289 | 0.331 | 0.083 | 0.141 | 0.141 | 0.016 | 0.09 | 0.34 | 0.12 | 0.13 | 0.11 | 0.15 |
| 1 | 223663505 | rs6690293 | 0.752 | 0.611 | 0.234 | 0.167 | 0.008 | 0.024 | 0.024 | 0.02 | 0.47 | 0.41 | 0.46 | 0.39 | 0.47 | 0.45 |
| 1 | 223664844 | rs17483288 | 1.495 | 0.704 | 0.112 | -0.186 | -0.034 | 0.108 | 0.108 | 0 | 0.17 | 0.01 | 0.09 | 0 | 0.15 | 0.03 |
| 1 | 223672283 | rs3922892 | 0.584 | 0.619 | 0.293 | 0.011 | 0.006 | 0.033 | 0.033 | 0.19 | 0.5 | 0.6 | 0.45 | 0.33 | 0.49 | 0.41 |
| 1 | 223674090 | rs6702233 | -0.584 | 0.543 | 0.318 | 0.102 | 0.086 | 0.105 | 0.105 | 1.131 | 0.06 | 0.35 | 0.11 | 0.08 | 0.06 | 0.14 |
| 1 | 223677665 | rs7549968 | 0.682 | 0.568 | 0.143 | 0.072 | 0.041 | 0.098 | 0.098 | 0 | 0.07 | 0.18 | 0.08 | 0.12 | 0.08 | 0.11 |
| 1 | 223682051 | rs4325124 | -0.248 | 0.595 | -0.521 | 0.072 | -0.065 | -0.08 | -0.08 | 0 | 0.38 | 0.13 | 0.44 | 0.56 | 0.42 | 0.42 |
| 1 | 223685954 | rs6667089 | 0.752 | 0.776 | -0.325 | 0.013 | 0.079 | 0.139 | 0.139 | 0 | 0.07 | 0.25 | 0.08 | 0.12 | 0.08 | 0.11 |
| 1 | 223689916 | rs12117655 | 0.144 | 0.843 | -0.383 | 0.326 | -0.005 | -0.009 | -0.009 | 0.411 | 0.16 | 0.12 | 0.22 | 0.29 | 0.2 | 0.27 |
| 1 | 223690077 | rs4421579 | -0.344 | 0.846 | -0.498 | 0.394 | -0.008 | -0.013 | -0.013 | 0.511 | 0.35 | 0.25 | 0.3 | 0.36 | 0.35 | 0.28 |
| 1 | 223693162 | rs17162209 | -0.046 | 0.851 | -0.309 | 0.593 | -0.002 | -0.004 | -0.004 | 0 | 0.3 | 0.25 | 0.29 | 0.36 | 0.3 | 0.26 |
| 1 | 223695145 | rs17542724 | 1.032 | 0.904 | -0.855 | 0.556 | -0.016 | -0.014 | -0.014 | 0 | 0.28 | 0.14 | 0.32 | 0.33 | 0.3 | 0.38 |
| 1 | 223697236 | rs10796478 | 0.048 | 0.547 | -0.95 | 0.684 | -0.002 | 0.006 | 0.006 | 0 | 0.61 | 0.61 | 0.65 | 0.63 | 0.66 | 0.71 |
| 1 | 223699609 | rs11261300 | 0.268 | 0.603 | -2.01 | 0.385 | -0.045 | -0.03 | -0.03 | 0.759 | 0.31 | 0.02 | 0.27 | 0.25 | 0.26 | 0.19 |
| 1 | 223701097 | rs12061221 | -0.165 | 0.884 | -0.869 | 0.499 | -0.003 | 0 | 0 | 0 | 0.25 | 0.17 | 0.25 | 0.28 | 0.3 | 0.25 |
| 1 | 223798234 | rs3890758 | -0.712 | 0.656 | -0.525 | 0.525 | -0.002 | 0.001 | 0.001 | 0 | 0.33 | 0.54 | 0.28 | 0.34 | 0.26 | 0.34 |
| 1 | 223806157 | rs35164209 | -0.397 | 0.721 | -0.9 | 0.063 | -0.005 | 0.035 | 0.035 | 0 | 0.23 | 0.16 | 0.2 | 0.11 | 0.2 | 0.05 |
| 1 | 223828623 | rs17483822 | -0.451 | 0.584 | -1.373 | -0.54 | -0.048 | 0.157 | 0.157 | 0 | 0.42 | 0.18 | 0.39 | 0.12 | 0.48 | 0.28 |
| 1 | 223828777 | rs12127418 | 0.33 | 0.575 | -1.069 | -0.198 | 0.019 | 0.022 | 0.022 | 0 | 0.17 | 0.24 | 0.18 | 0.25 | 0.19 | 0.28 |
| 1 | 223831591 | rs11584610 | 0.231 | 0.485 | -0.947 | -0.208 | -0.052 | 0.16 | 0.16 | 0.009 | 0.42 | 0.16 | 0.38 | 0.12 | 0.48 | 0.27 |
| 1 | 223832277 | rs9804135 | -0.378 | -0.127 | -1.299 | 0.366 | -0.007 | -0.008 | -0.008 | 0 | 0.6 | 0.7 | 0.58 | 0.43 | 0.59 | 0.5 |
| 1 | 223833290 | rs16841918 | 0.988 | -0.339 | -0.501 | 0.504 | -0.02 | -0.015 | -0.015 | 0 | 0.05 | 0.09 | 0.01 | 0.01 | 0.01 | 0.06 |
| 1 | 223834505 | rs1125346 | 1.476 | -0.352 | -0.542 | 0.5 | -0.026 | -0.09 | -0.09 | 0 | 0.23 | 0.04 | 0.23 | 0.32 | 0.22 | 0.22 |
| 1 | 223838950 | rs6677146 | 1.471 | -0.381 | -0.531 | 0.486 | -0.024 | -0.086 | -0.086 | 0 | 0.23 | 0.04 | 0.23 | 0.32 | 0.22 | 0.22 |
| 1 | 223839376 | rs1573980 | -0.526 | -0.336 | -0.253 | 0.497 | -0.053 | 0.125 | 0.125 | 0 | 0.18 | 0.53 | 0.2 | 0.31 | 0.11 | 0.23 |
| 1 | 223840617 | rs12132899 | 1.128 | -0.261 | -0.274 | 0.551 | -0.001 | -0.045 | -0.045 | 0.133 | 0.09 | 0.01 | 0.07 | 0.2 | 0.09 | 0.16 |
| 1 | 223841226 | rs1892079 | 0.813 | -0.198 | -0.561 | -0.009 | -0.007 | 0.028 | 0.028 | 0 | 0.32 | 0.23 | 0.34 | 0.11 | 0.34 | 0.18 |
| 1 | 223841682 | rs2096345 | 1.046 | -0.166 | -0.478 | 0.142 | 0.014 | 0.079 | 0.079 | 0 | 0.17 | 0.24 | 0.18 | 0.4 | 0.19 | 0.37 |
| 1 | 223841836 | rs2105405 | 1.17 | -0.165 | -0.478 | 0.138 | 0.016 | 0.023 | 0.023 | 0 | 0.16 | 0.24 | 0.18 | 0.24 | 0.19 | 0.28 |
| 1 | 223846564 | rs7516954 | 0.067 | -0.246 | -0.241 | 0.09 | 0.004 | 0.002 | 0.002 | 0 | 0.25 | 0.28 | 0.26 | 0.29 | 0.28 | 0.35 |
| 1 | 223847496 | rs1419308 | -0.206 | -0.276 | -0.329 | 0.024 | -0.007 | 0.067 | 0.067 | 0.05 | 0.52 | 0.72 | 0.61 | 0.71 | 0.51 | 0.57 |
| 1 | 223850733 | rs1109223 | -0.916 | -0.122 | 0.23 | -0.053 | -0.001 | 0.037 | 0.037 | 0 | 0.64 | 0.76 | 0.61 | 0.28 | 0.67 | 0.42 |
| 1 | 223853792 | rs10916022 | -0.028 | -0.271 | 0.508 | 0.003 | 0.002 | 0.003 | 0.003 | 0.093 | 0.48 | 0.41 | 0.52 | 0.36 | 0.56 | 0.46 |
| 1 | 223854684 | rs6692983 | -0.708 | -0.293 | 0.702 | -0.009 | -0.006 | 0 | 0 | 0 | 0.65 | 0.54 | 0.59 | 0.38 | 0.7 | 0.61 |
| 1 | 223857726 | rs10916078 | 0.136 | -0.416 | 0.744 | -0.055 | 0.002 | 0.011 | 0.011 | 0.796 | 0.45 | 0.35 | 0.4 | 0.3 | 0.55 | 0.4 |
| 1 | 223858133 | rs4653522 | -1.358 | -0.359 | 0.888 | -0.028 | -0.006 | -0.053 | -0.053 | 0.118 | 0.83 | 0.77 | 0.93 | 0.98 | 0.86 | 0.84 |
| 1 | 223859188 | rs1419310 | -1.311 | -0.356 | 0.915 | -0.015 | -0.011 | -0.038 | -0.038 | 0 | 0.9 | 0.77 | 0.91 | 0.96 | 0.91 | 0.94 |
| 1 | 223861401 | rs7547741 | -0.03 | -0.351 | 0.881 | -0.01 | 0.01 | 0.024 | 0.024 | 0.006 | 0.49 | 0.4 | 0.41 | 0.32 | 0.56 | 0.47 |
| 1 | 223869600 | rs11587370 | -0.03 | -0.351 | 0.881 | -0.01 | 0.009 | 0.022 | 0.022 | 0.006 | 0.49 | 0.4 | 0.41 | 0.32 | 0.56 | 0.47 |
| 1 | 223871057 | rs10495213 | 2.039 | -0.209 | 0.444 | 0.112 | -0.029 | -0.037 | -0.037 | 0 | 0.07 | 0 | 0.1 | 0.15 | 0.09 | 0.15 |
| 1 | 223872051 | rs12137025 | 1.47 | -0.249 | 0.503 | 0.082 | 0.003 | 0.011 | 0.011 | 0 | 0.08 | 0.03 | 0.08 | 0.05 | 0.09 | 0.06 |
| 1 | 223874213 | rs6662068 | 1.821 | -0.265 | 0.336 | 0.15 | -0.031 | -0.111 | -0.111 | 2.486 | 0.22 | 0.02 | 0.22 | 0.34 | 0.19 | 0.23 |
| 1 | 223878700 | rs17556570 | 1.216 | -0.212 | 0.466 | 0.071 | 0 | -0.019 | -0.019 | 0 | 0.16 | 0.12 | 0.06 | 0.02 | 0.14 | 0.15 |
| 1 | 223879083 | rs12568612 | 1.244 | -0.23 | 0.435 | 0.063 | 0.001 | -0.015 | -0.015 | 0.528 | 0.16 | 0.12 | 0.06 | 0.02 | 0.14 | 0.15 |

| | | | | | | | | | | | | | | | | |
|---|---|---|---|---|---|---|---|---|---|---|---|---|---|---|---|---|
| 1 | 223887164 | rs16842027 | 1.635 | -0.412 | 0.133 | 0.45 | -0.028 | -0.093 | -0.093 | 0 | 0.23 | 0.02 | 0.22 | 0.27 | 0.19 | 0.23 |
| 1 | 223894889 | rs7555491 | 0.596 | -0.394 | 0.804 | 0.617 | -0.011 | 0 | 0 | 0 | 0.17 | 0.32 | 0.27 | 0.19 | 0.14 | 0.16 |
| 1 | 223901545 | rs2186006 | 0.345 | -0.425 | 0.908 | 0.602 | -0.017 | -0.004 | -0.004 | 3.582 | 0.18 | 0.4 | 0.27 | 0.19 | 0.14 | 0.16 |
| 1 | 223903781 | rs10495215 | 0.871 | -0.599 | 0.804 | 0.463 | 0.005 | -0.024 | -0.024 | 0 | 0.09 | 0.13 | 0.05 | 0.01 | 0.11 | 0.04 |
| 1 | 223905833 | rs12080565 | 0.641 | -0.637 | 0.93 | 0.454 | -0.04 | 0.047 | 0.047 | 0 | 0.1 | 0.24 | 0.22 | 0.18 | 0.05 | 0.12 |
| 1 | 223906113 | rs6674492 | -1.333 | -0.971 | 0.909 | 0.446 | -0.027 | 0.073 | 0.073 | 0 | 0.28 | 0.44 | 0.34 | 0.42 | 0.24 | 0.22 |
| 1 | 223909655 | rs6685475 | 0.692 | -0.829 | 0.754 | 0.313 | 0.01 | -0.03 | -0.03 | 0 | 0.08 | 0.16 | 0.05 | 0.01 | 0.1 | 0.03 |
| 1 | 223912748 | rs1419305 | -0.44 | -0.954 | 1.106 | 0.781 | -0.018 | 0.002 | 0.002 | 0 | 0.19 | 0.36 | 0.09 | 0.18 | 0.14 | 0.16 |
| 1 | 223917907 | rs10916646 | 0.295 | -0.968 | 1.166 | 0.974 | -0.023 | 0.073 | 0.073 | 0 | 0.8 | 0.53 | 0.91 | 0.71 | 0.86 | 0.82 |
| 1 | 223924236 | rs1222145 | 0.768 | -1.14 | 0.997 | 0.906 | -0.027 | 0.028 | 0.028 | 0 | 0.73 | 0.53 | 0.83 | 0.69 | 0.77 | 0.78 |
| 1 | 223929086 | rs12754142 | 1.43 | -0.933 | -0.111 | 0.263 | -0.097 | 0.078 | 0.078 | 0 | 0.21 | 0.02 | 0.14 | 0.08 | 0.29 | 0.21 |
| 1 | 223931068 | rs12066704 | -0.484 | -1.101 | 0.586 | 0.52 | -0.005 | 0.023 | 0.023 | 0.824 | 0.14 | 0.64 | 0.27 | 0.19 | 0.1 | 0.05 |
| 1 | 223933079 | rs17598 | -0.484 | -1.101 | 0.586 | 0.52 | -0.005 | 0.023 | 0.023 | 0.824 | 0.14 | 0.64 | 0.27 | 0.19 | 0.1 | 0.05 |
| 1 | 223936168 | rs3767716 | 0.121 | -1.089 | 0.794 | 0.539 | 0.034 | 0.073 | 0.073 | 0 | 0.12 | 0.37 | 0.25 | 0.19 | 0.1 | 0.05 |
| 1 | 223938565 | rs1153968 | -1.351 | -1.276 | -0.605 | 0.429 | 0.055 | -0.059 | -0.059 | 0 | 0.48 | 0.88 | 0.44 | 0.44 | 0.33 | 0.37 |
| 1 | 223942295 | rs1153961 | -1.322 | -0.798 | 0.735 | 0.642 | 0.057 | 0.222 | 0.222 | 1.25 | 0.05 | 0.35 | 0.15 | 0.11 | 0.01 | 0.01 |
| 1 | 223951841 | rs751128 | -0.057 | -0.489 | 0.327 | 0.391 | -0.003 | -0.003 | -0.003 | 0 | 0.43 | 0.52 | 0.42 | 0.39 | 0.43 | 0.36 |
| 1 | 223953405 | rs1222154 | -0.646 | -0.547 | 0.007 | 0.353 | -0.027 | 0.016 | 0.016 | 0.153 | 0.54 | 0.83 | 0.62 | 0.6 | 0.52 | 0.42 |
| 1 | 223953470 | rs1222153 | -1.047 | 0.134 | 0.568 | 0.621 | 0.002 | -0.008 | -0.008 | 0 | 0.22 | 0.73 | 0.28 | 0.29 | 0.22 | 0.12 |
| 1 | 223958547 | rs1153952 | -1.031 | 0.028 | 0.71 | 0.685 | -0.006 | 0.011 | 0.011 | 0 | 0.22 | 0.71 | 0.27 | 0.3 | 0.18 | 0.13 |
| 1 | 223959796 | rs1153950 | -1.075 | 0.018 | 0.737 | 0.68 | -0.008 | 0.016 | 0.016 | 0 | 0.22 | 0.71 | 0.28 | 0.3 | 0.19 | 0.13 |
| 1 | 223960536 | rs1153947 | 0.881 | 0.008 | 0.841 | 0.698 | -0.014 | 0.095 | 0.095 | 0 | 0.1 | 0.27 | 0.2 | 0.2 | 0.09 | 0.08 |
| 1 | 223962929 | rs10961 | -0.676 | 0.117 | 0.515 | 0.602 | -0.046 | -0.031 | -0.031 | 0.008 | 0.41 | 0.88 | 0.48 | 0.47 | 0.42 | 0.33 |
| 1 | 223963468 | rs7331 | 1.369 | 0.148 | 0.57 | 0.593 | 0.027 | -0.038 | -0.038 | 0 | 0.09 | 0.22 | 0.05 | 0.02 | 0.09 | 0.04 |
| 1 | 223966391 | rs6678103 | 2.212 | 0.147 | 0.525 | 0.594 | -0.009 | 0.01 | 0.01 | 0 | 0.19 | 0.14 | 0.2 | 0.17 | 0.24 | 0.21 |
| 1 | 223967953 | rs17739 | -1.008 | 0.12 | 0.548 | 0.624 | -0.005 | 0.012 | 0.012 | 0 | 0.21 | 0.73 | 0.27 | 0.3 | 0.18 | 0.13 |
| 1 | 223979244 | rs1153937 | -0.338 | 0.095 | 0.262 | 0.638 | -0.04 | 0.176 | 0.176 | 0 | 0.14 | 0.53 | 0.23 | 0.28 | 0.1 | 0.08 |
| 1 | 223981970 | rs7517765 | 2.809 | 0.099 | -0.005 | 0.611 | -0.03 | -0.069 | -0.069 | 0.306 | 0.18 | 0.01 | 0.16 | 0.23 | 0.2 | 0.22 |
| 1 | 223989022 | rs16842257 | 1.459 | 0.139 | 0.022 | 0.6 | 0.031 | -0.035 | -0.035 | 0.397 | 0.08 | 0.21 | 0.05 | 0.02 | 0.09 | 0.04 |
| 1 | 223989966 | rs1153933 | 0.042 | 0.208 | -0.077 | 0.677 | -0.03 | 0.094 | 0.094 | 0 | 0.13 | 0.42 | 0.21 | 0.19 | 0.09 | 0.08 |
| 1 | 223993119 | rs1153930 | -0.075 | 0.208 | -0.078 | 0.677 | -0.034 | 0.086 | 0.086 | 0 | 0.13 | 0.42 | 0.21 | 0.19 | 0.09 | 0.08 |
| 1 | 224001624 | rs10915824 | 1.409 | 0.235 | -0.304 | 0.635 | -0.066 | -0.022 | -0.022 | 3.584 | 0.14 | 0.01 | 0.2 | 0.08 | 0.22 | 0.18 |
| 1 | 224002329 | rs6604725 | 2.731 | 0.273 | 0.065 | 0.794 | -0.038 | -0.072 | -0.072 | 0 | 0.17 | 0.01 | 0.15 | 0.23 | 0.19 | 0.22 |
| 1 | 224008913 | rs3738370 | 0.025 | 0.375 | 0.323 | 0.664 | -0.031 | 0.078 | 0.078 | 0 | 0.13 | 0.38 | 0.21 | 0.19 | 0.09 | 0.08 |
| 1 | 224021459 | rs2045733 | -0.843 | 0.474 | 0.188 | 0.677 | -0.037 | -0.022 | -0.022 | 0.059 | 0.4 | 0.83 | 0.51 | 0.48 | 0.43 | 0.33 |
| 1 | 224026137 | rs1890120 | -0.773 | 0.494 | 0.184 | 0.654 | -0.006 | 0.008 | 0.008 | 0 | 0.39 | 0.68 | 0.49 | 0.39 | 0.43 | 0.33 |
| 1 | 224035842 | rs745697 | 1.747 | 0.87 | 0.152 | 0.72 | -0.02 | -0.001 | -0.001 | 0 | 0.19 | 0.13 | 0.24 | 0.27 | 0.24 | 0.22 |
| 1 | 224041034 | rs10915880 | 1.953 | 1.025 | -0.233 | 0.73 | -0.037 | -0.014 | -0.014 | 0.263 | 0.09 | 0.01 | 0.09 | 0.05 | 0.14 | 0.15 |
| 1 | 224045479 | rs7535882 | 0.648 | 1.033 | 0.396 | 0.704 | 0.016 | -0.032 | -0.032 | 0 | 0.09 | 0.15 | 0.04 | 0.02 | 0.09 | 0.04 |
| 1 | 224046104 | rs16842321 | 0.564 | 0.778 | 0.526 | 1.059 | 0.003 | 0.097 | 0.097 | 0 | 0.07 | 0.18 | 0.18 | 0.19 | 0.07 | 0.07 |
| 1 | 224051439 | rs6604731 | -0.017 | 0.979 | 0.084 | 0.253 | 0.008 | -0.028 | -0.028 | 0 | 0.29 | 0.65 | 0.34 | 0.27 | 0.32 | 0.27 |
| 1 | 224058273 | rs10915917 | 0.428 | 1.073 | -0.872 | 0.245 | -0.001 | 0 | 0 | 0 | 0.26 | 0.68 | 0.28 | 0.29 | 0.25 | 0.23 |
| 1 | 224069384 | rs12073545 | 0.893 | 1.075 | -1.287 | 0.266 | -0.046 | -0.065 | -0.065 | 0.056 | 0.18 | 0.01 | 0.17 | 0.19 | 0.2 | 0.21 |
| 1 | 224070559 | rs1037952 | 0.468 | 1.215 | -0.275 | 0.118 | -0.002 | 0.004 | 0.004 | 0 | 0.05 | 0.12 | 0.07 | 0.08 | 0.07 | 0.06 |
| 1 | 224216822 | rs2488513 | 0.401 | 1.313 | -0.924 | -0.612 | 0.022 | -0.101 | -0.101 | 0 | 0.7 | 0.43 | 0.73 | 0.92 | 0.61 | 0.79 |
| 1 | 224230307 | rs10916197 | -0.444 | 0.958 | -2.171 | 0.219 | -0.086 | -0.015 | -0.015 | 0 | 0.35 | 0.08 | 0.47 | 0.68 | 0.54 | 0.5 |
| 1 | 224237655 | rs10916206 | -0.577 | 0.94 | -2.14 | 0.217 | -0.08 | -0.008 | -0.008 | 0 | 0.35 | 0.08 | 0.47 | 0.68 | 0.54 | 0.49 |
| 1 | 224237716 | rs4551564 | 0.002 | 1.72 | -2.364 | 0.826 | -0.101 | -0.116 | -0.116 | 0.035 | 0.22 | 0.07 | 0.4 | 0.47 | 0.4 | 0.36 |
| 1 | 224239031 | rs6604879 | 0.717 | 0.27 | -2.352 | -0.096 | 0.013 | 0.015 | 0.015 | 0 | 0.36 | 0.64 | 0.31 | 0.22 | 0.26 | 0.39 |
| 1 | 224240313 | rs10916209 | -0.005 | 0.578 | -2.125 | 0.108 | -0.046 | -0.06 | -0.06 | 0.318 | 0.13 | 0.01 | 0.07 | 0.21 | 0.15 | 0.15 |
| 1 | 224245754 | rs7552832 | 0.237 | 0.041 | -1.582 | -0.262 | -0.003 | 0.018 | 0.018 | 0.111 | 0.58 | 0.62 | 0.5 | 0.32 | 0.46 | 0.49 |
| 1 | 224246746 | rs6663154 | 0.619 | -0.374 | -1.462 | -0.302 | 0.044 | 0.062 | 0.062 | 0 | 0.24 | 0.12 | 0.2 | 0.08 | 0.18 | 0.07 |
| 1 | 224251666 | rs7517868 | -0.303 | 0.021 | -1.436 | 0.133 | 0.018 | 0.01 | 0.01 | 0 | 0.7 | 0.63 | 0.66 | 0.41 | 0.56 | 0.61 |
| 1 | 224251838 | rs7531519 | 0.819 | -0.197 | -0.634 | -0.072 | 0 | -0.006 | -0.006 | 0 | 0.32 | 0.33 | 0.3 | 0.23 | 0.31 | 0.42 |
| 1 | 224255729 | rs1492689 | 0.172 | -0.543 | -0.129 | 0.21 | -0.003 | -0.007 | -0.007 | 1.807 | 0.12 | 0.1 | 0.16 | 0.07 | 0.1 | 0.14 |
| 1 | 224255757 | rs7513964 | -0.531 | -0.191 | -0.31 | -0.089 | -0.003 | -0.016 | -0.016 | 0 | 0.41 | 0.53 | 0.49 | 0.73 | 0.41 | 0.35 |
| 1 | 224260824 | rs6604887 | -1.154 | -0.182 | -0.331 | -0.145 | -0.003 | 0.024 | 0.024 | 0 | 0.54 | 0.62 | 0.65 | 0.8 | 0.51 | 0.49 |
| 1 | 224262149 | rs7535721 | -1.154 | -0.182 | -0.331 | -0.145 | -0.003 | 0.024 | 0.024 | 0 | 0.54 | 0.62 | 0.65 | 0.8 | 0.51 | 0.49 |
| 1 | 224263933 | rs16848126 | 1.31 | -0.267 | -0.492 | -0.179 | 0.038 | 0.063 | 0.063 | 0 | 0.19 | 0.08 | 0.19 | 0.09 | 0.15 | 0.07 |
| 1 | 224265217 | rs7544572 | 0.481 | -0.285 | 0.337 | -0.166 | 0.016 | -0.013 | -0.013 | 1.487 | 0.68 | 0.49 | 0.55 | 0.29 | 0.64 | 0.59 |
| 1 | 224268954 | rs6683927 | 0.915 | -0.216 | 0.185 | -0.171 | -0.015 | 0.098 | 0.098 | 0 | 0.46 | 0.29 | 0.34 | 0.2 | 0.49 | 0.52 |
| 1 | 224272792 | rs2404849 | 0.699 | 0.261 | 0.533 | 0.344 | 0.037 | 0.023 | 0.023 | 0 | 0.88 | 0.75 | 0.79 | 0.82 | 0.86 | 0.92 |
| 1 | 224272942 | rs4653930 | 1.262 | 0.046 | 0.542 | 1.098 | -0.012 | -0.023 | -0.023 | 2.112 | 0.5 | 0.17 | 0.34 | 0.47 | 0.52 | 0.63 |
| 1 | 224273311 | rs2896987 | -0.962 | -0.112 | 0.909 | 1.474 | 0.007 | 0.012 | 0.012 | 0 | 0.23 | 0.24 | 0.39 | 0.36 | 0.27 | 0.27 |
| 1 | 224279663 | rs1891740 | -1.489 | -0.301 | 1.131 | 1.275 | 0.023 | 0.017 | 0.017 | 0 | 0.81 | 0.85 | 0.84 | 0.88 | 0.84 | 0.95 |
| 1 | 224283064 | rs1573185 | -1.556 | -0.163 | 1.219 | 1.476 | 0.006 | 0.023 | 0.023 | 0 | 0.28 | 0.59 | 0.44 | 0.42 | 0.28 | 0.27 |
| 1 | 224284375 | rs10916290 | -0.997 | -0.137 | 1.15 | 1.486 | 0.006 | 0.017 | 0.017 | 0 | 0.23 | 0.25 | 0.39 | 0.42 | 0.27 | 0.26 |
| 1 | 224314814 | rs4653564 | 1.147 | -0.274 | 1.64 | 1.321 | 0.026 | 0.042 | 0.042 | 0 | 0.52 | 0.29 | 0.42 | 0.4 | 0.57 | 0.62 |
| 1 | 224318758 | rs10157771 | 0.557 | -0.208 | 1.693 | 1.401 | 0.003 | -0.005 | -0.005 | 0 | 0.12 | 0.12 | 0.1 | 0.09 | 0.12 | 0.17 |

| | | | | | | | | | | | | | | | | |
|---|---|---|---|---|---|---|---|---|---|---|---|---|---|---|---|---|
| 1 | 224321637 | rs869945 | -1.06 | -0.275 | 1.695 | 1.313 | -0.001 | -0.002 | -0.002 | 1.319 | 0.43 | 0.42 | 0.54 | 0.59 | 0.43 | 0.37 |
| 1 | 224335422 | rs12122004 | 0.224 | -0.244 | 1.564 | 1.312 | 0.005 | 0.009 | 0.009 | 0 | 0.08 | 0.06 | 0.05 | 0.02 | 0.06 | 0.05 |
| 1 | 224339138 | rs10429876 | -0.973 | -0.306 | 1.603 | 1.256 | -0.001 | 0.004 | 0.004 | 0 | 0.24 | 0.21 | 0.37 | 0.48 | 0.26 | 0.27 |
| 1 | 224346959 | rs8328 | 1.096 | -0.306 | 1.284 | 1.172 | 0.029 | 0.042 | 0.042 | 0 | 0.52 | 0.29 | 0.41 | 0.4 | 0.57 | 0.62 |
| 1 | 224360933 | rs16844819 | -1.36 | -0.196 | 1.237 | 1.219 | -0.001 | -0.001 | -0.001 | 0.007 | 0.49 | 0.51 | 0.58 | 0.62 | 0.47 | 0.4 |
| 1 | 224361454 | rs16844823 | 0.961 | -0.27 | 0.998 | 1.178 | 0.024 | 0.018 | 0.018 | 0 | 0.2 | 0.16 | 0.17 | 0.12 | 0.17 | 0.1 |
| 1 | 224373608 | rs4653996 | 1.238 | -0.085 | 1.209 | 1.28 | -0.023 | 0.092 | 0.092 | 0 | 0.43 | 0.23 | 0.36 | 0.17 | 0.51 | 0.56 |
| 1 | 224380234 | rs908804 | -0.394 | -0.144 | 1.404 | 1.383 | 0.014 | 0.135 | 0.135 | 0 | 0.11 | 0.21 | 0.22 | 0.36 | 0.09 | 0.07 |
| 1 | 224381107 | rs908803 | -0.387 | -0.106 | 1.295 | 1.4 | 0.015 | 0.137 | 0.137 | 0 | 0.11 | 0.21 | 0.22 | 0.36 | 0.09 | 0.07 |
| 1 | 224387749 | rs12022755 | -0.205 | -0.043 | 1.319 | 1.396 | 0.007 | -0.045 | -0.045 | 0 | 0.38 | 0.23 | 0.35 | 0.49 | 0.38 | 0.33 |
| 1 | 224395782 | rs2176600 | 0.763 | 0.113 | 1.481 | 1.272 | 0.007 | 0.031 | 0.031 | 0 | 0.51 | 0.54 | 0.43 | 0.16 | 0.55 | 0.62 |
| 1 | 224396044 | rs2139155 | 1.301 | 0.064 | 1.637 | 1.239 | -0.036 | 0.133 | 0.133 | 0 | 0.43 | 0.18 | 0.35 | 0.15 | 0.5 | 0.58 |
| 1 | 224397220 | rs6685440 | 0.409 | 0.061 | 1.577 | 1.348 | 0.017 | 0.033 | 0.033 | 0 | 0.32 | 0.2 | 0.28 | 0.2 | 0.3 | 0.22 |
| 1 | 224397977 | rs4654001 | 0.759 | 0.083 | 1.519 | 1.381 | 0 | 0.044 | 0.044 | 0 | 0.05 | 0.04 | 0.05 | 0.29 | 0.05 | 0.08 |
| 1 | 224398639 | rs2011117 | 0.346 | 0.1 | 1.482 | 1.258 | 0.005 | 0.001 | 0.001 | 0 | 0.56 | 0.57 | 0.47 | 0.45 | 0.6 | 0.7 |
| 1 | 224401596 | rs6689187 | 0.996 | 0.142 | 1.132 | 0.717 | -0.032 | 0.14 | 0.14 | 0 | 0.43 | 0.18 | 0.31 | 0.12 | 0.51 | 0.58 |
| 1 | 224403769 | rs10495222 | 0.238 | 0.336 | 0.847 | 0.931 | 0.016 | 0.139 | 0.139 | 0 | 0.89 | 0.81 | 0.75 | 0.63 | 0.9 | 0.92 |
| 1 | 224411596 | rs10799541 | -0.054 | 0.31 | 0.983 | 0.916 | 0.017 | -0.021 | -0.021 | 0 | 0.38 | 0.35 | 0.38 | 0.51 | 0.35 | 0.32 |
| 1 | 224412157 | rs12134258 | -1.36 | 0.644 | 1.155 | 1.759 | 0.017 | 0.124 | 0.124 | 0 | 0.12 | 0.2 | 0.25 | 0.37 | 0.11 | 0.08 |
| 1 | 224426229 | rs6672211 | -0.825 | 0.374 | 1.124 | 1.47 | -0.019 | 0.008 | 0.008 | 0 | 0.08 | 0.2 | 0.05 | 0.06 | 0.04 | 0.03 |
| 1 | 224426825 | rs10916543 | 1.468 | 0.392 | 1.235 | 1.606 | 0.001 | -0.011 | -0.011 | 0 | 0.84 | 0.88 | 0.73 | 0.63 | 0.88 | 0.92 |
| 1 | 224427171 | rs10799551 | 0.718 | 0.654 | 1.225 | 1.745 | 0.015 | 0.123 | 0.123 | 0 | 0.88 | 0.79 | 0.74 | 0.63 | 0.89 | 0.91 |
| 1 | 224428261 | rs4654009 | 0.722 | 0.654 | 1.241 | 1.745 | 0.03 | 0.181 | 0.181 | 0 | 0.88 | 0.72 | 0.73 | 0.62 | 0.89 | 0.92 |
| 1 | 224429078 | rs6664230 | 1.361 | 0.642 | 0.882 | 1.727 | 0.027 | 0.177 | 0.177 | 0 | 0.82 | 0.55 | 0.68 | 0.61 | 0.85 | 0.89 |
| 1 | 224454099 | rs6682667 | -1.717 | 0.739 | 1.617 | 1.346 | 0.132 | 0.217 | 0.217 | 0 | 0.05 | 0.43 | 0.06 | 0.11 | 0.03 | 0.09 |
| 1 | 224476409 | rs10916581 | -0.212 | 0.701 | 0.883 | 1.956 | -0.019 | -0.085 | -0.085 | 0 | 0.12 | 0.01 | 0.2 | 0.41 | 0.13 | 0.14 |
| 1 | 224492746 | rs3754090 | 1.579 | 0.686 | 1.513 | 1.636 | 0.114 | 0.197 | 0.197 | 0.019 | 0.94 | 0.52 | 0.93 | 0.89 | 0.97 | 0.92 |
| 1 | 224495852 | rs7534447 | 1.579 | 0.686 | 1.515 | 1.636 | 0.116 | 0.199 | 0.199 | 0.019 | 0.94 | 0.53 | 0.93 | 0.89 | 0.97 | 0.92 |
| 1 | 224517842 | rs2896990 | 1.561 | 0.685 | 1.639 | 1.636 | 0.121 | 0.205 | 0.205 | 0.005 | 0.94 | 0.56 | 0.93 | 0.89 | 0.97 | 0.92 |
| 1 | 224531559 | rs7534101 | 1.514 | 0.677 | 1.642 | 1.632 | 0.117 | 0.2 | 0.2 | 0 | 0.94 | 0.58 | 0.93 | 0.89 | 0.97 | 0.92 |
| 1 | 224538940 | rs11579964 | 0.542 | 0.334 | 0.619 | 0.734 | -0.007 | 0.112 | 0.112 | 0 | 0.17 | 0.02 | 0.1 | 0 | 0.18 | 0.04 |
| 1 | 224544529 | rs7517754 | -0.519 | 0.939 | 0.248 | -0.317 | 0.006 | -0.086 | -0.086 | 0 | 0.77 | 0.36 | 0.81 | 0.99 | 0.75 | 0.89 |
| 1 | 224551593 | rs12128292 | 0.464 | 1.001 | -0.022 | -0.675 | 0.019 | -0.036 | -0.036 | 7.987 | 0.1 | 0.22 | 0.06 | 0.03 | 0.09 | 0.28 |
| 1 | 224566216 | rs11586729 | 1.02 | 1.031 | 0.077 | -0.695 | -0.002 | -0.032 | -0.032 | 0.03 | 0.2 | 0.22 | 0.13 | 0.01 | 0.24 | 0.11 |
| 1 | 224567004 | rs4654037 | -0.708 | 0.961 | 0.092 | -0.729 | 0.005 | -0.083 | -0.083 | 0 | 0.77 | 0.35 | 0.81 | 0.99 | 0.75 | 0.89 |
| 1 | 224569921 | rs1539207 | -0.751 | 0.961 | 0.1 | -0.729 | 0.007 | -0.088 | -0.088 | 0 | 0.77 | 0.35 | 0.81 | 0.99 | 0.75 | 0.89 |
| 1 | 224572573 | rs12740255 | 1.036 | 0.971 | 0.096 | -0.725 | -0.003 | -0.032 | -0.032 | 0.866 | 0.2 | 0.22 | 0.13 | 0.01 | 0.24 | 0.11 |
| 1 | 224575018 | rs17503938 | 1.021 | 0.972 | 0.096 | -0.724 | -0.003 | -0.028 | -0.028 | 0.015 | 0.2 | 0.22 | 0.13 | 0.01 | 0.24 | 0.11 |
| 1 | 224617323 | rs7526329 | 1.424 | 0.974 | -0.762 | -0.706 | 0.016 | 0.088 | 0.088 | 0 | 0.25 | 0.04 | 0.11 | 0.07 | 0.17 | 0.15 |
| 1 | 224625075 | rs10799573 | 0.094 | 1.071 | -1 | -0.463 | -0.056 | 0.188 | 0.188 | 0.083 | 0.66 | 0.13 | 0.75 | 0.95 | 0.66 | 0.61 |
| 1 | 224644549 | rs10799576 | 0.807 | 0.821 | -1.194 | 0.052 | 0.016 | -0.164 | -0.164 | 0.387 | 0.24 | 0.03 | 0.27 | 0.49 | 0.22 | 0.14 |
| 1 | 224659778 | rs2404785 | 0.533 | 1.55 | -0.425 | -0.153 | 0.028 | -0.053 | -0.053 | 0 | 0.88 | 0.7 | 0.9 | 0.97 | 0.86 | 0.72 |
| 1 | 224663434 | rs7539200 | 0.622 | 1.66 | -0.845 | -0.371 | -0.081 | 0.192 | 0.192 | 0 | 0.65 | 0.13 | 0.73 | 0.93 | 0.62 | 0.61 |
| 1 | 224682826 | rs1341868 | 0.848 | 1.564 | -1.08 | -0.5 | -0.099 | 0.212 | 0.212 | 0.359 | 0.57 | 0.12 | 0.67 | 0.85 | 0.53 | 0.51 |
| 1 | 224682884 | rs10916625 | -0.414 | 1.594 | -0.867 | -0.425 | 0.013 | -0.017 | -0.017 | 0 | 0.18 | 0.24 | 0.14 | 0.11 | 0.21 | 0.39 |
| 1 | 224687445 | rs1891243 | 0.665 | 1.771 | -0.354 | -0.148 | -0.053 | 0.039 | 0.039 | 0 | 0.66 | 0.26 | 0.57 | 0.77 | 0.57 | 0.61 |
| 1 | 224688598 | rs4653589 | 1.584 | 1.537 | 0.544 | 0.344 | 0.062 | 0.057 | 0.057 | 0 | 0.86 | 0.45 | 0.71 | 0.79 | 0.82 | 0.71 |
| 1 | 224696628 | rs712065 | -1.206 | 1.541 | 0.526 | 0.346 | 0.034 | 0.028 | 0.028 | 0 | 0.11 | 0.21 | 0.25 | 0.18 | 0.18 | 0.28 |
| 1 | 224699166 | rs712066 | -2.117 | 1.457 | 0.543 | 0.342 | 0.047 | 0.046 | 0.046 | 0 | 0.14 | 0.58 | 0.3 | 0.21 | 0.18 | 0.28 |
| 1 | 224704236 | rs6695773 | -1.857 | 1.401 | 0.842 | 0.278 | 0.069 | 0.057 | 0.057 | 0 | 0.13 | 0.48 | 0.3 | 0.2 | 0.18 | 0.27 |
| 1 | 224704913 | rs10916629 | -0.722 | 1.446 | 0.757 | 0.356 | 0.024 | 0.018 | 0.018 | 0 | 0.08 | 0.1 | 0.24 | 0.15 | 0.16 | 0.23 |
| 1 | 224705588 | rs6682205 | 0.416 | 1.802 | 0.43 | 0.124 | 0.001 | -0.016 | -0.016 | 0 | 0.11 | 0.13 | 0.08 | 0.02 | 0.14 | 0.07 |
| 1 | 224707177 | rs7550648 | -1.391 | 1.484 | 0.551 | 0.395 | 0.16 | 0.13 | 0.13 | 0 | 0.11 | 0.37 | 0.28 | 0.17 | 0.17 | 0.26 |
| 1 | 224713232 | rs10799582 | -1.04 | 0.931 | -0.068 | 0.586 | -0.004 | -0.004 | -0.004 | 0.288 | 0.14 | 0.14 | 0.24 | 0.25 | 0.18 | 0.38 |
| 1 | 224720286 | rs712074 | -1.121 | 0.868 | 0.072 | 0.578 | 0.022 | 0.103 | 0.103 | 1.194 | 0.16 | 0.52 | 0.26 | 0.25 | 0.18 | 0.38 |
| 1 | 224726060 | rs712082 | -1.356 | 0.838 | -0.219 | 0.595 | -0.004 | 0.056 | 0.056 | 0.5 | 0.17 | 0.6 | 0.26 | 0.29 | 0.18 | 0.4 |
| 1 | 224729933 | rs712086 | -0.509 | 0.497 | -1.531 | 0.005 | 0.015 | 0.195 | 0.195 | 0 | 0.52 | 0.81 | 0.61 | 0.77 | 0.46 | 0.66 |
| 1 | 224733066 | rs698279 | -0.58 | 0.655 | -1.199 | 0.489 | -0.017 | 0.014 | 0.014 | 0 | 0.91 | 1 | 0.94 | 0.94 | 0.89 | 0.94 |
| 1 | 224746577 | rs17571353 | -0.379 | 0.71 | -1.019 | 0.616 | 0 | -0.003 | -0.003 | 0 | 0.06 | 0.09 | 0.04 | 0.1 | 0.06 | 0.1 |
| 1 | 224752508 | rs1499294 | 0.203 | 0.447 | -1.403 | 0.763 | 0.012 | 0.017 | 0.017 | 0 | 0.34 | 0.14 | 0.33 | 0.28 | 0.29 | 0.26 |
| 1 | 224753137 | rs16841702 | -0.193 | 0.147 | -0.374 | 0.665 | -0.015 | -0.014 | -0.014 | 0 | 0.39 | 0.55 | 0.38 | 0.32 | 0.29 | 0.26 |
| 1 | 224755314 | rs12131542 | 0.945 | 0.468 | -0.952 | 0.259 | -0.151 | 0.252 | 0.252 | 0 | 0.37 | 0.05 | 0.32 | 0.04 | 0.45 | 0.21 |
| 1 | 224758560 | rs2088619 | -0.446 | 0.492 | -0.101 | 1.349 | -0.001 | 0.028 | 0.028 | 0.169 | 0.3 | 0.43 | 0.46 | 0.59 | 0.3 | 0.58 |
| 1 | 224768252 | rs6426142 | -0.716 | 0.592 | 0.075 | 1.462 | 0.089 | 0.275 | 0.275 | 0.007 | 0.08 | 0.31 | 0.13 | 0.29 | 0.11 | 0.3 |
| 1 | 224770763 | rs12044602 | -0.308 | 0.597 | 0.337 | 1.464 | 0.04 | 0.161 | 0.161 | 0 | 0.07 | 0.18 | 0.12 | 0.29 | 0.11 | 0.3 |
| 1 | 224772910 | rs4654049 | -0.39 | 0.556 | 0.204 | 1.243 | 0.001 | 0.034 | 0.034 | 0.306 | 0.29 | 0.45 | 0.46 | 0.6 | 0.3 | 0.58 |
| 1 | 224779653 | rs904506 | -0.912 | 0.934 | -0.111 | 1.253 | -0.023 | -0.02 | -0.02 | 0 | 0.17 | 0.08 | 0.11 | 0.2 | 0.19 | 0.13 |
| 1 | 224794913 | rs7535525 | 1.042 | 0.829 | -0.016 | 1.277 | -0.013 | -0.019 | -0.019 | 0.506 | 0.05 | 0 | 0.02 | 0.07 | 0.07 | 0.04 |
| 1 | 224805112 | rs1532557 | 0.396 | 0.63 | 0.33 | 1.835 | -0.012 | 0.02 | 0.02 | 0 | 0.24 | 0.32 | 0.36 | 0.34 | 0.19 | 0.28 |

| | | | | | | | | | | | | | | | | |
|---|---|---|---|---|---|---|---|---|---|---|---|---|---|---|---|---|
| 1 | 224806976 | rs9919286 | 0.347 | 0.795 | 0.119 | 1.86 | 0.003 | 0.027 | 0.027 | 0.006 | 0.05 | 0.07 | 0.09 | 0.19 | 0.11 | 0.29 |
| 1 | 224807561 | rs883115 | -1.881 | 0.754 | -0.027 | 1.729 | -0.025 | 0.176 | 0.176 | 0.235 | 0.5 | 0.77 | 0.6 | 0.77 | 0.5 | 0.73 |
| 1 | 224807973 | rs883116 | 0.563 | 0.559 | -0.1 | 1.742 | 0.012 | -0.057 | -0.057 | 0 | 0.23 | 0.13 | 0.34 | 0.36 | 0.19 | 0.28 |
| 1 | 224808134 | rs880496 | -0.784 | 0.607 | -0.03 | 1.54 | -0.02 | -0.017 | -0.017 | 0 | 0.16 | 0.08 | 0.11 | 0.18 | 0.19 | 0.12 |
| 1 | 224821550 | rs924362 | -1.117 | 0.55 | 0.222 | 1.657 | -0.003 | 0.003 | 0.003 | 0.056 | 0.41 | 0.47 | 0.48 | 0.57 | 0.39 | 0.46 |
| 1 | 224822482 | rs10799590 | -1.519 | 0.432 | 0.118 | 1.667 | 0.015 | 0.145 | 0.145 | 0 | 0.46 | 0.82 | 0.63 | 0.76 | 0.39 | 0.52 |
| 1 | 224824411 | rs12123347 | -2.002 | 0.454 | 0.393 | 1.998 | -0.001 | 0.012 | 0.012 | 0 | 0.24 | 0.66 | 0.29 | 0.38 | 0.2 | 0.26 |
| 1 | 224824529 | rs12145463 | -1.819 | 0.459 | 0.396 | 2 | -0.001 | 0.018 | 0.018 | 0 | 0.23 | 0.65 | 0.29 | 0.38 | 0.2 | 0.26 |
| 1 | 224832751 | rs12124222 | -0.085 | 0.449 | 0.36 | 2.062 | 0.002 | -0.024 | -0.024 | 0 | 0.05 | 0 | 0.12 | 0.15 | 0.03 | 0.09 |
| 1 | 224833490 | rs12065840 | 0.624 | 0.413 | 0.312 | 2.102 | 0.01 | -0.056 | -0.056 | 0.325 | 0.22 | 0.12 | 0.34 | 0.34 | 0.2 | 0.21 |
| 1 | 224835036 | rs12118224 | -1.336 | 0.436 | 0.574 | 2.121 | 0.006 | 0.029 | 0.029 | 0 | 0.18 | 0.24 | 0.22 | 0.33 | 0.19 | 0.19 |
| 1 | 224835372 | rs17513562 | -0.427 | 0.502 | 0.575 | 2.097 | 0.021 | 0.017 | 0.017 | 0 | 0.13 | 0.21 | 0.1 | 0.18 | 0.16 | 0.1 |
| 1 | 224836514 | rs12130499 | -1.733 | 0.473 | 0.625 | 2.049 | 0.013 | 0.06 | 0.06 | 0 | 0.45 | 0.66 | 0.62 | 0.72 | 0.4 | 0.47 |
| 1 | 224838018 | rs6672460 | -0.106 | 0.526 | 0.502 | 2.026 | 0.004 | -0.009 | -0.009 | 0 | 0.05 | 0.02 | 0.12 | 0.15 | 0.03 | 0.09 |
| 1 | 224842251 | rs298733 | -1.732 | 0.412 | 0.427 | 2.001 | -0.014 | 0.122 | 0.122 | 0.09 | 0.47 | 0.89 | 0.64 | 0.73 | 0.4 | 0.45 |
| 1 | 224844991 | rs298736 | -1.348 | 0.391 | 0.646 | 2.073 | 0.01 | 0.049 | 0.049 | 0 | 0.18 | 0.57 | 0.16 | 0.24 | 0.17 | 0.17 |
| 1 | 224848250 | rs1499282 | -0.508 | 0.392 | 0.753 | 2.083 | 0.018 | 0.026 | 0.026 | 0 | 0.14 | 0.26 | 0.11 | 0.19 | 0.17 | 0.1 |
| 1 | 224850846 | rs1369848 | -0.323 | 0.349 | 0.742 | 2.121 | -0.007 | 0.013 | 0.013 | 0 | 0.3 | 0.35 | 0.49 | 0.52 | 0.23 | 0.32 |
| 1 | 224854784 | rs1337538 | 0.498 | 0.368 | 0.631 | 2.253 | 0.012 | -0.018 | -0.018 | 0 | 0.23 | 0.18 | 0.35 | 0.34 | 0.2 | 0.22 |
| 1 | 224860443 | rs12133496 | -1.431 | 0.391 | 0.525 | 2.213 | 0.003 | 0.018 | 0.018 | 0 | 0.19 | 0.27 | 0.23 | 0.27 | 0.19 | 0.17 |
| 1 | 224863137 | rs12117838 | -0.084 | 0.394 | 0.456 | 2.224 | 0.007 | -0.02 | -0.02 | 0.088 | 0.05 | 0.01 | 0.12 | 0.16 | 0.03 | 0.07 |
| 1 | 224863140 | rs4653591 | 1.72 | 0.385 | 0.404 | 2.262 | -0.019 | 0.167 | 0.167 | 0.124 | 0.52 | 0.08 | 0.35 | 0.23 | 0.6 | 0.5 |
| 1 | 224864341 | rs6426147 | 0.514 | 0.388 | 0.548 | 2.302 | 0.001 | -0.002 | -0.002 | 2.081 | 0.23 | 0.22 | 0.36 | 0.37 | 0.2 | 0.22 |
| 1 | 224865644 | rs12136553 | -0.078 | 0.402 | 0.525 | 2.28 | 0.007 | -0.02 | -0.02 | 0 | 0.05 | 0.01 | 0.12 | 0.16 | 0.03 | 0.07 |
| 1 | 224872443 | rs3738364 | -1.664 | 0.464 | 0.618 | 2.324 | 0.004 | 0.049 | 0.049 | 0 | 0.21 | 0.41 | 0.25 | 0.31 | 0.2 | 0.2 |
| 1 | 224872473 | rs3738363 | -0.775 | 0.469 | 0.625 | 2.332 | 0.023 | 0.001 | 0.001 | 0 | 0.15 | 0.41 | 0.12 | 0.12 | 0.17 | 0.1 |
| 1 | 224874931 | rs10915695 | 0.512 | 0.458 | 0.609 | 2.331 | 0.001 | -0.002 | -0.002 | 0 | 0.23 | 0.22 | 0.35 | 0.37 | 0.2 | 0.22 |
| 1 | 224876002 | rs7549891 | -0.755 | 0.495 | 0.588 | 2.278 | 0.012 | 0.01 | 0.01 | 0 | 0.15 | 0.27 | 0.11 | 0.15 | 0.17 | 0.13 |
| 1 | 224879127 | rs7545574 | -1.556 | 0.684 | 0.489 | 2.079 | -0.001 | -0.002 | -0.002 | 0.044 | 0.44 | 0.64 | 0.6 | 0.65 | 0.4 | 0.4 |
| 1 | 224879577 | rs7543218 | -1.509 | 0.67 | 0.544 | 2.066 | -0.003 | -0.006 | -0.006 | 2.47 | 0.43 | 0.6 | 0.6 | 0.65 | 0.4 | 0.41 |
| 1 | 224880533 | rs6696394 | -0.699 | 0.784 | 0.118 | 2.163 | -0.002 | 0.028 | 0.028 | 0 | 0.06 | 0.07 | 0.13 | 0.15 | 0.04 | 0.11 |
| 1 | 224880653 | rs879363 | -0.597 | 0.687 | 0.116 | 2.163 | -0.005 | 0.025 | 0.025 | 0 | 0.06 | 0.06 | 0.13 | 0.15 | 0.03 | 0.11 |
| 1 | 224881828 | rs1436171 | 1.53 | 0.683 | 0.286 | 2.17 | 0.004 | 0.118 | 0.118 | 0.106 | 0.53 | 0.15 | 0.37 | 0.27 | 0.58 | 0.53 |
| 1 | 224881970 | rs4653593 | 0.55 | 0.668 | 0.379 | 2.234 | 0.047 | 0.085 | 0.085 | 0 | 0.83 | 0.56 | 0.87 | 0.78 | 0.83 | 0.87 |
| 1 | 224886068 | rs2897161 | 1.731 | 0.732 | 0.293 | 2.184 | -0.034 | 0.147 | 0.147 | 0.378 | 0.51 | 0.07 | 0.37 | 0.26 | 0.58 | 0.5 |
| 1 | 224886492 | rs9725216 | 0.452 | 0.828 | 0.382 | 2.207 | 0.058 | 0.08 | 0.08 | 0 | 0.84 | 0.56 | 0.88 | 0.81 | 0.83 | 0.87 |
| 1 | 224888421 | rs1465766 | 0.488 | 0.827 | 0.181 | 2.227 | 0.003 | -0.011 | -0.011 | 0.005 | 0.23 | 0.22 | 0.36 | 0.36 | 0.21 | 0.22 |
| 1 | 224890520 | rs1965776 | 0.867 | 0.872 | -0.127 | 1.772 | 0.001 | -0.02 | -0.02 | 0 | 0.58 | 0.18 | 0.5 | 0.42 | 0.62 | 0.61 |
| 1 | 224894095 | rs1369846 | 0.819 | 0.937 | -0.128 | 1.761 | 0 | -0.026 | -0.026 | 0 | 0.6 | 0.24 | 0.51 | 0.44 | 0.62 | 0.65 |
| 1 | 224900110 | rs892975 | -1.293 | 1.34 | 0.316 | 1.959 | 0.051 | 0.169 | 0.169 | 0 | 0.14 | 0.23 | 0.29 | 0.45 | 0.2 | 0.37 |
| 1 | 224903823 | rs6690131 | 0.098 | 1.328 | 0.674 | 1.789 | -0.002 | 0.001 | 0.001 | 0 | 0.26 | 0.28 | 0.32 | 0.34 | 0.23 | 0.31 |
| 1 | 224906158 | rs12741660 | -0.483 | 1.235 | 0.503 | 1.376 | -0.002 | 0.009 | 0.009 | 0 | 0.16 | 0.3 | 0.13 | 0.17 | 0.16 | 0.12 |
| 1 | 224906444 | rs1561109 | -1.171 | 1.023 | 0.422 | 1.278 | -0.004 | 0.019 | 0.019 | 0.033 | 0.42 | 0.57 | 0.63 | 0.79 | 0.43 | 0.68 |
| 1 | 224909092 | rs6426154 | -1.776 | 1.092 | 0.454 | 1.386 | 0.018 | 0.007 | 0.007 | 0 | 0.32 | 0.54 | 0.44 | 0.63 | 0.37 | 0.51 |
| 1 | 224910762 | rs12036311 | -0.976 | 0.972 | 0.191 | 1.184 | 0.035 | 0.048 | 0.048 | 0 | 0.23 | 0.48 | 0.21 | 0.29 | 0.25 | 0.3 |
| 1 | 224910935 | rs4653596 | -0.809 | 0.941 | 0.136 | 1.155 | 0 | 0.002 | 0.002 | 0 | 0.24 | 0.26 | 0.38 | 0.27 | 0.27 | 0.18 |
| 1 | 224911001 | rs892978 | 0.446 | 1.686 | 0.066 | 0.841 | 0.063 | -0.032 | -0.032 | 0 | 0.7 | 0.5 | 0.76 | 0.81 | 0.67 | 0.75 |
| 1 | 224911934 | rs2405523 | 1.646 | 1.186 | -0.051 | 1.189 | -0.017 | 0.021 | 0.021 | 0.926 | 0.43 | 0.31 | 0.35 | 0.38 | 0.48 | 0.43 |
| 1 | 224912930 | rs2012466 | -1.045 | 1.254 | -0.373 | 1.286 | 0.001 | -0.029 | -0.029 | 0.246 | 0.27 | 0.13 | 0.34 | 0.35 | 0.26 | 0.33 |
| 1 | 224915429 | rs1010735 | -1.196 | 1.25 | -0.252 | 1.285 | -0.002 | 0.008 | 0.008 | 0 | 0.28 | 0.24 | 0.34 | 0.35 | 0.26 | 0.33 |
| 1 | 224916592 | rs6660579 | -0.199 | 1.106 | -0.462 | 1.196 | 0.013 | 0.004 | 0.004 | 0 | 0.17 | 0.11 | 0.15 | 0.16 | 0.13 | 0.21 |
| 1 | 224916593 | rs16853130 | 0.044 | 1.11 | -0.533 | 0.864 | -0.014 | -0.028 | -0.028 | 4.157 | 0.28 | 0.52 | 0.31 | 0.22 | 0.26 | 0.23 |
| 1 | 224921487 | rs6426156 | -0.411 | 0.919 | -0.885 | 0.624 | -0.005 | -0.007 | -0.007 | 0.523 | 0.9 | 0.95 | 0.82 | 0.84 | 0.88 | 0.88 |
| 1 | 224925739 | rs7414300 | 1.219 | 0.852 | -1.022 | -0.483 | -0.001 | -0.063 | -0.063 | 0 | 0.55 | 0.51 | 0.32 | 0.14 | 0.56 | 0.38 |
| 1 | 224930810 | rs6674873 | -0.347 | 1.715 | -0.638 | 0.26 | 0.004 | -0.062 | -0.062 | 0 | 0.26 | 0.33 | 0.15 | 0.07 | 0.29 | 0.21 |
| 1 | 224936295 | rs10449246 | -0.758 | 1.16 | -0.762 | 1.429 | -0.019 | -0.09 | -0.09 | 1.397 | 0.16 | 0.02 | 0.15 | 0.31 | 0.15 | 0.22 |
| 1 | 224942850 | rs1436174 | 1.414 | 1.26 | -0.011 | 1.002 | -0.007 | 0.018 | 0.018 | 0.039 | 0.78 | 0.36 | 0.7 | 0.67 | 0.82 | 0.68 |
| 1 | 224945481 | rs2574877 | 0.339 | 1.294 | 0.092 | 1.021 | -0.004 | -0.016 | -0.016 | 0 | 0.87 | 0.96 | 0.77 | 0.76 | 0.88 | 0.79 |
| 1 | 224966644 | rs2574881 | 1.121 | 1.365 | 0.315 | 1.039 | 0 | 0.02 | 0.02 | 0 | 0.85 | 0.82 | 0.75 | 0.67 | 0.87 | 0.68 |
| 1 | 224971229 | rs2802559 | 0.112 | 1.358 | 0.217 | 1.053 | 0 | 0.003 | 0.003 | 0 | 0.89 | 0.93 | 0.78 | 0.76 | 0.93 | 0.79 |
| 1 | 224972014 | rs12033057 | -0.294 | 1.844 | -0.216 | 1.333 | -0.074 | -0.119 | -0.119 | 0 | 0.14 | 0.01 | 0.31 | 0.46 | 0.2 | 0.29 |
| 1 | 224987113 | rs2802745 | 1.837 | 1.654 | -0.018 | 0.82 | -0.005 | -0.005 | -0.005 | 0 | 0.74 | 0.36 | 0.68 | 0.68 | 0.78 | 0.55 |
| 1 | 224991534 | rs12081821 | -1.648 | 1.649 | 0.043 | 0.834 | 0.015 | -0.06 | -0.06 | 0 | 0.16 | 0.61 | 0.1 | 0.1 | 0.14 | 0.25 |
| 1 | 225000644 | rs1316883 | 0.322 | 1.675 | -0.079 | 0.841 | -0.01 | 0.023 | 0.023 | 0.15 | 0.07 | 0.01 | 0.03 | 0 | 0.08 | 0.14 |
| 1 | 225007842 | rs11590145 | 0.379 | 1.779 | -0.015 | 1.472 | -0.033 | -0.082 | -0.082 | 0 | 0.12 | 0 | 0.17 | 0.27 | 0.11 | 0.16 |
| 1 | 225008581 | rs11810181 | -0.897 | 1.631 | 0.205 | 0.958 | 0.073 | 0.015 | 0.015 | 0.383 | 0.07 | 0.46 | 0.06 | 0.02 | 0.06 | 0.04 |
| 1 | 225020648 | rs1936632 | -0.088 | 1.593 | -0.184 | 0.999 | 0 | -0.006 | -0.006 | 2.533 | 0.9 | 0.97 | 0.79 | 0.78 | 0.92 | 0.8 |
| 1 | 225026240 | rs3856143 | 1.597 | 1.672 | -0.262 | 0.323 | -0.005 | 0.019 | 0.019 | 0 | 0.53 | 0.59 | 0.34 | 0.19 | 0.5 | 0.35 |
| 1 | 225036472 | rs2662918 | -0.347 | 1.751 | -0.448 | 0.534 | -0.045 | -0.1 | -0.1 | 0.248 | 0.84 | 0.97 | 0.7 | 0.48 | 0.79 | 0.65 |

| | | | | | | | | | | | | | | | | |
|---|---|---|---|---|---|---|---|---|---|---|---|---|---|---|---|---|
| 1 | 225044284 | rs1892123 | 0.832 | 1.653 | -0.463 | 0.375 | -0.018 | 0.014 | 0.014 | 0 | 0.62 | 0.78 | 0.57 | 0.41 | 0.58 | 0.5 |
| 1 | 225061843 | rs16856829 | 0.033 | 1.712 | -0.019 | 0.798 | 0 | 0.007 | 0.007 | 0 | 0.08 | 0.04 | 0.21 | 0.22 | 0.08 | 0.15 |
| 1 | 225061975 | rs2802567 | -0.86 | 1.509 | 0.326 | 0.726 | 0.053 | 0.097 | 0.097 | 5.654 | 0.06 | 0.58 | 0.05 | 0.06 | 0 | 0.05 |
| 1 | 225062793 | rs2244113 | -1.52 | 1.66 | 0.201 | 0.666 | 0.021 | -0.002 | -0.002 | 0.055 | 0.1 | 0.63 | 0.09 | 0.08 | 0.07 | 0.09 |
| 1 | 225066930 | rs1892122 | -2.132 | 1.608 | 0.145 | 0.603 | -0.016 | -0.039 | -0.039 | 0 | 0.15 | 0.72 | 0.12 | 0.08 | 0.11 | 0.13 |
| 1 | 225067331 | rs2051158 | -2.037 | 1.702 | -0.007 | 0.727 | 0.002 | -0.002 | -0.002 | 0 | 0.22 | 0.8 | 0.33 | 0.29 | 0.19 | 0.34 |
| 1 | 225070168 | rs2254406 | -1.996 | 1.225 | -0.164 | 0.679 | 0.034 | -0.02 | -0.02 | 0.084 | 0.24 | 0.89 | 0.32 | 0.3 | 0.16 | 0.29 |
| 1 | 225090870 | rs1936635 | -0.808 | 0.997 | -0.194 | 0.587 | 0.001 | -0.017 | -0.017 | 0.011 | 0.16 | 0.04 | 0.25 | 0.24 | 0.16 | 0.21 |
| 1 | 225095329 | rs2501094 | -0.987 | 1.107 | 0.619 | 0.693 | 0.053 | 0.048 | 0.048 | 6.858 | 0.07 | 0.51 | 0.05 | 0.01 | 0.01 | 0.03 |
| 1 | 225106597 | rs2501082 | -1.718 | 1.163 | 0.527 | 0.759 | -0.006 | 0.072 | 0.072 | 2.287 | 0.15 | 0.55 | 0.27 | 0.25 | 0.12 | 0.2 |
| 1 | 225146177 | rs2128270 | -1.56 | 1.186 | 0.853 | 0.783 | 0.029 | 0.135 | 0.135 | 1.339 | 0.13 | 0.47 | 0.26 | 0.24 | 0.12 | 0.18 |
| 1 | 225146988 | rs2489352 | -2.02 | 1.105 | 1.043 | 0.826 | -0.012 | 0.063 | 0.063 | 1.931 | 0.15 | 0.66 | 0.27 | 0.29 | 0.12 | 0.22 |
| 1 | 225152505 | rs12023087 | -0.319 | 1.297 | 0.347 | 0.524 | -0.108 | -0.107 | -0.107 | 0 | 0.3 | 0.05 | 0.39 | 0.55 | 0.42 | 0.44 |
| 1 | 225156457 | rs16858926 | 1.122 | 1.344 | 0.28 | 0.857 | -0.02 | -0.068 | -0.068 | 0 | 0.1 | 0 | 0.17 | 0.3 | 0.1 | 0.14 |
| 1 | 225182795 | rs2489327 | -1.577 | 1.318 | 0.402 | 0.775 | -0.012 | 0.063 | 0.063 | 1.931 | 0.15 | 0.66 | 0.27 | 0.29 | 0.12 | 0.22 |
| 1 | 225191852 | rs2449285 | -1.694 | 1.445 | 0.252 | 0.681 | -0.022 | 0.031 | 0.031 | 0 | 0.94 | 1 | 0.97 | 1 | 0.94 | 0.99 |
| 1 | 225202821 | rs1506072 | 0.888 | 1.261 | 0.669 | 1.045 | 0.029 | 0.135 | 0.135 | 1.339 | 0.87 | 0.53 | 0.74 | 0.76 | 0.88 | 0.83 |
| 1 | 225206826 | rs2449276 | 0.694 | 1.251 | 0.685 | 1.048 | 0.022 | 0.197 | 0.197 | 1.172 | 0.91 | 0.56 | 0.78 | 0.78 | 0.96 | 0.88 |
| 1 | 225215085 | rs2449315 | 1.117 | 1.266 | 0.656 | 1.036 | 0.029 | 0.135 | 0.135 | 1.339 | 0.87 | 0.53 | 0.74 | 0.76 | 0.88 | 0.83 |
| 1 | 225227937 | rs2939363 | 1.541 | 1.252 | 0.613 | 1.039 | -0.012 | 0.062 | 0.062 | 1.832 | 0.85 | 0.34 | 0.73 | 0.71 | 0.88 | 0.78 |
| 1 | 225232264 | rs1506074 | 1.521 | 1.253 | 0.613 | 1.04 | -0.012 | 0.063 | 0.063 | 1.931 | 0.85 | 0.34 | 0.73 | 0.71 | 0.88 | 0.78 |
| 1 | 225234649 | rs12758848 | -0.795 | 1.312 | 0.521 | 1.002 | -0.108 | -0.107 | -0.107 | 0 | 0.3 | 0.05 | 0.39 | 0.55 | 0.42 | 0.44 |
| 1 | 225235080 | rs1506073 | 0.982 | 1.307 | 0.735 | 1.114 | -0.023 | 0.127 | 0.127 | 1.622 | 0.89 | 0.36 | 0.77 | 0.72 | 0.96 | 0.83 |
| 1 | 225268343 | rs3128651 | 0.966 | 1.306 | 0.735 | 1.106 | -0.022 | 0.132 | 0.132 | 1.427 | 0.89 | 0.37 | 0.77 | 0.73 | 0.96 | 0.83 |
| 1 | 225273248 | rs3128658 | 1 | 1.304 | 0.734 | 1.105 | -0.023 | 0.13 | 0.13 | 1.384 | 0.89 | 0.37 | 0.77 | 0.72 | 0.96 | 0.83 |
| 1 | 225287476 | rs4275419 | -0.809 | 1.297 | 0.97 | 1.099 | 0.06 | 0.106 | 0.106 | 3.19 | 0.06 | 0.53 | 0.04 | 0.06 | 0 | 0.04 |
| 1 | 225297453 | rs10915777 | 0.653 | 1.529 | 0.753 | 1.128 | -0.044 | -0.04 | -0.04 | 0 | 0.07 | 0.01 | 0.09 | 0.14 | 0.17 | 0.24 |
| 1 | 225302384 | rs10915779 | -1.742 | 1.08 | 0.915 | 1.082 | -0.01 | 0.071 | 0.071 | 1.665 | 0.15 | 0.65 | 0.27 | 0.29 | 0.12 | 0.22 |
| 1 | 225340241 | rs1842332 | 0.459 | 1.107 | 0.544 | 1.136 | -0.001 | 0.047 | 0.047 | 0 | 0.12 | 0.13 | 0.19 | 0.31 | 0.11 | 0.14 |
| 1 | 225341687 | rs16844377 | -0.56 | 1.128 | 1.243 | 1.22 | 0.021 | 0.152 | 0.152 | 0 | 0.08 | 0.27 | 0.2 | 0.22 | 0.05 | 0.12 |
| 1 | 225380088 | rs1572750 | -0.497 | 1.385 | 0.892 | 1.015 | -0.082 | -0.037 | -0.037 | 0.965 | 0.19 | 0.03 | 0.21 | 0.24 | 0.32 | 0.32 |
| 1 | 225410051 | rs649637 | -1.525 | 1.34 | 0.999 | 1.092 | -0.004 | 0.084 | 0.084 | 0 | 0.25 | 0.5 | 0.44 | 0.6 | 0.23 | 0.36 |
| 1 | 225418456 | rs650112 | -0.922 | 1.344 | 1.034 | 1.114 | -0.005 | 0.143 | 0.143 | 0 | 0.2 | 0.4 | 0.39 | 0.53 | 0.16 | 0.27 |
| 1 | 225419456 | rs664855 | -1.51 | 1.339 | 1.034 | 1.101 | -0.006 | 0.069 | 0.069 | 0.018 | 0.25 | 0.54 | 0.44 | 0.6 | 0.23 | 0.36 |
| 1 | 225427066 | rs584720 | -0.598 | 1.347 | 1.055 | 1.175 | -0.001 | 0.039 | 0.039 | 0 | 0.17 | 0.25 | 0.24 | 0.37 | 0.18 | 0.24 |
| 1 | 225429205 | rs12049495 | -1.052 | 1.093 | 0.734 | 0.795 | 0.016 | -0.025 | -0.025 | 0 | 0.33 | 0.42 | 0.29 | 0.24 | 0.39 | 0.45 |
| 1 | 225435891 | rs712060 | -0.655 | 1.233 | 0.96 | 1.194 | -0.001 | 0.015 | 0.015 | 0 | 0.17 | 0.21 | 0.24 | 0.37 | 0.18 | 0.24 |
| 1 | 225441678 | rs12759119 | 0.8 | 1.226 | 0.95 | 1.207 | -0.028 | -0.071 | -0.071 | 0 | 0.11 | 0 | 0.17 | 0.31 | 0.11 | 0.14 |
| 1 | 225443089 | rs638371 | -0.612 | 1.233 | 0.964 | 1.164 | -0.001 | 0.015 | 0.015 | 0 | 0.17 | 0.21 | 0.24 | 0.37 | 0.18 | 0.24 |
| 1 | 225449050 | rs12564045 | -0.728 | 1.325 | 0.665 | 1.145 | 0.019 | 0.146 | 0.146 | 0 | 0.08 | 0.26 | 0.2 | 0.22 | 0.05 | 0.12 |
| 1 | 225470144 | rs1553447 | 0.212 | 1.251 | 0.454 | 1.11 | 0 | 0.017 | 0.017 | 0 | 0.12 | 0.11 | 0.19 | 0.33 | 0.11 | 0.14 |
| 1 | 225474997 | rs11587822 | -0.112 | 1.266 | 0.006 | 1.133 | -0.003 | 0.031 | 0.031 | 0 | 0.13 | 0.15 | 0.2 | 0.33 | 0.11 | 0.14 |
| 1 | 225477546 | rs12042076 | -1.25 | 1.522 | -0.295 | 1.017 | -0.062 | -0.071 | -0.071 | 0 | 0.24 | 0.11 | 0.39 | 0.45 | 0.36 | 0.47 |
| 1 | 225493668 | rs12035153 | -0.043 | 1.536 | -0.166 | 1.078 | -0.019 | -0.019 | -0.019 | 0 | 0.08 | 0.03 | 0.11 | 0.18 | 0.19 | 0.21 |
| 1 | 225495656 | rs10495236 | -0.137 | 1.08 | -0.575 | 0.879 | -0.011 | -0.091 | -0.091 | 0.938 | 0.23 | 0.06 | 0.27 | 0.33 | 0.21 | 0.28 |
| 1 | 225495697 | rs7524527 | -1.158 | 1.055 | 0.225 | 0.851 | -0.006 | 0.004 | 0.004 | 0 | 0.34 | 0.53 | 0.53 | 0.62 | 0.32 | 0.38 |
| 1 | 225506427 | rs12403278 | 0.636 | 1.011 | 0.661 | 0.834 | 0.011 | 0.013 | 0.013 | 0.368 | 0.07 | 0.23 | 0.06 | 0.03 | 0.03 | 0.01 |
| 1 | 225508712 | rs648011 | -1.669 | 0.624 | 0.632 | 0.524 | -0.001 | -0.002 | -0.002 | 0.053 | 0.35 | 0.63 | 0.27 | 0.28 | 0.32 | 0.47 |
| 1 | 225510641 | rs1532771 | -1.038 | 0.819 | 0.609 | 0.688 | -0.005 | -0.017 | -0.017 | 0 | 0.35 | 0.34 | 0.41 | 0.48 | 0.37 | 0.56 |
| 1 | 225512083 | rs12032942 | 0.204 | 0.797 | 0.607 | 0.691 | -0.034 | -0.009 | -0.009 | 0.245 | 0.09 | 0.03 | 0.12 | 0.14 | 0.16 | 0.28 |
| 1 | 225512398 | rs6668011 | 0.605 | 0.805 | 0.693 | 0.685 | 0.018 | 0.015 | 0.015 | 0.461 | 0.07 | 0.25 | 0.06 | 0.03 | 0.04 | 0.01 |
| 1 | 225516861 | rs17521900 | 0.155 | 0.75 | 0.679 | 0.687 | -0.028 | -0.005 | -0.005 | 0 | 0.09 | 0.05 | 0.12 | 0.14 | 0.16 | 0.28 |
| 1 | 225520212 | rs12036489 | -1.597 | 0.716 | 0.652 | 0.655 | 0.008 | -0.032 | -0.032 | 0 | 0.22 | 0.36 | 0.21 | 0.18 | 0.25 | 0.36 |
| 1 | 225524656 | rs7521190 | -1.563 | 0.835 | 0.857 | 0.831 | 0.048 | 0.098 | 0.098 | 0 | 0.2 | 0.51 | 0.38 | 0.42 | 0.27 | 0.38 |
| 1 | 225528183 | rs3856145 | -1.618 | 0.723 | 0.842 | 0.799 | 0.001 | -0.008 | -0.008 | 0.151 | 0.42 | 0.64 | 0.47 | 0.51 | 0.4 | 0.57 |
| 1 | 225529840 | rs3913657 | -0.927 | 0.703 | 0.81 | 0.771 | 0.027 | -0.011 | -0.011 | 0.197 | 0.16 | 0.35 | 0.18 | 0.17 | 0.19 | 0.3 |
| 1 | 225533684 | rs6667999 | -1.616 | 0.703 | 0.827 | 0.772 | 0.001 | -0.006 | -0.006 | 1.698 | 0.42 | 0.64 | 0.47 | 0.51 | 0.4 | 0.57 |
| 1 | 225534219 | rs7527925 | -1.616 | 0.703 | 0.804 | 0.772 | 0.001 | -0.007 | -0.007 | 0 | 0.42 | 0.64 | 0.47 | 0.51 | 0.4 | 0.57 |
| 1 | 225534348 | rs7535953 | -1.626 | 0.693 | 0.8 | 0.767 | 0.001 | -0.007 | -0.007 | 0 | 0.42 | 0.64 | 0.47 | 0.51 | 0.4 | 0.57 |
| 1 | 225535308 | rs16844752 | -0.152 | 0.663 | 0.648 | 0.786 | -0.001 | 0.002 | 0.002 | 0 | 0.25 | 0.29 | 0.29 | 0.34 | 0.21 | 0.28 |
| 1 | 225538052 | rs4459070 | -0.585 | 0.678 | 0.624 | 0.791 | -0.006 | 0.009 | 0.009 | 0 | 0.3 | 0.3 | 0.33 | 0.37 | 0.24 | 0.29 |
| 1 | 225539363 | rs12408617 | 0.648 | 0.596 | 0.593 | 0.174 | 0.013 | 0.015 | 0.015 | 0.368 | 0.07 | 0.24 | 0.06 | 0.03 | 0.03 | 0.01 |
| 1 | 225540026 | rs3913654 | -0.577 | 0.616 | 0.619 | 0.197 | -0.003 | 0.043 | 0.043 | 0 | 0.3 | 0.51 | 0.49 | 0.59 | 0.29 | 0.36 |
| 1 | 225540100 | rs12040037 | -1.169 | 0.695 | 0.94 | 0.057 | 0.021 | -0.011 | -0.011 | 0.017 | 0.16 | 0.29 | 0.18 | 0.17 | 0.19 | 0.29 |
| 1 | 225562385 | rs10915816 | 1.13 | 0.527 | -0.004 | -0.179 | -0.008 | -0.067 | -0.067 | 0 | 0.81 | 0.36 | 0.86 | 0.97 | 0.84 | 0.68 |
| 1 | 225565015 | rs3856154 | 1.724 | 0.481 | -0.416 | -0.149 | 0.004 | 0.025 | 0.025 | 0.81 | 0.66 | 0.3 | 0.75 | 0.76 | 0.66 | 0.45 |
| 1 | 225569241 | rs950210 | 1.755 | 0.465 | -0.439 | -0.163 | 0.005 | 0.06 | 0.06 | 1.086 | 0.66 | 0.19 | 0.74 | 0.76 | 0.66 | 0.45 |
| 1 | 225582816 | rs3949347 | 2.057 | 0.35 | -0.616 | -0.329 | 0.007 | 0.101 | 0.101 | 2.449 | 0.61 | 0.17 | 0.71 | 0.74 | 0.62 | 0.4 |
| 1 | 225587463 | rs12073271 | -0.214 | 0.34 | -0.581 | -0.492 | 0.001 | 0.024 | 0.024 | 0 | 0.13 | 0.08 | 0.09 | 0.08 | 0.13 | 0.13 |

| | | | | | | | | | | | | | | | | |
|---|---|---|---|---|---|---|---|---|---|---|---|---|---|---|---|---|
| 1 | 225588533 | rs10915823 | 1.942 | 0.322 | -0.184 | -0.52 | 0.004 | 0.003 | 0.003 | 2.387 | 0.65 | 0.32 | 0.75 | 0.86 | 0.68 | 0.51 |
| 1 | 225589473 | rs7406 | -1.118 | 0.32 | -0.232 | -0.474 | -0.01 | -0.09 | -0.09 | 0 | 0.19 | 0.58 | 0.14 | 0.04 | 0.16 | 0.32 |
| 1 | 225598855 | rs17557867 | -0.254 | 0.294 | -0.525 | -0.513 | -0.03 | 0.03 | 0.03 | 0.07 | 0.1 | 0 | 0.07 | 0.03 | 0.12 | 0.27 |
| 1 | 225602259 | rs3766929 | 1.723 | 0.297 | -0.496 | -0.479 | -0.006 | -0.035 | -0.035 | 1.599 | 0.76 | 0.33 | 0.83 | 0.9 | 0.8 | 0.62 |
| 1 | 225607144 | rs2230419 | 1.151 | 0.332 | -0.423 | -0.463 | -0.011 | -0.07 | -0.07 | 0 | 0.81 | 0.36 | 0.86 | 0.96 | 0.84 | 0.69 |
| 1 | 225609763 | rs12410357 | -0.31 | 0.48 | -0.868 | -0.198 | -0.001 | 0.034 | 0.034 | 0 | 0.12 | 0.06 | 0.08 | 0.05 | 0.12 | 0.11 |
| 1 | 225610751 | rs6686405 | -0.778 | 0.43 | -2.028 | -0.114 | -0.084 | -0.082 | -0.082 | 0 | 0.56 | 0.89 | 0.49 | 0.57 | 0.55 | 0.53 |
| 1 | 225610861 | rs6678087 | -0.401 | 0.245 | -1.853 | -0.177 | 0.007 | -0.029 | -0.029 | 0.337 | 0.44 | 0.74 | 0.41 | 0.53 | 0.42 | 0.45 |
| 1 | 225611166 | rs1417522 | 0.072 | -0.257 | -2.403 | -0.828 | 0.041 | 0.086 | 0.086 | 0 | 0.63 | 0.79 | 0.61 | 0.85 | 0.66 | 0.72 |
| 1 | 225611845 | rs1340867 | 1.044 | 0.552 | -1.442 | -0.938 | 0.007 | -0.042 | -0.042 | 0.022 | 0.68 | 0.6 | 0.72 | 0.75 | 0.6 | 0.37 |
| 1 | 225613736 | rs16844878 | 0.777 | 0.391 | -1.663 | -0.878 | -0.028 | -0.083 | -0.083 | 0 | 0.29 | 0.08 | 0.29 | 0.41 | 0.33 | 0.2 |
| 1 | 225613976 | rs11588279 | 0.331 | 0.692 | -0.908 | -0.799 | 0.011 | -0.01 | -0.01 | 0 | 0.8 | 0.72 | 0.82 | 0.77 | 0.75 | 0.54 |
| 1 | 225618159 | rs2066233 | 1.353 | 0.665 | -0.703 | -0.813 | 0.004 | -0.047 | -0.047 | 0.258 | 0.7 | 0.5 | 0.75 | 0.75 | 0.65 | 0.42 |
| 1 | 225623076 | rs7539651 | -0.814 | 0.69 | -0.468 | -0.736 | 0.021 | -0.046 | -0.046 | 0.806 | 0.89 | 0.74 | 0.91 | 0.95 | 0.88 | 0.9 |
| 1 | 225624016 | rs4653637 | 1.567 | 0.725 | -0.702 | -0.673 | 0 | -0.032 | -0.032 | 0 | 0.7 | 0.55 | 0.65 | 0.7 | 0.64 | 0.41 |
| 1 | 225624635 | rs880434 | 0.204 | 0.773 | -0.798 | -0.649 | -0.022 | 0.032 | 0.032 | 0 | 0.06 | 0 | 0.05 | 0.02 | 0.09 | 0.11 |
| 1 | 225627509 | rs4653638 | -0.316 | 0.697 | -0.648 | -0.564 | 0.003 | 0.001 | 0.001 | 0 | 0.89 | 0.88 | 0.9 | 0.87 | 0.86 | 0.67 |
| 1 | 225631404 | rs1340865 | 0.322 | 1.245 | -0.548 | -0.583 | 0.051 | 0.026 | 0.026 | 0 | 0.87 | 0.74 | 0.87 | 0.82 | 0.83 | 0.64 |
| 1 | 225637606 | rs6426075 | 0.37 | 0.955 | -0.528 | -0.589 | 0.021 | 0.026 | 0.026 | 0 | 0.87 | 0.72 | 0.88 | 0.82 | 0.86 | 0.64 |
| 1 | 225641541 | rs10915829 | 0.76 | 1.002 | -0.6 | -0.597 | -0.001 | 0 | 0 | 0 | 0.11 | 0.09 | 0.17 | 0.1 | 0.13 | 0.11 |
| 1 | 225648044 | rs1340871 | 0.733 | 1.001 | -0.559 | -0.598 | -0.003 | -0.007 | -0.007 | 0.755 | 0.11 | 0.12 | 0.17 | 0.1 | 0.13 | 0.11 |
| 1 | 225653150 | rs10799312 | 0.731 | 0.957 | -0.567 | -0.631 | 0.012 | -0.003 | -0.003 | 0 | 0.84 | 0.76 | 0.86 | 0.8 | 0.82 | 0.57 |
| 1 | 225665284 | rs7515963 | 0.803 | 1.103 | -0.567 | -0.59 | 0.011 | -0.002 | -0.002 | 0 | 0.77 | 0.64 | 0.71 | 0.72 | 0.73 | 0.55 |
| 1 | 225668524 | rs10915840 | 1.944 | 0.964 | -1.12 | -0.609 | -0.082 | -0.003 | -0.003 | 0.004 | 0.31 | 0.01 | 0.29 | 0.16 | 0.28 | 0.11 |
| 1 | 225672596 | rs10799318 | 1.286 | 1.423 | -0.369 | -0.38 | 0.009 | -0.011 | -0.011 | 0 | 0.73 | 0.64 | 0.68 | 0.7 | 0.68 | 0.45 |
| 1 | 225675003 | rs3795443 | 0.096 | 1.465 | -0.471 | -0.391 | -0.004 | 0.009 | 0.009 | 0.019 | 0.13 | 0.08 | 0.19 | 0.12 | 0.16 | 0.21 |
| 1 | 225675459 | rs4653643 | 1.58 | 1.436 | -0.438 | -0.4 | 0.013 | -0.02 | -0.02 | 0 | 0.73 | 0.58 | 0.64 | 0.71 | 0.68 | 0.42 |
| 1 | 225676330 | rs954534 | 0.592 | 1.446 | -0.513 | -0.384 | -0.017 | -0.004 | -0.004 | 0 | 0.08 | 0.04 | 0.13 | 0.08 | 0.12 | 0.25 |
| 1 | 225680980 | rs1771316 | -0.229 | 1.425 | -0.448 | -0.386 | 0.001 | 0.025 | 0.025 | 0 | 0.95 | 0.87 | 0.97 | 0.91 | 0.96 | 0.91 |
| 1 | 225689665 | rs1891000 | 0.132 | 1.385 | -0.431 | -0.373 | 0.003 | 0.003 | 0.003 | 0 | 0.87 | 0.84 | 0.84 | 0.83 | 0.84 | 0.66 |
| 1 | 225693306 | rs16845000 | 0.904 | 1.409 | -0.45 | -0.371 | -0.016 | 0 | 0 | 0 | 0.09 | 0.02 | 0.16 | 0.1 | 0.11 | 0.1 |
| 1 | 225695170 | rs7514094 | 1.412 | 1.597 | -0.487 | -0.357 | 0.001 | 0.005 | 0.005 | 0 | 0.47 | 0.52 | 0.38 | 0.29 | 0.37 | 0.22 |
| 1 | 225699681 | rs2184778 | 0.688 | 1.487 | -0.368 | -0.274 | -0.001 | 0.005 | 0.005 | 0 | 0.1 | 0.07 | 0.16 | 0.1 | 0.12 | 0.12 |
| 1 | 225756293 | rs712062 | 1.03 | 1.393 | -0.34 | -0.114 | 0.011 | -0.021 | -0.021 | 0 | 0.72 | 0.57 | 0.64 | 0.7 | 0.68 | 0.42 |
| 1 | 225756751 | rs487591 | 0.601 | 1.419 | -0.763 | -0.104 | -0.017 | 0.008 | 0.008 | 0.041 | 0.74 | 0.85 | 0.66 | 0.7 | 0.69 | 0.44 |
| 1 | 225776831 | rs2657568 | 0.279 | 1.35 | -1.054 | -0.007 | -0.005 | 0.007 | 0.007 | 0 | 0.05 | 0.09 | 0.03 | 0.09 | 0.04 | 0.09 |
| 1 | 225781165 | rs16845098 | 0.62 | 1.462 | -1.407 | 0.15 | -0.048 | -0.128 | -0.128 | 0 | 0.22 | 0.02 | 0.23 | 0.41 | 0.29 | 0.18 |
| 1 | 225792814 | rs12038373 | 0.898 | 0.847 | -1.19 | 0.285 | -0.001 | 0 | 0 | 0 | 0.09 | 0.06 | 0.16 | 0.12 | 0.11 | 0.12 |
| 1 | 225797957 | rs2769689 | -1.579 | 0.677 | -1.22 | 0.158 | 0.028 | 0.001 | 0.001 | 0 | 0.63 | 0.76 | 0.72 | 0.72 | 0.74 | 0.78 |
| 1 | 225812911 | rs2639703 | -1.129 | 0.629 | -0.656 | 0.112 | 0 | 0.011 | 0.011 | 0.012 | 0.3 | 0.71 | 0.39 | 0.3 | 0.31 | 0.57 |
| 1 | 225824028 | rs2840967 | -0.996 | 0.693 | -0.627 | 0.142 | 0 | 0.009 | 0.009 | 0 | 0.29 | 0.71 | 0.39 | 0.3 | 0.31 | 0.58 |
| 1 | 225836677 | rs17503834 | 0.957 | 1.022 | -1.102 | 0.335 | -0.011 | 0.051 | 0.051 | 0 | 0.13 | 0.01 | 0.08 | 0.02 | 0.14 | 0.05 |
| 1 | 225842940 | rs2639712 | -1.132 | 1.483 | -0.442 | 0.944 | 0.022 | 0.021 | 0.021 | 0 | 0.28 | 0.43 | 0.37 | 0.35 | 0.34 | 0.57 |
| 1 | 225854495 | rs7555023 | -0.723 | 0.936 | -0.873 | 0.974 | -0.018 | -0.016 | -0.016 | 0.397 | 0.06 | 0 | 0.03 | 0.09 | 0.05 | 0.1 |
| 1 | 225859141 | rs2840945 | -0.784 | 0.529 | -0.508 | 1.183 | -0.014 | 0.037 | 0.037 | 0 | 0.28 | 0.33 | 0.29 | 0.42 | 0.2 | 0.26 |
| 1 | 225860552 | rs4653648 | -0.277 | 0.315 | 0.257 | 1.033 | -0.019 | -0.005 | -0.005 | 0 | 0.19 | 0.39 | 0.15 | 0.13 | 0.14 | 0.13 |
| 1 | 225863345 | rs7519220 | -0.944 | 0.737 | 0.064 | 1 | 0.003 | 0.009 | 0.009 | 0 | 0.28 | 0.45 | 0.21 | 0.26 | 0.29 | 0.27 |
| 1 | 225864622 | rs7365361 | -0.928 | 0.737 | 0.064 | 1 | 0.004 | 0.009 | 0.009 | 0 | 0.28 | 0.45 | 0.21 | 0.26 | 0.29 | 0.27 |
| 1 | 225865578 | rs2840956 | -1.788 | 0.803 | -0.07 | 0.65 | 0 | 0.102 | 0.102 | 0 | 0.51 | 0.7 | 0.59 | 0.83 | 0.48 | 0.52 |
| 1 | 225872632 | rs2769685 | -1.394 | 0.827 | 0.068 | 0.783 | 0.013 | 0.016 | 0.016 | 0 | 0.39 | 0.28 | 0.49 | 0.66 | 0.32 | 0.36 |
| 1 | 225873612 | rs2615060 | -1.025 | 0.809 | 0.088 | 0.736 | 0.015 | -0.012 | -0.012 | 1.778 | 0.32 | 0.16 | 0.44 | 0.65 | 0.25 | 0.32 |
| 1 | 225880082 | rs11799692 | 0.388 | 0.747 | 0.06 | 0.649 | -0.004 | -0.029 | -0.029 | 0 | 0.09 | 0.11 | 0.05 | 0.01 | 0.07 | 0.05 |
| 1 | 225880114 | rs1876083 | 0.388 | 0.677 | 0.049 | 0.83 | 0.001 | 0 | 0 | 0 | 0.06 | 0.06 | 0.03 | 0.06 | 0.03 | 0.07 |
| 1 | 225883399 | rs17504298 | 0.305 | 0.617 | 0.262 | 0.859 | -0.019 | -0.003 | -0.003 | 0.092 | 0.19 | 0.34 | 0.14 | 0.12 | 0.14 | 0.12 |
| 1 | 225885157 | rs902959 | -2.314 | 0.687 | 0.116 | 0.774 | -0.006 | 0.086 | 0.086 | 0 | 0.53 | 0.68 | 0.58 | 0.8 | 0.47 | 0.57 |
| 1 | 225886318 | rs943759 | -0.4 | 0.717 | -0.138 | 0.68 | -0.007 | 0.008 | 0.008 | 0.237 | 0.18 | 0.12 | 0.17 | 0.14 | 0.22 | 0.2 |
| 1 | 225889445 | rs2615066 | -2.062 | 0.686 | 0.076 | 0.779 | -0.001 | -0.007 | -0.007 | 0 | 0.35 | 0.4 | 0.44 | 0.75 | 0.33 | 0.41 |
| 1 | 225890232 | rs9943197 | 0.158 | 0.662 | 0.061 | 0.951 | -0.002 | 0.072 | 0.072 | 0 | 0.11 | 0.15 | 0.24 | 0.54 | 0.08 | 0.13 |
| 1 | 225892615 | rs2840964 | 0.281 | 0.667 | 0.063 | 0.926 | 0.001 | -0.001 | -0.001 | 0 | 0.07 | 0.07 | 0.03 | 0.07 | 0.03 | 0.08 |
| 1 | 225893396 | rs9943192 | 0.367 | 0.703 | -0.009 | 0.973 | -0.002 | 0.092 | 0.092 | 0 | 0.1 | 0.14 | 0.24 | 0.53 | 0.08 | 0.13 |
| 1 | 225894423 | rs6701838 | -1.234 | 0.754 | 0.033 | 0.865 | 0.001 | 0.005 | 0.005 | 0.596 | 0.28 | 0.32 | 0.41 | 0.68 | 0.3 | 0.33 |
| 1 | 225895403 | rs7512304 | -1.686 | 0.746 | 0.064 | 0.786 | -0.001 | -0.001 | -0.001 | 0 | 0.34 | 0.35 | 0.44 | 0.75 | 0.33 | 0.41 |
| 1 | 225901006 | rs10915864 | -0.176 | 1.644 | 0.087 | 1.3 | -0.013 | -0.016 | -0.016 | 0.426 | 0.19 | 0.18 | 0.23 | 0.26 | 0.21 | 0.16 |
| 1 | 225902582 | rs12044217 | -0.653 | 1.555 | 0.701 | 1.24 | 0.036 | 0 | 0 | 0 | 0.17 | 0.33 | 0.18 | 0.15 | 0.2 | 0.2 |
| 1 | 225903098 | rs12081218 | -0.116 | 1.555 | 0.604 | 1.179 | -0.017 | -0.016 | -0.016 | 0 | 0.21 | 0.18 | 0.24 | 0.26 | 0.23 | 0.18 |
| 1 | 225909711 | rs7525768 | -0.838 | 1.518 | 0.591 | 1.375 | -0.019 | -0.046 | -0.046 | 0 | 0.26 | 0.21 | 0.26 | 0.38 | 0.25 | 0.23 |
| 1 | 225909815 | rs6656088 | -0.244 | 1.528 | 0.668 | 1.302 | 0.056 | -0.006 | -0.006 | 0.367 | 0.08 | 0.32 | 0.06 | 0 | 0.07 | 0.03 |
| 1 | 225912620 | rs4653659 | 1.629 | 1.55 | 0.519 | 1.317 | -0.036 | 0.031 | 0.031 | 0.027 | 0.09 | 0.01 | 0.07 | 0.02 | 0.13 | 0.03 |
| 1 | 225921655 | rs4653662 | -0.783 | 1.327 | 0.584 | 1.454 | 0.011 | -0.004 | -0.004 | 0 | 0.29 | 0.51 | 0.42 | 0.26 | 0.3 | 0.22 |

| | | | | | | | | | | | | | | | | |
|---|---|---|---|---|---|---|---|---|---|---|---|---|---|---|---|---|
| 1 | 225928538 | rs10799322 | -1.824 | 1.287 | 0.513 | 1.404 | -0.02 | -0.007 | -0.007 | 0 | 0.44 | 0.68 | 0.57 | 0.54 | 0.44 | 0.42 |
| 1 | 225933475 | rs2615074 | -0.241 | 1.474 | 0.021 | 0.739 | -0.037 | -0.009 | -0.009 | 0.275 | 0.24 | 0.18 | 0.29 | 0.3 | 0.35 | 0.54 |
| 1 | 225937228 | rs2615071 | -1.278 | 1.428 | 0.21 | 0.752 | 0.004 | 0.006 | 0.006 | 0.003 | 0.48 | 0.57 | 0.51 | 0.39 | 0.51 | 0.63 |
| 1 | 225937585 | rs2615069 | 0.731 | 1.535 | -0.043 | 0.763 | -0.007 | 0.003 | 0.003 | 0.085 | 0.16 | 0.18 | 0.17 | 0.15 | 0.22 | 0.39 |
| 1 | 225941944 | rs12029847 | 0.991 | 1.517 | -0.232 | 0.746 | -0.037 | -0.039 | -0.039 | 0.039 | 0.08 | 0.01 | 0.11 | 0.14 | 0.12 | 0.16 |
| 1 | 225944165 | rs11588772 | 0.33 | 1 | 0.376 | 0.586 | 0.01 | -0.008 | -0.008 | 0 | 0.21 | 0.22 | 0.28 | 0.12 | 0.21 | 0.18 |
| 1 | 225944658 | rs2126345 | -0.968 | 0.065 | 0.691 | 0.586 | 0.025 | -0.002 | -0.002 | 1.198 | 0.58 | 0.68 | 0.65 | 0.54 | 0.65 | 0.83 |
| 1 | 225946690 | rs16845250 | -0.327 | 0.203 | 0.178 | 0.589 | -0.01 | 0.027 | 0.027 | 0 | 0.06 | 0.2 | 0.03 | 0.08 | 0.02 | 0.05 |
| 1 | 225949423 | rs7538185 | 0.036 | 0.386 | -0.202 | 0.54 | -0.001 | -0.028 | -0.028 | 0 | 0.06 | 0.09 | 0.04 | 0 | 0.07 | 0.03 |
| 1 | 225955919 | rs1557092 | -0.892 | 0.214 | -0.273 | 0.612 | 0.015 | -0.004 | -0.004 | 0 | 0.6 | 0.68 | 0.67 | 0.54 | 0.69 | 0.84 |
| 1 | 225958857 | rs6685014 | -0.303 | 0.405 | -0.524 | 0.686 | 0.018 | -0.012 | -0.012 | 0 | 0.46 | 0.36 | 0.59 | 0.47 | 0.57 | 0.75 |
| 1 | 225965803 | rs3738037 | -1.02 | 0.417 | -0.346 | 0.304 | 0.011 | -0.008 | -0.008 | 0 | 0.12 | 0.58 | 0.16 | 0.08 | 0.11 | 0.06 |
| 1 | 225967597 | rs6677132 | -0.97 | 0.344 | -1.167 | 0.133 | 0.001 | 0.008 | 0.008 | 0 | 0.36 | 0.66 | 0.37 | 0.2 | 0.34 | 0.25 |
| 1 | 225969031 | rs17571004 | 0.054 | 0.41 | -1.622 | 0.165 | -0.002 | -0.002 | -0.002 | 0 | 0.24 | 0.24 | 0.38 | 0.6 | 0.37 | 0.56 |
| 1 | 225971433 | rs12042925 | 1.274 | 0.972 | -1.785 | 0.665 | -0.063 | -0.062 | -0.062 | 0 | 0.1 | 0.01 | 0.21 | 0.33 | 0.2 | 0.3 |
| 1 | 225973777 | rs12568108 | 1.845 | 1.041 | -1.799 | 0.76 | -0.034 | -0.029 | -0.029 | 0 | 0.05 | 0.01 | 0.11 | 0.18 | 0.13 | 0.14 |
| 1 | 225974531 | rs4653433 | -1.163 | -0.028 | -2.156 | 0.058 | 0.016 | 0.07 | 0.07 | 0.456 | 0.52 | 0.39 | 0.65 | 0.8 | 0.58 | 0.77 |
| 1 | 225975271 | rs12031296 | 0.693 | 0.084 | -2.152 | 0.142 | 0.011 | 0.027 | 0.027 | 0.883 | 0.16 | 0.23 | 0.24 | 0.33 | 0.21 | 0.37 |
| 1 | 225978571 | rs12732891 | 1.034 | -0.393 | -2.293 | -0.072 | 0.086 | 0.01 | 0.01 | 0 | 0.22 | 0.01 | 0.15 | 0.2 | 0.09 | 0.19 |
| 1 | 225994515 | rs10799324 | -0.868 | -0.064 | -1.593 | -0.135 | 0 | -0.012 | -0.012 | 0 | 0.57 | 0.6 | 0.67 | 0.73 | 0.58 | 0.78 |
| 1 | 225995211 | rs4653436 | 0.616 | -0.044 | -2.444 | -0.159 | -0.045 | -0.021 | -0.021 | 0 | 0.28 | 0.13 | 0.26 | 0.23 | 0.33 | 0.21 |
| 1 | 225997070 | rs10737420 | -0.821 | -0.11 | -2.331 | -0.106 | -0.025 | 0.111 | 0.111 | 0.639 | 0.53 | 0.77 | 0.63 | 0.77 | 0.47 | 0.77 |
| 1 | 226004162 | rs17571067 | -0.097 | 0.133 | -2.242 | 0.107 | 0.003 | -0.002 | -0.002 | 0 | 0.32 | 0.43 | 0.34 | 0.31 | 0.4 | 0.36 |
| 1 | 226008101 | rs10753410 | 0.667 | -0.048 | -2.796 | 0.374 | 0.054 | -0.023 | -0.023 | 0.331 | 0.23 | 0.1 | 0.16 | 0.3 | 0.13 | 0.22 |
| 1 | 226012614 | rs2854451 | 0.825 | 0.11 | -3.074 | 0.458 | -0.047 | -0.089 | -0.089 | 0.238 | 0.2 | 0.04 | 0.25 | 0.33 | 0.25 | 0.38 |
| 1 | 226012686 | rs3753658 | 1.265 | 0.162 | -3.168 | 0.453 | -0.042 | -0.053 | -0.053 | 0.575 | 0.09 | 0.01 | 0.14 | 0.18 | 0.14 | 0.2 |
| 1 | 226012717 | rs3753659 | 1.265 | 0.162 | -3.168 | 0.453 | -0.042 | -0.053 | -0.053 | 0 | 0.09 | 0.01 | 0.14 | 0.18 | 0.14 | 0.2 |
| 1 | 226013041 | rs3738040 | 1.399 | 0.107 | -3.155 | 0.496 | 0.006 | -0.062 | -0.062 | 0 | 0.13 | 0 | 0.11 | 0.25 | 0.06 | 0.14 |
| 1 | 226013355 | rs1877724 | 0.874 | 0.234 | -3.097 | 0.536 | -0.047 | -0.089 | -0.089 | 0.072 | 0.19 | 0.04 | 0.25 | 0.33 | 0.25 | 0.38 |
| 1 | 226013388 | rs3738042 | 1.03 | 0.196 | -3.112 | 0.285 | -0.045 | 0.017 | 0.017 | 0 | 0.25 | 0.11 | 0.25 | 0.17 | 0.33 | 0.19 |
| 1 | 226014342 | rs3753661 | 0.866 | 0.008 | -2.827 | 0.347 | 0.019 | -0.055 | -0.055 | 0 | 0.17 | 0 | 0.11 | 0.24 | 0.08 | 0.19 |
| 1 | 226015017 | rs3766934 | 0.782 | 0.034 | -2.697 | 0.36 | 0.021 | -0.009 | -0.009 | 0 | 0.18 | 0.1 | 0.13 | 0.24 | 0.08 | 0.18 |
| 1 | 226015116 | rs2671272 | -1.703 | 0.023 | -2.627 | 0.188 | -0.003 | -0.027 | -0.027 | 0 | 0.74 | 0.39 | 0.78 | 0.92 | 0.79 | 0.96 |
| 1 | 226015299 | rs3738043 | 0.331 | 0.096 | -2.433 | 0.802 | 0.023 | -0.008 | -0.008 | 0 | 0.18 | 0.1 | 0.13 | 0.24 | 0.08 | 0.18 |
| 1 | 226015359 | rs2854456 | -0.183 | 0.972 | -2.165 | 1.062 | -0.047 | -0.089 | -0.089 | 0.675 | 0.19 | 0.04 | 0.25 | 0.33 | 0.25 | 0.38 |
| 1 | 226018027 | rs4149222 | 0.156 | 1.262 | -2.065 | 1.087 | -0.047 | -0.053 | -0.053 | 0 | 0.09 | 0.01 | 0.14 | 0.18 | 0.14 | 0.2 |
| 1 | 226018127 | rs2671270 | -0.178 | 1.022 | -2.19 | 1.053 | -0.032 | -0.089 | -0.089 | 0 | 0.19 | 0.03 | 0.25 | 0.34 | 0.25 | 0.38 |
| 1 | 226020988 | rs2740168 | -0.651 | 1.234 | -1.922 | 0.798 | -0.005 | -0.018 | -0.018 | 0 | 0.36 | 0.61 | 0.36 | 0.52 | 0.37 | 0.44 |
| 1 | 226024797 | rs2740170 | -1.53 | 0.786 | -2.601 | -0.06 | 0.001 | 0.082 | 0.082 | 3.445 | 0.75 | 0.83 | 0.86 | 0.96 | 0.78 | 0.86 |
| 1 | 226025528 | rs2740171 | -1.247 | 0.579 | -1.375 | 0.114 | -0.011 | -0.045 | -0.045 | 0 | 0.72 | 0.62 | 0.83 | 0.87 | 0.78 | 0.84 |
| 1 | 226025690 | rs2671267 | 0.239 | 0.571 | -1.034 | 0.272 | 0 | -0.001 | -0.001 | 9.389 | 0.13 | 0.13 | 0.07 | 0.12 | 0.1 | 0.23 |
| 1 | 226026406 | rs2234922 | -0.859 | 0.54 | -0.463 | 0.108 | 0.01 | -0.051 | -0.051 | 0 | 0.2 | 0.35 | 0.14 | 0.12 | 0.16 | 0.23 |
| 1 | 226027323 | rs4149226 | -0.829 | 0.803 | -1.054 | 0.379 | -0.018 | -0.019 | -0.019 | 0 | 0.42 | 0.72 | 0.51 | 0.57 | 0.53 | 0.46 |
| 1 | 226032229 | rs1051741 | 0.762 | 0.668 | -0.924 | 0.713 | 0 | -0.001 | -0.001 | 0 | 0.11 | 0.11 | 0.06 | 0.11 | 0.09 | 0.25 |
| 1 | 226032677 | rs4149228 | 1.317 | 0.636 | -0.845 | 0.834 | 0.005 | -0.006 | -0.006 | 0.689 | 0.06 | 0.04 | 0.08 | 0.16 | 0.05 | 0.09 |
| 1 | 226033969 | rs2740174 | 0.546 | 0.676 | -0.786 | 0.745 | 0 | 0 | 0 | 0 | 0.13 | 0.11 | 0.06 | 0.13 | 0.1 | 0.27 |
| 1 | 226044766 | rs9919303 | -0.326 | 0.45 | -0.698 | 0.569 | 0.005 | 0.069 | 0.069 | 0 | 0.29 | 0.55 | 0.31 | 0.52 | 0.26 | 0.47 |
| 1 | 226049648 | rs360080 | -1.559 | 0.484 | -0.72 | 0.434 | 0 | -0.014 | -0.014 | 0 | 0.95 | 0.93 | 0.96 | 1 | 0.97 | 0.99 |
| 1 | 226050609 | rs10158985 | 1.13 | 0.603 | -1.14 | 0.293 | -0.042 | 0.051 | 0.051 | 0 | 0.13 | 0.06 | 0.13 | 0.01 | 0.22 | 0.14 |
| 1 | 226055595 | rs2292564 | -0.297 | 0.503 | -0.838 | 0.529 | 0.008 | 0.082 | 0.082 | 0 | 0.28 | 0.52 | 0.31 | 0.52 | 0.26 | 0.47 |
| 1 | 226061658 | rs3820236 | 0.139 | 0.411 | -1.116 | 0.574 | -0.004 | -0.008 | -0.008 | 0.061 | 0.24 | 0.22 | 0.29 | 0.41 | 0.26 | 0.47 |
| 1 | 226065538 | rs360096 | 0.672 | 0.423 | -1.859 | 0.62 | 0.01 | -0.004 | -0.004 | 0.749 | 0.6 | 0.23 | 0.55 | 0.33 | 0.66 | 0.44 |
| 1 | 226065755 | rs360097 | 0.312 | 0.426 | -1.849 | 0.621 | 0.003 | -0.035 | -0.035 | 0 | 0.64 | 0.25 | 0.63 | 0.48 | 0.71 | 0.51 |
| 1 | 226067862 | rs360102 | 0.601 | 0.466 | -1.933 | 0.482 | 0.008 | 0 | 0 | 0 | 0.6 | 0.25 | 0.55 | 0.32 | 0.66 | 0.44 |
| 1 | 226071754 | rs3753668 | 0.445 | 0.533 | -1.795 | 0.544 | -0.022 | 0.076 | 0.076 | 0 | 0.08 | 0.28 | 0.1 | 0.24 | 0.05 | 0.09 |
| 1 | 226074373 | rs3766941 | 0.273 | 0.526 | -1.798 | 0.541 | -0.024 | 0.068 | 0.068 | 0 | 0.09 | 0.28 | 0.1 | 0.24 | 0.05 | 0.09 |
| 1 | 226077132 | rs360059 | 1.213 | 0.231 | -2.22 | 0.156 | -0.024 | 0.087 | 0.087 | 0.983 | 0.4 | 0.14 | 0.34 | 0.2 | 0.39 | 0.23 |
| 1 | 226077231 | rs1877726 | -0.304 | 0.228 | -2.259 | 0.207 | 0.005 | -0.018 | -0.018 | 0 | 0.28 | 0.34 | 0.3 | 0.35 | 0.32 | 0.29 |
| 1 | 226078915 | rs4653437 | -0.174 | 0.279 | -2.234 | 0.232 | 0.002 | -0.014 | -0.014 | 0.437 | 0.27 | 0.34 | 0.3 | 0.35 | 0.32 | 0.29 |
| 1 | 226082425 | rs1223245 | 1.065 | 0.372 | -2.221 | 0.181 | -0.02 | 0.094 | 0.094 | 0.978 | 0.4 | 0.14 | 0.34 | 0.2 | 0.39 | 0.23 |
| 1 | 226091206 | rs360073 | -0.526 | 0.291 | -1.727 | 0.439 | 0.007 | 0.025 | 0.025 | 0 | 0.75 | 0.51 | 0.71 | 0.68 | 0.77 | 0.71 |
| 1 | 226091735 | rs360071 | -0.052 | 0.241 | -1.535 | 0.558 | 0 | 0.001 | 0.001 | 0 | 0.58 | 0.51 | 0.58 | 0.56 | 0.65 | 0.43 |
| 1 | 226095048 | rs10915896 | 0.109 | 0.59 | -1.979 | 0.559 | -0.025 | 0.007 | 0.007 | 0 | 0.33 | 0.2 | 0.39 | 0.25 | 0.41 | 0.32 |
| 1 | 226107787 | rs14868 | 0.5 | 0.252 | -1.845 | 0.801 | -0.034 | -0.028 | -0.028 | 0 | 0.11 | 0.02 | 0.12 | 0.09 | 0.15 | 0.18 |
| 1 | 226123114 | rs12046656 | 0.085 | 0.077 | -1.717 | 0.684 | 0.004 | 0.015 | 0.015 | 0 | 0.18 | 0.07 | 0.12 | 0.17 | 0.15 | 0.23 |
| 1 | 226125617 | rs3007716 | 0.902 | -0.048 | -1.818 | 0.26 | -0.001 | 0.148 | 0.148 | 2.394 | 0.29 | 0.07 | 0.13 | 0.07 | 0.22 | 0.2 |
| 1 | 226131691 | rs2493163 | -1.123 | -0.053 | -0.307 | 1.508 | -0.034 | 0.062 | 0.062 | 0 | 0.27 | 0.39 | 0.33 | 0.45 | 0.19 | 0.31 |
| 1 | 226135132 | rs2038393 | -1.612 | 0.295 | 0.338 | 1.789 | 0.071 | 0.276 | 0.276 | 3.325 | 0.1 | 0.49 | 0.27 | 0.27 | 0.08 | 0.11 |
| 1 | 226144247 | rs2816336 | -1.313 | -0.257 | -0.012 | 1.038 | -0.036 | 0.118 | 0.118 | 1.04 | 0.12 | 0.46 | 0.08 | 0.22 | 0.04 | 0.12 |

| | | | | | | | | | | | | | | | | |
|---|---|---|---|---|---|---|---|---|---|---|---|---|---|---|---|---|
| 1 | 226149325 | rs12031056 | -0.381 | 0.04 | -1.033 | 0.69 | -0.026 | 0.002 | 0.002 | 0 | 0.15 | 0.01 | 0.14 | 0.08 | 0.15 | 0.1 |
| 1 | 226152996 | rs2749696 | -1.253 | -0.437 | 0.044 | 0.898 | -0.027 | 0.104 | 0.104 | 0.004 | 0.1 | 0.39 | 0.06 | 0.2 | 0.02 | 0.1 |
| 1 | 226156109 | rs12743653 | -0.462 | 0.457 | -0.86 | 1.099 | -0.136 | -0.123 | -0.123 | 0 | 0.32 | 0.07 | 0.4 | 0.51 | 0.45 | 0.41 |
| 1 | 226160511 | rs12728713 | -0.46 | 0.439 | -0.873 | 1.096 | -0.137 | -0.122 | -0.122 | 0 | 0.32 | 0.06 | 0.39 | 0.51 | 0.45 | 0.42 |
| 1 | 226160882 | rs2180421 | -0.709 | -0.96 | -1.224 | 0.141 | -0.004 | 0.048 | 0.048 | 0.445 | 0.59 | 0.77 | 0.58 | 0.74 | 0.59 | 0.66 |
| 1 | 226165299 | rs2755094 | -1.335 | -1.689 | -0.62 | -0.231 | -0.004 | -0.021 | -0.021 | 0.525 | 0.79 | 0.79 | 0.9 | 0.96 | 0.89 | 0.87 |
| 1 | 226166027 | rs2816330 | 0.504 | -1.726 | -0.097 | -0.185 | -0.002 | 0.005 | 0.005 | 0 | 0.4 | 0.39 | 0.55 | 0.32 | 0.42 | 0.36 |
| 1 | 226166555 | rs10915916 | 0.725 | -1.753 | -0.033 | -0.215 | -0.023 | 0.042 | 0.042 | 0 | 0.38 | 0.2 | 0.53 | 0.26 | 0.42 | 0.33 |
| 1 | 226169966 | rs2816326 | -0.493 | -1.824 | 0.357 | -0.214 | -0.072 | -0.055 | -0.055 | 0.91 | 0.73 | 0.57 | 0.87 | 0.84 | 0.87 | 0.78 |
| 1 | 226182455 | rs4653701 | 0.794 | -1.761 | 0.164 | -0.22 | -0.027 | 0.039 | 0.039 | 0 | 0.37 | 0.18 | 0.53 | 0.26 | 0.42 | 0.33 |
| 1 | 226184438 | rs1998273 | -1.28 | -1.801 | 0.31 | -0.261 | -0.019 | -0.044 | -0.044 | 0 | 0.79 | 0.74 | 0.89 | 0.96 | 0.89 | 0.87 |
| 1 | 226187899 | rs2755093 | -0.773 | -1.563 | 0.37 | -0.251 | -0.027 | 0.043 | 0.043 | 0.588 | 0.61 | 0.8 | 0.46 | 0.73 | 0.56 | 0.66 |
| 1 | 226195946 | rs11808474 | 1.012 | -1.586 | 0.687 | -0.301 | -0.028 | -0.035 | -0.035 | 0 | 0.21 | 0.29 | 0.11 | 0.11 | 0.12 | 0.15 |
| 1 | 226196624 | rs11586667 | 1.012 | -1.586 | 0.687 | -0.301 | -0.028 | -0.035 | -0.035 | 0.548 | 0.21 | 0.29 | 0.11 | 0.11 | 0.12 | 0.15 |
| 1 | 226199835 | rs2755095 | -0.001 | -1.186 | 0.791 | -0.348 | -0.046 | -0.053 | -0.053 | 0.237 | 0.28 | 0.58 | 0.17 | 0.19 | 0.16 | 0.25 |
| 1 | 226202334 | rs11580276 | 0.333 | -1.176 | 0.61 | -0.329 | -0.017 | -0.012 | -0.012 | 0 | 0.26 | 0.34 | 0.14 | 0.23 | 0.16 | 0.25 |
| 1 | 226203053 | rs11590226 | 1.04 | -1.177 | 0.609 | -0.333 | -0.028 | -0.035 | -0.035 | 2.66 | 0.21 | 0.29 | 0.11 | 0.11 | 0.12 | 0.15 |
| 1 | 226204746 | rs7554619 | 0.843 | -1.176 | 0.593 | -0.333 | -0.041 | -0.05 | -0.05 | 0.955 | 0.22 | 0.35 | 0.11 | 0.11 | 0.12 | 0.15 |
| 1 | 226214882 | rs6682659 | -0.609 | -0.799 | 0.566 | -0.081 | 0.029 | 0.027 | 0.027 | 0 | 0.65 | 0.59 | 0.68 | 0.49 | 0.57 | 0.61 |
| 1 | 226222657 | rs4653704 | -0.058 | -1.064 | 0.586 | -0.185 | 0.006 | -0.003 | -0.003 | 0 | 0.59 | 0.52 | 0.67 | 0.38 | 0.56 | 0.5 |
| 1 | 226228627 | rs12754943 | 0.354 | -1.084 | 0.128 | -0.612 | 0.038 | 0.041 | 0.041 | 0.207 | 0.39 | 0.3 | 0.36 | 0.24 | 0.34 | 0.35 |
| 1 | 226234883 | rs10915924 | -1.099 | -1.006 | 0.111 | -0.498 | 0.009 | 0.008 | 0.008 | 0 | 0.66 | 0.69 | 0.69 | 0.49 | 0.57 | 0.61 |
| 1 | 226239830 | rs6701318 | -1.099 | -1.006 | 0.111 | -0.506 | 0.009 | 0.008 | 0.008 | 0 | 0.66 | 0.69 | 0.69 | 0.49 | 0.57 | 0.61 |
| 1 | 226246771 | rs10753411 | 1.035 | -1.02 | 0.131 | -0.513 | -0.013 | -0.013 | -0.013 | 0 | 0.33 | 0.24 | 0.3 | 0.51 | 0.43 | 0.39 |
| 1 | 226257722 | rs4653440 | 2.012 | -1.155 | -0.027 | -0.502 | -0.007 | -0.066 | -0.066 | 1.802 | 0.11 | 0.01 | 0.24 | 0.18 | 0.14 | 0.04 |
| 1 | 226259807 | rs6700189 | 0.58 | -0.455 | -0.025 | -0.09 | 0.024 | 0.019 | 0.019 | 0 | 0.26 | 0.48 | 0.27 | 0.27 | 0.28 | 0.19 |
| 1 | 226266029 | rs4271181 | -0.211 | -0.464 | -0.295 | -0.083 | 0.002 | 0.008 | 0.008 | 0 | 0.7 | 0.42 | 0.71 | 0.64 | 0.71 | 0.72 |
| 1 | 226266440 | rs10915928 | 0.348 | -0.434 | -0.551 | -0.186 | 0.004 | 0.001 | 0.001 | 0 | 0.53 | 0.32 | 0.47 | 0.55 | 0.5 | 0.64 |
| 1 | 226276554 | rs12096737 | 1.208 | -0.283 | -0.368 | -0.053 | 0.019 | 0.014 | 0.014 | 0 | 0.21 | 0.24 | 0.24 | 0.26 | 0.28 | 0.16 |
| 1 | 226279050 | rs10915933 | 0.564 | -0.308 | -0.99 | -0.239 | -0.017 | -0.067 | -0.067 | 0 | 0.32 | 0.19 | 0.37 | 0.44 | 0.37 | 0.37 |
| 1 | 226285151 | rs1355391 | -0.262 | -0.366 | -1.032 | -0.288 | 0.002 | -0.029 | -0.029 | 0 | 0.63 | 0.34 | 0.54 | 0.49 | 0.66 | 0.62 |
| 1 | 226292385 | rs10915935 | -1.089 | -0.467 | -1.182 | -0.583 | 0.018 | 0.009 | 0.009 | 0 | 0.74 | 0.77 | 0.87 | 0.94 | 0.84 | 0.76 |
| 1 | 226292585 | rs11579988 | 0.146 | -0.413 | -1.355 | -0.67 | 0.015 | 0.099 | 0.099 | 0 | 0.32 | 0.15 | 0.16 | 0.12 | 0.25 | 0.31 |
| 1 | 226300488 | rs7515023 | 0.82 | -0.227 | -1.112 | -0.326 | -0.014 | -0.017 | -0.017 | 0 | 0.28 | 0.21 | 0.39 | 0.36 | 0.33 | 0.19 |
| 1 | 226313525 | rs1877155 | 0.122 | 0.057 | -1.136 | -0.266 | 0.008 | -0.014 | -0.014 | 0 | 0.34 | 0.29 | 0.38 | 0.54 | 0.33 | 0.38 |
| 1 | 226317841 | rs3887559 | 0.531 | 0.004 | -1.162 | 0.129 | 0.002 | -0.007 | -0.007 | 3.724 | 0.31 | 0.28 | 0.37 | 0.36 | 0.33 | 0.29 |
| 1 | 226319520 | rs1219671 | 0.08 | 0.003 | -1.108 | -0.044 | -0.001 | -0.001 | -0.001 | 1.059 | 0.57 | 0.58 | 0.55 | 0.47 | 0.55 | 0.59 |
| 1 | 226327068 | rs2666859 | 0.791 | 0.296 | -1.58 | -0.261 | 0.009 | 0.097 | 0.097 | 0 | 0.22 | 0.06 | 0.14 | 0.1 | 0.18 | 0.34 |
| 1 | 226332993 | rs1043641 | 0.992 | 0.335 | -1.582 | -0.278 | -0.006 | 0.044 | 0.044 | 0 | 0.2 | 0.08 | 0.22 | 0.08 | 0.19 | 0.08 |
| 1 | 226348355 | rs16860230 | 1.387 | 0.782 | -1.689 | 0.01 | -0.022 | -0.021 | -0.021 | 0.118 | 0.14 | 0.08 | 0.17 | 0.21 | 0.19 | 0.2 |
| 1 | 226350367 | rs1219663 | 1.074 | 0.713 | -1.571 | -0.031 | 0.013 | 0.026 | 0.026 | 0 | 0.24 | 0.17 | 0.16 | 0.15 | 0.19 | 0.4 |
| 1 | 226350383 | rs4653719 | -0.282 | 0.716 | -1.573 | -0.03 | -0.012 | 0.01 | 0.01 | 0 | 0.62 | 0.75 | 0.67 | 0.63 | 0.62 | 0.41 |
| 1 | 226352498 | rs2306120 | 0.326 | 0.572 | -1.435 | 0.168 | 0.002 | -0.009 | -0.009 | 0.753 | 0.57 | 0.62 | 0.64 | 0.45 | 0.61 | 0.34 |
| 1 | 226355373 | rs12402078 | 1.539 | 0.573 | -1.457 | 0.141 | -0.006 | 0.044 | 0.044 | 0 | 0.2 | 0.08 | 0.22 | 0.08 | 0.19 | 0.08 |
| 1 | 226360795 | rs9286998 | 0.824 | 0.518 | -1.379 | 0.178 | 0.002 | 0.009 | 0.009 | 0 | 0.37 | 0.53 | 0.42 | 0.37 | 0.42 | 0.26 |
| 1 | 226361285 | rs10915954 | 0.332 | 0.651 | -1.39 | 0.126 | 0.006 | 0.002 | 0.002 | 0 | 0.42 | 0.67 | 0.44 | 0.56 | 0.43 | 0.33 |
| 1 | 226363155 | rs10915955 | 1.332 | 0.773 | -1.337 | -0.019 | -0.034 | -0.031 | -0.031 | 0.258 | 0.13 | 0.05 | 0.17 | 0.21 | 0.19 | 0.19 |
| 1 | 226382922 | rs11584966 | 2.167 | 0.41 | -1.42 | -0.334 | 0.035 | 0.078 | 0.078 | 0 | 0.15 | 0.03 | 0.11 | 0.01 | 0.11 | 0.19 |
| 1 | 226385623 | rs10753415 | -0.006 | 0.539 | -1.349 | -0.29 | 0.018 | 0.032 | 0.032 | 0 | 0.52 | 0.6 | 0.59 | 0.65 | 0.61 | 0.44 |
| 1 | 226387618 | rs10495244 | 1.019 | 0.486 | -1.234 | -0.35 | -0.013 | -0.013 | -0.013 | 0 | 0.15 | 0.09 | 0.18 | 0.21 | 0.18 | 0.19 |
| 1 | 226389655 | rs10915958 | -0.302 | 0.484 | -1.19 | -0.369 | 0 | 0.001 | 0.001 | 0 | 0.64 | 0.66 | 0.66 | 0.73 | 0.68 | 0.57 |
| 1 | 226400759 | rs2927340 | 1.174 | 0.4 | -1.228 | -0.432 | 0.008 | 0.002 | 0.002 | 0 | 0.2 | 0.17 | 0.13 | 0.04 | 0.12 | 0.21 |
| 1 | 226401914 | rs11811952 | 0.995 | 0.672 | -1.248 | -0.275 | -0.009 | 0.023 | 0.023 | 0 | 0.25 | 0.09 | 0.26 | 0.21 | 0.24 | 0.28 |
| 1 | 226403979 | rs11810473 | 1.795 | 0.737 | -1.391 | 0.011 | -0.031 | -0.083 | -0.083 | 0 | 0.16 | 0.01 | 0.27 | 0.19 | 0.22 | 0.07 |
| 1 | 226404457 | rs3008192 | -0.872 | 0.805 | -0.75 | -0.578 | -0.003 | 0.02 | 0.02 | 0 | 0.62 | 0.52 | 0.62 | 0.51 | 0.61 | 0.73 |
| 1 | 226432521 | rs2889537 | -0.382 | 0.748 | -0.794 | -0.77 | 0.001 | 0.014 | 0.014 | 0 | 0.65 | 0.75 | 0.66 | 0.74 | 0.68 | 0.65 |
| 1 | 226440023 | rs7521958 | 1.18 | 0.814 | -0.817 | -0.91 | 0.001 | 0.01 | 0.01 | 0 | 0.24 | 0.17 | 0.25 | 0.2 | 0.23 | 0.21 |
| 1 | 226445528 | rs7367845 | 0.955 | 0.517 | -0.603 | -0.787 | 0 | 0.01 | 0.01 | 0 | 0.35 | 0.39 | 0.38 | 0.5 | 0.39 | 0.25 |
| 1 | 226473296 | rs10753418 | 0.747 | 0.495 | -0.591 | -0.779 | -0.001 | 0.023 | 0.023 | 0 | 0.37 | 0.47 | 0.38 | 0.5 | 0.39 | 0.27 |
| 1 | 226474132 | rs10799347 | -0.491 | 0.552 | -0.627 | -0.773 | -0.001 | 0.013 | 0.013 | 0 | 0.67 | 0.75 | 0.67 | 0.75 | 0.68 | 0.66 |
| 1 | 226474479 | rs10127776 | -0.258 | 0.705 | -0.637 | -0.712 | -0.007 | -0.013 | -0.013 | 0.237 | 0.63 | 0.57 | 0.65 | 0.71 | 0.68 | 0.58 |
| 1 | 226491656 | rs11802954 | 0.709 | 0.688 | -0.588 | -0.591 | -0.011 | 0.015 | 0.015 | 0 | 0.26 | 0.16 | 0.27 | 0.21 | 0.29 | 0.31 |
| 1 | 226498612 | rs2615762 | 0.98 | 0.463 | -0.43 | -0.286 | -0.014 | -0.003 | -0.003 | 0 | 0.06 | 0.15 | 0.04 | 0.04 | 0.02 | 0.04 |
| 1 | 226508645 | rs10915971 | 0.728 | 0.961 | -0.644 | -0.197 | -0.018 | -0.08 | -0.08 | 0 | 0.28 | 0.14 | 0.32 | 0.49 | 0.32 | 0.26 |
| 1 | 226521192 | rs10915975 | -0.478 | 0.725 | -1.142 | -0.904 | -0.008 | 0.033 | 0.033 | 0.053 | 0.54 | 0.43 | 0.48 | 0.26 | 0.5 | 0.55 |
| 1 | 226521979 | rs10915976 | -0.458 | 0.634 | -1.103 | -0.903 | -0.001 | -0.002 | -0.002 | 0 | 0.56 | 0.48 | 0.49 | 0.29 | 0.49 | 0.6 |
| 1 | 226523560 | rs11800726 | -0.337 | 0.838 | -1.014 | -0.313 | 0.001 | 0.006 | 0.006 | 0 | 0.34 | 0.32 | 0.33 | 0.29 | 0.3 | 0.39 |
| 1 | 226536645 | rs874583 | -0.481 | 0.982 | -1.244 | -0.863 | 0.019 | 0.13 | 0.13 | 0 | 0.54 | 0.7 | 0.65 | 0.9 | 0.57 | 0.67 |
| 1 | 226542049 | rs6426551 | -0.434 | 1.485 | -1.23 | 0.006 | -0.002 | 0.003 | 0.003 | 0 | 0.78 | 0.78 | 0.58 | 0.52 | 0.74 | 0.83 |

| | | | | | | | | | | | | | | | | |
|---|---|---|---|---|---|---|---|---|---|---|---|---|---|---|---|---|
| 1 | 226543909 | rs6675327 | 0.461 | 1.591 | -0.531 | -0.058 | 0.041 | 0.11 | 0.11 | 0 | 0.17 | 0.42 | 0.15 | 0.31 | 0.17 | 0.18 |
| 1 | 226544420 | rs12567614 | 0.621 | 1.491 | -0.475 | -0.097 | -0.003 | -0.006 | -0.006 | 1.294 | 0.6 | 0.36 | 0.44 | 0.15 | 0.57 | 0.56 |
| 1 | 226548554 | rs8679 | 1.571 | 1.04 | -1.146 | -0.56 | 0.025 | 0.174 | 0.174 | 0 | 0.31 | 0.03 | 0.19 | 0.07 | 0.23 | 0.25 |
| 1 | 226549498 | rs2271347 | 1.571 | 1.038 | -1.154 | -0.561 | 0.025 | 0.174 | 0.174 | 0 | 0.31 | 0.03 | 0.19 | 0.07 | 0.23 | 0.25 |
| 1 | 226550191 | rs6664761 | 0.447 | 0.969 | -0.535 | 0.003 | 0.028 | 0.081 | 0.081 | 0 | 0.17 | 0.33 | 0.14 | 0.31 | 0.17 | 0.18 |
| 1 | 226555302 | rs1136410 | 1.313 | 1.126 | -0.773 | 0.176 | -0.022 | -0.077 | -0.077 | 0 | 0.13 | 0.04 | 0.35 | 0.44 | 0.15 | 0.1 |
| 1 | 226562621 | rs3219104 | -1.234 | 1.108 | -0.715 | 0.168 | -0.001 | 0.001 | 0.001 | 0 | 0.86 | 0.89 | 0.64 | 0.56 | 0.85 | 0.9 |
| 1 | 226564691 | rs3219090 | 0.477 | 1.107 | -0.688 | 0.134 | 0.005 | -0.033 | -0.033 | 0 | 0.67 | 0.44 | 0.5 | 0.21 | 0.68 | 0.63 |
| 1 | 226566726 | rs907190 | -0.889 | 1.111 | -0.699 | 0.155 | 0 | 0.026 | 0.026 | 0 | 0.84 | 0.83 | 0.64 | 0.52 | 0.85 | 0.81 |
| 1 | 226568665 | rs1805410 | 0.894 | 1.342 | -0.676 | 0.112 | 0.001 | 0.021 | 0.021 | 0.27 | 0.15 | 0.08 | 0.09 | 0.03 | 0.15 | 0.25 |
| 1 | 226576296 | rs2293464 | 0.379 | 1.419 | -0.616 | 0.307 | 0.036 | 0.098 | 0.098 | 0 | 0.17 | 0.39 | 0.15 | 0.31 | 0.17 | 0.18 |
| 1 | 226578047 | rs1805411 | 0.768 | 1.135 | -1.203 | 0.404 | -0.003 | -0.014 | -0.014 | 0 | 0.13 | 0.09 | 0.35 | 0.44 | 0.15 | 0.1 |
| 1 | 226587442 | rs2793658 | 0.52 | 1.14 | -1.207 | 0.406 | -0.001 | 0.001 | 0.001 | 0 | 0.14 | 0.11 | 0.36 | 0.44 | 0.15 | 0.1 |
| 1 | 226610638 | rs10915989 | -0.167 | 1.219 | -1.54 | -0.057 | -0.04 | 0.087 | 0.087 | 0.004 | 0.21 | 0.06 | 0.13 | 0.04 | 0.23 | 0.28 |
| 1 | 226611363 | rs1527365 | 0.739 | 1.053 | -1.532 | 0.271 | -0.007 | -0.03 | -0.03 | 1.02 | 0.15 | 0.09 | 0.35 | 0.4 | 0.17 | 0.12 |
| 1 | 226612023 | rs12025226 | -0.223 | 0.99 | -1.462 | 0.058 | -0.017 | 0.081 | 0.081 | 0.008 | 0.2 | 0.05 | 0.11 | 0.04 | 0.2 | 0.26 |
| 1 | 226618382 | rs10915993 | -0.096 | 0.92 | -0.752 | 0.355 | 0.003 | 0.019 | 0.019 | 0 | 0.28 | 0.33 | 0.32 | 0.49 | 0.31 | 0.26 |
| 1 | 226621778 | rs1317646 | -0.042 | 0.808 | -0.122 | 0.294 | 0.006 | 0.079 | 0.079 | 0 | 0.27 | 0.42 | 0.3 | 0.46 | 0.28 | 0.24 |
| 1 | 226623766 | rs16845946 | 0.713 | 1.01 | -0.183 | -0.111 | 0.005 | -0.001 | -0.001 | 0.096 | 0.08 | 0.12 | 0.05 | 0.06 | 0.1 | 0.04 |
| 1 | 226624910 | rs2377311 | 0.041 | 0.669 | -0.194 | -0.465 | -0.002 | 0.042 | 0.042 | 0 | 0.53 | 0.39 | 0.32 | 0.13 | 0.5 | 0.62 |
| 1 | 226629656 | rs12118824 | -0.07 | 0.598 | -0.177 | -0.473 | -0.004 | 0.041 | 0.041 | 0 | 0.54 | 0.35 | 0.33 | 0.15 | 0.51 | 0.65 |
| 1 | 226631272 | rs10799352 | -0.039 | 0.578 | -0.015 | -0.483 | 0 | -0.012 | -0.012 | 0 | 0.55 | 0.48 | 0.34 | 0.15 | 0.51 | 0.65 |
| 1 | 226636060 | rs1341331 | -0.063 | 0.564 | -0.028 | -0.397 | 0 | -0.009 | -0.009 | 0.765 | 0.55 | 0.48 | 0.36 | 0.17 | 0.51 | 0.65 |
| 1 | 226636593 | rs4653741 | 0.322 | 0.07 | -0.169 | -0.738 | -0.046 | 0.204 | 0.204 | 0 | 0.45 | 0.14 | 0.28 | 0.11 | 0.47 | 0.48 |
| 1 | 226640376 | rs10495245 | -0.478 | 0.115 | -0.059 | -0.587 | -0.023 | 0.017 | 0.017 | 0 | 0.1 | 0.34 | 0.08 | 0.07 | 0.04 | 0.17 |
| 1 | 226642835 | rs1527362 | -0.079 | -0.202 | -0.667 | -1.098 | 0 | -0.012 | -0.012 | 0 | 0.29 | 0.24 | 0.14 | 0.1 | 0.23 | 0.32 |
| 1 | 226653478 | rs4653743 | -1.024 | -0.07 | -0.864 | -0.747 | -0.008 | -0.04 | -0.04 | 7.672 | 0.76 | 0.78 | 0.62 | 0.54 | 0.73 | 0.85 |
| 1 | 226654252 | rs2099380 | 0.935 | 0.007 | -0.95 | -0.206 | -0.052 | -0.067 | -0.067 | 0 | 0.1 | 0.02 | 0.08 | 0.27 | 0.12 | 0.04 |
| 1 | 226655011 | rs12033699 | -0.463 | -0.313 | -0.558 | 0.033 | -0.01 | 0.042 | 0.042 | 0 | 0.29 | 0.43 | 0.19 | 0.35 | 0.22 | 0.33 |
| 1 | 226665284 | rs10799355 | -0.571 | -0.014 | -0.552 | 0.148 | 0.002 | 0.007 | 0.007 | 0 | 0.62 | 0.52 | 0.34 | 0.43 | 0.55 | 0.66 |
| 1 | 226670795 | rs898834 | -0.836 | -0.001 | -0.754 | 0.238 | -0.011 | -0.031 | -0.031 | 0 | 0.77 | 0.77 | 0.6 | 0.58 | 0.71 | 0.84 |
| 1 | 226670999 | rs898833 | 1.003 | 0.139 | -0.412 | 0.107 | 0.003 | -0.018 | -0.018 | 2.153 | 0.21 | 0.31 | 0.28 | 0.19 | 0.22 | 0.25 |
| 1 | 226675933 | rs4653745 | -0.071 | -0.131 | -1.1 | 0.181 | -0.008 | 0.026 | 0.026 | 0 | 0.57 | 0.25 | 0.56 | 0.63 | 0.6 | 0.62 |
| 1 | 226676299 | rs11588657 | 0.761 | -0.023 | -1.2 | 0.278 | -0.009 | -0.018 | -0.018 | 0 | 0.11 | 0.05 | 0.06 | 0.31 | 0.1 | 0.09 |
| 1 | 226677818 | rs17519476 | 0.826 | -0.242 | -1.389 | 0.091 | -0.029 | 0.077 | 0.077 | 1.413 | 0.25 | 0.03 | 0.23 | 0.09 | 0.27 | 0.26 |
| 1 | 226678169 | rs688902 | -0.485 | -0.4 | -0.952 | 0.025 | 0 | 0 | 0 | 0.021 | 0.48 | 0.48 | 0.39 | 0.47 | 0.48 | 0.43 |
| 1 | 226683251 | rs12402021 | -0.176 | -0.356 | -1.175 | 0.038 | 0.011 | -0.04 | -0.04 | 0 | 0.38 | 0.24 | 0.28 | 0.44 | 0.35 | 0.47 |
| 1 | 226693542 | rs7524430 | 0.386 | -0.281 | -1.039 | 0.354 | -0.004 | 0.061 | 0.061 | 0 | 0.11 | 0.18 | 0.04 | 0.33 | 0.07 | 0.1 |
| 1 | 226694739 | rs1151815 | -0.649 | -0.139 | -1.418 | 0.004 | -0.058 | -0.018 | -0.018 | 0 | 0.6 | 0.8 | 0.47 | 0.56 | 0.51 | 0.61 |
| 1 | 226702300 | rs506128 | -1.138 | 0.235 | -1.259 | 0.299 | -0.016 | -0.042 | -0.042 | 8.956 | 0.81 | 0.91 | 0.75 | 0.74 | 0.77 | 0.87 |
| 1 | 226703467 | rs1287949 | -0.259 | 0.234 | -0.925 | 0.239 | 0.003 | 0.013 | 0.013 | 0 | 0.73 | 0.73 | 0.83 | 0.65 | 0.75 | 0.79 |
| 1 | 226704825 | rs11585765 | 1.33 | 0.08 | -1.037 | 0.43 | -0.013 | -0.041 | -0.041 | 0.87 | 0.17 | 0.11 | 0.3 | 0.31 | 0.21 | 0.08 |
| 1 | 226708398 | rs751684 | -0.127 | -0.396 | -0.583 | 0.108 | 0.005 | 0.012 | 0.012 | 0 | 0.27 | 0.32 | 0.21 | 0.38 | 0.28 | 0.32 |
| 1 | 226708481 | rs584297 | -1.197 | -0.779 | -0.968 | -0.208 | -0.093 | -0.105 | -0.105 | 0 | 0.6 | 0.91 | 0.48 | 0.53 | 0.57 | 0.74 |
| 1 | 226709011 | rs12043214 | 0.874 | -0.73 | -0.835 | -0.169 | -0.01 | 0.083 | 0.083 | 4.899 | 0.26 | 0.04 | 0.19 | 0.07 | 0.24 | 0.21 |
| 1 | 226709873 | rs11584545 | 1.067 | -0.329 | -0.246 | 0.248 | 0 | 0.006 | 0.006 | 0 | 0.17 | 0.13 | 0.14 | 0.31 | 0.2 | 0.18 |
| 1 | 226714204 | rs10916006 | -0.472 | -0.524 | 0.106 | 0.219 | 0.017 | 0.066 | 0.066 | 3.247 | 0.06 | 0.33 | 0.04 | 0.07 | 0.01 | 0.15 |
| 1 | 226717942 | rs7545563 | 0.12 | -0.207 | -0.516 | 0.262 | 0.006 | 0.002 | 0.002 | 0 | 0.28 | 0.32 | 0.22 | 0.37 | 0.34 | 0.33 |
| 1 | 226729492 | rs16846092 | 0.489 | 0.365 | -0.973 | 0.739 | -0.1 | -0.102 | -0.102 | 0 | 0.33 | 0.08 | 0.5 | 0.45 | 0.4 | 0.27 |
| 1 | 226731876 | rs4653451 | 0.203 | 0.455 | -0.619 | 0.706 | -0.024 | -0.043 | -0.043 | 0 | 0.38 | 0.25 | 0.52 | 0.53 | 0.41 | 0.41 |
| 1 | 226737202 | rs708741 | 0.162 | 0.37 | -0.101 | 0.838 | -0.003 | 0 | 0 | 0.208 | 0.91 | 0.94 | 0.88 | 0.95 | 0.88 | 0.87 |
| 1 | 226743163 | rs708745 | 0.246 | 0.366 | 0.017 | 0.836 | 0.003 | -0.001 | -0.001 | 0 | 0.91 | 0.87 | 0.87 | 0.95 | 0.88 | 0.86 |
| 1 | 226752447 | rs16846117 | -0.507 | 0.079 | 0.489 | 0.854 | -0.009 | 0.014 | 0.014 | 0 | 0.11 | 0.28 | 0.09 | 0.11 | 0.08 | 0.19 |
| 1 | 226755701 | rs672572 | -0.879 | 0.018 | 0.349 | 0.93 | -0.012 | 0.09 | 0.09 | 0.337 | 0.13 | 0.45 | 0.1 | 0.18 | 0.08 | 0.3 |
| 1 | 226760026 | rs12024134 | -0.971 | 0.302 | -0.445 | 1.343 | -0.001 | -0.006 | -0.006 | 0.175 | 0.14 | 0.16 | 0.19 | 0.28 | 0.16 | 0.38 |
| 1 | 226763678 | rs837444 | -0.167 | 0.576 | -0.802 | 1.065 | 0.002 | 0 | 0 | 2.274 | 0.4 | 0.58 | 0.67 | 0.52 | 0.49 | 0.46 |
| 1 | 226764155 | rs1151809 | -0.855 | -0.193 | -1.243 | 0.49 | -0.024 | 0.011 | 0.011 | 3.498 | 0.62 | 0.93 | 0.81 | 0.66 | 0.67 | 0.83 |
| 1 | 226764886 | rs6691873 | -0.438 | 0.006 | -1.037 | 0.614 | -0.01 | 0.035 | 0.035 | 0.818 | 0.07 | 0.2 | 0.06 | 0.12 | 0.07 | 0.25 |
| 1 | 226769529 | rs16846167 | 1.106 | -0.213 | -1.469 | 0.297 | 0.003 | -0.019 | -0.019 | 0 | 0.14 | 0.06 | 0.08 | 0.23 | 0.14 | 0.13 |
| 1 | 226769798 | rs589616 | 0.775 | -0.204 | -1.443 | 0.283 | -0.001 | -0.001 | -0.001 | 0 | 0.51 | 0.5 | 0.59 | 0.39 | 0.46 | 0.35 |
| 1 | 226772712 | rs837919 | -0.36 | -0.174 | -1.486 | 0.272 | -0.022 | -0.004 | -0.004 | 0.308 | 0.67 | 0.89 | 0.79 | 0.67 | 0.67 | 0.8 |
| 1 | 226772889 | rs12035052 | -0.16 | -0.194 | -1.448 | 0.27 | 0.009 | 0.048 | 0.048 | 0 | 0.16 | 0.29 | 0.19 | 0.28 | 0.21 | 0.45 |
| 1 | 226774009 | rs1151808 | -0.283 | 0.024 | -1.405 | 0.337 | -0.022 | -0.005 | -0.005 | 0 | 0.67 | 0.89 | 0.79 | 0.67 | 0.67 | 0.8 |
| 1 | 226774042 | rs865682 | -1.566 | 0.098 | -1.489 | 0.234 | -0.021 | 0.048 | 0.048 | 2.834 | 0.88 | 1 | 0.93 | 0.98 | 0.89 | 0.96 |
| 1 | 226778432 | rs528089 | -0.324 | 0.201 | -1.276 | 0.428 | -0.03 | 0.038 | 0.038 | 0 | 0.64 | 0.88 | 0.76 | 0.71 | 0.62 | 0.75 |
| 1 | 226778821 | rs2282426 | 0.99 | 0.125 | -1.189 | 0.456 | 0.001 | 0 | 0 | 1 | 0.41 | 0.45 | 0.54 | 0.39 | 0.39 | 0.32 |
| 1 | 226779449 | rs497911 | -0.628 | 0.3 | -1.027 | 0.616 | 0.036 | 0.153 | 0.153 | 0 | 0.15 | 0.41 | 0.21 | 0.33 | 0.21 | 0.45 |
| 1 | 226784188 | rs597429 | -0.887 | 0.31 | -0.87 | 0.352 | 0.01 | 0.025 | 0.025 | 0.057 | 0.53 | 0.68 | 0.45 | 0.62 | 0.56 | 0.65 |
| 1 | 226785742 | rs708754 | -0.24 | 0.411 | -0.815 | 0.475 | -0.001 | 0.001 | 0.001 | 0 | 0.42 | 0.39 | 0.29 | 0.33 | 0.4 | 0.39 |

| | | | | | | | | | | | | | | | | |
|---|---|---|---|---|---|---|---|---|---|---|---|---|---|---|---|---|
| 1 | 226790197 | rs12086463 | 0.264 | 0.507 | -0.659 | 0.595 | 0.077 | 0.225 | 0.225 | 0 | 0.11 | 0.44 | 0.18 | 0.28 | 0.15 | 0.25 |
| 1 | 226790403 | rs878149 | 1.499 | 0.77 | -1.035 | 0.562 | -0.003 | 0.084 | 0.084 | 0.133 | 0.14 | 0.28 | 0.31 | 0.29 | 0.13 | 0.13 |
| 1 | 226790621 | rs878148 | -0.089 | 1.161 | -0.672 | 0.795 | 0.002 | 0.014 | 0.014 | 0 | 0.34 | 0.16 | 0.19 | 0.31 | 0.29 | 0.23 |
| 1 | 226790951 | rs3820632 | 0.193 | 1.199 | -0.688 | 0.733 | 0.022 | 0.051 | 0.051 | 0 | 0.13 | 0.26 | 0.2 | 0.27 | 0.2 | 0.25 |
| 1 | 226794525 | rs1461920 | -0.395 | 1.046 | -1.26 | 0.861 | -0.002 | -0.016 | -0.016 | 0 | 0.39 | 0.24 | 0.23 | 0.41 | 0.35 | 0.42 |
| 1 | 226797733 | rs6697883 | -0.561 | 1.221 | -0.826 | 0.872 | 0.015 | 0.022 | 0.022 | 0 | 0.25 | 0.47 | 0.28 | 0.3 | 0.3 | 0.31 |
| 1 | 226807117 | rs2078211 | -0.146 | 1.39 | -1.379 | 1.185 | -0.001 | -0.003 | -0.003 | 0 | 0.15 | 0.13 | 0.1 | 0.23 | 0.15 | 0.24 |
| 1 | 226807512 | rs708757 | 0.35 | 2.141 | -1.677 | 1.267 | -0.064 | -0.047 | -0.047 | 0 | 0.74 | 0.84 | 0.69 | 0.62 | 0.56 | 0.65 |
| 1 | 226810146 | rs10916019 | -0.89 | 1.668 | -1.705 | 1.344 | -0.041 | -0.103 | -0.103 | 1.685 | 0.3 | 0.13 | 0.27 | 0.55 | 0.35 | 0.42 |
| 1 | 226812231 | rs7533764 | -0.28 | 1.562 | -1.253 | 1.234 | 0.002 | 0.002 | 0.002 | 0 | 0.09 | 0.12 | 0.06 | 0.14 | 0.08 | 0.15 |
| 1 | 226817467 | rs6696348 | 0.484 | 1.262 | -1.309 | 1.25 | 0.003 | -0.015 | -0.015 | 0.925 | 0.65 | 0.3 | 0.51 | 0.39 | 0.67 | 0.67 |
| 1 | 226817556 | rs697845 | -0.305 | 1.24 | -0.305 | 1.485 | -0.022 | -0.01 | -0.01 | 0.024 | 0.8 | 0.62 | 0.87 | 0.79 | 0.84 | 0.86 |
| 1 | 226817699 | rs697846 | -0.013 | 0.99 | -0.04 | 1.744 | 0.026 | 0.074 | 0.074 | 0 | 0.12 | 0.21 | 0.16 | 0.29 | 0.16 | 0.12 |
| 1 | 226819833 | rs1144840 | 0.259 | 0.995 | -0.241 | 1.654 | -0.003 | -0.004 | -0.004 | 0.016 | 0.1 | 0.08 | 0.07 | 0.15 | 0.09 | 0.14 |
| 1 | 226820500 | rs1050492 | -1.287 | 0.623 | -0.335 | 1.43 | 0.039 | -0.024 | -0.024 | 0 | 0.39 | 0.78 | 0.45 | 0.61 | 0.28 | 0.34 |
| 1 | 226821139 | rs3738723 | 0.009 | 0.539 | -0.177 | 1.526 | -0.062 | -0.007 | -0.007 | 0 | 0.22 | 0.4 | 0.14 | 0.25 | 0.1 | 0.15 |
| 1 | 226825341 | rs2236604 | 0.112 | 0.774 | 0.236 | 1.45 | -0.004 | -0.001 | -0.001 | 0.597 | 0.22 | 0.21 | 0.16 | 0.25 | 0.14 | 0.16 |
| 1 | 226828038 | rs1288934 | -1.138 | 0.958 | 0.178 | 1.53 | 0.002 | -0.002 | -0.002 | 0 | 0.35 | 0.66 | 0.4 | 0.47 | 0.24 | 0.39 |
| 1 | 226829990 | rs17522524 | 0.444 | 1.128 | -0.298 | 1.45 | -0.005 | -0.012 | -0.012 | 0 | 0.1 | 0.05 | 0.06 | 0.15 | 0.09 | 0.13 |
| 1 | 226833304 | rs1144836 | -1.796 | 0.963 | -0.606 | 1.138 | -0.104 | 0.051 | 0.051 | 4.364 | 0.55 | 0.98 | 0.58 | 0.77 | 0.42 | 0.49 |
| 1 | 226835214 | rs1144835 | -1.731 | 0.986 | -0.552 | 1.187 | -0.092 | 0.086 | 0.086 | 4.683 | 0.54 | 0.96 | 0.59 | 0.78 | 0.43 | 0.49 |
| 1 | 226836190 | rs4653762 | -1.34 | 1.101 | -0.22 | 1.321 | 0 | 0.003 | 0.003 | 1.314 | 0.49 | 0.51 | 0.55 | 0.76 | 0.43 | 0.49 |
| 1 | 226841347 | rs3820636 | -0.227 | 0.721 | -0.533 | 1.291 | 0.002 | 0 | 0 | 3.505 | 0.26 | 0.24 | 0.19 | 0.28 | 0.18 | 0.18 |
| 1 | 226851963 | rs3754390 | 1.096 | 0.764 | -1.584 | 0.887 | -0.042 | -0.061 | -0.061 | 0.886 | 0.3 | 0.01 | 0.26 | 0.2 | 0.26 | 0.38 |
| 1 | 226852155 | rs1288938 | 0.391 | 0.88 | -1.781 | 0.776 | -0.004 | 0.012 | 0.012 | 0 | 0.15 | 0.08 | 0.11 | 0.15 | 0.14 | 0.12 |
| 1 | 226859676 | rs12094617 | 0.412 | 0.857 | -1.138 | 0.738 | -0.006 | 0.003 | 0.003 | 1.368 | 0.24 | 0.32 | 0.2 | 0.3 | 0.22 | 0.18 |
| 1 | 226865254 | rs708766 | 0.311 | 0.866 | -0.911 | 0.634 | 0 | 0.002 | 0.002 | 0 | 0.18 | 0.17 | 0.12 | 0.15 | 0.15 | 0.15 |
| 1 | 226875552 | rs940659 | 0.486 | 0.817 | -0.84 | 0.835 | 0 | -0.005 | -0.005 | 0 | 0.22 | 0.17 | 0.19 | 0.3 | 0.22 | 0.17 |
| 1 | 226880081 | rs12404031 | 0.233 | 0.829 | -0.877 | 0.839 | -0.006 | 0.02 | 0.02 | 1.155 | 0.24 | 0.33 | 0.34 | 0.38 | 0.22 | 0.26 |
| 1 | 226883325 | rs697851 | 0.156 | 0.862 | -0.897 | 0.798 | -0.003 | -0.003 | -0.003 | 0.093 | 0.26 | 0.35 | 0.22 | 0.31 | 0.24 | 0.22 |
| 1 | 226884755 | rs1341283 | 0.238 | 0.808 | -0.844 | 0.791 | -0.01 | 0.02 | 0.02 | 1.552 | 0.25 | 0.38 | 0.35 | 0.38 | 0.22 | 0.26 |
| 1 | 226898992 | rs3768403 | 1.173 | 0.633 | -1.465 | 0.85 | -0.049 | -0.067 | -0.067 | 0 | 0.3 | 0.02 | 0.26 | 0.21 | 0.26 | 0.38 |
| 1 | 226900415 | rs3768405 | 1.395 | 0.466 | -1.627 | 0.586 | -0.062 | 0.03 | 0.03 | 0 | 0.21 | 0.01 | 0.1 | 0.07 | 0.2 | 0.28 |
| 1 | 226912891 | rs708772 | -0.423 | 0.599 | -1.435 | 0.69 | -0.033 | -0.019 | -0.019 | 0 | 0.37 | 0.21 | 0.38 | 0.42 | 0.42 | 0.31 |
| 1 | 226917898 | rs3768414 | -0.505 | 0.318 | -0.799 | 1.028 | -0.003 | 0.01 | 0.01 | 0 | 0.56 | 0.42 | 0.66 | 0.68 | 0.53 | 0.6 |
| 1 | 226920730 | rs708773 | -0.083 | 0.46 | -0.743 | 0.909 | -0.004 | 0.002 | 0.002 | 0.031 | 0.29 | 0.25 | 0.32 | 0.26 | 0.31 | 0.29 |
| 1 | 226923938 | rs6667260 | -0.642 | 0.47 | -0.401 | 0.579 | -0.001 | 0 | 0 | 0.143 | 0.57 | 0.61 | 0.66 | 0.68 | 0.53 | 0.6 |
| 1 | 226928864 | rs1110033 | -0.447 | 0.394 | -0.249 | 0.591 | 0 | 0.005 | 0.005 | 0.155 | 0.52 | 0.61 | 0.5 | 0.57 | 0.46 | 0.48 |
| 1 | 226933722 | rs708777 | -0.537 | 0.396 | -0.427 | 0.544 | -0.01 | 0.034 | 0.034 | 0.085 | 0.53 | 0.77 | 0.65 | 0.64 | 0.47 | 0.53 |
| 1 | 226947468 | rs7540797 | 1.046 | 0.343 | -0.591 | 0.844 | 0.004 | -0.014 | -0.014 | 0.596 | 0.25 | 0.15 | 0.19 | 0.35 | 0.22 | 0.2 |
| 1 | 226956181 | rs12729579 | 0.668 | 0.664 | -0.539 | 0.451 | -0.029 | 0.074 | 0.074 | 0 | 0.55 | 0.26 | 0.5 | 0.76 | 0.51 | 0.65 |
| 1 | 226959680 | rs10799366 | 0.575 | 0.654 | -0.557 | 0.418 | -0.055 | 0.109 | 0.109 | 0 | 0.55 | 0.17 | 0.5 | 0.76 | 0.51 | 0.65 |
| 1 | 226960025 | rs835673 | -0.9 | 0.732 | -0.295 | 0.246 | 0.003 | -0.039 | -0.039 | 0 | 0.83 | 0.84 | 0.84 | 1 | 0.77 | 0.94 |
| 1 | 226965174 | rs12033364 | 0.642 | 0.678 | -0.623 | 0.446 | -0.055 | 0.07 | 0.07 | 0 | 0.55 | 0.17 | 0.38 | 0.69 | 0.51 | 0.6 |
| 1 | 226965334 | rs4653468 | 0.732 | 0.67 | -0.519 | 0.381 | -0.026 | 0.072 | 0.072 | 0 | 0.55 | 0.27 | 0.51 | 0.76 | 0.51 | 0.65 |
| 1 | 226968814 | rs708761 | 0.98 | 0.634 | -0.737 | 0.363 | -0.039 | 0.061 | 0.061 | 0.615 | 0.22 | 0.08 | 0.11 | 0.06 | 0.25 | 0.31 |
| 1 | 226971029 | rs6684472 | -0.115 | 0.547 | -0.373 | 0.421 | 0.012 | -0.009 | -0.009 | 2.609 | 0.36 | 0.53 | 0.37 | 0.29 | 0.45 | 0.52 |
| 1 | 226972422 | rs835661 | -0.369 | 0.511 | -0.884 | 0.617 | -0.101 | -0.105 | -0.105 | 0.499 | 0.74 | 0.98 | 0.75 | 0.69 | 0.7 | 0.81 |
| 1 | 226973381 | rs10495250 | 0.666 | 0.172 | -0.15 | 0.526 | 0.011 | 0.019 | 0.019 | 0.068 | 0.2 | 0.41 | 0.22 | 0.19 | 0.21 | 0.25 |
| 1 | 226977944 | rs12738352 | -0.493 | 0.493 | -0.696 | 0.841 | -0.059 | 0 | 0 | 0.096 | 0.42 | 0.11 | 0.41 | 0.38 | 0.54 | 0.46 |
| 1 | 226999183 | rs6660716 | -0.069 | 0.629 | 0.118 | 1.027 | -0.003 | 0.005 | 0.005 | 0 | 0.21 | 0.24 | 0.21 | 0.28 | 0.16 | 0.15 |
| 1 | 227008502 | rs12405779 | 1.402 | 0.813 | -0.12 | 0.878 | -0.037 | 0.017 | 0.017 | 0 | 0.15 | 0.04 | 0.13 | 0.11 | 0.2 | 0.26 |
| 1 | 227013878 | rs6426548 | -1.018 | 0.581 | 0.227 | 0.861 | -0.001 | -0.001 | -0.001 | 1.315 | 0.41 | 0.58 | 0.5 | 0.61 | 0.4 | 0.57 |
| 1 | 227017031 | rs9660802 | 0.24 | 0.555 | 0.28 | 0.894 | 0.003 | -0.008 | -0.008 | 0 | 0.2 | 0.2 | 0.27 | 0.33 | 0.17 | 0.18 |
| 1 | 227047249 | rs1150897 | -1.033 | 0.574 | 0.212 | 0.872 | 0 | 0.002 | 0.002 | 0.063 | 0.41 | 0.59 | 0.5 | 0.61 | 0.4 | 0.58 |
| 1 | 227047256 | rs1150896 | 0.572 | 0.554 | -0.112 | 0.882 | -0.03 | -0.091 | -0.091 | 0 | 0.18 | 0.02 | 0.19 | 0.28 | 0.16 | 0.15 |
| 1 | 227059037 | rs1295645 | 0.536 | 0.547 | -0.114 | 0.879 | -0.029 | -0.091 | -0.091 | 0.942 | 0.18 | 0.02 | 0.19 | 0.28 | 0.17 | 0.15 |
| 1 | 227063671 | rs2073489 | -1.491 | 0.624 | -0.06 | 0.047 | 0.005 | 0.052 | 0.052 | 0.855 | 0.54 | 0.72 | 0.6 | 0.62 | 0.54 | 0.71 |
| 1 | 227069677 | rs11405 | -0.807 | 0.621 | 0.057 | 0.148 | 0.002 | -0.016 | -0.016 | 1.898 | 0.79 | 0.83 | 0.72 | 0.54 | 0.8 | 0.75 |
| 1 | 227069737 | rs6759 | 0.571 | 0.659 | -0.034 | 0.092 | 0.003 | 0.015 | 0.015 | 0.043 | 0.58 | 0.36 | 0.48 | 0.39 | 0.59 | 0.43 |
| 1 | 227070378 | rs1295640 | 0.578 | 0.663 | -0.033 | 0.094 | 0.003 | 0.015 | 0.015 | 0.049 | 0.58 | 0.36 | 0.48 | 0.39 | 0.59 | 0.43 |
| 1 | 227071525 | rs1046240 | 0.578 | 0.663 | -0.033 | 0.094 | 0.003 | 0.015 | 0.015 | 0.049 | 0.58 | 0.36 | 0.48 | 0.39 | 0.59 | 0.43 |
| 1 | 227075984 | rs1800680 | 1.377 | 0.824 | -0.318 | -0.086 | -0.039 | 0.046 | 0.046 | 0 | 0.1 | 0.02 | 0.1 | 0 | 0.18 | 0.05 |
| 1 | 227078592 | rs1800678 | 0.116 | 0.535 | -0.076 | 0.037 | -0.036 | 0.093 | 0.093 | 0 | 0.73 | 0.53 | 0.62 | 0.49 | 0.8 | 0.69 |
| 1 | 227084660 | rs2802268 | 1.25 | 0.458 | -0.396 | 0.275 | 0.02 | -0.001 | -0.001 | 0.014 | 0.34 | 0.25 | 0.3 | 0.38 | 0.27 | 0.22 |
| 1 | 227085462 | rs1800672 | 0.319 | 0.857 | 0.103 | 0.346 | 0.02 | -0.008 | -0.008 | 0 | 0.2 | 0.41 | 0.23 | 0.17 | 0.21 | 0.32 |
| 1 | 227088591 | rs12126925 | -0.237 | 1.131 | -0.58 | 0.12 | -0.007 | 0.006 | 0.006 | 0 | 0.13 | 0.09 | 0.09 | 0 | 0.16 | 0.06 |
| 1 | 227090500 | rs12405469 | 1.101 | 0.978 | -1.264 | -0.081 | -0.003 | 0.094 | 0.094 | 0 | 0.38 | 0.16 | 0.28 | 0.24 | 0.33 | 0.28 |
| 1 | 227092862 | rs6661550 | 0.612 | 1.184 | -0.879 | 0.587 | 0.002 | 0.004 | 0.004 | 0.023 | 0.21 | 0.2 | 0.26 | 0.3 | 0.25 | 0.42 |

| | | | | | | | | | | | | | | | |
|---|---|---|---|---|---|---|---|---|---|---|---|---|---|---|---|
| 1 | 227093321 | rs2313007 | -1.362 | 1.111 | -1.118 | 0.899 | -0.018 | -0.054 | -0.054 | 0.111 | 0.83 | 0.96 | 0.83 | 0.78 | 0.83 | 0.85 |
| 1 | 227097039 | rs1938386 | 0.234 | 1.07 | -0.841 | 0.776 | -0.015 | -0.014 | -0.014 | 3.16 | 0.61 | 0.73 | 0.57 | 0.48 | 0.58 | 0.43 |
| 1 | 227098588 | rs10916066 | -0.011 | 1.009 | -0.565 | 0.637 | 0.023 | -0.101 | -0.101 | 0 | 0.19 | 0.38 | 0.15 | 0.03 | 0.25 | 0.1 |
| 1 | 227103413 | rs12041000 | 1.188 | 0.722 | -0.854 | 1.014 | 0.01 | -0.013 | -0.013 | 0.311 | 0.4 | 0.14 | 0.39 | 0.43 | 0.34 | 0.29 |
| 1 | 227118912 | rs17319408 | -1.128 | 0.839 | -0.634 | 1.173 | -0.046 | 0.005 | 0.005 | 1.59 | 0.44 | 0.85 | 0.53 | 0.58 | 0.46 | 0.58 |
| 1 | 227124660 | rs6660932 | -0.915 | 0.695 | 0.169 | 1.144 | 0.039 | 0.035 | 0.035 | 1.311 | 0.09 | 0.4 | 0.13 | 0.07 | 0.1 | 0.1 |
| 1 | 227125997 | rs7513120 | -0.768 | 0.841 | -0.072 | 1.396 | -0.007 | -0.009 | -0.009 | 0 | 0.36 | 0.64 | 0.5 | 0.51 | 0.43 | 0.53 |
| 1 | 227127618 | rs2777840 | -0.334 | 0.309 | -0.534 | 1.069 | -0.017 | -0.013 | -0.013 | 0 | 0.45 | 0.75 | 0.5 | 0.51 | 0.49 | 0.55 |
| 1 | 227141493 | rs2802271 | -0.962 | 0.068 | -0.9 | 1.056 | 0.018 | -0.03 | -0.03 | 0.69 | 0.65 | 0.9 | 0.73 | 0.66 | 0.7 | 0.71 |
| 1 | 227152392 | rs12133428 | 1.715 | 0.293 | -1.043 | 1.123 | -0.036 | -0.015 | -0.015 | 1.458 | 0.11 | 0.02 | 0.15 | 0.16 | 0.15 | 0.38 |
| 1 | 227153930 | rs2154367 | 0.409 | 0.194 | -0.496 | 1.322 | 0.01 | 0.047 | 0.047 | 1.089 | 0.17 | 0.25 | 0.31 | 0.39 | 0.21 | 0.4 |
| 1 | 227162976 | rs2777843 | -1.365 | -0.421 | -1.018 | 0.907 | 0.031 | 0.015 | 0.015 | 0 | 0.82 | 0.93 | 0.87 | 0.88 | 0.87 | 0.81 |
| 1 | 227170768 | rs2256060 | -0.783 | -0.203 | -0.712 | 1.055 | 0.018 | 0.046 | 0.046 | 0 | 0.14 | 0.28 | 0.36 | 0.26 | 0.2 | 0.13 |
| 1 | 227173537 | rs3768420 | 1.819 | -0.114 | -1.029 | 0.494 | -0.034 | 0.018 | 0.018 | 0 | 0.22 | 0.06 | 0.16 | 0.15 | 0.22 | 0.13 |
| 1 | 227174210 | rs3738725 | 0.638 | -0.426 | -0.857 | 0.215 | -0.003 | 0.068 | 0.068 | 0 | 0.45 | 0.29 | 0.31 | 0.21 | 0.49 | 0.27 |
| 1 | 227174902 | rs8159 | 0.227 | -0.6 | -0.678 | 0.277 | 0.003 | 0.071 | 0.071 | 0 | 0.26 | 0.37 | 0.41 | 0.5 | 0.28 | 0.22 |
| 1 | 227175253 | rs10495274 | 0.221 | -0.689 | -0.639 | 0.208 | 0.007 | 0.011 | 0.011 | 0.148 | 0.58 | 0.36 | 0.42 | 0.34 | 0.61 | 0.39 |
| 1 | 227176511 | rs2802275 | -0.746 | -0.621 | -0.677 | 0.17 | -0.01 | -0.005 | -0.005 | 3.235 | 0.71 | 0.66 | 0.72 | 0.72 | 0.77 | 0.49 |
| 1 | 227178127 | rs3768425 | 1.035 | -0.554 | -0.833 | 0.142 | -0.017 | 0.04 | 0.04 | 0 | 0.41 | 0.29 | 0.29 | 0.21 | 0.44 | 0.22 |
| 1 | 227178700 | rs1045290 | 1.035 | -0.554 | -0.833 | 0.142 | -0.017 | 0.04 | 0.04 | 0 | 0.41 | 0.29 | 0.29 | 0.21 | 0.44 | 0.22 |
| 1 | 227178713 | rs1045289 | 1.035 | -0.554 | -0.833 | 0.142 | -0.017 | 0.04 | 0.04 | 0 | 0.41 | 0.29 | 0.29 | 0.21 | 0.44 | 0.22 |
| 1 | 227178948 | rs2989946 | -0.857 | -0.651 | -0.688 | 0.101 | 0.011 | 0.018 | 0.018 | 0 | 0.7 | 0.81 | 0.71 | 0.71 | 0.72 | 0.44 |
| 1 | 227179037 | rs1045286 | 1.062 | -0.659 | -0.774 | 0.092 | -0.017 | 0.04 | 0.04 | 0 | 0.41 | 0.29 | 0.29 | 0.21 | 0.44 | 0.22 |
| 1 | 227179054 | rs1045285 | 1.14 | -0.649 | -0.772 | 0.1 | -0.027 | 0.078 | 0.078 | 0 | 0.4 | 0.25 | 0.28 | 0.16 | 0.44 | 0.2 |
| 1 | 227179358 | rs2997438 | -1.651 | -0.709 | -0.824 | 0.116 | 0.027 | 0.023 | 0.023 | 0 | 0.81 | 0.9 | 0.87 | 0.89 | 0.87 | 0.82 |
| 1 | 227180858 | rs1801783 | 1.328 | -0.65 | -0.85 | 0.065 | -0.055 | 0.111 | 0.111 | 0.142 | 0.22 | 0.07 | 0.14 | 0 | 0.27 | 0.12 |
| 1 | 227180947 | rs3795445 | 0.707 | -0.663 | -0.82 | 0.121 | -0.005 | 0.097 | 0.097 | 0 | 0.45 | 0.27 | 0.31 | 0.16 | 0.49 | 0.24 |
| 1 | 227180981 | rs3795446 | 0.707 | -0.663 | -0.82 | 0.121 | -0.005 | 0.097 | 0.097 | 0 | 0.45 | 0.27 | 0.31 | 0.16 | 0.49 | 0.24 |
| 1 | 227181555 | rs1045252 | 0.983 | -0.509 | -0.693 | 0.234 | -0.018 | 0.041 | 0.041 | 0 | 0.41 | 0.28 | 0.29 | 0.21 | 0.44 | 0.22 |
| 1 | 227181838 | rs3795449 | 1.013 | -0.505 | -0.742 | 0.236 | -0.022 | 0.063 | 0.063 | 0 | 0.4 | 0.27 | 0.28 | 0.16 | 0.44 | 0.19 |
| 1 | 227181923 | rs3795451 | 1.021 | -0.51 | -0.744 | 0.22 | -0.022 | 0.046 | 0.046 | 0 | 0.4 | 0.27 | 0.29 | 0.21 | 0.44 | 0.22 |
| 1 | 227182033 | rs2802269 | -1.912 | -0.431 | -1.075 | 0.251 | 0.03 | 0.025 | 0.025 | 0 | 0.82 | 0.93 | 0.87 | 0.89 | 0.87 | 0.81 |
| 1 | 227182575 | rs1045247 | -1.966 | -0.443 | -1.076 | 0.21 | -0.037 | -0.014 | -0.014 | 0 | 0.86 | 0.98 | 0.84 | 0.83 | 0.83 | 0.61 |
| 1 | 227190456 | rs1298652 | -2.176 | -0.368 | -0.933 | 0.314 | 0.033 | 0.057 | 0.057 | 0 | 0.82 | 0.93 | 0.93 | 0.95 | 0.88 | 0.83 |
| 1 | 227192477 | rs3818694 | 0.264 | -0.079 | -0.581 | 0.627 | -0.005 | 0.007 | 0.007 | 0 | 0.22 | 0.33 | 0.25 | 0.27 | 0.21 | 0.2 |
| 1 | 227200100 | rs4653474 | 0.253 | -0.078 | -0.572 | 0.587 | -0.006 | 0.023 | 0.023 | 0 | 0.23 | 0.33 | 0.31 | 0.33 | 0.21 | 0.19 |
| 1 | 227204557 | rs17600747 | 1.798 | -0.067 | -0.608 | 0.372 | -0.016 | 0.014 | 0.014 | 0 | 0.22 | 0.09 | 0.17 | 0.15 | 0.22 | 0.13 |
| 1 | 227205331 | rs3795453 | 0.42 | -0.177 | -0.384 | 0.356 | -0.006 | 0.018 | 0.018 | 0 | 0.43 | 0.38 | 0.3 | 0.15 | 0.44 | 0.19 |
| 1 | 227205402 | rs3795454 | -0.045 | -0.2 | -0.393 | 0.392 | 0.004 | 0.051 | 0.051 | 0 | 0.25 | 0.38 | 0.47 | 0.55 | 0.29 | 0.21 |
| 1 | 227210820 | rs11580229 | 0.423 | -0.058 | -0.351 | 0.18 | -0.011 | 0.044 | 0.044 | 0 | 0.42 | 0.34 | 0.29 | 0.15 | 0.44 | 0.19 |
| 1 | 227212024 | rs10916077 | 0.717 | 0.204 | -0.177 | 0.568 | 0.02 | 0.014 | 0.014 | 0 | 0.09 | 0.24 | 0.15 | 0.09 | 0.1 | 0.05 |
| 1 | 227216662 | rs1929861 | -0.144 | 0.318 | -0.479 | 0.505 | 0.009 | 0.006 | 0.006 | 0 | 0.42 | 0.48 | 0.42 | 0.4 | 0.47 | 0.23 |
| 1 | 227216775 | rs1929860 | 0.798 | 0.375 | -0.499 | 0.446 | -0.006 | -0.006 | -0.006 | 3.093 | 0.24 | 0.19 | 0.33 | 0.4 | 0.33 | 0.19 |
| 1 | 227221715 | rs11808332 | 0.092 | 0.377 | -0.535 | 0.634 | -0.002 | 0.015 | 0.015 | 0 | 0.25 | 0.37 | 0.27 | 0.25 | 0.26 | 0.1 |
| 1 | 227223367 | rs10495272 | 2.139 | 0.39 | -0.742 | 0.511 | -0.038 | -0.022 | -0.022 | 0 | 0.11 | 0.02 | 0.14 | 0.17 | 0.15 | 0.35 |
| 1 | 227223736 | rs7521639 | -0.91 | 0.149 | -0.635 | 0.381 | -0.007 | 0.014 | 0.014 | 0 | 0.4 | 0.33 | 0.58 | 0.72 | 0.47 | 0.62 |
| 1 | 227228946 | rs1029063 | -0.834 | 0.195 | -0.697 | 0.475 | 0.006 | -0.001 | -0.001 | 0 | 0.47 | 0.58 | 0.53 | 0.49 | 0.45 | 0.29 |
| 1 | 227242247 | rs2150874 | 2.115 | 0.288 | -0.796 | 0.559 | -0.038 | -0.046 | -0.046 | 1.774 | 0.11 | 0.02 | 0.14 | 0.23 | 0.15 | 0.38 |
| 1 | 227280717 | rs4653789 | -1.2 | 0.123 | -0.507 | 0.509 | -0.006 | 0.022 | 0.022 | 0 | 0.71 | 0.65 | 0.67 | 0.58 | 0.74 | 0.78 |
| 1 | 227285635 | rs1304324 | 0.535 | 0.035 | -0.722 | 0.337 | 0.005 | 0.002 | 0.002 | 0 | 0.22 | 0.21 | 0.26 | 0.21 | 0.19 | 0.2 |
| 1 | 227290139 | rs533134 | -1.918 | 0.217 | -0.868 | 0.242 | 0.012 | 0.041 | 0.041 | 0 | 0.76 | 0.85 | 0.83 | 0.87 | 0.81 | 0.81 |
| 1 | 227304180 | rs498020 | -1.064 | 0.269 | -0.807 | 0.223 | -0.002 | 0.01 | 0.01 | 0.041 | 0.58 | 0.62 | 0.67 | 0.72 | 0.61 | 0.69 |
| 1 | 227306488 | rs2482453 | -1.311 | 0.236 | -1.13 | 0.159 | -0.009 | 0.061 | 0.061 | 0 | 0.72 | 0.82 | 0.78 | 0.87 | 0.73 | 0.8 |
| 1 | 227311310 | rs10495258 | 1.7 | 0.169 | -1.181 | 0.234 | -0.006 | 0.086 | 0.086 | 0.275 | 0.14 | 0.02 | 0.06 | 0 | 0.13 | 0.04 |
| 1 | 227318028 | rs17606040 | 1.438 | -0.026 | -1.236 | 0.395 | 0.032 | -0.016 | -0.016 | 0 | 0.14 | 0.04 | 0.11 | 0.16 | 0.1 | 0.13 |
| 1 | 227325759 | rs12134379 | 1.903 | 0.376 | -1.161 | 0.508 | -0.038 | -0.022 | -0.022 | 0 | 0.11 | 0.02 | 0.14 | 0.18 | 0.15 | 0.35 |
| 1 | 227348126 | rs11800062 | 1.009 | 0.082 | -0.619 | 0.089 | -0.026 | -0.096 | -0.096 | 1.018 | 0.18 | 0.3 | 0.09 | 0 | 0.13 | 0.04 |
| 1 | 227354535 | rs2175789 | -0.002 | 0.239 | -0.503 | 0.433 | 0.02 | 0.149 | 0.149 | 0.005 | 0.88 | 0.78 | 0.71 | 0.63 | 0.84 | 0.78 |
| 1 | 227356925 | rs12727754 | 1.664 | 0.449 | -0.712 | 0.449 | -0.012 | -0.011 | -0.011 | 0.033 | 0.05 | 0.04 | 0.13 | 0.08 | 0.08 | 0.08 |
| 1 | 227357475 | rs6426578 | 2.091 | 0.446 | -0.656 | 0.331 | -0.012 | 0.005 | 0.005 | 0.092 | 0.17 | 0.12 | 0.15 | 0.13 | 0.2 | 0.13 |
| 1 | 227371906 | rs12742066 | 0.65 | 0.163 | -0.637 | 0.272 | 0 | -0.001 | -0.001 | 0 | 0.23 | 0.22 | 0.26 | 0.25 | 0.19 | 0.21 |
| 1 | 227372471 | rs12035986 | 0.64 | 0.165 | -0.636 | 0.293 | 0 | -0.001 | -0.001 | 0 | 0.23 | 0.22 | 0.26 | 0.26 | 0.19 | 0.21 |
| 1 | 227374760 | rs12035998 | 1.784 | 0.17 | -0.581 | 0.294 | 0.012 | 0.043 | 0.043 | 0 | 0.24 | 0.15 | 0.17 | 0.13 | 0.19 | 0.18 |
| 1 | 227380754 | rs10916100 | -0.489 | 0.198 | -0.478 | 0.414 | -0.001 | 0.04 | 0.04 | 0 | 0.74 | 0.61 | 0.64 | 0.66 | 0.72 | 0.74 |
| 1 | 227383889 | rs6662583 | 1.047 | 0.217 | -0.527 | 0.388 | -0.004 | 0.009 | 0.009 | 0.059 | 0.42 | 0.37 | 0.33 | 0.34 | 0.39 | 0.33 |
| 1 | 227404082 | rs2813969 | -1.216 | 0.22 | -0.511 | 0.315 | -0.002 | 0.011 | 0.011 | 0.114 | 0.58 | 0.61 | 0.67 | 0.74 | 0.6 | 0.69 |
| 1 | 227405833 | rs2670452 | 0.26 | 0.258 | -0.475 | 0.4 | -0.001 | 0.032 | 0.032 | 0 | 0.26 | 0.38 | 0.37 | 0.34 | 0.28 | 0.26 |
| 1 | 227426205 | rs7519099 | 1.283 | 0.21 | -0.441 | 0.296 | -0.021 | -0.084 | -0.084 | 0.197 | 0.18 | 0.29 | 0.09 | 0 | 0.13 | 0.04 |
| 1 | 227450144 | rs12038272 | 2.43 | 0.082 | -0.836 | 0.202 | -0.009 | 0.008 | 0.008 | 0.02 | 0.17 | 0 | 0.13 | 0.13 | 0.14 | 0.14 |

| | | | | | | | | | | | | | | | | |
|---|---|---|---|---|---|---|---|---|---|---|---|---|---|---|---|---|
| 1 | 227452700 | rs1969230 | 0.307 | 0.163 | -0.637 | 0.289 | -0.001 | 0.032 | 0.032 | 0 | 0.26 | 0.38 | 0.36 | 0.34 | 0.28 | 0.26 |
| 1 | 227468945 | rs10916113 | 2.361 | 0.215 | -0.696 | 0.274 | -0.015 | -0.011 | -0.011 | 0.404 | 0.09 | 0.05 | 0.06 | 0.14 | 0.13 | 0.19 |
| 1 | 227477442 | rs7540055 | -0.074 | 0.187 | -0.577 | 0.198 | 0.002 | 0.002 | 0.002 | 0 | 0.57 | 0.48 | 0.49 | 0.53 | 0.51 | 0.61 |
| 1 | 227486433 | rs7519121 | -0.275 | 0.243 | -0.661 | 0.186 | 0.007 | 0.01 | 0.01 | 0 | 0.35 | 0.41 | 0.42 | 0.42 | 0.41 | 0.43 |
| 1 | 227494029 | rs7518813 | 2.355 | 0.248 | -0.672 | 0.181 | -0.015 | 0.011 | 0.011 | 0.085 | 0.09 | 0.03 | 0.05 | 0.08 | 0.13 | 0.17 |
| 1 | 227496438 | rs6660043 | 1.688 | 0.036 | -0.584 | 0.203 | 0.012 | 0.042 | 0.042 | 0 | 0.24 | 0.15 | 0.17 | 0.13 | 0.19 | 0.18 |
| 1 | 227499593 | rs11804709 | 0.036 | 0.116 | -0.4 | 0.478 | -0.001 | 0.031 | 0.031 | 0 | 0.26 | 0.38 | 0.36 | 0.34 | 0.28 | 0.26 |
| 1 | 227506948 | rs7549619 | 0.012 | 0.548 | -0.524 | 0.91 | 0.01 | 0.083 | 0.083 | 0 | 0.08 | 0.08 | 0.27 | 0.34 | 0.15 | 0.21 |
| 1 | 227513033 | rs10916122 | -0.327 | 0.268 | -0.494 | 0.517 | 0 | 0 | 0 | 0 | 0.27 | 0.25 | 0.29 | 0.27 | 0.25 | 0.27 |
| 1 | 227517737 | rs2953640 | -1.794 | 0.391 | -0.235 | 0.438 | -0.001 | 0 | 0 | 0 | 0.82 | 0.84 | 0.82 | 0.84 | 0.8 | 0.87 |
| 1 | 227518267 | rs4653828 | 1.262 | -0.085 | -0.353 | 0.295 | 0.002 | -0.044 | -0.044 | 0 | 0.29 | 0.19 | 0.41 | 0.45 | 0.28 | 0.32 |
| 1 | 227524891 | rs3127000 | 0.113 | -0.044 | -0.345 | 0.308 | -0.007 | 0.007 | 0.007 | 0 | 0.41 | 0.38 | 0.31 | 0.31 | 0.45 | 0.35 |
| 1 | 227529286 | rs3125123 | -0.019 | -0.111 | -0.387 | 0.232 | -0.003 | 0.006 | 0.006 | 0 | 0.43 | 0.4 | 0.32 | 0.34 | 0.46 | 0.4 |
| 1 | 227539205 | rs3014269 | -1.042 | -0.196 | -0.445 | 0.221 | -0.005 | 0.001 | 0.001 | 0 | 0.91 | 0.85 | 0.87 | 0.92 | 0.92 | 0.95 |
| 1 | 227542229 | rs4075196 | 0.635 | -0.096 | -0.567 | 0.045 | 0.005 | -0.02 | -0.02 | 0.043 | 0.42 | 0.26 | 0.52 | 0.44 | 0.42 | 0.37 |
| 1 | 227542811 | rs4075861 | 1.522 | -0.066 | -0.578 | 0.091 | -0.004 | -0.045 | -0.045 | 0.015 | 0.29 | 0.2 | 0.45 | 0.44 | 0.29 | 0.32 |
| 1 | 227547548 | rs3014252 | 0.235 | -0.237 | -0.547 | -0.115 | 0.022 | 0.051 | 0.051 | 0 | 0.65 | 0.77 | 0.72 | 0.8 | 0.68 | 0.73 |
| 1 | 227548499 | rs3014253 | 0.056 | -0.238 | -0.574 | -0.141 | 0.003 | 0.009 | 0.009 | 0 | 0.74 | 0.81 | 0.76 | 0.79 | 0.75 | 0.73 |
| 1 | 227552494 | rs4329477 | 1.455 | -0.315 | -0.247 | -0.044 | 0.007 | 0.023 | 0.023 | 0 | 0.13 | 0.09 | 0.07 | 0 | 0.13 | 0.05 |
| 1 | 227552917 | rs3014267 | 0.439 | -0.403 | -0.242 | -0.057 | 0.018 | 0.026 | 0.026 | 0 | 0.64 | 0.72 | 0.74 | 0.75 | 0.68 | 0.66 |
| 1 | 227554171 | rs2953655 | -0.366 | -0.33 | -0.026 | 0.133 | -0.033 | -0.002 | -0.002 | 0 | 0.79 | 0.7 | 0.83 | 0.8 | 0.86 | 0.8 |
| 1 | 227558822 | rs3014268 | 1.396 | 0.167 | 0.179 | 0.249 | 0.008 | 0.013 | 0.013 | 0.212 | 0.14 | 0.19 | 0.12 | 0.17 | 0.16 | 0.1 |
| 1 | 227563126 | rs10916131 | 0.339 | -0.02 | -0.005 | 0.198 | -0.008 | 0.002 | 0.002 | 0 | 0.16 | 0.17 | 0.18 | 0.22 | 0.14 | 0.23 |
| 1 | 227566134 | rs10916134 | 1.601 | 0.041 | 0.11 | 0.151 | 0.002 | 0.002 | 0.002 | 0.109 | 0.13 | 0.13 | 0.12 | 0.16 | 0.16 | 0.1 |
| 1 | 227576220 | rs7533453 | 0.076 | 0.02 | 0.279 | 0.095 | 0.013 | -0.007 | -0.007 | 0 | 0.23 | 0.45 | 0.18 | 0.2 | 0.22 | 0.19 |
| 1 | 227581017 | rs4606283 | 0.101 | -0.041 | 0.081 | 0.163 | -0.027 | 0.015 | 0.015 | 0 | 0.68 | 0.56 | 0.7 | 0.64 | 0.71 | 0.68 |
| 1 | 227582562 | rs6697269 | -0.866 | -0.045 | -0.139 | 0.232 | 0.009 | -0.006 | -0.006 | 0 | 0.85 | 0.88 | 0.83 | 0.8 | 0.88 | 0.78 |
| 1 | 227583800 | rs3014274 | 1.503 | 0.049 | -0.368 | 0.052 | -0.013 | -0.048 | -0.048 | 0 | 0.35 | 0.25 | 0.5 | 0.5 | 0.37 | 0.39 |
| 1 | 227584832 | rs3000787 | -0.885 | -0.355 | -0.278 | 0.306 | 0.017 | -0.011 | -0.011 | 0 | 0.85 | 0.9 | 0.76 | 0.8 | 0.88 | 0.79 |
| 1 | 227585983 | rs6670351 | 0.554 | -0.339 | -0.355 | 0.006 | 0.002 | 0.026 | 0.026 | 0 | 0.18 | 0.15 | 0.11 | 0.05 | 0.18 | 0.15 |
| 1 | 227586401 | rs6659701 | 0.252 | -0.286 | -0.483 | 0.091 | -0.022 | -0.003 | -0.003 | 0 | 0.62 | 0.7 | 0.51 | 0.44 | 0.57 | 0.59 |
| 1 | 227595966 | rs4601553 | 0.924 | -0.274 | -0.494 | 0.092 | 0.023 | 0.112 | 0.112 | 0 | 0.43 | 0.21 | 0.36 | 0.28 | 0.4 | 0.43 |
| 1 | 227598169 | rs3000784 | -0.765 | -0.39 | -0.13 | 0.234 | -0.038 | 0.011 | 0.011 | 0 | 0.82 | 0.74 | 0.75 | 0.81 | 0.87 | 0.79 |
| 1 | 227598446 | rs3014290 | -0.734 | -0.389 | -0.13 | 0.248 | -0.038 | 0.012 | 0.012 | 0 | 0.82 | 0.73 | 0.75 | 0.81 | 0.87 | 0.79 |
| 1 | 227599079 | rs3000790 | -0.16 | -0.457 | -0.28 | 0.076 | -0.057 | -0.033 | -0.033 | 0 | 0.38 | 0.23 | 0.33 | 0.41 | 0.44 | 0.31 |
| 1 | 227599174 | rs3000791 | -0.652 | -0.459 | -0.306 | 0.054 | -0.032 | 0.006 | 0.006 | 0 | 0.81 | 0.73 | 0.74 | 0.81 | 0.85 | 0.67 |
| 1 | 227603016 | rs10916142 | 1.919 | -0.239 | -0.306 | 0.179 | 0.012 | 0.007 | 0.007 | 0.914 | 0.1 | 0.11 | 0.11 | 0.14 | 0.15 | 0.1 |
| 1 | 227603207 | rs12025219 | 2.669 | -0.293 | -0.417 | 0.068 | 0.02 | 0.028 | 0.028 | 0 | 0.26 | 0.07 | 0.25 | 0.23 | 0.23 | 0.29 |
| 1 | 227603763 | rs3014294 | 0.247 | -0.162 | -0.313 | 0.141 | -0.044 | -0.043 | -0.043 | 3.902 | 0.27 | 0.18 | 0.27 | 0.36 | 0.33 | 0.21 |
| 1 | 227608904 | rs3000802 | 1.783 | 0.069 | -0.412 | 0.316 | -0.024 | -0.024 | -0.024 | 0 | 0.07 | 0 | 0.15 | 0.28 | 0.1 | 0.02 |
| 1 | 227610592 | rs6670816 | 1.926 | 0.075 | -0.278 | 0.161 | 0.001 | -0.001 | -0.001 | 0 | 0.11 | 0.12 | 0.07 | 0.04 | 0.11 | 0.09 |
| 1 | 227611295 | rs12028062 | 1.725 | 0.162 | -0.147 | 0.265 | 0.043 | 0.042 | 0.042 | 0.559 | 0.1 | 0.19 | 0.11 | 0.16 | 0.15 | 0.1 |
| 1 | 227617519 | rs6664147 | 1.013 | 0.107 | -0.426 | 0.097 | -0.04 | 0.036 | 0.036 | 0 | 0.2 | 0.13 | 0.12 | 0.08 | 0.23 | 0.19 |
| 1 | 227617761 | rs3014302 | -0.914 | -0.309 | -0.262 | -0.131 | 0.03 | 0.018 | 0.018 | 0 | 0.51 | 0.62 | 0.46 | 0.58 | 0.58 | 0.46 |
| 1 | 227618146 | rs7533662 | 0.518 | -0.253 | -0.259 | -0.048 | 0.007 | 0.008 | 0.008 | 0 | 0.57 | 0.4 | 0.65 | 0.75 | 0.63 | 0.58 |
| 1 | 227618683 | rs12125212 | 2.361 | -0.141 | -0.417 | -0.144 | -0.003 | 0.028 | 0.028 | 0 | 0.25 | 0.05 | 0.25 | 0.23 | 0.24 | 0.24 |
| 1 | 227618915 | rs16847813 | 0.682 | 0.095 | -0.269 | -0.11 | 0.018 | -0.004 | -0.004 | 0 | 0.21 | 0.28 | 0.17 | 0.2 | 0.26 | 0.19 |
| 1 | 227619332 | rs12126205 | 0.965 | -0.038 | -0.263 | -0.14 | -0.002 | -0.008 | -0.008 | 0 | 0.39 | 0.38 | 0.33 | 0.28 | 0.33 | 0.37 |
| 1 | 227622011 | rs7546172 | 0.585 | -0.004 | -0.482 | -0.02 | 0.011 | 0.014 | 0.014 | 0 | 0.48 | 0.44 | 0.49 | 0.39 | 0.41 | 0.53 |
| 1 | 227622222 | rs11586581 | 0.065 | 0.169 | -0.144 | 0.173 | 0.006 | 0.014 | 0.014 | 0.069 | 0.69 | 0.65 | 0.62 | 0.45 | 0.62 | 0.71 |
| 1 | 227624541 | rs10916146 | -0.292 | -0.308 | 0.072 | 0.658 | 0.002 | -0.084 | -0.084 | 0 | 0.81 | 0.93 | 0.75 | 0.61 | 0.8 | 0.81 |
| 1 | 227626662 | rs16847833 | 0.94 | 0.16 | 0.434 | 0.324 | -0.002 | -0.004 | -0.004 | 1.797 | 0.12 | 0.15 | 0.14 | 0.14 | 0.12 | 0.18 |
| 1 | 227636066 | rs903687 | 0.405 | 0.153 | 0.515 | 0.355 | -0.008 | 0.022 | 0.022 | 0 | 0.1 | 0.14 | 0.18 | 0.15 | 0.09 | 0.17 |
| 1 | 227637092 | rs2819516 | 0.542 | 0.088 | 0.578 | 0.335 | -0.027 | 0.018 | 0.018 | 0 | 0.76 | 0.57 | 0.66 | 0.69 | 0.78 | 0.59 |
| 1 | 227643154 | rs2819519 | 0.799 | 0.175 | 0.568 | 0.315 | -0.011 | 0.003 | 0.003 | 0 | 0.73 | 0.55 | 0.62 | 0.69 | 0.71 | 0.55 |
| 1 | 227654294 | rs1495855 | 0.005 | 0.115 | 0.424 | 0.39 | 0 | 0.006 | 0.006 | 0 | 0.12 | 0.13 | 0.2 | 0.16 | 0.15 | 0.22 |
| 1 | 227655222 | rs953537 | -0.765 | 0.364 | 0.211 | 0.146 | 0.002 | -0.017 | -0.017 | 0 | 0.32 | 0.38 | 0.26 | 0.2 | 0.31 | 0.37 |
| 1 | 227658753 | rs1587158 | -0.737 | 0.353 | 0.214 | 0.184 | -0.007 | -0.005 | -0.005 | 0.562 | 0.86 | 0.76 | 0.85 | 0.84 | 0.87 | 0.76 |
| 1 | 227668181 | rs2819522 | -0.232 | 0.428 | 0.196 | 0.019 | 0.012 | -0.043 | -0.043 | 0 | 0.8 | 0.69 | 0.87 | 0.91 | 0.81 | 0.76 |
| 1 | 227673227 | rs1390403 | -0.334 | 0.448 | 0.033 | 0.029 | 0.005 | -0.02 | -0.02 | 0.107 | 0.81 | 0.73 | 0.88 | 0.91 | 0.82 | 0.76 |
| 1 | 227677506 | rs903698 | 0.771 | 0.303 | -0.067 | 0.111 | -0.003 | 0.005 | 0.005 | 0 | 0.59 | 0.56 | 0.63 | 0.74 | 0.56 | 0.48 |
| 1 | 227689829 | rs2819493 | -1.109 | -0.023 | 0.001 | 0.462 | -0.001 | 0.008 | 0.008 | 0 | 0.37 | 0.62 | 0.41 | 0.52 | 0.34 | 0.4 |
| 1 | 227696594 | rs1495842 | 1.041 | -0.032 | 0.309 | 0.543 | 0.012 | 0.005 | 0.005 | 0 | 0.54 | 0.35 | 0.58 | 0.41 | 0.61 | 0.47 |
| 1 | 227697381 | rs2644140 | -1.244 | -0.054 | 0.608 | 0.541 | -0.004 | 0.003 | 0.003 | 0 | 0.44 | 0.52 | 0.4 | 0.56 | 0.38 | 0.47 |
| 1 | 227703397 | rs2644162 | -0.391 | -0.286 | 0.664 | 0.579 | 0.039 | -0.05 | -0.05 | 1.733 | 0.29 | 0.13 | 0.3 | 0.46 | 0.21 | 0.29 |
| 1 | 227707618 | rs2819506 | 1.593 | -0.363 | 0.601 | 0.691 | -0.003 | -0.011 | -0.011 | 0 | 0.06 | 0 | 0.11 | 0.06 | 0.06 | 0.07 |
| 1 | 227710381 | rs2453682 | -0.441 | -0.325 | 0.696 | 0.672 | 0.039 | -0.05 | -0.05 | 0.256 | 0.29 | 0.13 | 0.29 | 0.46 | 0.21 | 0.29 |
| 1 | 227714296 | rs2814084 | -0.472 | -0.338 | 0.915 | 0.67 | 0.025 | -0.028 | -0.028 | 0.021 | 0.29 | 0.18 | 0.3 | 0.46 | 0.21 | 0.29 |
| 1 | 227724401 | rs2819498 | -0.493 | -0.348 | 0.887 | 0.685 | 0.026 | -0.027 | -0.027 | 0 | 0.29 | 0.17 | 0.3 | 0.46 | 0.21 | 0.29 |

| | | | | | | | | | | | | | | | | |
|---|---|---|---|---|---|---|---|---|---|---|---|---|---|---|---|---|
| 1 | 227736363 | rs1495849 | -0.608 | -0.349 | 0.888 | 0.657 | 0.025 | -0.032 | -0.032 | 0 | 0.3 | 0.2 | 0.3 | 0.49 | 0.21 | 0.36 |
| 1 | 227736946 | rs10495276 | 1.456 | -0.305 | 0.872 | 0.627 | 0.016 | 0.007 | 0.007 | 0 | 0.53 | 0.32 | 0.58 | 0.41 | 0.61 | 0.47 |
| 1 | 227750068 | rs6696239 | 0.109 | -0.253 | 1.224 | 0.578 | 0.037 | -0.027 | -0.027 | 0.018 | 0.16 | 0.37 | 0.11 | 0.1 | 0.18 | 0.18 |
| 1 | 227753072 | rs4653856 | 2.234 | -0.259 | 1.022 | 0.566 | -0.035 | 0.104 | 0.104 | 0 | 0.38 | 0.12 | 0.29 | 0.17 | 0.43 | 0.25 |
| 1 | 227758072 | rs12025815 | 1.338 | -0.358 | 1.092 | 0.666 | 0.018 | 0.007 | 0.007 | 0 | 0.53 | 0.3 | 0.58 | 0.42 | 0.6 | 0.46 |
| 1 | 227772396 | rs3942992 | -0.328 | -0.349 | 1.156 | 0.63 | 0.066 | -0.037 | -0.037 | 0 | 0.83 | 0.54 | 0.88 | 0.9 | 0.82 | 0.82 |
| 1 | 227796748 | rs16847992 | -0.314 | -0.345 | 0.979 | 0.651 | 0.041 | -0.045 | -0.045 | 0 | 0.29 | 0.15 | 0.29 | 0.45 | 0.21 | 0.29 |
| 1 | 227798565 | rs10799428 | 0.401 | -0.299 | 1.085 | 0.666 | 0.013 | -0.011 | -0.011 | 2.463 | 0.15 | 0.25 | 0.1 | 0.1 | 0.18 | 0.18 |
| 1 | 227819490 | rs12070002 | -1.295 | -0.307 | 0.77 | 0.667 | 0.006 | -0.004 | -0.004 | 0.348 | 0.47 | 0.63 | 0.42 | 0.55 | 0.4 | 0.49 |
| 1 | 227819514 | rs12070036 | -1.295 | -0.307 | 0.77 | 0.667 | 0.006 | -0.004 | -0.004 | 3.681 | 0.47 | 0.63 | 0.42 | 0.55 | 0.4 | 0.49 |
| 1 | 227821255 | rs1810680 | -2.367 | -0.285 | 0.478 | 0.606 | -0.068 | 0.105 | 0.105 | 0 | 0.63 | 0.94 | 0.72 | 0.83 | 0.57 | 0.77 |
| 1 | 227834400 | rs6426468 | -2.363 | -0.276 | 0.539 | 0.637 | -0.057 | 0.1 | 0.1 | 0 | 0.63 | 0.92 | 0.72 | 0.82 | 0.57 | 0.77 |
| 1 | 227839384 | rs12096675 | -0.379 | -0.371 | 0.622 | 0.679 | 0.027 | -0.028 | -0.028 | 0 | 0.3 | 0.17 | 0.3 | 0.45 | 0.22 | 0.31 |
| 1 | 227862055 | rs1923821 | -0.318 | -0.379 | 0.729 | 0.675 | 0.01 | -0.011 | -0.011 | 0.017 | 0.3 | 0.21 | 0.3 | 0.45 | 0.22 | 0.31 |
| 1 | 227880275 | rs6671682 | 0.06 | -0.477 | 0.598 | 0.694 | 0.007 | -0.003 | -0.003 | 6.229 | 0.15 | 0.1 | 0.23 | 0.41 | 0.11 | 0.2 |
| 1 | 227885726 | rs1923822 | 0.035 | -0.371 | 0.8 | 0.664 | 0.001 | 0.035 | 0.035 | 0 | 0.16 | 0.28 | 0.29 | 0.24 | 0.17 | 0.22 |
| 1 | 227909907 | rs1475180 | 0.093 | -0.367 | 0.749 | 0.666 | 0.004 | 0.047 | 0.047 | 0 | 0.16 | 0.29 | 0.29 | 0.24 | 0.17 | 0.22 |
| 1 | 227912474 | rs12097239 | -0.147 | -0.294 | 0.828 | 0.579 | 0.017 | -0.019 | -0.019 | 0 | 0.22 | 0.37 | 0.22 | 0.16 | 0.25 | 0.25 |
| 1 | 227917645 | rs1570429 | 0.952 | -0.283 | 0.534 | 0.553 | 0 | -0.014 | -0.014 | 0 | 0.15 | 0.12 | 0.07 | 0.05 | 0.11 | 0.11 |
| 1 | 227918562 | rs6664307 | -0.217 | -0.247 | 0.701 | 0.57 | 0.029 | -0.032 | -0.032 | 0 | 0.23 | 0.43 | 0.22 | 0.16 | 0.25 | 0.22 |
| 1 | 227920061 | rs1060295 | -2.439 | -0.312 | 0.5 | 0.703 | -0.098 | 0.07 | 0.07 | 0.005 | 0.62 | 0.95 | 0.71 | 0.79 | 0.56 | 0.66 |
| 1 | 227920103 | rs3087908 | 0.349 | -0.287 | 0.614 | 0.82 | -0.003 | -0.011 | -0.011 | 0.27 | 0.13 | 0.14 | 0.27 | 0.21 | 0.15 | 0.14 |
| 1 | 227920447 | rs751748 | 0.349 | -0.287 | 0.614 | 0.82 | -0.003 | -0.011 | -0.011 | 1.272 | 0.13 | 0.14 | 0.27 | 0.21 | 0.15 | 0.14 |
| 1 | 227934606 | rs16848110 | 0.287 | -0.387 | 0.705 | 0.734 | -0.032 | -0.062 | -0.062 | 0 | 0.19 | 0.34 | 0.09 | 0.08 | 0.11 | 0.17 |
| 1 | 227934750 | rs16848112 | -0.074 | -0.463 | 0.745 | 0.801 | -0.033 | 0.085 | 0.085 | 3.072 | 0.1 | 0.17 | 0.14 | 0.35 | 0.05 | 0.12 |
| 1 | 227935444 | rs2236359 | -1.034 | -0.429 | 0.701 | 0.782 | 0.023 | 0.009 | 0.009 | 0 | 0.34 | 0.48 | 0.49 | 0.37 | 0.4 | 0.36 |
| 1 | 227935762 | rs2236358 | 0.579 | -0.409 | 0.617 | 0.806 | -0.019 | 0.079 | 0.079 | 1.027 | 0.08 | 0.1 | 0.13 | 0.35 | 0.05 | 0.12 |
| 1 | 227942842 | rs7533588 | -2.42 | -0.354 | 0.524 | 0.758 | -0.039 | 0.078 | 0.078 | 0 | 0.61 | 0.88 | 0.72 | 0.79 | 0.56 | 0.65 |
| 1 | 227943959 | rs12064596 | 0.45 | -0.229 | 0.401 | 0.902 | -0.026 | 0.106 | 0.106 | 1.516 | 0.08 | 0.14 | 0.13 | 0.35 | 0.05 | 0.12 |
| 1 | 227947434 | rs748405 | 0.925 | -0.013 | 0.15 | 0.385 | 0.004 | -0.001 | -0.001 | 0.117 | 0.16 | 0.1 | 0.07 | 0.05 | 0.11 | 0.11 |
| 1 | 227957591 | rs10916205 | -0.99 | 0.179 | 0.101 | 0.472 | 0.016 | 0.009 | 0.009 | 0 | 0.32 | 0.41 | 0.48 | 0.36 | 0.39 | 0.36 |
| 1 | 227959417 | rs6683757 | -1.013 | 0.309 | -0.039 | 0.548 | 0.005 | 0.003 | 0.003 | 0 | 0.3 | 0.34 | 0.46 | 0.36 | 0.36 | 0.32 |
| 1 | 227963491 | rs754224 | 0.428 | 0.481 | -0.306 | 0.682 | -0.031 | -0.053 | -0.053 | 0.394 | 0.11 | 0.03 | 0.26 | 0.2 | 0.14 | 0.14 |
| 1 | 227966581 | rs10799447 | -1.211 | 0.411 | -0.049 | 0.712 | 0 | 0.001 | 0.001 | 0 | 0.53 | 0.51 | 0.55 | 0.38 | 0.5 | 0.51 |
| 1 | 227995966 | rs4394649 | 0.019 | 1.074 | -0.201 | 1.312 | -0.04 | -0.044 | -0.044 | 0.111 | 0.07 | 0.01 | 0.24 | 0.19 | 0.13 | 0.13 |
| 1 | 228003247 | rs10916214 | 0.019 | 1.074 | -0.201 | 1.312 | -0.04 | -0.044 | -0.044 | 0 | 0.07 | 0.01 | 0.24 | 0.19 | 0.13 | 0.13 |
| 1 | 228006435 | rs10916216 | -0.609 | 0.803 | -0.124 | 0.915 | 0.004 | 0.014 | 0.014 | 0 | 0.27 | 0.27 | 0.18 | 0.12 | 0.25 | 0.18 |
| 1 | 228010782 | rs7410917 | 0.421 | 0.747 | -0.136 | 0.897 | 0.008 | 0.036 | 0.036 | 0 | 0.73 | 0.79 | 0.82 | 0.87 | 0.75 | 0.82 |
| 1 | 228017395 | rs4622126 | -0.55 | 0.825 | 0.075 | 0.837 | 0.024 | -0.012 | -0.012 | 0.052 | 0.1 | 0.23 | 0.08 | 0.07 | 0.14 | 0.11 |
| 1 | 228019842 | rs6426478 | 0.456 | 0.65 | 0.085 | 1.406 | 0.015 | -0.045 | -0.045 | 0 | 0.7 | 0.81 | 0.53 | 0.49 | 0.73 | 0.79 |
| 1 | 228024977 | rs6426481 | -0.234 | 0.617 | 0.127 | 1.406 | 0.019 | -0.015 | -0.015 | 1.567 | 0.77 | 0.83 | 0.78 | 0.68 | 0.86 | 0.92 |
| 1 | 228027310 | rs12565154 | -1.336 | 0.737 | 0.744 | 1.45 | 0.08 | 0.183 | 0.183 | 5.699 | 0.07 | 0.5 | 0.08 | 0.17 | 0.05 | 0.22 |
| 1 | 228029804 | rs12043266 | 1.531 | 0.554 | -0.092 | 1.314 | -0.019 | 0.137 | 0.137 | 2.936 | 0.49 | 0.09 | 0.36 | 0.25 | 0.54 | 0.49 |
| 1 | 228029955 | rs12129350 | 0.695 | 0.638 | 0.088 | 1.474 | 0.012 | 0.016 | 0.016 | 0 | 0.07 | 0.12 | 0.05 | 0.1 | 0.1 | 0.1 |
| 1 | 228037278 | rs11589239 | 1.507 | 0.637 | -0.164 | 1.369 | -0.018 | 0.139 | 0.139 | 3.035 | 0.5 | 0.09 | 0.36 | 0.25 | 0.55 | 0.49 |
| 1 | 228038399 | rs4233229 | 0.161 | 0.526 | 0.089 | 1.532 | -0.042 | -0.05 | -0.05 | 0.539 | 0.09 | 0.01 | 0.25 | 0.2 | 0.13 | 0.13 |
| 1 | 228045508 | rs2872243 | 0.198 | 0.18 | 0.276 | 1.524 | 0.025 | 0.002 | 0.002 | 0.614 | 0.7 | 0.79 | 0.74 | 0.69 | 0.8 | 0.9 |
| 1 | 228046401 | rs12135131 | -1.226 | 0.81 | 0.927 | 1.825 | 0.039 | 0.054 | 0.054 | 0 | 0.13 | 0.26 | 0.3 | 0.24 | 0.16 | 0.19 |
| 1 | 228050227 | rs12567590 | -0.649 | 0.451 | 0.54 | 1.955 | -0.001 | 0.072 | 0.072 | 0.731 | 0.1 | 0.17 | 0.16 | 0.26 | 0.1 | 0.19 |
| 1 | 228072809 | rs4074668 | -0.106 | 0.802 | 0.748 | 2.093 | 0 | 0 | 0 | 0 | 0.09 | 0.1 | 0.05 | 0.07 | 0.11 | 0.09 |
| 1 | 228079192 | rs10799454 | -0.098 | 0.543 | 0.715 | 1.94 | -0.017 | -0.017 | -0.017 | 0.048 | 0.15 | 0.24 | 0.17 | 0.15 | 0.12 | 0.2 |
| 1 | 228080785 | rs20488 | -1.697 | 0.285 | -0.257 | 1.849 | 0.019 | -0.043 | -0.043 | 0.015 | 0.32 | 0.83 | 0.31 | 0.43 | 0.25 | 0.36 |
| 1 | 228087561 | rs4653883 | -0.932 | 0.118 | -0.346 | 1.983 | -0.005 | 0.007 | 0.007 | 0.013 | 0.21 | 0.7 | 0.23 | 0.44 | 0.13 | 0.34 |
| 1 | 228098340 | rs4518905 | -0.75 | 0.771 | -0.881 | 1.979 | -0.002 | 0.026 | 0.026 | 0 | 0.11 | 0.08 | 0.22 | 0.4 | 0.14 | 0.11 |
| 1 | 228101119 | rs10916243 | 1.022 | 0.961 | -1.822 | 1.163 | -0.073 | -0.057 | -0.057 | 0 | 0.4 | 0.04 | 0.42 | 0.33 | 0.42 | 0.4 |
| 1 | 228103781 | rs1009658 | -0.009 | 0.507 | -1.769 | 0.543 | 0.013 | 0.049 | 0.049 | 0 | 0.5 | 0.66 | 0.53 | 0.64 | 0.53 | 0.4 |
| 1 | 228104375 | rs12748472 | 0.42 | 0.374 | -2.367 | 0.685 | -0.118 | -0.049 | -0.049 | 0 | 0.31 | 0.07 | 0.31 | 0.26 | 0.37 | 0.16 |
| 1 | 228106921 | rs12725747 | 1.066 | -0.051 | -2.57 | 0.329 | 0.012 | 0.021 | 0.021 | 7.538 | 0.14 | 0.08 | 0.07 | 0.09 | 0.09 | 0.15 |
| 1 | 228108703 | rs10916244 | 0.582 | 0.27 | -2.383 | 0.446 | -0.096 | -0.032 | -0.032 | 0 | 0.31 | 0.09 | 0.31 | 0.26 | 0.36 | 0.16 |
| 1 | 228110064 | rs10127943 | 0.142 | 0.355 | -1.833 | 0.432 | 0 | 0.015 | 0.015 | 0 | 0.33 | 0.56 | 0.35 | 0.33 | 0.32 | 0.26 |
| 1 | 228110734 | rs12046421 | 0.663 | -0.115 | -1.919 | -0.212 | 0 | 0.056 | 0.056 | 12.854 | 0.29 | 0.16 | 0.2 | 0.11 | 0.23 | 0.35 |
| 1 | 228115129 | rs3949294 | 1.541 | 0.078 | -1.674 | 0.068 | -0.002 | -0.006 | -0.006 | 0.038 | 0.08 | 0.04 | 0.06 | 0.1 | 0.07 | 0.12 |
| 1 | 228116831 | rs592736 | -0.488 | -0.328 | -0.985 | 0.458 | -0.005 | 0.002 | 0.002 | 0.236 | 0.45 | 0.66 | 0.55 | 0.39 | 0.48 | 0.42 |
| 1 | 228127227 | rs680997 | -1.746 | -0.317 | -0.732 | 0.416 | 0 | -0.008 | -0.008 | 0 | 0.92 | 0.91 | 0.96 | 0.96 | 0.92 | 0.85 |
| 1 | 228139672 | rs681239 | -1.417 | -0.162 | -0.622 | 0.797 | -0.006 | -0.044 | -0.044 | 0.49 | 0.65 | 0.89 | 0.65 | 0.61 | 0.67 | 0.55 |
| 1 | 228140460 | rs6665129 | -0.798 | -0.157 | -0.397 | 0.832 | -0.029 | 0.149 | 0.149 | 0.318 | 0.14 | 0.41 | 0.2 | 0.31 | 0.08 | 0.24 |
| 1 | 228148627 | rs11579417 | 1.821 | -0.077 | -0.662 | 0.626 | 0 | 0 | 0 | 0.04 | 0.08 | 0.05 | 0.05 | 0.09 | 0.07 | 0.11 |
| 1 | 228153917 | rs849898 | 0.609 | -0.07 | -0.85 | 0.54 | -0.148 | 0.113 | 0.113 | 0 | 0.3 | 0.05 | 0.32 | 0.08 | 0.39 | 0.17 |
| 1 | 228174528 | rs6426488 | 0.121 | -0.116 | -0.498 | 0.854 | -0.013 | 0.054 | 0.054 | 0.524 | 0.11 | 0.22 | 0.2 | 0.28 | 0.08 | 0.23 |

| | | | | | | | | | | | | | | | | |
|---|---|---|---|---|---|---|---|---|---|---|---|---|---|---|---|---|
| 1 | 228188545 | rs11589040 | 1.596 | -0.099 | -0.574 | 0.827 | -0.009 | -0.012 | -0.012 | 0.674 | 0.08 | 0.02 | 0.1 | 0.11 | 0.07 | 0.12 |
| 1 | 228193857 | rs964941 | 0.589 | 0.102 | -0.476 | 0.652 | -0.001 | 0 | 0 | 0.458 | 0.57 | 0.35 | 0.45 | 0.61 | 0.57 | 0.62 |
| 1 | 228206980 | rs3094913 | 0.687 | 0.079 | -0.461 | 0.627 | -0.017 | 0.018 | 0.018 | 1.924 | 0.55 | 0.16 | 0.43 | 0.61 | 0.57 | 0.61 |
| 1 | 228210179 | rs3094911 | -0.536 | 0.115 | -0.329 | 0.649 | 0.003 | 0 | 0 | 0 | 0.69 | 0.63 | 0.58 | 0.71 | 0.65 | 0.73 |
| 1 | 228216997 | rs708122 | 1.309 | 0.151 | -0.736 | 0.705 | -0.006 | -0.018 | -0.018 | 0 | 0.37 | 0.14 | 0.33 | 0.36 | 0.37 | 0.44 |
| 1 | 228219882 | rs10916258 | 1.073 | 0.22 | -0.591 | 0.763 | -0.005 | 0.006 | 0.006 | 2.669 | 0.2 | 0.23 | 0.2 | 0.24 | 0.2 | 0.16 |
| 1 | 228223036 | rs3121309 | -0.486 | 0.663 | -0.662 | 0.616 | -0.002 | -0.002 | -0.002 | 0.067 | 0.77 | 0.79 | 0.77 | 0.74 | 0.79 | 0.8 |
| 1 | 228224153 | rs1745423 | 0.233 | 0.427 | -0.543 | 0.915 | -0.014 | -0.009 | -0.009 | 0 | 0.67 | 0.76 | 0.62 | 0.64 | 0.63 | 0.67 |
| 1 | 228224824 | rs3121310 | -0.101 | 0.349 | -0.274 | 0.856 | 0.009 | 0.017 | 0.017 | 0 | 0.72 | 0.45 | 0.66 | 0.7 | 0.69 | 0.77 |
| 1 | 228227477 | rs7539664 | -0.725 | 0.327 | -0.191 | 0.983 | -0.015 | 0.084 | 0.084 | 0.864 | 0.13 | 0.42 | 0.17 | 0.23 | 0.08 | 0.15 |
| 1 | 228229067 | rs1034792 | 0.829 | 0.414 | -0.32 | 1.027 | 0 | 0.001 | 0.001 | 1.043 | 0.09 | 0.12 | 0.13 | 0.23 | 0.07 | 0.15 |
| 1 | 228239344 | rs882451 | 1.514 | 0.43 | -0.414 | 0.943 | -0.016 | -0.017 | -0.017 | 0 | 0.08 | 0.03 | 0.11 | 0.11 | 0.07 | 0.12 |
| 1 | 228247351 | rs752107 | -0.521 | 0.369 | -0.335 | 0.954 | -0.011 | 0.016 | 0.016 | 0 | 0.75 | 0.82 | 0.72 | 0.82 | 0.69 | 0.8 |
| 1 | 228260728 | rs1188977 | -0.039 | 0.282 | -0.133 | 1.114 | -0.002 | 0.144 | 0.144 | 3.366 | 0.09 | 0.37 | 0.15 | 0.22 | 0.06 | 0.09 |
| 1 | 228261291 | rs3738680 | 1.113 | 0.282 | -0.142 | 1.093 | 0.004 | -0.013 | -0.013 | 0 | 0.07 | 0.06 | 0.09 | 0.19 | 0.06 | 0.07 |
| 1 | 228268492 | rs3820624 | 1.236 | 0.303 | -0.212 | 1.104 | 0.004 | -0.011 | -0.011 | 0 | 0.07 | 0.05 | 0.09 | 0.19 | 0.06 | 0.07 |
| 1 | 228277690 | rs1188975 | -0.357 | 0.227 | -0.081 | 1.096 | -0.024 | 0.092 | 0.092 | 1.785 | 0.11 | 0.37 | 0.15 | 0.23 | 0.06 | 0.09 |
| 1 | 228279023 | rs1048503 | 1.131 | 0.257 | -0.188 | 1.091 | 0.004 | -0.01 | -0.01 | 0 | 0.07 | 0.07 | 0.09 | 0.19 | 0.06 | 0.07 |
| 1 | 228282695 | rs3768331 | 0.872 | 0.317 | -0.356 | 1.046 | -0.039 | 0.019 | 0.019 | 3.496 | 0.44 | 0.15 | 0.37 | 0.35 | 0.42 | 0.55 |
| 1 | 228294300 | rs12564763 | 0.533 | 0.273 | -0.11 | 0.974 | -0.006 | 0.006 | 0.006 | 0 | 0.63 | 0.41 | 0.6 | 0.61 | 0.64 | 0.71 |
| 1 | 228296745 | rs849749 | -0.545 | 0.307 | -0.03 | 0.974 | -0.003 | 0.004 | 0.004 | 0 | 0.74 | 0.78 | 0.73 | 0.82 | 0.7 | 0.8 |
| 1 | 228297613 | rs1620734 | 0.207 | 0.167 | -0.082 | 1.063 | -0.002 | 0.142 | 0.142 | 2.995 | 0.09 | 0.33 | 0.15 | 0.23 | 0.06 | 0.1 |
| 1 | 228303048 | rs10916266 | 1.227 | 0.037 | -0.18 | 1.115 | -0.004 | 0.013 | 0.013 | 3.808 | 0.07 | 0.08 | 0.07 | 0.2 | 0.04 | 0.06 |
| 1 | 228322826 | rs6426492 | 0.511 | 0.17 | -0.179 | 0.991 | -0.018 | 0.01 | 0.01 | 0 | 0.23 | 0.18 | 0.25 | 0.18 | 0.3 | 0.2 |
| 1 | 228326492 | rs11809443 | 1.117 | 0.25 | -0.093 | 1.172 | -0.001 | 0.002 | 0.002 | 1.01 | 0.19 | 0.23 | 0.25 | 0.25 | 0.21 | 0.16 |
| 1 | 228331040 | rs2313602 | 0.475 | 0.133 | -0.128 | 1.151 | -0.003 | 0.01 | 0.01 | 0 | 0.63 | 0.41 | 0.59 | 0.59 | 0.63 | 0.71 |
| 1 | 228333652 | rs11577976 | 1.417 | 0.094 | -0.123 | 1.163 | -0.002 | 0.013 | 0.013 | 4.376 | 0.07 | 0.07 | 0.07 | 0.2 | 0.04 | 0.06 |
| 1 | 228334277 | rs10916268 | 0.493 | 0.106 | -0.13 | 1.11 | -0.003 | 0.011 | 0.011 | 0 | 0.63 | 0.41 | 0.59 | 0.59 | 0.63 | 0.72 |
| 1 | 228344000 | rs6663726 | -0.294 | 0.083 | -0.086 | 1.04 | -0.038 | 0.146 | 0.146 | 3.854 | 0.11 | 0.4 | 0.13 | 0.24 | 0.04 | 0.07 |
| 1 | 228350774 | rs12134624 | 0.448 | 0.094 | -0.141 | 1.056 | -0.021 | 0.033 | 0.033 | 0 | 0.26 | 0.17 | 0.28 | 0.14 | 0.33 | 0.21 |
| 1 | 228359775 | rs2066237 | 0.447 | 0.113 | -0.147 | 1.071 | -0.003 | -0.001 | -0.001 | 0 | 0.63 | 0.42 | 0.59 | 0.62 | 0.63 | 0.72 |
| 1 | 228364605 | rs4653915 | -0.147 | 0.24 | -0.418 | 1.112 | -0.005 | 0.005 | 0.005 | 0 | 0.31 | 0.25 | 0.38 | 0.3 | 0.37 | 0.25 |
| 1 | 228372062 | rs1059778 | -0.261 | 0.046 | -0.397 | 1.13 | -0.006 | -0.008 | -0.008 | 0 | 0.32 | 0.28 | 0.38 | 0.39 | 0.37 | 0.28 |
| 1 | 228376627 | rs7539762 | 0.411 | 0.053 | -0.508 | 1.101 | -0.02 | 0.034 | 0.034 | 0 | 0.26 | 0.17 | 0.29 | 0.15 | 0.33 | 0.21 |
| 1 | 228377913 | rs12084338 | 0.263 | 0.057 | -0.521 | 1.108 | -0.014 | 0.015 | 0.015 | 0 | 0.27 | 0.2 | 0.29 | 0.21 | 0.33 | 0.22 |
| 1 | 228381518 | rs12120621 | 1.416 | 0.096 | -1.269 | 1.046 | -0.08 | 0 | 0 | 0 | 0.22 | 0.01 | 0.15 | 0.13 | 0.2 | 0.32 |
| 1 | 228383202 | rs12142165 | 0.719 | 0.225 | -0.956 | 1.044 | -0.021 | -0.02 | -0.02 | 0.764 | 0.43 | 0.14 | 0.32 | 0.37 | 0.38 | 0.56 |
| 1 | 228391808 | rs11800309 | 1.315 | 0.487 | -1.057 | 0.995 | -0.046 | -0.043 | -0.043 | 0.016 | 0.18 | 0.03 | 0.25 | 0.25 | 0.25 | 0.18 |
| 1 | 228402047 | rs1620111 | -0.28 | 0.277 | -0.675 | 0.874 | -0.004 | 0.012 | 0.012 | 0 | 0.68 | 0.68 | 0.64 | 0.82 | 0.64 | 0.8 |
| 1 | 228402121 | rs1771487 | -0.28 | 0.277 | -0.675 | 0.874 | -0.004 | 0.012 | 0.012 | 0 | 0.68 | 0.68 | 0.64 | 0.82 | 0.64 | 0.8 |
| 1 | 228406717 | rs10157526 | -0.496 | 0.187 | -0.397 | 1.1 | -0.007 | 0.148 | 0.148 | 2.689 | 0.05 | 0.24 | 0.02 | 0.18 | 0.01 | 0.05 |
| 1 | 228412308 | rs1757152 | -0.485 | 0.238 | -0.486 | 1.03 | -0.004 | -0.005 | -0.005 | 0.038 | 0.37 | 0.59 | 0.4 | 0.38 | 0.37 | 0.26 |
| 1 | 228424600 | rs1757139 | -0.452 | 0.217 | -0.429 | 1.055 | -0.004 | -0.003 | -0.003 | 0.075 | 0.37 | 0.6 | 0.4 | 0.39 | 0.36 | 0.26 |
| 1 | 228427193 | rs1771459 | -0.268 | 0.229 | -0.479 | 1.055 | -0.001 | -0.003 | -0.003 | 0 | 0.36 | 0.46 | 0.36 | 0.35 | 0.36 | 0.25 |
| 1 | 228428805 | rs6701173 | 0.898 | 0.323 | -0.566 | 1.01 | 0.009 | -0.001 | -0.001 | 3.138 | 0.22 | 0.25 | 0.29 | 0.25 | 0.28 | 0.2 |
| 1 | 228431930 | rs10916293 | -0.784 | 0.267 | -0.602 | 1.014 | 0.007 | -0.01 | -0.01 | 0.233 | 0.58 | 0.71 | 0.65 | 0.6 | 0.64 | 0.46 |
| 1 | 228433491 | rs1888681 | -0.784 | 0.314 | -0.66 | 0.945 | 0.007 | -0.01 | -0.01 | 0 | 0.58 | 0.71 | 0.65 | 0.6 | 0.64 | 0.46 |
| 1 | 228434395 | rs7517088 | -0.731 | 0.291 | -0.672 | 0.933 | 0.009 | -0.009 | -0.009 | 0 | 0.57 | 0.71 | 0.65 | 0.6 | 0.64 | 0.46 |
| 1 | 228442663 | rs4653544 | -1.361 | 0.157 | -0.944 | 1.023 | -0.059 | -0.104 | -0.104 | 0 | 0.74 | 0.98 | 0.8 | 0.69 | 0.73 | 0.63 |
| 1 | 228444741 | rs3795780 | -1.092 | 0.753 | -0.399 | 1.303 | 0.007 | -0.01 | -0.01 | 0.021 | 0.58 | 0.71 | 0.65 | 0.6 | 0.64 | 0.46 |
| 1 | 228460217 | rs424645 | 0.635 | 0.864 | -0.323 | 1.086 | 0.01 | -0.009 | -0.009 | 0 | 0.22 | 0.25 | 0.3 | 0.23 | 0.28 | 0.2 |
| 1 | 228464248 | rs1188721 | -1.397 | 0.868 | -0.397 | 1.329 | 0.009 | 0.029 | 0.029 | 2.67 | 0.6 | 0.85 | 0.7 | 0.75 | 0.66 | 0.46 |
| 1 | 228465346 | rs493945 | -0.214 | 0.821 | -0.262 | 1.46 | -0.004 | -0.007 | -0.007 | 0 | 0.33 | 0.36 | 0.36 | 0.38 | 0.36 | 0.19 |
| 1 | 228465370 | rs867599 | -1.163 | 0.651 | -0.69 | 0.877 | 0.017 | 0.04 | 0.04 | 0.417 | 0.56 | 0.75 | 0.7 | 0.74 | 0.67 | 0.46 |
| 1 | 228466750 | rs3795782 | -0.035 | 0.808 | -0.673 | 0.742 | -0.03 | 0.067 | 0.067 | 0 | 0.3 | 0.19 | 0.36 | 0.18 | 0.35 | 0.17 |
| 1 | 228473600 | rs1150907 | -0.037 | 0.31 | -0.554 | 0.731 | 0.032 | -0.053 | -0.053 | 0 | 0.34 | 0.47 | 0.42 | 0.25 | 0.48 | 0.27 |
| 1 | 228474817 | rs4653935 | 0.155 | 0.155 | -0.586 | 0.731 | 0.009 | 0.016 | 0.016 | 0.358 | 0.2 | 0.19 | 0.24 | 0.15 | 0.16 | 0.09 |
| 1 | 228491633 | rs11584418 | 2.05 | 0.14 | -0.62 | 0.721 | 0.004 | -0.005 | -0.005 | 0 | 0.05 | 0.04 | 0.06 | 0.1 | 0.03 | 0.04 |
| 1 | 228492044 | rs1150910 | 0.027 | 0.221 | -0.594 | 0.646 | 0.028 | -0.046 | -0.046 | 0 | 0.34 | 0.42 | 0.41 | 0.25 | 0.49 | 0.27 |
| 1 | 228494696 | rs4653939 | -0.578 | 0.284 | -0.842 | 0.824 | -0.013 | 0.053 | 0.053 | 0.084 | 0.24 | 0.37 | 0.29 | 0.5 | 0.19 | 0.17 |
| 1 | 228494790 | rs435776 | 0.329 | 0.248 | -0.884 | 0.77 | -0.113 | 0.057 | 0.057 | 0.207 | 0.3 | 0.06 | 0.38 | 0.25 | 0.48 | 0.27 |
| 1 | 228505204 | rs4653942 | -0.786 | 0.317 | -0.685 | 1.034 | -0.017 | 0.07 | 0.07 | 0.056 | 0.24 | 0.39 | 0.3 | 0.5 | 0.19 | 0.17 |
| 1 | 228525008 | rs3795809 | 0.876 | 0.459 | -0.821 | 0.921 | -0.032 | 0.037 | 0.037 | 0 | 0.13 | 0.02 | 0.17 | 0.06 | 0.13 | 0.06 |
| 1 | 228529129 | rs3795810 | 1.094 | 0.323 | -1.006 | 0.928 | 0.022 | -0.065 | -0.065 | 0.184 | 0.14 | 0.04 | 0.07 | 0.33 | 0.1 | 0.15 |
| 1 | 228532195 | rs417237 | -0.88 | 0.351 | -0.883 | 0.824 | 0 | -0.006 | -0.006 | 0 | 0.42 | 0.56 | 0.63 | 0.4 | 0.57 | 0.31 |
| 1 | 228538968 | rs3795811 | 1.359 | 0.689 | -1.441 | 0.864 | -0.008 | -0.056 | -0.056 | 0 | 0.13 | 0.05 | 0.05 | 0.31 | 0.13 | 0.15 |
| 1 | 228546242 | rs4653946 | 0.355 | 0.452 | -1.364 | 0.755 | -0.021 | -0.015 | -0.015 | 0 | 0.6 | 0.33 | 0.53 | 0.4 | 0.48 | 0.63 |
| 1 | 228547901 | rs12035900 | 1.068 | 0.434 | -1.339 | 0.702 | 0.01 | 0.025 | 0.025 | 6.768 | 0.4 | 0.11 | 0.31 | 0.27 | 0.33 | 0.54 |
| 1 | 228556788 | rs3738685 | 1.016 | 0.446 | -1.335 | 0.711 | -0.008 | 0.037 | 0.037 | 5.365 | 0.41 | 0.18 | 0.31 | 0.25 | 0.36 | 0.54 |

| | | | | | | | | | | | | | | | | |
|---|---|---|---|---|---|---|---|---|---|---|---|---|---|---|---|---|
| 1 | 228559994 | rs500049 | 0.247 | 0.635 | -1.155 | 1.142 | -0.022 | -0.018 | -0.018 | 0 | 0.66 | 0.71 | 0.39 | 0.56 | 0.5 | 0.69 |
| 1 | 228568159 | rs12760907 | 0.59 | 0.161 | -0.831 | 1.169 | -0.019 | 0.046 | 0.046 | 0.231 | 0.15 | 0.17 | 0.05 | 0.34 | 0.1 | 0.14 |
| 1 | 228568201 | rs12735877 | 1.117 | 0.204 | -1.048 | 1.134 | 0.021 | -0.052 | -0.052 | 0 | 0.13 | 0.04 | 0.04 | 0.31 | 0.1 | 0.14 |
| 1 | 228576253 | rs10916306 | -0.777 | 0.648 | -1.252 | 1.053 | -0.127 | -0.044 | -0.044 | 0 | 0.79 | 0.93 | 0.62 | 0.69 | 0.53 | 0.8 |
| 1 | 228579788 | rs4653951 | 1.211 | 0.256 | -1.333 | 0.871 | 0.005 | 0.008 | 0.008 | 0 | 0.1 | 0.03 | 0.12 | 0.06 | 0.06 | 0.07 |
| 1 | 228585562 | rs564309 | 0.819 | 0.059 | -1.024 | 1.008 | -0.023 | 0.043 | 0.043 | 0.269 | 0.15 | 0.14 | 0.04 | 0.31 | 0.1 | 0.14 |
| 1 | 228587435 | rs513467 | -0.046 | -0.171 | -0.945 | 0.949 | -0.055 | 0.062 | 0.062 | 5.714 | 0.2 | 0.27 | 0.13 | 0.38 | 0.11 | 0.16 |
| 1 | 228593222 | rs503391 | 0.691 | -0.155 | -0.979 | 0.952 | -0.033 | 0.06 | 0.06 | 0.154 | 0.16 | 0.19 | 0.05 | 0.31 | 0.1 | 0.14 |
| 1 | 228601638 | rs472594 | -1.26 | -0.239 | -1.122 | 0.745 | 0.094 | -0.042 | -0.042 | 0.039 | 0.38 | 0.85 | 0.33 | 0.44 | 0.21 | 0.23 |
| 1 | 228603029 | rs10799476 | 1.562 | -0.328 | -1.107 | 0.848 | 0.032 | 0.042 | 0.042 | 5.018 | 0.41 | 0.09 | 0.3 | 0.24 | 0.33 | 0.57 |
| 1 | 228606965 | rs493950 | -1.027 | -0.878 | -0.376 | 1.171 | 0.047 | -0.039 | -0.039 | 0 | 0.25 | 0.8 | 0.18 | 0.38 | 0.12 | 0.16 |
| 1 | 228612444 | rs1188473 | -1.131 | -0.705 | -0.156 | 1.192 | 0.019 | -0.06 | -0.06 | 0 | 0.87 | 0.96 | 0.96 | 0.7 | 0.89 | 0.86 |
| 1 | 228617493 | rs1188482 | -1.005 | -0.668 | -0.207 | 1.215 | 0.018 | -0.037 | -0.037 | 0.022 | 0.86 | 0.92 | 0.96 | 0.69 | 0.89 | 0.86 |
| 1 | 228622347 | rs4581279 | -0.343 | -0.678 | -0.152 | 1.208 | -0.036 | 0.077 | 0.077 | 0 | 0.83 | 0.73 | 0.91 | 0.67 | 0.89 | 0.85 |
| 1 | 228631641 | rs12131962 | 0.8 | -0.475 | -0.439 | 1.135 | 0.046 | -0.038 | -0.038 | 0 | 0.71 | 0.2 | 0.79 | 0.62 | 0.78 | 0.84 |
| 1 | 228633197 | rs16848614 | 0.738 | -0.491 | -0.643 | 0.949 | 0.009 | 0.015 | 0.015 | 0 | 0.11 | 0.04 | 0.12 | 0.07 | 0.06 | 0.07 |
| 1 | 228636824 | rs7529055 | -0.684 | -0.559 | -0.621 | 1.144 | -0.002 | -0.06 | -0.06 | 0 | 0.82 | 0.94 | 0.94 | 0.7 | 0.8 | 0.85 |
| 1 | 228641393 | rs7528331 | 0.269 | -0.622 | -0.307 | 1.148 | -0.014 | 0.046 | 0.046 | 17.728 | 0.09 | 0.54 | 0.12 | 0.08 | 0.02 | 0.01 |
| 1 | 228643447 | rs728727 | 1.65 | -0.062 | -0.63 | 1.333 | 0.003 | -0.002 | -0.002 | 0.603 | 0.54 | 0.11 | 0.58 | 0.36 | 0.6 | 0.72 |
| 1 | 228651011 | rs6426503 | -0.856 | -0.167 | -0.552 | 1.241 | -0.007 | -0.063 | -0.063 | 0 | 0.82 | 0.95 | 0.94 | 0.7 | 0.8 | 0.85 |
| 1 | 228651457 | rs1007686 | -0.823 | -0.166 | -0.551 | 1.242 | -0.006 | -0.063 | -0.063 | 0 | 0.82 | 0.94 | 0.94 | 0.7 | 0.8 | 0.85 |
| 1 | 228654823 | rs10916318 | 0.136 | -0.131 | -0.185 | 0.812 | -0.017 | 0.027 | 0.027 | 20.168 | 0.09 | 0.55 | 0.09 | 0.05 | 0.02 | 0.01 |
| 1 | 228660501 | rs10733022 | 0.713 | 0.015 | -0.607 | 0.924 | 0.003 | 0.003 | 0.003 | 1.665 | 0.59 | 0.6 | 0.65 | 0.31 | 0.63 | 0.72 |
| 1 | 228674518 | rs4233233 | 1.405 | -0.083 | -1.745 | 0.356 | -0.152 | -0.135 | -0.135 | 0.787 | 0.31 | 0.03 | 0.5 | 0.41 | 0.44 | 0.56 |
| 1 | 228681195 | rs3795816 | -0.303 | -0.505 | -1.53 | 0.075 | 0.012 | 0.207 | 0.207 | 0.219 | 0.19 | 0.45 | 0.24 | 0.51 | 0.17 | 0.21 |
| 1 | 228681749 | rs3795817 | 1.045 | -0.913 | -1.874 | -0.267 | -0.008 | 0.017 | 0.017 | 0 | 0.49 | 0.18 | 0.6 | 0.44 | 0.55 | 0.63 |
| 1 | 228685999 | rs3795818 | -0.341 | -0.871 | -1.668 | -0.462 | 0.006 | 0.044 | 0.044 | 0 | 0.68 | 0.7 | 0.84 | 0.95 | 0.72 | 0.84 |
| 1 | 228687796 | rs12072922 | 1.162 | -0.513 | -1.658 | -0.161 | -0.032 | 0.023 | 0.023 | 0 | 0.08 | 0.03 | 0.05 | 0.04 | 0.08 | 0.05 |
| 1 | 228689848 | rs13374918 | 1.014 | -0.91 | -1.464 | -0.025 | 0.002 | -0.01 | -0.01 | 0 | 0.18 | 0.16 | 0.11 | 0.03 | 0.11 | 0.08 |
| 1 | 228699477 | rs6665115 | 0.698 | -0.929 | -1.502 | 0.17 | 0.041 | 0.021 | 0.021 | 0 | 0.49 | 0.3 | 0.64 | 0.44 | 0.59 | 0.65 |
| 1 | 228703236 | rs12136157 | 0.59 | -1.223 | -1.873 | -0.222 | -0.042 | 0.061 | 0.061 | 0 | 0.21 | 0.02 | 0.1 | 0.06 | 0.2 | 0.24 |
| 1 | 228703289 | rs1925714 | -0.921 | -1.551 | -1.26 | -0.047 | -0.004 | -0.014 | -0.014 | 0.21 | 0.8 | 0.77 | 0.8 | 0.9 | 0.8 | 0.87 |
| 1 | 228707539 | rs1925716 | 2.11 | -1.819 | -1.329 | 0.273 | -0.099 | -0.097 | -0.097 | 0 | 0.3 | 0.12 | 0.41 | 0.49 | 0.42 | 0.46 |
| 1 | 228715780 | rs11588356 | 2.152 | -1.788 | -1.336 | 0.288 | -0.105 | -0.104 | -0.104 | 0 | 0.3 | 0.11 | 0.41 | 0.49 | 0.42 | 0.46 |
| 1 | 228731677 | rs744668 | -0.628 | -2.062 | -0.687 | 0.44 | -0.018 | -0.043 | -0.043 | 0.028 | 0.79 | 0.63 | 0.78 | 0.9 | 0.8 | 0.87 |
| 1 | 228741147 | rs7415895 | -0.757 | -1.975 | -0.912 | 0.496 | -0.006 | -0.015 | -0.015 | 0 | 0.81 | 0.76 | 0.8 | 0.91 | 0.81 | 0.88 |
| 1 | 228784907 | rs6672815 | -0.005 | -2.283 | -0.722 | 0.953 | -0.009 | 0.011 | 0.011 | 0.022 | 0.22 | 0.25 | 0.26 | 0.37 | 0.18 | 0.14 |
| 1 | 228796151 | rs6658934 | 0.241 | -2.262 | -0.892 | 0.951 | 0.024 | -0.044 | -0.044 | 0.632 | 0.21 | 0.13 | 0.25 | 0.36 | 0.17 | 0.13 |
| 1 | 228805855 | rs765070 | -1.894 | -2.73 | -1.138 | 0.873 | -0.061 | -0.042 | -0.042 | 0 | 0.66 | 0.89 | 0.56 | 0.63 | 0.45 | 0.51 |
| 1 | 228818506 | rs6692721 | -0.073 | -3.049 | -0.648 | 0.716 | -0.05 | -0.002 | -0.002 | 0 | 0.21 | 0.58 | 0.21 | 0.18 | 0.11 | 0.12 |
| 1 | 228818957 | rs10916354 | -2.169 | -3.098 | -1.117 | 0.779 | -0.121 | -0.109 | -0.109 | 0.174 | 0.66 | 0.96 | 0.56 | 0.62 | 0.45 | 0.51 |
| 1 | 228824972 | rs7526487 | -1.021 | -3.136 | -1.02 | 0.393 | 0.106 | 0.099 | 0.099 | 0 | 0.45 | 0.82 | 0.32 | 0.25 | 0.29 | 0.38 |
| 1 | 228829657 | rs12028277 | -0.038 | -3.157 | -0.563 | 0.25 | 0.007 | 0.022 | 0.022 | 0 | 0.25 | 0.16 | 0.13 | 0.07 | 0.2 | 0.31 |
| 1 | 228834637 | rs7555155 | -2.338 | -3.097 | -0.916 | 0.519 | -0.111 | -0.024 | -0.024 | 0 | 0.64 | 0.96 | 0.56 | 0.7 | 0.47 | 0.55 |
| 1 | 228866987 | rs2180162 | 0.083 | -3.17 | 0.122 | 0.282 | -0.066 | 0.074 | 0.074 | 0 | 0.17 | 0.37 | 0.22 | 0.24 | 0.09 | 0.1 |
| 1 | 228874049 | rs2295256 | -1.41 | -3.319 | 0.186 | 0.235 | 0.036 | 0.013 | 0.013 | 0 | 0.4 | 0.65 | 0.34 | 0.29 | 0.27 | 0.36 |
| 1 | 228878522 | rs4531263 | -2.125 | -3.42 | 0.16 | 0.211 | 0.047 | -0.001 | -0.001 | 0 | 0.57 | 0.81 | 0.44 | 0.36 | 0.38 | 0.4 |
| 1 | 228880021 | rs1062060 | 0.807 | -3.391 | 0.204 | 0.197 | -0.001 | 0.039 | 0.039 | 0 | 0.19 | 0.1 | 0.1 | 0.04 | 0.17 | 0.25 |
| 1 | 228880115 | rs6426514 | 0.854 | -3.406 | 0.229 | 0.191 | 0.009 | 0.01 | 0.01 | 0.083 | 0.14 | 0.13 | 0.09 | 0.03 | 0.1 | 0.02 |
| 1 | 228880466 | rs11580020 | 0.693 | -3.206 | 0.31 | 0.337 | -0.001 | 0.039 | 0.039 | 0 | 0.19 | 0.1 | 0.1 | 0.04 | 0.17 | 0.25 |
| 1 | 228881129 | rs13349 | -1.221 | -3.092 | 0.245 | 0.393 | 0.047 | 0.017 | 0.017 | 0 | 0.4 | 0.66 | 0.34 | 0.29 | 0.27 | 0.36 |
| 1 | 228884405 | rs10916364 | -1.081 | -3.031 | 0.252 | 0.445 | -0.022 | -0.006 | -0.006 | 3.067 | 0.38 | 0.5 | 0.32 | 0.29 | 0.27 | 0.36 |
| 1 | 228886880 | rs9660783 | -1.307 | -2.816 | 0.248 | 0.412 | 0.025 | 0.049 | 0.049 | 0 | 0.44 | 0.63 | 0.29 | 0.13 | 0.33 | 0.37 |
| 1 | 228887522 | rs4440831 | -1.016 | -2.79 | 0.252 | 0.465 | 0.005 | 0.004 | 0.004 | 0 | 0.42 | 0.58 | 0.29 | 0.13 | 0.33 | 0.36 |
| 1 | 228887667 | rs7535661 | -0.28 | -2.846 | 0.125 | 0.418 | -0.007 | -0.037 | -0.037 | 0.275 | 0.29 | 0.36 | 0.23 | 0.11 | 0.27 | 0.35 |
| 1 | 228891103 | rs6676800 | 1.451 | -2.675 | -0.519 | 0.489 | 0.025 | 0.006 | 0.006 | 0 | 0.45 | 0.23 | 0.5 | 0.64 | 0.58 | 0.55 |
| 1 | 228894575 | rs10916366 | -1.569 | -2.412 | -0.728 | 0.67 | 0.068 | 0 | 0 | 0 | 0.5 | 0.9 | 0.47 | 0.36 | 0.34 | 0.4 |
| 1 | 228903818 | rs728370 | 1.021 | -2.439 | -0.071 | 0.952 | 0.01 | -0.003 | -0.003 | 0.012 | 0.61 | 0.39 | 0.6 | 0.6 | 0.71 | 0.84 |
| 1 | 228904822 | rs11585991 | 0.348 | -2.577 | -0.158 | 0.83 | -0.004 | -0.016 | -0.016 | 0 | 0.23 | 0.26 | 0.14 | 0.07 | 0.17 | 0.32 |
| 1 | 228907642 | rs7518686 | 0.955 | -2.555 | -0.276 | 0.917 | -0.017 | 0.016 | 0.016 | 0.221 | 0.13 | 0.15 | 0.21 | 0.19 | 0.05 | 0.04 |
| 1 | 228909127 | rs6703541 | 1.704 | -2.395 | -0.718 | 0.92 | -0.038 | -0.088 | -0.088 | 0.031 | 0.39 | 0.13 | 0.47 | 0.53 | 0.55 | 0.52 |
| 1 | 228916634 | rs241324 | 0.86 | -2.448 | 0.049 | 1.287 | -0.004 | 0 | 0 | 0 | 0.71 | 0.7 | 0.76 | 0.63 | 0.82 | 0.87 |
| 1 | 228917051 | rs241323 | -0.326 | -1.991 | -0.455 | 0.977 | 0.019 | 0.007 | 0.007 | 0 | 0.84 | 0.87 | 0.93 | 0.85 | 0.89 | 0.93 |
| 1 | 228919202 | rs241315 | 0.818 | -1.553 | -0.094 | 1.171 | -0.005 | -0.002 | -0.002 | 0 | 0.76 | 0.72 | 0.74 | 0.77 | 0.82 | 0.92 |
| 1 | 228922262 | rs12067649 | -1.591 | -1.413 | 0.311 | 1.857 | -0.01 | 0.223 | 0.223 | 0 | 0.1 | 0.31 | 0.22 | 0.37 | 0.08 | 0.1 |
| 1 | 228924938 | rs171322 | 0.034 | -1.763 | -0.075 | 1.017 | -0.012 | 0 | 0 | 0.239 | 0.54 | 0.56 | 0.66 | 0.59 | 0.69 | 0.63 |
| 1 | 228927898 | rs127557 | 0.348 | -1.497 | -0.956 | 1.014 | 0.053 | -0.031 | -0.031 | 0 | 0.45 | 0.21 | 0.59 | 0.52 | 0.67 | 0.51 |
| 1 | 228937821 | rs241305 | 0.415 | -1.446 | -1.056 | 0.961 | 0.044 | -0.053 | -0.053 | 0 | 0.44 | 0.18 | 0.59 | 0.53 | 0.67 | 0.52 |
| 1 | 228943083 | rs2748081 | 0.089 | -1.505 | -0.434 | 0.59 | 0.034 | 0.041 | 0.041 | 0 | 0.5 | 0.61 | 0.73 | 0.73 | 0.6 | 0.61 |

| | | | | | | | | | | | | | | | | |
|---|---|---|---|---|---|---|---|---|---|---|---|---|---|---|---|---|
| 1 | 228943744 | rs2748084 | -0.655 | -1.813 | -0.205 | 0.087 | 0.006 | 0.014 | 0.014 | 0 | 0.35 | 0.33 | 0.31 | 0.15 | 0.33 | 0.21 |
| 1 | 228945692 | rs12749124 | -1.319 | -1.27 | 0.061 | 0.246 | 0.009 | -0.02 | -0.02 | 0 | 0.07 | 0.07 | 0.07 | 0 | 0.14 | 0.04 |
| 1 | 228949608 | rs241337 | -0.759 | -1.731 | 0.542 | 0.299 | 0.059 | 0.058 | 0.058 | 0 | 0.49 | 0.66 | 0.4 | 0.31 | 0.31 | 0.25 |
| 1 | 228962427 | rs241301 | -0.666 | -1.737 | 0.771 | 0.316 | 0.006 | 0.014 | 0.014 | 0.007 | 0.5 | 0.41 | 0.41 | 0.3 | 0.43 | 0.23 |
| 1 | 228968184 | rs2639757 | -1.715 | -1.842 | 0.967 | 0.419 | 0.007 | 0 | 0 | 0.084 | 0.05 | 0.33 | 0.05 | 0 | 0.01 | 0 |
| 1 | 228989201 | rs801109 | -0.6 | -2.442 | 0.843 | 0.427 | 0.031 | 0.017 | 0.017 | 1.636 | 0.46 | 0.59 | 0.33 | 0.38 | 0.28 | 0.24 |
| 1 | 228995219 | rs714349 | -0.689 | -2.215 | 0.917 | 0.499 | 0.013 | 0.016 | 0.016 | 0 | 0.49 | 0.4 | 0.33 | 0.17 | 0.37 | 0.23 |
| 1 | 228997835 | rs801114 | -1.264 | -2.155 | 0.262 | 0.622 | 0.029 | 0.015 | 0.015 | 1.628 | 0.57 | 0.84 | 0.42 | 0.4 | 0.38 | 0.28 |
| 1 | 229003739 | rs71074 | -0.55 | -2.05 | 0.389 | 0.88 | 0.09 | 0.028 | 0.028 | 0.534 | 0.48 | 0.71 | 0.36 | 0.43 | 0.32 | 0.19 |
| 1 | 229004119 | rs171326 | -1.023 | -2.021 | 0.381 | 0.944 | 0.03 | -0.021 | -0.021 | 0 | 0.55 | 0.79 | 0.44 | 0.49 | 0.4 | 0.28 |
| 1 | 229004264 | rs241362 | -2.645 | -2.027 | 0.58 | 0.974 | 0.011 | 0.073 | 0.073 | 0.02 | 0.12 | 0.3 | 0.15 | 0.28 | 0.11 | 0.11 |
| 1 | 229005123 | rs801116 | -0.64 | -2.069 | 0.545 | 1.001 | 0.103 | 0.013 | 0.013 | 0.022 | 0.5 | 0.71 | 0.38 | 0.49 | 0.32 | 0.24 |
| 1 | 229005833 | rs241359 | -2.403 | -1.508 | 0.362 | 1.26 | -0.019 | 0.134 | 0.134 | 0 | 0.06 | 0.23 | 0.08 | 0.28 | 0.02 | 0.07 |
| 1 | 229018916 | rs1341715 | -1.012 | -1.62 | 0.139 | 1.059 | 0.011 | -0.03 | -0.03 | 0 | 0.56 | 0.79 | 0.45 | 0.48 | 0.42 | 0.3 |
| 1 | 229027438 | rs6426525 | -0.74 | -1.605 | 0.224 | 0.97 | 0.03 | 0.075 | 0.075 | 1.705 | 0.52 | 0.66 | 0.39 | 0.17 | 0.39 | 0.24 |
| 1 | 229030081 | rs11804896 | -0.705 | -1.606 | 0.217 | 0.97 | 0.027 | 0.065 | 0.065 | 1.65 | 0.51 | 0.63 | 0.39 | 0.17 | 0.39 | 0.24 |
| 1 | 229038351 | rs7551503 | -0.967 | -1.813 | 0.143 | 1.171 | 0.106 | 0.049 | 0.049 | 0.448 | 0.47 | 0.81 | 0.39 | 0.41 | 0.33 | 0.23 |
| 1 | 229039470 | rs2236591 | -0.917 | -1.769 | 0.233 | 0.871 | 0.06 | 0.059 | 0.059 | 0.044 | 0.46 | 0.68 | 0.37 | 0.26 | 0.33 | 0.26 |
| 1 | 229041874 | rs12078733 | -0.921 | -1.607 | -0.167 | 0.734 | 0.03 | 0.045 | 0.045 | 0.053 | 0.44 | 0.62 | 0.35 | 0.18 | 0.33 | 0.23 |
| 1 | 229049984 | rs6426527 | -1.109 | -1.001 | -0.912 | 0.7 | 0.023 | 0.043 | 0.043 | 0.004 | 0.52 | 0.86 | 0.44 | 0.41 | 0.38 | 0.33 |
| 1 | 229052132 | rs2000163 | -1.184 | -1.036 | -0.897 | 0.681 | 0.017 | 0.039 | 0.039 | 0 | 0.53 | 0.83 | 0.39 | 0.4 | 0.39 | 0.37 |
| 1 | 229060300 | rs1891202 | -1.068 | -1.054 | -0.541 | 0.844 | 0.04 | 0.008 | 0.008 | 0.208 | 0.43 | 0.74 | 0.36 | 0.4 | 0.32 | 0.31 |
| 1 | 229060567 | rs1891203 | -0.124 | -1.288 | -0.545 | 0.655 | 0.007 | 0.069 | 0.069 | 0 | 0.27 | 0.11 | 0.1 | 0.07 | 0.2 | 0.17 |
| 1 | 229065083 | rs1389139 | -1.064 | -1.111 | -0.561 | 0.682 | 0.042 | 0.009 | 0.009 | 0.186 | 0.44 | 0.74 | 0.36 | 0.4 | 0.32 | 0.31 |
| 1 | 229073985 | rs2220741 | -1.299 | -0.945 | 0.383 | 1.316 | -0.028 | 0.102 | 0.102 | 0 | 0.22 | 0.59 | 0.33 | 0.49 | 0.15 | 0.2 |
| 1 | 229076157 | rs10916389 | 0.63 | -1.172 | 0.131 | 0.291 | -0.024 | -0.044 | -0.044 | 0 | 0.78 | 0.75 | 0.8 | 0.96 | 0.83 | 0.84 |
| 1 | 229076758 | rs710824 | 0.643 | -0.718 | 0.151 | 0.837 | -0.005 | -0.001 | -0.001 | 0 | 0.59 | 0.41 | 0.7 | 0.44 | 0.68 | 0.56 |
| 1 | 229076902 | rs12065469 | -1.073 | -0.63 | 0.244 | 1.445 | 0.005 | 0.149 | 0.149 | 0 | 0.05 | 0.12 | 0.03 | 0.28 | 0.04 | 0.1 |
| 1 | 229077631 | rs529443 | -0.296 | 0.06 | -0.115 | 1.229 | -0.003 | -0.069 | -0.069 | 0 | 0.38 | 0.13 | 0.28 | 0.38 | 0.34 | 0.47 |
| 1 | 229091091 | rs778374 | -0.325 | 0.061 | -0.079 | 0.666 | 0.014 | 0.021 | 0.021 | 0.607 | 0.8 | 0.69 | 0.77 | 0.71 | 0.74 | 0.78 |
| 1 | 229091234 | rs991304 | 1.068 | 0.122 | 0.002 | 0.769 | -0.016 | 0.012 | 0.012 | 0 | 0.35 | 0.19 | 0.35 | 0.36 | 0.35 | 0.31 |
| 1 | 229091575 | rs778373 | 0.478 | 0.105 | -0.085 | 0.748 | -0.046 | -0.093 | -0.093 | 0 | 0.36 | 0.03 | 0.26 | 0.31 | 0.34 | 0.46 |
| 1 | 229098232 | rs496663 | 0.426 | 0.197 | 0.211 | 0.955 | -0.002 | 0 | 0 | 0.071 | 0.42 | 0.4 | 0.49 | 0.45 | 0.41 | 0.33 |
| 1 | 229100829 | rs519791 | 1.077 | 0.315 | 0.229 | 0.736 | -0.014 | 0.013 | 0.013 | 0 | 0.34 | 0.25 | 0.45 | 0.32 | 0.36 | 0.28 |
| 1 | 229105145 | rs603404 | -0.199 | 0.179 | 0.485 | 0.744 | -0.009 | -0.008 | -0.008 | 2.056 | 0.63 | 0.56 | 0.63 | 0.39 | 0.68 | 0.56 |
| 1 | 229106195 | rs778381 | -0.539 | 0.081 | 0.569 | 0.88 | 0.015 | 0.089 | 0.089 | 0 | 0.13 | 0.3 | 0.13 | 0.29 | 0.18 | 0.18 |
| 1 | 229107279 | rs7513940 | -0.367 | -0.06 | 0.074 | 0.876 | -0.002 | -0.044 | -0.044 | 0 | 0.08 | 0.17 | 0.05 | 0 | 0.06 | 0.07 |
| 1 | 229109847 | rs10916394 | -1.182 | -0.181 | -0.303 | 1.163 | -0.036 | 0.079 | 0.079 | 4.61 | 0.16 | 0.64 | 0.1 | 0.37 | 0.08 | 0.17 |
| 1 | 229112170 | rs609384 | 0.504 | -0.001 | -1.677 | 0.912 | -0.029 | -0.077 | -0.077 | 0 | 0.37 | 0.03 | 0.48 | 0.43 | 0.33 | 0.37 |
| 1 | 229112223 | rs609447 | -0.415 | 0.246 | -1.597 | 0.565 | -0.081 | -0.007 | -0.007 | 0 | 0.62 | 0.86 | 0.67 | 0.74 | 0.53 | 0.6 |
| 1 | 229114198 | rs7529576 | -0.178 | -0.083 | -1.445 | -0.012 | 0.002 | -0.01 | -0.01 | 0 | 0.6 | 0.63 | 0.62 | 0.79 | 0.5 | 0.66 |
| 1 | 229114512 | rs710814 | -0.116 | -0.254 | -2.075 | -0.038 | 0.055 | 0.154 | 0.154 | 0.499 | 0.53 | 0.8 | 0.59 | 0.79 | 0.34 | 0.41 |
| 1 | 229114638 | rs778536 | -0.524 | 0.167 | -1.609 | 0.149 | -0.103 | 0.078 | 0.078 | 0.909 | 0.71 | 0.92 | 0.56 | 0.88 | 0.51 | 0.66 |
| 1 | 229118735 | rs10916399 | 0.122 | -0.338 | -1.346 | 0.68 | 0.049 | 0.014 | 0.014 | 0 | 0.51 | 0.18 | 0.32 | 0.47 | 0.39 | 0.5 |
| 1 | 229121521 | rs4654700 | -0.837 | -0.438 | -0.289 | 1.059 | -0.015 | 0.157 | 0.157 | 0 | 0.05 | 0.21 | 0.1 | 0.15 | 0 | 0.03 |
| 1 | 229126747 | rs710804 | 0.252 | 0.04 | -0.35 | 0.425 | -0.001 | -0.001 | -0.001 | 0 | 0.41 | 0.45 | 0.37 | 0.23 | 0.52 | 0.38 |
| 1 | 229128091 | rs778532 | 0.327 | 0.014 | -0.283 | 0.412 | -0.008 | 0.01 | 0.01 | 0.036 | 0.41 | 0.41 | 0.36 | 0.23 | 0.51 | 0.37 |
| 1 | 229132668 | rs729628 | 0.191 | -0.321 | -0.796 | 0.202 | 0.026 | 0.118 | 0.118 | 0.053 | 0.34 | 0.16 | 0.19 | 0.13 | 0.27 | 0.22 |
| 1 | 229132772 | rs602532 | 0.738 | -0.068 | -1.015 | 0.27 | -0.031 | 0.051 | 0.051 | 0.329 | 0.39 | 0.24 | 0.33 | 0.2 | 0.49 | 0.34 |
| 1 | 229135031 | rs12120173 | 0.106 | -0.379 | -1.512 | 0.108 | 0.014 | 0.075 | 0.075 | 0.264 | 0.33 | 0.08 | 0.19 | 0.18 | 0.28 | 0.2 |
| 1 | 229140452 | rs755482 | 0.224 | -0.083 | -0.919 | 0.248 | 0.002 | -0.043 | -0.043 | 0.368 | 0.37 | 0.47 | 0.29 | 0.2 | 0.33 | 0.26 |
| 1 | 229141933 | rs644850 | -0.887 | 0.151 | -1.515 | 0.747 | 0.001 | -0.058 | -0.058 | 0.186 | 0.83 | 0.96 | 0.8 | 0.76 | 0.85 | 0.85 |
| 1 | 229142111 | rs483375 | 0.836 | 0.187 | -1.491 | 0.194 | -0.085 | 0.021 | 0.021 | 0.843 | 0.36 | 0.06 | 0.38 | 0.27 | 0.44 | 0.3 |
| 1 | 229144461 | rs687199 | 0.267 | 0.385 | -0.541 | 1.008 | -0.007 | 0.045 | 0.045 | 0 | 0.76 | 0.7 | 0.69 | 0.5 | 0.79 | 0.58 |
| 1 | 229145883 | rs595950 | 0.214 | 0.444 | -0.837 | 1.037 | -0.001 | 0.012 | 0.012 | 0 | 0.76 | 0.74 | 0.69 | 0.49 | 0.79 | 0.58 |
| 1 | 229155165 | rs551755 | 0.244 | 0.863 | -1.246 | 0.261 | -0.017 | 0.044 | 0.044 | 0.01 | 0.64 | 0.82 | 0.74 | 0.78 | 0.62 | 0.79 |
| 1 | 229155711 | rs652363 | 0.269 | 1.119 | -0.55 | 0.189 | 0.011 | -0.014 | -0.014 | 1.296 | 0.69 | 0.57 | 0.75 | 0.77 | 0.66 | 0.81 |
| 1 | 229156248 | rs10916409 | -0.604 | 0.941 | -0.682 | 0.235 | 0.008 | 0.016 | 0.016 | 0 | 0.38 | 0.45 | 0.45 | 0.53 | 0.43 | 0.36 |
| 1 | 229157752 | rs1458762 | -0.117 | 1.132 | -1.235 | -0.315 | -0.03 | 0.092 | 0.092 | 0 | 0.17 | 0.01 | 0.22 | 0.01 | 0.18 | 0.26 |
| 1 | 229159643 | rs537250 | 0.412 | 1.382 | -0.634 | 0.331 | 0.009 | 0.016 | 0.016 | 0.027 | 0.72 | 0.64 | 0.71 | 0.66 | 0.68 | 0.8 |
| 1 | 229161379 | rs7528235 | 0.57 | 1.265 | -0.38 | 0.684 | 0.006 | 0.069 | 0.069 | 0 | 0.11 | 0.2 | 0.11 | 0.27 | 0.1 | 0.1 |
| 1 | 229164804 | rs567911 | 0.777 | 1.264 | 0.068 | 0.803 | 0.006 | 0.005 | 0.005 | 0 | 0.63 | 0.42 | 0.6 | 0.48 | 0.61 | 0.71 |
| 1 | 229168822 | rs7517398 | 0.955 | 0.875 | -0.413 | 0.584 | 0.012 | 0.046 | 0.046 | 0 | 0.47 | 0.34 | 0.47 | 0.28 | 0.46 | 0.46 |
| 1 | 229171102 | rs475862 | -1.095 | 1.188 | -0.112 | 1.051 | -0.001 | 0.014 | 0.014 | 0.29 | 0.85 | 0.89 | 0.88 | 0.95 | 0.85 | 0.96 |
| 1 | 229173352 | rs1870362 | 1.334 | 1.087 | -0.128 | 1.025 | 0.009 | 0.04 | 0.04 | 0 | 0.46 | 0.34 | 0.46 | 0.27 | 0.46 | 0.46 |
| 1 | 229181697 | rs666453 | 0.072 | 1.177 | -0.091 | 1.135 | -0.052 | -0.069 | -0.069 | 0.904 | 0.25 | 0.08 | 0.29 | 0.37 | 0.3 | 0.36 |
| 1 | 229182292 | rs7538164 | 0.995 | 1.138 | 0.112 | 1.145 | 0.009 | 0.094 | 0.094 | 0 | 0.11 | 0.21 | 0.12 | 0.26 | 0.1 | 0.11 |
| 1 | 229183555 | rs546206 | 1.149 | 1.148 | 0.022 | 1.099 | -0.005 | 0.007 | 0.007 | 0 | 0.17 | 0.06 | 0.15 | 0.16 | 0.15 | 0.25 |
| 1 | 229189441 | rs505557 | 0.351 | 0.818 | 0.191 | 1.099 | -0.004 | 0.05 | 0.05 | 0 | 0.73 | 0.53 | 0.74 | 0.65 | 0.76 | 0.85 |

| | | | | | | | | | | | | | | | | |
|---|---|---|---|---|---|---|---|---|---|---|---|---|---|---|---|---|
| 1 | 229191667 | rs491986 | -0.994 | 0.841 | 0.145 | 1.071 | 0 | 0.009 | 0.009 | 0.435 | 0.86 | 0.89 | 0.87 | 0.95 | 0.86 | 0.96 |
| 1 | 229196360 | rs4654715 | 0.549 | 0.747 | 0.57 | 1.054 | 0.003 | 0 | 0 | 0 | 0.58 | 0.55 | 0.58 | 0.58 | 0.56 | 0.58 |
| 1 | 229200011 | rs4654644 | 1.323 | 0.721 | 0.335 | 1.058 | 0.018 | 0.078 | 0.078 | 0 | 0.46 | 0.24 | 0.45 | 0.28 | 0.46 | 0.46 |
| 1 | 229200392 | rs652014 | -0.904 | 0.761 | 0.44 | 1.048 | 0.003 | -0.003 | -0.003 | 1.051 | 0.39 | 0.55 | 0.43 | 0.67 | 0.42 | 0.49 |
| 1 | 229202402 | rs778902 | -0.915 | 0.789 | 0.467 | 0.903 | 0 | -0.004 | -0.004 | 2.848 | 0.85 | 0.85 | 0.87 | 0.95 | 0.88 | 0.96 |
| 1 | 229203703 | rs4654717 | 0.369 | 0.744 | 0.461 | 1.15 | -0.001 | -0.001 | -0.001 | 0 | 0.58 | 0.55 | 0.58 | 0.58 | 0.58 | 0.59 |
| 1 | 229212018 | rs17451637 | 1.178 | 0.78 | 0.281 | 1.26 | 0.019 | 0.078 | 0.078 | 0 | 0.46 | 0.26 | 0.45 | 0.23 | 0.46 | 0.48 |
| 1 | 229216656 | rs12137005 | -0.094 | 0.898 | 0.372 | 1.742 | 0.013 | 0.094 | 0.094 | 0.019 | 0.14 | 0.27 | 0.14 | 0.36 | 0.13 | 0.12 |
| 1 | 229224746 | rs12139882 | -0.026 | 0.758 | 0.278 | 1.76 | 0.005 | 0.124 | 0.124 | 0.065 | 0.14 | 0.27 | 0.14 | 0.38 | 0.11 | 0.21 |
| 1 | 229227923 | rs10916422 | -0.734 | 0.719 | -0.027 | 1.87 | 0.003 | -0.023 | -0.023 | 0.691 | 0.18 | 0.14 | 0.2 | 0.59 | 0.15 | 0.33 |
| 1 | 229229775 | rs12089229 | 0.345 | 0.783 | 0.057 | 1.831 | -0.001 | 0.014 | 0.014 | 0.228 | 0.13 | 0.16 | 0.13 | 0.3 | 0.11 | 0.21 |
| 1 | 229231141 | rs619422 | -0.044 | 1.136 | -0.246 | 1.529 | -0.03 | -0.025 | -0.025 | 1.213 | 0.27 | 0.13 | 0.31 | 0.37 | 0.35 | 0.44 |
| 1 | 229233144 | rs724899 | -0.396 | 1.112 | 0.451 | 1.52 | 0.021 | 0.144 | 0.144 | 0.51 | 0.17 | 0.47 | 0.16 | 0.38 | 0.15 | 0.22 |
| 1 | 229258917 | rs4078437 | 1.299 | 0.846 | -0.35 | 1.085 | -0.027 | -0.019 | -0.019 | 0 | 0.61 | 0.18 | 0.53 | 0.52 | 0.59 | 0.56 |
| 1 | 229260643 | rs9283408 | 0.048 | 0.871 | -0.156 | 1.001 | -0.03 | -0.037 | -0.037 | 0 | 0.77 | 0.82 | 0.68 | 0.58 | 0.69 | 0.62 |
| 1 | 229262842 | rs10916424 | -0.404 | 0.412 | 0.41 | 0.757 | 0.007 | -0.086 | -0.086 | 1.868 | 0.13 | 0.46 | 0.14 | 0.01 | 0.1 | 0.02 |
| 1 | 229269540 | rs3936161 | -0.685 | 0.328 | 0.507 | 1.159 | -0.022 | 0.075 | 0.075 | 0 | 0.14 | 0.61 | 0.15 | 0.19 | 0.06 | 0.17 |
| 1 | 229272553 | rs12072775 | 0.512 | 0.322 | 0.595 | 1.156 | 0.001 | 0 | 0 | 0 | 0.09 | 0.07 | 0.11 | 0.19 | 0.05 | 0.17 |
| 1 | 229273375 | rs9699623 | 0.159 | 0.926 | 0.124 | 1.09 | -0.026 | -0.004 | -0.004 | 0.322 | 0.82 | 0.94 | 0.84 | 0.78 | 0.75 | 0.76 |
| 1 | 229277059 | rs3923210 | 2.016 | 0.061 | -0.476 | 1.151 | -0.051 | 0.111 | 0.111 | 0 | 0.46 | 0.12 | 0.46 | 0.28 | 0.51 | 0.4 |
| 1 | 229278349 | rs9287022 | -0.059 | 0.192 | -0.148 | 1.277 | -0.009 | 0.11 | 0.11 | 0 | 0.13 | 0.48 | 0.15 | 0.19 | 0.06 | 0.18 |
| 1 | 229281204 | rs11583419 | 1.33 | 0.199 | -1.019 | 0.672 | -0.097 | -0.058 | -0.058 | 0 | 0.18 | 0.03 | 0.2 | 0.2 | 0.3 | 0.19 |
| 1 | 229282076 | rs12064154 | 0.218 | 0.169 | -0.933 | 0.684 | -0.024 | 0.067 | 0.067 | 0 | 0.14 | 0.65 | 0.16 | 0.23 | 0.06 | 0.2 |
| 1 | 229282778 | rs11580448 | 1.288 | 0.346 | -1.37 | 0.747 | -0.097 | -0.06 | -0.06 | 0.122 | 0.18 | 0.03 | 0.2 | 0.2 | 0.3 | 0.2 |
| 1 | 229282932 | rs9660543 | -1.382 | 0.366 | -1.111 | 0.815 | -0.009 | -0.011 | -0.011 | 0.264 | 0.88 | 0.94 | 0.89 | 0.79 | 0.85 | 0.79 |
| 1 | 229285858 | rs9793739 | 1.028 | 0.272 | -1.039 | 0.763 | -0.005 | 0.041 | 0.041 | 0 | 0.09 | 0.17 | 0.12 | 0.2 | 0.05 | 0.18 |
| 1 | 229290907 | rs9701452 | 0.35 | 0.082 | -1.217 | 0.762 | 0.013 | -0.036 | -0.036 | 0.98 | 0.21 | 0.13 | 0.21 | 0.31 | 0.17 | 0.17 |
| 1 | 229291651 | rs9287024 | -0.488 | -0.218 | -1.184 | 0.348 | 0.063 | 0.162 | 0.162 | 0 | 0.52 | 0.2 | 0.57 | 0.71 | 0.63 | 0.55 |
| 1 | 229293357 | rs9435847 | 0.97 | -0.093 | -0.942 | 0.478 | -0.005 | -0.001 | -0.001 | 0 | 0.13 | 0.08 | 0.11 | 0.18 | 0.15 | 0.21 |
| 1 | 229300574 | rs342826 | -1.774 | -0.561 | -0.735 | 0.087 | 0.03 | 0.071 | 0.071 | 0 | 0.73 | 0.84 | 0.75 | 0.89 | 0.8 | 0.78 |
| 1 | 229301364 | rs12130319 | -0.316 | -0.413 | -0.798 | 0.257 | 0.041 | 0.073 | 0.073 | 0 | 0.51 | 0.32 | 0.52 | 0.68 | 0.62 | 0.52 |
| 1 | 229301909 | rs342824 | -0.556 | -0.417 | -0.781 | 0.254 | 0.031 | 0.074 | 0.074 | 0 | 0.55 | 0.74 | 0.56 | 0.68 | 0.62 | 0.52 |
| 1 | 229302836 | rs6693489 | 0.393 | -0.303 | -0.78 | 0.266 | -0.037 | -0.013 | -0.013 | 0.255 | 0.38 | 0.2 | 0.4 | 0.5 | 0.45 | 0.31 |
| 1 | 229304681 | rs400827 | -0.72 | -0.272 | -0.963 | -0.039 | 0.033 | 0.081 | 0.081 | 0 | 0.55 | 0.75 | 0.56 | 0.69 | 0.62 | 0.51 |
| 1 | 229305150 | rs431675 | -0.856 | 0.205 | -1.239 | 0.305 | -0.002 | -0.03 | -0.03 | 0 | 0.36 | 0.7 | 0.36 | 0.54 | 0.36 | 0.5 |
| 1 | 229305602 | rs365684 | 0.487 | 0.101 | -1.194 | 0.199 | 0.06 | 0.011 | 0.011 | 0 | 0.27 | 0.47 | 0.33 | 0.28 | 0.4 | 0.17 |
| 1 | 229307013 | rs449616 | -1.486 | -0.144 | -1.479 | -0.104 | 0.016 | 0.12 | 0.12 | 0 | 0.7 | 0.87 | 0.72 | 0.88 | 0.8 | 0.73 |
| 1 | 229307113 | rs424699 | -0.509 | -0.172 | -1.346 | 0.099 | 0.003 | -0.013 | -0.013 | 0 | 0.31 | 0.66 | 0.32 | 0.54 | 0.31 | 0.44 |
| 1 | 229307604 | rs416479 | -0.398 | -0.148 | -1.359 | 0.111 | 0 | 0.075 | 0.075 | 0 | 0.29 | 0.48 | 0.31 | 0.54 | 0.32 | 0.44 |
| 1 | 229316554 | rs12026629 | 0.06 | -0.659 | -1.626 | 0.193 | -0.04 | -0.057 | -0.057 | 0 | 0.37 | 0.06 | 0.32 | 0.35 | 0.39 | 0.31 |
| 1 | 229319552 | rs869676 | 0.041 | -0.66 | -1.653 | 0.203 | -0.054 | -0.077 | -0.077 | 0 | 0.38 | 0.06 | 0.32 | 0.35 | 0.39 | 0.31 |
| 1 | 229329912 | rs9435854 | -0.032 | -0.622 | -1.477 | 0.149 | -0.01 | -0.023 | -0.023 | 0 | 0.28 | 0.27 | 0.34 | 0.54 | 0.38 | 0.44 |
| 1 | 229331680 | rs9435856 | -0.148 | -0.597 | -1.456 | 0.159 | -0.009 | -0.024 | -0.024 | 0.042 | 0.3 | 0.28 | 0.34 | 0.54 | 0.38 | 0.44 |
| 1 | 229334294 | rs12023401 | 2.495 | -0.572 | -1.548 | 0.269 | 0.009 | -0.017 | -0.017 | 2.58 | 0.08 | 0.04 | 0.11 | 0.14 | 0.04 | 0.07 |
| 1 | 229334731 | rs4925498 | -0.625 | -0.594 | -1.403 | 0.235 | -0.001 | 0 | 0 | 0.056 | 0.37 | 0.33 | 0.46 | 0.67 | 0.42 | 0.5 |
| 1 | 229336116 | rs8179432 | 0.662 | -0.602 | -1.626 | 0.191 | -0.086 | -0.023 | -0.023 | 0.301 | 0.29 | 0.02 | 0.21 | 0.21 | 0.35 | 0.24 |
| 1 | 229339307 | rs10916446 | 0.689 | -0.629 | -2.094 | 0.218 | -0.087 | 0.006 | 0.006 | 0.367 | 0.29 | 0.02 | 0.16 | 0.18 | 0.34 | 0.24 |
| 1 | 229340480 | rs386978 | -1.568 | -0.247 | -1.977 | 0.193 | -0.006 | 0.115 | 0.115 | 0.016 | 0.71 | 0.88 | 0.71 | 0.89 | 0.78 | 0.75 |
| 1 | 229340752 | rs9435858 | -0.231 | 0.043 | -1.761 | 0.475 | -0.027 | -0.066 | -0.066 | 0.011 | 0.28 | 0.15 | 0.33 | 0.53 | 0.38 | 0.44 |
| 1 | 229350560 | rs342833 | -1.017 | -0.255 | -1.312 | 0.928 | 0.008 | 0.079 | 0.079 | 0 | 0.51 | 0.8 | 0.54 | 0.56 | 0.43 | 0.43 |
| 1 | 229351259 | rs342834 | 0.473 | -0.252 | -1.304 | 0.939 | 0 | -0.005 | -0.005 | 0 | 0.31 | 0.39 | 0.25 | 0.23 | 0.33 | 0.2 |
| 1 | 229352808 | rs12029314 | 0.205 | 0.104 | -1.469 | 1.24 | 0 | 0.019 | 0.019 | 0.873 | 0.13 | 0.22 | 0.17 | 0.14 | 0.09 | 0.07 |
| 1 | 229360981 | rs501082 | -0.627 | 0.144 | -2.108 | 0.8 | 0.037 | 0.027 | 0.027 | 0 | 0.43 | 0.79 | 0.48 | 0.55 | 0.35 | 0.38 |
| 1 | 229362876 | rs932927 | 0.813 | 1.13 | -1.911 | 1.063 | -0.055 | -0.074 | -0.074 | 0.395 | 0.16 | 0.01 | 0.11 | 0.33 | 0.2 | 0.32 |
| 1 | 229367115 | rs342818 | -0.581 | 0.633 | -1.099 | 0.823 | -0.002 | 0.007 | 0.007 | 0 | 0.31 | 0.53 | 0.25 | 0.26 | 0.3 | 0.22 |
| 1 | 229372113 | rs2941079 | -0.153 | 0.559 | -1.609 | 0.567 | -0.019 | -0.039 | -0.039 | 0.012 | 0.54 | 0.84 | 0.66 | 0.6 | 0.54 | 0.59 |
| 1 | 229372913 | rs9435863 | 0.159 | 0.744 | -1.125 | 0.533 | 0.001 | 0.007 | 0.007 | 0.658 | 0.15 | 0.17 | 0.19 | 0.19 | 0.16 | 0.08 |
| 1 | 229373197 | rs4925525 | 0.852 | 0.454 | -1.538 | 0.353 | -0.002 | -0.044 | -0.044 | 0 | 0.37 | 0.11 | 0.3 | 0.41 | 0.32 | 0.4 |
| 1 | 229376357 | rs440392 | -0.826 | 0.734 | -0.932 | 0.166 | 0.015 | -0.046 | -0.046 | 0.309 | 0.55 | 0.86 | 0.65 | 0.51 | 0.62 | 0.47 |
| 1 | 229377014 | rs9435865 | 0.553 | 0.885 | -0.908 | 0.219 | 0.001 | 0.003 | 0.003 | 0.086 | 0.14 | 0.15 | 0.2 | 0.18 | 0.17 | 0.08 |
| 1 | 229377038 | rs2036775 | 0.895 | 0.686 | -0.965 | -0.102 | -0.027 | 0.005 | 0.005 | 0 | 0.23 | 0.13 | 0.31 | 0.24 | 0.31 | 0.25 |
| 1 | 229379488 | rs12132864 | 0.386 | 0.615 | -0.456 | -0.135 | -0.002 | -0.029 | -0.029 | 0.929 | 0.14 | 0.24 | 0.1 | 0.08 | 0.14 | 0.15 |
| 1 | 229385256 | rs6695100 | -0.285 | 0.576 | -0.723 | 0.08 | -0.034 | -0.025 | -0.025 | 0.009 | 0.37 | 0.21 | 0.5 | 0.42 | 0.46 | 0.32 |
| 1 | 229391748 | rs12026082 | 0.542 | 0.802 | -0.762 | -0.009 | -0.041 | 0.007 | 0.007 | 0 | 0.23 | 0.09 | 0.32 | 0.24 | 0.31 | 0.25 |
| 1 | 229411709 | rs237749 | -0.777 | 0.77 | -0.326 | -0.201 | 0.081 | 0.011 | 0.011 | 0 | 0.5 | 0.72 | 0.65 | 0.49 | 0.67 | 0.47 |
| 1 | 229415637 | rs10737440 | 0.187 | 1.07 | 0.424 | -0.273 | -0.023 | -0.011 | -0.011 | 0 | 0.34 | 0.24 | 0.51 | 0.4 | 0.45 | 0.32 |
| 1 | 229416068 | rs10799523 | 0.181 | 1.066 | 0.504 | -0.27 | -0.021 | -0.011 | -0.011 | 0.259 | 0.34 | 0.25 | 0.52 | 0.4 | 0.45 | 0.35 |
| 1 | 229419421 | rs237756 | -1.154 | 1.029 | 0.562 | -0.294 | 0.076 | -0.004 | -0.004 | 0 | 0.53 | 0.77 | 0.67 | 0.49 | 0.71 | 0.48 |
| 1 | 229424118 | rs9660826 | 0.256 | 1.041 | 0.595 | -0.347 | 0.024 | -0.019 | -0.019 | 1.946 | 0.13 | 0.21 | 0.12 | 0.09 | 0.22 | 0.15 |

| | | | | | | | | | | | | | | | | |
|---|---|---|---|---|---|---|---|---|---|---|---|---|---|---|---|---|
| 1 | 229431916 | rs237777 | -0.132 | 1.041 | 0.595 | -0.347 | 0.046 | -0.037 | -0.037 | 0.569 | 0.15 | 0.33 | 0.13 | 0.09 | 0.22 | 0.15 |
| 1 | 229435892 | rs12048819 | 0.201 | 1.042 | 0.498 | -0.328 | -0.021 | -0.008 | -0.008 | 0.859 | 0.34 | 0.25 | 0.51 | 0.4 | 0.45 | 0.32 |
| 1 | 229436923 | rs12084468 | 0.253 | 1.042 | 0.498 | -0.34 | 0.024 | -0.019 | -0.019 | 0.147 | 0.13 | 0.21 | 0.12 | 0.09 | 0.22 | 0.15 |
| 1 | 229438828 | rs237787 | -0.806 | 1.043 | 0.498 | -0.339 | 0.08 | 0.012 | 0.012 | 0 | 0.5 | 0.72 | 0.65 | 0.49 | 0.67 | 0.47 |
| 1 | 229440190 | rs2069372 | 0.2 | 1.052 | 0.501 | -0.333 | -0.021 | -0.009 | -0.009 | 0.189 | 0.34 | 0.25 | 0.51 | 0.4 | 0.45 | 0.33 |
| 1 | 229448733 | rs10916463 | 0.214 | 1.077 | 0.326 | -0.36 | 0.023 | -0.02 | -0.02 | 0 | 0.13 | 0.21 | 0.12 | 0.09 | 0.22 | 0.15 |
| 1 | 229449017 | rs12132466 | 1.218 | 1.113 | 0.225 | -0.379 | -0.018 | 0.045 | 0.045 | 0 | 0.08 | 0.01 | 0.06 | 0 | 0.1 | 0.1 |
| 1 | 229458456 | rs1088 | 0.515 | 0.915 | 0.28 | -0.318 | 0.046 | -0.03 | -0.03 | 0 | 0.14 | 0.3 | 0.13 | 0.09 | 0.22 | 0.15 |
| 1 | 229458632 | rs237799 | 0.252 | 0.952 | -0.081 | -0.316 | 0.074 | -0.043 | -0.043 | 0 | 0.15 | 0.43 | 0.13 | 0.09 | 0.22 | 0.15 |
| 1 | 229461801 | rs237802 | -0.718 | 0.884 | -0.022 | -0.294 | 0.061 | 0.016 | 0.016 | 0 | 0.5 | 0.68 | 0.64 | 0.48 | 0.67 | 0.48 |
| 1 | 229464621 | rs237807 | -0.222 | 1.225 | -0.37 | 0.047 | 0.073 | -0.041 | -0.041 | 0.297 | 0.15 | 0.43 | 0.13 | 0.09 | 0.22 | 0.15 |
| 1 | 229468906 | rs164037 | -0.026 | 0.553 | -0.619 | 0.15 | 0.01 | 0.036 | 0.036 | 0 | 0.39 | 0.75 | 0.34 | 0.32 | 0.37 | 0.36 |
| 1 | 229471199 | rs237784 | -0.782 | 1.409 | -0.558 | 0.493 | 0.036 | -0.019 | -0.019 | 0 | 0.34 | 0.51 | 0.37 | 0.25 | 0.47 | 0.24 |
| 1 | 229475610 | rs238080 | -0.432 | 0.627 | -0.902 | 0.63 | 0 | 0 | 0 | 0 | 0.37 | 0.4 | 0.34 | 0.41 | 0.28 | 0.39 |
| 1 | 229479811 | rs238081 | -0.267 | 0.73 | -1.017 | 0.585 | 0.007 | -0.002 | -0.002 | 0.636 | 0.36 | 0.33 | 0.32 | 0.43 | 0.28 | 0.39 |
| 1 | 229483148 | rs238102 | -0.461 | 0.625 | -0.962 | 0.594 | 0.003 | 0.001 | 0.001 | 0.09 | 0.38 | 0.41 | 0.34 | 0.42 | 0.3 | 0.4 |
| 1 | 229499636 | rs238109 | -0.526 | 0.652 | -1.263 | 0.641 | 0.017 | -0.006 | -0.006 | 0 | 0.41 | 0.29 | 0.33 | 0.45 | 0.33 | 0.42 |
| 1 | 229508905 | rs238099 | -0.625 | 0.674 | -1.864 | -0.005 | -0.062 | 0.086 | 0.086 | 0 | 0.58 | 0.14 | 0.38 | 0.71 | 0.48 | 0.67 |
| 1 | 229519927 | rs7549589 | 0.205 | 0.608 | -1.398 | 0.287 | 0.016 | 0.056 | 0.056 | 0 | 0.56 | 0.72 | 0.65 | 0.27 | 0.64 | 0.41 |
| 1 | 229537513 | rs1002063 | -0.699 | 1.323 | -1.594 | 0.423 | -0.095 | 0.006 | 0.006 | 0 | 0.19 | 0.02 | 0.18 | 0.1 | 0.26 | 0.22 |
| 1 | 229545085 | rs867325 | -0.546 | 0.393 | -0.587 | 0.78 | -0.023 | 0.046 | 0.046 | 0 | 0.19 | 0.44 | 0.35 | 0.21 | 0.14 | 0.24 |
| 1 | 229556990 | rs598392 | -0.292 | -0.139 | -0.614 | 0.567 | -0.016 | -0.009 | -0.009 | 0.031 | 0.36 | 0.44 | 0.46 | 0.29 | 0.25 | 0.31 |
| 1 | 229564117 | rs728614 | 0.122 | -0.767 | -0.432 | 0.254 | -0.012 | -0.014 | -0.014 | 0 | 0.63 | 0.41 | 0.67 | 0.85 | 0.8 | 0.8 |
| 1 | 229565732 | rs499689 | -0.001 | -0.143 | -0.486 | 0.413 | -0.012 | 0.022 | 0.022 | 0.334 | 0.39 | 0.27 | 0.35 | 0.31 | 0.42 | 0.52 |
| 1 | 229566362 | rs526244 | 0.485 | -0.196 | -0.729 | 0.259 | 0 | -0.001 | -0.001 | 0 | 0.14 | 0.15 | 0.13 | 0.15 | 0.14 | 0.2 |
| 1 | 229571061 | rs506388 | -0.731 | -0.535 | -0.958 | 0.182 | -0.006 | -0.013 | -0.013 | 0.423 | 0.67 | 0.71 | 0.65 | 0.48 | 0.6 | 0.68 |
| 1 | 229575709 | rs10753458 | 0.783 | -0.795 | -0.36 | 0.304 | -0.039 | -0.055 | -0.055 | 0 | 0.28 | 0.51 | 0.3 | 0.15 | 0.18 | 0.14 |
| 1 | 229580221 | rs4338396 | 1.017 | -0.902 | 0.104 | 0.253 | -0.01 | -0.016 | -0.016 | 0 | 0.25 | 0.32 | 0.29 | 0.15 | 0.18 | 0.13 |
| 1 | 229594310 | rs10916480 | 1.316 | -0.356 | 0.047 | 0.087 | 0.006 | 0.189 | 0.189 | 0 | 0.13 | 0.26 | 0.25 | 0.53 | 0.13 | 0.21 |
| 1 | 229619516 | rs568916 | 0.6 | -0.247 | -0.245 | -0.344 | -0.011 | 0.015 | 0.015 | 0 | 0.21 | 0.07 | 0.15 | 0.16 | 0.18 | 0.27 |
| 1 | 229631734 | rs11805194 | 0.777 | -0.224 | 0.1 | -0.361 | 0.004 | 0.006 | 0.006 | 0.099 | 0.26 | 0.26 | 0.3 | 0.16 | 0.21 | 0.14 |
| 1 | 229635521 | rs10916495 | 0.917 | -0.226 | 0.129 | -0.362 | 0.012 | 0.024 | 0.024 | 0.092 | 0.25 | 0.14 | 0.29 | 0.16 | 0.2 | 0.14 |
| 1 | 229642652 | rs6671385 | 0.63 | -0.256 | 0.155 | -0.376 | 0.003 | 0.004 | 0.004 | 0 | 0.27 | 0.27 | 0.3 | 0.16 | 0.21 | 0.14 |
| 1 | 229656562 | rs10799533 | 0.843 | -0.23 | 0.182 | -0.344 | 0.009 | 0.013 | 0.013 | 0 | 0.27 | 0.2 | 0.3 | 0.16 | 0.2 | 0.15 |
| 1 | 229681006 | rs927204 | 1.778 | -0.261 | 0.088 | 0.172 | -0.01 | -0.028 | -0.028 | 0 | 0.1 | 0.07 | 0.22 | 0.31 | 0.13 | 0.17 |
| 1 | 229683264 | rs3738187 | -0.087 | -0.213 | 0.446 | -0.492 | -0.03 | -0.042 | -0.042 | 0 | 0.31 | 0.57 | 0.33 | 0.16 | 0.21 | 0.16 |
| 1 | 229693132 | rs12734075 | 0.99 | 0.475 | 0.102 | -0.484 | -0.071 | 0.061 | 0.061 | 0 | 0.1 | 0.01 | 0.11 | 0.01 | 0.24 | 0.06 |
| 1 | 229712340 | rs6587341 | 0.101 | -0.509 | 0.401 | 0.043 | -0.019 | -0.03 | -0.03 | 0 | 0.28 | 0.36 | 0.3 | 0.16 | 0.21 | 0.15 |
| 1 | 229718258 | rs12040438 | 0.103 | -0.116 | -0.225 | 0.168 | -0.008 | -0.017 | -0.017 | 0 | 0.27 | 0.37 | 0.31 | 0.17 | 0.24 | 0.19 |
| 1 | 229723016 | rs348334 | 0.381 | -0.636 | -0.314 | 0.8 | -0.035 | 0.042 | 0.042 | 0 | 0.77 | 0.74 | 0.71 | 0.6 | 0.89 | 0.88 |
| 1 | 229723828 | rs6690223 | 0.612 | -0.645 | -0.703 | -0.08 | 0.008 | 0.035 | 0.035 | 0 | 0.27 | 0.1 | 0.16 | 0.17 | 0.26 | 0.23 |
| 1 | 229724800 | rs6701445 | 0.193 | 0.537 | -0.334 | -0.397 | 0.027 | -0.043 | -0.043 | 0.567 | 0.84 | 0.7 | 0.68 | 0.94 | 0.79 | 0.78 |
| 1 | 229730452 | rs2295625 | 0.583 | 0.434 | -0.93 | -0.241 | -0.077 | 0.044 | 0.044 | 0.268 | 0.43 | 0.09 | 0.27 | 0.33 | 0.46 | 0.43 |
| 1 | 229733674 | rs1444726 | 0.739 | 0.035 | -0.379 | -0.096 | -0.028 | 0.189 | 0.189 | 0 | 0.11 | 0.21 | 0.24 | 0.43 | 0.06 | 0.07 |
| 1 | 229737680 | rs1816878 | 0.81 | 0.034 | -0.347 | -0.097 | -0.025 | 0.158 | 0.158 | 0 | 0.11 | 0.19 | 0.23 | 0.43 | 0.06 | 0.07 |
| 1 | 229738170 | rs3753886 | -0.176 | 0.443 | 0.1 | -0.188 | 0.004 | -0.023 | -0.023 | 0 | 0.57 | 0.49 | 0.54 | 0.76 | 0.52 | 0.57 |
| 1 | 229738236 | rs4925500 | -0.489 | 0.435 | 0.15 | -0.275 | -0.045 | 0.041 | 0.041 | 0 | 0.86 | 0.93 | 0.72 | 0.94 | 0.76 | 0.79 |
| 1 | 229739148 | rs10916525 | -0.106 | 0.475 | 0.04 | -0.232 | 0.001 | -0.01 | -0.01 | 0.297 | 0.56 | 0.41 | 0.54 | 0.76 | 0.51 | 0.57 |
| 1 | 229746019 | rs2273573 | -0.134 | 0.474 | 0.017 | -0.232 | 0.004 | -0.022 | -0.022 | 0 | 0.57 | 0.49 | 0.54 | 0.76 | 0.52 | 0.57 |
| 1 | 229750036 | rs7547607 | 1.623 | 0.444 | 0.182 | -0.263 | 0.003 | 0.017 | 0.017 | 0 | 0.26 | 0.13 | 0.15 | 0.18 | 0.24 | 0.22 |
| 1 | 229750343 | rs2273574 | 0.877 | 0.389 | -0.224 | -0.137 | -0.003 | -0.005 | -0.005 | 0 | 0.33 | 0.09 | 0.22 | 0.31 | 0.31 | 0.43 |
| 1 | 229772693 | rs3811473 | -0.617 | 0.451 | -0.078 | 0.042 | 0.004 | -0.02 | -0.02 | 0 | 0.57 | 0.48 | 0.54 | 0.76 | 0.52 | 0.57 |
| 1 | 229772932 | rs3811474 | 0.41 | 0.431 | -0.101 | 0.085 | -0.028 | 0.155 | 0.155 | 0 | 0.12 | 0.2 | 0.24 | 0.43 | 0.06 | 0.07 |
| 1 | 229785425 | rs6696449 | -0.038 | 0.528 | -0.288 | 0.027 | -0.002 | 0.053 | 0.053 | 0 | 0.45 | 0.24 | 0.3 | 0.33 | 0.45 | 0.5 |
| 1 | 229793082 | rs6686112 | 0.036 | 0.492 | -0.203 | -0.104 | -0.024 | 0.006 | 0.006 | 0.023 | 0.45 | 0.2 | 0.33 | 0.42 | 0.48 | 0.49 |
| 1 | 229795070 | rs1048306 | -0.764 | 0.448 | -0.106 | -0.071 | 0.002 | -0.028 | -0.028 | 0 | 0.58 | 0.51 | 0.58 | 0.83 | 0.55 | 0.56 |
| 1 | 229795330 | rs3820361 | -0.292 | 0.861 | -0.483 | -0.051 | -0.004 | 0.009 | 0.009 | 0.052 | 0.46 | 0.32 | 0.34 | 0.42 | 0.48 | 0.49 |
| 1 | 229798874 | rs12403112 | -0.282 | 0.878 | -0.662 | -0.043 | -0.007 | 0.012 | 0.012 | 0.069 | 0.46 | 0.28 | 0.34 | 0.42 | 0.48 | 0.49 |
| 1 | 229803654 | rs12738647 | -0.015 | 1.15 | -0.831 | -0.196 | -0.041 | 0.055 | 0.055 | 0 | 0.14 | 0.03 | 0.11 | 0.04 | 0.19 | 0.06 |
| 1 | 229805213 | rs12117146 | 0.166 | 1.06 | 0.256 | 0.206 | 0.012 | -0.012 | -0.012 | 0.008 | 0.92 | 0.85 | 0.93 | 0.93 | 0.87 | 0.85 |
| 1 | 229810806 | rs2891866 | 0.302 | 0.498 | 0.238 | -0.367 | 0.013 | 0.046 | 0.046 | 0 | 0.52 | 0.39 | 0.47 | 0.11 | 0.54 | 0.35 |
| 1 | 229822279 | rs12745210 | 0.347 | 0.386 | 0.26 | -0.429 | 0.034 | 0.236 | 0.236 | 0 | 0.49 | 0.18 | 0.45 | 0.11 | 0.54 | 0.35 |
| 1 | 229824672 | rs11583220 | 2.001 | 0.223 | 0.34 | -0.517 | -0.011 | -0.019 | -0.019 | 0 | 0.06 | 0.07 | 0.03 | 0.02 | 0.06 | 0.02 |
| 1 | 229827300 | rs4925464 | 0 | 0.344 | 0.568 | -1.184 | -0.005 | -0.019 | -0.019 | 0 | 0.41 | 0.57 | 0.46 | 0.82 | 0.34 | 0.49 |
| 1 | 229829737 | rs16850116 | 0.096 | 0.394 | 0.179 | -1.172 | 0.093 | 0.178 | 0.178 | 0.052 | 0.54 | 0.84 | 0.49 | 0.82 | 0.34 | 0.5 |
| 1 | 229835413 | rs7525615 | 0.375 | 0.314 | 0.383 | -1.023 | -0.007 | -0.025 | -0.025 | 0.225 | 0.32 | 0.35 | 0.42 | 0.81 | 0.31 | 0.51 |
| 1 | 229835854 | rs7518596 | 0.412 | 0.32 | 0.388 | -1.05 | -0.007 | -0.025 | -0.025 | 0 | 0.32 | 0.35 | 0.42 | 0.81 | 0.31 | 0.49 |
| 1 | 229840693 | rs7541744 | -0.106 | 0.498 | 0.093 | -0.996 | 0.001 | -0.007 | -0.007 | 4.723 | 0.42 | 0.45 | 0.47 | 0.81 | 0.39 | 0.5 |
| 1 | 229845799 | rs1931324 | -0.937 | 0.808 | -0.463 | -1.162 | -0.01 | 0.097 | 0.097 | 0.01 | 0.62 | 0.8 | 0.63 | 0.86 | 0.6 | 0.61 |

| | | | | | | | | | | | | | | | | |
|---|---|---|---|---|---|---|---|---|---|---|---|---|---|---|---|---|
| 1 | 229850656 | rs1889192 | -0.239 | 0.969 | 0.01 | -0.751 | 0.002 | -0.015 | -0.015 | 0 | 0.17 | 0.18 | 0.13 | 0.04 | 0.21 | 0.06 |
| 1 | 229861486 | rs1930903 | -0.379 | 1.005 | -0.215 | -0.413 | -0.016 | 0.028 | 0.028 | 0.221 | 0.57 | 0.67 | 0.49 | 0.3 | 0.49 | 0.42 |
| 1 | 229863007 | rs12742436 | 0.544 | 1.231 | -0.365 | -0.327 | -0.042 | 0.02 | 0.02 | 0.09 | 0.11 | 0.04 | 0.09 | 0.05 | 0.18 | 0.05 |
| 1 | 229867467 | rs914909 | -0.659 | 1.207 | -0.307 | 0.128 | -0.035 | -0.033 | -0.033 | 0.226 | 0.94 | 0.98 | 0.92 | 0.8 | 0.84 | 0.82 |
| 1 | 229874672 | rs2210590 | 0.446 | 0.786 | -0.279 | -0.215 | -0.042 | 0.052 | 0.052 | 0.637 | 0.19 | 0.06 | 0.13 | 0.05 | 0.25 | 0.1 |
| 1 | 229888337 | rs670054 | 0.605 | 0.364 | -0.219 | 0.166 | -0.007 | -0.007 | -0.007 | 0 | 0.31 | 0.32 | 0.32 | 0.25 | 0.24 | 0.26 |
| 1 | 229900511 | rs592228 | 0.464 | 0.784 | -0.068 | 0.646 | 0.004 | 0.008 | 0.008 | 1.028 | 0.9 | 0.85 | 0.9 | 0.81 | 0.84 | 0.81 |
| 1 | 229905386 | rs485056 | 0.464 | 0.784 | -0.047 | 0.654 | 0.008 | 0.014 | 0.014 | 1.028 | 0.9 | 0.84 | 0.9 | 0.81 | 0.84 | 0.81 |
| 1 | 229910737 | rs643009 | 0.919 | 0.721 | -0.088 | 0.478 | 0.002 | -0.009 | -0.009 | 0 | 0.62 | 0.45 | 0.7 | 0.73 | 0.57 | 0.64 |
| 1 | 229912382 | rs570701 | 0.847 | 0.816 | -0.21 | 1.199 | -0.005 | -0.001 | -0.001 | 0 | 0.83 | 0.81 | 0.8 | 0.64 | 0.75 | 0.76 |
| 1 | 229916660 | rs557208 | 0.336 | 0.147 | 0.539 | 1.069 | 0.036 | 0.037 | 0.037 | 0 | 0.7 | 0.57 | 0.72 | 0.61 | 0.56 | 0.67 |
| 1 | 229921927 | rs479531 | 0.138 | -0.075 | 1.017 | 0.977 | -0.072 | -0.075 | -0.075 | 0 | 0.73 | 0.93 | 0.76 | 0.61 | 0.57 | 0.68 |
| 1 | 229927798 | rs583645 | 0.586 | -0.037 | 1 | 1.013 | -0.002 | -0.002 | -0.002 | 0 | 0.68 | 0.37 | 0.71 | 0.61 | 0.56 | 0.68 |
| 1 | 229929602 | rs2780630 | 0.312 | 0.02 | 0.873 | 1.043 | 0.002 | 0.002 | 0.002 | 0 | 0.69 | 0.69 | 0.74 | 0.61 | 0.57 | 0.68 |
| 1 | 229939169 | rs657297 | 0.606 | -0.32 | 0.207 | 0.957 | 0.005 | -0.004 | -0.004 | 0 | 0.58 | 0.31 | 0.69 | 0.57 | 0.55 | 0.63 |
| 1 | 229958921 | rs662363 | 0.26 | -0.376 | 0.031 | 0.812 | 0.006 | 0.01 | 0.01 | 0 | 0.55 | 0.65 | 0.69 | 0.57 | 0.53 | 0.6 |
| 1 | 229974348 | rs687266 | 0.373 | -0.465 | 0.203 | 0.675 | 0 | 0.001 | 0.001 | 0 | 0.52 | 0.52 | 0.66 | 0.56 | 0.51 | 0.59 |
| 1 | 229981623 | rs528425 | 0.146 | -0.489 | 0.116 | 0.641 | -0.012 | 0.01 | 0.01 | 0 | 0.54 | 0.83 | 0.68 | 0.56 | 0.51 | 0.6 |
| 1 | 229983606 | rs1029119 | -1.569 | -0.676 | 1.111 | 0.52 | 0.048 | 0.044 | 0.044 | 0.939 | 0.06 | 0.47 | 0.05 | 0 | 0.01 | 0.01 |
| 1 | 229984347 | rs6587358 | -1.772 | -0.671 | 1.042 | 0.491 | -0.001 | -0.014 | -0.014 | 0 | 0.07 | 0.59 | 0.06 | 0 | 0.01 | 0.01 |
| 1 | 229984950 | rs16850412 | 0.541 | -0.792 | 0.499 | 0.442 | -0.02 | -0.04 | -0.04 | 0 | 0.16 | 0.01 | 0.12 | 0.16 | 0.15 | 0.09 |
| 1 | 229988711 | rs7548102 | -2.376 | -0.69 | 1.725 | 0.489 | -0.036 | -0.046 | -0.046 | 0 | 0.09 | 0.78 | 0.07 | 0 | 0.01 | 0.01 |
| 1 | 229993997 | rs6687252 | -1.634 | -0.677 | 1.965 | 0.495 | 0.059 | 0.043 | 0.043 | 0.762 | 0.06 | 0.45 | 0.04 | 0 | 0.01 | 0 |
| 1 | 229996754 | rs10495282 | -1.864 | -0.677 | 1.768 | 0.495 | 0.005 | -0.009 | -0.009 | 0.146 | 0.07 | 0.6 | 0.06 | 0 | 0.01 | 0 |
| 1 | 229998239 | rs12062782 | -2.302 | -0.683 | 1.68 | 0.492 | -0.027 | -0.039 | -0.039 | 0.609 | 0.08 | 0.7 | 0.07 | 0 | 0.01 | 0.01 |
| 1 | 230010403 | rs4087355 | -1.746 | -0.634 | 1.631 | 0.515 | 0.003 | -0.01 | -0.01 | 0 | 0.07 | 0.64 | 0.06 | 0 | 0.01 | 0 |
| 1 | 230018473 | rs2780624 | -1.025 | -0.379 | 0.649 | 0.406 | -0.111 | -0.003 | -0.003 | 0 | 0.81 | 0.99 | 0.71 | 0.9 | 0.69 | 0.86 |
| 1 | 230021629 | rs3853553 | -0.191 | -1.425 | 1.267 | 1.081 | 0.018 | -0.036 | -0.036 | 0 | 0.17 | 0 | 0.05 | 0.21 | 0.1 | 0.22 |
| 1 | 230051057 | rs6685413 | -2.225 | -0.755 | 2.569 | 1.228 | 0.049 | 0.04 | 0.04 | 0.279 | 0.06 | 0.42 | 0.04 | 0 | 0 | 0.01 |
| 1 | 230056781 | rs4925523 | -1.912 | -0.52 | 2.289 | 1.515 | 0 | 0.006 | 0.006 | 0.665 | 0.28 | 0.71 | 0.36 | 0.4 | 0.24 | 0.33 |
| 1 | 230074815 | rs7526242 | -2.339 | -0.364 | 2.867 | 1.466 | 0.081 | 0.064 | 0.064 | 0.179 | 0.06 | 0.49 | 0.04 | 0 | 0.01 | 0.01 |
| 1 | 230081298 | rs11578286 | -1.898 | -0.345 | 2.779 | 1.538 | -0.001 | 0 | 0 | 0 | 0.06 | 0.12 | 0.04 | 0.04 | 0.03 | 0.11 |
| 1 | 230088714 | rs11588095 | -3.532 | -0.391 | 3.075 | 1.53 | -0.008 | 0.003 | 0.003 | 0.026 | 0.1 | 0.57 | 0.07 | 0.04 | 0.03 | 0.12 |
| 1 | 230095259 | rs16841453 | -2.171 | -0.384 | 3.043 | 1.534 | 0.029 | 0.045 | 0.045 | 0 | 0.06 | 0.31 | 0.05 | 0.04 | 0.03 | 0.11 |
| 1 | 230104395 | rs7522922 | -1.67 | -0.354 | 2.477 | 1.548 | -0.003 | 0 | 0 | 0 | 0.05 | 0.12 | 0.04 | 0.04 | 0.03 | 0.11 |
| 1 | 230111146 | rs12029484 | -0.684 | -0.386 | 2.451 | 1.715 | -0.01 | -0.009 | -0.009 | 1.07 | 0.06 | 0.01 | 0.13 | 0.29 | 0.07 | 0.11 |
| 1 | 230121692 | rs11588464 | -2.428 | -0.391 | 2.561 | 1.501 | 0.002 | -0.03 | -0.03 | 0.687 | 0.08 | 0.32 | 0.05 | 0.01 | 0.03 | 0.12 |
| 1 | 230130480 | rs11122536 | -2.213 | -0.108 | 2.133 | 1.756 | 0.033 | 0.113 | 0.113 | 0 | 0.19 | 0.43 | 0.3 | 0.39 | 0.19 | 0.26 |
| 1 | 230131583 | rs11122542 | -1.704 | -0.36 | 1.833 | 1.285 | -0.002 | 0.002 | 0.002 | 1.697 | 0.13 | 0.12 | 0.06 | 0.06 | 0.12 | 0.11 |
| 1 | 230136986 | rs10779848 | -3.281 | -0.624 | 2.365 | 1.796 | 0.002 | -0.006 | -0.006 | 0 | 0.3 | 0.71 | 0.2 | 0.43 | 0.22 | 0.32 |
| 1 | 230138554 | rs9432181 | -2.287 | -0.728 | 3.097 | 1.87 | 0.023 | 0.331 | 0.331 | 0 | 0.1 | 0.4 | 0.12 | 0.36 | 0.09 | 0.14 |
| 1 | 230142626 | rs3811492 | -2.701 | -0.754 | 3.367 | 1.849 | -0.039 | -0.055 | -0.055 | 0 | 0.21 | 0.52 | 0.1 | 0.09 | 0.11 | 0.2 |
| 1 | 230143970 | rs12042904 | -1.359 | -0.714 | 3.339 | 1.909 | 0.023 | 0.253 | 0.253 | 0 | 0.07 | 0.24 | 0.09 | 0.34 | 0.09 | 0.09 |
| 1 | 230151884 | rs4628515 | -2.386 | -0.802 | 3.175 | 1.669 | -0.03 | -0.044 | -0.044 | 0 | 0.2 | 0.34 | 0.09 | 0.09 | 0.11 | 0.17 |
| 1 | 230152083 | rs4295887 | -2.469 | -0.803 | 3.167 | 1.669 | -0.039 | -0.057 | -0.057 | 0 | 0.21 | 0.39 | 0.09 | 0.09 | 0.11 | 0.17 |
| 1 | 230168963 | rs4847027 | -3.222 | -0.761 | 2.995 | 1.779 | 0 | -0.001 | -0.001 | 1.095 | 0.29 | 0.7 | 0.19 | 0.43 | 0.2 | 0.3 |
| 1 | 230180447 | rs877968 | -2.707 | -0.714 | 2.898 | 1.646 | 0.001 | -0.022 | -0.022 | 0 | 0.26 | 0.7 | 0.18 | 0.41 | 0.22 | 0.23 |
| 1 | 230186844 | rs7515331 | -1.556 | -0.777 | 2.893 | 1.652 | 0.015 | 0.127 | 0.127 | 0 | 0.09 | 0.13 | 0.08 | 0.35 | 0.11 | 0.12 |
| 1 | 230187526 | rs6663181 | 2.73 | -0.798 | 2.719 | 1.641 | 0.006 | -0.056 | -0.056 | 0 | 0.74 | 0.24 | 0.82 | 0.59 | 0.79 | 0.77 |
| 1 | 230188994 | rs12406877 | -1.192 | -0.834 | 2.806 | 1.633 | 0 | 0.01 | 0.01 | 0.174 | 0.12 | 0.09 | 0.04 | 0.06 | 0.1 | 0.09 |
| 1 | 230192848 | rs1467469 | 2.618 | -0.859 | 2.99 | 1.665 | -0.007 | 0.114 | 0.114 | 0 | 0.86 | 0.39 | 0.87 | 0.64 | 0.89 | 0.83 |
| 1 | 230195713 | rs3009715 | -2.953 | -0.868 | 2.584 | 1.66 | 0.002 | 0.203 | 0.203 | 0 | 0.14 | 0.38 | 0.13 | 0.34 | 0.11 | 0.21 |
| 1 | 230202028 | rs1321258 | -2.098 | -1.106 | 1.87 | 1.7 | 0.007 | 0.306 | 0.306 | 0 | 0.1 | 0.32 | 0.1 | 0.34 | 0.07 | 0.15 |
| 1 | 230207349 | rs7531967 | -3.164 | -1.256 | 0.517 | 1.615 | 0.01 | -0.015 | -0.015 | 0 | 0.2 | 0.8 | 0.16 | 0.35 | 0.09 | 0.26 |
| 1 | 230207406 | rs1474926 | 2.885 | -1.229 | 0.026 | 1.567 | 0.047 | -0.109 | -0.109 | 0 | 0.68 | 0.1 | 0.78 | 0.59 | 0.81 | 0.64 |
| 1 | 230208372 | rs2883025 | 0.863 | -0.662 | -0.075 | 1.728 | -0.019 | 0.009 | 0.009 | 0.038 | 0.9 | 0.96 | 0.94 | 0.95 | 0.9 | 0.9 |
| 1 | 230208852 | rs12046584 | -1.06 | -0.947 | -0.02 | 1.854 | -0.005 | 0.117 | 0.117 | 0 | 0.07 | 0.06 | 0.06 | 0.36 | 0.07 | 0.15 |
| 1 | 230209313 | rs13375918 | -1.254 | -0.821 | -0.247 | 0.916 | -0.011 | -0.024 | -0.024 | 1.071 | 0.05 | 0.12 | 0.01 | 0 | 0.02 | 0.03 |
| 1 | 230209790 | rs6684432 | 0.878 | -0.631 | -0.571 | 1.047 | -0.004 | 0.006 | 0.006 | 0.115 | 0.91 | 0.96 | 0.96 | 0.95 | 0.91 | 0.91 |
| 1 | 230211398 | rs6673349 | -1.294 | -1.389 | -0.722 | 0.949 | 0.029 | 0.009 | 0.009 | 0 | 0.23 | 0.81 | 0.27 | 0.13 | 0.09 | 0.21 |
| 1 | 230212533 | rs11122275 | -0.365 | -0.887 | -1.337 | 0.999 | -0.033 | -0.039 | -0.039 | 0 | 0.16 | 0.02 | 0.12 | 0.2 | 0.16 | 0.19 |
| 1 | 230214319 | rs910820 | 0.224 | -1.427 | -0.912 | 0.468 | -0.032 | -0.051 | -0.051 | 0.097 | 0.65 | 0.52 | 0.63 | 0.86 | 0.76 | 0.75 |
| 1 | 230230759 | rs7523343 | -0.658 | -1.238 | -1 | 1.486 | 0.002 | -0.004 | -0.004 | 0.314 | 0.25 | 0.2 | 0.17 | 0.39 | 0.21 | 0.23 |
| 1 | 230231546 | rs2057231 | -1.246 | -1.644 | -1.326 | 0.623 | -0.004 | 0 | 0 | 0 | 0.54 | 0.51 | 0.45 | 0.46 | 0.43 | 0.52 |
| 1 | 230237103 | rs11122316 | -1.244 | -1.915 | -1.352 | 0.908 | 0.002 | 0.01 | 0.01 | 0 | 0.57 | 0.32 | 0.49 | 0.57 | 0.44 | 0.6 |
| 1 | 230238507 | rs1998067 | 0.378 | -1.923 | -1.304 | 0.811 | 0.013 | 0.111 | 0.111 | 0 | 0.26 | 0.05 | 0.26 | 0.06 | 0.21 | 0.28 |
| 1 | 230238821 | rs12035297 | 1.141 | -0.402 | -1.53 | 0.779 | 0.07 | 0.024 | 0.024 | 0 | 0.18 | 0.35 | 0.22 | 0.26 | 0.26 | 0.21 |
| 1 | 230248739 | rs7550262 | 0.645 | -0.435 | -1.534 | 0.773 | 0.002 | -0.007 | -0.007 | 0 | 0.25 | 0.2 | 0.16 | 0.39 | 0.21 | 0.23 |
| 1 | 230250182 | rs754666 | -0.175 | -0.181 | -1.395 | 0.823 | 0.004 | 0.002 | 0.002 | 0 | 0.52 | 0.41 | 0.44 | 0.44 | 0.42 | 0.51 |

| | | | | | | | | | | | | | | | | |
|---|---|---|---|---|---|---|---|---|---|---|---|---|---|---|---|---|
| 1 | 230251823 | rs4846833 | -0.858 | -0.19 | -1.56 | 0.776 | -0.003 | -0.015 | -0.015 | 0 | 0.7 | 0.76 | 0.66 | 0.72 | 0.68 | 0.73 |
| 1 | 230263844 | rs2144297 | -0.258 | -0.208 | -1.532 | 0.744 | -0.001 | 0.03 | 0.03 | 0.325 | 0.48 | 0.65 | 0.54 | 0.67 | 0.47 | 0.5 |
| 1 | 230263959 | rs11122361 | 0.47 | -0.268 | -1.31 | 0.862 | -0.004 | 0.001 | 0.001 | 0 | 0.31 | 0.3 | 0.33 | 0.4 | 0.22 | 0.29 |
| 1 | 230265342 | rs7541570 | 0.957 | 0.368 | -1.071 | 0.955 | 0.071 | 0.018 | 0.018 | 0 | 0.16 | 0.34 | 0.22 | 0.25 | 0.24 | 0.2 |
| 1 | 230267209 | rs1884578 | -1.452 | 0.35 | -1.454 | 0.902 | -0.097 | 0.005 | 0.005 | 0 | 0.77 | 0.99 | 0.76 | 0.85 | 0.71 | 0.84 |
| 1 | 230272063 | rs1973943 | -0.381 | 0.118 | -1.355 | 0.781 | -0.003 | -0.004 | -0.004 | 0 | 0.53 | 0.66 | 0.44 | 0.45 | 0.51 | 0.52 |
| 1 | 230272925 | rs11122383 | -0.105 | 0.247 | -1.409 | 0.495 | 0.007 | -0.005 | -0.005 | 0 | 0.46 | 0.58 | 0.5 | 0.59 | 0.39 | 0.49 |
| 1 | 230279410 | rs4846904 | -0.296 | 0.291 | -1.123 | 0.578 | -0.002 | 0.03 | 0.03 | 0.076 | 0.8 | 0.77 | 0.84 | 0.34 | 0.8 | 0.76 |
| 1 | 230285056 | rs4846908 | 0.41 | 0.566 | -1.46 | -0.552 | 0 | 0.02 | 0.02 | 0 | 0.46 | 0.4 | 0.41 | 0.14 | 0.46 | 0.38 |
| 1 | 230287128 | rs1998064 | 0.798 | 0.463 | -1.297 | -0.179 | 0.003 | -0.02 | -0.02 | 0 | 0.22 | 0.25 | 0.15 | 0.06 | 0.25 | 0.16 |
| 1 | 230291868 | rs2352723 | -0.961 | -0.013 | -1.47 | 0.799 | -0.008 | -0.024 | -0.024 | 0 | 0.85 | 0.9 | 0.86 | 0.77 | 0.82 | 0.78 |
| 1 | 230294715 | rs4846913 | 0.507 | -0.069 | -1.937 | 0.068 | -0.083 | 0.174 | 0.174 | 0 | 0.45 | 0.06 | 0.54 | 0.2 | 0.6 | 0.4 |
| 1 | 230294916 | rs2144300 | 0.513 | -0.07 | -1.938 | 0.068 | -0.084 | 0.173 | 0.173 | 0 | 0.45 | 0.06 | 0.54 | 0.2 | 0.6 | 0.39 |
| 1 | 230297659 | rs2281719 | 0.636 | -0.024 | -1.916 | 0.165 | -0.071 | 0.131 | 0.131 | 0 | 0.46 | 0.06 | 0.54 | 0.26 | 0.61 | 0.41 |
| 1 | 230299949 | rs10779835 | 0.636 | -0.024 | -1.916 | 0.165 | -0.071 | 0.131 | 0.131 | 0 | 0.46 | 0.06 | 0.54 | 0.26 | 0.61 | 0.41 |
| 1 | 230301776 | rs2296065 | -1.081 | -0.025 | -1.616 | 0.121 | -0.001 | -0.008 | -0.008 | 0.101 | 0.86 | 0.85 | 0.92 | 0.96 | 0.84 | 0.85 |
| 1 | 230304988 | rs10489615 | 0.531 | -0.036 | -1.197 | 0.144 | 0.025 | 0.125 | 0.125 | 0 | 0.49 | 0.25 | 0.56 | 0.26 | 0.61 | 0.41 |
| 1 | 230308855 | rs11122458 | -0.385 | 0.162 | -1.248 | 0.857 | -0.01 | -0.037 | -0.037 | 0 | 0.26 | 0.13 | 0.3 | 0.31 | 0.2 | 0.4 |
| 1 | 230313437 | rs11122459 | -0.214 | 0.026 | -1.19 | 1.106 | 0.001 | 0.02 | 0.02 | 0.017 | 0.07 | 0.07 | 0.21 | 0.27 | 0.05 | 0.18 |
| 1 | 230313911 | rs678050 | -0.099 | -0.23 | -0.467 | 1.099 | 0.017 | 0.222 | 0.222 | 0 | 0.16 | 0.43 | 0.09 | 0.38 | 0.17 | 0.15 |
| 1 | 230324067 | rs611229 | -1.067 | -0.292 | -0.864 | 0.451 | 0.035 | 0.097 | 0.097 | 0 | 0.47 | 0.71 | 0.4 | 0.69 | 0.37 | 0.54 |
| 1 | 230327568 | rs10779837 | -0.972 | -0.009 | -0.339 | 1.23 | -0.007 | 0.083 | 0.083 | 0 | 0.17 | 0.25 | 0.22 | 0.53 | 0.14 | 0.37 |
| 1 | 230327766 | rs6678450 | -0.509 | 0.165 | -0.491 | 0.107 | -0.002 | 0.038 | 0.038 | 0 | 0.52 | 0.45 | 0.36 | 0.3 | 0.56 | 0.5 |
| 1 | 230384970 | rs1923950 | 0.441 | 0.305 | 0.296 | 0.331 | -0.002 | 0.007 | 0.007 | 0.045 | 0.15 | 0.1 | 0.07 | 0.09 | 0.15 | 0.13 |
| 1 | 230387283 | rs3789629 | -1.245 | -0.035 | 0.066 | -0.089 | 0.019 | 0.045 | 0.045 | 0 | 0.45 | 0.6 | 0.62 | 0.73 | 0.39 | 0.4 |
| 1 | 230390002 | rs901675 | -1.417 | 0.535 | -0.597 | -0.368 | -0.073 | 0.099 | 0.099 | 0 | 0.77 | 0.92 | 0.87 | 0.95 | 0.71 | 0.78 |
| 1 | 230398445 | rs3811485 | 0.092 | 0.163 | -0.21 | -0.578 | -0.021 | -0.047 | -0.047 | 0.487 | 0.15 | 0.25 | 0.14 | 0.04 | 0.12 | 0.14 |
| 1 | 230399410 | rs16851270 | 1.031 | 0.346 | -0.437 | -0.477 | 0.004 | 0.017 | 0.017 | 0 | 0.11 | 0.08 | 0.12 | 0.04 | 0.11 | 0.14 |
| 1 | 230408858 | rs11576324 | 0.393 | 0.315 | -0.648 | -1.417 | 0.003 | -0.008 | -0.008 | 0 | 0.44 | 0.38 | 0.35 | 0.08 | 0.45 | 0.43 |
| 1 | 230409058 | rs11588358 | 0.608 | 0.288 | -0.659 | -1.447 | 0.002 | -0.025 | -0.025 | 0 | 0.41 | 0.37 | 0.3 | 0.06 | 0.43 | 0.42 |
| 1 | 230409562 | rs17761459 | -0.08 | -0.41 | -0.33 | -0.956 | -0.001 | -0.006 | -0.006 | 0 | 0.17 | 0.18 | 0.08 | 0.06 | 0.15 | 0.12 |
| 1 | 230409624 | rs7533085 | 0.023 | -0.21 | -0.113 | -0.93 | 0.01 | 0.067 | 0.067 | 0.478 | 0.07 | 0.23 | 0.08 | 0.16 | 0.05 | 0.11 |
| 1 | 230410162 | rs12091838 | -0.49 | -0.38 | -0.451 | -1.249 | -0.017 | -0.043 | -0.043 | 0.15 | 0.2 | 0.25 | 0.15 | 0.04 | 0.14 | 0.06 |
| 1 | 230412797 | rs16851315 | 1.694 | -0.202 | -0.999 | -1.281 | 0.012 | 0.044 | 0.044 | 1.051 | 0.08 | 0.01 | 0.05 | 0 | 0.06 | 0.01 |
| 1 | 230415025 | rs6698963 | -1.203 | -0.133 | -1.631 | -1.94 | -0.093 | 0.286 | 0.286 | 0.057 | 0.57 | 0.89 | 0.66 | 0.95 | 0.46 | 0.59 |
| 1 | 230415244 | rs2273968 | 0.719 | 0.01 | -0.755 | -1.007 | 0.031 | 0.263 | 0.263 | 0 | 0.09 | 0.25 | 0.24 | 0.44 | 0.11 | 0.1 |
| 1 | 230415293 | rs2273967 | -0.041 | -0.199 | -0.419 | -1.15 | 0 | 0 | 0 | 0 | 0.3 | 0.3 | 0.17 | 0.18 | 0.23 | 0.36 |
| 1 | 230415924 | rs3213494 | 1.52 | -0.165 | -0.223 | -1.095 | 0.026 | 0.044 | 0.044 | 0.058 | 0.05 | 0.2 | 0.05 | 0.08 | 0.03 | 0.09 |
| 1 | 230416038 | rs3213495 | 0.657 | 0.001 | -0.579 | -1.05 | 0.086 | 0.298 | 0.298 | 0 | 0.08 | 0.25 | 0.25 | 0.44 | 0.12 | 0.11 |
| 1 | 230416744 | rs1043900 | 0.169 | -0.126 | -0.904 | -1.275 | -0.019 | 0.084 | 0.084 | 0 | 0.25 | 0.06 | 0.12 | 0.11 | 0.2 | 0.27 |
| 1 | 230417294 | rs1043908 | 0.666 | -0.338 | -0.685 | -1.396 | -0.004 | -0.029 | -0.029 | 0 | 0.15 | 0.19 | 0.08 | 0.05 | 0.12 | 0.14 |
| 1 | 230417371 | rs1043909 | 2.113 | -0.47 | -1.318 | -1.801 | -0.012 | -0.117 | -0.117 | 0 | 0.23 | 0.02 | 0.23 | 0.34 | 0.21 | 0.24 |
| 1 | 230417394 | rs3088075 | 0.267 | -0.451 | -0.821 | -1.816 | -0.008 | 0.231 | 0.231 | 0 | 0.14 | 0.37 | 0.26 | 0.47 | 0.1 | 0.11 |
| 1 | 230417408 | rs11620 | 0.196 | -0.459 | -0.878 | -1.906 | 0.005 | 0.001 | 0.001 | 0 | 0.52 | 0.57 | 0.45 | 0.51 | 0.44 | 0.5 |
| 1 | 230417999 | rs9431818 | -0.237 | -0.363 | -1.114 | -2.006 | -0.094 | 0.199 | 0.199 | 0 | 0.61 | 0.86 | 0.66 | 0.91 | 0.51 | 0.51 |
| 1 | 230419308 | rs12757317 | 2.326 | -0.504 | -1.084 | -1.931 | 0.017 | -0.062 | -0.062 | 0 | 0.24 | 0.07 | 0.23 | 0.34 | 0.21 | 0.24 |
| 1 | 230419344 | rs6680658 | 0.089 | -0.413 | -0.935 | -1.981 | 0.02 | 0.061 | 0.061 | 0 | 0.28 | 0.15 | 0.17 | 0.08 | 0.24 | 0.23 |
| 1 | 230420370 | rs4846948 | 0.034 | -0.589 | -0.646 | -1.859 | 0.002 | -0.013 | -0.013 | 0 | 0.55 | 0.53 | 0.59 | 0.85 | 0.43 | 0.46 |
| 1 | 230421477 | rs7511943 | 1.33 | -0.59 | -0.597 | -1.783 | 0.013 | 0.214 | 0.214 | 0 | 0.09 | 0.22 | 0.23 | 0.46 | 0.09 | 0.09 |
| 1 | 230421503 | rs6541309 | -0.6 | -0.503 | -0.799 | -1.809 | -0.058 | 0.201 | 0.201 | 0.074 | 0.62 | 0.85 | 0.66 | 0.92 | 0.47 | 0.49 |
| 1 | 230421722 | rs6687864 | 0.596 | -0.576 | -0.74 | -1.705 | -0.023 | -0.042 | -0.042 | 0 | 0.18 | 0.22 | 0.1 | 0.04 | 0.12 | 0.1 |
| 1 | 230422031 | rs17762687 | 1.578 | -0.301 | -0.74 | -1.491 | 0.01 | 0.019 | 0.019 | 0 | 0.09 | 0.01 | 0.04 | 0.04 | 0.06 | 0.09 |
| 1 | 230422210 | rs10864734 | 0.588 | -0.063 | -0.469 | -1.256 | 0.001 | -0.001 | -0.001 | 0 | 0.36 | 0.34 | 0.48 | 0.79 | 0.31 | 0.33 |
| 1 | 230423481 | rs11122478 | 0.253 | 0.306 | -0.146 | -0.842 | -0.001 | 0.192 | 0.192 | 0 | 0.13 | 0.29 | 0.25 | 0.46 | 0.1 | 0.09 |
| 1 | 230427840 | rs9793529 | -0.797 | 1.259 | -1.215 | -1.201 | -0.023 | 0.014 | 0.014 | 0.099 | 0.87 | 0.9 | 0.9 | 0.99 | 0.79 | 0.82 |
| 1 | 230428502 | rs17711945 | 0.606 | 0.561 | -1.224 | -1.197 | 0.007 | 0.023 | 0.023 | 0 | 0.19 | 0.12 | 0.11 | 0.07 | 0.16 | 0.21 |
| 1 | 230429143 | rs751645 | 0.7 | 0.586 | -1.271 | -1.158 | 0.009 | 0.026 | 0.026 | 0 | 0.19 | 0.09 | 0.13 | 0.1 | 0.16 | 0.21 |
| 1 | 230430380 | rs6666008 | 0.91 | 0.994 | -0.909 | -0.804 | 0.04 | -0.017 | -0.017 | 0 | 0.45 | 0.25 | 0.38 | 0.43 | 0.4 | 0.45 |
| 1 | 230431713 | rs11122484 | 0.714 | 0.577 | -0.517 | -0.773 | 0.052 | 0.043 | 0.043 | 0.231 | 0.06 | 0.29 | 0.04 | 0.06 | 0.04 | 0.04 |
| 1 | 230432227 | rs7529965 | -0.189 | 0.927 | -1.344 | -0.742 | -0.015 | 0.129 | 0.129 | 0 | 0.25 | 0.08 | 0.21 | 0.05 | 0.27 | 0.25 |
| 1 | 230435738 | rs2046614 | 0.397 | 1.168 | -0.794 | -0.356 | 0.018 | 0.015 | 0.015 | 0.449 | 0.62 | 0.48 | 0.54 | 0.52 | 0.55 | 0.54 |
| 1 | 230435991 | rs4846854 | 0.571 | 0.924 | -0.649 | -0.53 | -0.007 | 0.003 | 0.003 | 0 | 0.43 | 0.45 | 0.4 | 0.42 | 0.37 | 0.41 |
| 1 | 230436175 | rs4846857 | -0.349 | 1.627 | -0.874 | -0.736 | 0.013 | -0.026 | -0.026 | 0 | 0.9 | 0.85 | 0.94 | 0.99 | 0.83 | 0.87 |
| 1 | 230436577 | rs17712125 | -0.092 | 1.323 | -1.383 | -0.884 | -0.008 | 0.104 | 0.104 | 0 | 0.21 | 0.07 | 0.12 | 0.04 | 0.17 | 0.3 |
| 1 | 230436615 | rs11122486 | -0.162 | 1.904 | -0.943 | -0.129 | -0.006 | -0.006 | -0.006 | 0 | 0.28 | 0.26 | 0.56 | 0.61 | 0.38 | 0.28 |
| 1 | 230436932 | rs6690154 | -1.142 | 1.463 | -0.53 | -0.444 | -0.009 | -0.068 | -0.068 | 0.244 | 0.28 | 0.39 | 0.16 | 0.17 | 0.21 | 0.4 |
| 1 | 230437358 | rs901676 | 0.77 | 2.141 | -0.446 | 0.356 | 0.022 | 0.064 | 0.064 | 0 | 0.83 | 0.78 | 0.66 | 0.51 | 0.71 | 0.82 |
| 1 | 230437791 | rs730087 | 0.446 | 2.047 | -0.62 | -0.298 | -0.013 | 0.013 | 0.013 | 0 | 0.75 | 0.82 | 0.82 | 0.88 | 0.66 | 0.67 |
| 1 | 230437989 | rs9432168 | -0.159 | 2.061 | -0.642 | -0.257 | -0.021 | 0.018 | 0.018 | 0 | 0.89 | 0.94 | 0.91 | 0.96 | 0.77 | 0.87 |

| | | | | | | | | | | | | | | | | |
|---|---|---|---|---|---|---|---|---|---|---|---|---|---|---|---|---|
| 1 | 230438592 | rs4414046 | 0.845 | 1.441 | -0.457 | -0.276 | 0 | -0.007 | -0.007 | 0 | 0.61 | 0.64 | 0.68 | 0.77 | 0.56 | 0.51 |
| 1 | 230445193 | rs901668 | 1.441 | 1.21 | 0.174 | 0.107 | 0 | 0.003 | 0.003 | 0 | 0.65 | 0.36 | 0.54 | 0.38 | 0.69 | 0.55 |
| 1 | 230446556 | rs12133061 | -1.481 | 1.197 | 0.534 | 0.088 | 0.05 | 0.13 | 0.13 | 0 | 0.21 | 0.36 | 0.33 | 0.52 | 0.22 | 0.31 |
| 1 | 230447108 | rs11122490 | -1.367 | 1.054 | 0.43 | -0.015 | 0.073 | 0.183 | 0.183 | 0 | 0.19 | 0.37 | 0.33 | 0.5 | 0.21 | 0.25 |
| 1 | 230450968 | rs10779842 | 0.13 | 0.94 | 0.775 | -0.088 | -0.026 | -0.045 | -0.045 | 0 | 0.85 | 0.67 | 0.85 | 0.94 | 0.88 | 0.83 |
| 1 | 230452233 | rs11122493 | 2.118 | 0.948 | 0.478 | -0.139 | -0.017 | 0.07 | 0.07 | 0 | 0.53 | 0.21 | 0.44 | 0.31 | 0.58 | 0.46 |
| 1 | 230457347 | rs11122494 | -0.791 | 1.049 | 0.594 | 0.009 | 0.047 | 0.105 | 0.105 | 0 | 0.15 | 0.25 | 0.3 | 0.49 | 0.2 | 0.22 |
| 1 | 230458324 | rs12089782 | 0.723 | 1.005 | 0.636 | 0.016 | -0.002 | 0.038 | 0.038 | 0.021 | 0.06 | 0.27 | 0.05 | 0.07 | 0.03 | 0.1 |
| 1 | 230459926 | rs1565091 | 2.215 | 0.842 | 0.177 | -0.083 | -0.016 | 0.09 | 0.09 | 0 | 0.51 | 0.2 | 0.42 | 0.31 | 0.55 | 0.43 |
| 1 | 230461079 | rs2306813 | -0.365 | 1.16 | 0.218 | 0.339 | 0.036 | 0.131 | 0.131 | 0 | 0.1 | 0.15 | 0.26 | 0.49 | 0.16 | 0.15 |
| 1 | 230463537 | rs10864742 | 0.639 | 0.758 | -0.255 | -0.186 | -0.021 | -0.031 | -0.031 | 0.019 | 0.79 | 0.77 | 0.81 | 0.92 | 0.85 | 0.79 |
| 1 | 230467643 | rs12116691 | 0.316 | 1.185 | -0.768 | 0.1 | -0.023 | 0.043 | 0.043 | 0 | 0.44 | 0.16 | 0.39 | 0.28 | 0.48 | 0.44 |
| 1 | 230473545 | rs12726792 | 0.465 | 1.286 | -0.855 | -0.235 | 0.005 | 0.198 | 0.198 | 0 | 0.37 | 0.05 | 0.18 | 0.1 | 0.32 | 0.34 |
| 1 | 230474898 | rs10864745 | -0.245 | 0.951 | -0.489 | 0.229 | -0.029 | -0.032 | -0.032 | 0 | 0.44 | 0.16 | 0.4 | 0.42 | 0.48 | 0.45 |
| 1 | 230484673 | rs12142665 | -0.423 | 1.947 | -0.429 | 0.138 | -0.013 | 0.009 | 0.009 | 0 | 0.08 | 0.05 | 0.07 | 0.03 | 0.19 | 0.04 |
| 1 | 230484922 | rs721532 | -0.402 | 2.066 | -0.469 | 0.279 | -0.009 | -0.007 | -0.007 | 0 | 0.05 | 0.03 | 0.19 | 0.11 | 0.12 | 0.08 |
| 1 | 230489573 | rs6689340 | -0.897 | 1.447 | 0.732 | 0.022 | 0.052 | 0.048 | 0.048 | 1.086 | 0.05 | 0.42 | 0.04 | 0 | 0.01 | 0 |
| 1 | 230495588 | rs1545969 | 0.116 | 1.13 | -0.347 | -0.247 | 0.021 | 0.034 | 0.034 | 0 | 0.39 | 0.31 | 0.39 | 0.34 | 0.29 | 0.27 |
| 1 | 230496808 | rs1108707 | -0.789 | 1.551 | -0.082 | -0.37 | 0.069 | -0.007 | -0.007 | 0 | 0.07 | 0.38 | 0.07 | 0.01 | 0.05 | 0.09 |
| 1 | 230498579 | rs828480 | -0.459 | 1.286 | -0.26 | -0.507 | 0.025 | -0.017 | -0.017 | 1.851 | 0.62 | 0.26 | 0.55 | 0.44 | 0.67 | 0.49 |
| 1 | 230502591 | rs923302 | -0.131 | 1.38 | -0.247 | -0.576 | -0.079 | 0.02 | 0.02 | 0 | 0.22 | 0.05 | 0.15 | 0.12 | 0.34 | 0.22 |
| 1 | 230504498 | rs12405197 | 0.658 | 1.153 | -0.398 | -0.74 | -0.008 | 0.035 | 0.035 | 0 | 0.4 | 0.28 | 0.43 | 0.34 | 0.37 | 0.34 |
| 1 | 230504875 | rs2795050 | -0.915 | 1.19 | -0.236 | -0.705 | -0.026 | 0.017 | 0.017 | 0 | 0.65 | 0.44 | 0.6 | 0.46 | 0.72 | 0.57 |
| 1 | 230505870 | rs828436 | 1.056 | 1.532 | -0.386 | -0.766 | -0.055 | 0.001 | 0.001 | 0 | 0.08 | 0.02 | 0.09 | 0.03 | 0.24 | 0.05 |
| 1 | 230508313 | rs828453 | -1.42 | 1.255 | -0.229 | -0.696 | -0.006 | 0.007 | 0.007 | 0 | 0.69 | 0.74 | 0.63 | 0.46 | 0.72 | 0.57 |
| 1 | 230509756 | rs828459 | -1.392 | 1.247 | -0.26 | -0.7 | 0.003 | -0.004 | -0.004 | 0.073 | 0.69 | 0.75 | 0.63 | 0.45 | 0.72 | 0.57 |
| 1 | 230510977 | rs901669 | 0.823 | 1.186 | -0.618 | -0.764 | -0.068 | 0.012 | 0.012 | 0 | 0.37 | 0.02 | 0.4 | 0.32 | 0.33 | 0.27 |
| 1 | 230517993 | rs701170 | -0.159 | 1.18 | -0.747 | -0.734 | -0.106 | 0.009 | 0.009 | 0 | 0.22 | 0.03 | 0.14 | 0.12 | 0.34 | 0.22 |
| 1 | 230521031 | rs11806158 | 0.49 | 1.19 | -0.761 | -0.695 | -0.057 | -0.006 | -0.006 | 0.286 | 0.41 | 0.12 | 0.36 | 0.42 | 0.39 | 0.42 |
| 1 | 230521144 | rs828467 | 0.091 | 1.195 | -0.534 | -0.729 | 0.027 | -0.055 | -0.055 | 0 | 0.2 | 0.35 | 0.15 | 0.03 | 0.29 | 0.13 |
| 1 | 230524669 | rs828468 | -0.3 | 1.227 | -0.518 | -0.64 | 0.025 | -0.114 | -0.114 | 0.26 | 0.26 | 0.6 | 0.18 | 0.04 | 0.33 | 0.16 |
| 1 | 230527005 | rs828470 | -0.327 | 1.245 | -0.566 | -0.632 | 0.022 | -0.115 | -0.115 | 0 | 0.27 | 0.6 | 0.18 | 0.04 | 0.33 | 0.16 |
| 1 | 230527248 | rs17782420 | -0.315 | 0.736 | -0.813 | -0.195 | -0.001 | 0 | 0 | 0 | 0.17 | 0.13 | 0.11 | 0.1 | 0.14 | 0.21 |
| 1 | 230528430 | rs270855 | -0.005 | 0.988 | -1.146 | -0.081 | 0.024 | 0.054 | 0.054 | 0 | 0.49 | 0.41 | 0.42 | 0.35 | 0.57 | 0.37 |
| 1 | 230535548 | rs270852 | 0.019 | 0.939 | -1.149 | 0.536 | 0.001 | 0.006 | 0.006 | 0 | 0.78 | 0.8 | 0.77 | 0.75 | 0.74 | 0.74 |
| 1 | 230535966 | rs270853 | -0.425 | 1.506 | -1.938 | 0.168 | -0.1 | 0.016 | 0.016 | 2.276 | 0.21 | 0.02 | 0.16 | 0.12 | 0.31 | 0.1 |
| 1 | 230551909 | rs11122509 | -0.823 | 0.793 | -1.869 | 0.579 | -0.057 | 0.01 | 0.01 | 0 | 0.36 | 0.21 | 0.56 | 0.32 | 0.45 | 0.43 |
| 1 | 230560900 | rs12023508 | 0.476 | 0.328 | -1.932 | 1.167 | -0.002 | -0.084 | -0.084 | 0.047 | 0.14 | 0.01 | 0.07 | 0.31 | 0.08 | 0.09 |
| 1 | 230563455 | rs12031576 | 0.123 | 0.822 | -1.627 | 1.089 | 0.006 | 0.072 | 0.072 | 1.065 | 0.09 | 0.21 | 0.35 | 0.26 | 0.09 | 0.07 |
| 1 | 230594749 | rs11586533 | -0.275 | 1.019 | -1.329 | 0.765 | -0.005 | -0.002 | -0.002 | 0 | 0.25 | 0.22 | 0.49 | 0.36 | 0.4 | 0.2 |
| 1 | 230604127 | rs1418131 | 0.541 | 1.112 | -1.553 | 0.58 | -0.079 | 0.073 | 0.073 | 0 | 0.18 | 0.04 | 0.15 | 0.05 | 0.32 | 0.12 |
| 1 | 230605643 | rs6676503 | -0.381 | -0.073 | -1.088 | 0.46 | -0.036 | -0.005 | -0.005 | 0.757 | 0.33 | 0.51 | 0.24 | 0.28 | 0.2 | 0.44 |
| 1 | 230614110 | rs200162 | -0.141 | 0.263 | -1.525 | 0.487 | 0.074 | -0.002 | -0.002 | 0.183 | 0.46 | 0.2 | 0.64 | 0.54 | 0.64 | 0.49 |
| 1 | 230622001 | rs853021 | 1.023 | 0.666 | -1.552 | 0.524 | -0.013 | -0.008 | -0.008 | 0 | 0.22 | 0.08 | 0.16 | 0.19 | 0.24 | 0.29 |
| 1 | 230622709 | rs853022 | -1.189 | 0.255 | -1.336 | 0.376 | -0.062 | -0.021 | -0.021 | 0 | 0.67 | 0.51 | 0.75 | 0.78 | 0.76 | 0.66 |
| 1 | 230624272 | rs17599080 | 0.883 | 0.297 | -2.024 | -0.318 | -0.05 | 0.057 | 0.057 | 0.271 | 0.21 | 0.09 | 0.17 | 0.05 | 0.3 | 0.1 |
| 1 | 230624383 | rs12064727 | -0.371 | 0.29 | -1.935 | -0.287 | -0.007 | -0.021 | -0.021 | 0 | 0.21 | 0.19 | 0.09 | 0.29 | 0.19 | 0.23 |
| 1 | 230627480 | rs6674676 | 0.091 | 0.258 | -1.656 | -0.069 | 0 | 0.002 | 0.002 | 4.166 | 0.15 | 0.18 | 0.06 | 0.29 | 0.12 | 0.18 |
| 1 | 230628591 | rs701152 | 1.235 | 0.252 | -1.82 | -0.077 | -0.016 | -0.01 | -0.01 | 0 | 0.23 | 0.06 | 0.16 | 0.19 | 0.24 | 0.29 |
| 1 | 230635833 | rs1418134 | -0.85 | 0.411 | -1.53 | 0.031 | -0.006 | 0.001 | 0.001 | 0.548 | 0.39 | 0.39 | 0.25 | 0.34 | 0.44 | 0.29 |
| 1 | 230637961 | rs1341972 | 0.057 | 0.811 | -1.881 | -0.456 | -0.058 | 0.06 | 0.06 | 0.149 | 0.21 | 0.1 | 0.18 | 0.05 | 0.32 | 0.1 |
| 1 | 230639734 | rs853017 | 0.315 | -0.14 | -1.584 | 0.009 | -0.008 | -0.008 | -0.008 | 0 | 0.22 | 0.06 | 0.16 | 0.19 | 0.24 | 0.29 |
| 1 | 230641218 | rs7539779 | -0.512 | -0.325 | -1.016 | -0.069 | -0.005 | -0.022 | -0.022 | 0 | 0.19 | 0.19 | 0.1 | 0.05 | 0.13 | 0.09 |
| 1 | 230644983 | rs701153 | -0.968 | -0.4 | -0.839 | 0.031 | -0.002 | -0.002 | -0.002 | 0 | 0.3 | 0.3 | 0.15 | 0.19 | 0.22 | 0.26 |
| 1 | 230648705 | rs7555103 | 0.828 | -0.771 | -1.2 | 0.319 | -0.06 | -0.005 | -0.005 | 2.179 | 0.31 | 0.17 | 0.52 | 0.36 | 0.5 | 0.15 |
| 1 | 230654458 | rs6661286 | 0.054 | -1.133 | -0.496 | 0.605 | -0.044 | 0.002 | 0.002 | 0.738 | 0.71 | 0.51 | 0.45 | 0.26 | 0.78 | 0.59 |
| 1 | 230654589 | rs6661407 | 0.379 | -1.268 | -0.629 | 0.438 | -0.031 | -0.019 | -0.019 | 0 | 0.65 | 0.48 | 0.43 | 0.26 | 0.72 | 0.52 |
| 1 | 230655367 | rs2530360 | -1.129 | -1.439 | -0.476 | 1.118 | -0.001 | 0.026 | 0.026 | 0 | 0.89 | 0.91 | 0.57 | 0.56 | 0.89 | 0.83 |
| 1 | 230656763 | rs6671466 | 0.676 | -1.466 | -0.48 | 1.025 | -0.035 | 0.002 | 0.002 | 0.619 | 0.71 | 0.51 | 0.44 | 0.26 | 0.76 | 0.58 |
| 1 | 230668260 | rs853431 | 0.155 | -1.66 | -0.646 | 0.885 | 0.01 | 0.001 | 0.001 | 0 | 0.26 | 0.23 | 0.13 | 0.2 | 0.18 | 0.25 |
| 1 | 230673182 | rs853434 | -1.749 | -1.657 | -0.66 | 1.045 | 0.011 | -0.024 | -0.024 | 0.01 | 0.91 | 0.95 | 0.86 | 0.78 | 0.95 | 0.9 |
| 1 | 230674826 | rs701157 | -1.456 | -1.759 | -0.464 | 0.886 | 0.024 | 0.015 | 0.015 | 0.521 | 0.5 | 0.64 | 0.56 | 0.6 | 0.43 | 0.57 |
| 1 | 230676042 | rs3849295 | 0.171 | -1.519 | -0.619 | 1.015 | -0.003 | -0.002 | -0.002 | 0.01 | 0.09 | 0.09 | 0.31 | 0.22 | 0.13 | 0.12 |
| 1 | 230676674 | rs12042923 | 0.927 | -1.455 | -0.684 | 1.083 | 0.005 | 0.026 | 0.026 | 0 | 0.2 | 0.11 | 0.11 | 0.08 | 0.16 | 0.15 |
| 1 | 230681731 | rs853442 | -0.117 | -1.17 | -0.295 | 1.58 | -0.002 | 0.008 | 0.008 | 0.333 | 0.16 | 0.17 | 0.1 | 0.29 | 0.13 | 0.26 |
| 1 | 230682813 | rs853444 | -0.096 | -0.892 | -0.287 | 1.445 | -0.018 | 0.042 | 0.042 | 0 | 0.82 | 0.66 | 0.8 | 0.74 | 0.89 | 0.84 |
| 1 | 230683403 | rs6541320 | -0.109 | -0.254 | -0.707 | 1.251 | -0.002 | -0.004 | -0.004 | 0 | 0.13 | 0.1 | 0.29 | 0.39 | 0.18 | 0.26 |
| 1 | 230684176 | rs6541321 | -0.165 | -0.534 | -0.694 | 0.921 | 0.002 | -0.004 | -0.004 | 0 | 0.13 | 0.11 | 0.27 | 0.3 | 0.14 | 0.23 |
| 1 | 230685079 | rs4846979 | 0.35 | -0.693 | -0.576 | 0.471 | -0.002 | 0.003 | 0.003 | 0.923 | 0.42 | 0.4 | 0.53 | 0.35 | 0.56 | 0.21 |

| | | | | | | | | | | | | | | | | |
|---|---|---|---|---|---|---|---|---|---|---|---|---|---|---|---|---|
| 1 | 230685255 | rs701164 | 0.012 | -1.106 | -0.352 | -0.197 | -0.003 | -0.007 | -0.007 | 0.825 | 0.52 | 0.54 | 0.41 | 0.26 | 0.63 | 0.4 |
| 1 | 230685483 | rs16851934 | -0.167 | -0.906 | 0.46 | -0.257 | 0.042 | -0.036 | -0.036 | 0.034 | 0.07 | 0.21 | 0.05 | 0 | 0.09 | 0.05 |
| 1 | 230687711 | rs853452 | -0.672 | -1.473 | 0.355 | -0.037 | 0.029 | -0.055 | -0.055 | 0.485 | 0.26 | 0.15 | 0.13 | 0.37 | 0.19 | 0.31 |
| 1 | 230688466 | rs11122534 | -0.529 | -1.229 | 0.599 | -0.279 | 0.005 | -0.06 | -0.06 | 3.864 | 0.15 | 0.2 | 0.18 | 0.02 | 0.17 | 0.24 |
| 1 | 230695062 | rs11122535 | 0.509 | -1.257 | 0.583 | -0.249 | 0.023 | -0.034 | -0.034 | 0 | 0.08 | 0.18 | 0.14 | 0.01 | 0.09 | 0.15 |
| 1 | 230698513 | rs11580919 | 0.959 | -1.316 | 0.511 | -0.22 | -0.001 | -0.009 | -0.009 | 0 | 0.06 | 0.07 | 0.04 | 0 | 0.07 | 0.05 |
| 1 | 230700678 | rs1615876 | -0.049 | -1.826 | 0.198 | -0.111 | 0.022 | -0.095 | -0.095 | 0 | 0.16 | 0.04 | 0.1 | 0.39 | 0.13 | 0.27 |
| 1 | 230701565 | rs1327696 | -2.04 | -2.038 | 0.592 | -0.362 | -0.005 | 0.002 | 0.002 | 0 | 0.44 | 0.47 | 0.32 | 0.46 | 0.39 | 0.57 |
| 1 | 230704291 | rs11804015 | 1.425 | -2.098 | 0.66 | -0.38 | -0.019 | 0.009 | 0.009 | 0 | 0.59 | 0.51 | 0.71 | 0.55 | 0.67 | 0.45 |
| 1 | 230706868 | rs3761946 | 1.39 | -1.892 | 0.678 | -0.393 | 0.002 | -0.001 | -0.001 | 0 | 0.53 | 0.5 | 0.59 | 0.54 | 0.58 | 0.43 |
| 1 | 230710689 | rs3888630 | 1.439 | -1.879 | 1.034 | -0.384 | 0.045 | -0.001 | -0.001 | 0 | 0.5 | 0.33 | 0.56 | 0.55 | 0.57 | 0.4 |
| 1 | 230711435 | rs740492 | -0.287 | -1.857 | 0.234 | -0.374 | 0.027 | -0.082 | -0.082 | 0 | 0.2 | 0.08 | 0.12 | 0.4 | 0.17 | 0.28 |
| 1 | 230713942 | rs11122545 | -0.728 | -1.798 | 0.275 | -0.391 | 0 | -0.029 | -0.029 | 0 | 0.28 | 0.27 | 0.29 | 0.05 | 0.26 | 0.32 |
| 1 | 230715075 | rs1637941 | -1.616 | -1.677 | 0.204 | -0.271 | 0.003 | 0 | 0 | 0 | 0.49 | 0.48 | 0.42 | 0.45 | 0.43 | 0.62 |
| 1 | 230723481 | rs6691073 | 0.446 | -1.428 | -0.03 | -0.087 | -0.015 | 0.004 | 0.004 | 0 | 0.06 | 0.14 | 0.02 | 0.09 | 0.04 | 0.04 |
| 1 | 230725969 | rs11122548 | -0.109 | -1.417 | -0.247 | -0.236 | -0.041 | 0.093 | 0.093 | 0.055 | 0.23 | 0.06 | 0.23 | 0.03 | 0.27 | 0.24 |
| 1 | 230727362 | rs1410140 | -0.729 | -1.489 | -0.281 | -0.286 | -0.039 | 0.145 | 0.145 | 0 | 0.3 | 0.08 | 0.27 | 0.04 | 0.33 | 0.34 |
| 1 | 230728901 | rs200587 | 0.388 | -1.547 | -0.092 | -0.044 | 0.001 | 0.009 | 0.009 | 0.426 | 0.15 | 0.13 | 0.12 | 0.31 | 0.13 | 0.22 |
| 1 | 230730601 | rs200588 | -2.078 | -1.593 | 0.204 | 0.125 | -0.002 | 0.001 | 0.001 | 0 | 0.56 | 0.59 | 0.52 | 0.56 | 0.52 | 0.67 |
| 1 | 230732682 | rs2148581 | -1.091 | -1.557 | -0.04 | 0.085 | 0.001 | 0.06 | 0.06 | 0 | 0.36 | 0.26 | 0.29 | 0.13 | 0.37 | 0.38 |
| 1 | 230734649 | rs12091639 | -0.548 | -1.508 | -0.122 | -0.008 | -0.026 | 0.115 | 0.115 | 0 | 0.3 | 0.1 | 0.27 | 0.04 | 0.33 | 0.34 |
| 1 | 230735580 | rs11122552 | -0.93 | -1.443 | -0.055 | 0.072 | -0.018 | 0.103 | 0.103 | 0 | 0.34 | 0.11 | 0.36 | 0.16 | 0.34 | 0.41 |
| 1 | 230745894 | rs853463 | -1.674 | -1.267 | 0.002 | -0.032 | 0.003 | -0.002 | -0.002 | 0.034 | 0.58 | 0.58 | 0.53 | 0.56 | 0.51 | 0.67 |
| 1 | 230747290 | rs16852085 | 0.515 | -0.781 | -0.177 | 0.17 | -0.012 | -0.05 | -0.05 | 0 | 0.09 | 0.24 | 0.06 | 0 | 0.06 | 0.1 |
| 1 | 230749646 | rs853464 | 0.163 | -0.557 | -0.668 | 0.698 | 0.001 | -0.026 | -0.026 | 0.081 | 0.13 | 0.08 | 0.11 | 0.2 | 0.13 | 0.24 |
| 1 | 230754222 | rs9651127 | -0.846 | -0.612 | -0.263 | 0.638 | 0 | 0.03 | 0.03 | 0.116 | 0.32 | 0.44 | 0.38 | 0.39 | 0.32 | 0.31 |
| 1 | 230758605 | rs714214 | 0.645 | -0.273 | -0.648 | 0.535 | -0.053 | 0.017 | 0.017 | 0.206 | 0.08 | 0.01 | 0.21 | 0.06 | 0.17 | 0.2 |
| 1 | 230759310 | rs853469 | -0.148 | -0.788 | -0.312 | 0.649 | -0.023 | 0.025 | 0.025 | 1.324 | 0.2 | 0.33 | 0.16 | 0.26 | 0.15 | 0.28 |
| 1 | 230762648 | rs12029075 | 0.127 | -0.72 | -0.605 | 0.611 | -0.06 | 0.036 | 0.036 | 0 | 0.14 | 0.01 | 0.22 | 0.07 | 0.2 | 0.21 |
| 1 | 230766155 | rs12031571 | 0.276 | -0.867 | -0.13 | 0.878 | -0.003 | 0.049 | 0.049 | 0 | 0.1 | 0.12 | 0.11 | 0.23 | 0.08 | 0.06 |
| 1 | 230766411 | rs4846862 | 0 | -1.147 | -0.468 | 0.73 | -0.013 | 0.032 | 0.032 | 0.261 | 0.31 | 0.34 | 0.27 | 0.48 | 0.21 | 0.34 |
| 1 | 230768037 | rs4846997 | 0.638 | -0.542 | -0.678 | 0.524 | -0.015 | 0.009 | 0.009 | 0 | 0.47 | 0.27 | 0.46 | 0.44 | 0.51 | 0.39 |
| 1 | 230771642 | rs11122562 | 0.347 | -0.619 | -0.341 | 0.816 | -0.006 | 0.02 | 0.02 | 0 | 0.88 | 0.82 | 0.87 | 0.79 | 0.92 | 0.9 |
| 1 | 230773031 | rs927888 | -0.336 | -0.736 | -0.435 | 0.822 | -0.002 | 0.001 | 0.001 | 0 | 0.11 | 0.13 | 0.06 | 0.11 | 0.09 | 0.08 |
| 1 | 230774851 | rs1078197 | 0.551 | -0.696 | -0.179 | 0.887 | -0.013 | 0.045 | 0.045 | 0 | 0.87 | 0.72 | 0.86 | 0.79 | 0.91 | 0.9 |
| 1 | 230775676 | rs12032754 | 0.266 | -0.328 | -0.717 | 1.065 | -0.001 | 0.005 | 0.005 | 0 | 0.08 | 0.04 | 0.11 | 0.21 | 0.08 | 0.05 |
| 1 | 230775683 | rs16852125 | -0.569 | -0.538 | -0.932 | 0.577 | 0.002 | 0 | 0 | 0 | 0.14 | 0.14 | 0.07 | 0.11 | 0.1 | 0.08 |
| 1 | 230778879 | rs4846999 | -0.156 | 0.505 | -1.254 | 0.654 | -0.071 | -0.027 | -0.027 | 0 | 0.12 | 0.01 | 0.27 | 0.17 | 0.2 | 0.15 |
| 1 | 230779994 | rs1887490 | -1.063 | 0.11 | -1.513 | 0.424 | -0.048 | 0.01 | 0.01 | 0 | 0.22 | 0.08 | 0.34 | 0.22 | 0.25 | 0.22 |
| 1 | 230780571 | rs3827743 | -0.881 | 0.396 | -1.642 | -0.39 | -0.005 | -0.002 | -0.002 | 0 | 0.71 | 0.76 | 0.8 | 0.95 | 0.65 | 0.83 |
| 1 | 230781741 | rs4847000 | 1.487 | -0.371 | -1.293 | 0.066 | -0.009 | 0.029 | 0.029 | 0 | 0.26 | 0.33 | 0.22 | 0.45 | 0.19 | 0.4 |
| 1 | 230786968 | rs12405479 | 1.304 | -0.276 | -1.395 | 0.001 | 0.017 | 0.018 | 0.018 | 2.06 | 0.1 | 0.02 | 0.07 | 0.05 | 0.06 | 0.07 |
| 1 | 230788501 | rs6659928 | 1.787 | -0.282 | -1.489 | 0.011 | -0.023 | -0.004 | -0.004 | 0 | 0.05 | 0.02 | 0.08 | 0.05 | 0.16 | 0.04 |
| 1 | 230788550 | rs6700774 | -0.621 | -0.307 | -1.252 | -0.069 | -0.006 | 0.031 | 0.031 | 0.921 | 0.59 | 0.67 | 0.53 | 0.78 | 0.47 | 0.71 |
| 1 | 230795177 | rs1359866 | -0.89 | -0.307 | -1.052 | 0.029 | -0.008 | 0.079 | 0.079 | 0 | 0.62 | 0.77 | 0.74 | 0.9 | 0.6 | 0.76 |
| 1 | 230795641 | rs2225146 | -0.2 | -0.412 | -1.108 | 0.156 | 0.029 | 0.072 | 0.072 | 0 | 0.49 | 0.65 | 0.46 | 0.73 | 0.41 | 0.64 |
| 1 | 230798808 | rs2296802 | 1.824 | -0.411 | -1.223 | 0.457 | 0.031 | -0.042 | -0.042 | 0 | 0.09 | 0.01 | 0.14 | 0.18 | 0.03 | 0.08 |
| 1 | 230802015 | rs16852170 | 1.987 | -0.533 | -1.453 | 0.877 | -0.027 | -0.046 | -0.046 | 1.35 | 0.08 | 0.01 | 0.09 | 0.18 | 0.09 | 0.22 |
| 1 | 230804773 | rs2281951 | 0.677 | -0.396 | -0.848 | 0.582 | -0.001 | -0.004 | -0.004 | 0 | 0.15 | 0.17 | 0.23 | 0.23 | 0.19 | 0.12 |
| 1 | 230805786 | rs16852189 | 1.491 | -0.404 | -0.796 | 0.557 | -0.01 | 0.042 | 0.042 | 0.347 | 0.07 | 0.18 | 0.02 | 0.06 | 0.02 | 0.09 |
| 1 | 230808005 | rs10864766 | -0.034 | -0.654 | -0.963 | 0.371 | 0.029 | 0.057 | 0.057 | 0 | 0.49 | 0.41 | 0.44 | 0.7 | 0.4 | 0.56 |
| 1 | 230808744 | rs2148964 | 0.808 | -0.486 | -0.63 | 0.725 | 0.007 | -0.024 | -0.024 | 0.018 | 0.31 | 0.29 | 0.23 | 0.48 | 0.24 | 0.43 |
| 1 | 230809434 | rs3789643 | 0.465 | 0.129 | -0.254 | 0.968 | -0.004 | 0.053 | 0.053 | 0.104 | 0.07 | 0.2 | 0.02 | 0.06 | 0.02 | 0.1 |
| 1 | 230810362 | rs12041242 | 0.508 | 0.759 | -0.9 | 0.892 | -0.045 | -0.024 | -0.024 | 0.091 | 0.1 | 0.01 | 0.25 | 0.14 | 0.16 | 0.11 |
| 1 | 230815376 | rs943579 | 0.181 | 0.73 | -0.192 | 1.246 | 0.007 | 0.107 | 0.107 | 0 | 0.85 | 0.68 | 0.81 | 0.7 | 0.87 | 0.91 |
| 1 | 230815481 | rs2493144 | 0.391 | 0.689 | -0.545 | 0.505 | 0.003 | 0.067 | 0.067 | 0 | 0.52 | 0.29 | 0.35 | 0.24 | 0.54 | 0.49 |
| 1 | 230816269 | rs3789647 | 0.211 | 0.401 | -0.575 | 0.303 | -0.001 | -0.001 | -0.001 | 0 | 0.26 | 0.26 | 0.25 | 0.29 | 0.32 | 0.31 |
| 1 | 230816460 | rs3789649 | 0.02 | 0.091 | -0.182 | 0.034 | 0.022 | 0.128 | 0.128 | 0 | 0.18 | 0.32 | 0.44 | 0.59 | 0.21 | 0.31 |
| 1 | 230816557 | rs3789650 | 0.424 | 0.069 | -0.082 | 0.014 | 0.004 | 0.001 | 0.001 | 0 | 0.27 | 0.29 | 0.26 | 0.29 | 0.32 | 0.31 |
| 1 | 230817384 | rs2493142 | 0.561 | -0.247 | -0.114 | -0.178 | -0.024 | 0.021 | 0.021 | 0 | 0.65 | 0.56 | 0.51 | 0.39 | 0.75 | 0.64 |
| 1 | 230817859 | rs1467670 | -1.402 | -0.21 | -0.729 | -0.188 | 0.019 | 0.009 | 0.009 | 0.253 | 0.84 | 0.89 | 0.96 | 0.99 | 0.96 | 0.96 |
| 1 | 230818841 | rs2479131 | 1.533 | -0.155 | -0.768 | -0.167 | -0.07 | 0.117 | 0.117 | 0 | 0.37 | 0.09 | 0.23 | 0.1 | 0.43 | 0.33 |
| 1 | 230820021 | rs2479132 | 1.515 | -0.145 | -0.759 | -0.177 | -0.058 | 0.111 | 0.111 | 0 | 0.36 | 0.11 | 0.23 | 0.1 | 0.43 | 0.33 |
| 1 | 230821760 | rs11122571 | 0.414 | 0.198 | -0.443 | 0.082 | 0.01 | 0.002 | 0.002 | 0 | 0.26 | 0.34 | 0.26 | 0.29 | 0.32 | 0.31 |
| 1 | 230825427 | rs11122572 | -0.177 | -0.307 | -0.232 | 0.805 | 0.013 | 0.198 | 0.198 | 0 | 0.09 | 0.19 | 0.27 | 0.49 | 0.08 | 0.21 |
| 1 | 230828873 | rs3789662 | -0.458 | -0.309 | -0.268 | 0.725 | 0.01 | 0.066 | 0.066 | 0 | 0.18 | 0.23 | 0.44 | 0.59 | 0.21 | 0.32 |
| 1 | 230836702 | rs7536290 | -0.164 | -0.371 | -0.257 | 0.731 | 0.021 | 0.074 | 0.074 | 0 | 0.16 | 0.4 | 0.33 | 0.26 | 0.16 | 0.22 |
| 1 | 230838258 | rs2067853 | 1.074 | -0.366 | -1.087 | 0.39 | -0.024 | 0.102 | 0.102 | 0 | 0.33 | 0.07 | 0.19 | 0.11 | 0.34 | 0.29 |
| 1 | 230838331 | rs7079 | 1.074 | -0.366 | -1.087 | 0.39 | -0.024 | 0.102 | 0.102 | 0 | 0.33 | 0.07 | 0.19 | 0.11 | 0.34 | 0.29 |

| | | | | | | | | | | | | | | | | |
|---|---|---|---|---|---|---|---|---|---|---|---|---|---|---|---|---|
| 1 | 230841509 | rs2478523 | -0.979 | -2.02 | -0.717 | 0.753 | 0.108 | -0.019 | -0.019 | 0 | 0.51 | 0.88 | 0.54 | 0.52 | 0.34 | 0.54 |
| 1 | 230844121 | rs2478545 | -0.784 | -2.144 | -0.377 | 0.563 | -0.048 | -0.022 | -0.022 | 0 | 0.35 | 0.53 | 0.4 | 0.34 | 0.23 | 0.3 |
| 1 | 230844992 | rs6687360 | -0.748 | -2.227 | -0.388 | 0.61 | 0 | -0.004 | -0.004 | 0 | 0.37 | 0.59 | 0.46 | 0.68 | 0.36 | 0.5 |
| 1 | 230845794 | rs699 | -1.477 | -2.237 | -0.524 | 0.608 | -0.02 | 0.147 | 0.147 | 0 | 0.56 | 0.9 | 0.64 | 0.85 | 0.41 | 0.64 |
| 1 | 230848702 | rs2004776 | -0.135 | -2.196 | -0.336 | 0.699 | 0.01 | 0.092 | 0.092 | 0 | 0.26 | 0.49 | 0.33 | 0.58 | 0.23 | 0.38 |
| 1 | 230848831 | rs3889728 | -0.049 | -2.21 | -0.395 | 0.704 | -0.011 | -0.043 | -0.043 | 0 | 0.2 | 0.16 | 0.35 | 0.51 | 0.24 | 0.28 |
| 1 | 230849359 | rs2493134 | -1.47 | -2.278 | -0.499 | 0.518 | -0.011 | 0.151 | 0.151 | 0 | 0.56 | 0.89 | 0.64 | 0.85 | 0.41 | 0.64 |
| 1 | 230849799 | rs2148582 | -1.509 | -2.392 | -0.624 | 0.447 | -0.018 | 0.137 | 0.137 | 0 | 0.56 | 0.91 | 0.65 | 0.85 | 0.41 | 0.64 |
| 1 | 230849872 | rs5051 | -1.538 | -2.383 | -0.626 | 0.45 | -0.024 | 0.134 | 0.134 | 0 | 0.57 | 0.92 | 0.65 | 0.85 | 0.41 | 0.64 |
| 1 | 230850641 | rs2071406 | 0.487 | -2.337 | -0.43 | 0.49 | -0.075 | -0.019 | -0.019 | 0 | 0.18 | 0.34 | 0.17 | 0.16 | 0.05 | 0.13 |
| 1 | 230850940 | rs2071404 | 1.372 | -2.204 | -0.381 | 0.501 | -0.009 | 0.016 | 0.016 | 0 | 0.15 | 0.22 | 0.09 | 0.18 | 0.1 | 0.22 |
| 1 | 230852116 | rs2493137 | -1.178 | -2.164 | -0.346 | 0.552 | -0.002 | -0.001 | -0.001 | 0 | 0.41 | 0.62 | 0.54 | 0.67 | 0.31 | 0.42 |
| 1 | 230855071 | rs7549009 | -0.71 | -1.573 | 0.101 | 0.912 | 0.018 | 0.253 | 0.253 | 0.488 | 0.12 | 0.38 | 0.3 | 0.51 | 0.09 | 0.22 |
| 1 | 230857586 | rs7524283 | -0.435 | -1.553 | 0.04 | 0.917 | 0.014 | 0.203 | 0.203 | 0.163 | 0.11 | 0.29 | 0.28 | 0.51 | 0.09 | 0.22 |
| 1 | 230861493 | rs1326882 | 1.156 | -1.699 | 0.01 | 0.81 | 0.001 | -0.008 | -0.008 | 1.779 | 0.12 | 0.1 | 0.03 | 0.01 | 0.03 | 0.03 |
| 1 | 230861578 | rs7515609 | -0.269 | -1.67 | 0.047 | 0.854 | 0.007 | 0.147 | 0.147 | 0.076 | 0.11 | 0.23 | 0.28 | 0.51 | 0.09 | 0.22 |
| 1 | 230861589 | rs7551720 | -0.242 | -1.665 | 0 | 0.856 | 0.008 | 0.149 | 0.149 | 0.067 | 0.11 | 0.23 | 0.28 | 0.51 | 0.09 | 0.22 |
| 1 | 230862998 | rs1326889 | -1.322 | -2.185 | -0.62 | 0.312 | -0.177 | 0.113 | 0.113 | 0.452 | 0.72 | 0.96 | 0.67 | 0.92 | 0.49 | 0.78 |
| 1 | 230863396 | rs2478528 | -1.35 | -2.119 | -0.244 | 0.531 | 0.042 | 0.047 | 0.047 | 0 | 0.44 | 0.71 | 0.5 | 0.75 | 0.29 | 0.54 |
| 1 | 230863965 | rs2148579 | 1.035 | -2.039 | -0.214 | 0.586 | -0.013 | -0.025 | -0.025 | 0 | 0.13 | 0.12 | 0.04 | 0.02 | 0.04 | 0.04 |
| 1 | 230866219 | rs10864775 | -0.232 | -2.023 | -0.214 | 0.629 | -0.007 | 0.045 | 0.045 | 0 | 0.26 | 0.48 | 0.37 | 0.68 | 0.19 | 0.43 |
| 1 | 230867197 | rs2478520 | -0.864 | -1.96 | -0.31 | 0.653 | -0.019 | -0.006 | -0.006 | 0.62 | 0.38 | 0.57 | 0.41 | 0.69 | 0.23 | 0.47 |
| 1 | 230868063 | rs10864777 | 0.953 | -1.477 | -0.463 | 0.467 | -0.006 | 0.004 | 0.004 | 0 | 0.14 | 0.22 | 0.09 | 0.15 | 0.1 | 0.22 |
| 1 | 230872355 | rs6662726 | 0.516 | -1.452 | -0.72 | 0.443 | -0.035 | -0.007 | -0.007 | 0 | 0.21 | 0.35 | 0.11 | 0.2 | 0.11 | 0.24 |
| 1 | 230872906 | rs4259658 | 0.335 | -1.632 | -0.704 | 0.489 | -0.027 | 0.025 | 0.025 | 0.026 | 0.09 | 0.17 | 0.16 | 0.16 | 0.03 | 0.15 |
| 1 | 230877144 | rs16852515 | -0.164 | -1.758 | -0.898 | 0.377 | -0.065 | -0.024 | -0.024 | 1.107 | 0.24 | 0.39 | 0.27 | 0.21 | 0.11 | 0.23 |
| 1 | 230877400 | rs1317764 | -0.955 | -0.791 | -1.726 | 0.553 | -0.117 | 0.067 | 0.067 | 0.064 | 0.75 | 0.93 | 0.68 | 0.92 | 0.49 | 0.81 |
| 1 | 230878561 | rs2493151 | -0.25 | -0.88 | -1.687 | 0.425 | 0.099 | 0.032 | 0.032 | 0 | 0.45 | 0.74 | 0.38 | 0.41 | 0.22 | 0.47 |
| 1 | 230879952 | rs2478541 | -0.227 | 0.738 | -1.611 | 0.616 | -0.162 | -0.056 | -0.056 | 1.46 | 0.75 | 0.96 | 0.69 | 0.84 | 0.49 | 0.78 |
| 1 | 230880155 | rs6662197 | -0.782 | 0.252 | -0.579 | 0.581 | -0.022 | 0.018 | 0.018 | 0.014 | 0.09 | 0.2 | 0.16 | 0.14 | 0.04 | 0.15 |
| 1 | 230880755 | rs17787522 | -0.305 | 1.308 | -1.068 | 0.415 | -0.018 | 0.015 | 0.015 | 0 | 0.05 | 0.02 | 0.09 | 0.02 | 0.15 | 0.06 |
| 1 | 230881063 | rs2478540 | -0.719 | 0.729 | -1.395 | 0.198 | 0.029 | 0.062 | 0.062 | 0.047 | 0.45 | 0.75 | 0.3 | 0.24 | 0.36 | 0.57 |
| 1 | 230881658 | rs16852535 | -0.845 | 1.445 | -0.554 | 0.176 | 0.048 | -0.029 | -0.029 | 0 | 0.28 | 0.52 | 0.31 | 0.24 | 0.41 | 0.23 |
| 1 | 230882946 | rs3893225 | -1.233 | 1.456 | -0.776 | 0.189 | 0.036 | 0.007 | 0.007 | 0.433 | 0.47 | 0.64 | 0.6 | 0.45 | 0.71 | 0.39 |
| 1 | 230884363 | rs6541330 | -0.7 | 1.692 | -0.314 | 0.065 | 0.031 | -0.013 | -0.013 | 0 | 0.3 | 0.44 | 0.43 | 0.33 | 0.43 | 0.28 |
| 1 | 230885085 | rs17731204 | 0.844 | 1.719 | -0.553 | -0.046 | -0.083 | 0.02 | 0.02 | 0 | 0.12 | 0.03 | 0.28 | 0.11 | 0.28 | 0.07 |
| 1 | 230885713 | rs1316408 | 0.395 | 1.558 | -0.445 | -0.009 | 0.022 | -0.035 | -0.035 | 0 | 0.14 | 0.13 | 0.16 | 0.02 | 0.28 | 0.06 |
| 1 | 230885776 | rs1316409 | 0.313 | 0.741 | -0.815 | -0.377 | -0.05 | 0.062 | 0.062 | 0 | 0.25 | 0.14 | 0.38 | 0.16 | 0.34 | 0.24 |
| 1 | 230886191 | rs2493153 | -1.475 | 0.171 | -0.693 | -0.203 | 0.082 | -0.028 | -0.028 | 0.233 | 0.59 | 0.81 | 0.7 | 0.5 | 0.78 | 0.56 |
| 1 | 230891333 | rs2493146 | -0.621 | 0.562 | -0.103 | -0.088 | 0.024 | 0.017 | 0.017 | 0.298 | 0.37 | 0.58 | 0.54 | 0.48 | 0.48 | 0.43 |
| 1 | 230893608 | rs6698654 | -0.016 | 0.645 | 0.281 | -0.106 | 0.037 | 0.01 | 0.01 | 0 | 0.17 | 0.34 | 0.14 | 0.12 | 0.21 | 0.11 |
| 1 | 230894323 | rs1890349 | -1.32 | 0.559 | 0.408 | -0.049 | 0.077 | 0.004 | 0.004 | 0.075 | 0.5 | 0.72 | 0.67 | 0.51 | 0.69 | 0.48 |
| 1 | 230894365 | rs1890350 | -0.42 | 0.676 | 0.263 | -0.036 | 0.006 | -0.001 | -0.001 | 0 | 0.33 | 0.38 | 0.53 | 0.4 | 0.48 | 0.37 |
| 1 | 230895340 | rs28359608 | 0.865 | 0.078 | 0.266 | 0.076 | -0.002 | -0.007 | -0.007 | 0 | 0.16 | 0.17 | 0.06 | 0.08 | 0.15 | 0.28 |
| 1 | 230896093 | rs7515210 | 0.063 | -0.318 | 0.3 | -0.214 | -0.004 | -0.018 | -0.018 | 0 | 0.31 | 0.32 | 0.31 | 0.2 | 0.28 | 0.28 |
| 1 | 230896238 | rs11579420 | -0.563 | -0.919 | 0.132 | -0.444 | 0.001 | 0.009 | 0.009 | 0.232 | 0.42 | 0.68 | 0.44 | 0.3 | 0.57 | 0.49 |
| 1 | 230897463 | rs3790955 | -0.317 | -0.839 | 0.405 | 0.103 | 0.03 | 0.03 | 0.03 | 0.716 | 0.35 | 0.47 | 0.52 | 0.53 | 0.53 | 0.46 |
| 1 | 230897893 | rs7523730 | 1.82 | -0.775 | 0.036 | 0.045 | -0.085 | 0.012 | 0.012 | 0 | 0.12 | 0.02 | 0.3 | 0.12 | 0.34 | 0.08 |
| 1 | 230898955 | rs3790960 | -0.486 | -0.804 | 0.292 | 0.14 | -0.002 | -0.002 | -0.002 | 0.193 | 0.37 | 0.63 | 0.52 | 0.53 | 0.53 | 0.46 |
| 1 | 230900011 | rs4847011 | 0.93 | -0.492 | -0.017 | 0.15 | 0.011 | 0 | 0 | 0 | 0.15 | 0.19 | 0.3 | 0.2 | 0.34 | 0.13 |
| 1 | 230903006 | rs12027189 | 2.007 | -0.777 | -0.299 | 0.082 | 0.07 | -0.067 | -0.067 | 0.54 | 0.22 | 0.02 | 0.12 | 0.24 | 0.11 | 0.14 |
| 1 | 230904268 | rs872505 | -0.108 | -0.535 | 0.037 | -0.721 | -0.015 | 0.024 | 0.024 | 0 | 0.67 | 0.71 | 0.5 | 0.78 | 0.42 | 0.8 |
| 1 | 230904911 | rs11122593 | 1.327 | -0.183 | 0.135 | -0.41 | -0.093 | -0.005 | -0.005 | 0.005 | 0.13 | 0.02 | 0.29 | 0.12 | 0.34 | 0.08 |
| 1 | 230905723 | rs4847017 | 0.865 | -0.282 | 0.255 | -0.436 | 0.009 | 0 | 0 | 0 | 0.15 | 0.18 | 0.31 | 0.12 | 0.34 | 0.08 |
| 1 | 230906954 | rs6677009 | -1.141 | -0.445 | 0.461 | -0.54 | 0.006 | 0.002 | 0.002 | 0 | 0.62 | 0.69 | 0.63 | 0.41 | 0.71 | 0.72 |
| 1 | 230907231 | rs12404010 | 2.102 | -0.64 | 0.519 | -0.615 | 0.006 | 0.006 | 0.006 | 0 | 0.12 | 0.1 | 0.11 | 0.08 | 0.05 | 0.14 |
| 1 | 230908863 | rs3790973 | -0.503 | -0.617 | 0.45 | -0.622 | 0.001 | -0.002 | -0.002 | 0 | 0.48 | 0.51 | 0.32 | 0.28 | 0.36 | 0.64 |
| 1 | 230910339 | rs7517389 | 2.05 | -0.544 | 0.328 | -0.723 | 0.006 | 0.006 | 0.006 | 0 | 0.12 | 0.1 | 0.11 | 0.08 | 0.05 | 0.14 |
| 1 | 230913184 | rs11122597 | 0.509 | -0.467 | 0.404 | -0.716 | -0.012 | 0.01 | 0.01 | 0 | 0.25 | 0.18 | 0.41 | 0.2 | 0.39 | 0.22 |
| 1 | 230913434 | rs10779852 | -1.84 | -0.571 | 0.071 | -0.648 | 0.009 | -0.133 | -0.133 | 0.476 | 0.68 | 0.96 | 0.68 | 0.47 | 0.76 | 0.79 |
| 1 | 230914229 | rs12724252 | -0.555 | -0.617 | 0.582 | -0.501 | -0.016 | -0.048 | -0.048 | 0 | 0.38 | 0.55 | 0.22 | 0.19 | 0.32 | 0.5 |
| 1 | 230914706 | rs1933630 | 1.339 | -0.652 | 0.593 | -0.54 | -0.005 | 0.01 | 0.01 | 0 | 0.11 | 0.09 | 0.13 | 0.08 | 0.15 | 0.11 |
| 1 | 230914729 | rs1933631 | -0.2 | -0.459 | 0.717 | -0.433 | -0.003 | -0.016 | -0.016 | 0.655 | 0.36 | 0.39 | 0.21 | 0.19 | 0.32 | 0.5 |
| 1 | 230917513 | rs7553239 | 0.121 | -0.608 | 0.551 | -0.496 | -0.018 | -0.068 | -0.068 | 0.011 | 0.24 | 0.35 | 0.16 | 0.08 | 0.18 | 0.19 |
| 1 | 230923313 | rs12731961 | 0.18 | -0.762 | 0.355 | -0.766 | -0.007 | -0.041 | -0.041 | 0 | 0.24 | 0.32 | 0.15 | 0.08 | 0.18 | 0.19 |
| 1 | 230924376 | rs1933633 | 0.524 | -0.722 | -0.063 | -0.747 | 0.018 | 0.051 | 0.051 | 0.005 | 0.22 | 0.14 | 0.14 | 0.08 | 0.18 | 0.19 |
| 1 | 230924905 | rs12082061 | 0.18 | -0.287 | -0.075 | -0.504 | -0.033 | -0.076 | -0.076 | 1.4 | 0.36 | 0.48 | 0.27 | 0.16 | 0.23 | 0.33 |
| 1 | 230925464 | rs6661547 | 0.495 | 0.788 | 0.03 | 0.274 | 0.002 | 0.003 | 0.003 | 2.408 | 0.13 | 0.21 | 0.07 | 0.1 | 0.14 | 0.31 |
| 1 | 230928351 | rs2282321 | -0.483 | 1.405 | -0.239 | 0.083 | -0.009 | 0.004 | 0.004 | 1.516 | 0.23 | 0.16 | 0.33 | 0.17 | 0.39 | 0.3 |

| | | | | | | | | | | | | | | | | |
|---|---|---|---|---|---|---|---|---|---|---|---|---|---|---|---|---|
| 1 | 230929310 | rs3828128 | 0.944 | 1.767 | 0.338 | 0.069 | 0.023 | 0 | 0 | 0 | 0.08 | 0.07 | 0.26 | 0.11 | 0.25 | 0.03 |
| 1 | 230931615 | rs12084401 | 1.195 | 0.247 | -0.309 | -0.66 | 0.009 | 0 | 0 | 0 | 0.33 | 0.32 | 0.18 | 0.04 | 0.11 | 0.21 |
| 1 | 230932442 | rs4477283 | 0.384 | 0.163 | -0.059 | -0.599 | -0.024 | -0.034 | -0.034 | 0 | 0.47 | 0.48 | 0.24 | 0.12 | 0.23 | 0.48 |
| 1 | 230936770 | rs6667160 | 1.297 | 0.27 | 0.127 | -0.564 | -0.005 | -0.013 | -0.013 | 0 | 0.13 | 0.17 | 0.06 | 0.07 | 0.13 | 0.27 |
| 1 | 230940170 | rs10864783 | -0.484 | 0.604 | -0.052 | -0.621 | 0 | 0 | 0 | 0 | 0.68 | 0.67 | 0.67 | 0.34 | 0.68 | 0.65 |
| 1 | 230942815 | rs12740185 | 1.356 | 0.338 | 0.007 | -0.584 | 0.005 | -0.002 | -0.002 | 0 | 0.32 | 0.32 | 0.18 | 0.04 | 0.11 | 0.21 |
| 1 | 230947583 | rs12410698 | 0.804 | 0.827 | -0.282 | -0.38 | -0.015 | 0.014 | 0.014 | 0 | 0.14 | 0.09 | 0.16 | 0.13 | 0.18 | 0.14 |
| 1 | 230951172 | rs1202524 | -1.088 | 1.113 | -0.364 | -0.439 | -0.038 | -0.002 | -0.002 | 0 | 0.89 | 0.96 | 0.72 | 0.89 | 0.69 | 0.92 |
| 1 | 230953248 | rs1202526 | 0.213 | 0.926 | -0.385 | -0.114 | 0.005 | 0.001 | 0.001 | 0.179 | 0.41 | 0.36 | 0.37 | 0.61 | 0.37 | 0.49 |
| 1 | 230953983 | rs12062054 | 0.749 | 0.079 | -0.177 | -0.122 | -0.065 | -0.047 | -0.047 | 6.568 | 0.35 | 0.51 | 0.2 | 0.15 | 0.14 | 0.29 |
| 1 | 230959250 | rs16852740 | -0.036 | 1.153 | -0.527 | 0.505 | 0.038 | 0.016 | 0.016 | 0 | 0.17 | 0.29 | 0.19 | 0.2 | 0.24 | 0.21 |
| 1 | 230959298 | rs4847021 | 0.303 | 1.006 | -0.973 | 1.07 | -0.025 | -0.017 | -0.017 | 0 | 0.05 | 0.01 | 0.17 | 0.26 | 0.07 | 0.07 |
| 1 | 230966799 | rs1202543 | -0.333 | 0.947 | -0.906 | 0.653 | -0.008 | -0.006 | -0.006 | 0 | 0.27 | 0.24 | 0.5 | 0.48 | 0.34 | 0.24 |
| 1 | 230967445 | rs11122614 | 0.625 | -0.755 | -1.062 | -0.013 | 0.091 | 0.12 | 0.12 | 0.858 | 0.33 | 0.12 | 0.21 | 0.04 | 0.18 | 0.25 |
| 1 | 230974329 | rs6673201 | -0.42 | 0.099 | -0.209 | 0.866 | 0.07 | 0.211 | 0.211 | 0.365 | 0.06 | 0.44 | 0.05 | 0.07 | 0.02 | 0.05 |
| 1 | 230978726 | rs11122616 | -0.199 | -0.246 | -1.609 | 0.639 | 0.015 | -0.1 | -0.1 | 4.109 | 0.37 | 0.05 | 0.32 | 0.5 | 0.31 | 0.33 |
| 1 | 230981823 | rs2095769 | -0.008 | -0.1 | -1.381 | 1.19 | -0.021 | -0.099 | -0.099 | 0.938 | 0.19 | 0.03 | 0.12 | 0.4 | 0.18 | 0.22 |
| 1 | 230984594 | rs11805329 | 0.01 | -0.314 | -0.689 | 0.969 | 0.057 | 0.038 | 0.038 | 0.029 | 0.05 | 0.39 | 0.04 | 0 | 0.01 | 0.02 |
| 1 | 230988920 | rs2180478 | 0.022 | -0.3 | -0.818 | 1.061 | -0.01 | -0.08 | -0.08 | 0 | 0.19 | 0.05 | 0.12 | 0.4 | 0.17 | 0.22 |
| 1 | 230990550 | rs1202568 | -0.715 | 0.47 | -0.845 | 0.661 | 0.014 | 0.018 | 0.018 | 0 | 0.78 | 0.69 | 0.57 | 0.6 | 0.71 | 0.81 |
| 1 | 230993475 | rs12034355 | 0.655 | 0.462 | -0.902 | 0.66 | -0.011 | -0.094 | -0.094 | 0 | 0.17 | 0.03 | 0.11 | 0.4 | 0.13 | 0.21 |
| 1 | 230994593 | rs1999903 | 0.957 | 0.82 | -1.079 | 0.357 | -0.023 | 0.015 | 0.015 | 0 | 0.11 | 0.03 | 0.09 | 0.06 | 0.15 | 0.23 |
| 1 | 231000791 | rs1202580 | 0.621 | 0.694 | -0.855 | 0.317 | 0.01 | 0.014 | 0.014 | 0.411 | 0.61 | 0.55 | 0.46 | 0.48 | 0.5 | 0.54 |
| 1 | 231000867 | rs1202581 | -0.074 | 0.626 | -0.717 | -0.007 | 0.011 | 0.016 | 0.016 | 0 | 0.7 | 0.32 | 0.51 | 0.55 | 0.63 | 0.77 |
| 1 | 231002150 | rs1202584 | -0.462 | 0.674 | -0.813 | -0.005 | 0.005 | 0.003 | 0.003 | 0 | 0.74 | 0.7 | 0.56 | 0.61 | 0.66 | 0.79 |
| 1 | 231005504 | rs3811499 | -0.107 | 0.599 | -1.219 | 0.01 | -0.01 | -0.035 | -0.035 | 0.324 | 0.3 | 0.17 | 0.21 | 0.53 | 0.29 | 0.47 |
| 1 | 231008904 | rs1202589 | 1.062 | 0.087 | -1.35 | -0.288 | 0.053 | 0.173 | 0.173 | 0.249 | 0.4 | 0.13 | 0.3 | 0.03 | 0.34 | 0.3 |
| 1 | 231012625 | rs1202592 | 0.87 | 0.125 | -1.131 | -0.258 | 0.051 | 0.115 | 0.115 | 0 | 0.43 | 0.23 | 0.31 | 0.08 | 0.35 | 0.32 |
| 1 | 231018941 | rs9431872 | -0.252 | -0.115 | -1.044 | 0.349 | 0.028 | -0.021 | -0.021 | 0.16 | 0.26 | 0.19 | 0.34 | 0.33 | 0.17 | 0.26 |
| 1 | 231024653 | rs16852992 | -0.247 | -0.038 | -0.678 | -0.048 | -0.016 | -0.025 | -0.025 | 0 | 0.15 | 0.14 | 0.1 | 0.04 | 0.12 | 0.13 |
| 1 | 231028032 | rs6704527 | 0.493 | 0.047 | -0.427 | 0.274 | 0.006 | -0.017 | -0.017 | 0 | 0.22 | 0.17 | 0.11 | 0.38 | 0.17 | 0.25 |
| 1 | 231034396 | rs6673054 | 0.485 | 0.044 | -0.326 | 0.232 | 0.003 | -0.008 | -0.008 | 0 | 0.23 | 0.18 | 0.12 | 0.4 | 0.18 | 0.26 |
| 1 | 231034640 | rs10495301 | 0.26 | 1.241 | -0.724 | 0.084 | -0.082 | 0.006 | 0.006 | 0 | 0.14 | 0.02 | 0.25 | 0.15 | 0.26 | 0.28 |
| 1 | 231036505 | rs3762404 | -0.145 | 0.299 | -0.447 | -0.431 | -0.001 | -0.001 | -0.001 | 1.274 | 0.36 | 0.42 | 0.66 | 0.54 | 0.53 | 0.5 |
| 1 | 231041558 | rs7533188 | -1.085 | 0.222 | -0.463 | -0.461 | 0.04 | 0.016 | 0.016 | 0 | 0.5 | 0.61 | 0.78 | 0.58 | 0.67 | 0.61 |
| 1 | 231053063 | rs10864790 | -0.024 | 0.627 | -0.181 | -0.598 | 0 | 0 | 0 | 0 | 0.33 | 0.34 | 0.63 | 0.56 | 0.51 | 0.49 |
| 1 | 231054409 | rs6656765 | 0.702 | 0.567 | 0.007 | -0.651 | -0.004 | -0.023 | -0.023 | 0.017 | 0.12 | 0.13 | 0.11 | 0.03 | 0.13 | 0.11 |
| 1 | 231060736 | rs3748775 | 0.126 | 0.593 | -0.002 | -0.628 | 0.001 | 0.002 | 0.002 | 0 | 0.33 | 0.34 | 0.63 | 0.56 | 0.51 | 0.49 |
| 1 | 231066782 | rs11122623 | -0.348 | 0.548 | -0.237 | -0.6 | 0.006 | 0.014 | 0.014 | 0 | 0.38 | 0.57 | 0.66 | 0.56 | 0.54 | 0.5 |
| 1 | 231069814 | rs1890352 | -1.082 | 0.483 | -0.349 | -0.583 | 0.083 | 0.032 | 0.032 | 0 | 0.5 | 0.7 | 0.78 | 0.59 | 0.67 | 0.61 |
| 1 | 231078492 | rs12130934 | 0.562 | 0.687 | -0.273 | -0.479 | -0.004 | -0.023 | -0.023 | 0 | 0.12 | 0.13 | 0.11 | 0.04 | 0.13 | 0.11 |
| 1 | 231080298 | rs12135063 | 1.728 | 0.675 | -0.244 | -0.502 | 0.004 | -0.023 | -0.023 | 0.073 | 0.05 | 0.07 | 0.07 | 0 | 0.08 | 0.04 |
| 1 | 231081304 | rs6673085 | -0.331 | 0.623 | -0.285 | -0.624 | 0.008 | 0.017 | 0.017 | 0 | 0.38 | 0.55 | 0.66 | 0.56 | 0.54 | 0.5 |
| 1 | 231087075 | rs4846884 | -2.002 | -0.098 | -0.521 | -1.158 | 0.044 | 0.05 | 0.05 | 0.168 | 0.72 | 0.83 | 0.89 | 0.98 | 0.84 | 0.86 |
| 1 | 231090578 | rs6541240 | -0.314 | 1.241 | -0.458 | -0.808 | 0.007 | 0.008 | 0.008 | 0 | 0.83 | 0.87 | 0.55 | 0.64 | 0.66 | 0.81 |
| 1 | 231097042 | rs11122162 | 1.179 | 1.082 | -0.338 | -0.866 | 0 | -0.027 | -0.027 | 0 | 0.05 | 0.17 | 0.03 | 0 | 0.03 | 0.01 |
| 1 | 231098099 | rs12041215 | 0.353 | 1.145 | -0.553 | -0.867 | -0.022 | 0.01 | 0.01 | 0 | 0.26 | 0.14 | 0.28 | 0.24 | 0.31 | 0.41 |
| 1 | 231104225 | rs6541247 | -0.253 | 1.012 | -0.147 | -0.523 | 0.012 | 0.017 | 0.017 | 0 | 0.82 | 0.83 | 0.59 | 0.67 | 0.71 | 0.81 |
| 1 | 231104580 | rs7553253 | -0.54 | 1.036 | -0.142 | -0.52 | 0.01 | 0.014 | 0.014 | 0 | 0.83 | 0.85 | 0.59 | 0.66 | 0.71 | 0.83 |
| 1 | 231112794 | rs13374343 | 0.554 | 1.011 | -0.546 | -0.495 | 0.025 | 0.033 | 0.033 | 0 | 0.06 | 0.15 | 0.13 | 0.13 | 0.08 | 0.19 |
| 1 | 231113437 | rs11122173 | 1.561 | 0.852 | -0.628 | -0.714 | 0.025 | -0.009 | -0.009 | 0.131 | 0.58 | 0.3 | 0.23 | 0.51 | 0.34 | 0.49 |
| 1 | 231115887 | rs10864645 | 1.14 | 0.843 | -0.628 | -0.717 | 0.003 | -0.001 | -0.001 | 0 | 0.63 | 0.42 | 0.36 | 0.65 | 0.43 | 0.69 |
| 1 | 231118857 | rs11122177 | 1.325 | 0.799 | -0.495 | -0.728 | -0.002 | 0.003 | 0.003 | 0.068 | 0.61 | 0.43 | 0.25 | 0.51 | 0.36 | 0.51 |
| 1 | 231123051 | rs10864647 | 1.408 | 0.862 | -1.183 | -0.676 | 0.02 | -0.007 | -0.007 | 0.349 | 0.58 | 0.31 | 0.23 | 0.51 | 0.35 | 0.5 |
| 1 | 231124376 | rs10779805 | 1.408 | 0.845 | -1.225 | -0.676 | 0.025 | -0.008 | -0.008 | 0 | 0.58 | 0.3 | 0.23 | 0.51 | 0.35 | 0.5 |
| 1 | 231128319 | rs12732752 | 0.15 | 1.253 | -1.344 | -0.528 | -0.079 | 0.063 | 0.063 | 0 | 0.13 | 0.01 | 0.17 | 0.02 | 0.27 | 0.12 |
| 1 | 231134867 | rs11122186 | 1.044 | 0.159 | -1.097 | -0.308 | 0.009 | 0.002 | 0.002 | 0 | 0.16 | 0.12 | 0.06 | 0.12 | 0.11 | 0.15 |
| 1 | 231151239 | rs12133304 | 0.221 | 0.282 | -1.021 | -0.248 | 0.012 | 0.017 | 0.017 | 0 | 0.05 | 0.1 | 0.13 | 0.13 | 0.08 | 0.19 |
| 1 | 231153850 | rs11122192 | 0.942 | 0.196 | -1.319 | -0.328 | 0.004 | 0.002 | 0.002 | 0 | 0.16 | 0.14 | 0.06 | 0.12 | 0.11 | 0.15 |
| 1 | 231156029 | rs10864654 | -0.639 | 0.673 | -1.884 | -0.206 | -0.127 | -0.079 | -0.079 | 0 | 0.27 | 0.09 | 0.55 | 0.36 | 0.51 | 0.25 |
| 1 | 231156513 | rs4409638 | -0.327 | 0.221 | -1.699 | -0.701 | -0.004 | 0.088 | 0.088 | 0 | 0.41 | 0.29 | 0.3 | 0.11 | 0.45 | 0.25 |
| 1 | 231156743 | rs4304564 | 0.121 | 0.486 | -1.161 | -0.054 | 0.022 | 0.016 | 0.016 | 0 | 0.25 | 0.37 | 0.44 | 0.27 | 0.3 | 0.16 |
| 1 | 231158220 | rs11122193 | 0.495 | -0.404 | -1.141 | 0.254 | -0.007 | -0.008 | -0.008 | 0 | 0.19 | 0.23 | 0.07 | 0.12 | 0.12 | 0.21 |
| 1 | 231160646 | rs4658913 | -0.498 | -0.27 | -1.116 | 0.223 | 0.021 | -0.004 | -0.004 | 0 | 0.56 | 0.68 | 0.7 | 0.49 | 0.65 | 0.56 |
| 1 | 231162643 | rs4506448 | -0.025 | -1.241 | -0.622 | -0.202 | -0.09 | -0.102 | -0.102 | 0.636 | 0.31 | 0.47 | 0.16 | 0.13 | 0.11 | 0.34 |
| 1 | 231173379 | rs4378192 | -0.123 | -0.486 | -0.215 | 0.106 | -0.011 | 0.006 | 0.006 | 0 | 0.42 | 0.43 | 0.44 | 0.54 | 0.32 | 0.41 |
| 1 | 231174431 | rs4543780 | -0.066 | 1.327 | -0.157 | 0.14 | 0.043 | -0.023 | -0.023 | 0.01 | 0.14 | 0.22 | 0.16 | 0.07 | 0.3 | 0.1 |
| 1 | 231186593 | rs6686989 | 0.412 | 0.83 | 0.426 | 0.469 | 0.02 | 0.079 | 0.079 | 1.735 | 0.88 | 0.68 | 0.81 | 0.74 | 0.87 | 0.75 |
| 1 | 231189818 | rs6679453 | 0.754 | 0.279 | 0.197 | 0.041 | 0.012 | 0.008 | 0.008 | 0 | 0.39 | 0.31 | 0.32 | 0.31 | 0.29 | 0.37 |

| | | | | | | | | | | | | | | | | |
|---|---|---|---|---|---|---|---|---|---|---|---|---|---|---|---|---|
| 1 | 231194176 | rs12117901 | 0.232 | 0.134 | -0.028 | -0.168 | -0.023 | -0.01 | -0.01 | 0.203 | 0.37 | 0.28 | 0.43 | 0.42 | 0.5 | 0.29 |
| 1 | 231195342 | rs4658931 | 0.938 | 0.401 | -0.525 | 0.161 | -0.053 | -0.053 | -0.053 | 0 | 0.09 | 0.01 | 0.25 | 0.2 | 0.18 | 0.05 |
| 1 | 231199867 | rs11122213 | 0.46 | 0.168 | -0.426 | -0.21 | 0.025 | -0.031 | -0.031 | 0 | 0.31 | 0.19 | 0.27 | 0.41 | 0.23 | 0.26 |
| 1 | 231200830 | rs4399139 | -0.735 | 0.017 | -0.4 | -0.347 | -0.005 | -0.024 | -0.024 | 0.281 | 0.36 | 0.56 | 0.57 | 0.22 | 0.63 | 0.25 |
| 1 | 231204794 | rs10864665 | -0.357 | 0.467 | -0.185 | 0.06 | 0.005 | -0.006 | -0.006 | 0.182 | 0.57 | 0.66 | 0.57 | 0.45 | 0.63 | 0.44 |
| 1 | 231207771 | rs6674245 | -0.751 | 0.925 | 0.515 | -0.12 | 0.057 | -0.104 | -0.104 | 0 | 0.17 | 0.51 | 0.19 | 0 | 0.23 | 0.06 |
| 1 | 231211408 | rs3923542 | -0.216 | 0.885 | 0.765 | -0.115 | 0.003 | -0.057 | -0.057 | 0 | 0.12 | 0.22 | 0.12 | 0 | 0.14 | 0.05 |
| 1 | 231214726 | rs12405429 | -0.501 | 1.756 | 0.484 | 0.224 | -0.034 | -0.033 | -0.033 | 1.05 | 0.06 | 0.01 | 0.22 | 0.12 | 0.15 | 0.05 |
| 1 | 231220364 | rs4388693 | 1.511 | 1.295 | 0.714 | -0.176 | -0.022 | 0.023 | 0.023 | 1.445 | 0.7 | 0.26 | 0.55 | 0.87 | 0.58 | 0.75 |
| 1 | 231228173 | rs4322215 | 0.647 | 0.383 | -0.065 | 0.003 | 0.104 | -0.018 | -0.018 | 0.025 | 0.39 | 0.19 | 0.2 | 0.57 | 0.19 | 0.5 |
| 1 | 231232043 | rs12136287 | 0.27 | 0.636 | 0.193 | -0.011 | -0.002 | -0.071 | -0.071 | 0 | 0.15 | 0.26 | 0.13 | 0 | 0.16 | 0.09 |
| 1 | 231234360 | rs7518479 | -0.446 | 0.68 | 0.515 | 0.007 | 0 | -0.012 | -0.012 | 0.133 | 0.55 | 0.57 | 0.5 | 0.73 | 0.46 | 0.67 |
| 1 | 231234439 | rs7521117 | -1.555 | 0.692 | 0.414 | 0.028 | -0.126 | 0.007 | 0.007 | 0.278 | 0.71 | 0.97 | 0.64 | 0.74 | 0.62 | 0.76 |
| 1 | 231243211 | rs12091966 | -1.408 | 0.029 | 0.87 | 0.424 | 0.046 | -0.037 | -0.037 | 2.64 | 0.24 | 0.51 | 0.37 | 0.14 | 0.31 | 0.19 |
| 1 | 231243756 | rs12410954 | -1.17 | 0.263 | 0.516 | 0.638 | 0.011 | 0.007 | 0.007 | 0 | 0.08 | 0.12 | 0.22 | 0.13 | 0.15 | 0.1 |
| 1 | 231246212 | rs6697412 | -1.281 | -0.035 | 0.597 | 0.489 | 0.024 | 0.008 | 0.008 | 0.03 | 0.2 | 0.59 | 0.16 | 0.16 | 0.2 | 0.21 |
| 1 | 231254003 | rs10864669 | -0.703 | -0.151 | 0.215 | 0.289 | 0.002 | -0.022 | -0.022 | 0 | 0.57 | 0.51 | 0.54 | 0.67 | 0.54 | 0.68 |
| 1 | 231255113 | rs2883720 | 0.34 | -0.138 | 0.073 | 0.373 | 0.018 | -0.001 | -0.001 | 0 | 0.4 | 0.29 | 0.43 | 0.54 | 0.35 | 0.45 |
| 1 | 231260770 | rs2180275 | -0.477 | 0.063 | 0.456 | 0.472 | -0.005 | 0.058 | 0.058 | 0.049 | 0.54 | 0.66 | 0.58 | 0.76 | 0.5 | 0.62 |
| 1 | 231262020 | rs12028750 | -1.738 | -0.249 | 0.275 | 0.723 | -0.013 | -0.03 | -0.03 | 0 | 0.37 | 0.72 | 0.47 | 0.51 | 0.42 | 0.41 |
| 1 | 231262302 | rs12028816 | -0.927 | 0.004 | -0.276 | 0.978 | 0.013 | 0.036 | 0.036 | 0.116 | 0.24 | 0.28 | 0.4 | 0.51 | 0.35 | 0.29 |
| 1 | 231270636 | rs12023396 | -0.52 | -0.246 | -0.378 | 0.984 | -0.013 | -0.039 | -0.039 | 0 | 0.2 | 0.15 | 0.23 | 0.37 | 0.25 | 0.23 |
| 1 | 231273550 | rs9431646 | -0.075 | -0.466 | -0.329 | 0.878 | -0.011 | -0.003 | -0.003 | 0 | 0.44 | 0.53 | 0.23 | 0.34 | 0.31 | 0.5 |
| 1 | 231276007 | rs765265 | -0.034 | -0.752 | -0.276 | 0.908 | -0.008 | -0.002 | -0.002 | 0 | 0.44 | 0.46 | 0.23 | 0.33 | 0.31 | 0.51 |
| 1 | 231277763 | rs6684407 | -0.653 | -0.193 | -0.103 | 1.489 | -0.038 | 0.043 | 0.043 | 0 | 0.62 | 0.75 | 0.43 | 0.7 | 0.53 | 0.72 |
| 1 | 231282933 | rs9431652 | -0.536 | -0.152 | -0.427 | 1.276 | -0.007 | -0.001 | -0.001 | 0 | 0.6 | 0.61 | 0.35 | 0.34 | 0.52 | 0.59 |
| 1 | 231284033 | rs12138442 | 0.508 | -0.21 | -1.205 | 1.336 | -0.057 | -0.033 | -0.033 | 0 | 0.34 | 0.22 | 0.61 | 0.64 | 0.47 | 0.37 |
| 1 | 231284128 | rs6689582 | -0.477 | -0.616 | -1.526 | 1.312 | 0.065 | 0.089 | 0.089 | 0 | 0.57 | 0.86 | 0.77 | 0.79 | 0.68 | 0.58 |
| 1 | 231284626 | rs12032939 | -1.031 | -0.597 | -1.426 | 1.429 | -0.003 | 0.037 | 0.037 | 0 | 0.09 | 0.08 | 0.16 | 0.53 | 0.12 | 0.3 |
| 1 | 231285646 | rs1321131 | -0.203 | -0.617 | -1.303 | 1.067 | 0.046 | 0.049 | 0.049 | 0.042 | 0.48 | 0.76 | 0.67 | 0.37 | 0.59 | 0.31 |
| 1 | 231291359 | rs1321132 | 0.049 | 0.022 | -1.064 | 1.157 | -0.011 | -0.032 | -0.032 | 0 | 0.42 | 0.25 | 0.48 | 0.57 | 0.51 | 0.45 |
| 1 | 231302356 | rs9431947 | -0.231 | -0.783 | -0.331 | 0.666 | -0.024 | -0.071 | -0.071 | 0 | 0.17 | 0.25 | 0.09 | 0.01 | 0.11 | 0.12 |
| 1 | 231311867 | rs9431660 | -0.714 | -0.855 | -0.165 | 1.254 | -0.016 | 0.009 | 0.009 | 0 | 0.37 | 0.45 | 0.37 | 0.41 | 0.26 | 0.46 |
| 1 | 231313859 | rs1406869 | -1.262 | -0.978 | -0.28 | 1.214 | 0.057 | 0.013 | 0.013 | 0 | 0.47 | 0.91 | 0.55 | 0.61 | 0.34 | 0.57 |
| 1 | 231316174 | rs12048739 | -0.108 | -0.921 | 0.188 | 1.464 | 0.015 | 0.134 | 0.134 | 0 | 0.06 | 0.13 | 0.16 | 0.27 | 0.07 | 0.11 |
| 1 | 231317179 | rs4658871 | -0.559 | -0.927 | -0.05 | 1.415 | -0.008 | -0.002 | -0.002 | 5.945 | 0.37 | 0.43 | 0.37 | 0.33 | 0.26 | 0.46 |
| 1 | 231318678 | rs12032558 | -1.345 | -0.35 | 0.707 | 1.526 | 0.058 | 0.27 | 0.27 | 0 | 0.09 | 0.46 | 0.18 | 0.26 | 0.07 | 0.11 |
| 1 | 231323339 | rs16853942 | -1.701 | -0.229 | 1.018 | 0.889 | 0.032 | -0.04 | -0.04 | 0 | 0.12 | 0.4 | 0.09 | 0.08 | 0.11 | 0.17 |
| 1 | 231324483 | rs12063574 | -2.114 | -0.673 | 0.981 | 1.49 | -0.012 | 0.098 | 0.098 | 0 | 0.21 | 0.48 | 0.12 | 0.33 | 0.14 | 0.32 |
| 1 | 231326086 | rs1998027 | -2.215 | -1.095 | 0.15 | 1.467 | -0.02 | 0.083 | 0.083 | 0 | 0.51 | 0.97 | 0.6 | 0.69 | 0.41 | 0.64 |
| 1 | 231328889 | rs10864673 | -2.008 | -0.813 | 0.783 | 1.263 | -0.003 | 0.108 | 0.108 | 0.064 | 0.2 | 0.54 | 0.12 | 0.33 | 0.14 | 0.32 |
| 1 | 231337031 | rs11122252 | -1.443 | -0.562 | 0.812 | 1.077 | 0.061 | -0.03 | -0.03 | 0 | 0.12 | 0.47 | 0.1 | 0.07 | 0.11 | 0.17 |
| 1 | 231340433 | rs6700950 | -1.367 | -0.548 | 0.848 | 1.049 | 0.034 | -0.024 | -0.024 | 0 | 0.11 | 0.46 | 0.09 | 0.07 | 0.08 | 0.13 |
| 1 | 231341511 | rs6541261 | -2.23 | -0.681 | 0.893 | 1.193 | -0.03 | 0.045 | 0.045 | 0.114 | 0.21 | 0.62 | 0.13 | 0.3 | 0.12 | 0.3 |
| 1 | 231341569 | rs6679627 | -0.189 | -0.669 | 0.827 | 1.205 | -0.022 | 0.082 | 0.082 | 0.432 | 0.09 | 0.15 | 0.04 | 0.23 | 0.04 | 0.16 |
| 1 | 231342857 | rs2180273 | -2.432 | -0.598 | 0.789 | 1.194 | -0.012 | 0.029 | 0.029 | 0 | 0.24 | 0.6 | 0.14 | 0.3 | 0.16 | 0.36 |
| 1 | 231345300 | rs16841556 | -1.834 | -0.696 | 0.551 | 1.383 | 0.001 | -0.001 | -0.001 | 0.498 | 0.39 | 0.41 | 0.44 | 0.41 | 0.32 | 0.35 |
| 1 | 231346186 | rs4658872 | -0.576 | -0.704 | 0.478 | 1.287 | 0.006 | -0.028 | -0.028 | 0 | 0.26 | 0.12 | 0.39 | 0.35 | 0.25 | 0.27 |
| 1 | 231352767 | rs2281480 | -1.939 | -0.811 | 0.614 | 1.319 | -0.01 | 0 | 0 | 0 | 0.39 | 0.44 | 0.46 | 0.58 | 0.31 | 0.44 |
| 1 | 231353589 | rs4658907 | -0.581 | -0.805 | 0.576 | 1.352 | 0.005 | -0.032 | -0.032 | 0 | 0.26 | 0.12 | 0.4 | 0.35 | 0.27 | 0.27 |
| 1 | 231355070 | rs524331 | -1.964 | -0.732 | 0.589 | 1.277 | -0.005 | -0.001 | -0.001 | 0 | 0.42 | 0.47 | 0.46 | 0.57 | 0.35 | 0.5 |
| 1 | 231362923 | rs491182 | -1.322 | -0.301 | 0.648 | 1.282 | -0.001 | -0.001 | -0.001 | 0 | 0.28 | 0.29 | 0.42 | 0.35 | 0.29 | 0.3 |
| 1 | 231363000 | rs16854012 | 1.699 | -0.332 | 0.015 | 0.931 | -0.005 | -0.083 | -0.083 | 0.039 | 0.12 | 0.01 | 0.19 | 0.29 | 0.09 | 0.15 |
| 1 | 231365440 | rs551480 | -1.265 | 0.372 | 0.91 | 1.095 | -0.004 | 0.009 | 0.009 | 0 | 0.27 | 0.37 | 0.42 | 0.35 | 0.26 | 0.26 |
| 1 | 231368219 | rs11579396 | -1.104 | 0.501 | 0.622 | 1.285 | 0.002 | 0.009 | 0.009 | 0 | 0.16 | 0.18 | 0.19 | 0.26 | 0.19 | 0.19 |
| 1 | 231376806 | rs2274067 | 0.254 | 0.257 | 1.002 | 1.193 | -0.004 | 0.005 | 0.005 | 0 | 0.07 | 0.04 | 0.2 | 0.09 | 0.06 | 0.11 |
| 1 | 231377081 | rs518686 | -2.484 | 1.06 | 1.065 | 1.129 | 0.001 | 0.002 | 0.002 | 0 | 0.52 | 0.53 | 0.62 | 0.63 | 0.52 | 0.63 |
| 1 | 231389097 | rs503633 | -2.257 | 0.976 | 1.081 | 1.184 | -0.001 | -0.002 | -0.002 | 0.114 | 0.44 | 0.43 | 0.59 | 0.56 | 0.51 | 0.6 |
| 1 | 231393519 | rs3767761 | -2.84 | 0.988 | 1.003 | 1.132 | 0.01 | 0.018 | 0.018 | 0 | 0.54 | 0.7 | 0.63 | 0.63 | 0.53 | 0.64 |
| 1 | 231401204 | rs6702011 | -1.358 | 1.254 | 1.155 | 1.243 | 0.02 | -0.003 | -0.003 | 0.849 | 0.19 | 0.27 | 0.37 | 0.24 | 0.23 | 0.25 |
| 1 | 231406961 | rs539699 | -2.854 | 1.246 | 0.854 | 1.089 | 0.007 | 0.019 | 0.019 | 0 | 0.54 | 0.7 | 0.63 | 0.63 | 0.53 | 0.63 |
| 1 | 231408091 | rs11558492 | -0.47 | 1.236 | 0.627 | 1.133 | 0.004 | 0.036 | 0.036 | 0 | 0.2 | 0.07 | 0.17 | 0.1 | 0.21 | 0.16 |
| 1 | 231416547 | rs540479 | 1.966 | 1.134 | 0.602 | 1.054 | 0.007 | 0.014 | 0.014 | 0 | 0.47 | 0.36 | 0.38 | 0.37 | 0.46 | 0.37 |
| 1 | 231427220 | rs4658911 | -0.829 | 1.146 | 0.485 | 0.931 | 0.004 | -0.009 | -0.009 | 0.076 | 0.12 | 0.13 | 0.23 | 0.12 | 0.1 | 0.16 |
| 1 | 231441424 | rs203745 | 0.492 | 1.017 | 0.669 | 1.072 | -0.008 | 0.025 | 0.025 | 0 | 0.89 | 0.68 | 0.96 | 0.92 | 0.95 | 0.95 |
| 1 | 231445207 | rs12047385 | -1.09 | 1.77 | 0.173 | 1.288 | -0.012 | -0.024 | -0.024 | 0 | 0.06 | 0 | 0.08 | 0.11 | 0.07 | 0.06 |
| 1 | 231459478 | rs6541266 | 1.759 | 1.973 | 0.431 | 1.499 | -0.018 | -0.046 | -0.046 | 0.003 | 0.64 | 0.19 | 0.49 | 0.51 | 0.66 | 0.57 |
| 1 | 231470650 | rs10489609 | 1.777 | 1.931 | 0.955 | 1.549 | -0.008 | -0.022 | -0.022 | 0 | 0.64 | 0.27 | 0.5 | 0.52 | 0.66 | 0.56 |
| 1 | 231476789 | rs11122273 | 1.843 | 2.048 | 1.044 | 1.503 | -0.002 | -0.004 | -0.004 | 0 | 0.65 | 0.33 | 0.51 | 0.52 | 0.64 | 0.56 |

| | | | | | | | | | | | | | | | | |
|---|---|---|---|---|---|---|---|---|---|---|---|---|---|---|---|---|
| 1 | 231477442 | rs2355873 | 1.567 | 2.019 | 1.999 | 1.485 | 0.019 | 0.033 | 0.033 | 0 | 0.69 | 0.57 | 0.52 | 0.53 | 0.65 | 0.57 |
| 1 | 231487993 | rs203742 | 0.174 | 2.105 | 2.425 | 1.469 | 0.006 | -0.026 | -0.026 | 0.704 | 0.08 | 0.12 | 0.11 | 0 | 0.12 | 0.03 |
| 1 | 231488524 | rs2437150 | 2.396 | 2.08 | 2.555 | 1.462 | 0.012 | 0.027 | 0.027 | 0 | 0.66 | 0.44 | 0.52 | 0.52 | 0.65 | 0.55 |
| 1 | 231489353 | rs2749717 | 2.145 | 2.075 | 2.603 | 1.459 | 0.021 | 0.042 | 0.042 | 0 | 0.68 | 0.52 | 0.52 | 0.53 | 0.66 | 0.56 |
| 1 | 231498460 | rs1076804 | 2.562 | 2.087 | 2.545 | 1.465 | -0.006 | -0.017 | -0.017 | 2.301 | 0.65 | 0.26 | 0.52 | 0.53 | 0.66 | 0.57 |
| 1 | 231499236 | rs2009873 | 2.552 | 2.089 | 2.546 | 1.466 | -0.006 | -0.017 | -0.017 | 0 | 0.65 | 0.26 | 0.51 | 0.53 | 0.66 | 0.57 |
| 1 | 231520874 | rs973253 | 2.509 | 2.089 | 2.629 | 1.466 | -0.008 | -0.02 | -0.02 | 1.223 | 0.65 | 0.26 | 0.52 | 0.53 | 0.66 | 0.57 |
| 1 | 231521478 | rs2486736 | 2.058 | 2.088 | 2.654 | 1.466 | 0.021 | 0.039 | 0.039 | 0 | 0.69 | 0.57 | 0.53 | 0.53 | 0.66 | 0.57 |
| 1 | 231531627 | rs480902 | 2.066 | 2.111 | 2.636 | 1.464 | 0 | 0.049 | 0.049 | 0.237 | 0.73 | 0.39 | 0.63 | 0.53 | 0.78 | 0.6 |
| 1 | 231532312 | rs508618 | 0.716 | 2.105 | 2.918 | 1.623 | 0.016 | -0.008 | -0.008 | 0 | 0.86 | 0.81 | 0.7 | 0.84 | 0.83 | 0.79 |
| 1 | 231535584 | rs2486729 | 2.672 | 2.097 | 2.826 | 1.589 | -0.007 | -0.019 | -0.019 | 0 | 0.65 | 0.26 | 0.52 | 0.52 | 0.66 | 0.57 |
| 1 | 231547888 | rs2491417 | -0.898 | 2.124 | 2.912 | 1.635 | 0.036 | -0.034 | -0.034 | 0 | 0.11 | 0.26 | 0.12 | 0.04 | 0.13 | 0.07 |
| 1 | 231548480 | rs2808611 | -2.726 | 2.074 | 2.551 | 1.601 | 0.006 | 0.013 | 0.013 | 0 | 0.33 | 0.58 | 0.48 | 0.47 | 0.34 | 0.43 |
| 1 | 231550168 | rs2808614 | -2.843 | 2.068 | 2.218 | 1.59 | -0.018 | -0.04 | -0.04 | 0 | 0.35 | 0.8 | 0.49 | 0.48 | 0.35 | 0.44 |
| 1 | 231553787 | rs7542797 | -0.851 | 2.552 | 1.901 | 1.359 | -0.002 | 0.002 | 0.002 | 0 | 0.14 | 0.17 | 0.15 | 0.1 | 0.16 | 0.14 |
| 1 | 231559340 | rs7544596 | -0.415 | 2.157 | 2.073 | 1.322 | 0.015 | -0.051 | -0.051 | 0.551 | 0.09 | 0.19 | 0.11 | 0 | 0.12 | 0.03 |
| 1 | 231572255 | rs12407531 | -1.623 | 2.055 | 2.294 | 1.623 | 0.011 | 0.089 | 0.089 | 0.011 | 0.15 | 0.19 | 0.21 | 0.51 | 0.14 | 0.37 |
| 1 | 231579172 | rs1622146 | -1.212 | 2.189 | 2.586 | 1.435 | 0.075 | -0.116 | -0.116 | 0 | 0.19 | 0.54 | 0.38 | 0.03 | 0.25 | 0.1 |
| 1 | 231597964 | rs1614148 | -0.677 | 2.203 | 2.485 | 1.442 | 0.012 | -0.029 | -0.029 | 0 | 0.17 | 0.21 | 0.36 | 0.03 | 0.25 | 0.1 |
| 1 | 231608260 | rs16854388 | -1.993 | 1.993 | 2.566 | 1.564 | 0.027 | 0.202 | 0.202 | 0.375 | 0.17 | 0.39 | 0.22 | 0.51 | 0.14 | 0.37 |
| 1 | 231651391 | rs1769807 | -1.736 | 2.42 | 2.579 | 1.658 | 0.036 | 0.149 | 0.149 | 0.02 | 0.16 | 0.28 | 0.23 | 0.52 | 0.19 | 0.38 |
| 1 | 231654536 | rs11576167 | -0.693 | 2.05 | 2.604 | 1.555 | -0.007 | -0.001 | -0.001 | 0 | 0.18 | 0.16 | 0.35 | 0.04 | 0.24 | 0.1 |
| 1 | 231656418 | rs1765794 | -2.792 | 2.037 | 2.584 | 1.557 | -0.021 | -0.018 | -0.018 | 0 | 0.37 | 0.73 | 0.57 | 0.55 | 0.38 | 0.47 |
| 1 | 231658301 | rs1059595 | -1.659 | 2.007 | 2.407 | 1.635 | 0.017 | 0.15 | 0.15 | 0.017 | 0.16 | 0.28 | 0.2 | 0.52 | 0.14 | 0.38 |
| 1 | 231699474 | rs1655285 | -0.363 | 2.36 | 1.38 | 1.128 | -0.003 | 0.016 | 0.016 | 0 | 0.12 | 0.07 | 0.06 | 0 | 0.12 | 0.12 |
| 1 | 231707414 | rs7533573 | -2.338 | 2.448 | 1.733 | 1.749 | -0.007 | -0.003 | -0.003 | 1.04 | 0.3 | 0.69 | 0.55 | 0.37 | 0.34 | 0.32 |
| 1 | 231709404 | rs6668227 | -1.056 | 2.496 | 1.386 | 1.542 | -0.003 | -0.004 | -0.004 | 0.499 | 0.18 | 0.15 | 0.36 | 0.05 | 0.23 | 0.1 |
| 1 | 231711489 | rs1765782 | 1.537 | 2.418 | 1.869 | 1.802 | 0.024 | 0.008 | 0.008 | 0 | 0.72 | 0.38 | 0.45 | 0.63 | 0.66 | 0.68 |
| 1 | 231730121 | rs1765791 | 1.8 | 2.096 | 1.892 | 1.918 | -0.027 | -0.03 | -0.03 | 0 | 0.64 | 0.29 | 0.46 | 0.49 | 0.62 | 0.56 |
| 1 | 231742575 | rs4658879 | 0.964 | 2.225 | 2.04 | 2.142 | 0.042 | 0.161 | 0.161 | 0 | 0.84 | 0.62 | 0.63 | 0.55 | 0.8 | 0.81 |
| 1 | 231754600 | rs4603118 | 1.8 | 2.267 | 1.97 | 2.008 | 0.032 | 0.072 | 0.072 | 0 | 0.73 | 0.6 | 0.57 | 0.44 | 0.68 | 0.67 |
| 1 | 231758938 | rs10746510 | 2.13 | 2.244 | 2.58 | 2.017 | 0.014 | 0.239 | 0.239 | 0 | 0.88 | 0.48 | 0.91 | 0.75 | 0.92 | 0.85 |
| 1 | 231767890 | rs2082552 | -0.566 | 2.364 | 2.39 | 1.993 | -0.043 | -0.094 | -0.094 | 0 | 0.13 | 0.02 | 0.33 | 0.47 | 0.18 | 0.18 |
| 1 | 231768539 | rs751229 | -2.425 | 2.06 | 2.769 | 1.746 | 0.027 | -0.074 | -0.074 | 0 | 0.35 | 0.52 | 0.49 | 0.81 | 0.4 | 0.48 |
| 1 | 231771108 | rs823163 | -1.715 | 1.949 | 2.689 | 1.711 | 0.004 | -0.014 | -0.014 | 0 | 0.3 | 0.35 | 0.48 | 0.79 | 0.34 | 0.45 |
| 1 | 231794794 | rs1094653 | -2.286 | 2.344 | 2.528 | 1.585 | 0.017 | -0.057 | -0.057 | 3.275 | 0.36 | 0.53 | 0.5 | 0.8 | 0.41 | 0.49 |
| 1 | 231804093 | rs10864693 | -2.667 | 1.929 | 2.27 | 1.286 | -0.001 | -0.016 | -0.016 | 0 | 0.43 | 0.57 | 0.53 | 0.81 | 0.46 | 0.52 |
| 1 | 231808282 | rs11122319 | -2.025 | 1.944 | 2.083 | 1.275 | 0.005 | -0.019 | -0.019 | 4.44 | 0.35 | 0.39 | 0.5 | 0.8 | 0.41 | 0.49 |
| 1 | 231808385 | rs12030517 | -1.01 | 1.971 | 1.961 | 1.465 | -0.075 | -0.114 | -0.114 | 0.287 | 0.24 | 0.07 | 0.42 | 0.57 | 0.33 | 0.34 |
| 1 | 231813134 | rs6541281 | 2.116 | 1.644 | 1.616 | 0.836 | 0.03 | 0.176 | 0.176 | 0.097 | 0.52 | 0.22 | 0.41 | 0.18 | 0.46 | 0.44 |
| 1 | 231823662 | rs10429978 | -1.504 | 1.861 | 1.534 | 0.885 | 0.02 | 0.035 | 0.035 | 0.177 | 0.31 | 0.52 | 0.36 | 0.36 | 0.36 | 0.31 |
| 1 | 231826253 | rs12040259 | -1.625 | 1.747 | 1.458 | 0.712 | 0.01 | 0.017 | 0.017 | 0 | 0.32 | 0.55 | 0.37 | 0.34 | 0.36 | 0.3 |
| 1 | 231828076 | rs16854785 | -0.913 | 1.729 | 1.586 | 0.746 | 0.032 | 0.226 | 0.226 | 0.122 | 0.06 | 0.32 | 0.04 | 0.13 | 0.02 | 0.09 |
| 1 | 231828596 | rs1538974 | -1.182 | 1.804 | 1.149 | 0.762 | -0.017 | 0.016 | 0.016 | 0.097 | 0.28 | 0.23 | 0.34 | 0.22 | 0.34 | 0.22 |
| 1 | 231835610 | rs11122321 | -1.328 | 2.051 | 0.98 | 0.901 | -0.003 | -0.004 | -0.004 | 0.278 | 0.28 | 0.26 | 0.35 | 0.35 | 0.36 | 0.29 |
| 1 | 231835941 | rs11577035 | -0.906 | 2.093 | 0.415 | 0.827 | -0.047 | 0.012 | 0.012 | 0.019 | 0.24 | 0.12 | 0.33 | 0.22 | 0.33 | 0.2 |
| 1 | 231851279 | rs1934909 | 0.67 | 2.107 | 0.225 | 0.595 | -0.004 | -0.001 | -0.001 | 0 | 0.91 | 0.91 | 0.95 | 0.98 | 0.86 | 0.89 |
| 1 | 231859181 | rs11122324 | -1.18 | 1.463 | 0.444 | 0.937 | 0.02 | 0.025 | 0.025 | 0 | 0.31 | 0.43 | 0.37 | 0.35 | 0.37 | 0.28 |
| 1 | 231863204 | rs2487453 | 1.012 | 1.358 | 0.458 | 0.931 | 0.006 | 0.017 | 0.017 | 0 | 0.67 | 0.48 | 0.62 | 0.63 | 0.63 | 0.71 |
| 1 | 231864002 | rs2812385 | 1.256 | 2.221 | 0.216 | 0.704 | 0.025 | 0.026 | 0.026 | 0 | 0.68 | 0.46 | 0.56 | 0.59 | 0.63 | 0.66 |
| 1 | 231865376 | rs2812388 | 1.308 | 2.365 | 1.264 | 0.667 | 0.027 | 0.109 | 0.109 | 0.199 | 0.78 | 0.59 | 0.62 | 0.6 | 0.78 | 0.73 |
| 1 | 231866461 | rs2793102 | 1.166 | 2.463 | 1.243 | 0.667 | 0.012 | 0.073 | 0.073 | 0.039 | 0.79 | 0.67 | 0.64 | 0.6 | 0.8 | 0.73 |
| 1 | 231870689 | rs4658939 | -1.643 | 2.232 | 1.614 | 0.64 | 0.006 | 0.084 | 0.084 | 0.905 | 0.22 | 0.39 | 0.36 | 0.42 | 0.19 | 0.27 |
| 1 | 231873198 | rs12027635 | 0.579 | 0.974 | 0.243 | -0.501 | -0.044 | 0.151 | 0.151 | 0 | 0.26 | 0.03 | 0.09 | 0.05 | 0.24 | 0.13 |
| 1 | 231873511 | rs10864694 | -0.963 | 0.64 | 0.646 | -0.136 | -0.001 | 0.008 | 0.008 | 0.249 | 0.2 | 0.21 | 0.34 | 0.42 | 0.19 | 0.27 |
| 1 | 231876767 | rs4658883 | -0.709 | 0.613 | 0.663 | 0.081 | -0.01 | -0.04 | -0.04 | 0 | 0.18 | 0.09 | 0.32 | 0.32 | 0.18 | 0.22 |
| 1 | 231877073 | rs2793093 | 1.496 | 0.601 | 0.753 | -0.02 | -0.002 | 0.017 | 0.017 | 1.507 | 0.74 | 0.39 | 0.62 | 0.58 | 0.78 | 0.71 |
| 1 | 231879592 | rs10495309 | -1.135 | 0.678 | 0.581 | 0.054 | 0.004 | 0.076 | 0.076 | 0.735 | 0.21 | 0.35 | 0.35 | 0.42 | 0.19 | 0.27 |
| 1 | 231885217 | rs10864695 | -1.19 | 0.703 | 0.698 | 0.055 | -0.004 | 0.049 | 0.049 | 1.157 | 0.22 | 0.33 | 0.34 | 0.41 | 0.17 | 0.27 |
| 1 | 231894608 | rs2793091 | 1.004 | 0.438 | 0.037 | -0.046 | 0.051 | -0.017 | -0.017 | 0.185 | 0.56 | 0.35 | 0.57 | 0.55 | 0.72 | 0.65 |
| 1 | 231896868 | rs1538979 | 0.129 | 0.481 | 0.263 | 0.152 | -0.022 | -0.01 | -0.01 | 0 | 0.27 | 0.34 | 0.14 | 0.24 | 0.16 | 0.24 |
| 1 | 231897207 | rs17749164 | 1.423 | 1.111 | -0.533 | -0.423 | -0.056 | -0.069 | -0.069 | 0.094 | 0.28 | 0.15 | 0.42 | 0.4 | 0.41 | 0.28 |
| 1 | 231897303 | rs1538977 | -0.219 | 0.979 | -0.388 | -0.313 | -0.009 | -0.016 | -0.016 | 0 | 0.19 | 0.12 | 0.22 | 0.23 | 0.22 | 0.2 |
| 1 | 231899704 | rs2793086 | 0.337 | 0.551 | -0.446 | -0.356 | -0.02 | 0.033 | 0.033 | 0 | 0.15 | 0.08 | 0.09 | 0.02 | 0.19 | 0.11 |
| 1 | 231899929 | rs7529706 | 1.086 | 0.397 | 0.035 | -0.374 | 0.006 | 0.032 | 0.032 | 0 | 0.68 | 0.46 | 0.61 | 0.53 | 0.69 | 0.61 |
| 1 | 231906589 | rs2492367 | -0.016 | 0.19 | 0.26 | -0.375 | 0.004 | -0.003 | -0.003 | 0 | 0.13 | 0.21 | 0.07 | 0.14 | 0.11 | 0.13 |
| 1 | 231908347 | rs7534681 | 0.46 | 0.083 | -0.046 | -0.415 | 0.062 | -0.007 | -0.007 | 3.553 | 0.5 | 0.31 | 0.52 | 0.52 | 0.63 | 0.55 |
| 1 | 231908755 | rs12046794 | 0.799 | -0.108 | 0.193 | -0.665 | 0.011 | 0.012 | 0.012 | 0.046 | 0.27 | 0.21 | 0.15 | 0.11 | 0.16 | 0.18 |
| 1 | 231911011 | rs2812389 | -1.021 | -0.238 | 0.086 | -0.161 | -0.001 | 0 | 0 | 0 | 0.73 | 0.76 | 0.65 | 0.58 | 0.6 | 0.61 |

| | | | | | | | | | | | | | | | | |
|---|---|---|---|---|---|---|---|---|---|---|---|---|---|---|---|---|
| 1 | 231911598 | rs2812390 | -0.952 | -0.223 | -0.239 | -0.217 | 0.055 | 0.057 | 0.057 | 0 | 0.72 | 0.66 | 0.63 | 0.58 | 0.57 | 0.57 |
| 1 | 231911976 | rs2812391 | 0.386 | -0.718 | -0.531 | -0.467 | 0.076 | -0.004 | -0.004 | 0 | 0.42 | 0.22 | 0.39 | 0.46 | 0.26 | 0.41 |
| 1 | 231912592 | rs16854954 | -0.428 | -0.9 | -0.462 | -0.616 | -0.001 | -0.073 | -0.073 | 3.502 | 0.28 | 0.43 | 0.18 | 0.13 | 0.3 | 0.18 |
| 1 | 231912893 | rs16854957 | 0.322 | -0.971 | -0.589 | -0.578 | -0.004 | -0.059 | -0.059 | 0 | 0.17 | 0.28 | 0.1 | 0.02 | 0.18 | 0.1 |
| 1 | 231917877 | rs17766087 | 0.782 | -0.774 | -0.831 | -0.439 | -0.001 | -0.024 | -0.024 | 0 | 0.13 | 0.12 | 0.08 | 0.02 | 0.19 | 0.08 |
| 1 | 231922845 | rs11590192 | 0.979 | -0.967 | -0.768 | -0.252 | 0.042 | 0.029 | 0.029 | 0 | 0.32 | 0.4 | 0.54 | 0.41 | 0.47 | 0.45 |
| 1 | 231923345 | rs9803690 | 0.772 | -0.974 | -1.14 | -0.285 | 0.018 | 0.014 | 0.014 | 0 | 0.56 | 0.72 | 0.72 | 0.58 | 0.58 | 0.7 |
| 1 | 231928554 | rs12130935 | 0.193 | -1.283 | -1.125 | 0.365 | 0.062 | -0.076 | -0.076 | 0 | 0.19 | 0.04 | 0.12 | 0.28 | 0.1 | 0.14 |
| 1 | 231928935 | rs1322784 | 1.036 | -1.328 | -0.953 | 0.29 | 0.004 | -0.004 | -0.004 | 0 | 0.67 | 0.62 | 0.79 | 0.6 | 0.76 | 0.75 |
| 1 | 231929075 | rs1322783 | 0.445 | -1.033 | -0.796 | 0.454 | 0.023 | 0.004 | 0.004 | 0.013 | 0.9 | 0.81 | 0.92 | 0.88 | 0.87 | 0.92 |
| 1 | 231936150 | rs2255340 | 0.5 | -0.889 | -1.046 | 0.339 | 0.011 | -0.013 | -0.013 | 0 | 0.7 | 0.7 | 0.79 | 0.6 | 0.77 | 0.77 |
| 1 | 231945373 | rs2738875 | -0.496 | -1.27 | -1.052 | -0.146 | 0.017 | 0.013 | 0.013 | 0 | 0.79 | 0.84 | 0.85 | 0.86 | 0.9 | 0.76 |
| 1 | 231948646 | rs967244 | 0.144 | -1.181 | -1.244 | -0.336 | 0.03 | 0.026 | 0.026 | 0.217 | 0.72 | 0.81 | 0.8 | 0.82 | 0.84 | 0.72 |
| 1 | 231948821 | rs2759346 | 0.599 | -1.064 | -0.685 | -0.353 | 0.032 | 0.005 | 0.005 | 0 | 0.51 | 0.38 | 0.6 | 0.42 | 0.66 | 0.55 |
| 1 | 231953429 | rs7548597 | 0.284 | -1.151 | -0.402 | -0.421 | 0.023 | -0.039 | -0.039 | 0.105 | 0.14 | 0.26 | 0.13 | 0.04 | 0.17 | 0.08 |
| 1 | 231954101 | rs6675281 | 0.758 | -1.101 | -0.379 | -0.457 | 0.104 | -0.018 | -0.018 | 0 | 0.05 | 0.19 | 0.08 | 0 | 0.12 | 0.05 |
| 1 | 231954145 | rs12133766 | 1.292 | -1.1 | -0.439 | -0.448 | -0.008 | 0.02 | 0.02 | 0 | 0.09 | 0.01 | 0.04 | 0.03 | 0.05 | 0.03 |
| 1 | 231954427 | rs1535530 | 0.054 | -1.326 | -0.365 | -0.437 | -0.009 | -0.022 | -0.022 | 0 | 0.63 | 0.55 | 0.72 | 0.79 | 0.73 | 0.67 |
| 1 | 231954609 | rs1535529 | 1.307 | -0.624 | -0.383 | -0.811 | -0.002 | 0.004 | 0.004 | 0 | 0.2 | 0.17 | 0.21 | 0.38 | 0.26 | 0.25 |
| 1 | 231954682 | rs2295959 | 1.113 | -0.666 | -0.341 | -0.878 | -0.02 | 0.056 | 0.056 | 0 | 0.12 | 0.14 | 0.11 | 0.37 | 0.07 | 0.14 |
| 1 | 231960352 | rs2759329 | -0.104 | -0.622 | -0.405 | -0.912 | 0.001 | -0.002 | -0.002 | 0.988 | 0.57 | 0.55 | 0.67 | 0.81 | 0.66 | 0.65 |
| 1 | 231963601 | rs1000730 | 0.245 | -0.151 | -0.167 | -0.804 | 0.002 | -0.002 | -0.002 | 0.038 | 0.35 | 0.3 | 0.46 | 0.41 | 0.39 | 0.38 |
| 1 | 231971985 | rs12132978 | 0.045 | -0.02 | -0.166 | -0.826 | 0.002 | -0.019 | -0.019 | 0 | 0.2 | 0.2 | 0.17 | 0.04 | 0.24 | 0.11 |
| 1 | 231972026 | rs11122347 | -0.492 | -0.03 | -0.151 | -0.749 | 0 | 0.001 | 0.001 | 0 | 0.31 | 0.33 | 0.28 | 0.41 | 0.3 | 0.26 |
| 1 | 231972217 | rs7551537 | -0.829 | -0.1 | -0.077 | -0.712 | 0.006 | 0.002 | 0.002 | 0 | 0.57 | 0.67 | 0.64 | 0.45 | 0.64 | 0.52 |
| 1 | 231973922 | rs6663190 | -0.883 | -0.022 | -0.216 | -0.692 | -0.005 | -0.004 | -0.004 | 0 | 0.55 | 0.54 | 0.62 | 0.45 | 0.62 | 0.5 |
| 1 | 231974391 | rs12087793 | 0.595 | 0.218 | -0.164 | -0.147 | -0.003 | -0.012 | -0.012 | 0 | 0.23 | 0.19 | 0.36 | 0.37 | 0.28 | 0.29 |
| 1 | 231982271 | rs11588799 | -0.002 | 0.169 | -0.14 | -0.259 | -0.004 | -0.017 | -0.017 | 0 | 0.32 | 0.29 | 0.36 | 0.43 | 0.42 | 0.42 |
| 1 | 231983760 | rs12139880 | 0.897 | 0.03 | -0.62 | -0.568 | -0.049 | -0.084 | -0.084 | 1.241 | 0.25 | 0.05 | 0.23 | 0.41 | 0.25 | 0.24 |
| 1 | 231984517 | rs11122348 | 0.808 | -0.148 | -0.499 | -0.845 | 0.054 | 0.05 | 0.05 | 0.532 | 0.29 | 0.11 | 0.26 | 0.15 | 0.17 | 0.19 |
| 1 | 231985580 | rs4459095 | -0.395 | -0.019 | -0.205 | -0.834 | 0.016 | -0.03 | -0.03 | 0.22 | 0.93 | 0.84 | 0.93 | 0.99 | 0.88 | 0.9 |
| 1 | 231985633 | rs6677522 | -1.032 | -0.008 | -0.278 | -0.828 | -0.001 | -0.005 | -0.005 | 0 | 0.94 | 0.94 | 0.93 | 0.99 | 0.88 | 0.9 |
| 1 | 231987149 | rs11580219 | 0.275 | -0.202 | 0.045 | -0.919 | 0.014 | 0.018 | 0.018 | 0 | 0.51 | 0.41 | 0.61 | 0.83 | 0.64 | 0.64 |
| 1 | 231992097 | rs1079344 | 0.995 | 0.021 | 0.041 | -0.542 | -0.015 | -0.028 | -0.028 | 0 | 0.25 | 0.21 | 0.35 | 0.42 | 0.38 | 0.4 |
| 1 | 231993738 | rs12730369 | 0.729 | -0.248 | -0.005 | -0.629 | 0.044 | 0.04 | 0.04 | 1.215 | 0.28 | 0.11 | 0.26 | 0.15 | 0.17 | 0.19 |
| 1 | 231998863 | rs12757857 | -0.133 | -0.132 | -0.381 | -0.568 | -0.011 | 0.02 | 0.02 | 0 | 0.53 | 0.32 | 0.62 | 0.58 | 0.55 | 0.59 |
| 1 | 232002392 | rs3082 | 0.46 | 0.62 | -0.058 | -0.121 | -0.017 | -0.027 | -0.027 | 0.04 | 0.25 | 0.21 | 0.36 | 0.42 | 0.38 | 0.4 |
| 1 | 232003412 | rs1407601 | 1.095 | 0.246 | 0.324 | -0.179 | 0.046 | -0.047 | -0.047 | 0 | 0.91 | 0.69 | 0.9 | 0.99 | 0.86 | 0.9 |
| 1 | 232006959 | rs6698332 | -0.133 | 0.241 | -0.005 | -0.117 | 0.009 | -0.017 | -0.017 | 0 | 0.94 | 0.91 | 0.94 | 0.99 | 0.87 | 0.93 |
| 1 | 232007181 | rs9432023 | -0.27 | -0.103 | 0.007 | -0.662 | -0.002 | -0.037 | -0.037 | 0.13 | 0.43 | 0.5 | 0.35 | 0.16 | 0.37 | 0.36 |
| 1 | 232007219 | rs9432024 | -0.062 | -0.505 | 0.109 | -0.903 | 0.02 | -0.05 | -0.05 | 0.273 | 0.33 | 0.55 | 0.34 | 0.15 | 0.34 | 0.33 |
| 1 | 232014116 | rs9432040 | -0.641 | -0.567 | -0.077 | -0.328 | 0.001 | 0.001 | 0.001 | 0 | 0.55 | 0.55 | 0.52 | 0.45 | 0.47 | 0.52 |
| 1 | 232015040 | rs2356606 | 0.43 | -0.52 | -0.272 | -0.56 | 0.034 | 0.031 | 0.031 | 0 | 0.28 | 0.19 | 0.18 | 0.15 | 0.21 | 0.28 |
| 1 | 232019810 | rs4658889 | -0.635 | -0.315 | 0.173 | -0.356 | 0.053 | -0.033 | -0.033 | 0 | 0.92 | 0.77 | 0.93 | 0.99 | 0.87 | 0.93 |
| 1 | 232019859 | rs11122362 | 1.091 | -0.33 | 0.004 | -0.33 | 0.003 | -0.007 | -0.007 | 0 | 0.14 | 0.11 | 0.17 | 0.29 | 0.1 | 0.16 |
| 1 | 232021248 | rs9431714 | 0.003 | -0.256 | -0.088 | -0.367 | -0.003 | 0.058 | 0.058 | 0 | 0.41 | 0.28 | 0.33 | 0.18 | 0.37 | 0.36 |
| 1 | 232024081 | rs4658890 | 0.211 | -0.34 | -0.526 | -0.085 | 0.001 | -0.005 | -0.005 | 0 | 0.48 | 0.35 | 0.51 | 0.5 | 0.49 | 0.49 |
| 1 | 232025174 | rs1073179 | 0.947 | -0.324 | -0.576 | -0.123 | -0.002 | 0.005 | 0.005 | 0 | 0.31 | 0.29 | 0.28 | 0.14 | 0.36 | 0.31 |
| 1 | 232031815 | rs967433 | 0.362 | -0.866 | -0.736 | 0.021 | -0.002 | 0.005 | 0.005 | 0 | 0.54 | 0.44 | 0.47 | 0.63 | 0.53 | 0.61 |
| 1 | 232033478 | rs6541290 | 0.407 | -0.639 | -0.668 | 0.39 | -0.001 | -0.027 | -0.027 | 0 | 0.13 | 0.08 | 0.22 | 0.24 | 0.11 | 0.16 |
| 1 | 232036030 | rs821722 | -0.804 | -0.954 | -0.459 | -0.189 | 0.008 | 0.016 | 0.016 | 0 | 0.48 | 0.56 | 0.53 | 0.74 | 0.36 | 0.46 |
| 1 | 232036145 | rs860275 | -0.83 | -1.802 | 0.061 | 0.403 | -0.052 | 0.044 | 0.044 | 0 | 0.29 | 0.41 | 0.33 | 0.64 | 0.15 | 0.35 |
| 1 | 232050653 | rs10864702 | -0.754 | -1.312 | 0.089 | 0.417 | 0 | -0.002 | -0.002 | 0.145 | 0.25 | 0.19 | 0.29 | 0.69 | 0.13 | 0.3 |
| 1 | 232051185 | rs7541019 | 0.035 | -1.351 | -0.619 | 0.145 | 0.003 | 0.055 | 0.055 | 0 | 0.77 | 0.88 | 0.86 | 0.91 | 0.74 | 0.72 |
| 1 | 232054078 | rs11122374 | -1.072 | -0.957 | 0.299 | 0.77 | -0.052 | 0.019 | 0.019 | 0 | 0.27 | 0.5 | 0.32 | 0.69 | 0.13 | 0.3 |
| 1 | 232059805 | rs821717 | -0.022 | 0.25 | -0.015 | 0.521 | -0.04 | 0.044 | 0.044 | 0 | 0.84 | 0.92 | 0.9 | 0.9 | 0.77 | 0.88 |
| 1 | 232066277 | rs11584798 | -0.201 | 0.259 | -0.021 | 0.525 | -0.045 | 0.046 | 0.046 | 0 | 0.15 | 0.07 | 0.1 | 0.08 | 0.23 | 0.11 |
| 1 | 232067057 | rs821577 | -1.823 | 0.138 | 0.299 | 0.524 | 0.006 | 0.082 | 0.082 | 0 | 0.49 | 0.7 | 0.51 | 0.83 | 0.48 | 0.53 |
| 1 | 232073128 | rs701158 | -1.862 | 0.115 | 0.596 | 0.533 | 0.007 | 0.136 | 0.136 | 0 | 0.49 | 0.75 | 0.51 | 0.82 | 0.48 | 0.53 |
| 1 | 232074973 | rs872625 | -1.481 | -0.232 | 1.04 | 1.331 | -0.039 | -0.006 | -0.006 | 0 | 0.32 | 0.46 | 0.4 | 0.72 | 0.25 | 0.41 |
| 1 | 232078614 | rs4658966 | -0.975 | -0.001 | 0.465 | 1.515 | 0.007 | -0.01 | -0.01 | 1.076 | 0.22 | 0.18 | 0.32 | 0.71 | 0.16 | 0.28 |
| 1 | 232079126 | rs9431732 | -0.518 | -0.08 | 0.019 | 1.503 | -0.014 | -0.021 | -0.021 | 0 | 0.08 | 0.16 | 0.05 | 0.04 | 0.07 | 0.11 |
| 1 | 232085485 | rs16855691 | -0.077 | 0.301 | -0.352 | 2.478 | 0.024 | -0.018 | -0.018 | 1.996 | 0.07 | 0.01 | 0.09 | 0.29 | 0.03 | 0.11 |
| 1 | 232086153 | rs12137417 | -0.678 | 0.443 | -0.257 | 1.471 | 0.016 | -0.05 | -0.05 | 1.176 | 0.22 | 0.1 | 0.31 | 0.71 | 0.16 | 0.28 |
| 1 | 232090252 | rs7548491 | -1.287 | 2.115 | 0.376 | 1.686 | 0.058 | 0.058 | 0.058 | 1.351 | 0.19 | 0.32 | 0.29 | 0.3 | 0.27 | 0.26 |
| 1 | 232090493 | rs2772122 | 1.4 | 2.641 | 0.937 | 2.137 | 0.098 | 0.14 | 0.14 | 0 | 0.83 | 0.43 | 0.8 | 0.62 | 0.72 | 0.83 |
| 1 | 232090932 | rs821589 | -1.618 | 1.513 | 1.371 | 1.852 | -0.042 | 0.06 | 0.06 | 0.004 | 0.16 | 0.6 | 0.1 | 0.18 | 0.09 | 0.11 |
| 1 | 232092481 | rs701160 | -1.835 | 1.17 | 1.08 | 1.282 | -0.013 | -0.01 | -0.01 | 0 | 0.31 | 0.64 | 0.18 | 0.18 | 0.21 | 0.27 |
| 1 | 232096701 | rs6670775 | -1.4 | 1.299 | 1.652 | 1.253 | -0.006 | -0.024 | -0.024 | 0.187 | 0.23 | 0.31 | 0.11 | 0.09 | 0.21 | 0.26 |

| | | | | | | | | | | | | | | | | |
|---|---|---|---|---|---|---|---|---|---|---|---|---|---|---|---|---|
| 1 | 232099015 | rs12566923 | 0.254 | 0.889 | 1.027 | 1.793 | 0.005 | -0.03 | -0.03 | 4.147 | 0.08 | 0.03 | 0.08 | 0.21 | 0.05 | 0.06 |
| 1 | 232102264 | rs821597 | -2.239 | 0.984 | 1.226 | 1.674 | 0.008 | -0.069 | -0.069 | 0.199 | 0.43 | 0.87 | 0.29 | 0.43 | 0.36 | 0.47 |
| 1 | 232107814 | rs4658971 | -0.698 | 0.951 | 1.909 | 1.59 | -0.001 | 0.182 | 0.182 | 0 | 0.11 | 0.43 | 0.1 | 0.21 | 0.05 | 0.06 |
| 1 | 232121222 | rs821663 | -1.877 | 0.924 | 1.504 | 1.572 | 0 | -0.006 | -0.006 | 0.009 | 0.36 | 0.72 | 0.24 | 0.33 | 0.34 | 0.42 |
| 1 | 232125109 | rs701163 | -1.517 | 0.906 | 1.589 | 1.445 | 0 | -0.024 | -0.024 | 0 | 0.29 | 0.64 | 0.16 | 0.13 | 0.29 | 0.37 |
| 1 | 232143946 | rs821615 | -1.068 | 0.838 | 1.353 | 1.444 | 0.011 | -0.036 | -0.036 | 0 | 0.25 | 0.37 | 0.14 | 0.1 | 0.29 | 0.27 |
| 1 | 232147210 | rs16856152 | 0.651 | 0.883 | 1.274 | 1.646 | 0 | 0.009 | 0.009 | 0.083 | 0.07 | 0.05 | 0.1 | 0.2 | 0.05 | 0.08 |
| 1 | 232147251 | rs821628 | -1.12 | 0.889 | 1.206 | 1.593 | 0.008 | -0.022 | -0.022 | 0.177 | 0.25 | 0.35 | 0.14 | 0.14 | 0.28 | 0.37 |
| 1 | 232147568 | rs866055 | -1.635 | 0.854 | 0.485 | 1.611 | 0.01 | 0.043 | 0.043 | 0 | 0.35 | 0.83 | 0.19 | 0.19 | 0.32 | 0.31 |
| 1 | 232147600 | rs864752 | -1.886 | 0.888 | 0.228 | 1.534 | -0.005 | -0.09 | -0.09 | 2.633 | 0.43 | 0.91 | 0.29 | 0.42 | 0.36 | 0.48 |
| 1 | 232148135 | rs821631 | -1.585 | 0.557 | 0.229 | 1.691 | 0.017 | 0.056 | 0.056 | 0 | 0.43 | 0.74 | 0.51 | 0.63 | 0.39 | 0.51 |
| 1 | 232149520 | rs821634 | -1.11 | 0.084 | 0.437 | 1.765 | -0.009 | 0.032 | 0.032 | 0 | 0.35 | 0.51 | 0.43 | 0.42 | 0.33 | 0.4 |
| 1 | 232149843 | rs821639 | -0.872 | 0.208 | 0.36 | 1.507 | 0.002 | -0.012 | -0.012 | 0.607 | 0.3 | 0.36 | 0.3 | 0.17 | 0.32 | 0.3 |
| 1 | 232153793 | rs1475514 | 0.41 | 0.437 | -0.503 | 1.016 | -0.009 | 0.032 | 0.032 | 0 | 0.59 | 0.73 | 0.64 | 0.8 | 0.57 | 0.61 |
| 1 | 232154836 | rs928099 | 0.928 | 0.36 | -2.044 | 0.109 | -0.042 | 0.14 | 0.14 | 0 | 0.27 | 0.03 | 0.18 | 0.04 | 0.28 | 0.28 |
| 1 | 232156766 | rs7533169 | 0.665 | 0.49 | -1.41 | 0.799 | 0.001 | -0.006 | -0.006 | 0 | 0.18 | 0.23 | 0.15 | 0.14 | 0.2 | 0.13 |
| 1 | 232158679 | rs3524 | -0.263 | 0.579 | -2.001 | 0.823 | -0.049 | 0.065 | 0.065 | 0 | 0.55 | 0.9 | 0.56 | 0.7 | 0.52 | 0.56 |
| 1 | 232161683 | rs12088230 | -0.751 | 0.734 | -0.469 | 0.414 | 0.013 | -0.023 | -0.023 | 0.663 | 0.27 | 0.37 | 0.43 | 0.15 | 0.31 | 0.23 |
| 1 | 232168500 | rs9726024 | -0.607 | 0.017 | 0.204 | 0.273 | 0.005 | 0.009 | 0.009 | 0 | 0.39 | 0.34 | 0.26 | 0.27 | 0.37 | 0.32 |
| 1 | 232168867 | rs9729194 | 0.288 | 0.462 | 0.549 | 0.269 | -0.003 | -0.025 | -0.025 | 0 | 0.23 | 0.3 | 0.21 | 0.13 | 0.22 | 0.18 |
| 1 | 232174173 | rs9729179 | -0.542 | 0.345 | 0.582 | 0.637 | -0.007 | 0.004 | 0.004 | 0.156 | 0.32 | 0.54 | 0.31 | 0.32 | 0.29 | 0.28 |
| 1 | 232174775 | rs1411771 | -0.542 | 0.345 | 0.583 | 0.637 | -0.007 | 0.004 | 0.004 | 0.156 | 0.32 | 0.55 | 0.31 | 0.32 | 0.29 | 0.28 |
| 1 | 232176195 | rs980989 | 0.869 | 0.3 | 0.623 | 0.603 | -0.022 | 0.045 | 0.045 | 2.188 | 0.19 | 0.07 | 0.17 | 0.1 | 0.23 | 0.14 |
| 1 | 232176306 | rs9308481 | 0.268 | 0.328 | 0.698 | 0.621 | 0.001 | -0.017 | -0.017 | 0 | 0.24 | 0.27 | 0.22 | 0.14 | 0.25 | 0.19 |
| 1 | 232176987 | rs16856322 | -0.331 | 0.436 | 0.559 | 0.646 | 0 | 0.003 | 0.003 | 0.343 | 0.29 | 0.32 | 0.29 | 0.32 | 0.28 | 0.28 |
| 1 | 232177302 | rs1411775 | 1.136 | 0.362 | 0.011 | 0.548 | -0.015 | 0.009 | 0.009 | 0 | 0.11 | 0.03 | 0.06 | 0.07 | 0.12 | 0.14 |
| 1 | 232177487 | rs1411776 | 0.484 | 0.322 | 0.26 | 0.548 | -0.003 | 0.012 | 0.012 | 0 | 0.16 | 0.1 | 0.1 | 0.15 | 0.16 | 0.2 |
| 1 | 232180614 | rs11802446 | -0.452 | 0.264 | 0.583 | 0.838 | -0.003 | -0.008 | -0.008 | 0 | 0.28 | 0.22 | 0.29 | 0.32 | 0.33 | 0.23 |
| 1 | 232181815 | rs9308482 | -1.905 | 0.453 | 1.1 | 1.325 | 0.055 | 0.275 | 0.275 | 0.011 | 0.11 | 0.5 | 0.18 | 0.28 | 0.13 | 0.2 |
| 1 | 232191712 | rs16856377 | -1.658 | 0.349 | 1.175 | 1.231 | 0.086 | 0.35 | 0.35 | 1.443 | 0.06 | 0.43 | 0.1 | 0.19 | 0.03 | 0.06 |
| 1 | 232195369 | rs9727625 | -1.488 | 0.404 | 1.34 | 1.229 | 0.086 | 0.319 | 0.319 | 0.445 | 0.05 | 0.34 | 0.09 | 0.19 | 0.03 | 0.06 |
| 1 | 232200012 | rs2776625 | -0.137 | 0.345 | 0.707 | 0.447 | -0.005 | 0.056 | 0.056 | 0 | 0.36 | 0.22 | 0.27 | 0.2 | 0.39 | 0.38 |
| 1 | 232210903 | rs10864711 | 0.311 | -0.093 | 0.505 | 0.228 | -0.008 | -0.049 | -0.049 | 1.143 | 0.57 | 0.55 | 0.71 | 0.8 | 0.6 | 0.6 |
| 1 | 232218784 | rs9660187 | 0.361 | -0.056 | 0.504 | 0.433 | 0 | 0.001 | 0.001 | 0 | 0.55 | 0.48 | 0.64 | 0.6 | 0.58 | 0.58 |
| 1 | 232223436 | rs10864713 | 0.435 | 0.2 | 0.387 | 0.497 | 0.003 | -0.001 | -0.001 | 0 | 0.54 | 0.45 | 0.61 | 0.55 | 0.57 | 0.54 |
| 1 | 232226670 | rs9308484 | 1.047 | 0.135 | 0.334 | 0.612 | -0.032 | -0.02 | -0.02 | 1.935 | 0.31 | 0.15 | 0.43 | 0.41 | 0.34 | 0.39 |
| 1 | 232227786 | rs1411769 | -0.642 | -0.124 | 0.576 | 0.377 | -0.006 | 0.062 | 0.062 | 0 | 0.82 | 0.7 | 0.82 | 0.69 | 0.85 | 0.8 |
| 1 | 232227916 | rs1411768 | 0.005 | -0.264 | 0.624 | 0.318 | -0.028 | 0.036 | 0.036 | 0 | 0.77 | 0.64 | 0.82 | 0.69 | 0.85 | 0.8 |
| 1 | 232232335 | rs2026298 | 0.4 | -0.296 | 0.499 | 0.306 | -0.024 | 0.025 | 0.025 | 0 | 0.73 | 0.38 | 0.77 | 0.64 | 0.84 | 0.72 |
| 1 | 232236342 | rs9661282 | -1.336 | -0.202 | 0.253 | 0.251 | 0.007 | 0.01 | 0.01 | 0 | 0.86 | 0.91 | 0.92 | 0.89 | 0.88 | 0.82 |
| 1 | 232242925 | rs9730058 | 0.619 | -0.305 | -0.023 | 0.123 | -0.065 | 0.053 | 0.053 | 0 | 0.31 | 0.03 | 0.25 | 0.15 | 0.33 | 0.24 |
| 1 | 232246001 | rs9729778 | -0.779 | -0.269 | 0.633 | 0.392 | 0.091 | 0.342 | 0.342 | 0 | 0.07 | 0.4 | 0.11 | 0.22 | 0.04 | 0.03 |
| 1 | 232246418 | rs9308486 | -0.893 | -0.23 | 0.386 | 0.199 | -0.004 | 0.006 | 0.006 | 0.918 | 0.54 | 0.69 | 0.47 | 0.47 | 0.5 | 0.41 |
| 1 | 232249603 | rs11122415 | 2.35 | 0.147 | 0.072 | 0.605 | -0.001 | -0.027 | -0.027 | 0 | 0.14 | 0.04 | 0.22 | 0.25 | 0.09 | 0.09 |
| 1 | 232265261 | rs1160075 | 1.198 | 0.573 | 0.329 | 0.671 | -0.024 | -0.042 | -0.042 | 0 | 0.31 | 0.17 | 0.46 | 0.48 | 0.34 | 0.44 |
| 1 | 232266181 | rs17832313 | 1.193 | 0.523 | 0.606 | 0.58 | -0.003 | -0.013 | -0.013 | 0 | 0.15 | 0.22 | 0.11 | 0.12 | 0.15 | 0.17 |
| 1 | 232266922 | rs1888601 | -0.165 | 0.52 | 0.799 | 0.604 | 0.014 | 0.098 | 0.098 | 1.235 | 0.53 | 0.73 | 0.66 | 0.8 | 0.53 | 0.65 |
| 1 | 232269234 | rs17832385 | 1.623 | 0.501 | 0.66 | 0.55 | -0.036 | -0.02 | -0.02 | 0 | 0.07 | 0.01 | 0.08 | 0.07 | 0.14 | 0.06 |
| 1 | 232270985 | rs11122432 | 0.192 | 0.557 | 0.959 | 0.62 | 0.003 | 0.064 | 0.064 | 0 | 0.2 | 0.41 | 0.19 | 0.33 | 0.19 | 0.21 |
| 1 | 232271232 | rs10864720 | -0.202 | 0.549 | 0.801 | 0.607 | 0.01 | 0.124 | 0.124 | 1.286 | 0.53 | 0.77 | 0.66 | 0.79 | 0.53 | 0.65 |
| 1 | 232272707 | rs16856487 | -0.238 | 0.337 | 1.404 | 0.893 | 0.003 | 0.114 | 0.114 | 0 | 0.21 | 0.55 | 0.2 | 0.35 | 0.19 | 0.22 |
| 1 | 232277358 | rs12080088 | -0.205 | 0.27 | 1.288 | 0.706 | -0.003 | 0.104 | 0.104 | 0 | 0.21 | 0.52 | 0.2 | 0.34 | 0.19 | 0.21 |
| 1 | 232278540 | rs9662895 | -0.808 | -0.001 | 1.014 | 0.601 | -0.057 | 0.078 | 0.078 | 0.018 | 0.69 | 0.93 | 0.76 | 0.86 | 0.66 | 0.74 |
| 1 | 232280816 | rs12087592 | -0.376 | 0.202 | 1.273 | 0.831 | 0.066 | 0.278 | 0.278 | 0 | 0.06 | 0.35 | 0.09 | 0.22 | 0.04 | 0.04 |
| 1 | 232282491 | rs954745 | -0.574 | 0.488 | 1.037 | 0.724 | 0.018 | 0.075 | 0.075 | 0 | 0.25 | 0.57 | 0.25 | 0.4 | 0.27 | 0.26 |
| 1 | 232284603 | rs1041469 | -0.936 | 0.375 | 0.484 | 0.47 | -0.096 | 0.048 | 0.048 | 0 | 0.72 | 0.96 | 0.77 | 0.88 | 0.66 | 0.75 |
| 1 | 232287416 | rs12087019 | -0.245 | 0.2 | 0.895 | 0.433 | 0.002 | -0.045 | -0.045 | 0 | 0.23 | 0.52 | 0.18 | 0.18 | 0.24 | 0.24 |
| 1 | 232295535 | rs9793010 | 0.891 | 0.162 | 0.721 | 0.388 | -0.001 | -0.001 | -0.001 | 0 | 0.17 | 0.14 | 0.12 | 0.12 | 0.21 | 0.2 |
| 1 | 232296627 | rs12042307 | 1.083 | 0.179 | 0.569 | 0.582 | 0.004 | -0.008 | -0.008 | 0 | 0.4 | 0.24 | 0.56 | 0.56 | 0.41 | 0.5 |
| 1 | 232300841 | rs9659432 | -0.722 | 0.048 | 0.651 | 0.704 | -0.009 | 0.032 | 0.032 | 0 | 0.3 | 0.49 | 0.24 | 0.39 | 0.27 | 0.26 |
| 1 | 232301835 | rs11122437 | 1.245 | 0.143 | 0.4 | 0.733 | 0.004 | -0.026 | -0.026 | 0 | 0.41 | 0.15 | 0.52 | 0.47 | 0.41 | 0.48 |
| 1 | 232307419 | rs10779832 | 0.749 | 0.316 | 0.456 | 0.547 | 0.004 | 0.024 | 0.024 | 0 | 0.52 | 0.38 | 0.61 | 0.64 | 0.47 | 0.52 |
| 1 | 232309788 | rs10864723 | 0.665 | 0.375 | -0.039 | 0.575 | 0 | 0 | 0 | 0 | 0.53 | 0.49 | 0.62 | 0.64 | 0.48 | 0.53 |
| 1 | 232310039 | rs9308491 | -1.034 | 0.608 | -0.514 | 0.604 | -0.093 | 0.03 | 0.03 | 0 | 0.75 | 0.96 | 0.79 | 0.87 | 0.69 | 0.76 |
| 1 | 232311020 | rs9727900 | -0.625 | 0.2 | 0.455 | 0.876 | 0.005 | 0.018 | 0.018 | 0 | 0.2 | 0.36 | 0.16 | 0.24 | 0.21 | 0.22 |
| 1 | 232312344 | rs12754490 | -0.995 | 0.269 | 0.248 | 0.854 | 0.006 | 0.021 | 0.021 | 0 | 0.28 | 0.58 | 0.24 | 0.39 | 0.28 | 0.26 |
| 1 | 232312643 | rs12144995 | 0.414 | -0.79 | -0.036 | 0.607 | -0.004 | 0.001 | 0.001 | 0 | 0.63 | 0.53 | 0.69 | 0.62 | 0.65 | 0.64 |
| 1 | 232314505 | rs1571755 | -0.481 | -1.431 | 0.206 | -0.234 | 0.003 | -0.038 | -0.038 | 0 | 0.33 | 0.45 | 0.32 | 0.14 | 0.35 | 0.39 |
| 1 | 232315161 | rs11122445 | 0.772 | -1.885 | -0.002 | -0.598 | -0.004 | 0.043 | 0.043 | 0.001 | 0.46 | 0.33 | 0.56 | 0.82 | 0.5 | 0.32 |

| | | | | | | | | | | | | | | | | |
|---|---|---|---|---|---|---|---|---|---|---|---|---|---|---|---|---|
| 1 | 232316602 | rs1984027 | 1.396 | -1.896 | -0.051 | -0.245 | -0.063 | 0.073 | 0.073 | 0.193 | 0.36 | 0.16 | 0.45 | 0.17 | 0.46 | 0.25 |
| 1 | 232318791 | rs10864724 | 0.163 | -2.104 | 0.312 | -0.389 | 0.008 | 0.057 | 0.057 | 0 | 0.53 | 0.61 | 0.64 | 0.86 | 0.57 | 0.37 |
| 1 | 232319587 | rs2884064 | -1.084 | -2.165 | 0.35 | -0.109 | 0 | -0.025 | -0.025 | 5.377 | 0.78 | 0.81 | 0.88 | 0.97 | 0.86 | 0.7 |
| 1 | 232320938 | rs12746164 | 0.645 | -1.783 | 0.394 | 0.129 | -0.017 | 0.035 | 0.035 | 0 | 0.25 | 0.2 | 0.24 | 0.11 | 0.29 | 0.33 |
| 1 | 232325530 | rs12071555 | -0.499 | -1.716 | 0.608 | -0.228 | -0.014 | -0.034 | -0.034 | 0.001 | 0.11 | 0.28 | 0.09 | 0.04 | 0.09 | 0.05 |
| 1 | 232328401 | rs6666516 | 0.978 | -1.804 | 0.423 | -0.334 | 0.001 | 0.003 | 0.003 | 0 | 0.52 | 0.49 | 0.53 | 0.21 | 0.48 | 0.53 |
| 1 | 232330619 | rs10752796 | 0.589 | -1.23 | 0.331 | 0.201 | -0.003 | -0.017 | -0.017 | 1.447 | 0.63 | 0.59 | 0.61 | 0.5 | 0.58 | 0.6 |
| 1 | 232340539 | rs10910263 | 0.006 | -1.307 | 0.215 | 0.243 | -0.011 | -0.053 | -0.053 | 0 | 0.27 | 0.2 | 0.31 | 0.46 | 0.32 | 0.35 |
| 1 | 232349350 | rs7411832 | 1.014 | -1.328 | -0.283 | -0.123 | 0.002 | 0.05 | 0.05 | 0.69 | 0.43 | 0.28 | 0.53 | 0.23 | 0.54 | 0.27 |
| 1 | 232349755 | rs6605044 | -0.243 | -1.063 | -0.1 | -0.795 | -0.073 | -0.143 | -0.143 | 0 | 0.73 | 0.51 | 0.88 | 0.96 | 0.89 | 0.69 |
| 1 | 232356135 | rs7365975 | -0.973 | -0.047 | 0.5 | 0.314 | -0.016 | -0.066 | -0.066 | 0 | 0.3 | 0.19 | 0.34 | 0.46 | 0.35 | 0.37 |
| 1 | 232363015 | rs10910559 | -0.874 | 0.009 | -0.221 | 0.448 | -0.037 | -0.109 | -0.109 | 0 | 0.29 | 0.11 | 0.33 | 0.46 | 0.34 | 0.37 |
| 1 | 232363040 | rs9729080 | -0.848 | 0.05 | -0.23 | 0.469 | -0.004 | -0.045 | -0.045 | 0 | 0.3 | 0.21 | 0.34 | 0.46 | 0.34 | 0.37 |
| 1 | 232363541 | rs9725780 | -1.12 | 0 | -0.3 | 0.51 | -0.005 | -0.019 | -0.019 | 0 | 0.37 | 0.32 | 0.4 | 0.51 | 0.45 | 0.39 |
| 1 | 232379496 | rs9659337 | -0.504 | 0.153 | -0.065 | 1.167 | -0.046 | -0.071 | -0.071 | 1.144 | 0.11 | 0.01 | 0.24 | 0.37 | 0.14 | 0.17 |
| 1 | 232393417 | rs4596830 | -0.92 | -0.192 | -0.437 | 0.15 | 0.003 | 0.004 | 0.004 | 0 | 0.1 | 0.12 | 0.06 | 0.04 | 0.06 | 0.03 |
| 1 | 232394263 | rs4649384 | 0.257 | -0.421 | -1.466 | -0.102 | 0.018 | 0.089 | 0.089 | 0 | 0.55 | 0.9 | 0.46 | 0.28 | 0.56 | 0.46 |
| 1 | 232396598 | rs9286804 | -0.111 | -0.258 | -0.905 | 0.067 | 0.019 | 0.021 | 0.021 | 0 | 0.62 | 0.82 | 0.51 | 0.31 | 0.63 | 0.48 |
| 1 | 232403070 | rs9286807 | -0.532 | -0.652 | -0.522 | -0.383 | -0.038 | -0.025 | -0.025 | 0 | 0.12 | 0.66 | 0.16 | 0.05 | 0.03 | 0.05 |
| 1 | 232409847 | rs10910648 | -0.796 | -0.774 | -0.863 | -0.32 | -0.051 | 0.008 | 0.008 | 3.108 | 0.15 | 0.73 | 0.19 | 0.13 | 0.05 | 0.12 |
| 1 | 232411985 | rs4564083 | -0.106 | 0.376 | -1.609 | -0.126 | -0.095 | 0.11 | 0.11 | 0 | 0.61 | 0.93 | 0.47 | 0.18 | 0.53 | 0.43 |
| 1 | 232418114 | rs9660420 | -0.311 | 0.527 | -0.801 | 0.557 | -0.007 | 0.011 | 0.011 | 0 | 0.36 | 0.48 | 0.43 | 0.42 | 0.31 | 0.24 |
| 1 | 232418458 | rs12403985 | -0.5 | 0.582 | -1.035 | 0.282 | 0.004 | 0.002 | 0.002 | 0 | 0.15 | 0.1 | 0.19 | 0.02 | 0.16 | 0.1 |
| 1 | 232423863 | rs1338302 | -0.568 | 0.453 | -1.761 | 0.737 | 0.056 | 0.119 | 0.119 | 0 | 0.56 | 0.81 | 0.67 | 0.76 | 0.68 | 0.52 |
| 1 | 232429661 | rs9728263 | -0.277 | 1.46 | -0.935 | 1.174 | 0.017 | 0.004 | 0.004 | 0 | 0.21 | 0.35 | 0.24 | 0.21 | 0.24 | 0.24 |
| 1 | 232432558 | rs9728261 | -0.095 | 1.474 | -0.208 | 1.472 | -0.014 | 0.022 | 0.022 | 0 | 0.29 | 0.21 | 0.21 | 0.2 | 0.36 | 0.22 |
| 1 | 232436679 | rs12070718 | -1.164 | 1.657 | 0.071 | 1.467 | -0.007 | 0.04 | 0.04 | 0 | 0.44 | 0.68 | 0.54 | 0.7 | 0.54 | 0.44 |
| 1 | 232438081 | rs885444 | -0.691 | 1.613 | 0.562 | 1.694 | 0.004 | 0.084 | 0.084 | 0 | 0.13 | 0.22 | 0.29 | 0.39 | 0.17 | 0.2 |
| 1 | 232439116 | rs17778561 | -0.158 | 1.635 | 0.754 | 1.657 | 0.009 | 0.021 | 0.021 | 0 | 0.12 | 0.2 | 0.15 | 0.19 | 0.15 | 0.15 |
| 1 | 232441116 | rs12030593 | -0.636 | 1.522 | 0.705 | 1.698 | 0.012 | 0.137 | 0.137 | 0 | 0.13 | 0.25 | 0.3 | 0.39 | 0.17 | 0.2 |
| 1 | 232441842 | rs868585 | -0.876 | 2.025 | 0.604 | 1.448 | 0.029 | 0.07 | 0.07 | 0 | 0.3 | 0.47 | 0.43 | 0.51 | 0.41 | 0.34 |
| 1 | 232442213 | rs946130 | -1.53 | 1.342 | 0.361 | 1.076 | 0.051 | 0.072 | 0.072 | 0 | 0.49 | 0.67 | 0.62 | 0.72 | 0.65 | 0.48 |
| 1 | 232443051 | rs16856780 | -0.135 | 1.154 | 0.143 | 1.941 | 0.004 | 0.053 | 0.053 | 0 | 0.06 | 0.04 | 0.19 | 0.33 | 0.06 | 0.06 |
| 1 | 232447391 | rs1631842 | 0.397 | 1.027 | -0.537 | 0.889 | -0.025 | 0.094 | 0.094 | 0 | 0.44 | 0.22 | 0.35 | 0.21 | 0.44 | 0.46 |
| 1 | 232453032 | rs9424522 | 0.038 | 1.277 | 0.074 | 1.12 | 0.062 | 0.035 | 0.035 | 0 | 0.08 | 0.25 | 0.1 | 0.07 | 0.1 | 0.12 |
| 1 | 232455196 | rs12081587 | 0.561 | 0.92 | -0.341 | 1.036 | 0.013 | 0.009 | 0.009 | 0 | 0.13 | 0.08 | 0.17 | 0.1 | 0.09 | 0.08 |
| 1 | 232458444 | rs10495315 | 0.763 | 0.862 | -0.554 | 1.034 | 0.027 | 0.001 | 0.001 | 0 | 0.11 | 0.02 | 0.16 | 0.09 | 0.08 | 0.07 |
| 1 | 232461177 | rs12409282 | -0.309 | 1.416 | -0.269 | 1.156 | 0.053 | 0.15 | 0.15 | 0.008 | 0.11 | 0.17 | 0.25 | 0.39 | 0.25 | 0.2 |
| 1 | 232463715 | rs4649292 | -0.792 | 1 | -0.466 | 0.965 | -0.012 | 0.012 | 0.012 | 0 | 0.27 | 0.58 | 0.33 | 0.3 | 0.24 | 0.29 |
| 1 | 232466945 | rs3120760 | 0.723 | 0.683 | -0.829 | 0.555 | 0.027 | 0.061 | 0.061 | 0 | 0.42 | 0.27 | 0.33 | 0.21 | 0.37 | 0.4 |
| 1 | 232472932 | rs789367 | 0.036 | 0.335 | -0.841 | -0.13 | 0.005 | 0.056 | 0.056 | 0 | 0.59 | 0.75 | 0.75 | 0.85 | 0.66 | 0.7 |
| 1 | 232474875 | rs3120762 | -0.227 | 0.465 | -0.835 | -0.146 | -0.014 | 0.041 | 0.041 | 0 | 0.66 | 0.78 | 0.77 | 0.85 | 0.66 | 0.67 |
| 1 | 232476942 | rs585499 | 1.488 | 0.043 | -1.231 | 0.017 | -0.027 | -0.019 | -0.019 | 0 | 0.21 | 0.1 | 0.23 | 0.28 | 0.27 | 0.36 |
| 1 | 232477033 | rs10495317 | 1.486 | 0.084 | -0.772 | -0.108 | 0.049 | -0.029 | -0.029 | 1.662 | 0.06 | 0.16 | 0.07 | 0 | 0.15 | 0.11 |
| 1 | 232477143 | rs584639 | -0.408 | -0.116 | -0.776 | 0.27 | 0 | -0.003 | -0.003 | 0 | 0.57 | 0.68 | 0.53 | 0.49 | 0.43 | 0.5 |
| 1 | 232480575 | rs1933692 | -1.353 | -0.186 | -1.082 | 0.014 | 0.006 | -0.023 | -0.023 | 0 | 0.85 | 0.95 | 0.9 | 0.88 | 0.88 | 0.82 |
| 1 | 232480680 | rs12060562 | -0.429 | -0.062 | -0.563 | 0.435 | 0.01 | 0.014 | 0.014 | 0.565 | 0.42 | 0.6 | 0.36 | 0.27 | 0.31 | 0.25 |
| 1 | 232482539 | rs9659038 | -0.413 | -0.05 | -0.525 | 0.217 | 0.007 | 0.009 | 0.009 | 0.507 | 0.42 | 0.59 | 0.36 | 0.21 | 0.31 | 0.25 |
| 1 | 232484979 | rs9725734 | 1.251 | -0.365 | -0.391 | -0.069 | 0.033 | 0.02 | 0.02 | 1.906 | 0.21 | 0.14 | 0.07 | 0 | 0.1 | 0.04 |
| 1 | 232485359 | rs592522 | 1.938 | -0.241 | -0.562 | 0.049 | -0.076 | -0.033 | -0.033 | 0 | 0.19 | 0.01 | 0.2 | 0.16 | 0.22 | 0.25 |
| 1 | 232485906 | rs9660150 | -0.353 | -0.298 | -0.398 | -0.004 | -0.01 | -0.04 | -0.04 | 0.141 | 0.4 | 0.51 | 0.33 | 0.21 | 0.31 | 0.28 |
| 1 | 232486245 | rs9424427 | 1.922 | -0.154 | -0.733 | 0.095 | -0.076 | -0.024 | -0.024 | 0.229 | 0.19 | 0.01 | 0.19 | 0.15 | 0.22 | 0.24 |
| 1 | 232488827 | rs1766593 | -1.338 | -0.308 | -0.355 | 0.205 | -0.034 | 0.075 | 0.075 | 0.009 | 0.64 | 0.85 | 0.76 | 0.84 | 0.64 | 0.64 |
| 1 | 232489410 | rs9661059 | 0.82 | -0.178 | 0.042 | 0.39 | 0.001 | 0.001 | 0.001 | 0.126 | 0.22 | 0.21 | 0.26 | 0.49 | 0.15 | 0.19 |
| 1 | 232491315 | rs789360 | 0.693 | -0.13 | 0.082 | 0.126 | 0.046 | 0.033 | 0.033 | 0 | 0.29 | 0.45 | 0.48 | 0.58 | 0.43 | 0.39 |
| 1 | 232492037 | rs653891 | 0.05 | 0.057 | -0.104 | 0.349 | 0.004 | 0.049 | 0.049 | 0 | 0.5 | 0.62 | 0.57 | 0.64 | 0.57 | 0.56 |
| 1 | 232494124 | rs16856874 | 1.3 | 0.093 | -0.601 | 0.372 | 0.006 | -0.016 | -0.016 | 0 | 0.09 | 0.03 | 0.18 | 0.18 | 0.07 | 0.07 |
| 1 | 232498336 | rs789363 | -0.272 | 0.008 | -0.288 | 0.294 | 0 | 0.006 | 0.006 | 0.083 | 0.55 | 0.46 | 0.68 | 0.69 | 0.59 | 0.55 |
| 1 | 232498977 | rs17176041 | 1.149 | 0.252 | -0.681 | 0.002 | -0.008 | 0.027 | 0.027 | 0 | 0.07 | 0.02 | 0.07 | 0 | 0.06 | 0.02 |
| 1 | 232505957 | rs3120763 | 0.586 | 0.182 | -0.831 | 0.156 | -0.022 | 0.02 | 0.02 | 0 | 0.26 | 0.08 | 0.23 | 0.16 | 0.26 | 0.22 |
| 1 | 232507616 | rs3131811 | 0.048 | 0.134 | -0.65 | 0.342 | 0.003 | 0.022 | 0.022 | 0 | 0.25 | 0.6 | 0.31 | 0.57 | 0.3 | 0.3 |
| 1 | 232509380 | rs17176304 | 0.636 | 0.526 | -0.804 | -0.292 | 0 | -0.039 | -0.039 | 0 | 0.18 | 0.2 | 0.12 | 0 | 0.22 | 0.2 |
| 1 | 232512873 | rs3120768 | -0.1 | 0.522 | -0.667 | 0.23 | -0.003 | 0.01 | 0.01 | 0 | 0.28 | 0.18 | 0.25 | 0.2 | 0.29 | 0.29 |
| 1 | 232522748 | rs612191 | 0.16 | 0.066 | -0.596 | 0.465 | -0.002 | -0.007 | -0.007 | 0.093 | 0.34 | 0.36 | 0.36 | 0.27 | 0.3 | 0.38 |
| 1 | 232526512 | rs17176717 | 0.814 | -0.088 | -0.775 | 0.478 | 0.013 | 0.005 | 0.005 | 0 | 0.11 | 0.08 | 0.06 | 0 | 0.06 | 0.02 |
| 1 | 232527958 | rs2486022 | -0.098 | 0.362 | -0.304 | 0.684 | 0 | 0.002 | 0.002 | 0 | 0.39 | 0.55 | 0.5 | 0.69 | 0.41 | 0.48 |
| 1 | 232532734 | rs1890395 | 0.59 | 0.178 | -0.12 | 0.386 | 0.003 | 0.002 | 0.002 | 0 | 0.23 | 0.25 | 0.47 | 0.74 | 0.27 | 0.34 |
| 1 | 232533015 | rs600379 | -1.058 | -0.309 | -0.777 | -0.524 | -0.072 | 0.093 | 0.093 | 0.216 | 0.63 | 0.85 | 0.74 | 0.81 | 0.6 | 0.55 |
| 1 | 232534776 | rs586719 | 0.137 | -0.583 | -0.462 | -0.667 | -0.013 | -0.099 | -0.099 | 0.652 | 0.29 | 0.36 | 0.21 | 0 | 0.25 | 0.14 |

| | | | | | | | | | | | | | | | | |
|---|---|---|---|---|---|---|---|---|---|---|---|---|---|---|---|---|
| 1 | 232534957 | rs12043251 | 1.209 | -0.57 | -0.493 | -0.624 | 0.004 | 0.003 | 0.003 | 0 | 0.23 | 0.26 | 0.47 | 0.74 | 0.27 | 0.34 |
| 1 | 232541847 | rs12045182 | -0.128 | -0.452 | -0.744 | -0.313 | 0.01 | 0.04 | 0.04 | 0.008 | 0.5 | 0.6 | 0.65 | 0.74 | 0.47 | 0.47 |
| 1 | 232543280 | rs12145607 | 0.757 | 0.217 | -1.709 | -0.467 | -0.064 | 0.135 | 0.135 | 0 | 0.25 | 0.03 | 0.19 | 0.04 | 0.24 | 0.23 |
| 1 | 232549986 | rs1890315 | 0.614 | -0.227 | -1.121 | -0.086 | 0.01 | -0.003 | -0.003 | 0 | 0.12 | 0.12 | 0.06 | 0 | 0.07 | 0.02 |
| 1 | 232550642 | rs16857038 | 0.357 | 0.42 | -0.962 | 0.36 | 0.024 | 0.005 | 0.005 | 0 | 0.06 | 0.09 | 0.2 | 0.12 | 0.13 | 0.14 |
| 1 | 232552312 | rs6701214 | -0.626 | -0.038 | -1.322 | -0.076 | -0.042 | 0.057 | 0.057 | 0.567 | 0.62 | 0.75 | 0.75 | 0.78 | 0.56 | 0.51 |
| 1 | 232552461 | rs16857041 | -0.095 | -0.294 | -0.751 | 0.215 | -0.04 | 0.077 | 0.077 | 0.015 | 0.14 | 0.38 | 0.16 | 0.22 | 0.06 | 0.08 |
| 1 | 232558345 | rs6424228 | -0.791 | 0.089 | -1.394 | -0.137 | -0.045 | 0.06 | 0.06 | 0 | 0.63 | 0.79 | 0.77 | 0.79 | 0.58 | 0.53 |
| 1 | 232562732 | rs10910460 | 0.095 | 0.335 | -1.387 | -0.122 | 0.005 | -0.009 | -0.009 | 0 | 0.27 | 0.11 | 0.28 | 0.31 | 0.25 | 0.24 |
| 1 | 232563920 | rs7545528 | -0.719 | 1.152 | -0.742 | -0.173 | 0.016 | -0.077 | -0.077 | 0.436 | 0.14 | 0.24 | 0.19 | 0 | 0.19 | 0.1 |
| 1 | 232568041 | rs1547742 | 0.217 | 0.838 | -1.288 | 0.105 | -0.026 | 0.012 | 0.012 | 0 | 0.07 | 0.01 | 0.13 | 0.04 | 0.1 | 0.06 |
| 1 | 232568587 | rs12030201 | -0.674 | 0.807 | -0.796 | 0.609 | 0.014 | 0.123 | 0.123 | 0 | 0.17 | 0.25 | 0.33 | 0.46 | 0.2 | 0.32 |
| 1 | 232570049 | rs10910478 | 1.123 | 0.734 | -1.225 | -0.164 | -0.013 | 0.026 | 0.026 | 0 | 0.25 | 0.16 | 0.17 | 0.13 | 0.3 | 0.22 |
| 1 | 232570571 | rs6685115 | -0.482 | -0.486 | -1.062 | -0.534 | -0.011 | 0.006 | 0.006 | 0 | 0.36 | 0.6 | 0.55 | 0.55 | 0.31 | 0.4 |
| 1 | 232575539 | rs6664207 | -0.561 | -0.547 | -1.434 | -0.664 | -0.078 | 0.036 | 0.036 | 0 | 0.7 | 0.94 | 0.75 | 0.78 | 0.66 | 0.74 |
| 1 | 232576594 | rs1547740 | 0.67 | -0.426 | -1.022 | -0.466 | -0.006 | 0.007 | 0.007 | 0 | 0.07 | 0.08 | 0.2 | 0.16 | 0.03 | 0.18 |
| 1 | 232580116 | rs1536322 | -2.025 | -0.532 | -1.207 | -0.809 | -0.011 | 0.006 | 0.006 | 0 | 0.8 | 0.84 | 0.83 | 0.87 | 0.71 | 0.8 |
| 1 | 232581436 | rs12757779 | 1.176 | -0.743 | -1.1 | -0.442 | -0.026 | 0.022 | 0.022 | 0 | 0.1 | 0.04 | 0.17 | 0 | 0.19 | 0.1 |
| 1 | 232581667 | rs12723697 | 1.041 | -0.73 | -0.98 | -0.421 | -0.001 | -0.013 | -0.013 | 0 | 0.11 | 0.09 | 0.17 | 0 | 0.2 | 0.1 |
| 1 | 232583916 | rs17794366 | 0.313 | -0.816 | -1.159 | -0.554 | -0.011 | 0.026 | 0.026 | 0 | 0.12 | 0.05 | 0.16 | 0.05 | 0.14 | 0.1 |
| 1 | 232584061 | rs4649374 | -1.336 | -1.13 | -0.352 | -0.05 | 0 | 0.087 | 0.087 | 0 | 0.22 | 0.37 | 0.46 | 0.54 | 0.22 | 0.37 |
| 1 | 232590866 | rs2275302 | -0.941 | -1.115 | -0.287 | -0.141 | 0.006 | 0.086 | 0.086 | 0 | 0.21 | 0.32 | 0.46 | 0.55 | 0.23 | 0.37 |
| 1 | 232591925 | rs4649265 | -0.349 | -1.134 | -0.263 | -0.653 | -0.014 | 0.004 | 0.004 | 0 | 0.09 | 0.12 | 0.22 | 0.15 | 0.03 | 0.18 |
| 1 | 232595148 | rs12139442 | 0.682 | -1.096 | -0.257 | -0.698 | 0.036 | -0.058 | -0.058 | 0 | 0.11 | 0.17 | 0.17 | 0 | 0.19 | 0.1 |
| 1 | 232596129 | rs7523171 | 0.902 | -1.368 | -0.348 | -0.85 | 0.04 | 0.086 | 0.086 | 0 | 0.23 | 0.07 | 0.09 | 0.11 | 0.15 | 0.18 |
| 1 | 232599189 | rs10910527 | -1.215 | -0.232 | 0.596 | 0.156 | -0.028 | 0.047 | 0.047 | 0 | 0.1 | 0.17 | 0.29 | 0.7 | 0.04 | 0.28 |
| 1 | 232599944 | rs10797570 | -2.039 | -0.235 | 0.598 | 0.139 | -0.004 | 0.021 | 0.021 | 0 | 0.22 | 0.34 | 0.46 | 0.7 | 0.23 | 0.38 |
| 1 | 232601208 | rs7536385 | -1.242 | -0.102 | 0.635 | 0.202 | -0.028 | 0.047 | 0.047 | 0 | 0.1 | 0.17 | 0.29 | 0.7 | 0.04 | 0.28 |
| 1 | 232602117 | rs12117117 | 0.13 | 0.093 | 0.598 | 0.12 | 0.047 | -0.063 | -0.063 | 8.076 | 0.11 | 0.19 | 0.17 | 0 | 0.19 | 0.1 |
| 1 | 232604178 | rs11589920 | -0.318 | 0.093 | 0.305 | 0.313 | 0.036 | 0.146 | 0.146 | 0 | 0.12 | 0.24 | 0.25 | 0.56 | 0.2 | 0.21 |
| 1 | 232607986 | rs2225041 | -1.197 | 0.09 | -0.251 | -0.24 | -0.003 | 0.007 | 0.007 | 0 | 0.21 | 0.18 | 0.39 | 0.16 | 0.23 | 0.3 |
| 1 | 232608141 | rs1359630 | -1.153 | 0.091 | -0.267 | -0.239 | -0.005 | 0.01 | 0.01 | 0 | 0.2 | 0.15 | 0.39 | 0.16 | 0.23 | 0.3 |
| 1 | 232633312 | rs10495328 | -1.183 | 0.148 | -0.295 | -0.211 | -0.004 | 0.004 | 0.004 | 0 | 0.2 | 0.18 | 0.39 | 0.16 | 0.23 | 0.29 |
| 1 | 232638336 | rs12729264 | -0.205 | 0.332 | -0.101 | -0.142 | 0.012 | -0.024 | -0.024 | 0 | 0.06 | 0.1 | 0.13 | 0 | 0.11 | 0.07 |
| 1 | 232638409 | rs6674092 | -0.293 | -0.519 | 0.037 | -0.393 | 0.023 | 0.033 | 0.033 | 0 | 0.44 | 0.57 | 0.54 | 0.85 | 0.3 | 0.46 |
| 1 | 232638547 | rs6694142 | -0.187 | -0.51 | 0.068 | -0.388 | 0.001 | -0.011 | -0.011 | 0 | 0.43 | 0.52 | 0.54 | 0.85 | 0.29 | 0.46 |
| 1 | 232641589 | rs10910547 | 0.432 | -0.122 | -0.284 | -0.49 | -0.047 | 0.07 | 0.07 | 0 | 0.31 | 0.18 | 0.29 | 0.02 | 0.41 | 0.24 |
| 1 | 232642297 | rs7533227 | -0.795 | -0.57 | 0.857 | 0.728 | -0.013 | -0.016 | -0.016 | 0 | 0.41 | 0.44 | 0.45 | 0.33 | 0.28 | 0.38 |
| 1 | 232642837 | rs6672473 | -0.459 | -0.457 | 0.441 | 0.889 | 0.014 | 0.034 | 0.034 | 0 | 0.33 | 0.2 | 0.17 | 0.18 | 0.26 | 0.17 |
| 1 | 232650867 | rs4649383 | 0.735 | -1.57 | 0.554 | 1.207 | -0.023 | -0.006 | -0.006 | 0 | 0.62 | 0.46 | 0.64 | 0.33 | 0.8 | 0.61 |
| 1 | 232658363 | rs2357070 | 0.552 | -1.515 | 0.534 | 1.194 | -0.043 | -0.005 | -0.005 | 0 | 0.63 | 0.55 | 0.64 | 0.34 | 0.8 | 0.62 |
| 1 | 232671405 | rs12096229 | -0.811 | -1.438 | 0.09 | 0.964 | -0.011 | -0.011 | -0.011 | 0.024 | 0.36 | 0.6 | 0.21 | 0.17 | 0.15 | 0.24 |
| 1 | 232675032 | rs943140 | -0.452 | -0.933 | -0.169 | 0.855 | -0.001 | -0.001 | -0.001 | 0.995 | 0.27 | 0.26 | 0.17 | 0.3 | 0.17 | 0.15 |
| 1 | 232678397 | rs814216 | -1.025 | -0.389 | 0.403 | 0.968 | 0.011 | 0.079 | 0.079 | 1.529 | 0.09 | 0.51 | 0.04 | 0.11 | 0.02 | 0.11 |
| 1 | 232683980 | rs10495329 | -0.064 | -0.716 | -1.136 | 0.363 | 0.043 | 0.124 | 0.124 | 0.029 | 0.28 | 0.11 | 0.16 | 0.01 | 0.17 | 0.14 |
| 1 | 232684139 | rs789661 | -1.104 | -0.783 | -1.228 | -0.111 | -0.034 | 0.038 | 0.038 | 0 | 0.67 | 0.76 | 0.66 | 0.87 | 0.46 | 0.52 |
| 1 | 232684827 | rs789663 | 0.17 | -0.958 | -0.485 | 0.032 | -0.007 | 0.001 | 0.001 | 0.052 | 0.31 | 0.3 | 0.27 | 0.33 | 0.25 | 0.2 |
| 1 | 232686897 | rs789666 | -0.749 | -0.583 | -0.262 | 0.074 | 0.009 | -0.008 | -0.008 | 0 | 0.64 | 0.6 | 0.64 | 0.74 | 0.47 | 0.41 |
| 1 | 232688592 | rs10797578 | -0.774 | -0.566 | -0.729 | -0.456 | -0.07 | 0.105 | 0.105 | 0.538 | 0.65 | 0.82 | 0.64 | 0.89 | 0.46 | 0.58 |
| 1 | 232690086 | rs2243857 | -0.699 | -0.497 | -1.229 | -0.856 | -0.102 | 0.128 | 0.128 | 0.242 | 0.65 | 0.9 | 0.64 | 0.88 | 0.56 | 0.61 |
| 1 | 232692293 | rs10910557 | 0.115 | -1.012 | -1.092 | -0.796 | 0.013 | -0.044 | -0.044 | 0 | 0.27 | 0.16 | 0.3 | 0.57 | 0.19 | 0.35 |
| 1 | 232692908 | rs796876 | -1.159 | -1.048 | -0.789 | -0.863 | -0.06 | 0.092 | 0.092 | 0 | 0.65 | 0.86 | 0.59 | 0.87 | 0.56 | 0.62 |
| 1 | 232693851 | rs789651 | -0.969 | -1.145 | 0.335 | -0.564 | -0.016 | 0.365 | 0.365 | 0.054 | 0.1 | 0.35 | 0.23 | 0.56 | 0.03 | 0.15 |
| 1 | 232701166 | rs12069683 | -0.064 | -1.205 | 0.14 | -0.821 | -0.001 | 0 | 0 | 0 | 0.12 | 0.07 | 0.05 | 0.01 | 0.1 | 0.1 |
| 1 | 232702158 | rs2247872 | -1.205 | -1.258 | 0.433 | -0.72 | 0.009 | -0.008 | -0.008 | 0.051 | 0.41 | 0.61 | 0.37 | 0.57 | 0.29 | 0.45 |
| 1 | 232702788 | rs12072371 | -0.317 | -1.285 | 0.389 | -0.716 | 0.003 | 0.01 | 0.01 | 0 | 0.31 | 0.22 | 0.14 | 0.06 | 0.26 | 0.37 |
| 1 | 232703563 | rs789647 | -0.04 | -1.242 | 0.502 | -0.726 | -0.015 | -0.026 | -0.026 | 0 | 0.07 | 0.27 | 0.02 | 0 | 0.01 | 0 |
| 1 | 232706935 | rs2243422 | -1.191 | -1.218 | -0.059 | -0.704 | 0.029 | -0.017 | -0.017 | 0 | 0.42 | 0.7 | 0.38 | 0.57 | 0.29 | 0.46 |
| 1 | 232710264 | rs1285683 | -0.684 | -0.661 | -0.711 | -0.414 | 0.003 | -0.023 | -0.023 | 0 | 0.33 | 0.26 | 0.34 | 0.57 | 0.28 | 0.46 |
| 1 | 232717335 | rs2025724 | 0.458 | 0.451 | -0.801 | -0.199 | -0.001 | 0.004 | 0.004 | 0 | 0.24 | 0.23 | 0.28 | 0.17 | 0.25 | 0.29 |
| 1 | 232720492 | rs17191808 | 0.65 | 0.671 | -0.988 | -0.405 | -0.024 | 0.035 | 0.035 | 0.229 | 0.07 | 0.02 | 0.03 | 0.01 | 0.07 | 0.04 |
| 1 | 232721837 | rs11810061 | 0.512 | 0.438 | -0.56 | -0.692 | 0 | 0.055 | 0.055 | 0 | 0.53 | 0.33 | 0.58 | 0.75 | 0.55 | 0.55 |
| 1 | 232723174 | rs12753593 | 0.009 | 0.431 | -0.484 | -0.698 | 0.003 | -0.095 | -0.095 | 2.056 | 0.23 | 0.43 | 0.14 | 0.07 | 0.2 | 0.16 |
| 1 | 232724509 | rs1334401 | 0.938 | 0.416 | -0.302 | -0.687 | 0 | -0.01 | -0.01 | 1.294 | 0.24 | 0.35 | 0.28 | 0.17 | 0.25 | 0.29 |
| 1 | 232727151 | rs2210669 | -0.024 | 0.271 | -0.533 | -1.175 | -0.004 | -0.022 | -0.022 | 0 | 0.44 | 0.5 | 0.37 | 0.17 | 0.39 | 0.43 |
| 1 | 232727525 | rs11805493 | 1.051 | 0.263 | -0.498 | -1.281 | 0.007 | -0.011 | -0.011 | 0.351 | 0.2 | 0.17 | 0.1 | 0 | 0.15 | 0.16 |
| 1 | 232727572 | rs4649390 | -0.672 | 0.296 | -0.482 | -1.288 | 0.001 | -0.057 | -0.057 | 0 | 0.78 | 0.68 | 0.86 | 0.93 | 0.8 | 0.84 |
| 1 | 232727734 | rs4649391 | 0.303 | 0.441 | -0.843 | -1.039 | -0.006 | -0.083 | -0.083 | 1.376 | 0.56 | 0.5 | 0.75 | 0.92 | 0.64 | 0.63 |
| 1 | 232730293 | rs945084 | 1.165 | 0.649 | -1.297 | -0.46 | -0.017 | 0.026 | 0.026 | 0 | 0.31 | 0.15 | 0.31 | 0.27 | 0.37 | 0.32 |

| | | | | | | | | | | | | | | | | |
|---|---|---|---|---|---|---|---|---|---|---|---|---|---|---|---|---|
| 1 | 232735234 | rs4649274 | 1.166 | 0.664 | -1.162 | -0.44 | -0.006 | 0.002 | 0.002 | 0.496 | 0.21 | 0.14 | 0.2 | 0.16 | 0.23 | 0.26 |
| 1 | 232736249 | rs4545285 | -0.305 | 0.556 | -1.181 | -0.511 | 0.001 | 0.002 | 0.002 | 0 | 0.61 | 0.38 | 0.61 | 0.76 | 0.65 | 0.59 |
| 1 | 232736520 | rs16857753 | 0.921 | 0.409 | -1.736 | -0.759 | -0.036 | -0.018 | -0.018 | 0 | 0.35 | 0.13 | 0.34 | 0.34 | 0.41 | 0.36 |
| 1 | 232744422 | rs17805981 | 0.584 | 0.534 | -1.631 | -0.934 | -0.031 | 0.137 | 0.137 | 0 | 0.22 | 0.03 | 0.12 | 0 | 0.21 | 0.15 |
| 1 | 232746120 | rs3795367 | -0.17 | 0.637 | -0.692 | -0.1 | -0.02 | -0.094 | -0.094 | 0 | 0.22 | 0.45 | 0.18 | 0.08 | 0.16 | 0.14 |
| 1 | 232751245 | rs2039490 | 0.021 | 0.972 | -0.86 | 0.544 | 0.008 | 0.013 | 0.013 | 0 | 0.48 | 0.58 | 0.5 | 0.43 | 0.48 | 0.47 |
| 1 | 232757382 | rs7554230 | -1.002 | 1.266 | -0.41 | 1.035 | 0.007 | 0.008 | 0.008 | 0.58 | 0.3 | 0.43 | 0.48 | 0.59 | 0.34 | 0.42 |
| 1 | 232761116 | rs1970168 | -0.798 | 1.152 | -0.141 | 1.078 | -0.003 | 0.026 | 0.026 | 0 | 0.45 | 0.68 | 0.53 | 0.36 | 0.45 | 0.65 |
| 1 | 232768148 | rs6424243 | -1.241 | 1.425 | -0.056 | 1.486 | 0 | -0.001 | -0.001 | 0 | 0.39 | 0.39 | 0.46 | 0.41 | 0.44 | 0.44 |
| 1 | 232773478 | rs9424568 | -1.233 | 1.462 | -0.079 | 1.476 | -0.006 | -0.002 | -0.002 | 5.426 | 0.39 | 0.36 | 0.46 | 0.41 | 0.44 | 0.44 |
| 1 | 232779187 | rs6685037 | -1.133 | 1.469 | 0.027 | 1.477 | -0.02 | 0.065 | 0.065 | 0 | 0.38 | 0.24 | 0.37 | 0.19 | 0.4 | 0.41 |
| 1 | 232783851 | rs7537378 | -1.107 | 1.494 | 0.014 | 1.555 | 0.005 | 0.004 | 0.004 | 0 | 0.37 | 0.41 | 0.44 | 0.43 | 0.4 | 0.41 |
| 1 | 232785687 | rs9424572 | -0.128 | 1.481 | -0.23 | 1.556 | -0.014 | 0.016 | 0.016 | 0 | 0.26 | 0.19 | 0.28 | 0.19 | 0.3 | 0.33 |
| 1 | 232788317 | rs16857819 | -0.092 | 1.493 | -0.225 | 1.529 | 0.001 | -0.058 | -0.058 | 0 | 0.12 | 0.2 | 0.1 | 0 | 0.11 | 0.09 |
| 1 | 232795105 | rs4310395 | -1.329 | 1.406 | -0.227 | 1.558 | -0.001 | -0.001 | -0.001 | 0 | 0.39 | 0.39 | 0.45 | 0.43 | 0.43 | 0.45 |
| 1 | 232797904 | rs12568268 | -0.414 | 1.472 | -0.734 | 1.712 | 0.003 | 0.028 | 0.028 | 0 | 0.13 | 0.19 | 0.17 | 0.24 | 0.13 | 0.12 |
| 1 | 232803067 | rs17808035 | -0.293 | 1.204 | -1.078 | 0.764 | 0.001 | -0.017 | -0.017 | 0 | 0.14 | 0.15 | 0.11 | 0 | 0.13 | 0.08 |
| 1 | 232806859 | rs17808157 | -0.044 | 1.321 | -1.465 | 0.652 | -0.096 | -0.021 | -0.021 | 0 | 0.22 | 0.02 | 0.26 | 0.21 | 0.25 | 0.29 |
| 1 | 232810976 | rs2151874 | -0.141 | 1.054 | -0.745 | 0.531 | -0.005 | 0.011 | 0.011 | 0 | 0.42 | 0.6 | 0.46 | 0.69 | 0.49 | 0.56 |
| 1 | 232812003 | rs9424471 | -1.201 | 0.963 | -0.038 | 1.112 | 0.03 | 0.151 | 0.151 | 0 | 0.1 | 0.23 | 0.04 | 0.38 | 0.09 | 0.17 |
| 1 | 232812411 | rs10495334 | -0.275 | 0.373 | 0.245 | 0.432 | -0.018 | -0.035 | -0.035 | 1.438 | 0.1 | 0.25 | 0.09 | 0.07 | 0.08 | 0.03 |
| 1 | 232813002 | rs9424578 | -1.417 | 0.576 | 0.225 | 0.307 | 0.002 | 0.003 | 0.003 | 0 | 0.18 | 0.2 | 0.1 | 0.12 | 0.15 | 0.11 |
| 1 | 232813800 | rs7544399 | -0.411 | 0.532 | 0.204 | 0.328 | 0.006 | 0.005 | 0.005 | 0 | 0.1 | 0.11 | 0.07 | 0.06 | 0.08 | 0.04 |
| 1 | 232818269 | rs9424473 | -0.765 | 0.472 | -0.402 | 0.138 | 0.005 | -0.001 | -0.001 | 0 | 0.48 | 0.52 | 0.45 | 0.52 | 0.43 | 0.35 |
| 1 | 232819617 | rs945089 | 0.029 | 0.342 | -0.509 | 0.055 | -0.062 | -0.114 | -0.114 | 0 | 0.36 | 0.06 | 0.4 | 0.5 | 0.36 | 0.48 |
| 1 | 232824691 | rs10495335 | 0.041 | 0.408 | -0.492 | 0.015 | -0.045 | -0.084 | -0.084 | 0 | 0.34 | 0.08 | 0.31 | 0.48 | 0.33 | 0.47 |
| 1 | 232830616 | rs17808766 | 1.134 | 0.145 | -1.101 | -0.299 | -0.021 | -0.002 | -0.002 | 0 | 0.14 | 0 | 0.05 | 0.08 | 0.11 | 0.12 |
| 1 | 232831301 | rs10910594 | -0.656 | 0.347 | -0.797 | -0.247 | -0.007 | 0.003 | 0.003 | 0 | 0.5 | 0.68 | 0.49 | 0.55 | 0.45 | 0.58 |
| 1 | 232833416 | rs1413586 | 0.296 | 0.392 | -0.584 | -0.207 | -0.037 | -0.049 | -0.049 | 0 | 0.38 | 0.16 | 0.41 | 0.5 | 0.38 | 0.48 |
| 1 | 232840697 | rs9424584 | -0.837 | 0.043 | -0.631 | -0.548 | -0.03 | -0.093 | -0.093 | 4.527 | 0.21 | 0.42 | 0.13 | 0.1 | 0.13 | 0.16 |
| 1 | 232867727 | rs9660252 | 0.044 | 0.093 | -0.64 | -0.603 | -0.025 | -0.061 | -0.061 | 0 | 0.12 | 0.24 | 0.08 | 0.03 | 0.06 | 0.08 |
| 1 | 232869592 | rs10910604 | 0.044 | 0.093 | -0.64 | -0.582 | -0.025 | -0.032 | -0.032 | 0 | 0.12 | 0.24 | 0.08 | 0.08 | 0.06 | 0.09 |
| 1 | 232873994 | rs903790 | -0.307 | 0.27 | -0.529 | -0.485 | 0 | -0.033 | -0.033 | 0.929 | 0.9 | 0.82 | 0.95 | 0.98 | 0.93 | 0.94 |
| 1 | 232876993 | rs2795425 | 0.337 | 0.193 | -0.776 | -0.564 | -0.034 | -0.011 | -0.011 | 2.793 | 0.48 | 0.21 | 0.55 | 0.43 | 0.54 | 0.37 |
| 1 | 232878488 | rs1565999 | -0.669 | 0.35 | -0.225 | -0.161 | 0.026 | 0.022 | 0.022 | 0 | 0.08 | 0.28 | 0.06 | 0.08 | 0.04 | 0.1 |
| 1 | 232895254 | rs2644464 | 0.043 | 0.571 | -0.855 | -0.039 | 0.024 | -0.011 | -0.011 | 0 | 0.9 | 0.72 | 0.94 | 0.92 | 0.92 | 0.92 |
| 1 | 232896663 | rs2644466 | 0.073 | 0.575 | -0.853 | -0.037 | 0.021 | -0.014 | -0.014 | 0 | 0.9 | 0.71 | 0.94 | 0.92 | 0.92 | 0.92 |
| 1 | 232901272 | rs10910619 | 0.433 | 0.534 | -1.832 | -0.012 | -0.094 | -0.152 | -0.152 | 0.86 | 0.28 | 0.03 | 0.39 | 0.35 | 0.39 | 0.2 |
| 1 | 232908828 | rs2814548 | -1.372 | 0.307 | -1.062 | -0.187 | 0 | 0.005 | 0.005 | 0 | 0.34 | 0.31 | 0.26 | 0.29 | 0.31 | 0.22 |
| 1 | 232911400 | rs2814501 | -2.185 | 0.158 | -0.72 | 0.146 | 0.014 | 0.011 | 0.011 | 0 | 0.33 | 0.69 | 0.26 | 0.31 | 0.26 | 0.29 |
| 1 | 232916501 | rs6689943 | -0.679 | 0.165 | -0.675 | 0.142 | -0.002 | 0.021 | 0.021 | 0 | 0.14 | 0.18 | 0.15 | 0.2 | 0.13 | 0.14 |
| 1 | 232919919 | rs9424482 | -1.64 | 0.148 | -0.65 | -0.006 | -0.002 | -0.003 | -0.003 | 0.136 | 0.23 | 0.27 | 0.18 | 0.2 | 0.2 | 0.19 |
| 1 | 232930057 | rs2795467 | -2.304 | 0.204 | -0.843 | 0.036 | 0.019 | 0.023 | 0.023 | 0 | 0.34 | 0.72 | 0.27 | 0.28 | 0.26 | 0.29 |
| 1 | 232933687 | rs1948529 | -0.464 | 0.135 | -0.583 | 0.243 | -0.012 | 0.01 | 0.01 | 0 | 0.09 | 0.05 | 0.03 | 0.08 | 0.07 | 0.08 |
| 1 | 232937739 | rs1495986 | -2.029 | 0.158 | -0.546 | 0.311 | 0.016 | -0.004 | -0.004 | 0 | 0.3 | 0.74 | 0.29 | 0.33 | 0.22 | 0.23 |
| 1 | 232940447 | rs10910625 | -0.391 | 0.106 | -0.611 | 0.304 | -0.005 | 0.004 | 0.004 | 0 | 0.07 | 0.13 | 0.08 | 0.14 | 0.06 | 0.1 |
| 1 | 232940953 | rs10797593 | -0.417 | 0.1 | -0.653 | 0.302 | -0.006 | 0.013 | 0.013 | 0 | 0.07 | 0.19 | 0.08 | 0.14 | 0.06 | 0.1 |
| 1 | 232942251 | rs12048236 | -0.39 | 0.118 | -0.623 | 0.334 | 0 | 0.029 | 0.029 | 0 | 0.07 | 0.25 | 0.08 | 0.14 | 0.05 | 0.1 |
| 1 | 232944569 | rs1845789 | 1.606 | 0.184 | -0.765 | 0.273 | 0.004 | -0.001 | -0.001 | 0.014 | 0.67 | 0.17 | 0.67 | 0.67 | 0.74 | 0.72 |
| 1 | 232947295 | rs2357331 | -0.6 | 1.064 | -0.578 | 0.871 | -0.004 | 0.02 | 0.02 | 0 | 0.07 | 0.21 | 0.08 | 0.14 | 0.05 | 0.09 |
| 1 | 232949914 | rs2814530 | -0.934 | 1.321 | -1.065 | 0.82 | 0.013 | 0.013 | 0.013 | 0 | 0.14 | 0.17 | 0.16 | 0.19 | 0.16 | 0.16 |
| 1 | 232953612 | rs12025856 | -0.402 | 1.373 | -1.938 | 0.971 | -0.09 | -0.116 | -0.116 | 0.088 | 0.35 | 0.09 | 0.47 | 0.48 | 0.46 | 0.3 |
| 1 | 232962287 | rs2132410 | -1.246 | 1.062 | 0.28 | 0.592 | 0.027 | 0.015 | 0.015 | 0 | 0.05 | 0.24 | 0.13 | 0.02 | 0.02 | 0.06 |
| 1 | 232977131 | rs11804919 | -1.831 | 0.883 | 1.208 | 0.506 | 0.075 | 0.058 | 0.058 | 0 | 0.06 | 0.42 | 0.16 | 0.02 | 0.03 | 0.06 |
| 1 | 232982116 | rs2644441 | 2.106 | 0.984 | 1.143 | 0.848 | -0.001 | 0.001 | 0.001 | 0.032 | 0.88 | 0.21 | 0.74 | 0.86 | 0.91 | 0.85 |
| 1 | 232990257 | rs1874401 | -1.498 | 0.966 | 1.32 | 0.862 | 0.004 | 0.007 | 0.007 | 0 | 0.05 | 0.1 | 0.18 | 0.06 | 0.05 | 0.07 |
| 1 | 232995962 | rs1353434 | 2.153 | 1.214 | 1.064 | 0.914 | -0.001 | 0.002 | 0.002 | 0.026 | 0.85 | 0.25 | 0.65 | 0.81 | 0.88 | 0.81 |
| 1 | 233001571 | rs2753391 | -0.833 | 1.081 | -0.1 | 0.629 | -0.043 | 0.037 | 0.037 | 0 | 0.64 | 0.87 | 0.67 | 0.76 | 0.59 | 0.64 |
| 1 | 233005854 | rs2753393 | -2.41 | 0.71 | 0.312 | 0.768 | 0.001 | 0.003 | 0.003 | 0 | 0.19 | 0.75 | 0.29 | 0.14 | 0.14 | 0.29 |
| 1 | 233008104 | rs2753395 | -2.144 | 0.655 | 0.689 | 0.739 | 0.057 | -0.055 | -0.055 | 1.609 | 0.1 | 0.54 | 0.1 | 0 | 0.09 | 0.06 |
| 1 | 233010053 | rs765556 | -2.55 | 0.757 | 0.557 | 0.855 | 0 | -0.001 | -0.001 | 0.488 | 0.2 | 0.75 | 0.31 | 0.2 | 0.16 | 0.29 |
| 1 | 233013887 | rs4649415 | -2.128 | 0.728 | 0.922 | 0.834 | 0.036 | 0.094 | 0.094 | 0 | 0.06 | 0.39 | 0.14 | 0.09 | 0.02 | 0.12 |
| 1 | 233020124 | rs9424601 | -1.476 | 0.857 | 1.005 | 0.808 | 0.108 | 0.186 | 0.186 | 0.09 | 0.05 | 0.35 | 0.2 | 0.07 | 0.04 | 0.05 |
| 1 | 233020568 | rs9424602 | -2.468 | 0.668 | 0.938 | 0.656 | 0.014 | 0.001 | 0.001 | 0.157 | 0.13 | 0.61 | 0.25 | 0.08 | 0.09 | 0.09 |
| 1 | 233022055 | rs9424491 | -1.475 | 0.591 | 0.707 | 0.74 | 0.065 | 0.138 | 0.138 | 1.341 | 0.08 | 0.44 | 0.2 | 0.08 | 0.04 | 0.05 |
| 1 | 233023763 | rs9424606 | 0.136 | 0.608 | -0.507 | 0.721 | -0.027 | -0.046 | -0.046 | 0 | 0.31 | 0.21 | 0.32 | 0.48 | 0.35 | 0.29 |
| 1 | 233023930 | rs6697517 | -0.853 | 0.551 | -0.84 | 0.608 | -0.073 | -0.133 | -0.133 | 2.692 | 0.75 | 0.95 | 0.71 | 0.5 | 0.69 | 0.78 |
| 1 | 233027385 | rs6698720 | -2.324 | -0.64 | -0.375 | 0.685 | 0.016 | -0.003 | -0.003 | 0 | 0.24 | 0.78 | 0.29 | 0.19 | 0.13 | 0.28 |
| 1 | 233029452 | rs12024828 | -1.299 | -0.51 | -0.015 | 0.702 | -0.025 | 0.024 | 0.024 | 0 | 0.15 | 0.62 | 0.2 | 0.14 | 0.09 | 0.14 |

| | | | | | | | | | | | | | | | | |
|---|---|---|---|---|---|---|---|---|---|---|---|---|---|---|---|---|
| 1 | 233032683 | rs12403846 | -0.836 | -1.231 | -0.663 | 0.574 | 0.007 | 0.006 | 0.006 | 0 | 0.31 | 0.2 | 0.34 | 0.31 | 0.28 | 0.32 |
| 1 | 233040129 | rs16858453 | 1.011 | -1.554 | -0.677 | 0.612 | 0.007 | -0.007 | -0.007 | 0 | 0.12 | 0.08 | 0.13 | 0.16 | 0.08 | 0.04 |
| 1 | 233040790 | rs9424611 | 0.571 | -1.463 | -0.306 | 0.498 | -0.03 | 0.07 | 0.07 | 0 | 0.71 | 0.57 | 0.65 | 0.57 | 0.75 | 0.69 |
| 1 | 233040857 | rs9424612 | 1.081 | -1.425 | -0.779 | 0.48 | 0.008 | -0.008 | -0.008 | 0 | 0.61 | 0.36 | 0.54 | 0.53 | 0.67 | 0.61 |
| 1 | 233045682 | rs10797596 | 1.205 | -1.35 | -1.429 | 0.52 | 0.055 | -0.036 | -0.036 | 0.773 | 0.58 | 0.22 | 0.53 | 0.53 | 0.66 | 0.53 |
| 1 | 233052341 | rs2224402 | 0.992 | -1.527 | -1.709 | 0.481 | 0.079 | -0.05 | -0.05 | 0 | 0.6 | 0.13 | 0.57 | 0.65 | 0.73 | 0.61 |
| 1 | 233060511 | rs1555394 | -1.515 | -1.504 | -1.051 | 0.292 | -0.015 | -0.014 | -0.014 | 0.779 | 0.3 | 0.32 | 0.25 | 0.18 | 0.2 | 0.21 |
| 1 | 233061650 | rs4350151 | -0.546 | -1.687 | -1.068 | 0.196 | 0.036 | 0.039 | 0.039 | 0 | 0.13 | 0.06 | 0.04 | 0.01 | 0.04 | 0.07 |
| 1 | 233073561 | rs4649418 | -1.039 | -1.679 | -1.039 | 0.205 | 0.013 | 0.019 | 0.019 | 0 | 0.16 | 0.23 | 0.2 | 0.17 | 0.16 | 0.13 |
| 1 | 233084744 | rs3820124 | -2.235 | -1.733 | -1.062 | 0.264 | -0.014 | -0.012 | -0.012 | 0 | 0.31 | 0.32 | 0.15 | 0.18 | 0.2 | 0.21 |
| 1 | 233089728 | rs12744791 | 0.93 | -1.79 | -1.699 | 0.048 | -0.033 | -0.006 | -0.006 | 0 | 0.13 | 0 | 0.13 | 0.12 | 0.16 | 0.16 |
| 1 | 233093955 | rs6704443 | -2.709 | -1.824 | -2.188 | -0.046 | -0.164 | -0.167 | -0.167 | 0.487 | 0.66 | 0.98 | 0.55 | 0.44 | 0.59 | 0.65 |
| 1 | 233102792 | rs12035585 | -1.368 | -1.514 | -1.514 | 0.073 | -0.012 | -0.036 | -0.036 | 0.007 | 0.21 | 0.3 | 0.11 | 0.06 | 0.14 | 0.11 |
| 1 | 233113827 | rs1045158 | -1.801 | -1.263 | -1.356 | 0.219 | 0.013 | 0.001 | 0.001 | 0 | 0.21 | 0.32 | 0.15 | 0.18 | 0.21 | 0.14 |
| 1 | 233119951 | rs2885 | -1.232 | -1.174 | -1.336 | 0.255 | 0.027 | 0.008 | 0.008 | 0 | 0.17 | 0.23 | 0.13 | 0.16 | 0.2 | 0.12 |
| 1 | 233123883 | rs3766496 | -1.236 | -1.159 | -1.263 | 0.229 | 0.018 | 0.004 | 0.004 | 0 | 0.17 | 0.19 | 0.13 | 0.16 | 0.2 | 0.12 |
| 1 | 233127416 | rs743195 | -1.606 | -1.191 | -1.169 | 0.219 | -0.001 | -0.007 | -0.007 | 0 | 0.22 | 0.35 | 0.16 | 0.17 | 0.19 | 0.15 |
| 1 | 233130937 | rs10910644 | -1.327 | -1.123 | -1.145 | 0.106 | 0.005 | -0.025 | -0.025 | 0.054 | 0.22 | 0.31 | 0.17 | 0.12 | 0.22 | 0.13 |
| 1 | 233140890 | rs3766491 | 1.227 | -1.154 | -1.287 | 0.346 | -0.012 | -0.029 | -0.029 | 4.559 | 0.05 | 0 | 0.06 | 0.15 | 0.06 | 0.16 |
| 1 | 233144081 | rs6424262 | -1.835 | -1.55 | -0.577 | 0.451 | 0.043 | -0.003 | -0.003 | 0 | 0.45 | 0.71 | 0.36 | 0.54 | 0.33 | 0.31 |
| 1 | 233148271 | rs6702102 | -1.829 | -1.734 | -0.161 | 0.526 | 0.005 | -0.006 | -0.006 | 0 | 0.37 | 0.57 | 0.28 | 0.49 | 0.22 | 0.29 |
| 1 | 233148513 | rs3753423 | -1.03 | -1.67 | -0.029 | 0.453 | -0.038 | 0 | 0 | 0 | 0.24 | 0.39 | 0.17 | 0.22 | 0.1 | 0.18 |
| 1 | 233154981 | rs4649424 | 0.586 | -1.376 | 0.143 | 0.133 | -0.02 | -0.082 | -0.082 | 0 | 0.8 | 0.42 | 0.85 | 1 | 0.82 | 0.85 |
| 1 | 233157853 | rs17453089 | 0.44 | -1.415 | -0.092 | 0.199 | -0.002 | -0.025 | -0.025 | 0.293 | 0.13 | 0.16 | 0.09 | 0 | 0.13 | 0.11 |
| 1 | 233161822 | rs7541947 | -1.789 | -1.35 | -0.356 | 0.484 | 0.046 | -0.03 | -0.03 | 0 | 0.51 | 0.87 | 0.38 | 0.47 | 0.37 | 0.39 |
| 1 | 233165898 | rs12063911 | -2.133 | -0.703 | 0.768 | 0.538 | -0.013 | 0.015 | 0.015 | 0.536 | 0.23 | 0.55 | 0.19 | 0.29 | 0.21 | 0.18 |
| 1 | 233167058 | rs6657371 | -1.124 | -0.539 | 0.357 | 0.624 | -0.026 | -0.056 | -0.056 | 0 | 0.15 | 0.04 | 0.12 | 0.29 | 0.2 | 0.13 |
| 1 | 233171297 | rs6676202 | -1.864 | -0.712 | 1.035 | 0.394 | -0.047 | -0.099 | -0.099 | 0 | 0.14 | 0.48 | 0.11 | 0 | 0.08 | 0.08 |
| 1 | 233175761 | rs2273675 | 0.399 | -0.413 | 0.699 | 0.591 | -0.026 | -0.021 | -0.021 | 0.002 | 0.05 | 0 | 0.05 | 0.08 | 0.06 | 0.07 |
| 1 | 233176034 | rs2273673 | -1.398 | -0.495 | 1.278 | 0.532 | 0.013 | -0.003 | -0.003 | 0 | 0.1 | 0.22 | 0.06 | 0.08 | 0.07 | 0.09 |
| 1 | 233176090 | rs2273672 | -1.939 | -0.596 | 1.654 | 0.615 | -0.007 | 0.058 | 0.058 | 0.014 | 0.2 | 0.39 | 0.17 | 0.27 | 0.2 | 0.17 |
| 1 | 233178442 | rs3820116 | -1.691 | -0.609 | 1.737 | 0.62 | 0.015 | 0.097 | 0.097 | 0 | 0.14 | 0.38 | 0.11 | 0.27 | 0.13 | 0.11 |
| 1 | 233178910 | rs7541494 | -1.402 | -0.589 | 1.577 | 0.658 | 0 | -0.029 | -0.029 | 0 | 0.15 | 0.23 | 0.13 | 0.08 | 0.13 | 0.15 |
| 1 | 233188011 | rs10489576 | -0.444 | -0.553 | 0.882 | 0.664 | 0.004 | 0.048 | 0.048 | 0 | 0.1 | 0.15 | 0.09 | 0.27 | 0.12 | 0.08 |
| 1 | 233191368 | rs6424272 | -2.197 | -0.59 | 0.43 | 0.297 | 0.005 | 0.016 | 0.016 | 0.016 | 0.3 | 0.74 | 0.22 | 0.25 | 0.26 | 0.25 |
| 1 | 233193800 | rs6695322 | -1.557 | -0.239 | 0.539 | 0.576 | 0.016 | 0.019 | 0.019 | 0 | 0.21 | 0.49 | 0.14 | 0.25 | 0.18 | 0.17 |
| 1 | 233196709 | rs10797599 | -0.535 | -0.562 | -0.27 | 0.002 | 0.009 | 0.043 | 0.043 | 0 | 0.48 | 0.44 | 0.61 | 0.76 | 0.48 | 0.61 |
| 1 | 233197806 | rs2357427 | -1.532 | -0.592 | -0.249 | -0.18 | -0.004 | -0.016 | -0.016 | 0.938 | 0.22 | 0.68 | 0.19 | 0.17 | 0.2 | 0.1 |
| 1 | 233200867 | rs6424273 | -0.36 | -0.605 | 0.001 | -0.206 | 0.05 | -0.047 | -0.047 | 0 | 0.11 | 0.55 | 0.12 | 0.02 | 0.11 | 0.05 |
| 1 | 233204338 | rs1155739 | -1.692 | -0.228 | -0.535 | -0.638 | 0 | -0.003 | -0.003 | 0 | 0.35 | 0.68 | 0.25 | 0.23 | 0.32 | 0.16 |
| 1 | 233204882 | rs12085776 | -1.337 | -0.277 | -0.532 | -0.666 | 0.004 | -0.01 | -0.01 | 0 | 0.2 | 0.68 | 0.18 | 0.17 | 0.19 | 0.1 |
| 1 | 233205554 | rs11484500 | -0.464 | -0.23 | -0.512 | -0.66 | -0.005 | -0.146 | -0.146 | 1.04 | 0.23 | 0.56 | 0.17 | 0.02 | 0.2 | 0.06 |
| 1 | 233214528 | rs6424276 | -1.516 | -0.229 | -0.538 | -0.647 | -0.002 | -0.013 | -0.013 | 0 | 0.33 | 0.68 | 0.24 | 0.17 | 0.29 | 0.12 |
| 1 | 233214860 | rs16858621 | -0.239 | -0.218 | -0.502 | -0.619 | -0.001 | 0.006 | 0.006 | 0 | 0.1 | 0.1 | 0.06 | 0.15 | 0.09 | 0.06 |
| 1 | 233215885 | rs6693854 | -1.798 | -0.204 | -0.527 | -0.633 | 0.009 | -0.005 | -0.005 | 0 | 0.24 | 0.69 | 0.2 | 0.23 | 0.23 | 0.15 |
| 1 | 233217410 | rs6424278 | -0.472 | -0.225 | -0.344 | -0.62 | -0.007 | -0.122 | -0.122 | 0.035 | 0.23 | 0.51 | 0.17 | 0.01 | 0.2 | 0.06 |
| 1 | 233219627 | rs2273676 | -0.506 | -0.146 | -0.387 | -0.475 | 0.005 | 0.051 | 0.051 | 0 | 0.09 | 0.16 | 0.06 | 0.18 | 0.06 | 0.09 |
| 1 | 233220264 | rs16858637 | -0.166 | -0.102 | -0.407 | -0.498 | -0.003 | 0.021 | 0.021 | 0 | 0.1 | 0.15 | 0.07 | 0.14 | 0.07 | 0.05 |
| 1 | 233222823 | rs10752798 | -0.59 | -0.317 | -1.058 | -0.512 | 0 | -0.001 | -0.001 | 0 | 0.62 | 0.69 | 0.48 | 0.4 | 0.63 | 0.52 |
| 1 | 233224085 | rs12026554 | 1.233 | -0.639 | -1.476 | -0.052 | -0.035 | -0.091 | -0.091 | 0 | 0.14 | 0.03 | 0.03 | 0.36 | 0.11 | 0.15 |
| 1 | 233225848 | rs12059884 | 1.149 | -0.627 | -1.167 | -0.669 | 0.074 | -0.032 | -0.032 | 0 | 0.08 | 0.35 | 0.11 | 0 | 0.1 | 0.02 |
| 1 | 233226430 | rs2295746 | 0.927 | -0.861 | -1.968 | -0.571 | -0.076 | 0.127 | 0.127 | 0 | 0.42 | 0.11 | 0.28 | 0.2 | 0.42 | 0.4 |
| 1 | 233229848 | rs1005112 | -0.95 | -0.876 | -2.071 | -0.697 | -0.059 | 0.131 | 0.131 | 0 | 0.56 | 0.87 | 0.67 | 0.78 | 0.51 | 0.61 |
| 1 | 233236580 | rs1040852 | 0.333 | -0.707 | -1.887 | -0.471 | 0.003 | 0.012 | 0.012 | 0 | 0.16 | 0.17 | 0.12 | 0.25 | 0.16 | 0.12 |
| 1 | 233236731 | rs11580152 | 0.844 | -0.709 | -1.884 | -0.461 | -0.002 | 0.012 | 0.012 | 0 | 0.13 | 0.15 | 0.11 | 0.21 | 0.13 | 0.08 |
| 1 | 233237670 | rs16858662 | 1.089 | -0.713 | -1.872 | -0.454 | 0 | -0.01 | -0.01 | 0 | 0.13 | 0.08 | 0.1 | 0.21 | 0.13 | 0.08 |
| 1 | 233239255 | rs11580870 | 0.829 | -0.714 | -1.87 | -0.469 | -0.002 | 0.012 | 0.012 | 0 | 0.13 | 0.15 | 0.11 | 0.21 | 0.13 | 0.08 |
| 1 | 233241590 | rs6676695 | 0.335 | -0.701 | -1.873 | -0.467 | 0.003 | 0.012 | 0.012 | 0 | 0.16 | 0.17 | 0.12 | 0.25 | 0.16 | 0.12 |
| 1 | 233241677 | rs16858670 | 1.087 | -0.7 | -1.873 | -0.465 | 0 | -0.01 | -0.01 | 0 | 0.13 | 0.08 | 0.1 | 0.21 | 0.13 | 0.08 |
| 1 | 233242006 | rs10910660 | 0.821 | -1.044 | -1.831 | -0.6 | 0.005 | -0.062 | -0.062 | 0.279 | 0.26 | 0.41 | 0.4 | 0.14 | 0.26 | 0.35 |
| 1 | 233243187 | rs16858681 | 0.203 | -0.539 | -2.296 | -0.138 | 0.005 | 0.016 | 0.016 | 0 | 0.16 | 0.2 | 0.12 | 0.25 | 0.16 | 0.12 |
| 1 | 233256280 | rs717588 | -1.215 | -0.34 | -3.036 | -0.013 | -0.058 | 0.033 | 0.033 | 0 | 0.7 | 0.92 | 0.75 | 0.79 | 0.62 | 0.65 |
| 1 | 233262943 | rs16858694 | -0.024 | 0.083 | -2.203 | 0.394 | 0.014 | -0.019 | -0.019 | 0 | 0.18 | 0.15 | 0.15 | 0.41 | 0.1 | 0.18 |
| 1 | 233272372 | rs1016405 | -0.593 | 0.601 | -2.56 | 0.573 | -0.083 | -0.073 | -0.073 | 0 | 0.55 | 0.95 | 0.67 | 0.61 | 0.55 | 0.47 |
| 1 | 233273881 | rs6424279 | 0.457 | 0.642 | -2.363 | 0.758 | 0.005 | 0.071 | 0.071 | 0 | 0.11 | 0.14 | 0.15 | 0.4 | 0.1 | 0.11 |
| 1 | 233274249 | rs7522112 | -0.327 | 0.519 | -2.819 | 0.454 | -0.083 | -0.073 | -0.073 | 0 | 0.55 | 0.95 | 0.67 | 0.61 | 0.55 | 0.47 |
| 1 | 233279256 | rs6687497 | -0.516 | 0.53 | -2.186 | 0.206 | 0.035 | 0.109 | 0.109 | 0 | 0.48 | 0.81 | 0.39 | 0.71 | 0.41 | 0.38 |
| 1 | 233280927 | rs12118654 | 0.484 | 0.745 | -2.496 | -0.208 | -0.078 | 0.06 | 0.06 | 0 | 0.33 | 0.14 | 0.39 | 0.18 | 0.45 | 0.48 |
| 1 | 233281297 | rs2477859 | -0.102 | 0.498 | -1.666 | 0.207 | 0 | -0.005 | -0.005 | 0 | 0.39 | 0.37 | 0.42 | 0.45 | 0.35 | 0.31 |

| | | | | | | | | | | | | | | | | |
|---|---|---|---|---|---|---|---|---|---|---|---|---|---|---|---|---|
| 1 | 233282125 | rs12402073 | 1.008 | 0.073 | -1.697 | 0.416 | 0.008 | -0.021 | -0.021 | 0.542 | 0.19 | 0.05 | 0.16 | 0.24 | 0.15 | 0.15 |
| 1 | 233282323 | rs2477863 | 1.108 | 0.097 | -1.39 | 0.434 | -0.001 | 0 | 0 | 0 | 0.18 | 0.19 | 0.23 | 0.21 | 0.17 | 0.13 |
| 1 | 233284879 | rs6424282 | 0.246 | -0.249 | -0.994 | 0.529 | -0.033 | 0.01 | 0.01 | 0 | 0.3 | 0.59 | 0.22 | 0.39 | 0.24 | 0.24 |
| 1 | 233285897 | rs6685114 | -0.061 | 0.264 | -1.473 | 0.524 | 0.04 | -0.037 | -0.037 | 0 | 0.57 | 0.83 | 0.69 | 0.49 | 0.62 | 0.64 |
| 1 | 233292257 | rs2477868 | 0.024 | 0.15 | -0.909 | 0.47 | -0.001 | 0 | 0 | 0 | 0.6 | 0.6 | 0.51 | 0.41 | 0.63 | 0.69 |
| 1 | 233295282 | rs10910665 | 1.029 | 0.385 | -1.333 | 0.247 | -0.132 | 0.038 | 0.038 | 0 | 0.34 | 0.02 | 0.28 | 0.18 | 0.45 | 0.46 |
| 1 | 233301658 | rs9424523 | 0.079 | 0.543 | -0.465 | 0.691 | 0.006 | -0.063 | -0.063 | 0.096 | 0.64 | 0.85 | 0.56 | 0.46 | 0.66 | 0.73 |
| 1 | 233304042 | rs12036063 | -1.62 | 1.165 | 0.365 | 1.69 | 0.017 | 0.084 | 0.084 | 0 | 0.06 | 0.08 | 0.16 | 0.33 | 0.09 | 0.1 |
| 1 | 233316169 | rs4649440 | -0.807 | 1.132 | 0.469 | 1.524 | 0.02 | 0.066 | 0.066 | 0 | 0.2 | 0.42 | 0.29 | 0.26 | 0.16 | 0.16 |
| 1 | 233318181 | rs12407315 | 0.073 | 1.158 | 0.314 | 1.444 | -0.006 | -0.023 | -0.023 | 0.002 | 0.15 | 0.09 | 0.23 | 0.21 | 0.13 | 0.11 |
| 1 | 233332347 | rs10797444 | -1.509 | 0.961 | 0.598 | 1.47 | 0.021 | 0.11 | 0.11 | 0 | 0.24 | 0.44 | 0.43 | 0.59 | 0.21 | 0.2 |
| 1 | 233338691 | rs4649442 | -0.803 | 0.785 | 0.413 | 1.543 | -0.006 | -0.02 | -0.02 | 0.035 | 0.19 | 0.14 | 0.4 | 0.55 | 0.2 | 0.15 |
| 1 | 233342220 | rs16858853 | -0.049 | 0.558 | 0.17 | 0.867 | -0.001 | -0.018 | -0.018 | 0.748 | 0.14 | 0.11 | 0.23 | 0.21 | 0.12 | 0.11 |
| 1 | 233348885 | rs4649446 | -0.356 | 0.494 | 0.193 | 0.867 | -0.004 | 0.007 | 0.007 | 0 | 0.17 | 0.18 | 0.24 | 0.21 | 0.12 | 0.15 |
| 1 | 233355529 | rs12087818 | 0.161 | 0.696 | -0.167 | 0.075 | -0.005 | -0.014 | -0.014 | 0 | 0.3 | 0.32 | 0.23 | 0.19 | 0.27 | 0.36 |
| 1 | 233356416 | rs7516050 | 0.064 | 0.787 | -0.219 | 0.096 | 0 | -0.001 | -0.001 | 0 | 0.38 | 0.33 | 0.27 | 0.22 | 0.35 | 0.39 |
| 1 | 233357504 | rs701178 | -0.851 | 1.018 | -0.669 | 0.339 | -0.053 | 0.056 | 0.056 | 0 | 0.65 | 0.9 | 0.79 | 0.8 | 0.64 | 0.71 |
| 1 | 233357608 | rs701177 | -0.794 | 1.094 | -0.378 | 0.377 | -0.006 | 0.031 | 0.031 | 0 | 0.63 | 0.7 | 0.77 | 0.8 | 0.64 | 0.71 |
| 1 | 233362930 | rs701176 | -0.226 | 1.13 | -0.264 | 0.663 | 0.009 | 0.077 | 0.077 | 0 | 0.18 | 0.27 | 0.48 | 0.53 | 0.23 | 0.25 |
| 1 | 233368289 | rs701173 | -0.31 | 1.082 | -0.369 | -0.095 | -0.015 | 0.053 | 0.053 | 0 | 0.61 | 0.67 | 0.75 | 0.87 | 0.61 | 0.65 |
| 1 | 233372917 | rs12141456 | -0.323 | 1.345 | -0.202 | -0.094 | -0.018 | 0.045 | 0.045 | 0.138 | 0.15 | 0.07 | 0.09 | 0.01 | 0.15 | 0.13 |
| 1 | 233374589 | rs10752750 | -0.037 | 1.557 | 0.262 | 0.269 | 0.019 | 0.033 | 0.033 | 0 | 0.82 | 0.71 | 0.77 | 0.66 | 0.75 | 0.59 |
| 1 | 233375713 | rs11800619 | -0.848 | 1.205 | 0.944 | 0.125 | 0.045 | 0.017 | 0.017 | 0.809 | 0.05 | 0.31 | 0.03 | 0.01 | 0.02 | 0.05 |
| 1 | 233379363 | rs11810439 | -1.438 | 1.144 | 0.694 | 0.095 | 0.047 | 0.013 | 0.013 | 3.208 | 0.07 | 0.46 | 0.04 | 0.01 | 0.02 | 0.05 |
| 1 | 233387566 | rs7546115 | 0.844 | 1.3 | 0.129 | 0.381 | -0.021 | -0.013 | -0.013 | 1.075 | 0.67 | 0.68 | 0.68 | 0.58 | 0.6 | 0.45 |
| 1 | 233397267 | rs10489808 | -0.32 | 1.014 | 0.173 | 0.32 | -0.005 | 0.005 | 0.005 | 0 | 0.18 | 0.16 | 0.14 | 0.12 | 0.2 | 0.26 |
| 1 | 233398713 | rs1033325 | -1.362 | 0.969 | 0.037 | 0.207 | -0.021 | 0.051 | 0.051 | 0 | 0.33 | 0.21 | 0.23 | 0.16 | 0.34 | 0.46 |
| 1 | 233400424 | rs1033322 | -1.68 | 0.857 | 0.534 | 0.161 | 0 | -0.017 | -0.017 | 0.052 | 0.37 | 0.57 | 0.25 | 0.17 | 0.35 | 0.46 |
| 1 | 233402674 | rs10910122 | -1.234 | 0.892 | 0.206 | 0.23 | -0.016 | 0.049 | 0.049 | 0 | 0.32 | 0.21 | 0.22 | 0.14 | 0.32 | 0.42 |
| 1 | 233406552 | rs12126986 | -0.178 | 0.963 | -0.294 | 0.248 | -0.027 | 0.052 | 0.052 | 1.299 | 0.14 | 0.03 | 0.09 | 0.05 | 0.15 | 0.2 |
| 1 | 233411878 | rs6697791 | -0.367 | 0.947 | -0.1 | 0.234 | 0.004 | -0.022 | -0.022 | 0 | 0.21 | 0.45 | 0.16 | 0.12 | 0.2 | 0.26 |
| 1 | 233412104 | rs12130559 | -0.367 | 0.947 | -0.1 | 0.234 | 0.004 | -0.022 | -0.022 | 0 | 0.21 | 0.45 | 0.16 | 0.12 | 0.2 | 0.26 |
| 1 | 233415367 | rs6683734 | -1.56 | 0.939 | -1.253 | 0.207 | 0.004 | 0.015 | 0.015 | 0 | 0.45 | 0.78 | 0.3 | 0.36 | 0.4 | 0.64 |
| 1 | 233418194 | rs11588896 | -0.633 | 1.006 | -1.233 | 0.085 | -0.026 | 0.068 | 0.068 | 1.288 | 0.14 | 0.02 | 0.08 | 0.02 | 0.13 | 0.17 |
| 1 | 233428992 | rs6701999 | -0.794 | 1.103 | -1.225 | 0.533 | -0.019 | -0.019 | -0.019 | 0.04 | 0.44 | 0.29 | 0.39 | 0.45 | 0.42 | 0.63 |
| 1 | 233429825 | rs6667845 | -0.285 | 0.668 | -1.091 | 0.187 | -0.008 | -0.01 | -0.01 | 0.471 | 0.38 | 0.58 | 0.55 | 0.31 | 0.37 | 0.31 |
| 1 | 233433500 | rs6700744 | -0.019 | 1.331 | -1.619 | 0.344 | -0.046 | -0.068 | -0.068 | 0 | 0.24 | 0.09 | 0.17 | 0.4 | 0.29 | 0.27 |
| 1 | 233433909 | rs1294348 | -0.239 | 0.773 | -1.357 | -0.33 | -0.009 | -0.038 | -0.038 | 0.729 | 0.65 | 0.55 | 0.66 | 0.73 | 0.68 | 0.7 |
| 1 | 233436403 | rs1294341 | -0.145 | 0.734 | -1.4 | -0.351 | -0.004 | -0.013 | -0.013 | 0 | 0.64 | 0.48 | 0.65 | 0.72 | 0.68 | 0.7 |
| 1 | 233437866 | rs4649215 | 0.971 | 0.695 | -1.46 | -0.367 | -0.122 | -0.154 | -0.154 | 0 | 0.23 | 0.02 | 0.16 | 0.39 | 0.29 | 0.27 |
| 1 | 233438952 | rs1294338 | -0.546 | 0.671 | -1.21 | -0.521 | -0.017 | -0.024 | -0.024 | 0 | 0.73 | 0.67 | 0.72 | 0.79 | 0.78 | 0.82 |
| 1 | 233439271 | rs1294337 | 0.594 | 0.582 | -1.109 | -0.537 | 0.002 | 0.002 | 0.002 | 0 | 0.41 | 0.46 | 0.49 | 0.33 | 0.39 | 0.44 |
| 1 | 233448608 | rs1294330 | -0.04 | 0.639 | -1.034 | -0.555 | 0 | 0.001 | 0.001 | 0 | 0.51 | 0.57 | 0.56 | 0.4 | 0.49 | 0.56 |
| 1 | 233451096 | rs12745867 | 0.641 | 0.836 | -1.052 | -0.437 | -0.015 | 0.027 | 0.027 | 0 | 0.08 | 0.04 | 0.04 | 0 | 0.1 | 0.08 |
| 1 | 233464877 | rs1294313 | 0.559 | 0.542 | -0.647 | -0.826 | -0.002 | -0.054 | -0.054 | 0.537 | 0.15 | 0.27 | 0.12 | 0.01 | 0.15 | 0.05 |
| 1 | 233467618 | rs1294308 | -0.186 | 0.071 | -0.568 | -1.379 | -0.005 | -0.045 | -0.045 | 1.472 | 0.72 | 0.73 | 0.84 | 0.93 | 0.75 | 0.79 |
| 1 | 233468448 | rs1294303 | -0.362 | 0.045 | -0.467 | -1.413 | 0.004 | -0.012 | -0.012 | 0 | 0.74 | 0.8 | 0.87 | 1 | 0.76 | 0.83 |
| 1 | 233469561 | rs1294296 | -0.188 | 0.045 | -0.47 | -1.413 | 0.001 | -0.045 | -0.045 | 0 | 0.72 | 0.76 | 0.86 | 1 | 0.76 | 0.83 |
| 1 | 233475385 | rs7513538 | 0.568 | 0.477 | -0.658 | -1.275 | -0.012 | 0.019 | 0.019 | 0.335 | 0.37 | 0.55 | 0.63 | 0.61 | 0.33 | 0.52 |
| 1 | 233476263 | rs1294266 | 0.326 | 0.58 | -0.8 | -1.209 | -0.054 | -0.054 | -0.054 | 0 | 0.36 | 0.2 | 0.24 | 0.38 | 0.42 | 0.32 |
| 1 | 233484687 | rs10797448 | 0.193 | 0.631 | -0.715 | -1.123 | -0.001 | -0.004 | -0.004 | 0.465 | 0.39 | 0.62 | 0.65 | 0.62 | 0.41 | 0.52 |
| 1 | 233489499 | rs7539016 | 0.332 | 0.637 | -0.369 | -1.047 | 0.004 | 0.029 | 0.029 | 0.078 | 0.38 | 0.52 | 0.62 | 0.56 | 0.41 | 0.48 |
| 1 | 233490776 | rs4649220 | 1.02 | 0.408 | -0.215 | -1.112 | -0.007 | 0.015 | 0.015 | 0 | 0.27 | 0.38 | 0.42 | 0.4 | 0.24 | 0.36 |
| 1 | 233491437 | rs4649222 | -0.355 | 0.385 | -0.421 | -1.393 | 0.011 | -0.009 | -0.009 | 0 | 0.65 | 0.73 | 0.81 | 0.99 | 0.72 | 0.83 |
| 1 | 233495568 | rs10752755 | -0.267 | 0.331 | -0.446 | -1.468 | -0.02 | -0.137 | -0.137 | 0.136 | 0.6 | 0.54 | 0.74 | 0.99 | 0.66 | 0.82 |
| 1 | 233497374 | rs1294260 | 1.486 | 0.257 | -0.639 | -1.421 | 0.013 | 0.022 | 0.022 | 0.188 | 0.14 | 0.09 | 0.09 | 0 | 0.08 | 0.03 |
| 1 | 233498236 | rs12039989 | 1.59 | 0.173 | -0.562 | -1.423 | -0.001 | 0.002 | 0.002 | 0 | 0.23 | 0.28 | 0.35 | 0.33 | 0.19 | 0.29 |
| 1 | 233498944 | rs1294259 | 1.431 | 0.172 | -0.851 | -1.402 | 0.013 | 0.014 | 0.014 | 0.228 | 0.14 | 0.1 | 0.09 | 0 | 0.08 | 0.03 |
| 1 | 233505173 | rs16858997 | 0.373 | 0.654 | -0.693 | -1.121 | -0.01 | -0.027 | -0.027 | 0 | 0.31 | 0.27 | 0.55 | 0.55 | 0.36 | 0.48 |
| 1 | 233508564 | rs10737197 | 0.335 | 0.333 | -0.595 | -1.202 | 0.003 | 0.06 | 0.06 | 0 | 0.51 | 0.6 | 0.54 | 0.78 | 0.5 | 0.64 |
| 1 | 233509581 | rs35225245 | 1.11 | 0.648 | -0.742 | -0.722 | -0.033 | -0.03 | -0.03 | 0 | 0.09 | 0.01 | 0.05 | 0.1 | 0.12 | 0.11 |
| 1 | 233512313 | rs1294248 | 0.372 | 0.426 | -0.121 | -0.906 | 0.058 | -0.002 | -0.002 | 1.949 | 0.06 | 0.23 | 0.07 | 0 | 0.05 | 0.02 |
| 1 | 233514934 | rs3795375 | 1.118 | -0.22 | -0.386 | -1.064 | 0.062 | -0.025 | -0.025 | 1.787 | 0.3 | 0.19 | 0.34 | 0.34 | 0.17 | 0.31 |
| 1 | 233515071 | rs963982 | 1.089 | -0.198 | -0.269 | -1.057 | 0.024 | -0.009 | -0.009 | 0.15 | 0.31 | 0.24 | 0.34 | 0.34 | 0.17 | 0.31 |
| 1 | 233518998 | rs1294233 | 0.01 | 0.063 | -0.349 | -1.132 | 0.082 | 0.085 | 0.085 | 0 | 0.29 | 0.5 | 0.38 | 0.4 | 0.4 | 0.34 |
| 1 | 233519393 | rs1294231 | -0.452 | 0.021 | -0.144 | -1.154 | -0.012 | 0.104 | 0.104 | 0 | 0.6 | 0.8 | 0.73 | 0.75 | 0.58 | 0.66 |
| 1 | 233519825 | rs1294230 | -0.094 | 0.016 | -0.145 | -1.156 | 0.082 | 0.085 | 0.085 | 0 | 0.29 | 0.5 | 0.38 | 0.4 | 0.4 | 0.34 |
| 1 | 233520144 | rs1294229 | -0.118 | 0.017 | -0.141 | -1.155 | 0.077 | 0.079 | 0.079 | 0 | 0.29 | 0.51 | 0.38 | 0.4 | 0.4 | 0.34 |
| 1 | 233524111 | rs1294207 | -0.451 | 0.015 | -0.542 | -1.144 | -0.012 | 0.101 | 0.101 | 0 | 0.6 | 0.8 | 0.73 | 0.75 | 0.57 | 0.66 |

| | | | | | | | | | | | | | | | | |
|---|---|---|---|---|---|---|---|---|---|---|---|---|---|---|---|---|
| 1 | 233525193 | rs1582192 | 1.403 | -0.006 | -0.511 | -1.141 | 0.06 | -0.022 | -0.022 | 0.433 | 0.31 | 0.17 | 0.33 | 0.34 | 0.17 | 0.31 |
| 1 | 233526832 | rs1294187 | -0.348 | 0.165 | -0.509 | -1.102 | -0.013 | 0.103 | 0.103 | 0 | 0.6 | 0.8 | 0.73 | 0.75 | 0.57 | 0.66 |
| 1 | 233531179 | rs1743266 | -0.261 | 0.258 | -0.143 | -1.119 | 0.023 | 0.011 | 0.011 | 0 | 0.35 | 0.52 | 0.28 | 0.35 | 0.46 | 0.37 |
| 1 | 233533418 | rs10910156 | 0.561 | 0.266 | -0.256 | -1.334 | 0.004 | -0.035 | -0.035 | 0 | 0.17 | 0.25 | 0.09 | 0.04 | 0.17 | 0.12 |
| 1 | 233534615 | rs12132550 | 0.615 | 0.266 | -0.225 | -1.334 | -0.001 | -0.01 | -0.01 | 0.318 | 0.16 | 0.21 | 0.09 | 0.04 | 0.17 | 0.12 |
| 1 | 233538845 | rs12131991 | 0.81 | 0.274 | -0.238 | -1.372 | 0.03 | -0.014 | -0.014 | 0 | 0.07 | 0.12 | 0.05 | 0.02 | 0.13 | 0.06 |
| 1 | 233540281 | rs10910157 | -0.893 | 0.225 | -0.194 | -1.328 | 0.045 | -0.002 | -0.002 | 0.008 | 0.49 | 0.33 | 0.41 | 0.44 | 0.58 | 0.54 |
| 1 | 233551226 | rs6691808 | -0.879 | 0.521 | 0.352 | -1.112 | 0.084 | 0.007 | 0.007 | 0 | 0.94 | 0.75 | 0.94 | 1 | 0.94 | 0.98 |
| 1 | 233554135 | rs7539909 | 0.574 | 0.535 | -0.108 | -1.135 | -0.091 | -0.087 | -0.087 | 1.907 | 0.27 | 0.06 | 0.19 | 0.34 | 0.39 | 0.35 |
| 1 | 233554853 | rs6687087 | 0.476 | 0.499 | -0.057 | -1.164 | 0 | -0.015 | -0.015 | 0.22 | 0.74 | 0.28 | 0.71 | 0.9 | 0.77 | 0.82 |
| 1 | 233561654 | rs12137253 | 0.574 | 0.548 | 0.043 | -1.092 | -0.003 | 0.001 | 0.001 | 0 | 0.16 | 0.16 | 0.08 | 0.04 | 0.17 | 0.12 |
| 1 | 233563269 | rs10910167 | 0.404 | 0.608 | -0.153 | -0.818 | 0.065 | -0.009 | -0.009 | 0 | 0.48 | 0.27 | 0.53 | 0.56 | 0.38 | 0.47 |
| 1 | 233570725 | rs4649316 | 0.596 | 0.791 | -0.214 | -0.487 | 0.028 | -0.05 | -0.05 | 0 | 0.35 | 0.21 | 0.46 | 0.56 | 0.32 | 0.46 |
| 1 | 233579288 | rs10797455 | 0.05 | 0.951 | -0.462 | -0.064 | 0.014 | 0.004 | 0.004 | 0 | 0.94 | 0.84 | 0.92 | 0.94 | 0.94 | 0.93 |
| 1 | 233599529 | rs11578460 | -0.32 | 1.137 | -0.944 | 0.033 | -0.034 | 0.012 | 0.012 | 0 | 0.69 | 0.87 | 0.78 | 0.76 | 0.65 | 0.77 |
| 1 | 233601025 | rs12134099 | 0.306 | 1.549 | -0.446 | -0.264 | 0.052 | -0.035 | -0.035 | 0 | 0.07 | 0.18 | 0.05 | 0 | 0.11 | 0.05 |
| 1 | 233601813 | rs4430273 | -0.464 | 1.078 | -0.294 | -0.409 | 0.01 | -0.038 | -0.038 | 0.374 | 0.31 | 0.46 | 0.34 | 0.19 | 0.33 | 0.35 |
| 1 | 233608565 | rs12032895 | 0.612 | 1.272 | -0.37 | 0.034 | 0.001 | 0 | 0 | 0 | 0.51 | 0.51 | 0.5 | 0.6 | 0.35 | 0.5 |
| 1 | 233619908 | rs6687124 | 0.617 | 1.407 | -0.45 | 0.113 | -0.02 | -0.017 | -0.017 | 0.637 | 0.09 | 0.05 | 0.19 | 0.1 | 0.18 | 0.18 |
| 1 | 233620063 | rs6424203 | -0.608 | 1.512 | -0.435 | 0.035 | 0.003 | -0.012 | -0.012 | 0 | 0.94 | 0.92 | 0.95 | 1 | 0.94 | 0.98 |
| 1 | 233620588 | rs10797461 | 0.586 | 1.468 | -0.191 | 0.358 | 0.027 | 0.047 | 0.047 | 0.767 | 0.84 | 0.73 | 0.72 | 0.76 | 0.77 | 0.75 |
| 1 | 233621682 | rs10910185 | 0.371 | 1.48 | -0.474 | 0.387 | -0.009 | -0.013 | -0.013 | 0 | 0.85 | 0.86 | 0.73 | 0.76 | 0.77 | 0.76 |
| 1 | 233621704 | rs10910186 | 0.299 | 1.486 | -0.472 | 0.352 | -0.008 | -0.002 | -0.002 | 0 | 0.85 | 0.86 | 0.76 | 0.9 | 0.77 | 0.8 |
| 1 | 233621844 | rs10797463 | 0.605 | 1.195 | -0.283 | 0.261 | 0.04 | 0.002 | 0.002 | 0 | 0.5 | 0.4 | 0.48 | 0.53 | 0.36 | 0.49 |
| 1 | 233622037 | rs12740471 | 0.122 | 1.181 | -0.869 | 0.494 | -0.002 | 0.009 | 0.009 | 0 | 0.1 | 0.09 | 0.21 | 0.24 | 0.15 | 0.23 |
| 1 | 233622420 | rs12741135 | -0.243 | 0.942 | -1.249 | 0.265 | -0.03 | 0.003 | 0.003 | 0 | 0.41 | 0.18 | 0.29 | 0.33 | 0.46 | 0.42 |
| 1 | 233623658 | rs9645292 | -0.399 | 0.846 | -1.67 | 0.128 | 0.033 | 0.031 | 0.031 | 0 | 0.51 | 0.12 | 0.42 | 0.4 | 0.62 | 0.51 |
| 1 | 233624614 | rs4233060 | 0.176 | 0.677 | -1.256 | 0.313 | 0.022 | 0.02 | 0.02 | 0 | 0.48 | 0.68 | 0.54 | 0.58 | 0.38 | 0.53 |
| 1 | 233625039 | rs12086262 | 0.801 | 0.6 | -1.396 | 0.488 | -0.006 | 0.009 | 0.009 | 0 | 0.26 | 0.13 | 0.23 | 0.24 | 0.27 | 0.14 |
| 1 | 233628771 | rs4129547 | 0.042 | -0.119 | -1.222 | 0.63 | -0.013 | -0.012 | -0.012 | 0 | 0.55 | 0.52 | 0.47 | 0.32 | 0.67 | 0.48 |
| 1 | 233633819 | rs10797465 | -0.027 | -1.032 | -1.199 | 1.001 | -0.069 | 0.039 | 0.039 | 0 | 0.66 | 0.53 | 0.65 | 0.45 | 0.86 | 0.69 |
| 1 | 233636625 | rs12047167 | -0.104 | -0.499 | -1.192 | 0.965 | -0.037 | 0.031 | 0.031 | 1.32 | 0.64 | 0.52 | 0.79 | 0.56 | 0.76 | 0.67 |
| 1 | 233643110 | rs1557129 | -0.205 | 0.367 | -1.781 | 0.649 | -0.078 | -0.025 | -0.025 | 0 | 0.19 | 0.02 | 0.11 | 0.2 | 0.22 | 0.2 |
| 1 | 233647842 | rs10910201 | 0.62 | -0.243 | -1.808 | 0.486 | -0.01 | 0.004 | 0.004 | 2.556 | 0.22 | 0.02 | 0.09 | 0.13 | 0.17 | 0.1 |
| 1 | 233653024 | rs10910204 | 0.925 | -0.018 | -2 | 0.642 | -0.013 | -0.02 | -0.02 | 0 | 0.16 | 0.06 | 0.41 | 0.21 | 0.22 | 0.18 |
| 1 | 233655782 | rs780214 | -0.467 | -0.267 | -2.007 | 0.654 | 0.011 | -0.025 | -0.025 | 1.144 | 0.76 | 0.81 | 0.86 | 0.58 | 0.84 | 0.69 |
| 1 | 233659142 | rs7540816 | 1.536 | -0.277 | -1.499 | 0.3 | 0.01 | -0.091 | -0.091 | 0 | 0.24 | 0.38 | 0.16 | 0.1 | 0.24 | 0.25 |
| 1 | 233659290 | rs1497090 | 0.716 | -0.279 | -1.507 | 0.32 | 0.004 | -0.013 | -0.013 | 0 | 0.4 | 0.39 | 0.57 | 0.31 | 0.49 | 0.43 |
| 1 | 233661003 | rs780222 | 0.503 | -0.3 | -1.406 | 0.321 | -0.001 | 0.001 | 0.001 | 0 | 0.35 | 0.37 | 0.28 | 0.27 | 0.35 | 0.27 |
| 1 | 233663942 | rs2000179 | 0.255 | -0.295 | -1.429 | 0.226 | -0.008 | 0.018 | 0.018 | 0 | 0.46 | 0.63 | 0.68 | 0.39 | 0.55 | 0.48 |
| 1 | 233664671 | rs12726051 | 0.633 | -0.064 | -1.752 | 0.226 | -0.004 | -0.001 | -0.001 | 0.415 | 0.09 | 0.09 | 0.09 | 0.01 | 0.16 | 0.12 |
| 1 | 233668726 | rs780225 | 0.418 | -0.568 | -1.814 | 0.576 | -0.033 | 0.023 | 0.023 | 0.971 | 0.55 | 0.3 | 0.39 | 0.57 | 0.48 | 0.49 |
| 1 | 233668751 | rs780226 | 0.78 | -0.682 | -1.827 | 0.656 | 0.028 | -0.045 | -0.045 | 0 | 0.3 | 0.19 | 0.17 | 0.4 | 0.2 | 0.21 |
| 1 | 233670385 | rs4649328 | -0.281 | -0.523 | -1.877 | 0.482 | 0.076 | 0.04 | 0.04 | 0 | 0.53 | 0.31 | 0.71 | 0.63 | 0.69 | 0.68 |
| 1 | 233673389 | rs10797467 | 0.507 | -0.008 | -1.816 | 0.797 | -0.129 | -0.08 | -0.08 | 0 | 0.22 | 0.03 | 0.4 | 0.29 | 0.39 | 0.32 |
| 1 | 233674529 | rs780235 | -0.593 | -0.55 | -1.194 | 0.813 | -0.063 | 0.196 | 0.196 | 0 | 0.79 | 0.59 | 0.75 | 0.56 | 0.9 | 0.81 |
| 1 | 233674625 | rs1083873 | -1.72 | -0.532 | -1.198 | 0.852 | -0.033 | 0.098 | 0.098 | 0 | 0.88 | 0.7 | 0.81 | 0.83 | 0.94 | 0.93 |
| 1 | 233676333 | rs3845310 | 0.808 | -0.413 | -1.334 | 0.789 | -0.12 | -0.02 | -0.02 | 0 | 0.22 | 0.03 | 0.35 | 0.21 | 0.37 | 0.28 |
| 1 | 233677177 | rs6659880 | 0.853 | -0.523 | -1.205 | 0.645 | -0.001 | -0.002 | -0.002 | 0 | 0.07 | 0.12 | 0.05 | 0.05 | 0.06 | 0.12 |
| 1 | 233678785 | rs780237 | -0.755 | -0.385 | -1.292 | 0.683 | -0.003 | 0.021 | 0.021 | 0.006 | 0.74 | 0.72 | 0.51 | 0.64 | 0.71 | 0.78 |
| 1 | 233679176 | rs12731522 | 1.004 | -0.405 | -1.466 | 0.714 | -0.016 | -0.022 | -0.022 | 0 | 0.15 | 0.07 | 0.1 | 0.3 | 0.18 | 0.25 |
| 1 | 233680563 | rs780240 | -0.729 | -0.395 | -1.579 | 0.363 | -0.003 | 0.032 | 0.032 | 0.013 | 0.74 | 0.69 | 0.51 | 0.64 | 0.7 | 0.78 |
| 1 | 233684352 | rs12739347 | 1.433 | -0.268 | -2.623 | -0.55 | -0.074 | 0.085 | 0.085 | 0.02 | 0.22 | 0.01 | 0.12 | 0.07 | 0.25 | 0.24 |
| 1 | 233685391 | rs780242 | -0.209 | -0.39 | -1.841 | -0.021 | 0.005 | 0.052 | 0.052 | 0 | 0.27 | 0.19 | 0.19 | 0.19 | 0.24 | 0.24 |
| 1 | 233689959 | rs780231 | 0.619 | -0.247 | -1.858 | 0.051 | 0.005 | 0.026 | 0.026 | 0 | 0.2 | 0.17 | 0.15 | 0.14 | 0.18 | 0.12 |
| 1 | 233695394 | rs12740439 | 0.699 | -0.204 | -2.445 | 0.17 | -0.122 | 0.12 | 0.12 | 0 | 0.46 | 0.06 | 0.45 | 0.26 | 0.51 | 0.39 |
| 1 | 233696745 | rs780246 | 1.11 | -0.135 | -2.377 | 0.209 | -0.028 | 0.016 | 0.016 | 0 | 0.16 | 0.02 | 0.11 | 0.15 | 0.17 | 0.12 |
| 1 | 233700237 | rs4375204 | 0.037 | -0.068 | -1.771 | 0.286 | 0.014 | 0.023 | 0.023 | 0 | 0.93 | 0.8 | 0.95 | 0.94 | 0.95 | 0.86 |
| 1 | 233701296 | rs6660091 | 0.156 | 0.07 | -2.322 | 0.189 | -0.077 | 0.135 | 0.135 | 0 | 0.46 | 0.12 | 0.47 | 0.27 | 0.52 | 0.37 |
| 1 | 233711232 | rs904068 | -0.313 | 0.675 | -1.652 | 0.739 | -0.014 | -0.009 | -0.009 | 0 | 0.4 | 0.65 | 0.46 | 0.59 | 0.39 | 0.48 |
| 1 | 233714931 | rs4649335 | 0.176 | 0.589 | -2.22 | 0.604 | -0.022 | 0.016 | 0.016 | 0 | 0.33 | 0.13 | 0.26 | 0.25 | 0.27 | 0.45 |
| 1 | 233718296 | rs780249 | 0.704 | 0.345 | -2.492 | 0.308 | 0.05 | 0.084 | 0.084 | 0 | 0.37 | 0.07 | 0.21 | 0.21 | 0.27 | 0.22 |
| 1 | 233723370 | rs6666161 | -0.483 | 0.405 | -2.269 | 0.78 | 0.072 | -0.035 | -0.035 | 0 | 0.58 | 0.74 | 0.62 | 0.56 | 0.72 | 0.73 |
| 1 | 233723638 | rs6681321 | -0.059 | 0.544 | -1.634 | 0.989 | 0.06 | 0.097 | 0.097 | 0 | 0.09 | 0.5 | 0.11 | 0.21 | 0.1 | 0.15 |
| 1 | 233724765 | rs11801736 | -0.001 | 0.547 | -1.582 | 0.988 | 0.074 | 0.165 | 0.165 | 0.068 | 0.08 | 0.42 | 0.09 | 0.21 | 0.05 | 0.14 |
| 1 | 233727007 | rs12116631 | 1.851 | 0.665 | -1.767 | 0.948 | -0.061 | -0.035 | -0.035 | 0 | 0.12 | 0.02 | 0.13 | 0.16 | 0.27 | 0.11 |
| 1 | 233729446 | rs10910220 | 0.66 | 0.391 | -1.738 | 0.882 | -0.084 | -0.063 | -0.063 | 0.018 | 0.29 | 0.1 | 0.35 | 0.37 | 0.45 | 0.29 |
| 1 | 233729852 | rs7542878 | -0.744 | 0.473 | -0.668 | 1.112 | 0.079 | 0.286 | 0.286 | 0 | 0.11 | 0.35 | 0.25 | 0.39 | 0.12 | 0.18 |
| 1 | 233733327 | rs1160006 | 0.182 | 0.426 | -0.77 | 1.007 | 0.028 | 0.015 | 0.015 | 0 | 0.51 | 0.37 | 0.53 | 0.4 | 0.62 | 0.6 |

| | | | | | | | | | | | | | | | | |
|---|---|---|---|---|---|---|---|---|---|---|---|---|---|---|---|---|
| 1 | 233733628 | rs1160005 | -0.918 | 0.34 | -0.947 | 0.804 | 0.071 | 0.078 | 0.078 | 0 | 0.63 | 0.89 | 0.79 | 0.79 | 0.74 | 0.78 |
| 1 | 233735038 | rs16859269 | 0.11 | 0.624 | -0.678 | 1.027 | 0.061 | 0.109 | 0.109 | 0 | 0.06 | 0.24 | 0.06 | 0.16 | 0.04 | 0.13 |
| 1 | 233735468 | rs1773950 | -1.656 | 0.22 | -1.117 | 0.94 | 0.005 | -0.01 | -0.01 | 0.097 | 0.85 | 0.87 | 0.89 | 0.79 | 0.91 | 0.85 |
| 1 | 233736384 | rs10797474 | 1.179 | 0.381 | -1.419 | 1.116 | -0.069 | -0.064 | -0.064 | 0 | 0.16 | 0.04 | 0.14 | 0.29 | 0.3 | 0.16 |
| 1 | 233737019 | rs1773952 | -1.671 | 0.383 | -1.26 | 0.955 | -0.003 | 0.012 | 0.012 | 0 | 0.83 | 0.78 | 0.85 | 0.79 | 0.85 | 0.84 |
| 1 | 233738249 | rs2226114 | -0.256 | -0.235 | -0.888 | 1.045 | 0 | 0 | 0 | 0 | 0.37 | 0.35 | 0.37 | 0.33 | 0.47 | 0.33 |
| 1 | 233738342 | rs1693229 | 0.78 | -0.353 | -0.871 | 0.983 | -0.025 | -0.011 | -0.011 | 0 | 0.27 | 0.35 | 0.16 | 0.2 | 0.17 | 0.17 |
| 1 | 233738404 | rs12119367 | 1.065 | -0.22 | -1.108 | 1.021 | -0.005 | 0.025 | 0.025 | 0.099 | 0.19 | 0.04 | 0.2 | 0.08 | 0.17 | 0.17 |
| 1 | 233741894 | rs1693216 | 0.506 | -0.042 | -1.036 | 1.144 | 0.015 | -0.017 | -0.017 | 0 | 0.28 | 0.23 | 0.3 | 0.42 | 0.2 | 0.22 |
| 1 | 233742345 | rs1083877 | -1.064 | 0.022 | -1.046 | 1.246 | -0.004 | 0 | 0 | 0 | 0.66 | 0.7 | 0.67 | 0.7 | 0.67 | 0.55 |
| 1 | 233743103 | rs780256 | -0.217 | 0.528 | -1.082 | 1.27 | 0.01 | -0.012 | -0.012 | 0 | 0.28 | 0.24 | 0.29 | 0.42 | 0.2 | 0.22 |
| 1 | 233746259 | rs680341 | 0.157 | 1.019 | -2.387 | -0.05 | -0.071 | -0.033 | -0.033 | 3.333 | 0.15 | 0.04 | 0.14 | 0.25 | 0.3 | 0.17 |
| 1 | 233749021 | rs636811 | -0.029 | 0.092 | -2.118 | -0.189 | 0.031 | 0.116 | 0.116 | 0 | 0.22 | 0.04 | 0.1 | 0.04 | 0.17 | 0.07 |
| 1 | 233753991 | rs635698 | -0.18 | 1.245 | -1.777 | 0.219 | -0.092 | 0.015 | 0.015 | 0 | 0.79 | 0.95 | 0.79 | 0.8 | 0.64 | 0.77 |
| 1 | 233756065 | rs1039128 | 0.231 | 0.005 | -0.989 | 0.525 | -0.005 | 0.001 | 0.001 | 0 | 0.67 | 0.65 | 0.82 | 0.66 | 0.76 | 0.72 |
| 1 | 233757257 | rs701204 | -0.059 | 0.118 | -0.985 | 0.648 | -0.017 | -0.005 | -0.005 | 0.083 | 0.16 | 0.23 | 0.18 | 0.18 | 0.11 | 0.14 |
| 1 | 233758218 | rs701209 | -0.181 | 0.371 | -0.793 | 0.614 | 0.005 | 0.012 | 0.012 | 0 | 0.76 | 0.74 | 0.84 | 0.74 | 0.74 | 0.8 |
| 1 | 233759918 | rs701213 | -0.983 | 0.03 | -0.981 | 0.501 | 0.001 | 0 | 0 | 0 | 0.31 | 0.33 | 0.39 | 0.34 | 0.25 | 0.34 |
| 1 | 233759939 | rs701214 | -0.27 | 0.266 | -0.762 | 0.347 | -0.014 | -0.018 | -0.018 | 0.573 | 0.14 | 0.18 | 0.17 | 0.08 | 0.1 | 0.08 |
| 1 | 233765598 | rs10910229 | -1.157 | -0.185 | -0.215 | -0.122 | -0.067 | -0.084 | -0.084 | 0 | 0.14 | 0.61 | 0.08 | 0.01 | 0.03 | 0.06 |
| 1 | 233765691 | rs701220 | -1.31 | 0.1 | -1.304 | -0.521 | 0.026 | 0.136 | 0.136 | 0 | 0.39 | 0.8 | 0.27 | 0.16 | 0.31 | 0.27 |
| 1 | 233766695 | rs701223 | -1.275 | -0.035 | -1.331 | 0.346 | 0.066 | 0.007 | 0.007 | 0 | 0.38 | 0.8 | 0.27 | 0.35 | 0.25 | 0.3 |
| 1 | 233767059 | rs701226 | -1.275 | -0.035 | -1.331 | 0.346 | 0.066 | 0.007 | 0.007 | 0 | 0.38 | 0.8 | 0.27 | 0.35 | 0.25 | 0.3 |
| 1 | 233771247 | rs701229 | -1.091 | -0.193 | -0.928 | 0.865 | 0.003 | -0.012 | -0.012 | 0.263 | 0.3 | 0.72 | 0.44 | 0.45 | 0.28 | 0.32 |
| 1 | 233771619 | rs10910230 | -0.292 | -0.251 | -1.238 | 0.615 | -0.069 | 0.001 | 0.001 | 0.454 | 0.15 | 0.01 | 0.14 | 0.14 | 0.22 | 0.15 |
| 1 | 233771854 | rs12045321 | -0.682 | -0.219 | -1.063 | 0.594 | -0.035 | 0.042 | 0.042 | 0 | 0.19 | 0.07 | 0.18 | 0.14 | 0.26 | 0.15 |
| 1 | 233775512 | rs4649240 | 0.755 | -0.256 | -1.202 | 0.592 | -0.012 | -0.012 | -0.012 | 0 | 0.37 | 0.34 | 0.6 | 0.46 | 0.41 | 0.41 |
| 1 | 233780638 | rs7536764 | -1.097 | -0.634 | -0.3 | 0.372 | -0.003 | -0.007 | -0.007 | 0 | 0.33 | 0.67 | 0.4 | 0.21 | 0.3 | 0.19 |
| 1 | 233782052 | rs6668792 | -0.382 | -0.462 | 0.115 | 0.37 | 0.001 | 0.013 | 0.013 | 0 | 0.26 | 0.24 | 0.21 | 0.15 | 0.27 | 0.16 |
| 1 | 233784222 | rs6667354 | 0.768 | -0.555 | -0.438 | 0.319 | -0.089 | -0.08 | -0.08 | 0 | 0.23 | 0.11 | 0.44 | 0.52 | 0.39 | 0.48 |
| 1 | 233786170 | rs2282428 | -0.864 | -0.726 | 0.103 | 0.19 | 0.018 | 0.022 | 0.022 | 0 | 0.51 | 0.59 | 0.63 | 0.69 | 0.63 | 0.66 |
| 1 | 233788316 | rs12144885 | 0.906 | -0.306 | -0.326 | 0.46 | -0.081 | -0.088 | -0.088 | 0.103 | 0.22 | 0.11 | 0.44 | 0.39 | 0.37 | 0.36 |
| 1 | 233791469 | rs701232 | -0.672 | -0.217 | 0.105 | 0.546 | -0.004 | -0.001 | -0.001 | 0 | 0.43 | 0.4 | 0.52 | 0.46 | 0.52 | 0.44 |
| 1 | 233794459 | rs6692174 | 0.331 | -0.047 | -0.037 | 0.26 | -0.01 | 0.051 | 0.051 | 0.564 | 0.24 | 0.12 | 0.18 | 0.12 | 0.27 | 0.15 |
| 1 | 233796651 | rs780083 | -1.92 | -0.763 | -0.118 | 0.112 | -0.001 | 0 | 0 | 0 | 0.68 | 0.7 | 0.66 | 0.67 | 0.67 | 0.68 |
| 1 | 233800484 | rs630938 | -1.93 | -0.836 | -0.715 | -0.184 | 0.015 | 0.033 | 0.033 | 0 | 0.87 | 0.79 | 0.91 | 0.76 | 0.87 | 0.83 |
| 1 | 233800613 | rs1693210 | 0.296 | -0.915 | -0.744 | -0.292 | -0.007 | -0.016 | -0.016 | 0 | 0.35 | 0.29 | 0.45 | 0.52 | 0.34 | 0.5 |
| 1 | 233805172 | rs7552738 | 0.763 | -0.908 | -0.43 | -0.047 | -0.001 | -0.004 | -0.004 | 0 | 0.26 | 0.25 | 0.2 | 0.14 | 0.3 | 0.22 |
| 1 | 233805762 | rs7544073 | 1.077 | -0.904 | -0.57 | -0.107 | -0.014 | 0.058 | 0.058 | 1.521 | 0.23 | 0.12 | 0.19 | 0.05 | 0.29 | 0.15 |
| 1 | 233809111 | rs696867 | -0.111 | -1.109 | -0.576 | -0.234 | 0.001 | 0 | 0 | 0 | 0.56 | 0.55 | 0.53 | 0.52 | 0.49 | 0.52 |
| 1 | 233811744 | rs565657 | 0.339 | -1.17 | -0.333 | -0.262 | -0.001 | -0.004 | -0.004 | 0 | 0.34 | 0.34 | 0.25 | 0.21 | 0.35 | 0.26 |
| 1 | 233811877 | rs611394 | -1.481 | -1.123 | -0.342 | -0.191 | -0.001 | 0.034 | 0.034 | 0 | 0.78 | 0.69 | 0.72 | 0.65 | 0.79 | 0.74 |
| 1 | 233815034 | rs3893324 | 0.922 | -1.146 | -0.813 | 0.005 | 0.018 | 0.044 | 0.044 | 0 | 0.29 | 0.15 | 0.15 | 0.15 | 0.21 | 0.21 |
| 1 | 233816146 | rs517580 | 1.109 | -1.103 | -0.489 | -0.037 | 0.003 | -0.061 | -0.061 | 0 | 0.27 | 0.32 | 0.2 | 0.04 | 0.32 | 0.15 |
| 1 | 233816651 | rs4649345 | 1.388 | -1.079 | -0.851 | 0.39 | -0.017 | 0.022 | 0.022 | 0 | 0.08 | 0.12 | 0.05 | 0.17 | 0.03 | 0.1 |
| 1 | 233823925 | rs595191 | -0.265 | -0.818 | -0.929 | 0.387 | 0.002 | 0.002 | 0.002 | 0 | 0.51 | 0.71 | 0.5 | 0.56 | 0.53 | 0.51 |
| 1 | 233843795 | rs4142987 | 0.669 | -0.927 | -1.384 | 0.137 | 0.036 | -0.069 | -0.069 | 0 | 0.36 | 0.06 | 0.39 | 0.4 | 0.32 | 0.44 |
| 1 | 233846259 | rs12730679 | 1.251 | -0.945 | -1.122 | 0.009 | -0.004 | -0.003 | -0.003 | 0 | 0.18 | 0.23 | 0.1 | 0.15 | 0.14 | 0.16 |
| 1 | 233846876 | rs603932 | -2.619 | -0.905 | -1.164 | -0.033 | 0 | -0.006 | -0.006 | 0.389 | 0.94 | 0.96 | 0.98 | 0.92 | 0.95 | 0.91 |
| 1 | 233855973 | rs600400 | 1.429 | -0.864 | -0.655 | -0.113 | -0.052 | 0.06 | 0.06 | 0 | 0.25 | 0.13 | 0.22 | 0.04 | 0.37 | 0.18 |
| 1 | 233858751 | rs963666 | 1.496 | -1.108 | -1.001 | 0.049 | -0.023 | -0.083 | -0.083 | 0.377 | 0.29 | 0.01 | 0.36 | 0.34 | 0.29 | 0.36 |
| 1 | 233859008 | rs12144732 | 0.377 | -1.147 | -0.927 | 0.06 | 0.029 | -0.006 | -0.006 | 0 | 0.47 | 0.25 | 0.46 | 0.49 | 0.43 | 0.52 |
| 1 | 233859026 | rs12757689 | 1.16 | -0.668 | -0.549 | 0.08 | -0.004 | -0.004 | -0.004 | 0 | 0.19 | 0.23 | 0.1 | 0.15 | 0.14 | 0.17 |
| 1 | 233861559 | rs4595319 | 0.242 | -0.037 | -0.281 | 0.064 | 0.001 | -0.009 | -0.009 | 0.029 | 0.58 | 0.55 | 0.66 | 0.63 | 0.49 | 0.62 |
| 1 | 233869627 | rs12025703 | -0.22 | 0.252 | -0.203 | 0.221 | 0.016 | 0.102 | 0.102 | 0 | 0.21 | 0.44 | 0.16 | 0.38 | 0.18 | 0.28 |
| 1 | 233870153 | rs10157147 | 0.778 | 0.269 | 0.085 | 0.16 | 0.007 | 0.074 | 0.074 | 0 | 0.15 | 0.24 | 0.12 | 0.32 | 0.14 | 0.19 |
| 1 | 233871030 | rs651185 | 1.269 | 0.423 | -0.247 | 0.09 | -0.031 | 0.01 | 0.01 | 0 | 0.2 | 0.12 | 0.34 | 0.1 | 0.34 | 0.16 |
| 1 | 233879281 | rs651210 | -0.858 | 0.348 | -0.201 | -0.033 | -0.004 | -0.005 | -0.005 | 0 | 0.42 | 0.58 | 0.5 | 0.45 | 0.51 | 0.44 |
| 1 | 233887113 | rs6676367 | 0.243 | 0.521 | -0.07 | 0.037 | 0.025 | 0.079 | 0.079 | 0 | 0.19 | 0.33 | 0.14 | 0.37 | 0.18 | 0.26 |
| 1 | 233887989 | rs1603133 | 1.202 | -0.253 | -0.514 | -0.362 | 0.006 | -0.059 | -0.059 | 0.883 | 0.27 | 0.14 | 0.35 | 0.42 | 0.28 | 0.37 |
| 1 | 233893454 | rs1126292 | 2.029 | -0.548 | -0.517 | -0.58 | 0.009 | 0.011 | 0.011 | 0 | 0.23 | 0.24 | 0.12 | 0.11 | 0.16 | 0.17 |
| 1 | 233895488 | rs1389967 | -1.186 | -0.581 | -0.511 | -0.613 | -0.042 | 0.054 | 0.054 | 0 | 0.73 | 0.86 | 0.63 | 0.9 | 0.62 | 0.8 |
| 1 | 233895684 | rs12036656 | -0.272 | -0.505 | -0.277 | -0.446 | 0.033 | -0.001 | -0.001 | 0 | 0.46 | 0.72 | 0.28 | 0.49 | 0.34 | 0.43 |
| 1 | 233897291 | rs12037816 | 0.023 | -0.503 | -0.249 | -0.445 | 0.014 | 0.079 | 0.079 | 0 | 0.23 | 0.45 | 0.15 | 0.37 | 0.19 | 0.26 |
| 1 | 233897400 | rs1339430 | 1.673 | -0.473 | -0.308 | -0.439 | 0 | -0.001 | -0.001 | 0.124 | 0.24 | 0.27 | 0.13 | 0.11 | 0.16 | 0.17 |
| 1 | 233911242 | rs6671309 | 0.568 | -0.299 | -0.421 | -0.399 | 0.037 | 0.045 | 0.045 | 0 | 0.19 | 0.32 | 0.14 | 0.32 | 0.2 | 0.2 |
| 1 | 233917033 | rs7530423 | 1.379 | -0.621 | -0.516 | -0.433 | -0.022 | -0.05 | -0.05 | 0.77 | 0.25 | 0.32 | 0.13 | 0.09 | 0.16 | 0.17 |
| 1 | 233917201 | rs16859566 | 1.276 | -0.616 | -0.53 | -0.412 | -0.033 | -0.04 | -0.04 | 0.249 | 0.26 | 0.36 | 0.13 | 0.14 | 0.16 | 0.23 |
| 1 | 233921835 | rs12026196 | -0.252 | -0.453 | -0.826 | -0.323 | 0.042 | 0.006 | 0.006 | 0 | 0.47 | 0.78 | 0.29 | 0.47 | 0.34 | 0.42 |

| | | | | | | | | | | | | | | | | |
|---|---|---|---|---|---|---|---|---|---|---|---|---|---|---|---|---|
| 1 | 233926128 | rs960899 | -0.548 | -0.593 | -0.517 | -0.443 | -0.001 | 0 | 0 | 0.007 | 0.62 | 0.65 | 0.69 | 0.65 | 0.52 | 0.64 |
| 1 | 233927002 | rs10797484 | -0.277 | -0.682 | -0.355 | -0.46 | -0.005 | 0 | 0 | 0 | 0.39 | 0.43 | 0.57 | 0.56 | 0.35 | 0.47 |
| 1 | 233927721 | rs16859591 | 0.131 | -0.711 | -0.315 | -0.392 | 0.013 | -0.035 | -0.035 | 0 | 0.36 | 0.27 | 0.54 | 0.49 | 0.36 | 0.4 |
| 1 | 233928550 | rs11802935 | -0.854 | -0.584 | -0.186 | -0.5 | -0.029 | 0.005 | 0.005 | 0.196 | 0.65 | 0.78 | 0.73 | 0.65 | 0.56 | 0.66 |
| 1 | 233928628 | rs16859597 | 1.291 | -0.559 | -0.58 | -0.42 | -0.028 | -0.063 | -0.063 | 0 | 0.26 | 0.36 | 0.13 | 0.09 | 0.16 | 0.17 |
| 1 | 233928834 | rs16859600 | 1.1 | -0.635 | -0.64 | -0.355 | -0.002 | 0.001 | 0.001 | 0 | 0.09 | 0.13 | 0.2 | 0.06 | 0.08 | 0.04 |
| 1 | 233934559 | rs6661049 | 1.308 | -0.861 | -0.877 | -0.272 | -0.023 | -0.05 | -0.05 | 0 | 0.25 | 0.31 | 0.12 | 0.09 | 0.16 | 0.17 |
| 1 | 233937019 | rs12117523 | 0.298 | -1.043 | -1.025 | -0.392 | 0.004 | 0 | 0 | 0 | 0.46 | 0.44 | 0.44 | 0.59 | 0.39 | 0.57 |
| 1 | 233938537 | rs1494895 | 0.895 | -0.8 | -1.052 | -0.154 | -0.006 | -0.068 | -0.068 | 0 | 0.22 | 0.13 | 0.32 | 0.5 | 0.26 | 0.43 |
| 1 | 233938656 | rs1494896 | 0.895 | -0.8 | -1.052 | -0.154 | -0.006 | -0.068 | -0.068 | 0.538 | 0.22 | 0.13 | 0.32 | 0.5 | 0.26 | 0.43 |
| 1 | 233943028 | rs12068756 | 0.367 | -0.857 | -0.905 | -0.167 | -0.028 | 0.019 | 0.019 | 0.314 | 0.1 | 0.3 | 0.07 | 0.07 | 0.04 | 0.09 |
| 1 | 233944143 | rs4649260 | 0.783 | -0.909 | -1.04 | -0.177 | 0.001 | -0.016 | -0.016 | 0 | 0.23 | 0.19 | 0.33 | 0.44 | 0.26 | 0.36 |
| 1 | 233944248 | rs4649261 | -0.626 | -0.739 | -1.09 | -0.383 | 0.009 | 0.003 | 0.003 | 0 | 0.42 | 0.41 | 0.59 | 0.57 | 0.39 | 0.51 |
| 1 | 233947454 | rs7530809 | -0.758 | -0.524 | -1.084 | -0.459 | 0.016 | -0.004 | -0.004 | 0 | 0.65 | 0.63 | 0.71 | 0.65 | 0.59 | 0.73 |
| 1 | 233950276 | rs7547933 | 1.343 | -0.77 | -1.224 | -0.659 | 0.011 | 0.005 | 0.005 | 0 | 0.25 | 0.21 | 0.11 | 0.05 | 0.18 | 0.16 |
| 1 | 233951659 | rs1472283 | -0.664 | -0.37 | -1.368 | -0.452 | 0.009 | -0.059 | -0.059 | 0 | 0.69 | 0.82 | 0.59 | 0.49 | 0.68 | 0.47 |
| 1 | 233954283 | rs1494899 | 0.295 | -0.287 | -1.407 | -0.357 | 0.027 | 0.015 | 0.015 | 0 | 0.32 | 0.37 | 0.23 | 0.38 | 0.38 | 0.25 |
| 1 | 233955738 | rs2884385 | 0.881 | -0.624 | -1.951 | -0.48 | 0.041 | 0.099 | 0.099 | 0 | 0.26 | 0.12 | 0.1 | 0.01 | 0.18 | 0.16 |
| 1 | 233957343 | rs1948501 | -0.756 | -0.39 | -1.868 | 0.456 | -0.053 | -0.015 | -0.015 | 0.147 | 0.8 | 0.97 | 0.67 | 0.79 | 0.74 | 0.66 |
| 1 | 233957385 | rs1948502 | -0.328 | -0.28 | -1.857 | 0.55 | -0.064 | -0.054 | -0.054 | 0 | 0.77 | 0.94 | 0.63 | 0.69 | 0.7 | 0.54 |
| 1 | 233976729 | rs12086015 | 0.536 | -0.616 | -0.961 | 0.867 | -0.019 | 0.062 | 0.062 | 0 | 0.12 | 0.2 | 0.11 | 0.28 | 0.11 | 0.04 |
| 1 | 233980768 | rs16859717 | 0.482 | -0.418 | -1.106 | 0.629 | -0.003 | 0.003 | 0.003 | 0 | 0.08 | 0.12 | 0.21 | 0.08 | 0.08 | 0.03 |
| 1 | 233988285 | rs6679063 | -0.785 | -1.1 | -1.209 | 0.706 | 0.011 | -0.012 | -0.012 | 0 | 0.23 | 0.23 | 0.14 | 0.37 | 0.11 | 0.25 |
| 1 | 233991001 | rs10797493 | -1.057 | -1.102 | -1.228 | 0.367 | -0.001 | 0 | 0 | 0 | 0.68 | 0.67 | 0.41 | 0.66 | 0.64 | 0.59 |
| 1 | 233993871 | rs2088108 | -0.363 | -1.16 | -0.956 | 0.475 | -0.052 | -0.005 | -0.005 | 0 | 0.22 | 0.34 | 0.12 | 0.24 | 0.11 | 0.17 |
| 1 | 233997870 | rs923975 | 0.225 | -1.287 | -1.317 | 0.356 | 0.03 | 0.01 | 0.01 | 0 | 0.5 | 0.29 | 0.3 | 0.53 | 0.53 | 0.46 |
| 1 | 233998067 | rs923976 | 1.028 | -1.289 | -1.32 | 0.358 | -0.017 | 0.015 | 0.015 | 0 | 0.4 | 0.28 | 0.25 | 0.29 | 0.47 | 0.32 |
| 1 | 233999167 | rs16859749 | 0.032 | -1.222 | -1.052 | 0.386 | -0.026 | 0.002 | 0.002 | 0 | 0.15 | 0.24 | 0.1 | 0.13 | 0.07 | 0.11 |
| 1 | 233999944 | rs10910288 | -0.65 | -0.914 | -1.025 | 0.46 | -0.008 | -0.008 | -0.008 | 0.471 | 0.61 | 0.57 | 0.39 | 0.66 | 0.62 | 0.58 |
| 1 | 234009231 | rs1160305 | -0.946 | -1.015 | -1.498 | 0.258 | 0.014 | 0.048 | 0.048 | 0 | 0.46 | 0.69 | 0.64 | 0.66 | 0.45 | 0.49 |
| 1 | 234011119 | rs4649366 | 0.596 | -0.971 | -1.355 | 0.388 | 0.004 | 0.024 | 0.024 | 0 | 0.24 | 0.45 | 0.5 | 0.3 | 0.27 | 0.34 |
| 1 | 234016310 | rs16859773 | 1.233 | -0.976 | -1.405 | 0.384 | 0 | -0.001 | -0.001 | 0 | 0.18 | 0.14 | 0.1 | 0.22 | 0.15 | 0.05 |
| 1 | 234017719 | rs16859784 | 0.885 | -1.071 | -1.446 | 0.337 | 0.002 | -0.003 | -0.003 | 0 | 0.19 | 0.14 | 0.1 | 0.22 | 0.15 | 0.05 |
| 1 | 234017784 | rs4439408 | -0.422 | -1.102 | -1.482 | 0.251 | -0.018 | 0.059 | 0.059 | 0 | 0.71 | 0.6 | 0.46 | 0.47 | 0.72 | 0.55 |
| 1 | 234018653 | rs10910301 | -1.6 | -1.118 | -1.463 | 0.244 | -0.003 | 0.033 | 0.033 | 0 | 0.8 | 0.66 | 0.53 | 0.71 | 0.78 | 0.69 |
| 1 | 234020481 | rs4442431 | 1.283 | -1.085 | -1.585 | 0.21 | -0.008 | 0.099 | 0.099 | 0 | 0.45 | 0.3 | 0.31 | 0.11 | 0.5 | 0.37 |
| 1 | 234021539 | rs7539340 | 0.159 | -0.612 | -1.21 | 0.479 | -0.013 | 0.054 | 0.054 | 0 | 0.68 | 0.55 | 0.45 | 0.47 | 0.68 | 0.52 |
| 1 | 234036807 | rs4303127 | -0.15 | 0.135 | -1.107 | 0.341 | -0.002 | 0.001 | 0.001 | 0.114 | 0.41 | 0.4 | 0.26 | 0.3 | 0.43 | 0.44 |
| 1 | 234039857 | rs10910308 | -0.369 | -0.356 | -1.034 | 0.248 | 0.004 | 0.004 | 0.004 | 0.639 | 0.46 | 0.42 | 0.31 | 0.45 | 0.47 | 0.54 |
| 1 | 234046681 | rs6693898 | -0.025 | 0.15 | -0.709 | -0.707 | 0.033 | -0.094 | -0.094 | 0 | 0.7 | 0.58 | 0.78 | 0.94 | 0.67 | 0.78 |
| 1 | 234054014 | rs1937266 | -0.334 | -0.763 | -1.118 | -0.253 | 0.003 | -0.022 | -0.022 | 0 | 0.41 | 0.16 | 0.57 | 0.59 | 0.41 | 0.54 |
| 1 | 234063801 | rs11802061 | 0.469 | -1.016 | -0.368 | -0.274 | 0.032 | 0.028 | 0.028 | 0.185 | 0.49 | 0.24 | 0.3 | 0.38 | 0.54 | 0.45 |
| 1 | 234064302 | rs6702789 | 0.809 | -0.804 | -0.452 | -0.29 | -0.014 | -0.016 | -0.016 | 4.357 | 0.11 | 0.06 | 0.07 | 0.12 | 0.12 | 0.15 |
| 1 | 234065250 | rs11811551 | 1.436 | -0.781 | -0.493 | -0.276 | -0.032 | 0.046 | 0.046 | 0 | 0.37 | 0.17 | 0.22 | 0.26 | 0.41 | 0.3 |
| 1 | 234066333 | rs11576361 | -0.509 | -0.712 | -0.597 | -0.209 | 0.007 | 0.006 | 0.006 | 0 | 0.46 | 0.43 | 0.67 | 0.62 | 0.46 | 0.56 |
| 1 | 234068161 | rs4920239 | 0.214 | -0.716 | -1.419 | -0.158 | -0.007 | -0.086 | -0.086 | 0 | 0.33 | 0.1 | 0.55 | 0.47 | 0.34 | 0.42 |
| 1 | 234073691 | rs6667430 | -0.696 | -0.737 | -0.985 | -0.079 | -0.03 | -0.022 | -0.022 | 1.047 | 0.33 | 0.57 | 0.37 | 0.27 | 0.25 | 0.2 |
| 1 | 234075610 | rs7543553 | 0.2 | -0.958 | -1.053 | -0.475 | -0.056 | -0.114 | -0.114 | 1.275 | 0.17 | 0.34 | 0.08 | 0 | 0.09 | 0.02 |
| 1 | 234080599 | rs1892298 | 0.556 | -0.59 | -1.574 | -0.307 | -0.106 | -0.166 | -0.166 | 0.086 | 0.24 | 0.02 | 0.53 | 0.49 | 0.35 | 0.43 |
| 1 | 234080872 | rs4506523 | -0.032 | -0.808 | -0.777 | -0.506 | -0.05 | -0.048 | -0.048 | 0 | 0.23 | 0.44 | 0.12 | 0.1 | 0.15 | 0.13 |
| 1 | 234086848 | rs4339928 | -0.028 | 0.282 | -0.903 | -0.39 | -0.023 | 0.019 | 0.019 | 0 | 0.67 | 0.8 | 0.79 | 0.73 | 0.65 | 0.7 |
| 1 | 234095813 | rs4920264 | 0.352 | -0.336 | -0.247 | -0.594 | -0.001 | -0.039 | -0.039 | 0.311 | 0.46 | 0.48 | 0.34 | 0.09 | 0.53 | 0.36 |
| 1 | 234098168 | rs4330979 | -0.404 | -0.603 | -0.146 | -0.292 | -0.005 | 0.013 | 0.013 | 0 | 0.08 | 0.12 | 0.14 | 0.23 | 0.04 | 0.08 |
| 1 | 234099392 | rs9435588 | -0.164 | -0.754 | 0.2 | -0.382 | 0.02 | 0.083 | 0.083 | 0 | 0.46 | 0.26 | 0.52 | 0.67 | 0.44 | 0.49 |
| 1 | 234099819 | rs12049593 | 0.31 | -0.85 | 0.065 | 0.01 | -0.005 | -0.029 | -0.029 | 0.253 | 0.05 | 0 | 0.04 | 0.17 | 0.04 | 0.09 |
| 1 | 234101540 | rs2186104 | -0.249 | -1.151 | -0.057 | 0.028 | -0.001 | -0.012 | -0.012 | 0 | 0.27 | 0.23 | 0.34 | 0.62 | 0.27 | 0.39 |
| 1 | 234105211 | rs7556173 | 0.014 | -1.069 | -0.147 | 0.031 | -0.011 | -0.04 | -0.04 | 0 | 0.24 | 0.12 | 0.24 | 0.56 | 0.27 | 0.39 |
| 1 | 234107883 | rs11587514 | -1.113 | -1.132 | 0.231 | -0.052 | -0.006 | -0.02 | -0.02 | 0 | 0.38 | 0.59 | 0.42 | 0.8 | 0.32 | 0.55 |
| 1 | 234110781 | rs4348787 | -1.662 | -0.944 | 0.042 | -0.144 | -0.08 | 0.117 | 0.117 | 0 | 0.65 | 0.85 | 0.72 | 0.91 | 0.54 | 0.66 |
| 1 | 234110847 | rs12045592 | 0.034 | -0.346 | 0.461 | 0.546 | 0 | 0.023 | 0.023 | 0 | 0.21 | 0.23 | 0.22 | 0.39 | 0.23 | 0.3 |
| 1 | 234115914 | rs7541879 | -0.837 | -0.196 | 0.277 | 0.486 | 0.027 | 0.106 | 0.106 | 0 | 0.1 | 0.3 | 0.17 | 0.25 | 0.08 | 0.08 |
| 1 | 234119810 | rs12759054 | -0.302 | -0.01 | -0.524 | 0.475 | -0.01 | -0.05 | -0.05 | 0 | 0.25 | 0.15 | 0.18 | 0.38 | 0.28 | 0.37 |
| 1 | 234121014 | rs1937264 | 0.055 | -0.272 | -0.746 | 0.184 | 0.001 | -0.014 | -0.014 | 0.378 | 0.22 | 0.09 | 0.14 | 0.23 | 0.21 | 0.22 |
| 1 | 234131370 | rs1937261 | -0.553 | -0.267 | -0.162 | 0.117 | -0.003 | 0.014 | 0.014 | 0 | 0.28 | 0.33 | 0.33 | 0.49 | 0.25 | 0.25 |
| 1 | 234139581 | rs4623775 | -1.181 | -0.479 | -0.224 | 0.232 | -0.008 | 0.084 | 0.084 | 0 | 0.1 | 0.15 | 0.22 | 0.43 | 0.07 | 0.15 |
| 1 | 234142548 | rs9435539 | -0.535 | -0.51 | -0.688 | -0.002 | -0.107 | -0.174 | -0.174 | 0 | 0.79 | 0.99 | 0.66 | 0.58 | 0.69 | 0.65 |
| 1 | 234142710 | rs6662498 | 0.593 | -0.721 | -0.283 | -0.109 | -0.03 | -0.036 | -0.036 | 0 | 0.07 | 0.15 | 0.02 | 0 | 0.01 | 0.01 |
| 1 | 234142786 | rs4084672 | 1.812 | -0.979 | -0.389 | -0.403 | -0.099 | 0.21 | 0.21 | 0 | 0.37 | 0.09 | 0.24 | 0.08 | 0.44 | 0.32 |
| 1 | 234143175 | rs6586363 | -0.53 | -0.888 | -0.301 | -0.359 | -0.044 | -0.094 | -0.094 | 0 | 0.77 | 0.91 | 0.65 | 0.58 | 0.69 | 0.64 |

| | | | | | | | | | | | | | | | | |
|---|---|---|---|---|---|---|---|---|---|---|---|---|---|---|---|---|
| 1 | 234148558 | rs7548651 | -0.674 | -0.982 | -0.236 | -0.403 | -0.009 | 0.007 | 0.007 | 0 | 0.33 | 0.59 | 0.4 | 0.48 | 0.25 | 0.32 |
| 1 | 234154862 | rs1458597 | -1.174 | -0.999 | 0.285 | -0.389 | 0.016 | 0.201 | 0.201 | 0.064 | 0.05 | 0.25 | 0.05 | 0.16 | 0.02 | 0.1 |
| 1 | 234163840 | rs2358194 | 1.563 | -1.062 | 0.282 | -0.474 | -0.014 | 0.062 | 0.062 | 0 | 0.4 | 0.3 | 0.26 | 0.09 | 0.44 | 0.39 |
| 1 | 234164051 | rs4244367 | 1.099 | -1.048 | 0.557 | -0.452 | 0.002 | -0.019 | -0.019 | 0 | 0.46 | 0.53 | 0.46 | 0.34 | 0.48 | 0.45 |
| 1 | 234164519 | rs4920167 | 1.585 | -1.04 | 0.016 | -0.433 | -0.102 | 0.088 | 0.088 | 0 | 0.39 | 0.11 | 0.26 | 0.22 | 0.46 | 0.38 |
| 1 | 234166056 | rs4575150 | -0.718 | -0.623 | 0.137 | -0.255 | -0.097 | -0.163 | -0.163 | 0.062 | 0.79 | 0.97 | 0.68 | 0.57 | 0.72 | 0.66 |
| 1 | 234166129 | rs2199074 | -2.201 | -0.587 | 0.243 | -0.93 | -0.013 | 0.007 | 0.007 | 2.686 | 0.14 | 0.73 | 0.29 | 0.25 | 0.07 | 0.14 |
| 1 | 234170887 | rs1458587 | -1.946 | -0.58 | 0.248 | -0.926 | -0.015 | 0.02 | 0.02 | 2.831 | 0.13 | 0.7 | 0.28 | 0.25 | 0.07 | 0.14 |
| 1 | 234172161 | rs7516956 | -1.021 | -0.57 | 0.216 | -0.934 | 0.055 | 0.02 | 0.02 | 5.169 | 0.07 | 0.48 | 0.06 | 0 | 0.02 | 0.08 |
| 1 | 234177152 | rs12137461 | 0.527 | -0.511 | -0.112 | -0.903 | -0.022 | 0.06 | 0.06 | 0 | 0.2 | 0.01 | 0.11 | 0.07 | 0.18 | 0.12 |
| 1 | 234192686 | rs12118875 | 1.097 | -0.295 | -0.52 | -0.824 | -0.017 | 0.158 | 0.158 | 0 | 0.43 | 0.21 | 0.26 | 0.07 | 0.46 | 0.34 |
| 1 | 234202145 | rs4076949 | -0.474 | -0.326 | 0.108 | -0.304 | -0.031 | -0.103 | -0.103 | 0 | 0.27 | 0.51 | 0.19 | 0.07 | 0.2 | 0.19 |
| 1 | 234205662 | rs4261174 | 1.038 | 0.268 | 0.137 | 0.363 | 0.015 | 0.065 | 0.065 | 0 | 0.91 | 0.79 | 0.77 | 0.75 | 0.92 | 0.92 |
| 1 | 234210491 | rs16842515 | -0.707 | 0.934 | -0.124 | -0.215 | 0.033 | -0.042 | -0.042 | 0.11 | 0.23 | 0.32 | 0.36 | 0.18 | 0.29 | 0.24 |
| 1 | 234217852 | rs9435557 | -0.168 | -0.142 | -1.464 | -0.852 | 0.025 | 0.11 | 0.11 | 0 | 0.78 | 0.97 | 0.84 | 0.97 | 0.8 | 0.87 |
| 1 | 234221302 | rs4345854 | 0.17 | -0.333 | -1.55 | -1.316 | 0.036 | 0.064 | 0.064 | 0 | 0.77 | 0.94 | 0.85 | 0.98 | 0.82 | 0.9 |
| 1 | 234230388 | rs2045518 | -0.701 | -0.535 | -1.977 | -1.867 | 0.039 | 0.089 | 0.089 | 0.033 | 0.35 | 0.19 | 0.41 | 0.84 | 0.28 | 0.44 |
| 1 | 234232523 | rs12565924 | 0.042 | -0.466 | -1.577 | -2.625 | 0.003 | 0.02 | 0.02 | 0 | 0.47 | 0.43 | 0.49 | 0.83 | 0.54 | 0.62 |
| 1 | 234240974 | rs2045521 | 0.919 | -0.435 | -1.552 | -2.477 | -0.02 | -0.053 | -0.053 | 0 | 0.13 | 0.01 | 0.29 | 0.82 | 0.13 | 0.32 |
| 1 | 234250281 | rs10797527 | -0.026 | -0.121 | -1.613 | -2.901 | -0.055 | 0.069 | 0.069 | 0.541 | 0.32 | 0.16 | 0.41 | 0.83 | 0.4 | 0.56 |
| 1 | 234255370 | rs12140042 | 0.409 | -0.446 | -1.33 | -3.084 | 0.104 | 0.024 | 0.024 | 0.329 | 0.06 | 0.39 | 0.05 | 0 | 0.05 | 0.05 |
| 1 | 234255826 | rs16842564 | 1.093 | -0.512 | -1.512 | -3.166 | -0.017 | -0.048 | -0.048 | 0 | 0.09 | 0.26 | 0.05 | 0 | 0.04 | 0.05 |
| 1 | 234259825 | rs12059433 | 0.616 | -0.398 | -1.737 | -3.39 | -0.052 | 0.132 | 0.132 | 0 | 0.23 | 0.04 | 0.16 | 0.01 | 0.24 | 0.13 |
| 1 | 234260086 | rs6683325 | -1.411 | -0.377 | -1.711 | -3.274 | -0.004 | -0.022 | -0.022 | 0 | 0.64 | 0.46 | 0.59 | 0.8 | 0.69 | 0.73 |
| 1 | 234261466 | rs12145633 | 0.415 | -0.315 | -1.705 | -3.274 | 0.001 | -0.053 | -0.053 | 0.624 | 0.26 | 0.3 | 0.16 | 0.01 | 0.3 | 0.3 |
| 1 | 234261751 | rs11577949 | 0.085 | -0.328 | -1.453 | -3.287 | -0.024 | -0.002 | -0.002 | 0.442 | 0.23 | 0.38 | 0.31 | 0.78 | 0.17 | 0.37 |
| 1 | 234261839 | rs17512304 | 1.5 | -0.153 | -1.224 | -3.138 | -0.082 | 0.1 | 0.1 | 0.382 | 0.17 | 0.02 | 0.11 | 0.01 | 0.26 | 0.24 |
| 1 | 234263875 | rs6694267 | -0.505 | -0.253 | -0.887 | -2.944 | 0.014 | 0.131 | 0.131 | 0.047 | 0.49 | 0.28 | 0.32 | 0.02 | 0.56 | 0.4 |
| 1 | 234263897 | rs6699737 | -0.505 | -0.294 | -0.887 | -2.944 | 0.016 | 0.131 | 0.131 | 0.047 | 0.49 | 0.28 | 0.32 | 0.02 | 0.56 | 0.41 |
| 1 | 234264472 | rs2067795 | 0.379 | -0.391 | -0.377 | -2.853 | -0.016 | -0.046 | -0.046 | 0.114 | 0.08 | 0.25 | 0.05 | 0 | 0.04 | 0.06 |
| 1 | 234264555 | rs2067797 | -0.091 | -0.432 | -0.918 | -2.954 | -0.057 | 0.211 | 0.211 | 0.828 | 0.42 | 0.13 | 0.28 | 0.02 | 0.52 | 0.35 |
| 1 | 234269792 | rs10910359 | -0.345 | -0.602 | -0.871 | -2.918 | 0.004 | 0.109 | 0.109 | 0.878 | 0.45 | 0.31 | 0.31 | 0.03 | 0.55 | 0.42 |
| 1 | 234274658 | rs9435565 | -1.341 | -0.68 | -0.673 | -2.5 | -0.007 | 0.038 | 0.038 | 2.706 | 0.67 | 0.87 | 0.71 | 0.75 | 0.71 | 0.74 |
| 1 | 234276639 | rs7512377 | -0.438 | -0.343 | -0.221 | -2.942 | 0.032 | 0.104 | 0.104 | 0.737 | 0.5 | 0.37 | 0.34 | 0.03 | 0.6 | 0.42 |
| 1 | 234279644 | rs10910365 | 0.455 | -0.556 | -0.171 | -2.394 | -0.09 | 0.152 | 0.152 | 0.112 | 0.23 | 0.01 | 0.13 | 0.01 | 0.31 | 0.27 |
| 1 | 234293730 | rs6586372 | -0.974 | -0.702 | 0.883 | -1.784 | 0.003 | -0.007 | -0.007 | 1.248 | 0.08 | 0.54 | 0.06 | 0 | 0.01 | 0.04 |
| 1 | 234293963 | rs6586373 | -1.466 | -0.773 | 1.011 | -1.818 | -0.034 | -0.043 | -0.043 | 0.01 | 0.09 | 0.6 | 0.06 | 0 | 0.01 | 0.05 |
| 1 | 234296453 | rs6681198 | -0.977 | -0.692 | 0.998 | -1.779 | 0.003 | -0.007 | -0.007 | 0 | 0.08 | 0.54 | 0.06 | 0 | 0.01 | 0.04 |
| 1 | 234304728 | rs7522961 | -0.574 | -0.646 | 0.982 | -1.754 | 0.017 | 0.008 | 0.008 | 5.322 | 0.07 | 0.53 | 0.05 | 0 | 0 | 0.03 |
| 1 | 234310757 | rs4920216 | 0.342 | -0.868 | -0.732 | -2.425 | 0.059 | 0.248 | 0.248 | 0.785 | 0.61 | 0.11 | 0.81 | 0.99 | 0.72 | 0.68 |
| 1 | 234320769 | rs13328660 | -0.354 | -0.689 | -0.251 | -2.127 | 0.005 | 0.007 | 0.007 | 0.268 | 0.06 | 0.32 | 0.03 | 0.01 | 0.01 | 0.03 |
| 1 | 234344093 | rs4027079 | 0.76 | -0.838 | -1.485 | -2.507 | -0.074 | 0.172 | 0.172 | 0.12 | 0.38 | 0.12 | 0.39 | 0.14 | 0.5 | 0.31 |
| 1 | 234344295 | rs491673 | -0.276 | -0.805 | -1.324 | -2.578 | 0 | -0.075 | -0.075 | 0.004 | 0.63 | 0.67 | 0.76 | 0.99 | 0.59 | 0.63 |
| 1 | 234352349 | rs10910383 | -0.154 | -1.484 | -1.53 | -2.097 | 0.052 | -0.011 | -0.011 | 0 | 0.22 | 0.07 | 0.33 | 0.81 | 0.15 | 0.37 |
| 1 | 234352808 | rs521127 | -1.375 | -1.498 | -2.288 | -2.633 | 0.034 | 0.237 | 0.237 | 2.33 | 0.52 | 0.92 | 0.56 | 0.84 | 0.39 | 0.5 |
| 1 | 234352899 | rs4333882 | -0.611 | -1.267 | -2.311 | -2.598 | -0.047 | -0.054 | -0.054 | 0.016 | 0.33 | 0.42 | 0.37 | 0.83 | 0.23 | 0.42 |
| 1 | 234358216 | rs622815 | -0.611 | -0.567 | -3.012 | -2.061 | -0.009 | 0.138 | 0.138 | 0.134 | 0.28 | 0.87 | 0.22 | 0.01 | 0.23 | 0.12 |
| 1 | 234364285 | rs10910389 | -0.418 | -0.545 | -2.421 | -1.11 | 0.004 | -0.063 | -0.063 | 0 | 0.12 | 0.01 | 0.31 | 0.8 | 0.1 | 0.3 |
| 1 | 234370879 | rs604055 | -0.041 | -0.176 | -2.559 | -1.345 | -0.044 | 0.032 | 0.032 | 0 | 0.71 | 0.93 | 0.72 | 0.2 | 0.7 | 0.74 |
| 1 | 234371523 | rs4027081 | 0.78 | -0.014 | -2.293 | -1.435 | 0.002 | -0.012 | -0.012 | 0.375 | 0.35 | 0.58 | 0.48 | 0.16 | 0.42 | 0.32 |
| 1 | 234379913 | rs8179278 | 0.351 | 0.768 | -2.674 | -0.703 | -0.028 | -0.047 | -0.047 | 0 | 0.09 | 0.01 | 0.11 | 0.66 | 0.12 | 0.15 |
| 1 | 234382599 | rs12119646 | 0.317 | 0.766 | -2.675 | -0.719 | -0.028 | -0.049 | -0.049 | 0 | 0.09 | 0.01 | 0.11 | 0.68 | 0.12 | 0.16 |
| 1 | 234385050 | rs689345 | 0.4 | 0.7 | -2.764 | -1.245 | -0.073 | 0.195 | 0.195 | 2.265 | 0.56 | 0.85 | 0.62 | 0.14 | 0.52 | 0.56 |
| 1 | 234389425 | rs487890 | -0.503 | 1.25 | -2.279 | -0.783 | -0.03 | 0.058 | 0.058 | 0 | 0.77 | 0.91 | 0.8 | 0.87 | 0.77 | 0.89 |
| 1 | 234393208 | rs6695462 | -0.426 | 0.831 | -1.582 | -0.091 | 0.027 | 0.06 | 0.06 | 0 | 0.4 | 0.28 | 0.3 | 0.74 | 0.33 | 0.61 |
| 1 | 234405404 | rs7536858 | -1.005 | 2.098 | -0.993 | 0.715 | 0.053 | 0.047 | 0.047 | 0 | 0.07 | 0.14 | 0.12 | 0.23 | 0.17 | 0.13 |
| 1 | 234407996 | rs532495 | -0.47 | 2.017 | -0.981 | -0.814 | 0.001 | -0.039 | -0.039 | 0 | 0.63 | 0.69 | 0.8 | 1 | 0.65 | 0.88 |
| 1 | 234410567 | rs16842847 | -0.026 | 1.414 | -1.021 | -1.019 | 0.007 | 0.004 | 0.004 | 0 | 0.11 | 0.11 | 0.05 | 0.08 | 0.05 | 0.05 |
| 1 | 234411855 | rs12037730 | -0.014 | 1.416 | -1.021 | -1.018 | 0.006 | 0.003 | 0.003 | 0 | 0.1 | 0.11 | 0.05 | 0.08 | 0.05 | 0.05 |
| 1 | 234412292 | rs10910404 | -0.354 | 1.504 | -0.893 | -1.089 | 0.017 | 0 | 0 | 0 | 0.25 | 0.28 | 0.36 | 0.24 | 0.37 | 0.31 |
| 1 | 234412427 | rs10495345 | -0.52 | 1.43 | -0.736 | -1.118 | 0.004 | 0.005 | 0.005 | 0 | 0.27 | 0.24 | 0.29 | 0.17 | 0.26 | 0.23 |
| 1 | 234415806 | rs7552485 | -1.499 | 1.499 | -0.683 | -1.105 | 0 | -0.001 | -0.001 | 0 | 0.42 | 0.48 | 0.45 | 0.36 | 0.49 | 0.45 |
| 1 | 234417623 | rs1001737 | 0.596 | 1.305 | -0.523 | -1.027 | 0.002 | -0.018 | -0.018 | 0 | 0.17 | 0.24 | 0.25 | 0.09 | 0.19 | 0.18 |
| 1 | 234420787 | rs11586797 | 0.596 | 1.307 | -0.522 | -0.999 | 0.001 | -0.006 | -0.006 | 0 | 0.17 | 0.24 | 0.25 | 0.13 | 0.19 | 0.21 |
| 1 | 234421180 | rs7518644 | -0.417 | 1.299 | -0.537 | -1.013 | 0.003 | 0 | 0 | 0.11 | 0.25 | 0.34 | 0.3 | 0.25 | 0.27 | 0.32 |
| 1 | 234423165 | rs573822 | 0.36 | 1.423 | -0.477 | -1.17 | 0.006 | -0.009 | -0.009 | 2.291 | 0.72 | 0.6 | 0.67 | 0.75 | 0.68 | 0.68 |
| 1 | 234425287 | rs6698302 | -0.117 | 1.865 | -0.529 | -1.386 | -0.005 | 0.018 | 0.018 | 0 | 0.26 | 0.23 | 0.29 | 0.03 | 0.28 | 0.24 |
| 1 | 234425332 | rs10752781 | 1.072 | 1.462 | -0.588 | -1.61 | -0.001 | 0.008 | 0.008 | 0.056 | 0.47 | 0.39 | 0.28 | 0.15 | 0.47 | 0.23 |
| 1 | 234426240 | rs620939 | 0.665 | 1.884 | -0.369 | -1.39 | 0.025 | -0.005 | -0.005 | 0 | 0.84 | 0.73 | 0.83 | 0.84 | 0.78 | 0.89 |

| | | | | | | | | | | | | | | | | |
|---|---|---|---|---|---|---|---|---|---|---|---|---|---|---|---|---|
| 1 | 234426478 | rs480039 | -0.297 | 1.942 | -0.522 | -1.27 | -0.007 | 0.004 | 0.004 | 0.388 | 0.22 | 0.11 | 0.34 | 0.75 | 0.21 | 0.53 |
| 1 | 234427083 | rs12047575 | -0.082 | 1.454 | -0.423 | -0.922 | 0.001 | 0.036 | 0.036 | 0 | 0.37 | 0.27 | 0.48 | 0.19 | 0.38 | 0.39 |
| 1 | 234427534 | rs594349 | -1.121 | 1.029 | -0.38 | -0.664 | -0.002 | 0.002 | 0.002 | 0 | 0.65 | 0.63 | 0.71 | 0.42 | 0.69 | 0.61 |
| 1 | 234428358 | rs10910409 | 1.22 | 1.277 | -0.496 | -0.565 | -0.04 | 0.023 | 0.023 | 0 | 0.11 | 0.03 | 0.09 | 0.05 | 0.16 | 0.09 |
| 1 | 234428792 | rs686182 | -0.432 | 1.288 | -0.357 | -0.468 | 0.054 | 0.045 | 0.045 | 0 | 0.21 | 0.4 | 0.35 | 0.3 | 0.24 | 0.19 |
| 1 | 234429319 | rs683909 | -0.395 | 1.249 | -0.353 | -0.469 | 0.059 | -0.018 | -0.018 | 0 | 0.21 | 0.4 | 0.18 | 0.19 | 0.25 | 0.18 |
| 1 | 234429727 | rs1570105 | 0.593 | 1.053 | -0.333 | -0.352 | -0.014 | 0.007 | 0.007 | 0 | 0.37 | 0.41 | 0.26 | 0.58 | 0.29 | 0.37 |
| 1 | 234431320 | rs517527 | 0.007 | 1.145 | -0.173 | -0.306 | -0.004 | -0.017 | -0.017 | 0 | 0.45 | 0.46 | 0.33 | 0.8 | 0.39 | 0.48 |
| 1 | 234432004 | rs2281599 | 0.833 | 1.09 | -0.388 | -0.318 | 0.026 | -0.046 | -0.046 | 0.001 | 0.3 | 0.47 | 0.36 | 0.24 | 0.32 | 0.09 |
| 1 | 234432465 | rs628407 | -0.182 | 1.073 | -0.576 | -0.142 | -0.001 | 0.008 | 0.008 | 0 | 0.77 | 0.72 | 0.78 | 0.47 | 0.8 | 0.58 |
| 1 | 234432640 | rs484394 | -0.035 | 0.942 | -0.749 | -0.731 | 0.022 | 0.1 | 0.1 | 0 | 0.36 | 0.16 | 0.34 | 0.08 | 0.36 | 0.38 |
| 1 | 234433886 | rs561954 | -0.817 | 1.259 | -0.255 | -0.21 | 0.057 | -0.013 | -0.013 | 1.254 | 0.1 | 0.22 | 0.16 | 0.02 | 0.15 | 0.11 |
| 1 | 234434020 | rs12748153 | -0.078 | 1.056 | -0.932 | -0.534 | 0.012 | 0.027 | 0.027 | 0.013 | 0.33 | 0.17 | 0.41 | 0.21 | 0.34 | 0.26 |
| 1 | 234434120 | rs559987 | -0.398 | 1.521 | -1.001 | -0.23 | -0.032 | -0.021 | -0.021 | 0 | 0.8 | 0.89 | 0.71 | 0.69 | 0.72 | 0.73 |
| 1 | 234435865 | rs501664 | 0.405 | 1.466 | -0.635 | -0.237 | 0.031 | 0.031 | 0.031 | 0 | 0.74 | 0.69 | 0.46 | 0.6 | 0.65 | 0.56 |
| 1 | 234436023 | rs12136778 | 0.065 | 1.572 | -0.68 | -0.403 | 0.061 | -0.011 | -0.011 | 0 | 0.05 | 0.19 | 0.12 | 0 | 0.06 | 0.16 |
| 1 | 234438235 | rs642487 | -0.661 | 1.5 | -1.046 | -0.258 | -0.082 | -0.064 | -0.064 | 0.019 | 0.82 | 0.97 | 0.72 | 0.75 | 0.73 | 0.79 |
| 1 | 234449697 | rs594657 | 0.535 | 0.661 | -0.752 | -0.509 | 0.011 | 0.012 | 0.012 | 0.096 | 0.43 | 0.62 | 0.4 | 0.73 | 0.34 | 0.43 |
| 1 | 234449742 | rs6662048 | 0.542 | 0.431 | -0.893 | -0.688 | 0.001 | 0.001 | 0.001 | 0 | 0.59 | 0.56 | 0.41 | 0.59 | 0.62 | 0.51 |
| 1 | 234449994 | rs561159 | -0.586 | 0.844 | -0.559 | -0.716 | 0.014 | -0.045 | -0.045 | 0 | 0.34 | 0.46 | 0.36 | 0.16 | 0.39 | 0.31 |
| 1 | 234452263 | rs745858 | -0.338 | 0.604 | -0.195 | -0.275 | 0.02 | -0.019 | -0.019 | 0 | 0.06 | 0.16 | 0.08 | 0 | 0.04 | 0.12 |
| 1 | 234452980 | rs1407149 | -0.62 | 0.411 | -0.2 | -0.435 | -0.002 | -0.033 | -0.033 | 0 | 0.1 | 0.13 | 0.09 | 0.01 | 0.06 | 0.14 |
| 1 | 234455148 | rs519143 | -0.363 | 0.412 | -0.718 | -0.6 | -0.047 | 0.005 | 0.005 | 1.071 | 0.62 | 0.8 | 0.63 | 0.71 | 0.48 | 0.63 |
| 1 | 234457499 | rs474208 | 0.783 | -0.223 | -0.349 | -0.922 | 0.012 | -0.006 | -0.006 | 0 | 0.4 | 0.28 | 0.36 | 0.59 | 0.33 | 0.43 |
| 1 | 234459353 | rs10910416 | -0.913 | 0.329 | -0.469 | -0.975 | -0.001 | -0.062 | -0.062 | 0 | 0.12 | 0.24 | 0.11 | 0.01 | 0.07 | 0.15 |
| 1 | 234462088 | rs568614 | -0.129 | 0.173 | -1.245 | -0.995 | 0.018 | 0.022 | 0.022 | 2.276 | 0.53 | 0.67 | 0.48 | 0.29 | 0.44 | 0.3 |
| 1 | 234468324 | rs4128841 | 0.356 | 0.269 | -2.1 | -1.775 | -0.162 | 0.151 | 0.151 | 0.19 | 0.38 | 0.05 | 0.44 | 0.12 | 0.52 | 0.53 |
| 1 | 234468879 | rs4586012 | 0.898 | -0.293 | -1.996 | -2.2 | -0.002 | -0.051 | -0.051 | 0 | 0.22 | 0.23 | 0.13 | 0.01 | 0.21 | 0.09 |
| 1 | 234473914 | rs2991952 | 0.806 | 0.178 | -2.317 | -2.09 | -0.103 | 0.1 | 0.1 | 0 | 0.31 | 0.03 | 0.34 | 0.11 | 0.42 | 0.45 |
| 1 | 234475832 | rs2991953 | -0.04 | -0.038 | -1.587 | -1.841 | -0.017 | -0.021 | -0.021 | 0 | 0.75 | 0.62 | 0.71 | 0.17 | 0.84 | 0.67 |
| 1 | 234476398 | rs2262883 | -0.025 | 0.382 | -0.969 | -1.434 | -0.013 | -0.118 | -0.118 | 0.701 | 0.44 | 0.5 | 0.26 | 0.01 | 0.33 | 0.22 |
| 1 | 234481678 | rs3001707 | 0.436 | 1.014 | -0.836 | -0.276 | -0.001 | -0.001 | -0.001 | 0 | 0.7 | 0.7 | 0.65 | 0.31 | 0.67 | 0.56 |
| 1 | 234482040 | rs7512712 | 0.302 | 0.372 | -0.819 | -1.036 | 0.008 | 0.024 | 0.024 | 0 | 0.4 | 0.38 | 0.44 | 0.1 | 0.34 | 0.21 |
| 1 | 234490818 | rs2991956 | -1.169 | 0.075 | -0.428 | -0.539 | -0.017 | -0.033 | -0.033 | 0.055 | 0.36 | 0.52 | 0.31 | 0.3 | 0.35 | 0.34 |
| 1 | 234492059 | rs7536709 | -0.758 | 0.509 | -1.007 | -0.217 | -0.001 | 0 | 0 | 0 | 0.25 | 0.21 | 0.23 | 0.3 | 0.28 | 0.3 |
| 1 | 234493015 | rs2991957 | -0.635 | 0.528 | -1.298 | -0.153 | -0.004 | -0.007 | -0.007 | 0 | 0.23 | 0.14 | 0.31 | 0.31 | 0.26 | 0.3 |
| 1 | 234507977 | rs6586384 | -0.826 | 0.235 | -1.797 | -0.218 | -0.001 | 0.018 | 0.018 | 0.121 | 0.51 | 0.87 | 0.65 | 0.48 | 0.49 | 0.42 |
| 1 | 234508704 | rs12070750 | 0.559 | -0.18 | -1.603 | -0.412 | -0.01 | -0.013 | -0.013 | 0 | 0.08 | 0.09 | 0.03 | 0 | 0.05 | 0.08 |
| 1 | 234510262 | rs4300255 | -0.851 | -0.32 | -1.307 | -0.67 | 0.01 | -0.001 | -0.001 | 0 | 0.42 | 0.72 | 0.58 | 0.48 | 0.4 | 0.39 |
| 1 | 234513871 | rs2752 | -1.051 | -0.255 | -1.369 | -0.678 | -0.018 | 0.007 | 0.007 | 0.102 | 0.44 | 0.81 | 0.64 | 0.48 | 0.45 | 0.34 |
| 1 | 234515608 | rs6695813 | 0.537 | -0.242 | -1.332 | -0.659 | 0 | -0.015 | -0.015 | 0 | 0.08 | 0.06 | 0.08 | 0 | 0.09 | 0.03 |
| 1 | 234524045 | rs7527165 | -0.532 | -0.374 | -0.959 | -0.676 | -0.001 | -0.001 | -0.001 | 0 | 0.35 | 0.66 | 0.54 | 0.49 | 0.36 | 0.31 |
| 1 | 234525940 | rs6669190 | -1.2 | 0.012 | -0.856 | -0.707 | -0.038 | 0.01 | 0.01 | 0.414 | 0.44 | 0.8 | 0.64 | 0.5 | 0.48 | 0.35 |
| 1 | 234526868 | rs7524822 | -1.325 | -0.007 | -0.873 | -0.724 | -0.016 | 0.042 | 0.042 | 0 | 0.5 | 0.77 | 0.66 | 0.5 | 0.52 | 0.43 |
| 1 | 234529519 | rs1802871 | -0.546 | -0.426 | -0.875 | -0.734 | -0.067 | -0.033 | -0.033 | 0 | 0.17 | 0.54 | 0.18 | 0.09 | 0.09 | 0.13 |
| 1 | 234529552 | rs2175594 | -1.026 | -0.195 | -0.836 | -0.661 | -0.003 | 0 | 0 | 1.457 | 0.42 | 0.62 | 0.57 | 0.5 | 0.44 | 0.4 |
| 1 | 234529570 | rs2175593 | -0.98 | -0.249 | -0.835 | -0.671 | -0.002 | 0 | 0 | 0 | 0.41 | 0.61 | 0.57 | 0.49 | 0.43 | 0.4 |
| 1 | 234531009 | rs4027040 | -1.271 | -0.148 | -1.203 | -0.564 | -0.014 | 0.044 | 0.044 | 0 | 0.5 | 0.76 | 0.66 | 0.5 | 0.53 | 0.44 |
| 1 | 234537992 | rs12031728 | 0.027 | 0.088 | -1.355 | -0.437 | -0.033 | -0.045 | -0.045 | 0 | 0.13 | 0.05 | 0.29 | 0.38 | 0.2 | 0.2 |
| 1 | 234539157 | rs10910427 | -1.362 | 0.002 | -0.827 | -0.449 | 0.043 | 0.029 | 0.029 | 0 | 0.33 | 0.45 | 0.54 | 0.47 | 0.44 | 0.35 |
| 1 | 234540466 | rs3820605 | 0.174 | 0.045 | -0.752 | -0.313 | -0.014 | -0.023 | -0.023 | 0 | 0.14 | 0.07 | 0.25 | 0.31 | 0.18 | 0.18 |
| 1 | 234540547 | rs3768288 | -1.301 | -1.19 | -1.421 | -1.076 | 0.009 | 0.008 | 0.008 | 0.015 | 0.6 | 0.5 | 0.6 | 0.49 | 0.54 | 0.51 |
| 1 | 234541801 | rs12073170 | 0.636 | -0.82 | -1.246 | -0.978 | 0.01 | 0.005 | 0.005 | 0 | 0.07 | 0.15 | 0.17 | 0.09 | 0.09 | 0.08 |
| 1 | 234542225 | rs17517734 | 1.4 | -0.848 | -1.29 | -1.012 | -0.047 | -0.008 | -0.008 | 0 | 0.1 | 0.04 | 0.1 | 0.08 | 0.17 | 0.03 |
| 1 | 234551520 | rs10910428 | 1.183 | -1.108 | -1.258 | -1.111 | 0.046 | 0.036 | 0.036 | 4.051 | 0.19 | 0.09 | 0.04 | 0 | 0.06 | 0.07 |
| 1 | 234551603 | rs3754315 | 0.133 | -0.28 | -0.88 | -0.618 | -0.001 | 0 | 0 | 0 | 0.15 | 0.11 | 0.25 | 0.31 | 0.18 | 0.24 |
| 1 | 234552011 | rs3820604 | -0.73 | -0.321 | -0.829 | -0.616 | 0.02 | 0.027 | 0.027 | 0 | 0.22 | 0.3 | 0.43 | 0.39 | 0.28 | 0.32 |
| 1 | 234555310 | rs12035712 | -0.522 | -1.009 | -0.908 | -0.5 | -0.001 | -0.005 | -0.005 | 0 | 0.25 | 0.25 | 0.35 | 0.42 | 0.25 | 0.21 |
| 1 | 234557918 | rs750813 | -0.404 | -0.944 | -1.061 | -0.473 | -0.009 | -0.047 | -0.047 | 0 | 0.24 | 0.14 | 0.34 | 0.42 | 0.25 | 0.21 |
| 1 | 234563060 | rs443685 | 0.485 | -0.795 | -1.005 | -0.338 | 0.065 | 0.093 | 0.093 | 0.058 | 0.27 | 0.48 | 0.27 | 0.45 | 0.34 | 0.48 |
| 1 | 234565787 | rs915184 | 0.726 | -1.545 | -1.516 | -0.502 | 0.042 | -0.001 | -0.001 | 0 | 0.29 | 0.2 | 0.2 | 0.22 | 0.11 | 0.17 |
| 1 | 234569192 | rs3754311 | 0.726 | -1.545 | -1.516 | -0.502 | 0.042 | -0.001 | -0.001 | 0 | 0.29 | 0.2 | 0.2 | 0.22 | 0.11 | 0.17 |
| 1 | 234570685 | rs6682410 | 1.041 | -0.829 | -1.906 | -0.306 | -0.01 | 0.021 | 0.021 | 0 | 0.14 | 0.08 | 0.12 | 0 | 0.19 | 0.07 |
| 1 | 234581891 | rs3768286 | 0.639 | -0.399 | -1.485 | -0.126 | 0.002 | 0.002 | 0.002 | 0 | 0.28 | 0.27 | 0.38 | 0.3 | 0.29 | 0.25 |
| 1 | 234582651 | rs4920246 | 0.603 | -0.403 | -1.469 | -0.128 | -0.002 | -0.002 | -0.002 | 0 | 0.28 | 0.31 | 0.38 | 0.3 | 0.29 | 0.25 |
| 1 | 234584231 | rs2102596 | 0.655 | -0.353 | -1.7 | -0.113 | 0.006 | 0.003 | 0.003 | 0 | 0.28 | 0.26 | 0.37 | 0.3 | 0.29 | 0.25 |
| 1 | 234585003 | rs270524 | 0.808 | 0.128 | -1.483 | 0.253 | -0.087 | 0.001 | 0.001 | 0 | 0.29 | 0.47 | 0.25 | 0.22 | 0.12 | 0.14 |
| 1 | 234585790 | rs1619856 | 0.944 | 2.04 | -1.416 | 0.624 | 0.004 | -0.027 | -0.027 | 0 | 0.63 | 0.75 | 0.64 | 0.52 | 0.45 | 0.45 |
| 1 | 234587008 | rs4920247 | -0.714 | 1.652 | -1.438 | 1.484 | -0.071 | -0.093 | -0.093 | 0 | 0.2 | 0.08 | 0.19 | 0.45 | 0.28 | 0.45 |

| | | | | | | | | | | | | | | | | |
|---|---|---|---|---|---|---|---|---|---|---|---|---|---|---|---|---|
| 1 | 234601529 | rs3736838 | -0.887 | 0.556 | -0.859 | 1.295 | -0.012 | 0.066 | 0.066 | 0 | 0.07 | 0.11 | 0.21 | 0.26 | 0.04 | 0.27 |
| 1 | 234603386 | rs270502 | -0.627 | 1.122 | -0.926 | -0.287 | -0.031 | 0.034 | 0.034 | 0 | 0.2 | 0.1 | 0.14 | 0 | 0.31 | 0.07 |
| 1 | 234611332 | rs528774 | -0.957 | 0.049 | -0.68 | -0.999 | 0 | 0.007 | 0.007 | 0 | 0.41 | 0.38 | 0.32 | 0.09 | 0.41 | 0.22 |
| 1 | 234612995 | rs544322 | -1.011 | -0.55 | -0.144 | -1.115 | 0.016 | 0.115 | 0.115 | 0 | 0.5 | 0.67 | 0.42 | 0.18 | 0.48 | 0.3 |
| 1 | 234617092 | rs4442397 | -0.268 | -0.919 | 0.112 | -1.21 | -0.028 | -0.02 | -0.02 | 0.066 | 0.26 | 0.37 | 0.44 | 0.84 | 0.15 | 0.51 |
| 1 | 234618234 | rs512643 | -1.321 | -0.913 | -0.501 | -1.359 | -0.066 | 0.239 | 0.239 | 0 | 0.57 | 0.88 | 0.69 | 0.93 | 0.52 | 0.67 |
| 1 | 234626786 | rs271747 | -0.094 | -0.612 | -0.224 | -1.299 | -0.003 | -0.02 | -0.02 | 1.143 | 0.18 | 0.19 | 0.14 | 0 | 0.24 | 0.07 |
| 1 | 234627055 | rs271748 | -1.112 | -0.817 | -0.447 | -1.704 | -0.044 | 0.052 | 0.052 | 0 | 0.68 | 0.82 | 0.68 | 0.88 | 0.57 | 0.73 |
| 1 | 234633308 | rs3951276 | -0.242 | -1.941 | -0.286 | -2.133 | 0.021 | 0.01 | 0.01 | 0 | 0.42 | 0.37 | 0.21 | 0.05 | 0.34 | 0.22 |
| 1 | 234636107 | rs6667048 | -1.102 | -1.867 | -0.083 | -1.945 | -0.004 | 0.017 | 0.017 | 0.013 | 0.61 | 0.69 | 0.62 | 0.29 | 0.47 | 0.65 |
| 1 | 234636595 | rs2026044 | -0.801 | -1.752 | 0.362 | -1.792 | -0.005 | 0.003 | 0.003 | 0 | 0.19 | 0.37 | 0.42 | 0.22 | 0.16 | 0.41 |
| 1 | 234637347 | rs6586393 | -0.271 | -1.867 | -0.205 | -1.17 | -0.011 | 0.004 | 0.004 | 0 | 0.36 | 0.37 | 0.2 | 0.56 | 0.3 | 0.29 |
| 1 | 234637789 | rs16843362 | -1.168 | -1.299 | -0.234 | -1.017 | 0.024 | -0.025 | -0.025 | 0 | 0.06 | 0.2 | 0.06 | 0 | 0.05 | 0.01 |
| 1 | 234638534 | rs2793861 | 0.324 | -1.431 | -0.935 | -0.765 | -0.016 | -0.011 | -0.011 | 0.023 | 0.73 | 0.58 | 0.53 | 0.75 | 0.78 | 0.49 |
| 1 | 234648819 | rs6586395 | -1.101 | -1.021 | -1.52 | -0.525 | -0.001 | -0.014 | -0.014 | 0 | 0.14 | 0.82 | 0.15 | 0.01 | 0.09 | 0.09 |
| 1 | 234653716 | rs6673545 | -0.136 | -1.083 | -2.07 | -0.603 | 0.098 | 0.132 | 0.132 | 0.23 | 0.55 | 0.87 | 0.52 | 0.74 | 0.71 | 0.4 |
| 1 | 234656596 | rs696859 | -0.294 | -0.395 | -2.062 | -0.773 | -0.104 | 0.213 | 0.213 | 0.325 | 0.66 | 0.94 | 0.73 | 0.96 | 0.44 | 0.83 |
| 1 | 234658886 | rs523638 | 0.446 | -0.927 | -1.122 | 0.293 | 0.017 | -0.006 | -0.006 | 0 | 0.55 | 0.38 | 0.62 | 0.55 | 0.38 | 0.72 |
| 1 | 234662389 | rs271736 | -0.188 | -0.75 | -1.094 | -0.394 | -0.047 | 0.178 | 0.178 | 0 | 0.66 | 0.89 | 0.68 | 0.94 | 0.4 | 0.81 |
| 1 | 234664631 | rs271740 | 0.395 | -0.991 | 0.161 | 0.24 | 0.024 | 0.025 | 0.025 | 0 | 0.58 | 0.3 | 0.6 | 0.63 | 0.38 | 0.71 |
| 1 | 234669847 | rs271762 | 0.159 | -0.943 | 0.68 | -0.115 | -0.007 | -0.069 | -0.069 | 0 | 0.61 | 0.56 | 0.62 | 0.92 | 0.38 | 0.8 |
| 1 | 234674965 | rs271767 | 0.455 | -1.201 | 0.245 | -0.331 | 0.066 | 0.155 | 0.155 | 0 | 0.53 | 0.24 | 0.58 | 0.91 | 0.34 | 0.78 |
| 1 | 234680657 | rs271760 | 0.17 | -1.085 | 0.47 | 0.075 | 0.018 | 0.026 | 0.026 | 0 | 0.58 | 0.31 | 0.6 | 0.64 | 0.35 | 0.75 |
| 1 | 234686925 | rs4920258 | 0.54 | -1.282 | 0.211 | -0.093 | 0.096 | 0.018 | 0.018 | 0.072 | 0.52 | 0.15 | 0.59 | 0.61 | 0.33 | 0.68 |
| 1 | 234689712 | rs4920260 | 0.679 | -2.085 | -0.291 | -0.66 | 0.089 | -0.015 | -0.015 | 0 | 0.39 | 0.18 | 0.54 | 0.63 | 0.23 | 0.58 |
| 1 | 234693321 | rs10797553 | -0.755 | -0.366 | -0.383 | -1.036 | -0.072 | 0.137 | 0.137 | 2.686 | 0.7 | 0.89 | 0.79 | 0.98 | 0.6 | 0.91 |
| 1 | 234694041 | rs12142968 | 0.72 | -0.637 | -0.414 | -0.822 | 0.105 | -0.026 | -0.026 | 1.596 | 0.42 | 0.12 | 0.55 | 0.61 | 0.26 | 0.59 |
| 1 | 234699236 | rs7531368 | -1.51 | -0.62 | 0.004 | -0.653 | -0.022 | 0.129 | 0.129 | 0.043 | 0.09 | 0.2 | 0.04 | 0.34 | 0.04 | 0.12 |
| 1 | 234701860 | rs726570 | -1.084 | -0.653 | 0.257 | -0.949 | 0.09 | -0.042 | -0.042 | 2.527 | 0.6 | 0.9 | 0.44 | 0.39 | 0.74 | 0.42 |
| 1 | 234701910 | rs9970996 | -0.896 | -0.318 | 0.399 | -0.628 | -0.013 | 0.159 | 0.159 | 0 | 0.06 | 0.15 | 0.03 | 0.34 | 0.04 | 0.11 |
| 1 | 234704046 | rs6663049 | -0.872 | 0.234 | 0.55 | -0.725 | 0.066 | 0.127 | 0.127 | 0.428 | 0.16 | 0.35 | 0.16 | 0.37 | 0.27 | 0.25 |
| 1 | 234706724 | rs875268 | -0.087 | 0.112 | 0.1 | -1.731 | -0.006 | -0.158 | -0.158 | 0.368 | 0.37 | 0.51 | 0.22 | 0.02 | 0.36 | 0.12 |
| 1 | 234716506 | rs955613 | -0.751 | 0.032 | -1.1 | -1.227 | 0.073 | 0.025 | 0.025 | 0 | 0.6 | 0.92 | 0.44 | 0.3 | 0.73 | 0.41 |
| 1 | 234716593 | rs955612 | -0.658 | 1.33 | 0.037 | -0.637 | 0.223 | 0.202 | 0.202 | 0.015 | 0.13 | 0.45 | 0.17 | 0.27 | 0.28 | 0.22 |
| 1 | 234726012 | rs744487 | -0.347 | 0.767 | -0.748 | -0.935 | 0 | -0.002 | -0.002 | 0 | 0.31 | 0.26 | 0.22 | 0.22 | 0.32 | 0.2 |
| 1 | 234729643 | rs7532095 | -1.099 | 0.347 | -1.591 | -1.251 | -0.001 | -0.063 | -0.063 | 0 | 0.75 | 0.92 | 0.54 | 0.29 | 0.82 | 0.48 |
| 1 | 234738338 | rs6672925 | -0.909 | 0.333 | -1.591 | -1.341 | -0.005 | -0.033 | -0.033 | 0.827 | 0.73 | 0.87 | 0.55 | 0.27 | 0.77 | 0.45 |
| 1 | 234740141 | rs8823 | -0.918 | 0.33 | -1.604 | -1.352 | -0.005 | -0.03 | -0.03 | 0.129 | 0.73 | 0.87 | 0.55 | 0.26 | 0.77 | 0.45 |
| 1 | 234743237 | rs7543281 | 0.631 | 0.461 | -1.834 | -1.773 | -0.001 | 0.036 | 0.036 | 0 | 0.17 | 0.08 | 0.08 | 0.04 | 0.15 | 0.07 |
| 1 | 234744250 | rs8722 | 1.143 | 0.9 | -1.633 | -1.823 | -0.079 | -0.088 | -0.088 | 0.319 | 0.14 | 0.01 | 0.14 | 0.2 | 0.25 | 0.12 |
| 1 | 234752588 | rs1887917 | -1.257 | 0.676 | -1.491 | -1.72 | -0.013 | -0.04 | -0.04 | 0 | 0.74 | 0.88 | 0.55 | 0.27 | 0.78 | 0.45 |
| 1 | 234754829 | rs10797557 | -0.077 | 0.765 | -1.364 | -1.7 | 0.002 | 0.021 | 0.021 | 2.613 | 0.36 | 0.27 | 0.28 | 0.21 | 0.39 | 0.26 |
| 1 | 234756418 | rs3850718 | -0.06 | 0.812 | -1.698 | -1.678 | -0.013 | 0.026 | 0.026 | 1.957 | 0.35 | 0.17 | 0.28 | 0.21 | 0.4 | 0.26 |
| 1 | 234763145 | rs7536996 | -0.081 | 0.812 | -1.671 | -1.637 | -0.016 | 0.026 | 0.026 | 1.009 | 0.35 | 0.2 | 0.28 | 0.2 | 0.39 | 0.26 |
| 1 | 234764925 | rs530596 | -0.391 | 0.73 | -0.904 | -2.093 | 0.007 | -0.06 | -0.06 | 0 | 0.17 | 0.2 | 0.11 | 0.01 | 0.2 | 0.1 |
| 1 | 234764966 | rs643342 | -0.666 | 0.709 | -0.914 | -2.722 | -0.018 | 0.008 | 0.008 | 0 | 0.18 | 0.12 | 0.22 | 0.13 | 0.28 | 0.13 |
| 1 | 234765021 | rs16843708 | -0.961 | 0.529 | -0.417 | -3.099 | 0.004 | -0.017 | -0.017 | 0 | 0.07 | 0.22 | 0.04 | 0.03 | 0.05 | 0.01 |
| 1 | 234765256 | rs533483 | -1.17 | 0.623 | 0.053 | -3.375 | 0.046 | -0.154 | -0.154 | 0.124 | 0.2 | 0.43 | 0.13 | 0 | 0.23 | 0.07 |
| 1 | 234766016 | rs17520203 | 0.892 | 0.952 | -0.658 | -3.442 | -0.007 | 0.008 | 0.008 | 0 | 0.14 | 0.09 | 0.13 | 0.02 | 0.18 | 0.09 |
| 1 | 234766437 | rs565423 | -1.929 | 0.48 | -1.51 | -3.723 | -0.059 | -0.041 | -0.041 | 0 | 0.82 | 0.99 | 0.54 | 0.15 | 0.77 | 0.63 |
| 1 | 234769793 | rs9803801 | 0.771 | -0.479 | -1.013 | -3.802 | -0.017 | -0.013 | -0.013 | 0 | 0.13 | 0.16 | 0.04 | 0.03 | 0.05 | 0.11 |
| 1 | 234771816 | rs6683807 | 0.165 | -0.509 | -0.422 | -4.034 | 0.005 | -0.031 | -0.031 | 0 | 0.28 | 0.26 | 0.1 | 0.01 | 0.21 | 0.1 |
| 1 | 234775227 | rs16843723 | 0.692 | -0.242 | -0.414 | -3.9 | -0.004 | -0.009 | -0.009 | 0 | 0.08 | 0.14 | 0.07 | 0.06 | 0.06 | 0.13 |
| 1 | 234779059 | rs10910479 | -0.208 | -0.308 | -0.54 | -3.993 | -0.003 | -0.037 | -0.037 | 0 | 0.29 | 0.34 | 0.16 | 0.07 | 0.25 | 0.21 |
| 1 | 234782449 | rs12058613 | -0.612 | -0.098 | -0.739 | -4.026 | 0 | -0.032 | -0.032 | 0 | 0.43 | 0.45 | 0.27 | 0.09 | 0.42 | 0.24 |
| 1 | 234789114 | rs600511 | -0.757 | 0.003 | -0.796 | -4.074 | -0.006 | 0.096 | 0.096 | 0 | 0.45 | 0.68 | 0.32 | 0.07 | 0.45 | 0.37 |
| 1 | 234791721 | rs564212 | -1.02 | -0.023 | -0.752 | -4.008 | 0.021 | 0.085 | 0.085 | 0 | 0.57 | 0.7 | 0.47 | 0.13 | 0.62 | 0.4 |
| 1 | 234792524 | rs673283 | -0.659 | 0.814 | -0.367 | -3.414 | 0.04 | -0.084 | -0.084 | 0 | 0.21 | 0.39 | 0.23 | 0.06 | 0.26 | 0.3 |
| 1 | 234803619 | rs636705 | -0.973 | 1.519 | -0.595 | -3.313 | -0.001 | 0.046 | 0.046 | 0.008 | 0.61 | 0.7 | 0.46 | 0.17 | 0.57 | 0.37 |
| 1 | 234805957 | rs12071518 | 0.413 | 1.45 | -0.526 | -3.568 | 0.007 | 0.009 | 0.009 | 0 | 0.33 | 0.26 | 0.19 | 0.09 | 0.25 | 0.22 |
| 1 | 234808507 | rs605957 | -1.075 | 1.5 | -0.441 | -3.566 | -0.002 | -0.011 | -0.011 | 0 | 0.6 | 0.6 | 0.38 | 0.11 | 0.53 | 0.35 |
| 1 | 234811787 | rs3009952 | 0.369 | 0.852 | -0.432 | -3.398 | -0.029 | -0.064 | -0.064 | 0 | 0.23 | 0.38 | 0.15 | 0.07 | 0.16 | 0.1 |
| 1 | 234819885 | rs486650 | -0.331 | 0.93 | -0.426 | -3.452 | -0.001 | -0.016 | -0.016 | 0.173 | 0.41 | 0.53 | 0.39 | 0.16 | 0.43 | 0.3 |
| 1 | 234820827 | rs588965 | -1.14 | 0.89 | -0.478 | -3.491 | -0.014 | 0.044 | 0.044 | 0.043 | 0.59 | 0.68 | 0.45 | 0.17 | 0.51 | 0.37 |
| 1 | 234827319 | rs16843810 | 1.114 | 0.814 | -0.427 | -3.455 | 0.027 | 0.028 | 0.028 | 0 | 0.19 | 0.16 | 0.07 | 0.01 | 0.09 | 0.07 |
| 1 | 234827824 | rs541021 | -0.887 | 0.861 | -0.402 | -3.31 | -0.013 | 0.045 | 0.045 | 0.103 | 0.58 | 0.67 | 0.38 | 0.11 | 0.47 | 0.33 |
| 1 | 234832199 | rs624203 | -1.413 | 0.451 | -0.363 | -3.038 | -0.021 | 0.062 | 0.062 | 0.055 | 0.6 | 0.72 | 0.43 | 0.13 | 0.51 | 0.36 |
| 1 | 234833429 | rs10910487 | -0.116 | 0.384 | -0.434 | -3.056 | 0.018 | 0.014 | 0.014 | 0 | 0.32 | 0.22 | 0.15 | 0.06 | 0.19 | 0.2 |
| 1 | 234837359 | rs1753579 | -0.146 | 0.723 | -0.934 | -2.504 | 0.002 | 0.034 | 0.034 | 0 | 0.36 | 0.33 | 0.23 | 0.06 | 0.33 | 0.24 |

| | | | | | | | | | | | | | | | | |
|---|---|---|---|---|---|---|---|---|---|---|---|---|---|---|---|---|
| 1 | 234837437 | rs522209 | 0.03 | 0.983 | -1.412 | -2.318 | -0.033 | 0.144 | 0.144 | 0 | 0.33 | 0.13 | 0.21 | 0.06 | 0.31 | 0.24 |
| 1 | 234838565 | rs3888368 | -0.67 | -0.159 | -0.454 | -1.311 | -0.034 | -0.123 | -0.123 | 0.868 | 0.27 | 0.39 | 0.19 | 0.06 | 0.18 | 0.13 |
| 1 | 234840803 | rs487738 | 0.342 | 0.063 | -0.868 | -0.864 | 0.003 | -0.012 | -0.012 | 0 | 0.68 | 0.67 | 0.8 | 0.86 | 0.67 | 0.71 |
| 1 | 234846396 | rs6695664 | -0.748 | -1.074 | -0.872 | -0.854 | -0.025 | -0.022 | -0.022 | 0 | 0.22 | 0.29 | 0.14 | 0.07 | 0.16 | 0.2 |
| 1 | 234850303 | rs543837 | 0.93 | -1.405 | -1.279 | -0.763 | 0.01 | 0.003 | 0.003 | 0.029 | 0.41 | 0.73 | 0.6 | 0.7 | 0.35 | 0.5 |
| 1 | 234851184 | rs496708 | 1.332 | -1.676 | -0.754 | -0.739 | 0.003 | 0.01 | 0.01 | 0 | 0.34 | 0.42 | 0.56 | 0.67 | 0.35 | 0.45 |
| 1 | 234851396 | rs633269 | 0.852 | -1.572 | -1.407 | -0.763 | 0.002 | 0 | 0 | 0 | 0.4 | 0.63 | 0.59 | 0.7 | 0.35 | 0.5 |
| 1 | 234853059 | rs556107 | -0.299 | -1.471 | -1.783 | -0.803 | -0.096 | 0.114 | 0.114 | 0 | 0.62 | 0.88 | 0.72 | 0.83 | 0.52 | 0.71 |
| 1 | 234863602 | rs478425 | 0.624 | -1.405 | -2.105 | -0.147 | 0.016 | -0.045 | -0.045 | 0.324 | 0.42 | 0.79 | 0.6 | 0.61 | 0.35 | 0.48 |
| 1 | 234864214 | rs641154 | -0.04 | -1.307 | -2.552 | -0.307 | -0.102 | 0.051 | 0.051 | 1.169 | 0.58 | 0.94 | 0.71 | 0.74 | 0.49 | 0.66 |
| 1 | 234870917 | rs506367 | 1.542 | -1.331 | -2.455 | -0.186 | -0.075 | -0.125 | -0.125 | 0.251 | 0.27 | 0.06 | 0.46 | 0.57 | 0.3 | 0.41 |
| 1 | 234873046 | rs16843932 | 0.306 | -0.096 | -2.134 | 0.233 | -0.023 | -0.039 | -0.039 | 0.322 | 0.07 | 0 | 0.05 | 0.15 | 0.09 | 0.01 |
| 1 | 234873275 | rs822928 | -0.355 | -0.333 | -1.99 | 0.203 | -0.138 | 0.004 | 0.004 | 0.11 | 0.64 | 0.94 | 0.73 | 0.74 | 0.54 | 0.7 |
| 1 | 234875600 | rs4920270 | -0.625 | -0.267 | -0.79 | 0.447 | -0.003 | 0.001 | 0.001 | 0 | 0.15 | 0.15 | 0.12 | 0.12 | 0.13 | 0.17 |
| 1 | 234878441 | rs2587535 | 0.006 | -0.094 | -0.706 | -0.289 | 0.016 | 0.041 | 0.041 | 1.141 | 0.53 | 0.36 | 0.63 | 0.83 | 0.6 | 0.61 |
| 1 | 234878715 | rs2587536 | 0.227 | 0.091 | -0.692 | -0.036 | -0.003 | 0.002 | 0.002 | 0 | 0.57 | 0.52 | 0.47 | 0.58 | 0.55 | 0.63 |
| 1 | 234881190 | rs10910497 | -0.575 | 1.044 | -0.819 | -0.294 | -0.014 | 0.006 | 0.006 | 0 | 0.38 | 0.24 | 0.38 | 0.32 | 0.43 | 0.23 |
| 1 | 234883182 | rs12133803 | 1.106 | 0.815 | -0.552 | -0.694 | 0.001 | 0.085 | 0.085 | 0 | 0.5 | 0.31 | 0.5 | 0.69 | 0.52 | 0.71 |
| 1 | 234886224 | rs10910498 | 0.113 | 1.054 | -0.598 | -0.837 | -0.001 | -0.003 | -0.003 | 0 | 0.24 | 0.18 | 0.23 | 0.13 | 0.25 | 0.14 |
| 1 | 234892143 | rs3844068 | 0.258 | 0.965 | -0.992 | -0.835 | 0.003 | 0.025 | 0.025 | 0.173 | 0.64 | 0.34 | 0.65 | 0.89 | 0.67 | 0.79 |
| 1 | 234892797 | rs2587531 | 0.665 | 0.918 | -1.084 | -0.638 | -0.007 | 0.097 | 0.097 | 0 | 0.5 | 0.27 | 0.5 | 0.71 | 0.51 | 0.7 |
| 1 | 234893420 | rs2582650 | 0.23 | 0.474 | -0.691 | -1.344 | 0.012 | -0.08 | -0.08 | 0.634 | 0.73 | 0.57 | 0.77 | 0.89 | 0.71 | 0.8 |
| 1 | 234893452 | rs2786022 | 0.184 | -0.238 | -0.687 | -1.063 | -0.005 | -0.002 | -0.002 | 0 | 0.77 | 0.26 | 0.82 | 0.88 | 0.88 | 0.96 |
| 1 | 234896409 | rs3862042 | 0.774 | 0.659 | -1.279 | -1.319 | -0.03 | 0.044 | 0.044 | 0.036 | 0.12 | 0.05 | 0.09 | 0.05 | 0.17 | 0.33 |
| 1 | 234897243 | rs2786014 | -1.13 | 0.578 | -1.25 | -1.379 | -0.007 | 0.047 | 0.047 | 0.064 | 0.7 | 0.85 | 0.75 | 0.83 | 0.7 | 0.88 |
| 1 | 234898271 | rs11588636 | 0.372 | 0.663 | -0.591 | -1.455 | -0.003 | -0.046 | -0.046 | 0 | 0.18 | 0.34 | 0.09 | 0.05 | 0.17 | 0.09 |
| 1 | 234901940 | rs2587530 | 0.331 | 0.475 | -0.347 | -1.546 | -0.022 | 0.077 | 0.077 | 0 | 0.41 | 0.26 | 0.51 | 0.8 | 0.47 | 0.68 |
| 1 | 234903206 | rs2796172 | -1.398 | 0.581 | -0.434 | -1.095 | -0.002 | 0.009 | 0.009 | 0.515 | 0.39 | 0.63 | 0.53 | 0.66 | 0.41 | 0.41 |
| 1 | 234903771 | rs3844069 | 0.288 | 0.682 | -0.531 | -0.798 | -0.001 | -0.05 | -0.05 | 0 | 0.18 | 0.35 | 0.1 | 0.05 | 0.16 | 0.09 |
| 1 | 234903911 | rs12727896 | 0.211 | 0.957 | -1.139 | -0.369 | 0.001 | 0.052 | 0.052 | 0 | 0.31 | 0.16 | 0.19 | 0.19 | 0.29 | 0.47 |
| 1 | 234906829 | rs2439500 | -0.172 | 0.848 | -0.782 | 0.167 | 0.011 | 0.084 | 0.084 | 0 | 0.09 | 0.1 | 0.3 | 0.59 | 0.17 | 0.2 |
| 1 | 234908088 | rs12067652 | 0.043 | 1.006 | -0.984 | -0.361 | -0.023 | 0.068 | 0.068 | 0 | 0.36 | 0.21 | 0.22 | 0.2 | 0.36 | 0.45 |
| 1 | 234909298 | rs4626924 | -0.355 | 0.517 | -0.621 | -0.796 | -0.004 | -0.034 | -0.034 | 0.03 | 0.53 | 0.52 | 0.63 | 0.83 | 0.56 | 0.77 |
| 1 | 234956342 | rs3914503 | -0.858 | 0.448 | -0.241 | -0.233 | -0.012 | 0.008 | 0.008 | 0.314 | 0.18 | 0.43 | 0.16 | 0.17 | 0.14 | 0.24 |
| 1 | 234963968 | rs2802984 | 0.5 | 0.543 | 0.09 | -0.678 | 0.012 | -0.038 | -0.038 | 0 | 0.93 | 0.9 | 0.94 | 1 | 0.94 | 0.85 |
| 1 | 234966277 | rs2796160 | 1.329 | 0.675 | 0.159 | -0.714 | 0.049 | -0.037 | -0.037 | 0 | 0.91 | 0.7 | 0.9 | 1 | 0.92 | 0.84 |
| 1 | 234966498 | rs2786007 | 0.8 | 0.758 | -0.955 | -1.349 | -0.097 | 0.11 | 0.11 | 1.634 | 0.31 | 0.03 | 0.22 | 0.08 | 0.38 | 0.14 |
| 1 | 234974779 | rs2048791 | 0.812 | 0.583 | -0.167 | -0.766 | 0.004 | -0.031 | -0.031 | 0 | 0.93 | 0.91 | 0.94 | 1 | 0.93 | 0.85 |
| 1 | 234990504 | rs883278 | -1.614 | 0.533 | -0.113 | -0.852 | 0.029 | -0.07 | -0.07 | 0 | 0.11 | 0.3 | 0.11 | 0 | 0.09 | 0.15 |
| 1 | 234998587 | rs7538849 | -0.981 | 0.473 | 0.093 | -0.838 | 0.022 | -0.003 | -0.003 | 0 | 0.05 | 0.24 | 0.05 | 0 | 0.03 | 0.01 |
| 1 | 235001591 | rs6586430 | -0.992 | 0.404 | 0.056 | -1.018 | 0 | -0.018 | -0.018 | 0.2 | 0.16 | 0.18 | 0.09 | 0.02 | 0.13 | 0.17 |
| 1 | 235005122 | rs2490458 | -1.379 | 0.426 | -0.039 | -0.99 | -0.013 | -0.135 | -0.135 | 0 | 0.22 | 0.42 | 0.14 | 0.02 | 0.16 | 0.2 |
| 1 | 235008029 | rs12758963 | -0.076 | 0.783 | -0.323 | -0.208 | -0.001 | 0.017 | 0.017 | 0 | 0.25 | 0.35 | 0.44 | 0.69 | 0.23 | 0.37 |
| 1 | 235011530 | rs10797566 | -0.696 | 1.012 | -0.763 | -0.902 | 0.023 | 0.048 | 0.048 | 0.211 | 0.51 | 0.43 | 0.62 | 0.76 | 0.55 | 0.67 |
| 1 | 235011725 | rs10910522 | -0.478 | 1.4 | -0.731 | -0.802 | -0.002 | 0.012 | 0.012 | 0 | 0.43 | 0.39 | 0.57 | 0.65 | 0.4 | 0.59 |
| 1 | 235014035 | rs2802950 | -1.125 | 1.535 | -0.397 | -0.897 | 0.008 | 0.02 | 0.02 | 0 | 0.55 | 0.59 | 0.64 | 0.71 | 0.57 | 0.64 |
| 1 | 235015199 | rs2802955 | -1.785 | 0.865 | -0.99 | -1.124 | -0.026 | 0.148 | 0.148 | 0.44 | 0.68 | 0.97 | 0.85 | 0.94 | 0.67 | 0.82 |
| 1 | 235017637 | rs1515656 | -0.287 | 0.727 | -0.541 | -0.814 | 0.004 | 0 | 0 | 0 | 0.4 | 0.4 | 0.52 | 0.59 | 0.34 | 0.52 |
| 1 | 235017928 | rs1515658 | 0.229 | 0.263 | -0.595 | -0.266 | 0.007 | -0.027 | -0.027 | 0 | 0.28 | 0.26 | 0.45 | 0.48 | 0.24 | 0.34 |
| 1 | 235018206 | rs10495355 | 0.197 | 0.244 | -0.717 | -0.47 | -0.003 | 0.014 | 0.014 | 0 | 0.22 | 0.27 | 0.41 | 0.54 | 0.18 | 0.37 |
| 1 | 235019266 | rs16844140 | 0.143 | 0.375 | -0.264 | -0.818 | 0.017 | -0.034 | -0.034 | 0 | 0.08 | 0.3 | 0.07 | 0 | 0.05 | 0.01 |
| 1 | 235024001 | rs6664412 | 0.076 | -0.587 | -0.876 | -0.891 | 0.008 | 0.007 | 0.007 | 0 | 0.15 | 0.12 | 0.06 | 0.03 | 0.08 | 0.15 |
| 1 | 235025575 | rs16844145 | -0.413 | -0.51 | -0.652 | -0.515 | -0.012 | 0.002 | 0.002 | 0 | 0.18 | 0.22 | 0.09 | 0.24 | 0.11 | 0.19 |
| 1 | 235025603 | rs4920161 | 1.224 | 0.002 | -1.231 | -0.18 | -0.13 | -0.167 | -0.167 | 0 | 0.32 | 0.03 | 0.47 | 0.43 | 0.44 | 0.35 |
| 1 | 235030325 | rs2673968 | 0.396 | -0.402 | -0.326 | -0.8 | -0.041 | -0.019 | -0.019 | 0.096 | 0.72 | 0.51 | 0.79 | 0.73 | 0.81 | 0.75 |
| 1 | 235030542 | rs2673969 | 0.23 | -0.36 | -0.332 | -0.944 | -0.009 | -0.02 | -0.02 | 0.07 | 0.72 | 0.68 | 0.89 | 0.94 | 0.85 | 0.75 |
| 1 | 235032251 | rs2803003 | 0.82 | -0.165 | -0.217 | -0.984 | 0.004 | -0.008 | -0.008 | 0 | 0.09 | 0.04 | 0.06 | 0.03 | 0.14 | 0.09 |
| 1 | 235032864 | rs1399847 | -0.204 | -0.438 | -0.193 | -1.009 | -0.018 | -0.026 | -0.026 | 0.091 | 0.26 | 0.33 | 0.09 | 0.06 | 0.13 | 0.24 |
| 1 | 235035709 | rs2802998 | 0.469 | 0.44 | 0.111 | -0.501 | 0.001 | -0.008 | -0.008 | 1.289 | 0.09 | 0.04 | 0.06 | 0.03 | 0.11 | 0.08 |
| 1 | 235040835 | rs7537522 | -0.499 | 0.328 | 0.646 | -0.629 | -0.012 | -0.058 | -0.058 | 0.004 | 0.1 | 0.36 | 0.06 | 0 | 0.05 | 0.03 |
| 1 | 235048988 | rs2674005 | -0.479 | 0.393 | 0.386 | -0.684 | 0 | -0.013 | -0.013 | 0 | 0.13 | 0.09 | 0.08 | 0.03 | 0.15 | 0.12 |
| 1 | 235049575 | rs2802988 | 1.392 | 0.583 | -0.015 | -0.452 | -0.054 | -0.069 | -0.069 | 0 | 0.31 | 0.11 | 0.39 | 0.43 | 0.44 | 0.35 |
| 1 | 235049923 | rs2802987 | 1.282 | 0.473 | 0.383 | -0.444 | -0.01 | -0.021 | -0.021 | 0.208 | 0.75 | 0.52 | 0.75 | 0.73 | 0.79 | 0.82 |
| 1 | 235050804 | rs17526880 | 0.069 | 0.921 | -0.483 | -0.152 | -0.012 | 0.029 | 0.029 | 0 | 0.13 | 0.04 | 0.07 | 0.03 | 0.14 | 0.13 |
| 1 | 235051501 | rs908329 | 0.945 | 0.86 | -0.527 | -0.132 | -0.041 | -0.063 | -0.063 | 0.055 | 0.33 | 0.12 | 0.39 | 0.43 | 0.45 | 0.37 |
| 1 | 235053032 | rs12562007 | -0.206 | 1.019 | -0.534 | -0.228 | -0.015 | 0.032 | 0.032 | 0 | 0.13 | 0.04 | 0.07 | 0.03 | 0.15 | 0.13 |
| 1 | 235053101 | rs6671950 | -0.464 | 1.009 | -0.535 | -0.15 | -0.024 | -0.02 | -0.02 | 0 | 0.15 | 0.05 | 0.17 | 0.27 | 0.18 | 0.14 |
| 1 | 235054637 | rs7553292 | -0.488 | 0.81 | -0.008 | -0.534 | 0 | 0.001 | 0.001 | 0 | 0.53 | 0.55 | 0.42 | 0.43 | 0.46 | 0.57 |
| 1 | 235056447 | rs7542764 | 0.739 | 0.898 | 0.307 | -0.671 | 0.023 | -0.022 | -0.022 | 0 | 0.06 | 0.2 | 0.05 | 0.01 | 0.05 | 0.01 |

| | | | | | | | | | | | | | | | |
|---|---|---|---|---|---|---|---|---|---|---|---|---|---|---|---|
| 1 | 235056631 | rs2673978 | 0.007 | 0.744 | 0.591 | -0.812 | 0.01 | 0.026 | 0.026 | 0 | 0.52 | 0.47 | 0.58 | 0.68 | 0.59 | 0.44 |
| 1 | 235056669 | rs4920200 | 0.4 | 0.749 | 0.586 | -0.817 | 0.01 | 0.018 | 0.018 | 0 | 0.39 | 0.3 | 0.34 | 0.32 | 0.32 | 0.53 |
| 1 | 235056714 | rs2802944 | 0.032 | 0.757 | 0.603 | -0.695 | 0.008 | -0.051 | -0.051 | 0.679 | 0.91 | 0.77 | 0.92 | 1 | 0.92 | 0.97 |
| 1 | 235059175 | rs2673961 | -0.723 | 0.693 | 0.336 | -0.731 | 0.007 | 0.01 | 0.01 | 0 | 0.63 | 0.74 | 0.68 | 0.69 | 0.7 | 0.47 |
| 1 | 235059784 | rs4456126 | 0.885 | 0.743 | 0.065 | -0.732 | 0.002 | 0.002 | 0.002 | 0 | 0.32 | 0.23 | 0.28 | 0.32 | 0.27 | 0.44 |
| 1 | 235060439 | rs10797568 | 0.022 | 1.375 | 0.182 | -0.815 | -0.001 | -0.01 | -0.01 | 0 | 0.47 | 0.52 | 0.38 | 0.32 | 0.44 | 0.56 |
| 1 | 235062181 | rs908319 | 0.432 | 1.475 | 0.018 | -0.612 | 0.004 | 0.015 | 0.015 | 0.195 | 0.35 | 0.19 | 0.51 | 0.66 | 0.33 | 0.31 |
| 1 | 235062433 | rs938455 | 0.337 | 1.493 | 0.055 | -0.634 | -0.001 | 0.014 | 0.014 | 0 | 0.51 | 0.49 | 0.38 | 0.34 | 0.49 | 0.61 |
| 1 | 235063539 | rs6665064 | -1.064 | 0.761 | -0.112 | -0.514 | 0.013 | -0.019 | -0.019 | 0 | 0.18 | 0.38 | 0.13 | 0.14 | 0.21 | 0.07 |
| 1 | 235067148 | rs2674004 | 0.555 | 0.778 | -0.099 | -0.533 | 0.014 | -0.101 | -0.101 | 0.871 | 0.83 | 0.57 | 0.89 | 1 | 0.81 | 0.92 |
| 1 | 235068131 | rs2803016 | 1.124 | 1.15 | -0.444 | -0.402 | 0.012 | -0.033 | -0.033 | 0 | 0.27 | 0.1 | 0.44 | 0.52 | 0.21 | 0.26 |
| 1 | 235069505 | rs12123230 | -0.355 | 1.3 | -0.403 | -0.61 | 0.024 | -0.064 | -0.064 | 0 | 0.13 | 0.28 | 0.1 | 0 | 0.2 | 0.06 |
| 1 | 235072837 | rs7532165 | -0.808 | 0.844 | -0.747 | -0.349 | -0.003 | -0.004 | -0.004 | 0 | 0.11 | 0.1 | 0.1 | 0.13 | 0.11 | 0.05 |
| 1 | 235075929 | rs2673980 | 1.03 | 0.759 | -0.723 | -0.431 | 0.001 | 0.005 | 0.005 | 0 | 0.85 | 0.85 | 0.87 | 0.87 | 0.84 | 0.93 |
| 1 | 235076145 | rs1515665 | 0.668 | 0.368 | -0.65 | -0.403 | -0.011 | 0.012 | 0.012 | 0 | 0.3 | 0.31 | 0.33 | 0.54 | 0.22 | 0.31 |
| 1 | 235084066 | rs2802963 | -0.018 | 0.983 | -0.941 | -0.412 | 0.017 | 0.009 | 0.009 | 0.118 | 0.49 | 0.5 | 0.45 | 0.54 | 0.41 | 0.29 |
| 1 | 235094931 | rs2802928 | -0.141 | 0.92 | -1.729 | -1.431 | -0.008 | 0.103 | 0.103 | 0 | 0.64 | 0.86 | 0.83 | 1 | 0.65 | 0.86 |
| 1 | 235095183 | rs7531062 | 0.133 | 0.447 | -1.316 | -1.244 | 0.004 | 0.001 | 0.001 | 0 | 0.38 | 0.36 | 0.23 | 0.13 | 0.33 | 0.15 |
| 1 | 235095444 | rs2802929 | 0.511 | 0.733 | -1.366 | -1.203 | -0.008 | -0.017 | -0.017 | 0 | 0.38 | 0.51 | 0.51 | 0.74 | 0.34 | 0.45 |
| 1 | 235095905 | rs2674000 | -0.898 | 0.496 | -1.291 | -1.822 | -0.011 | 0.041 | 0.041 | 2.113 | 0.92 | 1 | 0.96 | 1 | 0.93 | 0.95 |
| 1 | 235097045 | rs10910536 | -0.901 | 0.407 | -1.445 | -1.891 | -0.033 | 0.025 | 0.025 | 0 | 0.14 | 0.03 | 0.11 | 0.07 | 0.17 | 0.1 |
| 1 | 235097172 | rs2244398 | 0.858 | -0.64 | -0.902 | -1.851 | 0.002 | 0.011 | 0.011 | 0 | 0.54 | 0.35 | 0.53 | 0.71 | 0.56 | 0.58 |
| 1 | 235100980 | rs2138290 | -0.54 | -1.882 | 0.129 | -2.218 | -0.032 | -0.064 | -0.064 | 0 | 0.23 | 0.28 | 0.14 | 0.03 | 0.11 | 0.12 |
| 1 | 235108308 | rs16844296 | -0.947 | -1.915 | 0.468 | -2.086 | -0.055 | -0.027 | -0.027 | 0 | 0.31 | 0.49 | 0.32 | 0.3 | 0.2 | 0.21 |
| 1 | 235110859 | rs3738622 | -1.168 | -1.924 | 0.161 | -2.09 | -0.014 | -0.007 | -0.007 | 0.416 | 0.33 | 0.57 | 0.32 | 0.29 | 0.2 | 0.22 |
| 1 | 235112899 | rs6586356 | -0.235 | -1.723 | -0.07 | -2.352 | -0.031 | 0.049 | 0.049 | 0 | 0.71 | 0.77 | 0.74 | 0.91 | 0.63 | 0.7 |
| 1 | 235114594 | rs10495358 | 1.719 | -1.72 | -0.333 | -2.327 | -0.015 | 0.053 | 0.053 | 0.856 | 0.1 | 0 | 0.1 | 0 | 0.08 | 0.07 |
| 1 | 235115509 | rs6586357 | 0.278 | -1.56 | -0.104 | -2.221 | 0.02 | -0.005 | -0.005 | 0 | 0.21 | 0.11 | 0.25 | 0.28 | 0.14 | 0.14 |
| 1 | 235117291 | rs7551616 | 0.151 | -1.515 | -0.033 | -2.872 | 0.016 | -0.079 | -0.079 | 0 | 0.63 | 0.49 | 0.76 | 0.92 | 0.55 | 0.65 |
| 1 | 235119040 | rs17600500 | 0.414 | -1.054 | -0.77 | -2.516 | -0.025 | 0.127 | 0.127 | 0 | 0.48 | 0.23 | 0.7 | 0.71 | 0.53 | 0.49 |
| 1 | 235119382 | rs6586361 | -0.754 | -0.824 | -0.253 | -1.933 | -0.002 | 0.107 | 0.107 | 0 | 0.55 | 0.72 | 0.33 | 0.2 | 0.44 | 0.43 |
| 1 | 235124951 | rs6684582 | 0.152 | -0.377 | -0.81 | -1.565 | 0.016 | 0.016 | 0.016 | 0.422 | 0.12 | 0.06 | 0.05 | 0.11 | 0.07 | 0.11 |
| 1 | 235126144 | rs7412124 | -0.526 | 0.299 | -0.423 | -1.289 | -0.001 | 0.008 | 0.008 | 0.358 | 0.36 | 0.36 | 0.25 | 0.08 | 0.35 | 0.31 |
| 1 | 235137472 | rs28758671 | 0.736 | 0.285 | -0.339 | -0.106 | -0.013 | 0.01 | 0.01 | 0 | 0.92 | 0.84 | 0.96 | 0.93 | 0.94 | 0.83 |
| 1 | 235152757 | rs9727449 | 0.548 | 0.638 | -0.742 | -0.43 | 0 | -0.032 | -0.032 | 0 | 0.8 | 0.8 | 0.85 | 1 | 0.79 | 0.98 |
| 1 | 235190683 | rs34360045 | -0.704 | 0.867 | -0.5 | 0.48 | 0.015 | 0.071 | 0.071 | 0 | 0.29 | 0.51 | 0.17 | 0.49 | 0.31 | 0.25 |
| 1 | 235248412 | rs9662936 | -0.25 | 0.3 | -1.048 | 0.351 | 0.033 | 0.022 | 0.022 | 0 | 0.61 | 0.72 | 0.47 | 0.32 | 0.69 | 0.36 |
| 1 | 235251384 | rs9659229 | -0.85 | -0.691 | -0.343 | -0.582 | -0.076 | -0.141 | -0.141 | 0 | 0.22 | 0.41 | 0.14 | 0.01 | 0.11 | 0.09 |
| 1 | 235254623 | rs9287215 | -0.793 | -1.099 | -1.021 | 0.367 | -0.062 | -0.085 | -0.085 | 0 | 0.69 | 0.99 | 0.71 | 0.73 | 0.73 | 0.49 |
| 1 | 235263162 | rs7544301 | 0.132 | -0.647 | -0.356 | 0.403 | -0.001 | 0.002 | 0.002 | 0 | 0.36 | 0.4 | 0.41 | 0.27 | 0.31 | 0.52 |
| 1 | 235267127 | rs11577354 | -0.202 | 0.337 | 0.407 | 0.592 | 0.001 | 0.001 | 0.001 | 0 | 0.47 | 0.55 | 0.5 | 0.38 | 0.52 | 0.58 |
| 1 | 235268017 | rs7554276 | -0.885 | 0.28 | 0.232 | 0.135 | 0.005 | -0.075 | -0.075 | 0 | 0.66 | 0.85 | 0.64 | 0.4 | 0.67 | 0.69 |
| 1 | 235270774 | rs4567344 | -1.96 | 0.263 | -0.277 | -0.012 | -0.044 | -0.167 | -0.167 | 0 | 0.79 | 0.99 | 0.69 | 0.41 | 0.76 | 0.74 |
| 1 | 235273179 | rs11301 | -0.323 | 0.136 | -0.281 | -0.071 | 0.009 | -0.068 | -0.068 | 0 | 0.65 | 0.87 | 0.64 | 0.39 | 0.67 | 0.67 |
| 1 | 235276289 | rs4551650 | 2.275 | 0.159 | -0.355 | -0.119 | -0.107 | -0.051 | -0.051 | 0 | 0.3 | 0.01 | 0.37 | 0.27 | 0.33 | 0.53 |
| 1 | 235279448 | rs4619036 | -0.287 | 0.121 | -0.217 | -0.152 | -0.006 | -0.036 | -0.036 | 0 | 0.19 | 0.26 | 0.13 | 0.02 | 0.15 | 0.09 |
| 1 | 235290922 | rs4329572 | -0.733 | 0.174 | -0.309 | -0.102 | -0.01 | -0.07 | -0.07 | 1.357 | 0.69 | 0.87 | 0.64 | 0.37 | 0.67 | 0.68 |
| 1 | 235294115 | rs7513381 | -0.396 | -0.114 | -0.326 | -0.228 | -0.006 | -0.039 | -0.039 | 0 | 0.16 | 0.25 | 0.12 | 0.02 | 0.15 | 0.08 |
| 1 | 235303150 | rs34003373 | -0.073 | -0.113 | -0.326 | -0.228 | -0.002 | -0.031 | -0.031 | 0 | 0.14 | 0.21 | 0.1 | 0.02 | 0.14 | 0.08 |
| 1 | 235315711 | rs2273741 | 2.105 | -0.154 | -0.379 | -0.109 | -0.105 | -0.004 | -0.004 | 0 | 0.31 | 0.02 | 0.37 | 0.23 | 0.33 | 0.53 |
| 1 | 235322000 | rs16832455 | -0.261 | -0.123 | -0.324 | -0.118 | -0.003 | -0.028 | -0.028 | 0 | 0.16 | 0.2 | 0.1 | 0.02 | 0.15 | 0.08 |
| 1 | 235324298 | rs2295815 | -0.421 | -0.102 | 0.207 | -0.145 | -0.018 | -0.071 | -0.071 | 0.997 | 0.16 | 0.4 | 0.12 | 0.02 | 0.12 | 0.08 |
| 1 | 235325570 | rs7530147 | 0.415 | -0.216 | 0.212 | -0.154 | -0.001 | 0.015 | 0.015 | 1.947 | 0.51 | 0.51 | 0.5 | 0.25 | 0.47 | 0.6 |
| 1 | 235327138 | rs12095653 | 0.444 | -0.24 | 0.178 | -0.233 | 0 | 0.032 | 0.032 | 2.253 | 0.5 | 0.45 | 0.49 | 0.25 | 0.47 | 0.6 |
| 1 | 235328341 | rs4660114 | 0.473 | -0.289 | 0.159 | -0.215 | 0.001 | 0.037 | 0.037 | 2.575 | 0.5 | 0.45 | 0.48 | 0.25 | 0.47 | 0.6 |
| 1 | 235375056 | rs16832469 | -0.056 | -0.349 | 0.193 | -0.241 | -0.007 | -0.026 | -0.026 | 0 | 0.13 | 0.22 | 0.08 | 0.02 | 0.11 | 0.06 |
| 1 | 235383188 | rs4659654 | 0.256 | -0.344 | 0.19 | -0.246 | 0 | 0.041 | 0.041 | 2.535 | 0.5 | 0.44 | 0.51 | 0.25 | 0.48 | 0.6 |
| 1 | 235384838 | rs12117563 | 0.056 | -0.88 | -0.617 | -0.367 | 0.015 | -0.08 | -0.08 | 0 | 0.19 | 0.06 | 0.21 | 0.61 | 0.14 | 0.3 |
| 1 | 235498059 | rs12730441 | 1.31 | -1.315 | -1.457 | -1.379 | -0.02 | 0.04 | 0.04 | 4.505 | 0.34 | 0.21 | 0.32 | 0.22 | 0.35 | 0.52 |
| 1 | 235499263 | rs10925482 | 1.32 | -1.27 | -1.446 | -1.367 | -0.019 | 0.04 | 0.04 | 4.423 | 0.34 | 0.21 | 0.32 | 0.23 | 0.35 | 0.53 |
| 1 | 235500649 | rs2803851 | -1.381 | -1.361 | -1.05 | -1.339 | -0.008 | 0.009 | 0.009 | 0 | 0.71 | 0.78 | 0.76 | 0.86 | 0.62 | 0.9 |
| 1 | 235508463 | rs10925503 | 3.763 | -1.071 | -1.205 | -0.864 | 0.012 | -0.058 | -0.058 | 0 | 0.1 | 0 | 0.27 | 0.59 | 0.09 | 0.27 |
| 1 | 235515534 | rs2789365 | -0.003 | -0.942 | -0.949 | -1.373 | 0 | 0.003 | 0.003 | 1.197 | 0.5 | 0.57 | 0.44 | 0.25 | 0.47 | 0.59 |
| 1 | 235523067 | rs7554192 | 0.895 | -0.834 | -1.317 | -1.472 | -0.005 | -0.021 | -0.021 | 0 | 0.17 | 0.24 | 0.09 | 0.02 | 0.16 | 0.08 |
| 1 | 235523555 | rs7516807 | -0.73 | -0.68 | -1.047 | -1.197 | -0.007 | 0.071 | 0.071 | 5.594 | 0.52 | 0.66 | 0.59 | 0.74 | 0.49 | 0.41 |
| 1 | 235525253 | rs6699315 | -0.471 | -0.62 | -1.083 | -1.148 | 0.002 | 0.055 | 0.055 | 3.735 | 0.5 | 0.59 | 0.58 | 0.74 | 0.48 | 0.41 |
| 1 | 235539020 | rs12745648 | 0.764 | -0.288 | -0.708 | -0.959 | 0.002 | -0.045 | -0.045 | 0 | 0.11 | 0.25 | 0.08 | 0.02 | 0.11 | 0.09 |
| 1 | 235575321 | rs4659520 | 0.028 | -0.867 | -0.554 | -0.496 | -0.002 | 0.001 | 0.001 | 0 | 0.58 | 0.61 | 0.63 | 0.63 | 0.55 | 0.47 |
| 1 | 235600129 | rs6429082 | 0.255 | -1.105 | -0.26 | -0.546 | 0 | -0.005 | -0.005 | 0 | 0.54 | 0.52 | 0.6 | 0.66 | 0.53 | 0.48 |

| | | | | | | | | | | | | | | | | |
|---|---|---|---|---|---|---|---|---|---|---|---|---|---|---|---|---|
| 1 | 235611093 | rs6429094 | 0.441 | -1.071 | -0.214 | -0.606 | -0.01 | -0.044 | -0.044 | 1.783 | 0.12 | 0.27 | 0.08 | 0.02 | 0.09 | 0.1 |
| 1 | 235612320 | rs704705 | -1.805 | -1.103 | -0.348 | -0.619 | -0.131 | -0.055 | -0.055 | 0.339 | 0.73 | 0.97 | 0.72 | 0.68 | 0.65 | 0.62 |
| 1 | 235621026 | rs704710 | -0.917 | -1.18 | -0.14 | -0.677 | -0.006 | 0.046 | 0.046 | 0 | 0.48 | 0.67 | 0.42 | 0.34 | 0.48 | 0.5 |
| 1 | 235628908 | rs1305556 | -1.05 | -1.27 | -0.518 | -0.487 | 0.001 | -0.015 | -0.015 | 0.368 | 0.56 | 0.57 | 0.71 | 0.84 | 0.51 | 0.78 |
| 1 | 235637421 | rs704716 | -1.045 | -1.266 | -0.516 | -0.485 | 0.001 | -0.015 | -0.015 | 0 | 0.56 | 0.58 | 0.71 | 0.84 | 0.51 | 0.78 |
| 1 | 235642359 | rs167092 | 1.482 | -1.324 | -0.493 | -0.488 | -0.049 | -0.003 | -0.003 | 0 | 0.28 | 0.12 | 0.29 | 0.32 | 0.35 | 0.38 |
| 1 | 235645079 | rs10925824 | -0.587 | -1.331 | -0.469 | -0.491 | -0.042 | 0.06 | 0.06 | 0 | 0.24 | 0.37 | 0.41 | 0.52 | 0.15 | 0.39 |
| 1 | 235647631 | rs291387 | -1.114 | -1.315 | -0.479 | -0.491 | 0 | -0.011 | -0.011 | 0 | 0.56 | 0.59 | 0.71 | 0.84 | 0.51 | 0.78 |
| 1 | 235652513 | rs291396 | -1.106 | -1.365 | -0.477 | -0.49 | 0 | -0.01 | -0.01 | 0 | 0.56 | 0.59 | 0.71 | 0.84 | 0.51 | 0.78 |
| 1 | 235661544 | rs696244 | -1.071 | -1.371 | -0.491 | -0.49 | 0 | -0.013 | -0.013 | 0.454 | 0.55 | 0.57 | 0.71 | 0.84 | 0.51 | 0.78 |
| 1 | 235664494 | rs4659905 | -0.666 | -1.222 | -0.838 | -0.387 | -0.047 | 0.067 | 0.067 | 0 | 0.24 | 0.38 | 0.41 | 0.52 | 0.15 | 0.39 |
| 1 | 235669988 | rs704722 | 1.489 | -1.023 | -1.222 | -0.436 | -0.136 | -0.059 | -0.059 | 0.009 | 0.27 | 0.03 | 0.28 | 0.33 | 0.35 | 0.38 |
| 1 | 235673555 | rs10925876 | 0.21 | -1.054 | -1.147 | -0.443 | 0.023 | -0.054 | -0.054 | 0.537 | 0.19 | 0.14 | 0.36 | 0.52 | 0.15 | 0.38 |
| 1 | 235674379 | rs291340 | 1.058 | -0.373 | -1.004 | -0.572 | -0.097 | -0.031 | -0.031 | 0 | 0.28 | 0.06 | 0.28 | 0.33 | 0.35 | 0.38 |
| 1 | 235688043 | rs291365 | 0.153 | -0.242 | -0.887 | -0.472 | -0.01 | 0.009 | 0.009 | 0 | 0.35 | 0.31 | 0.32 | 0.33 | 0.38 | 0.41 |
| 1 | 235705157 | rs2131921 | -1.067 | 0.069 | -1.325 | -0.943 | -0.135 | 0.094 | 0.094 | 0 | 0.72 | 0.98 | 0.81 | 0.88 | 0.65 | 0.62 |
| 1 | 235705673 | rs10802797 | -0.466 | -0.023 | -0.468 | -1.001 | 0.004 | -0.031 | -0.031 | 0 | 0.67 | 0.66 | 0.77 | 0.88 | 0.64 | 0.55 |
| 1 | 235708538 | rs12746334 | 0.475 | -0.387 | -0.823 | -1.037 | 0.003 | 0.003 | 0.003 | 0 | 0.14 | 0.13 | 0.14 | 0.04 | 0.08 | 0.1 |
| 1 | 235710276 | rs6429146 | 0.114 | -0.333 | -0.938 | -1.124 | 0.004 | 0.075 | 0.075 | 0 | 0.51 | 0.41 | 0.62 | 0.82 | 0.56 | 0.48 |
| 1 | 235714162 | rs12756717 | -0.853 | -0.077 | -0.515 | -0.444 | -0.027 | 0.029 | 0.029 | 0 | 0.28 | 0.57 | 0.39 | 0.68 | 0.2 | 0.34 |
| 1 | 235717340 | rs425581 | -0.295 | 1.029 | -0.812 | -0.669 | -0.038 | -0.018 | -0.018 | 0.084 | 0.17 | 0.1 | 0.14 | 0.17 | 0.34 | 0.23 |
| 1 | 235728896 | rs488618 | -0.516 | 0.214 | -0.679 | -1.456 | 0.001 | 0.015 | 0.015 | 0.824 | 0.46 | 0.54 | 0.7 | 0.8 | 0.58 | 0.6 |
| 1 | 235738515 | rs6674438 | 0.974 | 0.232 | -0.768 | -1.449 | 0.006 | 0.055 | 0.055 | 0 | 0.22 | 0.12 | 0.13 | 0 | 0.19 | 0.07 |
| 1 | 235743992 | rs2841873 | -1.199 | 0.287 | -1.428 | -1.363 | -0.085 | 0.111 | 0.111 | 0 | 0.65 | 0.97 | 0.79 | 0.85 | 0.66 | 0.64 |
| 1 | 235756737 | rs881070 | 0.158 | 0.015 | -0.773 | -1.243 | -0.033 | -0.091 | -0.091 | 0 | 0.42 | 0.53 | 0.23 | 0.06 | 0.27 | 0.18 |
| 1 | 235756821 | rs2774320 | -0.721 | 0.458 | -1.956 | -1.227 | -0.107 | 0.15 | 0.15 | 0 | 0.6 | 0.97 | 0.73 | 0.86 | 0.61 | 0.57 |
| 1 | 235760817 | rs10926189 | -0.477 | 0.775 | -1.222 | -0.508 | -0.006 | -0.003 | -0.003 | 0 | 0.2 | 0.19 | 0.12 | 0.2 | 0.28 | 0.13 |
| 1 | 235764497 | rs4659946 | 0.542 | 0.45 | -1.091 | -0.741 | 0.002 | -0.001 | -0.001 | 0 | 0.69 | 0.68 | 0.41 | 0.26 | 0.58 | 0.46 |
| 1 | 235766062 | rs12140087 | 0.267 | 0.731 | -1.472 | -0.451 | -0.029 | -0.013 | -0.013 | 4.14 | 0.16 | 0.1 | 0.09 | 0.17 | 0.25 | 0.16 |
| 1 | 235766162 | rs17547995 | 0.398 | 0.347 | -1.485 | -0.61 | 0.018 | 0.071 | 0.071 | 0.074 | 0.4 | 0.2 | 0.25 | 0.24 | 0.34 | 0.39 |
| 1 | 235768378 | rs7555113 | -0.473 | 0.943 | -1.043 | -0.265 | -0.041 | -0.011 | -0.011 | 2.766 | 0.18 | 0.09 | 0.1 | 0.18 | 0.28 | 0.17 |
| 1 | 235774836 | rs6429196 | -0.142 | -0.621 | -0.639 | -0.659 | 0.029 | 0.03 | 0.03 | 0.057 | 0.41 | 0.65 | 0.24 | 0.08 | 0.28 | 0.12 |
| 1 | 235779137 | rs10926264 | -1.006 | -1.138 | 0.553 | 0.806 | -0.064 | 0.167 | 0.167 | 0.467 | 0.18 | 0.34 | 0.35 | 0.57 | 0.06 | 0.24 |
| 1 | 235790922 | rs6429206 | -0.646 | -0.772 | -0.978 | 0.001 | 0.005 | 0.121 | 0.121 | 0.9 | 0.54 | 0.87 | 0.71 | 0.75 | 0.45 | 0.52 |
| 1 | 235798334 | rs6429207 | -0.439 | -0.912 | -0.536 | -0.367 | 0.002 | 0.055 | 0.055 | 0 | 0.64 | 0.72 | 0.62 | 0.85 | 0.62 | 0.71 |
| 1 | 235802387 | rs7554426 | 0.023 | -0.687 | -0.659 | -0.237 | 0.06 | 0.173 | 0.173 | 1.081 | 0.53 | 0.2 | 0.56 | 0.85 | 0.61 | 0.7 |
| 1 | 235814507 | rs10159416 | -1.118 | 0.193 | 0.057 | 0.179 | 0.011 | 0.047 | 0.047 | 0 | 0.17 | 0.26 | 0.28 | 0.27 | 0.2 | 0.31 |
| 1 | 235826583 | rs3754230 | -1.322 | 0.209 | 0.791 | -0.196 | 0.005 | 0.001 | 0.001 | 0 | 0.09 | 0.1 | 0.19 | 0.12 | 0.1 | 0.22 |
| 1 | 235827443 | rs3819013 | 0.05 | -0.114 | 0.032 | -1.604 | -0.026 | 0.184 | 0.184 | 0 | 0.43 | 0.22 | 0.3 | 0.04 | 0.56 | 0.13 |
| 1 | 235832555 | rs11810173 | 0.162 | 0.391 | -0.368 | -1.977 | -0.015 | 0.027 | 0.027 | 0 | 0.12 | 0.06 | 0.1 | 0.04 | 0.15 | 0.03 |
| 1 | 235849939 | rs7545788 | -0.651 | 0.038 | 0.263 | -2.141 | 0.009 | 0.12 | 0.12 | 0.249 | 0.49 | 0.76 | 0.4 | 0.04 | 0.56 | 0.16 |
| 1 | 235850916 | rs6696218 | 0.027 | 0.013 | 0.158 | -2.108 | -0.001 | 0.023 | 0.023 | 0 | 0.12 | 0.06 | 0.1 | 0.04 | 0.12 | 0.03 |
| 1 | 235883125 | rs12074371 | -1.005 | 0.273 | 0.283 | -2.584 | -0.013 | 0.007 | 0.007 | 0 | 0.33 | 0.69 | 0.32 | 0.07 | 0.43 | 0.18 |
| 1 | 235890096 | rs2208384 | -0.559 | 0.159 | 0.246 | -2.575 | -0.015 | 0.03 | 0.03 | 0 | 0.79 | 0.91 | 0.75 | 0.84 | 0.79 | 0.75 |
| 1 | 235893915 | rs2208383 | 0.271 | -0.433 | -0.482 | -3.386 | 0.013 | 0.1 | 0.1 | 0 | 0.39 | 0.15 | 0.32 | 0.1 | 0.38 | 0.09 |
| 1 | 235909485 | rs10926565 | 0.764 | -0.076 | 0.136 | -2.967 | 0.051 | -0.089 | -0.089 | 0 | 0.18 | 0.33 | 0.19 | 0.02 | 0.28 | 0.15 |
| 1 | 235910392 | rs6676149 | -0.91 | -0.1 | 0.119 | -2.981 | 0.034 | -0.11 | -0.11 | 0 | 0.32 | 0.49 | 0.3 | 0.07 | 0.43 | 0.18 |
| 1 | 235945294 | rs6696123 | -0.617 | 0.028 | 0.05 | -2.918 | 0.048 | -0.112 | -0.112 | 0 | 0.3 | 0.49 | 0.3 | 0.07 | 0.43 | 0.18 |
| 1 | 235971763 | rs3768066 | 0.525 | 0.072 | -0.156 | -2.92 | -0.016 | 0.027 | 0.027 | 0 | 0.12 | 0.06 | 0.1 | 0.04 | 0.15 | 0.02 |
| 1 | 235972435 | rs3820553 | -0.466 | -0.113 | -0.199 | -2.862 | 0.046 | -0.111 | -0.111 | 0 | 0.31 | 0.49 | 0.3 | 0.07 | 0.42 | 0.17 |
| 1 | 235972867 | rs1063128 | -0.466 | -0.113 | -0.199 | -2.862 | 0.046 | -0.111 | -0.111 | 0 | 0.31 | 0.49 | 0.3 | 0.07 | 0.43 | 0.17 |
| 1 | 235988566 | rs2474929 | 1.594 | -0.225 | -0.77 | -2.924 | 0.001 | 0.136 | 0.136 | 0.52 | 0.41 | 0.03 | 0.4 | 0.77 | 0.34 | 0.58 |
| 1 | 235994867 | rs3820555 | -0.667 | -0.199 | -0.483 | -3.015 | 0.011 | -0.051 | -0.051 | 0.953 | 0.33 | 0.63 | 0.31 | 0.07 | 0.43 | 0.17 |
| 1 | 235995280 | rs6703642 | -0.88 | -0.228 | -0.52 | -3.03 | -0.089 | 0.176 | 0.176 | 0.066 | 0.35 | 0.89 | 0.34 | 0.07 | 0.43 | 0.17 |
| 1 | 236039877 | rs9782955 | -0.484 | 0.064 | -0.349 | -2.772 | -0.069 | 0.051 | 0.051 | 0 | 0.77 | 0.98 | 0.75 | 0.88 | 0.77 | 0.76 |
| 1 | 236044714 | rs2385029 | -1.2 | -0.143 | -0.137 | -2.87 | -0.018 | -0.088 | -0.088 | 0 | 0.83 | 0.73 | 0.91 | 1 | 0.88 | 0.97 |
| 1 | 236051207 | rs4660123 | 0.946 | -0.371 | 0.006 | -3.048 | 0.027 | 0.025 | 0.025 | 0 | 0.48 | 0.62 | 0.47 | 0.9 | 0.33 | 0.64 |
| 1 | 236054172 | rs10754801 | -0.391 | 0.087 | 0.005 | -2.995 | 0.022 | -0.042 | -0.042 | 0.022 | 0.22 | 0.37 | 0.31 | 0.13 | 0.25 | 0.31 |
| 1 | 236055833 | rs6657655 | 0.156 | 0.288 | 0.082 | -2.976 | 0.082 | -0.127 | -0.127 | 0 | 0.18 | 0.51 | 0.2 | 0 | 0.27 | 0.07 |
| 1 | 236065239 | rs12404348 | 1.645 | 0.065 | -0.451 | -2.791 | 0.049 | 0.187 | 0.187 | 0.818 | 0.42 | 0.12 | 0.42 | 0.85 | 0.33 | 0.5 |
| 1 | 236066991 | rs10803141 | -0.295 | 0.159 | -0.166 | -2.678 | 0.046 | -0.055 | -0.055 | 0 | 0.22 | 0.37 | 0.32 | 0.13 | 0.27 | 0.31 |
| 1 | 236070001 | rs6691710 | -0.341 | 0.114 | -0.171 | -2.662 | 0.031 | -0.066 | -0.066 | 0 | 0.23 | 0.36 | 0.32 | 0.1 | 0.27 | 0.26 |
| 1 | 236077778 | rs4660126 | 1.196 | -0.477 | -0.343 | -2.334 | 0.039 | 0.071 | 0.071 | 0.339 | 0.42 | 0.29 | 0.42 | 0.83 | 0.3 | 0.49 |
| 1 | 236084124 | rs927729 | -0.118 | -0.448 | 0.31 | -1.628 | -0.013 | -0.078 | -0.078 | 0 | 0.82 | 0.72 | 0.9 | 0.98 | 0.84 | 0.91 |
| 1 | 236087324 | rs1885253 | 0.97 | -0.685 | 0.078 | -1.619 | -0.009 | -0.03 | -0.03 | 0.166 | 0.71 | 0.34 | 0.87 | 0.94 | 0.83 | 0.89 |
| 1 | 236095773 | rs10803154 | 1.042 | -0.043 | -0.178 | -1.508 | -0.001 | -0.013 | -0.013 | 0 | 0.69 | 0.7 | 0.86 | 0.88 | 0.74 | 0.88 |
| 1 | 236096171 | rs16832929 | 0.109 | 0.524 | -0.595 | -0.07 | -0.062 | -0.088 | -0.088 | 0.815 | 0.13 | 0.01 | 0.28 | 0.35 | 0.19 | 0.32 |
| 1 | 236097628 | rs12047081 | -0.971 | 0.28 | -0.001 | -0.017 | 0.005 | 0.006 | 0.006 | 0.126 | 0.21 | 0.18 | 0.09 | 0.12 | 0.15 | 0.08 |
| 1 | 236099394 | rs4445453 | -0.55 | 0.267 | -0.297 | -0.048 | -0.005 | 0.023 | 0.023 | 0.963 | 0.17 | 0.15 | 0.09 | 0.11 | 0.18 | 0.08 |

| | | | | | | | | | | | | | | | | |
|---|---|---|---|---|---|---|---|---|---|---|---|---|---|---|---|---|
| 1 | 236102226 | rs10803159 | -1.695 | 0.081 | 0.062 | 0.218 | -0.045 | 0.098 | 0.098 | 1.828 | 0.28 | 0.48 | 0.21 | 0.51 | 0.2 | 0.32 |
| 1 | 236103925 | rs10927073 | -0.59 | 0.066 | -0.273 | 0.155 | -0.002 | -0.05 | -0.05 | 0.298 | 0.19 | 0.13 | 0.17 | 0.48 | 0.19 | 0.29 |
| 1 | 236104822 | rs6669631 | -0.556 | 0.024 | -0.065 | 0.136 | -0.032 | -0.011 | -0.011 | 0.104 | 0.09 | 0.32 | 0.03 | 0.02 | 0.01 | 0.03 |
| 1 | 236105308 | rs4660131 | 1.393 | 0.756 | -0.992 | 0.424 | 0.005 | -0.048 | -0.048 | 1.52 | 0.71 | 0.77 | 0.77 | 0.51 | 0.7 | 0.68 |
| 1 | 236109373 | rs10927083 | -1.088 | 0.943 | -0.56 | 0.86 | 0 | -0.039 | -0.039 | 0 | 0.19 | 0.16 | 0.17 | 0.48 | 0.19 | 0.29 |
| 1 | 236135681 | rs766890 | -1.476 | 0.127 | -1.15 | 0.919 | 0.056 | -0.131 | -0.131 | 0 | 0.33 | 0.89 | 0.31 | 0.52 | 0.23 | 0.29 |
| 1 | 236140266 | rs1048253 | -1.409 | 0.007 | 0.162 | 1.288 | -0.054 | 0.092 | 0.092 | 0.857 | 0.24 | 0.55 | 0.28 | 0.61 | 0.14 | 0.28 |
| 1 | 236141174 | rs3213190 | -1.234 | -0.018 | 0.542 | 1.299 | -0.021 | 0.04 | 0.04 | 0.189 | 0.23 | 0.39 | 0.26 | 0.61 | 0.15 | 0.28 |
| 1 | 236144754 | rs3820558 | -1.082 | -0.152 | 0.682 | 1.315 | -0.003 | 0.003 | 0.003 | 0 | 0.22 | 0.26 | 0.25 | 0.63 | 0.14 | 0.27 |
| 1 | 236144951 | rs3738525 | -1.259 | -0.097 | 0.647 | 1.323 | -0.019 | 0.027 | 0.027 | 0.223 | 0.23 | 0.36 | 0.26 | 0.62 | 0.14 | 0.27 |
| 1 | 236150459 | rs4659618 | -0.502 | -0.3 | 0.323 | 1.002 | 0.006 | -0.013 | -0.013 | 0.127 | 0.78 | 0.87 | 0.83 | 0.73 | 0.78 | 0.62 |
| 1 | 236153161 | rs1570831 | -0.665 | -0.323 | 0.313 | 1.005 | -0.033 | 0.035 | 0.035 | 0.12 | 0.21 | 0.33 | 0.23 | 0.62 | 0.11 | 0.26 |
| 1 | 236154012 | rs1041111 | 1.486 | -0.309 | 0.38 | 0.955 | 0.02 | 0.028 | 0.028 | 4.977 | 0.52 | 0.33 | 0.56 | 0.11 | 0.64 | 0.35 |
| 1 | 236154365 | rs4660137 | -1.068 | -0.092 | -0.015 | 1.147 | -0.028 | 0.036 | 0.036 | 0.093 | 0.23 | 0.38 | 0.26 | 0.64 | 0.14 | 0.27 |
| 1 | 236155817 | rs4660139 | 0.035 | 0.454 | -0.35 | 1.546 | -0.003 | -0.032 | -0.032 | 0.016 | 0.75 | 0.81 | 0.77 | 0.49 | 0.74 | 0.6 |
| 1 | 236157277 | rs16833075 | -1.103 | 0.821 | -0.132 | 1.722 | -0.002 | 0.027 | 0.027 | 0 | 0.19 | 0.23 | 0.24 | 0.55 | 0.14 | 0.39 |
| 1 | 236166380 | rs9726880 | -1.21 | 1.978 | 0.342 | 1.107 | 0.147 | -0.012 | -0.012 | 0 | 0.15 | 0.37 | 0.19 | 0.1 | 0.36 | 0.22 |
| 1 | 236170665 | rs11590845 | -0.949 | 2.115 | 0.171 | 1.11 | -0.023 | 0.007 | 0.007 | 0.636 | 0.11 | 0.07 | 0.14 | 0.09 | 0.35 | 0.2 |
| 1 | 236175327 | rs3738531 | -0.194 | 0.956 | 0.07 | 1.676 | 0.005 | -0.005 | -0.005 | 0 | 0.13 | 0.11 | 0.31 | 0.26 | 0.09 | 0.27 |
| 1 | 236179042 | rs16833108 | -0.186 | 0.987 | 0.107 | 1.676 | 0.002 | 0.001 | 0.001 | 0.041 | 0.12 | 0.11 | 0.31 | 0.26 | 0.09 | 0.27 |
| 1 | 236182924 | rs12411075 | -0.462 | 0.907 | 0.108 | 1.664 | 0.027 | -0.05 | -0.05 | 0 | 0.14 | 0.05 | 0.31 | 0.26 | 0.1 | 0.26 |
| 1 | 236187365 | rs3738533 | -0.951 | 0.679 | 0.255 | 1.542 | -0.018 | 0.017 | 0.017 | 0.255 | 0.18 | 0.21 | 0.33 | 0.27 | 0.12 | 0.28 |
| 1 | 236192782 | rs3818238 | -0.96 | 1.135 | 0.153 | 2.02 | -0.004 | 0.321 | 0.321 | 0 | 0.07 | 0.25 | 0.13 | 0.41 | 0.04 | 0.23 |
| 1 | 236201609 | rs4659620 | -1.119 | 1.976 | -0.717 | 1.007 | -0.051 | 0.028 | 0.028 | 0 | 0.14 | 0.05 | 0.12 | 0.1 | 0.27 | 0.16 |
| 1 | 236203280 | rs3768085 | -0.569 | 1.4 | -0.903 | 1.203 | -0.004 | -0.034 | -0.034 | 1.512 | 0.08 | 0 | 0.1 | 0.11 | 0.08 | 0.06 |
| 1 | 236207687 | rs11583517 | -0.731 | 2.287 | -0.909 | 2.034 | -0.026 | -0.03 | -0.03 | 2.44 | 0.12 | 0.06 | 0.4 | 0.36 | 0.24 | 0.26 |
| 1 | 236209883 | rs927730 | -0.542 | 2.725 | -0.737 | 1.917 | 0.017 | 0.039 | 0.039 | 1.257 | 0.16 | 0.21 | 0.11 | 0.34 | 0.19 | 0.33 |
| 1 | 236212038 | rs6665008 | 0.11 | 2.853 | -0.918 | 1.712 | -0.014 | 0.023 | 0.023 | 0.647 | 0.1 | 0.03 | 0.07 | 0 | 0.11 | 0.09 |
| 1 | 236220574 | rs12409606 | -0.342 | 2.66 | -0.906 | 1.857 | -0.025 | -0.029 | -0.029 | 0 | 0.12 | 0.06 | 0.4 | 0.36 | 0.24 | 0.26 |
| 1 | 236245015 | rs2734815 | -1.362 | 1.697 | -1.912 | 0.266 | -0.048 | 0.142 | 0.142 | 0.918 | 0.71 | 0.99 | 0.88 | 0.96 | 0.75 | 0.86 |
| 1 | 236252880 | rs2753440 | -1.032 | 1.707 | -0.98 | 0.849 | 0.007 | 0 | 0 | 0 | 0.31 | 0.32 | 0.51 | 0.65 | 0.44 | 0.58 |
| 1 | 236263421 | rs17554571 | 0.09 | 0.948 | -0.988 | 0.607 | 0.006 | 0.04 | 0.04 | 0 | 0.22 | 0.12 | 0.17 | 0.16 | 0.2 | 0.25 |
| 1 | 236264213 | rs2734808 | -0.166 | 1.212 | -1.003 | 1.202 | -0.019 | -0.024 | -0.024 | 0 | 0.12 | 0.07 | 0.39 | 0.37 | 0.22 | 0.23 |
| 1 | 236265665 | rs16833290 | -0.199 | 0.805 | -0.355 | 0.907 | -0.011 | -0.015 | -0.015 | 0.056 | 0.22 | 0.3 | 0.14 | 0.18 | 0.18 | 0.15 |
| 1 | 236272711 | rs6682352 | -0.247 | 0.829 | -0.103 | 0.658 | 0.003 | -0.055 | -0.055 | 2.525 | 0.22 | 0.38 | 0.18 | 0.08 | 0.2 | 0.13 |
| 1 | 236274926 | rs7542662 | -1.138 | 1.018 | -0.288 | 1.033 | 0.003 | 0.012 | 0.012 | 0 | 0.33 | 0.38 | 0.49 | 0.44 | 0.39 | 0.36 |
| 1 | 236277789 | rs4659625 | -0.487 | 1.094 | -0.306 | 1.321 | 0.011 | 0.082 | 0.082 | 0 | 0.12 | 0.2 | 0.3 | 0.38 | 0.22 | 0.23 |
| 1 | 236281455 | rs6658832 | -0.712 | 0.499 | -0.321 | 0.42 | -0.002 | -0.005 | -0.005 | 0 | 0.48 | 0.54 | 0.55 | 0.69 | 0.45 | 0.59 |
| 1 | 236284300 | rs16833337 | 0.144 | 0.507 | -0.541 | -0.019 | 0.005 | 0.003 | 0.003 | 0 | 0.38 | 0.34 | 0.34 | 0.37 | 0.28 | 0.31 |
| 1 | 236285562 | rs10495370 | 0.34 | 0.539 | -0.517 | -0.249 | -0.015 | -0.022 | -0.022 | 0 | 0.1 | 0.18 | 0.06 | 0.06 | 0.06 | 0.04 |
| 1 | 236291032 | rs10924293 | -0.929 | 0.998 | -0.879 | -0.577 | -0.044 | 0.061 | 0.061 | 0 | 0.82 | 0.93 | 0.79 | 0.99 | 0.72 | 0.78 |
| 1 | 236291893 | rs16833356 | -0.032 | 0.534 | -0.294 | -0.485 | -0.028 | -0.012 | -0.012 | 0 | 0.12 | 0.29 | 0.07 | 0.13 | 0.07 | 0.09 |
| 1 | 236292483 | rs10754470 | 0.803 | 0.222 | -0.209 | -0.514 | 0 | 0.001 | 0.001 | 0.045 | 0.47 | 0.37 | 0.42 | 0.59 | 0.44 | 0.44 |
| 1 | 236292668 | rs10802244 | 0.239 | 0.34 | -0.529 | -0.81 | -0.013 | 0.01 | 0.01 | 0 | 0.64 | 0.63 | 0.65 | 0.87 | 0.47 | 0.66 |
| 1 | 236292851 | rs4659628 | 0.682 | 0.026 | -0.446 | -0.414 | 0.005 | -0.001 | -0.001 | 0 | 0.44 | 0.52 | 0.36 | 0.53 | 0.25 | 0.43 |
| 1 | 236293754 | rs7546012 | -0.145 | 0.61 | -0.396 | -0.648 | -0.006 | -0.006 | -0.006 | 0 | 0.74 | 0.71 | 0.78 | 0.91 | 0.57 | 0.73 |
| 1 | 236294825 | rs4659633 | 0.096 | 0.721 | -0.715 | -0.693 | -0.046 | 0.044 | 0.044 | 0 | 0.7 | 0.75 | 0.76 | 0.94 | 0.53 | 0.76 |
| 1 | 236295308 | rs12097024 | 1.163 | 0.264 | -0.591 | -0.456 | -0.018 | -0.016 | -0.016 | 0 | 0.08 | 0.15 | 0.04 | 0.05 | 0.04 | 0.03 |
| 1 | 236296322 | rs6689479 | 0.012 | -0.153 | -0.699 | -0.731 | -0.01 | -0.025 | -0.025 | 0.464 | 0.22 | 0.24 | 0.1 | 0.1 | 0.15 | 0.11 |
| 1 | 236296950 | rs12566852 | 0.45 | 0.121 | -0.707 | -0.122 | -0.014 | -0.016 | -0.016 | 3.391 | 0.24 | 0.17 | 0.35 | 0.27 | 0.27 | 0.27 |
| 1 | 236298158 | rs2025016 | 0.142 | 0.101 | -0.326 | 0.249 | -0.046 | 0.091 | 0.091 | 0 | 0.67 | 0.86 | 0.78 | 0.93 | 0.58 | 0.71 |
| 1 | 236300472 | rs10924330 | -1.131 | -0.023 | 0.039 | 0.962 | 0.012 | -0.015 | -0.015 | 0 | 0.43 | 0.67 | 0.54 | 0.49 | 0.43 | 0.39 |
| 1 | 236306825 | rs7543562 | 0.205 | -0.125 | 0.297 | 1.146 | 0.002 | 0.003 | 0.003 | 0 | 0.67 | 0.6 | 0.74 | 0.63 | 0.62 | 0.72 |
| 1 | 236307360 | rs10924341 | 1.011 | -0.173 | 0.321 | 1.163 | 0.011 | -0.015 | -0.015 | 0 | 0.53 | 0.27 | 0.44 | 0.5 | 0.51 | 0.6 |
| 1 | 236310024 | rs911271 | 0.55 | -0.123 | 0.165 | 0.9 | -0.001 | -0.002 | -0.002 | 0 | 0.59 | 0.63 | 0.51 | 0.79 | 0.52 | 0.69 |
| 1 | 236311886 | rs10802285 | -0.646 | 0.283 | 0.221 | 1.107 | -0.042 | 0.059 | 0.059 | 0 | 0.81 | 0.92 | 0.86 | 0.96 | 0.73 | 0.81 |
| 1 | 236312073 | rs16833387 | -1.586 | -0.374 | 1.301 | 2.159 | 0.013 | 0.344 | 0.344 | 2.665 | 0.06 | 0.34 | 0.07 | 0.31 | 0 | 0.08 |
| 1 | 236315070 | rs10924380 | -1.126 | -0.274 | 1.11 | 2.141 | -0.014 | 0.14 | 0.14 | 0.394 | 0.15 | 0.33 | 0.35 | 0.38 | 0.16 | 0.2 |
| 1 | 236327799 | rs10924453 | -2.165 | -0.264 | 1.09 | 2.102 | 0.035 | 0.325 | 0.325 | 1.5 | 0.07 | 0.5 | 0.09 | 0.18 | 0.01 | 0.09 |
| 1 | 236329513 | rs7537965 | -1.815 | -0.199 | 0.946 | 2.154 | 0.051 | 0.422 | 0.422 | 0.517 | 0.06 | 0.44 | 0.08 | 0.26 | 0.01 | 0.09 |
| 1 | 236335649 | rs4304570 | -0.351 | 0.021 | -0.075 | 1.694 | -0.007 | -0.034 | -0.034 | 0 | 0.11 | 0.03 | 0.26 | 0.18 | 0.14 | 0.15 |
| 1 | 236336829 | rs13374732 | -0.301 | 0.148 | 0.033 | 1.23 | 0.006 | -0.028 | -0.028 | 0 | 0.07 | 0.12 | 0.04 | 0.01 | 0.07 | 0.12 |
| 1 | 236338226 | rs4659635 | -0.035 | -0.19 | -0.459 | 1.382 | 0.002 | -0.004 | -0.004 | 0 | 0.18 | 0.16 | 0.22 | 0.28 | 0.15 | 0.16 |
| 1 | 236346213 | rs2385203 | -0.244 | 0.026 | -0.658 | 1.058 | -0.006 | -0.05 | -0.05 | 0.594 | 0.13 | 0.04 | 0.22 | 0.2 | 0.11 | 0.16 |
| 1 | 236357391 | rs10924691 | -1.054 | -0.358 | 0.416 | 0.928 | -0.028 | 0.018 | 0.018 | 0 | 0.18 | 0.29 | 0.24 | 0.23 | 0.11 | 0.2 |
| 1 | 236358092 | rs10924694 | -0.201 | -0.45 | 0.254 | 0.596 | 0.011 | 0.045 | 0.045 | 0.451 | 0.27 | 0.57 | 0.38 | 0.61 | 0.26 | 0.4 |
| 1 | 236370654 | rs1254205 | -1.133 | -0.465 | 0.494 | 0.108 | 0 | -0.033 | -0.033 | 0 | 0.43 | 0.52 | 0.61 | 0.83 | 0.41 | 0.49 |
| 1 | 236378709 | rs1110423 | -0.566 | -0.699 | 0.883 | -0.376 | -0.026 | 0.013 | 0.013 | 0.285 | 0.2 | 0.57 | 0.29 | 0.21 | 0.14 | 0.17 |
| 1 | 236378916 | rs1111667 | -0.577 | -0.705 | 0.947 | -0.38 | -0.03 | 0.017 | 0.017 | 0.058 | 0.2 | 0.54 | 0.29 | 0.21 | 0.14 | 0.17 |

| | | | | | | | | | | | | | | | | |
|---|---|---|---|---|---|---|---|---|---|---|---|---|---|---|---|---|
| 1 | 236379623 | rs2463193 | -0.325 | -0.831 | 1.164 | -0.511 | 0 | 0.002 | 0.002 | 0.385 | 0.28 | 0.31 | 0.35 | 0.68 | 0.26 | 0.36 |
| 1 | 236380611 | rs1726631 | -1.128 | -0.969 | 0.962 | -1.19 | -0.001 | -0.037 | -0.037 | 1.306 | 0.45 | 0.49 | 0.58 | 0.88 | 0.4 | 0.53 |
| 1 | 236384402 | rs1405623 | -1.221 | -0.991 | 0.945 | -1.169 | -0.001 | -0.028 | -0.028 | 0 | 0.44 | 0.48 | 0.58 | 0.88 | 0.4 | 0.53 |
| 1 | 236389999 | rs10924790 | -0.546 | -0.952 | 0.816 | -0.554 | -0.017 | 0.151 | 0.151 | 0 | 0.07 | 0.1 | 0.08 | 0.37 | 0.03 | 0.13 |
| 1 | 236394735 | rs12023793 | 0.25 | -0.957 | 0.945 | -0.576 | -0.011 | -0.015 | -0.015 | 0 | 0.16 | 0.25 | 0.21 | 0.12 | 0.12 | 0.21 |
| 1 | 236394926 | rs941238 | 0.788 | -0.992 | 0.928 | -0.607 | -0.001 | -0.036 | -0.036 | 0.008 | 0.6 | 0.22 | 0.44 | 0.49 | 0.63 | 0.6 |
| 1 | 236404391 | rs2695057 | 0.641 | -0.979 | 0.874 | -0.622 | -0.018 | -0.037 | -0.037 | 0.026 | 0.21 | 0.16 | 0.31 | 0.4 | 0.25 | 0.21 |
| 1 | 236407465 | rs1405628 | -0.256 | -0.985 | 0.91 | -0.618 | 0 | -0.001 | -0.001 | 1.777 | 0.27 | 0.3 | 0.39 | 0.77 | 0.28 | 0.32 |
| 1 | 236420985 | rs7527802 | 0.252 | -0.728 | 0.951 | -0.507 | -0.011 | -0.014 | -0.014 | 0 | 0.16 | 0.24 | 0.21 | 0.12 | 0.12 | 0.21 |
| 1 | 236426120 | rs1749582 | 0.441 | -0.617 | 0.689 | -0.507 | -0.012 | -0.026 | -0.026 | 1.61 | 0.21 | 0.19 | 0.31 | 0.4 | 0.25 | 0.21 |
| 1 | 236439916 | rs6662005 | -0.678 | -0.085 | 0.83 | -0.624 | 0.05 | 0.294 | 0.294 | 0.02 | 0.11 | 0.29 | 0.11 | 0.38 | 0.11 | 0.18 |
| 1 | 236441037 | rs1749553 | 0.537 | -0.453 | 0.328 | -1.472 | 0.003 | 0.072 | 0.072 | 0 | 0.54 | 0.33 | 0.43 | 0.14 | 0.48 | 0.46 |
| 1 | 236442464 | rs17566445 | 0.631 | -0.245 | 0.245 | -1.584 | -0.028 | 0.033 | 0.033 | 0.081 | 0.14 | 0.02 | 0.13 | 0.07 | 0.14 | 0.16 |
| 1 | 236451196 | rs557205 | 1.068 | -0.238 | 0.298 | -1.71 | 0.012 | 0.119 | 0.119 | 0 | 0.38 | 0.18 | 0.21 | 0.04 | 0.37 | 0.31 |
| 1 | 236465416 | rs17567019 | 1.068 | -0.62 | 0.632 | -1.411 | 0.002 | 0.13 | 0.13 | 0.005 | 0.43 | 0.2 | 0.27 | 0.04 | 0.44 | 0.35 |
| 1 | 236469925 | rs7524741 | 0.243 | -0.754 | 0.807 | -1.186 | -0.039 | 0.071 | 0.071 | 0 | 0.23 | 0.32 | 0.32 | 0.45 | 0.15 | 0.34 |
| 1 | 236474797 | rs7554436 | 0.958 | -0.669 | 0.79 | -1.216 | 0.005 | 0.033 | 0.033 | 0 | 0.46 | 0.38 | 0.29 | 0.04 | 0.46 | 0.35 |
| 1 | 236476805 | rs1562200 | 0.549 | -0.503 | 0.614 | -1.086 | -0.029 | 0.128 | 0.128 | 0.136 | 0.55 | 0.24 | 0.44 | 0.15 | 0.54 | 0.47 |
| 1 | 236480062 | rs9287214 | 0.365 | -0.427 | 0.582 | -1.125 | -0.015 | 0.049 | 0.049 | 0.652 | 0.57 | 0.34 | 0.44 | 0.14 | 0.54 | 0.48 |
| 1 | 236484653 | rs7521930 | 0.154 | -1.215 | 0.641 | -0.818 | -0.025 | 0.095 | 0.095 | 0 | 0.22 | 0.33 | 0.33 | 0.46 | 0.18 | 0.34 |
| 1 | 236494413 | rs16833586 | -0.36 | -1.902 | 0.709 | -0.836 | -0.036 | -0.025 | -0.025 | 0.626 | 0.15 | 0.29 | 0.19 | 0.1 | 0.08 | 0.13 |
| 1 | 236494471 | rs12040004 | -0.266 | -1.896 | 0.431 | -0.872 | -0.032 | 0.035 | 0.035 | 0 | 0.28 | 0.33 | 0.34 | 0.45 | 0.2 | 0.35 |
| 1 | 236506693 | rs688831 | -0.193 | -1.721 | 0.007 | -0.901 | -0.024 | -0.02 | -0.02 | 0 | 0.86 | 0.69 | 0.81 | 0.89 | 0.91 | 0.87 |
| 1 | 236506815 | rs10802528 | 0.793 | -1.802 | -0.039 | -1.032 | -0.002 | 0.022 | 0.022 | 0 | 0.47 | 0.37 | 0.32 | 0.07 | 0.52 | 0.41 |
| 1 | 236509101 | rs609516 | 0.823 | -1.856 | -0.006 | -0.939 | -0.034 | -0.039 | -0.039 | 3.459 | 0.75 | 0.41 | 0.75 | 0.86 | 0.85 | 0.79 |
| 1 | 236517110 | rs599110 | -0.594 | -1.681 | -0.185 | -0.865 | -0.003 | -0.007 | -0.007 | 0 | 0.86 | 0.88 | 0.82 | 0.89 | 0.88 | 0.85 |
| 1 | 236517124 | rs605998 | -1.319 | -1.12 | -0.034 | -0.37 | 0.018 | 0.053 | 0.053 | 0 | 0.5 | 0.6 | 0.55 | 0.85 | 0.43 | 0.54 |
| 1 | 236520701 | rs11577428 | 0.097 | -1.112 | -0.105 | -0.284 | -0.034 | -0.058 | -0.058 | 0.762 | 0.11 | 0.25 | 0.06 | 0.03 | 0.06 | 0.08 |
| 1 | 236521008 | rs10925109 | 0.245 | -0.372 | 0.041 | 0.383 | 0.026 | -0.052 | -0.052 | 0 | 0.18 | 0.11 | 0.28 | 0.42 | 0.12 | 0.29 |
| 1 | 236525568 | rs667325 | 0.663 | 0.615 | 0.852 | 1.011 | 0.044 | 0.029 | 0.029 | 0.1 | 0.92 | 0.75 | 0.82 | 0.9 | 0.92 | 0.86 |
| 1 | 236529878 | rs590470 | -0.745 | 0.485 | 0.276 | 1.057 | -0.01 | -0.023 | -0.023 | 2.14 | 0.22 | 0.19 | 0.2 | 0.39 | 0.26 | 0.19 |
| 1 | 236533247 | rs2463198 | -0.909 | 0.553 | -0.251 | 1.09 | 0.024 | 0.082 | 0.082 | 0.12 | 0.24 | 0.42 | 0.22 | 0.39 | 0.26 | 0.19 |
| 1 | 236546386 | rs2120959 | 0.332 | 0.028 | -1.046 | 0.793 | 0.044 | -0.009 | -0.009 | 0 | 0.55 | 0.17 | 0.61 | 0.45 | 0.65 | 0.52 |
| 1 | 236548122 | rs6695464 | 0.419 | -0.052 | 0.056 | 0.464 | 0.023 | -0.08 | -0.08 | 0 | 0.83 | 0.48 | 0.86 | 0.92 | 0.83 | 0.94 |
| 1 | 236552530 | rs4659660 | 0.17 | 0.013 | 0.684 | 0.43 | 0.038 | -0.013 | -0.013 | 0 | 0.92 | 0.79 | 0.92 | 0.92 | 0.87 | 0.94 |
| 1 | 236557412 | rs10802533 | 0.936 | -0.049 | 0.941 | 0.401 | 0.021 | -0.078 | -0.078 | 0 | 0.83 | 0.53 | 0.87 | 0.92 | 0.83 | 0.94 |
| 1 | 236557771 | rs966365 | 1.084 | -0.051 | 0.975 | 0.4 | 0.019 | -0.091 | -0.091 | 0 | 0.82 | 0.47 | 0.86 | 0.92 | 0.83 | 0.94 |
| 1 | 236557928 | rs966364 | 1.133 | -0.052 | 0.977 | 0.399 | 0.014 | -0.089 | -0.089 | 0 | 0.82 | 0.45 | 0.86 | 0.92 | 0.83 | 0.94 |
| 1 | 236560021 | rs12080914 | 0.78 | -0.637 | 0.214 | 0.773 | 0.037 | 0.004 | 0.004 | 2.057 | 0.48 | 0.24 | 0.43 | 0.37 | 0.58 | 0.51 |
| 1 | 236563287 | rs1999506 | 0.666 | -1.185 | 0.065 | 0.542 | 0.058 | 0.106 | 0.106 | 0 | 0.47 | 0.2 | 0.51 | 0.26 | 0.65 | 0.52 |
| 1 | 236567767 | rs593981 | 0.431 | -1.251 | 1.133 | 0.483 | 0.001 | 0 | 0 | 0 | 0.52 | 0.47 | 0.53 | 0.28 | 0.65 | 0.52 |
| 1 | 236568770 | rs608825 | 0.574 | -1.238 | 1.463 | 0.479 | 0.026 | 0.043 | 0.043 | 0 | 0.49 | 0.33 | 0.5 | 0.26 | 0.65 | 0.52 |
| 1 | 236573260 | rs645507 | -0.18 | -1.444 | 1.596 | 0.444 | 0 | 0.001 | 0.001 | 0 | 0.65 | 0.63 | 0.69 | 0.39 | 0.67 | 0.67 |
| 1 | 236574267 | rs630891 | -0.014 | -1.461 | 1.62 | 0.421 | -0.002 | 0.004 | 0.004 | 0 | 0.64 | 0.55 | 0.69 | 0.39 | 0.67 | 0.67 |
| 1 | 236575433 | rs16833645 | -0.733 | -1.417 | 1.005 | 0.405 | -0.004 | -0.002 | -0.002 | 0.575 | 0.18 | 0.19 | 0.22 | 0.13 | 0.09 | 0.16 |
| 1 | 236585417 | rs665504 | 0.135 | -1.167 | 0.742 | 0.42 | 0.012 | 0.016 | 0.016 | 0 | 0.6 | 0.25 | 0.64 | 0.35 | 0.67 | 0.63 |
| 1 | 236585542 | rs654862 | 0.135 | -1.167 | 0.742 | 0.42 | 0.012 | 0.016 | 0.016 | 0 | 0.6 | 0.24 | 0.64 | 0.35 | 0.67 | 0.63 |
| 1 | 236590816 | rs635487 | 0.823 | -0.974 | 0.716 | 0.368 | 0.002 | 0.068 | 0.068 | 0.008 | 0.45 | 0.27 | 0.46 | 0.26 | 0.58 | 0.47 |
| 1 | 236601735 | rs636877 | 0.459 | -1.236 | 1.099 | 0.502 | -0.065 | 0.093 | 0.093 | 2.582 | 0.79 | 0.45 | 0.71 | 0.37 | 0.9 | 0.82 |
| 1 | 236602721 | rs622391 | 0.174 | -1.246 | 1.178 | 0.68 | -0.059 | 0.169 | 0.169 | 0.424 | 0.81 | 0.63 | 0.73 | 0.41 | 0.9 | 0.8 |
| 1 | 236609529 | rs637489 | 0.721 | -1.042 | -0.227 | 0.142 | -0.014 | 0.191 | 0.191 | 0.001 | 0.45 | 0.09 | 0.44 | 0.11 | 0.63 | 0.43 |
| 1 | 236611959 | rs650429 | 0.574 | -1.072 | 0.179 | 0.226 | -0.041 | -0.031 | -0.031 | 0 | 0.87 | 0.54 | 0.92 | 0.95 | 0.95 | 0.93 |
| 1 | 236616134 | rs17582358 | -0.325 | -0.581 | -1.231 | -0.127 | 0.001 | -0.005 | -0.005 | 0 | 0.21 | 0.22 | 0.14 | 0.03 | 0.19 | 0.24 |
| 1 | 236618200 | rs1253618 | -0.028 | 0.25 | -1.974 | 0.531 | 0.003 | -0.005 | -0.005 | 0.31 | 0.33 | 0.28 | 0.42 | 0.43 | 0.34 | 0.25 |
| 1 | 236619529 | rs1253615 | 0.032 | 0.917 | -2.726 | 0.483 | -0.057 | -0.055 | -0.055 | 0 | 0.29 | 0.1 | 0.41 | 0.36 | 0.42 | 0.3 |
| 1 | 236621524 | rs7512191 | -0.443 | 1.292 | -2.562 | 0.14 | -0.013 | 0.042 | 0.042 | 0.122 | 0.33 | 0.2 | 0.34 | 0.14 | 0.41 | 0.27 |
| 1 | 236622590 | rs16833707 | 0.171 | 0.972 | -2.624 | 0.942 | -0.039 | -0.162 | -0.162 | 0 | 0.26 | 0.02 | 0.38 | 0.43 | 0.27 | 0.37 |
| 1 | 236623239 | rs12045819 | -0.547 | 1.096 | -1.599 | 0.43 | 0.001 | 0 | 0 | 0 | 0.11 | 0.13 | 0.1 | 0.08 | 0.14 | 0.13 |
| 1 | 236623318 | rs12035437 | -0.547 | 1.096 | -1.599 | 0.43 | 0.001 | 0 | 0 | 6.734 | 0.11 | 0.13 | 0.1 | 0.08 | 0.14 | 0.13 |
| 1 | 236631091 | rs1253601 | 0.265 | 1.003 | -2.208 | -0.238 | -0.051 | 0.039 | 0.039 | 0 | 0.55 | 0.14 | 0.52 | 0.6 | 0.53 | 0.53 |
| 1 | 236632983 | rs16833721 | -0.213 | 1.063 | -1.188 | -0.384 | 0.038 | -0.033 | -0.033 | 0.114 | 0.08 | 0.28 | 0.07 | 0 | 0.06 | 0.06 |
| 1 | 236634875 | rs11577180 | 0.123 | 0.718 | -2.325 | -1.227 | -0.033 | 0.143 | 0.143 | 0 | 0.28 | 0.03 | 0.19 | 0.03 | 0.22 | 0.21 |
| 1 | 236635341 | rs11587227 | -0.282 | 0.785 | -2.481 | -1.427 | -0.055 | 0.185 | 0.185 | 0 | 0.33 | 0.03 | 0.21 | 0.04 | 0.28 | 0.22 |
| 1 | 236636146 | rs585298 | -1.348 | 0.487 | -1.931 | -1.632 | 0.007 | -0.009 | -0.009 | 0 | 0.64 | 0.68 | 0.59 | 0.42 | 0.68 | 0.66 |
| 1 | 236636159 | rs585273 | -0.387 | 0.523 | -1.869 | -1.607 | -0.01 | 0.001 | 0.001 | 0.28 | 0.35 | 0.65 | 0.39 | 0.38 | 0.43 | 0.44 |
| 1 | 236644617 | rs653740 | -1.538 | 0.507 | -1.477 | -1.388 | 0.014 | -0.013 | -0.013 | 0.064 | 0.64 | 0.7 | 0.59 | 0.44 | 0.7 | 0.67 |
| 1 | 236645670 | rs604070 | 0.116 | 0.846 | -1.604 | -1.697 | -0.043 | 0.051 | 0.051 | 0 | 0.17 | 0.05 | 0.11 | 0.11 | 0.28 | 0.14 |
| 1 | 236653341 | rs12750497 | -0.159 | 0.596 | -1.269 | -1.718 | -0.013 | 0.085 | 0.085 | 0 | 0.34 | 0.23 | 0.26 | 0.05 | 0.32 | 0.31 |
| 1 | 236656457 | rs12753714 | -0.207 | 0.699 | -0.841 | -1.655 | -0.028 | 0.035 | 0.035 | 0 | 0.32 | 0.18 | 0.34 | 0.2 | 0.38 | 0.32 |

| | | | | | | | | | | | | | | | | |
|---|---|---|---|---|---|---|---|---|---|---|---|---|---|---|---|---|
| 1 | 236658679 | rs10489788 | -0.038 | 0.803 | -0.616 | -1.796 | -0.001 | 0.004 | 0.004 | 0 | 0.28 | 0.24 | 0.3 | 0.19 | 0.27 | 0.23 |
| 1 | 236659941 | rs10925137 | -1.029 | 0.505 | -0.221 | -1.739 | -0.004 | 0.025 | 0.025 | 0 | 0.6 | 0.73 | 0.52 | 0.36 | 0.58 | 0.58 |
| 1 | 236661920 | rs7513978 | -0.84 | 0.593 | 0.053 | -1.633 | 0 | -0.009 | -0.009 | 0 | 0.57 | 0.43 | 0.57 | 0.32 | 0.58 | 0.63 |
| 1 | 236666907 | rs10925145 | -0.815 | 0.104 | 0.152 | -1.68 | 0.012 | 0.04 | 0.04 | 0 | 0.54 | 0.38 | 0.56 | 0.23 | 0.61 | 0.51 |
| 1 | 236668845 | rs9727054 | -0.412 | 0.131 | -0.176 | -1.67 | 0 | 0.001 | 0.001 | 0 | 0.37 | 0.65 | 0.38 | 0.3 | 0.34 | 0.38 |
| 1 | 236672004 | rs6689881 | -1.086 | -0.195 | -0.848 | -1.915 | -0.027 | -0.073 | -0.073 | 0 | 0.36 | 0.48 | 0.32 | 0.19 | 0.3 | 0.37 |
| 1 | 236677586 | rs10754573 | -1.096 | 0.154 | -0.877 | -1.664 | 0.024 | 0.087 | 0.087 | 0 | 0.5 | 0.74 | 0.48 | 0.29 | 0.43 | 0.42 |
| 1 | 236680253 | rs16833739 | -1.066 | -0.083 | -0.828 | -1.501 | -0.021 | -0.092 | -0.092 | 0.313 | 0.2 | 0.33 | 0.08 | 0 | 0.14 | 0.13 |
| 1 | 236681990 | rs1266381 | -0.626 | 0.488 | -1.446 | -1.237 | -0.038 | 0.053 | 0.053 | 0 | 0.17 | 0.08 | 0.2 | 0.11 | 0.23 | 0.22 |
| 1 | 236682114 | rs10802546 | 0.032 | -0.528 | -1.29 | -0.799 | -0.013 | 0.002 | 0.002 | 0 | 0.36 | 0.52 | 0.43 | 0.63 | 0.28 | 0.48 |
| 1 | 236682248 | rs4659682 | 0.201 | -0.814 | -1.249 | -1.347 | -0.017 | -0.106 | -0.106 | 0.148 | 0.6 | 0.48 | 0.73 | 0.91 | 0.67 | 0.75 |
| 1 | 236682308 | rs10925157 | -0.258 | -1.163 | -1.525 | -0.198 | 0.047 | 0.047 | 0.047 | 0 | 0.33 | 0.19 | 0.39 | 0.22 | 0.22 | 0.43 |
| 1 | 236684794 | rs12041958 | -0.999 | -0.493 | -1.236 | 0.441 | 0.024 | 0.007 | 0.007 | 0 | 0.24 | 0.12 | 0.15 | 0.21 | 0.18 | 0.15 |
| 1 | 236685587 | rs2799426 | -1.008 | -0.253 | -0.755 | 0.673 | -0.002 | 0.001 | 0.001 | 0 | 0.11 | 0.12 | 0.09 | 0.13 | 0.07 | 0.09 |
| 1 | 236686053 | rs10925158 | -1.219 | -0.055 | -0.181 | 0.42 | 0.021 | -0.029 | -0.029 | 0.014 | 0.1 | 0.28 | 0.11 | 0.05 | 0.08 | 0.09 |
| 1 | 236687782 | rs3820564 | -1.449 | -0.791 | 0.453 | 0.743 | -0.025 | 0.066 | 0.066 | 0 | 0.09 | 0.27 | 0.04 | 0.11 | 0.03 | 0.04 |
| 1 | 236689568 | rs2251642 | 0.444 | -1.234 | -0.492 | 0.596 | 0.026 | -0.044 | -0.044 | 0 | 0.46 | 0.15 | 0.48 | 0.48 | 0.63 | 0.43 |
| 1 | 236699354 | rs16833777 | -1.299 | -1.536 | -0.44 | 1.027 | -0.032 | 0.05 | 0.05 | 0.209 | 0.09 | 0.22 | 0.12 | 0.18 | 0.03 | 0.07 |
| 1 | 236700823 | rs1041934 | 0.436 | -2.245 | 0.428 | 0.633 | 0.012 | -0.004 | -0.004 | 0 | 0.54 | 0.45 | 0.51 | 0.48 | 0.68 | 0.41 |
| 1 | 236701862 | rs2472124 | 0.436 | -2.245 | 0.428 | 0.633 | 0.012 | -0.004 | -0.004 | 0 | 0.54 | 0.45 | 0.51 | 0.48 | 0.68 | 0.41 |
| 1 | 236706278 | rs2273863 | -1.097 | -2.27 | 0.455 | 0.622 | -0.034 | 0.05 | 0.05 | 0.366 | 0.09 | 0.22 | 0.12 | 0.18 | 0.03 | 0.07 |
| 1 | 236707007 | rs2799410 | 1.083 | -2.348 | 0.887 | 0.606 | 0.026 | -0.034 | -0.034 | 1.086 | 0.44 | 0.22 | 0.48 | 0.48 | 0.64 | 0.4 |
| 1 | 236707816 | rs1124907 | 0.439 | -2.332 | 0.895 | 0.605 | 0.012 | -0.004 | -0.004 | 0 | 0.54 | 0.44 | 0.51 | 0.48 | 0.68 | 0.41 |
| 1 | 236709988 | rs2298098 | 0.567 | -2.655 | 1.218 | 0.616 | 0.003 | 0.09 | 0.09 | 0 | 0.07 | 0.14 | 0.11 | 0.28 | 0.06 | 0.34 |
| 1 | 236711323 | rs15701 | 0.905 | -2.637 | 1.068 | 0.627 | 0.012 | -0.004 | -0.004 | 0 | 0.54 | 0.44 | 0.51 | 0.48 | 0.68 | 0.41 |
| 1 | 236711618 | rs1041939 | -1.036 | -2.655 | 1.06 | 0.618 | -0.038 | 0.044 | 0.044 | 0.552 | 0.1 | 0.22 | 0.12 | 0.18 | 0.03 | 0.1 |
| 1 | 236711810 | rs2758997 | 0.192 | -2.746 | 0.977 | 0.565 | 0.002 | 0.001 | 0.001 | 0 | 0.65 | 0.69 | 0.76 | 0.69 | 0.74 | 0.55 |
| 1 | 236714064 | rs6989 | 1.08 | -2.737 | 0.982 | 0.556 | 0.011 | -0.004 | -0.004 | 0 | 0.54 | 0.43 | 0.51 | 0.48 | 0.68 | 0.41 |
| 1 | 236716959 | rs2275685 | 1.137 | -2.742 | 1.069 | 0.554 | 0.035 | -0.01 | -0.01 | 17.316 | 0.53 | 0.39 | 0.51 | 0.48 | 0.68 | 0.41 |
| 1 | 236717926 | rs2275687 | 1.099 | -2.742 | 1.069 | 0.554 | 0.033 | -0.011 | -0.011 | 0.656 | 0.54 | 0.39 | 0.51 | 0.48 | 0.68 | 0.41 |
| 1 | 236718620 | rs1126627 | 1.099 | -2.742 | 1.069 | 0.554 | 0.033 | -0.011 | -0.011 | 1.282 | 0.54 | 0.39 | 0.51 | 0.48 | 0.68 | 0.41 |
| 1 | 236721660 | rs2275689 | 0.208 | -2.756 | 1.145 | 0.591 | 0.003 | 0.002 | 0.002 | 0 | 0.65 | 0.7 | 0.76 | 0.68 | 0.74 | 0.51 |
| 1 | 236724605 | rs1077438 | 0.575 | -2.772 | 1.146 | 0.569 | -0.007 | -0.002 | -0.002 | 0.165 | 0.61 | 0.45 | 0.62 | 0.65 | 0.71 | 0.51 |
| 1 | 236725376 | rs766124 | 1.833 | -2.793 | 1.104 | 0.56 | -0.015 | -0.115 | -0.115 | 0.117 | 0.43 | 0.07 | 0.47 | 0.48 | 0.64 | 0.4 |
| 1 | 236728150 | rs2799434 | 1.464 | -2.784 | 1.227 | 0.572 | 0.088 | -0.047 | -0.047 | 0 | 0.51 | 0.13 | 0.5 | 0.48 | 0.68 | 0.41 |
| 1 | 236729956 | rs653737 | 0.464 | -2.958 | 1.101 | 0.458 | -0.031 | 0.122 | 0.122 | 0 | 0.24 | 0.01 | 0.1 | 0.04 | 0.19 | 0.14 |
| 1 | 236732776 | rs2794778 | 0.542 | -2.874 | 1.179 | 0.487 | -0.006 | 0 | 0 | 0 | 0.63 | 0.61 | 0.63 | 0.65 | 0.71 | 0.51 |
| 1 | 236742166 | rs16833958 | -1.24 | -2.819 | 0.911 | 0.505 | -0.02 | 0.022 | 0.022 | 0.233 | 0.08 | 0.14 | 0.11 | 0.18 | 0.03 | 0.05 |
| 1 | 236744712 | rs4518892 | 1.57 | -2.875 | 0.782 | 0.489 | 0.087 | -0.05 | -0.05 | 0.108 | 0.51 | 0.13 | 0.5 | 0.48 | 0.67 | 0.41 |
| 1 | 236746443 | rs2794768 | 1.201 | -2.89 | 0.91 | 0.49 | 0.044 | 0.028 | 0.028 | 0 | 0.55 | 0.3 | 0.59 | 0.65 | 0.68 | 0.45 |
| 1 | 236747412 | rs1891238 | 1.201 | -2.89 | 0.91 | 0.49 | 0.044 | 0.028 | 0.028 | 0 | 0.55 | 0.3 | 0.59 | 0.65 | 0.68 | 0.45 |
| 1 | 236747494 | rs16833979 | -0.418 | -2.928 | 0.876 | 0.462 | 0.008 | -0.006 | -0.006 | 0 | 0.1 | 0.07 | 0.11 | 0.18 | 0.05 | 0.05 |
| 1 | 236748570 | rs657629 | 0.025 | -3.333 | 0.761 | 0.213 | -0.007 | -0.107 | -0.107 | 0 | 0.18 | 0.34 | 0.1 | 0 | 0.15 | 0.05 |
| 1 | 236749626 | rs3738541 | -0.228 | -3.303 | 0.613 | 0.232 | 0.008 | -0.006 | -0.006 | 0 | 0.1 | 0.07 | 0.11 | 0.18 | 0.06 | 0.05 |
| 1 | 236749649 | rs2794763 | 2.63 | -3.345 | 0.577 | 0.2 | -0.068 | -0.164 | -0.164 | 0.377 | 0.42 | 0.03 | 0.46 | 0.47 | 0.62 | 0.39 |
| 1 | 236755814 | rs2794757 | 1.41 | -3.368 | 0.604 | 0.357 | 0.026 | 0.031 | 0.031 | 0 | 0.58 | 0.33 | 0.6 | 0.72 | 0.7 | 0.53 |
| 1 | 236758887 | rs2794751 | 1.41 | -3.368 | 0.604 | 0.336 | 0.026 | 0.033 | 0.033 | 0 | 0.58 | 0.33 | 0.6 | 0.72 | 0.7 | 0.53 |
| 1 | 236759825 | rs2853585 | 0.48 | -3.368 | 0.681 | 0.349 | -0.035 | 0.015 | 0.015 | 0 | 0.78 | 0.64 | 0.83 | 0.75 | 0.88 | 0.61 |
| 1 | 236761778 | rs2298099 | -0.65 | -3.321 | 0.106 | 0.37 | 0.015 | -0.017 | -0.017 | 0.085 | 0.12 | 0.07 | 0.11 | 0.25 | 0.06 | 0.11 |
| 1 | 236765676 | rs2853612 | 2.272 | -3.29 | 0.159 | 0.369 | 0.034 | -0.033 | -0.033 | 1.186 | 0.45 | 0.25 | 0.49 | 0.48 | 0.63 | 0.4 |
| 1 | 236767330 | rs10802551 | 1.739 | -3.286 | 0.252 | 0.346 | 0.046 | 0.071 | 0.071 | 0.028 | 0.53 | 0.33 | 0.72 | 0.75 | 0.69 | 0.54 |
| 1 | 236768404 | rs965399 | -1.461 | -3.093 | 0.027 | 0.643 | 0.003 | 0.005 | 0.005 | 0 | 0.09 | 0.08 | 0.23 | 0.28 | 0.06 | 0.14 |
| 1 | 236772113 | rs4477263 | -1.248 | -3.11 | -0.084 | 0.647 | 0.001 | 0.014 | 0.014 | 0 | 0.08 | 0.08 | 0.23 | 0.28 | 0.06 | 0.15 |
| 1 | 236772416 | rs630101 | 1.757 | -3.071 | -0.284 | 0.549 | 0.085 | -0.166 | -0.166 | 0 | 0.63 | 0.06 | 0.55 | 0.47 | 0.8 | 0.42 |
| 1 | 236796979 | rs4659695 | 1.375 | -2.477 | 0.38 | 0.403 | 0.029 | 0.062 | 0.062 | 0.314 | 0.59 | 0.15 | 0.71 | 0.71 | 0.73 | 0.51 |
| 1 | 236799281 | rs819648 | 0.954 | -2.654 | 0.959 | 0.457 | -0.001 | 0.007 | 0.007 | 0 | 0.74 | 0.3 | 0.64 | 0.7 | 0.77 | 0.63 |
| 1 | 236800698 | rs10495382 | 2.693 | -2.921 | 0.658 | 0.721 | -0.03 | -0.083 | -0.083 | 0.076 | 0.43 | 0.09 | 0.49 | 0.49 | 0.65 | 0.38 |
| 1 | 236800747 | rs663398 | -0.548 | -2.776 | 1.385 | 0.863 | -0.002 | -0.011 | -0.011 | 0 | 0.19 | 0.51 | 0.25 | 0.11 | 0.14 | 0.15 |
| 1 | 236802303 | rs490076 | 1.886 | -2.786 | 1.469 | 0.899 | -0.016 | 0.032 | 0.032 | 0.184 | 0.75 | 0.63 | 0.76 | 0.63 | 0.83 | 0.65 |
| 1 | 236803643 | rs819650 | 1.434 | -2.731 | 1.683 | 0.887 | 0.024 | 0.076 | 0.076 | 0.343 | 0.88 | 0.42 | 0.86 | 0.82 | 0.94 | 0.74 |
| 1 | 236803753 | rs10925178 | -0.563 | -2.659 | 1.305 | 0.923 | 0.018 | 0.051 | 0.051 | 0 | 0.07 | 0.16 | 0.11 | 0.18 | 0.06 | 0.24 |
| 1 | 236807017 | rs645206 | -1.607 | -2.743 | 1.092 | 0.892 | -0.034 | 0 | 0 | 0 | 0.2 | 0.33 | 0.18 | 0.2 | 0.12 | 0.13 |
| 1 | 236818787 | rs682757 | 0.768 | -3.25 | 0.491 | 0.223 | 0.006 | 0.003 | 0.003 | 0.012 | 0.58 | 0.42 | 0.62 | 0.64 | 0.77 | 0.49 |
| 1 | 236821847 | rs475488 | -1.323 | -3.416 | 0.73 | 0.186 | -0.001 | 0 | 0 | 0.012 | 0.22 | 0.24 | 0.29 | 0.22 | 0.14 | 0.19 |
| 1 | 236822322 | rs12140395 | 2.759 | -3.42 | 0.749 | 0.183 | -0.077 | -0.08 | -0.08 | 22.408 | 0.35 | 0.1 | 0.43 | 0.44 | 0.64 | 0.38 |
| 1 | 236829348 | rs10925193 | 0.891 | -3.544 | 0.821 | 0.099 | 0.005 | 0.013 | 0.013 | 0 | 0.11 | 0.15 | 0.19 | 0.11 | 0.09 | 0.12 |
| 1 | 236832848 | rs4659462 | 2.831 | -3.558 | 0.758 | 0.1 | -0.077 | -0.086 | -0.086 | 0 | 0.35 | 0.1 | 0.43 | 0.45 | 0.64 | 0.37 |
| 1 | 236836197 | rs1253245 | 0.264 | -3.577 | 0.797 | 0.093 | -0.023 | -0.022 | -0.022 | 0 | 0.7 | 0.58 | 0.81 | 0.76 | 0.87 | 0.62 |
| 1 | 236837395 | rs1036001 | 2.908 | -3.556 | 0.676 | 0.099 | -0.14 | -0.141 | -0.141 | 0.343 | 0.34 | 0.04 | 0.42 | 0.45 | 0.64 | 0.37 |

| | | | | | | | | | | | | | | | | |
|---|---|---|---|---|---|---|---|---|---|---|---|---|---|---|---|---|
| 1 | 236837527 | rs16834155 | 0.402 | -3.681 | 0.616 | 0.036 | -0.023 | -0.055 | -0.055 | 0.019 | 0.11 | 0 | 0.11 | 0.21 | 0.07 | 0.27 |
| 1 | 236842428 | rs617626 | -1.014 | -3.609 | 0.608 | 0.091 | 0.023 | 0.003 | 0.003 | 1.12 | 0.21 | 0.17 | 0.17 | 0.2 | 0.13 | 0.11 |
| 1 | 236848268 | rs4659700 | 2.918 | -2.626 | 0.241 | -0.005 | -0.16 | -0.137 | -0.137 | 0 | 0.32 | 0.04 | 0.35 | 0.45 | 0.53 | 0.36 |
| 1 | 236848751 | rs819666 | 0.45 | -2.476 | 0.64 | 0.084 | -0.007 | -0.019 | -0.019 | 0 | 0.68 | 0.57 | 0.74 | 0.76 | 0.76 | 0.61 |
| 1 | 236855172 | rs3893143 | 0.974 | -2.025 | 0.052 | 0.198 | -0.01 | 0.017 | 0.017 | 0.051 | 0.1 | 0.07 | 0.06 | 0.02 | 0.1 | 0.07 |
| 1 | 236861805 | rs6428966 | 0.614 | -2.245 | -0.045 | 0.354 | 0.015 | -0.042 | -0.042 | 0.897 | 0.9 | 0.96 | 0.86 | 0.79 | 0.92 | 0.9 |
| 1 | 236863831 | rs12736189 | 0.647 | -2.363 | 0.256 | 0.215 | -0.004 | -0.003 | -0.003 | 0.165 | 0.15 | 0.15 | 0.04 | 0.07 | 0.05 | 0.1 |
| 1 | 236864459 | rs3905125 | 2.763 | -2.598 | -0.273 | 0.19 | -0.158 | -0.134 | -0.134 | 0.312 | 0.38 | 0.04 | 0.39 | 0.46 | 0.54 | 0.37 |
| 1 | 236865568 | rs707216 | -0.413 | -2.747 | 0.014 | 0.139 | -0.002 | 0.014 | 0.014 | 0 | 0.17 | 0.17 | 0.09 | 0.13 | 0.19 | 0.14 |
| 1 | 236866836 | rs16834213 | 0.73 | -2.865 | 0.105 | 0.188 | 0.022 | 0.013 | 0.013 | 0 | 0.15 | 0.09 | 0.03 | 0.07 | 0.04 | 0.1 |
| 1 | 236867394 | rs707214 | 1.094 | -2.865 | 0.649 | 0.176 | 0.069 | 0.054 | 0.054 | 0 | 0.93 | 0.65 | 0.83 | 0.98 | 0.98 | 0.97 |
| 1 | 236872018 | rs707210 | 1.086 | -2.865 | 0.511 | 0.205 | -0.007 | -0.003 | -0.003 | 0 | 0.63 | 0.44 | 0.63 | 0.71 | 0.76 | 0.59 |
| 1 | 236879332 | rs1418071 | 0.485 | -3.159 | 0.173 | 0.135 | -0.045 | -0.029 | -0.029 | 1.155 | 0.75 | 0.63 | 0.79 | 0.89 | 0.88 | 0.84 |
| 1 | 236893599 | rs819639 | -1.95 | -3.115 | -0.114 | 0.394 | 0.043 | 0.008 | 0.008 | 0 | 0.5 | 0.6 | 0.35 | 0.56 | 0.31 | 0.53 |
| 1 | 236895000 | rs16834278 | 0.882 | -3.01 | -0.688 | 0.383 | 0.002 | 0.005 | 0.005 | 0 | 0.09 | 0.03 | 0.16 | 0.05 | 0.06 | 0.04 |
| 1 | 236896619 | rs2039744 | 0.183 | -3.125 | -0.572 | 0.362 | -0.001 | 0 | 0 | 0 | 0.18 | 0.19 | 0.16 | 0.2 | 0.13 | 0.22 |
| 1 | 236896660 | rs707205 | -0.741 | -3.252 | -0.639 | 0.302 | 0.01 | 0.003 | 0.003 | 0 | 0.26 | 0.21 | 0.18 | 0.22 | 0.19 | 0.29 |
| 1 | 236897376 | rs707204 | 0.458 | -3.333 | -0.43 | 0.33 | -0.027 | 0 | 0 | 0.478 | 0.69 | 0.56 | 0.79 | 0.73 | 0.82 | 0.67 |
| 1 | 236898035 | rs819643 | -0.403 | -2.968 | -0.966 | 0.366 | -0.027 | 0.028 | 0.028 | 0 | 0.92 | 0.99 | 0.96 | 0.98 | 0.9 | 0.96 |
| 1 | 236898163 | rs6428968 | -0.662 | -3.026 | -0.951 | 0.473 | 0.012 | -0.004 | -0.004 | 0 | 0.29 | 0.28 | 0.2 | 0.3 | 0.17 | 0.33 |
| 1 | 236898187 | rs7536332 | 1.389 | -3.057 | -1.021 | 0.419 | 0.022 | -0.005 | -0.005 | 0 | 0.1 | 0.07 | 0.03 | 0.07 | 0.04 | 0.07 |
| 1 | 236898395 | rs7536539 | 1.294 | -3.049 | -1.016 | 0.424 | 0.021 | -0.006 | -0.006 | 0 | 0.1 | 0.07 | 0.03 | 0.07 | 0.04 | 0.07 |
| 1 | 236899042 | rs2288600 | 0.027 | -3.05 | -1.087 | 0.423 | -0.007 | 0 | 0 | 0.511 | 0.19 | 0.22 | 0.17 | 0.19 | 0.13 | 0.25 |
| 1 | 236899343 | rs4659708 | 2.071 | -3.082 | -1.24 | 0.373 | -0.062 | -0.065 | -0.065 | 0 | 0.39 | 0.08 | 0.45 | 0.39 | 0.62 | 0.4 |
| 1 | 236902955 | rs7544174 | 1.459 | -2.964 | -1.117 | 0.319 | 0.04 | -0.038 | -0.038 | 0 | 0.48 | 0.12 | 0.48 | 0.47 | 0.65 | 0.5 |
| 1 | 236903791 | rs2385494 | 1.014 | -2.625 | -0.651 | 0.355 | 0.007 | 0.001 | 0.001 | 0 | 0.66 | 0.34 | 0.65 | 0.61 | 0.78 | 0.71 |
| 1 | 236907847 | rs3738543 | 0.132 | -2.454 | -0.724 | 0.372 | 0.027 | 0.018 | 0.018 | 0 | 0.15 | 0.08 | 0.06 | 0.07 | 0.08 | 0.09 |
| 1 | 236910863 | rs10802557 | -1.211 | -2.762 | -0.747 | 0.422 | 0.067 | -0.009 | -0.009 | 0.2 | 0.43 | 0.26 | 0.28 | 0.43 | 0.28 | 0.4 |
| 1 | 236911137 | rs10925211 | 0.442 | -2.872 | -0.556 | 0.592 | 0.031 | -0.007 | -0.007 | 0 | 0.15 | 0.07 | 0.18 | 0.11 | 0.06 | 0.14 |
| 1 | 236911343 | rs10802558 | -0.127 | -2.994 | -0.61 | 0.526 | 0.005 | 0 | 0 | 0 | 0.24 | 0.18 | 0.19 | 0.21 | 0.2 | 0.25 |
| 1 | 236912597 | rs2297955 | -0.574 | -3.605 | 0.343 | 0.541 | -0.02 | 0.001 | 0.001 | 0 | 0.22 | 0.24 | 0.21 | 0.24 | 0.08 | 0.17 |
| 1 | 236914576 | rs3738544 | 1.695 | -3.8 | 0.801 | 0.483 | 0.033 | 0.002 | 0.002 | 0 | 0.6 | 0.35 | 0.65 | 0.57 | 0.79 | 0.66 |
| 1 | 236915207 | rs4659713 | 0.606 | -3.656 | 0.679 | 0.542 | -0.04 | -0.001 | -0.001 | 0 | 0.74 | 0.67 | 0.77 | 0.76 | 0.85 | 0.88 |
| 1 | 236916612 | rs10802560 | -0.252 | -2.856 | 0.502 | 0.577 | -0.001 | -0.007 | -0.007 | 0 | 0.91 | 0.92 | 0.95 | 0.96 | 0.92 | 0.94 |
| 1 | 236924506 | rs2282366 | 0.899 | -3.042 | 0.579 | 0.693 | -0.015 | 0.007 | 0.007 | 0 | 0.83 | 0.79 | 0.87 | 0.74 | 0.89 | 0.73 |
| 1 | 236926445 | rs1803032 | 1.267 | -3.307 | 0.757 | 0.753 | -0.033 | 0.026 | 0.026 | 0 | 0.64 | 0.5 | 0.69 | 0.42 | 0.72 | 0.51 |
| 1 | 236926667 | rs7522310 | 2.102 | -3.589 | -0.206 | 0.567 | 0.049 | -0.109 | -0.109 | 0 | 0.46 | 0.03 | 0.59 | 0.42 | 0.65 | 0.48 |
| 1 | 236926873 | rs2153203 | -0.236 | -2.942 | 0.299 | 0.791 | 0.019 | -0.02 | -0.02 | 0.52 | 0.09 | 0.04 | 0.08 | 0.21 | 0.04 | 0.21 |
| 1 | 236928589 | rs7514673 | -0.102 | -2.554 | 0.498 | 0.682 | -0.009 | 0.016 | 0.016 | 0 | 0.19 | 0.28 | 0.18 | 0.2 | 0.17 | 0.21 |
| 1 | 236935938 | rs4659470 | -0.699 | -2.615 | 0.498 | 0.691 | -0.018 | -0.006 | -0.006 | 0.411 | 0.34 | 0.39 | 0.23 | 0.28 | 0.23 | 0.31 |
| 1 | 236938367 | rs6687802 | -0.771 | -2.647 | 0.558 | 0.762 | 0 | -0.001 | -0.001 | 3.8 | 0.36 | 0.67 | 0.28 | 0.41 | 0.23 | 0.33 |
| 1 | 236939395 | rs7546640 | -0.888 | -2.15 | 1.012 | 0.624 | -0.008 | -0.01 | -0.01 | 0 | 0.07 | 0.17 | 0.04 | 0.04 | 0.03 | 0.15 |
| 1 | 236946305 | rs10925228 | -0.008 | -1.954 | 0.507 | 0.667 | -0.001 | -0.003 | -0.003 | 0.407 | 0.92 | 0.91 | 0.96 | 0.94 | 0.94 | 0.95 |
| 1 | 236949304 | rs6658562 | 0.947 | -2.263 | 0.379 | 0.705 | -0.038 | 0 | 0 | 0.033 | 0.82 | 0.74 | 0.86 | 0.76 | 0.9 | 0.84 |
| 1 | 236971998 | rs7526063 | -1.277 | -2.006 | 0.58 | 0.733 | -0.008 | 0.009 | 0.009 | 0 | 0.05 | 0.14 | 0.04 | 0.05 | 0.03 | 0.09 |
| 1 | 236984562 | rs10925241 | -0.15 | -1.705 | 0.149 | 0.813 | 0.014 | 0 | 0 | 0.168 | 0.48 | 0.41 | 0.59 | 0.51 | 0.56 | 0.56 |
| 1 | 236996390 | rs883396 | 1.039 | -1.354 | -0.342 | 0.743 | -0.014 | -0.011 | -0.011 | 5.282 | 0.21 | 0.15 | 0.41 | 0.33 | 0.34 | 0.26 |
| 1 | 237006914 | rs4659727 | -0.297 | -1.913 | 0.074 | 0.277 | -0.04 | -0.062 | -0.062 | 0 | 0.26 | 0.34 | 0.18 | 0.1 | 0.17 | 0.32 |
| 1 | 237015628 | rs2275568 | 0.276 | -1.871 | 0.157 | 0.335 | 0.032 | 0.027 | 0.027 | 0 | 0.31 | 0.38 | 0.49 | 0.45 | 0.41 | 0.39 |
| 1 | 237034784 | rs4659736 | 0.987 | -1.988 | 0.364 | 0.583 | 0.016 | 0.054 | 0.054 | 0 | 0.17 | 0.03 | 0.12 | 0.07 | 0.16 | 0.16 |
| 1 | 237046160 | rs10925257 | -0.251 | -1.993 | 0.461 | 0.592 | -0.023 | -0.038 | -0.038 | 0.045 | 0.26 | 0.29 | 0.18 | 0.1 | 0.17 | 0.33 |
| 1 | 237048500 | rs1805087 | -0.19 | -1.989 | 0.462 | 0.594 | -0.023 | -0.039 | -0.039 | 0 | 0.25 | 0.28 | 0.18 | 0.11 | 0.17 | 0.32 |
| 1 | 237048676 | rs2275565 | -1.007 | -2.007 | 0.478 | 0.599 | -0.057 | -0.065 | -0.065 | 0 | 0.31 | 0.52 | 0.22 | 0.15 | 0.2 | 0.41 |
| 1 | 237050985 | rs3768147 | 1.04 | -1.933 | 0.42 | 0.737 | -0.028 | -0.027 | -0.027 | 0 | 0.25 | 0.16 | 0.45 | 0.43 | 0.39 | 0.3 |
| 1 | 237057608 | rs2297965 | -0.731 | -2.033 | 0.528 | 0.726 | -0.001 | -0.001 | -0.001 | 0 | 0.51 | 0.44 | 0.61 | 0.52 | 0.56 | 0.62 |
| 1 | 237060433 | rs3820571 | -1.527 | -2.046 | 0.333 | 0.702 | -0.018 | 0.116 | 0.116 | 2.334 | 0.59 | 0.91 | 0.68 | 0.78 | 0.58 | 0.73 |
| 1 | 237060850 | rs12022937 | -0.841 | -2.044 | 0.376 | 0.714 | 0.007 | 0.255 | 0.255 | 1.18 | 0.08 | 0.47 | 0.07 | 0.26 | 0.03 | 0.11 |
| 1 | 237061056 | rs2853522 | -0.729 | -2.048 | 0.365 | 0.711 | 0 | -0.001 | -0.001 | 0 | 0.51 | 0.44 | 0.61 | 0.53 | 0.56 | 0.62 |
| 1 | 237061097 | rs11799670 | 0.252 | -2.055 | 0.495 | 0.702 | -0.027 | -0.021 | -0.021 | 0 | 0.08 | 0.24 | 0.05 | 0.04 | 0.02 | 0.16 |
| 1 | 237062198 | rs2853523 | -1.832 | -2.255 | 0.373 | 0.599 | -0.086 | 0.06 | 0.06 | 0.75 | 0.63 | 0.95 | 0.69 | 0.78 | 0.59 | 0.74 |
| 1 | 237062305 | rs1050993 | -1.827 | -2.251 | 0.371 | 0.597 | -0.086 | 0.06 | 0.06 | 0.743 | 0.63 | 0.95 | 0.69 | 0.78 | 0.59 | 0.74 |
| 1 | 237062628 | rs1804742 | 0.417 | -2.262 | 0.395 | 0.583 | -0.025 | -0.044 | -0.044 | 0.02 | 0.22 | 0.29 | 0.17 | 0.1 | 0.16 | 0.27 |
| 1 | 237063748 | rs10925263 | -0.909 | -2.076 | 0.465 | 0.589 | -0.007 | 0.202 | 0.202 | 0.641 | 0.08 | 0.36 | 0.07 | 0.26 | 0.03 | 0.12 |
| 1 | 237077480 | rs4351629 | -1.488 | -2.089 | 0.322 | 0.538 | -0.029 | 0.078 | 0.078 | 0.696 | 0.63 | 0.84 | 0.68 | 0.78 | 0.59 | 0.73 |
| 1 | 237083719 | rs6428977 | -1.012 | -2.11 | 0.398 | 0.511 | -0.001 | 0.051 | 0.051 | 0.284 | 0.58 | 0.67 | 0.64 | 0.73 | 0.53 | 0.64 |
| 1 | 237089463 | rs11806999 | -0.184 | -2.071 | 0.449 | 0.537 | -0.026 | -0.019 | -0.019 | 0 | 0.08 | 0.24 | 0.05 | 0.03 | 0.02 | 0.16 |
| 1 | 237093381 | rs946936 | -0.79 | -1.905 | 0.093 | 0.379 | 0.018 | 0.009 | 0.009 | 1.538 | 0.56 | 0.61 | 0.65 | 0.57 | 0.58 | 0.71 |
| 1 | 237106237 | rs1252106 | -1.131 | -1.804 | -0.692 | 0.325 | -0.032 | -0.044 | -0.044 | 0 | 0.62 | 0.84 | 0.42 | 0.53 | 0.5 | 0.65 |
| 1 | 237110626 | rs7545717 | -0.297 | -2.24 | -0.052 | 0.137 | -0.068 | -0.029 | -0.029 | 0 | 0.14 | 0.46 | 0.09 | 0.08 | 0.05 | 0.15 |

| | | | | | | | | | | | | | | | | |
|---|---|---|---|---|---|---|---|---|---|---|---|---|---|---|---|---|
| 1 | 237110859 | rs4659482 | 1.104 | -2.51 | -0.381 | -0.024 | 0.016 | 0.037 | 0.037 | 0 | 0.16 | 0.05 | 0.07 | 0.04 | 0.11 | 0.16 |
| 1 | 237111188 | rs1252108 | -1.303 | -2.493 | -0.446 | 0.014 | -0.02 | -0.047 | -0.047 | 0 | 0.6 | 0.81 | 0.42 | 0.53 | 0.48 | 0.64 |
| 1 | 237111220 | rs1270486 | 1.207 | -2.252 | -0.534 | 0.268 | -0.076 | -0.02 | -0.02 | 0.568 | 0.24 | 0.03 | 0.21 | 0.21 | 0.3 | 0.28 |
| 1 | 237111682 | rs1252109 | 1.155 | -2.247 | -0.532 | 0.248 | -0.075 | -0.132 | -0.132 | 1.476 | 0.24 | 0.03 | 0.22 | 0.41 | 0.3 | 0.3 |
| 1 | 237114288 | rs16834612 | -0.092 | -2.173 | -0.196 | -0.297 | -0.067 | -0.021 | -0.021 | 0 | 0.13 | 0.38 | 0.09 | 0.08 | 0.05 | 0.15 |
| 1 | 237123771 | rs4659757 | 0.845 | -1.829 | -0.549 | -0.32 | 0.017 | 0.032 | 0.032 | 0 | 0.16 | 0.08 | 0.07 | 0.04 | 0.1 | 0.16 |
| 1 | 237126155 | rs10802577 | -1.169 | -1.876 | -0.528 | -0.179 | -0.016 | -0.01 | -0.01 | 0 | 0.75 | 0.85 | 0.79 | 0.74 | 0.7 | 0.72 |
| 1 | 237126385 | rs1417303 | -0.124 | -1.652 | -0.299 | -0.025 | 0.003 | 0.007 | 0.007 | 0 | 0.54 | 0.57 | 0.68 | 0.71 | 0.57 | 0.52 |
| 1 | 237129217 | rs6675175 | 0.348 | -0.317 | -1.378 | 0.021 | -0.046 | -0.049 | -0.049 | 4.435 | 0.43 | 0.12 | 0.53 | 0.55 | 0.5 | 0.45 |
| 1 | 237129335 | rs6683896 | 0.476 | -0.743 | -1.267 | 0.118 | 0.06 | 0.021 | 0.021 | 0 | 0.5 | 0.26 | 0.54 | 0.46 | 0.59 | 0.34 |
| 1 | 237130858 | rs4492605 | 0.237 | -0.004 | -0.967 | 0.511 | -0.003 | -0.003 | -0.003 | 0 | 0.78 | 0.8 | 0.67 | 0.65 | 0.66 | 0.61 |
| 1 | 237131515 | rs17609678 | -0.608 | -1.124 | -0.771 | -0.005 | 0.012 | 0.027 | 0.027 | 0 | 0.19 | 0.17 | 0.11 | 0.04 | 0.13 | 0.18 |
| 1 | 237132197 | rs6685619 | 1.122 | -1.363 | -0.395 | 0.159 | -0.007 | -0.008 | -0.008 | 0.2 | 0.6 | 0.57 | 0.56 | 0.66 | 0.65 | 0.52 |
| 1 | 237134311 | rs13374506 | -0.297 | -0.885 | -0.386 | 0.038 | -0.006 | 0.008 | 0.008 | 0 | 0.16 | 0.15 | 0.12 | 0.05 | 0.19 | 0.15 |
| 1 | 237137760 | rs6684983 | -0.786 | -1.09 | 0.024 | 0.506 | 0.014 | 0.038 | 0.038 | 0 | 0.2 | 0.42 | 0.16 | 0.2 | 0.21 | 0.23 |
| 1 | 237138582 | rs1252135 | -0.656 | -0.831 | -0.091 | 0.624 | 0.007 | -0.003 | -0.003 | 0 | 0.17 | 0.31 | 0.28 | 0.17 | 0.15 | 0.21 |
| 1 | 237140296 | rs1819088 | 1.112 | -1 | -1.475 | 0.56 | -0.157 | -0.136 | -0.136 | 0.165 | 0.41 | 0.02 | 0.44 | 0.54 | 0.53 | 0.47 |
| 1 | 237146261 | rs12133452 | 0.938 | -0.7 | -1.278 | 0.337 | -0.095 | -0.054 | -0.054 | 0.707 | 0.48 | 0.05 | 0.46 | 0.55 | 0.53 | 0.53 |
| 1 | 237148460 | rs11582059 | 0.376 | -0.693 | -0.872 | 0 | 0.024 | -0.036 | -0.036 | 0 | 0.09 | 0.15 | 0.09 | 0 | 0.16 | 0.11 |
| 1 | 237149323 | rs946933 | 0.551 | -0.987 | -0.212 | -0.214 | -0.004 | -0.097 | -0.097 | 0 | 0.67 | 0.63 | 0.78 | 0.98 | 0.7 | 0.78 |
| 1 | 237149479 | rs1729489 | -0.344 | -1.044 | 0.362 | -0.128 | 0.01 | 0.016 | 0.016 | 0.117 | 0.16 | 0.43 | 0.3 | 0.23 | 0.15 | 0.16 |
| 1 | 237151125 | rs1252139 | 0.571 | -1.037 | 0.356 | -0.127 | -0.005 | -0.127 | -0.127 | 0 | 0.67 | 0.59 | 0.78 | 0.98 | 0.7 | 0.78 |
| 1 | 237153527 | rs10925286 | 1.468 | -0.972 | -0.3 | -0.058 | -0.001 | 0.202 | 0.202 | 0 | 0.51 | 0.13 | 0.48 | 0.75 | 0.56 | 0.62 |
| 1 | 237153919 | rs1252145 | -0.076 | -1.191 | -0.55 | -0.118 | -0.001 | -0.033 | -0.033 | 0 | 0.77 | 0.8 | 0.88 | 0.98 | 0.86 | 0.88 |
| 1 | 237161467 | rs12047281 | 1.149 | -1.042 | -1.018 | 0.271 | -0.056 | -0.059 | -0.059 | 0 | 0.48 | 0.02 | 0.49 | 0.59 | 0.6 | 0.5 |
| 1 | 237165077 | rs1252160 | 0.129 | -0.944 | 0.003 | 0.187 | -0.008 | -0.091 | -0.091 | 0 | 0.82 | 0.72 | 0.89 | 0.98 | 0.86 | 0.89 |
| 1 | 237166773 | rs12410379 | -0.309 | -1.045 | -1.013 | 0.136 | -0.003 | -0.003 | -0.003 | 0 | 0.21 | 0.22 | 0.29 | 0.2 | 0.16 | 0.26 |
| 1 | 237167218 | rs4659762 | 0.697 | -0.69 | -1.692 | 0.316 | 0.008 | -0.094 | -0.094 | 0 | 0.54 | 0.03 | 0.51 | 0.55 | 0.67 | 0.53 |
| 1 | 237171386 | rs4659764 | -0.716 | 0.001 | -0.548 | 0.273 | -0.007 | -0.002 | -0.002 | 0 | 0.4 | 0.46 | 0.45 | 0.41 | 0.34 | 0.43 |
| 1 | 237174035 | rs10925289 | -1.302 | 0.131 | -0.064 | -0.083 | 0.009 | -0.063 | -0.063 | 6.984 | 0.11 | 0.29 | 0.09 | 0 | 0.08 | 0.02 |
| 1 | 237177572 | rs12724972 | -1.342 | -0.123 | 0.075 | -0.731 | -0.025 | -0.148 | -0.148 | 1.955 | 0.29 | 0.47 | 0.19 | 0.01 | 0.22 | 0.2 |
| 1 | 237189528 | rs7556491 | -1.524 | -0.368 | 0.267 | 0.412 | -0.022 | 0.032 | 0.032 | 2.524 | 0.24 | 0.61 | 0.19 | 0.39 | 0.13 | 0.17 |
| 1 | 237194523 | rs12239401 | -0.626 | 0.749 | -0.209 | 0.077 | -0.03 | 0.029 | 0.029 | 0 | 0.34 | 0.26 | 0.33 | 0.18 | 0.43 | 0.25 |
| 1 | 237194922 | rs10802580 | -0.844 | -0.221 | -1.294 | -0.312 | -0.008 | 0.101 | 0.101 | 0 | 0.71 | 0.98 | 0.67 | 0.9 | 0.74 | 0.6 |
| 1 | 237196106 | rs12032218 | 1.2 | -0.174 | -0.744 | 0.609 | 0.003 | -0.007 | -0.007 | 0 | 0.25 | 0.23 | 0.13 | 0.31 | 0.19 | 0.12 |
| 1 | 237196413 | rs12022449 | 0.107 | 1 | -0.745 | 0.921 | -0.062 | 0.032 | 0.032 | 1.661 | 0.28 | 0.08 | 0.24 | 0.19 | 0.35 | 0.17 |
| 1 | 237197508 | rs12403151 | -0.111 | 1.072 | 0.001 | 0.93 | 0.106 | 0.061 | 0.061 | 1.943 | 0.14 | 0.44 | 0.2 | 0.16 | 0.17 | 0.18 |
| 1 | 237199183 | rs11583513 | 0.042 | 0.985 | -0.214 | 0.626 | -0.058 | 0.047 | 0.047 | 3.345 | 0.29 | 0.08 | 0.24 | 0.19 | 0.36 | 0.17 |
| 1 | 237199775 | rs2385576 | -0.579 | 0.881 | 0.098 | 0.646 | -0.005 | -0.004 | -0.004 | 0 | 0.4 | 0.32 | 0.46 | 0.58 | 0.53 | 0.45 |
| 1 | 237200616 | rs4659766 | -0.39 | 0.834 | 0.327 | 0.435 | 0.035 | 0.019 | 0.019 | 0 | 0.36 | 0.45 | 0.37 | 0.42 | 0.45 | 0.37 |
| 1 | 237207189 | rs1881548 | 0.186 | 1.088 | -1.141 | 0.203 | -0.023 | -0.019 | -0.019 | 0 | 0.24 | 0.02 | 0.32 | 0.19 | 0.21 | 0.41 |
| 1 | 237207450 | rs12123449 | -0.049 | 1.416 | -0.858 | -0.029 | -0.024 | -0.003 | -0.003 | 0 | 0.1 | 0.05 | 0.16 | 0.09 | 0.18 | 0.17 |
| 1 | 237210003 | rs6678625 | 0.15 | 0.558 | -0.823 | -0.424 | 0.015 | 0.042 | 0.042 | 0 | 0.22 | 0.11 | 0.3 | 0.07 | 0.16 | 0.33 |
| 1 | 237212306 | rs10925298 | -1.712 | 1.243 | -0.27 | -0.051 | 0 | -0.023 | -0.023 | 0 | 0.36 | 0.56 | 0.29 | 0.19 | 0.38 | 0.25 |
| 1 | 237215263 | rs4659488 | -1.731 | 1.257 | -0.298 | -0.025 | 0.004 | -0.039 | -0.039 | 0 | 0.34 | 0.53 | 0.27 | 0.18 | 0.36 | 0.24 |
| 1 | 237215658 | rs1569188 | -0.901 | 1.843 | -0.413 | 0.056 | -0.067 | 0 | 0 | 0 | 0.12 | 0.03 | 0.16 | 0.07 | 0.25 | 0.1 |
| 1 | 237216900 | rs10925300 | -0.982 | 0.583 | 0.099 | 0.123 | -0.053 | -0.039 | -0.039 | 0.363 | 0.19 | 0.41 | 0.1 | 0.09 | 0.1 | 0.13 |
| 1 | 237227726 | rs16834780 | -0.663 | 1.543 | 0.096 | 0.446 | 0.051 | 0.027 | 0.027 | 0.315 | 0.11 | 0.27 | 0.13 | 0.17 | 0.2 | 0.11 |
| 1 | 237227916 | rs9428368 | 0.489 | 1.313 | -0.071 | 0.064 | 0.052 | -0.089 | -0.089 | 0.016 | 0.85 | 0.66 | 0.84 | 0.98 | 0.8 | 0.8 |
| 1 | 237235986 | rs10925304 | 0.067 | 0.758 | -1.583 | 0.764 | -0.053 | -0.13 | -0.13 | 0.801 | 0.28 | 0.04 | 0.29 | 0.48 | 0.3 | 0.45 |
| 1 | 237247063 | rs6684412 | 0.27 | -0.045 | -0.639 | 0.62 | -0.026 | -0.015 | -0.015 | 0 | 0.73 | 0.49 | 0.81 | 0.78 | 0.8 | 0.72 |
| 1 | 237248703 | rs6428989 | 0.712 | -0.321 | -0.258 | 0.861 | -0.024 | 0.057 | 0.057 | 0.816 | 0.87 | 0.49 | 0.85 | 0.82 | 0.94 | 0.93 |
| 1 | 237257650 | rs6428992 | 0.239 | 0.063 | -1.206 | 0.826 | -0.009 | -0.006 | -0.006 | 0 | 0.4 | 0.29 | 0.32 | 0.58 | 0.47 | 0.56 |
| 1 | 237261761 | rs7551249 | -0.223 | -0.099 | -1.2 | 0.722 | -0.031 | -0.014 | -0.014 | 0 | 0.88 | 0.42 | 0.79 | 0.93 | 0.96 | 0.93 |
| 1 | 237262184 | rs7551672 | -0.117 | -0.033 | -1.356 | 0.87 | -0.046 | 0.064 | 0.064 | 0.843 | 0.85 | 0.38 | 0.76 | 0.8 | 0.93 | 0.9 |
| 1 | 237262866 | rs12131303 | 0.046 | 0.299 | -2.451 | 0.57 | -0.019 | 0.132 | 0.132 | 0.344 | 0.42 | 0.1 | 0.43 | 0.13 | 0.44 | 0.31 |
| 1 | 237266263 | rs7511682 | 0.073 | 0.101 | -2.368 | 0.622 | -0.005 | 0.052 | 0.052 | 0 | 0.42 | 0.11 | 0.42 | 0.25 | 0.43 | 0.34 |
| 1 | 237269434 | rs6696837 | -0.168 | 0.287 | -2.585 | 0.851 | -0.133 | -0.113 | -0.113 | 0.014 | 0.61 | 0.94 | 0.62 | 0.43 | 0.51 | 0.49 |
| 1 | 237271632 | rs2485579 | 0.635 | -0.359 | -2.701 | 0.765 | 0.03 | 0.001 | 0.001 | 4.554 | 0.75 | 0.1 | 0.74 | 0.78 | 0.85 | 0.82 |
| 1 | 237272025 | rs1415714 | 0.971 | -0.13 | -1.917 | 0.927 | 0.01 | 0.009 | 0.009 | 2.013 | 0.73 | 0.64 | 0.72 | 0.67 | 0.72 | 0.68 |
| 1 | 237272139 | rs1980798 | 1.272 | -0.776 | -1.443 | 0.252 | 0.001 | 0.004 | 0.004 | 0 | 0.48 | 0.39 | 0.59 | 0.58 | 0.54 | 0.51 |
| 1 | 237272274 | rs17612827 | 0.002 | -0.622 | -1.42 | 0.084 | -0.006 | 0.003 | 0.003 | 0 | 0.12 | 0.08 | 0.2 | 0.09 | 0.2 | 0.15 |
| 1 | 237272384 | rs10925315 | -1.511 | -1.022 | -0.898 | 0.295 | 0.003 | 0.005 | 0.005 | 0 | 0.48 | 0.56 | 0.38 | 0.46 | 0.42 | 0.49 |
| 1 | 237272756 | rs12128519 | 1.063 | -1.349 | -0.578 | 0.414 | 0.004 | -0.004 | -0.004 | 0 | 0.61 | 0.35 | 0.78 | 0.48 | 0.73 | 0.63 |
| 1 | 237272800 | rs12125175 | 0.709 | -1.378 | -0.239 | 0.401 | -0.017 | 0.048 | 0.048 | 0 | 0.67 | 0.46 | 0.8 | 0.48 | 0.73 | 0.65 |
| 1 | 237272920 | rs17669728 | -0.729 | -1.379 | -0.348 | 0.248 | -0.001 | -0.007 | -0.007 | 1.226 | 0.22 | 0.19 | 0.22 | 0.09 | 0.21 | 0.2 |
| 1 | 237274019 | rs2485598 | -0.617 | -1.459 | -0.651 | 0.642 | -0.003 | 0.006 | 0.006 | 0 | 0.31 | 0.36 | 0.21 | 0.62 | 0.25 | 0.36 |
| 1 | 237276068 | rs2490390 | 0.334 | -1.323 | -0.865 | 0.678 | -0.026 | 0.023 | 0.023 | 0 | 0.69 | 0.53 | 0.78 | 0.36 | 0.79 | 0.61 |
| 1 | 237278149 | rs11810113 | 1.647 | -1.135 | -1.039 | 0.559 | -0.098 | 0.057 | 0.057 | 0 | 0.4 | 0.1 | 0.51 | 0.26 | 0.51 | 0.41 |

| | | | | | | | | | | | | | | | | |
|---|---|---|---|---|---|---|---|---|---|---|---|---|---|---|---|---|
| 1 | 237280629 | rs2485564 | 0.642 | -0.843 | -0.86 | 0.644 | 0 | 0 | 0 | 0 | 0.6 | 0.61 | 0.66 | 0.37 | 0.6 | 0.54 |
| 1 | 237282971 | rs2490385 | 0.703 | -1.031 | -1.715 | 0.617 | 0.101 | -0.083 | -0.083 | 0 | 0.48 | 0.08 | 0.61 | 0.41 | 0.71 | 0.5 |
| 1 | 237285607 | rs2485570 | 1.144 | -0.798 | -0.274 | 0.534 | 0.046 | 0.015 | 0.015 | 0 | 0.93 | 0.71 | 0.91 | 0.97 | 0.92 | 0.98 |
| 1 | 237285629 | rs6693720 | -0.73 | -1.548 | -0.027 | 0.247 | -0.013 | -0.02 | -0.02 | 0.036 | 0.18 | 0.21 | 0.1 | 0.12 | 0.11 | 0.19 |
| 1 | 237296231 | rs10737813 | 0.472 | -1.55 | -0.715 | 0.115 | 0.018 | 0.025 | 0.025 | 0 | 0.53 | 0.32 | 0.66 | 0.42 | 0.56 | 0.58 |
| 1 | 237299349 | rs1337797 | -0.415 | -1.401 | -0.26 | 0.546 | 0.002 | 0.029 | 0.029 | 0 | 0.68 | 0.59 | 0.73 | 0.6 | 0.66 | 0.75 |
| 1 | 237299708 | rs2490373 | -0.39 | -1.418 | -0.268 | 0.538 | 0.002 | 0.028 | 0.028 | 0 | 0.68 | 0.59 | 0.73 | 0.6 | 0.66 | 0.76 |
| 1 | 237303417 | rs2152885 | 1.445 | -1.497 | -0.551 | 0.489 | -0.066 | 0.046 | 0.046 | 0 | 0.18 | 0.03 | 0.17 | 0.08 | 0.24 | 0.11 |
| 1 | 237304812 | rs2490372 | 0.003 | -1.515 | -0.099 | 0.498 | 0 | 0.031 | 0.031 | 0 | 0.66 | 0.42 | 0.7 | 0.55 | 0.64 | 0.73 |
| 1 | 237307601 | rs6657602 | 0.461 | -1.485 | -0.242 | 0.388 | 0 | -0.029 | -0.029 | 0 | 0.09 | 0.12 | 0.07 | 0.03 | 0.07 | 0.16 |
| 1 | 237310235 | rs2490370 | -0.142 | -1.468 | -0.242 | 0.512 | 0.005 | -0.018 | -0.018 | 0 | 0.62 | 0.67 | 0.69 | 0.54 | 0.65 | 0.68 |
| 1 | 237313722 | rs1409052 | 1.061 | -1.324 | -0.948 | 0.311 | -0.152 | -0.005 | -0.005 | 0 | 0.38 | 0.04 | 0.53 | 0.3 | 0.5 | 0.36 |
| 1 | 237321914 | rs2490365 | -0.26 | -1.497 | 0.246 | 0.452 | -0.003 | 0.005 | 0.005 | 0 | 0.59 | 0.56 | 0.66 | 0.47 | 0.61 | 0.64 |
| 1 | 237322012 | rs1361116 | 0.368 | -1.533 | 0.532 | 0.434 | 0.023 | -0.003 | -0.003 | 0 | 0.53 | 0.37 | 0.63 | 0.46 | 0.6 | 0.61 |
| 1 | 237323484 | rs1415716 | 0.176 | -1.369 | 0.236 | 0.48 | 0.001 | 0.014 | 0.014 | 0 | 0.07 | 0.19 | 0.06 | 0.11 | 0.07 | 0.05 |
| 1 | 237333965 | rs884641 | 0.079 | -1.478 | 0.132 | 0.292 | -0.002 | -0.008 | -0.008 | 0.147 | 0.37 | 0.31 | 0.33 | 0.52 | 0.4 | 0.38 |
| 1 | 237337651 | rs2490356 | 1.046 | -1.707 | 0.048 | 0.182 | -0.056 | 0.057 | 0.057 | 0.72 | 0.38 | 0.19 | 0.53 | 0.29 | 0.49 | 0.39 |
| 1 | 237343092 | rs2490352 | 1.591 | -1.879 | -0.089 | 0.147 | -0.15 | 0.033 | 0.033 | 0.804 | 0.33 | 0.05 | 0.51 | 0.24 | 0.48 | 0.35 |
| 1 | 237343127 | rs2490351 | -0.022 | -1.892 | 0.4 | 0.243 | 0.007 | 0.013 | 0.013 | 0 | 0.53 | 0.57 | 0.63 | 0.37 | 0.57 | 0.51 |
| 1 | 237344644 | rs9730301 | -0.204 | -1.77 | 0.504 | 0.287 | -0.014 | -0.019 | -0.019 | 0 | 0.17 | 0.51 | 0.12 | 0.13 | 0.08 | 0.14 |
| 1 | 237349657 | rs6668069 | -0.147 | -1.845 | 0.331 | 0.269 | 0.002 | -0.02 | -0.02 | 1.115 | 0.4 | 0.29 | 0.33 | 0.5 | 0.39 | 0.4 |
| 1 | 237349738 | rs6683225 | -1.338 | -1.871 | 0.422 | 0.25 | -0.013 | 0.008 | 0.008 | 0.063 | 0.61 | 0.64 | 0.46 | 0.65 | 0.51 | 0.55 |
| 1 | 237351354 | rs11811696 | 0.244 | -1.166 | 0.778 | 0.616 | -0.017 | -0.042 | -0.042 | 0.838 | 0.16 | 0.2 | 0.09 | 0.03 | 0.11 | 0.06 |
| 1 | 237355844 | rs268787 | 0.384 | -1.207 | 0.832 | 0.651 | -0.01 | -0.004 | -0.004 | 0.171 | 0.59 | 0.46 | 0.66 | 0.42 | 0.61 | 0.53 |
| 1 | 237356775 | rs2211009 | 1.133 | -1.123 | 0.665 | 0.668 | -0.08 | -0.089 | -0.089 | 0 | 0.21 | 0.03 | 0.22 | 0.33 | 0.28 | 0.29 |
| 1 | 237357012 | rs1765887 | 0.615 | -1.124 | 0.802 | 0.729 | -0.09 | -0.065 | -0.065 | 0 | 0.78 | 0.48 | 0.88 | 0.9 | 0.87 | 0.8 |
| 1 | 237359988 | rs187910 | 0.566 | -0.909 | -0.025 | 0.739 | 0.026 | -0.007 | -0.007 | 0 | 0.52 | 0.27 | 0.6 | 0.51 | 0.56 | 0.49 |
| 1 | 237363378 | rs268790 | 1.184 | -0.903 | -0.186 | 0.672 | -0.026 | 0.001 | 0.001 | 0 | 0.43 | 0.21 | 0.53 | 0.44 | 0.48 | 0.36 |
| 1 | 237364582 | rs12401834 | -0.207 | -0.958 | -0.169 | 0.676 | 0 | 0.012 | 0.012 | 0 | 0.4 | 0.25 | 0.32 | 0.35 | 0.41 | 0.38 |
| 1 | 237369284 | rs637520 | 1.145 | -0.666 | -0.815 | 0.642 | -0.017 | 0.023 | 0.023 | 0 | 0.42 | 0.23 | 0.53 | 0.37 | 0.47 | 0.35 |
| 1 | 237373237 | rs628414 | 0.118 | -0.3 | -0.701 | 0.391 | -0.01 | -0.063 | -0.063 | 0 | 0.79 | 0.3 | 0.81 | 0.98 | 0.81 | 0.93 |
| 1 | 237374838 | rs2111736 | 1.049 | -0.284 | -0.913 | 0.535 | 0.008 | 0.023 | 0.023 | 0.072 | 0.65 | 0.25 | 0.72 | 0.76 | 0.7 | 0.67 |
| 1 | 237378155 | rs2152884 | 0.564 | 0.042 | -1.004 | 0.859 | -0.013 | -0.065 | -0.065 | 0 | 0.26 | 0.12 | 0.22 | 0.41 | 0.26 | 0.42 |
| 1 | 237379105 | rs6663087 | -0.539 | -0.062 | -0.796 | 0.543 | -0.01 | -0.002 | -0.002 | 0.907 | 0.39 | 0.28 | 0.3 | 0.58 | 0.37 | 0.57 |
| 1 | 237382689 | rs6661229 | -0.097 | 0.144 | -0.58 | 0.818 | -0.002 | -0.001 | -0.001 | 0.11 | 0.42 | 0.43 | 0.49 | 0.6 | 0.35 | 0.44 |
| 1 | 237383625 | rs870872 | 0.128 | 0.414 | -0.718 | 0.708 | -0.045 | 0.053 | 0.053 | 0 | 0.13 | 0.04 | 0.15 | 0.01 | 0.24 | 0.12 |
| 1 | 237390732 | rs6683160 | 0.22 | -0.142 | -0.354 | 1.348 | 0.003 | -0.053 | -0.053 | 2.383 | 0.33 | 0.13 | 0.4 | 0.46 | 0.33 | 0.36 |
| 1 | 237393277 | rs1362841 | 0.362 | -0.315 | -0.488 | 1.389 | -0.044 | -0.113 | -0.113 | 3.433 | 0.31 | 0.02 | 0.39 | 0.4 | 0.31 | 0.34 |
| 1 | 237395982 | rs10802592 | -0.681 | -0.185 | -0.306 | 1.312 | 0.001 | 0.002 | 0.002 | 0 | 0.44 | 0.44 | 0.51 | 0.66 | 0.38 | 0.5 |
| 1 | 237401055 | rs11587851 | 0.313 | 0.241 | -1.074 | 0.497 | -0.028 | -0.03 | -0.03 | 0 | 0.26 | 0.07 | 0.38 | 0.28 | 0.29 | 0.36 |
| 1 | 237406445 | rs666787 | 1.04 | 0.153 | 0.136 | 0.335 | 0.017 | -0.028 | -0.028 | 0 | 0.91 | 0.58 | 0.94 | 1 | 0.97 | 0.96 |
| 1 | 237411254 | rs10925349 | 0.147 | 0.375 | -0.412 | 0.5 | -0.002 | -0.003 | -0.003 | 0 | 0.43 | 0.49 | 0.54 | 0.32 | 0.39 | 0.44 |
| 1 | 237414375 | rs888439 | -0.244 | -0.062 | -1.092 | 0.318 | 0.039 | -0.014 | -0.014 | 0 | 0.25 | 0.08 | 0.12 | 0.27 | 0.16 | 0.25 |
| 1 | 237416049 | rs12133002 | -0.703 | 0.41 | -0.429 | -0.151 | 0.003 | -0.043 | -0.043 | 0 | 0.58 | 0.49 | 0.43 | 0.83 | 0.54 | 0.57 |
| 1 | 237419785 | rs10925354 | -0.493 | -0.337 | -0.368 | 0.081 | -0.009 | -0.016 | -0.016 | 3.569 | 0.42 | 0.44 | 0.29 | 0.7 | 0.33 | 0.37 |
| 1 | 237420086 | rs918241 | 0.977 | -0.464 | -0.725 | 0.231 | -0.029 | 0.066 | 0.066 | 0 | 0.42 | 0.23 | 0.55 | 0.31 | 0.44 | 0.47 |
| 1 | 237420681 | rs950964 | 0.265 | -0.645 | -0.906 | 0.513 | 0.006 | -0.009 | -0.009 | 0 | 0.19 | 0.17 | 0.21 | 0.54 | 0.14 | 0.23 |
| 1 | 237422989 | rs10495392 | -0.345 | -0.693 | -0.438 | 0.304 | -0.001 | -0.031 | -0.031 | 0 | 0.88 | 0.81 | 0.9 | 0.93 | 0.88 | 0.95 |
| 1 | 237423053 | rs961121 | 0.487 | -0.854 | -0.025 | 0.249 | -0.01 | -0.042 | -0.042 | 0 | 0.12 | 0.28 | 0.04 | 0.05 | 0.05 | 0.17 |
| 1 | 237427945 | rs2065985 | 1.898 | -0.736 | -0.601 | 0.263 | -0.141 | 0.139 | 0.139 | 0.097 | 0.38 | 0.02 | 0.49 | 0.14 | 0.42 | 0.43 |
| 1 | 237428520 | rs6429001 | -0.08 | -0.67 | -0.475 | 0.281 | 0.013 | -0.034 | -0.034 | 0.313 | 0.86 | 0.76 | 0.84 | 0.9 | 0.81 | 0.94 |
| 1 | 237430681 | rs7551532 | 0.476 | -0.704 | -0.377 | 0.329 | -0.009 | 0.022 | 0.022 | 0.073 | 0.69 | 0.57 | 0.62 | 0.38 | 0.68 | 0.73 |
| 1 | 237431505 | rs10495394 | 0.818 | -0.768 | -0.426 | 0.272 | -0.034 | -0.036 | -0.036 | 0 | 0.17 | 0.05 | 0.07 | 0.18 | 0.21 | 0.12 |
| 1 | 237432302 | rs12043216 | 0.677 | -0.604 | -0.388 | 0.343 | 0 | 0.01 | 0.01 | 0 | 0.17 | 0.18 | 0.22 | 0.54 | 0.14 | 0.22 |
| 1 | 237433239 | rs4659491 | 1.96 | -0.599 | -0.351 | 0.299 | -0.096 | 0.2 | 0.2 | 0.014 | 0.38 | 0.08 | 0.49 | 0.12 | 0.42 | 0.43 |
| 1 | 237433524 | rs2275288 | 0.485 | -0.516 | -0.347 | 0.414 | 0.012 | -0.022 | -0.022 | 0 | 0.19 | 0.14 | 0.21 | 0.52 | 0.14 | 0.22 |
| 1 | 237433625 | rs2275287 | -0.424 | -0.414 | -0.509 | 0.114 | 0.004 | 0.011 | 0.011 | 0 | 0.37 | 0.36 | 0.31 | 0.7 | 0.35 | 0.33 |
| 1 | 237434181 | rs7526874 | -0.657 | -0.343 | -0.68 | 0.121 | -0.017 | 0.015 | 0.015 | 0 | 0.9 | 0.92 | 0.88 | 1 | 0.83 | 0.99 |
| 1 | 237434910 | rs7519473 | 0.432 | -0.379 | -0.639 | 0.149 | -0.003 | -0.025 | -0.025 | 0 | 0.12 | 0.33 | 0.04 | 0.06 | 0.05 | 0.18 |
| 1 | 237435536 | rs2080679 | -1.294 | -0.359 | -0.846 | 0.074 | 0.047 | 0.158 | 0.158 | 0 | 0.5 | 0.75 | 0.36 | 0.77 | 0.4 | 0.52 |
| 1 | 237439204 | rs12049532 | 0.794 | -0.343 | -0.894 | 0.226 | 0.014 | -0.097 | -0.097 | 0.039 | 0.17 | 0.02 | 0.18 | 0.35 | 0.14 | 0.19 |
| 1 | 237444583 | rs7520389 | -0.334 | 0.273 | -0.54 | -0.487 | -0.021 | 0.016 | 0.016 | 0 | 0.89 | 0.92 | 0.87 | 1 | 0.8 | 0.98 |
| 1 | 237444888 | rs6429004 | -0.214 | -0.405 | -0.164 | -0.11 | 0.003 | -0.052 | -0.052 | 0.294 | 0.12 | 0.35 | 0.05 | 0.05 | 0.06 | 0.18 |
| 1 | 237446556 | rs7526759 | -0.469 | -0.022 | -1.117 | 0.123 | 0.068 | 0.148 | 0.148 | 0 | 0.5 | 0.75 | 0.34 | 0.76 | 0.36 | 0.5 |
| 1 | 237453992 | rs1345497 | 0.292 | 0.238 | -0.951 | 0.669 | -0.005 | 0.028 | 0.028 | 0.029 | 0.47 | 0.25 | 0.58 | 0.37 | 0.51 | 0.4 |
| 1 | 237455023 | rs7538075 | 0.032 | -0.193 | -1.09 | 0.416 | -0.052 | 0.082 | 0.082 | 0 | 0.34 | 0.07 | 0.24 | 0.18 | 0.33 | 0.37 |
| 1 | 237456093 | rs10802598 | 0.82 | -0.171 | -0.023 | 0.42 | 0.079 | 0.047 | 0.047 | 0.001 | 0.93 | 0.59 | 0.91 | 0.97 | 0.93 | 0.91 |
| 1 | 237460954 | rs11802494 | -0.065 | -0.384 | 0.171 | 0.739 | -0.001 | 0.156 | 0.156 | 0.896 | 0.07 | 0.28 | 0.05 | 0.2 | 0.03 | 0.03 |
| 1 | 237462985 | rs10754597 | 0.217 | -0.434 | -0.254 | 0.731 | -0.066 | 0.08 | 0.08 | 0 | 0.33 | 0.09 | 0.25 | 0.18 | 0.34 | 0.37 |

| | | | | | | | | | | | | | | | | |
|---|---|---|---|---|---|---|---|---|---|---|---|---|---|---|---|---|
| 1 | 237463399 | rs16835116 | -0.473 | -0.313 | -0.211 | 0.715 | -0.004 | -0.007 | -0.007 | 0 | 0.43 | 0.53 | 0.32 | 0.38 | 0.37 | 0.48 |
| 1 | 237471081 | rs10802600 | -0.532 | -0.305 | 0.022 | 0.724 | 0.002 | 0.008 | 0.008 | 0.007 | 0.45 | 0.6 | 0.34 | 0.42 | 0.4 | 0.56 |
| 1 | 237486945 | rs974892 | -0.579 | -0.121 | -0.402 | 0.798 | 0.015 | 0.023 | 0.023 | 1.011 | 0.45 | 0.75 | 0.35 | 0.39 | 0.37 | 0.46 |
| 1 | 237493362 | rs2808224 | -0.119 | -0.219 | -0.877 | 0.416 | -0.006 | 0.058 | 0.058 | 0 | 0.36 | 0.22 | 0.27 | 0.2 | 0.33 | 0.36 |
| 1 | 237498486 | rs2543038 | 0.08 | -0.594 | -0.927 | 0.597 | 0.021 | 0.019 | 0.019 | 0 | 0.55 | 0.27 | 0.71 | 0.59 | 0.64 | 0.28 |
| 1 | 237502215 | rs2808226 | -1.685 | -0.326 | 0.055 | 0.775 | 0.053 | 0.025 | 0.025 | 1.028 | 0.07 | 0.33 | 0.09 | 0.04 | 0.08 | 0.02 |
| 1 | 237504759 | rs10925374 | 0.055 | -0.347 | -0.418 | 1.153 | -0.004 | -0.026 | -0.026 | 0.731 | 0.11 | 0.07 | 0.17 | 0.34 | 0.13 | 0.07 |
| 1 | 237516073 | rs10802607 | -1.234 | -0.346 | -0.186 | 1.117 | 0.012 | 0.154 | 0.154 | 0.023 | 0.18 | 0.38 | 0.25 | 0.38 | 0.2 | 0.09 |
| 1 | 237525382 | rs4638118 | -1.474 | -0.435 | -0.371 | 0.987 | -0.001 | 0.006 | 0.006 | 0.044 | 0.26 | 0.31 | 0.31 | 0.32 | 0.34 | 0.1 |
| 1 | 237525415 | rs7514255 | -1.315 | -0.409 | -0.378 | 1 | 0 | 0.001 | 0.001 | 0.02 | 0.24 | 0.26 | 0.3 | 0.32 | 0.34 | 0.1 |
| 1 | 237530837 | rs7550611 | -0.659 | -1.215 | -0.559 | 0.446 | -0.014 | 0.014 | 0.014 | 0 | 0.74 | 0.82 | 0.84 | 0.83 | 0.72 | 0.56 |
| 1 | 237531393 | rs7551196 | -1.302 | -1.028 | -0.341 | 0.873 | -0.035 | 0.079 | 0.079 | 0.131 | 0.25 | 0.45 | 0.33 | 0.42 | 0.23 | 0.18 |
| 1 | 237536837 | rs10925387 | -0.64 | -0.987 | -0.322 | 0.792 | -0.02 | -0.029 | -0.029 | 0.124 | 0.13 | 0.33 | 0.09 | 0.06 | 0.1 | 0.1 |
| 1 | 237539667 | rs10925390 | 0.185 | -0.968 | -0.409 | 0.88 | -0.002 | 0.057 | 0.057 | 0 | 0.11 | 0.13 | 0.26 | 0.38 | 0.12 | 0.08 |
| 1 | 237540615 | rs10925391 | -1.459 | -0.846 | -0.417 | 0.857 | -0.032 | 0.094 | 0.094 | 0.256 | 0.27 | 0.55 | 0.34 | 0.43 | 0.24 | 0.19 |
| 1 | 237547162 | rs7511662 | -0.32 | -0.706 | -0.58 | 0.866 | 0.006 | 0.011 | 0.011 | 0 | 0.48 | 0.36 | 0.72 | 0.59 | 0.51 | 0.35 |
| 1 | 237547223 | rs17682073 | 0.237 | -0.81 | -0.657 | 0.919 | -0.104 | 0.159 | 0.159 | 0 | 0.4 | 0.07 | 0.48 | 0.22 | 0.4 | 0.27 |
| 1 | 237547355 | rs11583145 | 0.306 | -0.619 | -0.368 | 1.161 | -0.002 | 0.123 | 0.123 | 0 | 0.1 | 0.18 | 0.26 | 0.39 | 0.12 | 0.09 |
| 1 | 237547600 | rs4659786 | -0.708 | -0.658 | -0.418 | 0.969 | 0.001 | 0.002 | 0.002 | 0 | 0.53 | 0.55 | 0.75 | 0.65 | 0.57 | 0.43 |
| 1 | 237548941 | rs16835191 | -0.152 | -0.591 | -0.472 | 1.031 | -0.01 | 0.043 | 0.043 | 0.432 | 0.05 | 0.2 | 0.03 | 0.05 | 0.03 | 0.09 |
| 1 | 237552147 | rs10925399 | -0.458 | -0.165 | -0.549 | 1.022 | 0.009 | 0.018 | 0.018 | 0 | 0.53 | 0.42 | 0.73 | 0.65 | 0.57 | 0.43 |
| 1 | 237554538 | rs12078739 | -0.515 | 0.049 | -1.251 | 1.708 | 0.012 | -0.021 | -0.021 | 2.803 | 0.11 | 0.09 | 0.19 | 0.35 | 0.11 | 0.12 |
| 1 | 237557130 | rs10925401 | -0.116 | 0.542 | -1.298 | 0.96 | 0.009 | 0.026 | 0.026 | 0.375 | 0.48 | 0.3 | 0.64 | 0.68 | 0.48 | 0.65 |
| 1 | 237557302 | rs4393145 | 0.308 | 0.383 | -1.877 | 0.968 | -0.018 | -0.051 | -0.051 | 0 | 0.27 | 0.1 | 0.32 | 0.38 | 0.33 | 0.31 |
| 1 | 237557907 | rs12407312 | 0.145 | 0.021 | -1.851 | 1.207 | 0 | -0.079 | -0.079 | 0.434 | 0.17 | 0.03 | 0.28 | 0.32 | 0.15 | 0.27 |
| 1 | 237559453 | rs10925402 | -0.735 | -0.075 | -2.241 | 0.09 | 0.03 | 0.132 | 0.132 | 0 | 0.7 | 0.85 | 0.84 | 1 | 0.78 | 0.83 |
| 1 | 237559533 | rs6677656 | -1.04 | -0.179 | -1.088 | 0.128 | 0.021 | -0.004 | -0.004 | 0 | 0.08 | 0.38 | 0.05 | 0 | 0.04 | 0.07 |
| 1 | 237561361 | rs10925405 | 0.94 | 0.447 | -0.497 | 0.248 | 0.049 | 0.068 | 0.068 | 0 | 0.15 | 0.35 | 0.15 | 0.27 | 0.21 | 0.16 |
| 1 | 237572916 | rs10737816 | -0.524 | 0.497 | -0.399 | 0.1 | 0.006 | 0.001 | 0.001 | 1.034 | 0.26 | 0.71 | 0.23 | 0.28 | 0.3 | 0.21 |
| 1 | 237575133 | rs6673700 | -0.713 | 0.517 | -0.337 | 0.104 | 0.096 | -0.01 | -0.01 | 0.143 | 0.08 | 0.33 | 0.07 | 0 | 0.1 | 0.06 |
| 1 | 237576019 | rs12691536 | 0.605 | 0.535 | -0.395 | 0.121 | 0.023 | 0.037 | 0.037 | 0 | 0.18 | 0.38 | 0.16 | 0.28 | 0.2 | 0.15 |
| 1 | 237578208 | rs17631862 | 0.259 | 0.296 | -0.728 | -0.292 | 0.012 | 0.069 | 0.069 | 0.354 | 0.45 | 0.24 | 0.6 | 0.71 | 0.43 | 0.61 |
| 1 | 237578224 | rs16835219 | 0.588 | 0.335 | -0.739 | -0.283 | 0.044 | 0.028 | 0.028 | 0 | 0.19 | 0.38 | 0.18 | 0.28 | 0.27 | 0.15 |
| 1 | 237578555 | rs12145251 | 0.788 | 0.326 | -0.611 | -0.275 | 0.014 | 0.02 | 0.02 | 0 | 0.18 | 0.33 | 0.15 | 0.28 | 0.22 | 0.14 |
| 1 | 237578970 | rs11577402 | 1.034 | 0.049 | -0.828 | -0.144 | -0.016 | -0.047 | -0.047 | 0 | 0.3 | 0.13 | 0.34 | 0.41 | 0.36 | 0.4 |
| 1 | 237580606 | rs6673182 | -1.068 | 0.09 | -0.991 | -0.138 | 0.002 | 0.174 | 0.174 | 0 | 0.71 | 0.93 | 0.84 | 1 | 0.76 | 0.82 |
| 1 | 237586384 | rs16835237 | 0.924 | 0.058 | -0.893 | 0.16 | 0.007 | -0.043 | -0.043 | 0 | 0.14 | 0.06 | 0.27 | 0.31 | 0.1 | 0.17 |
| 1 | 237588487 | rs7525969 | -0.314 | 0.127 | -0.689 | 0.064 | 0.002 | 0.031 | 0.031 | 0 | 0.52 | 0.49 | 0.66 | 0.7 | 0.51 | 0.62 |
| 1 | 237589433 | rs6674675 | 0.725 | -0.058 | -0.888 | 0.47 | -0.002 | 0.01 | 0.01 | 0 | 0.31 | 0.22 | 0.35 | 0.21 | 0.39 | 0.31 |
| 1 | 237589565 | rs4659793 | -0.148 | -0.378 | -1.259 | 0.588 | 0.01 | -0.021 | -0.021 | 0.155 | 0.19 | 0.14 | 0.3 | 0.48 | 0.1 | 0.32 |
| 1 | 237593216 | rs12123446 | 0.332 | -0.367 | -1.188 | 0.637 | -0.002 | 0.013 | 0.013 | 0 | 0.15 | 0.17 | 0.29 | 0.28 | 0.11 | 0.17 |
| 1 | 237595465 | rs4376723 | -0.389 | -0.08 | -1.59 | -0.02 | 0.011 | 0.068 | 0.068 | 0 | 0.65 | 0.91 | 0.8 | 0.79 | 0.69 | 0.62 |
| 1 | 237597163 | rs12038715 | 0.554 | -0.156 | -1.552 | 0.064 | -0.061 | -0.052 | -0.052 | 0.104 | 0.42 | 0.17 | 0.4 | 0.47 | 0.48 | 0.39 |
| 1 | 237597309 | rs7537614 | -0.371 | -0.115 | -1.392 | -0.56 | 0.007 | 0.007 | 0.007 | 0.194 | 0.74 | 0.8 | 0.89 | 1 | 0.81 | 0.85 |
| 1 | 237599305 | rs10925415 | 0.47 | -0.316 | -0.73 | -0.319 | 0 | -0.001 | -0.001 | 0 | 0.57 | 0.44 | 0.51 | 0.7 | 0.61 | 0.62 |
| 1 | 237599446 | rs11803630 | 1.04 | 0.104 | -1.068 | -0.077 | -0.037 | -0.033 | -0.033 | 0.868 | 0.16 | 0.11 | 0.19 | 0.21 | 0.25 | 0.23 |
| 1 | 237600863 | rs7552202 | 0.452 | -0.526 | -0.541 | -0.452 | -0.004 | -0.009 | -0.009 | 0 | 0.58 | 0.46 | 0.54 | 0.7 | 0.67 | 0.63 |
| 1 | 237602666 | rs10925416 | 0.1 | -0.644 | -0.393 | -0.413 | 0.004 | 0.002 | 0.002 | 0 | 0.19 | 0.18 | 0.29 | 0.2 | 0.14 | 0.17 |
| 1 | 237604253 | rs6661949 | -0.508 | -0.306 | -0.716 | -0.351 | -0.032 | 0.13 | 0.13 | 0 | 0.8 | 0.98 | 0.9 | 0.99 | 0.82 | 0.83 |
| 1 | 237605048 | rs2998402 | -1.61 | -0.393 | 0.16 | 0.094 | -0.01 | 0.062 | 0.062 | 14.318 | 0.07 | 0.62 | 0.11 | 0.12 | 0.02 | 0.07 |
| 1 | 237612398 | rs2927932 | -1.41 | -0.549 | 0.534 | -0.045 | -0.003 | 0.063 | 0.063 | 0.994 | 0.07 | 0.34 | 0.04 | 0.07 | 0.01 | 0.03 |
| 1 | 237613571 | rs2086209 | 0.444 | -0.482 | -0.411 | -0.45 | 0.001 | 0.125 | 0.125 | 1.218 | 0.49 | 0.27 | 0.52 | 0.76 | 0.56 | 0.6 |
| 1 | 237613801 | rs4659796 | 1.008 | -0.388 | -0.631 | -0.269 | -0.019 | 0.003 | 0.003 | 0.665 | 0.41 | 0.25 | 0.48 | 0.55 | 0.52 | 0.4 |
| 1 | 237614434 | rs6429014 | 0.62 | -0.157 | -0.386 | -0.153 | 0.031 | 0.065 | 0.065 | 0.848 | 0.48 | 0.26 | 0.54 | 0.65 | 0.6 | 0.58 |
| 1 | 237617757 | rs3765097 | 0.663 | -0.149 | -0.379 | -0.224 | 0.039 | 0.118 | 0.118 | 1.009 | 0.5 | 0.28 | 0.54 | 0.74 | 0.6 | 0.6 |
| 1 | 237620049 | rs2045955 | 0.57 | -0.162 | -0.374 | -0.255 | -0.004 | 0.129 | 0.129 | 1.045 | 0.49 | 0.27 | 0.52 | 0.78 | 0.56 | 0.6 |
| 1 | 237621239 | rs1478912 | 0.541 | -0.183 | 0.468 | -0.275 | 0.001 | 0.028 | 0.028 | 0.47 | 0.51 | 0.5 | 0.55 | 0.78 | 0.56 | 0.6 |
| 1 | 237626405 | rs1382585 | 0.584 | -0.153 | 0.533 | -0.259 | -0.001 | 0.09 | 0.09 | 1.188 | 0.49 | 0.3 | 0.53 | 0.74 | 0.56 | 0.6 |
| 1 | 237627498 | rs2618679 | 0.207 | -0.403 | 0.211 | -0.504 | 0.001 | -0.001 | -0.001 | 0.774 | 0.25 | 0.3 | 0.27 | 0.16 | 0.18 | 0.19 |
| 1 | 237634946 | rs2618681 | -0.891 | -0.348 | -0.439 | -0.682 | -0.01 | 0.046 | 0.046 | 0 | 0.8 | 0.85 | 0.87 | 0.99 | 0.79 | 0.82 |
| 1 | 237636179 | rs10754604 | 0.623 | -0.316 | -0.315 | -0.337 | -0.029 | 0.098 | 0.098 | 0.826 | 0.47 | 0.18 | 0.51 | 0.7 | 0.56 | 0.57 |
| 1 | 237643234 | rs6429023 | -1.267 | -0.47 | -0.083 | -0.029 | -0.016 | 0.012 | 0.012 | 0.451 | 0.17 | 0.49 | 0.13 | 0.15 | 0.1 | 0.13 |
| 1 | 237647282 | rs12117452 | -0.593 | -0.547 | -1.079 | -0.046 | 0.02 | 0.013 | 0.013 | 0 | 0.31 | 0.19 | 0.23 | 0.31 | 0.24 | 0.41 |
| 1 | 237648221 | rs12060093 | -0.255 | -0.923 | -1.361 | -0.547 | 0.009 | 0.009 | 0.009 | 0.922 | 0.32 | 0.74 | 0.21 | 0.21 | 0.23 | 0.19 |
| 1 | 237648917 | rs6673954 | -0.138 | -1.518 | -1.772 | -0.796 | 0.045 | 0.06 | 0.06 | 0 | 0.45 | 0.26 | 0.29 | 0.33 | 0.31 | 0.46 |
| 1 | 237649065 | rs9428668 | 0.019 | -1.527 | -1.587 | -0.76 | 0.021 | 0.019 | 0.019 | 0 | 0.39 | 0.28 | 0.24 | 0.31 | 0.26 | 0.41 |
| 1 | 237651054 | rs12058655 | 0.275 | -1.69 | -1.234 | -0.78 | -0.028 | -0.054 | -0.054 | 0.111 | 0.28 | 0.52 | 0.18 | 0.16 | 0.22 | 0.14 |
| 1 | 237653255 | rs2997975 | 0.433 | -1.533 | -1.531 | -0.74 | 0.023 | 0.016 | 0.016 | 0 | 0.34 | 0.21 | 0.23 | 0.31 | 0.25 | 0.44 |
| 1 | 237654831 | rs2127155 | -1.444 | -0.781 | -1.599 | -0.358 | -0.168 | -0.169 | -0.169 | 0 | 0.72 | 0.97 | 0.52 | 0.55 | 0.55 | 0.61 |

| | | | | | | | | | | | | | | | | |
|---|---|---|---|---|---|---|---|---|---|---|---|---|---|---|---|---|
| 1 | 237658484 | rs2010045 | -0.783 | -1.203 | -1.098 | -0.543 | -0.043 | -0.031 | -0.031 | 0 | 0.65 | 0.73 | 0.48 | 0.46 | 0.52 | 0.64 |
| 1 | 237658615 | rs2010033 | 0.275 | -1.716 | -0.949 | -0.829 | 0.017 | 0.02 | 0.02 | 0 | 0.37 | 0.25 | 0.27 | 0.29 | 0.29 | 0.43 |
| 1 | 237658638 | rs2010032 | -1.048 | -1.467 | -1.018 | -0.751 | -0.047 | -0.031 | -0.031 | 0 | 0.66 | 0.74 | 0.49 | 0.47 | 0.52 | 0.67 |
| 1 | 237659306 | rs16835325 | 0.233 | -1.36 | -0.656 | -0.661 | -0.008 | 0.019 | 0.019 | 0.516 | 0.06 | 0.3 | 0.03 | 0.04 | 0.01 | 0.02 |
| 1 | 237663757 | rs2618698 | 0.393 | -1.66 | -0.73 | -0.865 | 0.003 | 0.003 | 0.003 | 1.268 | 0.52 | 0.43 | 0.61 | 0.63 | 0.67 | 0.52 |
| 1 | 237665859 | rs2618699 | -0.207 | -1.23 | -0.913 | -0.853 | 0.012 | 0.041 | 0.041 | 0 | 0.58 | 0.74 | 0.52 | 0.67 | 0.65 | 0.53 |
| 1 | 237670140 | rs2618702 | -0.565 | -1.29 | -1.063 | -0.888 | 0.006 | 0.068 | 0.068 | 0.09 | 0.63 | 0.86 | 0.62 | 0.73 | 0.68 | 0.54 |
| 1 | 237673871 | rs3766841 | 0.71 | -1.192 | -1.028 | -0.755 | -0.039 | -0.02 | -0.02 | 0 | 0.24 | 0.38 | 0.13 | 0.15 | 0.18 | 0.21 |
| 1 | 237677346 | rs2618687 | 1.149 | -0.673 | -1.351 | -0.553 | -0.139 | -0.146 | -0.146 | 0.527 | 0.24 | 0.02 | 0.34 | 0.49 | 0.43 | 0.28 |
| 1 | 237682895 | rs9633357 | -0.04 | -1.221 | -1.412 | -1.155 | 0.026 | 0.021 | 0.021 | 0 | 0.33 | 0.16 | 0.34 | 0.3 | 0.28 | 0.44 |
| 1 | 237685255 | rs2618693 | 0.113 | -1.292 | -0.909 | -1.216 | 0.001 | -0.001 | -0.001 | 0 | 0.4 | 0.39 | 0.5 | 0.6 | 0.51 | 0.31 |
| 1 | 237690817 | rs10737817 | 0.308 | -1.467 | -1.16 | -1.309 | 0.009 | 0.033 | 0.033 | 0 | 0.3 | 0.07 | 0.33 | 0.25 | 0.26 | 0.41 |
| 1 | 237690923 | rs2779360 | 0.097 | -1.481 | -0.95 | -1.336 | 0 | -0.003 | -0.003 | 0.75 | 0.42 | 0.42 | 0.51 | 0.62 | 0.53 | 0.32 |
| 1 | 237691284 | rs2618703 | 0.004 | -1.505 | -0.421 | -1.177 | -0.006 | -0.012 | -0.012 | 0.158 | 0.63 | 0.41 | 0.77 | 0.83 | 0.75 | 0.7 |
| 1 | 237693190 | rs2805446 | -0.843 | -1.083 | -0.237 | -1.06 | -0.019 | -0.053 | -0.053 | 3.109 | 0.74 | 0.68 | 0.86 | 0.9 | 0.8 | 0.67 |
| 1 | 237702709 | rs10925451 | -0.517 | -1.001 | -0.431 | -0.695 | 0.025 | 0.037 | 0.037 | 1.393 | 0.29 | 0.17 | 0.34 | 0.23 | 0.24 | 0.31 |
| 1 | 237703401 | rs2257096 | 0.709 | -0.428 | -0.322 | -0.352 | -0.018 | -0.02 | -0.02 | 0 | 0.25 | 0.2 | 0.36 | 0.5 | 0.43 | 0.26 |
| 1 | 237708177 | rs2618666 | 0.493 | -0.707 | 0.421 | -0.393 | 0.039 | 0.213 | 0.213 | 0.67 | 0.93 | 0.58 | 0.96 | 0.87 | 0.98 | 0.98 |
| 1 | 237710993 | rs2618663 | 1.224 | -0.816 | 0.435 | -0.471 | -0.034 | 0.066 | 0.066 | 1.94 | 0.87 | 0.49 | 0.94 | 0.87 | 0.96 | 0.97 |
| 1 | 237711457 | rs2618662 | 1.024 | -0.814 | 0.487 | -0.47 | -0.028 | 0.073 | 0.073 | 1.849 | 0.88 | 0.54 | 0.95 | 0.87 | 0.96 | 0.97 |
| 1 | 237714103 | rs2306238 | 0.464 | -0.832 | 0.239 | -0.563 | 0.006 | 0.002 | 0.002 | 0 | 0.23 | 0.05 | 0.3 | 0.22 | 0.21 | 0.3 |
| 1 | 237718716 | rs2805464 | 0.232 | -0.74 | 0.354 | -0.516 | -0.004 | -0.003 | -0.003 | 0 | 0.29 | 0.3 | 0.38 | 0.6 | 0.42 | 0.26 |
| 1 | 237721111 | rs2805468 | 0.772 | -0.861 | 0.273 | -0.588 | -0.001 | 0 | 0 | 0 | 0.61 | 0.39 | 0.6 | 0.65 | 0.71 | 0.67 |
| 1 | 237721988 | rs2805469 | 0.785 | -0.412 | 0.245 | -0.261 | -0.072 | -0.083 | -0.083 | 0 | 0.24 | 0.13 | 0.35 | 0.51 | 0.41 | 0.25 |
| 1 | 237723828 | rs2127148 | 0.344 | -0.707 | 0.666 | -0.268 | 0.014 | 0.046 | 0.046 | 0.354 | 0.94 | 0.7 | 0.98 | 0.96 | 0.98 | 0.98 |
| 1 | 237725545 | rs2779394 | 0.798 | -0.808 | 0.685 | -0.308 | -0.002 | 0.072 | 0.072 | 0.508 | 0.9 | 0.57 | 0.92 | 0.87 | 0.93 | 0.95 |
| 1 | 237725934 | rs722582 | 0.838 | -0.264 | 0.509 | -0.165 | 0.01 | 0.04 | 0.04 | 0 | 0.21 | 0.3 | 0.22 | 0.49 | 0.3 | 0.25 |
| 1 | 237727970 | rs1031862 | 0.691 | -0.331 | 0.369 | -0.205 | 0.003 | 0.025 | 0.025 | 0 | 0.63 | 0.24 | 0.67 | 0.27 | 0.62 | 0.68 |
| 1 | 237729040 | rs3766865 | 1.202 | -0.747 | 0.147 | -0.254 | 0.017 | -0.055 | -0.055 | 0.259 | 0.14 | 0 | 0.13 | 0.18 | 0.11 | 0.12 |
| 1 | 237735683 | rs2779408 | 0.432 | -0.344 | 0.218 | -0.15 | -0.003 | -0.003 | -0.003 | 0 | 0.66 | 0.37 | 0.74 | 0.26 | 0.7 | 0.7 |
| 1 | 237741484 | rs2618704 | -0.048 | -0.331 | -0.137 | -0.176 | -0.001 | 0 | 0 | 0 | 0.7 | 0.28 | 0.73 | 0.38 | 0.71 | 0.72 |
| 1 | 237741884 | rs2805407 | 0.778 | -0.151 | -0.469 | -0.347 | -0.088 | 0.005 | 0.005 | 3.803 | 0.14 | 0.02 | 0.33 | 0.09 | 0.23 | 0.15 |
| 1 | 237744972 | rs1842085 | -0.346 | -0.623 | 0 | -0.182 | -0.007 | 0.001 | 0.001 | 0 | 0.3 | 0.36 | 0.25 | 0.69 | 0.29 | 0.28 |
| 1 | 237758247 | rs1842088 | -0.113 | -0.613 | -0.724 | -0.275 | -0.114 | -0.008 | -0.008 | 0.342 | 0.36 | 0.14 | 0.55 | 0.31 | 0.49 | 0.32 |
| 1 | 237759914 | rs2990548 | 0.559 | -0.577 | -0.926 | -0.257 | -0.143 | -0.027 | -0.027 | 3.008 | 0.3 | 0.05 | 0.52 | 0.27 | 0.46 | 0.29 |
| 1 | 237760083 | rs1817410 | 0.019 | -0.28 | -0.608 | -0.1 | 0.007 | -0.011 | -0.011 | 0 | 0.58 | 0.48 | 0.42 | 0.69 | 0.5 | 0.65 |
| 1 | 237762445 | rs2127159 | 1.115 | 0.01 | -1.647 | -0.065 | 0.002 | 0.145 | 0.145 | 0 | 0.34 | 0.08 | 0.2 | 0.07 | 0.24 | 0.41 |
| 1 | 237765657 | rs3820572 | 0.312 | 0.356 | -1.268 | 0.674 | -0.009 | -0.056 | -0.056 | 0 | 0.19 | 0.1 | 0.17 | 0.48 | 0.22 | 0.22 |
| 1 | 237771576 | rs12116442 | 0.251 | 0.417 | -1.238 | 0.706 | -0.001 | -0.053 | -0.053 | 0 | 0.21 | 0.13 | 0.17 | 0.48 | 0.22 | 0.22 |
| 1 | 237773427 | rs1717783 | -0.585 | 0.509 | -0.829 | 0.631 | -0.03 | 0.008 | 0.008 | 0 | 0.33 | 0.19 | 0.52 | 0.29 | 0.4 | 0.28 |
| 1 | 237778084 | rs3766871 | -1.243 | 0.005 | 0.098 | 0.778 | 0.002 | 0.018 | 0.018 | 2.492 | 0.06 | 0.16 | 0.03 | 0.06 | 0.03 | 0.07 |
| 1 | 237778909 | rs596502 | 1.442 | -0.547 | 0.376 | 1.248 | 0.016 | -0.02 | -0.02 | 0 | 0.7 | 0.28 | 0.81 | 0.55 | 0.83 | 0.67 |
| 1 | 237779700 | rs10925474 | -1.251 | -0.413 | 0.901 | 1.154 | 0.022 | 0.043 | 0.043 | 4.209 | 0.07 | 0.28 | 0.04 | 0.06 | 0.03 | 0.09 |
| 1 | 237784313 | rs489088 | 1.658 | -0.582 | 0.405 | 1.242 | 0.022 | -0.097 | -0.097 | 0 | 0.65 | 0.12 | 0.71 | 0.56 | 0.73 | 0.67 |
| 1 | 237785217 | rs578741 | 0.621 | -0.584 | 1.297 | 1.259 | -0.01 | 0.008 | 0.008 | 0 | 0.78 | 0.7 | 0.86 | 0.7 | 0.84 | 0.79 |
| 1 | 237788786 | rs6683048 | -1.477 | -0.633 | 1.798 | 1.257 | 0.054 | 0.058 | 0.058 | 1.289 | 0.1 | 0.34 | 0.12 | 0.13 | 0.11 | 0.1 |
| 1 | 237789317 | rs512336 | 0.339 | -0.473 | 1.647 | 1.353 | -0.016 | -0.005 | -0.005 | 0 | 0.82 | 0.77 | 0.89 | 0.8 | 0.89 | 0.84 |
| 1 | 237790680 | rs939698 | 0.816 | -0.434 | 1.645 | 1.29 | -0.018 | 0.042 | 0.042 | 0 | 0.78 | 0.61 | 0.83 | 0.62 | 0.83 | 0.77 |
| 1 | 237793646 | rs678541 | 0.949 | -0.408 | 1.619 | 1.317 | -0.027 | 0.055 | 0.055 | 0 | 0.76 | 0.62 | 0.83 | 0.6 | 0.83 | 0.8 |
| 1 | 237794603 | rs1759123 | 1.388 | -0.413 | 1.403 | 1.345 | -0.01 | 0.02 | 0.02 | 0.209 | 0.7 | 0.37 | 0.75 | 0.58 | 0.74 | 0.76 |
| 1 | 237796237 | rs1891557 | -0.571 | 0.052 | 0.896 | 1.159 | -0.029 | -0.087 | -0.087 | 0.844 | 0.24 | 0.1 | 0.32 | 0.44 | 0.26 | 0.34 |
| 1 | 237796837 | rs10802626 | -1.077 | -0.012 | 1.5 | 1.227 | 0.021 | 0.065 | 0.065 | 0 | 0.29 | 0.53 | 0.38 | 0.48 | 0.32 | 0.38 |
| 1 | 237797082 | rs1967579 | -1.167 | -0.025 | 1.549 | 1.195 | 0.011 | 0.037 | 0.037 | 0 | 0.29 | 0.57 | 0.39 | 0.48 | 0.32 | 0.38 |
| 1 | 237800381 | rs578152 | 0.955 | -0.045 | 1.433 | 0.934 | 0.028 | 0.035 | 0.035 | 0 | 0.75 | 0.57 | 0.75 | 0.64 | 0.7 | 0.78 |
| 1 | 237803805 | rs1029088 | -1.163 | -0.134 | 1.432 | 0.784 | -0.026 | 0.004 | 0.004 | 0 | 0.32 | 0.38 | 0.35 | 0.59 | 0.24 | 0.52 |
| 1 | 237805301 | rs1475678 | 1.85 | -0.294 | 1.254 | 0.729 | 0.021 | 0.185 | 0.185 | 0 | 0.51 | 0.21 | 0.47 | 0.19 | 0.56 | 0.41 |
| 1 | 237806819 | rs11583646 | -0.553 | -0.333 | 1.196 | 0.839 | 0.002 | 0.022 | 0.022 | 0 | 0.09 | 0.22 | 0.08 | 0.13 | 0.1 | 0.15 |
| 1 | 237810013 | rs3766878 | -1.355 | -0.334 | 1.107 | 0.83 | -0.008 | 0.033 | 0.033 | 0 | 0.25 | 0.43 | 0.22 | 0.35 | 0.24 | 0.22 |
| 1 | 237812261 | rs12725455 | 1.803 | -0.367 | 1.151 | 0.816 | 0.028 | 0.201 | 0.201 | 0 | 0.51 | 0.21 | 0.46 | 0.18 | 0.56 | 0.41 |
| 1 | 237812498 | rs3766879 | -0.035 | -0.048 | 0.641 | 0.967 | -0.004 | 0.007 | 0.007 | 0 | 0.16 | 0.21 | 0.14 | 0.23 | 0.14 | 0.1 |
| 1 | 237812676 | rs490057 | -1.471 | 0.017 | 0.581 | 0.937 | -0.003 | -0.042 | -0.042 | 0 | 0.46 | 0.52 | 0.51 | 0.81 | 0.44 | 0.59 |
| 1 | 237814783 | rs684923 | -1.65 | 0.064 | 0.488 | 0.926 | -0.003 | -0.035 | -0.035 | 0 | 0.45 | 0.49 | 0.5 | 0.8 | 0.43 | 0.56 |
| 1 | 237815546 | rs622625 | -0.586 | 0.372 | -0.028 | 1.533 | -0.012 | -0.082 | -0.082 | 0.066 | 0.2 | 0.07 | 0.29 | 0.45 | 0.2 | 0.31 |
| 1 | 237826332 | rs1953076 | -1.456 | 0.379 | 0.806 | 1.013 | 0.054 | 0.131 | 0.131 | 2.502 | 0.06 | 0.38 | 0.04 | 0.08 | 0.01 | 0.13 |
| 1 | 237834701 | rs2012813 | -0.756 | 1.404 | 0.32 | 1.335 | 0.054 | 0.074 | 0.074 | 0 | 0.27 | 0.43 | 0.29 | 0.55 | 0.38 | 0.38 |
| 1 | 237844948 | rs1533772 | -1.516 | 2.293 | 0.479 | 1.666 | 0.036 | 0.043 | 0.043 | 0 | 0.27 | 0.39 | 0.36 | 0.32 | 0.37 | 0.43 |
| 1 | 237845083 | rs1464459 | 0.204 | 2.716 | -0.818 | 1.325 | -0.071 | -0.105 | -0.105 | 0 | 0.14 | 0.03 | 0.21 | 0.51 | 0.24 | 0.24 |
| 1 | 237855896 | rs12032093 | 1.322 | 1.753 | -1.336 | 0.422 | -0.071 | -0.044 | -0.044 | 0 | 0.57 | 0.09 | 0.54 | 0.64 | 0.54 | 0.5 |
| 1 | 237862587 | rs4427395 | 1.07 | 1.339 | -1.058 | 0.711 | -0.007 | -0.101 | -0.101 | 0 | 0.66 | 0.11 | 0.58 | 0.41 | 0.7 | 0.66 |

| | | | | | | | | | | | | | | | | |
|---|---|---|---|---|---|---|---|---|---|---|---|---|---|---|---|---|
| 1 | 237886146 | rs2090876 | -0.567 | 1.107 | -0.367 | 0.608 | -0.013 | -0.066 | -0.066 | 1.17 | 0.2 | 0.1 | 0.34 | 0.65 | 0.21 | 0.28 |
| 1 | 237896307 | rs12086956 | -0.444 | 0.979 | 0.65 | 0.614 | -0.03 | -0.042 | -0.042 | 1.346 | 0.08 | 0.27 | 0.04 | 0.01 | 0.03 | 0.12 |
| 1 | 237897844 | rs2685297 | 0.672 | -0.107 | -0.42 | -0.098 | -0.022 | -0.152 | -0.152 | 0.519 | 0.35 | 0.04 | 0.36 | 0.57 | 0.28 | 0.32 |
| 1 | 237901607 | rs10925501 | -0.389 | 0.236 | -0.002 | -0.059 | 0.034 | 0.024 | 0.024 | 0 | 0.49 | 0.68 | 0.51 | 0.67 | 0.39 | 0.45 |
| 1 | 237908911 | rs16835705 | 0.192 | 0.932 | -0.491 | -0.272 | -0.014 | -0.005 | -0.005 | 0.693 | 0.19 | 0.16 | 0.23 | 0.18 | 0.28 | 0.3 |
| 1 | 237911004 | rs2819770 | 1.125 | 0.934 | -0.694 | -0.349 | -0.029 | 0.125 | 0.125 | 0 | 0.27 | 0.05 | 0.22 | 0.04 | 0.28 | 0.17 |
| 1 | 237921596 | rs1464461 | 0.374 | 0.531 | -0.118 | -0.291 | 0.004 | 0.002 | 0.002 | 0 | 0.46 | 0.43 | 0.47 | 0.64 | 0.39 | 0.4 |
| 1 | 237923989 | rs6699085 | -1.268 | 0.511 | -0.097 | -0.289 | 0.001 | 0.003 | 0.003 | 0.01 | 0.1 | 0.11 | 0.09 | 0.12 | 0.13 | 0.16 |
| 1 | 237924112 | rs2056387 | 0.795 | 0.892 | 0.305 | -0.529 | -0.001 | -0.063 | -0.063 | 1.464 | 0.29 | 0.34 | 0.32 | 0.04 | 0.33 | 0.18 |
| 1 | 237925817 | rs1357150 | -0.362 | 0.855 | 0.467 | -0.468 | 0 | -0.003 | -0.003 | 0 | 0.51 | 0.47 | 0.49 | 0.21 | 0.57 | 0.48 |
| 1 | 237927340 | rs1949665 | -1.624 | 1.032 | 0.587 | -0.295 | -0.015 | -0.003 | -0.003 | 3.473 | 0.34 | 0.69 | 0.3 | 0.28 | 0.37 | 0.45 |
| 1 | 237928221 | rs10802632 | -1.102 | 0.475 | 0.474 | -0.522 | 0.037 | 0.03 | 0.03 | 0 | 0.56 | 0.74 | 0.6 | 0.3 | 0.67 | 0.5 |
| 1 | 237928320 | rs2253244 | -1.574 | 0.477 | 0.457 | -0.517 | 0.026 | -0.067 | -0.067 | 0 | 0.6 | 0.88 | 0.62 | 0.38 | 0.71 | 0.62 |
| 1 | 237929556 | rs2253083 | -1.609 | 0.479 | 0.532 | -0.475 | -0.02 | 0.025 | 0.025 | 0 | 0.08 | 0.2 | 0.04 | 0.08 | 0.05 | 0.16 |
| 1 | 237931094 | rs2250079 | -0.464 | 0.637 | 0.448 | -0.548 | -0.011 | 0.03 | 0.03 | 0.323 | 0.38 | 0.71 | 0.45 | 0.22 | 0.47 | 0.34 |
| 1 | 237931618 | rs2250042 | -0.079 | 0.973 | 0.322 | -0.542 | -0.024 | 0.012 | 0.012 | 0 | 0.22 | 0.1 | 0.16 | 0.16 | 0.22 | 0.3 |
| 1 | 237932989 | rs2249847 | -1.059 | 1.009 | 0.441 | -0.605 | -0.005 | -0.016 | -0.016 | 0.013 | 0.54 | 0.55 | 0.53 | 0.17 | 0.65 | 0.4 |
| 1 | 237938506 | rs2249179 | -0.598 | 1.184 | 0.495 | -0.739 | -0.002 | -0.137 | -0.137 | 0 | 0.26 | 0.45 | 0.16 | 0.03 | 0.21 | 0.2 |
| 1 | 237939116 | rs11584594 | 1.22 | 1.287 | -0.188 | -0.537 | -0.06 | 0.098 | 0.098 | 0 | 0.23 | 0.06 | 0.28 | 0.03 | 0.34 | 0.17 |
| 1 | 237945870 | rs12737847 | -0.955 | 0.941 | 0.14 | -0.255 | 0.01 | 0.058 | 0.058 | 0 | 0.06 | 0.17 | 0.04 | 0.08 | 0.04 | 0.07 |
| 1 | 237950431 | rs1773471 | 0.676 | 1.122 | -0.306 | -0.289 | 0.015 | 0.011 | 0.011 | 0 | 0.46 | 0.4 | 0.45 | 0.66 | 0.36 | 0.39 |
| 1 | 237951482 | rs16835818 | -0.923 | 0.941 | -0.057 | -0.583 | 0.005 | -0.022 | -0.022 | 0.032 | 0.28 | 0.58 | 0.2 | 0.12 | 0.28 | 0.2 |
| 1 | 237957161 | rs790901 | -0.33 | 1.012 | -0.076 | -0.827 | 0.035 | -0.133 | -0.133 | 0.279 | 0.71 | 0.57 | 0.8 | 0.95 | 0.65 | 0.89 |
| 1 | 237957678 | rs790899 | -0.078 | 0.975 | -0.104 | -0.851 | 0.026 | -0.141 | -0.141 | 0 | 0.67 | 0.55 | 0.79 | 0.94 | 0.63 | 0.86 |
| 1 | 237958093 | rs2790345 | 0.808 | 0.587 | -0.302 | -0.761 | 0 | -0.004 | -0.004 | 0 | 0.56 | 0.43 | 0.72 | 0.82 | 0.58 | 0.73 |
| 1 | 237965043 | rs2790347 | 0.591 | 0.583 | -0.346 | -0.14 | 0.012 | 0.023 | 0.023 | 0 | 0.52 | 0.4 | 0.41 | 0.63 | 0.4 | 0.47 |
| 1 | 237967937 | rs2794828 | 1.294 | 0.175 | -0.983 | 0.118 | -0.022 | -0.106 | -0.106 | 1.381 | 0.2 | 0.08 | 0.26 | 0.5 | 0.19 | 0.28 |
| 1 | 237973034 | rs790883 | 1.276 | 0.171 | -0.939 | 0.123 | -0.016 | -0.094 | -0.094 | 0 | 0.2 | 0.09 | 0.26 | 0.5 | 0.19 | 0.28 |
| 1 | 237974720 | rs813173 | -0.457 | 0.194 | -0.745 | -0.295 | 0.001 | 0.106 | 0.106 | 0 | 0.52 | 0.72 | 0.55 | 0.83 | 0.47 | 0.66 |
| 1 | 237979535 | rs2794834 | 1.371 | 0.209 | -0.739 | -0.22 | -0.033 | -0.128 | -0.128 | 0.127 | 0.2 | 0.05 | 0.26 | 0.55 | 0.19 | 0.29 |
| 1 | 237981846 | rs1891246 | -0.207 | 0.109 | -0.684 | -0.5 | 0.042 | 0.027 | 0.027 | 0 | 0.46 | 0.73 | 0.38 | 0.39 | 0.4 | 0.5 |
| 1 | 237990122 | rs2819742 | -1.126 | 0.294 | -1.697 | -0.573 | -0.151 | 0.177 | 0.177 | 0.595 | 0.68 | 0.98 | 0.66 | 0.94 | 0.6 | 0.83 |
| 1 | 237990337 | rs2819743 | 0.529 | 0.333 | -1.194 | -0.069 | 0.003 | -0.055 | -0.055 | 0 | 0.24 | 0.15 | 0.28 | 0.61 | 0.22 | 0.37 |
| 1 | 237993416 | rs6695129 | -0.681 | 0.013 | -0.972 | 0.019 | -0.042 | -0.022 | -0.022 | 0 | 0.63 | 0.75 | 0.43 | 0.39 | 0.51 | 0.59 |
| 1 | 237993724 | rs2275692 | 0.436 | 0.449 | -0.802 | 0.907 | -0.06 | 0.047 | 0.047 | 0 | 0.19 | 0.01 | 0.1 | 0.05 | 0.22 | 0.06 |
| 1 | 237997087 | rs12594 | -0.317 | 0.459 | -1.277 | 0.638 | -0.024 | 0.025 | 0.025 | 0 | 0.32 | 0.13 | 0.3 | 0.26 | 0.36 | 0.23 |
| 1 | 237997254 | rs16835904 | -0.398 | -0.115 | -0.255 | 0.956 | -0.034 | 0.074 | 0.074 | 0.003 | 0.16 | 0.38 | 0.11 | 0.26 | 0.08 | 0.3 |
| 1 | 238000883 | rs946585 | -0.564 | 0.187 | -0.974 | 0.578 | -0.04 | 0.026 | 0.026 | 2.133 | 0.73 | 0.95 | 0.73 | 0.77 | 0.69 | 0.87 |
| 1 | 238003342 | rs6700222 | 0.055 | 0.383 | -0.688 | 0.662 | 0.017 | -0.018 | -0.018 | 0 | 0.24 | 0.36 | 0.23 | 0.21 | 0.31 | 0.26 |
| 1 | 238003774 | rs6703045 | 0.719 | 0.269 | -0.88 | 0.913 | -0.004 | 0.014 | 0.014 | 0 | 0.41 | 0.17 | 0.41 | 0.3 | 0.35 | 0.39 |
| 1 | 238010537 | rs1891249 | -0.213 | 0.511 | -0.581 | -0.06 | -0.037 | 0.036 | 0.036 | 0.521 | 0.72 | 0.89 | 0.8 | 0.77 | 0.67 | 0.84 |
| 1 | 238015491 | rs2819750 | -1.024 | 0.284 | 0.498 | 0.096 | 0.005 | -0.02 | -0.02 | 0 | 0.34 | 0.54 | 0.25 | 0.23 | 0.3 | 0.25 |
| 1 | 238015523 | rs2819751 | -1.026 | 0.282 | 0.615 | 0.122 | 0.002 | -0.009 | -0.009 | 0 | 0.34 | 0.56 | 0.25 | 0.24 | 0.3 | 0.25 |
| 1 | 238017237 | rs707224 | -0.224 | 0.043 | 0.724 | 0.293 | 0.002 | -0.002 | -0.002 | 0 | 0.83 | 0.84 | 0.84 | 0.8 | 0.87 | 0.7 |
| 1 | 238026731 | rs2790356 | -0.293 | 0.067 | 1.127 | 0.313 | -0.017 | 0.008 | 0.008 | 0.273 | 0.18 | 0.26 | 0.11 | 0.2 | 0.12 | 0.14 |
| 1 | 238028623 | rs2790360 | 0.244 | 0.135 | 0.981 | 0.347 | -0.008 | 0.007 | 0.007 | 0.104 | 0.16 | 0.2 | 0.11 | 0.2 | 0.11 | 0.15 |
| 1 | 238028729 | rs2794847 | 0.352 | 0.136 | 0.971 | 0.347 | 0.009 | -0.016 | -0.016 | 0.094 | 0.15 | 0.08 | 0.1 | 0.2 | 0.11 | 0.15 |
| 1 | 238032494 | rs6696783 | 1.021 | -0.151 | 1.013 | 0.428 | -0.01 | -0.006 | -0.006 | 0 | 0.6 | 0.5 | 0.65 | 0.37 | 0.69 | 0.56 |
| 1 | 238032804 | rs1341873 | 0.25 | -0.229 | 0.68 | 0.412 | -0.006 | 0.006 | 0.006 | 0.074 | 0.16 | 0.22 | 0.11 | 0.2 | 0.11 | 0.15 |
| 1 | 238033066 | rs2794849 | 0.289 | -0.228 | 0.68 | 0.413 | -0.006 | 0.006 | 0.006 | 0.075 | 0.16 | 0.21 | 0.11 | 0.2 | 0.11 | 0.15 |
| 1 | 238034983 | rs2794852 | 0.289 | -0.228 | 0.732 | 0.413 | -0.009 | 0.012 | 0.012 | 0.075 | 0.16 | 0.25 | 0.11 | 0.2 | 0.11 | 0.15 |
| 1 | 238039652 | rs1339649 | -0.161 | -0.255 | 0.627 | 0.387 | -0.012 | 0.006 | 0.006 | 0 | 0.19 | 0.23 | 0.12 | 0.21 | 0.12 | 0.2 |
| 1 | 238043980 | rs946581 | 0.034 | -0.237 | 0.49 | 0.402 | -0.004 | 0.002 | 0.002 | 0.009 | 0.18 | 0.18 | 0.15 | 0.22 | 0.12 | 0.16 |
| 1 | 238045995 | rs2275694 | -0.189 | 0.098 | 0.894 | 0.505 | 0.043 | 0.165 | 0.165 | 0 | 0.18 | 0.37 | 0.19 | 0.43 | 0.21 | 0.25 |
| 1 | 238046603 | rs2487789 | -1.478 | -0.256 | 0.326 | 0.14 | 0.002 | 0.041 | 0.041 | 0 | 0.47 | 0.84 | 0.45 | 0.62 | 0.38 | 0.46 |
| 1 | 238052922 | rs2790365 | -1.151 | -0.304 | 0.271 | 0.096 | -0.002 | 0.021 | 0.021 | 0 | 0.41 | 0.86 | 0.4 | 0.63 | 0.34 | 0.57 |
| 1 | 238055618 | rs693420 | 0.626 | -0.296 | 0.272 | 0.165 | 0.007 | -0.014 | -0.014 | 1.094 | 0.16 | 0.15 | 0.15 | 0.22 | 0.12 | 0.16 |
| 1 | 238059704 | rs2636211 | -1.158 | -0.176 | -0.005 | 0.136 | -0.029 | 0.016 | 0.016 | 0 | 0.4 | 0.88 | 0.44 | 0.64 | 0.35 | 0.57 |
| 1 | 238059867 | rs487111 | 0.717 | -0.055 | 0.125 | 0.363 | 0.012 | -0.049 | -0.049 | 0.268 | 0.15 | 0.05 | 0.14 | 0.23 | 0.1 | 0.16 |
| 1 | 238064730 | rs12123002 | 1.055 | 0.106 | -0.212 | 0.008 | -0.047 | 0.039 | 0.039 | 0.665 | 0.39 | 0.14 | 0.43 | 0.34 | 0.49 | 0.27 |
| 1 | 238094343 | rs503027 | -0.811 | 0.308 | -0.092 | 0.534 | -0.004 | 0.069 | 0.069 | 0 | 0.27 | 0.57 | 0.31 | 0.44 | 0.27 | 0.43 |
| 1 | 238096886 | rs547984 | -0.613 | 0.496 | -0.679 | 0.06 | 0.02 | 0.008 | 0.008 | 0 | 0.53 | 0.74 | 0.52 | 0.51 | 0.58 | 0.62 |
| 1 | 238102080 | rs2451620 | -0.558 | 0.59 | -0.479 | -0.107 | -0.003 | 0.009 | 0.009 | 0 | 0.47 | 0.74 | 0.44 | 0.52 | 0.47 | 0.58 |
| 1 | 238102779 | rs2499606 | -0.341 | 0.571 | -0.456 | -0.114 | -0.003 | -0.001 | -0.001 | 0 | 0.43 | 0.61 | 0.41 | 0.52 | 0.46 | 0.57 |
| 1 | 238103501 | rs2499604 | -0.489 | 0.646 | -0.632 | -0.043 | -0.003 | 0.009 | 0.009 | 0 | 0.47 | 0.74 | 0.43 | 0.52 | 0.47 | 0.58 |
| 1 | 238126784 | rs2200958 | 0.125 | 1.037 | -0.466 | -0.567 | -0.011 | 0.018 | 0.018 | 0 | 0.15 | 0.12 | 0.18 | 0.03 | 0.19 | 0.13 |
| 1 | 238128633 | rs12134826 | 0.928 | 1.096 | -1.194 | -1.426 | -0.029 | 0.058 | 0.058 | 1.145 | 0.2 | 0.07 | 0.15 | 0 | 0.27 | 0.05 |
| 1 | 238154347 | rs2499566 | 0.807 | 0.558 | -0.312 | -0.367 | 0 | 0.009 | 0.009 | 0 | 0.51 | 0.62 | 0.46 | 0.51 | 0.51 | 0.36 |
| 1 | 238159999 | rs2487103 | -0.204 | 0.364 | -0.251 | -0.397 | 0.001 | -0.017 | -0.017 | 0.02 | 0.7 | 0.82 | 0.68 | 0.54 | 0.73 | 0.53 |

| | | | | | | | | | | | | | | | | |
|---|---|---|---|---|---|---|---|---|---|---|---|---|---|---|---|---|
| 1 | 238163813 | rs2487104 | 0.477 | 0.333 | -0.21 | -0.36 | -0.031 | -0.012 | -0.012 | 0 | 0.57 | 0.84 | 0.56 | 0.53 | 0.55 | 0.39 |
| 1 | 238179820 | rs1340456 | 0.85 | 0.236 | 0.409 | -0.414 | -0.001 | 0.003 | 0.003 | 0 | 0.49 | 0.43 | 0.44 | 0.52 | 0.52 | 0.36 |
| 1 | 238223495 | rs6684566 | 0.147 | -0.075 | 1.491 | 0.268 | -0.015 | -0.028 | -0.028 | 0 | 0.06 | 0.19 | 0.09 | 0.02 | 0.03 | 0.03 |
| 1 | 238224845 | rs16836211 | 0.193 | -0.074 | 1.492 | 0.268 | -0.014 | -0.027 | -0.027 | 0 | 0.06 | 0.19 | 0.09 | 0.02 | 0.03 | 0.03 |
| 1 | 238226188 | rs16836214 | 0.193 | -0.073 | 1.442 | 0.26 | -0.014 | -0.027 | -0.027 | 0 | 0.06 | 0.19 | 0.09 | 0.02 | 0.03 | 0.03 |
| 1 | 238234097 | rs2487084 | -0.245 | -0.053 | 1.229 | 0.21 | -0.004 | -0.05 | -0.05 | 0 | 0.3 | 0.12 | 0.31 | 0.45 | 0.27 | 0.48 |
| 1 | 238255771 | rs2487091 | 1.183 | -0.298 | 1.387 | 0.137 | -0.058 | 0.003 | 0.003 | 2.093 | 0.83 | 0.37 | 0.85 | 0.83 | 0.92 | 0.84 |
| 1 | 238270039 | rs1796942 | 1.291 | -0.289 | 1.495 | 0.311 | -0.058 | 0.054 | 0.054 | 0.936 | 0.87 | 0.36 | 0.87 | 0.81 | 0.95 | 0.85 |
| 1 | 238271681 | rs7525052 | 0.707 | -0.271 | 1.504 | 0.304 | -0.011 | -0.023 | -0.023 | 0.001 | 0.05 | 0.18 | 0.09 | 0.02 | 0.03 | 0.03 |
| 1 | 238271688 | rs7525053 | 0.481 | -0.272 | 1.519 | 0.303 | -0.009 | -0.023 | -0.023 | 0 | 0.06 | 0.24 | 0.09 | 0.02 | 0.03 | 0.03 |
| 1 | 238273454 | rs2487096 | 0.934 | -0.211 | 1.47 | 0.157 | 0.025 | 0.062 | 0.062 | 0 | 0.86 | 0.6 | 0.89 | 0.83 | 0.88 | 0.83 |
| 1 | 238273632 | rs2486217 | 1.816 | -0.215 | 1.379 | 0.154 | -0.02 | -0.006 | -0.006 | 0 | 0.8 | 0.34 | 0.8 | 0.81 | 0.85 | 0.81 |
| 1 | 238274479 | rs10802655 | -0.649 | -0.071 | 0.993 | 0.256 | 0.018 | 0.005 | 0.005 | 1.238 | 0.8 | 0.86 | 0.93 | 0.79 | 0.86 | 0.71 |
| 1 | 238277625 | rs16836356 | 0.057 | -0.2 | 1.135 | 0.227 | -0.016 | -0.015 | -0.015 | 0.782 | 0.07 | 0.16 | 0.09 | 0.06 | 0.04 | 0.16 |
| 1 | 238281405 | rs16836366 | -0.147 | -0.208 | 1.162 | 0.223 | -0.018 | -0.016 | -0.016 | 0 | 0.07 | 0.21 | 0.09 | 0.06 | 0.04 | 0.16 |
| 1 | 238291482 | rs17523257 | -1.02 | -0.21 | 1.314 | 0.333 | 0.01 | 0.013 | 0.013 | 0 | 0.14 | 0.21 | 0.27 | 0.2 | 0.14 | 0.16 |
| 1 | 238292209 | rs7555423 | -1.901 | -0.236 | 1.371 | 0.321 | -0.008 | -0.001 | -0.001 | 0.151 | 0.22 | 0.48 | 0.37 | 0.26 | 0.18 | 0.31 |
| 1 | 238293594 | rs7521808 | -1.25 | -0.214 | 1.303 | 0.333 | 0.018 | 0.03 | 0.03 | 0 | 0.16 | 0.3 | 0.33 | 0.26 | 0.15 | 0.27 |
| 1 | 238297568 | rs17523604 | -1.076 | -0.198 | 1.32 | 0.331 | 0.001 | 0.008 | 0.008 | 0 | 0.14 | 0.21 | 0.28 | 0.2 | 0.14 | 0.2 |
| 1 | 238298024 | rs12070012 | -1.796 | -0.204 | 1.345 | 0.327 | -0.003 | 0.002 | 0.002 | 0 | 0.2 | 0.35 | 0.35 | 0.26 | 0.18 | 0.31 |
| 1 | 238298452 | rs17523666 | -0.325 | -0.077 | 1.392 | 0.403 | 0.017 | 0.032 | 0.032 | 0 | 0.1 | 0.15 | 0.25 | 0.2 | 0.11 | 0.15 |
| 1 | 238300214 | rs10925573 | -0.47 | -0.084 | 1.254 | 0.37 | -0.005 | -0.014 | -0.014 | 0 | 0.06 | 0.14 | 0.08 | 0.06 | 0.04 | 0.12 |
| 1 | 238300737 | rs16836421 | -0.022 | -0.083 | 1.209 | 0.371 | -0.001 | -0.003 | -0.003 | 0 | 0.05 | 0.05 | 0.03 | 0.04 | 0.04 | 0.12 |
| 1 | 238301369 | rs7519935 | -1.405 | -0.147 | 1.601 | 0.34 | -0.004 | -0.019 | -0.019 | 0 | 0.08 | 0.24 | 0.09 | 0.06 | 0.05 | 0.12 |
| 1 | 238303362 | rs2066074 | 0.937 | -0.06 | 1.411 | 0.392 | 0.003 | -0.005 | -0.005 | 0.126 | 0.63 | 0.37 | 0.62 | 0.39 | 0.69 | 0.42 |
| 1 | 238303419 | rs2066073 | 1.291 | -0.053 | 1.366 | 0.397 | 0.016 | -0.007 | -0.007 | 0 | 0.58 | 0.27 | 0.55 | 0.38 | 0.66 | 0.41 |
| 1 | 238309715 | rs7533581 | -0.445 | -0.009 | 1.625 | 0.238 | 0.038 | 0.07 | 0.07 | 0 | 0.11 | 0.26 | 0.25 | 0.2 | 0.11 | 0.14 |
| 1 | 238310901 | rs1933980 | 1.025 | -0.055 | 1.716 | 0.27 | -0.001 | 0.002 | 0.002 | 0 | 0.62 | 0.57 | 0.59 | 0.45 | 0.68 | 0.53 |
| 1 | 238311680 | rs7547681 | 1.216 | -0.077 | 1.901 | 0.26 | -0.004 | 0.006 | 0.006 | 0 | 0.6 | 0.42 | 0.57 | 0.45 | 0.68 | 0.53 |
| 1 | 238312176 | rs10925576 | -0.886 | -0.144 | 1.771 | 0.292 | 0.004 | -0.015 | -0.015 | 0 | 0.79 | 0.87 | 0.9 | 0.76 | 0.83 | 0.69 |
| 1 | 238322892 | rs7555833 | -0.84 | -0.192 | 1.729 | 0.269 | -0.003 | -0.029 | -0.029 | 0.241 | 0.79 | 0.92 | 0.9 | 0.76 | 0.84 | 0.68 |
| 1 | 238326067 | rs6668206 | -0.324 | -0.194 | 1.793 | 0.274 | -0.004 | 0.005 | 0.005 | 0.612 | 0.76 | 0.7 | 0.89 | 0.66 | 0.83 | 0.67 |
| 1 | 238331394 | rs1933978 | -1.184 | -0.177 | 1.84 | 0.274 | 0.012 | 0.057 | 0.057 | 0 | 0.16 | 0.27 | 0.27 | 0.3 | 0.15 | 0.17 |
| 1 | 238332506 | rs12034052 | 1.573 | -0.236 | 1.793 | 0.237 | 0.024 | -0.062 | -0.062 | 1.169 | 0.56 | 0.16 | 0.55 | 0.42 | 0.66 | 0.4 |
| 1 | 238334315 | rs1933979 | -0.477 | -0.246 | 1.864 | 0.233 | -0.006 | -0.001 | -0.001 | 1.559 | 0.77 | 0.68 | 0.89 | 0.77 | 0.83 | 0.69 |
| 1 | 238334792 | rs12036196 | 1.57 | -0.226 | 1.608 | 0.226 | 0.03 | -0.044 | -0.044 | 2.66 | 0.56 | 0.17 | 0.55 | 0.41 | 0.66 | 0.4 |
| 1 | 238339110 | rs1572703 | -0.253 | -0.019 | 1.421 | 0.055 | -0.026 | 0.038 | 0.038 | 0 | 0.24 | 0.17 | 0.44 | 0.13 | 0.33 | 0.33 |
| 1 | 238340968 | rs6683290 | -0.986 | -0.028 | 1.52 | 0.054 | 0.003 | -0.021 | -0.021 | 0.021 | 0.3 | 0.64 | 0.53 | 0.18 | 0.35 | 0.45 |
| 1 | 238346577 | rs1338922 | -1.189 | -0.07 | 1.375 | 0.085 | -0.021 | -0.049 | -0.049 | 1.018 | 0.8 | 0.97 | 0.92 | 0.76 | 0.83 | 0.69 |
| 1 | 238351005 | rs7549399 | 1.499 | -0.124 | 1.46 | 0.099 | -0.009 | -0.007 | -0.007 | 0 | 0.46 | 0.23 | 0.37 | 0.48 | 0.47 | 0.23 |
| 1 | 238358586 | rs1351447 | -1.094 | -0.122 | 1.641 | 0.098 | 0.031 | 0.014 | 0.014 | 0 | 0.15 | 0.44 | 0.28 | 0.16 | 0.15 | 0.29 |
| 1 | 238366533 | rs518552 | 0.002 | -0.122 | 1.449 | 0.099 | -0.001 | 0 | 0 | 0 | 0.27 | 0.29 | 0.13 | 0.33 | 0.2 | 0.37 |
| 1 | 238375280 | rs10925586 | 1.475 | -0.042 | 1.474 | 0.084 | -0.013 | -0.011 | -0.011 | 0.512 | 0.46 | 0.21 | 0.37 | 0.48 | 0.48 | 0.23 |
| 1 | 238376059 | rs6668349 | 0.443 | -0.046 | 1.535 | 0.078 | 0.002 | 0.014 | 0.014 | 1.826 | 0.59 | 0.65 | 0.67 | 0.65 | 0.6 | 0.47 |
| 1 | 238376635 | rs10925587 | 1.47 | -0.003 | 1.21 | 0.103 | -0.016 | -0.014 | -0.014 | 0 | 0.45 | 0.21 | 0.37 | 0.48 | 0.47 | 0.22 |
| 1 | 238383296 | rs6697891 | -1.368 | 0.026 | 1.228 | -0.019 | 0 | 0.05 | 0.05 | 1.987 | 0.51 | 0.74 | 0.62 | 0.42 | 0.52 | 0.75 |
| 1 | 238386465 | rs17591250 | 1.29 | 0.106 | 0.967 | -0.049 | -0.035 | 0.046 | 0.046 | 0 | 0.13 | 0.04 | 0.19 | 0.01 | 0.2 | 0.16 |
| 1 | 238386642 | rs10737820 | -0.667 | 0.035 | 1.178 | -0.062 | 0 | -0.005 | -0.005 | 0 | 0.38 | 0.39 | 0.35 | 0.3 | 0.39 | 0.59 |
| 1 | 238388504 | rs722372 | 0.214 | -0.05 | 1.216 | 0.061 | -0.017 | 0 | 0 | 0 | 0.25 | 0.33 | 0.12 | 0.27 | 0.19 | 0.43 |
| 1 | 238391155 | rs7537266 | 0.398 | 0.047 | 1.037 | 0.181 | -0.001 | 0.002 | 0.002 | 0.914 | 0.25 | 0.24 | 0.14 | 0.33 | 0.24 | 0.47 |
| 1 | 238394485 | rs6657977 | 0.765 | -0.121 | 1.222 | 0.28 | -0.037 | 0.013 | 0.013 | 1.061 | 0.88 | 0.6 | 0.89 | 0.87 | 0.96 | 0.92 |
| 1 | 238395155 | rs10754629 | 1.856 | -0.134 | 0.651 | 0.225 | -0.033 | -0.034 | -0.034 | 0.206 | 0.38 | 0.14 | 0.38 | 0.32 | 0.42 | 0.18 |
| 1 | 238397598 | rs10802665 | -0.084 | 0.079 | 0.724 | 0.329 | -0.038 | -0.045 | -0.045 | 0 | 0.34 | 0.18 | 0.43 | 0.51 | 0.44 | 0.55 |
| 1 | 238403285 | rs4659846 | 0.532 | 0.094 | 0.972 | 0.426 | -0.014 | -0.009 | -0.009 | 0 | 0.6 | 0.53 | 0.78 | 0.62 | 0.67 | 0.58 |
| 1 | 238416905 | rs6684578 | 0.039 | 0.285 | 0.933 | 0.236 | -0.003 | -0.016 | -0.016 | 0.789 | 0.55 | 0.53 | 0.75 | 0.71 | 0.6 | 0.57 |
| 1 | 238418359 | rs9661467 | -0.529 | 0.217 | 1.211 | 0.182 | -0.007 | 0.032 | 0.032 | 1.594 | 0.62 | 0.79 | 0.81 | 0.71 | 0.64 | 0.58 |
| 1 | 238419413 | rs1481135 | 0.479 | 0.17 | 1.175 | 0.209 | 0.006 | 0.015 | 0.015 | 1.043 | 0.51 | 0.45 | 0.7 | 0.58 | 0.59 | 0.56 |
| 1 | 238420323 | rs1481151 | -0.6 | 0.273 | 0.769 | 0.224 | -0.002 | 0.033 | 0.033 | 1.96 | 0.63 | 0.8 | 0.81 | 0.71 | 0.66 | 0.58 |
| 1 | 238420949 | rs900006 | -0.378 | 0.295 | 0.777 | 0.21 | 0.006 | 0.011 | 0.011 | 2.265 | 0.6 | 0.73 | 0.76 | 0.58 | 0.65 | 0.57 |
| 1 | 238429670 | rs12049335 | 1.016 | 0.415 | 0.399 | 0.275 | -0.001 | -0.033 | -0.033 | 0 | 0.37 | 0.17 | 0.36 | 0.32 | 0.39 | 0.2 |
| 1 | 238432547 | rs9658992 | -0.781 | 0.46 | 0.987 | 0.28 | 0.027 | -0.042 | -0.042 | 0 | 0.09 | 0.32 | 0.07 | 0 | 0.07 | 0.04 |
| 1 | 238432884 | rs6672666 | -1.273 | 0.488 | 1.103 | 0.311 | 0.008 | 0.014 | 0.014 | 0 | 0.27 | 0.58 | 0.45 | 0.39 | 0.29 | 0.4 |
| 1 | 238432940 | rs259614 | 0.903 | 0.336 | 0.883 | 0.22 | 0.006 | -0.025 | -0.025 | 0 | 0.38 | 0.2 | 0.37 | 0.33 | 0.39 | 0.21 |
| 1 | 238434116 | rs16836628 | -0.094 | 0.333 | 0.852 | 0.343 | 0.004 | 0.014 | 0.014 | 0 | 0.18 | 0.27 | 0.38 | 0.39 | 0.22 | 0.36 |
| 1 | 238441600 | rs7536535 | 0.635 | 0.397 | 0.717 | 0.243 | -0.007 | -0.058 | -0.058 | 0.016 | 0.39 | 0.21 | 0.44 | 0.46 | 0.42 | 0.22 |
| 1 | 238441781 | rs259633 | -0.102 | 0.246 | 0.944 | 0.241 | 0.006 | 0.024 | 0.024 | 0.822 | 0.57 | 0.42 | 0.77 | 0.67 | 0.64 | 0.43 |
| 1 | 238446621 | rs259639 | -1.864 | 0.164 | 1.285 | 0.464 | 0.123 | 0.19 | 0.19 | 0 | 0.07 | 0.34 | 0.27 | 0.13 | 0.05 | 0.15 |
| 1 | 238446854 | rs259640 | 0.212 | -0.066 | 0.26 | 0.431 | 0.07 | 0.033 | 0.033 | 0.209 | 0.5 | 0.19 | 0.57 | 0.61 | 0.61 | 0.69 |
| 1 | 238451820 | rs259645 | 0.299 | -0.134 | 0.405 | 0.465 | 0.005 | 0.019 | 0.019 | 0.516 | 0.33 | 0.16 | 0.32 | 0.21 | 0.32 | 0.23 |

| | | | | | | | | | | | | | | | | |
|---|---|---|---|---|---|---|---|---|---|---|---|---|---|---|---|---|
| 1 | 238453671 | rs259649 | -0.489 | -0.257 | 1.047 | 0.555 | 0.021 | -0.011 | -0.011 | 0 | 0.42 | 0.7 | 0.38 | 0.44 | 0.35 | 0.31 |
| 1 | 238459434 | rs259654 | -1.183 | -0.231 | 0.966 | 0.554 | 0.007 | 0.202 | 0.202 | 2.07 | 0.09 | 0.57 | 0.07 | 0.24 | 0.03 | 0.08 |
| 1 | 238463077 | rs408420 | -0.403 | -0.159 | 0.532 | 0.652 | 0.037 | -0.018 | -0.018 | 0 | 0.43 | 0.81 | 0.39 | 0.43 | 0.35 | 0.31 |
| 1 | 238471995 | rs259576 | 0.675 | 0.25 | 0.145 | -0.794 | -0.015 | 0.065 | 0.065 | 0 | 0.45 | 0.26 | 0.38 | 0.82 | 0.44 | 0.48 |
| 1 | 238472891 | rs259573 | 0.861 | 0.038 | 0.067 | -1.162 | -0.004 | -0.02 | -0.02 | 0.535 | 0.62 | 0.37 | 0.49 | 0.82 | 0.62 | 0.57 |
| 1 | 238474457 | rs16836698 | -0.822 | -0.021 | 0.4 | -1.237 | -0.028 | -0.08 | -0.08 | 0 | 0.32 | 0.5 | 0.38 | 0.14 | 0.3 | 0.28 |
| 1 | 238476102 | rs12142622 | -0.071 | -0.123 | 0.506 | -1.254 | 0.006 | 0.002 | 0.002 | 0.396 | 0.13 | 0.09 | 0.11 | 0.01 | 0.15 | 0.02 |
| 1 | 238476967 | rs12746619 | -0.394 | -0.169 | 0.663 | -1.281 | -0.024 | -0.024 | -0.024 | 0 | 0.17 | 0.42 | 0.27 | 0.13 | 0.13 | 0.22 |
| 1 | 238477085 | rs16836704 | -0.484 | -0.155 | 0.472 | -1.389 | -0.006 | 0.049 | 0.049 | 0.776 | 0.3 | 0.18 | 0.21 | 0.01 | 0.33 | 0.12 |
| 1 | 238477178 | rs12748123 | 0.122 | -0.209 | 0.816 | -1.397 | -0.007 | -0.004 | -0.004 | 0 | 0.14 | 0.19 | 0.24 | 0.13 | 0.12 | 0.21 |
| 1 | 238477828 | rs370522 | 0.771 | -0.012 | 0.917 | -1.096 | -0.003 | 0.033 | 0.033 | 0 | 0.65 | 0.7 | 0.66 | 0.88 | 0.58 | 0.73 |
| 1 | 238479714 | rs194595 | 0.443 | -0.019 | 0.884 | -1.068 | -0.015 | 0.059 | 0.059 | 0.204 | 0.68 | 0.77 | 0.68 | 0.91 | 0.6 | 0.83 |
| 1 | 238485541 | rs1335975 | 1.169 | -0.508 | 1.143 | -0.995 | 0.014 | 0.033 | 0.033 | 0 | 0.63 | 0.3 | 0.67 | 0.88 | 0.66 | 0.73 |
| 1 | 238494665 | rs17528326 | 0.855 | -0.439 | 1.179 | -0.995 | -0.029 | 0.07 | 0.07 | 0.434 | 0.17 | 0.04 | 0.1 | 0 | 0.18 | 0.1 |
| 1 | 238502766 | rs9428720 | 0.777 | -0.403 | 1.299 | -0.962 | -0.005 | -0.034 | -0.034 | 0 | 0.67 | 0.58 | 0.69 | 0.88 | 0.66 | 0.74 |
| 1 | 238503593 | rs1335972 | 1.141 | -0.417 | 1.342 | -0.969 | 0.015 | 0.04 | 0.04 | 0 | 0.63 | 0.28 | 0.67 | 0.88 | 0.66 | 0.74 |
| 1 | 238504813 | rs7530131 | -0.759 | -0.381 | 1.18 | -0.96 | -0.009 | -0.075 | -0.075 | 0 | 0.3 | 0.37 | 0.2 | 0.03 | 0.27 | 0.22 |
| 1 | 238515411 | rs1342066 | -0.039 | -0.464 | 1.212 | -0.868 | -0.001 | -0.07 | -0.07 | 0.165 | 0.77 | 0.54 | 0.78 | 0.92 | 0.78 | 0.86 |
| 1 | 238518116 | rs12403783 | -0.655 | -0.687 | 1.031 | -0.943 | 0.003 | 0.003 | 0.003 | 0.075 | 0.21 | 0.2 | 0.28 | 0.15 | 0.17 | 0.24 |
| 1 | 238520256 | rs10925623 | 1.758 | -0.636 | 0.857 | -0.951 | 0.061 | 0.142 | 0.142 | 0.278 | 0.51 | 0.12 | 0.45 | 0.74 | 0.58 | 0.5 |
| 1 | 238522082 | rs10802677 | 1.763 | -0.16 | 1.037 | -0.585 | 0.01 | 0.069 | 0.069 | 3.349 | 0.88 | 0.45 | 0.83 | 0.89 | 0.89 | 0.85 |
| 1 | 238528098 | rs1414537 | 0.389 | -0.058 | 0.562 | -0.487 | -0.001 | -0.004 | -0.004 | 2.29 | 0.84 | 0.87 | 0.7 | 0.78 | 0.81 | 0.7 |
| 1 | 238529991 | rs7547699 | 0.958 | -0.325 | 0.787 | -0.611 | -0.006 | -0.011 | -0.011 | 0 | 0.64 | 0.5 | 0.55 | 0.72 | 0.65 | 0.53 |
| 1 | 238533500 | rs7526709 | 0.378 | -0.144 | 0.48 | -1.007 | 0.003 | -0.029 | -0.029 | 0 | 0.71 | 0.61 | 0.68 | 0.89 | 0.69 | 0.74 |
| 1 | 238534722 | rs7540190 | 0.027 | -0.192 | 0.546 | -1.032 | -0.014 | 0.035 | 0.035 | 0 | 0.74 | 0.78 | 0.7 | 0.89 | 0.71 | 0.75 |
| 1 | 238534934 | rs10737826 | 1.61 | -0.338 | 0.878 | -1.063 | 0.005 | 0.027 | 0.027 | 0 | 0.53 | 0.33 | 0.46 | 0.7 | 0.56 | 0.44 |
| 1 | 238535530 | rs4659870 | 0.024 | -0.167 | 0.656 | -1.009 | 0 | 0 | 0 | 0 | 0.13 | 0.14 | 0.19 | 0.13 | 0.12 | 0.2 |
| 1 | 238536274 | rs6679846 | -0.146 | -0.092 | 0.638 | -0.913 | -0.008 | 0.017 | 0.017 | 0 | 0.76 | 0.8 | 0.72 | 0.86 | 0.75 | 0.77 |
| 1 | 238536388 | rs10802680 | 1.016 | -0.217 | 0.747 | -0.996 | -0.004 | 0.016 | 0.016 | 0 | 0.6 | 0.6 | 0.51 | 0.76 | 0.58 | 0.54 |
| 1 | 238538758 | rs12137904 | 0.794 | 0.177 | 0.633 | -0.782 | 0.017 | -0.025 | -0.025 | 0 | 0.77 | 0.62 | 0.7 | 0.88 | 0.72 | 0.74 |
| 1 | 238539142 | rs12757566 | 0.214 | 0.133 | 0.482 | -0.594 | 0.002 | -0.005 | -0.005 | 0 | 0.11 | 0.13 | 0.22 | 0.15 | 0.1 | 0.2 |
| 1 | 238540156 | rs10754633 | 1.581 | 0.156 | 0.575 | -0.671 | 0.008 | -0.019 | -0.019 | 0 | 0.64 | 0.4 | 0.47 | 0.73 | 0.61 | 0.53 |
| 1 | 238540226 | rs7531928 | -0.857 | 0.05 | 0.624 | -0.636 | -0.015 | 0.016 | 0.016 | 0.627 | 0.14 | 0.38 | 0.26 | 0.15 | 0.11 | 0.21 |
| 1 | 238544006 | rs1021358 | 1.574 | 0.396 | 0.09 | -1.169 | -0.012 | 0.103 | 0.103 | 0 | 0.56 | 0.25 | 0.55 | 0.8 | 0.51 | 0.61 |
| 1 | 238549714 | rs6678429 | 0.819 | 0.511 | -0.297 | -1.217 | -0.099 | 0.055 | 0.055 | 0 | 0.73 | 0.88 | 0.66 | 0.87 | 0.59 | 0.7 |
| 1 | 238551688 | rs2171907 | 0.656 | 0.534 | -0.289 | -1.201 | -0.101 | 0.046 | 0.046 | 0 | 0.74 | 0.92 | 0.67 | 0.87 | 0.6 | 0.7 |
| 1 | 238554944 | rs12098216 | -0.145 | 0.09 | 0.461 | -0.983 | -0.041 | -0.055 | -0.055 | 0 | 0.1 | 0.16 | 0.04 | 0.01 | 0.06 | 0.02 |
| 1 | 238556742 | rs6690847 | 0.158 | 0.698 | 0.143 | -1.018 | -0.023 | -0.003 | -0.003 | 0 | 0.87 | 0.91 | 0.86 | 0.87 | 0.79 | 0.77 |
| 1 | 238558150 | rs10754634 | 0.395 | 0.661 | 0.405 | -1.101 | 0.007 | -0.022 | -0.022 | 0 | 0.86 | 0.77 | 0.88 | 0.93 | 0.83 | 0.87 |
| 1 | 238559372 | rs6429075 | 1.089 | 0.426 | 0.669 | -1.342 | 0.017 | -0.123 | -0.123 | 0.11 | 0.7 | 0.53 | 0.7 | 0.91 | 0.67 | 0.72 |
| 1 | 238559857 | rs923543 | 2.069 | -0.055 | 0.167 | -1.45 | 0.02 | 0.112 | 0.112 | 0 | 0.54 | 0.28 | 0.61 | 0.79 | 0.54 | 0.52 |
| 1 | 238560794 | rs10925637 | 1.091 | 0.484 | 0.126 | -1.362 | 0.006 | -0.065 | -0.065 | 0 | 0.7 | 0.61 | 0.71 | 0.91 | 0.67 | 0.72 |
| 1 | 238573691 | rs1021356 | -0.057 | 1.162 | -0.477 | -1.131 | -0.063 | 0.053 | 0.053 | 0 | 0.84 | 0.97 | 0.83 | 0.95 | 0.74 | 0.84 |
| 1 | 238574814 | rs6662339 | 1.228 | 0.518 | -0.377 | -1.415 | -0.087 | 0.13 | 0.13 | 0.043 | 0.72 | 0.93 | 0.78 | 0.95 | 0.64 | 0.75 |
| 1 | 238576412 | rs10925644 | -0.265 | 0.428 | -0.449 | -0.635 | -0.003 | 0.008 | 0.008 | 0 | 0.19 | 0.18 | 0.26 | 0.33 | 0.19 | 0.34 |
| 1 | 238576614 | rs12118091 | -0.725 | 0.653 | -0.497 | -0.649 | -0.061 | 0.054 | 0.054 | 0 | 0.84 | 0.95 | 0.83 | 0.95 | 0.75 | 0.84 |
| 1 | 238579228 | rs12131363 | 0.153 | 0.368 | -0.308 | -0.536 | -0.002 | 0.035 | 0.035 | 0 | 0.15 | 0.17 | 0.28 | 0.34 | 0.19 | 0.33 |
| 1 | 238582075 | rs12067134 | -0.152 | 0.139 | 0.106 | -0.773 | 0.004 | -0.062 | -0.062 | 0.068 | 0.1 | 0.18 | 0.05 | 0 | 0.09 | 0.08 |
| 1 | 238582365 | rs1418210 | 1.157 | -0.098 | -0.043 | -1.143 | -0.03 | 0.001 | 0.001 | 0 | 0.59 | 0.76 | 0.5 | 0.63 | 0.5 | 0.45 |
| 1 | 238582765 | rs12026222 | 2.141 | -0.179 | 0.071 | -1.146 | 0.003 | 0.012 | 0.012 | 0 | 0.43 | 0.31 | 0.4 | 0.63 | 0.4 | 0.37 |
| 1 | 238582836 | rs10925645 | 1.636 | -0.069 | 0.44 | -1.057 | 0.002 | -0.036 | -0.036 | 0.03 | 0.59 | 0.42 | 0.67 | 0.89 | 0.56 | 0.66 |
| 1 | 238588516 | rs10925648 | -1.149 | -0.251 | 0.543 | -0.961 | -0.052 | -0.089 | -0.089 | 0.149 | 0.13 | 0.32 | 0.07 | 0 | 0.05 | 0.04 |
| 1 | 238595161 | rs4288585 | 1.19 | 0.025 | 0.019 | -1.098 | 0 | -0.019 | -0.019 | 3.541 | 0.63 | 0.38 | 0.62 | 0.85 | 0.62 | 0.46 |
| 1 | 238598399 | rs1342078 | -1.099 | 0.45 | -0.279 | -0.99 | -0.029 | 0.025 | 0.025 | 0 | 0.86 | 0.91 | 0.89 | 0.99 | 0.76 | 0.84 |
| 1 | 238613141 | rs10925652 | 1.27 | -0.068 | -0.355 | -0.765 | -0.032 | 0.052 | 0.052 | 0 | 0.44 | 0.21 | 0.43 | 0.68 | 0.48 | 0.36 |
| 1 | 238622443 | rs10157478 | -0.742 | 0.173 | 0.157 | -0.559 | -0.033 | -0.064 | -0.064 | 0 | 0.11 | 0.3 | 0.05 | 0 | 0.04 | 0.04 |
| 1 | 238631540 | rs10925657 | 0.774 | 0.522 | -0.44 | -0.538 | -0.003 | 0.03 | 0.03 | 0.937 | 0.48 | 0.46 | 0.52 | 0.75 | 0.5 | 0.37 |
| 1 | 238636362 | rs7532779 | 0.057 | 0.189 | 0.054 | -0.75 | -0.001 | -0.02 | -0.02 | 1.937 | 0.6 | 0.49 | 0.67 | 0.84 | 0.63 | 0.41 |
| 1 | 238644100 | rs1054888 | -0.068 | 0.275 | -0.085 | -0.564 | -0.001 | -0.041 | -0.041 | 0 | 0.62 | 0.59 | 0.68 | 0.84 | 0.62 | 0.43 |
| 1 | 238645291 | rs12736761 | 0.16 | 0.404 | -0.464 | -0.282 | 0.012 | 0.097 | 0.097 | 0.276 | 0.52 | 0.37 | 0.65 | 0.8 | 0.57 | 0.45 |
| 1 | 238650133 | rs12035663 | -0.729 | 0.318 | -0.458 | 0.479 | 0.004 | 0.097 | 0.097 | 0 | 0.18 | 0.3 | 0.22 | 0.53 | 0.19 | 0.26 |
| 1 | 238654555 | rs6658308 | -0.779 | 0.646 | -1.029 | 0.348 | -0.052 | 0.008 | 0.008 | 0.376 | 0.72 | 0.85 | 0.68 | 0.78 | 0.64 | 0.55 |
| 1 | 238655389 | rs2392862 | 0.843 | -0.033 | -1.359 | -0.082 | -0.037 | 0.076 | 0.076 | 0.52 | 0.32 | 0.05 | 0.26 | 0.2 | 0.34 | 0.21 |
| 1 | 238669665 | rs6692597 | 0.358 | -0.536 | -0.053 | 0.155 | -0.063 | -0.095 | -0.095 | 0 | 0.16 | 0.29 | 0.16 | 0 | 0.05 | 0.07 |
| 1 | 238672156 | rs2893644 | -1.351 | -0.31 | -0.124 | -0.019 | -0.024 | 0.051 | 0.051 | 0 | 0.71 | 0.86 | 0.8 | 0.85 | 0.7 | 0.57 |
| 1 | 238672294 | rs6429079 | -1.135 | -0.338 | -0.244 | 0.348 | 0.041 | -0.027 | -0.027 | 4.424 | 0.38 | 0.75 | 0.39 | 0.57 | 0.29 | 0.31 |
| 1 | 238672931 | rs12722764 | 1.49 | -0.31 | -0.286 | 0.359 | -0.085 | -0.025 | -0.025 | 0 | 0.27 | 0.02 | 0.23 | 0.22 | 0.31 | 0.19 |
| 1 | 238676454 | rs6429080 | 0.123 | -0.287 | -0.165 | 0.397 | -0.034 | -0.103 | -0.103 | 0 | 0.19 | 0.3 | 0.2 | 0 | 0.12 | 0.09 |
| 1 | 238676995 | rs1327896 | -1.696 | -0.284 | -0.232 | 0.418 | 0.032 | 0.023 | 0.023 | 2.062 | 0.44 | 0.83 | 0.56 | 0.63 | 0.38 | 0.38 |

| | | | | | | | | | | | | | | | | |
|---|---|---|---|---|---|---|---|---|---|---|---|---|---|---|---|---|
| 1 | 238683538 | rs10495412 | -1.075 | -0.428 | -0.096 | 0.688 | 0.01 | 0.072 | 0.072 | 0 | 0.24 | 0.46 | 0.35 | 0.63 | 0.26 | 0.29 |
| 1 | 238684052 | rs10495415 | 0.541 | 0.011 | -0.379 | 0.371 | 0.004 | -0.032 | -0.032 | 0.007 | 0.06 | 0.1 | 0.06 | 0.01 | 0.1 | 0.1 |
| 1 | 238685207 | rs11807567 | 0.418 | -0.01 | -0.336 | 0.728 | 0 | 0.008 | 0.008 | 0 | 0.11 | 0.11 | 0.12 | 0.26 | 0.13 | 0.1 |
| 1 | 238689341 | rs17545913 | -0.05 | 0.158 | -0.381 | 0.828 | 0.002 | -0.002 | -0.002 | 0 | 0.09 | 0.15 | 0.19 | 0.08 | 0.12 | 0.16 |
| 1 | 238690302 | rs11810140 | 0.259 | 0.218 | -0.651 | 0.85 | -0.015 | 0.013 | 0.013 | 1.045 | 0.27 | 0.42 | 0.3 | 0.29 | 0.2 | 0.18 |
| 1 | 238693830 | rs6702645 | 0.61 | 0.516 | -1.341 | 0.807 | -0.116 | 0.004 | 0.004 | 0.105 | 0.3 | 0.02 | 0.28 | 0.21 | 0.38 | 0.22 |
| 1 | 238696036 | rs535043 | 0.166 | 0.374 | -0.121 | 1.093 | -0.002 | 0.009 | 0.009 | 0 | 0.66 | 0.61 | 0.75 | 0.59 | 0.7 | 0.56 |
| 1 | 238697492 | rs7550596 | -1.245 | 0.18 | 0.162 | 1.182 | -0.034 | 0.081 | 0.081 | 0 | 0.15 | 0.4 | 0.07 | 0.24 | 0.09 | 0.23 |
| 1 | 238698680 | rs523691 | 0.416 | 0.779 | -0.386 | 0.732 | -0.002 | 0.002 | 0.002 | 0.038 | 0.7 | 0.28 | 0.65 | 0.7 | 0.65 | 0.77 |
| 1 | 238699660 | rs10802704 | -1.333 | 0.284 | 0.178 | 1.496 | -0.018 | 0.128 | 0.128 | 0 | 0.09 | 0.22 | 0.1 | 0.21 | 0.04 | 0.08 |
| 1 | 238700272 | rs487974 | 0.061 | 0.478 | -0.372 | 1.249 | -0.002 | -0.014 | -0.014 | 0.136 | 0.3 | 0.26 | 0.43 | 0.48 | 0.34 | 0.32 |
| 1 | 238700707 | rs2820211 | -0.047 | 0.359 | -0.302 | 0.638 | 0 | -0.003 | -0.003 | 0 | 0.39 | 0.49 | 0.36 | 0.42 | 0.43 | 0.4 |
| 1 | 238705309 | rs515344 | 0.214 | -0.19 | -0.01 | 0.553 | -0.021 | 0 | 0 | 0 | 0.83 | 0.72 | 0.91 | 0.87 | 0.89 | 0.77 |
| 1 | 238705483 | rs12126899 | 0.658 | -0.175 | -0.108 | 0.541 | -0.003 | 0.032 | 0.032 | 0.344 | 0.35 | 0.18 | 0.25 | 0.31 | 0.38 | 0.34 |
| 1 | 238705853 | rs490745 | 1.039 | -0.303 | -0.574 | 0.45 | -0.164 | -0.134 | -0.134 | 2.139 | 0.28 | 0.02 | 0.4 | 0.37 | 0.39 | 0.38 |
| 1 | 238705929 | rs491556 | -0.332 | -0.301 | -0.213 | 0.286 | -0.014 | 0.019 | 0.019 | 0 | 0.87 | 0.72 | 0.92 | 0.87 | 0.9 | 0.79 |
| 1 | 238706721 | rs519899 | -0.055 | -0.453 | 0.301 | 0.159 | 0.033 | 0.029 | 0.029 | 0.029 | 0.51 | 0.35 | 0.64 | 0.56 | 0.55 | 0.57 |
| 1 | 238712017 | rs574280 | 0.303 | -0.38 | 0.226 | 0.145 | -0.012 | 0.005 | 0.005 | 0.184 | 0.45 | 0.23 | 0.59 | 0.54 | 0.49 | 0.45 |
| 1 | 238719031 | rs789375 | -0.318 | -0.196 | 0.407 | 0.102 | -0.003 | -0.006 | -0.006 | 0.432 | 0.54 | 0.47 | 0.64 | 0.59 | 0.53 | 0.58 |
| 1 | 238719158 | rs526612 | 0.458 | 0.022 | 0.143 | 0.08 | 0.006 | 0.01 | 0.01 | 0 | 0.07 | 0.1 | 0.11 | 0.08 | 0.09 | 0.05 |
| 1 | 238719720 | rs789099 | 0.201 | -0.117 | 0.195 | 0.073 | 0.01 | 0.024 | 0.024 | 0 | 0.48 | 0.34 | 0.61 | 0.55 | 0.52 | 0.48 |
| 1 | 238720444 | rs6429087 | 0.682 | 0.126 | -0.564 | 0.245 | -0.003 | 0.032 | 0.032 | 0.177 | 0.33 | 0.18 | 0.2 | 0.29 | 0.34 | 0.33 |
| 1 | 238724741 | rs10754644 | -0.019 | 0.141 | -0.52 | 0.269 | 0.005 | 0.014 | 0.014 | 0 | 0.42 | 0.34 | 0.28 | 0.38 | 0.43 | 0.45 |
| 1 | 238725567 | rs7516101 | 0.384 | 0.228 | -0.828 | -0.025 | -0.016 | 0.043 | 0.043 | 0 | 0.33 | 0.17 | 0.21 | 0.29 | 0.36 | 0.38 |
| 1 | 238725762 | rs560229 | -0.113 | 0.101 | -0.352 | 0.069 | -0.013 | -0.01 | -0.01 | 0.145 | 0.18 | 0.24 | 0.17 | 0.12 | 0.12 | 0.13 |
| 1 | 238726690 | rs1002726 | -0.821 | 0.564 | -0.662 | -0.075 | -0.01 | 0.014 | 0.014 | 0 | 0.29 | 0.2 | 0.27 | 0.23 | 0.33 | 0.27 |
| 1 | 238729070 | rs2078306 | -1.237 | 0.193 | -0.019 | -0.232 | -0.003 | -0.002 | -0.002 | 0 | 0.33 | 0.6 | 0.3 | 0.27 | 0.22 | 0.25 |
| 1 | 238729563 | rs7512016 | -0.583 | 0.102 | 0.251 | -0.143 | -0.016 | 0.048 | 0.048 | 0 | 0.06 | 0.19 | 0.07 | 0.11 | 0.01 | 0.06 |
| 1 | 238730264 | rs648701 | 1.138 | 0.226 | -0.146 | -0.42 | -0.002 | 0.135 | 0.135 | 1.654 | 0.4 | 0.19 | 0.28 | 0.14 | 0.4 | 0.33 |
| 1 | 238735153 | rs471748 | 0.379 | 0.082 | 0.309 | -0.016 | -0.008 | 0.012 | 0.012 | 0 | 0.94 | 0.73 | 0.91 | 0.95 | 0.99 | 0.94 |
| 1 | 238738542 | rs2653910 | 0.725 | 0.067 | 0.274 | -0.009 | -0.006 | 0.076 | 0.076 | 0 | 0.93 | 0.67 | 0.9 | 0.89 | 0.99 | 0.93 |
| 1 | 238741253 | rs624912 | 0.667 | 0.334 | -0.41 | 0.064 | 0 | 0 | 0 | 1.265 | 0.69 | 0.33 | 0.7 | 0.66 | 0.75 | 0.73 |
| 1 | 238747111 | rs482697 | -0.301 | 0.181 | -1.113 | 0.272 | 0.026 | -0.031 | -0.031 | 0.064 | 0.37 | 0.81 | 0.48 | 0.56 | 0.26 | 0.42 |
| 1 | 238749403 | rs652252 | -0.181 | 0.737 | -1.569 | 0.259 | -0.123 | -0.094 | -0.094 | 0 | 0.75 | 0.98 | 0.83 | 0.71 | 0.67 | 0.85 |
| 1 | 238751635 | rs683070 | 0.35 | 0.193 | -0.221 | 0.405 | 0.005 | -0.01 | -0.01 | 0 | 0.29 | 0.31 | 0.36 | 0.5 | 0.22 | 0.35 |
| 1 | 238758325 | rs585622 | 0.261 | 0.124 | 0.576 | 0.475 | -0.029 | 0.046 | 0.046 | 0 | 0.32 | 0.41 | 0.39 | 0.5 | 0.23 | 0.36 |
| 1 | 238759205 | rs526130 | -0.046 | 0.066 | 0.857 | 0.437 | 0 | -0.002 | -0.002 | 0 | 0.34 | 0.58 | 0.42 | 0.53 | 0.24 | 0.36 |
| 1 | 238768264 | rs572934 | -0.239 | 0.473 | -0.075 | 0.311 | -0.057 | -0.079 | -0.079 | 0 | 0.75 | 0.92 | 0.8 | 0.69 | 0.69 | 0.82 |
| 1 | 238771226 | rs7517092 | 0.178 | 0.964 | 0.211 | 0.052 | 0.003 | -0.062 | -0.062 | 0 | 0.17 | 0.2 | 0.13 | 0.01 | 0.17 | 0.03 |
| 1 | 238772150 | rs10802709 | 1.039 | 0.786 | 0.311 | 0.281 | -0.015 | -0.006 | -0.006 | 0 | 0.34 | 0.4 | 0.34 | 0.32 | 0.29 | 0.42 |
| 1 | 238773806 | rs2494076 | 0.019 | 0.789 | 0.285 | 0.264 | 0 | 0.002 | 0.002 | 0 | 0.46 | 0.47 | 0.45 | 0.36 | 0.42 | 0.36 |
| 1 | 238774832 | rs590734 | 1.089 | 0.749 | 0.17 | 0.312 | 0.006 | -0.009 | -0.009 | 0.628 | 0.29 | 0.27 | 0.34 | 0.36 | 0.25 | 0.34 |
| 1 | 238778872 | rs649812 | -0.791 | 0.827 | 0.099 | 0.35 | -0.05 | -0.102 | -0.102 | 0.463 | 0.81 | 0.98 | 0.85 | 0.7 | 0.77 | 0.82 |
| 1 | 238779890 | rs510798 | 1.297 | 0.761 | 0.153 | 0.313 | 0 | -0.025 | -0.025 | 0 | 0.28 | 0.18 | 0.31 | 0.35 | 0.26 | 0.36 |
| 1 | 238783484 | rs1502872 | -0.653 | 0.693 | 0.67 | 0.202 | -0.001 | 0.061 | 0.061 | 0 | 0.52 | 0.67 | 0.52 | 0.33 | 0.5 | 0.46 |
| 1 | 238783811 | rs6672862 | 0.907 | 0.899 | 0.812 | 0.12 | 0.009 | -0.002 | -0.002 | 0 | 0.31 | 0.24 | 0.35 | 0.33 | 0.25 | 0.38 |
| 1 | 238786017 | rs10754645 | 0.603 | 0.814 | 0.531 | -0.063 | 0.002 | -0.007 | -0.007 | 0 | 0.35 | 0.43 | 0.41 | 0.33 | 0.35 | 0.43 |
| 1 | 238786780 | rs12728263 | 0.304 | 0.845 | 0.219 | -0.084 | -0.001 | -0.022 | -0.022 | 0 | 0.16 | 0.11 | 0.11 | 0 | 0.16 | 0.03 |
| 1 | 238787614 | rs6681102 | -0.284 | 0.754 | 0.093 | 0.325 | 0.004 | 0.051 | 0.051 | 0 | 0.52 | 0.62 | 0.52 | 0.33 | 0.51 | 0.46 |
| 1 | 238811326 | rs10925719 | -0.174 | 0.692 | -0.265 | -0.031 | 0.003 | 0.008 | 0.008 | 0 | 0.53 | 0.42 | 0.54 | 0.78 | 0.6 | 0.61 |
| 1 | 238817430 | rs6678175 | -1.164 | 0.978 | -0.308 | 0.099 | 0.081 | 0.006 | 0.006 | 0 | 0.1 | 0.33 | 0.08 | 0.08 | 0.11 | 0.1 |
| 1 | 238818794 | rs16837283 | 1.707 | 0.756 | -0.823 | -0.21 | -0.008 | -0.059 | -0.059 | 0 | 0.12 | 0.01 | 0.12 | 0.2 | 0.1 | 0.25 |
| 1 | 238819523 | rs12726612 | 0.759 | 0.833 | -0.772 | 0.121 | -0.088 | -0.062 | -0.062 | 0 | 0.43 | 0.05 | 0.42 | 0.56 | 0.47 | 0.49 |
| 1 | 238827206 | rs7546888 | -1.127 | 0.912 | -0.436 | 0.188 | 0.056 | 0.067 | 0.067 | 0 | 0.1 | 0.25 | 0.1 | 0.23 | 0.14 | 0.15 |
| 1 | 238829636 | rs10925727 | -0.018 | 0.896 | -0.677 | 0.128 | 0.019 | 0.081 | 0.081 | 0 | 0.52 | 0.34 | 0.51 | 0.73 | 0.59 | 0.57 |
| 1 | 238837866 | rs2174076 | 0.404 | 0.555 | -0.97 | 0.124 | -0.05 | 0.029 | 0.029 | 0 | 0.51 | 0.11 | 0.48 | 0.56 | 0.54 | 0.52 |
| 1 | 238845573 | rs2653914 | -0.034 | 1.248 | -0.413 | 0.421 | 0.021 | 0.014 | 0.014 | 0 | 0.22 | 0.24 | 0.18 | 0.25 | 0.28 | 0.15 |
| 1 | 238849083 | rs2841377 | -0.41 | 0.794 | -0.364 | 0.093 | 0.014 | 0.038 | 0.038 | 0 | 0.6 | 0.67 | 0.59 | 0.74 | 0.66 | 0.64 |
| 1 | 238853437 | rs6698518 | 0.631 | 0.811 | -0.284 | 0.115 | 0.031 | -0.023 | -0.023 | 0 | 0.39 | 0.28 | 0.39 | 0.54 | 0.31 | 0.43 |
| 1 | 238855007 | rs10802716 | 0.691 | 0.804 | -0.287 | 0.152 | 0.028 | -0.014 | -0.014 | 0.011 | 0.38 | 0.26 | 0.38 | 0.46 | 0.31 | 0.43 |
| 1 | 238855427 | rs2841393 | -0.648 | 0.836 | -0.414 | -0.028 | -0.006 | -0.057 | -0.057 | 0.113 | 0.72 | 0.5 | 0.68 | 0.86 | 0.75 | 0.85 |
| 1 | 238860631 | rs6675135 | 0.941 | 0.695 | -0.864 | 0.139 | -0.001 | -0.139 | -0.139 | 0 | 0.35 | 0.02 | 0.35 | 0.44 | 0.33 | 0.43 |
| 1 | 238861548 | rs2202215 | 0.161 | 1.115 | -0.371 | 0.137 | 0.014 | 0.003 | 0.003 | 0 | 0.21 | 0.24 | 0.16 | 0.2 | 0.29 | 0.14 |
| 1 | 238874684 | rs10925739 | -0.05 | 1.131 | 0.32 | 0.032 | 0.008 | -0.009 | -0.009 | 0 | 0.6 | 0.51 | 0.52 | 0.61 | 0.61 | 0.62 |
| 1 | 238875754 | rs12735197 | 1.329 | 0.976 | -0.159 | -0.048 | -0.019 | -0.148 | -0.148 | 0.05 | 0.3 | 0.06 | 0.32 | 0.46 | 0.31 | 0.41 |
| 1 | 238886760 | rs10925740 | -0.368 | 1.098 | -0.09 | -0.078 | -0.041 | 0.063 | 0.063 | 0 | 0.61 | 0.79 | 0.49 | 0.74 | 0.58 | 0.61 |
| 1 | 238889951 | rs12744326 | 1.324 | 0.396 | -0.516 | 0.038 | -0.011 | -0.185 | -0.185 | 0.098 | 0.27 | 0.02 | 0.21 | 0.5 | 0.25 | 0.45 |
| 1 | 238890112 | rs17552002 | 0.315 | 0.799 | -0.377 | -0.212 | -0.031 | 0.036 | 0.036 | 0 | 0.3 | 0.19 | 0.22 | 0.16 | 0.38 | 0.16 |
| 1 | 238890738 | rs4601573 | 1.317 | 0.701 | -0.599 | -0.061 | -0.001 | -0.181 | -0.181 | 0 | 0.3 | 0.02 | 0.23 | 0.48 | 0.27 | 0.46 |

| | | | | | | | | | | | | | | | | |
|---|---|---|---|---|---|---|---|---|---|---|---|---|---|---|---|---|
| 1 | 238899903 | rs10925747 | -0.072 | 0.897 | 0.041 | -0.013 | 0.028 | 0.008 | 0.008 | 0 | 0.3 | 0.55 | 0.23 | 0.3 | 0.38 | 0.14 |
| 1 | 238903098 | rs12123186 | 1.179 | 0.523 | -0.631 | -0.525 | 0.007 | -0.046 | -0.046 | 0 | 0.16 | 0 | 0.18 | 0.2 | 0.11 | 0.24 |
| 1 | 238908229 | rs2091078 | 0.103 | 0.71 | -0.129 | -0.488 | 0.012 | -0.014 | -0.014 | 0 | 0.29 | 0.32 | 0.21 | 0.19 | 0.38 | 0.13 |
| 1 | 238911829 | rs2494766 | -1.185 | 0.368 | -0.178 | -0.522 | -0.006 | 0.003 | 0.003 | 0.162 | 0.69 | 0.58 | 0.68 | 0.4 | 0.71 | 0.55 |
| 1 | 238911955 | rs7531688 | -1.439 | 0.227 | -0.77 | -0.394 | 0.058 | -0.132 | -0.132 | 0.012 | 0.7 | 0.93 | 0.79 | 0.53 | 0.76 | 0.58 |
| 1 | 238914122 | rs2841406 | -0.334 | 0.75 | -0.677 | -0.616 | -0.015 | 0.015 | 0.015 | 0.096 | 0.31 | 0.42 | 0.34 | 0.4 | 0.24 | 0.34 |
| 1 | 238915864 | rs2841407 | -0.493 | 0.783 | -1.211 | -0.609 | -0.001 | 0 | 0 | 0 | 0.33 | 0.62 | 0.4 | 0.41 | 0.24 | 0.35 |
| 1 | 238918722 | rs10925754 | 0.902 | 1.588 | -1.374 | -1.328 | -0.096 | 0.1 | 0.1 | 0.295 | 0.19 | 0.03 | 0.16 | 0.02 | 0.29 | 0.07 |
| 1 | 238921247 | rs16837404 | 0.446 | 1.681 | -0.85 | -0.775 | 0.003 | -0.02 | -0.02 | 0 | 0.05 | 0.08 | 0.04 | 0 | 0.04 | 0.03 |
| 1 | 238921472 | rs12754272 | 0.63 | 1.775 | -1.042 | -0.72 | -0.051 | 0.033 | 0.033 | 0.565 | 0.15 | 0.04 | 0.24 | 0.07 | 0.27 | 0.1 |
| 1 | 238922048 | rs3011572 | -0.594 | 1.304 | -0.836 | -0.672 | -0.007 | 0.013 | 0.013 | 0 | 0.4 | 0.59 | 0.57 | 0.27 | 0.42 | 0.4 |
| 1 | 238926821 | rs12408234 | 0.514 | 1.345 | -0.718 | -0.778 | 0.007 | -0.038 | -0.038 | 0.887 | 0.16 | 0.19 | 0.21 | 0.03 | 0.15 | 0.05 |
| 1 | 238928481 | rs1357520 | -0.436 | 1.221 | -0.598 | -0.789 | -0.001 | -0.003 | -0.003 | 0.279 | 0.08 | 0.12 | 0.13 | 0.03 | 0.08 | 0.11 |
| 1 | 238930549 | rs16837416 | 0.173 | 0.838 | -0.033 | -0.518 | -0.005 | 0.089 | 0.089 | 0 | 0.05 | 0.21 | 0.02 | 0.13 | 0.01 | 0.04 |
| 1 | 238930859 | rs10925760 | -0.028 | 1.122 | -0.176 | -0.915 | -0.009 | -0.017 | -0.017 | 0 | 0.25 | 0.44 | 0.15 | 0.27 | 0.2 | 0.2 |
| 1 | 238931061 | rs2841417 | -0.117 | 1.312 | -1.089 | -1.36 | -0.023 | 0.097 | 0.097 | 0 | 0.28 | 0.16 | 0.34 | 0.05 | 0.32 | 0.24 |
| 1 | 238931277 | rs2689128 | 0.436 | 1.241 | -0.963 | -1.277 | -0.01 | 0.064 | 0.064 | 0 | 0.23 | 0.16 | 0.28 | 0.03 | 0.24 | 0.13 |
| 1 | 238942337 | rs7542176 | -0.367 | 0.728 | -0.266 | -1.232 | -0.009 | -0.007 | -0.007 | 0 | 0.33 | 0.34 | 0.29 | 0.27 | 0.24 | 0.33 |
| 1 | 238946546 | rs2841346 | -1.28 | 1.001 | -0.4 | -1.315 | 0.032 | 0.083 | 0.083 | 0 | 0.46 | 0.7 | 0.47 | 0.3 | 0.38 | 0.41 |
| 1 | 238947632 | rs4313384 | 2.011 | 1.416 | -0.596 | -1.269 | -0.051 | 0.04 | 0.04 | 0.229 | 0.12 | 0.01 | 0.11 | 0.03 | 0.13 | 0.05 |
| 1 | 238949005 | rs3011577 | -1.921 | 1.315 | -0.559 | -1.34 | -0.007 | 0.031 | 0.031 | 0.678 | 0.62 | 0.78 | 0.63 | 0.33 | 0.57 | 0.49 |
| 1 | 238952309 | rs6693951 | 0.659 | 1.142 | -0.384 | -0.869 | -0.006 | 0 | 0 | 0 | 0.19 | 0.2 | 0.14 | 0.25 | 0.12 | 0.22 |
| 1 | 238962668 | rs10925763 | -0.595 | 1.603 | 0.218 | -0.821 | 0.019 | -0.036 | -0.036 | 0 | 0.23 | 0.41 | 0.22 | 0.15 | 0.19 | 0.18 |
| 1 | 238962748 | rs12086208 | 0.882 | 1.674 | -0.035 | -0.762 | -0.014 | 0.027 | 0.027 | 0 | 0.14 | 0.09 | 0.12 | 0.03 | 0.13 | 0.05 |
| 1 | 238965663 | rs17619413 | -0.172 | 1.762 | -0.054 | -0.922 | -0.009 | 0.015 | 0.015 | 1.351 | 0.17 | 0.1 | 0.18 | 0.05 | 0.23 | 0.07 |
| 1 | 238972181 | rs2689160 | -1.971 | 1.504 | 0.203 | -1.026 | -0.036 | 0.074 | 0.074 | 1.817 | 0.41 | 0.74 | 0.43 | 0.19 | 0.44 | 0.22 |
| 1 | 238982199 | rs6664301 | -0.952 | 1.35 | 0.734 | -0.991 | 0.001 | 0.059 | 0.059 | 0 | 0.09 | 0.36 | 0.11 | 0.13 | 0.04 | 0.06 |
| 1 | 238985075 | rs2841375 | -0.429 | 1.498 | 0.604 | -1.047 | 0.075 | -0.124 | -0.124 | 0 | 0.21 | 0.35 | 0.23 | 0.01 | 0.28 | 0.09 |
| 1 | 238987220 | rs12406780 | 1.333 | 1.431 | 0.52 | -0.961 | -0.012 | -0.003 | -0.003 | 0 | 0.1 | 0.02 | 0.08 | 0.06 | 0.1 | 0.05 |
| 1 | 239001279 | rs1983433 | -2.081 | 1.423 | 0.486 | -1.002 | -0.037 | 0.078 | 0.078 | 1.762 | 0.4 | 0.75 | 0.43 | 0.19 | 0.43 | 0.22 |
| 1 | 239003522 | rs869035 | -0.789 | 1.432 | 0.621 | -0.997 | -0.014 | -0.001 | -0.001 | 0 | 0.19 | 0.39 | 0.2 | 0.18 | 0.15 | 0.13 |
| 1 | 239005080 | rs3001434 | 0.227 | 1.738 | 0.484 | -1.015 | 0.111 | -0.102 | -0.102 | 0 | 0.16 | 0.35 | 0.19 | 0.01 | 0.21 | 0.08 |
| 1 | 239021684 | rs2841385 | -1.47 | 1.656 | 0.436 | -1.037 | 0.044 | -0.097 | -0.097 | 0 | 0.34 | 0.43 | 0.35 | 0.08 | 0.42 | 0.19 |
| 1 | 239022848 | rs1523416 | 0.242 | 1.709 | 0.467 | -1.039 | 0.084 | -0.098 | -0.098 | 0.351 | 0.16 | 0.34 | 0.18 | 0.01 | 0.2 | 0.07 |
| 1 | 239023492 | rs1523417 | -1.465 | 1.629 | 0.432 | -1.027 | 0.011 | -0.045 | -0.045 | 0 | 0.35 | 0.55 | 0.37 | 0.08 | 0.42 | 0.19 |
| 1 | 239057106 | rs6674607 | -1.037 | 1.451 | 0.555 | -0.827 | 0.03 | -0.074 | -0.074 | 1.743 | 0.87 | 0.77 | 0.85 | 1 | 0.87 | 0.95 |
| 1 | 239058330 | rs6689380 | -0.576 | 1.452 | 0.402 | -0.909 | 0 | -0.009 | -0.009 | 0 | 0.08 | 0.07 | 0.08 | 0.02 | 0.11 | 0.07 |
| 1 | 239060960 | rs7551773 | 0.474 | 1.443 | 0.435 | -0.909 | 0.04 | -0.123 | -0.123 | 0 | 0.79 | 0.66 | 0.75 | 0.98 | 0.76 | 0.88 |
| 1 | 239063172 | rs6695351 | -0.141 | 1.435 | 0.368 | -0.89 | 0.026 | -0.102 | -0.102 | 0.836 | 0.18 | 0.29 | 0.2 | 0 | 0.2 | 0.07 |
| 1 | 239070768 | rs10495426 | 1.241 | 0.843 | 0.3 | -0.034 | -0.018 | -0.019 | -0.019 | 0 | 0.06 | 0.01 | 0.07 | 0.2 | 0.08 | 0.13 |
| 1 | 239072781 | rs9428765 | 0.811 | 0.597 | 0.227 | -0.333 | -0.012 | 0.011 | 0.011 | 0 | 0.12 | 0.02 | 0.17 | 0.05 | 0.14 | 0.1 |
| 1 | 239078410 | rs9428420 | 0.629 | 0.603 | 0.274 | -0.348 | -0.007 | 0.022 | 0.022 | 0 | 0.13 | 0.08 | 0.18 | 0.05 | 0.15 | 0.12 |
| 1 | 239080702 | rs1915254 | 1.323 | 0.581 | 0.617 | -0.458 | 0.014 | -0.087 | -0.087 | 1.309 | 0.73 | 0.56 | 0.64 | 0.93 | 0.68 | 0.81 |
| 1 | 239086183 | rs6689433 | 1.495 | 0.606 | 0.655 | -0.449 | 0.092 | -0.008 | -0.008 | 0.566 | 0.91 | 0.68 | 0.86 | 0.97 | 0.9 | 0.94 |
| 1 | 239094157 | rs9428779 | 0.551 | 0.546 | 0.59 | -0.445 | 0.005 | 0.013 | 0.013 | 0 | 0.13 | 0.13 | 0.18 | 0.05 | 0.14 | 0.1 |
| 1 | 239094695 | rs12087472 | -0.782 | 0.288 | 0.833 | -0.402 | -0.01 | 0.004 | 0.004 | 0 | 0.1 | 0.23 | 0.11 | 0.12 | 0.09 | 0.06 |
| 1 | 239094847 | rs17621459 | 0.626 | 0.288 | 0.829 | -0.402 | 0.005 | 0.013 | 0.013 | 0 | 0.13 | 0.12 | 0.18 | 0.05 | 0.14 | 0.1 |
| 1 | 239096030 | rs6429111 | -0.833 | 0.279 | 0.812 | -0.379 | -0.01 | 0.004 | 0.004 | 0.033 | 0.1 | 0.24 | 0.11 | 0.12 | 0.09 | 0.06 |
| 1 | 239097333 | rs10925783 | 1.521 | -0.081 | 0.131 | -0.55 | -0.007 | 0.005 | 0.005 | 0 | 0.17 | 0.01 | 0.13 | 0.12 | 0.12 | 0.18 |
| 1 | 239097408 | rs4659887 | 0.844 | 0.005 | 0.695 | -0.369 | 0.033 | 0.006 | 0.006 | 0.448 | 0.9 | 0.63 | 0.88 | 0.93 | 0.92 | 0.96 |
| 1 | 239109443 | rs12023560 | 1.614 | -0.05 | 0.571 | -0.331 | 0.009 | 0.026 | 0.026 | 0.151 | 0.67 | 0.25 | 0.57 | 0.8 | 0.67 | 0.79 |
| 1 | 239110043 | rs12752615 | 1.785 | -0.096 | 0.045 | -0.073 | -0.115 | -0.128 | -0.128 | 0.14 | 0.26 | 0.01 | 0.28 | 0.4 | 0.31 | 0.32 |
| 1 | 239110659 | rs6682752 | -0.203 | 0.114 | 1.123 | -0.141 | -0.003 | 0.043 | 0.043 | 0 | 0.06 | 0.26 | 0.04 | 0.06 | 0.01 | 0.02 |
| 1 | 239117641 | rs9428438 | 0.739 | 0.154 | 0.716 | -0.107 | -0.016 | -0.009 | -0.009 | 0 | 0.12 | 0.01 | 0.17 | 0.06 | 0.14 | 0.1 |
| 1 | 239120367 | rs12124650 | -0.716 | 0.151 | 0.87 | -0.118 | 0.001 | 0.002 | 0.002 | 0 | 0.19 | 0.23 | 0.27 | 0.13 | 0.22 | 0.13 |
| 1 | 239120492 | rs10732303 | 0.762 | 0.177 | 1.001 | -0.167 | 0.055 | 0.022 | 0.022 | 0.346 | 0.9 | 0.57 | 0.87 | 0.93 | 0.92 | 0.96 |
| 1 | 239124181 | rs2883158 | 2.006 | 0.165 | 0.902 | -0.186 | -0.003 | -0.009 | -0.009 | 1.166 | 0.78 | 0.28 | 0.73 | 0.86 | 0.81 | 0.89 |
| 1 | 239128572 | rs6656982 | 0.814 | 0.15 | 0.513 | -0.019 | -0.008 | -0.017 | -0.017 | 0 | 0.77 | 0.59 | 0.68 | 0.85 | 0.75 | 0.83 |
| 1 | 239143179 | rs1574985 | 1.359 | 0.089 | 0.189 | 0.174 | -0.005 | 0.003 | 0.003 | 0.505 | 0.66 | 0.1 | 0.53 | 0.77 | 0.67 | 0.8 |
| 1 | 239151401 | rs6699410 | -0.659 | 0.138 | 0.352 | 0.143 | 0.023 | 0.074 | 0.074 | 0 | 0.05 | 0.19 | 0.07 | 0.09 | 0.04 | 0.05 |
| 1 | 239156403 | rs16837628 | -1.23 | 0.156 | 0.401 | 0.144 | 0.053 | 0.076 | 0.076 | 0 | 0.06 | 0.24 | 0.08 | 0.09 | 0.07 | 0.05 |
| 1 | 239158189 | rs9428810 | -0.844 | 0.142 | 0.35 | -0.218 | -0.006 | -0.046 | -0.046 | 7.09 | 0.24 | 0.36 | 0.28 | 0.07 | 0.24 | 0.13 |
| 1 | 239159611 | rs1915278 | -0.352 | 0.121 | 0.257 | -0.184 | 0 | 0.002 | 0.002 | 0 | 0.19 | 0.2 | 0.22 | 0.07 | 0.2 | 0.12 |
| 1 | 239165770 | rs9428451 | -0.241 | 0.359 | -0.17 | -0.066 | -0.025 | 0.042 | 0.042 | 0.843 | 0.32 | 0.13 | 0.37 | 0.2 | 0.34 | 0.29 |
| 1 | 239166229 | rs9428452 | -0.08 | 0.335 | 0.028 | -0.266 | 0.004 | -0.003 | -0.003 | 0.243 | 0.13 | 0.13 | 0.22 | 0.06 | 0.2 | 0.12 |
| 1 | 239166691 | rs9428453 | 0.203 | 0.335 | 0.032 | -0.265 | -0.001 | -0.001 | -0.001 | 0 | 0.12 | 0.1 | 0.2 | 0.06 | 0.17 | 0.11 |
| 1 | 239170783 | rs12145112 | 0.991 | 0.219 | -0.293 | -0.387 | -0.027 | 0.023 | 0.023 | 0.855 | 0.15 | 0.01 | 0.08 | 0.09 | 0.16 | 0.16 |
| 1 | 239171877 | rs9428818 | 0.346 | 0.414 | 0.816 | 0.128 | 0.049 | 0.083 | 0.083 | 1.26 | 0.92 | 0.59 | 0.83 | 0.92 | 0.95 | 0.97 |
| 1 | 239176448 | rs10925792 | 1.325 | 0.402 | 0.714 | 0.369 | -0.005 | 0.054 | 0.054 | 0.168 | 0.49 | 0.27 | 0.32 | 0.41 | 0.51 | 0.5 |

| | | | | | | | | | | | | | | | | |
|---|---|---|---|---|---|---|---|---|---|---|---|---|---|---|---|---|
| 1 | 239177621 | rs1915247 | 0.064 | 0.41 | 0.654 | 0.382 | 0.008 | -0.056 | -0.056 | 0.092 | 0.3 | 0.18 | 0.29 | 0.46 | 0.23 | 0.36 |
| 1 | 239179054 | rs9428454 | 0.436 | 0.521 | 0.375 | 0.438 | -0.038 | -0.017 | -0.017 | 0 | 0.11 | 0.02 | 0.17 | 0.06 | 0.16 | 0.1 |
| 1 | 239184346 | rs12118040 | 1.37 | 0.529 | 0.432 | 0.408 | -0.021 | 0.062 | 0.062 | 0.273 | 0.49 | 0.18 | 0.32 | 0.41 | 0.51 | 0.5 |
| 1 | 239184393 | rs12129112 | 1.297 | 1.103 | 0.864 | 0.659 | 0.031 | -0.002 | -0.002 | 0 | 0.79 | 0.49 | 0.62 | 0.87 | 0.74 | 0.86 |
| 1 | 239184630 | rs17622574 | -0.35 | 0.753 | 0.383 | 0.798 | 0.001 | -0.004 | -0.004 | 0 | 0.12 | 0.12 | 0.2 | 0.04 | 0.17 | 0.11 |
| 1 | 239213531 | rs10925806 | 1.04 | 0.934 | -0.306 | 0.83 | -0.016 | 0.048 | 0.048 | 2.614 | 0.53 | 0.21 | 0.35 | 0.39 | 0.49 | 0.52 |
| 1 | 239214483 | rs1509701 | 1.46 | 0.713 | 0.284 | 0.617 | 0.024 | 0.036 | 0.036 | 0.323 | 0.85 | 0.66 | 0.72 | 0.86 | 0.83 | 0.89 |
| 1 | 239215774 | rs1396901 | 0.434 | 0.71 | -0.174 | 0.535 | -0.002 | 0.001 | 0.001 | 0 | 0.9 | 0.92 | 0.82 | 0.94 | 0.85 | 0.92 |
| 1 | 239221751 | rs1912230 | -1.027 | 0.144 | 0.33 | 0.734 | -0.016 | 0.114 | 0.114 | 0.27 | 0.1 | 0.37 | 0.21 | 0.14 | 0.07 | 0.07 |
| 1 | 239224547 | rs12041748 | -0.357 | 0.459 | -0.288 | 0.898 | -0.005 | -0.011 | -0.011 | 0.024 | 0.13 | 0.08 | 0.13 | 0.23 | 0.13 | 0.2 |
| 1 | 239227470 | rs7545552 | -0.627 | -0.404 | -0.392 | -0.12 | -0.016 | -0.034 | -0.034 | 0.241 | 0.31 | 0.31 | 0.4 | 0.12 | 0.22 | 0.33 |
| 1 | 239248743 | rs6682504 | -0.983 | -0.615 | 0.281 | -0.437 | -0.041 | -0.094 | -0.094 | 0 | 0.37 | 0.52 | 0.47 | 0.11 | 0.29 | 0.34 |
| 1 | 239249308 | rs12084592 | -0.564 | -0.612 | 0.282 | -0.405 | -0.024 | -0.043 | -0.043 | 0 | 0.31 | 0.37 | 0.4 | 0.11 | 0.22 | 0.28 |
| 1 | 239253108 | rs6429124 | -0.747 | -0.624 | 0.127 | -0.268 | -0.046 | -0.05 | -0.05 | 0 | 0.32 | 0.44 | 0.42 | 0.2 | 0.23 | 0.33 |
| 1 | 239254075 | rs1509689 | -0.585 | -0.629 | 0.146 | -0.27 | -0.002 | -0.001 | -0.001 | 0 | 0.3 | 0.27 | 0.4 | 0.2 | 0.22 | 0.29 |
| 1 | 239256802 | rs1588360 | -0.697 | -0.612 | -0.081 | -0.328 | -0.046 | -0.11 | -0.11 | 0.346 | 0.31 | 0.44 | 0.37 | 0.06 | 0.23 | 0.31 |
| 1 | 239262486 | rs16837758 | -0.469 | -0.611 | -0.348 | -0.219 | -0.033 | 0.02 | 0.02 | 0 | 0.23 | 0.37 | 0.26 | 0.24 | 0.16 | 0.31 |
| 1 | 239263935 | rs16837766 | -0.63 | -0.333 | -0.443 | -0.256 | -0.02 | 0.007 | 0.007 | 0 | 0.27 | 0.38 | 0.29 | 0.25 | 0.21 | 0.36 |
| 1 | 239265755 | rs11805939 | -1.595 | 0.315 | -1.431 | -0.053 | -0.058 | -0.11 | -0.11 | 0.786 | 0.68 | 0.97 | 0.76 | 0.6 | 0.69 | 0.7 |
| 1 | 239271778 | rs4659902 | -0.954 | 0.44 | -0.737 | 0.147 | 0.008 | 0.012 | 0.012 | 1.294 | 0.45 | 0.37 | 0.53 | 0.62 | 0.39 | 0.62 |
| 1 | 239272488 | rs12040128 | 0.12 | 0.395 | -0.996 | 0.131 | -0.036 | -0.087 | -0.087 | 0 | 0.21 | 0.03 | 0.23 | 0.33 | 0.2 | 0.3 |
| 1 | 239273035 | rs17640659 | 0.941 | 0.386 | -1.065 | 0.237 | -0.046 | -0.018 | -0.018 | 0 | 0.14 | 0.01 | 0.12 | 0.12 | 0.15 | 0.24 |
| 1 | 239275579 | rs10495440 | 0.554 | 0.492 | -0.894 | 0.145 | -0.045 | -0.106 | -0.106 | 0 | 0.21 | 0.01 | 0.23 | 0.33 | 0.2 | 0.3 |
| 1 | 239277139 | rs1567400 | -0.822 | 0.41 | -0.623 | 0.255 | -0.003 | -0.006 | -0.006 | 0 | 0.46 | 0.47 | 0.55 | 0.63 | 0.4 | 0.62 |
| 1 | 239278524 | rs17640830 | 1.063 | 0.033 | -0.736 | 0.049 | -0.008 | 0.041 | 0.041 | 0.468 | 0.24 | 0.18 | 0.17 | 0.07 | 0.24 | 0.09 |
| 1 | 239283303 | rs10495439 | -0.065 | 0.204 | -0.131 | 0.744 | -0.005 | 0.007 | 0.007 | 1.608 | 0.18 | 0.18 | 0.23 | 0.28 | 0.11 | 0.22 |
| 1 | 239289175 | rs16837808 | -0.076 | 0.286 | -0.397 | 0.783 | -0.026 | 0.045 | 0.045 | 2.131 | 0.19 | 0.28 | 0.23 | 0.27 | 0.12 | 0.21 |
| 1 | 239297423 | rs10495437 | -0.415 | 0.972 | -0.592 | 0.452 | 0.011 | -0.001 | -0.001 | 0 | 0.29 | 0.34 | 0.28 | 0.31 | 0.37 | 0.27 |
| 1 | 239305236 | rs1848751 | -0.78 | 1.003 | -0.515 | 0.602 | -0.011 | -0.005 | -0.005 | 0.083 | 0.25 | 0.21 | 0.38 | 0.26 | 0.28 | 0.33 |
| 1 | 239306319 | rs1858175 | -0.313 | 0.626 | -1.022 | -0.238 | -0.038 | 0.052 | 0.052 | 0 | 0.72 | 0.84 | 0.83 | 0.91 | 0.62 | 0.71 |
| 1 | 239306468 | rs6657123 | 1.079 | -0.466 | -0.687 | 0.18 | 0.035 | 0 | 0 | 0 | 0.34 | 0.21 | 0.31 | 0.34 | 0.25 | 0.22 |
| 1 | 239306871 | rs16837837 | -0.252 | 0.171 | -0.657 | -0.08 | 0.007 | -0.073 | -0.073 | 0 | 0.16 | 0.24 | 0.12 | 0.01 | 0.19 | 0.11 |
| 1 | 239309399 | rs17567797 | 0.945 | 0.254 | -0.712 | -0.023 | 0.015 | -0.037 | -0.037 | 0 | 0.08 | 0.1 | 0.05 | 0 | 0.13 | 0.04 |
| 1 | 239310416 | rs2354178 | -1.243 | 0.301 | -0.976 | -0.027 | -0.097 | 0.037 | 0.037 | 0 | 0.81 | 0.99 | 0.88 | 0.91 | 0.74 | 0.74 |
| 1 | 239311068 | rs7419201 | 0.774 | 0.427 | -1.225 | 0.066 | -0.074 | -0.061 | -0.061 | 0 | 0.14 | 0.03 | 0.31 | 0.24 | 0.19 | 0.22 |
| 1 | 239313573 | rs7541537 | -0.614 | -0.493 | -0.787 | 0.131 | -0.093 | -0.009 | -0.009 | 0 | 0.66 | 0.88 | 0.58 | 0.67 | 0.56 | 0.48 |
| 1 | 239316750 | rs2136521 | 0.632 | -0.922 | -0.476 | 0.142 | 0.006 | 0.003 | 0.003 | 0 | 0.47 | 0.41 | 0.41 | 0.4 | 0.34 | 0.34 |
| 1 | 239317135 | rs4659904 | -0.327 | -0.819 | -0.854 | 0.174 | -0.053 | 0.016 | 0.016 | 0 | 0.62 | 0.88 | 0.54 | 0.67 | 0.48 | 0.47 |
| 1 | 239320854 | rs12132104 | 1.217 | -1.584 | -1.143 | 0.096 | -0.006 | 0.003 | 0.003 | 0 | 0.2 | 0.17 | 0.27 | 0.23 | 0.17 | 0.17 |
| 1 | 239321213 | rs10925832 | 0.483 | -1.083 | -0.801 | 0.268 | 0.019 | 0.13 | 0.13 | 0.068 | 0.12 | 0.34 | 0.11 | 0.26 | 0.12 | 0.1 |
| 1 | 239323771 | rs9787221 | -0.028 | -1.158 | -1.817 | 0.018 | -0.008 | 0.017 | 0.017 | 0 | 0.27 | 0.65 | 0.22 | 0.33 | 0.24 | 0.24 |
| 1 | 239323855 | rs9787223 | -0.098 | -0.996 | -1.715 | 0.126 | -0.005 | 0.004 | 0.004 | 0 | 0.31 | 0.65 | 0.27 | 0.33 | 0.31 | 0.25 |
| 1 | 239327385 | rs1980004 | -0.065 | -1.144 | -1.306 | 0.103 | 0.023 | 0.004 | 0.004 | 0 | 0.35 | 0.24 | 0.3 | 0.34 | 0.23 | 0.23 |
| 1 | 239332114 | rs6668037 | 0.127 | -0.542 | -1.71 | 0.281 | -0.004 | -0.053 | -0.053 | 0.102 | 0.2 | 0.04 | 0.2 | 0.34 | 0.21 | 0.15 |
| 1 | 239338587 | rs1112903 | -0.656 | 0.416 | -2.292 | 0.014 | -0.079 | 0.005 | 0.005 | 0 | 0.36 | 0.12 | 0.35 | 0.41 | 0.46 | 0.38 |
| 1 | 239339558 | rs6686907 | 0.943 | 0.281 | -2.098 | -0.121 | 0 | -0.009 | -0.009 | 0.317 | 0.27 | 0.23 | 0.36 | 0.4 | 0.25 | 0.25 |
| 1 | 239340085 | rs17568414 | 1.026 | 0.147 | -1.835 | -0.518 | 0 | -0.031 | -0.031 | 0.521 | 0.15 | 0.14 | 0.16 | 0 | 0.17 | 0.16 |
| 1 | 239341528 | rs10925835 | 1.015 | -0.041 | -0.973 | -0.177 | 0.009 | 0.03 | 0.03 | 0 | 0.3 | 0.12 | 0.33 | 0.32 | 0.31 | 0.28 |
| 1 | 239341587 | rs10737832 | -1.151 | -0.2 | -0.615 | -0.197 | -0.04 | 0.029 | 0.029 | 0 | 0.81 | 0.64 | 0.88 | 0.8 | 0.88 | 0.77 |
| 1 | 239343728 | rs6703086 | -0.253 | -0.019 | -0.78 | -0.05 | 0 | 0.011 | 0.011 | 0.385 | 0.62 | 0.33 | 0.72 | 0.72 | 0.63 | 0.54 |
| 1 | 239344447 | rs12038672 | -0.059 | 0.422 | -0.402 | 0.383 | 0.002 | 0.009 | 0.009 | 0 | 0.32 | 0.39 | 0.4 | 0.39 | 0.31 | 0.26 |
| 1 | 239350728 | rs2664057 | -0.544 | -0.252 | -0.255 | 0.085 | 0.039 | 0.072 | 0.072 | 0.79 | 0.4 | 0.8 | 0.37 | 0.22 | 0.27 | 0.21 |
| 1 | 239352713 | rs17287547 | -0.015 | -0.552 | 0.464 | -0.076 | -0.011 | -0.004 | -0.004 | 0 | 0.19 | 0.28 | 0.21 | 0.17 | 0.13 | 0.1 |
| 1 | 239356155 | rs2039801 | 0.612 | -0.619 | 0.648 | -0.119 | 0.007 | -0.069 | -0.069 | 0 | 0.17 | 0.44 | 0.14 | 0.05 | 0.13 | 0.09 |
| 1 | 239359436 | rs10802757 | -0.018 | -0.585 | 0.634 | -0.063 | -0.006 | -0.002 | -0.002 | 0 | 0.19 | 0.22 | 0.2 | 0.17 | 0.13 | 0.1 |
| 1 | 239371117 | rs17642796 | -0.222 | -0.517 | 0.944 | -0.477 | 0.005 | 0.07 | 0.07 | 0.007 | 0.21 | 0.08 | 0.21 | 0.06 | 0.2 | 0.13 |
| 1 | 239377627 | rs7521497 | -0.528 | -0.579 | 0.995 | -0.51 | -0.003 | 0.083 | 0.083 | 0.011 | 0.33 | 0.17 | 0.31 | 0.1 | 0.31 | 0.22 |
| 1 | 239382027 | rs2790661 | -1.312 | -0.6 | 1.159 | -0.518 | 0.034 | 0.128 | 0.128 | 0.48 | 0.42 | 0.79 | 0.38 | 0.1 | 0.33 | 0.24 |
| 1 | 239383418 | rs16838036 | 1.015 | -0.332 | 0.727 | -0.175 | -0.031 | -0.049 | -0.049 | 0 | 0.15 | 0.07 | 0.23 | 0.42 | 0.22 | 0.23 |
| 1 | 239384288 | rs2105233 | -1.171 | -0.484 | 1.175 | -0.607 | -0.008 | -0.075 | -0.075 | 0 | 0.39 | 0.48 | 0.35 | 0.1 | 0.34 | 0.25 |
| 1 | 239397104 | rs1339738 | -1.906 | -0.577 | 0.896 | -0.434 | -0.024 | -0.044 | -0.044 | 0 | 0.89 | 1 | 0.95 | 0.87 | 0.9 | 0.9 |
| 1 | 239414759 | rs2790645 | -0.089 | -0.495 | 1.372 | -0.679 | -0.023 | -0.095 | -0.095 | 0 | 0.22 | 0.5 | 0.18 | 0.06 | 0.16 | 0.13 |
| 1 | 239423655 | rs2790634 | -0.02 | -0.497 | 1.384 | -0.592 | -0.02 | -0.055 | -0.055 | 0.002 | 0.22 | 0.46 | 0.17 | 0.11 | 0.16 | 0.13 |
| 1 | 239428730 | rs2790630 | -1.425 | -0.538 | 1.316 | -0.911 | 0.007 | 0.015 | 0.015 | 0.148 | 0.4 | 0.66 | 0.36 | 0.12 | 0.33 | 0.25 |
| 1 | 239430497 | rs9428826 | -0.055 | -0.458 | 1.191 | -0.819 | -0.022 | -0.091 | -0.091 | 0.095 | 0.22 | 0.49 | 0.17 | 0.06 | 0.16 | 0.15 |
| 1 | 239433828 | rs2790624 | 0.701 | -0.415 | 0.865 | -0.795 | 0 | -0.004 | -0.004 | 0 | 0.17 | 0.18 | 0.13 | 0.07 | 0.14 | 0.13 |
| 1 | 239436876 | rs2820040 | -0.045 | -0.451 | 0.931 | -0.819 | -0.016 | -0.066 | -0.066 | 0 | 0.22 | 0.41 | 0.17 | 0.06 | 0.17 | 0.15 |
| 1 | 239439085 | rs1416824 | -0.263 | -0.331 | 0.884 | -0.653 | -0.003 | -0.036 | -0.036 | 0 | 0.18 | 0.25 | 0.19 | 0.06 | 0.16 | 0.1 |
| 1 | 239446740 | rs2820050 | -1.145 | -0.389 | 0.688 | -0.66 | -0.014 | -0.054 | -0.054 | 0 | 0.26 | 0.7 | 0.25 | 0.07 | 0.19 | 0.12 |

| | | | | | | | | | | | | | | | | |
|---|---|---|---|---|---|---|---|---|---|---|---|---|---|---|---|---|
| 1 | 239453106 | rs16838192 | -0.202 | -0.194 | 0.216 | -0.535 | 0.004 | 0.026 | 0.026 | 0 | 0.17 | 0.15 | 0.18 | 0.07 | 0.17 | 0.1 |
| 1 | 239466347 | rs4568821 | -0.506 | -0.329 | -0.047 | -0.521 | 0.012 | 0.015 | 0.015 | 0.886 | 0.61 | 0.71 | 0.77 | 0.72 | 0.7 | 0.59 |
| 1 | 239468210 | rs10802760 | 0.459 | -0.309 | -0.167 | -0.516 | 0.018 | 0.04 | 0.04 | 1.757 | 0.37 | 0.1 | 0.21 | 0.28 | 0.3 | 0.41 |
| 1 | 239469734 | rs10925867 | 0.686 | 0.357 | -0.024 | -0.707 | 0.021 | -0.036 | -0.036 | 0 | 0.15 | 0.22 | 0.13 | 0.03 | 0.22 | 0.19 |
| 1 | 239473113 | rs10495435 | -0.002 | 0.4 | -0.094 | -0.786 | -0.001 | 0.001 | 0.001 | 0.326 | 0.42 | 0.46 | 0.62 | 0.69 | 0.48 | 0.39 |
| 1 | 239473965 | rs10495434 | 0.486 | 0.662 | -0.285 | -0.192 | 0.045 | 0.106 | 0.106 | 0 | 0.07 | 0.17 | 0.1 | 0.25 | 0.08 | 0.07 |
| 1 | 239474391 | rs17583977 | 1.459 | 0.444 | -0.695 | -0.611 | -0.004 | 0.005 | 0.005 | 0 | 0.1 | 0.07 | 0.14 | 0.03 | 0.1 | 0.04 |
| 1 | 239475761 | rs10495432 | 1.349 | 0.245 | -0.834 | -0.501 | -0.051 | -0.161 | -0.161 | 0 | 0.24 | 0.05 | 0.38 | 0.41 | 0.29 | 0.28 |
| 1 | 239475925 | rs10495431 | 0.266 | 0.51 | -0.5 | -0.644 | 0.01 | -0.031 | -0.031 | 0 | 0.15 | 0.2 | 0.12 | 0.03 | 0.2 | 0.19 |
| 1 | 239491288 | rs4659909 | -0.058 | -0.059 | -0.837 | -0.651 | -0.016 | 0.009 | 0.009 | 0 | 0.16 | 0.09 | 0.3 | 0.01 | 0.22 | 0.25 |
| 1 | 239491834 | rs6661899 | 0.973 | -0.251 | -0.949 | -0.536 | -0.001 | -0.001 | -0.001 | 0.02 | 0.66 | 0.39 | 0.59 | 0.69 | 0.72 | 0.64 |
| 1 | 239495889 | rs4390163 | 0.803 | -0.001 | -1.175 | -0.5 | -0.133 | 0.04 | 0.04 | 1.894 | 0.29 | 0.04 | 0.18 | 0.16 | 0.37 | 0.36 |
| 1 | 239498465 | rs4573503 | 1.173 | 0.069 | 0.218 | 0.097 | -0.002 | -0.003 | -0.003 | 0 | 0.69 | 0.42 | 0.62 | 0.69 | 0.74 | 0.64 |
| 1 | 239498915 | rs9428828 | 0.982 | 0.494 | 0.302 | 0.417 | 0.018 | 0.02 | 0.02 | 0 | 0.76 | 0.57 | 0.65 | 0.71 | 0.78 | 0.68 |
| 1 | 239510175 | rs7551236 | 0.108 | 0.822 | 0.31 | 0.341 | 0.051 | -0.06 | -0.06 | 0 | 0.75 | 0.54 | 0.8 | 0.84 | 0.73 | 0.76 |
| 1 | 239519066 | rs12036333 | 0.454 | 0.285 | -1.383 | 0.773 | -0.029 | -0.123 | -0.123 | 0 | 0.18 | 0.01 | 0.32 | 0.35 | 0.19 | 0.15 |
| 1 | 239519108 | rs6700381 | 0.502 | 0.495 | -0.551 | 0.458 | -0.001 | -0.004 | -0.004 | 0.748 | 0.72 | 0.32 | 0.72 | 0.6 | 0.74 | 0.66 |
| 1 | 239530742 | rs1317950 | -0.35 | -0.016 | -0.755 | 0.067 | -0.018 | 0.089 | 0.089 | 0 | 0.34 | 0.17 | 0.25 | 0.14 | 0.35 | 0.23 |
| 1 | 239535513 | rs4497194 | -1.971 | -0.572 | 0.821 | 0.862 | -0.022 | 0.062 | 0.062 | 0 | 0.15 | 0.35 | 0.11 | 0.26 | 0.1 | 0.16 |
| 1 | 239536074 | rs2354396 | -1.776 | -0.584 | 0.824 | 0.822 | -0.036 | 0.054 | 0.054 | 0.12 | 0.13 | 0.34 | 0.1 | 0.22 | 0.06 | 0.12 |
| 1 | 239538192 | rs11808381 | -1.525 | -0.53 | 0.493 | 0.87 | -0.024 | 0.028 | 0.028 | 0 | 0.11 | 0.19 | 0.08 | 0.22 | 0.06 | 0.08 |
| 1 | 239559348 | rs4130463 | -1.429 | -0.66 | 0.604 | 0.863 | -0.03 | 0.07 | 0.07 | 0 | 0.16 | 0.32 | 0.13 | 0.35 | 0.09 | 0.12 |
| 1 | 239564682 | rs4659544 | 1.458 | -0.765 | 0.739 | 0.802 | -0.057 | 0.093 | 0.093 | 0 | 0.81 | 0.47 | 0.85 | 0.68 | 0.91 | 0.84 |
| 1 | 239567065 | rs10925877 | -0.294 | -1.601 | -0.247 | 0.241 | 0.046 | -0.001 | -0.001 | 1.211 | 0.22 | 0.08 | 0.14 | 0.24 | 0.14 | 0.17 |
| 1 | 239570932 | rs12123857 | -0.177 | -0.835 | -0.53 | -0.464 | -0.02 | 0.018 | 0.018 | 0 | 0.19 | 0.18 | 0.15 | 0.1 | 0.27 | 0.19 |
| 1 | 239574661 | rs11577797 | 0.075 | -1.027 | -0.199 | -0.9 | -0.001 | -0.004 | -0.004 | 0.011 | 0.26 | 0.26 | 0.18 | 0.19 | 0.21 | 0.3 |
| 1 | 239575774 | rs6678395 | -0.092 | -1.075 | 0.158 | -0.853 | 0.01 | 0.015 | 0.015 | 0 | 0.27 | 0.2 | 0.17 | 0.26 | 0.21 | 0.31 |
| 1 | 239581298 | rs12137225 | -0.778 | -1.116 | 0.307 | -0.912 | -0.009 | 0.023 | 0.023 | 0 | 0.29 | 0.29 | 0.2 | 0.12 | 0.34 | 0.31 |
| 1 | 239597345 | rs6671410 | -1.134 | -1.562 | 0.352 | -1.168 | 0.013 | -0.002 | -0.002 | 0 | 0.66 | 0.64 | 0.4 | 0.39 | 0.58 | 0.68 |
| 1 | 239601307 | rs6676664 | -0.452 | -1.766 | 0.425 | -1.31 | -0.019 | -0.024 | -0.024 | 0 | 0.16 | 0.29 | 0.06 | 0.11 | 0.08 | 0.16 |
| 1 | 239601676 | rs6687984 | -1.085 | -1.649 | 0.335 | -1.402 | -0.002 | -0.015 | -0.015 | 0 | 0.33 | 0.44 | 0.2 | 0.21 | 0.32 | 0.37 |
| 1 | 239606858 | rs17645304 | 0.461 | -1.498 | 0.139 | -1.354 | -0.073 | -0.002 | -0.002 | 0 | 0.18 | 0.01 | 0.12 | 0.1 | 0.23 | 0.2 |
| 1 | 239612814 | rs12086449 | 0.384 | -1.371 | 0.53 | -1.215 | 0.014 | 0.029 | 0.029 | 2.345 | 0.41 | 0.35 | 0.26 | 0.26 | 0.33 | 0.4 |
| 1 | 239617831 | rs10925888 | 1.141 | -1.34 | 0.312 | -1.231 | -0.064 | -0.023 | -0.023 | 0.024 | 0.32 | 0.11 | 0.58 | 0.6 | 0.43 | 0.34 |
| 1 | 239622971 | rs12048639 | 0.859 | -1.311 | 0.159 | -1.206 | -0.015 | 0.087 | 0.087 | 1.271 | 0.34 | 0.02 | 0.2 | 0.21 | 0.27 | 0.29 |
| 1 | 239626368 | rs16838363 | 0.922 | -1.231 | 0.33 | -1.163 | -0.021 | -0.029 | -0.029 | 0 | 0.07 | 0.17 | 0.03 | 0.04 | 0.03 | 0.06 |
| 1 | 239626755 | rs10925889 | 0.633 | -1.117 | 0.067 | -1.061 | 0.041 | 0.106 | 0.106 | 1.07 | 0.37 | 0.15 | 0.23 | 0.21 | 0.28 | 0.36 |
| 1 | 239658863 | rs1796835 | -0.526 | -0.208 | 0.262 | -0.358 | 0.014 | 0 | 0 | 0 | 0.67 | 0.65 | 0.41 | 0.37 | 0.56 | 0.59 |
| 1 | 239676235 | rs1782349 | -0.741 | -0.033 | -0.036 | -0.188 | -0.001 | -0.015 | -0.015 | 0 | 0.08 | 0.17 | 0.07 | 0.07 | 0.08 | 0.11 |
| 1 | 239677004 | rs1782357 | 0.096 | -0.057 | -0.025 | -0.264 | 0.007 | -0.004 | -0.004 | 0 | 0.62 | 0.4 | 0.41 | 0.36 | 0.55 | 0.48 |
| 1 | 239683556 | rs12041334 | 0.847 | 0.405 | -0.68 | 0.325 | -0.12 | -0.164 | -0.164 | 0 | 0.22 | 0.01 | 0.39 | 0.49 | 0.35 | 0.25 |
| 1 | 239690844 | rs1218671 | 0.87 | 0.002 | -0.595 | -0.076 | 0.039 | 0.072 | 0.072 | 0.155 | 0.42 | 0.23 | 0.27 | 0.24 | 0.33 | 0.25 |
| 1 | 239701246 | rs16838444 | -0.109 | 0.551 | -0.031 | 0.239 | 0.02 | -0.019 | -0.019 | 0 | 0.06 | 0.14 | 0.12 | 0.01 | 0.07 | 0.11 |
| 1 | 239706595 | rs10925910 | 1.531 | 0.533 | 0.185 | 0.43 | 0.029 | -0.052 | -0.052 | 0 | 0.75 | 0.07 | 0.73 | 0.76 | 0.83 | 0.67 |
| 1 | 239713659 | rs10925915 | 0.253 | 1.005 | -0.04 | 0.372 | -0.109 | -0.097 | -0.097 | 0.181 | 0.27 | 0.08 | 0.46 | 0.58 | 0.45 | 0.4 |
| 1 | 239714774 | rs1992547 | -1.012 | 0.918 | 0.189 | 0.298 | -0.004 | -0.018 | -0.018 | 0 | 0.94 | 0.95 | 0.99 | 1 | 0.98 | 0.89 |
| 1 | 239719817 | rs12074530 | 0.185 | 1.093 | 0.294 | 0.374 | 0.032 | -0.016 | -0.016 | 0 | 0.07 | 0.14 | 0.14 | 0.01 | 0.1 | 0.1 |
| 1 | 239725150 | rs11804608 | -0.508 | 0.958 | 0.627 | 0.365 | 0.008 | -0.004 | -0.004 | 0.291 | 0.35 | 0.34 | 0.51 | 0.7 | 0.47 | 0.47 |
| 1 | 239725588 | rs6670728 | -0.243 | 1.165 | 0.314 | 0.388 | -0.017 | 0.008 | 0.008 | 0 | 0.89 | 0.68 | 0.94 | 0.88 | 0.93 | 0.83 |
| 1 | 239726247 | rs1984207 | 0.177 | 1.294 | -0.541 | 0.296 | -0.111 | -0.095 | -0.095 | 0 | 0.27 | 0.08 | 0.47 | 0.58 | 0.45 | 0.4 |
| 1 | 239743257 | rs12135445 | 0.269 | 1.315 | 0.029 | 0.743 | -0.056 | -0.039 | -0.039 | 0 | 0.09 | 0.01 | 0.11 | 0.12 | 0.18 | 0.17 |
| 1 | 239746732 | rs10127769 | 0.76 | 0.645 | 0.422 | 0.666 | -0.036 | 0.012 | 0.012 | 1.925 | 0.85 | 0.29 | 0.91 | 0.83 | 0.93 | 0.83 |
| 1 | 239760713 | rs10802779 | 0.603 | 0.663 | -0.046 | 0.674 | -0.031 | 0.06 | 0.06 | 1.091 | 0.86 | 0.48 | 0.93 | 0.83 | 0.93 | 0.83 |
| 1 | 239768800 | rs6429140 | 1.166 | 0.761 | -1.458 | 0.585 | -0.003 | -0.096 | -0.096 | 0 | 0.72 | 0.04 | 0.72 | 0.71 | 0.81 | 0.66 |
| 1 | 239774393 | rs17594385 | -0.223 | 1.244 | -1.058 | 0.404 | -0.027 | -0.02 | -0.02 | 0 | 0.06 | 0.01 | 0.09 | 0.09 | 0.14 | 0.09 |
| 1 | 239774670 | rs6685121 | -1.013 | 1.002 | -0.316 | -0.082 | -0.026 | -0.088 | -0.088 | 0 | 0.15 | 0.21 | 0.04 | 0.01 | 0.08 | 0.25 |
| 1 | 239775817 | rs6688669 | -1.475 | 1.255 | -0.554 | -0.157 | -0.008 | 0.004 | 0.004 | 0.156 | 0.3 | 0.74 | 0.28 | 0.26 | 0.35 | 0.22 |
| 1 | 239778660 | rs12080951 | -0.69 | 1.368 | 0.41 | -0.179 | 0.066 | -0.02 | -0.02 | 0 | 0.12 | 0.27 | 0.14 | 0.06 | 0.2 | 0.08 |
| 1 | 239781166 | rs6690612 | -1.247 | 1.313 | 0.732 | -0.122 | 0.102 | 0.086 | 0.086 | 0.593 | 0.14 | 0.48 | 0.21 | 0.18 | 0.21 | 0.11 |
| 1 | 239784089 | rs12070319 | -1.49 | 1.25 | 0.663 | -0.329 | 0.02 | -0.033 | -0.033 | 2.387 | 0.18 | 0.27 | 0.16 | 0.08 | 0.24 | 0.29 |
| 1 | 239784537 | rs2355228 | -0.532 | 1.279 | 0.672 | -0.323 | 0.033 | -0.014 | -0.014 | 0 | 0.11 | 0.21 | 0.12 | 0.07 | 0.18 | 0.08 |
| 1 | 239785165 | rs2355230 | -1.818 | 1.093 | -0.399 | -0.697 | -0.045 | 0.092 | 0.092 | 0.114 | 0.61 | 0.98 | 0.67 | 0.81 | 0.66 | 0.65 |
| 1 | 239794364 | rs726168 | -0.288 | 0.944 | -0.22 | -0.841 | -0.017 | -0.06 | -0.06 | 0.627 | 0.15 | 0.21 | 0.13 | 0.01 | 0.09 | 0.09 |
| 1 | 239794640 | rs12088787 | 0.768 | 0.93 | -0.323 | -0.842 | 0.018 | 0.051 | 0.051 | 2.698 | 0.1 | 0.03 | 0.03 | 0 | 0.07 | 0.07 |
| 1 | 239798412 | rs12037424 | 0.953 | 0.846 | -0.36 | -0.972 | -0.033 | -0.059 | -0.059 | 0.711 | 0.24 | 0.11 | 0.31 | 0.65 | 0.27 | 0.27 |
| 1 | 239807920 | rs1867263 | -1.549 | 0.773 | -0.167 | -1.024 | 0.004 | 0.023 | 0.023 | 0.26 | 0.59 | 0.68 | 0.77 | 0.86 | 0.61 | 0.67 |
| 1 | 239811709 | rs16832152 | 0.981 | 0.788 | -0.165 | -1.011 | -0.018 | -0.041 | -0.041 | 0.2 | 0.25 | 0.15 | 0.32 | 0.66 | 0.26 | 0.27 |
| 1 | 239812103 | rs6691263 | -0.703 | 0.777 | -0.059 | -1.027 | 0.009 | 0.002 | 0.002 | 0 | 0.19 | 0.25 | 0.32 | 0.2 | 0.24 | 0.3 |
| 1 | 239812538 | rs10925941 | -1.346 | 0.779 | -0.102 | -1.017 | 0.001 | -0.007 | -0.007 | 0 | 0.54 | 0.57 | 0.69 | 0.85 | 0.58 | 0.65 |

| | | | | | | | | | | | | | | | | |
|---|---|---|---|---|---|---|---|---|---|---|---|---|---|---|---|---|
| 1 | 239813953 | rs12090480 | -1.511 | 1.031 | 0.065 | -0.827 | 0.002 | -0.029 | -0.029 | 0 | 0.3 | 0.41 | 0.37 | 0.2 | 0.31 | 0.38 |
| 1 | 239819503 | rs4659550 | -1.36 | 0.879 | 0.171 | -1.012 | -0.002 | -0.042 | -0.042 | 0 | 0.57 | 0.6 | 0.78 | 0.86 | 0.6 | 0.66 |
| 1 | 239832680 | rs10802789 | 0.092 | 1.103 | -0.146 | -0.769 | -0.011 | 0.004 | 0.004 | 0.654 | 0.32 | 0.29 | 0.57 | 0.73 | 0.44 | 0.3 |
| 1 | 239833100 | rs10754677 | -0.479 | 1.092 | -0.162 | -0.78 | -0.022 | 0.023 | 0.023 | 2.475 | 0.39 | 0.32 | 0.6 | 0.74 | 0.5 | 0.5 |
| 1 | 239839305 | rs1867266 | 0.695 | 1.312 | -0.157 | -0.566 | -0.026 | 0.026 | 0.026 | 0 | 0.06 | 0.01 | 0.03 | 0.01 | 0.07 | 0.21 |
| 1 | 239846380 | rs6692711 | 1.467 | 0.858 | -0.55 | -0.906 | -0.022 | 0.128 | 0.128 | 0 | 0.27 | 0.06 | 0.13 | 0.05 | 0.25 | 0.29 |
| 1 | 239853105 | rs12406493 | 0.334 | 1.098 | 0.275 | -0.112 | 0.001 | 0 | 0 | 0 | 0.54 | 0.48 | 0.57 | 0.48 | 0.51 | 0.49 |
| 1 | 239857992 | rs4145784 | 0.008 | 1.137 | 0.477 | -0.225 | 0.003 | -0.03 | -0.03 | 0 | 0.9 | 0.88 | 0.87 | 1 | 0.89 | 0.95 |
| 1 | 239867143 | rs2278642 | 0.653 | 0.967 | 0.59 | -0.005 | 0.019 | 0.013 | 0.013 | 0 | 0.47 | 0.33 | 0.45 | 0.4 | 0.43 | 0.45 |
| 1 | 239870621 | rs10802794 | 0.273 | 1.051 | 0.4 | 0.105 | -0.002 | 0.003 | 0.003 | 0 | 0.53 | 0.78 | 0.57 | 0.4 | 0.45 | 0.47 |
| 1 | 239877721 | rs6663632 | 0.402 | 1.028 | 0.297 | 0.119 | 0.007 | 0.01 | 0.01 | 0 | 0.52 | 0.59 | 0.55 | 0.4 | 0.46 | 0.47 |
| 1 | 239879553 | rs1431718 | -0.101 | 1.098 | 0.046 | 0.168 | -0.003 | -0.025 | -0.025 | 0.216 | 0.61 | 0.71 | 0.62 | 0.49 | 0.6 | 0.68 |
| 1 | 239916015 | rs12124903 | -0.489 | 0.874 | -0.535 | 0.433 | -0.019 | -0.011 | -0.011 | 0 | 0.07 | 0.02 | 0.05 | 0.09 | 0.1 | 0.2 |
| 1 | 239917787 | rs12126146 | -0.346 | 0.889 | -0.537 | 0.44 | -0.011 | -0.006 | -0.006 | 0 | 0.06 | 0.02 | 0.04 | 0.09 | 0.1 | 0.2 |
| 1 | 239924343 | rs685475 | 1.315 | 0.588 | 0.192 | 0.663 | 0.022 | 0.098 | 0.098 | 0 | 0.85 | 0.43 | 0.79 | 0.76 | 0.87 | 0.7 |
| 1 | 239924408 | rs685550 | 1.456 | -0.323 | -0.381 | -0.105 | -0.03 | -0.023 | -0.023 | 1.227 | 0.67 | 0.51 | 0.74 | 0.76 | 0.76 | 0.63 |
| 1 | 239924486 | rs685960 | 0.943 | -0.447 | -0.167 | -0.229 | -0.04 | -0.097 | -0.097 | 0 | 0.82 | 0.56 | 0.89 | 0.96 | 0.9 | 0.83 |
| 1 | 239924805 | rs843030 | -0.89 | -0.11 | -0.347 | -0.478 | 0.001 | -0.022 | -0.022 | 0 | 0.11 | 0.16 | 0.1 | 0.01 | 0.12 | 0.05 |
| 1 | 239924870 | rs6703930 | 0.812 | 0.148 | -0.641 | 0.032 | -0.002 | -0.003 | -0.003 | 1.09 | 0.54 | 0.51 | 0.64 | 0.63 | 0.5 | 0.63 |
| 1 | 239925109 | rs7533134 | -0.659 | -0.312 | -1.266 | 0.328 | 0.025 | -0.015 | -0.015 | 0 | 0.35 | 0.16 | 0.36 | 0.57 | 0.28 | 0.23 |
| 1 | 239927009 | rs17657156 | -0.252 | 0.055 | -0.806 | 0.297 | -0.003 | 0.017 | 0.017 | 0 | 0.09 | 0.04 | 0.04 | 0 | 0.1 | 0.1 |
| 1 | 239931618 | rs532718 | 0.53 | -0.223 | -0.607 | 0.83 | 0.009 | -0.021 | -0.021 | 0 | 0.69 | 0.75 | 0.72 | 0.46 | 0.75 | 0.66 |
| 1 | 239934541 | rs2841037 | -0.826 | -0.334 | -0.814 | 1.022 | -0.008 | -0.024 | -0.024 | 1.683 | 0.86 | 0.92 | 0.83 | 0.85 | 0.84 | 0.75 |
| 1 | 239935351 | rs663927 | 0.808 | -0.369 | -0.311 | 0.968 | -0.014 | 0.014 | 0.014 | 0 | 0.68 | 0.63 | 0.71 | 0.43 | 0.75 | 0.56 |
| 1 | 239936582 | rs481036 | 1.044 | -0.383 | -0.504 | 1.022 | -0.01 | 0.003 | 0.003 | 0 | 0.65 | 0.43 | 0.69 | 0.43 | 0.75 | 0.49 |
| 1 | 239945157 | rs534615 | -0.533 | 0.194 | -0.852 | 0.936 | -0.002 | -0.004 | -0.004 | 0 | 0.85 | 0.87 | 0.82 | 0.87 | 0.83 | 0.75 |
| 1 | 239946076 | rs626694 | -0.566 | 0.179 | -0.859 | 0.929 | -0.001 | -0.003 | -0.003 | 0 | 0.85 | 0.87 | 0.82 | 0.87 | 0.83 | 0.75 |
| 1 | 239955676 | rs693948 | -0.094 | 0.389 | -0.766 | 0.864 | 0.013 | 0.041 | 0.041 | 0 | 0.4 | 0.72 | 0.43 | 0.21 | 0.37 | 0.4 |
| 1 | 239962002 | rs665159 | -0.831 | 0.409 | -1.296 | 1.064 | -0.018 | 0.049 | 0.049 | 0.648 | 0.53 | 0.87 | 0.53 | 0.6 | 0.46 | 0.55 |
| 1 | 239962686 | rs2790336 | -0.103 | 0.456 | -0.758 | 0.326 | 0.001 | 0.002 | 0.002 | 0 | 0.37 | 0.39 | 0.4 | 0.21 | 0.36 | 0.39 |
| 1 | 239970097 | rs12059546 | -0.788 | 0.574 | -0.447 | 0.494 | 0.001 | -0.001 | -0.001 | 0 | 0.23 | 0.57 | 0.25 | 0.32 | 0.19 | 0.21 |
| 1 | 239971919 | rs558438 | 0.328 | 0.496 | -1.277 | 0.602 | -0.003 | -0.041 | -0.041 | 0 | 0.23 | 0.1 | 0.12 | 0.32 | 0.23 | 0.18 |
| 1 | 239974006 | rs6690809 | 0.877 | 0.305 | -1.47 | 0.297 | -0.059 | -0.014 | -0.014 | 0 | 0.46 | 0.17 | 0.58 | 0.35 | 0.53 | 0.59 |
| 1 | 239979186 | rs7543259 | 0.227 | -0.054 | -0.839 | 0.225 | -0.003 | 0.006 | 0.006 | 0.594 | 0.31 | 0.28 | 0.15 | 0.47 | 0.27 | 0.27 |
| 1 | 239981643 | rs6429157 | -0.883 | -0.142 | -0.929 | 0.134 | -0.004 | 0.037 | 0.037 | 1.49 | 0.52 | 0.7 | 0.4 | 0.65 | 0.46 | 0.49 |
| 1 | 239982638 | rs1578180 | -0.021 | -0.052 | -0.548 | 0.146 | 0.002 | -0.007 | -0.007 | 0 | 0.19 | 0.23 | 0.22 | 0.18 | 0.18 | 0.22 |
| 1 | 239983913 | rs4523505 | 0.129 | -0.32 | -0.866 | 0.416 | 0.002 | -0.002 | -0.002 | 0.005 | 0.08 | 0.06 | 0.03 | 0.18 | 0.04 | 0.09 |
| 1 | 239984051 | rs10802807 | 1.559 | 0.113 | -1.515 | 0.245 | -0.057 | -0.075 | -0.075 | 2.759 | 0.16 | 0.01 | 0.24 | 0.2 | 0.19 | 0.11 |
| 1 | 239984925 | rs1934349 | -1.026 | 0.162 | -1.494 | -0.293 | -0.055 | 0.087 | 0.087 | 0 | 0.68 | 0.86 | 0.63 | 0.87 | 0.57 | 0.6 |
| 1 | 239985876 | rs12072181 | 0.273 | 0.254 | -1.029 | 0.136 | 0.005 | -0.007 | -0.007 | 0 | 0.3 | 0.25 | 0.15 | 0.33 | 0.26 | 0.21 |
| 1 | 239988849 | rs7536133 | 0.483 | 0.784 | -0.87 | 0.46 | -0.004 | -0.005 | -0.005 | 0 | 0.18 | 0.2 | 0.26 | 0.52 | 0.2 | 0.19 |
| 1 | 239989964 | rs589962 | 0.164 | 1.144 | -1.007 | -0.187 | -0.054 | 0.066 | 0.066 | 0.852 | 0.22 | 0.1 | 0.37 | 0.09 | 0.31 | 0.12 |
| 1 | 239992286 | rs621060 | 0.213 | 0.945 | -0.102 | -0.323 | 0.073 | -0.095 | -0.095 | 0 | 0.86 | 0.51 | 0.89 | 1 | 0.86 | 0.97 |
| 1 | 239994906 | rs685548 | -0.516 | 0.687 | -0.168 | -0.1 | 0.001 | 0.001 | 0.001 | 0 | 0.55 | 0.47 | 0.62 | 0.43 | 0.62 | 0.65 |
| 1 | 240002419 | rs1304352 | -1.027 | 0.869 | 0.602 | 0.008 | 0.082 | 0.033 | 0.033 | 0.004 | 0.08 | 0.55 | 0.07 | 0.07 | 0.04 | 0.04 |
| 1 | 240006785 | rs602117 | -0.305 | 0.563 | 0.118 | 0.085 | -0.002 | 0.017 | 0.017 | 0 | 0.49 | 0.39 | 0.46 | 0.45 | 0.51 | 0.6 |
| 1 | 240009037 | rs650751 | 0.525 | 0.391 | 0.665 | -0.246 | 0.007 | 0.003 | 0.003 | 4.494 | 0.92 | 0.49 | 0.92 | 1 | 1 | 0.97 |
| 1 | 240011753 | rs1594513 | 1.034 | 0.765 | 0.039 | -0.27 | -0.079 | 0.019 | 0.019 | 0 | 0.2 | 0.01 | 0.25 | 0.11 | 0.28 | 0.12 |
| 1 | 240015308 | rs10925994 | 1.203 | 0.902 | 0.436 | 0.539 | -0.031 | -0.006 | -0.006 | 0 | 0.07 | 0.01 | 0.11 | 0.34 | 0.1 | 0.09 |
| 1 | 240025977 | rs497576 | 0.666 | 0.57 | 0.5 | 0.043 | -0.016 | 0.008 | 0.008 | 0 | 0.46 | 0.17 | 0.46 | 0.54 | 0.57 | 0.56 |
| 1 | 240030399 | rs682355 | 0.673 | 0.686 | 0.555 | 0.078 | -0.026 | 0.013 | 0.013 | 0 | 0.46 | 0.19 | 0.46 | 0.54 | 0.54 | 0.57 |
| 1 | 240045908 | rs536477 | 0.643 | 0.457 | 0.921 | -0.086 | 0.02 | 0.068 | 0.068 | 0 | 0.54 | 0.21 | 0.58 | 0.62 | 0.6 | 0.58 |
| 1 | 240046468 | rs10926002 | -0.035 | 0.268 | 1.101 | -0.108 | -0.023 | -0.025 | -0.025 | 0 | 0.06 | 0.21 | 0.03 | 0 | 0 | 0.02 |
| 1 | 240048298 | rs2217533 | 0.098 | 0.403 | 0.806 | -0.157 | -0.009 | 0.012 | 0.012 | 0 | 0.1 | 0.08 | 0.06 | 0.07 | 0.1 | 0.05 |
| 1 | 240051340 | rs10495447 | 0.303 | 0.417 | 0.734 | -0.114 | 0 | -0.007 | -0.007 | 0.854 | 0.29 | 0.08 | 0.28 | 0.31 | 0.29 | 0.35 |
| 1 | 240055553 | rs10926004 | -0.732 | 0.578 | -0.252 | 0.136 | 0.019 | 0.053 | 0.053 | 0 | 0.49 | 0.66 | 0.41 | 0.38 | 0.43 | 0.43 |
| 1 | 240060328 | rs16839034 | -0.378 | 0.306 | -0.075 | 0.737 | -0.006 | 0.004 | 0.004 | 0.01 | 0.16 | 0.18 | 0.06 | 0.26 | 0.11 | 0.29 |
| 1 | 240061728 | rs16839045 | -0.815 | 0.839 | 0.106 | 0.756 | 0.057 | 0.046 | 0.046 | 0 | 0.05 | 0.18 | 0.07 | 0.07 | 0.07 | 0.03 |
| 1 | 240062123 | rs10926008 | -1.215 | 0.759 | -0.37 | 0.614 | 0.01 | -0.016 | -0.016 | 0.071 | 0.39 | 0.7 | 0.31 | 0.43 | 0.35 | 0.48 |
| 1 | 240063366 | rs16839051 | -1.874 | 0.89 | 0.41 | 0.68 | 0.046 | 0.137 | 0.137 | 0 | 0.08 | 0.27 | 0.09 | 0.18 | 0.11 | 0.08 |
| 1 | 240063699 | rs10926009 | -2.001 | 0.754 | 0.333 | 0.436 | 0.002 | 0.003 | 0.003 | 3.919 | 0.1 | 0.67 | 0.14 | 0.1 | 0.08 | 0.07 |
| 1 | 240063821 | rs4620530 | 0.578 | 0.953 | -1.135 | -0.178 | -0.061 | 0.14 | 0.14 | 0.326 | 0.57 | 0.08 | 0.53 | 0.77 | 0.58 | 0.69 |
| 1 | 240064538 | rs10399860 | -1.538 | 1.084 | -0.24 | 0.517 | 0 | 0.012 | 0.012 | 0.178 | 0.28 | 0.69 | 0.33 | 0.5 | 0.29 | 0.33 |
| 1 | 240065878 | rs12036109 | -1.507 | 1.185 | -0.503 | 0.2 | -0.001 | -0.004 | -0.004 | 0 | 0.44 | 0.59 | 0.49 | 0.49 | 0.45 | 0.39 |
| 1 | 240067260 | rs7520974 | -1.642 | 1.275 | -0.585 | 0.132 | -0.003 | -0.003 | -0.003 | 0 | 0.49 | 0.71 | 0.56 | 0.49 | 0.52 | 0.5 |
| 1 | 240070187 | rs6701181 | -1.508 | 1.315 | -1.022 | 0.148 | 0.006 | -0.011 | -0.011 | 0.469 | 0.45 | 0.69 | 0.47 | 0.49 | 0.43 | 0.39 |
| 1 | 240074981 | rs6682184 | -1.455 | 0.789 | -1.415 | 0.464 | 0 | -0.08 | -0.08 | 0.044 | 0.39 | 0.86 | 0.52 | 0.52 | 0.36 | 0.44 |
| 1 | 240076608 | rs10495449 | -0.494 | 0.781 | -1.196 | 0.572 | 0.019 | -0.01 | -0.01 | 0 | 0.18 | 0.66 | 0.25 | 0.14 | 0.19 | 0.22 |
| 1 | 240077023 | rs4431831 | -0.053 | 0.71 | -1.174 | 0.642 | -0.001 | 0.005 | 0.005 | 0 | 0.19 | 0.19 | 0.25 | 0.38 | 0.15 | 0.19 |

| | | | | | | | | | | | | | | | | |
|---|---|---|---|---|---|---|---|---|---|---|---|---|---|---|---|---|
| 1 | 240077309 | rs4659554 | -1.125 | 1.383 | -0.571 | 0.029 | 0.01 | -0.015 | -0.015 | 0.125 | 0.23 | 0.67 | 0.27 | 0.14 | 0.25 | 0.25 |
| 1 | 240091765 | rs6666471 | -0.446 | 0.955 | -0.763 | 0.018 | -0.002 | -0.007 | -0.007 | 3.067 | 0.43 | 0.55 | 0.48 | 0.61 | 0.41 | 0.53 |
| 1 | 240094975 | rs7531091 | -0.116 | 0.177 | -1.372 | -0.529 | 0.097 | 0.076 | 0.076 | 0 | 0.57 | 0.82 | 0.55 | 0.29 | 0.75 | 0.47 |
| 1 | 240095692 | rs10926018 | 0.152 | 0.537 | -0.859 | -0.572 | 0.015 | -0.008 | -0.008 | 0 | 0.06 | 0.11 | 0.03 | 0.04 | 0.07 | 0.06 |
| 1 | 240098415 | rs10081951 | -0.484 | -0.044 | -0.822 | -0.893 | 0.037 | 0.016 | 0.016 | 0 | 0.63 | 0.7 | 0.6 | 0.29 | 0.84 | 0.51 |
| 1 | 240102885 | rs17606620 | 0.533 | 1.321 | -0.714 | -1.373 | 0.044 | -0.044 | -0.044 | 0 | 0.12 | 0.17 | 0.23 | 0 | 0.31 | 0.08 |
| 1 | 240103855 | rs10926022 | 0.303 | 0.875 | -0.638 | -1.317 | -0.001 | 0.003 | 0.003 | 0 | 0.17 | 0.16 | 0.12 | 0.05 | 0.18 | 0.17 |
| 1 | 240107121 | rs12742834 | 0.51 | 0.863 | -0.793 | -1.227 | 0.006 | 0.036 | 0.036 | 0 | 0.3 | 0.2 | 0.2 | 0.13 | 0.31 | 0.21 |
| 1 | 240108543 | rs12563039 | -0.13 | 0.97 | -1.018 | -0.805 | 0 | 0.031 | 0.031 | 0 | 0.51 | 0.53 | 0.36 | 0.22 | 0.53 | 0.43 |
| 1 | 240111019 | rs10926023 | -0.321 | 0.602 | -1.396 | 0.376 | 0.003 | -0.058 | -0.058 | 0 | 0.21 | 0.13 | 0.36 | 0.44 | 0.19 | 0.17 |
| 1 | 240112465 | rs10926024 | -0.575 | 0.907 | -1.529 | 0.27 | -0.017 | 0.006 | 0.006 | 0 | 0.72 | 0.8 | 0.61 | 0.74 | 0.65 | 0.68 |
| 1 | 240112614 | rs10926025 | 0.472 | 0.501 | -1.862 | 0.183 | -0.044 | 0.139 | 0.139 | 0.4 | 0.45 | 0.12 | 0.32 | 0.27 | 0.47 | 0.38 |
| 1 | 240113107 | rs1538025 | 0.023 | 0.383 | -1.73 | 0.283 | -0.004 | 0.011 | 0.011 | 1.825 | 0.27 | 0.66 | 0.29 | 0.47 | 0.18 | 0.31 |
| 1 | 240113513 | rs12121905 | 1.095 | 0.511 | -2.295 | -0.163 | -0.014 | 0.135 | 0.135 | 0.028 | 0.21 | 0.01 | 0.06 | 0.02 | 0.15 | 0.16 |
| 1 | 240114540 | rs1471351 | 1.351 | 0.482 | -2.051 | 0.271 | -0.013 | -0.072 | -0.072 | 0 | 0.15 | 0.03 | 0.11 | 0.33 | 0.13 | 0.11 |
| 1 | 240117721 | rs10926030 | 1.679 | 0.456 | -2.079 | 0.034 | -0.003 | -0.004 | -0.004 | 0 | 0.16 | 0.17 | 0.22 | 0.24 | 0.23 | 0.17 |
| 1 | 240118265 | rs1984165 | -1.649 | 0.461 | -2.23 | 0.009 | -0.042 | -0.015 | -0.015 | 0 | 0.73 | 0.95 | 0.63 | 0.73 | 0.69 | 0.69 |
| 1 | 240119053 | rs6696393 | -0.612 | 0.387 | -1.181 | -0.141 | -0.006 | 0.025 | 0.025 | 0 | 0.53 | 0.6 | 0.49 | 0.4 | 0.52 | 0.58 |
| 1 | 240121383 | rs10926034 | 0.985 | 0.362 | -1.044 | -0.161 | 0.004 | 0.068 | 0.068 | 0.004 | 0.19 | 0.32 | 0.15 | 0.32 | 0.17 | 0.11 |
| 1 | 240121828 | rs7544436 | 0.224 | 0.288 | -0.963 | -0.317 | -0.027 | -0.068 | -0.068 | 0.366 | 0.29 | 0.41 | 0.21 | 0.16 | 0.19 | 0.38 |
| 1 | 240122305 | rs1903630 | -0.665 | 0.362 | -1.091 | -0.268 | 0.03 | -0.002 | -0.002 | 0 | 0.48 | 0.73 | 0.36 | 0.48 | 0.35 | 0.48 |
| 1 | 240124315 | rs4659556 | 0.796 | 0.52 | -0.967 | -0.132 | -0.01 | 0.01 | 0.01 | 0 | 0.24 | 0.19 | 0.27 | 0.2 | 0.33 | 0.21 |
| 1 | 240127293 | rs1124375 | 1.064 | 0.388 | -0.003 | -0.14 | 0.014 | 0.072 | 0.072 | 0.343 | 0.08 | 0.28 | 0.14 | 0.14 | 0.04 | 0.21 |
| 1 | 240130042 | rs12135917 | 1.101 | 0.353 | 0.415 | -0.183 | 0.004 | 0.061 | 0.061 | 0 | 0.18 | 0.3 | 0.15 | 0.32 | 0.17 | 0.1 |
| 1 | 240130088 | rs10926042 | 0.824 | 0.347 | 0.366 | -0.233 | 0.006 | -0.01 | -0.01 | 0 | 0.25 | 0.3 | 0.28 | 0.19 | 0.33 | 0.2 |
| 1 | 240131535 | rs1415278 | -1.018 | 0.328 | 0.333 | -0.228 | -0.017 | 0.104 | 0.104 | 0.177 | 0.51 | 0.92 | 0.57 | 0.65 | 0.54 | 0.52 |
| 1 | 240132039 | rs10926043 | 0.151 | 0.329 | 0.506 | -0.203 | -0.003 | 0.081 | 0.081 | 0.752 | 0.25 | 0.42 | 0.28 | 0.45 | 0.21 | 0.31 |
| 1 | 240133746 | rs12077571 | -0.127 | 0.355 | 0.488 | -0.282 | -0.001 | 0.002 | 0.002 | 0 | 0.32 | 0.68 | 0.42 | 0.33 | 0.38 | 0.42 |
| 1 | 240135971 | rs12133285 | -0.533 | 0.251 | 0.499 | -0.455 | -0.003 | -0.022 | -0.022 | 1.01 | 0.34 | 0.36 | 0.3 | 0.19 | 0.29 | 0.4 |
| 1 | 240136510 | rs16839215 | 1.412 | 0.236 | 0.62 | -0.426 | -0.004 | 0.03 | 0.03 | 0 | 0.07 | 0.16 | 0.18 | 0.14 | 0.04 | 0.19 |
| 1 | 240137682 | rs12743220 | 1.801 | 0.207 | 0.544 | -0.366 | -0.032 | -0.08 | -0.08 | 1.488 | 0.15 | 0.04 | 0.12 | 0.33 | 0.17 | 0.11 |
| 1 | 240137811 | rs10495451 | -0.305 | 0.205 | 0.664 | -0.394 | 0.006 | 0.027 | 0.027 | 0 | 0.34 | 0.28 | 0.29 | 0.19 | 0.28 | 0.4 |
| 1 | 240140563 | rs1381526 | -2.753 | 0.321 | 0.226 | -0.632 | -0.036 | -0.069 | -0.069 | 0.484 | 0.79 | 0.99 | 0.75 | 0.7 | 0.8 | 0.73 |
| 1 | 240140699 | rs12138402 | 1.782 | 0.31 | 0.206 | -0.657 | -0.024 | 0.071 | 0.071 | 0 | 0.19 | 0.01 | 0.06 | 0.05 | 0.14 | 0.16 |
| 1 | 240140986 | rs12119833 | 0.079 | 0.352 | 0.411 | -0.484 | -0.077 | -0.098 | -0.098 | 0 | 0.42 | 0.07 | 0.23 | 0.38 | 0.4 | 0.32 |
| 1 | 240141393 | rs11589497 | 1.441 | 0.413 | 0.463 | -0.498 | -0.001 | -0.013 | -0.013 | 0 | 0.09 | 0.05 | 0.1 | 0 | 0.11 | 0.05 |
| 1 | 240146525 | rs1415277 | -1.148 | 0.393 | 0.574 | -0.468 | 0 | -0.002 | -0.002 | 0 | 0.6 | 0.61 | 0.54 | 0.53 | 0.57 | 0.55 |
| 1 | 240147503 | rs1337088 | -2.27 | 0.476 | 0.395 | -0.458 | -0.035 | -0.068 | -0.068 | 0 | 0.79 | 0.98 | 0.75 | 0.7 | 0.8 | 0.73 |
| 1 | 240152077 | rs2169454 | -0.526 | 0.542 | 1.203 | -0.207 | 0.008 | -0.024 | -0.024 | 0 | 0.17 | 0.36 | 0.3 | 0.14 | 0.16 | 0.24 |
| 1 | 240157086 | rs7539606 | 0.731 | 0.583 | 0.585 | -0.195 | 0.002 | -0.03 | -0.03 | 0 | 0.09 | 0.08 | 0.12 | 0 | 0.12 | 0.05 |
| 1 | 240158598 | rs4659940 | 0.145 | 0.611 | 0.543 | -0.143 | 0 | -0.018 | -0.018 | 0 | 0.16 | 0.24 | 0.3 | 0.14 | 0.16 | 0.23 |
| 1 | 240164711 | rs6675551 | -0.027 | 0.655 | 0.33 | 0.018 | 0.005 | 0.094 | 0.094 | 0 | 0.22 | 0.4 | 0.12 | 0.51 | 0.19 | 0.25 |
| 1 | 240166887 | rs10802823 | -0.198 | 0.43 | -0.057 | -0.736 | 0.008 | 0.013 | 0.013 | 0 | 0.31 | 0.28 | 0.13 | 0.05 | 0.24 | 0.21 |
| 1 | 240167654 | rs7545703 | -0.252 | 0.428 | 0.121 | -0.688 | 0 | -0.001 | -0.001 | 0 | 0.35 | 0.34 | 0.24 | 0.21 | 0.36 | 0.37 |
| 1 | 240170796 | rs2065914 | -0.326 | 0.644 | 0.285 | -0.739 | 0.002 | -0.016 | -0.016 | 0 | 0.36 | 0.34 | 0.32 | 0.23 | 0.41 | 0.38 |
| 1 | 240172395 | rs6429174 | -0.837 | 0.303 | -0.187 | -0.541 | 0.004 | -0.012 | -0.012 | 0 | 0.59 | 0.81 | 0.65 | 0.55 | 0.66 | 0.44 |
| 1 | 240172647 | rs10926057 | -0.118 | 0.354 | -0.03 | -0.676 | -0.002 | 0.015 | 0.015 | 0 | 0.36 | 0.3 | 0.32 | 0.23 | 0.41 | 0.36 |
| 1 | 240173483 | rs16839314 | 1.378 | 0.343 | -0.046 | -0.415 | -0.006 | -0.047 | -0.047 | 0 | 0.17 | 0.07 | 0.18 | 0.33 | 0.22 | 0.21 |
| 1 | 240173998 | rs10926064 | -0.14 | 0.386 | 0.382 | -0.901 | -0.001 | 0 | 0 | 0 | 0.39 | 0.57 | 0.38 | 0.33 | 0.38 | 0.26 |
| 1 | 240174291 | rs922306 | -2.023 | 0.39 | 0.278 | -0.896 | 0.001 | -0.044 | -0.044 | 0 | 0.75 | 0.86 | 0.7 | 0.55 | 0.79 | 0.62 |
| 1 | 240175902 | rs16839331 | -0.195 | 0.398 | 0.198 | -0.864 | 0.001 | -0.02 | -0.02 | 0 | 0.37 | 0.39 | 0.32 | 0.22 | 0.41 | 0.36 |
| 1 | 240176633 | rs10802826 | 0.305 | 0.72 | 0.387 | -0.721 | -0.001 | 0.012 | 0.012 | 0 | 0.26 | 0.3 | 0.25 | 0.29 | 0.25 | 0.18 |
| 1 | 240177781 | rs922305 | -0.743 | 0.514 | 0.016 | -0.634 | -0.03 | 0.008 | 0.008 | 0 | 0.44 | 0.74 | 0.53 | 0.34 | 0.46 | 0.31 |
| 1 | 240181417 | rs2095629 | 1.197 | 0.835 | 0.175 | -0.477 | -0.009 | 0.007 | 0.007 | 0 | 0.16 | 0.07 | 0.17 | 0.15 | 0.22 | 0.18 |
| 1 | 240183949 | rs2066298 | -0.873 | 0.829 | 0.184 | -0.551 | -0.022 | -0.03 | -0.03 | 1.898 | 0.42 | 0.74 | 0.54 | 0.48 | 0.5 | 0.4 |
| 1 | 240184328 | rs12125242 | 0.319 | 0.708 | 0.055 | -0.681 | 0.001 | 0.057 | 0.057 | 0 | 0.25 | 0.12 | 0.17 | 0.05 | 0.25 | 0.21 |
| 1 | 240187633 | rs1891230 | -1.857 | 0.769 | 0.029 | -0.662 | -0.004 | -0.073 | -0.073 | 0 | 0.71 | 0.94 | 0.68 | 0.52 | 0.74 | 0.64 |
| 1 | 240188406 | rs16839364 | -0.38 | 0.761 | 0.554 | -0.559 | -0.008 | -0.03 | -0.03 | 0.198 | 0.32 | 0.42 | 0.23 | 0.2 | 0.29 | 0.32 |
| 1 | 240190181 | rs10926083 | 0.274 | 0.884 | 0.506 | -0.67 | 0.004 | 0.065 | 0.065 | 0 | 0.26 | 0.12 | 0.18 | 0.06 | 0.25 | 0.2 |
| 1 | 240190387 | rs922307 | 0.979 | 1.097 | 0.697 | -0.75 | 0.013 | 0.072 | 0.072 | 0 | 0.11 | 0.18 | 0.12 | 0.28 | 0.17 | 0.1 |
| 1 | 240195776 | rs12088486 | 0.265 | 1.084 | 0.42 | -1.141 | 0.005 | 0.045 | 0.045 | 0 | 0.26 | 0.14 | 0.18 | 0.06 | 0.25 | 0.2 |
| 1 | 240199952 | rs9887780 | 0.291 | 1.084 | 0.387 | -1.141 | 0.002 | 0.073 | 0.073 | 0 | 0.26 | 0.1 | 0.18 | 0.05 | 0.25 | 0.2 |
| 1 | 240204823 | rs12131924 | 0.954 | 1.086 | 0.326 | -1.178 | -0.013 | 0 | 0 | 0 | 0.1 | 0.01 | 0.03 | 0.05 | 0.08 | 0.11 |
| 1 | 240206467 | rs12134717 | 0.209 | 1.119 | 0.42 | -1.162 | 0.005 | 0.052 | 0.052 | 0 | 0.26 | 0.13 | 0.18 | 0.05 | 0.25 | 0.2 |
| 1 | 240206985 | rs10926087 | -0.562 | 1.091 | 0.489 | -1.226 | -0.024 | 0.13 | 0.13 | 0 | 0.42 | 0.19 | 0.36 | 0.09 | 0.45 | 0.32 |
| 1 | 240207152 | rs10926089 | -0.81 | 1.07 | 0.712 | -1.236 | 0.003 | -0.075 | -0.075 | 0 | 0.45 | 0.49 | 0.39 | 0.09 | 0.46 | 0.34 |
| 1 | 240213924 | rs6693552 | 1.037 | 1.223 | 0.271 | -0.833 | 0.001 | -0.055 | -0.055 | 0.535 | 0.18 | 0.17 | 0.16 | 0 | 0.22 | 0.09 |
| 1 | 240217385 | rs12140610 | 0.718 | 1.482 | 0.213 | -0.877 | -0.006 | 0.007 | 0.007 | 0 | 0.07 | 0.09 | 0.18 | 0.02 | 0.1 | 0.08 |
| 1 | 240218134 | rs12733694 | 1.091 | 1.278 | 0.305 | -0.855 | 0.002 | -0.068 | -0.068 | 0.071 | 0.18 | 0.21 | 0.16 | 0 | 0.22 | 0.09 |

| | | | | | | | | | | | | | | | | |
|---|---|---|---|---|---|---|---|---|---|---|---|---|---|---|---|---|
| 1 | 240219128 | rs12144492 | -1.324 | 1.248 | 0.156 | -0.604 | 0.015 | 0.006 | 0.006 | 0 | 0.49 | 0.7 | 0.65 | 0.38 | 0.55 | 0.4 |
| 1 | 240224754 | rs11579499 | -0.739 | 1.332 | 0.215 | -0.406 | -0.007 | 0 | 0 | 0 | 0.17 | 0.18 | 0.25 | 0.02 | 0.22 | 0.12 |
| 1 | 240225460 | rs6429178 | -1.338 | 1.24 | 0.213 | -0.189 | 0.041 | -0.059 | -0.059 | 0 | 0.67 | 0.81 | 0.85 | 0.41 | 0.78 | 0.5 |
| 1 | 240227047 | rs6701956 | -1.152 | 1.368 | 0.271 | 0.167 | 0.034 | -0.017 | -0.017 | 0.011 | 0.62 | 0.83 | 0.82 | 0.57 | 0.72 | 0.57 |
| 1 | 240227948 | rs10495457 | -0.433 | 1.184 | 0.214 | 0.003 | -0.001 | 0 | 0 | 0 | 0.43 | 0.41 | 0.67 | 0.32 | 0.5 | 0.31 |
| 1 | 240230726 | rs4659560 | -0.449 | 1.122 | -0.159 | 0.045 | -0.002 | 0.002 | 0.002 | 0.193 | 0.43 | 0.37 | 0.68 | 0.32 | 0.5 | 0.31 |
| 1 | 240236430 | rs6429180 | -1.62 | 1.253 | 0.409 | -0.179 | 0.004 | -0.018 | -0.018 | 0 | 0.32 | 0.66 | 0.36 | 0.17 | 0.34 | 0.24 |
| 1 | 240238025 | rs7515731 | -0.351 | 1.631 | -0.237 | -0.068 | -0.018 | 0.015 | 0.015 | 0 | 0.16 | 0.09 | 0.21 | 0.13 | 0.18 | 0.19 |
| 1 | 240240025 | rs4659562 | -1.236 | 1.07 | 0.161 | 0.192 | 0.001 | -0.01 | -0.01 | 3.132 | 0.14 | 0.17 | 0.14 | 0.08 | 0.15 | 0.05 |
| 1 | 240243920 | rs9803722 | -1.124 | 1.293 | 0.345 | 0.185 | 0.055 | -0.034 | -0.034 | 0.334 | 0.08 | 0.25 | 0.06 | 0.02 | 0.11 | 0.04 |
| 1 | 240253320 | rs1418049 | 0.409 | 1.348 | 0.054 | 0.704 | 0 | 0.011 | 0.011 | 0 | 0.75 | 0.73 | 0.75 | 0.45 | 0.79 | 0.75 |
| 1 | 240254828 | rs10754691 | 0.406 | 1.34 | 0.074 | 0.829 | -0.001 | 0.004 | 0.004 | 0 | 0.75 | 0.75 | 0.75 | 0.47 | 0.79 | 0.75 |
| 1 | 240259800 | rs2061632 | 0.857 | 1.003 | 0.19 | 0.249 | 0.001 | 0.044 | 0.044 | 0.026 | 0.58 | 0.23 | 0.4 | 0.34 | 0.63 | 0.52 |
| 1 | 240269323 | rs12062492 | 0.518 | 1.096 | 1.062 | 0.265 | 0.006 | -0.001 | -0.001 | 0 | 0.63 | 0.59 | 0.48 | 0.35 | 0.63 | 0.55 |
| 1 | 240280566 | rs6656902 | -1.28 | 1.371 | 1.29 | 0.237 | 0.047 | 0.008 | 0.008 | 0 | 0.05 | 0.2 | 0.05 | 0.03 | 0.09 | 0.07 |
| 1 | 240283081 | rs12145060 | 1.139 | 0.845 | 0.548 | -0.564 | -0.01 | 0.08 | 0.08 | 0 | 0.34 | 0.14 | 0.16 | 0.16 | 0.36 | 0.24 |
| 1 | 240286855 | rs12021945 | 1.22 | 0.854 | 0.278 | -0.668 | -0.072 | 0.083 | 0.083 | 0 | 0.33 | 0.03 | 0.15 | 0.16 | 0.36 | 0.24 |
| 1 | 240287470 | rs6660144 | 0.507 | 0.898 | 0.494 | -0.606 | 0.007 | -0.007 | -0.007 | 2.376 | 0.06 | 0.12 | 0.05 | 0.02 | 0.08 | 0.03 |
| 1 | 240288935 | rs10926135 | -0.054 | 0.566 | 0.653 | -0.797 | -0.001 | 0 | 0 | 0 | 0.57 | 0.57 | 0.42 | 0.34 | 0.57 | 0.49 |
| 1 | 240294563 | rs12562365 | 0.656 | 0.629 | 0.156 | -0.738 | 0.011 | -0.008 | -0.008 | 0.148 | 0.06 | 0.15 | 0.05 | 0.02 | 0.08 | 0.02 |
| 1 | 240296107 | rs6690640 | -0.352 | 0.6 | -0.391 | -0.81 | -0.026 | 0.007 | 0.007 | 0 | 0.63 | 0.84 | 0.47 | 0.34 | 0.62 | 0.52 |
| 1 | 240297030 | rs17669038 | 0.787 | 0.575 | -0.312 | -0.875 | 0.001 | 0 | 0 | 0 | 0.09 | 0.07 | 0.13 | 0.13 | 0.09 | 0.16 |
| 1 | 240297126 | rs6429184 | 0.221 | 0.574 | -0.284 | -0.879 | -0.013 | 0.083 | 0.083 | 0 | 0.54 | 0.75 | 0.33 | 0.21 | 0.53 | 0.37 |
| 1 | 240298817 | rs1982530 | 0.553 | 0.678 | -0.73 | -0.657 | 0.001 | 0 | 0 | 1.114 | 0.09 | 0.07 | 0.13 | 0.13 | 0.09 | 0.16 |
| 1 | 240299514 | rs10754692 | 0.655 | 0.606 | -0.865 | -0.688 | -0.048 | 0.094 | 0.094 | 0.47 | 0.4 | 0.06 | 0.23 | 0.19 | 0.4 | 0.25 |
| 1 | 240303322 | rs7537876 | 0.148 | 0.313 | -0.477 | -0.493 | 0.004 | -0.005 | -0.005 | 0.353 | 0.41 | 0.45 | 0.45 | 0.32 | 0.46 | 0.47 |
| 1 | 240305244 | rs10926138 | 0.059 | 0.145 | -0.181 | -0.547 | -0.004 | 0.006 | 0.006 | 1.81 | 0.43 | 0.64 | 0.47 | 0.32 | 0.47 | 0.44 |
| 1 | 240307048 | rs16848431 | 0.353 | -0.043 | -0.632 | -0.397 | 0.005 | 0.008 | 0.008 | 0 | 0.27 | 0.23 | 0.13 | 0.2 | 0.19 | 0.25 |
| 1 | 240307892 | rs1539200 | -0.24 | -0.047 | -0.748 | -0.363 | -0.042 | -0.096 | -0.096 | 0.569 | 0.71 | 0.92 | 0.58 | 0.52 | 0.66 | 0.69 |
| 1 | 240308200 | rs12127949 | -0.264 | -0.011 | -0.752 | -0.328 | -0.043 | -0.094 | -0.094 | 0 | 0.71 | 0.92 | 0.58 | 0.5 | 0.66 | 0.69 |
| 1 | 240310294 | rs17669357 | 0.494 | -0.245 | -0.652 | -0.229 | -0.02 | 0.014 | 0.014 | 0.259 | 0.58 | 0.22 | 0.48 | 0.44 | 0.52 | 0.65 |
| 1 | 240310698 | rs6667904 | 0.432 | -0.252 | -0.519 | -0.232 | -0.012 | 0.009 | 0.009 | 0.195 | 0.59 | 0.31 | 0.48 | 0.44 | 0.52 | 0.66 |
| 1 | 240311227 | rs12759640 | 0.266 | -0.291 | -0.249 | -0.108 | 0.009 | 0.023 | 0.023 | 0 | 0.06 | 0.13 | 0.05 | 0.07 | 0.06 | 0.07 |
| 1 | 240313215 | rs1350225 | 0.001 | -0.292 | -0.175 | -0.188 | 0.017 | 0.035 | 0.035 | 0 | 0.66 | 0.47 | 0.53 | 0.51 | 0.59 | 0.72 |
| 1 | 240314499 | rs10754693 | -0.114 | -0.289 | -0.192 | -0.186 | 0.014 | 0.028 | 0.028 | 0 | 0.67 | 0.55 | 0.54 | 0.51 | 0.59 | 0.72 |
| 1 | 240315515 | rs1891231 | 1.757 | -0.289 | -0.653 | -0.236 | -0.083 | 0.023 | 0.023 | 0.036 | 0.31 | 0.04 | 0.36 | 0.24 | 0.34 | 0.39 |
| 1 | 240324530 | rs10926149 | -0.681 | -0.346 | 0.116 | -0.224 | -0.023 | -0.018 | -0.018 | 0 | 0.35 | 0.54 | 0.18 | 0.29 | 0.24 | 0.33 |
| 1 | 240329312 | rs7525061 | 0.585 | -0.356 | 0.13 | -0.224 | 0.018 | 0.024 | 0.024 | 0.125 | 0.25 | 0.11 | 0.11 | 0.21 | 0.18 | 0.24 |
| 1 | 240336853 | rs6691735 | -0.548 | -0.354 | 0.217 | -0.221 | -0.03 | -0.023 | -0.023 | 0 | 0.33 | 0.49 | 0.17 | 0.29 | 0.24 | 0.31 |
| 1 | 240338827 | rs12723259 | -0.31 | -0.139 | 0.054 | -0.169 | 0.01 | 0.019 | 0.019 | 0 | 0.07 | 0.12 | 0.07 | 0.08 | 0.07 | 0.07 |
| 1 | 240339812 | rs7542728 | -1.882 | 0.097 | -0.335 | -0.337 | -0.029 | 0.031 | 0.031 | 0 | 0.94 | 1 | 0.89 | 1 | 0.92 | 0.94 |
| 1 | 240341268 | rs3765588 | -0.131 | -0.408 | 0.087 | -0.194 | -0.008 | -0.006 | -0.006 | 0 | 0.29 | 0.33 | 0.11 | 0.22 | 0.18 | 0.28 |
| 1 | 240345872 | rs7528413 | 0.22 | -0.332 | -0.187 | -0.157 | 0.015 | 0.01 | 0.01 | 0.086 | 0.27 | 0.19 | 0.11 | 0.22 | 0.18 | 0.26 |
| 1 | 240353592 | rs7544775 | -0.064 | -0.366 | -0.158 | -0.213 | 0.014 | 0.009 | 0.009 | 1.231 | 0.33 | 0.28 | 0.14 | 0.2 | 0.18 | 0.22 |
| 1 | 240354069 | rs2124724 | -0.168 | -0.193 | -0.375 | -0.232 | -0.044 | -0.051 | -0.051 | 0 | 0.67 | 0.83 | 0.48 | 0.36 | 0.58 | 0.65 |
| 1 | 240358485 | rs1037666 | -0.714 | -0.118 | -0.409 | -0.226 | -0.033 | -0.061 | -0.061 | 0 | 0.72 | 0.85 | 0.52 | 0.37 | 0.66 | 0.7 |
| 1 | 240360119 | rs12562593 | -0.675 | 0.109 | -0.36 | -0.298 | -0.009 | 0.006 | 0.006 | 0 | 0.07 | 0.05 | 0.07 | 0.03 | 0.14 | 0.06 |
| 1 | 240361422 | rs6660298 | 1.535 | -0.17 | 0.082 | -0.451 | -0.007 | 0.026 | 0.026 | 0 | 0.3 | 0.23 | 0.29 | 0.14 | 0.34 | 0.4 |
| 1 | 240363209 | rs1456660 | 0.139 | -0.688 | 0.377 | -0.338 | 0.008 | 0.001 | 0.001 | 1.047 | 0.36 | 0.33 | 0.15 | 0.29 | 0.19 | 0.26 |
| 1 | 240364187 | rs10926165 | -0.296 | -0.639 | 0.475 | -0.326 | -0.003 | -0.001 | -0.001 | 1.513 | 0.39 | 0.37 | 0.18 | 0.32 | 0.26 | 0.29 |
| 1 | 240373569 | rs10754695 | 0.011 | -1.215 | 0.236 | -0.477 | -0.017 | 0.001 | 0.001 | 0.581 | 0.33 | 0.37 | 0.19 | 0.3 | 0.23 | 0.23 |
| 1 | 240374809 | rs10926167 | 1.58 | -1.029 | -0.222 | -0.603 | -0.005 | -0.003 | -0.003 | 1.107 | 0.29 | 0.27 | 0.25 | 0.1 | 0.41 | 0.42 |
| 1 | 240382784 | rs6429189 | -1.214 | -1.089 | -0.311 | -0.595 | -0.042 | -0.088 | -0.088 | 0 | 0.76 | 0.91 | 0.5 | 0.42 | 0.71 | 0.77 |
| 1 | 240383334 | rs10495461 | 1.586 | -1.004 | -0.195 | -0.647 | -0.012 | 0.009 | 0.009 | 0 | 0.28 | 0.24 | 0.25 | 0.09 | 0.41 | 0.42 |
| 1 | 240388476 | rs7520065 | 1.521 | -0.862 | -0.136 | -0.573 | -0.014 | 0.015 | 0.015 | 0 | 0.3 | 0.24 | 0.24 | 0.1 | 0.41 | 0.42 |
| 1 | 240392809 | rs6689960 | 1.525 | -0.86 | -0.137 | -0.573 | -0.014 | 0.015 | 0.015 | 0 | 0.3 | 0.24 | 0.24 | 0.1 | 0.41 | 0.42 |
| 1 | 240394879 | rs12037713 | -0.227 | -1.043 | -0.352 | -0.552 | -0.022 | -0.042 | -0.042 | 0 | 0.66 | 0.8 | 0.45 | 0.42 | 0.64 | 0.68 |
| 1 | 240396790 | rs9651082 | 0.018 | -1.161 | -0.161 | -0.438 | 0.01 | -0.001 | -0.001 | 0 | 0.36 | 0.62 | 0.21 | 0.33 | 0.22 | 0.24 |
| 1 | 240396842 | rs7531340 | -1.32 | -1.011 | -0.329 | -0.344 | -0.039 | -0.09 | -0.09 | 0 | 0.78 | 0.9 | 0.51 | 0.44 | 0.73 | 0.78 |
| 1 | 240402395 | rs6429194 | -0.212 | -1.015 | 0.191 | -0.385 | -0.021 | -0.004 | -0.004 | 0.255 | 0.37 | 0.53 | 0.2 | 0.3 | 0.22 | 0.24 |
| 1 | 240403189 | rs1456645 | -0.803 | -0.897 | -0.045 | -0.341 | -0.008 | -0.036 | -0.036 | 0 | 0.71 | 0.81 | 0.49 | 0.42 | 0.7 | 0.72 |
| 1 | 240403456 | rs6698342 | -1.645 | -0.771 | -0.358 | -0.401 | -0.108 | 0.08 | 0.08 | 0 | 0.72 | 0.93 | 0.78 | 0.88 | 0.6 | 0.57 |
| 1 | 240404525 | rs7516886 | -1.662 | -0.9 | 0.092 | -0.243 | -0.058 | 0.037 | 0.037 | 0 | 0.72 | 0.87 | 0.78 | 0.84 | 0.61 | 0.61 |
| 1 | 240407163 | rs1456655 | -0.46 | -1.07 | 0.491 | -0.241 | 0 | 0 | 0 | 0.699 | 0.4 | 0.36 | 0.26 | 0.32 | 0.35 | 0.31 |
| 1 | 240408346 | rs2078095 | 0.431 | -1.247 | 0.387 | -0.323 | -0.004 | -0.024 | -0.024 | 0 | 0.27 | 0.07 | 0.17 | 0.25 | 0.27 | 0.24 |
| 1 | 240409592 | rs2008864 | 0.692 | -1.225 | 0.494 | -0.313 | -0.002 | 0.022 | 0.022 | 0.02 | 0.3 | 0.37 | 0.51 | 0.52 | 0.26 | 0.28 |
| 1 | 240409620 | rs1004076 | -0.026 | -1.336 | 0.596 | -0.31 | -0.007 | -0.001 | -0.001 | 0 | 0.3 | 0.34 | 0.19 | 0.25 | 0.28 | 0.24 |
| 1 | 240418702 | rs2124728 | 0.229 | -1.234 | 0.328 | -0.15 | -0.003 | -0.009 | -0.009 | 0.949 | 0.35 | 0.35 | 0.56 | 0.56 | 0.36 | 0.31 |
| 1 | 240420089 | rs897662 | -1.734 | -1.064 | 0.151 | -0.11 | -0.032 | 0.033 | 0.033 | 0 | 0.71 | 0.81 | 0.77 | 0.84 | 0.61 | 0.6 |

| | | | | | | | | | | | | | | | | |
|---|---|---|---|---|---|---|---|---|---|---|---|---|---|---|---|---|
| 1 | 240424857 | rs10926185 | -0.471 | -1.178 | 0.138 | -0.066 | -0.01 | -0.003 | -0.003 | 0.046 | 0.34 | 0.33 | 0.2 | 0.28 | 0.24 | 0.28 |
| 1 | 240427155 | rs716527 | -1.927 | -1.003 | -0.543 | -0.05 | -0.119 | 0.02 | 0.02 | 0 | 0.73 | 0.96 | 0.78 | 0.84 | 0.61 | 0.6 |
| 1 | 240428265 | rs10926188 | -1.71 | -0.96 | -0.337 | -0.026 | 0.012 | -0.014 | -0.014 | 0 | 0.71 | 0.67 | 0.77 | 0.84 | 0.61 | 0.6 |
| 1 | 240435787 | rs17626030 | 0.534 | -0.723 | -0.708 | 0.298 | 0.022 | -0.027 | -0.027 | 0 | 0.21 | 0.07 | 0.12 | 0.21 | 0.17 | 0.18 |
| 1 | 240437871 | rs6694063 | 0.097 | -0.133 | -0.145 | 0.526 | -0.001 | 0 | 0 | 0 | 0.08 | 0.09 | 0.05 | 0.1 | 0.07 | 0.08 |
| 1 | 240438307 | rs16839692 | 0.061 | -0.114 | -0.268 | 0.767 | -0.012 | -0.066 | -0.066 | 0 | 0.24 | 0.12 | 0.41 | 0.4 | 0.25 | 0.29 |
| 1 | 240442817 | rs10495462 | -0.723 | -0.171 | 0.167 | 0.592 | 0.003 | 0.004 | 0.004 | 0 | 0.36 | 0.41 | 0.58 | 0.64 | 0.4 | 0.32 |
| 1 | 240444169 | rs7531591 | -0.066 | -0.125 | 0.034 | 0.333 | -0.009 | 0.006 | 0.006 | 0 | 0.12 | 0.05 | 0.07 | 0.15 | 0.11 | 0.13 |
| 1 | 240444305 | rs2045349 | -0.353 | -0.524 | 0.51 | 0.223 | 0.003 | -0.018 | -0.018 | 2.454 | 0.31 | 0.23 | 0.51 | 0.42 | 0.29 | 0.34 |
| 1 | 240447774 | rs7519158 | -1.299 | -0.393 | 1.033 | 0.25 | 0.001 | -0.014 | -0.014 | 0.048 | 0.22 | 0.25 | 0.15 | 0.17 | 0.23 | 0.17 |
| 1 | 240449365 | rs11805911 | -1.821 | -0.452 | 1.341 | 0.212 | 0.003 | -0.044 | -0.044 | 0.534 | 0.24 | 0.65 | 0.18 | 0.17 | 0.23 | 0.16 |
| 1 | 240449759 | rs4659954 | 0.092 | -0.471 | 1.361 | 0.255 | -0.003 | -0.028 | -0.028 | 0 | 0.27 | 0.18 | 0.49 | 0.42 | 0.28 | 0.33 |
| 1 | 240453639 | rs6690616 | -0.113 | -0.363 | 1.447 | 0.163 | 0.021 | -0.041 | -0.041 | 0.043 | 0.1 | 0.2 | 0.07 | 0.02 | 0.12 | 0.03 |
| 1 | 240455395 | rs10802855 | -0.296 | -0.328 | 1.424 | 0.269 | -0.001 | 0.006 | 0.006 | 0 | 0.29 | 0.31 | 0.5 | 0.39 | 0.28 | 0.37 |
| 1 | 240456751 | rs6659900 | -1.979 | -0.383 | 1.086 | 0.154 | -0.078 | -0.102 | -0.102 | 0.209 | 0.54 | 0.95 | 0.67 | 0.58 | 0.52 | 0.53 |
| 1 | 240457973 | rs7526497 | -0.24 | -0.389 | 1.486 | 0.07 | 0.013 | -0.044 | -0.044 | 0 | 0.11 | 0.2 | 0.07 | 0.03 | 0.13 | 0.04 |
| 1 | 240459528 | rs10926203 | 0.359 | -0.379 | 1.284 | 0.303 | 0 | -0.004 | -0.004 | 0.556 | 0.22 | 0.17 | 0.36 | 0.27 | 0.21 | 0.23 |
| 1 | 240466440 | rs6699880 | -0.724 | -0.359 | 1.514 | 0.348 | 0.006 | -0.017 | -0.017 | 0 | 0.33 | 0.44 | 0.43 | 0.29 | 0.34 | 0.26 |
| 1 | 240473704 | rs1379074 | -0.806 | -0.309 | 1.336 | 0.406 | -0.004 | -0.018 | -0.018 | 1.555 | 0.16 | 0.09 | 0.18 | 0.2 | 0.14 | 0.22 |
| 1 | 240476158 | rs7545302 | -2.307 | -0.261 | 1.368 | 0.379 | 0 | 0.01 | 0.01 | 0 | 0.29 | 0.73 | 0.28 | 0.23 | 0.25 | 0.25 |
| 1 | 240479478 | rs4332348 | -1.055 | -0.311 | 1.406 | 0.365 | -0.003 | -0.019 | -0.019 | 0.011 | 0.38 | 0.29 | 0.55 | 0.46 | 0.36 | 0.44 |
| 1 | 240481401 | rs12023809 | 0.397 | -0.228 | 1.537 | 0.419 | -0.015 | 0.019 | 0.019 | 0 | 0.1 | 0 | 0.05 | 0.07 | 0.08 | 0.1 |
| 1 | 240483108 | rs7519584 | 0.22 | -0.226 | 1.609 | 0.443 | 0 | -0.001 | -0.001 | 0 | 0.24 | 0.2 | 0.37 | 0.26 | 0.23 | 0.23 |
| 1 | 240487547 | rs10495463 | -1.694 | -0.127 | 1.788 | 0.438 | -0.007 | 0.002 | 0.002 | 0 | 0.5 | 0.76 | 0.63 | 0.5 | 0.48 | 0.48 |
| 1 | 240489585 | rs6664970 | -0.102 | -0.077 | 2.009 | 0.426 | 0.034 | -0.044 | -0.044 | 0 | 0.1 | 0.27 | 0.06 | 0.03 | 0.12 | 0.03 |
| 1 | 240490119 | rs6679031 | -0.475 | -0.093 | 1.888 | 0.499 | -0.001 | 0.004 | 0.004 | 0 | 0.32 | 0.27 | 0.36 | 0.32 | 0.3 | 0.31 |
| 1 | 240492414 | rs6677726 | -1.238 | -0.079 | 1.947 | 0.529 | 0.021 | 0.116 | 0.116 | 0 | 0.07 | 0.23 | 0.21 | 0.15 | 0.06 | 0.13 |
| 1 | 240492734 | rs3795677 | -0.547 | -0.017 | 1.633 | 0.493 | 0 | -0.003 | -0.003 | 0 | 0.32 | 0.34 | 0.36 | 0.32 | 0.29 | 0.31 |
| 1 | 240496935 | rs4495690 | -0.853 | -0.025 | 1.392 | 0.45 | 0.018 | 0.021 | 0.021 | 0 | 0.36 | 0.53 | 0.55 | 0.43 | 0.39 | 0.37 |
| 1 | 240498229 | rs10926222 | -1.034 | 0.038 | 1.45 | 0.524 | -0.001 | 0.001 | 0.001 | 0 | 0.39 | 0.56 | 0.59 | 0.5 | 0.36 | 0.43 |
| 1 | 240499490 | rs7545070 | -0.81 | 0.006 | 1.456 | 0.534 | 0 | 0.012 | 0.012 | 0 | 0.37 | 0.53 | 0.58 | 0.49 | 0.35 | 0.44 |
| 1 | 240505407 | rs12407532 | -0.3 | 0.07 | 1.377 | 0.55 | 0.03 | -0.018 | -0.018 | 0.027 | 0.12 | 0.3 | 0.15 | 0.07 | 0.13 | 0.04 |
| 1 | 240507412 | rs4460624 | -1.707 | 0.009 | 0.963 | 0.543 | -0.071 | -0.073 | -0.073 | 2.828 | 0.52 | 0.92 | 0.65 | 0.5 | 0.49 | 0.5 |
| 1 | 240508458 | rs12130718 | -0.269 | 0.217 | 1.495 | 0.72 | -0.002 | -0.005 | -0.005 | 0 | 0.28 | 0.24 | 0.45 | 0.35 | 0.28 | 0.32 |
| 1 | 240510161 | rs7552602 | -0.452 | 0.271 | 1.11 | 0.766 | 0.006 | 0.032 | 0.032 | 0 | 0.29 | 0.41 | 0.46 | 0.35 | 0.29 | 0.34 |
| 1 | 240511965 | rs6658239 | 0.218 | 0.241 | 0.432 | 0.408 | -0.019 | 0.004 | 0.004 | 0 | 0.23 | 0.06 | 0.3 | 0.21 | 0.22 | 0.18 |
| 1 | 240519677 | rs9725819 | -1.18 | 0.429 | 0.855 | 0.491 | 0.011 | -0.033 | -0.033 | 0 | 0.14 | 0.31 | 0.14 | 0.06 | 0.14 | 0.08 |
| 1 | 240527029 | rs9660912 | -0.912 | 0.357 | 0.146 | 0.595 | 0 | -0.007 | -0.007 | 0 | 0.47 | 0.71 | 0.58 | 0.51 | 0.49 | 0.43 |
| 1 | 240527080 | rs12723298 | 0.356 | 0.459 | 0.214 | 0.45 | -0.003 | 0.008 | 0.008 | 0 | 0.23 | 0.14 | 0.32 | 0.19 | 0.24 | 0.19 |
| 1 | 240528954 | rs4412583 | -1.802 | 0.3 | -0.355 | 0.558 | -0.086 | 0.063 | 0.063 | 1.22 | 0.63 | 0.98 | 0.74 | 0.79 | 0.59 | 0.61 |
| 1 | 240530753 | rs6689621 | -0.637 | 0.274 | 0.33 | 0.748 | -0.004 | 0.001 | 0.001 | 0 | 0.37 | 0.6 | 0.5 | 0.36 | 0.38 | 0.29 |
| 1 | 240531291 | rs12064380 | -0.943 | 0.114 | 0.352 | 0.961 | 0.026 | 0.001 | 0.001 | 0 | 0.23 | 0.42 | 0.24 | 0.24 | 0.25 | 0.16 |
| 1 | 240532424 | rs12022143 | 0.057 | 0.133 | -0.479 | 1.082 | 0.015 | -0.032 | -0.032 | 0 | 0.13 | 0.09 | 0.12 | 0.35 | 0.09 | 0.19 |
| 1 | 240533352 | rs9725359 | -1.159 | 0.479 | -0.584 | 0.667 | 0.003 | 0.115 | 0.115 | 0 | 0.58 | 0.87 | 0.69 | 0.78 | 0.58 | 0.57 |
| 1 | 240538952 | rs9662927 | -1.008 | 0.213 | -0.333 | 0.874 | 0.012 | 0.035 | 0.035 | 0 | 0.47 | 0.75 | 0.54 | 0.65 | 0.43 | 0.51 |
| 1 | 240541011 | rs10802866 | -1.008 | 0.227 | -0.418 | 0.9 | 0.012 | 0.017 | 0.017 | 0 | 0.47 | 0.69 | 0.54 | 0.59 | 0.43 | 0.51 |
| 1 | 240547720 | rs12563220 | -0.196 | 0.296 | -0.869 | 0.894 | -0.015 | 0.012 | 0.012 | 0 | 0.08 | 0.03 | 0.06 | 0.06 | 0.1 | 0.1 |
| 1 | 240548111 | rs9661605 | -0.1 | 0.288 | -0.826 | 1.148 | -0.009 | -0.015 | -0.015 | 0.03 | 0.33 | 0.18 | 0.47 | 0.43 | 0.33 | 0.36 |
| 1 | 240550077 | rs1537850 | -0.112 | 0.296 | -0.807 | 1.052 | -0.009 | 0 | 0 | 0 | 0.33 | 0.19 | 0.47 | 0.38 | 0.34 | 0.35 |
| 1 | 240556931 | rs9659428 | -1.058 | 0.393 | -1 | 0.266 | 0.007 | 0.035 | 0.035 | 0 | 0.47 | 0.8 | 0.54 | 0.65 | 0.43 | 0.5 |
| 1 | 240561727 | rs9287233 | -0.532 | 0.59 | -1.256 | 0.724 | 0 | -0.014 | -0.014 | 0 | 0.34 | 0.27 | 0.47 | 0.58 | 0.33 | 0.37 |
| 1 | 240564513 | rs10802867 | -0.051 | 0.498 | -2.193 | -0.039 | -0.039 | 0.091 | 0.091 | 0.591 | 0.65 | 0.95 | 0.62 | 0.84 | 0.63 | 0.77 |
| 1 | 240565868 | rs4659971 | 0.568 | 0.205 | -1.985 | -0.466 | 0.008 | 0.022 | 0.022 | 0.056 | 0.57 | 0.38 | 0.59 | 0.23 | 0.65 | 0.61 |
| 1 | 240567903 | rs16839923 | 0.65 | 0.078 | -2.228 | 0.308 | -0.017 | -0.061 | -0.061 | 0 | 0.1 | 0.01 | 0.09 | 0.43 | 0.08 | 0.13 |
| 1 | 240570975 | rs17629839 | 0.999 | 0.737 | -1.564 | -0.064 | -0.03 | 0.029 | 0.029 | 0 | 0.06 | 0.01 | 0.11 | 0.01 | 0.1 | 0.19 |
| 1 | 240574210 | rs953247 | -0.022 | 0.624 | -1.16 | 0.047 | -0.002 | -0.039 | -0.039 | 0 | 0.2 | 0.12 | 0.19 | 0.59 | 0.16 | 0.22 |
| 1 | 240575168 | rs10926248 | 0.834 | 0.383 | -0.623 | -0.14 | 0.017 | 0.068 | 0.068 | 0 | 0.58 | 0.29 | 0.56 | 0.25 | 0.65 | 0.63 |
| 1 | 240586392 | rs1572417 | -1.028 | 0.406 | 0.078 | -0.163 | -0.002 | 0.004 | 0.004 | 0 | 0.82 | 0.75 | 0.78 | 0.82 | 0.8 | 0.85 |
| 1 | 240586695 | rs1414660 | -0.384 | 0.321 | 0.116 | -0.144 | -0.014 | 0.113 | 0.113 | 0 | 0.23 | 0.42 | 0.21 | 0.4 | 0.15 | 0.2 |
| 1 | 240586798 | rs1414659 | 0.03 | 0.122 | -0.2 | -0.638 | 0.001 | 0 | 0 | 0 | 0.45 | 0.57 | 0.32 | 0.43 | 0.39 | 0.39 |
| 1 | 240589117 | rs1414657 | 0.544 | 0.112 | -0.312 | -0.462 | 0.008 | 0.005 | 0.005 | 0 | 0.6 | 0.35 | 0.57 | 0.42 | 0.66 | 0.65 |
| 1 | 240596246 | rs9727462 | -1.013 | 0.214 | -0.135 | -0.332 | 0.014 | 0.04 | 0.04 | 0 | 0.84 | 0.66 | 0.79 | 0.82 | 0.82 | 0.87 |
| 1 | 240596675 | rs12145827 | 0.161 | 0.188 | -0.925 | -0.373 | -0.107 | -0.05 | -0.05 | 0 | 0.39 | 0.06 | 0.44 | 0.4 | 0.44 | 0.48 |
| 1 | 240598419 | rs12755409 | -0.845 | -0.284 | -0.063 | -0.469 | -0.009 | -0.02 | -0.02 | 0.048 | 0.34 | 0.57 | 0.32 | 0.2 | 0.24 | 0.27 |
| 1 | 240598887 | rs4659573 | 0.351 | -0.211 | -0.041 | -0.564 | -0.029 | 0.035 | 0.035 | 0 | 0.7 | 0.83 | 0.64 | 0.76 | 0.68 | 0.58 |
| 1 | 240600551 | rs9729181 | -0.184 | -0.238 | 0.619 | -0.414 | -0.021 | -0.019 | -0.019 | 0 | 0.09 | 0.34 | 0.06 | 0.01 | 0.02 | 0.08 |
| 1 | 240602059 | rs9728609 | 0.29 | -0.349 | 0.108 | -0.696 | -0.037 | 0.063 | 0.063 | 0 | 0.71 | 0.88 | 0.75 | 0.83 | 0.71 | 0.67 |
| 1 | 240603314 | rs2152556 | 1.349 | -0.503 | 0.31 | 0.112 | 0.002 | -0.001 | -0.001 | 0.357 | 0.55 | 0.56 | 0.64 | 0.52 | 0.63 | 0.5 |
| 1 | 240603534 | rs12139890 | -0.749 | -0.439 | 0.54 | 0.341 | -0.05 | 0.085 | 0.085 | 0 | 0.16 | 0.31 | 0.11 | 0.31 | 0.08 | 0.16 |

| | | | | | | | | | | | | | | | |
|---|---|---|---|---|---|---|---|---|---|---|---|---|---|---|---|
| 1 | 240605052 | rs12091371 | -0.92 | -0.456 | 0.526 | 0.343 | -0.062 | 0.13 | 0.13 | 0 | 0.16 | 0.48 | 0.13 | 0.31 | 0.07 | 0.16 |
| 1 | 240605412 | rs11577868 | 1.333 | -0.065 | 0.4 | 0.439 | -0.014 | 0.106 | 0.106 | 0.023 | 0.77 | 0.5 | 0.83 | 0.6 | 0.81 | 0.63 |
| 1 | 240605511 | rs11577878 | -1.588 | -0.147 | 0.123 | 0.013 | 0.002 | -0.003 | -0.003 | 0 | 0.38 | 0.62 | 0.35 | 0.44 | 0.31 | 0.31 |
| 1 | 240605623 | rs17682283 | -0.099 | 0.05 | -0.129 | -0.33 | 0.011 | 0.07 | 0.07 | 0 | 0.41 | 0.3 | 0.37 | 0.16 | 0.39 | 0.29 |
| 1 | 240608034 | rs2883924 | -0.771 | -0.082 | 0.337 | 0.092 | -0.023 | 0.1 | 0.1 | 0.035 | 0.19 | 0.46 | 0.17 | 0.32 | 0.14 | 0.17 |
| 1 | 240609633 | rs11576138 | 0.929 | 0.078 | 0.088 | 0.053 | 0.002 | 0.012 | 0.012 | 0.025 | 0.77 | 0.7 | 0.81 | 0.66 | 0.77 | 0.65 |
| 1 | 240610842 | rs12410581 | -0.109 | -0.067 | 0.075 | 0.059 | -0.002 | 0.008 | 0.008 | 0 | 0.76 | 0.72 | 0.82 | 0.63 | 0.77 | 0.68 |
| 1 | 240611115 | rs10926262 | 0.767 | 0.081 | 0.116 | -0.04 | -0.003 | 0 | 0 | 0 | 0.54 | 0.52 | 0.62 | 0.49 | 0.62 | 0.51 |
| 1 | 240611614 | rs12096679 | -0.368 | 0.2 | 0.213 | 0.049 | -0.005 | -0.007 | -0.007 | 0 | 0.22 | 0.52 | 0.31 | 0.2 | 0.22 | 0.15 |
| 1 | 240612857 | rs12064827 | 0.881 | 0.397 | -0.003 | -0.311 | 0.021 | -0.045 | -0.045 | 0 | 0.11 | 0.25 | 0.13 | 0.05 | 0.16 | 0.08 |
| 1 | 240612986 | rs11590081 | 1.243 | 0.255 | -0.429 | -0.385 | -0.062 | -0.044 | -0.044 | 0.863 | 0.42 | 0.16 | 0.46 | 0.42 | 0.49 | 0.33 |
| 1 | 240613707 | rs10802871 | 0.921 | 0.743 | 0.273 | -0.335 | 0.027 | -0.037 | -0.037 | 1.799 | 0.8 | 0.6 | 0.82 | 0.88 | 0.73 | 0.74 |
| 1 | 240613766 | rs10754705 | 0.414 | 0.566 | -0.107 | 0.101 | 0.006 | 0.001 | 0.001 | 0 | 0.91 | 0.86 | 0.95 | 0.93 | 0.89 | 0.81 |
| 1 | 240620131 | rs12403071 | 0.259 | 0.418 | 0.017 | 0.215 | 0.007 | 0.117 | 0.117 | 0 | 0.8 | 0.65 | 0.8 | 0.66 | 0.81 | 0.7 |
| 1 | 240620154 | rs17631686 | 0.347 | 0.179 | -0.882 | -0.665 | 0.033 | 0.089 | 0.089 | 0 | 0.36 | 0.19 | 0.24 | 0.12 | 0.31 | 0.28 |
| 1 | 240624682 | rs10926272 | -0.46 | 0.217 | -0.935 | -0.492 | -0.016 | -0.073 | -0.073 | 0 | 0.82 | 0.91 | 0.81 | 0.65 | 0.81 | 0.69 |
| 1 | 240626199 | rs10495470 | 0.28 | -0.031 | -1.01 | -0.638 | 0.025 | 0.018 | 0.018 | 0 | 0.13 | 0.04 | 0.1 | 0.1 | 0.1 | 0.13 |
| 1 | 240626698 | rs9659866 | 0.626 | 0.943 | -0.795 | -0.747 | 0.009 | 0.04 | 0.04 | 0 | 0.16 | 0.21 | 0.35 | 0.5 | 0.22 | 0.25 |
| 1 | 240628795 | rs7521428 | -0.003 | 0.538 | -0.495 | -0.876 | 0.002 | -0.02 | -0.02 | 2.234 | 0.4 | 0.56 | 0.24 | 0.1 | 0.31 | 0.28 |
| 1 | 240633273 | rs9287243 | -0.745 | 0.459 | -1.248 | -0.92 | -0.039 | -0.08 | -0.08 | 1.588 | 0.62 | 0.91 | 0.67 | 0.6 | 0.61 | 0.56 |
| 1 | 240637016 | rs4659973 | -0.686 | 0.522 | -1.185 | -1.017 | -0.02 | -0.076 | -0.076 | 0.202 | 0.61 | 0.89 | 0.67 | 0.58 | 0.61 | 0.54 |
| 1 | 240637571 | rs882869 | 0.614 | 0.49 | -1.203 | -0.983 | -0.086 | -0.107 | -0.107 | 0.832 | 0.21 | 0.09 | 0.41 | 0.49 | 0.33 | 0.29 |
| 1 | 240639558 | rs879081 | 0.232 | 0.479 | -0.964 | -0.993 | 0.008 | 0.014 | 0.014 | 0 | 0.24 | 0.31 | 0.43 | 0.48 | 0.33 | 0.28 |
| 1 | 240639611 | rs9729366 | 0.094 | -0.002 | -0.687 | -0.89 | -0.018 | -0.055 | -0.055 | 0.207 | 0.37 | 0.5 | 0.23 | 0.1 | 0.28 | 0.26 |
| 1 | 240641881 | rs9725597 | 0.349 | 0.408 | -0.42 | -0.927 | -0.02 | -0.114 | -0.114 | 0 | 0.3 | 0.36 | 0.19 | 0.02 | 0.24 | 0.21 |
| 1 | 240643167 | rs11578184 | -0.529 | 1.193 | -0.594 | 0.061 | -0.023 | -0.018 | -0.018 | 0 | 0.32 | 0.29 | 0.46 | 0.48 | 0.4 | 0.33 |
| 1 | 240643184 | rs9442211 | -0.716 | 1.213 | -0.946 | 0.099 | 0.027 | 0.02 | 0.02 | 0 | 0.34 | 0.49 | 0.48 | 0.48 | 0.4 | 0.33 |
| 1 | 240646538 | rs9287247 | -0.381 | 1.353 | -1.707 | 0.398 | -0.031 | -0.079 | -0.079 | 0.015 | 0.29 | 0.12 | 0.42 | 0.45 | 0.3 | 0.31 |
| 1 | 240646776 | rs9662136 | 0.055 | 1.1 | -2.287 | 0.251 | -0.058 | -0.122 | -0.122 | 0 | 0.57 | 0.95 | 0.59 | 0.58 | 0.44 | 0.57 |
| 1 | 240646941 | rs9661430 | 0.252 | 1.017 | -1.015 | -0.545 | 0.014 | -0.033 | -0.033 | 0 | 0.66 | 0.5 | 0.62 | 0.77 | 0.61 | 0.66 |
| 1 | 240647194 | rs9661447 | -1.244 | 0.558 | -0.23 | -0.498 | 0.067 | -0.047 | -0.047 | 0.288 | 0.1 | 0.46 | 0.1 | 0 | 0.09 | 0.06 |
| 1 | 240647894 | rs9661714 | -0.344 | 0.149 | -0.602 | -0.706 | -0.033 | -0.112 | -0.112 | 0 | 0.25 | 0.45 | 0.15 | 0.1 | 0.16 | 0.16 |
| 1 | 240647955 | rs11582667 | 0.391 | 1.297 | -1.67 | -0.165 | -0.105 | -0.092 | -0.092 | 0 | 0.17 | 0.04 | 0.44 | 0.43 | 0.4 | 0.23 |
| 1 | 240649079 | rs1414654 | -0.957 | 0.595 | -1.522 | -0.095 | -0.013 | -0.001 | -0.001 | 0 | 0.85 | 0.89 | 0.83 | 0.88 | 0.84 | 0.88 |
| 1 | 240652415 | rs16840250 | 1.05 | -0.68 | -1.867 | -0.272 | 0.089 | -0.141 | -0.141 | 0 | 0.23 | 0.05 | 0.15 | 0.33 | 0.11 | 0.3 |
| 1 | 240652947 | rs9697 | -0.7 | 0.183 | -1.265 | -0.247 | 0.003 | -0.001 | -0.001 | 0 | 0.84 | 0.82 | 0.83 | 0.88 | 0.83 | 0.88 |
| 1 | 240655330 | rs9287248 | 0.175 | 0.081 | -1.236 | -0.297 | -0.003 | 0.025 | 0.025 | 1.055 | 0.67 | 0.73 | 0.71 | 0.81 | 0.66 | 0.72 |
| 1 | 240658373 | rs10802881 | -0.573 | 0.103 | -1.283 | -0.088 | -0.004 | 0.001 | 0.001 | 0 | 0.76 | 0.82 | 0.66 | 0.79 | 0.7 | 0.73 |
| 1 | 240659736 | rs2152555 | 0.478 | 0.429 | -2.022 | 0.093 | 0.034 | -0.165 | -0.165 | 0 | 0.29 | 0.05 | 0.2 | 0.35 | 0.18 | 0.33 |
| 1 | 240660818 | rs12137158 | -0.412 | 1.166 | -1.023 | 0.337 | -0.031 | -0.001 | -0.001 | 0 | 0.34 | 0.27 | 0.45 | 0.46 | 0.47 | 0.31 |
| 1 | 240666156 | rs12136933 | -0.481 | 1.189 | -0.831 | 0.183 | -0.003 | 0.013 | 0.013 | 0 | 0.34 | 0.28 | 0.38 | 0.35 | 0.36 | 0.27 |
| 1 | 240666644 | rs16840259 | -0.857 | 1.134 | -0.485 | 1.203 | -0.002 | 0.13 | 0.13 | 0 | 0.13 | 0.22 | 0.31 | 0.47 | 0.13 | 0.24 |
| 1 | 240667002 | rs10465630 | -1.049 | 1.596 | -0.127 | 1.039 | 0.029 | 0.136 | 0.136 | 0.862 | 0.21 | 0.51 | 0.28 | 0.36 | 0.25 | 0.16 |
| 1 | 240667092 | rs10926286 | 0.374 | 1.644 | -0.418 | 0.699 | 0.006 | 0.048 | 0.048 | 0.017 | 0.74 | 0.65 | 0.55 | 0.42 | 0.71 | 0.56 |
| 1 | 240667391 | rs9428479 | 0.397 | 1.571 | -0.754 | 1.007 | -0.002 | -0.005 | -0.005 | 0 | 0.76 | 0.77 | 0.68 | 0.59 | 0.7 | 0.71 |
| 1 | 240667889 | rs951617 | 0.277 | 1.198 | -0.812 | 0.372 | -0.001 | -0.001 | -0.001 | 1.622 | 0.43 | 0.54 | 0.32 | 0.3 | 0.4 | 0.33 |
| 1 | 240672410 | rs11807703 | 0.047 | 1.348 | -0.887 | 0.136 | 0.006 | -0.038 | -0.038 | 0 | 0.1 | 0.15 | 0.08 | 0 | 0.13 | 0.01 |
| 1 | 240673104 | rs7545242 | -0.419 | 1.579 | -1.846 | 0.152 | -0.073 | -0.03 | -0.03 | 0.033 | 0.25 | 0.03 | 0.23 | 0.21 | 0.33 | 0.27 |
| 1 | 240674472 | rs2185285 | 0.127 | 0.742 | -1.516 | 0.079 | 0.005 | 0.009 | 0.009 | 0 | 0.48 | 0.46 | 0.46 | 0.39 | 0.43 | 0.35 |
| 1 | 240675199 | rs7540147 | -0.278 | 0.624 | -1.693 | 0.036 | 0.016 | 0.017 | 0.017 | 0 | 0.63 | 0.5 | 0.61 | 0.53 | 0.6 | 0.56 |
| 1 | 240678302 | rs6702434 | -0.967 | -0.507 | -1.797 | -0.249 | -0.025 | 0.061 | 0.061 | 0 | 0.59 | 0.73 | 0.57 | 0.83 | 0.53 | 0.69 |
| 1 | 240679626 | rs9428482 | 0.699 | -0.362 | -1.392 | 0.035 | -0.012 | -0.033 | -0.033 | 0.817 | 0.26 | 0.14 | 0.25 | 0.5 | 0.3 | 0.33 |
| 1 | 240680663 | rs9428483 | -0.691 | -0.515 | -1.18 | 0.025 | 0.005 | 0.009 | 0.009 | 0 | 0.42 | 0.35 | 0.41 | 0.71 | 0.38 | 0.49 |
| 1 | 240680789 | rs17685281 | -0.691 | -0.515 | -1.18 | 0.025 | 0.005 | 0.009 | 0.009 | 0 | 0.42 | 0.35 | 0.41 | 0.71 | 0.38 | 0.49 |
| 1 | 240680957 | rs16840284 | 0.106 | -0.797 | -1.381 | -0.145 | 0.03 | -0.017 | -0.017 | 0 | 0.15 | 0.09 | 0.15 | 0.22 | 0.08 | 0.17 |
| 1 | 240686578 | rs4659977 | 0.735 | 0.165 | -1.419 | -0.588 | -0.053 | 0.034 | 0.034 | 0 | 0.2 | 0.12 | 0.42 | 0.11 | 0.35 | 0.22 |
| 1 | 240690431 | rs12034261 | 0.164 | -0.373 | -1.233 | -0.364 | 0.006 | 0.001 | 0.001 | 0 | 0.3 | 0.3 | 0.21 | 0.3 | 0.34 | 0.33 |
| 1 | 240693711 | rs3748540 | 0.017 | -0.53 | -1.013 | -0.293 | -0.001 | -0.001 | -0.001 | 0 | 0.56 | 0.62 | 0.75 | 0.64 | 0.63 | 0.59 |
| 1 | 240694114 | rs3828024 | 0.299 | -1.048 | -1.053 | -0.18 | -0.003 | -0.001 | -0.001 | 0 | 0.51 | 0.46 | 0.58 | 0.39 | 0.63 | 0.52 |
| 1 | 240694163 | rs16840310 | 0.29 | -1.048 | -1.053 | -0.18 | -0.003 | -0.001 | -0.001 | 0 | 0.51 | 0.46 | 0.58 | 0.39 | 0.63 | 0.51 |
| 1 | 240694876 | rs6693656 | -1.082 | -0.654 | -1.254 | 0.266 | 0.023 | 0.037 | 0.037 | 0 | 0.87 | 0.97 | 0.94 | 0.95 | 0.93 | 0.9 |
| 1 | 240695255 | rs12129547 | 0.508 | -0.8 | -1.175 | 0.295 | 0.015 | -0.036 | -0.036 | 0 | 0.27 | 0.22 | 0.27 | 0.49 | 0.24 | 0.34 |
| 1 | 240698245 | rs4587552 | -0.172 | -0.325 | -0.949 | 0.789 | 0.018 | 0.023 | 0.023 | 0 | 0.86 | 0.93 | 0.93 | 0.95 | 0.93 | 0.9 |
| 1 | 240699814 | rs12027604 | -0.47 | 0.416 | -0.317 | 1.294 | 0 | 0.024 | 0.024 | 0 | 0.24 | 0.3 | 0.17 | 0.31 | 0.22 | 0.3 |
| 1 | 240706107 | rs10495471 | -0.744 | 0.337 | -0.852 | 1.582 | 0.004 | -0.033 | -0.033 | 0.056 | 0.13 | 0.05 | 0.14 | 0.26 | 0.14 | 0.12 |
| 1 | 240709136 | rs16840341 | -0.93 | 0.62 | -0.734 | 1.955 | -0.005 | -0.012 | -0.012 | 0 | 0.18 | 0.14 | 0.29 | 0.38 | 0.28 | 0.36 |
| 1 | 240712148 | rs12409520 | -0.674 | 0.852 | -0.16 | 2.158 | 0.035 | 0.05 | 0.05 | 0 | 0.06 | 0.21 | 0.16 | 0.1 | 0.09 | 0.05 |
| 1 | 240712898 | rs12069813 | -1.848 | -0.292 | -0.167 | 2.082 | 0.033 | -0.001 | -0.001 | 0.339 | 0.41 | 0.86 | 0.51 | 0.6 | 0.4 | 0.49 |
| 1 | 240714030 | rs10802887 | -1.202 | -0.677 | 0.391 | 2.088 | 0.01 | -0.029 | -0.029 | 0 | 0.29 | 0.22 | 0.31 | 0.46 | 0.3 | 0.44 |

| | | | | | | | | | | | | | | | | |
|---|---|---|---|---|---|---|---|---|---|---|---|---|---|---|---|---|
| 1 | 240718726 | rs10926298 | -1.089 | -0.668 | 0.337 | 2.158 | 0.009 | -0.044 | -0.044 | 0 | 0.28 | 0.14 | 0.32 | 0.4 | 0.32 | 0.37 |
| 1 | 240724339 | rs9428851 | -1.61 | -0.583 | 1.24 | 2.09 | 0.022 | 0.145 | 0.145 | 1.732 | 0.08 | 0.53 | 0.07 | 0.16 | 0.02 | 0.12 |
| 1 | 240733426 | rs4454537 | -1.031 | -0.612 | 0.899 | 2.27 | -0.005 | -0.089 | -0.089 | 1.532 | 0.27 | 0.07 | 0.3 | 0.38 | 0.31 | 0.37 |
| 1 | 240736355 | rs7543336 | -1.187 | -0.666 | 1.826 | 2.125 | -0.025 | 0.043 | 0.043 | 0 | 0.22 | 0.35 | 0.19 | 0.37 | 0.18 | 0.4 |
| 1 | 240740757 | rs10926303 | -2.018 | -0.79 | 1.226 | 2.037 | 0.006 | -0.004 | -0.004 | 0 | 0.29 | 0.65 | 0.26 | 0.34 | 0.2 | 0.36 |
| 1 | 240740946 | rs4453026 | -1.77 | -0.777 | 0.855 | 2.043 | -0.041 | 0.038 | 0.038 | 0 | 0.27 | 0.45 | 0.24 | 0.34 | 0.2 | 0.36 |
| 1 | 240742905 | rs7525142 | -2.367 | -0.693 | 0.338 | 2.081 | 0.051 | -0.171 | -0.171 | 0.092 | 0.34 | 0.95 | 0.29 | 0.52 | 0.22 | 0.48 |
| 1 | 240744031 | rs12117424 | -1.028 | -0.308 | 0.451 | 1.769 | 0.003 | -0.025 | -0.025 | 0 | 0.2 | 0.06 | 0.15 | 0.26 | 0.18 | 0.32 |
| 1 | 240744784 | rs10926304 | -1.404 | -0.248 | 0.359 | 1.607 | -0.032 | 0.061 | 0.061 | 0 | 0.25 | 0.42 | 0.22 | 0.36 | 0.2 | 0.37 |
| 1 | 240763283 | rs6429214 | 0.03 | 0.597 | -1.295 | 0.313 | -0.142 | 0.131 | 0.131 | 0.916 | 0.71 | 0.99 | 0.78 | 0.93 | 0.66 | 0.73 |
| 1 | 240765476 | rs1934346 | 0.442 | 0.512 | -2.117 | 0.236 | -0.066 | -0.057 | -0.057 | 0.541 | 0.23 | 0.02 | 0.25 | 0.23 | 0.27 | 0.33 |
| 1 | 240770788 | rs6686630 | -0.014 | 0.057 | -1.561 | 0.173 | -0.016 | 0.085 | 0.085 | 1.739 | 0.57 | 0.31 | 0.6 | 0.86 | 0.56 | 0.64 |
| 1 | 240770827 | rs12123578 | 0.564 | -0.063 | -1.368 | 0.407 | 0 | 0 | 0 | 0 | 0.06 | 0.07 | 0.06 | 0.02 | 0.08 | 0.06 |
| 1 | 240773666 | rs11578889 | 0.288 | -0.201 | -1.561 | 0.493 | 0 | -0.009 | -0.009 | 0 | 0.31 | 0.22 | 0.4 | 0.7 | 0.29 | 0.31 |
| 1 | 240776190 | rs2185282 | 0.431 | -0.219 | -1.906 | 0.491 | 0.017 | 0.026 | 0.026 | 3.843 | 0.18 | 0.07 | 0.11 | 0.14 | 0.16 | 0.07 |
| 1 | 240776222 | rs1934339 | 0.322 | 0.522 | -1.703 | 0.766 | -0.043 | -0.102 | -0.102 | 0.274 | 0.23 | 0.05 | 0.31 | 0.39 | 0.22 | 0.4 |
| 1 | 240778731 | rs7546054 | -0.083 | 0.415 | -0.521 | 0.285 | -0.014 | -0.069 | -0.069 | 0 | 0.3 | 0.55 | 0.24 | 0.13 | 0.28 | 0.27 |
| 1 | 240798089 | rs6429215 | 0.17 | 1.401 | -1.42 | 1.045 | -0.125 | -0.086 | -0.086 | 0.065 | 0.72 | 0.95 | 0.43 | 0.31 | 0.61 | 0.51 |
| 1 | 240799543 | rs4659986 | -0.56 | 0.553 | -0.976 | 1.069 | 0 | -0.004 | -0.004 | 0 | 0.37 | 0.66 | 0.31 | 0.36 | 0.31 | 0.41 |
| 1 | 240800342 | rs7545099 | -1.174 | 0.052 | -0.049 | 1.288 | 0.022 | 0.166 | 0.166 | 1.372 | 0.07 | 0.3 | 0.13 | 0.23 | 0.04 | 0.15 |
| 1 | 240803170 | rs6691111 | -0.561 | 0.063 | -0.581 | 0.821 | -0.005 | 0.001 | 0.001 | 0 | 0.16 | 0.16 | 0.18 | 0.12 | 0.14 | 0.18 |
| 1 | 240814351 | rs6429216 | 0.108 | 0.171 | -1.022 | 0.31 | 0.003 | 0.034 | 0.034 | 0 | 0.49 | 0.64 | 0.47 | 0.41 | 0.49 | 0.48 |
| 1 | 240816220 | rs4348710 | 0.243 | 0.479 | -1.507 | -0.174 | -0.013 | 0.041 | 0.041 | 0 | 0.1 | 0.02 | 0.06 | 0.03 | 0.1 | 0.09 |
| 1 | 240819499 | rs7526951 | -0.561 | 0.224 | -1.677 | -0.23 | -0.083 | 0.083 | 0.083 | 0 | 0.76 | 0.91 | 0.79 | 0.88 | 0.67 | 0.78 |
| 1 | 240821761 | rs7513948 | -0.045 | -0.014 | -1.223 | -0.505 | -0.001 | -0.018 | -0.018 | 0 | 0.6 | 0.4 | 0.72 | 0.87 | 0.62 | 0.71 |
| 1 | 240827275 | rs12035526 | 0.496 | 0.22 | -1.738 | 0.339 | -0.063 | -0.101 | -0.101 | 0.225 | 0.23 | 0.02 | 0.28 | 0.32 | 0.26 | 0.18 |
| 1 | 240828396 | rs12079180 | -0.021 | 0.072 | -1.221 | 0.396 | 0 | 0.01 | 0.01 | 0 | 0.49 | 0.34 | 0.5 | 0.55 | 0.53 | 0.38 |
| 1 | 240835685 | rs4659995 | 0.132 | 0.27 | -1.264 | 0.517 | -0.009 | -0.007 | -0.007 | 0.373 | 0.54 | 0.68 | 0.64 | 0.53 | 0.58 | 0.51 |
| 1 | 240842076 | rs11583286 | -0.642 | 0.633 | -0.65 | 0.54 | 0.008 | -0.01 | -0.01 | 0.434 | 0.28 | 0.49 | 0.22 | 0.25 | 0.3 | 0.31 |
| 1 | 240845574 | rs10926328 | 0.31 | 0.331 | -0.958 | 0.539 | -0.002 | 0.006 | 0.006 | 0 | 0.53 | 0.58 | 0.57 | 0.38 | 0.51 | 0.46 |
| 1 | 240845654 | rs10926329 | 0.741 | -0.049 | -1.485 | 0.236 | -0.049 | -0.019 | -0.019 | 0.026 | 0.42 | 0.12 | 0.53 | 0.38 | 0.49 | 0.41 |
| 1 | 240846124 | rs4659996 | 0.25 | 0.118 | -1.339 | 0.523 | -0.013 | 0.038 | 0.038 | 0 | 0.52 | 0.75 | 0.6 | 0.39 | 0.51 | 0.45 |
| 1 | 240852596 | rs10802901 | 0.186 | 0.255 | -1.585 | 0.539 | -0.015 | 0.089 | 0.089 | 0 | 0.47 | 0.26 | 0.45 | 0.26 | 0.53 | 0.43 |
| 1 | 240856523 | rs6696418 | 0.186 | 0.171 | -1.536 | 0.809 | 0.027 | 0.002 | 0.002 | 0.647 | 0.52 | 0.27 | 0.55 | 0.42 | 0.59 | 0.53 |
| 1 | 240866465 | rs16840492 | -0.861 | -0.516 | -1.059 | 0.766 | 0.048 | 0.003 | 0.003 | 0 | 0.47 | 0.68 | 0.44 | 0.55 | 0.32 | 0.51 |
| 1 | 240867476 | rs16840493 | -0.489 | -0.294 | -0.454 | 1.244 | -0.018 | 0.024 | 0.024 | 2.856 | 0.07 | 0.16 | 0.17 | 0.14 | 0.04 | 0.05 |
| 1 | 240870238 | rs7536260 | -0.701 | -0.31 | -0.293 | 1.228 | -0.017 | 0.034 | 0.034 | 3.074 | 0.08 | 0.22 | 0.19 | 0.14 | 0.04 | 0.07 |
| 1 | 240871487 | rs16840501 | -0.902 | -0.24 | -0.747 | 1.363 | 0.001 | -0.023 | -0.023 | 0 | 0.1 | 0.02 | 0.05 | 0.14 | 0.06 | 0.07 |
| 1 | 240879700 | rs6683045 | -1.051 | -0.035 | -0.316 | 1.514 | 0.016 | 0.096 | 0.096 | 0 | 0.25 | 0.49 | 0.26 | 0.42 | 0.26 | 0.36 |
| 1 | 240882543 | rs10802903 | 0.427 | 0.434 | -1.601 | 1.383 | -0.011 | -0.084 | -0.084 | 0 | 0.58 | 0.1 | 0.68 | 0.53 | 0.66 | 0.69 |
| 1 | 240886646 | rs7518789 | -1.319 | 0.442 | -0.599 | 1.725 | 0.007 | 0.042 | 0.042 | 0 | 0.07 | 0.17 | 0.05 | 0.15 | 0.06 | 0.08 |
| 1 | 240887547 | rs12753782 | -0.429 | 0.464 | -0.798 | 1.932 | -0.031 | -0.048 | -0.048 | 0 | 0.16 | 0.04 | 0.27 | 0.31 | 0.22 | 0.29 |
| 1 | 240888800 | rs4625268 | -1.292 | 0.424 | -0.751 | 1.873 | -0.007 | -0.02 | -0.02 | 0 | 0.23 | 0.18 | 0.33 | 0.45 | 0.28 | 0.38 |
| 1 | 240891609 | rs12076411 | -1.341 | 0.355 | -0.666 | 1.765 | -0.001 | 0.009 | 0.009 | 0 | 0.21 | 0.73 | 0.27 | 0.36 | 0.18 | 0.22 |
| 1 | 240893967 | rs947667 | -0.586 | 0.265 | -0.609 | 1.512 | -0.006 | 0.023 | 0.023 | 0.067 | 0.32 | 0.59 | 0.4 | 0.58 | 0.31 | 0.47 |
| 1 | 240900515 | rs2051032 | 0.162 | 0.624 | -1.662 | 1.087 | -0.107 | 0.012 | 0.012 | 0 | 0.38 | 0.07 | 0.52 | 0.35 | 0.45 | 0.43 |
| 1 | 240901194 | rs11802621 | 0.689 | 0.205 | -1.982 | 1.154 | 0.006 | -0.053 | -0.053 | 0 | 0.26 | 0.07 | 0.16 | 0.28 | 0.25 | 0.38 |
| 1 | 240901337 | rs2066379 | -0.697 | 0.138 | -1.434 | 1.283 | -0.006 | 0.029 | 0.029 | 0 | 0.71 | 0.81 | 0.82 | 0.89 | 0.72 | 0.85 |
| 1 | 240901508 | rs2066380 | -0.527 | 0.211 | -0.869 | 1.153 | -0.004 | 0.008 | 0.008 | 0 | 0.68 | 0.51 | 0.72 | 0.68 | 0.71 | 0.82 |
| 1 | 240913885 | rs12565039 | 0.085 | 0.624 | -1.422 | 0.828 | -0.131 | 0.025 | 0.025 | 0 | 0.39 | 0.08 | 0.56 | 0.33 | 0.49 | 0.45 |
| 1 | 240917525 | rs7542845 | -0.292 | 0.324 | -0.805 | 0.974 | -0.001 | 0.003 | 0.003 | 0 | 0.69 | 0.7 | 0.64 | 0.54 | 0.74 | 0.65 |
| 1 | 240921065 | rs12133167 | 0.031 | 0.334 | -1.047 | 0.45 | -0.008 | 0.027 | 0.027 | 0.315 | 0.1 | 0.04 | 0.09 | 0.03 | 0.11 | 0.11 |
| 1 | 240932039 | rs1418479 | -0.581 | 0.008 | -0.439 | 0.517 | -0.005 | 0.011 | 0.011 | 0 | 0.69 | 0.65 | 0.65 | 0.59 | 0.78 | 0.62 |
| 1 | 240933445 | rs6677808 | -0.228 | -0.314 | 0.131 | 0.431 | 0.009 | 0.082 | 0.082 | 0.521 | 0.07 | 0.3 | 0.05 | 0.1 | 0.04 | 0.01 |
| 1 | 240934837 | rs4568818 | 1.096 | 0.04 | -0.726 | -0.167 | -0.142 | -0.079 | -0.079 | 0 | 0.22 | 0.02 | 0.34 | 0.32 | 0.4 | 0.32 |
| 1 | 240937073 | rs10926352 | 0.188 | -0.373 | -0.361 | -0.085 | 0.038 | 0.021 | 0.021 | 0.83 | 0.54 | 0.33 | 0.56 | 0.47 | 0.68 | 0.53 |
| 1 | 240939391 | rs12757054 | 0.923 | -0.296 | -0.461 | -0.011 | -0.002 | 0.027 | 0.027 | 0.305 | 0.08 | 0.03 | 0.04 | 0.03 | 0.07 | 0.08 |
| 1 | 240943100 | rs10926354 | -1.176 | -0.178 | -0.246 | -0.068 | 0.011 | -0.067 | -0.067 | 0 | 0.69 | 0.83 | 0.65 | 0.59 | 0.76 | 0.62 |
| 1 | 240954544 | rs10495475 | 0.276 | -0.322 | 0.048 | -0.164 | 0.073 | 0.057 | 0.057 | 0.34 | 0.5 | 0.16 | 0.53 | 0.47 | 0.65 | 0.49 |
| 1 | 240956849 | rs1342445 | -0.486 | -0.271 | 0.244 | -0.167 | 0.015 | -0.03 | -0.03 | 0.204 | 0.6 | 0.74 | 0.6 | 0.57 | 0.68 | 0.52 |
| 1 | 240957393 | rs1832905 | 0.446 | -0.37 | 0.321 | -0.168 | 0.072 | 0.046 | 0.046 | 0.294 | 0.5 | 0.21 | 0.53 | 0.47 | 0.65 | 0.49 |
| 1 | 240957471 | rs1970533 | -1.199 | -0.381 | 0.34 | -0.181 | 0.01 | -0.078 | -0.078 | 1.29 | 0.69 | 0.86 | 0.64 | 0.6 | 0.76 | 0.61 |
| 1 | 240960073 | rs6429223 | -0.291 | -0.392 | 0.275 | -0.187 | 0.004 | -0.004 | -0.004 | 0 | 0.59 | 0.61 | 0.59 | 0.57 | 0.69 | 0.51 |
| 1 | 240961900 | rs3901167 | 0.993 | -0.264 | 0.166 | -0.133 | -0.002 | 0.032 | 0.032 | 0.176 | 0.08 | 0.04 | 0.04 | 0.02 | 0.08 | 0.08 |
| 1 | 240962372 | rs3932692 | 0.331 | -0.272 | 0.17 | -0.129 | 0.067 | 0.042 | 0.042 | 0.01 | 0.51 | 0.21 | 0.53 | 0.47 | 0.65 | 0.49 |
| 1 | 240967427 | rs6679827 | -0.824 | -0.255 | 0.643 | -0.047 | -0.012 | 0.024 | 0.024 | 0 | 0.66 | 0.64 | 0.64 | 0.59 | 0.76 | 0.59 |
| 1 | 240975634 | rs12239695 | -0.89 | -0.239 | 0.482 | -0.066 | 0.012 | -0.032 | -0.032 | 0 | 0.67 | 0.79 | 0.64 | 0.59 | 0.76 | 0.61 |
| 1 | 240977252 | rs1361748 | 0.523 | 0.007 | 0.343 | -0.255 | -0.024 | 0.017 | 0.017 | 0.089 | 0.46 | 0.18 | 0.43 | 0.47 | 0.55 | 0.42 |
| 1 | 240977379 | rs1342447 | -1.244 | 0.052 | 0.303 | -0.222 | 0.012 | -0.029 | -0.029 | 0 | 0.68 | 0.8 | 0.65 | 0.59 | 0.77 | 0.61 |

| | | | | | | | | | | | | | | | | |
|---|---|---|---|---|---|---|---|---|---|---|---|---|---|---|---|---|
| 1 | 240977964 | rs12744536 | -0.571 | 0.198 | -0.025 | -0.138 | 0.017 | 0 | 0 | 0 | 0.6 | 0.38 | 0.58 | 0.49 | 0.71 | 0.57 |
| 1 | 240979962 | rs3820259 | -0.249 | 0.632 | 0.11 | -0.027 | -0.001 | 0.002 | 0.002 | 0 | 0.17 | 0.17 | 0.19 | 0.12 | 0.2 | 0.17 |
| 1 | 240986292 | rs10926365 | -0.639 | 0.19 | 0.185 | 0.273 | 0.008 | 0.002 | 0.002 | 0 | 0.5 | 0.58 | 0.51 | 0.53 | 0.57 | 0.52 |
| 1 | 240987634 | rs13376172 | 0.364 | 0.435 | -0.327 | 0.264 | -0.068 | -0.034 | -0.034 | 0 | 0.34 | 0.17 | 0.44 | 0.47 | 0.51 | 0.45 |
| 1 | 241000790 | rs12042312 | -0.101 | 0.455 | -0.66 | 0.247 | -0.041 | -0.054 | -0.054 | 0 | 0.43 | 0.16 | 0.48 | 0.5 | 0.54 | 0.49 |
| 1 | 241003220 | rs796786 | -0.198 | 0.505 | -0.122 | 0.386 | 0.01 | -0.001 | -0.001 | 0.954 | 0.52 | 0.65 | 0.64 | 0.56 | 0.6 | 0.52 |
| 1 | 241006953 | rs10495476 | 1.04 | 0.595 | 0.494 | 0.792 | -0.012 | 0.014 | 0.014 | 1.513 | 0.68 | 0.56 | 0.73 | 0.62 | 0.72 | 0.63 |
| 1 | 241015766 | rs6429226 | 0.79 | 0.997 | 0.723 | 0.794 | 0.034 | 0.12 | 0.12 | 2.258 | 0.79 | 0.51 | 0.73 | 0.59 | 0.78 | 0.71 |
| 1 | 241016184 | rs1674746 | 0.409 | 1.123 | -0.192 | 0.853 | -0.046 | -0.081 | -0.081 | 0.018 | 0.82 | 0.93 | 0.76 | 0.59 | 0.79 | 0.71 |
| 1 | 241016693 | rs1380304 | 0.7 | 1.14 | -0.13 | 0.857 | 0.009 | 0.033 | 0.033 | 2.827 | 0.8 | 0.73 | 0.74 | 0.58 | 0.78 | 0.68 |
| 1 | 241027058 | rs2995378 | 0.632 | 1.226 | -0.416 | 0.741 | -0.012 | -0.017 | -0.017 | 0.091 | 0.79 | 0.8 | 0.7 | 0.62 | 0.72 | 0.68 |
| 1 | 241027742 | rs410331 | 0.143 | 1.256 | -0.758 | 0.507 | -0.105 | -0.083 | -0.083 | 0.116 | 0.82 | 0.99 | 0.68 | 0.68 | 0.71 | 0.7 |
| 1 | 241034693 | rs166370 | 0.002 | 0.499 | -1.011 | -0.248 | 0.105 | 0.143 | 0.143 | 0.983 | 0.49 | 0.71 | 0.22 | 0.16 | 0.31 | 0.29 |
| 1 | 241039187 | rs4660010 | 0.475 | 0.095 | -0.658 | -0.704 | 0.004 | -0.008 | -0.008 | 0 | 0.64 | 0.41 | 0.56 | 0.64 | 0.6 | 0.63 |
| 1 | 241041334 | rs12129550 | 0.745 | -0.154 | -0.826 | -0.947 | 0.021 | 0.069 | 0.069 | 0 | 0.29 | 0.15 | 0.13 | 0.14 | 0.24 | 0.21 |
| 1 | 241042131 | rs2796742 | 0.453 | -0.165 | -0.31 | -0.976 | -0.009 | 0.015 | 0.015 | 0 | 0.46 | 0.36 | 0.52 | 0.53 | 0.5 | 0.51 |
| 1 | 241047490 | rs3856237 | 0.406 | -0.218 | 0.037 | -0.965 | -0.014 | 0.025 | 0.025 | 0 | 0.47 | 0.27 | 0.51 | 0.53 | 0.5 | 0.52 |
| 1 | 241048722 | rs12069867 | 1.189 | -0.291 | 0.169 | -1.035 | -0.033 | -0.049 | -0.049 | 0.46 | 0.33 | 0.15 | 0.44 | 0.51 | 0.39 | 0.46 |
| 1 | 241049361 | rs2995381 | -0.765 | -0.088 | 0.31 | -0.904 | 0.015 | 0.014 | 0.014 | 0.007 | 0.82 | 0.83 | 0.68 | 0.69 | 0.74 | 0.74 |
| 1 | 241050375 | rs17698283 | 0.402 | -0.286 | 0.007 | -0.94 | 0.006 | 0.011 | 0.011 | 0.334 | 0.13 | 0.11 | 0.08 | 0.02 | 0.12 | 0.06 |
| 1 | 241058174 | rs12751419 | -0.325 | -0.023 | -0.173 | -1.04 | 0 | 0.051 | 0.051 | 0.053 | 0.17 | 0.11 | 0.1 | 0 | 0.16 | 0.08 |
| 1 | 241065033 | rs17378711 | -0.359 | -0.035 | -0.179 | -1.046 | 0.001 | 0.053 | 0.053 | 0.046 | 0.18 | 0.11 | 0.1 | 0 | 0.16 | 0.08 |
| 1 | 241067471 | rs261804 | 0.072 | -0.156 | -0.141 | -1.122 | 0.001 | 0.042 | 0.042 | 0 | 0.5 | 0.28 | 0.45 | 0.46 | 0.48 | 0.44 |
| 1 | 241068852 | rs390488 | -1.15 | -0.182 | 0.403 | -0.829 | -0.013 | -0.025 | -0.025 | 0 | 0.7 | 0.74 | 0.57 | 0.49 | 0.65 | 0.52 |
| 1 | 241070371 | rs261827 | -0.818 | -0.129 | 0.196 | -0.905 | 0 | 0.001 | 0.001 | 0 | 0.68 | 0.38 | 0.56 | 0.45 | 0.66 | 0.52 |
| 1 | 241078810 | rs191735 | -0.381 | -0.331 | 0.17 | -0.904 | 0 | 0.001 | 0.001 | 0 | 0.62 | 0.4 | 0.55 | 0.45 | 0.62 | 0.51 |
| 1 | 241087708 | rs422256 | -0.995 | -0.547 | -0.453 | -0.992 | -0.055 | -0.091 | -0.091 | 0.087 | 0.66 | 0.84 | 0.57 | 0.51 | 0.61 | 0.5 |
| 1 | 241090153 | rs195778 | 0.312 | -0.69 | -0.699 | -1.178 | -0.128 | -0.053 | -0.053 | 0 | 0.32 | 0.03 | 0.39 | 0.32 | 0.42 | 0.33 |
| 1 | 241095576 | rs261861 | -0.429 | -0.721 | -0.026 | -1.023 | -0.022 | -0.008 | -0.008 | 0.434 | 0.61 | 0.65 | 0.51 | 0.47 | 0.53 | 0.49 |
| 1 | 241100122 | rs261857 | 0.223 | -0.765 | 0.554 | -1.146 | -0.002 | 0.047 | 0.047 | 0 | 0.44 | 0.26 | 0.44 | 0.25 | 0.46 | 0.39 |
| 1 | 241105515 | rs6697598 | -0.869 | -0.76 | 0.763 | -0.922 | 0.002 | -0.018 | -0.018 | 0 | 0.1 | 0.37 | 0.05 | 0.02 | 0.04 | 0.02 |
| 1 | 241107061 | rs12038552 | 0.867 | -0.641 | -0.295 | -0.868 | -0.099 | -0.019 | -0.019 | 0 | 0.36 | 0.09 | 0.46 | 0.69 | 0.41 | 0.55 |
| 1 | 241110180 | rs261796 | 1.713 | -0.399 | 0.558 | -0.887 | 0.021 | -0.015 | -0.015 | 0 | 0.9 | 0.58 | 0.93 | 0.96 | 0.92 | 0.87 |
| 1 | 241111278 | rs261798 | -0.056 | -0.399 | 0.162 | -0.797 | -0.008 | -0.005 | -0.005 | 0 | 0.95 | 0.9 | 0.99 | 0.99 | 0.99 | 0.98 |
| 1 | 241117850 | rs431188 | 1.517 | -0.451 | 0.151 | -0.7 | -0.006 | -0.001 | -0.001 | 0 | 0.86 | 0.77 | 0.93 | 0.9 | 0.91 | 0.83 |
| 1 | 241119341 | rs2502436 | 1.428 | -0.423 | 0.161 | -0.687 | -0.007 | -0.001 | -0.001 | 0 | 0.86 | 0.77 | 0.93 | 0.9 | 0.91 | 0.83 |
| 1 | 241125990 | rs3912106 | -0.134 | -0.323 | 0.191 | -0.84 | -0.002 | 0.011 | 0.011 | 0.753 | 0.43 | 0.4 | 0.42 | 0.28 | 0.46 | 0.49 |
| 1 | 241126234 | rs3912107 | 0.199 | -0.329 | 0.196 | -0.839 | -0.001 | 0.007 | 0.007 | 0.168 | 0.38 | 0.36 | 0.39 | 0.24 | 0.41 | 0.36 |
| 1 | 241131301 | rs12119557 | -0.059 | -0.271 | 0.122 | -0.672 | 0.004 | 0.085 | 0.085 | 0 | 0.44 | 0.26 | 0.41 | 0.26 | 0.43 | 0.39 |
| 1 | 241133108 | rs6429230 | 0.334 | -0.359 | 0.18 | -0.766 | -0.003 | -0.022 | -0.022 | 0 | 0.82 | 0.38 | 0.88 | 0.91 | 0.85 | 0.88 |
| 1 | 241133704 | rs11586002 | 0.985 | -0.338 | 0.097 | -0.799 | -0.052 | -0.035 | -0.035 | 0.079 | 0.36 | 0.12 | 0.47 | 0.65 | 0.41 | 0.48 |
| 1 | 241134342 | rs7532910 | -0.671 | -0.335 | 0.613 | -0.837 | 0.004 | -0.094 | -0.094 | 0 | 0.86 | 0.71 | 0.92 | 1 | 0.9 | 0.98 |
| 1 | 241135110 | rs6682084 | 0.108 | -0.315 | 0.042 | -0.819 | -0.015 | 0.087 | 0.087 | 0 | 0.47 | 0.29 | 0.43 | 0.3 | 0.49 | 0.51 |
| 1 | 241139151 | rs4660017 | -0.137 | -0.161 | 0.19 | -0.73 | -0.013 | -0.103 | -0.103 | 0 | 0.84 | 0.47 | 0.9 | 1 | 0.9 | 0.99 |
| 1 | 241145243 | rs3911618 | 0.661 | -0.054 | -0.699 | -0.615 | -0.055 | 0.028 | 0.028 | 0 | 0.37 | 0.12 | 0.47 | 0.69 | 0.42 | 0.48 |
| 1 | 241148960 | rs10802917 | 0.495 | -0.084 | -0.675 | -0.599 | -0.046 | 0.016 | 0.016 | 0 | 0.39 | 0.18 | 0.51 | 0.67 | 0.47 | 0.6 |
| 1 | 241151974 | rs1478251 | -0.615 | -0.05 | -0.616 | -0.683 | 0.001 | -0.01 | -0.01 | 0 | 0.86 | 0.91 | 0.93 | 1 | 0.89 | 0.98 |
| 1 | 241152090 | rs9786932 | 0.088 | -0.072 | -0.47 | -0.678 | 0.008 | 0.028 | 0.028 | 0 | 0.47 | 0.56 | 0.41 | 0.28 | 0.42 | 0.34 |
| 1 | 241152827 | rs6673612 | -0.331 | 0.144 | -0.691 | -0.418 | 0 | -0.01 | -0.01 | 0.018 | 0.87 | 0.92 | 0.93 | 1 | 0.89 | 0.98 |
| 1 | 241158335 | rs2678785 | 0.412 | 0.062 | -1.255 | -0.569 | -0.081 | -0.02 | -0.02 | 0 | 0.39 | 0.1 | 0.49 | 0.62 | 0.47 | 0.51 |
| 1 | 241159338 | rs6657011 | 0.034 | -0.116 | -0.561 | -0.54 | 0 | 0 | 0 | 0 | 0.45 | 0.56 | 0.44 | 0.41 | 0.46 | 0.53 |
| 1 | 241161151 | rs10926391 | 0.197 | -0.291 | -0.846 | -0.816 | -0.001 | 0.018 | 0.018 | 0 | 0.45 | 0.42 | 0.43 | 0.64 | 0.47 | 0.6 |
| 1 | 241161648 | rs575226 | 0.218 | -0.353 | -1.627 | -0.754 | -0.007 | 0.027 | 0.027 | 0 | 0.38 | 0.2 | 0.45 | 0.34 | 0.38 | 0.36 |
| 1 | 241163924 | rs624331 | -0.168 | -0.24 | -2.207 | -0.78 | -0.007 | 0.063 | 0.063 | 0 | 0.21 | 0.1 | 0.15 | 0.06 | 0.2 | 0.12 |
| 1 | 241164526 | rs2502448 | 0.74 | -0.29 | -2.364 | -0.597 | -0.03 | -0.039 | -0.039 | 0.887 | 0.34 | 0.14 | 0.36 | 0.58 | 0.4 | 0.54 |
| 1 | 241167010 | rs785539 | -0.219 | -0.122 | -2.238 | -0.789 | -0.009 | 0.072 | 0.072 | 0 | 0.22 | 0.09 | 0.16 | 0.05 | 0.2 | 0.15 |
| 1 | 241168080 | rs6680767 | 0.05 | -0.381 | -2.148 | -0.635 | -0.053 | 0.05 | 0.05 | 0 | 0.36 | 0.14 | 0.39 | 0.31 | 0.4 | 0.33 |
| 1 | 241168535 | rs10802919 | -0.31 | -0.398 | -2.013 | -0.789 | -0.003 | 0.042 | 0.042 | 0.365 | 0.4 | 0.29 | 0.4 | 0.31 | 0.4 | 0.34 |
| 1 | 241170150 | rs12026885 | -0.082 | -0.825 | -2.137 | -0.692 | 0.018 | 0.004 | 0.004 | 0.268 | 0.2 | 0.14 | 0.21 | 0.22 | 0.14 | 0.22 |
| 1 | 241172790 | rs599578 | 1.282 | -0.656 | -2.463 | -0.557 | -0.054 | -0.036 | -0.036 | 0 | 0.44 | 0.09 | 0.42 | 0.55 | 0.45 | 0.48 |
| 1 | 241174285 | rs639108 | 1.241 | -0.518 | -2.405 | -0.501 | -0.064 | -0.071 | -0.071 | 0.357 | 0.45 | 0.08 | 0.42 | 0.52 | 0.45 | 0.48 |
| 1 | 241176871 | rs546547 | -1.307 | -0.482 | -1.709 | -0.887 | 0.01 | 0.056 | 0.056 | 2.116 | 0.31 | 0.13 | 0.28 | 0.2 | 0.26 | 0.26 |
| 1 | 241178272 | rs10802921 | -0.696 | -0.143 | -1.31 | -0.465 | -0.006 | 0.004 | 0.004 | 0.187 | 0.2 | 0.1 | 0.22 | 0.19 | 0.19 | 0.22 |
| 1 | 241180198 | rs530479 | 0.84 | -0.714 | -0.501 | -0.89 | -0.023 | -0.097 | -0.097 | 0 | 0.82 | 0.66 | 0.89 | 1 | 0.88 | 0.91 |
| 1 | 241183190 | rs12068733 | -1.073 | -1.446 | -0.28 | -1.097 | -0.033 | -0.03 | -0.03 | 0 | 0.24 | 0.33 | 0.17 | 0.12 | 0.13 | 0.16 |
| 1 | 241183612 | rs601675 | -2.1 | -2.419 | -0.636 | -1.343 | 0.157 | 0.136 | 0.136 | 0.083 | 0.47 | 0.87 | 0.35 | 0.21 | 0.22 | 0.36 |
| 1 | 241183893 | rs602676 | -0.66 | -2.261 | 0.256 | -1.668 | -0.028 | -0.094 | -0.094 | 0.154 | 0.16 | 0.26 | 0.11 | 0 | 0.08 | 0.02 |
| 1 | 241185048 | rs6663803 | -0.951 | -2.21 | 0.704 | -1.574 | -0.058 | -0.056 | -0.056 | 0 | 0.26 | 0.41 | 0.18 | 0.12 | 0.13 | 0.16 |
| 1 | 241185050 | rs10926393 | 0.071 | -2.167 | 0.591 | -1.527 | 0.006 | 0.001 | 0.001 | 0 | 0.13 | 0.08 | 0.12 | 0.09 | 0.07 | 0.08 |

| | | | | | | | | | | | | | | | | |
|---|---|---|---|---|---|---|---|---|---|---|---|---|---|---|---|---|
| 1 | 241186056 | rs523361 | -1.41 | -1.876 | 0.058 | -2.117 | -0.005 | 0.074 | 0.074 | 0 | 0.59 | 0.68 | 0.43 | 0.12 | 0.44 | 0.34 |
| 1 | 241186585 | rs662023 | -1.16 | -1.906 | 0.186 | -2.131 | 0 | 0.013 | 0.013 | 0 | 0.57 | 0.53 | 0.42 | 0.12 | 0.44 | 0.34 |
| 1 | 241186891 | rs10802923 | -1.343 | -1.906 | 0.185 | -2.131 | -0.004 | 0.073 | 0.073 | 0 | 0.58 | 0.68 | 0.43 | 0.12 | 0.44 | 0.34 |
| 1 | 241188093 | rs472519 | -1.137 | -1.994 | 0.185 | -2.13 | 0.002 | 0.015 | 0.015 | 0 | 0.56 | 0.53 | 0.42 | 0.12 | 0.43 | 0.34 |
| 1 | 241191352 | rs616935 | -1.306 | -1.884 | 0.083 | -2.121 | -0.003 | 0.077 | 0.077 | 0 | 0.58 | 0.68 | 0.43 | 0.12 | 0.44 | 0.35 |
| 1 | 241196335 | rs785544 | -1.126 | -1.814 | 0.066 | -2.014 | 0.002 | 0.041 | 0.041 | 0 | 0.56 | 0.58 | 0.46 | 0.14 | 0.44 | 0.32 |
| 1 | 241198268 | rs596695 | -0.944 | -1.668 | 0.469 | -2.044 | 0.01 | 0.013 | 0.013 | 0 | 0.49 | 0.48 | 0.39 | 0.12 | 0.4 | 0.33 |
| 1 | 241198832 | rs583951 | 0.591 | -2.008 | 0.753 | -2.094 | 0.024 | 0.035 | 0.035 | 0.99 | 0.54 | 0.39 | 0.62 | 0.86 | 0.66 | 0.75 |
| 1 | 241199956 | rs2500241 | 0.68 | -1.283 | 1.071 | -2.242 | 0.01 | -0.119 | -0.119 | 0 | 0.69 | 0.55 | 0.76 | 0.96 | 0.68 | 0.81 |
| 1 | 241199995 | rs538423 | 1.919 | -0.984 | 0.786 | -1.948 | -0.162 | -0.098 | -0.098 | 0 | 0.3 | 0.02 | 0.42 | 0.69 | 0.41 | 0.41 |
| 1 | 241201934 | rs2459950 | 1.69 | 0.388 | 2.147 | -1.662 | 0.002 | -0.093 | -0.093 | 0.349 | 0.65 | 0.39 | 0.73 | 0.96 | 0.65 | 0.81 |
| 1 | 241202525 | rs11590400 | -0.578 | 0.393 | 1.892 | -1.679 | -0.013 | 0.011 | 0.011 | 0.152 | 0.2 | 0.1 | 0.14 | 0.01 | 0.25 | 0.14 |
| 1 | 241205433 | rs12059322 | -0.889 | 0.36 | 1.881 | -1.642 | 0.002 | -0.028 | -0.028 | 0.023 | 0.07 | 0.17 | 0.06 | 0.02 | 0.06 | 0.04 |
| 1 | 241210870 | rs10926408 | -2.019 | 0.436 | 1.923 | -1.639 | 0.008 | -0.123 | -0.123 | 0 | 0.34 | 0.5 | 0.25 | 0.04 | 0.35 | 0.19 |
| 1 | 241224878 | rs3002682 | 1.527 | 0.796 | 2.138 | -1.247 | 0.018 | -0.083 | -0.083 | 0.425 | 0.67 | 0.47 | 0.76 | 0.84 | 0.66 | 0.81 |
| 1 | 241240132 | rs12751978 | -1.593 | 0.592 | 2.462 | -0.959 | -0.002 | -0.095 | -0.095 | 0.064 | 0.23 | 0.32 | 0.17 | 0.02 | 0.21 | 0.09 |
| 1 | 241242460 | rs1878729 | -2.777 | 0.728 | 1.849 | -0.885 | -0.073 | 0.093 | 0.093 | 0 | 0.41 | 0.91 | 0.36 | 0.3 | 0.45 | 0.32 |
| 1 | 241242629 | rs2678778 | -0.431 | 0.899 | 2.081 | -0.927 | 0.022 | 0.015 | 0.015 | 0 | 0.11 | 0.21 | 0.14 | 0.13 | 0.16 | 0.17 |
| 1 | 241247504 | rs2815844 | -0.683 | 0.901 | 2.036 | -1.176 | 0.051 | -0.012 | -0.012 | 0 | 0.12 | 0.36 | 0.15 | 0.09 | 0.16 | 0.17 |
| 1 | 241249273 | rs2500246 | -1.306 | 0.714 | 1.757 | -1.208 | 0.001 | -0.101 | -0.101 | 0 | 0.2 | 0.32 | 0.11 | 0.01 | 0.18 | 0.09 |
| 1 | 241254147 | rs2502426 | -1.258 | 1.094 | 1.604 | -1.025 | 0.022 | -0.139 | -0.139 | 0 | 0.19 | 0.5 | 0.13 | 0.01 | 0.18 | 0.09 |
| 1 | 241257765 | rs2492988 | -1.119 | 0.905 | 1.155 | -0.955 | 0.041 | -0.093 | -0.093 | 0.572 | 0.22 | 0.53 | 0.2 | 0.09 | 0.24 | 0.14 |
| 1 | 241258422 | rs2678780 | -1.716 | 0.905 | 0.053 | -0.86 | -0.039 | 0.101 | 0.101 | 0 | 0.36 | 0.84 | 0.35 | 0.19 | 0.39 | 0.33 |
| 1 | 241260733 | rs2502432 | -1.284 | 1.01 | 0.582 | -0.648 | 0.042 | -0.071 | -0.071 | 0 | 0.2 | 0.39 | 0.17 | 0.09 | 0.24 | 0.14 |
| 1 | 241264216 | rs12086317 | -1.199 | 0.536 | 0.287 | -0.718 | -0.003 | -0.046 | -0.046 | 0 | 0.17 | 0.18 | 0.16 | 0.02 | 0.15 | 0.1 |
| 1 | 241274518 | rs7521857 | -1.71 | 1.151 | 0.147 | -0.266 | 0 | -0.005 | -0.005 | 0 | 0.13 | 0.83 | 0.19 | 0.09 | 0.15 | 0.05 |
| 1 | 241280303 | rs12024588 | -0.266 | 0.563 | 0.165 | -0.286 | -0.01 | -0.006 | -0.006 | 0 | 0.23 | 0.24 | 0.16 | 0.18 | 0.19 | 0.15 |
| 1 | 241281193 | rs4339853 | -1.355 | 0.932 | 0.949 | 0.902 | 0.028 | 0.194 | 0.194 | 1.34 | 0.08 | 0.33 | 0.13 | 0.28 | 0.05 | 0.09 |
| 1 | 241286525 | rs7550909 | -1.311 | 0.811 | 1.07 | 0.426 | 0.026 | 0.04 | 0.04 | 0.514 | 0.19 | 0.4 | 0.2 | 0.21 | 0.22 | 0.12 |
| 1 | 241288878 | rs4131804 | -1.116 | 0.329 | 0.968 | 0.24 | 0.031 | -0.05 | -0.05 | 0.542 | 0.35 | 0.83 | 0.32 | 0.38 | 0.25 | 0.26 |
| 1 | 241290401 | rs6685517 | -0.741 | 0.305 | 1.072 | 0.3 | 0.01 | 0.034 | 0.034 | 0 | 0.07 | 0.15 | 0.13 | 0.17 | 0.06 | 0.08 |
| 1 | 241290674 | rs6696557 | -1.089 | 0.744 | 0.986 | 0.224 | -0.001 | -0.038 | -0.038 | 0 | 0.36 | 0.89 | 0.34 | 0.41 | 0.28 | 0.26 |
| 1 | 241292292 | rs12067659 | -1.91 | 0.708 | 1.147 | 0.209 | 0.074 | 0.12 | 0.12 | 2.974 | 0.05 | 0.48 | 0.03 | 0.07 | 0 | 0 |
| 1 | 241294032 | rs6429241 | -1.83 | 0.83 | 0.531 | 1.335 | -0.084 | 0.075 | 0.075 | 2.132 | 0.49 | 0.97 | 0.54 | 0.69 | 0.46 | 0.39 |
| 1 | 241304206 | rs4660032 | -1.609 | 0.605 | 0.69 | 1.424 | -0.035 | 0.136 | 0.136 | 3.492 | 0.11 | 0.61 | 0.13 | 0.25 | 0.02 | 0.05 |
| 1 | 241319268 | rs7543441 | -1.382 | 0.779 | 0.413 | 1.148 | 0.041 | 0.048 | 0.048 | 0.946 | 0.46 | 0.78 | 0.49 | 0.63 | 0.41 | 0.36 |
| 1 | 241319347 | rs12041617 | 0.463 | 0.877 | 0.101 | 1.122 | -0.009 | -0.007 | -0.007 | 0 | 0.22 | 0.23 | 0.16 | 0.16 | 0.19 | 0.18 |
| 1 | 241320367 | rs10465632 | -1.274 | 1.014 | -0.653 | 0.776 | 0.011 | 0.068 | 0.068 | 2.59 | 0.48 | 0.95 | 0.51 | 0.7 | 0.41 | 0.36 |
| 1 | 241326426 | rs6665888 | -0.777 | 1.945 | -0.733 | 0.532 | -0.014 | 0.051 | 0.051 | 0 | 0.43 | 0.77 | 0.52 | 0.68 | 0.38 | 0.45 |
| 1 | 241326594 | rs12567677 | 1.104 | 1.499 | -1.439 | 0.336 | -0.044 | 0.079 | 0.079 | 0.437 | 0.41 | 0.04 | 0.39 | 0.28 | 0.43 | 0.45 |
| 1 | 241326886 | rs6675503 | 0.194 | 1.085 | -0.949 | 0.234 | -0.003 | -0.04 | -0.04 | 0 | 0.12 | 0.13 | 0.07 | 0 | 0.12 | 0.02 |
| 1 | 241328227 | rs10802928 | 0.408 | 1.087 | -2.261 | 0.027 | -0.082 | -0.048 | -0.048 | 0 | 0.31 | 0.04 | 0.34 | 0.3 | 0.32 | 0.41 |
| 1 | 241329258 | rs4660034 | -0.185 | 1.007 | -1.963 | 0.274 | 0.014 | -0.012 | -0.012 | 0 | 0.66 | 0.31 | 0.69 | 0.64 | 0.71 | 0.68 |
| 1 | 241330394 | rs4660036 | -0.058 | 0.585 | -2.538 | -0.293 | 0.011 | 0.031 | 0.031 | 3.148 | 0.43 | 0.29 | 0.4 | 0.58 | 0.39 | 0.33 |
| 1 | 241331883 | rs12126298 | 0.779 | 0.559 | -2.512 | -0.097 | 0.002 | -0.025 | -0.025 | 0.326 | 0.26 | 0.17 | 0.27 | 0.36 | 0.25 | 0.28 |
| 1 | 241334777 | rs6429244 | -1.617 | 0.866 | -2.555 | -0.34 | -0.062 | 0.101 | 0.101 | 0 | 0.82 | 0.97 | 0.84 | 0.99 | 0.72 | 0.87 |
| 1 | 241334960 | rs6429246 | 0.5 | 0.359 | -1.593 | -0.059 | 0.003 | 0.05 | 0.05 | 1.744 | 0.12 | 0.39 | 0.13 | 0.2 | 0.09 | 0.27 |
| 1 | 241338558 | rs10926423 | 1.648 | 0.359 | -1.881 | 0.015 | -0.004 | -0.06 | -0.06 | 0.02 | 0.16 | 0.07 | 0.13 | 0.23 | 0.14 | 0.05 |
| 1 | 241339851 | rs11578627 | 0.737 | 0.148 | -1.875 | -0.605 | 0.011 | 0.089 | 0.089 | 0 | 0.52 | 0.31 | 0.48 | 0.17 | 0.43 | 0.29 |
| 1 | 241340415 | rs6677639 | 0.804 | -0.074 | -1.732 | -0.721 | -0.047 | -0.176 | -0.176 | 0 | 0.32 | 0.45 | 0.2 | 0 | 0.19 | 0.09 |
| 1 | 241342330 | rs4072897 | -0.056 | 0.48 | -1.185 | 0.222 | 0.03 | 0.179 | 0.179 | 0 | 0.84 | 0.6 | 0.83 | 0.6 | 0.81 | 0.67 |
| 1 | 241345433 | rs10802929 | -0.52 | 0.491 | -1.738 | -0.078 | -0.016 | 0.018 | 0.018 | 0.412 | 0.37 | 0.32 | 0.4 | 0.63 | 0.44 | 0.41 |
| 1 | 241353778 | rs12127470 | 0.389 | -0.053 | -1.623 | 0 | 0.007 | -0.01 | -0.01 | 0 | 0.71 | 0.74 | 0.73 | 0.41 | 0.73 | 0.42 |
| 1 | 241356618 | rs12025118 | -0.162 | -0.111 | -1.587 | 0.204 | -0.003 | -0.003 | -0.003 | 1.484 | 0.81 | 0.76 | 0.87 | 0.82 | 0.83 | 0.66 |
| 1 | 241357782 | rs11802564 | 1.254 | -0.079 | -1.648 | 0.324 | -0.007 | -0.061 | -0.061 | 0 | 0.1 | 0 | 0.14 | 0.4 | 0.1 | 0.24 |
| 1 | 241361793 | rs1915872 | 1.868 | -0.188 | -2.365 | -0.306 | -0.071 | -0.012 | -0.012 | 0 | 0.21 | 0.02 | 0.32 | 0.19 | 0.23 | 0.19 |
| 1 | 241363241 | rs1915877 | 1.249 | -0.578 | -2.312 | -0.877 | -0.041 | -0.137 | -0.137 | 0.042 | 0.25 | 0.56 | 0.13 | 0 | 0.14 | 0.05 |
| 1 | 241364184 | rs2815873 | -0.466 | -0.337 | -2.114 | -0.615 | -0.018 | -0.014 | -0.014 | 0 | 0.8 | 0.66 | 0.86 | 0.82 | 0.83 | 0.67 |
| 1 | 241365280 | rs10802934 | 0.83 | 0.239 | -2.035 | -0.277 | -0.061 | -0.023 | -0.023 | 0 | 0.24 | 0.07 | 0.3 | 0.26 | 0.36 | 0.19 |
| 1 | 241367721 | rs2177113 | 0.998 | 0.281 | -1.888 | -0.07 | -0.001 | 0.014 | 0.014 | 0 | 0.08 | 0.04 | 0.11 | 0.25 | 0.09 | 0.05 |
| 1 | 241370019 | rs12722977 | 1.038 | 0.709 | -1.948 | -0.241 | -0.011 | 0 | 0 | 0 | 0.1 | 0.08 | 0.21 | 0.15 | 0.12 | 0.08 |
| 1 | 241372527 | rs4659592 | 0.617 | 0.779 | -2.119 | -0.362 | -0.051 | -0.094 | -0.094 | 0 | 0.29 | 0.15 | 0.33 | 0.42 | 0.39 | 0.4 |
| 1 | 241374115 | rs2686228 | 0.609 | 0.228 | -1.795 | -0.701 | -0.023 | -0.008 | -0.008 | 0 | 0.3 | 0.41 | 0.22 | 0.24 | 0.22 | 0.2 |
| 1 | 241375848 | rs2177116 | 0.66 | 0.212 | -1.797 | -0.711 | -0.031 | -0.02 | -0.02 | 0.264 | 0.29 | 0.45 | 0.22 | 0.21 | 0.2 | 0.18 |
| 1 | 241376147 | rs4400595 | -1.527 | 0.12 | -1.896 | -0.849 | -0.024 | -0.055 | -0.055 | 0 | 0.75 | 0.56 | 0.82 | 0.84 | 0.79 | 0.73 |
| 1 | 241376512 | rs2686226 | -1.956 | 0.121 | -1.741 | -0.849 | 0.002 | 0.004 | 0.004 | 0 | 0.77 | 0.78 | 0.84 | 0.84 | 0.79 | 0.73 |
| 1 | 241378929 | rs1879745 | 0.639 | 0.229 | -1.736 | -0.706 | -0.024 | -0.018 | -0.018 | 0.051 | 0.3 | 0.42 | 0.22 | 0.21 | 0.22 | 0.2 |
| 1 | 241381794 | rs4660045 | 1.915 | 0.331 | -1.902 | -0.676 | -0.026 | -0.076 | -0.076 | 0 | 0.11 | 0.01 | 0.14 | 0.4 | 0.14 | 0.27 |
| 1 | 241383503 | rs2815864 | -0.992 | 0.413 | -1.831 | -0.892 | -0.001 | -0.013 | -0.013 | 0 | 0.58 | 0.56 | 0.63 | 0.83 | 0.53 | 0.55 |

| | | | | | | | | | | | | | | | | |
|---|---|---|---|---|---|---|---|---|---|---|---|---|---|---|---|---|
| 1 | 241384522 | rs11584669 | 0.356 | 0.403 | -1.605 | -0.899 | -0.025 | -0.026 | -0.026 | 0 | 0.31 | 0.42 | 0.22 | 0.22 | 0.23 | 0.19 |
| 1 | 241384805 | rs6682206 | 2.243 | 0.781 | -1.826 | -0.945 | -0.025 | 0.016 | 0.016 | 0 | 0.16 | 0.09 | 0.2 | 0.01 | 0.26 | 0.14 |
| 1 | 241388129 | rs6662036 | 2.369 | 0.749 | -1.829 | -0.931 | -0.023 | 0.016 | 0.016 | 0 | 0.16 | 0.08 | 0.2 | 0.01 | 0.25 | 0.14 |
| 1 | 241388393 | rs2341688 | 0.897 | 0.46 | -1.726 | -0.532 | -0.015 | -0.018 | -0.018 | 0.283 | 0.15 | 0.06 | 0.13 | 0.22 | 0.14 | 0.13 |
| 1 | 241391530 | rs12140345 | 0.995 | 0.209 | -1.502 | -0.483 | 0.013 | -0.033 | -0.033 | 0 | 0.14 | 0.04 | 0.11 | 0.2 | 0.09 | 0.12 |
| 1 | 241392925 | rs2686231 | 1.356 | 0.209 | -1.032 | -0.837 | -0.005 | -0.104 | -0.104 | 2.12 | 0.15 | 0.33 | 0.1 | 0 | 0.12 | 0.06 |
| 1 | 241394361 | rs2815858 | -0.345 | 0.213 | -0.822 | -0.407 | 0.003 | 0.048 | 0.048 | 0 | 0.75 | 0.67 | 0.68 | 0.54 | 0.72 | 0.6 |
| 1 | 241397662 | rs2815851 | 0.252 | 0.087 | -1.062 | -0.631 | -0.005 | -0.005 | -0.005 | 0.065 | 0.43 | 0.46 | 0.35 | 0.34 | 0.34 | 0.43 |
| 1 | 241400004 | rs2686240 | 0.461 | 0.239 | -1.02 | -0.805 | -0.002 | 0.002 | 0.002 | 0 | 0.45 | 0.45 | 0.34 | 0.38 | 0.49 | 0.47 |
| 1 | 241401205 | rs11589937 | 1.805 | 0.169 | -1.097 | -0.651 | 0.002 | 0.006 | 0.006 | 0.31 | 0.15 | 0.08 | 0.13 | 0.15 | 0.12 | 0.07 |
| 1 | 241401288 | rs2815834 | 0.825 | 0.112 | -0.899 | -0.704 | 0.002 | -0.012 | -0.012 | 0 | 0.26 | 0.21 | 0.14 | 0.33 | 0.23 | 0.34 |
| 1 | 241403497 | rs4359023 | 0.039 | 0.132 | -0.985 | -0.474 | 0.002 | -0.001 | -0.001 | 0.151 | 0.29 | 0.17 | 0.26 | 0.36 | 0.26 | 0.2 |
| 1 | 241406762 | rs10926438 | 0.624 | 0.46 | -1.174 | -0.414 | -0.031 | -0.125 | -0.125 | 0.005 | 0.21 | 0.09 | 0.18 | 0.53 | 0.24 | 0.3 |
| 1 | 241408548 | rs9428501 | -1.636 | 0.505 | -0.853 | -0.559 | 0.006 | 0.007 | 0.007 | 0 | 0.73 | 0.72 | 0.63 | 0.71 | 0.62 | 0.63 |
| 1 | 241409847 | rs1879744 | -0.057 | 0.298 | -0.836 | -0.495 | -0.003 | 0.001 | 0.001 | 0 | 0.44 | 0.45 | 0.45 | 0.49 | 0.36 | 0.36 |
| 1 | 241413586 | rs12402337 | 2.593 | 0.42 | -0.912 | -0.425 | 0.001 | 0.004 | 0.004 | 0 | 0.14 | 0.1 | 0.21 | 0.02 | 0.12 | 0.09 |
| 1 | 241420962 | rs12404897 | 1.407 | 0.78 | -0.696 | -0.325 | 0.015 | 0.02 | 0.02 | 0 | 0.25 | 0.32 | 0.33 | 0.3 | 0.37 | 0.37 |
| 1 | 241421855 | rs10926442 | 1.628 | 0.445 | -0.997 | -0.374 | 0.003 | 0.006 | 0.006 | 0.309 | 0.15 | 0.1 | 0.14 | 0.14 | 0.11 | 0.08 |
| 1 | 241421911 | rs10926443 | 1.297 | 0.443 | -1.076 | -0.372 | -0.007 | -0.014 | -0.014 | 0 | 0.13 | 0.09 | 0.14 | 0.22 | 0.14 | 0.12 |
| 1 | 241428109 | rs10926446 | -0.979 | 0.27 | -0.889 | -0.441 | 0.001 | -0.02 | -0.02 | 0 | 0.74 | 0.81 | 0.84 | 0.67 | 0.75 | 0.72 |
| 1 | 241428657 | rs7511717 | 1.663 | 0.317 | -0.672 | -0.5 | 0.002 | 0.038 | 0.038 | 0.203 | 0.25 | 0.39 | 0.34 | 0.28 | 0.26 | 0.39 |
| 1 | 241433465 | rs1996806 | 0.372 | 0.257 | -0.805 | -0.61 | 0.004 | -0.023 | -0.023 | 0 | 0.28 | 0.2 | 0.21 | 0.46 | 0.24 | 0.28 |
| 1 | 241433847 | rs4660051 | -0.384 | 0.235 | -0.833 | -0.652 | 0.012 | 0.039 | 0.039 | 0.08 | 0.55 | 0.3 | 0.56 | 0.73 | 0.6 | 0.53 |
| 1 | 241437308 | rs7555768 | -2.271 | 0.333 | -1.006 | -0.686 | -0.001 | -0.041 | -0.041 | 0 | 0.84 | 0.8 | 0.9 | 1 | 0.86 | 0.92 |
| 1 | 241439295 | rs10442664 | 3.291 | 0.411 | -1.226 | -0.714 | -0.064 | 0.033 | 0.033 | 0 | 0.14 | 0.01 | 0.19 | 0.06 | 0.22 | 0.12 |
| 1 | 241439672 | rs4659596 | 0.481 | -0.005 | -1.04 | -0.651 | -0.033 | -0.027 | -0.027 | 0 | 0.39 | 0.21 | 0.38 | 0.59 | 0.46 | 0.42 |
| 1 | 241440418 | rs7520347 | 2.187 | 0.024 | -1.078 | -0.576 | -0.025 | -0.061 | -0.061 | 0 | 0.1 | 0 | 0.06 | 0.33 | 0.11 | 0.18 |
| 1 | 241442019 | rs10802943 | -0.668 | -0.054 | -0.971 | -0.641 | 0.003 | 0.009 | 0.009 | 0 | 0.58 | 0.38 | 0.56 | 0.73 | 0.6 | 0.52 |
| 1 | 241445752 | rs3964281 | 0.902 | 0.023 | -1.087 | -0.617 | -0.032 | -0.06 | -0.06 | 0.964 | 0.33 | 0.15 | 0.31 | 0.58 | 0.41 | 0.4 |
| 1 | 241447310 | rs10802944 | 1.09 | 0 | -0.952 | -0.652 | 0.001 | 0.011 | 0.011 | 0.256 | 0.31 | 0.43 | 0.41 | 0.29 | 0.32 | 0.42 |
| 1 | 241449703 | rs1114839 | 1.317 | 0.021 | -1.164 | -1.001 | -0.034 | 0.023 | 0.023 | 0 | 0.27 | 0.12 | 0.31 | 0.26 | 0.35 | 0.24 |
| 1 | 241450347 | rs11584771 | 1.721 | 0.004 | -1.134 | -1.016 | 0.001 | 0.011 | 0.011 | 0.41 | 0.16 | 0.08 | 0.13 | 0.13 | 0.13 | 0.09 |
| 1 | 241451891 | rs12097296 | 1.173 | 0.177 | -1.19 | -0.853 | -0.034 | 0.023 | 0.023 | 0 | 0.27 | 0.13 | 0.31 | 0.26 | 0.35 | 0.24 |
| 1 | 241452500 | rs12090898 | 1.953 | 0.174 | -1.414 | -0.75 | -0.06 | -0.016 | -0.016 | 0 | 0.21 | 0.05 | 0.25 | 0.26 | 0.3 | 0.21 |
| 1 | 241453482 | rs10802947 | 1.179 | -0.314 | -1.009 | -0.199 | -0.105 | -0.002 | -0.002 | 0 | 0.37 | 0.09 | 0.37 | 0.37 | 0.47 | 0.42 |
| 1 | 241455756 | rs10737856 | 0.602 | -0.404 | -0.02 | -0.408 | -0.01 | -0.075 | -0.075 | 0 | 0.65 | 0.57 | 0.72 | 0.87 | 0.74 | 0.82 |
| 1 | 241458102 | rs6429256 | -0.356 | -0.337 | 0.255 | 0.102 | -0.009 | -0.025 | -0.025 | 0 | 0.19 | 0.24 | 0.17 | 0.14 | 0.13 | 0.1 |
| 1 | 241458939 | rs9782891 | -0.556 | -0.703 | 0.23 | 0.368 | -0.005 | 0.035 | 0.035 | 0 | 0.37 | 0.21 | 0.42 | 0.27 | 0.42 | 0.24 |
| 1 | 241459462 | rs7355007 | 0.452 | -0.7 | 0.811 | 0.064 | -0.024 | -0.131 | -0.131 | 0 | 0.73 | 0.47 | 0.81 | 0.98 | 0.77 | 0.89 |
| 1 | 241461507 | rs12062959 | -0.745 | -0.523 | 0.757 | 1.099 | -0.03 | 0.085 | 0.085 | 0.028 | 0.06 | 0.19 | 0.1 | 0.15 | 0.01 | 0.21 |
| 1 | 241467163 | rs7539874 | -0.663 | -0.385 | -0.276 | 0.747 | -0.042 | 0.028 | 0.028 | 0 | 0.28 | 0.47 | 0.22 | 0.38 | 0.18 | 0.36 |
| 1 | 241469725 | rs10926458 | 0.503 | 0.34 | -0.847 | 0.863 | 0.013 | -0.041 | -0.041 | 0 | 0.22 | 0.14 | 0.31 | 0.5 | 0.17 | 0.14 |
| 1 | 241470634 | rs10802948 | 0.432 | 1.125 | -0.423 | 0.307 | -0.001 | 0 | 0 | 0 | 0.8 | 0.82 | 0.86 | 0.9 | 0.81 | 0.84 |
| 1 | 241475718 | rs11589951 | -1.127 | 1.291 | -0.292 | 0.768 | 0.01 | 0.014 | 0.014 | 0 | 0.08 | 0.13 | 0.23 | 0.14 | 0.14 | 0.24 |
| 1 | 241483770 | rs1341468 | -1.237 | 1.015 | -0.075 | 0.481 | 0.012 | -0.004 | -0.004 | 0.908 | 0.36 | 0.62 | 0.42 | 0.26 | 0.4 | 0.4 |
| 1 | 241494578 | rs7512603 | -0.999 | 0.577 | 0.222 | 0.679 | 0 | -0.001 | -0.001 | 0.391 | 0.26 | 0.27 | 0.16 | 0.25 | 0.19 | 0.19 |
| 1 | 241499174 | rs11803606 | -1.061 | 0.172 | 0.363 | 0.672 | -0.044 | 0.021 | 0.021 | 0 | 0.27 | 0.56 | 0.16 | 0.25 | 0.15 | 0.21 |
| 1 | 241502909 | rs12128314 | -0.99 | 0.736 | -0.504 | 0.887 | -0.017 | -0.022 | -0.022 | 0 | 0.06 | 0.02 | 0.19 | 0.16 | 0.1 | 0.32 |
| 1 | 241504787 | rs10802949 | -1.075 | 0.423 | -0.516 | 0.402 | -0.002 | -0.001 | -0.001 | 5.836 | 0.33 | 0.36 | 0.38 | 0.28 | 0.37 | 0.36 |
| 1 | 241505994 | rs10737860 | 0.103 | -1.08 | -0.085 | -0.236 | -0.04 | -0.043 | -0.043 | 0 | 0.7 | 0.5 | 0.82 | 0.84 | 0.82 | 0.85 |
| 1 | 241507247 | rs10802951 | -0.705 | -1.444 | -0.059 | -0.406 | -0.02 | -0.026 | -0.026 | 1.955 | 0.12 | 0.3 | 0.21 | 0.06 | 0.07 | 0.3 |
| 1 | 241509795 | rs10737861 | 0.085 | -1.407 | 0.089 | -0.339 | -0.01 | -0.011 | -0.011 | 0.614 | 0.69 | 0.34 | 0.82 | 0.85 | 0.82 | 0.85 |
| 1 | 241519317 | rs6677041 | 0.894 | -0.959 | 0.551 | -0.302 | -0.006 | -0.036 | -0.036 | 0 | 0.86 | 0.73 | 0.82 | 0.94 | 0.89 | 0.75 |
| 1 | 241522325 | rs12038803 | 0.869 | -1.446 | 0.781 | -0.374 | -0.016 | -0.02 | -0.02 | 0 | 0.64 | 0.44 | 0.7 | 0.81 | 0.75 | 0.68 |
| 1 | 241522475 | rs6429264 | -1.044 | -1.39 | 1.168 | -0.283 | -0.016 | -0.002 | -0.002 | 0.234 | 0.14 | 0.21 | 0.19 | 0.13 | 0.11 | 0.19 |
| 1 | 241525507 | rs4660071 | -0.082 | -1.136 | 1.157 | -0.349 | -0.004 | -0.014 | -0.014 | 0 | 0.1 | 0.15 | 0.18 | 0.09 | 0.12 | 0.21 |
| 1 | 241525522 | rs1341463 | 0.91 | -1.426 | 0.963 | -0.086 | -0.03 | 0.015 | 0.015 | 1.976 | 0.68 | 0.51 | 0.75 | 0.76 | 0.74 | 0.72 |
| 1 | 241526489 | rs10926468 | 0.622 | -1.424 | 0.788 | 0.021 | -0.018 | 0.009 | 0.009 | 5.339 | 0.73 | 0.33 | 0.83 | 0.76 | 0.84 | 0.87 |
| 1 | 241533392 | rs10926472 | 0.385 | -1.431 | 0.722 | 0.038 | -0.027 | 0.017 | 0.017 | 1.749 | 0.76 | 0.34 | 0.83 | 0.76 | 0.85 | 0.88 |
| 1 | 241535092 | rs4660075 | 0.914 | -0.852 | -0.016 | 0.294 | 0.012 | -0.015 | -0.015 | 0 | 0.58 | 0.27 | 0.55 | 0.49 | 0.6 | 0.81 |
| 1 | 241535414 | rs9651084 | 0.049 | -0.611 | -0.286 | 0.552 | -0.018 | 0.012 | 0.012 | 0 | 0.2 | 0.1 | 0.25 | 0.24 | 0.24 | 0.09 |
| 1 | 241537354 | rs6429269 | 0.402 | -0.595 | -0.031 | 0.84 | 0.002 | 0.007 | 0.007 | 0.231 | 0.14 | 0.12 | 0.15 | 0.26 | 0.14 | 0.08 |
| 1 | 241539109 | rs1592072 | 0.663 | -0.441 | -0.107 | 0.471 | -0.001 | -0.001 | -0.001 | 1.425 | 0.61 | 0.39 | 0.61 | 0.59 | 0.6 | 0.78 |
| 1 | 241539802 | rs6699695 | -0.418 | -0.36 | -0.441 | 0.521 | -0.001 | -0.001 | -0.001 | 0 | 0.33 | 0.27 | 0.33 | 0.43 | 0.29 | 0.28 |
| 1 | 241542880 | rs12140273 | 1.303 | 0.189 | -1.104 | 0.268 | -0.124 | -0.022 | -0.022 | 0 | 0.26 | 0.01 | 0.27 | 0.23 | 0.32 | 0.28 |
| 1 | 241546751 | rs2341938 | 0.001 | -0.039 | -0.373 | -0.014 | -0.001 | 0.013 | 0.013 | 0 | 0.51 | 0.56 | 0.52 | 0.62 | 0.47 | 0.61 |
| 1 | 241550350 | rs1557078 | -0.313 | -0.02 | -0.492 | -0.013 | -0.042 | 0.011 | 0.011 | 0 | 0.54 | 0.81 | 0.55 | 0.62 | 0.48 | 0.62 |
| 1 | 241554603 | rs10926476 | -0.031 | 0.2 | -0.251 | 0.025 | -0.017 | -0.009 | -0.009 | 2.595 | 0.44 | 0.16 | 0.38 | 0.4 | 0.48 | 0.37 |
| 1 | 241557595 | rs1341455 | 0.779 | 0.259 | -0.239 | 0.107 | -0.045 | -0.065 | -0.065 | 0 | 0.33 | 0.09 | 0.34 | 0.4 | 0.38 | 0.31 |

| | | | | | | | | | | | | | | | | |
|---|---|---|---|---|---|---|---|---|---|---|---|---|---|---|---|---|
| 1 | 241561050 | rs10802960 | 0.563 | 0.279 | -0.036 | 0.112 | 0 | 0 | 0 | 0 | 0.35 | 0.32 | 0.35 | 0.4 | 0.39 | 0.34 |
| 1 | 241566725 | rs1557079 | -0.207 | 0.029 | 0.192 | 0.173 | -0.023 | 0.003 | 0.003 | 0 | 0.82 | 0.6 | 0.83 | 0.83 | 0.85 | 0.79 |
| 1 | 241569364 | rs1935095 | -0.635 | 0.037 | 0.073 | 0.177 | -0.014 | 0.001 | 0.001 | 0 | 0.84 | 0.73 | 0.84 | 0.84 | 0.86 | 0.81 |
| 1 | 241570018 | rs2341942 | 0.897 | 0.382 | -0.481 | 0.157 | -0.029 | -0.038 | -0.038 | 0 | 0.32 | 0.09 | 0.3 | 0.36 | 0.36 | 0.31 |
| 1 | 241574867 | rs7529232 | 0.756 | 0.181 | -0.316 | 0.349 | -0.033 | -0.008 | -0.008 | 0.499 | 0.4 | 0.09 | 0.37 | 0.36 | 0.39 | 0.33 |
| 1 | 241575771 | rs9428874 | -0.544 | 0.113 | -0.005 | 0.331 | 0.005 | 0.004 | 0.004 | 0 | 0.59 | 0.5 | 0.54 | 0.53 | 0.54 | 0.56 |
| 1 | 241576287 | rs1341448 | 0.827 | 0.127 | -0.001 | 0.338 | -0.033 | -0.008 | -0.008 | 0 | 0.4 | 0.08 | 0.37 | 0.36 | 0.39 | 0.33 |
| 1 | 241576302 | rs1341449 | -0.365 | 0.269 | -0.066 | 0.4 | 0.008 | 0.006 | 0.006 | 0 | 0.58 | 0.45 | 0.53 | 0.53 | 0.54 | 0.55 |
| 1 | 241577094 | rs1341452 | -0.45 | 0.345 | -0.163 | 0.437 | 0.005 | 0.004 | 0.004 | 0 | 0.59 | 0.5 | 0.54 | 0.53 | 0.54 | 0.56 |
| 1 | 241581283 | rs3014560 | -0.058 | 0.407 | -0.428 | 0.527 | -0.01 | 0.002 | 0.002 | 0 | 0.16 | 0.17 | 0.16 | 0.16 | 0.14 | 0.2 |
| 1 | 241582668 | rs3014559 | -0.076 | 0.5 | -0.524 | 0.506 | 0.002 | 0.002 | 0.002 | 0.184 | 0.62 | 0.54 | 0.55 | 0.57 | 0.55 | 0.57 |
| 1 | 241583418 | rs10926483 | 0.013 | -0.055 | -1.068 | -0.299 | -0.009 | 0.129 | 0.129 | 0 | 0.52 | 0.31 | 0.6 | 0.84 | 0.48 | 0.69 |
| 1 | 241585499 | rs1539098 | -0.454 | -0.781 | -0.533 | -0.519 | -0.019 | 0.154 | 0.154 | 1.081 | 0.52 | 0.79 | 0.55 | 0.81 | 0.45 | 0.7 |
| 1 | 241585802 | rs1539099 | 0.619 | -1.092 | -0.463 | -0.077 | 0.008 | 0.011 | 0.011 | 1.195 | 0.3 | 0.59 | 0.31 | 0.41 | 0.27 | 0.43 |
| 1 | 241586834 | rs1544190 | -0.468 | -1.001 | -0.463 | 0.178 | 0.015 | -0.097 | -0.097 | 0.961 | 0.2 | 0.09 | 0.23 | 0.42 | 0.18 | 0.27 |
| 1 | 241596981 | rs3014552 | 0.177 | -0.965 | 0.12 | -0.028 | 0.009 | 0.007 | 0.007 | 0 | 0.32 | 0.55 | 0.41 | 0.4 | 0.26 | 0.39 |
| 1 | 241603322 | rs12184326 | 1.41 | -1.03 | 0.012 | -0.322 | -0.019 | 0.048 | 0.048 | 0 | 0.29 | 0.13 | 0.23 | 0.15 | 0.36 | 0.25 |
| 1 | 241603554 | rs2994968 | 1.345 | -1.161 | -0.146 | -0.247 | -0.11 | -0.018 | -0.018 | 0 | 0.24 | 0.01 | 0.33 | 0.24 | 0.22 | 0.25 |
| 1 | 241608403 | rs1578488 | 0.502 | -1.287 | 1.218 | 0.158 | -0.019 | 0.017 | 0.017 | 1.054 | 0.69 | 0.38 | 0.69 | 0.56 | 0.74 | 0.81 |
| 1 | 241611462 | rs4660089 | 1.416 | -1.183 | 0.9 | 0.234 | -0.025 | 0.039 | 0.039 | 0 | 0.27 | 0.12 | 0.23 | 0.15 | 0.36 | 0.25 |
| 1 | 241615673 | rs1578486 | -0.184 | -1.068 | 1.072 | 0.342 | -0.008 | 0.044 | 0.044 | 0.296 | 0.78 | 0.72 | 0.72 | 0.58 | 0.76 | 0.92 |
| 1 | 241625692 | rs10802965 | 0.232 | -0.967 | 1.011 | 0.272 | 0 | -0.003 | -0.003 | 0 | 0.43 | 0.41 | 0.43 | 0.38 | 0.36 | 0.58 |
| 1 | 241625863 | rs7540470 | 0.682 | -0.853 | 0.845 | 0.225 | -0.002 | -0.04 | -0.04 | 0 | 0.16 | 0.24 | 0.13 | 0.07 | 0.19 | 0.06 |
| 1 | 241633413 | rs2994978 | 0.079 | -0.801 | 0.647 | 0.279 | -0.007 | 0 | 0 | 0.307 | 0.43 | 0.3 | 0.44 | 0.42 | 0.36 | 0.61 |
| 1 | 241637536 | rs10926494 | 0.908 | -0.353 | 0.616 | 0.086 | -0.026 | 0.033 | 0.033 | 2.349 | 0.3 | 0.16 | 0.35 | 0.19 | 0.29 | 0.54 |
| 1 | 241639401 | rs10754728 | 0.294 | -0.59 | 1.036 | 0.242 | -0.008 | -0.015 | -0.015 | 0 | 0.41 | 0.49 | 0.32 | 0.34 | 0.41 | 0.33 |
| 1 | 241646566 | rs2994984 | 1.11 | -0.69 | 0.928 | 0.247 | -0.006 | 0.025 | 0.025 | 1.709 | 0.3 | 0.19 | 0.35 | 0.19 | 0.28 | 0.55 |
| 1 | 241650163 | rs10157091 | 0.638 | -0.675 | 0.731 | 0.319 | -0.001 | -0.009 | -0.009 | 0.693 | 0.34 | 0.35 | 0.36 | 0.24 | 0.28 | 0.56 |
| 1 | 241659359 | rs947306 | 1.239 | -0.829 | 0.703 | 0.236 | 0.018 | -0.012 | -0.012 | 0.199 | 0.28 | 0.31 | 0.26 | 0.2 | 0.39 | 0.24 |
| 1 | 241667244 | rs4659607 | -0.501 | -0.825 | 0.684 | 0.241 | -0.001 | -0.044 | -0.044 | 0 | 0.74 | 0.8 | 0.69 | 0.57 | 0.7 | 0.89 |
| 1 | 241671638 | rs11802956 | 0.113 | -0.779 | 0.601 | 0.256 | 0 | -0.012 | -0.012 | 0 | 0.41 | 0.45 | 0.33 | 0.33 | 0.43 | 0.33 |
| 1 | 241678668 | rs2992644 | 1.228 | -2.027 | 1.552 | 0.878 | 0.003 | -0.002 | -0.002 | 0 | 0.61 | 0.57 | 0.61 | 0.43 | 0.67 | 0.79 |
| 1 | 241684831 | rs6678986 | -0.042 | -2.032 | 1.079 | 0.897 | -0.008 | -0.002 | -0.002 | 0 | 0.12 | 0.12 | 0.03 | 0.04 | 0.03 | 0.09 |
| 1 | 241690554 | rs6678766 | 0.312 | -2.132 | 1.145 | 0.919 | -0.001 | 0.014 | 0.014 | 0 | 0.73 | 0.64 | 0.67 | 0.58 | 0.71 | 0.86 |
| 1 | 241694414 | rs10926507 | 1.663 | -2.133 | 0.451 | 0.638 | 0.097 | -0.02 | -0.02 | 4.287 | 0.54 | 0.06 | 0.58 | 0.4 | 0.67 | 0.75 |
| 1 | 241694949 | rs10926508 | -0.114 | -1.275 | 0.809 | -0.273 | -0.024 | 0.002 | 0.002 | 0.228 | 0.11 | 0.11 | 0.06 | 0.14 | 0.03 | 0.1 |
| 1 | 241695792 | rs3819976 | -0.468 | -1.272 | 0.799 | -0.271 | -0.045 | 0.004 | 0.004 | 0.133 | 0.12 | 0.18 | 0.07 | 0.14 | 0.03 | 0.1 |
| 1 | 241697819 | rs6429278 | -0.391 | -1.22 | 0.477 | -0.246 | -0.058 | 0.008 | 0.008 | 0.31 | 0.13 | 0.19 | 0.07 | 0.14 | 0.03 | 0.1 |
| 1 | 241700371 | rs2992641 | -0.882 | -0.869 | -0.078 | -0.109 | -0.023 | -0.03 | -0.03 | 0 | 0.35 | 0.54 | 0.25 | 0.23 | 0.28 | 0.2 |
| 1 | 241703089 | rs1932440 | 0.975 | -0.613 | 0.05 | 0.07 | -0.027 | -0.012 | -0.012 | 0.108 | 0.84 | 0.42 | 0.85 | 0.86 | 0.91 | 0.88 |
| 1 | 241703480 | rs12030573 | 1.079 | -0.634 | 0.213 | -0.545 | -0.007 | -0.142 | -0.142 | 0 | 0.74 | 0.55 | 0.84 | 0.96 | 0.76 | 0.91 |
| 1 | 241704726 | rs2050512 | -0.847 | -0.81 | -0.345 | -0.004 | -0.037 | 0.027 | 0.027 | 0.754 | 0.27 | 0.52 | 0.37 | 0.57 | 0.15 | 0.19 |
| 1 | 241705226 | rs2050515 | 0.52 | -0.647 | -0.684 | 1.032 | -0.03 | -0.003 | -0.003 | 0.585 | 0.06 | 0 | 0.15 | 0.3 | 0.09 | 0.09 |
| 1 | 241706007 | rs6677357 | -1.116 | -0.893 | -0.811 | 0.1 | 0.009 | 0.005 | 0.005 | 0.005 | 0.37 | 0.62 | 0.44 | 0.62 | 0.26 | 0.28 |
| 1 | 241706533 | rs2050516 | 0.718 | -0.217 | -1.667 | -0.434 | -0.088 | 0.075 | 0.075 | 0 | 0.26 | 0.09 | 0.23 | 0.14 | 0.42 | 0.21 |
| 1 | 241707102 | rs11587924 | 0.469 | -0.231 | -1.605 | -0.441 | -0.065 | 0.068 | 0.068 | 0.03 | 0.28 | 0.14 | 0.23 | 0.14 | 0.42 | 0.21 |
| 1 | 241711324 | rs2050505 | 0.048 | -0.418 | -2.059 | -0.451 | -0.162 | 0.071 | 0.071 | 0.007 | 0.34 | 0.03 | 0.27 | 0.24 | 0.44 | 0.42 |
| 1 | 241711701 | rs11808201 | -0.928 | -0.645 | -1.485 | -0.251 | 0.011 | 0.014 | 0.014 | 0 | 0.25 | 0.23 | 0.39 | 0.2 | 0.22 | 0.29 |
| 1 | 241715422 | rs12079041 | -0.979 | 0.303 | -0.685 | -0.286 | 0.056 | -0.035 | -0.035 | 0 | 0.07 | 0.26 | 0.04 | 0 | 0.07 | 0.03 |
| 1 | 241720543 | rs3007738 | -0.262 | -0.994 | -0.266 | -0.546 | -0.069 | -0.028 | -0.028 | 0 | 0.82 | 0.64 | 0.75 | 0.84 | 0.94 | 0.88 |
| 1 | 241720559 | rs3007737 | 0.594 | -0.501 | -1.14 | -0.127 | 0.038 | 0.005 | 0.005 | 0.341 | 0.49 | 0.22 | 0.45 | 0.43 | 0.59 | 0.44 |
| 1 | 241724922 | rs3753214 | 1.361 | 0.054 | -1.403 | 0.354 | -0.004 | 0.026 | 0.026 | 1.523 | 0.06 | 0.04 | 0.09 | 0.19 | 0.09 | 0.05 |
| 1 | 241727309 | rs12139931 | 0.793 | 0.175 | -1.357 | 0.537 | -0.015 | -0.024 | -0.024 | 0 | 0.28 | 0.15 | 0.24 | 0.35 | 0.32 | 0.35 |
| 1 | 241730054 | rs2275163 | 0.529 | 0.54 | -1.995 | 0.199 | -0.03 | -0.048 | -0.048 | 4.975 | 0.28 | 0.16 | 0.31 | 0.38 | 0.35 | 0.23 |
| 1 | 241731095 | rs4660103 | 1.082 | 0.418 | -1.952 | 0.128 | -0.008 | 0.019 | 0.019 | 1.347 | 0.23 | 0.16 | 0.21 | 0.19 | 0.26 | 0.18 |
| 1 | 241747838 | rs648708 | 0.006 | 0.398 | -1.914 | 0.492 | -0.019 | 0.025 | 0.025 | 0 | 0.3 | 0.35 | 0.36 | 0.45 | 0.2 | 0.32 |
| 1 | 241755348 | rs1053230 | 1.455 | 0.433 | -2.048 | 0.514 | -0.073 | 0.071 | 0.071 | 0 | 0.2 | 0.01 | 0.13 | 0.05 | 0.22 | 0.21 |
| 1 | 241755567 | rs1053221 | 1.018 | 0.429 | -1.411 | 1.249 | -0.055 | -0.077 | -0.077 | 0 | 0.12 | 0.01 | 0.1 | 0.26 | 0.14 | 0.2 |
| 1 | 241755925 | rs640718 | 0.781 | 0.383 | -1.398 | 1.228 | -0.072 | -0.004 | -0.004 | 0 | 0.18 | 0.3 | 0.26 | 0.19 | 0.06 | 0.12 |
| 1 | 241758302 | rs588246 | 0.791 | 0.665 | -1.112 | 1.36 | -0.047 | 0.033 | 0.033 | 0.46 | 0.14 | 0.3 | 0.26 | 0.19 | 0.05 | 0.09 |
| 1 | 241758631 | rs1053183 | 0.157 | 0.858 | -0.994 | 1.005 | -0.013 | 0.047 | 0.047 | 0 | 0.27 | 0.35 | 0.36 | 0.45 | 0.19 | 0.29 |
| 1 | 241761661 | rs650833 | -0.995 | 0.946 | -1.08 | 0.822 | 0 | 0.016 | 0.016 | 0 | 0.94 | 0.91 | 0.88 | 0.85 | 0.87 | 0.91 |
| 1 | 241766551 | rs3753216 | 0.212 | 0.778 | -1.055 | 0.035 | 0.007 | -0.031 | -0.031 | 0 | 0.66 | 0.54 | 0.51 | 0.27 | 0.68 | 0.61 |
| 1 | 241768617 | rs616593 | 0.279 | 0.897 | -1.009 | -0.339 | 0.009 | -0.089 | -0.089 | 0.094 | 0.34 | 0.44 | 0.22 | 0.09 | 0.32 | 0.23 |
| 1 | 241768630 | rs646355 | -0.166 | 1.387 | -1.573 | 0.149 | -0.088 | 0.031 | 0.031 | 0.662 | 0.79 | 0.98 | 0.79 | 0.87 | 0.68 | 0.81 |
| 1 | 241772421 | rs613032 | -0.254 | 1.203 | -0.522 | 1.103 | 0.031 | 0.125 | 0.125 | 0 | 0.09 | 0.18 | 0.06 | 0.27 | 0.1 | 0.09 |
| 1 | 241772881 | rs3765813 | -0.193 | 1.583 | -0.022 | 1.667 | 0.054 | 0.068 | 0.068 | 0 | 0.15 | 0.29 | 0.26 | 0.33 | 0.23 | 0.26 |
| 1 | 241773027 | rs3765814 | -0.087 | -0.036 | -0.286 | 0.859 | -0.035 | -0.003 | -0.003 | 0.52 | 0.25 | 0.35 | 0.36 | 0.27 | 0.16 | 0.2 |
| 1 | 241776501 | rs672477 | 0.058 | -0.661 | -0.584 | 0.905 | 0.02 | -0.019 | -0.019 | 0 | 0.25 | 0.18 | 0.12 | 0.34 | 0.2 | 0.2 |

| | | | | | | | | | | | | | | | | |
|---|---|---|---|---|---|---|---|---|---|---|---|---|---|---|---|---|
| 1 | 241780207 | rs6660991 | 0.024 | -0.418 | -0.37 | 0.665 | -0.019 | -0.027 | -0.027 | 2.683 | 0.12 | 0.16 | 0.07 | 0.05 | 0.06 | 0.1 |
| 1 | 241794231 | rs3765821 | -0.368 | 0.076 | -0.002 | 0.777 | 0.007 | 0.019 | 0.019 | 0 | 0.54 | 0.36 | 0.65 | 0.69 | 0.58 | 0.63 |
| 1 | 241796284 | rs676750 | -1.515 | 0.052 | -0.165 | 0.78 | -0.046 | 0.003 | 0.003 | 0.223 | 0.7 | 0.89 | 0.76 | 0.76 | 0.66 | 0.77 |
| 1 | 241798631 | rs3737604 | -1.612 | 0.488 | 0.952 | 1.459 | -0.007 | 0.02 | 0.02 | 0 | 0.31 | 0.23 | 0.26 | 0.22 | 0.32 | 0.37 |
| 1 | 241800324 | rs3819981 | -0.59 | 0.597 | 0.828 | 1.489 | -0.025 | 0.005 | 0.005 | 0 | 0.19 | 0.1 | 0.19 | 0.18 | 0.25 | 0.26 |
| 1 | 241804866 | rs12141693 | -0.638 | 0.527 | 0.783 | 1.288 | 0.008 | 0.043 | 0.043 | 0.332 | 0.12 | 0.02 | 0.06 | 0.05 | 0.08 | 0.11 |
| 1 | 241808411 | rs6656762 | -0.525 | 0.683 | 0.668 | 1.641 | 0.009 | -0.016 | -0.016 | 0.773 | 0.12 | 0.1 | 0.08 | 0.2 | 0.07 | 0.1 |
| 1 | 241809586 | rs663814 | -1.995 | 0.794 | 0.714 | 1.862 | 0.003 | 0.004 | 0.004 | 0 | 0.34 | 0.59 | 0.35 | 0.66 | 0.36 | 0.45 |
| 1 | 241809865 | rs6663216 | -1.808 | 0.792 | 0.778 | 1.861 | 0.004 | 0.004 | 0.004 | 0 | 0.33 | 0.39 | 0.34 | 0.65 | 0.35 | 0.45 |
| 1 | 241809896 | rs6663223 | -0.385 | 0.803 | 0.787 | 1.918 | 0.022 | 0.064 | 0.064 | 0 | 0.18 | 0.25 | 0.22 | 0.44 | 0.24 | 0.33 |
| 1 | 241811131 | rs1537802 | 1.768 | 0.842 | 0.664 | 1.864 | 0.003 | -0.003 | -0.003 | 0.037 | 0.62 | 0.4 | 0.55 | 0.34 | 0.57 | 0.54 |
| 1 | 241813771 | rs10802975 | 0.913 | 0.914 | 0.216 | 1.823 | -0.002 | 0.002 | 0.002 | 0 | 0.82 | 0.77 | 0.78 | 0.78 | 0.82 | 0.87 |
| 1 | 241813906 | rs7411883 | -0.988 | 0.79 | -0.382 | 1.347 | -0.001 | -0.002 | -0.002 | 0 | 0.33 | 0.34 | 0.39 | 0.59 | 0.38 | 0.42 |
| 1 | 241818980 | rs12723036 | -1.34 | 0.393 | -0.042 | 1.379 | -0.007 | 0.027 | 0.027 | 0 | 0.06 | 0.12 | 0.07 | 0.12 | 0.05 | 0.16 |
| 1 | 241821181 | rs6429284 | 1.131 | -0.089 | 0.192 | 1.663 | -0.01 | 0.019 | 0.019 | 0 | 0.75 | 0.7 | 0.83 | 0.63 | 0.83 | 0.72 |
| 1 | 241822182 | rs7416113 | -0.304 | -0.285 | 0.23 | 1.232 | 0 | 0.004 | 0.004 | 0 | 0.25 | 0.24 | 0.4 | 0.52 | 0.25 | 0.33 |
| 1 | 241826698 | rs6682476 | 1.231 | -0.484 | 0.288 | 1.065 | 0.019 | 0.038 | 0.038 | 0 | 0.58 | 0.37 | 0.46 | 0.12 | 0.65 | 0.51 |
| 1 | 241829586 | rs12564480 | -0.705 | -0.258 | 0.511 | 1.257 | -0.003 | 0.017 | 0.017 | 0 | 0.15 | 0.23 | 0.12 | 0.25 | 0.12 | 0.14 |
| 1 | 241830876 | rs6429288 | 1.696 | -0.215 | 0.476 | 1.167 | 0.015 | 0.1 | 0.1 | 0 | 0.48 | 0.34 | 0.42 | 0.12 | 0.56 | 0.38 |
| 1 | 241833287 | rs7513188 | -0.211 | -0.165 | 0.44 | 1.623 | -0.006 | 0.068 | 0.068 | 0 | 0.8 | 0.71 | 0.72 | 0.6 | 0.82 | 0.71 |
| 1 | 241836977 | rs6661311 | 0.902 | -0.001 | 0.184 | 0.94 | 0.003 | 0.029 | 0.029 | 0 | 0.58 | 0.37 | 0.57 | 0.26 | 0.63 | 0.45 |
| 1 | 241841958 | rs7419051 | -1.043 | 0.251 | -0.135 | 1.181 | -0.013 | -0.035 | -0.035 | 0.105 | 0.11 | 0.06 | 0.2 | 0.19 | 0.13 | 0.08 |
| 1 | 241849047 | rs7527828 | 1.078 | 0.042 | 0.104 | 0.376 | -0.001 | -0.034 | -0.034 | 0 | 0.53 | 0.56 | 0.46 | 0.12 | 0.59 | 0.37 |
| 1 | 241849430 | rs7414398 | -1.061 | -0.198 | 0.298 | 1.516 | 0.005 | -0.012 | -0.012 | 0 | 0.18 | 0.14 | 0.29 | 0.37 | 0.16 | 0.26 |
| 1 | 241849572 | rs7516595 | 1.4 | -0.076 | 0.989 | 1.755 | 0.048 | 0.208 | 0.208 | 0 | 0.82 | 0.56 | 0.62 | 0.63 | 0.81 | 0.87 |
| 1 | 241849779 | rs6429290 | 0.919 | -0.315 | 0.529 | 1.613 | -0.015 | 0.009 | 0.009 | 0 | 0.72 | 0.65 | 0.6 | 0.38 | 0.76 | 0.71 |
| 1 | 241851745 | rs12133073 | 0.132 | -0.031 | -0.254 | 0.671 | -0.007 | 0.036 | 0.036 | 0.356 | 0.53 | 0.68 | 0.47 | 0.4 | 0.49 | 0.55 |
| 1 | 241854641 | rs10926534 | 1.524 | 0.149 | -0.304 | 0.575 | -0.005 | 0.044 | 0.044 | 0 | 0.31 | 0.17 | 0.28 | 0.17 | 0.32 | 0.29 |
| 1 | 241854809 | rs9428886 | 0.244 | -0.038 | -0.462 | 0.679 | -0.031 | 0.007 | 0.007 | 0 | 0.15 | 0.29 | 0.09 | 0.18 | 0.1 | 0.22 |
| 1 | 241855644 | rs7418366 | 0.498 | 0.255 | -1.263 | 0.779 | -0.001 | 0.003 | 0.003 | 0 | 0.1 | 0.08 | 0.13 | 0.22 | 0.07 | 0.13 |
| 1 | 241862049 | rs10926538 | -0.545 | 0.587 | -1.486 | 0.678 | -0.075 | -0.108 | -0.108 | 0 | 0.68 | 0.9 | 0.54 | 0.55 | 0.6 | 0.78 |
| 1 | 241865219 | rs12117802 | 0.484 | 0.698 | -1.056 | 0.239 | -0.014 | 0.065 | 0.065 | 0 | 0.3 | 0.11 | 0.27 | 0.15 | 0.32 | 0.28 |
| 1 | 241868349 | rs9428890 | -0.057 | 1.241 | -0.427 | 0.244 | 0.043 | -0.039 | -0.039 | 0 | 0.1 | 0.19 | 0.13 | 0.01 | 0.17 | 0.03 |
| 1 | 241869867 | rs7417070 | -0.533 | 1.24 | -0.823 | -0.096 | -0.013 | 0.027 | 0.027 | 0 | 0.07 | 0.02 | 0.11 | 0.02 | 0.06 | 0.12 |
| 1 | 241875326 | rs12092919 | -1.071 | 1.112 | 0.168 | -0.145 | 0.08 | -0.078 | -0.078 | 0 | 0.15 | 0.39 | 0.24 | 0.02 | 0.23 | 0.13 |
| 1 | 241875533 | rs10802980 | -0.828 | -0.032 | -0.011 | 0.463 | 0.025 | -0.042 | -0.042 | 1.336 | 0.78 | 0.89 | 0.76 | 0.48 | 0.81 | 0.7 |
| 1 | 241878353 | rs7418763 | 0.401 | -0.162 | -0.186 | 0.338 | 0.057 | 0.1 | 0.1 | 0 | 0.5 | 0.34 | 0.57 | 0.19 | 0.55 | 0.46 |
| 1 | 241881362 | rs10926545 | 0.065 | -0.103 | -0.198 | 0.334 | 0.023 | 0.031 | 0.031 | 0.019 | 0.11 | 0.26 | 0.22 | 0.09 | 0.11 | 0.16 |
| 1 | 241881940 | rs10802982 | -1.623 | -0.249 | -0.301 | 0.576 | 0.043 | -0.054 | -0.054 | 0 | 0.8 | 0.95 | 0.77 | 0.52 | 0.83 | 0.75 |
| 1 | 241882665 | rs12725593 | 0.439 | -0.353 | 0.037 | 0.137 | -0.009 | -0.026 | -0.026 | 0 | 0.23 | 0.13 | 0.16 | 0.29 | 0.24 | 0.23 |
| 1 | 241885307 | rs7519667 | -1.09 | -0.449 | 0.498 | 0.06 | -0.029 | 0.043 | 0.043 | 1.48 | 0.76 | 0.72 | 0.74 | 0.47 | 0.8 | 0.7 |
| 1 | 241891040 | rs6690359 | -1.025 | 0.44 | 0.258 | 0.129 | 0.001 | 0.003 | 0.003 | 0 | 0.39 | 0.4 | 0.48 | 0.4 | 0.4 | 0.5 |
| 1 | 241892235 | rs9782914 | -1.021 | 0.364 | 0.483 | 0.222 | 0 | 0.033 | 0.033 | 0 | 0.17 | 0.39 | 0.17 | 0.21 | 0.14 | 0.15 |
| 1 | 241897079 | rs7367532 | -0.641 | 0.451 | 0.038 | 0.115 | 0 | 0.021 | 0.021 | 0 | 0.11 | 0.11 | 0.16 | 0.28 | 0.1 | 0.16 |
| 1 | 241897899 | rs10926556 | -0.693 | 0.444 | 0.518 | 0.111 | 0.009 | 0.15 | 0.15 | 0 | 0.11 | 0.26 | 0.17 | 0.28 | 0.1 | 0.16 |
| 1 | 241899337 | rs10926558 | -0.574 | 0.132 | 0.516 | -0.275 | -0.018 | -0.024 | -0.024 | 0.378 | 0.3 | 0.34 | 0.31 | 0.22 | 0.2 | 0.35 |
| 1 | 241901183 | rs6695661 | -1.434 | 0.462 | 0.604 | -0.207 | 0 | -0.011 | -0.011 | 0 | 0.35 | 0.32 | 0.44 | 0.51 | 0.31 | 0.49 |
| 1 | 241907691 | rs7418599 | -1.359 | 0.499 | 0.438 | -0.221 | -0.032 | -0.087 | -0.087 | 1.897 | 0.34 | 0.12 | 0.42 | 0.51 | 0.31 | 0.48 |
| 1 | 241907817 | rs10802990 | -1.624 | 1.054 | 0.431 | -0.471 | -0.027 | -0.009 | -0.009 | 0.463 | 0.45 | 0.82 | 0.57 | 0.51 | 0.47 | 0.5 |
| 1 | 241914626 | rs7530797 | -1.4 | 0.839 | 0.296 | -0.587 | 0.04 | -0.041 | -0.041 | 0.959 | 0.71 | 0.86 | 0.71 | 0.56 | 0.75 | 0.65 |
| 1 | 241925878 | rs7551270 | -1.281 | 0.844 | 0.485 | -0.585 | -0.004 | 0.001 | 0.001 | 0 | 0.7 | 0.7 | 0.71 | 0.56 | 0.75 | 0.65 |
| 1 | 241926236 | rs11587279 | -0.269 | 1.215 | -0.225 | -0.565 | -0.028 | 0.04 | 0.04 | 0 | 0.12 | 0.05 | 0.16 | 0.03 | 0.15 | 0.14 |
| 1 | 241931654 | rs10926569 | -0.636 | 1.096 | -0.152 | -0.536 | 0.005 | -0.017 | -0.017 | 0.033 | 0.59 | 0.18 | 0.57 | 0.56 | 0.59 | 0.63 |
| 1 | 241934757 | rs12091522 | 1.182 | 1.128 | 0.052 | -0.617 | 0.084 | -0.043 | -0.043 | 0 | 0.09 | 0.29 | 0.13 | 0 | 0.16 | 0.02 |
| 1 | 241940024 | rs7514880 | 1.217 | 1.128 | 0.048 | -0.616 | 0.058 | -0.042 | -0.042 | 0 | 0.09 | 0.22 | 0.12 | 0 | 0.16 | 0.02 |
| 1 | 241945917 | rs4658506 | 0.763 | 1.195 | -0.034 | -0.591 | -0.032 | -0.037 | -0.037 | 0 | 0.06 | 0 | 0.18 | 0.16 | 0.09 | 0.15 |
| 1 | 241946294 | rs952084 | 1.409 | 1.105 | 0.051 | -0.595 | 0.036 | -0.036 | -0.036 | 0 | 0.29 | 0.13 | 0.28 | 0.43 | 0.25 | 0.35 |
| 1 | 241951329 | rs12074374 | 1.283 | 1.131 | 0.038 | -0.64 | 0.058 | -0.042 | -0.042 | 0 | 0.09 | 0.22 | 0.13 | 0.01 | 0.16 | 0.02 |
| 1 | 241951703 | rs7524442 | -0.429 | 1.132 | -0.036 | -0.614 | -0.009 | 0.003 | 0.003 | 0.369 | 0.54 | 0.32 | 0.39 | 0.41 | 0.5 | 0.48 |
| 1 | 241958977 | rs6673963 | -0.851 | 1.092 | -0.034 | -0.607 | 0.003 | -0.001 | -0.001 | 0 | 0.61 | 0.42 | 0.6 | 0.57 | 0.6 | 0.63 |
| 1 | 241960669 | rs1342866 | -1.307 | 1.206 | 0.07 | -0.589 | 0.135 | -0.038 | -0.038 | 0 | 0.9 | 0.56 | 0.87 | 1 | 0.85 | 0.98 |
| 1 | 241963501 | rs4658647 | -0.304 | 1.083 | -0.254 | -0.75 | -0.003 | 0.076 | 0.076 | 0 | 0.26 | 0.16 | 0.21 | 0.09 | 0.25 | 0.28 |
| 1 | 241972287 | rs9428895 | 0.263 | 0.533 | -0.057 | -0.424 | 0.041 | -0.002 | -0.002 | 0.056 | 0.82 | 0.4 | 0.68 | 0.84 | 0.76 | 0.83 |
| 1 | 241978738 | rs12565340 | -0.136 | 0.543 | -0.11 | -0.302 | 0.03 | 0.06 | 0.06 | 0 | 0.08 | 0.17 | 0.19 | 0.16 | 0.09 | 0.17 |
| 1 | 241979567 | rs9428896 | -1.317 | 0.527 | -0.725 | -0.395 | 0.019 | 0.066 | 0.066 | 0 | 0.85 | 0.74 | 0.82 | 0.72 | 0.81 | 0.93 |
| 1 | 241979706 | rs12081499 | -0.108 | 0.51 | -1.398 | -0.47 | -0.056 | 0.269 | 0.269 | 0 | 0.48 | 0.19 | 0.33 | 0.13 | 0.48 | 0.4 |
| 1 | 241980222 | rs1342872 | -0.104 | 0.524 | -1.394 | -0.467 | -0.052 | 0.269 | 0.269 | 0 | 0.48 | 0.19 | 0.34 | 0.13 | 0.49 | 0.4 |
| 1 | 241984595 | rs10489352 | 0.962 | 0.448 | -1.532 | -0.322 | 0.023 | 0.032 | 0.032 | 0 | 0.13 | 0.07 | 0.07 | 0.07 | 0.07 | 0.11 |
| 1 | 241993831 | rs10489350 | -0.006 | 0.881 | -1.039 | 0.246 | -0.042 | -0.02 | -0.02 | 0 | 0.12 | 0.03 | 0.22 | 0.14 | 0.16 | 0.19 |

| | | | | | | | | | | | | | | | | |
|---|---|---|---|---|---|---|---|---|---|---|---|---|---|---|---|---|
| 1 | 241994183 | rs12038477 | 0.708 | 0.846 | -1.099 | 0.271 | 0.03 | -0.04 | -0.04 | 0 | 0.29 | 0.12 | 0.29 | 0.45 | 0.26 | 0.36 |
| 1 | 241997104 | rs6691782 | 0.106 | 1.001 | -1.42 | 0.202 | -0.009 | 0.007 | 0.007 | 0 | 0.41 | 0.4 | 0.53 | 0.59 | 0.4 | 0.54 |
| 1 | 242000788 | rs9287256 | -0.071 | 1.071 | -2.336 | -0.461 | -0.034 | 0.176 | 0.176 | 0.026 | 0.41 | 0.19 | 0.29 | 0.13 | 0.4 | 0.38 |
| 1 | 242009307 | rs12118937 | -0.967 | 1.371 | -1.907 | 0.283 | -0.072 | -0.028 | -0.028 | 0 | 0.15 | 0.02 | 0.24 | 0.14 | 0.2 | 0.2 |
| 1 | 242010116 | rs3754093 | -0.641 | 0.188 | -1.752 | 0.54 | 0.005 | -0.003 | -0.003 | 0 | 0.33 | 0.29 | 0.31 | 0.4 | 0.17 | 0.39 |
| 1 | 242011244 | rs1776181 | -0.073 | 0.21 | -1.797 | -0.064 | 0.012 | 0.052 | 0.052 | 0.675 | 0.49 | 0.65 | 0.51 | 0.72 | 0.52 | 0.49 |
| 1 | 242011406 | rs1776179 | -0.308 | -0.124 | -2.052 | -0.532 | 0.003 | 0.001 | 0.001 | 0 | 0.65 | 0.66 | 0.6 | 0.86 | 0.69 | 0.69 |
| 1 | 242012149 | rs1776177 | 0.757 | -0.373 | -1.5 | -0.974 | 0.012 | -0.016 | -0.016 | 0 | 0.4 | 0.47 | 0.46 | 0.71 | 0.46 | 0.41 |
| 1 | 242012385 | rs4149852 | 0.82 | -0.444 | -1.734 | -1.127 | -0.027 | -0.026 | -0.026 | 0 | 0.06 | 0.01 | 0.21 | 0.09 | 0.09 | 0.1 |
| 1 | 242012660 | rs2073490 | 0.097 | -0.591 | -1.227 | -1.219 | -0.048 | 0 | 0 | 0 | 0.21 | 0.02 | 0.12 | 0.14 | 0.2 | 0.25 |
| 1 | 242016838 | rs4149868 | 1.066 | -0.573 | -0.83 | -1.097 | -0.014 | -0.019 | -0.019 | 0 | 0.05 | 0.14 | 0.02 | 0.01 | 0.02 | 0.02 |
| 1 | 242022130 | rs2526699 | -0.109 | -0.608 | -0.867 | -1.214 | 0.021 | 0.165 | 0.165 | 2.138 | 0.54 | 0.86 | 0.7 | 0.81 | 0.57 | 0.63 |
| 1 | 242025098 | rs1776133 | 0.056 | -0.584 | -0.871 | -1.203 | 0.025 | 0.083 | 0.083 | 1.43 | 0.52 | 0.72 | 0.7 | 0.8 | 0.57 | 0.53 |
| 1 | 242027363 | rs1776139 | 0.902 | -0.539 | -0.961 | -1.181 | -0.01 | 0.01 | 0.01 | 0.828 | 0.4 | 0.39 | 0.46 | 0.7 | 0.46 | 0.42 |
| 1 | 242027786 | rs2797604 | 0.928 | -0.727 | -1.027 | -0.463 | -0.091 | 0.042 | 0.042 | 1.195 | 0.36 | 0.16 | 0.29 | 0.28 | 0.48 | 0.43 |
| 1 | 242028027 | rs1635509 | -0.329 | -0.547 | -0.963 | -0.705 | 0.015 | 0.04 | 0.04 | 1.259 | 0.51 | 0.62 | 0.69 | 0.8 | 0.57 | 0.53 |
| 1 | 242030151 | rs735943 | -0.329 | -0.547 | -0.931 | -0.705 | 0.015 | 0.04 | 0.04 | 1.259 | 0.51 | 0.62 | 0.69 | 0.8 | 0.57 | 0.53 |
| 1 | 242035382 | rs4149963 | 0.71 | -0.345 | -1.226 | -0.568 | -0.027 | -0.026 | -0.026 | 1.529 | 0.06 | 0.01 | 0.21 | 0.09 | 0.09 | 0.1 |
| 1 | 242038713 | rs1776140 | 0.01 | 0.205 | -1.043 | -0.96 | -0.01 | 0.009 | 0.009 | 1.098 | 0.4 | 0.41 | 0.45 | 0.69 | 0.46 | 0.42 |
| 1 | 242042301 | rs1047840 | -0.505 | -0.167 | -0.818 | -1.077 | -0.003 | -0.036 | -0.036 | 0.71 | 0.38 | 0.48 | 0.39 | 0.23 | 0.37 | 0.26 |
| 1 | 242042545 | rs1776148 | -0.312 | -0.448 | -0.642 | -1.278 | 0.014 | 0.03 | 0.03 | 0 | 0.6 | 0.69 | 0.69 | 0.81 | 0.65 | 0.57 |
| 1 | 242042768 | rs4149982 | -0.088 | -0.493 | -0.749 | -0.604 | 0.012 | -0.015 | -0.015 | 0.106 | 0.14 | 0.1 | 0.08 | 0.24 | 0.07 | 0.18 |
| 1 | 242048674 | rs9350 | 0.446 | -0.075 | -0.802 | -0.56 | -0.002 | 0.037 | 0.037 | 0.057 | 0.18 | 0.18 | 0.32 | 0.47 | 0.15 | 0.18 |
| 1 | 242050736 | rs4150005 | 0.461 | -0.074 | -0.801 | -0.554 | -0.002 | 0.04 | 0.04 | 0.031 | 0.18 | 0.18 | 0.32 | 0.47 | 0.15 | 0.18 |
| 1 | 242057540 | rs7521516 | 0.889 | -0.173 | -1.045 | -1.011 | -0.003 | 0.106 | 0.106 | 0 | 0.25 | 0.09 | 0.16 | 0.02 | 0.26 | 0.24 |
| 1 | 242059221 | rs4658535 | 0.012 | -0.126 | -0.925 | -0.838 | -0.012 | 0.018 | 0.018 | 1.882 | 0.25 | 0.3 | 0.34 | 0.6 | 0.18 | 0.25 |
| 1 | 242063452 | rs851785 | -0.069 | 0.254 | -0.847 | -0.481 | -0.001 | 0 | 0 | 2.906 | 0.36 | 0.37 | 0.34 | 0.4 | 0.39 | 0.49 |
| 1 | 242064151 | rs1776170 | -0.764 | 0.384 | -1.029 | -0.351 | -0.027 | -0.061 | -0.061 | 3.03 | 0.66 | 0.54 | 0.75 | 0.87 | 0.72 | 0.68 |
| 1 | 242064633 | rs7551606 | 0.502 | 0.433 | -1.17 | -0.32 | -0.011 | -0.039 | -0.039 | 0 | 0.35 | 0.21 | 0.42 | 0.49 | 0.34 | 0.27 |
| 1 | 242065332 | rs1635516 | -0.128 | 0.166 | -0.82 | -0.014 | -0.018 | -0.044 | -0.044 | 0.108 | 0.67 | 0.6 | 0.76 | 0.87 | 0.72 | 0.68 |
| 1 | 242070731 | rs12568965 | -0.437 | 0.616 | -1.045 | 0.614 | -0.003 | -0.014 | -0.014 | 0 | 0.18 | 0.15 | 0.18 | 0.27 | 0.19 | 0.26 |
| 1 | 242071602 | rs6681581 | -0.524 | -0.198 | -0.714 | 0.164 | -0.037 | -0.035 | -0.035 | 0 | 0.31 | 0.5 | 0.35 | 0.19 | 0.27 | 0.23 |
| 1 | 242073301 | rs12139416 | 0.245 | 0.063 | -0.933 | -0.419 | 0 | -0.002 | -0.002 | 0.002 | 0.5 | 0.49 | 0.65 | 0.77 | 0.46 | 0.48 |
| 1 | 242074809 | rs6660916 | 0.223 | -0.024 | -0.915 | -0.318 | 0 | 0.002 | 0.002 | 0.063 | 0.51 | 0.46 | 0.66 | 0.72 | 0.48 | 0.48 |
| 1 | 242080853 | rs12722924 | 0.242 | 0.172 | -1.44 | -0.699 | -0.028 | 0.049 | 0.049 | 0 | 0.2 | 0.06 | 0.13 | 0.13 | 0.22 | 0.13 |
| 1 | 242081840 | rs6666851 | -0.241 | -0.249 | -1.24 | -0.887 | -0.004 | -0.013 | -0.013 | 0 | 0.11 | 0.17 | 0.07 | 0.03 | 0.08 | 0.12 |
| 1 | 242081875 | rs6664298 | -0.583 | 0.09 | -1.217 | -0.746 | -0.016 | 0.027 | 0.027 | 0.106 | 0.42 | 0.31 | 0.29 | 0.32 | 0.46 | 0.51 |
| 1 | 242086079 | rs12057998 | 0.142 | 0.304 | -1.08 | -0.212 | -0.002 | 0.032 | 0.032 | 0 | 0.3 | 0.41 | 0.32 | 0.42 | 0.28 | 0.18 |
| 1 | 242086319 | rs6689502 | -0.943 | 0.018 | -0.629 | 0.189 | 0.005 | 0.048 | 0.048 | 0 | 0.93 | 0.86 | 0.85 | 0.84 | 0.93 | 0.88 |
| 1 | 242087206 | rs6699609 | 1.537 | -0.221 | -1.003 | -0.256 | -0.002 | -0.038 | -0.038 | 0.003 | 0.22 | 0.22 | 0.16 | 0.04 | 0.24 | 0.15 |
| 1 | 242087644 | rs4658565 | 0.035 | 0.061 | -1.359 | 0.519 | 0.003 | 0.005 | 0.005 | 0 | 0.49 | 0.42 | 0.45 | 0.45 | 0.5 | 0.38 |
| 1 | 242087718 | rs12060816 | 0.267 | -0.315 | -1.349 | 0.706 | -0.01 | -0.025 | -0.025 | 0.394 | 0.09 | 0.04 | 0.09 | 0.16 | 0.07 | 0.13 |
| 1 | 242089795 | rs7516206 | 0.629 | -0.561 | -1.339 | 0.246 | 0.013 | 0.029 | 0.029 | 0.26 | 0.13 | 0.03 | 0.05 | 0 | 0.09 | 0.05 |
| 1 | 242090414 | rs10803002 | -0.53 | -1.291 | -1.235 | 0.706 | -0.033 | 0.009 | 0.009 | 0 | 0.72 | 0.35 | 0.65 | 0.66 | 0.86 | 0.7 |
| 1 | 242094314 | rs4658573 | -0.343 | -0.604 | -1.161 | 0.376 | 0.053 | 0.054 | 0.054 | 0 | 0.59 | 0.32 | 0.68 | 0.72 | 0.69 | 0.71 |
| 1 | 242096551 | rs3844252 | -1.342 | -0.499 | -0.692 | 0.372 | -0.016 | -0.006 | -0.006 | 0 | 0.89 | 0.85 | 0.94 | 0.88 | 0.92 | 0.88 |
| 1 | 242098415 | rs3845563 | -0.112 | 0.264 | -1.011 | 0.328 | -0.021 | -0.011 | -0.011 | 0.103 | 0.37 | 0.26 | 0.34 | 0.39 | 0.43 | 0.5 |
| 1 | 242106296 | rs11581005 | -0.521 | 0.197 | -0.906 | 0.317 | 0 | 0.001 | 0.001 | 0 | 0.51 | 0.54 | 0.43 | 0.39 | 0.55 | 0.56 |
| 1 | 242109010 | rs28674674 | -0.245 | 0.63 | -1.227 | 0.108 | -0.001 | -0.003 | -0.003 | 0 | 0.26 | 0.27 | 0.24 | 0.2 | 0.24 | 0.36 |
| 1 | 242121796 | rs3851304 | -0.006 | 0.752 | -1.586 | -0.063 | -0.011 | 0.024 | 0.024 | 1.227 | 0.36 | 0.26 | 0.32 | 0.23 | 0.37 | 0.28 |
| 1 | 242127836 | rs1040264 | -0.338 | 0.478 | -1.507 | 0.298 | -0.029 | 0.049 | 0.049 | 0 | 0.69 | 0.41 | 0.71 | 0.42 | 0.77 | 0.73 |
| 1 | 242140841 | rs28477715 | 0.052 | 1.18 | -2.029 | -0.069 | -0.031 | 0.02 | 0.02 | 0 | 0.14 | 0.02 | 0.09 | 0.07 | 0.13 | 0.07 |
| 1 | 242148832 | rs2102911 | -0.468 | 1.244 | -2.355 | -0.408 | -0.068 | 0.13 | 0.13 | 0 | 0.3 | 0.04 | 0.19 | 0.07 | 0.31 | 0.2 |
| 1 | 242150334 | rs28373959 | -0.723 | 1.534 | -2.536 | -0.55 | -0.139 | 0.17 | 0.17 | 0 | 0.37 | 0.06 | 0.26 | 0.13 | 0.46 | 0.33 |
| 1 | 242158192 | rs28757855 | -0.057 | 1.02 | -1.729 | 0.263 | 0.004 | 0.003 | 0.003 | 0.162 | 0.42 | 0.55 | 0.48 | 0.45 | 0.53 | 0.58 |
| 1 | 242162874 | rs28534947 | -0.108 | 0.852 | -1.659 | 0.233 | 0.001 | 0.001 | 0.001 | 0 | 0.43 | 0.52 | 0.59 | 0.53 | 0.58 | 0.68 |
| 1 | 242171251 | rs28407556 | 0.007 | 0.351 | -1.482 | -0.623 | -0.022 | -0.075 | -0.075 | 0 | 0.65 | 0.63 | 0.82 | 0.93 | 0.77 | 0.77 |
| 1 | 242184078 | rs28586974 | -0.278 | 0.797 | -2.242 | -0.692 | -0.079 | 0.256 | 0.256 | 0 | 0.46 | 0.1 | 0.3 | 0.13 | 0.53 | 0.41 |
| 1 | 242195928 | rs28593183 | -0.782 | 1.327 | -1.285 | 0.377 | 0.024 | 0.083 | 0.083 | 0.361 | 0.21 | 0.32 | 0.26 | 0.58 | 0.24 | 0.39 |
| 1 | 242198204 | rs871243 | -0.17 | 1.26 | -1.625 | 0.154 | -0.001 | 0.009 | 0.009 | 1.97 | 0.13 | 0.08 | 0.15 | 0.06 | 0.13 | 0.11 |
| 1 | 242200884 | rs6673338 | -0.162 | 0.838 | -1.57 | 0.241 | 0.005 | -0.002 | -0.002 | 0 | 0.37 | 0.67 | 0.47 | 0.59 | 0.34 | 0.54 |
| 1 | 242201247 | rs10926604 | 0.634 | 0.801 | -2.074 | -0.376 | -0.028 | 0.083 | 0.083 | 0 | 0.32 | 0.16 | 0.19 | 0.14 | 0.32 | 0.28 |
| 1 | 242203795 | rs4641297 | -0.333 | 0.732 | -1.855 | 0.026 | 0.001 | -0.001 | -0.001 | 1.233 | 0.66 | 0.7 | 0.69 | 0.7 | 0.73 | 0.81 |
| 1 | 242208009 | rs10754748 | 0.1 | 0.746 | -1.507 | 0.184 | 0.005 | -0.002 | -0.002 | 0 | 0.53 | 0.58 | 0.53 | 0.56 | 0.63 | 0.65 |
| 1 | 242222470 | rs12042710 | 0.219 | 0.88 | -1.255 | 0.334 | 0 | 0 | 0 | 0 | 0.23 | 0.23 | 0.25 | 0.27 | 0.28 | 0.29 |
| 1 | 242223343 | rs10158939 | 0.51 | 0.769 | -0.988 | 0.182 | 0 | -0.003 | -0.003 | 0 | 0.27 | 0.2 | 0.33 | 0.3 | 0.32 | 0.39 |
| 1 | 242233314 | rs9428536 | -0.424 | 0.586 | -1.036 | 0.117 | -0.009 | 0.007 | 0.007 | 0 | 0.66 | 0.78 | 0.58 | 0.7 | 0.53 | 0.62 |
| 1 | 242238751 | rs7528333 | 1.136 | 0.225 | -0.691 | -0.295 | -0.008 | -0.037 | -0.037 | 0.557 | 0.1 | 0.19 | 0.09 | 0.01 | 0.1 | 0.05 |
| 1 | 242243178 | rs10803015 | 1.158 | -0.292 | -1.037 | -0.609 | 0.046 | 0.058 | 0.058 | 0 | 0.31 | 0.2 | 0.18 | 0.13 | 0.21 | 0.27 |

| | | | | | | | | | | | | | | | | |
|---|---|---|---|---|---|---|---|---|---|---|---|---|---|---|---|---|
| 1 | 242243325 | rs6681101 | -0.129 | 0.2 | -0.7 | -0.228 | 0.003 | 0.01 | 0.01 | 0 | 0.56 | 0.65 | 0.38 | 0.27 | 0.42 | 0.37 |
| 1 | 242249484 | rs6429332 | -0.071 | 0.225 | -0.657 | -0.416 | 0.006 | 0.013 | 0.013 | 0.06 | 0.55 | 0.64 | 0.39 | 0.25 | 0.41 | 0.39 |
| 1 | 242251696 | rs10926618 | -0.174 | 0.216 | -0.655 | -0.558 | 0.009 | 0.024 | 0.024 | 0.074 | 0.54 | 0.65 | 0.39 | 0.23 | 0.41 | 0.37 |
| 1 | 242253024 | rs7523793 | 0.488 | 0.137 | -0.608 | -0.543 | -0.003 | -0.015 | -0.015 | 0 | 0.41 | 0.45 | 0.31 | 0.22 | 0.34 | 0.33 |
| 1 | 242263834 | rs12135876 | 1.258 | 0.011 | -0.829 | -0.489 | 0.033 | 0.069 | 0.069 | 0 | 0.28 | 0.17 | 0.22 | 0.11 | 0.21 | 0.25 |
| 1 | 242265419 | rs12129551 | 1.258 | 0.011 | -0.828 | -0.489 | 0.032 | 0.064 | 0.064 | 0 | 0.28 | 0.18 | 0.22 | 0.11 | 0.21 | 0.25 |
| 1 | 242266434 | rs2164422 | -0.806 | 0.824 | -0.644 | 0.258 | -0.007 | -0.016 | -0.016 | 0 | 0.72 | 0.83 | 0.68 | 0.67 | 0.64 | 0.63 |
| 1 | 242270341 | rs4274045 | -0.545 | 0.83 | -0.515 | -0.067 | -0.001 | 0.015 | 0.015 | 0.584 | 0.45 | 0.59 | 0.59 | 0.75 | 0.48 | 0.59 |
| 1 | 242274908 | rs10803016 | -0.515 | 0.948 | -0.547 | 0.316 | 0.003 | 0.001 | 0.001 | 0.073 | 0.44 | 0.53 | 0.57 | 0.57 | 0.45 | 0.52 |
| 1 | 242275356 | rs10926621 | 1.042 | 0.58 | -0.905 | -0.019 | 0.017 | 0.027 | 0.027 | 0 | 0.15 | 0.08 | 0.07 | 0.09 | 0.11 | 0.19 |
| 1 | 242279403 | rs12129770 | 1.087 | 0.545 | -0.624 | 0.004 | -0.017 | -0.011 | -0.011 | 0 | 0.14 | 0.19 | 0.06 | 0.1 | 0.08 | 0.19 |
| 1 | 242281455 | rs1435951 | 1.477 | 0.817 | -1.27 | 0.335 | -0.046 | -0.076 | -0.076 | 0.609 | 0.12 | 0.01 | 0.21 | 0.33 | 0.11 | 0.2 |
| 1 | 242281976 | rs2800757 | -0.794 | 1.252 | -0.769 | 0.106 | 0.042 | -0.021 | -0.021 | 0.034 | 0.82 | 0.7 | 0.82 | 0.86 | 0.73 | 0.94 |
| 1 | 242283539 | rs2800756 | -1.718 | 0.81 | -1.445 | 0.208 | -0.054 | -0.027 | -0.027 | 0.697 | 0.87 | 0.99 | 0.87 | 0.86 | 0.79 | 0.97 |
| 1 | 242284412 | rs6672510 | -0.163 | 0.379 | -1.382 | 0.135 | 0.042 | 0.058 | 0.058 | 0 | 0.47 | 0.68 | 0.34 | 0.26 | 0.34 | 0.44 |
| 1 | 242287684 | rs10926623 | 1.098 | 0.392 | -1.456 | 0.193 | 0.006 | 0.009 | 0.009 | 0 | 0.13 | 0.1 | 0.07 | 0.1 | 0.1 | 0.18 |
| 1 | 242292280 | rs2252440 | 0.342 | 0.621 | -1.56 | 0.174 | -0.009 | -0.015 | -0.015 | 0 | 0.39 | 0.24 | 0.52 | 0.59 | 0.41 | 0.52 |
| 1 | 242292690 | rs1898152 | -0.887 | 1.108 | -1.656 | 0.075 | -0.099 | 0.018 | 0.018 | 0 | 0.82 | 0.99 | 0.84 | 0.88 | 0.73 | 0.95 |
| 1 | 242293918 | rs7541781 | 1.497 | 0.295 | -1.069 | 0.235 | -0.001 | -0.004 | -0.004 | 0 | 0.17 | 0.19 | 0.16 | 0.07 | 0.15 | 0.17 |
| 1 | 242297423 | rs1550125 | 0.459 | 0.599 | -1.265 | 0.594 | -0.03 | -0.019 | -0.019 | 0.819 | 0.25 | 0.12 | 0.3 | 0.27 | 0.33 | 0.27 |
| 1 | 242301657 | rs1978790 | -0.458 | 0.047 | -1.113 | 0.26 | 0.02 | 0.036 | 0.036 | 0 | 0.55 | 0.34 | 0.65 | 0.76 | 0.56 | 0.71 |
| 1 | 242301917 | rs2840609 | -1.372 | -0.165 | -1.53 | -0.04 | -0.062 | 0.052 | 0.052 | 0 | 0.85 | 0.99 | 0.84 | 0.94 | 0.76 | 0.97 |
| 1 | 242302144 | rs7538792 | 0.801 | -0.268 | -0.779 | 0.201 | 0.027 | 0 | 0 | 0.605 | 0.27 | 0.36 | 0.31 | 0.23 | 0.34 | 0.29 |
| 1 | 242304648 | rs892929 | 1.206 | -0.393 | -0.978 | 0.282 | 0.002 | -0.028 | -0.028 | 0 | 0.14 | 0.09 | 0.28 | 0.44 | 0.09 | 0.21 |
| 1 | 242306890 | rs2636257 | 0.189 | -0.475 | -0.915 | 0.241 | -0.018 | 0.016 | 0.016 | 0.083 | 0.55 | 0.78 | 0.47 | 0.35 | 0.54 | 0.5 |
| 1 | 242310046 | rs10926624 | 0.615 | -0.364 | -0.645 | 0.392 | 0.028 | -0.006 | -0.006 | 8.104 | 0.29 | 0.39 | 0.31 | 0.23 | 0.34 | 0.32 |
| 1 | 242322529 | rs316840 | -0.123 | -0.705 | -0.943 | 0.463 | -0.004 | -0.024 | -0.024 | 0 | 0.4 | 0.74 | 0.39 | 0.41 | 0.41 | 0.39 |
| 1 | 242322870 | rs737133 | 1.084 | -0.605 | -0.873 | 0.628 | -0.042 | -0.029 | -0.029 | 2.852 | 0.25 | 0.1 | 0.3 | 0.29 | 0.34 | 0.31 |
| 1 | 242327576 | rs12037915 | 1.48 | -0.65 | -0.823 | 0.709 | -0.033 | -0.08 | -0.08 | 0 | 0.12 | 0.01 | 0.27 | 0.36 | 0.09 | 0.17 |
| 1 | 242328222 | rs695021 | -1.477 | -0.631 | -0.995 | 0.502 | -0.065 | 0.061 | 0.061 | 0 | 0.84 | 0.99 | 0.83 | 0.95 | 0.76 | 0.96 |
| 1 | 242329791 | rs1393299 | 0.049 | -0.748 | -0.71 | 0.467 | -0.006 | 0.01 | 0.01 | 0 | 0.42 | 0.6 | 0.61 | 0.7 | 0.44 | 0.52 |
| 1 | 242329814 | rs316896 | 0.354 | -0.841 | -0.618 | 0.517 | 0.009 | 0.011 | 0.011 | 0 | 0.42 | 0.38 | 0.22 | 0.25 | 0.32 | 0.43 |
| 1 | 242332277 | rs10926631 | 1.164 | -0.606 | -0.36 | 0.84 | -0.001 | -0.021 | -0.021 | 0 | 0.13 | 0.09 | 0.29 | 0.44 | 0.09 | 0.18 |
| 1 | 242342504 | rs316873 | 1.101 | -0.621 | -0.422 | 0.907 | 0 | 0.005 | 0.005 | 0 | 0.13 | 0.1 | 0.28 | 0.44 | 0.09 | 0.21 |
| 1 | 242343502 | rs10926640 | 2.457 | -0.847 | -0.558 | 0.65 | 0.006 | 0.049 | 0.049 | 0 | 0.16 | 0.06 | 0.08 | 0 | 0.14 | 0.12 |
| 1 | 242343622 | rs316868 | 1.198 | -0.783 | -0.511 | 0.781 | 0 | 0.013 | 0.013 | 0 | 0.13 | 0.1 | 0.28 | 0.5 | 0.09 | 0.21 |
| 1 | 242344395 | rs11809579 | 1.09 | -0.712 | -0.502 | 0.629 | 0.045 | 0.015 | 0.015 | 0 | 0.23 | 0.37 | 0.28 | 0.22 | 0.29 | 0.3 |
| 1 | 242345289 | rs2013655 | 1.608 | -0.941 | -0.429 | 0.493 | -0.017 | 0.002 | 0.002 | 0 | 0.11 | 0.17 | 0.08 | 0.12 | 0.07 | 0.06 |
| 1 | 242345428 | rs725601 | 0.114 | -0.857 | -0.614 | 0.41 | 0.001 | -0.001 | -0.001 | 0.108 | 0.49 | 0.45 | 0.47 | 0.54 | 0.39 | 0.58 |
| 1 | 242347587 | rs12135329 | 0.947 | -0.837 | -0.727 | 0.184 | 0.015 | 0.101 | 0.101 | 0 | 0.35 | 0.15 | 0.18 | 0.11 | 0.3 | 0.37 |
| 1 | 242347608 | rs316833 | -1.194 | -0.548 | -0.824 | 0.409 | -0.01 | 0.018 | 0.018 | 0 | 0.83 | 0.86 | 0.82 | 0.95 | 0.76 | 0.94 |
| 1 | 242350414 | rs316837 | -1.461 | -0.493 | -1.051 | 0.436 | -0.07 | 0.059 | 0.059 | 0 | 0.84 | 0.99 | 0.83 | 0.94 | 0.76 | 0.96 |
| 1 | 242350620 | rs189779 | 1.438 | -0.684 | -0.945 | 0.827 | -0.001 | -0.001 | -0.001 | 0 | 0.12 | 0.1 | 0.28 | 0.3 | 0.09 | 0.19 |
| 1 | 242357455 | rs427498 | -1.528 | -0.286 | -1.068 | 0.28 | -0.068 | 0.047 | 0.047 | 0 | 0.86 | 0.99 | 0.84 | 0.94 | 0.77 | 0.97 |
| 1 | 242359678 | rs10803021 | 0.361 | -0.401 | -0.854 | 0.641 | 0.028 | -0.054 | -0.054 | 0 | 0.24 | 0.19 | 0.35 | 0.5 | 0.16 | 0.24 |
| 1 | 242360179 | rs2036408 | 0.494 | 0.015 | -0.628 | 0.483 | 0.014 | -0.002 | -0.002 | 0 | 0.38 | 0.26 | 0.54 | 0.63 | 0.32 | 0.37 |
| 1 | 242365657 | rs11577736 | 0.715 | -0.113 | -0.786 | 0.139 | -0.006 | 0.041 | 0.041 | 0 | 0.19 | 0.06 | 0.1 | 0.11 | 0.17 | 0.26 |
| 1 | 242366282 | rs2653165 | -0.852 | 0.068 | -0.3 | 0.131 | -0.009 | 0.063 | 0.063 | 0 | 0.68 | 0.77 | 0.74 | 0.94 | 0.62 | 0.85 |
| 1 | 242368250 | rs12098208 | 1.314 | 0.552 | 0.05 | 0.748 | -0.006 | 0.005 | 0.005 | 0 | 0.09 | 0.14 | 0.09 | 0.12 | 0.09 | 0.07 |
| 1 | 242375779 | rs6429339 | -0.928 | 0.742 | 0.045 | 0.604 | 0.013 | 0.002 | 0.002 | 0 | 0.87 | 0.83 | 0.85 | 0.89 | 0.77 | 0.92 |
| 1 | 242380057 | rs1604506 | 0.413 | 0.351 | -0.3 | 0.558 | 0.037 | 0.083 | 0.083 | 0 | 0.39 | 0.15 | 0.32 | 0.2 | 0.31 | 0.38 |
| 1 | 242380948 | rs10926652 | 0.492 | 0.333 | -0.114 | 0.656 | 0 | -0.011 | -0.011 | 0 | 0.72 | 0.24 | 0.71 | 0.67 | 0.67 | 0.81 |
| 1 | 242381784 | rs1847097 | 0.178 | 0.057 | -0.009 | 0.618 | -0.009 | 0.003 | 0.003 | 0 | 0.42 | 0.39 | 0.5 | 0.64 | 0.43 | 0.52 |
| 1 | 242384733 | rs1609951 | 0.655 | 0.258 | -0.37 | 0.669 | 0.008 | -0.031 | -0.031 | 1.132 | 0.36 | 0.29 | 0.42 | 0.55 | 0.28 | 0.38 |
| 1 | 242385038 | rs10737874 | 0.655 | 0.258 | -0.37 | 0.669 | 0.008 | -0.031 | -0.031 | 0 | 0.36 | 0.29 | 0.42 | 0.55 | 0.28 | 0.38 |
| 1 | 242389687 | rs2033665 | 0.012 | 0.292 | -0.13 | 0.679 | 0.031 | 0.015 | 0.015 | 0 | 0.52 | 0.41 | 0.43 | 0.36 | 0.41 | 0.47 |
| 1 | 242393905 | rs434200 | -1.466 | 0.445 | -0.475 | 0.708 | -0.066 | 0.005 | 0.005 | 0 | 0.88 | 0.98 | 0.86 | 0.92 | 0.77 | 0.95 |
| 1 | 242396830 | rs10803028 | 0.659 | 0.189 | -0.582 | 0.504 | -0.03 | -0.021 | -0.021 | 0 | 0.4 | 0.15 | 0.39 | 0.48 | 0.37 | 0.54 |
| 1 | 242398159 | rs400838 | -0.516 | 0.25 | -0.667 | 0.604 | -0.009 | -0.012 | -0.012 | 1.224 | 0.79 | 0.82 | 0.75 | 0.75 | 0.69 | 0.88 |
| 1 | 242398403 | rs390956 | -0.307 | 0.216 | -0.668 | 0.605 | -0.006 | -0.024 | -0.024 | 1.234 | 0.78 | 0.8 | 0.75 | 0.7 | 0.69 | 0.84 |
| 1 | 242404260 | rs425246 | 1.255 | 0.173 | -0.468 | 0.441 | -0.025 | 0.093 | 0.093 | 0.733 | 0.1 | 0.37 | 0.12 | 0.17 | 0.07 | 0.09 |
| 1 | 242408299 | rs12040225 | 1.527 | 0.514 | -0.883 | 0.406 | -0.022 | -0.042 | -0.042 | 0 | 0.26 | 0.09 | 0.31 | 0.38 | 0.21 | 0.28 |
| 1 | 242409100 | rs404323 | -0.304 | -0.307 | -0.503 | 0.153 | -0.031 | -0.001 | -0.001 | 0.103 | 0.73 | 0.44 | 0.8 | 0.74 | 0.77 | 0.67 |
| 1 | 242411114 | rs316901 | -0.246 | -0.018 | -0.576 | 0.13 | -0.036 | -0.018 | -0.018 | 0 | 0.73 | 0.58 | 0.73 | 0.86 | 0.82 | 0.84 |
| 1 | 242416045 | rs2574672 | -0.854 | 0.699 | -0.886 | 0.321 | -0.016 | 0.021 | 0.021 | 0 | 0.13 | 0.07 | 0.12 | 0.08 | 0.2 | 0.06 |
| 1 | 242419232 | rs316916 | -1.248 | 0.713 | -0.839 | 0.43 | -0.024 | 0.024 | 0.024 | 0 | 0.1 | 0.04 | 0.1 | 0.06 | 0.18 | 0.07 |
| 1 | 242424870 | rs12033351 | 0.123 | 0.067 | -1.145 | -0.097 | -0.001 | 0.173 | 0.173 | 0 | 0.37 | 0.1 | 0.27 | 0.1 | 0.33 | 0.31 |
| 1 | 242427162 | rs3913529 | -0.063 | -0.243 | -0.147 | 0.386 | -0.01 | -0.019 | -0.019 | 0 | 0.41 | 0.47 | 0.3 | 0.25 | 0.33 | 0.37 |
| 1 | 242453104 | rs4123234 | 1.1 | -0.769 | 0.547 | 1.454 | -0.008 | 0.006 | 0.006 | 0 | 0.72 | 0.3 | 0.61 | 0.36 | 0.8 | 0.68 |

| | | | | | | | | | | | | | | | | |
|---|---|---|---|---|---|---|---|---|---|---|---|---|---|---|---|---|
| 1 | 242465655 | rs4039095 | 0.633 | -0.409 | 1.27 | 1.765 | 0.031 | 0.326 | 0.326 | 0.167 | 0.89 | 0.58 | 0.84 | 0.48 | 0.89 | 0.73 |
| 1 | 242477255 | rs3863747 | -1.938 | -0.437 | 0.946 | 1.328 | -0.032 | 0.058 | 0.058 | 0.146 | 0.29 | 0.56 | 0.22 | 0.42 | 0.23 | 0.31 |
| 1 | 242531619 | rs1439524 | 1.464 | -0.616 | 1.124 | 0.612 | -0.053 | -0.051 | -0.051 | 0 | 0.8 | 0.6 | 0.86 | 0.86 | 0.9 | 0.9 |
| 1 | 242531654 | rs1439523 | 1.099 | -0.64 | 0.933 | 0.603 | -0.035 | 0.002 | 0.002 | 0.068 | 0.89 | 0.66 | 0.92 | 0.89 | 0.98 | 0.94 |
| 1 | 242535075 | rs2880314 | -1.001 | -0.523 | 0.566 | 0.647 | 0.007 | 0.01 | 0.01 | 1.399 | 0.11 | 0.1 | 0.1 | 0.09 | 0.08 | 0.07 |
| 1 | 242537612 | rs10754763 | 1.721 | -0.459 | 0.694 | 0.782 | -0.031 | -0.006 | -0.006 | 0 | 0.8 | 0.66 | 0.79 | 0.8 | 0.88 | 0.89 |
| 1 | 242537723 | rs10803040 | 0.644 | -0.32 | 0.741 | 0.846 | -0.01 | 0.043 | 0.043 | 0 | 0.91 | 0.78 | 0.9 | 0.89 | 0.96 | 0.96 |
| 1 | 242539072 | rs6694290 | 1.268 | -0.304 | 0.628 | 0.861 | -0.023 | 0.009 | 0.009 | 0.271 | 0.82 | 0.67 | 0.8 | 0.8 | 0.89 | 0.89 |
| 1 | 242547329 | rs6691318 | -0.494 | -0.047 | -0.472 | -0.181 | -0.006 | 0.158 | 0.158 | 0.55 | 0.47 | 0.24 | 0.48 | 0.09 | 0.43 | 0.34 |
| 1 | 242547880 | rs12097629 | -0.522 | 0.123 | -0.164 | -0.254 | 0.01 | 0.01 | 0.01 | 0 | 0.1 | 0.19 | 0.2 | 0.08 | 0.1 | 0.04 |
| 1 | 242547948 | rs12097667 | -1.009 | 0.114 | -0.138 | -0.202 | 0.001 | 0.035 | 0.035 | 0 | 0.12 | 0.2 | 0.31 | 0.18 | 0.11 | 0.05 |
| 1 | 242551565 | rs1342361 | 0.086 | 0.041 | 0.527 | 0.296 | 0.016 | 0.131 | 0.131 | 0 | 0.91 | 0.75 | 0.91 | 0.8 | 0.93 | 0.77 |
| 1 | 242551698 | rs1342362 | 0.527 | 0.022 | 0.603 | 0.29 | 0.004 | 0.207 | 0.207 | 0 | 0.89 | 0.65 | 0.8 | 0.7 | 0.92 | 0.75 |
| 1 | 242553664 | rs2654861 | 0.926 | -0.075 | 0.614 | 0.244 | -0.002 | -0.001 | -0.001 | 0 | 0.42 | 0.44 | 0.32 | 0.6 | 0.49 | 0.43 |
| 1 | 242554564 | rs2654862 | -1.157 | -0.075 | 0.612 | 0.244 | -0.008 | -0.006 | -0.006 | 0 | 0.59 | 0.65 | 0.69 | 0.4 | 0.5 | 0.57 |
| 1 | 242555949 | rs2796135 | -1.118 | -0.037 | 0.598 | 0.144 | 0.016 | 0.13 | 0.13 | 0 | 0.16 | 0.39 | 0.25 | 0.28 | 0.15 | 0.24 |
| 1 | 242562721 | rs2809985 | -0.458 | 0.004 | 0.293 | 0.145 | 0.003 | 0.011 | 0.011 | 0 | 0.48 | 0.56 | 0.39 | 0.22 | 0.4 | 0.52 |
| 1 | 242565280 | rs2810006 | 0.087 | 0.013 | -0.007 | -0.205 | 0.001 | 0.035 | 0.035 | 0 | 0.27 | 0.18 | 0.28 | 0.1 | 0.26 | 0.3 |
| 1 | 242565475 | rs2810007 | -1.792 | 0.144 | -0.297 | -0.031 | -0.023 | -0.071 | -0.071 | 0 | 0.71 | 0.85 | 0.73 | 0.47 | 0.63 | 0.7 |
| 1 | 242566034 | rs924774 | -0.829 | -0.025 | 0.6 | -0.467 | -0.03 | -0.049 | -0.049 | 0 | 0.34 | 0.46 | 0.19 | 0.19 | 0.2 | 0.22 |
| 1 | 242566422 | rs963319 | -0.285 | -0.19 | 0.477 | -0.617 | -0.005 | -0.006 | -0.006 | 0 | 0.19 | 0.23 | 0.08 | 0.11 | 0.06 | 0.07 |
| 1 | 242572649 | rs10803042 | -1.628 | -0.022 | 0.491 | -0.339 | 0.003 | -0.003 | -0.003 | 0 | 0.41 | 0.6 | 0.44 | 0.42 | 0.34 | 0.37 |
| 1 | 242578411 | rs7545843 | -1.774 | -0.049 | 0.497 | -0.359 | 0.027 | -0.012 | -0.012 | 0.786 | 0.43 | 0.75 | 0.45 | 0.42 | 0.33 | 0.39 |
| 1 | 242578674 | rs7532175 | -1.774 | -0.049 | 0.497 | -0.359 | 0.027 | -0.012 | -0.012 | 0 | 0.43 | 0.75 | 0.45 | 0.42 | 0.33 | 0.39 |
| 1 | 242580359 | rs2810017 | -1.936 | -0.174 | 0.17 | -0.638 | -0.025 | -0.067 | -0.067 | 0 | 0.67 | 0.87 | 0.73 | 0.42 | 0.64 | 0.6 |
| 1 | 242585078 | rs12130625 | 0.149 | -0.2 | 0.19 | -0.663 | 0.025 | 0.019 | 0.019 | 0 | 0.17 | 0.05 | 0.07 | 0.09 | 0.06 | 0.06 |
| 1 | 242587281 | rs2580232 | 0.937 | -0.07 | 0.254 | -0.718 | -0.025 | 0.088 | 0.088 | 0 | 0.26 | 0.1 | 0.28 | 0.01 | 0.31 | 0.21 |
| 1 | 242591769 | rs2580231 | 0.954 | -0.072 | 0.251 | -0.719 | -0.025 | 0.087 | 0.087 | 0 | 0.26 | 0.11 | 0.28 | 0.01 | 0.31 | 0.22 |
| 1 | 242593858 | rs1394060 | 0.843 | -0.071 | 0.352 | -0.719 | -0.002 | -0.025 | -0.025 | 0 | 0.26 | 0.24 | 0.29 | 0.01 | 0.31 | 0.23 |
| 1 | 242593873 | rs1503794 | -0.493 | -0.082 | 0.337 | -0.696 | 0.012 | 0.083 | 0.083 | 0 | 0.44 | 0.29 | 0.36 | 0.1 | 0.37 | 0.29 |
| 1 | 242599000 | rs4339844 | -0.504 | 0.12 | 0.792 | -0.807 | -0.007 | -0.002 | -0.002 | 0 | 0.2 | 0.55 | 0.14 | 0.21 | 0.19 | 0.27 |
| 1 | 242600263 | rs2654881 | -1.315 | 0.084 | 0.367 | -0.878 | -0.046 | 0.059 | 0.059 | 0 | 0.54 | 0.89 | 0.66 | 0.35 | 0.59 | 0.56 |
| 1 | 242600367 | rs6668435 | 0.049 | 0.219 | 0.799 | -0.744 | -0.007 | -0.039 | -0.039 | 0 | 0.15 | 0.31 | 0.08 | 0.11 | 0.13 | 0.16 |
| 1 | 242603059 | rs2055536 | -0.386 | -0.007 | 0.42 | -0.874 | 0.006 | -0.02 | -0.02 | 0 | 0.38 | 0.49 | 0.51 | 0.24 | 0.45 | 0.36 |
| 1 | 242603873 | rs2810032 | 0.054 | 0.013 | 0.286 | -0.877 | -0.039 | 0.085 | 0.085 | 0 | 0.33 | 0.18 | 0.39 | 0.11 | 0.44 | 0.3 |
| 1 | 242606388 | rs2810034 | -0.119 | -0.013 | 0.364 | -1.023 | 0.001 | -0.015 | -0.015 | 2.878 | 0.34 | 0.38 | 0.45 | 0.16 | 0.38 | 0.22 |
| 1 | 242607583 | rs2654882 | -1.058 | 0.023 | -0.161 | -1.051 | -0.055 | 0.159 | 0.159 | 0.247 | 0.51 | 0.87 | 0.69 | 0.29 | 0.53 | 0.4 |
| 1 | 242613284 | rs2654870 | 0.109 | 0.335 | 0.059 | -0.938 | -0.041 | 0.122 | 0.122 | 0 | 0.29 | 0.12 | 0.34 | 0.01 | 0.37 | 0.16 |
| 1 | 242616988 | rs10926742 | -0.026 | 0.351 | 0.244 | -0.92 | 0.002 | -0.069 | -0.069 | 0 | 0.31 | 0.33 | 0.35 | 0.02 | 0.37 | 0.18 |
| 1 | 242618940 | rs1473576 | -0.511 | 0.534 | -0.12 | -0.8 | 0.004 | 0.01 | 0.01 | 0.212 | 0.35 | 0.67 | 0.58 | 0.24 | 0.39 | 0.26 |
| 1 | 242619825 | rs10926744 | -0.379 | 0.584 | -0.366 | -0.941 | -0.001 | 0.011 | 0.011 | 0 | 0.4 | 0.35 | 0.58 | 0.17 | 0.44 | 0.25 |
| 1 | 242620466 | rs6700801 | -0.814 | 0.745 | -0.683 | -0.412 | -0.032 | -0.041 | -0.041 | 1.494 | 0.12 | 0.02 | 0.07 | 0.12 | 0.14 | 0.23 |
| 1 | 242627009 | rs11589847 | -0.924 | 0.805 | -0.68 | -0.607 | 0.018 | 0.033 | 0.033 | 0 | 0.35 | 0.78 | 0.52 | 0.24 | 0.32 | 0.45 |
| 1 | 242627059 | rs12751376 | -0.113 | 0.719 | -0.598 | -0.653 | -0.003 | 0.045 | 0.045 | 0 | 0.25 | 0.16 | 0.39 | 0.11 | 0.26 | 0.34 |
| 1 | 242628042 | rs4658484 | -1.618 | 0.843 | -0.928 | -0.684 | 0.019 | 0.04 | 0.04 | 0 | 0.54 | 0.83 | 0.75 | 0.4 | 0.54 | 0.54 |
| 1 | 242628122 | rs4658486 | -1.836 | 0.792 | -1.051 | -0.715 | 0.013 | -0.023 | -0.023 | 0 | 0.59 | 0.87 | 0.78 | 0.44 | 0.61 | 0.57 |
| 1 | 242630652 | rs2134686 | -1.542 | 0.79 | -0.898 | -0.724 | 0.013 | 0.033 | 0.033 | 0 | 0.55 | 0.85 | 0.75 | 0.4 | 0.54 | 0.55 |
| 1 | 242633259 | rs6429357 | -1.38 | 0.764 | -0.779 | -0.76 | 0.031 | 0.052 | 0.052 | 0 | 0.53 | 0.82 | 0.75 | 0.4 | 0.54 | 0.54 |
| 1 | 242633878 | rs10926748 | 0.377 | 0.773 | -0.703 | -0.69 | -0.002 | -0.004 | -0.004 | 0 | 0.06 | 0.07 | 0.04 | 0.04 | 0.07 | 0.03 |
| 1 | 242635682 | rs872216 | 0.477 | 1.059 | -0.964 | -0.718 | -0.063 | 0.014 | 0.014 | 2.049 | 0.17 | 0.02 | 0.2 | 0.15 | 0.16 | 0.08 |
| 1 | 242641923 | rs1548160 | -1.833 | 0.955 | -0.909 | -0.932 | 0.005 | 0.005 | 0.005 | 0 | 0.6 | 0.87 | 0.68 | 0.38 | 0.61 | 0.57 |
| 1 | 242652356 | rs7526731 | -1.224 | 0.945 | -0.481 | -0.885 | 0.004 | 0.059 | 0.059 | 0.841 | 0.43 | 0.76 | 0.57 | 0.28 | 0.44 | 0.48 |
| 1 | 242653512 | rs10803047 | 0.109 | 1.026 | -0.477 | -0.814 | -0.016 | 0.044 | 0.044 | 0 | 0.27 | 0.21 | 0.42 | 0.1 | 0.32 | 0.35 |
| 1 | 242654482 | rs6429360 | 0.045 | 0.656 | 0.149 | -0.433 | 0.053 | 0.159 | 0.159 | 0 | 0.08 | 0.45 | 0.11 | 0.14 | 0.05 | 0.1 |
| 1 | 242656412 | rs10926753 | -0.964 | 0.823 | -0.433 | -0.896 | 0.018 | 0.044 | 0.044 | 0 | 0.52 | 0.41 | 0.58 | 0.24 | 0.56 | 0.5 |
| 1 | 242656745 | rs7539893 | -0.57 | 1.168 | -0.803 | -0.998 | -0.018 | 0.054 | 0.054 | 0 | 0.32 | 0.23 | 0.36 | 0.1 | 0.38 | 0.4 |
| 1 | 242657895 | rs9919289 | 0.569 | 0.817 | -0.512 | -0.57 | -0.008 | 0.003 | 0.003 | 0 | 0.35 | 0.41 | 0.25 | 0.63 | 0.34 | 0.39 |
| 1 | 242658747 | rs1875757 | 0.137 | 1.255 | -0.429 | -0.764 | -0.001 | -0.001 | -0.001 | 0.017 | 0.47 | 0.48 | 0.31 | 0.63 | 0.46 | 0.44 |
| 1 | 242662040 | rs10926756 | 0.038 | 1.396 | -0.409 | -1.034 | 0.007 | 0.033 | 0.033 | 0 | 0.35 | 0.22 | 0.44 | 0.23 | 0.33 | 0.25 |
| 1 | 242663386 | rs905581 | 0.715 | 1.377 | -0.201 | -1 | -0.002 | 0.002 | 0.002 | 0 | 0.44 | 0.2 | 0.28 | 0.62 | 0.43 | 0.43 |
| 1 | 242663902 | rs2809988 | -0.67 | 1.426 | -0.063 | -0.926 | -0.002 | -0.001 | -0.001 | 0 | 0.84 | 0.83 | 0.76 | 0.87 | 0.82 | 0.74 |
| 1 | 242669380 | rs2654856 | -0.854 | 1.151 | 0.231 | -1.153 | -0.005 | -0.094 | -0.094 | 0.289 | 0.27 | 0.55 | 0.18 | 0.05 | 0.27 | 0.18 |
| 1 | 242673425 | rs2809992 | -1.577 | 1.161 | -0.243 | -1.165 | -0.023 | 0.062 | 0.062 | 0 | 0.59 | 0.84 | 0.62 | 0.3 | 0.6 | 0.42 |
| 1 | 242673570 | rs2654892 | -0.623 | 1.479 | 0.13 | -0.893 | -0.003 | -0.061 | -0.061 | 0.263 | 0.24 | 0.29 | 0.16 | 0.03 | 0.22 | 0.14 |
| 1 | 242675029 | rs2654894 | -1.417 | 1.575 | -0.415 | -0.651 | -0.018 | 0.062 | 0.062 | 0 | 0.56 | 0.84 | 0.6 | 0.32 | 0.57 | 0.46 |
| 1 | 242675158 | rs12140143 | 0.048 | 1.855 | -0.033 | -0.648 | -0.002 | 0.002 | 0.002 | 0 | 0.21 | 0.19 | 0.25 | 0.15 | 0.21 | 0.15 |
| 1 | 242677578 | rs2654897 | -1.021 | 1.573 | -0.261 | -0.86 | 0 | -0.014 | -0.014 | 0.056 | 0.33 | 0.33 | 0.33 | 0.11 | 0.36 | 0.26 |
| 1 | 242682532 | rs2810005 | -1.587 | 1.706 | -0.665 | -0.66 | -0.004 | 0.078 | 0.078 | 0 | 0.57 | 0.91 | 0.57 | 0.28 | 0.6 | 0.51 |
| 1 | 242687390 | rs2810008 | -1.162 | 1.804 | -0.01 | -0.263 | -0.008 | -0.033 | -0.033 | 0 | 0.32 | 0.37 | 0.31 | 0.16 | 0.33 | 0.26 |

| | | | | | | | | | | | | | | | | |
|---|---|---|---|---|---|---|---|---|---|---|---|---|---|---|---|---|
| 1 | 242690382 | rs2810011 | 0.707 | 2.306 | -0.044 | -0.343 | 0.05 | -0.048 | -0.048 | 0 | 0.68 | 0.43 | 0.7 | 0.82 | 0.64 | 0.75 |
| 1 | 242690935 | rs12067469 | -0.625 | 2.225 | 0.084 | 0.28 | 0.011 | 0.041 | 0.041 | 0 | 0.13 | 0.24 | 0.16 | 0.19 | 0.17 | 0.14 |
| 1 | 242694809 | rs11576273 | -1.144 | 2.092 | -0.615 | 0.359 | -0.007 | -0.002 | -0.002 | 0 | 0.26 | 0.1 | 0.22 | 0.21 | 0.29 | 0.22 |
| 1 | 242695639 | rs7551349 | -0.311 | 2.087 | -0.092 | 0.958 | 0.021 | 0.003 | 0.003 | 0 | 0.16 | 0.25 | 0.18 | 0.15 | 0.18 | 0.09 |
| 1 | 242696547 | rs6677308 | -0.956 | 1.344 | -0.36 | 1.018 | -0.006 | 0.027 | 0.027 | 0 | 0.12 | 0.15 | 0.13 | 0.21 | 0.14 | 0.1 |
| 1 | 242700803 | rs2343121 | 0.792 | 2.39 | -0.142 | 1.315 | 0.085 | 0.079 | 0.079 | 0.063 | 0.87 | 0.78 | 0.75 | 0.74 | 0.73 | 0.83 |
| 1 | 242701567 | rs11804678 | -0.403 | 1.716 | -0.543 | 0.605 | 0.014 | 0.026 | 0.026 | 0 | 0.14 | 0.08 | 0.07 | 0.03 | 0.1 | 0.12 |
| 1 | 242706443 | rs12403816 | 1.129 | 1.273 | -0.68 | 0.43 | -0.001 | -0.001 | -0.001 | 0 | 0.54 | 0.59 | 0.56 | 0.53 | 0.51 | 0.58 |
| 1 | 242708729 | rs6689437 | 0.388 | 0.383 | -1.254 | -0.221 | -0.036 | -0.068 | -0.068 | 0 | 0.23 | 0.65 | 0.14 | 0.06 | 0.1 | 0.18 |
| 1 | 242712951 | rs10458443 | 0.098 | 1.133 | -1.535 | -0.044 | -0.074 | -0.073 | -0.073 | 3.315 | 0.12 | 0.01 | 0.22 | 0.21 | 0.23 | 0.15 |
| 1 | 242718333 | rs12405135 | 0.063 | 0.45 | -1.572 | -0.445 | -0.001 | -0.044 | -0.044 | 5.734 | 0.11 | 0.01 | 0.13 | 0.16 | 0.08 | 0.25 |
| 1 | 242723279 | rs4618952 | -0.642 | 0.65 | -0.971 | -0.403 | 0.023 | 0.017 | 0.017 | 0 | 0.21 | 0.29 | 0.42 | 0.31 | 0.31 | 0.29 |
| 1 | 242726365 | rs4658825 | 0.974 | 0.405 | -1.188 | -0.634 | -0.082 | 0.035 | 0.035 | 0.205 | 0.54 | 0.11 | 0.43 | 0.66 | 0.53 | 0.53 |
| 1 | 242727483 | rs1343747 | -0.481 | 0.431 | -1.068 | -0.605 | 0.03 | 0.02 | 0.02 | 0 | 0.21 | 0.3 | 0.42 | 0.31 | 0.32 | 0.29 |
| 1 | 242733044 | rs7523794 | 1.675 | 0.034 | -0.507 | -0.802 | -0.041 | -0.024 | -0.024 | 0.009 | 0.12 | 0.27 | 0.06 | 0.05 | 0.05 | 0.11 |
| 1 | 242738254 | rs1318463 | 0.73 | 0.07 | -0.426 | -0.831 | 0.033 | -0.007 | -0.007 | 1.33 | 0.7 | 0.51 | 0.53 | 0.7 | 0.63 | 0.65 |
| 1 | 242739083 | rs10926773 | -0.483 | 0.266 | -0.666 | -1.073 | 0.011 | 0.006 | 0.006 | 0 | 0.23 | 0.27 | 0.42 | 0.31 | 0.33 | 0.29 |
| 1 | 242740810 | rs6658662 | 1.661 | 0.172 | -0.762 | -0.824 | -0.079 | -0.069 | -0.069 | 0 | 0.47 | 0.08 | 0.4 | 0.46 | 0.49 | 0.38 |
| 1 | 242741023 | rs1938341 | 1.356 | 0.487 | -0.565 | -0.748 | 0.003 | 0.003 | 0.003 | 0.617 | 0.6 | 0.45 | 0.48 | 0.51 | 0.54 | 0.49 |
| 1 | 242746708 | rs4658827 | 1.282 | 0.271 | -0.695 | -0.81 | -0.012 | -0.004 | -0.004 | 0.041 | 0.58 | 0.39 | 0.45 | 0.54 | 0.51 | 0.5 |
| 1 | 242748068 | rs1938329 | 0.283 | 0.181 | -0.732 | -0.857 | -0.001 | -0.003 | -0.003 | 0.811 | 0.78 | 0.76 | 0.84 | 0.8 | 0.79 | 0.75 |
| 1 | 242764180 | rs6429365 | 0.884 | 0.52 | -0.652 | -0.684 | -0.003 | 0.005 | 0.005 | 0.087 | 0.64 | 0.41 | 0.49 | 0.69 | 0.58 | 0.64 |
| 1 | 242765473 | rs7527225 | -0.167 | 0.814 | -0.453 | -0.388 | 0.021 | 0.033 | 0.033 | 0 | 0.18 | 0.23 | 0.38 | 0.29 | 0.26 | 0.26 |
| 1 | 242767005 | rs6429366 | -1.044 | 0.642 | -0.718 | -0.366 | -0.002 | -0.001 | -0.001 | 0 | 0.55 | 0.54 | 0.59 | 0.54 | 0.56 | 0.62 |
| 1 | 242767678 | rs7523607 | -0.325 | 0.382 | -0.921 | -0.975 | -0.003 | -0.096 | -0.096 | 0 | 0.76 | 0.58 | 0.86 | 0.97 | 0.78 | 0.86 |
| 1 | 242767951 | rs4658489 | 0.638 | 0.396 | -1.139 | -0.611 | -0.032 | -0.034 | -0.034 | 0.907 | 0.46 | 0.21 | 0.46 | 0.48 | 0.46 | 0.4 |
| 1 | 242768780 | rs2343126 | -0.638 | 0.709 | -0.479 | -0.352 | 0.01 | 0.011 | 0.011 | 0 | 0.2 | 0.22 | 0.26 | 0.29 | 0.28 | 0.27 |
| 1 | 242769507 | rs11577288 | -0.041 | 0.894 | -0.544 | -0.325 | -0.031 | -0.032 | -0.032 | 0.021 | 0.12 | 0.06 | 0.21 | 0.2 | 0.19 | 0.18 |
| 1 | 242777990 | rs7549010 | 1 | 0.81 | -0.256 | -0.7 | -0.001 | -0.022 | -0.022 | 1.168 | 0.11 | 0.16 | 0.07 | 0.03 | 0.08 | 0.12 |
| 1 | 242789505 | rs12729887 | 1.304 | 0.701 | -0.909 | -1.109 | -0.033 | 0.107 | 0.107 | 0.039 | 0.25 | 0.05 | 0.22 | 0.08 | 0.24 | 0.25 |
| 1 | 242791710 | rs1316440 | 0.152 | 0.587 | -0.655 | -1.343 | -0.001 | -0.038 | -0.038 | 0.375 | 0.23 | 0.3 | 0.13 | 0.06 | 0.21 | 0.22 |
| 1 | 242795726 | rs6697901 | 1.523 | 0.622 | -0.525 | -1.269 | -0.01 | 0.09 | 0.09 | 0.053 | 0.26 | 0.09 | 0.23 | 0.09 | 0.24 | 0.25 |
| 1 | 242796014 | rs10926796 | 1.56 | 0.635 | -0.571 | -1.266 | -0.033 | 0.107 | 0.107 | 0.064 | 0.25 | 0.05 | 0.22 | 0.08 | 0.24 | 0.25 |
| 1 | 242798972 | rs4658840 | 0.592 | 0.604 | -0.284 | -1.286 | -0.061 | 0.156 | 0.156 | 2.529 | 0.36 | 0.07 | 0.28 | 0.11 | 0.36 | 0.35 |
| 1 | 242800080 | rs10926799 | 1.522 | 0.514 | 0.184 | -1.132 | 0.003 | -0.031 | -0.031 | 0 | 0.3 | 0.19 | 0.25 | 0.5 | 0.27 | 0.35 |
| 1 | 242805757 | rs1413216 | 1.758 | 0.552 | 0.199 | -1.075 | -0.028 | 0.02 | 0.02 | 5.366 | 0.1 | 0.02 | 0.06 | 0.02 | 0.12 | 0.1 |
| 1 | 242805821 | rs1413214 | 0.608 | 0.509 | 0.256 | -1.068 | -0.027 | -0.037 | -0.037 | 0 | 0.18 | 0.13 | 0.35 | 0.34 | 0.26 | 0.19 |
| 1 | 242806621 | rs6668917 | -1.085 | 0.337 | 0.407 | -1.007 | -0.017 | 0.037 | 0.037 | 0.248 | 0.67 | 0.61 | 0.73 | 0.5 | 0.74 | 0.63 |
| 1 | 242807999 | rs10926804 | 1.725 | 0.301 | 0.156 | -1.051 | -0.004 | -0.083 | -0.083 | 0 | 0.27 | 0.08 | 0.23 | 0.49 | 0.24 | 0.34 |
| 1 | 242808496 | rs10803065 | 1.546 | 0.523 | 0.18 | -0.944 | 0 | -0.002 | -0.002 | 0 | 0.3 | 0.26 | 0.25 | 0.49 | 0.24 | 0.36 |
| 1 | 242808798 | rs10803066 | 1.316 | 0.246 | 0.195 | -0.923 | -0.001 | -0.011 | -0.011 | 0 | 0.32 | 0.26 | 0.26 | 0.49 | 0.28 | 0.41 |
| 1 | 242811594 | rs2184185 | -0.244 | 0.357 | 0.173 | -0.886 | 0.021 | 0.012 | 0.012 | 0 | 0.53 | 0.32 | 0.66 | 0.49 | 0.6 | 0.52 |
| 1 | 242812831 | rs12042953 | 0.524 | 0.546 | -0.091 | -0.847 | -0.035 | -0.047 | -0.047 | 0.951 | 0.18 | 0.13 | 0.38 | 0.37 | 0.28 | 0.2 |
| 1 | 242819974 | rs1333696 | -1.391 | 0.358 | 0.008 | -0.847 | -0.01 | 0.052 | 0.052 | 0.167 | 0.66 | 0.58 | 0.72 | 0.53 | 0.71 | 0.62 |
| 1 | 242820837 | rs10159343 | 1.245 | 0.03 | -0.681 | -0.806 | -0.001 | 0.087 | 0.087 | 0.412 | 0.24 | 0.08 | 0.22 | 0.09 | 0.23 | 0.22 |
| 1 | 242827332 | rs11580100 | 0.374 | 0.059 | -0.878 | -0.69 | 0.033 | 0.118 | 0.118 | 0.336 | 0.51 | 0.3 | 0.63 | 0.84 | 0.58 | 0.58 |
| 1 | 242828277 | rs7526164 | 1.373 | 0.082 | -1.24 | -0.422 | 0.006 | 0.008 | 0.008 | 2.383 | 0.09 | 0.06 | 0.03 | 0.02 | 0.07 | 0.09 |
| 1 | 242836169 | rs12691546 | 0.537 | 0.14 | -1.137 | -0.338 | 0.001 | 0.069 | 0.069 | 0.535 | 0.3 | 0.17 | 0.25 | 0.13 | 0.3 | 0.31 |
| 1 | 242840908 | rs10926819 | 0.801 | 0.44 | -1.321 | -0.082 | -0.011 | -0.045 | -0.045 | 0 | 0.34 | 0.18 | 0.27 | 0.47 | 0.33 | 0.4 |
| 1 | 242840987 | rs914949 | -0.313 | 0.855 | -1.122 | 0.225 | -0.003 | -0.004 | -0.004 | 0.174 | 0.28 | 0.29 | 0.42 | 0.4 | 0.34 | 0.28 |
| 1 | 242842300 | rs5020631 | 0.393 | 0.474 | -1.421 | -0.109 | -0.048 | 0.073 | 0.073 | 0.149 | 0.22 | 0.03 | 0.21 | 0.09 | 0.23 | 0.22 |
| 1 | 242855782 | rs12127991 | 0.687 | 0.569 | -0.097 | 0.262 | 0.013 | -0.009 | -0.009 | 0 | 0.78 | 0.5 | 0.58 | 0.81 | 0.77 | 0.74 |
| 1 | 242856566 | rs6429370 | 0.287 | 0.478 | 0.199 | 0.229 | 0.003 | 0 | 0 | 0 | 0.84 | 0.78 | 0.61 | 0.83 | 0.77 | 0.77 |
| 1 | 242856854 | rs10926825 | 1.594 | 0.168 | 0.127 | 0.746 | 0.016 | -0.018 | -0.018 | 0 | 0.61 | 0.27 | 0.41 | 0.45 | 0.65 | 0.57 |
| 1 | 242856905 | rs6659331 | -0.399 | 0.191 | -0.018 | 0.893 | -0.015 | 0.032 | 0.032 | 0 | 0.18 | 0.22 | 0.11 | 0.33 | 0.12 | 0.16 |
| 1 | 242872032 | rs1413202 | -1.257 | 0.319 | 0.293 | 0.688 | -0.003 | 0.04 | 0.04 | 0 | 0.3 | 0.55 | 0.49 | 0.44 | 0.31 | 0.42 |
| 1 | 242872549 | rs11584716 | 0.229 | 0.241 | -0.301 | 0.138 | 0.002 | 0.018 | 0.018 | 2.33 | 0.24 | 0.16 | 0.12 | 0.04 | 0.2 | 0.15 |
| 1 | 242872709 | rs11586761 | 0.229 | 0.241 | -0.301 | 0.138 | 0.002 | 0.018 | 0.018 | 2.33 | 0.24 | 0.16 | 0.12 | 0.04 | 0.2 | 0.15 |
| 1 | 242873595 | rs10926828 | 1.555 | -0.005 | -0.554 | -0.038 | -0.137 | -0.164 | -0.164 | 0 | 0.39 | 0.03 | 0.32 | 0.51 | 0.44 | 0.4 |
| 1 | 242873955 | rs6657754 | -0.349 | -0.015 | -0.126 | -0.015 | -0.014 | 0.03 | 0.03 | 0.06 | 0.22 | 0.62 | 0.27 | 0.32 | 0.17 | 0.22 |
| 1 | 242874434 | rs12561770 | 0.106 | 0.056 | -0.346 | 0.025 | -0.001 | 0.003 | 0.003 | 0 | 0.18 | 0.21 | 0.23 | 0.33 | 0.17 | 0.2 |
| 1 | 242874959 | rs10803075 | -0.733 | 0.199 | -0.283 | 0.124 | -0.001 | -0.002 | -0.002 | 0.328 | 0.43 | 0.59 | 0.37 | 0.37 | 0.37 | 0.37 |
| 1 | 242876052 | rs12060035 | 0.148 | 0.374 | -0.681 | -0.196 | -0.002 | 0.035 | 0.035 | 0 | 0.25 | 0.17 | 0.14 | 0.06 | 0.23 | 0.1 |
| 1 | 242878998 | rs6690236 | -0.626 | 0.553 | -0.819 | -0.014 | 0.017 | 0.079 | 0.079 | 0 | 0.46 | 0.67 | 0.49 | 0.13 | 0.43 | 0.38 |
| 1 | 242881445 | rs10803076 | 0.555 | 0.734 | -1.141 | 0.648 | -0.048 | -0.07 | -0.07 | 0.332 | 0.22 | 0.05 | 0.2 | 0.33 | 0.22 | 0.22 |
| 1 | 242881598 | rs12756245 | 0.233 | 0.512 | -0.617 | 0.58 | 0 | 0.001 | 0.001 | 0 | 0.6 | 0.42 | 0.52 | 0.49 | 0.61 | 0.59 |
| 1 | 242884679 | rs1879158 | 0.06 | 0.309 | -0.246 | 0.564 | -0.003 | -0.016 | -0.016 | 0.372 | 0.29 | 0.25 | 0.27 | 0.5 | 0.27 | 0.39 |
| 1 | 242890160 | rs10926833 | -0.145 | 0.109 | -0.093 | 0.485 | -0.001 | 0.007 | 0.007 | 0 | 0.33 | 0.39 | 0.28 | 0.5 | 0.27 | 0.42 |
| 1 | 242901577 | rs12184253 | -0.655 | -0.25 | -0.264 | -0.282 | -0.006 | -0.063 | -0.063 | 0 | 0.19 | 0.27 | 0.1 | 0 | 0.14 | 0.06 |

| | | | | | | | | | | | | | | | | |
|---|---|---|---|---|---|---|---|---|---|---|---|---|---|---|---|---|
| 1 | 242907976 | rs12075764 | -0.48 | -0.196 | -0.687 | -0.237 | 0.003 | -0.002 | -0.002 | 0 | 0.17 | 0.19 | 0.09 | 0 | 0.14 | 0.06 |
| 1 | 242911065 | rs2027047 | -0.762 | -0.314 | -0.398 | 0.296 | -0.03 | 0.033 | 0.033 | 0 | 0.07 | 0.12 | 0.06 | 0.14 | 0.04 | 0.28 |
| 1 | 242915189 | rs1007097 | 1.157 | -0.492 | -0.326 | 0.258 | -0.009 | 0.008 | 0.008 | 0.326 | 0.65 | 0.45 | 0.68 | 0.52 | 0.7 | 0.4 |
| 1 | 242915247 | rs1556559 | -0.121 | -0.35 | 0.021 | 0.302 | 0.075 | -0.025 | -0.025 | 0 | 0.94 | 0.78 | 0.95 | 1 | 0.92 | 0.98 |
| 1 | 242916149 | rs10803081 | 0.477 | -0.384 | 0.051 | 0.289 | 0.063 | -0.052 | -0.052 | 0 | 0.91 | 0.72 | 0.92 | 0.99 | 0.9 | 0.92 |
| 1 | 242919558 | rs12407826 | 0.027 | -0.448 | -0.011 | 0.449 | -0.013 | 0.116 | 0.116 | 0.07 | 0.05 | 0.12 | 0.08 | 0.26 | 0.03 | 0.11 |
| 1 | 242919916 | rs1855368 | 1.885 | -0.509 | 0.009 | 0.429 | -0.005 | 0.098 | 0.098 | 0.211 | 0.8 | 0.62 | 0.79 | 0.63 | 0.82 | 0.65 |
| 1 | 242920580 | rs10803083 | 1.181 | -0.356 | -0.026 | 0.467 | 0.01 | -0.024 | -0.024 | 0 | 0.85 | 0.74 | 0.87 | 0.92 | 0.86 | 0.76 |
| 1 | 242921641 | rs10436949 | 0.914 | -0.294 | -0.781 | 0.428 | -0.057 | -0.116 | -0.116 | 0 | 0.28 | 0.04 | 0.4 | 0.36 | 0.32 | 0.2 |
| 1 | 242927370 | rs1591217 | -0.217 | -0.428 | -0.429 | 0.327 | 0.012 | -0.012 | -0.012 | 0 | 0.59 | 0.52 | 0.54 | 0.35 | 0.58 | 0.66 |
| 1 | 242928602 | rs7527876 | 0.435 | -0.328 | -0.472 | 0.364 | 0.008 | -0.006 | -0.006 | 0.074 | 0.15 | 0.07 | 0.11 | 0.19 | 0.15 | 0.42 |
| 1 | 242928876 | rs7514112 | -0.506 | -0.427 | -0.509 | 0.552 | -0.03 | -0.023 | -0.023 | 0 | 0.53 | 0.17 | 0.54 | 0.52 | 0.51 | 0.67 |
| 1 | 242931513 | rs10926848 | 0.48 | -0.373 | -0.317 | 0.585 | -0.049 | -0.034 | -0.034 | 0 | 0.38 | 0.09 | 0.43 | 0.35 | 0.36 | 0.25 |
| 1 | 242941556 | rs4658858 | -1.219 | -0.423 | -0.147 | 0.546 | 0.007 | 0.044 | 0.044 | 0 | 0.7 | 0.59 | 0.59 | 0.57 | 0.71 | 0.73 |
| 1 | 242943269 | rs12085924 | -0.419 | -0.375 | -0.415 | 0.653 | -0.016 | 0.099 | 0.099 | 0 | 0.23 | 0.03 | 0.13 | 0.1 | 0.18 | 0.22 |
| 1 | 242946169 | rs9943068 | -0.123 | -0.263 | -0.244 | 0.853 | -0.046 | -0.065 | -0.065 | 1.343 | 0.47 | 0.09 | 0.43 | 0.52 | 0.42 | 0.49 |
| 1 | 242952078 | rs914951 | -0.949 | -0.033 | -0.182 | 0.471 | -0.002 | -0.021 | -0.021 | 0 | 0.59 | 0.57 | 0.61 | 0.84 | 0.58 | 0.81 |
| 1 | 242959355 | rs4630115 | 0.782 | -0.036 | -0.301 | 0.699 | -0.021 | -0.009 | -0.009 | 0.081 | 0.34 | 0.13 | 0.37 | 0.31 | 0.32 | 0.25 |
| 1 | 242959776 | rs2780784 | -1.058 | -0.03 | -0.175 | 0.686 | -0.001 | -0.011 | -0.011 | 0 | 0.59 | 0.62 | 0.62 | 0.84 | 0.58 | 0.81 |
| 1 | 242964870 | rs12126980 | 0.5 | 0.219 | -0.208 | 0.947 | -0.017 | -0.009 | -0.009 | 0 | 0.34 | 0.13 | 0.37 | 0.3 | 0.31 | 0.24 |
| 1 | 242965985 | rs1889220 | -1.267 | 0.297 | -0.077 | 1.102 | 0.059 | 0.298 | 0.298 | 0 | 0.09 | 0.27 | 0.15 | 0.43 | 0.15 | 0.31 |
| 1 | 242969964 | rs6661201 | 0.444 | 0.228 | -0.446 | 0.809 | -0.023 | -0.013 | -0.013 | 0 | 0.34 | 0.11 | 0.37 | 0.3 | 0.31 | 0.24 |
| 1 | 242977333 | rs2780790 | -0.787 | 0.31 | -0.284 | 0.886 | 0.003 | -0.003 | -0.003 | 0.878 | 0.23 | 0.26 | 0.19 | 0.28 | 0.25 | 0.46 |
| 1 | 242989157 | rs2210612 | -0.438 | 0.124 | -0.177 | 1.093 | -0.023 | 0.108 | 0.108 | 0.429 | 0.2 | 0.3 | 0.21 | 0.52 | 0.18 | 0.51 |
| 1 | 243000992 | rs4658860 | -0.327 | 0.235 | -0.137 | 0.891 | -0.01 | -0.003 | -0.003 | 0 | 0.5 | 0.76 | 0.53 | 0.53 | 0.47 | 0.52 |
| 1 | 243003907 | rs914942 | -0.261 | -0.04 | 0.34 | 0.704 | 0 | 0.061 | 0.061 | 0 | 0.48 | 0.59 | 0.58 | 0.81 | 0.45 | 0.65 |
| 1 | 243004424 | rs10926883 | 0.131 | -0.101 | 0.649 | 0.881 | -0.001 | 0.002 | 0.002 | 0 | 0.43 | 0.39 | 0.46 | 0.34 | 0.39 | 0.32 |
| 1 | 243004669 | rs12407970 | -0.954 | -0.101 | 0.648 | 0.882 | 0.004 | 0.419 | 0.419 | 0 | 0.05 | 0.2 | 0.12 | 0.47 | 0.05 | 0.33 |
| 1 | 243005376 | rs6429381 | -0.442 | -0.112 | 0.72 | 0.872 | 0.003 | 0.154 | 0.154 | 0 | 0.49 | 0.71 | 0.58 | 0.81 | 0.44 | 0.65 |
| 1 | 243005986 | rs1333691 | 0.11 | -0.046 | 0.703 | 0.893 | -0.001 | 0.002 | 0.002 | 0 | 0.43 | 0.39 | 0.46 | 0.34 | 0.39 | 0.32 |
| 1 | 243027814 | rs10926898 | -0.173 | 0.116 | 0.436 | 0.39 | -0.001 | 0.047 | 0.047 | 0 | 0.48 | 0.58 | 0.57 | 0.8 | 0.44 | 0.64 |
| 1 | 243028780 | rs12022794 | -0.854 | 0.181 | 0.441 | 0.509 | 0.039 | 0.207 | 0.207 | 0 | 0.07 | 0.31 | 0.12 | 0.27 | 0.08 | 0.15 |
| 1 | 243033446 | rs4535989 | 0.282 | 0.212 | -0.533 | 0.255 | -0.038 | -0.066 | -0.066 | 0.112 | 0.4 | 0.1 | 0.46 | 0.53 | 0.4 | 0.5 |
| 1 | 243034750 | rs4233436 | 0.243 | 0.284 | -0.548 | 0.265 | -0.007 | -0.027 | -0.027 | 0 | 0.41 | 0.17 | 0.46 | 0.53 | 0.39 | 0.49 |
| 1 | 243040970 | rs12032023 | 0.357 | 0.205 | -0.56 | 0.287 | 0 | -0.029 | -0.029 | 0 | 0.38 | 0.16 | 0.44 | 0.53 | 0.35 | 0.49 |
| 1 | 243041047 | rs4658504 | -0.307 | 0.299 | -0.834 | -0.386 | -0.062 | 0.159 | 0.159 | 0 | 0.64 | 0.89 | 0.69 | 0.89 | 0.59 | 0.89 |
| 1 | 243053057 | rs4278360 | 0.024 | -0.133 | -0.811 | 0.526 | 0.024 | -0.078 | -0.078 | 0 | 0.2 | 0.09 | 0.32 | 0.54 | 0.17 | 0.19 |
| 1 | 243056401 | rs4658507 | 0.633 | 0.469 | -1.474 | -0.331 | -0.109 | -0.022 | -0.022 | 0.138 | 0.29 | 0.09 | 0.21 | 0.25 | 0.43 | 0.12 |
| 1 | 243056785 | rs2491828 | -0.287 | 0.083 | -1.596 | -0.932 | -0.062 | 0.124 | 0.124 | 0 | 0.81 | 0.99 | 0.78 | 0.99 | 0.76 | 0.76 |
| 1 | 243065369 | rs12077950 | -0.282 | -0.428 | -0.042 | -0.654 | -0.044 | -0.08 | -0.08 | 0 | 0.14 | 0.63 | 0.09 | 0.03 | 0.05 | 0.16 |
| 1 | 243071916 | rs2502350 | -0.247 | -0.492 | 0.134 | -0.659 | -0.05 | -0.077 | -0.077 | 0 | 0.13 | 0.56 | 0.09 | 0.03 | 0.04 | 0.17 |
| 1 | 243078995 | rs2502308 | -0.575 | -0.085 | -0.441 | -0.847 | 0.009 | -0.043 | -0.043 | 0 | 0.87 | 0.86 | 0.79 | 0.99 | 0.79 | 0.81 |
| 1 | 243080523 | rs6679417 | 0.028 | 0.174 | -0.404 | -0.878 | 0.028 | -0.067 | -0.067 | 0.273 | 0.12 | 0.44 | 0.08 | 0.03 | 0.09 | 0.21 |
| 1 | 243080833 | rs11576569 | 0.855 | 0.516 | -2.066 | -0.683 | -0.07 | 0.008 | 0.008 | 0 | 0.28 | 0.09 | 0.2 | 0.21 | 0.36 | 0.12 |
| 1 | 243086195 | rs2502313 | 0.181 | -0.199 | -1.839 | 0.118 | 0.031 | -0.093 | -0.093 | 0.428 | 0.25 | 0.13 | 0.36 | 0.56 | 0.21 | 0.15 |
| 1 | 243087106 | rs3003542 | -0.208 | 0.036 | -1.73 | -0.176 | 0.043 | 0.045 | 0.045 | 0 | 0.4 | 0.17 | 0.49 | 0.71 | 0.33 | 0.43 |
| 1 | 243102869 | rs2998676 | 0.095 | 0.252 | -1.042 | 0.249 | 0.003 | 0 | 0 | 0 | 0.52 | 0.54 | 0.5 | 0.5 | 0.53 | 0.5 |
| 1 | 243119242 | rs12741729 | -0.215 | -0.009 | -1.133 | 0.013 | 0.017 | 0.125 | 0.125 | 0 | 0.43 | 0.2 | 0.22 | 0.12 | 0.4 | 0.58 |
| 1 | 243140736 | rs2907176 | 0.743 | 0.373 | -1.027 | -0.003 | -0.022 | 0.029 | 0.029 | 0 | 0.23 | 0.18 | 0.19 | 0.07 | 0.28 | 0.1 |
| 1 | 243147031 | rs12039141 | -0.21 | 0.09 | -0.463 | 0.237 | 0.002 | 0.012 | 0.012 | 0.001 | 0.7 | 0.29 | 0.62 | 0.51 | 0.7 | 0.84 |
| 1 | 243181599 | rs4098386 | 0.047 | -0.416 | -0.226 | 0.463 | 0.025 | 0.012 | 0.012 | 0 | 0.4 | 0.24 | 0.25 | 0.33 | 0.34 | 0.44 |
| 1 | 243277476 | rs4658537 | -0.354 | 0.067 | 0.365 | 0.75 | 0.017 | 0.078 | 0.078 | 0.059 | 0.07 | 0.13 | 0.2 | 0.26 | 0.12 | 0.13 |
| 1 | 243355204 | rs10926961 | 0.174 | -0.554 | 1.071 | -0.347 | 0.013 | -0.004 | -0.004 | 0 | 0.36 | 0.17 | 0.23 | 0.3 | 0.34 | 0.44 |
| 1 | 243380513 | rs7518350 | -0.479 | -0.732 | 1.222 | -0.536 | 0.037 | 0.042 | 0.042 | 0 | 0.45 | 0.33 | 0.27 | 0.29 | 0.38 | 0.57 |
| 1 | 243385307 | rs6662118 | -1.008 | -0.766 | 1.495 | -0.559 | 0.055 | 0.068 | 0.068 | 0 | 0.51 | 0.69 | 0.32 | 0.29 | 0.39 | 0.57 |
| 1 | 243391131 | rs4658547 | -0.463 | -0.684 | 1.557 | -0.548 | 0.031 | 0.005 | 0.005 | 0 | 0.93 | 0.79 | 0.87 | 0.95 | 0.9 | 0.9 |
| 1 | 243397782 | rs10926975 | 0.184 | -1.02 | 1.166 | -0.933 | 0.001 | -0.003 | -0.003 | 0 | 0.27 | 0.24 | 0.2 | 0.28 | 0.26 | 0.41 |
| 1 | 243410208 | rs6662991 | -0.748 | -0.772 | 1.325 | -0.73 | 0.051 | 0.06 | 0.06 | 0 | 0.44 | 0.21 | 0.26 | 0.29 | 0.38 | 0.55 |
| 1 | 243412936 | rs4658552 | -0.924 | -0.762 | 1.23 | -0.726 | 0.007 | 0.007 | 0.007 | 0 | 0.48 | 0.55 | 0.3 | 0.29 | 0.38 | 0.57 |
| 1 | 243440352 | rs12026068 | 1.057 | -0.653 | 1.098 | -1.239 | -0.031 | -0.021 | -0.021 | 0 | 0.05 | 0.01 | 0.12 | 0.06 | 0.1 | 0.1 |
| 1 | 243449881 | rs3818802 | 1.633 | -0.639 | 1.108 | -1.179 | -0.037 | 0.108 | 0.108 | 0 | 0.42 | 0.09 | 0.57 | 0.74 | 0.54 | 0.34 |
| 1 | 243454014 | rs10803133 | 1.633 | -0.639 | 1.108 | -1.179 | -0.037 | 0.108 | 0.108 | 0 | 0.42 | 0.09 | 0.57 | 0.74 | 0.54 | 0.34 |
| 1 | 243454083 | rs10754800 | 0.678 | 0.064 | 1.607 | -0.802 | 0.051 | 0.005 | 0.005 | 0.074 | 0.89 | 0.6 | 0.85 | 0.94 | 0.89 | 0.9 |
| 1 | 243462153 | rs10926984 | 0.558 | 0.295 | 1.498 | -0.696 | -0.033 | -0.022 | -0.022 | 0 | 0.05 | 0.01 | 0.14 | 0.06 | 0.12 | 0.1 |
| 1 | 243470453 | rs2932634 | 1.119 | 0.187 | 1.642 | -0.749 | 0.007 | 0.029 | 0.029 | 0 | 0.53 | 0.34 | 0.61 | 0.74 | 0.55 | 0.42 |
| 1 | 243483368 | rs10926989 | -0.577 | 0.149 | 1.547 | -0.759 | 0.031 | 0.107 | 0.107 | 0 | 0.37 | 0.14 | 0.21 | 0.19 | 0.32 | 0.44 |
| 1 | 243492643 | rs2490400 | 0.829 | 0.449 | 1.285 | -0.817 | 0.091 | -0.005 | -0.005 | 0 | 0.87 | 0.54 | 0.72 | 0.93 | 0.8 | 0.8 |
| 1 | 243498312 | rs2802723 | 1.179 | 0.498 | 0.844 | -0.742 | 0.035 | 0.133 | 0.133 | 0 | 0.51 | 0.18 | 0.6 | 0.74 | 0.56 | 0.39 |
| 1 | 243498533 | rs2802722 | 1.031 | 0.598 | 0.901 | -0.744 | 0.014 | 0.06 | 0.06 | 0 | 0.53 | 0.3 | 0.62 | 0.74 | 0.55 | 0.38 |

| | | | | | | | | | | | | | | | | |
|---|---|---|---|---|---|---|---|---|---|---|---|---|---|---|---|---|
| 1 | 243501047 | rs2636319 | 1.148 | 0.565 | 0.631 | -0.727 | 0.032 | 0.128 | 0.128 | 0 | 0.51 | 0.19 | 0.59 | 0.74 | 0.56 | 0.39 |
| 1 | 243505841 | rs4658558 | 1.328 | 0.814 | 0.243 | -1.007 | -0.002 | 0.182 | 0.182 | 0 | 0.5 | 0.14 | 0.47 | 0.81 | 0.45 | 0.35 |
| 1 | 243513047 | rs10754803 | 1.504 | 0.637 | 0.363 | -0.724 | -0.067 | 0.172 | 0.172 | 0 | 0.48 | 0.07 | 0.56 | 0.81 | 0.48 | 0.4 |
| 1 | 243518151 | rs2802728 | 1.37 | 0.537 | 1.024 | -0.721 | -0.033 | -0.039 | -0.039 | 0 | 0.86 | 0.37 | 0.87 | 0.91 | 0.9 | 0.84 |
| 1 | 243522906 | rs2490393 | 0.098 | 0.665 | 0.841 | -0.714 | 0.003 | 0 | 0 | 0 | 0.7 | 0.77 | 0.84 | 0.97 | 0.73 | 0.67 |
| 1 | 243546954 | rs12239876 | -0.714 | 0.786 | 0.651 | -0.447 | -0.005 | -0.009 | -0.009 | 0 | 0.1 | 0.14 | 0.09 | 0.08 | 0.1 | 0.14 |
| 1 | 243577313 | rs4658568 | -0.776 | 0.806 | 0.971 | -0.403 | 0.011 | 0.004 | 0.004 | 0 | 0.1 | 0.27 | 0.1 | 0.08 | 0.1 | 0.15 |
| 1 | 243579112 | rs10927011 | 1.692 | 0.746 | 0.681 | -0.526 | -0.032 | 0.165 | 0.165 | 0 | 0.51 | 0.09 | 0.54 | 0.78 | 0.5 | 0.42 |
| 1 | 243582314 | rs2039838 | -0.862 | 0.818 | 0.969 | -0.263 | 0.001 | -0.008 | -0.008 | 0 | 0.1 | 0.22 | 0.1 | 0.08 | 0.1 | 0.15 |
| 1 | 243590990 | rs6429425 | -0.319 | 1.143 | 0.76 | -0.02 | -0.004 | -0.003 | -0.003 | 0 | 0.06 | 0.08 | 0.15 | 0.08 | 0.12 | 0.1 |
| 1 | 243593660 | rs12049318 | -1.207 | 1.137 | 0.972 | -0.049 | 0.021 | 0.011 | 0.011 | 0.013 | 0.09 | 0.23 | 0.14 | 0.08 | 0.14 | 0.1 |
| 1 | 243603132 | rs6685276 | -1.441 | 1.18 | 0.886 | -0.063 | 0.02 | 0.007 | 0.007 | 0 | 0.09 | 0.21 | 0.14 | 0.08 | 0.16 | 0.1 |
| 1 | 243608642 | rs6702894 | -2.05 | 1.167 | 0.92 | -0.057 | 0.024 | 0.019 | 0.019 | 0.086 | 0.13 | 0.24 | 0.18 | 0.16 | 0.21 | 0.24 |
| 1 | 243609927 | rs3006925 | 0.273 | 0.767 | 0.977 | -0.388 | 0.003 | 0.007 | 0.007 | 0 | 0.74 | 0.76 | 0.81 | 0.95 | 0.78 | 0.67 |
| 1 | 243613748 | rs11800122 | -1.792 | 1.273 | 1.229 | -0.12 | 0.032 | 0.009 | 0.009 | 0 | 0.11 | 0.27 | 0.14 | 0.09 | 0.16 | 0.11 |
| 1 | 243615589 | rs10803143 | -1.449 | 1.119 | 1.633 | -0.275 | 0.027 | -0.048 | -0.048 | 0 | 0.22 | 0.36 | 0.21 | 0.09 | 0.26 | 0.18 |
| 1 | 243617092 | rs12028447 | -0.221 | 1.135 | 1.597 | -0.261 | 0.015 | 0.022 | 0.022 | 0.265 | 0.06 | 0.15 | 0.11 | 0.09 | 0.1 | 0.1 |
| 1 | 243617188 | rs11810833 | -0.352 | 1.071 | 1.35 | -0.247 | 0.003 | -0.044 | -0.044 | 0.424 | 0.15 | 0.16 | 0.1 | 0 | 0.16 | 0.08 |
| 1 | 243639218 | rs10927025 | 1.862 | 0.644 | 1.022 | 0.089 | 0.015 | 0.102 | 0.102 | 0 | 0.59 | 0.18 | 0.7 | 0.78 | 0.61 | 0.44 |
| 1 | 243641420 | rs2994339 | -1.459 | 0.903 | 1.362 | 0.254 | 0.041 | 0.006 | 0.006 | 1.114 | 0.14 | 0.28 | 0.14 | 0.12 | 0.19 | 0.14 |
| 1 | 243641971 | rs3006917 | -1.365 | 0.914 | 1.404 | 0.26 | 0.041 | 0.007 | 0.007 | 0.374 | 0.13 | 0.23 | 0.13 | 0.12 | 0.19 | 0.14 |
| 1 | 243644497 | rs1008173 | -1.783 | 0.822 | 1.419 | 1.183 | 0.058 | 0.19 | 0.19 | 0.511 | 0.16 | 0.38 | 0.21 | 0.41 | 0.21 | 0.17 |
| 1 | 243647921 | rs2994329 | -1.808 | 0.898 | 1.421 | 1.262 | 0.065 | 0.185 | 0.185 | 1.693 | 0.16 | 0.4 | 0.22 | 0.37 | 0.2 | 0.19 |
| 1 | 243654278 | rs884328 | -1.583 | 1.182 | 0.794 | 1.187 | -0.06 | -0.077 | -0.077 | 3.175 | 0.15 | 0.03 | 0.25 | 0.36 | 0.23 | 0.22 |
| 1 | 243657399 | rs3006927 | 0.712 | 0.845 | 0.889 | 1.123 | 0.001 | 0.001 | 0.001 | 0 | 0.82 | 0.83 | 0.9 | 0.8 | 0.84 | 0.61 |
| 1 | 243658030 | rs3006928 | 0.414 | 0.915 | 0.801 | 1.117 | 0.009 | 0.014 | 0.014 | 0 | 0.84 | 0.96 | 0.93 | 0.89 | 0.85 | 0.67 |
| 1 | 243662027 | rs9428576 | -2.499 | 1.49 | 1.025 | 1.428 | -0.016 | -0.008 | -0.008 | 0 | 0.35 | 0.73 | 0.39 | 0.36 | 0.42 | 0.33 |
| 1 | 243662773 | rs10927029 | -0.58 | 1.33 | 1.588 | 1.489 | 0.015 | 0.012 | 0.012 | 0 | 0.13 | 0.18 | 0.17 | 0.11 | 0.17 | 0.11 |
| 1 | 243664857 | rs1058304 | -2.581 | 0.099 | 1.69 | 1.221 | 0.001 | -0.001 | -0.001 | 0 | 0.26 | 0.7 | 0.3 | 0.27 | 0.26 | 0.23 |
| 1 | 243673099 | rs12045585 | -1.981 | 0.103 | 1.613 | 1.424 | -0.006 | -0.006 | -0.006 | 0.003 | 0.09 | 0.07 | 0.15 | 0.25 | 0.13 | 0.18 |
| 1 | 243674387 | rs12140414 | -1.975 | 0.098 | 1.883 | 1.338 | 0.051 | 0.069 | 0.069 | 0.185 | 0.1 | 0.22 | 0.17 | 0.26 | 0.18 | 0.21 |
| 1 | 243678389 | rs6704286 | 1.282 | -0.153 | 2.004 | 1.426 | -0.013 | -0.003 | -0.003 | 0 | 0.8 | 0.79 | 0.9 | 0.84 | 0.86 | 0.69 |
| 1 | 243703982 | rs10927035 | 3.017 | -0.13 | 1.693 | 1.355 | 0.034 | -0.081 | -0.081 | 2.653 | 0.58 | 0.09 | 0.53 | 0.5 | 0.63 | 0.45 |
| 1 | 243710407 | rs1121276 | -2.45 | -0.467 | 1.628 | 1.345 | -0.006 | 0.009 | 0.009 | 0 | 0.27 | 0.15 | 0.42 | 0.19 | 0.34 | 0.37 |
| 1 | 243723214 | rs12144559 | 2.826 | -0.863 | 2.157 | 1.174 | 0.065 | -0.098 | -0.098 | 0 | 0.59 | 0.1 | 0.56 | 0.5 | 0.67 | 0.45 |
| 1 | 243731693 | rs10927040 | -2.129 | -0.876 | 2.553 | 1.189 | 0.002 | 0.019 | 0.019 | 0 | 0.18 | 0.32 | 0.25 | 0.24 | 0.2 | 0.24 |
| 1 | 243768906 | rs3856231 | 2.897 | -0.956 | 2.475 | 1.192 | 0.068 | -0.079 | -0.079 | 0 | 0.6 | 0.12 | 0.56 | 0.5 | 0.67 | 0.45 |
| 1 | 243797799 | rs320332 | 1.51 | -0.934 | 2.454 | 1.146 | -0.017 | 0.008 | 0.008 | 0 | 0.77 | 0.74 | 0.81 | 0.73 | 0.83 | 0.68 |
| 1 | 243801190 | rs10927046 | -1.11 | -1.343 | 2.456 | 1.242 | -0.005 | -0.002 | -0.002 | 0.175 | 0.08 | 0.06 | 0.14 | 0.24 | 0.15 | 0.2 |
| 1 | 243804158 | rs10754807 | -2.503 | -1.384 | 2.634 | 1.223 | -0.003 | 0.028 | 0.028 | 0.526 | 0.19 | 0.49 | 0.27 | 0.23 | 0.2 | 0.23 |
| 1 | 243846167 | rs10803152 | -1.958 | -1.365 | 2.439 | 1.17 | 0.035 | 0.101 | 0.101 | 0.123 | 0.12 | 0.31 | 0.22 | 0.31 | 0.16 | 0.22 |
| 1 | 243846271 | rs4658588 | 2.924 | -1.434 | 2.311 | 1.158 | 0.07 | -0.071 | -0.071 | 0.443 | 0.59 | 0.13 | 0.56 | 0.42 | 0.66 | 0.47 |
| 1 | 243853830 | rs6686591 | 1.334 | -1.331 | 2.426 | 1.202 | -0.061 | 0.039 | 0.039 | 0 | 0.78 | 0.57 | 0.81 | 0.73 | 0.86 | 0.69 |
| 1 | 243892532 | rs9782883 | -2.228 | -1.323 | 2.419 | 1.186 | 0.002 | 0.077 | 0.077 | 0.167 | 0.19 | 0.47 | 0.26 | 0.34 | 0.2 | 0.24 |
| 1 | 243894782 | rs10803155 | 1.098 | -1.302 | 2.395 | 1.2 | -0.013 | 0.005 | 0.005 | 0 | 0.79 | 0.77 | 0.83 | 0.75 | 0.86 | 0.7 |
| 1 | 243900640 | rs4553169 | -2.347 | -1.286 | 2.268 | 1.188 | -0.007 | 0.033 | 0.033 | 0.503 | 0.19 | 0.63 | 0.27 | 0.32 | 0.2 | 0.24 |
| 1 | 243919773 | rs971285 | 2.91 | -1.273 | 1.919 | 1.179 | 0.056 | -0.105 | -0.105 | 0.053 | 0.59 | 0.08 | 0.55 | 0.43 | 0.66 | 0.43 |
| 1 | 243940076 | rs12048930 | -1.875 | -1.215 | 2.004 | 1.186 | 0.005 | 0.029 | 0.029 | 0.011 | 0.17 | 0.3 | 0.24 | 0.32 | 0.2 | 0.24 |
| 1 | 243954407 | rs12047209 | -0.593 | -1.258 | 1.61 | 1.203 | -0.008 | 0.003 | 0.003 | 0 | 0.08 | 0.05 | 0.13 | 0.31 | 0.15 | 0.21 |
| 1 | 243957150 | rs10733129 | 0.21 | -1.261 | 1.456 | 1.172 | 0.007 | -0.051 | -0.051 | 0 | 0.86 | 1 | 0.87 | 0.86 | 0.88 | 0.79 |
| 1 | 243976837 | rs6703013 | 0.779 | -1.173 | 1.644 | 0.978 | -0.06 | 0.029 | 0.029 | 0 | 0.78 | 0.58 | 0.81 | 0.76 | 0.86 | 0.7 |
| 1 | 243995041 | rs10157763 | 1.371 | -0.942 | 1.147 | 0.353 | 0.043 | -0.121 | -0.121 | 0 | 0.59 | 0.06 | 0.55 | 0.42 | 0.66 | 0.46 |
| 1 | 244015993 | rs4430311 | 0.796 | -0.86 | 0.655 | 0.428 | -0.001 | 0.002 | 0.002 | 0.095 | 0.65 | 0.38 | 0.64 | 0.44 | 0.65 | 0.49 |
| 1 | 244023667 | rs3008657 | 1.052 | -0.653 | -0.12 | 0.492 | 0.071 | -0.069 | -0.069 | 0 | 0.57 | 0.13 | 0.54 | 0.42 | 0.63 | 0.46 |
| 1 | 244034805 | rs12083813 | -1.031 | -0.758 | 0.619 | -0.16 | -0.006 | -0.011 | -0.011 | 0 | 0.44 | 0.52 | 0.39 | 0.71 | 0.4 | 0.57 |
| 1 | 244048656 | rs4658597 | -1.167 | -0.652 | 0.599 | -0.221 | 0.021 | 0.072 | 0.072 | 0.287 | 0.48 | 0.71 | 0.42 | 0.72 | 0.46 | 0.59 |
| 1 | 244055001 | rs12090270 | -0.838 | -0.737 | 0.765 | -0.259 | -0.008 | -0.014 | -0.014 | 0 | 0.43 | 0.47 | 0.37 | 0.72 | 0.39 | 0.6 |
| 1 | 244060651 | rs10803168 | -0.642 | -0.777 | 0.408 | -0.278 | 0.015 | 0.031 | 0.031 | 0 | 0.41 | 0.26 | 0.34 | 0.72 | 0.39 | 0.6 |
| 1 | 244061500 | rs4658405 | -1.392 | -0.823 | 0.308 | -0.3 | 0.04 | 0.116 | 0.116 | 0.902 | 0.5 | 0.76 | 0.42 | 0.73 | 0.45 | 0.61 |
| 1 | 244073630 | rs10927090 | -0.809 | -0.578 | -0.193 | -0.172 | 0.011 | 0.024 | 0.024 | 0 | 0.42 | 0.3 | 0.34 | 0.71 | 0.39 | 0.58 |
| 1 | 244096047 | rs559080 | 0.33 | -0.881 | -0.546 | -0.368 | -0.021 | -0.013 | -0.013 | 0.135 | 0.58 | 0.51 | 0.66 | 0.32 | 0.68 | 0.44 |
| 1 | 244096256 | rs471567 | 0.994 | -0.994 | -0.828 | -0.605 | -0.043 | 0.026 | 0.026 | 0 | 0.42 | 0.06 | 0.58 | 0.29 | 0.59 | 0.39 |
| 1 | 244100097 | rs482261 | 0.365 | -1.014 | -0.718 | -0.634 | 0.022 | 0.023 | 0.023 | 0 | 0.52 | 0.43 | 0.6 | 0.29 | 0.65 | 0.43 |
| 1 | 244100402 | rs524489 | 1.008 | -0.868 | -0.934 | -0.559 | -0.057 | 0.02 | 0.02 | 0 | 0.42 | 0.05 | 0.55 | 0.29 | 0.57 | 0.38 |
| 1 | 244102151 | rs543626 | 0.75 | -0.627 | -0.496 | -0.16 | 0.007 | 0.007 | 0.007 | 0 | 0.57 | 0.37 | 0.61 | 0.3 | 0.66 | 0.46 |
| 1 | 244106113 | rs990794 | -0.715 | -0.562 | -0.686 | 0.134 | -0.009 | -0.003 | -0.003 | 0 | 0.35 | 0.36 | 0.35 | 0.7 | 0.26 | 0.5 |
| 1 | 244112606 | rs472276 | -0.632 | 0.16 | -0.929 | 0.317 | -0.001 | -0.004 | -0.004 | 0 | 0.44 | 0.46 | 0.49 | 0.6 | 0.39 | 0.62 |
| 1 | 244121341 | rs500202 | -0.504 | 0.331 | -1.704 | 0.119 | 0.003 | 0.101 | 0.101 | 3.362 | 0.48 | 0.8 | 0.61 | 0.28 | 0.51 | 0.6 |
| 1 | 244123391 | rs484459 | 0.067 | 0.577 | -2.207 | -0.074 | -0.109 | -0.022 | -0.022 | 0 | 0.23 | 0.01 | 0.32 | 0.18 | 0.29 | 0.37 |

| | | | | | | | | | | | | | | | | |
|---|---|---|---|---|---|---|---|---|---|---|---|---|---|---|---|---|
| 1 | 244138990 | rs10927094 | 1.564 | 0.26 | -2.735 | 0.486 | -0.11 | -0.113 | -0.113 | 0 | 0.31 | 0.02 | 0.25 | 0.39 | 0.37 | 0.23 |
| 1 | 244139478 | rs960091 | 1.164 | 0.119 | -2.499 | -0.007 | -0.014 | 0.016 | 0.016 | 0.568 | 0.13 | 0.07 | 0.21 | 0.05 | 0.17 | 0.19 |
| 1 | 244140051 | rs1223870 | -1.008 | -0.009 | -1.402 | -0.038 | -0.001 | 0.005 | 0.005 | 2.648 | 0.45 | 0.54 | 0.54 | 0.23 | 0.51 | 0.59 |
| 1 | 244140225 | rs10803171 | 0.867 | -0.279 | -1.071 | -0.443 | 0.003 | -0.005 | -0.005 | 0 | 0.13 | 0.15 | 0.21 | 0.07 | 0.17 | 0.19 |
| 1 | 244147535 | rs505706 | -0.647 | -0.565 | -0.528 | -0.78 | 0.008 | -0.105 | -0.105 | 0.405 | 0.28 | 0.38 | 0.33 | 0.08 | 0.31 | 0.28 |
| 1 | 244150083 | rs472967 | 0.94 | -0.614 | -0.982 | -0.962 | -0.016 | 0.238 | 0.238 | 0 | 0.45 | 0.15 | 0.44 | 0.08 | 0.54 | 0.3 |
| 1 | 244155726 | rs4658609 | 0.809 | -1.062 | -0.918 | -0.866 | -0.089 | 0.038 | 0.038 | 0 | 0.25 | 0.1 | 0.32 | 0.15 | 0.32 | 0.34 |
| 1 | 244155946 | rs7511723 | -0.228 | -1.037 | -0.378 | -0.682 | -0.022 | 0.031 | 0.031 | 0 | 0.19 | 0.34 | 0.28 | 0.72 | 0.14 | 0.29 |
| 1 | 244160904 | rs536441 | -1.268 | -1.051 | -0.402 | -0.617 | -0.001 | -0.045 | -0.045 | 0.547 | 0.62 | 0.49 | 0.74 | 0.74 | 0.64 | 0.73 |
| 1 | 244161728 | rs564887 | 1.209 | -1.08 | -0.507 | -0.383 | -0.038 | -0.011 | -0.011 | 0 | 0.17 | 0.03 | 0.17 | 0.12 | 0.2 | 0.19 |
| 1 | 244161982 | rs1711347 | -0.103 | -0.926 | -0.359 | -0.501 | -0.01 | 0.008 | 0.008 | 0.35 | 0.38 | 0.28 | 0.48 | 0.26 | 0.47 | 0.5 |
| 1 | 244164997 | rs12119867 | -0.423 | -0.904 | -0.388 | -0.595 | 0.022 | -0.042 | -0.042 | 1.357 | 0.24 | 0.21 | 0.27 | 0.45 | 0.17 | 0.22 |
| 1 | 244165607 | rs692981 | 0.189 | -0.867 | -0.395 | -0.66 | -0.064 | 0.016 | 0.016 | 0.221 | 0.35 | 0.16 | 0.44 | 0.29 | 0.45 | 0.5 |
| 1 | 244167131 | rs1009375 | 2.392 | -0.894 | -0.49 | -0.699 | -0.04 | 0.157 | 0.157 | 1.508 | 0.29 | 0.04 | 0.18 | 0.05 | 0.3 | 0.09 |
| 1 | 244167596 | rs6673663 | -1.202 | -0.893 | -0.183 | -0.545 | -0.032 | 0.006 | 0.006 | 0 | 0.33 | 0.56 | 0.35 | 0.66 | 0.25 | 0.4 |
| 1 | 244168955 | rs12118536 | 0.817 | -0.805 | -0.703 | -0.545 | -0.002 | 0.02 | 0.02 | 0.211 | 0.11 | 0.08 | 0.06 | 0 | 0.11 | 0.05 |
| 1 | 244169914 | rs510280 | 1.156 | -0.789 | -0.725 | -0.555 | -0.013 | -0.002 | -0.002 | 0.509 | 0.17 | 0.07 | 0.17 | 0.15 | 0.21 | 0.19 |
| 1 | 244171167 | rs12066849 | 0.635 | -1.008 | -0.449 | -0.302 | -0.003 | 0.023 | 0.023 | 0.246 | 0.11 | 0.08 | 0.06 | 0 | 0.11 | 0.05 |
| 1 | 244171992 | rs6429443 | 0.12 | -0.773 | -0.289 | -0.133 | 0.037 | 0.106 | 0.106 | 0.404 | 0.09 | 0.35 | 0.08 | 0.21 | 0.08 | 0.18 |
| 1 | 244173426 | rs12122824 | -0.735 | -0.715 | -0.73 | -0.974 | -0.004 | 0 | 0 | 0 | 0.34 | 0.65 | 0.37 | 0.66 | 0.25 | 0.4 |
| 1 | 244176424 | rs476141 | 0.052 | -0.292 | -1.423 | -1.349 | -0.053 | 0.058 | 0.058 | 0.011 | 0.35 | 0.17 | 0.44 | 0.21 | 0.46 | 0.52 |
| 1 | 244182973 | rs12041799 | -0.444 | -0.541 | -0.983 | -1.152 | 0 | -0.001 | -0.001 | 0 | 0.22 | 0.24 | 0.31 | 0.74 | 0.18 | 0.32 |
| 1 | 244184508 | rs12131509 | -0.444 | -0.541 | -0.974 | -1.147 | 0 | 0 | 0 | 0.024 | 0.22 | 0.24 | 0.31 | 0.74 | 0.18 | 0.32 |
| 1 | 244186620 | rs12131048 | -0.191 | -0.599 | -0.975 | -1.418 | -0.006 | -0.005 | -0.005 | 0 | 0.29 | 0.36 | 0.34 | 0.75 | 0.22 | 0.26 |
| 1 | 244187600 | rs1000543 | -0.917 | -0.317 | -0.994 | -1.36 | -0.116 | 0.107 | 0.107 | 0 | 0.62 | 0.93 | 0.59 | 0.83 | 0.51 | 0.51 |
| 1 | 244193100 | rs2454222 | -0.71 | -0.393 | -0.955 | -1.399 | -0.09 | 0.066 | 0.066 | 0 | 0.6 | 0.91 | 0.56 | 0.77 | 0.5 | 0.5 |
| 1 | 244194842 | rs2454221 | -0.858 | -0.216 | -0.925 | -1.316 | -0.113 | 0.046 | 0.046 | 0 | 0.62 | 0.93 | 0.57 | 0.77 | 0.5 | 0.5 |
| 1 | 244197500 | rs2039789 | -0.5 | 0.301 | -1.25 | -0.578 | 0.012 | 0.06 | 0.06 | 0.017 | 0.42 | 0.26 | 0.46 | 0.76 | 0.38 | 0.36 |
| 1 | 244209836 | rs35083998 | -0.036 | 0.72 | -1.424 | -0.207 | -0.019 | 0.038 | 0.038 | 0.011 | 0.07 | 0 | 0.04 | 0 | 0.06 | 0.04 |
| 1 | 244215864 | rs1389622 | -0.549 | 1.246 | -1.725 | -0.533 | -0.033 | 0.037 | 0.037 | 0 | 0.17 | 0.06 | 0.27 | 0.07 | 0.23 | 0.27 |
| 1 | 244221036 | rs11590416 | 0.645 | 0.425 | -2.418 | -1.207 | 0.003 | 0.136 | 0.136 | 0.062 | 0.21 | 0.01 | 0.12 | 0 | 0.19 | 0.1 |
| 1 | 244225402 | rs1565824 | -0.099 | 0.51 | -2.255 | -0.618 | 0.002 | -0.009 | -0.009 | 0 | 0.54 | 0.31 | 0.51 | 0.55 | 0.51 | 0.28 |
| 1 | 244226640 | rs3123390 | -0.322 | 1.081 | -1.847 | -0.036 | -0.078 | -0.086 | -0.086 | 0 | 0.86 | 0.98 | 0.87 | 0.7 | 0.8 | 0.77 |
| 1 | 244228699 | rs2353097 | 0.247 | 0.768 | -1.23 | 0.138 | -0.002 | -0.017 | -0.017 | 0.001 | 0.31 | 0.23 | 0.29 | 0.44 | 0.29 | 0.17 |
| 1 | 244229199 | rs10927118 | -0.312 | 0.693 | -1.432 | 0.082 | -0.007 | -0.001 | -0.001 | 0 | 0.39 | 0.56 | 0.33 | 0.44 | 0.31 | 0.21 |
| 1 | 244232976 | rs2486538 | 0.343 | 0.909 | -1.389 | 0.021 | 0.013 | 0.058 | 0.058 | 0 | 0.49 | 0.29 | 0.53 | 0.26 | 0.5 | 0.58 |
| 1 | 244238657 | rs2486541 | 0.038 | 0.595 | -1.357 | 0.116 | -0.014 | -0.006 | -0.006 | 0 | 0.67 | 0.82 | 0.71 | 0.62 | 0.66 | 0.8 |
| 1 | 244244542 | rs2454228 | 0.767 | 0.834 | -0.785 | 0.296 | -0.002 | -0.001 | -0.001 | 0 | 0.63 | 0.39 | 0.5 | 0.65 | 0.57 | 0.52 |
| 1 | 244249487 | rs10927123 | -0.597 | 0.49 | -0.791 | 0.503 | 0.003 | -0.011 | -0.011 | 0 | 0.26 | 0.21 | 0.33 | 0.26 | 0.24 | 0.35 |
| 1 | 244250317 | rs2454238 | 1.25 | 0.363 | -0.787 | 0.401 | 0.009 | 0.036 | 0.036 | 0.636 | 0.5 | 0.19 | 0.48 | 0.6 | 0.54 | 0.45 |
| 1 | 244256339 | rs4658409 | -0.587 | -0.008 | -0.49 | 0.493 | -0.013 | -0.01 | -0.01 | 0 | 0.24 | 0.19 | 0.34 | 0.24 | 0.28 | 0.35 |
| 1 | 244256594 | rs2453200 | 0.629 | 0.358 | -0.722 | 0.645 | 0.002 | 0.001 | 0.001 | 0.174 | 0.63 | 0.32 | 0.59 | 0.74 | 0.7 | 0.58 |
| 1 | 244260475 | rs4658613 | 0.725 | 0.437 | -0.61 | 1.226 | -0.003 | 0.01 | 0.01 | 0.199 | 0.79 | 0.78 | 0.88 | 0.71 | 0.8 | 0.79 |
| 1 | 244268142 | rs6663169 | -1.033 | 0.217 | 0.28 | 1.564 | -0.002 | 0.145 | 0.145 | 0.154 | 0.19 | 0.48 | 0.21 | 0.36 | 0.15 | 0.23 |
| 1 | 244268866 | rs12727340 | -1.242 | 0.071 | 0.94 | 1.744 | 0.032 | 0.006 | 0.006 | 1.526 | 0.12 | 0.33 | 0.18 | 0.11 | 0.12 | 0.23 |
| 1 | 244271051 | rs2500499 | 0.414 | -0.433 | -0.153 | 1.379 | -0.02 | 0.021 | 0.021 | 0 | 0.21 | 0.02 | 0.23 | 0.13 | 0.18 | 0.12 |
| 1 | 244272338 | rs2883247 | -0.664 | -0.364 | 0.126 | 1.38 | 0.015 | -0.003 | -0.003 | 0 | 0.12 | 0.27 | 0.17 | 0.1 | 0.12 | 0.23 |
| 1 | 244275252 | rs12042441 | -0.139 | -0.376 | 0.131 | 1.381 | 0.013 | -0.019 | -0.019 | 0 | 0.1 | 0.19 | 0.09 | 0.03 | 0.11 | 0.09 |
| 1 | 244280081 | rs10803175 | -1.409 | -0.337 | 0.474 | 1.398 | -0.008 | 0.1 | 0.1 | 0.598 | 0.2 | 0.35 | 0.33 | 0.43 | 0.16 | 0.36 |
| 1 | 244287517 | rs10927136 | 0.083 | -0.848 | 0.7 | 1.19 | 0.024 | 0.031 | 0.031 | 0 | 0.67 | 0.72 | 0.71 | 0.82 | 0.77 | 0.76 |
| 1 | 244288473 | rs10927137 | 1.544 | -0.957 | 1.253 | 1.164 | 0.029 | 0.012 | 0.012 | 0 | 0.46 | 0.23 | 0.39 | 0.38 | 0.62 | 0.41 |
| 1 | 244288525 | rs6695842 | -0.051 | -0.935 | 0.927 | 1.161 | 0.045 | 0.067 | 0.067 | 0 | 0.68 | 0.85 | 0.72 | 0.82 | 0.77 | 0.76 |
| 1 | 244298163 | rs12402376 | 0.272 | -0.946 | 0.981 | 1.223 | 0.007 | 0.004 | 0.004 | 0 | 0.08 | 0.05 | 0.14 | 0.33 | 0.05 | 0.12 |
| 1 | 244301114 | rs6696761 | 0.278 | -0.95 | 1.22 | 1.187 | 0.01 | -0.01 | -0.01 | 0 | 0.09 | 0.13 | 0.09 | 0.04 | 0.11 | 0.08 |
| 1 | 244302030 | rs6429449 | 0.699 | -1.052 | 0.973 | 1.206 | 0.032 | -0.014 | -0.014 | 0 | 0.59 | 0.65 | 0.57 | 0.49 | 0.73 | 0.64 |
| 1 | 244303822 | rs4333836 | -0.001 | -0.985 | 0.991 | 1.11 | -0.023 | -0.018 | -0.018 | 0 | 0.69 | 0.61 | 0.63 | 0.75 | 0.79 | 0.62 |
| 1 | 244306471 | rs7541167 | -0.191 | -0.948 | 0.218 | 1.075 | 0.031 | 0.046 | 0.046 | 0 | 0.72 | 0.87 | 0.73 | 0.82 | 0.79 | 0.77 |
| 1 | 244307139 | rs4658411 | -0.18 | -0.947 | 0.208 | 1.075 | 0.029 | 0.04 | 0.04 | 0 | 0.72 | 0.85 | 0.73 | 0.82 | 0.79 | 0.77 |
| 1 | 244309299 | rs6695226 | 0.292 | -0.86 | 0.218 | 1.285 | 0.036 | -0.048 | -0.048 | 0 | 0.64 | 0.78 | 0.6 | 0.46 | 0.75 | 0.64 |
| 1 | 244309780 | rs10927144 | 1.298 | 0.172 | -0.44 | 0.792 | -0.113 | 0.086 | 0.086 | 0.187 | 0.41 | 0.04 | 0.33 | 0.21 | 0.48 | 0.22 |
| 1 | 244314476 | rs913992 | -0.111 | 0.429 | -0.186 | 0.659 | -0.005 | -0.012 | -0.012 | 0 | 0.73 | 0.75 | 0.67 | 0.62 | 0.69 | 0.73 |
| 1 | 244314860 | rs12131999 | -0.994 | 0.291 | -0.467 | 1.01 | 0 | -0.071 | -0.071 | 0 | 0.15 | 0.07 | 0.27 | 0.41 | 0.17 | 0.31 |
| 1 | 244314952 | rs2183478 | -0.377 | 0.424 | -0.432 | 1.041 | -0.018 | -0.017 | -0.017 | 0 | 0.06 | 0.04 | 0.12 | 0.15 | 0.16 | 0.18 |
| 1 | 244315180 | rs913994 | 0.61 | 0.406 | -0.034 | 0.721 | 0.004 | 0.012 | 0.012 | 0.042 | 0.5 | 0.4 | 0.53 | 0.45 | 0.53 | 0.44 |
| 1 | 244315274 | rs877884 | -0.624 | 0.175 | -0.692 | -0.095 | 0.013 | 0.028 | 0.028 | 0 | 0.72 | 0.82 | 0.76 | 0.82 | 0.79 | 0.78 |
| 1 | 244315918 | rs12080155 | 0.255 | 0.702 | -0.996 | -0.206 | 0.002 | -0.031 | -0.031 | 0 | 0.06 | 0.14 | 0.06 | 0 | 0.04 | 0.06 |
| 1 | 244316323 | rs12076800 | 1.036 | 0.967 | -1.642 | -0.017 | -0.009 | 0.109 | 0.109 | 0.197 | 0.36 | 0.17 | 0.25 | 0.02 | 0.39 | 0.12 |
| 1 | 244316996 | rs6691729 | -1.029 | 1.632 | -1.222 | 1.224 | 0.004 | 0.013 | 0.013 | 1.259 | 0.2 | 0.25 | 0.27 | 0.53 | 0.33 | 0.44 |
| 1 | 244319340 | rs10803181 | -0.329 | 1.282 | -1.844 | 0.493 | 0.024 | 0.046 | 0.046 | 0 | 0.5 | 0.72 | 0.54 | 0.63 | 0.42 | 0.55 |

| | | | | | | | | | | | | | | | | |
|---|---|---|---|---|---|---|---|---|---|---|---|---|---|---|---|---|
| 1 | 244319585 | rs12035147 | 0.03 | 1.332 | -2.062 | -0.266 | -0.06 | 0.11 | 0.11 | 1.197 | 0.64 | 0.86 | 0.72 | 0.87 | 0.46 | 0.84 |
| 1 | 244322387 | rs4658413 | -0.54 | 0.556 | -1.087 | -0.092 | -0.02 | -0.101 | -0.101 | 0 | 0.21 | 0.48 | 0.13 | 0.07 | 0.17 | 0.23 |
| 1 | 244328091 | rs6429451 | -0.292 | 0.218 | -1.861 | 0.871 | 0.012 | -0.078 | -0.078 | 4.07 | 0.84 | 0.96 | 0.76 | 0.61 | 0.87 | 0.79 |
| 1 | 244334507 | rs12087666 | -0.402 | 0.602 | -1.714 | 0.955 | -0.002 | 0.008 | 0.008 | 0.447 | 0.42 | 0.71 | 0.48 | 0.32 | 0.44 | 0.41 |
| 1 | 244337988 | rs11587569 | -0.67 | 0.464 | -1.665 | 1.12 | -0.033 | -0.039 | -0.039 | 1.371 | 0.16 | 0.04 | 0.14 | 0.16 | 0.16 | 0.19 |
| 1 | 244340598 | rs11588708 | -0.767 | 0.441 | -1.474 | 1.238 | -0.007 | -0.025 | -0.025 | 0 | 0.17 | 0.1 | 0.15 | 0.19 | 0.16 | 0.2 |
| 1 | 244343876 | rs6696289 | -1.827 | 0.024 | -0.814 | 1.231 | -0.013 | 0.012 | 0.012 | 1.673 | 0.34 | 0.59 | 0.25 | 0.62 | 0.24 | 0.56 |
| 1 | 244345666 | rs6685563 | -0.165 | 0.023 | -0.614 | 1.201 | -0.038 | -0.026 | -0.026 | 0.043 | 0.1 | 0.38 | 0.06 | 0.03 | 0.02 | 0.06 |
| 1 | 244351608 | rs6681086 | -1.535 | 0.272 | -1.015 | 1.194 | -0.009 | 0.01 | 0.01 | 1.307 | 0.36 | 0.57 | 0.26 | 0.6 | 0.28 | 0.55 |
| 1 | 244353555 | rs11577259 | -0.842 | 0.962 | -1.494 | 0.738 | -0.006 | -0.01 | -0.01 | 0.268 | 0.17 | 0.07 | 0.11 | 0.15 | 0.18 | 0.2 |
| 1 | 244357987 | rs10803182 | -0.492 | 1.069 | -1.902 | 0.279 | -0.057 | 0.057 | 0.057 | 0 | 0.26 | 0.03 | 0.2 | 0.1 | 0.32 | 0.21 |
| 1 | 244358043 | rs9659599 | -0.781 | 1.499 | -1.34 | 0.177 | -0.008 | 0.014 | 0.014 | 0 | 0.28 | 0.19 | 0.23 | 0.1 | 0.36 | 0.21 |
| 1 | 244358190 | rs1556922 | -0.848 | 2.153 | -2.01 | 0.206 | -0.118 | 0.054 | 0.054 | 0 | 0.33 | 0.03 | 0.31 | 0.16 | 0.55 | 0.26 |
| 1 | 244358308 | rs7535810 | -0.412 | 0.977 | -1.345 | 0.791 | -0.028 | 0.055 | 0.055 | 0.056 | 0.31 | 0.44 | 0.37 | 0.57 | 0.24 | 0.51 |
| 1 | 244359621 | rs1411613 | -0.369 | 1.219 | -1.814 | 0.82 | -0.041 | -0.06 | -0.06 | 0 | 0.16 | 0.04 | 0.12 | 0.24 | 0.18 | 0.2 |
| 1 | 244366307 | rs7544355 | 0.604 | -0.271 | -2.248 | 0.618 | 0.061 | -0.082 | -0.082 | 0.14 | 0.29 | 0.14 | 0.3 | 0.48 | 0.2 | 0.39 |
| 1 | 244369276 | rs12032278 | -0.295 | -0.19 | -2.203 | 0.392 | 0.002 | -0.003 | -0.003 | 0.038 | 0.06 | 0.02 | 0.03 | 0.13 | 0.03 | 0.05 |
| 1 | 244370950 | rs6659442 | -0.215 | 0.454 | -2.343 | -0.079 | -0.01 | 0.012 | 0.012 | 0 | 0.18 | 0.1 | 0.15 | 0.14 | 0.2 | 0.18 |
| 1 | 244372986 | rs6684323 | -0.257 | 0.406 | -1.923 | -0.105 | 0.002 | -0.007 | -0.007 | 0 | 0.18 | 0.18 | 0.17 | 0.13 | 0.2 | 0.18 |
| 1 | 244373559 | rs12139580 | 0.656 | 0.381 | -2.193 | -0.167 | -0.017 | -0.061 | -0.061 | 0 | 0.14 | 0.01 | 0.1 | 0.21 | 0.13 | 0.17 |
| 1 | 244374051 | rs945652 | -0.701 | 0.616 | -2.074 | -0.934 | -0.056 | 0.071 | 0.071 | 0 | 0.77 | 0.9 | 0.77 | 0.95 | 0.65 | 0.84 |
| 1 | 244377774 | rs10737892 | -0.405 | 0.451 | -1.781 | -0.959 | -0.036 | 0.042 | 0.042 | 0 | 0.74 | 0.82 | 0.74 | 0.95 | 0.63 | 0.84 |
| 1 | 244377850 | rs7522710 | -0.405 | 0.497 | -1.777 | -0.896 | -0.041 | 0.05 | 0.05 | 0 | 0.74 | 0.83 | 0.74 | 0.96 | 0.63 | 0.84 |
| 1 | 244377938 | rs10803183 | 0.457 | 0.226 | -1.681 | -0.847 | 0.007 | 0.036 | 0.036 | 0 | 0.56 | 0.63 | 0.58 | 0.81 | 0.44 | 0.63 |
| 1 | 244380551 | rs4634891 | -0.167 | 0.146 | -1.836 | -0.773 | -0.046 | 0.094 | 0.094 | 0.018 | 0.37 | 0.15 | 0.41 | 0.21 | 0.54 | 0.3 |
| 1 | 244380895 | rs12726392 | -0.145 | 0.61 | -1.865 | -0.807 | -0.085 | 0.067 | 0.067 | 0.009 | 0.35 | 0.15 | 0.4 | 0.23 | 0.49 | 0.3 |
| 1 | 244382369 | rs12135811 | 0.78 | -0.135 | -1.482 | -0.784 | 0.077 | 0.077 | 0.077 | 0.059 | 0.46 | 0.21 | 0.48 | 0.74 | 0.34 | 0.55 |
| 1 | 244382698 | rs10927166 | 1.401 | -0.109 | -1.5 | -0.753 | 0.064 | -0.033 | -0.033 | 0 | 0.34 | 0.15 | 0.33 | 0.56 | 0.18 | 0.4 |
| 1 | 244383838 | rs4658616 | 2.033 | 0.194 | -1.637 | -0.681 | -0.027 | -0.036 | -0.036 | 0 | 0.11 | 0.01 | 0.14 | 0.18 | 0.15 | 0.15 |
| 1 | 244386176 | rs6686011 | -0.627 | 0.245 | -0.798 | -0.999 | 0.04 | -0.062 | -0.062 | 0 | 0.75 | 0.63 | 0.7 | 0.94 | 0.6 | 0.79 |
| 1 | 244386214 | rs6662316 | 1.658 | -0.633 | -1.084 | -0.843 | 0.019 | -0.004 | -0.004 | 0 | 0.24 | 0.18 | 0.1 | 0.01 | 0.1 | 0.15 |
| 1 | 244389589 | rs12048859 | -0.307 | 0.682 | -0.911 | 0.167 | -0.006 | 0.013 | 0.013 | 0 | 0.15 | 0.19 | 0.1 | 0.19 | 0.12 | 0.13 |
| 1 | 244390564 | rs10429924 | -0.317 | 0.948 | -0.792 | 0.831 | 0.004 | -0.031 | -0.031 | 0 | 0.11 | 0.04 | 0.2 | 0.37 | 0.09 | 0.24 |
| 1 | 244391618 | rs6692542 | 0.392 | 1.718 | -0.464 | -0.155 | 0.001 | -0.007 | -0.007 | 0 | 0.78 | 0.81 | 0.74 | 0.93 | 0.63 | 0.78 |
| 1 | 244391742 | rs6664367 | 1.015 | 1.109 | -0.085 | 0.523 | 0.01 | 0.039 | 0.039 | 0 | 0.93 | 0.88 | 0.95 | 0.85 | 0.91 | 0.84 |
| 1 | 244392845 | rs879065 | -0.345 | 1.272 | -0.335 | 0.078 | -0.03 | 0.055 | 0.055 | 4.482 | 0.37 | 0.25 | 0.34 | 0.16 | 0.41 | 0.33 |
| 1 | 244401066 | rs7548336 | -0.069 | 0.735 | 0.636 | 1.048 | 0.023 | 0.046 | 0.046 | 0 | 0.51 | 0.33 | 0.65 | 0.68 | 0.61 | 0.61 |
| 1 | 244404834 | rs7555626 | -0.222 | 0.901 | 0.754 | 0.986 | 0.002 | 0.021 | 0.021 | 0 | 0.55 | 0.66 | 0.66 | 0.71 | 0.63 | 0.71 |
| 1 | 244406046 | rs879725 | -0.302 | 0.941 | 0.754 | 0.993 | 0.002 | 0.009 | 0.009 | 0 | 0.54 | 0.58 | 0.65 | 0.71 | 0.62 | 0.71 |
| 1 | 244437407 | rs622169 | -0.796 | 1.61 | 0.688 | 0.988 | -0.13 | 0.18 | 0.18 | 0.12 | 0.33 | 0.08 | 0.32 | 0.03 | 0.46 | 0.43 |
| 1 | 244448353 | rs515513 | -0.905 | 1.608 | 1.028 | 1.005 | -0.039 | 0.093 | 0.093 | 0 | 0.37 | 0.25 | 0.34 | 0.03 | 0.48 | 0.44 |
| 1 | 244452911 | rs651529 | -1.019 | 1.715 | 1.023 | 1.3 | -0.003 | 0.006 | 0.006 | 0 | 0.42 | 0.59 | 0.53 | 0.2 | 0.53 | 0.55 |
| 1 | 244459783 | rs678004 | -0.982 | 1.761 | 0.803 | 1.089 | 0.025 | -0.015 | -0.015 | 0 | 0.37 | 0.49 | 0.4 | 0.32 | 0.51 | 0.52 |
| 1 | 244464197 | rs7539318 | -1.565 | 1.824 | 1.353 | 1.243 | 0.016 | 0.039 | 0.039 | 0 | 0.23 | 0.66 | 0.26 | 0.33 | 0.23 | 0.34 |
| 1 | 244480389 | rs3120118 | -1.51 | 1.763 | 0.912 | 0.903 | 0.026 | -0.004 | -0.004 | 1.058 | 0.05 | 0.14 | 0.02 | 0.03 | 0.05 | 0.07 |
| 1 | 244490369 | rs602620 | -0.871 | 1.697 | 0.526 | 0.482 | 0.011 | -0.038 | -0.038 | 0 | 0.4 | 0.56 | 0.32 | 0.22 | 0.44 | 0.42 |
| 1 | 244496482 | rs559798 | 0.388 | 2.29 | 0.314 | 1.395 | -0.001 | 0.012 | 0.012 | 0 | 0.91 | 0.9 | 0.84 | 0.75 | 0.85 | 0.89 |
| 1 | 244497186 | rs3102455 | 1.119 | 2.599 | 0.814 | 1.392 | 0.074 | 0.114 | 0.114 | 0 | 0.77 | 0.55 | 0.73 | 0.56 | 0.68 | 0.76 |
| 1 | 244507556 | rs6663981 | -1.11 | 2.141 | 0.42 | 1.367 | 0.002 | 0.029 | 0.029 | 0 | 0.12 | 0.2 | 0.11 | 0.23 | 0.14 | 0.28 |
| 1 | 244507827 | rs4658623 | -0.591 | 2.263 | 0.098 | 1.38 | -0.015 | -0.011 | -0.011 | 0 | 0.07 | 0.06 | 0.16 | 0.2 | 0.17 | 0.09 |
| 1 | 244516334 | rs10927202 | -0.797 | 1.443 | 0.092 | 0.405 | -0.018 | -0.065 | -0.065 | 0 | 0.35 | 0.42 | 0.46 | 0.82 | 0.31 | 0.56 |
| 1 | 244522070 | rs4658417 | 0.718 | 0.909 | -0.085 | -0.589 | -0.024 | 0.112 | 0.112 | 0 | 0.45 | 0.29 | 0.34 | 0.14 | 0.47 | 0.29 |
| 1 | 244524345 | rs7547047 | 0.639 | 0.231 | 0.393 | 0.011 | -0.018 | -0.006 | -0.006 | 0.009 | 0.7 | 0.55 | 0.81 | 0.73 | 0.76 | 0.78 |
| 1 | 244528127 | rs9782996 | -0.205 | 0.33 | 0.373 | 0.078 | 0.007 | -0.038 | -0.038 | 0 | 0.78 | 0.66 | 0.85 | 0.91 | 0.77 | 0.83 |
| 1 | 244531679 | rs10927219 | 0.197 | 0.248 | 0.384 | 0.265 | -0.009 | 0.131 | 0.131 | 0.012 | 0.22 | 0.37 | 0.43 | 0.51 | 0.24 | 0.38 |
| 1 | 244532816 | rs11580053 | 1.295 | -0.248 | -0.588 | -0.294 | -0.108 | 0.195 | 0.195 | 0 | 0.38 | 0.08 | 0.35 | 0.13 | 0.49 | 0.28 |
| 1 | 244532937 | rs3123705 | -0.264 | -0.337 | -0.241 | 0.037 | -0.015 | 0.055 | 0.055 | 0 | 0.74 | 0.67 | 0.56 | 0.48 | 0.75 | 0.59 |
| 1 | 244534781 | rs3101456 | -0.719 | -0.231 | -0.299 | -0.415 | -0.022 | -0.052 | -0.052 | 0 | 0.41 | 0.47 | 0.54 | 0.8 | 0.34 | 0.57 |
| 1 | 244539868 | rs3102458 | -0.538 | 0.229 | -0.685 | -0.075 | -0.003 | 0.002 | 0.002 | 0 | 0.23 | 0.15 | 0.22 | 0.21 | 0.24 | 0.27 |
| 1 | 244544441 | rs3123711 | -0.978 | 0.191 | -0.645 | 0.021 | 0.008 | 0.015 | 0.015 | 3.532 | 0.38 | 0.56 | 0.63 | 0.43 | 0.43 | 0.65 |
| 1 | 244544458 | rs3102463 | -0.688 | 0.003 | -1.195 | -0.523 | -0.019 | 0.047 | 0.047 | 0 | 0.65 | 0.74 | 0.68 | 0.83 | 0.59 | 0.76 |
| 1 | 244544818 | rs3123710 | -0.708 | -0.523 | 0.375 | -0.312 | 0.011 | 0.02 | 0.02 | 0 | 0.53 | 0.6 | 0.52 | 0.27 | 0.58 | 0.67 |
| 1 | 244545766 | rs4658636 | -0.944 | -0.059 | 0.428 | -0.173 | 0.015 | 0.011 | 0.011 | 0.508 | 0.55 | 0.8 | 0.67 | 0.59 | 0.56 | 0.52 |
| 1 | 244562587 | rs12730102 | 1.528 | 0.212 | 1.045 | -0.318 | 0.002 | -0.042 | -0.042 | 0.263 | 0.16 | 0.18 | 0.11 | 0 | 0.17 | 0.06 |
| 1 | 244566343 | rs12404529 | -0.251 | 0.171 | 1.16 | 0.04 | 0.013 | 0.099 | 0.099 | 0 | 0.07 | 0.23 | 0.23 | 0.15 | 0.06 | 0.13 |
| 1 | 244569746 | rs11580674 | -0.266 | 0.095 | 1.195 | 0.069 | 0 | 0 | 0 | 0 | 0.48 | 0.54 | 0.45 | 0.46 | 0.49 | 0.39 |
| 1 | 244570385 | rs12084013 | -0.74 | 0.027 | 1.108 | 0.051 | -0.009 | 0.042 | 0.042 | 0 | 0.12 | 0.28 | 0.24 | 0.15 | 0.09 | 0.15 |
| 1 | 244573030 | rs2306084 | 0.309 | 0.059 | 1.129 | 0.075 | 0 | 0.006 | 0.006 | 0 | 0.06 | 0.05 | 0.21 | 0.14 | 0.06 | 0.13 |
| 1 | 244573756 | rs6429464 | 0.143 | 0.163 | 1.103 | -0.021 | -0.001 | -0.027 | -0.027 | 1.25 | 0.42 | 0.39 | 0.44 | 0.06 | 0.53 | 0.32 |

| | | | | | | | | | | | | | | | | |
|---|---|---|---|---|---|---|---|---|---|---|---|---|---|---|---|---|
| 1 | 244576717 | rs3102460 | -1.236 | 0.257 | 0.916 | 0.114 | -0.005 | 0 | 0 | 0 | 0.68 | 0.67 | 0.66 | 0.69 | 0.76 | 0.72 |
| 1 | 244578462 | rs12087973 | -0.011 | 0.294 | 0.927 | 0.146 | 0.009 | 0.084 | 0.084 | 0 | 0.07 | 0.2 | 0.23 | 0.15 | 0.06 | 0.13 |
| 1 | 244579519 | rs3123696 | -0.054 | 0.401 | 0.416 | 0.266 | -0.001 | 0.013 | 0.013 | 0 | 0.2 | 0.2 | 0.2 | 0.63 | 0.19 | 0.38 |
| 1 | 244580348 | rs3123697 | -2.042 | 0.531 | -0.266 | -0.325 | 0.035 | 0.031 | 0.031 | 1.442 | 0.76 | 0.9 | 0.89 | 0.84 | 0.81 | 0.84 |
| 1 | 244599858 | rs3127464 | 1.546 | 0.29 | -0.079 | -0.095 | 0.01 | 0.043 | 0.043 | 0.551 | 0.28 | 0.13 | 0.18 | 0.16 | 0.28 | 0.16 |
| 1 | 244601152 | rs3127465 | -0.471 | 0.522 | -0.171 | 0.205 | -0.011 | -0.036 | -0.036 | 0 | 0.84 | 0.95 | 0.85 | 0.77 | 0.83 | 0.67 |
| 1 | 244618005 | rs3127468 | 0.149 | 0.274 | 0.72 | -0.049 | 0 | -0.012 | -0.012 | 0 | 0.48 | 0.53 | 0.32 | 0.18 | 0.47 | 0.2 |
| 1 | 244624337 | rs10927237 | 0.542 | 0.675 | 0.701 | -0.354 | 0.02 | -0.108 | -0.108 | 0.688 | 0.19 | 0.37 | 0.15 | 0.01 | 0.21 | 0.06 |
| 1 | 244624887 | rs3124058 | -0.804 | 0.665 | 0.579 | -0.118 | -0.001 | 0 | 0 | 0.239 | 0.64 | 0.69 | 0.62 | 0.74 | 0.64 | 0.46 |
| 1 | 244631821 | rs3003207 | 0.1 | 0.557 | 0.562 | -0.037 | 0.016 | 0.045 | 0.045 | 0 | 0.43 | 0.33 | 0.47 | 0.75 | 0.41 | 0.35 |
| 1 | 244635604 | rs3124064 | -0.882 | 0.588 | 0.414 | -0.008 | -0.028 | 0.047 | 0.047 | 1.436 | 0.64 | 0.84 | 0.63 | 0.76 | 0.62 | 0.41 |
| 1 | 244636995 | rs12068598 | 0.352 | 0.731 | 0.364 | 0.109 | 0.024 | -0.112 | -0.112 | 0 | 0.19 | 0.39 | 0.16 | 0.01 | 0.21 | 0.06 |
| 1 | 244652123 | rs3003301 | 0.489 | 0.69 | 0.451 | 1.127 | 0 | 0.005 | 0.005 | 2.221 | 0.48 | 0.59 | 0.37 | 0.46 | 0.51 | 0.29 |
| 1 | 244652199 | rs4658642 | -1.243 | 0.603 | 0.808 | 1.835 | -0.017 | 0.104 | 0.104 | 0 | 0.13 | 0.23 | 0.25 | 0.29 | 0.08 | 0.11 |
| 1 | 244660547 | rs3006024 | -1.61 | 1.358 | 1.774 | 1.258 | 0.014 | -0.012 | -0.012 | 0.012 | 0.22 | 0.6 | 0.21 | 0.19 | 0.24 | 0.29 |
| 1 | 244661477 | rs3006029 | -1.157 | 1.361 | 2.074 | 1.208 | 0.078 | -0.011 | -0.011 | 0 | 0.08 | 0.36 | 0.06 | 0 | 0.07 | 0.07 |
| 1 | 244689029 | rs3005928 | -2.365 | 1.471 | 2.065 | 1.188 | 0.01 | -0.02 | -0.02 | 0 | 0.22 | 0.59 | 0.13 | 0.18 | 0.21 | 0.27 |
| 1 | 244713825 | rs3003222 | 0.944 | 1.313 | 1.961 | 1.13 | 0.082 | -0.005 | -0.005 | 0 | 0.93 | 0.66 | 0.94 | 1 | 0.93 | 0.93 |
| 1 | 244724092 | rs7544794 | -2.182 | 1.162 | 2.168 | 2.216 | 0.045 | 0.388 | 0.388 | 0.015 | 0.1 | 0.43 | 0.24 | 0.38 | 0.08 | 0.17 |
| 1 | 244735858 | rs10927267 | -2.205 | 1.201 | 2.233 | 2.212 | 0.042 | 0.363 | 0.363 | 0.036 | 0.1 | 0.43 | 0.25 | 0.37 | 0.08 | 0.17 |
| 1 | 244738592 | rs7517558 | -1.839 | 1.23 | 2.256 | 2.227 | 0.05 | 0.37 | 0.37 | 0 | 0.08 | 0.39 | 0.24 | 0.37 | 0.08 | 0.17 |
| 1 | 244747150 | rs6429487 | -1.813 | 1.234 | 2.274 | 2.228 | 0.048 | 0.361 | 0.361 | 0 | 0.08 | 0.38 | 0.24 | 0.37 | 0.08 | 0.17 |
| 1 | 244750094 | rs6429491 | -2.232 | 1.185 | 2.232 | 2.241 | 0.027 | 0.268 | 0.268 | 0 | 0.09 | 0.33 | 0.23 | 0.37 | 0.08 | 0.15 |
| 1 | 244777718 | rs10803228 | 2.14 | 1.652 | 2.39 | 2.063 | 0.011 | 0.025 | 0.025 | 0 | 0.74 | 0.4 | 0.6 | 0.58 | 0.69 | 0.7 |
| 1 | 244782870 | rs6670863 | 2.184 | 1.488 | 2.494 | 2.035 | 0.036 | 0.075 | 0.075 | 0 | 0.75 | 0.48 | 0.61 | 0.57 | 0.7 | 0.7 |
| 1 | 244785995 | rs10465638 | -2.047 | 1.32 | 2.68 | 1.923 | -0.002 | 0.005 | 0.005 | 0 | 0.17 | 0.35 | 0.17 | 0.16 | 0.14 | 0.21 |
| 1 | 244792069 | rs3000691 | 1.41 | 1.294 | 2.497 | 1.956 | 0 | -0.001 | -0.001 | 0 | 0.86 | 0.83 | 0.85 | 0.85 | 0.86 | 0.8 |
| 1 | 244797942 | rs2653151 | 1.091 | 1.249 | 2.42 | 1.906 | 0 | -0.001 | -0.001 | 0 | 0.89 | 0.85 | 0.91 | 0.91 | 0.91 | 0.86 |
| 1 | 244801873 | rs6688788 | 1.932 | 1.235 | 2.654 | 1.962 | 0.086 | 0.121 | 0.121 | 0 | 0.89 | 0.6 | 0.76 | 0.74 | 0.84 | 0.91 |
| 1 | 244802142 | rs6676542 | 2.611 | 1.196 | 2.696 | 1.908 | 0.025 | 0.029 | 0.029 | 0 | 0.76 | 0.46 | 0.66 | 0.65 | 0.74 | 0.77 |
| 1 | 244802705 | rs10754842 | 2.252 | 1.239 | 2.513 | 1.896 | 0.037 | 0.064 | 0.064 | 0 | 0.79 | 0.59 | 0.68 | 0.61 | 0.75 | 0.77 |
| 1 | 244809396 | rs2653142 | 2.304 | 1.17 | 2.446 | 1.771 | -0.027 | -0.044 | -0.044 | 0 | 0.91 | 0.71 | 0.97 | 1 | 0.97 | 0.97 |
| 1 | 244809662 | rs1995391 | 2.974 | 1.232 | 2.644 | 1.884 | -0.006 | 0.028 | 0.028 | 0 | 0.73 | 0.46 | 0.65 | 0.62 | 0.74 | 0.73 |
| 1 | 244809936 | rs987179 | -2.231 | 1.397 | 1.895 | 1.964 | 0.005 | 0.014 | 0.014 | 0 | 0.18 | 0.22 | 0.31 | 0.38 | 0.23 | 0.23 |
| 1 | 244813471 | rs989284 | -2.586 | 1.398 | 1.815 | 2.01 | 0.002 | 0.013 | 0.013 | 0 | 0.2 | 0.26 | 0.3 | 0.38 | 0.23 | 0.22 |
| 1 | 244815439 | rs2653158 | -1.99 | 1.357 | 1.501 | 1.97 | 0.028 | 0.025 | 0.025 | 0 | 0.22 | 0.43 | 0.28 | 0.28 | 0.23 | 0.17 |
| 1 | 244822569 | rs2819018 | -2.256 | 1.881 | 0.554 | 2.521 | -0.077 | -0.103 | -0.103 | 0.18 | 0.14 | 0.03 | 0.28 | 0.41 | 0.25 | 0.16 |
| 1 | 244824192 | rs2819017 | -1.465 | 1.678 | 0.549 | 2.342 | -0.018 | -0.038 | -0.038 | 2.095 | 0.08 | 0.01 | 0.06 | 0.08 | 0.1 | 0.04 |
| 1 | 244829280 | rs12404604 | 1.877 | 1.539 | 0.772 | 2.599 | 0.003 | 0.033 | 0.033 | 0 | 0.86 | 0.84 | 0.76 | 0.61 | 0.8 | 0.9 |
| 1 | 244838822 | rs12122462 | 2.118 | 1.26 | 1.007 | 2.361 | -0.002 | 0.016 | 0.016 | 0 | 0.72 | 0.34 | 0.69 | 0.62 | 0.75 | 0.83 |
| 1 | 244854821 | rs4658649 | 1.255 | 0.986 | -0.243 | 1.809 | -0.083 | -0.013 | -0.013 | 0 | 0.28 | 0.03 | 0.37 | 0.2 | 0.28 | 0.16 |
| 1 | 244858017 | rs11589363 | 0.67 | 0.243 | -0.058 | 1.711 | -0.002 | -0.001 | -0.001 | 0 | 0.26 | 0.24 | 0.22 | 0.3 | 0.26 | 0.35 |
| 1 | 244862210 | rs12057937 | -0.72 | 0.235 | -0.093 | 1.698 | 0.064 | 0.086 | 0.086 | 0.02 | 0.42 | 0.87 | 0.29 | 0.3 | 0.32 | 0.42 |
| 1 | 244868689 | rs12060258 | 0.086 | 0.228 | -0.078 | 1.76 | -0.024 | -0.012 | -0.012 | 0 | 0.3 | 0.39 | 0.24 | 0.3 | 0.26 | 0.36 |
| 1 | 244869479 | rs2242448 | 0.118 | 0.229 | -0.078 | 1.755 | -0.024 | 0.007 | 0.007 | 0 | 0.31 | 0.39 | 0.24 | 0.34 | 0.27 | 0.36 |
| 1 | 244873070 | rs6694009 | 1.039 | 0.402 | -0.244 | 1.385 | 0.014 | 0.039 | 0.039 | 0 | 0.1 | 0.03 | 0.02 | 0 | 0.05 | 0.07 |
| 1 | 244873684 | rs10927298 | 0.78 | 0.46 | -0.131 | 1.562 | -0.011 | 0.072 | 0.072 | 0.142 | 0.48 | 0.28 | 0.49 | 0.31 | 0.49 | 0.47 |
| 1 | 244874837 | rs2365966 | -0.254 | 0.832 | -0.456 | 1.348 | -0.005 | -0.005 | -0.005 | 1.596 | 0.74 | 0.75 | 0.81 | 0.76 | 0.75 | 0.66 |
| 1 | 244877955 | rs6695943 | -0.594 | 0.951 | -0.383 | 1.3 | 0.008 | 0.06 | 0.06 | 0 | 0.22 | 0.36 | 0.18 | 0.34 | 0.23 | 0.16 |
| 1 | 244883815 | rs10803241 | 0.355 | 0.736 | -0.997 | 0.86 | 0.005 | 0.019 | 0.019 | 0 | 0.59 | 0.71 | 0.71 | 0.66 | 0.6 | 0.54 |
| 1 | 244885285 | rs10927306 | -0.06 | 0.316 | -0.557 | 0.271 | 0.003 | 0.011 | 0.011 | 0 | 0.55 | 0.59 | 0.74 | 0.65 | 0.59 | 0.63 |
| 1 | 244886134 | rs6429500 | 0.285 | 0.384 | -0.589 | 0.459 | -0.019 | 0.004 | 0.004 | 0 | 0.44 | 0.34 | 0.58 | 0.4 | 0.5 | 0.45 |
| 1 | 244887026 | rs11577325 | 0.268 | 0.173 | -0.605 | -0.057 | 0.01 | -0.004 | -0.004 | 0 | 0.47 | 0.26 | 0.67 | 0.54 | 0.54 | 0.55 |
| 1 | 244893103 | rs4658653 | 0.27 | 0.617 | -0.464 | -0.143 | 0.022 | 0.007 | 0.007 | 0 | 0.29 | 0.36 | 0.58 | 0.66 | 0.39 | 0.48 |
| 1 | 244894273 | rs1566661 | 0.209 | 0.154 | -0.1 | 0.132 | 0.007 | 0.005 | 0.005 | 0 | 0.41 | 0.47 | 0.63 | 0.52 | 0.52 | 0.52 |
| 1 | 244902468 | rs6699271 | 0.458 | -0.313 | -0.282 | 0.018 | -0.008 | -0.006 | -0.006 | 0 | 0.39 | 0.32 | 0.62 | 0.56 | 0.53 | 0.52 |
| 1 | 244909440 | rs6663759 | -0.698 | -0.636 | -0.652 | -0.149 | -0.049 | -0.029 | -0.029 | 0 | 0.67 | 0.98 | 0.77 | 0.71 | 0.7 | 0.67 |
| 1 | 244915579 | rs2134268 | 0.209 | -0.393 | -0.155 | 0.075 | 0.002 | 0.002 | 0.002 | 0.372 | 0.52 | 0.49 | 0.69 | 0.67 | 0.68 | 0.58 |
| 1 | 244916619 | rs7534801 | -0.423 | -0.233 | -1.274 | 0.181 | 0.072 | 0.049 | 0.049 | 3.004 | 0.61 | 0.9 | 0.76 | 0.71 | 0.76 | 0.66 |
| 1 | 244920820 | rs2134271 | 0.123 | 0.04 | -1.05 | -0.055 | 0.009 | 0.069 | 0.069 | 3.334 | 0.46 | 0.7 | 0.65 | 0.71 | 0.59 | 0.61 |
| 1 | 244921530 | rs2365958 | -0.605 | 0.132 | -1.161 | -0.023 | 0.054 | 0.072 | 0.072 | 3.745 | 0.61 | 0.74 | 0.74 | 0.8 | 0.77 | 0.7 |
| 1 | 244921577 | rs6697884 | -0.369 | 0.66 | -0.843 | 0.541 | 0.002 | 0.015 | 0.015 | 2.994 | 0.45 | 0.56 | 0.64 | 0.71 | 0.58 | 0.61 |
| 1 | 244922115 | rs4658662 | 0.082 | 0.922 | -0.608 | 0.767 | 0.032 | -0.012 | -0.012 | 6.163 | 0.36 | 0.46 | 0.6 | 0.67 | 0.49 | 0.48 |
| 1 | 244924975 | rs2047762 | -1.247 | 0.781 | -0.611 | 1.161 | -0.004 | -0.005 | -0.005 | 0 | 0.07 | 0.11 | 0.04 | 0.05 | 0.08 | 0.18 |
| 1 | 244930726 | rs2088239 | -0.859 | 0.86 | -0.864 | 1.579 | -0.009 | -0.029 | -0.029 | 0 | 0.26 | 0.18 | 0.47 | 0.63 | 0.27 | 0.44 |
| 1 | 244934673 | rs9661248 | -1.452 | 0.27 | -0.409 | 1.686 | 0 | 0.007 | 0.007 | 0 | 0.23 | 0.71 | 0.24 | 0.44 | 0.19 | 0.21 |
| 1 | 244938498 | rs9727042 | -1.273 | -0.243 | -0.079 | 0.914 | -0.027 | -0.022 | -0.022 | 0 | 0.21 | 0.57 | 0.15 | 0.12 | 0.15 | 0.1 |
| 1 | 244952372 | rs9730373 | -0.973 | -0.986 | 0.381 | 1.278 | -0.018 | 0.023 | 0.023 | 0.044 | 0.29 | 0.6 | 0.42 | 0.51 | 0.15 | 0.33 |
| 1 | 244953565 | rs9658889 | -1.417 | -1.24 | 0.729 | 1.074 | -0.022 | -0.024 | -0.024 | 0.26 | 0.13 | 0.31 | 0.08 | 0.05 | 0.1 | 0.07 |

| | | | | | | | | | | | | | | | | |
|---|---|---|---|---|---|---|---|---|---|---|---|---|---|---|---|---|
| 1 | 244957186 | rs9662967 | -1.052 | -1.207 | 0.195 | 1.068 | 0.002 | -0.003 | -0.003 | 0 | 0.37 | 0.63 | 0.53 | 0.59 | 0.3 | 0.46 |
| 1 | 244964713 | rs751715 | 0.622 | -1.983 | 0.411 | 0.678 | -0.069 | 0.015 | 0.015 | 0 | 0.68 | 0.47 | 0.58 | 0.38 | 0.83 | 0.6 |
| 1 | 244965588 | rs2365687 | 0.43 | -1.819 | 0.002 | 0.672 | -0.03 | 0.008 | 0.008 | 0 | 0.69 | 0.64 | 0.59 | 0.39 | 0.84 | 0.61 |
| 1 | 244967771 | rs2365685 | 0.437 | -1.212 | -0.733 | -0.05 | 0.082 | 0.035 | 0.035 | 0 | 0.53 | 0.33 | 0.66 | 0.66 | 0.66 | 0.55 |
| 1 | 244997953 | rs10927330 | -1.401 | -0.814 | -0.2 | -0.66 | -0.024 | -0.107 | -0.107 | 0.323 | 0.32 | 0.42 | 0.14 | 0.01 | 0.22 | 0.15 |
| 1 | 245000918 | rs12139064 | -1.117 | -0.822 | -0.447 | -0.59 | -0.004 | -0.048 | -0.048 | 0 | 0.27 | 0.28 | 0.11 | 0.01 | 0.17 | 0.13 |
| 1 | 245005713 | rs12095737 | -1.024 | -0.759 | -0.718 | -0.559 | 0.022 | 0.01 | 0.01 | 2.969 | 0.26 | 0.15 | 0.1 | 0.01 | 0.17 | 0.13 |
| 1 | 245005828 | rs12070200 | -1.024 | -0.759 | -0.718 | -0.548 | 0.022 | 0.013 | 0.013 | 2.969 | 0.26 | 0.15 | 0.1 | 0.01 | 0.17 | 0.13 |
| 1 | 245012509 | rs12146095 | -1.335 | -0.871 | -0.623 | -0.536 | 0.018 | -0.002 | -0.002 | 0 | 0.27 | 0.18 | 0.11 | 0.01 | 0.19 | 0.15 |
| 1 | 245013807 | rs1495946 | -0.881 | -0.889 | -0.728 | -0.506 | 0.012 | -0.007 | -0.007 | 0.013 | 0.23 | 0.15 | 0.09 | 0.01 | 0.16 | 0.13 |
| 1 | 245018624 | rs10927336 | -0.881 | -0.889 | -0.728 | -0.506 | 0.012 | -0.007 | -0.007 | 0 | 0.23 | 0.15 | 0.09 | 0.01 | 0.16 | 0.12 |
| 1 | 245020285 | rs1874393 | -1.031 | -0.815 | -0.692 | -0.46 | 0.019 | 0 | 0 | 3.009 | 0.27 | 0.17 | 0.11 | 0.01 | 0.19 | 0.14 |
| 1 | 245042312 | rs4658673 | -1.107 | -0.033 | -0.789 | 0.454 | -0.014 | -0.026 | -0.026 | 0 | 0.15 | 0.1 | 0.19 | 0.22 | 0.21 | 0.18 |
| 1 | 245051886 | rs12566094 | -1.852 | -0.398 | -0.501 | 1.036 | 0.001 | -0.006 | -0.006 | 0 | 0.21 | 0.21 | 0.24 | 0.3 | 0.25 | 0.31 |
| 1 | 245052662 | rs13375200 | -1.089 | -0.871 | 0.684 | 0.733 | 0.076 | 0.179 | 0.179 | 2.031 | 0.06 | 0.44 | 0.04 | 0.09 | 0.01 | 0.02 |
| 1 | 245054094 | rs10927345 | -1.195 | -0.906 | 0.632 | 0.676 | 0 | -0.002 | -0.002 | 0.571 | 0.17 | 0.23 | 0.2 | 0.14 | 0.2 | 0.17 |
| 1 | 245062907 | rs7529728 | -2.32 | -1.05 | 1.563 | 0.969 | -0.02 | 0.172 | 0.172 | 0 | 0.16 | 0.55 | 0.18 | 0.34 | 0.13 | 0.26 |
| 1 | 245067198 | rs9730519 | -1.75 | -1.396 | 1.426 | 0.615 | -0.018 | -0.001 | -0.001 | 0 | 0.22 | 0.31 | 0.16 | 0.21 | 0.17 | 0.07 |
| 1 | 245084254 | rs2172955 | -2.289 | -1.71 | 1.738 | 0.647 | -0.031 | 0.016 | 0.016 | 0 | 0.35 | 0.56 | 0.3 | 0.42 | 0.26 | 0.37 |
| 1 | 245089333 | rs1979569 | -1.902 | -1.696 | 1.774 | 0.553 | 0.032 | -0.028 | -0.028 | 0.012 | 0.29 | 0.16 | 0.22 | 0.36 | 0.25 | 0.36 |
| 1 | 245093639 | rs6684661 | -2.13 | -1.743 | 1.75 | 0.504 | -0.032 | 0.005 | 0.005 | 0 | 0.36 | 0.56 | 0.25 | 0.39 | 0.26 | 0.36 |
| 1 | 245099878 | rs10927348 | -0.777 | -0.474 | 0.792 | 0.065 | -0.006 | -0.001 | -0.001 | 0 | 0.33 | 0.33 | 0.27 | 0.4 | 0.41 | 0.37 |
| 1 | 245102866 | rs6676378 | -1.505 | -0.102 | 0.376 | -0.28 | 0.025 | 0.075 | 0.075 | 0 | 0.5 | 0.76 | 0.4 | 0.4 | 0.52 | 0.39 |
| 1 | 245108240 | rs6429511 | -1.671 | -0.54 | 0.857 | -0.909 | -0.034 | -0.149 | -0.149 | 2.843 | 0.24 | 0.44 | 0.16 | 0 | 0.18 | 0.04 |
| 1 | 245109861 | rs12747042 | -1.362 | -0.637 | 0.264 | -0.947 | 0.023 | 0.037 | 0.037 | 0.057 | 0.2 | 0.11 | 0.1 | 0 | 0.15 | 0.03 |
| 1 | 245116281 | rs7524543 | -0.814 | -0.544 | 0.229 | -0.971 | -0.013 | -0.022 | -0.022 | 0.018 | 0.39 | 0.42 | 0.28 | 0.18 | 0.3 | 0.21 |
| 1 | 245121226 | rs7540416 | -1.364 | -0.827 | -0.188 | -1.389 | 0.096 | 0.194 | 0.194 | 2.487 | 0.51 | 0.72 | 0.26 | 0.1 | 0.35 | 0.21 |
| 1 | 245121498 | rs12082673 | -1.377 | -0.91 | 0.028 | -1.379 | 0.056 | 0.113 | 0.113 | 2.418 | 0.5 | 0.69 | 0.27 | 0.13 | 0.37 | 0.22 |
| 1 | 245123655 | rs6660957 | -1.359 | -0.873 | 0.079 | -1.147 | -0.05 | -0.101 | -0.101 | 0.625 | 0.21 | 0.33 | 0.09 | 0 | 0.12 | 0.05 |
| 1 | 245130641 | rs6700087 | 0.356 | 0.011 | -0.745 | -0.966 | 0.008 | 0.124 | 0.124 | 1.963 | 0.48 | 0.19 | 0.42 | 0.72 | 0.58 | 0.64 |
| 1 | 245135714 | rs11590498 | 0.306 | 0.184 | -0.328 | -1.078 | 0.022 | 0.087 | 0.087 | 2.108 | 0.52 | 0.29 | 0.64 | 0.73 | 0.58 | 0.67 |
| 1 | 245142256 | rs4658680 | -0.121 | 0.112 | 0.043 | -1.269 | 0.002 | -0.068 | -0.068 | 0 | 0.59 | 0.41 | 0.75 | 0.99 | 0.69 | 0.88 |
| 1 | 245144712 | rs1021282 | 0.413 | 0.105 | 0.061 | -1.27 | 0.01 | 0.04 | 0.04 | 1.775 | 0.53 | 0.37 | 0.65 | 0.73 | 0.59 | 0.67 |
| 1 | 245148734 | rs12403396 | 0.508 | 0.099 | 0.051 | -1.273 | 0.022 | 0.085 | 0.085 | 1.729 | 0.52 | 0.29 | 0.65 | 0.73 | 0.59 | 0.67 |
| 1 | 245156511 | rs4658683 | -0.392 | 0.293 | 0.24 | -1.408 | -0.028 | -0.132 | -0.132 | 0 | 0.63 | 0.46 | 0.79 | 1 | 0.76 | 0.95 |
| 1 | 245168038 | rs7552510 | 0.037 | 0.325 | 0.421 | -1.361 | 0.007 | 0.068 | 0.068 | 0 | 0.05 | 0.09 | 0.11 | 0.27 | 0.11 | 0.2 |
| 1 | 245170495 | rs6684005 | 1.614 | 1.025 | -0.133 | -0.573 | -0.049 | -0.075 | -0.075 | 0.138 | 0.11 | 0.01 | 0.13 | 0.43 | 0.13 | 0.15 |
| 1 | 245173460 | rs10803251 | -1.891 | 0.461 | 0.697 | -0.921 | 0.057 | 0.099 | 0.099 | 0 | 0.14 | 0.34 | 0.16 | 0.27 | 0.18 | 0.28 |
| 1 | 245179188 | rs1196366 | -0.623 | 0.266 | 1.29 | -0.783 | 0.019 | 0.081 | 0.081 | 4.147 | 0.63 | 0.81 | 0.75 | 0.87 | 0.66 | 0.84 |
| 1 | 245179212 | rs6696296 | -0.979 | 0.282 | 1.2 | -0.775 | 0.029 | 0.042 | 0.042 | 0.063 | 0.09 | 0.24 | 0.09 | 0.15 | 0.07 | 0.16 |
| 1 | 245179960 | rs10927369 | 0.667 | 0.278 | 1.06 | -0.777 | 0.02 | 0.127 | 0.127 | 3.008 | 0.48 | 0.17 | 0.63 | 0.73 | 0.59 | 0.64 |
| 1 | 245190615 | rs493092 | 1.118 | 0.375 | 1.143 | -0.55 | 0.056 | 0.054 | 0.054 | 0.181 | 0.87 | 0.55 | 0.89 | 0.88 | 0.88 | 0.84 |
| 1 | 245191095 | rs600025 | 0.144 | 0.233 | 1.303 | -0.9 | 0.02 | 0.1 | 0.1 | 5.637 | 0.41 | 0.23 | 0.26 | 0.15 | 0.39 | 0.18 |
| 1 | 245191893 | rs586013 | -0.245 | 0.184 | 1.306 | -0.892 | -0.01 | -0.016 | -0.016 | 0.274 | 0.45 | 0.5 | 0.31 | 0.29 | 0.38 | 0.27 |
| 1 | 245205890 | rs12728471 | 0.219 | 0.251 | 0.666 | -0.87 | 0.068 | 0.103 | 0.103 | 5.061 | 0.52 | 0.23 | 0.67 | 0.71 | 0.61 | 0.7 |
| 1 | 245209134 | rs4020082 | 0.054 | 0.252 | 0.75 | -0.87 | 0.036 | 0.053 | 0.053 | 5.313 | 0.54 | 0.33 | 0.68 | 0.71 | 0.62 | 0.73 |
| 1 | 245219054 | rs611185 | -0.011 | 0.279 | 0.901 | -0.902 | 0.02 | 0.101 | 0.101 | 5.768 | 0.46 | 0.29 | 0.28 | 0.17 | 0.44 | 0.26 |
| 1 | 245223064 | rs576872 | -0.574 | 0.374 | 0.938 | -0.936 | -0.007 | 0.076 | 0.076 | 3.861 | 0.53 | 0.38 | 0.32 | 0.18 | 0.51 | 0.35 |
| 1 | 245228886 | rs6692143 | 1.033 | 0.361 | 0.946 | -0.92 | -0.026 | 0.026 | 0.026 | 3.675 | 0.41 | 0.25 | 0.61 | 0.68 | 0.48 | 0.53 |
| 1 | 245238442 | rs6668039 | -0.074 | 0.41 | 0.959 | -0.902 | 0.013 | -0.031 | -0.031 | 0 | 0.07 | 0.13 | 0.05 | 0 | 0.07 | 0.09 |
| 1 | 245238995 | rs4658690 | 1.073 | 0.417 | 1.011 | -0.847 | -0.025 | 0.019 | 0.019 | 2.454 | 0.4 | 0.24 | 0.61 | 0.65 | 0.48 | 0.54 |
| 1 | 245245378 | rs10927386 | -0.474 | 0.043 | 0.816 | -0.631 | 0.024 | 0.115 | 0.115 | 3.882 | 0.48 | 0.29 | 0.29 | 0.19 | 0.45 | 0.29 |
| 1 | 245245402 | rs10927387 | -0.438 | 0.049 | 0.756 | -0.628 | 0.023 | 0.123 | 0.123 | 4.241 | 0.48 | 0.25 | 0.29 | 0.19 | 0.45 | 0.29 |
| 1 | 245268412 | rs4658698 | 0.135 | 0.341 | 0.443 | -0.469 | 0.002 | -0.002 | -0.002 | 0.68 | 0.91 | 0.89 | 0.96 | 0.91 | 0.88 | 0.89 |
| 1 | 245271418 | rs2886208 | 0.462 | -0.397 | 0.785 | -0.816 | 0.041 | 0.06 | 0.06 | 4.038 | 0.52 | 0.36 | 0.71 | 0.69 | 0.67 | 0.76 |
| 1 | 245273390 | rs5015334 | 0.444 | -0.292 | 1.156 | -0.8 | 0.064 | 0.072 | 0.072 | 3.244 | 0.53 | 0.34 | 0.72 | 0.7 | 0.7 | 0.76 |
| 1 | 245276494 | rs7538816 | 0.273 | -0.455 | 1.315 | -0.907 | 0.044 | 0.047 | 0.047 | 6.756 | 0.33 | 0.22 | 0.17 | 0.01 | 0.21 | 0.05 |
| 1 | 245281347 | rs4072543 | 0.905 | -0.125 | 1.193 | -1.22 | 0.049 | 0.102 | 0.102 | 0 | 0.93 | 0.7 | 0.96 | 0.87 | 0.94 | 0.92 |
| 1 | 245281678 | rs7531729 | 0.26 | -0.129 | 0.641 | -1.196 | 0.044 | 0.084 | 0.084 | 3.456 | 0.53 | 0.34 | 0.74 | 0.86 | 0.67 | 0.85 |
| 1 | 245283910 | rs6693918 | -0.345 | 0.119 | 0.671 | -0.904 | 0 | 0.012 | 0.012 | 2.554 | 0.47 | 0.41 | 0.52 | 0.21 | 0.44 | 0.25 |
| 1 | 245284307 | rs6694326 | 1.033 | -0.415 | 1.016 | -0.656 | -0.023 | -0.087 | -0.087 | 0 | 0.82 | 0.72 | 0.91 | 1 | 0.92 | 0.94 |
| 1 | 245288239 | rs945953 | -2.064 | 0.067 | 1.254 | 0.129 | -0.007 | 0.03 | 0.03 | 0 | 0.13 | 0.34 | 0.37 | 0.13 | 0.09 | 0.14 |
| 1 | 245288493 | rs6672147 | -2.468 | -0.026 | 1.176 | 0.233 | -0.033 | 0.019 | 0.019 | 0 | 0.21 | 0.48 | 0.49 | 0.28 | 0.16 | 0.29 |
| 1 | 245290111 | rs4658702 | -2.125 | 0.059 | 0.939 | 0.303 | -0.022 | 0.028 | 0.028 | 0 | 0.15 | 0.37 | 0.4 | 0.19 | 0.11 | 0.25 |
| 1 | 245302155 | rs12404246 | -1.539 | 0.187 | 0.81 | 0.39 | 0.008 | 0.125 | 0.125 | 0 | 0.07 | 0.25 | 0.15 | 0.18 | 0.04 | 0.07 |
| 1 | 245303909 | rs4489540 | -2.312 | 0.086 | 1.071 | 0.189 | -0.017 | 0.013 | 0.013 | 0 | 0.15 | 0.24 | 0.22 | 0.21 | 0.11 | 0.13 |
| 1 | 245305530 | rs7549529 | -2.125 | 0.431 | 0.804 | -0.557 | -0.007 | -0.104 | -0.104 | 0 | 0.29 | 0.42 | 0.27 | 0.1 | 0.24 | 0.2 |
| 1 | 245306896 | rs4083908 | -1.791 | 0.517 | 0.826 | -0.488 | -0.002 | -0.071 | -0.071 | 0 | 0.26 | 0.27 | 0.19 | 0.04 | 0.23 | 0.18 |
| 1 | 245307406 | rs2365084 | -0.879 | 0.568 | 0.794 | -0.469 | -0.001 | -0.007 | -0.007 | 1.429 | 0.1 | 0.08 | 0.11 | 0.03 | 0.09 | 0.04 |

| | | | | | | | | | | | | | | | | |
|---|---|---|---|---|---|---|---|---|---|---|---|---|---|---|---|---|
| 1 | 245312162 | rs4658437 | 0.58 | 0.784 | 0.39 | -0.414 | 0.01 | 0.011 | 0.011 | 0.167 | 0.32 | 0.54 | 0.42 | 0.35 | 0.37 | 0.2 |
| 1 | 245312295 | rs4658439 | -0.234 | 0.751 | -0.079 | -0.455 | -0.001 | 0.001 | 0.001 | 0.576 | 0.42 | 0.64 | 0.54 | 0.42 | 0.47 | 0.34 |
| 1 | 245316219 | rs6676019 | 0.764 | 0.828 | -0.428 | -0.189 | 0.003 | -0.012 | -0.012 | 0 | 0.42 | 0.11 | 0.32 | 0.55 | 0.4 | 0.64 |
| 1 | 245318408 | rs6671940 | -0.878 | 0.956 | 0.238 | -0.016 | -0.013 | 0.025 | 0.025 | 0.302 | 0.1 | 0.06 | 0.05 | 0 | 0.11 | 0.13 |
| 1 | 245328085 | rs12039117 | 1.124 | 0.576 | 0.58 | 0.339 | 0.005 | -0.013 | -0.013 | 0 | 0.87 | 0.57 | 0.86 | 0.91 | 0.92 | 0.95 |
| 1 | 245329981 | rs945951 | 1.106 | 0.622 | 0.233 | 0.33 | -0.009 | -0.03 | -0.03 | 0.093 | 0.86 | 0.64 | 0.87 | 0.92 | 0.92 | 0.95 |
| 1 | 245335643 | rs6656693 | -1.046 | 1.096 | -0.559 | 0.956 | -0.011 | -0.022 | -0.022 | 2.144 | 0.3 | 0.75 | 0.25 | 0.47 | 0.3 | 0.29 |
| 1 | 245336796 | rs4658714 | 0.687 | 1.009 | -0.606 | 0.978 | -0.014 | -0.007 | -0.007 | 0 | 0.83 | 0.87 | 0.84 | 0.74 | 0.78 | 0.71 |
| 1 | 245341153 | rs6668478 | -0.938 | -0.085 | 0.254 | 1.362 | -0.034 | 0.173 | 0.173 | 0 | 0.14 | 0.4 | 0.05 | 0.31 | 0.08 | 0.12 |
| 1 | 245341817 | rs6661783 | -1.203 | -0.029 | 0.365 | 1.421 | 0.011 | -0.054 | -0.054 | 0 | 0.14 | 0.46 | 0.08 | 0.09 | 0.11 | 0.11 |
| 1 | 245341997 | rs6701995 | -1.315 | -0.127 | 0.396 | 1.335 | -0.027 | 0.211 | 0.211 | 0 | 0.17 | 0.47 | 0.07 | 0.4 | 0.12 | 0.19 |
| 1 | 245344629 | rs3795475 | 0.691 | -0.449 | -0.087 | -0.202 | 0.02 | 0.108 | 0.108 | 0 | 0.45 | 0.29 | 0.53 | 0.17 | 0.58 | 0.4 |
| 1 | 245345109 | rs6429530 | 0.401 | -1.033 | 0.928 | -0.252 | -0.038 | -0.104 | -0.104 | 2.214 | 0.82 | 0.7 | 0.89 | 1 | 0.89 | 0.93 |
| 1 | 245348263 | rs10737770 | 0.498 | -1.096 | 1.148 | -0.244 | -0.053 | -0.125 | -0.125 | 2.198 | 0.82 | 0.6 | 0.89 | 1 | 0.89 | 0.93 |
| 1 | 245348828 | rs10924112 | 1.029 | -1.388 | 0.756 | -0.164 | -0.062 | -0.075 | -0.075 | 0.41 | 0.69 | 0.54 | 0.8 | 0.89 | 0.82 | 0.74 |
| 1 | 245350885 | rs7537950 | 1.127 | -1.372 | 0.358 | -0.124 | -0.022 | -0.027 | -0.027 | 0.751 | 0.68 | 0.57 | 0.8 | 0.88 | 0.81 | 0.72 |
| 1 | 245352307 | rs4658446 | 0.226 | -0.892 | 0.333 | 0.307 | -0.031 | -0.008 | -0.008 | 0.261 | 0.84 | 0.72 | 0.89 | 0.88 | 0.9 | 0.78 |
| 1 | 245358619 | rs6671796 | 0.591 | -0.398 | -0.334 | -0.023 | 0.017 | 0.09 | 0.09 | 0.418 | 0.49 | 0.69 | 0.53 | 0.25 | 0.52 | 0.44 |
| 1 | 245363423 | rs6428904 | -0.606 | -0.37 | 0.068 | 0.49 | 0 | -0.001 | -0.001 | 0 | 0.21 | 0.19 | 0.11 | 0.2 | 0.18 | 0.13 |
| 1 | 245366662 | rs10802198 | 1.507 | -0.644 | 0.367 | 0.287 | 0.022 | 0.085 | 0.085 | 0.125 | 0.55 | 0.28 | 0.59 | 0.28 | 0.58 | 0.49 |
| 1 | 245367423 | rs6697031 | 0.587 | -0.812 | 0.952 | 0.289 | -0.032 | -0.047 | -0.047 | 0 | 0.82 | 0.5 | 0.89 | 0.93 | 0.87 | 0.89 |
| 1 | 245367534 | rs6684360 | 1.923 | -0.795 | 0.872 | 0.272 | 0.019 | 0.084 | 0.084 | 0.113 | 0.55 | 0.29 | 0.59 | 0.27 | 0.57 | 0.49 |
| 1 | 245371026 | rs7527554 | 0.119 | -0.784 | 0.941 | 0.299 | -0.003 | -0.005 | -0.005 | 0 | 0.85 | 0.81 | 0.93 | 0.94 | 0.88 | 0.89 |
| 1 | 245371412 | rs6662127 | -0.722 | -0.788 | 0.551 | 1.214 | 0 | 0.005 | 0.005 | 0 | 0.22 | 0.23 | 0.24 | 0.57 | 0.24 | 0.3 |
| 1 | 245371492 | rs12129648 | 1.402 | -0.889 | -0.006 | 1.113 | -0.016 | -0.045 | -0.045 | 6.483 | 0.57 | 0.03 | 0.63 | 0.38 | 0.64 | 0.59 |
| 1 | 245382117 | rs4497254 | 1.26 | -0.651 | 0.836 | 0.873 | 0.004 | 0.035 | 0.035 | 1.1 | 0.62 | 0.22 | 0.65 | 0.3 | 0.64 | 0.6 |
| 1 | 245388149 | rs10924125 | -1.208 | -0.971 | 0.743 | 0.583 | 0.013 | 0.024 | 0.024 | 0.069 | 0.41 | 0.25 | 0.33 | 0.6 | 0.39 | 0.41 |
| 1 | 245400227 | rs7513943 | -1.386 | -0.478 | 1.118 | 0.553 | -0.002 | -0.008 | -0.008 | 0 | 0.4 | 0.57 | 0.36 | 0.61 | 0.38 | 0.4 |
| 1 | 245400309 | rs7547227 | -1.317 | -0.476 | 1.267 | 0.535 | -0.002 | -0.016 | -0.016 | 0 | 0.39 | 0.51 | 0.36 | 0.71 | 0.38 | 0.4 |
| 1 | 245403490 | rs2941285 | 1.337 | -0.587 | 1.105 | 0.463 | 0.017 | 0.09 | 0.09 | 0.361 | 0.55 | 0.22 | 0.6 | 0.29 | 0.58 | 0.57 |
| 1 | 245413424 | rs10924134 | -1.016 | 0.143 | 1.325 | 0.705 | -0.008 | 0.024 | 0.024 | 0 | 0.37 | 0.49 | 0.48 | 0.55 | 0.31 | 0.42 |
| 1 | 245413516 | rs7554953 | -0.628 | 0.466 | 1.254 | 0.54 | 0.002 | -0.012 | -0.012 | 0.022 | 0.16 | 0.19 | 0.16 | 0.07 | 0.17 | 0.07 |
| 1 | 245417562 | rs6692147 | -0.999 | 0.412 | 1.28 | 0.534 | 0.011 | 0.01 | 0.01 | 0 | 0.17 | 0.36 | 0.17 | 0.14 | 0.18 | 0.08 |
| 1 | 245419312 | rs907015 | -1.486 | 0.922 | 1.456 | 0.53 | -0.002 | 0.008 | 0.008 | 0 | 0.47 | 0.6 | 0.61 | 0.61 | 0.47 | 0.5 |
| 1 | 245419426 | rs12075902 | -0.519 | 0.859 | 1.453 | 0.529 | 0.001 | -0.009 | -0.009 | 0 | 0.35 | 0.36 | 0.46 | 0.54 | 0.3 | 0.43 |
| 1 | 245419947 | rs6677893 | -2.167 | 0.849 | 0.946 | 0.498 | -0.035 | 0.074 | 0.074 | 0 | 0.55 | 0.85 | 0.65 | 0.68 | 0.52 | 0.5 |
| 1 | 245420757 | rs4658733 | 0.591 | 0.89 | 0.769 | 1.272 | -0.054 | -0.076 | -0.076 | 0.014 | 0.14 | 0.01 | 0.29 | 0.22 | 0.13 | 0.15 |
| 1 | 245422577 | rs6428906 | -2.145 | 0.86 | 0.87 | 0.992 | -0.017 | 0.034 | 0.034 | 0 | 0.56 | 0.81 | 0.66 | 0.63 | 0.53 | 0.53 |
| 1 | 245423334 | rs7524902 | -1.71 | 0.867 | 0.854 | 0.962 | -0.009 | 0.039 | 0.039 | 0 | 0.48 | 0.82 | 0.56 | 0.65 | 0.46 | 0.49 |
| 1 | 245423526 | rs6428909 | -0.924 | 0.876 | 1.104 | 0.949 | 0 | -0.001 | -0.001 | 0 | 0.4 | 0.59 | 0.5 | 0.64 | 0.39 | 0.48 |
| 1 | 245423939 | rs2292319 | 0.6 | 0.903 | 0.963 | 0.957 | -0.009 | -0.006 | -0.006 | 0 | 0.1 | 0.02 | 0.11 | 0.06 | 0.1 | 0.04 |
| 1 | 245426659 | rs4658734 | -2.223 | 0.889 | 0.955 | 0.998 | -0.026 | 0.053 | 0.053 | 0 | 0.57 | 0.84 | 0.67 | 0.69 | 0.56 | 0.52 |
| 1 | 245428802 | rs12740199 | 0.242 | 0.762 | 0.69 | 0.719 | -0.001 | 0.003 | 0.003 | 0 | 0.07 | 0.05 | 0.04 | 0 | 0.07 | 0.02 |
| 1 | 245430915 | rs12732812 | -0.821 | 0.738 | 0.672 | 0.711 | -0.006 | 0.033 | 0.033 | 0 | 0.18 | 0.07 | 0.15 | 0.05 | 0.17 | 0.06 |
| 1 | 245431939 | rs7513602 | -1.447 | 1.12 | 0.692 | 0.773 | -0.001 | 0.001 | 0.001 | 0 | 0.47 | 0.57 | 0.56 | 0.61 | 0.48 | 0.53 |
| 1 | 245434303 | rs2364340 | -2.247 | 0.905 | 0.454 | 0.452 | 0 | 0.026 | 0.026 | 0 | 0.61 | 0.75 | 0.69 | 0.72 | 0.61 | 0.57 |
| 1 | 245441251 | rs9660396 | -1.033 | 1.07 | 0.416 | 0.655 | 0.002 | -0.009 | -0.009 | 0.123 | 0.2 | 0.26 | 0.18 | 0.13 | 0.22 | 0.06 |
| 1 | 245446721 | rs6428910 | -2.119 | 0.907 | 0.695 | 0.498 | 0.001 | 0.021 | 0.021 | 0 | 0.6 | 0.73 | 0.71 | 0.73 | 0.59 | 0.56 |
| 1 | 245446812 | rs7544367 | -2.065 | 0.912 | 0.7 | 0.497 | -0.001 | 0 | 0 | 0 | 0.59 | 0.65 | 0.69 | 0.72 | 0.56 | 0.56 |
| 1 | 245447738 | rs4658739 | -1.267 | 0.908 | 0.656 | 0.48 | 0 | 0.013 | 0.013 | 0 | 0.5 | 0.49 | 0.63 | 0.67 | 0.48 | 0.54 |
| 1 | 245448886 | rs7528364 | -2.216 | 0.892 | 0.589 | 0.541 | -0.002 | 0.012 | 0.012 | 0 | 0.61 | 0.71 | 0.7 | 0.72 | 0.58 | 0.58 |
| 1 | 245451992 | rs10924157 | -1.279 | 0.877 | 0.554 | 0.541 | 0 | 0.008 | 0.008 | 0 | 0.51 | 0.5 | 0.63 | 0.67 | 0.48 | 0.54 |
| 1 | 245453417 | rs12748466 | -0.333 | 0.71 | 0.546 | 0.544 | 0.001 | -0.039 | -0.039 | 0.356 | 0.1 | 0.17 | 0.07 | 0.02 | 0.11 | 0.02 |
| 1 | 245456075 | rs12084148 | -1.392 | 0.727 | 0.536 | 0.542 | -0.002 | 0.029 | 0.029 | 0 | 0.52 | 0.63 | 0.64 | 0.68 | 0.5 | 0.54 |
| 1 | 245459768 | rs4020773 | -2.407 | 0.707 | 0.441 | 0.535 | -0.02 | 0.034 | 0.034 | 0 | 0.63 | 0.84 | 0.72 | 0.73 | 0.61 | 0.58 |
| 1 | 245459950 | rs12239562 | -1.215 | 0.681 | 0.434 | 0.578 | -0.011 | -0.015 | -0.015 | 0 | 0.2 | 0.31 | 0.17 | 0.11 | 0.19 | 0.06 |
| 1 | 245461759 | rs12043715 | -0.192 | 0.693 | 0.166 | 0.581 | -0.001 | -0.004 | -0.004 | 0 | 0.1 | 0.11 | 0.06 | 0.05 | 0.07 | 0.02 |
| 1 | 245461981 | rs2364339 | 2.568 | 0.582 | -0.324 | 0.417 | -0.105 | -0.001 | -0.001 | 0 | 0.31 | 0.01 | 0.24 | 0.19 | 0.35 | 0.33 |
| 1 | 245462248 | rs6665809 | -0.568 | 0.513 | 0.312 | 0.645 | -0.002 | 0.003 | 0.003 | 0 | 0.38 | 0.43 | 0.5 | 0.61 | 0.33 | 0.5 |
| 1 | 245470001 | rs12120571 | 0.281 | 0.556 | 0.355 | 0.546 | -0.001 | -0.008 | -0.008 | 0 | 0.22 | 0.23 | 0.17 | 0.43 | 0.2 | 0.34 |
| 1 | 245470154 | rs10802209 | -0.815 | 0.47 | 0.179 | 0.541 | -0.01 | 0.001 | 0.001 | 0 | 0.33 | 0.49 | 0.5 | 0.33 | 0.33 | 0.19 |
| 1 | 245474078 | rs907014 | -1.389 | 0.435 | -0.35 | 0.278 | -0.007 | 0.015 | 0.015 | 0 | 0.63 | 0.71 | 0.71 | 0.76 | 0.6 | 0.61 |
| 1 | 245475743 | rs12738350 | -0.768 | 0.302 | 0.055 | 0.718 | -0.002 | 0.041 | 0.041 | 0 | 0.08 | 0.01 | 0.06 | 0 | 0.08 | 0.04 |
| 1 | 245477121 | rs11811491 | -0.856 | 0.297 | 0.054 | 0.718 | -0.016 | 0.059 | 0.059 | 0 | 0.11 | 0.02 | 0.07 | 0 | 0.13 | 0.06 |
| 1 | 245480950 | rs10924169 | -1.026 | 0.172 | 0.803 | 1.237 | -0.01 | 0.011 | 0.011 | 0 | 0.12 | 0.25 | 0.29 | 0.16 | 0.08 | 0.12 |
| 1 | 245481432 | rs4658748 | -1.236 | 0.273 | 0.336 | 1.278 | -0.006 | 0.005 | 0.005 | 0 | 0.48 | 0.75 | 0.63 | 0.53 | 0.5 | 0.56 |
| 1 | 245486835 | rs6428914 | -1.237 | 0.293 | 0.219 | 1.171 | -0.002 | 0 | 0 | 0 | 0.49 | 0.76 | 0.61 | 0.49 | 0.5 | 0.55 |
| 1 | 245490640 | rs10924173 | -1.602 | -0.365 | 0.787 | 1.764 | 0.016 | -0.051 | -0.051 | 0 | 0.13 | 0.08 | 0.32 | 0.32 | 0.08 | 0.15 |
| 1 | 245491406 | rs4658753 | -2.412 | -0.533 | 1.227 | 1.437 | -0.016 | -0.061 | -0.061 | 0 | 0.13 | 0.7 | 0.11 | 0.01 | 0.09 | 0.03 |

| | | | | | | | | | | | | | | | |
|---|---|---|---|---|---|---|---|---|---|---|---|---|---|---|---|
| 1 | 245498162 | rs11803433 | -1.756 | -0.516 | 0.994 | 1.883 | -0.025 | 0.123 | 0.123 | 0 | 0.18 | 0.58 | 0.38 | 0.37 | 0.1 | 0.21 |
| 1 | 245511851 | rs12072874 | -1.737 | -0.749 | 0.827 | 1.746 | -0.071 | 0.094 | 0.094 | 0.048 | 0.17 | 0.59 | 0.15 | 0.29 | 0.05 | 0.1 |
| 1 | 245520118 | rs4256853 | -1.529 | -0.522 | 0.847 | 1.92 | -0.042 | 0.16 | 0.16 | 3.141 | 0.09 | 0.67 | 0.14 | 0.24 | 0.02 | 0.02 |
| 1 | 245528780 | rs1173847 | -0.396 | -0.423 | -0.74 | 1.575 | 0.027 | -0.008 | -0.008 | 0 | 0.51 | 0.89 | 0.61 | 0.55 | 0.44 | 0.48 |
| 1 | 245534952 | rs10924184 | -0.705 | -0.846 | -0.1 | 1.867 | -0.016 | 0.035 | 0.035 | 0 | 0.11 | 0.15 | 0.05 | 0.38 | 0.04 | 0.16 |
| 1 | 245535862 | rs1173842 | 0.413 | -0.77 | -0.297 | 1.151 | 0.006 | 0.071 | 0.071 | 0 | 0.47 | 0.32 | 0.6 | 0.19 | 0.54 | 0.55 |
| 1 | 245537353 | rs551022 | -0.695 | -0.852 | -0.757 | 0.818 | -0.04 | -0.099 | -0.099 | 0 | 0.7 | 0.88 | 0.72 | 0.58 | 0.66 | 0.72 |
| 1 | 245538064 | rs482106 | -0.635 | -0.761 | -0.753 | 0.834 | -0.048 | -0.097 | -0.097 | 0.958 | 0.71 | 0.88 | 0.72 | 0.59 | 0.65 | 0.73 |
| 1 | 245542859 | rs1174002 | 0.636 | -0.934 | -0.653 | 0.59 | 0.021 | 0.114 | 0.114 | 0 | 0.5 | 0.81 | 0.65 | 0.28 | 0.55 | 0.49 |
| 1 | 245543358 | rs819896 | -1.01 | -1.039 | -0.284 | 0.631 | 0.021 | -0.009 | -0.009 | 1.503 | 0.21 | 0.16 | 0.11 | 0.31 | 0.11 | 0.18 |
| 1 | 245550394 | rs1173663 | 0.213 | -0.845 | -0.054 | 0.268 | 0.014 | 0.039 | 0.039 | 0 | 0.56 | 0.7 | 0.66 | 0.31 | 0.56 | 0.54 |
| 1 | 245551682 | rs819883 | 0.326 | -0.652 | -0.249 | 0.201 | 0.002 | -0.032 | -0.032 | 0 | 0.14 | 0.06 | 0.07 | 0.29 | 0.1 | 0.2 |
| 1 | 245552025 | rs491725 | 0.689 | -0.621 | 0.73 | 0.15 | 0.012 | 0.029 | 0.029 | 0 | 0.5 | 0.53 | 0.63 | 0.31 | 0.54 | 0.54 |
| 1 | 245555664 | rs819876 | 0.926 | -0.262 | 0.565 | 0.129 | -0.054 | 0.106 | 0.106 | 0.553 | 0.49 | 0.1 | 0.56 | 0.39 | 0.52 | 0.69 |
| 1 | 245556857 | rs12141949 | -0.092 | -0.281 | 0.891 | 0.133 | 0.003 | 0.176 | 0.176 | 1.467 | 0.12 | 0.48 | 0.12 | 0.23 | 0.06 | 0.03 |
| 1 | 245557294 | rs1173837 | -0.698 | -0.272 | 0.819 | 0.132 | -0.001 | -0.004 | -0.004 | 0 | 0.7 | 0.68 | 0.72 | 0.62 | 0.65 | 0.74 |
| 1 | 245558828 | rs1173829 | -0.698 | -0.272 | 0.819 | 0.132 | -0.001 | -0.004 | -0.004 | 0 | 0.7 | 0.68 | 0.72 | 0.63 | 0.65 | 0.74 |
| 1 | 245560145 | rs11579379 | -1.126 | -0.462 | 0.896 | 0.189 | -0.017 | 0.024 | 0.024 | 1.558 | 0.21 | 0.57 | 0.16 | 0.23 | 0.12 | 0.05 |
| 1 | 245563382 | rs7545405 | -0.171 | -0.48 | 0.911 | 0.096 | -0.003 | -0.055 | -0.055 | 0.35 | 0.1 | 0.29 | 0.05 | 0 | 0.05 | 0.02 |
| 1 | 245563442 | rs496529 | -0.659 | 0.064 | 0.511 | 0.15 | -0.06 | -0.087 | -0.087 | 0 | 0.72 | 0.9 | 0.75 | 0.63 | 0.64 | 0.74 |
| 1 | 245566986 | rs1704427 | 0.117 | 0.004 | 0.674 | 0.005 | 0.002 | -0.006 | -0.006 | 0.302 | 0.62 | 0.57 | 0.62 | 0.39 | 0.58 | 0.72 |
| 1 | 245572481 | rs4658769 | -0.85 | -0.009 | 0.644 | 0.031 | -0.055 | 0.03 | 0.03 | 2.898 | 0.22 | 0.56 | 0.17 | 0.23 | 0.12 | 0.05 |
| 1 | 245572552 | rs676292 | -0.587 | 0.36 | 0.081 | 0.141 | -0.076 | -0.111 | -0.111 | 0.042 | 0.73 | 0.9 | 0.73 | 0.62 | 0.64 | 0.77 |
| 1 | 245574714 | rs1069232 | -0.372 | 0.66 | 0.974 | -0.819 | 0.016 | 0.016 | 0.016 | 0 | 0.73 | 0.62 | 0.56 | 0.43 | 0.69 | 0.75 |
| 1 | 245575003 | rs1173642 | 0.077 | 0.97 | 1.323 | -1.345 | -0.016 | 0.009 | 0.009 | 0 | 0.66 | 0.36 | 0.43 | 0.33 | 0.55 | 0.67 |
| 1 | 245575183 | rs1173644 | 0.007 | 0.975 | 1.88 | -1.362 | 0.044 | -0.026 | -0.026 | 0 | 0.66 | 0.48 | 0.45 | 0.33 | 0.56 | 0.67 |
| 1 | 245575250 | rs1173645 | 0.105 | 0.953 | 1.893 | -1.415 | -0.008 | 0.005 | 0.005 | 0 | 0.65 | 0.34 | 0.44 | 0.31 | 0.56 | 0.68 |
| 1 | 245575879 | rs479631 | -0.061 | 0.983 | 1.96 | -1.4 | 0.013 | -0.01 | -0.01 | 0.024 | 0.67 | 0.44 | 0.44 | 0.31 | 0.55 | 0.68 |
| 1 | 245576050 | rs1093962 | -0.638 | 1.07 | 1.988 | -1.382 | 0.147 | -0.003 | -0.003 | 0 | 0.06 | 0.3 | 0.09 | 0 | 0.13 | 0.01 |
| 1 | 245576457 | rs1093961 | -0.662 | 0.994 | 1.688 | -1.367 | -0.002 | 0 | 0 | 0 | 0.73 | 0.77 | 0.54 | 0.32 | 0.68 | 0.67 |
| 1 | 245577304 | rs2795071 | -0.556 | 1.034 | 1.31 | -1.35 | 0.019 | -0.011 | -0.011 | 0 | 0.72 | 0.63 | 0.53 | 0.32 | 0.68 | 0.67 |
| 1 | 245578185 | rs6701279 | -0.382 | 0.824 | 1.033 | -1.502 | 0.011 | -0.04 | -0.04 | 0 | 0.11 | 0.18 | 0.11 | 0 | 0.17 | 0.03 |
| 1 | 245578692 | rs1173650 | 0.935 | 0.292 | 0.142 | -2.003 | -0.01 | 0.053 | 0.053 | 0 | 0.29 | 0.17 | 0.29 | 0.11 | 0.32 | 0.41 |
| 1 | 245581534 | rs1148919 | 0.302 | 0.888 | 0.3 | -1.493 | -0.001 | -0.003 | -0.003 | 0 | 0.59 | 0.42 | 0.42 | 0.3 | 0.51 | 0.63 |
| 1 | 245582111 | rs1111813 | -0.465 | 0.873 | -0.065 | -1.106 | 0.008 | -0.006 | -0.006 | 0.248 | 0.72 | 0.71 | 0.53 | 0.32 | 0.68 | 0.66 |
| 1 | 245583654 | rs10924197 | 0.153 | 0.851 | -0.023 | -1.42 | 0.007 | -0.087 | -0.087 | 0 | 0.68 | 0.47 | 0.76 | 0.9 | 0.68 | 0.67 |
| 1 | 245586529 | rs1773825 | -1.044 | 0.69 | 0.481 | -0.925 | 0.047 | -0.019 | -0.019 | 2.508 | 0.08 | 0.29 | 0.14 | 0.01 | 0.08 | 0.03 |
| 1 | 245589159 | rs2971998 | -0.837 | 0.71 | 0.374 | -0.916 | 0.025 | -0.03 | -0.03 | 0 | 0.07 | 0.2 | 0.13 | 0.01 | 0.08 | 0.03 |
| 1 | 245594106 | rs1756921 | -0.802 | 0.684 | 0.19 | -1.348 | -0.002 | -0.071 | -0.071 | 0 | 0.36 | 0.47 | 0.3 | 0.2 | 0.29 | 0.36 |
| 1 | 245594168 | rs1687098 | -0.306 | 1.005 | 0.259 | -1.137 | 0.066 | -0.023 | -0.023 | 0 | 0.1 | 0.18 | 0.08 | 0.08 | 0.13 | 0.16 |
| 1 | 245595999 | rs538165 | 0.287 | 0.581 | -0.009 | -1.372 | 0 | -0.041 | -0.041 | 0 | 0.9 | 0.84 | 0.91 | 1 | 0.88 | 0.98 |
| 1 | 245599511 | rs1173849 | 0.233 | 0.232 | -0.397 | -1.131 | 0.002 | -0.004 | -0.004 | 0 | 0.16 | 0.12 | 0.12 | 0.22 | 0.15 | 0.19 |
| 1 | 245600808 | rs704649 | 0.678 | -0.025 | -0.39 | -1.148 | 0.037 | 0.045 | 0.045 | 0 | 0.55 | 0.34 | 0.63 | 0.63 | 0.61 | 0.64 |
| 1 | 245608819 | rs1093937 | -0.647 | 0.145 | -0.295 | -0.919 | -0.01 | -0.027 | -0.027 | 0 | 0.95 | 0.92 | 0.96 | 1 | 0.95 | 0.99 |
| 1 | 245611389 | rs819744 | -0.647 | 0.145 | -0.303 | -0.919 | -0.01 | -0.026 | -0.026 | 0 | 0.95 | 0.92 | 0.96 | 1 | 0.95 | 0.99 |
| 1 | 245612896 | rs12094249 | 0.842 | 0.503 | -1.163 | -0.425 | -0.013 | -0.048 | -0.048 | 0.042 | 0.29 | 0.07 | 0.28 | 0.28 | 0.27 | 0.27 |
| 1 | 245613630 | rs1126065 | 0.012 | 0.891 | -0.36 | -0.242 | 0.001 | 0.016 | 0.016 | 0 | 0.23 | 0.28 | 0.3 | 0.33 | 0.2 | 0.28 |
| 1 | 245616799 | rs10924200 | 1.225 | 0.974 | -0.84 | -0.06 | -0.001 | -0.08 | -0.08 | 0 | 0.12 | 0.02 | 0.18 | 0.27 | 0.08 | 0.16 |
| 1 | 245623061 | rs1173656 | 0.112 | 0.981 | -0.617 | -0.257 | 0.001 | 0.012 | 0.012 | 0 | 0.19 | 0.26 | 0.23 | 0.28 | 0.22 | 0.2 |
| 1 | 245625652 | rs669314 | 0.64 | 0.834 | -0.652 | -0.243 | -0.008 | 0.001 | 0.001 | 0 | 0.66 | 0.71 | 0.63 | 0.66 | 0.63 | 0.64 |
| 1 | 245630414 | rs523744 | -0.07 | 0.648 | -0.909 | 0.028 | -0.014 | -0.047 | -0.047 | 0 | 0.8 | 0.88 | 0.73 | 0.66 | 0.8 | 0.71 |
| 1 | 245633002 | rs1093920 | -0.281 | 0.769 | -0.888 | 0.037 | -0.027 | -0.056 | -0.056 | 0 | 0.81 | 0.9 | 0.73 | 0.66 | 0.8 | 0.71 |
| 1 | 245634069 | rs12078625 | 0.426 | 0.839 | -0.613 | -0.005 | 0.004 | -0.004 | -0.004 | 0.095 | 0.06 | 0.08 | 0.07 | 0.05 | 0.11 | 0.12 |
| 1 | 245634267 | rs1173660 | -0.274 | 0.83 | -0.529 | -0.009 | 0.023 | -0.018 | -0.018 | 0 | 0.93 | 0.86 | 0.92 | 0.95 | 0.89 | 0.88 |
| 1 | 245636059 | rs1538472 | -0.492 | 0.627 | -0.396 | -0.156 | -0.004 | 0.001 | 0.001 | 0 | 0.46 | 0.43 | 0.52 | 0.56 | 0.51 | 0.46 |
| 1 | 245636915 | rs6428922 | 0.532 | 0.077 | 0 | 0.528 | -0.02 | 0.002 | 0.002 | 0 | 0.75 | 0.66 | 0.64 | 0.76 | 0.76 | 0.78 |
| 1 | 245637039 | rs6428923 | -0.664 | 0.179 | -0.12 | 1.205 | -0.004 | 0.059 | 0.059 | 0 | 0.08 | 0.11 | 0.06 | 0.23 | 0.05 | 0.16 |
| 1 | 245641147 | rs6687098 | 0.482 | 0.067 | -0.343 | 0.979 | 0.01 | -0.014 | -0.014 | 0 | 0.12 | 0.07 | 0.08 | 0.14 | 0.06 | 0.1 |
| 1 | 245642219 | rs1538471 | 1.707 | 0.38 | -0.09 | 1.114 | -0.001 | 0.02 | 0.02 | 0.066 | 0.54 | 0.36 | 0.5 | 0.41 | 0.52 | 0.53 |
| 1 | 245645242 | rs10924209 | -1.062 | 0.119 | 0.222 | 1.259 | -0.027 | 0.042 | 0.042 | 0 | 0.23 | 0.51 | 0.29 | 0.3 | 0.19 | 0.2 |
| 1 | 245647726 | rs7523252 | -0.869 | 0.316 | -0.302 | 0.78 | -0.012 | 0.048 | 0.048 | 0.124 | 0.3 | 0.5 | 0.24 | 0.35 | 0.26 | 0.23 |
| 1 | 245649651 | rs1111815 | 0.976 | 0.59 | -0.401 | 0.108 | -0.001 | 0.006 | 0.006 | 0 | 0.66 | 0.29 | 0.71 | 0.65 | 0.65 | 0.62 |
| 1 | 245649841 | rs12089406 | -0.647 | 1.15 | 0.121 | -0.294 | 0.039 | -0.018 | -0.018 | 0.537 | 0.13 | 0.22 | 0.12 | 0.08 | 0.15 | 0.24 |
| 1 | 245653992 | rs1538467 | -0.029 | 0.84 | -0.174 | -0.442 | 0.001 | 0.004 | 0.004 | 0.805 | 0.48 | 0.46 | 0.63 | 0.8 | 0.49 | 0.59 |
| 1 | 245654625 | rs2363902 | 0.321 | 0.878 | -0.506 | -0.748 | -0.054 | 0.226 | 0.226 | 1.295 | 0.43 | 0.08 | 0.28 | 0.1 | 0.45 | 0.37 |
| 1 | 245655241 | rs1556874 | 1.219 | 0.544 | 0.253 | 0.014 | 0.071 | 0.047 | 0.047 | 0.857 | 0.92 | 0.64 | 0.93 | 0.94 | 0.95 | 0.96 |
| 1 | 245668132 | rs6428926 | -0.445 | 0.39 | -0.659 | -0.333 | -0.001 | -0.016 | -0.016 | 0.269 | 0.32 | 0.32 | 0.29 | 0.1 | 0.23 | 0.14 |
| 1 | 245681650 | rs6662614 | -0.561 | 0.583 | -1.294 | -0.414 | 0.003 | 0.126 | 0.126 | 0 | 0.34 | 0.13 | 0.29 | 0.07 | 0.31 | 0.16 |
| 1 | 245689787 | rs6698390 | -0.296 | 0.875 | -0.833 | 0.272 | 0 | 0.004 | 0.004 | 0 | 0.2 | 0.18 | 0.2 | 0.42 | 0.2 | 0.34 |

| | | | | | | | | | | | | | | | | |
|---|---|---|---|---|---|---|---|---|---|---|---|---|---|---|---|---|
| 1 | 245697260 | rs10924235 | -0.468 | 0.943 | -1.425 | 0.307 | -0.06 | -0.165 | -0.165 | 0 | 0.2 | 0.02 | 0.32 | 0.43 | 0.25 | 0.39 |
| 1 | 245699085 | rs10802219 | 0.203 | 0.892 | -1.217 | -1.008 | -0.004 | 0.078 | 0.078 | 0 | 0.53 | 0.72 | 0.36 | 0.02 | 0.48 | 0.45 |
| 1 | 245703284 | rs7536825 | -0.043 | 0.808 | -1.022 | -0.023 | 0.001 | 0 | 0 | 0 | 0.53 | 0.47 | 0.43 | 0.49 | 0.56 | 0.42 |
| 1 | 245703819 | rs7551348 | -0.247 | 0.466 | -1.082 | 0.225 | -0.005 | -0.005 | -0.005 | 0.209 | 0.42 | 0.45 | 0.39 | 0.51 | 0.45 | 0.51 |
| 1 | 245703990 | rs9919234 | -0.034 | 0.282 | -0.905 | 0.31 | -0.001 | 0.001 | 0.001 | 0.197 | 0.61 | 0.68 | 0.55 | 0.64 | 0.58 | 0.63 |
| 1 | 245711056 | rs12133360 | 0.172 | 0.234 | -1.079 | 0.304 | -0.046 | 0.02 | 0.02 | 0 | 0.34 | 0.09 | 0.27 | 0.32 | 0.36 | 0.19 |
| 1 | 245711179 | rs10802220 | 0.046 | 0.112 | -1.019 | 0.482 | 0 | 0.032 | 0.032 | 0 | 0.37 | 0.47 | 0.48 | 0.48 | 0.39 | 0.49 |
| 1 | 245715272 | rs1771503 | -0.1 | 0.329 | -0.84 | 0.556 | 0.063 | 0.019 | 0.019 | 0 | 0.05 | 0.31 | 0.03 | 0.01 | 0.03 | 0.08 |
| 1 | 245717009 | rs947106 | 0.245 | 0.177 | -1.418 | 0.909 | -0.048 | 0.096 | 0.096 | 0 | 0.56 | 0.89 | 0.55 | 0.72 | 0.55 | 0.49 |
| 1 | 245717698 | rs1621632 | -0.582 | 0.22 | -0.792 | 1.061 | -0.009 | -0.012 | -0.012 | 0 | 0.77 | 0.89 | 0.79 | 0.79 | 0.78 | 0.73 |
| 1 | 245720262 | rs9628692 | -0.048 | 0.382 | -0.361 | 0.627 | 0.003 | 0.041 | 0.041 | 0.063 | 0.67 | 0.49 | 0.59 | 0.49 | 0.68 | 0.63 |
| 1 | 245722181 | rs1715780 | 0.051 | 0.303 | -0.293 | 0.118 | -0.004 | -0.033 | -0.033 | 0 | 0.24 | 0.31 | 0.25 | 0.13 | 0.28 | 0.3 |
| 1 | 245725714 | rs6687026 | -0.487 | 0.166 | 0.133 | 1.116 | 0.003 | 0.192 | 0.192 | 0 | 0.11 | 0.27 | 0.26 | 0.57 | 0.1 | 0.18 |
| 1 | 245727474 | rs10924243 | 0.082 | -0.192 | -0.045 | -0.38 | 0.012 | 0.016 | 0.016 | 0 | 0.62 | 0.35 | 0.71 | 0.85 | 0.73 | 0.62 |
| 1 | 245727532 | rs1715778 | -0.757 | -0.466 | 0.711 | 0.145 | 0.091 | 0.232 | 0.232 | 0 | 0.07 | 0.41 | 0.22 | 0.2 | 0.07 | 0.16 |
| 1 | 245728083 | rs1771492 | -0.882 | -0.638 | -0.112 | -0.726 | 0.025 | 0.11 | 0.11 | 0.075 | 0.43 | 0.81 | 0.47 | 0.28 | 0.34 | 0.38 |
| 1 | 245730311 | rs6690829 | 0.06 | -0.606 | 0.05 | -0.775 | -0.002 | -0.039 | -0.039 | 0.013 | 0.3 | 0.28 | 0.22 | 0.12 | 0.24 | 0.22 |
| 1 | 245733608 | rs10924245 | 0.307 | -0.574 | -0.208 | -0.773 | -0.006 | -0.011 | -0.011 | 0.763 | 0.08 | 0.19 | 0.11 | 0.08 | 0.06 | 0.11 |
| 1 | 245734311 | rs1616300 | -1.656 | -0.399 | -0.972 | -1.056 | -0.069 | 0.104 | 0.104 | 0.032 | 0.7 | 0.95 | 0.73 | 0.88 | 0.68 | 0.64 |
| 1 | 245737772 | rs6699551 | 0.383 | -1.105 | -0.189 | -0.891 | 0.014 | -0.002 | -0.002 | 0 | 0.16 | 0.15 | 0.07 | 0 | 0.09 | 0.03 |
| 1 | 245743584 | rs6663746 | -0.611 | -0.912 | 0.005 | -0.615 | 0 | -0.007 | -0.007 | 0 | 0.45 | 0.53 | 0.4 | 0.27 | 0.34 | 0.3 |
| 1 | 245745337 | rs1620343 | -1.013 | -0.596 | -0.765 | -0.369 | 0.014 | 0.115 | 0.115 | 0 | 0.54 | 0.83 | 0.51 | 0.37 | 0.39 | 0.39 |
| 1 | 245753173 | rs7550851 | -0.481 | -0.685 | 0.11 | -0.138 | -0.033 | -0.045 | -0.045 | 0.887 | 0.17 | 0.26 | 0.07 | 0.15 | 0.09 | 0.08 |
| 1 | 245762071 | rs1771528 | -1.306 | -0.161 | 0.38 | 0.422 | 0.017 | 0.004 | 0.004 | 0 | 0.49 | 0.62 | 0.42 | 0.48 | 0.47 | 0.46 |
| 1 | 245762190 | rs1771527 | -0.87 | -0.239 | 0.583 | 0.474 | -0.004 | 0 | 0 | 0 | 0.39 | 0.45 | 0.34 | 0.38 | 0.38 | 0.39 |
| 1 | 245762525 | rs1572893 | -0.59 | -0.216 | 0.434 | 0.485 | 0.007 | -0.016 | -0.016 | 0 | 0.36 | 0.23 | 0.32 | 0.38 | 0.38 | 0.38 |
| 1 | 245766651 | rs947103 | -0.745 | -0.252 | 0.359 | 0.47 | -0.009 | -0.03 | -0.03 | 0 | 0.34 | 0.19 | 0.24 | 0.45 | 0.39 | 0.37 |
| 1 | 245767748 | rs12406586 | -0.769 | -0.436 | 0.565 | 1.013 | -0.022 | 0.034 | 0.034 | 0.042 | 0.15 | 0.34 | 0.34 | 0.23 | 0.1 | 0.16 |
| 1 | 245767834 | rs12401628 | -0.676 | -0.339 | 0.086 | 1.106 | -0.01 | 0.013 | 0.013 | 0 | 0.14 | 0.19 | 0.33 | 0.23 | 0.11 | 0.16 |
| 1 | 245770642 | rs10924249 | 0.269 | -0.348 | -0.445 | 1.137 | 0.009 | -0.085 | -0.085 | 0 | 0.2 | 0.06 | 0.12 | 0.33 | 0.21 | 0.23 |
| 1 | 245772529 | rs7555351 | -0.021 | -0.347 | -0.415 | 1.13 | -0.001 | 0.006 | 0.006 | 0 | 0.08 | 0.09 | 0.22 | 0.2 | 0.08 | 0.08 |
| 1 | 245773041 | rs12041565 | 0.304 | -0.262 | -0.508 | 1.143 | 0.014 | -0.086 | -0.086 | 0.041 | 0.2 | 0.05 | 0.12 | 0.33 | 0.2 | 0.23 |
| 1 | 245773223 | rs1771523 | -1.28 | -0.215 | 0.023 | 0.439 | 0.024 | 0.023 | 0.023 | 0 | 0.49 | 0.61 | 0.56 | 0.65 | 0.42 | 0.51 |
| 1 | 245774993 | rs10754458 | -1.045 | -0.374 | -0.058 | 0.553 | -0.01 | -0.004 | -0.004 | 0 | 0.44 | 0.45 | 0.5 | 0.65 | 0.36 | 0.48 |
| 1 | 245778984 | rs1771522 | -0.957 | 0.615 | 0.095 | 0.562 | 0.008 | 0.019 | 0.019 | 0.839 | 0.35 | 0.46 | 0.35 | 0.42 | 0.38 | 0.44 |
| 1 | 245779716 | rs1715789 | -1.214 | 0.459 | 0.123 | 0.159 | 0.009 | -0.007 | -0.007 | 0.1 | 0.54 | 0.57 | 0.72 | 0.89 | 0.7 | 0.69 |
| 1 | 245785508 | rs10754459 | -0.04 | 0.71 | 0.25 | 0.29 | -0.004 | -0.006 | -0.006 | 0 | 0.2 | 0.16 | 0.17 | 0.4 | 0.23 | 0.33 |
| 1 | 245790762 | rs1715815 | -0.172 | 0.674 | 0.484 | 0.531 | -0.025 | -0.057 | -0.057 | 0 | 0.29 | 0.17 | 0.29 | 0.37 | 0.38 | 0.22 |
| 1 | 245804664 | rs10924270 | -1.011 | 0.601 | 0.883 | 0.372 | -0.014 | -0.024 | -0.024 | 0.466 | 0.39 | 0.69 | 0.45 | 0.43 | 0.48 | 0.39 |
| 1 | 245806987 | rs12084726 | -0.556 | 0.356 | 1.174 | 0.221 | -0.001 | -0.019 | -0.019 | 3.499 | 0.18 | 0.25 | 0.12 | 0.02 | 0.18 | 0.06 |
| 1 | 245807983 | rs6663144 | -1.258 | 0.693 | 1.051 | 0.236 | 0.001 | -0.011 | -0.011 | 0.079 | 0.48 | 0.69 | 0.46 | 0.45 | 0.53 | 0.42 |
| 1 | 245808095 | rs10802234 | -0.898 | 0.936 | 1.245 | 0.332 | 0.05 | 0.03 | 0.03 | 0 | 0.05 | 0.2 | 0.12 | 0.07 | 0.08 | 0.14 |
| 1 | 245811646 | rs10158765 | -0.245 | 0.799 | 1.146 | 0.57 | 0.003 | -0.031 | -0.031 | 0 | 0.31 | 0.43 | 0.33 | 0.25 | 0.35 | 0.34 |
| 1 | 245813813 | rs6701020 | 0.202 | 1.176 | 1.063 | 0.615 | -0.003 | 0.023 | 0.023 | 0 | 0.52 | 0.33 | 0.49 | 0.35 | 0.55 | 0.28 |
| 1 | 245816347 | rs7527246 | -2.001 | 1.195 | 1.66 | 0.67 | 0.071 | 0.076 | 0.076 | 0.054 | 0.09 | 0.51 | 0.19 | 0.1 | 0.09 | 0.19 |
| 1 | 245817499 | rs4658792 | 0.615 | 1.201 | 0.781 | 0.586 | -0.079 | 0.151 | 0.151 | 0 | 0.47 | 0.03 | 0.4 | 0.72 | 0.47 | 0.5 |
| 1 | 245818715 | rs6428942 | -1.247 | 1.129 | 1.031 | 0.673 | -0.004 | 0.001 | 0.001 | 0.212 | 0.13 | 0.09 | 0.2 | 0.51 | 0.14 | 0.45 |
| 1 | 245818820 | rs12410304 | -0.287 | 1.125 | 0.868 | 0.678 | -0.02 | -0.053 | -0.053 | 0 | 0.1 | 0.02 | 0.11 | 0.46 | 0.09 | 0.34 |
| 1 | 245819198 | rs6682995 | -0.007 | 1.199 | 1.322 | 0.581 | -0.003 | -0.007 | -0.007 | 0 | 0.57 | 0.67 | 0.59 | 0.35 | 0.6 | 0.4 |
| 1 | 245819228 | rs12746314 | 1.23 | 1.175 | 1.147 | 0.599 | 0 | -0.007 | -0.007 | 0 | 0.4 | 0.13 | 0.38 | 0.29 | 0.42 | 0.24 |
| 1 | 245826358 | rs10924294 | -1.535 | 1.151 | 1.138 | 0.181 | 0.008 | -0.042 | -0.042 | 0 | 0.17 | 0.62 | 0.22 | 0.06 | 0.18 | 0.17 |
| 1 | 245826977 | rs1538961 | -0.584 | 1.174 | 1.134 | 0.077 | 0 | 0.016 | 0.016 | 0 | 0.66 | 0.65 | 0.7 | 0.81 | 0.68 | 0.73 |
| 1 | 245835059 | rs10924299 | -0.124 | 1.086 | 0.917 | 0.296 | -0.028 | 0.102 | 0.102 | 0 | 0.58 | 0.77 | 0.61 | 0.8 | 0.56 | 0.68 |
| 1 | 245837801 | rs6674082 | 0.01 | 1.21 | 0.649 | 0.322 | 0 | -0.028 | -0.028 | 0 | 0.57 | 0.54 | 0.69 | 0.81 | 0.56 | 0.68 |
| 1 | 245840409 | rs2282381 | -1.234 | 1.032 | 0.477 | 1.359 | -0.001 | 0.084 | 0.084 | 0 | 0.13 | 0.15 | 0.27 | 0.49 | 0.12 | 0.42 |
| 1 | 245846319 | rs4658466 | 0.331 | 1.421 | 0.588 | 1.059 | 0.007 | 0.023 | 0.023 | 0 | 0.68 | 0.57 | 0.72 | 0.55 | 0.73 | 0.49 |
| 1 | 245847034 | rs10924303 | -0.269 | 1.656 | 0.632 | 1.262 | -0.017 | -0.002 | -0.002 | 0 | 0.06 | 0.02 | 0.1 | 0.06 | 0.2 | 0.04 |
| 1 | 245850095 | rs871840 | -1.086 | 1.391 | 0.897 | 0.901 | 0.027 | -0.087 | -0.087 | 0 | 0.16 | 0.23 | 0.16 | 0.02 | 0.24 | 0.05 |
| 1 | 245866511 | rs3737918 | -1.457 | 0.606 | 1.012 | 1.402 | -0.001 | 0.067 | 0.067 | 0.204 | 0.29 | 0.55 | 0.36 | 0.55 | 0.27 | 0.49 |
| 1 | 245868363 | rs2000101 | -1.278 | 0.185 | 1.151 | 1.46 | -0.01 | 0.032 | 0.032 | 0 | 0.26 | 0.38 | 0.31 | 0.42 | 0.24 | 0.53 |
| 1 | 245873836 | rs1538959 | -1.345 | 0.102 | 1.126 | 1.468 | -0.002 | 0.075 | 0.075 | 0.357 | 0.25 | 0.55 | 0.3 | 0.45 | 0.21 | 0.45 |
| 1 | 245882414 | rs7524184 | -1.076 | 0.179 | 1.056 | 1.387 | -0.011 | 0.101 | 0.101 | 0.259 | 0.23 | 0.41 | 0.29 | 0.42 | 0.17 | 0.37 |
| 1 | 245884152 | rs10802251 | -1.308 | 0.16 | 1.023 | 1.352 | 0 | 0.095 | 0.095 | 0.118 | 0.25 | 0.44 | 0.29 | 0.48 | 0.21 | 0.41 |
| 1 | 245901545 | rs10802263 | -1.641 | -0.151 | 0.98 | 0.944 | -0.014 | -0.003 | -0.003 | 0 | 0.45 | 0.47 | 0.39 | 0.45 | 0.29 | 0.47 |
| 1 | 245904087 | rs10802264 | -1.612 | -0.111 | 1.151 | 0.955 | 0.013 | 0 | 0 | 0 | 0.47 | 0.58 | 0.39 | 0.51 | 0.33 | 0.5 |
| 1 | 245904391 | rs10802265 | -1.606 | -0.123 | 1.131 | 0.952 | -0.015 | -0.002 | -0.002 | 0.023 | 0.45 | 0.47 | 0.38 | 0.44 | 0.28 | 0.46 |
| 1 | 245907373 | rs12121171 | -0.792 | -0.48 | 1.716 | 0.599 | -0.041 | -0.14 | -0.14 | 0 | 0.19 | 0.5 | 0.11 | 0 | 0.1 | 0.07 |
| 1 | 245908008 | rs2791359 | -1.03 | -0.477 | 1.492 | 0.623 | -0.016 | -0.099 | -0.099 | 0 | 0.2 | 0.5 | 0.1 | 0.07 | 0.14 | 0.1 |
| 1 | 245911746 | rs2791362 | -1.022 | -0.497 | 1.505 | 0.679 | -0.015 | -0.035 | -0.035 | 0 | 0.2 | 0.48 | 0.25 | 0.16 | 0.15 | 0.1 |

| | | | | | | | | | | | | | | | | |
|---|---|---|---|---|---|---|---|---|---|---|---|---|---|---|---|---|
| 1 | 245912840 | rs8025 | -1.102 | -0.447 | 1.421 | 0.752 | -0.018 | 0.025 | 0.025 | 0 | 0.31 | 0.32 | 0.34 | 0.47 | 0.22 | 0.39 |
| 1 | 245914284 | rs2362932 | -1.535 | -0.446 | 1.399 | 0.745 | -0.015 | 0.036 | 0.036 | 0 | 0.35 | 0.47 | 0.35 | 0.55 | 0.27 | 0.43 |
| 1 | 245920863 | rs2791372 | -0.913 | -0.45 | 1.279 | 0.86 | -0.039 | 0.087 | 0.087 | 0 | 0.28 | 0.42 | 0.3 | 0.47 | 0.19 | 0.38 |
| 1 | 245922680 | rs2791376 | -1.726 | -0.766 | 1.052 | 0.789 | -0.007 | 0 | 0 | 0 | 0.46 | 0.55 | 0.51 | 0.51 | 0.26 | 0.44 |
| 1 | 245924864 | rs12045474 | -1.164 | -0.62 | 0.66 | 0.836 | 0.013 | -0.028 | -0.028 | 0 | 0.3 | 0.18 | 0.44 | 0.64 | 0.24 | 0.43 |
| 1 | 245926038 | rs12088988 | -1.717 | -0.701 | 0.592 | 0.596 | 0.031 | -0.021 | -0.021 | 4.612 | 0.34 | 0.19 | 0.42 | 0.46 | 0.22 | 0.31 |
| 1 | 245931249 | rs1316384 | 0.047 | -0.745 | 0.416 | 0.306 | 0.002 | -0.004 | -0.004 | 0 | 0.11 | 0.08 | 0.17 | 0 | 0.08 | 0.05 |
| 1 | 245932939 | rs2362928 | -1.496 | -1.04 | 0.29 | 0.5 | 0.077 | -0.014 | -0.014 | 0.332 | 0.29 | 0.12 | 0.25 | 0.32 | 0.14 | 0.24 |
| 1 | 245936738 | rs2817494 | -1.655 | -1.253 | -0.085 | 0.088 | 0.103 | 0.162 | 0.162 | 0 | 0.41 | 0.84 | 0.44 | 0.88 | 0.24 | 0.49 |
| 1 | 245939067 | rs12567813 | -0.063 | -1.252 | 0.606 | 0.202 | -0.001 | 0.076 | 0.076 | 0.008 | 0.18 | 0.29 | 0.16 | 0.49 | 0.16 | 0.28 |
| 1 | 245940014 | rs2817487 | -2.216 | -1.2 | 0.386 | 0.083 | 0.105 | 0.281 | 0.281 | 0 | 0.52 | 0.91 | 0.59 | 0.88 | 0.32 | 0.54 |
| 1 | 245940334 | rs6687221 | -0.931 | -0.194 | 0.34 | 0 | -0.002 | 0 | 0 | 0 | 0.18 | 0.17 | 0.4 | 0.13 | 0.22 | 0.21 |
| 1 | 245944439 | rs2262933 | -1.456 | -0.726 | 0.378 | 0.022 | 0.089 | 0.137 | 0.137 | 0.014 | 0.41 | 0.88 | 0.29 | 0.78 | 0.25 | 0.48 |
| 1 | 245946320 | rs1317002 | -0.955 | -0.138 | 1.163 | 0.269 | -0.016 | 0.09 | 0.09 | 0 | 0.25 | 0.45 | 0.18 | 0.5 | 0.2 | 0.3 |
| 1 | 245946830 | rs1317297 | -0.529 | 0.488 | 1.346 | -0.221 | 0 | -0.022 | -0.022 | 0 | 0.12 | 0.09 | 0.16 | 0 | 0.07 | 0.05 |
| 1 | 245950642 | rs7555819 | -2.087 | 0.795 | 1.336 | -0.063 | 0 | -0.005 | -0.005 | 3.588 | 0.22 | 0.74 | 0.31 | 0.2 | 0.22 | 0.19 |
| 1 | 245953416 | rs2791384 | -1.378 | 0.937 | 0.777 | 0.237 | 0.052 | 0.037 | 0.037 | 0 | 0.15 | 0.5 | 0.29 | 0.19 | 0.16 | 0.08 |
| 1 | 245959395 | rs2791391 | -1.201 | 1.442 | 0.243 | -0.092 | 0.026 | -0.133 | -0.133 | 0 | 0.17 | 0.46 | 0.11 | 0 | 0.15 | 0.12 |
| 1 | 245964761 | rs2791396 | -1.56 | 1.36 | 0.123 | -0.035 | 0.004 | -0.083 | -0.083 | 0.641 | 0.21 | 0.44 | 0.17 | 0.1 | 0.18 | 0.17 |
| 1 | 245965551 | rs2791398 | -0.034 | 1.99 | -0.216 | -0.168 | -0.049 | 0.001 | 0.001 | 0.062 | 0.91 | 0.99 | 0.83 | 0.9 | 0.86 | 0.93 |
| 1 | 245966242 | rs2791399 | -1.23 | 1.169 | 0.139 | -0.131 | 0.024 | -0.132 | -0.132 | 0 | 0.17 | 0.38 | 0.1 | 0 | 0.15 | 0.11 |
| 1 | 245966754 | rs2791400 | -0.363 | 0.534 | -1.241 | 0.34 | 0.026 | 0.022 | 0.022 | 0.026 | 0.56 | 0.91 | 0.7 | 0.39 | 0.67 | 0.51 |
| 1 | 245969011 | rs2817510 | 1.002 | 0.788 | -0.246 | 1.26 | -0.025 | -0.037 | -0.037 | 0 | 0.89 | 0.97 | 0.77 | 0.82 | 0.88 | 0.81 |
| 1 | 245970462 | rs12038196 | -1.541 | 0.692 | -0.49 | 0.954 | -0.018 | 0.039 | 0.039 | 0 | 0.18 | 0.02 | 0.2 | 0.1 | 0.15 | 0.18 |
| 1 | 245970550 | rs4654285 | -1.233 | 0.396 | 0.237 | 1.864 | -0.009 | 0.048 | 0.048 | 0 | 0.06 | 0.13 | 0.13 | 0.2 | 0.04 | 0.15 |
| 1 | 245973612 | rs2069207 | 1.954 | 0.593 | 0.819 | 2.103 | 0.004 | 0.009 | 0.009 | 2.104 | 0.65 | 0.57 | 0.48 | 0.46 | 0.61 | 0.45 |
| 1 | 245973740 | rs2069206 | 1.706 | -0.555 | 1.861 | 2.194 | -0.005 | 0.02 | 0.02 | 0 | 0.81 | 0.59 | 0.66 | 0.81 | 0.84 | 0.74 |
| 1 | 245977467 | rs10802268 | 2.502 | -0.541 | 2.198 | 2.56 | -0.001 | 0.002 | 0.002 | 0.035 | 0.74 | 0.28 | 0.54 | 0.56 | 0.81 | 0.62 |
| 1 | 245979357 | rs12031535 | -1.015 | -0.513 | 1.931 | 2.571 | -0.025 | -0.055 | -0.055 | 0 | 0.12 | 0.04 | 0.23 | 0.2 | 0.11 | 0.2 |
| 1 | 245986815 | rs12030297 | -1.842 | -0.519 | 2.481 | 2.575 | -0.005 | 0.021 | 0.021 | 0 | 0.15 | 0.23 | 0.27 | 0.26 | 0.12 | 0.24 |
| 1 | 245989865 | rs6700401 | 1.997 | -0.531 | 2.945 | 2.553 | 0.038 | 0.189 | 0.189 | 0.827 | 0.9 | 0.57 | 0.83 | 0.82 | 0.94 | 0.86 |
| 1 | 245992363 | rs4654127 | 2.34 | -0.545 | 3.035 | 2.6 | -0.008 | 0.104 | 0.104 | 0.041 | 0.78 | 0.45 | 0.58 | 0.63 | 0.81 | 0.66 |
| 1 | 245992462 | rs12407509 | -1.031 | -0.531 | 2.894 | 2.542 | -0.01 | -0.032 | -0.032 | 0 | 0.12 | 0.07 | 0.24 | 0.18 | 0.13 | 0.2 |
| 1 | 245994035 | rs7514038 | 2.201 | -0.482 | 3.122 | 2.567 | 0.024 | 0.266 | 0.266 | 3.95 | 0.9 | 0.48 | 0.69 | 0.68 | 0.94 | 0.85 |
| 1 | 245995920 | rs2153409 | 2.357 | -0.49 | 3.112 | 2.489 | 0.004 | 0.19 | 0.19 | 4.662 | 0.89 | 0.43 | 0.68 | 0.69 | 0.94 | 0.85 |
| 1 | 245996629 | rs10754474 | 2.57 | -0.418 | 3.131 | 2.581 | -0.003 | 0.108 | 0.108 | 0.156 | 0.76 | 0.47 | 0.56 | 0.5 | 0.8 | 0.62 |
| 1 | 246000684 | rs12038930 | -1.395 | -0.338 | 2.509 | 2.53 | -0.022 | -0.031 | -0.031 | 0 | 0.14 | 0.06 | 0.13 | 0.19 | 0.14 | 0.23 |
| 1 | 246003615 | rs12137129 | -1.193 | -0.551 | 2.072 | 2.465 | -0.01 | -0.067 | -0.067 | 0.645 | 0.11 | 0.01 | 0.26 | 0.28 | 0.08 | 0.15 |
| 1 | 246003671 | rs9660719 | -0.795 | -1.054 | 2.011 | 2.221 | -0.031 | 0.101 | 0.101 | 0 | 0.16 | 0.23 | 0.07 | 0.4 | 0.1 | 0.28 |
| 1 | 246004257 | rs2362913 | 1.976 | -0.897 | 1.267 | 1.64 | 0.051 | 0.177 | 0.177 | 0 | 0.55 | 0.25 | 0.5 | 0.13 | 0.67 | 0.33 |
| 1 | 246014333 | rs10924361 | -1.702 | -0.71 | 2.199 | 2.886 | -0.05 | 0.176 | 0.176 | 0 | 0.17 | 0.35 | 0.27 | 0.59 | 0.1 | 0.24 |
| 1 | 246014758 | rs9729771 | -1.859 | -0.633 | 2.571 | 2.944 | 0.007 | 0.037 | 0.037 | 0 | 0.15 | 0.31 | 0.18 | 0.16 | 0.09 | 0.22 |
| 1 | 246015491 | rs9729891 | -1.9 | -0.788 | 2.591 | 2.948 | -0.001 | 0.042 | 0.042 | 0 | 0.15 | 0.31 | 0.19 | 0.16 | 0.09 | 0.23 |
| 1 | 246022159 | rs2185369 | 1.551 | -1.001 | 2.768 | 3.073 | -0.014 | 0.034 | 0.034 | 0 | 0.82 | 0.75 | 0.73 | 0.37 | 0.88 | 0.64 |
| 1 | 246027410 | rs10924366 | -0.912 | -1.002 | 2.474 | 3.205 | 0.011 | 0.012 | 0.012 | 0 | 0.08 | 0.03 | 0.11 | 0.46 | 0.07 | 0.16 |
| 1 | 246030183 | rs12038045 | 2.756 | -0.944 | 2.052 | 2.479 | 0.029 | 0.036 | 0.036 | 0 | 0.6 | 0.31 | 0.52 | 0.19 | 0.72 | 0.39 |
| 1 | 246038016 | rs12123988 | -2.219 | -0.492 | 1.967 | 2.516 | -0.002 | 0.001 | 0.001 | 0 | 0.15 | 0.14 | 0.16 | 0.17 | 0.11 | 0.32 |
| 1 | 246053770 | rs13374895 | -1.723 | -0.604 | 1.969 | 2.512 | -0.006 | 0.001 | 0.001 | 0 | 0.12 | 0.13 | 0.09 | 0.13 | 0.07 | 0.21 |
| 1 | 246058909 | rs4654068 | -1.886 | -0.812 | 2.195 | 2.49 | -0.03 | 0.034 | 0.034 | 0 | 0.11 | 0.17 | 0.09 | 0.17 | 0.07 | 0.28 |
| 1 | 246063859 | rs2362255 | 1.628 | -0.2 | 2.497 | 2.886 | 0.039 | 0.191 | 0.191 | 0 | 0.87 | 0.62 | 0.83 | 0.63 | 0.86 | 0.79 |
| 1 | 246083889 | rs10924388 | -1.734 | -0.34 | 2.317 | 2.824 | -0.028 | 0.107 | 0.107 | 0 | 0.13 | 0.28 | 0.2 | 0.28 | 0.08 | 0.15 |
| 1 | 246090818 | rs2275312 | 1.201 | -0.415 | 2.03 | 2.391 | 0.006 | 0.132 | 0.132 | 0 | 0.88 | 0.81 | 0.86 | 0.48 | 0.88 | 0.79 |
| 1 | 246097238 | rs1933525 | 1.25 | -0.405 | 2.176 | 2.377 | 0.023 | 0.248 | 0.248 | 0 | 0.88 | 0.66 | 0.86 | 0.5 | 0.89 | 0.8 |
| 1 | 246097984 | rs7355113 | -1.179 | -0.358 | 1.617 | 2.401 | 0.004 | 0.095 | 0.095 | 0 | 0.11 | 0.13 | 0.13 | 0.51 | 0.11 | 0.21 |
| 1 | 246110573 | rs10802300 | 1.967 | -0.641 | 0.688 | 2.133 | 0.056 | -0.101 | -0.101 | 0 | 0.7 | 0.09 | 0.74 | 0.47 | 0.81 | 0.64 |
| 1 | 246113980 | rs12093398 | -1.859 | -0.807 | 1.204 | 1.288 | -0.08 | -0.042 | -0.042 | 0.014 | 0.18 | 0.42 | 0.08 | 0.08 | 0.08 | 0.17 |
| 1 | 246117191 | rs4142898 | 1.211 | -0.752 | 1.126 | 1.245 | -0.065 | -0.043 | -0.043 | 0.226 | 0.84 | 0.66 | 0.94 | 0.92 | 0.93 | 0.85 |
| 1 | 246118050 | rs6695279 | 0.969 | -0.02 | 0.617 | 1.236 | 0 | -0.003 | -0.003 | 0 | 0.59 | 0.57 | 0.62 | 0.44 | 0.65 | 0.42 |
| 1 | 246123475 | rs4233248 | -1.041 | -0.135 | 1.077 | 0.856 | -0.018 | -0.082 | -0.082 | 0 | 0.25 | 0.41 | 0.23 | 0.13 | 0.18 | 0.28 |
| 1 | 246125283 | rs1544149 | 1.632 | -0.624 | 0.762 | 0.919 | 0.051 | 0.018 | 0.018 | 0 | 0.52 | 0.34 | 0.6 | 0.44 | 0.66 | 0.4 |
| 1 | 246125818 | rs9659850 | -0.933 | -0.227 | 1.155 | 0.993 | 0.119 | 0.392 | 0.392 | 1.406 | 0.07 | 0.36 | 0.2 | 0.4 | 0.11 | 0.22 |
| 1 | 246127777 | rs2153028 | -0.157 | -0.302 | 1.242 | 0.875 | -0.021 | -0.029 | -0.029 | 0.021 | 0.16 | 0.17 | 0.16 | 0.1 | 0.08 | 0.22 |
| 1 | 246128858 | rs10924414 | -0.317 | -0.269 | 1.096 | 0.935 | 0.041 | 0.196 | 0.196 | 0.222 | 0.06 | 0.19 | 0.19 | 0.4 | 0.11 | 0.22 |
| 1 | 246130872 | rs2066028 | -1.1 | -0.494 | 1.005 | 1.294 | -0.001 | 0.003 | 0.003 | 0 | 0.27 | 0.33 | 0.34 | 0.61 | 0.26 | 0.38 |
| 1 | 246136553 | rs11576666 | -1.103 | -0.498 | 1.088 | 1.292 | -0.003 | 0.013 | 0.013 | 0 | 0.27 | 0.32 | 0.35 | 0.61 | 0.26 | 0.38 |
| 1 | 246142250 | rs10802305 | -1.491 | -0.615 | 0.973 | 1.172 | 0.005 | -0.012 | -0.012 | 0.201 | 0.29 | 0.29 | 0.34 | 0.62 | 0.26 | 0.38 |
| 1 | 246146328 | rs1538294 | 1.668 | -0.428 | 1.382 | 1.47 | 0.039 | -0.01 | -0.01 | 0.168 | 0.68 | 0.23 | 0.71 | 0.67 | 0.8 | 0.64 |
| 1 | 246148791 | rs4654180 | -0.651 | -0.436 | 1.406 | 1.436 | 0.021 | 0.023 | 0.023 | 0 | 0.18 | 0.06 | 0.15 | 0.1 | 0.09 | 0.23 |
| 1 | 246152175 | rs13373879 | -0.929 | -0.437 | 1.438 | 1.435 | -0.019 | -0.024 | -0.024 | 0 | 0.19 | 0.18 | 0.17 | 0.1 | 0.09 | 0.23 |

| | | | | | | | | | | | | | | | | |
|---|---|---|---|---|---|---|---|---|---|---|---|---|---|---|---|---|
| 1 | 246152478 | rs10802308 | 0.84 | -0.331 | 1.51 | 1.525 | 0.028 | 0.132 | 0.132 | 0 | 0.88 | 0.6 | 0.89 | 0.77 | 0.89 | 0.88 |
| 1 | 246154179 | rs6426242 | -0.373 | -0.307 | 1.033 | 1.619 | 0.002 | 0.101 | 0.101 | 0 | 0.06 | 0.07 | 0.18 | 0.43 | 0.1 | 0.22 |
| 1 | 246156599 | rs7548294 | -0.642 | -0.457 | 1.058 | 1.336 | 0.039 | 0.045 | 0.045 | 0.019 | 0.17 | 0.02 | 0.14 | 0.11 | 0.09 | 0.23 |
| 1 | 246158895 | rs10802315 | 1.251 | 0.088 | 1.229 | 1.744 | 0.054 | 0.206 | 0.206 | 0 | 0.88 | 0.51 | 0.89 | 0.77 | 0.89 | 0.88 |
| 1 | 246170087 | rs946145 | -1.625 | 0.334 | 0.202 | 1.61 | -0.017 | -0.034 | -0.034 | 0 | 0.16 | 0.11 | 0.23 | 0.41 | 0.19 | 0.2 |
| 1 | 246203214 | rs12084862 | -1.911 | -0.007 | 0.385 | 0.782 | 0.025 | 0.037 | 0.037 | 0.671 | 0.36 | 0.67 | 0.38 | 0.82 | 0.27 | 0.5 |
| 1 | 246204285 | rs9660598 | -1.979 | 0.009 | 0.444 | 0.784 | 0.005 | 0.024 | 0.024 | 0 | 0.36 | 0.67 | 0.41 | 0.8 | 0.34 | 0.49 |
| 1 | 246223120 | rs6693295 | -0.905 | -0.033 | 1.438 | 1.259 | -0.023 | 0.096 | 0.096 | 0.059 | 0.22 | 0.37 | 0.33 | 0.41 | 0.18 | 0.3 |
| 1 | 246226566 | rs4445429 | 2.154 | -0.133 | 1.623 | 1.151 | -0.016 | -0.09 | -0.09 | 1.526 | 0.62 | 0.55 | 0.63 | 0.05 | 0.72 | 0.38 |
| 1 | 246228225 | rs6426271 | 1.129 | -0.081 | 1.541 | 1.224 | -0.002 | 0.006 | 0.006 | 0.28 | 0.85 | 0.86 | 0.92 | 0.6 | 0.84 | 0.85 |
| 1 | 246242855 | rs2788014 | -1.648 | -0.277 | 1.428 | 0.974 | 0.021 | -0.034 | -0.034 | 0 | 0.28 | 0.76 | 0.42 | 0.42 | 0.23 | 0.39 |
| 1 | 246242908 | rs2788015 | -1.004 | -0.285 | 1.609 | 0.97 | -0.021 | 0.031 | 0.031 | 0.018 | 0.23 | 0.27 | 0.33 | 0.42 | 0.14 | 0.34 |
| 1 | 246243336 | rs10924497 | -1.925 | -0.211 | 1.613 | 0.99 | 0.111 | 0.003 | 0.003 | 0.94 | 0.08 | 0.51 | 0.14 | 0 | 0.12 | 0.1 |
| 1 | 246251363 | rs2787991 | 1.773 | -0.131 | 1.253 | 0.859 | -0.001 | 0.026 | 0.026 | 0.022 | 0.8 | 0.79 | 0.9 | 0.47 | 0.81 | 0.67 |
| 1 | 246261526 | rs7514524 | -1.765 | -0.253 | 1.238 | 0.82 | 0.018 | -0.022 | -0.022 | 0 | 0.3 | 0.72 | 0.37 | 0.56 | 0.21 | 0.53 |
| 1 | 246266221 | rs4654087 | 0.605 | -0.307 | 1.215 | 0.84 | -0.002 | 0.018 | 0.018 | 0 | 0.86 | 0.86 | 0.92 | 0.62 | 0.83 | 0.84 |
| 1 | 246270251 | rs4654197 | 1.965 | -0.326 | 1.561 | 0.768 | -0.021 | -0.103 | -0.103 | 0 | 0.61 | 0.55 | 0.62 | 0.05 | 0.71 | 0.37 |
| 1 | 246278250 | rs12084046 | -0.602 | -0.341 | 1.747 | 0.884 | -0.014 | 0.012 | 0.012 | 0 | 0.21 | 0.22 | 0.3 | 0.42 | 0.1 | 0.3 |
| 1 | 246280508 | rs7553693 | -2.113 | -0.305 | 1.689 | 0.775 | 0.188 | 0.075 | 0.075 | 3.066 | 0.05 | 0.45 | 0.08 | 0 | 0.09 | 0.05 |
| 1 | 246282263 | rs12404212 | -2.184 | -0.304 | 1.468 | 0.797 | 0.051 | 0.06 | 0.06 | 0.616 | 0.07 | 0.31 | 0.07 | 0.14 | 0.1 | 0.22 |
| 1 | 246284662 | rs6426280 | 1.925 | -0.384 | 1.303 | 0.729 | 0.013 | 0.039 | 0.039 | 0 | 0.61 | 0.32 | 0.52 | 0.17 | 0.65 | 0.53 |
| 1 | 246289111 | rs12031436 | -1.306 | -0.271 | 1.324 | 0.812 | -0.007 | 0.056 | 0.056 | 0.028 | 0.05 | 0.17 | 0.02 | 0.15 | 0.02 | 0.22 |
| 1 | 246292470 | rs10924512 | 1.021 | -0.198 | 1.293 | 0.975 | 0.007 | 0.028 | 0.028 | 0.112 | 0.84 | 0.8 | 0.88 | 0.62 | 0.75 | 0.84 |
| 1 | 246292814 | rs10924513 | -0.855 | -0.455 | 1.384 | 0.975 | -0.065 | 0.117 | 0.117 | 0 | 0.21 | 0.37 | 0.34 | 0.45 | 0.1 | 0.3 |
| 1 | 246308279 | rs3845512 | 0.494 | -0.199 | 1.014 | 1.393 | 0.012 | 0.063 | 0.063 | 0 | 0.88 | 0.82 | 0.91 | 0.39 | 0.85 | 0.73 |
| 1 | 246310585 | rs6426282 | -1.796 | -0.418 | 1.017 | 1.513 | -0.012 | 0.001 | 0.001 | 0 | 0.26 | 0.63 | 0.37 | 0.35 | 0.15 | 0.42 |
| 1 | 246310748 | rs6692380 | -1.493 | 0.043 | 0.979 | 1.674 | -0.002 | 0.001 | 0.001 | 0.339 | 0.08 | 0.06 | 0.06 | 0.27 | 0.09 | 0.26 |
| 1 | 246313723 | rs4607842 | -1.07 | -0.179 | 1.089 | 1.541 | -0.025 | -0.037 | -0.037 | 0 | 0.19 | 0.43 | 0.33 | 0.15 | 0.17 | 0.17 |
| 1 | 246329014 | rs3845516 | -2.095 | -0.195 | 0.954 | 1.627 | -0.007 | 0 | 0 | 0 | 0.28 | 0.62 | 0.39 | 0.36 | 0.23 | 0.43 |
| 1 | 246341231 | rs7531193 | -0.201 | 0.149 | 0.842 | 1.768 | 0.014 | 0.06 | 0.06 | 0 | 0.9 | 0.85 | 0.94 | 0.69 | 0.84 | 0.87 |
| 1 | 246344768 | rs10802343 | -0.794 | -0.062 | 0.922 | 1.643 | -0.065 | -0.025 | -0.025 | 0 | 0.18 | 0.35 | 0.32 | 0.15 | 0.09 | 0.18 |
| 1 | 246346297 | rs12406235 | -0.812 | -0.126 | 0.746 | 1.648 | 0.018 | -0.042 | -0.042 | 0.27 | 0.08 | 0.01 | 0.04 | 0.26 | 0.04 | 0.19 |
| 1 | 246347153 | rs6680215 | -2.036 | -0.115 | 0.87 | 1.624 | -0.024 | 0.01 | 0.01 | 0 | 0.27 | 0.54 | 0.37 | 0.36 | 0.23 | 0.43 |
| 1 | 246353934 | rs12038162 | 0.033 | -0.323 | -0.358 | 1.511 | 0.042 | -0.042 | -0.042 | 0 | 0.18 | 0.1 | 0.06 | 0.24 | 0.1 | 0.23 |
| 1 | 246355668 | rs10924537 | 0.726 | 0.073 | -0.294 | 1.163 | -0.013 | 0.018 | 0.018 | 0 | 0.66 | 0.53 | 0.8 | 0.64 | 0.7 | 0.69 |
| 1 | 246364256 | rs12406327 | -0.943 | 0.514 | -0.628 | 1.514 | -0.007 | -0.027 | -0.027 | 0 | 0.16 | 0.13 | 0.35 | 0.36 | 0.18 | 0.23 |
| 1 | 246364811 | rs12079361 | -1.495 | 0.02 | 0.31 | 1.836 | 0.001 | 0.291 | 0.291 | 2.098 | 0.08 | 0.44 | 0.04 | 0.25 | 0.01 | 0.17 |
| 1 | 246374869 | rs7340039 | -1.674 | 0.073 | 0.092 | 1.882 | -0.013 | 0.102 | 0.102 | 0.064 | 0.08 | 0.2 | 0.05 | 0.24 | 0.04 | 0.18 |
| 1 | 246375856 | rs11585648 | -1.057 | 0.072 | 0.08 | 1.848 | -0.005 | -0.029 | -0.029 | 0 | 0.06 | 0.13 | 0.05 | 0 | 0.04 | 0.01 |
| 1 | 246376182 | rs4654220 | -0.284 | 0.408 | -0.962 | 1.084 | -0.033 | -0.061 | -0.061 | 1.118 | 0.39 | 0.14 | 0.59 | 0.44 | 0.45 | 0.36 |
| 1 | 246391677 | rs4654096 | -0.367 | 0.489 | -0.717 | 1.175 | -0.001 | 0.056 | 0.056 | 0.331 | 0.16 | 0.19 | 0.33 | 0.6 | 0.14 | 0.39 |
| 1 | 246410112 | rs1361409 | -1.32 | 0.72 | -0.793 | 1.094 | -0.007 | 0.058 | 0.058 | 0.872 | 0.22 | 0.33 | 0.37 | 0.6 | 0.18 | 0.4 |
| 1 | 246414404 | rs10924625 | -0.343 | 0.716 | -0.778 | 1.088 | -0.001 | 0.057 | 0.057 | 0.331 | 0.16 | 0.19 | 0.33 | 0.6 | 0.14 | 0.39 |
| 1 | 246425984 | rs1770011 | -1.319 | 0.717 | -0.777 | 1.09 | -0.007 | 0.06 | 0.06 | 0.872 | 0.22 | 0.33 | 0.37 | 0.6 | 0.18 | 0.4 |
| 1 | 246429058 | rs12760011 | -0.836 | 1.397 | -0.983 | 1.122 | 0.009 | 0.031 | 0.031 | 0.511 | 0.18 | 0.2 | 0.37 | 0.61 | 0.22 | 0.45 |
| 1 | 246435928 | rs12042203 | -0.008 | 0.782 | -1.29 | 1.15 | -0.01 | -0.094 | -0.094 | 0 | 0.14 | 0.02 | 0.09 | 0.42 | 0.13 | 0.35 |
| 1 | 246441421 | rs12404422 | 0.194 | 0.718 | -1.568 | 1.202 | -0.029 | -0.089 | -0.089 | 0.118 | 0.17 | 0.03 | 0.26 | 0.31 | 0.16 | 0.23 |
| 1 | 246443337 | rs10754506 | -1.161 | 0.725 | -1.364 | 1.062 | 0.003 | 0.005 | 0.005 | 0.887 | 0.32 | 0.22 | 0.37 | 0.73 | 0.29 | 0.57 |
| 1 | 246444111 | rs1621854 | -1.856 | 0.444 | -2.104 | 0.266 | -0.114 | 0.044 | 0.044 | 0 | 0.8 | 0.99 | 0.81 | 0.9 | 0.7 | 0.81 |
| 1 | 246457214 | rs7516664 | 1.447 | 0.277 | -2.069 | 0.221 | -0.062 | 0.027 | 0.027 | 0 | 0.19 | 0.03 | 0.24 | 0.08 | 0.28 | 0.14 |
| 1 | 246465955 | rs1148716 | -1.361 | 0.072 | -1.471 | 0.345 | -0.008 | -0.031 | -0.031 | 0.114 | 0.39 | 0.55 | 0.49 | 0.83 | 0.34 | 0.62 |
| 1 | 246468500 | rs12410831 | 0.989 | 0.173 | -1.6 | 0.49 | -0.002 | -0.087 | -0.087 | 0 | 0.16 | 0.02 | 0.24 | 0.32 | 0.12 | 0.22 |
| 1 | 246475061 | rs12048180 | 0.62 | 0.211 | -1.516 | 0.72 | -0.001 | -0.041 | -0.041 | 0.83 | 0.14 | 0.07 | 0.09 | 0.37 | 0.14 | 0.34 |
| 1 | 246479095 | rs10924688 | 0.721 | 0.197 | -1.536 | 0.717 | -0.021 | -0.086 | -0.086 | 0 | 0.13 | 0.02 | 0.09 | 0.38 | 0.14 | 0.34 |
| 1 | 246483783 | rs10924690 | 0.814 | 0.196 | -1.335 | 0.695 | 0.034 | 0.026 | 0.026 | 0 | 0.12 | 0.23 | 0.12 | 0.18 | 0.18 | 0.15 |
| 1 | 246483863 | rs1148724 | 0.513 | 0.161 | -1.47 | 0.665 | 0.009 | -0.08 | -0.08 | 0 | 0.18 | 0.04 | 0.24 | 0.32 | 0.13 | 0.22 |
| 1 | 246495633 | rs6659024 | 0.632 | 0.216 | -1.386 | 0.689 | 0.005 | 0.005 | 0.005 | 0.499 | 0.66 | 0.71 | 0.63 | 0.27 | 0.69 | 0.43 |
| 1 | 246512218 | rs10924702 | -0.625 | 0.487 | -1.156 | 0.899 | -0.006 | 0.101 | 0.101 | 0 | 0.19 | 0.33 | 0.2 | 0.51 | 0.18 | 0.39 |
| 1 | 246521189 | rs10924704 | 1.107 | 0.542 | -1.226 | 0.888 | -0.036 | -0.038 | -0.038 | 0 | 0.11 | 0.01 | 0.08 | 0.18 | 0.15 | 0.16 |
| 1 | 246534452 | rs10802397 | -0.394 | 0.704 | -1.159 | 0.933 | -0.002 | 0.078 | 0.078 | 0 | 0.18 | 0.28 | 0.19 | 0.51 | 0.18 | 0.4 |
| 1 | 246550861 | rs1339965 | 0.327 | 0.384 | -0.914 | 0.612 | -0.003 | 0.012 | 0.012 | 0 | 0.46 | 0.31 | 0.51 | 0.4 | 0.44 | 0.44 |
| 1 | 246556279 | rs10802398 | 0.086 | 0.33 | -0.907 | 0.601 | -0.003 | -0.005 | -0.005 | 0.098 | 0.3 | 0.3 | 0.32 | 0.47 | 0.4 | 0.44 |
| 1 | 246556508 | rs7555661 | 1.451 | 0.458 | -0.998 | 0.292 | -0.02 | 0.03 | 0.03 | 0 | 0.23 | 0.15 | 0.22 | 0.09 | 0.28 | 0.2 |
| 1 | 246557566 | rs960227 | 0.078 | 0.501 | -0.915 | 0.482 | -0.003 | -0.005 | -0.005 | 0 | 0.3 | 0.3 | 0.33 | 0.48 | 0.4 | 0.44 |
| 1 | 246567550 | rs6665003 | 0.21 | 0.332 | -0.754 | 0.217 | 0.005 | 0.053 | 0.053 | 0 | 0.5 | 0.3 | 0.58 | 0.32 | 0.49 | 0.48 |
| 1 | 246568701 | rs6678204 | 0.066 | 0.296 | -0.576 | 0.449 | 0.015 | 0.347 | 0.347 | 1.143 | 0.06 | 0.31 | 0.07 | 0.32 | 0.03 | 0.2 |
| 1 | 246576661 | rs4654110 | 0.947 | 0.256 | -0.784 | 0.306 | -0.049 | -0.066 | -0.066 | 0 | 0.25 | 0.01 | 0.34 | 0.25 | 0.22 | 0.27 |
| 1 | 246580718 | rs6673291 | 0.699 | 0.611 | -0.472 | 0.471 | -0.003 | -0.003 | -0.003 | 0 | 0.23 | 0.22 | 0.28 | 0.35 | 0.36 | 0.26 |
| 1 | 246586841 | rs10924727 | 0.029 | 0.59 | -0.479 | 0.386 | 0.009 | 0.003 | 0.003 | 0 | 0.28 | 0.34 | 0.35 | 0.67 | 0.43 | 0.47 |

| | | | | | | | | | | | | | | | | |
|---|---|---|---|---|---|---|---|---|---|---|---|---|---|---|---|---|
| 1 | 246590002 | rs11800474 | 1.196 | 0.633 | -0.684 | 0.195 | -0.015 | 0.035 | 0.035 | 0 | 0.22 | 0.13 | 0.21 | 0.07 | 0.27 | 0.22 |
| 1 | 246603657 | rs4654253 | 0.133 | 0.671 | -0.426 | 0.91 | 0.012 | 0.032 | 0.032 | 0 | 0.73 | 0.63 | 0.79 | 0.62 | 0.64 | 0.74 |
| 1 | 246606143 | rs2340780 | -0.144 | 0.09 | -0.544 | 0.833 | -0.039 | 0.015 | 0.015 | 0.358 | 0.72 | 0.52 | 0.77 | 0.68 | 0.84 | 0.73 |
| 1 | 246606218 | rs10754517 | -0.028 | 0.19 | -0.557 | 0.862 | -0.027 | 0.02 | 0.02 | 0.159 | 0.71 | 0.5 | 0.76 | 0.67 | 0.81 | 0.73 |
| 1 | 246613039 | rs7518504 | 0.022 | -0.403 | -1.097 | 0.768 | 0.03 | -0.028 | -0.028 | 1.159 | 0.32 | 0.14 | 0.15 | 0.3 | 0.27 | 0.19 |
| 1 | 246618280 | rs11802785 | 0.443 | -0.187 | -1.393 | -0.323 | -0.007 | 0.033 | 0.033 | 0.432 | 0.25 | 0.17 | 0.19 | 0 | 0.24 | 0.21 |
| 1 | 246618880 | rs10924734 | 1.392 | 0.051 | -1.66 | 0.008 | -0.019 | -0.013 | -0.013 | 0 | 0.1 | 0.05 | 0.25 | 0.47 | 0.15 | 0.21 |
| 1 | 246618945 | rs7550853 | 1.108 | -0.056 | -1.259 | -0.2 | 0.019 | -0.012 | -0.012 | 0 | 0.33 | 0.21 | 0.17 | 0.3 | 0.29 | 0.21 |
| 1 | 246622814 | rs4654261 | -0.197 | -0.061 | -0.309 | -0.202 | -0.001 | -0.001 | -0.001 | 0 | 0.62 | 0.58 | 0.42 | 0.32 | 0.55 | 0.42 |
| 1 | 246632947 | rs10924739 | 0.136 | -0.11 | -0.02 | -0.251 | -0.001 | 0 | 0 | 0 | 0.58 | 0.42 | 0.37 | 0.3 | 0.53 | 0.41 |
| 1 | 246645797 | rs10802412 | 1.137 | -0.446 | -0.294 | -0.336 | -0.101 | 0.007 | 0.007 | 0 | 0.22 | 0.02 | 0.29 | 0.17 | 0.28 | 0.25 |
| 1 | 246646270 | rs7537318 | 1.064 | -0.775 | -0.085 | -0.29 | 0.021 | -0.041 | -0.041 | 1.252 | 0.34 | 0.14 | 0.18 | 0.31 | 0.33 | 0.19 |
| 1 | 246656303 | rs10924745 | 1.016 | -0.788 | 0.334 | -0.244 | 0.018 | -0.024 | -0.024 | 0 | 0.34 | 0.18 | 0.18 | 0.31 | 0.33 | 0.2 |
| 1 | 246657606 | rs7517207 | -0.553 | -1.733 | 0.179 | -0.507 | -0.04 | 0.068 | 0.068 | 0 | 0.25 | 0.52 | 0.37 | 0.36 | 0.21 | 0.19 |
| 1 | 246661236 | rs11800820 | 0.624 | -1.596 | 0.016 | -1.316 | 0.024 | 0.074 | 0.074 | 0 | 0.43 | 0.31 | 0.25 | 0.06 | 0.36 | 0.27 |
| 1 | 246664442 | rs6668455 | 2.843 | -1.646 | -0.046 | -1.331 | 0.02 | 0.022 | 0.022 | 4.033 | 0.17 | 0.1 | 0.07 | 0.05 | 0.11 | 0.04 |
| 1 | 246664479 | rs10924754 | -0.34 | -1.673 | 0.156 | -1.362 | 0 | -0.033 | -0.033 | 0 | 0.51 | 0.52 | 0.69 | 0.9 | 0.63 | 0.56 |
| 1 | 246665393 | rs10802415 | -0.34 | -1.655 | 0.156 | -1.362 | 0 | -0.033 | -0.033 | 0 | 0.51 | 0.52 | 0.69 | 0.9 | 0.63 | 0.56 |
| 1 | 246666882 | rs10924756 | -0.432 | -1.718 | 0.273 | -1.337 | -0.022 | -0.006 | -0.006 | 0.352 | 0.59 | 0.53 | 0.74 | 0.72 | 0.69 | 0.66 |
| 1 | 246668230 | rs4654117 | -0.559 | -1.513 | 0.304 | -1.274 | 0 | 0.019 | 0.019 | 0.412 | 0.45 | 0.62 | 0.33 | 0.18 | 0.46 | 0.37 |
| 1 | 246668544 | rs4654274 | 0.771 | -1.429 | -0.123 | -1.3 | -0.005 | 0.008 | 0.008 | 0.495 | 0.27 | 0.24 | 0.19 | 0 | 0.28 | 0.24 |
| 1 | 246676039 | rs12122591 | -0.24 | -1.262 | 0.101 | -0.862 | -0.009 | 0.049 | 0.049 | 0 | 0.21 | 0.46 | 0.15 | 0.18 | 0.21 | 0.16 |
| 1 | 246686111 | rs4654279 | -1.059 | -1 | -0.39 | -1.109 | -0.006 | -0.044 | -0.044 | 0 | 0.66 | 0.62 | 0.73 | 0.91 | 0.71 | 0.75 |
| 1 | 246687586 | rs3129586 | -0.08 | -1.1 | -0.448 | -1.144 | 0.007 | -0.014 | -0.014 | 0 | 0.58 | 0.52 | 0.38 | 0.24 | 0.55 | 0.35 |
| 1 | 246693055 | rs12026838 | 0.842 | -0.978 | -0.847 | -0.875 | 0.004 | -0.069 | -0.069 | 0 | 0.15 | 0.04 | 0.11 | 0.22 | 0.14 | 0.14 |
| 1 | 246694480 | rs10924771 | -0.87 | -0.848 | -0.65 | -1.207 | 0.002 | -0.017 | -0.017 | 1.932 | 0.61 | 0.4 | 0.7 | 0.95 | 0.65 | 0.89 |
| 1 | 246696596 | rs10924772 | 0.128 | -0.81 | -0.602 | -1.138 | -0.034 | 0.005 | 0.005 | 0 | 0.42 | 0.19 | 0.55 | 0.66 | 0.5 | 0.58 |
| 1 | 246697698 | rs2105134 | -0.248 | -0.446 | 0.102 | -1.563 | 0.011 | -0.016 | -0.016 | 0.327 | 0.95 | 0.8 | 0.95 | 0.99 | 0.96 | 1 |
| 1 | 246698653 | rs3120693 | 0.724 | -0.522 | -0.352 | -1.607 | -0.003 | -0.047 | -0.047 | 0 | 0.9 | 0.43 | 0.92 | 0.99 | 0.95 | 0.99 |
| 1 | 246701819 | rs10924774 | 1.257 | 0.237 | -1.282 | -1.462 | -0.017 | 0.007 | 0.007 | 0 | 0.13 | 0.08 | 0.23 | 0.1 | 0.21 | 0.15 |
| 1 | 246704563 | rs764243 | 1.296 | -0.274 | -1.366 | -1.649 | -0.031 | 0.163 | 0.163 | 2.993 | 0.28 | 0.04 | 0.2 | 0 | 0.28 | 0.1 |
| 1 | 246705277 | rs3129567 | 0.813 | -0.266 | -1.17 | -1.654 | 0.006 | 0.02 | 0.02 | 1.003 | 0.33 | 0.18 | 0.22 | 0 | 0.31 | 0.11 |
| 1 | 246707858 | rs3129568 | 1.089 | -0.19 | -1.114 | -1.559 | 0.003 | 0.006 | 0.006 | 2.179 | 0.3 | 0.17 | 0.22 | 0 | 0.28 | 0.1 |
| 1 | 246716604 | rs4654291 | 0.626 | 0.265 | -1.099 | -1.467 | -0.023 | 0.069 | 0.069 | 0.124 | 0.41 | 0.25 | 0.45 | 0.12 | 0.46 | 0.28 |
| 1 | 246718977 | rs7536768 | 0.348 | -0.085 | -1.048 | -1.157 | -0.019 | 0.04 | 0.04 | 0.582 | 0.26 | 0.36 | 0.34 | 0.65 | 0.24 | 0.5 |
| 1 | 246725264 | rs10924779 | 0.812 | 0.319 | -1.708 | -1.554 | -0.178 | 0.182 | 0.182 | 1.334 | 0.37 | 0.03 | 0.42 | 0.12 | 0.44 | 0.28 |
| 1 | 246733870 | rs12062784 | 0.857 | 0.571 | -1.562 | -1.469 | -0.081 | 0.095 | 0.095 | 2.178 | 0.35 | 0.1 | 0.41 | 0.17 | 0.37 | 0.23 |
| 1 | 246746101 | rs4531261 | 0.865 | 0.444 | -0.769 | -1.389 | 0.001 | -0.024 | -0.024 | 0.03 | 0.57 | 0.48 | 0.54 | 0.18 | 0.55 | 0.35 |
| 1 | 246747586 | rs4494115 | -0.314 | 0.253 | -0.312 | -0.993 | -0.003 | -0.002 | -0.002 | 0 | 0.31 | 0.32 | 0.35 | 0.68 | 0.28 | 0.6 |
| 1 | 246755124 | rs6702823 | -0.827 | 0.234 | -0.421 | -0.838 | -0.009 | 0 | 0 | 1.961 | 0.89 | 0.8 | 0.92 | 0.89 | 0.91 | 0.94 |
| 1 | 246757449 | rs6663956 | -0.979 | 0.252 | -0.971 | -0.829 | 0.006 | -0.002 | -0.002 | 0 | 0.9 | 0.92 | 0.94 | 0.89 | 0.9 | 0.94 |
| 1 | 246766435 | rs6426312 | -0.862 | 0.265 | -0.964 | -0.824 | 0.009 | 0.002 | 0.002 | 0.185 | 0.89 | 0.91 | 0.93 | 0.89 | 0.9 | 0.94 |
| 1 | 246780619 | rs3124075 | 1.138 | 0.239 | -1.038 | -1.122 | 0.038 | 0.13 | 0.13 | 1.247 | 0.51 | 0.31 | 0.52 | 0.21 | 0.56 | 0.32 |
| 1 | 246793880 | rs7518651 | -0.952 | 0.279 | -1.425 | -1.002 | 0.026 | 0.063 | 0.063 | 5.861 | 0.42 | 0.75 | 0.43 | 0.7 | 0.37 | 0.63 |
| 1 | 246811696 | rs11802065 | 0.162 | 0.343 | -1.009 | -0.98 | 0.001 | -0.003 | -0.003 | 1.422 | 0.25 | 0.21 | 0.33 | 0.7 | 0.25 | 0.53 |
| 1 | 246814806 | rs12126135 | -0.238 | 0.645 | -0.273 | -0.567 | 0.007 | -0.083 | -0.083 | 0.024 | 0.13 | 0.49 | 0.08 | 0 | 0.09 | 0.08 |
| 1 | 246815596 | rs10924791 | -0.052 | 1.408 | -1.022 | -0.587 | -0.031 | 0.021 | 0.021 | 0 | 0.17 | 0.1 | 0.27 | 0.14 | 0.23 | 0.17 |
| 1 | 246818634 | rs3124089 | 0.898 | 1.313 | -1.1 | -0.303 | -0.018 | 0.109 | 0.109 | 0 | 0.55 | 0.19 | 0.56 | 0.71 | 0.55 | 0.65 |
| 1 | 246823857 | rs7534740 | -0.069 | 1.159 | -0.59 | 0.36 | 0.008 | 0.012 | 0.012 | 0 | 0.18 | 0.26 | 0.39 | 0.35 | 0.24 | 0.25 |
| 1 | 246829616 | rs4130648 | -1.221 | 1.015 | -0.578 | 0.449 | 0.003 | 0.002 | 0.002 | 0 | 0.52 | 0.61 | 0.62 | 0.7 | 0.62 | 0.59 |
| 1 | 246829873 | rs3795472 | -1.226 | 0.976 | -0.571 | 0.48 | 0.001 | 0.001 | 0.001 | 0 | 0.52 | 0.61 | 0.62 | 0.69 | 0.62 | 0.59 |
| 1 | 246830293 | rs3795471 | -1.411 | 0.954 | -1.104 | 0.463 | 0.03 | 0.026 | 0.026 | 0.331 | 0.55 | 0.94 | 0.65 | 0.7 | 0.62 | 0.59 |
| 1 | 246832757 | rs9426254 | -0.282 | 1.105 | -0.789 | 0.137 | 0 | 0.027 | 0.027 | 0.003 | 0.84 | 0.91 | 0.95 | 1 | 0.88 | 0.89 |
| 1 | 246837340 | rs10924800 | 0.003 | 1.242 | -0.497 | 0.367 | 0.02 | 0.012 | 0.012 | 1.178 | 0.37 | 0.39 | 0.45 | 0.44 | 0.43 | 0.55 |
| 1 | 246840223 | rs6683786 | -0.63 | 1.63 | 0.476 | 0.834 | 0.031 | 0.103 | 0.103 | 0 | 0.16 | 0.62 | 0.31 | 0.29 | 0.17 | 0.21 |
| 1 | 246841405 | rs12567575 | 1.111 | 1.641 | 0.537 | 0.425 | 0.001 | -0.013 | -0.013 | 0 | 0.6 | 0.44 | 0.6 | 0.76 | 0.52 | 0.48 |
| 1 | 246843070 | rs11588722 | 0.637 | 1.439 | 0.219 | 0.662 | -0.046 | 0.054 | 0.054 | 0 | 0.14 | 0.03 | 0.2 | 0.06 | 0.2 | 0.21 |
| 1 | 246843315 | rs4072061 | -0.964 | 1.36 | 0.308 | 0.468 | -0.012 | 0.006 | 0.006 | 0.093 | 0.33 | 0.33 | 0.43 | 0.23 | 0.47 | 0.42 |
| 1 | 246844066 | rs12045922 | -0.374 | 1.162 | 0.319 | 0.68 | 0.076 | 0.004 | 0.004 | 0 | 0.21 | 0.44 | 0.24 | 0.17 | 0.27 | 0.2 |
| 1 | 246854862 | rs10802432 | 1.134 | 0.592 | -0.656 | 0.664 | 0.025 | 0.065 | 0.065 | 0.038 | 0.44 | 0.05 | 0.29 | 0.36 | 0.34 | 0.39 |
| 1 | 246859182 | rs10802434 | -0.645 | 0.661 | 0.067 | 0.529 | 0.009 | -0.009 | -0.009 | 0.026 | 0.32 | 0.45 | 0.43 | 0.26 | 0.36 | 0.41 |
| 1 | 246870293 | rs7532928 | -0.931 | 0.068 | 0.224 | 0.538 | 0.001 | -0.002 | -0.002 | 0 | 0.49 | 0.52 | 0.65 | 0.77 | 0.59 | 0.57 |
| 1 | 246874875 | rs10802440 | -0.663 | 0.095 | 0.34 | 0.813 | 0.003 | 0.012 | 0.012 | 0 | 0.34 | 0.53 | 0.41 | 0.45 | 0.36 | 0.39 |
| 1 | 246886265 | rs6697433 | -0.606 | 0.223 | 0.604 | 0.952 | 0.057 | 0.082 | 0.082 | 0 | 0.1 | 0.3 | 0.18 | 0.18 | 0.09 | 0.1 |
| 1 | 246888053 | rs10754532 | 0.908 | 0.023 | 0.347 | 0.712 | 0 | 0.006 | 0.006 | 0 | 0.48 | 0.4 | 0.38 | 0.24 | 0.47 | 0.49 |
| 1 | 246891269 | rs3935011 | 0.691 | -0.004 | 0.463 | 0.726 | 0 | -0.002 | -0.002 | 0.003 | 0.52 | 0.46 | 0.52 | 0.41 | 0.49 | 0.51 |
| 1 | 246893948 | rs6657743 | 0.298 | 0.048 | 0.678 | 0.895 | -0.031 | -0.037 | -0.037 | 5.886 | 0.2 | 0.09 | 0.25 | 0.32 | 0.25 | 0.19 |
| 1 | 246898134 | rs7526237 | -0.278 | 0.095 | 0.847 | 0.782 | 0.035 | 0.077 | 0.077 | 0 | 0.07 | 0.21 | 0.17 | 0.18 | 0.07 | 0.09 |
| 1 | 246907206 | rs7513496 | -0.892 | 0.064 | 0.852 | 0.767 | 0.098 | 0.169 | 0.169 | 0 | 0.09 | 0.43 | 0.18 | 0.17 | 0.07 | 0.09 |

| | | | | | | | | | | | | | | | | |
|---|---|---|---|---|---|---|---|---|---|---|---|---|---|---|---|---|
| 1 | 246928540 | rs10924820 | 1.188 | -0.176 | 0.484 | 0.594 | -0.043 | 0.09 | 0.09 | 0 | 0.41 | 0.07 | 0.34 | 0.23 | 0.42 | 0.47 |
| 1 | 246929018 | rs4926440 | -0.141 | -0.203 | 0.745 | 0.633 | 0.065 | 0.196 | 0.196 | 0 | 0.07 | 0.37 | 0.19 | 0.18 | 0.03 | 0.05 |
| 1 | 246930874 | rs1127149 | 0.242 | -0.2 | 0.746 | 0.634 | 0.081 | 0.217 | 0.217 | 0 | 0.06 | 0.33 | 0.18 | 0.18 | 0.03 | 0.05 |
| 1 | 246931242 | rs10127783 | 1.183 | -0.218 | 0.619 | 0.608 | -0.035 | 0.094 | 0.094 | 0 | 0.42 | 0.09 | 0.33 | 0.23 | 0.42 | 0.46 |
| 1 | 246931824 | rs10924824 | 1.276 | -0.215 | 0.633 | 0.609 | -0.045 | 0.088 | 0.088 | 0 | 0.41 | 0.07 | 0.33 | 0.23 | 0.42 | 0.47 |
| 1 | 246934957 | rs10924826 | 0.343 | -1.05 | 0.115 | 0.533 | 0.033 | 0.085 | 0.085 | 3.435 | 0.53 | 0.2 | 0.54 | 0.7 | 0.68 | 0.66 |
| 1 | 246935011 | rs6699332 | 0.716 | -0.213 | 0.878 | 0.771 | 0 | 0.036 | 0.036 | 0 | 0.05 | 0.11 | 0.16 | 0.18 | 0.03 | 0.05 |
| 1 | 246939943 | rs6685364 | 1.094 | -0.208 | 0.879 | 0.756 | -0.011 | 0.075 | 0.075 | 0 | 0.39 | 0.09 | 0.32 | 0.23 | 0.38 | 0.42 |
| 1 | 246940442 | rs3795462 | 0.369 | -0.106 | 0.825 | 0.827 | -0.001 | -0.001 | -0.001 | 0 | 0.49 | 0.5 | 0.52 | 0.4 | 0.45 | 0.51 |
| 1 | 246941990 | rs4926456 | 0.013 | 0.142 | 0.661 | 1.08 | -0.002 | 0.038 | 0.038 | 0 | 0.05 | 0.11 | 0.16 | 0.17 | 0.03 | 0.05 |
| 1 | 246942107 | rs4926443 | 0.013 | 0.142 | 0.661 | 1.08 | -0.002 | 0.038 | 0.038 | 0 | 0.05 | 0.11 | 0.16 | 0.17 | 0.03 | 0.05 |
| 1 | 246954166 | rs6694274 | -0.299 | 0.318 | -0.004 | 1.077 | 0.005 | 0.024 | 0.024 | 0 | 0.17 | 0.21 | 0.19 | 0.31 | 0.17 | 0.2 |
| 1 | 246960463 | rs7551836 | 0.459 | 0.165 | -0.01 | 1.006 | -0.008 | 0.058 | 0.058 | 0.151 | 0.92 | 0.78 | 0.83 | 0.83 | 0.97 | 0.94 |
| 1 | 246961246 | rs4926461 | 0.401 | -0.009 | -0.156 | 0.662 | -0.002 | -0.006 | -0.006 | 0.628 | 0.66 | 0.62 | 0.6 | 0.69 | 0.62 | 0.79 |
| 1 | 246961396 | rs4926446 | 1.263 | -0.04 | -1.011 | 0.466 | -0.043 | 0.004 | 0.004 | 0 | 0.39 | 0.09 | 0.32 | 0.32 | 0.36 | 0.42 |
| 1 | 246963019 | rs3935608 | 0.059 | 0.172 | -0.558 | 0.232 | -0.002 | 0.031 | 0.031 | 0 | 0.26 | 0.15 | 0.23 | 0.13 | 0.34 | 0.17 |
| 1 | 246968455 | rs4926465 | 0.207 | -0.033 | -0.115 | 0.17 | -0.006 | -0.047 | -0.047 | 0.12 | 0.64 | 0.56 | 0.72 | 0.85 | 0.6 | 0.71 |
| 1 | 246969804 | rs4926469 | 0.03 | -0.011 | -0.437 | 0.211 | 0 | -0.003 | -0.003 | 0.045 | 0.71 | 0.68 | 0.77 | 0.84 | 0.65 | 0.73 |
| 1 | 246973094 | rs11584835 | -0.27 | -0.072 | -0.457 | 0.166 | -0.009 | -0.074 | -0.074 | 0.04 | 0.3 | 0.48 | 0.23 | 0.13 | 0.35 | 0.27 |
| 1 | 246978504 | rs10924843 | -0.27 | 0.115 | -1.781 | 0.461 | -0.006 | 0 | 0 | 0 | 0.35 | 0.17 | 0.37 | 0.34 | 0.42 | 0.38 |
| 1 | 246988179 | rs12038733 | 0.531 | 0.035 | -2.088 | 0.737 | -0.037 | -0.071 | -0.071 | 0.023 | 0.37 | 0.04 | 0.31 | 0.4 | 0.3 | 0.33 |
| 1 | 246992698 | rs10924848 | -0.129 | 0.211 | -1.626 | 0.805 | 0.01 | 0.004 | 0.004 | 0 | 0.44 | 0.39 | 0.42 | 0.59 | 0.36 | 0.36 |
| 1 | 246992842 | rs10924849 | -0.864 | 0.341 | -2.334 | 1.147 | -0.01 | -0.07 | -0.07 | 0.367 | 0.24 | 0.1 | 0.38 | 0.35 | 0.24 | 0.24 |
| 1 | 246993684 | rs1110068 | -1.059 | 0.64 | -3.305 | 0.936 | -0.095 | -0.04 | -0.04 | 0 | 0.6 | 0.98 | 0.75 | 0.66 | 0.6 | 0.58 |
| 1 | 246994906 | rs10924852 | -0.706 | 0.655 | -2.696 | 0.665 | 0.001 | -0.012 | -0.012 | 0.494 | 0.35 | 0.72 | 0.52 | 0.43 | 0.34 | 0.37 |
| 1 | 247006314 | rs12401878 | 0.505 | 0.739 | -2.449 | 0.709 | 0.005 | 0 | 0 | 2.405 | 0.19 | 0.22 | 0.34 | 0.19 | 0.22 | 0.17 |
| 1 | 247008439 | rs7549408 | 0.101 | 0.814 | -1.711 | 0.433 | 0.105 | -0.005 | -0.005 | 0.033 | 0.09 | 0.47 | 0.09 | 0.01 | 0.09 | 0.14 |
| 1 | 247027779 | rs1691251 | -1.401 | 0.697 | -2.443 | 0.472 | -0.124 | -0.012 | -0.012 | 0 | 0.53 | 0.98 | 0.71 | 0.59 | 0.54 | 0.52 |
| 1 | 247028789 | rs10924861 | 0.157 | 0.425 | -2.104 | 0.817 | -0.021 | -0.063 | -0.063 | 0 | 0.22 | 0.08 | 0.4 | 0.36 | 0.24 | 0.2 |
| 1 | 247031972 | rs4971287 | 0.907 | 0.152 | -2.502 | 0.451 | -0.061 | -0.114 | -0.114 | 0.053 | 0.28 | 0.01 | 0.14 | 0.34 | 0.29 | 0.45 |
| 1 | 247035708 | rs10924862 | 0.324 | -0.218 | -0.893 | 0.383 | -0.009 | 0 | 0 | 0.279 | 0.21 | 0.11 | 0.27 | 0.19 | 0.21 | 0.16 |
| 1 | 247039948 | rs2067916 | 0.279 | -0.225 | -0.537 | 0.379 | 0 | -0.001 | -0.001 | 0.142 | 0.21 | 0.21 | 0.28 | 0.19 | 0.21 | 0.16 |
| 1 | 247044692 | rs2642984 | -0.764 | -0.496 | -0.154 | 0.317 | -0.025 | 0.02 | 0.02 | 0 | 0.31 | 0.56 | 0.31 | 0.35 | 0.28 | 0.36 |
| 1 | 247046778 | rs10924863 | 0.087 | -0.513 | -0.174 | 0.373 | -0.002 | -0.035 | -0.035 | 0 | 0.23 | 0.13 | 0.28 | 0.33 | 0.23 | 0.2 |
| 1 | 247055427 | rs2800219 | -1.931 | -0.562 | 0.196 | 0.275 | -0.022 | -0.011 | -0.011 | 1.83 | 0.53 | 0.93 | 0.62 | 0.54 | 0.5 | 0.52 |
| 1 | 247061466 | rs12023594 | 0.572 | -0.497 | -0.107 | 0.308 | -0.015 | -0.004 | -0.004 | 0 | 0.2 | 0.09 | 0.27 | 0.19 | 0.21 | 0.16 |
| 1 | 247065472 | rs1691255 | -1.783 | -0.502 | -0.076 | 0.308 | 0.009 | 0.013 | 0.013 | 0 | 0.52 | 0.73 | 0.59 | 0.54 | 0.5 | 0.52 |
| 1 | 247069232 | rs7529090 | -0.755 | -0.533 | 0.26 | 0.3 | 0.004 | -0.006 | -0.006 | 0.166 | 0.29 | 0.28 | 0.3 | 0.35 | 0.28 | 0.36 |
| 1 | 247081383 | rs1039010 | -2.244 | -0.6 | -0.624 | 0.284 | -0.168 | -0.092 | -0.092 | 0 | 0.61 | 0.98 | 0.63 | 0.62 | 0.52 | 0.54 |
| 1 | 247084953 | rs10924867 | 0.24 | -0.27 | -0.051 | 0.296 | -0.001 | -0.001 | -0.001 | 0.762 | 0.2 | 0.22 | 0.26 | 0.19 | 0.2 | 0.16 |
| 1 | 247098587 | rs4971278 | -1.094 | -0.18 | -0.128 | 0.172 | -0.001 | -0.044 | -0.044 | 0 | 0.38 | 0.2 | 0.46 | 0.52 | 0.38 | 0.36 |
| 1 | 247106126 | rs1691234 | -2.066 | -0.28 | 0.017 | 0.139 | 0.002 | 0.002 | 0.002 | 0 | 0.57 | 0.5 | 0.6 | 0.56 | 0.52 | 0.38 |
| 1 | 247109168 | rs6426191 | -0.873 | -0.332 | -0.042 | 0.126 | 0.002 | 0.008 | 0.008 | 0 | 0.39 | 0.32 | 0.4 | 0.33 | 0.38 | 0.21 |
| 1 | 247109267 | rs2642983 | -1.816 | -0.176 | -0.124 | -0.101 | 0.001 | 0 | 0 | 0 | 0.56 | 0.48 | 0.59 | 0.56 | 0.52 | 0.38 |
| 1 | 247111062 | rs1779962 | -2.071 | -0.204 | -0.035 | -0.117 | -0.019 | -0.019 | -0.019 | 0 | 0.58 | 0.67 | 0.61 | 0.56 | 0.54 | 0.4 |
| 1 | 247112320 | rs1691246 | -1.365 | -0.097 | -0.758 | -0.129 | 0.002 | 0.003 | 0.003 | 0 | 0.49 | 0.45 | 0.57 | 0.47 | 0.51 | 0.36 |
| 1 | 247116302 | rs10924874 | -0.685 | 0.185 | -1.823 | -0.274 | -0.041 | -0.092 | -0.092 | 0 | 0.35 | 0.06 | 0.45 | 0.44 | 0.35 | 0.34 |
| 1 | 247125355 | rs11579381 | -0.71 | 0.129 | -1.802 | -0.266 | -0.039 | -0.132 | -0.132 | 0 | 0.36 | 0.06 | 0.44 | 0.48 | 0.35 | 0.35 |
| 1 | 247129945 | rs1779985 | -2.165 | -0.182 | -1.872 | -0.826 | -0.162 | -0.172 | -0.172 | 0 | 0.63 | 0.97 | 0.64 | 0.54 | 0.54 | 0.55 |
| 1 | 247130577 | rs4971282 | -1.008 | -0.193 | -1.356 | -0.59 | 0.018 | -0.064 | -0.064 | 0.021 | 0.37 | 0.13 | 0.43 | 0.44 | 0.33 | 0.33 |
| 1 | 247139343 | rs6673828 | -0.932 | -0.621 | -0.71 | -1.679 | -0.026 | -0.049 | -0.049 | 0 | 0.29 | 0.5 | 0.27 | 0.1 | 0.18 | 0.19 |
| 1 | 247140240 | rs2642963 | -1.463 | -0.18 | -0.832 | -2.099 | 0.028 | 0.136 | 0.136 | 0.424 | 0.43 | 0.78 | 0.29 | 0.16 | 0.37 | 0.35 |
| 1 | 247142239 | rs7555643 | 0.991 | -0.286 | -0.697 | -2.326 | -0.01 | 0.289 | 0.289 | 0 | 0.57 | 0.09 | 0.72 | 0.97 | 0.64 | 0.61 |
| 1 | 247145690 | rs4506496 | 0.862 | -0.342 | -0.02 | -2.354 | 0.011 | -0.006 | -0.006 | 0 | 0.6 | 0.32 | 0.75 | 0.97 | 0.66 | 0.62 |
| 1 | 247149996 | rs4971310 | 0.97 | -0.488 | -0.317 | -2.329 | 0.01 | 0.181 | 0.181 | 0 | 0.57 | 0.16 | 0.72 | 0.97 | 0.64 | 0.63 |
| 1 | 247150740 | rs3749427 | 0.982 | -0.481 | -0.314 | -2.326 | 0.011 | 0.182 | 0.182 | 0 | 0.57 | 0.16 | 0.72 | 0.97 | 0.64 | 0.63 |
| 1 | 247151557 | rs2642992 | 0.76 | -0.396 | -0.047 | -2.308 | 0.003 | 0.064 | 0.064 | 0 | 0.59 | 0.29 | 0.74 | 0.97 | 0.64 | 0.63 |
| 1 | 247155176 | rs6668864 | 0.668 | -0.49 | -0.265 | -2.383 | -0.003 | 0.006 | 0.006 | 0 | 0.65 | 0.29 | 0.77 | 0.98 | 0.66 | 0.62 |
| 1 | 247156066 | rs2642978 | -0.962 | -0.616 | -0.154 | -2.417 | 0.001 | -0.009 | -0.009 | 0 | 0.88 | 0.89 | 0.96 | 0.99 | 0.89 | 0.81 |
| 1 | 247162658 | rs2642973 | 0.5 | -0.631 | -0.149 | -2.43 | 0.007 | 0.018 | 0.018 | 0 | 0.79 | 0.83 | 0.93 | 0.97 | 0.83 | 0.62 |
| 1 | 247167077 | rs7513728 | 1.239 | -0.586 | 0.025 | -2.447 | 0.003 | -0.007 | -0.007 | 0 | 0.59 | 0.34 | 0.76 | 0.97 | 0.64 | 0.58 |
| 1 | 247172257 | rs2453407 | -1.564 | -0.645 | -0.066 | -2.427 | 0.008 | 0.071 | 0.071 | 0 | 0.41 | 0.72 | 0.24 | 0.03 | 0.35 | 0.41 |
| 1 | 247173682 | rs12091227 | -0.001 | -0.595 | -0.064 | -2.355 | 0.022 | -0.144 | -0.144 | 0 | 0.19 | 0.49 | 0.17 | 0 | 0.2 | 0.04 |
| 1 | 247179076 | rs28540229 | 0.777 | -0.544 | -0.843 | -2.254 | -0.009 | 0.054 | 0.054 | 0 | 0.11 | 0.02 | 0.02 | 0.01 | 0.1 | 0.19 |
| 1 | 247180121 | rs2642966 | -1.387 | -0.444 | -0.894 | -2.333 | -0.035 | 0.274 | 0.274 | 0 | 0.43 | 0.87 | 0.27 | 0.03 | 0.38 | 0.42 |
| 1 | 247196539 | rs2255481 | -1.693 | -0.451 | -1.274 | -2.386 | -0.069 | 0.218 | 0.218 | 0 | 0.58 | 0.91 | 0.36 | 0.14 | 0.5 | 0.53 |
| 1 | 247200639 | rs12141826 | -0.123 | -0.684 | -1.336 | -2.491 | -0.003 | 0.033 | 0.033 | 0 | 0.19 | 0.08 | 0.12 | 0.13 | 0.17 | 0.27 |
| 1 | 247203606 | rs7547915 | -0.763 | -0.643 | -0.295 | -2.408 | -0.005 | -0.147 | -0.147 | 1.327 | 0.37 | 0.5 | 0.21 | 0.01 | 0.31 | 0.26 |
| 1 | 247207285 | rs12116484 | 1.121 | -0.599 | -0.499 | -2.328 | -0.027 | 0.064 | 0.064 | 0 | 0.11 | 0.01 | 0.02 | 0.01 | 0.1 | 0.18 |

| | | | | | | | | | | | | | | | | |
|---|---|---|---|---|---|---|---|---|---|---|---|---|---|---|---|---|
| 1 | 247226344 | rs10924899 | 0.858 | -0.608 | -0.097 | -2.333 | -0.002 | -0.037 | -0.037 | 0 | 0.12 | 0.14 | 0.03 | 0.01 | 0.1 | 0.18 |
| 1 | 247236374 | rs4925612 | -0.31 | -0.616 | 0.059 | -2.366 | -0.01 | -0.181 | -0.181 | 0 | 0.26 | 0.47 | 0.19 | 0 | 0.21 | 0.09 |
| 1 | 247236563 | rs4925614 | -0.25 | -0.589 | -0.044 | -2.353 | -0.004 | -0.135 | -0.135 | 0 | 0.25 | 0.38 | 0.18 | 0 | 0.21 | 0.09 |
| 1 | 247241668 | rs6701757 | -2.009 | -0.569 | -0.857 | -1.684 | -0.11 | -0.116 | -0.116 | 0 | 0.67 | 0.97 | 0.44 | 0.44 | 0.57 | 0.62 |
| 1 | 247251482 | rs6669242 | -1.7 | -1.075 | -0.204 | -2.008 | 0.069 | 0.275 | 0.275 | 0 | 0.5 | 0.84 | 0.26 | 0.13 | 0.35 | 0.48 |
| 1 | 247252836 | rs12144733 | -1.257 | -1.107 | 0.385 | -2.033 | -0.006 | -0.027 | -0.027 | 0.164 | 0.34 | 0.61 | 0.22 | 0.13 | 0.26 | 0.33 |
| 1 | 247253566 | rs2890103 | -1.277 | -1.111 | 0.378 | -2.035 | -0.004 | -0.022 | -0.022 | 1.692 | 0.34 | 0.61 | 0.22 | 0.13 | 0.26 | 0.32 |
| 1 | 247261502 | rs10924910 | -0.375 | -1.097 | 0.724 | -2.048 | -0.007 | -0.028 | -0.028 | 0 | 0.2 | 0.29 | 0.17 | 0.12 | 0.21 | 0.26 |
| 1 | 247262625 | rs6699898 | -1.126 | -1.116 | 0.846 | -2.061 | 0.004 | 0.005 | 0.005 | 0 | 0.34 | 0.29 | 0.19 | 0.13 | 0.31 | 0.44 |
| 1 | 247267668 | rs2270616 | -1.457 | -1.02 | 0.938 | -2.01 | 0.008 | 0.005 | 0.005 | 0.06 | 0.47 | 0.39 | 0.34 | 0.13 | 0.39 | 0.52 |
| 1 | 247272298 | rs10924919 | -1.33 | -0.85 | 0.897 | -1.926 | 0.001 | -0.014 | -0.014 | 0.576 | 0.45 | 0.41 | 0.32 | 0.13 | 0.39 | 0.5 |
| 1 | 247276394 | rs6426220 | -1.695 | -0.964 | 0.769 | -1.936 | 0.03 | 0.096 | 0.096 | 0 | 0.51 | 0.72 | 0.41 | 0.14 | 0.39 | 0.53 |
| 1 | 247283320 | rs10924924 | -1.653 | -1.132 | 0.673 | -1.598 | -0.009 | -0.034 | -0.034 | 0 | 0.35 | 0.61 | 0.32 | 0.12 | 0.28 | 0.34 |
| 1 | 247305873 | rs12093759 | -0.828 | -1.139 | 0.417 | -1.612 | 0.023 | 0.065 | 0.065 | 0.013 | 0.24 | 0.1 | 0.14 | 0.13 | 0.2 | 0.31 |
| 1 | 247306833 | rs3844080 | -2.069 | -1.219 | 0.206 | -1.713 | 0.061 | 0.258 | 0.258 | 0 | 0.47 | 0.9 | 0.39 | 0.13 | 0.35 | 0.53 |
| 1 | 247308691 | rs7521596 | -1.589 | -1.233 | 0.53 | -1.703 | -0.005 | -0.027 | -0.027 | 0.917 | 0.38 | 0.62 | 0.22 | 0.13 | 0.27 | 0.51 |
| 1 | 247313462 | rs6663210 | -0.127 | -0.903 | 0.667 | -1.416 | 0.034 | -0.035 | -0.035 | 0 | 0.08 | 0.29 | 0.17 | 0 | 0.08 | 0.02 |
| 1 | 247319489 | rs3767857 | -0.147 | -0.856 | 0.272 | -1.396 | -0.011 | 0.003 | 0.003 | 0 | 0.09 | 0.06 | 0.1 | 0.09 | 0.15 | 0.1 |
| 1 | 247320868 | rs11806327 | -0.694 | -0.731 | 0.057 | -1.347 | 0.004 | -0.047 | -0.047 | 0 | 0.09 | 0.2 | 0.17 | 0 | 0.08 | 0.02 |
| 1 | 247323260 | rs9793102 | -1.173 | -0.825 | -0.624 | -1.327 | 0.007 | 0.004 | 0.004 | 0 | 0.36 | 0.64 | 0.31 | 0.13 | 0.28 | 0.33 |
| 1 | 247336726 | rs1609840 | -0.331 | -0.43 | -0.532 | -0.533 | -0.009 | 0 | 0 | 0 | 0.4 | 0.57 | 0.22 | 0.43 | 0.27 | 0.57 |
| 1 | 247338187 | rs10802474 | -0.515 | -0.356 | -0.513 | -0.066 | 0.013 | 0.004 | 0.004 | 0 | 0.49 | 0.62 | 0.31 | 0.53 | 0.41 | 0.64 |
| 1 | 247338427 | rs3850818 | -0.457 | -0.317 | -0.957 | -0.177 | 0.006 | 0.009 | 0.009 | 0 | 0.51 | 0.79 | 0.34 | 0.44 | 0.42 | 0.65 |
| 1 | 247338764 | rs10924933 | -0.379 | -0.59 | -1.211 | -0.009 | 0.041 | 0.042 | 0.042 | 0 | 0.51 | 0.77 | 0.37 | 0.42 | 0.41 | 0.68 |
| 1 | 247339768 | rs10924936 | -0.641 | -0.427 | -1.641 | -0.093 | -0.037 | -0.102 | -0.102 | 0 | 0.57 | 0.9 | 0.44 | 0.48 | 0.42 | 0.71 |
| 1 | 247340756 | rs6659817 | -0.471 | -0.243 | -1.508 | -0.005 | 0.02 | -0.038 | -0.038 | 0 | 0.53 | 0.79 | 0.43 | 0.48 | 0.42 | 0.65 |
| 1 | 247340784 | rs6684931 | -0.487 | -0.237 | -1.612 | -0.003 | 0.012 | -0.051 | -0.051 | 0.327 | 0.53 | 0.81 | 0.43 | 0.48 | 0.42 | 0.65 |
| 1 | 247346189 | rs12039124 | 0.481 | -0.202 | -0.647 | -0.454 | -0.024 | -0.116 | -0.116 | 0 | 0.8 | 0.64 | 0.88 | 0.99 | 0.83 | 0.91 |
| 1 | 247347971 | rs6684181 | 0.367 | -0.163 | -1.818 | -0.28 | -0.048 | -0.059 | -0.059 | 0.575 | 0.25 | 0 | 0.13 | 0.2 | 0.19 | 0.35 |
| 1 | 247352621 | rs10754538 | 1.196 | 0.304 | -2.464 | -0.285 | -0.075 | -0.128 | -0.128 | 0 | 0.32 | 0.06 | 0.51 | 0.59 | 0.38 | 0.3 |
| 1 | 247353330 | rs4351686 | 0.759 | 0.378 | -1.598 | -0.334 | 0.009 | 0.003 | 0.003 | 0 | 0.27 | 0.16 | 0.15 | 0.2 | 0.18 | 0.47 |
| 1 | 247355029 | rs12131404 | -0.554 | 0.024 | -0.614 | -0.564 | -0.012 | -0.003 | -0.003 | 0 | 0.44 | 0.6 | 0.36 | 0.4 | 0.45 | 0.62 |
| 1 | 247360368 | rs7551512 | -0.087 | 0.13 | -0.189 | -0.485 | 0.08 | 0.036 | 0.036 | 0 | 0.17 | 0.36 | 0.21 | 0.2 | 0.27 | 0.14 |
| 1 | 247364063 | rs17720872 | 0.419 | 0.144 | -0.132 | -0.303 | -0.008 | -0.013 | -0.013 | 0.253 | 0.06 | 0.04 | 0.08 | 0.12 | 0.11 | 0.08 |
| 1 | 247366124 | rs11806853 | 2.089 | -0.002 | 0.114 | -0.385 | -0.029 | -0.03 | -0.03 | 0 | 0.06 | 0.13 | 0.02 | 0 | 0 | 0 |
| 1 | 247366435 | rs11811599 | -0.231 | -0.157 | 0.095 | -0.4 | 0.016 | 0 | 0 | 0 | 0.4 | 0.65 | 0.27 | 0.33 | 0.29 | 0.46 |
| 1 | 247371918 | rs7555638 | -0.469 | -0.154 | -0.014 | -0.769 | -0.044 | 0.051 | 0.051 | 0 | 0.79 | 0.92 | 0.8 | 0.91 | 0.71 | 0.86 |
| 1 | 247372163 | rs17269923 | 0.202 | -0.295 | 0.237 | -0.424 | -0.001 | -0.004 | -0.004 | 0.153 | 0.34 | 0.33 | 0.17 | 0.17 | 0.18 | 0.39 |
| 1 | 247393655 | rs6693807 | 0.47 | -0.326 | 0.157 | -0.341 | -0.001 | 0 | 0 | 0 | 0.29 | 0.29 | 0.23 | 0.3 | 0.28 | 0.36 |
| 1 | 247400259 | rs10924968 | 0.799 | -0.493 | -0.124 | 0.198 | 0.033 | -0.014 | -0.014 | 1.254 | 0.27 | 0.11 | 0.15 | 0.25 | 0.17 | 0.28 |
| 1 | 247401645 | rs2793289 | 0.189 | -0.598 | -0.212 | 0.128 | 0.051 | -0.003 | -0.003 | 0 | 0.58 | 0.32 | 0.73 | 0.63 | 0.73 | 0.52 |
| 1 | 247404911 | rs12118988 | -0.226 | -0.147 | -0.621 | -0.465 | -0.01 | 0.01 | 0.01 | 0 | 0.32 | 0.28 | 0.3 | 0.09 | 0.35 | 0.15 |
| 1 | 247412899 | rs1778538 | 0.354 | -0.37 | -0.245 | -0.367 | -0.046 | 0 | 0 | 0.28 | 0.83 | 0.51 | 0.89 | 0.84 | 0.9 | 0.85 |
| 1 | 247413084 | rs1771942 | -0.327 | -0.309 | -0.772 | -0.75 | 0.01 | 0.005 | 0.005 | 0 | 0.48 | 0.46 | 0.72 | 0.62 | 0.64 | 0.55 |
| 1 | 247415879 | rs1614597 | 1.268 | 0.103 | -0.79 | -0.432 | -0.083 | -0.142 | -0.142 | 2.328 | 0.17 | 0.01 | 0.49 | 0.53 | 0.3 | 0.38 |
| 1 | 247419664 | rs1778541 | 0.473 | -0.162 | -0.381 | -0.596 | -0.042 | -0.008 | -0.008 | 0 | 0.82 | 0.52 | 0.89 | 0.84 | 0.89 | 0.85 |
| 1 | 247436234 | rs2890342 | -0.217 | 0.066 | -0.419 | -1.259 | 0.003 | -0.095 | -0.095 | 0.01 | 0.4 | 0.49 | 0.27 | 0.06 | 0.39 | 0.2 |
| 1 | 247455719 | rs1192831 | 1.278 | 0.577 | -0.833 | -1.6 | -0.01 | -0.002 | -0.002 | 0.033 | 0.13 | 0.08 | 0.35 | 0.52 | 0.19 | 0.37 |
| 1 | 247455896 | rs1192832 | 0.871 | -0.119 | -0.881 | -1.586 | -0.034 | -0.048 | -0.048 | 0 | 0.18 | 0.09 | 0.5 | 0.52 | 0.32 | 0.4 |
| 1 | 247459414 | rs1057996 | -0.608 | -0.362 | -0.274 | -1.674 | 0.057 | 0.027 | 0.027 | 0.227 | 0.94 | 0.71 | 0.97 | 1 | 0.99 | 0.92 |
| 1 | 247460727 | rs3850825 | 0.947 | -0.954 | -0.717 | -2.104 | 0.057 | -0.026 | -0.026 | 0 | 0.38 | 0.21 | 0.22 | 0.41 | 0.3 | 0.33 |
| 1 | 247462701 | rs1616299 | 1.056 | -0.732 | -0.76 | -1.982 | 0.006 | 0.024 | 0.024 | 0.06 | 0.18 | 0.19 | 0.51 | 0.52 | 0.33 | 0.41 |
| 1 | 247470703 | rs4925618 | 1.905 | -0.551 | -1.389 | -2.223 | -0.038 | 0.056 | 0.056 | 0 | 0.1 | 0.01 | 0.06 | 0 | 0.13 | 0.04 |
| 1 | 247475532 | rs11581163 | -0.239 | -0.946 | -1.241 | -2.137 | -0.021 | 0.114 | 0.114 | 0.016 | 0.4 | 0.26 | 0.24 | 0.06 | 0.39 | 0.2 |
| 1 | 247476616 | rs1778549 | 1.173 | -0.341 | -1.49 | -1.772 | -0.067 | -0.094 | -0.094 | 0 | 0.17 | 0.05 | 0.5 | 0.52 | 0.33 | 0.4 |
| 1 | 247490707 | rs6690864 | 1.469 | -0.619 | -1.598 | -1.782 | 0.034 | -0.079 | -0.079 | 0 | 0.3 | 0.09 | 0.2 | 0.42 | 0.27 | 0.31 |
| 1 | 247491143 | rs10924987 | 1.331 | -0.56 | -1.529 | -1.76 | 0.022 | -0.038 | -0.038 | 0 | 0.32 | 0.19 | 0.22 | 0.45 | 0.29 | 0.34 |
| 1 | 247500650 | rs10924989 | 0.825 | -0.363 | -1.505 | -1.361 | -0.112 | -0.153 | -0.153 | 9.016 | 0.18 | 0.01 | 0.49 | 0.51 | 0.33 | 0.42 |
| 1 | 247503507 | rs10737800 | 0.248 | -1.557 | -0.839 | -2.202 | -0.048 | -0.104 | -0.104 | 0 | 0.83 | 0.44 | 0.9 | 0.99 | 0.93 | 0.89 |
| 1 | 247507887 | rs11585647 | 0.4 | -1.841 | -0.972 | -2.243 | -0.026 | -0.029 | -0.029 | 0.207 | 0.71 | 0.34 | 0.79 | 0.9 | 0.9 | 0.87 |
| 1 | 247507908 | rs10924990 | 0.109 | -1.234 | -1.379 | -2.013 | -0.09 | -0.04 | -0.04 | 0 | 0.32 | 0.11 | 0.59 | 0.68 | 0.54 | 0.57 |
| 1 | 247511781 | rs4925626 | 0.602 | -1.904 | -1.082 | -1.852 | 0.042 | 0.035 | 0.035 | 2.229 | 0.17 | 0.09 | 0.05 | 0 | 0.04 | 0.02 |
| 1 | 247512163 | rs28450758 | 0.848 | -1.318 | -0.971 | -1.641 | -0.014 | -0.001 | -0.001 | 0 | 0.08 | 0.05 | 0.05 | 0.07 | 0.09 | 0.11 |
| 1 | 247513049 | rs6702432 | 0.644 | -1.209 | -0.461 | -1.641 | -0.011 | -0.002 | -0.002 | 0 | 0.33 | 0.29 | 0.6 | 0.7 | 0.57 | 0.55 |
| 1 | 247519614 | rs4925539 | 0.026 | -0.951 | -0.211 | -1.292 | 0.03 | 0.059 | 0.059 | 0 | 0.52 | 0.66 | 0.78 | 0.73 | 0.63 | 0.6 |
| 1 | 247519784 | rs2000127 | 0.233 | -0.004 | 0.236 | -0.394 | -0.004 | -0.003 | -0.003 | 0 | 0.74 | 0.43 | 0.83 | 0.8 | 0.78 | 0.84 |
| 1 | 247535535 | rs10399826 | -0.437 | -0.248 | -0.25 | -0.808 | -0.044 | -0.165 | -0.165 | 0 | 0.23 | 0.51 | 0.15 | 0.01 | 0.15 | 0.1 |
| 1 | 247536083 | rs10399672 | -0.164 | -0.159 | -1.306 | -0.782 | -0.001 | -0.002 | -0.002 | 0.027 | 0.51 | 0.52 | 0.41 | 0.67 | 0.44 | 0.49 |
| 1 | 247537126 | rs10925000 | 0.771 | 0.677 | -1.698 | -0.981 | -0.004 | 0.002 | 0.002 | 0 | 0.17 | 0.12 | 0.33 | 0.16 | 0.26 | 0.36 |

| | | | | | | | | | | | | | | | | |
|---|---|---|---|---|---|---|---|---|---|---|---|---|---|---|---|---|
| 1 | 247537146 | rs12043526 | 0.706 | 0.29 | -2.512 | -0.829 | -0.055 | -0.092 | -0.092 | 0 | 0.34 | 0.02 | 0.28 | 0.56 | 0.32 | 0.31 |
| 1 | 247540313 | rs10925002 | -0.245 | 0.259 | -2.225 | -1.076 | -0.049 | 0.144 | 0.144 | 0.257 | 0.63 | 0.18 | 0.51 | 0.94 | 0.59 | 0.68 |
| 1 | 247542052 | rs12130139 | 0.59 | 0.562 | -2.537 | -1.035 | -0.079 | 0.089 | 0.089 | 0.479 | 0.5 | 0.02 | 0.49 | 0.63 | 0.54 | 0.58 |
| 1 | 247550331 | rs7542425 | -0.18 | -0.112 | -2.027 | -0.864 | 0.009 | -0.028 | -0.028 | 0 | 0.42 | 0.54 | 0.45 | 0.24 | 0.45 | 0.53 |
| 1 | 247552084 | rs10925006 | -0.06 | -0.292 | -1.339 | -0.886 | -0.025 | -0.004 | -0.004 | 0 | 0.69 | 0.55 | 0.72 | 0.64 | 0.73 | 0.73 |
| 1 | 247552549 | rs6426244 | 0.274 | -0.281 | -1.337 | -0.958 | -0.016 | -0.02 | -0.02 | 0 | 0.64 | 0.54 | 0.67 | 0.64 | 0.64 | 0.72 |
| 1 | 247554949 | rs2386551 | -0.214 | -0.539 | -1.29 | -0.433 | 0.002 | 0.034 | 0.034 | 0 | 0.82 | 0.64 | 0.84 | 0.68 | 0.79 | 0.73 |
| 1 | 247559156 | rs10802494 | 0.166 | -0.337 | -2.183 | -0.355 | 0.003 | 0.028 | 0.028 | 0.2 | 0.59 | 0.08 | 0.65 | 0.63 | 0.64 | 0.66 |
| 1 | 247559367 | rs12122076 | 0.17 | -0.306 | -2.206 | -0.354 | -0.003 | 0.026 | 0.026 | 0.262 | 0.59 | 0.07 | 0.65 | 0.63 | 0.64 | 0.65 |
| 1 | 247560979 | rs10802495 | 0.337 | -0.218 | -2.068 | -0.29 | 0.012 | 0.072 | 0.072 | 0 | 0.53 | 0.21 | 0.53 | 0.59 | 0.55 | 0.65 |
| 1 | 247561708 | rs10925009 | 0.757 | -0.166 | -2.445 | -0.234 | -0.118 | 0.002 | 0.002 | 0.097 | 0.44 | 0.03 | 0.49 | 0.61 | 0.52 | 0.55 |
| 1 | 247561724 | rs10925010 | 0.74 | -0.088 | -2.407 | -0.205 | -0.083 | 0.031 | 0.031 | 0.018 | 0.45 | 0.07 | 0.49 | 0.61 | 0.52 | 0.55 |
| 1 | 247565953 | rs4508048 | 0.296 | -0.405 | -1.917 | 0.3 | 0.045 | -0.024 | -0.024 | 2.272 | 0.61 | 0.17 | 0.66 | 0.55 | 0.67 | 0.71 |
| 1 | 247575768 | rs7529058 | 0.488 | -0.39 | -1.781 | 0.483 | 0.037 | 0.019 | 0.019 | 0 | 0.52 | 0.26 | 0.58 | 0.45 | 0.54 | 0.6 |
| 1 | 247577079 | rs10925013 | -0.045 | -1.177 | -1.44 | -0.094 | -0.004 | -0.025 | -0.025 | 0 | 0.84 | 0.82 | 0.93 | 0.99 | 0.92 | 0.77 |
| 1 | 247578441 | rs2027432 | 0.365 | -1.396 | -1.7 | -0.06 | 0.026 | 0.046 | 0.046 | 0 | 0.74 | 0.84 | 0.88 | 0.93 | 0.83 | 0.74 |
| 1 | 247579435 | rs3738448 | 0.718 | -1.52 | -1.935 | 0.234 | 0.003 | -0.048 | -0.048 | 0 | 0.09 | 0.01 | 0.07 | 0.13 | 0.05 | 0.04 |
| 1 | 247580568 | rs4925648 | 0.61 | -1.432 | -1.609 | 0.617 | 0.007 | 0.041 | 0.041 | 0 | 0.06 | 0.11 | 0.23 | 0.23 | 0.11 | 0.18 |
| 1 | 247581209 | rs12137901 | -0.047 | -1.868 | -0.914 | 0.406 | 0.005 | 0.098 | 0.098 | 0 | 0.17 | 0.36 | 0.31 | 0.31 | 0.23 | 0.21 |
| 1 | 247584075 | rs4925650 | 1.974 | -1.767 | -1.063 | 0.161 | -0.098 | -0.07 | -0.07 | 0 | 0.33 | 0.1 | 0.31 | 0.48 | 0.44 | 0.45 |
| 1 | 247586336 | rs3806265 | -1.252 | -1.774 | -0.632 | 0.197 | -0.017 | 0.016 | 0.016 | 0 | 0.4 | 0.54 | 0.55 | 0.46 | 0.33 | 0.43 |
| 1 | 247587408 | rs7525979 | 0.067 | -1.737 | -0.335 | 0.062 | -0.024 | 0.024 | 0.024 | 0 | 0.11 | 0.23 | 0.09 | 0.13 | 0.05 | 0.04 |
| 1 | 247587531 | rs4925543 | -1.744 | -1.719 | -0.374 | 0.074 | -0.014 | -0.028 | -0.028 | 0 | 0.92 | 0.84 | 0.95 | 1 | 0.96 | 0.88 |
| 1 | 247588960 | rs3738447 | -0.081 | -1.431 | -0.414 | 0.236 | -0.023 | 0.035 | 0.035 | 0 | 0.11 | 0.27 | 0.11 | 0.13 | 0.06 | 0.03 |
| 1 | 247594890 | rs12564791 | 0.03 | -1.369 | -0.64 | 0.188 | -0.001 | 0 | 0 | 0 | 0.21 | 0.18 | 0.1 | 0.15 | 0.14 | 0.06 |
| 1 | 247596057 | rs4925654 | 0.058 | -1.29 | -0.55 | 0.184 | -0.01 | -0.007 | -0.007 | 0 | 0.75 | 0.74 | 0.87 | 0.85 | 0.79 | 0.9 |
| 1 | 247599070 | rs4612666 | 0.552 | -1.033 | -0.263 | 0.46 | -0.025 | 0.058 | 0.058 | 0.911 | 0.71 | 0.43 | 0.62 | 0.55 | 0.76 | 0.65 |
| 1 | 247599232 | rs10754557 | 0.576 | -1.109 | -0.479 | 0.172 | 0.035 | 0.002 | 0.002 | 0.767 | 0.62 | 0.17 | 0.46 | 0.66 | 0.62 | 0.6 |
| 1 | 247602968 | rs4925546 | -0.554 | -0.528 | 0.043 | -0.251 | 0.036 | 0.032 | 0.032 | 0 | 0.51 | 0.62 | 0.64 | 0.62 | 0.65 | 0.62 |
| 1 | 247603463 | rs4925659 | 0.363 | 0.038 | -0.058 | -0.202 | -0.073 | -0.084 | -0.084 | 2.417 | 0.28 | 0.11 | 0.51 | 0.51 | 0.4 | 0.5 |
| 1 | 247607052 | rs10159239 | -0.547 | 0.104 | -0.087 | -0.456 | 0.036 | 0.026 | 0.026 | 1.001 | 0.5 | 0.65 | 0.62 | 0.57 | 0.51 | 0.62 |
| 1 | 247607642 | rs12130711 | 0.483 | 0.192 | -0.386 | -0.505 | 0.058 | 0.013 | 0.013 | 1.271 | 0.46 | 0.21 | 0.24 | 0.42 | 0.45 | 0.37 |
| 1 | 247609328 | rs12565738 | 0.436 | 0.21 | -0.175 | -0.509 | -0.014 | -0.065 | -0.065 | 0 | 0.2 | 0.35 | 0.12 | 0.07 | 0.1 | 0.12 |
| 1 | 247612562 | rs10925027 | -0.279 | 0.9 | -0.116 | -0.131 | 0.048 | 0.041 | 0.041 | 0 | 0.69 | 0.52 | 0.44 | 0.47 | 0.6 | 0.49 |
| 1 | 247614386 | rs12070953 | -0.646 | 0.424 | -0.362 | -0.555 | -0.005 | -0.053 | -0.053 | 0 | 0.15 | 0.2 | 0.18 | 0 | 0.14 | 0.04 |
| 1 | 247614407 | rs12065526 | -0.646 | 0.393 | -0.362 | -0.555 | -0.006 | -0.053 | -0.053 | 0.902 | 0.15 | 0.2 | 0.18 | 0 | 0.13 | 0.04 |
| 1 | 247614553 | rs12086048 | -0.643 | 0.39 | -0.363 | -0.556 | -0.006 | -0.053 | -0.053 | 0.361 | 0.15 | 0.19 | 0.18 | 0 | 0.13 | 0.04 |
| 1 | 247615512 | rs6673459 | -0.969 | 0.533 | -0.226 | -0.745 | 0.003 | -0.056 | -0.056 | 0 | 0.29 | 0.37 | 0.22 | 0.09 | 0.28 | 0.2 |
| 1 | 247617036 | rs4353135 | -0.755 | 0.19 | -0.884 | -0.513 | 0.064 | -0.013 | -0.013 | 0 | 0.6 | 0.79 | 0.74 | 0.59 | 0.67 | 0.69 |
| 1 | 247619310 | rs9988572 | 0.262 | 0.201 | -0.093 | -0.607 | -0.021 | -0.088 | -0.088 | 0 | 0.86 | 0.63 | 0.83 | 1 | 0.91 | 0.96 |
| 1 | 247622874 | rs4925671 | 0.494 | 0.187 | -0.071 | -0.685 | -0.002 | -0.011 | -0.011 | 0 | 0.71 | 0.67 | 0.81 | 0.84 | 0.72 | 0.78 |
| 1 | 247622940 | rs10737808 | -0.079 | 0.121 | 0.23 | -0.645 | 0.002 | 0.001 | 0.001 | 0 | 0.5 | 0.49 | 0.56 | 0.57 | 0.55 | 0.66 |
| 1 | 247652126 | rs11577815 | 0.694 | 0.574 | 0.382 | -0.959 | -0.047 | 0.008 | 0.008 | 0 | 0.12 | 0.04 | 0.06 | 0.08 | 0.17 | 0.15 |
| 1 | 247652603 | rs10754563 | -0.173 | 0.324 | 1.165 | -1.079 | 0.021 | -0.023 | -0.023 | 0 | 0.85 | 0.76 | 0.92 | 0.92 | 0.83 | 0.85 |
| 1 | 247653412 | rs10802516 | 0.222 | 0.728 | 1.019 | -1.262 | -0.001 | -0.005 | -0.005 | 2.012 | 0.5 | 0.47 | 0.69 | 0.74 | 0.53 | 0.56 |
| 1 | 247654993 | rs10925061 | 0.959 | 0.735 | 1.012 | -0.913 | -0.009 | -0.01 | -0.01 | 0 | 0.35 | 0.45 | 0.25 | 0.27 | 0.32 | 0.26 |
| 1 | 247657548 | rs11577737 | 0.59 | 0.757 | 0.546 | -0.971 | -0.047 | -0.012 | -0.012 | 0.992 | 0.16 | 0.04 | 0.06 | 0.1 | 0.17 | 0.18 |
| 1 | 247666165 | rs4472787 | 1.583 | 0.471 | 0.53 | -0.786 | -0.004 | 0.024 | 0.024 | 0 | 0.19 | 0.11 | 0.1 | 0.13 | 0.17 | 0.22 |
| 1 | 247689532 | rs1881797 | -0.247 | 0.377 | 0.844 | -0.837 | -0.027 | -0.111 | -0.111 | 0 | 0.82 | 0.42 | 0.76 | 0.97 | 0.84 | 0.93 |
| 1 | 247690417 | rs12404733 | 0.523 | 0.436 | 0.665 | -0.821 | -0.006 | -0.066 | -0.066 | 0.107 | 0.16 | 0.26 | 0.22 | 0.03 | 0.17 | 0.07 |
| 1 | 247696047 | rs10802521 | -0.889 | 0.242 | 0.204 | -0.814 | -0.002 | -0.004 | -0.004 | 0 | 0.74 | 0.73 | 0.85 | 0.8 | 0.78 | 0.71 |
| 1 | 247697724 | rs6426255 | -0.279 | 0.757 | -0.057 | -0.942 | 0.005 | -0.045 | -0.045 | 0 | 0.25 | 0.48 | 0.26 | 0.12 | 0.29 | 0.13 |
| 1 | 247701607 | rs10925073 | 0.587 | 0.758 | -1.178 | -0.472 | 0.009 | -0.022 | -0.022 | 0 | 0.35 | 0.11 | 0.22 | 0.32 | 0.34 | 0.29 |
| 1 | 247702886 | rs10925076 | -0.351 | 1.171 | -0.721 | -0.042 | -0.071 | -0.131 | -0.131 | 0.282 | 0.82 | 0.97 | 0.61 | 0.53 | 0.82 | 0.66 |
| 1 | 247710624 | rs1105488 | -0.275 | 0.503 | -0.855 | 0.018 | 0 | -0.003 | -0.003 | 0 | 0.28 | 0.67 | 0.24 | 0.41 | 0.31 | 0.27 |
| 1 | 247710867 | rs1105489 | -0.288 | 0.487 | -0.766 | 0.02 | -0.002 | 0.011 | 0.011 | 0 | 0.28 | 0.64 | 0.24 | 0.41 | 0.31 | 0.28 |
| 1 | 247715687 | rs4925559 | 0.08 | -0.001 | -2.2 | -1.379 | 0.047 | 0.118 | 0.118 | 1.772 | 0.53 | 0.27 | 0.68 | 0.74 | 0.63 | 0.71 |
| 1 | 247715998 | rs6426257 | -1.55 | 0.037 | -2.295 | -1.443 | -0.001 | 0.092 | 0.092 | 1.014 | 0.73 | 0.94 | 0.79 | 0.85 | 0.8 | 0.85 |
| 1 | 247719377 | rs3862182 | -0.054 | 0.258 | -1.921 | -1.228 | 0.015 | -0.062 | -0.062 | 0 | 0.11 | 0.65 | 0.1 | 0.01 | 0.12 | 0.06 |
| 1 | 247723670 | rs3844081 | 0.688 | 0.73 | -2.687 | -1.467 | -0.141 | -0.026 | -0.026 | 0 | 0.28 | 0.04 | 0.51 | 0.72 | 0.4 | 0.54 |
| 1 | 247728684 | rs1176064 | -0.633 | 0.326 | -2.015 | -1.927 | 0.001 | -0.017 | -0.017 | 0 | 0.63 | 0.43 | 0.65 | 0.78 | 0.6 | 0.75 |
| 1 | 247739213 | rs12402077 | 1.719 | 0.33 | -2.265 | -2.023 | -0.034 | 0.03 | 0.03 | 0.007 | 0.17 | 0.01 | 0.19 | 0.07 | 0.19 | 0.09 |
| 1 | 247740764 | rs12405268 | 1.719 | 0.33 | -2.265 | -2.023 | -0.034 | 0.03 | 0.03 | 0.007 | 0.17 | 0.01 | 0.19 | 0.07 | 0.19 | 0.09 |
| 1 | 247742577 | rs12075454 | 0.096 | 0.077 | -1.923 | -1.829 | -0.011 | -0.026 | -0.026 | 0 | 0.08 | 0.12 | 0.04 | 0.01 | 0.05 | 0.1 |
| 1 | 247745227 | rs10925083 | 1.111 | 0.245 | -2.357 | -1.992 | -0.118 | 0.05 | 0.05 | 0 | 0.33 | 0.06 | 0.51 | 0.76 | 0.41 | 0.49 |
| 1 | 247745313 | rs10802523 | 1.041 | 0.264 | -2.156 | -1.983 | -0.049 | 0.063 | 0.063 | 0 | 0.34 | 0.16 | 0.52 | 0.76 | 0.41 | 0.49 |
| 1 | 247748235 | rs1176046 | 0.138 | 0.186 | -1.764 | -1.649 | 0.001 | 0.009 | 0.009 | 0 | 0.56 | 0.42 | 0.64 | 0.78 | 0.58 | 0.67 |
| 1 | 247749572 | rs7539699 | -0.141 | 0.002 | -1.43 | -1.548 | 0.006 | 0.019 | 0.019 | 0.005 | 0.39 | 0.3 | 0.31 | 0.21 | 0.37 | 0.25 |
| 1 | 247752161 | rs10925085 | 0.034 | 0.259 | -2.004 | -0.66 | -0.05 | -0.149 | -0.149 | 0.786 | 0.36 | 0.06 | 0.35 | 0.52 | 0.41 | 0.28 |

| | | | | | | | | | | | | | | | | |
|---|---|---|---|---|---|---|---|---|---|---|---|---|---|---|---|---|
| 1 | 247752367 | rs869111 | -0.12 | 0.011 | -1.666 | -1.449 | -0.009 | -0.004 | -0.004 | 0.003 | 0.44 | 0.31 | 0.34 | 0.44 | 0.52 | 0.34 |
| 1 | 247760810 | rs6676750 | 0.041 | 0.037 | -1.654 | -1.444 | -0.011 | -0.004 | -0.004 | 0 | 0.42 | 0.3 | 0.35 | 0.44 | 0.5 | 0.31 |
| 1 | 247762249 | rs7537011 | 1.737 | -0.033 | -1.466 | -1.532 | 0.014 | 0.019 | 0.019 | 0.316 | 0.17 | 0.12 | 0.19 | 0.06 | 0.15 | 0.07 |
| 1 | 247762328 | rs7523061 | -0.515 | -0.132 | -1.045 | -1.44 | -0.033 | -0.056 | -0.056 | 0 | 0.63 | 0.79 | 0.57 | 0.5 | 0.65 | 0.39 |
| 1 | 247763857 | rs4925689 | 0.278 | -0.046 | -0.417 | -1.411 | 0.002 | 0 | 0 | 0 | 0.51 | 0.39 | 0.49 | 0.5 | 0.53 | 0.34 |
| 1 | 247778888 | rs6671837 | 1.117 | 0.575 | -0.421 | -1.11 | -0.019 | 0.028 | 0.028 | 0 | 0.06 | 0.01 | 0.04 | 0 | 0.11 | 0.03 |
| 1 | 247781569 | rs1144815 | -0.494 | 0.438 | -0.348 | -1.173 | 0 | -0.006 | -0.006 | 0 | 0.52 | 0.49 | 0.63 | 0.77 | 0.55 | 0.61 |
| 1 | 247783890 | rs1144814 | 0.614 | 0.401 | 0.036 | -0.463 | 0 | 0.013 | 0.013 | 0 | 0.22 | 0.2 | 0.23 | 0.49 | 0.24 | 0.19 |
| 1 | 247789226 | rs1176018 | -0.329 | 0.185 | 0.039 | -0.585 | 0.001 | 0.004 | 0.004 | 0 | 0.44 | 0.35 | 0.34 | 0.65 | 0.39 | 0.38 |
| 1 | 247815357 | rs1391483 | 0.772 | -0.597 | -0.286 | -1.106 | -0.006 | 0 | 0 | 0 | 0.36 | 0.3 | 0.62 | 0.61 | 0.42 | 0.42 |
| 1 | 247828797 | rs28394701 | 0.974 | -0.63 | -0.36 | -1.198 | -0.018 | 0.069 | 0.069 | 0.104 | 0.29 | 0.13 | 0.2 | 0.18 | 0.31 | 0.24 |
| 1 | 247833196 | rs1151643 | 0.75 | -0.353 | -0.324 | -1.092 | -0.007 | 0.001 | 0.001 | 0 | 0.36 | 0.3 | 0.62 | 0.6 | 0.42 | 0.43 |
| 1 | 247834126 | rs28723634 | 0.398 | -0.367 | -0.26 | -1.11 | 0.015 | 0.025 | 0.025 | 0.73 | 0.43 | 0.3 | 0.24 | 0.38 | 0.35 | 0.37 |
| 1 | 247835352 | rs1151641 | -0.87 | -0.355 | -0.278 | -1.11 | 0.008 | -0.08 | -0.08 | 0.088 | 0.85 | 0.6 | 0.9 | 0.98 | 0.82 | 0.83 |
| 1 | 247835674 | rs28446289 | 0.276 | -0.22 | -1.194 | -0.99 | 0.016 | 0.027 | 0.027 | 0.578 | 0.43 | 0.3 | 0.24 | 0.38 | 0.35 | 0.37 |
| 1 | 247835950 | rs1151640 | 0.085 | 0.283 | -1.27 | -0.124 | -0.071 | -0.02 | -0.02 | 0 | 0.33 | 0.13 | 0.59 | 0.6 | 0.42 | 0.43 |
| 1 | 247840784 | rs28620741 | 0.474 | -0.019 | -0.914 | 0.096 | -0.008 | 0.001 | 0.001 | 0 | 0.13 | 0.12 | 0.04 | 0.18 | 0.04 | 0.13 |
| 1 | 247841859 | rs1151633 | -0.355 | 0.342 | -0.645 | -0.113 | 0.006 | -0.01 | -0.01 | 1.839 | 0.43 | 0.5 | 0.68 | 0.6 | 0.46 | 0.46 |
| 1 | 247844562 | rs1176002 | -0.034 | 0.425 | -1.066 | -0.129 | -0.067 | -0.016 | -0.016 | 0 | 0.34 | 0.13 | 0.6 | 0.6 | 0.42 | 0.43 |
| 1 | 247845256 | rs1176001 | -0.09 | 0.41 | -0.761 | -0.136 | -0.035 | -0.003 | -0.003 | 0 | 0.35 | 0.21 | 0.61 | 0.6 | 0.42 | 0.43 |
| 1 | 247857274 | rs7555595 | -0.173 | 0.35 | -0.791 | -0.381 | -0.011 | 0.037 | 0.037 | 0 | 0.09 | 0.03 | 0.07 | 0 | 0.07 | 0.04 |
| 1 | 247861516 | rs6679472 | -0.236 | -0.487 | -0.655 | -1.146 | -0.019 | -0.015 | -0.015 | 0.66 | 0.32 | 0.58 | 0.13 | 0.23 | 0.2 | 0.25 |
| 1 | 247862621 | rs7526880 | -1.012 | -0.597 | -0.44 | -1.2 | 0.062 | 0.104 | 0.104 | 1.01 | 0.42 | 0.79 | 0.22 | 0.24 | 0.28 | 0.29 |
| 1 | 247865309 | rs12095040 | 0.045 | -0.883 | 0.12 | -1.361 | -0.02 | -0.095 | -0.095 | 0 | 0.19 | 0.36 | 0.07 | 0.02 | 0.16 | 0.17 |
| 1 | 247865773 | rs3862188 | -1.422 | -0.893 | 0.004 | -1.36 | 0.047 | 0.119 | 0.119 | 1.075 | 0.43 | 0.93 | 0.24 | 0.24 | 0.28 | 0.3 |
| 1 | 247868113 | rs880143 | -1.428 | -0.904 | 0 | -1.365 | 0.048 | 0.119 | 0.119 | 1.065 | 0.43 | 0.93 | 0.24 | 0.24 | 0.28 | 0.3 |
| 1 | 247874551 | rs12091656 | 0.291 | -1.033 | 0.319 | -1.375 | -0.049 | 0.03 | 0.03 | 1.94 | 0.19 | 0.43 | 0.08 | 0.23 | 0.07 | 0.16 |
| 1 | 247875415 | rs2282316 | -1.071 | -1.08 | 0.066 | -1.75 | 0.018 | 0.024 | 0.024 | 1.229 | 0.35 | 0.69 | 0.2 | 0.24 | 0.21 | 0.26 |
| 1 | 247881348 | rs6587397 | 0.279 | -1.12 | 0.13 | -1.715 | 0.018 | -0.025 | -0.025 | 0.237 | 0.08 | 0.35 | 0.07 | 0 | 0.04 | 0.06 |
| 1 | 247896549 | rs1416117 | -0.204 | -1.188 | 1 | -1.736 | -0.052 | -0.012 | -0.012 | 0 | 0.24 | 0.58 | 0.13 | 0.22 | 0.09 | 0.2 |
| 1 | 247901567 | rs7552632 | 0.793 | -1.109 | 1.086 | -1.695 | -0.009 | -0.033 | -0.033 | 0 | 0.07 | 0.14 | 0.04 | 0 | 0.04 | 0.06 |
| 1 | 247910283 | rs841164 | -1.734 | -1.466 | 0.85 | -1.94 | 0.057 | 0.184 | 0.184 | 0.249 | 0.51 | 0.94 | 0.28 | 0.24 | 0.32 | 0.36 |
| 1 | 247910460 | rs841163 | -0.504 | -1.358 | 1.019 | -1.814 | -0.016 | -0.042 | -0.042 | 0 | 0.11 | 0.46 | 0.07 | 0.02 | 0.05 | 0.12 |
| 1 | 247913766 | rs841158 | -0.66 | -1.499 | 1.004 | -1.84 | -0.006 | -0.004 | -0.004 | 0 | 0.29 | 0.65 | 0.18 | 0.24 | 0.15 | 0.24 |
| 1 | 247918781 | rs2640034 | -0.172 | -1.624 | 0.728 | -2.198 | 0.005 | -0.02 | -0.02 | 0.089 | 0.8 | 0.8 | 0.81 | 1 | 0.84 | 0.83 |
| 1 | 247919340 | rs10788766 | 0.536 | -1.398 | 1.081 | -1.952 | 0.018 | -0.007 | -0.007 | 4.087 | 0.92 | 0.59 | 0.93 | 0.99 | 0.96 | 0.88 |
| 1 | 247921352 | rs1552813 | 0.552 | -1.398 | 1.108 | -1.952 | 0.021 | -0.005 | -0.005 | 0 | 0.92 | 0.58 | 0.92 | 0.99 | 0.96 | 0.88 |
| 1 | 247925390 | rs10788785 | -0.344 | -1.377 | 0.39 | -1.4 | -0.013 | 0.045 | 0.045 | 0 | 0.2 | 0.34 | 0.17 | 0.34 | 0.18 | 0.43 |
| 1 | 247929510 | rs10888379 | 0.798 | -1.17 | -0.334 | -2.31 | 0.012 | 0.089 | 0.089 | 0 | 0.66 | 0.17 | 0.65 | 0.98 | 0.71 | 0.77 |
| 1 | 247931319 | rs6685883 | 0.578 | -1.129 | -0.151 | -2.22 | 0.003 | -0.018 | -0.018 | 0 | 0.81 | 0.18 | 0.82 | 0.99 | 0.87 | 0.9 |
| 1 | 247937019 | rs12747565 | 2.086 | -0.093 | -0.423 | -2.039 | -0.089 | 0.004 | 0.004 | 0.335 | 0.24 | 0.02 | 0.34 | 0.13 | 0.31 | 0.19 |
| 1 | 247941192 | rs6694222 | 1.291 | -0.604 | 0.421 | -1.964 | 0.02 | 0.067 | 0.067 | 0.931 | 0.73 | 0.15 | 0.81 | 0.98 | 0.82 | 0.79 |
| 1 | 247952900 | rs2101702 | 0.388 | -0.724 | 0.93 | -1.506 | 0.03 | 0.036 | 0.036 | 0.342 | 0.64 | 0.2 | 0.77 | 0.71 | 0.79 | 0.75 |
| 1 | 247963276 | rs10788720 | 0.567 | -0.525 | 1.889 | -1.44 | -0.033 | -0.073 | -0.073 | 0 | 0.88 | 0.47 | 0.9 | 0.99 | 0.96 | 0.94 |
| 1 | 247966718 | rs10749629 | 1.395 | -0.536 | 1.631 | -1.502 | 0.013 | 0 | 0 | 1.905 | 0.78 | 0.17 | 0.82 | 0.99 | 0.88 | 0.89 |
| 1 | 247972002 | rs7529313 | -0.586 | -0.517 | 1.799 | -1.492 | 0.043 | -0.068 | -0.068 | 0 | 0.89 | 0.53 | 0.9 | 1 | 0.92 | 0.99 |
| 1 | 247975655 | rs7541715 | 1.314 | -0.529 | 1.764 | -1.5 | -0.001 | -0.039 | -0.039 | 0 | 0.78 | 0.23 | 0.82 | 0.99 | 0.88 | 0.93 |
| 1 | 247977420 | rs10736372 | 1.314 | -0.529 | 1.764 | -1.5 | -0.001 | -0.039 | -0.039 | 1.832 | 0.78 | 0.23 | 0.82 | 0.99 | 0.88 | 0.93 |
| 1 | 247978996 | rs10888249 | -0.578 | -0.44 | 1.655 | -1.441 | 0.043 | -0.067 | -0.067 | 0.304 | 0.89 | 0.54 | 0.9 | 1 | 0.92 | 0.99 |
| 1 | 247986129 | rs12130569 | -0.957 | -0.377 | 1.406 | -1.639 | -0.008 | -0.132 | -0.132 | 0 | 0.25 | 0.38 | 0.24 | 0.02 | 0.23 | 0.22 |
| 1 | 247991609 | rs4925724 | -0.022 | -0.415 | 1.305 | -1.668 | -0.024 | 0.077 | 0.077 | 0.098 | 0.18 | 0.04 | 0.16 | 0.01 | 0.21 | 0.15 |
| 1 | 247993295 | rs1389228 | 1.004 | -0.327 | 1.483 | -1.667 | 0 | -0.033 | -0.033 | 0.193 | 0.8 | 0.19 | 0.82 | 0.99 | 0.88 | 0.89 |
| 1 | 247993701 | rs12133205 | 0.299 | -0.31 | 1.427 | -1.652 | -0.024 | 0.062 | 0.062 | 0.514 | 0.16 | 0.04 | 0.16 | 0 | 0.19 | 0.13 |
| 1 | 247995497 | rs11807791 | -0.655 | -0.306 | 1.654 | -1.621 | -0.012 | -0.04 | -0.04 | 0.806 | 0.1 | 0.35 | 0.09 | 0.02 | 0.04 | 0.09 |
| 1 | 247997613 | rs10888252 | 1.739 | -0.276 | 1.197 | -1.653 | 0.014 | 0.134 | 0.134 | 1.539 | 0.63 | 0.13 | 0.66 | 0.98 | 0.68 | 0.76 |
| 1 | 248000141 | rs8179377 | 1.243 | -0.067 | 1.708 | -1.248 | -0.009 | -0.092 | -0.092 | 0 | 0.72 | 0.35 | 0.73 | 0.98 | 0.76 | 0.78 |
| 1 | 248004296 | rs6681483 | -1.375 | 0.066 | 1.709 | -0.953 | 0.028 | -0.072 | -0.072 | 0.09 | 0.12 | 0.36 | 0.13 | 0 | 0.1 | 0.07 |
| 1 | 248004775 | rs10888256 | 0.743 | 0.093 | 1.33 | -0.94 | 0.012 | -0.055 | -0.055 | 0 | 0.89 | 0.78 | 0.88 | 1 | 0.9 | 0.93 |
| 1 | 248012296 | rs7530535 | 1.805 | -0.193 | 1.123 | -0.641 | 0.011 | 0.021 | 0.021 | 1.557 | 0.69 | 0.23 | 0.68 | 0.79 | 0.74 | 0.78 |
| 1 | 248014871 | rs10888260 | 1.064 | -0.031 | 1.766 | -0.656 | 0.067 | -0.039 | -0.039 | 0 | 0.9 | 0.56 | 0.87 | 1 | 0.9 | 0.94 |
| 1 | 248016894 | rs12096473 | -0.292 | -0.039 | 1.799 | -0.63 | 0.06 | 0.012 | 0.012 | 0 | 0.05 | 0.3 | 0.09 | 0 | 0.05 | 0.06 |
| 1 | 248018426 | rs7513527 | -0.248 | -0.094 | 1.688 | -0.672 | 0.012 | -0.088 | -0.088 | 0 | 0.14 | 0.29 | 0.17 | 0 | 0.15 | 0.11 |
| 1 | 248019808 | rs4925742 | 0.776 | -0.137 | 1.441 | -0.504 | 0.001 | 0.102 | 0.102 | 0 | 0.91 | 0.73 | 0.93 | 0.8 | 0.94 | 0.98 |
| 1 | 248021747 | rs12045264 | 1.88 | -0.104 | 0.732 | -0.837 | 0.018 | 0.078 | 0.078 | 3.586 | 0.62 | 0.16 | 0.63 | 0.8 | 0.63 | 0.75 |
| 1 | 248023966 | rs3737277 | -1.454 | -0.161 | 1.001 | -0.726 | -0.002 | -0.008 | -0.008 | 0 | 0.27 | 0.33 | 0.27 | 0.19 | 0.27 | 0.2 |
| 1 | 248027152 | rs4925572 | -1.769 | -0.19 | 1.057 | -0.709 | -0.01 | -0.034 | -0.034 | 2.605 | 0.31 | 0.59 | 0.3 | 0.2 | 0.31 | 0.2 |
| 1 | 248032168 | rs6657427 | -2.132 | 0.161 | 0.714 | -0.267 | 0.002 | -0.003 | -0.003 | 0.446 | 0.34 | 0.62 | 0.44 | 0.32 | 0.36 | 0.35 |
| 1 | 248035778 | rs12404622 | -0.883 | 0.316 | 0.318 | -0.108 | 0.005 | 0.037 | 0.037 | 0 | 0.13 | 0.17 | 0.23 | 0.31 | 0.17 | 0.21 |
| 1 | 248039713 | rs3811445 | -1.909 | 0.449 | -0.086 | 0.322 | 0 | 0 | 0 | 0 | 0.52 | 0.53 | 0.71 | 0.54 | 0.59 | 0.61 |

| | | | | | | | | | | | | | | | | |
|---|---|---|---|---|---|---|---|---|---|---|---|---|---|---|---|---|
| 1 | 248040018 | rs4925574 | -1.372 | 0.723 | -0.366 | 0.531 | -0.028 | -0.068 | -0.068 | 0 | 0.22 | 0.07 | 0.36 | 0.31 | 0.26 | 0.31 |
| 1 | 248040104 | rs4925753 | -0.218 | 0.774 | -0.3 | 0.613 | -0.003 | -0.005 | -0.005 | 0 | 0.07 | 0.05 | 0.21 | 0.13 | 0.1 | 0.16 |
| 1 | 248043646 | rs11204537 | -1.077 | 0.554 | -0.128 | 0.407 | -0.007 | -0.004 | -0.004 | 0 | 0.19 | 0.26 | 0.18 | 0.18 | 0.17 | 0.16 |
| 1 | 248046272 | rs11204538 | 0.988 | 0.591 | -0.111 | 0.217 | -0.001 | -0.003 | -0.003 | 0.578 | 0.61 | 0.62 | 0.5 | 0.78 | 0.52 | 0.6 |
| 1 | 248058265 | rs6587386 | 0.72 | 0.396 | -0.305 | 1.052 | -0.003 | -0.004 | -0.004 | 0.083 | 0.72 | 0.72 | 0.61 | 0.53 | 0.65 | 0.59 |
| 1 | 248059423 | rs10888267 | 1.156 | -0.523 | -0.676 | 0.935 | -0.028 | -0.032 | -0.032 | 0 | 0.44 | 0.14 | 0.35 | 0.57 | 0.53 | 0.42 |
| 1 | 248059476 | rs12139390 | 1.618 | -0.54 | 1.036 | 1.884 | -0.011 | 0.041 | 0.041 | 0 | 0.77 | 0.26 | 0.6 | 0.58 | 0.8 | 0.75 |
| 1 | 248059712 | rs11204546 | 2.181 | -0.409 | 1.379 | 1.772 | 0.007 | -0.023 | -0.023 | 0 | 0.65 | 0.26 | 0.46 | 0.59 | 0.67 | 0.48 |
| 1 | 248059856 | rs10888268 | 2.088 | -0.402 | 1.382 | 1.742 | 0.008 | -0.022 | -0.022 | 0 | 0.64 | 0.26 | 0.45 | 0.59 | 0.67 | 0.47 |
| 1 | 248060541 | rs6587388 | -1.347 | -0.03 | 2.246 | 1.971 | 0.004 | 0.106 | 0.106 | 0 | 0.17 | 0.32 | 0.32 | 0.39 | 0.19 | 0.25 |
| 1 | 248069912 | rs11204553 | 1.895 | -0.047 | 2.362 | 1.676 | -0.021 | -0.092 | -0.092 | 0.646 | 0.78 | 0.47 | 0.74 | 0.96 | 0.84 | 0.68 |
| 1 | 248071583 | rs6670590 | 1.206 | 0.01 | 2.717 | 1.703 | 0.01 | -0.066 | -0.066 | 0 | 0.84 | 0.64 | 0.79 | 0.97 | 0.86 | 0.74 |
| 1 | 248072026 | rs6587399 | -0.736 | 0.035 | 2.584 | 1.72 | -0.01 | -0.024 | -0.024 | 0 | 0.06 | 0.19 | 0.05 | 0 | 0.02 | 0.05 |
| 1 | 248073488 | rs4259697 | -1.405 | 0.084 | 2.704 | 1.745 | -0.003 | -0.072 | -0.072 | 0 | 0.19 | 0.31 | 0.24 | 0.03 | 0.16 | 0.3 |
| 1 | 248074978 | rs11204555 | 1.652 | -0.011 | 2.213 | 2.011 | -0.008 | -0.006 | -0.006 | 0 | 0.67 | 0.46 | 0.57 | 0.76 | 0.71 | 0.49 |
| 1 | 248076269 | rs11204557 | -1.122 | -0.362 | 2.094 | 2.264 | 0.021 | -0.08 | -0.08 | 0.929 | 0.23 | 0.04 | 0.24 | 0.26 | 0.15 | 0.46 |
| 1 | 248079963 | rs10888274 | -1.733 | -0.413 | 2.662 | 2.181 | -0.037 | 0.061 | 0.061 | 0.646 | 0.28 | 0.47 | 0.28 | 0.35 | 0.2 | 0.46 |
| 1 | 248082848 | rs10888277 | 1.653 | -0.393 | 2.539 | 2.337 | 0.013 | -0.008 | -0.008 | 0 | 0.68 | 0.35 | 0.69 | 0.45 | 0.79 | 0.51 |
| 1 | 248083564 | rs12046496 | -1.607 | -0.363 | 2.46 | 2.351 | -0.011 | 0.013 | 0.013 | 0.812 | 0.28 | 0.3 | 0.27 | 0.55 | 0.2 | 0.49 |
| 1 | 248085784 | rs4925775 | 1.397 | -0.352 | 2.264 | 2.388 | -0.01 | 0.011 | 0.011 | 0.98 | 0.72 | 0.68 | 0.72 | 0.45 | 0.79 | 0.51 |
| 1 | 248090251 | rs12094001 | -0.844 | -0.336 | 1.245 | 2.459 | 0.017 | -0.052 | -0.052 | 0.696 | 0.22 | 0.1 | 0.24 | 0.33 | 0.17 | 0.43 |
| 1 | 248094461 | rs11204565 | 1.483 | -0.313 | 1.16 | 2.04 | 0.003 | -0.003 | -0.003 | 0 | 0.68 | 0.36 | 0.7 | 0.45 | 0.79 | 0.51 |
| 1 | 248095483 | rs7525640 | 1.559 | -0.311 | 1.161 | 2.041 | 0.005 | -0.004 | -0.004 | 0 | 0.67 | 0.36 | 0.7 | 0.45 | 0.79 | 0.51 |
| 1 | 248111359 | rs1416874 | 1.936 | -0.329 | 0.814 | 2.034 | -0.023 | 0.042 | 0.042 | 5.5 | 0.9 | 0.3 | 0.93 | 0.88 | 0.95 | 0.95 |
| 1 | 248115436 | rs2184995 | 1.936 | -0.329 | 0.816 | 2.034 | -0.021 | 0.051 | 0.051 | 5.5 | 0.9 | 0.31 | 0.93 | 0.88 | 0.95 | 0.95 |
| 1 | 248117416 | rs10888287 | 1.498 | -0.336 | 0.926 | 2.041 | 0.026 | 0.113 | 0.113 | 6.081 | 0.94 | 0.38 | 0.94 | 0.92 | 0.99 | 0.97 |
| 1 | 248125297 | rs2065975 | 2.297 | -0.336 | 0.887 | 2.036 | -0.02 | 0.054 | 0.054 | 5.355 | 0.9 | 0.32 | 0.93 | 0.88 | 0.95 | 0.95 |
| 1 | 248126669 | rs10788748 | 2.553 | -0.361 | 0.848 | 2.024 | -0.029 | 0.037 | 0.037 | 4.237 | 0.89 | 0.31 | 0.93 | 0.88 | 0.95 | 0.95 |
| 1 | 248128929 | rs6664332 | -1.317 | -0.117 | 0.585 | 2.031 | 0.002 | -0.022 | -0.022 | 0 | 0.33 | 0.29 | 0.36 | 0.46 | 0.33 | 0.44 |
| 1 | 248129240 | rs4478844 | 1.35 | -0.069 | -0.271 | 1.819 | 0.025 | -0.043 | -0.043 | 0 | 0.58 | 0.09 | 0.57 | 0.46 | 0.65 | 0.53 |
| 1 | 248131036 | rs4925798 | 0.977 | 0.278 | 0.392 | 2.068 | -0.006 | 0.006 | 0.006 | 0 | 0.63 | 0.15 | 0.66 | 0.38 | 0.63 | 0.62 |
| 1 | 248131388 | rs7556514 | -2.898 | 0.12 | 2.182 | 2.628 | 0.07 | 0.379 | 0.379 | 0 | 0.08 | 0.46 | 0.11 | 0.27 | 0.07 | 0.06 |
| 1 | 248138217 | rs4451578 | 2.234 | 0.259 | 1.925 | 2.349 | 0.055 | 0.236 | 0.236 | 1.75 | 0.92 | 0.56 | 0.89 | 0.8 | 0.95 | 0.98 |
| 1 | 248145644 | rs6695603 | -2.707 | -0.077 | 2.412 | 2.557 | -0.019 | 0.075 | 0.075 | 1.998 | 0.16 | 0.73 | 0.12 | 0.32 | 0.12 | 0.12 |
| 1 | 248148188 | rs11576203 | -2.986 | -0.02 | 2.22 | 2.404 | -0.003 | 0.009 | 0.009 | 3.663 | 0.2 | 0.79 | 0.14 | 0.32 | 0.16 | 0.14 |
| 1 | 248178332 | rs11802312 | -2.154 | 0.001 | 2.153 | 2.444 | -0.003 | 0.026 | 0.026 | 0 | 0.06 | 0.12 | 0.04 | 0.16 | 0.05 | 0.07 |
| 1 | 248191707 | rs7511998 | -3.017 | 0.041 | 2.403 | 2.443 | 0.001 | 0.028 | 0.028 | 0.812 | 0.15 | 0.25 | 0.1 | 0.3 | 0.15 | 0.14 |
| 1 | 248220954 | rs12043704 | -0.935 | -0.164 | 1.925 | 2.83 | 0.023 | -0.036 | -0.036 | 0 | 0.1 | 0.02 | 0.07 | 0.36 | 0.05 | 0.21 |
| 1 | 248224216 | rs6658227 | -2.886 | -0.271 | 2.794 | 2.657 | 0.008 | 0.061 | 0.061 | 0.654 | 0.14 | 0.3 | 0.1 | 0.25 | 0.14 | 0.08 |
| 1 | 248227201 | rs7527770 | 3.204 | -0.391 | 2.714 | 2.625 | 0.015 | -0.036 | -0.036 | 0 | 0.8 | 0.09 | 0.85 | 0.75 | 0.85 | 0.92 |
| 1 | 248249895 | rs11204589 | 3.363 | -0.401 | 3.202 | 2.614 | 0.005 | -0.011 | -0.011 | 0 | 0.81 | 0.17 | 0.86 | 0.75 | 0.85 | 0.92 |
| 1 | 248255265 | rs12037329 | -2.718 | -0.389 | 3.437 | 2.65 | 0.065 | 0.123 | 0.123 | 0.584 | 0.07 | 0.43 | 0.05 | 0.09 | 0.06 | 0.05 |
| 1 | 248256924 | rs7525316 | -1.749 | -0.348 | 3.564 | 2.66 | 0.003 | 0.022 | 0.022 | 0 | 0.09 | 0.11 | 0.06 | 0.17 | 0.08 | 0.03 |
| 1 | 248264018 | rs10749643 | 3.167 | -0.594 | 3.244 | 2.472 | 0.003 | -0.006 | -0.006 | 2.491 | 0.76 | 0.21 | 0.84 | 0.72 | 0.8 | 0.86 |
| 1 | 248266555 | rs6669478 | 2.801 | -0.609 | 3.489 | 2.578 | 0.016 | 0.055 | 0.055 | 1.752 | 0.89 | 0.43 | 0.92 | 0.9 | 0.91 | 0.95 |
| 1 | 248279484 | rs11204592 | 3.069 | -0.435 | 3.224 | 2.702 | 0.025 | -0.044 | -0.044 | 0 | 0.67 | 0.08 | 0.74 | 0.39 | 0.74 | 0.68 |
| 1 | 248286356 | rs9435872 | -1.968 | -0.092 | 3.599 | 2.156 | 0 | 0.098 | 0.098 | 0.768 | 0.1 | 0.34 | 0.07 | 0.18 | 0.07 | 0.04 |
| 1 | 248290903 | rs12043363 | -1.168 | -0.092 | 3.643 | 2.232 | 0 | 0.009 | 0.009 | 0.004 | 0.06 | 0.06 | 0.02 | 0.1 | 0.06 | 0.05 |
| 1 | 248297531 | rs3001300 | -1.975 | -0.076 | 3.629 | 2.154 | 0 | 0.098 | 0.098 | 0.768 | 0.1 | 0.34 | 0.07 | 0.18 | 0.07 | 0.04 |
| 1 | 248316406 | rs7553764 | -3.275 | -0.043 | 3.241 | 2.128 | -0.02 | 0.063 | 0.063 | 0.188 | 0.18 | 0.63 | 0.11 | 0.26 | 0.14 | 0.08 |
| 1 | 248322479 | rs11804999 | -3.502 | -0.031 | 3.144 | 2.131 | -0.011 | 0.049 | 0.049 | 1.131 | 0.19 | 0.65 | 0.12 | 0.27 | 0.16 | 0.08 |
| 1 | 248322906 | rs9435885 | -1.528 | -0.006 | 3.018 | 2.155 | 0.002 | 0.085 | 0.085 | 0.017 | 0.09 | 0.23 | 0.06 | 0.18 | 0.07 | 0.03 |
| 1 | 248323242 | rs6692816 | -1.268 | -0.001 | 2.9 | 2.154 | 0 | 0.004 | 0.004 | 0 | 0.06 | 0.06 | 0.02 | 0.08 | 0.06 | 0.04 |
| 1 | 248329013 | rs9435886 | -1.638 | -0.003 | 2.915 | 1.817 | 0.002 | 0.085 | 0.085 | 0.012 | 0.09 | 0.23 | 0.06 | 0.18 | 0.07 | 0.03 |
| 1 | 248340600 | rs3001306 | -2.412 | 0.004 | 2.174 | 1.812 | -0.007 | 0.058 | 0.058 | 3.365 | 0.13 | 0.64 | 0.11 | 0.19 | 0.09 | 0.04 |
| 1 | 248342577 | rs6683114 | 1.219 | 0.076 | 1.883 | 1.29 | 0.007 | -0.061 | -0.061 | 1.85 | 0.89 | 0.73 | 0.91 | 1 | 0.91 | 0.98 |
| 1 | 248356143 | rs12025124 | -1.202 | 0.682 | 2.338 | 1.979 | 0.031 | 0.155 | 0.155 | 0 | 0.16 | 0.34 | 0.2 | 0.36 | 0.15 | 0.33 |
| 1 | 248360026 | rs7548525 | 0.84 | 0.78 | 2.235 | 1.778 | 0.001 | -0.001 | -0.001 | 1.73 | 0.76 | 0.74 | 0.78 | 0.79 | 0.76 | 0.93 |
| 1 | 248362339 | rs7550643 | 1.904 | 0.734 | 2.254 | 1.764 | -0.005 | -0.001 | -0.001 | 2.832 | 0.6 | 0.41 | 0.51 | 0.44 | 0.61 | 0.6 |
| 1 | 248363421 | rs12142371 | -0.702 | 0.792 | 2.058 | 1.8 | 0.027 | 0.137 | 0.137 | 0 | 0.15 | 0.3 | 0.27 | 0.35 | 0.15 | 0.33 |
| 1 | 248376070 | rs6587439 | 0.606 | 0.606 | 1.698 | 1.664 | -0.039 | -0.012 | -0.012 | 0 | 0.78 | 0.94 | 0.8 | 0.8 | 0.77 | 0.92 |
| 1 | 248401116 | rs4347239 | 1.155 | 0.477 | 2.57 | 1.35 | 0.013 | -0.019 | -0.019 | 0 | 0.72 | 0.43 | 0.75 | 0.8 | 0.74 | 0.91 |
| 1 | 248402758 | rs6666148 | 1.155 | 0.479 | 2.909 | 1.35 | 0.013 | -0.017 | -0.017 | 0.055 | 0.72 | 0.43 | 0.75 | 0.8 | 0.73 | 0.91 |
| 1 | 248411889 | rs4453078 | 0.738 | 0.442 | 2.956 | 1.463 | 0.043 | 0.019 | 0.019 | 0 | 0.78 | 0.67 | 0.75 | 0.73 | 0.71 | 0.89 |
| 1 | 248415662 | rs6587440 | 0.817 | 0.451 | 2.897 | 1.349 | 0.045 | -0.015 | -0.015 | 0 | 0.77 | 0.62 | 0.76 | 0.83 | 0.72 | 0.89 |
| 1 | 248426484 | rs4517383 | 0.358 | 0.382 | 2.896 | 1.332 | 0.01 | -0.003 | -0.003 | 0 | 0.81 | 0.82 | 0.79 | 0.86 | 0.72 | 0.9 |
| 1 | 248428227 | rs10888337 | 0.66 | 0.339 | 2.886 | 1.312 | 0.032 | -0.015 | -0.015 | 0.074 | 0.79 | 0.68 | 0.77 | 0.86 | 0.72 | 0.9 |
| 1 | 248429719 | rs10749646 | 0.272 | 0.283 | 2.699 | 1.423 | -0.053 | 0.006 | 0.006 | 0 | 0.81 | 0.93 | 0.79 | 0.86 | 0.72 | 0.9 |
| 1 | 248435831 | rs7521689 | 0.19 | 0.315 | 2.713 | 1.438 | -0.052 | 0.005 | 0.005 | 0 | 0.81 | 0.93 | 0.79 | 0.86 | 0.72 | 0.9 |

| | | | | | | | | | | | | | | | | |
|---|---|---|---|---|---|---|---|---|---|---|---|---|---|---|---|---|
| 1 | 248437568 | rs4586010 | 0.301 | 0.287 | 2.747 | 1.439 | 0.021 | -0.005 | -0.005 | 0 | 0.81 | 0.78 | 0.78 | 0.86 | 0.73 | 0.9 |
| 1 | 248455512 | rs10888354 | -0.806 | 0.356 | 2.104 | 1.476 | -0.031 | -0.113 | -0.113 | 0.115 | 0.2 | 0.01 | 0.25 | 0.36 | 0.17 | 0.35 |
| 1 | 248464330 | rs4430369 | 0.408 | 0.403 | 2.252 | 1.339 | -0.024 | 0.009 | 0.009 | 0 | 0.8 | 0.89 | 0.78 | 0.86 | 0.72 | 0.9 |
| 1 | 248464871 | rs4372295 | 0.307 | 0.412 | 2.398 | 1.343 | 0.036 | -0.01 | -0.01 | 0 | 0.81 | 0.75 | 0.77 | 0.86 | 0.72 | 0.9 |
| 1 | 248471222 | rs4244183 | 0.244 | 0.416 | 2.304 | 1.345 | 0.022 | -0.006 | -0.006 | 0.971 | 0.81 | 0.79 | 0.78 | 0.86 | 0.72 | 0.9 |
| 1 | 248475798 | rs4310471 | 0.383 | 0.413 | 2.294 | 1.342 | -0.019 | 0.009 | 0.009 | 0 | 0.8 | 0.86 | 0.77 | 0.86 | 0.71 | 0.9 |
| 1 | 248486913 | rs4244185 | 0.416 | 0.423 | 2.335 | 1.342 | 0.019 | -0.009 | -0.009 | 0 | 0.8 | 0.77 | 0.77 | 0.86 | 0.71 | 0.9 |
| 1 | 248487016 | rs4451579 | -0.804 | 0.354 | 2.289 | 1.457 | -0.035 | -0.125 | -0.125 | 0 | 0.21 | 0.01 | 0.25 | 0.36 | 0.17 | 0.36 |
| 1 | 248487638 | rs7555310 | 0.42 | 0.407 | 2.341 | 1.232 | 0.02 | -0.007 | -0.007 | 0.01 | 0.8 | 0.77 | 0.77 | 0.86 | 0.71 | 0.9 |
| 1 | 248493511 | rs4244187 | 0.385 | 0.409 | 2.134 | 1.196 | -0.019 | 0.009 | 0.009 | 0 | 0.8 | 0.87 | 0.77 | 0.86 | 0.71 | 0.9 |
| 1 | 248496863 | rs4481887 | 0.189 | 0.424 | 2.128 | 1.214 | -0.023 | 0.006 | 0.006 | 0 | 0.81 | 0.9 | 0.78 | 0.86 | 0.72 | 0.9 |
| 1 | 248509182 | rs1339992 | 0.84 | 0.42 | 2.559 | 1.212 | 0.059 | -0.036 | -0.036 | 0.24 | 0.77 | 0.46 | 0.74 | 0.86 | 0.71 | 0.89 |
| 1 | 248509495 | rs1339990 | 0.84 | 0.42 | 2.559 | 1.212 | 0.059 | -0.036 | -0.036 | 0 | 0.77 | 0.46 | 0.75 | 0.86 | 0.71 | 0.89 |
| 1 | 248519821 | rs1538703 | 0.987 | 0.366 | 2.196 | 1.354 | 0.043 | 0.001 | 0.001 | 0 | 0.73 | 0.44 | 0.7 | 0.77 | 0.65 | 0.81 |
| 1 | 248521032 | rs946761 | 0.779 | 0.317 | 1.894 | 1.319 | 0.027 | -0.001 | -0.001 | 0 | 0.74 | 0.58 | 0.72 | 0.78 | 0.66 | 0.81 |
| 1 | 248528981 | rs6669587 | -0.825 | 0.215 | 1.59 | 1.303 | -0.001 | 0.03 | 0.03 | 0 | 0.22 | 0.24 | 0.27 | 0.43 | 0.18 | 0.36 |
| 1 | 248533285 | rs6421453 | 0.379 | 0.378 | 1.621 | 1.347 | -0.057 | -0.037 | -0.037 | 0 | 0.77 | 0.95 | 0.75 | 0.77 | 0.67 | 0.8 |
| 1 | 248550894 | rs7418152 | -1.209 | 0.302 | 1.705 | 1.439 | -0.004 | 0.126 | 0.126 | 0 | 0.23 | 0.52 | 0.32 | 0.44 | 0.18 | 0.37 |
| 1 | 248552478 | rs1538706 | -0.769 | 0.43 | 0.705 | 1.501 | 0.001 | 0.037 | 0.037 | 0 | 0.2 | 0.22 | 0.29 | 0.44 | 0.18 | 0.37 |
| 1 | 248562691 | rs28571660 | -0.769 | 0.43 | 0.617 | 1.516 | 0.001 | 0.03 | 0.03 | 0 | 0.2 | 0.22 | 0.29 | 0.42 | 0.18 | 0.37 |
| 1 | 248568202 | rs28524986 | -0.792 | 0.431 | 0.623 | 1.477 | 0.002 | 0.044 | 0.044 | 0 | 0.2 | 0.25 | 0.29 | 0.44 | 0.18 | 0.37 |
| 1 | 248577691 | rs6587472 | -1.009 | 0.443 | 0.546 | 1.566 | 0.002 | 0.134 | 0.134 | 0 | 0.22 | 0.58 | 0.32 | 0.44 | 0.18 | 0.36 |
| 2 | 295255 | rs6548221 | -0.79 | 0.649 | -0.01 | -0.416 | 0.01 | 0.019 | 0.019 | 0.407 | 0.69 | 0.77 | 0.57 | 0.85 | 0.76 | 0.69 |
| 2 | 309480 | rs6717613 | -0.596 | 1.03 | -0.771 | 0.874 | -0.004 | 0.003 | 0.003 | 0.147 | 0.35 | 0.69 | 0.41 | 0.32 | 0.38 | 0.24 |
| 2 | 312992 | rs9309730 | 0.724 | 0.393 | -0.256 | 1.264 | -0.053 | 0.142 | 0.142 | 0.018 | 0.79 | 0.63 | 0.85 | 0.49 | 0.88 | 0.8 |
| 2 | 320574 | rs6719944 | 0.317 | 1.386 | -0.866 | 0.627 | -0.05 | 0.028 | 0.028 | 0 | 0.62 | 0.84 | 0.68 | 0.77 | 0.59 | 0.66 |
| 2 | 333665 | rs2941874 | 0.322 | 1.329 | -0.423 | 0.448 | 0.025 | -0.022 | -0.022 | 0 | 0.62 | 0.55 | 0.66 | 0.78 | 0.56 | 0.67 |
| 2 | 343517 | rs6749467 | -0.602 | 0.41 | -0.459 | -0.25 | 0 | -0.008 | -0.008 | 0 | 0.44 | 0.57 | 0.54 | 0.79 | 0.44 | 0.61 |
| 2 | 343588 | rs6548226 | -0.566 | -0.246 | -0.487 | -0.574 | -0.016 | 0.083 | 0.083 | 0.388 | 0.47 | 0.24 | 0.55 | 0.7 | 0.48 | 0.45 |
| 2 | 345703 | rs6548227 | -0.228 | -0.042 | 0.076 | -0.29 | -0.018 | 0.019 | 0.019 | 0.161 | 0.09 | 0.26 | 0.07 | 0.11 | 0.06 | 0.19 |
| 2 | 348470 | rs12472764 | -0.95 | -0.346 | -0.286 | -0.633 | -0.005 | 0.004 | 0.004 | 0.147 | 0.57 | 0.61 | 0.59 | 0.8 | 0.51 | 0.66 |
| 2 | 354241 | rs1858308 | 0.143 | -0.203 | -0.423 | 0.191 | -0.007 | -0.042 | -0.042 | 0 | 0.29 | 0.19 | 0.42 | 0.39 | 0.28 | 0.25 |
| 2 | 355197 | rs6730397 | -0.034 | -0.301 | 0.136 | -0.253 | -0.01 | -0.024 | -0.024 | 1.489 | 0.05 | 0.17 | 0.03 | 0 | 0.03 | 0.04 |
| 2 | 355800 | rs6740249 | -1.363 | -0.602 | -0.116 | -0.56 | -0.113 | 0.154 | 0.154 | 1.648 | 0.59 | 0.92 | 0.61 | 0.8 | 0.49 | 0.63 |
| 2 | 358768 | rs7581663 | 0.092 | -0.632 | 0.2 | -0.5 | -0.011 | -0.025 | -0.025 | 0 | 0.05 | 0.19 | 0.03 | 0 | 0.03 | 0.04 |
| 2 | 366410 | rs958877 | -1.279 | -0.66 | -0.289 | -0.523 | -0.104 | 0.157 | 0.157 | 0.911 | 0.58 | 0.92 | 0.6 | 0.8 | 0.48 | 0.61 |
| 2 | 367736 | rs7595480 | -0.367 | -0.65 | -0.019 | -0.54 | -0.018 | 0.103 | 0.103 | 0.401 | 0.43 | 0.29 | 0.51 | 0.79 | 0.41 | 0.54 |
| 2 | 371148 | rs11693451 | -0.781 | -0.483 | -0.032 | -0.443 | -0.004 | 0.091 | 0.091 | 0.047 | 0.52 | 0.72 | 0.57 | 0.79 | 0.44 | 0.59 |
| 2 | 382409 | rs11686764 | -0.752 | 0.041 | -0.657 | -0.447 | 0.018 | 0.14 | 0.14 | 0.254 | 0.54 | 0.77 | 0.52 | 0.82 | 0.36 | 0.59 |
| 2 | 383458 | rs4374407 | -0.145 | 0.549 | -0.553 | 0.06 | -0.031 | -0.011 | -0.011 | 0 | 0.07 | 0.01 | 0.04 | 0.08 | 0.1 | 0.14 |
| 2 | 384236 | rs7565976 | -0.383 | 0.37 | -0.288 | 0.42 | 0.012 | 0 | 0 | 0 | 0.39 | 0.63 | 0.48 | 0.33 | 0.3 | 0.35 |
| 2 | 391781 | rs12475610 | -0.466 | 0.298 | -0.353 | 0.569 | 0.042 | -0.021 | -0.021 | 0 | 0.42 | 0.68 | 0.47 | 0.46 | 0.32 | 0.48 |
| 2 | 406981 | rs4630798 | -1.021 | 1.527 | 0.166 | 1.066 | 0.058 | 0.047 | 0.047 | 0 | 0.2 | 0.3 | 0.37 | 0.29 | 0.31 | 0.3 |
| 2 | 419464 | rs6726633 | -0.899 | 1.098 | 0.35 | 1.238 | 0.014 | 0.183 | 0.183 | 0.09 | 0.05 | 0.13 | 0.06 | 0.24 | 0.08 | 0.06 |
| 2 | 429880 | rs10460598 | -0.931 | 1.091 | 0.345 | 1.255 | 0.011 | 0.179 | 0.179 | 0.108 | 0.05 | 0.13 | 0.06 | 0.25 | 0.08 | 0.06 |
| 2 | 437664 | rs2685230 | -1.373 | 1.593 | 0.909 | 1.25 | 0.034 | 0.021 | 0.021 | 0 | 0.28 | 0.52 | 0.37 | 0.36 | 0.35 | 0.34 |
| 2 | 440230 | rs7585330 | -1.174 | 1.549 | 0.775 | 1.246 | -0.057 | -0.036 | -0.036 | 1.431 | 0.24 | 0.1 | 0.33 | 0.32 | 0.35 | 0.33 |
| 2 | 443281 | rs4854307 | -1.68 | 1.245 | 1.342 | 1.405 | -0.005 | 0.235 | 0.235 | 4.292 | 0.08 | 0.23 | 0.21 | 0.34 | 0.07 | 0.07 |
| 2 | 449487 | rs2667214 | -2.303 | 1.595 | 1.37 | 1.658 | 0.063 | 0.172 | 0.172 | 0 | 0.22 | 0.45 | 0.41 | 0.45 | 0.29 | 0.38 |
| 2 | 451581 | rs2724856 | -2.051 | 1.31 | 1.633 | 1.572 | 0.061 | 0.185 | 0.185 | 1.326 | 0.07 | 0.41 | 0.15 | 0.14 | 0.04 | 0.04 |
| 2 | 451832 | rs17041942 | -1.159 | 0.91 | 1.35 | 1.281 | -0.017 | 0.013 | 0.013 | 0 | 0.09 | 0.15 | 0.13 | 0.1 | 0.05 | 0.13 |
| 2 | 451914 | rs2667203 | -1.636 | 1.094 | 1.74 | 1.497 | 0.058 | 0.146 | 0.146 | 0.648 | 0.06 | 0.36 | 0.13 | 0.11 | 0.03 | 0.04 |
| 2 | 454860 | rs1511324 | 0.043 | 0.12 | 0.825 | 1.211 | -0.002 | 0 | 0 | 0 | 0.37 | 0.39 | 0.27 | 0.29 | 0.25 | 0.34 |
| 2 | 455683 | rs6729656 | -0.543 | 0.114 | 0.747 | 1.647 | -0.005 | 0.079 | 0.079 | 0 | 0.09 | 0.13 | 0.07 | 0.37 | 0.1 | 0.08 |
| 2 | 465300 | rs868559 | -1.256 | 0.448 | 0.311 | 1.473 | 0.057 | 0.08 | 0.08 | 0 | 0.23 | 0.42 | 0.4 | 0.35 | 0.3 | 0.34 |
| 2 | 467148 | rs2724860 | -1.582 | 0.996 | 0.015 | 2.161 | -0.178 | -0.079 | -0.079 | 0 | 0.63 | 0.98 | 0.71 | 0.7 | 0.59 | 0.71 |
| 2 | 467580 | rs10209725 | -0.937 | 0.809 | 0.459 | 1.866 | -0.033 | 0.172 | 0.172 | 0.414 | 0.14 | 0.37 | 0.12 | 0.32 | 0.11 | 0.12 |
| 2 | 469494 | rs12469725 | 0.348 | 1.018 | -0.279 | 1.428 | -0.01 | 0.013 | 0.013 | 0 | 0.09 | 0.05 | 0.28 | 0.03 | 0.19 | 0.22 |
| 2 | 471514 | rs2724861 | -1.059 | 0.268 | -0.554 | 1.196 | 0.007 | 0.033 | 0.033 | 3.045 | 0.46 | 0.64 | 0.61 | 0.68 | 0.45 | 0.64 |
| 2 | 477364 | rs2724863 | -0.564 | 0.099 | -0.624 | 1.544 | -0.008 | -0.052 | -0.052 | 0.153 | 0.32 | 0.29 | 0.54 | 0.51 | 0.31 | 0.49 |
| 2 | 481504 | rs2724913 | -1.008 | -0.232 | -0.602 | 0.822 | 0.016 | -0.07 | -0.07 | 0.32 | 0.63 | 0.56 | 0.74 | 0.88 | 0.55 | 0.68 |
| 2 | 483114 | rs4854313 | -0.196 | -0.619 | -1.747 | 0.899 | -0.049 | -0.069 | -0.069 | 0 | 0.32 | 0.15 | 0.44 | 0.55 | 0.36 | 0.46 |
| 2 | 487043 | rs2667215 | -1.335 | -0.335 | -1.399 | 0.629 | -0.058 | 0.099 | 0.099 | 0 | 0.67 | 0.88 | 0.77 | 0.91 | 0.57 | 0.68 |
| 2 | 499823 | rs2685217 | -0.462 | -0.457 | -0.596 | 1.249 | -0.009 | 0.021 | 0.021 | 0 | 0.31 | 0.6 | 0.21 | 0.45 | 0.25 | 0.29 |
| 2 | 502023 | rs4854316 | -0.608 | -0.389 | -0.566 | 0.737 | -0.021 | -0.003 | -0.003 | 0 | 0.16 | 0.11 | 0.34 | 0.16 | 0.19 | 0.24 |
| 2 | 504243 | rs11891417 | 0.711 | -0.259 | -0.455 | 0.826 | 0.022 | 0.009 | 0.009 | 0 | 0.06 | 0.09 | 0.23 | 0.14 | 0.14 | 0.17 |
| 2 | 504767 | rs17041279 | 0.812 | -0.586 | -0.295 | 1.052 | -0.015 | 0.045 | 0.045 | 0 | 0.17 | 0.19 | 0.29 | 0.61 | 0.11 | 0.23 |
| 2 | 506733 | rs11127475 | -0.297 | -1.156 | -0.107 | 0.518 | -0.002 | -0.006 | -0.006 | 0.273 | 0.17 | 0.17 | 0.29 | 0.28 | 0.18 | 0.23 |
| 2 | 510975 | rs12465676 | 1.316 | -1.189 | -0.256 | 0.598 | 0.013 | -0.108 | -0.108 | 0 | 0.16 | 0.01 | 0.25 | 0.44 | 0.11 | 0.23 |

| | | | | | | | | | | | | | | | | |
|---|---|---|---|---|---|---|---|---|---|---|---|---|---|---|---|---|
| 2 | 515593 | rs2724929 | -1.679 | -1.617 | -0.339 | 0.206 | 0.051 | 0.041 | 0.041 | 2.751 | 0.37 | 0.83 | 0.39 | 0.31 | 0.28 | 0.31 |
| 2 | 516565 | rs13006698 | 1.512 | -1.525 | -0.54 | 0.034 | -0.036 | -0.029 | -0.029 | 0.189 | 0.17 | 0.04 | 0.15 | 0.16 | 0.2 | 0.19 |
| 2 | 522617 | rs4854318 | -0.508 | -1.356 | -0.239 | 0.208 | 0.01 | 0.009 | 0.009 | 0 | 0.33 | 0.18 | 0.54 | 0.74 | 0.3 | 0.46 |
| 2 | 522921 | rs2724855 | -1.854 | -1.38 | -0.185 | 0.194 | 0.03 | 0.045 | 0.045 | 0.271 | 0.51 | 0.66 | 0.63 | 0.77 | 0.4 | 0.55 |
| 2 | 523613 | rs10865547 | 1.521 | -1.392 | -0.277 | 0.351 | -0.006 | -0.118 | -0.118 | 0 | 0.15 | 0 | 0.24 | 0.46 | 0.11 | 0.22 |
| 2 | 524774 | rs1317448 | 0.707 | -1.396 | 0.197 | -0.131 | -0.021 | -0.04 | -0.04 | 0 | 0.12 | 0.16 | 0.05 | 0.01 | 0.09 | 0.08 |
| 2 | 535043 | rs2974256 | -0.447 | -1.375 | 0.019 | -0.308 | -0.019 | -0.04 | -0.04 | 0 | 0.31 | 0.52 | 0.23 | 0.17 | 0.29 | 0.24 |
| 2 | 548121 | rs2685261 | -2.107 | -1.229 | -0.216 | -0.394 | -0.062 | 0.094 | 0.094 | 0 | 0.66 | 0.86 | 0.75 | 0.86 | 0.59 | 0.7 |
| 2 | 553264 | rs2685244 | -2.138 | -1.071 | -0.444 | -0.442 | -0.077 | 0.158 | 0.158 | 0 | 0.65 | 0.91 | 0.8 | 0.93 | 0.6 | 0.74 |
| 2 | 554109 | rs13411762 | 0.799 | -1.065 | -0.166 | -0.457 | -0.012 | -0.037 | -0.037 | 0.821 | 0.1 | 0.13 | 0.04 | 0 | 0.08 | 0.08 |
| 2 | 557590 | rs2685268 | -1.689 | -1.055 | -0.522 | -0.314 | -0.058 | 0.106 | 0.106 | 0 | 0.6 | 0.84 | 0.71 | 0.8 | 0.57 | 0.67 |
| 2 | 559913 | rs17042018 | 1.33 | -1.057 | -0.661 | -0.06 | -0.005 | -0.132 | -0.132 | 3.191 | 0.16 | 0 | 0.24 | 0.47 | 0.11 | 0.23 |
| 2 | 569139 | rs2724918 | -0.847 | -1.104 | -0.155 | -0.301 | -0.079 | 0.002 | 0.002 | 0.682 | 0.18 | 0.62 | 0.17 | 0.17 | 0.07 | 0.1 |
| 2 | 573207 | rs2683968 | -1.883 | -0.992 | -0.435 | -0.41 | -0.082 | 0.168 | 0.168 | 0 | 0.63 | 0.88 | 0.79 | 0.93 | 0.58 | 0.67 |
| 2 | 581414 | rs17042043 | 0.8 | -0.591 | -0.63 | -0.132 | -0.024 | -0.046 | -0.046 | 0.323 | 0.14 | 0.01 | 0.1 | 0.2 | 0.16 | 0.19 |
| 2 | 584523 | rs6712649 | -1.059 | -0.346 | -0.171 | -0.215 | -0.004 | 0.018 | 0.018 | 0 | 0.56 | 0.42 | 0.67 | 0.79 | 0.54 | 0.61 |
| 2 | 587649 | rs17042065 | -0.249 | -0.106 | -0.348 | 0.03 | -0.006 | -0.011 | -0.011 | 0 | 0.11 | 0.14 | 0.08 | 0.04 | 0.08 | 0.05 |
| 2 | 596532 | rs12714409 | 0.12 | -0.13 | -0.237 | 0.26 | -0.042 | 0.194 | 0.194 | 0 | 0.21 | 0.46 | 0.28 | 0.49 | 0.13 | 0.26 |
| 2 | 596629 | rs4854267 | -0.692 | 0.402 | -0.857 | -0.55 | -0.016 | 0.061 | 0.061 | 0 | 0.58 | 0.69 | 0.73 | 0.93 | 0.56 | 0.68 |
| 2 | 597557 | rs17784455 | -0.475 | 0.758 | -0.792 | 0.185 | -0.032 | -0.009 | -0.009 | 0 | 0.06 | 0.02 | 0.09 | 0.05 | 0.16 | 0.06 |
| 2 | 601905 | rs2867105 | -0.293 | 0.548 | -0.732 | -0.06 | 0 | 0.01 | 0.01 | 0 | 0.83 | 0.87 | 0.85 | 0.91 | 0.83 | 0.73 |
| 2 | 602036 | rs12996547 | -0.314 | 0.244 | -0.535 | -0.71 | -0.001 | -0.073 | -0.073 | 0.798 | 0.31 | 0.52 | 0.27 | 0.11 | 0.36 | 0.25 |
| 2 | 602475 | rs1316802 | 1.076 | 0.556 | -0.766 | -0.112 | -0.017 | -0.072 | -0.072 | 0 | 0.18 | 0.08 | 0.33 | 0.47 | 0.17 | 0.26 |
| 2 | 603079 | rs4241319 | 0.939 | 0.251 | -0.578 | -0.322 | -0.021 | 0.052 | 0.052 | 0.213 | 0.49 | 0.21 | 0.37 | 0.41 | 0.49 | 0.41 |
| 2 | 607049 | rs2903497 | 0.252 | 0.182 | -0.451 | -0.365 | -0.005 | 0.006 | 0.006 | 0 | 0.11 | 0.1 | 0.09 | 0.05 | 0.16 | 0.11 |
| 2 | 607107 | rs2867133 | -0.179 | 0.165 | -0.138 | -0.374 | -0.001 | -0.003 | -0.003 | 0 | 0.64 | 0.68 | 0.52 | 0.46 | 0.66 | 0.51 |
| 2 | 608411 | rs12714411 | -0.599 | 0.315 | -0.224 | -0.218 | 0.014 | -0.043 | -0.043 | 0 | 0.15 | 0.47 | 0.15 | 0.05 | 0.17 | 0.11 |
| 2 | 610088 | rs2903496 | 0.874 | 0.382 | -0.998 | 0.102 | -0.059 | -0.145 | -0.145 | 6.228 | 0.18 | 0.01 | 0.32 | 0.45 | 0.17 | 0.26 |
| 2 | 613782 | rs2966402 | 0.184 | 0.28 | -0.373 | 0.008 | 0 | -0.001 | -0.001 | 0 | 0.6 | 0.62 | 0.49 | 0.42 | 0.66 | 0.54 |
| 2 | 619653 | rs10153797 | 0.601 | 0.326 | -0.874 | -0.161 | -0.007 | 0.027 | 0.027 | 0 | 0.16 | 0.11 | 0.13 | 0.06 | 0.19 | 0.15 |
| 2 | 622827 | rs2867125 | -0.909 | 0.318 | -0.826 | -0.182 | -0.001 | 0.028 | 0.028 | 0.02 | 0.83 | 0.9 | 0.86 | 0.91 | 0.83 | 0.81 |
| 2 | 623588 | rs12468153 | 1.098 | 0.358 | -0.964 | -0.062 | -0.025 | -0.089 | -0.089 | 0 | 0.17 | 0.05 | 0.33 | 0.5 | 0.17 | 0.26 |
| 2 | 629914 | rs6732471 | -0.993 | 0.352 | -0.869 | -0.055 | -0.005 | 0.035 | 0.035 | 0 | 0.83 | 0.9 | 0.86 | 0.91 | 0.83 | 0.87 |
| 2 | 638144 | rs4854344 | -0.956 | 0.34 | -0.289 | -0.061 | 0 | -0.02 | -0.02 | 0 | 0.83 | 0.79 | 0.86 | 0.91 | 0.83 | 0.87 |
| 2 | 644953 | rs7561317 | -0.789 | 0.322 | -0.273 | -0.076 | -0.003 | -0.024 | -0.024 | 0 | 0.82 | 0.76 | 0.85 | 0.91 | 0.83 | 0.87 |
| 2 | 647760 | rs4854348 | -0.662 | 0.236 | 0.459 | -0.113 | 0.002 | -0.066 | -0.066 | 0 | 0.81 | 0.59 | 0.84 | 0.91 | 0.83 | 0.87 |
| 2 | 648527 | rs4640417 | 0.015 | 0.019 | 0.402 | -0.077 | -0.005 | -0.035 | -0.035 | 0 | 0.37 | 0.22 | 0.48 | 0.53 | 0.36 | 0.46 |
| 2 | 651641 | rs9784046 | 0.336 | 0.271 | -0.041 | -0.082 | 0 | -0.001 | -0.001 | 0 | 0.09 | 0.07 | 0.1 | 0.05 | 0.18 | 0.16 |
| 2 | 652797 | rs6720683 | 0.731 | 0.129 | 0.066 | -0.072 | -0.039 | 0.018 | 0.018 | 0.134 | 0.43 | 0.16 | 0.36 | 0.37 | 0.52 | 0.39 |
| 2 | 653195 | rs13396935 | 0.488 | 0.289 | 0.258 | 0.069 | -0.001 | -0.022 | -0.022 | 0 | 0.17 | 0.24 | 0.15 | 0.09 | 0.17 | 0.15 |
| 2 | 654472 | rs6728479 | -0.744 | 0.171 | 0.292 | 1.044 | -0.014 | 0.111 | 0.111 | 0.226 | 0.11 | 0.21 | 0.23 | 0.35 | 0.07 | 0.18 |
| 2 | 654685 | rs11689974 | -0.214 | 0.335 | 0.509 | 1.183 | 0.012 | 0.095 | 0.095 | 3.536 | 0.8 | 0.49 | 0.77 | 0.64 | 0.85 | 0.69 |
| 2 | 657569 | rs7574078 | -0.423 | 0.088 | 0.368 | 0.968 | -0.001 | 0.033 | 0.033 | 6.056 | 0.32 | 0.38 | 0.45 | 0.55 | 0.29 | 0.57 |
| 2 | 657652 | rs4854350 | 1.379 | 0.156 | -0.354 | 0.643 | -0.111 | -0.091 | -0.091 | 1.318 | 0.28 | 0.02 | 0.25 | 0.32 | 0.33 | 0.18 |
| 2 | 658937 | rs6548243 | -0.401 | -0.079 | 0.262 | 0.902 | -0.003 | 0.038 | 0.038 | 5.521 | 0.32 | 0.42 | 0.46 | 0.55 | 0.29 | 0.57 |
| 2 | 659958 | rs7592497 | -0.267 | -0.023 | 0.072 | 0.897 | -0.001 | -0.003 | -0.003 | 0.02 | 0.3 | 0.24 | 0.46 | 0.56 | 0.28 | 0.57 |
| 2 | 661188 | rs17042314 | -0.142 | -0.195 | 0.492 | 0.469 | -0.026 | -0.084 | -0.084 | 0.384 | 0.16 | 0.31 | 0.1 | 0.02 | 0.11 | 0.12 |
| 2 | 661645 | rs4854351 | -0.011 | -0.259 | 0.798 | 0.679 | -0.007 | 0.032 | 0.032 | 0 | 0.88 | 0.75 | 0.78 | 0.84 | 0.86 | 0.79 |
| 2 | 666343 | rs17729386 | 1.314 | 0.028 | 0.183 | 0.717 | -0.039 | -0.05 | -0.05 | 0.741 | 0.31 | 0.14 | 0.21 | 0.36 | 0.3 | 0.18 |
| 2 | 668618 | rs17729501 | 0.593 | -0.045 | 0.283 | 0.635 | 0.013 | 0.022 | 0.022 | 0 | 0.13 | 0.09 | 0.07 | 0.02 | 0.1 | 0.12 |
| 2 | 668685 | rs17042334 | 0.492 | -0.028 | 0.252 | 0.673 | -0.011 | -0.042 | -0.042 | 0 | 0.11 | 0.02 | 0.2 | 0.16 | 0.14 | 0.21 |
| 2 | 674297 | rs4241323 | 0.072 | -0.067 | 0.602 | 0.653 | -0.013 | 0.027 | 0.027 | 0 | 0.86 | 0.71 | 0.78 | 0.84 | 0.86 | 0.8 |
| 2 | 674963 | rs12990777 | 0.756 | -0.079 | 0.38 | 0.639 | -0.03 | -0.027 | -0.027 | 0 | 0.72 | 0.48 | 0.69 | 0.83 | 0.76 | 0.68 |
| 2 | 675831 | rs2293084 | 0.274 | 0.075 | 0.201 | 0.707 | -0.034 | -0.018 | -0.018 | 0 | 0.4 | 0.23 | 0.47 | 0.46 | 0.46 | 0.5 |
| 2 | 679179 | rs1320333 | 0.103 | 0.182 | 0.231 | 0.555 | -0.003 | 0.006 | 0.006 | 0 | 0.1 | 0.09 | 0.05 | 0.02 | 0.1 | 0.09 |
| 2 | 681739 | rs1879524 | 0.056 | 0.152 | 0.6 | 0.531 | 0.003 | 0.003 | 0.003 | 0 | 0.87 | 0.85 | 0.77 | 0.87 | 0.84 | 0.78 |
| 2 | 682363 | rs2867116 | 0.095 | 0.07 | 0.611 | 0.459 | -0.001 | 0.002 | 0.002 | 0 | 0.11 | 0.08 | 0.09 | 0.02 | 0.12 | 0.1 |
| 2 | 686675 | rs2867117 | 1.304 | 0.029 | 0.714 | 0.446 | -0.006 | -0.035 | -0.035 | 0 | 0.73 | 0.64 | 0.67 | 0.85 | 0.72 | 0.67 |
| 2 | 692784 | rs1320334 | 1.096 | -0.136 | 0.4 | 0.465 | -0.017 | -0.001 | -0.001 | 0 | 0.46 | 0.27 | 0.43 | 0.57 | 0.43 | 0.44 |
| 2 | 696458 | rs2293085 | 0.944 | -0.183 | 0.543 | 0.713 | -0.043 | -0.041 | -0.041 | 0 | 0.39 | 0.16 | 0.43 | 0.36 | 0.42 | 0.36 |
| 2 | 697499 | rs2867134 | 0.289 | -0.139 | 0.773 | 0.654 | 0.007 | 0.016 | 0.016 | 0 | 0.6 | 0.4 | 0.68 | 0.72 | 0.59 | 0.66 |
| 2 | 705636 | rs6707477 | 0.392 | -0.469 | 0.727 | 0.67 | 0.028 | 0.039 | 0.039 | 0.135 | 0.59 | 0.34 | 0.68 | 0.72 | 0.61 | 0.65 |
| 2 | 706400 | rs2867137 | 0.268 | -0.365 | 0.516 | 0.644 | 0.004 | 0.004 | 0.004 | 0 | 0.11 | 0.08 | 0.08 | 0.02 | 0.1 | 0.12 |
| 2 | 709119 | rs4854272 | -1.136 | -0.285 | 0.649 | 0.744 | -0.004 | 0.044 | 0.044 | 0.061 | 0.78 | 0.71 | 0.76 | 0.59 | 0.83 | 0.7 |
| 2 | 710539 | rs10208465 | -0.499 | -0.359 | 0.316 | 0.559 | -0.02 | -0.048 | -0.048 | 0 | 0.14 | 0.27 | 0.09 | 0.02 | 0.12 | 0.13 |
| 2 | 711622 | rs4378835 | 0.513 | -0.117 | -0.107 | 0.787 | 0.001 | -0.005 | -0.005 | 0.058 | 0.25 | 0.14 | 0.26 | 0.27 | 0.29 | 0.23 |
| 2 | 716367 | rs13001660 | -0.311 | -0.15 | -0.051 | 0.808 | 0.001 | 0.021 | 0.021 | 0 | 0.49 | 0.31 | 0.5 | 0.6 | 0.47 | 0.57 |
| 2 | 719889 | rs6711931 | 1.33 | -0.17 | -0.011 | 0.943 | -0.002 | -0.001 | -0.001 | 0.064 | 0.21 | 0.21 | 0.26 | 0.28 | 0.2 | 0.28 |
| 2 | 724368 | rs4549126 | -0.695 | -0.222 | 0.141 | 0.815 | -0.001 | -0.003 | -0.003 | 0.277 | 0.58 | 0.4 | 0.6 | 0.56 | 0.62 | 0.64 |

| | | | | | | | | | | | | | | | | |
|---|---|---|---|---|---|---|---|---|---|---|---|---|---|---|---|---|
| 2 | 729552 | rs6741835 | -0.591 | -0.251 | 0.237 | 0.82 | -0.003 | -0.002 | -0.002 | 0 | 0.57 | 0.35 | 0.6 | 0.59 | 0.62 | 0.64 |
| 2 | 729948 | rs7580986 | -0.7 | -0.236 | 0.181 | 0.828 | 0 | 0 | 0 | 0 | 0.58 | 0.42 | 0.61 | 0.6 | 0.63 | 0.64 |
| 2 | 741003 | rs4854364 | -0.148 | -0.683 | 0.442 | 0.731 | -0.014 | -0.005 | -0.005 | 0 | 0.34 | 0.5 | 0.27 | 0.34 | 0.25 | 0.34 |
| 2 | 746124 | rs4854366 | 1.037 | -0.666 | 0.666 | 0.747 | 0.015 | 0.02 | 0.02 | 0.159 | 0.24 | 0.31 | 0.24 | 0.32 | 0.24 | 0.33 |
| 2 | 746192 | rs4854367 | -0.34 | -0.809 | 0.556 | 0.711 | 0.007 | -0.005 | -0.005 | 0 | 0.36 | 0.62 | 0.41 | 0.39 | 0.3 | 0.54 |
| 2 | 752530 | rs4854369 | -0.753 | -0.479 | -0.254 | 0.683 | 0.003 | 0.107 | 0.107 | 4.378 | 0.53 | 0.75 | 0.62 | 0.8 | 0.57 | 0.74 |
| 2 | 762741 | rs7423929 | 0.942 | -0.108 | -0.782 | 0.717 | 0 | 0.001 | 0.001 | 0.082 | 0.24 | 0.22 | 0.25 | 0.32 | 0.25 | 0.34 |
| 2 | 769289 | rs4271795 | 0.866 | -0.064 | -0.769 | 0.74 | 0.001 | 0.003 | 0.003 | 0.105 | 0.25 | 0.27 | 0.25 | 0.32 | 0.25 | 0.34 |
| 2 | 772153 | rs6756708 | 0.893 | -0.023 | -0.916 | 0.715 | 0.01 | 0.018 | 0.018 | 0.043 | 0.25 | 0.35 | 0.31 | 0.32 | 0.25 | 0.33 |
| 2 | 773278 | rs4380275 | -0.896 | 0.126 | -1.567 | 0.65 | -0.051 | 0.001 | 0.001 | 2.749 | 0.65 | 0.89 | 0.78 | 0.74 | 0.61 | 0.81 |
| 2 | 776119 | rs4443027 | 0.684 | -0.25 | -0.852 | 0.553 | 0.007 | 0.005 | 0.005 | 0 | 0.57 | 0.39 | 0.51 | 0.57 | 0.61 | 0.53 |
| 2 | 776287 | rs4854379 | -0.845 | -0.282 | -0.56 | 0.621 | -0.006 | -0.001 | -0.001 | 0 | 0.41 | 0.53 | 0.48 | 0.44 | 0.39 | 0.47 |
| 2 | 776605 | rs4854278 | 0.369 | -0.687 | -0.907 | 0.407 | -0.017 | -0.009 | -0.009 | 0 | 0.73 | 0.31 | 0.76 | 0.8 | 0.8 | 0.64 |
| 2 | 777156 | rs4375898 | -1.043 | -1.069 | -0.557 | 0.175 | -0.012 | 0 | 0 | 0 | 0.28 | 0.31 | 0.31 | 0.33 | 0.18 | 0.36 |
| 2 | 777252 | rs6714163 | 1.232 | -0.916 | -1.132 | 0.259 | 0.052 | 0.029 | 0.029 | 0 | 0.53 | 0.28 | 0.53 | 0.59 | 0.63 | 0.52 |
| 2 | 782581 | rs10183412 | -0.38 | -0.456 | -1.607 | -0.078 | -0.005 | 0.124 | 0.124 | 0 | 0.28 | 0.1 | 0.23 | 0.12 | 0.27 | 0.37 |
| 2 | 788807 | rs6746997 | -0.515 | -0.169 | -0.758 | 0.508 | 0.002 | 0.016 | 0.016 | 0 | 0.15 | 0.3 | 0.26 | 0.25 | 0.18 | 0.19 |
| 2 | 796058 | rs12987038 | 0.427 | -0.041 | -0.472 | 0.482 | -0.012 | -0.007 | -0.007 | 0 | 0.67 | 0.41 | 0.83 | 0.74 | 0.76 | 0.88 |
| 2 | 796546 | rs6737221 | -0.201 | 0.064 | -0.353 | 0.658 | -0.004 | 0.006 | 0.006 | 2.083 | 0.26 | 0.27 | 0.13 | 0.31 | 0.21 | 0.1 |
| 2 | 820721 | rs12467306 | 1.056 | -0.305 | -0.392 | 0.945 | 0.025 | -0.036 | -0.036 | 2.616 | 0.71 | 0.23 | 0.71 | 0.46 | 0.75 | 0.77 |
| 2 | 822116 | rs6757749 | -0.518 | -0.179 | -0.151 | 0.859 | 0.001 | 0.003 | 0.003 | 0 | 0.23 | 0.23 | 0.15 | 0.19 | 0.24 | 0.15 |
| 2 | 825721 | rs4358155 | -0.76 | -0.16 | 0.345 | 0.941 | 0 | 0.01 | 0.01 | 0 | 0.9 | 0.89 | 0.81 | 0.79 | 0.86 | 0.84 |
| 2 | 826069 | rs4589796 | 0.692 | 0.261 | -0.048 | 0.687 | 0.007 | 0.063 | 0.063 | 0 | 0.72 | 0.4 | 0.58 | 0.52 | 0.68 | 0.76 |
| 2 | 826392 | rs4854279 | 1.142 | 0.302 | -0.164 | 0.698 | -0.006 | -0.015 | -0.015 | 0.512 | 0.09 | 0.04 | 0.19 | 0.21 | 0.14 | 0.16 |
| 2 | 831480 | rs11127524 | 1.537 | 0.214 | -0.235 | 0.585 | -0.02 | 0.08 | 0.08 | 0.015 | 0.49 | 0.16 | 0.43 | 0.34 | 0.43 | 0.62 |
| 2 | 832145 | rs12714424 | -0.158 | 0.194 | 0.205 | 0.68 | 0.007 | 0.145 | 0.145 | 0 | 0.85 | 0.74 | 0.77 | 0.53 | 0.81 | 0.8 |
| 2 | 836821 | rs12465993 | -0.23 | 0.115 | 0.166 | 0.568 | 0.002 | 0.005 | 0.005 | 0 | 0.24 | 0.23 | 0.25 | 0.18 | 0.25 | 0.14 |
| 2 | 837120 | rs10175009 | -1.225 | 0.277 | 0.88 | 1.125 | 0.021 | 0.211 | 0.211 | 0 | 0.94 | 0.79 | 0.96 | 0.76 | 0.95 | 0.97 |
| 2 | 837567 | rs4366938 | -0.187 | 0.142 | 0.533 | 0.495 | 0.003 | 0.007 | 0.007 | 0 | 0.24 | 0.23 | 0.15 | 0.19 | 0.25 | 0.14 |
| 2 | 839425 | rs13397770 | 1.009 | 0.148 | 0.58 | 0.507 | -0.003 | -0.005 | -0.005 | 0 | 0.61 | 0.52 | 0.62 | 0.34 | 0.56 | 0.66 |
| 2 | 839984 | rs4632380 | -1.158 | 0.079 | 0.323 | 0.461 | -0.002 | -0.004 | -0.004 | 0 | 0.9 | 0.89 | 0.81 | 0.8 | 0.86 | 0.83 |
| 2 | 846164 | rs4533500 | 1.563 | 0.094 | 0.346 | 0.423 | -0.011 | 0.046 | 0.046 | 0.006 | 0.49 | 0.17 | 0.43 | 0.38 | 0.43 | 0.62 |
| 2 | 851144 | rs11682609 | 0.935 | 0.166 | 0.273 | 0.638 | 0.011 | 0.195 | 0.195 | 0 | 0.05 | 0.15 | 0.04 | 0.26 | 0.05 | 0.04 |
| 2 | 851775 | rs10179370 | -0.182 | 0.164 | 0.254 | 0.628 | 0.005 | 0.121 | 0.121 | 0 | 0.85 | 0.75 | 0.76 | 0.53 | 0.81 | 0.79 |
| 2 | 857713 | rs12714428 | 0.576 | -0.062 | -0.439 | -0.181 | -0.054 | 0.104 | 0.104 | 0.038 | 0.29 | 0.13 | 0.24 | 0 | 0.33 | 0.08 |
| 2 | 858062 | rs6548270 | -0.344 | -0.142 | -0.458 | -0.04 | -0.019 | -0.042 | -0.042 | 0 | 0.28 | 0.16 | 0.35 | 0.57 | 0.39 | 0.32 |
| 2 | 861841 | rs4992454 | -0.31 | -0.133 | -0.315 | -0.833 | -0.002 | -0.003 | -0.003 | 0.474 | 0.28 | 0.23 | 0.27 | 0.37 | 0.38 | 0.27 |
| 2 | 865861 | rs4854288 | 0.632 | -0.139 | -0.193 | -0.771 | 0.019 | 0.098 | 0.098 | 0 | 0.14 | 0.22 | 0.15 | 0.31 | 0.17 | 0.15 |
| 2 | 867187 | rs4263142 | -0.117 | -0.224 | -0.463 | -1.015 | -0.044 | 0.152 | 0.152 | 0 | 0.58 | 0.76 | 0.62 | 0.94 | 0.46 | 0.8 |
| 2 | 868450 | rs6760982 | 0.955 | -0.35 | -0.381 | -1.03 | 0.014 | -0.003 | -0.003 | 0.416 | 0.39 | 0.25 | 0.45 | 0.63 | 0.29 | 0.62 |
| 2 | 872859 | rs4854414 | -0.915 | -0.383 | -0.655 | -1.376 | 0.021 | 0.015 | 0.015 | 0 | 0.68 | 0.58 | 0.54 | 0.35 | 0.64 | 0.61 |
| 2 | 873102 | rs12997230 | 1.147 | -0.03 | -0.789 | -1.32 | -0.044 | 0.077 | 0.077 | 0.016 | 0.26 | 0.11 | 0.25 | 0 | 0.33 | 0.07 |
| 2 | 882064 | rs4854417 | 1.943 | -0.101 | -0.473 | -0.849 | 0.02 | 0.1 | 0.1 | 0 | 0.16 | 0.22 | 0.32 | 0.58 | 0.16 | 0.24 |
| 2 | 884441 | rs7421139 | -0.712 | -0.833 | -0.861 | -1.079 | 0.02 | 0.018 | 0.018 | 0.578 | 0.78 | 0.8 | 0.87 | 0.96 | 0.87 | 0.65 |
| 2 | 896473 | rs6419706 | -0.072 | -0.691 | -0.784 | -1.088 | 0.046 | 0.153 | 0.153 | 0 | 0.61 | 0.79 | 0.75 | 0.9 | 0.66 | 0.53 |
| 2 | 903583 | rs4854424 | 0.141 | -0.665 | -0.41 | -1.05 | -0.001 | -0.027 | -0.027 | 0.493 | 0.56 | 0.55 | 0.72 | 0.84 | 0.63 | 0.49 |
| 2 | 904394 | rs4241338 | 0.226 | -0.646 | -0.697 | -1.036 | -0.014 | -0.013 | -0.013 | 0 | 0.57 | 0.47 | 0.64 | 0.68 | 0.65 | 0.62 |
| 2 | 910327 | rs6548281 | 0.316 | -0.811 | -0.754 | -0.996 | -0.04 | 0.05 | 0.05 | 0.341 | 0.24 | 0.38 | 0.17 | 0.31 | 0.16 | 0.31 |
| 2 | 914015 | rs4854428 | -0.546 | -0.518 | -1.339 | -1.575 | 0.001 | -0.026 | -0.026 | 0.009 | 0.81 | 0.8 | 0.81 | 1 | 0.82 | 0.95 |
| 2 | 914564 | rs4854431 | -0.091 | -0.281 | -1.28 | -1.482 | 0.025 | 0.087 | 0.087 | 0.088 | 0.61 | 0.71 | 0.68 | 0.94 | 0.6 | 0.82 |
| 2 | 918533 | rs7420683 | -0.151 | -0.246 | -1.344 | -1.464 | 0.024 | 0.095 | 0.095 | 0.213 | 0.61 | 0.75 | 0.68 | 0.93 | 0.6 | 0.82 |
| 2 | 929469 | rs13415540 | 0.276 | 0.4 | -1.214 | -1.61 | -0.004 | -0.026 | -0.026 | 0 | 0.27 | 0.24 | 0.2 | 0.01 | 0.28 | 0.21 |
| 2 | 933573 | rs4971358 | -0.331 | -0.079 | -1.267 | -2.067 | 0 | -0.02 | -0.02 | 0 | 0.48 | 0.58 | 0.5 | 0.14 | 0.5 | 0.46 |
| 2 | 934215 | rs4971337 | 1.267 | -0.357 | -1.135 | -2.097 | 0 | -0.004 | -0.004 | 0.177 | 0.11 | 0.11 | 0.26 | 0.04 | 0.09 | 0.26 |
| 2 | 943250 | rs4971346 | -0.082 | -0.376 | -1.148 | -2.163 | -0.025 | -0.078 | -0.078 | 0 | 0.35 | 0.5 | 0.42 | 0.12 | 0.29 | 0.35 |
| 2 | 949196 | rs7559853 | -0.145 | -0.376 | -1.108 | -2.164 | -0.03 | -0.093 | -0.093 | 0 | 0.35 | 0.51 | 0.42 | 0.11 | 0.29 | 0.35 |
| 2 | 950291 | rs6759670 | -1.275 | -0.443 | -1.749 | -1.725 | -0.05 | -0.136 | -0.136 | 3.152 | 0.72 | 0.96 | 0.68 | 0.45 | 0.74 | 0.6 |
| 2 | 954306 | rs10178601 | 0.012 | -0.748 | -1.059 | -2.109 | -0.004 | -0.027 | -0.027 | 0 | 0.32 | 0.34 | 0.4 | 0.12 | 0.29 | 0.34 |
| 2 | 963326 | rs6548284 | -0.282 | -0.856 | -0.937 | -2.129 | -0.014 | -0.086 | -0.086 | 0.072 | 0.35 | 0.46 | 0.25 | 0.06 | 0.35 | 0.12 |
| 2 | 963334 | rs6548285 | -0.282 | -0.856 | -0.937 | -2.129 | -0.014 | -0.086 | -0.086 | 1.005 | 0.35 | 0.46 | 0.25 | 0.06 | 0.35 | 0.12 |
| 2 | 970417 | rs7355580 | 0.14 | -0.685 | -1.05 | -2.045 | -0.002 | -0.028 | -0.028 | 0.942 | 0.31 | 0.32 | 0.23 | 0.06 | 0.34 | 0.13 |
| 2 | 973232 | rs6741262 | 0.136 | -0.663 | -1.035 | -2.034 | -0.003 | -0.041 | -0.041 | 0.162 | 0.31 | 0.34 | 0.24 | 0.06 | 0.34 | 0.13 |
| 2 | 974720 | rs4524155 | -0.333 | -0.657 | -0.997 | -2.053 | -0.001 | -0.029 | -0.029 | 0 | 0.4 | 0.38 | 0.5 | 0.08 | 0.42 | 0.38 |
| 2 | 1001060 | rs7598261 | -0.483 | -0.775 | -1.131 | -2.101 | 0.003 | 0.031 | 0.031 | 0 | 0.55 | 0.41 | 0.6 | 0.1 | 0.5 | 0.51 |
| 2 | 1003737 | rs11675840 | -0.001 | -0.846 | -1.263 | -2.209 | 0.04 | 0.087 | 0.087 | 0 | 0.46 | 0.31 | 0.38 | 0.07 | 0.42 | 0.26 |
| 2 | 1004716 | rs12104941 | 0.003 | -0.823 | -1.579 | -2.24 | 0.019 | 0.16 | 0.16 | 0.159 | 0.35 | 0.16 | 0.46 | 0.06 | 0.34 | 0.43 |
| 2 | 1010474 | rs6706944 | 0.076 | -0.84 | -1.356 | -2.334 | 0.034 | 0.136 | 0.136 | 0 | 0.45 | 0.24 | 0.37 | 0.06 | 0.46 | 0.24 |
| 2 | 1010713 | rs6707252 | -1.062 | -0.627 | -1.304 | -2.159 | 0.006 | -0.013 | -0.013 | 0 | 0.78 | 0.83 | 0.8 | 0.32 | 0.84 | 0.75 |
| 2 | 1013294 | rs11894899 | 2.154 | -0.892 | -1.5 | -2.293 | -0.027 | 0.031 | 0.031 | 1.285 | 0.12 | 0.03 | 0.12 | 0.04 | 0.12 | 0.06 |

| | | | | | | | | | | | | | | | | |
|---|---|---|---|---|---|---|---|---|---|---|---|---|---|---|---|---|
| 2 | 1014757 | rs7574097 | 1.39 | -0.973 | -1.505 | -2.368 | 0.005 | 0.121 | 0.121 | 0 | 0.28 | 0.11 | 0.24 | 0.02 | 0.31 | 0.18 |
| 2 | 1014837 | rs7574207 | 0.3 | -0.969 | -1.448 | -2.354 | 0.034 | 0.133 | 0.133 | 0 | 0.45 | 0.25 | 0.37 | 0.07 | 0.46 | 0.24 |
| 2 | 1018983 | rs7589320 | 2.978 | -0.842 | -1.344 | -2.232 | -0.022 | -0.024 | -0.024 | 0 | 0.05 | 0.11 | 0.01 | 0 | 0.03 | 0 |
| 2 | 1019512 | rs7580579 | 0.469 | -0.323 | -1.153 | -2.195 | 0.026 | -0.02 | -0.02 | 0 | 0.34 | 0.45 | 0.29 | 0.25 | 0.4 | 0.34 |
| 2 | 1019753 | rs9309723 | -1.253 | -0.269 | -1.033 | -2.026 | -0.065 | -0.101 | -0.101 | 1.048 | 0.81 | 0.99 | 0.82 | 0.31 | 0.82 | 0.74 |
| 2 | 1020741 | rs7603971 | 1.08 | -0.12 | -0.582 | -1.753 | 0.063 | -0.001 | -0.001 | 0.515 | 0.22 | 0.42 | 0.18 | 0.21 | 0.28 | 0.29 |
| 2 | 1027081 | rs10207582 | -0.335 | -0.342 | -0.637 | -1.864 | 0.008 | -0.029 | -0.029 | 0 | 0.38 | 0.58 | 0.45 | 0.26 | 0.38 | 0.55 |
| 2 | 1027830 | rs4971336 | -0.129 | -0.154 | -0.555 | -1.782 | 0.005 | -0.011 | -0.011 | 0 | 0.34 | 0.65 | 0.45 | 0.25 | 0.36 | 0.49 |
| 2 | 1033530 | rs10779977 | 0.323 | -0.04 | -0.547 | -1.725 | 0.023 | -0.015 | -0.015 | 0 | 0.3 | 0.42 | 0.42 | 0.25 | 0.35 | 0.49 |
| 2 | 1037783 | rs4971329 | 0.75 | 0.222 | -0.261 | -1.552 | 0.016 | -0.007 | -0.007 | 0 | 0.24 | 0.38 | 0.18 | 0.21 | 0.28 | 0.3 |
| 2 | 1043352 | rs4246558 | -0.647 | -0.027 | -0.076 | -0.44 | 0 | 0.004 | 0.004 | 0 | 0.83 | 0.89 | 0.85 | 0.48 | 0.87 | 0.78 |
| 2 | 1044617 | rs4488692 | -0.359 | -0.061 | 0.15 | -0.436 | -0.006 | 0.067 | 0.067 | 0 | 0.12 | 0.3 | 0.09 | 0.18 | 0.09 | 0.11 |
| 2 | 1046996 | rs9309724 | 0.257 | -0.047 | -0.025 | -0.584 | -0.002 | -0.042 | -0.042 | 0 | 0.16 | 0.23 | 0.15 | 0.04 | 0.14 | 0.06 |
| 2 | 1048359 | rs4581927 | -0.431 | -0.009 | 0.085 | -0.355 | -0.007 | 0.045 | 0.045 | 0 | 0.12 | 0.23 | 0.09 | 0.2 | 0.09 | 0.12 |
| 2 | 1052961 | rs6734393 | 0.562 | -0.034 | 0.124 | -0.469 | -0.008 | 0.001 | 0.001 | 0 | 0.6 | 0.51 | 0.4 | 0.39 | 0.66 | 0.58 |
| 2 | 1061182 | rs35801395 | 0.928 | 0.216 | -0.142 | -0.481 | 0 | 0.002 | 0.002 | 0.071 | 0.19 | 0.16 | 0.1 | 0.21 | 0.18 | 0.21 |
| 2 | 1063504 | rs34787379 | 0.946 | 0.216 | -0.141 | -0.481 | 0 | 0.002 | 0.002 | 0.061 | 0.18 | 0.16 | 0.1 | 0.21 | 0.18 | 0.21 |
| 2 | 1064700 | rs7426276 | 1.116 | 0.252 | -0.265 | -0.412 | -0.046 | -0.024 | -0.024 | 0 | 0.08 | 0.01 | 0.27 | 0.51 | 0.12 | 0.14 |
| 2 | 1064808 | rs6751259 | 0.866 | -0.099 | -0.033 | -0.384 | 0 | 0.002 | 0.002 | 0.061 | 0.18 | 0.16 | 0.1 | 0.22 | 0.18 | 0.21 |
| 2 | 1069949 | rs4586665 | 0.843 | -0.3 | 0.233 | -0.44 | 0.012 | 0.019 | 0.019 | 0 | 0.52 | 0.47 | 0.59 | 0.69 | 0.62 | 0.53 |
| 2 | 1076684 | rs4971371 | 0.882 | -0.276 | 0.135 | -0.474 | 0.014 | 0.015 | 0.015 | 0 | 0.51 | 0.43 | 0.59 | 0.64 | 0.62 | 0.51 |
| 2 | 1079320 | rs4971432 | 0.014 | -0.62 | 0.154 | -0.404 | -0.029 | 0.019 | 0.019 | 1.164 | 0.73 | 0.56 | 0.78 | 0.68 | 0.8 | 0.58 |
| 2 | 1081594 | rs10865542 | -1.064 | -0.479 | 0.163 | -0.351 | 0.002 | 0.04 | 0.04 | 0.01 | 0.82 | 0.77 | 0.9 | 0.72 | 0.84 | 0.8 |
| 2 | 1091101 | rs6717701 | -1.47 | -0.457 | 0.42 | -0.001 | -0.001 | 0.002 | 0.002 | 0 | 0.6 | 0.65 | 0.68 | 0.71 | 0.54 | 0.65 |
| 2 | 1092195 | rs4971439 | -1.533 | 0.039 | 0.798 | 0.376 | 0.005 | 0.021 | 0.021 | 0 | 0.54 | 0.71 | 0.65 | 0.66 | 0.52 | 0.66 |
| 2 | 1096540 | rs7606829 | -0.095 | -0.008 | 0.821 | 0.293 | -0.006 | -0.025 | -0.025 | 0.652 | 0.16 | 0.21 | 0.19 | 0.03 | 0.15 | 0.09 |
| 2 | 1097714 | rs4366936 | 0.435 | -0.051 | 0.595 | 0.403 | 0.022 | -0.019 | -0.019 | 0 | 0.07 | 0.01 | 0.1 | 0.07 | 0.03 | 0.25 |
| 2 | 1097795 | rs6716119 | 1.553 | -0.029 | 0.663 | 0.428 | -0.003 | 0.039 | 0.039 | 0 | 0.45 | 0.18 | 0.34 | 0.35 | 0.49 | 0.34 |
| 2 | 1098891 | rs10180028 | -1.61 | 0.017 | 0.599 | 0.451 | 0.008 | 0.034 | 0.034 | 0 | 0.53 | 0.7 | 0.66 | 0.66 | 0.51 | 0.66 |
| 2 | 1101127 | rs12469446 | 0.026 | 0.144 | 0.48 | 0.454 | 0.007 | 0.012 | 0.012 | 0 | 0.3 | 0.32 | 0.36 | 0.55 | 0.33 | 0.32 |
| 2 | 1116648 | rs4971463 | -0.586 | 0.045 | 0.455 | 0.635 | 0 | 0 | 0 | 1.814 | 0.45 | 0.52 | 0.54 | 0.58 | 0.49 | 0.44 |
| 2 | 1124416 | rs7580578 | -0.983 | 0.295 | -0.041 | 0.575 | 0.01 | 0.056 | 0.056 | 0 | 0.53 | 0.73 | 0.67 | 0.73 | 0.53 | 0.69 |
| 2 | 1148426 | rs4305308 | 1.311 | -0.03 | 1.491 | 1.411 | -0.035 | 0.043 | 0.043 | 0.12 | 0.87 | 0.57 | 0.81 | 0.86 | 0.92 | 0.79 |
| 2 | 1150095 | rs7579535 | 0.128 | 0.413 | 0.839 | 1.8 | 0.046 | 0.16 | 0.16 | 0 | 0.92 | 0.85 | 0.72 | 0.56 | 0.87 | 0.81 |
| 2 | 1152434 | rs4246561 | 1.368 | 0.169 | 0.606 | 1.717 | -0.071 | 0.009 | 0.009 | 1.037 | 0.82 | 0.56 | 0.86 | 0.8 | 0.95 | 0.7 |
| 2 | 1154664 | rs4467306 | 0.685 | 0.222 | 0.59 | 1.454 | 0.015 | 0.164 | 0.164 | 0 | 0.81 | 0.58 | 0.69 | 0.45 | 0.85 | 0.66 |
| 2 | 1157068 | rs4971397 | -1.142 | 0.274 | 0.277 | 1.296 | -0.004 | 0.071 | 0.071 | 0 | 0.27 | 0.53 | 0.44 | 0.62 | 0.2 | 0.54 |
| 2 | 1164595 | rs4453719 | -0.95 | -0.138 | -0.411 | 0.845 | 0.028 | 0.126 | 0.126 | 0 | 0.49 | 0.81 | 0.55 | 0.74 | 0.46 | 0.54 |
| 2 | 1168171 | rs4583504 | 0.247 | -0.275 | -0.437 | 0.804 | -0.024 | -0.053 | -0.053 | 0 | 0.16 | 0.13 | 0.4 | 0.43 | 0.25 | 0.24 |
| 2 | 1173285 | rs6714337 | -0.143 | -0.449 | 0.352 | 0.641 | -0.028 | -0.078 | -0.078 | 6.861 | 0.24 | 0.41 | 0.1 | 0.1 | 0.17 | 0.16 |
| 2 | 1183846 | rs4971407 | 0.362 | -0.347 | 0.35 | 0.71 | -0.04 | -0.072 | -0.072 | 0 | 0.15 | 0.07 | 0.39 | 0.43 | 0.25 | 0.24 |
| 2 | 1186496 | rs4270381 | -0.72 | -0.373 | 0.482 | 0.707 | -0.006 | -0.019 | -0.019 | 0 | 0.23 | 0.23 | 0.45 | 0.63 | 0.29 | 0.34 |
| 2 | 1188495 | rs4312525 | -0.371 | -0.6 | 0.75 | 0.737 | -0.011 | 0.11 | 0.11 | 0 | 0.08 | 0.19 | 0.05 | 0.21 | 0.04 | 0.1 |
| 2 | 1204796 | rs6751090 | -0.737 | -0.675 | 0.728 | 0.527 | -0.089 | -0.067 | -0.067 | 0 | 0.19 | 0.48 | 0.07 | 0.1 | 0.06 | 0.19 |
| 2 | 1216394 | rs4971418 | -1.093 | -0.831 | -0.258 | 0.162 | 0.099 | -0.04 | -0.04 | 5.448 | 0.43 | 0.76 | 0.3 | 0.5 | 0.31 | 0.51 |
| 2 | 1228487 | rs6548209 | -1.173 | -0.782 | 0.022 | -0.092 | 0.008 | 0.003 | 0.003 | 0 | 0.36 | 0.65 | 0.22 | 0.34 | 0.24 | 0.4 |
| 2 | 1235479 | rs28678499 | -1.173 | -0.782 | -0.039 | -0.095 | 0.006 | 0.003 | 0.003 | 0 | 0.36 | 0.65 | 0.22 | 0.34 | 0.24 | 0.41 |
| 2 | 1244060 | rs35847752 | 0.648 | -0.616 | -0.298 | -0.083 | -0.012 | 0.064 | 0.064 | 0.091 | 0.13 | 0.04 | 0.04 | 0.02 | 0.12 | 0.17 |
| 2 | 1253328 | rs34539029 | -0.718 | -0.741 | 0.304 | 0.06 | -0.062 | 0.039 | 0.039 | 0.34 | 0.24 | 0.42 | 0.17 | 0.3 | 0.12 | 0.22 |
| 2 | 1262839 | rs9286350 | -0.75 | -0.733 | 0.33 | 0.063 | -0.062 | 0.035 | 0.035 | 0.902 | 0.24 | 0.43 | 0.18 | 0.3 | 0.12 | 0.23 |
| 2 | 1263891 | rs9712145 | -0.747 | -0.741 | 0.323 | 0.06 | -0.063 | 0.037 | 0.037 | 3.446 | 0.24 | 0.45 | 0.18 | 0.3 | 0.13 | 0.23 |
| 2 | 1282121 | rs9750959 | -0.485 | -0.711 | 0.395 | 0.093 | -0.027 | 0.026 | 0.026 | 0 | 0.21 | 0.26 | 0.16 | 0.31 | 0.12 | 0.23 |
| 2 | 1283031 | rs11674954 | 0.412 | -0.267 | -0.164 | 0.185 | -0.039 | -0.081 | -0.081 | 0 | 0.17 | 0.07 | 0.35 | 0.49 | 0.25 | 0.26 |
| 2 | 1288106 | rs6754505 | 0.164 | -0.487 | 0.364 | -0.154 | -0.001 | -0.009 | -0.009 | 0 | 0.6 | 0.54 | 0.43 | 0.33 | 0.52 | 0.58 |
| 2 | 1296139 | rs11676238 | 0.244 | -0.507 | 0.346 | -0.167 | -0.001 | -0.027 | -0.027 | 1.607 | 0.27 | 0.19 | 0.45 | 0.55 | 0.3 | 0.33 |
| 2 | 1297615 | rs9753172 | -1.131 | -0.713 | -0.206 | -0.321 | -0.087 | 0.029 | 0.029 | 0 | 0.6 | 0.84 | 0.67 | 0.72 | 0.63 | 0.7 |
| 2 | 1298915 | rs9677798 | -0.996 | -0.728 | -0.002 | -0.329 | -0.022 | 0.036 | 0.036 | 0.082 | 0.58 | 0.69 | 0.66 | 0.72 | 0.63 | 0.69 |
| 2 | 1309956 | rs4293594 | -1.033 | -0.623 | -0.084 | -0.408 | -0.029 | 0.002 | 0.002 | 0.12 | 0.58 | 0.75 | 0.66 | 0.64 | 0.64 | 0.7 |
| 2 | 1311157 | rs13021490 | 0.82 | -0.63 | -0.106 | -0.4 | 0.009 | 0.049 | 0.049 | 0.306 | 0.16 | 0.06 | 0.07 | 0.07 | 0.15 | 0.28 |
| 2 | 1314780 | rs4641984 | 0.854 | -0.55 | -0.136 | -0.367 | -0.011 | -0.03 | -0.03 | 0.648 | 0.06 | 0 | 0.13 | 0.21 | 0.04 | 0.11 |
| 2 | 1316373 | rs9677943 | 0.668 | -0.45 | 0.198 | -0.514 | -0.028 | -0.005 | -0.005 | 0 | 0.45 | 0.24 | 0.48 | 0.41 | 0.4 | 0.36 |
| 2 | 1318413 | rs13011951 | 0.5 | -0.172 | -0.057 | -0.66 | 0.003 | 0.02 | 0.02 | 0 | 0.14 | 0.08 | 0.09 | 0.03 | 0.09 | 0.11 |
| 2 | 1320143 | rs4927635 | -0.082 | -0.342 | 0.136 | -0.523 | -0.001 | 0 | 0 | 0.936 | 0.61 | 0.45 | 0.53 | 0.44 | 0.54 | 0.59 |
| 2 | 1327698 | rs12718455 | -0.058 | -0.272 | -0.319 | -0.442 | -0.001 | 0 | 0 | 2.358 | 0.59 | 0.44 | 0.56 | 0.45 | 0.54 | 0.6 |
| 2 | 1350207 | rs9677202 | -0.074 | 0.309 | -0.623 | -0.579 | 0 | 0.081 | 0.081 | 1.309 | 0.61 | 0.77 | 0.6 | 0.86 | 0.59 | 0.55 |
| 2 | 1359621 | rs9749785 | 0.813 | 0.276 | -0.258 | -0.399 | -0.001 | 0.005 | 0.005 | 0.063 | 0.46 | 0.37 | 0.43 | 0.42 | 0.45 | 0.5 |
| 2 | 1361922 | rs9677878 | -0.328 | 0.177 | -0.123 | -0.454 | -0.005 | -0.068 | -0.068 | 0 | 0.73 | 0.66 | 0.75 | 0.95 | 0.74 | 0.8 |
| 2 | 1361992 | rs4927600 | -0.328 | 0.177 | -0.123 | -0.454 | -0.005 | -0.068 | -0.068 | 0 | 0.73 | 0.66 | 0.75 | 0.95 | 0.74 | 0.8 |
| 2 | 1367511 | rs11894739 | -0.04 | 0.262 | 0.535 | 0.136 | -0.012 | -0.054 | -0.054 | 0 | 0.26 | 0.19 | 0.31 | 0.53 | 0.3 | 0.31 |

| | | | | | | | | | | | | | | | | |
|---|---|---|---|---|---|---|---|---|---|---|---|---|---|---|---|---|
| 2 | 1372758 | rs10198003 | -1.079 | 0.172 | 0.545 | 0.033 | 0.02 | 0.022 | 0.022 | 0.732 | 0.05 | 0.3 | 0.06 | 0.01 | 0.01 | 0.02 |
| 2 | 1376705 | rs4490233 | -0.575 | 0.431 | -0.283 | 0.378 | -0.002 | 0.013 | 0.013 | 0.304 | 0.31 | 0.61 | 0.34 | 0.49 | 0.31 | 0.32 |
| 2 | 1377042 | rs13423589 | 0.165 | 0.704 | -0.903 | 0.271 | -0.023 | 0.04 | 0.04 | 0.707 | 0.14 | 0.03 | 0.11 | 0.06 | 0.15 | 0.16 |
| 2 | 1378060 | rs4927602 | -0.343 | 0.552 | -1.253 | 0.562 | 0.036 | 0.032 | 0.032 | 0.26 | 0.51 | 0.8 | 0.58 | 0.6 | 0.52 | 0.52 |
| 2 | 1378943 | rs9326161 | -0.931 | 0.573 | 0.141 | 0.418 | 0.054 | -0.005 | -0.005 | 0.153 | 0.08 | 0.38 | 0.05 | 0 | 0.04 | 0.03 |
| 2 | 1378969 | rs4076290 | 0.396 | 0.623 | -0.469 | 0.297 | 0.015 | 0.217 | 0.217 | 0.809 | 0.5 | 0.16 | 0.59 | 0.23 | 0.6 | 0.42 |
| 2 | 1384725 | rs11897977 | -1.821 | 0.482 | 0.116 | 1.073 | -0.031 | 0.066 | 0.066 | 2.965 | 0.16 | 0.7 | 0.15 | 0.31 | 0.08 | 0.2 |
| 2 | 1397415 | rs10153889 | -0.631 | 0.547 | 0.423 | 0.999 | 0.016 | -0.04 | -0.04 | 0 | 0.08 | 0.33 | 0.04 | 0 | 0.05 | 0.04 |
| 2 | 1398300 | rs1996207 | 0.936 | 0.561 | 0.408 | 1.224 | -0.055 | -0.081 | -0.081 | 0.459 | 0.32 | 0.06 | 0.24 | 0.4 | 0.31 | 0.3 |
| 2 | 1401781 | rs938326 | -0.328 | 0.733 | 0.649 | 1.274 | 0.018 | 0.035 | 0.035 | 0 | 0.5 | 0.63 | 0.64 | 0.61 | 0.53 | 0.65 |
| 2 | 1404495 | rs11211644 | 0.352 | 0.763 | 0.551 | 1.29 | -0.036 | 0.085 | 0.085 | 0.02 | 0.41 | 0.22 | 0.52 | 0.25 | 0.49 | 0.39 |
| 2 | 1407078 | rs1546588 | -0.142 | 1.181 | 0.869 | 1.479 | 0.047 | 0.176 | 0.176 | 0.108 | 0.07 | 0.14 | 0.17 | 0.33 | 0.08 | 0.05 |
| 2 | 1407628 | rs11675342 | -0.142 | 1.115 | 0.852 | 1.344 | -0.002 | 0.049 | 0.049 | 0 | 0.43 | 0.3 | 0.47 | 0.32 | 0.46 | 0.59 |
| 2 | 1407815 | rs11675434 | 0.141 | 1.09 | 0.832 | 1.331 | -0.011 | 0.033 | 0.033 | 0 | 0.4 | 0.29 | 0.46 | 0.32 | 0.43 | 0.45 |
| 2 | 1411883 | rs13400534 | -0.644 | 1.247 | 1.076 | 1.355 | 0.015 | 0.047 | 0.047 | 0 | 0.51 | 0.64 | 0.66 | 0.66 | 0.54 | 0.65 |
| 2 | 1412230 | rs11682968 | -0.635 | 1.215 | 1.076 | 1.354 | 0.017 | 0.048 | 0.048 | 0 | 0.51 | 0.64 | 0.66 | 0.66 | 0.54 | 0.66 |
| 2 | 1417199 | rs2071402 | 0.141 | 1.255 | 1.246 | 1.334 | -0.003 | 0.038 | 0.038 | 0.008 | 0.41 | 0.32 | 0.47 | 0.25 | 0.44 | 0.53 |
| 2 | 1417244 | rs2071403 | -0.531 | 1.125 | 1.024 | 1.14 | 0.02 | 0.054 | 0.054 | 0 | 0.51 | 0.65 | 0.66 | 0.67 | 0.54 | 0.66 |
| 2 | 1423283 | rs10193983 | -1.179 | 1.446 | 1.604 | 1.191 | 0.059 | 0.026 | 0.026 | 0.089 | 0.1 | 0.25 | 0.17 | 0.09 | 0.13 | 0.06 |
| 2 | 1423500 | rs1967512 | -0.722 | 1.413 | 1.282 | 1.051 | 0.019 | -0.01 | -0.01 | 0 | 0.34 | 0.48 | 0.48 | 0.31 | 0.37 | 0.3 |
| 2 | 1426346 | rs2048726 | -1.249 | 1.406 | 0.936 | 0.975 | -0.03 | 0.008 | 0.008 | 0 | 0.41 | 0.76 | 0.57 | 0.39 | 0.42 | 0.59 |
| 2 | 1426621 | rs2276702 | 0.82 | 1.315 | 1.39 | 1.133 | 0.009 | 0.006 | 0.006 | 0 | 0.61 | 0.46 | 0.48 | 0.55 | 0.56 | 0.56 |
| 2 | 1427107 | rs9678469 | 0.594 | 1.229 | 1.099 | 1.161 | 0.036 | 0.011 | 0.011 | 0 | 0.87 | 0.8 | 0.8 | 0.84 | 0.79 | 0.91 |
| 2 | 1431590 | rs10519477 | 0.144 | 1.188 | 0.929 | 1.162 | -0.004 | -0.012 | -0.012 | 0 | 0.25 | 0.31 | 0.32 | 0.19 | 0.24 | 0.27 |
| 2 | 1431799 | rs13424221 | 1.102 | 1.199 | 0.483 | 1.231 | -0.015 | -0.01 | -0.01 | 0 | 0.55 | 0.32 | 0.4 | 0.52 | 0.52 | 0.38 |
| 2 | 1435296 | rs4927606 | -0.098 | 0.511 | 0.185 | 0.876 | 0.003 | 0.034 | 0.034 | 0.033 | 0.28 | 0.22 | 0.35 | 0.16 | 0.25 | 0.43 |
| 2 | 1441178 | rs1514684 | -0.268 | 0.462 | 0.452 | 0.86 | 0.002 | 0.007 | 0.007 | 0.008 | 0.29 | 0.3 | 0.36 | 0.16 | 0.25 | 0.43 |
| 2 | 1441305 | rs1514682 | -0.526 | 0.355 | 1.028 | 0.962 | -0.015 | -0.012 | -0.012 | 0 | 0.32 | 0.51 | 0.37 | 0.27 | 0.25 | 0.55 |
| 2 | 1441549 | rs9751407 | 0.841 | 0.834 | 1.722 | 1.177 | 0.028 | 0.005 | 0.005 | 0.001 | 0.87 | 0.72 | 0.91 | 0.89 | 0.86 | 0.93 |
| 2 | 1449966 | rs7602332 | 0.914 | 0.826 | 1.172 | 1.312 | -0.004 | 0.013 | 0.013 | 0.009 | 0.85 | 0.75 | 0.87 | 0.81 | 0.87 | 0.93 |
| 2 | 1451697 | rs10204515 | -0.422 | 0.653 | 0.661 | 1.126 | -0.005 | -0.014 | -0.014 | 0 | 0.43 | 0.5 | 0.4 | 0.29 | 0.43 | 0.64 |
| 2 | 1453528 | rs4927608 | -0.537 | 0.653 | 0.753 | 1.116 | -0.005 | -0.01 | -0.01 | 0.303 | 0.44 | 0.51 | 0.4 | 0.29 | 0.42 | 0.64 |
| 2 | 1458001 | rs2070882 | 1.21 | 0.698 | 0.776 | 1.175 | -0.016 | -0.026 | -0.026 | 0 | 0.42 | 0.3 | 0.48 | 0.53 | 0.44 | 0.29 |
| 2 | 1459461 | rs6706775 | 0.451 | 0.639 | 1.223 | 1.041 | -0.006 | -0.026 | -0.026 | 4.978 | 0.57 | 0.49 | 0.58 | 0.78 | 0.57 | 0.46 |
| 2 | 1460004 | rs4927611 | -0.072 | 1.534 | 2.156 | 1.357 | -0.003 | -0.012 | -0.012 | 5.043 | 0.36 | 0.39 | 0.37 | 0.18 | 0.37 | 0.52 |
| 2 | 1460140 | rs4927612 | -0.035 | 0.779 | 1.62 | 1.082 | -0.001 | -0.006 | -0.006 | 0 | 0.35 | 0.38 | 0.37 | 0.18 | 0.37 | 0.51 |
| 2 | 1460220 | rs11694726 | 0.608 | 0.815 | 1.555 | 1.104 | 0.007 | 0.024 | 0.024 | 0 | 0.06 | 0.1 | 0.08 | 0.14 | 0.07 | 0.04 |
| 2 | 1462885 | rs4927616 | -1.164 | 0.795 | 1.387 | 1.017 | -0.004 | -0.015 | -0.015 | 0 | 0.92 | 0.88 | 0.95 | 0.96 | 0.94 | 0.97 |
| 2 | 1463412 | rs6732480 | 0.316 | 0.882 | 1.395 | 1.066 | -0.006 | -0.026 | -0.026 | 0 | 0.57 | 0.49 | 0.59 | 0.78 | 0.57 | 0.46 |
| 2 | 1467210 | rs6723319 | -1.069 | 0.934 | 1.369 | 1.233 | 0.004 | 0.047 | 0.047 | 0 | 0.16 | 0.29 | 0.11 | 0.25 | 0.13 | 0.16 |
| 2 | 1474131 | rs11211654 | 0.537 | 0.948 | 1.388 | 1.201 | 0.013 | 0.039 | 0.039 | 0 | 0.06 | 0.13 | 0.08 | 0.14 | 0.07 | 0.04 |
| 2 | 1476213 | rs13431646 | 0.008 | 0.855 | 1.687 | 1.025 | 0.02 | -0.027 | -0.027 | 0 | 0.06 | 0.21 | 0.06 | 0 | 0.03 | 0.17 |
| 2 | 1476991 | rs10201842 | 1.547 | 0.747 | 1.527 | 1.039 | -0.009 | -0.014 | -0.014 | 0 | 0.42 | 0.33 | 0.48 | 0.53 | 0.44 | 0.3 |
| 2 | 1482940 | rs6588678 | -0.028 | 0.658 | 1.193 | 1.081 | 0.007 | 0.014 | 0.014 | 0 | 0.66 | 0.85 | 0.66 | 0.74 | 0.6 | 0.64 |
| 2 | 1494373 | rs10196514 | -1.266 | 0.906 | 1.878 | 1.327 | -0.01 | 0.02 | 0.02 | 0.074 | 0.12 | 0.33 | 0.04 | 0.09 | 0.06 | 0.13 |
| 2 | 1495800 | rs2048722 | -1.892 | 0.714 | 1.865 | 1.329 | -0.003 | -0.016 | -0.016 | 0 | 0.47 | 0.71 | 0.46 | 0.5 | 0.46 | 0.65 |
| 2 | 1497831 | rs10189329 | -0.479 | 0.69 | 2.091 | 1.314 | 0.005 | 0.032 | 0.032 | 0 | 0.1 | 0.25 | 0.04 | 0.09 | 0.06 | 0.12 |
| 2 | 1499728 | rs2276704 | 0.071 | 0.701 | 2.276 | 1.385 | -0.002 | -0.003 | -0.003 | 0 | 0.24 | 0.21 | 0.29 | 0.27 | 0.31 | 0.32 |
| 2 | 1499927 | rs732609 | -1.496 | 0.682 | 2.346 | 1.374 | 0 | 0.001 | 0.001 | 0.272 | 0.39 | 0.65 | 0.44 | 0.41 | 0.4 | 0.53 |
| 2 | 1502757 | rs3755551 | 0.05 | 0.737 | 1.678 | 1.355 | -0.002 | -0.004 | -0.004 | 0 | 0.24 | 0.21 | 0.29 | 0.27 | 0.31 | 0.32 |
| 2 | 1506142 | rs938330 | -0.567 | 0.692 | 1.67 | 1.354 | 0 | -0.002 | -0.002 | 1.317 | 0.3 | 0.34 | 0.35 | 0.27 | 0.34 | 0.49 |
| 2 | 1508685 | rs4927621 | -1.25 | 0.693 | 1.526 | 1.342 | 0.001 | 0.009 | 0.009 | 0.455 | 0.41 | 0.55 | 0.45 | 0.5 | 0.44 | 0.65 |
| 2 | 1516365 | rs12465127 | 1.337 | 1.152 | 0.962 | 1.508 | 0.013 | 0.009 | 0.009 | 0.094 | 0.44 | 0.34 | 0.24 | 0.35 | 0.34 | 0.37 |
| 2 | 1516680 | rs4927624 | -1.822 | 1.148 | 1.118 | 1.439 | 0.041 | 0.088 | 0.088 | 0 | 0.5 | 0.82 | 0.67 | 0.63 | 0.57 | 0.69 |
| 2 | 1516787 | rs13398180 | -0.347 | 1.297 | 1.284 | 1.073 | -0.004 | 0.001 | 0.001 | 0 | 0.19 | 0.18 | 0.23 | 0.15 | 0.23 | 0.1 |
| 2 | 1518112 | rs11896517 | -1.265 | 1.353 | 1.515 | 1.165 | 0.094 | 0.065 | 0.065 | 0.582 | 0.12 | 0.36 | 0.19 | 0.17 | 0.16 | 0.17 |
| 2 | 1519064 | rs17732233 | 0.052 | 1.33 | 1.215 | 1.148 | 0 | 0.002 | 0.002 | 0.01 | 0.25 | 0.14 | 0.28 | 0.24 | 0.31 | 0.32 |
| 2 | 1520676 | rs1126799 | -1.661 | 1.339 | 1.209 | 1.019 | 0.011 | 0.03 | 0.03 | 0.077 | 0.51 | 0.83 | 0.6 | 0.58 | 0.54 | 0.69 |
| 2 | 1523751 | rs2048727 | -0.234 | 1.413 | 1.274 | 0.715 | 0.002 | -0.004 | -0.004 | 0.306 | 0.33 | 0.19 | 0.34 | 0.33 | 0.36 | 0.39 |
| 2 | 1525529 | rs4927625 | -1.039 | 1.369 | 1.456 | 0.734 | 0.004 | -0.009 | -0.009 | 0 | 0.43 | 0.46 | 0.55 | 0.37 | 0.52 | 0.59 |
| 2 | 1547230 | rs6605278 | -1.737 | 1.594 | 1.571 | 0.828 | 0.097 | 0.025 | 0.025 | 3.403 | 0.17 | 0.5 | 0.26 | 0.18 | 0.2 | 0.29 |
| 2 | 1552841 | rs10206900 | -1.727 | 1.575 | 1.684 | 0.839 | 0.119 | 0.041 | 0.041 | 2.523 | 0.17 | 0.56 | 0.26 | 0.18 | 0.2 | 0.29 |
| 2 | 1553393 | rs10192959 | -0.476 | 1.529 | 1.302 | 0.902 | 0.065 | -0.014 | -0.014 | 0.501 | 0.07 | 0.21 | 0.16 | 0.05 | 0.14 | 0.06 |
| 2 | 1556330 | rs13414537 | 0.01 | 1.541 | 1.195 | 0.908 | 0.038 | -0.008 | -0.008 | 0 | 0.06 | 0.15 | 0.15 | 0.05 | 0.14 | 0.06 |
| 2 | 1563833 | rs7583169 | 1.262 | 1.486 | 0.687 | 0.891 | 0.006 | -0.041 | -0.041 | 0.294 | 0.48 | 0.06 | 0.38 | 0.47 | 0.43 | 0.3 |
| 2 | 1565555 | rs7584264 | -0.296 | 1.559 | 1.033 | 0.888 | 0.161 | 0.023 | 0.023 | 0.018 | 0.07 | 0.35 | 0.18 | 0.05 | 0.14 | 0.06 |
| 2 | 1580250 | rs6721243 | -1.465 | 1.523 | 0.905 | 0.944 | 0.132 | 0.119 | 0.119 | 1.533 | 0.12 | 0.47 | 0.21 | 0.18 | 0.17 | 0.15 |
| 2 | 1580504 | rs6720874 | -0.307 | 1.62 | 0.715 | 0.721 | 0.162 | 0.011 | 0.011 | 0.568 | 0.08 | 0.36 | 0.18 | 0.05 | 0.15 | 0.06 |
| 2 | 1582242 | rs7564265 | 1.04 | 1.418 | 0.264 | 0.72 | 0.02 | 0.006 | 0.006 | 0.013 | 0.49 | 0.37 | 0.39 | 0.47 | 0.43 | 0.31 |

| | | | | | | | | | | | | | | | | |
|---|---|---|---|---|---|---|---|---|---|---|---|---|---|---|---|---|
| 2 | 1584345 | rs28532847 | -0.95 | 1.47 | 0.09 | 0.65 | 0.09 | -0.036 | -0.036 | 0 | 0.11 | 0.3 | 0.19 | 0.05 | 0.17 | 0.1 |
| 2 | 1619121 | rs10183727 | 0.983 | 0.468 | -1.172 | 0.658 | 0.037 | 0 | 0 | 0 | 0.49 | 0.21 | 0.66 | 0.57 | 0.57 | 0.6 |
| 2 | 1620721 | rs12614899 | -0.308 | 0.29 | -1.132 | 0.517 | -0.042 | -0.066 | -0.066 | 0 | 0.2 | 0.59 | 0.14 | 0.09 | 0.1 | 0.19 |
| 2 | 1621608 | rs10779972 | 0.203 | 0.553 | -1.091 | 0.799 | -0.002 | -0.002 | -0.002 | 0 | 0.63 | 0.39 | 0.74 | 0.73 | 0.75 | 0.74 |
| 2 | 1622140 | rs9752413 | 0.249 | 1.055 | -1.132 | 0.994 | -0.023 | -0.028 | -0.028 | 0 | 0.12 | 0.04 | 0.07 | 0.15 | 0.18 | 0.11 |
| 2 | 1625121 | rs9309692 | 0.455 | 0.727 | -0.951 | 0.913 | 0.001 | -0.003 | -0.003 | 0 | 0.38 | 0.62 | 0.41 | 0.47 | 0.36 | 0.51 |
| 2 | 1625269 | rs9679409 | -0.231 | 0.953 | -0.797 | 0.559 | -0.007 | -0.017 | -0.017 | 0 | 0.37 | 0.49 | 0.42 | 0.37 | 0.33 | 0.46 |
| 2 | 1626392 | rs9752743 | -0.716 | 1.061 | -1.05 | 0.566 | -0.021 | -0.021 | -0.021 | 0 | 0.53 | 0.78 | 0.54 | 0.49 | 0.54 | 0.55 |
| 2 | 1627730 | rs10196165 | 0.971 | 1.85 | -0.02 | 0.797 | -0.001 | -0.004 | -0.004 | 0 | 0.41 | 0.4 | 0.4 | 0.34 | 0.42 | 0.33 |
| 2 | 1627779 | rs11904364 | -0.534 | 1.835 | -0.046 | 0.571 | -0.001 | -0.003 | -0.003 | 0.05 | 0.41 | 0.65 | 0.54 | 0.44 | 0.42 | 0.52 |
| 2 | 1629108 | rs9711675 | 0.811 | 2.417 | 0.516 | 0.767 | -0.005 | 0.024 | 0.024 | 0 | 0.57 | 0.25 | 0.53 | 0.68 | 0.48 | 0.52 |
| 2 | 1637201 | rs4240 | 0.633 | 2.296 | 0.987 | 0.643 | -0.011 | -0.007 | -0.007 | 0.129 | 0.62 | 0.29 | 0.45 | 0.6 | 0.59 | 0.57 |
| 2 | 1637648 | rs1054241 | 0.534 | 2.277 | 1.143 | 0.618 | -0.014 | -0.01 | -0.01 | 0.011 | 0.64 | 0.28 | 0.45 | 0.6 | 0.61 | 0.6 |
| 2 | 1639060 | rs2241458 | -0.207 | 2.289 | 1.043 | 0.64 | -0.002 | 0.008 | 0.008 | 0 | 0.29 | 0.29 | 0.42 | 0.39 | 0.25 | 0.4 |
| 2 | 1640170 | rs12714344 | 0.537 | 2.122 | 0.992 | 0.321 | 0.124 | 0.021 | 0.021 | 0.175 | 0.93 | 0.58 | 0.93 | 1 | 0.91 | 0.98 |
| 2 | 1641991 | rs12714333 | 0.172 | 2.131 | 0.987 | 0.326 | 0.104 | 0.012 | 0.012 | 0 | 0.94 | 0.65 | 0.94 | 1 | 0.91 | 0.98 |
| 2 | 1642070 | rs12714332 | 1.121 | 2.114 | 0.945 | 0.324 | 0.156 | -0.024 | -0.024 | 0 | 0.91 | 0.52 | 0.86 | 0.99 | 0.84 | 0.97 |
| 2 | 1643456 | rs7575438 | 1.061 | 2.085 | 0.809 | 0.321 | 0.004 | 0.004 | 0.004 | 0 | 0.37 | 0.35 | 0.36 | 0.31 | 0.34 | 0.33 |
| 2 | 1647171 | rs6730800 | 0.222 | 1.833 | 0.161 | 0.391 | 0.069 | -0.019 | -0.019 | 0 | 0.94 | 0.77 | 0.96 | 1 | 0.89 | 0.98 |
| 2 | 1652660 | rs3811613 | 1.102 | 1.916 | -0.155 | 0.398 | 0.048 | 0.018 | 0.018 | 0 | 0.34 | 0.12 | 0.27 | 0.28 | 0.28 | 0.31 |
| 2 | 1664654 | rs17841813 | 0.243 | 2.19 | 0.046 | 0.334 | 0.002 | -0.038 | -0.038 | 0 | 0.77 | 0.73 | 0.82 | 0.98 | 0.69 | 0.87 |
| 2 | 1675299 | rs7590885 | -0.262 | 2.355 | -0.617 | 0.805 | 0.053 | 0.08 | 0.08 | 0 | 0.2 | 0.42 | 0.42 | 0.32 | 0.23 | 0.37 |
| 2 | 1676388 | rs1544846 | 0.618 | 1.839 | -1.884 | 0.963 | 0.002 | 0.01 | 0.01 | 0 | 0.5 | 0.07 | 0.36 | 0.54 | 0.43 | 0.47 |
| 2 | 1679912 | rs11127327 | -0.262 | 1.724 | -0.884 | 1.177 | 0.027 | 0.032 | 0.032 | 0.13 | 0.26 | 0.34 | 0.4 | 0.31 | 0.4 | 0.32 |
| 2 | 1685800 | rs11892515 | -0.489 | 1.228 | -1.847 | 0.884 | -0.042 | -0.08 | -0.08 | 0 | 0.59 | 0.93 | 0.68 | 0.52 | 0.61 | 0.58 |
| 2 | 1688065 | rs2288184 | -0.651 | 1.139 | -0.421 | 0.704 | 0.032 | -0.036 | -0.036 | 1.215 | 0.1 | 0.36 | 0.12 | 0.04 | 0.09 | 0.07 |
| 2 | 1693035 | rs7580177 | -0.979 | 1.42 | -0.23 | 1.278 | 0.011 | 0.034 | 0.034 | 0 | 0.22 | 0.32 | 0.24 | 0.32 | 0.26 | 0.3 |
| 2 | 1696016 | rs2288183 | -0.269 | 0.503 | -0.173 | 0.389 | -0.013 | 0.017 | 0.017 | 0.158 | 0.67 | 0.58 | 0.72 | 0.35 | 0.67 | 0.69 |
| 2 | 1697488 | rs2241452 | 0.511 | 0.481 | -0.168 | 1.048 | 0.001 | 0.079 | 0.079 | 0 | 0.94 | 0.93 | 0.91 | 0.69 | 0.94 | 0.93 |
| 2 | 1700986 | rs890054 | 0.932 | 0.422 | 0.355 | 1.025 | 0.036 | 0.229 | 0.229 | 3.804 | 0.93 | 0.72 | 0.82 | 0.69 | 0.93 | 0.9 |
| 2 | 1705569 | rs1863136 | 0.543 | 0.43 | 0.796 | 1.109 | 0 | -0.001 | -0.001 | 0 | 0.52 | 0.57 | 0.61 | 0.59 | 0.47 | 0.47 |
| 2 | 1708502 | rs7588729 | -1.058 | 0.497 | 0.76 | 1.021 | -0.008 | 0.021 | 0.021 | 0 | 0.52 | 0.8 | 0.46 | 0.36 | 0.53 | 0.57 |
| 2 | 1726761 | rs6735777 | -0.073 | 0.482 | 0.518 | 1.304 | -0.05 | -0.101 | -0.101 | 0.894 | 0.14 | 0.01 | 0.15 | 0.27 | 0.15 | 0.29 |
| 2 | 1727930 | rs4853843 | -0.056 | -0.038 | 0.653 | 1.129 | -0.032 | 0.002 | 0.002 | 0 | 0.67 | 0.49 | 0.79 | 0.63 | 0.69 | 0.8 |
| 2 | 1731405 | rs4853842 | 1.354 | 0.026 | 0.585 | 0.834 | -0.049 | 0.213 | 0.213 | 0 | 0.43 | 0.16 | 0.5 | 0.05 | 0.44 | 0.39 |
| 2 | 1735480 | rs6714537 | -0.166 | -0.17 | 0.91 | 1.388 | 0.002 | 0.071 | 0.071 | 0.021 | 0.14 | 0.17 | 0.11 | 0.43 | 0.12 | 0.07 |
| 2 | 1735964 | rs11127323 | 1.284 | -0.235 | 0.586 | 1.159 | -0.058 | 0.073 | 0.073 | 0 | 0.44 | 0.19 | 0.58 | 0.33 | 0.44 | 0.38 |
| 2 | 1742043 | rs4853841 | -0.154 | -0.321 | 1.091 | 0.882 | -0.013 | -0.086 | -0.086 | 0.729 | 0.72 | 0.61 | 0.81 | 0.94 | 0.68 | 0.81 |
| 2 | 1744932 | rs6740357 | -0.979 | -0.454 | 0.981 | 1.45 | -0.009 | 0.04 | 0.04 | 0.01 | 0.2 | 0.22 | 0.13 | 0.38 | 0.13 | 0.08 |
| 2 | 1745326 | rs4853759 | 0.982 | -0.739 | 1.193 | 1.263 | -0.007 | 0.01 | 0.01 | 0 | 0.62 | 0.62 | 0.73 | 0.52 | 0.64 | 0.67 |
| 2 | 1752229 | rs2382558 | -1.744 | -0.379 | 1.264 | 1.521 | 0.012 | 0.046 | 0.046 | 0 | 0.26 | 0.56 | 0.29 | 0.56 | 0.25 | 0.36 |
| 2 | 1755898 | rs13416067 | -1.235 | -0.357 | 1.274 | 1.442 | 0.005 | 0.029 | 0.029 | 0.115 | 0.22 | 0.31 | 0.23 | 0.55 | 0.2 | 0.34 |
| 2 | 1761496 | rs6756670 | -2.35 | -0.462 | 0.99 | 1.333 | 0.002 | -0.003 | -0.003 | 0 | 0.31 | 0.68 | 0.3 | 0.57 | 0.26 | 0.39 |
| 2 | 1761661 | rs6706107 | -2.18 | -0.191 | 0.906 | 1.466 | 0.006 | 0.023 | 0.023 | 0 | 0.28 | 0.53 | 0.29 | 0.57 | 0.26 | 0.39 |
| 2 | 1771963 | rs13403634 | -0.623 | -0.102 | 0.775 | 1.745 | 0.01 | 0.089 | 0.089 | 0 | 0.17 | 0.27 | 0.14 | 0.38 | 0.15 | 0.12 |
| 2 | 1773347 | rs2059412 | -0.54 | -0.148 | 0.918 | 1.737 | 0.011 | 0.196 | 0.196 | 0 | 0.07 | 0.27 | 0.07 | 0.18 | 0.03 | 0.25 |
| 2 | 1773400 | rs718002 | -0.403 | -0.147 | 0.909 | 1.737 | 0.006 | 0.178 | 0.178 | 0 | 0.07 | 0.24 | 0.06 | 0.18 | 0.03 | 0.25 |
| 2 | 1773830 | rs12475777 | -1.098 | -0.134 | 0.665 | 1.868 | 0.019 | 0.063 | 0.063 | 0 | 0.2 | 0.3 | 0.11 | 0.31 | 0.19 | 0.12 |
| 2 | 1775648 | rs10048625 | -0.216 | -0.071 | 0.456 | 1.86 | 0.024 | 0.114 | 0.114 | 0 | 0.14 | 0.24 | 0.07 | 0.32 | 0.14 | 0.09 |
| 2 | 1778558 | rs1124200 | -0.758 | -0.049 | -0.349 | 1.308 | 0.002 | 0.026 | 0.026 | 0.069 | 0.27 | 0.59 | 0.17 | 0.5 | 0.25 | 0.37 |
| 2 | 1779523 | rs6726699 | 0.441 | -0.088 | -0.905 | 1.198 | 0.02 | 0.001 | 0.001 | 0 | 0.78 | 0.94 | 0.81 | 0.77 | 0.77 | 0.79 |
| 2 | 1781022 | rs12617500 | -1.641 | 0.232 | -0.334 | 1.572 | -0.003 | 0.068 | 0.068 | 0 | 0.12 | 0.19 | 0.11 | 0.62 | 0.1 | 0.23 |
| 2 | 1786200 | rs6759577 | 0.855 | 0.924 | 0.137 | 1.039 | 0 | -0.01 | -0.01 | 1.901 | 0.68 | 0.59 | 0.62 | 0.29 | 0.66 | 0.67 |
| 2 | 1787933 | rs6751762 | -0.037 | 1.024 | 0.089 | 1.03 | -0.003 | -0.018 | -0.018 | 0 | 0.78 | 0.81 | 0.76 | 0.44 | 0.72 | 0.83 |
| 2 | 1788847 | rs10519486 | -1.592 | 1.105 | 0.639 | 0.64 | 0.033 | -0.014 | -0.014 | 0 | 0.23 | 0.46 | 0.22 | 0.16 | 0.25 | 0.29 |
| 2 | 1793809 | rs6742365 | -1.299 | 1.07 | 1.102 | 0.627 | 0.006 | -0.013 | -0.013 | 0 | 0.16 | 0.25 | 0.15 | 0.08 | 0.16 | 0.21 |
| 2 | 1794410 | rs17338616 | 1.531 | 1.008 | 0.771 | 0.625 | -0.008 | 0.082 | 0.082 | 0 | 0.53 | 0.26 | 0.52 | 0.29 | 0.46 | 0.53 |
| 2 | 1799691 | rs7588942 | 1.002 | 0.988 | 0.715 | 0.514 | 0 | -0.002 | -0.002 | 0 | 0.6 | 0.49 | 0.6 | 0.37 | 0.58 | 0.64 |
| 2 | 1802922 | rs7563788 | 0.3 | 1.1 | 0.561 | 0.485 | 0.007 | -0.03 | -0.03 | 0 | 0.07 | 0.1 | 0.04 | 0 | 0.08 | 0.03 |
| 2 | 1805430 | rs2288178 | -0.715 | 1.071 | 0.191 | 0.674 | 0.018 | -0.027 | -0.027 | 0 | 0.09 | 0 | 0.08 | 0.08 | 0.05 | 0.11 |
| 2 | 1805489 | rs2288179 | -0.715 | 1.071 | 0.191 | 0.674 | 0.018 | -0.027 | -0.027 | 0 | 0.09 | 0.01 | 0.08 | 0.08 | 0.05 | 0.11 |
| 2 | 1806093 | rs10200829 | -0.987 | 1.129 | 0.238 | 0.681 | -0.003 | -0.02 | -0.02 | 1.954 | 0.11 | 0.06 | 0.12 | 0.14 | 0.1 | 0.18 |
| 2 | 1806879 | rs13006826 | 0.17 | 1.145 | 0.393 | 0.687 | 0.007 | 0.04 | 0.04 | 0 | 0.71 | 0.61 | 0.69 | 0.45 | 0.64 | 0.79 |
| 2 | 1810830 | rs11684279 | 0.853 | 1.059 | 0.072 | 0.45 | 0.001 | -0.004 | -0.004 | 0.092 | 0.58 | 0.41 | 0.55 | 0.31 | 0.52 | 0.58 |
| 2 | 1824122 | rs6755207 | -0.993 | 1.081 | 0.074 | 0.773 | 0.022 | 0.032 | 0.032 | 0 | 0.09 | 0.31 | 0.11 | 0.06 | 0.07 | 0.15 |
| 2 | 1828774 | rs4072464 | -0.731 | 0.909 | -1.782 | 0.088 | -0.028 | -0.003 | -0.003 | 0 | 0.59 | 0.98 | 0.68 | 0.7 | 0.62 | 0.6 |
| 2 | 1842774 | rs6728613 | -1.252 | 1.183 | -0.921 | 0.622 | -0.029 | -0.009 | -0.009 | 0 | 0.35 | 0.83 | 0.35 | 0.66 | 0.37 | 0.38 |
| 2 | 1855717 | rs7421372 | 1.05 | 1.28 | -0.639 | 0.667 | -0.031 | 0.016 | 0.016 | 0 | 0.63 | 0.27 | 0.64 | 0.3 | 0.54 | 0.6 |
| 2 | 1860120 | rs4571084 | -1.081 | 1.449 | 0.804 | 1.47 | 0.052 | 0.096 | 0.096 | 0.055 | 0.12 | 0.36 | 0.11 | 0.24 | 0.15 | 0.1 |

| | | | | | | | | | | | | | | | | |
|---|---|---|---|---|---|---|---|---|---|---|---|---|---|---|---|---|
| 2 | 1867240 | rs11674865 | -0.333 | 1.628 | 0.751 | 1.338 | 0.006 | -0.022 | -0.022 | 0 | 0.05 | 0.05 | 0.03 | 0 | 0.09 | 0.02 |
| 2 | 1871083 | rs11904794 | -1.263 | 1.805 | 0.937 | 1.425 | 0.117 | 0.216 | 0.216 | 0 | 0.07 | 0.31 | 0.09 | 0.23 | 0.1 | 0.1 |
| 2 | 1875792 | rs4853754 | -1.64 | 1.779 | 1.375 | 1.731 | 0 | 0.044 | 0.044 | 0 | 0.23 | 0.67 | 0.23 | 0.43 | 0.21 | 0.21 |
| 2 | 1877998 | rs4305302 | -1.573 | 1.79 | 1.691 | 1.737 | 0.008 | 0.087 | 0.087 | 0 | 0.22 | 0.6 | 0.22 | 0.43 | 0.21 | 0.21 |
| 2 | 1881459 | rs4557004 | -2.22 | 1.808 | 2.012 | 1.757 | 0.003 | 0.039 | 0.039 | 0 | 0.28 | 0.62 | 0.26 | 0.46 | 0.26 | 0.27 |
| 2 | 1884069 | rs6747509 | -2.191 | 1.837 | 2.399 | 2.011 | -0.005 | -0.009 | -0.009 | 0 | 0.16 | 0.39 | 0.14 | 0.13 | 0.12 | 0.15 |
| 2 | 1887049 | rs11127306 | -1.068 | 1.871 | 2.258 | 2.03 | 0.018 | 0.109 | 0.109 | 0 | 0.11 | 0.22 | 0.12 | 0.34 | 0.12 | 0.1 |
| 2 | 1907446 | rs2195903 | -3.1 | 1.854 | 2.43 | 1.955 | 0.009 | 0.022 | 0.022 | 0.247 | 0.24 | 0.67 | 0.2 | 0.37 | 0.27 | 0.21 |
| 2 | 1911871 | rs17039158 | -1.834 | 1.832 | 2.431 | 1.903 | -0.005 | -0.025 | -0.025 | 0 | 0.08 | 0.08 | 0.04 | 0 | 0.05 | 0.03 |
| 2 | 1912759 | rs12471985 | -3.896 | 1.683 | 2.285 | 2.051 | -0.029 | -0.011 | -0.011 | 0.045 | 0.35 | 0.86 | 0.34 | 0.66 | 0.37 | 0.3 |
| 2 | 1915297 | rs1368233 | -3.425 | 1.712 | 2.266 | 2.361 | 0.003 | 0.017 | 0.017 | 0 | 0.28 | 0.69 | 0.27 | 0.48 | 0.32 | 0.27 |
| 2 | 1915399 | rs12988893 | -1.163 | 1.472 | 2.025 | 2.105 | 0.001 | -0.007 | -0.007 | 0 | 0.08 | 0.06 | 0.04 | 0 | 0.06 | 0.03 |
| 2 | 1919847 | rs1432539 | -1.769 | 1.666 | 2.724 | 2.286 | 0.037 | 0.127 | 0.127 | 0 | 0.1 | 0.21 | 0.12 | 0.35 | 0.13 | 0.15 |
| 2 | 1924666 | rs13404264 | -1.606 | 1.721 | 2.858 | 2.361 | 0.003 | 0.041 | 0.041 | 0 | 0.1 | 0.14 | 0.15 | 0.27 | 0.1 | 0.09 |
| 2 | 1925993 | rs2241685 | -1.43 | 1.708 | 2.723 | 2.35 | 0.019 | 0.142 | 0.142 | 0 | 0.09 | 0.21 | 0.11 | 0.35 | 0.1 | 0.13 |
| 2 | 1940723 | rs1864881 | -3.755 | 1.768 | 2.708 | 2.226 | -0.011 | -0.01 | -0.01 | 2.915 | 0.33 | 0.76 | 0.34 | 0.64 | 0.37 | 0.37 |
| 2 | 1946914 | rs3748988 | -3.69 | 1.773 | 2.766 | 2.228 | -0.004 | -0.004 | -0.004 | 0.257 | 0.32 | 0.71 | 0.33 | 0.64 | 0.37 | 0.37 |
| 2 | 1946968 | rs3748989 | -0.952 | 1.765 | 2.694 | 2.289 | 0.023 | 0.123 | 0.123 | 0 | 0.07 | 0.14 | 0.11 | 0.34 | 0.09 | 0.12 |
| 2 | 1950858 | rs13405738 | -2.764 | 1.749 | 2.672 | 2.487 | 0 | 0.01 | 0.01 | 0 | 0.14 | 0.18 | 0.12 | 0.28 | 0.14 | 0.21 |
| 2 | 1951254 | rs7569044 | -2.821 | 1.715 | 2.828 | 2.627 | -0.044 | 0.048 | 0.048 | 0.012 | 0.17 | 0.58 | 0.13 | 0.29 | 0.07 | 0.14 |
| 2 | 1951389 | rs11678039 | -2.968 | 1.746 | 2.711 | 2.569 | 0.004 | 0.01 | 0.01 | 0 | 0.19 | 0.22 | 0.2 | 0.3 | 0.26 | 0.25 |
| 2 | 1952576 | rs919718 | -1.037 | 1.727 | 2.686 | 2.578 | 0.029 | 0.126 | 0.126 | 0 | 0.07 | 0.16 | 0.11 | 0.34 | 0.09 | 0.12 |
| 2 | 1964906 | rs10185662 | -3.417 | 1.676 | 2.599 | 2.316 | 0.021 | 0.082 | 0.082 | 1.08 | 0.22 | 0.53 | 0.21 | 0.3 | 0.22 | 0.22 |
| 2 | 1969755 | rs4853816 | -3.489 | 1.433 | 2.69 | 2.215 | 0.009 | 0.069 | 0.069 | 0 | 0.2 | 0.46 | 0.2 | 0.27 | 0.17 | 0.2 |
| 2 | 1970226 | rs11127298 | -2.47 | 1.439 | 2.361 | 1.778 | 0.016 | -0.048 | -0.048 | 0 | 0.16 | 0.45 | 0.12 | 0.06 | 0.14 | 0.09 |
| 2 | 1970675 | rs6751072 | -1.972 | 1.319 | 2.174 | 1.881 | -0.033 | -0.022 | -0.022 | 0 | 0.16 | 0.33 | 0.13 | 0.06 | 0.08 | 0.17 |
| 2 | 1980644 | rs11888121 | -2.218 | 1.29 | 2.122 | 1.937 | -0.015 | 0.018 | 0.018 | 0.471 | 0.18 | 0.48 | 0.16 | 0.13 | 0.1 | 0.18 |
| 2 | 1981503 | rs6721182 | -2.42 | 1.544 | 2.125 | 2.207 | 0.022 | 0.067 | 0.067 | 0 | 0.15 | 0.29 | 0.15 | 0.34 | 0.17 | 0.18 |
| 2 | 1983619 | rs7561518 | -3.73 | 1.674 | 2.004 | 2.179 | -0.01 | -0.036 | -0.036 | 0 | 0.34 | 0.78 | 0.32 | 0.48 | 0.3 | 0.35 |
| 2 | 1987338 | rs7569308 | -3.595 | 1.564 | 2.015 | 2.183 | 0.003 | -0.034 | -0.034 | 0.196 | 0.33 | 0.78 | 0.3 | 0.48 | 0.27 | 0.34 |
| 2 | 1989289 | rs4853812 | -3.329 | 1.582 | 2.083 | 2.34 | 0.006 | 0.045 | 0.045 | 0 | 0.28 | 0.58 | 0.28 | 0.54 | 0.26 | 0.3 |
| 2 | 1991561 | rs17039243 | -1.416 | 1.6 | 2.143 | 2.359 | 0.016 | 0.06 | 0.06 | 0.001 | 0.1 | 0.11 | 0.13 | 0.4 | 0.15 | 0.15 |
| 2 | 1992624 | rs891878 | 0.593 | 1.41 | 0.81 | 1.62 | -0.001 | -0.001 | -0.001 | 0 | 0.48 | 0.48 | 0.41 | 0.62 | 0.53 | 0.6 |
| 2 | 1993347 | rs7608969 | 1.759 | 1.383 | 0.518 | 1.377 | -0.016 | 0.036 | 0.036 | 0.065 | 0.35 | 0.2 | 0.26 | 0.22 | 0.37 | 0.45 |
| 2 | 1998275 | rs13382326 | -0.162 | 1.059 | 0.414 | 1.186 | -0.023 | 0.002 | 0.002 | 1.025 | 0.11 | 0.2 | 0.11 | 0.1 | 0.06 | 0.03 |
| 2 | 2027348 | rs1978703 | -0.518 | 1.395 | -0.732 | 1.087 | -0.084 | -0.072 | -0.072 | 0 | 0.71 | 0.98 | 0.57 | 0.74 | 0.65 | 0.77 |
| 2 | 2043349 | rs891877 | -0.212 | 1.791 | 0.536 | 1.279 | 0.009 | 0.046 | 0.046 | 0 | 0.09 | 0.17 | 0.08 | 0.09 | 0.06 | 0.14 |
| 2 | 2047397 | rs6759709 | -1.873 | 1.807 | 0.81 | 1.211 | 0.033 | 0.007 | 0.007 | 0.898 | 0.23 | 0.37 | 0.18 | 0.14 | 0.26 | 0.21 |
| 2 | 2071493 | rs11685526 | -1.353 | 2.034 | 0.685 | 1.665 | 0.014 | 0.07 | 0.07 | 0 | 0.09 | 0.15 | 0.1 | 0.31 | 0.11 | 0.18 |
| 2 | 2090280 | rs1614012 | -3.517 | 1.975 | 0.493 | 1.681 | -0.051 | -0.08 | -0.08 | 0 | 0.35 | 0.87 | 0.27 | 0.45 | 0.36 | 0.39 |
| 2 | 2091619 | rs1421614 | -2.086 | 1.994 | 0.644 | 1.432 | 0.004 | -0.003 | -0.003 | 0 | 0.26 | 0.71 | 0.15 | 0.14 | 0.25 | 0.2 |
| 2 | 2102687 | rs6755567 | -1.934 | 2.371 | 0.496 | 1.636 | 0.044 | 0.015 | 0.015 | 0.296 | 0.22 | 0.41 | 0.12 | 0.14 | 0.25 | 0.21 |
| 2 | 2116870 | rs751150 | -0.811 | 2.475 | 0.285 | 1.691 | 0.002 | 0.009 | 0.009 | 0 | 0.37 | 0.4 | 0.37 | 0.45 | 0.36 | 0.57 |
| 2 | 2124706 | rs17039339 | -1.627 | 2.453 | 0.656 | 1.897 | 0.02 | 0.039 | 0.039 | 0 | 0.08 | 0.13 | 0.09 | 0.2 | 0.11 | 0.14 |
| 2 | 2128585 | rs1620250 | 0.358 | 2.55 | 0.492 | 1.952 | -0.037 | -0.032 | -0.032 | 0 | 0.85 | 0.99 | 0.86 | 0.84 | 0.83 | 0.87 |
| 2 | 2135309 | rs17247373 | 0.89 | 2.427 | 0.746 | 1.871 | -0.003 | -0.029 | -0.029 | 0 | 0.06 | 0.02 | 0.01 | 0.08 | 0.04 | 0.06 |
| 2 | 2154853 | rs2042142 | -2.579 | 2.445 | 0.832 | 1.8 | 0.022 | 0.031 | 0.031 | 0.006 | 0.3 | 0.55 | 0.2 | 0.4 | 0.32 | 0.36 |
| 2 | 2163076 | rs1465389 | 0.895 | 2.392 | 0.66 | 1.48 | 0.018 | 0 | 0 | 0 | 0.87 | 0.69 | 0.9 | 0.88 | 0.86 | 0.86 |
| 2 | 2165519 | rs11687473 | -0.297 | 2.406 | 0.568 | 1.455 | 0.03 | 0.046 | 0.046 | 0 | 0.05 | 0.1 | 0.08 | 0.17 | 0.1 | 0.08 |
| 2 | 2170631 | rs17039361 | -2.088 | 2.421 | 0.605 | 1.226 | 0.025 | 0.008 | 0.008 | 0.234 | 0.3 | 0.54 | 0.2 | 0.27 | 0.34 | 0.34 |
| 2 | 2178658 | rs11674222 | -0.754 | 2.037 | 0.682 | 1.024 | 0.038 | 0.042 | 0.042 | 0 | 0.06 | 0.16 | 0.09 | 0.13 | 0.09 | 0.07 |
| 2 | 2192750 | rs6709211 | -0.397 | 1.933 | 0.587 | 1.027 | 0.098 | 0.053 | 0.053 | 0 | 0.06 | 0.18 | 0.1 | 0.12 | 0.14 | 0.06 |
| 2 | 2197239 | rs17039365 | -0.433 | 1.939 | 0.511 | 1.026 | 0.108 | 0.054 | 0.054 | 0.01 | 0.08 | 0.28 | 0.11 | 0.12 | 0.14 | 0.06 |
| 2 | 2199705 | rs7596980 | -0.947 | 1.868 | 0.32 | 1.027 | 0.153 | 0.084 | 0.084 | 0.989 | 0.1 | 0.44 | 0.12 | 0.12 | 0.14 | 0.06 |
| 2 | 2204863 | rs12471456 | -1.444 | 1.867 | 0.187 | 1.025 | 0.127 | 0.025 | 0.025 | 0.779 | 0.12 | 0.53 | 0.14 | 0.12 | 0.2 | 0.08 |
| 2 | 2210909 | rs11887279 | -0.907 | 1.601 | -0.673 | 0.921 | -0.003 | -0.003 | -0.003 | 0 | 0.23 | 0.28 | 0.1 | 0.13 | 0.2 | 0.26 |
| 2 | 2211665 | rs7587635 | -0.089 | 1.969 | -0.775 | 0.974 | 0.097 | 0.022 | 0.022 | 0.117 | 0.08 | 0.25 | 0.11 | 0.1 | 0.14 | 0.06 |
| 2 | 2214115 | rs1990855 | -0.299 | 1.684 | -1.38 | 1.018 | 0.053 | -0.039 | -0.039 | 0 | 0.23 | 0.45 | 0.18 | 0.15 | 0.27 | 0.28 |
| 2 | 2214468 | rs7582528 | -0.749 | 1.785 | -1.797 | 1.217 | -0.055 | -0.072 | -0.072 | 0.238 | 0.86 | 0.99 | 0.86 | 0.76 | 0.78 | 0.92 |
| 2 | 2215203 | rs13012468 | -0.259 | 1.43 | -1.32 | 0.756 | 0.003 | -0.024 | -0.024 | 0 | 0.14 | 0.01 | 0.04 | 0.09 | 0.08 | 0.09 |
| 2 | 2222530 | rs17039395 | -0.442 | 1.547 | -1.189 | 0.753 | 0.056 | -0.041 | -0.041 | 0 | 0.23 | 0.46 | 0.18 | 0.15 | 0.27 | 0.28 |
| 2 | 2222968 | rs17247310 | -0.96 | 1.509 | -1.092 | 0.857 | 0.007 | -0.025 | -0.025 | 1.408 | 0.38 | 0.49 | 0.24 | 0.24 | 0.37 | 0.38 |
| 2 | 2227388 | rs11127371 | -0.322 | 1.387 | -1.719 | 0.666 | -0.013 | 0.012 | 0.012 | 0 | 0.27 | 0.14 | 0.13 | 0.19 | 0.23 | 0.31 |
| 2 | 2227707 | rs10865533 | -0.28 | 0.8 | -1.535 | 0.142 | 0.006 | 0.041 | 0.041 | 0 | 0.69 | 0.79 | 0.58 | 0.8 | 0.7 | 0.67 |
| 2 | 2228072 | rs888627 | -0.206 | 0.816 | -1.423 | 0.203 | 0.005 | 0.015 | 0.015 | 0 | 0.68 | 0.73 | 0.57 | 0.78 | 0.7 | 0.67 |
| 2 | 2233492 | rs10184190 | -0.537 | 0.82 | -1.539 | -0.108 | -0.008 | 0.038 | 0.038 | 0 | 0.29 | 0.17 | 0.16 | 0.14 | 0.37 | 0.3 |
| 2 | 2236281 | rs4853946 | -0.346 | 0.544 | -1.483 | -0.571 | 0.007 | 0.078 | 0.078 | 0 | 0.54 | 0.68 | 0.59 | 0.78 | 0.47 | 0.7 |
| 2 | 2236799 | rs17247253 | -0.799 | 0.365 | -1.546 | -0.429 | -0.001 | -0.001 | -0.001 | 0 | 0.13 | 0.11 | 0.25 | 0.17 | 0.14 | 0.37 |
| 2 | 2241344 | rs1862112 | -0.242 | 0.349 | -1.237 | -0.355 | 0.023 | 0.04 | 0.04 | 0 | 0.49 | 0.61 | 0.37 | 0.65 | 0.42 | 0.46 |

| | | | | | | | | | | | | | | | | |
|---|---|---|---|---|---|---|---|---|---|---|---|---|---|---|---|---|
| 2 | 2244388 | rs7582260 | -0.498 | 0.149 | -0.664 | -0.431 | -0.001 | 0.01 | 0.01 | 0 | 0.53 | 0.53 | 0.46 | 0.82 | 0.51 | 0.63 |
| 2 | 2249041 | rs2008419 | -0.914 | -0.113 | -0.034 | 0.006 | -0.016 | 0.021 | 0.021 | 0.05 | 0.28 | 0.4 | 0.29 | 0.35 | 0.27 | 0.29 |
| 2 | 2249052 | rs2008412 | -1.344 | -0.298 | 0.142 | -0.336 | -0.001 | -0.003 | -0.003 | 0 | 0.35 | 0.62 | 0.39 | 0.77 | 0.31 | 0.47 |
| 2 | 2249553 | rs9808093 | -0.498 | -0.07 | 0.147 | -0.455 | -0.001 | -0.001 | -0.001 | 0.162 | 0.2 | 0.16 | 0.11 | 0.1 | 0.26 | 0.24 |
| 2 | 2250426 | rs1421612 | -0.683 | -0.145 | 0.628 | -0.464 | 0.018 | -0.018 | -0.018 | 0 | 0.09 | 0.24 | 0.05 | 0.02 | 0.08 | 0.15 |
| 2 | 2256417 | rs11127366 | 0.517 | -0.028 | 0.414 | -0.848 | 0 | 0 | 0 | 0 | 0.26 | 0.3 | 0.38 | 0.74 | 0.24 | 0.25 |
| 2 | 2257857 | rs10201099 | 0.836 | 0.06 | 0.045 | -0.961 | -0.003 | 0.01 | 0.01 | 0.515 | 0.13 | 0.05 | 0.06 | 0.08 | 0.15 | 0.1 |
| 2 | 2259540 | rs4473412 | -0.454 | -0.202 | -0.346 | -1.28 | 0.002 | -0.003 | -0.003 | 0 | 0.44 | 0.46 | 0.45 | 0.83 | 0.38 | 0.58 |
| 2 | 2263952 | rs7600426 | -0.453 | -0.185 | -0.516 | -1.27 | 0.003 | 0.003 | 0.003 | 0.057 | 0.44 | 0.44 | 0.45 | 0.83 | 0.38 | 0.58 |
| 2 | 2268526 | rs12470297 | -0.809 | 0.138 | -0.994 | -1.223 | -0.016 | 0.029 | 0.029 | 0.063 | 0.6 | 0.7 | 0.52 | 0.91 | 0.54 | 0.69 |
| 2 | 2283434 | rs7568465 | 0.052 | 0.175 | -1.126 | -1.185 | 0.006 | 0.026 | 0.026 | 0 | 0.44 | 0.6 | 0.45 | 0.85 | 0.39 | 0.47 |
| 2 | 2284083 | rs10495485 | -0.603 | 0.125 | -1.402 | -0.508 | -0.006 | 0.019 | 0.019 | 0 | 0.15 | 0.04 | 0.28 | 0.07 | 0.15 | 0.21 |
| 2 | 2284303 | rs10519519 | -0.603 | 0.125 | -1.402 | -0.508 | -0.006 | 0.019 | 0.019 | 0 | 0.15 | 0.04 | 0.28 | 0.07 | 0.15 | 0.21 |
| 2 | 2285829 | rs7588195 | -0.216 | 0.587 | -1.028 | -0.522 | -0.006 | 0.037 | 0.037 | 0.129 | 0.44 | 0.67 | 0.4 | 0.1 | 0.44 | 0.28 |
| 2 | 2294426 | rs10190125 | -0.952 | 0.205 | -0.709 | -0.606 | 0.003 | 0 | 0 | 0 | 0.38 | 0.35 | 0.22 | 0.17 | 0.32 | 0.5 |
| 2 | 2305125 | rs6710116 | -0.864 | 0.242 | -0.586 | -0.667 | 0.004 | -0.006 | -0.006 | 0 | 0.36 | 0.34 | 0.19 | 0.08 | 0.27 | 0.47 |
| 2 | 2314997 | rs7581403 | -0.985 | 0.01 | 0.02 | 0.296 | -0.014 | 0.01 | 0.01 | 0.104 | 0.37 | 0.49 | 0.25 | 0.44 | 0.27 | 0.48 |
| 2 | 2315612 | rs7588535 | -0.545 | -0.033 | -0.296 | -0.076 | -0.016 | 0.071 | 0.071 | 0 | 0.23 | 0.14 | 0.16 | 0.06 | 0.24 | 0.3 |
| 2 | 2327295 | rs6715366 | -0.569 | 0.016 | -0.421 | -0.052 | -0.011 | 0.057 | 0.057 | 0 | 0.24 | 0.16 | 0.16 | 0.06 | 0.24 | 0.31 |
| 2 | 2336685 | rs13419681 | -1.528 | -0.263 | -0.4 | -0.255 | 0.012 | 0.009 | 0.009 | 0 | 0.38 | 0.36 | 0.21 | 0.08 | 0.27 | 0.53 |
| 2 | 2352566 | rs12479155 | -1.309 | -0.028 | -1.042 | 0.113 | -0.06 | -0.1 | -0.1 | 0 | 0.34 | 0.06 | 0.29 | 0.41 | 0.36 | 0.39 |
| 2 | 2368632 | rs10197525 | -0.374 | 0.169 | -0.945 | 0.259 | -0.051 | 0.049 | 0.049 | 0 | 0.21 | 0.02 | 0.15 | 0.1 | 0.26 | 0.28 |
| 2 | 2371324 | rs3942245 | -1.204 | 0.014 | -0.976 | 0.267 | -0.004 | -0.06 | -0.06 | 0.236 | 0.2 | 0.08 | 0.18 | 0.36 | 0.16 | 0.31 |
| 2 | 2372181 | rs6759198 | -2.511 | -0.406 | 0.071 | 0.873 | 0.006 | 0.019 | 0.019 | 0.505 | 0.5 | 0.44 | 0.61 | 0.72 | 0.44 | 0.73 |
| 2 | 2381297 | rs12612846 | -1.932 | -0.279 | -0.036 | 0.799 | -0.001 | -0.002 | -0.002 | 0 | 0.37 | 0.35 | 0.56 | 0.72 | 0.36 | 0.47 |
| 2 | 2381613 | rs11884676 | -0.156 | -0.272 | 0.553 | 0.721 | 0.015 | -0.037 | -0.037 | 0.575 | 0.07 | 0.18 | 0.03 | 0 | 0.07 | 0.06 |
| 2 | 2382311 | rs10084179 | -0.208 | -0.237 | 0.267 | 0.82 | -0.054 | 0.03 | 0.03 | 0 | 0.17 | 0.01 | 0.14 | 0.09 | 0.25 | 0.21 |
| 2 | 2388352 | rs13389796 | -1.863 | -0.297 | 0.783 | 0.765 | -0.001 | -0.001 | -0.001 | 0.411 | 0.36 | 0.5 | 0.47 | 0.36 | 0.38 | 0.58 |
| 2 | 2388613 | rs749285 | -1.299 | -0.337 | 0.885 | 0.894 | -0.011 | 0.025 | 0.025 | 0 | 0.3 | 0.41 | 0.44 | 0.36 | 0.31 | 0.52 |
| 2 | 2391725 | rs713347 | -2.251 | -0.342 | 0.772 | 0.836 | -0.003 | -0.01 | -0.01 | 0 | 0.45 | 0.51 | 0.6 | 0.73 | 0.4 | 0.64 |
| 2 | 2396203 | rs6729986 | -0.7 | -0.462 | 0.669 | 0.986 | 0.015 | -0.035 | -0.035 | 0 | 0.16 | 0.13 | 0.16 | 0.38 | 0.09 | 0.12 |
| 2 | 2397565 | rs11901302 | -1.809 | -0.607 | 0.978 | 0.755 | -0.035 | -0.005 | -0.005 | 0.38 | 0.33 | 0.44 | 0.31 | 0.68 | 0.24 | 0.57 |
| 2 | 2398899 | rs17039644 | -1.034 | -0.436 | 0.623 | 1.07 | 0.005 | -0.055 | -0.055 | 0 | 0.2 | 0.07 | 0.22 | 0.29 | 0.18 | 0.25 |
| 2 | 2399187 | rs10199854 | -0.279 | -0.317 | 0.787 | 0.942 | 0.008 | -0.03 | -0.03 | 0 | 0.12 | 0.14 | 0.08 | 0.01 | 0.18 | 0.12 |
| 2 | 2399331 | rs10192243 | -0.279 | -0.317 | 0.787 | 0.942 | 0.008 | -0.03 | -0.03 | 0 | 0.12 | 0.14 | 0.08 | 0.01 | 0.18 | 0.12 |
| 2 | 2403308 | rs2668827 | -1.161 | -0.31 | 1.063 | 1.105 | -0.028 | 0.23 | 0.23 | 0 | 0.15 | 0.4 | 0.34 | 0.43 | 0.12 | 0.33 |
| 2 | 2403590 | rs1567675 | -1.764 | -0.233 | 1.061 | 1.071 | 0 | 0 | 0 | 0 | 0.36 | 0.34 | 0.33 | 0.31 | 0.36 | 0.3 |
| 2 | 2404930 | rs4381808 | -0.559 | -0.679 | 0.961 | 1.539 | 0.023 | -0.068 | -0.068 | 0 | 0.14 | 0.05 | 0.17 | 0.28 | 0.12 | 0.1 |
| 2 | 2409704 | rs2551212 | -1.19 | -0.684 | 1.066 | 1.543 | -0.025 | 0.164 | 0.164 | 0 | 0.14 | 0.26 | 0.32 | 0.43 | 0.12 | 0.33 |
| 2 | 2413449 | rs6719038 | -0.847 | -0.662 | 0.894 | 1.519 | 0.006 | -0.008 | -0.008 | 0 | 0.16 | 0.16 | 0.18 | 0.27 | 0.12 | 0.1 |
| 2 | 2415390 | rs6752827 | -0.533 | -0.617 | 0.567 | 1.464 | 0.001 | -0.091 | -0.091 | 0.003 | 0.14 | 0.01 | 0.14 | 0.25 | 0.12 | 0.1 |
| 2 | 2417721 | rs6741513 | 1.277 | -0.14 | 0.878 | 1.716 | -0.031 | 0.277 | 0.277 | 0 | 0.85 | 0.6 | 0.64 | 0.53 | 0.86 | 0.67 |
| 2 | 2424495 | rs1606310 | 0.393 | -0.021 | 0.221 | 0.888 | -0.017 | -0.017 | -0.017 | 0 | 0.8 | 0.74 | 0.86 | 0.89 | 0.83 | 0.76 |
| 2 | 2428072 | rs1355608 | -1.406 | 0.18 | -0.248 | 1.12 | 0.031 | -0.033 | -0.033 | 0 | 0.3 | 0.82 | 0.25 | 0.59 | 0.21 | 0.28 |
| 2 | 2437074 | rs7577250 | -1.507 | 0.195 | 0.184 | 0.956 | 0.001 | -0.002 | -0.002 | 1.825 | 0.3 | 0.68 | 0.22 | 0.53 | 0.19 | 0.28 |
| 2 | 2439727 | rs7563098 | -1.446 | 0.17 | 0.191 | 0.942 | 0.018 | -0.034 | -0.034 | 0 | 0.3 | 0.75 | 0.23 | 0.52 | 0.19 | 0.34 |
| 2 | 2440670 | rs12472646 | -0.34 | 0.485 | -0.162 | 0.637 | 0.009 | -0.045 | -0.045 | 0 | 0.13 | 0.18 | 0.16 | 0 | 0.16 | 0.13 |
| 2 | 2443235 | rs1606312 | 1.735 | 0.539 | -0.594 | 0.593 | -0.001 | -0.023 | -0.023 | 0 | 0.54 | 0.16 | 0.58 | 0.46 | 0.6 | 0.59 |
| 2 | 2447965 | rs4853940 | 0.026 | 0.586 | -0.671 | 0.308 | -0.032 | 0.055 | 0.055 | 0 | 0.89 | 0.98 | 0.91 | 1 | 0.87 | 0.92 |
| 2 | 2448088 | rs4853939 | 0.563 | 0.513 | -0.609 | 0.719 | -0.019 | -0.036 | -0.036 | 0 | 0.08 | 0.02 | 0.06 | 0.18 | 0.1 | 0.08 |
| 2 | 2451437 | rs1858409 | 1.677 | 0.375 | 0.233 | 0.639 | 0 | -0.01 | -0.01 | 0 | 0.58 | 0.45 | 0.62 | 0.74 | 0.61 | 0.61 |
| 2 | 2456024 | rs844011 | -1.487 | 0.422 | 0.408 | 0.448 | -0.001 | -0.134 | -0.134 | 0 | 0.33 | 0.45 | 0.31 | 0.02 | 0.29 | 0.31 |
| 2 | 2460597 | rs823543 | -2.174 | 0.671 | -0.255 | 0.703 | -0.024 | 0.03 | 0.03 | 0.89 | 0.49 | 0.86 | 0.45 | 0.59 | 0.44 | 0.39 |
| 2 | 2464401 | rs17338303 | 0.228 | 0.844 | -0.461 | 0.424 | -0.028 | 0.052 | 0.052 | 0 | 0.11 | 0.02 | 0.14 | 0.01 | 0.12 | 0.1 |
| 2 | 2464583 | rs823546 | -1.697 | 0.771 | 0.168 | 0.596 | -0.008 | 0.014 | 0.014 | 0 | 0.42 | 0.41 | 0.36 | 0.25 | 0.42 | 0.35 |
| 2 | 2467867 | rs823552 | -1.155 | 0.884 | 0.099 | 0.313 | -0.018 | 0.057 | 0.057 | 0.677 | 0.33 | 0.19 | 0.27 | 0.02 | 0.34 | 0.28 |
| 2 | 2468250 | rs4853934 | 0.383 | 0.669 | 0.09 | 0.579 | -0.014 | -0.026 | -0.026 | 0 | 0.1 | 0.06 | 0.08 | 0.2 | 0.13 | 0.09 |
| 2 | 2469769 | rs2137318 | -1.224 | 0.495 | 0.748 | 0.81 | 0.031 | 0.378 | 0.378 | 3.522 | 0.07 | 0.43 | 0.09 | 0.28 | 0.01 | 0.04 |
| 2 | 2472155 | rs1706871 | -2.337 | 0.45 | 0.464 | 0.504 | -0.031 | -0.03 | -0.03 | 0.305 | 0.53 | 0.86 | 0.48 | 0.5 | 0.47 | 0.45 |
| 2 | 2472331 | rs1706870 | 0.026 | 0.384 | 0.44 | 0.47 | -0.035 | 0.069 | 0.069 | 0 | 0.14 | 0.04 | 0.11 | 0 | 0.16 | 0.08 |
| 2 | 2472608 | rs12617042 | -0.271 | 0.314 | 0.733 | 0.655 | -0.001 | -0.002 | -0.002 | 2.192 | 0.14 | 0.14 | 0.1 | 0.21 | 0.14 | 0.14 |
| 2 | 2472901 | rs12615680 | -0.137 | 0.315 | 0.685 | 0.656 | -0.008 | -0.024 | -0.024 | 0 | 0.13 | 0.08 | 0.1 | 0.21 | 0.14 | 0.11 |
| 2 | 2473647 | rs13013201 | -1.205 | 0.305 | 0.67 | 0.607 | -0.018 | 0.16 | 0.16 | 0.011 | 0.2 | 0.56 | 0.19 | 0.49 | 0.15 | 0.18 |
| 2 | 2475208 | rs6706645 | -1.936 | 0.38 | 0.232 | 0.485 | 0.007 | -0.014 | -0.014 | 2.765 | 0.27 | 0.69 | 0.22 | 0.49 | 0.19 | 0.29 |
| 2 | 2478424 | rs6760886 | -1.933 | 0.507 | 0.237 | 0.48 | 0.004 | -0.009 | -0.009 | 0 | 0.27 | 0.68 | 0.23 | 0.5 | 0.19 | 0.28 |
| 2 | 2485163 | rs1024143 | -1.638 | 0.354 | 0.355 | 0.558 | -0.062 | 0.051 | 0.051 | 6.499 | 0.17 | 0.57 | 0.14 | 0.29 | 0.06 | 0.2 |
| 2 | 2489141 | rs17039844 | -0.056 | 0.389 | 0.356 | 0.544 | 0.001 | -0.002 | -0.002 | 0 | 0.06 | 0.04 | 0.06 | 0.02 | 0.04 | 0.12 |
| 2 | 2493770 | rs823538 | -2.263 | 0.447 | 0.144 | 0.412 | -0.043 | -0.043 | -0.043 | 0.16 | 0.55 | 0.85 | 0.48 | 0.51 | 0.53 | 0.5 |
| 2 | 2495732 | rs6728339 | -1.995 | 0.111 | 0.658 | 0.556 | -0.054 | 0.033 | 0.033 | 7.178 | 0.18 | 0.6 | 0.14 | 0.29 | 0.06 | 0.2 |

| | | | | | | | | | | | | | | | | |
|---|---|---|---|---|---|---|---|---|---|---|---|---|---|---|---|---|
| 2 | 2501833 | rs6755430 | -0.932 | 0.312 | 0.721 | 0.731 | -0.038 | 0.184 | 0.184 | 2.783 | 0.1 | 0.36 | 0.06 | 0.26 | 0.02 | 0.08 |
| 2 | 2504572 | rs12053164 | -1.017 | 0.341 | 0.567 | 0.772 | -0.031 | 0.011 | 0.011 | 0 | 0.2 | 0.45 | 0.12 | 0.19 | 0.13 | 0.14 |
| 2 | 2504833 | rs6707223 | 0.469 | 1 | 0.214 | 0.508 | -0.043 | 0.058 | 0.058 | 1.341 | 0.12 | 0.02 | 0.16 | 0.01 | 0.19 | 0.12 |
| 2 | 2504891 | rs6712933 | -0.688 | 0.924 | 0.721 | 0.525 | 0.014 | -0.061 | -0.061 | 0 | 0.23 | 0.32 | 0.24 | 0.06 | 0.27 | 0.2 |
| 2 | 2511874 | rs2063548 | -1.086 | 1.105 | 0.607 | 0.968 | 0.042 | 0.306 | 0.306 | 0.072 | 0.07 | 0.26 | 0.08 | 0.38 | 0.08 | 0.1 |
| 2 | 2512660 | rs823557 | -1.834 | 1.112 | -0.503 | 0.696 | -0.055 | -0.111 | -0.111 | 0 | 0.84 | 0.99 | 0.66 | 0.62 | 0.82 | 0.75 |
| 2 | 2516992 | rs1877440 | -0.067 | 0.846 | 0.38 | 0.591 | 0.029 | 0.24 | 0.24 | 0 | 0.1 | 0.35 | 0.06 | 0.22 | 0.07 | 0.06 |
| 2 | 2517775 | rs1706862 | -1.175 | 0.828 | 0.068 | 0.386 | -0.003 | 0.011 | 0.011 | 0.597 | 0.44 | 0.43 | 0.41 | 0.37 | 0.47 | 0.46 |
| 2 | 2518821 | rs7587997 | -1.177 | 0.816 | 0.092 | 0.38 | -0.003 | 0.012 | 0.012 | 0.608 | 0.44 | 0.43 | 0.42 | 0.37 | 0.47 | 0.46 |
| 2 | 2519513 | rs10199521 | -0.235 | 0.839 | -0.078 | 0.356 | -0.001 | 0.002 | 0.002 | 0 | 0.22 | 0.28 | 0.18 | 0.36 | 0.2 | 0.27 |
| 2 | 2519911 | rs1706864 | -1.974 | 0.895 | -0.331 | 0.368 | -0.022 | -0.007 | -0.007 | 0 | 0.54 | 0.81 | 0.48 | 0.59 | 0.54 | 0.51 |
| 2 | 2521301 | rs2668843 | -0.165 | 1.028 | -0.486 | 0.028 | -0.009 | 0.007 | 0.007 | 0 | 0.19 | 0.11 | 0.16 | 0.22 | 0.19 | 0.24 |
| 2 | 2522028 | rs908356 | -0.687 | 0.927 | -0.055 | 0.042 | 0.013 | -0.049 | -0.049 | 0 | 0.28 | 0.41 | 0.32 | 0.16 | 0.32 | 0.27 |
| 2 | 2526178 | rs10865530 | -0.85 | 0.649 | -0.536 | 0.281 | 0.001 | 0.007 | 0.007 | 0.685 | 0.49 | 0.51 | 0.52 | 0.6 | 0.5 | 0.43 |
| 2 | 2527547 | rs6760668 | -0.052 | 0.782 | -1.467 | 0.148 | -0.02 | 0.052 | 0.052 | 0.185 | 0.27 | 0.15 | 0.17 | 0.11 | 0.29 | 0.1 |
| 2 | 2527641 | rs6720427 | -0.081 | 0.785 | -1.279 | 0.15 | -0.004 | 0.008 | 0.008 | 0.012 | 0.27 | 0.23 | 0.17 | 0.11 | 0.29 | 0.1 |
| 2 | 2528351 | rs13004261 | -0.089 | 0.72 | -0.988 | 0.482 | 0.021 | -0.025 | -0.025 | 2.663 | 0.05 | 0.13 | 0.05 | 0 | 0.09 | 0.03 |
| 2 | 2529122 | rs4853930 | -0.168 | 0.333 | -1.349 | 0.768 | 0.009 | -0.055 | -0.055 | 0 | 0.19 | 0.11 | 0.34 | 0.37 | 0.21 | 0.28 |
| 2 | 2533203 | rs4853800 | 0.058 | 0.479 | -1.107 | -0.181 | 0.013 | 0.03 | 0.03 | 0.03 | 0.55 | 0.42 | 0.56 | 0.85 | 0.46 | 0.62 |
| 2 | 2536915 | rs12618856 | 0.292 | 0.893 | -1.721 | 0.267 | -0.073 | -0.082 | -0.082 | 0.114 | 0.25 | 0.04 | 0.24 | 0.29 | 0.28 | 0.41 |
| 2 | 2537804 | rs6721113 | 0.41 | 0.444 | -1.732 | -0.851 | -0.001 | 0.046 | 0.046 | 0.011 | 0.25 | 0.13 | 0.14 | 0 | 0.25 | 0.11 |
| 2 | 2538202 | rs11900612 | 0.525 | 0.717 | -1.553 | -0.554 | 0.015 | 0.03 | 0.03 | 0.6 | 0.06 | 0.14 | 0.04 | 0.11 | 0.06 | 0.06 |
| 2 | 2540292 | rs2037148 | 0.001 | 0.332 | -1.465 | -0.617 | 0.053 | 0.087 | 0.087 | 1.601 | 0.5 | 0.36 | 0.7 | 0.69 | 0.61 | 0.65 |
| 2 | 2540386 | rs2037147 | 0.467 | 0.274 | -1.288 | -0.657 | -0.035 | -0.03 | -0.03 | 0 | 0.39 | 0.21 | 0.61 | 0.54 | 0.48 | 0.56 |
| 2 | 2540775 | rs1980145 | -1.6 | 0.136 | -1.582 | -0.621 | -0.003 | -0.095 | -0.095 | 0 | 0.82 | 0.99 | 0.87 | 0.69 | 0.87 | 0.77 |
| 2 | 2545078 | rs7568276 | -0.149 | 0.323 | -0.685 | -1.231 | 0.003 | -0.058 | -0.058 | 0 | 0.34 | 0.54 | 0.2 | 0.12 | 0.32 | 0.17 |
| 2 | 2548524 | rs4853924 | 0.753 | 0.27 | -1.283 | -1.18 | -0.084 | 0.013 | 0.013 | 0.117 | 0.31 | 0.12 | 0.6 | 0.71 | 0.46 | 0.56 |
| 2 | 2549317 | rs6758216 | -1.433 | 0.135 | -1.4 | -1.304 | -0.037 | 0.019 | 0.019 | 0.291 | 0.77 | 0.97 | 0.85 | 0.81 | 0.8 | 0.73 |
| 2 | 2550499 | rs10182442 | 0.573 | 0.157 | -0.725 | -1.295 | 0.008 | 0.004 | 0.004 | 0 | 0.25 | 0.15 | 0.12 | 0 | 0.2 | 0.11 |
| 2 | 2554399 | rs908355 | -0.13 | 0.457 | -0.658 | -0.814 | -0.058 | 0.07 | 0.07 | 0.154 | 0.4 | 0.19 | 0.67 | 0.74 | 0.54 | 0.61 |
| 2 | 2564422 | rs11689408 | -0.151 | 0.492 | -0.576 | -0.791 | -0.039 | 0.058 | 0.058 | 0.347 | 0.4 | 0.26 | 0.67 | 0.74 | 0.54 | 0.61 |
| 2 | 2571142 | rs869759 | -0.904 | 0.08 | 0.614 | -0.818 | -0.032 | -0.104 | -0.104 | 1.573 | 0.14 | 0.48 | 0.07 | 0 | 0.07 | 0.04 |
| 2 | 2574659 | rs17040031 | -1.289 | 0.109 | 0.547 | -0.893 | -0.028 | -0.091 | -0.091 | 0.014 | 0.14 | 0.57 | 0.11 | 0.03 | 0.07 | 0.03 |
| 2 | 2574682 | rs10192523 | -1.515 | 0.091 | 0.451 | -0.914 | -0.022 | -0.085 | -0.085 | 5.501 | 0.21 | 0.64 | 0.13 | 0.03 | 0.12 | 0.06 |
| 2 | 2580483 | rs6754007 | -1.218 | 0.164 | 0.84 | -0.602 | -0.011 | -0.088 | -0.088 | 0 | 0.14 | 0.45 | 0.11 | 0.03 | 0.07 | 0.03 |
| 2 | 2580875 | rs6548093 | -2.057 | 0.181 | 0.497 | -0.763 | 0.014 | 0.038 | 0.038 | 0.658 | 0.3 | 0.75 | 0.26 | 0.16 | 0.25 | 0.15 |
| 2 | 2580965 | rs6548092 | -1.618 | 0.201 | 0.58 | -0.692 | -0.01 | -0.021 | -0.021 | 0 | 0.25 | 0.62 | 0.24 | 0.16 | 0.21 | 0.15 |
| 2 | 2582266 | rs867535 | -1.097 | 0.449 | 0.228 | -0.863 | 0.001 | 0.014 | 0.014 | 0.004 | 0.38 | 0.64 | 0.27 | 0.24 | 0.36 | 0.25 |
| 2 | 2583212 | rs6742158 | -0.184 | 0.519 | 0.531 | -0.777 | 0.009 | -0.005 | -0.005 | 0 | 0.25 | 0.41 | 0.2 | 0.22 | 0.27 | 0.19 |
| 2 | 2583517 | rs4853793 | -1.041 | 0.71 | -0.507 | -0.872 | -0.024 | -0.032 | -0.032 | 0.048 | 0.85 | 0.95 | 0.78 | 0.8 | 0.77 | 0.68 |
| 2 | 2585323 | rs6746066 | -1.133 | 0.571 | 0.051 | -1.464 | -0.001 | -0.077 | -0.077 | 0 | 0.33 | 0.55 | 0.16 | 0.01 | 0.32 | 0.13 |
| 2 | 2585729 | rs6548091 | -1.491 | 0.737 | -0.099 | -1.591 | -0.001 | -0.026 | -0.026 | 0.665 | 0.41 | 0.56 | 0.21 | 0.02 | 0.4 | 0.17 |
| 2 | 2586698 | rs17246978 | 0.075 | 0.727 | 0.245 | -1.6 | 0.002 | -0.039 | -0.039 | 0 | 0.22 | 0.19 | 0.1 | 0.01 | 0.24 | 0.09 |
| 2 | 2590381 | rs2598592 | 1.284 | 0.698 | 0.079 | -1.625 | 0 | -0.026 | -0.026 | 0.265 | 0.58 | 0.45 | 0.79 | 0.99 | 0.6 | 0.83 |
| 2 | 2592143 | rs17719904 | -1.195 | 0.432 | 0.238 | -1.028 | -0.005 | -0.083 | -0.083 | 0 | 0.18 | 0.26 | 0.1 | 0 | 0.16 | 0.07 |
| 2 | 2594614 | rs12995456 | 0.082 | 0.635 | -0.146 | -0.638 | 0 | -0.037 | -0.037 | 0.876 | 0.07 | 0.14 | 0.03 | 0 | 0.06 | 0.04 |
| 2 | 2603338 | rs1451198 | 0.254 | 0.785 | -0.78 | -0.738 | -0.032 | 0.051 | 0.051 | 0 | 0.77 | 0.93 | 0.82 | 0.85 | 0.72 | 0.86 |
| 2 | 2604016 | rs7586549 | 0.17 | 0.343 | -1.416 | -0.575 | -0.009 | 0.079 | 0.079 | 0 | 0.53 | 0.86 | 0.39 | 0.34 | 0.49 | 0.59 |
| 2 | 2604923 | rs11690425 | 0.183 | 0.19 | -0.42 | -0.69 | 0.013 | -0.024 | -0.024 | 0 | 0.1 | 0.29 | 0.05 | 0.02 | 0.08 | 0.04 |
| 2 | 2605585 | rs7565295 | 0.526 | 0.151 | 0.368 | -0.808 | 0.001 | -0.004 | -0.004 | 0 | 0.44 | 0.47 | 0.34 | 0.33 | 0.42 | 0.54 |
| 2 | 2605757 | rs7556672 | -0.346 | 0.563 | 0.482 | -1.049 | 0.025 | -0.06 | -0.06 | 2.015 | 0.28 | 0.53 | 0.26 | 0.13 | 0.31 | 0.36 |
| 2 | 2607024 | rs12714359 | 0.197 | 0.534 | 0.385 | -1.092 | -0.013 | 0.061 | 0.061 | 0 | 0.59 | 0.77 | 0.61 | 0.71 | 0.47 | 0.55 |
| 2 | 2608294 | rs7591027 | 1.203 | 0.508 | 0.468 | -1.102 | 0.028 | 0.044 | 0.044 | 0.338 | 0.47 | 0.42 | 0.55 | 0.69 | 0.38 | 0.48 |
| 2 | 2608466 | rs17496843 | 1.21 | 0.577 | 0.501 | -1.093 | -0.087 | 0.009 | 0.009 | 0 | 0.17 | 0.01 | 0.2 | 0.13 | 0.24 | 0.31 |
| 2 | 2608702 | rs7598374 | 1.525 | 0.47 | 0.603 | -1.044 | 0.004 | -0.035 | -0.035 | 0 | 0.06 | 0.14 | 0.03 | 0 | 0.06 | 0.05 |
| 2 | 2608936 | rs4853787 | 0.766 | 0.46 | 0.564 | -0.997 | 0.011 | 0.017 | 0.017 | 0.34 | 0.5 | 0.56 | 0.56 | 0.71 | 0.4 | 0.5 |
| 2 | 2612152 | rs4853901 | 0.905 | 0.461 | 0.649 | -0.982 | 0.027 | 0.041 | 0.041 | 0.375 | 0.47 | 0.43 | 0.55 | 0.69 | 0.38 | 0.48 |
| 2 | 2613267 | rs4853896 | 0.744 | 0.482 | 0.515 | -0.976 | 0.01 | 0.015 | 0.015 | 0.36 | 0.51 | 0.57 | 0.56 | 0.71 | 0.4 | 0.5 |
| 2 | 2624398 | rs7566479 | 0.699 | 0.733 | 0.402 | -0.852 | 0 | 0 | 0 | 0 | 0.54 | 0.52 | 0.57 | 0.7 | 0.43 | 0.55 |
| 2 | 2624754 | rs12475114 | 0.48 | 0.938 | 0.498 | -0.849 | 0.031 | 0.016 | 0.016 | 0 | 0.19 | 0.26 | 0.27 | 0.26 | 0.26 | 0.34 |
| 2 | 2624846 | rs7586516 | 0.521 | 0.8 | 0.199 | -0.855 | 0.013 | -0.023 | -0.023 | 0 | 0.61 | 0.54 | 0.6 | 0.7 | 0.48 | 0.56 |
| 2 | 2625248 | rs1020530 | 0.387 | 0.939 | 0.281 | -0.751 | 0.028 | -0.023 | -0.023 | 0.227 | 0.64 | 0.52 | 0.61 | 0.69 | 0.53 | 0.57 |
| 2 | 2626800 | rs2385306 | 0.086 | 0.684 | 0.568 | -0.422 | 0.01 | 0.015 | 0.015 | 0 | 0.22 | 0.36 | 0.27 | 0.26 | 0.25 | 0.37 |
| 2 | 2627251 | rs1376427 | -0.943 | 0.688 | 0.396 | -0.547 | -0.001 | 0 | 0 | 0 | 0.34 | 0.6 | 0.32 | 0.26 | 0.33 | 0.4 |
| 2 | 2629446 | rs12620686 | -0.663 | 0.541 | 0.518 | -0.462 | 0 | -0.01 | -0.01 | 0 | 0.16 | 0.29 | 0.12 | 0.14 | 0.15 | 0.14 |
| 2 | 2631443 | rs1451199 | -0.667 | 0.545 | 0.497 | -0.215 | 0 | 0.03 | 0.03 | 0 | 0.16 | 0.3 | 0.13 | 0.2 | 0.15 | 0.14 |
| 2 | 2632709 | rs11685339 | 1.101 | 0.627 | 0.118 | -0.284 | 0.01 | -0.013 | -0.013 | 0 | 0.17 | 0.21 | 0.21 | 0.11 | 0.23 | 0.29 |
| 2 | 2633258 | rs13031108 | 0.657 | 0.692 | -0.286 | -0.295 | 0.004 | -0.034 | -0.034 | 1.983 | 0.09 | 0.1 | 0.04 | 0 | 0.09 | 0.03 |
| 2 | 2633388 | rs7600331 | 1.181 | 0.304 | -0.665 | -0.17 | 0.027 | -0.082 | -0.082 | 0 | 0.3 | 0.13 | 0.46 | 0.45 | 0.26 | 0.26 |

| | | | | | | | | | | | | | | | |
|---|---|---|---|---|---|---|---|---|---|---|---|---|---|---|---|
| 2 | 2633681 | rs6548079 | -0.777 | 0.473 | -0.807 | -0.175 | -0.014 | 0.042 | 0.042 | 0 | 0.65 | 0.75 | 0.81 | 0.77 | 0.65 | 0.7 |
| 2 | 2637735 | rs7576292 | -0.13 | 0.618 | -1.005 | 0.106 | 0.011 | 0.017 | 0.017 | 0 | 0.32 | 0.51 | 0.32 | 0.38 | 0.37 | 0.46 |
| 2 | 2641899 | rs1076061 | -0.085 | 0.668 | -1.012 | 0.079 | 0.012 | 0.014 | 0.014 | 0 | 0.32 | 0.5 | 0.31 | 0.34 | 0.36 | 0.43 |
| 2 | 2643225 | rs1551128 | 0.49 | 0.447 | -0.906 | 0.299 | 0.009 | -0.014 | -0.014 | 0 | 0.3 | 0.24 | 0.49 | 0.46 | 0.28 | 0.27 |
| 2 | 2644566 | rs1545457 | 0.103 | 0.968 | -0.784 | -0.036 | -0.003 | -0.017 | -0.017 | 0 | 0.87 | 0.87 | 0.91 | 1 | 0.85 | 0.97 |
| 2 | 2644858 | rs17040193 | -1.01 | 0.394 | -0.638 | 0.271 | -0.027 | 0.004 | 0.004 | 0 | 0.24 | 0.35 | 0.14 | 0.22 | 0.16 | 0.21 |
| 2 | 2646590 | rs1868849 | 0.005 | 0.892 | -1.261 | 0.05 | -0.037 | 0.014 | 0.014 | 0 | 0.89 | 0.98 | 0.94 | 0.95 | 0.86 | 0.91 |
| 2 | 2649905 | rs4853888 | -0.479 | 0.873 | -1.085 | 0.544 | 0.001 | 0.021 | 0.021 | 0 | 0.14 | 0.16 | 0.14 | 0.26 | 0.17 | 0.24 |
| 2 | 2650131 | rs4853887 | 0.533 | 0.818 | -1.378 | -0.043 | -0.018 | 0.048 | 0.048 | 0 | 0.84 | 0.91 | 0.9 | 0.95 | 0.82 | 0.91 |
| 2 | 2654542 | rs719874 | 0.889 | 0.744 | -1.082 | -0.164 | -0.001 | -0.001 | -0.001 | 0 | 0.55 | 0.47 | 0.63 | 0.34 | 0.53 | 0.37 |
| 2 | 2654731 | rs719875 | 0.34 | 0.573 | -0.904 | -0.308 | 0.005 | 0.013 | 0.013 | 0 | 0.09 | 0.07 | 0.04 | 0 | 0.06 | 0.05 |
| 2 | 2658100 | rs1466155 | -0.032 | 0.628 | -0.296 | -0.046 | -0.001 | -0.001 | -0.001 | 0 | 0.88 | 0.91 | 0.92 | 0.96 | 0.87 | 0.96 |
| 2 | 2659311 | rs7581110 | 0.053 | 0.629 | -0.413 | 0 | -0.004 | 0.049 | 0.049 | 0.379 | 0.15 | 0.22 | 0.13 | 0.45 | 0.13 | 0.35 |
| 2 | 2662292 | rs7575112 | 0.715 | 0.693 | -0.359 | -0.014 | 0.005 | -0.027 | -0.027 | 0 | 0.84 | 0.7 | 0.88 | 0.93 | 0.84 | 0.94 |
| 2 | 2665294 | rs1349454 | 1.202 | 0.62 | -0.787 | -0.066 | -0.005 | 0.033 | 0.033 | 1.607 | 0.66 | 0.47 | 0.74 | 0.48 | 0.67 | 0.55 |
| 2 | 2666773 | rs896227 | -0.232 | 0.549 | -1.199 | -0.106 | -0.019 | 0.046 | 0.046 | 5.557 | 0.86 | 0.97 | 0.92 | 0.96 | 0.85 | 0.97 |
| 2 | 2666966 | rs10202240 | 0.228 | 0.758 | -1.118 | -0.498 | -0.004 | 0.015 | 0.015 | 0.267 | 0.43 | 0.34 | 0.27 | 0.4 | 0.41 | 0.56 |
| 2 | 2667339 | rs896226 | 0.307 | 0.676 | -0.752 | -0.362 | 0.007 | -0.035 | -0.035 | 0 | 0.82 | 0.68 | 0.86 | 0.91 | 0.79 | 0.92 |
| 2 | 2670178 | rs6713893 | 0.646 | 0.874 | -1.477 | -0.379 | -0.026 | 0.064 | 0.064 | 0 | 0.2 | 0.06 | 0.12 | 0 | 0.26 | 0.15 |
| 2 | 2678138 | rs3829839 | -0.388 | 0.58 | -0.994 | 0.539 | 0.001 | 0 | 0 | 0.98 | 0.2 | 0.21 | 0.16 | 0.39 | 0.17 | 0.18 |
| 2 | 2679481 | rs10184017 | 0.142 | 1.064 | -0.397 | 0.466 | 0.013 | 0.023 | 0.023 | 0 | 0.7 | 0.58 | 0.53 | 0.7 | 0.71 | 0.8 |
| 2 | 2685581 | rs1868852 | 0.349 | 0.819 | -0.617 | 0.136 | -0.001 | -0.012 | -0.012 | 0.399 | 0.85 | 0.87 | 0.86 | 0.98 | 0.81 | 0.96 |
| 2 | 2686579 | rs11694900 | -0.219 | 1.209 | -0.178 | 0.212 | 0.019 | 0.001 | 0.001 | 0 | 0.24 | 0.24 | 0.25 | 0.24 | 0.28 | 0.18 |
| 2 | 2687911 | rs1470738 | 0.883 | 0.955 | -0.192 | 0.109 | 0.008 | -0.015 | -0.015 | 0 | 0.6 | 0.6 | 0.6 | 0.75 | 0.53 | 0.77 |
| 2 | 2694190 | rs1551130 | 0.075 | 1.052 | -0.662 | 0.353 | -0.046 | -0.058 | -0.058 | 0 | 0.19 | 0.1 | 0.2 | 0.3 | 0.23 | 0.16 |
| 2 | 2696828 | rs13428254 | -0.365 | 1.156 | -0.331 | 0.002 | 0.002 | -0.006 | -0.006 | 0 | 0.17 | 0.14 | 0.13 | 0.02 | 0.19 | 0.03 |
| 2 | 2701665 | rs1993081 | 0.151 | 0.916 | -0.16 | 0.056 | 0.016 | -0.054 | -0.054 | 0 | 0.75 | 0.64 | 0.66 | 0.88 | 0.72 | 0.7 |
| 2 | 2705929 | rs1993082 | -0.358 | 0.879 | -0.615 | 0.14 | 0 | -0.002 | -0.002 | 0 | 0.79 | 0.76 | 0.71 | 0.85 | 0.78 | 0.73 |
| 2 | 2709646 | rs734614 | 0.242 | 0.679 | -0.567 | 0.588 | -0.002 | 0 | 0 | 0 | 0.7 | 0.66 | 0.57 | 0.63 | 0.73 | 0.61 |
| 2 | 2712943 | rs7589592 | -0.088 | 0.296 | -0.625 | 0.836 | -0.006 | -0.01 | -0.01 | 0 | 0.42 | 0.38 | 0.44 | 0.5 | 0.44 | 0.29 |
| 2 | 2714308 | rs17495097 | 1.027 | 0.603 | -1.646 | 0.07 | -0.04 | 0.073 | 0.073 | 0 | 0.3 | 0.12 | 0.18 | 0.15 | 0.35 | 0.22 |
| 2 | 2715340 | rs17040276 | 0.129 | 0.298 | -1.112 | 0.469 | 0.008 | -0.003 | -0.003 | 0 | 0.74 | 0.76 | 0.62 | 0.67 | 0.79 | 0.64 |
| 2 | 2718705 | rs17040287 | -0.387 | 0.574 | -1.076 | 0.194 | 0.001 | 0.003 | 0.003 | 8.742 | 0.11 | 0.06 | 0.05 | 0 | 0.1 | 0.02 |
| 2 | 2720083 | rs10168424 | 0.41 | 0.978 | -0.783 | -0.266 | 0.016 | -0.067 | -0.067 | 0 | 0.67 | 0.56 | 0.77 | 0.88 | 0.63 | 0.84 |
| 2 | 2720103 | rs2385311 | 0.49 | 1.017 | -1.549 | 0.067 | 0.009 | -0.001 | -0.001 | 0 | 0.47 | 0.42 | 0.54 | 0.49 | 0.59 | 0.59 |
| 2 | 2721958 | rs10206131 | 0.323 | 0.814 | -1.92 | -0.238 | 0.003 | -0.007 | -0.007 | 0.08 | 0.17 | 0.09 | 0.24 | 0.22 | 0.15 | 0.22 |
| 2 | 2722262 | rs10205815 | -0.578 | 0.51 | -1.714 | 0.412 | -0.003 | 0.003 | 0.003 | 0 | 0.28 | 0.26 | 0.3 | 0.45 | 0.22 | 0.32 |
| 2 | 2722295 | rs6719442 | 1.554 | 0.298 | -1.959 | 0.113 | -0.012 | 0.015 | 0.015 | 0 | 0.27 | 0.21 | 0.21 | 0 | 0.29 | 0.15 |
| 2 | 2723155 | rs1026208 | -1.079 | 0.196 | -1.291 | 0.364 | -0.01 | 0.065 | 0.065 | 0 | 0.91 | 0.86 | 0.86 | 0.77 | 0.94 | 0.75 |
| 2 | 2729381 | rs11127338 | 0.204 | 0.213 | -1.395 | 0.283 | -0.001 | -0.001 | -0.001 | 0 | 0.36 | 0.38 | 0.35 | 0.35 | 0.43 | 0.29 |
| 2 | 2730803 | rs11688980 | 0.14 | 0.188 | -1.607 | 0.365 | 0.012 | -0.049 | -0.049 | 0 | 0.19 | 0.1 | 0.27 | 0.39 | 0.17 | 0.43 |
| 2 | 2732877 | rs896232 | 0.605 | -0.127 | -1.8 | -0.319 | -0.007 | 0.031 | 0.031 | 0 | 0.37 | 0.22 | 0.29 | 0.28 | 0.37 | 0.22 |
| 2 | 2741289 | rs1451196 | -0.528 | -0.125 | -1.62 | -0.785 | -0.005 | -0.014 | -0.014 | 0 | 0.64 | 0.45 | 0.65 | 0.73 | 0.69 | 0.84 |
| 2 | 2741446 | rs1451197 | 0.149 | 0.011 | -1.684 | -0.58 | 0.007 | 0.034 | 0.034 | 0.597 | 0.51 | 0.36 | 0.53 | 0.38 | 0.45 | 0.5 |
| 2 | 2742629 | rs2122728 | 0.881 | -0.049 | -1.728 | -0.688 | 0.007 | 0.011 | 0.011 | 0.019 | 0.29 | 0.21 | 0.27 | 0.27 | 0.27 | 0.12 |
| 2 | 2742713 | rs11682663 | 2.229 | 0.235 | -1.574 | -0.703 | 0.003 | -0.035 | -0.035 | 0 | 0.07 | 0.12 | 0.04 | 0 | 0.07 | 0.06 |
| 2 | 2749520 | rs2053906 | -1.805 | 0.446 | -1.341 | -0.57 | 0.058 | -0.016 | -0.016 | 0 | 0.92 | 0.65 | 0.92 | 1 | 0.96 | 0.97 |
| 2 | 2754703 | rs17038881 | -0.98 | 0.355 | -1.448 | -0.639 | -0.013 | -0.069 | -0.069 | 0 | 0.73 | 0.59 | 0.68 | 0.85 | 0.79 | 0.85 |
| 2 | 2755205 | rs11675796 | 1.077 | 0.136 | -1.708 | -0.567 | 0.016 | -0.005 | -0.005 | 0 | 0.25 | 0.16 | 0.23 | 0.26 | 0.19 | 0.19 |
| 2 | 2755337 | rs13025008 | 0.337 | 0.412 | -1.521 | -0.439 | -0.001 | -0.002 | -0.002 | 0 | 0.34 | 0.33 | 0.38 | 0.45 | 0.43 | 0.57 |
| 2 | 2755448 | rs12714352 | -0.526 | 0.536 | -1.714 | -0.466 | -0.011 | 0.007 | 0.007 | 0 | 0.77 | 0.68 | 0.72 | 0.74 | 0.8 | 0.84 |
| 2 | 2761141 | rs2043544 | 2.098 | 0.619 | -1.99 | -0.278 | -0.002 | -0.044 | -0.044 | 0 | 0.1 | 0.04 | 0.17 | 0.26 | 0.09 | 0.12 |
| 2 | 2763051 | rs12620694 | 1.94 | 0.773 | -1.873 | -0.191 | -0.002 | -0.007 | -0.007 | 0 | 0.2 | 0.19 | 0.22 | 0.26 | 0.19 | 0.12 |
| 2 | 2763249 | rs10203375 | 0.233 | 0.669 | -2.298 | 0.284 | 0.015 | 0.043 | 0.043 | 0.051 | 0.37 | 0.24 | 0.33 | 0.15 | 0.28 | 0.19 |
| 2 | 2764201 | rs1031348 | 0.067 | 0.582 | -1.538 | 0.964 | 0.012 | 0.039 | 0.039 | 0.283 | 0.14 | 0.18 | 0.14 | 0.31 | 0.25 | 0.45 |
| 2 | 2765664 | rs744321 | 0.736 | 0.353 | -1.403 | 0.304 | -0.008 | -0.046 | -0.046 | 0 | 0.17 | 0.16 | 0.07 | 0 | 0.1 | 0.07 |
| 2 | 2765887 | rs730344 | -0.66 | 1.246 | -1.38 | 1.233 | -0.013 | -0.018 | -0.018 | 0 | 0.3 | 0.26 | 0.35 | 0.57 | 0.42 | 0.58 |
| 2 | 2770201 | rs12619156 | -0.633 | 0.788 | -1.21 | 1.616 | -0.029 | -0.06 | -0.06 | 0 | 0.1 | 0.02 | 0.11 | 0.54 | 0.14 | 0.4 |
| 2 | 2775695 | rs11686290 | 0.342 | 0.388 | -0.798 | -0.111 | 0.002 | -0.018 | -0.018 | 0 | 0.46 | 0.37 | 0.36 | 0.03 | 0.44 | 0.29 |
| 2 | 2776734 | rs2082832 | -0.242 | 0.29 | -0.191 | 0.57 | -0.032 | 0.06 | 0.06 | 0 | 0.28 | 0.5 | 0.39 | 0.39 | 0.18 | 0.18 |
| 2 | 2777618 | rs875601 | -0.238 | 0.398 | 0.123 | 0.37 | 0.031 | 0.044 | 0.044 | 0 | 0.06 | 0.2 | 0.12 | 0.06 | 0.05 | 0.12 |
| 2 | 2778016 | rs2385315 | -0.329 | -0.314 | -0.25 | 0.109 | -0.001 | 0.01 | 0.01 | 0 | 0.51 | 0.64 | 0.58 | 0.38 | 0.5 | 0.49 |
| 2 | 2778929 | rs10495490 | 0.337 | -0.217 | -0.715 | 0.11 | -0.018 | -0.002 | -0.002 | 0 | 0.38 | 0.1 | 0.27 | 0.3 | 0.36 | 0.31 |
| 2 | 2787149 | rs1469813 | 0.368 | -0.255 | -0.158 | 0.324 | -0.024 | 0.042 | 0.042 | 0 | 0.76 | 0.63 | 0.64 | 0.34 | 0.8 | 0.6 |
| 2 | 2794222 | rs1865581 | -0.29 | -0.135 | -0.438 | 0.456 | 0.014 | 0.031 | 0.031 | 0 | 0.51 | 0.54 | 0.6 | 0.26 | 0.59 | 0.48 |
| 2 | 2795206 | rs11695235 | 1.016 | 0.005 | -0.986 | 0.27 | -0.059 | -0.077 | -0.077 | 0.562 | 0.11 | 0.01 | 0.16 | 0.31 | 0.2 | 0.2 |
| 2 | 2795866 | rs17441173 | 1.452 | 0.006 | -0.659 | -0.447 | 0.014 | -0.069 | -0.069 | 0.234 | 0.25 | 0.22 | 0.2 | 0.01 | 0.35 | 0.17 |
| 2 | 2797454 | rs13017799 | 0.601 | 0.049 | -0.966 | -0.412 | -0.003 | -0.001 | -0.001 | 0 | 0.35 | 0.27 | 0.3 | 0.18 | 0.43 | 0.25 |
| 2 | 2797712 | rs2385324 | 1.035 | 0.58 | -0.817 | 0.628 | -0.001 | -0.02 | -0.02 | 0 | 0.83 | 0.85 | 0.68 | 0.67 | 0.81 | 0.53 |

| | | | | | | | | | | | | | | | | |
|---|---|---|---|---|---|---|---|---|---|---|---|---|---|---|---|---|
| 2 | 2799469 | rs7569763 | -0.003 | 0.614 | -0.819 | 0.734 | 0.007 | -0.014 | -0.014 | 0 | 0.85 | 0.86 | 0.88 | 0.74 | 0.89 | 0.66 |
| 2 | 2802431 | rs6758587 | -0.256 | 0.835 | -0.235 | 1.033 | 0.004 | 0.092 | 0.092 | 0.085 | 0.15 | 0.24 | 0.23 | 0.5 | 0.2 | 0.22 |
| 2 | 2802619 | rs6707221 | 0.964 | 0.837 | -0.293 | 1.018 | -0.003 | -0.022 | -0.022 | 0 | 0.84 | 0.88 | 0.68 | 0.68 | 0.81 | 0.54 |
| 2 | 2811664 | rs934555 | 0.212 | 0.776 | -0.302 | 1.049 | -0.001 | 0.017 | 0.017 | 0 | 0.9 | 0.89 | 0.78 | 0.71 | 0.88 | 0.71 |
| 2 | 2812016 | rs2385327 | -1.035 | 0.881 | -0.361 | 1.017 | -0.002 | 0.002 | 0.002 | 2.09 | 0.23 | 0.25 | 0.35 | 0.53 | 0.33 | 0.4 |
| 2 | 2813636 | rs1435549 | -0.419 | 0.932 | -0.58 | 0.983 | -0.033 | 0.017 | 0.017 | 0.464 | 0.08 | 0.03 | 0.12 | 0.03 | 0.13 | 0.19 |
| 2 | 2818117 | rs934556 | 1.142 | 0.252 | -0.182 | 1.168 | -0.007 | 0.123 | 0.123 | 0 | 0.88 | 0.71 | 0.77 | 0.71 | 0.88 | 0.71 |
| 2 | 2827345 | rs823221 | 0.822 | 0.374 | -0.942 | 1.229 | -0.006 | 0.039 | 0.039 | 0 | 0.89 | 0.83 | 0.78 | 0.71 | 0.88 | 0.72 |
| 2 | 2831484 | rs823231 | 0.17 | 0.696 | -1.988 | 0.703 | -0.053 | 0.049 | 0.049 | 1.469 | 0.53 | 0.13 | 0.6 | 0.59 | 0.45 | 0.55 |
| 2 | 2832206 | rs709276 | -0.551 | 1.179 | -0.903 | 0.562 | 0.077 | 0.05 | 0.05 | 0 | 0.86 | 0.67 | 0.81 | 0.81 | 0.79 | 0.84 |
| 2 | 2833031 | rs6755122 | -0.134 | 1.074 | -0.428 | 0.433 | -0.027 | -0.05 | -0.05 | 0 | 0.29 | 0.47 | 0.15 | 0.22 | 0.24 | 0.27 |
| 2 | 2836286 | rs4853769 | 0.838 | 1.147 | -1.022 | -0.055 | -0.012 | 0.066 | 0.066 | 2.237 | 0.53 | 0.15 | 0.62 | 0.6 | 0.47 | 0.58 |
| 2 | 2836350 | rs823234 | 0.634 | 1.068 | -0.31 | -0.181 | -0.004 | -0.022 | -0.022 | 0 | 0.14 | 0.25 | 0.1 | 0.14 | 0.13 | 0.18 |
| 2 | 2838270 | rs10200599 | 1.282 | 0.356 | -1.393 | 0.222 | -0.004 | -0.043 | -0.043 | 0 | 0.31 | 0.07 | 0.28 | 0.37 | 0.32 | 0.43 |
| 2 | 2839594 | rs935630 | 0.508 | 0.327 | -1.374 | 0.195 | -0.016 | 0.064 | 0.064 | 1.545 | 0.48 | 0.12 | 0.61 | 0.59 | 0.46 | 0.55 |
| 2 | 2842914 | rs823237 | 0.45 | -0.441 | -0.87 | 0.27 | -0.039 | 0.06 | 0.06 | 0.155 | 0.73 | 0.38 | 0.6 | 0.59 | 0.78 | 0.78 |
| 2 | 2842932 | rs10189124 | 0.5 | -0.267 | -0.078 | 0.443 | -0.004 | -0.019 | -0.019 | 0 | 0.11 | 0.23 | 0.08 | 0.08 | 0.11 | 0.07 |
| 2 | 2843308 | rs13403149 | 0.627 | -0.367 | -0.288 | 0.266 | 0.051 | 0.037 | 0.037 | 1.251 | 0.45 | 0.15 | 0.35 | 0.37 | 0.39 | 0.49 |
| 2 | 2845963 | rs13401708 | 0.125 | -0.361 | -0.271 | 0.273 | 0.024 | 0.019 | 0.019 | 0.251 | 0.53 | 0.14 | 0.41 | 0.45 | 0.49 | 0.56 |
| 2 | 2848102 | rs823243 | 0.091 | -0.36 | 0.234 | 0.305 | 0.016 | 0.395 | 0.395 | 0 | 0.93 | 0.63 | 0.71 | 0.67 | 0.96 | 0.89 |
| 2 | 2849167 | rs823246 | 0.565 | -0.33 | 0.16 | 0.332 | -0.037 | 0.055 | 0.055 | 0 | 0.36 | 0.22 | 0.29 | 0.23 | 0.45 | 0.31 |
| 2 | 2849451 | rs12988342 | -0.261 | -0.419 | 0.268 | 0.319 | 0.002 | 0.002 | 0.002 | 0 | 0.45 | 0.67 | 0.6 | 0.55 | 0.51 | 0.42 |
| 2 | 2850540 | rs823247 | 0.141 | -0.333 | 0.107 | 0.113 | -0.001 | -0.002 | -0.002 | 0 | 0.39 | 0.45 | 0.32 | 0.23 | 0.47 | 0.31 |
| 2 | 2850880 | rs11678833 | 0.341 | -0.372 | 0.299 | 0.32 | -0.009 | -0.021 | -0.021 | 0 | 0.11 | 0.24 | 0.07 | 0.08 | 0.1 | 0.08 |
| 2 | 2864210 | rs1656313 | -0.051 | -0.459 | 0.021 | 0.562 | 0.025 | 0.014 | 0.014 | 0 | 0.52 | 0.62 | 0.53 | 0.63 | 0.39 | 0.66 |
| 2 | 2866812 | rs4853857 | 0.178 | -0.469 | -0.173 | 0.562 | -0.002 | 0.035 | 0.035 | 0 | 0.44 | 0.12 | 0.37 | 0.3 | 0.59 | 0.28 |
| 2 | 2871305 | rs1656318 | 0.161 | -0.5 | 0.078 | 0.54 | 0.01 | 0.007 | 0.007 | 0 | 0.5 | 0.47 | 0.52 | 0.63 | 0.39 | 0.66 |
| 2 | 2876145 | rs3748974 | 0.159 | -0.396 | -0.859 | 0.504 | -0.006 | 0.231 | 0.231 | 0 | 0.44 | 0.14 | 0.33 | 0.11 | 0.59 | 0.27 |
| 2 | 2877949 | rs13404653 | 0.651 | -0.712 | -1.705 | 0.833 | 0.067 | -0.035 | -0.035 | 0 | 0.32 | 0.04 | 0.33 | 0.29 | 0.21 | 0.33 |
| 2 | 2878583 | rs7601893 | 0.66 | 0.161 | -1.455 | 0.443 | -0.07 | 0.035 | 0.035 | 0 | 0.19 | 0.09 | 0.2 | 0.09 | 0.32 | 0.19 |
| 2 | 2879779 | rs1656325 | -0.293 | -1.936 | -1.473 | 0.288 | 0.055 | 0.009 | 0.009 | 0 | 0.64 | 0.27 | 0.73 | 0.67 | 0.79 | 0.73 |
| 2 | 2884918 | rs7598986 | 0.529 | -1.861 | -1.603 | 0.055 | -0.068 | 0.04 | 0.04 | 0.11 | 0.28 | 0.17 | 0.28 | 0.12 | 0.46 | 0.27 |
| 2 | 2886795 | rs13014742 | 1.021 | -1.955 | -0.972 | 0.031 | -0.086 | 0.046 | 0.046 | 0 | 0.25 | 0.1 | 0.21 | 0.1 | 0.39 | 0.18 |
| 2 | 2887102 | rs823191 | 0.22 | -1.379 | -0.387 | 0.577 | -0.005 | -0.126 | -0.126 | 0.141 | 0.34 | 0.06 | 0.45 | 0.58 | 0.31 | 0.46 |
| 2 | 2894549 | rs10211532 | 0.466 | -1.186 | 0.194 | 0.29 | 0.025 | -0.017 | -0.017 | 0 | 0.05 | 0.13 | 0.09 | 0.02 | 0.08 | 0.11 |
| 2 | 2895193 | rs17040523 | 1.673 | -1.23 | -0.357 | 0.361 | -0.03 | -0.009 | -0.009 | 0.167 | 0.06 | 0 | 0.27 | 0.31 | 0.11 | 0.22 |
| 2 | 2896238 | rs4853851 | 0.379 | -1.501 | -0.246 | -0.187 | 0.034 | 0.128 | 0.128 | 0 | 0.5 | 0.26 | 0.33 | 0.2 | 0.56 | 0.39 |
| 2 | 2899178 | rs823208 | 0.497 | -1.131 | 0.462 | 0.297 | 0.002 | 0.265 | 0.265 | 2.812 | 0.92 | 0.49 | 0.89 | 0.79 | 0.98 | 0.97 |
| 2 | 2899681 | rs4853848 | 0.648 | -1.642 | -0.387 | -0.435 | 0.067 | 0.082 | 0.082 | 0.176 | 0.54 | 0.13 | 0.68 | 0.72 | 0.68 | 0.65 |
| 2 | 2901761 | rs17486311 | 1.3 | -1.848 | -0.143 | -0.34 | -0.005 | -0.054 | -0.054 | 0.012 | 0.14 | 0.01 | 0.09 | 0.23 | 0.1 | 0.13 |
| 2 | 2908338 | rs4594476 | 0.462 | -2.016 | 0.542 | -1.037 | 0.001 | -0.044 | -0.044 | 0 | 0.27 | 0.29 | 0.13 | 0.03 | 0.19 | 0.16 |
| 2 | 2912305 | rs1729917 | 0.215 | -1.642 | 0.858 | -0.672 | 0.025 | 0.228 | 0.228 | 2.034 | 0.92 | 0.66 | 0.89 | 0.79 | 0.94 | 0.94 |
| 2 | 2915622 | rs1667045 | 0.595 | -1.539 | 0.629 | -0.566 | -0.011 | -0.008 | -0.008 | 3.554 | 0.1 | 0.2 | 0.08 | 0.05 | 0.07 | 0.13 |
| 2 | 2916955 | rs12714318 | 0.83 | -1.596 | 0.293 | -0.588 | -0.043 | 0.055 | 0.055 | 1.051 | 0.82 | 0.39 | 0.82 | 0.74 | 0.91 | 0.83 |
| 2 | 2926013 | rs17491951 | 1.418 | -1.094 | -0.114 | -0.71 | -0.138 | 0.036 | 0.036 | 0 | 0.25 | 0.02 | 0.18 | 0.13 | 0.36 | 0.12 |
| 2 | 2928690 | rs1729888 | 0.109 | -1.61 | 1.186 | -0.43 | 0.004 | -0.004 | -0.004 | 0 | 0.09 | 0.24 | 0.09 | 0.05 | 0.09 | 0.13 |
| 2 | 2934355 | rs1463146 | 0.116 | -1.587 | 0.919 | -0.395 | 0.02 | 0.008 | 0.008 | 0 | 0.1 | 0.34 | 0.1 | 0.05 | 0.09 | 0.13 |
| 2 | 2935910 | rs4853954 | 1.129 | -1.569 | -0.398 | -0.57 | 0.087 | 0.08 | 0.08 | 0.078 | 0.54 | 0.15 | 0.7 | 0.72 | 0.68 | 0.65 |
| 2 | 2942376 | rs7594094 | -0.566 | -1.368 | 0.227 | 0.012 | -0.078 | -0.15 | -0.15 | 0 | 0.23 | 0.45 | 0.07 | 0.01 | 0.09 | 0.18 |
| 2 | 2942896 | rs4853951 | 0.707 | -0.176 | 0.391 | 1.009 | -0.011 | 0.126 | 0.126 | 0.018 | 0.88 | 0.69 | 0.85 | 0.74 | 0.9 | 0.84 |
| 2 | 2945455 | rs1667060 | 0.979 | -0.403 | 0.382 | 0.304 | -0.044 | -0.075 | -0.075 | 0 | 0.87 | 0.7 | 0.95 | 1 | 0.96 | 0.97 |
| 2 | 2946046 | rs4853948 | -0.376 | -0.312 | 0.028 | 0.531 | 0.007 | -0.007 | -0.007 | 0.318 | 0.33 | 0.31 | 0.43 | 0.36 | 0.27 | 0.43 |
| 2 | 2953815 | rs1667032 | 0.345 | 0.234 | 0.232 | 0.042 | 0.02 | 0.005 | 0.005 | 0.192 | 0.71 | 0.69 | 0.74 | 0.67 | 0.61 | 0.72 |
| 2 | 2953847 | rs1667033 | -0.441 | 0.573 | 0.212 | 0.098 | -0.021 | 0.026 | 0.026 | 0.008 | 0.82 | 0.92 | 0.84 | 0.92 | 0.72 | 0.91 |
| 2 | 2956085 | rs1729896 | 0.177 | 0.583 | 0.273 | -0.046 | -0.001 | -0.002 | -0.002 | 0 | 0.35 | 0.32 | 0.54 | 0.53 | 0.39 | 0.44 |
| 2 | 2963057 | rs1729898 | -0.57 | 0.789 | 0.009 | 0.06 | -0.027 | 0.031 | 0.031 | 0 | 0.82 | 0.93 | 0.84 | 0.92 | 0.72 | 0.91 |
| 2 | 2964695 | rs1729901 | 0.036 | 1.004 | 0.425 | 0.058 | 0.045 | 0 | 0 | 1.91 | 0.86 | 0.73 | 0.85 | 0.87 | 0.75 | 0.91 |
| 2 | 2970150 | rs1729907 | 0.078 | 0.754 | 0.299 | -0.226 | -0.054 | 0.008 | 0.008 | 0 | 0.45 | 0.14 | 0.63 | 0.55 | 0.48 | 0.45 |
| 2 | 2974127 | rs1729911 | 0.8 | 0.683 | 0.429 | -0.273 | 0.002 | 0.001 | 0.001 | 3.077 | 0.75 | 0.36 | 0.8 | 0.69 | 0.69 | 0.69 |
| 2 | 2975483 | rs1375223 | -0.324 | 0.557 | -0.022 | -0.282 | -0.007 | 0.024 | 0.024 | 0 | 0.74 | 0.78 | 0.83 | 0.85 | 0.7 | 0.88 |
| 2 | 2977450 | rs1667043 | 0.65 | 0.527 | -0.388 | -0.117 | -0.046 | -0.032 | -0.032 | 0 | 0.08 | 0.01 | 0.1 | 0.1 | 0.2 | 0.11 |
| 2 | 2978747 | rs1624522 | -0.041 | 0.459 | -0.148 | -0.146 | 0.007 | -0.008 | -0.008 | 0 | 0.63 | 0.49 | 0.72 | 0.61 | 0.58 | 0.68 |
| 2 | 2978905 | rs17437842 | 1.518 | 0.405 | -0.324 | -0.188 | -0.012 | 0.128 | 0.128 | 0 | 0.27 | 0.01 | 0.14 | 0.08 | 0.18 | 0.26 |
| 2 | 2979113 | rs1729916 | 0.276 | 0.412 | -0.222 | -0.184 | -0.008 | -0.008 | -0.008 | 0 | 0.33 | 0.29 | 0.57 | 0.53 | 0.4 | 0.42 |
| 2 | 2984615 | rs1618617 | 0.136 | 0.44 | 0.016 | -0.04 | 0.001 | -0.004 | -0.004 | 0.004 | 0.62 | 0.45 | 0.72 | 0.61 | 0.59 | 0.68 |
| 2 | 3003010 | rs7591035 | 0.418 | 0.486 | -0.34 | 0.596 | -0.004 | 0.025 | 0.025 | 0 | 0.25 | 0.26 | 0.47 | 0.43 | 0.21 | 0.32 |
| 2 | 3004998 | rs10210870 | -0.481 | 0.618 | -0.304 | -0.045 | 0.01 | -0.04 | -0.04 | 0.071 | 0.24 | 0.29 | 0.19 | 0.07 | 0.27 | 0.2 |
| 2 | 3005457 | rs17437576 | 0.711 | 0.478 | -1.105 | -0.302 | -0.032 | 0.09 | 0.09 | 0 | 0.25 | 0.01 | 0.18 | 0.09 | 0.23 | 0.22 |
| 2 | 3007972 | rs7592135 | -0.357 | 0.333 | -0.479 | 0.012 | -0.001 | -0.004 | -0.004 | 0 | 0.39 | 0.38 | 0.31 | 0.34 | 0.39 | 0.43 |

| | | | | | | | | | | | | | | | | |
|---|---|---|---|---|---|---|---|---|---|---|---|---|---|---|---|---|
| 2 | 3008413 | rs1881659 | 0.344 | 0.653 | 0.074 | 0.367 | 0.033 | 0.058 | 0.058 | 0 | 0.77 | 0.62 | 0.77 | 0.66 | 0.72 | 0.63 |
| 2 | 3012290 | rs2031028 | 1.551 | 0.593 | -0.84 | 0.596 | -0.07 | 0.071 | 0.071 | 2.752 | 0.47 | 0.06 | 0.39 | 0.29 | 0.46 | 0.38 |
| 2 | 3012316 | rs17040704 | 0.221 | 0.497 | -0.066 | 0.904 | -0.005 | -0.012 | -0.012 | 0 | 0.12 | 0.09 | 0.11 | 0.27 | 0.13 | 0.14 |
| 2 | 3012861 | rs12479004 | 1.528 | 0.498 | -0.102 | 0.88 | -0.037 | -0.001 | -0.001 | 0 | 0.57 | 0.19 | 0.46 | 0.37 | 0.58 | 0.45 |
| 2 | 3012947 | rs12470135 | 1.505 | 0.536 | -0.068 | 0.899 | -0.032 | 0.001 | 0.001 | 0.502 | 0.57 | 0.21 | 0.46 | 0.37 | 0.58 | 0.45 |
| 2 | 3013207 | rs12623874 | 0.322 | 0.639 | -0.189 | 0.943 | -0.091 | -0.112 | -0.112 | 1.346 | 0.16 | 0.02 | 0.36 | 0.35 | 0.25 | 0.36 |
| 2 | 3025336 | rs17040721 | -0.45 | 0.827 | 0.416 | 1.09 | 0.011 | 0.034 | 0.034 | 0 | 0.28 | 0.46 | 0.42 | 0.34 | 0.28 | 0.41 |
| 2 | 3027789 | rs17040726 | -0.349 | 0.865 | 0.485 | 1.124 | 0.007 | 0.012 | 0.012 | 0 | 0.28 | 0.41 | 0.42 | 0.29 | 0.28 | 0.4 |
| 2 | 3029042 | rs17040731 | -0.23 | 0.969 | -0.193 | 1.13 | 0.014 | 0.069 | 0.069 | 0 | 0.13 | 0.24 | 0.12 | 0.27 | 0.13 | 0.13 |
| 2 | 3037092 | rs6709949 | -0.637 | 0.943 | -0.53 | 0.841 | 0.003 | 0.008 | 0.008 | 0 | 0.27 | 0.28 | 0.4 | 0.34 | 0.29 | 0.41 |
| 2 | 3062055 | rs13022539 | -0.686 | 0.955 | -0.661 | 0.8 | -0.005 | -0.001 | -0.001 | 0 | 0.43 | 0.57 | 0.61 | 0.57 | 0.52 | 0.51 |
| 2 | 3062238 | rs12714369 | -0.34 | 0.654 | 0.005 | 0.647 | 0.001 | 0 | 0 | 2.127 | 0.32 | 0.27 | 0.57 | 0.55 | 0.47 | 0.42 |
| 2 | 3062748 | rs13007553 | -1.179 | -0.096 | -0.636 | 0.107 | -0.023 | 0.123 | 0.123 | 0 | 0.66 | 0.87 | 0.74 | 0.86 | 0.65 | 0.66 |
| 2 | 3063237 | rs13004913 | 0.767 | 0.468 | -0.525 | 0.432 | -0.023 | -0.024 | -0.024 | 2.348 | 0.18 | 0.16 | 0.51 | 0.41 | 0.32 | 0.31 |
| 2 | 3066698 | rs6605287 | -1.597 | 0.326 | -0.484 | 0.134 | -0.03 | 0.086 | 0.086 | 0 | 0.67 | 0.83 | 0.74 | 0.85 | 0.64 | 0.64 |
| 2 | 3073183 | rs7583267 | 0.293 | 0.295 | -0.289 | 0.369 | 0 | 0.002 | 0.002 | 0.593 | 0.22 | 0.26 | 0.48 | 0.38 | 0.29 | 0.28 |
| 2 | 3078181 | rs6744756 | -1.242 | 0.237 | -0.135 | 0.229 | -0.04 | 0.071 | 0.071 | 0 | 0.6 | 0.76 | 0.69 | 0.73 | 0.51 | 0.56 |
| 2 | 3079021 | rs11127390 | -0.959 | -0.004 | 0.057 | 0.05 | 0.006 | -0.036 | -0.036 | 0.005 | 0.37 | 0.41 | 0.61 | 0.74 | 0.43 | 0.51 |
| 2 | 3080161 | rs7419318 | -1.823 | 0.079 | 0.009 | 0.065 | -0.009 | 0.049 | 0.049 | 0 | 0.66 | 0.74 | 0.73 | 0.85 | 0.64 | 0.65 |
| 2 | 3082104 | rs11127398 | -0.561 | 0.019 | 0.161 | 0.113 | 0.002 | 0.005 | 0.005 | 0 | 0.48 | 0.42 | 0.6 | 0.49 | 0.5 | 0.41 |
| 2 | 3082663 | rs12614860 | -0.501 | -0.006 | -0.215 | 0.258 | 0.011 | -0.02 | -0.02 | 0.076 | 0.15 | 0.11 | 0.1 | 0.34 | 0.11 | 0.2 |
| 2 | 3083625 | rs7371392 | -0.868 | 0.057 | 0.079 | 0.184 | 0.008 | -0.003 | -0.003 | 0.99 | 0.43 | 0.4 | 0.22 | 0.45 | 0.33 | 0.35 |
| 2 | 3084950 | rs6754030 | -1.166 | 0.169 | -0.116 | 0.308 | -0.002 | -0.014 | -0.014 | 0 | 0.22 | 0.19 | 0.16 | 0.45 | 0.25 | 0.29 |
| 2 | 3085996 | rs12464090 | -0.385 | 0.104 | -0.4 | 0.325 | 0.019 | -0.036 | -0.036 | 0 | 0.14 | 0.09 | 0.11 | 0.34 | 0.11 | 0.2 |
| 2 | 3086833 | rs10169682 | -1.045 | 0.153 | -0.447 | -0.071 | 0.004 | 0.069 | 0.069 | 0 | 0.59 | 0.69 | 0.72 | 0.83 | 0.61 | 0.57 |
| 2 | 3092604 | rs7604183 | -1.598 | 0.399 | -0.757 | 0.098 | -0.029 | 0.063 | 0.063 | 0 | 0.68 | 0.84 | 0.75 | 0.83 | 0.64 | 0.64 |
| 2 | 3101073 | rs992905 | -0.129 | 1.342 | -0.981 | 0.095 | -0.047 | -0.007 | -0.007 | 0 | 0.07 | 0.02 | 0.09 | 0.05 | 0.21 | 0.05 |
| 2 | 3101683 | rs12472632 | 0.224 | 0.793 | -0.969 | 0.256 | 0.001 | -0.016 | -0.016 | 0 | 0.17 | 0.15 | 0.46 | 0.43 | 0.16 | 0.32 |
| 2 | 3101773 | rs767885 | 1.111 | 0.558 | -0.933 | -0.132 | 0.022 | 0.042 | 0.042 | 0.345 | 0.22 | 0.1 | 0.05 | 0 | 0.12 | 0.09 |
| 2 | 3103097 | rs4438515 | -0.703 | 0.568 | -0.874 | 0.078 | 0.013 | -0.053 | -0.053 | 0.158 | 0.28 | 0.17 | 0.5 | 0.43 | 0.25 | 0.42 |
| 2 | 3105711 | rs744334 | -1.335 | 0.79 | -0.438 | -0.338 | -0.013 | 0.02 | 0.02 | 0.598 | 0.39 | 0.37 | 0.68 | 0.78 | 0.54 | 0.53 |
| 2 | 3106689 | rs11127438 | -0.869 | 1.29 | -0.507 | -0.508 | 0.002 | -0.013 | -0.013 | 0.005 | 0.63 | 0.64 | 0.76 | 0.78 | 0.56 | 0.62 |
| 2 | 3107035 | rs4854190 | 0.658 | 1.422 | -0.693 | 0.03 | -0.05 | -0.059 | -0.059 | 1.152 | 0.09 | 0.01 | 0.12 | 0.26 | 0.16 | 0.12 |
| 2 | 3107942 | rs1551296 | -0.835 | 1.506 | -0.605 | -0.09 | -0.01 | -0.003 | -0.003 | 0.059 | 0.94 | 0.97 | 0.93 | 0.94 | 0.86 | 0.95 |
| 2 | 3107978 | rs6707574 | -0.931 | 1.551 | -0.577 | -0.153 | 0.001 | -0.013 | -0.013 | 0 | 0.63 | 0.63 | 0.77 | 0.78 | 0.57 | 0.62 |
| 2 | 3108215 | rs1973049 | 1.399 | 1.413 | -0.533 | -0.133 | -0.003 | -0.028 | -0.028 | 0 | 0.31 | 0.34 | 0.17 | 0.16 | 0.28 | 0.33 |
| 2 | 3109259 | rs17040814 | -0.386 | 1.128 | -0.624 | -0.162 | -0.013 | -0.035 | -0.035 | 1.565 | 0.09 | 0.18 | 0.06 | 0.01 | 0.05 | 0.04 |
| 2 | 3109628 | rs7564506 | -0.317 | 1.142 | -1.321 | -0.573 | 0 | 0.002 | 0.002 | 0 | 0.45 | 0.55 | 0.67 | 0.77 | 0.42 | 0.48 |
| 2 | 3111589 | rs11690361 | 0.017 | 1.119 | -1.026 | 0.052 | -0.008 | 0.004 | 0.004 | 0 | 0.45 | 0.43 | 0.35 | 0.44 | 0.49 | 0.45 |
| 2 | 3113371 | rs7585606 | -0.739 | 0.85 | -1.32 | 0.415 | 0.007 | 0.049 | 0.049 | 0 | 0.69 | 0.82 | 0.83 | 0.83 | 0.71 | 0.68 |
| 2 | 3113710 | rs10206572 | 0.073 | 1.031 | -0.961 | 1.069 | -0.002 | 0.031 | 0.031 | 0 | 0.13 | 0.13 | 0.33 | 0.39 | 0.11 | 0.28 |
| 2 | 3114755 | rs7592329 | 0.342 | 1.738 | -0.982 | 0.623 | -0.016 | 0.024 | 0.024 | 0 | 0.25 | 0.17 | 0.17 | 0.18 | 0.3 | 0.31 |
| 2 | 3116598 | rs13023918 | -1.395 | 1.506 | -0.508 | 1.094 | -0.007 | -0.002 | -0.002 | 0 | 0.1 | 0.07 | 0.18 | 0.12 | 0.17 | 0.11 |
| 2 | 3117247 | rs6761955 | 1.066 | 1.308 | 0.031 | 1.404 | 0.034 | 0.137 | 0.137 | 0 | 0.79 | 0.58 | 0.67 | 0.57 | 0.74 | 0.76 |
| 2 | 3120184 | rs4854192 | 0.285 | 1.294 | -0.3 | 1.254 | 0.001 | 0.002 | 0.002 | 0 | 0.79 | 0.81 | 0.76 | 0.75 | 0.69 | 0.69 |
| 2 | 3122870 | rs6754432 | 0.222 | 1.32 | 0.184 | 1.288 | 0.016 | -0.032 | -0.032 | 0.05 | 0.8 | 0.71 | 0.81 | 0.88 | 0.73 | 0.88 |
| 2 | 3123061 | rs6747687 | 1.257 | 0.913 | -0.458 | 1.368 | 0.004 | -0.035 | -0.035 | 0.024 | 0.59 | 0.24 | 0.59 | 0.57 | 0.57 | 0.58 |
| 2 | 3125571 | rs965273 | 1.132 | 0.983 | -0.236 | 1.497 | 0.006 | -0.003 | -0.003 | 0 | 0.67 | 0.23 | 0.68 | 0.69 | 0.65 | 0.84 |
| 2 | 3127370 | rs7565101 | 0.275 | 0.868 | -0.238 | 1.251 | -0.008 | 0.001 | 0.001 | 0.04 | 0.76 | 0.82 | 0.61 | 0.81 | 0.61 | 0.66 |
| 2 | 3127643 | rs7559077 | 0.699 | 0.708 | -0.88 | 0.845 | -0.019 | 0.028 | 0.028 | 0.389 | 0.64 | 0.24 | 0.51 | 0.75 | 0.54 | 0.79 |
| 2 | 3128923 | rs1078821 | 0.548 | 0.204 | -0.889 | 0.844 | 0 | -0.001 | -0.001 | 0 | 0.74 | 0.27 | 0.79 | 0.59 | 0.76 | 0.77 |
| 2 | 3129088 | rs1078819 | -0.143 | -0.448 | -1.216 | 0.132 | 0.091 | 0.113 | 0.113 | 1.763 | 0.47 | 0.2 | 0.69 | 0.72 | 0.7 | 0.62 |
| 2 | 3131096 | rs6548160 | 0.461 | -0.339 | -0.664 | 0.468 | -0.029 | -0.015 | -0.015 | 0 | 0.74 | 0.65 | 0.87 | 0.84 | 0.81 | 0.72 |
| 2 | 3135165 | rs4854194 | 0.277 | -0.088 | -1.359 | 0.192 | -0.023 | 0.113 | 0.113 | 2.137 | 0.44 | 0.19 | 0.57 | 0.24 | 0.51 | 0.45 |
| 2 | 3135690 | rs7569266 | -0.456 | -0.338 | -0.186 | 0.173 | -0.025 | -0.038 | -0.038 | 1.856 | 0.16 | 0.5 | 0.1 | 0.06 | 0.1 | 0.27 |
| 2 | 3149987 | rs6548165 | 0.403 | -0.371 | 1.444 | 0.926 | 0.023 | 0.004 | 0.004 | 2.756 | 0.82 | 0.49 | 0.83 | 0.83 | 0.8 | 0.83 |
| 2 | 3151657 | rs11675522 | -1.332 | -0.702 | 1.349 | 1.46 | 0.009 | 0.022 | 0.022 | 2.595 | 0.33 | 0.54 | 0.35 | 0.53 | 0.43 | 0.44 |
| 2 | 3152466 | rs2887312 | 0.384 | -0.692 | 1.342 | 1.645 | 0.019 | 0.109 | 0.109 | 0 | 0.83 | 0.75 | 0.82 | 0.51 | 0.78 | 0.74 |
| 2 | 3153867 | rs1551298 | 0.254 | -0.658 | 1.168 | 1.63 | 0.046 | 0.197 | 0.197 | 0 | 0.84 | 0.7 | 0.82 | 0.54 | 0.78 | 0.74 |
| 2 | 3163738 | rs6730025 | 0.854 | -1.023 | 0.303 | 0.944 | -0.019 | 0 | 0 | 0 | 0.62 | 0.53 | 0.77 | 0.41 | 0.67 | 0.54 |
| 2 | 3172438 | rs13021132 | 0.84 | 0.097 | -0.091 | 0.498 | -0.001 | -0.003 | -0.003 | 0.143 | 0.61 | 0.38 | 0.58 | 0.26 | 0.51 | 0.46 |
| 2 | 3172635 | rs10495516 | 1.031 | 0.249 | -1.431 | 0.244 | -0.187 | -0.024 | -0.024 | 0 | 0.34 | 0.03 | 0.55 | 0.28 | 0.44 | 0.3 |
| 2 | 3172653 | rs10495515 | 0.276 | 0.78 | -0.891 | 0.276 | -0.012 | 0.018 | 0.018 | 0.627 | 0.79 | 0.85 | 0.81 | 0.89 | 0.71 | 0.73 |
| 2 | 3173024 | rs4854131 | 0.744 | 0.323 | -0.186 | 0.477 | -0.001 | 0.029 | 0.029 | 0.745 | 0.72 | 0.48 | 0.76 | 0.41 | 0.72 | 0.66 |
| 2 | 3175376 | rs7578217 | 0.851 | 0.281 | 0.097 | 0.918 | 0.005 | 0.145 | 0.145 | 0 | 0.93 | 0.88 | 0.88 | 0.48 | 0.91 | 0.75 |
| 2 | 3175992 | rs10208086 | 1.262 | 1.162 | 0.499 | 0.559 | 0.02 | -0.023 | -0.023 | 0 | 0.62 | 0.52 | 0.57 | 0.31 | 0.49 | 0.5 |
| 2 | 3180126 | rs7591433 | 0.786 | 1.245 | 0.36 | 0.638 | 0.008 | 0.168 | 0.168 | 0 | 0.93 | 0.9 | 0.88 | 0.49 | 0.91 | 0.74 |
| 2 | 3187698 | rs11682390 | 1.037 | 1.244 | 0.464 | 0.638 | 0.01 | 0.197 | 0.197 | 0 | 0.92 | 0.86 | 0.88 | 0.49 | 0.91 | 0.74 |
| 2 | 3190384 | rs2887311 | 1.772 | 1.216 | 0.726 | 0.648 | 0.025 | 0.251 | 0.251 | 0 | 0.91 | 0.67 | 0.87 | 0.42 | 0.91 | 0.74 |

| | | | | | | | | | | | | | | | | |
|---|---|---|---|---|---|---|---|---|---|---|---|---|---|---|---|---|
| 2 | 3190969 | rs4402798 | 1.772 | 1.216 | 0.738 | 0.639 | 0.026 | 0.271 | 0.271 | 0 | 0.91 | 0.67 | 0.87 | 0.43 | 0.91 | 0.74 |
| 2 | 3192798 | rs6836 | 1.745 | 1.218 | 0.731 | 0.64 | 0.026 | 0.269 | 0.269 | 0 | 0.91 | 0.67 | 0.87 | 0.43 | 0.91 | 0.74 |
| 2 | 3193040 | rs17682 | -0.5 | 1.23 | 0.298 | 0.542 | -0.133 | 0.055 | 0.055 | 0 | 0.25 | 0.03 | 0.25 | 0.15 | 0.39 | 0.22 |
| 2 | 3201417 | rs7563563 | 1.205 | 1.284 | 0.632 | 0.704 | 0.016 | 0.197 | 0.197 | 0 | 0.92 | 0.85 | 0.88 | 0.48 | 0.9 | 0.76 |
| 2 | 3204480 | rs7585437 | 1.331 | 1.307 | 1.265 | 0.351 | -0.003 | 0.001 | 0.001 | 0 | 0.63 | 0.41 | 0.59 | 0.27 | 0.47 | 0.53 |
| 2 | 3212247 | rs7584823 | 0.95 | 1.482 | 1.705 | 0.177 | 0.005 | -0.007 | -0.007 | 0 | 0.68 | 0.63 | 0.7 | 0.78 | 0.55 | 0.72 |
| 2 | 3213822 | rs4854106 | 1.115 | 1.413 | 1.448 | 0.242 | 0.01 | 0.02 | 0.02 | 0.038 | 0.94 | 0.82 | 0.97 | 0.98 | 0.95 | 0.93 |
| 2 | 3220709 | rs7595382 | 1.239 | 1.38 | 1.556 | 0.275 | 0.005 | -0.003 | -0.003 | 0 | 0.63 | 0.39 | 0.64 | 0.74 | 0.48 | 0.69 |
| 2 | 3238675 | rs6548133 | 1.189 | 1.234 | 1.202 | 0.32 | 0.018 | 0.042 | 0.042 | 0.045 | 0.94 | 0.76 | 0.97 | 0.97 | 0.96 | 0.93 |
| 2 | 3243238 | rs2694103 | 1.525 | 1.221 | 1.372 | 0.311 | 0.042 | 0.065 | 0.065 | 0.402 | 0.94 | 0.69 | 0.96 | 0.97 | 0.96 | 0.93 |
| 2 | 3243694 | rs2602762 | 1.143 | 1.227 | 1.269 | 0.314 | 0.021 | 0.041 | 0.041 | 0.014 | 0.95 | 0.76 | 0.97 | 0.97 | 0.96 | 0.93 |
| 2 | 3250248 | rs1869416 | 1.057 | 1.578 | 1.141 | 0.181 | 0.013 | -0.005 | -0.005 | 0 | 0.68 | 0.63 | 0.67 | 0.76 | 0.49 | 0.69 |
| 2 | 3258174 | rs2694093 | 0.564 | 1.686 | 1.15 | 0.183 | 0.018 | -0.014 | -0.014 | 0.068 | 0.74 | 0.69 | 0.72 | 0.8 | 0.61 | 0.8 |
| 2 | 3302962 | rs4546035 | -0.544 | 1.601 | 0.678 | 0.355 | 0.001 | -0.001 | -0.001 | 0 | 0.17 | 0.2 | 0.2 | 0.2 | 0.26 | 0.17 |
| 2 | 3309800 | rs10186914 | -1.339 | 1.541 | 0.723 | 0.38 | 0.033 | 0.006 | 0.006 | 0.746 | 0.23 | 0.34 | 0.27 | 0.25 | 0.38 | 0.25 |
| 2 | 3311400 | rs4854198 | -1.045 | 0.27 | -0.015 | -0.567 | 0.015 | 0.071 | 0.071 | 0.037 | 0.38 | 0.23 | 0.2 | 0.11 | 0.28 | 0.27 |
| 2 | 3313428 | rs6749474 | -0.138 | 0.452 | -0.141 | -0.388 | -0.007 | -0.002 | -0.002 | 0 | 0.11 | 0.12 | 0.13 | 0.15 | 0.19 | 0.17 |
| 2 | 3313543 | rs7606684 | 0.116 | -0.601 | -0.146 | -0.785 | -0.023 | -0.053 | -0.053 | 0 | 0.18 | 0.15 | 0.05 | 0.01 | 0.05 | 0.09 |
| 2 | 3320502 | rs12621987 | -1.32 | -0.303 | -0.429 | -0.366 | 0.029 | 0.12 | 0.12 | 0 | 0.5 | 0.65 | 0.33 | 0.82 | 0.42 | 0.55 |
| 2 | 3325148 | rs4854150 | -1.171 | -0.66 | -0.399 | -0.376 | 0.024 | 0.034 | 0.034 | 0 | 0.51 | 0.54 | 0.32 | 0.76 | 0.36 | 0.51 |
| 2 | 3326045 | rs6548153 | -0.662 | -0.75 | -0.178 | -0.136 | -0.019 | -0.004 | -0.004 | 0 | 0.36 | 0.53 | 0.17 | 0.62 | 0.18 | 0.41 |
| 2 | 3329409 | rs7570426 | 0.16 | -0.983 | -0.338 | -0.178 | -0.109 | 0.076 | 0.076 | 0.471 | 0.26 | 0.41 | 0.12 | 0.57 | 0.08 | 0.26 |
| 2 | 3340094 | rs1111657 | -0.227 | -0.791 | -0.256 | -0.121 | -0.099 | 0.024 | 0.024 | 0 | 0.32 | 0.42 | 0.13 | 0.57 | 0.11 | 0.36 |
| 2 | 3342009 | rs3811617 | -1.527 | -0.702 | -0.516 | -0.232 | 0.033 | 0.185 | 0.185 | 0 | 0.52 | 0.76 | 0.34 | 0.8 | 0.42 | 0.55 |
| 2 | 3348220 | rs11688647 | -0.717 | -0.702 | -0.486 | 0.616 | 0.058 | -0.043 | -0.043 | 0 | 0.37 | 0.71 | 0.19 | 0.39 | 0.2 | 0.41 |
| 2 | 3356345 | rs13421307 | -0.157 | -0.768 | -0.058 | 0.654 | -0.042 | 0.032 | 0.032 | 0 | 0.31 | 0.39 | 0.15 | 0.43 | 0.16 | 0.38 |
| 2 | 3357930 | rs11127416 | 0.082 | -0.63 | 0 | 0.632 | -0.066 | 0.053 | 0.053 | 0.092 | 0.67 | 0.5 | 0.84 | 0.57 | 0.83 | 0.59 |
| 2 | 3360352 | rs2324584 | 0.021 | -0.598 | -0.007 | 0.661 | -0.055 | 0.045 | 0.045 | 0.047 | 0.68 | 0.52 | 0.83 | 0.58 | 0.83 | 0.58 |
| 2 | 3363111 | rs4854179 | -0.071 | 0.143 | -0.234 | 0.847 | -0.072 | -0.041 | -0.041 | 0 | 0.13 | 0.01 | 0.16 | 0.17 | 0.22 | 0.11 |
| 2 | 3380568 | rs12464175 | -0.003 | 0.183 | -0.304 | 0.933 | -0.022 | -0.017 | -0.017 | 0 | 0.12 | 0.08 | 0.16 | 0.19 | 0.2 | 0.11 |
| 2 | 3382118 | rs6548154 | -0.066 | -0.306 | -0.383 | 0.6 | 0.017 | 0.017 | 0.017 | 1.217 | 0.47 | 0.42 | 0.7 | 0.35 | 0.63 | 0.47 |
| 2 | 3382440 | rs7601188 | 0.685 | -0.076 | -0.192 | 0.75 | -0.002 | 0 | 0 | 0.011 | 0.34 | 0.33 | 0.54 | 0.28 | 0.43 | 0.37 |
| 2 | 3383741 | rs12992865 | 0.572 | -0.076 | -0.198 | 0.837 | -0.002 | -0.002 | -0.002 | 0 | 0.35 | 0.34 | 0.54 | 0.48 | 0.43 | 0.38 |
| 2 | 3387126 | rs9808246 | 0.131 | 0.553 | 0.204 | 1.098 | 0.008 | 0.016 | 0.016 | 0.574 | 0.34 | 0.42 | 0.48 | 0.45 | 0.35 | 0.42 |
| 2 | 3395383 | rs11692425 | -0.679 | 0.441 | 0.241 | 0.792 | -0.006 | 0 | 0 | 0 | 0.71 | 0.77 | 0.5 | 0.68 | 0.62 | 0.68 |
| 2 | 3402591 | rs10165099 | -0.641 | 0.45 | 0.339 | 0.787 | 0.015 | 0 | 0 | 1.982 | 0.71 | 0.7 | 0.5 | 0.67 | 0.62 | 0.69 |
| 2 | 3403361 | rs6738313 | -1.14 | 0.637 | 0.238 | 0.727 | -0.045 | -0.031 | -0.031 | 0 | 0.75 | 0.85 | 0.56 | 0.65 | 0.68 | 0.7 |
| 2 | 3411923 | rs4627605 | -0.614 | 0.175 | 0.059 | 0.955 | -0.028 | -0.032 | -0.032 | 0 | 0.67 | 0.78 | 0.53 | 0.45 | 0.62 | 0.63 |
| 2 | 3434286 | rs12989971 | 0.365 | -0.388 | -0.007 | 0.543 | 0.007 | 0.082 | 0.082 | 0 | 0.22 | 0.32 | 0.24 | 0.38 | 0.27 | 0.46 |
| 2 | 3435668 | rs7595325 | 0.699 | -0.783 | -0.472 | 0.013 | 0.059 | 0.068 | 0.068 | 1.419 | 0.27 | 0.14 | 0.11 | 0.01 | 0.14 | 0.05 |
| 2 | 3447827 | rs4482520 | -1.007 | -0.329 | -0.592 | 0.377 | -0.022 | 0.006 | 0.006 | 0.084 | 0.69 | 0.77 | 0.53 | 0.66 | 0.62 | 0.68 |
| 2 | 3456506 | rs10207406 | -0.321 | -1.364 | 0.13 | 0.133 | -0.05 | -0.041 | -0.041 | 0 | 0.32 | 0.45 | 0.14 | 0.2 | 0.15 | 0.11 |
| 2 | 3458400 | rs6730178 | -0.272 | -1.276 | 0.18 | 0.313 | -0.048 | -0.013 | -0.013 | 0 | 0.24 | 0.63 | 0.14 | 0.22 | 0.11 | 0.07 |
| 2 | 3463978 | rs10182502 | -1.19 | -0.959 | 0.45 | 0.13 | 0.015 | 0.011 | 0.011 | 3.358 | 0.06 | 0.41 | 0.03 | 0.01 | 0.01 | 0 |
| 2 | 3464810 | rs7585799 | -0.212 | -0.722 | -0.04 | 0.163 | 0.001 | 0.005 | 0.005 | 0 | 0.46 | 0.43 | 0.34 | 0.27 | 0.48 | 0.51 |
| 2 | 3466385 | rs10198978 | -1.498 | -0.512 | 0.449 | 0.36 | 0.023 | 0.077 | 0.077 | 6.068 | 0.07 | 0.46 | 0.05 | 0.08 | 0.01 | 0.06 |
| 2 | 3470254 | rs11677515 | -0.641 | -0.743 | -0.268 | 0.394 | 0.037 | -0.004 | -0.004 | 0 | 0.49 | 0.75 | 0.36 | 0.46 | 0.45 | 0.5 |
| 2 | 3470416 | rs4352260 | 0.264 | -0.862 | 0.433 | 0.25 | -0.009 | 0.012 | 0.012 | 0 | 0.32 | 0.4 | 0.22 | 0.35 | 0.29 | 0.29 |
| 2 | 3471146 | rs11127428 | -1.094 | -0.965 | 0.326 | 0.343 | 0.042 | 0.007 | 0.007 | 0 | 0.49 | 0.71 | 0.37 | 0.46 | 0.43 | 0.52 |
| 2 | 3472358 | rs12994580 | -0.126 | -0.991 | 0.447 | 0.314 | 0 | 0.017 | 0.017 | 0 | 0.13 | 0.09 | 0.11 | 0.08 | 0.14 | 0.21 |
| 2 | 3472665 | rs6548164 | -0.984 | -0.867 | 0.362 | 0.323 | 0.044 | 0.001 | 0.001 | 0.037 | 0.51 | 0.79 | 0.36 | 0.46 | 0.43 | 0.5 |
| 2 | 3475389 | rs12714377 | -0.659 | -0.964 | 0.478 | 0.298 | -0.012 | -0.042 | -0.042 | 0.058 | 0.16 | 0.29 | 0.12 | 0.08 | 0.14 | 0.22 |
| 2 | 3477014 | rs13429373 | 0.447 | -1.004 | 0.746 | 0.296 | 0.005 | -0.016 | -0.016 | 0 | 0.32 | 0.26 | 0.23 | 0.38 | 0.28 | 0.29 |
| 2 | 3478452 | rs10178877 | -1.6 | -1.046 | 0.227 | 0.01 | -0.119 | -0.06 | -0.06 | 0.164 | 0.64 | 0.92 | 0.43 | 0.53 | 0.51 | 0.58 |
| 2 | 3479737 | rs7591761 | 0.061 | -1.054 | 0.399 | -0.003 | -0.013 | -0.036 | -0.036 | 0 | 0.15 | 0.27 | 0.08 | 0.06 | 0.09 | 0.06 |
| 2 | 3479959 | rs10188054 | -0.204 | -1.074 | 0.467 | -0.019 | -0.006 | -0.063 | -0.063 | 1.934 | 0.14 | 0.22 | 0.1 | 0.01 | 0.13 | 0.11 |
| 2 | 3480366 | rs10188449 | -0.564 | -1.035 | 0.526 | 0.041 | 0.018 | 0.003 | 0.003 | 0 | 0.48 | 0.6 | 0.35 | 0.47 | 0.42 | 0.51 |
| 2 | 3480563 | rs7592700 | -1.508 | -1.035 | 0.499 | 0.042 | -0.101 | -0.044 | -0.044 | 0.327 | 0.63 | 0.89 | 0.43 | 0.53 | 0.51 | 0.58 |
| 2 | 3481659 | rs4321399 | -0.602 | -1.12 | 0.743 | 0.334 | 0.005 | -0.002 | -0.002 | 1.27 | 0.45 | 0.4 | 0.34 | 0.47 | 0.42 | 0.51 |
| 2 | 3483205 | rs6767 | -0.821 | -1.074 | 0.512 | 0.357 | 0.018 | 0.003 | 0.003 | 0.003 | 0.48 | 0.6 | 0.35 | 0.47 | 0.42 | 0.51 |
| 2 | 3489932 | rs12714380 | -1.198 | -1.13 | 0.329 | 0.191 | -0.038 | -0.097 | -0.097 | 2.925 | 0.19 | 0.53 | 0.14 | 0.06 | 0.14 | 0.14 |
| 2 | 3491772 | rs9750503 | 0.494 | -1.093 | -0.14 | 0.298 | -0.004 | 0.009 | 0.009 | 0.011 | 0.31 | 0.31 | 0.23 | 0.38 | 0.26 | 0.29 |
| 2 | 3494374 | rs9752321 | -0.582 | -1.19 | -0.289 | 0.26 | 0.014 | 0.009 | 0.009 | 0 | 0.45 | 0.35 | 0.28 | 0.36 | 0.4 | 0.42 |
| 2 | 3494562 | rs9752760 | -0.304 | -1.072 | -0.342 | 0.334 | 0.012 | 0.012 | 0.012 | 0 | 0.42 | 0.34 | 0.24 | 0.3 | 0.33 | 0.36 |
| 2 | 3495078 | rs9309717 | -0.687 | -1.143 | -0.781 | 0.261 | 0.089 | -0.108 | -0.108 | 0 | 0.43 | 0.91 | 0.25 | 0.5 | 0.29 | 0.42 |
| 2 | 3497167 | rs9678657 | -1.575 | -1.056 | 0.238 | -0.002 | -0.032 | 0.1 | 0.1 | 2.854 | 0.1 | 0.48 | 0.05 | 0.14 | 0.03 | 0.13 |
| 2 | 3498883 | rs12620727 | 0.722 | -1.421 | -0.207 | -0.26 | 0.003 | 0.002 | 0.002 | 0 | 0.23 | 0.18 | 0.27 | 0.21 | 0.16 | 0.16 |
| 2 | 3502035 | rs1130333 | -0.031 | -1.436 | -0.317 | -0.303 | -0.008 | -0.004 | -0.004 | 4.657 | 0.3 | 0.68 | 0.32 | 0.22 | 0.17 | 0.2 |
| 2 | 3502055 | rs1130319 | -0.031 | -1.436 | -0.317 | -0.303 | -0.008 | -0.004 | -0.004 | 0 | 0.3 | 0.68 | 0.32 | 0.22 | 0.17 | 0.2 |

| | | | | | | | | | | | | | | | | |
|---|---|---|---|---|---|---|---|---|---|---|---|---|---|---|---|---|
| 2 | 3502398 | rs1130165 | -0.031 | -1.375 | -0.317 | -0.303 | -0.008 | -0.004 | -0.004 | 0.359 | 0.3 | 0.68 | 0.32 | 0.22 | 0.17 | 0.2 |
| 2 | 3502625 | rs1130127 | -0.007 | -1.405 | -0.326 | -0.311 | -0.009 | -0.005 | -0.005 | 0 | 0.29 | 0.68 | 0.32 | 0.22 | 0.17 | 0.2 |
| 2 | 3504687 | rs9950 | -1.246 | -1.139 | -0.795 | -0.277 | 0.074 | -0.016 | -0.016 | 0.085 | 0.5 | 0.89 | 0.45 | 0.45 | 0.39 | 0.34 |
| 2 | 3505732 | rs13392776 | -1.452 | -1.115 | -0.788 | -0.591 | 0.07 | 0.044 | 0.044 | 0.669 | 0.5 | 0.89 | 0.43 | 0.43 | 0.39 | 0.34 |
| 2 | 3508131 | rs11680305 | -1.471 | -1.121 | -0.832 | -0.594 | 0.043 | -0.016 | -0.016 | 1.979 | 0.52 | 0.91 | 0.44 | 0.46 | 0.39 | 0.42 |
| 2 | 3508994 | rs9798096 | 0.608 | -1.082 | -0.757 | -0.562 | 0 | -0.003 | -0.003 | 0.602 | 0.14 | 0.08 | 0.08 | 0.13 | 0.15 | 0.11 |
| 2 | 3514549 | rs7586092 | -1.442 | -1.064 | -0.819 | -0.625 | 0.042 | -0.017 | -0.017 | 0 | 0.52 | 0.91 | 0.44 | 0.46 | 0.39 | 0.42 |
| 2 | 3519739 | rs6422709 | -0.255 | -0.694 | -0.419 | -0.415 | -0.001 | 0 | 0 | 0 | 0.2 | 0.21 | 0.14 | 0.22 | 0.22 | 0.14 |
| 2 | 3526603 | rs10205909 | -0.106 | 0.196 | -0.41 | -1.717 | -0.002 | -0.077 | -0.077 | 0.014 | 0.63 | 0.48 | 0.65 | 0.81 | 0.65 | 0.73 |
| 2 | 3579585 | rs1129241 | 0.251 | 0.4 | -0.692 | -1.77 | 0.005 | 0.031 | 0.031 | 0.808 | 0.16 | 0.1 | 0.1 | 0 | 0.12 | 0.05 |
| 2 | 3586529 | rs10197283 | -1.554 | 0.962 | -0.928 | -2.119 | -0.003 | 0.049 | 0.049 | 1.074 | 0.57 | 0.79 | 0.52 | 0.41 | 0.57 | 0.49 |
| 2 | 3586769 | rs6742649 | -0.889 | 1.123 | -0.526 | -2.221 | 0 | -0.002 | -0.002 | 0 | 0.39 | 0.62 | 0.23 | 0.24 | 0.35 | 0.31 |
| 2 | 3595834 | rs13002108 | 0.345 | 1.102 | -0.545 | -2.23 | 0.001 | -0.002 | -0.002 | 0 | 0.1 | 0.08 | 0.27 | 0.17 | 0.19 | 0.17 |
| 2 | 3597974 | rs10186193 | -0.882 | 1.037 | -0.559 | -2.22 | 0 | -0.004 | -0.004 | 0 | 0.39 | 0.62 | 0.23 | 0.24 | 0.35 | 0.31 |
| 2 | 3599700 | rs7563960 | -2.099 | 0.892 | -0.932 | -2.218 | -0.021 | -0.008 | -0.008 | 6.489 | 0.64 | 0.88 | 0.56 | 0.42 | 0.66 | 0.54 |
| 2 | 3603340 | rs7607888 | -0.521 | 0.784 | -0.327 | -2.23 | -0.04 | -0.129 | -0.129 | 0 | 0.3 | 0.46 | 0.16 | 0.07 | 0.19 | 0.24 |
| 2 | 3605468 | rs12711976 | 0.411 | 1.303 | -0.321 | -2.255 | 0.001 | -0.013 | -0.013 | 0 | 0.69 | 0.68 | 0.59 | 0.81 | 0.63 | 0.72 |
| 2 | 3607238 | rs6716194 | 0.317 | 1.447 | -0.647 | -2.149 | 0 | -0.001 | -0.001 | 1.168 | 0.1 | 0.09 | 0.27 | 0.17 | 0.19 | 0.17 |
| 2 | 3608756 | rs4583476 | 2 | 0.962 | -1.138 | -2.286 | 0.01 | -0.014 | -0.014 | 0 | 0.35 | 0.1 | 0.4 | 0.58 | 0.3 | 0.47 |
| 2 | 3612844 | rs4133261 | -0.99 | 1.624 | -0.133 | -1.946 | 0.048 | 0.001 | 0.001 | 0 | 0.29 | 0.51 | 0.4 | 0.37 | 0.45 | 0.3 |
| 2 | 3615822 | rs6542658 | -0.76 | 1.338 | -0.607 | -1.866 | -0.019 | 0.051 | 0.051 | 0.537 | 0.84 | 0.91 | 0.88 | 1 | 0.83 | 0.95 |
| 2 | 3623540 | rs3087873 | -0.139 | 1.457 | -0.469 | -1.804 | -0.003 | -0.005 | -0.005 | 0 | 0.77 | 0.75 | 0.84 | 0.99 | 0.76 | 0.87 |
| 2 | 3628383 | rs2071639 | -0.151 | 1.799 | -0.04 | -1.082 | -0.004 | 0.016 | 0.016 | 0 | 0.11 | 0.2 | 0.05 | 0.18 | 0.11 | 0.07 |
| 2 | 3629178 | rs10203413 | -0.77 | 1.921 | 0.024 | -1.144 | 0.001 | -0.006 | -0.006 | 0 | 0.18 | 0.29 | 0.11 | 0.18 | 0.22 | 0.12 |
| 2 | 3630569 | rs4850031 | -0.526 | 2.416 | -0.322 | -1.115 | -0.009 | -0.007 | -0.007 | 0 | 0.09 | 0.07 | 0.28 | 0.19 | 0.19 | 0.17 |
| 2 | 3631802 | rs7608995 | -1.082 | 2.435 | -0.002 | -1.197 | 0.064 | 0.011 | 0.011 | 0 | 0.11 | 0.22 | 0.32 | 0.18 | 0.25 | 0.18 |
| 2 | 3633917 | rs11684944 | -1.044 | 2.281 | -0.514 | -1.78 | -0.029 | 0.063 | 0.063 | 1.266 | 0.33 | 0.76 | 0.22 | 0.16 | 0.35 | 0.25 |
| 2 | 3634753 | rs4850047 | -1.162 | 1.857 | -1.179 | -1.819 | 0.022 | -0.048 | -0.048 | 0.301 | 0.8 | 0.96 | 0.79 | 0.25 | 0.84 | 0.56 |
| 2 | 3642165 | rs4849953 | 1.26 | 1.984 | -1.019 | -1.893 | 0.047 | 0.112 | 0.112 | 1.322 | 0.41 | 0.26 | 0.38 | 0.08 | 0.33 | 0.22 |
| 2 | 3642361 | rs3820897 | -0.983 | 1.8 | -1.166 | -1.676 | 0.008 | -0.015 | -0.015 | 0 | 0.77 | 0.96 | 0.76 | 0.25 | 0.78 | 0.51 |
| 2 | 3645429 | rs13017757 | -0.311 | 1.506 | -0.416 | -1.477 | 0.037 | -0.058 | -0.058 | 0 | 0.23 | 0.5 | 0.29 | 0.17 | 0.25 | 0.21 |
| 2 | 3647117 | rs11902670 | -0.549 | 1.001 | -0.696 | -0.788 | 0.005 | -0.036 | -0.036 | 0 | 0.09 | 0.1 | 0.19 | 0 | 0.1 | 0.11 |
| 2 | 3648396 | rs13004318 | 0.19 | 0.904 | -0.886 | -0.633 | -0.042 | 0.037 | 0.037 | 1.234 | 0.66 | 0.74 | 0.51 | 0.23 | 0.54 | 0.37 |
| 2 | 3658521 | rs13022258 | 0.48 | 0.244 | 0.005 | 0.076 | 0.074 | 0.066 | 0.066 | 0 | 0.79 | 0.54 | 0.8 | 0.7 | 0.71 | 0.91 |
| 2 | 3658674 | rs13000184 | 0.494 | 0.239 | 0.002 | 0.074 | 0.073 | 0.065 | 0.065 | 0 | 0.79 | 0.54 | 0.8 | 0.7 | 0.71 | 0.91 |
| 2 | 3658724 | rs17017737 | -0.745 | -0.143 | 0.123 | -0.167 | -0.024 | -0.041 | -0.041 | 0 | 0.09 | 0.22 | 0.03 | 0.01 | 0.03 | 0.11 |
| 2 | 3660208 | rs12468922 | 0.775 | -0.241 | 0.4 | -0.224 | 0.037 | 0.028 | 0.028 | 0 | 0.75 | 0.48 | 0.8 | 0.7 | 0.71 | 0.9 |
| 2 | 3660752 | rs3811531 | 1.241 | -0.557 | 0.071 | -0.346 | 0.004 | 0.027 | 0.027 | 3.769 | 0.62 | 0.2 | 0.75 | 0.69 | 0.67 | 0.74 |
| 2 | 3660856 | rs7567724 | -0.249 | -0.698 | -0.259 | -0.528 | 0.008 | 0.011 | 0.011 | 0 | 0.14 | 0.11 | 0.07 | 0.02 | 0.11 | 0.09 |
| 2 | 3668126 | rs2872168 | 0.951 | 0.042 | 0.056 | 0.228 | 0.017 | 0.01 | 0.01 | 2.956 | 0.72 | 0.52 | 0.78 | 0.7 | 0.7 | 0.85 |
| 2 | 3673648 | rs10170348 | -0.62 | 0.855 | -0.52 | 0.921 | -0.022 | 0.011 | 0.011 | 0 | 0.06 | 0.01 | 0.07 | 0.31 | 0.16 | 0.06 |
| 2 | 3677022 | rs731034 | -0.802 | 0.844 | -0.454 | 0.948 | 0.005 | 0.044 | 0.044 | 0 | 0.07 | 0.09 | 0.08 | 0.32 | 0.16 | 0.06 |
| 2 | 3682981 | rs17017778 | -0.288 | -0.301 | -0.58 | 1.254 | 0.003 | -0.056 | -0.056 | 2.691 | 0.1 | 0.02 | 0.09 | 0.33 | 0.09 | 0.07 |
| 2 | 3684342 | rs10183413 | 0.607 | -0.147 | -0.453 | 0.439 | -0.008 | 0.018 | 0.018 | 0 | 0.84 | 0.89 | 0.76 | 0.88 | 0.81 | 0.89 |
| 2 | 3684912 | rs3811529 | 1.795 | -0.711 | -0.264 | 0.116 | 0.047 | 0.079 | 0.079 | 2.904 | 0.66 | 0.2 | 0.67 | 0.86 | 0.78 | 0.76 |
| 2 | 3685046 | rs3811528 | 0.419 | -0.708 | -0.295 | 0.12 | -0.009 | 0.02 | 0.02 | 0 | 0.84 | 0.89 | 0.76 | 0.88 | 0.81 | 0.89 |
| 2 | 3686001 | rs11123567 | -1.076 | -0.724 | 0.285 | 0.187 | -0.073 | -0.081 | -0.081 | 0.392 | 0.13 | 0.63 | 0.07 | 0 | 0.02 | 0.08 |
| 2 | 3691548 | rs7567833 | -1.577 | -0.725 | 0.027 | 0.184 | -0.011 | -0.022 | -0.022 | 0 | 0.18 | 0.74 | 0.09 | 0.02 | 0.03 | 0.12 |
| 2 | 3694994 | rs13011199 | 1.776 | -0.531 | 0.32 | 0.504 | 0.029 | -0.018 | -0.018 | 0 | 0.61 | 0.26 | 0.75 | 0.6 | 0.7 | 0.74 |
| 2 | 3696636 | rs7589966 | 0.804 | -0.488 | 0.558 | 0.744 | 0.002 | -0.017 | -0.017 | 0 | 0.8 | 0.87 | 0.85 | 0.72 | 0.79 | 0.85 |
| 2 | 3698371 | rs13031619 | 0.889 | -0.503 | 0.803 | 0.737 | 0.001 | -0.005 | -0.005 | 0 | 0.79 | 0.84 | 0.85 | 0.72 | 0.79 | 0.85 |
| 2 | 3701972 | rs11894441 | -2.026 | -0.714 | 2.031 | 0.312 | -0.029 | -0.055 | -0.055 | 0 | 0.34 | 0.5 | 0.2 | 0.16 | 0.26 | 0.19 |
| 2 | 3705712 | rs9308790 | -1.769 | -0.868 | 2.435 | 0.365 | -0.015 | 0.102 | 0.102 | 3.499 | 0.19 | 0.34 | 0.14 | 0.31 | 0.16 | 0.07 |
| 2 | 3706055 | rs4404281 | -2.525 | -0.881 | 1.921 | 0.302 | 0.051 | -0.043 | -0.043 | 0.015 | 0.4 | 0.91 | 0.28 | 0.33 | 0.33 | 0.24 |
| 2 | 3706812 | rs13425985 | -1.194 | -0.835 | 1.993 | 0.34 | 0.035 | 0.197 | 0.197 | 0 | 0.05 | 0.21 | 0.06 | 0.17 | 0.06 | 0.01 |
| 2 | 3707423 | rs6754459 | -2.227 | -0.856 | 1.773 | 0.312 | 0.045 | 0.066 | 0.066 | 2.089 | 0.35 | 0.75 | 0.23 | 0.16 | 0.26 | 0.22 |
| 2 | 3707775 | rs10208384 | -0.888 | -0.891 | 1.601 | 0.386 | -0.019 | -0.065 | -0.065 | 2.67 | 0.17 | 0.26 | 0.12 | 0.02 | 0.15 | 0.13 |
| 2 | 3709108 | rs1965732 | -2.287 | -0.731 | 0.709 | 0.458 | 0.067 | -0.022 | -0.022 | 11.69 | 0.39 | 0.82 | 0.27 | 0.33 | 0.31 | 0.24 |
| 2 | 3714129 | rs10173297 | -1.583 | -0.422 | 1.44 | 0.786 | -0.02 | 0.116 | 0.116 | 0.826 | 0.15 | 0.36 | 0.1 | 0.31 | 0.12 | 0.06 |
| 2 | 3714261 | rs7558370 | -0.422 | -0.088 | 1.298 | 0.763 | -0.001 | 0.008 | 0.008 | 2.654 | 0.09 | 0.1 | 0.06 | 0.12 | 0.07 | 0.04 |
| 2 | 3715015 | rs13395090 | -1.933 | 0.124 | 0.935 | 0.288 | 0 | 0.023 | 0.023 | 0.359 | 0.42 | 0.67 | 0.31 | 0.31 | 0.44 | 0.28 |
| 2 | 3718227 | rs7576701 | -0.865 | 0.333 | 0.938 | 0.464 | 0.005 | -0.05 | -0.05 | 0 | 0.19 | 0.31 | 0.13 | 0.05 | 0.21 | 0.16 |
| 2 | 3718692 | rs13409104 | -1.866 | 0.311 | 1.058 | 0.53 | -0.003 | 0.005 | 0.005 | 0.263 | 0.34 | 0.55 | 0.23 | 0.3 | 0.33 | 0.22 |
| 2 | 3722494 | rs13409287 | -0.972 | 0.372 | 0.789 | 0.81 | -0.007 | 0.04 | 0.04 | 0.353 | 0.15 | 0.23 | 0.1 | 0.25 | 0.12 | 0.06 |
| 2 | 3726163 | rs2043080 | -1.812 | 0.204 | 0.576 | 0.519 | 0.022 | 0.037 | 0.037 | 0 | 0.49 | 0.7 | 0.31 | 0.38 | 0.48 | 0.27 |
| 2 | 3727000 | rs34616879 | -0.379 | 0.248 | 0.72 | 0.493 | -0.007 | 0.033 | 0.033 | 0 | 0.22 | 0.06 | 0.11 | 0.12 | 0.23 | 0.14 |
| 2 | 3729254 | rs13426642 | -1.42 | 0.023 | 1.603 | 0.52 | 0.011 | 0.013 | 0.013 | 0 | 0.14 | 0.35 | 0.13 | 0.13 | 0.17 | 0.1 |
| 2 | 3744728 | rs2303902 | -1.555 | 0.083 | 1.833 | 0.598 | 0.003 | 0.011 | 0.011 | 0 | 0.4 | 0.35 | 0.22 | 0.24 | 0.37 | 0.23 |
| 2 | 3748971 | rs6730148 | -1.268 | 0.07 | 2.095 | 0.649 | 0.022 | 0.062 | 0.062 | 0.057 | 0.06 | 0.26 | 0.08 | 0.09 | 0.07 | 0.03 |

| | | | | | | | | | | | | | | | | |
|---|---|---|---|---|---|---|---|---|---|---|---|---|---|---|---|---|
| 2 | 3749546 | rs11682163 | -2.107 | 0.244 | 1.954 | 0.675 | 0.015 | 0.029 | 0.029 | 0 | 0.47 | 0.71 | 0.31 | 0.33 | 0.45 | 0.27 |
| 2 | 3753153 | rs13030620 | -0.979 | 0.781 | 1.78 | 0.708 | -0.084 | 0.073 | 0.073 | 0.163 | 0.3 | 0.07 | 0.29 | 0.13 | 0.4 | 0.22 |
| 2 | 3754240 | rs7575946 | -2.249 | 0.798 | 2.267 | 0.927 | -0.006 | 0.012 | 0.012 | 0 | 0.46 | 0.65 | 0.38 | 0.36 | 0.47 | 0.26 |
| 2 | 3755153 | rs2163263 | -1.152 | 0.784 | 2.464 | 0.842 | -0.045 | 0.084 | 0.084 | 2.155 | 0.16 | 0.56 | 0.1 | 0.23 | 0.07 | 0.04 |
| 2 | 3756073 | rs17505481 | -1.873 | 0.793 | 2.354 | 0.809 | -0.001 | -0.004 | -0.004 | 0.546 | 0.42 | 0.38 | 0.36 | 0.23 | 0.45 | 0.23 |
| 2 | 3758225 | rs12711968 | -1.686 | 0.79 | 1.819 | 0.808 | -0.028 | 0.063 | 0.063 | 0.018 | 0.41 | 0.18 | 0.34 | 0.23 | 0.45 | 0.23 |
| 2 | 3759830 | rs6542651 | -1.607 | 0.38 | 1.818 | 1.366 | -0.044 | 0.046 | 0.046 | 4.337 | 0.21 | 0.69 | 0.16 | 0.41 | 0.08 | 0.05 |
| 2 | 3763411 | rs13019382 | -0.636 | 0.679 | 1.458 | 1.1 | -0.029 | 0.038 | 0.038 | 0 | 0.24 | 0.1 | 0.22 | 0.11 | 0.29 | 0.17 |
| 2 | 3764796 | rs12612303 | -0.667 | 0.613 | 1.299 | 1.55 | -0.044 | -0.019 | -0.019 | 0.093 | 0.23 | 0.07 | 0.23 | 0.29 | 0.33 | 0.23 |
| 2 | 3765506 | rs11886426 | -2.159 | 0.564 | 1.6 | 1.313 | -0.005 | -0.003 | -0.003 | 0 | 0.45 | 0.62 | 0.39 | 0.48 | 0.5 | 0.29 |
| 2 | 3767991 | rs7558456 | -1.05 | 0.678 | 1.488 | 1.468 | 0.033 | 0.075 | 0.075 | 0 | 0.27 | 0.5 | 0.23 | 0.39 | 0.3 | 0.23 |
| 2 | 3772338 | rs2603148 | 1.189 | 1.728 | 0.642 | 1.355 | -0.011 | -0.015 | -0.015 | 0 | 0.64 | 0.31 | 0.67 | 0.48 | 0.52 | 0.69 |
| 2 | 3775086 | rs357942 | 1.267 | 1.593 | 0.447 | 1.413 | -0.011 | -0.03 | -0.03 | 0.987 | 0.64 | 0.31 | 0.55 | 0.45 | 0.56 | 0.71 |
| 2 | 3779094 | rs9308791 | 0.782 | 1.374 | 0.542 | 1.245 | 0.002 | 0.01 | 0.01 | 0.045 | 0.92 | 0.86 | 0.81 | 0.82 | 0.85 | 0.92 |
| 2 | 3780595 | rs9308792 | -1.45 | 1.413 | 0.505 | 1.364 | 0 | -0.013 | -0.013 | 0 | 0.42 | 0.65 | 0.38 | 0.51 | 0.52 | 0.3 |
| 2 | 3780896 | rs357953 | 1.764 | 1.312 | 0.346 | 1.111 | 0.103 | 0.099 | 0.099 | 0 | 0.51 | 0.23 | 0.43 | 0.31 | 0.33 | 0.63 |
| 2 | 3782675 | rs357956 | 1.087 | 1.577 | 0.772 | 0.256 | 0.015 | -0.126 | -0.126 | 1.134 | 0.66 | 0.56 | 0.81 | 0.98 | 0.64 | 0.87 |
| 2 | 3783688 | rs357957 | 2.024 | 1.594 | 0.675 | 0.244 | 0.028 | 0.049 | 0.049 | 0 | 0.51 | 0.38 | 0.66 | 0.74 | 0.41 | 0.62 |
| 2 | 3783746 | rs10171874 | -1.155 | 1.675 | 0.733 | 0.306 | 0.019 | 0.01 | 0.01 | 0 | 0.15 | 0.19 | 0.13 | 0.24 | 0.21 | 0.25 |
| 2 | 3785545 | rs7563964 | -1.644 | 1.615 | 0.346 | 0.265 | 0.037 | 0.01 | 0.01 | 0 | 0.19 | 0.44 | 0.14 | 0.23 | 0.21 | 0.24 |
| 2 | 3789383 | rs2249380 | 0.537 | 1.754 | -0.899 | 0.305 | -0.096 | 0.136 | 0.136 | 0 | 0.7 | 0.89 | 0.78 | 0.98 | 0.59 | 0.87 |
| 2 | 3792231 | rs17446920 | -1.084 | 1.686 | -0.74 | 0.746 | -0.021 | -0.006 | -0.006 | 0.016 | 0.14 | 0.04 | 0.1 | 0.15 | 0.16 | 0.21 |
| 2 | 3794065 | rs357977 | 0.141 | 1.687 | -0.987 | 0.304 | -0.033 | 0.04 | 0.04 | 0 | 0.74 | 0.85 | 0.86 | 0.86 | 0.71 | 0.85 |
| 2 | 3794523 | rs357976 | 1.581 | 1.094 | -1.412 | 0.259 | 0.047 | 0.032 | 0.032 | 0 | 0.41 | 0.18 | 0.5 | 0.33 | 0.39 | 0.67 |
| 2 | 3798725 | rs357973 | -0.19 | 1.246 | -0.564 | 0.092 | -0.001 | -0.001 | -0.001 | 0 | 0.89 | 0.88 | 0.89 | 0.95 | 0.9 | 0.91 |
| 2 | 3800383 | rs357969 | 1.149 | 1.107 | -0.673 | 0.231 | -0.017 | 0.141 | 0.141 | 0 | 0.53 | 0.17 | 0.59 | 0.68 | 0.55 | 0.72 |
| 2 | 3801681 | rs357967 | 0.845 | 1.199 | -0.054 | 0.228 | 0.001 | -0.013 | -0.013 | 0 | 0.58 | 0.46 | 0.67 | 0.73 | 0.57 | 0.75 |
| 2 | 3811167 | rs10200015 | -0.052 | 1.245 | 0.39 | 0.486 | 0.01 | 0.043 | 0.043 | 0 | 0.17 | 0.35 | 0.13 | 0.22 | 0.14 | 0.08 |
| 2 | 3812716 | rs13010824 | 1.375 | 0.66 | -0.155 | 0.215 | 0.03 | 0.031 | 0.031 | 0 | 0.41 | 0.22 | 0.36 | 0.36 | 0.34 | 0.58 |
| 2 | 3813083 | rs6761958 | -0.103 | 0.952 | -0.078 | 0.058 | 0.017 | -0.029 | -0.029 | 0 | 0.73 | 0.56 | 0.81 | 0.77 | 0.72 | 0.84 |
| 2 | 3814614 | rs4849926 | -0.311 | 1.111 | -0.176 | 0.018 | -0.024 | -0.068 | -0.068 | 0 | 0.3 | 0.15 | 0.44 | 0.42 | 0.38 | 0.25 |
| 2 | 3814727 | rs4849989 | 1.627 | 1.077 | -0.236 | -0.015 | 0.02 | 0.04 | 0.04 | 0 | 0.41 | 0.24 | 0.36 | 0.35 | 0.34 | 0.59 |
| 2 | 3815250 | rs10180257 | 0.004 | 1.118 | 0.316 | 0.017 | 0.019 | -0.008 | -0.008 | 1.061 | 0.72 | 0.64 | 0.79 | 0.7 | 0.68 | 0.81 |
| 2 | 3817635 | rs6720903 | 1.267 | 0.886 | 0.44 | 0.117 | 0.025 | 0.034 | 0.034 | 0 | 0.42 | 0.27 | 0.37 | 0.36 | 0.34 | 0.59 |
| 2 | 3818623 | rs7559461 | 0.688 | 0.941 | 0.847 | 0.186 | -0.002 | 0.004 | 0.004 | 0 | 0.56 | 0.43 | 0.52 | 0.64 | 0.48 | 0.69 |
| 2 | 3820113 | rs882418 | 0.81 | 0.167 | 0.454 | -0.182 | -0.008 | 0.054 | 0.054 | 0.013 | 0.33 | 0.22 | 0.32 | 0.09 | 0.36 | 0.43 |
| 2 | 3820193 | rs12463627 | 1.864 | 0.167 | 0.382 | -0.137 | -0.025 | -0.092 | -0.092 | 0 | 0.13 | 0 | 0.15 | 0.29 | 0.14 | 0.1 |
| 2 | 3820763 | rs6751953 | 0.528 | 0.128 | 0.423 | -0.164 | 0.011 | -0.005 | -0.005 | 0 | 0.36 | 0.31 | 0.3 | 0.55 | 0.26 | 0.32 |
| 2 | 3824135 | rs10195488 | 0.599 | 0.196 | 0.339 | -0.163 | -0.003 | -0.016 | -0.016 | 0 | 0.34 | 0.27 | 0.38 | 0.38 | 0.36 | 0.47 |
| 2 | 3826597 | rs13035992 | 0.614 | 0.197 | 0.341 | -0.163 | -0.003 | -0.017 | -0.017 | 0 | 0.34 | 0.27 | 0.37 | 0.38 | 0.36 | 0.47 |
| 2 | 3828093 | rs17018009 | 0.686 | 0.185 | 0.205 | -0.049 | 0.048 | -0.078 | -0.078 | 0 | 0.35 | 0.09 | 0.28 | 0.55 | 0.26 | 0.31 |
| 2 | 3829098 | rs17018014 | 0.672 | 0.176 | 0.339 | -0.063 | 0.037 | -0.032 | -0.032 | 0 | 0.35 | 0.19 | 0.29 | 0.55 | 0.27 | 0.31 |
| 2 | 3830906 | rs11687254 | 1.983 | 0.238 | 0.076 | -0.128 | 0.001 | -0.013 | -0.013 | 0 | 0.21 | 0.26 | 0.23 | 0.1 | 0.22 | 0.37 |
| 2 | 3832358 | rs1466227 | -0.834 | 0.269 | 0.125 | -0.233 | -0.012 | 0.045 | 0.045 | 0 | 0.71 | 0.81 | 0.73 | 0.92 | 0.68 | 0.78 |
| 2 | 3836887 | rs11886335 | 0.338 | 0.377 | 0.255 | -0.109 | 0.091 | 0.1 | 0.1 | 0 | 0.13 | 0.36 | 0.21 | 0.27 | 0.19 | 0.09 |
| 2 | 3837303 | rs11886534 | -0.415 | 0.363 | 0.222 | -0.156 | -0.001 | -0.01 | -0.01 | 0 | 0.48 | 0.52 | 0.5 | 0.82 | 0.46 | 0.41 |
| 2 | 3838080 | rs17018062 | 1.875 | 0.15 | 0.201 | -0.105 | 0.035 | -0.016 | -0.016 | 0.138 | 0.22 | 0.13 | 0.13 | 0.26 | 0.12 | 0.21 |
| 2 | 3838226 | rs13396069 | 0.255 | 0.17 | 0.18 | -0.103 | -0.013 | 0.042 | 0.042 | 0 | 0.34 | 0.49 | 0.34 | 0.54 | 0.3 | 0.3 |
| 2 | 3839093 | rs17018070 | 1.887 | 0.181 | -0.171 | -0.092 | 0.034 | -0.016 | -0.016 | 0 | 0.21 | 0.13 | 0.13 | 0.26 | 0.11 | 0.21 |
| 2 | 3839131 | rs17018072 | 0.741 | 0.193 | -0.343 | -0.091 | 0.024 | 0.032 | 0.032 | 0 | 0.11 | 0.21 | 0.2 | 0.27 | 0.19 | 0.09 |
| 2 | 3841136 | rs9789761 | 1.993 | 0.207 | -1.043 | -0.414 | 0.039 | -0.05 | -0.05 | 0 | 0.2 | 0.05 | 0.13 | 0.26 | 0.11 | 0.21 |
| 2 | 3841699 | rs12466058 | -0.093 | 0.437 | -0.899 | -0.497 | 0.011 | 0.049 | 0.049 | 0.064 | 0.48 | 0.62 | 0.49 | 0.82 | 0.44 | 0.39 |
| 2 | 3848532 | rs1454242 | 1.902 | 0.463 | -0.966 | -0.528 | -0.007 | 0.025 | 0.025 | 0 | 0.21 | 0.17 | 0.22 | 0.1 | 0.22 | 0.37 |
| 2 | 3849806 | rs10174064 | 1.89 | 0.468 | -0.976 | -0.526 | -0.006 | 0.021 | 0.021 | 0 | 0.21 | 0.17 | 0.22 | 0.1 | 0.22 | 0.37 |
| 2 | 3851381 | rs7566217 | -0.197 | 0.653 | -0.908 | -0.096 | -0.011 | 0.036 | 0.036 | 0.016 | 0.69 | 0.8 | 0.72 | 0.91 | 0.66 | 0.76 |
| 2 | 3854501 | rs10195076 | -0.011 | 0.664 | -0.879 | -0.056 | -0.002 | -0.001 | -0.001 | 0 | 0.7 | 0.73 | 0.74 | 0.91 | 0.69 | 0.79 |
| 2 | 3855896 | rs1111281 | -0.226 | 0.832 | -1.47 | 0.089 | -0.026 | 0.059 | 0.059 | 0 | 0.73 | 0.87 | 0.74 | 0.91 | 0.68 | 0.79 |
| 2 | 3873845 | rs12052433 | 0.621 | 0.225 | -1.222 | 0.822 | -0.008 | -0.083 | -0.083 | 0 | 0.14 | 0.04 | 0.15 | 0.4 | 0.14 | 0.11 |
| 2 | 3874580 | rs12711980 | 0.473 | 0.249 | -0.867 | -0.056 | -0.001 | -0.006 | -0.006 | 0 | 0.52 | 0.47 | 0.44 | 0.28 | 0.5 | 0.68 |
| 2 | 3874837 | rs6748936 | -0.033 | -0.679 | -0.881 | -0.588 | -0.029 | -0.047 | -0.047 | 0 | 0.44 | 0.44 | 0.29 | 0.15 | 0.28 | 0.4 |
| 2 | 3877393 | rs1013972 | -0.353 | -0.08 | -0.541 | -0.139 | 0.002 | 0 | 0 | 0 | 0.47 | 0.41 | 0.43 | 0.58 | 0.48 | 0.52 |
| 2 | 3877557 | rs1563173 | -0.705 | 0.114 | -0.319 | -0.408 | 0 | -0.002 | -0.002 | 0 | 0.55 | 0.58 | 0.6 | 0.67 | 0.61 | 0.56 |
| 2 | 3877747 | rs17018121 | 0.963 | 0.269 | -0.11 | -0.889 | 0.001 | -0.021 | -0.021 | 0 | 0.16 | 0.2 | 0.19 | 0.12 | 0.19 | 0.32 |
| 2 | 3880340 | rs1454237 | -0.194 | 0.172 | -0.275 | -0.854 | 0.001 | -0.049 | -0.049 | 0 | 0.34 | 0.37 | 0.3 | 0.14 | 0.38 | 0.41 |
| 2 | 3880436 | rs1454236 | -0.267 | 0.173 | -0.436 | -0.854 | -0.001 | -0.092 | -0.092 | 1.023 | 0.36 | 0.51 | 0.3 | 0.14 | 0.38 | 0.41 |
| 2 | 3881424 | rs10495492 | 0.239 | 0.394 | 0.005 | -0.35 | 0.002 | 0.032 | 0.032 | 0 | 0.7 | 0.56 | 0.55 | 0.51 | 0.68 | 0.81 |
| 2 | 3882061 | rs2034732 | -0.13 | 0.36 | -0.097 | -1.107 | 0.002 | -0.004 | -0.004 | 0 | 0.54 | 0.41 | 0.56 | 0.17 | 0.6 | 0.6 |
| 2 | 3882855 | rs11693058 | 0.417 | 0.296 | -0.087 | -1.139 | 0 | 0.009 | 0.009 | 0 | 0.44 | 0.39 | 0.4 | 0.17 | 0.48 | 0.52 |
| 2 | 3883127 | rs12467066 | 0.931 | 0.817 | -0.052 | -0.683 | 0.004 | -0.01 | -0.01 | 0 | 0.06 | 0.06 | 0.16 | 0 | 0.11 | 0.05 |

| | | | | | | | | | | | | | | | | |
|---|---|---|---|---|---|---|---|---|---|---|---|---|---|---|---|---|
| 2 | 3890350 | rs4849930 | 0.852 | 0.633 | -0.016 | -0.869 | 0.007 | 0.007 | 0.007 | 0.422 | 0.32 | 0.28 | 0.29 | 0.15 | 0.27 | 0.44 |
| 2 | 3890627 | rs12612077 | 1.896 | 0.553 | -0.237 | -0.591 | -0.005 | -0.07 | -0.07 | 0 | 0.11 | 0.01 | 0.17 | 0.43 | 0.1 | 0.1 |
| 2 | 3891773 | rs1563171 | 0.382 | 0.624 | -0.126 | -0.724 | 0.007 | -0.057 | -0.057 | 0.124 | 0.29 | 0.16 | 0.26 | 0.44 | 0.28 | 0.19 |
| 2 | 3891792 | rs897067 | -1.67 | 0.66 | -0.054 | -0.743 | -0.004 | -0.002 | -0.002 | 0 | 0.93 | 0.96 | 0.89 | 0.92 | 0.89 | 0.97 |
| 2 | 3892049 | rs897065 | -1.255 | 0.549 | 0.102 | -0.722 | 0.006 | -0.007 | -0.007 | 0.551 | 0.89 | 0.88 | 0.85 | 0.91 | 0.85 | 0.95 |
| 2 | 3893207 | rs17462475 | 1.257 | 0.598 | 0.082 | -0.719 | -0.001 | 0.007 | 0.007 | 0 | 0.18 | 0.14 | 0.09 | 0.01 | 0.18 | 0.1 |
| 2 | 3894559 | rs968796 | -0.442 | 0.33 | 0.279 | -0.364 | -0.001 | -0.001 | -0.001 | 0.165 | 0.59 | 0.59 | 0.59 | 0.48 | 0.57 | 0.75 |
| 2 | 3898745 | rs1454232 | 0.691 | 0.42 | -0.077 | -0.332 | -0.056 | -0.029 | -0.029 | 0.011 | 0.25 | 0.11 | 0.28 | 0.33 | 0.3 | 0.31 |
| 2 | 3902926 | rs4849996 | 0.892 | 0.336 | 0.027 | -0.392 | 0.008 | 0.014 | 0.014 | 0.463 | 0.31 | 0.25 | 0.29 | 0.15 | 0.27 | 0.44 |
| 2 | 3904446 | rs6748872 | -0.992 | 0.394 | 0.042 | -0.388 | -0.003 | 0.004 | 0.004 | 0.012 | 0.71 | 0.75 | 0.77 | 0.9 | 0.67 | 0.85 |
| 2 | 3904709 | rs1454229 | 1.361 | 0.543 | -0.513 | -0.298 | -0.018 | -0.033 | -0.033 | 0 | 0.19 | 0.1 | 0.12 | 0.32 | 0.19 | 0.27 |
| 2 | 3905742 | rs10495494 | 1.333 | 0.062 | -0.372 | -0.095 | -0.009 | 0.104 | 0.104 | 0.02 | 0.12 | 0.15 | 0.18 | 0.42 | 0.1 | 0.1 |
| 2 | 3905982 | rs722087 | -0.108 | 0.062 | -0.432 | -0.102 | 0 | 0 | 0 | 0.371 | 0.44 | 0.39 | 0.47 | 0.58 | 0.37 | 0.54 |
| 2 | 3909150 | rs4849997 | -0.027 | 0.445 | -0.55 | 0.595 | 0 | 0.001 | 0.001 | 0.415 | 0.57 | 0.41 | 0.57 | 0.48 | 0.57 | 0.75 |
| 2 | 3912820 | rs4849999 | 0.624 | 0.359 | -0.49 | 0.926 | -0.004 | 0.015 | 0.015 | 0 | 0.93 | 0.91 | 0.94 | 0.83 | 0.92 | 0.82 |
| 2 | 3913856 | rs12185617 | -0.8 | -0.022 | -0.225 | 0.501 | -0.028 | -0.03 | -0.03 | 1.731 | 0.19 | 0.29 | 0.11 | 0.12 | 0.14 | 0.25 |
| 2 | 3914743 | rs754013 | -0.894 | 0.441 | 0.069 | 1.199 | 0.037 | 0.155 | 0.155 | 0.486 | 0.15 | 0.34 | 0.3 | 0.31 | 0.17 | 0.25 |
| 2 | 3915801 | rs763792 | 0.015 | 0.78 | -1.075 | 0.036 | -0.103 | 0.156 | 0.156 | 0.1 | 0.27 | 0.04 | 0.28 | 0.04 | 0.34 | 0.16 |
| 2 | 3916161 | rs968098 | 0.371 | 0.17 | -0.707 | 0.233 | -0.007 | 0.027 | 0.027 | 0 | 0.29 | 0.33 | 0.16 | 0.42 | 0.21 | 0.39 |
| 2 | 3918124 | rs4850000 | -0.434 | -0.024 | -0.095 | -0.128 | -0.004 | -0.001 | -0.001 | 0 | 0.63 | 0.63 | 0.65 | 0.87 | 0.57 | 0.78 |
| 2 | 3920685 | rs17509530 | 0.339 | -0.093 | -0.283 | -0.341 | -0.001 | -0.007 | -0.007 | 0 | 0.08 | 0.07 | 0.05 | 0 | 0.11 | 0.1 |
| 2 | 3921326 | rs766904 | 1.246 | -0.149 | -0.237 | -0.373 | -0.002 | 0.001 | 0.001 | 0 | 0.07 | 0.09 | 0.06 | 0.09 | 0.07 | 0.06 |
| 2 | 3922214 | rs10184902 | 0.059 | -0.219 | 0.075 | -0.423 | -0.001 | -0.03 | -0.03 | 0.424 | 0.54 | 0.42 | 0.59 | 0.87 | 0.46 | 0.69 |
| 2 | 3923606 | rs10186254 | 0.232 | -0.328 | 0.024 | -0.491 | 0 | -0.001 | -0.001 | 0 | 0.52 | 0.4 | 0.58 | 0.87 | 0.45 | 0.66 |
| 2 | 3924073 | rs11123620 | 1.187 | -0.204 | -0.319 | -0.429 | -0.085 | 0.162 | 0.162 | 0 | 0.28 | 0.04 | 0.27 | 0.04 | 0.33 | 0.15 |
| 2 | 3924842 | rs2313468 | -0.494 | -0.183 | 0.019 | -0.134 | -0.001 | 0 | 0 | 0 | 0.14 | 0.14 | 0.12 | 0.1 | 0.18 | 0.16 |
| 2 | 3927175 | rs12620133 | 0.89 | -0.495 | -0.32 | -0.2 | -0.003 | -0.046 | -0.046 | 2.159 | 0.24 | 0.11 | 0.13 | 0.35 | 0.19 | 0.39 |
| 2 | 3928737 | rs7591808 | -0.287 | -0.817 | -0.438 | -0.118 | -0.001 | -0.003 | -0.003 | 0 | 0.47 | 0.45 | 0.44 | 0.63 | 0.49 | 0.65 |
| 2 | 3947654 | rs883258 | 0.161 | -0.528 | -0.589 | 0.04 | -0.012 | 0.1 | 0.1 | 0.238 | 0.12 | 0.23 | 0.24 | 0.36 | 0.12 | 0.13 |
| 2 | 3948987 | rs2047120 | 0.721 | -0.45 | -1.065 | -0.14 | -0.014 | 0.089 | 0.089 | 1.29 | 0.36 | 0.23 | 0.31 | 0.04 | 0.37 | 0.18 |
| 2 | 3952385 | rs12472158 | -0.304 | -0.132 | -0.598 | 0.855 | 0.001 | 0.007 | 0.007 | 0 | 0.19 | 0.18 | 0.26 | 0.3 | 0.21 | 0.25 |
| 2 | 3959927 | rs13398385 | -0.193 | 0.156 | -0.469 | 0.935 | 0.01 | 0.024 | 0.024 | 0 | 0.15 | 0.18 | 0.25 | 0.28 | 0.2 | 0.22 |
| 2 | 3963865 | rs6735949 | -0.067 | 0.261 | -0.507 | 0.863 | -0.003 | 0.004 | 0.004 | 0 | 0.45 | 0.66 | 0.6 | 0.38 | 0.54 | 0.35 |
| 2 | 3972403 | rs2313472 | -0.736 | 0.513 | 0.081 | 1.439 | 0.035 | 0.214 | 0.214 | 0 | 0.16 | 0.34 | 0.28 | 0.49 | 0.21 | 0.24 |
| 2 | 3974358 | rs2029730 | -0.956 | 0.11 | 0.082 | 1.197 | -0.019 | 0.089 | 0.089 | 0 | 0.12 | 0.27 | 0.28 | 0.34 | 0.07 | 0.09 |
| 2 | 3991471 | rs4241163 | -0.725 | -0.415 | -0.129 | 0.181 | 0.008 | 0.004 | 0.004 | 1.293 | 0.43 | 0.59 | 0.42 | 0.36 | 0.31 | 0.27 |
| 2 | 3994038 | rs4850014 | -1.2 | 0.305 | -0.903 | -0.421 | -0.041 | -0.059 | -0.059 | 0 | 0.7 | 0.89 | 0.69 | 0.38 | 0.61 | 0.54 |
| 2 | 3996533 | rs1494053 | -1.108 | 0.289 | -0.887 | -0.429 | -0.034 | -0.051 | -0.051 | 8.14 | 0.7 | 0.87 | 0.68 | 0.38 | 0.61 | 0.54 |
| 2 | 4006424 | rs12464708 | -0.212 | 0.033 | -0.347 | -0.874 | -0.01 | -0.005 | -0.005 | 0.043 | 0.32 | 0.23 | 0.49 | 0.34 | 0.35 | 0.34 |
| 2 | 4006656 | rs6542664 | -0.727 | -0.339 | -0.274 | -0.777 | -0.009 | 0.005 | 0.005 | 0 | 0.24 | 0.25 | 0.33 | 0.33 | 0.16 | 0.21 |
| 2 | 4010128 | rs11901335 | 0.498 | -0.032 | -0.383 | -1.084 | 0.018 | 0.015 | 0.015 | 0 | 0.25 | 0.18 | 0.12 | 0.02 | 0.18 | 0.11 |
| 2 | 4011444 | rs6745246 | 0.252 | 0.318 | -0.157 | -0.972 | 0.019 | 0.004 | 0.004 | 0.005 | 0.25 | 0.35 | 0.46 | 0.25 | 0.29 | 0.21 |
| 2 | 4014285 | rs923889 | -0.008 | 0.184 | -0.24 | -1.132 | -0.011 | -0.081 | -0.081 | 1.051 | 0.34 | 0.37 | 0.19 | 0.06 | 0.27 | 0.26 |
| 2 | 4014874 | rs953648 | -0.476 | 0.549 | -0.039 | -0.832 | -0.042 | 0.027 | 0.027 | 0 | 0.61 | 0.88 | 0.66 | 0.3 | 0.57 | 0.46 |
| 2 | 4018661 | rs4850022 | 0.15 | 1.276 | 0.319 | -0.876 | -0.042 | 0.099 | 0.099 | 0 | 0.24 | 0.06 | 0.15 | 0 | 0.29 | 0.09 |
| 2 | 4018730 | rs4849938 | -0.306 | 1.491 | 0.272 | -0.932 | -0.098 | 0.135 | 0.135 | 0 | 0.28 | 0.06 | 0.19 | 0 | 0.37 | 0.12 |
| 2 | 4030608 | rs1494055 | -0.806 | 1.014 | 1.315 | -0.957 | 0.021 | -0.136 | -0.136 | 0 | 0.34 | 0.53 | 0.22 | 0.01 | 0.4 | 0.17 |
| 2 | 4035184 | rs1389583 | -1.982 | 0.851 | 1.578 | -0.473 | 0.02 | 0.395 | 0.395 | 0 | 0.07 | 0.45 | 0.22 | 0.26 | 0.01 | 0.05 |
| 2 | 4045175 | rs4410308 | -1.013 | 0.951 | 1.373 | -0.641 | 0.011 | -0.014 | -0.014 | 0 | 0.38 | 0.55 | 0.42 | 0.27 | 0.41 | 0.23 |
| 2 | 4049789 | rs6542668 | -2.108 | 1.072 | 1.698 | -0.727 | -0.026 | 0.067 | 0.067 | 0 | 0.18 | 0.41 | 0.28 | 0.23 | 0.14 | 0.17 |
| 2 | 4050728 | rs6542670 | -1.482 | 1.088 | 1.601 | -0.797 | 0.002 | -0.013 | -0.013 | 0 | 0.44 | 0.54 | 0.45 | 0.24 | 0.49 | 0.29 |
| 2 | 4054980 | rs13416585 | 0.406 | 0.94 | 1.273 | -0.983 | -0.031 | 0.131 | 0.131 | 0 | 0.64 | 0.84 | 0.76 | 0.99 | 0.64 | 0.81 |
| 2 | 4057069 | rs2196650 | 0.831 | 0.803 | 1.704 | -0.779 | 0.003 | -0.017 | -0.017 | 0 | 0.6 | 0.41 | 0.61 | 0.81 | 0.57 | 0.79 |
| 2 | 4057728 | rs4850024 | 0.896 | 0.815 | 1.734 | -0.773 | 0.012 | -0.045 | -0.045 | 4.592 | 0.6 | 0.45 | 0.61 | 0.82 | 0.57 | 0.79 |
| 2 | 4069041 | rs934757 | -0.661 | 0.92 | 1.484 | -0.778 | -0.018 | 0.069 | 0.069 | 0.323 | 0.75 | 0.9 | 0.89 | 1 | 0.78 | 0.95 |
| 2 | 4071892 | rs1370514 | 0.468 | 0.894 | 1.588 | -0.775 | -0.001 | -0.048 | -0.048 | 0 | 0.64 | 0.54 | 0.59 | 0.83 | 0.68 | 0.85 |
| 2 | 4086161 | rs10166392 | 0.823 | 0.846 | 1.534 | -0.747 | 0.001 | -0.01 | -0.01 | 0.16 | 0.58 | 0.54 | 0.53 | 0.81 | 0.54 | 0.75 |
| 2 | 4092059 | rs17510511 | 0.461 | 0.837 | 1.306 | -0.734 | -0.094 | 0.164 | 0.164 | 0 | 0.26 | 0.04 | 0.13 | 0 | 0.31 | 0.11 |
| 2 | 4102330 | rs7420856 | 0.837 | 0.798 | 1.509 | -0.622 | 0.004 | -0.023 | -0.023 | 0 | 0.58 | 0.47 | 0.53 | 0.79 | 0.57 | 0.76 |
| 2 | 4130779 | rs10194645 | 1.639 | 0.837 | 1.255 | -0.446 | -0.024 | -0.004 | -0.004 | 0 | 0.44 | 0.22 | 0.44 | 0.56 | 0.49 | 0.64 |
| 2 | 4133051 | rs7584928 | -1.223 | 1.148 | 1.208 | -0.944 | -0.002 | 0 | 0 | 0.001 | 0.44 | 0.42 | 0.48 | 0.19 | 0.48 | 0.27 |
| 2 | 4135272 | rs12478345 | -0.522 | 1.154 | 1.179 | -0.959 | 0 | 0.02 | 0.02 | 0 | 0.09 | 0.13 | 0.3 | 0.16 | 0.09 | 0.09 |
| 2 | 4136145 | rs1024027 | -0.672 | 1.179 | 1.155 | -0.991 | -0.008 | 0.036 | 0.036 | 0.161 | 0.37 | 0.3 | 0.4 | 0.18 | 0.38 | 0.19 |
| 2 | 4137992 | rs6759055 | -1.026 | 1.067 | 0.732 | -0.994 | 0 | 0.006 | 0.006 | 0 | 0.13 | 0.09 | 0.11 | 0.04 | 0.15 | 0.13 |
| 2 | 4140685 | rs1546130 | -0.221 | 1.185 | 1.014 | -0.838 | -0.001 | 0.006 | 0.006 | 0.005 | 0.29 | 0.26 | 0.35 | 0.15 | 0.24 | 0.11 |
| 2 | 4140801 | rs1546131 | -0.276 | 1.188 | 0.933 | -0.837 | 0 | -0.023 | -0.023 | 0.188 | 0.3 | 0.34 | 0.35 | 0.15 | 0.24 | 0.11 |
| 2 | 4142259 | rs1545136 | -0.299 | 1.092 | 0.277 | -0.657 | -0.005 | 0.004 | 0.004 | 0 | 0.92 | 0.95 | 0.96 | 0.99 | 0.89 | 0.96 |
| 2 | 4146859 | rs6753345 | -0.841 | 1.053 | -0.155 | -0.381 | 0.067 | 0.124 | 0.124 | 0 | 0.12 | 0.41 | 0.31 | 0.18 | 0.12 | 0.1 |
| 2 | 4147979 | rs13033221 | 0.012 | 0.807 | -1.304 | -0.655 | -0.038 | 0.156 | 0.156 | 0.15 | 0.24 | 0.02 | 0.11 | 0 | 0.2 | 0.08 |

| | | | | | | | | | | | | | | | | |
|---|---|---|---|---|---|---|---|---|---|---|---|---|---|---|---|---|
| 2 | 4148784 | rs6761374 | 0.245 | 0.945 | -0.708 | -0.232 | -0.001 | -0.001 | -0.001 | 0 | 0.59 | 0.57 | 0.53 | 0.62 | 0.62 | 0.74 |
| 2 | 4152928 | rs1347684 | -0.432 | 1.062 | -0.655 | -1.845 | 0.002 | -0.049 | -0.049 | 0.244 | 0.58 | 0.48 | 0.42 | 0.15 | 0.6 | 0.45 |
| 2 | 4154552 | rs9751543 | -0.199 | 1.016 | -0.491 | -1.856 | -0.004 | -0.035 | -0.035 | 0.226 | 0.4 | 0.56 | 0.34 | 0.15 | 0.38 | 0.43 |
| 2 | 4170619 | rs1437700 | 0.022 | 1.05 | -1.32 | -1.804 | -0.033 | 0.125 | 0.125 | 0 | 0.33 | 0.06 | 0.29 | 0.14 | 0.34 | 0.37 |
| 2 | 4173729 | rs1437701 | -0.885 | 1.336 | -1.441 | -1.934 | 0 | 0.131 | 0.131 | 0 | 0.48 | 0.28 | 0.42 | 0.16 | 0.55 | 0.51 |
| 2 | 4176692 | rs7601231 | -0.727 | 1.162 | -0.502 | -1.532 | 0.019 | -0.064 | -0.064 | 0 | 0.15 | 0.23 | 0.14 | 0 | 0.2 | 0.14 |
| 2 | 4186566 | rs4850056 | -0.829 | 0.987 | -0.667 | -1.747 | 0 | -0.07 | -0.07 | 0.163 | 0.33 | 0.3 | 0.26 | 0.01 | 0.37 | 0.29 |
| 2 | 4187268 | rs17316964 | 0.095 | 1.008 | -0.991 | -1.812 | -0.019 | 0.032 | 0.032 | 0 | 0.15 | 0.05 | 0.12 | 0 | 0.18 | 0.09 |
| 2 | 4188698 | rs6714273 | 0.24 | 0.578 | -1.052 | -1.539 | -0.001 | -0.014 | -0.014 | 0 | 0.4 | 0.65 | 0.46 | 0.52 | 0.38 | 0.46 |
| 2 | 4191699 | rs7556893 | -1.149 | 0.617 | -1.45 | -1.627 | 0.047 | -0.003 | -0.003 | 1.458 | 0.54 | 0.82 | 0.6 | 0.43 | 0.6 | 0.68 |
| 2 | 4193107 | rs10180291 | -1.007 | 0.704 | -1.016 | -1.207 | -0.004 | -0.035 | -0.035 | 0 | 0.56 | 0.76 | 0.57 | 0.54 | 0.54 | 0.55 |
| 2 | 4196243 | rs10190553 | -0.489 | 1.179 | 0.14 | -0.501 | 0.018 | 0.062 | 0.062 | 0 | 0.26 | 0.55 | 0.39 | 0.44 | 0.28 | 0.41 |
| 2 | 4224068 | rs1175861 | 1.009 | 0.807 | 0.485 | -1.669 | -0.002 | -0.064 | -0.064 | 2.282 | 0.59 | 0.4 | 0.71 | 0.99 | 0.6 | 0.64 |
| 2 | 4228747 | rs17018722 | -0.098 | 0.875 | 0.712 | -1.81 | -0.001 | -0.025 | -0.025 | 0.3 | 0.13 | 0.13 | 0.12 | 0 | 0.17 | 0.19 |
| 2 | 4230420 | rs2577189 | 1.21 | 0.86 | 0.785 | -1.805 | 0.001 | -0.093 | -0.093 | 0 | 0.6 | 0.43 | 0.71 | 0.99 | 0.6 | 0.64 |
| 2 | 4231211 | rs17018741 | -0.347 | 0.509 | 1.222 | -1.442 | 0.047 | -0.068 | -0.068 | 0 | 0.11 | 0.27 | 0.11 | 0 | 0.13 | 0.16 |
| 2 | 4232904 | rs2577190 | 1.271 | -0.328 | 1.602 | -1.406 | -0.033 | -0.116 | -0.116 | 0.122 | 0.78 | 0.42 | 0.88 | 0.99 | 0.86 | 0.91 |
| 2 | 4235543 | rs2577193 | -0.139 | -0.349 | 1.833 | -1.326 | -0.011 | -0.036 | -0.036 | 0.108 | 0.88 | 0.8 | 0.93 | 0.99 | 0.92 | 0.93 |
| 2 | 4237884 | rs2679461 | 1.373 | -0.385 | 2.015 | -1.348 | -0.053 | -0.132 | -0.132 | 0.026 | 0.79 | 0.46 | 0.89 | 0.99 | 0.88 | 0.86 |
| 2 | 4245520 | rs1898518 | 0.589 | -0.338 | 2.128 | -1.353 | 0 | -0.019 | -0.019 | 2.352 | 0.86 | 0.85 | 0.91 | 0.99 | 0.87 | 0.87 |
| 2 | 4246080 | rs2577202 | -0.361 | -0.337 | 2.136 | -1.337 | -0.005 | -0.02 | -0.02 | 0.003 | 0.9 | 0.88 | 0.94 | 0.99 | 0.92 | 0.93 |
| 2 | 4246462 | rs2118380 | 1.816 | -0.407 | 2.116 | -1.44 | -0.028 | -0.108 | -0.108 | 0.1 | 0.71 | 0.42 | 0.86 | 0.99 | 0.81 | 0.85 |
| 2 | 4247411 | rs11683373 | -1.579 | -0.39 | 2.051 | -1.425 | -0.029 | -0.119 | -0.119 | 0 | 0.17 | 0.42 | 0.08 | 0 | 0.11 | 0.08 |
| 2 | 4253421 | rs6718311 | -0.425 | -0.327 | 1.688 | -1.353 | 0 | -0.033 | -0.033 | 0 | 0.06 | 0.13 | 0.03 | 0 | 0.05 | 0.06 |
| 2 | 4257313 | rs1464233 | -1.02 | -0.366 | 1.698 | -1.32 | -0.033 | -0.089 | -0.089 | 0 | 0.2 | 0.39 | 0.11 | 0.04 | 0.13 | 0.14 |
| 2 | 4267783 | rs11900881 | -0.735 | -0.349 | 1.326 | -1.286 | -0.023 | -0.062 | -0.062 | 0 | 0.19 | 0.33 | 0.11 | 0.03 | 0.13 | 0.14 |
| 2 | 4270392 | rs6542690 | -0.397 | -0.373 | 1.089 | -1.303 | -0.025 | -0.063 | -0.063 | 0 | 0.11 | 0.24 | 0.05 | 0 | 0.05 | 0.02 |
| 2 | 4275204 | rs6755583 | -0.88 | -0.212 | 0.676 | -1.144 | -0.029 | -0.075 | -0.075 | 0.061 | 0.2 | 0.37 | 0.1 | 0.03 | 0.13 | 0.14 |
| 2 | 4275332 | rs6706825 | -0.837 | -0.204 | 0.475 | -1.14 | -0.022 | -0.06 | -0.06 | 0 | 0.2 | 0.3 | 0.1 | 0.03 | 0.13 | 0.14 |
| 2 | 4278838 | rs6542691 | -0.84 | 0.315 | -0.626 | -0.397 | 0.002 | -0.026 | -0.026 | 0 | 0.55 | 0.8 | 0.56 | 0.45 | 0.56 | 0.48 |
| 2 | 4279290 | rs7574256 | -1.1 | -0.185 | 0.31 | -0.677 | -0.062 | -0.15 | -0.15 | 0.009 | 0.22 | 0.51 | 0.12 | 0.02 | 0.14 | 0.13 |
| 2 | 4279882 | rs17018879 | -0.105 | 0.028 | 0.579 | -0.565 | 0.007 | -0.04 | -0.04 | 0.653 | 0.07 | 0.17 | 0.05 | 0 | 0.07 | 0.02 |
| 2 | 4279955 | rs7573313 | -0.49 | -0.067 | -0.255 | -0.388 | -0.101 | 0.076 | 0.076 | 0 | 0.6 | 0.89 | 0.37 | 0.32 | 0.45 | 0.61 |
| 2 | 4279962 | rs13421136 | -0.161 | -0.06 | 0.467 | -0.317 | 0.026 | -0.041 | -0.041 | 0.011 | 0.09 | 0.3 | 0.06 | 0 | 0.08 | 0.02 |
| 2 | 4280385 | rs13424039 | -0.797 | -0.08 | -0.207 | 0.192 | -0.097 | -0.118 | -0.118 | 0.056 | 0.73 | 0.92 | 0.54 | 0.48 | 0.6 | 0.78 |
| 2 | 4282214 | rs17018886 | 0.288 | -0.128 | -0.139 | 0.212 | -0.008 | -0.059 | -0.059 | 0.455 | 0.1 | 0.24 | 0.05 | 0 | 0.09 | 0.02 |
| 2 | 4283339 | rs2314221 | -0.518 | -0.196 | -0.395 | 0.876 | -0.044 | 0.118 | 0.118 | 0 | 0.15 | 0.54 | 0.08 | 0.25 | 0.09 | 0.03 |
| 2 | 4285500 | rs7595190 | -1.193 | -0.163 | -0.385 | 1.12 | -0.012 | -0.002 | -0.002 | 1.398 | 0.21 | 0.66 | 0.11 | 0.2 | 0.14 | 0.05 |
| 2 | 4287408 | rs6754487 | -0.475 | -0.181 | -0.338 | 1.185 | -0.044 | 0.056 | 0.056 | 0 | 0.15 | 0.54 | 0.08 | 0.2 | 0.09 | 0.03 |
| 2 | 4288725 | rs4849957 | 0.453 | -0.05 | -0.414 | 1.173 | -0.007 | -0.004 | -0.004 | 0.341 | 0.55 | 0.4 | 0.46 | 0.46 | 0.48 | 0.73 |
| 2 | 4289810 | rs7601152 | -0.671 | -0.28 | 0.578 | 1.329 | -0.026 | 0.055 | 0.055 | 0 | 0.16 | 0.44 | 0.09 | 0.2 | 0.12 | 0.05 |
| 2 | 4290661 | rs7419807 | -0.921 | -0.182 | -0.49 | 1.154 | 0.007 | -0.013 | -0.013 | 0 | 0.31 | 0.68 | 0.25 | 0.39 | 0.26 | 0.17 |
| 2 | 4294129 | rs7606147 | -1.19 | -0.373 | -0.189 | 0.969 | -0.018 | -0.002 | -0.002 | 0 | 0.26 | 0.68 | 0.16 | 0.23 | 0.15 | 0.24 |
| 2 | 4294293 | rs7572261 | -1.159 | -0.45 | 0.512 | 0.911 | -0.055 | 0.013 | 0.013 | 1.769 | 0.23 | 0.59 | 0.14 | 0.23 | 0.12 | 0.24 |
| 2 | 4294389 | rs1105749 | -1.493 | -0.342 | 0.325 | 1.412 | -0.032 | 0.059 | 0.059 | 1.012 | 0.27 | 0.61 | 0.25 | 0.45 | 0.21 | 0.26 |
| 2 | 4296217 | rs7576476 | -1.85 | -0.388 | 0.406 | 1.413 | -0.002 | 0.002 | 0.002 | 1.288 | 0.29 | 0.67 | 0.27 | 0.44 | 0.24 | 0.26 |
| 2 | 4297644 | rs7567558 | -1.225 | -0.413 | 1.134 | 1.244 | -0.049 | -0.029 | -0.029 | 0 | 0.2 | 0.49 | 0.14 | 0.15 | 0.12 | 0.1 |
| 2 | 4302786 | rs7577104 | -1.266 | -0.128 | 0.902 | 1.389 | 0.001 | 0.001 | 0.001 | 0.05 | 0.1 | 0.15 | 0.11 | 0.09 | 0.11 | 0.08 |
| 2 | 4307217 | rs6743559 | -1.376 | 0.23 | 0.591 | 0.972 | -0.002 | 0.003 | 0.003 | 1.184 | 0.25 | 0.24 | 0.19 | 0.16 | 0.3 | 0.19 |
| 2 | 4311020 | rs6542697 | -1.777 | 0.179 | 0.653 | 1.164 | 0.002 | 0.035 | 0.035 | 0.488 | 0.32 | 0.53 | 0.31 | 0.45 | 0.37 | 0.31 |
| 2 | 4317473 | rs10179510 | -1.061 | 0.334 | -0.062 | 0.605 | -0.001 | -0.001 | -0.001 | 0.096 | 0.42 | 0.58 | 0.42 | 0.44 | 0.45 | 0.43 |
| 2 | 4318976 | rs11686135 | -0.629 | 0.299 | -0.208 | 0.229 | -0.002 | -0.011 | -0.011 | 0.028 | 0.13 | 0.09 | 0.07 | 0 | 0.13 | 0.08 |
| 2 | 4323188 | rs6708052 | -0.875 | 0.133 | 0.102 | 0.119 | -0.005 | -0.01 | -0.01 | 0 | 0.23 | 0.32 | 0.15 | 0.15 | 0.16 | 0.36 |
| 2 | 4325172 | rs6716364 | -1.407 | -0.297 | -0.156 | -0.351 | 0.029 | 0.09 | 0.09 | 0.068 | 0.53 | 0.78 | 0.3 | 0.3 | 0.37 | 0.5 |
| 2 | 4327104 | rs11123636 | 0.933 | 0.066 | -0.608 | 0.001 | 0.016 | 0.01 | 0.01 | 4.814 | 0.16 | 0.17 | 0.35 | 0.2 | 0.2 | 0.2 |
| 2 | 4327339 | rs7586198 | 0.904 | -0.261 | -0.366 | -0.31 | -0.005 | -0.02 | -0.02 | 0 | 0.22 | 0.24 | 0.14 | 0.14 | 0.2 | 0.07 |
| 2 | 4335063 | rs7589994 | -1.823 | -0.385 | -0.38 | -0.249 | -0.036 | -0.06 | -0.06 | 0 | 0.65 | 0.83 | 0.61 | 0.5 | 0.55 | 0.64 |
| 2 | 4338128 | rs11123638 | -2.085 | -0.376 | -0.578 | -0.225 | -0.095 | -0.141 | -0.141 | 0 | 0.68 | 0.92 | 0.63 | 0.51 | 0.59 | 0.74 |
| 2 | 4345324 | rs7594822 | -0.335 | -0.77 | -0.321 | -0.371 | 0.003 | -0.001 | -0.001 | 0 | 0.37 | 0.41 | 0.53 | 0.35 | 0.4 | 0.6 |
| 2 | 4348672 | rs938767 | -0.194 | -0.905 | -0.156 | -0.446 | -0.057 | -0.065 | -0.065 | 0.436 | 0.31 | 0.46 | 0.09 | 0.16 | 0.19 | 0.16 |
| 2 | 4349814 | rs2139164 | 0.458 | -1.065 | 0.021 | -0.529 | 0.001 | 0.019 | 0.019 | 0.092 | 0.15 | 0.11 | 0.12 | 0.02 | 0.13 | 0.24 |
| 2 | 4351869 | rs17019005 | -0.101 | -0.947 | 0.441 | -0.385 | -0.024 | 0.095 | 0.095 | 1.136 | 0.1 | 0.36 | 0.02 | 0.15 | 0.01 | 0.09 |
| 2 | 4353141 | rs4585039 | 0.758 | -0.833 | -0.168 | -0.377 | 0.017 | 0.016 | 0.016 | 0 | 0.2 | 0.08 | 0.06 | 0 | 0.15 | 0.04 |
| 2 | 4371080 | rs12053357 | 0.069 | -0.731 | 0.021 | -0.199 | -0.03 | 0.019 | 0.019 | 0.014 | 0.09 | 0.15 | 0.01 | 0.15 | 0.01 | 0.09 |
| 2 | 4377723 | rs11123646 | 0.21 | -0.765 | -0.107 | -0.383 | -0.001 | -0.016 | -0.016 | 3.642 | 0.27 | 0.18 | 0.39 | 0.34 | 0.25 | 0.25 |
| 2 | 4378203 | rs12465466 | -1.108 | -0.872 | 0.362 | -0.493 | 0.006 | 0.013 | 0.013 | 0.057 | 0.5 | 0.42 | 0.45 | 0.33 | 0.4 | 0.38 |
| 2 | 4382168 | rs6750318 | -2.109 | -0.788 | 0.25 | -0.458 | -0.013 | -0.022 | -0.022 | 0 | 0.67 | 0.69 | 0.6 | 0.52 | 0.55 | 0.73 |
| 2 | 4394440 | rs35704616 | 0.695 | -0.783 | -0.038 | -0.497 | 0.017 | 0.023 | 0.023 | 0 | 0.19 | 0.09 | 0.06 | 0 | 0.15 | 0.04 |
| 2 | 4396305 | rs11897918 | -0.146 | -0.399 | 0.302 | 0.054 | -0.027 | -0.045 | -0.045 | 0.299 | 0.08 | 0.19 | 0.01 | 0 | 0.01 | 0.08 |

| | | | | | | | | | | | | | | | | |
|---|---|---|---|---|---|---|---|---|---|---|---|---|---|---|---|---|
| 2 | 4402182 | rs6542705 | -0.787 | -0.176 | -0.179 | 0.37 | 0 | -0.003 | -0.003 | 0 | 0.19 | 0.15 | 0.22 | 0.23 | 0.16 | 0.39 |
| 2 | 4404065 | rs12476119 | -0.641 | -0.556 | -0.232 | 0.214 | -0.002 | -0.003 | -0.003 | 0 | 0.61 | 0.6 | 0.6 | 0.55 | 0.56 | 0.71 |
| 2 | 4409059 | rs9808493 | -0.929 | -1.207 | -0.015 | 0.058 | 0.007 | 0.007 | 0.007 | 5.284 | 0.33 | 0.68 | 0.14 | 0.22 | 0.21 | 0.33 |
| 2 | 4409380 | rs17019063 | 1.032 | -1.244 | -0.112 | 0.248 | 0.01 | 0.02 | 0.02 | 0 | 0.07 | 0.13 | 0.16 | 0.13 | 0.07 | 0.13 |
| 2 | 4409558 | rs6542706 | -1.363 | -1.167 | -0.158 | 0.281 | 0.046 | 0.039 | 0.039 | 1.453 | 0.41 | 0.84 | 0.35 | 0.37 | 0.3 | 0.5 |
| 2 | 4409755 | rs6542707 | -1.091 | -1.123 | 0.656 | 0.477 | -0.058 | -0.091 | -0.091 | 0.344 | 0.17 | 0.5 | 0.04 | 0.05 | 0.07 | 0.28 |
| 2 | 4418917 | rs6717653 | -0.277 | -0.872 | 0.441 | 0.788 | 0.013 | 0.026 | 0.026 | 0.19 | 0.09 | 0.17 | 0.22 | 0.17 | 0.1 | 0.2 |
| 2 | 4419477 | rs4131040 | -1.205 | -1.167 | 0.899 | 0.646 | -0.01 | -0.042 | -0.042 | 0 | 0.14 | 0.41 | 0.08 | 0.05 | 0.07 | 0.24 |
| 2 | 4419844 | rs1318482 | -1.151 | -1.098 | 0.914 | 0.675 | 0.043 | 0.005 | 0.005 | 0 | 0.09 | 0.33 | 0.08 | 0.05 | 0.07 | 0.24 |
| 2 | 4420858 | rs12712001 | -0.008 | -1.265 | 0.904 | 0.699 | 0.029 | 0.022 | 0.022 | 0 | 0.06 | 0.24 | 0.03 | 0.05 | 0.05 | 0.17 |
| 2 | 4424713 | rs1377583 | 0.536 | -1.766 | -0.137 | 0.903 | 0.028 | -0.047 | -0.047 | 0 | 0.26 | 0.14 | 0.11 | 0.47 | 0.15 | 0.27 |
| 2 | 4438142 | rs1840194 | 1.164 | -1.062 | 0.171 | 0.866 | -0.022 | -0.022 | -0.022 | 0.157 | 0.83 | 0.45 | 0.93 | 0.86 | 0.89 | 0.77 |
| 2 | 4449762 | rs630845 | -0.141 | -1.034 | -0.197 | 0.849 | 0.055 | 0.005 | 0.005 | 0.254 | 0.94 | 0.69 | 0.95 | 0.95 | 0.91 | 0.88 |
| 2 | 4449917 | rs619773 | 0.648 | -0.961 | -0.473 | 0.807 | 0.015 | -0.015 | -0.015 | 1.314 | 0.36 | 0.23 | 0.2 | 0.42 | 0.3 | 0.43 |
| 2 | 4451884 | rs10165034 | 0.568 | -0.904 | -0.461 | 0.756 | 0.054 | 0.001 | 0.001 | 0 | 0.05 | 0.23 | 0.03 | 0.02 | 0.08 | 0.12 |
| 2 | 4451952 | rs590049 | 1.026 | -0.901 | -0.661 | 0.792 | -0.026 | 0.019 | 0.019 | 0.973 | 0.76 | 0.35 | 0.73 | 0.66 | 0.84 | 0.71 |
| 2 | 4456055 | rs2602629 | -0.841 | -0.217 | -1 | 0.891 | -0.062 | 0.065 | 0.065 | 0.23 | 0.62 | 0.81 | 0.48 | 0.75 | 0.48 | 0.8 |
| 2 | 4465553 | rs4572604 | -1.334 | 0.087 | -0.572 | 1.002 | 0.004 | 0.11 | 0.11 | 0 | 0.21 | 0.54 | 0.29 | 0.39 | 0.2 | 0.28 |
| 2 | 4469132 | rs9973340 | 0.208 | 0.353 | -0.331 | 0.28 | -0.061 | 0.116 | 0.116 | 0.325 | 0.39 | 0.11 | 0.5 | 0.22 | 0.54 | 0.29 |
| 2 | 4470836 | rs686173 | 0.297 | 0.414 | -0.406 | 0.336 | 0.051 | 0.035 | 0.035 | 0 | 0.48 | 0.32 | 0.4 | 0.57 | 0.35 | 0.54 |
| 2 | 4471138 | rs2677912 | 0.168 | 0.554 | -0.809 | 0.503 | 0 | -0.001 | -0.001 | 0 | 0.55 | 0.56 | 0.46 | 0.6 | 0.44 | 0.58 |
| 2 | 4474196 | rs634273 | -0.126 | 0.756 | -1.702 | 0.171 | -0.027 | 0.123 | 0.123 | 0 | 0.6 | 0.87 | 0.47 | 0.78 | 0.43 | 0.71 |
| 2 | 4476883 | rs587913 | 0.84 | 2.332 | -0.806 | 0.202 | 0.129 | -0.022 | -0.022 | 5.416 | 0.92 | 0.78 | 0.8 | 0.97 | 0.77 | 0.88 |
| 2 | 4479479 | rs2314703 | 0.186 | 1.442 | -1.085 | 0.134 | -0.004 | -0.001 | -0.001 | 0.5 | 0.48 | 0.49 | 0.38 | 0.38 | 0.31 | 0.44 |
| 2 | 4485192 | rs608974 | 0.14 | 1.022 | -1.118 | -0.609 | 0.017 | 0.038 | 0.038 | 0.563 | 0.35 | 0.26 | 0.33 | 0.17 | 0.24 | 0.33 |
| 2 | 4489752 | rs17019119 | 0.434 | 0.939 | -1.256 | -0.712 | -0.022 | 0.05 | 0.05 | 0.907 | 0.31 | 0.01 | 0.29 | 0.16 | 0.22 | 0.33 |
| 2 | 4492173 | rs7596238 | -0.218 | 0.579 | -1.481 | -0.353 | -0.023 | 0.003 | 0.003 | 0 | 0.13 | 0.27 | 0.09 | 0.1 | 0.1 | 0.06 |
| 2 | 4493374 | rs633168 | -1.401 | 0.399 | -1.121 | -0.402 | 0.007 | 0.028 | 0.028 | 0.111 | 0.77 | 0.68 | 0.75 | 0.4 | 0.73 | 0.67 |
| 2 | 4494928 | rs11895724 | 0.023 | 0.446 | -1.116 | -0.274 | -0.026 | 0.02 | 0.02 | 0 | 0.1 | 0.28 | 0.06 | 0.1 | 0.07 | 0.05 |
| 2 | 4495174 | rs651418 | 0.438 | 0.361 | -0.894 | -0.501 | 0.037 | 0.197 | 0.197 | 0.176 | 0.5 | 0.18 | 0.46 | 0.17 | 0.42 | 0.41 |
| 2 | 4499533 | rs2602628 | -0.549 | 0.236 | -0.607 | -0.282 | -0.011 | 0.02 | 0.02 | 0 | 0.68 | 0.58 | 0.71 | 0.39 | 0.74 | 0.64 |
| 2 | 4509942 | rs606243 | -0.498 | 0.18 | -0.623 | -0.226 | -0.011 | 0.021 | 0.021 | 0 | 0.68 | 0.58 | 0.69 | 0.4 | 0.73 | 0.65 |
| 2 | 4514506 | rs17619675 | 0.542 | -0.254 | -0.613 | -0.762 | 0.013 | 0.003 | 0.003 | 0.183 | 0.21 | 0.13 | 0.11 | 0 | 0.17 | 0.06 |
| 2 | 4516625 | rs589804 | -0.377 | 0.636 | -0.685 | -0.476 | 0 | 0.007 | 0.007 | 0.119 | 0.56 | 0.44 | 0.45 | 0.26 | 0.53 | 0.43 |
| 2 | 4517511 | rs648196 | 1.403 | 0.564 | -1.011 | -0.617 | -0.04 | -0.046 | -0.046 | 0 | 0.24 | 0.01 | 0.25 | 0.21 | 0.21 | 0.32 |
| 2 | 4524510 | rs616251 | -0.94 | 0.673 | -0.776 | -0.321 | -0.005 | 0.012 | 0.012 | 0.002 | 0.61 | 0.51 | 0.56 | 0.45 | 0.62 | 0.63 |
| 2 | 4540319 | rs642267 | 0.952 | 0.686 | -0.876 | -0.245 | 0.012 | 0.009 | 0.009 | 0 | 0.26 | 0.11 | 0.25 | 0.2 | 0.21 | 0.32 |
| 2 | 4548958 | rs671518 | -1.296 | 0.888 | -0.646 | -0.019 | 0.008 | -0.008 | -0.008 | 0.017 | 0.59 | 0.61 | 0.55 | 0.45 | 0.59 | 0.58 |
| 2 | 4549436 | rs673811 | -1.299 | 1.237 | -0.646 | -0.021 | 0.008 | -0.008 | -0.008 | 0 | 0.59 | 0.61 | 0.55 | 0.45 | 0.59 | 0.58 |
| 2 | 4557059 | rs591265 | -0.094 | 1.503 | -1.047 | -0.025 | 0.028 | 0.113 | 0.113 | 0 | 0.44 | 0.21 | 0.35 | 0.21 | 0.38 | 0.37 |
| 2 | 4557832 | rs11682901 | 0.963 | 1.111 | -0.779 | -0.111 | 0.027 | -0.028 | -0.028 | 0 | 0.05 | 0.15 | 0.06 | 0 | 0.08 | 0.01 |
| 2 | 4559812 | rs7605831 | 1.42 | 0.784 | -1.335 | 0.159 | 0.017 | 0.056 | 0.056 | 0 | 0.17 | 0.05 | 0.06 | 0 | 0.1 | 0.05 |
| 2 | 4561186 | rs13012375 | -0.127 | 0.724 | -0.746 | 0.896 | 0 | 0 | 0 | 0 | 0.65 | 0.69 | 0.58 | 0.54 | 0.6 | 0.44 |
| 2 | 4579907 | rs1453598 | -0.113 | 0.149 | -1.02 | 0.67 | -0.007 | 0.018 | 0.018 | 0 | 0.66 | 0.73 | 0.59 | 0.67 | 0.62 | 0.71 |
| 2 | 4581232 | rs1670528 | -1.112 | -0.718 | -1.028 | 1.003 | -0.011 | -0.08 | -0.08 | 0 | 0.8 | 0.89 | 0.53 | 0.49 | 0.77 | 0.74 |
| 2 | 4581932 | rs6542609 | -0.288 | -1.131 | -0.013 | 0.546 | -0.024 | -0.055 | -0.055 | 0 | 0.1 | 0.38 | 0.08 | 0 | 0.09 | 0.02 |
| 2 | 4583204 | rs1564331 | -0.645 | -1.02 | -0.114 | 0.462 | -0.005 | -0.078 | -0.078 | 0 | 0.12 | 0.38 | 0.1 | 0 | 0.13 | 0.04 |
| 2 | 4584797 | rs6724821 | -0.777 | -1.367 | 0.039 | 0.239 | -0.067 | -0.129 | -0.129 | 0 | 0.16 | 0.5 | 0.11 | 0 | 0.11 | 0.02 |
| 2 | 4587751 | rs4849963 | 0.799 | -1.402 | -0.029 | 0.396 | 0.003 | 0.018 | 0.018 | 0 | 0.11 | 0.13 | 0.13 | 0.21 | 0.13 | 0.25 |
| 2 | 4591164 | rs7603898 | 1.044 | -1.344 | 0.035 | 0.341 | -0.008 | -0.006 | -0.006 | 0 | 0.1 | 0.06 | 0.13 | 0.25 | 0.13 | 0.25 |
| 2 | 4592041 | rs10495502 | 1.471 | -1.753 | -0.213 | 0.009 | -0.098 | 0.158 | 0.158 | 5.285 | 0.34 | 0.07 | 0.21 | 0.11 | 0.35 | 0.15 |
| 2 | 4595048 | rs10177804 | -1.289 | -1.779 | 0.033 | 0.158 | 0.01 | 0 | 0 | 0 | 0.43 | 0.59 | 0.32 | 0.5 | 0.38 | 0.58 |
| 2 | 4597061 | rs9308783 | -1.41 | -1.78 | -0.013 | 0.16 | 0.022 | 0.006 | 0.006 | 0 | 0.45 | 0.65 | 0.34 | 0.5 | 0.42 | 0.6 |
| 2 | 4601476 | rs10495504 | -0.622 | -1.442 | 0.47 | 0.294 | -0.024 | -0.035 | -0.035 | 0 | 0.7 | 0.81 | 0.43 | 0.35 | 0.64 | 0.5 |
| 2 | 4602607 | rs6712089 | -0.925 | -1.479 | 0.682 | 0.27 | -0.025 | -0.008 | -0.008 | 0 | 0.31 | 0.56 | 0.2 | 0.24 | 0.28 | 0.35 |
| 2 | 4603577 | rs10174999 | -0.13 | -1.488 | 0.54 | 0.295 | -0.009 | 0.004 | 0.004 | 0 | 0.23 | 0.28 | 0.15 | 0.24 | 0.22 | 0.33 |
| 2 | 4606117 | rs17019420 | 1.56 | -1.429 | 0.398 | 0.185 | -0.002 | -0.015 | -0.015 | 0.012 | 0.06 | 0.07 | 0.05 | 0 | 0.09 | 0.02 |
| 2 | 4606754 | rs1904321 | 0.101 | -1.329 | 0.283 | 0.313 | -0.041 | 0.004 | 0.004 | 0 | 0.25 | 0.41 | 0.16 | 0.22 | 0.22 | 0.33 |
| 2 | 4608698 | rs1382743 | -1.803 | -1.308 | 0.187 | 0.373 | -0.008 | -0.037 | -0.037 | 0.625 | 0.8 | 0.85 | 0.56 | 0.61 | 0.77 | 0.74 |
| 2 | 4609193 | rs17564063 | 1.239 | -1.306 | 0.234 | 0.321 | -0.093 | 0.151 | 0.151 | 5.58 | 0.36 | 0.07 | 0.22 | 0.15 | 0.36 | 0.14 |
| 2 | 4613258 | rs10864993 | 0.264 | -1.257 | 0.38 | 0.365 | 0.025 | 0.136 | 0.136 | 0 | 0.12 | 0.29 | 0.15 | 0.25 | 0.12 | 0.25 |
| 2 | 4613367 | rs10864994 | 0.264 | -1.257 | 0.38 | 0.365 | 0.025 | 0.136 | 0.136 | 0 | 0.12 | 0.29 | 0.15 | 0.25 | 0.12 | 0.25 |
| 2 | 4617571 | rs1585756 | -0.792 | -1.416 | 0.319 | 0.284 | 0.004 | 0.056 | 0.056 | 0 | 0.22 | 0.4 | 0.18 | 0.25 | 0.19 | 0.28 |
| 2 | 4620195 | rs4849917 | 0.337 | -1.411 | 0.317 | 0.289 | 0.026 | 0.138 | 0.138 | 0 | 0.12 | 0.29 | 0.15 | 0.25 | 0.12 | 0.26 |
| 2 | 4626527 | rs6751518 | -0.707 | -1.406 | 0.215 | 0.295 | 0.004 | 0.056 | 0.056 | 0 | 0.22 | 0.4 | 0.18 | 0.25 | 0.19 | 0.28 |
| 2 | 4630194 | rs6542610 | -0.124 | -1.386 | -0.095 | 0.277 | 0.008 | 0.003 | 0.003 | 2.271 | 0.55 | 0.53 | 0.4 | 0.4 | 0.55 | 0.42 |
| 2 | 4638414 | rs12711951 | 0.704 | -1.096 | -0.284 | 0.391 | -0.005 | 0.001 | 0.001 | 0 | 0.24 | 0.3 | 0.18 | 0.2 | 0.25 | 0.3 |
| 2 | 4641479 | rs1479237 | 0.885 | -0.767 | -0.452 | 0.668 | -0.01 | -0.016 | -0.016 | 0 | 0.1 | 0.07 | 0.13 | 0.27 | 0.13 | 0.25 |
| 2 | 4641827 | rs10196371 | -1.484 | -0.838 | -0.469 | 0.657 | -0.007 | -0.037 | -0.037 | 0.922 | 0.79 | 0.86 | 0.56 | 0.62 | 0.77 | 0.72 |

| | | | | | | | | | | | | | | | | |
|---|---|---|---|---|---|---|---|---|---|---|---|---|---|---|---|---|
| 2 | 4645034 | rs10206503 | -0.874 | -0.267 | -0.178 | 0.739 | 0.001 | 0 | 0 | 0 | 0.49 | 0.45 | 0.37 | 0.41 | 0.53 | 0.45 |
| 2 | 4650896 | rs10172284 | -0.762 | -0.27 | -0.167 | 0.743 | 0.01 | 0.004 | 0.004 | 0 | 0.48 | 0.36 | 0.36 | 0.41 | 0.52 | 0.45 |
| 2 | 4651792 | rs11673833 | -0.196 | -0.197 | -0.716 | 0.542 | 0.005 | -0.045 | -0.045 | 0 | 0.38 | 0.41 | 0.24 | 0.14 | 0.37 | 0.16 |
| 2 | 4653262 | rs17019490 | -0.324 | -0.306 | -1.657 | 0.761 | -0.028 | -0.071 | -0.071 | 0.218 | 0.44 | 0.05 | 0.35 | 0.43 | 0.52 | 0.45 |
| 2 | 4653622 | rs8179839 | -0.327 | -0.291 | -1.714 | 0.763 | -0.031 | -0.074 | -0.074 | 0 | 0.44 | 0.1 | 0.36 | 0.42 | 0.52 | 0.46 |
| 2 | 4654356 | rs17019492 | 0.159 | -1.578 | -0.649 | 1.036 | -0.013 | 0.032 | 0.032 | 0.043 | 0.22 | 0.58 | 0.14 | 0.19 | 0.18 | 0.2 |
| 2 | 4654488 | rs10177009 | -0.465 | -1.763 | -1.384 | 0.156 | 0.046 | 0.049 | 0.049 | 0 | 0.43 | 0.23 | 0.17 | 0.68 | 0.32 | 0.48 |
| 2 | 4659821 | rs13006169 | 0.136 | -1.72 | -0.861 | 0.287 | 0.011 | -0.018 | -0.018 | 0 | 0.24 | 0.19 | 0.11 | 0.25 | 0.18 | 0.19 |
| 2 | 4662656 | rs13405591 | -0.011 | -1.982 | -0.667 | 0.445 | 0.006 | -0.03 | -0.03 | 3.147 | 0.27 | 0.16 | 0.12 | 0.54 | 0.24 | 0.4 |
| 2 | 4666383 | rs12623182 | 0.938 | -1.956 | -0.781 | 0.465 | -0.011 | -0.079 | -0.079 | 0 | 0.19 | 0.04 | 0.08 | 0.25 | 0.16 | 0.2 |
| 2 | 4666664 | rs7581129 | -0.089 | -1.976 | 0.002 | 0.326 | -0.008 | 0.062 | 0.062 | 0 | 0.1 | 0.12 | 0.04 | 0.28 | 0.07 | 0.21 |
| 2 | 4669324 | rs6760967 | 0.057 | -1.883 | -0.142 | -0.345 | -0.005 | -0.012 | -0.012 | 0 | 0.15 | 0.25 | 0.09 | 0.1 | 0.1 | 0.13 |
| 2 | 4677204 | rs10165964 | -1.13 | -2.366 | -0.317 | -0.451 | 0.007 | 0 | 0 | 0 | 0.31 | 0.32 | 0.14 | 0.35 | 0.2 | 0.38 |
| 2 | 4677905 | rs826027 | -1.816 | -2.576 | -0.577 | -0.556 | -0.013 | -0.1 | -0.1 | 0 | 0.79 | 0.98 | 0.9 | 0.74 | 0.82 | 0.77 |
| 2 | 4680024 | rs9808088 | 0.147 | -2.438 | -0.317 | -0.479 | -0.014 | -0.027 | -0.027 | 0 | 0.16 | 0.3 | 0.09 | 0.1 | 0.1 | 0.13 |
| 2 | 4684425 | rs826021 | -1.763 | -2.776 | 0.256 | -0.643 | 0.078 | 0.025 | 0.025 | 1.279 | 0.39 | 0.75 | 0.16 | 0.35 | 0.2 | 0.37 |
| 2 | 4684505 | rs826022 | -0.055 | -2.676 | 0.621 | -0.597 | -0.025 | -0.051 | -0.051 | 0 | 0.18 | 0.46 | 0.09 | 0.1 | 0.1 | 0.13 |
| 2 | 4686524 | rs2121221 | 0.332 | -2.566 | 0.107 | -0.502 | 0.02 | -0.037 | -0.037 | 0 | 0.14 | 0.04 | 0.05 | 0.25 | 0.09 | 0.23 |
| 2 | 4705220 | rs2872782 | 1.261 | -2.089 | -0.325 | -0.949 | -0.124 | 0.102 | 0.102 | 0 | 0.21 | 0.01 | 0.19 | 0.03 | 0.34 | 0.15 |
| 2 | 4705263 | rs699549 | -1.71 | -3.379 | 0.137 | -0.565 | -0.011 | -0.09 | -0.09 | 0 | 0.77 | 0.98 | 0.89 | 0.75 | 0.8 | 0.76 |
| 2 | 4712182 | rs826013 | 0.229 | -3.473 | 0.708 | -0.598 | 0.004 | -0.002 | -0.002 | 0 | 0.55 | 0.5 | 0.81 | 0.53 | 0.7 | 0.5 |
| 2 | 4717515 | rs11123571 | 1.184 | -3.507 | 0.764 | -0.603 | -0.003 | -0.023 | -0.023 | 1.308 | 0.38 | 0.16 | 0.72 | 0.43 | 0.61 | 0.38 |
| 2 | 4718647 | rs13392225 | 1.44 | -3.493 | 0.688 | -0.556 | -0.016 | -0.034 | -0.034 | 5.367 | 0.06 | 0 | 0.04 | 0.23 | 0.07 | 0.24 |
| 2 | 4726231 | rs285545 | 0.287 | -3.541 | 1.013 | -0.605 | 0.002 | -0.002 | -0.002 | 0 | 0.56 | 0.61 | 0.82 | 0.53 | 0.71 | 0.51 |
| 2 | 4730927 | rs4849968 | 1.147 | -3.585 | 1.226 | -0.649 | 0.009 | -0.001 | -0.001 | 0 | 0.39 | 0.3 | 0.7 | 0.42 | 0.6 | 0.38 |
| 2 | 4730954 | rs10176368 | 0.011 | -3.272 | 1.243 | -0.581 | -0.02 | -0.022 | -0.022 | 0 | 0.06 | 0.2 | 0.01 | 0 | 0.01 | 0.01 |
| 2 | 4731202 | rs17019650 | 0.049 | -3.276 | 1.205 | -0.578 | 0.018 | -0.027 | -0.027 | 0 | 0.13 | 0.05 | 0.05 | 0.23 | 0.09 | 0.23 |
| 2 | 4734663 | rs7608994 | 0.232 | -3.277 | 1.043 | -0.609 | 0.014 | 0.02 | 0.02 | 0 | 0.15 | 0.09 | 0.1 | 0.11 | 0.1 | 0.14 |
| 2 | 4734920 | rs10495507 | -1.235 | -3.132 | 1.352 | -0.285 | -0.06 | 0.008 | 0.008 | 0 | 0.2 | 0.33 | 0.07 | 0.23 | 0.1 | 0.25 |
| 2 | 4744787 | rs1446145 | -1.212 | -2.742 | 1.625 | -0.717 | -0.018 | -0.021 | -0.021 | 0 | 0.08 | 0.31 | 0.02 | 0 | 0.01 | 0.01 |
| 2 | 4747638 | rs10202915 | -1.356 | -2.726 | 1.275 | -0.708 | -0.013 | -0.018 | -0.018 | 0 | 0.09 | 0.42 | 0.03 | 0 | 0.01 | 0.01 |
| 2 | 4753673 | rs1374128 | -0.116 | -2.268 | -0.274 | -0.465 | -0.01 | -0.003 | -0.003 | 0 | 0.66 | 0.67 | 0.86 | 0.77 | 0.77 | 0.73 |
| 2 | 4753961 | rs1374127 | -0.151 | -1.883 | -0.148 | -0.294 | -0.001 | 0 | 0 | 0 | 0.67 | 0.7 | 0.84 | 0.77 | 0.74 | 0.71 |
| 2 | 4756013 | rs10165056 | 0.208 | -1.435 | -0.089 | 0.12 | -0.007 | -0.002 | -0.002 | 0 | 0.56 | 0.44 | 0.82 | 0.67 | 0.72 | 0.67 |
| 2 | 4767683 | rs13419691 | -1.016 | -0.886 | -0.26 | 0.073 | -0.033 | -0.029 | -0.029 | 0 | 0.22 | 0.26 | 0.06 | 0.09 | 0.09 | 0.08 |
| 2 | 4767895 | rs11695408 | -1.288 | -0.363 | -0.318 | 0.032 | -0.004 | -0.009 | -0.009 | 0.042 | 0.19 | 0.21 | 0.13 | 0.11 | 0.16 | 0.09 |
| 2 | 4770187 | rs1838770 | -0.538 | 0.063 | -0.364 | 0.173 | -0.007 | -0.004 | -0.004 | 0 | 0.07 | 0.03 | 0.14 | 0.1 | 0.1 | 0.08 |
| 2 | 4774401 | rs2044429 | 0.537 | -0.034 | 0.127 | 0.559 | -0.004 | 0.033 | 0.033 | 0 | 0.91 | 0.87 | 0.96 | 0.82 | 0.93 | 0.82 |
| 2 | 4776763 | rs12467016 | 0.339 | -0.062 | 0.057 | 0.376 | -0.006 | 0.011 | 0.011 | 4.753 | 0.29 | 0.2 | 0.45 | 0.27 | 0.35 | 0.26 |
| 2 | 4778512 | rs11123574 | 1.886 | -0.649 | 0.92 | 0.455 | 0.011 | -0.001 | -0.001 | 0 | 0.71 | 0.23 | 0.76 | 0.7 | 0.8 | 0.7 |
| 2 | 4799191 | rs957504 | -0.49 | -0.474 | 1.172 | 0.577 | -0.006 | 0.065 | 0.065 | 0 | 0.08 | 0.14 | 0.03 | 0.22 | 0.06 | 0.21 |
| 2 | 4806334 | rs880767 | -0.494 | -0.468 | 1.077 | 0.58 | -0.004 | 0.022 | 0.022 | 0 | 0.08 | 0.13 | 0.03 | 0.22 | 0.06 | 0.21 |
| 2 | 4807546 | rs13405551 | -1.682 | -0.483 | 1.198 | 0.543 | 0.015 | 0.047 | 0.047 | 0 | 0.2 | 0.33 | 0.21 | 0.29 | 0.19 | 0.3 |
| 2 | 4808027 | rs825937 | -1.654 | -0.507 | 1.216 | 0.497 | -0.002 | -0.047 | -0.047 | 0 | 0.2 | 0.58 | 0.24 | 0.08 | 0.16 | 0.1 |
| 2 | 4818625 | rs825914 | -1.869 | -0.153 | 1.243 | 0.663 | -0.001 | -0.001 | -0.001 | 0 | 0.25 | 0.7 | 0.31 | 0.29 | 0.22 | 0.3 |
| 2 | 4819159 | rs17019820 | -1.03 | 0.141 | 1.213 | 0.349 | 0.03 | -0.015 | -0.015 | 0 | 0.13 | 0.23 | 0.24 | 0.08 | 0.15 | 0.08 |
| 2 | 4821442 | rs825923 | -0.694 | -0.433 | 0.355 | 0.821 | -0.032 | 0.039 | 0.039 | 2.776 | 0.24 | 0.41 | 0.33 | 0.68 | 0.15 | 0.47 |
| 2 | 4821951 | rs7604940 | -0.091 | -0.427 | -0.269 | -0.725 | -0.003 | -0.096 | -0.096 | 0.149 | 0.41 | 0.53 | 0.22 | 0 | 0.38 | 0.11 |
| 2 | 4822438 | rs13395076 | 0.212 | -0.785 | -0.943 | -0.36 | 0.035 | -0.013 | -0.013 | 0 | 0.16 | 0.04 | 0.14 | 0.2 | 0.12 | 0.3 |
| 2 | 4822972 | rs1446137 | -0.831 | 0.403 | -0.47 | -0.833 | 0.006 | -0.007 | -0.007 | 0 | 0.34 | 0.29 | 0.31 | 0.34 | 0.42 | 0.43 |
| 2 | 4823625 | rs9646941 | 0.184 | 0.205 | -0.289 | -1.055 | -0.008 | 0.003 | 0.003 | 0 | 0.22 | 0.17 | 0.18 | 0.14 | 0.33 | 0.18 |
| 2 | 4824622 | rs12616546 | -1.152 | -0.365 | -0.537 | -1.036 | 0.001 | -0.003 | -0.003 | 0 | 0.65 | 0.6 | 0.65 | 0.49 | 0.67 | 0.73 |
| 2 | 4825362 | rs13431987 | 1.834 | -0.361 | -0.64 | -1.033 | 0.018 | -0.028 | -0.028 | 0 | 0.16 | 0.04 | 0.14 | 0.25 | 0.15 | 0.29 |
| 2 | 4825542 | rs17581422 | 1.341 | -0.492 | -0.406 | -1.202 | -0.002 | 0.016 | 0.016 | 0 | 0.24 | 0.14 | 0.31 | 0.15 | 0.19 | 0.26 |
| 2 | 4827559 | rs6710557 | -0.069 | -0.517 | -0.333 | -1.196 | 0.006 | 0.065 | 0.065 | 0 | 0.43 | 0.28 | 0.47 | 0.25 | 0.5 | 0.41 |
| 2 | 4830190 | rs2165183 | -0.329 | 0.083 | -0.259 | -0.758 | -0.003 | 0.007 | 0.007 | 0 | 0.45 | 0.34 | 0.36 | 0.39 | 0.47 | 0.49 |
| 2 | 4830542 | rs10185569 | 0.39 | 0.078 | -0.257 | -0.777 | 0.002 | -0.001 | -0.001 | 0 | 0.36 | 0.31 | 0.32 | 0.39 | 0.42 | 0.44 |
| 2 | 4832338 | rs12711955 | -2.009 | -0.326 | -0.292 | -0.781 | 0.028 | 0.025 | 0.025 | 0 | 0.87 | 0.94 | 0.92 | 0.96 | 0.94 | 0.9 |
| 2 | 4833848 | rs1446147 | -0.765 | -0.273 | -0.023 | -0.77 | 0.002 | -0.011 | -0.011 | 0 | 0.66 | 0.65 | 0.66 | 0.55 | 0.64 | 0.74 |
| 2 | 4838853 | rs13425189 | 1.073 | -0.22 | -0.118 | -0.832 | 0.002 | -0.004 | -0.004 | 0 | 0.15 | 0.14 | 0.15 | 0.05 | 0.26 | 0.17 |
| 2 | 4840214 | rs7570834 | 1.845 | -0.286 | -0.133 | -0.777 | -0.007 | -0.003 | -0.003 | 0.056 | 0.2 | 0.31 | 0.29 | 0.16 | 0.16 | 0.24 |
| 2 | 4854428 | rs10207322 | 0.509 | -0.374 | -0.275 | -0.784 | 0.019 | 0.001 | 0.001 | 0 | 0.32 | 0.19 | 0.23 | 0.34 | 0.23 | 0.35 |
| 2 | 4865185 | rs1866608 | 0.554 | -0.458 | -0.195 | -0.783 | 0.027 | 0.003 | 0.003 | 0 | 0.32 | 0.15 | 0.27 | 0.33 | 0.23 | 0.35 |
| 2 | 4866246 | rs11691826 | 2.014 | -0.339 | -0.144 | -0.78 | -0.001 | -0.001 | -0.001 | 0 | 0.19 | 0.22 | 0.3 | 0.17 | 0.18 | 0.25 |
| 2 | 4869564 | rs7569982 | 0.072 | 0.225 | -0.206 | -0.781 | 0.021 | -0.053 | -0.053 | 0 | 0.19 | 0.29 | 0.14 | 0.05 | 0.2 | 0.14 |
| 2 | 4870574 | rs12619133 | 1.591 | 0.192 | -0.415 | -0.703 | 0.04 | -0.052 | -0.052 | 0 | 0.23 | 0.04 | 0.22 | 0.32 | 0.18 | 0.3 |
| 2 | 4874946 | rs13427718 | 1.497 | 0.377 | -0.28 | -0.803 | 0.029 | -0.02 | -0.02 | 0.296 | 0.11 | 0.15 | 0.11 | 0.05 | 0.18 | 0.13 |
| 2 | 4882104 | rs1837447 | -1.111 | -0.072 | -0.677 | -0.849 | -0.017 | 0.056 | 0.056 | 0 | 0.8 | 0.66 | 0.82 | 0.58 | 0.87 | 0.78 |
| 2 | 4882122 | rs1821990 | 0.072 | -0.04 | -0.674 | -0.838 | 0.005 | -0.013 | -0.013 | 0 | 0.37 | 0.57 | 0.43 | 0.23 | 0.39 | 0.4 |

| | | | | | | | | | | | | | | | | |
|---|---|---|---|---|---|---|---|---|---|---|---|---|---|---|---|---|
| 2 | 4887124 | rs2028368 | -0.556 | -0.02 | -0.732 | -0.814 | -0.004 | -0.007 | -0.007 | 0 | 0.65 | 0.63 | 0.69 | 0.58 | 0.6 | 0.72 |
| 2 | 4893566 | rs12621974 | 1.535 | -0.114 | -0.708 | -0.765 | 0.059 | -0.05 | -0.05 | 0.021 | 0.26 | 0.07 | 0.25 | 0.34 | 0.2 | 0.32 |
| 2 | 4893723 | rs6542638 | 0.181 | 0.073 | -0.545 | -0.868 | 0.001 | 0.002 | 0.002 | 0 | 0.57 | 0.46 | 0.66 | 0.56 | 0.56 | 0.69 |
| 2 | 4896970 | rs1465875 | -0.108 | 0.082 | -0.622 | -0.916 | -0.001 | 0 | 0 | 0 | 0.58 | 0.54 | 0.61 | 0.51 | 0.42 | 0.6 |
| 2 | 4897117 | rs1866611 | 0.45 | 0.086 | -0.589 | -0.884 | 0.01 | -0.001 | -0.001 | 0 | 0.48 | 0.53 | 0.57 | 0.51 | 0.37 | 0.55 |
| 2 | 4900640 | rs10176555 | 0.694 | 0.081 | -0.586 | -0.883 | 0 | -0.001 | -0.001 | 0.033 | 0.47 | 0.52 | 0.57 | 0.51 | 0.37 | 0.55 |
| 2 | 4901791 | rs13383006 | 1.386 | 0.165 | -0.509 | -0.872 | 0.014 | -0.013 | -0.013 | 0 | 0.11 | 0.13 | 0.1 | 0.05 | 0.18 | 0.12 |
| 2 | 4907067 | rs17020172 | 1.583 | -0.126 | -0.468 | -0.781 | 0.004 | -0.001 | -0.001 | 0.853 | 0.21 | 0.44 | 0.32 | 0.17 | 0.18 | 0.24 |
| 2 | 4916939 | rs10495513 | 1.498 | -0.106 | -0.628 | -0.771 | 0 | 0 | 0 | 0 | 0.22 | 0.23 | 0.32 | 0.17 | 0.2 | 0.27 |
| 2 | 4917761 | rs1439829 | -0.701 | -0.241 | -0.635 | -0.839 | 0.013 | 0.033 | 0.033 | 0 | 0.68 | 0.45 | 0.7 | 0.56 | 0.59 | 0.72 |
| 2 | 4918229 | rs1439828 | 0.229 | -0.302 | -0.669 | -0.869 | -0.008 | 0.004 | 0.004 | 0 | 0.19 | 0.14 | 0.12 | 0.05 | 0.2 | 0.14 |
| 2 | 4924044 | rs7602114 | 0.552 | -0.366 | -0.713 | -1.013 | 0.043 | -0.011 | -0.011 | 0.036 | 0.47 | 0.32 | 0.57 | 0.51 | 0.39 | 0.58 |
| 2 | 4925630 | rs10177436 | 0.31 | -0.38 | -0.714 | -1.015 | 0.048 | 0 | 0 | 2.151 | 0.5 | 0.36 | 0.57 | 0.51 | 0.39 | 0.58 |
| 2 | 4930330 | rs6542644 | 0.133 | -0.328 | -0.604 | -1.991 | -0.001 | -0.006 | -0.006 | 0.031 | 0.42 | 0.57 | 0.47 | 0.17 | 0.48 | 0.34 |
| 2 | 4933355 | rs1371479 | -0.402 | -0.196 | -0.503 | -1.909 | -0.005 | -0.007 | -0.007 | 0 | 0.65 | 0.52 | 0.53 | 0.7 | 0.69 | 0.61 |
| 2 | 4938567 | rs1371488 | -1.303 | 0.041 | -0.427 | -1.671 | -0.013 | 0.022 | 0.022 | 0 | 0.84 | 0.7 | 0.83 | 0.79 | 0.88 | 0.82 |
| 2 | 4943820 | rs13016616 | 1.774 | 0.354 | -0.716 | -2.331 | -0.047 | 0.055 | 0.055 | 0 | 0.16 | 0.05 | 0.13 | 0 | 0.26 | 0.07 |
| 2 | 4946245 | rs6542647 | 0.471 | 0.254 | -0.387 | -2.191 | 0.049 | -0.049 | -0.049 | 0 | 0.14 | 0.28 | 0.11 | 0.05 | 0.18 | 0.14 |
| 2 | 4948300 | rs1345001 | 1.211 | -0.324 | -1.132 | -2.343 | 0.048 | -0.055 | -0.055 | 0 | 0.3 | 0.08 | 0.35 | 0.65 | 0.23 | 0.33 |
| 2 | 4948472 | rs1345000 | 1.316 | -0.282 | -1.111 | -2.319 | 0.042 | -0.057 | -0.057 | 0.042 | 0.29 | 0.07 | 0.35 | 0.65 | 0.23 | 0.33 |
| 2 | 4948693 | rs1344999 | -0.215 | -0.268 | -0.938 | -2.339 | 0.086 | 0.089 | 0.089 | 0 | 0.5 | 0.82 | 0.55 | 0.34 | 0.63 | 0.54 |
| 2 | 4949915 | rs6542649 | -0.216 | -0.264 | -1.003 | -2.337 | 0.086 | 0.089 | 0.089 | 0 | 0.5 | 0.81 | 0.55 | 0.34 | 0.63 | 0.54 |
| 2 | 4951850 | rs7577926 | 1.246 | 0.076 | -0.521 | -2.18 | 0.017 | 0.009 | 0.009 | 0.266 | 0.16 | 0.31 | 0.25 | 0.15 | 0.17 | 0.22 |
| 2 | 4952078 | rs4355130 | -0.19 | 0.105 | -0.822 | -2.087 | -0.01 | -0.068 | -0.068 | 0.622 | 0.77 | 0.71 | 0.87 | 0.99 | 0.85 | 0.86 |
| 2 | 4952545 | rs13393948 | 0.383 | 0.646 | -1.196 | -1.619 | -0.018 | 0.018 | 0.018 | 0 | 0.24 | 0.14 | 0.15 | 0 | 0.28 | 0.12 |
| 2 | 4957189 | rs4144195 | 0.616 | 0.283 | -1.357 | -1.523 | -0.002 | 0.015 | 0.015 | 0 | 0.28 | 0.2 | 0.42 | 0.77 | 0.31 | 0.46 |
| 2 | 4957412 | rs6712088 | -0.077 | 0.026 | -1.455 | -1.798 | 0.01 | 0.043 | 0.043 | 0 | 0.37 | 0.28 | 0.45 | 0.77 | 0.33 | 0.47 |
| 2 | 4958832 | rs9989821 | 1.788 | 0.259 | -1.529 | -2.008 | -0.026 | 0.038 | 0.038 | 0 | 0.15 | 0.06 | 0.12 | 0 | 0.22 | 0.1 |
| 2 | 4958988 | rs2620437 | -0.866 | -0.115 | -1.414 | -2.25 | 0.008 | 0.023 | 0.023 | 0 | 0.64 | 0.71 | 0.54 | 0.22 | 0.63 | 0.46 |
| 2 | 4960266 | rs7588463 | 1.545 | 0.043 | -1.477 | -2.47 | -0.028 | 0.068 | 0.068 | 0 | 0.2 | 0.09 | 0.15 | 0.06 | 0.22 | 0.14 |
| 2 | 4960962 | rs11123600 | 0.38 | 0.047 | -1.162 | -2.445 | 0.037 | -0.067 | -0.067 | 0 | 0.32 | 0.49 | 0.36 | 0.16 | 0.39 | 0.3 |
| 2 | 4964038 | rs722148 | 1.41 | 0.059 | -1.171 | -2.256 | 0.001 | 0.059 | 0.059 | 0 | 0.05 | 0.12 | 0.07 | 0.12 | 0.07 | 0.1 |
| 2 | 4964465 | rs721087 | 0.115 | 0.051 | -1.196 | -2.373 | 0.001 | 0.036 | 0.036 | 0 | 0.5 | 0.59 | 0.61 | 0.82 | 0.42 | 0.56 |
| 2 | 4965676 | rs736687 | -0.813 | 0.119 | -1.307 | -2.31 | 0 | -0.039 | -0.039 | 1.063 | 0.77 | 0.8 | 0.83 | 0.99 | 0.72 | 0.83 |
| 2 | 4967202 | rs2966429 | -0.152 | 0.345 | -1.236 | -2.173 | 0.006 | -0.031 | -0.031 | 0 | 0.66 | 0.73 | 0.8 | 0.99 | 0.69 | 0.81 |
| 2 | 4973779 | rs13016249 | 0.158 | 0.789 | -1.029 | -1.594 | -0.023 | 0.017 | 0.017 | 0 | 0.32 | 0.25 | 0.28 | 0.05 | 0.41 | 0.23 |
| 2 | 4974397 | rs12463956 | -0.258 | 1.096 | -0.926 | -1.138 | -0.023 | 0.026 | 0.026 | 0 | 0.32 | 0.26 | 0.38 | 0.13 | 0.46 | 0.33 |
| 2 | 4974780 | rs12464103 | 0.142 | 1.195 | -0.744 | -0.805 | 0.015 | 0.007 | 0.007 | 0.438 | 0.12 | 0.2 | 0.19 | 0.12 | 0.14 | 0.15 |
| 2 | 4975953 | rs6422705 | 0.038 | 0.894 | -1.075 | -0.446 | -0.01 | 0.012 | 0.012 | 0 | 0.91 | 0.82 | 0.92 | 0.86 | 0.92 | 0.88 |
| 2 | 4976727 | rs7591716 | 0.252 | 1.203 | -1.321 | -0.615 | -0.02 | 0.041 | 0.041 | 0.473 | 0.27 | 0.17 | 0.26 | 0.13 | 0.33 | 0.19 |
| 2 | 4981785 | rs12621874 | 0.384 | 1.237 | -0.925 | -0.1 | -0.003 | 0.006 | 0.006 | 0.414 | 0.68 | 0.73 | 0.63 | 0.75 | 0.66 | 0.72 |
| 2 | 4983682 | rs11123605 | 0.094 | 1.379 | -1.053 | 0.148 | -0.006 | -0.006 | -0.006 | 0.007 | 0.18 | 0.22 | 0.33 | 0.63 | 0.2 | 0.39 |
| 2 | 4985756 | rs6422704 | -0.852 | 1.416 | -1.186 | 0.292 | -0.004 | 0 | 0 | 0 | 0.85 | 0.89 | 0.84 | 0.82 | 0.83 | 0.83 |
| 2 | 4985980 | rs17020354 | -0.051 | 1.18 | -0.904 | 0.885 | -0.012 | 0.009 | 0.009 | 3.907 | 0.15 | 0.04 | 0.2 | 0.08 | 0.17 | 0.12 |
| 2 | 4986983 | rs2315629 | -1.074 | 1.174 | -0.702 | 0.792 | 0.009 | -0.012 | -0.012 | 0.219 | 0.34 | 0.44 | 0.54 | 0.71 | 0.37 | 0.5 |
| 2 | 4989067 | rs12052647 | -0.946 | 0.998 | -1.21 | 1.682 | -0.06 | -0.158 | -0.158 | 0 | 0.18 | 0.01 | 0.18 | 0.52 | 0.18 | 0.43 |
| 2 | 4992822 | rs17020407 | -1.377 | 0.93 | -0.463 | 2.138 | 0.014 | 0.178 | 0.178 | 0 | 0.07 | 0.13 | 0.14 | 0.49 | 0.1 | 0.32 |
| 2 | 5173840 | rs8179663 | -1.224 | -0.391 | 0.819 | 1.86 | -0.067 | 0.106 | 0.106 | 0 | 0.24 | 0.43 | 0.23 | 0.53 | 0.1 | 0.43 |
| 2 | 5173889 | rs8179639 | -1.224 | -0.391 | 0.819 | 1.86 | -0.067 | 0.106 | 0.106 | 0 | 0.24 | 0.42 | 0.23 | 0.53 | 0.1 | 0.43 |
| 2 | 5180567 | rs10184695 | -1.075 | -0.439 | 1.028 | 2.212 | -0.034 | 0.009 | 0.009 | 0 | 0.21 | 0.29 | 0.17 | 0.33 | 0.07 | 0.39 |
| 2 | 5181373 | rs4552162 | -1.075 | -0.439 | 1.03 | 2.212 | -0.032 | 0.008 | 0.008 | 0 | 0.21 | 0.29 | 0.17 | 0.33 | 0.08 | 0.39 |
| 2 | 5181658 | rs13393547 | -1.46 | -0.437 | 1.069 | 2.222 | -0.063 | 0.102 | 0.102 | 0 | 0.23 | 0.42 | 0.23 | 0.53 | 0.1 | 0.43 |
| 2 | 5196712 | rs4452110 | -0.894 | -0.349 | 0.418 | 1.956 | 0.019 | 0.039 | 0.039 | 0 | 0.12 | 0.01 | 0.07 | 0.04 | 0.06 | 0.15 |
| 2 | 5196902 | rs11690345 | 1.546 | -0.023 | 0.523 | 1.561 | 0.037 | -0.042 | -0.042 | 1.911 | 0.62 | 0.18 | 0.49 | 0.47 | 0.72 | 0.43 |
| 2 | 5215885 | rs12613788 | 1.128 | 0.067 | -0.787 | 0.563 | -0.107 | -0.091 | -0.091 | 4.719 | 0.43 | 0.02 | 0.25 | 0.42 | 0.4 | 0.19 |
| 2 | 5217992 | rs6749133 | -0.711 | -0.085 | 0.421 | 0.555 | 0.03 | 0.047 | 0.047 | 0.818 | 0.18 | 0.6 | 0.37 | 0.21 | 0.22 | 0.19 |
| 2 | 5219219 | rs16863310 | -0.01 | -0.187 | 0.788 | 0.449 | -0.033 | -0.046 | -0.046 | 0 | 0.11 | 0.24 | 0.15 | 0.03 | 0.04 | 0.16 |
| 2 | 5224268 | rs3943477 | -1.287 | -0.125 | 1.187 | 0.468 | 0.006 | 0.122 | 0.122 | 0.175 | 0.24 | 0.48 | 0.44 | 0.53 | 0.27 | 0.42 |
| 2 | 5226543 | rs1154756 | 0.637 | -0.092 | 1.089 | 0.089 | -0.009 | 0.008 | 0.008 | 1.454 | 0.22 | 0.15 | 0.16 | 0.01 | 0.29 | 0.24 |
| 2 | 5226989 | rs1453780 | -1.033 | -0.143 | 1.296 | 0.269 | 0.007 | 0.035 | 0.035 | 0 | 0.19 | 0.28 | 0.37 | 0.33 | 0.24 | 0.38 |
| 2 | 5232834 | rs1453778 | -0.355 | -0.043 | 1.084 | -0.023 | 0.057 | -0.081 | -0.081 | 0.199 | 0.14 | 0.36 | 0.29 | 0 | 0.2 | 0.18 |
| 2 | 5238812 | rs731436 | 0.681 | -0.174 | -0.081 | -0.597 | 0.014 | 0.014 | 0.014 | 0.274 | 0.28 | 0.19 | 0.25 | 0.03 | 0.19 | 0.17 |
| 2 | 5239929 | rs13015498 | 0.261 | 0.187 | -0.063 | -0.245 | -0.012 | -0.011 | -0.011 | 0 | 0.25 | 0.2 | 0.23 | 0.36 | 0.35 | 0.45 |
| 2 | 5240144 | rs13403240 | -0.294 | 0.541 | -0.45 | -0.435 | -0.002 | -0.003 | -0.003 | 1.384 | 0.18 | 0.17 | 0.38 | 0.36 | 0.28 | 0.36 |
| 2 | 5241229 | rs10181839 | -0.435 | 0.432 | -0.466 | -0.41 | -0.062 | -0.008 | -0.008 | 0 | 0.34 | 0.18 | 0.5 | 0.38 | 0.53 | 0.58 |
| 2 | 5249937 | rs4668780 | 1.327 | 0.137 | -0.659 | -0.358 | -0.039 | -0.103 | -0.103 | 4.196 | 0.29 | 0.07 | 0.16 | 0.41 | 0.27 | 0.21 |
| 2 | 5256677 | rs6732278 | 1.02 | 0.048 | -0.227 | -0.425 | -0.031 | -0.132 | -0.132 | 0 | 0.29 | 0.42 | 0.25 | 0.03 | 0.18 | 0.16 |
| 2 | 5256734 | rs11685553 | 0.96 | 0.062 | -0.248 | -0.343 | 0.003 | -0.015 | -0.015 | 0 | 0.33 | 0.26 | 0.17 | 0.41 | 0.28 | 0.21 |
| 2 | 5257356 | rs1869410 | 0.936 | 0.543 | -1.012 | -0.893 | -0.089 | 0.094 | 0.094 | 4.553 | 0.18 | 0.04 | 0.15 | 0.01 | 0.29 | 0.23 |

| | | | | | | | | | | | | | | | | |
|---|---|---|---|---|---|---|---|---|---|---|---|---|---|---|---|---|
| 2 | 5260481 | rs11889165 | 0.452 | 0.455 | -0.625 | -0.763 | -0.023 | -0.103 | -0.103 | 0 | 0.31 | 0.38 | 0.22 | 0.03 | 0.18 | 0.16 |
| 2 | 5261410 | rs6432420 | 0.214 | 0.81 | -0.687 | -0.916 | 0.002 | -0.108 | -0.108 | 0 | 0.36 | 0.39 | 0.29 | 0.02 | 0.34 | 0.26 |
| 2 | 5266006 | rs7572452 | 0.808 | 0.829 | -1.56 | -0.293 | -0.042 | -0.136 | -0.136 | 0 | 0.28 | 0.05 | 0.15 | 0.45 | 0.29 | 0.23 |
| 2 | 5266939 | rs17330726 | 0.061 | 0.796 | -1.261 | -0.172 | -0.006 | 0.033 | 0.033 | 1.47 | 0.45 | 0.3 | 0.41 | 0.38 | 0.43 | 0.55 |
| 2 | 5269417 | rs1453798 | 0.439 | 1.087 | -1.135 | -0.836 | -0.003 | 0.072 | 0.072 | 0 | 0.55 | 0.33 | 0.46 | 0.82 | 0.45 | 0.59 |
| 2 | 5271928 | rs4670034 | 0.773 | 1.346 | -0.384 | -0.658 | 0.061 | -0.043 | -0.043 | 0.306 | 0.79 | 0.55 | 0.8 | 0.88 | 0.76 | 0.75 |
| 2 | 5283993 | rs7577322 | -0.006 | 0.081 | -0.764 | -1.291 | -0.026 | -0.067 | -0.067 | 0 | 0.61 | 0.55 | 0.56 | 0.15 | 0.69 | 0.58 |
| 2 | 5286577 | rs4407219 | 0.169 | 0.198 | -0.58 | -1.443 | 0.01 | 0.009 | 0.009 | 0 | 0.57 | 0.62 | 0.52 | 0.15 | 0.63 | 0.54 |
| 2 | 5287827 | rs9679610 | 0.518 | 0.463 | -0.992 | -1.888 | 0 | -0.029 | -0.029 | 1.181 | 0.43 | 0.56 | 0.27 | 0.14 | 0.39 | 0.38 |
| 2 | 5288644 | rs4668488 | 0.011 | 0.597 | -1.234 | -1.746 | 0.015 | 0.033 | 0.033 | 0 | 0.34 | 0.26 | 0.4 | 0.81 | 0.32 | 0.39 |
| 2 | 5292766 | rs17272087 | 2.144 | 0.436 | -1.696 | -2.02 | -0.066 | 0.079 | 0.079 | 0 | 0.23 | 0.01 | 0.16 | 0.1 | 0.25 | 0.21 |
| 2 | 5293171 | rs17331611 | 0.992 | 0.333 | -1.173 | -1.601 | -0.013 | 0.032 | 0.032 | 0.312 | 0.06 | 0.01 | 0.05 | 0 | 0.06 | 0.05 |
| 2 | 5293622 | rs6740049 | 1.085 | 0.167 | -1.242 | -0.958 | 0.011 | -0.047 | -0.047 | 0 | 0.17 | 0.12 | 0.16 | 0.42 | 0.17 | 0.14 |
| 2 | 5293922 | rs16863430 | 0.037 | 0.503 | -0.798 | -0.511 | -0.006 | 0.031 | 0.031 | 0 | 0.17 | 0.16 | 0.23 | 0.39 | 0.15 | 0.25 |
| 2 | 5294800 | rs7593216 | 0.058 | 0.544 | -1.022 | -0.587 | 0.006 | -0.023 | -0.023 | 0.139 | 0.08 | 0.13 | 0.07 | 0.01 | 0.1 | 0.08 |
| 2 | 5300664 | rs10929427 | -0.071 | 0.333 | -1.2 | -0.3 | 0.003 | -0.006 | -0.006 | 0 | 0.17 | 0.11 | 0.23 | 0.37 | 0.15 | 0.25 |
| 2 | 5304015 | rs13421030 | 0.518 | 0.281 | -1.13 | -1.237 | 0.013 | 0.042 | 0.042 | 1.223 | 0.28 | 0.14 | 0.21 | 0.16 | 0.25 | 0.3 |
| 2 | 5325906 | rs10929445 | 0.564 | 0.918 | -0.445 | -1.419 | 0.001 | -0.061 | -0.061 | 0.12 | 0.65 | 0.35 | 0.59 | 0.95 | 0.61 | 0.68 |
| 2 | 5336076 | rs7604248 | -0.352 | 0.863 | -0.362 | -1.427 | -0.022 | -0.003 | -0.003 | 0.329 | 0.28 | 0.28 | 0.35 | 0.04 | 0.39 | 0.25 |
| 2 | 5340502 | rs6718709 | 0.503 | 0.883 | -0.866 | -1.369 | -0.036 | 0.063 | 0.063 | 0 | 0.62 | 0.25 | 0.52 | 0.95 | 0.55 | 0.61 |
| 2 | 5345278 | rs13399059 | -0.8 | 0.91 | -0.64 | -0.622 | -0.002 | 0 | 0 | 0 | 0.14 | 0.12 | 0.17 | 0.07 | 0.14 | 0.12 |
| 2 | 5345732 | rs4462764 | -0.377 | 0.635 | -0.522 | 0.122 | -0.007 | 0.015 | 0.015 | 0.293 | 0.38 | 0.53 | 0.4 | 0.42 | 0.34 | 0.29 |
| 2 | 5349911 | rs7425501 | -0.271 | 0.351 | -0.243 | -0.331 | -0.01 | -0.013 | -0.013 | 0 | 0.32 | 0.42 | 0.39 | 0.7 | 0.29 | 0.53 |
| 2 | 5357276 | rs1453791 | 0.455 | 0.308 | -0.605 | -0.491 | 0.014 | -0.049 | -0.049 | 0 | 0.17 | 0.03 | 0.12 | 0.32 | 0.14 | 0.17 |
| 2 | 5361397 | rs10181393 | -0.619 | 0.818 | 0.716 | -0.689 | -0.024 | 0.039 | 0.039 | 0 | 0.43 | 0.27 | 0.42 | 0.65 | 0.52 | 0.63 |
| 2 | 5371547 | rs17273872 | -0.138 | 0.923 | 0.95 | -0.771 | -0.012 | -0.003 | -0.003 | 0 | 0.32 | 0.24 | 0.36 | 0.65 | 0.38 | 0.52 |
| 2 | 5372402 | rs13013629 | 0.907 | 0.907 | 1.115 | -0.792 | -0.041 | 0.057 | 0.057 | 0 | 0.1 | 0.01 | 0.05 | 0 | 0.14 | 0.06 |
| 2 | 5376583 | rs2123733 | 0.589 | 0.48 | 1.443 | -0.836 | -0.004 | 0.061 | 0.061 | 1.778 | 0.58 | 0.13 | 0.6 | 0.31 | 0.61 | 0.42 |
| 2 | 5389987 | rs16863472 | 0.625 | 0.46 | 1.333 | -0.78 | -0.061 | -0.094 | -0.094 | 0.485 | 0.22 | 0.05 | 0.29 | 0.6 | 0.28 | 0.41 |
| 2 | 5395873 | rs6756390 | -0.844 | 0.417 | 2.249 | -0.818 | 0.006 | -0.005 | -0.005 | 0 | 0.36 | 0.48 | 0.39 | 0.64 | 0.39 | 0.56 |
| 2 | 5407073 | rs10432395 | -0.597 | 0.508 | 2.275 | -0.735 | 0.003 | 0.001 | 0.001 | 0 | 0.33 | 0.33 | 0.37 | 0.67 | 0.38 | 0.5 |
| 2 | 5411779 | rs4669198 | 0.781 | 0.501 | 1.984 | -0.705 | -0.003 | -0.016 | -0.016 | 0.829 | 0.07 | 0.09 | 0.02 | 0.04 | 0.05 | 0.12 |
| 2 | 5418756 | rs2198493 | 0.259 | 0.514 | 2.101 | -0.685 | 0.012 | 0.004 | 0.004 | 0 | 0.67 | 0.61 | 0.62 | 0.33 | 0.62 | 0.5 |
| 2 | 5422373 | rs4349322 | -0.614 | 0.524 | 2.094 | -0.68 | 0.012 | 0.002 | 0.002 | 0 | 0.35 | 0.44 | 0.38 | 0.67 | 0.38 | 0.5 |
| 2 | 5426351 | rs10929492 | 0.316 | 0.561 | 1.876 | -0.623 | 0.014 | 0.009 | 0.009 | 0 | 0.64 | 0.49 | 0.62 | 0.36 | 0.62 | 0.49 |
| 2 | 5427628 | rs7592529 | -0.479 | 0.579 | 1.606 | -0.595 | -0.005 | -0.004 | -0.004 | 0 | 0.33 | 0.29 | 0.37 | 0.64 | 0.38 | 0.51 |
| 2 | 5428308 | rs13388472 | -0.093 | 0.596 | 1.449 | -0.6 | -0.009 | -0.017 | -0.017 | 0 | 0.3 | 0.2 | 0.32 | 0.64 | 0.33 | 0.48 |
| 2 | 5434679 | rs7561631 | 0.308 | 0.545 | 1.206 | -0.569 | 0.009 | 0.008 | 0.008 | 0 | 0.66 | 0.6 | 0.63 | 0.36 | 0.62 | 0.5 |
| 2 | 5436975 | rs13010129 | 0.733 | 0.547 | 0.679 | -0.878 | -0.001 | 0.004 | 0.004 | 0 | 0.54 | 0.37 | 0.61 | 0.3 | 0.56 | 0.49 |
| 2 | 5437543 | rs13004938 | 0.369 | 0.781 | 0.012 | -0.872 | -0.028 | 0.028 | 0.028 | 0 | 0.14 | 0.1 | 0.11 | 0 | 0.21 | 0.09 |
| 2 | 5439411 | rs4669209 | -0.085 | 0.561 | 0.366 | -0.65 | 0.003 | -0.003 | -0.003 | 0 | 0.27 | 0.25 | 0.27 | 0.7 | 0.24 | 0.43 |
| 2 | 5449582 | rs10803691 | 0.217 | 0.429 | -0.138 | -0.783 | -0.043 | -0.048 | -0.048 | 0 | 0.24 | 0.01 | 0.29 | 0.71 | 0.24 | 0.43 |
| 2 | 5451280 | rs6754266 | 0.798 | 0.339 | 0.341 | -1.315 | -0.048 | 0.144 | 0.144 | 0 | 0.47 | 0.16 | 0.52 | 0.29 | 0.49 | 0.47 |
| 2 | 5455479 | rs1404253 | 0.702 | 0.309 | 0.822 | -1.321 | -0.01 | 0.095 | 0.095 | 0 | 0.48 | 0.29 | 0.52 | 0.29 | 0.49 | 0.48 |
| 2 | 5464026 | rs10206851 | -0.431 | 0.085 | 1.096 | -1.529 | 0.006 | -0.097 | -0.097 | 0 | 0.74 | 0.67 | 0.82 | 1 | 0.71 | 0.9 |
| 2 | 5464044 | rs12477790 | 0.746 | 0.08 | 1.16 | -1.518 | -0.014 | 0.123 | 0.123 | 0 | 0.48 | 0.25 | 0.51 | 0.24 | 0.49 | 0.47 |
| 2 | 5464761 | rs10190732 | -0.294 | 0.092 | 1.536 | -1.132 | -0.023 | -0.017 | -0.017 | 0.197 | 0.28 | 0.37 | 0.3 | 0.76 | 0.22 | 0.43 |
| 2 | 5464904 | rs11900728 | 1.457 | 0.008 | 0.492 | -1.542 | -0.026 | 0.081 | 0.081 | 0 | 0.24 | 0.14 | 0.31 | 0.03 | 0.26 | 0.17 |
| 2 | 5465154 | rs9287696 | -0.605 | 0.146 | 0.405 | -1.418 | -0.005 | 0.077 | 0.077 | 0 | 0.58 | 0.72 | 0.68 | 0.79 | 0.56 | 0.61 |
| 2 | 5467135 | rs10432402 | 0.61 | 0.207 | 0.054 | -1.077 | -0.008 | 0.009 | 0.009 | 0 | 0.21 | 0.25 | 0.28 | 0.75 | 0.2 | 0.4 |
| 2 | 5467894 | rs1357752 | 0.417 | 0.123 | -0.258 | -1.14 | 0.002 | -0.079 | -0.079 | 0.108 | 0.36 | 0.48 | 0.42 | 0.03 | 0.37 | 0.19 |
| 2 | 5469833 | rs6734151 | -0.496 | 0.182 | -1.528 | -1.019 | -0.008 | 0.106 | 0.106 | 0 | 0.57 | 0.78 | 0.7 | 0.78 | 0.57 | 0.59 |
| 2 | 5487097 | rs4668565 | 0.361 | 0.315 | -1.408 | -0.658 | 0.011 | -0.01 | -0.01 | 0.063 | 0.2 | 0.09 | 0.28 | 0.78 | 0.19 | 0.4 |
| 2 | 5487627 | rs10164517 | -0.507 | 0.38 | -1.512 | -0.014 | -0.016 | -0.025 | -0.025 | 0.074 | 0.55 | 0.84 | 0.66 | 0.54 | 0.59 | 0.57 |
| 2 | 5488174 | rs2349682 | -0.11 | 0.513 | -1.173 | 0.737 | 0.012 | -0.036 | -0.036 | 0 | 0.18 | 0.13 | 0.22 | 0.52 | 0.18 | 0.42 |
| 2 | 5488456 | rs6727285 | -1.179 | 1.15 | -0.547 | -0.431 | 0.032 | -0.099 | -0.099 | 0.258 | 0.27 | 0.59 | 0.22 | 0.05 | 0.34 | 0.32 |
| 2 | 5488519 | rs10929516 | -1.505 | 0.506 | -0.431 | -0.244 | -0.003 | -0.014 | -0.014 | 0.045 | 0.21 | 0.74 | 0.14 | 0.06 | 0.18 | 0.19 |
| 2 | 5494048 | rs955146 | 0.773 | 0.636 | -0.901 | 0.137 | -0.063 | -0.155 | -0.155 | 0.167 | 0.23 | 0.01 | 0.42 | 0.49 | 0.27 | 0.55 |
| 2 | 5495567 | rs10177362 | -0.543 | 0.55 | 0.04 | 0.096 | -0.004 | -0.007 | -0.007 | 0 | 0.09 | 0.4 | 0.07 | 0.02 | 0.06 | 0.02 |
| 2 | 5495963 | rs11677035 | -0.446 | 0.719 | -0.262 | -0.137 | -0.029 | -0.023 | -0.023 | 0 | 0.41 | 0.22 | 0.55 | 0.49 | 0.46 | 0.62 |
| 2 | 5498227 | rs10929518 | -0.341 | 1.053 | 0.106 | 0.246 | -0.002 | 0.007 | 0.007 | 0 | 0.33 | 0.4 | 0.51 | 0.52 | 0.36 | 0.58 |
| 2 | 5503200 | rs1404257 | -0.325 | 0.945 | -0.182 | 0.153 | 0.008 | 0.027 | 0.027 | 0 | 0.34 | 0.49 | 0.55 | 0.56 | 0.42 | 0.57 |
| 2 | 5508545 | rs1524490 | -0.622 | 0.559 | -0.4 | 0.124 | 0.027 | -0.031 | -0.031 | 0 | 0.07 | 0.13 | 0.07 | 0.01 | 0.09 | 0.03 |
| 2 | 5510754 | rs700952 | 0.615 | 0.511 | -0.817 | 0.104 | 0.001 | -0.032 | -0.032 | 0.377 | 0.83 | 0.2 | 0.82 | 0.96 | 0.82 | 0.93 |
| 2 | 5513490 | rs792049 | 0.224 | 0.289 | -1.472 | 0.206 | -0.03 | -0.074 | -0.074 | 0 | 0.42 | 0.08 | 0.3 | 0.41 | 0.36 | 0.27 |
| 2 | 5515545 | rs10170790 | -0.334 | 0.566 | -0.902 | -0.271 | 0.005 | 0.022 | 0.022 | 0 | 0.49 | 0.58 | 0.61 | 0.28 | 0.56 | 0.58 |
| 2 | 5517283 | rs6720039 | 0.021 | 0.601 | -1.826 | -0.026 | -0.077 | 0.075 | 0.075 | 1.847 | 0.46 | 0.08 | 0.53 | 0.27 | 0.52 | 0.56 |
| 2 | 5529462 | rs12466138 | 0.642 | -0.045 | -0.434 | 0.009 | -0.035 | -0.042 | -0.042 | 5.123 | 0.83 | 0.72 | 0.9 | 0.95 | 0.92 | 0.87 |
| 2 | 5531532 | rs792106 | -0.015 | 0.115 | -1.075 | 0.135 | -0.081 | -0.066 | -0.066 | 0 | 0.43 | 0.14 | 0.39 | 0.5 | 0.47 | 0.35 |

| | | | | | | | | | | | | | | | | |
|---|---|---|---|---|---|---|---|---|---|---|---|---|---|---|---|---|
| 2 | 5535233 | rs792111 | -0.647 | -0.01 | -1.222 | -0.271 | 0.001 | -0.045 | -0.045 | 0.116 | 0.65 | 0.86 | 0.56 | 0.6 | 0.65 | 0.53 |
| 2 | 5540419 | rs6431957 | -0.628 | 0.073 | -1.251 | -0.329 | 0.019 | -0.035 | -0.035 | 0 | 0.63 | 0.83 | 0.58 | 0.6 | 0.67 | 0.53 |
| 2 | 5542127 | rs792048 | 0.324 | 0.246 | -1.278 | -0.293 | -0.001 | -0.045 | -0.045 | 0 | 0.46 | 0.15 | 0.43 | 0.5 | 0.54 | 0.38 |
| 2 | 5546290 | rs1658999 | 0.109 | 0.884 | -1.795 | -0.722 | -0.008 | -0.043 | -0.043 | 0 | 0.4 | 0.22 | 0.45 | 0.52 | 0.39 | 0.37 |
| 2 | 5546998 | rs1722373 | -0.729 | 0.997 | -1.851 | -0.551 | -0.03 | 0.036 | 0.036 | 0.523 | 0.9 | 0.93 | 0.91 | 1 | 0.84 | 0.93 |
| 2 | 5554540 | rs12466470 | 0.528 | 0.748 | -2.419 | -0.757 | 0.035 | -0.076 | -0.076 | 0 | 0.35 | 0.07 | 0.45 | 0.4 | 0.28 | 0.4 |
| 2 | 5559013 | rs1524483 | 0.466 | 0.755 | -2.415 | -0.746 | 0.045 | -0.07 | -0.07 | 0.021 | 0.36 | 0.07 | 0.45 | 0.39 | 0.27 | 0.41 |
| 2 | 5560403 | rs17291890 | -0.145 | 1.209 | -2.078 | -0.65 | -0.036 | 0.057 | 0.057 | 0.315 | 0.19 | 0.12 | 0.12 | 0 | 0.23 | 0.13 |
| 2 | 5564064 | rs700954 | 0.102 | 1.323 | -1.901 | -0.12 | -0.054 | -0.079 | -0.079 | 0.672 | 0.25 | 0.12 | 0.3 | 0.58 | 0.33 | 0.3 |
| 2 | 5571360 | rs13412303 | -0.125 | 1.123 | -2.142 | -0.047 | -0.043 | -0.018 | -0.018 | 1.998 | 0.4 | 0.14 | 0.38 | 0.41 | 0.42 | 0.39 |
| 2 | 5573939 | rs10495519 | -0.452 | 0.68 | -1.238 | -0.078 | 0.004 | 0.021 | 0.021 | 0 | 0.16 | 0.09 | 0.08 | 0.03 | 0.17 | 0.12 |
| 2 | 5576475 | rs792088 | 0.492 | 1.015 | -0.6 | 0.022 | -0.004 | -0.03 | -0.03 | 0 | 0.4 | 0.16 | 0.5 | 0.5 | 0.37 | 0.53 |
| 2 | 5578212 | rs792084 | 0.596 | 1.002 | -0.345 | 0.034 | -0.004 | -0.036 | -0.036 | 0 | 0.4 | 0.2 | 0.52 | 0.5 | 0.38 | 0.56 |
| 2 | 5579420 | rs2462394 | 0.4 | 1.007 | -0.071 | 0.119 | -0.005 | 0 | 0 | 0 | 0.44 | 0.34 | 0.6 | 0.41 | 0.47 | 0.59 |
| 2 | 5586877 | rs10495520 | -0.635 | 0.816 | 0.017 | 0.026 | 0.002 | -0.036 | -0.036 | 0 | 0.09 | 0.14 | 0.1 | 0.03 | 0.12 | 0.06 |
| 2 | 5590257 | rs813779 | 0.643 | 0.71 | -0.308 | -0.318 | -0.008 | -0.065 | -0.065 | 0 | 0.78 | 0.32 | 0.78 | 0.97 | 0.78 | 0.9 |
| 2 | 5591550 | rs969768 | 0.362 | 0.771 | -0.983 | -0.01 | 0.007 | -0.015 | -0.015 | 0.453 | 0.42 | 0.28 | 0.44 | 0.41 | 0.37 | 0.62 |
| 2 | 5593978 | rs4668614 | -0.271 | 0.747 | -0.636 | -0.717 | 0.002 | -0.013 | -0.013 | 0 | 0.51 | 0.44 | 0.67 | 0.87 | 0.54 | 0.73 |
| 2 | 5594768 | rs792071 | 0.015 | 0.675 | 0.535 | -0.257 | 0.072 | 0.117 | 0.117 | 3.035 | 0.89 | 0.62 | 0.93 | 0.86 | 0.9 | 0.88 |
| 2 | 5600812 | rs792068 | 0.413 | 0.621 | 0.109 | -0.362 | -0.002 | -0.003 | -0.003 | 0 | 0.68 | 0.3 | 0.76 | 0.7 | 0.69 | 0.72 |
| 2 | 5602284 | rs17293732 | 0.461 | 0.516 | 0.354 | -0.405 | -0.003 | -0.004 | -0.004 | 0 | 0.18 | 0.12 | 0.15 | 0.2 | 0.17 | 0.19 |
| 2 | 5603942 | rs792095 | 1.343 | 0.448 | -0.493 | -0.594 | 0.032 | 0.135 | 0.135 | 0.3 | 0.39 | 0.17 | 0.29 | 0.15 | 0.34 | 0.25 |
| 2 | 5607349 | rs10180670 | 0.597 | 0.507 | 0.016 | -0.522 | 0.015 | 0.023 | 0.023 | 1.952 | 0.19 | 0.3 | 0.17 | 0.19 | 0.17 | 0.19 |
| 2 | 5610080 | rs10929583 | 0.433 | 0.482 | -0.61 | -0.52 | -0.032 | -0.065 | -0.065 | 0 | 0.3 | 0.13 | 0.47 | 0.55 | 0.36 | 0.47 |
| 2 | 5611610 | rs792040 | 1.35 | 0.455 | -0.493 | -0.555 | 0.006 | 0.09 | 0.09 | 0.37 | 0.39 | 0.27 | 0.28 | 0.15 | 0.39 | 0.26 |
| 2 | 5611779 | rs10194520 | -0.728 | 0.566 | -0.931 | -0.338 | -0.022 | 0.007 | 0.007 | 2.126 | 0.39 | 0.17 | 0.52 | 0.66 | 0.44 | 0.55 |
| 2 | 5612893 | rs17360019 | 0.609 | 0.003 | -0.797 | -0.116 | 0.013 | -0.019 | -0.019 | 1.899 | 0.08 | 0.03 | 0.04 | 0.1 | 0.08 | 0.08 |
| 2 | 5615785 | rs6725308 | 0.607 | -0.039 | -0.846 | -0.446 | 0.096 | 0.175 | 0.175 | 0 | 0.5 | 0.19 | 0.29 | 0.22 | 0.4 | 0.47 |
| 2 | 5617681 | rs6432010 | 0.636 | 0.258 | -0.401 | -0.115 | 0.001 | 0.014 | 0.014 | 0.733 | 0.2 | 0.2 | 0.26 | 0.58 | 0.26 | 0.31 |
| 2 | 5621715 | rs4600622 | 0.645 | -0.112 | -0.27 | -0.201 | 0.003 | -0.004 | -0.004 | 2.795 | 0.66 | 0.59 | 0.52 | 0.71 | 0.65 | 0.63 |
| 2 | 5623047 | rs17294604 | -0.073 | -0.919 | -0.226 | 0.068 | 0.013 | 0.004 | 0.004 | 0.122 | 0.11 | 0.03 | 0.04 | 0.1 | 0.1 | 0.14 |
| 2 | 5623567 | rs4669393 | 0.401 | -1.165 | -0.377 | -0.096 | 0.014 | 0.005 | 0.005 | 0.381 | 0.19 | 0.15 | 0.12 | 0.15 | 0.09 | 0.29 |
| 2 | 5625500 | rs1370937 | 0.86 | -1.109 | -0.303 | 0.063 | 0.067 | 0.059 | 0.059 | 0 | 0.55 | 0.25 | 0.77 | 0.75 | 0.71 | 0.53 |
| 2 | 5633637 | rs13414559 | 0.13 | -1.019 | -0.556 | 0.368 | 0.042 | 0.002 | 0.002 | 0.213 | 0.26 | 0.19 | 0.15 | 0.24 | 0.14 | 0.3 |
| 2 | 5634436 | rs17360555 | -0.288 | -0.783 | -0.291 | 0.504 | 0 | 0 | 0 | 0 | 0.13 | 0.1 | 0.05 | 0.1 | 0.15 | 0.14 |
| 2 | 5635976 | rs16863845 | 0.727 | -0.457 | -0.717 | 0.631 | -0.067 | -0.089 | -0.089 | 0.629 | 0.13 | 0.01 | 0.18 | 0.46 | 0.17 | 0.2 |
| 2 | 5636444 | rs7561204 | 0.105 | -0.852 | -0.077 | 0.296 | 0.028 | 0.011 | 0.011 | 0 | 0.28 | 0.15 | 0.19 | 0.24 | 0.22 | 0.33 |
| 2 | 5643937 | rs1438628 | 1.074 | -0.987 | -0.091 | 0.233 | 0.086 | 0.036 | 0.036 | 0 | 0.49 | 0.07 | 0.72 | 0.65 | 0.6 | 0.48 |
| 2 | 5645041 | rs4613246 | -0.418 | -0.996 | 0.167 | 0.257 | -0.007 | -0.01 | -0.01 | 0 | 0.14 | 0.18 | 0.06 | 0.1 | 0.17 | 0.15 |
| 2 | 5645610 | rs1866364 | -1.237 | -1.013 | 0.283 | 0.302 | 0.072 | 0.048 | 0.048 | 0 | 0.47 | 0.71 | 0.27 | 0.34 | 0.39 | 0.49 |
| 2 | 5646522 | rs12619163 | -0.069 | -0.805 | 0.311 | 0.42 | 0.005 | 0.001 | 0.001 | 0 | 0.31 | 0.29 | 0.19 | 0.27 | 0.22 | 0.34 |
| 2 | 5650768 | rs4669467 | -0.393 | -0.853 | 0.23 | 0.529 | -0.001 | 0.005 | 0.005 | 0.979 | 0.12 | 0.07 | 0.13 | 0.24 | 0.14 | 0.21 |
| 2 | 5651525 | rs6737885 | 0.426 | -0.991 | 0.317 | 0.391 | -0.01 | -0.002 | -0.002 | 0 | 0.24 | 0.24 | 0.07 | 0.21 | 0.12 | 0.22 |
| 2 | 5656809 | rs4669474 | -0.768 | -0.903 | 0.09 | 0.414 | -0.007 | 0.014 | 0.014 | 0 | 0.13 | 0.18 | 0.14 | 0.2 | 0.16 | 0.22 |
| 2 | 5662656 | rs34440296 | -0.171 | -0.872 | -0.022 | 0.373 | 0 | -0.002 | -0.002 | 0 | 0.1 | 0.07 | 0.13 | 0.2 | 0.15 | 0.19 |
| 2 | 5662812 | rs7566447 | -0.1 | -1.117 | -0.04 | 0.3 | -0.008 | -0.001 | -0.001 | 0 | 0.7 | 0.33 | 0.9 | 0.78 | 0.86 | 0.71 |
| 2 | 5666520 | rs10929629 | 0.859 | -1.106 | -0.8 | 0.289 | -0.009 | -0.001 | -0.001 | 0.151 | 0.8 | 0.27 | 0.83 | 0.8 | 0.84 | 0.79 |
| 2 | 5672063 | rs10194381 | 0.468 | -0.87 | -1.203 | 0.349 | 0.12 | 0.005 | 0.005 | 0.683 | 0.57 | 0.19 | 0.77 | 0.58 | 0.71 | 0.57 |
| 2 | 5680539 | rs7575439 | 0.069 | -0.277 | -1.428 | 0.451 | -0.07 | -0.096 | -0.096 | 0.407 | 0.72 | 0.89 | 0.66 | 0.57 | 0.65 | 0.59 |
| 2 | 5688981 | rs2882274 | 0.785 | -0.066 | -2.386 | 0.623 | -0.086 | 0.101 | 0.101 | 0.688 | 0.39 | 0.11 | 0.51 | 0.21 | 0.48 | 0.3 |
| 2 | 5689697 | rs1561208 | -0.704 | 0.074 | -2.373 | 0.699 | -0.03 | -0.065 | -0.065 | 0.028 | 0.79 | 0.87 | 0.71 | 0.58 | 0.76 | 0.7 |
| 2 | 5693834 | rs10495521 | -0.054 | -0.389 | -1.72 | 0.799 | 0 | 0.007 | 0.007 | 0.046 | 0.09 | 0.1 | 0.1 | 0.16 | 0.15 | 0.15 |
| 2 | 5693857 | rs10185641 | -0.492 | -0.425 | -2.295 | 0.706 | 0.085 | 0.016 | 0.016 | 0.698 | 0.4 | 0.77 | 0.2 | 0.37 | 0.28 | 0.4 |
| 2 | 5700470 | rs13036193 | -0.421 | 0.602 | -1.874 | 0.643 | -0.044 | 0.073 | 0.073 | 0 | 0.19 | 0.01 | 0.1 | 0.07 | 0.21 | 0.16 |
| 2 | 5703784 | rs10929661 | 0.366 | 0.802 | -1.579 | 0.968 | -0.002 | 0.018 | 0.018 | 0 | 0.61 | 0.27 | 0.51 | 0.61 | 0.63 | 0.65 |
| 2 | 5712665 | rs17362697 | -0.55 | 1.176 | -1.4 | 0.988 | -0.043 | -0.004 | -0.004 | 0.244 | 0.12 | 0.01 | 0.1 | 0.13 | 0.17 | 0.11 |
| 2 | 5723564 | rs10168340 | -1.117 | 1.606 | -0.923 | 0.362 | 0.01 | -0.018 | -0.018 | 0.116 | 0.32 | 0.34 | 0.32 | 0.24 | 0.46 | 0.24 |
| 2 | 5725248 | rs13007983 | 0.192 | 1.609 | -0.891 | 0.357 | -0.022 | 0.022 | 0.022 | 0 | 0.17 | 0.12 | 0.17 | 0 | 0.25 | 0.05 |
| 2 | 5729629 | rs17363466 | -0.146 | 1.467 | -0.865 | 0.407 | -0.007 | 0.006 | 0.006 | 0 | 0.12 | 0.06 | 0.11 | 0.14 | 0.18 | 0.1 |
| 2 | 5730428 | rs11895508 | 0.954 | 1.36 | -1.559 | -0.518 | -0.033 | 0.09 | 0.09 | 0 | 0.16 | 0.01 | 0.11 | 0.02 | 0.16 | 0.17 |
| 2 | 5733323 | rs7557343 | -1.188 | 1.162 | -0.85 | 0.242 | 0.008 | -0.001 | -0.001 | 0.838 | 0.53 | 0.56 | 0.42 | 0.45 | 0.63 | 0.37 |
| 2 | 5747177 | rs16863992 | 0.03 | 0.872 | -0.124 | 0.984 | -0.001 | 0.001 | 0.001 | 0.865 | 0.18 | 0.18 | 0.12 | 0.26 | 0.15 | 0.16 |
| 2 | 5747267 | rs11673923 | -0.79 | 0.8 | 0.115 | 0.671 | 0.026 | 0.019 | 0.019 | 0.019 | 0.16 | 0.32 | 0.17 | 0.19 | 0.22 | 0.17 |
| 2 | 5754189 | rs1821797 | -0.824 | 0.751 | -0.37 | 0.715 | 0.003 | 0.01 | 0.01 | 0 | 0.17 | 0.24 | 0.18 | 0.21 | 0.23 | 0.17 |
| 2 | 5798242 | rs16864032 | -1.327 | 0.638 | -0.253 | 0.523 | 0.001 | -0.023 | -0.023 | 0 | 0.09 | 0.17 | 0.05 | 0.03 | 0.06 | 0.02 |
| 2 | 5802276 | rs13416623 | -1.054 | 0.644 | -0.234 | 0.526 | 0.001 | -0.012 | -0.012 | 0.281 | 0.08 | 0.11 | 0.05 | 0.03 | 0.06 | 0.02 |
| 2 | 5802628 | rs11685276 | -0.393 | 0.415 | -0.301 | 0.802 | -0.005 | 0.018 | 0.018 | 0 | 0.22 | 0.3 | 0.2 | 0.38 | 0.17 | 0.35 |
| 2 | 5804497 | rs4433972 | -1.305 | 0.389 | -0.256 | 0.724 | -0.002 | 0.002 | 0.002 | 0 | 0.34 | 0.66 | 0.28 | 0.41 | 0.29 | 0.42 |
| 2 | 5810657 | rs10188973 | -0.741 | 0.841 | -0.538 | 0.506 | -0.01 | -0.004 | -0.004 | 0.056 | 0.06 | 0.05 | 0.04 | 0.05 | 0.06 | 0.02 |

| | | | | | | | | | | | | | | | | |
|---|---|---|---|---|---|---|---|---|---|---|---|---|---|---|---|---|
| 2 | 5817238 | rs6727490 | -0.149 | 0.907 | -0.633 | -0.268 | -0.006 | -0.007 | -0.007 | 0.46 | 0.64 | 0.68 | 0.7 | 0.92 | 0.6 | 0.74 |
| 2 | 5819725 | rs16864048 | -0.026 | 0.719 | -0.983 | 0.449 | 0.014 | -0.023 | -0.023 | 0 | 0.09 | 0.01 | 0.17 | 0.31 | 0.09 | 0.13 |
| 2 | 5821748 | rs4547512 | -0.32 | 0.846 | -0.766 | 0.051 | -0.022 | 0.038 | 0.038 | 0 | 0.65 | 0.74 | 0.71 | 0.92 | 0.6 | 0.74 |
| 2 | 5829114 | rs12052773 | 0.146 | 1.093 | -0.607 | -0.123 | -0.023 | 0.035 | 0.035 | 0 | 0.63 | 0.71 | 0.52 | 0.81 | 0.56 | 0.65 |
| 2 | 5830599 | rs3922853 | -1.249 | 0.641 | 0.248 | -0.324 | -0.008 | -0.09 | -0.09 | 0 | 0.22 | 0.6 | 0.33 | 0.04 | 0.17 | 0.19 |
| 2 | 5838030 | rs4371338 | -1.237 | 0.512 | 0.442 | -0.078 | -0.006 | 0.022 | 0.022 | 0 | 0.36 | 0.47 | 0.57 | 0.48 | 0.37 | 0.4 |
| 2 | 5839686 | rs13419910 | -1.506 | 0.508 | 0.611 | -0.083 | 0.007 | -0.015 | -0.015 | 0 | 0.38 | 0.63 | 0.58 | 0.48 | 0.37 | 0.39 |
| 2 | 5840461 | rs6729137 | 0.031 | 0.391 | 0.706 | -0.131 | -0.007 | -0.002 | -0.002 | 0 | 0.08 | 0.08 | 0.09 | 0.13 | 0.07 | 0.08 |
| 2 | 5847365 | rs12476710 | -0.16 | 0.578 | 0.632 | -0.246 | -0.002 | -0.003 | -0.003 | 0 | 0.09 | 0.08 | 0.23 | 0.01 | 0.09 | 0.17 |
| 2 | 5847675 | rs4322814 | -1.477 | 0.543 | 1.061 | -0.297 | 0.001 | -0.047 | -0.047 | 0.689 | 0.19 | 0.34 | 0.34 | 0.14 | 0.19 | 0.26 |
| 2 | 5850764 | rs16855588 | -0.519 | 0.704 | 0.93 | -0.312 | 0.021 | -0.044 | -0.044 | 0 | 0.11 | 0.16 | 0.25 | 0.01 | 0.15 | 0.19 |
| 2 | 5852483 | rs7604153 | 0.191 | 0.648 | 0.9 | -0.299 | -0.009 | -0.002 | -0.002 | 0 | 0.08 | 0.1 | 0.09 | 0.13 | 0.07 | 0.09 |
| 2 | 5855300 | rs7596062 | -1.713 | 0.709 | 0.936 | -0.276 | -0.002 | -0.002 | -0.002 | 0 | 0.42 | 0.51 | 0.59 | 0.48 | 0.49 | 0.44 |
| 2 | 5859603 | rs2722553 | -1.068 | 0.691 | 1.127 | -0.3 | 0.03 | -0.04 | -0.04 | 0 | 0.09 | 0.37 | 0.05 | 0 | 0.06 | 0.01 |
| 2 | 5859691 | rs2564003 | -1.522 | 0.721 | 1.006 | -0.302 | -0.001 | 0 | 0 | 0 | 0.36 | 0.58 | 0.51 | 0.35 | 0.42 | 0.36 |
| 2 | 5860471 | rs1864262 | -1.589 | 0.701 | 0.452 | -0.127 | 0.001 | 0.01 | 0.01 | 0 | 0.37 | 0.4 | 0.56 | 0.48 | 0.43 | 0.43 |
| 2 | 5863154 | rs2722557 | -1.577 | 0.741 | 0.295 | -0.084 | 0.004 | 0.023 | 0.023 | 0 | 0.37 | 0.45 | 0.56 | 0.48 | 0.43 | 0.43 |
| 2 | 5863289 | rs7569783 | 0.576 | 1.076 | -0.546 | 0.089 | -0.009 | -0.047 | -0.047 | 0 | 0.16 | 0.07 | 0.21 | 0.34 | 0.2 | 0.16 |
| 2 | 5867787 | rs12478711 | -0.458 | 1.126 | -0.509 | 0.088 | -0.035 | -0.038 | -0.038 | 0 | 0.26 | 0.13 | 0.46 | 0.35 | 0.36 | 0.35 |
| 2 | 5869348 | rs2722572 | 0.308 | 0.313 | -0.56 | -0.059 | -0.007 | -0.007 | -0.007 | 0 | 0.31 | 0.36 | 0.34 | 0.27 | 0.27 | 0.32 |
| 2 | 5869890 | rs2564045 | -0.766 | -0.021 | -0.559 | -0.118 | 0.034 | -0.009 | -0.009 | 0 | 0.52 | 0.78 | 0.49 | 0.53 | 0.44 | 0.44 |
| 2 | 5875648 | rs16864085 | -0.132 | -0.31 | -0.449 | -0.204 | 0.022 | 0.002 | 0.002 | 0 | 0.2 | 0.11 | 0.11 | 0.22 | 0.15 | 0.15 |
| 2 | 5876302 | rs2252087 | 0.835 | -0.264 | -0.356 | -0.186 | -0.004 | 0.003 | 0.003 | 0 | 0.29 | 0.37 | 0.36 | 0.31 | 0.27 | 0.29 |
| 2 | 5878277 | rs2564050 | -1.136 | -0.236 | -0.652 | -0.224 | -0.031 | -0.051 | -0.051 | 0.092 | 0.58 | 0.85 | 0.52 | 0.53 | 0.45 | 0.53 |
| 2 | 5890171 | rs1366733 | 1.086 | -0.134 | -0.247 | -0.187 | -0.004 | 0.015 | 0.015 | 0 | 0.26 | 0.36 | 0.38 | 0.3 | 0.26 | 0.32 |
| 2 | 5891097 | rs13391560 | 0.906 | 0.065 | -1.079 | -0.215 | -0.028 | -0.057 | -0.057 | 0 | 0.3 | 0.1 | 0.24 | 0.31 | 0.33 | 0.2 |
| 2 | 5891437 | rs2722602 | 0.262 | -0.005 | -0.864 | -0.28 | -0.001 | 0.001 | 0.001 | 1.676 | 0.58 | 0.59 | 0.62 | 0.61 | 0.59 | 0.52 |
| 2 | 5892202 | rs11679426 | 0.415 | 0.004 | -1.264 | -0.317 | -0.027 | -0.01 | -0.01 | 0 | 0.15 | 0.01 | 0.11 | 0.14 | 0.13 | 0.31 |
| 2 | 5892291 | rs11679471 | 0.359 | 0.031 | -1.227 | -0.322 | -0.01 | 0.006 | 0.006 | 0.984 | 0.15 | 0.04 | 0.11 | 0.15 | 0.13 | 0.31 |
| 2 | 5892514 | rs16864121 | 1.456 | -0.17 | -1.03 | -0.224 | -0.003 | 0.026 | 0.026 | 0 | 0.07 | 0.02 | 0.02 | 0 | 0.05 | 0.08 |
| 2 | 5892564 | rs16864123 | 0.234 | -0.231 | -1.009 | -0.197 | 0.016 | -0.037 | -0.037 | 0.252 | 0.39 | 0.17 | 0.3 | 0.52 | 0.37 | 0.2 |
| 2 | 5893301 | rs1594102 | 0.109 | -0.195 | -0.739 | 0.138 | -0.004 | 0.038 | 0.038 | 0 | 0.11 | 0.15 | 0.12 | 0.25 | 0.08 | 0.13 |
| 2 | 5896537 | rs11888581 | -1.082 | -0.386 | -0.714 | 0.2 | -0.021 | 0.118 | 0.118 | 0 | 0.13 | 0.24 | 0.16 | 0.47 | 0.07 | 0.11 |
| 2 | 5897864 | rs16864140 | -0.079 | -0.316 | -0.802 | 0.265 | -0.008 | 0.079 | 0.079 | 0 | 0.06 | 0.09 | 0.09 | 0.26 | 0.03 | 0.05 |
| 2 | 5898727 | rs16864143 | 1.154 | -0.427 | -0.978 | 0.001 | -0.006 | -0.052 | -0.052 | 1.23 | 0.29 | 0.14 | 0.21 | 0.31 | 0.3 | 0.14 |
| 2 | 5899644 | rs16864146 | -0.012 | -0.369 | -0.845 | -0.041 | -0.009 | 0.095 | 0.095 | 0 | 0.06 | 0.11 | 0.1 | 0.26 | 0.04 | 0.05 |
| 2 | 5901174 | rs11892518 | 1.052 | -0.233 | -0.928 | -0.051 | -0.005 | -0.054 | -0.054 | 0 | 0.29 | 0.13 | 0.22 | 0.32 | 0.3 | 0.14 |
| 2 | 5903697 | rs1429244 | -0.668 | -0.234 | -1.11 | -0.395 | -0.053 | 0.057 | 0.057 | 2.214 | 0.73 | 0.88 | 0.75 | 0.81 | 0.66 | 0.6 |
| 2 | 5904390 | rs12619507 | -0.167 | -0.405 | -0.912 | -0.48 | -0.009 | 0.004 | 0.004 | 0 | 0.14 | 0.2 | 0.1 | 0.21 | 0.1 | 0.18 |
| 2 | 5907263 | rs1366746 | 1.039 | -0.459 | -0.854 | -0.553 | -0.012 | -0.097 | -0.097 | 1.188 | 0.28 | 0.57 | 0.36 | 0.02 | 0.25 | 0.24 |
| 2 | 5908151 | rs1560246 | 0.208 | -0.449 | -0.894 | -0.43 | -0.062 | 0.04 | 0.04 | 7.654 | 0.6 | 0.84 | 0.66 | 0.6 | 0.57 | 0.42 |
| 2 | 5917248 | rs10929818 | 0.447 | -0.434 | -0.902 | -0.436 | 0 | -0.039 | -0.039 | 0 | 0.4 | 0.18 | 0.29 | 0.53 | 0.38 | 0.29 |
| 2 | 5920176 | rs4668769 | 0.534 | -0.483 | -0.859 | -0.396 | 0.003 | 0.001 | 0.001 | 0.888 | 0.32 | 0.65 | 0.45 | 0.28 | 0.28 | 0.31 |
| 2 | 5921095 | rs16864193 | 1.379 | -0.249 | -1.015 | -0.657 | -0.004 | 0.022 | 0.022 | 0.086 | 0.07 | 0.02 | 0.02 | 0 | 0.06 | 0.12 |
| 2 | 5923637 | rs2564075 | 0.064 | -0.143 | -0.494 | -0.632 | 0.007 | 0.018 | 0.018 | 0 | 0.42 | 0.6 | 0.46 | 0.23 | 0.37 | 0.43 |
| 2 | 5925857 | rs1560238 | 0.619 | -0.06 | -0.422 | -0.348 | 0.007 | -0.002 | -0.002 | 0 | 0.32 | 0.28 | 0.3 | 0.58 | 0.32 | 0.18 |
| 2 | 5926156 | rs16864200 | 0.767 | -0.063 | -0.419 | -0.35 | 0.006 | -0.007 | -0.007 | 0 | 0.31 | 0.23 | 0.29 | 0.58 | 0.32 | 0.18 |
| 2 | 5928093 | rs16864209 | 0.638 | -0.04 | -0.434 | -0.338 | 0.007 | -0.002 | -0.002 | 0.473 | 0.32 | 0.28 | 0.3 | 0.58 | 0.32 | 0.18 |
| 2 | 5942145 | rs2722583 | 0.193 | 0.417 | -0.234 | -0.351 | -0.006 | -0.041 | -0.041 | 0 | 0.08 | 0.14 | 0.03 | 0 | 0.06 | 0.11 |
| 2 | 5945721 | rs1109509 | 0.766 | 0.21 | -0.21 | -0.712 | -0.014 | -0.134 | -0.134 | 0 | 0.33 | 0.54 | 0.32 | 0.02 | 0.27 | 0.26 |
| 2 | 5946313 | rs2722591 | 0.162 | 0.284 | -0.719 | -0.262 | -0.141 | 0.054 | 0.054 | 3.742 | 0.68 | 0.97 | 0.7 | 0.8 | 0.6 | 0.49 |
| 2 | 5950473 | rs4668786 | -0.581 | 0.285 | -0.215 | -0.165 | -0.048 | 0 | 0 | 0 | 0.81 | 0.94 | 0.72 | 0.83 | 0.73 | 0.67 |
| 2 | 5950557 | rs6741254 | 0.533 | -0.038 | -0.046 | -0.003 | 0.004 | 0.004 | 0.004 | 0.03 | 0.35 | 0.61 | 0.33 | 0.27 | 0.21 | 0.29 |
| 2 | 5958708 | rs1429238 | -1.105 | 0.139 | -0.226 | -0.008 | -0.057 | 0.006 | 0.006 | 0 | 0.79 | 0.94 | 0.71 | 0.83 | 0.7 | 0.67 |
| 2 | 5958997 | rs10929838 | 1.902 | 0.134 | -0.367 | 0.306 | -0.1 | -0.072 | -0.072 | 0 | 0.24 | 0.05 | 0.2 | 0.25 | 0.35 | 0.15 |
| 2 | 5959288 | rs12614917 | 0.411 | 0.12 | -0.255 | 0.012 | -0.107 | -0.034 | -0.034 | 0 | 0.4 | 0.12 | 0.35 | 0.56 | 0.48 | 0.38 |
| 2 | 5961696 | rs1429231 | -0.187 | 0.069 | -0.235 | -0.045 | 0.1 | 0.141 | 0.141 | 0 | 0.46 | 0.83 | 0.37 | 0.27 | 0.28 | 0.4 |
| 2 | 5965388 | rs11691578 | 1.165 | 0.16 | -0.087 | -0.037 | -0.043 | -0.135 | -0.135 | 0 | 0.28 | 0.42 | 0.26 | 0.02 | 0.18 | 0.22 |
| 2 | 5966207 | rs1429229 | 0.161 | 0.385 | -0.166 | 0.083 | 0.027 | 0.038 | 0.038 | 0 | 0.4 | 0.7 | 0.43 | 0.29 | 0.28 | 0.32 |
| 2 | 5966363 | rs1366740 | 0.616 | 0.091 | -0.087 | 0.144 | -0.03 | -0.029 | -0.029 | 0.494 | 0.35 | 0.51 | 0.33 | 0.26 | 0.22 | 0.28 |
| 2 | 5968922 | rs995949 | 0.421 | 0.1 | -0.132 | 0.149 | 0.019 | 0.018 | 0.018 | 0 | 0.37 | 0.69 | 0.35 | 0.26 | 0.22 | 0.28 |
| 2 | 5971934 | rs6732445 | -0.218 | 0.242 | -0.032 | 0.292 | -0.002 | 0.001 | 0.001 | 0 | 0.44 | 0.44 | 0.38 | 0.56 | 0.48 | 0.39 |
| 2 | 5973504 | rs2709447 | -0.15 | 0.283 | -0.069 | 0.31 | 0.007 | -0.006 | -0.006 | 0.544 | 0.43 | 0.52 | 0.39 | 0.57 | 0.48 | 0.39 |
| 2 | 5973886 | rs2563999 | 0.572 | 0.337 | -0.47 | 0.25 | -0.005 | -0.047 | -0.047 | 0 | 0.08 | 0.2 | 0.03 | 0 | 0.05 | 0.11 |
| 2 | 5977648 | rs13405548 | 0.508 | 0.345 | -0.591 | 0.362 | -0.091 | -0.048 | -0.048 | 0.007 | 0.34 | 0.1 | 0.35 | 0.58 | 0.44 | 0.33 |
| 2 | 5978992 | rs2709460 | -0.994 | 0.213 | -0.647 | 0.263 | -0.016 | 0.015 | 0.015 | 0 | 0.79 | 0.88 | 0.71 | 0.85 | 0.7 | 0.71 |
| 2 | 5979192 | rs4669905 | 0.414 | 0.405 | -0.574 | 0.444 | -0.076 | -0.032 | -0.032 | 0 | 0.35 | 0.12 | 0.35 | 0.58 | 0.44 | 0.35 |
| 2 | 5983436 | rs2709458 | 0.27 | 0.356 | -0.59 | 0.685 | 0.001 | 0.076 | 0.076 | 0 | 0.12 | 0.24 | 0.15 | 0.32 | 0.09 | 0.16 |
| 2 | 5987178 | rs10495525 | 1.198 | 0.623 | -0.772 | 0.766 | -0.012 | -0.004 | -0.004 | 0 | 0.26 | 0.17 | 0.2 | 0.25 | 0.33 | 0.23 |

| | | | | | | | | | | | | | | | | |
|---|---|---|---|---|---|---|---|---|---|---|---|---|---|---|---|---|
| 2 | 5988639 | rs2564010 | -0.032 | 0.538 | -1.014 | 0.769 | 0 | -0.002 | -0.002 | 0 | 0.45 | 0.58 | 0.39 | 0.55 | 0.47 | 0.44 |
| 2 | 5989380 | rs7593997 | -0.308 | -0.025 | -0.831 | 0.545 | -0.012 | -0.019 | -0.019 | 0.934 | 0.63 | 0.67 | 0.55 | 0.52 | 0.58 | 0.56 |
| 2 | 5989536 | rs11690032 | -0.227 | -0.568 | -0.717 | 0.322 | -0.031 | -0.075 | -0.075 | 0.375 | 0.24 | 0.27 | 0.09 | 0 | 0.14 | 0.12 |
| 2 | 5991798 | rs2709468 | -0.79 | 0.209 | -1.111 | 1.771 | 0 | 0.045 | 0.045 | 0 | 0.08 | 0.05 | 0.21 | 0.46 | 0.08 | 0.15 |
| 2 | 5991969 | rs2163025 | -0.649 | -0.118 | -2.256 | 0.709 | 0.049 | 0.202 | 0.202 | 0 | 0.49 | 0.91 | 0.49 | 0.77 | 0.39 | 0.45 |
| 2 | 5994603 | rs10929853 | 0.21 | 0.076 | -2.337 | 0.471 | -0.053 | -0.044 | -0.044 | 0 | 0.32 | 0.1 | 0.19 | 0.32 | 0.34 | 0.26 |
| 2 | 6002657 | rs12471085 | -0.322 | -0.283 | -1.822 | 0.896 | -0.012 | 0 | 0 | 0 | 0.07 | 0.01 | 0.19 | 0.49 | 0.07 | 0.19 |
| 2 | 6003646 | rs6750041 | -0.283 | -0.648 | -0.944 | -0.031 | -0.039 | -0.062 | -0.062 | 0.356 | 0.14 | 0.16 | 0.04 | 0 | 0.05 | 0.04 |
| 2 | 6005793 | rs11692731 | -1.269 | 0.115 | -0.321 | 0.101 | 0.021 | -0.05 | -0.05 | 2.835 | 0.09 | 0.28 | 0.07 | 0 | 0.09 | 0.03 |
| 2 | 6008030 | rs891251 | -1.372 | 0.106 | -0.179 | 0.097 | 0.03 | -0.051 | -0.051 | 0 | 0.09 | 0.32 | 0.08 | 0 | 0.09 | 0.03 |
| 2 | 6008310 | rs891250 | -0.996 | 0.136 | -0.209 | 0.112 | 0.024 | -0.037 | -0.037 | 0 | 0.07 | 0.18 | 0.06 | 0 | 0.09 | 0.03 |
| 2 | 6010100 | rs11899694 | -1.79 | -0.2 | 0.093 | 0.162 | -0.005 | -0.012 | -0.012 | 0.755 | 0.06 | 0.32 | 0.03 | 0 | 0 | 0 |
| 2 | 6010700 | rs2059912 | -0.333 | -0.903 | 0.042 | -0.36 | -0.051 | -0.084 | -0.084 | 0 | 0.2 | 0.32 | 0.12 | 0 | 0.07 | 0.05 |
| 2 | 6013787 | rs2709443 | -0.761 | -0.898 | 0.128 | -0.354 | -0.038 | 0.041 | 0.041 | 4.69 | 0.6 | 0.8 | 0.5 | 0.64 | 0.49 | 0.38 |
| 2 | 6016600 | rs1429225 | 0.317 | -1.081 | 0.083 | -0.08 | -0.006 | -0.001 | -0.001 | 0 | 0.38 | 0.27 | 0.21 | 0.32 | 0.36 | 0.24 |
| 2 | 6020296 | rs2709440 | -0.859 | -0.959 | 0.307 | -0.063 | -0.032 | 0.038 | 0.038 | 4.69 | 0.6 | 0.77 | 0.49 | 0.64 | 0.48 | 0.38 |
| 2 | 6021463 | rs2564033 | 0.916 | -1.045 | 0.973 | -0.026 | -0.035 | -0.059 | -0.059 | 0 | 0.16 | 0.16 | 0.09 | 0 | 0.07 | 0.04 |
| 2 | 6026244 | rs2564036 | -0.886 | -0.814 | 1.05 | -0.096 | -0.015 | 0.016 | 0.016 | 5.139 | 0.62 | 0.75 | 0.5 | 0.64 | 0.51 | 0.39 |
| 2 | 6029084 | rs6760889 | 0.401 | -0.799 | 1.191 | -0.045 | -0.008 | 0 | 0 | 0 | 0.38 | 0.34 | 0.23 | 0.32 | 0.39 | 0.25 |
| 2 | 6038606 | rs2464989 | 0.046 | -0.759 | 1.005 | 0.015 | -0.058 | 0.052 | 0.052 | 5.004 | 0.76 | 0.57 | 0.72 | 0.68 | 0.87 | 0.86 |
| 2 | 6039177 | rs16864477 | 0.996 | -0.79 | 0.448 | -0.496 | 0.008 | -0.007 | -0.007 | 0.051 | 0.16 | 0.08 | 0.08 | 0 | 0.07 | 0.04 |
| 2 | 6040913 | rs17373058 | 0.12 | -0.749 | 0.447 | -0.47 | 0.012 | 0.088 | 0.088 | 0 | 0.5 | 0.27 | 0.53 | 0.36 | 0.5 | 0.59 |
| 2 | 6041853 | rs4455137 | -0.009 | -0.108 | 1.013 | -0.257 | -0.016 | 0.034 | 0.034 | 1.79 | 0.58 | 0.6 | 0.49 | 0.67 | 0.51 | 0.38 |
| 2 | 6042843 | rs7587957 | 0.025 | -0.175 | 0.985 | -0.294 | -0.003 | 0.003 | 0.003 | 2.463 | 0.61 | 0.35 | 0.48 | 0.67 | 0.52 | 0.42 |
| 2 | 6046983 | rs10200963 | -0.922 | 0.15 | 0.576 | -0.366 | 0.067 | -0.006 | -0.006 | 0 | 0.05 | 0.25 | 0.06 | 0 | 0.06 | 0.02 |
| 2 | 6047944 | rs4668815 | -0.386 | 0.122 | -0.269 | -0.215 | -0.061 | 0.037 | 0.037 | 5.999 | 0.65 | 0.85 | 0.54 | 0.69 | 0.56 | 0.42 |
| 2 | 6049411 | rs2350571 | -0.017 | 0.063 | -0.104 | -0.069 | -0.004 | 0.003 | 0.003 | 1.284 | 0.62 | 0.36 | 0.48 | 0.71 | 0.51 | 0.44 |
| 2 | 6056656 | rs4433977 | 0.748 | -0.265 | -0.341 | 0.434 | 0.072 | 0.2 | 0.2 | 0 | 0.5 | 0.14 | 0.41 | 0.24 | 0.66 | 0.52 |
| 2 | 6057847 | rs10495528 | -0.003 | -0.36 | -0.119 | 0.337 | -0.02 | 0.043 | 0.043 | 0 | 0.3 | 0.39 | 0.32 | 0.45 | 0.19 | 0.3 |
| 2 | 6060989 | rs4293540 | 0.071 | -0.077 | -0.223 | 0.408 | 0.002 | 0.014 | 0.014 | 0 | 0.91 | 0.96 | 0.95 | 1 | 0.92 | 0.98 |
| 2 | 6066942 | rs7565497 | 0.685 | -0.15 | 0.104 | 0.562 | -0.018 | 0.036 | 0.036 | 0 | 0.79 | 0.63 | 0.64 | 0.71 | 0.83 | 0.8 |
| 2 | 6070009 | rs12469824 | -0.125 | 0.58 | -0.804 | 0.212 | -0.034 | 0.045 | 0.045 | 0 | 0.13 | 0.04 | 0.23 | 0.02 | 0.19 | 0.06 |
| 2 | 6072230 | rs4485544 | 0.29 | 0.085 | -1.371 | 0.719 | 0.01 | -0.111 | -0.111 | 0.078 | 0.15 | 0.01 | 0.06 | 0.39 | 0.1 | 0.23 |
| 2 | 6073955 | rs16864520 | -0.098 | -0.196 | -0.738 | 0.622 | 0.004 | -0.014 | -0.014 | 0 | 0.17 | 0.15 | 0.32 | 0.3 | 0.16 | 0.21 |
| 2 | 6075682 | rs4669966 | 1.245 | -0.512 | -0.865 | 0.365 | 0.017 | 0.101 | 0.101 | 0 | 0.47 | 0.14 | 0.34 | 0.31 | 0.56 | 0.5 |
| 2 | 6084916 | rs10495532 | 0.055 | -0.651 | -0.206 | 0.493 | -0.038 | 0.105 | 0.105 | 0.053 | 0.17 | 0.34 | 0.07 | 0.38 | 0.08 | 0.18 |
| 2 | 6085200 | rs13432692 | 0.533 | -0.329 | -0.404 | 0.165 | 0.003 | -0.006 | -0.006 | 0.737 | 0.6 | 0.47 | 0.36 | 0.64 | 0.54 | 0.59 |
| 2 | 6086333 | rs13425571 | 0.709 | -0.298 | -0.401 | 0.222 | 0.012 | -0.016 | -0.016 | 0 | 0.15 | 0.13 | 0.05 | 0.38 | 0.08 | 0.18 |
| 2 | 6087070 | rs5018486 | 1.072 | -0.055 | -0.557 | 0.063 | -0.071 | 0.093 | 0.093 | 0 | 0.44 | 0.14 | 0.3 | 0.26 | 0.47 | 0.41 |
| 2 | 6088403 | rs12692524 | 0.225 | 0.638 | -0.894 | 0.201 | -0.043 | 0.051 | 0.051 | 0 | 0.13 | 0.03 | 0.23 | 0.02 | 0.2 | 0.07 |
| 2 | 6088751 | rs16864545 | 0.598 | 0.266 | -1.354 | 0.173 | -0.045 | -0.037 | -0.037 | 0.138 | 0.27 | 0.15 | 0.25 | 0.33 | 0.4 | 0.37 |
| 2 | 6091222 | rs11674680 | 0.858 | 0.217 | -2.116 | -0.274 | -0.011 | -0.019 | -0.019 | 0 | 0.34 | 0.04 | 0.31 | 0.33 | 0.27 | 0.38 |
| 2 | 6092674 | rs1806463 | 0.631 | 0.491 | -1.487 | -0.373 | 0.012 | -0.018 | -0.018 | 0 | 0.13 | 0.17 | 0.21 | 0.03 | 0.19 | 0.06 |
| 2 | 6111604 | rs4383308 | 0.958 | 0.514 | -1.289 | -0.37 | 0.028 | -0.016 | -0.016 | 0.509 | 0.07 | 0.17 | 0.14 | 0.02 | 0.08 | 0.03 |
| 2 | 6112516 | rs11691189 | 0.465 | 0.428 | -1.39 | -0.413 | 0.008 | -0.017 | -0.017 | 0 | 0.08 | 0.14 | 0.14 | 0.02 | 0.08 | 0.03 |
| 2 | 6112702 | rs13015403 | 0.756 | 0.409 | -1.854 | -0.46 | -0.135 | -0.079 | -0.079 | 4.022 | 0.3 | 0.03 | 0.29 | 0.27 | 0.46 | 0.31 |
| 2 | 6114008 | rs10198477 | 0.752 | 0.143 | -1.73 | -0.551 | 0.009 | 0.048 | 0.048 | 0 | 0.42 | 0.25 | 0.46 | 0.35 | 0.34 | 0.41 |
| 2 | 6117147 | rs4669989 | -0.312 | -0.16 | -0.816 | -0.514 | -0.013 | 0.034 | 0.034 | 0 | 0.25 | 0.36 | 0.3 | 0.38 | 0.2 | 0.29 |
| 2 | 6119019 | rs2609178 | -0.53 | 0.148 | -0.922 | -0.172 | -0.095 | -0.073 | -0.073 | 0 | 0.64 | 0.89 | 0.57 | 0.62 | 0.43 | 0.63 |
| 2 | 6119270 | rs12623061 | 0.682 | 0.01 | -0.722 | -0.184 | -0.008 | 0.13 | 0.13 | 0 | 0.12 | 0.27 | 0.14 | 0.37 | 0.08 | 0.21 |
| 2 | 6120589 | rs1880800 | -0.414 | 0.369 | -1.052 | -0.325 | -0.113 | -0.027 | -0.027 | 0.056 | 0.66 | 0.89 | 0.58 | 0.7 | 0.49 | 0.67 |
| 2 | 6123492 | rs2709437 | 0.092 | 0.298 | -0.33 | -0.21 | 0.012 | -0.009 | -0.009 | 0 | 0.62 | 0.62 | 0.5 | 0.7 | 0.53 | 0.65 |
| 2 | 6125021 | rs1079417 | -0.219 | 0.042 | 0.383 | -0.098 | 0.025 | 0.013 | 0.013 | 0 | 0.67 | 0.57 | 0.57 | 0.62 | 0.6 | 0.68 |
| 2 | 6134940 | rs1405262 | 0.651 | -1.295 | 0.532 | 0.169 | -0.01 | 0.068 | 0.068 | 0 | 0.1 | 0.16 | 0.15 | 0.37 | 0.09 | 0.31 |
| 2 | 6141286 | rs941009 | -1.391 | -1.672 | 0.384 | -0.201 | 0.057 | -0.014 | -0.014 | 0 | 0.37 | 0.76 | 0.4 | 0.39 | 0.27 | 0.39 |
| 2 | 6146154 | rs4670020 | 0.474 | -1.72 | 0.805 | -0.17 | 0.006 | 0.006 | 0.006 | 0 | 0.17 | 0.08 | 0.15 | 0 | 0.16 | 0.05 |
| 2 | 6152005 | rs2609195 | -1.02 | -1.732 | 0.584 | -0.048 | -0.05 | -0.006 | -0.006 | 0 | 0.68 | 0.81 | 0.53 | 0.67 | 0.5 | 0.66 |
| 2 | 6152133 | rs2693833 | -1.147 | -1.731 | 0.467 | -0.048 | -0.131 | -0.06 | -0.06 | 0.088 | 0.69 | 0.95 | 0.54 | 0.67 | 0.5 | 0.66 |
| 2 | 6159981 | rs12468070 | 0.746 | -1.943 | 1.164 | -0.011 | 0.039 | -0.006 | -0.006 | 0.239 | 0.41 | 0.3 | 0.33 | 0.39 | 0.3 | 0.49 |
| 2 | 6160127 | rs2609187 | -0.002 | -1.958 | 1.411 | -0.151 | -0.052 | 0.034 | 0.034 | 0 | 0.22 | 0.36 | 0.19 | 0.27 | 0.17 | 0.16 |
| 2 | 6162581 | rs12997510 | 0.992 | -1.808 | 1.488 | 0.37 | 0.022 | 0.054 | 0.054 | 0 | 0.28 | 0.2 | 0.14 | 0.02 | 0.19 | 0.16 |
| 2 | 6162992 | rs1918046 | 0.248 | -1.871 | 2.002 | 0.435 | -0.034 | -0.015 | -0.015 | 0 | 0.34 | 0.57 | 0.19 | 0.29 | 0.22 | 0.23 |
| 2 | 6163005 | rs1918045 | 1.073 | -1.86 | 1.862 | 0.436 | 0.023 | 0.075 | 0.075 | 0 | 0.28 | 0.17 | 0.14 | 0.02 | 0.19 | 0.16 |
| 2 | 6165923 | rs2693822 | 0.38 | -1.895 | 1.87 | 0.448 | -0.028 | -0.01 | -0.01 | 0.485 | 0.33 | 0.51 | 0.19 | 0.29 | 0.22 | 0.23 |
| 2 | 6169727 | rs7566764 | 0.793 | -1.845 | 1.58 | 0.529 | 0 | 0 | 0 | 0 | 0.3 | 0.33 | 0.12 | 0.28 | 0.21 | 0.23 |
| 2 | 6178547 | rs2257197 | 0.406 | -1.833 | 1.727 | 0.473 | -0.023 | -0.022 | -0.022 | 0 | 0.34 | 0.6 | 0.18 | 0.22 | 0.22 | 0.24 |
| 2 | 6185634 | rs2609101 | 0.21 | -1.864 | 1.459 | 0.454 | 0.058 | 0.057 | 0.057 | 1.643 | 0.36 | 0.8 | 0.2 | 0.21 | 0.22 | 0.23 |
| 2 | 6195196 | rs6718476 | -0.189 | -1.972 | 1.54 | 1.009 | 0.012 | 0.005 | 0.005 | 0.04 | 0.12 | 0.08 | 0.21 | 0.42 | 0.08 | 0.29 |
| 2 | 6195847 | rs1949914 | -0.803 | -1.934 | 1.727 | 0.933 | 0.106 | 0.079 | 0.079 | 0.466 | 0.49 | 0.79 | 0.42 | 0.64 | 0.32 | 0.53 |

| | | | | | | | | | | | | | | | | |
|---|---|---|---|---|---|---|---|---|---|---|---|---|---|---|---|---|
| 2 | 6200387 | rs13014761 | 0.845 | -1.976 | 2.207 | 0.893 | 0.011 | 0.012 | 0.012 | 0.324 | 0.3 | 0.25 | 0.17 | 0.02 | 0.22 | 0.16 |
| 2 | 6204312 | rs12992673 | 0.452 | -1.906 | 2.284 | 0.95 | -0.005 | -0.007 | -0.007 | 0 | 0.34 | 0.4 | 0.19 | 0.22 | 0.24 | 0.24 |
| 2 | 6204508 | rs2693818 | -1.007 | -1.994 | 2.251 | 1.057 | -0.045 | 0.255 | 0.255 | 0 | 0.15 | 0.36 | 0.23 | 0.42 | 0.1 | 0.29 |
| 2 | 6215483 | rs1818962 | -0.096 | -1.837 | 1.831 | 0.203 | 0.008 | -0.025 | -0.025 | 0.532 | 0.42 | 0.39 | 0.26 | 0.02 | 0.31 | 0.24 |
| 2 | 6215499 | rs2693862 | -1.022 | -1.924 | 1.771 | 0.562 | 0.024 | 0.052 | 0.052 | 0 | 0.54 | 0.67 | 0.5 | 0.26 | 0.4 | 0.48 |
| 2 | 6217962 | rs2693865 | 0.312 | -1.839 | 1.988 | 0.648 | -0.014 | -0.088 | -0.088 | 2.844 | 0.34 | 0.59 | 0.21 | 0.02 | 0.22 | 0.22 |
| 2 | 6218776 | rs16864678 | -0.043 | -1.801 | 1.844 | 0.71 | 0.015 | -0.016 | -0.016 | 3.173 | 0.1 | 0.07 | 0.22 | 0.24 | 0.08 | 0.22 |
| 2 | 6221936 | rs2693864 | -0.574 | -1.741 | 1.844 | 0.902 | 0.062 | 0.027 | 0.027 | 0.071 | 0.47 | 0.73 | 0.47 | 0.61 | 0.33 | 0.49 |
| 2 | 6231750 | rs2609095 | -0.731 | -1.813 | 1.328 | 0.594 | 0.107 | 0.029 | 0.029 | 0.443 | 0.48 | 0.88 | 0.48 | 0.61 | 0.34 | 0.49 |
| 2 | 6233146 | rs7585444 | 0.007 | -1.815 | 1.75 | 0.44 | -0.014 | 0.05 | 0.05 | 0 | 0.1 | 0.14 | 0.23 | 0.27 | 0.09 | 0.2 |
| 2 | 6233660 | rs2693872 | 0.679 | -1.859 | 1.787 | 0.231 | 0.011 | -0.007 | -0.007 | 0.652 | 0.31 | 0.28 | 0.21 | 0.01 | 0.22 | 0.22 |
| 2 | 6235208 | rs2249724 | -0.488 | -1.84 | 1.588 | 0.318 | 0.072 | 0.079 | 0.079 | 0.296 | 0.46 | 0.76 | 0.49 | 0.25 | 0.32 | 0.41 |
| 2 | 6246405 | rs6719357 | -0.551 | -1.671 | 1.837 | 0.533 | -0.021 | 0.149 | 0.149 | 0 | 0.1 | 0.23 | 0.05 | 0.4 | 0.06 | 0.1 |
| 2 | 6247449 | rs9798084 | 0.422 | -1.671 | 1.309 | 0.261 | 0.009 | 0.03 | 0.03 | 0.236 | 0.34 | 0.25 | 0.26 | 0.01 | 0.28 | 0.22 |
| 2 | 6254752 | rs2693889 | -1.38 | -1.289 | 1.549 | 0.759 | -0.043 | 0.185 | 0.185 | 0.262 | 0.14 | 0.57 | 0.07 | 0.4 | 0.06 | 0.1 |
| 2 | 6265868 | rs2609129 | -0.096 | -1.26 | 1.6 | 0.71 | -0.013 | -0.041 | -0.041 | 0.344 | 0.08 | 0.25 | 0.03 | 0 | 0.03 | 0.02 |
| 2 | 6271312 | rs1526374 | -0.562 | -1.149 | 0.701 | 0.983 | 0.013 | -0.031 | -0.031 | 2.381 | 0.12 | 0.08 | 0.26 | 0.23 | 0.12 | 0.21 |
| 2 | 6273208 | rs1358135 | -1.042 | -1.313 | 0.102 | 0.143 | -0.081 | -0.03 | -0.03 | 0 | 0.58 | 0.96 | 0.59 | 0.64 | 0.46 | 0.53 |
| 2 | 6290953 | rs10495534 | -1.722 | -1.055 | 1.422 | 1.248 | -0.031 | 0.155 | 0.155 | 0 | 0.17 | 0.41 | 0.3 | 0.62 | 0.15 | 0.29 |
| 2 | 6297760 | rs12622790 | -1.535 | -0.927 | 1.099 | 0.504 | -0.017 | 0.141 | 0.141 | 0 | 0.12 | 0.32 | 0.25 | 0.22 | 0.09 | 0.23 |
| 2 | 6304703 | rs7585073 | -1.072 | -0.854 | 0.406 | 0.54 | 0 | 0.007 | 0.007 | 0 | 0.1 | 0.07 | 0.23 | 0.22 | 0.08 | 0.23 |
| 2 | 6304780 | rs7587995 | -1.072 | -0.854 | 0.406 | 0.54 | 0 | 0.007 | 0.007 | 0 | 0.1 | 0.07 | 0.23 | 0.22 | 0.08 | 0.23 |
| 2 | 6307291 | rs4669042 | -0.968 | -0.879 | 0.274 | 0.506 | 0.006 | -0.006 | -0.006 | 0.028 | 0.08 | 0.06 | 0.22 | 0.23 | 0.08 | 0.22 |
| 2 | 6307401 | rs872460 | -0.779 | -0.819 | 0.301 | 0.536 | 0.009 | -0.013 | -0.013 | 0 | 0.09 | 0.05 | 0.23 | 0.23 | 0.09 | 0.23 |
| 2 | 6310017 | rs16864841 | -1.184 | -0.697 | 0.583 | -0.009 | -0.002 | -0.007 | -0.007 | 0 | 0.07 | 0.12 | 0.11 | 0.08 | 0.07 | 0.13 |
| 2 | 6310880 | rs7566793 | -1.23 | -1.105 | 0.202 | -0.322 | 0.004 | 0.003 | 0.003 | 0 | 0.23 | 0.19 | 0.31 | 0.19 | 0.18 | 0.26 |
| 2 | 6318487 | rs13397476 | -0.743 | -1.223 | -0.063 | -0.531 | 0.017 | 0.02 | 0.02 | 0.362 | 0.19 | 0.07 | 0.24 | 0.11 | 0.15 | 0.14 |
| 2 | 6325751 | rs10178048 | -1.442 | -1.313 | 0.698 | -0.554 | -0.018 | -0.01 | -0.01 | 0.996 | 0.1 | 0.21 | 0.11 | 0.08 | 0.06 | 0.12 |
| 2 | 6329241 | rs10173194 | -1.397 | -1.282 | 0.738 | -0.558 | -0.012 | -0.009 | -0.009 | 0 | 0.1 | 0.15 | 0.11 | 0.08 | 0.07 | 0.12 |
| 2 | 6329601 | rs12463667 | 0.069 | -1.295 | 0.963 | -0.581 | 0.002 | 0.001 | 0.001 | 0 | 0.14 | 0.1 | 0.16 | 0.11 | 0.12 | 0.16 |
| 2 | 6330746 | rs1562624 | -0.042 | -1.257 | 0.935 | -1.151 | 0 | 0.009 | 0.009 | 0.191 | 0.2 | 0.13 | 0.13 | 0.01 | 0.24 | 0.16 |
| 2 | 6333379 | rs10439238 | -1.791 | -1.373 | 1.181 | -1.181 | 0.015 | 0.033 | 0.033 | 0.051 | 0.46 | 0.6 | 0.42 | 0.19 | 0.42 | 0.43 |
| 2 | 6334684 | rs9287668 | -1.637 | -1.345 | 1.077 | -1.287 | 0.001 | -0.006 | -0.006 | 0 | 0.45 | 0.56 | 0.42 | 0.18 | 0.43 | 0.44 |
| 2 | 6341885 | rs12692305 | -0.301 | -1.063 | 1.378 | -1.142 | -0.001 | -0.045 | -0.045 | 0 | 0.2 | 0.2 | 0.13 | 0.01 | 0.23 | 0.16 |
| 2 | 6342769 | rs6710611 | -1.683 | -1.468 | 1.027 | -1.585 | 0.009 | 0.057 | 0.057 | 0.708 | 0.53 | 0.66 | 0.49 | 0.17 | 0.48 | 0.5 |
| 2 | 6344766 | rs4396686 | -1.376 | -1.48 | 0.928 | -1.609 | 0.02 | 0.111 | 0.111 | 0.655 | 0.48 | 0.3 | 0.46 | 0.17 | 0.48 | 0.5 |
| 2 | 6351862 | rs1015230 | -0.759 | -1.522 | 0.87 | -1.61 | -0.008 | -0.022 | -0.022 | 0.365 | 0.34 | 0.4 | 0.37 | 0.16 | 0.25 | 0.39 |
| 2 | 6351943 | rs1448572 | -0.128 | -1.442 | 0.803 | -1.62 | -0.011 | -0.112 | -0.112 | 0 | 0.22 | 0.37 | 0.14 | 0.01 | 0.23 | 0.15 |
| 2 | 6352799 | rs10204418 | -0.021 | -1.435 | 0.697 | -1.617 | -0.007 | -0.087 | -0.087 | 0 | 0.21 | 0.29 | 0.13 | 0.01 | 0.23 | 0.15 |
| 2 | 6354992 | rs2044651 | 0.101 | -0.97 | 0.243 | -1.102 | 0.002 | 0.052 | 0.052 | 0 | 0.21 | 0.05 | 0.23 | 0.09 | 0.17 | 0.25 |
| 2 | 6358830 | rs16864928 | -0.53 | -0.614 | 0.329 | -0.826 | -0.001 | 0.001 | 0.001 | 0 | 0.07 | 0.07 | 0.1 | 0.06 | 0.08 | 0.14 |
| 2 | 6361101 | rs13389679 | -0.902 | -0.618 | 0.346 | -0.716 | -0.001 | 0.009 | 0.009 | 0 | 0.1 | 0.28 | 0.12 | 0.09 | 0.08 | 0.14 |
| 2 | 6363082 | rs2166927 | -0.605 | -0.077 | -0.413 | -0.818 | 0.012 | 0.061 | 0.061 | 0.19 | 0.41 | 0.77 | 0.44 | 0.23 | 0.33 | 0.46 |
| 2 | 6363474 | rs17377128 | -1.04 | 0.378 | -0.248 | -0.656 | -0.009 | -0.003 | -0.003 | 0 | 0.14 | 0.09 | 0.12 | 0 | 0.23 | 0.13 |
| 2 | 6365649 | rs13388785 | -1.434 | 0.289 | 0.767 | -0.637 | 0.054 | -0.034 | -0.034 | 0.441 | 0.09 | 0.4 | 0.07 | 0 | 0.07 | 0.04 |
| 2 | 6366150 | rs2351104 | 0.834 | 0.281 | 0.395 | -0.663 | 0.002 | -0.017 | -0.017 | 0.027 | 0.68 | 0.61 | 0.67 | 0.82 | 0.65 | 0.74 |
| 2 | 6366715 | rs4444491 | 0.42 | 0.231 | 0.583 | -0.754 | -0.004 | 0.01 | 0.01 | 0 | 0.78 | 0.87 | 0.74 | 0.84 | 0.76 | 0.78 |
| 2 | 6367089 | rs7592788 | -0.854 | 0.224 | 0.508 | -0.742 | 0.012 | -0.027 | -0.027 | 0 | 0.08 | 0.2 | 0.06 | 0.01 | 0.11 | 0.08 |
| 2 | 6368302 | rs12478200 | 0.15 | 0.485 | -0.25 | -0.053 | -0.004 | -0.024 | -0.024 | 0.003 | 0.16 | 0.1 | 0.28 | 0.38 | 0.18 | 0.25 |
| 2 | 6368894 | rs7586450 | -1.741 | 0.653 | 0.245 | -0.24 | 0 | -0.002 | -0.002 | 0 | 0.34 | 0.57 | 0.4 | 0.39 | 0.34 | 0.39 |
| 2 | 6374104 | rs6743687 | -0.067 | 0.77 | -0.163 | -0.148 | 0.01 | -0.023 | -0.023 | 0.007 | 0.07 | 0.12 | 0.06 | 0.01 | 0.11 | 0.08 |
| 2 | 6375910 | rs13384785 | 0.997 | 0.71 | -0.208 | -0.176 | -0.006 | 0.093 | 0.093 | 0 | 0.6 | 0.22 | 0.61 | 0.83 | 0.59 | 0.64 |
| 2 | 6380352 | rs10191492 | 1.563 | 0.652 | -0.359 | -0.09 | -0.013 | -0.053 | -0.053 | 0 | 0.44 | 0.15 | 0.35 | 0.45 | 0.4 | 0.38 |
| 2 | 6380948 | rs10519722 | 1.14 | 0.609 | -0.462 | 0.483 | 0.023 | 0.005 | 0.005 | 0 | 0.53 | 0.14 | 0.55 | 0.56 | 0.57 | 0.53 |
| 2 | 6384728 | rs1562617 | -0.829 | 0.87 | 0.179 | 1.189 | -0.008 | -0.007 | -0.007 | 0.025 | 0.11 | 0.12 | 0.27 | 0.27 | 0.19 | 0.21 |
| 2 | 6388637 | rs11685714 | -1.059 | 0.851 | 0.676 | 1.208 | 0.03 | 0.049 | 0.049 | 0 | 0.12 | 0.24 | 0.29 | 0.27 | 0.19 | 0.21 |
| 2 | 6389676 | rs7567289 | 0.255 | 0.991 | 0.394 | 1.067 | -0.014 | -0.009 | -0.009 | 0 | 0.83 | 0.91 | 0.76 | 0.84 | 0.78 | 0.78 |
| 2 | 6389792 | rs13003180 | 1.251 | 0.958 | 0.615 | 1.088 | -0.001 | 0.001 | 0.001 | 0 | 0.72 | 0.7 | 0.63 | 0.64 | 0.68 | 0.66 |
| 2 | 6392335 | rs921229 | 0.931 | 0.89 | 0.661 | 0.959 | -0.004 | -0.002 | -0.002 | 0.293 | 0.79 | 0.73 | 0.81 | 0.8 | 0.83 | 0.7 |
| 2 | 6396424 | rs2351100 | -0.262 | 0.933 | 0.688 | 1.026 | -0.004 | 0.029 | 0.029 | 0 | 0.11 | 0.21 | 0.13 | 0.2 | 0.1 | 0.12 |
| 2 | 6398203 | rs10193208 | 1.49 | 0.926 | 1.465 | 1.457 | 0.014 | 0.066 | 0.066 | 0 | 0.76 | 0.47 | 0.63 | 0.64 | 0.73 | 0.64 |
| 2 | 6402296 | rs4538171 | 1.533 | 0.902 | 1.575 | 1.435 | 0.006 | 0.036 | 0.036 | 0 | 0.76 | 0.4 | 0.62 | 0.64 | 0.73 | 0.64 |
| 2 | 6413931 | rs6431789 | -0.192 | 0.916 | 1.135 | 1.231 | 0.002 | 0.006 | 0.006 | 0 | 0.9 | 0.93 | 0.94 | 1 | 0.93 | 0.83 |
| 2 | 6415415 | rs10199875 | -0.728 | 0.793 | 0.821 | 1.24 | -0.005 | -0.005 | -0.005 | 0 | 0.12 | 0.05 | 0.19 | 0.15 | 0.15 | 0.09 |
| 2 | 6421045 | rs12623864 | 1.978 | 0.63 | 0.852 | 1.45 | -0.011 | -0.036 | -0.036 | 6.075 | 0.62 | 0.21 | 0.42 | 0.49 | 0.57 | 0.55 |
| 2 | 6421388 | rs12329052 | 1.601 | 0.644 | 1.092 | 1.385 | -0.011 | -0.013 | -0.013 | 0 | 0.7 | 0.52 | 0.67 | 0.76 | 0.69 | 0.65 |
| 2 | 6424508 | rs13391757 | -0.823 | 0.809 | 0.475 | 1.435 | -0.005 | -0.009 | -0.009 | 0.292 | 0.11 | 0.05 | 0.19 | 0.15 | 0.15 | 0.09 |
| 2 | 6434587 | rs4337447 | -1.373 | 0.798 | 1.316 | 1.544 | 0.068 | 0.235 | 0.235 | 0 | 0.1 | 0.36 | 0.27 | 0.27 | 0.13 | 0.11 |
| 2 | 6440370 | rs751394 | 0.607 | 1.155 | 0.852 | 1.131 | -0.012 | -0.029 | -0.029 | 0 | 0.72 | 0.78 | 0.59 | 0.45 | 0.65 | 0.69 |

| | | | | | | | | | | | | | | | | |
|---|---|---|---|---|---|---|---|---|---|---|---|---|---|---|---|---|
| 2 | 6443216 | rs6752376 | 1.342 | 1.406 | 1.07 | 0.6 | 0.012 | 0.021 | 0.021 | 0 | 0.51 | 0.38 | 0.41 | 0.34 | 0.42 | 0.51 |
| 2 | 6444393 | rs1816975 | 0.809 | 1.528 | 0.859 | 0.582 | -0.016 | 0.022 | 0.022 | 0.063 | 0.57 | 0.66 | 0.47 | 0.35 | 0.49 | 0.52 |
| 2 | 6449896 | rs1562622 | 1.678 | 1.074 | -0.261 | 0.237 | 0.006 | 0.032 | 0.032 | 0 | 0.42 | 0.18 | 0.37 | 0.35 | 0.4 | 0.38 |
| 2 | 6450098 | rs1562621 | 1.143 | 0.837 | -0.119 | 0.063 | -0.016 | 0.041 | 0.041 | 1.127 | 0.54 | 0.22 | 0.42 | 0.38 | 0.48 | 0.47 |
| 2 | 6461421 | rs7591996 | -0.748 | 0.555 | -0.599 | -0.563 | 0.022 | 0.042 | 0.042 | 1.789 | 0.55 | 0.84 | 0.54 | 0.64 | 0.46 | 0.51 |
| 2 | 6465033 | rs10929424 | 1.036 | 0.625 | -0.257 | -0.591 | -0.018 | 0.005 | 0.005 | 0.448 | 0.37 | 0.24 | 0.37 | 0.35 | 0.33 | 0.43 |
| 2 | 6466163 | rs6751561 | 1.01 | 0.52 | -0.015 | -0.672 | -0.001 | -0.005 | -0.005 | 0 | 0.11 | 0.09 | 0.07 | 0.09 | 0.11 | 0.1 |
| 2 | 6474334 | rs7568761 | 1.291 | 0.545 | -0.163 | -0.676 | -0.042 | -0.004 | -0.004 | 0 | 0.37 | 0.14 | 0.36 | 0.35 | 0.34 | 0.44 |
| 2 | 6478428 | rs4669060 | -0.994 | 0.69 | 0.075 | -0.663 | 0.025 | -0.028 | -0.028 | 0 | 0.93 | 0.89 | 0.92 | 1 | 0.81 | 0.94 |
| 2 | 6478838 | rs13403411 | 0.556 | 0.496 | 0.428 | -0.739 | 0.028 | -0.027 | -0.027 | 0 | 0.37 | 0.2 | 0.42 | 0.54 | 0.32 | 0.4 |
| 2 | 6485953 | rs12995389 | 0.469 | 0.538 | 0.469 | -0.763 | -0.006 | 0.031 | 0.031 | 0 | 0.48 | 0.23 | 0.43 | 0.43 | 0.44 | 0.51 |
| 2 | 6486246 | rs16865045 | 1.142 | 0.561 | 0.493 | -0.759 | -0.001 | -0.003 | -0.003 | 0 | 0.11 | 0.09 | 0.08 | 0.09 | 0.11 | 0.07 |
| 2 | 6489121 | rs16865056 | 1.047 | 0.555 | 0.712 | -0.762 | -0.001 | -0.008 | -0.008 | 0 | 0.11 | 0.12 | 0.08 | 0.09 | 0.11 | 0.07 |
| 2 | 6492395 | rs726342 | 1.44 | 0.542 | 0.582 | -0.787 | -0.074 | -0.015 | -0.015 | 0.548 | 0.37 | 0.1 | 0.35 | 0.34 | 0.33 | 0.43 |
| 2 | 6494761 | rs9287675 | -0.909 | 0.442 | 0.999 | -0.81 | 0.006 | 0.032 | 0.032 | 0.333 | 0.55 | 0.68 | 0.55 | 0.65 | 0.48 | 0.5 |
| 2 | 6502497 | rs10205734 | -0.1 | 0.426 | 1.069 | -0.802 | -0.014 | -0.005 | -0.005 | 0 | 0.43 | 0.5 | 0.48 | 0.56 | 0.36 | 0.43 |
| 2 | 6503318 | rs11694771 | -0.698 | 0.382 | 0.851 | -0.806 | 0.01 | 0.037 | 0.037 | 0.37 | 0.53 | 0.61 | 0.56 | 0.65 | 0.47 | 0.5 |
| 2 | 6503855 | rs11679546 | -0.876 | 0.503 | 0.472 | -0.747 | -0.007 | 0.056 | 0.056 | 0.895 | 0.56 | 0.8 | 0.57 | 0.65 | 0.47 | 0.5 |
| 2 | 6522333 | rs6739653 | -0.831 | 0.726 | 0.541 | -0.715 | 0.032 | -0.029 | -0.029 | 0 | 0.94 | 0.9 | 0.92 | 1 | 0.8 | 0.94 |
| 2 | 6523425 | rs9989754 | -0.653 | 0.581 | 0.847 | -0.933 | -0.013 | -0.079 | -0.079 | 0.142 | 0.2 | 0.48 | 0.16 | 0.09 | 0.19 | 0.12 |
| 2 | 6523796 | rs10445712 | 1.507 | 0.49 | 0.477 | -0.958 | -0.069 | -0.012 | -0.012 | 0 | 0.37 | 0.1 | 0.34 | 0.35 | 0.33 | 0.43 |
| 2 | 6523994 | rs10208273 | 0.608 | 0.397 | 0.584 | -0.867 | 0.026 | -0.018 | -0.018 | 0.019 | 0.35 | 0.21 | 0.4 | 0.55 | 0.29 | 0.39 |
| 2 | 6529635 | rs1564031 | -0.409 | 0.353 | 0.963 | -0.919 | 0.021 | 0.026 | 0.026 | 0 | 0.47 | 0.34 | 0.49 | 0.65 | 0.39 | 0.46 |
| 2 | 6533291 | rs16865111 | 1.266 | 0.295 | 0.909 | -0.855 | -0.003 | -0.009 | -0.009 | 0 | 0.11 | 0.08 | 0.07 | 0.09 | 0.1 | 0.06 |
| 2 | 6533326 | rs6713599 | 0.603 | 0.392 | 1.175 | -0.899 | 0 | -0.006 | -0.006 | 0.062 | 0.47 | 0.59 | 0.44 | 0.35 | 0.41 | 0.48 |
| 2 | 6536892 | rs4669066 | 0.251 | 0.477 | 1.064 | -0.831 | -0.001 | -0.002 | -0.002 | 0 | 0.51 | 0.48 | 0.45 | 0.44 | 0.46 | 0.52 |
| 2 | 6541599 | rs13030822 | 1.337 | 0.677 | 0.606 | -0.825 | -0.067 | -0.025 | -0.025 | 0.293 | 0.37 | 0.07 | 0.33 | 0.35 | 0.32 | 0.44 |
| 2 | 6548066 | rs16865126 | 0.576 | 0.618 | 0.881 | -0.759 | -0.008 | -0.024 | -0.024 | 0 | 0.12 | 0.21 | 0.08 | 0.09 | 0.1 | 0.07 |
| 2 | 6549520 | rs7577880 | 1.081 | 0.396 | 0.865 | -0.786 | -0.008 | 0.018 | 0.018 | 0.019 | 0.4 | 0.3 | 0.4 | 0.35 | 0.35 | 0.47 |
| 2 | 6550266 | rs1462060 | 0.478 | 0.199 | 1.137 | -0.813 | -0.027 | -0.015 | -0.015 | 0 | 0.11 | 0.27 | 0.08 | 0.09 | 0.08 | 0.07 |
| 2 | 6551857 | rs6723180 | -0.442 | 0.147 | 0.575 | -1.217 | 0 | -0.036 | -0.036 | 0 | 0.4 | 0.56 | 0.32 | 0.16 | 0.49 | 0.21 |
| 2 | 6553800 | rs4594412 | 0.591 | -0.009 | 0.245 | -1.097 | 0.002 | -0.049 | -0.049 | 0.891 | 0.25 | 0.12 | 0.27 | 0.49 | 0.19 | 0.32 |
| 2 | 6555732 | rs17380176 | 1.243 | -0.047 | 0.354 | -1.161 | 0.001 | 0.054 | 0.054 | 0 | 0.26 | 0.2 | 0.19 | 0.06 | 0.27 | 0.09 |
| 2 | 6561446 | rs7577180 | 0.7 | -0.533 | 0.413 | -0.694 | -0.025 | -0.013 | -0.013 | 0 | 0.1 | 0.24 | 0.08 | 0.1 | 0.08 | 0.07 |
| 2 | 6566079 | rs730632 | 0.314 | -0.358 | 0.155 | -0.603 | 0.017 | -0.044 | -0.044 | 0 | 0.25 | 0.13 | 0.27 | 0.5 | 0.15 | 0.27 |
| 2 | 6566543 | rs384180 | -0.514 | -0.063 | 0.763 | -0.343 | -0.013 | -0.004 | -0.004 | 0 | 0.9 | 0.68 | 0.91 | 0.92 | 0.92 | 0.92 |
| 2 | 6569684 | rs12619623 | 0.194 | -0.099 | -0.082 | -0.463 | -0.009 | 0.009 | 0.009 | 3.655 | 0.58 | 0.24 | 0.45 | 0.58 | 0.43 | 0.4 |
| 2 | 6573786 | rs10208426 | -0.212 | 0.191 | 0.126 | -0.27 | 0.029 | 0 | 0 | 0.751 | 0.65 | 0.54 | 0.54 | 0.64 | 0.5 | 0.44 |
| 2 | 6576150 | rs17380246 | 0.855 | 0.347 | 0.085 | -0.402 | 0.014 | 0.079 | 0.079 | 0 | 0.33 | 0.15 | 0.21 | 0 | 0.31 | 0.1 |
| 2 | 6576255 | rs16865157 | -0.083 | 0.573 | -0.252 | 0.18 | -0.003 | -0.023 | -0.023 | 0 | 0.2 | 0.17 | 0.33 | 0.43 | 0.19 | 0.22 |
| 2 | 6577458 | rs306158 | -0.901 | 0.418 | 0.076 | -0.897 | -0.022 | -0.164 | -0.164 | 0 | 0.63 | 0.51 | 0.75 | 1 | 0.69 | 0.88 |
| 2 | 6580745 | rs10495536 | -0.156 | 1.092 | -0.522 | -0.416 | -0.008 | 0.02 | 0.02 | 0.272 | 0.18 | 0.11 | 0.14 | 0.04 | 0.16 | 0.16 |
| 2 | 6581866 | rs13003560 | 0.411 | 1.439 | -0.258 | -0.24 | -0.019 | 0.041 | 0.041 | 0.076 | 0.37 | 0.27 | 0.57 | 0.85 | 0.47 | 0.61 |
| 2 | 6583763 | rs4669070 | -0.433 | 1.903 | 0.003 | 1.198 | 0 | -0.014 | -0.014 | 0 | 0.05 | 0.01 | 0.03 | 0.11 | 0.06 | 0.09 |
| 2 | 6585051 | rs7558641 | -0.963 | 1.898 | 0.048 | 1.229 | -0.001 | -0.001 | -0.001 | 0 | 0.23 | 0.2 | 0.18 | 0.15 | 0.22 | 0.25 |
| 2 | 6593923 | rs16865170 | -1.335 | 1.452 | 0.985 | 1.784 | 0.094 | 0.05 | 0.05 | 0 | 0.09 | 0.35 | 0.17 | 0.1 | 0.13 | 0.1 |
| 2 | 6594973 | rs1585535 | -0.731 | 1.236 | 0.032 | 1.487 | -0.031 | 0.007 | 0.007 | 0 | 0.35 | 0.29 | 0.5 | 0.61 | 0.44 | 0.58 |
| 2 | 6600177 | rs931007 | -0.69 | 1.096 | 0.291 | 1.771 | -0.034 | -0.037 | -0.037 | 0 | 0.32 | 0.23 | 0.57 | 0.47 | 0.51 | 0.57 |
| 2 | 6619062 | rs167378 | 1.195 | 0.634 | 1.854 | 1.524 | -0.006 | 0.04 | 0.04 | 0.115 | 0.87 | 0.52 | 0.89 | 0.85 | 0.91 | 0.88 |
| 2 | 6624325 | rs381770 | 0.883 | 0.664 | 1.66 | 1.531 | -0.008 | 0.016 | 0.016 | 0 | 0.9 | 0.8 | 0.92 | 0.85 | 0.91 | 0.89 |
| 2 | 6627807 | rs306196 | 0.662 | 0.648 | 1.31 | 1.553 | 0.004 | -0.009 | -0.009 | 0 | 0.91 | 0.94 | 0.92 | 0.85 | 0.91 | 0.91 |
| 2 | 6641549 | rs4668498 | -0.98 | 0.496 | 0.993 | 1.647 | -0.016 | -0.037 | -0.037 | 1.667 | 0.38 | 0.23 | 0.55 | 0.48 | 0.53 | 0.58 |
| 2 | 6643906 | rs413642 | -0.867 | 0.366 | 0.633 | 1.664 | -0.043 | -0.086 | -0.086 | 2.104 | 0.37 | 0.12 | 0.54 | 0.49 | 0.54 | 0.58 |
| 2 | 6646177 | rs306184 | -1.51 | 0.919 | 1.059 | 1.336 | -0.007 | -0.036 | -0.036 | 1.155 | 0.36 | 0.54 | 0.24 | 0.19 | 0.37 | 0.27 |
| 2 | 6646250 | rs306183 | -0.046 | 0.919 | 0.985 | 1.302 | -0.021 | 0.057 | 0.057 | 0 | 0.22 | 0.1 | 0.13 | 0.04 | 0.28 | 0.15 |
| 2 | 6646953 | rs306182 | -0.132 | 0.936 | 0.993 | 1.282 | -0.016 | 0.054 | 0.054 | 0 | 0.23 | 0.12 | 0.13 | 0.03 | 0.28 | 0.15 |
| 2 | 6650482 | rs1462063 | 1.298 | 0.937 | 1.053 | 1.363 | 0.011 | 0.029 | 0.029 | 0 | 0.6 | 0.28 | 0.75 | 0.73 | 0.62 | 0.73 |
| 2 | 6650842 | rs16865273 | -1.24 | 0.993 | 0.317 | 1.389 | 0.009 | -0.015 | -0.015 | 0 | 0.12 | 0.08 | 0.07 | 0.2 | 0.11 | 0.1 |
| 2 | 6651312 | rs16865276 | -1.24 | 0.993 | 0.317 | 1.389 | 0.009 | -0.015 | -0.015 | 0 | 0.12 | 0.08 | 0.07 | 0.2 | 0.11 | 0.1 |
| 2 | 6652490 | rs11696092 | 1.854 | 1.026 | 0.279 | 1.499 | -0.003 | -0.004 | -0.004 | 0.069 | 0.47 | 0.39 | 0.33 | 0.34 | 0.35 | 0.3 |
| 2 | 6652942 | rs16865282 | -0.682 | 0.838 | -0.385 | 1.853 | -0.087 | -0.095 | -0.095 | 0 | 0.14 | 0.02 | 0.43 | 0.42 | 0.26 | 0.44 |
| 2 | 6653510 | rs12162425 | -1.315 | 0.237 | -0.072 | 0.984 | 0.002 | -0.03 | -0.03 | 0 | 0.1 | 0.01 | 0.07 | 0.17 | 0.09 | 0.1 |
| 2 | 6655781 | rs16865292 | -0.323 | 0.24 | 0.26 | 0.861 | 0.001 | -0.009 | -0.009 | 0 | 0.06 | 0.08 | 0.04 | 0.04 | 0.03 | 0.06 |
| 2 | 6657578 | rs16865299 | -1.774 | 0.399 | 0.788 | 1.131 | 0.036 | 0.067 | 0.067 | 0 | 0.27 | 0.34 | 0.55 | 0.46 | 0.38 | 0.57 |
| 2 | 6657760 | rs16865301 | -0.189 | 0.402 | 0.756 | 1.058 | -0.002 | -0.001 | -0.001 | 0 | 0.13 | 0.14 | 0.12 | 0.27 | 0.15 | 0.11 |
| 2 | 6659744 | rs1379465 | 0.54 | -0.1 | 0.575 | 1.052 | 0.038 | -0.047 | -0.047 | 0 | 0.92 | 0.98 | 0.98 | 0.78 | 0.98 | 0.96 |
| 2 | 6660497 | rs17381941 | -0.643 | 0.243 | 0.764 | 1.113 | -0.012 | 0.025 | 0.025 | 0.061 | 0.08 | 0.04 | 0.08 | 0 | 0.14 | 0.08 |
| 2 | 6665456 | rs3919811 | -0.496 | 0.245 | 0.846 | 1.157 | 0.043 | 0.163 | 0.163 | 0 | 0.13 | 0.23 | 0.42 | 0.42 | 0.2 | 0.43 |
| 2 | 6666613 | rs6752421 | -1.243 | -0.266 | 0.837 | 0.857 | 0.001 | -0.023 | -0.023 | 0.203 | 0.14 | 0.18 | 0.11 | 0.04 | 0.18 | 0.15 |

| | | | | | | | | | | | | | | | | |
|---|---|---|---|---|---|---|---|---|---|---|---|---|---|---|---|---|
| 2 | 6679793 | rs4669075 | 1.059 | -0.849 | 0.361 | 0.057 | -0.053 | -0.065 | -0.065 | 0.11 | 0.68 | 0.43 | 0.85 | 0.83 | 0.81 | 0.85 |
| 2 | 6683070 | rs6431809 | -0.369 | -0.405 | -0.183 | 0.039 | -0.01 | -0.04 | -0.04 | 0.295 | 0.16 | 0.18 | 0.05 | 0.01 | 0.07 | 0.08 |
| 2 | 6692014 | rs10202071 | 0.48 | -0.24 | -0.75 | -0.083 | -0.048 | 0.055 | 0.055 | 0 | 0.35 | 0.19 | 0.27 | 0.16 | 0.42 | 0.34 |
| 2 | 6694519 | rs10495537 | 0.434 | -0.736 | -1.311 | 0.361 | -0.02 | -0.095 | -0.095 | 0.366 | 0.33 | 0.05 | 0.26 | 0.44 | 0.3 | 0.38 |
| 2 | 6695161 | rs1870131 | 0.665 | -0.78 | -1.018 | 0.514 | 0.008 | 0.01 | 0.01 | 0 | 0.05 | 0.06 | 0.08 | 0.08 | 0.08 | 0.06 |
| 2 | 6699717 | rs12185687 | -0.051 | -0.974 | -0.184 | 0.485 | 0.021 | 0.038 | 0.038 | 1.634 | 0.05 | 0.29 | 0.03 | 0.03 | 0.01 | 0.03 |
| 2 | 6700873 | rs6748722 | -0.153 | -0.758 | -0.112 | 0.501 | -0.003 | 0.001 | 0.001 | 0.424 | 0.44 | 0.51 | 0.37 | 0.55 | 0.39 | 0.47 |
| 2 | 6703291 | rs4669076 | 0.981 | -0.323 | -0.541 | 0.485 | -0.116 | 0.091 | 0.091 | 0.011 | 0.33 | 0.08 | 0.27 | 0.16 | 0.42 | 0.34 |
| 2 | 6704579 | rs16865340 | -0.409 | 0.096 | 0.227 | 0.967 | 0.035 | 0.039 | 0.039 | 0 | 0.06 | 0.15 | 0.1 | 0.08 | 0.08 | 0.06 |
| 2 | 6706459 | rs1159673 | 0.365 | -0.18 | 0.153 | 0.994 | -0.008 | 0.026 | 0.026 | 0 | 0.81 | 0.73 | 0.66 | 0.69 | 0.81 | 0.81 |
| 2 | 6707387 | rs16865345 | -0.915 | -0.064 | 0.667 | 0.741 | 0.003 | -0.006 | -0.006 | 0 | 0.19 | 0.35 | 0.26 | 0.16 | 0.26 | 0.2 |
| 2 | 6711467 | rs2219379 | 0.938 | -0.47 | 1.055 | 0.366 | -0.006 | -0.022 | -0.022 | 0 | 0.73 | 0.67 | 0.73 | 0.85 | 0.72 | 0.75 |
| 2 | 6711631 | rs4669077 | 0.783 | -0.345 | 0.143 | 0.047 | -0.033 | 0.071 | 0.071 | 0 | 0.41 | 0.14 | 0.25 | 0.28 | 0.41 | 0.35 |
| 2 | 6717572 | rs6746495 | 0.471 | 0.209 | 0.401 | -0.738 | 0 | -0.104 | -0.104 | 0.001 | 0.35 | 0.44 | 0.3 | 0.03 | 0.32 | 0.33 |
| 2 | 6720602 | rs11891157 | 0.468 | 0.21 | 0.526 | -0.738 | 0 | -0.053 | -0.053 | 0.146 | 0.35 | 0.4 | 0.3 | 0.03 | 0.32 | 0.33 |
| 2 | 6723895 | rs1563613 | -0.362 | 0.283 | 0.515 | -0.573 | 0 | -0.016 | -0.016 | 2.479 | 0.6 | 0.41 | 0.67 | 0.85 | 0.64 | 0.65 |
| 2 | 6725121 | rs16865372 | 1.45 | 0.294 | 0.355 | -0.244 | 0.021 | -0.081 | -0.081 | 0 | 0.23 | 0.11 | 0.25 | 0.51 | 0.21 | 0.34 |
| 2 | 6727808 | rs7560214 | 0.33 | 0.313 | 0.065 | -0.337 | 0 | -0.015 | -0.015 | 0.191 | 0.46 | 0.53 | 0.39 | 0.25 | 0.44 | 0.45 |
| 2 | 6729707 | rs17803843 | 0.482 | 0.417 | 0.186 | -0.406 | -0.018 | -0.017 | -0.017 | 0.157 | 0.18 | 0.09 | 0.14 | 0.21 | 0.22 | 0.11 |
| 2 | 6730225 | rs1457515 | 1.76 | 0.412 | 0.174 | -0.422 | 0.008 | -0.143 | -0.143 | 0.634 | 0.22 | 0.03 | 0.24 | 0.52 | 0.21 | 0.34 |
| 2 | 6731740 | rs12988213 | 1.699 | 0.425 | 0.177 | -0.416 | 0.02 | -0.112 | -0.112 | 0 | 0.23 | 0.09 | 0.24 | 0.52 | 0.21 | 0.34 |
| 2 | 6735155 | rs13383761 | 0.973 | 0.435 | 0.238 | -0.431 | 0.01 | -0.035 | -0.035 | 0 | 0.1 | 0.18 | 0.21 | 0.02 | 0.14 | 0.12 |
| 2 | 6746052 | rs11675022 | 0.662 | 0.44 | 0.031 | -0.385 | 0 | -0.011 | -0.011 | 0 | 0.43 | 0.45 | 0.37 | 0.14 | 0.4 | 0.42 |
| 2 | 6757157 | rs755411 | 1.385 | 0.428 | 0.051 | -0.358 | -0.04 | -0.033 | -0.033 | 0.502 | 0.55 | 0.19 | 0.53 | 0.56 | 0.53 | 0.65 |
| 2 | 6758957 | rs4668504 | 0.716 | 0.463 | 0.353 | -0.047 | -0.007 | 0.06 | 0.06 | 0 | 0.84 | 0.49 | 0.88 | 0.8 | 0.88 | 0.88 |
| 2 | 6767108 | rs10495541 | 0.189 | 0.41 | 0.352 | -0.033 | -0.001 | 0.024 | 0.024 | 0 | 0.07 | 0.06 | 0.07 | 0.21 | 0.07 | 0.07 |
| 2 | 6769988 | rs1563948 | 0.343 | 0.248 | 0.372 | 0.024 | -0.015 | 0.028 | 0.028 | 0 | 0.85 | 0.58 | 0.91 | 0.9 | 0.91 | 0.89 |
| 2 | 6771256 | rs10495540 | 1.351 | 0.194 | 0.317 | -0.022 | 0.022 | -0.082 | -0.082 | 0 | 0.23 | 0.11 | 0.24 | 0.51 | 0.21 | 0.35 |
| 2 | 6773920 | rs11694697 | -0.367 | 0.568 | 0.944 | -0.364 | -0.008 | 0.019 | 0.019 | 0 | 0.12 | 0.27 | 0.08 | 0.1 | 0.09 | 0.11 |
| 2 | 6777829 | rs17815605 | 0.857 | 0.569 | 0.568 | -0.385 | -0.007 | 0.004 | 0.004 | 0 | 0.12 | 0.07 | 0.07 | 0 | 0.15 | 0.04 |
| 2 | 6777992 | rs11692570 | -0.716 | 0.353 | 0.664 | 0.001 | -0.002 | 0.036 | 0.036 | 0 | 0.13 | 0.37 | 0.08 | 0.1 | 0.09 | 0.11 |
| 2 | 6779277 | rs2219613 | 0.692 | 0.376 | 0.468 | 0.013 | -0.015 | 0.028 | 0.028 | 0 | 0.85 | 0.58 | 0.91 | 0.9 | 0.91 | 0.89 |
| 2 | 6787890 | rs930976 | 0.823 | 1.034 | -0.148 | -0.409 | -0.008 | -0.069 | -0.069 | 0.121 | 0.24 | 0.31 | 0.21 | 0.04 | 0.2 | 0.26 |
| 2 | 6788641 | rs16865459 | 1.259 | 0.804 | -0.809 | -0.733 | 0.05 | -0.048 | -0.048 | 1.419 | 0.26 | 0.07 | 0.26 | 0.66 | 0.17 | 0.38 |
| 2 | 6790038 | rs10495538 | 1.41 | 1.323 | -0.884 | -1.16 | -0.012 | 0.019 | 0.019 | 0 | 0.12 | 0.07 | 0.1 | 0 | 0.18 | 0.04 |
| 2 | 6793933 | rs4436930 | 0.725 | 1.669 | -0.954 | -1.329 | -0.014 | 0.047 | 0.047 | 0 | 0.19 | 0.08 | 0.23 | 0.05 | 0.21 | 0.15 |
| 2 | 6795143 | rs12620029 | 1.481 | 1.399 | -0.829 | -1.013 | 0.048 | -0.044 | -0.044 | 0.329 | 0.25 | 0.08 | 0.26 | 0.7 | 0.17 | 0.37 |
| 2 | 6795330 | rs10172433 | 0.491 | 1.465 | -0.69 | -0.991 | 0 | -0.027 | -0.027 | 0.385 | 0.43 | 0.41 | 0.45 | 0.09 | 0.4 | 0.41 |
| 2 | 6803109 | rs380019 | 0.265 | 1.532 | -0.838 | -0.884 | -0.001 | 0.03 | 0.03 | 0 | 0.38 | 0.16 | 0.39 | 0.69 | 0.34 | 0.52 |
| 2 | 6810021 | rs308011 | 0.352 | 1.463 | -0.852 | -1.069 | -0.019 | 0.035 | 0.035 | 0.261 | 0.14 | 0.08 | 0.16 | 0.04 | 0.18 | 0.16 |
| 2 | 6825279 | rs672463 | 0.02 | 1.534 | -0.721 | -1.038 | -0.002 | -0.074 | -0.074 | 0 | 0.43 | 0.48 | 0.41 | 0.88 | 0.43 | 0.43 |
| 2 | 6833940 | rs2351935 | 0.305 | 1.033 | -1.138 | -0.829 | -0.003 | -0.064 | -0.064 | 0 | 0.44 | 0.49 | 0.45 | 0.09 | 0.41 | 0.4 |
| 2 | 6834951 | rs10929447 | 0.605 | 0.576 | -1.639 | 0.174 | 0.036 | -0.076 | -0.076 | 0 | 0.25 | 0.08 | 0.25 | 0.63 | 0.17 | 0.37 |
| 2 | 6835876 | rs7582948 | -0.507 | 1.281 | -1.194 | -0.22 | -0.028 | 0.043 | 0.043 | 0.164 | 0.34 | 0.23 | 0.37 | 0.13 | 0.43 | 0.26 |
| 2 | 6836359 | rs308003 | 0.753 | 1.1 | -0.659 | 0.131 | 0.033 | 0.007 | 0.007 | 0.028 | 0.77 | 0.66 | 0.63 | 0.79 | 0.71 | 0.8 |
| 2 | 6839127 | rs11694230 | -0.643 | -0.375 | -1.498 | 0.015 | -0.011 | 0.011 | 0.011 | 0.06 | 0.36 | 0.1 | 0.37 | 0.73 | 0.37 | 0.63 |
| 2 | 6841330 | rs307973 | -1.564 | -1.036 | -1.606 | -0.485 | -0.066 | 0.149 | 0.149 | 0 | 0.62 | 0.82 | 0.68 | 0.91 | 0.54 | 0.83 |
| 2 | 6851976 | rs307992 | 0.292 | -1.101 | -0.368 | -0.595 | -0.006 | -0.086 | -0.086 | 0 | 0.16 | 0.27 | 0.23 | 0 | 0.12 | 0.1 |
| 2 | 6852941 | rs13382813 | -0.333 | -1.256 | -0.077 | -0.506 | 0.034 | 0.201 | 0.201 | 0 | 0.08 | 0.38 | 0.07 | 0.15 | 0.05 | 0.08 |
| 2 | 6856893 | rs16865531 | -0.213 | -1.259 | -0.041 | -0.506 | 0.045 | 0.261 | 0.261 | 0 | 0.08 | 0.39 | 0.07 | 0.17 | 0.05 | 0.09 |
| 2 | 6866483 | rs13413734 | -0.25 | -1.261 | 0.584 | -0.508 | 0.022 | 0.202 | 0.202 | 0 | 0.07 | 0.31 | 0.06 | 0.17 | 0.05 | 0.08 |
| 2 | 6866987 | rs12620748 | -0.158 | -1.517 | 0.046 | -0.689 | -0.011 | 0.061 | 0.061 | 0 | 0.36 | 0.15 | 0.38 | 0.8 | 0.37 | 0.62 |
| 2 | 6869496 | rs308020 | -1.992 | -1.557 | -0.161 | -0.789 | -0.174 | 0.248 | 0.248 | 0 | 0.64 | 0.95 | 0.68 | 0.93 | 0.52 | 0.81 |
| 2 | 6870151 | rs11687816 | 0.357 | -1.041 | 0.125 | -0.55 | -0.022 | 0.024 | 0.024 | 0.256 | 0.09 | 0.04 | 0.07 | 0.01 | 0.13 | 0.14 |
| 2 | 6871849 | rs1870744 | 0.332 | -1.04 | 0.079 | -0.55 | -0.034 | 0.04 | 0.04 | 0.334 | 0.09 | 0.02 | 0.07 | 0.01 | 0.13 | 0.15 |
| 2 | 6873356 | rs308022 | -0.029 | -1.082 | 0.436 | -0.488 | -0.002 | -0.121 | -0.121 | 0.007 | 0.16 | 0.37 | 0.24 | 0 | 0.12 | 0.1 |
| 2 | 6874452 | rs308027 | 0.132 | -1.049 | 0.047 | -0.47 | -0.006 | -0.089 | -0.089 | 1.882 | 0.16 | 0.27 | 0.23 | 0 | 0.12 | 0.1 |
| 2 | 6886617 | rs954845 | -0.997 | -1.078 | -0.112 | 0.186 | 0 | 0 | 0 | 3.443 | 0.44 | 0.42 | 0.54 | 0.43 | 0.4 | 0.5 |
| 2 | 6898693 | rs11690692 | -0.229 | -1.172 | -0.71 | -0.444 | -0.022 | -0.02 | -0.02 | 0 | 0.21 | 0.27 | 0.16 | 0.12 | 0.15 | 0.15 |
| 2 | 6904791 | rs950370 | -0.068 | -0.833 | -1.075 | -0.52 | -0.01 | -0.021 | -0.021 | 0.311 | 0.13 | 0.17 | 0.14 | 0.07 | 0.09 | 0.19 |
| 2 | 6908119 | rs4668506 | -0.788 | -0.399 | -1.916 | -0.792 | -0.077 | 0.215 | 0.215 | 0 | 0.61 | 0.91 | 0.7 | 0.93 | 0.5 | 0.79 |
| 2 | 6914714 | rs1006422 | -0.756 | -0.415 | -1.218 | 0.473 | -0.001 | -0.001 | -0.001 | 0 | 0.14 | 0.11 | 0.3 | 0.67 | 0.16 | 0.33 |
| 2 | 6922184 | rs6760979 | -0.885 | -0.401 | -1.451 | -0.134 | -0.005 | 0.096 | 0.096 | 0 | 0.55 | 0.84 | 0.6 | 0.8 | 0.43 | 0.68 |
| 2 | 6924167 | rs1566982 | -0.098 | -0.225 | -1.114 | -0.195 | -0.026 | -0.112 | -0.112 | 2.807 | 0.23 | 0.37 | 0.11 | 0.03 | 0.14 | 0.25 |
| 2 | 6929318 | rs4669100 | -0.551 | 0.236 | -1.629 | 0.711 | -0.017 | -0.063 | -0.063 | 1.2 | 0.26 | 0.07 | 0.44 | 0.65 | 0.28 | 0.42 |
| 2 | 6934071 | rs10495543 | -1.041 | 0.358 | -1.215 | 1.346 | 0 | 0.044 | 0.044 | 0 | 0.08 | 0.12 | 0.31 | 0.65 | 0.13 | 0.23 |
| 2 | 6935239 | rs905968 | -0.502 | -0.116 | -1.18 | 0.092 | 0.001 | 0.003 | 0.003 | 0.023 | 0.54 | 0.6 | 0.66 | 0.68 | 0.48 | 0.59 |
| 2 | 6936438 | rs12466123 | 0.22 | -0.678 | -0.983 | 0.506 | 0.006 | -0.011 | -0.011 | 0 | 0.28 | 0.27 | 0.28 | 0.36 | 0.24 | 0.3 |
| 2 | 6937310 | rs2174519 | -0.44 | -0.697 | -0.675 | 0.54 | 0.085 | 0.014 | 0.014 | 0.035 | 0.52 | 0.77 | 0.41 | 0.38 | 0.38 | 0.47 |

| | | | | | | | | | | | | | | | | |
|---|---|---|---|---|---|---|---|---|---|---|---|---|---|---|---|---|
| 2 | 6937593 | rs10200467 | 0.371 | -0.271 | -1.142 | 0.217 | -0.007 | -0.027 | -0.027 | 3.46 | 0.23 | 0.19 | 0.37 | 0.37 | 0.25 | 0.26 |
| 2 | 6937744 | rs6742876 | 0.103 | -0.953 | -0.635 | 0.089 | 0.035 | -0.005 | -0.005 | 0 | 0.43 | 0.27 | 0.36 | 0.38 | 0.37 | 0.33 |
| 2 | 6937849 | rs6742989 | -0.285 | -0.997 | -0.167 | 0.068 | 0.026 | 0.007 | 0.007 | 0.131 | 0.48 | 0.4 | 0.38 | 0.38 | 0.38 | 0.34 |
| 2 | 6941288 | rs7600900 | -0.214 | -0.655 | -0.284 | 0.24 | 0.04 | 0.008 | 0.008 | 0 | 0.47 | 0.63 | 0.38 | 0.38 | 0.38 | 0.46 |
| 2 | 6944753 | rs771175 | -0.133 | 0.165 | -0.224 | 0.071 | 0.008 | -0.017 | -0.017 | 0.713 | 0.39 | 0.39 | 0.38 | 0.35 | 0.45 | 0.37 |
| 2 | 6946945 | rs6431831 | -0.156 | -0.134 | 0.407 | -0.44 | -0.002 | 0.002 | 0.002 | 0 | 0.6 | 0.42 | 0.6 | 0.67 | 0.56 | 0.66 |
| 2 | 6947587 | rs1876105 | -0.326 | -0.05 | 0.226 | -0.442 | -0.005 | 0.007 | 0.007 | 0 | 0.62 | 0.7 | 0.62 | 0.67 | 0.55 | 0.66 |
| 2 | 6948715 | rs771177 | -1.323 | 0.022 | -0.21 | -0.453 | 0 | 0.029 | 0.029 | 0 | 0.94 | 0.99 | 0.97 | 1 | 0.94 | 0.99 |
| 2 | 6951030 | rs1077583 | 0.41 | -0.346 | -0.257 | -0.588 | -0.014 | 0.142 | 0.142 | 0 | 0.52 | 0.28 | 0.55 | 0.8 | 0.5 | 0.62 |
| 2 | 6951833 | rs9789516 | 0.237 | -0.529 | -0.463 | -0.502 | 0.01 | 0.081 | 0.081 | 0 | 0.42 | 0.22 | 0.48 | 0.76 | 0.35 | 0.55 |
| 2 | 6954392 | rs12053057 | 0.727 | -0.527 | -0.987 | -0.086 | -0.037 | -0.052 | -0.052 | 0.083 | 0.14 | 0.03 | 0.09 | 0.2 | 0.14 | 0.17 |
| 2 | 6954913 | rs7560241 | -1.103 | -1.142 | -1.491 | -1.053 | 0.049 | 0.181 | 0.181 | 0.226 | 0.51 | 0.89 | 0.43 | 0.77 | 0.35 | 0.53 |
| 2 | 6956119 | rs16865632 | -0.142 | -1.113 | -0.343 | -0.969 | 0.026 | 0.017 | 0.017 | 1.647 | 0.06 | 0.31 | 0.04 | 0 | 0.02 | 0 |
| 2 | 6956133 | rs10495544 | -0.145 | -1.167 | -0.171 | -1.004 | 0.007 | 0.022 | 0.022 | 0 | 0.36 | 0.27 | 0.34 | 0.77 | 0.28 | 0.49 |
| 2 | 6958384 | rs6431834 | -0.141 | -1.134 | -0.615 | -1.031 | -0.043 | -0.147 | -0.147 | 1.812 | 0.19 | 0.39 | 0.09 | 0.01 | 0.09 | 0.12 |
| 2 | 6960928 | rs700940 | 1.208 | -0.895 | -1.68 | -0.931 | -0.084 | 0.119 | 0.119 | 0 | 0.4 | 0.13 | 0.5 | 0.22 | 0.55 | 0.45 |
| 2 | 6965430 | rs6734387 | -0.22 | -0.731 | -1.263 | -0.467 | -0.024 | 0.012 | 0.012 | 0 | 0.3 | 0.08 | 0.27 | 0.73 | 0.3 | 0.42 |
| 2 | 6966583 | rs771171 | 0.358 | -0.43 | -0.71 | -0.495 | -0.006 | -0.068 | -0.068 | 0 | 0.87 | 0.77 | 0.9 | 1 | 0.84 | 0.92 |
| 2 | 6967495 | rs7575062 | -0.397 | -0.595 | -0.823 | -0.029 | 0.002 | -0.003 | -0.003 | 0 | 0.29 | 0.28 | 0.24 | 0.66 | 0.28 | 0.37 |
| 2 | 6967943 | rs771172 | -1.314 | -0.124 | -0.422 | -0.174 | -0.008 | -0.035 | -0.035 | 0.003 | 0.19 | 0.23 | 0.26 | 0.02 | 0.13 | 0.16 |
| 2 | 6970375 | rs2712032 | -0.823 | 0.354 | -0.728 | 0.531 | -0.031 | -0.045 | -0.045 | 0 | 0.19 | 0.09 | 0.13 | 0.25 | 0.25 | 0.32 |
| 2 | 6976190 | rs2250846 | 1.73 | -0.189 | 0.455 | 0.48 | -0.021 | 0.073 | 0.073 | 0.517 | 0.9 | 0.55 | 0.93 | 0.91 | 0.97 | 0.96 |
| 2 | 6982261 | rs6758745 | 0.915 | -0.179 | 0.41 | 0.496 | -0.009 | -0.01 | -0.01 | 0 | 0.94 | 0.84 | 0.97 | 0.98 | 0.98 | 0.98 |
| 2 | 6984091 | rs2197829 | 1.034 | 0.175 | 0.154 | 0.489 | -0.004 | -0.048 | -0.048 | 0.401 | 0.78 | 0.47 | 0.84 | 0.9 | 0.8 | 0.9 |
| 2 | 6988669 | rs13308 | 1.093 | 0.175 | 0.145 | 0.47 | -0.015 | -0.066 | -0.066 | 0.593 | 0.76 | 0.46 | 0.84 | 0.9 | 0.8 | 0.85 |
| 2 | 6990020 | rs10495545 | -0.451 | 0.115 | -0.467 | 1.403 | -0.016 | -0.068 | -0.068 | 0.412 | 0.12 | 0.02 | 0.1 | 0.27 | 0.12 | 0.04 |
| 2 | 7002410 | rs4669107 | 1.283 | 0.004 | 0.809 | 0.185 | 0.006 | 0.003 | 0.003 | 1.594 | 0.94 | 0.64 | 0.97 | 1 | 1 | 1 |
| 2 | 7024874 | rs16865704 | -0.901 | -0.273 | 0.637 | 0.058 | -0.013 | -0.011 | -0.011 | 0.07 | 0.06 | 0.08 | 0.01 | 0.02 | 0.01 | 0.11 |
| 2 | 7028438 | rs4669111 | 1.553 | -0.543 | 1.147 | -0.198 | -0.033 | -0.038 | -0.038 | 0 | 0.76 | 0.54 | 0.83 | 0.86 | 0.81 | 0.72 |
| 2 | 7030541 | rs2305258 | -0.129 | -0.522 | 1.069 | -0.198 | -0.008 | -0.004 | -0.004 | 0 | 0.09 | 0.04 | 0.05 | 0.1 | 0.15 | 0.05 |
| 2 | 7032618 | rs16865717 | 1.579 | -0.715 | 0.638 | -0.498 | -0.026 | -0.038 | -0.038 | 0 | 0.63 | 0.42 | 0.75 | 0.76 | 0.72 | 0.58 |
| 2 | 7036529 | rs4669113 | -0.716 | 0.216 | 1.137 | -0.546 | 0.019 | -0.029 | -0.029 | 0.214 | 0.07 | 0.09 | 0.04 | 0.02 | 0.08 | 0.13 |
| 2 | 7036621 | rs4669114 | -0.716 | 0.216 | 1.137 | -0.546 | 0.019 | -0.029 | -0.029 | 0 | 0.07 | 0.09 | 0.04 | 0.02 | 0.08 | 0.13 |
| 2 | 7036678 | rs7570384 | 1.347 | 0.367 | 1.219 | -0.56 | 0.006 | -0.031 | -0.031 | 0 | 0.69 | 0.63 | 0.8 | 0.78 | 0.65 | 0.67 |
| 2 | 7036756 | rs10495546 | -0.69 | 0.395 | 1.092 | -0.536 | 0.019 | -0.029 | -0.029 | 0 | 0.07 | 0.09 | 0.04 | 0.02 | 0.08 | 0.13 |
| 2 | 7036811 | rs6431837 | 1.223 | 0.342 | 0.902 | -0.485 | 0.003 | -0.041 | -0.041 | 0.124 | 0.63 | 0.51 | 0.72 | 0.75 | 0.62 | 0.56 |
| 2 | 7048009 | rs4668516 | -0.314 | 0.636 | 0.106 | -1.009 | -0.013 | 0.031 | 0.031 | 0 | 0.79 | 0.88 | 0.79 | 0.98 | 0.76 | 0.84 |
| 2 | 7048241 | rs10153810 | 0.581 | 0.277 | -0.472 | -1.496 | -0.037 | 0.049 | 0.049 | 0 | 0.14 | 0.07 | 0.13 | 0.03 | 0.21 | 0.17 |
| 2 | 7048776 | rs10171510 | 0.546 | 0.262 | -0.48 | -1.504 | -0.037 | 0.051 | 0.051 | 0 | 0.15 | 0.07 | 0.13 | 0.03 | 0.21 | 0.17 |
| 2 | 7050887 | rs3922041 | -0.007 | -0.305 | -0.754 | -1.415 | -0.003 | -0.001 | -0.001 | 0 | 0.32 | 0.59 | 0.29 | 0.69 | 0.26 | 0.38 |
| 2 | 7056183 | rs3806609 | -0.315 | -0.681 | -0.501 | -1.24 | 0.01 | 0.067 | 0.067 | 0 | 0.46 | 0.69 | 0.43 | 0.71 | 0.47 | 0.56 |
| 2 | 7060629 | rs771274 | -0.433 | -1.082 | 0.104 | -0.919 | -0.051 | 0.168 | 0.168 | 0.071 | 0.86 | 0.66 | 0.8 | 0.42 | 0.92 | 0.82 |
| 2 | 7061691 | rs4669119 | 0.682 | -1.074 | 0.207 | -1 | -0.057 | 0.07 | 0.07 | 0 | 0.28 | 0.09 | 0.18 | 0.13 | 0.36 | 0.38 |
| 2 | 7063149 | rs13397645 | 0.687 | -1.105 | 0.245 | -0.88 | -0.042 | 0.084 | 0.084 | 0 | 0.33 | 0.14 | 0.39 | 0.14 | 0.38 | 0.39 |
| 2 | 7063401 | rs771283 | 0.108 | -1.107 | 0.348 | -0.886 | -0.001 | -0.01 | -0.01 | 0.149 | 0.53 | 0.48 | 0.4 | 0.28 | 0.54 | 0.42 |
| 2 | 7068024 | rs11694597 | 0.78 | -0.925 | 0.253 | -0.739 | 0 | 0.002 | 0.002 | 0 | 0.33 | 0.28 | 0.19 | 0.32 | 0.3 | 0.27 |
| 2 | 7068927 | rs720388 | -0.761 | -1.109 | -0.038 | -0.654 | -0.03 | 0.224 | 0.224 | 0 | 0.88 | 0.7 | 0.75 | 0.47 | 0.92 | 0.82 |
| 2 | 7072935 | rs10167072 | 0.886 | -1.001 | -0.322 | -0.644 | 0.002 | -0.01 | -0.01 | 0 | 0.32 | 0.14 | 0.18 | 0.34 | 0.29 | 0.26 |
| 2 | 7075439 | rs771300 | 0.916 | -0.845 | -0.852 | -0.656 | -0.041 | 0.09 | 0.09 | 0.002 | 0.19 | 0.05 | 0.15 | 0.01 | 0.23 | 0.1 |
| 2 | 7075768 | rs11683539 | -0.17 | -1.003 | -0.087 | 0.034 | -0.019 | 0.044 | 0.044 | 0 | 0.34 | 0.26 | 0.42 | 0.14 | 0.4 | 0.39 |
| 2 | 7080792 | rs1797557 | 0.652 | -1.482 | 0.501 | 1.056 | -0.034 | 0.15 | 0.15 | 0 | 0.86 | 0.76 | 0.75 | 0.46 | 0.92 | 0.8 |
| 2 | 7081787 | rs771306 | 1.344 | -1.199 | 0.83 | 0.891 | -0.006 | 0.11 | 0.11 | 1.04 | 0.89 | 0.52 | 0.93 | 0.86 | 0.96 | 0.95 |
| 2 | 7085191 | rs771311 | -1.226 | -1.907 | 0.766 | 1.621 | -0.063 | 0.094 | 0.094 | 0 | 0.23 | 0.53 | 0.2 | 0.42 | 0.11 | 0.28 |
| 2 | 7088624 | rs716896 | 0.77 | -2.868 | -0.052 | 1.523 | 0.078 | 0.019 | 0.019 | 0.642 | 0.6 | 0.3 | 0.79 | 0.39 | 0.84 | 0.6 |
| 2 | 7090819 | rs10192053 | 1.342 | -2.931 | -0.561 | 1.473 | 0.174 | 0.151 | 0.151 | 1.25 | 0.54 | 0.14 | 0.67 | 0.27 | 0.77 | 0.53 |
| 2 | 7093715 | rs771313 | 0.364 | -2.302 | -0.122 | 1.392 | -0.002 | -0.001 | -0.001 | 0 | 0.73 | 0.7 | 0.84 | 0.7 | 0.86 | 0.77 |
| 2 | 7096989 | rs3772019 | -0.732 | -1.864 | -0.543 | 1.397 | -0.007 | 0.005 | 0.005 | 0 | 0.1 | 0.13 | 0.15 | 0.13 | 0.09 | 0.08 |
| 2 | 7097523 | rs771315 | -0.037 | -1.589 | -0.942 | 1.223 | 0.018 | -0.079 | -0.079 | 0.083 | 0.82 | 0.99 | 0.89 | 0.76 | 0.89 | 0.83 |
| 2 | 7104166 | rs771320 | 0.548 | -1.31 | -0.83 | 0.508 | 0.023 | 0.019 | 0.019 | 0 | 0.52 | 0.47 | 0.68 | 0.36 | 0.63 | 0.42 |
| 2 | 7106364 | rs771323 | 0.107 | -1.538 | -1.495 | 0.211 | 0.04 | 0.081 | 0.081 | 0.624 | 0.74 | 0.99 | 0.9 | 0.9 | 0.81 | 0.78 |
| 2 | 7108212 | rs4669122 | 0.082 | -0.928 | -1.293 | -0.265 | -0.002 | -0.014 | -0.014 | 0 | 0.57 | 0.41 | 0.43 | 0.33 | 0.54 | 0.41 |
| 2 | 7111737 | rs3772003 | 0.504 | -1.012 | -2.119 | -0.948 | -0.112 | 0.22 | 0.22 | 0 | 0.38 | 0.03 | 0.21 | 0.09 | 0.37 | 0.31 |
| 2 | 7111899 | rs3772002 | 0.859 | -1.073 | -1.54 | -1.001 | -0.005 | 0.02 | 0.02 | 0.017 | 0.14 | 0.16 | 0.19 | 0.24 | 0.14 | 0.07 |
| 2 | 7114174 | rs771253 | 0.54 | -1.406 | -1.2 | -1.142 | 0.002 | -0.03 | -0.03 | 0 | 0.3 | 0.41 | 0.27 | 0.06 | 0.38 | 0.16 |
| 2 | 7116077 | rs4669123 | 0.209 | -1.011 | -0.97 | -1.1 | 0.002 | 0.03 | 0.03 | 0 | 0.54 | 0.32 | 0.41 | 0.32 | 0.51 | 0.38 |
| 2 | 7125756 | rs771261 | 0.095 | -0.842 | -0.735 | -1.27 | 0.003 | -0.006 | -0.006 | 0 | 0.76 | 0.77 | 0.67 | 0.38 | 0.82 | 0.5 |
| 2 | 7126569 | rs700943 | 0.751 | -0.905 | -0.767 | -1.492 | 0.005 | -0.029 | -0.029 | 0 | 0.23 | 0.35 | 0.25 | 0.05 | 0.32 | 0.13 |
| 2 | 7126676 | rs932196 | 1.464 | -0.98 | -1.077 | -1.644 | -0.074 | 0.089 | 0.089 | 0 | 0.14 | 0.01 | 0.18 | 0 | 0.26 | 0.11 |
| 2 | 7129248 | rs13019196 | 1.505 | -0.584 | -1.059 | -1.753 | -0.004 | 0.105 | 0.105 | 0 | 0.35 | 0.16 | 0.2 | 0.09 | 0.31 | 0.3 |

| | | | | | | | | | | | | | | | | |
|---|---|---|---|---|---|---|---|---|---|---|---|---|---|---|---|---|
| 2 | 7132067 | rs771268 | -0.384 | -0.573 | -1.09 | -1.699 | -0.004 | 0.004 | 0.004 | 0 | 0.41 | 0.61 | 0.47 | 0.29 | 0.51 | 0.2 |
| 2 | 7132884 | rs11679967 | 1.474 | -0.617 | -1.194 | -1.753 | -0.024 | 0.165 | 0.165 | 0 | 0.36 | 0.11 | 0.2 | 0.1 | 0.31 | 0.34 |
| 2 | 7134144 | rs771272 | -1.162 | -0.786 | -0.928 | -1.742 | 0.009 | 0.07 | 0.07 | 0.028 | 0.48 | 0.75 | 0.61 | 0.68 | 0.49 | 0.62 |
| 2 | 7137067 | rs364891 | -1.432 | -0.83 | -1.003 | -1.765 | -0.002 | -0.002 | -0.002 | 0 | 0.82 | 0.87 | 0.8 | 0.78 | 0.8 | 0.94 |
| 2 | 7137435 | rs10495548 | 0.49 | -0.915 | -0.872 | -1.79 | 0.02 | -0.06 | -0.06 | 0 | 0.23 | 0.41 | 0.26 | 0.06 | 0.32 | 0.11 |
| 2 | 7146214 | rs309318 | -1.264 | -0.785 | -0.72 | -1.597 | 0.004 | 0.006 | 0.006 | 0 | 0.82 | 0.79 | 0.79 | 0.78 | 0.8 | 0.94 |
| 2 | 7150330 | rs447721 | 0.572 | -0.91 | -0.823 | -1.745 | 0 | -0.006 | -0.006 | 0.005 | 0.57 | 0.45 | 0.45 | 0.14 | 0.62 | 0.45 |
| 2 | 7150654 | rs381214 | 0.687 | -1.066 | -0.786 | -1.864 | -0.005 | -0.036 | -0.036 | 0 | 0.55 | 0.5 | 0.45 | 0.14 | 0.62 | 0.44 |
| 2 | 7151074 | rs412098 | -0.397 | -1.139 | -0.827 | -1.592 | 0.001 | -0.016 | -0.016 | 0.284 | 0.71 | 0.6 | 0.67 | 0.19 | 0.72 | 0.66 |
| 2 | 7153253 | rs6720046 | -0.431 | -1.11 | -0.848 | -1.417 | -0.013 | 0.034 | 0.034 | 0 | 0.25 | 0.31 | 0.15 | 0.56 | 0.17 | 0.49 |
| 2 | 7156947 | rs309311 | 0.217 | -1.104 | -0.532 | -1.545 | -0.004 | 0.02 | 0.02 | 0 | 0.56 | 0.6 | 0.58 | 0.76 | 0.47 | 0.76 |
| 2 | 7158425 | rs3771964 | 0.655 | -1.036 | -0.728 | -1.373 | -0.014 | 0.153 | 0.153 | 0 | 0.13 | 0.22 | 0.13 | 0.55 | 0.09 | 0.41 |
| 2 | 7159485 | rs309309 | -1.574 | -1.019 | -0.868 | -1.4 | -0.011 | -0.013 | -0.013 | 0.018 | 0.83 | 0.89 | 0.81 | 0.77 | 0.8 | 0.93 |
| 2 | 7161314 | rs3755503 | 0.591 | -1.346 | -0.701 | -1.096 | -0.01 | 0.115 | 0.115 | 0 | 0.13 | 0.17 | 0.12 | 0.55 | 0.09 | 0.41 |
| 2 | 7161646 | rs419970 | -0.272 | -1.431 | -0.771 | -1.213 | -0.016 | 0.038 | 0.038 | 1.53 | 0.56 | 0.28 | 0.52 | 0.72 | 0.47 | 0.81 |
| 2 | 7163141 | rs451763 | 1.28 | -1.665 | -1.063 | -1.245 | -0.032 | 0.102 | 0.102 | 0.33 | 0.33 | 0.07 | 0.38 | 0.17 | 0.32 | 0.33 |
| 2 | 7164946 | rs447031 | 0.634 | -1.413 | -0.919 | -1.11 | 0.015 | 0.19 | 0.19 | 2.11 | 0.43 | 0.1 | 0.39 | 0.17 | 0.35 | 0.4 |
| 2 | 7166507 | rs377617 | 0.628 | -1.37 | -0.979 | -0.833 | 0.003 | 0.195 | 0.195 | 2.066 | 0.43 | 0.09 | 0.39 | 0.17 | 0.35 | 0.4 |
| 2 | 7171605 | rs367107 | -0.216 | -1.362 | -0.648 | -0.771 | -0.006 | 0.039 | 0.039 | 0.731 | 0.57 | 0.3 | 0.51 | 0.72 | 0.44 | 0.81 |
| 2 | 7175132 | rs433983 | 1.636 | -1.82 | -1.13 | -1.16 | -0.057 | 0.019 | 0.019 | 0 | 0.23 | 0.04 | 0.34 | 0.17 | 0.27 | 0.26 |
| 2 | 7177700 | rs996508 | 0.815 | -1.877 | -1.076 | -1.174 | -0.01 | 0.109 | 0.109 | 0 | 0.13 | 0.17 | 0.12 | 0.55 | 0.09 | 0.41 |
| 2 | 7181486 | rs309304 | -1.004 | -1.937 | -0.745 | -1.363 | 0.023 | -0.018 | -0.018 | 0 | 0.62 | 0.46 | 0.6 | 0.72 | 0.54 | 0.89 |
| 2 | 7181560 | rs309303 | 0.897 | -1.871 | -0.823 | -1.407 | -0.004 | 0.006 | 0.006 | 0 | 0.29 | 0.26 | 0.43 | 0.16 | 0.37 | 0.34 |
| 2 | 7182919 | rs17668045 | 1.234 | -1.915 | -0.75 | -1.318 | 0.049 | 0.123 | 0.123 | 1.505 | 0.2 | 0.04 | 0.04 | 0 | 0.08 | 0.14 |
| 2 | 7188104 | rs16865879 | 0.31 | -1.808 | -0.576 | -1.222 | 0.009 | -0.005 | -0.005 | 0 | 0.08 | 0.15 | 0.1 | 0.06 | 0.11 | 0.1 |
| 2 | 7188435 | rs309294 | 0.249 | -1.984 | -0.647 | -0.63 | 0.015 | 0.146 | 0.146 | 1.628 | 0.48 | 0.16 | 0.46 | 0.29 | 0.4 | 0.56 |
| 2 | 7189397 | rs309292 | 0.234 | -1.977 | -0.622 | -0.627 | 0.021 | 0.141 | 0.141 | 1.683 | 0.49 | 0.18 | 0.46 | 0.29 | 0.4 | 0.58 |
| 2 | 7190175 | rs6431843 | 0.528 | -1.962 | -0.451 | -0.505 | -0.034 | 0.248 | 0.248 | 0 | 0.09 | 0.24 | 0.04 | 0.45 | 0.03 | 0.24 |
| 2 | 7196439 | rs309281 | 1.463 | -2.056 | -0.642 | -0.624 | -0.069 | -0.021 | -0.021 | 0.804 | 0.27 | 0.04 | 0.42 | 0.29 | 0.32 | 0.44 |
| 2 | 7197199 | rs309278 | -0.391 | -2.069 | -0.49 | -0.657 | 0 | -0.018 | -0.018 | 0 | 0.45 | 0.48 | 0.6 | 0.8 | 0.48 | 0.79 |
| 2 | 7197780 | rs6431844 | 1.098 | -2.014 | -0.475 | -0.52 | -0.025 | 0.212 | 0.212 | 0 | 0.07 | 0.16 | 0.06 | 0.44 | 0.03 | 0.25 |
| 2 | 7200864 | rs309273 | -0.493 | -2.074 | -0.581 | -0.593 | -0.008 | 0.014 | 0.014 | 2.466 | 0.58 | 0.33 | 0.57 | 0.36 | 0.53 | 0.67 |
| 2 | 7202496 | rs309271 | 1.499 | -1.975 | -0.839 | -0.589 | -0.064 | -0.019 | -0.019 | 2.188 | 0.27 | 0.05 | 0.42 | 0.29 | 0.31 | 0.44 |
| 2 | 7205349 | rs35366716 | -0.463 | -1.798 | -0.618 | -0.365 | -0.008 | 0.249 | 0.249 | 0 | 0.16 | 0.38 | 0.18 | 0.51 | 0.17 | 0.34 |
| 2 | 7209885 | rs7605141 | -0.282 | -1.638 | -0.525 | -0.253 | -0.021 | 0.012 | 0.012 | 0 | 0.37 | 0.39 | 0.22 | 0.49 | 0.24 | 0.48 |
| 2 | 7216231 | rs10445713 | 1.358 | -1.345 | -0.422 | -0.25 | -0.087 | 0.034 | 0.034 | 0.373 | 0.25 | 0.12 | 0.29 | 0.15 | 0.39 | 0.07 |
| 2 | 7216352 | rs4669130 | 1.263 | -1.352 | -0.246 | -0.254 | -0.042 | 0.025 | 0.025 | 0.217 | 0.26 | 0.18 | 0.29 | 0.15 | 0.39 | 0.07 |
| 2 | 7217012 | rs7558716 | 0.525 | -1.163 | -0.167 | 0.054 | 0.012 | -0.003 | -0.003 | 0.092 | 0.1 | 0.14 | 0.12 | 0.07 | 0.18 | 0.07 |
| 2 | 7217472 | rs16865916 | 0.758 | -1.493 | -0.372 | 0.322 | 0.008 | -0.014 | -0.014 | 0 | 0.23 | 0.18 | 0.08 | 0.43 | 0.09 | 0.36 |
| 2 | 7217823 | rs1901765 | -1.077 | -1.624 | -0.113 | 0.248 | 0 | 0.001 | 0.001 | 0.178 | 0.68 | 0.72 | 0.52 | 0.71 | 0.7 | 0.53 |
| 2 | 7219384 | rs1453122 | -0.062 | -1.444 | -0.283 | 0.26 | -0.006 | -0.007 | -0.007 | 0 | 0.45 | 0.53 | 0.44 | 0.28 | 0.61 | 0.18 |
| 2 | 7231233 | rs10182281 | -1.31 | -1.37 | -0.524 | 0.278 | -0.031 | -0.008 | -0.008 | 0 | 0.72 | 0.95 | 0.54 | 0.71 | 0.7 | 0.54 |
| 2 | 7231665 | rs7567367 | -0.157 | -1.463 | -0.225 | 0.274 | -0.006 | -0.009 | -0.009 | 2.209 | 0.46 | 0.57 | 0.41 | 0.28 | 0.6 | 0.17 |
| 2 | 7232478 | rs4669131 | -0.193 | -1.46 | -0.218 | 0.281 | -0.006 | -0.008 | -0.008 | 0.334 | 0.47 | 0.57 | 0.42 | 0.28 | 0.61 | 0.18 |
| 2 | 7234095 | rs4668520 | 1.281 | -1.325 | -0.487 | 0.211 | -0.106 | -0.006 | -0.006 | 0 | 0.27 | 0.13 | 0.29 | 0.21 | 0.41 | 0.09 |
| 2 | 7242813 | rs6712003 | -1.286 | -1.333 | -0.138 | 0.225 | 0 | 0.009 | 0.009 | 0.229 | 0.7 | 0.78 | 0.49 | 0.73 | 0.69 | 0.52 |
| 2 | 7244036 | rs6722663 | -0.176 | -1.482 | 0.128 | 0.252 | -0.009 | -0.009 | -0.009 | 0 | 0.46 | 0.53 | 0.42 | 0.3 | 0.61 | 0.18 |
| 2 | 7244329 | rs4668521 | -1.421 | -1.461 | -0.021 | 0.261 | -0.002 | 0.013 | 0.013 | 0.192 | 0.71 | 0.82 | 0.5 | 0.73 | 0.69 | 0.53 |
| 2 | 7246745 | rs17624427 | 1.412 | -1.295 | -0.467 | 0.367 | -0.094 | 0.006 | 0.006 | 0.047 | 0.28 | 0.14 | 0.3 | 0.2 | 0.41 | 0.09 |
| 2 | 7248419 | rs6431850 | 0.279 | -1.33 | -0.352 | 0.393 | -0.002 | -0.013 | -0.013 | 0 | 0.17 | 0.25 | 0.11 | 0.1 | 0.2 | 0.09 |
| 2 | 7248699 | rs6742329 | 0.462 | -0.97 | -0.783 | 0.658 | -0.014 | 0.003 | 0.003 | 0 | 0.24 | 0.23 | 0.08 | 0.43 | 0.09 | 0.35 |
| 2 | 7249578 | rs6746186 | 0.442 | -0.946 | -0.868 | 0.67 | -0.026 | 0.013 | 0.013 | 0 | 0.24 | 0.25 | 0.07 | 0.43 | 0.09 | 0.35 |
| 2 | 7250562 | rs2352400 | 0 | -0.309 | -0.631 | 0.489 | 0.006 | 0.025 | 0.025 | 0 | 0.52 | 0.4 | 0.38 | 0.63 | 0.49 | 0.45 |
| 2 | 7252209 | rs4669133 | -0.595 | 0.002 | -0.521 | 0.559 | 0.005 | -0.004 | -0.004 | 0.46 | 0.57 | 0.57 | 0.46 | 0.6 | 0.64 | 0.47 |
| 2 | 7256134 | rs12104459 | 0.415 | 0.555 | -0.377 | 0.578 | 0.019 | 0.021 | 0.021 | 0 | 0.16 | 0.22 | 0.12 | 0.29 | 0.18 | 0.28 |
| 2 | 7256592 | rs1453129 | -0.127 | 0.576 | -0.455 | 0.52 | -0.01 | 0.016 | 0.016 | 0.302 | 0.4 | 0.34 | 0.34 | 0.29 | 0.47 | 0.19 |
| 2 | 7258204 | rs4669138 | -0.403 | 0.558 | -0.58 | 0.513 | 0.001 | 0.019 | 0.019 | 0.9 | 0.44 | 0.26 | 0.38 | 0.42 | 0.51 | 0.37 |
| 2 | 7260133 | rs4669139 | -1.116 | 0.54 | -0.559 | 0.553 | 0.001 | -0.001 | -0.001 | 0.192 | 0.57 | 0.54 | 0.48 | 0.63 | 0.65 | 0.47 |
| 2 | 7262081 | rs10221761 | -1.12 | 0.537 | -0.56 | 0.551 | 0.001 | -0.001 | -0.001 | 0.011 | 0.57 | 0.54 | 0.48 | 0.63 | 0.65 | 0.47 |
| 2 | 7262101 | rs10221834 | 0.112 | 0.64 | -0.753 | 0.251 | -0.003 | 0.002 | 0.002 | 0.097 | 0.12 | 0.21 | 0.07 | 0.16 | 0.14 | 0.11 |
| 2 | 7262831 | rs16865965 | 0.027 | 0.642 | -0.83 | 0.252 | -0.002 | 0.004 | 0.004 | 0.247 | 0.13 | 0.26 | 0.08 | 0.16 | 0.14 | 0.11 |
| 2 | 7267159 | rs13018029 | 0.207 | 0.967 | -1.419 | 0.24 | -0.077 | 0.052 | 0.052 | 3.23 | 0.27 | 0.08 | 0.26 | 0.14 | 0.32 | 0.08 |
| 2 | 7278187 | rs6748226 | -0.544 | 0.624 | -1.116 | -0.16 | 0 | -0.012 | -0.012 | 0.798 | 0.29 | 0.26 | 0.19 | 0.12 | 0.3 | 0.3 |
| 2 | 7284883 | rs4669141 | -0.465 | 0.646 | -1.268 | -0.125 | -0.003 | 0.057 | 0.057 | 0 | 0.28 | 0.15 | 0.19 | 0.16 | 0.3 | 0.3 |
| 2 | 7285635 | rs10929468 | 0.315 | 0.638 | -1.301 | -0.127 | -0.011 | 0.022 | 0.022 | 0 | 0.17 | 0.08 | 0.12 | 0.16 | 0.22 | 0.21 |
| 2 | 7292558 | rs13387376 | 0.177 | 0.613 | -1.282 | -0.13 | 0 | 0.013 | 0.013 | 0 | 0.18 | 0.13 | 0.12 | 0.16 | 0.21 | 0.22 |
| 2 | 7295543 | rs11685708 | 0.066 | 0.599 | -1.138 | -0.137 | -0.002 | -0.01 | -0.01 | 0 | 0.18 | 0.2 | 0.12 | 0.15 | 0.21 | 0.24 |
| 2 | 7301432 | rs1917681 | -1.069 | 0.724 | -1.746 | 0.066 | 0.068 | -0.051 | -0.051 | 0 | 0.55 | 0.91 | 0.46 | 0.62 | 0.65 | 0.47 |
| 2 | 7302999 | rs930767 | -0.816 | 1.127 | -1.084 | 0.256 | 0 | 0.001 | 0.001 | 0.006 | 0.51 | 0.52 | 0.37 | 0.36 | 0.51 | 0.36 |

| | | | | | | | | | | | | | | | | |
|---|---|---|---|---|---|---|---|---|---|---|---|---|---|---|---|---|
| 2 | 7304656 | rs16865980 | 1.169 | 1.131 | -1.177 | 0.27 | 0.002 | 0.001 | 0.001 | 1.579 | 0.21 | 0.23 | 0.17 | 0.2 | 0.22 | 0.06 |
| 2 | 7311632 | rs10495552 | -0.776 | 1.179 | -1.33 | 0.29 | 0 | -0.002 | -0.002 | 0 | 0.51 | 0.5 | 0.36 | 0.36 | 0.51 | 0.36 |
| 2 | 7313755 | rs9287684 | 0.433 | 0.855 | -1.062 | 0.535 | -0.001 | -0.006 | -0.006 | 0.171 | 0.18 | 0.2 | 0.12 | 0.16 | 0.21 | 0.24 |
| 2 | 7314628 | rs16865987 | 1.122 | 0.695 | -1.663 | 0.567 | -0.007 | -0.125 | -0.125 | 0 | 0.23 | 0.01 | 0.44 | 0.28 | 0.23 | 0.3 |
| 2 | 7320746 | rs7588701 | -0.154 | 1.142 | -0.752 | 0.468 | 0.136 | 0.065 | 0.065 | 0 | 0.16 | 0.41 | 0.17 | 0.27 | 0.23 | 0.17 |
| 2 | 7322437 | rs12151441 | -0.463 | -0.05 | -1.363 | 0.166 | 0.014 | 0.037 | 0.037 | 0.212 | 0.26 | 0.14 | 0.15 | 0.19 | 0.24 | 0.29 |
| 2 | 7323039 | rs12623068 | -0.543 | -0.308 | -1.077 | 0.542 | -0.016 | 0.013 | 0.013 | 0.203 | 0.31 | 0.4 | 0.24 | 0.44 | 0.24 | 0.38 |
| 2 | 7325615 | rs7596634 | -0.142 | 0.859 | -1.799 | -0.344 | -0.121 | 0.121 | 0.121 | 0 | 0.25 | 0.04 | 0.22 | 0.07 | 0.37 | 0.15 |
| 2 | 7329760 | rs6706981 | 0.563 | 0.803 | -1.562 | 0.221 | -0.043 | -0.049 | -0.049 | 0 | 0.65 | 0.78 | 0.58 | 0.51 | 0.51 | 0.73 |
| 2 | 7330487 | rs6755106 | -0.186 | 0.277 | -1.863 | -0.394 | 0.096 | -0.022 | -0.022 | 0 | 0.51 | 0.17 | 0.6 | 0.44 | 0.62 | 0.52 |
| 2 | 7331801 | rs7607300 | -0.358 | 0.197 | -1.6 | -0.54 | 0.075 | -0.006 | -0.006 | 0 | 0.53 | 0.25 | 0.67 | 0.48 | 0.63 | 0.53 |
| 2 | 7334656 | rs1917679 | -0.819 | 0.119 | -0.934 | -0.576 | 0.02 | 0.243 | 0.243 | 0 | 0.91 | 0.77 | 0.88 | 0.61 | 0.9 | 0.91 |
| 2 | 7335598 | rs6739227 | 0.939 | -0.084 | -0.156 | -0.693 | 0.006 | -0.007 | -0.007 | 1.076 | 0.36 | 0.31 | 0.19 | 0.13 | 0.27 | 0.34 |
| 2 | 7337945 | rs17672973 | 0.462 | 0.864 | -0.491 | -0.825 | 0.086 | -0.028 | -0.028 | 0.253 | 0.05 | 0.13 | 0.06 | 0 | 0.15 | 0.04 |
| 2 | 7338199 | rs17672986 | 0.201 | 0.823 | -0.548 | -0.82 | -0.056 | -0.034 | -0.034 | 0 | 0.47 | 0.13 | 0.61 | 0.47 | 0.48 | 0.47 |
| 2 | 7339394 | rs7603211 | 1.144 | 0.736 | -0.045 | -0.859 | 0.04 | 0.118 | 0.118 | 0 | 0.34 | 0.14 | 0.17 | 0.12 | 0.26 | 0.32 |
| 2 | 7340107 | rs6431861 | 0.254 | 1.186 | -0.207 | -0.534 | 0.08 | 0.152 | 0.152 | 0 | 0.86 | 0.71 | 0.82 | 0.61 | 0.75 | 0.88 |
| 2 | 7340453 | rs12998506 | 0.169 | 0.848 | -0.274 | -0.616 | 0.005 | -0.05 | -0.05 | 0.044 | 0.59 | 0.53 | 0.39 | 0.19 | 0.5 | 0.52 |
| 2 | 7344495 | rs4669153 | 0.686 | 1.179 | -0.649 | -0.562 | 0.004 | -0.019 | -0.019 | 2.907 | 0.27 | 0.24 | 0.43 | 0.42 | 0.22 | 0.33 |
| 2 | 7349867 | rs17679036 | 0.543 | 1.038 | -0.783 | -0.858 | 0.01 | -0.025 | -0.025 | 0 | 0.14 | 0.19 | 0.16 | 0.06 | 0.18 | 0.14 |
| 2 | 7350493 | rs4669155 | -0.02 | 1.164 | -0.622 | -0.747 | 0 | 0 | 0 | 0 | 0.37 | 0.35 | 0.58 | 0.47 | 0.39 | 0.43 |
| 2 | 7350916 | rs4669156 | 1 | 1.088 | -0.635 | -0.734 | 0.005 | -0.041 | -0.041 | 2.553 | 0.26 | 0.17 | 0.43 | 0.42 | 0.22 | 0.33 |
| 2 | 7351271 | rs4669157 | -0.569 | 0.952 | -0.597 | -0.783 | -0.002 | 0 | 0 | 1.347 | 0.47 | 0.47 | 0.66 | 0.48 | 0.57 | 0.58 |
| 2 | 7356055 | rs1980339 | -0.832 | 0.891 | -0.574 | -0.756 | 0 | 0.048 | 0.048 | 0 | 0.91 | 0.89 | 0.86 | 0.62 | 0.88 | 0.92 |
| 2 | 7361280 | rs11682769 | 0.804 | 0.879 | -0.506 | -0.849 | -0.021 | -0.074 | -0.074 | 0 | 0.44 | 0.47 | 0.2 | 0.14 | 0.32 | 0.35 |
| 2 | 7364691 | rs870863 | -0.544 | 0.908 | -0.565 | -0.825 | 0.019 | 0 | 0 | 0.771 | 0.46 | 0.39 | 0.65 | 0.48 | 0.57 | 0.57 |
| 2 | 7367462 | rs1525492 | 0.946 | 1.073 | -0.639 | -0.771 | 0.004 | -0.072 | -0.072 | 0 | 0.26 | 0.12 | 0.41 | 0.42 | 0.22 | 0.33 |
| 2 | 7370196 | rs952810 | 0.017 | 1.001 | -0.615 | -0.757 | -0.051 | -0.054 | -0.054 | 0 | 0.35 | 0.21 | 0.49 | 0.42 | 0.4 | 0.47 |
| 2 | 7382776 | rs17628484 | 0.477 | 0.726 | -0.727 | -0.883 | 0.003 | -0.043 | -0.043 | 0.013 | 0.18 | 0.24 | 0.19 | 0.07 | 0.19 | 0.15 |
| 2 | 7383183 | rs2056635 | 0.687 | 0.725 | -0.837 | -0.883 | -0.017 | 0.051 | 0.051 | 0 | 0.17 | 0.07 | 0.17 | 0.07 | 0.19 | 0.15 |
| 2 | 7383538 | rs4669169 | 0.952 | 0.74 | -1.005 | -0.916 | -0.011 | -0.16 | -0.16 | 0 | 0.27 | 0.03 | 0.4 | 0.41 | 0.22 | 0.33 |
| 2 | 7387447 | rs2140855 | 0.866 | 1.447 | -0.266 | -0.641 | 0.019 | 0.065 | 0.065 | 0.44 | 0.7 | 0.49 | 0.63 | 0.53 | 0.63 | 0.71 |
| 2 | 7392102 | rs7591725 | 1.074 | 1.292 | -0.346 | -0.626 | 0.009 | 0.013 | 0.013 | 1.239 | 0.63 | 0.53 | 0.63 | 0.53 | 0.45 | 0.61 |
| 2 | 7396577 | rs7599262 | -0.689 | 1.344 | -0.349 | -0.692 | 0.046 | -0.083 | -0.083 | 0 | 0.27 | 0.36 | 0.26 | 0.07 | 0.38 | 0.25 |
| 2 | 7404954 | rs6431869 | 1.075 | 1.307 | -0.66 | -0.623 | 0.001 | 0.001 | 0.001 | 1.085 | 0.66 | 0.67 | 0.64 | 0.51 | 0.45 | 0.67 |
| 2 | 7405579 | rs876997 | 0.41 | 1.18 | -0.167 | -0.202 | 0.029 | 0.191 | 0.191 | 0.405 | 0.88 | 0.7 | 0.87 | 0.58 | 0.83 | 0.88 |
| 2 | 7405900 | rs10495553 | -0.178 | 1.249 | -0.441 | -0.18 | 0.014 | 0.104 | 0.104 | 0 | 0.09 | 0.15 | 0.12 | 0.39 | 0.15 | 0.1 |
| 2 | 7406067 | rs959724 | 0.871 | 1.35 | -0.563 | -0.321 | 0.052 | 0.038 | 0.038 | 0.752 | 0.78 | 0.68 | 0.79 | 0.62 | 0.64 | 0.74 |
| 2 | 7406992 | rs17629193 | 0.264 | 0.732 | -1.437 | -0.934 | -0.003 | 0.152 | 0.152 | 0 | 0.3 | 0.07 | 0.23 | 0.06 | 0.26 | 0.17 |
| 2 | 7407734 | rs4669172 | 0.651 | 0.676 | -1.771 | -0.732 | -0.047 | -0.134 | -0.134 | 0 | 0.16 | 0.01 | 0.42 | 0.42 | 0.22 | 0.27 |
| 2 | 7407863 | rs16866054 | 0.025 | 0.965 | -0.817 | -0.862 | 0.031 | 0.156 | 0.156 | 0.041 | 0.13 | 0.3 | 0.15 | 0.4 | 0.15 | 0.16 |
| 2 | 7415677 | rs4669173 | 0.374 | 0.503 | -0.73 | -1.107 | 0.039 | -0.015 | -0.015 | 0 | 0.4 | 0.22 | 0.52 | 0.54 | 0.36 | 0.51 |
| 2 | 7416777 | rs13007576 | 0.392 | 0.497 | -0.568 | -1.185 | 0.002 | 0.071 | 0.071 | 0 | 0.2 | 0.33 | 0.18 | 0.4 | 0.2 | 0.2 |
| 2 | 7417009 | rs11898813 | -0.752 | 0.458 | -0.579 | -1.197 | 0.004 | -0.062 | -0.062 | 0 | 0.61 | 0.56 | 0.7 | 0.94 | 0.56 | 0.72 |
| 2 | 7424153 | rs4668535 | -0.177 | 0.791 | -0.18 | -0.828 | -0.012 | 0.102 | 0.102 | 0 | 0.23 | 0.51 | 0.2 | 0.4 | 0.2 | 0.22 |
| 2 | 7431316 | rs17630005 | 0.76 | 1.47 | -0.16 | -1.087 | 0.11 | -0.031 | -0.031 | 2.235 | 0.06 | 0.11 | 0.07 | 0 | 0.18 | 0.08 |
| 2 | 7431380 | rs10191135 | 0.063 | 0.708 | -0.121 | -1.157 | 0.02 | -0.02 | -0.02 | 0 | 0.35 | 0.27 | 0.37 | 0.53 | 0.3 | 0.39 |
| 2 | 7434265 | rs919507 | -0.675 | 0.646 | -0.19 | -1.188 | 0 | 0.003 | 0.003 | 0 | 0.72 | 0.72 | 0.63 | 0.59 | 0.7 | 0.67 |
| 2 | 7437247 | rs10182503 | -0.685 | 0.651 | -0.197 | -1.186 | 0 | 0.004 | 0.004 | 0 | 0.72 | 0.71 | 0.64 | 0.59 | 0.7 | 0.67 |
| 2 | 7439390 | rs17681533 | 0.914 | 0.581 | -0.439 | -1.316 | -0.063 | 0.141 | 0.141 | 0.092 | 0.38 | 0.16 | 0.26 | 0.06 | 0.43 | 0.29 |
| 2 | 7442667 | rs12617706 | 0.234 | 0.559 | -0.39 | -1.287 | -0.002 | 0.003 | 0.003 | 0 | 0.32 | 0.35 | 0.36 | 0.53 | 0.27 | 0.35 |
| 2 | 7445627 | rs4669177 | 0.91 | 0.622 | -0.474 | -1.261 | -0.128 | 0.222 | 0.222 | 0.158 | 0.37 | 0.06 | 0.25 | 0.06 | 0.44 | 0.25 |
| 2 | 7454360 | rs1430289 | -0.179 | 0.851 | -0.379 | -1.15 | -0.008 | 0.016 | 0.016 | 0 | 0.37 | 0.44 | 0.41 | 0.53 | 0.35 | 0.37 |
| 2 | 7456361 | rs6750297 | 0.682 | 0.677 | -0.491 | -1.167 | -0.04 | 0.101 | 0.101 | 0 | 0.39 | 0.24 | 0.26 | 0.06 | 0.43 | 0.25 |
| 2 | 7460260 | rs10179297 | -0.297 | 0.176 | -0.871 | -0.963 | -0.011 | -0.008 | -0.008 | 0 | 0.39 | 0.36 | 0.45 | 0.54 | 0.47 | 0.44 |
| 2 | 7461967 | rs10495554 | -0.435 | 0.722 | -0.834 | -0.693 | 0 | -0.014 | -0.014 | 0.309 | 0.74 | 0.76 | 0.66 | 0.6 | 0.73 | 0.66 |
| 2 | 7463251 | rs10495555 | -0.489 | 1.377 | -0.694 | -1.144 | 0.005 | -0.027 | -0.027 | 0.237 | 0.22 | 0.22 | 0.18 | 0.02 | 0.36 | 0.16 |
| 2 | 7477186 | rs11682970 | -0.388 | 1.082 | -0.805 | -0.472 | -0.003 | -0.015 | -0.015 | 0 | 0.2 | 0.2 | 0.29 | 0.62 | 0.19 | 0.3 |
| 2 | 7484783 | rs4669183 | -0.376 | 0.9 | -0.744 | -1.597 | 0.006 | -0.067 | -0.067 | 2.626 | 0.57 | 0.55 | 0.65 | 0.8 | 0.51 | 0.49 |
| 2 | 7488859 | rs734271 | -0.021 | 0.706 | -0.627 | -1.542 | -0.002 | -0.004 | -0.004 | 0 | 0.23 | 0.34 | 0.35 | 0.73 | 0.19 | 0.31 |
| 2 | 7508833 | rs13005109 | 1.034 | 0.869 | -0.972 | -1.519 | 0.014 | 0.043 | 0.043 | 0 | 0.34 | 0.24 | 0.3 | 0.11 | 0.3 | 0.18 |
| 2 | 7517231 | rs13429103 | 0.508 | 1.081 | -1.034 | -1.321 | 0.095 | -0.055 | -0.055 | 0.787 | 0.09 | 0.24 | 0.08 | 0 | 0.17 | 0.14 |
| 2 | 7519326 | rs10929482 | -0.26 | 0.467 | -0.961 | -1.074 | -0.002 | -0.001 | -0.001 | 0 | 0.23 | 0.3 | 0.35 | 0.74 | 0.19 | 0.29 |
| 2 | 7520388 | rs10929483 | -0.307 | 0.409 | -1.443 | -1.366 | -0.012 | 0.038 | 0.038 | 0 | 0.6 | 0.73 | 0.75 | 0.92 | 0.54 | 0.56 |
| 2 | 7521310 | rs4669192 | 0.572 | 0.272 | -2.118 | -1.235 | -0.139 | 0.223 | 0.223 | 0 | 0.41 | 0.03 | 0.38 | 0.1 | 0.46 | 0.25 |
| 2 | 7522315 | rs2028381 | -0.569 | 0.13 | -1.572 | -1.092 | -0.051 | 0.147 | 0.147 | 0.46 | 0.61 | 0.83 | 0.68 | 0.91 | 0.51 | 0.51 |
| 2 | 7522349 | rs2028380 | -0.744 | -0.034 | -0.485 | -0.822 | -0.008 | -0.094 | -0.094 | 0.243 | 0.14 | 0.34 | 0.06 | 0 | 0.09 | 0.05 |
| 2 | 7522717 | rs4669194 | 0.799 | -0.11 | -0.116 | 0.443 | 0.011 | 0.032 | 0.032 | 0 | 0.36 | 0.41 | 0.36 | 0.43 | 0.34 | 0.29 |
| 2 | 7523040 | rs16866174 | -0.893 | -0.087 | 0.997 | 1.772 | -0.002 | 0.04 | 0.04 | 0 | 0.11 | 0.19 | 0.08 | 0.33 | 0.1 | 0.18 |

| | | | | | | | | | | | | | | | | |
|---|---|---|---|---|---|---|---|---|---|---|---|---|---|---|---|---|
| 2 | 7523858 | rs4146108 | -0.916 | -0.1 | 0.917 | 2.179 | 0.039 | 0.321 | 0.321 | 0.042 | 0.08 | 0.3 | 0.09 | 0.29 | 0.04 | 0.09 |
| 2 | 7525664 | rs7582300 | -1.633 | -0.079 | 0.213 | 1.671 | 0.035 | -0.026 | -0.026 | 0.1 | 0.31 | 0.81 | 0.22 | 0.63 | 0.21 | 0.33 |
| 2 | 7533435 | rs6756247 | 0.277 | 0.565 | 0.518 | 2.186 | 0 | -0.004 | -0.004 | 2.059 | 0.64 | 0.43 | 0.55 | 0.5 | 0.63 | 0.49 |
| 2 | 7537486 | rs2163256 | 0.209 | 0.552 | 0.719 | 2.179 | 0.002 | 0.053 | 0.053 | 0.261 | 0.66 | 0.57 | 0.56 | 0.5 | 0.63 | 0.51 |
| 2 | 7538773 | rs11897984 | -0.433 | 0.555 | 0.802 | 2.462 | 0.004 | -0.024 | -0.024 | 0 | 0.73 | 0.69 | 0.76 | 0.81 | 0.69 | 0.6 |
| 2 | 7541865 | rs10929484 | 0.52 | 0.519 | 0.831 | 2.439 | 0.008 | -0.033 | -0.033 | 0 | 0.63 | 0.3 | 0.54 | 0.5 | 0.63 | 0.5 |
| 2 | 7544326 | rs16855398 | -0.635 | 0.493 | 0.762 | 2.512 | -0.008 | 0 | 0 | 0 | 0.06 | 0 | 0.19 | 0.33 | 0.06 | 0.14 |
| 2 | 7552684 | rs1583118 | 0.781 | 0.564 | 0.347 | 2.651 | -0.01 | 0.039 | 0.039 | 0.056 | 0.57 | 0.29 | 0.48 | 0.33 | 0.52 | 0.4 |
| 2 | 7554698 | rs3910068 | 0.853 | 0.565 | 0.306 | 2.651 | -0.009 | 0.045 | 0.045 | 0.002 | 0.56 | 0.28 | 0.47 | 0.33 | 0.52 | 0.38 |
| 2 | 7554772 | rs3910069 | 0.385 | 0.704 | -0.186 | 2.556 | -0.001 | -0.002 | -0.002 | 0 | 0.62 | 0.69 | 0.55 | 0.51 | 0.55 | 0.47 |
| 2 | 7560605 | rs1430296 | 0.41 | 0.709 | -0.242 | 2.26 | 0.001 | 0 | 0 | 0.045 | 0.62 | 0.67 | 0.54 | 0.62 | 0.56 | 0.48 |
| 2 | 7561886 | rs1367272 | 1.673 | 0.496 | -0.305 | 2.659 | -0.004 | 0.017 | 0.017 | 0.168 | 0.87 | 0.87 | 0.73 | 0.52 | 0.83 | 0.73 |
| 2 | 7564293 | rs8179776 | 1.77 | 0.065 | -0.198 | 2.439 | -0.039 | 0.151 | 0.151 | 0.511 | 0.77 | 0.52 | 0.72 | 0.57 | 0.82 | 0.63 |
| 2 | 7564768 | rs6732885 | -0.367 | 0.706 | -0.319 | 2.07 | 0 | -0.006 | -0.006 | 0 | 0.44 | 0.47 | 0.45 | 0.34 | 0.43 | 0.44 |
| 2 | 7566026 | rs1367273 | 1.274 | 0.606 | 0.104 | 2.588 | 0 | -0.016 | -0.016 | 0 | 0.92 | 0.98 | 0.8 | 0.82 | 0.92 | 0.81 |
| 2 | 7567667 | rs16866239 | -0.733 | 0.676 | 0.389 | 2.267 | -0.004 | -0.033 | -0.033 | 0 | 0.06 | 0.18 | 0.03 | 0 | 0.03 | 0.02 |
| 2 | 7568539 | rs3811606 | 1.031 | 1.05 | -0.096 | 2.771 | -0.01 | 0.039 | 0.039 | 0 | 0.93 | 0.97 | 0.77 | 0.68 | 0.87 | 0.82 |
| 2 | 7574792 | rs4668543 | -1.165 | 0.691 | 0.323 | 2.412 | 0.021 | 0.215 | 0.215 | 0 | 0.06 | 0.16 | 0.07 | 0.38 | 0.07 | 0.23 |
| 2 | 7575903 | rs16866241 | -1.165 | 0.691 | 0.323 | 2.412 | 0.021 | 0.215 | 0.215 | 0 | 0.06 | 0.16 | 0.07 | 0.38 | 0.07 | 0.23 |
| 2 | 7579103 | rs732078 | -2.146 | 0.771 | 0.604 | 2.396 | 0.02 | 0.126 | 0.126 | 0.166 | 0.19 | 0.34 | 0.27 | 0.57 | 0.2 | 0.47 |
| 2 | 7581220 | rs10181784 | -0.543 | 1.017 | 0.269 | 2.349 | -0.038 | -0.031 | -0.031 | 0.627 | 0.06 | 0.01 | 0.05 | 0.13 | 0.11 | 0.21 |
| 2 | 7584668 | rs16866255 | -1.631 | 0.163 | 0.728 | 2.321 | 0.004 | 0.124 | 0.124 | 0.158 | 0.09 | 0.17 | 0.09 | 0.37 | 0.12 | 0.24 |
| 2 | 7598525 | rs6728724 | -1.287 | -0.778 | 1.462 | 1.437 | 0.023 | -0.013 | -0.013 | 0.409 | 0.4 | 0.66 | 0.54 | 0.51 | 0.32 | 0.45 |
| 2 | 7609558 | rs13409750 | -0.811 | -1.031 | 1.949 | 1.297 | -0.042 | -0.076 | -0.076 | 0.514 | 0.29 | 0.55 | 0.27 | 0.09 | 0.2 | 0.18 |
| 2 | 7610700 | rs7567384 | -0.818 | -1.029 | 1.951 | 1.298 | -0.042 | -0.076 | -0.076 | 0.078 | 0.29 | 0.55 | 0.27 | 0.09 | 0.2 | 0.18 |
| 2 | 7614263 | rs12105729 | -1.183 | -1.067 | 2.058 | 1.284 | -0.03 | 0.238 | 0.238 | 0 | 0.17 | 0.51 | 0.33 | 0.44 | 0.14 | 0.32 |
| 2 | 7620637 | rs1439975 | -1.681 | -1.066 | 1.592 | 1.265 | 0.07 | -0.013 | -0.013 | 0.017 | 0.43 | 0.76 | 0.57 | 0.52 | 0.33 | 0.47 |
| 2 | 7631459 | rs10495557 | -1.383 | -0.891 | 1.583 | 1.192 | -0.01 | 0.01 | 0.01 | 0 | 0.37 | 0.45 | 0.39 | 0.47 | 0.32 | 0.45 |
| 2 | 7636919 | rs11890644 | -0.491 | -0.812 | 1.327 | 1.19 | -0.001 | 0.076 | 0.076 | 0 | 0.13 | 0.19 | 0.3 | 0.43 | 0.15 | 0.29 |
| 2 | 7640278 | rs6752396 | -0.587 | -0.869 | 1.309 | 0.487 | -0.036 | -0.103 | -0.103 | 0.174 | 0.26 | 0.36 | 0.11 | 0.03 | 0.16 | 0.17 |
| 2 | 7642484 | rs6706159 | -0.541 | -0.878 | 1.244 | 0.482 | -0.023 | -0.079 | -0.079 | 3.332 | 0.25 | 0.32 | 0.11 | 0.03 | 0.16 | 0.17 |
| 2 | 7644366 | rs10191864 | -1.449 | -1.026 | 0.834 | 0.3 | 0.072 | 0.122 | 0.122 | 0.229 | 0.44 | 0.8 | 0.23 | 0.18 | 0.31 | 0.45 |
| 2 | 7647007 | rs1561574 | -0.686 | -1.227 | 1.605 | 0.181 | -0.053 | -0.104 | -0.104 | 0 | 0.23 | 0.47 | 0.12 | 0.06 | 0.13 | 0.11 |
| 2 | 7648655 | rs1439964 | -0.6 | -1.284 | 1.71 | 0.148 | -0.003 | -0.01 | -0.01 | 0 | 0.15 | 0.21 | 0.08 | 0.12 | 0.16 | 0.23 |
| 2 | 7654144 | rs1829483 | -0.334 | -1.369 | 1.807 | 0.104 | -0.043 | -0.104 | -0.104 | 2.79 | 0.2 | 0.46 | 0.11 | 0.04 | 0.12 | 0.08 |
| 2 | 7656440 | rs6431885 | -2.409 | -1.57 | 1.393 | 0.557 | 0.053 | 0.062 | 0.062 | 0.769 | 0.52 | 0.91 | 0.51 | 0.71 | 0.42 | 0.63 |
| 2 | 7657807 | rs10519830 | -0.218 | -1.358 | 1.435 | 0.608 | 0.036 | 0.058 | 0.058 | 0 | 0.06 | 0.17 | 0.04 | 0.12 | 0.11 | 0.2 |
| 2 | 7659087 | rs7604664 | -1.314 | -1.35 | 0.918 | 0.665 | -0.006 | 0.015 | 0.015 | 1.314 | 0.26 | 0.27 | 0.36 | 0.55 | 0.19 | 0.34 |
| 2 | 7663990 | rs16866330 | -0.026 | -1.018 | 0.571 | 1.084 | -0.009 | -0.056 | -0.056 | 0.017 | 0.12 | 0.02 | 0.29 | 0.4 | 0.14 | 0.29 |
| 2 | 7664359 | rs13394510 | -0.696 | -0.849 | 0.847 | 0.862 | 0.006 | 0.002 | 0.002 | 0 | 0.24 | 0.48 | 0.13 | 0.28 | 0.25 | 0.29 |
| 2 | 7665560 | rs2172114 | -1.425 | -0.896 | 0.109 | 0.555 | 0.023 | 0.021 | 0.021 | 0.398 | 0.43 | 0.83 | 0.47 | 0.71 | 0.39 | 0.62 |
| 2 | 7667932 | rs11691367 | -0.964 | -0.569 | 0.604 | -0.038 | -0.005 | -0.052 | -0.052 | 3.97 | 0.35 | 0.47 | 0.16 | 0.13 | 0.31 | 0.33 |
| 2 | 7675435 | rs1850909 | -0.623 | -0.611 | 0.774 | -0.053 | -0.001 | 0.013 | 0.013 | 0 | 0.31 | 0.3 | 0.15 | 0.13 | 0.31 | 0.33 |
| 2 | 7682371 | rs979290 | -1.262 | -0.676 | 1.135 | -0.017 | -0.001 | -0.012 | -0.012 | 0 | 0.37 | 0.65 | 0.2 | 0.18 | 0.33 | 0.37 |
| 2 | 7693796 | rs11676805 | -0.801 | -0.801 | 1.049 | -0.354 | -0.003 | -0.028 | -0.028 | 0 | 0.38 | 0.6 | 0.19 | 0.12 | 0.36 | 0.32 |
| 2 | 7700390 | rs6749863 | -0.726 | -0.85 | 0.923 | -0.436 | 0.007 | 0.024 | 0.024 | 0 | 0.42 | 0.67 | 0.21 | 0.13 | 0.38 | 0.33 |
| 2 | 7702036 | rs7589284 | -0.446 | -0.885 | 0.882 | -0.454 | 0 | 0.014 | 0.014 | 0 | 0.37 | 0.33 | 0.18 | 0.13 | 0.38 | 0.33 |
| 2 | 7710141 | rs880236 | -1.295 | -0.948 | 0.457 | -0.116 | 0.015 | 0.062 | 0.062 | 0 | 0.5 | 0.73 | 0.44 | 0.34 | 0.45 | 0.45 |
| 2 | 7716244 | rs2049072 | 0.397 | -0.836 | 0.364 | -0.074 | -0.025 | -0.041 | -0.041 | 6.481 | 0.07 | 0.01 | 0.23 | 0.21 | 0.07 | 0.11 |
| 2 | 7751923 | rs1879195 | -0.843 | -0.976 | 0.551 | -0.275 | 0.009 | 0.029 | 0.029 | 0 | 0.5 | 0.64 | 0.43 | 0.35 | 0.45 | 0.42 |
| 2 | 7763995 | rs270824 | -1.184 | -0.979 | 0.451 | -0.258 | -0.004 | 0.034 | 0.034 | 0 | 0.54 | 0.75 | 0.46 | 0.38 | 0.47 | 0.46 |
| 2 | 7765144 | rs270823 | -1.767 | -0.863 | -0.056 | -0.558 | -0.084 | 0.187 | 0.187 | 0 | 0.64 | 0.96 | 0.73 | 0.9 | 0.59 | 0.78 |
| 2 | 7775304 | rs270843 | -1.063 | -0.717 | 0.173 | -0.369 | 0.008 | 0.012 | 0.012 | 0 | 0.49 | 0.64 | 0.42 | 0.4 | 0.45 | 0.44 |
| 2 | 7807245 | rs12615861 | -0.726 | -0.457 | -0.963 | -0.363 | -0.004 | 0.031 | 0.031 | 0 | 0.44 | 0.24 | 0.39 | 0.35 | 0.46 | 0.44 |
| 2 | 7809505 | rs7597526 | 0.467 | -0.508 | -0.768 | -1.045 | -0.022 | -0.131 | -0.131 | 0 | 0.61 | 0.52 | 0.82 | 0.97 | 0.69 | 0.8 |
| 2 | 7812716 | rs11682611 | -0.868 | -0.506 | -0.812 | -0.547 | 0.005 | -0.024 | -0.024 | 0 | 0.14 | 0.17 | 0.08 | 0.04 | 0.14 | 0.15 |
| 2 | 7813271 | rs7573161 | -1.279 | -0.806 | -1.619 | -0.501 | -0.09 | 0.025 | 0.025 | 0 | 0.6 | 0.87 | 0.51 | 0.73 | 0.5 | 0.8 |
| 2 | 7817636 | rs10182315 | -0.658 | -0.703 | -0.755 | -0.403 | 0.016 | -0.056 | -0.056 | 0 | 0.13 | 0.25 | 0.09 | 0.02 | 0.13 | 0.14 |
| 2 | 7826501 | rs810067 | 0.982 | -1.301 | -0.825 | -0.07 | 0.021 | 0.116 | 0.116 | 0.055 | 0.51 | 0.11 | 0.44 | 0.24 | 0.62 | 0.37 |
| 2 | 7829406 | rs812811 | 0.127 | -1.835 | -0.039 | -0.085 | -0.001 | 0.005 | 0.005 | 0 | 0.77 | 0.74 | 0.58 | 0.25 | 0.81 | 0.52 |
| 2 | 7833261 | rs11676467 | -0.489 | -1.97 | 0.716 | -0.215 | -0.019 | -0.116 | -0.116 | 0 | 0.19 | 0.38 | 0.07 | 0.01 | 0.15 | 0.13 |
| 2 | 7833821 | rs772436 | 1.134 | -1.879 | 0.31 | -0.128 | 0.035 | 0.078 | 0.078 | 0 | 0.55 | 0.18 | 0.47 | 0.24 | 0.65 | 0.38 |
| 2 | 7838257 | rs772433 | 1.313 | -1.911 | 0.213 | -0.059 | 0.05 | 0.115 | 0.115 | 0 | 0.52 | 0.16 | 0.44 | 0.23 | 0.65 | 0.38 |
| 2 | 7849665 | rs772442 | 1.534 | -2.163 | -0.02 | -0.017 | -0.011 | 0.093 | 0.093 | 0 | 0.43 | 0.08 | 0.4 | 0.21 | 0.62 | 0.32 |
| 2 | 7856399 | rs2948927 | 1.309 | -2.216 | 1.343 | 0.346 | -0.039 | -0.037 | -0.037 | 0 | 0.78 | 0.42 | 0.87 | 0.86 | 0.89 | 0.77 |
| 2 | 7862742 | rs2948926 | 1.612 | -2.281 | 0.931 | 0.333 | 0.092 | 0.129 | 0.129 | 1.968 | 0.64 | 0.09 | 0.81 | 0.84 | 0.8 | 0.7 |
| 2 | 7871194 | rs772458 | 1.245 | -1.959 | 1.8 | 0.56 | -0.05 | -0.023 | -0.023 | 0 | 0.82 | 0.56 | 0.89 | 0.86 | 0.91 | 0.84 |
| 2 | 7877131 | rs1656168 | 1.604 | -1.913 | 1.304 | 0.788 | 0.043 | 0.075 | 0.075 | 1.996 | 0.65 | 0.07 | 0.68 | 0.77 | 0.79 | 0.65 |
| 2 | 7879097 | rs700946 | 0.667 | -1.931 | 2.09 | 0.815 | 0.015 | -0.06 | -0.06 | 0.036 | 0.84 | 0.52 | 0.88 | 0.94 | 0.87 | 0.86 |

| | | | | | | | | | | | | | | | | |
|---|---|---|---|---|---|---|---|---|---|---|---|---|---|---|---|---|
| 2 | 7882930 | rs2948925 | 0.103 | -1.921 | 1.341 | 1.199 | 0.011 | -0.104 | -0.104 | 0.966 | 0.2 | 0.05 | 0.28 | 0.5 | 0.17 | 0.3 |
| 2 | 7883907 | rs12692341 | 2.221 | -1.913 | 1.242 | 1.193 | -0.024 | 0.044 | 0.044 | 0.274 | 0.46 | 0.05 | 0.4 | 0.27 | 0.63 | 0.33 |
| 2 | 7886879 | rs812810 | -0.125 | -1.914 | 1.419 | 1.185 | -0.002 | -0.015 | -0.015 | 0 | 0.88 | 0.84 | 0.93 | 0.95 | 0.9 | 0.91 |
| 2 | 7893850 | rs13395925 | 2.125 | -1.881 | 1.352 | 0.989 | -0.024 | 0.044 | 0.044 | 0.274 | 0.46 | 0.05 | 0.41 | 0.27 | 0.63 | 0.33 |
| 2 | 7897634 | rs772428 | -1.085 | -1.872 | 1.47 | 0.999 | 0.012 | -0.038 | -0.038 | 1.425 | 0.34 | 0.2 | 0.36 | 0.55 | 0.3 | 0.45 |
| 2 | 7907486 | rs6757465 | -0.425 | -2.13 | 1.682 | 0.543 | -0.049 | 0.072 | 0.072 | 0.813 | 0.13 | 0.43 | 0.2 | 0.18 | 0.03 | 0.14 |
| 2 | 7912594 | rs4669210 | -0.342 | -1.772 | 0.877 | 0.178 | 0.008 | -0.032 | -0.032 | 0 | 0.06 | 0.17 | 0.03 | 0 | 0.05 | 0.09 |
| 2 | 7921970 | rs6745856 | -0.767 | -1.787 | 0.766 | 0.151 | 0.003 | -0.152 | -0.152 | 0.632 | 0.2 | 0.51 | 0.13 | 0.01 | 0.17 | 0.22 |
| 2 | 7924833 | rs6708659 | 0.159 | -1.782 | 0.674 | 0.151 | 0.02 | -0.053 | -0.053 | 0 | 0.1 | 0.27 | 0.07 | 0.01 | 0.08 | 0.14 |
| 2 | 7928168 | rs6431896 | 0.345 | -1.775 | 0.64 | 0.154 | -0.007 | -0.059 | -0.059 | 0.311 | 0.1 | 0.21 | 0.05 | 0 | 0.09 | 0.08 |
| 2 | 7930884 | rs6760840 | 1.256 | -1.476 | -0.615 | 0.479 | 0.049 | 0.149 | 0.149 | 0.594 | 0.46 | 0.16 | 0.37 | 0.2 | 0.63 | 0.31 |
| 2 | 7943978 | rs10803690 | 0.152 | -1.496 | -0.944 | 0.755 | -0.017 | -0.064 | -0.064 | 0 | 0.15 | 0.01 | 0.22 | 0.25 | 0.15 | 0.25 |
| 2 | 7946856 | rs9287693 | -0.126 | -0.776 | -0.394 | 0.307 | 0.007 | 0.007 | 0.007 | 0.017 | 0.57 | 0.46 | 0.45 | 0.21 | 0.71 | 0.45 |
| 2 | 7949693 | rs7561052 | 0.123 | -0.216 | -1.477 | 0.707 | 0.028 | 0.156 | 0.156 | 0.736 | 0.46 | 0.09 | 0.36 | 0.19 | 0.63 | 0.31 |
| 2 | 7958572 | rs16866463 | -1.323 | 0.312 | -0.19 | 1.282 | 0.011 | 0.022 | 0.022 | 0 | 0.12 | 0.26 | 0.06 | 0.17 | 0.11 | 0.2 |
| 2 | 7961407 | rs6751195 | -1.466 | 0.113 | -0.415 | 1.727 | -0.018 | 0.093 | 0.093 | 0 | 0.05 | 0.1 | 0.3 | 0.26 | 0.01 | 0.14 |
| 2 | 7962632 | rs798440 | -1.363 | 0.398 | -0.739 | 1.366 | -0.006 | 0.005 | 0.005 | 0 | 0.12 | 0.11 | 0.12 | 0.11 | 0.12 | 0.12 |
| 2 | 7966977 | rs7601867 | -1.498 | 0.194 | -0.161 | 0.823 | -0.009 | -0.072 | -0.072 | 1.119 | 0.14 | 0.29 | 0.11 | 0 | 0.1 | 0.1 |
| 2 | 7968275 | rs798443 | 1.739 | -0.625 | -0.834 | 1.418 | 0.061 | -0.076 | -0.076 | 0 | 0.68 | 0.06 | 0.67 | 0.63 | 0.82 | 0.67 |
| 2 | 7971295 | rs798444 | 0.39 | -0.623 | 0.203 | 1.572 | -0.013 | 0.046 | 0.046 | 0 | 0.88 | 0.84 | 0.85 | 0.73 | 0.92 | 0.82 |
| 2 | 7973726 | rs6750299 | -1.253 | -0.643 | 0.88 | 1.539 | -0.013 | -0.032 | -0.032 | 0 | 0.15 | 0.33 | 0.13 | 0.09 | 0.09 | 0.1 |
| 2 | 7974860 | rs798433 | 1.621 | -0.681 | 1.942 | 1.561 | -0.037 | 0.07 | 0.07 | 1.153 | 0.82 | 0.41 | 0.8 | 0.65 | 0.91 | 0.72 |
| 2 | 7978996 | rs798420 | 1.156 | -0.479 | 0.402 | 1.25 | -0.069 | -0.031 | -0.031 | 0 | 0.34 | 0.04 | 0.23 | 0.29 | 0.41 | 0.4 |
| 2 | 7983193 | rs798425 | 0.019 | -0.407 | 1.191 | 1.326 | -0.015 | 0.056 | 0.056 | 0 | 0.88 | 0.82 | 0.86 | 0.73 | 0.92 | 0.82 |
| 2 | 7983713 | rs798428 | 0.063 | -0.407 | 1.274 | 1.326 | -0.019 | 0.089 | 0.089 | 0 | 0.87 | 0.78 | 0.85 | 0.73 | 0.92 | 0.82 |
| 2 | 7988726 | rs6732533 | 1.082 | -0.411 | 1.453 | 1.324 | -0.035 | 0.042 | 0.042 | 1.128 | 0.82 | 0.39 | 0.8 | 0.73 | 0.91 | 0.74 |
| 2 | 7988851 | rs10183806 | 0.944 | -0.401 | 1.397 | 1.284 | -0.04 | 0.075 | 0.075 | 0.36 | 0.83 | 0.46 | 0.81 | 0.73 | 0.91 | 0.74 |
| 2 | 7989723 | rs1030051 | 1.474 | -0.517 | 1.363 | 1.181 | -0.041 | 0.032 | 0.032 | 0 | 0.73 | 0.55 | 0.73 | 0.67 | 0.83 | 0.76 |
| 2 | 7989914 | rs1030052 | 1.756 | -0.294 | 0.695 | 1.123 | 0.033 | -0.041 | -0.041 | 0.041 | 0.63 | 0.14 | 0.61 | 0.59 | 0.71 | 0.51 |
| 2 | 7990214 | rs12472063 | 1.937 | 0.124 | 0.627 | 0.695 | 0.015 | 0.037 | 0.037 | 1.114 | 0.57 | 0.09 | 0.6 | 0.67 | 0.64 | 0.57 |
| 2 | 7992985 | rs2216174 | 0.096 | -0.155 | 0.585 | 0.728 | -0.037 | 0.01 | 0.01 | 0 | 0.9 | 0.99 | 0.94 | 0.96 | 0.86 | 0.95 |
| 2 | 7993391 | rs17430926 | -0.48 | -0.138 | 0.652 | 0.694 | -0.014 | 0.037 | 0.037 | 0 | 0.08 | 0.01 | 0.07 | 0 | 0.06 | 0.02 |
| 2 | 7997419 | rs1861500 | 0.965 | -0.11 | 1.262 | 0.665 | -0.007 | 0.002 | 0.002 | 1.254 | 0.7 | 0.56 | 0.72 | 0.7 | 0.7 | 0.65 |
| 2 | 8001770 | rs9287694 | 2.257 | -0.216 | 1.376 | 0.646 | 0.031 | 0.032 | 0.032 | 1.833 | 0.54 | 0.1 | 0.59 | 0.66 | 0.63 | 0.55 |
| 2 | 8003686 | rs10168890 | 2.291 | -0.216 | 1.435 | 0.646 | 0.026 | 0.026 | 0.026 | 1.946 | 0.54 | 0.07 | 0.58 | 0.66 | 0.62 | 0.55 |
| 2 | 8004700 | rs2041874 | 1.748 | -0.212 | 1.46 | 0.649 | -0.006 | -0.005 | -0.005 | 0 | 0.59 | 0.45 | 0.63 | 0.67 | 0.63 | 0.55 |
| 2 | 8010162 | rs968964 | 2.171 | -0.202 | 1.269 | 0.654 | 0.035 | 0.034 | 0.034 | 1.481 | 0.55 | 0.17 | 0.59 | 0.66 | 0.63 | 0.55 |
| 2 | 8020302 | rs6736951 | 1.608 | -0.224 | 1.305 | 0.677 | -0.007 | -0.002 | -0.002 | 0.084 | 0.61 | 0.48 | 0.64 | 0.66 | 0.63 | 0.65 |
| 2 | 8027020 | rs2302672 | 0.636 | -0.198 | 1.268 | 0.644 | 0.001 | -0.006 | -0.006 | 0 | 0.72 | 0.75 | 0.8 | 0.66 | 0.72 | 0.67 |
| 2 | 8028725 | rs968763 | 1.752 | -0.255 | 1.336 | 0.607 | -0.002 | -0.002 | -0.002 | 0.411 | 0.59 | 0.4 | 0.67 | 0.64 | 0.64 | 0.57 |
| 2 | 8037331 | rs16866524 | -2.561 | 0.132 | 1.103 | -0.163 | -0.026 | -0.082 | -0.082 | 0.024 | 0.14 | 0.55 | 0.15 | 0.03 | 0.08 | 0.09 |
| 2 | 8038518 | rs7564066 | 1.785 | 0.592 | 0.228 | -0.04 | -0.073 | 0.055 | 0.055 | 1.703 | 0.54 | 0.06 | 0.51 | 0.29 | 0.57 | 0.44 |
| 2 | 8039194 | rs10929507 | 1.752 | 0.076 | 0.35 | -0.388 | 0.001 | 0.113 | 0.113 | 0.786 | 0.54 | 0.05 | 0.48 | 0.24 | 0.64 | 0.41 |
| 2 | 8039425 | rs13389085 | 1.434 | -0.276 | 0.498 | -0.502 | 0.006 | 0.18 | 0.18 | 0.011 | 0.48 | 0.1 | 0.56 | 0.19 | 0.67 | 0.35 |
| 2 | 8044396 | rs11691520 | 1.08 | -0.415 | 1.813 | 0.252 | -0.009 | -0.002 | -0.002 | 5.789 | 0.67 | 0.35 | 0.66 | 0.3 | 0.76 | 0.57 |
| 2 | 8045187 | rs1861506 | 1.26 | -0.529 | 1.235 | 0.255 | 0.01 | -0.054 | -0.054 | 0.397 | 0.9 | 0.67 | 0.89 | 1 | 0.94 | 0.91 |
| 2 | 8046782 | rs10189216 | 0.947 | -0.54 | 1.063 | 0.264 | 0.029 | 0.011 | 0.011 | 0.756 | 0.66 | 0.24 | 0.73 | 0.27 | 0.78 | 0.57 |
| 2 | 8048559 | rs4669221 | 0.993 | -0.401 | -0.573 | 0.348 | 0.044 | 0.012 | 0.012 | 0.596 | 0.65 | 0.21 | 0.74 | 0.29 | 0.78 | 0.57 |
| 2 | 8050508 | rs6739973 | 0.965 | -0.304 | -0.935 | 0.474 | 0.039 | 0.05 | 0.05 | 0.37 | 0.58 | 0.12 | 0.65 | 0.29 | 0.72 | 0.48 |
| 2 | 8051734 | rs888132 | 0.973 | -0.313 | -0.924 | 0.088 | 0.034 | 0.113 | 0.113 | 0.305 | 0.58 | 0.13 | 0.61 | 0.23 | 0.71 | 0.49 |
| 2 | 8060544 | rs6759088 | -1.457 | -0.272 | 0.02 | 0.05 | -0.003 | -0.022 | -0.022 | 0.054 | 0.15 | 0.19 | 0.09 | 0.12 | 0.14 | 0.13 |
| 2 | 8063121 | rs12469588 | 2.1 | -0.575 | 0.086 | -0.204 | 0.031 | 0.075 | 0.075 | 1.207 | 0.7 | 0.17 | 0.81 | 0.84 | 0.8 | 0.79 |
| 2 | 8065966 | rs10191104 | 1.741 | -0.628 | 0.575 | -0.233 | -0.015 | -0.041 | -0.041 | 0 | 0.77 | 0.32 | 0.84 | 0.88 | 0.85 | 0.86 |
| 2 | 8070689 | rs10170110 | 0.633 | -0.754 | 0.616 | -0.196 | -0.004 | -0.022 | -0.022 | 0 | 0.91 | 0.86 | 0.97 | 0.96 | 0.95 | 0.92 |
| 2 | 8072360 | rs10187145 | 2.125 | -0.814 | 0.538 | -0.232 | 0.061 | 0.134 | 0.134 | 2.437 | 0.68 | 0.11 | 0.81 | 0.84 | 0.82 | 0.79 |
| 2 | 8072756 | rs10164530 | 2.01 | -0.805 | 0.724 | -0.228 | 0.039 | 0.065 | 0.065 | 1.662 | 0.69 | 0.21 | 0.82 | 0.84 | 0.82 | 0.8 |
| 2 | 8086574 | rs11894196 | -1.569 | -0.826 | 0.949 | -0.227 | -0.032 | -0.065 | -0.065 | 0 | 0.19 | 0.35 | 0.11 | 0.12 | 0.13 | 0.13 |
| 2 | 8095158 | rs41009 | 1.285 | -0.522 | -0.853 | 0.06 | 0.037 | 0.067 | 0.067 | 2.058 | 0.74 | 0.12 | 0.84 | 0.83 | 0.85 | 0.8 |
| 2 | 8096549 | rs13024465 | -1.177 | -0.182 | -0.493 | 0.402 | -0.006 | -0.009 | -0.009 | 0 | 0.16 | 0.28 | 0.13 | 0.1 | 0.12 | 0.36 |
| 2 | 8096829 | rs193925 | 0.947 | -0.019 | -0.855 | 0.964 | 0.005 | -0.013 | -0.013 | 0 | 0.65 | 0.3 | 0.7 | 0.48 | 0.71 | 0.57 |
| 2 | 8098622 | rs40996 | 0.893 | -0.257 | -1.532 | 1.037 | 0.05 | -0.072 | -0.072 | 0.672 | 0.62 | 0.1 | 0.71 | 0.36 | 0.72 | 0.51 |
| 2 | 8099581 | rs999520 | 0.495 | 0.104 | -2.283 | 0.945 | -0.016 | 0.026 | 0.026 | 1.186 | 0.57 | 0.09 | 0.68 | 0.39 | 0.66 | 0.55 |
| 2 | 8099955 | rs2111073 | -0.259 | 0.36 | -1.281 | 1.122 | 0.016 | 0.017 | 0.017 | 0 | 0.27 | 0.43 | 0.21 | 0.34 | 0.34 | 0.21 |
| 2 | 8100477 | rs1477375 | -0.684 | 0.315 | -1.228 | 1.139 | 0.006 | -0.003 | -0.003 | 0 | 0.52 | 0.61 | 0.54 | 0.53 | 0.57 | 0.57 |
| 2 | 8101004 | rs4669228 | 0.325 | 0.327 | -0.991 | 1.027 | 0.005 | -0.005 | -0.005 | 0 | 0.22 | 0.25 | 0.29 | 0.16 | 0.21 | 0.29 |
| 2 | 8101337 | rs7597307 | 0.691 | -0.353 | -0.563 | 1.037 | -0.032 | 0.029 | 0.029 | 1.32 | 0.16 | 0.33 | 0.1 | 0.17 | 0.11 | 0.06 |
| 2 | 8102347 | rs12468177 | -0.18 | -0.305 | -0.539 | 1.12 | 0.005 | 0.003 | 0.003 | 0 | 0.45 | 0.41 | 0.49 | 0.47 | 0.55 | 0.53 |
| 2 | 8104849 | rs6753101 | -0.7 | -0.354 | 0.151 | 1.241 | 0.001 | -0.002 | -0.002 | 0 | 0.44 | 0.57 | 0.3 | 0.5 | 0.46 | 0.32 |
| 2 | 8107435 | rs193931 | 0.4 | -0.363 | -0.2 | 1.004 | -0.002 | -0.003 | -0.003 | 0 | 0.56 | 0.5 | 0.66 | 0.35 | 0.55 | 0.67 |

| | | | | | | | | | | | | | | | |
|---|---|---|---|---|---|---|---|---|---|---|---|---|---|---|---|
| 2 | 8111231 | rs41005 | 0.402 | -0.358 | -0.349 | 1.008 | -0.003 | -0.005 | -0.005 | 0.516 | 0.56 | 0.52 | 0.66 | 0.35 | 0.54 | 0.67 |
| 2 | 8113921 | rs41002 | 0.544 | -0.269 | -0.974 | 1.037 | 0.006 | 0.015 | 0.015 | 0.453 | 0.53 | 0.37 | 0.64 | 0.35 | 0.53 | 0.67 |
| 2 | 8117416 | rs10469612 | -0.66 | -0.792 | -0.572 | 0.999 | -0.048 | 0 | 0 | 1.362 | 0.24 | 0.55 | 0.18 | 0.27 | 0.14 | 0.1 |
| 2 | 8118123 | rs13021734 | 0.31 | -0.604 | -1.79 | 0.87 | 0.027 | -0.059 | -0.059 | 0 | 0.68 | 0.1 | 0.71 | 0.65 | 0.74 | 0.65 |
| 2 | 8118171 | rs13021915 | 0.577 | -0.435 | -1.127 | 0.572 | 0.1 | 0.129 | 0.129 | 0.014 | 0.53 | 0.18 | 0.54 | 0.68 | 0.75 | 0.74 |
| 2 | 8123746 | rs4669232 | -1.092 | -0.116 | 0.99 | 0.978 | -0.022 | 0.014 | 0.014 | 1.044 | 0.14 | 0.25 | 0.12 | 0.21 | 0.08 | 0.05 |
| 2 | 8128866 | rs1423712 | -2.122 | 0.208 | 1.693 | 1.248 | 0.027 | 0.11 | 0.11 | 3.071 | 0.1 | 0.53 | 0.13 | 0.22 | 0.07 | 0.04 |
| 2 | 8143969 | rs13405482 | -2.377 | 0.096 | 2.286 | 1.296 | -0.008 | 0.054 | 0.054 | 1.432 | 0.12 | 0.53 | 0.09 | 0.14 | 0.05 | 0.06 |
| 2 | 8145998 | rs4669238 | -1.973 | 0.438 | 2.065 | 1.779 | -0.003 | -0.035 | -0.035 | 0 | 0.31 | 0.69 | 0.29 | 0.54 | 0.3 | 0.24 |
| 2 | 8148765 | rs13426351 | -1.315 | 0.527 | 1.968 | 1.733 | 0 | -0.001 | -0.001 | 0.073 | 0.07 | 0.07 | 0.06 | 0.09 | 0.05 | 0.06 |
| 2 | 8149516 | rs6431916 | -1.555 | 0.536 | 2.126 | 1.737 | 0.014 | 0.019 | 0.019 | 0 | 0.07 | 0.23 | 0.07 | 0.09 | 0.05 | 0.06 |
| 2 | 8149627 | rs1364058 | -0.83 | 0.458 | 2.039 | 1.982 | -0.003 | 0.001 | 0.001 | 0.052 | 0.19 | 0.14 | 0.17 | 0.41 | 0.25 | 0.17 |
| 2 | 8159804 | rs7602105 | -1.865 | 0.238 | 2.095 | 1.57 | 0.02 | 0.219 | 0.219 | 4.031 | 0.08 | 0.51 | 0.08 | 0.18 | 0.02 | 0.05 |
| 2 | 8163722 | rs7597251 | -0.873 | 0.805 | 0.252 | 1.629 | -0.007 | -0.013 | -0.013 | 0 | 0.22 | 0.2 | 0.23 | 0.49 | 0.28 | 0.28 |
| 2 | 8165519 | rs6728934 | -1.561 | 0.037 | -0.26 | 1.413 | 0.015 | 0.025 | 0.025 | 0 | 0.35 | 0.25 | 0.28 | 0.67 | 0.29 | 0.35 |
| 2 | 8166036 | rs6757233 | -1.046 | 0.085 | -0.269 | 1.483 | -0.006 | -0.045 | -0.045 | 0 | 0.26 | 0.17 | 0.25 | 0.47 | 0.27 | 0.29 |
| 2 | 8178222 | rs6734986 | -1.453 | 0.401 | 0.761 | 0.994 | 0.021 | -0.015 | -0.015 | 0.009 | 0.08 | 0.14 | 0.22 | 0.05 | 0.12 | 0.03 |
| 2 | 8180189 | rs10929510 | 0.044 | 0.867 | 0.226 | 1.383 | -0.099 | -0.121 | -0.121 | 0 | 0.84 | 0.99 | 0.83 | 0.63 | 0.73 | 0.69 |
| 2 | 8189345 | rs6747840 | 0.982 | 0.806 | 0.486 | 1.171 | -0.024 | -0.055 | -0.055 | 0.792 | 0.73 | 0.8 | 0.79 | 0.53 | 0.68 | 0.58 |
| 2 | 8192868 | rs963825 | 1.068 | 0.827 | -0.352 | 0.19 | -0.019 | 0.14 | 0.14 | 0.151 | 0.32 | 0.12 | 0.3 | 0.05 | 0.32 | 0.2 |
| 2 | 8201262 | rs6718111 | 0.079 | 0.696 | -0.041 | 0.258 | -0.001 | 0.072 | 0.072 | 0.117 | 0.46 | 0.74 | 0.58 | 0.27 | 0.46 | 0.27 |
| 2 | 8207090 | rs6431922 | -0.035 | 1.266 | 0.189 | 0.271 | -0.018 | -0.001 | -0.001 | 0 | 0.53 | 0.79 | 0.53 | 0.57 | 0.53 | 0.48 |
| 2 | 8212939 | rs4668561 | 0.135 | 1.084 | 0.44 | 0.23 | -0.002 | -0.001 | -0.001 | 0 | 0.53 | 0.55 | 0.49 | 0.57 | 0.46 | 0.52 |
| 2 | 8216404 | rs1025053 | -0.114 | 1.184 | -0.305 | 0.21 | -0.012 | -0.013 | -0.013 | 0.936 | 0.56 | 0.76 | 0.52 | 0.58 | 0.51 | 0.53 |
| 2 | 8216735 | rs2052573 | -0.615 | 1.53 | -0.263 | 0.4 | -0.003 | 0.009 | 0.009 | 0.153 | 0.84 | 0.89 | 0.81 | 0.63 | 0.72 | 0.71 |
| 2 | 8227802 | rs6759347 | -0.254 | 0.753 | -0.105 | 0.43 | 0.014 | 0.052 | 0.052 | 0 | 0.49 | 0.59 | 0.5 | 0.38 | 0.47 | 0.48 |
| 2 | 8228731 | rs2355056 | -0.017 | 0.522 | -0.279 | 0.5 | -0.001 | -0.004 | -0.004 | 0 | 0.47 | 0.42 | 0.49 | 0.37 | 0.44 | 0.44 |
| 2 | 8230798 | rs6717377 | -0.017 | 0.283 | -0.827 | 0.612 | 0.041 | -0.012 | -0.012 | 0 | 0.48 | 0.25 | 0.48 | 0.49 | 0.58 | 0.48 |
| 2 | 8235239 | rs6712604 | -0.795 | -0.164 | 0.51 | 0.728 | 0.007 | -0.002 | -0.002 | 0 | 0.2 | 0.46 | 0.11 | 0.18 | 0.17 | 0.13 |
| 2 | 8239925 | rs16866597 | -0.54 | -0.212 | 0.688 | 0.752 | 0.016 | -0.009 | -0.009 | 0.037 | 0.16 | 0.25 | 0.08 | 0.11 | 0.16 | 0.12 |
| 2 | 8245597 | rs11673723 | -1.776 | -0.743 | 0.927 | 0.474 | -0.008 | -0.017 | -0.017 | 2.151 | 0.27 | 0.65 | 0.14 | 0.11 | 0.18 | 0.15 |
| 2 | 8259185 | rs11685783 | -1.251 | -0.777 | 1.024 | 0.468 | -0.001 | -0.001 | -0.001 | 0 | 0.22 | 0.26 | 0.1 | 0.18 | 0.16 | 0.16 |
| 2 | 8266416 | rs17317684 | -1.127 | -0.772 | 0.964 | 0.452 | 0.002 | 0.011 | 0.011 | 0.885 | 0.22 | 0.22 | 0.09 | 0.11 | 0.16 | 0.15 |
| 2 | 8272952 | rs181129 | 0.813 | -1.181 | 0.225 | 0.665 | 0.024 | -0.035 | -0.035 | 0 | 0.67 | 0.28 | 0.42 | 0.47 | 0.75 | 0.61 |
| 2 | 8282704 | rs12463875 | -0.259 | -1.304 | 0.436 | 0.992 | -0.007 | -0.05 | -0.05 | 0 | 0.09 | 0.01 | 0.08 | 0.18 | 0.09 | 0.13 |
| 2 | 8283740 | rs6735900 | 0.049 | -1.571 | 0.486 | 0.646 | 0.018 | -0.057 | -0.057 | 1.825 | 0.17 | 0.33 | 0.08 | 0.04 | 0.14 | 0.12 |
| 2 | 8284866 | rs181133 | 1.413 | -1.653 | 0.044 | 0.68 | 0.058 | -0.039 | -0.039 | 0 | 0.65 | 0.2 | 0.42 | 0.4 | 0.75 | 0.59 |
| 2 | 8285148 | rs181134 | 2.083 | -1.616 | -0.045 | 0.577 | 0.049 | 0.116 | 0.116 | 0.614 | 0.5 | 0.18 | 0.37 | 0.29 | 0.62 | 0.46 |
| 2 | 8287950 | rs7591430 | 0.47 | -1.808 | -0.104 | 0.437 | 0.002 | 0.004 | 0.004 | 0 | 0.62 | 0.7 | 0.38 | 0.34 | 0.6 | 0.64 |
| 2 | 8290719 | rs11695288 | -0.439 | -1.956 | 0.537 | 0.659 | -0.007 | -0.098 | -0.098 | 0 | 0.2 | 0.5 | 0.11 | 0.03 | 0.13 | 0.15 |
| 2 | 8294778 | rs181136 | -1.869 | -1.906 | -0.145 | 0.719 | -0.055 | 0.031 | 0.031 | 0.241 | 0.58 | 0.91 | 0.7 | 0.67 | 0.45 | 0.53 |
| 2 | 8310994 | rs1981456 | -2.155 | -2.107 | 0.002 | 0.879 | -0.087 | 0.071 | 0.071 | 0.667 | 0.59 | 0.97 | 0.68 | 0.74 | 0.47 | 0.49 |
| 2 | 8311011 | rs4669244 | -1.1 | -2.089 | 1.164 | 1.614 | -0.032 | 0.019 | 0.019 | 1.327 | 0.31 | 0.49 | 0.53 | 0.38 | 0.29 | 0.24 |
| 2 | 8311232 | rs6718040 | -1.194 | -2.075 | 1.456 | 1.622 | -0.019 | 0.185 | 0.185 | 0.005 | 0.07 | 0.19 | 0.04 | 0.31 | 0.03 | 0.1 |
| 2 | 8319260 | rs11891372 | -1.647 | -2.081 | 1.417 | 1.596 | 0.018 | 0.01 | 0.01 | 0.057 | 0.36 | 0.7 | 0.56 | 0.7 | 0.32 | 0.34 |
| 2 | 8325774 | rs10194137 | -0.263 | -1.917 | 1.748 | 1.828 | -0.008 | -0.041 | -0.041 | 0 | 0.18 | 0.08 | 0.48 | 0.37 | 0.24 | 0.24 |
| 2 | 8338845 | rs11684514 | -0.472 | -1.864 | 1.973 | 1.396 | 0.008 | -0.011 | -0.011 | 0.037 | 0.16 | 0.14 | 0.07 | 0.31 | 0.1 | 0.1 |
| 2 | 8349845 | rs6431930 | -1.186 | -1.894 | 2.252 | 1.387 | -0.065 | 0.116 | 0.116 | 0.696 | 0.2 | 0.53 | 0.11 | 0.32 | 0.1 | 0.12 |
| 2 | 8359262 | rs11684085 | -0.821 | -1.756 | 1.817 | 1.471 | -0.029 | 0.068 | 0.068 | 1.067 | 0.17 | 0.25 | 0.09 | 0.31 | 0.1 | 0.11 |
| 2 | 8360731 | rs6725841 | -1.569 | -1.645 | 1.274 | 1.499 | -0.033 | 0.013 | 0.013 | 1.163 | 0.25 | 0.69 | 0.15 | 0.31 | 0.12 | 0.14 |
| 2 | 8363269 | rs16866627 | -1.635 | -0.912 | 0.782 | 0.983 | -0.037 | -0.045 | -0.045 | 0 | 0.09 | 0.3 | 0.06 | 0 | 0.03 | 0.05 |
| 2 | 8371388 | rs6720719 | 0.083 | -0.419 | 0.065 | 1.647 | -0.005 | 0.011 | 0.011 | 0 | 0.85 | 0.87 | 0.51 | 0.55 | 0.73 | 0.8 |
| 2 | 8373397 | rs11904059 | 1.528 | -1.065 | 0.502 | 1.475 | -0.011 | -0.004 | -0.004 | 0 | 0.65 | 0.51 | 0.45 | 0.42 | 0.66 | 0.62 |
| 2 | 8374968 | rs16866633 | -0.913 | -1.029 | 0.556 | 1.46 | 0.014 | -0.001 | -0.001 | 0 | 0.16 | 0.13 | 0.05 | 0.13 | 0.07 | 0.18 |
| 2 | 8376228 | rs238620 | 1.131 | -1.013 | 0.83 | 1.489 | 0 | 0.146 | 0.146 | 1.868 | 0.76 | 0.48 | 0.46 | 0.59 | 0.72 | 0.78 |
| 2 | 8383579 | rs171152 | -0.02 | -0.731 | 0.34 | 1.656 | 0.004 | 0.046 | 0.046 | 0 | 0.83 | 0.82 | 0.51 | 0.58 | 0.73 | 0.82 |
| 2 | 8391187 | rs238627 | -0.058 | -0.715 | 0.311 | 1.696 | 0.006 | 0.053 | 0.053 | 0 | 0.84 | 0.81 | 0.51 | 0.6 | 0.73 | 0.83 |
| 2 | 8400057 | rs238618 | 1.413 | -0.932 | 0.431 | 1.583 | -0.01 | 0.038 | 0.038 | 0.695 | 0.66 | 0.45 | 0.47 | 0.52 | 0.7 | 0.65 |
| 2 | 8400895 | rs10490219 | -0.501 | -0.695 | -0.274 | 1.688 | -0.002 | 0 | 0 | 0 | 0.14 | 0.2 | 0.04 | 0.16 | 0.05 | 0.16 |
| 2 | 8404110 | rs238617 | 0.303 | -0.247 | -0.544 | 1.565 | 0.008 | 0.036 | 0.036 | 0 | 0.83 | 0.78 | 0.52 | 0.67 | 0.75 | 0.81 |
| 2 | 8409113 | rs238630 | 0.096 | -0.367 | -1.494 | 1.201 | 0.019 | 0.026 | 0.026 | 0.072 | 0.17 | 0.1 | 0.06 | 0.09 | 0.09 | 0.05 |
| 2 | 8409494 | rs238631 | 0.232 | 0.434 | -1.155 | 1.697 | -0.031 | -0.053 | -0.053 | 2.014 | 0.85 | 0.94 | 0.55 | 0.68 | 0.75 | 0.84 |
| 2 | 8423057 | rs17494521 | -0.461 | 0.628 | -1.259 | 1.266 | -0.032 | -0.036 | -0.036 | 0.398 | 0.4 | 0.2 | 0.48 | 0.55 | 0.42 | 0.43 |
| 2 | 8424313 | rs7595804 | -0.86 | 0.446 | 0.099 | 1.379 | 0.069 | 0.162 | 0.162 | 3.475 | 0.08 | 0.51 | 0.09 | 0.1 | 0.07 | 0.11 |
| 2 | 8436099 | rs346830 | -0.918 | 0.036 | 0.384 | 1.639 | -0.041 | 0.085 | 0.085 | 0.031 | 0.25 | 0.49 | 0.44 | 0.42 | 0.16 | 0.26 |
| 2 | 8437675 | rs2562011 | -0.949 | 0.299 | 1.141 | 1.634 | -0.011 | 0.081 | 0.081 | 0 | 0.23 | 0.44 | 0.47 | 0.4 | 0.16 | 0.26 |
| 2 | 8439905 | rs10206877 | -0.441 | 0.777 | 0.499 | 1.464 | -0.015 | -0.018 | -0.018 | 0 | 0.3 | 0.28 | 0.55 | 0.44 | 0.38 | 0.37 |
| 2 | 8440346 | rs10207297 | -0.332 | 0.765 | 0.575 | 1.459 | -0.021 | -0.026 | -0.026 | 0 | 0.29 | 0.25 | 0.55 | 0.44 | 0.38 | 0.37 |
| 2 | 8440718 | rs3097288 | -0.448 | 0.299 | 0.29 | 1.146 | 0.014 | -0.005 | -0.005 | 1.475 | 0.69 | 0.76 | 0.8 | 0.63 | 0.77 | 0.69 |

| | | | | | | | | | | | | | | | | |
|---|---|---|---|---|---|---|---|---|---|---|---|---|---|---|---|---|
| 2 | 8442248 | rs10174949 | 1.095 | 0.424 | 0.094 | 1.339 | -0.07 | -0.049 | -0.049 | 2.548 | 0.21 | 0.06 | 0.27 | 0.3 | 0.31 | 0.26 |
| 2 | 8446377 | rs4315494 | 0.192 | 0.032 | 0.962 | 1.368 | -0.018 | 0.02 | 0.02 | 0 | 0.14 | 0.24 | 0.11 | 0.25 | 0.09 | 0.04 |
| 2 | 8446695 | rs658243 | 1.597 | 0.094 | 1.094 | 1.289 | -0.041 | 0.068 | 0.068 | 0 | 0.34 | 0.1 | 0.2 | 0.18 | 0.36 | 0.31 |
| 2 | 8446866 | rs11887635 | 0.276 | 0.059 | 1.245 | 1.273 | -0.025 | 0.024 | 0.024 | 0 | 0.15 | 0.29 | 0.13 | 0.25 | 0.1 | 0.04 |
| 2 | 8447876 | rs3102937 | 0.663 | 0.055 | 1.396 | 1.309 | 0.004 | -0.004 | -0.004 | 0 | 0.44 | 0.42 | 0.49 | 0.34 | 0.46 | 0.41 |
| 2 | 8447909 | rs642501 | -0.25 | 0.058 | 1.336 | 1.311 | -0.001 | -0.003 | -0.003 | 1.382 | 0.57 | 0.54 | 0.61 | 0.59 | 0.55 | 0.46 |
| 2 | 8447979 | rs690249 | 0.841 | 0.075 | 1.176 | 1.305 | -0.026 | 0.009 | 0.009 | 0 | 0.42 | 0.26 | 0.48 | 0.34 | 0.46 | 0.41 |
| 2 | 8450533 | rs3111415 | -1.696 | 0.062 | 0.892 | 1.305 | 0.015 | 0.003 | 0.003 | 0 | 0.83 | 0.97 | 0.91 | 0.87 | 0.86 | 0.73 |
| 2 | 8451479 | rs439614 | -1.424 | 0.11 | 0.935 | 1.423 | -0.019 | 0.074 | 0.074 | 0 | 0.49 | 0.86 | 0.71 | 0.66 | 0.5 | 0.41 |
| 2 | 8451498 | rs13395467 | 1.087 | 0.48 | 1.039 | 1.443 | -0.065 | -0.034 | -0.034 | 0.429 | 0.21 | 0.08 | 0.27 | 0.28 | 0.32 | 0.26 |
| 2 | 8452497 | rs3102960 | -1.187 | 0.923 | 0.883 | 1.121 | -0.032 | 0.013 | 0.013 | 0 | 0.81 | 0.97 | 0.89 | 0.87 | 0.79 | 0.72 |
| 2 | 8453175 | rs12470607 | 0.437 | 0.569 | 0.947 | 1.157 | -0.005 | -0.003 | -0.003 | 0 | 0.07 | 0.06 | 0.27 | 0.15 | 0.07 | 0.11 |
| 2 | 8456003 | rs404236 | -1.128 | 0.547 | 1.165 | 1.144 | -0.027 | -0.016 | -0.016 | 0.049 | 0.81 | 0.76 | 0.89 | 0.87 | 0.85 | 0.72 |
| 2 | 8457497 | rs184706 | -0.096 | 0.579 | 1.009 | 1.324 | -0.003 | -0.001 | -0.001 | 0.023 | 0.56 | 0.63 | 0.54 | 0.55 | 0.48 | 0.47 |
| 2 | 8458707 | rs16866676 | 0.48 | 0.595 | 0.477 | 1.349 | 0 | -0.003 | -0.003 | 0 | 0.07 | 0.05 | 0.27 | 0.15 | 0.06 | 0.11 |
| 2 | 8459919 | rs2630745 | 0.744 | 0.595 | 0.468 | 1.224 | 0.025 | 0.033 | 0.033 | 0 | 0.45 | 0.31 | 0.24 | 0.39 | 0.38 | 0.35 |
| 2 | 8475614 | rs3097278 | -1.154 | 0.529 | 0.19 | 1.225 | 0.018 | 0.01 | 0.01 | 0 | 0.81 | 0.96 | 0.9 | 0.87 | 0.85 | 0.73 |
| 2 | 8477175 | rs3111422 | -0.557 | 0.776 | 0.143 | 1.088 | -0.01 | -0.011 | -0.011 | 2.593 | 0.37 | 0.67 | 0.68 | 0.48 | 0.46 | 0.4 |
| 2 | 8480734 | rs424620 | 0.862 | 0.815 | -0.466 | 0.927 | -0.053 | -0.047 | -0.047 | 0 | 0.21 | 0.14 | 0.33 | 0.34 | 0.32 | 0.24 |
| 2 | 8486659 | rs3111424 | -1.175 | 0.71 | -0.859 | 0.934 | 0.014 | 0.012 | 0.012 | 0 | 0.81 | 0.98 | 0.9 | 0.88 | 0.85 | 0.74 |
| 2 | 8487296 | rs9287698 | 0.704 | 0.924 | -0.679 | 0.957 | -0.084 | -0.065 | -0.065 | 0 | 0.22 | 0.09 | 0.32 | 0.37 | 0.34 | 0.24 |
| 2 | 8501165 | rs3097284 | -0.705 | 0.308 | -0.674 | 0.188 | 0.038 | 0.055 | 0.055 | 0 | 0.78 | 0.97 | 0.89 | 0.89 | 0.84 | 0.76 |
| 2 | 8501908 | rs3102943 | -0.813 | 0.213 | -0.324 | 0.054 | -0.016 | -0.003 | -0.003 | 0 | 0.88 | 0.84 | 0.95 | 0.91 | 0.95 | 0.83 |
| 2 | 8507486 | rs3102941 | 0.716 | 0.102 | -0.153 | -0.163 | 0.006 | 0.046 | 0.046 | 1.304 | 0.52 | 0.33 | 0.29 | 0.33 | 0.49 | 0.36 |
| 2 | 8508080 | rs12623235 | 0.216 | 0.182 | 0.088 | -0.069 | 0.023 | 0.01 | 0.01 | 0.441 | 0.28 | 0.35 | 0.59 | 0.56 | 0.38 | 0.4 |
| 2 | 8512104 | rs16866692 | -0.938 | 0.165 | 0.428 | -0.088 | -0.007 | -0.012 | -0.012 | 0.061 | 0.09 | 0.21 | 0.08 | 0.08 | 0.08 | 0.06 |
| 2 | 8516939 | rs4669256 | 0.616 | -0.061 | -0.122 | -0.33 | 0.016 | -0.026 | -0.026 | 0 | 0.41 | 0.27 | 0.65 | 0.5 | 0.56 | 0.55 |
| 2 | 8520621 | rs12613635 | 0.638 | 0.057 | -0.501 | -0.214 | 0.02 | -0.011 | -0.011 | 0.175 | 0.35 | 0.22 | 0.21 | 0.39 | 0.31 | 0.18 |
| 2 | 8522443 | rs13384783 | 1.589 | 0.069 | -0.677 | -0.242 | 0.017 | -0.05 | -0.05 | 0 | 0.23 | 0.08 | 0.13 | 0.39 | 0.21 | 0.15 |
| 2 | 8522651 | rs12616139 | -0.413 | 0.729 | -0.609 | -0.123 | 0.002 | 0.003 | 0.003 | 0.406 | 0.5 | 0.43 | 0.57 | 0.58 | 0.54 | 0.36 |
| 2 | 8524541 | rs4669261 | 0.131 | 0.803 | -0.38 | 0.144 | -0.006 | -0.002 | -0.002 | 0 | 0.1 | 0.08 | 0.32 | 0.11 | 0.18 | 0.14 |
| 2 | 8529486 | rs7557740 | -1.076 | 0.749 | -0.483 | 0.129 | 0.011 | -0.005 | -0.005 | 0.426 | 0.2 | 0.23 | 0.38 | 0.2 | 0.26 | 0.18 |
| 2 | 8531810 | rs12476361 | -0.861 | 0.445 | -0.558 | 0.151 | 0.012 | -0.006 | -0.006 | 2.68 | 0.19 | 0.24 | 0.38 | 0.2 | 0.25 | 0.18 |
| 2 | 8537949 | rs6759998 | 0.586 | 0.873 | -1.285 | 0.231 | -0.001 | 0.009 | 0.009 | 1.066 | 0.61 | 0.29 | 0.52 | 0.73 | 0.62 | 0.48 |
| 2 | 8542955 | rs4668568 | -0.358 | 1.196 | -2.168 | 0.478 | -0.014 | -0.01 | -0.01 | 0 | 0.67 | 0.75 | 0.74 | 0.55 | 0.58 | 0.71 |
| 2 | 8543151 | rs7587352 | -0.612 | 1.238 | -2.17 | 0.481 | -0.003 | -0.007 | -0.007 | 0 | 0.7 | 0.79 | 0.82 | 0.6 | 0.68 | 0.83 |
| 2 | 8545555 | rs4233860 | -0.808 | 1.202 | -2.616 | 0.602 | -0.015 | -0.028 | -0.028 | 0 | 0.72 | 0.87 | 0.82 | 0.6 | 0.68 | 0.83 |
| 2 | 8548997 | rs901350 | -0.097 | 1.093 | -2.643 | 0.94 | -0.047 | -0.145 | -0.145 | 0.004 | 0.37 | 0.03 | 0.35 | 0.44 | 0.35 | 0.4 |
| 2 | 8556513 | rs436385 | -1.126 | 1.347 | -1.609 | 0.929 | -0.002 | -0.003 | -0.003 | 0 | 0.05 | 0.06 | 0.08 | 0.12 | 0.11 | 0.05 |
| 2 | 8561739 | rs1872603 | 0.58 | 0.988 | -1.91 | 0.895 | 0 | -0.012 | -0.012 | 0 | 0.75 | 0.84 | 0.79 | 0.66 | 0.76 | 0.8 |
| 2 | 8563326 | rs950968 | 0.476 | 0.914 | -2.601 | 0.842 | 0.003 | -0.003 | -0.003 | 0 | 0.47 | 0.78 | 0.64 | 0.45 | 0.54 | 0.71 |
| 2 | 8564462 | rs35406633 | 0.261 | 0.919 | -2.588 | 0.873 | 0.009 | -0.006 | -0.006 | 0 | 0.23 | 0.14 | 0.14 | 0.22 | 0.22 | 0.11 |
| 2 | 8566687 | rs403081 | 0.564 | 0.877 | -2.611 | 0.966 | -0.001 | -0.017 | -0.017 | 0 | 0.08 | 0.05 | 0.08 | 0.18 | 0.08 | 0.08 |
| 2 | 8566885 | rs4669273 | -0.1 | 0.981 | -3.025 | 0.827 | -0.051 | -0.048 | -0.048 | 0.874 | 0.42 | 0.15 | 0.39 | 0.52 | 0.47 | 0.61 |
| 2 | 8567632 | rs2655470 | 0.451 | 0.403 | -2.276 | 0.549 | -0.001 | -0.016 | -0.016 | 0 | 0.08 | 0.04 | 0.08 | 0.15 | 0.08 | 0.08 |
| 2 | 8571139 | rs2655450 | -0.54 | 0.311 | -2.098 | 0.409 | 0.003 | -0.014 | -0.014 | 0 | 0.32 | 0.22 | 0.24 | 0.39 | 0.3 | 0.2 |
| 2 | 8571477 | rs2630720 | -0.812 | 0.356 | -1.466 | 0.381 | 0.002 | -0.001 | -0.001 | 0.027 | 0.41 | 0.62 | 0.31 | 0.44 | 0.33 | 0.24 |
| 2 | 8572964 | rs379131 | -0.521 | 0.926 | -1.386 | 0.506 | -0.015 | -0.024 | -0.024 | 0 | 0.09 | 0.04 | 0.1 | 0.21 | 0.11 | 0.09 |
| 2 | 8577685 | rs453322 | 0.611 | 0.574 | -0.935 | -0.585 | -0.009 | -0.035 | -0.035 | 0.231 | 0.75 | 0.71 | 0.71 | 0.86 | 0.78 | 0.76 |
| 2 | 8579149 | rs2002685 | -0.589 | 0.642 | -1.21 | -0.6 | -0.026 | 0.024 | 0.024 | 0.465 | 0.14 | 0.03 | 0.14 | 0.1 | 0.17 | 0.12 |
| 2 | 8580884 | rs2655455 | -0.101 | 0.561 | -1.004 | -1.599 | 0.017 | 0.068 | 0.068 | 0 | 0.46 | 0.67 | 0.51 | 0.85 | 0.46 | 0.62 |
| 2 | 8582292 | rs420609 | -0.044 | 0.174 | -0.756 | -1.98 | 0.006 | 0.011 | 0.011 | 0.287 | 0.58 | 0.38 | 0.67 | 0.84 | 0.63 | 0.69 |
| 2 | 8590041 | rs1504240 | -0.213 | 0.385 | -0.089 | -2.094 | 0 | -0.012 | -0.012 | 0 | 0.52 | 0.51 | 0.48 | 0.87 | 0.56 | 0.64 |
| 2 | 8592437 | rs10427278 | -0.228 | 0.43 | -0.063 | -2.072 | -0.001 | 0.006 | 0.006 | 0.531 | 0.52 | 0.53 | 0.48 | 0.86 | 0.57 | 0.64 |
| 2 | 8592500 | rs10929523 | -0.263 | 0.412 | -0.338 | -2.081 | 0.002 | 0.045 | 0.045 | 0.112 | 0.5 | 0.4 | 0.47 | 0.86 | 0.57 | 0.64 |
| 2 | 8602278 | rs6431940 | 0.151 | 0.326 | -0.769 | -2.056 | -0.04 | 0.232 | 0.232 | 0.483 | 0.47 | 0.21 | 0.5 | 0.13 | 0.46 | 0.31 |
| 2 | 8605131 | rs1875812 | 0.332 | 0.316 | -0.918 | -2.273 | -0.001 | -0.008 | -0.008 | 0 | 0.15 | 0.15 | 0.11 | 0.08 | 0.14 | 0.1 |
| 2 | 8608271 | rs12474952 | -0.165 | 0.177 | -0.612 | -2.356 | 0.004 | -0.002 | -0.002 | 0.012 | 0.55 | 0.46 | 0.74 | 0.8 | 0.64 | 0.7 |
| 2 | 8609001 | rs4668572 | -0.045 | -0.132 | -0.539 | -2.283 | -0.012 | 0.011 | 0.011 | 0 | 0.34 | 0.33 | 0.43 | 0.85 | 0.44 | 0.55 |
| 2 | 8609400 | rs10929527 | 0.784 | -0.025 | -1.131 | -2.219 | -0.03 | 0.153 | 0.153 | 0 | 0.33 | 0.09 | 0.41 | 0.05 | 0.32 | 0.2 |
| 2 | 8609639 | rs6727210 | -0.177 | -0.329 | -0.473 | -1.434 | -0.035 | -0.061 | -0.061 | 0 | 0.7 | 0.61 | 0.85 | 0.9 | 0.77 | 0.8 |
| 2 | 8621162 | rs6733104 | 0.114 | -0.073 | -0.724 | -0.496 | -0.004 | 0.01 | 0.01 | 0 | 0.75 | 0.83 | 0.73 | 0.18 | 0.73 | 0.62 |
| 2 | 8630749 | rs12328987 | -0.9 | -0.58 | -0.488 | -0.202 | 0.042 | 0.012 | 0.012 | 0 | 0.1 | 0 | 0.13 | 0.03 | 0.01 | 0.09 |
| 2 | 8631081 | rs10192647 | -1.459 | -0.536 | 0.182 | -0.17 | -0.032 | 0.006 | 0.006 | 0 | 0.12 | 0.26 | 0.17 | 0.09 | 0.03 | 0.22 |
| 2 | 8631462 | rs6431942 | -1.912 | -0.699 | 0.58 | -0.259 | -0.061 | -0.008 | -0.008 | 0 | 0.16 | 0.6 | 0.22 | 0.09 | 0.04 | 0.21 |
| 2 | 8632587 | rs4668574 | -0.865 | -0.577 | -0.332 | -0.236 | -0.029 | -0.014 | -0.014 | 0 | 0.1 | 0.14 | 0.14 | 0.03 | 0.01 | 0.11 |
| 2 | 8644700 | rs4669280 | -0.825 | -0.319 | -0.541 | -0.003 | -0.009 | 0.001 | 0.001 | 0 | 0.25 | 0.64 | 0.48 | 0.3 | 0.2 | 0.26 |
| 2 | 8654581 | rs876753 | 0.335 | 0.287 | -1.816 | 0.32 | -0.164 | -0.178 | -0.178 | 0 | 0.3 | 0.02 | 0.31 | 0.54 | 0.43 | 0.6 |
| 2 | 8657302 | rs6431945 | -0.086 | 0.071 | -1.689 | 0.538 | -0.068 | -0.1 | -0.1 | 0 | 0.8 | 0.96 | 0.84 | 0.62 | 0.71 | 0.82 |

| | | | | | | | | | | | | | | | | |
|---|---|---|---|---|---|---|---|---|---|---|---|---|---|---|---|---|
| 2 | 8673584 | rs2356132 | 0.007 | -1.029 | -1.273 | -0.289 | 0.053 | 0.067 | 0.067 | 0 | 0.73 | 0.84 | 0.73 | 0.96 | 0.87 | 0.83 |
| 2 | 8673927 | rs1532791 | 0.775 | -0.658 | -0.536 | 0.083 | 0.009 | 0.008 | 0.008 | 0 | 0.48 | 0.41 | 0.44 | 0.5 | 0.54 | 0.6 |
| 2 | 8677844 | rs6431949 | 0.465 | -0.602 | -0.58 | 0.118 | 0.029 | -0.016 | -0.016 | 0 | 0.54 | 0.75 | 0.49 | 0.54 | 0.6 | 0.65 |
| 2 | 8680931 | rs6740252 | 0.525 | -0.789 | 0.1 | 0.335 | -0.027 | 0.001 | 0.001 | 0 | 0.67 | 0.46 | 0.71 | 0.41 | 0.76 | 0.62 |
| 2 | 8684493 | rs1007832 | 0.539 | -0.89 | -0.163 | 0.312 | 0.037 | 0.009 | 0.009 | 0 | 0.64 | 0.28 | 0.54 | 0.45 | 0.74 | 0.63 |
| 2 | 8691159 | rs345903 | -1.302 | -0.803 | -0.335 | 0.094 | 0.051 | 0.064 | 0.064 | 0.306 | 0.48 | 0.74 | 0.5 | 0.61 | 0.35 | 0.41 |
| 2 | 8694850 | rs4399693 | 0.783 | -0.83 | -0.178 | 0.095 | 0.026 | -0.006 | -0.006 | 0 | 0.57 | 0.28 | 0.62 | 0.47 | 0.67 | 0.63 |
| 2 | 8695654 | rs2883914 | 0.951 | -1.438 | -0.367 | -0.09 | 0.069 | -0.023 | -0.023 | 0 | 0.55 | 0.26 | 0.62 | 0.51 | 0.72 | 0.68 |
| 2 | 8712608 | rs11695862 | 1.576 | -0.887 | -0.922 | -0.393 | -0.116 | -0.041 | -0.041 | 0.364 | 0.42 | 0.05 | 0.41 | 0.44 | 0.51 | 0.59 |
| 2 | 8718430 | rs11904418 | -0.604 | -0.68 | -0.145 | -0.346 | 0 | 0.006 | 0.006 | 0 | 0.19 | 0.16 | 0.36 | 0.62 | 0.25 | 0.23 |
| 2 | 8718961 | rs11679703 | -0.133 | -0.695 | -0.168 | -0.331 | -0.001 | 0.007 | 0.007 | 0 | 0.16 | 0.13 | 0.34 | 0.62 | 0.23 | 0.22 |
| 2 | 8719699 | rs6744749 | 0.459 | -1.399 | 0.327 | -0.747 | -0.03 | -0.159 | -0.159 | 0 | 0.69 | 0.57 | 0.86 | 1 | 0.78 | 0.81 |
| 2 | 8728607 | rs4668578 | 1.284 | -1.403 | 0.776 | -0.087 | -0.027 | 0.002 | 0.002 | 0 | 0.69 | 0.45 | 0.6 | 0.36 | 0.78 | 0.73 |
| 2 | 8728909 | rs3856442 | -1.055 | -1.275 | 0.758 | -0.004 | -0.019 | 0.022 | 0.022 | 0.014 | 0.23 | 0.29 | 0.35 | 0.64 | 0.18 | 0.25 |
| 2 | 8739461 | rs4669299 | 0.605 | -1.146 | 0.611 | -0.278 | 0.016 | 0.012 | 0.012 | 0.22 | 0.94 | 0.63 | 0.96 | 1 | 0.99 | 1 |
| 2 | 8750103 | rs13399561 | -0.839 | -1.432 | -0.505 | -0.67 | 0.007 | -0.022 | -0.022 | 0 | 0.24 | 0.25 | 0.09 | 0 | 0.16 | 0.15 |
| 2 | 8750266 | rs3856447 | 1.705 | -1.405 | -1.397 | -0.59 | -0.089 | -0.097 | -0.097 | 3.128 | 0.43 | 0.02 | 0.49 | 0.44 | 0.61 | 0.59 |
| 2 | 8750673 | rs3856448 | 1.409 | -1.464 | -1.293 | -0.686 | 0.019 | 0.313 | 0.313 | 0 | 0.47 | 0.16 | 0.65 | 0.96 | 0.64 | 0.73 |
| 2 | 8752881 | rs7423613 | -0.047 | -1.404 | -1.183 | -0.613 | 0.006 | -0.009 | -0.009 | 0 | 0.71 | 0.37 | 0.73 | 0.96 | 0.8 | 0.88 |
| 2 | 8756583 | rs13428852 | -0.421 | -1.028 | -0.504 | 0.185 | 0.001 | -0.016 | -0.016 | 0 | 0.16 | 0.18 | 0.05 | 0 | 0.11 | 0.13 |
| 2 | 8761162 | rs3856450 | -1.097 | -0.885 | -0.439 | 0.442 | -0.012 | 0.015 | 0.015 | 0 | 0.3 | 0.41 | 0.26 | 0.55 | 0.18 | 0.29 |
| 2 | 8770564 | rs4669318 | 1.022 | -0.39 | -0.321 | 0.082 | -0.011 | -0.06 | -0.06 | 0.589 | 0.9 | 0.78 | 0.93 | 1 | 0.92 | 0.97 |
| 2 | 8775493 | rs1344807 | 0.231 | -0.337 | -1.522 | 0.129 | 0.047 | 0.067 | 0.067 | 0.005 | 0.46 | 0.15 | 0.3 | 0.36 | 0.35 | 0.29 |
| 2 | 8776102 | rs3927971 | -0.44 | -0.349 | -1.362 | -0.059 | 0.001 | -0.004 | -0.004 | 0 | 0.43 | 0.44 | 0.63 | 0.4 | 0.53 | 0.52 |
| 2 | 8779824 | rs7572345 | -0.867 | -0.319 | -1.855 | 0.111 | -0.059 | -0.055 | -0.055 | 0.013 | 0.64 | 0.88 | 0.68 | 0.63 | 0.6 | 0.66 |
| 2 | 8786568 | rs3856451 | 0.391 | -0.435 | -1.764 | -0.069 | -0.075 | -0.084 | -0.084 | 1.249 | 0.27 | 0.14 | 0.54 | 0.45 | 0.42 | 0.31 |
| 2 | 8788439 | rs4669322 | -1.056 | -0.61 | -1.622 | -0.201 | -0.007 | 0.008 | 0.008 | 0 | 0.58 | 0.81 | 0.69 | 0.42 | 0.59 | 0.55 |
| 2 | 8790973 | rs3889716 | -1.474 | -1.157 | 0.086 | -0.719 | -0.008 | -0.034 | -0.034 | 0.397 | 0.25 | 0.33 | 0.11 | 0.07 | 0.14 | 0.26 |
| 2 | 8794240 | rs3856456 | 1.627 | -1.364 | 0.839 | -0.923 | -0.02 | 0.023 | 0.023 | 0 | 0.86 | 0.6 | 0.93 | 0.82 | 0.95 | 0.8 |
| 2 | 8797869 | rs3856460 | 0.831 | -1.555 | 1.067 | -1.143 | -0.029 | -0.121 | -0.121 | 1.914 | 0.78 | 0.58 | 0.86 | 0.98 | 0.85 | 0.8 |
| 2 | 8798656 | rs7606528 | -1.52 | -1.717 | 1.307 | -1.054 | -0.053 | -0.017 | -0.017 | 0 | 0.17 | 0.41 | 0.07 | 0.19 | 0.05 | 0.2 |
| 2 | 8800807 | rs4669325 | 1.766 | -1.694 | 1.291 | -1.096 | -0.038 | -0.092 | -0.092 | 1.89 | 0.87 | 0.47 | 0.93 | 1 | 0.94 | 0.97 |
| 2 | 8807250 | rs7573522 | 1.893 | -1.644 | 0.996 | -1.069 | -0.039 | -0.085 | -0.085 | 0 | 0.86 | 0.34 | 0.91 | 1 | 0.94 | 0.96 |
| 2 | 8811354 | rs1054765 | 2.088 | -1.856 | 0.286 | -1.274 | 0.124 | 0.209 | 0.209 | 0.021 | 0.59 | 0.07 | 0.79 | 0.83 | 0.8 | 0.59 |
| 2 | 8814245 | rs7567871 | -2.236 | -1.657 | 0.728 | -1.144 | -0.038 | -0.08 | -0.08 | 0.14 | 0.14 | 0.71 | 0.1 | 0 | 0.06 | 0.04 |
| 2 | 8817803 | rs1434123 | 1.856 | -1.634 | 0.368 | -1.153 | -0.016 | -0.052 | -0.052 | 0 | 0.85 | 0.24 | 0.9 | 1 | 0.93 | 0.95 |
| 2 | 8826750 | rs4669330 | 1.73 | -1.434 | -0.02 | -1.03 | 0.087 | 0.11 | 0.11 | 0.001 | 0.61 | 0.14 | 0.73 | 0.74 | 0.76 | 0.61 |
| 2 | 8834580 | rs10200926 | 1.209 | -1.5 | 1.008 | -1.006 | -0.073 | 0.034 | 0.034 | 0 | 0.84 | 0.56 | 0.9 | 0.77 | 0.97 | 0.86 |
| 2 | 8837017 | rs6709598 | 1.364 | -1.451 | 1.254 | -1.207 | -0.032 | -0.08 | -0.08 | 0 | 0.73 | 0.51 | 0.8 | 0.86 | 0.81 | 0.75 |
| 2 | 8842714 | rs4668588 | 1.839 | -1.547 | 0.903 | -1.409 | 0.058 | 0.092 | 0.092 | 0 | 0.62 | 0.23 | 0.73 | 0.87 | 0.74 | 0.66 |
| 2 | 8843347 | rs6728858 | -0.911 | -1.544 | 1.374 | -1.428 | 0 | -0.082 | -0.082 | 0.351 | 0.18 | 0.3 | 0.13 | 0 | 0.2 | 0.21 |
| 2 | 8845429 | rs6723945 | -0.27 | -1.464 | 1.79 | -1.153 | -0.045 | -0.014 | -0.014 | 0 | 0.1 | 0.29 | 0.1 | 0.12 | 0.01 | 0.06 |
| 2 | 8847321 | rs7593804 | 0.715 | -1.285 | 1.175 | -1.182 | -0.031 | -0.056 | -0.056 | 0 | 0.9 | 0.81 | 0.96 | 0.99 | 0.95 | 0.93 |
| 2 | 8852462 | rs13010395 | -0.377 | -0.81 | 0.381 | -1.292 | -0.012 | 0.016 | 0.016 | 0 | 0.23 | 0.2 | 0.27 | 0.14 | 0.34 | 0.29 |
| 2 | 8858736 | rs9789497 | 1.326 | -0.983 | -0.171 | -1.199 | -0.013 | 0.071 | 0.071 | 0.388 | 0.47 | 0.09 | 0.53 | 0.62 | 0.56 | 0.56 |
| 2 | 8865152 | rs7606661 | -0.893 | -1.046 | 0.574 | -1.201 | 0.016 | 0.008 | 0.008 | 1.43 | 0.05 | 0.31 | 0.04 | 0 | 0.01 | 0 |
| 2 | 8871342 | rs1044280 | 0.336 | -1.102 | 0.954 | -1.24 | 0.006 | -0.004 | -0.004 | 0 | 0.22 | 0.24 | 0.24 | 0.16 | 0.29 | 0.24 |
| 2 | 8887180 | rs3351 | -0.175 | -1.211 | 0.884 | -1.209 | 0.045 | 0.031 | 0.031 | 0 | 0.22 | 0.1 | 0.16 | 0.18 | 0.13 | 0.18 |
| 2 | 8893432 | rs2118471 | -0.06 | -1.223 | 0.974 | -1.212 | 0.02 | 0.018 | 0.018 | 0 | 0.79 | 0.86 | 0.85 | 0.83 | 0.87 | 0.82 |
| 2 | 8948794 | rs8179714 | -0.395 | -0.83 | 0.816 | -1.272 | 0.02 | 0.033 | 0.033 | 0 | 0.26 | 0.15 | 0.18 | 0.18 | 0.19 | 0.23 |
| 2 | 8949070 | rs7587928 | 0.008 | -0.732 | 0.324 | -1.277 | 0.01 | -0.012 | -0.012 | 0 | 0.24 | 0.33 | 0.26 | 0.19 | 0.29 | 0.27 |
| 2 | 8953644 | rs17631263 | 0.183 | -0.73 | 0.285 | -1.279 | -0.003 | 0.001 | 0.001 | 0 | 0.23 | 0.26 | 0.26 | 0.17 | 0.29 | 0.27 |
| 2 | 8960809 | rs13390283 | 1.666 | -0.642 | -0.575 | -1.234 | -0.082 | 0.023 | 0.023 | 0.055 | 0.43 | 0.08 | 0.51 | 0.62 | 0.51 | 0.5 |
| 2 | 8967745 | rs12053486 | 1.832 | -0.825 | -0.819 | -1.475 | -0.078 | -0.011 | -0.011 | 0 | 0.38 | 0.08 | 0.48 | 0.61 | 0.47 | 0.5 |
| 2 | 8968557 | rs4669342 | 1.596 | -0.84 | -0.596 | -1.306 | -0.084 | 0.03 | 0.03 | 0.055 | 0.43 | 0.08 | 0.51 | 0.62 | 0.51 | 0.51 |
| 2 | 8971962 | rs7578535 | -0.44 | -0.059 | -0.283 | -0.853 | -0.019 | 0.004 | 0.004 | 0 | 0.25 | 0.16 | 0.26 | 0.21 | 0.3 | 0.28 |
| 2 | 8976403 | rs10173234 | -0.862 | 0.339 | -0.549 | -0.664 | -0.02 | -0.021 | -0.021 | 0 | 0.05 | 0.02 | 0.14 | 0.19 | 0.11 | 0.1 |
| 2 | 8977770 | rs3749007 | -2.086 | 0.194 | -0.442 | -0.754 | 0.008 | 0.03 | 0.03 | 0.718 | 0.13 | 0.15 | 0.26 | 0.33 | 0.17 | 0.16 |
| 2 | 8978676 | rs6757496 | -0.492 | -0.287 | -0.986 | -0.865 | 0.006 | 0.02 | 0.02 | 0 | 0.44 | 0.65 | 0.44 | 0.35 | 0.42 | 0.37 |
| 2 | 8980346 | rs13008591 | -1.2 | -0.135 | -0.866 | -0.2 | 0.004 | 0.021 | 0.021 | 0.02 | 0.06 | 0.09 | 0.14 | 0.16 | 0.1 | 0.09 |
| 2 | 8981020 | rs11894892 | -0.844 | -0.363 | -1.177 | -0.821 | 0.025 | 0.057 | 0.057 | 0.084 | 0.21 | 0.07 | 0.09 | 0.01 | 0.16 | 0.17 |
| 2 | 8982094 | rs6431966 | 0.821 | -0.327 | -0.666 | -0.638 | -0.009 | -0.033 | -0.033 | 0 | 0.88 | 0.84 | 0.94 | 0.99 | 0.92 | 0.81 |
| 2 | 8982985 | rs6716851 | 0.627 | -0.159 | -0.406 | -0.566 | 0.002 | 0.007 | 0.007 | 0.233 | 0.57 | 0.44 | 0.74 | 0.65 | 0.63 | 0.55 |
| 2 | 8986124 | rs16866817 | -0.108 | -0.441 | -0.594 | -0.948 | -0.036 | 0.097 | 0.097 | 0.367 | 0.21 | 0.06 | 0.08 | 0.05 | 0.21 | 0.13 |
| 2 | 8990247 | rs13396431 | -1.474 | -0.319 | 0.693 | -1.113 | 0.004 | -0.104 | -0.104 | 0 | 0.32 | 0.55 | 0.18 | 0.06 | 0.29 | 0.28 |
| 2 | 8992153 | rs6759797 | -1.824 | -0.337 | 1.406 | -1.114 | 0.111 | 0.044 | 0.044 | 2.743 | 0.06 | 0.42 | 0.06 | 0 | 0.05 | 0.11 |
| 2 | 8993505 | rs3811602 | -0.792 | -0.344 | 1.366 | -1.128 | -0.021 | 0.083 | 0.083 | 0.653 | 0.25 | 0.1 | 0.11 | 0.06 | 0.24 | 0.16 |
| 2 | 8996326 | rs1550255 | -2.308 | -0.364 | 1.534 | -1.206 | 0.008 | -0.095 | -0.095 | 0.12 | 0.17 | 0.6 | 0.14 | 0.01 | 0.15 | 0.21 |
| 2 | 9001492 | rs1550253 | -1.732 | -0.488 | 1.902 | -1.239 | 0.007 | -0.078 | -0.078 | 0 | 0.13 | 0.32 | 0.08 | 0.01 | 0.1 | 0.1 |

| | | | | | | | | | | | | | | | | |
|---|---|---|---|---|---|---|---|---|---|---|---|---|---|---|---|---|
| 2 | 9001909 | rs16866828 | -1.209 | -0.473 | 1.968 | -1.232 | 0 | -0.041 | -0.041 | 0 | 0.11 | 0.18 | 0.07 | 0.01 | 0.1 | 0.1 |
| 2 | 9007718 | rs16866835 | 0.429 | -0.47 | 1.943 | -1.232 | -0.019 | 0.034 | 0.034 | 0 | 0.07 | 0.01 | 0.02 | 0 | 0.06 | 0.05 |
| 2 | 9008904 | rs10194749 | -2.366 | -0.467 | 1.99 | -0.95 | -0.002 | -0.04 | -0.04 | 0 | 0.18 | 0.61 | 0.14 | 0.06 | 0.14 | 0.21 |
| 2 | 9017034 | rs998967 | -2.377 | -0.659 | 1.903 | -0.969 | 0.012 | 0.005 | 0.005 | 0.087 | 0.33 | 0.71 | 0.25 | 0.27 | 0.25 | 0.55 |
| 2 | 9060219 | rs7605431 | -1.081 | -0.561 | 1.871 | -0.914 | 0.001 | -0.047 | -0.047 | 0 | 0.11 | 0.19 | 0.07 | 0.01 | 0.1 | 0.1 |
| 2 | 9064191 | rs13415968 | -1.138 | -0.561 | 1.879 | -0.915 | 0.003 | -0.059 | -0.059 | 0.019 | 0.11 | 0.21 | 0.07 | 0.01 | 0.1 | 0.1 |
| 2 | 9082822 | rs10192594 | -0.142 | -0.577 | 1.847 | -0.956 | -0.001 | 0.01 | 0.01 | 0.106 | 0.83 | 0.91 | 0.92 | 0.96 | 0.84 | 0.89 |
| 2 | 9118739 | rs7585370 | -1.194 | -0.537 | 1.765 | -1.025 | 0.002 | -0.06 | -0.06 | 0 | 0.12 | 0.2 | 0.07 | 0.01 | 0.1 | 0.1 |
| 2 | 9127208 | rs13006592 | -0.088 | -0.334 | 1.623 | -1 | -0.036 | -0.063 | -0.063 | 0 | 0.13 | 0.01 | 0.12 | 0.2 | 0.16 | 0.33 |
| 2 | 9130182 | rs1512290 | 0.102 | -0.387 | 1.681 | -1.003 | 0 | -0.018 | -0.018 | 0.354 | 0.07 | 0.07 | 0.03 | 0.01 | 0.07 | 0.08 |
| 2 | 9165288 | rs1473101 | 1.94 | -0.467 | 1.786 | -1.068 | 0 | -0.001 | -0.001 | 0.735 | 0.56 | 0.52 | 0.74 | 0.75 | 0.6 | 0.47 |
| 2 | 9185559 | rs6431973 | 1.509 | -0.466 | 1.76 | -0.95 | -0.007 | -0.028 | -0.028 | 0.42 | 0.65 | 0.63 | 0.78 | 0.75 | 0.68 | 0.54 |
| 2 | 9186516 | rs6706366 | 1.293 | -0.395 | 1.693 | -0.915 | -0.001 | -0.002 | -0.002 | 0 | 0.67 | 0.74 | 0.8 | 0.75 | 0.68 | 0.55 |
| 2 | 9191332 | rs1006387 | 1.083 | -0.59 | 1.684 | -1.027 | -0.003 | -0.107 | -0.107 | 0 | 0.77 | 0.48 | 0.86 | 0.93 | 0.81 | 0.8 |
| 2 | 9205549 | rs1913002 | 0.246 | -0.599 | 1.285 | -1.116 | -0.006 | -0.116 | -0.116 | 2.187 | 0.83 | 0.67 | 0.91 | 0.99 | 0.85 | 0.9 |
| 2 | 9207028 | rs4668598 | -0.767 | -0.759 | 1.334 | -0.83 | 0.006 | -0.01 | -0.01 | 0 | 0.16 | 0.14 | 0.1 | 0.24 | 0.11 | 0.31 |
| 2 | 9212749 | rs1543082 | 0.49 | -0.685 | 1.329 | -0.821 | 0.002 | -0.093 | -0.093 | 0 | 0.8 | 0.68 | 0.88 | 0.95 | 0.81 | 0.78 |
| 2 | 9219822 | rs12471187 | -0.96 | -0.699 | 1.185 | -0.999 | -0.018 | 0.056 | 0.056 | 0 | 0.16 | 0.34 | 0.11 | 0.24 | 0.1 | 0.28 |
| 2 | 9227762 | rs16866884 | -1.353 | -0.801 | 0.45 | -1.127 | 0.022 | -0.006 | -0.006 | 0.708 | 0.27 | 0.11 | 0.23 | 0.23 | 0.2 | 0.37 |
| 2 | 9234386 | rs4668599 | 0.887 | -1.095 | 0.68 | -1.706 | -0.008 | -0.071 | -0.071 | 0.218 | 0.74 | 0.73 | 0.89 | 1 | 0.79 | 0.8 |
| 2 | 9234484 | rs4668601 | -0.188 | -1.11 | 0.669 | -1.595 | 0.006 | -0.009 | -0.009 | 0 | 0.1 | 0.08 | 0.16 | 0.21 | 0.06 | 0.25 |
| 2 | 9235586 | rs6740488 | -0.206 | -1.1 | 1.142 | -1.63 | 0.009 | -0.023 | -0.023 | 0 | 0.16 | 0.41 | 0.13 | 0.1 | 0.15 | 0.19 |
| 2 | 9237190 | rs6735385 | -0.923 | -1.119 | 1.333 | -1.949 | -0.019 | -0.107 | -0.107 | 1.281 | 0.23 | 0.42 | 0.12 | 0.06 | 0.19 | 0.26 |
| 2 | 9242770 | rs10754952 | 1.461 | -1.29 | 1.161 | -2.111 | 0.001 | -0.026 | -0.026 | 0 | 0.62 | 0.4 | 0.79 | 0.93 | 0.69 | 0.62 |
| 2 | 9244409 | rs11688561 | 2.975 | -1.62 | 0.906 | -2.236 | -0.062 | 0.104 | 0.104 | 0.663 | 0.42 | 0.07 | 0.57 | 0.72 | 0.57 | 0.33 |
| 2 | 9244458 | rs12474118 | -0.215 | -1.629 | 1.009 | -2.227 | 0.02 | -0.011 | -0.011 | 0 | 0.16 | 0.08 | 0.18 | 0.21 | 0.11 | 0.33 |
| 2 | 9245951 | rs874878 | 2.025 | -1.615 | 1.136 | -2.237 | 0.088 | 0.128 | 0.128 | 0.079 | 0.57 | 0.17 | 0.66 | 0.78 | 0.73 | 0.53 |
| 2 | 9247025 | rs16866900 | 0.127 | -1.492 | 1.127 | -2.185 | -0.007 | 0.023 | 0.023 | 0 | 0.15 | 0.08 | 0.09 | 0.07 | 0.15 | 0.2 |
| 2 | 9247782 | rs328632 | 0.386 | -1.444 | 1.474 | -2.149 | -0.019 | -0.122 | -0.122 | 0 | 0.8 | 0.63 | 0.89 | 1 | 0.84 | 0.87 |
| 2 | 9250038 | rs1109670 | -1.435 | -1.461 | 1.257 | -1.895 | 0.012 | 0.016 | 0.016 | 0 | 0.35 | 0.3 | 0.31 | 0.29 | 0.27 | 0.54 |
| 2 | 9253906 | rs13423661 | 1.703 | -0.692 | 1.168 | -2.252 | -0.031 | 0.238 | 0.238 | 0.162 | 0.46 | 0.11 | 0.47 | 0.86 | 0.54 | 0.61 |
| 2 | 9259598 | rs9636252 | 1.702 | -0.647 | 2.012 | -1.954 | 0.077 | 0.213 | 0.213 | 0.198 | 0.56 | 0.1 | 0.57 | 0.84 | 0.68 | 0.66 |
| 2 | 9262177 | rs4669361 | 0.522 | -0.629 | 1.906 | -1.946 | 0.008 | 0.001 | 0.001 | 0.851 | 0.07 | 0.04 | 0.09 | 0 | 0.05 | 0.06 |
| 2 | 9263555 | rs12475939 | -0.153 | -0.6 | 1.928 | -1.928 | -0.016 | 0.118 | 0.118 | 0 | 0.22 | 0.03 | 0.11 | 0 | 0.2 | 0.17 |
| 2 | 9267131 | rs10194786 | 1.359 | -0.605 | 1.982 | -1.931 | 0.039 | 0.098 | 0.098 | 0 | 0.63 | 0.18 | 0.67 | 0.84 | 0.72 | 0.71 |
| 2 | 9277612 | rs2175334 | 0.135 | -0.438 | 1.613 | -1.948 | -0.046 | 0.149 | 0.149 | 0.018 | 0.76 | 0.97 | 0.87 | 1 | 0.76 | 0.82 |
| 2 | 9279631 | rs1543042 | -1.648 | -0.406 | 2.008 | -1.863 | 0.048 | 0.138 | 0.138 | 0 | 0.07 | 0.4 | 0.14 | 0.16 | 0.06 | 0.1 |
| 2 | 9281704 | rs6736097 | 1.58 | -0.473 | 1.121 | -1.937 | 0.066 | 0.22 | 0.22 | 0.082 | 0.55 | 0.11 | 0.65 | 0.83 | 0.66 | 0.64 |
| 2 | 9285525 | rs4669367 | 1.512 | -0.631 | 1.112 | -1.919 | 0.041 | 0.224 | 0.224 | 0.188 | 0.56 | 0.12 | 0.73 | 0.84 | 0.63 | 0.65 |
| 2 | 9288363 | rs4668606 | 0.683 | -0.668 | 1.474 | -1.929 | -0.006 | 0.058 | 0.058 | 0 | 0.7 | 0.84 | 0.82 | 0.84 | 0.68 | 0.7 |
| 2 | 9290191 | rs1467079 | -1.403 | -0.67 | 2.131 | -1.859 | -0.06 | 0.037 | 0.037 | 0 | 0.14 | 0.51 | 0.08 | 0.15 | 0.08 | 0.18 |
| 2 | 9290324 | rs1344185 | -2.198 | -0.668 | 1.634 | -1.85 | -0.002 | 0.003 | 0.003 | 0.988 | 0.2 | 0.79 | 0.11 | 0.15 | 0.09 | 0.19 |
| 2 | 9294188 | rs2356976 | -0.922 | -0.565 | 1.39 | -1.877 | 0 | -0.013 | -0.013 | 0 | 0.3 | 0.37 | 0.18 | 0.15 | 0.32 | 0.29 |
| 2 | 9302037 | rs4669374 | 1.457 | -0.569 | 0.461 | -1.927 | 0.049 | 0.237 | 0.237 | 0 | 0.55 | 0.11 | 0.63 | 0.84 | 0.63 | 0.61 |
| 2 | 9308148 | rs907165 | -1.643 | -0.157 | 0.774 | -1.699 | -0.008 | 0.122 | 0.122 | 0.451 | 0.52 | 0.7 | 0.49 | 0.24 | 0.49 | 0.63 |
| 2 | 9310604 | rs6431983 | -0.521 | -0.228 | 1.196 | -1.686 | -0.028 | -0.02 | -0.02 | 0 | 0.23 | 0.47 | 0.13 | 0.19 | 0.19 | 0.25 |
| 2 | 9321834 | rs17626269 | -0.061 | -0.294 | 0.38 | -1.716 | -0.056 | 0.14 | 0.14 | 0 | 0.23 | 0.04 | 0.13 | 0.01 | 0.26 | 0.17 |
| 2 | 9324513 | rs12692379 | -0.571 | -0.046 | 0.983 | -1.48 | -0.002 | -0.001 | -0.001 | 0 | 0.39 | 0.34 | 0.26 | 0.21 | 0.43 | 0.39 |
| 2 | 9324811 | rs6431986 | 0.93 | 0.147 | 0.555 | -1.284 | -0.019 | 0.042 | 0.042 | 0.087 | 0.43 | 0.27 | 0.47 | 0.61 | 0.45 | 0.33 |
| 2 | 9328776 | rs7556775 | -0.877 | 0.172 | 1.335 | -1.306 | 0.027 | -0.117 | -0.117 | 0 | 0.21 | 0.39 | 0.16 | 0.04 | 0.24 | 0.19 |
| 2 | 9333231 | rs16866956 | -0.864 | 0.402 | 0.857 | -1.116 | -0.014 | -0.031 | -0.031 | 0.135 | 0.15 | 0.37 | 0.08 | 0.11 | 0.1 | 0.12 |
| 2 | 9340227 | rs6743148 | -0.917 | 0.503 | -0.111 | -1.035 | 0.001 | 0.065 | 0.065 | 0 | 0.4 | 0.71 | 0.28 | 0.05 | 0.4 | 0.28 |
| 2 | 9344570 | rs6431990 | -0.684 | 0.637 | -0.563 | -0.873 | -0.001 | -0.112 | -0.112 | 0 | 0.37 | 0.49 | 0.25 | 0.05 | 0.38 | 0.26 |
| 2 | 9352584 | rs12615983 | -0.346 | 0.407 | -0.832 | -0.215 | 0.003 | 0.003 | 0.003 | 0 | 0.43 | 0.38 | 0.41 | 0.69 | 0.33 | 0.5 |
| 2 | 9352648 | rs12622238 | -0.187 | 0.387 | -0.869 | -1.47 | 0.004 | 0.011 | 0.011 | 0 | 0.52 | 0.56 | 0.7 | 0.91 | 0.46 | 0.67 |
| 2 | 9355318 | rs10166616 | -0.084 | 0.418 | -1.434 | -2.075 | -0.091 | 0.085 | 0.085 | 0.563 | 0.63 | 0.9 | 0.48 | 0.2 | 0.58 | 0.48 |
| 2 | 9356117 | rs2666205 | -0.81 | 0.522 | -1.504 | -2.27 | -0.013 | 0.047 | 0.047 | 4.907 | 0.91 | 0.99 | 0.95 | 1 | 0.91 | 0.98 |
| 2 | 9362985 | rs2136842 | -0.061 | 0.058 | -1.532 | -2.337 | -0.023 | 0.097 | 0.097 | 0.741 | 0.6 | 0.84 | 0.46 | 0.17 | 0.49 | 0.45 |
| 2 | 9380338 | rs6758205 | 0.347 | -0.394 | -1.442 | -2.165 | -0.014 | 0.008 | 0.008 | 0 | 0.18 | 0.08 | 0.09 | 0.12 | 0.21 | 0.24 |
| 2 | 9382343 | rs17685287 | 1.202 | -0.93 | -1.232 | -2.744 | -0.007 | 0.047 | 0.047 | 0.92 | 0.23 | 0.08 | 0.15 | 0.03 | 0.25 | 0.12 |
| 2 | 9391628 | rs2666212 | 0.026 | -1.206 | 0.712 | -2.893 | -0.039 | -0.059 | -0.059 | 1.941 | 0.32 | 0.57 | 0.27 | 0.14 | 0.24 | 0.31 |
| 2 | 9391691 | rs2356779 | 1.211 | -1.204 | 0.768 | -2.927 | -0.001 | -0.018 | -0.018 | 0.948 | 0.23 | 0.22 | 0.15 | 0.03 | 0.24 | 0.11 |
| 2 | 9397003 | rs1510796 | -0.523 | -1.008 | 0.522 | -2.805 | -0.006 | 0.098 | 0.098 | 0.276 | 0.56 | 0.8 | 0.44 | 0.17 | 0.49 | 0.43 |
| 2 | 9409543 | rs7602301 | 0.363 | -0.665 | 0.816 | -3.041 | -0.015 | -0.122 | -0.122 | 0 | 0.25 | 0.36 | 0.15 | 0 | 0.22 | 0.21 |
| 2 | 9413213 | rs10179702 | 0.166 | -0.328 | 1.366 | -2.641 | 0.033 | 0.004 | 0.004 | 0 | 0.08 | 0.22 | 0.14 | 0.06 | 0.11 | 0.07 |
| 2 | 9416090 | rs7605582 | 0.208 | -0.455 | 1.557 | -2.193 | -0.006 | -0.038 | -0.038 | 0 | 0.62 | 0.46 | 0.69 | 0.82 | 0.65 | 0.74 |
| 2 | 9417334 | rs10439230 | -0.041 | -0.196 | 1.131 | -1.994 | 0.011 | 0.005 | 0.005 | 0 | 0.08 | 0.16 | 0.13 | 0.08 | 0.11 | 0.07 |
| 2 | 9419106 | rs10192751 | 0.177 | -0.196 | 1.078 | -1.993 | 0.008 | 0.003 | 0.003 | 0 | 0.08 | 0.13 | 0.13 | 0.07 | 0.11 | 0.07 |
| 2 | 9419134 | rs10192759 | 0.192 | -0.194 | 1.073 | -2.02 | 0.006 | 0.001 | 0.001 | 0 | 0.08 | 0.12 | 0.13 | 0.07 | 0.11 | 0.07 |

| | | | | | | | | | | | | | | | | |
|---|---|---|---|---|---|---|---|---|---|---|---|---|---|---|---|---|
| 2 | 9422526 | rs10192441 | -0.387 | -0.322 | 1.256 | -2.335 | 0 | -0.029 | -0.029 | 0.717 | 0.36 | 0.42 | 0.23 | 0.14 | 0.36 | 0.24 |
| 2 | 9423931 | rs11689743 | 1.394 | -0.354 | 1.185 | -2.35 | -0.012 | 0.044 | 0.044 | 0.236 | 0.23 | 0.12 | 0.16 | 0.03 | 0.25 | 0.12 |
| 2 | 9434811 | rs4610024 | 0.508 | -0.579 | 1.206 | -2.359 | 0.002 | 0.166 | 0.166 | 1.537 | 0.58 | 0.11 | 0.73 | 0.85 | 0.63 | 0.73 |
| 2 | 9439432 | rs6743955 | -0.291 | -0.421 | 1.344 | -2.287 | 0.001 | -0.033 | -0.033 | 0 | 0.35 | 0.42 | 0.23 | 0.14 | 0.36 | 0.25 |
| 2 | 9444014 | rs6736437 | 0.146 | -0.432 | 1.453 | -2.292 | -0.006 | 0.008 | 0.008 | 0.005 | 0.31 | 0.25 | 0.21 | 0.08 | 0.36 | 0.19 |
| 2 | 9459406 | rs4627537 | 0.128 | -0.366 | 1.106 | -2.264 | -0.015 | 0.046 | 0.046 | 0.196 | 0.31 | 0.22 | 0.2 | 0.08 | 0.36 | 0.19 |
| 2 | 9462949 | rs10210653 | -0.67 | -0.414 | 1.503 | -2.279 | 0.003 | 0.027 | 0.027 | 0.589 | 0.39 | 0.68 | 0.26 | 0.15 | 0.37 | 0.28 |
| 2 | 9465687 | rs10181704 | -0.505 | -0.348 | 1.199 | -2.245 | -0.001 | -0.017 | -0.017 | 0 | 0.38 | 0.6 | 0.24 | 0.14 | 0.36 | 0.26 |
| 2 | 9467815 | rs7569295 | 0.042 | -0.205 | 0.454 | -2.223 | -0.034 | 0.108 | 0.108 | 0.038 | 0.32 | 0.18 | 0.2 | 0.08 | 0.36 | 0.2 |
| 2 | 9496291 | rs2709591 | -1.116 | -0.12 | 0.263 | -2.528 | -0.122 | 0.109 | 0.109 | 1.596 | 0.67 | 0.94 | 0.53 | 0.14 | 0.58 | 0.54 |
| 2 | 9499377 | rs2709576 | -0.246 | -0.353 | 0.965 | -2.409 | -0.005 | -0.033 | -0.033 | 0.032 | 0.39 | 0.58 | 0.36 | 0.14 | 0.3 | 0.25 |
| 2 | 9513850 | rs4599081 | -1.606 | -0.003 | 1.703 | -1.809 | 0.058 | -0.06 | -0.06 | 0 | 0.12 | 0.53 | 0.09 | 0.03 | 0.11 | 0.08 |
| 2 | 9514573 | rs2709580 | -2.015 | -0.041 | 1.366 | -1.814 | 0 | 0 | 0 | 0 | 0.17 | 0.8 | 0.11 | 0.12 | 0.11 | 0.18 |
| 2 | 9543221 | rs11674625 | -1.717 | 0.08 | 1.108 | -1.744 | -0.004 | -0.007 | -0.007 | 0.003 | 0.16 | 0.71 | 0.14 | 0.13 | 0.11 | 0.19 |
| 2 | 9546030 | rs10593 | 1.235 | 0.178 | 0.981 | -1.651 | 0.051 | 0.045 | 0.045 | 0 | 0.87 | 0.6 | 0.87 | 0.86 | 0.87 | 0.79 |
| 2 | 9549753 | rs10169262 | 0.63 | -0.189 | 0.456 | -2.35 | 0.003 | -0.037 | -0.037 | 0 | 0.3 | 0.25 | 0.14 | 0.02 | 0.19 | 0.08 |
| 2 | 9554255 | rs10208358 | -1.34 | -0.091 | -0.226 | -2.783 | -0.159 | 0.044 | 0.044 | 0.732 | 0.73 | 0.98 | 0.45 | 0.15 | 0.59 | 0.51 |
| 2 | 9560346 | rs4436931 | 0.84 | -0.053 | 0.222 | -2.728 | 0.015 | 0.015 | 0.015 | 3.291 | 0.31 | 0.22 | 0.15 | 0.02 | 0.2 | 0.08 |
| 2 | 9588091 | rs10176656 | 0.929 | -0.034 | 0.116 | -2.722 | 0.021 | 0.131 | 0.131 | 4.331 | 0.29 | 0.13 | 0.13 | 0.02 | 0.19 | 0.07 |
| 2 | 9619204 | rs7561588 | -0.1 | 0.055 | 0.459 | -2.483 | 0.018 | 0.001 | 0.001 | 0 | 0.06 | 0.13 | 0.06 | 0.07 | 0.07 | 0.09 |
| 2 | 9647091 | rs1056204 | 0.705 | -0.096 | 0.084 | -2.775 | -0.027 | 0.142 | 0.142 | 1.388 | 0.29 | 0.07 | 0.18 | 0 | 0.32 | 0.23 |
| 2 | 9661775 | rs17590882 | 0.631 | -0.328 | 0.337 | -2.616 | 0.029 | 0.059 | 0.059 | 1.959 | 0.3 | 0.2 | 0.15 | 0.02 | 0.19 | 0.1 |
| 2 | 9681797 | rs1880439 | 0.029 | 0.047 | 0.196 | -2.638 | 0.007 | -0.019 | -0.019 | 1.264 | 0.12 | 0.13 | 0.09 | 0.07 | 0.16 | 0.1 |
| 2 | 9683696 | rs10179642 | -0.31 | 0.041 | 0.265 | -2.64 | 0.013 | -0.033 | -0.033 | 1.048 | 0.13 | 0.18 | 0.1 | 0.07 | 0.16 | 0.1 |
| 2 | 9697171 | rs11684747 | 0.405 | 0.085 | -0.057 | -2.421 | 0.029 | 0.046 | 0.046 | 2.496 | 0.31 | 0.2 | 0.15 | 0.02 | 0.19 | 0.1 |
| 2 | 9703226 | rs10929588 | -0.633 | 0.076 | 0.002 | -2.36 | 0.026 | 0.066 | 0.066 | 3.665 | 0.42 | 0.67 | 0.27 | 0.17 | 0.29 | 0.22 |
| 2 | 9703439 | rs10929589 | -1.473 | 0.152 | 0.644 | -2.126 | 0.051 | 0.061 | 0.061 | 2.71 | 0.12 | 0.46 | 0.12 | 0.13 | 0.11 | 0.13 |
| 2 | 9721896 | rs6432018 | -0.864 | 0.213 | -0.116 | -2.689 | -0.051 | 0.175 | 0.175 | 0.272 | 0.64 | 0.86 | 0.41 | 0.06 | 0.59 | 0.36 |
| 2 | 9736066 | rs10495571 | 1.266 | 0.169 | 0.077 | -2.922 | -0.005 | 0.115 | 0.115 | 1.074 | 0.23 | 0.04 | 0.12 | 0.01 | 0.18 | 0.1 |
| 2 | 9748518 | rs16867073 | -0.155 | 0.217 | 0.467 | -2.886 | -0.004 | -0.004 | -0.004 | 0 | 0.15 | 0.31 | 0.13 | 0.11 | 0.13 | 0.09 |
| 2 | 9750609 | rs7608161 | 0.355 | 0.192 | 0.506 | -3.007 | -0.001 | -0.091 | -0.091 | 0.582 | 0.29 | 0.37 | 0.2 | 0.01 | 0.29 | 0.18 |
| 2 | 9751478 | rs7597961 | -0.305 | -0.29 | 0.698 | -2.427 | -0.002 | -0.009 | -0.009 | 0 | 0.15 | 0.33 | 0.1 | 0.1 | 0.13 | 0.08 |
| 2 | 9754439 | rs17362839 | 0.785 | -0.211 | -0.587 | -2.549 | 0.003 | 0.101 | 0.101 | 0.73 | 0.23 | 0.08 | 0.12 | 0.01 | 0.18 | 0.1 |
| 2 | 9758621 | rs6432025 | -0.515 | 0.015 | -0.551 | -2.289 | 0.051 | 0.235 | 0.235 | 2.64 | 0.44 | 0.84 | 0.26 | 0.06 | 0.31 | 0.2 |
| 2 | 9770050 | rs13417081 | -0.123 | 0.351 | 0.036 | -1.209 | -0.008 | -0.023 | -0.023 | 0 | 0.62 | 0.51 | 0.48 | 0.23 | 0.64 | 0.53 |
| 2 | 9773586 | rs3762535 | -0.612 | 0.099 | -0.211 | -1.419 | -0.088 | 0.116 | 0.116 | 0 | 0.62 | 0.86 | 0.37 | 0.17 | 0.52 | 0.48 |
| 2 | 9774331 | rs6715748 | 0.481 | -0.082 | 0.127 | -1.607 | 0.004 | 0.043 | 0.043 | 0.194 | 0.09 | 0.2 | 0.06 | 0.16 | 0.08 | 0.07 |
| 2 | 9777229 | rs4668626 | -0.37 | -0.406 | 0.055 | -1.97 | 0.035 | 0.162 | 0.162 | 0.127 | 0.48 | 0.77 | 0.3 | 0.16 | 0.4 | 0.33 |
| 2 | 9780744 | rs1534043 | 0.526 | -0.411 | 0.423 | -2.014 | 0.002 | -0.052 | -0.052 | 0 | 0.36 | 0.34 | 0.2 | 0.01 | 0.29 | 0.26 |
| 2 | 9781094 | rs10427360 | -0.336 | -0.453 | -0.479 | -1.959 | 0.027 | 0.169 | 0.169 | 0.137 | 0.48 | 0.78 | 0.29 | 0.16 | 0.4 | 0.33 |
| 2 | 9782006 | rs10929592 | 0.102 | -0.113 | -0.506 | -1.663 | 0 | -0.023 | -0.023 | 0.276 | 0.14 | 0.13 | 0.08 | 0.01 | 0.12 | 0.15 |
| 2 | 9783635 | rs884427 | 0.657 | -0.103 | -0.642 | -1.508 | -0.002 | 0.013 | 0.013 | 0 | 0.08 | 0.08 | 0.05 | 0.16 | 0.07 | 0.06 |
| 2 | 9786695 | rs17453871 | -0.207 | -0.171 | -0.798 | -1.92 | -0.024 | 0.206 | 0.206 | 0.056 | 0.43 | 0.17 | 0.25 | 0.01 | 0.41 | 0.4 |
| 2 | 9787768 | rs6747234 | -0.054 | -0.213 | -0.248 | -1.678 | -0.006 | -0.111 | -0.111 | 0.015 | 0.36 | 0.49 | 0.24 | 0.04 | 0.33 | 0.25 |
| 2 | 9792908 | rs16867105 | -0.218 | -0.453 | -0.905 | -1.333 | 0.011 | 0.009 | 0.009 | 0 | 0.3 | 0.2 | 0.18 | 0.21 | 0.23 | 0.2 |
| 2 | 9795757 | rs7608996 | -0.945 | -0.677 | -1.284 | -1.495 | 0.061 | 0.064 | 0.064 | 0.04 | 0.4 | 0.75 | 0.22 | 0.22 | 0.27 | 0.21 |
| 2 | 9796791 | rs951629 | -1.112 | -0.88 | -1.809 | -1.814 | -0.096 | 0.028 | 0.028 | 0 | 0.64 | 0.89 | 0.36 | 0.26 | 0.46 | 0.49 |
| 2 | 9797196 | rs10189576 | -0.428 | -0.909 | -0.919 | -1.243 | -0.045 | -0.081 | -0.081 | 0 | 0.12 | 0.58 | 0.06 | 0.01 | 0.06 | 0.03 |
| 2 | 9797994 | rs4669407 | -0.237 | -0.315 | -1.69 | -0.926 | 0.009 | -0.024 | -0.024 | 0 | 0.25 | 0.03 | 0.13 | 0.2 | 0.18 | 0.18 |
| 2 | 9805923 | rs4669413 | -0.143 | 0.488 | -1.78 | -0.718 | -0.044 | -0.011 | -0.011 | 0 | 0.31 | 0.24 | 0.63 | 0.69 | 0.44 | 0.43 |
| 2 | 9806527 | rs2357258 | -0.795 | 0.029 | -2.172 | -1.278 | 0.065 | 0.104 | 0.104 | 0.589 | 0.49 | 0.3 | 0.73 | 0.8 | 0.65 | 0.53 |
| 2 | 9807984 | rs6713719 | 0.81 | -0.305 | -1.703 | -1.427 | -0.016 | 0.009 | 0.009 | 0 | 0.13 | 0.17 | 0.09 | 0.15 | 0.08 | 0.14 |
| 2 | 9809611 | rs11677837 | 0.427 | 0.008 | -1.692 | -1.273 | -0.01 | 0.037 | 0.037 | 0 | 0.17 | 0.06 | 0.11 | 0.11 | 0.21 | 0.1 |
| 2 | 9814374 | rs10179015 | 0.515 | -0.066 | -1.702 | -1.377 | -0.053 | -0.018 | -0.018 | 0 | 0.3 | 0.19 | 0.63 | 0.7 | 0.42 | 0.43 |
| 2 | 9814481 | rs6734784 | 0.472 | -0.085 | -1.123 | -1.366 | 0 | -0.012 | -0.012 | 0 | 0.17 | 0.18 | 0.12 | 0.1 | 0.21 | 0.1 |
| 2 | 9815766 | rs16867116 | 0.959 | -0.607 | -0.647 | -1.413 | 0.036 | 0.011 | 0.011 | 1.563 | 0.06 | 0.34 | 0.03 | 0 | 0.02 | 0.04 |
| 2 | 9815939 | rs9287712 | -0.004 | -0.543 | -1.029 | -1.33 | 0.025 | 0.046 | 0.046 | 0.524 | 0.46 | 0.36 | 0.75 | 0.8 | 0.62 | 0.49 |
| 2 | 9823199 | rs13422487 | -0.303 | 0.181 | -1.209 | -1.037 | -0.006 | 0.013 | 0.013 | 0 | 0.35 | 0.25 | 0.5 | 0.24 | 0.37 | 0.31 |
| 2 | 9829235 | rs6713691 | -0.328 | -0.106 | -0.767 | -1.049 | -0.027 | 0.013 | 0.013 | 0.149 | 0.22 | 0.48 | 0.17 | 0.19 | 0.16 | 0.22 |
| 2 | 9830260 | rs13028921 | 0.437 | 0.016 | -1.713 | -1.542 | -0.026 | 0.051 | 0.051 | 0 | 0.4 | 0.12 | 0.69 | 0.73 | 0.57 | 0.49 |
| 2 | 9831821 | rs10188732 | -0.241 | 0.115 | -0.758 | -1.533 | 0.009 | -0.011 | -0.011 | 1.165 | 0.27 | 0.47 | 0.26 | 0.24 | 0.28 | 0.29 |
| 2 | 9833224 | rs4668631 | 1.086 | 0.291 | -1.35 | -1.628 | -0.01 | 0.027 | 0.027 | 0 | 0.22 | 0.16 | 0.33 | 0.07 | 0.28 | 0.09 |
| 2 | 9833804 | rs10495567 | -1.185 | -0.073 | -1.754 | -1.868 | 0.041 | 0.145 | 0.145 | 0.008 | 0.78 | 1 | 0.91 | 1 | 0.86 | 0.76 |
| 2 | 9837863 | rs12992481 | 1.66 | 0.189 | -1.644 | -1.769 | -0.033 | -0.025 | -0.025 | 0 | 0.08 | 0.04 | 0.08 | 0.12 | 0.13 | 0.15 |
| 2 | 9837880 | rs13426484 | 0.45 | -0.592 | -1.164 | -1.997 | 0.002 | -0.045 | -0.045 | 0.034 | 0.27 | 0.31 | 0.34 | 0.07 | 0.3 | 0.09 |
| 2 | 9840416 | rs4668634 | -0.338 | -0.249 | -0.813 | -1.748 | 0.024 | 0.066 | 0.066 | 0 | 0.55 | 0.71 | 0.59 | 0.93 | 0.59 | 0.69 |
| 2 | 9840603 | rs4669424 | -0.469 | -0.223 | -1.223 | -1.721 | 0.031 | 0.169 | 0.169 | 0 | 0.57 | 0.82 | 0.59 | 0.93 | 0.6 | 0.69 |
| 2 | 9844178 | rs11680429 | 0.045 | -0.432 | -0.155 | -0.889 | -0.004 | -0.002 | -0.002 | 0 | 0.59 | 0.62 | 0.73 | 0.33 | 0.67 | 0.48 |
| 2 | 9844958 | rs6735280 | 0.141 | -0.575 | 0.094 | -0.922 | 0.01 | 0.005 | 0.005 | 0 | 0.58 | 0.41 | 0.67 | 0.33 | 0.69 | 0.47 |

| | | | | | | | | | | | | | | | | |
|---|---|---|---|---|---|---|---|---|---|---|---|---|---|---|---|---|
| 2 | 9849528 | rs12615047 | 1.158 | 0.01 | 0.209 | -0.908 | 0.046 | 0.004 | 0.004 | 0 | 0.1 | 0.17 | 0.09 | 0.09 | 0.14 | 0.09 |
| 2 | 9852110 | rs11685771 | 0.446 | -0.25 | 0.123 | -0.898 | 0.008 | 0.142 | 0.142 | 0 | 0.52 | 0.21 | 0.64 | 0.22 | 0.58 | 0.38 |
| 2 | 9855737 | rs4605345 | 1.346 | -0.12 | -0.189 | -1.014 | -0.058 | 0.018 | 0.018 | 0 | 0.3 | 0.06 | 0.54 | 0.21 | 0.38 | 0.26 |
| 2 | 9865547 | rs10929604 | 0.871 | -0.233 | 1.013 | -0.406 | 0.041 | 0.003 | 0.003 | 0 | 0.11 | 0.19 | 0.1 | 0.1 | 0.14 | 0.09 |
| 2 | 9869267 | rs4669444 | 0.859 | -0.317 | 1.078 | -0.52 | -0.015 | 0.153 | 0.153 | 0 | 0.52 | 0.19 | 0.63 | 0.22 | 0.56 | 0.42 |
| 2 | 9869850 | rs11901380 | 0.365 | -1.312 | 2.221 | -0.2 | -0.006 | -0.002 | -0.002 | 0 | 0.63 | 0.56 | 0.75 | 0.32 | 0.71 | 0.47 |
| 2 | 9870631 | rs7592044 | 0.253 | -1.334 | 2.313 | -0.011 | 0.021 | 0.004 | 0.004 | 0 | 0.64 | 0.72 | 0.76 | 0.32 | 0.71 | 0.47 |
| 2 | 9890152 | rs6754662 | 0.433 | -1.16 | 2.625 | 0.283 | -0.014 | 0.006 | 0.006 | 0 | 0.68 | 0.59 | 0.77 | 0.32 | 0.73 | 0.56 |
| 2 | 9890804 | rs4668644 | 0.57 | -1.173 | 2.564 | 0.28 | -0.019 | -0.002 | -0.002 | 0.697 | 0.66 | 0.49 | 0.76 | 0.31 | 0.73 | 0.57 |
| 2 | 9892899 | rs10189991 | -0.611 | -1.212 | 2.64 | 0.785 | -0.018 | 0.06 | 0.06 | 0 | 0.19 | 0.39 | 0.17 | 0.41 | 0.16 | 0.17 |
| 2 | 9893138 | rs10178017 | 0.923 | -1.208 | 2.624 | 0.797 | -0.008 | 0 | 0 | 0 | 0.66 | 0.43 | 0.76 | 0.33 | 0.72 | 0.58 |
| 2 | 9893454 | rs13412895 | -0.681 | -1.192 | 2.613 | 0.79 | -0.029 | 0.075 | 0.075 | 0.072 | 0.2 | 0.47 | 0.18 | 0.41 | 0.16 | 0.17 |
| 2 | 9895018 | rs1533388 | 0.11 | -0.913 | 2.06 | 0.833 | -0.037 | 0.098 | 0.098 | 0 | 0.8 | 0.5 | 0.9 | 0.73 | 0.88 | 0.75 |
| 2 | 9899127 | rs16867150 | 0.321 | -0.859 | 1.431 | 0.875 | -0.006 | -0.045 | -0.045 | 3.698 | 0.13 | 0.07 | 0.15 | 0.39 | 0.15 | 0.13 |
| 2 | 9900476 | rs12329266 | -0.242 | -0.82 | 1.615 | 0.894 | -0.015 | 0.068 | 0.068 | 0 | 0.17 | 0.36 | 0.16 | 0.38 | 0.15 | 0.16 |
| 2 | 9902056 | rs6709608 | -0.077 | -0.614 | 0.874 | 0.882 | -0.011 | 0.045 | 0.045 | 0 | 0.82 | 0.73 | 0.91 | 0.73 | 0.87 | 0.74 |
| 2 | 9906145 | rs10200632 | 0.481 | -0.608 | 0.066 | 0.661 | -0.041 | 0.009 | 0.009 | 0 | 0.74 | 0.45 | 0.87 | 0.83 | 0.83 | 0.73 |
| 2 | 9906471 | rs13030983 | 0.416 | -0.236 | -0.795 | 0.335 | 0.026 | 0.007 | 0.007 | 0.096 | 0.6 | 0.29 | 0.68 | 0.34 | 0.69 | 0.57 |
| 2 | 9908010 | rs12995890 | -0.047 | 0.47 | -0.949 | -0.389 | 0 | -0.04 | -0.04 | 0.058 | 0.59 | 0.49 | 0.64 | 0.92 | 0.59 | 0.74 |
| 2 | 9908651 | rs10929612 | -0.457 | 0.565 | -1.548 | -0.084 | 0.024 | -0.045 | -0.045 | 0 | 0.65 | 0.79 | 0.63 | 0.47 | 0.67 | 0.62 |
| 2 | 9908901 | rs4669453 | -0.078 | 1.033 | -1.916 | -0.209 | -0.008 | -0.001 | -0.001 | 0 | 0.28 | 0.26 | 0.39 | 0.3 | 0.33 | 0.29 |
| 2 | 9909114 | rs4669454 | -0.847 | 0.793 | -2.273 | -0.212 | 0.044 | -0.101 | -0.101 | 0 | 0.71 | 0.94 | 0.68 | 0.53 | 0.76 | 0.64 |
| 2 | 9912089 | rs11676697 | 0.586 | 1.346 | -1.951 | -0.277 | -0.033 | 0.052 | 0.052 | 0 | 0.1 | 0.02 | 0.06 | 0.01 | 0.14 | 0.15 |
| 2 | 9912829 | rs871820 | -0.753 | 1.275 | -2.072 | -0.308 | -0.126 | 0.028 | 0.028 | 3.331 | 0.42 | 0.06 | 0.41 | 0.26 | 0.46 | 0.51 |
| 2 | 9913006 | rs10184938 | 0.629 | 1.438 | -1.911 | -0.386 | -0.055 | 0.077 | 0.077 | 0.897 | 0.16 | 0.04 | 0.09 | 0.01 | 0.21 | 0.21 |
| 2 | 9913515 | rs13395893 | -0.166 | 1.092 | -1.584 | -0.293 | -0.016 | 0.018 | 0.018 | 3.319 | 0.19 | 0.64 | 0.18 | 0.26 | 0.15 | 0.16 |
| 2 | 9914468 | rs11674369 | -0.398 | 1.153 | -1.774 | -0.295 | -0.069 | 0.016 | 0.016 | 7.815 | 0.36 | 0.15 | 0.39 | 0.24 | 0.44 | 0.41 |
| 2 | 9914751 | rs11674480 | -0.889 | 0.86 | -1.421 | 0.112 | 0.04 | -0.05 | -0.05 | 0 | 0.56 | 0.89 | 0.62 | 0.52 | 0.59 | 0.58 |
| 2 | 9922283 | rs4669458 | -0.026 | 1.009 | -1.08 | 0.264 | 0.026 | 0.05 | 0.05 | 0.156 | 0.2 | 0.37 | 0.31 | 0.24 | 0.22 | 0.2 |
| 2 | 9924387 | rs2357266 | -0.706 | 0.505 | -1.609 | 0.427 | 0.028 | -0.028 | -0.028 | 0 | 0.62 | 0.79 | 0.66 | 0.52 | 0.67 | 0.59 |
| 2 | 9931094 | rs17363477 | -0.406 | 0.794 | -1.443 | 0.38 | -0.003 | -0.006 | -0.006 | 0.781 | 0.19 | 0.12 | 0.29 | 0.16 | 0.19 | 0.2 |
| 2 | 9933956 | rs4669463 | -0.133 | 0.124 | -1.153 | -0.511 | -0.019 | -0.088 | -0.088 | 0 | 0.71 | 0.66 | 0.84 | 0.97 | 0.75 | 0.8 |
| 2 | 9935529 | rs13421149 | 0.467 | 0.564 | -1.008 | 0.316 | 0 | -0.004 | -0.004 | 0 | 0.29 | 0.28 | 0.3 | 0.4 | 0.3 | 0.33 |
| 2 | 9938881 | rs7592934 | -0.012 | 1.064 | -0.768 | 0.633 | 0.035 | 0.069 | 0.069 | 0 | 0.07 | 0.14 | 0.27 | 0.18 | 0.13 | 0.12 |
| 2 | 9940897 | rs6739059 | -0.364 | 0.818 | -1.179 | 0.372 | -0.007 | 0.03 | 0.03 | 0 | 0.59 | 0.67 | 0.7 | 0.72 | 0.57 | 0.71 |
| 2 | 9942159 | rs908293 | 1.074 | 0.634 | -1.496 | 0.376 | 0.007 | 0.023 | 0.023 | 0 | 0.11 | 0 | 0.06 | 0.02 | 0.08 | 0.19 |
| 2 | 9946213 | rs10929616 | -0.781 | 0.744 | -1.74 | 0.37 | -0.017 | -0.028 | -0.028 | 0 | 0.67 | 0.9 | 0.61 | 0.6 | 0.66 | 0.63 |
| 2 | 9948107 | rs4669468 | 0.51 | 1.429 | -1.427 | 0.467 | -0.022 | -0.023 | -0.023 | 0.538 | 0.13 | 0.08 | 0.1 | 0.27 | 0.21 | 0.16 |
| 2 | 9948404 | rs6734693 | -0.839 | 0.591 | -1.008 | 0.054 | -0.001 | -0.002 | -0.002 | 0 | 0.45 | 0.65 | 0.37 | 0.44 | 0.51 | 0.49 |
| 2 | 9949464 | rs16867175 | 0.237 | 0.274 | -0.778 | -0.133 | -0.01 | -0.029 | -0.029 | 0.151 | 0.14 | 0.23 | 0.1 | 0.02 | 0.1 | 0.18 |
| 2 | 9951970 | rs7568887 | 0.251 | -0.316 | -1.092 | -0.038 | -0.007 | -0.001 | -0.001 | 0.051 | 0.47 | 0.45 | 0.4 | 0.43 | 0.35 | 0.32 |
| 2 | 9952199 | rs7559852 | -0.033 | 0.291 | -1.049 | -0.139 | 0 | 0 | 0 | 0 | 0.53 | 0.52 | 0.48 | 0.46 | 0.43 | 0.36 |
| 2 | 9952744 | rs1913680 | -1.733 | 0.455 | -1.431 | 0.025 | -0.006 | 0.018 | 0.018 | 0 | 0.91 | 0.96 | 0.93 | 0.95 | 0.92 | 0.86 |
| 2 | 9953949 | rs908295 | 1.744 | 1.057 | -1.555 | 0.005 | -0.041 | 0.029 | 0.029 | 0 | 0.12 | 0.02 | 0.17 | 0.04 | 0.15 | 0.17 |
| 2 | 9954768 | rs908296 | 0.427 | 0.083 | -1.534 | -0.314 | 0.002 | -0.004 | -0.004 | 0 | 0.31 | 0.28 | 0.35 | 0.43 | 0.25 | 0.24 |
| 2 | 9956981 | rs16867179 | 1.64 | 0.111 | -1.635 | -0.202 | 0.006 | 0.004 | 0.004 | 0 | 0.13 | 0.06 | 0.09 | 0.09 | 0.13 | 0.1 |
| 2 | 9957097 | rs10803704 | -0.184 | 0.332 | -1.353 | -0.175 | 0 | 0 | 0 | 0 | 0.46 | 0.59 | 0.36 | 0.44 | 0.51 | 0.45 |
| 2 | 9958771 | rs4533450 | -1.069 | -0.198 | -1.317 | -0.566 | -0.084 | 0.099 | 0.099 | 0 | 0.71 | 0.92 | 0.66 | 0.89 | 0.64 | 0.72 |
| 2 | 9964947 | rs16867184 | 1.63 | -0.689 | -0.493 | -0.271 | 0.002 | -0.002 | -0.002 | 2.574 | 0.08 | 0.07 | 0.03 | 0.17 | 0.06 | 0.15 |
| 2 | 9971366 | rs287980 | 0.058 | -0.991 | -0.19 | -0.423 | -0.035 | 0.003 | 0.003 | 0 | 0.73 | 0.48 | 0.83 | 0.75 | 0.78 | 0.8 |
| 2 | 9972442 | rs287982 | 0.214 | -1.097 | -0.15 | -0.475 | -0.045 | -0.002 | -0.002 | 0 | 0.71 | 0.49 | 0.83 | 0.75 | 0.78 | 0.8 |
| 2 | 9975557 | rs7568330 | 0.268 | -1.035 | -0.374 | -0.465 | -0.009 | 0.001 | 0.001 | 0 | 0.16 | 0.2 | 0.09 | 0.17 | 0.12 | 0.16 |
| 2 | 9983686 | rs2303914 | -0.098 | -0.978 | -0.795 | -0.526 | -0.005 | 0.027 | 0.027 | 0 | 0.46 | 0.41 | 0.59 | 0.68 | 0.52 | 0.55 |
| 2 | 9985606 | rs287986 | 1.97 | -0.862 | -0.914 | -0.504 | -0.052 | 0.107 | 0.107 | 0 | 0.24 | 0.07 | 0.23 | 0.05 | 0.25 | 0.24 |
| 2 | 9988299 | rs16867193 | 1.536 | -0.76 | -0.917 | -0.15 | -0.04 | -0.044 | -0.044 | 0.132 | 0.08 | 0.01 | 0.24 | 0.28 | 0.13 | 0.16 |
| 2 | 9988348 | rs1469544 | 1.013 | -0.852 | -1.116 | -0.094 | -0.071 | -0.098 | -0.098 | 0 | 0.15 | 0.01 | 0.18 | 0.3 | 0.21 | 0.23 |
| 2 | 9989442 | rs16867195 | 0.487 | -0.889 | -1.05 | -0.11 | -0.037 | -0.047 | -0.047 | 0.084 | 0.2 | 0.08 | 0.23 | 0.3 | 0.27 | 0.24 |
| 2 | 9991867 | rs287992 | 2.401 | -0.96 | -1.117 | -0.169 | -0.098 | 0.126 | 0.126 | 0 | 0.25 | 0.01 | 0.25 | 0.05 | 0.3 | 0.25 |
| 2 | 9996156 | rs1014509 | -0.006 | -0.168 | -0.185 | 0.545 | -0.037 | -0.047 | -0.047 | 0.268 | 0.2 | 0.08 | 0.23 | 0.3 | 0.27 | 0.24 |
| 2 | 10017166 | rs16867210 | -0.4 | -0.204 | -0.078 | 0.516 | -0.002 | 0.002 | 0.002 | 0 | 0.29 | 0.4 | 0.39 | 0.39 | 0.26 | 0.3 |
| 2 | 10033277 | rs420122 | 2.031 | -0.151 | -0.475 | 0.525 | -0.061 | 0.099 | 0.099 | 0 | 0.35 | 0.07 | 0.31 | 0.14 | 0.38 | 0.27 |
| 2 | 10040351 | rs1022312 | -0.533 | -0.154 | -0.054 | 0.718 | -0.009 | 0.002 | 0.002 | 0 | 0.3 | 0.5 | 0.4 | 0.38 | 0.28 | 0.32 |
| 2 | 10045014 | rs400917 | -0.897 | -0.134 | -0.035 | 0.761 | -0.033 | 0.066 | 0.066 | 0 | 0.88 | 0.66 | 0.94 | 0.83 | 0.93 | 0.83 |
| 2 | 10047496 | rs16867226 | 0.989 | -0.075 | -0.121 | 0.794 | 0.067 | -0.037 | -0.037 | 1.964 | 0.55 | 0.13 | 0.54 | 0.45 | 0.66 | 0.51 |
| 2 | 10050947 | rs1054561 | 0.951 | -0.079 | -0.123 | 0.792 | 0.069 | -0.036 | -0.036 | 0 | 0.55 | 0.13 | 0.54 | 0.45 | 0.66 | 0.51 |
| 2 | 10059097 | rs2303921 | -1.293 | -0.037 | -0.079 | 0.792 | -0.005 | 0.001 | 0.001 | 0 | 0.46 | 0.57 | 0.48 | 0.55 | 0.4 | 0.49 |
| 2 | 10059770 | rs1820965 | 1.014 | -0.142 | -0.925 | 1.001 | 0.074 | -0.024 | -0.024 | 0.295 | 0.55 | 0.13 | 0.54 | 0.48 | 0.67 | 0.52 |
| 2 | 10059844 | rs16867245 | 1.065 | -0.271 | -0.907 | 1.023 | -0.04 | -0.044 | -0.044 | 0 | 0.08 | 0.01 | 0.24 | 0.23 | 0.13 | 0.16 |
| 2 | 10060587 | rs4668651 | -0.21 | -0.189 | -0.794 | 0.846 | 0.004 | 0.038 | 0.038 | 0 | 0.12 | 0.15 | 0.22 | 0.28 | 0.16 | 0.21 |

| | | | | | | | | | | | | | | | | |
|---|---|---|---|---|---|---|---|---|---|---|---|---|---|---|---|---|
| 2 | 10060980 | rs4668652 | 0.233 | -0.546 | -0.474 | 0.68 | -0.02 | 0.027 | 0.027 | 0.028 | 0.74 | 0.56 | 0.63 | 0.66 | 0.79 | 0.72 |
| 2 | 10061592 | rs2357391 | 0.5 | -0.026 | -1.679 | 0.525 | 0.064 | -0.058 | -0.058 | 0 | 0.53 | 0.09 | 0.53 | 0.43 | 0.64 | 0.5 |
| 2 | 10077293 | rs750872 | 0.352 | 0.737 | -0.902 | 1.273 | -0.006 | 0.026 | 0.026 | 1.372 | 0.76 | 0.31 | 0.68 | 0.54 | 0.79 | 0.65 |
| 2 | 10079684 | rs2288643 | 0.597 | 0.991 | -1.239 | 0.677 | -0.011 | 0.105 | 0.105 | 0 | 0.53 | 0.25 | 0.41 | 0.25 | 0.51 | 0.43 |
| 2 | 10083009 | rs4669507 | -0.637 | 0.896 | -0.625 | 1.127 | 0.002 | 0.04 | 0.04 | 0.773 | 0.29 | 0.42 | 0.38 | 0.48 | 0.3 | 0.4 |
| 2 | 10085722 | rs1109037 | -0.269 | 0.787 | -0.795 | 0.95 | 0.001 | 0.001 | 0.001 | 0.019 | 0.43 | 0.42 | 0.49 | 0.44 | 0.46 | 0.4 |
| 2 | 10089722 | rs3762536 | -0.947 | 0.354 | -0.538 | 1.069 | -0.014 | 0.014 | 0.014 | 0.004 | 0.13 | 0.21 | 0.34 | 0.22 | 0.13 | 0.09 |
| 2 | 10090572 | rs1864539 | -1.575 | 0.407 | -0.181 | 1.397 | -0.015 | 0.044 | 0.044 | 0.928 | 0.21 | 0.35 | 0.4 | 0.34 | 0.23 | 0.13 |
| 2 | 10095185 | rs16867251 | -0.673 | 0.762 | 0.532 | 0.625 | 0.037 | -0.123 | -0.123 | 0 | 0.18 | 0.44 | 0.1 | 0.01 | 0.15 | 0.07 |
| 2 | 10098227 | rs6708858 | -1.07 | 0.209 | -0.469 | -0.078 | -0.006 | 0.111 | 0.111 | 0.774 | 0.52 | 0.77 | 0.38 | 0.26 | 0.45 | 0.31 |
| 2 | 10099635 | rs3791765 | -0.092 | 0.183 | -0.42 | -0.107 | 0.005 | 0.007 | 0.007 | 0 | 0.33 | 0.3 | 0.28 | 0.25 | 0.3 | 0.24 |
| 2 | 10101468 | rs16867256 | 0.043 | 0.179 | -0.355 | -0.132 | 0.036 | -0.122 | -0.122 | 0.001 | 0.18 | 0.44 | 0.11 | 0.01 | 0.15 | 0.07 |
| 2 | 10103414 | rs11902457 | -0.007 | 0.137 | 0.45 | -0.187 | -0.006 | -0.024 | -0.024 | 0 | 0.07 | 0.22 | 0.03 | 0.01 | 0.02 | 0.01 |
| 2 | 10108037 | rs7575106 | 0.93 | 0.264 | 0.608 | -0.217 | 0.005 | -0.03 | -0.03 | 0 | 0.08 | 0.1 | 0.04 | 0 | 0.08 | 0.05 |
| 2 | 10110517 | rs2163336 | -1.013 | 0.188 | 0.882 | -0.34 | 0.015 | 0.039 | 0.039 | 2.036 | 0.45 | 0.65 | 0.29 | 0.15 | 0.33 | 0.26 |
| 2 | 10110666 | rs2116016 | -2.01 | 0.163 | 0.828 | -0.167 | -0.027 | -0.038 | -0.038 | 0 | 0.6 | 0.8 | 0.7 | 0.5 | 0.58 | 0.39 |
| 2 | 10111219 | rs11883961 | 0.431 | 0.066 | 0.943 | -0.246 | 0.005 | 0.003 | 0.003 | 0.369 | 0.27 | 0.27 | 0.2 | 0.14 | 0.18 | 0.19 |
| 2 | 10117868 | rs11902236 | -0.877 | -0.041 | 0.777 | -0.407 | 0.014 | 0.018 | 0.018 | 0.379 | 0.4 | 0.65 | 0.23 | 0.15 | 0.29 | 0.22 |
| 2 | 10120605 | rs2011516 | -0.954 | 0.044 | 0.867 | -0.263 | 0.002 | -0.002 | -0.002 | 0 | 0.24 | 0.29 | 0.41 | 0.26 | 0.27 | 0.16 |
| 2 | 10124656 | rs731453 | -0.458 | 0.067 | 0.878 | -0.243 | 0.002 | 0.004 | 0.004 | 0 | 0.2 | 0.21 | 0.39 | 0.26 | 0.23 | 0.14 |
| 2 | 10126330 | rs2303920 | -0.517 | 0.042 | 0.933 | -0.255 | 0.002 | 0.008 | 0.008 | 0 | 0.21 | 0.31 | 0.4 | 0.26 | 0.23 | 0.14 |
| 2 | 10126416 | rs16867274 | -1.386 | -0.027 | 0.726 | -0.313 | 0.033 | 0.06 | 0.06 | 0 | 0.49 | 0.69 | 0.62 | 0.4 | 0.44 | 0.34 |
| 2 | 10131122 | rs1367389 | -0.473 | 0.029 | 0.637 | -0.255 | 0.002 | 0.004 | 0.004 | 0 | 0.2 | 0.21 | 0.39 | 0.26 | 0.23 | 0.14 |
| 2 | 10133284 | rs12618788 | 0.416 | -0.041 | 0.913 | -0.263 | -0.006 | 0.049 | 0.049 | 0.119 | 0.12 | 0.2 | 0.35 | 0.26 | 0.15 | 0.09 |
| 2 | 10133820 | rs6750565 | 0.901 | -0.083 | 0.807 | -0.321 | 0.016 | 0.031 | 0.031 | 0.58 | 0.23 | 0.13 | 0.19 | 0.14 | 0.18 | 0.19 |
| 2 | 10136592 | rs16867277 | 1.239 | -0.024 | 0.736 | -0.313 | 0.005 | -0.028 | -0.028 | 0 | 0.08 | 0.08 | 0.05 | 0 | 0.08 | 0.05 |
| 2 | 10136936 | rs6705493 | -0.313 | -0.07 | 0.815 | -0.316 | -0.022 | -0.006 | -0.006 | 2.601 | 0.37 | 0.43 | 0.55 | 0.4 | 0.33 | 0.28 |
| 2 | 10137119 | rs2357394 | -2.018 | 0.397 | 0.141 | -0.077 | -0.017 | -0.003 | -0.003 | 0.738 | 0.73 | 0.89 | 0.85 | 0.79 | 0.76 | 0.68 |
| 2 | 10140981 | rs934303 | 0.073 | 0.268 | 0.391 | -0.263 | -0.021 | -0.006 | -0.006 | 0.019 | 0.37 | 0.43 | 0.55 | 0.4 | 0.33 | 0.28 |
| 2 | 10141191 | rs1465638 | -0.093 | 0.335 | 0.095 | -0.237 | -0.014 | -0.005 | -0.005 | 0 | 0.38 | 0.55 | 0.56 | 0.4 | 0.33 | 0.28 |
| 2 | 10142073 | rs1052835 | -0.96 | 0.319 | 0.107 | -0.258 | 0.041 | 0.074 | 0.074 | 0 | 0.5 | 0.73 | 0.62 | 0.41 | 0.44 | 0.34 |
| 2 | 10142302 | rs11246 | -0.105 | 0.356 | 0.056 | -0.219 | -0.014 | -0.005 | -0.005 | 0 | 0.38 | 0.55 | 0.56 | 0.4 | 0.33 | 0.28 |
| 2 | 10145348 | rs1469541 | 1.157 | 0.471 | -0.029 | -0.182 | 0.002 | -0.019 | -0.019 | 0 | 0.09 | 0.07 | 0.05 | 0 | 0.12 | 0.06 |
| 2 | 10156725 | rs11686388 | -0.647 | 0.22 | 0.071 | 0.141 | -0.015 | 0.035 | 0.035 | 0.024 | 0.25 | 0.52 | 0.34 | 0.31 | 0.28 | 0.15 |
| 2 | 10159679 | rs6716186 | 0.448 | 0.346 | 0.016 | 0.212 | -0.005 | 0.032 | 0.032 | 0 | 0.72 | 0.47 | 0.63 | 0.65 | 0.67 | 0.72 |
| 2 | 10162037 | rs4073397 | 0.477 | 0.772 | -0.43 | 0.174 | 0.013 | 0.03 | 0.03 | 0 | 0.28 | 0.34 | 0.33 | 0.52 | 0.32 | 0.41 |
| 2 | 10163328 | rs10803714 | -0.159 | 0.432 | -0.771 | 0.588 | -0.002 | -0.004 | -0.004 | 0 | 0.8 | 0.85 | 0.67 | 0.68 | 0.7 | 0.83 |
| 2 | 10167212 | rs35454789 | 0.038 | 0.277 | -0.976 | 0.325 | 0.006 | 0.002 | 0.002 | 0 | 0.56 | 0.38 | 0.52 | 0.57 | 0.55 | 0.62 |
| 2 | 10175442 | rs6727702 | 0.208 | 0.872 | -1.114 | 0.066 | -0.007 | 0.005 | 0.005 | 0 | 0.21 | 0.21 | 0.18 | 0.14 | 0.23 | 0.24 |
| 2 | 10177158 | rs6747078 | -0.132 | -0.206 | -1.45 | -0.736 | 0.063 | 0.098 | 0.098 | 0 | 0.42 | 0.73 | 0.22 | 0.19 | 0.27 | 0.14 |
| 2 | 10178236 | rs4396680 | -0.471 | 0.451 | -1.693 | -0.542 | -0.052 | -0.056 | -0.056 | 0 | 0.9 | 0.99 | 0.89 | 0.83 | 0.82 | 0.84 |
| 2 | 10182496 | rs4669520 | 0.067 | 0.294 | -1.168 | -0.614 | 0.003 | 0.004 | 0.004 | 0 | 0.87 | 0.81 | 0.86 | 0.83 | 0.82 | 0.85 |
| 2 | 10193178 | rs4444493 | 1.191 | -0.161 | -1.511 | -1.03 | -0.001 | 0.085 | 0.085 | 0 | 0.15 | 0.02 | 0.07 | 0 | 0.12 | 0.07 |
| 2 | 10194192 | rs4669522 | 0.005 | -0.13 | -1.336 | -1.005 | 0.002 | 0.003 | 0.003 | 0 | 0.86 | 0.8 | 0.87 | 0.83 | 0.82 | 0.85 |
| 2 | 10194487 | rs7632 | 0.894 | -0.605 | -1.027 | -1.398 | -0.003 | 0.005 | 0.005 | 0 | 0.43 | 0.56 | 0.65 | 0.66 | 0.54 | 0.55 |
| 2 | 10197615 | rs10208406 | -0.411 | -0.432 | -0.811 | -1.294 | 0.008 | 0.011 | 0.011 | 0 | 0.87 | 0.8 | 0.87 | 0.83 | 0.82 | 0.85 |
| 2 | 10198113 | rs6717452 | 0.52 | -0.603 | -0.489 | -1.421 | -0.002 | -0.013 | -0.013 | 0.353 | 0.53 | 0.47 | 0.72 | 0.83 | 0.59 | 0.72 |
| 2 | 10201514 | rs10171924 | -0.163 | -0.293 | -0.414 | -1.058 | 0.022 | 0.031 | 0.031 | 0 | 0.86 | 0.72 | 0.87 | 0.83 | 0.82 | 0.85 |
| 2 | 10201829 | rs7596501 | 0.1 | -0.421 | -0.585 | -1.151 | 0.003 | 0.024 | 0.024 | 0 | 0.69 | 0.74 | 0.8 | 0.83 | 0.71 | 0.79 |
| 2 | 10206686 | rs12466448 | 0.796 | -0.279 | -0.616 | -0.926 | -0.001 | 0.028 | 0.028 | 0 | 0.22 | 0.27 | 0.43 | 0.6 | 0.31 | 0.32 |
| 2 | 10209993 | rs4233874 | -0.09 | -0.403 | -0.396 | -1.031 | -0.003 | 0 | 0 | 0 | 0.4 | 0.4 | 0.5 | 0.59 | 0.42 | 0.38 |
| 2 | 10210244 | rs6432057 | -0.264 | -0.065 | -0.488 | -1.193 | -0.002 | 0.001 | 0.001 | 0 | 0.65 | 0.65 | 0.69 | 0.77 | 0.69 | 0.66 |
| 2 | 10217952 | rs4583435 | 0.928 | 0.21 | -0.79 | -0.859 | 0.008 | -0.001 | -0.001 | 0 | 0.15 | 0.12 | 0.23 | 0.15 | 0.24 | 0.25 |
| 2 | 10218946 | rs12622986 | 0.733 | -0.723 | -0.441 | -1.075 | 0.001 | 0.008 | 0.008 | 0 | 0.2 | 0.27 | 0.13 | 0.18 | 0.18 | 0.24 |
| 2 | 10227261 | rs12622575 | 0.47 | -0.793 | -0.848 | -1.198 | -0.005 | -0.002 | -0.002 | 0 | 0.35 | 0.32 | 0.65 | 0.65 | 0.53 | 0.49 |
| 2 | 10229911 | rs11680378 | 0.352 | -0.748 | -0.858 | -1.115 | -0.022 | -0.007 | -0.007 | 0 | 0.35 | 0.25 | 0.64 | 0.66 | 0.53 | 0.46 |
| 2 | 10234179 | rs7587317 | -0.007 | -0.932 | -0.467 | -0.678 | -0.018 | -0.083 | -0.083 | 0 | 0.78 | 0.7 | 0.88 | 1 | 0.84 | 0.96 |
| 2 | 10240175 | rs11897268 | -0.307 | -0.838 | -0.934 | -0.367 | 0.031 | 0.031 | 0.031 | 0 | 0.25 | 0.2 | 0.14 | 0.11 | 0.17 | 0.2 |
| 2 | 10241915 | rs4233877 | 0.503 | -0.528 | -0.38 | 0.14 | -0.013 | 0.034 | 0.034 | 1.258 | 0.85 | 0.76 | 0.88 | 0.81 | 0.84 | 0.76 |
| 2 | 10259201 | rs7585544 | -0.199 | -0.768 | -0.465 | -0.687 | -0.026 | -0.112 | -0.112 | 0.266 | 0.27 | 0.34 | 0.15 | 0.01 | 0.2 | 0.09 |
| 2 | 10261486 | rs7582263 | -0.009 | -0.783 | -0.566 | -0.695 | 0.031 | 0.093 | 0.093 | 0.747 | 0.26 | 0.08 | 0.13 | 0.01 | 0.2 | 0.09 |
| 2 | 10262920 | rs1130609 | 0.19 | -0.063 | -0.887 | -0.22 | -0.067 | -0.089 | -0.089 | 0 | 0.77 | 0.95 | 0.55 | 0.34 | 0.73 | 0.56 |
| 2 | 10266563 | rs6759180 | -0.114 | -0.02 | -0.753 | -0.231 | -0.05 | -0.065 | -0.065 | 0 | 0.81 | 0.94 | 0.55 | 0.34 | 0.72 | 0.57 |
| 2 | 10268798 | rs4668664 | -0.174 | -0.15 | -0.768 | -0.212 | -0.043 | -0.065 | -0.065 | 0 | 0.81 | 0.94 | 0.56 | 0.34 | 0.73 | 0.58 |
| 2 | 10271196 | rs1138729 | 0.446 | -0.293 | -0.867 | -0.328 | 0.021 | 0.026 | 0.026 | 1.423 | 0.23 | 0.15 | 0.12 | 0.01 | 0.16 | 0.1 |
| 2 | 10272673 | rs4309551 | 0.913 | -0.388 | -0.463 | -0.373 | 0.007 | 0.023 | 0.023 | 0.732 | 0.55 | 0.39 | 0.42 | 0.33 | 0.57 | 0.47 |
| 2 | 10275746 | rs2241113 | 1.336 | 0.057 | -0.315 | 0.311 | -0.001 | -0.015 | -0.015 | 0 | 0.67 | 0.51 | 0.46 | 0.29 | 0.67 | 0.49 |
| 2 | 10281514 | rs17313542 | -0.301 | 0.328 | -1.135 | 0.456 | 0 | 0.032 | 0.032 | 0 | 0.11 | 0.01 | 0.07 | 0.05 | 0.07 | 0.08 |
| 2 | 10286466 | rs6749325 | 0.492 | 0.38 | -0.91 | 0.515 | 0.002 | 0.014 | 0.014 | 0 | 0.64 | 0.72 | 0.46 | 0.29 | 0.65 | 0.49 |

| | | | | | | | | | | | | | | | | |
|---|---|---|---|---|---|---|---|---|---|---|---|---|---|---|---|---|
| 2 | 10298766 | rs16855895 | -0.845 | 0.682 | -0.623 | 1.657 | 0.013 | 0.146 | 0.146 | 0 | 0.08 | 0.15 | 0.27 | 0.47 | 0.1 | 0.28 |
| 2 | 10303224 | rs16855903 | -0.998 | 0.949 | -0.648 | 1.996 | 0.019 | 0.144 | 0.144 | 0.053 | 0.09 | 0.14 | 0.28 | 0.51 | 0.14 | 0.24 |
| 2 | 10305928 | rs17313689 | 0.099 | 0.953 | -1.3 | 1.665 | -0.01 | -0.032 | -0.032 | 3.803 | 0.16 | 0.06 | 0.14 | 0.21 | 0.14 | 0.36 |
| 2 | 10310957 | rs16855939 | 0.013 | 0.366 | -1.281 | 1.312 | 0.023 | 0.022 | 0.022 | 0 | 0.13 | 0.06 | 0.07 | 0.02 | 0.04 | 0.06 |
| 2 | 10312826 | rs10929635 | 0.634 | 0.429 | -0.74 | 1.085 | -0.006 | -0.003 | -0.003 | 0 | 0.63 | 0.45 | 0.49 | 0.36 | 0.66 | 0.62 |
| 2 | 10314596 | rs1003187 | 0.626 | 0.226 | -0.743 | 0.896 | 0.031 | -0.041 | -0.041 | 0 | 0.63 | 0.31 | 0.53 | 0.48 | 0.69 | 0.67 |
| 2 | 10330025 | rs12621615 | -0.175 | 0.249 | -0.969 | 0.753 | 0 | 0.002 | 0.002 | 0 | 0.46 | 0.36 | 0.41 | 0.5 | 0.52 | 0.44 |
| 2 | 10332539 | rs13392520 | 0.313 | 0.259 | -0.742 | 0.916 | 0.021 | -0.024 | -0.024 | 0.654 | 0.39 | 0.36 | 0.36 | 0.41 | 0.31 | 0.52 |
| 2 | 10334833 | rs10191873 | -0.346 | 0.921 | -0.419 | 1.257 | 0.041 | 0.183 | 0.183 | 0 | 0.14 | 0.35 | 0.19 | 0.32 | 0.17 | 0.26 |
| 2 | 10336946 | rs6745337 | 0.239 | 1.236 | -1.374 | 0.542 | -0.058 | 0.034 | 0.034 | 0 | 0.17 | 0.06 | 0.25 | 0.1 | 0.22 | 0.1 |
| 2 | 10341468 | rs2357468 | 0.07 | 0.731 | -1.65 | 0.424 | -0.021 | 0.099 | 0.099 | 0.931 | 0.54 | 0.18 | 0.47 | 0.31 | 0.49 | 0.34 |
| 2 | 10344465 | rs6741014 | 0.158 | 0.707 | -1.179 | 0.424 | -0.001 | 0.014 | 0.014 | 0 | 0.31 | 0.6 | 0.35 | 0.55 | 0.32 | 0.54 |
| 2 | 10347635 | rs887962 | -2.095 | 0.764 | -1.319 | 0.212 | -0.002 | -0.001 | -0.001 | 0 | 0.91 | 0.94 | 0.92 | 1 | 0.86 | 0.97 |
| 2 | 10349858 | rs17267630 | 0.864 | 0.514 | -1.872 | 0.045 | -0.059 | 0.011 | 0.011 | 0 | 0.31 | 0.07 | 0.33 | 0.24 | 0.32 | 0.21 |
| 2 | 10357818 | rs10207325 | -0.155 | 0.353 | -0.807 | 0.118 | 0.001 | 0.016 | 0.016 | 0 | 0.24 | 0.57 | 0.25 | 0.29 | 0.26 | 0.33 |
| 2 | 10359536 | rs16855961 | 0.626 | 0.371 | -0.615 | 0.167 | -0.001 | 0.005 | 0.005 | 0 | 0.07 | 0.1 | 0.09 | 0.1 | 0.06 | 0.09 |
| 2 | 10361023 | rs11676328 | 0.462 | 0.218 | -0.603 | -0.082 | 0.008 | 0.031 | 0.031 | 0 | 0.62 | 0.32 | 0.67 | 0.71 | 0.6 | 0.65 |
| 2 | 10362170 | rs1861234 | -2.32 | 0.304 | -0.524 | -0.075 | -0.001 | -0.009 | -0.009 | 0.055 | 0.9 | 0.92 | 0.91 | 1 | 0.86 | 0.97 |
| 2 | 10364272 | rs16855971 | 0.635 | 0.282 | 0.011 | 0.041 | 0.014 | 0.003 | 0.003 | 0 | 0.17 | 0.33 | 0.16 | 0.19 | 0.2 | 0.23 |
| 2 | 10368321 | rs1006866 | 0.76 | 0.432 | -0.792 | -0.265 | -0.003 | 0.111 | 0.111 | 0 | 0.51 | 0.2 | 0.6 | 0.7 | 0.51 | 0.6 |
| 2 | 10368606 | rs7587651 | 1.21 | 0.436 | -1.162 | -0.411 | -0.046 | -0.041 | -0.041 | 0 | 0.36 | 0.12 | 0.52 | 0.62 | 0.36 | 0.48 |
| 2 | 10368871 | rs7601478 | -1.394 | 0.389 | -0.774 | -0.295 | -0.006 | -0.003 | -0.003 | 0.054 | 0.81 | 0.87 | 0.86 | 1 | 0.78 | 0.93 |
| 2 | 10369975 | rs16855984 | -0.635 | 0.465 | -0.252 | 0.271 | -0.002 | -0.001 | -0.001 | 0.173 | 0.3 | 0.67 | 0.26 | 0.3 | 0.26 | 0.33 |
| 2 | 10371990 | rs6432079 | 0.155 | 0.794 | 0.335 | 0.48 | 0.089 | 0.256 | 0.256 | 0.724 | 0.05 | 0.33 | 0.09 | 0.19 | 0.04 | 0.19 |
| 2 | 10372061 | rs16855996 | 0.694 | 1.128 | -0.246 | 0.878 | 0.052 | 0.167 | 0.167 | 0 | 0.06 | 0.12 | 0.18 | 0.37 | 0.12 | 0.25 |
| 2 | 10376784 | rs759340 | 0.096 | 0.689 | -0.158 | 0.518 | 0.016 | 0.001 | 0.001 | 0.05 | 0.19 | 0.52 | 0.17 | 0.18 | 0.2 | 0.25 |
| 2 | 10377112 | rs16856013 | 0.642 | 0.408 | -0.851 | 0.388 | 0.025 | 0.014 | 0.014 | 0 | 0.29 | 0.12 | 0.15 | 0.22 | 0.21 | 0.27 |
| 2 | 10377834 | rs10929641 | 0.815 | 0.516 | -0.903 | 0.512 | 0.001 | 0.136 | 0.136 | 0 | 0.09 | 0.12 | 0.18 | 0.44 | 0.08 | 0.22 |
| 2 | 10378227 | rs741260 | -0.38 | 0.412 | -0.602 | 0.16 | 0.008 | 0.005 | 0.005 | 0 | 0.42 | 0.64 | 0.38 | 0.61 | 0.29 | 0.51 |
| 2 | 10380209 | rs759341 | 0.15 | 0.774 | -0.67 | -0.089 | 0.001 | -0.035 | -0.035 | 0 | 0.32 | 0.29 | 0.24 | 0.16 | 0.27 | 0.41 |
| 2 | 10381434 | rs1468780 | 1.594 | 0.625 | -1.077 | 0.302 | 0 | 0.005 | 0.005 | 0 | 0.12 | 0.07 | 0.08 | 0.12 | 0.12 | 0.06 |
| 2 | 10381996 | rs10929642 | 0.205 | 0.632 | -0.785 | 0.613 | 0.003 | 0.017 | 0.017 | 0 | 0.46 | 0.34 | 0.56 | 0.71 | 0.39 | 0.67 |
| 2 | 10385097 | rs10929646 | -0.358 | 0.722 | -0.779 | 1.023 | 0.001 | 0.007 | 0.007 | 0 | 0.29 | 0.3 | 0.35 | 0.45 | 0.37 | 0.28 |
| 2 | 10394082 | rs887958 | -0.069 | 0.611 | -1.546 | 0.81 | -0.01 | 0.066 | 0.066 | 0 | 0.6 | 0.81 | 0.55 | 0.68 | 0.6 | 0.48 |
| 2 | 10394490 | rs11695911 | 0.374 | 0.946 | -1.103 | 0.977 | 0.121 | 0.153 | 0.153 | 2.526 | 0.07 | 0.3 | 0.17 | 0.15 | 0.12 | 0.04 |
| 2 | 10394558 | rs887959 | -0.698 | 0.908 | -2.023 | 0.733 | 0.003 | 0.03 | 0.03 | 0 | 0.76 | 0.96 | 0.81 | 0.85 | 0.74 | 0.74 |
| 2 | 10395868 | rs759343 | 0.125 | 1.149 | -1.95 | 0.744 | -0.037 | -0.014 | -0.014 | 0 | 0.47 | 0.16 | 0.49 | 0.59 | 0.44 | 0.56 |
| 2 | 10395944 | rs12476240 | 0.287 | 1.217 | -1.96 | 0.693 | -0.035 | -0.023 | -0.023 | 0 | 0.42 | 0.14 | 0.3 | 0.41 | 0.4 | 0.35 |
| 2 | 10402632 | rs7556858 | 0.925 | 0.687 | -1.667 | 0.04 | 0.01 | 0.049 | 0.049 | 5.562 | 0.3 | 0.14 | 0.33 | 0.18 | 0.27 | 0.21 |
| 2 | 10402763 | rs7597744 | 0.016 | 0.42 | -1.356 | 0.239 | -0.005 | 0.008 | 0.008 | 1.238 | 0.65 | 0.57 | 0.75 | 0.59 | 0.69 | 0.66 |
| 2 | 10404367 | rs16856070 | 0.045 | 0.363 | -1.833 | 0.066 | 0.034 | 0.088 | 0.088 | 0 | 0.27 | 0.09 | 0.12 | 0.1 | 0.2 | 0.17 |
| 2 | 10407935 | rs2110638 | -0.162 | 0.955 | -1.188 | 0.539 | -0.005 | -0.005 | -0.005 | 2.635 | 0.81 | 0.82 | 0.75 | 0.7 | 0.77 | 0.68 |
| 2 | 10408604 | rs6707453 | -0.155 | 1.245 | -1.022 | 0.518 | -0.015 | 0.003 | 0.003 | 0.062 | 0.78 | 0.87 | 0.8 | 0.76 | 0.74 | 0.61 |
| 2 | 10408886 | rs4669561 | 0.044 | 0.872 | -0.579 | 0.976 | 0.025 | 0.103 | 0.103 | 0 | 0.2 | 0.3 | 0.31 | 0.42 | 0.25 | 0.23 |
| 2 | 10413431 | rs17379538 | 0.585 | 0.739 | -1.078 | 0.283 | -0.025 | 0.092 | 0.092 | 0.015 | 0.41 | 0.2 | 0.47 | 0.21 | 0.45 | 0.34 |
| 2 | 10414143 | rs6432083 | -1.02 | 0.527 | 0.058 | 0.414 | 0.004 | 0.054 | 0.054 | 0.52 | 0.16 | 0.45 | 0.16 | 0.17 | 0.12 | 0.05 |
| 2 | 10422342 | rs4669564 | 0.437 | 0.742 | -0.455 | 0.287 | -0.005 | -0.011 | -0.011 | 0 | 0.66 | 0.6 | 0.83 | 0.78 | 0.68 | 0.66 |
| 2 | 10424886 | rs6432085 | 0.071 | 0.825 | -0.932 | 0.12 | 0.008 | -0.003 | -0.003 | 0 | 0.68 | 0.81 | 0.79 | 0.69 | 0.72 | 0.51 |
| 2 | 10427767 | rs10929654 | 0.307 | 1.176 | -0.504 | 0.377 | 0.042 | 0.025 | 0.025 | 0 | 0.09 | 0.13 | 0.1 | 0.16 | 0.14 | 0.09 |
| 2 | 10433946 | rs2357541 | -0.209 | 0.034 | -0.957 | 0.163 | 0.026 | -0.032 | -0.032 | 0 | 0.25 | 0.11 | 0.2 | 0.4 | 0.15 | 0.11 |
| 2 | 10437333 | rs1574463 | 0.184 | 0.503 | -0.639 | 0.069 | -0.004 | -0.025 | -0.025 | 0.374 | 0.09 | 0.12 | 0.03 | 0 | 0.05 | 0.04 |
| 2 | 10438621 | rs4668676 | 0.765 | 0.902 | -0.587 | 0.466 | 0 | 0 | 0 | 0.016 | 0.61 | 0.59 | 0.67 | 0.4 | 0.61 | 0.56 |
| 2 | 10440987 | rs2884208 | 0.28 | 1.258 | -1.585 | 0.673 | -0.032 | -0.066 | -0.066 | 0.762 | 0.71 | 0.86 | 0.74 | 0.42 | 0.69 | 0.61 |
| 2 | 10444031 | rs13023336 | -0.891 | 1.011 | -0.937 | 0.299 | -0.007 | 0.01 | 0.01 | 0 | 0.1 | 0.09 | 0.08 | 0.03 | 0.12 | 0.07 |
| 2 | 10450398 | rs6734475 | -1.469 | 0.512 | 0.337 | 0.628 | 0.029 | 0.115 | 0.115 | 4.222 | 0.06 | 0.35 | 0.04 | 0.1 | 0.03 | 0.03 |
| 2 | 10451165 | rs16856098 | 0.393 | 0.142 | 0.113 | -0.248 | -0.015 | -0.043 | -0.043 | 0 | 0.6 | 0.5 | 0.71 | 0.83 | 0.61 | 0.75 |
| 2 | 10453849 | rs2160189 | -0.069 | 0.088 | 0.206 | -0.276 | 0.004 | 0.157 | 0.157 | 0.117 | 0.66 | 0.92 | 0.77 | 0.92 | 0.63 | 0.79 |
| 2 | 10454622 | rs12052519 | 0.442 | 0.016 | 0.232 | -0.309 | -0.008 | -0.027 | -0.027 | 0 | 0.59 | 0.44 | 0.71 | 0.81 | 0.61 | 0.75 |
| 2 | 10462852 | rs11694489 | 0.252 | -0.106 | 0.254 | -0.475 | -0.009 | -0.021 | -0.021 | 0.018 | 0.12 | 0.27 | 0.07 | 0.06 | 0.1 | 0.03 |
| 2 | 10467171 | rs2014889 | -0.579 | -0.156 | -0.1 | -0.523 | -0.003 | 0.092 | 0.092 | 0 | 0.78 | 0.95 | 0.82 | 0.99 | 0.73 | 0.82 |
| 2 | 10467823 | rs16856119 | 0.273 | -0.17 | 0.28 | -0.026 | 0.009 | 0.009 | 0.009 | 1.467 | 0.11 | 0.03 | 0.06 | 0.07 | 0.1 | 0.03 |
| 2 | 10468296 | rs1003652 | -1.268 | -0.256 | 0.259 | -0.026 | -0.056 | 0.015 | 0.015 | 1.471 | 0.19 | 0.51 | 0.11 | 0.18 | 0.12 | 0.06 |
| 2 | 10469516 | rs1003653 | -0.36 | 0.107 | 0.497 | 1.023 | -0.01 | 0.134 | 0.134 | 0.005 | 0.14 | 0.25 | 0.25 | 0.37 | 0.11 | 0.06 |
| 2 | 10475308 | rs4669572 | -0.263 | 0.842 | -0.033 | 0.261 | -0.017 | 0.017 | 0.017 | 0 | 0.72 | 0.83 | 0.65 | 0.8 | 0.62 | 0.78 |
| 2 | 10476532 | rs6745569 | -0.202 | 0.654 | 0.572 | -0.027 | 0.013 | -0.051 | -0.051 | 0.063 | 0.17 | 0.4 | 0.11 | 0.06 | 0.15 | 0.25 |
| 2 | 10478422 | rs1558628 | -0.273 | 0.652 | 0.621 | -0.125 | 0.019 | -0.098 | -0.098 | 0 | 0.17 | 0.43 | 0.11 | 0.03 | 0.15 | 0.25 |
| 2 | 10478936 | rs4669573 | -0.298 | 0.269 | -0.328 | 0.049 | 0.015 | 0.06 | 0.06 | 0 | 0.42 | 0.74 | 0.29 | 0.23 | 0.36 | 0.43 |
| 2 | 10480201 | rs6758633 | 0.177 | 0.525 | -0.372 | 0.145 | 0.022 | -0.024 | -0.024 | 0 | 0.09 | 0.15 | 0.08 | 0.03 | 0.14 | 0.25 |
| 2 | 10485409 | rs10197851 | -0.328 | 0.367 | -0.804 | 0.021 | 0.019 | 0.127 | 0.127 | 0 | 0.47 | 0.8 | 0.33 | 0.22 | 0.4 | 0.6 |

| | | | | | | | | | | | | | | | | |
|---|---|---|---|---|---|---|---|---|---|---|---|---|---|---|---|---|
| 2 | 10489435 | rs17434144 | 0.642 | 0.116 | -1.124 | -0.117 | 0.056 | 0.045 | 0.045 | 0 | 0.46 | 0.23 | 0.43 | 0.36 | 0.39 | 0.62 |
| 2 | 10491985 | rs12990794 | 0.689 | 0.918 | -1.029 | -0.021 | 0.003 | 0 | 0 | 0.54 | 0.09 | 0.05 | 0.12 | 0.16 | 0.18 | 0.1 |
| 2 | 10495928 | rs1476986 | -0.222 | 1.082 | -0.679 | -0.304 | -0.006 | -0.018 | -0.018 | 0 | 0.74 | 0.76 | 0.84 | 0.99 | 0.71 | 0.82 |
| 2 | 10495974 | rs1476985 | -0.405 | 1.083 | -0.602 | -0.216 | 0.018 | 0.031 | 0.031 | 0 | 0.28 | 0.51 | 0.49 | 0.68 | 0.31 | 0.21 |
| 2 | 10496935 | rs12621622 | 0.121 | 1.106 | -0.414 | -0.67 | 0.009 | -0.091 | -0.091 | 1.504 | 0.71 | 0.62 | 0.82 | 0.98 | 0.71 | 0.76 |
| 2 | 10496944 | rs12618307 | 0.812 | 1.081 | -0.383 | -0.628 | -0.01 | 0.025 | 0.025 | 0 | 0.6 | 0.33 | 0.68 | 0.81 | 0.52 | 0.66 |
| 2 | 10501681 | rs2884210 | -0.014 | 1.252 | -0.522 | -0.646 | 0.01 | -0.052 | -0.052 | 0.748 | 0.78 | 0.71 | 0.85 | 0.98 | 0.73 | 0.75 |
| 2 | 10502733 | rs11677361 | -0.887 | 0.435 | -0.312 | -0.148 | 0.008 | -0.002 | -0.002 | 0.065 | 0.31 | 0.55 | 0.5 | 0.7 | 0.32 | 0.23 |
| 2 | 10507096 | rs7578524 | 1.091 | 0.009 | -0.46 | -0.249 | 0.011 | 0.023 | 0.023 | 0.468 | 0.12 | 0.06 | 0.07 | 0 | 0.1 | 0.05 |
| 2 | 10510040 | rs6755271 | 0.68 | -0.009 | -0.584 | -0.48 | -0.003 | 0.005 | 0.005 | 0 | 0.55 | 0.44 | 0.45 | 0.32 | 0.53 | 0.45 |
| 2 | 10512382 | rs7563637 | 0.954 | 0.186 | -1.013 | -0.362 | 0.011 | 0.033 | 0.033 | 0 | 0.5 | 0.6 | 0.32 | 0.31 | 0.43 | 0.4 |
| 2 | 10512580 | rs7563835 | -0.78 | 0.226 | -1.09 | -0.39 | -0.086 | 0.12 | 0.12 | 0 | 0.74 | 0.95 | 0.73 | 0.95 | 0.64 | 0.61 |
| 2 | 10514296 | rs2041741 | -1.145 | -0.231 | -0.662 | -0.001 | -0.027 | 0.323 | 0.323 | 7.859 | 0.12 | 0.64 | 0.2 | 0.47 | 0.03 | 0.07 |
| 2 | 10516429 | rs9967756 | 0.253 | -0.185 | -0.742 | -0.017 | 0.006 | 0.103 | 0.103 | 0 | 0.54 | 0.77 | 0.55 | 0.77 | 0.44 | 0.48 |
| 2 | 10517308 | rs13392890 | -0.164 | -0.147 | -0.817 | 0.065 | -0.018 | 0.121 | 0.121 | 0 | 0.14 | 0.28 | 0.19 | 0.48 | 0.06 | 0.2 |
| 2 | 10517835 | rs7560149 | -0.455 | -0.165 | -0.936 | -0.017 | -0.066 | 0.095 | 0.095 | 0 | 0.63 | 0.86 | 0.62 | 0.8 | 0.52 | 0.49 |
| 2 | 10521126 | rs6721270 | 1.366 | 0.214 | -0.621 | 0.344 | -0.032 | 0.007 | 0.007 | 0 | 0.32 | 0.1 | 0.32 | 0.3 | 0.37 | 0.27 |
| 2 | 10521719 | rs1420107 | -0.389 | 0.23 | -0.678 | 0.776 | -0.017 | 0.086 | 0.086 | 3.195 | 0.19 | 0.29 | 0.24 | 0.48 | 0.12 | 0.12 |
| 2 | 10524780 | rs10198351 | 0.342 | 0.618 | -0.724 | 0.463 | -0.003 | 0.022 | 0.022 | 0.309 | 0.56 | 0.58 | 0.48 | 0.3 | 0.55 | 0.33 |
| 2 | 10527833 | rs12476436 | -0.723 | 0.619 | -0.051 | 1.137 | -0.008 | 0.002 | 0.002 | 0.209 | 0.2 | 0.29 | 0.32 | 0.79 | 0.17 | 0.2 |
| 2 | 10530325 | rs11888704 | -1.28 | 1.091 | 0.83 | 1.814 | 0.065 | 0.251 | 0.251 | 2.532 | 0.09 | 0.36 | 0.25 | 0.31 | 0.07 | 0.05 |
| 2 | 10532869 | rs7582990 | -0.794 | 1.313 | 0.434 | 1.554 | -0.004 | 0.008 | 0.008 | 0.374 | 0.42 | 0.64 | 0.54 | 0.4 | 0.55 | 0.69 |
| 2 | 10533921 | rs7583465 | -0.453 | 2.23 | 1.085 | 1.525 | 0.011 | -0.005 | -0.005 | 0 | 0.3 | 0.38 | 0.45 | 0.33 | 0.38 | 0.53 |
| 2 | 10534375 | rs17434886 | -0.276 | 2.225 | 1.205 | 1.43 | 0.033 | -0.005 | -0.005 | 0 | 0.11 | 0.2 | 0.08 | 0.07 | 0.18 | 0.16 |
| 2 | 10537018 | rs2270302 | -0.983 | 2.2 | 1.606 | 1.588 | 0.015 | -0.003 | -0.003 | 0 | 0.36 | 0.41 | 0.39 | 0.32 | 0.49 | 0.67 |
| 2 | 10537462 | rs11676661 | -1.404 | 2.039 | 1.891 | 1.578 | 0.001 | -0.011 | -0.011 | 0 | 0.42 | 0.53 | 0.42 | 0.32 | 0.56 | 0.7 |
| 2 | 10537914 | rs3771147 | 0.116 | 2.545 | 1.463 | 1.538 | -0.006 | -0.004 | -0.004 | 0 | 0.14 | 0.12 | 0.26 | 0.2 | 0.25 | 0.28 |
| 2 | 10540386 | rs4668685 | -1.573 | 2.59 | 1.699 | 2.103 | -0.006 | -0.005 | -0.005 | 0.608 | 0.32 | 0.35 | 0.44 | 0.59 | 0.42 | 0.57 |
| 2 | 10541712 | rs9917245 | -2.125 | 2.19 | 1.607 | 1.402 | 0.01 | 0.032 | 0.032 | 0.065 | 0.47 | 0.64 | 0.44 | 0.32 | 0.56 | 0.69 |
| 2 | 10544640 | rs3771138 | -0.321 | 2.345 | 1.538 | 1.462 | -0.03 | -0.016 | -0.016 | 0 | 0.18 | 0.07 | 0.28 | 0.23 | 0.28 | 0.47 |
| 2 | 10546236 | rs3755264 | -0.359 | 2.331 | 1.532 | 1.455 | -0.003 | -0.002 | -0.002 | 0 | 0.18 | 0.15 | 0.29 | 0.23 | 0.28 | 0.47 |
| 2 | 10548577 | rs876614 | -1.03 | 2.322 | 1.682 | 1.419 | 0.039 | -0.011 | -0.011 | 0 | 0.25 | 0.42 | 0.32 | 0.23 | 0.32 | 0.48 |
| 2 | 10548826 | rs3771130 | -1.79 | 2.398 | 1.655 | 1.438 | 0.031 | -0.013 | -0.013 | 0 | 0.36 | 0.46 | 0.4 | 0.31 | 0.51 | 0.66 |
| 2 | 10549357 | rs7579181 | -1.748 | 1.923 | 1.829 | 1.245 | 0.009 | -0.069 | -0.069 | 0.056 | 0.28 | 0.45 | 0.15 | 0.08 | 0.27 | 0.24 |
| 2 | 10553898 | rs3755261 | -1.02 | 1.938 | 1.598 | 1.268 | -0.004 | 0.005 | 0.005 | 0 | 0.21 | 0.21 | 0.14 | 0.08 | 0.23 | 0.21 |
| 2 | 10555214 | rs3821199 | -1.32 | 1.88 | 1.528 | 1.719 | 0.005 | 0.002 | 0.002 | 0 | 0.22 | 0.25 | 0.16 | 0.2 | 0.23 | 0.21 |
| 2 | 10555776 | rs3771122 | -1.82 | 2.106 | 1.334 | 1.867 | -0.004 | 0.019 | 0.019 | 0 | 0.07 | 0.03 | 0.22 | 0.69 | 0.07 | 0.24 |
| 2 | 10557093 | rs13383253 | -1.179 | 2.068 | 1.519 | 1.572 | 0.027 | -0.059 | -0.059 | 0 | 0.22 | 0.59 | 0.16 | 0.08 | 0.21 | 0.2 |
| 2 | 10561383 | rs6432096 | -2.256 | 1.543 | 0.988 | 1.523 | 0.051 | 0.111 | 0.111 | 0.605 | 0.5 | 0.76 | 0.43 | 0.68 | 0.56 | 0.68 |
| 2 | 10563913 | rs1974676 | -1.68 | 1.678 | 1.106 | 1.596 | 0.005 | -0.02 | -0.02 | 0.773 | 0.4 | 0.47 | 0.29 | 0.17 | 0.51 | 0.56 |
| 2 | 10566033 | rs3755259 | -0.793 | 1.711 | 0.912 | 1.538 | 0.004 | -0.018 | -0.018 | 0 | 0.19 | 0.17 | 0.16 | 0.04 | 0.28 | 0.2 |
| 2 | 10566190 | rs3755258 | -0.781 | 1.711 | 0.912 | 1.538 | 0.004 | -0.018 | -0.018 | 0 | 0.19 | 0.17 | 0.16 | 0.04 | 0.28 | 0.2 |
| 2 | 10566414 | rs3755256 | 0.448 | 1.829 | 0.519 | 1.593 | -0.032 | 0.026 | 0.026 | 0 | 0.11 | 0.05 | 0.12 | 0.04 | 0.24 | 0.17 |
| 2 | 10567895 | rs1465323 | -1.278 | 1.579 | 0.768 | 1.74 | 0.012 | 0.006 | 0.006 | 1.231 | 0.31 | 0.41 | 0.33 | 0.64 | 0.37 | 0.49 |
| 2 | 10573198 | rs11694911 | -1.343 | 0.945 | 0.409 | 2.067 | 0.006 | -0.029 | -0.029 | 0 | 0.16 | 0.08 | 0.12 | 0.36 | 0.13 | 0.2 |
| 2 | 10580967 | rs1049500 | -1.822 | 0.649 | 1.315 | 2.048 | -0.034 | 0.195 | 0.195 | 0.209 | 0.11 | 0.3 | 0.13 | 0.57 | 0.05 | 0.25 |
| 2 | 10583160 | rs1405948 | -0.189 | 0.997 | 0.746 | 1.499 | -0.001 | 0.021 | 0.021 | 0.027 | 0.06 | 0.08 | 0.23 | 0.19 | 0.07 | 0.11 |
| 2 | 10595578 | rs6432097 | -0.53 | 0.996 | 0.363 | 0.82 | 0.002 | -0.059 | -0.059 | 1.271 | 0.23 | 0.41 | 0.17 | 0.06 | 0.24 | 0.37 |
| 2 | 10599845 | rs10803719 | -1.228 | 1.236 | -0.364 | 0.702 | 0.006 | 0.019 | 0.019 | 0.081 | 0.55 | 0.68 | 0.5 | 0.33 | 0.59 | 0.7 |
| 2 | 10607465 | rs818161 | -0.322 | 1.212 | -0.607 | 0.962 | -0.023 | 0.001 | 0.001 | 0 | 0.21 | 0.17 | 0.12 | 0.26 | 0.26 | 0.25 |
| 2 | 10612091 | rs818169 | -0.723 | 1.28 | -0.35 | 0.582 | -0.001 | -0.002 | -0.002 | 0.018 | 0.27 | 0.28 | 0.17 | 0.11 | 0.29 | 0.37 |
| 2 | 10612630 | rs818167 | -1.421 | 1.094 | -0.799 | 0.322 | -0.021 | 0.145 | 0.145 | 0.541 | 0.55 | 0.84 | 0.46 | 0.81 | 0.57 | 0.87 |
| 2 | 10613163 | rs10177521 | -0.367 | 0.864 | -0.223 | 0.982 | -0.017 | 0.15 | 0.15 | 0 | 0.11 | 0.22 | 0.16 | 0.5 | 0.09 | 0.22 |
| 2 | 10613284 | rs818165 | -1.049 | 0.704 | -0.373 | 0.45 | 0.027 | 0.014 | 0.014 | 0 | 0.42 | 0.77 | 0.35 | 0.61 | 0.37 | 0.6 |
| 2 | 10615346 | rs6755985 | -1.163 | 0.601 | -0.386 | 0.11 | 0.012 | 0.067 | 0.067 | 1.292 | 0.5 | 0.7 | 0.52 | 0.73 | 0.55 | 0.65 |
| 2 | 10615835 | rs16856271 | -0.222 | 0.504 | -0.637 | 0.099 | -0.019 | -0.008 | -0.008 | 0.932 | 0.11 | 0.13 | 0.04 | 0.08 | 0.06 | 0.11 |
| 2 | 10616214 | rs7591360 | 0.351 | 0.926 | -0.918 | 0.024 | -0.049 | 0.025 | 0.025 | 0.909 | 0.2 | 0.13 | 0.29 | 0.18 | 0.32 | 0.29 |
| 2 | 10617598 | rs1728149 | -0.844 | 0.182 | -0.812 | -0.316 | -0.009 | -0.047 | -0.047 | 0.54 | 0.63 | 0.6 | 0.55 | 0.78 | 0.72 | 0.8 |
| 2 | 10617864 | rs1728147 | -0.6 | -0.385 | -1.322 | -0.292 | 0.056 | 0.137 | 0.137 | 0 | 0.51 | 0.71 | 0.53 | 0.79 | 0.39 | 0.62 |
| 2 | 10618025 | rs2430424 | -0.376 | -0.205 | -0.948 | 0.564 | -0.033 | 0.041 | 0.041 | 0 | 0.29 | 0.44 | 0.21 | 0.52 | 0.24 | 0.32 |
| 2 | 10619702 | rs11890785 | -0.282 | 0.323 | -1.455 | 0.673 | -0.014 | -0.015 | -0.015 | 0 | 0.09 | 0.05 | 0.1 | 0.14 | 0.12 | 0.16 |
| 2 | 10620167 | rs2430425 | 1.074 | -0.312 | -1.214 | 1.083 | -0.006 | 0.019 | 0.019 | 0.835 | 0.78 | 0.29 | 0.77 | 0.58 | 0.78 | 0.7 |
| 2 | 10627544 | rs818182 | 0.644 | -0.04 | -2.122 | 0.352 | 0.005 | 0.108 | 0.108 | 2.142 | 0.44 | 0.15 | 0.37 | 0.23 | 0.43 | 0.26 |
| 2 | 10627860 | rs1641129 | 0.403 | -0.068 | -1.978 | 0.523 | 0.014 | -0.018 | -0.018 | 0.553 | 0.11 | 0.06 | 0.14 | 0.22 | 0.07 | 0.12 |
| 2 | 10637325 | rs818148 | 0.25 | 0.477 | -1.73 | 0.61 | -0.002 | -0.008 | -0.008 | 0 | 0.7 | 0.71 | 0.62 | 0.51 | 0.68 | 0.6 |
| 2 | 10643327 | rs818155 | 0.631 | 0.542 | -1.587 | -0.186 | 0.001 | 0.076 | 0.076 | 0.479 | 0.46 | 0.28 | 0.37 | 0.22 | 0.45 | 0.23 |
| 2 | 10648857 | rs818185 | -0.053 | 0.623 | -1.513 | -0.078 | -0.001 | -0.003 | -0.003 | 0 | 0.64 | 0.36 | 0.49 | 0.41 | 0.62 | 0.48 |
| 2 | 10649626 | rs818184 | 1.049 | 0.639 | -1.768 | -0.565 | -0.035 | 0.114 | 0.114 | 0.096 | 0.42 | 0.09 | 0.33 | 0.2 | 0.41 | 0.21 |
| 2 | 10650739 | rs7609435 | -0.049 | 0.831 | -0.903 | -1.29 | 0.005 | -0.046 | -0.046 | 0 | 0.59 | 0.51 | 0.5 | 0.16 | 0.56 | 0.45 |

| | | | | | | | | | | | | | | | | |
|---|---|---|---|---|---|---|---|---|---|---|---|---|---|---|---|---|
| 2 | 10654259 | rs6710665 | -0.666 | 0.687 | -1.044 | -1.024 | 0.006 | -0.043 | -0.043 | 0 | 0.7 | 0.6 | 0.59 | 0.19 | 0.69 | 0.59 |
| 2 | 10656284 | rs10185605 | -0.165 | 0.895 | -1.096 | -0.901 | 0.009 | 0.001 | 0.001 | 0 | 0.11 | 0.22 | 0.2 | 0.1 | 0.12 | 0.18 |
| 2 | 10660472 | rs12475194 | 0.157 | 0.798 | -1.966 | -0.856 | -0.018 | 0.093 | 0.093 | 2.11 | 0.21 | 0.06 | 0.12 | 0.07 | 0.2 | 0.27 |
| 2 | 10661047 | rs1405950 | -0.672 | 0.835 | -1.983 | -0.854 | 0 | -0.011 | -0.011 | 0 | 0.28 | 0.25 | 0.17 | 0.07 | 0.27 | 0.32 |
| 2 | 10662366 | rs12466300 | -0.003 | 0.531 | -2.252 | -0.648 | 0.006 | 0.025 | 0.025 | 0.66 | 0.22 | 0.17 | 0.13 | 0.1 | 0.19 | 0.27 |
| 2 | 10663183 | rs11887760 | 0.089 | 0.704 | -2.513 | -0.378 | -0.115 | 0.184 | 0.184 | 3.044 | 0.39 | 0.08 | 0.44 | 0.11 | 0.48 | 0.31 |
| 2 | 10665979 | rs16856345 | -0.473 | 0.264 | -1.989 | 0.444 | 0.025 | -0.034 | -0.034 | 0 | 0.31 | 0.22 | 0.23 | 0.47 | 0.26 | 0.43 |
| 2 | 10666469 | rs13005655 | 0.394 | 0.446 | -1.871 | -0.59 | 0.005 | -0.013 | -0.013 | 0 | 0.44 | 0.56 | 0.37 | 0.05 | 0.42 | 0.29 |
| 2 | 10667035 | rs16856347 | 0.061 | 1.167 | -2.151 | 0.293 | -0.066 | -0.038 | -0.038 | 0 | 0.23 | 0.08 | 0.33 | 0.28 | 0.28 | 0.25 |
| 2 | 10671640 | rs2430428 | -0.325 | 0.449 | -2.25 | 0.367 | 0 | -0.007 | -0.007 | 0.023 | 0.42 | 0.24 | 0.44 | 0.4 | 0.38 | 0.38 |
| 2 | 10672033 | rs1641147 | -0.824 | 0.297 | -1.484 | 0.26 | -0.01 | -0.013 | -0.013 | 0 | 0.6 | 0.66 | 0.53 | 0.42 | 0.51 | 0.59 |
| 2 | 10673359 | rs1641149 | -0.91 | 0.333 | -1.445 | 0.26 | -0.005 | -0.01 | -0.01 | 0 | 0.66 | 0.67 | 0.55 | 0.43 | 0.59 | 0.61 |
| 2 | 10674949 | rs1728143 | 0.37 | 0.031 | -1.311 | 0.383 | -0.004 | 0.003 | 0.003 | 0 | 0.1 | 0.09 | 0.03 | 0.14 | 0.04 | 0.08 |
| 2 | 10682245 | rs1527884 | 0.963 | 0.24 | -1.529 | 0.608 | -0.023 | -0.058 | -0.058 | 0 | 0.32 | 0.1 | 0.33 | 0.43 | 0.36 | 0.21 |
| 2 | 10682449 | rs12613865 | 0.452 | 0.195 | -1.675 | 0.415 | -0.049 | 0.001 | 0.001 | 0 | 0.29 | 0.12 | 0.39 | 0.26 | 0.34 | 0.28 |
| 2 | 10684391 | rs4669588 | -0.164 | -0.067 | -0.934 | 0.63 | -0.007 | 0.067 | 0.067 | 0.004 | 0.87 | 0.82 | 0.85 | 0.73 | 0.92 | 0.75 |
| 2 | 10685243 | rs13024441 | 0.707 | 0.155 | -1.053 | 0.481 | -0.019 | -0.024 | -0.024 | 0 | 0.37 | 0.16 | 0.43 | 0.43 | 0.43 | 0.25 |
| 2 | 10689608 | rs10194050 | -0.137 | -0.365 | -0.552 | 0.567 | -0.019 | -0.004 | -0.004 | 0 | 0.12 | 0.18 | 0.04 | 0.12 | 0.04 | 0.09 |
| 2 | 10692600 | rs11883723 | 0.126 | -0.342 | -0.513 | 0.427 | 0.021 | 0.005 | 0.005 | 1.262 | 0.49 | 0.62 | 0.38 | 0.49 | 0.37 | 0.4 |
| 2 | 10692766 | rs11883805 | -0.473 | -0.532 | -0.345 | 0.071 | 0.032 | 0.027 | 0.027 | 0.321 | 0.51 | 0.42 | 0.57 | 0.62 | 0.6 | 0.56 |
| 2 | 10693478 | rs1918705 | 0.022 | -0.244 | 0.218 | 0.012 | -0.014 | -0.012 | -0.012 | 0 | 0.63 | 0.5 | 0.68 | 0.71 | 0.76 | 0.49 |
| 2 | 10696877 | rs2357650 | 0.478 | -0.271 | 0.182 | 0.016 | -0.005 | 0.006 | 0.006 | 0 | 0.33 | 0.31 | 0.34 | 0.2 | 0.42 | 0.3 |
| 2 | 10698364 | rs16856412 | 1.31 | -0.399 | 0.058 | 0.096 | -0.006 | 0.022 | 0.022 | 0 | 0.25 | 0.19 | 0.24 | 0.15 | 0.29 | 0.14 |
| 2 | 10709816 | rs759400 | -0.397 | -0.56 | 0.291 | 0.271 | 0.03 | 0.024 | 0.024 | 0.203 | 0.67 | 0.81 | 0.75 | 0.75 | 0.79 | 0.52 |
| 2 | 10717282 | rs10929676 | 0.391 | -0.427 | 0.666 | 0.274 | 0.029 | 0.035 | 0.035 | 0.004 | 0.57 | 0.31 | 0.43 | 0.32 | 0.68 | 0.42 |
| 2 | 10717806 | rs2287059 | 1.673 | -0.365 | 0.337 | 0.302 | -0.079 | 0.021 | 0.021 | 0 | 0.24 | 0.01 | 0.18 | 0.13 | 0.27 | 0.12 |
| 2 | 10747229 | rs6744848 | 0.784 | -0.268 | 0.731 | 0.494 | -0.003 | 0.198 | 0.198 | 0 | 0.05 | 0.12 | 0.15 | 0.39 | 0.05 | 0.08 |
| 2 | 10758859 | rs6714413 | -0.254 | -0.275 | 0.502 | 0.377 | -0.034 | -0.004 | -0.004 | 0 | 0.41 | 0.77 | 0.56 | 0.62 | 0.49 | 0.39 |
| 2 | 10768598 | rs6733161 | -0.683 | -0.186 | 0.704 | 0.504 | 0.049 | 0.386 | 0.386 | 0.001 | 0.09 | 0.37 | 0.31 | 0.43 | 0.12 | 0.1 |
| 2 | 10784491 | rs3732111 | -0.521 | -0.148 | 0.626 | 0.495 | 0.034 | 0.286 | 0.286 | 0 | 0.08 | 0.3 | 0.3 | 0.42 | 0.12 | 0.1 |
| 2 | 10789249 | rs744809 | 0.822 | -0.213 | 0.423 | 0.291 | -0.031 | 0.04 | 0.04 | 0.104 | 0.29 | 0.17 | 0.22 | 0.19 | 0.35 | 0.28 |
| 2 | 10796181 | rs1023588 | 0.384 | -0.235 | 0.644 | 0.422 | 0.026 | 0.3 | 0.3 | 0.125 | 0.9 | 0.66 | 0.69 | 0.57 | 0.88 | 0.88 |
| 2 | 10807748 | rs1012580 | 1.689 | -0.329 | 0.394 | 0.271 | -0.085 | 0.036 | 0.036 | 0.014 | 0.25 | 0.01 | 0.19 | 0.12 | 0.28 | 0.12 |
| 2 | 10808204 | rs2357656 | 0.013 | -0.286 | 0.529 | 0.237 | -0.01 | -0.012 | -0.012 | 0 | 0.37 | 0.4 | 0.28 | 0.25 | 0.28 | 0.51 |
| 2 | 10815833 | rs9287721 | 1.133 | -0.238 | 0.438 | 0.259 | -0.033 | 0.04 | 0.04 | 0.018 | 0.28 | 0.14 | 0.21 | 0.19 | 0.32 | 0.25 |
| 2 | 10827157 | rs11684157 | -0.598 | -0.601 | 0.528 | 0.274 | 0.007 | 0.008 | 0.008 | 0.481 | 0.63 | 0.52 | 0.47 | 0.38 | 0.56 | 0.64 |
| 2 | 10830744 | rs17364568 | -0.073 | -0.823 | 0.332 | 0.285 | 0.022 | 0.035 | 0.035 | 0.025 | 0.35 | 0.16 | 0.25 | 0.24 | 0.26 | 0.5 |
| 2 | 10840338 | rs12614934 | 0.245 | -0.735 | 0.4 | -0.071 | 0.03 | 0.02 | 0.02 | 0.171 | 0.56 | 0.38 | 0.57 | 0.35 | 0.72 | 0.41 |
| 2 | 10846202 | rs12993133 | -1.139 | -0.596 | 0.699 | -0.043 | -0.004 | -0.068 | -0.068 | 0 | 0.73 | 0.49 | 0.79 | 0.87 | 0.75 | 0.89 |
| 2 | 10849552 | rs13396733 | -0.111 | -0.757 | 0.56 | -0.242 | -0.041 | 0.039 | 0.039 | 0 | 0.31 | 0.45 | 0.43 | 0.61 | 0.18 | 0.35 |
| 2 | 10850929 | rs7422405 | -0.248 | -0.572 | 0.108 | -0.066 | -0.001 | 0.003 | 0.003 | 0.718 | 0.56 | 0.57 | 0.62 | 0.7 | 0.42 | 0.46 |
| 2 | 10857763 | rs6736664 | 0.796 | -0.174 | -0.459 | 0.186 | -0.01 | 0.075 | 0.075 | 0 | 0.28 | 0.13 | 0.19 | 0.08 | 0.27 | 0.12 |
| 2 | 10862188 | rs12468286 | 0.484 | 0.023 | -0.491 | 0.399 | -0.048 | 0.006 | 0.006 | 0.303 | 0.29 | 0.18 | 0.31 | 0.26 | 0.44 | 0.34 |
| 2 | 10874512 | rs4669613 | -0.477 | -0.193 | -0.602 | 0.452 | -0.009 | -0.007 | -0.007 | 0.119 | 0.34 | 0.34 | 0.33 | 0.24 | 0.22 | 0.46 |
| 2 | 10886098 | rs4669621 | 0.163 | 0.179 | -0.729 | 0.469 | -0.01 | -0.009 | -0.009 | 0 | 0.57 | 0.57 | 0.76 | 0.66 | 0.64 | 0.54 |
| 2 | 10887075 | rs10192975 | -0.546 | -0.094 | -0.502 | 0.483 | 0.003 | 0.086 | 0.086 | 0 | 0.87 | 0.72 | 0.92 | 0.73 | 0.9 | 0.87 |
| 2 | 10889788 | rs10929698 | 0.15 | -0.561 | -1.181 | 0.316 | 0.039 | 0.112 | 0.112 | 0.606 | 0.48 | 0.2 | 0.47 | 0.3 | 0.62 | 0.64 |
| 2 | 10890205 | rs2884288 | -0.839 | -0.655 | -0.725 | 0.251 | -0.027 | 0.045 | 0.045 | 0 | 0.71 | 0.47 | 0.58 | 0.63 | 0.79 | 0.79 |
| 2 | 10899269 | rs1198859 | 0.348 | -0.186 | -0.584 | -0.03 | 0 | 0.005 | 0.005 | 0 | 0.36 | 0.61 | 0.39 | 0.54 | 0.36 | 0.41 |
| 2 | 10901492 | rs4539765 | -0.061 | -0.17 | -0.968 | -0.05 | -0.041 | 0.148 | 0.148 | 0 | 0.61 | 0.85 | 0.72 | 0.9 | 0.57 | 0.62 |
| 2 | 10903412 | rs1198872 | 0.331 | -0.018 | 0.12 | 0.124 | 0.002 | 0.03 | 0.03 | 0 | 0.33 | 0.43 | 0.36 | 0.54 | 0.34 | 0.38 |
| 2 | 10908159 | rs1198844 | -0.302 | 0.572 | 0.482 | -0.074 | 0.047 | -0.04 | -0.04 | 0.559 | 0.85 | 0.74 | 0.84 | 0.95 | 0.79 | 0.94 |
| 2 | 10910851 | rs6757005 | -0.322 | 0.138 | 0.122 | -0.068 | -0.019 | -0.001 | -0.001 | 0 | 0.37 | 0.31 | 0.43 | 0.33 | 0.46 | 0.31 |
| 2 | 10910877 | rs12464402 | 0.459 | 0.234 | 0.235 | -0.459 | 0.035 | -0.034 | -0.034 | 0.484 | 0.15 | 0.26 | 0.15 | 0.05 | 0.21 | 0.07 |
| 2 | 10911552 | rs17455547 | -0.005 | 0.067 | -0.036 | -0.61 | -0.007 | 0.005 | 0.005 | 0 | 0.12 | 0.08 | 0.11 | 0.01 | 0.18 | 0.23 |
| 2 | 10916670 | rs3856465 | -0.646 | -0.197 | 0.567 | -0.584 | 0.007 | -0.039 | -0.039 | 0 | 0.26 | 0.33 | 0.2 | 0.07 | 0.32 | 0.24 |
| 2 | 10916693 | rs1734440 | 0.662 | -0.115 | 0.587 | -0.204 | 0.026 | 0.008 | 0.008 | 0.016 | 0.41 | 0.28 | 0.37 | 0.4 | 0.29 | 0.4 |
| 2 | 10922516 | rs1686482 | 0.241 | -0.31 | 0.423 | -0.117 | -0.028 | 0.005 | 0.005 | 0 | 0.5 | 0.14 | 0.48 | 0.45 | 0.48 | 0.45 |
| 2 | 10923666 | rs1060011 | 2.214 | -0.32 | 0.576 | -0.141 | -0.016 | -0.024 | -0.024 | 0 | 0.05 | 0.06 | 0.01 | 0 | 0.01 | 0.02 |
| 2 | 10923700 | rs1060005 | 2.214 | -0.32 | 0.576 | -0.141 | -0.016 | -0.024 | -0.024 | 0 | 0.05 | 0.06 | 0.01 | 0 | 0.01 | 0.02 |
| 2 | 10924524 | rs1198873 | -0.511 | -0.285 | 0.807 | -0.109 | -0.001 | 0.003 | 0.003 | 0 | 0.58 | 0.55 | 0.55 | 0.47 | 0.59 | 0.63 |
| 2 | 10925406 | rs6756520 | -1.109 | -0.168 | 0.896 | -0.065 | 0.069 | -0.009 | -0.009 | 0.05 | 0.08 | 0.41 | 0.08 | 0.02 | 0.11 | 0.19 |
| 2 | 10926767 | rs17455708 | 0.678 | 0.195 | 0.706 | -0.103 | -0.068 | 0.045 | 0.045 | 0 | 0.19 | 0.01 | 0.15 | 0.06 | 0.21 | 0.07 |
| 2 | 10930903 | rs4807 | 1.215 | 0.205 | 0.195 | -0.077 | 0.014 | -0.043 | -0.043 | 0 | 0.31 | 0.11 | 0.34 | 0.39 | 0.27 | 0.38 |
| 2 | 10933249 | rs4799 | 0.157 | 0.247 | 0.21 | -0.089 | -0.021 | 0.006 | 0.006 | 0 | 0.51 | 0.16 | 0.48 | 0.46 | 0.48 | 0.46 |
| 2 | 10936092 | rs1734340 | 0.612 | 0.219 | 0.196 | -0.118 | 0.005 | -0.012 | -0.012 | 0 | 0.37 | 0.19 | 0.39 | 0.41 | 0.38 | 0.57 |
| 2 | 10937037 | rs1734343 | 0.171 | 0.279 | 0.322 | -0.034 | -0.02 | 0.008 | 0.008 | 0 | 0.5 | 0.16 | 0.47 | 0.45 | 0.47 | 0.45 |
| 2 | 10942687 | rs4758 | -0.294 | 0.296 | 0.338 | -0.006 | 0 | -0.021 | -0.021 | 0 | 0.56 | 0.2 | 0.52 | 0.46 | 0.58 | 0.64 |
| 2 | 10956867 | rs10197595 | 0.312 | 0.327 | 0.82 | -0.247 | 0.074 | -0.004 | -0.004 | 0.803 | 0.06 | 0.25 | 0.07 | 0.01 | 0.11 | 0.17 |

| | | | | | | | | | | | | | | | | |
|---|---|---|---|---|---|---|---|---|---|---|---|---|---|---|---|---|
| 2 | 10958235 | rs12463454 | 0.848 | 0.361 | 0.327 | -0.327 | -0.063 | 0.038 | 0.038 | 0 | 0.17 | 0.01 | 0.13 | 0.06 | 0.19 | 0.05 |
| 2 | 10963420 | rs1626087 | -0.835 | 0.26 | 0.551 | -0.291 | -0.001 | -0.007 | -0.007 | 0 | 0.63 | 0.6 | 0.68 | 0.81 | 0.64 | 0.72 |
| 2 | 10963527 | rs1106184 | -1.408 | 0.206 | -0.151 | -0.322 | -0.063 | 0.059 | 0.059 | 0 | 0.7 | 0.88 | 0.7 | 0.81 | 0.65 | 0.74 |
| 2 | 10963570 | rs1106185 | 0.624 | 0.264 | -0.203 | -0.209 | 0.019 | 0.036 | 0.036 | 0 | 0.38 | 0.24 | 0.48 | 0.75 | 0.34 | 0.5 |
| 2 | 10969328 | rs1734444 | -0.106 | 0.327 | -0.632 | -0.066 | -0.029 | 0.155 | 0.155 | 0 | 0.55 | 0.14 | 0.59 | 0.8 | 0.53 | 0.56 |
| 2 | 10970124 | rs17365477 | 0.602 | 0.373 | -0.133 | 0.29 | -0.011 | 0.064 | 0.064 | 0 | 0.44 | 0.37 | 0.41 | 0.24 | 0.5 | 0.3 |
| 2 | 10970473 | rs1686484 | 0.233 | 0.245 | -0.346 | 0.971 | 0.002 | -0.002 | -0.002 | 0.066 | 0.35 | 0.33 | 0.41 | 0.44 | 0.28 | 0.41 |
| 2 | 10971017 | rs4669632 | -0.362 | 0.374 | -0.412 | 1.193 | -0.009 | 0.112 | 0.112 | 0 | 0.09 | 0.14 | 0.21 | 0.4 | 0.08 | 0.15 |
| 2 | 10976790 | rs4669633 | -0.343 | 0.6 | -0.846 | 0.348 | -0.009 | 0.026 | 0.026 | 0 | 0.45 | 0.29 | 0.44 | 0.58 | 0.5 | 0.62 |
| 2 | 10980653 | rs1623958 | 1.024 | 0.689 | -1.069 | -0.241 | 0.013 | -0.002 | -0.002 | 0 | 0.3 | 0.11 | 0.32 | 0.34 | 0.23 | 0.29 |
| 2 | 10984698 | rs1734383 | 0.528 | 0.265 | -0.309 | -0.258 | 0.003 | 0.035 | 0.035 | 0.956 | 0.7 | 0.37 | 0.54 | 0.49 | 0.65 | 0.63 |
| 2 | 10985364 | rs1734389 | 0.427 | 0.254 | -0.401 | -0.263 | 0.012 | 0.067 | 0.067 | 0.125 | 0.71 | 0.48 | 0.56 | 0.49 | 0.66 | 0.63 |
| 2 | 10986626 | rs1686468 | -0.226 | 0.656 | -0.438 | -0.293 | -0.01 | -0.004 | -0.004 | 0 | 0.62 | 0.49 | 0.64 | 0.4 | 0.68 | 0.54 |
| 2 | 10988588 | rs9287724 | 0.222 | 0.768 | -0.543 | -0.25 | 0.011 | -0.022 | -0.022 | 0 | 0.1 | 0.16 | 0.16 | 0.03 | 0.16 | 0.15 |
| 2 | 10990594 | rs2969891 | -0.073 | 0.525 | -0.516 | -0.07 | -0.01 | -0.004 | -0.004 | 0.134 | 0.62 | 0.5 | 0.63 | 0.41 | 0.68 | 0.54 |
| 2 | 10993273 | rs11692302 | 0.198 | 0.601 | -1.186 | -0.292 | -0.003 | 0.098 | 0.098 | 0 | 0.44 | 0.3 | 0.36 | 0.18 | 0.43 | 0.32 |
| 2 | 10993949 | rs1198855 | -0.463 | 0.475 | -1.158 | 0.057 | 0.009 | -0.011 | -0.011 | 0 | 0.82 | 0.84 | 0.75 | 0.53 | 0.86 | 0.71 |
| 2 | 10994937 | rs1198851 | 0.723 | 1.19 | -1.094 | 0.282 | -0.07 | -0.097 | -0.097 | 0 | 0.27 | 0.06 | 0.29 | 0.33 | 0.28 | 0.25 |
| 2 | 11000883 | rs1734337 | -0.188 | 1.056 | -1.003 | 0.368 | 0.003 | -0.004 | -0.004 | 0 | 0.39 | 0.51 | 0.56 | 0.68 | 0.39 | 0.35 |
| 2 | 11001132 | rs7586507 | 0.09 | 1.008 | -1.293 | 0.219 | -0.012 | 0.027 | 0.027 | 0.02 | 0.36 | 0.31 | 0.27 | 0.17 | 0.43 | 0.39 |
| 2 | 11001859 | rs16856756 | 0.508 | -0.215 | -1.48 | 0.339 | 0.02 | 0.043 | 0.043 | 0.025 | 0.28 | 0.19 | 0.18 | 0.13 | 0.23 | 0.23 |
| 2 | 11004489 | rs1318011 | 0.28 | -0.328 | -1.316 | 0.733 | 0 | 0.001 | 0.001 | 0 | 0.35 | 0.35 | 0.24 | 0.16 | 0.31 | 0.24 |
| 2 | 11004803 | rs1997523 | -0.493 | 0.356 | -0.884 | 1.229 | -0.012 | -0.008 | -0.008 | 0 | 0.43 | 0.65 | 0.51 | 0.51 | 0.51 | 0.55 |
| 2 | 11011192 | rs10178826 | -1.008 | -0.002 | -0.495 | 1.861 | -0.018 | 0.122 | 0.122 | 0 | 0.06 | 0.15 | 0.06 | 0.33 | 0.02 | 0.2 |
| 2 | 11016542 | rs2178740 | 0.468 | -0.316 | -0.495 | 1.304 | -0.001 | 0 | 0 | 0 | 0.55 | 0.41 | 0.67 | 0.51 | 0.63 | 0.41 |
| 2 | 11017815 | rs2357880 | 0.239 | -0.621 | 0.012 | 1.458 | 0.007 | 0.011 | 0.011 | 0 | 0.67 | 0.64 | 0.77 | 0.71 | 0.7 | 0.6 |
| 2 | 11018680 | rs1317981 | 0.238 | -0.68 | 0.191 | 1.534 | 0.004 | 0.004 | 0.004 | 0 | 0.67 | 0.64 | 0.7 | 0.67 | 0.69 | 0.6 |
| 2 | 11019047 | rs1574182 | 0.152 | -0.594 | 0.182 | 1.587 | 0.007 | 0.004 | 0.004 | 0 | 0.68 | 0.67 | 0.71 | 0.66 | 0.72 | 0.63 |
| 2 | 11021849 | rs4669639 | -1.297 | -0.972 | 0.291 | 1.501 | -0.035 | 0.061 | 0.061 | 0.709 | 0.16 | 0.33 | 0.14 | 0.21 | 0.11 | 0.24 |
| 2 | 11023436 | rs12471848 | 1.394 | -1.084 | -0.431 | 0.982 | -0.066 | -0.061 | -0.061 | 0.065 | 0.44 | 0.09 | 0.53 | 0.47 | 0.54 | 0.36 |
| 2 | 11024358 | rs1922717 | -0.707 | -1.484 | -0.214 | 1.084 | 0.001 | -0.001 | -0.001 | 0.202 | 0.86 | 0.84 | 0.8 | 0.85 | 0.88 | 0.77 |
| 2 | 11024620 | rs1317768 | -1.296 | -1.711 | 1.007 | 0.919 | -0.035 | -0.071 | -0.071 | 0 | 0.14 | 0.42 | 0.09 | 0.02 | 0.06 | 0.03 |
| 2 | 11025664 | rs4365435 | -2.295 | -1.911 | 1.002 | 0.862 | 0.047 | -0.033 | -0.033 | 0 | 0.34 | 0.83 | 0.35 | 0.4 | 0.2 | 0.26 |
| 2 | 11025705 | rs7576288 | -0.615 | -1.909 | 1.188 | 0.912 | 0.01 | 0.14 | 0.14 | 0.667 | 0.06 | 0.17 | 0.11 | 0.22 | 0.04 | 0.1 |
| 2 | 11025731 | rs2357883 | -1.16 | -1.922 | 0.99 | 0.96 | 0.002 | -0.026 | -0.026 | 0 | 0.87 | 0.96 | 0.81 | 0.84 | 0.88 | 0.8 |
| 2 | 11026710 | rs7570842 | -2.367 | -1.929 | 1.057 | 0.934 | 0.043 | -0.03 | -0.03 | 0 | 0.34 | 0.8 | 0.35 | 0.4 | 0.2 | 0.27 |
| 2 | 11031430 | rs1317761 | 2.228 | -1.956 | 1.052 | 0.992 | 0.09 | 0.006 | 0.006 | 1.411 | 0.53 | 0.11 | 0.38 | 0.43 | 0.69 | 0.53 |
| 2 | 11031548 | rs7586618 | -0.657 | -1.994 | 1.596 | 0.968 | -0.021 | 0.001 | 0.001 | 0 | 0.13 | 0.29 | 0.09 | 0.16 | 0.07 | 0.15 |
| 2 | 11031788 | rs1882646 | -2.072 | -2.015 | 1.702 | 0.957 | 0.011 | 0.011 | 0.011 | 1.035 | 0.27 | 0.76 | 0.17 | 0.18 | 0.14 | 0.19 |
| 2 | 11033225 | rs12622387 | -1.783 | -2.001 | 2.257 | 0.991 | -0.039 | 0.124 | 0.124 | 0.043 | 0.23 | 0.49 | 0.27 | 0.39 | 0.15 | 0.23 |
| 2 | 11033567 | rs4233891 | 0.846 | -1.975 | 1.853 | 0.862 | 0.002 | -0.001 | -0.001 | 0 | 0.72 | 0.68 | 0.64 | 0.66 | 0.8 | 0.63 |
| 2 | 11038833 | rs13387419 | -1.944 | -1.713 | 1.764 | 0.62 | -0.043 | -0.109 | -0.109 | 0.112 | 0.2 | 0.62 | 0.13 | 0.02 | 0.11 | 0.13 |
| 2 | 11039291 | rs7605799 | -1.45 | -1.626 | 1.591 | 0.811 | -0.011 | -0.034 | -0.034 | 0.076 | 0.11 | 0.41 | 0.08 | 0.04 | 0.04 | 0.1 |
| 2 | 11046638 | rs2091817 | 1.157 | -1.386 | 0.852 | 0.888 | 0.03 | -0.037 | -0.037 | 0 | 0.61 | 0.27 | 0.39 | 0.52 | 0.69 | 0.6 |
| 2 | 11055298 | rs17365678 | 0.37 | -0.933 | 0.443 | 0.364 | -0.005 | 0.023 | 0.023 | 0 | 0.09 | 0.03 | 0.03 | 0 | 0.08 | 0.15 |
| 2 | 11055328 | rs12623956 | -0.131 | -0.884 | 0.898 | 1.013 | 0.028 | 0.16 | 0.16 | 0.18 | 0.05 | 0.14 | 0.19 | 0.2 | 0.06 | 0.07 |
| 2 | 11056828 | rs13425611 | -0.852 | -1.013 | 1.023 | 1.015 | -0.025 | -0.011 | -0.011 | 0 | 0.27 | 0.37 | 0.19 | 0.3 | 0.18 | 0.29 |
| 2 | 11059730 | rs4668701 | -1.188 | -1.074 | 0.466 | 0.729 | 0.002 | 0.04 | 0.04 | 0 | 0.87 | 0.98 | 0.85 | 0.97 | 0.85 | 0.81 |
| 2 | 11061353 | rs921605 | 2.36 | -1.002 | 0.681 | 0.762 | -0.062 | -0.037 | -0.037 | 0.09 | 0.43 | 0.13 | 0.41 | 0.46 | 0.52 | 0.29 |
| 2 | 11064348 | rs4669647 | 0.105 | -1.231 | 1.261 | 0.78 | 0.003 | -0.02 | -0.02 | 0 | 0.8 | 0.66 | 0.75 | 0.87 | 0.79 | 0.75 |
| 2 | 11066699 | rs1453084 | 0.425 | -1.332 | 1.139 | 0.971 | -0.054 | 0.174 | 0.174 | 0.623 | 0.78 | 0.54 | 0.63 | 0.49 | 0.88 | 0.71 |
| 2 | 11072956 | rs1467993 | 1.695 | -1.264 | 0.75 | 0.923 | 0.021 | 0.055 | 0.055 | 0.176 | 0.48 | 0.28 | 0.35 | 0.36 | 0.57 | 0.31 |
| 2 | 11073209 | rs1453087 | -0.511 | -0.863 | -0.31 | 0.452 | -0.012 | 0.041 | 0.041 | 0 | 0.26 | 0.08 | 0.25 | 0.14 | 0.27 | 0.4 |
| 2 | 11075317 | rs12692423 | -0.726 | -0.802 | -0.261 | 0.456 | -0.016 | 0.04 | 0.04 | 0 | 0.28 | 0.18 | 0.27 | 0.13 | 0.3 | 0.39 |
| 2 | 11081549 | rs6758252 | 0.608 | -2.118 | -0.582 | 0.392 | 0.116 | 0.073 | 0.073 | 1.02 | 0.55 | 0.22 | 0.64 | 0.72 | 0.77 | 0.71 |
| 2 | 11081916 | rs4669652 | 0.207 | -2.527 | 0.09 | 0.174 | -0.028 | -0.016 | -0.016 | 0 | 0.71 | 0.69 | 0.85 | 0.86 | 0.84 | 0.76 |
| 2 | 11083259 | rs4669653 | 1.96 | -2.749 | -0.347 | 0.222 | 0.01 | -0.053 | -0.053 | 0 | 0.38 | 0.12 | 0.51 | 0.56 | 0.62 | 0.39 |
| 2 | 11083409 | rs6727014 | 0.096 | -2.798 | -0.164 | 0.131 | -0.01 | -0.041 | -0.041 | 0 | 0.15 | 0.18 | 0.06 | 0.01 | 0.09 | 0.16 |
| 2 | 11083550 | rs1869288 | -0.67 | -2.811 | -0.356 | 0.146 | -0.002 | 0.001 | 0.001 | 0 | 0.78 | 0.79 | 0.9 | 0.88 | 0.88 | 0.8 |
| 2 | 11084495 | rs12620687 | -0.607 | -2.403 | -0.013 | 0.204 | -0.001 | 0.001 | 0.001 | 0.128 | 0.1 | 0.14 | 0.15 | 0.17 | 0.07 | 0.09 |
| 2 | 11085744 | rs4442966 | -1.64 | -2.265 | -0.224 | 0.253 | 0.023 | 0.017 | 0.017 | 0.126 | 0.39 | 0.67 | 0.39 | 0.32 | 0.26 | 0.41 |
| 2 | 11085940 | rs11680593 | -1.595 | -2.241 | -0.276 | 0.265 | 0 | 0 | 0 | 0.058 | 0.39 | 0.64 | 0.38 | 0.32 | 0.26 | 0.41 |
| 2 | 11087314 | rs13386117 | -0.014 | -2.032 | -0.446 | 0.444 | 0.02 | -0.004 | -0.004 | 0 | 0.16 | 0.07 | 0.13 | 0.18 | 0.05 | 0.21 |
| 2 | 11087429 | rs9677245 | -0.79 | -2.279 | 0.01 | 0.333 | -0.022 | 0.025 | 0.025 | 0.903 | 0.23 | 0.5 | 0.14 | 0.23 | 0.19 | 0.28 |
| 2 | 11091780 | rs6730326 | -0.848 | -2 | -0.307 | 0.204 | -0.002 | 0 | 0 | 0 | 0.3 | 0.31 | 0.25 | 0.31 | 0.25 | 0.27 |
| 2 | 11092613 | rs10199787 | -2.107 | -1.588 | -0.214 | 0.263 | -0.022 | 0.002 | 0.002 | 0 | 0.62 | 0.76 | 0.6 | 0.62 | 0.41 | 0.64 |
| 2 | 11093401 | rs10202936 | -1.74 | -0.833 | 0.063 | 0.843 | -0.003 | 0 | 0 | 0 | 0.39 | 0.39 | 0.5 | 0.6 | 0.31 | 0.51 |
| 2 | 11093430 | rs10202946 | -0.967 | -0.58 | -0.523 | 1.088 | 0.002 | -0.006 | -0.006 | 0 | 0.25 | 0.21 | 0.41 | 0.56 | 0.19 | 0.36 |
| 2 | 11099093 | rs6432153 | -0.672 | 0.027 | -0.55 | 0.784 | -0.005 | -0.01 | -0.01 | 0 | 0.27 | 0.2 | 0.37 | 0.39 | 0.27 | 0.34 |

| | | | | | | | | | | | | | | | | |
|---|---|---|---|---|---|---|---|---|---|---|---|---|---|---|---|---|
| 2 | 11100051 | rs6741453 | -0.854 | -0.199 | -0.678 | 0.499 | 0.02 | 0.001 | 0.001 | 0 | 0.51 | 0.62 | 0.57 | 0.5 | 0.39 | 0.55 |
| 2 | 11103159 | rs6432155 | 0.305 | 0.249 | -0.597 | 0.037 | 0 | -0.001 | -0.001 | 0 | 0.46 | 0.53 | 0.36 | 0.52 | 0.59 | 0.48 |
| 2 | 11103409 | rs13424719 | -0.091 | 0.19 | -0.504 | -0.202 | -0.002 | 0.013 | 0.013 | 0 | 0.35 | 0.37 | 0.55 | 0.44 | 0.32 | 0.48 |
| 2 | 11103496 | rs4669654 | 0.445 | 0.13 | -0.117 | -0.456 | -0.016 | -0.066 | -0.066 | 0 | 0.79 | 0.71 | 0.88 | 0.95 | 0.87 | 0.86 |
| 2 | 11107245 | rs6748518 | 0.691 | 0.24 | -0.22 | -0.479 | -0.022 | -0.114 | -0.114 | 1.525 | 0.77 | 0.62 | 0.84 | 0.98 | 0.81 | 0.91 |
| 2 | 11113531 | rs4669658 | -0.679 | 0.524 | -1.02 | 0.013 | 0.007 | -0.029 | -0.029 | 0.058 | 0.47 | 0.82 | 0.64 | 0.54 | 0.4 | 0.58 |
| 2 | 11113859 | rs6744989 | -0.463 | 0.422 | -0.464 | 0.225 | -0.014 | 0.019 | 0.019 | 0 | 0.23 | 0.58 | 0.35 | 0.31 | 0.16 | 0.34 |
| 2 | 11117817 | rs4143555 | -0.384 | 0.311 | 0.602 | 0.421 | -0.033 | 0.112 | 0.112 | 0 | 0.16 | 0.37 | 0.39 | 0.36 | 0.06 | 0.2 |
| 2 | 11118277 | rs6432157 | -1.373 | 0.643 | 1.002 | 0.323 | -0.005 | 0.013 | 0.013 | 0 | 0.37 | 0.71 | 0.38 | 0.28 | 0.39 | 0.44 |
| 2 | 11118410 | rs6432158 | -1.954 | 0.651 | 0.949 | 0.329 | -0.018 | -0.031 | -0.031 | 0 | 0.47 | 0.82 | 0.65 | 0.54 | 0.43 | 0.58 |
| 2 | 11118836 | rs13390165 | -0.967 | 0.595 | 1.007 | 0.315 | -0.037 | 0.068 | 0.068 | 0 | 0.16 | 0.43 | 0.31 | 0.26 | 0.06 | 0.17 |
| 2 | 11120801 | rs896757 | -1.388 | 0.555 | 1.258 | 0.247 | -0.006 | 0.075 | 0.075 | 0 | 0.21 | 0.42 | 0.42 | 0.41 | 0.15 | 0.36 |
| 2 | 11121074 | rs896754 | -1.773 | 0.513 | 1.174 | 0.157 | 0.002 | 0.019 | 0.019 | 0 | 0.4 | 0.51 | 0.57 | 0.53 | 0.36 | 0.55 |
| 2 | 11123515 | rs10495577 | -1.396 | 0.51 | 1.05 | 0.206 | -0.007 | 0.058 | 0.058 | 0.014 | 0.21 | 0.38 | 0.42 | 0.41 | 0.15 | 0.36 |
| 2 | 11126753 | rs754798 | -1.973 | 0.544 | 0.86 | 0.127 | -0.014 | -0.035 | -0.035 | 0 | 0.47 | 0.79 | 0.64 | 0.53 | 0.42 | 0.58 |
| 2 | 11127705 | rs7573335 | -0.759 | 0.451 | 1.05 | 0.156 | 0.016 | -0.017 | -0.017 | 0 | 0.1 | 0.27 | 0.05 | 0.05 | 0.1 | 0.2 |
| 2 | 11131612 | rs2357885 | -0.852 | 1.043 | 0.853 | 0.492 | 0.008 | 0.045 | 0.045 | 0 | 0.33 | 0.38 | 0.45 | 0.46 | 0.37 | 0.46 |
| 2 | 11132304 | rs6739736 | -1.077 | 0.232 | -0.196 | -0.059 | -0.01 | -0.025 | -0.025 | 0 | 0.64 | 0.73 | 0.63 | 0.55 | 0.65 | 0.8 |
| 2 | 11134604 | rs4668703 | -0.945 | 0.555 | -0.229 | -0.061 | 0.005 | -0.032 | -0.032 | 0.12 | 0.65 | 0.74 | 0.64 | 0.53 | 0.68 | 0.81 |
| 2 | 11142721 | rs1901753 | -0.276 | 0.728 | 0.401 | 0.253 | 0.041 | -0.009 | -0.009 | 0 | 0.31 | 0.4 | 0.28 | 0.23 | 0.46 | 0.44 |
| 2 | 11144390 | rs16857004 | 0.174 | 0.37 | 0.244 | 0.425 | -0.025 | 0.071 | 0.071 | 0 | 0.07 | 0.07 | 0.2 | 0.28 | 0.01 | 0.13 |
| 2 | 11147253 | rs11682194 | -0.467 | 0.518 | 0.13 | 0.282 | 0.002 | -0.007 | -0.007 | 0 | 0.1 | 0.05 | 0.23 | 0.21 | 0.07 | 0.12 |
| 2 | 11147985 | rs16857022 | -0.062 | 0.759 | 0.543 | 0.218 | 0.055 | 0.036 | 0.036 | 0 | 0.06 | 0.2 | 0.04 | 0.05 | 0.08 | 0.15 |
| 2 | 11150378 | rs10520062 | -1.472 | 0.386 | 0.843 | 0.001 | 0.012 | -0.026 | -0.026 | 0.155 | 0.14 | 0.52 | 0.08 | 0.06 | 0.12 | 0.18 |
| 2 | 11151597 | rs735206 | -0.675 | 0.388 | 0.215 | -0.134 | 0.031 | 0.032 | 0.032 | 0 | 0.49 | 0.65 | 0.6 | 0.55 | 0.64 | 0.61 |
| 2 | 11152503 | rs4668705 | 0.117 | 0.615 | 0.27 | -0.112 | -0.002 | -0.007 | -0.007 | 0 | 0.44 | 0.47 | 0.44 | 0.23 | 0.41 | 0.35 |
| 2 | 11154270 | rs12472523 | 0.689 | 0.601 | -0.326 | -0.128 | 0.02 | 0.049 | 0.049 | 0.804 | 0.37 | 0.16 | 0.37 | 0.23 | 0.33 | 0.33 |
| 2 | 11154991 | rs4668706 | 0.547 | 0.532 | -0.265 | -0.158 | -0.016 | -0.015 | -0.015 | 0 | 0.3 | 0.25 | 0.27 | 0.37 | 0.41 | 0.47 |
| 2 | 11158072 | rs6733287 | -0.448 | 0.484 | -0.205 | -0.249 | -0.003 | -0.003 | -0.003 | 0 | 0.46 | 0.51 | 0.4 | 0.38 | 0.59 | 0.54 |
| 2 | 11159080 | rs6432163 | -1.465 | 0.653 | -0.808 | 0.149 | -0.017 | -0.086 | -0.086 | 0 | 0.87 | 0.98 | 0.79 | 0.6 | 0.92 | 0.87 |
| 2 | 11160768 | rs12053436 | 0.399 | 0.818 | -0.572 | 0.079 | -0.043 | -0.048 | -0.048 | 0 | 0.31 | 0.13 | 0.26 | 0.38 | 0.42 | 0.47 |
| 2 | 11161544 | rs11903722 | 0.016 | 0.801 | -0.617 | 0.023 | -0.059 | -0.04 | -0.04 | 0 | 0.34 | 0.15 | 0.28 | 0.38 | 0.47 | 0.48 |
| 2 | 11162280 | rs10176051 | 0.177 | 0.872 | -0.037 | -0.251 | -0.003 | 0.015 | 0.015 | 0 | 0.53 | 0.63 | 0.4 | 0.19 | 0.5 | 0.31 |
| 2 | 11162735 | rs1349722 | 0.341 | 1.014 | -0.268 | -0.113 | -0.016 | -0.021 | -0.021 | 2.625 | 0.3 | 0.2 | 0.33 | 0.32 | 0.42 | 0.37 |
| 2 | 11164636 | rs4668708 | 0.793 | 0.933 | -0.442 | -0.093 | -0.023 | -0.075 | -0.075 | 1.695 | 0.26 | 0.09 | 0.28 | 0.33 | 0.33 | 0.36 |
| 2 | 11165291 | rs2357887 | 0.758 | 0.433 | -0.582 | -0.567 | 0.042 | 0.034 | 0.034 | 1.021 | 0.51 | 0.2 | 0.35 | 0.33 | 0.45 | 0.48 |
| 2 | 11175395 | rs11689191 | 0.082 | 0.607 | -0.263 | -0.591 | 0.013 | 0.006 | 0.006 | 0.593 | 0.66 | 0.38 | 0.53 | 0.51 | 0.62 | 0.59 |
| 2 | 11180113 | rs4669682 | -0.939 | 0.683 | 0.15 | -0.586 | 0.01 | 0.048 | 0.048 | 0 | 0.78 | 0.59 | 0.63 | 0.57 | 0.79 | 0.77 |
| 2 | 11182136 | rs6432164 | -1.285 | 0.67 | -0.155 | -0.589 | -0.028 | -0.058 | -0.058 | 4.655 | 0.81 | 0.87 | 0.66 | 0.57 | 0.8 | 0.77 |
| 2 | 11183443 | rs4668711 | -1.325 | 0.682 | -0.398 | -0.583 | -0.035 | -0.067 | -0.067 | 0 | 0.81 | 0.91 | 0.66 | 0.57 | 0.8 | 0.77 |
| 2 | 11184914 | rs2357888 | 0.504 | 0.696 | -0.388 | -0.603 | 0 | -0.001 | -0.001 | 0 | 0.6 | 0.39 | 0.47 | 0.5 | 0.5 | 0.56 |
| 2 | 11188257 | rs10168758 | 1.786 | 0.629 | -0.828 | -0.474 | -0.002 | -0.017 | -0.017 | 3.067 | 0.09 | 0.05 | 0.14 | 0.18 | 0.1 | 0.09 |
| 2 | 11189849 | rs11677588 | 0.887 | 0.477 | -0.907 | -0.613 | 0.05 | 0.053 | 0.053 | 0.993 | 0.33 | 0.15 | 0.21 | 0.18 | 0.21 | 0.2 |
| 2 | 11195795 | rs2100369 | -0.723 | 0.801 | -0.535 | -0.524 | -0.034 | -0.074 | -0.074 | 0.024 | 0.69 | 0.85 | 0.54 | 0.44 | 0.68 | 0.73 |
| 2 | 11200347 | rs7564899 | 0.123 | 0.89 | -0.053 | -0.577 | 0.007 | 0.008 | 0.008 | 0 | 0.59 | 0.46 | 0.43 | 0.4 | 0.53 | 0.58 |
| 2 | 11205359 | rs10197275 | -0.833 | 1.108 | 0.107 | -0.458 | 0.107 | -0.017 | -0.017 | 0 | 0.11 | 0.35 | 0.11 | 0.04 | 0.15 | 0.15 |
| 2 | 11205421 | rs16857117 | -0.685 | 1.156 | 0.111 | -0.459 | 0.086 | 0.044 | 0.044 | 0.125 | 0.07 | 0.34 | 0.05 | 0.04 | 0.06 | 0.13 |
| 2 | 11210087 | rs11892303 | 0.613 | 1.274 | -0.846 | -0.441 | -0.032 | 0.026 | 0.026 | 0 | 0.05 | 0.01 | 0.06 | 0 | 0.11 | 0.02 |
| 2 | 11212039 | rs7577930 | -0.123 | 0.808 | -0.79 | -0.595 | -0.01 | -0.02 | -0.02 | 0 | 0.6 | 0.69 | 0.5 | 0.4 | 0.61 | 0.59 |
| 2 | 11213936 | rs4669685 | 0.661 | 0.647 | -0.61 | -0.416 | -0.006 | 0.064 | 0.064 | 0 | 0.25 | 0.34 | 0.28 | 0.37 | 0.29 | 0.38 |
| 2 | 11215734 | rs6432168 | -0.985 | 0.624 | -1.38 | -0.663 | 0.005 | -0.017 | -0.017 | 0 | 0.52 | 0.8 | 0.46 | 0.42 | 0.57 | 0.65 |
| 2 | 11220562 | rs6739054 | 0.079 | 0.509 | -1.626 | -0.387 | 0.002 | -0.094 | -0.094 | 0 | 0.31 | 0.08 | 0.37 | 0.4 | 0.32 | 0.45 |
| 2 | 11223628 | rs1073201 | 0.488 | 0.504 | -1.094 | -0.834 | 0 | 0.009 | 0.009 | 0 | 0.48 | 0.47 | 0.5 | 0.54 | 0.49 | 0.56 |
| 2 | 11223962 | rs13019468 | 1.12 | 0.706 | -1.043 | -0.781 | -0.015 | 0.021 | 0.021 | 0 | 0.05 | 0.02 | 0.04 | 0 | 0.08 | 0.02 |
| 2 | 11226507 | rs4668712 | 0.843 | 0.922 | -0.806 | -0.812 | 0.01 | -0.048 | -0.048 | 0.914 | 0.82 | 0.73 | 0.88 | 0.92 | 0.85 | 0.76 |
| 2 | 11229946 | rs4669687 | -0.102 | 0.744 | -0.851 | -0.826 | -0.004 | -0.03 | -0.03 | 0 | 0.89 | 0.76 | 0.93 | 0.94 | 0.95 | 0.8 |
| 2 | 11233323 | rs4669688 | -0.147 | 0.87 | -1.146 | -0.864 | -0.001 | 0 | 0 | 0 | 0.62 | 0.64 | 0.58 | 0.55 | 0.62 | 0.59 |
| 2 | 11237119 | rs7568379 | 0.33 | 0.825 | -1.832 | -1.001 | 0.025 | 0.064 | 0.064 | 0.028 | 0.47 | 0.26 | 0.48 | 0.55 | 0.47 | 0.51 |
| 2 | 11237705 | rs11678376 | 0.374 | 1.502 | -1.406 | -0.61 | -0.02 | 0.019 | 0.019 | 0 | 0.06 | 0.03 | 0.05 | 0 | 0.11 | 0.01 |
| 2 | 11242204 | rs908275 | 1.009 | 1.44 | -1.823 | -0.677 | 0.018 | -0.042 | -0.042 | 0.166 | 0.47 | 0.14 | 0.4 | 0.54 | 0.4 | 0.43 |
| 2 | 11243387 | rs866995 | -0.467 | 2.096 | -2.173 | -1.017 | -0.083 | -0.11 | -0.11 | 0 | 0.26 | 0.06 | 0.38 | 0.5 | 0.34 | 0.39 |
| 2 | 11248242 | rs12478033 | 2.059 | 1.351 | -1.994 | -0.924 | -0.015 | -0.043 | -0.043 | 0.651 | 0.37 | 0.08 | 0.35 | 0.41 | 0.33 | 0.32 |
| 2 | 11250697 | rs10929716 | 0.056 | 1.527 | -1.649 | -0.909 | -0.015 | -0.055 | -0.055 | 0 | 0.25 | 0.14 | 0.36 | 0.47 | 0.29 | 0.37 |
| 2 | 11257494 | rs9287725 | 1.761 | 1.538 | -1.556 | -1.006 | -0.04 | 0.031 | 0.031 | 0.054 | 0.45 | 0.16 | 0.42 | 0.42 | 0.47 | 0.34 |
| 2 | 11257505 | rs16857178 | -0.5 | 1.182 | -0.355 | -0.78 | 0.038 | 0.023 | 0.023 | 5.739 | 0.07 | 0.44 | 0.05 | 0.01 | 0.02 | 0.06 |
| 2 | 11257726 | rs6731491 | 1.188 | 1.228 | -0.783 | -0.759 | 0.006 | 0.014 | 0.014 | 1.563 | 0.7 | 0.3 | 0.78 | 0.89 | 0.75 | 0.7 |
| 2 | 11259067 | rs6750442 | -1.107 | 1.524 | -0.776 | -0.859 | -0.001 | -0.008 | -0.008 | 2.899 | 0.37 | 0.23 | 0.41 | 0.55 | 0.38 | 0.4 |
| 2 | 11259605 | rs6739182 | -1.705 | 1.436 | -0.236 | -0.839 | 0.001 | 0.012 | 0.012 | 0 | 0.48 | 0.56 | 0.49 | 0.57 | 0.48 | 0.45 |
| 2 | 11262066 | rs12614177 | -0.406 | 1.07 | -0.55 | -0.508 | 0.017 | -0.054 | -0.054 | 0 | 0.27 | 0.15 | 0.34 | 0.49 | 0.23 | 0.38 |

| | | | | | | | | | | | | | | | | |
|---|---|---|---|---|---|---|---|---|---|---|---|---|---|---|---|---|
| 2 | 11263948 | rs4669694 | 0.375 | 1.362 | 0.297 | -0.803 | 0.009 | -0.058 | -0.058 | 0.043 | 0.88 | 0.84 | 0.89 | 1 | 0.87 | 0.88 |
| 2 | 11264444 | rs6752604 | 0.891 | 1.333 | 0.757 | -0.865 | 0.009 | -0.071 | -0.071 | 0 | 0.86 | 0.76 | 0.84 | 1 | 0.84 | 0.88 |
| 2 | 11264486 | rs6737910 | -1.398 | 1.471 | 1.094 | -0.953 | 0.028 | -0.091 | -0.091 | 0 | 0.13 | 0.48 | 0.1 | 0 | 0.09 | 0.02 |
| 2 | 11264830 | rs7560331 | 1.703 | 1.631 | 0.493 | -1.065 | 0.044 | -0.163 | -0.163 | 1.596 | 0.7 | 0.5 | 0.73 | 1 | 0.65 | 0.86 |
| 2 | 11265972 | rs7608117 | 1.693 | 1.489 | 0.151 | -0.535 | 0.022 | -0.003 | -0.003 | 0 | 0.51 | 0.24 | 0.5 | 0.55 | 0.49 | 0.51 |
| 2 | 11272576 | rs10495579 | -0.386 | 1.614 | -0.547 | -0.668 | -0.048 | 0.041 | 0.041 | 0 | 0.1 | 0.03 | 0.07 | 0 | 0.21 | 0.03 |
| 2 | 11272784 | rs17366322 | 2.041 | 1.647 | -0.644 | -0.602 | -0.051 | -0.106 | -0.106 | 0.612 | 0.47 | 0.03 | 0.46 | 0.53 | 0.46 | 0.52 |
| 2 | 11273757 | rs6733410 | -1.128 | 1.231 | 0.247 | -0.227 | 0.006 | 0.048 | 0.048 | 4.162 | 0.08 | 0.57 | 0.05 | 0.04 | 0.02 | 0.01 |
| 2 | 11273813 | rs6751137 | -0.246 | 1.267 | -0.016 | -0.279 | -0.021 | 0.026 | 0.026 | 0 | 0.93 | 0.98 | 0.89 | 1 | 0.9 | 0.94 |
| 2 | 11283965 | rs6432178 | -1.323 | 1.019 | 0.092 | -0.195 | -0.013 | 0.026 | 0.026 | 6.145 | 0.08 | 0.61 | 0.06 | 0.04 | 0.02 | 0.01 |
| 2 | 11284415 | rs6713473 | -1.81 | 0.997 | 0.097 | -0.207 | -0.037 | -0.002 | -0.002 | 0.075 | 0.1 | 0.65 | 0.06 | 0.04 | 0.02 | 0.01 |
| 2 | 11284768 | rs6717030 | -1.316 | 0.998 | 0.098 | -0.207 | -0.015 | 0.024 | 0.024 | 6.145 | 0.08 | 0.62 | 0.06 | 0.04 | 0.02 | 0.01 |
| 2 | 11284866 | rs6756515 | -1.316 | 0.998 | 0.098 | -0.207 | -0.015 | 0.024 | 0.024 | 6.145 | 0.08 | 0.62 | 0.06 | 0.04 | 0.02 | 0.01 |
| 2 | 11285016 | rs6717388 | -1.316 | 0.998 | 0.098 | -0.194 | -0.014 | 0.03 | 0.03 | 6.145 | 0.08 | 0.61 | 0.06 | 0.05 | 0.02 | 0.01 |
| 2 | 11285693 | rs908276 | -1.316 | 0.998 | 0.094 | -0.213 | -0.015 | 0.024 | 0.024 | 6.145 | 0.08 | 0.62 | 0.06 | 0.04 | 0.02 | 0.01 |
| 2 | 11286090 | rs11903207 | -1.489 | 1.007 | 0.002 | -0.208 | -0.037 | -0.002 | -0.002 | 0.239 | 0.1 | 0.65 | 0.06 | 0.04 | 0.02 | 0.01 |
| 2 | 11288577 | rs12995564 | -0.503 | 1.127 | -0.495 | -0.237 | 0.012 | -0.047 | -0.047 | 0.23 | 0.28 | 0.16 | 0.31 | 0.47 | 0.24 | 0.39 |
| 2 | 11292605 | rs4669695 | -0.493 | 1.177 | -0.611 | -0.211 | 0.016 | -0.054 | -0.054 | 0.02 | 0.28 | 0.16 | 0.31 | 0.47 | 0.24 | 0.38 |
| 2 | 11293336 | rs7581322 | 1.447 | 1.238 | -0.938 | -0.077 | 0.017 | -0.048 | -0.048 | 0 | 0.45 | 0.09 | 0.43 | 0.51 | 0.43 | 0.5 |
| 2 | 11300732 | rs2271622 | 1.379 | 1.201 | -0.444 | 0.134 | 0.024 | 0.015 | 0.015 | 0.022 | 0.48 | 0.13 | 0.45 | 0.47 | 0.47 | 0.49 |
| 2 | 11300869 | rs6432182 | 0.05 | 1.178 | -0.547 | -0.049 | -0.028 | 0.048 | 0.048 | 0.077 | 0.9 | 0.99 | 0.88 | 1 | 0.88 | 0.88 |
| 2 | 11317226 | rs13018466 | 1.369 | 0.865 | -0.192 | 0.024 | -0.006 | 0.014 | 0.014 | 0 | 0.53 | 0.22 | 0.49 | 0.45 | 0.57 | 0.5 |
| 2 | 11319660 | rs921322 | 1.825 | 0.837 | -0.22 | 0.003 | -0.016 | 0.013 | 0.013 | 0.544 | 0.46 | 0.13 | 0.55 | 0.39 | 0.46 | 0.48 |
| 2 | 11320877 | rs8996 | 1.785 | 1.011 | -0.242 | -0.009 | -0.012 | 0.012 | 0.012 | 0.079 | 0.45 | 0.16 | 0.55 | 0.39 | 0.46 | 0.48 |
| 2 | 11323276 | rs978906 | 1.785 | 1.011 | -0.242 | -0.009 | -0.012 | 0.012 | 0.012 | 0.079 | 0.45 | 0.18 | 0.55 | 0.39 | 0.46 | 0.48 |
| 2 | 11353200 | rs7598032 | 1.327 | 0.997 | -0.101 | -0.017 | 0.006 | 0.03 | 0.03 | 0 | 0.5 | 0.26 | 0.58 | 0.45 | 0.55 | 0.49 |
| 2 | 11356190 | rs726843 | 1.888 | 0.985 | -0.133 | -0.015 | -0.047 | -0.014 | -0.014 | 0 | 0.44 | 0.09 | 0.54 | 0.39 | 0.46 | 0.43 |
| 2 | 11382829 | rs1868585 | -0.312 | 0.949 | 0.835 | -0.058 | -0.015 | -0.069 | -0.069 | 0.062 | 0.28 | 0.09 | 0.23 | 0.55 | 0.29 | 0.4 |
| 2 | 11385834 | rs6716817 | 1.997 | 0.96 | 0.842 | -0.08 | 0.008 | 0.015 | 0.015 | 0.019 | 0.48 | 0.31 | 0.58 | 0.39 | 0.53 | 0.48 |
| 2 | 11398736 | rs6760802 | 0.547 | 0.944 | 1.362 | -0.105 | -0.001 | -0.073 | -0.073 | 0 | 0.85 | 0.69 | 0.86 | 1 | 0.89 | 0.9 |
| 2 | 11401165 | rs6759490 | 1.443 | 0.95 | 1.351 | -0.106 | 0.009 | -0.088 | -0.088 | 0 | 0.8 | 0.55 | 0.83 | 0.94 | 0.8 | 0.89 |
| 2 | 11401538 | rs4669700 | -0.201 | 0.936 | 0.931 | -0.087 | 0.032 | -0.062 | -0.062 | 0.033 | 0.27 | 0.1 | 0.21 | 0.55 | 0.21 | 0.4 |
| 2 | 11411483 | rs10929728 | -0.79 | 0.888 | 0.907 | -0.084 | -0.001 | -0.012 | -0.012 | 0 | 0.33 | 0.25 | 0.27 | 0.55 | 0.34 | 0.41 |
| 2 | 11451984 | rs10203916 | 1.362 | 0.901 | 0.872 | -0.117 | -0.015 | -0.121 | -0.121 | 0.208 | 0.81 | 0.47 | 0.84 | 0.94 | 0.87 | 0.89 |
| 2 | 11465522 | rs753770 | 1.346 | 0.903 | 0.873 | -0.116 | -0.014 | -0.12 | -0.12 | 2.101 | 0.81 | 0.47 | 0.84 | 0.94 | 0.87 | 0.89 |
| 2 | 11489531 | rs7581184 | -1.228 | 0.932 | 0.691 | 0.281 | -0.002 | -0.005 | -0.005 | 0.02 | 0.37 | 0.34 | 0.23 | 0.46 | 0.41 | 0.43 |
| 2 | 11494852 | rs6432187 | -0.634 | 0.79 | 0.231 | 0.319 | 0.007 | -0.071 | -0.071 | 0 | 0.31 | 0.14 | 0.18 | 0.52 | 0.28 | 0.42 |
| 2 | 11499032 | rs6432188 | 0.03 | 0.996 | 0.371 | 0.157 | -0.013 | -0.003 | -0.003 | 0 | 0.89 | 0.91 | 0.81 | 0.9 | 0.83 | 0.93 |
| 2 | 11502656 | rs6705170 | 1.19 | 0.811 | 0.461 | 0.265 | 0 | 0 | 0 | 0 | 0.54 | 0.53 | 0.63 | 0.49 | 0.55 | 0.51 |
| 2 | 11503351 | rs7424263 | -1.019 | 0.216 | 0.456 | -0.036 | 0.023 | -0.087 | -0.087 | 0 | 0.17 | 0.32 | 0.18 | 0 | 0.21 | 0.08 |
| 2 | 11505213 | rs4533449 | 0.328 | 0.069 | 0.251 | 0.324 | -0.011 | -0.006 | -0.006 | 0 | 0.64 | 0.59 | 0.74 | 0.63 | 0.73 | 0.57 |
| 2 | 11509634 | rs4668723 | 0.653 | 0.227 | -0.745 | 0.499 | 0.041 | 0.078 | 0.078 | 0.117 | 0.54 | 0.19 | 0.66 | 0.6 | 0.63 | 0.56 |
| 2 | 11526716 | rs12466910 | 0.016 | 0 | -0.705 | 0.885 | 0.002 | 0.001 | 0.001 | 0 | 0.46 | 0.48 | 0.53 | 0.54 | 0.45 | 0.45 |
| 2 | 11529510 | rs11683268 | 0.3 | -0.177 | -1.077 | 1.255 | -0.068 | -0.066 | -0.066 | 0.017 | 0.39 | 0.03 | 0.38 | 0.37 | 0.37 | 0.42 |
| 2 | 11530369 | rs4669724 | 0.051 | -0.303 | -1.134 | 0.65 | -0.036 | -0.007 | -0.007 | 0 | 0.78 | 0.46 | 0.88 | 0.84 | 0.83 | 0.87 |
| 2 | 11531731 | rs6761608 | 0.013 | 0.213 | -1.326 | 0.839 | -0.057 | 0.033 | 0.033 | 0 | 0.36 | 0.16 | 0.47 | 0.28 | 0.45 | 0.43 |
| 2 | 11535523 | rs4260216 | 0.057 | -0.257 | -1.082 | 0.899 | 0 | -0.001 | -0.001 | 0 | 0.63 | 0.38 | 0.6 | 0.48 | 0.71 | 0.62 |
| 2 | 11540150 | rs12619687 | 0.45 | -0.297 | -1.487 | 1.08 | 0.02 | -0.065 | -0.065 | 0 | 0.22 | 0.06 | 0.12 | 0.39 | 0.17 | 0.2 |
| 2 | 11540489 | rs4669727 | 0.483 | 0.418 | -0.716 | 0.379 | 0.011 | -0.035 | -0.035 | 0.834 | 0.86 | 0.79 | 0.88 | 0.95 | 0.84 | 0.93 |
| 2 | 11546173 | rs10929735 | 1.126 | 0.121 | -0.63 | 0.299 | 0.014 | 0.018 | 0.018 | 1.286 | 0.72 | 0.23 | 0.78 | 0.79 | 0.81 | 0.9 |
| 2 | 11546354 | rs13418118 | -0.614 | 0.246 | -0.586 | -0.157 | -0.011 | 0.033 | 0.033 | 0 | 0.18 | 0.1 | 0.13 | 0.05 | 0.17 | 0.08 |
| 2 | 11548607 | rs7580173 | 0.031 | 0.783 | -0.443 | -0.481 | -0.025 | 0.058 | 0.058 | 0 | 0.38 | 0.25 | 0.55 | 0.73 | 0.4 | 0.55 |
| 2 | 11562630 | rs6708001 | 0.107 | 0.797 | -0.023 | -0.215 | 0.023 | 0.031 | 0.031 | 0 | 0.37 | 0.25 | 0.53 | 0.64 | 0.35 | 0.48 |
| 2 | 11572815 | rs11687681 | -0.278 | 0.743 | 0.864 | -0.213 | 0.002 | 0.002 | 0.002 | 0 | 0.44 | 0.63 | 0.58 | 0.62 | 0.38 | 0.47 |
| 2 | 11579054 | rs11686138 | -0.206 | 0.764 | 0.907 | -0.193 | 0.001 | 0.001 | 0.001 | 0 | 0.45 | 0.62 | 0.58 | 0.61 | 0.39 | 0.51 |
| 2 | 11580569 | rs13414905 | 0.424 | 1.2 | 0.916 | -0.349 | 0.023 | -0.024 | -0.024 | 0.029 | 0.14 | 0.14 | 0.17 | 0.04 | 0.21 | 0.09 |
| 2 | 11589362 | rs4292073 | -0.16 | 1.211 | 0.511 | -0.312 | -0.012 | -0.012 | -0.012 | 0 | 0.45 | 0.56 | 0.57 | 0.61 | 0.39 | 0.48 |
| 2 | 11610082 | rs6724463 | -0.537 | 0.557 | -0.122 | -0.019 | -0.024 | 0.034 | 0.034 | 0 | 0.6 | 0.92 | 0.73 | 0.64 | 0.58 | 0.61 |
| 2 | 11618069 | rs1561014 | -0.208 | 0.596 | -0.144 | -0.489 | 0.049 | 0.201 | 0.201 | 0.044 | 0.54 | 0.21 | 0.59 | 0.81 | 0.62 | 0.8 |
| 2 | 11619674 | rs4669730 | 0.111 | 0.106 | 0.638 | -0.292 | -0.002 | 0.005 | 0.005 | 0 | 0.46 | 0.35 | 0.6 | 0.7 | 0.49 | 0.66 |
| 2 | 11627410 | rs7601812 | 0.087 | 0.189 | 0.558 | -0.271 | -0.008 | 0.035 | 0.035 | 0.274 | 0.46 | 0.34 | 0.56 | 0.68 | 0.48 | 0.64 |
| 2 | 11652795 | rs9287731 | -0.235 | -0.046 | 0.809 | -0.438 | 0.003 | 0.131 | 0.131 | 0 | 0.49 | 0.75 | 0.61 | 0.75 | 0.49 | 0.61 |
| 2 | 11654532 | rs13430148 | -1.541 | 0.839 | 1.142 | -0.279 | 0.077 | -0.065 | -0.065 | 1.293 | 0.11 | 0.26 | 0.09 | 0.01 | 0.14 | 0.08 |
| 2 | 11654588 | rs13395155 | -1.609 | 0.83 | 1.052 | -0.284 | 0.095 | -0.069 | -0.069 | 0 | 0.11 | 0.37 | 0.1 | 0.01 | 0.14 | 0.08 |
| 2 | 11655487 | rs13407457 | -1.608 | 0.807 | 1.054 | -0.32 | 0.09 | -0.069 | -0.069 | 0.625 | 0.11 | 0.37 | 0.1 | 0.01 | 0.14 | 0.08 |
| 2 | 11664349 | rs7582396 | -0.363 | 0.125 | 0.487 | -0.361 | 0.002 | 0.001 | 0.001 | 0.527 | 0.61 | 0.61 | 0.66 | 0.74 | 0.7 | 0.85 |
| 2 | 11668593 | rs7597563 | 0.851 | -0.094 | -0.222 | -0.08 | -0.099 | -0.141 | -0.141 | 1.753 | 0.38 | 0.03 | 0.51 | 0.52 | 0.55 | 0.66 |
| 2 | 11674295 | rs11685032 | 0.878 | -0.597 | 0.301 | 0.042 | -0.096 | -0.165 | -0.165 | 3.003 | 0.38 | 0.03 | 0.52 | 0.49 | 0.56 | 0.66 |
| 2 | 11675472 | rs16857592 | -1.066 | -0.701 | 0.548 | 0.035 | -0.042 | 0.003 | 0.003 | 0.064 | 0.23 | 0.58 | 0.14 | 0.22 | 0.15 | 0.19 |

| | | | | | | | | | | | | | | | | |
|---|---|---|---|---|---|---|---|---|---|---|---|---|---|---|---|---|
| 2 | 11681438 | rs1865574 | -0.652 | -0.674 | 0.724 | -0.123 | -0.005 | -0.007 | -0.007 | 0 | 0.62 | 0.6 | 0.65 | 0.71 | 0.69 | 0.85 |
| 2 | 11689048 | rs6728463 | -1.404 | -0.655 | 1.244 | 0.063 | -0.017 | 0.068 | 0.068 | 1.108 | 0.14 | 0.49 | 0.1 | 0.16 | 0.08 | 0.07 |
| 2 | 11690990 | rs6718650 | -0.527 | -0.661 | 1.258 | 0.057 | -0.004 | -0.014 | -0.014 | 1.931 | 0.09 | 0.11 | 0.06 | 0.03 | 0.07 | 0.05 |
| 2 | 11692106 | rs6737520 | -1.809 | -0.792 | 1.124 | -0.012 | -0.024 | 0.077 | 0.077 | 1.881 | 0.08 | 0.28 | 0.05 | 0.13 | 0.03 | 0.02 |
| 2 | 11692236 | rs17600933 | 0.707 | -0.824 | 0.21 | -0.124 | 0.001 | 0.064 | 0.064 | 2.876 | 0.43 | 0.13 | 0.57 | 0.67 | 0.57 | 0.81 |
| 2 | 11692664 | rs4669749 | -0.891 | -0.857 | -0.014 | -0.235 | 0.009 | 0.041 | 0.041 | 0.368 | 0.65 | 0.75 | 0.68 | 0.83 | 0.66 | 0.88 |
| 2 | 11694006 | rs6432212 | -1.333 | -0.906 | -0.328 | -0.124 | -0.019 | 0.008 | 0.008 | 5.642 | 0.2 | 0.68 | 0.15 | 0.23 | 0.11 | 0.27 |
| 2 | 11702661 | rs10929757 | 0.98 | -0.943 | -0.902 | -0.107 | -0.1 | -0.119 | -0.119 | 1.273 | 0.41 | 0.03 | 0.51 | 0.55 | 0.54 | 0.59 |
| 2 | 11702960 | rs6432216 | -1.614 | -1.089 | -0.773 | -0.075 | 0.018 | -0.002 | -0.002 | 0.008 | 0.24 | 0.8 | 0.18 | 0.23 | 0.13 | 0.27 |
| 2 | 11710716 | rs6432220 | -0.444 | -0.776 | -0.949 | 0 | 0.022 | 0.062 | 0.062 | 0.45 | 0.65 | 0.94 | 0.71 | 0.76 | 0.67 | 0.86 |
| 2 | 11717429 | rs7570979 | -0.495 | -0.772 | -1.312 | 0.075 | -0.025 | 0.041 | 0.041 | 1.155 | 0.67 | 0.96 | 0.65 | 0.73 | 0.67 | 0.86 |
| 2 | 11718858 | rs7578132 | 0.587 | -0.527 | -1.219 | -0.276 | -0.024 | 0.051 | 0.051 | 0.268 | 0.45 | 0.18 | 0.47 | 0.62 | 0.52 | 0.58 |
| 2 | 11721535 | rs11674184 | -0.278 | -0.792 | -0.705 | -0.397 | 0.022 | 0.016 | 0.016 | 0 | 0.45 | 0.21 | 0.47 | 0.4 | 0.44 | 0.4 |
| 2 | 11727087 | rs17529680 | 0.725 | -0.721 | -1.094 | -0.319 | -0.058 | 0.003 | 0.003 | 0.017 | 0.31 | 0.01 | 0.38 | 0.23 | 0.33 | 0.15 |
| 2 | 11731774 | rs2358040 | -1.033 | -0.358 | -0.799 | -0.155 | -0.106 | -0.083 | -0.083 | 3.978 | 0.62 | 0.9 | 0.59 | 0.47 | 0.52 | 0.47 |
| 2 | 11731846 | rs2884374 | -1.033 | -0.358 | -0.799 | -0.155 | -0.106 | -0.083 | -0.083 | 0 | 0.62 | 0.9 | 0.59 | 0.47 | 0.52 | 0.47 |
| 2 | 11748406 | rs12476319 | 0.267 | -0.387 | 0.096 | -0.302 | -0.001 | -0.004 | -0.004 | 0 | 0.42 | 0.41 | 0.45 | 0.7 | 0.47 | 0.72 |
| 2 | 11753697 | rs3828298 | -0.772 | -0.862 | -0.41 | 0.05 | 0.004 | 0.004 | 0.004 | 1.354 | 0.3 | 0.64 | 0.19 | 0.18 | 0.25 | 0.32 |
| 2 | 11756260 | rs7584842 | -0.026 | -0.479 | -0.232 | 0.321 | 0.017 | 0.04 | 0.04 | 0 | 0.5 | 0.6 | 0.42 | 0.29 | 0.45 | 0.24 |
| 2 | 11756429 | rs2304400 | 0.078 | 0.02 | -0.988 | 0.313 | -0.075 | -0.106 | -0.106 | 0.936 | 0.39 | 0.09 | 0.4 | 0.51 | 0.48 | 0.47 |
| 2 | 11762550 | rs33969665 | -0.788 | 0.024 | -0.18 | 0.739 | -0.009 | 0.01 | 0.01 | 0 | 0.31 | 0.42 | 0.26 | 0.38 | 0.26 | 0.43 |
| 2 | 11768872 | rs6718783 | -0.636 | -0.063 | 0.628 | 0.552 | 0.015 | 0.016 | 0.016 | 0 | 0.08 | 0.25 | 0.11 | 0.07 | 0.04 | 0.12 |
| 2 | 11774324 | rs2304402 | -0.32 | -0.251 | -0.132 | 0.163 | -0.001 | 0.016 | 0.016 | 0 | 0.53 | 0.42 | 0.36 | 0.3 | 0.48 | 0.45 |
| 2 | 11775032 | rs10178511 | -0.087 | -0.482 | -0.15 | -0.024 | -0.002 | -0.014 | -0.014 | 1.19 | 0.12 | 0.29 | 0.09 | 0.08 | 0.1 | 0.14 |
| 2 | 11775137 | rs13016287 | 1 | -0.528 | -0.688 | -0.204 | 0.016 | 0.027 | 0.027 | 2.254 | 0.28 | 0.03 | 0.16 | 0.16 | 0.22 | 0.17 |
| 2 | 11777757 | rs3749064 | 0.273 | -0.889 | -0.911 | 0.249 | 0.089 | -0.02 | -0.02 | 0 | 0.39 | 0.13 | 0.24 | 0.41 | 0.28 | 0.35 |
| 2 | 11779603 | rs13406323 | -0.937 | -0.454 | -0.744 | 0.46 | 0.012 | 0.013 | 0.013 | 0 | 0.63 | 0.44 | 0.44 | 0.56 | 0.55 | 0.64 |
| 2 | 11783775 | rs4516403 | 0.375 | -0.432 | -1.106 | 0.583 | 0.03 | -0.045 | -0.045 | 0.076 | 0.09 | 0.01 | 0.07 | 0.23 | 0.06 | 0.18 |
| 2 | 11786832 | rs13001573 | -0.216 | -1.004 | -0.786 | 0.272 | 0.063 | 0.046 | 0.046 | 2.205 | 0.51 | 0.38 | 0.72 | 0.64 | 0.68 | 0.61 |
| 2 | 11787239 | rs7567094 | -0.682 | -0.874 | -0.522 | 0.317 | -0.001 | 0.001 | 0.001 | 0.25 | 0.2 | 0.18 | 0.14 | 0.17 | 0.19 | 0.3 |
| 2 | 11791754 | rs4473360 | 0.742 | -1.163 | -0.293 | 0.573 | 0.031 | 0.009 | 0.009 | 1.443 | 0.5 | 0.38 | 0.65 | 0.47 | 0.64 | 0.61 |
| 2 | 11792857 | rs11690767 | 0.085 | -0.423 | -0.074 | 1.146 | -0.004 | -0.014 | -0.014 | 0 | 0.06 | 0.04 | 0.11 | 0.08 | 0.06 | 0.03 |
| 2 | 11793393 | rs4401188 | 0.18 | -0.531 | 0.383 | 1.109 | -0.02 | 0.022 | 0.022 | 0.765 | 0.8 | 0.71 | 0.89 | 0.76 | 0.89 | 0.78 |
| 2 | 11793770 | rs6725504 | 1.001 | -0.237 | 0.219 | 1.186 | -0.014 | 0.016 | 0.016 | 0.069 | 0.66 | 0.43 | 0.78 | 0.58 | 0.72 | 0.65 |
| 2 | 11794196 | rs13389925 | -0.488 | 0.47 | 0.523 | 0.95 | 0.021 | 0.001 | 0.001 | 0 | 0.16 | 0.41 | 0.12 | 0.18 | 0.18 | 0.15 |
| 2 | 11796361 | rs12692444 | 0.544 | 0.658 | -0.143 | 1.007 | 0.004 | 0.033 | 0.033 | 0 | 0.51 | 0.31 | 0.55 | 0.41 | 0.51 | 0.64 |
| 2 | 11796774 | rs10210917 | 0.491 | 0.656 | -0.065 | 0.991 | 0.004 | 0.026 | 0.026 | 0 | 0.52 | 0.37 | 0.56 | 0.4 | 0.51 | 0.64 |
| 2 | 11811286 | rs7567183 | -0.15 | 0.847 | -0.121 | 1.118 | -0.004 | 0.001 | 0.001 | 0 | 0.41 | 0.4 | 0.26 | 0.47 | 0.34 | 0.33 |
| 2 | 11815492 | rs4358094 | 0.315 | 0.607 | -0.968 | 0.852 | 0.009 | 0.035 | 0.035 | 0.003 | 0.5 | 0.32 | 0.54 | 0.4 | 0.51 | 0.65 |
| 2 | 11823585 | rs6432226 | -0.668 | 0.593 | -0.793 | 1.021 | 0.001 | -0.008 | -0.008 | 0 | 0.34 | 0.32 | 0.29 | 0.54 | 0.3 | 0.4 |
| 2 | 11834635 | rs7606137 | -1.089 | 0.381 | 0.24 | 0.72 | 0.062 | 0.047 | 0.047 | 4.873 | 0.05 | 0.49 | 0.05 | 0.01 | 0.02 | 0.02 |
| 2 | 11838141 | rs4296403 | -2.136 | 0.318 | 0.047 | 1.256 | -0.007 | 0.053 | 0.053 | 4.984 | 0.15 | 0.68 | 0.13 | 0.3 | 0.14 | 0.09 |
| 2 | 11839631 | rs4233899 | 0.671 | 0.202 | -0.303 | 0.547 | 0.029 | 0.076 | 0.076 | 0 | 0.53 | 0.34 | 0.65 | 0.8 | 0.64 | 0.63 |
| 2 | 11843277 | rs4669772 | 1.27 | 0.549 | -0.426 | 0.66 | -0.068 | -0.047 | -0.047 | 1.066 | 0.44 | 0.15 | 0.53 | 0.53 | 0.51 | 0.56 |
| 2 | 11844839 | rs3923671 | -0.192 | 0.26 | 0.319 | 0.542 | 0.011 | 0.02 | 0.02 | 0 | 0.31 | 0.23 | 0.27 | 0.14 | 0.22 | 0.3 |
| 2 | 11853913 | rs4614906 | 0.302 | 0.228 | -0.087 | 0.517 | -0.038 | 0.005 | 0.005 | 0 | 0.25 | 0.01 | 0.23 | 0.14 | 0.21 | 0.27 |
| 2 | 11854012 | rs10182021 | -0.783 | 0.301 | 0.356 | 0.549 | 0.016 | 0.073 | 0.073 | 0.207 | 0.11 | 0.23 | 0.1 | 0.27 | 0.14 | 0.09 |
| 2 | 11855489 | rs10803740 | 1.54 | 0.717 | 0.17 | 0.907 | -0.001 | -0.028 | -0.028 | 0 | 0.72 | 0.21 | 0.79 | 0.6 | 0.75 | 0.85 |
| 2 | 11857546 | rs7599540 | -0.715 | 0.587 | 0.523 | 0.909 | 0.039 | 0.114 | 0.114 | 0 | 0.08 | 0.22 | 0.1 | 0.25 | 0.12 | 0.05 |
| 2 | 11857590 | rs4669777 | -0.666 | 0.519 | 0.373 | 0.885 | 0 | -0.032 | -0.032 | 0.47 | 0.33 | 0.25 | 0.35 | 0.46 | 0.32 | 0.3 |
| 2 | 11861920 | rs4076903 | 1.724 | 0.347 | -0.055 | 0.588 | -0.005 | -0.026 | -0.026 | 0 | 0.66 | 0.13 | 0.75 | 0.63 | 0.68 | 0.78 |
| 2 | 11864946 | rs13421506 | -0.729 | 0.153 | 0.373 | 0.116 | -0.019 | -0.043 | -0.043 | 0 | 0.11 | 0.18 | 0.04 | 0.01 | 0.07 | 0.08 |
| 2 | 11865652 | rs6749129 | -1.941 | 0.306 | -0.198 | 0.219 | -0.122 | -0.068 | -0.068 | 0 | 0.59 | 0.94 | 0.39 | 0.41 | 0.48 | 0.41 |
| 2 | 11870305 | rs7599768 | -1.173 | 0.357 | 0.612 | -0.288 | -0.006 | -0.086 | -0.086 | 0 | 0.21 | 0.31 | 0.14 | 0.07 | 0.2 | 0.16 |
| 2 | 11879105 | rs10185333 | -0.647 | 0.705 | -0.437 | -0.749 | -0.078 | 0.138 | 0.138 | 0 | 0.43 | 0.08 | 0.33 | 0.22 | 0.38 | 0.29 |
| 2 | 11880969 | rs10191952 | -1.148 | 0.768 | 0.196 | -0.806 | -0.001 | -0.001 | -0.001 | 0 | 0.52 | 0.53 | 0.4 | 0.22 | 0.42 | 0.34 |
| 2 | 11881039 | rs10191977 | -1.148 | 0.768 | 0.196 | -0.806 | -0.001 | -0.001 | -0.001 | 0 | 0.52 | 0.53 | 0.4 | 0.22 | 0.42 | 0.34 |
| 2 | 11882480 | rs893350 | -1.207 | 0.596 | 0.474 | -0.679 | 0.053 | -0.009 | -0.009 | 0.729 | 0.08 | 0.42 | 0.06 | 0 | 0.04 | 0.05 |
| 2 | 11882761 | rs893346 | -1.681 | 0.545 | 0.226 | -0.718 | -0.017 | -0.061 | -0.061 | 0 | 0.11 | 0.68 | 0.09 | 0 | 0.07 | 0.06 |
| 2 | 11889385 | rs4076055 | -1.565 | 0.347 | -0.99 | -1.25 | -0.047 | 0.099 | 0.099 | 1.788 | 0.63 | 0.94 | 0.41 | 0.19 | 0.56 | 0.35 |
| 2 | 11896975 | rs7595221 | -1.393 | 0.28 | -0.705 | -1.273 | 0.004 | 0.073 | 0.073 | 1.541 | 0.61 | 0.83 | 0.4 | 0.19 | 0.55 | 0.34 |
| 2 | 11898931 | rs6726960 | -0.546 | 0.327 | -0.751 | -1.38 | -0.001 | -0.102 | -0.102 | 0.014 | 0.23 | 0.71 | 0.17 | 0 | 0.21 | 0.12 |
| 2 | 11917278 | rs6717910 | -0.435 | 0.318 | -1.218 | -1.246 | -0.048 | 0.119 | 0.119 | 0 | 0.39 | 0.11 | 0.24 | 0.19 | 0.34 | 0.23 |
| 2 | 11922054 | rs11695702 | 0.05 | 0.327 | -1.063 | -1.358 | 0.025 | -0.073 | -0.073 | 0 | 0.28 | 0.42 | 0.22 | 0.06 | 0.29 | 0.23 |
| 2 | 11924293 | rs3762583 | 0.445 | 0.193 | -2.325 | -1.425 | -0.036 | -0.011 | -0.011 | 0 | 0.16 | 0.01 | 0.09 | 0.14 | 0.15 | 0.07 |
| 2 | 11924950 | rs3795974 | -1.962 | -0.009 | -2.488 | -1.426 | 0.012 | 0.013 | 0.013 | 0.204 | 0.65 | 0.8 | 0.52 | 0.25 | 0.6 | 0.41 |
| 2 | 11925782 | rs11695081 | 0.289 | 0.53 | -2.085 | -1.418 | 0.001 | 0.013 | 0.013 | 0 | 0.17 | 0.12 | 0.1 | 0.11 | 0.15 | 0.07 |
| 2 | 11930279 | rs2358126 | -1.129 | 0.311 | -2.726 | -1.538 | -0.045 | 0.109 | 0.109 | 0.134 | 0.42 | 0.74 | 0.33 | 0.19 | 0.45 | 0.33 |
| 2 | 11936798 | rs2577264 | -1.45 | 0.041 | -3.023 | -1.377 | 0.003 | 0.17 | 0.17 | 0.738 | 0.58 | 0.95 | 0.41 | 0.14 | 0.57 | 0.39 |

| | | | | | | | | | | | | | | | | |
|---|---|---|---|---|---|---|---|---|---|---|---|---|---|---|---|---|
| 2 | 11940171 | rs11690125 | -0.37 | -0.22 | -1.972 | -0.881 | -0.033 | 0.025 | 0.025 | 0 | 0.16 | 0.01 | 0.09 | 0.09 | 0.15 | 0.07 |
| 2 | 11959905 | rs2716609 | -0.234 | -1.07 | -1.374 | -0.768 | -0.042 | -0.026 | -0.026 | 0 | 0.34 | 0.61 | 0.24 | 0.16 | 0.14 | 0.13 |
| 2 | 11965814 | rs1050800 | -0.106 | 0.043 | -1.559 | -0.35 | -0.031 | -0.055 | -0.055 | 0 | 0.11 | 0.02 | 0.13 | 0.22 | 0.14 | 0.11 |
| 2 | 11972490 | rs2577256 | 0.182 | 0.295 | -1.338 | -1.313 | -0.009 | 0.027 | 0.027 | 0.269 | 0.58 | 0.71 | 0.38 | 0.18 | 0.49 | 0.23 |
| 2 | 11980639 | rs6709884 | -0.412 | 0.195 | -0.934 | -1.659 | -0.005 | -0.073 | -0.073 | 0 | 0.37 | 0.51 | 0.38 | 0.87 | 0.35 | 0.61 |
| 2 | 11983283 | rs1012566 | -0.424 | 0.491 | -0.788 | -1.963 | -0.001 | -0.017 | -0.017 | 0 | 0.38 | 0.63 | 0.4 | 0.82 | 0.37 | 0.59 |
| 2 | 11989805 | rs7575492 | -0.83 | 0.375 | -0.434 | -1.94 | -0.004 | -0.066 | -0.066 | 0.138 | 0.3 | 0.33 | 0.19 | 0.04 | 0.23 | 0.23 |
| 2 | 11991593 | rs6735995 | -0.179 | 0.338 | -0.751 | -1.716 | -0.009 | -0.048 | -0.048 | 5.663 | 0.2 | 0.19 | 0.09 | 0 | 0.12 | 0.19 |
| 2 | 11992128 | rs10495585 | -0.899 | 1.199 | -0.878 | -1.132 | 0.016 | -0.031 | -0.031 | 0.79 | 0.16 | 0.17 | 0.16 | 0.02 | 0.23 | 0.09 |
| 2 | 11995686 | rs6740809 | -0.555 | 0.875 | -1.223 | -1.253 | 0.001 | 0.085 | 0.085 | 2.897 | 0.38 | 0.23 | 0.28 | 0.09 | 0.32 | 0.32 |
| 2 | 11996330 | rs2577268 | 0.436 | 0.414 | -1.178 | -1.427 | 0.001 | -0.01 | -0.01 | 0 | 0.09 | 0.09 | 0.04 | 0 | 0.04 | 0.01 |
| 2 | 12003659 | rs10193112 | -0.6 | 0.032 | -0.646 | -1.469 | -0.01 | -0.07 | -0.07 | 0.132 | 0.25 | 0.54 | 0.17 | 0.14 | 0.21 | 0.15 |
| 2 | 12006615 | rs953071 | -0.079 | 0.52 | -0.839 | -1.614 | -0.001 | 0.052 | 0.052 | 0 | 0.59 | 0.78 | 0.61 | 0.78 | 0.61 | 0.6 |
| 2 | 12008798 | rs10929771 | -0.697 | 0.577 | -0.581 | -1.041 | 0 | 0.076 | 0.076 | 0.653 | 0.5 | 0.72 | 0.42 | 0.26 | 0.42 | 0.4 |
| 2 | 12009981 | rs13409643 | 0.238 | 0.471 | -0.12 | -1.104 | -0.002 | -0.008 | -0.008 | 0 | 0.19 | 0.25 | 0.12 | 0.13 | 0.16 | 0.15 |
| 2 | 12015763 | rs2716628 | 2.058 | 0.86 | -0.528 | -1.188 | -0.005 | -0.009 | -0.009 | 0 | 0.26 | 0.14 | 0.42 | 0.72 | 0.33 | 0.41 |
| 2 | 12017371 | rs893341 | -0.124 | 1.064 | -0.754 | -1.129 | -0.004 | 0 | 0 | 0 | 0.87 | 0.88 | 0.88 | 0.86 | 0.83 | 0.85 |
| 2 | 12019150 | rs11894033 | -0.472 | 1.122 | -0.586 | -0.957 | 0.014 | -0.002 | -0.002 | 0.24 | 0.15 | 0.25 | 0.18 | 0.15 | 0.11 | 0.11 |
| 2 | 12028293 | rs4669783 | 1.161 | 0.965 | -1.054 | -1.007 | -0.048 | 0.04 | 0.04 | 0.851 | 0.24 | 0.01 | 0.2 | 0.08 | 0.2 | 0.22 |
| 2 | 12029246 | rs7576131 | -1.356 | 1.067 | -0.401 | -0.823 | -0.002 | 0.002 | 0.002 | 0 | 0.33 | 0.3 | 0.27 | 0.16 | 0.31 | 0.22 |
| 2 | 12031541 | rs17546105 | 0.172 | 1.259 | -0.71 | -0.495 | -0.028 | 0.036 | 0.036 | 0 | 0.07 | 0.01 | 0.04 | 0 | 0.09 | 0.06 |
| 2 | 12037235 | rs4446054 | -1.171 | 1.017 | -0.381 | -0.135 | -0.001 | -0.001 | -0.001 | 0.144 | 0.31 | 0.33 | 0.33 | 0.33 | 0.34 | 0.31 |
| 2 | 12037492 | rs4027132 | -1.336 | 0.981 | -0.495 | -0.233 | 0.001 | 0.008 | 0.008 | 0 | 0.59 | 0.57 | 0.6 | 0.45 | 0.55 | 0.54 |
| 2 | 12037951 | rs4233904 | -1.595 | 0.816 | -0.419 | -0.157 | -0.01 | 0.023 | 0.023 | 0 | 0.66 | 0.62 | 0.64 | 0.46 | 0.66 | 0.56 |
| 2 | 12037968 | rs4669784 | 0.363 | 0.79 | -0.393 | -0.17 | -0.006 | 0.048 | 0.048 | 0 | 0.35 | 0.26 | 0.29 | 0.11 | 0.32 | 0.24 |
| 2 | 12038123 | rs4296404 | -1.627 | 0.921 | -0.258 | 0.4 | 0.004 | 0.006 | 0.006 | 0 | 0.32 | 0.38 | 0.35 | 0.35 | 0.34 | 0.32 |
| 2 | 12038710 | rs10183830 | -0.825 | 1.11 | -0.608 | 0.559 | -0.03 | -0.015 | -0.015 | 0 | 0.17 | 0.11 | 0.2 | 0.19 | 0.21 | 0.17 |
| 2 | 12040592 | rs2581094 | 0.396 | 1.102 | -0.484 | 0.477 | 0.02 | 0.098 | 0.098 | 0 | 0.95 | 0.84 | 0.9 | 0.86 | 0.92 | 0.89 |
| 2 | 12041594 | rs16857996 | -0.679 | 1.151 | -0.68 | 0.399 | -0.028 | -0.007 | -0.007 | 0 | 0.16 | 0.12 | 0.21 | 0.18 | 0.22 | 0.16 |
| 2 | 12044820 | rs2716651 | 0.542 | 1.332 | -0.583 | 0.663 | 0.041 | 0.124 | 0.124 | 0 | 0.94 | 0.84 | 0.86 | 0.83 | 0.88 | 0.86 |
| 2 | 12045457 | rs10204395 | -0.911 | 1.084 | -1.207 | 0.218 | -0.017 | 0.016 | 0.016 | 0 | 0.24 | 0.16 | 0.23 | 0.19 | 0.25 | 0.17 |
| 2 | 12056323 | rs966391 | -0.179 | 0.694 | -1.774 | 0.037 | -0.011 | 0.039 | 0.039 | 0.973 | 0.75 | 0.99 | 0.76 | 0.88 | 0.79 | 0.73 |
| 2 | 12057524 | rs2581087 | 0.576 | 0.822 | -0.996 | 0.058 | 0.002 | 0.01 | 0.01 | 0.15 | 0.61 | 0.36 | 0.54 | 0.75 | 0.67 | 0.64 |
| 2 | 12068261 | rs17546884 | -0.318 | 0.785 | -0.471 | -0.087 | -0.003 | 0.01 | 0.01 | 0 | 0.09 | 0.08 | 0.06 | 0.01 | 0.11 | 0.08 |
| 2 | 12069382 | rs2581084 | 0.915 | 1.014 | -0.975 | 0.095 | -0.01 | -0.003 | -0.003 | 0 | 0.51 | 0.15 | 0.54 | 0.59 | 0.49 | 0.58 |
| 2 | 12071398 | rs16858030 | -1.105 | 1.243 | -0.259 | 0.171 | 0.006 | 0.007 | 0.007 | 0.433 | 0.26 | 0.68 | 0.27 | 0.3 | 0.26 | 0.21 |
| 2 | 12073058 | rs13386620 | -1.063 | 1.204 | -0.115 | 0.181 | 0.05 | 0.025 | 0.025 | 0.716 | 0.14 | 0.57 | 0.19 | 0.14 | 0.14 | 0.1 |
| 2 | 12074453 | rs7590488 | 0.852 | 1.042 | -0.453 | 0.281 | 0.001 | -0.008 | -0.008 | 0 | 0.12 | 0.1 | 0.1 | 0.16 | 0.13 | 0.1 |
| 2 | 12078838 | rs13025593 | 1.269 | 0.788 | -0.719 | 0.229 | -0.014 | -0.074 | -0.074 | 0.022 | 0.38 | 0.11 | 0.38 | 0.55 | 0.39 | 0.45 |
| 2 | 12081365 | rs2581081 | 0.155 | 0.74 | -0.022 | 0.344 | -0.005 | 0 | 0 | 0.116 | 0.59 | 0.46 | 0.58 | 0.63 | 0.59 | 0.6 |
| 2 | 12081764 | rs12468031 | -0.291 | 0.811 | 0.021 | 0.282 | -0.001 | -0.02 | -0.02 | 0 | 0.08 | 0.12 | 0.05 | 0.01 | 0.1 | 0.01 |
| 2 | 12083477 | rs9287739 | 0.275 | 0.76 | 0.393 | 0.372 | -0.002 | 0 | 0 | 0 | 0.62 | 0.65 | 0.55 | 0.64 | 0.64 | 0.56 |
| 2 | 12084726 | rs6716648 | 0.006 | 0.82 | 0.15 | 0.405 | -0.019 | -0.012 | -0.012 | 0 | 0.13 | 0.09 | 0.16 | 0.16 | 0.14 | 0.18 |
| 2 | 12092323 | rs10197022 | 1.779 | 0.783 | 0.275 | 0.382 | -0.012 | -0.029 | -0.029 | 0.04 | 0.37 | 0.21 | 0.37 | 0.47 | 0.39 | 0.41 |
| 2 | 12100863 | rs2165221 | 0.15 | 0.747 | 0.42 | 0.378 | -0.001 | 0 | 0 | 2.511 | 0.59 | 0.51 | 0.58 | 0.63 | 0.61 | 0.64 |
| 2 | 12103269 | rs11891008 | 1.55 | 0.748 | 0.029 | 0.42 | -0.035 | -0.036 | -0.036 | 0 | 0.39 | 0.16 | 0.38 | 0.46 | 0.42 | 0.45 |
| 2 | 12104445 | rs2255293 | -1.271 | 0.74 | 0.091 | 0.398 | 0.011 | -0.016 | -0.016 | 0 | 0.77 | 0.93 | 0.75 | 0.8 | 0.81 | 0.75 |
| 2 | 12108100 | rs2581064 | 0.091 | 0.742 | -0.108 | 0.304 | -0.024 | 0.011 | 0.011 | 0 | 0.13 | 0.08 | 0.12 | 0.1 | 0.15 | 0.18 |
| 2 | 12111495 | rs1595809 | -0.869 | 0.796 | 0.066 | 0.224 | -0.002 | 0 | 0 | 0 | 0.29 | 0.34 | 0.27 | 0.27 | 0.32 | 0.28 |
| 2 | 12116884 | rs7575078 | 1.46 | 0.818 | 0.219 | 0.212 | 0 | -0.015 | -0.015 | 0 | 0.42 | 0.23 | 0.44 | 0.58 | 0.44 | 0.46 |
| 2 | 12118454 | rs2716644 | -0.21 | 0.985 | -0.264 | 0.206 | -0.008 | 0.006 | 0.006 | 0 | 0.14 | 0.14 | 0.13 | 0.1 | 0.15 | 0.18 |
| 2 | 12119806 | rs2716643 | -0.967 | 0.957 | -0.284 | 0.288 | -0.001 | 0 | 0 | 0 | 0.29 | 0.35 | 0.27 | 0.2 | 0.31 | 0.28 |
| 2 | 12123584 | rs2716629 | 0.342 | 0.85 | -0.191 | 0.397 | -0.001 | 0.011 | 0.011 | 0.018 | 0.16 | 0.25 | 0.13 | 0.17 | 0.17 | 0.1 |
| 2 | 12123760 | rs13011455 | 1.407 | 0.879 | -0.957 | 0.481 | -0.06 | -0.16 | -0.16 | 0.831 | 0.35 | 0.03 | 0.38 | 0.55 | 0.35 | 0.45 |
| 2 | 12124162 | rs2581053 | -1.135 | 0.732 | -0.757 | 0.135 | -0.008 | 0.052 | 0.052 | 0 | 0.37 | 0.75 | 0.3 | 0.23 | 0.41 | 0.21 |
| 2 | 12128492 | rs10166800 | -0.472 | 1.088 | -0.081 | -0.048 | 0.021 | -0.011 | -0.011 | 0 | 0.17 | 0.37 | 0.34 | 0.13 | 0.17 | 0.09 |
| 2 | 12129050 | rs1530145 | -2.113 | 1.526 | -0.462 | -0.4 | -0.01 | 0.04 | 0.04 | 0.021 | 0.43 | 0.81 | 0.6 | 0.32 | 0.52 | 0.24 |
| 2 | 12129329 | rs6432244 | -1.73 | 1.715 | -0.298 | -0.45 | -0.018 | 0.019 | 0.019 | 0.127 | 0.32 | 0.75 | 0.51 | 0.19 | 0.4 | 0.14 |
| 2 | 12129369 | rs4477888 | -1.497 | 1.738 | -0.294 | -0.475 | -0.006 | 0.007 | 0.007 | 0.229 | 0.28 | 0.74 | 0.47 | 0.13 | 0.33 | 0.13 |
| 2 | 12132638 | rs1036668 | -0.62 | 1.023 | -0.688 | -0.37 | -0.001 | -0.001 | -0.001 | 0 | 0.73 | 0.72 | 0.72 | 0.76 | 0.76 | 0.77 |
| 2 | 12134346 | rs12613478 | 0.258 | 0.995 | -0.663 | -0.371 | -0.001 | -0.001 | -0.001 | 0.908 | 0.58 | 0.42 | 0.41 | 0.65 | 0.6 | 0.67 |
| 2 | 12138656 | rs16858133 | 0.328 | 0.834 | -0.192 | -0.359 | 0.037 | -0.046 | -0.046 | 0 | 0.08 | 0.22 | 0.12 | 0 | 0.15 | 0.03 |
| 2 | 12139070 | rs4669793 | -1.551 | 0.703 | -0.704 | -0.329 | 0.011 | -0.021 | -0.021 | 0 | 0.81 | 0.94 | 0.78 | 0.83 | 0.83 | 0.81 |
| 2 | 12139219 | rs4669794 | 1.908 | 0.803 | -0.934 | -0.263 | -0.039 | -0.073 | -0.073 | 0.658 | 0.4 | 0.05 | 0.21 | 0.46 | 0.36 | 0.51 |
| 2 | 12143553 | rs1550398 | -1.331 | 0.709 | -0.637 | -0.151 | 0.016 | -0.009 | -0.009 | 0 | 0.79 | 0.87 | 0.78 | 0.8 | 0.83 | 0.79 |
| 2 | 12146012 | rs11894474 | 0.085 | 0.816 | -0.292 | -0.137 | 0.003 | 0.093 | 0.093 | 0.351 | 0.15 | 0.5 | 0.13 | 0.22 | 0.12 | 0.09 |
| 2 | 12146345 | rs7565113 | -0.428 | 0.974 | -0.592 | -0.332 | 0.026 | -0.025 | -0.025 | 0 | 0.21 | 0.26 | 0.42 | 0.1 | 0.33 | 0.19 |
| 2 | 12146869 | rs11897150 | 0.073 | 0.98 | -0.686 | -0.336 | -0.006 | 0.008 | 0.008 | 2.675 | 0.62 | 0.7 | 0.38 | 0.73 | 0.55 | 0.65 |
| 2 | 12147336 | rs12692451 | -0.034 | 0.851 | -0.507 | -0.323 | -0.014 | -0.012 | -0.012 | 0.97 | 0.65 | 0.46 | 0.47 | 0.73 | 0.69 | 0.69 |

| | | | | | | | | | | | | | | | | |
|---|---|---|---|---|---|---|---|---|---|---|---|---|---|---|---|---|
| 2 | 12150821 | rs12618497 | 0.816 | 0.633 | -1.161 | -0.485 | -0.009 | 0 | 0 | 0.445 | 0.48 | 0.14 | 0.52 | 0.57 | 0.52 | 0.66 |
| 2 | 12151365 | rs6744092 | -0.483 | 0.102 | -0.875 | -0.572 | -0.014 | -0.007 | -0.007 | 0 | 0.72 | 0.27 | 0.75 | 0.8 | 0.83 | 0.78 |
| 2 | 12162113 | rs10929777 | 0.407 | -0.256 | -0.998 | -0.541 | 0.085 | -0.021 | -0.021 | 0 | 0.61 | 0.21 | 0.67 | 0.59 | 0.73 | 0.76 |
| 2 | 12164827 | rs12471113 | 0.041 | -0.283 | -0.976 | -0.556 | 0.061 | -0.023 | -0.023 | 0 | 0.64 | 0.22 | 0.69 | 0.62 | 0.75 | 0.77 |
| 2 | 12166432 | rs35921314 | 0.957 | 0.059 | -1.043 | -0.592 | -0.021 | 0.019 | 0.019 | 0 | 0.09 | 0.07 | 0.18 | 0.02 | 0.2 | 0.04 |
| 2 | 12168075 | rs12471545 | 1.041 | 0.022 | -1.098 | -0.591 | 0.056 | 0.009 | 0.009 | 0.663 | 0.53 | 0.14 | 0.56 | 0.6 | 0.57 | 0.73 |
| 2 | 12171525 | rs4387758 | 2.27 | -0.209 | -1.516 | -0.907 | -0.04 | 0.154 | 0.154 | 0.031 | 0.25 | 0.03 | 0.1 | 0.04 | 0.25 | 0.09 |
| 2 | 12173301 | rs4600620 | 0.02 | 0.194 | -0.683 | -0.556 | 0.027 | -0.004 | -0.004 | 0 | 0.62 | 0.33 | 0.7 | 0.63 | 0.73 | 0.76 |
| 2 | 12174330 | rs11687540 | 1.184 | 0.269 | -0.695 | -0.432 | -0.022 | -0.002 | -0.002 | 0 | 0.18 | 0.1 | 0.42 | 0.14 | 0.23 | 0.24 |
| 2 | 12177253 | rs10495587 | 1.188 | 0.344 | -0.844 | -0.459 | -0.03 | 0.027 | 0.027 | 0 | 0.09 | 0.06 | 0.18 | 0.02 | 0.19 | 0.04 |
| 2 | 12177316 | rs7585074 | -1.627 | -0.358 | -0.382 | -0.421 | -0.031 | 0.004 | 0.004 | 0 | 0.82 | 0.76 | 0.82 | 0.83 | 0.87 | 0.86 |
| 2 | 12180914 | rs2165220 | -1.085 | -0.172 | -0.509 | -0.292 | -0.069 | -0.002 | -0.002 | 0 | 0.78 | 0.55 | 0.78 | 0.82 | 0.84 | 0.84 |
| 2 | 12181825 | rs11690527 | 1.488 | 0.143 | -1.188 | -0.249 | -0.021 | -0.125 | -0.125 | 0 | 0.36 | 0.04 | 0.18 | 0.5 | 0.3 | 0.45 |
| 2 | 12183610 | rs10803742 | -0.013 | -0.148 | -0.613 | -0.071 | -0.009 | 0.002 | 0.002 | 0.031 | 0.65 | 0.38 | 0.71 | 0.63 | 0.73 | 0.77 |
| 2 | 12185325 | rs10929782 | 0.203 | -0.089 | -0.988 | -0.077 | 0.079 | -0.027 | -0.027 | 0 | 0.61 | 0.19 | 0.69 | 0.58 | 0.73 | 0.76 |
| 2 | 12186335 | rs893787 | -1 | 0.531 | -0.501 | -0.284 | -0.003 | 0.003 | 0.003 | 0.202 | 0.4 | 0.7 | 0.4 | 0.32 | 0.46 | 0.38 |
| 2 | 12186432 | rs736447 | -0.366 | 0.587 | -0.284 | -0.335 | -0.003 | 0.007 | 0.007 | 0 | 0.22 | 0.56 | 0.23 | 0.22 | 0.24 | 0.1 |
| 2 | 12191492 | rs4669800 | -1.148 | 0.72 | -0.569 | -0.454 | -0.001 | 0.005 | 0.005 | 0 | 0.75 | 0.81 | 0.82 | 0.87 | 0.74 | 0.84 |
| 2 | 12193065 | rs17549909 | -0.348 | 0.873 | -0.736 | -0.229 | 0.008 | 0.001 | 0.001 | 0.051 | 0.3 | 0.36 | 0.27 | 0.31 | 0.38 | 0.37 |
| 2 | 12193343 | rs11893707 | 0.881 | 0.962 | -0.995 | -0.119 | -0.003 | 0 | 0 | 0 | 0.1 | 0.08 | 0.11 | 0.2 | 0.15 | 0.07 |
| 2 | 12195376 | rs3856469 | -0.838 | 0.799 | -0.971 | -0.242 | -0.014 | -0.037 | -0.037 | 0 | 0.74 | 0.7 | 0.83 | 0.88 | 0.77 | 0.85 |
| 2 | 12198305 | rs4669803 | 0.312 | 0.921 | -1.712 | 0.01 | -0.033 | 0.031 | 0.031 | 0.644 | 0.18 | 0.08 | 0.16 | 0.1 | 0.23 | 0.26 |
| 2 | 12200752 | rs4669806 | 0.535 | 0.423 | -2.086 | 0.21 | 0.008 | -0.148 | -0.148 | 0.063 | 0.32 | 0.04 | 0.44 | 0.52 | 0.25 | 0.46 |
| 2 | 12207377 | rs10495589 | 0.034 | 0.659 | -1.444 | 0.029 | -0.028 | 0.032 | 0.032 | 0.125 | 0.69 | 0.82 | 0.56 | 0.83 | 0.61 | 0.63 |
| 2 | 12208211 | rs13382821 | 0.711 | 0.477 | -0.625 | 0.467 | 0.033 | 0.085 | 0.085 | 0 | 0.1 | 0.26 | 0.08 | 0.21 | 0.12 | 0.05 |
| 2 | 12212445 | rs16858228 | 0.82 | 0.113 | -1.038 | 0.138 | 0.026 | -0.002 | -0.002 | 0.149 | 0.08 | 0 | 0.13 | 0.05 | 0.02 | 0.06 |
| 2 | 12214108 | rs6729317 | 0.656 | 0.088 | -1.64 | 0.103 | -0.062 | -0.06 | -0.06 | 0 | 0.35 | 0.06 | 0.41 | 0.38 | 0.4 | 0.51 |
| 2 | 12215603 | rs7569619 | 1.314 | -0.063 | -1.118 | 0.18 | 0.005 | 0.062 | 0.062 | 0.357 | 0.12 | 0.23 | 0.06 | 0.3 | 0.13 | 0.05 |
| 2 | 12216408 | rs3845756 | 1.126 | -0.212 | -1.08 | -0.085 | 0.011 | 0.015 | 0.015 | 1.013 | 0.22 | 0.09 | 0.13 | 0.16 | 0.2 | 0.12 |
| 2 | 12219836 | rs6753131 | -0.854 | -0.023 | -0.13 | -0.142 | -0.008 | 0.011 | 0.011 | 0 | 0.75 | 0.84 | 0.65 | 0.85 | 0.72 | 0.68 |
| 2 | 12220993 | rs6432257 | 0.354 | -0.085 | -0.056 | -0.115 | -0.003 | 0.022 | 0.022 | 0 | 0.32 | 0.49 | 0.3 | 0.48 | 0.33 | 0.21 |
| 2 | 12221688 | rs10803743 | -0.392 | 0.045 | 0.401 | -0.31 | 0.006 | -0.047 | -0.047 | 0 | 0.74 | 0.65 | 0.64 | 0.84 | 0.75 | 0.66 |
| 2 | 12225930 | rs10166582 | -0.163 | 0.011 | 0.316 | -0.322 | 0 | -0.014 | -0.014 | 0 | 0.72 | 0.41 | 0.63 | 0.84 | 0.75 | 0.68 |
| 2 | 12228696 | rs6707418 | -0.674 | 0.025 | 0.45 | -0.31 | 0 | -0.004 | -0.004 | 0 | 0.76 | 0.8 | 0.67 | 0.85 | 0.76 | 0.68 |
| 2 | 12229926 | rs6432261 | -0.292 | 0.087 | 0.046 | -0.28 | 0.003 | -0.048 | -0.048 | 0 | 0.73 | 0.5 | 0.64 | 0.85 | 0.75 | 0.68 |
| 2 | 12235711 | rs12611533 | 0.58 | 0.083 | -0.704 | 0.19 | -0.002 | 0.009 | 0.009 | 0 | 0.32 | 0.11 | 0.29 | 0.28 | 0.3 | 0.44 |
| 2 | 12236493 | rs10495591 | 1.069 | 0.083 | -0.476 | 0.418 | -0.002 | 0.11 | 0.11 | 1.418 | 0.14 | 0.31 | 0.09 | 0.34 | 0.13 | 0.08 |
| 2 | 12237248 | rs6732455 | -0.352 | 0.036 | -0.553 | 0.263 | 0.002 | 0.002 | 0.002 | 0.132 | 0.48 | 0.55 | 0.39 | 0.61 | 0.42 | 0.51 |
| 2 | 12241022 | rs11695912 | 0.821 | 0.122 | -1.205 | -0.089 | -0.027 | 0.034 | 0.034 | 0 | 0.19 | 0.02 | 0.09 | 0.07 | 0.17 | 0.05 |
| 2 | 12242095 | rs12998142 | 0.781 | 0.114 | -1.209 | -0.093 | -0.026 | 0.035 | 0.035 | 0 | 0.2 | 0.01 | 0.09 | 0.07 | 0.17 | 0.1 |
| 2 | 12243960 | rs6738871 | 0.874 | 0.121 | -1.019 | -0.086 | 0.001 | 0.006 | 0.006 | 0 | 0.18 | 0.11 | 0.11 | 0.07 | 0.17 | 0.1 |
| 2 | 12244434 | rs13404506 | 0.297 | 0.662 | -0.972 | 0.29 | -0.027 | -0.036 | -0.036 | 0 | 0.2 | 0.09 | 0.26 | 0.29 | 0.26 | 0.35 |
| 2 | 12248484 | rs10495592 | 0.401 | 0.679 | -0.985 | 0.357 | -0.025 | -0.044 | -0.044 | 0 | 0.2 | 0.08 | 0.24 | 0.33 | 0.25 | 0.41 |
| 2 | 12248589 | rs1470013 | 0.302 | 0.511 | -0.293 | -0.031 | -0.006 | -0.025 | -0.025 | 0 | 0.73 | 0.68 | 0.67 | 0.88 | 0.79 | 0.67 |
| 2 | 12252350 | rs17551536 | -0.198 | 0.98 | -0.18 | -0.055 | 0.014 | -0.083 | -0.083 | 0 | 0.18 | 0.27 | 0.11 | 0.01 | 0.16 | 0.21 |
| 2 | 12261370 | rs17551690 | 0.576 | 0.932 | -0.301 | -0.094 | -0.014 | 0.016 | 0.016 | 0 | 0.11 | 0.07 | 0.05 | 0.01 | 0.13 | 0.12 |
| 2 | 12261650 | rs13416258 | -0.608 | 0.969 | -0.364 | 0.155 | -0.002 | 0 | 0 | 0 | 0.44 | 0.44 | 0.54 | 0.4 | 0.45 | 0.56 |
| 2 | 12262421 | rs10201732 | -0.406 | 1.114 | -0.776 | 0.077 | -0.027 | 0.005 | 0.005 | 0 | 0.86 | 0.96 | 0.84 | 0.91 | 0.81 | 0.82 |
| 2 | 12265210 | rs7601706 | 0.388 | 1.219 | -0.908 | -0.053 | -0.027 | 0.034 | 0.034 | 0 | 0.81 | 0.92 | 0.81 | 0.91 | 0.74 | 0.8 |
| 2 | 12266205 | rs2033808 | 1.125 | 1.224 | -0.846 | 0.181 | -0.002 | 0.002 | 0.002 | 0 | 0.24 | 0.3 | 0.42 | 0.3 | 0.21 | 0.39 |
| 2 | 12269747 | rs16858314 | 1.776 | 1.273 | -0.949 | 0.169 | 0.033 | -0.102 | -0.102 | 0 | 0.29 | 0.08 | 0.18 | 0.53 | 0.24 | 0.15 |
| 2 | 12272871 | rs10201223 | 0.686 | 1.31 | -1.036 | 0.371 | 0.002 | -0.01 | -0.01 | 0 | 0.3 | 0.34 | 0.48 | 0.44 | 0.27 | 0.42 |
| 2 | 12272949 | rs10164741 | 1.004 | 1.315 | -0.832 | 0.195 | 0.007 | 0.105 | 0.105 | 0 | 0.57 | 0.76 | 0.63 | 0.83 | 0.45 | 0.55 |
| 2 | 12273632 | rs11901591 | -0.78 | 1.433 | -0.765 | 0.341 | -0.005 | 0.002 | 0.002 | 0 | 0.56 | 0.59 | 0.67 | 0.45 | 0.57 | 0.66 |
| 2 | 12274022 | rs1344935 | -0.78 | 1.433 | -0.765 | 0.341 | -0.005 | 0.002 | 0.002 | 0 | 0.56 | 0.59 | 0.67 | 0.45 | 0.57 | 0.65 |
| 2 | 12279879 | rs916522 | -0.327 | 1.639 | -0.761 | -0.077 | -0.031 | 0.039 | 0.039 | 0 | 0.24 | 0.18 | 0.18 | 0.08 | 0.3 | 0.25 |
| 2 | 12284801 | rs934857 | -0.409 | 1.45 | -0.591 | 0.459 | 0.005 | 0.004 | 0.004 | 0 | 0.46 | 0.58 | 0.39 | 0.48 | 0.47 | 0.36 |
| 2 | 12285268 | rs16858333 | 0.94 | 1.276 | -0.784 | 0.673 | 0.035 | -0.018 | -0.018 | 0.143 | 0.27 | 0.15 | 0.16 | 0.38 | 0.21 | 0.11 |
| 2 | 12288187 | rs10204156 | -0.145 | 1.664 | -0.616 | 0.77 | 0.004 | 0.006 | 0.006 | 0 | 0.5 | 0.66 | 0.57 | 0.51 | 0.42 | 0.71 |
| 2 | 12291656 | rs1971516 | -0.45 | 1.456 | -0.288 | 0.209 | 0.035 | 0.024 | 0.024 | 0.401 | 0.06 | 0.2 | 0.06 | 0.06 | 0.06 | 0.08 |
| 2 | 12293125 | rs7558848 | 0.2 | 1.474 | -0.62 | 0.117 | -0.008 | -0.079 | -0.079 | 0 | 0.39 | 0.59 | 0.44 | 0.07 | 0.32 | 0.23 |
| 2 | 12294875 | rs6760764 | -0.354 | 1.482 | -1.027 | 0.598 | 0.007 | 0.011 | 0.011 | 0 | 0.55 | 0.76 | 0.63 | 0.58 | 0.53 | 0.75 |
| 2 | 12304387 | rs6432266 | -0.035 | 1.557 | -0.588 | 0.649 | -0.001 | -0.001 | -0.001 | 0 | 0.51 | 0.56 | 0.61 | 0.5 | 0.46 | 0.69 |
| 2 | 12304753 | rs3954862 | -0.206 | 1.467 | -1.015 | 0.416 | -0.027 | 0.015 | 0.015 | 0 | 0.86 | 0.92 | 0.83 | 0.95 | 0.82 | 0.93 |
| 2 | 12307810 | rs12617169 | 0.027 | 1.55 | -0.887 | 0.462 | 0.003 | -0.01 | -0.01 | 0 | 0.53 | 0.72 | 0.64 | 0.5 | 0.5 | 0.7 |
| 2 | 12310078 | rs16858361 | 0.196 | 1.462 | -0.625 | 0.418 | -0.001 | -0.001 | -0.001 | 0 | 0.49 | 0.52 | 0.6 | 0.49 | 0.45 | 0.68 |
| 2 | 12310905 | rs12692470 | 0.29 | 1.464 | -0.627 | 0.419 | 0.003 | 0.004 | 0.004 | 0 | 0.5 | 0.61 | 0.61 | 0.49 | 0.45 | 0.69 |
| 2 | 12314198 | rs13394440 | -0.638 | 1.429 | -0.737 | 0.365 | 0.012 | -0.01 | -0.01 | 0 | 0.08 | 0.16 | 0.05 | 0.04 | 0.1 | 0.12 |
| 2 | 12314960 | rs10929791 | -0.18 | 1.404 | -0.709 | 0.365 | -0.006 | -0.004 | -0.004 | 0 | 0.07 | 0.21 | 0.03 | 0.04 | 0.05 | 0.1 |

| | | | | | | | | | | | | | | | |
|---|---|---|---|---|---|---|---|---|---|---|---|---|---|---|---|
| 2 | 12316968 | rs10167975 | -0.362 | 1.412 | -1.09 | 0.351 | -0.061 | -0.098 | -0.098 | 0 | 0.59 | 0.92 | 0.67 | 0.54 | 0.55 | 0.79 |
| 2 | 12320631 | rs4669822 | 0.884 | 1.426 | -1.142 | 0.196 | -0.001 | -0.112 | -0.112 | 1.73 | 0.38 | 0.55 | 0.46 | 0.05 | 0.36 | 0.22 |
| 2 | 12321069 | rs4669825 | -0.16 | 1.446 | -1.078 | 0.262 | -0.016 | -0.012 | -0.012 | 0 | 0.56 | 0.75 | 0.64 | 0.53 | 0.51 | 0.77 |
| 2 | 12322109 | rs12996777 | 1.353 | 1.498 | -1.44 | 0.327 | -0.006 | -0.092 | -0.092 | 0 | 0.27 | 0.04 | 0.16 | 0.41 | 0.26 | 0.15 |
| 2 | 12323369 | rs10803744 | -0.408 | 1.652 | -1.446 | 0.245 | -0.04 | -0.027 | -0.027 | 0 | 0.93 | 0.99 | 0.87 | 0.95 | 0.87 | 0.94 |
| 2 | 12326036 | rs6755003 | 0.071 | 1.349 | -1.66 | 0.175 | -0.014 | -0.094 | -0.094 | 0.095 | 0.4 | 0.06 | 0.21 | 0.46 | 0.39 | 0.25 |
| 2 | 12327974 | rs6432268 | 0.435 | 1.365 | -1.475 | 0.103 | 0.023 | 0.168 | 0.168 | 0 | 0.45 | 0.78 | 0.28 | 0.05 | 0.37 | 0.22 |
| 2 | 12331260 | rs10205430 | 0.339 | 1.345 | -1.477 | 0.088 | 0.007 | -0.033 | -0.033 | 0.433 | 0.09 | 0.14 | 0.07 | 0.01 | 0.09 | 0.02 |
| 2 | 12331619 | rs10195126 | 0.163 | 1.343 | -1.534 | 0.08 | -0.02 | 0.168 | 0.168 | 0.528 | 0.42 | 0.08 | 0.53 | 0.89 | 0.43 | 0.7 |
| 2 | 12332543 | rs4669826 | -0.464 | 1.331 | -1.495 | 0.091 | -0.004 | 0.059 | 0.059 | 0 | 0.52 | 0.32 | 0.54 | 0.9 | 0.52 | 0.72 |
| 2 | 12334573 | rs12477218 | -0.007 | 1.342 | -1.531 | 0.15 | -0.005 | -0.028 | -0.028 | 5.192 | 0.43 | 0.19 | 0.25 | 0.47 | 0.44 | 0.27 |
| 2 | 12335122 | rs12465732 | 1.175 | 1.329 | -1.531 | 0.153 | -0.001 | -0.045 | -0.045 | 0 | 0.11 | 0.03 | 0.29 | 0.43 | 0.09 | 0.46 |
| 2 | 12335338 | rs1225214 | 0.134 | 1.425 | -1.54 | 0.145 | -0.01 | 0.001 | 0.001 | 0 | 0.53 | 0.8 | 0.63 | 0.49 | 0.45 | 0.67 |
| 2 | 12336415 | rs2602615 | 0.113 | 1.442 | -1.54 | 0.154 | -0.03 | -0.021 | -0.021 | 0.017 | 0.94 | 0.97 | 0.84 | 0.95 | 0.85 | 0.93 |
| 2 | 12337977 | rs10202729 | 0.838 | 1.38 | -1.412 | 0.167 | -0.006 | 0.068 | 0.068 | 0 | 0.12 | 0.19 | 0.3 | 0.43 | 0.09 | 0.46 |
| 2 | 12339043 | rs6432269 | 0.849 | 1.393 | -1.4 | 0.003 | -0.005 | -0.089 | -0.089 | 0 | 0.41 | 0.55 | 0.32 | 0.06 | 0.36 | 0.22 |
| 2 | 12339714 | rs11686692 | 1.634 | 1.298 | -1.441 | 0.047 | -0.004 | 0.033 | 0.033 | 0 | 0.06 | 0.01 | 0.03 | 0 | 0.05 | 0.01 |
| 2 | 12342726 | rs6432270 | 0.422 | 1.277 | -1.449 | 0.014 | -0.004 | 0.014 | 0.014 | 0 | 0.46 | 0.6 | 0.38 | 0.1 | 0.47 | 0.28 |
| 2 | 12344474 | rs1869381 | 0.502 | 1.154 | -1.649 | 0.08 | -0.016 | 0.077 | 0.077 | 0.008 | 0.37 | 0.09 | 0.44 | 0.85 | 0.37 | 0.6 |
| 2 | 12349389 | rs2380422 | 0.857 | 0.908 | -1.677 | -0.37 | 0.007 | 0.019 | 0.019 | 0 | 0.18 | 0.11 | 0.06 | 0.02 | 0.12 | 0.11 |
| 2 | 12354664 | rs1542327 | -0.102 | 1.062 | -1.93 | -0.012 | 0.008 | -0.021 | -0.021 | 0 | 0.82 | 0.86 | 0.68 | 0.53 | 0.86 | 0.47 |
| 2 | 12354856 | rs1542328 | -0.394 | 1.075 | -1.788 | -0.026 | -0.007 | 0.001 | 0.001 | 0.095 | 0.93 | 0.88 | 0.97 | 0.96 | 0.95 | 0.92 |
| 2 | 12361873 | rs16858479 | 0.352 | 1.148 | -1.798 | -0.354 | -0.01 | -0.118 | -0.118 | 0 | 0.37 | 0.51 | 0.27 | 0.05 | 0.3 | 0.26 |
| 2 | 12362221 | rs10929792 | -0.517 | 1.474 | -1.924 | -0.339 | 0.007 | 0 | 0 | 0 | 0.57 | 0.54 | 0.6 | 0.47 | 0.49 | 0.72 |
| 2 | 12365264 | rs12692473 | 0.751 | 1.61 | -1.938 | -0.323 | 0.028 | 0.029 | 0.029 | 0 | 0.65 | 0.57 | 0.43 | 0.47 | 0.56 | 0.37 |
| 2 | 12366908 | rs1453488 | -0.715 | 1.311 | -2.064 | -0.215 | 0.006 | -0.049 | -0.049 | 0.011 | 0.79 | 0.89 | 0.66 | 0.54 | 0.82 | 0.53 |
| 2 | 12367698 | rs12692474 | 1.629 | 1.3 | -2.462 | -0.23 | 0.021 | -0.098 | -0.098 | 0 | 0.28 | 0.05 | 0.15 | 0.43 | 0.26 | 0.11 |
| 2 | 12369870 | rs4669834 | 0.582 | 1.43 | -2.31 | -0.082 | -0.031 | -0.128 | -0.128 | 0 | 0.2 | 0.01 | 0.32 | 0.45 | 0.18 | 0.47 |
| 2 | 12371945 | rs7587168 | 0.478 | 1.047 | -2.285 | -0.268 | 0.023 | 0.003 | 0.003 | 0 | 0.48 | 0.61 | 0.25 | 0.46 | 0.38 | 0.26 |
| 2 | 12372412 | rs10197467 | -0.171 | 1.333 | -2.296 | -0.167 | -0.038 | -0.024 | -0.024 | 0.098 | 0.59 | 0.83 | 0.39 | 0.51 | 0.51 | 0.42 |
| 2 | 12372510 | rs10197576 | 1.013 | 0.927 | -2.7 | -0.04 | 0.05 | -0.077 | -0.077 | 0.234 | 0.36 | 0.13 | 0.31 | 0.52 | 0.31 | 0.21 |
| 2 | 12372623 | rs4669835 | -0.03 | 0.93 | -2.452 | -0.319 | 0 | -0.043 | -0.043 | 0 | 0.73 | 0.38 | 0.82 | 0.96 | 0.78 | 0.79 |
| 2 | 12373840 | rs11673803 | 0.507 | 0.806 | -2.771 | 0.199 | -0.06 | -0.149 | -0.149 | 0.164 | 0.22 | 0.05 | 0.36 | 0.58 | 0.22 | 0.47 |
| 2 | 12375804 | rs12613308 | 0.818 | 0.956 | -2.74 | 0.123 | -0.037 | 0.051 | 0.051 | 0.152 | 0.51 | 0.1 | 0.36 | 0.37 | 0.52 | 0.36 |
| 2 | 12381269 | rs13006265 | 0.603 | 0.552 | -2.819 | 0.222 | -0.037 | 0.026 | 0.026 | 0.502 | 0.53 | 0.1 | 0.4 | 0.38 | 0.57 | 0.37 |
| 2 | 12397435 | rs1020605 | 0.142 | 0.544 | -2.163 | 0.316 | 0.01 | 0.011 | 0.011 | 0 | 0.95 | 0.87 | 0.89 | 0.94 | 0.91 | 0.93 |
| 2 | 12399050 | rs2380426 | 0.487 | 0.272 | -2.465 | 0.344 | 0.029 | -0.003 | -0.003 | 0.61 | 0.56 | 0.16 | 0.45 | 0.39 | 0.63 | 0.41 |
| 2 | 12401923 | rs1901851 | 0.032 | 0.069 | -2.332 | 0.252 | 0.038 | -0.048 | -0.048 | 0 | 0.63 | 0.23 | 0.58 | 0.47 | 0.76 | 0.48 |
| 2 | 12404697 | rs1120601 | 0.434 | -0.116 | -2.637 | -0.792 | 0.038 | 0.09 | 0.09 | 0 | 0.31 | 0.12 | 0.26 | 0.2 | 0.25 | 0.17 |
| 2 | 12404849 | rs10192295 | 0.028 | -0.712 | -2.079 | -1.303 | 0.002 | 0.004 | 0.004 | 0.089 | 0.5 | 0.43 | 0.52 | 0.74 | 0.46 | 0.7 |
| 2 | 12404974 | rs6711476 | -1.157 | -0.122 | -2.146 | -1.421 | 0.03 | -0.042 | -0.042 | 0 | 0.8 | 0.71 | 0.76 | 0.93 | 0.64 | 0.92 |
| 2 | 12406113 | rs1453497 | 0.509 | -0.551 | -1.962 | -1.2 | -0.003 | -0.017 | -0.017 | 0.097 | 0.38 | 0.43 | 0.45 | 0.62 | 0.38 | 0.61 |
| 2 | 12409041 | rs12989835 | 1.202 | -0.542 | -2.037 | -1.317 | -0.015 | 0.023 | 0.023 | 0 | 0.12 | 0.05 | 0.08 | 0.04 | 0.13 | 0.07 |
| 2 | 12409153 | rs10205200 | -0.633 | -0.331 | -1.822 | -1.117 | -0.031 | 0.054 | 0.054 | 0.095 | 0.68 | 0.8 | 0.7 | 0.86 | 0.56 | 0.83 |
| 2 | 12411048 | rs10164492 | 0.564 | -0.309 | -2.04 | -1.037 | -0.005 | -0.016 | -0.016 | 0 | 0.39 | 0.44 | 0.47 | 0.63 | 0.41 | 0.61 |
| 2 | 12413298 | rs17462600 | 0.471 | -0.189 | -2.199 | -0.991 | -0.113 | 0.059 | 0.059 | 0.229 | 0.31 | 0.05 | 0.29 | 0.15 | 0.43 | 0.17 |
| 2 | 12415940 | rs7567643 | -1.419 | -0.403 | -1.871 | -0.877 | 0.034 | 0.018 | 0.018 | 0 | 0.81 | 0.91 | 0.82 | 0.81 | 0.9 | 0.84 |
| 2 | 12418070 | rs10754960 | -0.088 | 0.142 | -1.425 | -0.606 | -0.003 | -0.012 | -0.012 | 1.367 | 0.41 | 0.51 | 0.49 | 0.66 | 0.41 | 0.61 |
| 2 | 12421708 | rs11681455 | -0.349 | 0.337 | -1.755 | -0.553 | -0.083 | 0.005 | 0.005 | 0.166 | 0.25 | 0.02 | 0.29 | 0.15 | 0.29 | 0.17 |
| 2 | 12431440 | rs723504 | 1.202 | -0.214 | -1.534 | -0.839 | 0.018 | 0.086 | 0.086 | 0.693 | 0.56 | 0.22 | 0.54 | 0.79 | 0.62 | 0.65 |
| 2 | 12437828 | rs16858619 | 0.406 | -0.214 | -0.759 | -0.712 | -0.026 | 0.005 | 0.005 | 0 | 0.07 | 0.12 | 0.05 | 0.04 | 0.01 | 0.08 |
| 2 | 12444332 | rs4669838 | 0.408 | -0.259 | -0.479 | -0.799 | -0.002 | -0.045 | -0.045 | 0.662 | 0.72 | 0.64 | 0.69 | 0.88 | 0.71 | 0.84 |
| 2 | 12445505 | rs6733870 | 0.142 | -0.221 | -0.649 | -0.8 | -0.002 | 0.023 | 0.023 | 0.184 | 0.73 | 0.79 | 0.7 | 0.89 | 0.72 | 0.84 |
| 2 | 12449224 | rs16858628 | -0.279 | -0.099 | -0.619 | -0.683 | -0.034 | -0.002 | -0.002 | 0.171 | 0.08 | 0.17 | 0.06 | 0.04 | 0.01 | 0.08 |
| 2 | 12453068 | rs11684242 | -0.093 | -0.015 | -0.826 | -0.757 | -0.047 | 0.039 | 0.039 | 0.236 | 0.23 | 0.08 | 0.3 | 0.12 | 0.31 | 0.17 |
| 2 | 12456170 | rs12463456 | 0.901 | -0.27 | 0.053 | -0.515 | -0.029 | 0.024 | 0.024 | 0.391 | 0.9 | 0.61 | 0.92 | 0.96 | 0.98 | 0.93 |
| 2 | 12480360 | rs6432279 | 0.138 | -0.187 | 0.1 | -0.515 | -0.023 | -0.003 | -0.003 | 0.225 | 0.92 | 0.83 | 0.94 | 0.96 | 0.97 | 0.94 |
| 2 | 12481154 | rs6717093 | 0.737 | -0.222 | 0.278 | -0.532 | -0.02 | 0.014 | 0.014 | 2.289 | 0.9 | 0.65 | 0.92 | 0.96 | 0.97 | 0.94 |
| 2 | 12488813 | rs12612952 | 0.813 | -0.241 | -0.677 | -0.524 | 0 | -0.046 | -0.046 | 0 | 0.23 | 0.13 | 0.18 | 0.59 | 0.18 | 0.51 |
| 2 | 12499715 | rs6432285 | 0.435 | -0.119 | -0.687 | -0.945 | -0.019 | -0.045 | -0.045 | 0.138 | 0.71 | 0.61 | 0.68 | 0.86 | 0.78 | 0.75 |
| 2 | 12509020 | rs6708491 | 0.184 | 0.211 | -0.445 | -0.651 | -0.009 | 0 | 0 | 1.033 | 0.91 | 0.81 | 0.94 | 0.94 | 0.94 | 0.94 |
| 2 | 12511957 | rs11690119 | 0.233 | 0.229 | -0.437 | -0.642 | -0.009 | 0.001 | 0.001 | 1.464 | 0.91 | 0.81 | 0.94 | 0.94 | 0.94 | 0.94 |
| 2 | 12515567 | rs2890415 | 1.346 | 0.226 | -0.638 | -0.741 | -0.016 | -0.02 | -0.02 | 0.22 | 0.84 | 0.59 | 0.88 | 0.91 | 0.9 | 0.92 |
| 2 | 12520854 | rs13030874 | -0.237 | 0.627 | -1.607 | -0.806 | -0.048 | -0.011 | -0.011 | 1.006 | 0.17 | 0.04 | 0.16 | 0.14 | 0.26 | 0.16 |
| 2 | 12523721 | rs7590007 | -0.013 | 0.008 | -1.678 | -1.149 | 0.019 | 0.039 | 0.039 | 0.432 | 0.29 | 0.72 | 0.32 | 0.09 | 0.21 | 0.22 |
| 2 | 12531197 | rs7563729 | -1.006 | 0.147 | -2.049 | -1.063 | -0.072 | 0.103 | 0.103 | 0 | 0.55 | 0.87 | 0.54 | 0.27 | 0.51 | 0.42 |
| 2 | 12539288 | rs16858807 | -0.601 | 0.054 | -1.159 | -0.99 | 0 | -0.004 | -0.004 | 0 | 0.25 | 0.2 | 0.18 | 0.15 | 0.33 | 0.18 |
| 2 | 12548283 | rs11676011 | 0.382 | -0.131 | -1.013 | -0.613 | -0.001 | -0.001 | -0.001 | 0 | 0.36 | 0.45 | 0.51 | 0.38 | 0.4 | 0.29 |
| 2 | 12549176 | rs13022361 | -0.242 | -0.136 | -0.899 | -0.866 | -0.006 | 0.027 | 0.027 | 0 | 0.43 | 0.29 | 0.27 | 0.15 | 0.49 | 0.32 |

| | | | | | | | | | | | | | | | | |
|---|---|---|---|---|---|---|---|---|---|---|---|---|---|---|---|---|
| 2 | 12553838 | rs11677203 | 0.059 | 0.125 | -0.568 | -0.745 | -0.003 | 0.001 | 0.001 | 0 | 0.21 | 0.18 | 0.17 | 0.14 | 0.3 | 0.17 |
| 2 | 12558059 | rs6742536 | 0.945 | 0.014 | -0.666 | -0.716 | 0.002 | -0.003 | -0.003 | 0 | 0.27 | 0.38 | 0.33 | 0.2 | 0.35 | 0.11 |
| 2 | 12559864 | rs873715 | 0.4 | 0.141 | -0.769 | -0.435 | -0.08 | 0.028 | 0.028 | 0 | 0.15 | 0.01 | 0.12 | 0.08 | 0.24 | 0.12 |
| 2 | 12567278 | rs6728483 | -0.202 | 0.066 | -0.764 | -0.44 | 0.026 | -0.035 | -0.035 | 0 | 0.23 | 0.33 | 0.18 | 0.14 | 0.31 | 0.18 |
| 2 | 12567937 | rs13022088 | 0.401 | 0.427 | -1.23 | -0.24 | 0.022 | -0.005 | -0.005 | 0.476 | 0.17 | 0.19 | 0.15 | 0.15 | 0.29 | 0.16 |
| 2 | 12568903 | rs16858854 | 0.84 | 0.066 | -1.596 | 0.004 | -0.013 | -0.003 | -0.003 | 0.664 | 0.09 | 0 | 0.07 | 0.09 | 0.08 | 0.03 |
| 2 | 12568996 | rs10929808 | -0.644 | 0.172 | -1.194 | 0.262 | 0.008 | -0.066 | -0.066 | 0.001 | 0.81 | 0.91 | 0.81 | 0.52 | 0.9 | 0.61 |
| 2 | 12575134 | rs1534432 | -0.06 | 0.12 | -0.947 | -0.589 | 0 | 0.026 | 0.026 | 0 | 0.32 | 0.27 | 0.22 | 0.18 | 0.29 | 0.35 |
| 2 | 12576670 | rs4669845 | 1.146 | 0.153 | -0.861 | -0.938 | -0.005 | 0.021 | 0.021 | 0 | 0.25 | 0.17 | 0.44 | 0.1 | 0.32 | 0.08 |
| 2 | 12579179 | rs10211429 | -0.129 | -0.019 | -0.457 | -1.215 | 0 | -0.044 | -0.044 | 1.421 | 0.32 | 0.56 | 0.24 | 0.25 | 0.33 | 0.33 |
| 2 | 12580132 | rs1527988 | -0.911 | -0.193 | -0.272 | -0.901 | -0.003 | -0.007 | -0.007 | 0 | 0.76 | 0.83 | 0.77 | 0.38 | 0.8 | 0.61 |
| 2 | 12581267 | rs6733711 | 0.1 | -0.2 | -0.291 | -0.91 | -0.001 | -0.007 | -0.007 | 0 | 0.58 | 0.44 | 0.66 | 0.27 | 0.61 | 0.45 |
| 2 | 12585080 | rs12692481 | 0.637 | -0.074 | -0.314 | -0.854 | -0.009 | -0.054 | -0.054 | 0 | 0.18 | 0.39 | 0.12 | 0.1 | 0.2 | 0.16 |
| 2 | 12585727 | rs10176670 | -0.268 | -0.091 | -0.41 | -0.894 | 0.006 | 0.057 | 0.057 | 0 | 0.49 | 0.65 | 0.34 | 0.27 | 0.48 | 0.51 |
| 2 | 12589208 | rs949981 | 1.2 | 0.009 | -1.031 | -0.842 | -0.009 | 0.036 | 0.036 | 0 | 0.25 | 0.16 | 0.43 | 0.1 | 0.32 | 0.07 |
| 2 | 12591069 | rs10210780 | 0.584 | -0.05 | -0.615 | -0.557 | -0.001 | -0.013 | -0.013 | 0 | 0.15 | 0.15 | 0.1 | 0.1 | 0.2 | 0.15 |
| 2 | 12591695 | rs1898605 | 0.14 | 0.097 | -1.245 | -0.43 | 0 | -0.001 | -0.001 | 0 | 0.33 | 0.29 | 0.21 | 0.18 | 0.28 | 0.36 |
| 2 | 12598397 | rs6432294 | 0.295 | -0.023 | -0.823 | 0.251 | -0.026 | -0.022 | -0.022 | 0.014 | 0.77 | 0.38 | 0.85 | 0.83 | 0.8 | 0.81 |
| 2 | 12606015 | rs12052996 | -0.139 | 0.48 | -1.487 | 0.71 | -0.017 | -0.042 | -0.042 | 0 | 0.23 | 0.09 | 0.19 | 0.27 | 0.19 | 0.08 |
| 2 | 12609639 | rs10203605 | -0.578 | 0.427 | -0.835 | 1.332 | -0.001 | 0.138 | 0.138 | 0 | 0.12 | 0.24 | 0.22 | 0.43 | 0.1 | 0.36 |
| 2 | 12610405 | rs7565747 | 0.244 | 0.373 | -0.877 | 0.588 | 0 | -0.011 | -0.011 | 0 | 0.28 | 0.28 | 0.24 | 0.19 | 0.31 | 0.24 |
| 2 | 12611100 | rs7592226 | 0.002 | 0.532 | -0.814 | 0.87 | -0.002 | 0.133 | 0.133 | 0 | 0.88 | 0.75 | 0.77 | 0.58 | 0.9 | 0.64 |
| 2 | 12613216 | rs7587865 | -0.244 | 0.218 | -1.392 | 0.006 | -0.008 | 0.041 | 0.041 | 0.012 | 0.48 | 0.72 | 0.64 | 0.62 | 0.49 | 0.52 |
| 2 | 12617507 | rs1011575 | 0.77 | -0.107 | -1.619 | -0.038 | 0.031 | -0.076 | -0.076 | 0.266 | 0.2 | 0.05 | 0.09 | 0.36 | 0.17 | 0.23 |
| 2 | 12617959 | rs6740022 | 0.207 | 0.093 | -1.551 | -0.176 | 0.023 | 0.091 | 0.091 | 0 | 0.53 | 0.69 | 0.5 | 0.22 | 0.57 | 0.54 |
| 2 | 12618597 | rs1997249 | 0.093 | 0.009 | -1.493 | -0.235 | 0.034 | 0.034 | 0.034 | 0 | 0.54 | 0.7 | 0.69 | 0.65 | 0.62 | 0.7 |
| 2 | 12619109 | rs1547587 | -0.058 | 0.084 | -0.961 | 0.109 | -0.006 | 0.003 | 0.003 | 0.007 | 0.7 | 0.68 | 0.78 | 0.65 | 0.79 | 0.86 |
| 2 | 12621757 | rs1527986 | -0.438 | 0.455 | -0.75 | -0.087 | -0.001 | -0.003 | -0.003 | 0.098 | 0.41 | 0.59 | 0.38 | 0.5 | 0.41 | 0.36 |
| 2 | 12624618 | rs12614015 | -0.223 | -0.248 | -0.735 | -0.292 | 0.012 | 0.013 | 0.013 | 0 | 0.45 | 0.43 | 0.43 | 0.77 | 0.37 | 0.43 |
| 2 | 12625033 | rs6432302 | -0.079 | -0.457 | -0.426 | -0.201 | -0.008 | -0.005 | -0.005 | 0.001 | 0.42 | 0.49 | 0.23 | 0.29 | 0.32 | 0.25 |
| 2 | 12625491 | rs4377309 | 0.157 | -0.619 | 0.182 | -0.611 | -0.042 | -0.11 | -0.11 | 0 | 0.78 | 0.42 | 0.85 | 0.96 | 0.84 | 0.9 |
| 2 | 12625737 | rs6760700 | 0.322 | 0 | 0.123 | -0.856 | 0.005 | -0.039 | -0.039 | 2.817 | 0.25 | 0.27 | 0.28 | 0.05 | 0.32 | 0.21 |
| 2 | 12625997 | rs11679693 | 1.191 | -0.316 | 0.66 | -0.586 | 0.027 | -0.031 | -0.031 | 0 | 0.93 | 0.72 | 0.89 | 1 | 0.94 | 0.95 |
| 2 | 12626376 | rs11889524 | 0.354 | -0.688 | 0.058 | -0.769 | -0.075 | 0.158 | 0.158 | 0 | 0.38 | 0.15 | 0.32 | 0.16 | 0.45 | 0.46 |
| 2 | 12627522 | rs2196700 | 0.948 | -0.289 | 0.566 | 0.197 | 0.007 | 0.266 | 0.266 | 0.328 | 0.88 | 0.68 | 0.66 | 0.56 | 0.88 | 0.72 |
| 2 | 12631916 | rs10929817 | 0.957 | -0.736 | 0.584 | -0.071 | -0.006 | -0.007 | -0.007 | 0.037 | 0.38 | 0.18 | 0.28 | 0.34 | 0.41 | 0.23 |
| 2 | 12642746 | rs1569120 | -0.655 | -0.556 | 0.907 | -0.376 | -0.002 | 0.014 | 0.014 | 0 | 0.56 | 0.63 | 0.63 | 0.84 | 0.58 | 0.55 |
| 2 | 12644429 | rs16858992 | 0.963 | -0.448 | 0.804 | 0.113 | -0.004 | 0.279 | 0.279 | 0 | 0.06 | 0.13 | 0.12 | 0.36 | 0.05 | 0.25 |
| 2 | 12644736 | rs1534430 | 1.073 | -0.508 | 0.585 | 0.07 | -0.018 | -0.04 | -0.04 | 0.531 | 0.39 | 0.18 | 0.39 | 0.48 | 0.43 | 0.25 |
| 2 | 12646409 | rs7594033 | -0.749 | -0.087 | 0.565 | 0.461 | 0.036 | 0.206 | 0.206 | 0 | 0.12 | 0.28 | 0.22 | 0.37 | 0.14 | 0.32 |
| 2 | 12651822 | rs11904698 | 0.242 | -0.222 | -0.005 | -0.014 | -0.034 | 0.115 | 0.115 | 0.515 | 0.41 | 0.19 | 0.34 | 0.19 | 0.42 | 0.43 |
| 2 | 12652931 | rs10169963 | -0.501 | -0.438 | 0.377 | 0.183 | 0.002 | 0.002 | 0.002 | 0 | 0.51 | 0.43 | 0.59 | 0.61 | 0.53 | 0.68 |
| 2 | 12655145 | rs4669855 | -0.63 | -0.314 | 0.601 | 0.244 | 0 | 0 | 0 | 0 | 0.54 | 0.52 | 0.6 | 0.61 | 0.53 | 0.69 |
| 2 | 12657754 | rs7606568 | 0.367 | -0.306 | 0.282 | 0.224 | -0.001 | 0.079 | 0.079 | 0 | 0.12 | 0.18 | 0.24 | 0.38 | 0.12 | 0.27 |
| 2 | 12657896 | rs13021445 | 0.494 | -0.349 | 0.169 | 0.19 | 0 | -0.002 | -0.002 | 1.724 | 0.44 | 0.39 | 0.39 | 0.43 | 0.47 | 0.31 |
| 2 | 12660466 | rs10188470 | 0.055 | -0.01 | -0.606 | 0.001 | -0.031 | 0.113 | 0.113 | 0.179 | 0.42 | 0.21 | 0.36 | 0.19 | 0.43 | 0.45 |
| 2 | 12673361 | rs6712833 | 0.123 | 0.283 | -0.76 | -0.106 | -0.007 | 0.022 | 0.022 | 0 | 0.33 | 0.23 | 0.23 | 0.06 | 0.33 | 0.24 |
| 2 | 12674125 | rs10495600 | 0.51 | -0.275 | -0.528 | -0.548 | -0.015 | -0.032 | -0.032 | 0 | 0.22 | 0.27 | 0.18 | 0.13 | 0.15 | 0.29 |
| 2 | 12675092 | rs779921 | -0.419 | 0.559 | -1.004 | 0.121 | -0.022 | -0.021 | -0.021 | 0 | 0.37 | 0.24 | 0.41 | 0.44 | 0.42 | 0.51 |
| 2 | 12677035 | rs779925 | -0.39 | 0.584 | -1.47 | 0.209 | -0.055 | -0.089 | -0.089 | 0 | 0.35 | 0.07 | 0.37 | 0.46 | 0.38 | 0.53 |
| 2 | 12679433 | rs6432310 | -0.489 | 0.187 | -0.897 | -0.272 | 0 | -0.001 | -0.001 | 0 | 0.59 | 0.42 | 0.63 | 0.78 | 0.62 | 0.8 |
| 2 | 12683997 | rs779919 | -0.014 | 0.149 | -1.216 | -0.121 | 0.009 | 0.025 | 0.025 | 0 | 0.67 | 0.88 | 0.75 | 0.8 | 0.71 | 0.81 |
| 2 | 12720311 | rs6432315 | -0.581 | 0.872 | -1.484 | 0.395 | 0.001 | 0.014 | 0.014 | 0 | 0.57 | 0.63 | 0.48 | 0.37 | 0.6 | 0.62 |
| 2 | 12720452 | rs4669862 | -0.505 | 0.578 | -1.749 | 1.325 | -0.022 | 0.012 | 0.012 | 0.381 | 0.07 | 0.01 | 0.19 | 0.39 | 0.07 | 0.2 |
| 2 | 12720673 | rs12466724 | -0.607 | 0.509 | -1.411 | 0.189 | 0.011 | 0.014 | 0.014 | 0 | 0.25 | 0.18 | 0.2 | 0.19 | 0.24 | 0.22 |
| 2 | 12725251 | rs17465326 | -0.771 | 1.122 | -1.105 | -0.147 | -0.003 | -0.025 | -0.025 | 0 | 0.24 | 0.2 | 0.19 | 0.02 | 0.37 | 0.3 |
| 2 | 12725389 | rs6432316 | 0.844 | 0.416 | -1.117 | 0.979 | 0.001 | 0.03 | 0.03 | 0 | 0.9 | 0.9 | 0.76 | 0.36 | 0.93 | 0.8 |
| 2 | 12726543 | rs1401702 | -0.919 | -0.166 | -0.926 | 1.15 | -0.012 | 0.026 | 0.026 | 0 | 0.16 | 0.22 | 0.29 | 0.68 | 0.1 | 0.26 |
| 2 | 12727616 | rs4668771 | 0.366 | -0.274 | -0.297 | 0.446 | 0.02 | -0.039 | -0.039 | 0 | 0.81 | 0.59 | 0.78 | 0.89 | 0.82 | 0.79 |
| 2 | 12732821 | rs13023811 | 0.148 | -0.106 | -0.654 | 0.103 | -0.006 | -0.009 | -0.009 | 0.51 | 0.08 | 0.06 | 0.06 | 0.13 | 0.09 | 0.08 |
| 2 | 12735727 | rs11675322 | 0.166 | -0.15 | -0.511 | 0.081 | 0.001 | 0.003 | 0.003 | 0 | 0.08 | 0.09 | 0.07 | 0.13 | 0.09 | 0.08 |
| 2 | 12736138 | rs7599581 | 0.606 | 0.042 | -0.485 | -0.339 | -0.013 | 0.112 | 0.112 | 0 | 0.86 | 0.71 | 0.74 | 0.35 | 0.91 | 0.79 |
| 2 | 12736826 | rs1533652 | -0.903 | 0.059 | -0.4 | -0.355 | -0.005 | 0.05 | 0.05 | 0 | 0.94 | 0.96 | 0.78 | 0.35 | 0.99 | 0.85 |
| 2 | 12741317 | rs1519543 | -0.374 | 0.043 | -0.329 | -0.363 | -0.027 | 0.235 | 0.235 | 0 | 0.93 | 0.78 | 0.76 | 0.35 | 0.99 | 0.85 |
| 2 | 12741369 | rs2139380 | 1.252 | 0.024 | -0.337 | -0.414 | -0.026 | -0.017 | -0.017 | 0 | 0.76 | 0.58 | 0.65 | 0.19 | 0.83 | 0.66 |
| 2 | 12742090 | rs2139381 | 1.83 | -0.978 | -1.389 | -1.146 | -0.162 | 0.223 | 0.223 | 0 | 0.36 | 0.04 | 0.39 | 0.07 | 0.53 | 0.33 |
| 2 | 12745303 | rs939219 | -1.483 | -1.063 | -1.399 | -1.224 | -0.025 | 0.099 | 0.099 | 0 | 0.57 | 0.87 | 0.39 | 0.27 | 0.43 | 0.5 |
| 2 | 12745420 | rs939221 | -1.519 | -1.081 | -1.428 | -1.096 | -0.037 | 0.084 | 0.084 | 0 | 0.57 | 0.9 | 0.39 | 0.27 | 0.43 | 0.49 |
| 2 | 12754657 | rs6432320 | -0.148 | -1.457 | -0.46 | -1.261 | -0.006 | -0.014 | -0.014 | 0.068 | 0.25 | 0.32 | 0.22 | 0.11 | 0.2 | 0.21 |

| | | | | | | | | | | | | | | | | |
|---|---|---|---|---|---|---|---|---|---|---|---|---|---|---|---|---|
| 2 | 12763243 | rs7568831 | -1.115 | -1.578 | 0.085 | -1.171 | -0.008 | 0.006 | 0.006 | 0.369 | 0.29 | 0.63 | 0.34 | 0.66 | 0.19 | 0.39 |
| 2 | 12764164 | rs11900869 | -0.697 | -1.604 | 0.445 | -1.211 | -0.016 | -0.133 | -0.133 | 1.327 | 0.24 | 0.38 | 0.11 | 0.03 | 0.17 | 0.22 |
| 2 | 12772713 | rs13021800 | 0.007 | -1.789 | 0.243 | -1.281 | 0.002 | -0.001 | -0.001 | 0.137 | 0.14 | 0.12 | 0.1 | 0.15 | 0.11 | 0.13 |
| 2 | 12776335 | rs11690210 | -0.996 | -1.945 | 0.367 | -1.362 | -0.036 | -0.034 | -0.034 | 1.409 | 0.28 | 0.42 | 0.33 | 0.78 | 0.15 | 0.33 |
| 2 | 12778345 | rs6432321 | -0.636 | -1.978 | 0.213 | -1.524 | 0 | 0.005 | 0.005 | 0 | 0.83 | 0.87 | 0.81 | 0.9 | 0.8 | 0.8 |
| 2 | 12779813 | rs11684315 | -0.271 | -1.962 | 0.129 | -1.619 | 0.016 | 0.009 | 0.009 | 0 | 0.23 | 0.2 | 0.12 | 0.16 | 0.13 | 0.15 |
| 2 | 12781453 | rs1401697 | 1.058 | -1.787 | 0.052 | -1.79 | -0.004 | -0.058 | -0.058 | 0.533 | 0.55 | 0.45 | 0.47 | 0.12 | 0.66 | 0.48 |
| 2 | 12787237 | rs559703 | -0.218 | -2.171 | 0.007 | -1.216 | -0.015 | 0.011 | 0.011 | 0 | 0.77 | 0.69 | 0.7 | 0.32 | 0.89 | 0.71 |
| 2 | 12787913 | rs651254 | -0.787 | -2.185 | -0.169 | -1.277 | 0.008 | 0.003 | 0.003 | 0 | 0.29 | 0.2 | 0.25 | 0.23 | 0.25 | 0.3 |
| 2 | 12790328 | rs1000053 | 0.022 | -2.185 | -0.299 | -1.16 | 0.015 | -0.006 | -0.006 | 0 | 0.16 | 0.11 | 0.08 | 0.16 | 0.09 | 0.11 |
| 2 | 12796124 | rs12614329 | 0.495 | -2.043 | -0.17 | -1.181 | -0.005 | 0.001 | 0.001 | 0 | 0.11 | 0.13 | 0.07 | 0.14 | 0.07 | 0.1 |
| 2 | 12805227 | rs10490550 | -1.56 | -1.072 | -0.778 | -1.76 | -0.015 | 0.076 | 0.076 | 0.083 | 0.62 | 0.73 | 0.48 | 0.86 | 0.45 | 0.64 |
| 2 | 12807185 | rs515602 | 1.662 | -1.026 | -0.723 | -1.306 | -0.019 | -0.02 | -0.02 | 0 | 0.06 | 0 | 0.14 | 0.07 | 0.11 | 0.15 |
| 2 | 12809444 | rs10490547 | 0.051 | -1.292 | -0.43 | -1.367 | -0.027 | -0.012 | -0.012 | 0 | 0.32 | 0.37 | 0.34 | 0.71 | 0.21 | 0.37 |
| 2 | 12819623 | rs11889770 | -0.121 | 0.366 | -0.49 | -1.217 | 0.006 | -0.001 | -0.001 | 2.773 | 0.7 | 0.64 | 0.51 | 0.74 | 0.55 | 0.65 |
| 2 | 12821349 | rs890073 | -0.561 | 0.195 | -0.072 | -1.265 | -0.002 | -0.043 | -0.043 | 0 | 0.56 | 0.57 | 0.6 | 0.84 | 0.44 | 0.56 |
| 2 | 12826964 | rs504581 | 0.127 | 0.442 | -0.339 | -1.392 | 0.01 | 0.025 | 0.025 | 0 | 0.48 | 0.42 | 0.46 | 0.84 | 0.38 | 0.53 |
| 2 | 12828973 | rs625842 | -0.702 | 0.405 | -0.439 | -1.429 | -0.005 | -0.003 | -0.003 | 0 | 0.6 | 0.61 | 0.53 | 0.84 | 0.49 | 0.67 |
| 2 | 12833928 | rs501596 | -0.044 | 0.297 | -0.342 | -1.303 | 0.013 | 0.031 | 0.031 | 0.011 | 0.49 | 0.56 | 0.46 | 0.84 | 0.38 | 0.53 |
| 2 | 12845368 | rs11902450 | -0.177 | 0.241 | -0.345 | -0.715 | -0.023 | -0.076 | -0.076 | 0 | 0.12 | 0.28 | 0.07 | 0 | 0.1 | 0.13 |
| 2 | 12862905 | rs6719434 | 1.235 | 0.932 | -1.299 | -0.722 | -0.111 | -0.037 | -0.037 | 0 | 0.21 | 0.01 | 0.21 | 0.19 | 0.28 | 0.15 |
| 2 | 12864766 | rs2288192 | 0.663 | 0.553 | -1.955 | -1.042 | -0.006 | -0.075 | -0.075 | 0 | 0.21 | 0.06 | 0.27 | 0.63 | 0.22 | 0.37 |
| 2 | 12867504 | rs890069 | -0.779 | -0.277 | -1.115 | -1.76 | 0.014 | -0.039 | -0.039 | 0.045 | 0.7 | 0.57 | 0.61 | 0.82 | 0.66 | 0.65 |
| 2 | 12867780 | rs17465002 | 0.648 | -0.357 | -1.413 | -1.806 | -0.088 | 0.164 | 0.164 | 1.963 | 0.43 | 0.07 | 0.48 | 0.82 | 0.5 | 0.51 |
| 2 | 12874437 | rs7609362 | 0.073 | -0.504 | -0.704 | -1.883 | -0.048 | -0.142 | -0.142 | 0.632 | 0.28 | 0.6 | 0.14 | 0 | 0.16 | 0.15 |
| 2 | 12876468 | rs1863175 | 0.347 | -0.346 | -0.114 | -2.201 | -0.018 | -0.112 | -0.112 | 0 | 0.15 | 0.51 | 0.11 | 0.02 | 0.11 | 0.06 |
| 2 | 12878668 | rs2042566 | 1.047 | -0.236 | -1.013 | -2.139 | -0.07 | 0.028 | 0.028 | 0 | 0.35 | 0.08 | 0.4 | 0.73 | 0.4 | 0.51 |
| 2 | 12881526 | rs1176 | 2.09 | 0.1 | -1.049 | -2.143 | -0.047 | 0.041 | 0.041 | 0 | 0.07 | 0.01 | 0.1 | 0 | 0.16 | 0.03 |
| 2 | 12888761 | rs6722342 | 0.625 | -0.177 | -0.699 | -2.075 | 0.006 | 0.019 | 0.019 | 0.2 | 0.07 | 0.13 | 0.04 | 0.1 | 0.09 | 0.08 |
| 2 | 12890650 | rs7596623 | 0.447 | -0.189 | -0.672 | -2.081 | 0.005 | 0.018 | 0.018 | 0.319 | 0.07 | 0.13 | 0.04 | 0.1 | 0.09 | 0.09 |
| 2 | 12890860 | rs2113818 | -0.119 | -0.387 | -0.689 | -2.284 | -0.006 | 0.094 | 0.094 | 0.097 | 0.49 | 0.29 | 0.47 | 0.83 | 0.52 | 0.62 |
| 2 | 12892929 | rs2059428 | -0.744 | -0.005 | -0.664 | -2.012 | 0.008 | -0.018 | -0.018 | 0 | 0.7 | 0.68 | 0.6 | 0.84 | 0.65 | 0.67 |
| 2 | 12896713 | rs12616457 | 0.892 | -0.058 | -1.03 | -1.938 | -0.059 | 0.039 | 0.039 | 0 | 0.33 | 0.06 | 0.39 | 0.75 | 0.37 | 0.5 |
| 2 | 12901531 | rs4668785 | 0.175 | -0.278 | -0.482 | -2.176 | -0.032 | 0.151 | 0.151 | 0.01 | 0.45 | 0.1 | 0.47 | 0.77 | 0.48 | 0.55 |
| 2 | 12901718 | rs10191541 | -0.552 | -0.157 | -0.137 | -2.29 | -0.016 | -0.066 | -0.066 | 0 | 0.24 | 0.54 | 0.18 | 0.11 | 0.22 | 0.14 |
| 2 | 12905490 | rs973977 | -1.092 | -0.221 | -0.143 | -2.325 | -0.004 | -0.003 | -0.003 | 0.095 | 0.69 | 0.7 | 0.59 | 0.86 | 0.62 | 0.66 |
| 2 | 12907151 | rs13413220 | 0.375 | -0.225 | 0.049 | -2.287 | 0.003 | 0.02 | 0.02 | 0.632 | 0.08 | 0.19 | 0.04 | 0.1 | 0.09 | 0.09 |
| 2 | 12910084 | rs890074 | -0.299 | -0.242 | -0.037 | -2.246 | -0.007 | 0.025 | 0.025 | 0 | 0.57 | 0.64 | 0.5 | 0.85 | 0.51 | 0.61 |
| 2 | 12912547 | rs13000767 | 2.048 | -0.261 | -0.15 | -2.231 | 0.005 | -0.011 | -0.011 | 0 | 0.13 | 0.12 | 0.05 | 0 | 0.04 | 0.02 |
| 2 | 12922923 | rs11674847 | 2.028 | 0.041 | -0.48 | -2.337 | -0.05 | 0.05 | 0.05 | 0 | 0.22 | 0.01 | 0.1 | 0.07 | 0.23 | 0.14 |
| 2 | 12936286 | rs7557984 | -0.033 | -0.111 | -0.119 | -2.198 | -0.033 | -0.062 | -0.062 | 0 | 0.21 | 0.49 | 0.15 | 0.1 | 0.18 | 0.12 |
| 2 | 12942042 | rs2194724 | 0.66 | -0.004 | -0.3 | -2.165 | -0.042 | 0.06 | 0.06 | 0 | 0.37 | 0.05 | 0.39 | 0.76 | 0.39 | 0.54 |
| 2 | 12951390 | rs6733784 | -1.995 | 0.101 | -0.737 | -2.272 | -0.048 | 0.005 | 0.005 | 0 | 0.83 | 0.98 | 0.74 | 0.87 | 0.83 | 0.84 |
| 2 | 12961220 | rs6710360 | 0.047 | 0.115 | -0.334 | -2.158 | 0.02 | 0.064 | 0.064 | 1.286 | 0.09 | 0.39 | 0.06 | 0.09 | 0.07 | 0.07 |
| 2 | 12962315 | rs10191751 | -0.854 | 0.185 | -0.339 | -2.211 | -0.002 | -0.006 | -0.006 | 0 | 0.69 | 0.67 | 0.61 | 0.87 | 0.66 | 0.65 |
| 2 | 12962818 | rs17465721 | 1.248 | 0.548 | -0.225 | -1.611 | -0.061 | 0.03 | 0.03 | 0.005 | 0.28 | 0.05 | 0.12 | 0.14 | 0.32 | 0.21 |
| 2 | 12964428 | rs10186351 | -0.677 | 0.462 | 0.164 | -2.095 | -0.01 | 0.012 | 0.012 | 0 | 0.59 | 0.62 | 0.52 | 0.85 | 0.54 | 0.64 |
| 2 | 12964510 | rs17465833 | 1.41 | 0.505 | 0.078 | -2.076 | -0.062 | 0.049 | 0.049 | 0 | 0.1 | 0.01 | 0.17 | 0.01 | 0.2 | 0.15 |
| 2 | 12968900 | rs10180496 | -0.056 | -0.059 | 0.244 | -1.973 | 0.02 | 0.057 | 0.057 | 0.909 | 0.08 | 0.32 | 0.05 | 0.09 | 0.07 | 0.07 |
| 2 | 12970820 | rs13415579 | 0.036 | -0.059 | 0.234 | -1.973 | 0.02 | 0.057 | 0.057 | 0.757 | 0.08 | 0.32 | 0.05 | 0.09 | 0.07 | 0.07 |
| 2 | 12974244 | rs4669886 | 0.128 | 0.004 | -0.018 | -1.911 | 0.014 | 0.172 | 0.172 | 0 | 0.47 | 0.11 | 0.43 | 0.77 | 0.44 | 0.56 |
| 2 | 12992430 | rs13396447 | -0.195 | -0.1 | 0.262 | -1.741 | 0.09 | 0.048 | 0.048 | 0.947 | 0.1 | 0.45 | 0.1 | 0.09 | 0.12 | 0.08 |
| 2 | 12996275 | rs1521244 | 0.869 | -0.108 | 0.122 | -1.821 | -0.04 | 0.033 | 0.033 | 0 | 0.31 | 0.08 | 0.37 | 0.77 | 0.37 | 0.5 |
| 2 | 13002998 | rs1357060 | -0.054 | 0.777 | 0.618 | -1.282 | -0.002 | 0.061 | 0.061 | 0 | 0.54 | 0.64 | 0.51 | 0.85 | 0.51 | 0.64 |
| 2 | 13015428 | rs13425286 | 0.027 | 0.973 | 0.253 | -1.13 | -0.004 | 0.043 | 0.043 | 0 | 0.53 | 0.62 | 0.52 | 0.79 | 0.53 | 0.55 |
| 2 | 13024831 | rs1879543 | -0.003 | 1.008 | -0.455 | -1.141 | -0.017 | 0.135 | 0.135 | 0 | 0.53 | 0.72 | 0.52 | 0.79 | 0.53 | 0.57 |
| 2 | 13028547 | rs6739633 | 0.86 | 1.004 | -0.458 | -1.126 | 0.014 | 0.083 | 0.083 | 0 | 0.42 | 0.12 | 0.41 | 0.7 | 0.42 | 0.48 |
| 2 | 13035974 | rs1357061 | -0.594 | 1.339 | -0.234 | -0.88 | 0.099 | 0.071 | 0.071 | 1.734 | 0.1 | 0.5 | 0.1 | 0.09 | 0.1 | 0.05 |
| 2 | 13037343 | rs10495605 | 0.238 | 1.196 | -0.426 | -0.698 | 0.105 | 0.13 | 0.13 | 0.058 | 0.06 | 0.36 | 0.07 | 0.09 | 0.07 | 0.04 |
| 2 | 13050694 | rs711868 | -0.34 | 1.08 | -0.974 | -0.758 | 0.017 | 0.031 | 0.031 | 0 | 0.39 | 0.69 | 0.36 | 0.29 | 0.31 | 0.27 |
| 2 | 13052225 | rs779357 | -0.39 | 0.679 | -0.736 | -0.273 | 0.009 | 0.043 | 0.043 | 0 | 0.53 | 0.64 | 0.55 | 0.3 | 0.54 | 0.44 |
| 2 | 13055358 | rs16859504 | -1.088 | 0.663 | 0.231 | 0.601 | 0.073 | 0.128 | 0.128 | 0.089 | 0.07 | 0.4 | 0.07 | 0.08 | 0.07 | 0.04 |
| 2 | 13069079 | rs7607052 | -0.065 | 0.79 | -0.975 | 0.642 | 0.001 | 0.003 | 0.003 | 0 | 0.5 | 0.66 | 0.34 | 0.43 | 0.51 | 0.34 |
| 2 | 13070276 | rs779374 | -0.094 | 0.68 | -1.898 | -0.727 | -0.004 | 0.227 | 0.227 | 0.11 | 0.56 | 0.9 | 0.72 | 0.95 | 0.56 | 0.58 |
| 2 | 13070917 | rs7587118 | 0.283 | 0.662 | -1.205 | -0.909 | 0.014 | -0.031 | -0.031 | 0 | 0.1 | 0.19 | 0.08 | 0.04 | 0.12 | 0.13 |
| 2 | 13075444 | rs779376 | 0.277 | 0.85 | -1.279 | -0.66 | -0.005 | -0.009 | -0.009 | 1.856 | 0.16 | 0.15 | 0.16 | 0.2 | 0.22 | 0.15 |
| 2 | 13080607 | rs779379 | -0.901 | -0.033 | -1.765 | -1.246 | 0.01 | 0.074 | 0.074 | 0.979 | 0.85 | 0.97 | 0.93 | 1 | 0.88 | 0.84 |
| 2 | 13081694 | rs779380 | 0.809 | 0.114 | -1.442 | -1.105 | 0.003 | 0.005 | 0.005 | 0.019 | 0.12 | 0.18 | 0.14 | 0.16 | 0.19 | 0.14 |
| 2 | 13085166 | rs779388 | 0.236 | 0.004 | -0.933 | -0.896 | 0.004 | 0.026 | 0.026 | 0 | 0.45 | 0.43 | 0.46 | 0.72 | 0.57 | 0.43 |

| | | | | | | | | | | | | | | | | |
|---|---|---|---|---|---|---|---|---|---|---|---|---|---|---|---|---|
| 2 | 13088380 | rs17381690 | 1.347 | 0.128 | -0.96 | -0.856 | -0.058 | 0.095 | 0.095 | 0 | 0.27 | 0.11 | 0.16 | 0.03 | 0.36 | 0.14 |
| 2 | 13089226 | rs711870 | -1.074 | -0.073 | -1.021 | -0.974 | 0.067 | 0.173 | 0.173 | 0.262 | 0.64 | 0.92 | 0.73 | 0.99 | 0.74 | 0.67 |
| 2 | 13089732 | rs779392 | -0.46 | -0.088 | -0.833 | -0.96 | 0.017 | 0.025 | 0.025 | 0.084 | 0.56 | 0.67 | 0.55 | 0.77 | 0.68 | 0.56 |
| 2 | 13093444 | rs7582647 | 0.663 | -0.158 | -0.495 | -0.964 | 0.001 | 0.018 | 0.018 | 0 | 0.09 | 0.13 | 0.28 | 0.22 | 0.09 | 0.16 |
| 2 | 13096218 | rs779403 | 0.416 | -0.127 | -0.359 | -0.882 | 0.005 | 0.022 | 0.022 | 0 | 0.17 | 0.26 | 0.29 | 0.69 | 0.2 | 0.29 |
| 2 | 13102613 | rs779406 | -1.032 | -0.147 | -0.353 | -0.926 | 0.057 | 0.092 | 0.092 | 1.025 | 0.66 | 0.86 | 0.83 | 0.99 | 0.78 | 0.74 |
| 2 | 13102991 | rs779407 | 1.135 | -0.014 | -0.457 | -0.79 | 0.004 | 0.017 | 0.017 | 0.017 | 0.13 | 0.19 | 0.14 | 0.21 | 0.18 | 0.14 |
| 2 | 13108251 | rs12994713 | 0.588 | 0.004 | -0.466 | -0.799 | -0.036 | 0.037 | 0.037 | 0 | 0.37 | 0.27 | 0.45 | 0.26 | 0.46 | 0.31 |
| 2 | 13109389 | rs1402545 | -0.08 | -0.022 | -0.419 | -0.811 | 0.003 | 0.005 | 0.005 | 0 | 0.47 | 0.48 | 0.53 | 0.3 | 0.58 | 0.44 |
| 2 | 13111388 | rs779350 | -0.145 | -0.092 | -0.473 | -0.594 | 0 | 0 | 0 | 0 | 0.52 | 0.55 | 0.61 | 0.46 | 0.65 | 0.46 |
| 2 | 13118395 | rs779345 | -0.71 | 0.216 | -0.589 | -0.471 | 0.007 | -0.11 | -0.11 | 1.721 | 0.76 | 0.68 | 0.83 | 0.99 | 0.77 | 0.84 |
| 2 | 13118520 | rs13415994 | 0.176 | 0.306 | -0.39 | 0.223 | 0 | 0.063 | 0.063 | 4.972 | 0.11 | 0.14 | 0.36 | 0.4 | 0.1 | 0.16 |
| 2 | 13118651 | rs11688528 | 0.697 | 0.351 | -0.951 | 0.345 | -0.049 | -0.055 | -0.055 | 0.03 | 0.35 | 0.16 | 0.22 | 0.35 | 0.37 | 0.46 |
| 2 | 13120011 | rs983491 | 0.733 | 0.376 | -1.091 | 0.359 | -0.068 | -0.074 | -0.074 | 0 | 0.34 | 0.11 | 0.21 | 0.35 | 0.37 | 0.46 |
| 2 | 13120212 | rs12991972 | 0.508 | 0.017 | -0.37 | 0.69 | -0.02 | 0.05 | 0.05 | 0.011 | 0.14 | 0 | 0.1 | 0.05 | 0.15 | 0.06 |
| 2 | 13121525 | rs4669900 | -0.587 | 0.089 | -0.153 | 0.834 | 0.002 | 0.226 | 0.226 | 2.322 | 0.15 | 0.47 | 0.39 | 0.42 | 0.11 | 0.2 |
| 2 | 13126278 | rs7590964 | -0.554 | 0.204 | -1.179 | -0.07 | -0.026 | -0.053 | -0.053 | 0 | 0.33 | 0.54 | 0.35 | 0.19 | 0.3 | 0.27 |
| 2 | 13133086 | rs2380458 | -0.021 | 0.535 | -1.51 | 0.106 | 0.002 | 0.028 | 0.028 | 0 | 0.42 | 0.34 | 0.56 | 0.71 | 0.38 | 0.65 |
| 2 | 13133155 | rs12476708 | 0.171 | 0.502 | -1.803 | -0.158 | -0.027 | -0.016 | -0.016 | 0.403 | 0.24 | 0.1 | 0.22 | 0.26 | 0.28 | 0.15 |
| 2 | 13134803 | rs12692488 | -0.447 | 0.463 | -1.944 | 0.252 | -0.005 | -0.019 | -0.019 | 0 | 0.4 | 0.78 | 0.4 | 0.45 | 0.38 | 0.21 |
| 2 | 13137013 | rs2138500 | 0.505 | 0.56 | -2.091 | 0.43 | -0.002 | -0.004 | -0.004 | 0 | 0.17 | 0.12 | 0.36 | 0.16 | 0.17 | 0.2 |
| 2 | 13140080 | rs1516289 | -0.442 | 0.501 | -1.972 | 0.443 | -0.038 | -0.017 | -0.017 | 0 | 0.56 | 0.87 | 0.74 | 0.6 | 0.53 | 0.39 |
| 2 | 13148064 | rs9784164 | -0.84 | 0.357 | -0.7 | 0.708 | -0.017 | 0.004 | 0.004 | 0 | 0.33 | 0.29 | 0.36 | 0.32 | 0.4 | 0.26 |
| 2 | 13149840 | rs10929842 | 0.206 | 0.563 | -0.701 | 0.668 | -0.007 | -0.004 | -0.004 | 0 | 0.14 | 0.12 | 0.15 | 0.17 | 0.17 | 0.12 |
| 2 | 13154596 | rs16859628 | -0.319 | 0.407 | -0.285 | 0.664 | 0.007 | 0.031 | 0.031 | 0 | 0.1 | 0.29 | 0.2 | 0.17 | 0.08 | 0.04 |
| 2 | 13162309 | rs938495 | -0.627 | 0.116 | 0.598 | 0.518 | -0.004 | 0.06 | 0.06 | 0.054 | 0.17 | 0.59 | 0.24 | 0.23 | 0.15 | 0.12 |
| 2 | 13166369 | rs953006 | 0.755 | 0.11 | 0.36 | 0.504 | -0.002 | 0.001 | 0.001 | 0 | 0.07 | 0.04 | 0.11 | 0.06 | 0.08 | 0.03 |
| 2 | 13167258 | rs4668795 | -1.381 | 0.118 | 0.454 | 0.504 | -0.004 | -0.002 | -0.002 | 0.132 | 0.24 | 0.66 | 0.35 | 0.27 | 0.24 | 0.15 |
| 2 | 13168232 | rs1818595 | -1.954 | -0.08 | -0.155 | 0.534 | -0.076 | -0.079 | -0.079 | 0 | 0.45 | 0.92 | 0.52 | 0.51 | 0.43 | 0.31 |
| 2 | 13197725 | rs11688153 | 0.356 | -0.28 | 0.558 | 0.576 | -0.004 | 0.002 | 0.002 | 0 | 0.11 | 0.14 | 0.12 | 0.11 | 0.09 | 0.13 |
| 2 | 13202216 | rs11674766 | 0.159 | -0.294 | 0.701 | 0.569 | -0.007 | 0.005 | 0.005 | 0 | 0.12 | 0.22 | 0.13 | 0.11 | 0.09 | 0.13 |
| 2 | 13204511 | rs1913995 | -1.424 | -0.567 | 1.049 | 0.913 | -0.017 | 0.018 | 0.018 | 0 | 0.26 | 0.66 | 0.36 | 0.37 | 0.18 | 0.21 |
| 2 | 13209268 | rs12477177 | 0.933 | -0.595 | 0.695 | 0.997 | -0.052 | -0.049 | -0.049 | 0 | 0.12 | 0.01 | 0.13 | 0.15 | 0.15 | 0.09 |
| 2 | 13230033 | rs13427923 | -0.856 | -0.833 | 1.325 | 0.978 | 0.004 | -0.011 | -0.011 | 0 | 0.2 | 0.14 | 0.25 | 0.26 | 0.17 | 0.18 |
| 2 | 13238258 | rs7587457 | -2.456 | -0.842 | 1.532 | 0.954 | 0.004 | -0.004 | -0.004 | 0 | 0.29 | 0.75 | 0.39 | 0.36 | 0.19 | 0.22 |
| 2 | 13244117 | rs9808277 | -2.491 | -0.872 | 1.492 | 1.043 | 0.005 | -0.005 | -0.005 | 0 | 0.3 | 0.75 | 0.43 | 0.37 | 0.2 | 0.24 |
| 2 | 13248391 | rs12612960 | 0.421 | -0.852 | 1.544 | 1.021 | -0.002 | 0 | 0 | 0 | 0.11 | 0.13 | 0.12 | 0.12 | 0.09 | 0.14 |
| 2 | 13248752 | rs6760545 | -1.571 | -1.252 | 1.348 | 0.852 | -0.01 | -0.001 | -0.001 | 0.571 | 0.26 | 0.75 | 0.27 | 0.23 | 0.11 | 0.18 |
| 2 | 13257628 | rs7568997 | -0.069 | -1.037 | 1.275 | 1.041 | -0.009 | 0.07 | 0.07 | 0 | 0.12 | 0.3 | 0.21 | 0.24 | 0.07 | 0.16 |
| 2 | 13282236 | rs931237 | -1.555 | -1.067 | 1.218 | 0.821 | -0.021 | 0.009 | 0.009 | 0.203 | 0.31 | 0.61 | 0.4 | 0.38 | 0.19 | 0.24 |
| 2 | 13295833 | rs16859903 | -0.334 | -0.936 | 0.594 | 0.852 | -0.015 | -0.007 | -0.007 | 0 | 0.18 | 0.2 | 0.18 | 0.14 | 0.11 | 0.08 |
| 2 | 13298930 | rs10187895 | -0.194 | -0.932 | 0.531 | 0.854 | -0.006 | -0.003 | -0.003 | 0 | 0.17 | 0.12 | 0.17 | 0.14 | 0.11 | 0.08 |
| 2 | 13302734 | rs12614088 | 1.111 | -0.844 | 0.264 | 0.888 | 0.001 | -0.011 | -0.011 | 0.002 | 0.09 | 0.05 | 0.1 | 0.13 | 0.06 | 0.13 |
| 2 | 13321120 | rs753822 | -1.217 | -0.86 | 0.376 | 0.872 | -0.044 | 0.03 | 0.03 | 0.202 | 0.31 | 0.49 | 0.39 | 0.39 | 0.19 | 0.24 |
| 2 | 13321498 | rs7604358 | -0.864 | -0.821 | 0.344 | 0.893 | -0.027 | -0.002 | -0.002 | 0 | 0.28 | 0.33 | 0.28 | 0.27 | 0.17 | 0.21 |
| 2 | 13323221 | rs6719067 | -0.977 | -0.763 | 0.141 | 0.787 | -0.042 | 0.036 | 0.036 | 0.053 | 0.31 | 0.41 | 0.38 | 0.39 | 0.18 | 0.24 |
| 2 | 13330848 | rs6742358 | 0.798 | -0.335 | -0.515 | 0.571 | -0.064 | -0.036 | -0.036 | 0.964 | 0.14 | 0.01 | 0.16 | 0.15 | 0.2 | 0.11 |
| 2 | 13333326 | rs748615 | 0.232 | -0.952 | 0.377 | 0.217 | -0.047 | 0.016 | 0.016 | 5.005 | 0.1 | 0.21 | 0.08 | 0.12 | 0.03 | 0.05 |
| 2 | 13337188 | rs13030841 | 0.195 | -1.121 | 0.031 | -0.388 | 0 | -0.022 | -0.022 | 0 | 0.34 | 0.6 | 0.33 | 0.14 | 0.31 | 0.24 |
| 2 | 13338190 | rs1871957 | -0.016 | -1.14 | -0.072 | -0.348 | -0.035 | 0.023 | 0.023 | 1.638 | 0.18 | 0.26 | 0.18 | 0.25 | 0.09 | 0.22 |
| 2 | 13345954 | rs13392510 | 0.7 | -1.181 | 0.521 | -0.454 | 0.007 | 0.012 | 0.012 | 0 | 0.1 | 0.14 | 0.18 | 0.13 | 0.1 | 0.05 |
| 2 | 13366874 | rs2046266 | -1.162 | -1.212 | -0.03 | -0.497 | -0.02 | 0.024 | 0.024 | 0.453 | 0.54 | 0.87 | 0.49 | 0.39 | 0.39 | 0.45 |
| 2 | 13368225 | rs13403000 | 0.054 | -1.161 | 0.027 | -0.46 | 0.001 | -0.033 | -0.033 | 0.133 | 0.35 | 0.57 | 0.31 | 0.21 | 0.31 | 0.24 |
| 2 | 13372691 | rs10209739 | -1.099 | -1.189 | 0.109 | -0.466 | 0.028 | -0.019 | -0.019 | 0 | 0.31 | 0.75 | 0.38 | 0.39 | 0.18 | 0.26 |
| 2 | 13380032 | rs9287761 | -1.169 | -1.19 | 0.109 | -0.467 | 0.039 | -0.024 | -0.024 | 6.308 | 0.32 | 0.79 | 0.38 | 0.39 | 0.18 | 0.26 |
| 2 | 13384186 | rs9678154 | 0.306 | -1.173 | 0.177 | -0.457 | 0.036 | 0.075 | 0.075 | 0 | 0.12 | 0.43 | 0.21 | 0.2 | 0.1 | 0.05 |
| 2 | 13392313 | rs13400473 | 0.892 | -1.155 | -0.013 | -0.459 | -0.043 | 0.001 | 0.001 | 5.648 | 0.09 | 0.18 | 0.07 | 0.05 | 0.03 | 0.04 |
| 2 | 13394398 | rs412783 | 1.301 | -1.158 | 0.056 | -0.485 | -0.001 | 0.001 | 0.001 | 0 | 0.09 | 0.08 | 0.1 | 0.13 | 0.06 | 0.17 |
| 2 | 13394400 | rs7608314 | 1.597 | -1.148 | 0.001 | -0.501 | -0.021 | 0.07 | 0.07 | 0 | 0.21 | 0.08 | 0.1 | 0.01 | 0.21 | 0.19 |
| 2 | 13396424 | rs389710 | -0.339 | -1.11 | 0.026 | -0.471 | -0.002 | -0.007 | -0.007 | 1.887 | 0.4 | 0.38 | 0.28 | 0.19 | 0.3 | 0.4 |
| 2 | 13403476 | rs397822 | 0.449 | -0.402 | 0.815 | 0.349 | -0.004 | 0.077 | 0.077 | 0 | 0.1 | 0.2 | 0.16 | 0.18 | 0.06 | 0.18 |
| 2 | 13413820 | rs6737979 | -0.981 | -0.469 | 0.656 | 0.29 | -0.013 | 0.043 | 0.043 | 3.258 | 0.16 | 0.67 | 0.27 | 0.25 | 0.1 | 0.06 |
| 2 | 13416593 | rs2380487 | -0.46 | -0.457 | 0.768 | 0.156 | 0.036 | 0.074 | 0.074 | 0 | 0.13 | 0.51 | 0.21 | 0.2 | 0.1 | 0.05 |
| 2 | 13416658 | rs453359 | -1.016 | -0.548 | 0.191 | -0.016 | -0.106 | 0.088 | 0.088 | 0 | 0.81 | 0.97 | 0.73 | 0.97 | 0.64 | 0.8 |
| 2 | 13423903 | rs10495612 | -1.446 | -1.434 | 1.203 | 0.709 | -0.028 | -0.018 | -0.018 | 0 | 0.22 | 0.66 | 0.23 | 0.2 | 0.12 | 0.09 |
| 2 | 13457796 | rs2890439 | -1.842 | -1.413 | 1.116 | 0.817 | 0.04 | -0.019 | -0.019 | 0 | 0.32 | 0.8 | 0.37 | 0.37 | 0.17 | 0.22 |
| 2 | 13466618 | rs7592566 | -1.515 | -0.984 | 0.989 | 0.371 | 0 | 0.031 | 0.031 | 1.528 | 0.15 | 0.62 | 0.21 | 0.2 | 0.1 | 0.06 |
| 2 | 13486778 | rs9287763 | -0.86 | -1.049 | 0.765 | 0.471 | -0.023 | 0.018 | 0.018 | 1.215 | 0.18 | 0.22 | 0.18 | 0.27 | 0.13 | 0.16 |
| 2 | 13490911 | rs901593 | -0.665 | -1.155 | 1.505 | 0.462 | -0.017 | 0.016 | 0.016 | 1.384 | 0.16 | 0.19 | 0.17 | 0.27 | 0.12 | 0.15 |

| | | | | | | | | | | | | | | | | |
|---|---|---|---|---|---|---|---|---|---|---|---|---|---|---|---|---|
| 2 | 13521116 | rs12692499 | -0.63 | -1.249 | 1.39 | 0.39 | -0.02 | 0.006 | 0.006 | 0.819 | 0.21 | 0.23 | 0.18 | 0.27 | 0.13 | 0.15 |
| 2 | 13524913 | rs1843802 | -0.731 | -1.272 | 1.676 | 0.378 | -0.039 | 0.012 | 0.012 | 0.784 | 0.22 | 0.31 | 0.19 | 0.27 | 0.13 | 0.15 |
| 2 | 13536772 | rs311998 | -1.376 | -1.387 | 1.914 | 0.321 | -0.007 | 0 | 0 | 0.379 | 0.27 | 0.7 | 0.22 | 0.27 | 0.14 | 0.15 |
| 2 | 13565136 | rs4290631 | -0.711 | -1.434 | 1.969 | -0.082 | -0.096 | -0.154 | -0.154 | 0.098 | 0.21 | 0.53 | 0.08 | 0 | 0.1 | 0.1 |
| 2 | 13574479 | rs4541235 | -0.697 | -1.429 | 1.971 | -0.066 | -0.096 | -0.148 | -0.148 | 0 | 0.21 | 0.53 | 0.08 | 0 | 0.1 | 0.1 |
| 2 | 13576289 | rs1594385 | -1.495 | -1.342 | 1.07 | -0.68 | -0.194 | 0.205 | 0.205 | 1.423 | 0.69 | 0.98 | 0.61 | 0.97 | 0.48 | 0.74 |
| 2 | 13578484 | rs1835395 | -0.18 | -1.253 | 2.11 | 0.099 | -0.006 | -0.032 | -0.032 | 0.799 | 0.17 | 0.19 | 0.05 | 0 | 0.1 | 0.1 |
| 2 | 13578736 | rs6414058 | -1.135 | -1.247 | 1.925 | 0.102 | -0.029 | -0.063 | -0.063 | 0 | 0.23 | 0.74 | 0.1 | 0 | 0.1 | 0.1 |
| 2 | 13585999 | rs10495613 | -1.053 | -1.249 | 1.924 | 0.101 | -0.029 | -0.064 | -0.064 | 0 | 0.22 | 0.74 | 0.1 | 0 | 0.1 | 0.09 |
| 2 | 13587240 | rs16860308 | -0.553 | -1.058 | 1.338 | 0.2 | -0.063 | -0.111 | -0.111 | 0.746 | 0.19 | 0.39 | 0.07 | 0 | 0.1 | 0.1 |
| 2 | 13599799 | rs13417970 | 0.327 | -0.837 | 0.251 | 0.844 | -0.01 | -0.068 | -0.068 | 0 | 0.22 | 0.05 | 0.23 | 0.41 | 0.21 | 0.29 |
| 2 | 13606441 | rs10166475 | 0.444 | -0.917 | 0.279 | 0.838 | -0.008 | -0.049 | -0.049 | 0 | 0.22 | 0.07 | 0.28 | 0.4 | 0.21 | 0.29 |
| 2 | 13614307 | rs10495614 | -0.265 | -0.86 | 0.285 | 1.054 | 0 | -0.001 | -0.001 | 0 | 0.28 | 0.68 | 0.33 | 0.39 | 0.21 | 0.34 |
| 2 | 13619589 | rs1996957 | -1.02 | -0.693 | -0.407 | 0.945 | 0.051 | 0.035 | 0.035 | 0.076 | 0.47 | 0.68 | 0.42 | 0.64 | 0.33 | 0.53 |
| 2 | 13621702 | rs2380510 | 0.009 | -0.404 | -0.592 | 1.149 | 0 | 0.003 | 0.003 | 0 | 0.24 | 0.21 | 0.28 | 0.39 | 0.21 | 0.32 |
| 2 | 13623160 | rs10197142 | -0.684 | -0.069 | -1.115 | 1.017 | 0.02 | 0.054 | 0.054 | 2.022 | 0.51 | 0.89 | 0.48 | 0.65 | 0.44 | 0.56 |
| 2 | 13625370 | rs480881 | -1.337 | -0.446 | -0.531 | 2.164 | 0.004 | 0.009 | 0.009 | 0.535 | 0.09 | 0.05 | 0.05 | 0.35 | 0.07 | 0.19 |
| 2 | 13626114 | rs511773 | -1.996 | -0.304 | -0.584 | 2.283 | -0.005 | -0.016 | -0.016 | 0.012 | 0.13 | 0.07 | 0.09 | 0.4 | 0.16 | 0.21 |
| 2 | 13627258 | rs2380508 | 2.046 | -0.009 | 0.13 | 2.365 | 0.011 | 0.046 | 0.046 | 1.279 | 0.75 | 0.66 | 0.75 | 0.47 | 0.74 | 0.58 |
| 2 | 13627507 | rs12997126 | 1.777 | 0.046 | 0.107 | 2.419 | -0.002 | 0.14 | 0.14 | 0.344 | 0.84 | 0.67 | 0.89 | 0.61 | 0.86 | 0.74 |
| 2 | 13628120 | rs13022060 | 2.632 | -0.759 | -0.087 | 2.078 | -0.004 | 0.037 | 0.037 | 0.631 | 0.57 | 0.11 | 0.63 | 0.35 | 0.61 | 0.5 |
| 2 | 13634635 | rs16860443 | -0.56 | -0.761 | 0.192 | 2.069 | -0.004 | -0.005 | -0.005 | 0 | 0.09 | 0.09 | 0.15 | 0.15 | 0.12 | 0.15 |
| 2 | 13638298 | rs11695059 | 2.552 | -0.764 | 0.484 | 1.94 | 0.017 | 0.039 | 0.039 | 0.411 | 0.57 | 0.17 | 0.64 | 0.35 | 0.61 | 0.51 |
| 2 | 13649918 | rs494516 | 0.258 | -0.752 | 0.431 | 1.953 | 0.045 | -0.015 | -0.015 | 1.919 | 0.94 | 0.76 | 0.96 | 1 | 0.93 | 0.92 |
| 2 | 13652009 | rs10929862 | 2.246 | -0.777 | 0.807 | 1.96 | -0.002 | -0.002 | -0.002 | 0 | 0.68 | 0.34 | 0.69 | 0.41 | 0.64 | 0.57 |
| 2 | 13652424 | rs12998776 | 2.544 | -0.789 | 0.853 | 1.955 | 0.009 | 0.033 | 0.033 | 0.017 | 0.58 | 0.22 | 0.65 | 0.35 | 0.61 | 0.51 |
| 2 | 13652731 | rs12999397 | 2.175 | -0.681 | 1.034 | 1.996 | 0.008 | 0.009 | 0.009 | 2.44 | 0.69 | 0.4 | 0.69 | 0.41 | 0.64 | 0.57 |
| 2 | 13654269 | rs2163324 | 1.893 | -1.09 | 1.413 | 2.277 | 0.001 | 0.125 | 0.125 | 1.429 | 0.79 | 0.49 | 0.77 | 0.46 | 0.81 | 0.62 |
| 2 | 13660465 | rs13018069 | 2.62 | -1.218 | 1.344 | 2.701 | 0.045 | -0.024 | -0.024 | 0.188 | 0.66 | 0.2 | 0.8 | 0.61 | 0.73 | 0.63 |
| 2 | 13662904 | rs7583247 | 1.691 | -1.148 | 1.484 | 2.636 | -0.001 | 0.136 | 0.136 | 0.113 | 0.84 | 0.68 | 0.89 | 0.61 | 0.86 | 0.75 |
| 2 | 13663715 | rs7579086 | -1.053 | -1.067 | 1.242 | 2.223 | 0.001 | -0.003 | -0.003 | 0.055 | 0.07 | 0.14 | 0.05 | 0.05 | 0.09 | 0.03 |
| 2 | 13674575 | rs2177463 | 1.488 | -1.205 | 1.17 | 2.32 | -0.049 | 0.007 | 0.007 | 0 | 0.77 | 0.45 | 0.88 | 0.74 | 0.88 | 0.79 |
| 2 | 13675599 | rs2177462 | 2.192 | -1.207 | 0.83 | 2.423 | 0.049 | -0.051 | -0.051 | 1.152 | 0.66 | 0.21 | 0.62 | 0.53 | 0.74 | 0.63 |
| 2 | 13676862 | rs2140525 | 1.479 | -1.242 | 1.362 | 2.636 | -0.006 | 0.301 | 0.301 | 0.406 | 0.87 | 0.49 | 0.87 | 0.59 | 0.92 | 0.74 |
| 2 | 13677244 | rs12052341 | -1.415 | -1.387 | 1.471 | 2.359 | 0.013 | 0.124 | 0.124 | 0 | 0.1 | 0.26 | 0.26 | 0.24 | 0.08 | 0.26 |
| 2 | 13677300 | rs12052345 | 0.185 | -1.277 | 1.225 | 1.909 | 0.009 | 0.062 | 0.062 | 0 | 0.81 | 0.72 | 0.82 | 0.66 | 0.82 | 0.73 |
| 2 | 13681063 | rs7604694 | -0.404 | -1.49 | 0.634 | 0.742 | 0.001 | 0.012 | 0.012 | 0.256 | 0.14 | 0.12 | 0.09 | 0.06 | 0.16 | 0.05 |
| 2 | 13683034 | rs7600167 | 0.351 | -1.647 | 0.543 | 0.613 | -0.007 | -0.007 | -0.007 | 1.093 | 0.62 | 0.58 | 0.67 | 0.25 | 0.73 | 0.33 |
| 2 | 13684367 | rs12618935 | 0.849 | -1.283 | 0.087 | 0.751 | 0.006 | 0.034 | 0.034 | 0.985 | 0.47 | 0.34 | 0.56 | 0.19 | 0.57 | 0.28 |
| 2 | 13685707 | rs4668809 | 1.121 | -1.228 | -0.147 | -0.065 | -0.036 | 0.248 | 0.248 | 1.813 | 0.43 | 0.1 | 0.45 | 0.04 | 0.55 | 0.21 |
| 2 | 13687188 | rs1608539 | 0.831 | -1.187 | -0.125 | 0.076 | 0.012 | 0.016 | 0.016 | 0.956 | 0.47 | 0.34 | 0.49 | 0.05 | 0.58 | 0.27 |
| 2 | 13691129 | rs6432397 | -1.166 | -1.579 | 0.275 | 1.02 | -0.024 | 0.011 | 0.011 | 0.065 | 0.23 | 0.63 | 0.32 | 0.28 | 0.19 | 0.18 |
| 2 | 13698137 | rs4669937 | 0.813 | -2.219 | 0.425 | 1.154 | -0.073 | 0.033 | 0.033 | 0 | 0.77 | 0.71 | 0.75 | 0.35 | 0.93 | 0.52 |
| 2 | 13699487 | rs1524007 | 0.863 | -2.214 | 0.407 | 1.145 | -0.101 | 0.05 | 0.05 | 0 | 0.77 | 0.64 | 0.77 | 0.34 | 0.93 | 0.54 |
| 2 | 13716541 | rs10929869 | 0.613 | -2.154 | 0.212 | 1.168 | -0.033 | 0.012 | 0.012 | 0 | 0.79 | 0.78 | 0.79 | 0.36 | 0.92 | 0.53 |
| 2 | 13717866 | rs4668812 | -0.588 | -1.886 | 0.13 | 1.407 | 0.008 | -0.032 | -0.032 | 0.479 | 0.07 | 0.01 | 0.21 | 0.19 | 0.04 | 0.11 |
| 2 | 13719632 | rs6713918 | -1.12 | -1.959 | 1.075 | 1.512 | -0.002 | -0.036 | -0.036 | 0 | 0.17 | 0.44 | 0.2 | 0.09 | 0.17 | 0.24 |
| 2 | 13722345 | rs16860573 | -2.031 | -1.964 | 1.162 | 1.5 | -0.029 | 0.03 | 0.03 | 0.072 | 0.24 | 0.51 | 0.4 | 0.29 | 0.21 | 0.35 |
| 2 | 13723317 | rs7600795 | 0.796 | -2.192 | 1.302 | 2.361 | -0.032 | 0.229 | 0.229 | 0 | 0.89 | 0.7 | 0.93 | 0.65 | 0.95 | 0.86 |
| 2 | 13726242 | rs11678087 | -0.413 | -2.196 | 1.114 | 2.41 | 0.008 | -0.032 | -0.032 | 0.296 | 0.07 | 0.01 | 0.2 | 0.21 | 0.04 | 0.11 |
| 2 | 13729211 | rs1403967 | 2.013 | -2.217 | 1.133 | 2.274 | 0.053 | -0.077 | -0.077 | 0 | 0.72 | 0.13 | 0.7 | 0.58 | 0.78 | 0.63 |
| 2 | 13736861 | rs2221783 | -0.704 | -2.106 | 1.263 | 2.284 | 0.001 | 0.013 | 0.013 | 0 | 0.07 | 0.06 | 0.2 | 0.24 | 0.03 | 0.11 |
| 2 | 13741861 | rs6760320 | -1.364 | -2.066 | 1.802 | 1.76 | 0.016 | 0.013 | 0.013 | 9.333 | 0.05 | 0.3 | 0.03 | 0 | 0 | 0.01 |
| 2 | 13742696 | rs16860626 | -0.897 | -1.866 | 1.536 | 1.881 | -0.001 | -0.004 | -0.004 | 0 | 0.14 | 0.18 | 0.18 | 0.11 | 0.17 | 0.25 |
| 2 | 13745887 | rs2136851 | 1.212 | -1.678 | 1.333 | 2.028 | -0.025 | 0.147 | 0.147 | 0 | 0.85 | 0.75 | 0.73 | 0.43 | 0.91 | 0.76 |
| 2 | 13778370 | rs10754962 | 1.388 | -1.586 | 1.717 | 2.107 | -0.032 | 0.242 | 0.242 | 0 | 0.83 | 0.55 | 0.72 | 0.42 | 0.9 | 0.78 |
| 2 | 13792156 | rs1510830 | 2.076 | -2.249 | 1.528 | 1.683 | 0.07 | 0.014 | 0.014 | 0.775 | 0.62 | 0.15 | 0.58 | 0.33 | 0.73 | 0.48 |
| 2 | 13792992 | rs1510832 | -0.498 | -2.294 | 1.635 | 1.666 | -0.001 | -0.007 | -0.007 | 0.142 | 0.11 | 0.15 | 0.09 | 0.02 | 0.14 | 0.11 |
| 2 | 13795489 | rs6432404 | 0.855 | -2.304 | 2.189 | 1.682 | -0.015 | 0.102 | 0.102 | 0 | 0.85 | 0.76 | 0.73 | 0.42 | 0.89 | 0.77 |
| 2 | 13799285 | rs13402033 | -1.781 | -2.261 | 2.391 | 1.596 | -0.029 | -0.046 | -0.046 | 3.104 | 0.21 | 0.36 | 0.14 | 0.09 | 0.17 | 0.29 |
| 2 | 13806941 | rs2890449 | -0.702 | -2.365 | 2.019 | 1.642 | -0.007 | 0.025 | 0.025 | 0 | 0.12 | 0.07 | 0.11 | 0.05 | 0.15 | 0.14 |
| 2 | 13810106 | rs6723675 | 1.266 | -2.823 | 2.418 | 1.663 | -0.053 | 0.239 | 0.239 | 0 | 0.83 | 0.62 | 0.85 | 0.46 | 0.91 | 0.78 |
| 2 | 13815205 | rs907168 | 2.019 | -2.711 | 2.355 | 1.681 | 0.002 | -0.004 | -0.004 | 0 | 0.72 | 0.29 | 0.72 | 0.42 | 0.76 | 0.66 |
| 2 | 13817699 | rs2380525 | -1.71 | -2.598 | 1.829 | 1.623 | 0.005 | -0.013 | -0.013 | 0 | 0.19 | 0.2 | 0.27 | 0.37 | 0.17 | 0.23 |
| 2 | 13819064 | rs2089784 | -1.785 | -2.597 | 1.788 | 1.624 | -0.009 | 0.032 | 0.032 | 0 | 0.19 | 0.28 | 0.28 | 0.37 | 0.17 | 0.23 |
| 2 | 13892092 | rs1588477 | -0.725 | -2.46 | 1.24 | 1.77 | 0.015 | 0.011 | 0.011 | 0 | 0.07 | 0.03 | 0.16 | 0.35 | 0.03 | 0.11 |
| 2 | 13902858 | rs17302935 | 1.621 | -3.041 | 0.121 | 1.081 | 0.039 | 0.109 | 0.109 | 0 | 0.43 | 0.11 | 0.56 | 0.24 | 0.6 | 0.32 |
| 2 | 13903204 | rs12623139 | 1.614 | -3.061 | 0.111 | 1.073 | 0.04 | 0.107 | 0.107 | 0 | 0.43 | 0.11 | 0.56 | 0.24 | 0.6 | 0.32 |
| 2 | 13907633 | rs721173 | -0.274 | -3.123 | 0.093 | 1.62 | 0.019 | -0.015 | -0.015 | 0 | 0.08 | 0.02 | 0.06 | 0.31 | 0.04 | 0.09 |

| | | | | | | | | | | | | | | | | |
|---|---|---|---|---|---|---|---|---|---|---|---|---|---|---|---|---|
| 2 | 13907832 | rs6432412 | -0.191 | -3.28 | 0.608 | 1.095 | 0.002 | -0.011 | -0.011 | 0 | 0.76 | 0.78 | 0.82 | 0.96 | 0.75 | 0.83 |
| 2 | 13911161 | rs6744168 | -0.962 | -2.959 | 1.274 | 1.464 | -0.054 | 0.108 | 0.108 | 0.004 | 0.21 | 0.4 | 0.17 | 0.38 | 0.11 | 0.42 |
| 2 | 13916789 | rs1446548 | -1.44 | -2.922 | 0.958 | 1.483 | -0.069 | 0.075 | 0.075 | 1.474 | 0.24 | 0.57 | 0.18 | 0.38 | 0.11 | 0.41 |
| 2 | 13926932 | rs12471894 | -0.086 | -1.807 | 0.1 | 0.831 | -0.017 | 0.008 | 0.008 | 0.398 | 0.35 | 0.3 | 0.4 | 0.24 | 0.37 | 0.44 |
| 2 | 13931667 | rs6725485 | -0.423 | -2.104 | -0.167 | 0.172 | 0.001 | -0.058 | -0.058 | 1.067 | 0.72 | 0.59 | 0.66 | 0.84 | 0.69 | 0.81 |
| 2 | 13934384 | rs2380538 | 0.14 | -2.183 | -0.2 | -0.028 | -0.009 | -0.054 | -0.054 | 0 | 0.64 | 0.49 | 0.62 | 0.82 | 0.65 | 0.66 |
| 2 | 13942346 | rs2034264 | 0.367 | -2.681 | -0.584 | 0.44 | -0.031 | 0.083 | 0.083 | 0 | 0.12 | 0.2 | 0.11 | 0.4 | 0.04 | 0.13 |
| 2 | 13943821 | rs1348801 | -1.091 | -2.753 | -0.092 | 0.347 | -0.006 | 0.026 | 0.026 | 0 | 0.75 | 0.81 | 0.69 | 0.84 | 0.69 | 0.81 |
| 2 | 13945596 | rs11683451 | 1.95 | -2.696 | -0.81 | 0.066 | 0.016 | 0.038 | 0.038 | 0 | 0.19 | 0.1 | 0.13 | 0.07 | 0.19 | 0.08 |
| 2 | 13950758 | rs1900295 | -1.215 | -2.43 | -0.709 | 0.116 | 0.002 | -0.035 | -0.035 | 0.724 | 0.74 | 0.69 | 0.67 | 0.84 | 0.69 | 0.81 |
| 2 | 13951524 | rs6432417 | 0.76 | -2.421 | -0.845 | 0.14 | 0.07 | -0.005 | -0.005 | 0.119 | 0.5 | 0.21 | 0.51 | 0.42 | 0.61 | 0.53 |
| 2 | 13952333 | rs12053008 | -0.634 | -1.947 | -0.775 | 0.346 | -0.023 | 0.038 | 0.038 | 0.037 | 0.22 | 0.37 | 0.18 | 0.41 | 0.14 | 0.29 |
| 2 | 13952619 | rs2053730 | -0.728 | -1.827 | -0.828 | -0.026 | -0.001 | -0.012 | -0.012 | 6.574 | 0.55 | 0.48 | 0.53 | 0.77 | 0.5 | 0.73 |
| 2 | 13953494 | rs10194191 | -1.348 | -1.278 | -1.363 | 0.153 | 0.09 | 0.083 | 0.083 | 0.185 | 0.44 | 0.87 | 0.43 | 0.75 | 0.28 | 0.55 |
| 2 | 13970627 | rs1446540 | -0.556 | -1.501 | -0.648 | 0.558 | -0.1 | 0.14 | 0.14 | 0.598 | 0.23 | 0.49 | 0.09 | 0.47 | 0.08 | 0.2 |
| 2 | 13973964 | rs6714585 | -0.505 | -1.143 | -0.597 | 0.554 | -0.052 | 0.135 | 0.135 | 0.971 | 0.23 | 0.49 | 0.11 | 0.47 | 0.14 | 0.21 |
| 2 | 13975917 | rs1349841 | -1.307 | -1.474 | -0.813 | 0.144 | 0.032 | 0.121 | 0.121 | 0.161 | 0.52 | 0.77 | 0.52 | 0.8 | 0.46 | 0.44 |
| 2 | 13978355 | rs4669956 | -1.763 | -0.251 | -0.923 | 0.007 | -0.039 | 0.096 | 0.096 | 0.236 | 0.59 | 0.85 | 0.63 | 0.83 | 0.52 | 0.65 |
| 2 | 13979062 | rs2380541 | 0.308 | -0.303 | -0.619 | 0.352 | -0.035 | 0.023 | 0.023 | 0.851 | 0.21 | 0.4 | 0.1 | 0.27 | 0.13 | 0.17 |
| 2 | 13981991 | rs17310819 | 1.835 | -0.436 | -0.693 | 0.265 | 0.004 | 0.029 | 0.029 | 0.615 | 0.11 | 0.04 | 0.05 | 0 | 0.09 | 0.03 |
| 2 | 13983167 | rs1813530 | 1.146 | -0.409 | -0.442 | 0.324 | -0.023 | 0.102 | 0.102 | 0 | 0.1 | 0.23 | 0.05 | 0.27 | 0.04 | 0.16 |
| 2 | 13994015 | rs16855675 | 1.27 | -0.397 | -0.958 | 0.059 | -0.022 | -0.062 | -0.062 | 0 | 0.25 | 0.04 | 0.36 | 0.33 | 0.29 | 0.21 |
| 2 | 13994643 | rs6753171 | 0.997 | -0.33 | -0.844 | 0.076 | 0.007 | -0.004 | -0.004 | 0.495 | 0.06 | 0.11 | 0.11 | 0.02 | 0.06 | 0.12 |
| 2 | 13998923 | rs12995842 | 1.919 | -0.292 | -1.161 | -0.166 | -0.007 | -0.007 | -0.007 | 0.451 | 0.19 | 0.1 | 0.2 | 0.2 | 0.21 | 0.16 |
| 2 | 14001382 | rs10199692 | -1.232 | -0.55 | -0.784 | 0.209 | 0.023 | 0.093 | 0.093 | 0.509 | 0.52 | 0.74 | 0.52 | 0.76 | 0.47 | 0.53 |
| 2 | 14005264 | rs16860891 | 0.674 | -0.144 | -0.808 | 0.502 | -0.041 | -0.061 | -0.061 | 1.724 | 0.26 | 0.04 | 0.38 | 0.34 | 0.33 | 0.33 |
| 2 | 14009046 | rs1947429 | -0.665 | 0.053 | -0.014 | 0.862 | -0.03 | 0.153 | 0.153 | 0.477 | 0.19 | 0.44 | 0.11 | 0.43 | 0.13 | 0.19 |
| 2 | 14017309 | rs9287766 | -1.296 | 0.442 | 0.015 | 1.07 | 0.006 | 0.11 | 0.11 | 0.639 | 0.21 | 0.62 | 0.21 | 0.46 | 0.13 | 0.4 |
| 2 | 14021058 | rs10495622 | -0.829 | 0.431 | 0.427 | 1.199 | -0.007 | 0.136 | 0.136 | 0 | 0.14 | 0.37 | 0.08 | 0.26 | 0.07 | 0.25 |
| 2 | 14031175 | rs16860924 | 0.059 | 0.434 | 0.424 | 1.216 | -0.024 | 0.139 | 0.139 | 0 | 0.1 | 0.27 | 0.05 | 0.26 | 0.03 | 0.13 |
| 2 | 14034822 | rs2890456 | -1.255 | 0.371 | -0.03 | 0.912 | -0.021 | 0.061 | 0.061 | 2.738 | 0.64 | 0.85 | 0.69 | 0.82 | 0.6 | 0.71 |
| 2 | 14035554 | rs7581691 | -0.642 | 0.562 | -0.35 | 1.017 | 0.083 | 0.093 | 0.093 | 1.797 | 0.11 | 0.34 | 0.16 | 0.19 | 0.1 | 0.27 |
| 2 | 14040663 | rs6432423 | -0.377 | 0.518 | -0.619 | 1.079 | 0 | 0 | 0 | 0 | 0.54 | 0.58 | 0.64 | 0.55 | 0.57 | 0.59 |
| 2 | 14044883 | rs11682654 | -0.121 | 0.448 | -0.6 | 1.006 | -0.001 | 0.004 | 0.004 | 2.007 | 0.52 | 0.42 | 0.56 | 0.63 | 0.54 | 0.56 |
| 2 | 14053239 | rs11900054 | -0.013 | 0.526 | -1.239 | 1.126 | 0.004 | 0.016 | 0.016 | 0.172 | 0.07 | 0.14 | 0.11 | 0.19 | 0.06 | 0.15 |
| 2 | 14059586 | rs921692 | 0.134 | 0.389 | -1.539 | 0.88 | -0.024 | 0.029 | 0.029 | 0.541 | 0.4 | 0.16 | 0.47 | 0.36 | 0.46 | 0.3 |
| 2 | 14063359 | rs10206108 | -1.059 | 0.498 | -0.766 | 1.295 | 0 | 0.008 | 0.008 | 0.113 | 0.27 | 0.68 | 0.3 | 0.34 | 0.25 | 0.2 |
| 2 | 14078213 | rs10929893 | -1.046 | 0.549 | -0.796 | 1.31 | 0.002 | 0.016 | 0.016 | 0.327 | 0.27 | 0.65 | 0.31 | 0.34 | 0.27 | 0.19 |
| 2 | 14078931 | rs1546669 | -0.033 | 0.332 | -1.192 | 1.674 | 0.002 | -0.019 | -0.019 | 0 | 0.05 | 0.01 | 0.02 | 0.19 | 0.03 | 0.04 |
| 2 | 14081267 | rs452005 | -0.325 | 0.277 | -1.429 | 1.593 | -0.031 | -0.131 | -0.131 | 0 | 0.17 | 0.01 | 0.29 | 0.43 | 0.18 | 0.34 |
| 2 | 14081905 | rs91166 | -0.201 | 0.186 | -0.652 | 1.081 | -0.001 | 0 | 0 | 0 | 0.41 | 0.39 | 0.64 | 0.4 | 0.49 | 0.46 |
| 2 | 14082251 | rs747387 | -1.25 | -0.891 | 0.52 | 0.833 | -0.023 | 0.064 | 0.064 | 0.003 | 0.2 | 0.59 | 0.14 | 0.31 | 0.16 | 0.15 |
| 2 | 14082777 | rs7597867 | -1.676 | -1.103 | 0.848 | 0.648 | -0.073 | 0.153 | 0.153 | 1.759 | 0.16 | 0.47 | 0.08 | 0.28 | 0.06 | 0.1 |
| 2 | 14093277 | rs16861009 | -1.254 | -1.013 | 0.445 | 0.621 | -0.017 | 0.036 | 0.036 | 0.137 | 0.13 | 0.18 | 0.05 | 0.29 | 0.07 | 0.13 |
| 2 | 14097616 | rs13416408 | -1.633 | -1.133 | 0.701 | 0.24 | -0.063 | -0.043 | -0.043 | 0 | 0.19 | 0.57 | 0.09 | 0.15 | 0.07 | 0.12 |
| 2 | 14111157 | rs17313662 | -0.403 | -1.023 | 1.08 | 0.395 | -0.01 | 0.025 | 0.025 | 0 | 0.05 | 0.15 | 0.04 | 0.17 | 0.04 | 0.08 |
| 2 | 14116991 | rs10495620 | -2.585 | -1.278 | 1.183 | 0.595 | -0.005 | 0 | 0 | 0 | 0.2 | 0.77 | 0.1 | 0.4 | 0.08 | 0.16 |
| 2 | 14120389 | rs13385255 | -2.129 | -1.181 | 1.545 | 0.592 | -0.021 | 0.287 | 0.287 | 0.728 | 0.12 | 0.45 | 0.07 | 0.39 | 0.08 | 0.16 |
| 2 | 14137805 | rs483394 | 0.019 | -1.133 | 0.642 | -0.145 | -0.005 | 0.013 | 0.013 | 0 | 0.85 | 0.93 | 0.91 | 0.99 | 0.85 | 0.95 |
| 2 | 14147318 | rs7596094 | -0.408 | -1.661 | 0.871 | -0.617 | -0.038 | -0.133 | -0.133 | 0.044 | 0.19 | 0.5 | 0.1 | 0.01 | 0.12 | 0.03 |
| 2 | 14147448 | rs7596212 | -1.804 | -1.873 | 0.488 | -0.498 | 0.08 | -0.009 | -0.009 | 0 | 0.33 | 0.85 | 0.17 | 0.36 | 0.18 | 0.2 |
| 2 | 14154731 | rs6432431 | -0.644 | -1.634 | 0.708 | -0.379 | -0.054 | 0.039 | 0.039 | 0.76 | 0.21 | 0.46 | 0.11 | 0.21 | 0.14 | 0.11 |
| 2 | 14156600 | rs1878831 | -0.579 | -1.599 | 0.584 | -0.464 | -0.045 | -0.118 | -0.118 | 0 | 0.2 | 0.55 | 0.11 | 0.01 | 0.12 | 0.02 |
| 2 | 14162065 | rs391769 | 1.127 | -1.06 | -0.583 | -0.835 | -0.156 | -0.141 | -0.141 | 0 | 0.26 | 0.02 | 0.54 | 0.56 | 0.41 | 0.44 |
| 2 | 14169341 | rs13422102 | -1.543 | -2.203 | 0.384 | -0.607 | -0.002 | 0.274 | 0.274 | 5.816 | 0.08 | 0.47 | 0.06 | 0.19 | 0.01 | 0.11 |
| 2 | 14170424 | rs13425805 | -1.521 | -2.254 | 0.4 | -0.607 | -0.007 | 0.274 | 0.274 | 5.544 | 0.08 | 0.47 | 0.06 | 0.19 | 0.01 | 0.11 |
| 2 | 14194649 | rs11900823 | -1.491 | -2.401 | 0.359 | -0.947 | -0.073 | 0.035 | 0.035 | 3.885 | 0.15 | 0.56 | 0.07 | 0.13 | 0.03 | 0.11 |
| 2 | 14213045 | rs17315655 | -0.005 | -2.858 | -0.502 | -1.415 | 0.036 | 0.163 | 0.163 | 0.149 | 0.33 | 0.08 | 0.23 | 0.12 | 0.31 | 0.26 |
| 2 | 14218094 | rs360488 | 0.007 | -2.766 | 0.591 | -0.824 | -0.012 | 0.065 | 0.065 | 0.924 | 0.78 | 0.94 | 0.86 | 0.93 | 0.78 | 0.9 |
| 2 | 14229959 | rs360472 | 1.248 | -2.694 | 0.617 | -0.748 | 0.024 | 0.023 | 0.023 | 1.723 | 0.54 | 0.42 | 0.71 | 0.68 | 0.68 | 0.7 |
| 2 | 14233052 | rs360468 | 1.338 | -2.675 | 0.692 | -0.748 | 0.03 | 0.029 | 0.029 | 0.574 | 0.52 | 0.41 | 0.7 | 0.68 | 0.68 | 0.68 |
| 2 | 14235849 | rs6729531 | -0.641 | -2.435 | 0.94 | -0.498 | 0.008 | 0.008 | 0.008 | 4.463 | 0.25 | 0.23 | 0.14 | 0.07 | 0.18 | 0.09 |
| 2 | 14236023 | rs6729808 | -0.564 | -2.421 | 0.871 | -0.492 | 0.013 | 0.032 | 0.032 | 4.705 | 0.24 | 0.17 | 0.14 | 0.07 | 0.18 | 0.09 |
| 2 | 14247375 | rs7425639 | -0.427 | -2.441 | 0.894 | -0.501 | 0.013 | 0.086 | 0.086 | 4.598 | 0.23 | 0.08 | 0.13 | 0.07 | 0.18 | 0.09 |
| 2 | 14255029 | rs7571786 | -0.722 | -2.49 | 1.202 | -0.487 | -0.018 | -0.065 | -0.065 | 0 | 0.26 | 0.36 | 0.16 | 0.08 | 0.18 | 0.09 |
| 2 | 14255041 | rs7583932 | -1.295 | -2.548 | 1.088 | -0.481 | -0.034 | -0.034 | -0.034 | 0.915 | 0.31 | 0.57 | 0.18 | 0.21 | 0.19 | 0.24 |
| 2 | 14258368 | rs6432444 | -0.689 | -2.425 | 0.573 | -0.353 | -0.016 | -0.005 | -0.005 | 0 | 0.14 | 0.21 | 0.11 | 0.13 | 0.11 | 0.09 |
| 2 | 14271246 | rs10174530 | -0.937 | -1.91 | 0.234 | -0.237 | 0.017 | 0.017 | 0.017 | 3.187 | 0.24 | 0.16 | 0.13 | 0.2 | 0.19 | 0.22 |
| 2 | 14286717 | rs6711956 | -1.152 | -1.831 | 0.42 | -0.299 | -0.044 | -0.125 | -0.125 | 0.542 | 0.27 | 0.48 | 0.17 | 0.07 | 0.19 | 0.22 |

| | | | | | | | | | | | | | | | | |
|---|---|---|---|---|---|---|---|---|---|---|---|---|---|---|---|---|
| 2 | 14286928 | rs7572670 | -1.235 | -1.847 | 0.375 | -0.212 | -0.052 | -0.048 | -0.048 | 0 | 0.28 | 0.49 | 0.17 | 0.21 | 0.18 | 0.31 |
| 2 | 14291599 | rs7355592 | -0.024 | -1.818 | 0.021 | -0.233 | -0.001 | 0 | 0 | 0 | 0.09 | 0.09 | 0.08 | 0.06 | 0.11 | 0.07 |
| 2 | 14303639 | rs16861267 | -0.483 | -1.831 | 0.176 | -0.213 | -0.038 | -0.011 | -0.011 | 0 | 0.17 | 0.31 | 0.08 | 0.14 | 0.08 | 0.24 |
| 2 | 14303716 | rs13384764 | -0.868 | -1.729 | 0.077 | 0.048 | -0.032 | -0.01 | -0.01 | 0 | 0.27 | 0.41 | 0.22 | 0.26 | 0.19 | 0.31 |
| 2 | 14306016 | rs2380560 | -0.17 | -1.635 | -0.078 | 0.482 | -0.017 | -0.016 | -0.016 | 0 | 0.77 | 0.63 | 0.64 | 0.78 | 0.79 | 0.84 |
| 2 | 14307383 | rs10495632 | 0.456 | -2.195 | -0.005 | 0.654 | 0.017 | -0.023 | -0.023 | 0 | 0.1 | 0.07 | 0.05 | 0.17 | 0.06 | 0.18 |
| 2 | 14307530 | rs1839206 | 0.642 | -2.214 | 0.006 | 0.686 | -0.027 | 0.01 | 0.01 | 0.007 | 0.66 | 0.56 | 0.59 | 0.61 | 0.73 | 0.66 |
| 2 | 14309625 | rs12692515 | -0.427 | -2.2 | 0.062 | 0.673 | -0.015 | -0.014 | -0.014 | 0 | 0.77 | 0.65 | 0.64 | 0.78 | 0.79 | 0.84 |
| 2 | 14312862 | rs10495633 | 1.289 | -2.711 | -0.502 | 0.778 | 0.053 | 0.032 | 0.032 | 3.649 | 0.45 | 0.09 | 0.43 | 0.38 | 0.63 | 0.48 |
| 2 | 14316914 | rs1448971 | -0.063 | -2.645 | -0.472 | 0.842 | -0.042 | 0.035 | 0.035 | 0 | 0.2 | 0.32 | 0.13 | 0.24 | 0.1 | 0.18 |
| 2 | 14317807 | rs10193126 | 0.849 | -2.319 | -0.615 | 0.838 | 0.06 | 0.046 | 0.046 | 1.605 | 0.47 | 0.26 | 0.46 | 0.38 | 0.62 | 0.48 |
| 2 | 14319301 | rs13027107 | -0.189 | -2.119 | -0.216 | 1.034 | -0.04 | 0.078 | 0.078 | 0 | 0.13 | 0.29 | 0.08 | 0.2 | 0.05 | 0.13 |
| 2 | 14319838 | rs6706080 | -1.073 | -2.372 | -0.34 | 1.109 | -0.042 | 0.045 | 0.045 | 0.116 | 0.23 | 0.35 | 0.13 | 0.36 | 0.11 | 0.31 |
| 2 | 14325551 | rs12052673 | 0.199 | -2.616 | -0.901 | 0.687 | 0.046 | -0.017 | -0.017 | 0 | 0.21 | 0.12 | 0.12 | 0.2 | 0.1 | 0.12 |
| 2 | 14327042 | rs11693062 | -0.79 | -2.83 | -0.788 | 0.344 | 0.044 | 0.043 | 0.043 | 0.836 | 0.49 | 0.57 | 0.29 | 0.22 | 0.29 | 0.29 |
| 2 | 14330209 | rs11885245 | -0.802 | -2.792 | -1.225 | 0.642 | 0.128 | -0.007 | -0.007 | 0 | 0.51 | 0.74 | 0.31 | 0.39 | 0.3 | 0.39 |
| 2 | 14332239 | rs1003981 | -0.802 | -2.792 | -1.225 | 0.633 | 0.128 | -0.007 | -0.007 | 1.002 | 0.51 | 0.75 | 0.31 | 0.39 | 0.3 | 0.39 |
| 2 | 14334195 | rs7571070 | -0.58 | -2.396 | -1.062 | 0.294 | 0.039 | 0.061 | 0.061 | 0.941 | 0.41 | 0.71 | 0.25 | 0.16 | 0.3 | 0.36 |
| 2 | 14346642 | rs10929904 | -0.719 | -0.924 | -0.942 | 0.765 | -0.015 | -0.013 | -0.013 | 0 | 0.83 | 0.85 | 0.82 | 0.77 | 0.77 | 0.69 |
| 2 | 14356926 | rs7580932 | -0.158 | -1.357 | -0.016 | 0.731 | -0.017 | 0 | 0 | 1.932 | 0.19 | 0.23 | 0.28 | 0.2 | 0.11 | 0.08 |
| 2 | 14361674 | rs16861458 | 1.201 | -1.27 | -0.011 | 0.629 | -0.005 | 0.006 | 0.006 | 0 | 0.35 | 0.4 | 0.26 | 0.21 | 0.46 | 0.25 |
| 2 | 14364007 | rs1947203 | 1.201 | -1.27 | -0.011 | 0.608 | -0.005 | 0.005 | 0.005 | 0 | 0.35 | 0.4 | 0.26 | 0.19 | 0.46 | 0.25 |
| 2 | 14366111 | rs753194 | 0.358 | -1.336 | 0.346 | 0.547 | 0.002 | 0.011 | 0.011 | 0.045 | 0.5 | 0.58 | 0.56 | 0.38 | 0.59 | 0.42 |
| 2 | 14366849 | rs1839211 | 1.266 | -1.292 | 0.444 | 0.551 | 0 | -0.01 | -0.01 | 0 | 0.37 | 0.42 | 0.26 | 0.19 | 0.46 | 0.3 |
| 2 | 14371461 | rs12469265 | -0.289 | -1.384 | 0.67 | 0.656 | -0.07 | 0.006 | 0.006 | 2.914 | 0.19 | 0.38 | 0.26 | 0.2 | 0.06 | 0.07 |
| 2 | 14374195 | rs6719569 | 0.245 | -1.38 | 0.583 | 0.854 | -0.035 | 0.005 | 0.005 | 1.599 | 0.17 | 0.22 | 0.24 | 0.2 | 0.05 | 0.07 |
| 2 | 14393459 | rs12986520 | 1.107 | -1.241 | 0.287 | 0.663 | 0.006 | 0.048 | 0.048 | 0 | 0.47 | 0.38 | 0.32 | 0.33 | 0.48 | 0.34 |
| 2 | 14400474 | rs12476448 | 1.187 | -1.419 | 0.042 | 0.649 | -0.019 | 0.097 | 0.097 | 0 | 0.46 | 0.24 | 0.32 | 0.32 | 0.51 | 0.35 |
| 2 | 14408629 | rs1448979 | -0.873 | -1.353 | 0.037 | 0.784 | 0.007 | 0.024 | 0.024 | 0 | 0.8 | 0.78 | 0.66 | 0.55 | 0.72 | 0.57 |
| 2 | 14410954 | rs13408357 | 1.501 | -1.588 | -0.101 | 0.711 | 0.028 | 0.013 | 0.013 | 0.058 | 0.17 | 0.05 | 0.08 | 0.13 | 0.1 | 0.12 |
| 2 | 14411651 | rs2197844 | -0.824 | -1.539 | 0.026 | 0.937 | 0.052 | -0.013 | -0.013 | 0.038 | 0.39 | 0.74 | 0.51 | 0.56 | 0.2 | 0.27 |
| 2 | 14413397 | rs1823309 | 0.345 | -1.406 | 0.224 | 0.979 | -0.016 | -0.017 | -0.017 | 0 | 0.64 | 0.69 | 0.57 | 0.54 | 0.56 | 0.42 |
| 2 | 14422091 | rs1900854 | -0.116 | -1.411 | 0.167 | 0.867 | -0.019 | 0.009 | 0.009 | 0.031 | 0.69 | 0.78 | 0.73 | 0.75 | 0.61 | 0.54 |
| 2 | 14427728 | rs2380591 | 0.4 | -1.378 | -0.074 | 0.814 | -0.034 | -0.038 | -0.038 | 0 | 0.65 | 0.74 | 0.58 | 0.54 | 0.56 | 0.42 |
| 2 | 14433542 | rs11684967 | 1.988 | -1.24 | -0.28 | 0.745 | -0.098 | 0.085 | 0.085 | 0.017 | 0.31 | 0.05 | 0.22 | 0.18 | 0.41 | 0.27 |
| 2 | 14437473 | rs9287778 | 0.56 | -1.062 | 0.273 | 1.063 | 0.032 | 0.036 | 0.036 | 2.167 | 0.13 | 0.05 | 0.02 | 0.01 | 0.04 | 0.03 |
| 2 | 14454304 | rs6432451 | -0.401 | -1.001 | 0.607 | 1.195 | 0.018 | 0.009 | 0.009 | 0 | 0.8 | 0.68 | 0.79 | 0.76 | 0.76 | 0.72 |
| 2 | 14455500 | rs6735064 | -0.247 | -1.128 | 0.913 | 1.03 | -0.033 | 0.008 | 0.008 | 0.759 | 0.16 | 0.2 | 0.24 | 0.21 | 0.05 | 0.07 |
| 2 | 14471739 | rs2380595 | -0.285 | -1.013 | 0.847 | 1.092 | -0.016 | 0.02 | 0.02 | 0.36 | 0.14 | 0.29 | 0.18 | 0.2 | 0.08 | 0.15 |
| 2 | 14480276 | rs10170192 | 1.341 | -1.007 | 0.524 | 1.227 | 0.04 | -0.019 | -0.019 | 0 | 0.15 | 0.06 | 0.07 | 0.16 | 0.05 | 0.07 |
| 2 | 14482894 | rs4322816 | 0.424 | -0.861 | 0.97 | 1.186 | 0.031 | 0.093 | 0.093 | 0.278 | 0.75 | 0.39 | 0.58 | 0.56 | 0.74 | 0.69 |
| 2 | 14491367 | rs1400028 | -0.61 | -0.774 | 0.396 | 1.179 | -0.032 | -0.031 | -0.031 | 5.432 | 0.83 | 0.87 | 0.81 | 0.75 | 0.76 | 0.73 |
| 2 | 14508416 | rs17344674 | 1.579 | -0.851 | -0.051 | 0.81 | -0.064 | 0.171 | 0.171 | 1.324 | 0.39 | 0.08 | 0.27 | 0.18 | 0.45 | 0.34 |
| 2 | 14512194 | rs13411127 | 0.558 | 0.217 | 1.098 | 1.366 | -0.015 | -0.027 | -0.027 | 0 | 0.08 | 0.1 | 0.02 | 0.01 | 0.03 | 0.02 |
| 2 | 14512212 | rs7562383 | -0.075 | 0.191 | 1.618 | 1.495 | 0.029 | 0.022 | 0.022 | 0 | 0.82 | 0.71 | 0.8 | 0.78 | 0.76 | 0.73 |
| 2 | 14515498 | rs13422220 | -0.194 | 0.234 | 1.652 | 1.555 | 0.003 | 0.042 | 0.042 | 0 | 0.1 | 0.15 | 0.17 | 0.21 | 0.09 | 0.14 |
| 2 | 14515733 | rs1515935 | 0.609 | 0.242 | 1.748 | 1.629 | 0.034 | 0.098 | 0.098 | 0 | 0.76 | 0.57 | 0.59 | 0.56 | 0.74 | 0.69 |
| 2 | 14528945 | rs925782 | 0.765 | 0.21 | 1.19 | 1.584 | 0.011 | 0.04 | 0.04 | 0.586 | 0.74 | 0.31 | 0.58 | 0.56 | 0.74 | 0.69 |
| 2 | 14530593 | rs13387341 | -0.504 | 0.034 | 0.897 | 1.371 | 0.008 | 0.013 | 0.013 | 0 | 0.17 | 0.11 | 0.09 | 0.03 | 0.18 | 0.15 |
| 2 | 14541972 | rs1515931 | -1.315 | 0.074 | 1.231 | 1.585 | -0.028 | 0.038 | 0.038 | 0.199 | 0.29 | 0.42 | 0.45 | 0.43 | 0.25 | 0.33 |
| 2 | 14542524 | rs2090305 | 0.303 | 0.302 | 1.108 | 1.732 | 0.079 | 0.114 | 0.114 | 0 | 0.81 | 0.64 | 0.66 | 0.59 | 0.7 | 0.6 |
| 2 | 14546780 | rs6748459 | 1.554 | 0.1 | 0.945 | 1.37 | 0.002 | -0.007 | -0.007 | 0 | 0.55 | 0.31 | 0.49 | 0.55 | 0.55 | 0.52 |
| 2 | 14555406 | rs1400030 | -0.407 | 0.464 | 1.28 | 1.593 | -0.012 | 0.002 | 0.002 | 0.317 | 0.18 | 0.21 | 0.27 | 0.22 | 0.18 | 0.17 |
| 2 | 14555574 | rs4668827 | -0.443 | 0.461 | 1.285 | 1.592 | -0.011 | 0.002 | 0.002 | 0.382 | 0.18 | 0.22 | 0.27 | 0.22 | 0.18 | 0.17 |
| 2 | 14558592 | rs2199850 | -0.263 | 0.461 | 1.341 | 1.592 | 0.003 | -0.002 | -0.002 | 0 | 0.17 | 0.13 | 0.27 | 0.22 | 0.18 | 0.17 |
| 2 | 14567453 | rs13426290 | -0.643 | 0.447 | 1.434 | 1.584 | -0.02 | 0.002 | 0.002 | 0.951 | 0.19 | 0.31 | 0.28 | 0.22 | 0.18 | 0.18 |
| 2 | 14568877 | rs6752334 | 1.316 | 0.461 | 1.414 | 1.65 | 0.02 | 0.027 | 0.027 | 0.003 | 0.65 | 0.49 | 0.53 | 0.56 | 0.57 | 0.55 |
| 2 | 14572871 | rs1913909 | 0.985 | 0.803 | 0.923 | 1.638 | 0.072 | -0.001 | -0.001 | 0 | 0.75 | 0.53 | 0.77 | 0.79 | 0.66 | 0.66 |
| 2 | 14576248 | rs2380609 | 0.16 | 0.821 | 0.741 | 1.646 | 0.077 | 0.024 | 0.024 | 0 | 0.84 | 0.71 | 0.78 | 0.8 | 0.67 | 0.76 |
| 2 | 14579016 | rs4416191 | 0.855 | 0.868 | 0.614 | 1.673 | 0.07 | -0.001 | -0.001 | 0 | 0.75 | 0.55 | 0.76 | 0.79 | 0.66 | 0.66 |
| 2 | 14583841 | rs4669988 | -0.709 | 0.932 | 0.572 | 1.885 | -0.002 | -0.014 | -0.014 | 0.596 | 0.19 | 0.15 | 0.42 | 0.46 | 0.24 | 0.28 |
| 2 | 14588024 | rs1400025 | 1.701 | 0.783 | 0.41 | 1.644 | -0.002 | -0.011 | -0.011 | 0 | 0.57 | 0.46 | 0.35 | 0.33 | 0.42 | 0.38 |
| 2 | 14588290 | rs7573188 | 1.709 | 0.77 | 0.41 | 1.643 | -0.002 | -0.011 | -0.011 | 0 | 0.57 | 0.46 | 0.35 | 0.33 | 0.42 | 0.38 |
| 2 | 14591055 | rs12329145 | -0.213 | 0.696 | 0.02 | 1.774 | 0.014 | -0.014 | -0.014 | 0.038 | 0.16 | 0.09 | 0.27 | 0.22 | 0.18 | 0.17 |
| 2 | 14592465 | rs993678 | 1.105 | 1.006 | -0.022 | 1.466 | 0.001 | -0.006 | -0.006 | 0 | 0.58 | 0.44 | 0.36 | 0.33 | 0.44 | 0.41 |
| 2 | 14595204 | rs4233908 | 0.21 | 1.912 | -0.416 | 1.79 | -0.051 | -0.053 | -0.053 | 0 | 0.91 | 0.98 | 0.76 | 0.82 | 0.69 | 0.8 |
| 2 | 14610061 | rs2865931 | 0.545 | 1.315 | -0.305 | 1.594 | -0.021 | -0.02 | -0.02 | 0.311 | 0.59 | 0.75 | 0.54 | 0.52 | 0.51 | 0.51 |
| 2 | 14618691 | rs9287779 | 0.762 | 1.303 | -0.209 | 1.565 | 0.004 | 0.063 | 0.063 | 0 | 0.57 | 0.7 | 0.43 | 0.24 | 0.46 | 0.39 |
| 2 | 14627914 | rs10169019 | 0.813 | 1.298 | -0.244 | 1.563 | 0.002 | 0.011 | 0.011 | 0 | 0.57 | 0.62 | 0.36 | 0.24 | 0.46 | 0.39 |

| | | | | | | | | | | | | | | | | |
|---|---|---|---|---|---|---|---|---|---|---|---|---|---|---|---|---|
| 2 | 14632564 | rs6760986 | 0.545 | 1.288 | -0.246 | 1.58 | -0.035 | -0.033 | -0.033 | 0 | 0.6 | 0.82 | 0.55 | 0.52 | 0.51 | 0.5 |
| 2 | 14648430 | rs6725529 | 0.771 | 1.077 | -0.591 | 0.604 | -0.001 | -0.015 | -0.015 | 0 | 0.53 | 0.5 | 0.33 | 0.23 | 0.43 | 0.39 |
| 2 | 14651181 | rs4669992 | 0.841 | 1.473 | -0.691 | 1.348 | -0.021 | -0.01 | -0.01 | 0 | 0.74 | 0.76 | 0.53 | 0.35 | 0.56 | 0.69 |
| 2 | 14651888 | rs10929915 | 1.037 | 1.138 | -0.615 | 1.316 | -0.014 | 0.006 | 0.006 | 0 | 0.69 | 0.71 | 0.73 | 0.79 | 0.59 | 0.64 |
| 2 | 14652352 | rs10203476 | 1.11 | 1.215 | -0.759 | 1.722 | -0.004 | -0.015 | -0.015 | 0.136 | 0.79 | 0.85 | 0.65 | 0.53 | 0.77 | 0.72 |
| 2 | 14653650 | rs7565739 | -1.481 | 1.021 | -1.007 | 0.991 | -0.041 | -0.035 | -0.035 | 0 | 0.4 | 0.76 | 0.63 | 0.54 | 0.48 | 0.47 |
| 2 | 14653846 | rs10195150 | -1.844 | 0.823 | -1.072 | 0.842 | -0.009 | 0.033 | 0.033 | 3.438 | 0.48 | 0.86 | 0.56 | 0.58 | 0.52 | 0.48 |
| 2 | 14654594 | rs10427335 | -1.518 | 1.119 | -1.193 | 0.809 | -0.006 | -0.025 | -0.025 | 0 | 0.41 | 0.78 | 0.49 | 0.53 | 0.38 | 0.39 |
| 2 | 14657186 | rs4669995 | 0.163 | 1.075 | -0.918 | 0.842 | -0.019 | 0.013 | 0.013 | 0.598 | 0.16 | 0.1 | 0.29 | 0.11 | 0.2 | 0.16 |
| 2 | 14659139 | rs6735488 | -1.571 | 1.122 | -0.548 | 0.895 | 0.001 | 0.01 | 0.01 | 0.805 | 0.32 | 0.62 | 0.38 | 0.47 | 0.34 | 0.32 |
| 2 | 14664859 | rs11895101 | -2.177 | 1.123 | -0.682 | 0.802 | 0.012 | 0.02 | 0.02 | 2.795 | 0.47 | 0.81 | 0.68 | 0.58 | 0.55 | 0.5 |
| 2 | 14666507 | rs6432463 | -2.221 | 1.114 | -0.754 | 0.784 | 0.003 | 0.017 | 0.017 | 3.387 | 0.47 | 0.87 | 0.68 | 0.58 | 0.55 | 0.5 |
| 2 | 14673583 | rs11893043 | -0.787 | 1.262 | -0.532 | -0.122 | -0.074 | 0.048 | 0.048 | 0 | 0.22 | 0.08 | 0.31 | 0.14 | 0.35 | 0.2 |
| 2 | 14678285 | rs7598113 | -1.855 | 0.485 | 0.205 | -0.502 | 0.024 | 0.074 | 0.074 | 0 | 0.49 | 0.68 | 0.54 | 0.24 | 0.55 | 0.46 |
| 2 | 14679266 | rs2049681 | 0.079 | 0.612 | 0.294 | -0.515 | 0.002 | -0.01 | -0.01 | 0 | 0.08 | 0.06 | 0.09 | 0.01 | 0.17 | 0.1 |
| 2 | 14681774 | rs2675888 | -2.092 | 0.543 | 0.448 | -0.527 | 0.039 | 0.105 | 0.105 | 0 | 0.52 | 0.76 | 0.56 | 0.26 | 0.58 | 0.46 |
| 2 | 14684454 | rs10181495 | -0.386 | 0.511 | 0.696 | -0.585 | -0.04 | 0.032 | 0.032 | 0.126 | 0.28 | 0.17 | 0.37 | 0.18 | 0.33 | 0.24 |
| 2 | 14684465 | rs2696895 | -1.829 | 0.493 | 0.889 | -0.595 | -0.003 | 0.061 | 0.061 | 0.074 | 0.45 | 0.69 | 0.47 | 0.25 | 0.43 | 0.36 |
| 2 | 14686152 | rs4670003 | -0.548 | 0.493 | 1.023 | -0.595 | -0.006 | 0.005 | 0.005 | 0.975 | 0.31 | 0.32 | 0.38 | 0.2 | 0.34 | 0.24 |
| 2 | 14687525 | rs16861832 | 0.136 | 0.493 | 1.077 | -0.577 | -0.012 | -0.014 | -0.014 | 1.249 | 0.11 | 0.11 | 0.05 | 0.05 | 0.07 | 0.11 |
| 2 | 14688476 | rs10177323 | -0.301 | 0.623 | 1.004 | -0.543 | -0.016 | 0.014 | 0.014 | 0.138 | 0.28 | 0.26 | 0.36 | 0.18 | 0.31 | 0.24 |
| 2 | 14691035 | rs13419352 | -1.034 | 0.54 | 0.926 | -0.543 | 0.007 | -0.013 | -0.013 | 0 | 0.34 | 0.42 | 0.41 | 0.21 | 0.36 | 0.25 |
| 2 | 14691935 | rs6724180 | -2.38 | 0.264 | 0.529 | -0.592 | 0 | 0.074 | 0.074 | 0.027 | 0.59 | 0.87 | 0.58 | 0.26 | 0.59 | 0.47 |
| 2 | 14692761 | rs4670006 | -0.137 | 0.551 | 0.356 | -0.345 | -0.016 | -0.017 | -0.017 | 0 | 0.16 | 0.16 | 0.06 | 0.05 | 0.07 | 0.13 |
| 2 | 14692896 | rs2675895 | -1.223 | 0.568 | 0.344 | -0.367 | -0.012 | -0.101 | -0.101 | 0 | 0.26 | 0.46 | 0.18 | 0.08 | 0.26 | 0.24 |
| 2 | 14693546 | rs6710377 | -0.959 | 0.629 | -0.005 | 0.187 | 0.001 | 0.016 | 0.016 | 0 | 0.42 | 0.39 | 0.43 | 0.23 | 0.38 | 0.36 |
| 2 | 14702349 | rs10181834 | -0.33 | 0.327 | -0.097 | 0.338 | 0.006 | 0.014 | 0.014 | 0 | 0.2 | 0.15 | 0.18 | 0.09 | 0.17 | 0.24 |
| 2 | 14702424 | rs4668834 | 0.641 | 0.354 | 0.433 | 0.385 | 0.003 | 0.006 | 0.006 | 0 | 0.88 | 0.8 | 0.75 | 0.88 | 0.91 | 0.82 |
| 2 | 14705390 | rs2675899 | 1.379 | 0.302 | 0.481 | 0.376 | -0.017 | -0.029 | -0.029 | 0 | 0.66 | 0.59 | 0.56 | 0.77 | 0.73 | 0.57 |
| 2 | 14708545 | rs6734939 | -0.248 | 0.347 | 0.343 | 0.397 | 0.009 | 0.04 | 0.04 | 0 | 0.21 | 0.06 | 0.17 | 0.11 | 0.18 | 0.25 |
| 2 | 14709611 | rs2380617 | -1.257 | 0.374 | 0.565 | 0.428 | 0.011 | 0.015 | 0.015 | 0 | 0.13 | 0.32 | 0.26 | 0.12 | 0.09 | 0.18 |
| 2 | 14711241 | rs13414125 | -1.518 | 0.393 | 0.557 | 0.436 | -0.016 | -0.029 | -0.029 | 0 | 0.32 | 0.42 | 0.43 | 0.23 | 0.26 | 0.42 |
| 2 | 14712421 | rs17348835 | 0.196 | 0.429 | -0.051 | 0.328 | -0.034 | 0.028 | 0.028 | 0 | 0.17 | 0.01 | 0.16 | 0.09 | 0.17 | 0.24 |
| 2 | 14712895 | rs13420936 | -0.337 | 0.436 | -0.019 | 0.329 | 0.003 | 0.005 | 0.005 | 0 | 0.21 | 0.21 | 0.18 | 0.11 | 0.17 | 0.24 |
| 2 | 14735509 | rs1434978 | -0.624 | 0.549 | -0.626 | 0 | -0.001 | -0.026 | -0.026 | 0 | 0.21 | 0.76 | 0.18 | 0.07 | 0.21 | 0.18 |
| 2 | 14743474 | rs16861918 | -0.109 | 0.665 | -0.629 | 0.237 | -0.014 | -0.011 | -0.011 | 0 | 0.08 | 0.01 | 0.17 | 0.09 | 0.09 | 0.18 |
| 2 | 14745146 | rs2015245 | -0.847 | 0.355 | -0.491 | 0.714 | -0.012 | -0.048 | -0.048 | 0 | 0.41 | 0.83 | 0.5 | 0.51 | 0.42 | 0.48 |
| 2 | 14765200 | rs11681353 | -0.727 | -0.049 | 0.12 | 0.436 | 0 | 0.031 | 0.031 | 1.761 | 0.31 | 0.2 | 0.22 | 0.14 | 0.3 | 0.33 |
| 2 | 14766820 | rs4670011 | -1.503 | -0.342 | 0.273 | 0.638 | 0.015 | -0.002 | -0.002 | 0 | 0.42 | 0.89 | 0.39 | 0.35 | 0.36 | 0.39 |
| 2 | 14768496 | rs6704919 | -1.148 | -0.303 | 1.76 | 0.638 | 0.043 | 0.179 | 0.179 | 0.378 | 0.08 | 0.34 | 0.16 | 0.2 | 0.06 | 0.07 |
| 2 | 14784871 | rs10183223 | -0.516 | -0.32 | 2.022 | 0.504 | 0.001 | 0.006 | 0.006 | 0.338 | 0.28 | 0.26 | 0.19 | 0.13 | 0.25 | 0.32 |
| 2 | 14796606 | rs11690472 | -0.678 | -0.252 | 1.529 | 0.532 | -0.016 | -0.046 | -0.046 | 0.079 | 0.31 | 0.46 | 0.2 | 0.13 | 0.25 | 0.32 |
| 2 | 14797165 | rs10202947 | -0.993 | -0.308 | 0.898 | 0.508 | -0.001 | 0 | 0 | 0 | 0.36 | 0.64 | 0.35 | 0.37 | 0.31 | 0.4 |
| 2 | 14800749 | rs765786 | -1.221 | -0.264 | 0.127 | 0.468 | 0.023 | 0.006 | 0.006 | 2.387 | 0.41 | 0.86 | 0.36 | 0.34 | 0.35 | 0.4 |
| 2 | 14805647 | rs1019999 | -1.416 | -0.456 | -0.098 | 0.407 | 0.022 | -0.011 | -0.011 | 0 | 0.46 | 0.92 | 0.47 | 0.38 | 0.37 | 0.37 |
| 2 | 14815508 | rs4670016 | -0.936 | -0.553 | 0.866 | 0.266 | 0.007 | -0.005 | -0.005 | 1.886 | 0.38 | 0.3 | 0.39 | 0.38 | 0.33 | 0.37 |
| 2 | 14867040 | rs1897912 | -1.742 | -0.894 | 0.777 | 0.208 | 0.043 | -0.003 | -0.003 | 0.912 | 0.51 | 0.9 | 0.47 | 0.4 | 0.38 | 0.49 |
| 2 | 14873608 | rs2714235 | -1.058 | -1.144 | 1.523 | 0.107 | 0.012 | -0.018 | -0.018 | 0 | 0.39 | 0.19 | 0.38 | 0.4 | 0.33 | 0.47 |
| 2 | 14877094 | rs2714240 | -1.697 | -1.115 | 2.09 | 0.123 | 0.021 | 0.009 | 0.009 | 3.548 | 0.47 | 0.6 | 0.45 | 0.4 | 0.38 | 0.49 |
| 2 | 14879690 | rs2571626 | -0.947 | -0.82 | 2.364 | 0.299 | -0.005 | 0.009 | 0.009 | 0.294 | 0.37 | 0.36 | 0.41 | 0.44 | 0.33 | 0.4 |
| 2 | 14881995 | rs7607607 | -0.604 | -0.957 | 2.523 | 0.225 | -0.01 | 0.066 | 0.066 | 0 | 0.28 | 0.51 | 0.39 | 0.38 | 0.26 | 0.3 |
| 2 | 14891882 | rs2010753 | -2.204 | -0.836 | 2.48 | 0.179 | 0.027 | -0.011 | -0.011 | 0 | 0.47 | 0.84 | 0.49 | 0.42 | 0.4 | 0.42 |
| 2 | 14892816 | rs2571631 | -1.559 | -0.842 | 2.375 | 0.069 | -0.012 | -0.036 | -0.036 | 0 | 0.17 | 0.25 | 0.06 | 0.04 | 0.09 | 0.12 |
| 2 | 14898514 | rs1434969 | -0.883 | -0.763 | 2.209 | 0.174 | -0.008 | 0.054 | 0.054 | 0 | 0.3 | 0.51 | 0.37 | 0.38 | 0.26 | 0.32 |
| 2 | 14900902 | rs1529833 | -2.037 | -0.773 | 2.071 | 0.174 | 0.008 | -0.018 | -0.018 | 0 | 0.46 | 0.89 | 0.48 | 0.39 | 0.4 | 0.42 |
| 2 | 14907998 | rs2705845 | -1.619 | -0.467 | 2.002 | 0.223 | 0.005 | -0.003 | -0.003 | 0.176 | 0.41 | 0.63 | 0.43 | 0.41 | 0.34 | 0.42 |
| 2 | 14919425 | rs2571637 | -0.937 | -0.515 | 1.979 | 0.093 | -0.007 | -0.039 | -0.039 | 0 | 0.11 | 0.14 | 0.05 | 0 | 0.07 | 0.05 |
| 2 | 14945300 | rs16862033 | -0.869 | -0.331 | 2.135 | 0.134 | 0.001 | 0.036 | 0.036 | 0 | 0.31 | 0.52 | 0.39 | 0.39 | 0.31 | 0.4 |
| 2 | 14952185 | rs6724694 | -0.928 | -0.346 | 2.146 | 0.127 | -0.001 | 0.022 | 0.022 | 0 | 0.32 | 0.55 | 0.39 | 0.39 | 0.31 | 0.4 |
| 2 | 14958858 | rs2705839 | -0.464 | -0.342 | 2.113 | 0.075 | -0.001 | -0.018 | -0.018 | 0.072 | 0.09 | 0.08 | 0.03 | 0 | 0.06 | 0.05 |
| 2 | 14964657 | rs2679440 | -2.115 | -0.382 | 2.177 | 0.135 | -0.014 | -0.019 | -0.019 | 0 | 0.47 | 0.88 | 0.49 | 0.4 | 0.43 | 0.45 |
| 2 | 14971042 | rs10211519 | -1.101 | -0.255 | 2.735 | 0.221 | 0.001 | -0.007 | -0.007 | 0.294 | 0.33 | 0.68 | 0.41 | 0.4 | 0.31 | 0.39 |
| 2 | 14985079 | rs2679421 | -2.652 | -0.331 | 2.989 | 0.151 | -0.01 | -0.131 | -0.131 | 1.709 | 0.17 | 0.53 | 0.11 | 0 | 0.12 | 0.08 |
| 2 | 15002033 | rs11889848 | -1.532 | -0.317 | 3.014 | 0.176 | 0.026 | 0.003 | 0.003 | 0.891 | 0.06 | 0.32 | 0.03 | 0 | 0.01 | 0.02 |
| 2 | 15004748 | rs1157907 | -0.406 | -0.251 | 1.877 | 0.209 | -0.02 | -0.056 | -0.056 | 0 | 0.26 | 0.13 | 0.36 | 0.4 | 0.3 | 0.36 |
| 2 | 15005388 | rs11884086 | -2.54 | -0.325 | 1.702 | 0.063 | -0.01 | -0.082 | -0.082 | 0.142 | 0.19 | 0.65 | 0.13 | 0 | 0.16 | 0.16 |
| 2 | 15006960 | rs1878847 | -2.462 | -0.308 | 1.232 | 0.102 | -0.019 | -0.065 | -0.065 | 0.869 | 0.14 | 0.7 | 0.12 | 0.01 | 0.1 | 0.11 |
| 2 | 15007094 | rs7423486 | -1.809 | -0.183 | 0.454 | 0.327 | -0.07 | -0.048 | -0.048 | 0 | 0.45 | 0.92 | 0.48 | 0.39 | 0.44 | 0.43 |
| 2 | 15022751 | rs7577316 | -1.188 | -0.281 | 1.171 | -0.049 | -0.003 | -0.05 | -0.05 | 2.531 | 0.09 | 0.22 | 0.05 | 0 | 0.06 | 0.04 |

| | | | | | | | | | | | | | | | | |
|---|---|---|---|---|---|---|---|---|---|---|---|---|---|---|---|---|
| 2 | 15025094 | rs4670032 | -0.179 | -0.368 | 1.073 | 0.071 | -0.035 | -0.027 | -0.027 | 0 | 0.22 | 0.08 | 0.21 | 0.28 | 0.26 | 0.27 |
| 2 | 15026865 | rs17367196 | -0.551 | -0.368 | 1.171 | 0.063 | -0.002 | -0.026 | -0.026 | 0 | 0.07 | 0.09 | 0.04 | 0 | 0.06 | 0.04 |
| 2 | 15027047 | rs6432484 | -2.351 | -0.531 | 1.743 | -0.057 | -0.014 | -0.124 | -0.124 | 1.134 | 0.19 | 0.51 | 0.11 | 0.01 | 0.15 | 0.16 |
| 2 | 15029661 | rs6730959 | -2.048 | -0.535 | 1.589 | 0 | 0.003 | 0.079 | 0.079 | 0 | 0.44 | 0.8 | 0.35 | 0.3 | 0.41 | 0.43 |
| 2 | 15029971 | rs6734284 | -1.822 | -0.492 | 2.107 | -0.038 | 0.015 | -0.018 | -0.018 | 6.262 | 0.09 | 0.39 | 0.05 | 0.01 | 0.05 | 0.11 |
| 2 | 15036358 | rs1518793 | -2.47 | -0.467 | 1.796 | 0.2 | -0.01 | -0.01 | -0.01 | 0 | 0.2 | 0.73 | 0.12 | 0.16 | 0.14 | 0.15 |
| 2 | 15041961 | rs7606159 | -0.212 | -0.724 | 1.228 | -0.328 | -0.002 | -0.009 | -0.009 | 2.067 | 0.24 | 0.22 | 0.19 | 0.06 | 0.26 | 0.16 |
| 2 | 15046146 | rs875199 | -1.619 | -0.459 | 1.454 | -0.196 | 0.002 | 0.012 | 0.012 | 0 | 0.26 | 0.62 | 0.21 | 0.36 | 0.26 | 0.31 |
| 2 | 15046881 | rs11896467 | -1.324 | -0.468 | 2.076 | -0.201 | 0.009 | 0.022 | 0.022 | 0 | 0.23 | 0.27 | 0.19 | 0.36 | 0.25 | 0.31 |
| 2 | 15048635 | rs6432491 | -2.016 | -0.49 | 1.469 | -0.206 | -0.002 | 0.071 | 0.071 | 0 | 0.5 | 0.75 | 0.4 | 0.42 | 0.52 | 0.46 |
| 2 | 15058898 | rs7577131 | -0.612 | -1.611 | 0.828 | -0.894 | 0.022 | 0.037 | 0.037 | 0 | 0.47 | 0.45 | 0.23 | 0.22 | 0.34 | 0.31 |
| 2 | 15059183 | rs17367970 | 0.317 | -1.798 | 0.676 | -1.089 | 0.032 | 0.045 | 0.045 | 0.579 | 0.27 | 0.18 | 0.12 | 0.07 | 0.14 | 0.17 |
| 2 | 15059250 | rs7577600 | 0.846 | -1.812 | 0.701 | -1.099 | 0.011 | -0.025 | -0.025 | 0 | 0.05 | 0.11 | 0.04 | 0 | 0.1 | 0.01 |
| 2 | 15061867 | rs11885902 | -0.54 | -1.815 | 0.695 | -1.079 | 0.042 | 0.057 | 0.057 | 0 | 0.42 | 0.34 | 0.19 | 0.22 | 0.24 | 0.29 |
| 2 | 15064363 | rs10176960 | 1.129 | -1.797 | 0.686 | -1.066 | 0.012 | 0.013 | 0.013 | 0 | 0.15 | 0.09 | 0.06 | 0.15 | 0.1 | 0.13 |
| 2 | 15066145 | rs1568283 | -0.006 | -1.827 | 0.758 | -1.051 | -0.007 | 0.002 | 0.002 | 0 | 0.2 | 0.3 | 0.19 | 0.18 | 0.21 | 0.22 |
| 2 | 15067543 | rs4303691 | -0.021 | -1.828 | 0.721 | -1.051 | -0.007 | 0.002 | 0.002 | 0 | 0.2 | 0.3 | 0.19 | 0.18 | 0.21 | 0.22 |
| 2 | 15068315 | rs10200717 | -0.878 | -1.753 | 0.364 | -1.295 | 0.021 | 0.035 | 0.035 | 0 | 0.47 | 0.45 | 0.23 | 0.22 | 0.34 | 0.31 |
| 2 | 15069050 | rs1010957 | 1.301 | -1.727 | 0.421 | -1.246 | 0.002 | 0.002 | 0.002 | 0 | 0.15 | 0.17 | 0.07 | 0.15 | 0.1 | 0.13 |
| 2 | 15071076 | rs4670044 | 1.04 | -1.748 | 0.292 | -1.259 | 0.006 | 0.008 | 0.008 | 0 | 0.2 | 0.18 | 0.11 | 0.07 | 0.13 | 0.16 |
| 2 | 15071980 | rs12692539 | 0.837 | -1.571 | 0.294 | -1.171 | 0.01 | -0.024 | -0.024 | 0 | 0.05 | 0.11 | 0.04 | 0 | 0.1 | 0.01 |
| 2 | 15074706 | rs11887847 | 0.512 | -1.573 | -0.077 | -1.055 | -0.029 | -0.031 | -0.031 | 0 | 0.18 | 0.34 | 0.08 | 0.15 | 0.1 | 0.13 |
| 2 | 15086877 | rs13411807 | 0.552 | -1.572 | -0.09 | -1.054 | -0.021 | -0.023 | -0.023 | 0 | 0.18 | 0.29 | 0.08 | 0.15 | 0.1 | 0.13 |
| 2 | 15088235 | rs793831 | 1.934 | -1.57 | -0.496 | -1.039 | -0.163 | -0.071 | -0.071 | 0 | 0.31 | 0.05 | 0.56 | 0.62 | 0.45 | 0.47 |
| 2 | 15091566 | rs17368670 | 0.186 | -0.99 | 0.151 | -0.598 | 0 | -0.003 | -0.003 | 0.058 | 0.14 | 0.21 | 0.15 | 0.11 | 0.16 | 0.19 |
| 2 | 15098291 | rs16862183 | 0.008 | -0.78 | -0.877 | -0.471 | -0.027 | -0.03 | -0.03 | 0.741 | 0.18 | 0.32 | 0.1 | 0.15 | 0.1 | 0.13 |
| 2 | 15099176 | rs17368979 | 0.59 | -0.505 | -2.028 | -0.367 | 0.014 | 0.016 | 0.016 | 0 | 0.15 | 0.09 | 0.07 | 0.13 | 0.1 | 0.12 |
| 2 | 15099605 | rs793827 | -0.406 | 0.058 | -1.937 | -0.039 | -0.005 | 0.029 | 0.029 | 0 | 0.79 | 0.91 | 0.87 | 0.89 | 0.79 | 0.82 |
| 2 | 15105026 | rs1594420 | 0.35 | 0.271 | -2.046 | 0.278 | -0.108 | -0.014 | -0.014 | 0 | 0.4 | 0.08 | 0.58 | 0.63 | 0.48 | 0.46 |
| 2 | 15106804 | rs4670047 | -0.718 | -0.639 | -1.304 | 0.295 | -0.025 | 0.003 | 0.003 | 0 | 0.32 | 0.54 | 0.31 | 0.35 | 0.18 | 0.17 |
| 2 | 15109381 | rs6748204 | -0.71 | -1.088 | 0.14 | 0.288 | -0.051 | -0.006 | -0.006 | 0 | 0.29 | 0.58 | 0.27 | 0.24 | 0.1 | 0.17 |
| 2 | 15110323 | rs7587023 | -1.567 | -1.054 | -0.283 | -0.27 | 0.072 | 0.064 | 0.064 | 0.062 | 0.48 | 0.72 | 0.42 | 0.34 | 0.35 | 0.23 |
| 2 | 15114088 | rs7595667 | -1.566 | -0.952 | -0.239 | -0.267 | 0.062 | 0.057 | 0.057 | 0 | 0.48 | 0.7 | 0.44 | 0.35 | 0.39 | 0.24 |
| 2 | 15121323 | rs12620296 | 0.053 | -1.018 | 0.455 | -0.336 | 0.018 | 0.014 | 0.014 | 0 | 0.25 | 0.19 | 0.12 | 0.16 | 0.1 | 0.15 |
| 2 | 15130998 | rs10198232 | 0 | -1.077 | 0.588 | -0.48 | 0.026 | 0.019 | 0.019 | 0 | 0.25 | 0.19 | 0.11 | 0.15 | 0.1 | 0.15 |
| 2 | 15133451 | rs10495643 | -0.314 | -0.942 | 0.452 | -0.843 | -0.068 | 0.106 | 0.106 | 0 | 0.17 | 0.02 | 0.15 | 0 | 0.28 | 0.07 |
| 2 | 15138286 | rs6432496 | -1.247 | -0.69 | 0.554 | -1.094 | 0.017 | 0.005 | 0.005 | 0 | 0.48 | 0.41 | 0.28 | 0.01 | 0.43 | 0.33 |
| 2 | 15148093 | rs7607769 | -1.343 | -0.565 | 0.629 | -1.066 | 0.002 | 0.012 | 0.012 | 0 | 0.4 | 0.32 | 0.27 | 0 | 0.41 | 0.29 |
| 2 | 15148338 | rs6748326 | -1.239 | -0.512 | 0.558 | -1.003 | -0.002 | -0.078 | -0.078 | 0 | 0.41 | 0.48 | 0.28 | 0.11 | 0.41 | 0.35 |
| 2 | 15162318 | rs4668845 | -0.491 | 0.336 | 0.56 | -0.698 | 0.031 | -0.003 | -0.003 | 0 | 0.22 | 0.33 | 0.22 | 0.19 | 0.32 | 0.36 |
| 2 | 15167324 | rs11889589 | 0.238 | -0.189 | 0.726 | -1.078 | 0.018 | 0.046 | 0.046 | 0.114 | 0.66 | 0.8 | 0.86 | 1 | 0.73 | 0.75 |
| 2 | 15171773 | rs934315 | 0.122 | 0.13 | 0.694 | -1.158 | 0.011 | 0.144 | 0.144 | 0.126 | 0.68 | 0.92 | 0.86 | 1 | 0.73 | 0.75 |
| 2 | 15172249 | rs1367484 | 0.435 | -0.006 | 1.103 | -1.08 | -0.005 | -0.033 | -0.033 | 0.439 | 0.63 | 0.57 | 0.83 | 0.88 | 0.73 | 0.72 |
| 2 | 15179746 | rs16862250 | -0.111 | -0.385 | 0.942 | -1.196 | -0.002 | -0.039 | -0.039 | 0.382 | 0.23 | 0.38 | 0.19 | 0.12 | 0.24 | 0.26 |
| 2 | 15180592 | rs12619056 | -0.086 | -0.464 | 0.593 | -1.25 | 0.047 | 0.138 | 0.138 | 0.166 | 0.67 | 0.9 | 0.87 | 0.99 | 0.77 | 0.69 |
| 2 | 15181133 | rs6720338 | -0.637 | -0.44 | 1.065 | -1.232 | 0.017 | 0.002 | 0.002 | 1.196 | 0.3 | 0.46 | 0.24 | 0.29 | 0.35 | 0.42 |
| 2 | 15186976 | rs1430805 | 2.065 | -0.443 | 0.495 | -0.974 | -0.098 | -0.095 | -0.095 | 0.086 | 0.33 | 0.06 | 0.45 | 0.62 | 0.42 | 0.27 |
| 2 | 15189758 | rs2033352 | -1.578 | -0.256 | 1.145 | -0.533 | -0.013 | -0.027 | -0.027 | 2.431 | 0.36 | 0.84 | 0.42 | 0.38 | 0.36 | 0.42 |
| 2 | 15189846 | rs2033351 | -1.368 | -0.239 | 1.534 | -0.651 | -0.004 | -0.034 | -0.034 | 0.172 | 0.38 | 0.6 | 0.27 | 0.12 | 0.35 | 0.33 |
| 2 | 15194559 | rs1430804 | -0.967 | 0.273 | 2.118 | -0.12 | -0.005 | -0.064 | -0.064 | 0 | 0.3 | 0.46 | 0.23 | 0.11 | 0.31 | 0.25 |
| 2 | 15200653 | rs4668867 | -1.5 | 0.261 | 1.52 | 0.003 | -0.017 | -0.039 | -0.039 | 1.52 | 0.38 | 0.85 | 0.43 | 0.39 | 0.38 | 0.42 |
| 2 | 15208275 | rs1178207 | 0.893 | 0.339 | 1.278 | -0.076 | 0.004 | -0.004 | -0.004 | 0 | 0.64 | 0.42 | 0.73 | 0.7 | 0.63 | 0.59 |
| 2 | 15214955 | rs6714740 | -0.885 | 0.116 | 1.087 | 0.448 | -0.001 | 0.004 | 0.004 | 0.351 | 0.26 | 0.34 | 0.18 | 0.33 | 0.24 | 0.32 |
| 2 | 15229630 | rs13391879 | -0.6 | 0.975 | 0.881 | 0.536 | 0.057 | 0.03 | 0.03 | 0.806 | 0.05 | 0.24 | 0.1 | 0.06 | 0.09 | 0.04 |
| 2 | 15230603 | rs10211655 | -1.051 | 0.67 | 0.895 | 0.774 | -0.002 | 0.014 | 0.014 | 0.045 | 0.2 | 0.26 | 0.14 | 0.27 | 0.19 | 0.36 |
| 2 | 15231280 | rs12464454 | -0.962 | 0.671 | 0.72 | 0.475 | -0.001 | -0.002 | -0.002 | 0.213 | 0.32 | 0.33 | 0.19 | 0.27 | 0.29 | 0.39 |
| 2 | 15232825 | rs10171235 | 0.942 | 0.601 | 1.317 | 0.633 | 0.048 | 0.083 | 0.083 | 0 | 0.9 | 0.66 | 0.79 | 0.86 | 0.9 | 0.94 |
| 2 | 15242150 | rs1178184 | 1.206 | 0.26 | 1.231 | 0.731 | 0.017 | 0.016 | 0.016 | 0 | 0.57 | 0.27 | 0.72 | 0.56 | 0.62 | 0.54 |
| 2 | 15250422 | rs12623344 | -0.636 | 0.234 | 1.116 | 0.693 | 0.003 | -0.002 | -0.002 | 0 | 0.95 | 0.98 | 0.98 | 1 | 0.96 | 0.96 |
| 2 | 15256606 | rs1990851 | -0.844 | 0.24 | 1.643 | 0.708 | -0.016 | -0.007 | -0.007 | 0.395 | 0.34 | 0.4 | 0.22 | 0.37 | 0.29 | 0.4 |
| 2 | 15262385 | rs13025761 | 1.434 | 0.391 | 1.156 | 0.701 | -0.016 | -0.055 | -0.055 | 0 | 0.54 | 0.08 | 0.56 | 0.48 | 0.57 | 0.55 |
| 2 | 15265510 | rs2216536 | -1.004 | 0.258 | 1.379 | 0.672 | 0.002 | -0.001 | -0.001 | 0.106 | 0.36 | 0.62 | 0.39 | 0.43 | 0.33 | 0.41 |
| 2 | 15268841 | rs10180512 | -0.827 | 0.274 | 1.454 | 0.687 | 0 | -0.009 | -0.009 | 3.546 | 0.34 | 0.37 | 0.36 | 0.13 | 0.32 | 0.37 |
| 2 | 15270541 | rs4668873 | 0.634 | 0.42 | 1.247 | 1.119 | -0.008 | 0.112 | 0.112 | 0 | 0.92 | 0.84 | 0.96 | 0.7 | 0.95 | 0.92 |
| 2 | 15274793 | rs2080804 | -0.347 | 0.287 | 0.882 | 0.59 | 0.018 | -0.033 | -0.033 | 0 | 0.25 | 0.38 | 0.22 | 0.14 | 0.3 | 0.27 |
| 2 | 15276188 | rs1862107 | -0.442 | 0.243 | 0.905 | 1.231 | 0.026 | 0.02 | 0.02 | 0 | 0.27 | 0.47 | 0.23 | 0.28 | 0.3 | 0.28 |
| 2 | 15277831 | rs16862376 | -0.422 | 0.23 | 0.921 | 1.363 | 0.027 | -0.02 | -0.02 | 0.137 | 0.27 | 0.47 | 0.22 | 0.19 | 0.3 | 0.28 |
| 2 | 15278561 | rs11688502 | 1.608 | -0.312 | 0.439 | 1.296 | 0.009 | 0.033 | 0.033 | 0.894 | 0.47 | 0.2 | 0.43 | 0.39 | 0.53 | 0.34 |
| 2 | 15280132 | rs6714977 | 1.084 | -0.196 | 0.411 | 1.451 | -0.022 | 0.053 | 0.053 | 0 | 0.74 | 0.61 | 0.65 | 0.54 | 0.78 | 0.62 |

| | | | | | | | | | | | | | | | | |
|---|---|---|---|---|---|---|---|---|---|---|---|---|---|---|---|---|
| 2 | 15281353 | rs12469371 | -0.85 | -0.535 | -0.163 | 1.658 | 0.024 | -0.037 | -0.037 | 0 | 0.16 | 0.09 | 0.25 | 0.46 | 0.12 | 0.34 |
| 2 | 15282571 | rs2193908 | -0.703 | -0.499 | -0.142 | 1.828 | 0.011 | 0.019 | 0.019 | 0 | 0.09 | 0.06 | 0.23 | 0.44 | 0.05 | 0.22 |
| 2 | 15284746 | rs10192667 | -1.079 | -0.748 | -0.002 | 1.392 | 0.01 | -0.006 | -0.006 | 0 | 0.4 | 0.67 | 0.46 | 0.59 | 0.33 | 0.52 |
| 2 | 15293757 | rs7578396 | -0.788 | -0.376 | 0.393 | 1.761 | -0.01 | 0.049 | 0.049 | 0 | 0.16 | 0.27 | 0.26 | 0.25 | 0.12 | 0.23 |
| 2 | 15294683 | rs12466427 | -0.828 | -1.196 | -0.103 | 1.503 | 0.008 | 0.009 | 0.009 | 0.581 | 0.54 | 0.67 | 0.6 | 0.55 | 0.45 | 0.42 |
| 2 | 15295760 | rs6431679 | -1.561 | -1.387 | -0.466 | 1.485 | -0.095 | 0.018 | 0.018 | 0.576 | 0.67 | 0.9 | 0.67 | 0.75 | 0.51 | 0.68 |
| 2 | 15297306 | rs4294976 | -0.622 | -1.437 | -0.321 | 1.343 | 0.048 | 0.018 | 0.018 | 2.129 | 0.5 | 0.61 | 0.54 | 0.56 | 0.34 | 0.4 |
| 2 | 15299860 | rs4668877 | 0.28 | -1.297 | -0.238 | 1.362 | 0 | -0.001 | -0.001 | 0 | 0.33 | 0.34 | 0.3 | 0.33 | 0.24 | 0.18 |
| 2 | 15303754 | rs4668879 | -0.457 | -1.216 | -0.045 | 1.696 | 0.021 | 0.167 | 0.167 | 0 | 0.16 | 0.27 | 0.28 | 0.52 | 0.17 | 0.34 |
| 2 | 15303781 | rs6732271 | 0.224 | -1.629 | 0.019 | 1.514 | -0.015 | -0.002 | -0.002 | 0.003 | 0.34 | 0.38 | 0.17 | 0.23 | 0.22 | 0.15 |
| 2 | 15303803 | rs4668880 | -0.073 | -1.498 | 0.051 | 1.628 | -0.032 | 0.214 | 0.214 | 0 | 0.13 | 0.25 | 0.23 | 0.51 | 0.07 | 0.35 |
| 2 | 15305054 | rs7586087 | 0.747 | -0.921 | -0.426 | 1.418 | -0.051 | -0.024 | -0.024 | 0 | 0.11 | 0.01 | 0.11 | 0.09 | 0.15 | 0.13 |
| 2 | 15305407 | rs10221671 | -0.846 | -1.665 | 0.113 | 1.634 | 0.062 | 0.073 | 0.073 | 0.201 | 0.51 | 0.67 | 0.44 | 0.72 | 0.37 | 0.49 |
| 2 | 15305738 | rs10754948 | -0.11 | -1.752 | 0.13 | 1.515 | -0.03 | 0.006 | 0.006 | 2.756 | 0.37 | 0.5 | 0.22 | 0.35 | 0.26 | 0.28 |
| 2 | 15309714 | rs4668436 | -0.404 | -1.39 | -0.264 | 1.814 | 0.011 | 0.124 | 0.124 | 0 | 0.07 | 0.13 | 0.29 | 0.34 | 0.04 | 0.09 |
| 2 | 15310820 | rs10179288 | 0.356 | -1.642 | -0.273 | 1.636 | -0.018 | -0.036 | -0.036 | 0.5 | 0.33 | 0.45 | 0.19 | 0.18 | 0.26 | 0.2 |
| 2 | 15311954 | rs1469017 | -2.009 | -1.756 | -0.562 | 1.645 | -0.085 | 0.087 | 0.087 | 1.983 | 0.63 | 0.88 | 0.65 | 0.84 | 0.45 | 0.67 |
| 2 | 15322577 | rs6730450 | -1.351 | -1.706 | -0.458 | 1.728 | -0.043 | 0.076 | 0.076 | 0 | 0.25 | 0.4 | 0.44 | 0.64 | 0.15 | 0.44 |
| 2 | 15326834 | rs2302940 | -0.286 | -1.73 | -0.481 | 1.789 | 0.026 | -0.018 | -0.018 | 0 | 0.11 | 0.05 | 0.09 | 0.11 | 0.08 | 0.12 |
| 2 | 15334251 | rs4668882 | -0.678 | -1.658 | -0.404 | 1.822 | -0.01 | 0.157 | 0.157 | 0 | 0.15 | 0.27 | 0.33 | 0.53 | 0.09 | 0.33 |
| 2 | 15351651 | rs13007383 | -1.573 | -1.639 | -0.553 | 1.682 | -0.019 | 0.067 | 0.067 | 0 | 0.19 | 0.26 | 0.39 | 0.45 | 0.11 | 0.21 |
| 2 | 15370718 | rs7568947 | -1.999 | -1.696 | -0.707 | 1.502 | -0.115 | 0.1 | 0.1 | 2.073 | 0.65 | 0.94 | 0.65 | 0.86 | 0.45 | 0.68 |
| 2 | 15372562 | rs6710817 | -0.305 | -1.617 | -0.558 | 1.525 | -0.005 | -0.001 | -0.001 | 1.548 | 0.12 | 0.14 | 0.1 | 0.1 | 0.06 | 0.12 |
| 2 | 15372731 | rs10929352 | 0.479 | -1.732 | -0.554 | 1.371 | -0.013 | -0.016 | -0.016 | 0.082 | 0.28 | 0.43 | 0.16 | 0.18 | 0.2 | 0.2 |
| 2 | 15378381 | rs3764922 | -0.202 | -1.357 | -0.899 | 1.446 | -0.001 | 0.022 | 0.022 | 0 | 0.13 | 0.13 | 0.31 | 0.34 | 0.1 | 0.1 |
| 2 | 15382331 | rs4668441 | -0.968 | -1.37 | -0.913 | 1.317 | -0.007 | 0.009 | 0.009 | 0 | 0.26 | 0.26 | 0.42 | 0.43 | 0.18 | 0.23 |
| 2 | 15384392 | rs7601870 | -1.361 | -0.858 | -1.278 | 1.154 | -0.113 | 0.023 | 0.023 | 2.764 | 0.63 | 0.95 | 0.64 | 0.79 | 0.42 | 0.65 |
| 2 | 15385484 | rs9287655 | -0.742 | -0.834 | -0.611 | 0.804 | 0.004 | 0.003 | 0.003 | 1.05 | 0.55 | 0.64 | 0.34 | 0.58 | 0.41 | 0.5 |
| 2 | 15400577 | rs6711181 | -1.3 | -0.772 | -0.41 | -0.17 | -0.112 | 0.145 | 0.145 | 3.279 | 0.61 | 0.93 | 0.56 | 0.86 | 0.44 | 0.62 |
| 2 | 15409565 | rs735447 | 0.313 | -0.967 | 0.037 | -0.214 | 0.01 | 0.01 | 0.01 | 1.796 | 0.33 | 0.33 | 0.15 | 0.17 | 0.21 | 0.11 |
| 2 | 15410130 | rs4465744 | -1.425 | -0.934 | -0.163 | -0.222 | -0.049 | 0.108 | 0.108 | 1.663 | 0.61 | 0.84 | 0.56 | 0.86 | 0.46 | 0.62 |
| 2 | 15428438 | rs9287656 | -0.054 | -0.861 | -0.027 | -0.127 | -0.027 | -0.017 | -0.017 | 0 | 0.37 | 0.62 | 0.24 | 0.25 | 0.23 | 0.21 |
| 2 | 15434967 | rs7598652 | -1.447 | -0.611 | -0.268 | -0.181 | -0.031 | 0.072 | 0.072 | 1.142 | 0.62 | 0.79 | 0.56 | 0.86 | 0.45 | 0.63 |
| 2 | 15451774 | rs10173491 | -0.06 | -0.306 | -0.497 | 0.138 | -0.008 | -0.002 | -0.002 | 0.078 | 0.35 | 0.4 | 0.21 | 0.29 | 0.22 | 0.12 |
| 2 | 15455991 | rs759806 | -0.385 | 0.396 | -0.635 | 0.681 | -0.042 | -0.069 | -0.069 | 0 | 0.18 | 0.06 | 0.29 | 0.6 | 0.18 | 0.39 |
| 2 | 15471002 | rs6759219 | -0.361 | 0.371 | -0.197 | 0.29 | -0.021 | 0.031 | 0.031 | 3.118 | 0.42 | 0.71 | 0.52 | 0.26 | 0.45 | 0.36 |
| 2 | 15496708 | rs3731997 | 0.601 | 0.113 | 2.007 | 0.517 | -0.002 | -0.001 | -0.001 | 0 | 0.11 | 0.13 | 0.05 | 0.1 | 0.06 | 0.01 |
| 2 | 15505716 | rs13015492 | 1.399 | -0.12 | 1.374 | 0.313 | 0.035 | 0.133 | 0.133 | 0.714 | 0.56 | 0.13 | 0.57 | 0.2 | 0.69 | 0.42 |
| 2 | 15514982 | rs2049720 | 1.149 | -0.09 | 1.544 | 0.256 | 0.013 | 0.039 | 0.039 | 0.225 | 0.59 | 0.35 | 0.6 | 0.18 | 0.69 | 0.52 |
| 2 | 15548390 | rs2031011 | 1.36 | -0.022 | 1.357 | 0.222 | 0.083 | 0.142 | 0.142 | 1.277 | 0.57 | 0.17 | 0.59 | 0.16 | 0.73 | 0.43 |
| 2 | 15564799 | rs6748427 | 1.174 | -0.039 | 1.545 | 0.267 | -0.001 | -0.01 | -0.01 | 0 | 0.59 | 0.45 | 0.6 | 0.17 | 0.69 | 0.52 |
| 2 | 15576750 | rs1473178 | 1.19 | -0.045 | 1.532 | 0.267 | 0.004 | 0.008 | 0.008 | 0.025 | 0.59 | 0.4 | 0.6 | 0.17 | 0.69 | 0.51 |
| 2 | 15614404 | rs13014906 | 1.112 | -0.06 | 1.66 | 0.283 | -0.008 | -0.03 | -0.03 | 0.15 | 0.61 | 0.55 | 0.62 | 0.22 | 0.69 | 0.53 |
| 2 | 15615910 | rs1990754 | 1.165 | -0.05 | 1.663 | 0.288 | -0.011 | -0.045 | -0.045 | 0 | 0.59 | 0.52 | 0.6 | 0.17 | 0.69 | 0.51 |
| 2 | 15616215 | rs2111449 | 0.554 | -0.023 | 1.706 | 0.339 | 0 | 0.036 | 0.036 | 0.234 | 0.73 | 0.58 | 0.67 | 0.34 | 0.76 | 0.55 |
| 2 | 15618954 | rs6710729 | 0.018 | 0.112 | 1.872 | 0.406 | -0.036 | -0.013 | -0.013 | 0 | 0.17 | 0.31 | 0.08 | 0.11 | 0.06 | 0.01 |
| 2 | 15672071 | rs4668921 | 0.355 | -0.224 | 1.67 | 0.122 | 0.005 | 0.011 | 0.011 | 0 | 0.59 | 0.62 | 0.63 | 0.22 | 0.69 | 0.54 |
| 2 | 15674389 | rs13002869 | 0.355 | -0.224 | 1.67 | 0.122 | 0.005 | 0.011 | 0.011 | 0 | 0.59 | 0.62 | 0.63 | 0.22 | 0.69 | 0.54 |
| 2 | 15674686 | rs13029846 | 0.741 | -0.145 | 1.415 | -0.047 | 0.048 | 0.146 | 0.146 | 0.63 | 0.55 | 0.21 | 0.58 | 0.16 | 0.69 | 0.42 |
| 2 | 15678368 | rs7595512 | 0.438 | -0.131 | 1.27 | -0.039 | -0.006 | -0.024 | -0.024 | 0 | 0.58 | 0.56 | 0.6 | 0.17 | 0.69 | 0.51 |
| 2 | 15720031 | rs7591019 | 0.613 | 0.109 | -0.649 | 0.084 | 0.041 | 0.103 | 0.103 | 0.38 | 0.55 | 0.27 | 0.58 | 0.16 | 0.69 | 0.43 |
| 2 | 15722380 | rs7601397 | -0.927 | -0.133 | 0.181 | 0.892 | -0.051 | 0.07 | 0.07 | 0 | 0.18 | 0.47 | 0.15 | 0.29 | 0.14 | 0.37 |
| 2 | 15722503 | rs13391772 | 0.065 | -0.318 | -0.251 | 0.366 | 0.024 | 0.043 | 0.043 | 0 | 0.49 | 0.63 | 0.52 | 0.25 | 0.57 | 0.35 |
| 2 | 15724206 | rs2098581 | -1.033 | -0.096 | -0.516 | 0.169 | 0.018 | 0.146 | 0.146 | 3.841 | 0.58 | 0.83 | 0.65 | 0.87 | 0.59 | 0.8 |
| 2 | 15725243 | rs10206789 | 0.554 | 0.528 | -0.623 | 0.008 | -0.025 | 0.036 | 0.036 | 0.197 | 0.21 | 0.11 | 0.26 | 0.09 | 0.26 | 0.25 |
| 2 | 15726041 | rs13425614 | 0.554 | 0.516 | -0.609 | 0.008 | -0.024 | 0.034 | 0.034 | 0.197 | 0.21 | 0.12 | 0.26 | 0.09 | 0.26 | 0.25 |
| 2 | 15729583 | rs3755130 | -1.114 | 0.479 | -0.301 | 0.68 | -0.01 | 0.065 | 0.065 | 0.293 | 0.45 | 0.71 | 0.49 | 0.29 | 0.42 | 0.55 |
| 2 | 15729820 | rs3755132 | -0.313 | 0.484 | -0.281 | 0.695 | -0.026 | -0.011 | -0.011 | 0.471 | 0.23 | 0.59 | 0.23 | 0.19 | 0.15 | 0.3 |
| 2 | 15731583 | rs3755133 | -0.313 | 0.484 | -0.221 | 0.695 | -0.027 | -0.011 | -0.011 | 0 | 0.23 | 0.59 | 0.23 | 0.19 | 0.16 | 0.3 |
| 2 | 15731859 | rs10153729 | -1.863 | 0.477 | -0.27 | 0.692 | 0.021 | 0.168 | 0.168 | 3.583 | 0.57 | 0.85 | 0.65 | 0.87 | 0.58 | 0.79 |
| 2 | 15735648 | rs2302929 | 0.235 | 0.461 | -0.26 | 1.392 | -0.024 | -0.023 | -0.023 | 0 | 0.1 | 0.03 | 0.14 | 0.37 | 0.16 | 0.2 |
| 2 | 15758114 | rs4668944 | -1.772 | 0.467 | -0.284 | 1.332 | 0.026 | 0.176 | 0.176 | 3.085 | 0.56 | 0.84 | 0.62 | 0.87 | 0.55 | 0.79 |
| 2 | 15772865 | rs976016 | -0.468 | 0.413 | 0.05 | 1.367 | -0.032 | 0.113 | 0.113 | 0 | 0.24 | 0.51 | 0.24 | 0.4 | 0.15 | 0.33 |
| 2 | 15781195 | rs4668951 | -0.101 | 0.535 | 0.442 | 1.191 | -0.003 | 0.002 | 0.002 | 0 | 0.19 | 0.22 | 0.19 | 0.19 | 0.15 | 0.3 |
| 2 | 15783051 | rs2287637 | 0.332 | 0.617 | 0.289 | 1.171 | -0.025 | 0.037 | 0.037 | 0.083 | 0.21 | 0.12 | 0.25 | 0.09 | 0.26 | 0.24 |
| 2 | 15793904 | rs807597 | -0.881 | 0.337 | 0.613 | 1.62 | 0.017 | 0.101 | 0.101 | 0.262 | 0.13 | 0.24 | 0.17 | 0.38 | 0.16 | 0.23 |
| 2 | 15799651 | rs753383 | -1.051 | 0.707 | 0.824 | 1.423 | 0.048 | 0.073 | 0.073 | 1.68 | 0.29 | 0.42 | 0.37 | 0.5 | 0.37 | 0.41 |
| 2 | 15801028 | rs6735541 | -0.087 | 0.735 | 0.888 | 1.444 | 0.051 | -0.079 | -0.079 | 0 | 0.19 | 0.37 | 0.24 | 0.08 | 0.22 | 0.24 |
| 2 | 15804456 | rs807576 | -0.118 | 0.787 | 0.73 | 1.517 | -0.017 | -0.006 | -0.006 | 0 | 0.09 | 0.05 | 0.15 | 0.42 | 0.15 | 0.2 |

| | | | | | | | | | | | | | | | | |
|---|---|---|---|---|---|---|---|---|---|---|---|---|---|---|---|---|
| 2 | 15807070 | rs12479010 | 0.414 | 0.683 | 0.709 | 1.166 | -0.031 | 0.041 | 0.041 | 0 | 0.18 | 0.07 | 0.19 | 0.08 | 0.22 | 0.21 |
| 2 | 15807637 | rs807573 | -1.582 | 0.515 | 0.931 | 1.078 | -0.003 | -0.001 | -0.001 | 0 | 0.39 | 0.39 | 0.45 | 0.6 | 0.44 | 0.41 |
| 2 | 15808935 | rs876718 | -0.006 | 0.296 | 0.797 | 1 | -0.011 | 0.03 | 0.03 | 0 | 0.14 | 0.23 | 0.2 | 0.23 | 0.09 | 0.3 |
| 2 | 15809109 | rs807571 | -0.727 | 0.238 | 0.712 | 0.993 | 0.007 | 0.071 | 0.071 | 0 | 0.19 | 0.29 | 0.23 | 0.52 | 0.21 | 0.2 |
| 2 | 15811833 | rs10177349 | 0.724 | 0.196 | 0.64 | 0.978 | -0.006 | 0.002 | 0.002 | 0 | 0.18 | 0.14 | 0.19 | 0.07 | 0.22 | 0.2 |
| 2 | 15812261 | rs7579979 | -1.285 | 0.315 | 0.68 | 1.026 | -0.001 | -0.001 | -0.001 | 3.651 | 0.4 | 0.61 | 0.44 | 0.6 | 0.4 | 0.42 |
| 2 | 15812393 | rs807570 | -2.307 | 0.294 | 0.352 | 1.012 | -0.01 | 0.176 | 0.176 | 0.21 | 0.55 | 0.86 | 0.66 | 0.83 | 0.53 | 0.73 |
| 2 | 15812891 | rs8179514 | -1.001 | 0.519 | 0.221 | 1.251 | -0.004 | 0.123 | 0.123 | 0.006 | 0.19 | 0.36 | 0.24 | 0.52 | 0.17 | 0.19 |
| 2 | 15814651 | rs7597353 | -0.223 | 0.281 | -0.303 | 0.68 | 0.001 | -0.021 | -0.021 | 0 | 0.22 | 0.24 | 0.21 | 0.07 | 0.23 | 0.2 |
| 2 | 15825003 | rs11693212 | -0.257 | -0.27 | -0.845 | 1.246 | 0.019 | -0.065 | -0.065 | 1.136 | 0.11 | 0.01 | 0.1 | 0.27 | 0.09 | 0.24 |
| 2 | 15825988 | rs6743299 | 1.383 | -0.139 | -0.677 | 1.029 | -0.006 | 0.078 | 0.078 | 0.208 | 0.47 | 0.3 | 0.44 | 0.2 | 0.51 | 0.23 |
| 2 | 15826700 | rs6733775 | -0.426 | 0.679 | -0.377 | 1.713 | 0.005 | 0.127 | 0.127 | 0 | 0.1 | 0.19 | 0.19 | 0.28 | 0.08 | 0.1 |
| 2 | 15827634 | rs6431722 | 0.166 | 0.544 | -0.556 | 1.053 | -0.016 | 0.002 | 0.002 | 0.515 | 0.77 | 0.81 | 0.76 | 0.77 | 0.7 | 0.6 |
| 2 | 15833099 | rs6743428 | -0.321 | -0.362 | -0.51 | 1.335 | 0.038 | -0.048 | -0.048 | 0.167 | 0.13 | 0.03 | 0.05 | 0.24 | 0.05 | 0.27 |
| 2 | 15833184 | rs1123392 | 0.04 | -0.185 | -0.434 | 1.484 | -0.01 | -0.003 | -0.003 | 0.004 | 0.78 | 0.81 | 0.85 | 0.78 | 0.72 | 0.69 |
| 2 | 15833720 | rs6754090 | -0.356 | -0.729 | 0.103 | 1.902 | -0.021 | 0.104 | 0.104 | 0 | 0.09 | 0.21 | 0.15 | 0.27 | 0.05 | 0.08 |
| 2 | 15834052 | rs6754458 | -0.866 | -0.744 | 0.123 | 1.851 | -0.008 | 0.01 | 0.01 | 0.049 | 0.21 | 0.24 | 0.2 | 0.51 | 0.1 | 0.32 |
| 2 | 15834309 | rs11682665 | -1.093 | -0.908 | 0.174 | 1.776 | -0.004 | 0 | 0 | 0 | 0.27 | 0.29 | 0.22 | 0.53 | 0.15 | 0.34 |
| 2 | 15835039 | rs7606345 | 0.024 | -0.284 | 0.529 | 1.926 | 0.019 | 0.005 | 0.005 | 0 | 0.78 | 0.73 | 0.84 | 0.77 | 0.71 | 0.71 |
| 2 | 15839424 | rs4668960 | 1.089 | -0.382 | 0.826 | 1.966 | 0.013 | 0.036 | 0.036 | 0 | 0.68 | 0.5 | 0.68 | 0.49 | 0.63 | 0.63 |
| 2 | 15842835 | rs6733488 | 0.053 | -0.251 | 1.198 | 1.901 | 0.02 | 0 | 0 | 0 | 0.78 | 0.7 | 0.83 | 0.76 | 0.7 | 0.7 |
| 2 | 15842888 | rs6720135 | 0.751 | -0.402 | 1.184 | 1.549 | 0.011 | 0.008 | 0.008 | 0 | 0.6 | 0.53 | 0.56 | 0.47 | 0.57 | 0.53 |
| 2 | 15843840 | rs1862101 | 1.513 | -0.556 | 0.939 | 1.536 | -0.043 | 0.27 | 0.27 | 0 | 0.51 | 0.13 | 0.41 | 0.15 | 0.51 | 0.47 |
| 2 | 15847172 | rs16862941 | -0.44 | -0.439 | 0.901 | 1.547 | -0.004 | 0.047 | 0.047 | 0 | 0.1 | 0.14 | 0.14 | 0.32 | 0.08 | 0.06 |
| 2 | 15850680 | rs12618251 | 0.842 | -0.465 | 0.95 | 1.647 | -0.007 | -0.044 | -0.044 | 0 | 0.16 | 0.04 | 0.22 | 0.22 | 0.13 | 0.08 |
| 2 | 15850849 | rs12618280 | 0.758 | -0.452 | 0.984 | 1.601 | -0.013 | -0.017 | -0.017 | 0 | 0.6 | 0.34 | 0.54 | 0.47 | 0.58 | 0.54 |
| 2 | 15852847 | rs16862954 | -0.547 | -0.252 | 0.815 | 1.724 | -0.004 | 0.026 | 0.026 | 0 | 0.1 | 0.14 | 0.14 | 0.24 | 0.08 | 0.06 |
| 2 | 15854600 | rs1421608 | -0.089 | -0.194 | 1.051 | 1.44 | 0.04 | 0.019 | 0.019 | 0 | 0.81 | 0.67 | 0.85 | 0.75 | 0.73 | 0.7 |
| 2 | 15856372 | rs6746441 | 1.36 | -0.288 | 0.764 | 1.03 | -0.004 | 0.144 | 0.144 | 0 | 0.51 | 0.23 | 0.49 | 0.25 | 0.49 | 0.54 |
| 2 | 15867762 | rs4668964 | -0.239 | -0.19 | 0.425 | 1.258 | -0.001 | 0 | 0 | 0.124 | 0.22 | 0.24 | 0.31 | 0.2 | 0.2 | 0.11 |
| 2 | 15870242 | rs7572151 | 0.162 | 0.159 | 0.687 | 1.236 | 0.021 | -0.004 | -0.004 | 0 | 0.83 | 0.76 | 0.86 | 0.85 | 0.74 | 0.72 |
| 2 | 15871493 | rs880565 | 0.865 | -0.279 | 1.389 | 1.405 | 0.019 | -0.016 | -0.016 | 0 | 0.62 | 0.48 | 0.56 | 0.33 | 0.57 | 0.6 |
| 2 | 15872266 | rs6431733 | -0.384 | -0.608 | 1.107 | 1.057 | -0.047 | -0.015 | -0.015 | 0 | 0.22 | 0.33 | 0.11 | 0.21 | 0.11 | 0.38 |
| 2 | 15872327 | rs2080799 | 0.584 | -0.493 | 0.847 | 1.292 | -0.009 | -0.034 | -0.034 | 1.645 | 0.17 | 0.06 | 0.27 | 0.2 | 0.17 | 0.08 |
| 2 | 15880014 | rs7571026 | -0.26 | -0.088 | 0.653 | 1.36 | 0.002 | 0.009 | 0.009 | 0 | 0.79 | 0.77 | 0.83 | 0.55 | 0.69 | 0.67 |
| 2 | 15880499 | rs4668967 | 0.904 | -0.024 | 0.149 | 1.398 | -0.014 | -0.031 | -0.031 | 0 | 0.15 | 0.07 | 0.25 | 0.22 | 0.18 | 0.07 |
| 2 | 15884555 | rs10186235 | 0.797 | -0.079 | 0.2 | 1.317 | 0.002 | 0.003 | 0.003 | 0 | 0.09 | 0.18 | 0.08 | 0.09 | 0.08 | 0.15 |
| 2 | 15885286 | rs4668968 | -0.853 | 0.037 | 0.192 | 1.319 | 0.008 | -0.008 | -0.008 | 0.975 | 0.36 | 0.3 | 0.35 | 0.41 | 0.3 | 0.47 |
| 2 | 15887726 | rs10929383 | 1.302 | 0.054 | -0.197 | 1.392 | 0.003 | 0.076 | 0.076 | 0 | 0.41 | 0.29 | 0.45 | 0.13 | 0.41 | 0.22 |
| 2 | 15889025 | rs12623348 | 0.988 | 0.338 | -0.276 | 1.492 | 0.009 | 0.003 | 0.003 | 0 | 0.47 | 0.45 | 0.5 | 0.44 | 0.44 | 0.27 |
| 2 | 15889419 | rs751310 | 0.681 | 0.457 | 0.047 | 1.277 | 0.007 | -0.012 | -0.012 | 0.094 | 0.6 | 0.51 | 0.73 | 0.63 | 0.58 | 0.34 |
| 2 | 15893633 | rs10198578 | -1.332 | 0.662 | 0.199 | 1.734 | 0.01 | 0.118 | 0.118 | 0 | 0.07 | 0.21 | 0.07 | 0.2 | 0.06 | 0.06 |
| 2 | 15893891 | rs6705499 | 1.209 | 0.689 | -0.448 | 1.548 | 0.016 | 0.104 | 0.104 | 0.598 | 0.49 | 0.16 | 0.48 | 0.34 | 0.42 | 0.29 |
| 2 | 15895707 | rs1345564 | 0.464 | 0.727 | 0.365 | 1.449 | 0.008 | 0.005 | 0.005 | 0 | 0.58 | 0.55 | 0.61 | 0.54 | 0.51 | 0.34 |
| 2 | 15896271 | rs6431735 | 0.269 | 0.479 | 0.964 | 1.639 | -0.049 | 0.001 | 0.001 | 0 | 0.63 | 0.86 | 0.65 | 0.64 | 0.53 | 0.43 |
| 2 | 15896356 | rs7567922 | -1.566 | 0.253 | 0.875 | 1.679 | 0.045 | 0.194 | 0.194 | 0 | 0.09 | 0.39 | 0.12 | 0.21 | 0.08 | 0.05 |
| 2 | 15898212 | rs13417353 | -1.36 | 0.307 | 0.637 | 1.68 | 0.014 | 0.085 | 0.085 | 0 | 0.08 | 0.22 | 0.1 | 0.21 | 0.08 | 0.06 |
| 2 | 15899188 | rs7576557 | 1.541 | 0.368 | 0.47 | 1.558 | -0.024 | 0.104 | 0.104 | 1.758 | 0.49 | 0.11 | 0.47 | 0.33 | 0.46 | 0.3 |
| 2 | 15900480 | rs7583242 | 0.784 | 0.542 | 0.402 | 1.65 | 0.002 | 0.002 | 0.002 | 0.05 | 0.71 | 0.7 | 0.68 | 0.56 | 0.6 | 0.67 |
| 2 | 15904734 | rs6728750 | 0.323 | 0.588 | 0.314 | 1.709 | -0.032 | -0.023 | -0.023 | 0 | 0.75 | 0.82 | 0.8 | 0.66 | 0.64 | 0.7 |
| 2 | 15905332 | rs16863012 | -0.188 | 0.433 | 0.451 | 1.726 | 0.005 | -0.007 | -0.007 | 0 | 0.16 | 0.1 | 0.13 | 0.24 | 0.11 | 0.29 |
| 2 | 15906778 | rs2544535 | 1.069 | 0.378 | 0.53 | 1.684 | 0.009 | -0.003 | -0.003 | 0 | 0.59 | 0.54 | 0.67 | 0.62 | 0.53 | 0.43 |
| 2 | 15909600 | rs6706165 | 1.358 | 0.398 | 0.957 | 1.73 | -0.011 | 0.008 | 0.008 | 0.484 | 0.56 | 0.39 | 0.6 | 0.42 | 0.51 | 0.41 |
| 2 | 15916430 | rs1981806 | 1.757 | 0.411 | 0.056 | 1.497 | 0.008 | 0.117 | 0.117 | 2.685 | 0.49 | 0.17 | 0.56 | 0.32 | 0.46 | 0.3 |
| 2 | 15917514 | rs11895482 | 1.72 | 0.497 | 0.046 | 1.483 | 0.004 | 0.126 | 0.126 | 2.787 | 0.5 | 0.17 | 0.57 | 0.32 | 0.47 | 0.29 |
| 2 | 15922288 | rs2693019 | -0.349 | 0.612 | 0.16 | 1.469 | -0.033 | 0.017 | 0.017 | 0 | 0.8 | 0.91 | 0.87 | 0.83 | 0.69 | 0.76 |
| 2 | 15922934 | rs12052505 | 1.9 | 0.562 | -0.046 | 1.387 | 0.007 | 0.119 | 0.119 | 2.687 | 0.49 | 0.18 | 0.56 | 0.32 | 0.47 | 0.29 |
| 2 | 15923441 | rs1363058 | 0.234 | 0.429 | 0.127 | 1.364 | -0.139 | 0.04 | 0.04 | 0 | 0.76 | 0.98 | 0.78 | 0.83 | 0.63 | 0.7 |
| 2 | 15926168 | rs2544527 | -0.098 | 0.053 | 0.447 | 1.71 | -0.103 | -0.011 | -0.011 | 0 | 0.78 | 0.99 | 0.85 | 0.79 | 0.71 | 0.7 |
| 2 | 15929276 | rs4668989 | -1.322 | -0.285 | 1.062 | 1.882 | -0.033 | 0.194 | 0.194 | 0 | 0.12 | 0.23 | 0.21 | 0.41 | 0.06 | 0.2 |
| 2 | 15930012 | rs10929389 | 1.599 | -0.255 | 0.367 | 1.522 | -0.028 | 0.118 | 0.118 | 2.387 | 0.53 | 0.13 | 0.52 | 0.28 | 0.57 | 0.37 |
| 2 | 15931387 | rs2380674 | -1.198 | -0.401 | 1.034 | 1.787 | 0 | -0.003 | -0.003 | 0 | 0.42 | 0.51 | 0.34 | 0.6 | 0.37 | 0.53 |
| 2 | 15931402 | rs2380675 | -1.075 | -0.41 | 1.05 | 1.878 | 0.002 | -0.011 | -0.011 | 0 | 0.41 | 0.51 | 0.32 | 0.34 | 0.37 | 0.46 |
| 2 | 15931514 | rs3954898 | -1.434 | -0.461 | 1.075 | 1.359 | 0.002 | -0.012 | -0.012 | 0.998 | 0.5 | 0.53 | 0.54 | 0.85 | 0.43 | 0.67 |
| 2 | 15934372 | rs12998452 | 1.304 | -0.432 | 1.126 | 1.329 | 0.007 | 0.132 | 0.132 | 0.025 | 0.49 | 0.34 | 0.45 | 0.14 | 0.57 | 0.33 |
| 2 | 15935706 | rs1121915 | -0.7 | -0.297 | 1.146 | 1.341 | 0.001 | -0.019 | -0.019 | 0 | 0.07 | 0.06 | 0.03 | 0 | 0.06 | 0.04 |
| 2 | 15935838 | rs1121916 | 1.915 | -0.352 | 1.378 | 1.36 | 0.007 | -0.036 | -0.036 | 1.21 | 0.86 | 0.63 | 0.92 | 0.91 | 0.91 | 0.95 |
| 2 | 15937754 | rs2380677 | -0.726 | -0.429 | 0.919 | 1.52 | -0.023 | -0.095 | -0.095 | 0 | 0.39 | 0.14 | 0.46 | 0.45 | 0.37 | 0.54 |
| 2 | 15938544 | rs16863041 | -1.253 | -0.098 | 1.527 | 1.432 | 0.026 | -0.054 | -0.054 | 0 | 0.14 | 0.32 | 0.13 | 0.04 | 0.15 | 0.08 |

| | | | | | | | | | | | | | | | | |
|---|---|---|---|---|---|---|---|---|---|---|---|---|---|---|---|---|
| 2 | 15940948 | rs11894034 | -0.683 | -0.121 | 1.288 | 1.479 | 0.005 | -0.043 | -0.043 | 0 | 0.12 | 0.33 | 0.13 | 0.06 | 0.11 | 0.07 |
| 2 | 15941752 | rs7565696 | -0.304 | -0.052 | 0.684 | 1.596 | -0.017 | -0.07 | -0.07 | 0 | 0.33 | 0.16 | 0.41 | 0.48 | 0.33 | 0.49 |
| 2 | 15945626 | rs2380680 | -1.813 | -0.075 | 0.294 | 1.467 | -0.038 | 0.142 | 0.142 | 0 | 0.56 | 0.85 | 0.64 | 0.84 | 0.49 | 0.71 |
| 2 | 15947105 | rs7604003 | -0.393 | -0.014 | -0.342 | 1.473 | -0.02 | -0.075 | -0.075 | 0.301 | 0.34 | 0.16 | 0.41 | 0.48 | 0.33 | 0.5 |
| 2 | 15949434 | rs9678924 | 0.95 | 0.031 | -0.116 | 1.335 | 0.002 | 0.249 | 0.249 | 0 | 0.83 | 0.46 | 0.65 | 0.52 | 0.85 | 0.73 |
| 2 | 15950899 | rs12476664 | 1.694 | 0.134 | -1.256 | 1.279 | -0.115 | 0.009 | 0.009 | 0 | 0.4 | 0.05 | 0.36 | 0.31 | 0.49 | 0.29 |
| 2 | 15952057 | rs11687741 | -0.283 | -0.085 | -0.953 | 1.354 | -0.005 | 0.077 | 0.077 | 0.402 | 0.33 | 0.11 | 0.25 | 0.2 | 0.3 | 0.4 |
| 2 | 15957255 | rs4669001 | 1.309 | -0.243 | -0.015 | 1.913 | -0.007 | 0.019 | 0.019 | 2.714 | 0.81 | 0.26 | 0.8 | 0.71 | 0.86 | 0.79 |
| 2 | 15963882 | rs2380699 | 1.338 | -0.369 | 0.272 | 0.982 | -0.021 | -0.048 | -0.048 | 2.543 | 0.86 | 0.73 | 0.9 | 0.94 | 0.93 | 0.88 |
| 2 | 15963991 | rs7575008 | -0.462 | -0.511 | -0.14 | 0.989 | 0.02 | 0.012 | 0.012 | 0 | 0.24 | 0.13 | 0.24 | 0.21 | 0.18 | 0.37 |
| 2 | 15964882 | rs4668468 | 0.731 | -0.463 | 0.322 | 0.841 | -0.014 | -0.036 | -0.036 | 0 | 0.67 | 0.63 | 0.7 | 0.83 | 0.75 | 0.65 |
| 2 | 15965628 | rs7595922 | -1.271 | -0.228 | 0.204 | 1.353 | -0.008 | 0.008 | 0.008 | 0 | 0.39 | 0.48 | 0.39 | 0.44 | 0.35 | 0.54 |
| 2 | 15975615 | rs12613665 | 1.047 | -0.805 | -0.404 | 1.497 | 0.076 | -0.03 | -0.03 | 0 | 0.63 | 0.19 | 0.67 | 0.47 | 0.75 | 0.46 |
| 2 | 15979519 | rs12612477 | 0.98 | -0.782 | -0.15 | 1.525 | 0.021 | -0.01 | -0.01 | 0 | 0.64 | 0.33 | 0.69 | 0.47 | 0.75 | 0.45 |
| 2 | 15982423 | rs11680265 | -0.814 | -0.465 | -0.521 | 1.23 | 0.018 | -0.029 | -0.029 | 0.946 | 0.11 | 0.04 | 0.11 | 0.16 | 0.13 | 0.19 |
| 2 | 15983501 | rs7604985 | 0.931 | -0.691 | -0.448 | 0.836 | -0.004 | -0.013 | -0.013 | 3.152 | 0.91 | 0.91 | 0.95 | 0.97 | 0.95 | 0.96 |
| 2 | 15983519 | rs7564455 | -1.361 | -0.05 | -0.213 | 0.868 | 0.008 | 0.004 | 0.004 | 0 | 0.09 | 0.12 | 0.11 | 0.13 | 0.15 | 0.18 |
| 2 | 15984215 | rs4490149 | 0.11 | -0.103 | -1.683 | 0.141 | 0.002 | 0.133 | 0.133 | 1.227 | 0.52 | 0.04 | 0.47 | 0.25 | 0.57 | 0.51 |
| 2 | 15987278 | rs6726347 | -0.799 | -0.084 | -1.473 | -0.137 | 0.003 | -0.004 | -0.004 | 0 | 0.7 | 0.72 | 0.51 | 0.41 | 0.7 | 0.63 |
| 2 | 15998357 | rs2380684 | 0.67 | -0.062 | -0.917 | -0.481 | -0.026 | 0.084 | 0.084 | 0.009 | 0.43 | 0.29 | 0.29 | 0.22 | 0.42 | 0.3 |
| 2 | 16004423 | rs2890495 | 1.008 | 0.909 | -0.529 | -0.515 | -0.056 | 0.129 | 0.129 | 1.508 | 0.4 | 0.06 | 0.23 | 0.22 | 0.36 | 0.31 |
| 2 | 16011511 | rs7576743 | -0.961 | 0.913 | 0.168 | -0.526 | 0.08 | -0.024 | -0.024 | 2.765 | 0.09 | 0.29 | 0.12 | 0.04 | 0.13 | 0.13 |
| 2 | 16012630 | rs2380707 | -0.16 | 0.716 | 0.035 | -0.715 | -0.014 | -0.04 | -0.04 | 0 | 0.81 | 0.73 | 0.87 | 0.91 | 0.85 | 0.82 |
| 2 | 16015937 | rs1896495 | 0.105 | 0.893 | -0.349 | -0.413 | -0.006 | 0.003 | 0.003 | 0 | 0.06 | 0.03 | 0.07 | 0.04 | 0.12 | 0.13 |
| 2 | 16023245 | rs2559063 | 1.031 | 0.853 | -0.268 | -0.441 | 0.053 | -0.033 | -0.033 | 0 | 0.9 | 0.72 | 0.89 | 0.96 | 0.87 | 0.87 |
| 2 | 16025538 | rs10929398 | 1.209 | 0.983 | -1.348 | -0.817 | -0.046 | 0.068 | 0.068 | 0.878 | 0.31 | 0.05 | 0.18 | 0.17 | 0.28 | 0.27 |
| 2 | 16029453 | rs2559069 | 0.013 | 0.975 | -0.744 | -0.573 | 0.002 | 0.009 | 0.009 | 0.921 | 0.47 | 0.49 | 0.63 | 0.75 | 0.52 | 0.58 |
| 2 | 16030034 | rs2693013 | -0.013 | 0.798 | -0.781 | -0.66 | 0.018 | 0.112 | 0.112 | 0.493 | 0.51 | 0.64 | 0.65 | 0.76 | 0.52 | 0.59 |
| 2 | 16030060 | rs7578895 | 1.078 | 0.886 | -0.841 | -0.699 | -0.06 | 0.094 | 0.094 | 2.537 | 0.35 | 0.05 | 0.22 | 0.2 | 0.32 | 0.27 |
| 2 | 16035256 | rs10495647 | -0.154 | 0.931 | -0.814 | -0.567 | -0.002 | 0.002 | 0.002 | 0 | 0.1 | 0.08 | 0.06 | 0.17 | 0.05 | 0.22 |
| 2 | 16036368 | rs11676479 | 0.664 | 0.974 | -0.897 | -0.617 | 0.003 | 0.134 | 0.134 | 2.188 | 0.39 | 0.23 | 0.26 | 0.18 | 0.33 | 0.27 |
| 2 | 16044792 | rs7594782 | 0.214 | 0.849 | -0.829 | -0.473 | -0.001 | -0.002 | -0.002 | 0.19 | 0.44 | 0.48 | 0.6 | 0.65 | 0.5 | 0.46 |
| 2 | 16059928 | rs11691519 | -0.293 | 0.979 | -1.056 | -0.418 | 0.005 | 0.069 | 0.069 | 0 | 0.52 | 0.39 | 0.32 | 0.26 | 0.5 | 0.55 |
| 2 | 16060483 | rs12476446 | 0.802 | 0.797 | -1.288 | -0.508 | -0.07 | 0.094 | 0.094 | 0 | 0.35 | 0.07 | 0.22 | 0.15 | 0.35 | 0.29 |
| 2 | 16060860 | rs13018245 | 0.802 | 0.766 | -1.288 | -0.508 | -0.067 | 0.094 | 0.094 | 0 | 0.35 | 0.07 | 0.22 | 0.15 | 0.35 | 0.29 |
| 2 | 16062413 | rs1863803 | 0.47 | 1.223 | -0.981 | -0.142 | 0.015 | 0.086 | 0.086 | 0 | 0.43 | 0.3 | 0.26 | 0.19 | 0.38 | 0.41 |
| 2 | 16072049 | rs12692281 | -0.124 | 1.281 | -0.658 | 0.08 | 0 | 0.025 | 0.025 | 0 | 0.49 | 0.48 | 0.29 | 0.23 | 0.47 | 0.55 |
| 2 | 16079754 | rs9653226 | -0.127 | 0.957 | -0.798 | 0.575 | 0.019 | 0.009 | 0.009 | 0 | 0.57 | 0.38 | 0.38 | 0.58 | 0.61 | 0.59 |
| 2 | 16106797 | rs6734021 | -0.623 | 2.311 | -0.394 | 0.814 | 0.044 | -0.048 | -0.048 | 1.171 | 0.13 | 0.19 | 0.17 | 0.01 | 0.27 | 0.11 |
| 2 | 16108123 | rs13431879 | -0.077 | 2.21 | -0.411 | 1.269 | 0.015 | 0.055 | 0.055 | 0 | 0.15 | 0.2 | 0.33 | 0.4 | 0.23 | 0.21 |
| 2 | 16112814 | rs16863159 | -0.018 | 1.932 | -0.585 | 1.183 | 0.005 | 0.241 | 0.241 | 1.688 | 0.13 | 0.49 | 0.27 | 0.28 | 0.09 | 0.13 |
| 2 | 16112828 | rs6716344 | 0.672 | 2.178 | -0.772 | 1.096 | 0.084 | 0.026 | 0.026 | 0 | 0.72 | 0.56 | 0.51 | 0.63 | 0.51 | 0.76 |
| 2 | 16113723 | rs6741005 | -1.138 | 1.538 | -0.427 | 1.57 | 0.011 | 0.117 | 0.117 | 0.272 | 0.23 | 0.5 | 0.32 | 0.34 | 0.22 | 0.25 |
| 2 | 16114295 | rs16855322 | -0.383 | 1.98 | -0.952 | 1.338 | 0.012 | 0.037 | 0.037 | 0 | 0.14 | 0.18 | 0.3 | 0.36 | 0.2 | 0.18 |
| 2 | 16115180 | rs6745287 | -0.82 | 1.247 | -1.188 | 0.599 | -0.007 | 0.007 | 0.007 | 0.055 | 0.12 | 0.09 | 0.07 | 0.07 | 0.13 | 0.12 |
| 2 | 16121864 | rs11685620 | -0.332 | 1.039 | -1.138 | 0.564 | 0 | 0.009 | 0.009 | 0 | 0.46 | 0.42 | 0.56 | 0.36 | 0.53 | 0.48 |
| 2 | 16128823 | rs6751458 | -0.285 | 1.032 | -1.49 | 0.426 | -0.041 | -0.003 | -0.003 | 1.587 | 0.32 | 0.17 | 0.33 | 0.36 | 0.53 | 0.28 |
| 2 | 16133863 | rs17467679 | -0.051 | 1.468 | -1.618 | 0.395 | -0.079 | -0.046 | -0.046 | 0.007 | 0.26 | 0.1 | 0.23 | 0.36 | 0.38 | 0.28 |
| 2 | 16136922 | rs9287666 | -0.597 | 0.671 | -1.397 | 0.363 | 0.005 | 0.036 | 0.036 | 0 | 0.43 | 0.41 | 0.64 | 0.81 | 0.55 | 0.55 |
| 2 | 16137611 | rs2114802 | -0.661 | -0.398 | -1.32 | -0.163 | -0.045 | -0.12 | -0.12 | 0 | 0.78 | 0.68 | 0.89 | 1 | 0.87 | 0.9 |
| 2 | 16137840 | rs6709176 | 0.076 | -0.412 | -1.418 | -0.192 | -0.019 | 0.034 | 0.034 | 1.058 | 0.72 | 0.68 | 0.55 | 0.55 | 0.78 | 0.64 |
| 2 | 16141257 | rs11677992 | -0.212 | -0.455 | -1.356 | -0.321 | 0.013 | 0.166 | 0.166 | 0.239 | 0.2 | 0.38 | 0.36 | 0.48 | 0.18 | 0.29 |
| 2 | 16142703 | rs13396055 | 0.925 | -0.55 | -1.268 | -0.394 | 0.013 | 0.036 | 0.036 | 2.257 | 0.31 | 0.24 | 0.24 | 0.13 | 0.3 | 0.32 |
| 2 | 16145517 | rs6748658 | 0.476 | -0.288 | -1.234 | -0.368 | 0.007 | 0.003 | 0.003 | 0 | 0.32 | 0.35 | 0.31 | 0.39 | 0.39 | 0.29 |
| 2 | 16149780 | rs1365966 | -0.203 | -0.211 | -1.683 | -0.068 | 0.003 | 0.033 | 0.033 | 0.556 | 0.46 | 0.22 | 0.54 | 0.61 | 0.48 | 0.62 |
| 2 | 16150760 | rs13008990 | 0.436 | -0.144 | -1.432 | 0.041 | -0.027 | 0.051 | 0.051 | 0 | 0.09 | 0 | 0.03 | 0 | 0.08 | 0.03 |
| 2 | 16162079 | rs6735335 | 0.336 | -0.712 | -0.967 | 0.473 | -0.007 | 0.002 | 0.002 | 0 | 0.6 | 0.51 | 0.52 | 0.48 | 0.68 | 0.43 |
| 2 | 16166988 | rs6431749 | 0.089 | -0.909 | -0.534 | 0.147 | -0.014 | -0.017 | -0.017 | 0 | 0.84 | 0.68 | 0.7 | 0.9 | 0.85 | 0.75 |
| 2 | 16169371 | rs6716719 | 0.52 | -0.981 | -0.604 | 0.361 | -0.019 | -0.006 | -0.006 | 0 | 0.64 | 0.48 | 0.61 | 0.69 | 0.71 | 0.55 |
| 2 | 16170523 | rs16863206 | 0.028 | -1.115 | -0.1 | -0.083 | -0.003 | -0.045 | -0.045 | 0.205 | 0.1 | 0.32 | 0.06 | 0 | 0.03 | 0.01 |
| 2 | 16176894 | rs10194143 | -0.647 | -1.342 | -0.194 | -0.135 | -0.056 | -0.052 | -0.052 | 0 | 0.26 | 0.47 | 0.18 | 0.14 | 0.14 | 0.18 |
| 2 | 16182802 | rs6727571 | 0.128 | -0.211 | -0.553 | 0.128 | 0 | 0 | 0 | 0 | 0.73 | 0.69 | 0.64 | 0.8 | 0.74 | 0.72 |
| 2 | 16183815 | rs7588005 | -0.027 | -0.145 | -0.718 | 0.151 | -0.005 | 0.005 | 0.005 | 0 | 0.75 | 0.78 | 0.65 | 0.81 | 0.74 | 0.75 |
| 2 | 16187902 | rs4669031 | 0.512 | -0.29 | -1.318 | 0.553 | 0.02 | -0.034 | -0.034 | 0 | 0.28 | 0.16 | 0.33 | 0.41 | 0.26 | 0.43 |
| 2 | 16187988 | rs4669033 | 0.079 | 0.756 | -1.113 | 0.315 | -0.017 | 0.011 | 0.011 | 0 | 0.74 | 0.79 | 0.77 | 0.82 | 0.71 | 0.72 |
| 2 | 16190882 | rs934059 | -0.178 | 1.214 | -0.844 | 0.372 | 0.01 | -0.005 | -0.005 | 0 | 0.73 | 0.75 | 0.62 | 0.78 | 0.63 | 0.84 |
| 2 | 16192664 | rs12612245 | -0.184 | 1.666 | -0.671 | 1.215 | 0.01 | 0.037 | 0.037 | 0 | 0.08 | 0.1 | 0.16 | 0.32 | 0.14 | 0.12 |
| 2 | 16194104 | rs10186232 | -0.257 | 1.12 | -1.146 | 0.657 | -0.027 | 0.047 | 0.047 | 0 | 0.59 | 0.81 | 0.58 | 0.73 | 0.5 | 0.68 |
| 2 | 16195030 | rs16863222 | 0.777 | 0.526 | -1.139 | 0.634 | 0.039 | -0.021 | -0.021 | 0 | 0.41 | 0.14 | 0.29 | 0.37 | 0.33 | 0.54 |

| | | | | | | | | | | | | | | | | |
|---|---|---|---|---|---|---|---|---|---|---|---|---|---|---|---|---|
| 2 | 16200946 | rs7605722 | 0.961 | 0.653 | -1.168 | 0.411 | 0.016 | -0.047 | -0.047 | 0 | 0.33 | 0.22 | 0.25 | 0.44 | 0.27 | 0.5 |
| 2 | 16202336 | rs2114797 | 0.579 | 0.907 | -1.196 | 0.525 | 0.003 | 0.028 | 0.028 | 0 | 0.08 | 0.09 | 0.16 | 0.32 | 0.14 | 0.14 |
| 2 | 16205386 | rs11885671 | -1.073 | 0.615 | -1.158 | 0.274 | -0.033 | 0.049 | 0.049 | 0 | 0.67 | 0.85 | 0.61 | 0.85 | 0.57 | 0.79 |
| 2 | 16209473 | rs1427682 | -1.056 | 0.492 | -0.446 | 0.292 | -0.003 | 0.008 | 0.008 | 0 | 0.29 | 0.43 | 0.29 | 0.39 | 0.27 | 0.27 |
| 2 | 16214286 | rs4073262 | -0.012 | 0.892 | -0.624 | 0.47 | -0.005 | -0.005 | -0.005 | 0 | 0.42 | 0.27 | 0.5 | 0.48 | 0.56 | 0.35 |
| 2 | 16228929 | rs4669037 | -0.735 | 0.683 | -0.87 | 0.587 | 0.006 | -0.004 | -0.004 | 0 | 0.59 | 0.47 | 0.67 | 0.55 | 0.64 | 0.47 |
| 2 | 16230808 | rs4669039 | -0.712 | 0.699 | -1.132 | 0.61 | 0.013 | -0.006 | -0.006 | 0 | 0.58 | 0.45 | 0.67 | 0.55 | 0.64 | 0.47 |
| 2 | 16241361 | rs4668485 | 0.722 | 0.977 | -0.692 | 0.954 | 0.023 | 0.071 | 0.071 | 0.176 | 0.81 | 0.58 | 0.74 | 0.63 | 0.79 | 0.87 |
| 2 | 16251998 | rs7370296 | 0.128 | 0.784 | -1.866 | 0.267 | -0.065 | 0.151 | 0.151 | 2.953 | 0.48 | 0.14 | 0.41 | 0.78 | 0.51 | 0.7 |
| 2 | 16252603 | rs28666066 | -0.324 | -0.296 | -1.157 | -0.147 | -0.003 | -0.034 | -0.034 | 0.066 | 0.2 | 0.21 | 0.24 | 0.03 | 0.23 | 0.19 |
| 2 | 16252614 | rs28549395 | 0.142 | 0.312 | -1.122 | 0.403 | 0.008 | 0.007 | 0.007 | 0 | 0.42 | 0.32 | 0.52 | 0.41 | 0.55 | 0.36 |
| 2 | 16358744 | rs4047462 | -0.547 | 0.483 | -0.815 | 0.23 | -0.006 | -0.014 | -0.014 | 0.926 | 0.09 | 0.1 | 0.11 | 0.05 | 0.07 | 0.06 |
| 2 | 16361503 | rs7422511 | -0.186 | 0.559 | -0.914 | 1.039 | -0.003 | 0.007 | 0.007 | 0 | 0.09 | 0.02 | 0.16 | 0.5 | 0.11 | 0.34 |
| 2 | 16361798 | rs11096635 | -0.216 | 0.534 | -0.83 | 1.026 | 0 | 0.028 | 0.028 | 0 | 0.1 | 0.03 | 0.23 | 0.5 | 0.11 | 0.34 |
| 2 | 16366130 | rs7559122 | 0.313 | 0.249 | -0.749 | -0.469 | 0.044 | 0.108 | 0.108 | 0 | 0.5 | 0.34 | 0.52 | 0.23 | 0.61 | 0.38 |
| 2 | 16367712 | rs870720 | 0.538 | 1.058 | -1.011 | -0.85 | 0.043 | 0.104 | 0.104 | 0 | 0.42 | 0.19 | 0.4 | 0.07 | 0.37 | 0.29 |
| 2 | 16369728 | rs7560874 | -0.148 | 1.024 | -1.064 | -0.808 | 0.036 | 0.034 | 0.034 | 0 | 0.63 | 0.84 | 0.73 | 0.32 | 0.75 | 0.46 |
| 2 | 16372364 | rs12710645 | -0.07 | 1.457 | -0.686 | -0.365 | 0.079 | 0.002 | 0.002 | 0.009 | 0.06 | 0.1 | 0.13 | 0.07 | 0.23 | 0.07 |
| 2 | 16377083 | rs11681625 | 0.157 | 1.436 | -0.577 | -0.353 | 0.123 | 0.01 | 0.01 | 0 | 0.05 | 0.09 | 0.13 | 0.08 | 0.24 | 0.06 |
| 2 | 16378972 | rs1567965 | 0.196 | -0.352 | -0.43 | -0.828 | 0.023 | 0.048 | 0.048 | 0 | 0.52 | 0.44 | 0.5 | 0.22 | 0.42 | 0.41 |
| 2 | 16381942 | rs10201439 | -1.124 | -1.137 | -0.198 | -0.898 | -0.01 | -0.042 | -0.042 | 0 | 0.29 | 0.59 | 0.43 | 0.11 | 0.19 | 0.31 |
| 2 | 16388004 | rs340729 | 1.077 | -1.63 | -0.484 | -1.403 | 0.047 | 0.085 | 0.085 | 0 | 0.55 | 0.35 | 0.48 | 0.93 | 0.72 | 0.64 |
| 2 | 16390699 | rs340724 | 1.287 | -1.611 | -0.528 | -1.409 | 0.078 | 0.196 | 0.196 | 0 | 0.53 | 0.26 | 0.47 | 0.93 | 0.72 | 0.62 |
| 2 | 16393486 | rs17383430 | -0.742 | -1.688 | -0.622 | -1.48 | 0.018 | 0.066 | 0.066 | 0 | 0.18 | 0.08 | 0.08 | 0.03 | 0.1 | 0.17 |
| 2 | 16399845 | rs687584 | 0.748 | -1.751 | -0.529 | -1.515 | 0.009 | -0.001 | -0.001 | 0.034 | 0.8 | 0.86 | 0.91 | 1 | 0.88 | 0.82 |
| 2 | 16403550 | rs340718 | 1.277 | -1.743 | 0.28 | -1.569 | -0.018 | -0.126 | -0.126 | 0.493 | 0.64 | 0.53 | 0.54 | 0.94 | 0.75 | 0.69 |
| 2 | 16406949 | rs664510 | 1.786 | -2.043 | -0.352 | -0.572 | 0.081 | 0.035 | 0.035 | 0 | 0.47 | 0.24 | 0.44 | 0.6 | 0.7 | 0.43 |
| 2 | 16408829 | rs585810 | 1.203 | -1.845 | 0.204 | -0.347 | -0.027 | -0.001 | -0.001 | 0.051 | 0.6 | 0.49 | 0.5 | 0.6 | 0.74 | 0.56 |
| 2 | 16410452 | rs1429410 | -0.815 | -1.484 | -0.037 | -0.437 | -0.041 | -0.134 | -0.134 | 0 | 0.24 | 0.53 | 0.15 | 0.02 | 0.15 | 0.19 |
| 2 | 16417425 | rs7564195 | -0.784 | -1.077 | -0.443 | 0.634 | 0.001 | 0.017 | 0.017 | 0.07 | 0.1 | 0.06 | 0.09 | 0.26 | 0.07 | 0.26 |
| 2 | 16419751 | rs4832659 | -0.806 | -1.133 | -0.423 | 0.778 | 0.003 | 0.011 | 0.011 | 0 | 0.09 | 0.06 | 0.06 | 0.23 | 0.06 | 0.28 |
| 2 | 16420456 | rs17468049 | -0.605 | -1.835 | -0.399 | 0.059 | 0.008 | -0.017 | -0.017 | 0 | 0.19 | 0.18 | 0.07 | 0.01 | 0.08 | 0.13 |
| 2 | 16430070 | rs2163063 | 0.287 | -1.986 | -0.345 | 0.089 | 0.012 | 0.009 | 0.009 | 0 | 0.11 | 0.08 | 0.05 | 0 | 0.07 | 0.11 |
| 2 | 16431781 | rs16982164 | -0.714 | -2.215 | 0.265 | -0.073 | -0.029 | -0.09 | -0.09 | 0.44 | 0.17 | 0.27 | 0.09 | 0 | 0.09 | 0.14 |
| 2 | 16433908 | rs7569508 | 0.702 | -1.833 | 0.645 | -0.231 | -0.025 | -0.025 | -0.025 | 0 | 0.22 | 0.14 | 0.28 | 0.57 | 0.36 | 0.36 |
| 2 | 16435236 | rs16982176 | -0.653 | -2.049 | 0.79 | -0.213 | -0.052 | -0.107 | -0.107 | 0.445 | 0.2 | 0.37 | 0.14 | 0.02 | 0.09 | 0.19 |
| 2 | 16438076 | rs340738 | -1.289 | -2.265 | 1.06 | -0.243 | -0.083 | 0.029 | 0.029 | 0 | 0.27 | 0.45 | 0.23 | 0.28 | 0.11 | 0.45 |
| 2 | 16439187 | rs9306904 | -0.553 | -2.187 | 0.987 | -0.672 | -0.06 | -0.131 | -0.131 | 0 | 0.21 | 0.45 | 0.1 | 0.01 | 0.09 | 0.19 |
| 2 | 16443492 | rs999518 | -0.4 | -2.231 | 0.939 | -0.703 | -0.049 | -0.101 | -0.101 | 0 | 0.19 | 0.41 | 0.09 | 0.01 | 0.09 | 0.17 |
| 2 | 16445487 | rs11685981 | 0.814 | -1.837 | 0.609 | -0.615 | -0.111 | -0.108 | -0.108 | 0.247 | 0.19 | 0.04 | 0.3 | 0.64 | 0.33 | 0.37 |
| 2 | 16446833 | rs340749 | -1.694 | -2.522 | 0.466 | -1.005 | -0.039 | 0.137 | 0.137 | 0 | 0.59 | 0.83 | 0.56 | 0.93 | 0.48 | 0.86 |
| 2 | 16451390 | rs340743 | -0.25 | -2.791 | 0.681 | -0.854 | -0.032 | 0.129 | 0.129 | 0 | 0.09 | 0.16 | 0.14 | 0.29 | 0.03 | 0.28 |
| 2 | 16452047 | rs12465207 | -0.378 | -2.799 | 0.714 | -0.929 | -0.028 | -0.091 | -0.091 | 0 | 0.29 | 0.34 | 0.11 | 0.01 | 0.13 | 0.21 |
| 2 | 16452715 | rs340747 | -1.23 | -2.66 | 0.282 | -0.813 | -0.021 | -0.002 | -0.002 | 0 | 0.39 | 0.64 | 0.26 | 0.31 | 0.16 | 0.51 |
| 2 | 16459536 | rs2333979 | -0.575 | -2.147 | -0.051 | -1.232 | -0.003 | 0.12 | 0.12 | 0 | 0.51 | 0.68 | 0.52 | 0.83 | 0.46 | 0.79 |
| 2 | 16459653 | rs340767 | -0.584 | -2.137 | -0.057 | -1.217 | -0.005 | 0.129 | 0.129 | 0 | 0.51 | 0.69 | 0.53 | 0.83 | 0.46 | 0.79 |
| 2 | 16463362 | rs340772 | -1.317 | -2.039 | -0.284 | -1.32 | -0.061 | 0.09 | 0.09 | 0 | 0.69 | 0.83 | 0.61 | 0.91 | 0.55 | 0.87 |
| 2 | 16464227 | rs733555 | 0.967 | -2.378 | -0.11 | -1.026 | -0.006 | -0.006 | -0.006 | 0 | 0.3 | 0.24 | 0.32 | 0.56 | 0.37 | 0.47 |
| 2 | 16467187 | rs11894986 | 1.002 | -2.614 | 0.121 | -1.373 | -0.016 | -0.031 | -0.031 | 0 | 0.21 | 0.24 | 0.12 | 0.04 | 0.12 | 0.22 |
| 2 | 16469396 | rs1366835 | 1.081 | -2.55 | 0.06 | -1.345 | 0 | 0 | 0 | 1.015 | 0.3 | 0.27 | 0.32 | 0.61 | 0.37 | 0.48 |
| 2 | 16469662 | rs340739 | -0.111 | -2.456 | -0.074 | -1.315 | -0.003 | 0.048 | 0.048 | 0 | 0.52 | 0.66 | 0.47 | 0.65 | 0.5 | 0.7 |
| 2 | 16469823 | rs340740 | -1.144 | -2.041 | -0.378 | -1.24 | -0.038 | 0.05 | 0.05 | 0 | 0.68 | 0.79 | 0.61 | 0.85 | 0.55 | 0.88 |
| 2 | 16493067 | rs4832490 | 0.852 | -1.852 | -0.781 | -1.037 | -0.036 | -0.036 | -0.036 | 0.425 | 0.29 | 0.16 | 0.31 | 0.57 | 0.38 | 0.46 |
| 2 | 16498117 | rs340786 | -0.521 | -1.323 | -0.993 | -1.738 | -0.034 | 0.059 | 0.059 | 0 | 0.65 | 0.77 | 0.61 | 0.91 | 0.53 | 0.84 |
| 2 | 16502338 | rs340781 | -0.533 | -2.365 | -0.717 | -1.67 | 0 | -0.006 | -0.006 | 0 | 0.76 | 0.78 | 0.72 | 0.93 | 0.76 | 0.88 |
| 2 | 16509408 | rs340778 | 0.42 | -2.812 | -0.618 | -1.879 | -0.012 | -0.02 | -0.02 | 0 | 0.23 | 0.23 | 0.14 | 0.05 | 0.14 | 0.24 |
| 2 | 16514787 | rs186875 | -0.205 | -2.549 | -1.037 | -1.786 | -0.003 | -0.031 | -0.031 | 0.018 | 0.71 | 0.61 | 0.62 | 0.87 | 0.73 | 0.84 |
| 2 | 16527624 | rs1430041 | -0.805 | -2.521 | -1.309 | -1.814 | -0.007 | 0.045 | 0.045 | 0.312 | 0.76 | 0.78 | 0.66 | 0.93 | 0.75 | 0.89 |
| 2 | 16531549 | rs7355377 | -0.095 | -2.934 | -1.209 | -1.868 | -0.047 | -0.05 | -0.05 | 0 | 0.27 | 0.56 | 0.2 | 0.11 | 0.16 | 0.29 |
| 2 | 16541456 | rs11690983 | -0.31 | -3.055 | -1.35 | -1.95 | 0.031 | 0.119 | 0.119 | 0.049 | 0.7 | 0.89 | 0.67 | 0.92 | 0.77 | 0.84 |
| 2 | 16543233 | rs10495650 | 0.177 | -2.219 | -0.964 | -1.541 | -0.053 | -0.051 | -0.051 | 0.425 | 0.23 | 0.36 | 0.16 | 0.11 | 0.14 | 0.27 |
| 2 | 16544056 | rs919432 | 0.572 | -2.183 | -1.012 | -1.516 | 0.066 | 0.144 | 0.144 | 0.075 | 0.49 | 0.23 | 0.47 | 0.82 | 0.61 | 0.6 |
| 2 | 16547545 | rs13389373 | -0.437 | -1.56 | -0.956 | -0.569 | -0.052 | -0.067 | -0.067 | 0.063 | 0.25 | 0.5 | 0.2 | 0.11 | 0.18 | 0.25 |
| 2 | 16551383 | rs12710671 | 0.587 | -1.3 | -1.861 | -0.426 | 0.071 | 0.164 | 0.164 | 0 | 0.46 | 0.12 | 0.51 | 0.2 | 0.63 | 0.52 |
| 2 | 16556128 | rs1367160 | -0.172 | -1.429 | -1.881 | 0.088 | 0.001 | -0.037 | -0.037 | 0 | 0.11 | 0.03 | 0.22 | 0.66 | 0.1 | 0.27 |
| 2 | 16556650 | rs12465117 | 0.912 | -1.708 | -2.104 | -0.119 | 0.093 | 0.218 | 0.218 | 0 | 0.49 | 0.11 | 0.55 | 0.17 | 0.64 | 0.52 |
| 2 | 16557140 | rs1156704 | -0.45 | -1.862 | -2.051 | -0.284 | -0.004 | -0.002 | -0.002 | 0 | 0.92 | 0.93 | 0.94 | 1 | 0.9 | 0.95 |
| 2 | 16559585 | rs6531084 | 0.002 | -1.816 | -2.062 | -0.233 | -0.001 | -0.04 | -0.04 | 0 | 0.11 | 0.03 | 0.22 | 0.67 | 0.1 | 0.26 |
| 2 | 16560645 | rs10196096 | 1.029 | -1.75 | -2.076 | -0.67 | 0.098 | 0.224 | 0.224 | 0 | 0.5 | 0.11 | 0.55 | 0.17 | 0.64 | 0.51 |

| | | | | | | | | | | | | | | | | |
|---|---|---|---|---|---|---|---|---|---|---|---|---|---|---|---|---|
| 2 | 16568383 | rs2340759 | 0.766 | -1.753 | -1.75 | -0.72 | 0.051 | 0.091 | 0.091 | 0 | 0.62 | 0.74 | 0.75 | 0.92 | 0.74 | 0.69 |
| 2 | 16568845 | rs983249 | 0.737 | -0.991 | -1.146 | -0.029 | 0.011 | 0.016 | 0.016 | 0 | 0.86 | 0.88 | 0.87 | 0.96 | 0.91 | 0.81 |
| 2 | 16585720 | rs1430060 | -1.241 | -1.018 | 0.486 | 0.084 | -0.005 | 0.004 | 0.004 | 4.353 | 0.11 | 0.32 | 0.07 | 0.07 | 0.04 | 0.07 |
| 2 | 16590965 | rs1367158 | 1.355 | -1.604 | 0.059 | -0.257 | 0.085 | 0.149 | 0.149 | 0.448 | 0.55 | 0.17 | 0.4 | 0.17 | 0.7 | 0.47 |
| 2 | 16598521 | rs11884584 | 0.78 | -1.807 | 0.828 | 0.048 | -0.012 | -0.051 | -0.051 | 0 | 0.78 | 0.45 | 0.67 | 0.88 | 0.84 | 0.84 |
| 2 | 16600041 | rs2115849 | 0.532 | -1.809 | 0.899 | 0.047 | -0.005 | -0.034 | -0.034 | 0 | 0.79 | 0.64 | 0.68 | 0.88 | 0.84 | 0.84 |
| 2 | 16605110 | rs1316661 | 1.574 | -1.782 | 0.777 | -0.11 | 0.032 | 0.071 | 0.071 | 0.432 | 0.64 | 0.19 | 0.72 | 0.8 | 0.74 | 0.74 |
| 2 | 16606596 | rs4832591 | 0.941 | -1.907 | 1.293 | -0.365 | 0.008 | -0.13 | -0.13 | 0 | 0.78 | 0.57 | 0.74 | 0.99 | 0.8 | 0.82 |
| 2 | 16607101 | rs11096549 | 2.466 | -2.049 | 1.242 | 0.028 | 0.065 | 0.095 | 0.095 | 1.667 | 0.57 | 0.17 | 0.47 | 0.21 | 0.69 | 0.48 |
| 2 | 16609693 | rs2115850 | 0.763 | -1.945 | 1.331 | -0.185 | 0.002 | -0.068 | -0.068 | 0 | 0.79 | 0.71 | 0.75 | 0.99 | 0.8 | 0.82 |
| 2 | 16617781 | rs6531114 | 0.877 | -1.165 | 1.211 | -0.222 | 0.051 | -0.124 | -0.124 | 0 | 0.78 | 0.61 | 0.72 | 1 | 0.74 | 0.8 |
| 2 | 16620803 | rs6759839 | 1.846 | -1.248 | 1.137 | -0.247 | -0.007 | 0.085 | 0.085 | 1.257 | 0.64 | 0.2 | 0.63 | 0.93 | 0.66 | 0.74 |
| 2 | 16630784 | rs4832615 | 1.48 | -1.161 | 1.249 | -0.263 | -0.03 | 0.045 | 0.045 | 0 | 0.77 | 0.83 | 0.7 | 0.95 | 0.74 | 0.81 |
| 2 | 16631488 | rs6728955 | 1.314 | -1.153 | 1.23 | -0.264 | -0.088 | 0.108 | 0.108 | 0 | 0.78 | 0.95 | 0.72 | 0.95 | 0.74 | 0.81 |
| 2 | 16635110 | rs13399375 | -0.52 | -1.213 | 1.724 | -0.237 | 0.055 | 0.036 | 0.036 | 0 | 0.05 | 0.36 | 0.04 | 0 | 0.01 | 0.03 |
| 2 | 16638478 | rs6531144 | 1.929 | -1.599 | 1.443 | -0.056 | -0.002 | 0.021 | 0.021 | 0.204 | 0.66 | 0.74 | 0.55 | 0.26 | 0.71 | 0.54 |
| 2 | 16639566 | rs1346751 | 1.306 | -1.562 | 1.388 | -0.101 | -0.068 | 0.141 | 0.141 | 0 | 0.74 | 0.93 | 0.65 | 0.95 | 0.74 | 0.8 |
| 2 | 16644134 | rs16982422 | 0.131 | -1.578 | 1.407 | 0.068 | 0.002 | 0.053 | 0.053 | 0 | 0.05 | 0.06 | 0.08 | 0.68 | 0.03 | 0.17 |
| 2 | 16648888 | rs4832633 | 1.823 | -1.609 | 1.534 | -0.088 | -0.001 | -0.077 | -0.077 | 0 | 0.68 | 0.42 | 0.61 | 0.93 | 0.73 | 0.76 |
| 2 | 16653570 | rs10175438 | -1.001 | -1.587 | 1.617 | -0.086 | 0.064 | 0.036 | 0.036 | 0.241 | 0.06 | 0.51 | 0.04 | 0 | 0.01 | 0.04 |
| 2 | 16664193 | rs2340761 | 1.881 | -1.602 | 1.508 | -0.1 | -0.018 | 0.077 | 0.077 | 0.168 | 0.64 | 0.82 | 0.53 | 0.21 | 0.7 | 0.51 |
| 2 | 16668723 | rs7566990 | 1.539 | -1.609 | 1.56 | -0.107 | -0.065 | 0.141 | 0.141 | 0 | 0.71 | 0.91 | 0.62 | 0.94 | 0.73 | 0.77 |
| 2 | 16670420 | rs4832644 | -1.234 | -1.687 | 1.503 | -0.098 | 0.004 | 0.013 | 0.013 | 0 | 0.08 | 0.07 | 0.08 | 0.73 | 0.04 | 0.19 |
| 2 | 16684487 | rs1430058 | 2.555 | -1.424 | 1.462 | -0.155 | 0.017 | 0.132 | 0.132 | 2.89 | 0.54 | 0.17 | 0.44 | 0.21 | 0.62 | 0.48 |
| 2 | 16686128 | rs12463697 | 1.996 | -1.42 | 1.484 | -0.277 | -0.002 | -0.095 | -0.095 | 0 | 0.62 | 0.47 | 0.6 | 0.93 | 0.66 | 0.63 |
| 2 | 16688759 | rs4240228 | 1.586 | -1.198 | 1.285 | -0.172 | -0.001 | -0.018 | -0.018 | 0.793 | 0.7 | 0.66 | 0.67 | 0.95 | 0.73 | 0.75 |
| 2 | 16702165 | rs4347841 | 1.004 | -1.122 | 1.362 | -0.107 | 0.015 | -0.083 | -0.083 | 0 | 0.78 | 0.71 | 0.7 | 1 | 0.75 | 0.81 |
| 2 | 16703441 | rs7576783 | 1.422 | -1.133 | 1.353 | -0.102 | 0.005 | -0.036 | -0.036 | 0 | 0.74 | 0.65 | 0.68 | 0.94 | 0.74 | 0.78 |
| 2 | 16708548 | rs4832465 | -2.272 | -1.126 | 1.51 | -0.052 | 0 | -0.004 | -0.004 | 0 | 0.34 | 0.65 | 0.5 | 0.74 | 0.29 | 0.37 |
| 2 | 16710565 | rs4832468 | -2.628 | -1.176 | 1.507 | -0.073 | 0.004 | 0.02 | 0.02 | 1.136 | 0.38 | 0.74 | 0.5 | 0.74 | 0.3 | 0.45 |
| 2 | 16726739 | rs4832657 | -1.833 | -1.093 | 1.539 | -0.042 | 0.011 | 0.018 | 0.018 | 0 | 0.3 | 0.58 | 0.48 | 0.67 | 0.29 | 0.33 |
| 2 | 16733054 | rs10172734 | -2.2 | -1.074 | 1.541 | 0.02 | -0.001 | 0.001 | 0.001 | 0.009 | 0.34 | 0.69 | 0.5 | 0.72 | 0.29 | 0.37 |
| 2 | 16733412 | rs10182313 | -1.472 | -0.76 | 1.563 | 0.266 | 0.013 | 0.31 | 0.31 | 0 | 0.1 | 0.36 | 0.18 | 0.66 | 0.05 | 0.15 |
| 2 | 16738867 | rs12989232 | -0.515 | -0.671 | 0.997 | -0.573 | -0.015 | 0.019 | 0.019 | 0 | 0.22 | 0.16 | 0.3 | 0.13 | 0.23 | 0.19 |
| 2 | 16741442 | rs7586823 | -2.331 | -0.556 | 0.696 | -0.893 | -0.016 | 0.16 | 0.16 | 0.185 | 0.49 | 0.88 | 0.49 | 0.24 | 0.4 | 0.46 |
| 2 | 16744902 | rs11096634 | 0.35 | -0.619 | 1.121 | -0.929 | 0.006 | -0.054 | -0.054 | 0 | 0.09 | 0.23 | 0.08 | 0 | 0.08 | 0.03 |
| 2 | 16751119 | rs10206736 | 0.247 | -0.636 | 0.772 | -0.933 | -0.01 | 0.014 | 0.014 | 0 | 0.09 | 0.13 | 0.06 | 0.1 | 0.04 | 0.09 |
| 2 | 16752999 | rs11676612 | -0.986 | -0.724 | 0.641 | -1.004 | -0.008 | 0.028 | 0.028 | 0 | 0.29 | 0.24 | 0.33 | 0.13 | 0.27 | 0.22 |
| 2 | 16764536 | rs10169258 | -2.312 | -0.634 | 0.509 | -1.423 | 0.007 | 0.262 | 0.262 | 0.149 | 0.49 | 0.87 | 0.49 | 0.13 | 0.38 | 0.45 |
| 2 | 16770592 | rs10171652 | -0.222 | -0.487 | 0.706 | -0.843 | -0.012 | 0.012 | 0.012 | 0 | 0.09 | 0.14 | 0.06 | 0.1 | 0.04 | 0.09 |
| 2 | 16770723 | rs10176532 | -1.429 | -0.499 | 0.718 | -0.85 | -0.017 | -0.032 | -0.032 | 0 | 0.23 | 0.31 | 0.15 | 0.11 | 0.16 | 0.15 |
| 2 | 16773622 | rs11893415 | -0.717 | 0.003 | -0.147 | -0.928 | -0.027 | -0.119 | -0.119 | 3.011 | 0.17 | 0.42 | 0.11 | 0.01 | 0.11 | 0.18 |
| 2 | 16784998 | rs7425164 | -0.581 | 1.266 | -1.177 | -0.782 | -0.07 | 0.07 | 0.07 | 0 | 0.22 | 0.06 | 0.35 | 0.02 | 0.35 | 0.21 |
| 2 | 16790749 | rs4832469 | -0.996 | 0.942 | -1.038 | -0.591 | 0.002 | 0.001 | 0.001 | 0 | 0.16 | 0.2 | 0.34 | 0.85 | 0.16 | 0.25 |
| 2 | 16791265 | rs6743717 | -0.112 | 0.614 | -1.102 | -1.62 | -0.019 | -0.054 | -0.054 | 0 | 0.29 | 0.44 | 0.28 | 0.08 | 0.21 | 0.3 |
| 2 | 16795553 | rs17643147 | 1.078 | 1.111 | -2.017 | -1.998 | -0.138 | 0.161 | 0.161 | 0 | 0.24 | 0.02 | 0.22 | 0 | 0.39 | 0.21 |
| 2 | 16796596 | rs888575 | -0.885 | 0.464 | -1.16 | -1.361 | -0.001 | -0.001 | -0.001 | 0 | 0.17 | 0.16 | 0.33 | 0.86 | 0.15 | 0.2 |
| 2 | 16797214 | rs16982572 | -0.632 | 0.844 | -1.275 | -0.816 | -0.021 | -0.013 | -0.013 | 0 | 0.07 | 0.03 | 0.22 | 0.75 | 0.12 | 0.15 |
| 2 | 16798475 | rs4832660 | -0.022 | 0.621 | -1.27 | -1.21 | -0.035 | 0.007 | 0.007 | 0 | 0.7 | 0.78 | 0.69 | 0.79 | 0.62 | 0.63 |
| 2 | 16799241 | rs10167880 | 0.65 | 0.11 | -1.083 | -1.604 | 0.013 | 0.069 | 0.069 | 0 | 0.5 | 0.34 | 0.28 | 0.1 | 0.47 | 0.46 |
| 2 | 16803169 | rs10188832 | -0.668 | -0.147 | -1.029 | -0.926 | -0.002 | 0.048 | 0.048 | 0 | 0.22 | 0.31 | 0.33 | 0.71 | 0.21 | 0.28 |
| 2 | 16803285 | rs2880359 | -1.307 | -0.295 | -1.501 | -1.355 | -0.088 | 0.24 | 0.24 | 0 | 0.66 | 0.97 | 0.78 | 0.99 | 0.62 | 0.65 |
| 2 | 16805353 | rs1024925 | -0.028 | -0.026 | -0.912 | -1.015 | 0.003 | -0.003 | -0.003 | 0 | 0.37 | 0.61 | 0.42 | 0.28 | 0.38 | 0.35 |
| 2 | 16807318 | rs11897275 | -0.096 | -0.004 | -0.881 | -1.01 | 0.012 | -0.073 | -0.073 | 0.408 | 0.15 | 0.33 | 0.12 | 0.06 | 0.14 | 0.09 |
| 2 | 16808970 | rs12052781 | -0.311 | 0.093 | -0.89 | -0.581 | 0.014 | -0.007 | -0.007 | 0 | 0.36 | 0.5 | 0.41 | 0.29 | 0.39 | 0.34 |
| 2 | 16809949 | rs2342437 | -0.428 | 0.271 | -1.839 | -0.12 | -0.089 | -0.029 | -0.029 | 0 | 0.39 | 0.09 | 0.39 | 0.36 | 0.43 | 0.39 |
| 2 | 16817049 | rs7597205 | -0.656 | -0.34 | -0.86 | -0.426 | -0.023 | -0.072 | -0.072 | 0 | 0.16 | 0.24 | 0.13 | 0.01 | 0.06 | 0.17 |
| 2 | 16817937 | rs2193776 | -0.08 | -0.752 | -1.623 | -0.756 | 0.007 | 0.029 | 0.029 | 0 | 0.25 | 0.12 | 0.14 | 0.13 | 0.21 | 0.12 |
| 2 | 16820305 | rs1722418 | -1.533 | -1.343 | -2.071 | -0.943 | 0.004 | 0.313 | 0.313 | 0 | 0.55 | 0.95 | 0.58 | 0.96 | 0.38 | 0.55 |
| 2 | 16821938 | rs16982630 | -1.002 | -1.381 | -1.201 | -0.128 | -0.011 | -0.004 | -0.004 | 0 | 0.46 | 0.51 | 0.3 | 0.36 | 0.3 | 0.38 |
| 2 | 16827463 | rs10514636 | 0.159 | -1.436 | -0.351 | -0.264 | -0.053 | -0.069 | -0.069 | 0 | 0.11 | 0.31 | 0.03 | 0 | 0.02 | 0.1 |
| 2 | 16829430 | rs16982644 | -0.347 | -1.878 | -0.599 | -0.356 | 0.011 | 0.008 | 0.008 | 0 | 0.09 | 0.08 | 0.05 | 0.06 | 0.04 | 0.07 |
| 2 | 16829764 | rs940532 | 0.215 | -1.841 | -0.208 | -0.461 | 0.003 | 0.003 | 0.003 | 0.962 | 0.2 | 0.16 | 0.1 | 0.13 | 0.13 | 0.07 |
| 2 | 16836063 | rs10198624 | -0.176 | -1.864 | 0.126 | -0.609 | -0.064 | -0.1 | -0.1 | 0 | 0.15 | 0.4 | 0.04 | 0 | 0.03 | 0.09 |
| 2 | 16838500 | rs1024924 | -0.614 | -2.046 | 0.431 | -0.755 | -0.025 | -0.06 | -0.06 | 0 | 0.19 | 0.24 | 0.11 | 0.04 | 0.07 | 0.22 |
| 2 | 16838966 | rs1024922 | -1.171 | -2.097 | 0.845 | -0.748 | -0.071 | -0.096 | -0.096 | 7.529 | 0.25 | 0.46 | 0.17 | 0.11 | 0.1 | 0.28 |
| 2 | 16840722 | rs1404292 | 0.403 | -0.743 | 0.364 | -0.346 | 0.023 | 0.02 | 0.02 | 0 | 0.08 | 0.09 | 0.09 | 0.17 | 0.18 | 0.07 |
| 2 | 16841933 | rs809538 | -1.205 | -0.66 | 0.085 | -0.151 | -0.029 | 0.047 | 0.047 | 0 | 0.94 | 1 | 0.72 | 0.45 | 0.89 | 0.82 |
| 2 | 16847569 | rs810064 | 1.438 | -1.035 | 0.402 | -0.468 | -0.024 | -0.049 | -0.049 | 0.421 | 0.12 | 0.16 | 0.05 | 0 | 0.04 | 0.01 |

| | | | | | | | | | | | | | | | | |
|---|---|---|---|---|---|---|---|---|---|---|---|---|---|---|---|---|
| 2 | 16850621 | rs1404293 | -1.684 | -1.393 | 0.162 | -0.762 | 0.161 | 0.164 | 0.164 | 0.021 | 0.52 | 0.83 | 0.3 | 0.22 | 0.22 | 0.35 |
| 2 | 16854462 | rs1990834 | 1.419 | -1.322 | 0.552 | -0.705 | -0.04 | -0.071 | -0.071 | 0 | 0.13 | 0.26 | 0.06 | 0 | 0.04 | 0.01 |
| 2 | 16854602 | rs2002992 | -1.496 | -1.325 | 0.352 | -0.704 | 0.153 | 0.149 | 0.149 | 0 | 0.48 | 0.79 | 0.28 | 0.22 | 0.21 | 0.34 |
| 2 | 16858380 | rs1722423 | -1.98 | -1.148 | 0.116 | -0.644 | -0.097 | -0.036 | -0.036 | 0 | 0.56 | 0.97 | 0.38 | 0.38 | 0.39 | 0.41 |
| 2 | 16866013 | rs1643212 | -1.56 | -1.295 | 0.225 | -0.74 | 0.187 | 0.178 | 0.178 | 0 | 0.48 | 0.89 | 0.29 | 0.23 | 0.21 | 0.34 |
| 2 | 16869031 | rs1722405 | -0.002 | -1.257 | 0.89 | -0.766 | -0.055 | -0.136 | -0.136 | 0.029 | 0.21 | 0.57 | 0.19 | 0.02 | 0.08 | 0.11 |
| 2 | 16869319 | rs724237 | 1.217 | -1.259 | 0.964 | -0.764 | -0.044 | -0.076 | -0.076 | 0.684 | 0.14 | 0.29 | 0.06 | 0 | 0.04 | 0.01 |
| 2 | 16872118 | rs6725232 | 0.312 | -1.209 | 0.808 | -0.716 | -0.001 | -0.013 | -0.013 | 0.063 | 0.06 | 0.05 | 0.09 | 0.02 | 0.03 | 0.1 |
| 2 | 16885881 | rs16982680 | 0.923 | -1.339 | 0.757 | -0.701 | -0.012 | -0.001 | -0.001 | 0.613 | 0.05 | 0.09 | 0.04 | 0.06 | 0.03 | 0.06 |
| 2 | 16889001 | rs10205624 | 1.085 | -1.451 | 0.857 | -0.675 | -0.007 | 0.007 | 0.007 | 0 | 0.1 | 0.14 | 0.06 | 0.13 | 0.07 | 0.07 |
| 2 | 16889372 | rs810066 | -1.256 | -1.593 | 0.679 | -0.752 | 0.143 | 0.071 | 0.071 | 1.921 | 0.35 | 0.94 | 0.21 | 0.21 | 0.16 | 0.16 |
| 2 | 16892189 | rs798379 | 1.666 | -1.591 | 0.892 | -0.754 | 0.001 | -0.015 | -0.015 | 0 | 0.12 | 0.18 | 0.06 | 0 | 0.04 | 0.01 |
| 2 | 16893593 | rs1643196 | 0.448 | -1.552 | 0.884 | -0.733 | -0.048 | -0.056 | -0.056 | 0.01 | 0.19 | 0.35 | 0.1 | 0.06 | 0.08 | 0.07 |
| 2 | 16893949 | rs1722412 | -0.504 | -1.556 | 0.898 | -0.734 | -0.04 | -0.039 | -0.039 | 0.536 | 0.24 | 0.69 | 0.14 | 0.09 | 0.09 | 0.09 |
| 2 | 16896756 | rs798365 | -0.367 | -1.082 | 0.607 | -0.643 | -0.069 | -0.049 | -0.049 | 2.714 | 0.3 | 0.58 | 0.17 | 0.19 | 0.16 | 0.14 |
| 2 | 16896822 | rs17645454 | 0.244 | -0.463 | 0.376 | -0.482 | -0.039 | -0.03 | -0.03 | 0 | 0.09 | 0.03 | 0.16 | 0.16 | 0.21 | 0.15 |
| 2 | 16904254 | rs6716519 | 1.221 | -0.583 | 0.308 | -0.557 | -0.001 | 0.009 | 0.009 | 1.618 | 0.45 | 0.34 | 0.4 | 0.11 | 0.56 | 0.44 |
| 2 | 16926228 | rs1862059 | 0.356 | -0.729 | 0.26 | -0.539 | -0.019 | -0.01 | -0.01 | 0 | 0.21 | 0.22 | 0.1 | 0.17 | 0.12 | 0.15 |
| 2 | 16926263 | rs1643189 | -1.938 | -0.396 | -0.577 | -0.828 | -0.122 | -0.078 | -0.078 | 1.361 | 0.57 | 0.98 | 0.52 | 0.44 | 0.52 | 0.56 |
| 2 | 16926284 | rs1862058 | 0.686 | -0.372 | -0.54 | -1.107 | -0.02 | -0.01 | -0.01 | 0 | 0.21 | 0.21 | 0.1 | 0.17 | 0.12 | 0.16 |
| 2 | 16928655 | rs1722409 | -0.348 | -0.103 | -0.199 | -1.036 | 0.015 | -0.019 | -0.019 | 0 | 0.26 | 0.39 | 0.21 | 0.18 | 0.28 | 0.22 |
| 2 | 16940144 | rs488541 | 0.364 | -0.056 | -0.436 | -1.027 | -0.031 | 0.007 | 0.007 | 0 | 0.19 | 0.17 | 0.17 | 0.18 | 0.25 | 0.18 |
| 2 | 16940157 | rs2287518 | 0.771 | -0.173 | -0.525 | -0.962 | -0.03 | 0.014 | 0.014 | 0 | 0.07 | 0.01 | 0.05 | 0.03 | 0.13 | 0.18 |
| 2 | 16943302 | rs6531262 | -0.004 | -0.387 | -0.21 | -1.061 | -0.025 | -0.047 | -0.047 | 0 | 0.21 | 0.73 | 0.14 | 0.03 | 0.08 | 0.06 |
| 2 | 16943918 | rs6531263 | -1.326 | 0.362 | -0.276 | -0.932 | 0.026 | 0.09 | 0.09 | 0.977 | 0.59 | 0.84 | 0.33 | 0.29 | 0.38 | 0.41 |
| 2 | 16944727 | rs12476939 | 0.318 | 0.683 | -0.241 | -0.856 | 0.007 | -0.002 | -0.002 | 0.113 | 0.1 | 0.09 | 0.12 | 0.18 | 0.2 | 0.17 |
| 2 | 16944775 | rs6531266 | -0.08 | 0.332 | -0.013 | -0.952 | 0 | 0.01 | 0.01 | 0.988 | 0.41 | 0.36 | 0.27 | 0.38 | 0.38 | 0.34 |
| 2 | 16944785 | rs6531267 | -1.701 | 0.343 | -0.323 | -0.699 | -0.099 | -0.096 | -0.096 | 0.231 | 0.68 | 0.95 | 0.46 | 0.47 | 0.58 | 0.59 |
| 2 | 16949426 | rs16982747 | 0.263 | 0.099 | 0.226 | -0.635 | -0.024 | -0.025 | -0.025 | 0.07 | 0.09 | 0.18 | 0.06 | 0.05 | 0.02 | 0.01 |
| 2 | 16959153 | rs12710751 | -0.116 | 0.614 | -0.114 | -0.599 | -0.006 | -0.019 | -0.019 | 0 | 0.45 | 0.48 | 0.32 | 0.24 | 0.37 | 0.37 |
| 2 | 16960303 | rs1024913 | -0.814 | 0.777 | -0.068 | -0.569 | -0.001 | 0.002 | 0.002 | 0.612 | 0.58 | 0.56 | 0.39 | 0.42 | 0.57 | 0.58 |
| 2 | 16960560 | rs2111873 | -1.097 | 0.261 | -0.374 | -0.23 | 0.028 | 0.006 | 0.006 | 0 | 0.17 | 0.35 | 0.1 | 0.17 | 0.2 | 0.23 |
| 2 | 16966630 | rs534990 | -0.419 | -0.221 | -0.937 | -0.739 | 0.002 | 0.001 | 0.001 | 0 | 0.11 | 0.09 | 0.08 | 0.05 | 0.06 | 0.08 |
| 2 | 16971986 | rs2341997 | -0.369 | -0.136 | -1.414 | -0.81 | -0.024 | 0.104 | 0.104 | 0.14 | 0.53 | 0.83 | 0.41 | 0.31 | 0.45 | 0.4 |
| 2 | 16981878 | rs2042078 | -0.44 | -0.144 | -0.215 | -0.6 | 0.018 | 0.023 | 0.023 | 0.007 | 0.57 | 0.67 | 0.46 | 0.35 | 0.58 | 0.45 |
| 2 | 16990911 | rs2341991 | -0.254 | -0.158 | 0.01 | -0.608 | -0.003 | -0.004 | -0.004 | 0 | 0.54 | 0.56 | 0.45 | 0.36 | 0.57 | 0.44 |
| 2 | 16991253 | rs558912 | -0.395 | -0.094 | -0.029 | -0.549 | 0.016 | 0.021 | 0.021 | 0 | 0.57 | 0.66 | 0.46 | 0.36 | 0.58 | 0.45 |
| 2 | 16994907 | rs6717502 | -0.526 | -0.146 | -0.078 | -0.556 | -0.012 | -0.012 | -0.012 | 0.84 | 0.6 | 0.59 | 0.45 | 0.38 | 0.59 | 0.51 |
| 2 | 16998604 | rs16982805 | -0.288 | -0.42 | 0.197 | -0.2 | -0.021 | -0.003 | -0.003 | 4.155 | 0.11 | 0.18 | 0.09 | 0.1 | 0.04 | 0.04 |
| 2 | 17009381 | rs6531270 | -1.11 | -0.858 | 0.324 | -0.348 | -0.011 | -0.003 | -0.003 | 0 | 0.33 | 0.63 | 0.54 | 0.74 | 0.18 | 0.31 |
| 2 | 17011894 | rs11096700 | -0.488 | -0.913 | -0.235 | -0.453 | -0.006 | -0.011 | -0.011 | 0 | 0.24 | 0.27 | 0.17 | 0.1 | 0.12 | 0.07 |
| 2 | 17031396 | rs713395 | 0.437 | -0.005 | -0.617 | -0.37 | -0.097 | -0.025 | -0.025 | 0.019 | 0.19 | 0.01 | 0.11 | 0.13 | 0.26 | 0.25 |
| 2 | 17033807 | rs7422674 | -0.612 | -0.761 | 0.124 | 0.01 | 0.012 | 0.027 | 0.027 | 0 | 0.23 | 0.18 | 0.16 | 0.13 | 0.17 | 0.1 |
| 2 | 17034032 | rs11096701 | -0.005 | -0.654 | 0.318 | 0.238 | -0.001 | 0.106 | 0.106 | 0 | 0.08 | 0.13 | 0.32 | 0.55 | 0.07 | 0.24 |
| 2 | 17043703 | rs16982855 | -1.615 | -0.785 | 0.472 | -0.15 | -0.018 | -0.024 | -0.024 | 0 | 0.35 | 0.47 | 0.5 | 0.75 | 0.25 | 0.33 |
| 2 | 17048151 | rs2193698 | -1.676 | -0.756 | 0.542 | -0.136 | -0.022 | -0.028 | -0.028 | 0.448 | 0.36 | 0.58 | 0.52 | 0.75 | 0.25 | 0.33 |
| 2 | 17062078 | rs10210697 | -1.698 | -0.757 | 0.541 | -0.14 | -0.022 | -0.027 | -0.027 | 0 | 0.36 | 0.57 | 0.52 | 0.75 | 0.25 | 0.33 |
| 2 | 17062275 | rs995406 | -1.698 | -0.757 | 0.545 | -0.14 | -0.023 | -0.028 | -0.028 | 0 | 0.36 | 0.56 | 0.52 | 0.75 | 0.25 | 0.34 |
| 2 | 17064165 | rs7563111 | -1.668 | -0.757 | 0.544 | -0.14 | -0.026 | -0.032 | -0.032 | 0 | 0.36 | 0.55 | 0.51 | 0.75 | 0.25 | 0.33 |
| 2 | 17079437 | rs12615806 | -1.8 | -0.715 | 0.361 | -0.148 | 0.003 | 0.003 | 0.003 | 0.151 | 0.41 | 0.63 | 0.55 | 0.73 | 0.29 | 0.35 |
| 2 | 17108634 | rs10490431 | 0.612 | -0.263 | 0.277 | 0.223 | 0.011 | 0.035 | 0.035 | 0 | 0.1 | 0.03 | 0.06 | 0 | 0.06 | 0.02 |
| 2 | 17118296 | rs1019401 | -0.505 | -0.206 | 0.088 | 0.334 | -0.027 | 0.092 | 0.092 | 0 | 0.19 | 0.39 | 0.4 | 0.61 | 0.13 | 0.26 |
| 2 | 17119478 | rs17649532 | -0.624 | 0.24 | -0.549 | -0.331 | 0 | 0.014 | 0.014 | 0 | 0.15 | 0.1 | 0.13 | 0.12 | 0.14 | 0.09 |
| 2 | 17156703 | rs2341504 | 0.134 | -0.248 | -0.084 | -0.897 | 0.013 | 0.024 | 0.024 | 0 | 0.52 | 0.59 | 0.74 | 0.76 | 0.6 | 0.51 |
| 2 | 17164488 | rs7557152 | -0.2 | -0.632 | -0.177 | -1.39 | 0.022 | 0.025 | 0.025 | 0 | 0.35 | 0.27 | 0.15 | 0.22 | 0.21 | 0.39 |
| 2 | 17165823 | rs7603644 | 0.948 | -0.747 | -0.379 | -1.564 | -0.002 | 0.029 | 0.029 | 0 | 0.43 | 0.3 | 0.38 | 0.17 | 0.5 | 0.31 |
| 2 | 17167209 | rs13023785 | 0.511 | -0.623 | -0.251 | -1.367 | -0.012 | 0.009 | 0.009 | 0 | 0.14 | 0.09 | 0.11 | 0.01 | 0.2 | 0.1 |
| 2 | 17170129 | rs2341505 | -0.613 | -1.191 | 0.09 | -1.595 | -0.079 | -0.065 | -0.065 | 0 | 0.32 | 0.5 | 0.46 | 0.8 | 0.18 | 0.51 |
| 2 | 17171039 | rs11888257 | -0.455 | -1.224 | 0.111 | -1.61 | -0.06 | -0.047 | -0.047 | 0 | 0.29 | 0.45 | 0.44 | 0.8 | 0.15 | 0.48 |
| 2 | 17181389 | rs883144 | -0.679 | -1.208 | 0.067 | -1.603 | -0.073 | -0.065 | -0.065 | 0 | 0.32 | 0.52 | 0.46 | 0.8 | 0.18 | 0.51 |
| 2 | 17182831 | rs1515987 | -0.679 | -1.196 | 0.06 | -1.643 | -0.073 | -0.067 | -0.067 | 0 | 0.32 | 0.52 | 0.46 | 0.8 | 0.18 | 0.51 |
| 2 | 17209850 | rs1112297 | -0.49 | -0.287 | -0.273 | -1.695 | -0.002 | 0.005 | 0.005 | 0.34 | 0.33 | 0.28 | 0.39 | 0.6 | 0.37 | 0.36 |
| 2 | 17216459 | rs1515966 | 0.115 | -0.253 | -0.427 | -1.718 | -0.006 | 0.02 | 0.02 | 0.176 | 0.27 | 0.18 | 0.18 | 0.01 | 0.3 | 0.14 |
| 2 | 17221404 | rs4538201 | 0.034 | -0.159 | -0.893 | -1.614 | -0.004 | 0 | 0 | 0.324 | 0.3 | 0.19 | 0.35 | 0.6 | 0.34 | 0.35 |
| 2 | 17223128 | rs6746364 | 0.015 | -0.143 | -0.933 | -1.606 | -0.002 | 0.001 | 0.001 | 0.36 | 0.3 | 0.21 | 0.36 | 0.6 | 0.34 | 0.35 |
| 2 | 17225905 | rs1515968 | -1.001 | 0.34 | -1.34 | -2.078 | 0.005 | 0.005 | 0.005 | 0.61 | 0.57 | 0.42 | 0.71 | 0.79 | 0.68 | 0.71 |
| 2 | 17229231 | rs11676781 | -0.201 | 0.256 | -1.343 | -1.254 | -0.004 | 0 | 0 | 0 | 0.29 | 0.18 | 0.35 | 0.6 | 0.32 | 0.34 |
| 2 | 17232378 | rs1515972 | -0.419 | 0.494 | -1.27 | -1.052 | 0.001 | 0.012 | 0.012 | 2.146 | 0.39 | 0.3 | 0.4 | 0.6 | 0.41 | 0.35 |
| 2 | 17247134 | rs2342546 | 0.239 | 0.92 | -1.735 | -1.539 | -0.04 | 0.097 | 0.097 | 0 | 0.77 | 0.91 | 0.86 | 1 | 0.74 | 0.91 |

| | | | | | | | | | | | | | | | | |
|---|---|---|---|---|---|---|---|---|---|---|---|---|---|---|---|---|
| 2 | 17249333 | rs1400042 | 0.284 | 1.215 | -1.701 | -1.243 | -0.001 | -0.001 | -0.001 | 0.766 | 0.2 | 0.17 | 0.2 | 0.14 | 0.22 | 0.33 |
| 2 | 17261089 | rs2342547 | 0.79 | 1.02 | -1.347 | -1.331 | -0.006 | 0.044 | 0.044 | 0 | 0.56 | 0.64 | 0.65 | 0.86 | 0.53 | 0.58 |
| 2 | 17271863 | rs10177601 | 0.324 | 1.289 | -0.859 | -1.288 | -0.008 | 0.009 | 0.009 | 0 | 0.71 | 0.77 | 0.7 | 0.87 | 0.65 | 0.83 |
| 2 | 17282048 | rs7572654 | 0.117 | 0.84 | -0.864 | -1.956 | 0.005 | 0.036 | 0.036 | 0 | 0.5 | 0.38 | 0.59 | 0.84 | 0.55 | 0.61 |
| 2 | 17305810 | rs13006099 | 0.33 | 0.714 | -1.006 | -2.395 | -0.035 | 0.078 | 0.078 | 0 | 0.22 | 0.13 | 0.23 | 0.05 | 0.24 | 0.2 |
| 2 | 17306653 | rs11687542 | 1.018 | 0.254 | -0.997 | -2.534 | -0.026 | 0.11 | 0.11 | 0.146 | 0.44 | 0.19 | 0.28 | 0.06 | 0.47 | 0.39 |
| 2 | 17307199 | rs1515965 | -0.398 | 0.19 | -0.635 | -2.566 | 0.009 | -0.069 | -0.069 | 0 | 0.76 | 0.62 | 0.75 | 0.95 | 0.76 | 0.8 |
| 2 | 17307871 | rs17386061 | 0.461 | 0.266 | -0.807 | -2.564 | 0.009 | 0.026 | 0.026 | 0 | 0.24 | 0.12 | 0.17 | 0.11 | 0.21 | 0.18 |
| 2 | 17340388 | rs11096723 | -0.075 | 0.175 | -0.825 | -2.643 | -0.004 | 0.047 | 0.047 | 0 | 0.3 | 0.22 | 0.54 | 0.93 | 0.34 | 0.47 |
| 2 | 17347701 | rs13002956 | 0.363 | 0.485 | -0.534 | -2.314 | 0.004 | 0.08 | 0.08 | 0 | 0.23 | 0.11 | 0.1 | 0.02 | 0.2 | 0.14 |
| 2 | 17355811 | rs13022263 | 1.059 | 0.314 | -0.506 | -2.234 | 0.033 | 0.053 | 0.053 | 0 | 0.13 | 0.02 | 0.05 | 0 | 0.07 | 0.09 |
| 2 | 17359410 | rs11893703 | -0.886 | 0.549 | 0.037 | -2.277 | 0.008 | -0.038 | -0.038 | 0 | 0.15 | 0.23 | 0.17 | 0.01 | 0.18 | 0.09 |
| 2 | 17368202 | rs1812211 | -1.161 | 0.52 | 0.56 | -2.184 | 0.026 | 0.028 | 0.028 | 0 | 0.17 | 0.54 | 0.26 | 0.14 | 0.17 | 0.13 |
| 2 | 17388200 | rs13031329 | -0.646 | 0.42 | 0.457 | -2.218 | -0.004 | -0.002 | -0.002 | 0 | 0.13 | 0.15 | 0.14 | 0.01 | 0.15 | 0.09 |
| 2 | 17408334 | rs885302 | -0.438 | 0.558 | 0.557 | -2.219 | 0.004 | -0.023 | -0.023 | 5.688 | 0.12 | 0.15 | 0.11 | 0.01 | 0.16 | 0.09 |
| 2 | 17415773 | rs287305 | -0.376 | 0.163 | 0.061 | -2.805 | 0.025 | -0.004 | -0.004 | 0 | 0.75 | 0.85 | 0.6 | 0.22 | 0.76 | 0.68 |
| 2 | 17416353 | rs7602077 | 1.509 | 0.356 | 0.118 | -2.993 | -0.036 | 0.111 | 0.111 | 0 | 0.27 | 0.08 | 0.14 | 0.06 | 0.25 | 0.38 |
| 2 | 17426034 | rs1983376 | 2.012 | 0.425 | 0.103 | -3.025 | -0.022 | 0.035 | 0.035 | 0 | 0.11 | 0.05 | 0.07 | 0.02 | 0.16 | 0.06 |
| 2 | 17432318 | rs17477667 | 0.747 | 0.084 | 0.331 | -2.832 | -0.011 | 0.047 | 0.047 | 0 | 0.41 | 0.22 | 0.26 | 0.02 | 0.43 | 0.18 |
| 2 | 17432556 | rs13383210 | 0.884 | 0.15 | 0.22 | -2.74 | -0.004 | -0.025 | -0.025 | 0 | 0.3 | 0.41 | 0.28 | 0.19 | 0.25 | 0.42 |
| 2 | 17433256 | rs6709439 | -0.187 | 0.245 | -0.786 | -2.655 | 0.023 | 0.142 | 0.142 | 0.194 | 0.56 | 0.78 | 0.48 | 0.16 | 0.6 | 0.31 |
| 2 | 17435001 | rs6720417 | 1.109 | 0.314 | -0.875 | -2.615 | -0.037 | 0.119 | 0.119 | 0 | 0.27 | 0.06 | 0.18 | 0.06 | 0.24 | 0.38 |
| 2 | 17444053 | rs13418365 | -0.067 | 0.436 | -0.751 | -2.58 | 0.006 | -0.019 | -0.019 | 1.187 | 0.1 | 0.12 | 0.1 | 0.02 | 0.15 | 0.1 |
| 2 | 17444904 | rs7585992 | 0.175 | 0.457 | -0.836 | -2.655 | -0.037 | 0.143 | 0.143 | 0 | 0.37 | 0.15 | 0.27 | 0.07 | 0.38 | 0.48 |
| 2 | 17445026 | rs7572210 | 0.708 | 0.487 | -0.843 | -2.669 | -0.004 | 0.1 | 0.1 | 0 | 0.32 | 0.15 | 0.2 | 0.06 | 0.27 | 0.39 |
| 2 | 17447130 | rs1607653 | -0.215 | 0.486 | -0.826 | -2.633 | -0.003 | -0.004 | -0.004 | 0.098 | 0.1 | 0.09 | 0.1 | 0.02 | 0.15 | 0.1 |
| 2 | 17451733 | rs13020891 | 0.287 | 0.348 | -0.665 | -2.527 | -0.02 | 0.105 | 0.105 | 0 | 0.38 | 0.21 | 0.28 | 0.07 | 0.38 | 0.47 |
| 2 | 17453209 | rs1519966 | -0.308 | 0.363 | -0.624 | -1.982 | 0.003 | -0.01 | -0.01 | 0 | 0.85 | 0.89 | 0.65 | 0.24 | 0.83 | 0.7 |
| 2 | 17453968 | rs7558682 | -0.26 | 0.363 | -0.599 | -1.982 | -0.008 | 0.027 | 0.027 | 0 | 0.84 | 0.81 | 0.65 | 0.24 | 0.83 | 0.69 |
| 2 | 17466012 | rs1976809 | -0.085 | 0.255 | -0.841 | -2.422 | 0.008 | 0.049 | 0.049 | 0 | 0.4 | 0.64 | 0.23 | 0.06 | 0.35 | 0.48 |
| 2 | 17467210 | rs1568385 | 0.908 | 0.209 | -0.941 | -2.576 | -0.012 | 0.17 | 0.17 | 0 | 0.28 | 0.05 | 0.1 | 0.02 | 0.23 | 0.37 |
| 2 | 17469243 | rs12988998 | -0.636 | 0.247 | -0.417 | -2.445 | 0.042 | -0.04 | -0.04 | 1.252 | 0.12 | 0.52 | 0.12 | 0.04 | 0.12 | 0.1 |
| 2 | 17470470 | rs10205002 | -0.497 | 0.36 | -0.403 | -2.434 | 0.021 | -0.025 | -0.025 | 0 | 0.11 | 0.52 | 0.09 | 0.04 | 0.08 | 0.1 |
| 2 | 17486497 | rs1401882 | 0.494 | 0.266 | -0.047 | -2.464 | -0.027 | -0.123 | -0.123 | 0.767 | 0.31 | 0.46 | 0.2 | 0.07 | 0.23 | 0.41 |
| 2 | 17487397 | rs10199328 | 0.138 | 0.736 | -0.474 | -2.242 | 0.001 | 0.035 | 0.035 | 0 | 0.52 | 0.39 | 0.37 | 0.05 | 0.51 | 0.28 |
| 2 | 17497137 | rs1519973 | 1.152 | 0.831 | -0.549 | -2.32 | -0.006 | 0.154 | 0.154 | 0 | 0.27 | 0.05 | 0.1 | 0.02 | 0.23 | 0.37 |
| 2 | 17506784 | rs4832491 | -0.747 | 0.974 | -0.694 | -2.119 | -0.049 | -0.044 | -0.044 | 0 | 0.91 | 0.99 | 0.89 | 0.91 | 0.87 | 0.89 |
| 2 | 17518050 | rs4502415 | 0.761 | 0.892 | -0.491 | -2.468 | -0.024 | -0.115 | -0.115 | 0.578 | 0.31 | 0.47 | 0.21 | 0.07 | 0.23 | 0.4 |
| 2 | 17530815 | rs7602088 | 0.584 | 0.567 | -1.312 | -2.757 | -0.027 | 0.066 | 0.066 | 2.419 | 0.27 | 0.2 | 0.24 | 0.02 | 0.31 | 0.08 |
| 2 | 17557962 | rs1401705 | -0.153 | 0.482 | -1.083 | -2.864 | -0.008 | -0.027 | -0.027 | 0 | 0.42 | 0.37 | 0.33 | 0.02 | 0.45 | 0.23 |
| 2 | 17561931 | rs6740930 | 0.891 | -0.127 | -1.673 | -3.03 | 0.042 | 0.101 | 0.101 | 0 | 0.24 | 0.07 | 0.07 | 0.01 | 0.18 | 0.23 |
| 2 | 17564763 | rs16983552 | -0.374 | -0.473 | -0.849 | -3.176 | -0.021 | -0.02 | -0.02 | 0.418 | 0.11 | 0.32 | 0.18 | 0.09 | 0.05 | 0.08 |
| 2 | 17568267 | rs871458 | 0.722 | -0.128 | -0.978 | -3.662 | 0.006 | -0.122 | -0.122 | 0.098 | 0.43 | 0.45 | 0.35 | 0.04 | 0.46 | 0.26 |
| 2 | 17581491 | rs932212 | 1.497 | -0.268 | -1.004 | -3.549 | 0.044 | 0.117 | 0.117 | 0 | 0.26 | 0.07 | 0.09 | 0.01 | 0.2 | 0.23 |
| 2 | 17582097 | rs11681567 | 0.232 | -0.205 | -0.971 | -3.508 | -0.004 | -0.036 | -0.036 | 0.133 | 0.38 | 0.39 | 0.28 | 0.1 | 0.25 | 0.31 |
| 2 | 17621418 | rs6737193 | -0.058 | 0.429 | -0.315 | -3.656 | -0.002 | -0.085 | -0.085 | 4.212 | 0.44 | 0.5 | 0.39 | 0.09 | 0.4 | 0.17 |
| 2 | 17626455 | rs11688504 | 0.93 | 0.331 | -0.395 | -3.918 | -0.058 | 0.269 | 0.269 | 0 | 0.36 | 0.02 | 0.16 | 0.02 | 0.34 | 0.42 |
| 2 | 17636816 | rs951375 | 0.585 | 0.29 | -0.036 | -3.875 | 0.008 | 0.025 | 0.025 | 0 | 0.06 | 0.23 | 0.05 | 0.09 | 0.04 | 0.05 |
| 2 | 17655480 | rs394874 | -0.437 | 0.21 | -0.229 | -3.997 | 0.005 | 0.125 | 0.125 | 0 | 0.59 | 0.23 | 0.66 | 0.9 | 0.65 | 0.8 |
| 2 | 17674793 | rs13020775 | 1.052 | 0.504 | -0.189 | -4.042 | -0.014 | 0.079 | 0.079 | 0 | 0.32 | 0.14 | 0.19 | 0.01 | 0.32 | 0.09 |
| 2 | 17683231 | rs17380121 | 2.492 | 0.496 | -0.68 | -4.119 | -0.012 | 0.084 | 0.084 | 0 | 0.14 | 0.01 | 0.05 | 0.01 | 0.11 | 0.03 |
| 2 | 17690666 | rs11689517 | 0.63 | 0.518 | -0.483 | -4.01 | -0.012 | 0.019 | 0.019 | 0 | 0.33 | 0.21 | 0.22 | 0.01 | 0.34 | 0.09 |
| 2 | 17698678 | rs665312 | 0.102 | 0.5 | -0.679 | -3.984 | -0.006 | 0.198 | 0.198 | 0 | 0.58 | 0.16 | 0.66 | 0.88 | 0.65 | 0.8 |
| 2 | 17701234 | rs11885715 | 0.592 | 0.547 | -0.642 | -3.969 | -0.001 | -0.06 | -0.06 | 1.874 | 0.33 | 0.29 | 0.22 | 0.01 | 0.34 | 0.09 |
| 2 | 17720922 | rs7596935 | 1.5 | 0.54 | -1.042 | -3.972 | -0.043 | 0.253 | 0.253 | 0 | 0.34 | 0.02 | 0.13 | 0.02 | 0.31 | 0.36 |
| 2 | 17754316 | rs1996610 | 0.12 | 0.538 | -1.089 | -3.768 | -0.02 | 0.232 | 0.232 | 0 | 0.57 | 0.16 | 0.65 | 0.99 | 0.62 | 0.83 |
| 2 | 17768968 | rs16983681 | 0.404 | 0.605 | -0.461 | -3.316 | 0.005 | 0.031 | 0.031 | 0 | 0.07 | 0.24 | 0.06 | 0.11 | 0.05 | 0.05 |
| 2 | 17770095 | rs1519473 | 0.187 | 0.702 | -0.483 | -3.306 | 0.016 | 0.029 | 0.029 | 0 | 0.08 | 0.24 | 0.06 | 0.13 | 0.07 | 0.06 |
| 2 | 17773524 | rs2680827 | 0.056 | 0.823 | -0.619 | -3.493 | -0.001 | -0.034 | -0.034 | 1.085 | 0.65 | 0.37 | 0.7 | 0.99 | 0.69 | 0.88 |
| 2 | 17774701 | rs4510229 | 0.136 | 0.823 | -0.532 | -3.367 | 0.013 | 0.033 | 0.033 | 0 | 0.08 | 0.24 | 0.06 | 0.13 | 0.07 | 0.06 |
| 2 | 17790513 | rs7567064 | 0.309 | 0.952 | -0.481 | -3.337 | 0.013 | -0.096 | -0.096 | 0.304 | 0.3 | 0.42 | 0.26 | 0.05 | 0.31 | 0.07 |
| 2 | 17822675 | rs2114352 | 1.263 | 0.786 | -1.37 | -3.328 | -0.108 | 0.222 | 0.222 | 0 | 0.4 | 0.02 | 0.17 | 0.1 | 0.39 | 0.41 |
| 2 | 17830028 | rs2018091 | 2.373 | 0.613 | -0.761 | -3.078 | 0.008 | 0.055 | 0.055 | 0 | 0.1 | 0 | 0.02 | 0 | 0.06 | 0.01 |
| 2 | 17834754 | rs2555099 | -0.455 | 0.786 | 0.034 | -3.15 | 0.001 | -0.111 | -0.111 | 0.652 | 0.33 | 0.54 | 0.23 | 0.04 | 0.33 | 0.1 |
| 2 | 17840090 | rs1991315 | -1.285 | 0.781 | 0.305 | -3.162 | -0.001 | -0.011 | -0.011 | 0.002 | 0.42 | 0.64 | 0.3 | 0.08 | 0.42 | 0.17 |
| 2 | 17846162 | rs2555107 | -0.61 | 0.732 | 0.588 | -3.188 | -0.004 | -0.143 | -0.143 | 0 | 0.34 | 0.52 | 0.22 | 0.01 | 0.33 | 0.08 |
| 2 | 17851241 | rs17315242 | 1.584 | 0.8 | 0.284 | -3.199 | -0.032 | 0.1 | 0.1 | 0 | 0.17 | 0.01 | 0.1 | 0.01 | 0.14 | 0.03 |
| 2 | 17884495 | rs1065381 | -0.202 | 0.779 | 0.526 | -3.074 | -0.001 | -0.073 | -0.073 | 0.325 | 0.33 | 0.35 | 0.22 | 0.01 | 0.32 | 0.08 |
| 2 | 17886901 | rs7571131 | -0.233 | 0.78 | 0.507 | -3.073 | -0.001 | -0.084 | -0.084 | 0 | 0.33 | 0.38 | 0.22 | 0.01 | 0.32 | 0.08 |

| | | | | | | | | | | | | | | | | |
|---|---|---|---|---|---|---|---|---|---|---|---|---|---|---|---|---|
| 2 | 17888457 | rs6734462 | -1.528 | 0.747 | 0.423 | -3.096 | -0.009 | 0.306 | 0.306 | 0 | 0.49 | 0.87 | 0.36 | 0.08 | 0.44 | 0.19 |
| 2 | 17889705 | rs9306855 | -1.528 | 0.747 | 0.423 | -3.096 | -0.009 | 0.306 | 0.306 | 0 | 0.49 | 0.87 | 0.36 | 0.08 | 0.44 | 0.19 |
| 2 | 17927571 | rs16983818 | -0.41 | 0.605 | 0.925 | -2.608 | 0 | -0.05 | -0.05 | 0 | 0.13 | 0.29 | 0.12 | 0.03 | 0.11 | 0.1 |
| 2 | 17930195 | rs6761104 | -0.55 | 0.605 | 0.961 | -2.608 | 0.015 | -0.025 | -0.025 | 0 | 0.14 | 0.41 | 0.12 | 0.06 | 0.11 | 0.11 |
| 2 | 17934162 | rs7561505 | 0.279 | 0.781 | 1.027 | -2.79 | -0.014 | 0.022 | 0.022 | 0 | 0.33 | 0.24 | 0.22 | 0.01 | 0.36 | 0.1 |
| 2 | 17935780 | rs6531053 | -1.074 | 0.769 | 1.07 | -2.796 | -0.048 | 0.268 | 0.268 | 0 | 0.48 | 0.85 | 0.36 | 0.08 | 0.48 | 0.21 |
| 2 | 17938501 | rs2615051 | 0.823 | 0.766 | 1.068 | -2.788 | -0.047 | 0.269 | 0.269 | 0 | 0.52 | 0.16 | 0.64 | 0.92 | 0.52 | 0.79 |
| 2 | 17942775 | rs1812152 | -0.499 | 0.766 | 1.087 | -2.788 | -0.018 | 0.038 | 0.038 | 0 | 0.67 | 0.77 | 0.78 | 0.99 | 0.64 | 0.9 |
| 2 | 17943015 | rs2615047 | -0.499 | 0.766 | 1 | -2.788 | -0.014 | 0.022 | 0.022 | 0 | 0.67 | 0.76 | 0.78 | 0.99 | 0.64 | 0.9 |
| 2 | 17946243 | rs16981869 | -0.264 | 0.78 | 0.851 | -2.781 | -0.002 | -0.046 | -0.046 | 0 | 0.13 | 0.24 | 0.12 | 0.03 | 0.11 | 0.1 |
| 2 | 17948649 | rs7574887 | -0.264 | 0.78 | 0.781 | -2.781 | 0 | -0.048 | -0.048 | 0 | 0.13 | 0.27 | 0.12 | 0.03 | 0.11 | 0.1 |
| 2 | 17953560 | rs300174 | -0.372 | 0.784 | 0.689 | -2.792 | 0.008 | -0.122 | -0.122 | 0 | 0.66 | 0.57 | 0.76 | 0.99 | 0.64 | 0.9 |
| 2 | 17954647 | rs300178 | -0.368 | 0.792 | 0.693 | -2.788 | 0.008 | -0.121 | -0.121 | 0 | 0.66 | 0.57 | 0.76 | 0.99 | 0.64 | 0.9 |
| 2 | 17962450 | rs300168 | 0.77 | 0.791 | 0.693 | -2.788 | -0.048 | 0.27 | 0.27 | 0 | 0.52 | 0.15 | 0.64 | 0.92 | 0.52 | 0.79 |
| 2 | 17962518 | rs300169 | -0.289 | 0.667 | 0.678 | -2.77 | 0.008 | -0.121 | -0.121 | 0.024 | 0.66 | 0.57 | 0.76 | 0.99 | 0.64 | 0.9 |
| 2 | 17966421 | rs300172 | -0.307 | 0.629 | 0.684 | -2.764 | 0.01 | -0.122 | -0.122 | 2.599 | 0.66 | 0.57 | 0.76 | 0.99 | 0.64 | 0.9 |
| 2 | 17981399 | rs300161 | -0.23 | 0.571 | 0.325 | -2.638 | 0.013 | -0.113 | -0.113 | 0 | 0.66 | 0.48 | 0.77 | 0.96 | 0.63 | 0.86 |
| 2 | 17981684 | rs300159 | 0.537 | 0.615 | -0.214 | -2.6 | -0.02 | 0.262 | 0.262 | 0 | 0.53 | 0.17 | 0.67 | 0.95 | 0.56 | 0.8 |
| 2 | 17991974 | rs765396 | -0.132 | 0.898 | -0.28 | -2.167 | 0.004 | -0.115 | -0.115 | 0.07 | 0.62 | 0.51 | 0.74 | 0.96 | 0.61 | 0.87 |
| 2 | 18008390 | rs6713888 | -0.243 | 1.15 | -0.457 | -1.751 | 0.018 | -0.025 | -0.025 | 0 | 0.07 | 0.16 | 0.1 | 0.04 | 0.13 | 0.02 |
| 2 | 18009796 | rs9306859 | 0.217 | 1.159 | -1.176 | -1.924 | -0.02 | 0.074 | 0.074 | 0 | 0.15 | 0.04 | 0.05 | 0 | 0.14 | 0.1 |
| 2 | 18016788 | rs4302214 | 0.311 | 0.814 | -1.004 | -2.205 | 0 | 0.089 | 0.089 | 0 | 0.35 | 0.21 | 0.25 | 0.03 | 0.34 | 0.16 |
| 2 | 18027520 | rs11096489 | 0.162 | 0.608 | -0.592 | -1.748 | -0.001 | -0.027 | -0.027 | 0.363 | 0.07 | 0.07 | 0.04 | 0 | 0.06 | 0.06 |
| 2 | 18031386 | rs16983978 | -1.006 | 0.499 | -0.575 | -1.807 | -0.011 | -0.046 | -0.046 | 0 | 0.11 | 0.15 | 0.07 | 0 | 0.07 | 0.1 |
| 2 | 18033770 | rs6531058 | -0.136 | 0.566 | -0.481 | -0.406 | -0.001 | -0.002 | -0.002 | 0 | 0.54 | 0.51 | 0.69 | 0.62 | 0.54 | 0.72 |
| 2 | 18033926 | rs7596542 | 0.278 | 0.707 | -0.726 | -0.139 | -0.003 | 0.073 | 0.073 | 0 | 0.91 | 0.86 | 0.94 | 0.71 | 0.91 | 0.88 |
| 2 | 18036252 | rs4832399 | 0.256 | 0.745 | -0.586 | -0.138 | 0.007 | 0.134 | 0.134 | 0 | 0.9 | 0.79 | 0.93 | 0.72 | 0.89 | 0.83 |
| 2 | 18037140 | rs12479002 | 0.182 | 0.779 | -0.633 | -0.34 | 0 | 0.001 | 0.001 | 0 | 0.49 | 0.46 | 0.65 | 0.64 | 0.48 | 0.57 |
| 2 | 18043777 | rs10180853 | 0.256 | 0.834 | -1.052 | -0.283 | -0.009 | 0.029 | 0.029 | 0 | 0.87 | 0.89 | 0.93 | 0.99 | 0.88 | 0.88 |
| 2 | 18046715 | rs12614549 | 0.29 | 0.958 | -1.757 | -0.279 | -0.055 | -0.109 | -0.109 | 0 | 0.24 | 0.04 | 0.16 | 0.31 | 0.26 | 0.21 |
| 2 | 18050100 | rs17388142 | 0.07 | 1.061 | -1.073 | -0.025 | 0.031 | 0.015 | 0.015 | 0 | 0.1 | 0.2 | 0.17 | 0.14 | 0.18 | 0.07 |
| 2 | 18056447 | rs9306861 | 1.171 | 1.086 | -1.001 | -0.04 | 0.037 | 0.01 | 0.01 | 0 | 0.06 | 0.11 | 0.08 | 0.07 | 0.16 | 0.04 |
| 2 | 18063035 | rs6531064 | 0.408 | 0.714 | -1.143 | -0.165 | -0.001 | 0.014 | 0.014 | 0 | 0.29 | 0.29 | 0.51 | 0.59 | 0.31 | 0.55 |
| 2 | 18077007 | rs4832514 | 1.483 | 0.52 | -1.373 | -0.251 | 0.029 | -0.047 | -0.047 | 0 | 0.22 | 0.09 | 0.34 | 0.46 | 0.15 | 0.48 |
| 2 | 18082049 | rs10184119 | 0.015 | 0.177 | -1.528 | -1.024 | 0.001 | -0.094 | -0.094 | 0 | 0.41 | 0.39 | 0.34 | 0.01 | 0.5 | 0.2 |
| 2 | 18083764 | rs7557530 | -0.123 | 0.641 | -2.037 | -1.185 | -0.034 | 0.072 | 0.072 | 0.785 | 0.73 | 0.81 | 0.82 | 0.95 | 0.69 | 0.82 |
| 2 | 18083850 | rs7583266 | -0.048 | 0.782 | -2.207 | -1.365 | -0.042 | 0.092 | 0.092 | 0.02 | 0.69 | 0.8 | 0.76 | 0.96 | 0.65 | 0.83 |
| 2 | 18083997 | rs11680994 | 0.995 | -0.294 | -1.826 | -0.841 | 0.033 | -0.045 | -0.045 | 0 | 0.27 | 0.09 | 0.44 | 0.47 | 0.2 | 0.53 |
| 2 | 18098857 | rs1564004 | 1.795 | -0.041 | -1.75 | -0.491 | -0.051 | -0.14 | -0.14 | 0 | 0.31 | 0.03 | 0.23 | 0.36 | 0.29 | 0.21 |
| 2 | 18099832 | rs4553827 | 0.398 | -0.136 | -1.213 | -0.495 | 0.05 | -0.037 | -0.037 | 2.516 | 0.64 | 0.24 | 0.52 | 0.52 | 0.74 | 0.44 |
| 2 | 18099919 | rs1870822 | -0.663 | 0.162 | -0.724 | -0.372 | 0.019 | -0.006 | -0.006 | 0.128 | 0.94 | 0.83 | 0.93 | 0.99 | 0.94 | 0.96 |
| 2 | 18102223 | rs1461949 | -0.939 | 0.054 | -0.749 | -0.409 | 0.018 | -0.042 | -0.042 | 0.262 | 0.76 | 0.74 | 0.8 | 0.98 | 0.61 | 0.87 |
| 2 | 18113508 | rs3747516 | 0.729 | -0.318 | -0.205 | -0.428 | -0.022 | 0.082 | 0.082 | 0 | 0.19 | 0.33 | 0.38 | 0.47 | 0.16 | 0.45 |
| 2 | 18113623 | rs4832524 | -1.058 | -0.329 | -0.221 | -0.434 | 0.02 | -0.043 | -0.043 | 0 | 0.77 | 0.74 | 0.81 | 0.98 | 0.61 | 0.87 |
| 2 | 18113789 | rs3747515 | 1.457 | -0.359 | -0.844 | -0.367 | -0.039 | -0.096 | -0.096 | 0 | 0.36 | 0.1 | 0.28 | 0.37 | 0.34 | 0.26 |
| 2 | 18115241 | rs6712976 | 1.259 | -0.044 | -0.319 | 0.171 | 0.016 | -0.023 | -0.023 | 0 | 0.14 | 0.07 | 0.32 | 0.39 | 0.1 | 0.43 |
| 2 | 18116319 | rs1461943 | 0.602 | 0.026 | -0.301 | 0.084 | -0.02 | 0.076 | 0.076 | 0 | 0.19 | 0.3 | 0.38 | 0.47 | 0.16 | 0.45 |
| 2 | 18122198 | rs1471342 | 0.774 | -0.029 | -0.651 | 0.158 | -0.038 | -0.096 | -0.096 | 4.474 | 0.42 | 0.06 | 0.26 | 0.35 | 0.37 | 0.31 |
| 2 | 18123412 | rs4832404 | 0.227 | 0.235 | -0.119 | 0.208 | 0.007 | -0.02 | -0.02 | 0 | 0.37 | 0.36 | 0.43 | 0.16 | 0.5 | 0.29 |
| 2 | 18135019 | rs6711333 | -0.566 | 0.108 | -0.585 | 0.199 | -0.022 | -0.019 | -0.019 | 0 | 0.62 | 0.32 | 0.43 | 0.52 | 0.49 | 0.48 |
| 2 | 18139878 | rs6706996 | 0.026 | -0.337 | -1.203 | 0.222 | -0.018 | -0.039 | -0.039 | 0.317 | 0.25 | 0.34 | 0.19 | 0.16 | 0.19 | 0.22 |
| 2 | 18141228 | rs10495673 | -0.192 | -0.104 | -1.311 | 0.662 | 0.008 | -0.08 | -0.08 | 0 | 0.39 | 0.05 | 0.24 | 0.35 | 0.33 | 0.27 |
| 2 | 18142133 | rs11903301 | -0.141 | -0.03 | -1.27 | 0.704 | 0.04 | -0.016 | -0.016 | 0.454 | 0.41 | 0.15 | 0.25 | 0.35 | 0.33 | 0.27 |
| 2 | 18147893 | rs1806644 | -0.545 | 0.086 | -0.848 | 0.551 | 0 | -0.004 | -0.004 | 0 | 0.37 | 0.32 | 0.46 | 0.77 | 0.42 | 0.59 |
| 2 | 18148175 | rs10186459 | -0.367 | -0.004 | -0.887 | 0.425 | -0.001 | -0.012 | -0.012 | 0 | 0.33 | 0.33 | 0.44 | 0.77 | 0.38 | 0.59 |
| 2 | 18150570 | rs10170651 | -0.57 | 0.164 | -0.84 | 1.004 | 0.12 | 0.114 | 0.114 | 0 | 0.08 | 0.26 | 0.15 | 0.17 | 0.17 | 0.1 |
| 2 | 18160897 | rs2952821 | -1.288 | -0.867 | -0.286 | 1.015 | 0.023 | 0.008 | 0.008 | 0 | 0.48 | 0.61 | 0.51 | 0.55 | 0.45 | 0.4 |
| 2 | 18171812 | rs2732936 | -1.427 | -1.07 | -0.382 | 0.113 | -0.02 | 0.003 | 0.003 | 0 | 0.63 | 0.72 | 0.72 | 0.97 | 0.54 | 0.8 |
| 2 | 18185590 | rs12470501 | -0.153 | -1.264 | -0.486 | 0.302 | -0.01 | -0.004 | -0.004 | 0 | 0.31 | 0.17 | 0.21 | 0.29 | 0.3 | 0.3 |
| 2 | 18189056 | rs12052676 | 1.201 | -1.137 | -0.468 | -0.067 | -0.014 | -0.019 | -0.019 | 0.473 | 0.16 | 0.01 | 0.05 | 0.12 | 0.11 | 0.16 |
| 2 | 18189420 | rs2732922 | -0.39 | -1.055 | -0.2 | -0.124 | 0.022 | 0.133 | 0.133 | 0.818 | 0.05 | 0.34 | 0.18 | 0.09 | 0.01 | 0.07 |
| 2 | 18192841 | rs2710017 | -1.254 | -0.843 | -0.223 | -0.307 | -0.011 | -0.033 | -0.033 | 0 | 0.64 | 0.68 | 0.71 | 0.97 | 0.56 | 0.81 |
| 2 | 18206413 | rs11886084 | -0.221 | -0.569 | -0.458 | 0.058 | 0.003 | -0.021 | -0.021 | 0 | 0.37 | 0.28 | 0.35 | 0.47 | 0.41 | 0.31 |
| 2 | 18209764 | rs2345097 | -0.536 | -0.513 | -0.229 | -0.564 | -0.001 | -0.071 | -0.071 | 0 | 0.61 | 0.6 | 0.71 | 0.97 | 0.49 | 0.77 |
| 2 | 18213422 | rs2957566 | 0.188 | -0.363 | 0.381 | -0.44 | 0.002 | -0.071 | -0.071 | 0 | 0.11 | 0.28 | 0.06 | 0 | 0.09 | 0.03 |
| 2 | 18214457 | rs2710005 | -0.235 | -0.666 | 0.436 | -0.243 | -0.018 | -0.007 | -0.007 | 0.133 | 0.26 | 0.4 | 0.26 | 0.21 | 0.19 | 0.15 |
| 2 | 18216417 | rs2345088 | 1.497 | -0.126 | -0.089 | 0.018 | -0.056 | -0.08 | -0.08 | 0 | 0.12 | 0.01 | 0.12 | 0.28 | 0.18 | 0.13 |
| 2 | 18217217 | rs2256213 | -0.646 | -0.121 | -0.175 | -0.289 | 0.013 | 0.034 | 0.034 | 1.359 | 0.5 | 0.73 | 0.63 | 0.6 | 0.47 | 0.65 |
| 2 | 18218755 | rs2732933 | -1.419 | -0.214 | -0.23 | -0.416 | -0.013 | 0.142 | 0.142 | 2.153 | 0.63 | 0.86 | 0.82 | 0.98 | 0.66 | 0.8 |

| | | | | | | | | | | | | | | | | |
|---|---|---|---|---|---|---|---|---|---|---|---|---|---|---|---|---|
| 2 | 18227025 | rs4832532 | -0.417 | -0.271 | 0.088 | -0.355 | 0.001 | -0.029 | -0.029 | 0 | 0.51 | 0.49 | 0.69 | 0.86 | 0.56 | 0.68 |
| 2 | 18230779 | rs2345089 | 0.24 | -0.227 | 0.078 | -0.124 | -0.017 | -0.044 | -0.044 | 0 | 0.28 | 0.17 | 0.35 | 0.51 | 0.3 | 0.31 |
| 2 | 18232621 | rs4073911 | 0.932 | -0.265 | 0.074 | -0.045 | 0.008 | 0.032 | 0.032 | 0.195 | 0.24 | 0.37 | 0.35 | 0.37 | 0.26 | 0.44 |
| 2 | 18233942 | rs10198155 | 1.344 | -0.278 | -0.069 | -0.078 | 0 | -0.008 | -0.008 | 0 | 0.21 | 0.21 | 0.22 | 0.09 | 0.25 | 0.18 |
| 2 | 18234446 | rs16984239 | 1.668 | -0.254 | -0.422 | 0.083 | -0.046 | -0.071 | -0.071 | 0 | 0.13 | 0.02 | 0.14 | 0.28 | 0.2 | 0.12 |
| 2 | 18242052 | rs12477900 | 0.043 | -0.744 | 0.298 | 0.455 | -0.014 | -0.039 | -0.039 | 0 | 0.15 | 0.25 | 0.11 | 0.09 | 0.11 | 0.08 |
| 2 | 18243534 | rs4075511 | -1.627 | -0.687 | -0.507 | 0.375 | -0.032 | 0.036 | 0.036 | 0.611 | 0.54 | 0.85 | 0.68 | 0.7 | 0.47 | 0.71 |
| 2 | 18244718 | rs7559591 | 0.685 | -0.78 | -0.502 | 0.608 | 0.04 | -0.08 | -0.08 | 0 | 0.16 | 0.03 | 0.21 | 0.31 | 0.07 | 0.15 |
| 2 | 18245160 | rs11892741 | 0.453 | -0.256 | 0.294 | 0.259 | -0.007 | 0.013 | 0.013 | 1.622 | 0.21 | 0.15 | 0.24 | 0.09 | 0.27 | 0.19 |
| 2 | 18260937 | rs10194386 | -1.226 | -0.171 | 0.193 | 0.095 | -0.001 | -0.032 | -0.032 | 0.295 | 0.38 | 0.63 | 0.35 | 0.1 | 0.38 | 0.34 |
| 2 | 18265521 | rs7580025 | -1.209 | -0.172 | 0.239 | 0.097 | -0.001 | -0.042 | -0.042 | 0 | 0.37 | 0.61 | 0.35 | 0.1 | 0.38 | 0.34 |
| 2 | 18265610 | rs7593736 | -1.129 | -0.168 | 0.151 | 0.098 | -0.001 | -0.035 | -0.035 | 1.377 | 0.37 | 0.61 | 0.35 | 0.11 | 0.38 | 0.31 |
| 2 | 18270135 | rs7578543 | -0.47 | -0.104 | -0.071 | 0.336 | -0.025 | 0.032 | 0.032 | 0.483 | 0.19 | 0.53 | 0.23 | 0.26 | 0.13 | 0.34 |
| 2 | 18274609 | rs4832414 | -1.067 | -0.088 | -1.093 | 0.245 | 0.005 | 0.051 | 0.051 | 1.064 | 0.42 | 0.85 | 0.5 | 0.36 | 0.39 | 0.6 |
| 2 | 18275380 | rs7570141 | -1.086 | -1.074 | -0.624 | 0.823 | -0.004 | 0.055 | 0.055 | 0 | 0.14 | 0.17 | 0.13 | 0.41 | 0.11 | 0.18 |
| 2 | 18276905 | rs17388897 | -0.459 | -1.359 | -0.938 | 0.047 | -0.007 | 0.048 | 0.048 | 0 | 0.31 | 0.23 | 0.3 | 0.07 | 0.33 | 0.22 |
| 2 | 18278512 | rs3866196 | -1.32 | -1.675 | -0.2 | 0.153 | 0 | 0 | 0 | 0 | 0.44 | 0.6 | 0.55 | 0.35 | 0.38 | 0.43 |
| 2 | 18280088 | rs2345102 | -0.577 | -1.585 | -0.21 | 0.206 | -0.001 | -0.031 | -0.031 | 0.1 | 0.33 | 0.36 | 0.3 | 0.09 | 0.33 | 0.24 |
| 2 | 18280641 | rs7560092 | 0.336 | -1.365 | -0.42 | 0.448 | 0.013 | -0.011 | -0.011 | 0 | 0.09 | 0.07 | 0.22 | 0.26 | 0.05 | 0.2 |
| 2 | 18282609 | rs6726628 | -0.813 | -1.165 | -0.394 | 0.098 | 0.007 | 0.022 | 0.022 | 0 | 0.39 | 0.27 | 0.49 | 0.31 | 0.38 | 0.42 |
| 2 | 18282657 | rs6713642 | -0.972 | -1.665 | 0.418 | 0.196 | -0.028 | 0.125 | 0.125 | 0.138 | 0.16 | 0.3 | 0.21 | 0.52 | 0.11 | 0.35 |
| 2 | 18283243 | rs9306867 | -0.733 | -1.778 | 0.142 | -1.022 | 0.019 | 0.032 | 0.032 | 0 | 0.51 | 0.45 | 0.64 | 0.8 | 0.41 | 0.69 |
| 2 | 18283354 | rs9284577 | -0.575 | -1.88 | 0.25 | -0.908 | 0.02 | 0.028 | 0.028 | 0.074 | 0.43 | 0.37 | 0.6 | 0.79 | 0.35 | 0.68 |
| 2 | 18285777 | rs6738939 | -1.101 | -1.492 | 0.216 | -0.797 | 0.008 | 0.033 | 0.033 | 0 | 0.54 | 0.63 | 0.71 | 0.83 | 0.41 | 0.75 |
| 2 | 18286852 | rs6419085 | -1.365 | -1.389 | 0.24 | -0.764 | -0.02 | 0.13 | 0.13 | 0 | 0.59 | 0.8 | 0.73 | 0.93 | 0.44 | 0.82 |
| 2 | 18288226 | rs6716074 | 0.326 | -1.359 | 0.443 | -0.026 | -0.01 | 0.028 | 0.028 | 0.048 | 0.18 | 0.2 | 0.33 | 0.34 | 0.15 | 0.25 |
| 2 | 18288633 | rs9973591 | -0.273 | -1.366 | 0.695 | -0.139 | -0.024 | 0.085 | 0.085 | 0.139 | 0.22 | 0.51 | 0.4 | 0.33 | 0.21 | 0.26 |
| 2 | 18288716 | rs11887971 | -0.846 | -1.436 | 0.738 | -0.13 | 0.003 | -0.045 | -0.045 | 0.036 | 0.17 | 0.43 | 0.14 | 0.09 | 0.14 | 0.2 |
| 2 | 18292201 | rs2345493 | 0.034 | -1.496 | 0.214 | -0.176 | -0.004 | 0.021 | 0.021 | 0.059 | 0.21 | 0.31 | 0.36 | 0.32 | 0.22 | 0.24 |
| 2 | 18293827 | rs11900434 | -1.204 | -1.684 | 0.095 | -0.433 | 0.004 | -0.01 | -0.01 | 0 | 0.52 | 0.59 | 0.7 | 0.81 | 0.41 | 0.73 |
| 2 | 18299134 | rs12473016 | -1.877 | -1.684 | -0.296 | -0.51 | -0.099 | 0.202 | 0.202 | 0 | 0.62 | 0.95 | 0.74 | 0.83 | 0.47 | 0.77 |
| 2 | 18301858 | rs4295006 | 0.946 | -1.713 | 0.155 | -0.027 | 0.014 | 0.021 | 0.021 | 0.193 | 0.14 | 0.06 | 0.11 | 0.06 | 0.09 | 0.12 |
| 2 | 18314546 | rs12471785 | -1.57 | -1.741 | 0.179 | -0.038 | 0.044 | 0.009 | 0.009 | 0.686 | 0.39 | 0.72 | 0.43 | 0.37 | 0.27 | 0.47 |
| 2 | 18315407 | rs10169347 | 0.433 | -1.629 | 0.229 | 0.041 | 0.009 | 0.149 | 0.149 | 0 | 0.06 | 0.14 | 0.19 | 0.24 | 0.06 | 0.15 |
| 2 | 18316158 | rs11684345 | 0.845 | -1.625 | -0.019 | 0.027 | 0.013 | 0.006 | 0.006 | 0 | 0.09 | 0.04 | 0.05 | 0.07 | 0.05 | 0.14 |
| 2 | 18327703 | rs4832543 | 0.891 | -1.586 | -0.112 | 0.046 | 0 | -0.15 | -0.15 | 0.617 | 0.18 | 0.01 | 0.25 | 0.46 | 0.12 | 0.3 |
| 2 | 18330524 | rs13416608 | -1.609 | -1.579 | 0.023 | 0.019 | 0.011 | 0.089 | 0.089 | 0 | 0.56 | 0.72 | 0.71 | 0.83 | 0.42 | 0.77 |
| 2 | 18332162 | rs10166150 | 0.005 | -1.706 | 0.212 | 0.158 | 0.038 | 0.184 | 0.184 | 0.02 | 0.07 | 0.22 | 0.2 | 0.23 | 0.08 | 0.15 |
| 2 | 18336818 | rs2881165 | -0.608 | -2.085 | -0.342 | 0.132 | 0.015 | 0.017 | 0.017 | 0.234 | 0.25 | 0.21 | 0.14 | 0.13 | 0.14 | 0.25 |
| 2 | 18338029 | rs6709406 | 0.212 | -1.583 | -0.555 | 0.093 | -0.033 | -0.04 | -0.04 | 1.243 | 0.17 | 0.26 | 0.1 | 0.12 | 0.09 | 0.16 |
| 2 | 18339926 | rs7568386 | -1.437 | -1.582 | -0.6 | 0.065 | 0.042 | 0.103 | 0.103 | 0 | 0.54 | 0.69 | 0.63 | 0.89 | 0.35 | 0.74 |
| 2 | 18339985 | rs896608 | -1.535 | -1.031 | 0.606 | 1.039 | -0.023 | 0.007 | 0.007 | 1.011 | 0.25 | 0.62 | 0.3 | 0.31 | 0.16 | 0.3 |
| 2 | 18340920 | rs2345505 | -0.827 | -0.711 | -0.374 | 0.404 | -0.034 | -0.041 | -0.041 | 0.128 | 0.18 | 0.26 | 0.11 | 0.12 | 0.1 | 0.16 |
| 2 | 18351537 | rs2011027 | -0.58 | -0.561 | -0.763 | 0.853 | 0.015 | -0.007 | -0.007 | 0 | 0.36 | 0.34 | 0.44 | 0.61 | 0.25 | 0.54 |
| 2 | 18355149 | rs11690190 | 1.544 | -0.939 | 0.771 | 2.161 | -0.013 | 0.21 | 0.21 | 1.294 | 0.86 | 0.4 | 0.82 | 0.65 | 0.92 | 0.72 |
| 2 | 18356051 | rs1377081 | -1.408 | -0.878 | 1.437 | 2.09 | 0.047 | 0.118 | 0.118 | 0.401 | 0.06 | 0.26 | 0.08 | 0.11 | 0.03 | 0.15 |
| 2 | 18360228 | rs1993163 | 1.684 | -1.087 | 1.316 | 2.35 | -0.024 | 0.088 | 0.088 | 0.015 | 0.82 | 0.59 | 0.77 | 0.33 | 0.86 | 0.71 |
| 2 | 18365934 | rs10495681 | -1.127 | -1.08 | 1.587 | 2.488 | -0.001 | 0.179 | 0.179 | 0 | 0.06 | 0.24 | 0.1 | 0.21 | 0.04 | 0.15 |
| 2 | 18371951 | rs7570727 | 1.689 | -1.104 | 1.644 | 2.217 | -0.022 | 0.067 | 0.067 | 0.058 | 0.82 | 0.61 | 0.78 | 0.31 | 0.86 | 0.7 |
| 2 | 18375769 | rs2123216 | 2.372 | -0.956 | 1.548 | 2.446 | 0.003 | 0.063 | 0.063 | 0 | 0.75 | 0.59 | 0.77 | 0.47 | 0.8 | 0.65 |
| 2 | 18377604 | rs6726366 | -1.606 | -0.957 | 1.419 | 2.516 | -0.009 | 0.139 | 0.139 | 0 | 0.13 | 0.22 | 0.15 | 0.42 | 0.11 | 0.16 |
| 2 | 18390215 | rs2345518 | 1.363 | -0.882 | 1.474 | 1.869 | 0.017 | 0.052 | 0.052 | 0 | 0.94 | 0.88 | 0.93 | 0.89 | 0.96 | 0.84 |
| 2 | 18391296 | rs2001954 | 1.445 | -1.031 | 1.831 | 1.791 | 0.021 | 0.016 | 0.016 | 0.166 | 0.85 | 0.46 | 0.88 | 0.89 | 0.9 | 0.81 |
| 2 | 18399929 | rs7563158 | 1.361 | -1.902 | 1.804 | 1.099 | 0.03 | 0.034 | 0.034 | 2.445 | 0.65 | 0.23 | 0.72 | 0.73 | 0.8 | 0.6 |
| 2 | 18402432 | rs1452425 | 1.358 | -2.08 | 1.826 | 1.044 | -0.023 | 0.001 | 0.001 | 1.156 | 0.75 | 0.42 | 0.78 | 0.72 | 0.84 | 0.64 |
| 2 | 18403518 | rs4832564 | 1.765 | -2.12 | 1.701 | 1.103 | 0.053 | 0.028 | 0.028 | 2.806 | 0.64 | 0.18 | 0.68 | 0.67 | 0.8 | 0.6 |
| 2 | 18404520 | rs13390392 | -0.775 | -1.981 | 1.631 | 1.13 | -0.013 | -0.016 | -0.016 | 0.041 | 0.07 | 0.16 | 0.02 | 0.02 | 0.03 | 0.04 |
| 2 | 18427836 | rs1452414 | 0.916 | -1.899 | 1.984 | 1.192 | 0.025 | 0.23 | 0.23 | 0.266 | 0.94 | 0.68 | 0.85 | 0.75 | 0.98 | 0.78 |
| 2 | 18429449 | rs11096525 | 0.645 | -1.898 | 2.089 | 1.192 | -0.021 | 0.122 | 0.122 | 0 | 0.95 | 0.83 | 0.87 | 0.76 | 0.99 | 0.81 |
| 2 | 18430121 | rs1356318 | 1.87 | -1.933 | 1.999 | 1.165 | -0.018 | 0.12 | 0.12 | 0.653 | 0.9 | 0.66 | 0.84 | 0.75 | 0.97 | 0.74 |
| 2 | 18432631 | rs1452429 | 1.937 | -1.942 | 1.993 | 1.163 | -0.027 | 0.112 | 0.112 | 1.079 | 0.9 | 0.67 | 0.84 | 0.75 | 0.97 | 0.77 |
| 2 | 18437963 | rs16984540 | -0.474 | -2.024 | 1.933 | 1.12 | -0.002 | 0.001 | 0.001 | 0 | 0.11 | 0.12 | 0.18 | 0.24 | 0.06 | 0.2 |
| 2 | 18439946 | rs7422512 | -2.28 | -3.093 | 1.405 | 0.374 | 0.052 | 0.141 | 0.141 | 0.062 | 0.55 | 0.77 | 0.58 | 0.91 | 0.31 | 0.75 |
| 2 | 18441064 | rs13420705 | -0.147 | -3.129 | 1.757 | 0.41 | -0.001 | -0.003 | -0.003 | 0 | 0.07 | 0.26 | 0.04 | 0.02 | 0.03 | 0.04 |
| 2 | 18445474 | rs16984571 | -0.682 | -3.173 | 1.624 | 0.439 | -0.005 | 0.006 | 0.006 | 0 | 0.16 | 0.19 | 0.21 | 0.25 | 0.11 | 0.25 |
| 2 | 18445728 | rs12476695 | -0.175 | -3.107 | 1.731 | 0.879 | 0.015 | -0.048 | -0.048 | 0.001 | 0.28 | 0.15 | 0.3 | 0.53 | 0.2 | 0.45 |
| 2 | 18447504 | rs6744889 | -0.902 | -2.984 | 1.621 | 0.698 | -0.004 | 0.004 | 0.004 | 0 | 0.17 | 0.22 | 0.22 | 0.25 | 0.13 | 0.22 |
| 2 | 18448477 | rs10195586 | 0.081 | -2.929 | 1.609 | 0.684 | -0.013 | -0.014 | -0.014 | 0 | 0.07 | 0.15 | 0.02 | 0.02 | 0.03 | 0.04 |
| 2 | 18468568 | rs2345450 | -2.364 | -2.888 | 1.582 | 0.565 | 0.007 | 0.029 | 0.029 | 0 | 0.55 | 0.63 | 0.45 | 0.66 | 0.34 | 0.55 |

| | | | | | | | | | | | | | | | | |
|---|---|---|---|---|---|---|---|---|---|---|---|---|---|---|---|---|
| 2 | 18469418 | rs11096527 | -0.981 | -2.893 | 1.497 | 0.579 | 0.014 | 0.002 | 0.002 | 0 | 0.38 | 0.29 | 0.36 | 0.59 | 0.24 | 0.49 |
| 2 | 18486802 | rs10205974 | -1.772 | -2.84 | 1.527 | 0.582 | -0.001 | -0.016 | -0.016 | 0 | 0.12 | 0.17 | 0.17 | 0.03 | 0.11 | 0.07 |
| 2 | 18489316 | rs1516903 | -2.463 | -2.999 | 1.533 | 0.485 | 0 | 0.027 | 0.027 | 0 | 0.57 | 0.65 | 0.5 | 0.86 | 0.37 | 0.79 |
| 2 | 18489476 | rs1516902 | -0.412 | -2.906 | 1.331 | 0.636 | 0.009 | -0.005 | -0.005 | 0 | 0.32 | 0.24 | 0.27 | 0.62 | 0.21 | 0.48 |
| 2 | 18496694 | rs12478663 | -2.019 | -2.904 | 0.814 | 0.576 | 0.011 | 0.068 | 0.068 | 0 | 0.53 | 0.63 | 0.52 | 0.82 | 0.38 | 0.75 |
| 2 | 18497448 | rs1516919 | -0.865 | -2.857 | 0.954 | 0.045 | -0.006 | -0.014 | -0.014 | 0.095 | 0.37 | 0.38 | 0.43 | 0.81 | 0.27 | 0.73 |
| 2 | 18497843 | rs17389100 | -0.043 | -2.473 | 0.942 | 0.147 | 0.023 | -0.024 | -0.024 | 0 | 0.05 | 0.11 | 0.04 | 0 | 0.07 | 0.01 |
| 2 | 18498682 | rs7608319 | 1.324 | -2.627 | 1.183 | 0.125 | 0.005 | -0.044 | -0.044 | 0 | 0.91 | 0.61 | 0.94 | 0.99 | 0.93 | 0.96 |
| 2 | 18501368 | rs1516916 | -0.456 | -2.557 | 0.735 | 0.234 | 0.009 | -0.049 | -0.049 | 0.293 | 0.28 | 0.16 | 0.32 | 0.55 | 0.22 | 0.44 |
| 2 | 18516059 | rs7588548 | -0.718 | -2.047 | 0.605 | 0.004 | -0.025 | -0.107 | -0.107 | 0.021 | 0.17 | 0.33 | 0.1 | 0.01 | 0.12 | 0.05 |
| 2 | 18519421 | rs17677983 | 0.061 | -0.573 | -0.731 | -0.359 | -0.023 | 0.088 | 0.088 | 0 | 0.27 | 0.05 | 0.2 | 0.11 | 0.27 | 0.24 |
| 2 | 18519734 | rs11689614 | 1.909 | -0.107 | -1.53 | -0.831 | -0.06 | 0.069 | 0.069 | 0 | 0.26 | 0.09 | 0.21 | 0.1 | 0.34 | 0.12 |
| 2 | 18519857 | rs13032026 | 1.495 | -0.311 | -1.068 | -0.618 | -0.015 | 0.038 | 0.038 | 0 | 0.31 | 0.21 | 0.24 | 0.1 | 0.35 | 0.15 |
| 2 | 18520021 | rs16984751 | -0.91 | -0.413 | -0.554 | 0.281 | 0 | -0.002 | -0.002 | 0 | 0.53 | 0.5 | 0.47 | 0.41 | 0.49 | 0.45 |
| 2 | 18521402 | rs2203648 | 1.235 | -0.504 | -0.663 | 0.724 | -0.009 | -0.007 | -0.007 | 0.882 | 0.13 | 0.05 | 0.16 | 0.14 | 0.13 | 0.17 |
| 2 | 18522633 | rs10196021 | 1.915 | -0.433 | -0.56 | 0.638 | -0.011 | 0.029 | 0.029 | 0 | 0.31 | 0.22 | 0.24 | 0.14 | 0.35 | 0.15 |
| 2 | 18522991 | rs17678232 | 0.152 | -0.487 | -0.529 | 0.446 | -0.004 | 0.163 | 0.163 | 0 | 0.39 | 0.11 | 0.25 | 0.11 | 0.36 | 0.27 |
| 2 | 18528678 | rs1038699 | -1.038 | -0.252 | -0.096 | 0.583 | -0.001 | 0 | 0 | 0 | 0.55 | 0.56 | 0.48 | 0.42 | 0.5 | 0.45 |
| 2 | 18531499 | rs1516913 | 1.885 | -0.181 | -0.317 | 0.771 | -0.006 | 0.007 | 0.007 | 0 | 0.31 | 0.24 | 0.24 | 0.19 | 0.34 | 0.15 |
| 2 | 18533874 | rs16984782 | -0.324 | -0.365 | -0.484 | 0.756 | -0.002 | -0.008 | -0.008 | 0.316 | 0.41 | 0.32 | 0.46 | 0.42 | 0.4 | 0.39 |
| 2 | 18535266 | rs17740738 | 0.048 | -0.335 | -0.77 | 0.696 | -0.02 | 0.104 | 0.104 | 0 | 0.32 | 0.12 | 0.28 | 0.11 | 0.32 | 0.28 |
| 2 | 18538201 | rs7575143 | 0.007 | -0.15 | -0.854 | 0.422 | -0.083 | 0.128 | 0.128 | 0 | 0.34 | 0.05 | 0.37 | 0.19 | 0.38 | 0.34 |
| 2 | 18539036 | rs7562205 | -0.486 | -0.083 | -0.748 | 0.492 | -0.015 | 0.005 | 0.005 | 0 | 0.4 | 0.24 | 0.42 | 0.34 | 0.4 | 0.33 |
| 2 | 18539232 | rs12619043 | -0.681 | -0.057 | -0.59 | 0.606 | 0.024 | 0.013 | 0.013 | 0 | 0.51 | 0.24 | 0.56 | 0.47 | 0.52 | 0.47 |
| 2 | 18544218 | rs1356315 | 0.853 | 0.03 | -0.033 | 0.964 | -0.001 | 0.021 | 0.021 | 0 | 0.11 | 0.09 | 0.16 | 0.29 | 0.12 | 0.13 |
| 2 | 18548744 | rs6726747 | -0.549 | -0.07 | 0.832 | 0.778 | 0.024 | 0.212 | 0.212 | 0.115 | 0.18 | 0.47 | 0.22 | 0.43 | 0.15 | 0.15 |
| 2 | 18550057 | rs1400452 | -0.533 | -0.201 | 1.033 | 0.632 | 0.003 | 0.173 | 0.173 | 0 | 0.2 | 0.41 | 0.26 | 0.57 | 0.16 | 0.41 |
| 2 | 18551410 | rs12622302 | 0.352 | -0.169 | 0.78 | 0.718 | 0.005 | 0.173 | 0.173 | 0.464 | 0.13 | 0.28 | 0.17 | 0.34 | 0.1 | 0.14 |
| 2 | 18554766 | rs4832432 | 0.992 | 0.064 | 0.045 | 0.269 | -0.039 | 0.032 | 0.032 | 0 | 0.12 | 0.02 | 0.15 | 0.07 | 0.15 | 0.16 |
| 2 | 18557563 | rs12611619 | 0.04 | -0.065 | 0.109 | -0.005 | 0.003 | -0.028 | -0.028 | 0 | 0.6 | 0.44 | 0.65 | 0.68 | 0.56 | 0.52 |
| 2 | 18559177 | rs2037221 | 1.375 | -0.152 | -0.395 | -0.153 | 0.006 | 0.024 | 0.024 | 0 | 0.32 | 0.16 | 0.29 | 0.22 | 0.3 | 0.28 |
| 2 | 18563358 | rs11674256 | -0.187 | -0.195 | -0.003 | -0.44 | -0.004 | -0.091 | -0.091 | 0.395 | 0.63 | 0.49 | 0.72 | 0.94 | 0.64 | 0.81 |
| 2 | 18563567 | rs6760471 | 1.106 | -0.067 | -0.106 | -0.219 | -0.008 | -0.012 | -0.012 | 0.009 | 0.13 | 0.06 | 0.16 | 0.17 | 0.14 | 0.19 |
| 2 | 18565167 | rs2345834 | -0.44 | -0.317 | 0.688 | -0.232 | 0.002 | -0.083 | -0.083 | 1.556 | 0.68 | 0.6 | 0.73 | 0.91 | 0.66 | 0.88 |
| 2 | 18566435 | rs4832574 | 0.412 | -0.354 | 0.734 | -0.194 | 0.003 | 0.009 | 0.009 | 0 | 0.52 | 0.45 | 0.55 | 0.38 | 0.55 | 0.54 |
| 2 | 18574952 | rs2345835 | 0.541 | -0.389 | 0.882 | -0.123 | -0.001 | 0.019 | 0.019 | 0.038 | 0.53 | 0.38 | 0.6 | 0.56 | 0.53 | 0.48 |
| 2 | 18582725 | rs969030 | -1.339 | -0.36 | 0.664 | -0.235 | -0.005 | 0.032 | 0.032 | 0.371 | 0.75 | 0.86 | 0.85 | 0.97 | 0.73 | 0.9 |
| 2 | 18583761 | rs4832581 | 1.432 | -0.364 | 0.464 | -0.149 | -0.028 | 0.004 | 0.004 | 0 | 0.35 | 0.1 | 0.37 | 0.27 | 0.38 | 0.31 |
| 2 | 18585657 | rs17690910 | 1.855 | -0.27 | 0.649 | 0.105 | -0.001 | 0 | 0 | 0.001 | 0.08 | 0.05 | 0.14 | 0.07 | 0.11 | 0.16 |
| 2 | 18586015 | rs776 | 0.057 | -0.385 | 0.99 | 0.202 | 0.017 | 0.031 | 0.031 | 0.306 | 0.21 | 0.47 | 0.23 | 0.31 | 0.2 | 0.42 |
| 2 | 18591751 | rs10167380 | 1.088 | -0.378 | 0.982 | 0.176 | 0.011 | -0.034 | -0.034 | 1.301 | 0.14 | 0.23 | 0.18 | 0.07 | 0.15 | 0.17 |
| 2 | 18609404 | rs13026592 | 1.009 | -0.454 | 0.763 | 0.093 | 0.005 | 0.004 | 0.004 | 0 | 0.42 | 0.32 | 0.44 | 0.37 | 0.44 | 0.32 |
| 2 | 18610429 | rs1489672 | -0.06 | -0.655 | 0.216 | -0.023 | 0.025 | 0.035 | 0.035 | 0 | 0.66 | 0.49 | 0.62 | 0.45 | 0.59 | 0.49 |
| 2 | 18610473 | rs1489671 | 1.674 | -0.109 | 0.112 | 0.223 | -0.047 | 0.014 | 0.014 | 0.421 | 0.33 | 0.05 | 0.29 | 0.21 | 0.35 | 0.27 |
| 2 | 18612284 | rs4832589 | -0.376 | 0.161 | 0.238 | 0.178 | -0.008 | 0.001 | 0.001 | 0.004 | 0.64 | 0.8 | 0.68 | 0.68 | 0.64 | 0.74 |
| 2 | 18616479 | rs6531100 | 0.832 | 0.481 | 0.102 | 0.072 | -0.038 | 0.039 | 0.039 | 0 | 0.13 | 0.04 | 0.17 | 0.07 | 0.15 | 0.17 |
| 2 | 18619517 | rs6757023 | -0.061 | 0.202 | 0.309 | 0.342 | -0.001 | 0.017 | 0.017 | 0.002 | 0.19 | 0.12 | 0.11 | 0.02 | 0.19 | 0.07 |
| 2 | 18637640 | rs2704631 | 0.122 | 0.248 | 0.411 | 1.133 | -0.001 | 0.003 | 0.003 | 0.326 | 0.92 | 0.94 | 0.89 | 0.94 | 0.91 | 0.96 |
| 2 | 18638057 | rs3912722 | -0.787 | 0.024 | 0.421 | 1.144 | -0.027 | 0.023 | 0.023 | 0 | 0.08 | 0.28 | 0.05 | 0.05 | 0.01 | 0.15 |
| 2 | 18646773 | rs10168276 | -1.234 | 0.348 | 0.207 | 1.14 | -0.002 | 0.073 | 0.073 | 0.41 | 0.13 | 0.25 | 0.16 | 0.28 | 0.12 | 0.28 |
| 2 | 18647868 | rs4832437 | -0.458 | 0.468 | -0.343 | 0.856 | -0.006 | 0.013 | 0.013 | 0 | 0.32 | 0.43 | 0.33 | 0.41 | 0.33 | 0.45 |
| 2 | 18648350 | rs11687269 | 0.138 | 0.745 | -0.751 | 0.314 | 0.025 | 0.094 | 0.094 | 0 | 0.52 | 0.33 | 0.51 | 0.18 | 0.57 | 0.51 |
| 2 | 18648687 | rs7589253 | -0.828 | 1.36 | -0.211 | 0.897 | 0.017 | 0.036 | 0.036 | 0 | 0.19 | 0.27 | 0.32 | 0.41 | 0.24 | 0.4 |
| 2 | 18652507 | rs11096539 | -0.698 | 1.212 | -0.058 | 0.336 | 0.002 | -0.025 | -0.025 | 0 | 0.06 | 0.1 | 0.04 | 0 | 0.04 | 0.08 |
| 2 | 18652804 | rs12473747 | 1.741 | 0.988 | 0.556 | -0.074 | -0.008 | -0.076 | -0.076 | 0 | 0.81 | 0.31 | 0.8 | 0.97 | 0.86 | 0.81 |
| 2 | 18653880 | rs2704625 | -0.082 | 0.983 | 0.521 | -0.042 | -0.01 | 0.024 | 0.024 | 0 | 0.92 | 0.97 | 0.91 | 0.97 | 0.91 | 0.88 |
| 2 | 18666310 | rs10174485 | -0.759 | 0.934 | 1.254 | 0.001 | 0.001 | -0.038 | -0.038 | 0 | 0.07 | 0.16 | 0.03 | 0 | 0.04 | 0.08 |
| 2 | 18668035 | rs6531105 | -1.777 | 0.911 | 1.51 | -0.01 | 0.013 | -0.058 | -0.058 | 0 | 0.11 | 0.57 | 0.09 | 0 | 0.04 | 0.08 |
| 2 | 18668523 | rs16985140 | -0.892 | 0.926 | 1.495 | -0.003 | 0.003 | -0.045 | -0.045 | 1.712 | 0.08 | 0.2 | 0.04 | 0 | 0.04 | 0.08 |
| 2 | 18673234 | rs2704619 | 0.284 | 1.269 | -0.821 | 0.209 | -0.019 | 0.005 | 0.005 | 0 | 0.58 | 0.8 | 0.63 | 0.63 | 0.56 | 0.46 |
| 2 | 18680065 | rs2345722 | 0.078 | 1.327 | -1.456 | -0.173 | -0.063 | 0.095 | 0.095 | 0 | 0.72 | 0.94 | 0.77 | 0.88 | 0.67 | 0.66 |
| 2 | 18681809 | rs6745120 | 0.022 | 1.957 | -1.606 | -0.056 | -0.048 | -0.055 | -0.055 | 0 | 0.11 | 0.03 | 0.36 | 0.67 | 0.19 | 0.25 |
| 2 | 18687534 | rs4119150 | -0.833 | 1.609 | -1.409 | -0.569 | -0.023 | 0.029 | 0.029 | 0 | 0.37 | 0.2 | 0.33 | 0.24 | 0.39 | 0.45 |
| 2 | 18693427 | rs7606752 | -0.022 | 1.409 | -1.454 | -0.849 | -0.048 | 0.079 | 0.079 | 0 | 0.24 | 0.06 | 0.19 | 0.09 | 0.28 | 0.27 |
| 2 | 18694885 | rs9808434 | -0.72 | 1.565 | -0.64 | -1.227 | 0.002 | 0 | 0 | 0 | 0.28 | 0.34 | 0.54 | 0.73 | 0.34 | 0.45 |
| 2 | 18695083 | rs9808120 | -1.182 | 1.518 | -0.809 | -1.492 | 0.002 | 0.011 | 0.011 | 0.172 | 0.36 | 0.31 | 0.51 | 0.82 | 0.39 | 0.67 |
| 2 | 18695114 | rs11096540 | -1.327 | 1.528 | -0.863 | -1.512 | 0.002 | 0.008 | 0.008 | 0.04 | 0.38 | 0.34 | 0.51 | 0.83 | 0.39 | 0.67 |
| 2 | 18695331 | rs9677265 | -0.518 | 1.602 | -0.946 | -1.533 | -0.058 | 0.047 | 0.047 | 0.021 | 0.29 | 0.08 | 0.5 | 0.84 | 0.36 | 0.64 |
| 2 | 18697952 | rs16985176 | -0.726 | 1.425 | 0.064 | -1.237 | 0.006 | -0.01 | -0.01 | 0.47 | 0.15 | 0.22 | 0.18 | 0.1 | 0.15 | 0.09 |

| | | | | | | | | | | | | | | | | |
|---|---|---|---|---|---|---|---|---|---|---|---|---|---|---|---|---|
| 2 | 18702313 | rs13430465 | 0.441 | 1.385 | -0.004 | -0.967 | 0.001 | 0.084 | 0.084 | 0 | 0.07 | 0.09 | 0.25 | 0.66 | 0.07 | 0.08 |
| 2 | 18704988 | rs13403095 | 0.312 | 1.105 | 0.218 | -1.581 | -0.01 | -0.084 | -0.084 | 0.07 | 0.21 | 0.3 | 0.11 | 0 | 0.15 | 0.1 |
| 2 | 18705277 | rs7565008 | 0.259 | 1.137 | 0.094 | -1.636 | -0.004 | -0.001 | -0.001 | 0.765 | 0.09 | 0.11 | 0.11 | 0.11 | 0.08 | 0.08 |
| 2 | 18708240 | rs11679389 | 0.007 | 1.363 | 0.028 | -1.96 | -0.06 | 0.043 | 0.043 | 0 | 0.31 | 0.15 | 0.36 | 0.22 | 0.44 | 0.61 |
| 2 | 18709585 | rs4832440 | 0.398 | 1.544 | 0.125 | -1.989 | -0.023 | 0.022 | 0.022 | 0 | 0.26 | 0.14 | 0.22 | 0.2 | 0.28 | 0.57 |
| 2 | 18709614 | rs4832599 | -0.25 | 1.52 | 0.149 | -2.001 | -0.001 | 0.002 | 0.002 | 0 | 0.13 | 0.14 | 0.16 | 0.15 | 0.11 | 0.14 |
| 2 | 18710497 | rs13031613 | -0.529 | 1.485 | 0.598 | -2.018 | 0.007 | -0.018 | -0.018 | 0 | 0.36 | 0.43 | 0.35 | 0.31 | 0.36 | 0.65 |
| 2 | 18712365 | rs17694878 | -0.051 | 1.484 | 0.274 | -1.901 | 0.022 | -0.03 | -0.03 | 0 | 0.06 | 0.09 | 0.14 | 0.01 | 0.15 | 0.04 |
| 2 | 18713455 | rs4832602 | -1.482 | 1.373 | -0.007 | -2.064 | -0.001 | 0.03 | 0.03 | 0 | 0.64 | 0.92 | 0.62 | 0.3 | 0.67 | 0.78 |
| 2 | 18714547 | rs11694160 | 0.38 | 1.406 | 0.008 | -2.047 | -0.042 | 0.028 | 0.028 | 0 | 0.25 | 0.1 | 0.22 | 0.19 | 0.28 | 0.56 |
| 2 | 18716201 | rs10191044 | -0.463 | 1.351 | 0.258 | -2.085 | 0.013 | 0.084 | 0.084 | 0 | 0.13 | 0.32 | 0.3 | 0.72 | 0.09 | 0.16 |
| 2 | 18717487 | rs11687867 | 1.1 | 1.309 | 0.078 | -2.047 | -0.035 | 0.034 | 0.034 | 0 | 0.2 | 0.08 | 0.17 | 0.14 | 0.25 | 0.48 |
| 2 | 18719405 | rs4528759 | -0.127 | 1.236 | 0.176 | -2.089 | -0.009 | -0.102 | -0.102 | 0.529 | 0.36 | 0.47 | 0.36 | 0.11 | 0.36 | 0.22 |
| 2 | 18722637 | rs4832606 | 0.159 | 1.14 | 0.03 | -1.692 | 0.044 | 0.05 | 0.05 | 0 | 0.06 | 0.26 | 0.05 | 0.05 | 0.03 | 0.06 |
| 2 | 18726434 | rs1987115 | -1 | 1.167 | -0.034 | -1.725 | -0.027 | 0.044 | 0.044 | 0 | 0.62 | 0.9 | 0.6 | 0.28 | 0.63 | 0.75 |
| 2 | 18727332 | rs10210151 | -0.342 | 1.056 | 0.675 | -0.941 | 0.035 | 0.046 | 0.046 | 0 | 0.06 | 0.26 | 0.05 | 0.05 | 0.03 | 0.06 |
| 2 | 18727980 | rs4035898 | 0.1 | 1.149 | -0.259 | -0.834 | 0.007 | 0.061 | 0.061 | 0 | 0.08 | 0.11 | 0.26 | 0.69 | 0.09 | 0.11 |
| 2 | 18734072 | rs16985278 | -0.688 | 0.797 | -0.108 | -1.271 | -0.007 | -0.048 | -0.048 | 0.935 | 0.29 | 0.66 | 0.28 | 0.05 | 0.24 | 0.16 |
| 2 | 18734872 | rs13403096 | -0.456 | 0.924 | -0.444 | -1.181 | 0.031 | -0.009 | -0.009 | 0 | 0.17 | 0.3 | 0.2 | 0.12 | 0.16 | 0.12 |
| 2 | 18735164 | rs16985284 | -0.014 | 0.39 | -0.716 | -1.005 | 0.024 | 0.013 | 0.013 | 0.034 | 0.21 | 0.12 | 0.09 | 0 | 0.13 | 0.08 |
| 2 | 18737788 | rs2346135 | -0.305 | 1.004 | -0.544 | -0.669 | 0.011 | -0.039 | -0.039 | 0.154 | 0.33 | 0.39 | 0.36 | 0.15 | 0.37 | 0.59 |
| 2 | 18738545 | rs2346136 | -0.365 | 0.987 | -0.546 | -0.671 | 0.012 | -0.046 | -0.046 | 1.458 | 0.33 | 0.4 | 0.37 | 0.15 | 0.37 | 0.59 |
| 2 | 18742258 | rs7587775 | -0.297 | 0.924 | -0.532 | -0.092 | -0.001 | 0 | 0 | 0 | 0.43 | 0.59 | 0.56 | 0.59 | 0.4 | 0.28 |
| 2 | 18744758 | rs3747660 | -1.385 | 0.556 | 0.596 | -0.049 | 0.098 | 0.166 | 0.166 | 4.068 | 0.06 | 0.46 | 0.17 | 0.08 | 0.05 | 0.02 |
| 2 | 18762337 | rs2346137 | 0.187 | 0.354 | -0.456 | -0.714 | -0.037 | 0.021 | 0.021 | 0 | 0.39 | 0.24 | 0.33 | 0.39 | 0.44 | 0.67 |
| 2 | 18767384 | rs11096544 | 1.082 | 0.321 | -0.547 | -0.814 | -0.122 | -0.037 | -0.037 | 0 | 0.2 | 0.01 | 0.17 | 0.18 | 0.3 | 0.17 |
| 2 | 18768850 | rs16985306 | -0.085 | -0.773 | -0.972 | -1.251 | 0.044 | 0.047 | 0.047 | 0 | 0.36 | 0.66 | 0.32 | 0.12 | 0.2 | 0.18 |
| 2 | 18778577 | rs11889898 | -0.47 | -0.822 | -0.598 | -1.15 | 0.026 | 0.079 | 0.079 | 6.592 | 0.07 | 0.55 | 0.18 | 0.09 | 0.05 | 0.02 |
| 2 | 18787470 | rs2252736 | 0.411 | -0.691 | -1.129 | -1.31 | 0.066 | 0.081 | 0.081 | 0 | 0.52 | 0.31 | 0.41 | 0.39 | 0.61 | 0.73 |
| 2 | 18794610 | rs17701746 | 0.341 | -0.912 | -1.114 | -1.425 | 0.045 | 0.056 | 0.056 | 0.301 | 0.29 | 0.15 | 0.15 | 0.04 | 0.16 | 0.14 |
| 2 | 18795283 | rs2256536 | -0.659 | -0.818 | -0.943 | -1.105 | 0.013 | 0.183 | 0.183 | 0.642 | 0.82 | 0.51 | 0.71 | 0.45 | 0.82 | 0.91 |
| 2 | 18802573 | rs2253665 | 0.608 | -0.824 | -0.962 | -1.116 | 0.054 | 0.085 | 0.085 | 0.859 | 0.52 | 0.33 | 0.42 | 0.34 | 0.61 | 0.75 |
| 2 | 18803872 | rs17752832 | 0.334 | -0.762 | -1.196 | -1.056 | 0.039 | 0.051 | 0.051 | 0.127 | 0.3 | 0.16 | 0.29 | 0.11 | 0.21 | 0.15 |
| 2 | 18811566 | rs16985353 | 1.625 | -0.649 | -1.195 | -0.892 | -0.05 | -0.083 | -0.083 | 1.871 | 0.12 | 0.02 | 0.26 | 0.52 | 0.17 | 0.08 |
| 2 | 18830980 | rs12185564 | 0.742 | -1.297 | -1.305 | -0.827 | 0.041 | 0.025 | 0.025 | 0 | 0.17 | 0.05 | 0.18 | 0.08 | 0.11 | 0.07 |
| 2 | 18832346 | rs12053481 | 1.625 | -1.171 | -1.281 | -0.309 | -0.048 | -0.088 | -0.088 | 0 | 0.13 | 0.02 | 0.27 | 0.47 | 0.17 | 0.08 |
| 2 | 18834651 | rs1445130 | 0.66 | -1.493 | -1.126 | -0.586 | -0.008 | -0.02 | -0.02 | 0 | 0.14 | 0.15 | 0.11 | 0.04 | 0.12 | 0.08 |
| 2 | 18835760 | rs4832444 | -0.436 | -1.56 | -0.751 | -0.603 | -0.025 | -0.014 | -0.014 | 0 | 0.29 | 0.31 | 0.27 | 0.15 | 0.2 | 0.16 |
| 2 | 18836829 | rs12989389 | -0.491 | -1.561 | -0.733 | -0.491 | -0.012 | -0.006 | -0.006 | 0.016 | 0.29 | 0.28 | 0.28 | 0.15 | 0.22 | 0.16 |
| 2 | 18839194 | rs13016448 | 0.669 | -1.58 | -0.784 | -0.58 | -0.008 | -0.014 | -0.014 | 0 | 0.14 | 0.15 | 0.11 | 0.04 | 0.12 | 0.08 |
| 2 | 18843057 | rs1978995 | 0.184 | -1.621 | -0.65 | -0.599 | -0.008 | -0.004 | -0.004 | 0 | 0.71 | 0.71 | 0.72 | 0.88 | 0.8 | 0.85 |
| 2 | 18847154 | rs4605374 | 1.421 | -1.596 | -0.8 | 0.068 | -0.055 | -0.095 | -0.095 | 0 | 0.14 | 0.02 | 0.29 | 0.37 | 0.19 | 0.08 |
| 2 | 18851257 | rs2881318 | 1.507 | -1.516 | -0.786 | -0.293 | -0.057 | -0.03 | -0.03 | 0.116 | 0.41 | 0.21 | 0.34 | 0.51 | 0.5 | 0.7 |
| 2 | 18851635 | rs6734865 | 0.837 | -1.413 | -0.654 | -0.394 | -0.007 | -0.01 | -0.01 | 1.073 | 0.14 | 0.13 | 0.1 | 0.04 | 0.09 | 0.07 |
| 2 | 18866226 | rs16985416 | 0.437 | -1.222 | -0.599 | -0.294 | 0.007 | 0.005 | 0.005 | 0.38 | 0.57 | 0.6 | 0.43 | 0.4 | 0.6 | 0.73 |
| 2 | 18868188 | rs4614937 | 0.183 | -1.167 | -0.434 | -0.18 | -0.008 | 0.003 | 0.003 | 2.24 | 0.66 | 0.38 | 0.52 | 0.45 | 0.71 | 0.8 |
| 2 | 18868661 | rs11096553 | -0.329 | -0.786 | -0.176 | 0.071 | 0.032 | 0.084 | 0.084 | 0 | 0.28 | 0.41 | 0.42 | 0.51 | 0.31 | 0.19 |
| 2 | 18882100 | rs6745716 | -0.797 | -1.011 | -0.053 | -0.188 | 0.009 | 0.002 | 0.002 | 0 | 0.41 | 0.46 | 0.58 | 0.53 | 0.48 | 0.24 |
| 2 | 18884786 | rs16985453 | -0.649 | -1.011 | -0.101 | -0.188 | -0.005 | -0.002 | -0.002 | 0.707 | 0.41 | 0.36 | 0.57 | 0.54 | 0.48 | 0.24 |
| 2 | 18897492 | rs7605697 | -0.766 | -1.661 | -0.58 | -0.706 | 0.075 | 0.071 | 0.071 | 1.656 | 0.71 | 0.85 | 0.88 | 0.87 | 0.85 | 0.56 |
| 2 | 18901864 | rs6723603 | -1.073 | -1.309 | -0.276 | -0.422 | 0.034 | 0.028 | 0.028 | 1.704 | 0.78 | 0.85 | 0.89 | 0.86 | 0.87 | 0.6 |
| 2 | 18903448 | rs16985487 | -0.039 | -1.239 | 0.07 | -0.491 | 0.002 | -0.045 | -0.045 | 0.55 | 0.25 | 0.38 | 0.28 | 0.09 | 0.26 | 0.13 |
| 2 | 18907874 | rs2346563 | 0.466 | -1.187 | -0.062 | -0.445 | -0.001 | -0.015 | -0.015 | 0.057 | 0.1 | 0.13 | 0.08 | 0.01 | 0.11 | 0.05 |
| 2 | 18914751 | rs4832616 | -1.124 | -1.241 | -0.159 | -0.446 | 0.026 | 0.023 | 0.023 | 1.579 | 0.78 | 0.84 | 0.89 | 0.86 | 0.87 | 0.6 |
| 2 | 18921391 | rs10495693 | 1.1 | -1.124 | -0.217 | -0.433 | 0.004 | 0.019 | 0.019 | 0 | 0.07 | 0.03 | 0.05 | 0.01 | 0.06 | 0.04 |
| 2 | 18924471 | rs10173922 | 0.214 | -1.068 | 0.028 | -0.271 | -0.003 | 0.003 | 0.003 | 2.09 | 0.58 | 0.45 | 0.54 | 0.45 | 0.58 | 0.47 |
| 2 | 18925494 | rs7558736 | 1.206 | -1.045 | 0.019 | -0.31 | 0.01 | 0.015 | 0.015 | 0.675 | 0.51 | 0.42 | 0.48 | 0.37 | 0.5 | 0.45 |
| 2 | 18943278 | rs4129693 | 0.227 | -0.825 | 0.238 | -0.321 | -0.001 | -0.035 | -0.035 | 0 | 0.21 | 0.22 | 0.23 | 0.01 | 0.21 | 0.12 |
| 2 | 18947315 | rs11885014 | -0.443 | -0.863 | 0.46 | -0.331 | -0.008 | -0.053 | -0.053 | 0 | 0.26 | 0.58 | 0.28 | 0.07 | 0.24 | 0.12 |
| 2 | 18948187 | rs7587939 | -0.384 | -0.847 | 0.48 | -0.324 | -0.006 | -0.086 | -0.086 | 0 | 0.25 | 0.46 | 0.27 | 0.07 | 0.25 | 0.12 |
| 2 | 18954914 | rs3890826 | -0.861 | -0.84 | 0.265 | -0.183 | -0.016 | -0.022 | -0.022 | 0 | 0.37 | 0.26 | 0.54 | 0.49 | 0.43 | 0.24 |
| 2 | 18958882 | rs6738546 | 1.285 | -0.802 | 0.284 | -0.198 | -0.002 | -0.012 | -0.012 | 0.19 | 0.13 | 0.1 | 0.18 | 0 | 0.14 | 0.07 |
| 2 | 18971405 | rs4371357 | 0.099 | -0.805 | 0.36 | -0.216 | -0.001 | -0.08 | -0.08 | 0.373 | 0.22 | 0.31 | 0.25 | 0.01 | 0.22 | 0.11 |
| 2 | 18973451 | rs11892969 | 1.716 | -0.871 | 0.112 | -0.174 | -0.023 | -0.016 | -0.016 | 0.073 | 0.38 | 0.2 | 0.3 | 0.36 | 0.36 | 0.33 |
| 2 | 18974696 | rs6721356 | -1.106 | -0.926 | 0.109 | -0.18 | 0.024 | 0.017 | 0.017 | 0.725 | 0.78 | 0.86 | 0.88 | 0.85 | 0.84 | 0.6 |
| 2 | 18978316 | rs4416215 | -0.687 | -0.915 | 0.174 | -0.159 | 0.016 | 0.027 | 0.027 | 0 | 0.37 | 0.48 | 0.55 | 0.5 | 0.43 | 0.25 |
| 2 | 18984638 | rs6740293 | 1.759 | -0.903 | 0.117 | -0.169 | -0.028 | -0.017 | -0.017 | 0 | 0.38 | 0.2 | 0.3 | 0.35 | 0.36 | 0.37 |
| 2 | 18993027 | rs6531119 | 1.639 | -0.91 | -0.086 | 0.065 | -0.038 | -0.054 | -0.054 | 2.164 | 0.13 | 0.06 | 0.28 | 0.43 | 0.21 | 0.13 |
| 2 | 18995463 | rs13011363 | 0.501 | -1.01 | 0.059 | -0.206 | 0.004 | -0.023 | -0.023 | 0 | 0.17 | 0.23 | 0.21 | 0.06 | 0.18 | 0.08 |

| | | | | | | | | | | | | | | | | |
|---|---|---|---|---|---|---|---|---|---|---|---|---|---|---|---|---|
| 2 | 18996307 | rs11886548 | -0.075 | -0.992 | -0.074 | -0.177 | -0.009 | -0.065 | -0.065 | 0.145 | 0.23 | 0.42 | 0.27 | 0.07 | 0.22 | 0.11 |
| 2 | 18997162 | rs4608518 | -0.53 | -0.957 | -0.368 | 0.07 | 0.024 | 0.019 | 0.019 | 0.709 | 0.78 | 0.86 | 0.89 | 0.87 | 0.84 | 0.63 |
| 2 | 19000674 | rs4240223 | 0.628 | -0.125 | -0.467 | 0.301 | 0.003 | 0 | 0 | 0 | 0.37 | 0.4 | 0.38 | 0.36 | 0.44 | 0.23 |
| 2 | 19006214 | rs13000922 | 0.456 | 0.136 | -0.881 | 0.682 | -0.005 | 0.011 | 0.011 | 0 | 0.15 | 0.15 | 0.32 | 0.53 | 0.23 | 0.19 |
| 2 | 19012808 | rs4240224 | -0.021 | -1.304 | -0.576 | -0.365 | -0.053 | -0.085 | -0.085 | 0 | 0.68 | 0.54 | 0.85 | 0.86 | 0.81 | 0.5 |
| 2 | 19013114 | rs4240225 | -0.354 | -1.218 | -0.71 | -0.271 | 0.009 | 0.056 | 0.056 | 0 | 0.17 | 0.07 | 0.21 | 0.01 | 0.15 | 0.04 |
| 2 | 19026974 | rs6531122 | -1.341 | -1.755 | 0.228 | 0.527 | -0.051 | 0.084 | 0.084 | 2.137 | 0.2 | 0.55 | 0.19 | 0.37 | 0.11 | 0.38 |
| 2 | 19029816 | rs6758285 | -1.563 | -2.333 | 0.542 | 0.348 | -0.039 | 0.013 | 0.013 | 0.002 | 0.35 | 0.42 | 0.28 | 0.6 | 0.17 | 0.61 |
| 2 | 19033928 | rs6720271 | -1.867 | -2.243 | 0.458 | 0.348 | 0.024 | -0.005 | -0.005 | 0 | 0.38 | 0.64 | 0.31 | 0.6 | 0.2 | 0.62 |
| 2 | 19051404 | rs4398278 | -1.763 | -2.284 | 0.339 | 0.207 | 0.037 | 0 | 0 | 0.446 | 0.38 | 0.66 | 0.32 | 0.63 | 0.19 | 0.62 |
| 2 | 19066057 | rs7562720 | -2.009 | -2.687 | -0.05 | -0.277 | 0.047 | 0.12 | 0.12 | 0 | 0.56 | 0.73 | 0.48 | 0.99 | 0.32 | 0.82 |
| 2 | 19066741 | rs12621971 | -0.699 | -2.642 | 0.366 | 0.171 | -0.005 | 0.028 | 0.028 | 4.163 | 0.16 | 0.19 | 0.16 | 0.41 | 0.1 | 0.34 |
| 2 | 19072345 | rs13390982 | 0.281 | -2.515 | 0.538 | 0.283 | 0.004 | 0.002 | 0.002 | 0 | 0.14 | 0.12 | 0.12 | 0.37 | 0.08 | 0.17 |
| 2 | 19074654 | rs7570667 | 1.194 | -2.587 | 1.013 | 0.097 | -0.006 | 0.001 | 0.001 | 0 | 0.65 | 0.63 | 0.74 | 0.42 | 0.8 | 0.46 |
| 2 | 19075909 | rs4482482 | 0.017 | -2.599 | 1.063 | 0.129 | -0.015 | -0.008 | -0.008 | 3.735 | 0.78 | 0.74 | 0.85 | 0.83 | 0.91 | 0.77 |
| 2 | 19077800 | rs4263113 | 0.435 | -2.442 | 1.056 | 0.033 | 0.018 | -0.019 | -0.019 | 1.418 | 0.66 | 0.73 | 0.74 | 0.46 | 0.83 | 0.61 |
| 2 | 19082713 | rs6531132 | -1.057 | -2.3 | 1.602 | -0.007 | -0.009 | 0.145 | 0.145 | 0 | 0.15 | 0.32 | 0.11 | 0.39 | 0.12 | 0.31 |
| 2 | 19083453 | rs3900754 | 1.293 | -2.427 | 1.31 | -0.181 | 0.053 | 0.081 | 0.081 | 0 | 0.49 | 0.65 | 0.63 | 0.07 | 0.7 | 0.31 |
| 2 | 19085355 | rs7603714 | 1.32 | -2.419 | 1.285 | -0.204 | 0.037 | 0.028 | 0.028 | 0 | 0.49 | 0.61 | 0.62 | 0.05 | 0.7 | 0.28 |
| 2 | 19087571 | rs11096563 | -1.294 | -2.482 | 1.52 | -0.097 | -0.031 | 0.18 | 0.18 | 0.05 | 0.17 | 0.57 | 0.13 | 0.38 | 0.11 | 0.29 |
| 2 | 19096394 | rs12464512 | 1.682 | -2.491 | 1.37 | -0.148 | 0.081 | 0.264 | 0.264 | 0.174 | 0.45 | 0.12 | 0.59 | 0.04 | 0.71 | 0.28 |
| 2 | 19110370 | rs6531134 | -1.08 | -2.34 | 1.654 | -0.008 | -0.019 | 0.003 | 0.003 | 0 | 0.4 | 0.42 | 0.29 | 0.58 | 0.19 | 0.42 |
| 2 | 19126513 | rs4435408 | 0.787 | -2.345 | 1.2 | -0.024 | 0.006 | -0.003 | -0.003 | 4.162 | 0.59 | 0.39 | 0.67 | 0.42 | 0.8 | 0.57 |
| 2 | 19131180 | rs6710040 | 0.478 | -1.773 | 0.94 | 0.288 | 0.01 | -0.013 | -0.013 | 0 | 0.69 | 0.74 | 0.73 | 0.43 | 0.82 | 0.58 |
| 2 | 19140546 | rs1317019 | 0.487 | -1.458 | 0.97 | 0.48 | -0.019 | 0.031 | 0.031 | 0 | 0.7 | 0.65 | 0.76 | 0.52 | 0.8 | 0.69 |
| 2 | 19142858 | rs16985720 | -0.413 | -1.344 | 0.685 | 0.556 | 0.006 | -0.003 | -0.003 | 0 | 0.1 | 0.06 | 0.07 | 0.38 | 0.08 | 0.28 |
| 2 | 19155925 | rs6531140 | 0.357 | -1.313 | 0.966 | 0.52 | 0.003 | -0.006 | -0.006 | 0 | 0.69 | 0.71 | 0.75 | 0.54 | 0.8 | 0.69 |
| 2 | 19161901 | rs7580276 | 0.375 | -1.326 | 0.85 | 0.435 | 0.007 | -0.017 | -0.017 | 1.178 | 0.7 | 0.75 | 0.75 | 0.54 | 0.8 | 0.7 |
| 2 | 19166928 | rs2347005 | 1.026 | -1.098 | 0.195 | -0.491 | 0.029 | 0.072 | 0.072 | 0.044 | 0.54 | 0.33 | 0.62 | 0.13 | 0.68 | 0.41 |
| 2 | 19168314 | rs12476467 | 1.035 | -1.081 | 0.15 | -0.49 | 0.021 | 0.045 | 0.045 | 0.278 | 0.54 | 0.37 | 0.62 | 0.13 | 0.68 | 0.41 |
| 2 | 19172488 | rs16985798 | 0.98 | -1.068 | 0.099 | -0.473 | 0.014 | 0.024 | 0.024 | 0.111 | 0.55 | 0.39 | 0.62 | 0.13 | 0.68 | 0.41 |
| 2 | 19184369 | rs16985823 | -0.889 | -0.498 | -0.036 | -0.036 | -0.014 | 0.002 | 0.002 | 0 | 0.09 | 0.04 | 0.06 | 0.1 | 0.11 | 0.23 |
| 2 | 19184996 | rs884895 | -1.536 | -0.862 | 0.81 | -0.064 | -0.034 | -0.025 | -0.025 | 0 | 0.2 | 0.55 | 0.16 | 0.17 | 0.15 | 0.32 |
| 2 | 19229771 | rs10776480 | 2.221 | -0.777 | 1.317 | -0.294 | -0.013 | -0.069 | -0.069 | 0 | 0.8 | 0.43 | 0.81 | 0.9 | 0.85 | 0.69 |
| 2 | 19229882 | rs17756190 | -0.994 | -0.751 | 1.256 | -0.282 | -0.001 | 0.009 | 0.009 | 0 | 0.07 | 0.06 | 0.05 | 0.03 | 0.07 | 0.05 |
| 2 | 19232805 | rs17756262 | -1.75 | -0.749 | 1.355 | -0.282 | -0.002 | 0.002 | 0.002 | 0 | 0.1 | 0.12 | 0.09 | 0.08 | 0.12 | 0.22 |
| 2 | 19244727 | rs4666376 | -2.793 | -0.733 | 1.646 | -0.287 | 0 | -0.055 | -0.055 | 0 | 0.18 | 0.41 | 0.23 | 0.1 | 0.15 | 0.31 |
| 2 | 19253232 | rs7559321 | 2.716 | -0.832 | 1.749 | -0.341 | -0.012 | -0.06 | -0.06 | 0 | 0.8 | 0.4 | 0.74 | 0.9 | 0.84 | 0.69 |
| 2 | 19262669 | rs2346567 | -2.008 | -0.776 | 1.342 | -0.277 | -0.005 | -0.033 | -0.033 | 0 | 0.11 | 0.22 | 0.16 | 0.03 | 0.09 | 0.12 |
| 2 | 19266789 | rs11096580 | 0.577 | -0.801 | 0.529 | -0.681 | 0.028 | 0.038 | 0.038 | 0 | 0.48 | 0.4 | 0.33 | 0.71 | 0.35 | 0.44 |
| 2 | 19271689 | rs17757050 | 0.064 | -0.772 | 0.481 | -0.672 | -0.002 | 0.003 | 0.003 | 0 | 0.05 | 0.05 | 0.04 | 0.02 | 0.06 | 0.04 |
| 2 | 19272190 | rs1574072 | -1.997 | -0.828 | 0.752 | -0.734 | -0.013 | -0.053 | -0.053 | 0 | 0.14 | 0.28 | 0.19 | 0.03 | 0.11 | 0.13 |
| 2 | 19273243 | rs7564073 | -2.122 | -0.607 | 0.722 | -0.816 | -0.01 | 0.007 | 0.007 | 0 | 0.19 | 0.31 | 0.31 | 0.21 | 0.17 | 0.2 |
| 2 | 19277788 | rs6715010 | 0.002 | -0.562 | 0.601 | -0.927 | -0.035 | 0.06 | 0.06 | 0 | 0.37 | 0.23 | 0.45 | 0.04 | 0.51 | 0.23 |
| 2 | 19279238 | rs10208636 | 1.071 | -0.613 | 0.702 | -0.928 | 0.05 | 0.075 | 0.075 | 0 | 0.46 | 0.29 | 0.32 | 0.71 | 0.35 | 0.44 |
| 2 | 19284343 | rs11895316 | 0.663 | -0.562 | 0.819 | -0.915 | 0 | -0.004 | -0.004 | 0.057 | 0.52 | 0.51 | 0.36 | 0.71 | 0.37 | 0.46 |
| 2 | 19286097 | rs11694107 | 0.586 | -0.515 | 0.811 | -0.907 | -0.039 | 0.043 | 0.043 | 0 | 0.31 | 0.17 | 0.41 | 0.02 | 0.44 | 0.19 |
| 2 | 19286391 | rs10207804 | 0.609 | -0.526 | 0.876 | -0.909 | 0.007 | 0.024 | 0.024 | 0 | 0.53 | 0.43 | 0.37 | 0.73 | 0.42 | 0.48 |
| 2 | 19286943 | rs4666451 | 0.493 | -0.493 | 0.835 | -0.905 | -0.025 | 0.013 | 0.013 | 0 | 0.31 | 0.2 | 0.41 | 0.02 | 0.44 | 0.19 |
| 2 | 19287515 | rs4422126 | 0.346 | -0.476 | 0.638 | -0.821 | 0.012 | 0.062 | 0.062 | 0 | 0.89 | 0.74 | 0.8 | 0.74 | 0.88 | 0.68 |
| 2 | 19295604 | rs4666460 | -0.26 | -0.46 | 0.67 | -0.922 | 0.02 | -0.004 | -0.004 | 0 | 0.93 | 0.76 | 0.93 | 0.94 | 0.94 | 0.75 |
| 2 | 19298727 | rs4574071 | 0.216 | -0.391 | 0.309 | -0.882 | -0.005 | -0.038 | -0.038 | 0.036 | 0.32 | 0.26 | 0.42 | 0.02 | 0.44 | 0.19 |
| 2 | 19301126 | rs4480951 | -0.486 | -0.508 | 0.05 | -0.639 | -0.001 | 0.021 | 0.021 | 0 | 0.43 | 0.36 | 0.59 | 0.23 | 0.57 | 0.33 |
| 2 | 19301822 | rs4233760 | -0.559 | -0.49 | -0.041 | -0.26 | 0.046 | 0.025 | 0.025 | 0 | 0.51 | 0.35 | 0.65 | 0.36 | 0.63 | 0.51 |
| 2 | 19302161 | rs6531152 | 0.202 | -1.264 | -0.052 | -0.796 | -0.015 | -0.011 | -0.011 | 0.65 | 0.73 | 0.67 | 0.64 | 0.69 | 0.81 | 0.78 |
| 2 | 19310223 | rs6722781 | 0.107 | -1.186 | -0.209 | -0.712 | 0.08 | 0.115 | 0.115 | 0.058 | 0.48 | 0.14 | 0.32 | 0.32 | 0.32 | 0.55 |
| 2 | 19311718 | rs12710695 | 0.088 | -1.144 | 0.297 | -0.406 | -0.003 | 0.001 | 0.001 | 0 | 0.44 | 0.48 | 0.56 | 0.54 | 0.62 | 0.38 |
| 2 | 19323073 | rs4358083 | 0.345 | -1.21 | 0.398 | -0.534 | -0.001 | -0.002 | -0.002 | 0 | 0.38 | 0.4 | 0.58 | 0.6 | 0.6 | 0.35 |
| 2 | 19328111 | rs4666323 | -0.183 | -1.603 | 0.121 | -0.704 | -0.032 | -0.011 | -0.011 | 0.065 | 0.58 | 0.52 | 0.74 | 0.69 | 0.78 | 0.72 |
| 2 | 19331518 | rs4666464 | -0.139 | -1.624 | 0.331 | -0.714 | -0.008 | -0.001 | -0.001 | 0 | 0.58 | 0.43 | 0.74 | 0.69 | 0.78 | 0.72 |
| 2 | 19336049 | rs16985956 | 1.352 | -1.354 | 0.547 | -0.562 | -0.013 | -0.024 | -0.024 | 0 | 0.05 | 0.1 | 0.02 | 0 | 0.02 | 0.02 |
| 2 | 19341657 | rs12464722 | 1.452 | -1.296 | 0.128 | -0.509 | -0.02 | -0.025 | -0.025 | 0.639 | 0.05 | 0.01 | 0.08 | 0.14 | 0.06 | 0.05 |
| 2 | 19354741 | rs12105891 | -0.122 | -1.21 | 0.265 | -0.662 | 0.002 | -0.014 | -0.014 | 0 | 0.33 | 0.33 | 0.4 | 0.24 | 0.33 | 0.45 |
| 2 | 19358024 | rs2028175 | 0.848 | -1.515 | 0.373 | -0.784 | 0.047 | 0.041 | 0.041 | 0.054 | 0.32 | 0.22 | 0.14 | 0.17 | 0.15 | 0.22 |
| 2 | 19368078 | rs2118381 | 0.028 | -1.469 | 0.426 | -0.926 | 0.003 | 0.004 | 0.004 | 0.154 | 0.54 | 0.44 | 0.73 | 0.7 | 0.77 | 0.72 |
| 2 | 19369318 | rs851339 | -0.403 | -1.013 | 0.569 | -0.709 | -0.019 | -0.036 | -0.036 | 0 | 0.92 | 0.68 | 0.95 | 1 | 0.98 | 0.98 |
| 2 | 19376903 | rs16981939 | 0.463 | -1.065 | 0.33 | -0.892 | -0.003 | -0.004 | -0.004 | 0 | 0.41 | 0.42 | 0.66 | 0.65 | 0.66 | 0.41 |
| 2 | 19378518 | rs851374 | -0.377 | -1.126 | 0.099 | -0.934 | -0.003 | -0.029 | -0.029 | 0 | 0.88 | 0.82 | 0.92 | 0.96 | 0.91 | 0.94 |
| 2 | 19389299 | rs851348 | 1.072 | -1.429 | -0.062 | -1.074 | 0.101 | -0.005 | -0.005 | 0.12 | 0.32 | 0.06 | 0.17 | 0.26 | 0.16 | 0.24 |

| | | | | | | | | | | | | | | | | |
|---|---|---|---|---|---|---|---|---|---|---|---|---|---|---|---|---|
| 2 | 19400895 | rs1527485 | 0.951 | -1.418 | 0.184 | -1.136 | -0.004 | -0.056 | -0.056 | 0.371 | 0.14 | 0.25 | 0.09 | 0.05 | 0.09 | 0.31 |
| 2 | 19405595 | rs851329 | 2.104 | -1.797 | -0.178 | -1.375 | 0.037 | 0.122 | 0.122 | 0.065 | 0.21 | 0.02 | 0.06 | 0 | 0.13 | 0.13 |
| 2 | 19409407 | rs16986058 | 1.164 | -1.714 | -0.086 | -1.317 | -0.002 | -0.06 | -0.06 | 0 | 0.14 | 0.26 | 0.09 | 0.05 | 0.09 | 0.3 |
| 2 | 19420316 | rs851320 | 0.246 | -1.718 | -0.099 | -1.33 | -0.014 | -0.11 | -0.11 | 0 | 0.74 | 0.54 | 0.83 | 0.91 | 0.82 | 0.63 |
| 2 | 19420375 | rs10169702 | 0.818 | -1.645 | -0.094 | -1.256 | 0.002 | 0.003 | 0.003 | 0 | 0.41 | 0.4 | 0.66 | 0.65 | 0.66 | 0.4 |
| 2 | 19427429 | rs851422 | -0.469 | -1.64 | -0.118 | -1.255 | 0.016 | 0.016 | 0.016 | 0 | 0.53 | 0.43 | 0.31 | 0.35 | 0.32 | 0.58 |
| 2 | 19434656 | rs13403560 | 0.749 | -1.266 | -0.668 | -1.138 | 0.006 | -0.01 | -0.01 | 0 | 0.2 | 0.25 | 0.3 | 0.2 | 0.22 | 0.13 |
| 2 | 19435346 | rs13034680 | 2.472 | -1.227 | -0.853 | -1.147 | -0.063 | -0.076 | -0.076 | 0 | 0.2 | 0.09 | 0.34 | 0.44 | 0.43 | 0.27 |
| 2 | 19441388 | rs1561271 | 0.711 | -1.331 | -0.843 | -1.168 | 0 | 0 | 0 | 0 | 0.41 | 0.44 | 0.65 | 0.63 | 0.64 | 0.39 |
| 2 | 19443735 | rs851408 | 1.654 | -0.93 | -1.587 | -0.841 | 0.015 | 0.111 | 0.111 | 0 | 0.21 | 0.04 | 0.07 | 0 | 0.16 | 0.13 |
| 2 | 19447101 | rs851400 | 0.033 | -0.841 | -1.292 | -0.604 | 0.066 | 0.019 | 0.019 | 0 | 0.39 | 0.22 | 0.24 | 0.33 | 0.25 | 0.29 |
| 2 | 19455852 | rs851387 | 0.55 | -0.998 | -1.427 | -0.478 | 0.101 | 0.001 | 0.001 | 0.059 | 0.33 | 0.14 | 0.18 | 0.28 | 0.17 | 0.24 |
| 2 | 19457189 | rs1527493 | 0.616 | 0.245 | -0.72 | -0.109 | -0.009 | 0.034 | 0.034 | 0 | 0.11 | 0.03 | 0.18 | 0.05 | 0.12 | 0.09 |
| 2 | 19465659 | rs1530002 | 0 | -0.075 | -0.136 | -0.426 | 0.008 | 0.045 | 0.045 | 0 | 0.31 | 0.16 | 0.25 | 0.12 | 0.22 | 0.38 |
| 2 | 19473075 | rs851357 | -0.827 | -0.157 | -0.105 | -0.469 | -0.042 | -0.049 | -0.049 | 0.001 | 0.7 | 0.75 | 0.62 | 0.55 | 0.56 | 0.66 |
| 2 | 19474112 | rs851356 | 0.531 | -0.242 | -0.06 | -0.504 | -0.022 | -0.008 | -0.008 | 0 | 0.37 | 0.44 | 0.26 | 0.29 | 0.27 | 0.24 |
| 2 | 19474892 | rs851353 | 0.211 | -0.246 | -0.201 | -0.506 | -0.004 | 0 | 0 | 0.289 | 0.39 | 0.57 | 0.38 | 0.43 | 0.33 | 0.27 |
| 2 | 19475331 | rs851352 | 0.728 | -0.272 | -0.108 | -0.519 | 0.01 | 0.003 | 0.003 | 0.913 | 0.34 | 0.29 | 0.25 | 0.29 | 0.27 | 0.24 |
| 2 | 19475649 | rs851351 | 0.344 | -0.167 | -0.858 | -0.522 | 0.012 | -0.003 | -0.003 | 2.014 | 0.41 | 0.62 | 0.38 | 0.43 | 0.33 | 0.27 |
| 2 | 19476851 | rs851350 | -1.81 | 0.122 | -1.15 | -0.519 | -0.081 | -0.113 | -0.113 | 0.002 | 0.79 | 0.9 | 0.69 | 0.57 | 0.64 | 0.71 |
| 2 | 19481841 | rs2118384 | 0.397 | 0.034 | -1.057 | -0.6 | 0.009 | 0.084 | 0.084 | 0 | 0.31 | 0.12 | 0.25 | 0.16 | 0.23 | 0.38 |
| 2 | 19483101 | rs851436 | 0.364 | 0.002 | -1.027 | -0.569 | 0.016 | 0 | 0 | 0.247 | 0.41 | 0.65 | 0.39 | 0.38 | 0.33 | 0.28 |
| 2 | 19486628 | rs1965786 | -0.333 | 0.592 | -1.066 | -0.143 | -0.06 | -0.061 | -0.061 | 0 | 0.71 | 0.79 | 0.61 | 0.52 | 0.55 | 0.65 |
| 2 | 19487589 | rs10209041 | -0.042 | 0.58 | -0.608 | -0.353 | 0.005 | -0.013 | -0.013 | 0 | 0.07 | 0.1 | 0.05 | 0.02 | 0.08 | 0.05 |
| 2 | 19489081 | rs703323 | 0.732 | 0.085 | -0.487 | -0.837 | -0.011 | -0.068 | -0.068 | 1.471 | 0.18 | 0.22 | 0.08 | 0 | 0.12 | 0.09 |
| 2 | 19490759 | rs6707044 | -0.044 | 0.821 | -0.913 | -0.15 | 0 | 0.036 | 0.036 | 0.01 | 0.13 | 0.09 | 0.34 | 0.48 | 0.2 | 0.35 |
| 2 | 19492579 | rs1727177 | -1.553 | 0.79 | -1.57 | -0.719 | -0.054 | 0.165 | 0.165 | 0 | 0.7 | 0.89 | 0.84 | 0.99 | 0.7 | 0.87 |
| 2 | 19495989 | rs825537 | 0.461 | 0.544 | -1.447 | -0.7 | -0.016 | -0.012 | -0.012 | 0 | 0.33 | 0.36 | 0.17 | 0.25 | 0.25 | 0.31 |
| 2 | 19497218 | rs875974 | 0.955 | 0.514 | -1.32 | -0.717 | -0.002 | -0.002 | -0.002 | 0.077 | 0.21 | 0.16 | 0.3 | 0.26 | 0.25 | 0.21 |
| 2 | 19497993 | rs10514626 | 1.466 | 0.12 | -1.315 | -0.661 | -0.011 | -0.04 | -0.04 | 1.177 | 0.07 | 0.01 | 0.06 | 0.25 | 0.09 | 0.18 |
| 2 | 19500695 | rs703316 | 0.424 | 0.221 | -1.109 | -0.545 | 0.031 | 0.06 | 0.06 | 0 | 0.51 | 0.31 | 0.46 | 0.28 | 0.54 | 0.35 |
| 2 | 19502877 | rs703313 | -0.517 | 0.188 | -1.029 | -0.562 | -0.035 | -0.028 | -0.028 | 0.345 | 0.65 | 0.55 | 0.82 | 0.75 | 0.74 | 0.69 |
| 2 | 19524172 | rs6725667 | 0.532 | 0.405 | -0.869 | -0.475 | 0.012 | 0.142 | 0.142 | 0 | 0.13 | 0.24 | 0.33 | 0.47 | 0.17 | 0.23 |
| 2 | 19531545 | rs703308 | -0.759 | 0.517 | -1.217 | -0.678 | -0.001 | -0.004 | -0.004 | 0 | 0.64 | 0.64 | 0.75 | 0.71 | 0.65 | 0.66 |
| 2 | 19546795 | rs12710706 | -1.285 | 0.465 | -0.527 | 0.298 | -0.002 | -0.016 | -0.016 | 0 | 0.37 | 0.5 | 0.61 | 0.7 | 0.4 | 0.55 |
| 2 | 19549981 | rs13011502 | 0.1 | 0.502 | -0.555 | 0.215 | 0 | 0 | 0 | 0 | 0.18 | 0.14 | 0.25 | 0.2 | 0.19 | 0.18 |
| 2 | 19550625 | rs9677277 | -0.896 | 0.473 | -0.695 | -0.3 | -0.001 | -0.025 | -0.025 | 0 | 0.38 | 0.5 | 0.61 | 0.72 | 0.4 | 0.55 |
| 2 | 19552389 | rs851066 | 0.344 | 0.193 | -0.847 | -0.578 | 0 | -0.028 | -0.028 | 2.496 | 0.85 | 0.84 | 0.94 | 1 | 0.92 | 0.83 |
| 2 | 19562083 | rs1978996 | -0.038 | 0.545 | -1.055 | -0.085 | -0.042 | -0.057 | -0.057 | 0 | 0.09 | 0.01 | 0.31 | 0.73 | 0.17 | 0.31 |
| 2 | 19563477 | rs13033424 | -0.557 | 0.63 | -0.818 | -0.48 | 0.002 | 0.004 | 0.004 | 0 | 0.12 | 0.13 | 0.2 | 0.1 | 0.17 | 0.12 |
| 2 | 19602083 | rs11685576 | -0.648 | 0.375 | -0.646 | -0.754 | -0.013 | -0.004 | -0.004 | 3.137 | 0.16 | 0.24 | 0.08 | 0.11 | 0.11 | 0.14 |
| 2 | 19604309 | rs12620067 | 0.347 | 0.657 | -0.9 | -0.648 | -0.049 | -0.06 | -0.06 | 0 | 0.09 | 0.01 | 0.32 | 0.73 | 0.18 | 0.28 |
| 2 | 19617302 | rs2166219 | -1.677 | 0.385 | -0.355 | -0.904 | 0.002 | -0.013 | -0.013 | 0 | 0.35 | 0.44 | 0.57 | 0.79 | 0.39 | 0.45 |
| 2 | 19625116 | rs13387360 | 0.205 | 0.419 | -0.428 | -0.997 | -0.004 | -0.036 | -0.036 | 0 | 0.11 | 0.21 | 0.06 | 0.02 | 0.07 | 0.06 |
| 2 | 19625636 | rs2198161 | -1.311 | 0.306 | -0.921 | -1.305 | -0.001 | -0.022 | -0.022 | 0 | 0.52 | 0.54 | 0.65 | 0.88 | 0.51 | 0.62 |
| 2 | 19625670 | rs13023304 | 0.952 | 0.563 | -1.007 | -1.15 | -0.012 | 0.008 | 0.008 | 0 | 0.06 | 0.02 | 0.16 | 0.05 | 0.1 | 0.04 |
| 2 | 19625818 | rs703297 | -1.112 | 0.276 | -0.921 | -1.18 | -0.001 | 0.012 | 0.012 | 0 | 0.5 | 0.54 | 0.64 | 0.88 | 0.51 | 0.62 |
| 2 | 19646090 | rs17442753 | -0.011 | -0.053 | -1.533 | -1.203 | 0.051 | 0.112 | 0.112 | 0.199 | 0.4 | 0.1 | 0.18 | 0.2 | 0.27 | 0.39 |
| 2 | 19646310 | rs11692444 | 0.565 | -0.028 | -0.988 | -1.563 | -0.003 | 0.008 | 0.008 | 0 | 0.36 | 0.33 | 0.38 | 0.05 | 0.45 | 0.28 |
| 2 | 19650271 | rs7596217 | 0.952 | 0.191 | -0.957 | -1.241 | -0.017 | 0.002 | 0.002 | 0 | 0.08 | 0.04 | 0.15 | 0.08 | 0.12 | 0.1 |
| 2 | 19652154 | rs703281 | -0.424 | -0.55 | -1.127 | -1.793 | 0.035 | 0.061 | 0.061 | 0.014 | 0.66 | 0.8 | 0.86 | 0.82 | 0.75 | 0.63 |
| 2 | 19653669 | rs10084260 | 0.123 | -0.643 | -0.558 | -1.759 | 0 | -0.004 | -0.004 | 0 | 0.54 | 0.58 | 0.66 | 0.74 | 0.62 | 0.52 |
| 2 | 19657569 | rs824475 | -0.36 | -0.569 | -0.738 | -1.398 | -0.021 | -0.041 | -0.041 | 0 | 0.9 | 0.84 | 0.96 | 1 | 0.94 | 0.93 |
| 2 | 19663338 | rs824466 | -0.404 | -0.412 | -0.839 | -1 | -0.008 | 0.105 | 0.105 | 0.038 | 0.14 | 0.32 | 0.27 | 0.67 | 0.12 | 0.23 |
| 2 | 19672181 | rs11689050 | 0.386 | -0.465 | -1.769 | -1.23 | -0.044 | 0.111 | 0.111 | 0 | 0.2 | 0.01 | 0.09 | 0.01 | 0.17 | 0.1 |
| 2 | 19673874 | rs12991182 | 0.956 | 0.191 | -1.675 | -1.188 | -0.018 | 0.01 | 0.01 | 0 | 0.09 | 0.05 | 0.2 | 0.08 | 0.15 | 0.1 |
| 2 | 19674224 | rs1820588 | 0.209 | 0 | -1.741 | -1.359 | 0.015 | 0.058 | 0.058 | 0.048 | 0.33 | 0.09 | 0.16 | 0.2 | 0.26 | 0.4 |
| 2 | 19675939 | rs997676 | 0.893 | 0.227 | -1.631 | -1.177 | -0.016 | 0.004 | 0.004 | 0 | 0.24 | 0.06 | 0.11 | 0.2 | 0.2 | 0.32 |
| 2 | 19680042 | rs11096592 | 0.895 | 0.496 | -0.917 | -1.08 | 0.002 | -0.03 | -0.03 | 1.674 | 0.26 | 0.27 | 0.3 | 0.07 | 0.32 | 0.14 |
| 2 | 19686824 | rs12995711 | 0.646 | 0.844 | -1.162 | -0.268 | 0.001 | -0.039 | -0.039 | 0.441 | 0.33 | 0.1 | 0.28 | 0.45 | 0.3 | 0.43 |
| 2 | 19689308 | rs824446 | 0.052 | 1.05 | -0.833 | -1.132 | 0.002 | 0.019 | 0.019 | 0 | 0.49 | 0.41 | 0.48 | 0.15 | 0.5 | 0.3 |
| 2 | 19690015 | rs824447 | -0.029 | 1.299 | -0.659 | -0.738 | 0.068 | -0.095 | -0.095 | 0 | 0.86 | 0.63 | 0.86 | 1 | 0.82 | 0.89 |
| 2 | 19692292 | rs824453 | -0.076 | 0.86 | -1.563 | -0.512 | -0.006 | 0.028 | 0.028 | 0 | 0.16 | 0.07 | 0.13 | 0.03 | 0.16 | 0.09 |
| 2 | 19692404 | rs2347283 | -0.013 | 0.854 | -1.62 | -0.581 | -0.006 | 0.073 | 0.073 | 0 | 0.34 | 0.16 | 0.33 | 0.12 | 0.37 | 0.21 |
| 2 | 19693078 | rs4621133 | 0.266 | 1.031 | -1.729 | -0.273 | -0.006 | 0.035 | 0.035 | 0.163 | 0.6 | 0.26 | 0.49 | 0.35 | 0.61 | 0.56 |
| 2 | 19693360 | rs2881591 | -0.318 | 0.827 | -1.6 | -0.564 | -0.024 | 0.001 | 0.001 | 0.434 | 0.38 | 0.28 | 0.42 | 0.55 | 0.46 | 0.45 |
| 2 | 19693784 | rs824454 | 0.133 | 0.439 | -0.943 | -1.521 | 0.002 | -0.022 | -0.022 | 1.11 | 0.52 | 0.56 | 0.48 | 0.88 | 0.44 | 0.7 |
| 2 | 19694200 | rs6705213 | -0.267 | 0.374 | -0.494 | -2.14 | 0.007 | 0.012 | 0.012 | 0 | 0.48 | 0.55 | 0.5 | 0.11 | 0.5 | 0.34 |
| 2 | 19694526 | rs13383201 | 0.331 | 0.289 | -0.297 | -2.27 | 0.012 | -0.028 | -0.028 | 0 | 0.19 | 0.26 | 0.26 | 0.08 | 0.17 | 0.13 |

| | | | | | | | | | | | | | | | | |
|---|---|---|---|---|---|---|---|---|---|---|---|---|---|---|---|---|
| 2 | 19694875 | rs6706023 | -0.985 | 0.315 | -0.34 | -2.26 | -0.002 | -0.019 | -0.019 | 0 | 0.58 | 0.57 | 0.61 | 0.13 | 0.65 | 0.4 |
| 2 | 19696456 | rs17758655 | 1.478 | 0.465 | -0.389 | -2.145 | -0.026 | 0.025 | 0.025 | 0 | 0.09 | 0.02 | 0.1 | 0.04 | 0.11 | 0.08 |
| 2 | 19697223 | rs17707800 | 0.958 | 0.359 | -0.235 | -2.213 | -0.028 | 0.04 | 0.04 | 0 | 0.27 | 0.17 | 0.22 | 0.03 | 0.33 | 0.21 |
| 2 | 19699023 | rs1649361 | -0.187 | -0.115 | -0.51 | -2.215 | 0.028 | 0.113 | 0.113 | 0 | 0.49 | 0.22 | 0.42 | 0.74 | 0.56 | 0.52 |
| 2 | 19699611 | rs1718641 | -0.786 | -0.078 | -0.515 | -2.137 | -0.005 | 0.023 | 0.023 | 0 | 0.39 | 0.32 | 0.45 | 0.79 | 0.39 | 0.44 |
| 2 | 19705084 | rs4666491 | 2.452 | -0.05 | -0.577 | -1.866 | -0.001 | 0 | 0 | 1.096 | 0.07 | 0.05 | 0.15 | 0.04 | 0.04 | 0.04 |
| 2 | 19705959 | rs1851411 | 0.697 | -0.122 | -0.483 | -1.925 | 0.028 | 0.038 | 0.038 | 0 | 0.32 | 0.24 | 0.3 | 0.18 | 0.28 | 0.35 |
| 2 | 19708876 | rs1718651 | 0.315 | 0.049 | -0.546 | -2.191 | -0.008 | -0.004 | -0.004 | 0 | 0.2 | 0.16 | 0.14 | 0.01 | 0.27 | 0.17 |
| 2 | 19713604 | rs12478269 | 1.512 | -0.035 | -1.056 | -2.462 | -0.099 | 0.12 | 0.12 | 0 | 0.23 | 0.02 | 0.2 | 0.04 | 0.29 | 0.2 |
| 2 | 19716396 | rs11096597 | 1.429 | -0.338 | -1.162 | -2.441 | 0.046 | -0.073 | -0.073 | 2.982 | 0.15 | 0.02 | 0.11 | 0.7 | 0.08 | 0.26 |
| 2 | 19722914 | rs1579204 | 1.982 | -0.249 | -0.839 | -2.511 | -0.023 | 0.015 | 0.015 | 0 | 0.08 | 0.02 | 0.15 | 0.06 | 0.11 | 0.09 |
| 2 | 19723280 | rs10208877 | 1.982 | -0.249 | -0.839 | -2.511 | -0.023 | 0.015 | 0.015 | 0 | 0.08 | 0.02 | 0.15 | 0.06 | 0.11 | 0.09 |
| 2 | 19723613 | rs10174214 | -0.346 | -0.31 | 0.045 | -2.617 | 0.014 | -0.081 | -0.081 | 0 | 0.72 | 0.57 | 0.69 | 0.95 | 0.68 | 0.79 |
| 2 | 19724103 | rs13407212 | -0.094 | -0.386 | 0.189 | -2.63 | 0.006 | 0.015 | 0.015 | 0.596 | 0.33 | 0.27 | 0.45 | 0.8 | 0.24 | 0.39 |
| 2 | 19729561 | rs17759637 | 1.655 | -0.377 | 0.08 | -2.562 | 0.007 | 0.015 | 0.015 | 0.114 | 0.22 | 0.18 | 0.18 | 0.16 | 0.2 | 0.24 |
| 2 | 19729725 | rs934012 | -0.438 | -0.162 | 0.285 | -2.334 | -0.003 | -0.005 | -0.005 | 0 | 0.73 | 0.69 | 0.69 | 0.95 | 0.68 | 0.78 |
| 2 | 19732698 | rs9973758 | 0.028 | -0.255 | 0.363 | -2.164 | -0.022 | -0.007 | -0.007 | 0 | 0.27 | 0.37 | 0.29 | 0.76 | 0.18 | 0.34 |
| 2 | 19741318 | rs1649387 | -1.184 | -0.265 | 0.191 | -2.203 | 0.003 | 0.058 | 0.058 | 0.406 | 0.58 | 0.65 | 0.65 | 0.8 | 0.58 | 0.59 |
| 2 | 19743086 | rs1522490 | 0.917 | -0.369 | 0.408 | -2.225 | -0.057 | 0.059 | 0.059 | 0.012 | 0.17 | 0.27 | 0.14 | 0.7 | 0.08 | 0.25 |
| 2 | 19743763 | rs17760223 | 0.118 | -0.355 | 0.402 | -2.244 | -0.005 | 0.008 | 0.008 | 0.625 | 0.59 | 0.57 | 0.48 | 0.9 | 0.5 | 0.66 |
| 2 | 19749304 | rs6712764 | 0.441 | -0.423 | 0.561 | -2.033 | -0.076 | 0.075 | 0.075 | 0 | 0.19 | 0.35 | 0.14 | 0.7 | 0.08 | 0.27 |
| 2 | 19749554 | rs6757410 | 0.304 | -0.128 | 0.408 | -2.071 | -0.001 | -0.009 | -0.009 | 0 | 0.56 | 0.5 | 0.57 | 0.24 | 0.57 | 0.48 |
| 2 | 19750744 | rs824484 | -0.704 | 0.269 | 0.6 | -1.818 | 0.075 | -0.023 | -0.023 | 0.022 | 0.88 | 0.53 | 0.82 | 0.94 | 0.89 | 0.92 |
| 2 | 19755159 | rs1522482 | 1.48 | -0.05 | 0.359 | -2.058 | -0.007 | -0.004 | -0.004 | 0.141 | 0.18 | 0.2 | 0.14 | 0.16 | 0.13 | 0.19 |
| 2 | 19759365 | rs1365930 | -0.389 | -0.051 | 0.364 | -2.101 | -0.011 | -0.028 | -0.028 | 0 | 0.7 | 0.38 | 0.68 | 0.78 | 0.75 | 0.73 |
| 2 | 19759954 | rs6705393 | 1.396 | 0.02 | -0.261 | -2.36 | -0.041 | 0.089 | 0.089 | 0 | 0.24 | 0.11 | 0.2 | 0.04 | 0.29 | 0.2 |
| 2 | 19760041 | rs1863776 | 1.069 | -0.078 | -0.354 | -2.416 | 0.02 | -0.002 | -0.002 | 1.264 | 0.26 | 0.06 | 0.33 | 0.73 | 0.2 | 0.34 |
| 2 | 19763760 | rs1427538 | 1.249 | 0.003 | -0.368 | -2.328 | -0.017 | 0.041 | 0.041 | 0.488 | 0.16 | 0.06 | 0.11 | 0.02 | 0.23 | 0.18 |
| 2 | 19763888 | rs6728887 | 0.675 | -0.018 | -0.301 | -2.32 | -0.001 | 0.095 | 0.095 | 0.264 | 0.41 | 0.25 | 0.34 | 0.2 | 0.42 | 0.39 |
| 2 | 19766789 | rs6740759 | 0.998 | -0.138 | 0.136 | -1.921 | 0.024 | -0.023 | -0.023 | 0 | 0.11 | 0.23 | 0.17 | 0.06 | 0.11 | 0.08 |
| 2 | 19770970 | rs13022722 | -0.082 | -0.128 | -0.389 | -1.849 | 0.002 | 0.015 | 0.015 | 0.298 | 0.58 | 0.38 | 0.46 | 0.22 | 0.64 | 0.57 |
| 2 | 19773144 | rs17686199 | 0.512 | -0.054 | -0.689 | -1.849 | -0.073 | 0.168 | 0.168 | 1.611 | 0.39 | 0.14 | 0.31 | 0.06 | 0.51 | 0.38 |
| 2 | 19774351 | rs11890596 | 0.686 | 0.023 | -0.158 | -1.016 | -0.003 | -0.001 | -0.001 | 2.612 | 0.18 | 0.18 | 0.14 | 0.16 | 0.13 | 0.19 |
| 2 | 19775396 | rs1015378 | -0.667 | 1.127 | 0.174 | -1.04 | 0.079 | -0.048 | -0.048 | 0.708 | 0.14 | 0.32 | 0.22 | 0.05 | 0.19 | 0.08 |
| 2 | 19775673 | rs1469366 | 0.047 | 0.183 | 0.302 | -0.513 | -0.048 | 0.113 | 0.113 | 2.609 | 0.82 | 0.43 | 0.78 | 0.63 | 0.9 | 0.84 |
| 2 | 19779372 | rs1820584 | 0.157 | 0.415 | -0.345 | -1.085 | 0 | -0.009 | -0.009 | 0 | 0.54 | 0.52 | 0.43 | 0.29 | 0.46 | 0.38 |
| 2 | 19780684 | rs12105038 | 0.423 | 0.473 | -0.788 | -1.759 | 0.003 | 0.035 | 0.035 | 0 | 0.33 | 0.26 | 0.26 | 0.01 | 0.34 | 0.17 |
| 2 | 19780732 | rs17686549 | 0.205 | 1.092 | -0.511 | -0.926 | -0.067 | -0.031 | -0.031 | 0.242 | 0.13 | 0.02 | 0.22 | 0.14 | 0.26 | 0.19 |
| 2 | 19791294 | rs12710719 | 0.877 | 0.878 | -0.316 | -1.178 | 0.027 | 0.179 | 0.179 | 0 | 0.47 | 0.27 | 0.42 | 0.17 | 0.45 | 0.43 |
| 2 | 19803695 | rs4666507 | -0.176 | 0.585 | 0.255 | -1.11 | -0.002 | -0.002 | -0.002 | 0.072 | 0.22 | 0.21 | 0.36 | 0.8 | 0.18 | 0.41 |
| 2 | 19808099 | rs4145607 | -1.002 | 0.708 | 0.486 | -1.213 | 0.027 | -0.11 | -0.11 | 0 | 0.8 | 0.63 | 0.88 | 0.99 | 0.8 | 0.89 |
| 2 | 19808582 | rs4666339 | 1.252 | 0.742 | 0.34 | -1.096 | -0.036 | -0.029 | -0.029 | 0.034 | 0.06 | 0.01 | 0.15 | 0.08 | 0.1 | 0.11 |
| 2 | 19809750 | rs7586843 | 0.02 | 0.63 | 0.502 | -1.154 | -0.021 | 0.007 | 0.007 | 0 | 0.2 | 0.23 | 0.25 | 0.74 | 0.12 | 0.32 |
| 2 | 19809777 | rs1427550 | 0.718 | 0.688 | 0.466 | -1.205 | 0.008 | 0.017 | 0.017 | 0 | 0.6 | 0.4 | 0.63 | 0.25 | 0.68 | 0.56 |
| 2 | 19811385 | rs7596636 | 0.725 | 0.707 | 0.645 | -1.19 | -0.022 | -0.049 | -0.049 | 0 | 0.72 | 0.51 | 0.58 | 0.19 | 0.78 | 0.57 |
| 2 | 19811522 | rs6531176 | -0.809 | 0.806 | 0.662 | -1.192 | -0.001 | 0.002 | 0.002 | 0 | 0.83 | 0.84 | 0.77 | 0.28 | 0.88 | 0.67 |
| 2 | 19812104 | rs11696024 | -0.034 | 0.807 | 0.663 | -1.192 | -0.017 | 0.012 | 0.012 | 0 | 0.77 | 0.67 | 0.72 | 0.27 | 0.83 | 0.65 |
| 2 | 19826777 | rs11687591 | 0.854 | 0.813 | 0.667 | -1.195 | 0.017 | -0.065 | -0.065 | 0.063 | 0.2 | 0.27 | 0.11 | 0.02 | 0.2 | 0.11 |
| 2 | 19834713 | rs4666524 | 0.282 | 0.837 | 0.727 | -1.163 | 0 | -0.024 | -0.024 | 0 | 0.63 | 0.59 | 0.66 | 0.26 | 0.68 | 0.56 |
| 2 | 19835526 | rs6755311 | 1.086 | 0.843 | 0.664 | -1.167 | -0.004 | 0.058 | 0.058 | 0 | 0.54 | 0.39 | 0.49 | 0.17 | 0.58 | 0.46 |
| 2 | 19842306 | rs13432997 | -1.086 | 0.793 | 0.872 | -1.14 | 0.035 | 0.04 | 0.04 | 0 | 0.12 | 0.52 | 0.19 | 0.1 | 0.1 | 0.1 |
| 2 | 19843041 | rs1026293 | 1.175 | 0.812 | 0.574 | -1.187 | -0.011 | 0.202 | 0.202 | 0 | 0.52 | 0.2 | 0.47 | 0.17 | 0.58 | 0.45 |
| 2 | 19845159 | rs7567542 | 0.872 | 0.805 | 0.509 | -1.228 | 0 | -0.031 | -0.031 | 0 | 0.52 | 0.54 | 0.49 | 0.18 | 0.56 | 0.46 |
| 2 | 19855082 | rs6725609 | 0.184 | 0.856 | 0.923 | -0.783 | -0.011 | -0.007 | -0.007 | 0 | 0.8 | 0.69 | 0.67 | 0.19 | 0.84 | 0.63 |
| 2 | 19860539 | rs10173007 | 1.55 | 0.915 | 1.008 | -0.943 | -0.025 | 0.184 | 0.184 | 0 | 0.54 | 0.22 | 0.47 | 0.07 | 0.57 | 0.44 |
| 2 | 19861003 | rs17554501 | 1.445 | 0.955 | 0.911 | -0.795 | -0.009 | -0.039 | -0.039 | 0 | 0.1 | 0.01 | 0.08 | 0.12 | 0.08 | 0.07 |
| 2 | 19865377 | rs4666533 | 1.232 | 0.948 | 1.065 | -0.799 | 0.003 | -0.005 | -0.005 | 0 | 0.11 | 0.08 | 0.09 | 0.12 | 0.08 | 0.07 |
| 2 | 19870143 | rs17619443 | 1.419 | 0.967 | 1.029 | -0.79 | 0.001 | -0.003 | -0.003 | 0 | 0.1 | 0.06 | 0.09 | 0.12 | 0.08 | 0.08 |
| 2 | 19870341 | rs4666343 | -0.897 | 1.042 | 1.077 | -0.823 | 0.039 | -0.028 | -0.028 | 0.358 | 0.25 | 0.43 | 0.2 | 0.15 | 0.26 | 0.19 |
| 2 | 19876126 | rs6744666 | 0.605 | 1.048 | 1.147 | -0.817 | 0.025 | -0.03 | -0.03 | 0 | 0.69 | 0.52 | 0.67 | 0.24 | 0.71 | 0.52 |
| 2 | 19878194 | rs16986826 | 1.149 | 0.974 | 0.818 | -0.978 | -0.013 | -0.062 | -0.062 | 0 | 0.65 | 0.49 | 0.57 | 0.1 | 0.72 | 0.51 |
| 2 | 19878796 | rs6531190 | -0.664 | 0.943 | 0.87 | -0.936 | -0.013 | 0.043 | 0.043 | 0 | 0.84 | 0.75 | 0.79 | 0.26 | 0.88 | 0.64 |
| 2 | 19878883 | rs6531191 | 1.785 | 0.966 | 0.685 | -0.965 | 0 | 0.199 | 0.199 | 0 | 0.5 | 0.26 | 0.45 | 0.07 | 0.54 | 0.39 |
| 2 | 19881081 | rs724822 | 1.744 | 1.083 | 0.547 | -0.809 | 0.014 | 0.193 | 0.193 | 0 | 0.5 | 0.25 | 0.44 | 0.09 | 0.51 | 0.42 |
| 2 | 19883495 | rs2198368 | 0.372 | 0.617 | 0.467 | -0.433 | -0.002 | 0.007 | 0.007 | 0 | 0.76 | 0.7 | 0.72 | 0.26 | 0.77 | 0.6 |
| 2 | 19885907 | rs9973954 | 0.89 | 0.67 | 0.085 | -0.657 | -0.012 | -0.073 | -0.073 | 0.351 | 0.66 | 0.55 | 0.56 | 0.11 | 0.69 | 0.54 |
| 2 | 19897377 | rs1551261 | 1.311 | 0.705 | -0.082 | -0.667 | 0.013 | 0.047 | 0.047 | 0.099 | 0.51 | 0.41 | 0.44 | 0.12 | 0.44 | 0.37 |
| 2 | 19901073 | rs1453205 | 0.944 | 0.858 | 0.115 | -0.175 | -0.004 | 0.004 | 0.004 | 0.042 | 0.59 | 0.42 | 0.6 | 0.32 | 0.54 | 0.45 |
| 2 | 19903658 | rs6733236 | -0.264 | 1.003 | 0.299 | -0.252 | -0.001 | 0.006 | 0.006 | 0.699 | 0.84 | 0.84 | 0.69 | 0.34 | 0.84 | 0.61 |

| | | | | | | | | | | | | | | | | |
|---|---|---|---|---|---|---|---|---|---|---|---|---|---|---|---|---|
| 2 | 19906545 | rs7593979 | -0.613 | 1.033 | 0.365 | -0.363 | 0.048 | -0.13 | -0.13 | 0 | 0.25 | 0.42 | 0.18 | 0.02 | 0.32 | 0.16 |
| 2 | 19908539 | rs17620259 | -0.312 | 1.018 | 0.234 | -0.324 | 0.011 | -0.041 | -0.041 | 0.032 | 0.2 | 0.23 | 0.17 | 0.02 | 0.29 | 0.15 |
| 2 | 19915438 | rs4572555 | 1.123 | 0.882 | -0.038 | -0.226 | -0.005 | 0.004 | 0.004 | 0.113 | 0.59 | 0.42 | 0.5 | 0.33 | 0.52 | 0.38 |
| 2 | 19915600 | rs2167796 | -0.248 | 0.903 | -0.111 | -0.204 | -0.007 | 0.006 | 0.006 | 0 | 0.16 | 0.14 | 0.12 | 0.04 | 0.22 | 0.14 |
| 2 | 19918242 | rs2347857 | 0.829 | 0.227 | -0.026 | 0.092 | 0.014 | 0.013 | 0.013 | 2.712 | 0.63 | 0.49 | 0.7 | 0.54 | 0.6 | 0.45 |
| 2 | 19926886 | rs13029325 | 0.429 | 0.125 | 0.02 | 0.367 | 0.019 | 0.053 | 0.053 | 0 | 0.74 | 0.47 | 0.69 | 0.63 | 0.68 | 0.59 |
| 2 | 19927019 | rs13034588 | -0.078 | 0.531 | 0.071 | 0.153 | -0.006 | -0.018 | -0.018 | 0 | 0.14 | 0.09 | 0.11 | 0.2 | 0.15 | 0.17 |
| 2 | 19933592 | rs896802 | 0.072 | 0.13 | -0.707 | -0.103 | 0.002 | -0.016 | -0.016 | 0 | 0.34 | 0.3 | 0.34 | 0.62 | 0.34 | 0.51 |
| 2 | 19934280 | rs1078150 | 1.941 | 0.241 | -1.549 | -0.345 | -0.054 | 0.095 | 0.095 | 0 | 0.22 | 0.05 | 0.2 | 0.03 | 0.28 | 0.08 |
| 2 | 19939783 | rs7571810 | -0.182 | 0.417 | -0.823 | 0.081 | 0.017 | 0.018 | 0.018 | 0 | 0.81 | 0.55 | 0.84 | 0.82 | 0.81 | 0.84 |
| 2 | 19941190 | rs10196450 | 0.75 | 0.552 | -1.795 | -0.124 | -0.037 | 0.132 | 0.132 | 0.319 | 0.42 | 0.09 | 0.49 | 0.22 | 0.43 | 0.24 |
| 2 | 19948614 | rs7558226 | -0.013 | 1.304 | -1.317 | 0.213 | -0.072 | 0.015 | 0.015 | 0 | 0.11 | 0.02 | 0.09 | 0.03 | 0.18 | 0.14 |
| 2 | 19948931 | rs4666562 | -0.363 | 0.881 | -0.461 | -0.022 | -0.004 | -0.052 | -0.052 | 0 | 0.16 | 0.32 | 0.18 | 0.06 | 0.15 | 0.13 |
| 2 | 19948952 | rs4666563 | -0.852 | 0.348 | -0.432 | 0.237 | 0 | -0.01 | -0.01 | 0 | 0.61 | 0.62 | 0.49 | 0.58 | 0.53 | 0.71 |
| 2 | 19950193 | rs13002741 | 1.08 | 0.176 | -0.343 | 0.191 | 0.061 | -0.086 | -0.086 | 0.018 | 0.3 | 0.09 | 0.19 | 0.48 | 0.16 | 0.42 |
| 2 | 19951502 | rs12710728 | 0.049 | 0.342 | 0.51 | 0.13 | 0.021 | -0.052 | -0.052 | 0.286 | 0.86 | 0.54 | 0.81 | 0.93 | 0.86 | 0.89 |
| 2 | 19954038 | rs11683563 | -0.423 | 0.387 | 0.597 | -0.003 | -0.01 | 0.02 | 0.02 | 0 | 0.58 | 0.35 | 0.55 | 0.9 | 0.44 | 0.85 |
| 2 | 19955245 | rs2123535 | 0.176 | 0.348 | 0.453 | 0.147 | -0.001 | 0 | 0 | 0 | 0.4 | 0.4 | 0.49 | 0.44 | 0.49 | 0.28 |
| 2 | 19958008 | rs1453202 | -0.736 | 0.444 | 0.398 | 0.757 | -0.014 | 0.137 | 0.137 | 0 | 0.1 | 0.17 | 0.23 | 0.58 | 0.07 | 0.27 |
| 2 | 19961624 | rs7559401 | 0.24 | 0.599 | 0.063 | 0.83 | -0.091 | 0.061 | 0.061 | 0 | 0.15 | 0.01 | 0.09 | 0.03 | 0.21 | 0.22 |
| 2 | 19966874 | rs1584784 | -0.535 | 0.438 | 0.551 | 0.935 | 0.038 | -0.007 | -0.007 | 0 | 0.89 | 0.67 | 0.83 | 0.93 | 0.84 | 0.94 |
| 2 | 19967944 | rs7601531 | -0.055 | 0.179 | 0.608 | 0.898 | 0.026 | -0.008 | -0.008 | 0.586 | 0.56 | 0.31 | 0.48 | 0.61 | 0.56 | 0.68 |
| 2 | 19973210 | rs12710729 | 0.857 | 0.31 | 0.097 | 0.862 | -0.016 | 0.154 | 0.154 | 0.63 | 0.32 | 0.08 | 0.26 | 0.03 | 0.37 | 0.13 |
| 2 | 19975732 | rs11681722 | -0.62 | 0.658 | 1.087 | 1.979 | -0.087 | 0.068 | 0.068 | 0 | 0.16 | 0.02 | 0.09 | 0.03 | 0.2 | 0.21 |
| 2 | 19976506 | rs896809 | -1.302 | 0.675 | 1.276 | 2.064 | 0.031 | 0.084 | 0.084 | 0.572 | 0.26 | 0.41 | 0.34 | 0.39 | 0.27 | 0.41 |
| 2 | 19984546 | rs9679438 | 1.841 | 0.14 | 1.42 | 2.536 | -0.012 | 0.271 | 0.271 | 1.051 | 0.91 | 0.67 | 0.79 | 0.61 | 0.96 | 0.84 |
| 2 | 19987440 | rs1204005 | -2.591 | 0.156 | 2.032 | 2.691 | 0.018 | 0.395 | 0.395 | 2.343 | 0.1 | 0.48 | 0.22 | 0.35 | 0.04 | 0.16 |
| 2 | 19988877 | rs10201889 | -1.366 | 0.181 | 1.969 | 2.685 | -0.004 | -0.005 | -0.005 | 0.185 | 0.21 | 0.64 | 0.21 | 0.16 | 0.16 | 0.17 |
| 2 | 19989521 | rs7604613 | -2.135 | 0.166 | 1.488 | 2.638 | 0.013 | -0.049 | -0.049 | 0 | 0.28 | 0.85 | 0.4 | 0.53 | 0.19 | 0.3 |
| 2 | 19992957 | rs1437864 | -0.933 | 0.188 | 0.628 | 1.654 | -0.083 | 0.149 | 0.149 | 0 | 0.7 | 0.93 | 0.69 | 0.92 | 0.56 | 0.81 |
| 2 | 19994998 | rs16987112 | -0.894 | 0.324 | 0.569 | 1.87 | -0.009 | -0.021 | -0.021 | 0 | 0.07 | 0.05 | 0.03 | 0.09 | 0.07 | 0.03 |
| 2 | 19997343 | rs10495699 | -0.034 | 0.13 | 1.105 | 2.076 | -0.008 | 0.03 | 0.03 | 0 | 0.56 | 0.66 | 0.61 | 0.63 | 0.48 | 0.64 |
| 2 | 20001026 | rs11096619 | -0.797 | 0.181 | 1.635 | 2.087 | 0.046 | 0.05 | 0.05 | 0 | 0.13 | 0.27 | 0.17 | 0.16 | 0.16 | 0.13 |
| 2 | 20009261 | rs993832 | -1.332 | -0.83 | 2.266 | 1.907 | -0.061 | 0.064 | 0.064 | 0.675 | 0.14 | 0.23 | 0.12 | 0.23 | 0.03 | 0.07 |
| 2 | 20009456 | rs1437860 | -1.934 | -0.847 | 2.451 | 1.889 | -0.105 | 0.216 | 0.216 | 0.673 | 0.18 | 0.45 | 0.16 | 0.46 | 0.05 | 0.16 |
| 2 | 20009943 | rs1437861 | -0.788 | -0.86 | 2.148 | 1.691 | -0.059 | 0.166 | 0.166 | 0 | 0.62 | 0.87 | 0.62 | 0.93 | 0.42 | 0.7 |
| 2 | 20025978 | rs6727158 | -0.815 | -0.854 | 2.097 | 1.655 | -0.076 | 0.187 | 0.187 | 0 | 0.62 | 0.88 | 0.63 | 0.93 | 0.44 | 0.7 |
| 2 | 20027663 | rs10184446 | 0.554 | -0.79 | 2.151 | 1.733 | -0.002 | -0.006 | -0.006 | 0 | 0.43 | 0.37 | 0.45 | 0.47 | 0.37 | 0.54 |
| 2 | 20042817 | rs6732272 | 0.413 | -0.902 | 1.742 | 1.57 | 0.001 | -0.007 | -0.007 | 0.492 | 0.44 | 0.43 | 0.47 | 0.69 | 0.4 | 0.63 |
| 2 | 20045737 | rs16987155 | -0.69 | -0.889 | 0.871 | 1.584 | -0.008 | 0.006 | 0.006 | 0.021 | 0.12 | 0.11 | 0.12 | 0.24 | 0.03 | 0.07 |
| 2 | 20046065 | rs11096623 | 0.8 | -0.887 | 0.797 | 1.609 | -0.017 | -0.028 | -0.028 | 0.04 | 0.41 | 0.26 | 0.43 | 0.48 | 0.36 | 0.52 |
| 2 | 20048018 | rs10200763 | -0.044 | -0.643 | 0.71 | 1.602 | -0.001 | 0.023 | 0.023 | 0 | 0.53 | 0.44 | 0.56 | 0.74 | 0.4 | 0.6 |
| 2 | 20050028 | rs893396 | -0.542 | -0.559 | -0.269 | 1.133 | -0.095 | 0.213 | 0.213 | 0 | 0.6 | 0.89 | 0.62 | 0.94 | 0.45 | 0.69 |
| 2 | 20054057 | rs10184466 | -1.518 | -1 | 0.965 | 2.269 | -0.079 | 0.127 | 0.127 | 1.105 | 0.19 | 0.6 | 0.18 | 0.45 | 0.08 | 0.18 |
| 2 | 20054392 | rs10184780 | -1.546 | -0.992 | 0.963 | 2.29 | -0.076 | 0.108 | 0.108 | 0.888 | 0.19 | 0.6 | 0.18 | 0.51 | 0.08 | 0.18 |
| 2 | 20059323 | rs7559602 | -1.28 | -0.892 | 0.38 | 1.896 | -0.105 | 0.216 | 0.216 | 0.051 | 0.18 | 0.49 | 0.15 | 0.39 | 0.07 | 0.17 |
| 2 | 20063586 | rs11096624 | 0.183 | -1.115 | -0.313 | 1.328 | -0.003 | -0.001 | -0.001 | 0 | 0.58 | 0.59 | 0.76 | 0.58 | 0.74 | 0.64 |
| 2 | 20064528 | rs7566585 | 0.551 | -0.15 | -0.183 | 1.3 | -0.004 | 0.005 | 0.005 | 0 | 0.44 | 0.39 | 0.6 | 0.36 | 0.54 | 0.44 |
| 2 | 20067295 | rs11687529 | 0.593 | -0.459 | -0.314 | 1.329 | -0.007 | 0.104 | 0.104 | 0.422 | 0.85 | 0.59 | 0.92 | 0.76 | 0.89 | 0.93 |
| 2 | 20069380 | rs2276607 | 0.626 | -0.176 | -0.581 | 1.053 | -0.004 | 0.058 | 0.058 | 0 | 0.16 | 0.29 | 0.26 | 0.3 | 0.13 | 0.25 |
| 2 | 20069414 | rs2276606 | -0.54 | -0.167 | -0.832 | 1.034 | 0.001 | -0.002 | -0.002 | 0 | 0.54 | 0.62 | 0.39 | 0.6 | 0.44 | 0.57 |
| 2 | 20073860 | rs11096626 | -0.286 | -0.014 | -0.963 | 0.742 | 0.001 | 0 | 0 | 0 | 0.49 | 0.48 | 0.32 | 0.4 | 0.38 | 0.43 |
| 2 | 20074271 | rs7595902 | 0.399 | -0.116 | -0.997 | 0.692 | -0.009 | -0.036 | -0.036 | 1.824 | 0.38 | 0.06 | 0.27 | 0.37 | 0.31 | 0.38 |
| 2 | 20076454 | rs3731669 | -0.161 | 0.23 | -0.839 | 0.87 | 0.009 | 0.078 | 0.078 | 0 | 0.77 | 0.65 | 0.66 | 0.5 | 0.78 | 0.62 |
| 2 | 20078571 | rs13002069 | 0.862 | 0.486 | -1.301 | 0.455 | -0.033 | 0.054 | 0.054 | 0 | 0.31 | 0.17 | 0.35 | 0.11 | 0.42 | 0.19 |
| 2 | 20081474 | rs17689262 | 0.213 | 0.127 | -1.439 | 0.328 | -0.008 | 0.053 | 0.053 | 0.789 | 0.16 | 0.03 | 0.11 | 0.05 | 0.15 | 0.07 |
| 2 | 20081742 | rs7590829 | -0.993 | 0.164 | -1.979 | -0.148 | -0.093 | 0.1 | 0.1 | 0 | 0.65 | 0.89 | 0.6 | 0.89 | 0.48 | 0.76 |
| 2 | 20082891 | rs4468781 | 0.466 | -0.314 | -1.341 | 0.519 | -0.011 | 0.012 | 0.012 | 0 | 0.15 | 0.19 | 0.14 | 0.22 | 0.08 | 0.17 |
| 2 | 20084133 | rs10204428 | 0.722 | -0.646 | -0.427 | 0.717 | -0.032 | -0.01 | -0.01 | 0.003 | 0.19 | 0.41 | 0.13 | 0.19 | 0.1 | 0.24 |
| 2 | 20084387 | rs10168217 | 0.095 | -0.661 | -0.139 | 0.974 | -0.006 | 0.062 | 0.062 | 1.038 | 0.24 | 0.44 | 0.21 | 0.41 | 0.19 | 0.4 |
| 2 | 20086166 | rs12466802 | 0.524 | -0.674 | -0.188 | 0.834 | -0.008 | -0.018 | -0.018 | 0 | 0.17 | 0.1 | 0.21 | 0.25 | 0.19 | 0.18 |
| 2 | 20086462 | rs13432782 | 0.053 | -0.684 | -0.156 | 0.821 | -0.014 | 0.068 | 0.068 | 1.108 | 0.25 | 0.51 | 0.21 | 0.41 | 0.19 | 0.4 |
| 2 | 20086610 | rs2278525 | -1.502 | -0.632 | -0.327 | 0.783 | -0.035 | 0.121 | 0.121 | 0 | 0.56 | 0.8 | 0.57 | 0.88 | 0.47 | 0.75 |
| 2 | 20087809 | rs10856791 | -1.485 | -0.631 | -0.328 | 0.784 | -0.034 | 0.12 | 0.12 | 0 | 0.56 | 0.79 | 0.57 | 0.88 | 0.47 | 0.75 |
| 2 | 20087984 | rs16987205 | 0.416 | -0.678 | -0.346 | 0.855 | -0.008 | -0.018 | -0.018 | 0.104 | 0.17 | 0.1 | 0.21 | 0.25 | 0.19 | 0.18 |
| 2 | 20090739 | rs16987206 | 0.004 | -0.901 | -0.012 | 0.799 | 0.038 | -0.029 | -0.029 | 0 | 0.3 | 0.19 | 0.23 | 0.4 | 0.18 | 0.38 |
| 2 | 20095690 | rs1816647 | -2.274 | -0.746 | 0.299 | 0.612 | -0.055 | 0.108 | 0.108 | 0 | 0.64 | 0.89 | 0.64 | 0.88 | 0.55 | 0.79 |
| 2 | 20096936 | rs7563410 | 0.888 | -0.85 | 0.25 | 0.607 | 0.022 | 0.011 | 0.011 | 0.725 | 0.19 | 0.09 | 0.12 | 0.17 | 0.11 | 0.23 |
| 2 | 20098208 | rs16987218 | 1.217 | -0.838 | 0.264 | 0.622 | 0.014 | -0.025 | -0.025 | 0 | 0.11 | 0.06 | 0.11 | 0.23 | 0.07 | 0.16 |

| | | | | | | | | | | | | | | | | |
|---|---|---|---|---|---|---|---|---|---|---|---|---|---|---|---|---|
| 2 | 20101717 | rs2304588 | 1.123 | -0.9 | 0.291 | 0.613 | 0.021 | -0.002 | -0.002 | 0 | 0.18 | 0.02 | 0.12 | 0.17 | 0.11 | 0.23 |
| 2 | 20101878 | rs1370585 | 0.245 | -0.887 | 0.392 | 0.647 | -0.006 | -0.019 | -0.019 | 0.597 | 0.2 | 0.11 | 0.21 | 0.22 | 0.22 | 0.19 |
| 2 | 20109061 | rs10175486 | -0.133 | -0.883 | 0.366 | 0.714 | 0.033 | -0.026 | -0.026 | 0 | 0.3 | 0.2 | 0.23 | 0.41 | 0.18 | 0.4 |
| 2 | 20112269 | rs10197890 | 0.508 | -0.998 | 0.478 | 0.565 | 0.065 | -0.004 | -0.004 | 0 | 0.06 | 0.23 | 0.05 | 0.03 | 0.07 | 0.03 |
| 2 | 20112796 | rs957614 | 0.16 | -1.001 | 0.655 | 0.643 | -0.002 | 0.007 | 0.007 | 0 | 0.21 | 0.26 | 0.23 | 0.23 | 0.22 | 0.19 |
| 2 | 20113088 | rs1056233 | 0.072 | -1.003 | 0.654 | 0.642 | -0.003 | 0.006 | 0.006 | 0 | 0.22 | 0.26 | 0.23 | 0.23 | 0.22 | 0.19 |
| 2 | 20113221 | rs6748924 | 0.513 | -0.962 | 0.417 | 0.648 | 0.065 | -0.004 | -0.004 | 0 | 0.06 | 0.23 | 0.05 | 0.03 | 0.07 | 0.03 |
| 2 | 20131079 | rs1191778 | 2.615 | -0.981 | 0.236 | 0.639 | -0.059 | 0.097 | 0.097 | 0 | 0.35 | 0.11 | 0.36 | 0.12 | 0.44 | 0.21 |
| 2 | 20135283 | rs6741091 | -1.198 | -0.732 | 0.262 | 0.749 | 0.004 | 0.004 | 0.004 | 0 | 0.5 | 0.47 | 0.46 | 0.62 | 0.4 | 0.6 |
| 2 | 20173255 | rs10153717 | -2.077 | -0.733 | -0.383 | 0.735 | -0.012 | 0.085 | 0.085 | 0 | 0.6 | 0.78 | 0.62 | 0.88 | 0.55 | 0.78 |
| 2 | 20189015 | rs1060742 | 1.106 | -0.785 | -0.337 | 0.713 | 0.003 | -0.022 | -0.022 | 1.809 | 0.17 | 0 | 0.12 | 0.17 | 0.11 | 0.22 |
| 2 | 20189089 | rs3731663 | -1.647 | -0.752 | -0.285 | 0.743 | -0.023 | 0.025 | 0.025 | 0 | 0.56 | 0.77 | 0.51 | 0.65 | 0.47 | 0.62 |
| 2 | 20190683 | rs721933 | 0.068 | -0.632 | -0.471 | 0.831 | -0.007 | -0.022 | -0.022 | 0 | 0.19 | 0.12 | 0.21 | 0.23 | 0.22 | 0.19 |
| 2 | 20199089 | rs3795965 | 0.988 | -0.585 | -0.454 | 0.618 | 0.015 | -0.023 | -0.023 | 0.325 | 0.11 | 0.06 | 0.11 | 0.21 | 0.07 | 0.17 |
| 2 | 20202811 | rs11681705 | 0.365 | -0.375 | -0.351 | 0.36 | -0.008 | 0.034 | 0.034 | 0.128 | 0.17 | 0.08 | 0.19 | 0.04 | 0.19 | 0.17 |
| 2 | 20202929 | rs28452699 | -0.075 | -0.447 | -0.259 | 0.324 | 0 | 0.017 | 0.017 | 0.108 | 0.19 | 0.14 | 0.2 | 0.04 | 0.2 | 0.18 |
| 2 | 20205680 | rs28401180 | -0.982 | -0.595 | -0.077 | 0.343 | 0.009 | 0.007 | 0.007 | 0 | 0.51 | 0.53 | 0.46 | 0.62 | 0.4 | 0.59 |
| 2 | 20207619 | rs11694716 | -1.85 | -0.586 | -0.086 | 0.191 | -0.008 | 0.084 | 0.084 | 0 | 0.61 | 0.78 | 0.69 | 0.89 | 0.59 | 0.82 |
| 2 | 20208412 | rs10856792 | -0.309 | -0.846 | 0.078 | 0.735 | -0.045 | 0.023 | 0.023 | 0 | 0.32 | 0.42 | 0.23 | 0.48 | 0.19 | 0.41 |
| 2 | 20218658 | rs12478941 | 0.083 | -0.62 | -0.554 | 0.788 | 0.048 | -0.063 | -0.063 | 0.604 | 0.28 | 0.09 | 0.14 | 0.42 | 0.16 | 0.44 |
| 2 | 20224999 | rs1191812 | 0.921 | -0.567 | -0.373 | 0.683 | 0.014 | 0.075 | 0.075 | 0.019 | 0.78 | 0.38 | 0.67 | 0.59 | 0.78 | 0.75 |
| 2 | 20227258 | rs1191818 | 1.264 | -0.532 | -0.376 | 0.634 | 0.038 | 0.137 | 0.137 | 0 | 0.5 | 0.24 | 0.52 | 0.17 | 0.62 | 0.28 |
| 2 | 20231348 | rs2704616 | 1.264 | -0.532 | -0.376 | 0.634 | 0.038 | 0.137 | 0.137 | 0 | 0.5 | 0.24 | 0.52 | 0.17 | 0.62 | 0.28 |
| 2 | 20232287 | rs1191825 | 1.264 | -0.532 | -0.376 | 0.634 | 0.039 | 0.144 | 0.144 | 0 | 0.5 | 0.23 | 0.52 | 0.17 | 0.62 | 0.28 |
| 2 | 20233135 | rs1191824 | 1.196 | -0.52 | -0.364 | 0.732 | 0.035 | 0.09 | 0.09 | 0 | 0.51 | 0.24 | 0.58 | 0.23 | 0.63 | 0.29 |
| 2 | 20236330 | rs1147116 | 0.213 | -0.528 | 0.097 | 0.85 | -0.035 | 0.015 | 0.015 | 0 | 0.63 | 0.45 | 0.8 | 0.49 | 0.8 | 0.5 |
| 2 | 20240668 | rs3731673 | 0.694 | -0.208 | -0.321 | 0.962 | -0.049 | -0.038 | -0.038 | 0.018 | 0.08 | 0.01 | 0.2 | 0.31 | 0.17 | 0.21 |
| 2 | 20242416 | rs13010075 | -0.943 | -0.601 | -0.294 | 0.574 | 0.001 | 0.076 | 0.076 | 0 | 0.51 | 0.31 | 0.56 | 0.75 | 0.48 | 0.76 |
| 2 | 20252014 | rs1513830 | 0.76 | -0.415 | -0.341 | 0.752 | -0.05 | -0.04 | -0.04 | 0.02 | 0.08 | 0.01 | 0.2 | 0.3 | 0.18 | 0.21 |
| 2 | 20252695 | rs6705765 | -0.573 | -0.755 | 0.028 | 0.422 | -0.001 | -0.009 | -0.009 | 0 | 0.37 | 0.36 | 0.43 | 0.78 | 0.33 | 0.68 |
| 2 | 20253754 | rs12476858 | -0.513 | -0.756 | 0.023 | 0.421 | 0 | 0.006 | 0.006 | 0.547 | 0.36 | 0.33 | 0.42 | 0.78 | 0.33 | 0.68 |
| 2 | 20263404 | rs988318 | -1.235 | -0.782 | 0.116 | 0.343 | -0.005 | -0.001 | -0.001 | 0 | 0.4 | 0.62 | 0.48 | 0.43 | 0.45 | 0.42 |
| 2 | 20270027 | rs2126551 | -0.83 | -0.66 | -0.005 | 0.446 | 0.012 | 0.008 | 0.008 | 0 | 0.53 | 0.61 | 0.53 | 0.59 | 0.42 | 0.61 |
| 2 | 20274303 | rs1513833 | -1.568 | -0.752 | 0.492 | 0.587 | 0.025 | 0.15 | 0.15 | 3.134 | 0.08 | 0.49 | 0.12 | 0.11 | 0.04 | 0.05 |
| 2 | 20275591 | rs7585694 | -1.815 | -0.907 | -0.073 | 0.314 | -0.047 | 0.09 | 0.09 | 0.618 | 0.68 | 0.89 | 0.74 | 0.9 | 0.64 | 0.85 |
| 2 | 20277748 | rs878983 | 0.352 | -0.212 | 0.225 | 0.733 | -0.06 | -0.056 | -0.056 | 0 | 0.09 | 0.01 | 0.16 | 0.32 | 0.2 | 0.23 |
| 2 | 20279912 | rs1028052 | -1.22 | -0.405 | 0.696 | 0.643 | 0.05 | 0.173 | 0.173 | 0.803 | 0.06 | 0.39 | 0.11 | 0.11 | 0.04 | 0.05 |
| 2 | 20292618 | rs10199614 | -0.956 | -0.124 | 0.692 | 0.718 | 0.126 | 0.166 | 0.166 | 0 | 0.16 | 0.52 | 0.22 | 0.37 | 0.23 | 0.26 |
| 2 | 20297379 | rs6531204 | -0.258 | -0.104 | 0.333 | 0.705 | 0.064 | 0.076 | 0.076 | 0 | 0.12 | 0.2 | 0.18 | 0.32 | 0.2 | 0.24 |
| 2 | 20300673 | rs907865 | -1.327 | -0.537 | -0.239 | 0.29 | -0.004 | 0.196 | 0.196 | 0.854 | 0.5 | 0.83 | 0.58 | 0.84 | 0.43 | 0.71 |
| 2 | 20312438 | rs7578149 | -0.377 | -0.203 | -0.113 | 0.521 | -0.003 | 0.003 | 0.003 | 0 | 0.6 | 0.47 | 0.47 | 0.46 | 0.43 | 0.63 |
| 2 | 20322122 | rs6717177 | 0.298 | -0.557 | -0.337 | 0.09 | 0.044 | 0.101 | 0.101 | 0 | 0.46 | 0.28 | 0.34 | 0.25 | 0.38 | 0.35 |
| 2 | 20323678 | rs6531208 | -1.526 | -0.274 | -0.567 | -0.129 | -0.042 | 0.072 | 0.072 | 0 | 0.77 | 0.92 | 0.82 | 0.95 | 0.69 | 0.88 |
| 2 | 20329353 | rs6753830 | 0.183 | -0.107 | -0.088 | 0.055 | 0.009 | -0.017 | -0.017 | 0 | 0.6 | 0.58 | 0.62 | 0.69 | 0.52 | 0.48 |
| 2 | 20331338 | rs975951 | -0.104 | -0.16 | 0.104 | 0.034 | -0.017 | 0.016 | 0.016 | 0.08 | 0.63 | 0.7 | 0.63 | 0.69 | 0.52 | 0.48 |
| 2 | 20334723 | rs6531209 | -1.412 | -0.138 | -0.133 | -0.206 | -0.023 | 0.029 | 0.029 | 0 | 0.78 | 0.83 | 0.82 | 0.96 | 0.71 | 0.86 |
| 2 | 20336237 | rs17652287 | 0.805 | -0.313 | -0.683 | -0.406 | 0.02 | 0.114 | 0.114 | 0.618 | 0.39 | 0.2 | 0.27 | 0.13 | 0.34 | 0.23 |
| 2 | 20338020 | rs6531212 | -0.74 | -0.223 | -0.831 | -0.381 | -0.038 | 0.085 | 0.085 | 0 | 0.71 | 0.85 | 0.72 | 0.93 | 0.64 | 0.81 |
| 2 | 20338288 | rs4666363 | 1.434 | -0.34 | -0.736 | 0.077 | 0.008 | 0.005 | 0.005 | 0 | 0.07 | 0 | 0.19 | 0.39 | 0.05 | 0.16 |
| 2 | 20341401 | rs11681098 | 0.9 | -0.587 | -0.936 | -0.454 | -0.027 | 0.175 | 0.175 | 0.151 | 0.33 | 0.09 | 0.25 | 0.07 | 0.34 | 0.21 |
| 2 | 20346312 | rs16987405 | 0.86 | -0.222 | 0.099 | -0.059 | -0.009 | 0.302 | 0.302 | 0 | 0.08 | 0.23 | 0.2 | 0.39 | 0.05 | 0.14 |
| 2 | 20346651 | rs981396 | 0.232 | -0.569 | 0.471 | -0.377 | 0.022 | 0.05 | 0.05 | 0.018 | 0.48 | 0.38 | 0.39 | 0.17 | 0.41 | 0.32 |
| 2 | 20347824 | rs4666369 | -0.392 | -0.304 | 0.697 | -0.302 | 0.003 | 0.032 | 0.032 | 0 | 0.19 | 0.26 | 0.37 | 0.64 | 0.16 | 0.47 |
| 2 | 20349026 | rs16987411 | 0.465 | -0.258 | 0.605 | -0.275 | 0.014 | 0.032 | 0.032 | 0 | 0.12 | 0.17 | 0.17 | 0.26 | 0.11 | 0.32 |
| 2 | 20350783 | rs10495703 | 0.296 | -0.332 | 0.706 | -0.352 | 0.003 | 0.052 | 0.052 | 0 | 0.5 | 0.35 | 0.45 | 0.39 | 0.47 | 0.57 |
| 2 | 20353356 | rs1398950 | 0.728 | -0.24 | 0.446 | -0.376 | 0.018 | 0.058 | 0.058 | 0.225 | 0.43 | 0.3 | 0.31 | 0.12 | 0.36 | 0.26 |
| 2 | 20356606 | rs7583442 | 1.287 | 0.011 | 0.463 | -0.076 | -0.013 | 0.172 | 0.172 | 0 | 0.08 | 0.13 | 0.2 | 0.4 | 0.05 | 0.14 |
| 2 | 20356944 | rs7577335 | -0.108 | 0.052 | 0.312 | -0.275 | 0.006 | -0.011 | -0.011 | 0 | 0.57 | 0.48 | 0.55 | 0.41 | 0.51 | 0.63 |
| 2 | 20368196 | rs11894335 | -0.95 | 0.259 | 0.089 | -0.43 | -0.061 | 0.099 | 0.099 | 0.176 | 0.67 | 0.88 | 0.74 | 0.87 | 0.58 | 0.73 |
| 2 | 20368519 | rs1473886 | 0.136 | 0.177 | 0.258 | -0.377 | 0 | 0.001 | 0.001 | 0 | 0.52 | 0.52 | 0.5 | 0.41 | 0.46 | 0.58 |
| 2 | 20370868 | rs907869 | -1.049 | 0.179 | 0.222 | -0.411 | -0.07 | 0.109 | 0.109 | 0.065 | 0.67 | 0.9 | 0.75 | 0.87 | 0.58 | 0.73 |
| 2 | 20370926 | rs868933 | 1.319 | -0.152 | 0.083 | -0.859 | 0.041 | 0.06 | 0.06 | 0 | 0.28 | 0.17 | 0.22 | 0.12 | 0.23 | 0.13 |
| 2 | 20371167 | rs907868 | -0.28 | 0.108 | 0.338 | -0.583 | -0.009 | 0.022 | 0.022 | 0 | 0.56 | 0.67 | 0.51 | 0.39 | 0.46 | 0.59 |
| 2 | 20372414 | rs11692531 | 1.137 | 0.216 | 0.225 | -0.556 | -0.006 | 0 | 0 | 0 | 0.07 | 0.06 | 0.15 | 0.26 | 0.12 | 0.3 |
| 2 | 20373372 | rs11687133 | 0.354 | 0.134 | 0.326 | -0.62 | 0.011 | 0.004 | 0.004 | 0 | 0.45 | 0.42 | 0.35 | 0.14 | 0.33 | 0.27 |
| 2 | 20374158 | rs11096641 | -0.203 | 0.083 | 0.225 | -0.57 | -0.002 | 0.002 | 0.002 | 0 | 0.55 | 0.61 | 0.51 | 0.4 | 0.45 | 0.57 |
| 2 | 20377070 | rs6531217 | -0.696 | 0.236 | 0.096 | -0.61 | 0.032 | 0.019 | 0.019 | 0 | 0.23 | 0.46 | 0.23 | 0.28 | 0.21 | 0.4 |
| 2 | 20378023 | rs4462755 | -0.547 | 0.832 | 0.029 | -0.724 | 0.064 | 0.012 | 0.012 | 0 | 0.22 | 0.36 | 0.29 | 0.29 | 0.34 | 0.41 |
| 2 | 20378882 | rs13428318 | 0.866 | 0.754 | -0.084 | -0.793 | -0.002 | 0.015 | 0.015 | 0.17 | 0.32 | 0.22 | 0.21 | 0.17 | 0.33 | 0.29 |

| | | | | | | | | | | | | | | | | |
|---|---|---|---|---|---|---|---|---|---|---|---|---|---|---|---|---|
| 2 | 20381830 | rs3924843 | -0.597 | 0.999 | -0.201 | -0.612 | 0.086 | 0.017 | 0.017 | 0 | 0.23 | 0.41 | 0.3 | 0.3 | 0.34 | 0.42 |
| 2 | 20384552 | rs10856796 | -0.744 | 0.698 | -0.704 | -0.728 | 0.003 | 0.015 | 0.015 | 0.252 | 0.43 | 0.66 | 0.42 | 0.38 | 0.53 | 0.6 |
| 2 | 20386339 | rs4358082 | -0.137 | 1.108 | -0.387 | -0.788 | 0.045 | 0.031 | 0.031 | 0.253 | 0.18 | 0.32 | 0.22 | 0.3 | 0.19 | 0.42 |
| 2 | 20388484 | rs6531222 | 0.104 | 0.648 | -1.313 | -0.681 | 0.057 | 0.127 | 0.127 | 0.408 | 0.5 | 0.3 | 0.38 | 0.2 | 0.38 | 0.27 |
| 2 | 20388982 | rs7581159 | 0.8 | 0.181 | -1.107 | -0.606 | 0 | 0.021 | 0.021 | 0 | 0.16 | 0.07 | 0.09 | 0.09 | 0.14 | 0.12 |
| 2 | 20391853 | rs2881926 | 0.102 | 0.6 | -1.016 | -0.37 | -0.015 | 0.015 | 0.015 | 0 | 0.57 | 0.29 | 0.53 | 0.49 | 0.51 | 0.57 |
| 2 | 20396122 | rs6749689 | -0.183 | 0.614 | -0.546 | 0.021 | -0.001 | -0.001 | -0.001 | 0 | 0.47 | 0.51 | 0.55 | 0.51 | 0.56 | 0.58 |
| 2 | 20406245 | rs3771254 | -1.305 | 0.708 | -0.416 | -0.215 | 0.003 | -0.034 | -0.034 | 0 | 0.1 | 0.19 | 0.07 | 0.01 | 0.1 | 0.07 |
| 2 | 20409322 | rs2015110 | 0.739 | 0.487 | -1.131 | -0.891 | 0.008 | 0.145 | 0.145 | 0 | 0.32 | 0.09 | 0.24 | 0.05 | 0.31 | 0.18 |
| 2 | 20412227 | rs3771243 | 0.172 | -0.074 | -0.337 | -0.764 | 0.023 | 0.032 | 0.032 | 0 | 0.5 | 0.44 | 0.34 | 0.18 | 0.39 | 0.37 |
| 2 | 20420732 | rs4666298 | 0.699 | 1.073 | 0.036 | -0.085 | 0.088 | 0.102 | 0.102 | 0 | 0.06 | 0.16 | 0.21 | 0.27 | 0.14 | 0.19 |
| 2 | 20427191 | rs7585590 | 0.21 | 0.815 | 0.205 | -0.37 | 0.015 | 0.018 | 0.018 | 0 | 0.49 | 0.44 | 0.33 | 0.18 | 0.39 | 0.36 |
| 2 | 20428737 | rs10495704 | 0.843 | 0.669 | 0.386 | -0.459 | 0.041 | 0.098 | 0.098 | 0 | 0.41 | 0.19 | 0.29 | 0.17 | 0.36 | 0.32 |
| 2 | 20430049 | rs12373772 | 0.202 | 0.663 | 0.576 | -0.465 | 0.029 | 0.05 | 0.05 | 0 | 0.48 | 0.37 | 0.32 | 0.18 | 0.39 | 0.36 |
| 2 | 20445639 | rs2110410 | 1.334 | 0.549 | 0.443 | -0.517 | 0.004 | 0.157 | 0.157 | 0 | 0.35 | 0.07 | 0.26 | 0.08 | 0.33 | 0.27 |
| 2 | 20447610 | rs6744963 | -0.923 | 0.562 | 1.119 | -0.494 | -0.019 | -0.048 | -0.048 | 0.071 | 0.09 | 0.28 | 0.04 | 0 | 0.04 | 0.04 |
| 2 | 20448561 | rs1043419 | 0.828 | 0.644 | 0.417 | -0.842 | -0.053 | 0.051 | 0.051 | 0 | 0.17 | 0.01 | 0.11 | 0.06 | 0.18 | 0.2 |
| 2 | 20449174 | rs1043400 | 1.522 | 0.619 | 0.569 | -0.749 | 0.037 | 0.11 | 0.11 | 0 | 0.41 | 0.17 | 0.29 | 0.2 | 0.35 | 0.31 |
| 2 | 20467929 | rs4666299 | 0.456 | 0.732 | 0.801 | -0.889 | -0.005 | -0.001 | -0.001 | 0 | 0.23 | 0.23 | 0.46 | 0.74 | 0.28 | 0.42 |
| 2 | 20472847 | rs10164749 | -1.696 | 0.992 | 1.051 | -0.91 | 0.022 | -0.028 | -0.028 | 0 | 0.94 | 0.89 | 0.92 | 1 | 0.86 | 0.99 |
| 2 | 20475446 | rs10856797 | 1.035 | 0.853 | 0.949 | -0.921 | 0.041 | 0.082 | 0.082 | 0 | 0.49 | 0.34 | 0.32 | 0.21 | 0.39 | 0.36 |
| 2 | 20479467 | rs7606633 | -0.648 | 0.811 | 0.998 | -0.99 | -0.051 | 0.081 | 0.081 | 0.074 | 0.78 | 0.95 | 0.86 | 0.94 | 0.75 | 0.79 |
| 2 | 20491321 | rs11896241 | 1.732 | 0.644 | 1.034 | -0.98 | 0.037 | 0.11 | 0.11 | 0 | 0.4 | 0.13 | 0.28 | 0.2 | 0.33 | 0.32 |
| 2 | 20502198 | rs6748274 | 1.819 | 0.651 | 0.909 | -0.96 | 0.034 | 0.118 | 0.118 | 0 | 0.4 | 0.12 | 0.29 | 0.2 | 0.35 | 0.32 |
| 2 | 20520878 | rs6531230 | -0.367 | 0.734 | 0.7 | -0.896 | -0.007 | 0.007 | 0.007 | 0.298 | 0.78 | 0.83 | 0.85 | 0.95 | 0.75 | 0.79 |
| 2 | 20521954 | rs7565347 | -0.312 | 0.8 | 0.261 | -0.856 | -0.02 | -0.042 | -0.042 | 0 | 0.08 | 0.16 | 0.02 | 0 | 0.04 | 0.04 |
| 2 | 20529948 | rs10153828 | 0.132 | 0.86 | -0.274 | -0.826 | 0.007 | 0.004 | 0.004 | 0 | 0.06 | 0.02 | 0.02 | 0 | 0.04 | 0.04 |
| 2 | 20536055 | rs6706872 | -0.521 | 1.03 | -0.229 | -0.658 | -0.052 | 0.073 | 0.073 | 0 | 0.8 | 0.95 | 0.86 | 0.94 | 0.75 | 0.79 |
| 2 | 20558421 | rs4666382 | -0.13 | 1.117 | 0.123 | -0.323 | 0.009 | -0.002 | -0.002 | 0 | 0.24 | 0.29 | 0.46 | 0.75 | 0.28 | 0.42 |
| 2 | 20581058 | rs343161 | 0.621 | 0.747 | -0.254 | -0.473 | -0.002 | 0.044 | 0.044 | 0.123 | 0.56 | 0.3 | 0.55 | 0.72 | 0.54 | 0.56 |
| 2 | 20590635 | rs6723605 | -0.046 | 0.759 | -0.223 | -0.04 | 0 | 0.012 | 0.012 | 0 | 0.79 | 0.77 | 0.68 | 0.48 | 0.76 | 0.75 |
| 2 | 20592798 | rs343155 | -0.24 | 0.744 | -0.437 | 0.023 | -0.006 | -0.087 | -0.087 | 0 | 0.29 | 0.14 | 0.28 | 0.47 | 0.27 | 0.42 |
| 2 | 20595958 | rs343151 | -0.712 | 0.952 | -0.06 | -0.127 | 0.053 | 0.028 | 0.028 | 0 | 0.86 | 0.75 | 0.7 | 0.75 | 0.77 | 0.8 |
| 2 | 20596551 | rs343150 | 0.397 | 0.817 | -0.284 | -0.152 | -0.041 | -0.156 | -0.156 | 0.157 | 0.24 | 0.03 | 0.25 | 0.48 | 0.24 | 0.38 |
| 2 | 20600312 | rs6742129 | -0.219 | 0.597 | 0.148 | -0.201 | -0.024 | -0.006 | -0.006 | 0 | 0.22 | 0.6 | 0.17 | 0.19 | 0.14 | 0.07 |
| 2 | 20601563 | rs343144 | -0.091 | 0.659 | 0.095 | -0.245 | 0.043 | 0.013 | 0.013 | 0 | 0.82 | 0.73 | 0.68 | 0.75 | 0.73 | 0.77 |
| 2 | 20602382 | rs13431644 | 0.01 | 0.629 | 0.338 | -0.249 | -0.026 | 0.006 | 0.006 | 0.649 | 0.21 | 0.45 | 0.16 | 0.19 | 0.15 | 0.07 |
| 2 | 20603229 | rs343142 | -0.593 | 0.932 | 0.294 | -0.175 | 0.035 | 0.016 | 0.016 | 0 | 0.87 | 0.82 | 0.71 | 0.76 | 0.77 | 0.81 |
| 2 | 20605249 | rs16987610 | -0.359 | 0.873 | 0.208 | -0.259 | -0.007 | -0.034 | -0.034 | 0 | 0.07 | 0.14 | 0.04 | 0 | 0.04 | 0.05 |
| 2 | 20609954 | rs953927 | -0.591 | 0.873 | 0.146 | -0.197 | 0.069 | 0.012 | 0.012 | 0 | 0.21 | 0.41 | 0.33 | 0.24 | 0.27 | 0.25 |
| 2 | 20616033 | rs6710171 | 0.74 | 0.822 | 0.061 | -0.669 | -0.002 | -0.024 | -0.024 | 0 | 0.44 | 0.44 | 0.4 | 0.15 | 0.44 | 0.35 |
| 2 | 20618441 | rs2128833 | -0.362 | 1.244 | 0.063 | -0.357 | 0.027 | -0.018 | -0.018 | 0 | 0.14 | 0.18 | 0.12 | 0.1 | 0.21 | 0.2 |
| 2 | 20620093 | rs11096663 | -0.75 | 0.685 | -0.047 | -0.339 | -0.003 | -0.012 | -0.012 | 0 | 0.4 | 0.56 | 0.56 | 0.67 | 0.36 | 0.57 |
| 2 | 20623085 | rs10186941 | -0.814 | 0.54 | 0.406 | -0.414 | 0.009 | 0.02 | 0.02 | 0 | 0.42 | 0.73 | 0.59 | 0.67 | 0.36 | 0.59 |
| 2 | 20624426 | rs6721647 | -0.687 | -0.643 | 0.052 | -0.897 | 0.012 | 0.011 | 0.011 | 0.988 | 0.49 | 0.54 | 0.26 | 0.15 | 0.4 | 0.18 |
| 2 | 20624972 | rs4643494 | 1.134 | -0.58 | -0.086 | -0.698 | -0.022 | 0.125 | 0.125 | 0 | 0.14 | 0.2 | 0.33 | 0.41 | 0.1 | 0.3 |
| 2 | 20625174 | rs6743033 | 0.607 | -0.701 | 0.087 | -0.757 | 0.001 | -0.014 | -0.014 | 0 | 0.29 | 0.21 | 0.13 | 0.07 | 0.24 | 0.07 |
| 2 | 20629723 | rs967151 | 0.207 | -0.476 | 0.159 | -0.165 | 0.001 | 0.009 | 0.009 | 0 | 0.16 | 0.13 | 0.12 | 0.09 | 0.16 | 0.11 |
| 2 | 20630272 | rs872284 | 0.347 | -0.55 | 0.232 | -0.023 | -0.04 | 0.183 | 0.183 | 0 | 0.14 | 0.3 | 0.27 | 0.41 | 0.09 | 0.29 |
| 2 | 20636133 | rs342101 | -0.543 | -0.304 | 0.151 | -0.981 | -0.018 | -0.046 | -0.046 | 0 | 0.19 | 0.38 | 0.13 | 0.09 | 0.13 | 0.12 |
| 2 | 20648211 | rs1062292 | 0.241 | -0.32 | -0.913 | -1.551 | -0.059 | 0.147 | 0.147 | 0.491 | 0.41 | 0.21 | 0.64 | 0.85 | 0.48 | 0.72 |
| 2 | 20649490 | rs7579125 | 0.84 | -0.657 | -0.637 | -1.654 | -0.019 | -0.044 | -0.044 | 0 | 0.1 | 0.16 | 0.07 | 0 | 0.06 | 0.04 |
| 2 | 20650965 | rs342054 | -0.612 | -0.598 | -0.651 | -1.826 | 0.02 | 0.097 | 0.097 | 0.111 | 0.6 | 0.76 | 0.81 | 0.91 | 0.65 | 0.79 |
| 2 | 20652507 | rs342056 | -0.863 | -0.792 | -0.81 | -1.979 | -0.01 | 0.033 | 0.033 | 0.05 | 0.75 | 0.81 | 0.85 | 0.91 | 0.71 | 0.84 |
| 2 | 20653928 | rs4666396 | 0.275 | -0.811 | -0.897 | -2.039 | -0.023 | 0.222 | 0.222 | 0 | 0.56 | 0.14 | 0.66 | 0.89 | 0.54 | 0.78 |
| 2 | 20654782 | rs342105 | -0.563 | -0.998 | -0.781 | -2.03 | -0.043 | 0.154 | 0.154 | 0.226 | 0.64 | 0.88 | 0.79 | 0.92 | 0.6 | 0.8 |
| 2 | 20655947 | rs1453051 | 0.043 | -1.019 | -0.084 | -2.019 | 0.004 | -0.045 | -0.045 | 0.175 | 0.57 | 0.54 | 0.68 | 0.86 | 0.5 | 0.76 |
| 2 | 20656478 | rs342107 | -0.309 | -0.838 | -0.967 | -1.934 | -0.052 | 0.106 | 0.106 | 0.035 | 0.62 | 0.81 | 0.72 | 0.86 | 0.52 | 0.76 |
| 2 | 20665887 | rs13021908 | 0.624 | -0.879 | -1.253 | -1.93 | -0.109 | 0.16 | 0.16 | 1.859 | 0.39 | 0.11 | 0.61 | 0.84 | 0.45 | 0.7 |
| 2 | 20668253 | rs342119 | -0.474 | -0.85 | -1.208 | -1.941 | -0.042 | 0.166 | 0.166 | 0.16 | 0.63 | 0.84 | 0.79 | 0.92 | 0.6 | 0.8 |
| 2 | 20675366 | rs11898154 | 0.151 | -0.158 | -0.755 | -1.489 | -0.04 | 0.077 | 0.077 | 2.347 | 0.39 | 0.21 | 0.63 | 0.84 | 0.45 | 0.68 |
| 2 | 20682303 | rs12997261 | 0.179 | -0.187 | -1.385 | -1.161 | -0.094 | 0.105 | 0.105 | 0.645 | 0.37 | 0.13 | 0.63 | 0.83 | 0.44 | 0.66 |
| 2 | 20682493 | rs13016649 | -0.02 | -0.112 | -1.462 | -1.025 | -0.119 | 0.084 | 0.084 | 0 | 0.39 | 0.11 | 0.63 | 0.81 | 0.46 | 0.57 |
| 2 | 20685471 | rs342070 | -0.64 | 0.363 | -0.962 | -0.442 | -0.003 | -0.004 | -0.004 | 0 | 0.74 | 0.77 | 0.69 | 0.38 | 0.75 | 0.53 |
| 2 | 20688941 | rs931257 | 0.475 | -0.008 | -0.92 | -0.925 | -0.015 | -0.03 | -0.03 | 0 | 0.36 | 0.58 | 0.22 | 0.13 | 0.24 | 0.29 |
| 2 | 20690143 | rs2046335 | -0.215 | 0.272 | -1.08 | -1.007 | 0.025 | 0.044 | 0.044 | 0.627 | 0.52 | 0.67 | 0.55 | 0.32 | 0.44 | 0.36 |
| 2 | 20695707 | rs4666412 | 0.993 | 0.282 | -1.071 | -0.985 | -0.008 | -0.011 | -0.011 | 0.107 | 0.16 | 0.1 | 0.35 | 0.19 | 0.2 | 0.07 |
| 2 | 20696241 | rs16987690 | 0.68 | 0.079 | -1.099 | -1.054 | -0.066 | -0.1 | -0.1 | 1.851 | 0.34 | 0.51 | 0.15 | 0.1 | 0.18 | 0.22 |
| 2 | 20696256 | rs342096 | 0.545 | 0.787 | -0.971 | -0.71 | -0.001 | -0.015 | -0.015 | 0 | 0.24 | 0.23 | 0.15 | 0.06 | 0.31 | 0.18 |

| | | | | | | | | | | | | | | | | |
|---|---|---|---|---|---|---|---|---|---|---|---|---|---|---|---|---|
| 2 | 20696907 | rs342095 | -0.909 | 0.79 | -0.831 | -0.664 | 0.004 | -0.017 | -0.017 | 0 | 0.43 | 0.51 | 0.56 | 0.27 | 0.57 | 0.31 |
| 2 | 20697925 | rs342092 | -0.055 | 0.846 | -0.961 | -0.578 | -0.001 | -0.011 | -0.011 | 0 | 0.25 | 0.23 | 0.15 | 0.06 | 0.31 | 0.17 |
| 2 | 20700786 | rs961496 | -0.235 | 0.955 | -0.953 | 0.078 | 0.005 | 0.016 | 0.016 | 0 | 0.38 | 0.37 | 0.38 | 0.77 | 0.31 | 0.63 |
| 2 | 20702032 | rs12162347 | -0.712 | 2.229 | -1.018 | 1.061 | -0.076 | -0.075 | -0.075 | 0 | 0.11 | 0.01 | 0.23 | 0.34 | 0.23 | 0.07 |
| 2 | 20703869 | rs12710739 | -0.277 | 1.91 | -0.289 | 0.286 | 0.006 | -0.089 | -0.089 | 0.336 | 0.3 | 0.38 | 0.18 | 0.1 | 0.27 | 0.21 |
| 2 | 20704614 | rs12997448 | 0.117 | 1.858 | -0.209 | 0.276 | -0.041 | 0.1 | 0.1 | 0 | 0.26 | 0.06 | 0.16 | 0.1 | 0.26 | 0.2 |
| 2 | 20705910 | rs342076 | 0.356 | 1.494 | 0.538 | 0.222 | -0.001 | -0.054 | -0.054 | 1.01 | 0.59 | 0.53 | 0.73 | 0.8 | 0.67 | 0.69 |
| 2 | 20706742 | rs342075 | 1.333 | 1.405 | 0.343 | 0.225 | 0.079 | 0.137 | 0.137 | 0 | 0.39 | 0.16 | 0.19 | 0.14 | 0.31 | 0.28 |
| 2 | 20707384 | rs2084920 | -0.382 | 1.54 | 0.364 | 0.616 | 0.001 | 0.018 | 0.018 | 0 | 0.22 | 0.24 | 0.39 | 0.43 | 0.21 | 0.43 |
| 2 | 20708719 | rs12618114 | -0.746 | 1.435 | 0.557 | 0.658 | 0.01 | 0.002 | 0.002 | 0 | 0.49 | 0.4 | 0.47 | 0.52 | 0.52 | 0.38 |
| 2 | 20709694 | rs342138 | 0.145 | 1.713 | 1.064 | 1.549 | 0.071 | 0.203 | 0.203 | 0 | 0.91 | 0.76 | 0.69 | 0.67 | 0.85 | 0.67 |
| 2 | 20711656 | rs11096667 | -0.539 | 1.274 | -0.136 | 1.273 | -0.023 | -0.005 | -0.005 | 0 | 0.08 | 0.03 | 0.23 | 0.31 | 0.2 | 0.08 |
| 2 | 20712253 | rs921598 | -0.277 | 0.22 | -0.41 | 0.471 | -0.031 | -0.007 | -0.007 | 0.915 | 0.12 | 0.17 | 0.03 | 0.09 | 0.05 | 0.09 |
| 2 | 20714027 | rs6728695 | -0.186 | 0.101 | -1.342 | -0.131 | 0.023 | 0.076 | 0.076 | 2.909 | 0.31 | 0.17 | 0.13 | 0.13 | 0.22 | 0.21 |
| 2 | 20714385 | rs342132 | -0.427 | 0.482 | -1.818 | 0.383 | -0.031 | -0.027 | -0.027 | 0.114 | 0.37 | 0.13 | 0.26 | 0.4 | 0.35 | 0.31 |
| 2 | 20715656 | rs342130 | 0.081 | 0.311 | -0.935 | 0.156 | 0.005 | -0.028 | -0.028 | 0 | 0.38 | 0.44 | 0.71 | 0.74 | 0.56 | 0.43 |
| 2 | 20716123 | rs342129 | -0.132 | 0.096 | -0.663 | 0.806 | 0.006 | 0.016 | 0.016 | 1.608 | 0.43 | 0.3 | 0.49 | 0.36 | 0.57 | 0.44 |
| 2 | 20719131 | rs7568428 | -0.391 | -0.384 | -0.214 | 0.356 | 0.026 | 0.061 | 0.061 | 0.518 | 0.48 | 0.66 | 0.75 | 0.76 | 0.63 | 0.57 |
| 2 | 20720237 | rs4666415 | -0.119 | 0.17 | -0.025 | 0.414 | 0.014 | 0.009 | 0.009 | 0 | 0.3 | 0.38 | 0.5 | 0.56 | 0.4 | 0.41 |
| 2 | 20720724 | rs7572838 | 0.617 | 0.3 | -0.071 | 0.288 | 0.014 | 0.005 | 0.005 | 0.06 | 0.18 | 0.28 | 0.26 | 0.21 | 0.23 | 0.16 |
| 2 | 20724694 | rs11899825 | -0.329 | 0.28 | -0.183 | 0.568 | -0.017 | -0.013 | -0.013 | 0 | 0.39 | 0.29 | 0.55 | 0.49 | 0.53 | 0.41 |
| 2 | 20728089 | rs7579437 | 0.039 | -0.048 | 0.022 | 0.121 | -0.026 | -0.092 | -0.092 | 0.842 | 0.78 | 0.47 | 0.87 | 0.96 | 0.84 | 0.84 |
| 2 | 20728931 | rs2289083 | -0.141 | -0.033 | 0.271 | 0.127 | 0.033 | 0.001 | 0.001 | 0.036 | 0.08 | 0.23 | 0.06 | 0.04 | 0.07 | 0.1 |
| 2 | 20729597 | rs2289084 | -0.848 | -0.12 | 0.158 | 0.153 | -0.002 | -0.01 | -0.01 | 0 | 0.16 | 0.37 | 0.15 | 0.13 | 0.12 | 0.28 |
| 2 | 20732911 | rs12710741 | -1.014 | -0.191 | 0.066 | 0.085 | -0.03 | -0.083 | -0.083 | 0.444 | 0.87 | 0.74 | 0.93 | 1 | 0.91 | 0.98 |
| 2 | 20734842 | rs11096668 | 0.78 | -0.134 | -0.218 | 0.259 | -0.041 | -0.048 | -0.048 | 0.923 | 0.22 | 0.09 | 0.35 | 0.29 | 0.3 | 0.32 |
| 2 | 20738876 | rs11900205 | 0.843 | -0.251 | -0.045 | 0.201 | -0.023 | -0.066 | -0.066 | 0 | 0.13 | 0.21 | 0.07 | 0 | 0.09 | 0.06 |
| 2 | 20743716 | rs11096672 | 1.114 | -0.319 | -0.555 | 0.293 | 0.035 | -0.026 | -0.026 | 0.145 | 0.37 | 0.14 | 0.29 | 0.42 | 0.31 | 0.31 |
| 2 | 20743917 | rs6725427 | 0.028 | -0.341 | -0.195 | 0.321 | 0.024 | -0.035 | -0.035 | 0.407 | 0.08 | 0.27 | 0.04 | 0 | 0.06 | 0.01 |
| 2 | 20744067 | rs6728591 | 0.188 | -0.242 | -0.407 | 0.372 | -0.003 | 0.002 | 0.002 | 0 | 0.59 | 0.49 | 0.51 | 0.62 | 0.58 | 0.52 |
| 2 | 20746932 | rs10183220 | 0.678 | -0.046 | -1.396 | 0.55 | 0.007 | 0.009 | 0.009 | 0 | 0.51 | 0.2 | 0.46 | 0.6 | 0.52 | 0.47 |
| 2 | 20747671 | rs12052803 | 0.311 | 0.515 | -1.481 | 1.105 | -0.042 | -0.071 | -0.071 | 0 | 0.21 | 0.08 | 0.33 | 0.32 | 0.29 | 0.34 |
| 2 | 20748457 | rs10495706 | 0.466 | 0.242 | -1.699 | 0.89 | -0.061 | -0.157 | -0.157 | 0.042 | 0.25 | 0.03 | 0.28 | 0.44 | 0.3 | 0.28 |
| 2 | 20749369 | rs4666419 | 0.02 | 0.573 | -0.624 | 0.529 | 0.076 | 0.015 | 0.015 | 0.652 | 0.22 | 0.5 | 0.3 | 0.28 | 0.24 | 0.33 |
| 2 | 20749739 | rs10495707 | 0.814 | 0.063 | -1.26 | 0.237 | -0.003 | -0.028 | -0.028 | 0 | 0.3 | 0.12 | 0.42 | 0.37 | 0.3 | 0.38 |
| 2 | 20751009 | rs2348868 | 0.932 | -0.191 | -1.254 | -0.094 | 0.025 | -0.006 | -0.006 | 0 | 0.27 | 0.17 | 0.1 | 0.01 | 0.17 | 0.04 |
| 2 | 20751163 | rs4666420 | 0.019 | 1.2 | -0.492 | 0.877 | 0.009 | 0.014 | 0.014 | 1.563 | 0.21 | 0.24 | 0.28 | 0.56 | 0.31 | 0.21 |
| 2 | 20752152 | rs2082873 | -1.285 | 1.216 | -0.25 | 0.929 | 0 | -0.005 | -0.005 | 0 | 0.4 | 0.59 | 0.46 | 0.63 | 0.53 | 0.58 |
| 2 | 20752478 | rs17656011 | 0.1 | 1.214 | -0.333 | 0.543 | 0.043 | -0.034 | -0.034 | 0 | 0.19 | 0.35 | 0.19 | 0.11 | 0.24 | 0.36 |
| 2 | 20752612 | rs11096674 | -0.635 | 0.301 | -1.003 | 0.342 | -0.001 | 0 | 0 | 4.694 | 0.4 | 0.61 | 0.46 | 0.67 | 0.52 | 0.58 |
| 2 | 20756900 | rs2060916 | 0.449 | -0.53 | -1.306 | 0.337 | 0.035 | 0.013 | 0.013 | 0.18 | 0.74 | 0.2 | 0.81 | 0.77 | 0.87 | 0.66 |
| 2 | 20760365 | rs6717824 | 0.055 | -0.19 | -1.529 | 0.309 | -0.059 | -0.037 | -0.037 | 0 | 0.15 | 0.01 | 0.21 | 0.17 | 0.22 | 0.13 |
| 2 | 20762613 | rs1435849 | -0.182 | -0.505 | -0.624 | 0.256 | -0.009 | 0.015 | 0.015 | 0.224 | 0.86 | 0.74 | 0.9 | 0.81 | 0.89 | 0.74 |
| 2 | 20763911 | rs16987836 | 0.846 | -0.614 | -1.853 | 0.257 | 0.002 | -0.128 | -0.128 | 0 | 0.36 | 0.02 | 0.43 | 0.38 | 0.28 | 0.33 |
| 2 | 20765963 | rs13006062 | -0.605 | -0.785 | -1.036 | -0.045 | 0.001 | 0.001 | 0.001 | 0 | 0.43 | 0.45 | 0.47 | 0.33 | 0.43 | 0.3 |
| 2 | 20768247 | rs10186749 | -0.642 | -1.041 | -0.205 | -0.164 | -0.021 | -0.057 | -0.057 | 0 | 0.87 | 0.82 | 0.94 | 1 | 0.94 | 0.96 |
| 2 | 20777149 | rs10166792 | -0.168 | -1.212 | 0.314 | -0.083 | -0.087 | -0.039 | -0.039 | 0 | 0.8 | 0.56 | 0.89 | 0.86 | 0.94 | 0.8 |
| 2 | 20781537 | rs4666431 | 0.512 | -1.344 | 0.488 | -0.245 | -0.056 | -0.052 | -0.052 | 0 | 0.73 | 0.57 | 0.84 | 0.84 | 0.83 | 0.7 |
| 2 | 20784873 | rs11694519 | 1.506 | -1.402 | -0.23 | -0.062 | 0.012 | -0.061 | -0.061 | 0 | 0.31 | 0.03 | 0.31 | 0.32 | 0.19 | 0.27 |
| 2 | 20789958 | rs766474 | 0.002 | -1.277 | 0.66 | -0.317 | 0.024 | 0.015 | 0.015 | 0.691 | 0.2 | 0.11 | 0.1 | 0.15 | 0.09 | 0.21 |
| 2 | 20791125 | rs921599 | 1.167 | -1.292 | 0.591 | -0.369 | 0.084 | 0.083 | 0.083 | 0.023 | 0.63 | 0.2 | 0.77 | 0.81 | 0.77 | 0.62 |
| 2 | 20794079 | rs7595132 | -0.336 | -1.25 | 0.642 | -0.285 | -0.008 | -0.016 | -0.016 | 0 | 0.12 | 0.19 | 0.1 | 0.05 | 0.12 | 0.15 |
| 2 | 20797032 | rs13031724 | 0.757 | -1.122 | -0.297 | -0.22 | 0.043 | 0.039 | 0.039 | 0.128 | 0.65 | 0.28 | 0.78 | 0.81 | 0.79 | 0.62 |
| 2 | 20804471 | rs1109689 | 0.333 | 0.033 | 0.076 | 0.187 | -0.003 | -0.023 | -0.023 | 0.085 | 0.05 | 0.11 | 0.03 | 0 | 0.07 | 0.01 |
| 2 | 20814371 | rs13027694 | -0.105 | 1.427 | -0.573 | 0.475 | -0.056 | -0.003 | -0.003 | 0 | 0.16 | 0.08 | 0.17 | 0.15 | 0.31 | 0.12 |
| 2 | 20814715 | rs6728166 | 0.211 | 1.287 | -0.641 | 0.202 | -0.002 | 0.031 | 0.031 | 0 | 0.21 | 0.14 | 0.15 | 0.09 | 0.21 | 0.19 |
| 2 | 20818286 | rs3821217 | -0.261 | 1.227 | -0.147 | 0.172 | -0.001 | -0.024 | -0.024 | 0.69 | 0.79 | 0.74 | 0.84 | 0.91 | 0.79 | 0.81 |
| 2 | 20818458 | rs3732151 | 1.82 | 1.092 | 0.392 | 0.074 | 0.091 | 0.032 | 0.032 | 5.086 | 0.48 | 0.24 | 0.39 | 0.6 | 0.28 | 0.46 |
| 2 | 20818764 | rs3732149 | 0.627 | 1.046 | 0.507 | 0.048 | -0.009 | 0.024 | 0.024 | 0 | 0.18 | 0.12 | 0.14 | 0.09 | 0.2 | 0.19 |
| 2 | 20819291 | rs870309 | -0.276 | 1.083 | 0.799 | 0.03 | -0.013 | 0 | 0 | 1.428 | 0.2 | 0.19 | 0.27 | 0.18 | 0.37 | 0.18 |
| 2 | 20821354 | rs6721558 | -0.17 | 1.021 | 0.942 | 0.112 | -0.004 | -0.031 | -0.031 | 0 | 0.78 | 0.68 | 0.84 | 0.85 | 0.79 | 0.81 |
| 2 | 20824559 | rs2305457 | 2.171 | 0.851 | 0.755 | 0.039 | 0.073 | -0.125 | -0.125 | 0 | 0.43 | 0.03 | 0.37 | 0.49 | 0.26 | 0.33 |
| 2 | 20824708 | rs10191266 | 1.606 | 1.264 | 1.107 | -0.184 | -0.012 | -0.006 | -0.006 | 0 | 0.55 | 0.46 | 0.45 | 0.6 | 0.35 | 0.49 |
| 2 | 20826748 | rs4456661 | 1.347 | 1.314 | 1.029 | -0.19 | 0.032 | -0.024 | -0.024 | 0.411 | 0.7 | 0.39 | 0.65 | 0.82 | 0.56 | 0.79 |
| 2 | 20827198 | rs6531249 | 0.131 | 1.36 | 1.11 | -0.167 | 0.099 | 0.001 | 0.001 | 0.229 | 0.16 | 0.36 | 0.26 | 0.15 | 0.32 | 0.13 |
| 2 | 20828748 | rs934601 | 1.228 | 1.221 | 0.73 | -0.155 | 0.023 | 0.039 | 0.039 | 2.164 | 0.54 | 0.38 | 0.52 | 0.73 | 0.36 | 0.61 |
| 2 | 20837469 | rs1865678 | 1.867 | 1.421 | 0.285 | 0.037 | 0.085 | 0.009 | 0.009 | 3.963 | 0.47 | 0.24 | 0.31 | 0.57 | 0.28 | 0.46 |
| 2 | 20837719 | rs11680086 | 0.092 | 1.481 | 0.259 | 0.023 | 0.01 | 0 | 0 | 0 | 0.22 | 0.26 | 0.27 | 0.22 | 0.26 | 0.34 |
| 2 | 20838318 | rs3796064 | 1.951 | 1.465 | 0.126 | 0.046 | 0.095 | 0.003 | 0.003 | 3.887 | 0.47 | 0.23 | 0.31 | 0.56 | 0.28 | 0.45 |

| | | | | | | | | | | | | | | | | |
|---|---|---|---|---|---|---|---|---|---|---|---|---|---|---|---|---|
| 2 | 20839003 | rs10166174 | -1.268 | 2.21 | -0.028 | 0.512 | 0.017 | -0.008 | -0.008 | 0 | 0.29 | 0.47 | 0.4 | 0.25 | 0.34 | 0.36 |
| 2 | 20839346 | rs7586298 | -0.016 | 2.387 | -0.986 | 0.713 | 0.004 | -0.021 | -0.021 | 0.443 | 0.18 | 0.21 | 0.15 | 0.1 | 0.2 | 0.2 |
| 2 | 20841182 | rs1344836 | -0.712 | 1.313 | -0.945 | 0.541 | -0.006 | -0.004 | -0.004 | 0 | 0.32 | 0.29 | 0.5 | 0.39 | 0.53 | 0.42 |
| 2 | 20845273 | rs2304423 | 0.391 | 1.042 | -1.235 | 0.219 | -0.014 | 0.011 | 0.011 | 0 | 0.68 | 0.79 | 0.73 | 0.71 | 0.66 | 0.83 |
| 2 | 20855176 | rs4666453 | 0.585 | 0.791 | -0.954 | 0.143 | 0.014 | 0.014 | 0.014 | 0 | 0.46 | 0.37 | 0.56 | 0.57 | 0.43 | 0.52 |
| 2 | 20855911 | rs12615616 | -0.264 | 0.505 | -0.853 | 0.091 | -0.016 | 0.003 | 0.003 | 0 | 0.41 | 0.28 | 0.32 | 0.43 | 0.42 | 0.36 |
| 2 | 20863286 | rs1013363 | 0.584 | 0.699 | -1.175 | 0.12 | -0.006 | 0.011 | 0.011 | 0 | 0.47 | 0.43 | 0.62 | 0.57 | 0.52 | 0.55 |
| 2 | 20868179 | rs7589372 | 0.023 | 0.743 | -1.744 | 0.009 | -0.027 | 0.06 | 0.06 | 0.076 | 0.69 | 0.81 | 0.83 | 0.84 | 0.69 | 0.91 |
| 2 | 20876386 | rs746776 | -0.681 | -0.241 | -1.577 | 0.606 | 0.016 | -0.012 | -0.012 | 0 | 0.28 | 0.72 | 0.29 | 0.36 | 0.17 | 0.42 |
| 2 | 20876760 | rs11678116 | 0.19 | 0.285 | -2.259 | 0.418 | -0.043 | -0.019 | -0.019 | 0 | 0.57 | 0.18 | 0.54 | 0.46 | 0.56 | 0.69 |
| 2 | 20878406 | rs3072 | 0.706 | 0.555 | -2.742 | 0.205 | 0.028 | -0.103 | -0.103 | 0 | 0.46 | 0.07 | 0.3 | 0.48 | 0.37 | 0.4 |
| 2 | 20878719 | rs9679195 | -0.241 | 0.666 | -2.03 | 0.288 | -0.009 | 0.093 | 0.093 | 0 | 0.89 | 0.81 | 0.84 | 0.7 | 0.9 | 0.88 |
| 2 | 20878820 | rs7255 | -0.547 | 0.278 | -1.094 | 0.038 | 0.001 | 0.015 | 0.015 | 0 | 0.43 | 0.4 | 0.52 | 0.22 | 0.53 | 0.43 |
| 2 | 20880345 | rs11903198 | -0.178 | 0.981 | -1.159 | -0.288 | -0.002 | -0.013 | -0.013 | 0 | 0.3 | 0.32 | 0.3 | 0.1 | 0.34 | 0.3 |
| 2 | 20882056 | rs13394027 | 2.245 | 0.681 | -1.469 | -0.333 | 0.037 | -0.178 | -0.178 | 0.608 | 0.3 | 0.02 | 0.2 | 0.45 | 0.23 | 0.32 |
| 2 | 20882949 | rs11675383 | -0.589 | 1.306 | -0.942 | -0.157 | 0.003 | -0.043 | -0.043 | 0.225 | 0.4 | 0.46 | 0.33 | 0.11 | 0.46 | 0.33 |
| 2 | 20883216 | rs10170771 | 1.978 | 1.169 | -1.053 | -0.152 | 0.07 | -0.177 | -0.177 | 0.063 | 0.35 | 0.02 | 0.27 | 0.45 | 0.25 | 0.36 |
| 2 | 20883953 | rs11693547 | 0.866 | 1.199 | -1.015 | -0.137 | -0.009 | 0 | 0 | 0 | 0.12 | 0.09 | 0.21 | 0.14 | 0.18 | 0.13 |
| 2 | 20884973 | rs9085 | 1.048 | 1.558 | -1.207 | -0.337 | 0.039 | -0.004 | -0.004 | 0 | 0.52 | 0.26 | 0.37 | 0.52 | 0.41 | 0.47 |
| 2 | 20888021 | rs6531253 | 0.1 | 1.57 | -1.11 | -0.339 | 0.007 | -0.038 | -0.038 | 0 | 0.17 | 0.22 | 0.1 | 0.06 | 0.16 | 0.12 |
| 2 | 20888419 | rs13423093 | 1.181 | 1.455 | -0.463 | -0.337 | 0.054 | -0.028 | -0.028 | 0.006 | 0.5 | 0.18 | 0.35 | 0.52 | 0.4 | 0.47 |
| 2 | 20889460 | rs16987985 | -0.525 | 1.804 | 0.344 | 0.105 | -0.001 | -0.041 | -0.041 | 0.314 | 0.18 | 0.23 | 0.09 | 0.07 | 0.14 | 0.17 |
| 2 | 20889714 | rs340595 | -0.612 | 1.924 | 0.479 | 0.154 | 0.057 | 0.131 | 0.131 | 0 | 0.23 | 0.44 | 0.39 | 0.42 | 0.29 | 0.25 |
| 2 | 20889754 | rs340596 | -1.369 | 1.707 | 0.011 | 0.198 | -0.024 | -0.018 | -0.018 | 2.981 | 0.41 | 0.71 | 0.48 | 0.49 | 0.42 | 0.42 |
| 2 | 20891799 | rs340598 | -1.383 | 1.976 | 0.606 | 0.639 | 0.013 | 0.002 | 0.002 | 0 | 0.37 | 0.6 | 0.5 | 0.37 | 0.43 | 0.38 |
| 2 | 20892006 | rs340600 | -0.83 | 2.261 | 0.757 | 0.493 | -0.01 | -0.07 | -0.07 | 0 | 0.23 | 0.54 | 0.25 | 0.08 | 0.18 | 0.14 |
| 2 | 20897345 | rs4971525 | -1.483 | 2.203 | 1.282 | 0.603 | 0.042 | 0.082 | 0.082 | 0 | 0.26 | 0.59 | 0.41 | 0.49 | 0.34 | 0.38 |
| 2 | 20897536 | rs4971548 | -1.078 | 2.372 | 1.444 | 0.804 | 0 | 0.06 | 0.06 | 0 | 0.08 | 0.16 | 0.08 | 0.26 | 0.1 | 0.18 |
| 2 | 20906599 | rs11674108 | -0.868 | 2.422 | 1.434 | 0.761 | 0.007 | 0.017 | 0.017 | 0 | 0.11 | 0.11 | 0.34 | 0.35 | 0.26 | 0.3 |
| 2 | 20924380 | rs17664438 | -0.768 | 2.413 | 1.443 | 0.757 | 0.003 | 0.012 | 0.012 | 0.018 | 0.11 | 0.09 | 0.34 | 0.35 | 0.26 | 0.3 |
| 2 | 20926462 | rs7576149 | -0.39 | 2.33 | 1.638 | 0.565 | 0.003 | -0.096 | -0.096 | 0.003 | 0.17 | 0.32 | 0.12 | 0 | 0.16 | 0.06 |
| 2 | 20927997 | rs6729843 | -0.845 | 2.234 | 1.649 | 0.745 | 0.057 | 0.163 | 0.163 | 0 | 0.08 | 0.4 | 0.07 | 0.13 | 0.06 | 0.06 |
| 2 | 20936516 | rs564125 | 2.059 | 2.605 | 1.03 | 0.666 | -0.042 | -0.067 | -0.067 | 0.002 | 0.6 | 0.11 | 0.44 | 0.51 | 0.46 | 0.62 |
| 2 | 20943742 | rs2164720 | -0.454 | 2.688 | 1.628 | 0.669 | -0.003 | -0.008 | -0.008 | 0 | 0.14 | 0.2 | 0.11 | 0.12 | 0.15 | 0.08 |
| 2 | 20945762 | rs4971547 | 0.157 | 2.747 | 1.659 | 0.736 | 0.085 | 0.082 | 0.082 | 0.614 | 0.08 | 0.09 | 0.3 | 0.24 | 0.22 | 0.13 |
| 2 | 20947197 | rs667529 | 1.876 | 2.716 | 1.731 | 0.726 | -0.021 | -0.035 | -0.035 | 0 | 0.63 | 0.22 | 0.45 | 0.5 | 0.46 | 0.68 |
| 2 | 20951332 | rs619562 | 1.132 | 2.697 | 1.858 | 0.719 | 0.007 | -0.04 | -0.04 | 0 | 0.74 | 0.51 | 0.77 | 0.87 | 0.71 | 0.86 |
| 2 | 20953188 | rs2625569 | 1.423 | 2.717 | 1.692 | 0.712 | 0.003 | -0.008 | -0.008 | 0 | 0.71 | 0.31 | 0.75 | 0.75 | 0.67 | 0.81 |
| 2 | 20962670 | rs17721572 | -0.558 | 2.512 | 1.633 | 0.756 | 0.117 | 0.116 | 0.116 | 1.022 | 0.1 | 0.18 | 0.31 | 0.25 | 0.22 | 0.13 |
| 2 | 20970053 | rs661929 | 1.118 | 1.645 | 1.191 | 0.228 | 0.003 | 0 | 0 | 0 | 0.49 | 0.46 | 0.41 | 0.5 | 0.3 | 0.59 |
| 2 | 20982391 | rs601325 | -0.903 | 1.723 | 1.933 | 0.309 | 0.018 | 0.054 | 0.054 | 0 | 0.11 | 0.28 | 0.08 | 0.14 | 0.09 | 0.08 |
| 2 | 21004251 | rs16988072 | 0.243 | 1.381 | 1.206 | 0.328 | -0.011 | -0.024 | -0.024 | 0 | 0.05 | 0.08 | 0.08 | 0 | 0.03 | 0.02 |
| 2 | 21007613 | rs560872 | -2.038 | 1.292 | 1.359 | 0.294 | 0.006 | 0.003 | 0.003 | 1.037 | 0.2 | 0.81 | 0.2 | 0.15 | 0.13 | 0.14 |
| 2 | 21015763 | rs492399 | 0.967 | 1.043 | 1.58 | 0.228 | -0.007 | -0.039 | -0.039 | 0.07 | 0.9 | 0.47 | 0.87 | 1 | 0.96 | 0.98 |
| 2 | 21026745 | rs7600933 | 1.161 | 1.066 | 1.134 | 0.239 | -0.021 | -0.051 | -0.051 | 0 | 0.9 | 0.43 | 0.87 | 1 | 0.96 | 0.93 |
| 2 | 21045744 | rs4971531 | -0.653 | 1.167 | 0.343 | 0.289 | 0.014 | -0.016 | -0.016 | 2.009 | 0.08 | 0.43 | 0.1 | 0 | 0.04 | 0.02 |
| 2 | 21047295 | rs6709100 | -1.159 | 1.52 | -1.192 | 0.488 | -0.061 | -0.028 | -0.028 | 0 | 0.45 | 0.86 | 0.55 | 0.41 | 0.47 | 0.44 |
| 2 | 21052728 | rs4246577 | -1.384 | 0.968 | -1.25 | 0.494 | -0.032 | 0.041 | 0.041 | 0 | 0.5 | 0.81 | 0.56 | 0.37 | 0.48 | 0.52 |
| 2 | 21053915 | rs1898432 | -0.257 | 0.874 | -0.642 | 0.373 | -0.005 | -0.002 | -0.002 | 0 | 0.16 | 0.17 | 0.15 | 0.14 | 0.13 | 0.15 |
| 2 | 21055386 | rs7587472 | 0.001 | 0.862 | 0.119 | 0.266 | 0.007 | -0.03 | -0.03 | 0 | 0.07 | 0.21 | 0.06 | 0 | 0.05 | 0.02 |
| 2 | 21059930 | rs10495711 | -0.121 | 0.952 | -0.086 | 0.256 | -0.036 | -0.02 | -0.02 | 4.822 | 0.14 | 0.06 | 0.21 | 0.15 | 0.22 | 0.18 |
| 2 | 21061934 | rs7571978 | -2.1 | 0.836 | 0.006 | 0.289 | -0.012 | 0.021 | 0.021 | 0 | 0.41 | 0.78 | 0.47 | 0.3 | 0.41 | 0.41 |
| 2 | 21067060 | rs6744750 | -1.139 | 0.83 | 0.279 | 0.203 | -0.016 | -0.015 | -0.015 | 0 | 0.25 | 0.64 | 0.25 | 0.15 | 0.18 | 0.18 |
| 2 | 21069265 | rs4971534 | -0.249 | 1.008 | 0.058 | 0.774 | -0.034 | -0.04 | -0.04 | 0 | 0.15 | 0.06 | 0.23 | 0.24 | 0.23 | 0.19 |
| 2 | 21082489 | rs6531255 | -1.826 | 0.761 | 0.192 | 0.674 | -0.012 | 0.05 | 0.05 | 0 | 0.49 | 0.68 | 0.58 | 0.58 | 0.5 | 0.53 |
| 2 | 21084332 | rs4971536 | -1.467 | 0.893 | 0.333 | 0.786 | 0.007 | 0.004 | 0.004 | 0 | 0.4 | 0.5 | 0.51 | 0.53 | 0.42 | 0.5 |
| 2 | 21086693 | rs1002755 | -1.29 | 0.368 | 0.505 | 0.161 | 0.006 | -0.096 | -0.096 | 0 | 0.17 | 0.44 | 0.13 | 0.02 | 0.15 | 0.15 |
| 2 | 21091802 | rs13396936 | -1.701 | 0.203 | 0.452 | 0.355 | -0.025 | -0.003 | -0.003 | 0 | 0.23 | 0.48 | 0.17 | 0.2 | 0.17 | 0.35 |
| 2 | 21098831 | rs7587653 | -1.497 | 0.242 | 0.289 | 0.38 | -0.015 | 0.005 | 0.005 | 0 | 0.22 | 0.39 | 0.16 | 0.2 | 0.17 | 0.33 |
| 2 | 21101382 | rs6722139 | -0.087 | -0.37 | -0.18 | 0.046 | -0.037 | -0.061 | -0.061 | 0 | 0.25 | 0.42 | 0.14 | 0.15 | 0.17 | 0.1 |
| 2 | 21101745 | rs7598775 | -1.052 | 0.164 | -0.397 | 0.352 | -0.013 | 0.026 | 0.026 | 0 | 0.12 | 0.2 | 0.08 | 0.18 | 0.07 | 0.22 |
| 2 | 21111763 | rs35364714 | -0.91 | 0.715 | -0.986 | -0.239 | -0.053 | 0.083 | 0.083 | 0 | 0.18 | 0.04 | 0.13 | 0.02 | 0.25 | 0.17 |
| 2 | 21118983 | rs12994068 | -0.652 | 0.059 | -0.936 | -0.683 | 0.047 | 0.07 | 0.07 | 1.119 | 0.39 | 0.82 | 0.36 | 0.18 | 0.27 | 0.12 |
| 2 | 21121130 | rs4971538 | 0.112 | 0.059 | -1.029 | -0.755 | -0.014 | 0.006 | 0.006 | 0 | 0.09 | 0.06 | 0.18 | 0.05 | 0.11 | 0.06 |
| 2 | 21123661 | rs7588415 | -0.075 | -0.102 | -0.923 | -0.817 | 0.055 | 0.054 | 0.054 | 3.951 | 0.31 | 0.79 | 0.18 | 0.13 | 0.16 | 0.06 |
| 2 | 21131683 | rs12998321 | 0.711 | -0.162 | -0.771 | -0.936 | -0.007 | -0.05 | -0.05 | 0 | 0.14 | 0.21 | 0.09 | 0 | 0.11 | 0.02 |
| 2 | 21134040 | rs10188503 | -0.289 | -0.106 | -0.794 | -0.977 | 0 | -0.03 | -0.03 | 0 | 0.21 | 0.21 | 0.15 | 0 | 0.2 | 0.06 |
| 2 | 21135577 | rs4341893 | -0.525 | 0.747 | -1.544 | -0.877 | -0.031 | -0.012 | -0.012 | 0.054 | 0.8 | 0.88 | 0.72 | 0.75 | 0.7 | 0.65 |
| 2 | 21136327 | rs12710748 | -1.068 | 0.607 | -1.293 | -1.47 | -0.007 | 0.003 | 0.003 | 0 | 0.35 | 0.34 | 0.4 | 0.06 | 0.44 | 0.18 |

| | | | | | | | | | | | | | | | | |
|---|---|---|---|---|---|---|---|---|---|---|---|---|---|---|---|---|
| 2 | 21137425 | rs4643493 | 0.944 | 0.7 | -1.413 | -1.405 | -0.028 | 0.03 | 0.03 | 0 | 0.1 | 0.02 | 0.11 | 0.02 | 0.2 | 0.09 |
| 2 | 21140033 | rs11693076 | -1.582 | 0.418 | -1.576 | -1.462 | 0.02 | 0.182 | 0.182 | 0.009 | 0.49 | 0.85 | 0.48 | 0.2 | 0.46 | 0.22 |
| 2 | 21140540 | rs11096689 | -1.705 | 0.595 | -1.739 | -1.511 | -0.071 | -0.006 | -0.006 | 0.241 | 0.89 | 0.99 | 0.76 | 0.89 | 0.76 | 0.83 |
| 2 | 21141815 | rs4533439 | -1.233 | 0.395 | -1.34 | -1.516 | 0.017 | 0.137 | 0.137 | 0 | 0.46 | 0.73 | 0.34 | 0.16 | 0.43 | 0.21 |
| 2 | 21141832 | rs13414987 | -0.536 | 0.331 | -1.299 | -1.489 | 0.031 | 0.047 | 0.047 | 0.669 | 0.36 | 0.72 | 0.22 | 0.14 | 0.22 | 0.1 |
| 2 | 21143329 | rs4507059 | -0.631 | 0.416 | -0.548 | -1.485 | 0.009 | -0.071 | -0.071 | 0 | 0.29 | 0.57 | 0.37 | 0.1 | 0.35 | 0.15 |
| 2 | 21143982 | rs4426495 | 1.171 | 0.222 | -0.495 | -1.565 | 0.01 | 0.019 | 0.019 | 0.335 | 0.57 | 0.54 | 0.4 | 0.84 | 0.37 | 0.61 |
| 2 | 21146521 | rs6732011 | 1.107 | 0.225 | -0.538 | -1.576 | -0.006 | -0.032 | -0.032 | 0 | 0.58 | 0.49 | 0.4 | 0.84 | 0.37 | 0.62 |
| 2 | 21184545 | rs1344063 | 0.678 | 0.636 | -0.351 | -1.436 | 0.08 | -0.055 | -0.055 | 0 | 0.73 | 0.45 | 0.48 | 0.85 | 0.57 | 0.75 |
| 2 | 21184764 | rs2337382 | 0.529 | 0.642 | -0.275 | -1.429 | 0.077 | -0.093 | -0.093 | 2.652 | 0.75 | 0.54 | 0.61 | 0.89 | 0.57 | 0.76 |
| 2 | 21184847 | rs17397667 | -0.902 | 0.683 | -0.22 | -1.449 | -0.014 | 0.012 | 0.012 | 0.71 | 0.87 | 0.88 | 0.75 | 0.94 | 0.77 | 0.85 |
| 2 | 21186891 | rs4665639 | 0.782 | 0.731 | -0.208 | -1.453 | 0.089 | -0.053 | -0.053 | 0.971 | 0.74 | 0.45 | 0.48 | 0.85 | 0.56 | 0.75 |
| 2 | 21193946 | rs6728178 | 2.177 | 0.383 | -0.354 | -1.654 | 0.028 | 0.006 | 0.006 | 0 | 0.33 | 0.18 | 0.26 | 0.7 | 0.24 | 0.57 |
| 2 | 21196112 | rs10495712 | -0.793 | 0.44 | -0.51 | -1.638 | -0.018 | 0.017 | 0.017 | 0.34 | 0.86 | 0.86 | 0.75 | 0.94 | 0.76 | 0.85 |
| 2 | 21197159 | rs11685356 | 0.941 | 0.351 | -0.494 | -1.598 | -0.002 | -0.003 | -0.003 | 0 | 0.22 | 0.19 | 0.15 | 0 | 0.23 | 0.07 |
| 2 | 21205563 | rs10164442 | -0.344 | 0.445 | -0.425 | -1.52 | 0.017 | 0.058 | 0.058 | 0 | 0.41 | 0.27 | 0.21 | 0.15 | 0.32 | 0.17 |
| 2 | 21219001 | rs6725189 | 0.348 | 0.342 | -0.495 | -1.34 | -0.011 | 0.003 | 0.003 | 0 | 0.12 | 0.11 | 0.26 | 0.07 | 0.24 | 0.1 |
| 2 | 21225281 | rs1042034 | -2.075 | 0 | -0.621 | -1.502 | 0.04 | -0.001 | -0.001 | 4.795 | 0.68 | 0.88 | 0.75 | 0.28 | 0.78 | 0.42 |
| 2 | 21225753 | rs1042031 | 0.933 | 0.097 | -0.132 | -1.469 | 0.031 | -0.023 | -0.023 | 0 | 0.1 | 0.16 | 0.12 | 0.04 | 0.21 | 0.09 |
| 2 | 21227331 | rs12720851 | 2.162 | -0.137 | -0.233 | -1.589 | -0.031 | -0.036 | -0.036 | 0 | 0.07 | 0.13 | 0.02 | 0 | 0.01 | 0 |
| 2 | 21228827 | rs1801701 | 2.421 | 0.48 | -0.613 | -1.658 | -0.028 | 0.026 | 0.026 | 0.108 | 0.05 | 0.01 | 0.1 | 0 | 0.09 | 0.02 |
| 2 | 21231524 | rs676210 | 1.958 | 0.013 | -0.529 | -1.61 | 0.039 | -0.002 | -0.002 | 0 | 0.32 | 0.12 | 0.25 | 0.72 | 0.22 | 0.57 |
| 2 | 21232195 | rs693 | 0.32 | 0.002 | -0.503 | -1.732 | -0.051 | 0.073 | 0.073 | 0 | 0.33 | 0.21 | 0.38 | 0.06 | 0.44 | 0.22 |
| 2 | 21237544 | rs673548 | 1.81 | -0.164 | -0.654 | -1.505 | 0.014 | 0.002 | 0.002 | 0 | 0.32 | 0.19 | 0.26 | 0.71 | 0.22 | 0.57 |
| 2 | 21237786 | rs2854725 | 1.384 | -0.093 | -0.85 | -1.391 | 0.003 | -0.025 | -0.025 | 0 | 0.13 | 0.04 | 0.06 | 0.15 | 0.09 | 0.1 |
| 2 | 21240364 | rs11126598 | -1.765 | 0.031 | -0.919 | -1.491 | 0.034 | 0.02 | 0.02 | 0 | 0.65 | 0.87 | 0.71 | 0.27 | 0.73 | 0.42 |
| 2 | 21241505 | rs12713956 | 0.892 | 0.254 | -0.657 | -1.409 | -0.001 | -0.008 | -0.008 | 0 | 0.15 | 0.19 | 0.21 | 0.04 | 0.12 | 0.11 |
| 2 | 21244358 | rs11676704 | 2.333 | 0.135 | -1.196 | -0.654 | -0.032 | -0.063 | -0.063 | 1.385 | 0.11 | 0.01 | 0.12 | 0.37 | 0.15 | 0.26 |
| 2 | 21250914 | rs679899 | 1.141 | 0.209 | -0.771 | -1.392 | -0.038 | 0.177 | 0.177 | 0 | 0.42 | 0.13 | 0.39 | 0.86 | 0.47 | 0.65 |
| 2 | 21259562 | rs1469513 | 0.113 | 0.051 | -0.485 | -1.574 | -0.032 | 0.045 | 0.045 | 0 | 0.3 | 0.22 | 0.34 | 0.1 | 0.4 | 0.18 |
| 2 | 21259612 | rs520354 | 1.2 | 0.22 | -0.605 | -1.486 | -0.031 | 0.176 | 0.176 | 0 | 0.43 | 0.14 | 0.4 | 0.87 | 0.48 | 0.67 |
| 2 | 21260601 | rs550619 | -0.674 | 0.628 | 0.279 | -0.868 | -0.009 | -0.025 | -0.025 | 0.057 | 0.84 | 0.81 | 0.79 | 0.96 | 0.89 | 0.88 |
| 2 | 21263900 | rs1367117 | -0.149 | 1.02 | -0.207 | -0.951 | -0.057 | 0.033 | 0.033 | 0 | 0.17 | 0.08 | 0.29 | 0.12 | 0.3 | 0.13 |
| 2 | 21267782 | rs512535 | 1.353 | 1.032 | 0.017 | -0.987 | -0.029 | 0.071 | 0.071 | 0.011 | 0.6 | 0.19 | 0.51 | 0.81 | 0.51 | 0.73 |
| 2 | 21270554 | rs585967 | -0.597 | 1.22 | 0.443 | -0.931 | 0.012 | -0.103 | -0.103 | 0.45 | 0.83 | 0.71 | 0.87 | 1 | 0.81 | 0.88 |
| 2 | 21271323 | rs1713222 | -0.549 | 1.221 | 0.4 | -0.931 | 0.008 | -0.096 | -0.096 | 0.039 | 0.82 | 0.71 | 0.87 | 1 | 0.81 | 0.88 |
| 2 | 21273490 | rs7575840 | -0.193 | 1.499 | -0.053 | -0.766 | -0.039 | 0.016 | 0.016 | 0.022 | 0.15 | 0.12 | 0.29 | 0.12 | 0.32 | 0.13 |
| 2 | 21277922 | rs6548010 | -0.469 | 1.426 | -0.076 | -0.804 | -0.038 | 0.02 | 0.02 | 0.085 | 0.16 | 0.12 | 0.29 | 0.12 | 0.32 | 0.14 |
| 2 | 21288321 | rs562338 | -0.295 | 1.292 | 0.132 | -0.927 | 0 | -0.076 | -0.076 | 0.258 | 0.79 | 0.35 | 0.82 | 0.98 | 0.8 | 0.87 |
| 2 | 21293914 | rs571468 | 1.791 | 0.965 | -0.109 | -0.71 | -0.057 | 0.06 | 0.06 | 0.259 | 0.54 | 0.08 | 0.5 | 0.72 | 0.46 | 0.72 |
| 2 | 21294975 | rs541041 | -0.506 | 0.806 | 0.071 | -0.717 | 0.004 | -0.092 | -0.092 | 0 | 0.8 | 0.4 | 0.82 | 0.99 | 0.8 | 0.88 |
| 2 | 21305587 | rs11886946 | 1.269 | 0.747 | 0.116 | -0.545 | -0.021 | 0.025 | 0.025 | 7.603 | 0.08 | 0.09 | 0.02 | 0.14 | 0.01 | 0 |
| 2 | 21320516 | rs2337383 | -0.159 | 1.058 | -0.095 | -0.646 | -0.033 | 0.021 | 0.021 | 0 | 0.16 | 0.11 | 0.29 | 0.12 | 0.29 | 0.14 |
| 2 | 21327635 | rs97458 | 0.209 | 0.945 | -0.005 | -0.811 | 0.024 | -0.063 | -0.063 | 0 | 0.71 | 0.53 | 0.74 | 0.86 | 0.66 | 0.76 |
| 2 | 21337403 | rs312951 | 2.221 | 0.873 | -0.149 | -0.823 | 0.019 | -0.066 | -0.066 | 0 | 0.42 | 0.07 | 0.4 | 0.6 | 0.34 | 0.6 |
| 2 | 21360583 | rs312966 | 1.386 | 0.742 | 0.025 | -0.739 | 0.015 | 0.03 | 0.03 | 0.507 | 0.55 | 0.4 | 0.45 | 0.74 | 0.37 | 0.63 |
| 2 | 21383717 | rs4560142 | -0.847 | 0.845 | 0.171 | -0.799 | 0.051 | -0.133 | -0.133 | 0 | 0.81 | 0.59 | 0.83 | 0.99 | 0.77 | 0.87 |
| 2 | 21385538 | rs575905 | 1.979 | 1.03 | -0.234 | -0.744 | -0.031 | -0.057 | -0.057 | 0.034 | 0.46 | 0.07 | 0.45 | 0.59 | 0.42 | 0.6 |
| 2 | 21386078 | rs12471982 | -0.252 | 0.756 | 0.246 | -0.707 | -0.001 | 0 | 0 | 0.042 | 0.16 | 0.19 | 0.11 | 0.13 | 0.14 | 0.13 |
| 2 | 21394682 | rs11687850 | 0.875 | 0.751 | 0.195 | -0.67 | -0.013 | -0.018 | -0.018 | 0 | 0.1 | 0.02 | 0.08 | 0.12 | 0.1 | 0.1 |
| 2 | 21397182 | rs506585 | -0.583 | 0.75 | 0.647 | -0.857 | 0.039 | -0.112 | -0.112 | 0.919 | 0.81 | 0.64 | 0.83 | 0.99 | 0.77 | 0.86 |
| 2 | 21405790 | rs567071 | 0.164 | 0.804 | 0.636 | -0.881 | 0.04 | -0.156 | -0.156 | 0 | 0.75 | 0.52 | 0.79 | 0.98 | 0.73 | 0.84 |
| 2 | 21414142 | rs503662 | 0.493 | 0.772 | 0.627 | -0.897 | 0.002 | -0.063 | -0.063 | 0.444 | 0.72 | 0.29 | 0.77 | 0.98 | 0.72 | 0.83 |
| 2 | 21420358 | rs1878512 | -0.126 | 0.651 | 0.681 | -0.924 | 0.003 | -0.134 | -0.134 | 1.839 | 0.23 | 0.57 | 0.18 | 0.01 | 0.23 | 0.15 |
| 2 | 21420828 | rs10206521 | -2.24 | 0.676 | 0.212 | -0.814 | -0.022 | -0.041 | -0.041 | 0.023 | 0.54 | 0.87 | 0.48 | 0.42 | 0.55 | 0.39 |
| 2 | 21423989 | rs10169543 | 0.343 | 0.493 | 0.565 | -0.799 | 0.016 | -0.106 | -0.106 | 1.546 | 0.2 | 0.31 | 0.15 | 0.01 | 0.23 | 0.16 |
| 2 | 21429198 | rs312050 | -1.746 | 0.41 | 0.269 | -0.776 | -0.052 | 0.103 | 0.103 | 0 | 0.46 | 0.9 | 0.4 | 0.26 | 0.42 | 0.28 |
| 2 | 21438863 | rs436941 | -1.799 | 0.442 | 0.436 | -0.784 | -0.016 | 0.09 | 0.09 | 0 | 0.45 | 0.83 | 0.39 | 0.26 | 0.42 | 0.28 |
| 2 | 21445569 | rs7605304 | -0.426 | 0.397 | 0.557 | -0.571 | -0.008 | 0.048 | 0.048 | 0.582 | 0.2 | 0.42 | 0.2 | 0.25 | 0.18 | 0.1 |
| 2 | 21449987 | rs949790 | -0.202 | 0.371 | 0.627 | -0.508 | 0.007 | 0.019 | 0.019 | 0.443 | 0.22 | 0.35 | 0.22 | 0.25 | 0.22 | 0.12 |
| 2 | 21451176 | rs925864 | -0.26 | 0.224 | 0.245 | -0.468 | -0.001 | 0.001 | 0.001 | 0.016 | 0.12 | 0.12 | 0.09 | 0.17 | 0.11 | 0.12 |
| 2 | 21451827 | rs386397 | -0.041 | 0.264 | 0.1 | -0.934 | -0.014 | 0.024 | 0.024 | 0 | 0.74 | 0.79 | 0.78 | 0.93 | 0.71 | 0.86 |
| 2 | 21453211 | rs10198972 | 0.675 | 0.043 | 0.426 | -0.941 | -0.026 | 0.006 | 0.006 | 0 | 0.16 | 0.23 | 0.07 | 0.14 | 0.06 | 0.15 |
| 2 | 21454424 | rs7566040 | 0.432 | -0.021 | 0.473 | -0.879 | -0.011 | 0.019 | 0.019 | 0.111 | 0.64 | 0.69 | 0.69 | 0.71 | 0.61 | 0.72 |
| 2 | 21455941 | rs950661 | 1.712 | 0.144 | 0.232 | -0.906 | -0.001 | -0.008 | -0.008 | 0.694 | 0.39 | 0.31 | 0.55 | 0.57 | 0.44 | 0.53 |
| 2 | 21457859 | rs4665632 | 1.836 | 0.161 | -0.417 | -0.893 | -0.015 | -0.043 | -0.043 | 0 | 0.38 | 0.16 | 0.54 | 0.57 | 0.44 | 0.53 |
| 2 | 21460827 | rs312028 | -0.412 | 0.211 | -0.8 | -0.928 | -0.017 | 0.071 | 0.071 | 0.641 | 0.77 | 0.88 | 0.87 | 0.94 | 0.79 | 0.83 |
| 2 | 21462096 | rs12615034 | 1.611 | -0.044 | -0.884 | -0.708 | -0.019 | -0.088 | -0.088 | 1.886 | 0.36 | 0.09 | 0.48 | 0.58 | 0.38 | 0.51 |
| 2 | 21469336 | rs13398007 | -0.037 | 0.2 | -0.425 | -0.691 | -0.007 | -0.047 | -0.047 | 0 | 0.21 | 0.31 | 0.12 | 0.05 | 0.16 | 0.15 |

| | | | | | | | | | | | | | | | | |
|---|---|---|---|---|---|---|---|---|---|---|---|---|---|---|---|---|
| 2 | 21473385 | rs11689154 | 1.244 | -0.124 | -1.17 | -0.525 | 0.007 | -0.004 | -0.004 | 0 | 0.34 | 0.12 | 0.47 | 0.67 | 0.35 | 0.51 |
| 2 | 21473751 | rs10197693 | 1.244 | -0.124 | -1.17 | -0.525 | 0.007 | -0.004 | -0.004 | 0 | 0.34 | 0.12 | 0.47 | 0.67 | 0.35 | 0.51 |
| 2 | 21475265 | rs4665243 | 1.169 | 0.043 | -0.957 | -0.307 | 0.007 | -0.019 | -0.019 | 0 | 0.41 | 0.14 | 0.47 | 0.61 | 0.39 | 0.58 |
| 2 | 21479786 | rs6733564 | -1.165 | -0.601 | -0.399 | -0.293 | 0.049 | -0.004 | -0.004 | 0 | 0.35 | 0.75 | 0.29 | 0.32 | 0.2 | 0.23 |
| 2 | 21480075 | rs13007394 | 0.008 | -0.383 | -0.417 | -0.267 | 0.001 | -0.01 | -0.01 | 0 | 0.1 | 0.06 | 0.1 | 0.15 | 0.1 | 0.11 |
| 2 | 21481950 | rs10746529 | 0.134 | -0.407 | 0.092 | -0.217 | -0.003 | 0.052 | 0.052 | 0 | 0.93 | 0.82 | 0.94 | 0.93 | 0.96 | 0.93 |
| 2 | 21482292 | rs10195618 | 1.21 | -0.588 | 0.179 | -0.357 | -0.012 | 0.004 | 0.004 | 0 | 0.87 | 0.75 | 0.87 | 0.93 | 0.89 | 0.9 |
| 2 | 21484754 | rs870638 | -1.899 | -0.68 | -0.114 | -0.532 | 0.082 | 0.016 | 0.016 | 4.386 | 0.42 | 0.8 | 0.33 | 0.3 | 0.27 | 0.26 |
| 2 | 21487118 | rs11897368 | -0.732 | -0.798 | 0.192 | -0.584 | -0.034 | -0.019 | -0.019 | 0 | 0.17 | 0.53 | 0.1 | 0.07 | 0.09 | 0.06 |
| 2 | 21489083 | rs6714773 | -2.114 | -0.786 | 0.119 | -0.58 | 0.111 | 0.01 | 0.01 | 4.44 | 0.43 | 0.84 | 0.34 | 0.3 | 0.25 | 0.27 |
| 2 | 21492840 | rs13415405 | -0.113 | -0.776 | 0.338 | -0.537 | -0.038 | 0.064 | 0.064 | 0.029 | 0.11 | 0.48 | 0.07 | 0.07 | 0.04 | 0.04 |
| 2 | 21494476 | rs870223 | 0.673 | -0.733 | 0.263 | -0.567 | -0.005 | 0.001 | 0.001 | 1.377 | 0.77 | 0.74 | 0.83 | 0.77 | 0.84 | 0.87 |
| 2 | 21496859 | rs4665252 | 1.793 | -0.588 | -0.307 | -0.48 | 0.098 | -0.002 | -0.002 | 0 | 0.55 | 0.11 | 0.65 | 0.68 | 0.72 | 0.73 |
| 2 | 21498067 | rs875919 | -0.659 | -0.565 | -0.349 | -0.405 | 0.015 | -0.037 | -0.037 | 0.223 | 0.21 | 0.1 | 0.18 | 0.26 | 0.17 | 0.14 |
| 2 | 21501344 | rs4665662 | 1.963 | -0.203 | -0.436 | -0.516 | 0.023 | 0.045 | 0.045 | 3.619 | 0.5 | 0.12 | 0.59 | 0.67 | 0.59 | 0.71 |
| 2 | 21504377 | rs7598331 | 0.163 | -0.556 | -0.148 | -0.433 | 0 | -0.001 | -0.001 | 0 | 0.09 | 0.05 | 0.09 | 0.1 | 0.1 | 0.09 |
| 2 | 21505938 | rs1877059 | 1.398 | -0.704 | -0.072 | -0.517 | 0.007 | 0.004 | 0.004 | 0 | 0.71 | 0.72 | 0.75 | 0.8 | 0.82 | 0.78 |
| 2 | 21507912 | rs4665670 | 0.806 | -0.723 | -0.122 | -0.5 | 0.026 | -0.001 | -0.001 | 0 | 0.77 | 0.83 | 0.8 | 0.81 | 0.86 | 0.85 |
| 2 | 21515070 | rs10171175 | -0.076 | -0.736 | 0.027 | -0.522 | -0.015 | 0.014 | 0.014 | 0 | 0.07 | 0.16 | 0.06 | 0.07 | 0.04 | 0.07 |
| 2 | 21515580 | rs10185624 | -0.076 | -0.736 | 0.027 | -0.522 | -0.015 | 0.014 | 0.014 | 0 | 0.07 | 0.16 | 0.06 | 0.07 | 0.04 | 0.07 |
| 2 | 21520172 | rs2196165 | 0.16 | -0.794 | 0.577 | -0.626 | -0.017 | -0.072 | -0.072 | 0.011 | 0.12 | 0.28 | 0.06 | 0 | 0.06 | 0.04 |
| 2 | 21520243 | rs219509 | -0.169 | 0.139 | 0.284 | -0.209 | -0.035 | 0.02 | 0.02 | 0 | 0.84 | 0.92 | 0.84 | 0.87 | 0.72 | 0.79 |
| 2 | 21524829 | rs391070 | 0.753 | 0.139 | 0.294 | -0.212 | -0.019 | 0.002 | 0.002 | 0.016 | 0.78 | 0.81 | 0.79 | 0.8 | 0.69 | 0.72 |
| 2 | 21527036 | rs6739502 | 1.998 | 0.319 | 0.222 | -0.272 | 0 | -0.002 | -0.002 | 0 | 0.54 | 0.45 | 0.59 | 0.72 | 0.5 | 0.59 |
| 2 | 21534836 | rs219542 | -0.63 | 0.051 | 0.042 | 0.194 | 0.001 | 0.002 | 0.002 | 1.987 | 0.14 | 0.15 | 0.1 | 0.09 | 0.06 | 0.1 |
| 2 | 21535673 | rs219543 | 1.681 | 0.087 | -0.024 | 0.2 | -0.001 | 0.01 | 0.01 | 0 | 0.64 | 0.65 | 0.69 | 0.72 | 0.62 | 0.63 |
| 2 | 21536873 | rs13427673 | -0.416 | 0.145 | 0.121 | 0.294 | -0.016 | 0.015 | 0.015 | 0 | 0.07 | 0.16 | 0.06 | 0.07 | 0.04 | 0.07 |
| 2 | 21537081 | rs219545 | -1.204 | 0.179 | 0.095 | 0.305 | -0.015 | -0.001 | -0.001 | 0 | 0.21 | 0.29 | 0.15 | 0.15 | 0.1 | 0.16 |
| 2 | 21538826 | rs219549 | 0.916 | 1.158 | 0.238 | 0.516 | 0.012 | -0.004 | -0.004 | 0 | 0.82 | 0.78 | 0.77 | 0.85 | 0.7 | 0.77 |
| 2 | 21539585 | rs219550 | -1.484 | -0.062 | 0.798 | 0.855 | -0.052 | 0.098 | 0.098 | 0 | 0.2 | 0.44 | 0.19 | 0.29 | 0.12 | 0.29 |
| 2 | 21540399 | rs7605009 | -1.34 | 0.424 | 0.812 | 0.462 | 0.014 | -0.011 | -0.011 | 0 | 0.14 | 0.41 | 0.12 | 0.08 | 0.1 | 0.12 |
| 2 | 21551222 | rs930265 | 0.882 | 0.795 | 0.392 | 0.673 | -0.008 | -0.001 | -0.001 | 0.133 | 0.77 | 0.78 | 0.68 | 0.77 | 0.68 | 0.84 |
| 2 | 21577743 | rs219553 | -1.612 | -0.084 | 0.227 | 0.517 | 0.017 | 0.016 | 0.016 | 0.841 | 0.3 | 0.77 | 0.28 | 0.25 | 0.22 | 0.23 |
| 2 | 21578241 | rs219555 | -1.016 | -0.058 | 1.293 | 0.527 | 0.023 | -0.004 | -0.004 | 0 | 0.2 | 0.48 | 0.25 | 0.17 | 0.2 | 0.13 |
| 2 | 21578382 | rs219556 | -1.594 | 0.012 | 0.289 | 0.162 | 0.001 | 0.064 | 0.064 | 0.067 | 0.43 | 0.77 | 0.37 | 0.31 | 0.38 | 0.33 |
| 2 | 21579130 | rs219559 | 0.262 | -0.181 | 0.505 | 0.467 | 0.006 | -0.008 | -0.008 | 0.139 | 0.74 | 0.6 | 0.78 | 0.73 | 0.71 | 0.77 |
| 2 | 21580112 | rs219562 | -1.237 | -0.698 | 0.464 | 0.773 | 0 | 0.008 | 0.008 | 0 | 0.35 | 0.33 | 0.27 | 0.25 | 0.33 | 0.34 |
| 2 | 21581183 | rs10865318 | -0.756 | -0.753 | 0.249 | 0.777 | -0.002 | 0.029 | 0.029 | 0 | 0.3 | 0.25 | 0.19 | 0.13 | 0.29 | 0.26 |
| 2 | 21581464 | rs635141 | -1.776 | -0.805 | 0.057 | 0.727 | 0.001 | 0.146 | 0.146 | 0.008 | 0.4 | 0.92 | 0.31 | 0.19 | 0.31 | 0.3 |
| 2 | 21586117 | rs219513 | -0.719 | -0.743 | 0.158 | 0.83 | -0.007 | 0.009 | 0.009 | 0 | 0.3 | 0.22 | 0.24 | 0.24 | 0.3 | 0.27 |
| 2 | 21593045 | rs219518 | -1.156 | -0.695 | -0.266 | 0.989 | -0.012 | -0.045 | -0.045 | 0 | 0.41 | 0.91 | 0.34 | 0.37 | 0.35 | 0.38 |
| 2 | 21600366 | rs7601920 | -1.19 | -0.745 | -0.22 | 1 | 0.003 | -0.039 | -0.039 | 0 | 0.41 | 0.91 | 0.33 | 0.39 | 0.32 | 0.39 |
| 2 | 21605125 | rs7584918 | -1.078 | -0.721 | -0.146 | 0.956 | 0.009 | -0.031 | -0.031 | 0 | 0.4 | 0.88 | 0.33 | 0.39 | 0.33 | 0.37 |
| 2 | 21609898 | rs6546015 | -1.145 | -0.412 | 0.576 | 1.041 | 0.046 | -0.018 | -0.018 | 2.047 | 0.45 | 0.77 | 0.34 | 0.4 | 0.33 | 0.37 |
| 2 | 21620269 | rs736442 | -1.14 | -0.35 | 1.61 | 1.177 | 0.029 | -0.095 | -0.095 | 0 | 0.44 | 0.92 | 0.35 | 0.43 | 0.33 | 0.37 |
| 2 | 21638968 | rs4557007 | -0.749 | -0.362 | 1.203 | 1.122 | 0.009 | 0.002 | 0.002 | 0 | 0.38 | 0.3 | 0.28 | 0.34 | 0.33 | 0.31 |
| 2 | 21647952 | rs7568314 | -1.245 | -0.324 | 1.241 | 0.991 | 0.042 | -0.074 | -0.074 | 0.017 | 0.44 | 0.88 | 0.35 | 0.41 | 0.33 | 0.37 |
| 2 | 21653983 | rs11693815 | -0.849 | -0.291 | 0.477 | 1.005 | 0.016 | 0.014 | 0.014 | 0 | 0.39 | 0.28 | 0.27 | 0.31 | 0.33 | 0.3 |
| 2 | 21683285 | rs6749957 | -1.03 | -0.024 | -0.516 | 0.77 | 0.054 | -0.073 | -0.073 | 0 | 0.46 | 0.89 | 0.35 | 0.41 | 0.33 | 0.36 |
| 2 | 21690814 | rs13424136 | -0.606 | -0.295 | 1.352 | 0.512 | -0.061 | -0.043 | -0.043 | 0 | 0.16 | 0.51 | 0.07 | 0.07 | 0.06 | 0.09 |
| 2 | 21692897 | rs890081 | -0.717 | -0.363 | 1.254 | 0.479 | -0.038 | -0.028 | -0.028 | 0 | 0.15 | 0.31 | 0.07 | 0.07 | 0.06 | 0.09 |
| 2 | 21693244 | rs890082 | -0.412 | -0.363 | 1.155 | 0.48 | 0.019 | 0.018 | 0.018 | 0 | 0.13 | 0.1 | 0.04 | 0.05 | 0.05 | 0.09 |
| 2 | 21694451 | rs11897825 | -1.76 | 0.432 | 0.814 | 0.408 | -0.093 | 0 | 0 | 0 | 0.41 | 0.91 | 0.37 | 0.34 | 0.42 | 0.4 |
| 2 | 21699789 | rs2081409 | -0.453 | -0.27 | 1.631 | 0.466 | -0.007 | -0.007 | -0.007 | 1.639 | 0.14 | 0.15 | 0.04 | 0.05 | 0.05 | 0.09 |
| 2 | 21701941 | rs3922224 | -1.148 | -0.272 | 1.717 | 0.473 | -0.076 | -0.056 | -0.056 | 0.016 | 0.18 | 0.55 | 0.08 | 0.12 | 0.05 | 0.1 |
| 2 | 21706105 | rs2194758 | -2.709 | 0.29 | 2.038 | 0.665 | 0.074 | 0.094 | 0.094 | 1.71 | 0.05 | 0.53 | 0.05 | 0.04 | 0.01 | 0 |
| 2 | 21822998 | rs1991126 | 2.169 | 0.619 | 2.301 | 0.763 | 0.125 | 0.229 | 0.229 | 0.076 | 0.94 | 0.54 | 0.92 | 0.84 | 0.96 | 0.95 |
| 2 | 21863733 | rs7595261 | -2.779 | 0.471 | 2.71 | 0.384 | 0.024 | 0.016 | 0.016 | 3.7 | 0.08 | 0.58 | 0.08 | 0.03 | 0.03 | 0 |
| 2 | 21864717 | rs2006198 | -2.779 | 0.471 | 2.775 | 0.384 | 0.019 | 0.011 | 0.011 | 3.7 | 0.08 | 0.59 | 0.08 | 0.03 | 0.03 | 0 |
| 2 | 21889579 | rs6547818 | 1.095 | 0.703 | 1.685 | -0.145 | -0.054 | 0.07 | 0.07 | 0.639 | 0.6 | 0.16 | 0.63 | 0.79 | 0.53 | 0.71 |
| 2 | 21896380 | rs17042870 | -2.153 | 0.633 | 2.798 | -0.082 | 0.068 | 0.064 | 0.064 | 2.699 | 0.07 | 0.39 | 0.07 | 0.04 | 0.03 | 0.01 |
| 2 | 21915774 | rs17042879 | -2.52 | 0.62 | 2.742 | -0.125 | 0.027 | 0.014 | 0.014 | 5.066 | 0.08 | 0.6 | 0.09 | 0.04 | 0.03 | 0.01 |
| 2 | 21922209 | rs17042892 | -2.008 | 0.595 | 2.559 | -0.422 | -0.006 | -0.047 | -0.047 | 0 | 0.13 | 0.58 | 0.13 | 0.04 | 0.09 | 0.04 |
| 2 | 21927375 | rs17042897 | -2.551 | 0.538 | 2.492 | -0.541 | -0.02 | -0.062 | -0.062 | 3.579 | 0.14 | 0.39 | 0.11 | 0.04 | 0.1 | 0.06 |
| 2 | 21934414 | rs1901346 | -2.057 | 0.524 | 2.593 | -0.538 | -0.022 | -0.045 | -0.045 | 0.878 | 0.12 | 0.31 | 0.1 | 0.04 | 0.09 | 0.06 |
| 2 | 21935603 | rs7597407 | -2.218 | 0.603 | 2.576 | -0.535 | -0.016 | -0.034 | -0.034 | 0 | 0.13 | 0.31 | 0.11 | 0.06 | 0.1 | 0.11 |
| 2 | 21938062 | rs9636422 | -1.725 | 0.586 | 2.601 | -0.532 | -0.031 | -0.033 | -0.033 | 0.859 | 0.1 | 0.28 | 0.09 | 0.04 | 0.07 | 0.05 |
| 2 | 21940844 | rs11891720 | -1.727 | 0.571 | 2.684 | -0.539 | -0.031 | -0.033 | -0.033 | 0 | 0.1 | 0.28 | 0.09 | 0.04 | 0.07 | 0.05 |
| 2 | 21942735 | rs11683091 | -0.065 | 0.732 | 2.125 | -0.697 | -0.071 | 0.048 | 0.048 | 0 | 0.21 | 0.03 | 0.19 | 0.08 | 0.32 | 0.15 |

| | | | | | | | | | | | | | | | | |
|---|---|---|---|---|---|---|---|---|---|---|---|---|---|---|---|---|
| 2 | 21943231 | rs7568769 | -1.155 | 0.557 | 2.238 | -0.867 | 0.001 | 0 | 0 | 0 | 0.08 | 0.15 | 0.06 | 0.06 | 0.07 | 0.08 |
| 2 | 21967993 | rs10495721 | -1.359 | 0.792 | 2.131 | -0.943 | -0.012 | -0.013 | -0.013 | 0 | 0.09 | 0.17 | 0.08 | 0.04 | 0.07 | 0.04 |
| 2 | 22005145 | rs11689185 | -1.237 | 0.889 | 2.054 | -0.941 | 0 | -0.001 | -0.001 | 0 | 0.09 | 0.13 | 0.08 | 0.04 | 0.07 | 0.04 |
| 2 | 22007094 | rs7587610 | -1.255 | 0.935 | 2.263 | -1.026 | 0.003 | -0.015 | -0.015 | 0 | 0.34 | 0.61 | 0.31 | 0.13 | 0.42 | 0.31 |
| 2 | 22011916 | rs2338557 | -1.222 | 0.896 | 2.331 | -1 | 0.007 | -0.022 | -0.022 | 0 | 0.34 | 0.59 | 0.31 | 0.14 | 0.41 | 0.32 |
| 2 | 22021330 | rs11691878 | -1.029 | 0.845 | 2.273 | -1.094 | 0.005 | -0.031 | -0.031 | 0 | 0.32 | 0.6 | 0.29 | 0.13 | 0.38 | 0.28 |
| 2 | 22033527 | rs10495722 | -1.004 | 0.873 | 2.543 | -1.126 | 0.011 | -0.058 | -0.058 | 0 | 0.32 | 0.52 | 0.28 | 0.12 | 0.37 | 0.28 |
| 2 | 22036018 | rs992996 | -1.102 | 0.874 | 2.52 | -1.126 | 0.003 | -0.026 | -0.026 | 0.524 | 0.33 | 0.61 | 0.29 | 0.12 | 0.37 | 0.29 |
| 2 | 22056486 | rs1022377 | -1.118 | 0.897 | 2.118 | -1.498 | 0 | -0.015 | -0.015 | 0 | 0.42 | 0.57 | 0.37 | 0.13 | 0.49 | 0.32 |
| 2 | 22079770 | rs1519491 | -0.422 | 0.77 | 2.014 | -1.679 | -0.039 | 0.162 | 0.162 | 0 | 0.32 | 0.07 | 0.27 | 0.06 | 0.41 | 0.21 |
| 2 | 22080649 | rs1401644 | -1.922 | 0.352 | 1.651 | -1.94 | -0.026 | 0.097 | 0.097 | 0.948 | 0.6 | 0.82 | 0.56 | 0.18 | 0.63 | 0.44 |
| 2 | 22098922 | rs10200675 | -1.224 | 0.338 | 1.558 | -1.972 | 0.007 | 0.072 | 0.072 | 0 | 0.49 | 0.65 | 0.38 | 0.1 | 0.52 | 0.36 |
| 2 | 22113056 | rs17674649 | -0.561 | 0.297 | 1.353 | -1.903 | 0.001 | -0.057 | -0.057 | 0.002 | 0.42 | 0.5 | 0.35 | 0.11 | 0.49 | 0.33 |
| 2 | 22113775 | rs2176696 | -0.718 | 0.295 | 1.224 | -1.904 | 0 | -0.004 | -0.004 | 0 | 0.43 | 0.61 | 0.37 | 0.11 | 0.5 | 0.34 |
| 2 | 22119666 | rs1574838 | -0.388 | 0.291 | 1.142 | -1.894 | -0.036 | 0.168 | 0.168 | 0.018 | 0.4 | 0.14 | 0.33 | 0.1 | 0.5 | 0.33 |
| 2 | 22125367 | rs4666291 | -0.079 | 0.403 | 0.889 | -1.821 | -0.001 | 0 | 0 | 0.207 | 0.1 | 0.1 | 0.16 | 0.08 | 0.1 | 0.06 |
| 2 | 22170066 | rs13401585 | -0.664 | 0.333 | 0.349 | -1.923 | -0.001 | -0.027 | -0.027 | 0.561 | 0.39 | 0.58 | 0.33 | 0.07 | 0.48 | 0.34 |
| 2 | 22186722 | rs2203879 | -1.628 | 0.644 | 0.649 | -1.186 | 0.009 | 0.003 | 0.003 | 0 | 0.08 | 0.08 | 0.02 | 0 | 0.03 | 0.05 |
| 2 | 22187647 | rs1123302 | -1.205 | 0.721 | 0.73 | -1.345 | -0.004 | -0.002 | -0.002 | 0 | 0.37 | 0.7 | 0.38 | 0.13 | 0.5 | 0.36 |
| 2 | 22189988 | rs329777 | -1.933 | 0.178 | 0.959 | -1.281 | -0.009 | -0.052 | -0.052 | 0 | 0.09 | 0.28 | 0.05 | 0 | 0.04 | 0.04 |
| 2 | 22227379 | rs1004750 | -1.881 | 0.143 | 0.666 | -1.299 | 0.001 | -0.015 | -0.015 | 0.151 | 0.1 | 0.12 | 0.05 | 0 | 0.04 | 0.06 |
| 2 | 22237848 | rs2338775 | -1.423 | 0.157 | 0.636 | -1.276 | 0.006 | -0.003 | -0.003 | 0 | 0.08 | 0.08 | 0.04 | 0 | 0.04 | 0.06 |
| 2 | 22302013 | rs328539 | -1.282 | 0.303 | -0.217 | -1.276 | -0.054 | 0.17 | 0.17 | 1.431 | 0.49 | 0.88 | 0.49 | 0.24 | 0.52 | 0.39 |
| 2 | 22325795 | rs388065 | -2.465 | -0.125 | 0.962 | -1.179 | -0.031 | -0.088 | -0.088 | 0.371 | 0.13 | 0.53 | 0.08 | 0 | 0.06 | 0.08 |
| 2 | 22335940 | rs7561156 | -2.005 | -0.045 | 0.789 | -1.154 | -0.028 | -0.081 | -0.081 | 0 | 0.16 | 0.26 | 0.08 | 0 | 0.08 | 0.09 |
| 2 | 22348539 | rs7601596 | -1.704 | -0.213 | 0.846 | -1.203 | -0.035 | -0.073 | -0.073 | 0 | 0.13 | 0.27 | 0.07 | 0 | 0.05 | 0.06 |
| 2 | 22363413 | rs6735395 | -2.018 | -0.215 | 0.946 | -1.204 | -0.042 | -0.112 | -0.112 | 0.167 | 0.17 | 0.42 | 0.1 | 0 | 0.09 | 0.09 |
| 2 | 22363508 | rs6722493 | -2.082 | -0.218 | 0.924 | -1.206 | -0.047 | -0.125 | -0.125 | 0 | 0.17 | 0.47 | 0.1 | 0 | 0.09 | 0.09 |
| 2 | 22397135 | rs7601985 | -1.861 | -0.154 | 0.503 | -1.168 | -0.035 | -0.099 | -0.099 | 1.773 | 0.16 | 0.37 | 0.1 | 0 | 0.08 | 0.09 |
| 2 | 22407334 | rs2176472 | -1.82 | -0.171 | 0.197 | -1.177 | -0.048 | -0.116 | -0.116 | 1.952 | 0.19 | 0.41 | 0.1 | 0 | 0.08 | 0.11 |
| 2 | 22426074 | rs1607267 | -1.059 | 0.263 | -0.759 | -1.007 | 0.002 | 0.02 | 0.02 | 0.014 | 0.23 | 0.15 | 0.1 | 0 | 0.16 | 0.1 |
| 2 | 22430795 | rs1554266 | 0.496 | -0.035 | -0.598 | -1.227 | 0.009 | 0.056 | 0.056 | 0 | 0.48 | 0.41 | 0.64 | 0.91 | 0.46 | 0.64 |
| 2 | 22439514 | rs10193722 | 0.515 | -0.141 | -0.122 | -1.298 | 0.013 | 0.102 | 0.102 | 0 | 0.48 | 0.35 | 0.64 | 0.91 | 0.46 | 0.64 |
| 2 | 22443343 | rs4362588 | -0.521 | -0.225 | 0.205 | -1.632 | -0.018 | 0.011 | 0.011 | 0 | 0.76 | 0.78 | 0.8 | 0.94 | 0.6 | 0.82 |
| 2 | 22476860 | rs11904182 | -1.286 | -0.744 | 1.017 | -1.03 | -0.03 | 0.056 | 0.056 | 0 | 0.11 | 0.29 | 0.18 | 0.15 | 0.04 | 0.08 |
| 2 | 22477818 | rs6748263 | -0.383 | -1.112 | 0.702 | -1.437 | -0.114 | 0.122 | 0.122 | 0 | 0.73 | 0.92 | 0.76 | 0.94 | 0.55 | 0.79 |
| 2 | 22492399 | rs10181985 | -0.381 | -1.107 | 0.733 | -1.435 | -0.096 | 0.105 | 0.105 | 0 | 0.73 | 0.89 | 0.76 | 0.94 | 0.55 | 0.79 |
| 2 | 22498479 | rs4589792 | 1.022 | -1.366 | 1.414 | -0.627 | -0.087 | -0.014 | -0.014 | 0 | 0.54 | 0.07 | 0.39 | 0.63 | 0.49 | 0.66 |
| 2 | 22500090 | rs8179610 | 0.793 | -1.359 | 1.572 | -0.558 | -0.023 | 0.07 | 0.07 | 0 | 0.57 | 0.28 | 0.52 | 0.76 | 0.49 | 0.69 |
| 2 | 22502937 | rs11898839 | -0.109 | -1.341 | 1.919 | -0.533 | -0.024 | 0.024 | 0.024 | 0 | 0.07 | 0.15 | 0.11 | 0.11 | 0.02 | 0.07 |
| 2 | 22521853 | rs7425354 | -0.876 | -1.18 | 1.898 | -0.668 | -0.084 | 0.02 | 0.02 | 1.102 | 0.8 | 0.92 | 0.74 | 0.87 | 0.6 | 0.79 |
| 2 | 22524305 | rs6725884 | -0.18 | -1.186 | 1.966 | -0.655 | -0.028 | 0.025 | 0.025 | 0 | 0.07 | 0.19 | 0.12 | 0.11 | 0.02 | 0.07 |
| 2 | 22525053 | rs7425479 | -1.186 | -0.889 | 2.094 | -0.618 | 0.081 | 0.077 | 0.077 | 0 | 0.05 | 0.42 | 0.04 | 0.01 | 0.01 | 0.02 |
| 2 | 22537264 | rs4665177 | 0.665 | -0.896 | 1.482 | -0.606 | -0.007 | 0.004 | 0.004 | 0 | 0.63 | 0.34 | 0.63 | 0.86 | 0.5 | 0.75 |
| 2 | 22539355 | rs7585241 | -0.085 | -0.551 | 1.259 | -0.317 | 0.046 | 0.011 | 0.011 | 1.55 | 0.24 | 0.34 | 0.39 | 0.25 | 0.41 | 0.24 |
| 2 | 22546852 | rs5015511 | 0.425 | -0.775 | 0.844 | -0.481 | 0.023 | -0.051 | -0.051 | 0 | 0.64 | 0.46 | 0.66 | 0.87 | 0.53 | 0.73 |
| 2 | 22548774 | rs6741228 | -0.754 | -1.293 | 0.889 | -0.753 | -0.001 | 0.004 | 0.004 | 0 | 0.46 | 0.51 | 0.51 | 0.31 | 0.54 | 0.46 |
| 2 | 22551105 | rs12992463 | 0.041 | -1.37 | 1.273 | -0.897 | -0.041 | 0.055 | 0.055 | 0.14 | 0.31 | 0.16 | 0.31 | 0.12 | 0.46 | 0.4 |
| 2 | 22553712 | rs6743484 | -0.837 | -2.037 | 0.873 | -1.294 | -0.052 | -0.053 | -0.053 | 0 | 0.29 | 0.39 | 0.25 | 0.14 | 0.15 | 0.13 |
| 2 | 22558609 | rs4558612 | -0.377 | -1.782 | 0.925 | -1.072 | 0.039 | -0.059 | -0.059 | 0 | 0.64 | 0.46 | 0.63 | 0.84 | 0.48 | 0.6 |
| 2 | 22568159 | rs17391760 | -0.174 | -1.961 | 1.12 | -1.193 | -0.039 | -0.01 | -0.01 | 3.854 | 0.17 | 0.21 | 0.17 | 0.14 | 0.07 | 0.11 |
| 2 | 22568473 | rs17043657 | 0.608 | -1.753 | 0.403 | -1.083 | -0.009 | -0.043 | -0.043 | 0 | 0.12 | 0.18 | 0.08 | 0 | 0.08 | 0.02 |
| 2 | 22570074 | rs1463979 | 1.151 | -1.529 | -0.023 | -0.905 | -0.092 | 0.028 | 0.028 | 0.324 | 0.36 | 0.06 | 0.4 | 0.73 | 0.38 | 0.48 |
| 2 | 22576100 | rs1605834 | -0.788 | -1.801 | 0.772 | -1.008 | -0.055 | -0.06 | -0.06 | 0 | 0.3 | 0.53 | 0.28 | 0.16 | 0.18 | 0.08 |
| 2 | 22576329 | rs13418342 | -0.539 | -1.822 | 0.543 | -0.969 | -0.07 | 0.042 | 0.042 | 0.349 | 0.7 | 0.88 | 0.67 | 0.8 | 0.57 | 0.61 |
| 2 | 22588525 | rs1355178 | 1.686 | -1.909 | 0.513 | -1.153 | -0.043 | -0.016 | -0.016 | 0.958 | 0.33 | 0.13 | 0.35 | 0.63 | 0.36 | 0.44 |
| 2 | 22589842 | rs10865106 | -0.026 | -1.908 | 0.726 | -1.163 | 0.011 | -0.042 | -0.042 | 0.018 | 0.62 | 0.55 | 0.66 | 0.87 | 0.53 | 0.55 |
| 2 | 22598926 | rs2339518 | -0.537 | -1.97 | 0.686 | -1.148 | -0.039 | 0.047 | 0.047 | 0.53 | 0.68 | 0.77 | 0.69 | 0.86 | 0.55 | 0.65 |
| 2 | 22603707 | rs2136377 | 1.75 | -2.126 | 0.392 | -1.094 | -0.123 | -0.092 | -0.092 | 0 | 0.32 | 0.02 | 0.35 | 0.61 | 0.37 | 0.44 |
| 2 | 22605630 | rs7569443 | -0.016 | -2.291 | 1.456 | -0.976 | -0.03 | -0.063 | -0.063 | 6.642 | 0.17 | 0.27 | 0.08 | 0.03 | 0.09 | 0.04 |
| 2 | 22612638 | rs1396725 | -1.44 | -2.312 | 1.861 | -0.978 | -0.043 | -0.042 | -0.042 | 0 | 0.35 | 0.54 | 0.28 | 0.17 | 0.17 | 0.18 |
| 2 | 22616801 | rs1509361 | -1.496 | -2.099 | 1.812 | -0.97 | 0.002 | 0.005 | 0.005 | 8.724 | 0.36 | 0.64 | 0.33 | 0.25 | 0.19 | 0.21 |
| 2 | 22618537 | rs6749647 | -0.979 | -2.071 | 1.752 | -0.961 | -0.055 | -0.031 | -0.031 | 0.391 | 0.31 | 0.43 | 0.31 | 0.24 | 0.18 | 0.11 |
| 2 | 22622168 | rs4563252 | 0.095 | -1.638 | 1.023 | -0.97 | 0.001 | 0.036 | 0.036 | 0 | 0.49 | 0.58 | 0.37 | 0.3 | 0.56 | 0.44 |
| 2 | 22622627 | rs9308966 | 1.017 | -1.63 | 1.248 | -0.775 | -0.031 | 0.079 | 0.079 | 0 | 0.41 | 0.26 | 0.29 | 0.14 | 0.51 | 0.28 |
| 2 | 22627847 | rs6543938 | -1.189 | -1.676 | 1.128 | -0.721 | -0.026 | 0.042 | 0.042 | 0 | 0.08 | 0.23 | 0.07 | 0.11 | 0.05 | 0.16 |
| 2 | 22636854 | rs4077557 | 0.387 | -1.656 | 1.01 | -0.471 | -0.003 | 0.028 | 0.028 | 0 | 0.08 | 0.09 | 0.17 | 0.19 | 0.08 | 0.04 |
| 2 | 22650830 | rs7596646 | -0.995 | -1.412 | 0.902 | -0.564 | -0.011 | -0.011 | -0.011 | 0 | 0.68 | 0.76 | 0.58 | 0.41 | 0.7 | 0.51 |
| 2 | 22652833 | rs11901977 | 0.17 | -1.404 | 0.887 | -0.661 | -0.002 | 0.062 | 0.062 | 0 | 0.49 | 0.62 | 0.37 | 0.17 | 0.57 | 0.44 |

| | | | | | | | | | | | | | | | | |
|---|---|---|---|---|---|---|---|---|---|---|---|---|---|---|---|---|
| 2 | 22665048 | rs1876766 | 0.266 | -1.476 | 1.066 | -0.573 | 0.002 | 0.058 | 0.058 | 0 | 0.49 | 0.61 | 0.41 | 0.18 | 0.58 | 0.45 |
| 2 | 22685090 | rs7559638 | 1.195 | -1.512 | 1.014 | -0.59 | -0.021 | 0.047 | 0.047 | 0 | 0.4 | 0.28 | 0.32 | 0.1 | 0.52 | 0.28 |
| 2 | 22688188 | rs1463982 | -1.08 | -1.561 | 0.526 | -0.548 | -0.085 | -0.107 | -0.107 | 0.287 | 0.69 | 0.97 | 0.65 | 0.41 | 0.71 | 0.53 |
| 2 | 22688256 | rs4533498 | 0.222 | -1.505 | 0.886 | -0.468 | 0.001 | -0.001 | -0.001 | 0 | 0.18 | 0.14 | 0.21 | 0.22 | 0.13 | 0.07 |
| 2 | 22692241 | rs6751273 | 0.5 | -1.367 | 0.987 | -0.312 | 0.005 | 0.063 | 0.063 | 0 | 0.07 | 0.11 | 0.18 | 0.21 | 0.08 | 0.05 |
| 2 | 22704855 | rs2339516 | 0.944 | -1.343 | 0.961 | -0.287 | 0.004 | 0.055 | 0.055 | 0 | 0.06 | 0.09 | 0.15 | 0.2 | 0.08 | 0.04 |
| 2 | 22712071 | rs17043898 | -0.62 | -1.344 | 0.972 | -0.387 | -0.017 | 0.034 | 0.034 | 0 | 0.06 | 0.14 | 0.07 | 0.09 | 0.05 | 0.17 |
| 2 | 22712247 | rs1876770 | -0.66 | -1.345 | 0.972 | -0.387 | -0.018 | 0.033 | 0.033 | 0 | 0.06 | 0.14 | 0.07 | 0.09 | 0.05 | 0.17 |
| 2 | 22736987 | rs17043947 | -0.75 | -1.307 | 0.74 | -0.347 | -0.004 | 0.062 | 0.062 | 0 | 0.05 | 0.23 | 0.07 | 0.09 | 0.05 | 0.16 |
| 2 | 22739528 | rs1509387 | 0.322 | -1.274 | 0.364 | -0.505 | 0.01 | 0.1 | 0.1 | 0 | 0.49 | 0.66 | 0.41 | 0.18 | 0.59 | 0.47 |
| 2 | 22748324 | rs17043976 | 0.587 | -1.172 | 0.065 | -0.052 | 0.004 | 0.073 | 0.073 | 0 | 0.07 | 0.11 | 0.17 | 0.22 | 0.08 | 0.04 |
| 2 | 22753225 | rs6544021 | -0.482 | -1.294 | -0.012 | -0.417 | -0.031 | 0.114 | 0.114 | 1.86 | 0.63 | 0.88 | 0.48 | 0.21 | 0.64 | 0.51 |
| 2 | 22756232 | rs4988826 | -0.612 | -1.285 | 0.183 | -0.263 | 0 | 0.011 | 0.011 | 0 | 0.45 | 0.44 | 0.6 | 0.74 | 0.44 | 0.49 |
| 2 | 22759343 | rs4665181 | -0.38 | -0.765 | 0.15 | 0.358 | -0.076 | -0.1 | -0.1 | 0.383 | 0.62 | 0.95 | 0.62 | 0.44 | 0.66 | 0.57 |
| 2 | 22763794 | rs2019046 | 1.237 | -1.058 | 1.04 | 0.146 | -0.009 | 0.023 | 0.023 | 0 | 0.44 | 0.34 | 0.33 | 0.18 | 0.51 | 0.36 |
| 2 | 22765580 | rs900928 | -0.656 | -1.05 | 0.794 | 0.166 | 0.007 | 0.026 | 0.026 | 0 | 0.12 | 0.27 | 0.23 | 0.17 | 0.13 | 0.06 |
| 2 | 22766554 | rs937930 | 0.056 | -1.009 | 0.231 | 0.184 | -0.034 | -0.025 | -0.025 | 0 | 0.38 | 0.1 | 0.4 | 0.6 | 0.34 | 0.48 |
| 2 | 22776833 | rs729108 | 1.082 | -1.065 | 0.666 | 0.223 | -0.022 | 0.137 | 0.137 | 0 | 0.44 | 0.21 | 0.31 | 0.09 | 0.52 | 0.32 |
| 2 | 22778763 | rs6748330 | 1.11 | -1.063 | 0.672 | 0.222 | -0.028 | 0.147 | 0.147 | 0 | 0.44 | 0.2 | 0.31 | 0.09 | 0.52 | 0.32 |
| 2 | 22779339 | rs2220153 | -1.128 | -1.086 | 1.204 | 0.247 | -0.003 | 0.021 | 0.021 | 0 | 0.53 | 0.63 | 0.64 | 0.82 | 0.47 | 0.58 |
| 2 | 22780029 | rs4146005 | 0.072 | -1.101 | 1.161 | 0.372 | 0.001 | 0.014 | 0.014 | 0.023 | 0.38 | 0.21 | 0.4 | 0.59 | 0.34 | 0.43 |
| 2 | 22790891 | rs1471767 | -1.14 | -0.998 | 0.867 | 0.305 | -0.001 | 0.013 | 0.013 | 0 | 0.53 | 0.62 | 0.64 | 0.79 | 0.47 | 0.57 |
| 2 | 22810978 | rs1601287 | -0.976 | -0.586 | 0.747 | 0.567 | 0.002 | 0.017 | 0.017 | 0 | 0.12 | 0.2 | 0.22 | 0.17 | 0.12 | 0.07 |
| 2 | 22821061 | rs11124579 | 0.721 | -0.608 | 0.627 | 0.467 | -0.002 | -0.005 | -0.005 | 0 | 0.88 | 0.83 | 0.78 | 0.88 | 0.84 | 0.93 |
| 2 | 22823750 | rs1997325 | 0.73 | -0.578 | 0.824 | 0.476 | 0.003 | -0.004 | -0.004 | 0.339 | 0.88 | 0.77 | 0.77 | 0.87 | 0.84 | 0.93 |
| 2 | 22826752 | rs936751 | -1.723 | -0.718 | 0.735 | 0.401 | -0.014 | -0.02 | -0.02 | 0 | 0.21 | 0.6 | 0.3 | 0.19 | 0.21 | 0.13 |
| 2 | 22834065 | rs2220152 | -0.522 | -1.101 | 0.232 | 0.415 | -0.001 | -0.001 | -0.001 | 0.101 | 0.63 | 0.66 | 0.55 | 0.43 | 0.68 | 0.57 |
| 2 | 22843449 | rs10495730 | 1.174 | -0.876 | -0.158 | 0.376 | -0.116 | 0.093 | 0.093 | 0 | 0.42 | 0.06 | 0.26 | 0.23 | 0.48 | 0.43 |
| 2 | 22844400 | rs12986993 | 1.284 | -0.947 | -0.166 | 0.364 | -0.132 | 0.083 | 0.083 | 0 | 0.4 | 0.06 | 0.26 | 0.21 | 0.48 | 0.39 |
| 2 | 22848683 | rs1484680 | 0.613 | -0.953 | 1.654 | 0.811 | 0 | 0.006 | 0.006 | 0 | 0.53 | 0.42 | 0.38 | 0.37 | 0.56 | 0.53 |
| 2 | 22853294 | rs7604930 | -0.134 | -0.873 | 1.308 | 0.855 | 0 | -0.001 | -0.001 | 0 | 0.09 | 0.12 | 0.06 | 0.09 | 0.08 | 0.1 |
| 2 | 22862384 | rs1881552 | -1.224 | -0.956 | 1.636 | 0.785 | 0.003 | 0.001 | 0.001 | 0 | 0.47 | 0.41 | 0.49 | 0.51 | 0.45 | 0.42 |
| 2 | 22886365 | rs11687600 | -1.642 | -1.024 | 1.456 | 0.708 | -0.002 | -0.003 | -0.003 | 0.63 | 0.15 | 0.15 | 0.25 | 0.13 | 0.16 | 0.08 |
| 2 | 22892638 | rs10495729 | 1.663 | -1.043 | 1.661 | 0.731 | -0.007 | 0.013 | 0.013 | 0 | 0.44 | 0.32 | 0.31 | 0.37 | 0.46 | 0.53 |
| 2 | 22900121 | rs10495728 | 0.5 | -1.036 | 0.958 | 0.764 | -0.051 | -0.092 | -0.092 | 0.004 | 0.28 | 0.07 | 0.35 | 0.4 | 0.3 | 0.29 |
| 2 | 22903486 | rs1534608 | -1.654 | -0.808 | 1.222 | 0.776 | -0.014 | -0.027 | -0.027 | 0 | 0.18 | 0.28 | 0.13 | 0.09 | 0.16 | 0.11 |
| 2 | 22907216 | rs10495727 | 0.639 | -0.841 | 1.06 | 0.943 | -0.062 | -0.1 | -0.1 | 0 | 0.26 | 0.03 | 0.31 | 0.38 | 0.29 | 0.29 |
| 2 | 22909155 | rs4505562 | -1.838 | -0.861 | 1.295 | 0.693 | -0.001 | -0.002 | -0.002 | 0.023 | 0.52 | 0.51 | 0.64 | 0.61 | 0.52 | 0.47 |
| 2 | 22915864 | rs1385272 | -2.033 | -0.896 | 1.424 | 0.728 | -0.026 | 0.039 | 0.039 | 0 | 0.19 | 0.4 | 0.17 | 0.25 | 0.17 | 0.18 |
| 2 | 22922022 | rs1872325 | -1.005 | -0.832 | 1.31 | 0.679 | -0.003 | -0.008 | -0.008 | 0.064 | 0.42 | 0.42 | 0.6 | 0.69 | 0.41 | 0.43 |
| 2 | 22931540 | rs7558407 | -1.033 | -0.894 | 1.073 | 0.423 | -0.018 | 0.015 | 0.015 | 0 | 0.1 | 0.14 | 0.2 | 0.21 | 0.04 | 0.14 |
| 2 | 22936495 | rs6747932 | -1.661 | -0.771 | 0.998 | 0.306 | -0.02 | 0.097 | 0.097 | 0 | 0.55 | 0.72 | 0.6 | 0.86 | 0.51 | 0.6 |
| 2 | 22940016 | rs1528777 | -0.516 | -0.514 | 0.563 | 0.47 | -0.032 | -0.06 | -0.06 | 0.18 | 0.38 | 0.17 | 0.28 | 0.5 | 0.4 | 0.39 |
| 2 | 22960628 | rs1919401 | -0.87 | -0.35 | 0.791 | 0.418 | -0.008 | 0.009 | 0.009 | 0 | 0.42 | 0.35 | 0.36 | 0.63 | 0.46 | 0.42 |
| 2 | 22961072 | rs12712618 | 0.135 | -0.411 | 0.756 | 0.438 | -0.004 | -0.007 | -0.007 | 0 | 0.66 | 0.69 | 0.71 | 0.46 | 0.65 | 0.59 |
| 2 | 22967228 | rs12469094 | 0.058 | -0.381 | 0.768 | 0.449 | -0.006 | -0.013 | -0.013 | 2.639 | 0.67 | 0.73 | 0.72 | 0.46 | 0.66 | 0.65 |
| 2 | 22969224 | rs1528779 | 1.575 | -0.459 | 0.835 | 0.373 | -0.007 | 0.076 | 0.076 | 0 | 0.46 | 0.32 | 0.39 | 0.13 | 0.51 | 0.41 |
| 2 | 22973119 | rs6733405 | 1.158 | -0.256 | 1.083 | 0.678 | 0.001 | 0.012 | 0.012 | 0 | 0.58 | 0.37 | 0.48 | 0.24 | 0.6 | 0.52 |
| 2 | 22973348 | rs11900261 | -1.356 | -0.077 | 1.152 | 0.889 | -0.027 | 0.037 | 0.037 | 0 | 0.1 | 0.19 | 0.19 | 0.22 | 0.04 | 0.14 |
| 2 | 22979281 | rs13385210 | -2.299 | -0.089 | 0.974 | 0.911 | -0.048 | 0.063 | 0.063 | 0 | 0.14 | 0.31 | 0.24 | 0.25 | 0.08 | 0.11 |
| 2 | 22980155 | rs7601387 | -1.035 | -0.06 | 0.986 | 0.925 | -0.025 | 0.094 | 0.094 | 0 | 0.08 | 0.18 | 0.2 | 0.22 | 0.04 | 0.09 |
| 2 | 22981898 | rs12053564 | -1.052 | 0.039 | 0.73 | 0.952 | 0.025 | -0.018 | -0.018 | 0 | 0.34 | 0.51 | 0.49 | 0.73 | 0.39 | 0.5 |
| 2 | 22984176 | rs2339564 | -2.05 | -0.635 | 1.285 | 1.104 | -0.02 | 0.162 | 0.162 | 0 | 0.13 | 0.45 | 0.25 | 0.29 | 0.08 | 0.1 |
| 2 | 23003253 | rs10195540 | -2.591 | -0.587 | 0.922 | 1.107 | -0.016 | 0.025 | 0.025 | 0.016 | 0.24 | 0.69 | 0.35 | 0.47 | 0.17 | 0.18 |
| 2 | 23016634 | rs10195794 | -1.568 | -0.287 | 0.614 | 0.815 | -0.014 | 0.016 | 0.016 | 0.197 | 0.12 | 0.18 | 0.11 | 0.18 | 0.1 | 0.08 |
| 2 | 23018729 | rs7567950 | -1.434 | -0.162 | 0.791 | 0.62 | 0.031 | 0.033 | 0.033 | 0 | 0.06 | 0.31 | 0.04 | 0.07 | 0.04 | 0.01 |
| 2 | 23020557 | rs4665518 | -1.771 | -0.508 | 0.307 | 0.607 | 0.009 | -0.01 | -0.01 | 0 | 0.27 | 0.76 | 0.36 | 0.44 | 0.19 | 0.19 |
| 2 | 23022235 | rs12621242 | 0.743 | -0.318 | 0.125 | 0.787 | 0.003 | -0.032 | -0.032 | 0 | 0.06 | 0.01 | 0.18 | 0.24 | 0.04 | 0.07 |
| 2 | 23024906 | rs13425940 | -0.928 | 0.344 | 0.004 | 0.195 | -0.016 | 0.047 | 0.047 | 0.095 | 0.38 | 0.3 | 0.47 | 0.84 | 0.41 | 0.54 |
| 2 | 23040066 | rs7582711 | -0.773 | 0.161 | 0.047 | 0.305 | -0.03 | 0.056 | 0.056 | 0 | 0.14 | 0.29 | 0.23 | 0.3 | 0.08 | 0.09 |
| 2 | 23041318 | rs4665523 | -1.71 | 0.276 | 0.323 | 0.211 | -0.019 | 0.043 | 0.043 | 0.065 | 0.26 | 0.6 | 0.33 | 0.44 | 0.2 | 0.18 |
| 2 | 23044808 | rs7605946 | -1.635 | 0.269 | 0.365 | 0.2 | -0.01 | 0.029 | 0.029 | 0.077 | 0.25 | 0.66 | 0.33 | 0.44 | 0.19 | 0.18 |
| 2 | 23045279 | rs4665183 | -0.333 | 0.271 | 0.298 | 0.201 | -0.023 | 0.07 | 0.07 | 0 | 0.12 | 0.29 | 0.22 | 0.3 | 0.08 | 0.08 |
| 2 | 23050423 | rs6742401 | -1.874 | 0.33 | 0.195 | 0.069 | 0.016 | 0.08 | 0.08 | 0 | 0.55 | 0.75 | 0.63 | 0.99 | 0.56 | 0.65 |
| 2 | 23050710 | rs4487114 | -0.13 | 0.486 | -0.005 | 0.168 | -0.12 | -0.097 | -0.097 | 1.021 | 0.3 | 0.04 | 0.27 | 0.59 | 0.38 | 0.48 |
| 2 | 23056602 | rs4665524 | -0.381 | 0.194 | 0.884 | 0.754 | -0.017 | 0.103 | 0.103 | 0 | 0.11 | 0.3 | 0.22 | 0.31 | 0.08 | 0.09 |
| 2 | 23058877 | rs4594478 | -1.936 | 0.219 | 0.496 | 0.792 | -0.01 | 0.013 | 0.013 | 0.014 | 0.27 | 0.63 | 0.36 | 0.38 | 0.23 | 0.19 |
| 2 | 23064073 | rs4665525 | -1.81 | 0.373 | 0.335 | 0.822 | -0.01 | 0.019 | 0.019 | 0 | 0.27 | 0.58 | 0.35 | 0.36 | 0.24 | 0.2 |
| 2 | 23067410 | rs6544251 | -1.014 | 0.503 | -0.294 | 0.37 | 0.004 | -0.019 | -0.019 | 0 | 0.11 | 0.17 | 0.08 | 0.06 | 0.11 | 0.09 |

| | | | | | | | | | | | | | | | | |
|---|---|---|---|---|---|---|---|---|---|---|---|---|---|---|---|---|
| 2 | 23069479 | rs6716353 | -0.925 | 0.482 | -0.496 | 0.587 | -0.002 | 0.002 | 0.002 | 0.336 | 0.36 | 0.68 | 0.4 | 0.48 | 0.32 | 0.24 |
| 2 | 23078177 | rs4665527 | -0.89 | 0.668 | -1.027 | -0.265 | -0.014 | 0.088 | 0.088 | 0 | 0.74 | 0.86 | 0.71 | 0.95 | 0.69 | 0.69 |
| 2 | 23083158 | rs6544272 | -0.142 | 0.684 | -0.907 | -0.541 | 0.001 | 0.019 | 0.019 | 0 | 0.57 | 0.35 | 0.58 | 0.78 | 0.54 | 0.62 |
| 2 | 23091376 | rs2001229 | -0.371 | 0.654 | -1.083 | -0.443 | 0.013 | 0.031 | 0.031 | 0 | 0.59 | 0.27 | 0.62 | 0.72 | 0.6 | 0.54 |
| 2 | 23098341 | rs2001427 | 0.005 | 0.555 | -1.806 | -0.66 | 0.004 | 0.139 | 0.139 | 0 | 0.52 | 0.09 | 0.51 | 0.74 | 0.5 | 0.55 |
| 2 | 23102550 | rs3738876 | -0.507 | 0.721 | -1.652 | -0.646 | -0.018 | 0.062 | 0.062 | 0 | 0.61 | 0.2 | 0.47 | 0.79 | 0.52 | 0.59 |
| 2 | 23106030 | rs2879555 | 0.006 | 0.561 | -2.116 | -0.947 | 0.008 | 0.092 | 0.092 | 0 | 0.45 | 0.18 | 0.33 | 0.75 | 0.42 | 0.48 |
| 2 | 23108116 | rs17321444 | -0.302 | 0.959 | -1.91 | -0.256 | -0.005 | 0.054 | 0.054 | 0 | 0.16 | 0.05 | 0.15 | 0.03 | 0.11 | 0.1 |
| 2 | 23114577 | rs9789418 | -0.324 | 1.306 | -1.913 | 0.455 | -0.072 | -0.111 | -0.111 | 0 | 0.17 | 0.01 | 0.19 | 0.58 | 0.21 | 0.27 |
| 2 | 23115536 | rs2001795 | -0.183 | 0.894 | -1.414 | -0.326 | 0.003 | 0.015 | 0.015 | 0.13 | 0.48 | 0.57 | 0.56 | 0.26 | 0.56 | 0.52 |
| 2 | 23130359 | rs10184453 | -0.374 | -0.065 | -1.044 | -1.016 | -0.058 | -0.075 | -0.075 | 3.027 | 0.36 | 0.48 | 0.16 | 0.17 | 0.19 | 0.29 |
| 2 | 23135466 | rs2339703 | -0.335 | -0.122 | -0.844 | -0.988 | -0.006 | -0.015 | -0.015 | 0 | 0.36 | 0.35 | 0.15 | 0.17 | 0.19 | 0.29 |
| 2 | 23148411 | rs1838899 | 0.045 | -0.381 | -0.981 | -1.294 | 0.056 | 0.138 | 0.138 | 0 | 0.27 | 0.04 | 0.11 | 0.04 | 0.18 | 0.22 |
| 2 | 23149442 | rs1542190 | -0.4 | -0.282 | -0.716 | -1.194 | 0.018 | 0.014 | 0.014 | 0 | 0.31 | 0.23 | 0.14 | 0.08 | 0.22 | 0.23 |
| 2 | 23153079 | rs17720928 | -0.425 | 0.209 | -0.796 | -1.318 | 0.003 | 0.105 | 0.105 | 0 | 0.32 | 0.14 | 0.17 | 0.06 | 0.31 | 0.22 |
| 2 | 23155213 | rs10206536 | -1.103 | 0.15 | -0.56 | -1.347 | -0.011 | -0.091 | -0.091 | 0.385 | 0.38 | 0.45 | 0.2 | 0.06 | 0.32 | 0.29 |
| 2 | 23155862 | rs7586848 | -0.724 | 0.188 | -0.629 | -1.328 | 0.009 | 0.026 | 0.026 | 0 | 0.34 | 0.29 | 0.18 | 0.06 | 0.31 | 0.22 |
| 2 | 23165621 | rs10175523 | -1.431 | 0.201 | -0.391 | -1.062 | -0.011 | -0.014 | -0.014 | 0.676 | 0.06 | 0.29 | 0.02 | 0 | 0 | 0.08 |
| 2 | 23166176 | rs2681017 | 0.357 | 0.293 | -0.637 | -1.725 | 0.003 | 0.039 | 0.039 | 0 | 0.54 | 0.35 | 0.72 | 0.93 | 0.57 | 0.62 |
| 2 | 23166973 | rs1446869 | -0.762 | 0.491 | -1.157 | -0.899 | 0.04 | 0.018 | 0.018 | 0.241 | 0.54 | 0.79 | 0.59 | 0.4 | 0.58 | 0.63 |
| 2 | 23173111 | rs7580662 | -0.167 | 1.001 | -0.806 | -0.738 | -0.01 | -0.005 | -0.005 | 0 | 0.27 | 0.26 | 0.29 | 0.29 | 0.31 | 0.35 |
| 2 | 23174479 | rs6726919 | -1.09 | 1.178 | -0.547 | -0.892 | -0.004 | -0.001 | -0.001 | 0 | 0.42 | 0.65 | 0.53 | 0.39 | 0.49 | 0.53 |
| 2 | 23174557 | rs6756361 | -0.036 | 0.997 | -0.395 | -0.799 | 0.058 | -0.003 | -0.003 | 5.036 | 0.15 | 0.46 | 0.25 | 0.11 | 0.18 | 0.19 |
| 2 | 23175970 | rs713514 | -0.036 | 0.997 | -0.459 | -0.799 | 0.053 | -0.005 | -0.005 | 0 | 0.15 | 0.44 | 0.25 | 0.11 | 0.18 | 0.19 |
| 2 | 23176757 | rs7572707 | -1.067 | 0.855 | -1.217 | -0.652 | 0.061 | 0.001 | 0.001 | 0.275 | 0.54 | 0.81 | 0.63 | 0.41 | 0.62 | 0.62 |
| 2 | 23179531 | rs6737109 | -0.907 | 0.967 | -1.166 | -0.978 | 0.009 | 0.05 | 0.05 | 0 | 0.45 | 0.62 | 0.47 | 0.21 | 0.43 | 0.45 |
| 2 | 23182078 | rs17321693 | -0.272 | 1.242 | -0.87 | -1.103 | -0.023 | 0.028 | 0.028 | 3.448 | 0.31 | 0.18 | 0.23 | 0.16 | 0.37 | 0.27 |
| 2 | 23182262 | rs1348864 | -1.903 | 1.304 | -1.128 | -1.101 | 0.018 | -0.004 | -0.004 | 0 | 0.63 | 0.66 | 0.63 | 0.38 | 0.67 | 0.64 |
| 2 | 23182754 | rs12614422 | -0.385 | 1.393 | -0.549 | -0.802 | 0.001 | 0.006 | 0.006 | 0 | 0.27 | 0.24 | 0.37 | 0.22 | 0.27 | 0.36 |
| 2 | 23183222 | rs17393683 | 0.261 | 1.757 | -0.726 | -0.68 | -0.031 | 0.002 | 0.002 | 0.002 | 0.18 | 0.07 | 0.15 | 0.16 | 0.23 | 0.18 |
| 2 | 23184327 | rs4665184 | -0.693 | 1.341 | -0.993 | -0.612 | -0.05 | 0.034 | 0.034 | 0.689 | 0.19 | 0.07 | 0.31 | 0.12 | 0.25 | 0.24 |
| 2 | 23185972 | rs2592781 | 0.682 | 1.422 | -0.311 | -0.188 | -0.009 | 0.006 | 0.006 | 0 | 0.58 | 0.38 | 0.54 | 0.67 | 0.53 | 0.66 |
| 2 | 23186695 | rs2592779 | 1.83 | 1.265 | -0.948 | -0.165 | 0.024 | -0.175 | -0.175 | 0 | 0.36 | 0.04 | 0.33 | 0.56 | 0.3 | 0.43 |
| 2 | 23187504 | rs2681019 | 0.647 | 1.604 | 0.722 | 0.463 | -0.028 | 0.018 | 0.018 | 0 | 0.62 | 0.74 | 0.58 | 0.73 | 0.54 | 0.67 |
| 2 | 23192909 | rs2592774 | 0.407 | 1.708 | 0.918 | 0.466 | 0.035 | 0.049 | 0.049 | 0.066 | 0.84 | 0.64 | 0.73 | 0.83 | 0.85 | 0.85 |
| 2 | 23196362 | rs2592768 | 1.151 | 1.745 | 1.023 | 0.419 | -0.001 | 0 | 0 | 0 | 0.57 | 0.46 | 0.48 | 0.63 | 0.48 | 0.64 |
| 2 | 23196895 | rs2681010 | 1.016 | 1.763 | 0.65 | 0.182 | 0.013 | -0.014 | -0.014 | 0 | 0.6 | 0.49 | 0.56 | 0.74 | 0.54 | 0.67 |
| 2 | 23197435 | rs12474212 | -0.253 | 1.843 | 0.47 | 0.311 | -0.057 | 0.072 | 0.072 | 0 | 0.24 | 0.06 | 0.17 | 0.1 | 0.31 | 0.18 |
| 2 | 23199583 | rs6707660 | 0.016 | 2.078 | 0.589 | 0.462 | -0.015 | 0.015 | 0.015 | 0 | 0.2 | 0.12 | 0.17 | 0.11 | 0.28 | 0.16 |
| 2 | 23215201 | rs2592762 | 1.023 | 1.998 | -0.097 | 0.82 | -0.047 | -0.054 | -0.054 | 0 | 0.55 | 0.14 | 0.45 | 0.59 | 0.51 | 0.61 |
| 2 | 23216262 | rs1446877 | -1.094 | 1.616 | 0.833 | 1.423 | 0.015 | 0.047 | 0.047 | 0 | 0.19 | 0.71 | 0.36 | 0.32 | 0.2 | 0.21 |
| 2 | 23223292 | rs1446881 | -0.596 | 1.124 | 0.099 | 0.784 | -0.004 | -0.005 | -0.005 | 0.125 | 0.59 | 0.52 | 0.68 | 0.6 | 0.63 | 0.56 |
| 2 | 23225050 | rs6745196 | -0.275 | 1.169 | -0.022 | 0.72 | 0.021 | -0.036 | -0.036 | 0 | 0.3 | 0.4 | 0.26 | 0.19 | 0.36 | 0.38 |
| 2 | 23226264 | rs11676189 | -1.009 | 1.129 | -0.012 | 0.73 | 0 | 0 | 0 | 0 | 0.58 | 0.56 | 0.67 | 0.6 | 0.6 | 0.56 |
| 2 | 23227422 | rs4665534 | 0.037 | 0.751 | 0.078 | 1.162 | 0.013 | -0.051 | -0.051 | 0.223 | 0.29 | 0.16 | 0.42 | 0.41 | 0.25 | 0.17 |
| 2 | 23228356 | rs6718240 | -0.674 | 0.855 | -0.079 | 1.054 | -0.012 | 0.005 | 0.005 | 0 | 0.4 | 0.46 | 0.52 | 0.54 | 0.33 | 0.45 |
| 2 | 23230226 | rs2592759 | 0.385 | 1.216 | 0.006 | 0.97 | 0 | 0 | 0 | 0.453 | 0.9 | 0.89 | 0.87 | 0.89 | 0.9 | 0.71 |
| 2 | 23230861 | rs13402936 | -1.195 | 1.254 | 0.583 | 1.06 | 0.049 | 0.032 | 0.032 | 0.067 | 0.11 | 0.36 | 0.1 | 0.15 | 0.12 | 0.15 |
| 2 | 23231141 | rs2681033 | 1.201 | 1.207 | 0.639 | 1.298 | 0.08 | 0.182 | 0.182 | 0 | 0.94 | 0.73 | 0.93 | 0.86 | 0.94 | 0.85 |
| 2 | 23233599 | rs2681034 | -0.357 | 1.61 | -0.318 | 1.618 | -0.01 | -0.006 | -0.006 | 0 | 0.16 | 0.14 | 0.28 | 0.42 | 0.25 | 0.17 |
| 2 | 23238004 | rs10495737 | 0.492 | 0.955 | -1.437 | 0.452 | -0.059 | 0.257 | 0.257 | 0.145 | 0.46 | 0.08 | 0.3 | 0.14 | 0.51 | 0.54 |
| 2 | 23238328 | rs2879542 | 0.471 | 0.928 | -1.413 | 0.439 | -0.047 | 0.25 | 0.25 | 0.155 | 0.47 | 0.1 | 0.3 | 0.14 | 0.51 | 0.54 |
| 2 | 23239848 | rs1867914 | 0.885 | 0.582 | -1.824 | 0.655 | -0.007 | -0.116 | -0.116 | 0 | 0.22 | 0.01 | 0.27 | 0.34 | 0.17 | 0.25 |
| 2 | 23241481 | rs2121450 | 0.72 | 0.673 | -1.617 | 0.445 | -0.102 | 0.285 | 0.285 | 0.238 | 0.46 | 0.05 | 0.3 | 0.13 | 0.51 | 0.56 |
| 2 | 23247881 | rs12466137 | 0.32 | 0.688 | -0.768 | 0.603 | -0.001 | -0.007 | -0.007 | 0 | 0.7 | 0.3 | 0.64 | 0.55 | 0.67 | 0.73 |
| 2 | 23252410 | rs1470513 | 0.374 | 0.677 | -0.576 | 0.592 | 0 | 0.003 | 0.003 | 0 | 0.71 | 0.3 | 0.64 | 0.55 | 0.68 | 0.82 |
| 2 | 23254448 | rs12467905 | 0.712 | 0.79 | -0.874 | 0.429 | -0.025 | 0.171 | 0.171 | 0.097 | 0.47 | 0.21 | 0.33 | 0.21 | 0.5 | 0.57 |
| 2 | 23257855 | rs1900374 | 0.79 | 0.776 | -0.878 | 0.428 | -0.051 | 0.207 | 0.207 | 0.188 | 0.46 | 0.17 | 0.33 | 0.2 | 0.5 | 0.51 |
| 2 | 23260036 | rs750044 | 0.836 | 0.741 | -0.784 | 0.411 | -0.073 | 0.223 | 0.223 | 0.072 | 0.46 | 0.15 | 0.33 | 0.2 | 0.5 | 0.47 |
| 2 | 23262942 | rs1004586 | 0.773 | 0.558 | -0.753 | 0.548 | -0.011 | -0.129 | -0.129 | 0 | 0.23 | 0.02 | 0.29 | 0.33 | 0.18 | 0.25 |
| 2 | 23263220 | rs11903039 | 0.03 | 0.443 | 0.375 | 0.446 | -0.011 | 0.033 | 0.033 | 0.057 | 0.11 | 0.59 | 0.11 | 0.13 | 0.05 | 0.03 |
| 2 | 23269996 | rs7593862 | 0.887 | 0.556 | -0.257 | -0.193 | 0.008 | 0.027 | 0.027 | 0 | 0.3 | 0.17 | 0.2 | 0.08 | 0.29 | 0.31 |
| 2 | 23273852 | rs4143808 | 1.014 | 0.739 | -0.901 | -0.046 | -0.03 | -0.035 | -0.035 | 0 | 0.12 | 0.06 | 0.09 | 0.19 | 0.15 | 0.12 |
| 2 | 23279149 | rs13033261 | 1.061 | 0.753 | -0.905 | -0.071 | -0.037 | 0.182 | 0.182 | 0 | 0.29 | 0.03 | 0.18 | 0.07 | 0.28 | 0.31 |
| 2 | 23281048 | rs4665195 | -0.668 | 1.342 | -0.48 | 0.319 | -0.033 | -0.043 | -0.043 | 1.57 | 0.81 | 0.88 | 0.65 | 0.59 | 0.69 | 0.81 |
| 2 | 23282771 | rs11689205 | 1.041 | 1.018 | -0.557 | 0.672 | -0.013 | 0.011 | 0.011 | 0 | 0.06 | 0.01 | 0.19 | 0.35 | 0.08 | 0.07 |
| 2 | 23295124 | rs10188372 | 0.035 | 0.343 | 0.128 | 0.225 | -0.009 | -0.002 | -0.002 | 1.427 | 0.36 | 0.42 | 0.37 | 0.33 | 0.27 | 0.39 |
| 2 | 23297148 | rs4665543 | 0.33 | 0.335 | 0.259 | 0.282 | 0.018 | -0.011 | -0.011 | 1.182 | 0.32 | 0.3 | 0.35 | 0.36 | 0.22 | 0.31 |
| 2 | 23298851 | rs1562753 | 1.109 | 0.306 | 0.014 | 0.096 | -0.017 | -0.099 | -0.099 | 0 | 0.23 | 0.07 | 0.3 | 0.38 | 0.19 | 0.3 |

| | | | | | | | | | | | | | | | | |
|---|---|---|---|---|---|---|---|---|---|---|---|---|---|---|---|---|
| 2 | 23300031 | rs12994268 | 0.547 | 0.052 | -0.137 | -0.448 | -0.027 | 0.104 | 0.104 | 0 | 0.35 | 0.15 | 0.22 | 0.08 | 0.41 | 0.37 |
| 2 | 23300822 | rs1446875 | -0.598 | 0.033 | 0.094 | -0.406 | -0.006 | -0.016 | -0.016 | 0.101 | 0.7 | 0.75 | 0.6 | 0.47 | 0.67 | 0.69 |
| 2 | 23314185 | rs4665548 | 1.576 | 0.023 | -0.031 | -0.418 | -0.005 | -0.075 | -0.075 | 0 | 0.22 | 0.1 | 0.3 | 0.38 | 0.19 | 0.3 |
| 2 | 23325109 | rs12617874 | 1.077 | -0.22 | -0.669 | -0.608 | -0.103 | 0.151 | 0.151 | 0.143 | 0.27 | 0.04 | 0.19 | 0.07 | 0.38 | 0.34 |
| 2 | 23325818 | rs1867916 | -0.509 | 0.044 | -0.467 | -0.197 | -0.01 | -0.029 | -0.029 | 0 | 0.69 | 0.75 | 0.59 | 0.51 | 0.67 | 0.67 |
| 2 | 23326056 | rs2044766 | -0.528 | 0.015 | -0.49 | -0.211 | -0.028 | -0.065 | -0.065 | 0 | 0.71 | 0.83 | 0.6 | 0.51 | 0.67 | 0.67 |
| 2 | 23328503 | rs1446872 | 1.05 | 0.256 | -1.028 | 0.02 | -0.016 | -0.125 | -0.125 | 0 | 0.25 | 0.07 | 0.32 | 0.43 | 0.22 | 0.28 |
| 2 | 23329669 | rs6734383 | 1.105 | 0.112 | -0.465 | -0.191 | -0.007 | -0.022 | -0.022 | 0.137 | 0.07 | 0.12 | 0.03 | 0 | 0.04 | 0.01 |
| 2 | 23334779 | rs4430896 | -0.519 | 0.403 | -0.469 | 0.13 | -0.001 | 0.027 | 0.027 | 0 | 0.47 | 0.55 | 0.34 | 0.3 | 0.5 | 0.51 |
| 2 | 23344587 | rs17667100 | -0.309 | 0.632 | -0.332 | 0.084 | 0.003 | -0.024 | -0.024 | 0 | 0.23 | 0.3 | 0.19 | 0.12 | 0.3 | 0.28 |
| 2 | 23346547 | rs4665552 | -0.649 | 0.744 | -0.415 | -0.02 | -0.002 | 0.002 | 0.002 | 0.207 | 0.69 | 0.71 | 0.79 | 0.79 | 0.71 | 0.8 |
| 2 | 23347247 | rs12468314 | 0.945 | 0.907 | -0.318 | -0.01 | -0.003 | 0.002 | 0.002 | 0.528 | 0.14 | 0.16 | 0.13 | 0.08 | 0.2 | 0.25 |
| 2 | 23350490 | rs4665554 | 0.241 | 0.8 | -0.186 | -0.018 | 0 | 0.001 | 0.001 | 0 | 0.52 | 0.5 | 0.64 | 0.7 | 0.46 | 0.54 |
| 2 | 23351731 | rs1470671 | 0.361 | 0.237 | -0.425 | 0.361 | -0.005 | 0.007 | 0.007 | 0 | 0.25 | 0.25 | 0.43 | 0.45 | 0.19 | 0.15 |
| 2 | 23353834 | rs6731656 | -0.441 | 0.586 | 0.227 | 0.499 | 0 | 0 | 0 | 0.038 | 0.45 | 0.51 | 0.36 | 0.38 | 0.5 | 0.66 |
| 2 | 23359101 | rs1530758 | 0.362 | 1.005 | -0.234 | 0.263 | -0.075 | -0.038 | -0.038 | 0 | 0.26 | 0.04 | 0.19 | 0.23 | 0.29 | 0.37 |
| 2 | 23364711 | rs1449965 | -0.408 | 0.874 | 0.546 | 0.525 | 0.023 | 0.024 | 0.024 | 0.109 | 0.25 | 0.43 | 0.33 | 0.33 | 0.31 | 0.33 |
| 2 | 23365209 | rs6736482 | -1.354 | 0.736 | 0.154 | 0.484 | 0 | -0.012 | -0.012 | 0 | 0.52 | 0.85 | 0.59 | 0.55 | 0.61 | 0.73 |
| 2 | 23370317 | rs10197888 | -0.508 | 0.801 | 0.361 | 0.664 | -0.004 | 0.038 | 0.038 | 0 | 0.1 | 0.16 | 0.19 | 0.25 | 0.09 | 0.09 |
| 2 | 23382331 | rs1839415 | 0.636 | 1.03 | 0.53 | 0.532 | 0.008 | -0.009 | -0.009 | 0.045 | 0.15 | 0.21 | 0.13 | 0.09 | 0.22 | 0.25 |
| 2 | 23383825 | rs13012579 | -0.232 | 0.862 | 0.836 | 0.494 | 0.005 | -0.014 | -0.014 | 0.6 | 0.3 | 0.25 | 0.35 | 0.46 | 0.29 | 0.43 |
| 2 | 23393713 | rs1597407 | -0.552 | 0.858 | 1.4 | 0.519 | -0.005 | 0.049 | 0.049 | 0 | 0.33 | 0.46 | 0.4 | 0.5 | 0.35 | 0.45 |
| 2 | 23402150 | rs6716929 | -1.516 | 0.868 | 1.272 | 0.459 | -0.002 | 0.005 | 0.005 | 2.424 | 0.52 | 0.86 | 0.57 | 0.58 | 0.61 | 0.7 |
| 2 | 23402458 | rs6744878 | -0.025 | 0.815 | 1.516 | 0.515 | 0 | -0.003 | -0.003 | 1.159 | 0.28 | 0.31 | 0.23 | 0.24 | 0.32 | 0.41 |
| 2 | 23411774 | rs17673557 | 0.524 | 0.949 | 1.462 | 0.549 | 0.002 | -0.005 | -0.005 | 0 | 0.16 | 0.2 | 0.15 | 0.08 | 0.25 | 0.25 |
| 2 | 23414651 | rs1449984 | 0.211 | 0.961 | 1.454 | 0.554 | -0.001 | 0 | 0 | 0.005 | 0.18 | 0.22 | 0.18 | 0.11 | 0.32 | 0.28 |
| 2 | 23419038 | rs4665202 | -0.727 | 0.682 | 1.58 | 0.545 | -0.014 | 0.152 | 0.152 | 0.06 | 0.1 | 0.35 | 0.19 | 0.26 | 0.05 | 0.07 |
| 2 | 23428461 | rs17673828 | 0.397 | 0.736 | 0.962 | 0.458 | -0.004 | -0.007 | -0.007 | 0 | 0.25 | 0.05 | 0.2 | 0.2 | 0.26 | 0.33 |
| 2 | 23431574 | rs4665567 | 0.356 | 0.782 | 1.171 | 0.458 | -0.012 | 0.02 | 0.02 | 0.085 | 0.59 | 0.67 | 0.67 | 0.72 | 0.51 | 0.4 |
| 2 | 23438337 | rs6719952 | 1.332 | 0.742 | 0.85 | 0.449 | 0.008 | 0.01 | 0.01 | 1.728 | 0.46 | 0.15 | 0.42 | 0.42 | 0.4 | 0.33 |
| 2 | 23438532 | rs12712887 | 0.217 | 0.887 | 0.659 | 0.62 | -0.009 | -0.035 | -0.035 | 2.156 | 0.26 | 0.09 | 0.35 | 0.33 | 0.3 | 0.37 |
| 2 | 23442158 | rs1449968 | 0.002 | 1.07 | 1.019 | 0.397 | 0.038 | 0.005 | 0.005 | 0.226 | 0.78 | 0.62 | 0.78 | 0.73 | 0.65 | 0.67 |
| 2 | 23443266 | rs4665203 | -0.388 | 0.729 | 0.974 | -0.143 | -0.024 | -0.015 | -0.015 | 1.103 | 0.27 | 0.35 | 0.35 | 0.21 | 0.26 | 0.31 |
| 2 | 23444212 | rs7589562 | 0.014 | 0.894 | 0.201 | -0.335 | 0.012 | 0.007 | 0.007 | 0 | 0.3 | 0.13 | 0.36 | 0.22 | 0.3 | 0.34 |
| 2 | 23444263 | rs7598792 | -0.588 | 0.703 | 0.266 | -0.202 | -0.001 | -0.002 | -0.002 | 0.137 | 0.53 | 0.47 | 0.52 | 0.32 | 0.58 | 0.69 |
| 2 | 23445415 | rs11677047 | 0.784 | 0.443 | -0.605 | -0.145 | -0.014 | -0.005 | -0.005 | 0.22 | 0.4 | 0.11 | 0.42 | 0.66 | 0.35 | 0.27 |
| 2 | 23446664 | rs10166879 | 0.441 | 0.561 | -0.112 | -0.243 | 0.013 | 0.037 | 0.037 | 0 | 0.45 | 0.34 | 0.46 | 0.71 | 0.4 | 0.37 |
| 2 | 23453834 | rs13394434 | -1.008 | 0.652 | 0.517 | -0.069 | 0.033 | 0.052 | 0.052 | 1.338 | 0.08 | 0.31 | 0.04 | 0.1 | 0.06 | 0.11 |
| 2 | 23455498 | rs11883447 | 0.463 | 0.632 | 0.534 | -0.084 | -0.016 | -0.028 | -0.028 | 0.243 | 0.05 | 0.15 | 0.01 | 0 | 0.02 | 0.04 |
| 2 | 23458669 | rs11685427 | 0.518 | 0.918 | 0.129 | -0.405 | -0.033 | 0.037 | 0.037 | 0.143 | 0.29 | 0.09 | 0.22 | 0.2 | 0.34 | 0.21 |
| 2 | 23468307 | rs1037301 | -0.946 | 0.755 | 0.504 | -0.839 | -0.011 | -0.027 | -0.027 | 0 | 0.59 | 0.5 | 0.44 | 0.23 | 0.68 | 0.64 |
| 2 | 23468850 | rs7570968 | 0.883 | 0.721 | 0.349 | -0.835 | 0.021 | 0.013 | 0.013 | 0 | 0.38 | 0.28 | 0.54 | 0.67 | 0.29 | 0.29 |
| 2 | 23494356 | rs2577734 | -0.568 | 0.817 | 0.438 | -0.806 | 0.008 | 0.02 | 0.02 | 0 | 0.51 | 0.44 | 0.38 | 0.23 | 0.59 | 0.61 |
| 2 | 23499042 | rs4665578 | 0.723 | 0.839 | 0.68 | -0.76 | 0.035 | 0.062 | 0.062 | 0 | 0.44 | 0.19 | 0.56 | 0.67 | 0.37 | 0.32 |
| 2 | 23500116 | rs12621867 | 0.667 | 0.825 | 0.732 | -0.757 | 0.029 | 0.044 | 0.044 | 0 | 0.45 | 0.28 | 0.57 | 0.67 | 0.37 | 0.32 |
| 2 | 23507017 | rs2577732 | -0.339 | 0.974 | 0.873 | -0.761 | 0.03 | -0.075 | -0.075 | 0 | 0.27 | 0.5 | 0.19 | 0.13 | 0.29 | 0.48 |
| 2 | 23507416 | rs1025703 | 0.069 | 0.974 | 0.856 | -0.763 | 0.004 | -0.062 | -0.062 | 0 | 0.24 | 0.32 | 0.16 | 0.03 | 0.25 | 0.41 |
| 2 | 23512029 | rs1438134 | 0.168 | 0.957 | 0.765 | -0.758 | 0.021 | -0.066 | -0.066 | 0.383 | 0.73 | 0.45 | 0.8 | 0.87 | 0.71 | 0.52 |
| 2 | 23512957 | rs4665582 | -0.082 | 0.963 | 0.74 | -0.755 | 0.016 | -0.048 | -0.048 | 0 | 0.74 | 0.62 | 0.82 | 0.87 | 0.72 | 0.53 |
| 2 | 23516743 | rs4665585 | 0.177 | 0.958 | 0.541 | -0.756 | 0.012 | -0.045 | -0.045 | 0.01 | 0.73 | 0.4 | 0.79 | 0.87 | 0.71 | 0.52 |
| 2 | 23532507 | rs12478810 | -0.069 | 0.853 | -0.584 | -0.393 | -0.033 | 0.021 | 0.021 | 1.581 | 0.64 | 0.22 | 0.56 | 0.68 | 0.58 | 0.7 |
| 2 | 23536135 | rs7587361 | -0.091 | 0.309 | -0.265 | -0.098 | -0.011 | -0.019 | -0.019 | 0 | 0.41 | 0.54 | 0.24 | 0.25 | 0.34 | 0.32 |
| 2 | 23538632 | rs1002159 | -0.629 | -0.113 | 0.243 | 0.781 | -0.007 | 0.026 | 0.026 | 0 | 0.77 | 0.71 | 0.84 | 0.62 | 0.78 | 0.88 |
| 2 | 23538843 | rs1002158 | -0.143 | -0.099 | 0.645 | 0.653 | -0.012 | 0.043 | 0.043 | 0 | 0.72 | 0.55 | 0.81 | 0.62 | 0.74 | 0.86 |
| 2 | 23539934 | rs1025702 | -0.165 | -0.209 | -0.04 | 0.431 | 0.003 | -0.014 | -0.014 | 0.962 | 0.41 | 0.68 | 0.6 | 0.53 | 0.38 | 0.39 |
| 2 | 23544124 | rs893425 | 1.01 | -0.182 | 0.437 | 0.698 | -0.008 | -0.003 | -0.003 | 0 | 0.25 | 0.45 | 0.32 | 0.25 | 0.23 | 0.24 |
| 2 | 23546661 | rs755422 | 0.046 | -0.154 | 0.453 | 0.654 | 0.002 | 0.004 | 0.004 | 0.164 | 0.47 | 0.54 | 0.47 | 0.61 | 0.44 | 0.37 |
| 2 | 23549628 | rs4665588 | 0.503 | -0.102 | 0.275 | 0.738 | -0.004 | -0.011 | -0.011 | 0.016 | 0.15 | 0.12 | 0.27 | 0.31 | 0.14 | 0.15 |
| 2 | 23550244 | rs10495742 | 1.63 | -0.258 | 0.365 | 0.744 | -0.012 | 0.008 | 0.008 | 1.613 | 0.22 | 0.08 | 0.13 | 0.18 | 0.22 | 0.19 |
| 2 | 23558014 | rs2723124 | -1.076 | -0.217 | 0.597 | 0.855 | 0.001 | -0.005 | -0.005 | 0 | 0.77 | 0.77 | 0.84 | 0.65 | 0.78 | 0.88 |
| 2 | 23565258 | rs1465832 | 1.182 | -0.191 | 0.376 | 0.684 | -0.001 | 0.011 | 0.011 | 0 | 0.1 | 0.02 | 0.19 | 0.08 | 0.07 | 0.13 |
| 2 | 23569877 | rs2723094 | 1.035 | -0.221 | 0.983 | 0.505 | -0.015 | 0.073 | 0.073 | 0 | 0.33 | 0.21 | 0.21 | 0.08 | 0.36 | 0.44 |
| 2 | 23571601 | rs6745559 | 0.076 | -0.227 | 1.292 | 0.779 | 0.033 | 0.192 | 0.192 | 0 | 0.08 | 0.16 | 0.26 | 0.36 | 0.11 | 0.11 |
| 2 | 23572734 | rs2577710 | 0.323 | -0.252 | 1.267 | 0.733 | 0.004 | 0 | 0 | 0 | 0.42 | 0.51 | 0.49 | 0.43 | 0.47 | 0.54 |
| 2 | 23572757 | rs2723140 | -1.628 | -0.295 | 1.306 | 0.767 | -0.008 | 0.034 | 0.034 | 0 | 0.8 | 0.85 | 0.86 | 0.95 | 0.77 | 0.9 |
| 2 | 23577270 | rs12995589 | -0.026 | -0.308 | 1.12 | 0.773 | 0.009 | -0.011 | -0.011 | 0 | 0.38 | 0.34 | 0.37 | 0.52 | 0.3 | 0.36 |
| 2 | 23577934 | rs2723137 | 0.914 | -0.228 | 1.219 | 0.352 | -0.008 | 0.033 | 0.033 | 0 | 0.33 | 0.27 | 0.22 | 0.09 | 0.36 | 0.44 |
| 2 | 23581171 | rs726512 | 1.386 | 0.106 | 0.713 | 0.632 | -0.045 | 0.057 | 0.057 | 0 | 0.17 | 0.05 | 0.12 | 0.06 | 0.22 | 0.09 |
| 2 | 23581558 | rs6749822 | 1.282 | 0.205 | 1.011 | 0.679 | 0.001 | -0.019 | -0.019 | 0 | 0.2 | 0.18 | 0.13 | 0.05 | 0.23 | 0.1 |

| | | | | | | | | | | | | | | | | |
|---|---|---|---|---|---|---|---|---|---|---|---|---|---|---|---|---|
| 2 | 23582314 | rs12053097 | -0.542 | 0.277 | 0.824 | 0.62 | 0 | 0.019 | 0.019 | 0 | 0.55 | 0.53 | 0.56 | 0.83 | 0.57 | 0.44 |
| 2 | 23582890 | rs12613333 | 0.447 | -0.227 | 0.778 | 0.96 | -0.002 | 0.025 | 0.025 | 0 | 0.24 | 0.32 | 0.17 | 0.29 | 0.24 | 0.13 |
| 2 | 23582983 | rs7566157 | 0.422 | -0.228 | 0.667 | 0.92 | 0 | 0.046 | 0.046 | 0 | 0.24 | 0.36 | 0.2 | 0.32 | 0.25 | 0.14 |
| 2 | 23584565 | rs875077 | 0.166 | -0.348 | 0.498 | 0.455 | 0.012 | 0.056 | 0.056 | 0 | 0.3 | 0.5 | 0.33 | 0.49 | 0.31 | 0.35 |
| 2 | 23584807 | rs2723129 | 0.045 | -0.265 | 0.039 | 0.379 | 0.003 | 0.001 | 0.001 | 0 | 0.06 | 0.07 | 0.05 | 0.05 | 0.1 | 0.03 |
| 2 | 23585786 | rs13427504 | 1.046 | -0.606 | -0.378 | 0.125 | -0.068 | 0.108 | 0.108 | 0 | 0.3 | 0.07 | 0.22 | 0.11 | 0.33 | 0.38 |
| 2 | 23585867 | rs12465302 | -0.247 | -0.661 | 0.062 | 0.53 | -0.006 | 0.031 | 0.031 | 0 | 0.14 | 0.19 | 0.42 | 0.35 | 0.1 | 0.16 |
| 2 | 23592145 | rs2577704 | -0.042 | -0.465 | 0.435 | 0.401 | 0.024 | 0.014 | 0.014 | 0 | 0.06 | 0.15 | 0.06 | 0.05 | 0.1 | 0.03 |
| 2 | 23602298 | rs1561322 | -0.057 | -0.427 | -0.373 | 0.385 | -0.014 | -0.043 | -0.043 | 0 | 0.38 | 0.23 | 0.54 | 0.56 | 0.36 | 0.47 |
| 2 | 23603909 | rs2723119 | -0.266 | -0.257 | -0.646 | 0.385 | -0.005 | -0.002 | -0.002 | 0 | 0.07 | 0.05 | 0.05 | 0.05 | 0.11 | 0.03 |
| 2 | 23606177 | rs12997821 | 0.098 | -0.306 | -0.989 | 0.186 | -0.052 | -0.006 | -0.006 | 0 | 0.35 | 0.08 | 0.35 | 0.27 | 0.3 | 0.43 |
| 2 | 23610838 | rs11125011 | 0.621 | -0.531 | -0.714 | -0.02 | -0.01 | 0.027 | 0.027 | 0 | 0.29 | 0.21 | 0.22 | 0.11 | 0.32 | 0.38 |
| 2 | 23620246 | rs920271 | -1.083 | -0.512 | -0.628 | 0.318 | 0.011 | -0.011 | -0.011 | 0 | 0.76 | 0.85 | 0.8 | 0.75 | 0.74 | 0.87 |
| 2 | 23621521 | rs7560798 | 0.806 | -0.447 | -0.798 | 0.03 | -0.007 | 0.008 | 0.008 | 0 | 0.23 | 0.19 | 0.2 | 0.12 | 0.27 | 0.32 |
| 2 | 23623283 | rs1530046 | -0.415 | -0.277 | -0.934 | 0.053 | -0.014 | -0.02 | -0.02 | 0 | 0.06 | 0.18 | 0.04 | 0.01 | 0.03 | 0.02 |
| 2 | 23627416 | rs4665209 | 0.646 | -0.117 | -1.439 | 0.157 | -0.122 | -0.084 | -0.084 | 0 | 0.28 | 0.02 | 0.34 | 0.3 | 0.3 | 0.37 |
| 2 | 23639980 | rs7572683 | -0.593 | -0.109 | -0.087 | 0.095 | -0.004 | -0.049 | -0.049 | 0.072 | 0.09 | 0.28 | 0.04 | 0 | 0.05 | 0.04 |
| 2 | 23652025 | rs729616 | -1.331 | -0.27 | -0.909 | 0.063 | -0.042 | 0.016 | 0.016 | 0.091 | 0.53 | 0.9 | 0.48 | 0.43 | 0.44 | 0.5 |
| 2 | 23658616 | rs873543 | -0.842 | -0.344 | -0.809 | 0.037 | -0.007 | -0.008 | -0.008 | 0 | 0.07 | 0.1 | 0.04 | 0.03 | 0.05 | 0.02 |
| 2 | 23664914 | rs4665211 | -0.945 | -0.375 | -0.37 | -0.043 | 0.003 | 0.012 | 0.012 | 0 | 0.44 | 0.67 | 0.45 | 0.42 | 0.39 | 0.49 |
| 2 | 23671725 | rs2164866 | -1.363 | -0.372 | -0.38 | -0.041 | 0.007 | 0.039 | 0.039 | 0.031 | 0.51 | 0.77 | 0.48 | 0.43 | 0.44 | 0.5 |
| 2 | 23672718 | rs11885576 | -0.494 | -0.402 | 0.015 | 0 | 0.028 | 0.208 | 0.208 | 0 | 0.1 | 0.38 | 0.21 | 0.29 | 0.07 | 0.08 |
| 2 | 23676071 | rs6544858 | -1.562 | -0.408 | 0.02 | -0.004 | -0.014 | 0.118 | 0.118 | 0 | 0.16 | 0.46 | 0.24 | 0.31 | 0.12 | 0.1 |
| 2 | 23676505 | rs6544861 | -1.541 | -0.394 | -0.033 | 0.05 | 0.011 | 0.23 | 0.23 | 0.978 | 0.1 | 0.41 | 0.22 | 0.31 | 0.08 | 0.06 |
| 2 | 23676916 | rs11689283 | -0.71 | -0.383 | -0.113 | 0.007 | -0.003 | -0.009 | -0.009 | 0 | 0.07 | 0.08 | 0.03 | 0.01 | 0.05 | 0.02 |
| 2 | 23677053 | rs13417938 | -0.358 | -0.348 | -0.351 | 0.001 | 0.001 | 0.008 | 0.008 | 0 | 0.35 | 0.31 | 0.24 | 0.13 | 0.32 | 0.41 |
| 2 | 23684388 | rs4665594 | -1.73 | -0.097 | -0.198 | 0.644 | 0.028 | 0.233 | 0.233 | 2.796 | 0.09 | 0.39 | 0.22 | 0.3 | 0.08 | 0.05 |
| 2 | 23689034 | rs1530045 | -0.531 | 0.31 | -0.481 | 0.367 | -0.001 | 0.013 | 0.013 | 0 | 0.28 | 0.33 | 0.37 | 0.39 | 0.29 | 0.35 |
| 2 | 23692679 | rs744109 | -0.675 | 0.137 | -1.113 | 0.687 | 0.002 | -0.003 | -0.003 | 0 | 0.34 | 0.61 | 0.35 | 0.41 | 0.25 | 0.37 |
| 2 | 23700603 | rs10178369 | -0.667 | 0.226 | -1.149 | 0.736 | 0.002 | -0.002 | -0.002 | 0.074 | 0.35 | 0.6 | 0.36 | 0.41 | 0.26 | 0.37 |
| 2 | 23705389 | rs13011734 | 0.49 | 0.682 | -2.032 | 0.754 | 0.005 | 0.021 | 0.021 | 0 | 0.5 | 0.08 | 0.46 | 0.41 | 0.41 | 0.37 |
| 2 | 23705676 | rs1465677 | -0.001 | 1.544 | -1.042 | 0.715 | 0.072 | -0.025 | -0.025 | 0 | 0.85 | 0.69 | 0.82 | 0.91 | 0.74 | 0.79 |
| 2 | 23707766 | rs6712599 | 0.167 | 1.729 | -0.829 | 0.663 | 0.044 | -0.018 | -0.018 | 0.286 | 0.86 | 0.75 | 0.84 | 0.91 | 0.79 | 0.81 |
| 2 | 23710164 | rs4665218 | 0.388 | 1.792 | -0.372 | 1.184 | 0.004 | 0.059 | 0.059 | 0 | 0.07 | 0.12 | 0.18 | 0.25 | 0.11 | 0.22 |
| 2 | 23710200 | rs10208318 | 0.388 | 1.792 | -0.372 | 1.184 | 0.004 | 0.059 | 0.059 | 0 | 0.07 | 0.12 | 0.18 | 0.25 | 0.11 | 0.22 |
| 2 | 23711427 | rs1368080 | 0.644 | 1.844 | -0.273 | 1.149 | 0.045 | 0.058 | 0.058 | 0 | 0.79 | 0.67 | 0.67 | 0.67 | 0.68 | 0.56 |
| 2 | 23711564 | rs10490750 | -0.027 | 1.612 | 0.124 | 0.94 | -0.005 | -0.008 | -0.008 | 0 | 0.26 | 0.4 | 0.18 | 0.24 | 0.23 | 0.19 |
| 2 | 23713107 | rs7601971 | 0.931 | 1.404 | -0.243 | 0.637 | 0.017 | 0.039 | 0.039 | 0 | 0.52 | 0.26 | 0.54 | 0.59 | 0.48 | 0.47 |
| 2 | 23716694 | rs6742202 | -0.524 | 1.169 | -0.234 | 0.66 | 0.003 | 0.023 | 0.023 | 0 | 0.21 | 0.28 | 0.38 | 0.25 | 0.22 | 0.36 |
| 2 | 23719023 | rs1344751 | 1.183 | 0.298 | -0.798 | 0.291 | -0.007 | -0.061 | -0.061 | 0 | 0.31 | 0.13 | 0.33 | 0.46 | 0.3 | 0.24 |
| 2 | 23720804 | rs1653752 | 0.255 | 0.333 | -0.418 | -0.354 | -0.001 | -0.024 | -0.024 | 0 | 0.35 | 0.34 | 0.25 | 0.08 | 0.43 | 0.34 |
| 2 | 23721872 | rs1709330 | 0.202 | 0.355 | -0.42 | -0.365 | 0.013 | -0.059 | -0.059 | 0 | 0.36 | 0.39 | 0.26 | 0.08 | 0.45 | 0.34 |
| 2 | 23724991 | rs1653763 | -0.418 | 0.314 | -0.201 | -0.437 | -0.005 | 0.014 | 0.014 | 0 | 0.44 | 0.6 | 0.31 | 0.1 | 0.52 | 0.45 |
| 2 | 23729232 | rs1653765 | 0.053 | 0.308 | -0.259 | -0.411 | 0.013 | -0.074 | -0.074 | 0 | 0.38 | 0.43 | 0.26 | 0.08 | 0.44 | 0.35 |
| 2 | 23733367 | rs1653769 | -0.009 | -0.296 | -0.06 | 0.064 | -0.004 | 0.002 | 0.002 | 0 | 0.73 | 0.72 | 0.64 | 0.65 | 0.79 | 0.69 |
| 2 | 23736076 | rs6722403 | 0.968 | -0.236 | -0.405 | 0.062 | -0.013 | 0.071 | 0.071 | 0.227 | 0.2 | 0.29 | 0.27 | 0.55 | 0.13 | 0.25 |
| 2 | 23741410 | rs12712975 | 0.483 | -0.102 | -0.039 | 0.252 | -0.008 | 0.002 | 0.002 | 0 | 0.66 | 0.62 | 0.58 | 0.64 | 0.71 | 0.65 |
| 2 | 23754948 | rs748860 | 0.595 | 0.518 | -0.272 | 0.338 | 0.001 | -0.001 | -0.001 | 0 | 0.55 | 0.42 | 0.5 | 0.6 | 0.59 | 0.52 |
| 2 | 23755080 | rs748861 | 0.381 | 0.495 | -0.511 | 0.376 | 0.001 | -0.001 | -0.001 | 0.136 | 0.57 | 0.59 | 0.53 | 0.6 | 0.61 | 0.53 |
| 2 | 23759508 | rs1709317 | 0.402 | 0.613 | -1.148 | -0.158 | -0.008 | 0.128 | 0.128 | 0 | 0.66 | 0.86 | 0.8 | 0.89 | 0.65 | 0.78 |
| 2 | 23759607 | rs1709318 | 0.402 | 0.613 | -1.148 | -0.158 | -0.008 | 0.128 | 0.128 | 0 | 0.66 | 0.86 | 0.8 | 0.89 | 0.65 | 0.78 |
| 2 | 23763576 | rs6745031 | -0.786 | 0.851 | -0.778 | -0.19 | 0.004 | -0.007 | -0.007 | 0 | 0.33 | 0.57 | 0.36 | 0.72 | 0.35 | 0.42 |
| 2 | 23764107 | rs1709296 | -0.89 | 0.608 | -0.465 | -0.287 | -0.009 | -0.029 | -0.029 | 0 | 0.08 | 0.18 | 0.05 | 0.02 | 0.06 | 0.08 |
| 2 | 23773262 | rs10179169 | -0.415 | 0.609 | 0.545 | 0.047 | 0.062 | 0.259 | 0.259 | 0.581 | 0.05 | 0.32 | 0.27 | 0.21 | 0.05 | 0.11 |
| 2 | 23773948 | rs17445645 | -0.726 | 1.128 | 0.241 | -0.349 | -0.022 | 0.04 | 0.04 | 0.608 | 0.2 | 0.12 | 0.17 | 0.04 | 0.32 | 0.19 |
| 2 | 23774752 | rs4665609 | 0.069 | 0.929 | 0.46 | -0.401 | -0.003 | 0.002 | 0.002 | 0 | 0.6 | 0.6 | 0.57 | 0.32 | 0.56 | 0.59 |
| 2 | 23781218 | rs4665612 | -0.403 | 0.974 | 0.67 | -0.233 | 0.076 | 0.266 | 0.266 | 0.263 | 0.06 | 0.34 | 0.27 | 0.21 | 0.05 | 0.12 |
| 2 | 23783753 | rs1709339 | -0.133 | 1.101 | 0.734 | -0.424 | 0.004 | -0.029 | -0.029 | 0 | 0.73 | 0.55 | 0.6 | 0.79 | 0.74 | 0.75 |
| 2 | 23787723 | rs6544949 | -0.442 | 1.11 | 0.681 | -0.401 | 0.055 | 0.242 | 0.242 | 0.123 | 0.07 | 0.3 | 0.31 | 0.23 | 0.06 | 0.12 |
| 2 | 23791431 | rs13035193 | 0.279 | 1.084 | 0.491 | -0.464 | -0.003 | 0.036 | 0.036 | 0 | 0.69 | 0.2 | 0.53 | 0.77 | 0.72 | 0.7 |
| 2 | 23791950 | rs1653721 | -1.024 | 1.061 | 0.492 | -0.451 | -0.019 | 0.071 | 0.071 | 0.056 | 0.8 | 0.89 | 0.88 | 1 | 0.8 | 0.87 |
| 2 | 23798079 | rs874868 | 0.068 | 1.058 | 0.578 | -0.434 | 0.006 | 0.01 | 0.01 | 0.119 | 0.72 | 0.28 | 0.57 | 0.77 | 0.77 | 0.76 |
| 2 | 23806335 | rs1653725 | -0.986 | 1.1 | 0.134 | -0.564 | -0.024 | 0.088 | 0.088 | 0.233 | 0.8 | 0.9 | 0.88 | 1 | 0.8 | 0.88 |
| 2 | 23817398 | rs606374 | 0.038 | 1.114 | 0.625 | -0.326 | 0 | -0.01 | -0.01 | 0.029 | 0.7 | 0.58 | 0.57 | 0.72 | 0.71 | 0.75 |
| 2 | 23821742 | rs1030281 | 0.163 | 1.112 | 0.482 | -0.333 | -0.007 | -0.015 | -0.015 | 0 | 0.68 | 0.46 | 0.56 | 0.71 | 0.71 | 0.75 |
| 2 | 23825044 | rs17045693 | 0.226 | 1.295 | 0.137 | -0.25 | -0.019 | -0.039 | -0.039 | 0.966 | 0.17 | 0.14 | 0.27 | 0.58 | 0.19 | 0.29 |
| 2 | 23829145 | rs583168 | 0.904 | 1.547 | 0.023 | -0.348 | 0.01 | 0.013 | 0.013 | 0 | 0.47 | 0.53 | 0.51 | 0.37 | 0.39 | 0.43 |
| 2 | 23838148 | rs6714495 | -0.117 | 1.539 | -0.01 | -0.26 | 0.082 | 0.34 | 0.34 | 0.02 | 0.07 | 0.34 | 0.31 | 0.3 | 0.08 | 0.12 |
| 2 | 23842690 | rs605750 | 1.396 | 1.508 | -1.002 | -1.088 | 0.027 | 0.125 | 0.125 | 0 | 0.39 | 0.17 | 0.19 | 0.09 | 0.32 | 0.26 |

| | | | | | | | | | | | | | | | | |
|---|---|---|---|---|---|---|---|---|---|---|---|---|---|---|---|---|
| 2 | 23847980 | rs577293 | -0.183 | 1.434 | -1.12 | -0.637 | 0.005 | -0.014 | -0.014 | 0.026 | 0.74 | 0.76 | 0.87 | 1 | 0.79 | 0.86 |
| 2 | 23854896 | rs1548020 | -0.857 | 1.504 | -1.13 | 0.231 | -0.056 | 0.017 | 0.017 | 0 | 0.12 | 0.02 | 0.1 | 0.04 | 0.21 | 0.13 |
| 2 | 23859550 | rs11688819 | -0.834 | 1.265 | -0.81 | 0.325 | 0.003 | -0.005 | -0.005 | 0 | 0.14 | 0.26 | 0.09 | 0.09 | 0.16 | 0.13 |
| 2 | 23860732 | rs689184 | 0.516 | 1.224 | -1.455 | 0.615 | -0.002 | -0.042 | -0.042 | 0.075 | 0.54 | 0.46 | 0.69 | 0.9 | 0.48 | 0.63 |
| 2 | 23862487 | rs543012 | 0.528 | 1.316 | -1.368 | 0.399 | -0.001 | -0.005 | -0.005 | 0.006 | 0.52 | 0.44 | 0.36 | 0.43 | 0.48 | 0.55 |
| 2 | 23872765 | rs7581165 | -0.288 | 1.434 | -2.225 | 0.719 | -0.117 | -0.156 | -0.156 | 0.058 | 0.27 | 0.06 | 0.48 | 0.44 | 0.37 | 0.43 |
| 2 | 23875431 | rs6721623 | -0.098 | 0.943 | -2.306 | 0.074 | 0.001 | -0.015 | -0.015 | 3.598 | 0.28 | 0.21 | 0.29 | 0.39 | 0.31 | 0.31 |
| 2 | 23877927 | rs3795932 | 0.235 | 0.762 | -2.835 | -0.227 | -0.115 | -0.142 | -0.142 | 0 | 0.35 | 0.02 | 0.51 | 0.52 | 0.41 | 0.45 |
| 2 | 23885168 | rs889866 | -1.66 | -0.109 | -2.014 | -1.046 | -0.053 | -0.129 | -0.129 | 0 | 0.82 | 0.44 | 0.9 | 0.99 | 0.92 | 0.95 |
| 2 | 23887437 | rs1991083 | 0.132 | -0.06 | -2.341 | -1.453 | 0.036 | 0.056 | 0.056 | 0 | 0.56 | 0.32 | 0.76 | 0.93 | 0.67 | 0.73 |
| 2 | 23889614 | rs1007195 | -1.551 | 0.02 | -2.284 | -1.429 | -0.053 | -0.138 | -0.138 | 0.389 | 0.79 | 0.44 | 0.88 | 0.99 | 0.89 | 0.92 |
| 2 | 23891433 | rs737665 | -0.773 | 0.303 | -2.399 | -1.522 | -0.009 | -0.091 | -0.091 | 0 | 0.72 | 0.35 | 0.82 | 0.99 | 0.79 | 0.84 |
| 2 | 23891551 | rs6715626 | -0.897 | 0.211 | -2.391 | -1.505 | -0.027 | -0.104 | -0.104 | 0 | 0.73 | 0.38 | 0.83 | 0.99 | 0.84 | 0.84 |
| 2 | 23891866 | rs13397566 | 0.85 | 0.409 | -2.393 | -1.204 | 0 | 0.005 | 0.005 | 0 | 0.11 | 0.13 | 0.07 | 0.19 | 0.13 | 0.13 |
| 2 | 23898966 | rs11125163 | 1.034 | 0.573 | -2.568 | -1.108 | -0.141 | -0.069 | -0.069 | 0.499 | 0.34 | 0.02 | 0.6 | 0.6 | 0.41 | 0.46 |
| 2 | 23913985 | rs2551350 | -2.454 | 0.427 | -1.375 | -1.364 | -0.036 | -0.1 | -0.1 | 0 | 0.86 | 0.62 | 0.92 | 1 | 0.94 | 0.98 |
| 2 | 23917721 | rs2551353 | -0.645 | 0.37 | -1.508 | -1.568 | -0.038 | -0.169 | -0.169 | 0 | 0.75 | 0.5 | 0.86 | 1 | 0.82 | 0.85 |
| 2 | 23919375 | rs3795942 | 0.525 | 0.709 | -1.705 | -1.708 | 0.015 | 0.065 | 0.065 | 0 | 0.5 | 0.37 | 0.7 | 0.78 | 0.54 | 0.58 |
| 2 | 23920545 | rs3795944 | 0.618 | 0.708 | -1.718 | -1.709 | 0.009 | 0.036 | 0.036 | 0 | 0.51 | 0.41 | 0.7 | 0.79 | 0.55 | 0.59 |
| 2 | 23926659 | rs3795948 | 0.943 | 0.723 | -1.712 | -1.699 | -0.007 | 0.022 | 0.022 | 0 | 0.44 | 0.39 | 0.67 | 0.78 | 0.46 | 0.55 |
| 2 | 23929979 | rs17045819 | -0.001 | 0.719 | -1.698 | -1.675 | -0.013 | 0.065 | 0.065 | 0 | 0.12 | 0.02 | 0.06 | 0 | 0.12 | 0.13 |
| 2 | 23930714 | rs2712098 | -2.605 | 0.72 | -1.568 | -1.581 | -0.037 | -0.109 | -0.109 | 3.849 | 0.86 | 0.52 | 0.92 | 1 | 0.94 | 0.98 |
| 2 | 23932461 | rs2712102 | 0.813 | 0.936 | -1.648 | -1.558 | -0.043 | -0.025 | -0.025 | 1.695 | 0.11 | 0.03 | 0.1 | 0.16 | 0.2 | 0.16 |
| 2 | 23947589 | rs12478744 | 0.004 | 0.963 | -1.607 | -1.517 | 0 | -0.005 | -0.005 | 1.016 | 0.58 | 0.43 | 0.73 | 0.78 | 0.59 | 0.68 |
| 2 | 23948953 | rs7608623 | 0.01 | 0.955 | -1.618 | -1.54 | 0 | -0.002 | -0.002 | 1.705 | 0.58 | 0.43 | 0.73 | 0.78 | 0.59 | 0.68 |
| 2 | 23956799 | rs2042481 | 0.916 | 1.077 | -1.634 | -1.685 | 0 | 0.043 | 0.043 | 0 | 0.4 | 0.26 | 0.62 | 0.77 | 0.39 | 0.52 |
| 2 | 23973603 | rs2712065 | 0.238 | 0.792 | -1.406 | -1.966 | 0.005 | 0.007 | 0.007 | 0 | 0.24 | 0.22 | 0.14 | 0.07 | 0.2 | 0.21 |
| 2 | 23973799 | rs6751857 | 1.304 | 0.905 | -1.422 | -1.973 | -0.013 | 0.044 | 0.044 | 0 | 0.4 | 0.27 | 0.64 | 0.78 | 0.42 | 0.52 |
| 2 | 23979729 | rs2551325 | 0.287 | 0.908 | -1.416 | -1.953 | 0.009 | 0.08 | 0.08 | 0 | 0.24 | 0.12 | 0.13 | 0.07 | 0.2 | 0.21 |
| 2 | 23982378 | rs2551326 | 0.076 | 0.907 | -1.384 | -1.951 | 0.013 | 0.07 | 0.07 | 0 | 0.25 | 0.14 | 0.14 | 0.07 | 0.2 | 0.21 |
| 2 | 24039779 | rs2339923 | 1.248 | 0.905 | -1.339 | -1.96 | -0.006 | 0.021 | 0.021 | 0 | 0.41 | 0.31 | 0.64 | 0.78 | 0.43 | 0.52 |
| 2 | 24056668 | rs17761971 | 1.511 | 0.963 | -1.405 | -2.023 | -0.035 | -0.028 | -0.028 | 0 | 0.08 | 0.02 | 0.08 | 0.15 | 0.18 | 0.13 |
| 2 | 24066193 | rs4233703 | 1.341 | 0.959 | -1.387 | -1.934 | -0.013 | 0.044 | 0.044 | 0 | 0.4 | 0.27 | 0.62 | 0.76 | 0.43 | 0.47 |
| 2 | 24071427 | rs7588286 | 0.858 | 0.996 | -1.394 | -1.908 | 0.003 | 0.077 | 0.077 | 0 | 0.2 | 0.08 | 0.09 | 0.06 | 0.16 | 0.16 |
| 2 | 24113354 | rs6731333 | -0.091 | 1.011 | -1.384 | -1.863 | 0.006 | 0.011 | 0.011 | 0 | 0.26 | 0.23 | 0.14 | 0.07 | 0.23 | 0.22 |
| 2 | 24154192 | rs11674248 | 0.634 | 0.947 | -1.413 | -1.912 | 0.015 | 0.071 | 0.071 | 0 | 0.51 | 0.33 | 0.71 | 0.78 | 0.55 | 0.64 |
| 2 | 24162018 | rs7572949 | 0.634 | 1 | -1.414 | -1.912 | 0.013 | 0.068 | 0.068 | 0 | 0.51 | 0.33 | 0.71 | 0.78 | 0.55 | 0.64 |
| 2 | 24163121 | rs6729507 | -1.47 | 0.996 | -1.386 | -1.906 | -0.004 | -0.044 | -0.044 | 1.15 | 0.77 | 0.63 | 0.85 | 0.85 | 0.77 | 0.84 |
| 2 | 24181159 | rs12616678 | 0.15 | 0.95 | -1.366 | -1.902 | 0.002 | -0.004 | -0.004 | 0 | 0.25 | 0.24 | 0.14 | 0.07 | 0.21 | 0.21 |
| 2 | 24231982 | rs12475277 | -1.423 | 1.467 | -1.398 | -1.913 | -0.01 | -0.049 | -0.049 | 0 | 0.76 | 0.56 | 0.86 | 0.85 | 0.78 | 0.84 |
| 2 | 24234440 | rs925229 | 1.278 | 1.401 | -1.491 | -2.039 | 0.005 | -0.031 | -0.031 | 0 | 0.39 | 0.43 | 0.66 | 0.77 | 0.41 | 0.45 |
| 2 | 24235704 | rs6545222 | -1.394 | 1.328 | -1.373 | -1.949 | -0.009 | -0.036 | -0.036 | 2.924 | 0.76 | 0.59 | 0.86 | 0.84 | 0.77 | 0.79 |
| 2 | 24243127 | rs2136264 | -0.142 | 1.301 | -1.456 | -2.05 | -0.011 | -0.072 | -0.072 | 0 | 0.64 | 0.56 | 0.8 | 0.88 | 0.68 | 0.68 |
| 2 | 24246175 | rs12989842 | -1.478 | 1.334 | -1.41 | -1.96 | -0.006 | -0.054 | -0.054 | 0.026 | 0.77 | 0.59 | 0.87 | 0.89 | 0.77 | 0.83 |
| 2 | 24246679 | rs925228 | -1.504 | 1.331 | -1.458 | -1.991 | -0.005 | -0.054 | -0.054 | 0 | 0.77 | 0.63 | 0.87 | 0.9 | 0.77 | 0.83 |
| 2 | 24251787 | rs2339940 | 0.804 | 1.175 | -1.539 | -2.051 | 0.001 | -0.031 | -0.031 | 0 | 0.46 | 0.5 | 0.7 | 0.83 | 0.52 | 0.48 |
| 2 | 24252436 | rs3204 | 1.263 | 1.222 | -1.538 | -2.047 | 0 | -0.017 | -0.017 | 0 | 0.41 | 0.39 | 0.66 | 0.83 | 0.44 | 0.45 |
| 2 | 24256071 | rs17763463 | -0.427 | 1.093 | -1.342 | -1.813 | 0.005 | 0.038 | 0.038 | 0 | 0.12 | 0.05 | 0.07 | 0.01 | 0.1 | 0.16 |
| 2 | 24262060 | rs3731620 | 0.539 | 1.077 | -1.308 | -1.758 | -0.016 | 0.017 | 0.017 | 0 | 0.1 | 0.08 | 0.05 | 0.04 | 0.14 | 0.18 |
| 2 | 24266198 | rs4270307 | 1.635 | 0.829 | -1.213 | -1.991 | -0.049 | -0.131 | -0.131 | 0 | 0.2 | 0.35 | 0.11 | 0.01 | 0.11 | 0.09 |
| 2 | 24272396 | rs13413450 | 0.686 | 0.831 | -1.215 | -1.99 | -0.016 | 0.017 | 0.017 | 0 | 0.1 | 0.08 | 0.05 | 0.04 | 0.14 | 0.18 |
| 2 | 24275306 | rs10185680 | -0.87 | 0.573 | -1.157 | -2.184 | 0.004 | 0.006 | 0.006 | 0 | 0.48 | 0.55 | 0.3 | 0.1 | 0.47 | 0.48 |
| 2 | 24276736 | rs11125393 | 0.521 | 0.583 | -1.104 | -2.143 | -0.019 | -0.129 | -0.129 | 0.115 | 0.3 | 0.45 | 0.17 | 0.05 | 0.25 | 0.26 |
| 2 | 24278841 | rs7568163 | 0.452 | 0.579 | -1.128 | -2.136 | -0.02 | -0.118 | -0.118 | 0.067 | 0.3 | 0.43 | 0.17 | 0.05 | 0.25 | 0.26 |
| 2 | 24281484 | rs11902184 | 1.695 | 0.542 | -1.111 | -2.124 | -0.05 | -0.131 | -0.131 | 0 | 0.2 | 0.35 | 0.11 | 0.01 | 0.11 | 0.08 |
| 2 | 24288775 | rs2160943 | -1.331 | 0.496 | -1.144 | -1.886 | 0.002 | -0.04 | -0.04 | 0 | 0.35 | 0.53 | 0.23 | 0.15 | 0.37 | 0.44 |
| 2 | 24289487 | rs11895257 | 0.047 | 0.488 | -1.136 | -1.888 | 0.028 | 0.007 | 0.007 | 1.798 | 0.07 | 0.37 | 0.05 | 0.01 | 0.02 | 0.05 |
| 2 | 24293756 | rs10189243 | -1.368 | 0.489 | -1.138 | -1.89 | 0.002 | -0.04 | -0.04 | 0 | 0.35 | 0.53 | 0.23 | 0.15 | 0.37 | 0.44 |
| 2 | 24296100 | rs11890987 | 0.945 | 0.509 | -1.141 | -1.887 | -0.014 | 0.015 | 0.015 | 0 | 0.09 | 0.08 | 0.05 | 0.04 | 0.14 | 0.17 |
| 2 | 24297527 | rs13416994 | 0.975 | 0.517 | -0.994 | -1.884 | -0.021 | 0.018 | 0.018 | 0 | 0.09 | 0.06 | 0.05 | 0.04 | 0.14 | 0.17 |
| 2 | 24300563 | rs10209238 | 0.975 | 0.517 | -0.995 | -1.884 | -0.021 | 0.018 | 0.018 | 0 | 0.09 | 0.07 | 0.05 | 0.04 | 0.14 | 0.17 |
| 2 | 24302246 | rs2303287 | 0.879 | 0.511 | -1.002 | -1.888 | -0.059 | -0.009 | -0.009 | 0.842 | 0.17 | 0.01 | 0.13 | 0.11 | 0.22 | 0.22 |
| 2 | 24305512 | rs4149372 | -0.495 | 0.531 | -0.986 | -1.879 | -0.029 | 0.036 | 0.036 | 0 | 0.28 | 0.16 | 0.18 | 0.14 | 0.36 | 0.39 |
| 2 | 24335554 | rs13028767 | 0.939 | 0.539 | -0.981 | -1.867 | -0.014 | 0.013 | 0.013 | 0 | 0.09 | 0.08 | 0.05 | 0.04 | 0.14 | 0.17 |
| 2 | 24337594 | rs10865262 | 0.808 | 0.438 | -1.016 | -1.859 | -0.006 | 0.068 | 0.068 | 0 | 0.48 | 0.65 | 0.67 | 0.78 | 0.46 | 0.46 |
| 2 | 24337861 | rs10200481 | 0.973 | 0.543 | -1.265 | -1.804 | -0.014 | 0.015 | 0.015 | 0 | 0.09 | 0.08 | 0.05 | 0.04 | 0.14 | 0.17 |
| 2 | 24342532 | rs1134516 | 0.092 | 0.541 | -1.268 | -1.806 | -0.003 | 0.001 | 0.001 | 0 | 0.13 | 0.13 | 0.08 | 0.05 | 0.16 | 0.22 |
| 2 | 24344914 | rs13006081 | 0.965 | 0.56 | -1.279 | -1.789 | -0.018 | 0.018 | 0.018 | 0 | 0.09 | 0.07 | 0.05 | 0.04 | 0.14 | 0.17 |
| 2 | 24352398 | rs10197527 | -0.565 | 0.387 | -1.215 | -1.724 | -0.014 | 0.032 | 0.032 | 0 | 0.31 | 0.21 | 0.21 | 0.15 | 0.37 | 0.44 |

| | | | | | | | | | | | | | | | | |
|---|---|---|---|---|---|---|---|---|---|---|---|---|---|---|---|---|
| 2 | 24358652 | rs13035774 | 0.815 | 0.655 | -1.236 | -1.449 | -0.015 | 0.054 | 0.054 | 0 | 0.44 | 0.31 | 0.65 | 0.8 | 0.46 | 0.46 |
| 2 | 24369019 | rs6754391 | 0.961 | 1.047 | -0.619 | -0.868 | 0.018 | -0.019 | -0.019 | 0 | 0.67 | 0.45 | 0.76 | 0.78 | 0.59 | 0.59 |
| 2 | 24369674 | rs11693860 | 0.346 | 0.857 | -0.265 | -0.671 | 0.008 | 0.043 | 0.043 | 1.728 | 0.9 | 0.59 | 0.93 | 0.92 | 0.94 | 0.95 |
| 2 | 24380715 | rs4665665 | 1.181 | 0.723 | -0.332 | -0.749 | -0.036 | -0.017 | -0.017 | 0.014 | 0.85 | 0.54 | 0.91 | 0.91 | 0.92 | 0.91 |
| 2 | 24391129 | rs6723257 | 1.424 | 0.821 | -0.442 | -0.828 | 0.009 | -0.013 | -0.013 | 0 | 0.68 | 0.4 | 0.77 | 0.78 | 0.62 | 0.6 |
| 2 | 24392618 | rs10199334 | 0.279 | 0.835 | -0.496 | -0.825 | -0.061 | -0.013 | -0.013 | 0 | 0.17 | 0.09 | 0.14 | 0.13 | 0.3 | 0.31 |
| 2 | 24397572 | rs11688445 | 1.034 | 0.791 | -0.34 | -0.805 | -0.03 | -0.025 | -0.025 | 0.008 | 0.86 | 0.58 | 0.91 | 0.92 | 0.92 | 0.91 |
| 2 | 24398281 | rs2042467 | 0.156 | 0.843 | -0.329 | -0.767 | 0.005 | 0.025 | 0.025 | 1.303 | 0.9 | 0.62 | 0.94 | 0.93 | 0.94 | 0.96 |
| 2 | 24413298 | rs2288073 | -0.118 | 0.953 | -0.651 | -0.69 | -0.033 | -0.007 | -0.007 | 0.807 | 0.19 | 0.14 | 0.15 | 0.15 | 0.31 | 0.32 |
| 2 | 24425620 | rs12052954 | 0.609 | 0.975 | -0.723 | -0.675 | -0.041 | -0.018 | -0.018 | 0 | 0.15 | 0.08 | 0.14 | 0.12 | 0.26 | 0.2 |
| 2 | 24431184 | rs2303291 | 0.518 | 0.923 | -0.805 | -0.7 | -0.078 | -0.048 | -0.048 | 0.002 | 0.15 | 0.02 | 0.13 | 0.12 | 0.26 | 0.2 |
| 2 | 24432211 | rs2303293 | 0.313 | 0.788 | -0.345 | -0.82 | -0.018 | -0.011 | -0.011 | 0.08 | 0.87 | 0.68 | 0.93 | 0.92 | 0.91 | 0.93 |
| 2 | 24432620 | rs2303294 | 1.314 | 0.737 | -0.266 | -0.871 | 0.005 | -0.014 | -0.014 | 0 | 0.69 | 0.58 | 0.76 | 0.79 | 0.63 | 0.69 |
| 2 | 24432839 | rs2303296 | 0.78 | 0.75 | -0.284 | -0.861 | -0.077 | -0.049 | -0.049 | 0.102 | 0.14 | 0.02 | 0.14 | 0.12 | 0.26 | 0.2 |
| 2 | 24434424 | rs2111882 | 0.803 | 0.743 | -0.279 | -0.864 | 0.007 | -0.003 | -0.003 | 0 | 0.73 | 0.64 | 0.79 | 0.8 | 0.65 | 0.73 |
| 2 | 24439048 | rs3731625 | -0.032 | 0.703 | -0.274 | -0.864 | 0.003 | -0.001 | -0.001 | 0.013 | 0.1 | 0.04 | 0.05 | 0.08 | 0.09 | 0.07 |
| 2 | 24442949 | rs1477220 | 0.005 | 0.71 | -0.271 | -0.861 | 0.002 | -0.001 | -0.001 | 1.42 | 0.1 | 0.04 | 0.05 | 0.08 | 0.09 | 0.07 |
| 2 | 24443271 | rs2543662 | -0.367 | 0.747 | -0.123 | -0.866 | -0.006 | 0 | 0 | 0 | 0.82 | 0.85 | 0.83 | 0.87 | 0.72 | 0.76 |
| 2 | 24443330 | rs1203 | 1.301 | 0.736 | -0.076 | -0.871 | 0.004 | -0.014 | -0.014 | 0.667 | 0.69 | 0.58 | 0.76 | 0.79 | 0.63 | 0.69 |
| 2 | 24443505 | rs2543663 | -0.452 | 0.723 | -0.074 | -0.87 | -0.019 | -0.016 | -0.016 | 0 | 0.13 | 0.27 | 0.07 | 0.08 | 0.09 | 0.07 |
| 2 | 24452180 | rs7589373 | 1.142 | 0.724 | -0.076 | -0.876 | -0.035 | -0.044 | -0.044 | 0 | 0.83 | 0.65 | 0.9 | 0.92 | 0.89 | 0.88 |
| 2 | 24455830 | rs1559472 | 2.003 | -0.054 | -0.627 | -0.863 | -0.049 | -0.037 | -0.037 | 0 | 0.43 | 0.14 | 0.65 | 0.56 | 0.49 | 0.42 |
| 2 | 24461334 | rs17046192 | 2.072 | -0.084 | -0.868 | -0.877 | -0.155 | -0.134 | -0.134 | 0.025 | 0.42 | 0.02 | 0.64 | 0.56 | 0.49 | 0.43 |
| 2 | 24468132 | rs4665675 | 1.657 | 0.56 | 0.091 | -0.878 | 0.005 | -0.012 | -0.012 | 1.042 | 0.65 | 0.43 | 0.74 | 0.78 | 0.58 | 0.58 |
| 2 | 24502554 | rs17789375 | 0.388 | 0.132 | 0.451 | -0.503 | -0.009 | -0.004 | -0.004 | 0.013 | 0.15 | 0.12 | 0.14 | 0.12 | 0.26 | 0.2 |
| 2 | 24509370 | rs17733015 | 0.482 | 0.133 | 0.445 | -0.502 | -0.032 | -0.017 | -0.017 | 0 | 0.14 | 0.07 | 0.14 | 0.12 | 0.26 | 0.2 |
| 2 | 24514996 | rs10495747 | -1.45 | 0.148 | 0.392 | -0.399 | -0.015 | -0.026 | -0.026 | 0.163 | 0.15 | 0.27 | 0.07 | 0.07 | 0.12 | 0.07 |
| 2 | 24524958 | rs7603997 | -1.116 | 0.069 | 0.404 | -0.374 | 0.009 | -0.004 | -0.004 | 0 | 0.28 | 0.39 | 0.21 | 0.19 | 0.35 | 0.27 |
| 2 | 24540591 | rs2551122 | -0.975 | -0.076 | 0.421 | -0.327 | -0.017 | -0.02 | -0.02 | 0.344 | 0.13 | 0.3 | 0.07 | 0.07 | 0.09 | 0.07 |
| 2 | 24587652 | rs2702080 | -0.696 | -0.071 | 0.273 | -0.308 | -0.009 | -0.013 | -0.013 | 0 | 0.12 | 0.2 | 0.06 | 0.07 | 0.09 | 0.07 |
| 2 | 24619883 | rs12470303 | 0.829 | 0.005 | 0.181 | -0.422 | 0 | -0.008 | -0.008 | 0.044 | 0.66 | 0.36 | 0.75 | 0.78 | 0.61 | 0.63 |
| 2 | 24632122 | rs7595372 | -0.879 | 0.086 | 0.333 | -0.069 | 0.002 | -0.007 | -0.007 | 0 | 0.39 | 0.49 | 0.32 | 0.6 | 0.41 | 0.47 |
| 2 | 24646213 | rs11125582 | -0.726 | 0.002 | 0.323 | -0.064 | -0.004 | 0.004 | 0.004 | 0 | 0.38 | 0.28 | 0.3 | 0.61 | 0.41 | 0.47 |
| 2 | 24652464 | rs2702050 | -1.045 | -0.095 | 0.233 | -0.132 | 0 | 0.001 | 0.001 | 0 | 0.45 | 0.53 | 0.28 | 0.61 | 0.45 | 0.54 |
| 2 | 24655725 | rs10189542 | -0.072 | -0.364 | 0.274 | 0.077 | -0.048 | 0.137 | 0.137 | 0 | 0.17 | 0.32 | 0.3 | 0.49 | 0.08 | 0.21 |
| 2 | 24658162 | rs2012222 | -0.308 | -0.193 | 0.027 | -0.17 | -0.025 | 0.059 | 0.059 | 0 | 0.36 | 0.22 | 0.23 | 0.12 | 0.44 | 0.38 |
| 2 | 24658794 | rs1545257 | -0.765 | 0.126 | -0.252 | -0.257 | 0.01 | 0.042 | 0.042 | 0 | 0.5 | 0.64 | 0.54 | 0.62 | 0.49 | 0.56 |
| 2 | 24662502 | rs2702056 | -0.821 | 0.332 | -0.479 | -0.289 | -0.008 | 0.083 | 0.083 | 0 | 0.54 | 0.7 | 0.37 | 0.82 | 0.5 | 0.68 |
| 2 | 24664347 | rs2584923 | -0.893 | 0.305 | -0.454 | -0.269 | -0.012 | 0.074 | 0.074 | 0 | 0.55 | 0.7 | 0.38 | 0.81 | 0.53 | 0.69 |
| 2 | 24672179 | rs13399755 | 0.202 | 0.374 | -1.425 | -0.2 | -0.011 | 0.081 | 0.081 | 0 | 0.15 | 0.05 | 0.09 | 0.02 | 0.15 | 0.07 |
| 2 | 24674318 | rs11675098 | 0.165 | 0.173 | -0.715 | 0.323 | -0.016 | 0.196 | 0.196 | 0 | 0.16 | 0.35 | 0.13 | 0.51 | 0.1 | 0.19 |
| 2 | 24674385 | rs2584937 | -0.108 | 0.161 | -0.825 | -0.615 | -0.018 | 0.005 | 0.005 | 0 | 0.74 | 0.63 | 0.84 | 0.3 | 0.82 | 0.67 |
| 2 | 24677866 | rs10180416 | 0.928 | 0.375 | -1.978 | -0.987 | -0.121 | 0.229 | 0.229 | 0 | 0.45 | 0.06 | 0.62 | 0.17 | 0.48 | 0.31 |
| 2 | 24680249 | rs1031092 | -0.368 | 0.373 | -1.749 | -0.99 | -0.008 | 0.001 | 0.001 | 0 | 0.75 | 0.66 | 0.85 | 0.28 | 0.81 | 0.67 |
| 2 | 24681375 | rs10201521 | 0.365 | 0.659 | -2.699 | -0.713 | -0.128 | 0.234 | 0.234 | 0 | 0.46 | 0.06 | 0.62 | 0.17 | 0.49 | 0.32 |
| 2 | 24686441 | rs6756785 | -0.489 | 0.8 | -1.955 | -0.514 | -0.022 | 0.068 | 0.068 | 0 | 0.26 | 0.09 | 0.4 | 0.1 | 0.29 | 0.28 |
| 2 | 24689512 | rs2702068 | 0.453 | 0.529 | -0.951 | -0.71 | -0.013 | -0.078 | -0.078 | 0 | 0.79 | 0.59 | 0.84 | 0.89 | 0.84 | 0.75 |
| 2 | 24692010 | rs1031089 | 0.232 | 0.444 | -0.388 | -0.802 | -0.004 | -0.117 | -0.117 | 0 | 0.84 | 0.57 | 0.9 | 1 | 0.89 | 0.83 |
| 2 | 24702158 | rs4665701 | 0.804 | 0.748 | -1.461 | -1.294 | -0.023 | 0.212 | 0.212 | 0 | 0.5 | 0.14 | 0.65 | 0.76 | 0.51 | 0.62 |
| 2 | 24724553 | rs9677360 | 0.791 | 0.725 | -1.604 | -1.001 | 0.022 | 0.061 | 0.061 | 0.036 | 0.66 | 0.29 | 0.79 | 0.87 | 0.76 | 0.74 |
| 2 | 24724785 | rs1822300 | 1.318 | 0.746 | -1.651 | -0.976 | -0.064 | 0.211 | 0.211 | 0.047 | 0.5 | 0.09 | 0.66 | 0.76 | 0.54 | 0.62 |
| 2 | 24725464 | rs11125627 | 0.09 | 0.754 | -1.638 | -0.964 | -0.015 | -0.018 | -0.018 | 1.193 | 0.22 | 0.23 | 0.17 | 0.3 | 0.27 | 0.17 |
| 2 | 24739410 | rs1545255 | 0.53 | 0.724 | -1.59 | -0.989 | 0.023 | 0.088 | 0.088 | 0.005 | 0.67 | 0.25 | 0.78 | 0.87 | 0.76 | 0.74 |
| 2 | 24769815 | rs11125662 | -0.139 | 0.583 | -1.63 | -1.031 | -0.056 | 0.049 | 0.049 | 0 | 0.25 | 0.14 | 0.18 | 0.14 | 0.31 | 0.13 |
| 2 | 24784168 | rs7598617 | 1.522 | 0.504 | -1.327 | -1.042 | 0 | 0.036 | 0.036 | 0 | 0.48 | 0.19 | 0.64 | 0.57 | 0.5 | 0.56 |
| 2 | 24799847 | rs12474894 | 2.586 | 0.32 | -2.157 | -1.521 | -0.05 | 0.057 | 0.057 | 0 | 0.18 | 0.01 | 0.29 | 0.05 | 0.18 | 0.12 |
| 2 | 24807533 | rs11688048 | -1.549 | 0.292 | -0.724 | -1.305 | 0 | -0.06 | -0.06 | 0 | 0.88 | 0.84 | 0.94 | 1 | 0.9 | 0.83 |
| 2 | 24810255 | rs995647 | -1.088 | 0.304 | -0.758 | -1.299 | 0.006 | -0.102 | -0.102 | 0 | 0.86 | 0.63 | 0.92 | 1 | 0.9 | 0.83 |
| 2 | 24846815 | rs11677500 | 1.067 | 0.503 | -1.066 | -1.224 | -0.019 | 0.013 | 0.013 | 0.308 | 0.54 | 0.21 | 0.67 | 0.57 | 0.55 | 0.57 |
| 2 | 24856759 | rs11894248 | 0.777 | 0.176 | -1.132 | -1.298 | -0.018 | 0.021 | 0.021 | 0 | 0.2 | 0.15 | 0.13 | 0.11 | 0.23 | 0.15 |
| 2 | 24878211 | rs6708615 | 1.489 | 0.142 | -1.155 | -1.291 | -0.059 | 0.007 | 0.007 | 0 | 0.46 | 0.07 | 0.62 | 0.57 | 0.5 | 0.57 |
| 2 | 24879449 | rs7561414 | -0.848 | 0.067 | -1.002 | -1.275 | -0.015 | -0.107 | -0.107 | 0.027 | 0.84 | 0.52 | 0.91 | 1 | 0.9 | 0.83 |
| 2 | 24881666 | rs17046446 | 0.308 | 0.074 | -1.08 | -1.243 | 0.012 | 0.002 | 0.002 | 0 | 0.08 | 0.03 | 0.04 | 0.1 | 0.04 | 0.08 |
| 2 | 24902072 | rs11693687 | 1.254 | 0.235 | -1.182 | -1.333 | -0.053 | -0.019 | -0.019 | 1.396 | 0.14 | 0.03 | 0.11 | 0.12 | 0.2 | 0.1 |
| 2 | 24914125 | rs2119120 | -1.176 | 0.222 | -1.053 | -1.353 | 0.002 | -0.1 | -0.1 | 0 | 0.86 | 0.63 | 0.92 | 1 | 0.9 | 0.83 |
| 2 | 24915342 | rs6749833 | -0.62 | 0.215 | -1.187 | -1.36 | -0.007 | 0.019 | 0.019 | 0 | 0.29 | 0.19 | 0.18 | 0.24 | 0.28 | 0.22 |
| 2 | 24925294 | rs11125751 | -0.074 | 0.255 | -1.003 | -1.248 | -0.017 | 0.004 | 0.004 | 0 | 0.8 | 0.48 | 0.87 | 0.81 | 0.85 | 0.77 |
| 2 | 24956027 | rs4640343 | 1.397 | 0.391 | -1.098 | -1.254 | -0.029 | 0.022 | 0.022 | 0.383 | 0.07 | 0 | 0.05 | 0.03 | 0.08 | 0.01 |
| 2 | 24994570 | rs9309308 | 0.583 | 0.405 | -1.119 | -1.304 | -0.024 | 0.017 | 0.017 | 0 | 0.18 | 0.09 | 0.12 | 0.12 | 0.22 | 0.12 |

| | | | | | | | | | | | | | | | | |
|---|---|---|---|---|---|---|---|---|---|---|---|---|---|---|---|---|
| 2 | 25009966 | rs12713393 | -0.676 | 0.328 | -0.878 | -1.28 | -0.02 | -0.104 | -0.104 | 0 | 0.84 | 0.49 | 0.9 | 1 | 0.9 | 0.84 |
| 2 | 25010850 | rs6750317 | 0.718 | 0.342 | -1.146 | -1.229 | 0.004 | 0.002 | 0.002 | 0 | 0.06 | 0.03 | 0.04 | 0.1 | 0.04 | 0.08 |
| 2 | 25013062 | rs10177595 | 1.453 | 0.159 | -1.363 | -1.384 | -0.004 | 0.032 | 0.032 | 0 | 0.46 | 0.15 | 0.6 | 0.58 | 0.49 | 0.56 |
| 2 | 25014764 | rs6545729 | 0.558 | 0.09 | -1.269 | -1.536 | 0.002 | 0.009 | 0.009 | 0.105 | 0.59 | 0.42 | 0.72 | 0.81 | 0.63 | 0.61 |
| 2 | 25022598 | rs1550116 | 0.067 | 0.097 | -1.261 | -1.52 | -0.014 | -0.018 | -0.018 | 0 | 0.13 | 0.11 | 0.11 | 0.21 | 0.15 | 0.09 |
| 2 | 25029603 | rs1550111 | 1.521 | 0.078 | -1.309 | -1.525 | -0.023 | 0.05 | 0.05 | 0 | 0.45 | 0.12 | 0.6 | 0.61 | 0.49 | 0.56 |
| 2 | 25032054 | rs6720845 | 0.91 | 0.082 | -1.288 | -1.525 | 0.003 | 0.004 | 0.004 | 0 | 0.06 | 0.09 | 0.04 | 0.07 | 0.04 | 0.06 |
| 2 | 25032151 | rs10208038 | 1.695 | 0.078 | -1.281 | -1.53 | -0.023 | 0.033 | 0.033 | 0 | 0.43 | 0.17 | 0.6 | 0.61 | 0.49 | 0.54 |
| 2 | 25037626 | rs11125797 | 0.634 | 0.216 | -1.162 | -1.482 | -0.029 | 0.02 | 0.02 | 0 | 0.19 | 0.07 | 0.15 | 0.11 | 0.23 | 0.13 |
| 2 | 25038268 | rs2082881 | 0.751 | 0.218 | -1.132 | -1.476 | 0.002 | 0.007 | 0.007 | 0 | 0.68 | 0.36 | 0.79 | 0.79 | 0.75 | 0.73 |
| 2 | 25040082 | rs2278483 | 0.722 | 0.233 | -1.088 | -1.458 | 0 | -0.001 | -0.001 | 0 | 0.18 | 0.2 | 0.15 | 0.11 | 0.23 | 0.1 |
| 2 | 25041620 | rs1550115 | 0.6 | 0.34 | -1.043 | -1.399 | 0 | -0.024 | -0.024 | 0.067 | 0.7 | 0.48 | 0.8 | 0.79 | 0.75 | 0.73 |
| 2 | 25044957 | rs17799872 | 1.488 | 0.399 | -1.113 | -1.303 | -0.004 | 0.005 | 0.005 | 0.393 | 0.07 | 0.06 | 0.06 | 0.04 | 0.08 | 0.01 |
| 2 | 25046090 | rs1127568 | -0.083 | 0.387 | -1.123 | -1.302 | -0.001 | -0.044 | -0.044 | 0 | 0.65 | 0.53 | 0.77 | 0.88 | 0.67 | 0.69 |
| 2 | 25046355 | rs4665273 | 0.692 | 0.431 | -1.224 | -1.276 | 0.002 | 0.044 | 0.044 | 0 | 0.56 | 0.35 | 0.7 | 0.71 | 0.6 | 0.61 |
| 2 | 25057263 | rs2278485 | 1.35 | 0.319 | -1.531 | -1.301 | -0.098 | -0.002 | -0.002 | 0.072 | 0.41 | 0.08 | 0.59 | 0.62 | 0.49 | 0.46 |
| 2 | 25063142 | rs934608 | -0.341 | 0.51 | -1.184 | -1.371 | -0.004 | -0.053 | -0.053 | 0.005 | 0.36 | 0.43 | 0.25 | 0.12 | 0.35 | 0.33 |
| 2 | 25064193 | rs2241759 | 0.923 | 0.591 | -1.45 | -1.198 | 0.003 | 0.079 | 0.079 | 0 | 0.47 | 0.36 | 0.44 | 0.74 | 0.48 | 0.56 |
| 2 | 25070645 | rs1344840 | 0.46 | 0.707 | -1.677 | -0.94 | -0.045 | 0.008 | 0.008 | 0.003 | 0.2 | 0.04 | 0.16 | 0.12 | 0.26 | 0.13 |
| 2 | 25074086 | rs11678026 | -0.294 | 0.628 | -1.44 | -0.355 | -0.032 | -0.005 | -0.005 | 0.413 | 0.14 | 0.03 | 0.08 | 0.1 | 0.14 | 0.12 |
| 2 | 25076126 | rs13387729 | -1.679 | 0.561 | -1.799 | 0.023 | 0.006 | -0.026 | -0.026 | 0 | 0.47 | 0.89 | 0.33 | 0.48 | 0.4 | 0.48 |
| 2 | 25077991 | rs6760328 | -0.612 | 0.546 | -0.759 | 0.048 | -0.004 | 0.034 | 0.034 | 0 | 0.29 | 0.35 | 0.18 | 0.37 | 0.28 | 0.4 |
| 2 | 25082926 | rs11687089 | -1.563 | 0.654 | -0.989 | 0.081 | 0.004 | -0.032 | -0.032 | 0.97 | 0.43 | 0.8 | 0.3 | 0.48 | 0.39 | 0.44 |
| 2 | 25097644 | rs13407913 | -1.978 | 0.61 | -1.01 | 0.08 | -0.009 | -0.024 | -0.024 | 0.493 | 0.47 | 0.88 | 0.34 | 0.48 | 0.43 | 0.48 |
| 2 | 25109302 | rs6545790 | -1.839 | 0.707 | -0.958 | -0.021 | -0.015 | 0.01 | 0.01 | 0.862 | 0.46 | 0.85 | 0.34 | 0.43 | 0.42 | 0.48 |
| 2 | 25120086 | rs6752483 | -1.775 | 0.761 | -0.954 | 0.005 | -0.024 | 0.001 | 0.001 | 1.093 | 0.46 | 0.86 | 0.35 | 0.42 | 0.43 | 0.46 |
| 2 | 25126046 | rs6712986 | -0.461 | 0.692 | -0.398 | -0.046 | -0.005 | -0.101 | -0.101 | 0 | 0.17 | 0.51 | 0.13 | 0.05 | 0.12 | 0.08 |
| 2 | 25131736 | rs13382979 | -0.673 | 0.77 | -0.412 | 0.034 | 0.007 | 0.033 | 0.033 | 0 | 0.29 | 0.34 | 0.22 | 0.39 | 0.31 | 0.38 |
| 2 | 25138987 | rs7593130 | -1.178 | 0.863 | -0.526 | 0.041 | 0.006 | 0.05 | 0.05 | 0.201 | 0.23 | 0.55 | 0.23 | 0.38 | 0.26 | 0.24 |
| 2 | 25141538 | rs11676272 | -1.809 | 1.055 | -1.054 | 0.045 | -0.075 | -0.017 | -0.017 | 0 | 0.47 | 0.91 | 0.39 | 0.43 | 0.46 | 0.47 |
| 2 | 25153986 | rs6726261 | -1.112 | 0.964 | -0.379 | 0.255 | -0.001 | 0.005 | 0.005 | 0.204 | 0.37 | 0.69 | 0.25 | 0.3 | 0.34 | 0.44 |
| 2 | 25158281 | rs713587 | -1.754 | 1.021 | -0.731 | 0.341 | -0.083 | -0.049 | -0.049 | 0 | 0.46 | 0.9 | 0.38 | 0.48 | 0.47 | 0.48 |
| 2 | 25160743 | rs7566051 | 0.569 | 1.083 | -0.215 | 0.448 | 0.025 | 0.048 | 0.048 | 0 | 0.1 | 0.21 | 0.13 | 0.18 | 0.12 | 0.03 |
| 2 | 25165825 | rs17046742 | 1.023 | 0.947 | 0.074 | 0.221 | -0.004 | -0.037 | -0.037 | 0.218 | 0.15 | 0.19 | 0.06 | 0 | 0.1 | 0.22 |
| 2 | 25168477 | rs17046751 | -0.611 | 1.06 | 0.75 | 0.115 | 0.032 | -0.039 | -0.039 | 0.107 | 0.09 | 0.39 | 0.05 | 0 | 0.05 | 0.08 |
| 2 | 25169200 | rs1172294 | -2.054 | 1.184 | 0.553 | 0.215 | -0.106 | -0.079 | -0.079 | 4.579 | 0.45 | 0.95 | 0.38 | 0.49 | 0.47 | 0.51 |
| 2 | 25190127 | rs6749438 | -0.71 | 1.118 | 0.922 | 0.296 | 0.038 | -0.036 | -0.036 | 0 | 0.09 | 0.4 | 0.05 | 0 | 0.05 | 0.08 |
| 2 | 25276284 | rs6733301 | 1.102 | 1.024 | 0.589 | 0.163 | -0.003 | -0.041 | -0.041 | 0 | 0.14 | 0.2 | 0.06 | 0 | 0.1 | 0.22 |
| 2 | 25302237 | rs17744519 | 0.837 | 1.151 | 0.438 | 0.06 | -0.016 | 0.002 | 0.002 | 0 | 0.18 | 0.16 | 0.12 | 0.01 | 0.23 | 0.3 |
| 2 | 25305504 | rs551573 | 1.181 | 0.63 | -0.273 | -0.224 | 0.003 | 0.182 | 0.182 | 0 | 0.46 | 0.09 | 0.57 | 0.78 | 0.45 | 0.53 |
| 2 | 25305756 | rs1530016 | 1.19 | 0.621 | -0.308 | 0.32 | -0.038 | -0.104 | -0.104 | 0 | 0.45 | 0.04 | 0.54 | 0.51 | 0.45 | 0.44 |
| 2 | 25314186 | rs2918630 | 0.447 | 0.52 | 0.24 | 0.19 | 0.001 | 0.015 | 0.015 | 0 | 0.54 | 0.48 | 0.61 | 0.79 | 0.49 | 0.6 |
| 2 | 25316045 | rs10495751 | -0.176 | 0.394 | 0.43 | 0.31 | 0.053 | 0.009 | 0.009 | 0 | 0.07 | 0.38 | 0.03 | 0 | 0.04 | 0.06 |
| 2 | 25327771 | rs6742730 | -0.062 | -0.309 | -0.071 | -0.012 | 0.05 | 0.001 | 0.001 | 0 | 0.07 | 0.35 | 0.03 | 0 | 0.03 | 0.06 |
| 2 | 25329898 | rs12613265 | 0.924 | -0.523 | -1.121 | -0.12 | 0.005 | -0.082 | -0.082 | 0.585 | 0.15 | 0.03 | 0.4 | 0.51 | 0.15 | 0.31 |
| 2 | 25339658 | rs4665284 | 0.498 | -0.574 | -1.221 | -0.044 | -0.004 | -0.115 | -0.115 | 0 | 0.22 | 0.06 | 0.45 | 0.52 | 0.26 | 0.36 |
| 2 | 25341838 | rs12992354 | 0.143 | -0.408 | -1.326 | 0.466 | -0.041 | -0.089 | -0.089 | 0 | 0.25 | 0.05 | 0.49 | 0.32 | 0.29 | 0.33 |
| 2 | 25342914 | rs7575363 | 0.389 | -0.871 | -0.815 | 0.002 | -0.002 | -0.028 | -0.028 | 0 | 0.29 | 0.25 | 0.27 | 0.48 | 0.3 | 0.17 |
| 2 | 25350321 | rs7596747 | 0.675 | -0.91 | -0.464 | 0.112 | 0.001 | -0.002 | -0.002 | 0 | 0.19 | 0.13 | 0.44 | 0.22 | 0.21 | 0.29 |
| 2 | 25359254 | rs10202934 | 0.075 | -0.964 | -0.259 | -0.049 | -0.001 | 0.001 | 0.001 | 0.002 | 0.23 | 0.23 | 0.48 | 0.59 | 0.26 | 0.53 |
| 2 | 25359422 | rs3731631 | 0.473 | -0.88 | -0.217 | -0.104 | 0 | 0 | 0 | 4.917 | 0.29 | 0.32 | 0.3 | 0.39 | 0.3 | 0.15 |
| 2 | 25362194 | rs6718083 | 0.11 | -1.161 | -0.355 | -0.366 | -0.022 | 0.003 | 0.003 | 1.237 | 0.31 | 0.38 | 0.44 | 0.3 | 0.26 | 0.35 |
| 2 | 25362515 | rs4665765 | -1.542 | -1.108 | -0.52 | -0.407 | -0.074 | 0.146 | 0.146 | 0 | 0.64 | 0.87 | 0.8 | 0.99 | 0.6 | 0.74 |
| 2 | 25373298 | rs13428823 | 0.543 | -1.068 | -0.538 | -0.366 | -0.009 | -0.015 | -0.015 | 0 | 0.57 | 0.55 | 0.5 | 0.7 | 0.62 | 0.6 |
| 2 | 25374912 | rs17046887 | 2.244 | -1.04 | -0.261 | -0.686 | -0.028 | -0.051 | -0.051 | 0 | 0.1 | 0.15 | 0.01 | 0 | 0.04 | 0.06 |
| 2 | 25377176 | rs2164808 | -1.405 | -1.039 | -0.747 | -0.806 | -0.101 | 0.204 | 0.204 | 0 | 0.64 | 0.89 | 0.8 | 0.99 | 0.6 | 0.74 |
| 2 | 25377329 | rs2118404 | -0.099 | -1.098 | -0.557 | -0.667 | -0.014 | 0.002 | 0.002 | 1.432 | 0.31 | 0.34 | 0.46 | 0.59 | 0.27 | 0.45 |
| 2 | 25378372 | rs7589318 | 2.255 | -0.999 | -1.097 | -0.651 | -0.048 | 0.105 | 0.105 | 0 | 0.26 | 0.11 | 0.16 | 0.01 | 0.27 | 0.21 |
| 2 | 25380573 | rs1866146 | -0.12 | -1.051 | -1.007 | -0.538 | -0.032 | -0.033 | -0.033 | 0.005 | 0.69 | 0.92 | 0.52 | 0.41 | 0.65 | 0.54 |
| 2 | 25384833 | rs6713532 | -0.436 | -1.063 | -0.516 | -0.56 | -0.014 | 0 | 0 | 1.415 | 0.33 | 0.52 | 0.5 | 0.59 | 0.27 | 0.46 |
| 2 | 25385485 | rs6545975 | -0.049 | -0.955 | -0.469 | -0.521 | 0 | 0.001 | 0.001 | 0.269 | 0.61 | 0.54 | 0.67 | 0.6 | 0.61 | 0.7 |
| 2 | 25385638 | rs7565427 | -1.445 | -0.801 | -0.49 | -0.973 | -0.003 | -0.01 | -0.01 | 0 | 0.89 | 0.89 | 0.95 | 1 | 0.88 | 0.96 |
| 2 | 25386064 | rs7565877 | 0.502 | -0.668 | -0.197 | -0.635 | -0.012 | -0.075 | -0.075 | 0 | 0.17 | 0.42 | 0.12 | 0.04 | 0.13 | 0.21 |
| 2 | 25389224 | rs934778 | 0.504 | -0.386 | -0.484 | -0.392 | -0.049 | 0.081 | 0.081 | 0 | 0.25 | 0.06 | 0.26 | 0.12 | 0.31 | 0.24 |
| 2 | 25394650 | rs6545976 | -0.177 | -0.557 | 0.085 | -0.214 | -0.033 | 0.022 | 0.022 | 0 | 0.13 | 0.5 | 0.07 | 0.1 | 0.06 | 0.19 |
| 2 | 25402280 | rs874401 | 0.564 | -0.262 | -0.605 | -0.424 | -0.002 | -0.002 | -0.002 | 0 | 0.22 | 0.21 | 0.43 | 0.2 | 0.2 | 0.17 |
| 2 | 25412032 | rs17046917 | 0.84 | -0.114 | -0.582 | -0.35 | -0.003 | -0.002 | -0.002 | 0 | 0.2 | 0.22 | 0.42 | 0.2 | 0.2 | 0.14 |
| 2 | 25448050 | rs4665783 | 0.827 | -0.031 | -0.838 | -0.332 | 0.004 | 0 | 0 | 0 | 0.2 | 0.15 | 0.41 | 0.21 | 0.2 | 0.15 |
| 2 | 25455389 | rs752208 | 0.385 | 0.034 | -0.485 | -0.408 | 0.001 | -0.069 | -0.069 | 0 | 0.22 | 0.3 | 0.2 | 0.05 | 0.23 | 0.08 |

| | | | | | | | | | | | | | | | | |
|---|---|---|---|---|---|---|---|---|---|---|---|---|---|---|---|---|
| 2 | 25458379 | rs10084238 | 0.201 | 0.295 | -1.123 | -0.096 | 0.004 | 0.118 | 0.118 | 0.066 | 0.53 | 0.11 | 0.53 | 0.29 | 0.58 | 0.42 |
| 2 | 25458546 | rs2304429 | 0.208 | 0.296 | -1.137 | -0.089 | 0 | 0.115 | 0.115 | 0.098 | 0.52 | 0.1 | 0.53 | 0.31 | 0.58 | 0.43 |
| 2 | 25466888 | rs2289093 | -0.366 | 0.287 | -0.551 | 0.102 | -0.015 | 0.018 | 0.018 | 0.054 | 0.67 | 0.47 | 0.69 | 0.62 | 0.74 | 0.55 |
| 2 | 25482883 | rs7594432 | 0.107 | 0.377 | -0.797 | -0.043 | -0.002 | 0.011 | 0.011 | 0 | 0.42 | 0.53 | 0.57 | 0.53 | 0.43 | 0.34 |
| 2 | 25482970 | rs11694842 | 1.07 | 0.458 | -1.28 | -0.042 | -0.112 | 0.09 | 0.09 | 0.628 | 0.26 | 0.01 | 0.14 | 0.09 | 0.31 | 0.24 |
| 2 | 25483121 | rs10460566 | -0.49 | 0.563 | -0.809 | 0.276 | -0.006 | 0.012 | 0.012 | 0.018 | 0.69 | 0.68 | 0.72 | 0.62 | 0.74 | 0.57 |
| 2 | 25485735 | rs11887120 | -0.468 | 0.536 | -1.013 | 0.475 | 0 | 0 | 0 | 0 | 0.43 | 0.51 | 0.54 | 0.51 | 0.41 | 0.34 |
| 2 | 25487658 | rs6546045 | 0.006 | 0.622 | -1.577 | 0.129 | -0.004 | 0.067 | 0.067 | 0 | 0.69 | 0.88 | 0.62 | 0.78 | 0.68 | 0.76 |
| 2 | 25491016 | rs7586294 | -0.267 | 0.817 | -1.262 | 0.178 | 0.006 | -0.008 | -0.008 | 3.849 | 0.42 | 0.47 | 0.39 | 0.35 | 0.55 | 0.38 |
| 2 | 25512438 | rs12999687 | -0.261 | 0.889 | -1.261 | -0.271 | 0 | 0.01 | 0.01 | 0 | 0.53 | 0.51 | 0.37 | 0.23 | 0.5 | 0.41 |
| 2 | 25518470 | rs13401241 | -0.104 | 1.432 | -1.19 | -0.219 | 0 | 0.001 | 0.001 | 0 | 0.51 | 0.46 | 0.38 | 0.27 | 0.54 | 0.38 |
| 2 | 25524944 | rs7581217 | -0.242 | 0.538 | -1.258 | -0.252 | -0.002 | 0.019 | 0.019 | 0.439 | 0.65 | 0.62 | 0.51 | 0.57 | 0.63 | 0.7 |
| 2 | 25530028 | rs749130 | 0.108 | 0.005 | -0.907 | -0.333 | 0 | 0.013 | 0.013 | 0 | 0.47 | 0.58 | 0.41 | 0.3 | 0.54 | 0.37 |
| 2 | 25531350 | rs6711622 | 0.938 | 0.018 | -0.764 | -0.366 | -0.002 | 0.022 | 0.022 | 0 | 0.37 | 0.25 | 0.3 | 0.2 | 0.4 | 0.24 |
| 2 | 25532969 | rs13036246 | 0.342 | 0.015 | -0.952 | -0.515 | -0.019 | 0.026 | 0.026 | 0 | 0.39 | 0.36 | 0.31 | 0.22 | 0.47 | 0.37 |
| 2 | 25539357 | rs6722613 | -0.225 | 0.305 | -0.904 | -0.153 | -0.003 | 0.019 | 0.019 | 0 | 0.57 | 0.65 | 0.43 | 0.28 | 0.57 | 0.46 |
| 2 | 25548902 | rs7575625 | 0.999 | 0.023 | -0.696 | -0.139 | 0.01 | -0.025 | -0.025 | 0 | 0.38 | 0.43 | 0.35 | 0.21 | 0.43 | 0.25 |
| 2 | 25550995 | rs12987326 | 0.116 | 0.21 | -0.633 | 0.179 | 0.005 | 0.008 | 0.008 | 0 | 0.56 | 0.58 | 0.47 | 0.31 | 0.6 | 0.48 |
| 2 | 25565907 | rs1550117 | -1.216 | -0.432 | -1.052 | 0.005 | 0.006 | -0.024 | -0.024 | 0 | 0.9 | 0.97 | 0.75 | 0.78 | 0.93 | 0.94 |
| 2 | 25568821 | rs7560488 | 0.749 | -0.487 | -0.832 | -0.171 | 0.003 | 0.004 | 0.004 | 0 | 0.44 | 0.39 | 0.33 | 0.15 | 0.41 | 0.33 |
| 2 | 25584582 | rs10182590 | -0.072 | -0.141 | -0.621 | -0.522 | -0.009 | -0.023 | -0.023 | 7.207 | 0.15 | 0.18 | 0.07 | 0.05 | 0.11 | 0.22 |
| 2 | 25585084 | rs934615 | -0.918 | -0.152 | -0.821 | -0.441 | 0.009 | -0.023 | -0.023 | 0 | 0.89 | 0.96 | 0.75 | 0.77 | 0.93 | 0.94 |
| 2 | 25587851 | rs6705138 | 0.833 | -0.031 | -0.86 | -0.482 | 0.007 | -0.006 | -0.006 | 1.113 | 0.57 | 0.46 | 0.53 | 0.55 | 0.56 | 0.5 |
| 2 | 25589466 | rs10196664 | -0.444 | -0.106 | -0.96 | -0.473 | 0.001 | -0.002 | -0.002 | 0 | 0.14 | 0.11 | 0.06 | 0.06 | 0.11 | 0.22 |
| 2 | 25595194 | rs1010658 | 0.02 | -0.311 | -0.911 | -0.568 | 0.005 | 0.005 | 0.005 | 0.873 | 0.62 | 0.63 | 0.46 | 0.25 | 0.68 | 0.67 |
| 2 | 25597681 | rs730072 | 0.023 | 0.066 | -0.984 | -0.682 | -0.002 | 0.003 | 0.003 | 0 | 0.43 | 0.39 | 0.43 | 0.64 | 0.49 | 0.58 |
| 2 | 25602310 | rs2304427 | 0.42 | -0.122 | -0.787 | -1.019 | -0.006 | -0.075 | -0.075 | 1.599 | 0.35 | 0.33 | 0.22 | 0.07 | 0.32 | 0.38 |
| 2 | 25613146 | rs7577599 | 0.324 | -0.303 | -0.286 | -0.342 | 0.03 | 0.126 | 0.126 | 2.007 | 0.17 | 0.4 | 0.28 | 0.55 | 0.2 | 0.2 |
| 2 | 25621530 | rs7571570 | -0.631 | -0.592 | -1.213 | -0.524 | 0.03 | 0.002 | 0.002 | 0 | 0.36 | 0.71 | 0.36 | 0.56 | 0.29 | 0.37 |
| 2 | 25633242 | rs10180663 | -0.624 | -0.385 | -1.614 | -0.674 | 0.033 | 0.003 | 0.003 | 0.085 | 0.37 | 0.84 | 0.36 | 0.56 | 0.33 | 0.31 |
| 2 | 25643944 | rs17745923 | 1.125 | 0.194 | -2.147 | -1.3 | -0.064 | 0.107 | 0.107 | 0.037 | 0.21 | 0.04 | 0.18 | 0.01 | 0.26 | 0.23 |
| 2 | 25655446 | rs2292225 | -0.08 | -0.129 | -1.645 | -1.804 | -0.002 | -0.038 | -0.038 | 0.078 | 0.57 | 0.46 | 0.43 | 0.08 | 0.63 | 0.57 |
| 2 | 25655772 | rs7583475 | -0.83 | 0.369 | -2.044 | -2.656 | 0.002 | -0.089 | -0.089 | 0.059 | 0.34 | 0.33 | 0.21 | 0.01 | 0.35 | 0.27 |
| 2 | 25658808 | rs2384234 | 2.048 | 0.691 | -2.048 | -2.183 | -0.035 | 0.053 | 0.053 | 0.748 | 0.17 | 0.08 | 0.17 | 0.01 | 0.24 | 0.23 |
| 2 | 25659244 | rs6746082 | 0.241 | 0.622 | -1.827 | -1.862 | 0.047 | 0.094 | 0.094 | 0.365 | 0.2 | 0.51 | 0.28 | 0.6 | 0.21 | 0.23 |
| 2 | 25662578 | rs10210057 | -0.627 | 0.214 | -2.023 | -1.992 | 0 | -0.001 | -0.001 | 0 | 0.82 | 0.84 | 0.72 | 0.72 | 0.86 | 0.81 |
| 2 | 25663991 | rs1006752 | 1.724 | 0.425 | -2.156 | -1.909 | -0.047 | -0.065 | -0.065 | 1.121 | 0.11 | 0.01 | 0.18 | 0.57 | 0.14 | 0.18 |
| 2 | 25668692 | rs6546181 | -1.963 | 0.357 | -2.074 | -1.916 | -0.006 | -0.055 | -0.055 | 0.156 | 0.88 | 0.99 | 0.76 | 0.73 | 0.91 | 0.88 |
| 2 | 25671806 | rs6546183 | 0.569 | 0.387 | -1.762 | -1.744 | 0.004 | 0.115 | 0.115 | 0.35 | 0.18 | 0.5 | 0.25 | 0.61 | 0.14 | 0.2 |
| 2 | 25675695 | rs10495753 | 1.153 | 0.495 | -2.5 | -1.816 | -0.045 | 0.062 | 0.062 | 0 | 0.24 | 0.05 | 0.18 | 0.11 | 0.26 | 0.22 |
| 2 | 25683447 | rs6546199 | -1.328 | 0.643 | -2.22 | -1.283 | -0.08 | -0.011 | -0.011 | 0.308 | 0.78 | 0.97 | 0.76 | 0.79 | 0.69 | 0.67 |
| 2 | 25705428 | rs6751254 | 0.314 | 0.415 | -1.923 | -0.956 | -0.003 | 0.13 | 0.13 | 0.637 | 0.52 | 0.26 | 0.31 | 0.26 | 0.49 | 0.43 |
| 2 | 25714916 | rs1507705 | -0.257 | 0.447 | -0.627 | -0.478 | -0.014 | 0.096 | 0.096 | 2.545 | 0.25 | 0.5 | 0.43 | 0.53 | 0.19 | 0.22 |
| 2 | 25727910 | rs9309386 | 0.606 | 0.768 | -0.819 | -0.036 | 0.005 | 0.087 | 0.087 | 0 | 0.38 | 0.11 | 0.39 | 0.27 | 0.33 | 0.33 |
| 2 | 25730439 | rs517403 | -0.082 | 1.257 | 0.37 | 0.572 | 0.07 | 0.193 | 0.193 | 0 | 0.19 | 0.4 | 0.3 | 0.45 | 0.24 | 0.21 |
| 2 | 25755848 | rs936012 | -0.246 | 1.232 | 0.709 | 0.61 | 0.065 | 0.19 | 0.19 | 0.044 | 0.19 | 0.55 | 0.3 | 0.45 | 0.23 | 0.23 |
| 2 | 25759913 | rs476731 | 0.352 | 1.309 | 0.501 | 0.65 | 0.069 | 0.187 | 0.187 | 0 | 0.16 | 0.32 | 0.27 | 0.45 | 0.23 | 0.23 |
| 2 | 25807106 | rs11126114 | 0.18 | 0.345 | -0.32 | 0.093 | 0.033 | 0.031 | 0.031 | 0 | 0.14 | 0.02 | 0.06 | 0.03 | 0.11 | 0.08 |
| 2 | 25821689 | rs1377780 | 0.391 | 0.973 | 0.203 | 0.214 | 0.001 | -0.021 | -0.021 | 0.674 | 0.54 | 0.5 | 0.49 | 0.27 | 0.52 | 0.44 |
| 2 | 25829201 | rs12613835 | 0.428 | 1.498 | -0.241 | 0.432 | -0.08 | -0.006 | -0.006 | 0.382 | 0.24 | 0.07 | 0.26 | 0.22 | 0.33 | 0.29 |
| 2 | 25837156 | rs11678268 | -0.615 | 1.465 | 0.043 | 0.611 | -0.023 | 0.014 | 0.014 | 0 | 0.37 | 0.26 | 0.49 | 0.73 | 0.49 | 0.46 |
| 2 | 25845467 | rs6546314 | 0.958 | 1.353 | 0.063 | 0.599 | 0.006 | 0.077 | 0.077 | 0.482 | 0.12 | 0.12 | 0.23 | 0.5 | 0.16 | 0.16 |
| 2 | 25853737 | rs12613340 | -0.624 | 1.277 | 0.144 | 0.355 | -0.025 | 0.022 | 0.022 | 0 | 0.38 | 0.24 | 0.5 | 0.73 | 0.49 | 0.44 |
| 2 | 25867203 | rs858648 | -0.896 | 1.201 | 0.418 | 0.345 | 0.005 | -0.025 | -0.025 | 0 | 0.42 | 0.49 | 0.53 | 0.76 | 0.5 | 0.49 |
| 2 | 25873082 | rs17047116 | 0.874 | 1.267 | -0.383 | 0.534 | 0.001 | 0.051 | 0.051 | 4.124 | 0.13 | 0.11 | 0.24 | 0.53 | 0.15 | 0.17 |
| 2 | 25894556 | rs17391976 | -0.003 | 1.371 | -1.063 | -0.383 | -0.047 | 0.014 | 0.014 | 0.133 | 0.25 | 0.12 | 0.26 | 0.23 | 0.33 | 0.27 |
| 2 | 25899768 | rs7604239 | -0.057 | 0.542 | -0.84 | -1.082 | 0.011 | 0.027 | 0.027 | 1.457 | 0.61 | 0.65 | 0.53 | 0.23 | 0.67 | 0.66 |
| 2 | 25911304 | rs11892454 | 0.021 | 0.953 | -0.577 | -1.465 | 0.005 | -0.063 | -0.063 | 0.469 | 0.25 | 0.41 | 0.25 | 0.14 | 0.24 | 0.28 |
| 2 | 25915721 | rs10182758 | -0.538 | 0.404 | -0.207 | -1.718 | -0.016 | -0.037 | -0.037 | 0.003 | 0.63 | 0.44 | 0.54 | 0.27 | 0.68 | 0.67 |
| 2 | 25915978 | rs12713638 | 0.018 | 0.543 | -0.038 | -1.888 | 0.008 | 0.031 | 0.031 | 0.355 | 0.55 | 0.33 | 0.47 | 0.26 | 0.62 | 0.55 |
| 2 | 25920167 | rs35324955 | 1.19 | 0.344 | -0.078 | -1.837 | 0 | -0.004 | -0.004 | 0 | 0.06 | 0.05 | 0.04 | 0 | 0.04 | 0.07 |
| 2 | 25921254 | rs10199771 | 0.226 | 0.339 | 0.017 | -1.859 | -0.005 | 0.031 | 0.031 | 0.383 | 0.46 | 0.36 | 0.37 | 0.13 | 0.5 | 0.51 |
| 2 | 25948600 | rs890682 | -0.09 | 0.625 | 0.373 | -1.727 | 0.017 | -0.042 | -0.042 | 0.003 | 0.21 | 0.34 | 0.24 | 0.09 | 0.25 | 0.21 |
| 2 | 25959477 | rs13004845 | -0.767 | 0.587 | 0.589 | -1.913 | 0.023 | -0.071 | -0.071 | 0 | 0.27 | 0.42 | 0.27 | 0.1 | 0.31 | 0.28 |
| 2 | 25961583 | rs10153560 | -0.435 | 0.633 | 0.681 | -1.928 | 0.013 | -0.047 | -0.047 | 0 | 0.24 | 0.32 | 0.25 | 0.1 | 0.28 | 0.25 |
| 2 | 25969729 | rs6747116 | -0.659 | 0.174 | 0.559 | -2.363 | -0.01 | -0.014 | -0.014 | 0 | 0.79 | 0.73 | 0.57 | 0.14 | 0.8 | 0.73 |
| 2 | 25981213 | rs2052826 | 1.307 | 0.13 | 0.476 | -2.38 | -0.024 | 0.252 | 0.252 | 0 | 0.49 | 0.16 | 0.32 | 0.11 | 0.49 | 0.44 |
| 2 | 25981272 | rs6546452 | 1.134 | 0.196 | 0.581 | -2.386 | -0.006 | 0.006 | 0.006 | 0.014 | 0.16 | 0.13 | 0.19 | 0.1 | 0.22 | 0.18 |
| 2 | 26003230 | rs7605347 | -0.119 | 0.182 | 0.623 | -2.381 | 0.003 | -0.018 | -0.018 | 0 | 0.23 | 0.23 | 0.22 | 0.1 | 0.28 | 0.25 |

| | | | | | | | | | | | | | | | | |
|---|---|---|---|---|---|---|---|---|---|---|---|---|---|---|---|---|
| 2 | 26013180 | rs11685550 | 0.489 | 0.259 | 0.404 | -2.625 | 0.004 | -0.025 | -0.025 | 1.518 | 0.69 | 0.49 | 0.47 | 0.21 | 0.64 | 0.68 |
| 2 | 26029485 | rs10210497 | -0.236 | 0.039 | 0.598 | -2.826 | 0.017 | -0.058 | -0.058 | 0 | 0.24 | 0.33 | 0.25 | 0.1 | 0.28 | 0.25 |
| 2 | 26053166 | rs10168718 | -0.587 | 0.044 | 0.575 | -2.829 | 0.024 | -0.074 | -0.074 | 0.085 | 0.27 | 0.42 | 0.27 | 0.1 | 0.31 | 0.28 |
| 2 | 26075213 | rs11894773 | 0.001 | -0.022 | 0.59 | -2.766 | -0.024 | -0.028 | -0.028 | 0.143 | 0.76 | 0.63 | 0.56 | 0.15 | 0.8 | 0.72 |
| 2 | 26088769 | rs6760135 | 0.931 | 0.141 | 0.313 | -2.638 | -0.004 | 0.003 | 0.003 | 0.014 | 0.16 | 0.13 | 0.19 | 0.1 | 0.22 | 0.18 |
| 2 | 26112833 | rs2138390 | 1.008 | -0.386 | 0.735 | -2.419 | -0.004 | -0.022 | -0.022 | 0.492 | 0.64 | 0.38 | 0.48 | 0.14 | 0.71 | 0.71 |
| 2 | 26113120 | rs11895615 | 1.207 | -0.401 | 0.775 | -2.432 | 0.016 | 0.025 | 0.025 | 0 | 0.62 | 0.31 | 0.46 | 0.14 | 0.69 | 0.69 |
| 2 | 26117455 | rs13033978 | -0.684 | -0.415 | 0.923 | -2.428 | -0.013 | -0.045 | -0.045 | 0 | 0.12 | 0.19 | 0.06 | 0 | 0.05 | 0.08 |
| 2 | 26149502 | rs1061485 | 1.04 | -0.321 | 0.721 | -2.386 | -0.025 | -0.074 | -0.074 | 0 | 0.68 | 0.52 | 0.5 | 0.14 | 0.72 | 0.73 |
| 2 | 26150589 | rs6738270 | 0.252 | -0.238 | 0.574 | -2.259 | -0.014 | -0.038 | -0.038 | 0.639 | 0.77 | 0.66 | 0.55 | 0.14 | 0.77 | 0.78 |
| 2 | 26150773 | rs6728684 | 0.213 | -0.211 | 0.588 | -2.246 | -0.013 | -0.038 | -0.038 | 0.298 | 0.77 | 0.65 | 0.55 | 0.14 | 0.77 | 0.78 |
| 2 | 26156655 | rs2384298 | 1.092 | -0.415 | -0.42 | -2.842 | 0.073 | 0.183 | 0.183 | 0 | 0.56 | 0.22 | 0.47 | 0.1 | 0.7 | 0.57 |
| 2 | 26159940 | rs6546620 | 0.616 | -0.382 | 0.2 | -2.685 | -0.028 | -0.086 | -0.086 | 0 | 0.7 | 0.6 | 0.56 | 0.1 | 0.79 | 0.65 |
| 2 | 26174103 | rs6546642 | -0.649 | -0.213 | 0.356 | -2.809 | -0.017 | -0.14 | -0.14 | 0.022 | 0.23 | 0.36 | 0.13 | 0 | 0.15 | 0.12 |
| 2 | 26177198 | rs17047211 | -0.615 | -0.214 | 0.358 | -2.809 | -0.01 | -0.095 | -0.095 | 0.556 | 0.23 | 0.27 | 0.12 | 0 | 0.15 | 0.12 |
| 2 | 26183142 | rs12994424 | -1.115 | -0.106 | 0.433 | -2.909 | 0.012 | -0.049 | -0.049 | 0 | 0.1 | 0.22 | 0.06 | 0 | 0.09 | 0.08 |
| 2 | 26190923 | rs7586869 | -0.915 | 0.156 | -0.463 | -3.5 | -0.004 | 0.129 | 0.129 | 0 | 0.6 | 0.75 | 0.39 | 0.08 | 0.57 | 0.4 |
| 2 | 26191296 | rs7603456 | -0.477 | 0.198 | -0.438 | -3.589 | 0.008 | -0.168 | -0.168 | 0 | 0.27 | 0.48 | 0.17 | 0 | 0.21 | 0.17 |
| 2 | 26196925 | rs12105041 | -1.51 | 0.261 | -0.852 | -3.62 | -0.094 | 0.154 | 0.154 | 0 | 0.72 | 0.98 | 0.78 | 0.94 | 0.66 | 0.64 |
| 2 | 26199220 | rs7563440 | 0.456 | -0.351 | -0.303 | -3.369 | -0.001 | -0.08 | -0.08 | 0.046 | 0.15 | 0.2 | 0.11 | 0 | 0.11 | 0.08 |
| 2 | 26199725 | rs2384317 | -0.052 | -0.102 | -0.717 | -3.022 | 0.051 | -0.043 | -0.043 | 0 | 0.08 | 0.21 | 0.06 | 0 | 0.1 | 0.1 |
| 2 | 26202280 | rs6706373 | -0.614 | -0.17 | -1.713 | -3.173 | -0.122 | 0.276 | 0.276 | 0 | 0.6 | 0.94 | 0.36 | 0.06 | 0.47 | 0.41 |
| 2 | 26206255 | rs2384319 | -0.145 | -0.658 | -1.052 | -2.795 | 0.024 | 0.033 | 0.033 | 0 | 0.17 | 0.08 | 0.41 | 0.92 | 0.11 | 0.25 |
| 2 | 26206843 | rs6719890 | -1.154 | -0.541 | -0.89 | -2.888 | 0.032 | 0.067 | 0.067 | 0 | 0.43 | 0.67 | 0.59 | 0.92 | 0.28 | 0.42 |
| 2 | 26207421 | rs12713741 | -1.221 | -0.592 | -1.134 | -2.89 | 0.06 | 0.185 | 0.185 | 0 | 0.44 | 0.77 | 0.59 | 0.92 | 0.28 | 0.42 |
| 2 | 26214015 | rs6745174 | 0.787 | -0.425 | -0.495 | -2.538 | -0.012 | -0.096 | -0.096 | 0.099 | 0.81 | 0.76 | 0.94 | 1 | 0.89 | 0.83 |
| 2 | 26223547 | rs12990137 | 0.852 | -0.503 | -1.185 | -2.946 | -0.019 | 0.258 | 0.258 | 0.973 | 0.43 | 0.16 | 0.33 | 0.01 | 0.58 | 0.45 |
| 2 | 26226280 | rs1992300 | 0.835 | -0.151 | -0.041 | -2.128 | -0.012 | -0.094 | -0.094 | 0 | 0.82 | 0.61 | 0.92 | 0.97 | 0.89 | 0.85 |
| 2 | 26251361 | rs4665809 | -0.2 | -0.452 | -0.551 | -2.21 | -0.002 | -0.015 | -0.015 | 0 | 0.72 | 0.72 | 0.81 | 0.89 | 0.79 | 0.78 |
| 2 | 26255263 | rs6749855 | -0.412 | -0.578 | -0.207 | -2.273 | 0.018 | 0.061 | 0.061 | 0.423 | 0.75 | 0.86 | 0.8 | 0.9 | 0.79 | 0.79 |
| 2 | 26272219 | rs10170359 | -0.45 | -0.585 | -0.123 | -2.239 | 0.014 | 0.045 | 0.045 | 0.382 | 0.75 | 0.86 | 0.81 | 0.89 | 0.79 | 0.78 |
| 2 | 26281239 | rs11677219 | 1.593 | -0.666 | -1.159 | -3.131 | -0.186 | 0.214 | 0.214 | 1.797 | 0.3 | 0.02 | 0.27 | 0.03 | 0.5 | 0.38 |
| 2 | 26307140 | rs12471809 | -0.39 | -0.562 | -0.704 | -2.821 | -0.004 | -0.017 | -0.017 | 0 | 0.72 | 0.71 | 0.81 | 0.89 | 0.79 | 0.77 |
| 2 | 26313406 | rs4665304 | -0.097 | -0.645 | -0.168 | -2.678 | -0.034 | -0.081 | -0.081 | 0.286 | 0.7 | 0.58 | 0.79 | 0.89 | 0.79 | 0.73 |
| 2 | 26321526 | rs11678157 | -0.204 | -0.636 | -0.192 | -2.674 | -0.02 | -0.053 | -0.053 | 0 | 0.71 | 0.64 | 0.8 | 0.89 | 0.79 | 0.72 |
| 2 | 26322040 | rs9332419 | 1.222 | -0.989 | 0.138 | -2.95 | 0.012 | 0.148 | 0.148 | 0.74 | 0.45 | 0.24 | 0.37 | 0.03 | 0.59 | 0.45 |
| 2 | 26324631 | rs750450 | -0.249 | -0.69 | 0.14 | -2.472 | 0.016 | 0.032 | 0.032 | 1.243 | 0.73 | 0.81 | 0.82 | 0.89 | 0.79 | 0.77 |
| 2 | 26344787 | rs11126375 | -0.212 | -0.687 | 0.135 | -2.471 | -0.008 | -0.025 | -0.025 | 0 | 0.72 | 0.69 | 0.81 | 0.89 | 0.79 | 0.78 |
| 2 | 26392741 | rs7564397 | 0.083 | -0.486 | 0.187 | -2.262 | 0.007 | 0.014 | 0.014 | 0.05 | 0.72 | 0.77 | 0.79 | 0.79 | 0.75 | 0.66 |
| 2 | 26409950 | rs892447 | 0.14 | -0.43 | -0.064 | -2.356 | -0.036 | 0.116 | 0.116 | 0.034 | 0.71 | 0.97 | 0.79 | 0.86 | 0.72 | 0.76 |
| 2 | 26410214 | rs1019972 | 1.046 | -0.435 | 0.218 | -2.257 | 0.005 | 0.027 | 0.027 | 0 | 0.06 | 0.01 | 0.03 | 0 | 0.05 | 0.01 |
| 2 | 26411894 | rs2241739 | 0.288 | -0.199 | 1.084 | -1.973 | -0.01 | -0.036 | -0.036 | 0 | 0.75 | 0.73 | 0.8 | 0.85 | 0.78 | 0.76 |
| 2 | 26466975 | rs3806515 | -0.161 | -0.403 | 0.58 | -1.695 | -0.001 | 0.065 | 0.065 | 0.038 | 0.23 | 0.04 | 0.16 | 0.15 | 0.21 | 0.22 |
| 2 | 26477650 | rs2033317 | -0.304 | -0.409 | 0.667 | -1.698 | 0.014 | 0.039 | 0.039 | 0 | 0.24 | 0.13 | 0.18 | 0.15 | 0.21 | 0.23 |
| 2 | 26497541 | rs3792007 | -0.15 | -0.407 | 0.606 | -1.697 | -0.022 | 0.052 | 0.052 | 0.015 | 0.23 | 0.02 | 0.16 | 0.15 | 0.21 | 0.22 |
| 2 | 26522041 | rs13021661 | 0.5 | -0.268 | 0.358 | -1.585 | -0.023 | 0.065 | 0.065 | 0.003 | 0.17 | 0.02 | 0.14 | 0.09 | 0.19 | 0.2 |
| 2 | 26524254 | rs7603119 | -0.07 | -0.255 | 0.491 | -1.581 | -0.018 | 0.042 | 0.042 | 0 | 0.21 | 0.07 | 0.44 | 0.87 | 0.25 | 0.26 |
| 2 | 26532491 | rs2007556 | 0.395 | -0.297 | 0.843 | -0.912 | 0 | 0.002 | 0.002 | 1.215 | 0.67 | 0.72 | 0.75 | 0.75 | 0.72 | 0.62 |
| 2 | 26535363 | rs11126434 | 0.789 | -0.324 | 0.864 | -0.931 | -0.005 | -0.009 | -0.009 | 0 | 0.62 | 0.48 | 0.69 | 0.71 | 0.69 | 0.59 |
| 2 | 26536694 | rs2052937 | 0.714 | -0.45 | 0.415 | -0.946 | 0.006 | 0.009 | 0.009 | 0.343 | 0.61 | 0.7 | 0.71 | 0.67 | 0.69 | 0.55 |
| 2 | 26554063 | rs1897115 | 1.489 | -0.441 | 0.696 | -0.518 | -0.022 | -0.001 | -0.001 | 0 | 0.72 | 0.41 | 0.61 | 0.31 | 0.84 | 0.74 |
| 2 | 26568503 | rs11900180 | -0.521 | -0.447 | 1.504 | -0.427 | -0.046 | -0.088 | -0.088 | 1.817 | 0.13 | 0.4 | 0.09 | 0 | 0.03 | 0.03 |
| 2 | 26573452 | rs7601607 | 0.049 | -0.469 | 1.081 | -0.878 | -0.061 | -0.017 | -0.017 | 0.239 | 0.29 | 0.07 | 0.3 | 0.29 | 0.37 | 0.32 |
| 2 | 26575986 | rs11898247 | 1.517 | -0.498 | 1.277 | -0.965 | -0.005 | 0.133 | 0.133 | 0 | 0.48 | 0.3 | 0.32 | 0.02 | 0.49 | 0.48 |
| 2 | 26576282 | rs7592979 | 0.528 | -0.472 | 1.29 | -0.936 | -0.017 | -0.051 | -0.051 | 0 | 0.1 | 0.38 | 0.08 | 0 | 0.02 | 0.02 |
| 2 | 26592429 | rs6546949 | 0.63 | -0.352 | 1.114 | -0.881 | -0.037 | 0.072 | 0.072 | 0 | 0.61 | 0.77 | 0.41 | 0.02 | 0.53 | 0.52 |
| 2 | 26602262 | rs11678646 | 0.006 | -0.319 | 0.914 | -0.801 | -0.074 | -0.012 | -0.012 | 0 | 0.29 | 0.06 | 0.29 | 0.25 | 0.36 | 0.32 |
| 2 | 26606821 | rs7565734 | 1.802 | -0.532 | 1.139 | -0.941 | -0.035 | 0.203 | 0.203 | 0 | 0.45 | 0.2 | 0.31 | 0.02 | 0.5 | 0.46 |
| 2 | 26613329 | rs7588538 | 0.1 | -0.499 | 1.164 | -0.822 | -0.074 | -0.013 | -0.013 | 0 | 0.28 | 0.06 | 0.29 | 0.25 | 0.36 | 0.32 |
| 2 | 26618543 | rs7092 | 1.702 | -0.529 | 1.198 | -1.035 | -0.037 | 0.207 | 0.207 | 0 | 0.46 | 0.2 | 0.31 | 0.02 | 0.5 | 0.46 |
| 2 | 26623160 | rs6713849 | 0.927 | -0.522 | 1.493 | -0.585 | -0.06 | -0.071 | -0.071 | 0 | 0.82 | 0.46 | 0.88 | 0.89 | 0.94 | 0.94 |
| 2 | 26632944 | rs6729941 | -0.599 | -0.595 | 1.418 | -0.544 | -0.058 | -0.069 | -0.069 | 0.15 | 0.15 | 0.38 | 0.05 | 0.06 | 0.04 | 0.03 |
| 2 | 26638322 | rs6722027 | 1.208 | -0.561 | 1.243 | -0.453 | -0.046 | 0.027 | 0.027 | 0 | 0.78 | 0.54 | 0.66 | 0.33 | 0.88 | 0.79 |
| 2 | 26638983 | rs13409838 | -0.662 | -0.587 | 1.4 | -0.455 | 0.063 | 0.059 | 0.059 | 0 | 0.06 | 0.34 | 0.06 | 0.04 | 0.03 | 0.02 |
| 2 | 26640781 | rs4491689 | 0.413 | -0.571 | 0.918 | -0.443 | 0.012 | -0.02 | -0.02 | 0 | 0.84 | 0.88 | 0.72 | 0.36 | 0.91 | 0.8 |
| 2 | 26653457 | rs4450561 | 1.532 | -0.503 | 0.313 | -0.873 | 0.033 | 0.218 | 0.218 | 0.051 | 0.47 | 0.17 | 0.35 | 0.02 | 0.58 | 0.44 |
| 2 | 26664327 | rs11126483 | 1.29 | -0.361 | 0.302 | -0.369 | -0.02 | -0.002 | -0.002 | 0 | 0.75 | 0.34 | 0.64 | 0.26 | 0.88 | 0.78 |
| 2 | 26667130 | rs3795958 | 0.464 | -0.389 | 0.03 | -0.402 | -0.021 | -0.074 | -0.074 | 1.004 | 0.15 | 0.01 | 0.19 | 0.24 | 0.19 | 0.23 |
| 2 | 26671544 | rs4072407 | 1.59 | 0.055 | 0.64 | 0.059 | 0 | -0.001 | -0.001 | 0 | 0.72 | 0.31 | 0.6 | 0.28 | 0.82 | 0.74 |

| | | | | | | | | | | | | | | | | |
|---|---|---|---|---|---|---|---|---|---|---|---|---|---|---|---|---|
| 2 | 26676395 | rs12623642 | -0.55 | 0.328 | 0.281 | 0.576 | -0.007 | 0.057 | 0.057 | 0 | 0.05 | 0.03 | 0.27 | 0.59 | 0.07 | 0.09 |
| 2 | 26677335 | rs17005349 | -1.275 | 0.007 | 0.874 | 0.031 | -0.015 | -0.056 | -0.056 | 0.51 | 0.17 | 0.34 | 0.11 | 0.08 | 0.09 | 0.11 |
| 2 | 26678496 | rs11126491 | -1.039 | 0.204 | 0.966 | 0.403 | -0.001 | -0.001 | -0.001 | 0.717 | 0.36 | 0.6 | 0.58 | 0.68 | 0.35 | 0.46 |
| 2 | 26679993 | rs10211527 | -2.384 | 0.393 | 1.476 | 0.298 | -0.017 | -0.077 | -0.077 | 0 | 0.14 | 0.7 | 0.13 | 0 | 0.09 | 0.07 |
| 2 | 26687761 | rs17005371 | -1.685 | 0.292 | 1.705 | 0.609 | 0.022 | 0.018 | 0.018 | 0.393 | 0.07 | 0.54 | 0.08 | 0 | 0.01 | 0.02 |
| 2 | 26704468 | rs4417678 | -0.802 | 0.941 | 0.012 | 0.899 | -0.066 | -0.082 | -0.082 | 0 | 0.18 | 0.04 | 0.25 | 0.34 | 0.27 | 0.34 |
| 2 | 26710421 | rs3739075 | -1.94 | 1.102 | 0.394 | 0.603 | 0.001 | -0.037 | -0.037 | 0.005 | 0.41 | 0.43 | 0.42 | 0.91 | 0.48 | 0.59 |
| 2 | 26711829 | rs12052886 | -0.115 | 1.196 | 0.23 | 0.853 | -0.082 | -0.09 | -0.09 | 4.641 | 0.15 | 0.01 | 0.23 | 0.31 | 0.26 | 0.33 |
| 2 | 26717244 | rs4665858 | -1.605 | 1.137 | 0.791 | 0.593 | 0.003 | -0.032 | -0.032 | 0 | 0.34 | 0.35 | 0.37 | 0.89 | 0.42 | 0.49 |
| 2 | 26719612 | rs11688361 | -1.095 | 1.806 | 0.889 | 0.761 | -0.055 | -0.046 | -0.046 | 0.265 | 0.25 | 0.12 | 0.33 | 0.39 | 0.4 | 0.39 |
| 2 | 26720434 | rs17392390 | -0.922 | 1.801 | 0.885 | 0.758 | -0.093 | -0.079 | -0.079 | 0 | 0.23 | 0.07 | 0.33 | 0.39 | 0.4 | 0.39 |
| 2 | 26722125 | rs7559985 | -1.333 | 1.555 | 0.938 | 0.654 | -0.005 | 0.022 | 0.022 | 0 | 0.24 | 0.7 | 0.18 | 0.5 | 0.19 | 0.19 |
| 2 | 26725045 | rs7592534 | 0.005 | 1.612 | 0.982 | 0.618 | -0.003 | -0.057 | -0.057 | 0.228 | 0.12 | 0.21 | 0.08 | 0 | 0.1 | 0.07 |
| 2 | 26727694 | rs1554481 | -0.998 | 1.645 | 0.671 | 0.631 | -0.041 | -0.033 | -0.033 | 0.22 | 0.26 | 0.14 | 0.34 | 0.39 | 0.4 | 0.4 |
| 2 | 26728179 | rs7590084 | -1.514 | 1.411 | 1.024 | 0.831 | -0.019 | 0.139 | 0.139 | 0 | 0.23 | 0.44 | 0.16 | 0.5 | 0.19 | 0.19 |
| 2 | 26731244 | rs13029647 | 1.113 | 1.88 | 1.022 | 0.593 | 0.012 | -0.032 | -0.032 | 0 | 0.7 | 0.54 | 0.66 | 0.18 | 0.61 | 0.53 |
| 2 | 26732732 | rs13006991 | 1.968 | 1.633 | 1.17 | 0.452 | 0.066 | 0.183 | 0.183 | 0.362 | 0.49 | 0.24 | 0.49 | 0.12 | 0.39 | 0.36 |
| 2 | 26733677 | rs7582117 | -1.519 | 1.437 | 0.988 | 0.838 | -0.024 | 0.216 | 0.216 | 0.05 | 0.16 | 0.39 | 0.11 | 0.49 | 0.12 | 0.19 |
| 2 | 26737900 | rs7604720 | 0.855 | 1.812 | 0.326 | 0.674 | -0.007 | -0.002 | -0.002 | 0 | 0.62 | 0.41 | 0.57 | 0.4 | 0.48 | 0.5 |
| 2 | 26740624 | rs4665332 | 0.959 | 1.149 | 0.004 | 0.62 | 0.015 | 0.005 | 0.005 | 0 | 0.53 | 0.34 | 0.51 | 0.44 | 0.42 | 0.5 |
| 2 | 26741551 | rs7592040 | -0.095 | 1.361 | 0.362 | 0.437 | -0.038 | -0.054 | -0.054 | 0 | 0.17 | 0.08 | 0.24 | 0.56 | 0.3 | 0.36 |
| 2 | 26742511 | rs6746918 | 1.157 | 1.002 | 0.61 | 0.232 | 0.001 | -0.002 | -0.002 | 0.168 | 0.6 | 0.47 | 0.6 | 0.39 | 0.54 | 0.43 |
| 2 | 26744437 | rs17320550 | -0.259 | 1.109 | -0.187 | 0.261 | -0.105 | -0.118 | -0.118 | 2.6 | 0.2 | 0.03 | 0.25 | 0.55 | 0.35 | 0.38 |
| 2 | 26744856 | rs12621611 | -0.496 | 0.803 | 0.383 | 0.179 | -0.016 | -0.073 | -0.073 | 0 | 0.19 | 0.3 | 0.14 | 0.01 | 0.11 | 0.15 |
| 2 | 26747562 | rs939818 | -0.53 | 0.773 | 0.404 | 0.171 | -0.02 | -0.079 | -0.079 | 0.021 | 0.2 | 0.33 | 0.14 | 0.01 | 0.11 | 0.15 |
| 2 | 26748504 | rs13025681 | 1.334 | 0.684 | 0.017 | 0.069 | 0.023 | -0.024 | -0.024 | 0 | 0.71 | 0.1 | 0.8 | 0.67 | 0.78 | 0.77 |
| 2 | 26751425 | rs7583853 | -0.404 | 0.626 | 0.111 | -0.544 | -0.018 | -0.119 | -0.119 | 0 | 0.21 | 0.35 | 0.14 | 0 | 0.14 | 0.16 |
| 2 | 26755551 | rs4665871 | 1.034 | 0.643 | -0.769 | -0.528 | -0.039 | 0.136 | 0.136 | 2.112 | 0.58 | 0.78 | 0.62 | 0.17 | 0.46 | 0.41 |
| 2 | 26757705 | rs1522099 | 0.134 | 0.978 | -0.884 | -0.189 | -0.042 | -0.017 | -0.017 | 0 | 0.81 | 0.93 | 0.75 | 0.28 | 0.66 | 0.62 |
| 2 | 26761165 | rs1522101 | 0.708 | -0.269 | -0.894 | -0.364 | 0.076 | 0.112 | 0.112 | 1.78 | 0.64 | 0.17 | 0.79 | 0.87 | 0.78 | 0.75 |
| 2 | 26777597 | rs4665879 | 0.471 | -0.324 | -0.077 | -0.276 | -0.034 | -0.038 | -0.038 | 0.086 | 0.77 | 0.45 | 0.8 | 0.84 | 0.83 | 0.82 |
| 2 | 26778413 | rs10167513 | -0.984 | -0.421 | 0.374 | -0.364 | -0.008 | -0.011 | -0.011 | 0 | 0.34 | 0.55 | 0.21 | 0.26 | 0.23 | 0.24 |
| 2 | 26781765 | rs7556908 | 1.15 | -0.051 | -0.352 | -0.968 | -0.021 | 0.13 | 0.13 | 0.015 | 0.39 | 0.18 | 0.51 | 0.13 | 0.37 | 0.33 |
| 2 | 26791454 | rs939815 | 0.284 | -0.01 | 0.5 | -0.762 | -0.019 | -0.071 | -0.071 | 0 | 0.64 | 0.45 | 0.78 | 0.91 | 0.75 | 0.76 |
| 2 | 26799031 | rs2272464 | 0.218 | 0.272 | 0.101 | -0.509 | -0.006 | -0.011 | -0.011 | 0 | 0.24 | 0.18 | 0.23 | 0.62 | 0.34 | 0.45 |
| 2 | 26803391 | rs2280513 | 0.74 | 0.098 | 0.27 | -0.615 | -0.002 | -0.001 | -0.001 | 0 | 0.12 | 0.11 | 0.06 | 0.09 | 0.05 | 0.06 |
| 2 | 26804122 | rs744539 | -0.889 | 0.097 | 0.601 | -0.402 | -0.008 | 0.03 | 0.03 | 0 | 0.56 | 0.63 | 0.43 | 0.76 | 0.55 | 0.66 |
| 2 | 26804143 | rs744540 | -1.066 | 1.157 | 0.532 | -0.501 | 0.009 | 0.023 | 0.023 | 0 | 0.59 | 0.66 | 0.49 | 0.78 | 0.62 | 0.69 |
| 2 | 26804247 | rs935172 | -0.652 | 0.717 | 0.28 | 0.111 | 0 | 0.001 | 0.001 | 0 | 0.51 | 0.54 | 0.63 | 0.69 | 0.5 | 0.6 |
| 2 | 26818006 | rs1148962 | 0.028 | 0.108 | 0.447 | -0.31 | -0.019 | -0.088 | -0.088 | 0 | 0.84 | 0.73 | 0.91 | 1 | 0.89 | 0.92 |
| 2 | 26821836 | rs1148967 | 0.905 | -0.266 | 0.523 | -0.626 | 0.025 | 0.004 | 0.004 | 0.385 | 0.7 | 0.26 | 0.84 | 0.99 | 0.85 | 0.79 |
| 2 | 26830665 | rs11685352 | -0.823 | -0.229 | 0.722 | -0.689 | -0.003 | -0.035 | -0.035 | 0 | 0.27 | 0.26 | 0.38 | 0.05 | 0.18 | 0.21 |
| 2 | 26832205 | rs12613651 | 0.83 | 0.293 | 0.181 | -0.353 | -0.035 | -0.035 | -0.035 | 0 | 0.06 | 0 | 0.06 | 0.11 | 0.12 | 0.08 |
| 2 | 26832666 | rs12474099 | 0.201 | 0.168 | 0.53 | -0.633 | 0.001 | -0.004 | -0.004 | 0.534 | 0.13 | 0.13 | 0.29 | 0.04 | 0.07 | 0.13 |
| 2 | 26834387 | rs1000835 | 0.477 | -0.172 | 0.248 | -0.345 | -0.004 | -0.032 | -0.032 | 0 | 0.67 | 0.79 | 0.75 | 0.54 | 0.65 | 0.57 |
| 2 | 26837395 | rs1731240 | 1.749 | -0.355 | 0.08 | -0.165 | -0.003 | 0.156 | 0.156 | 0 | 0.46 | 0.25 | 0.4 | 0.14 | 0.55 | 0.42 |
| 2 | 26840492 | rs6727852 | 0.021 | 0.056 | 0.311 | -0.184 | -0.006 | -0.013 | -0.013 | 0 | 0.19 | 0.12 | 0.18 | 0.46 | 0.25 | 0.35 |
| 2 | 26854065 | rs7563543 | 0.747 | -0.121 | -0.57 | -1.808 | -0.033 | 0.057 | 0.057 | 0 | 0.34 | 0.23 | 0.28 | 0.04 | 0.4 | 0.34 |
| 2 | 26858305 | rs10495759 | -1.34 | 0.328 | 0.33 | -1.239 | -0.008 | -0.005 | -0.005 | 0 | 0.08 | 0.16 | 0.27 | 0.06 | 0.04 | 0.1 |
| 2 | 26865646 | rs1275928 | 0.701 | 0.301 | 0.118 | -1.226 | 0.012 | 0.191 | 0.191 | 0 | 0.53 | 0.25 | 0.59 | 0.1 | 0.54 | 0.45 |
| 2 | 26871233 | rs1275931 | 0.547 | 0.953 | 0.751 | -0.871 | -0.003 | -0.065 | -0.065 | 0 | 0.55 | 0.54 | 0.65 | 0.1 | 0.58 | 0.39 |
| 2 | 26873672 | rs752472 | 0.795 | 0.883 | 0.468 | -0.706 | 0.006 | 0.105 | 0.105 | 0.021 | 0.51 | 0.25 | 0.59 | 0.3 | 0.54 | 0.38 |
| 2 | 26876429 | rs1731253 | 0.638 | 0.936 | 0.398 | -0.656 | 0.006 | 0.072 | 0.072 | 0 | 0.52 | 0.67 | 0.57 | 0.33 | 0.46 | 0.37 |
| 2 | 26887425 | rs1731229 | 0.349 | 0.607 | 0.5 | -1.008 | -0.009 | -0.103 | -0.103 | 0 | 0.63 | 0.47 | 0.64 | 0.1 | 0.63 | 0.45 |
| 2 | 26892720 | rs1275962 | 0.228 | 0.755 | -0.235 | -0.932 | -0.007 | -0.1 | -0.1 | 0.244 | 0.64 | 0.49 | 0.65 | 0.1 | 0.62 | 0.49 |
| 2 | 26894589 | rs1009221 | 0.309 | 0.706 | -0.726 | -0.594 | 0.001 | -0.016 | -0.016 | 0 | 0.56 | 0.43 | 0.63 | 0.18 | 0.5 | 0.4 |
| 2 | 26897501 | rs1275957 | 0.497 | 0.238 | -1.491 | -0.821 | 0.032 | 0.181 | 0.181 | 0.022 | 0.54 | 0.11 | 0.68 | 0.21 | 0.58 | 0.47 |
| 2 | 26901215 | rs6707973 | -0.338 | 0.268 | -0.143 | -0.64 | 0.09 | 0.073 | 0.073 | 0 | 0.06 | 0.43 | 0.06 | 0.06 | 0.05 | 0.12 |
| 2 | 26912079 | rs1275986 | 0.603 | 0.194 | -0.936 | -0.866 | 0.005 | 0.105 | 0.105 | 0 | 0.57 | 0.2 | 0.71 | 0.23 | 0.59 | 0.56 |
| 2 | 26916969 | rs1275980 | 0.758 | 0.154 | -1.136 | -0.866 | -0.003 | 0.134 | 0.134 | 0.287 | 0.55 | 0.14 | 0.7 | 0.24 | 0.59 | 0.56 |
| 2 | 26928811 | rs11126666 | -0.279 | 0.24 | -0.868 | -0.613 | -0.017 | -0.017 | -0.017 | 0.031 | 0.24 | 0.17 | 0.17 | 0.69 | 0.27 | 0.28 |
| 2 | 26938182 | rs1662987 | 0.033 | 0.262 | -0.51 | -0.317 | 0.005 | 0.005 | 0.005 | 0 | 0.78 | 0.69 | 0.8 | 0.72 | 0.76 | 0.77 |
| 2 | 26940234 | rs1662988 | 0.005 | 0.04 | -0.793 | -0.632 | -0.015 | 0.027 | 0.027 | 0 | 0.32 | 0.55 | 0.25 | 0.53 | 0.28 | 0.41 |
| 2 | 26942256 | rs2384463 | 0.8 | -0.121 | -0.907 | -0.948 | -0.029 | 0.117 | 0.117 | 0 | 0.43 | 0.18 | 0.56 | 0.2 | 0.5 | 0.33 |
| 2 | 26952998 | rs11126678 | 0.635 | -0.293 | -0.991 | -0.939 | -0.025 | 0.002 | 0.002 | 0 | 0.21 | 0.07 | 0.16 | 0.25 | 0.22 | 0.22 |
| 2 | 26953354 | rs1731260 | -1.026 | -0.391 | -0.834 | -0.989 | -0.018 | 0.107 | 0.107 | 0 | 0.57 | 0.79 | 0.45 | 0.8 | 0.5 | 0.68 |
| 2 | 26954991 | rs2891519 | 0.521 | -0.374 | -0.675 | -0.899 | 0 | 0.003 | 0.003 | 0 | 0.21 | 0.17 | 0.18 | 0.25 | 0.22 | 0.22 |
| 2 | 26961857 | rs1275935 | -0.712 | -0.293 | -0.165 | -1.24 | -0.011 | 0.033 | 0.033 | 0 | 0.62 | 0.29 | 0.46 | 0.84 | 0.59 | 0.72 |
| 2 | 26964015 | rs870535 | 0.649 | -0.324 | 0.177 | -1.257 | -0.044 | 0.061 | 0.061 | 0 | 0.35 | 0.15 | 0.48 | 0.16 | 0.41 | 0.28 |

| | | | | | | | | | | | | | | | | |
|---|---|---|---|---|---|---|---|---|---|---|---|---|---|---|---|---|
| 2 | 26990507 | rs828259 | 0.372 | -0.343 | 1.004 | -1.287 | -0.015 | 0.039 | 0.039 | 0 | 0.38 | 0.73 | 0.54 | 0.16 | 0.41 | 0.28 |
| 2 | 27003021 | rs10186142 | 0.201 | -0.453 | 0.774 | -1.345 | 0.007 | -0.048 | -0.048 | 5.916 | 0.33 | 0.47 | 0.27 | 0.3 | 0.33 | 0.26 |
| 2 | 27008442 | rs3806517 | 0.109 | -0.456 | 0.658 | -1.347 | 0.001 | -0.021 | -0.021 | 0 | 0.34 | 0.6 | 0.3 | 0.3 | 0.33 | 0.26 |
| 2 | 27008458 | rs3806518 | -0.564 | -0.443 | 0.139 | -1.336 | 0.036 | -0.063 | -0.063 | 0 | 0.69 | 0.9 | 0.78 | 0.46 | 0.74 | 0.54 |
| 2 | 27009478 | rs12467466 | 1.757 | -0.207 | -0.473 | -1.454 | -0.085 | -0.06 | -0.06 | 0 | 0.14 | 0.01 | 0.39 | 0.16 | 0.23 | 0.13 |
| 2 | 27014980 | rs12992046 | 0.706 | -0.355 | -0.064 | -1.252 | -0.047 | 0.047 | 0.047 | 0 | 0.3 | 0.12 | 0.24 | 0.29 | 0.33 | 0.27 |
| 2 | 27024478 | rs2060390 | -0.51 | -0.387 | 0.118 | -1.061 | 0.02 | -0.021 | -0.021 | 0 | 0.68 | 0.75 | 0.77 | 0.47 | 0.74 | 0.55 |
| 2 | 27028992 | rs714447 | -0.153 | -0.285 | 0.04 | -1.05 | 0.024 | -0.015 | -0.015 | 0.198 | 0.63 | 0.36 | 0.74 | 0.47 | 0.71 | 0.55 |
| 2 | 27033604 | rs999087 | -1.085 | -0.26 | -0.115 | -1.14 | -0.037 | 0.089 | 0.089 | 0 | 0.59 | 0.88 | 0.65 | 0.2 | 0.56 | 0.39 |
| 2 | 27044227 | rs7606808 | -0.602 | -0.28 | 0.404 | -1.196 | 0.002 | 0.015 | 0.015 | 0 | 0.53 | 0.42 | 0.57 | 0.19 | 0.53 | 0.36 |
| 2 | 27058451 | rs4513249 | 0.397 | -0.314 | -0.328 | -1.252 | -0.121 | 0.022 | 0.022 | 0 | 0.32 | 0.08 | 0.46 | 0.19 | 0.4 | 0.27 |
| 2 | 27064577 | rs6547275 | -1.305 | -0.346 | -0.098 | -1.233 | -0.026 | 0.087 | 0.087 | 0 | 0.6 | 0.84 | 0.64 | 0.19 | 0.55 | 0.39 |
| 2 | 27069555 | rs934493 | -1.364 | -0.456 | 0.527 | -1.031 | 0.031 | 0.007 | 0.007 | 3.142 | 0.06 | 0.38 | 0.07 | 0 | 0.02 | 0.02 |
| 2 | 27070362 | rs6547284 | -1.364 | -0.456 | 0.527 | -1.031 | 0.031 | 0.007 | 0.007 | 3.142 | 0.06 | 0.38 | 0.06 | 0 | 0.02 | 0.02 |
| 2 | 27083489 | rs6756245 | -0.16 | -0.302 | 0.263 | -1 | -0.015 | 0.03 | 0.03 | 0 | 0.37 | 0.37 | 0.5 | 0.15 | 0.39 | 0.31 |
| 2 | 27101281 | rs7596707 | 0.321 | -0.776 | 0.227 | -0.796 | 0.025 | 0.041 | 0.041 | 0 | 0.18 | 0.09 | 0.07 | 0 | 0.1 | 0.09 |
| 2 | 27128529 | rs509047 | -0.8 | -0.422 | 1.181 | 0.046 | 0.003 | 0.005 | 0.005 | 0 | 0.38 | 0.52 | 0.57 | 0.29 | 0.37 | 0.28 |
| 2 | 27141515 | rs9309559 | -0.73 | -0.451 | 1.418 | 0.041 | -0.001 | -0.003 | -0.003 | 0 | 0.37 | 0.4 | 0.56 | 0.3 | 0.37 | 0.28 |
| 2 | 27152874 | rs1371614 | -0.452 | -0.5 | 1.704 | -0.108 | -0.001 | -0.01 | -0.01 | 0 | 0.27 | 0.36 | 0.39 | 0.15 | 0.24 | 0.2 |
| 2 | 27155848 | rs3820821 | 0.867 | -0.493 | 1.689 | -0.031 | 0 | -0.001 | -0.001 | 0 | 0.6 | 0.58 | 0.44 | 0.69 | 0.62 | 0.67 |
| 2 | 27157788 | rs920435 | -1.203 | -0.662 | 1.694 | -0.115 | 0.008 | -0.01 | -0.01 | 0.863 | 0.42 | 0.28 | 0.55 | 0.31 | 0.39 | 0.35 |
| 2 | 27163044 | rs3739085 | 0.901 | -0.32 | 1.823 | 0.009 | -0.001 | 0 | 0 | 0.605 | 0.6 | 0.57 | 0.45 | 0.69 | 0.62 | 0.68 |
| 2 | 27169490 | rs3769137 | 0.981 | -0.112 | 2.022 | 0.134 | -0.001 | 0.003 | 0.003 | 0.016 | 0.62 | 0.53 | 0.48 | 0.71 | 0.63 | 0.68 |
| 2 | 27169842 | rs11126784 | 1.097 | -0.126 | 2.239 | 0.138 | -0.001 | 0.001 | 0.001 | 0.105 | 0.61 | 0.42 | 0.46 | 0.71 | 0.63 | 0.68 |
| 2 | 27170982 | rs13017554 | 1.175 | -0.149 | 2.563 | 0.103 | 0.005 | 0.001 | 0.001 | 0.661 | 0.59 | 0.27 | 0.44 | 0.71 | 0.63 | 0.68 |
| 2 | 27171245 | rs10181727 | -1.159 | -0.149 | 2.682 | 0.104 | -0.006 | -0.012 | -0.012 | 1.157 | 0.36 | 0.27 | 0.49 | 0.29 | 0.36 | 0.32 |
| 2 | 27184446 | rs2891534 | 1.1 | -0.134 | 2.751 | -0.115 | 0.002 | 0.005 | 0.005 | 0.454 | 0.6 | 0.35 | 0.47 | 0.77 | 0.63 | 0.68 |
| 2 | 27187814 | rs1440042 | 1.209 | -0.158 | 2.756 | -0.126 | 0.008 | 0.021 | 0.021 | 1.012 | 0.59 | 0.25 | 0.44 | 0.77 | 0.62 | 0.68 |
| 2 | 27191473 | rs7564363 | 0.65 | -0.234 | 2.686 | -0.199 | -0.015 | 0.022 | 0.022 | 0.017 | 0.78 | 0.54 | 0.77 | 0.84 | 0.82 | 0.83 |
| 2 | 27194013 | rs4572551 | 0.97 | -0.08 | 2.83 | -0.07 | -0.001 | -0.001 | -0.001 | 0 | 0.6 | 0.38 | 0.45 | 0.77 | 0.62 | 0.68 |
| 2 | 27201857 | rs893790 | -1.088 | -0.039 | 2.837 | -0.081 | -0.004 | 0.007 | 0.007 | 0.001 | 0.38 | 0.32 | 0.52 | 0.23 | 0.38 | 0.32 |
| 2 | 27204650 | rs13020526 | 0.987 | -0.036 | 2.726 | -0.079 | 0.003 | 0.01 | 0.01 | 0.664 | 0.59 | 0.32 | 0.45 | 0.77 | 0.62 | 0.68 |
| 2 | 27212526 | rs920433 | 0.887 | 0.018 | 2.095 | -0.053 | 0 | -0.009 | -0.009 | 2.48 | 0.61 | 0.5 | 0.47 | 0.77 | 0.62 | 0.68 |
| 2 | 27250603 | rs11681562 | 0.822 | -0.023 | 1.955 | -0.035 | -0.002 | -0.006 | -0.006 | 0 | 0.62 | 0.54 | 0.48 | 0.77 | 0.63 | 0.68 |
| 2 | 27260469 | rs1124649 | -1.028 | -0.011 | 1.952 | -0.069 | -0.007 | -0.05 | -0.05 | 0.017 | 0.36 | 0.45 | 0.4 | 0.07 | 0.31 | 0.28 |
| 2 | 27263122 | rs1866654 | 1.097 | -0.023 | 2.123 | -0.066 | 0.007 | 0.012 | 0.012 | 1.612 | 0.59 | 0.33 | 0.45 | 0.77 | 0.62 | 0.69 |
| 2 | 27273128 | rs11126836 | 1.061 | -0.033 | 2.173 | -0.071 | 0.009 | 0.016 | 0.016 | 1.605 | 0.59 | 0.31 | 0.45 | 0.77 | 0.62 | 0.69 |
| 2 | 27290525 | rs11681145 | 0.702 | -0.027 | 2.097 | -0.08 | 0.003 | -0.009 | -0.009 | 0 | 0.63 | 0.36 | 0.58 | 0.93 | 0.69 | 0.72 |
| 2 | 27293150 | rs1053609 | 0.975 | -0.018 | 1.845 | -0.082 | 0.012 | 0.02 | 0.02 | 1.706 | 0.58 | 0.29 | 0.45 | 0.77 | 0.62 | 0.69 |
| 2 | 27302561 | rs2011616 | -1.368 | -0.085 | 0.985 | -0.135 | 0.012 | 0.038 | 0.038 | 0.01 | 0.44 | 0.27 | 0.52 | 0.23 | 0.39 | 0.35 |
| 2 | 27314882 | rs12714092 | -0.764 | 0.076 | 1.041 | -0.016 | -0.026 | -0.042 | -0.042 | 0.165 | 0.34 | 0.13 | 0.47 | 0.23 | 0.36 | 0.3 |
| 2 | 27315252 | rs2304681 | -1.019 | 0.045 | 1.152 | 0.01 | -0.002 | 0.003 | 0.003 | 0.021 | 0.37 | 0.3 | 0.5 | 0.23 | 0.38 | 0.3 |
| 2 | 27323385 | rs1131375 | -0.917 | 0.082 | 0.451 | 0.044 | -0.024 | -0.02 | -0.02 | 0.169 | 0.36 | 0.16 | 0.48 | 0.24 | 0.37 | 0.31 |
| 2 | 27335586 | rs1866559 | -0.889 | 0.155 | 0.483 | 0.114 | -0.025 | -0.027 | -0.027 | 0 | 0.36 | 0.17 | 0.49 | 0.24 | 0.38 | 0.31 |
| 2 | 27345484 | rs7605526 | -0.884 | 0.162 | 0.303 | 0.131 | -0.004 | -0.005 | -0.005 | 0.611 | 0.36 | 0.29 | 0.49 | 0.23 | 0.37 | 0.29 |
| 2 | 27348029 | rs4665947 | -0.905 | 0.126 | 0.155 | 0.207 | -0.013 | -0.015 | -0.015 | 2.573 | 0.38 | 0.24 | 0.52 | 0.3 | 0.36 | 0.33 |
| 2 | 27351279 | rs1866561 | 0.657 | 0.069 | 0.731 | 0.168 | 0.001 | 0.002 | 0.002 | 0.038 | 0.6 | 0.62 | 0.5 | 0.71 | 0.64 | 0.67 |
| 2 | 27355991 | rs2289358 | -0.347 | -0.015 | 0.534 | 0.114 | 0.006 | -0.003 | -0.003 | 0 | 0.24 | 0.27 | 0.18 | 0.17 | 0.16 | 0.15 |
| 2 | 27356278 | rs2289359 | -0.334 | 0.016 | 0.564 | 0.14 | 0.004 | -0.003 | -0.003 | 0 | 0.23 | 0.28 | 0.18 | 0.17 | 0.17 | 0.15 |
| 2 | 27361246 | rs2304680 | -0.449 | -0.123 | 1.336 | 0.189 | -0.035 | 0.001 | 0.001 | 1.339 | 0.26 | 0.52 | 0.2 | 0.17 | 0.16 | 0.15 |
| 2 | 27362371 | rs11691585 | -0.473 | -0.138 | 1.404 | 0.19 | -0.034 | 0.003 | 0.003 | 1.568 | 0.26 | 0.57 | 0.21 | 0.18 | 0.17 | 0.15 |
| 2 | 27366408 | rs1659711 | 1.243 | 0.317 | 1.024 | -0.107 | -0.011 | 0.023 | 0.023 | 0.546 | 0.44 | 0.22 | 0.38 | 0.7 | 0.51 | 0.61 |
| 2 | 27370970 | rs1275513 | 1.252 | 0.299 | 1.015 | -0.116 | -0.011 | 0.023 | 0.023 | 0.614 | 0.44 | 0.22 | 0.38 | 0.7 | 0.51 | 0.61 |
| 2 | 27374009 | rs1275509 | 0.664 | 0.558 | 0.551 | -0.009 | -0.004 | 0.018 | 0.018 | 0.603 | 0.55 | 0.3 | 0.44 | 0.7 | 0.59 | 0.66 |
| 2 | 27384579 | rs7588910 | 0.322 | 0.592 | 0.457 | 0.026 | 0.002 | -0.005 | -0.005 | 0 | 0.59 | 0.34 | 0.44 | 0.6 | 0.62 | 0.66 |
| 2 | 27385275 | rs11883596 | -0.772 | 0.26 | 0.84 | -0.443 | -0.004 | -0.106 | -0.106 | 0 | 0.17 | 0.31 | 0.1 | 0 | 0.11 | 0.08 |
| 2 | 27386799 | rs1275539 | 1.053 | 0.034 | 0.052 | -0.48 | 0.043 | 0.18 | 0.18 | 0.572 | 0.55 | 0.09 | 0.52 | 0.77 | 0.66 | 0.64 |
| 2 | 27393030 | rs1659689 | 0.916 | -0.386 | 0.427 | -0.719 | 0.046 | 0.129 | 0.129 | 0.023 | 0.58 | 0.19 | 0.53 | 0.87 | 0.7 | 0.61 |
| 2 | 27419282 | rs1275501 | 1.001 | -0.463 | 0.401 | -0.733 | 0.071 | 0.171 | 0.171 | 0.345 | 0.57 | 0.16 | 0.53 | 0.85 | 0.71 | 0.59 |
| 2 | 27420690 | rs1659685 | 1.001 | -0.463 | 0.408 | -0.733 | 0.07 | 0.167 | 0.167 | 0.345 | 0.57 | 0.16 | 0.53 | 0.85 | 0.71 | 0.59 |
| 2 | 27422605 | rs7081 | 0.935 | -0.571 | 1.376 | -0.815 | -0.019 | -0.078 | -0.078 | 0 | 0.88 | 0.43 | 0.9 | 1 | 0.93 | 0.94 |
| 2 | 27424636 | rs1395 | 1.33 | -0.885 | 1.301 | -0.972 | 0.086 | 0.191 | 0.191 | 0.146 | 0.55 | 0.16 | 0.52 | 0.85 | 0.69 | 0.55 |
| 2 | 27426362 | rs2580761 | -0.207 | -0.801 | 1.335 | -0.935 | 0.005 | 0.005 | 0.005 | 0 | 0.31 | 0.27 | 0.37 | 0.15 | 0.22 | 0.35 |
| 2 | 27428702 | rs1275525 | 0.867 | -0.798 | 1.367 | -0.931 | -0.019 | -0.078 | -0.078 | 0 | 0.88 | 0.43 | 0.9 | 1 | 0.93 | 0.94 |
| 2 | 27430781 | rs1275528 | 1.136 | -0.801 | 1.347 | -0.935 | 0.07 | 0.167 | 0.167 | 0.345 | 0.57 | 0.16 | 0.53 | 0.85 | 0.71 | 0.59 |
| 2 | 27432642 | rs6547496 | 0.305 | -0.813 | 1.737 | -0.942 | -0.002 | -0.015 | -0.015 | 0 | 0.07 | 0.07 | 0.04 | 0 | 0.05 | 0.06 |
| 2 | 27435374 | rs11608 | 1.21 | -0.769 | 1.733 | -0.951 | 0.063 | 0.168 | 0.168 | 0.412 | 0.57 | 0.17 | 0.53 | 0.85 | 0.7 | 0.58 |
| 2 | 27438710 | rs11126918 | -0.198 | -0.792 | 1.756 | -0.919 | 0.005 | 0.006 | 0.006 | 0 | 0.3 | 0.27 | 0.37 | 0.15 | 0.22 | 0.35 |
| 2 | 27439751 | rs7437 | 0.181 | -0.796 | 2.094 | -0.93 | -0.006 | -0.024 | -0.024 | 0.105 | 0.07 | 0.11 | 0.04 | 0 | 0.05 | 0.06 |

| | | | | | | | | | | | | | | | | |
|---|---|---|---|---|---|---|---|---|---|---|---|---|---|---|---|---|
| 2 | 27441397 | rs1275538 | 1.167 | -0.794 | 2.069 | -0.929 | 0.07 | 0.167 | 0.167 | 0.37 | 0.57 | 0.16 | 0.53 | 0.85 | 0.71 | 0.59 |
| 2 | 27443196 | rs4665958 | -1.479 | -0.794 | 2.069 | -0.929 | 0.07 | 0.167 | 0.167 | 0.37 | 0.43 | 0.84 | 0.47 | 0.15 | 0.29 | 0.41 |
| 2 | 27455281 | rs6747264 | 0.212 | -0.777 | 2.062 | -0.906 | -0.003 | -0.017 | -0.017 | 0 | 0.07 | 0.09 | 0.04 | 0 | 0.05 | 0.07 |
| 2 | 27455726 | rs13399758 | -1.712 | -0.789 | 2.136 | -0.913 | -0.031 | -0.081 | -0.081 | 0.111 | 0.13 | 0.58 | 0.1 | 0 | 0.06 | 0.07 |
| 2 | 27471025 | rs1561535 | -1.494 | -0.786 | 1.921 | -0.934 | 0.082 | 0.189 | 0.189 | 0.547 | 0.43 | 0.87 | 0.47 | 0.15 | 0.28 | 0.41 |
| 2 | 27478231 | rs13390228 | -1.512 | -0.688 | 2.099 | -0.847 | 0.091 | 0.074 | 0.074 | 1.531 | 0.05 | 0.48 | 0.06 | 0 | 0.01 | 0 |
| 2 | 27504312 | rs1992291 | -0.757 | -0.824 | 2.248 | -0.916 | 0.016 | 0.026 | 0.026 | 0.206 | 0.34 | 0.73 | 0.43 | 0.16 | 0.22 | 0.35 |
| 2 | 27508710 | rs11686131 | -0.06 | -0.835 | 2.274 | -0.922 | 0.003 | 0.003 | 0.003 | 0 | 0.28 | 0.27 | 0.36 | 0.16 | 0.2 | 0.35 |
| 2 | 27519254 | rs13404446 | -1.287 | -0.861 | 2.344 | -0.945 | 0.066 | 0.156 | 0.156 | 0.438 | 0.41 | 0.86 | 0.46 | 0.16 | 0.25 | 0.41 |
| 2 | 27531665 | rs6731600 | 0.295 | -0.83 | 2.275 | -0.927 | -0.008 | -0.025 | -0.025 | 0 | 0.07 | 0.1 | 0.03 | 0 | 0.03 | 0.06 |
| 2 | 27542041 | rs1975384 | -0.069 | -0.773 | 2.233 | -0.879 | 0.003 | 0.003 | 0.003 | 0 | 0.28 | 0.26 | 0.36 | 0.16 | 0.2 | 0.35 |
| 2 | 27550967 | rs1049817 | -0.656 | 0.241 | 1.483 | -1.229 | -0.008 | 0.066 | 0.066 | 0.264 | 0.4 | 0.74 | 0.48 | 0.16 | 0.38 | 0.4 |
| 2 | 27558252 | rs11684134 | -0.744 | -0.249 | 1.402 | -1.195 | -0.001 | 0 | 0 | 0.238 | 0.44 | 0.47 | 0.48 | 0.23 | 0.42 | 0.46 |
| 2 | 27568565 | rs10205219 | -0.49 | -0.079 | 1.543 | -1.198 | -0.003 | 0.006 | 0.006 | 0.169 | 0.39 | 0.67 | 0.47 | 0.16 | 0.38 | 0.4 |
| 2 | 27574953 | rs4665969 | -0.25 | -0.081 | 1.376 | -1.199 | -0.001 | -0.01 | -0.01 | 4.512 | 0.37 | 0.41 | 0.45 | 0.16 | 0.38 | 0.4 |
| 2 | 27583273 | rs6716894 | -0.532 | -0.097 | 1.5 | -1.225 | 0.003 | -0.058 | -0.058 | 0 | 0.39 | 0.51 | 0.46 | 0.17 | 0.37 | 0.4 |
| 2 | 27584666 | rs7586601 | -1.237 | -0.238 | 1.488 | -1.348 | -0.006 | 0.24 | 0.24 | 1.076 | 0.49 | 0.85 | 0.52 | 0.17 | 0.41 | 0.46 |
| 2 | 27586155 | rs11695549 | 0.066 | -0.239 | 1.619 | -1.348 | -0.021 | 0.063 | 0.063 | 0.199 | 0.34 | 0.26 | 0.43 | 0.16 | 0.36 | 0.4 |
| 2 | 27589810 | rs2280737 | -0.452 | -0.212 | 1.537 | -1.326 | 0.003 | -0.058 | -0.058 | 1.873 | 0.39 | 0.51 | 0.46 | 0.17 | 0.37 | 0.4 |
| 2 | 27592423 | rs7602534 | -0.223 | -0.272 | 1.575 | -1.352 | 0.008 | -0.069 | -0.069 | 0 | 0.37 | 0.49 | 0.47 | 0.16 | 0.36 | 0.4 |
| 2 | 27599132 | rs12475426 | -0.401 | -0.292 | 1.727 | -1.389 | -0.006 | 0.067 | 0.067 | 0.181 | 0.4 | 0.76 | 0.49 | 0.17 | 0.38 | 0.44 |
| 2 | 27600239 | rs13472 | -0.005 | -0.271 | 1.766 | -1.368 | -0.005 | 0.011 | 0.011 | 0.047 | 0.36 | 0.37 | 0.45 | 0.17 | 0.37 | 0.43 |
| 2 | 27604279 | rs4582 | -0.36 | -0.274 | 1.597 | -1.369 | -0.005 | 0.066 | 0.066 | 0.323 | 0.4 | 0.76 | 0.48 | 0.17 | 0.37 | 0.44 |
| 2 | 27613617 | rs11891554 | 0.464 | -0.284 | 1.597 | -1.358 | -0.015 | -0.042 | -0.042 | 0 | 0.1 | 0.14 | 0.04 | 0 | 0.04 | 0.06 |
| 2 | 27635463 | rs1728918 | -1.37 | 0.113 | 0.871 | -0.926 | -0.052 | -0.077 | -0.077 | 0 | 0.75 | 0.97 | 0.75 | 0.77 | 0.73 | 0.86 |
| 2 | 27635582 | rs1060525 | -0.256 | 0.511 | 1.388 | -0.976 | -0.006 | 0.064 | 0.064 | 0.392 | 0.4 | 0.76 | 0.49 | 0.17 | 0.37 | 0.44 |
| 2 | 27642734 | rs11675428 | 2.575 | 0.728 | 1.197 | -0.972 | -0.03 | 0.027 | 0.027 | 0 | 0.05 | 0.01 | 0.05 | 0 | 0.16 | 0.05 |
| 2 | 27663416 | rs1260342 | 0.106 | 0.358 | 1.574 | -0.806 | -0.005 | 0.047 | 0.047 | 0.476 | 0.39 | 0.73 | 0.48 | 0.17 | 0.38 | 0.43 |
| 2 | 27667297 | rs4803 | 0.096 | 0.358 | 1.659 | -0.806 | -0.005 | 0.048 | 0.048 | 0.459 | 0.39 | 0.73 | 0.48 | 0.17 | 0.37 | 0.44 |
| 2 | 27670307 | rs11126999 | 0.503 | 0.355 | 1.758 | -0.808 | -0.005 | 0.012 | 0.012 | 0.032 | 0.36 | 0.37 | 0.45 | 0.17 | 0.37 | 0.43 |
| 2 | 27716494 | rs704795 | 0.116 | 0.371 | 1.481 | -0.799 | -0.001 | 0 | 0 | 0.41 | 0.39 | 0.67 | 0.48 | 0.17 | 0.38 | 0.44 |
| 2 | 27718474 | rs780090 | -2.434 | 0.547 | 0.992 | -0.858 | -0.03 | 0.035 | 0.035 | 0 | 0.93 | 0.99 | 0.97 | 1 | 0.91 | 0.99 |
| 2 | 27722416 | rs1260320 | 0.269 | 0.488 | 1.319 | -0.78 | 0 | -0.026 | -0.026 | 2.84 | 0.38 | 0.61 | 0.48 | 0.16 | 0.38 | 0.44 |
| 2 | 27729480 | rs2293571 | 0.551 | 0.502 | 1.208 | -0.762 | 0 | -0.019 | -0.019 | 0.111 | 0.36 | 0.4 | 0.46 | 0.17 | 0.37 | 0.48 |
| 2 | 27730817 | rs8179219 | 1.042 | 0.476 | 0.875 | -0.791 | -0.009 | -0.03 | -0.03 | 0 | 0.09 | 0.1 | 0.04 | 0 | 0.04 | 0.05 |
| 2 | 27730940 | rs1260326 | -0.954 | 0.456 | 0.375 | -0.209 | -0.037 | -0.021 | -0.021 | 0 | 0.58 | 0.91 | 0.64 | 0.52 | 0.59 | 0.8 |
| 2 | 27731212 | rs3817588 | 0.955 | 0.901 | 0.463 | -0.056 | -0.051 | -0.068 | -0.068 | 0 | 0.1 | 0.01 | 0.12 | 0.34 | 0.21 | 0.26 |
| 2 | 27741237 | rs780094 | -0.944 | 0.608 | 0.545 | -0.085 | -0.021 | -0.02 | -0.02 | 0 | 0.59 | 0.87 | 0.64 | 0.52 | 0.59 | 0.8 |
| 2 | 27742603 | rs780093 | -1.026 | 0.612 | 0.525 | -0.104 | -0.034 | -0.032 | -0.032 | 0 | 0.6 | 0.91 | 0.64 | 0.52 | 0.59 | 0.8 |
| 2 | 27743154 | rs780092 | 0.035 | 0.666 | 1.095 | -0.069 | 0.003 | 0.066 | 0.066 | 0 | 0.13 | 0.24 | 0.14 | 0.34 | 0.17 | 0.25 |
| 2 | 27743423 | rs11681351 | 0.84 | 0.652 | 1.35 | -0.384 | -0.011 | 0.031 | 0.031 | 0.069 | 0.35 | 0.29 | 0.45 | 0.18 | 0.37 | 0.49 |
| 2 | 27773648 | rs2384626 | -0.261 | 0.641 | 1.466 | -0.147 | 0.005 | 0.012 | 0.012 | 0 | 0.51 | 0.4 | 0.66 | 0.85 | 0.48 | 0.51 |
| 2 | 27783801 | rs4665382 | 1.341 | 0.591 | 1.318 | 0.1 | -0.021 | -0.165 | -0.165 | 2.12 | 0.3 | 0.02 | 0.51 | 0.51 | 0.26 | 0.17 |
| 2 | 27801418 | rs1919126 | -0.926 | 0.567 | 1.537 | 0.078 | -0.001 | 0.103 | 0.103 | 0 | 0.55 | 0.74 | 0.69 | 0.85 | 0.5 | 0.51 |
| 2 | 27801493 | rs1919127 | 1.317 | 0.561 | 1.584 | 0.078 | 0.017 | -0.048 | -0.048 | 0 | 0.3 | 0.17 | 0.52 | 0.51 | 0.26 | 0.17 |
| 2 | 27802805 | rs3811644 | 0.147 | 0.571 | 1.61 | 0.088 | -0.011 | -0.052 | -0.052 | 0 | 0.18 | 0.05 | 0.11 | 0.34 | 0.21 | 0.33 |
| 2 | 27808154 | rs6734059 | -0.139 | 0.589 | 1.587 | 0.084 | -0.001 | 0.021 | 0.021 | 0 | 0.2 | 0.22 | 0.13 | 0.34 | 0.22 | 0.33 |
| 2 | 27812252 | rs6760250 | 1.239 | 0.578 | 1.519 | 0.073 | 0.019 | -0.048 | -0.048 | 0.513 | 0.31 | 0.17 | 0.52 | 0.51 | 0.26 | 0.17 |
| 2 | 27827196 | rs2141372 | -0.757 | 0.627 | 1.465 | 0.106 | -0.015 | 0.049 | 0.049 | 0.184 | 0.24 | 0.54 | 0.15 | 0.34 | 0.22 | 0.33 |
| 2 | 27838058 | rs11127071 | -0.867 | 0.685 | 1.283 | 0.068 | 0 | 0.094 | 0.094 | 0 | 0.54 | 0.71 | 0.69 | 0.85 | 0.5 | 0.51 |
| 2 | 27844601 | rs1881396 | -0.258 | 0.677 | 1.305 | -0.064 | 0 | 0.001 | 0.001 | 0 | 0.2 | 0.17 | 0.13 | 0.33 | 0.22 | 0.33 |
| 2 | 27846603 | rs1528404 | -0.723 | 0.799 | 1.063 | -0.813 | -0.003 | 0.103 | 0.103 | 0 | 0.54 | 0.72 | 0.68 | 0.9 | 0.5 | 0.51 |
| 2 | 27851120 | rs12615690 | -0.652 | 0.782 | 1.127 | -0.809 | 0.003 | 0.049 | 0.049 | 0 | 0.54 | 0.65 | 0.67 | 0.9 | 0.48 | 0.52 |
| 2 | 27855239 | rs4666005 | -0.414 | 0.754 | 1.097 | -0.818 | 0 | -0.039 | -0.039 | 0 | 0.53 | 0.46 | 0.66 | 0.9 | 0.48 | 0.52 |
| 2 | 27857517 | rs1528402 | -0.524 | 0.781 | 1.006 | -0.813 | 0 | -0.041 | -0.041 | 0.232 | 0.53 | 0.53 | 0.68 | 0.9 | 0.5 | 0.52 |
| 2 | 27861707 | rs11692692 | 0.515 | 0.699 | 0.889 | -0.865 | -0.013 | 0.206 | 0.206 | 0.065 | 0.45 | 0.14 | 0.32 | 0.08 | 0.53 | 0.5 |
| 2 | 27862397 | rs7564625 | 0.4 | 0.795 | 0.957 | -0.851 | 0.008 | 0.087 | 0.087 | 0 | 0.47 | 0.29 | 0.35 | 0.08 | 0.53 | 0.51 |
| 2 | 27874563 | rs7595739 | 0.036 | 0.686 | 1.329 | -0.874 | 0.016 | 0.026 | 0.026 | 0 | 0.5 | 0.63 | 0.38 | 0.08 | 0.55 | 0.51 |
| 2 | 27878686 | rs4666010 | 0.012 | 0.604 | 1.271 | -0.588 | 0 | -0.015 | -0.015 | 0 | 0.2 | 0.13 | 0.43 | 0.58 | 0.23 | 0.34 |
| 2 | 27887034 | rs9678851 | 0.558 | 0.643 | 0.867 | -0.677 | 0.012 | 0.117 | 0.117 | 0 | 0.47 | 0.25 | 0.35 | 0.08 | 0.54 | 0.51 |
| 2 | 27895073 | rs2178198 | 2.086 | 0.983 | 0.198 | -0.627 | -0.037 | 0.038 | 0.038 | 0 | 0.08 | 0.02 | 0.06 | 0 | 0.13 | 0.01 |
| 2 | 27910706 | rs13023094 | 1.216 | 0.492 | 0.309 | -0.397 | -0.025 | -0.06 | -0.06 | 0 | 0.27 | 0.02 | 0.17 | 0.33 | 0.22 | 0.15 |
| 2 | 27928797 | rs13030973 | 1.003 | 0.531 | -0.383 | -0.409 | -0.01 | -0.036 | -0.036 | 0 | 0.28 | 0.03 | 0.18 | 0.33 | 0.23 | 0.15 |
| 2 | 27930837 | rs4665385 | 0.351 | 0.809 | 0.289 | 0.181 | -0.033 | -0.069 | -0.069 | 0 | 0.12 | 0.01 | 0.4 | 0.59 | 0.18 | 0.27 |
| 2 | 27951658 | rs867282 | 0.895 | 0.75 | 0.82 | 0.097 | -0.007 | 0.046 | 0.046 | 0.37 | 0.76 | 0.48 | 0.55 | 0.41 | 0.76 | 0.69 |
| 2 | 27975394 | rs6547796 | 1.212 | 0.775 | 0.873 | -0.02 | -0.002 | 0 | 0 | 0 | 0.72 | 0.32 | 0.52 | 0.41 | 0.73 | 0.67 |
| 2 | 27988821 | rs13013484 | 1.376 | 0.752 | 1.015 | -0.078 | -0.003 | -0.002 | -0.002 | 0 | 0.7 | 0.36 | 0.5 | 0.41 | 0.71 | 0.67 |
| 2 | 27995931 | rs3792252 | 0.987 | 0.873 | 1.276 | -0.069 | 0.006 | 0.018 | 0.018 | 0.04 | 0.73 | 0.51 | 0.54 | 0.41 | 0.73 | 0.67 |
| 2 | 28002002 | rs3792253 | -0.706 | 0.696 | 1.135 | -0.035 | -0.002 | -0.009 | -0.009 | 0 | 0.19 | 0.17 | 0.41 | 0.59 | 0.22 | 0.32 |

| | | | | | | | | | | | | | | | | |
|---|---|---|---|---|---|---|---|---|---|---|---|---|---|---|---|---|
| 2 | 28019367 | rs4666015 | -1.012 | 0.613 | 1.16 | -0.031 | 0.001 | 0.035 | 0.035 | 0 | 0.22 | 0.39 | 0.44 | 0.6 | 0.23 | 0.32 |
| 2 | 28055881 | rs4666021 | -0.028 | 0.691 | 0.95 | -0.208 | 0.01 | 0.038 | 0.038 | 0 | 0.13 | 0.16 | 0.39 | 0.63 | 0.2 | 0.24 |
| 2 | 28062396 | rs4361084 | -1.031 | 0.761 | 0.151 | -0.829 | -0.072 | 0.122 | 0.122 | 0 | 0.8 | 0.97 | 0.84 | 1 | 0.74 | 0.95 |
| 2 | 28090059 | rs10171517 | 0.858 | 0.557 | 0.639 | -0.226 | -0.001 | 0.08 | 0.08 | 0.172 | 0.77 | 0.38 | 0.84 | 0.55 | 0.77 | 0.71 |
| 2 | 28090820 | rs898034 | 0.833 | 0.561 | 0.642 | -0.224 | 0 | 0.082 | 0.082 | 0.1 | 0.77 | 0.39 | 0.84 | 0.55 | 0.77 | 0.71 |
| 2 | 28118134 | rs7606935 | -0.247 | 0.556 | 0.754 | -0.347 | -0.024 | -0.054 | -0.054 | 0 | 0.11 | 0.21 | 0.05 | 0 | 0.06 | 0.1 |
| 2 | 28144067 | rs1870325 | 0.405 | 0.615 | 0.659 | -0.3 | 0.018 | 0.051 | 0.051 | 0 | 0.8 | 0.6 | 0.58 | 0.38 | 0.77 | 0.7 |
| 2 | 28151354 | rs10185472 | 0.405 | 0.615 | 0.66 | -0.3 | 0.017 | 0.05 | 0.05 | 0 | 0.8 | 0.6 | 0.58 | 0.38 | 0.77 | 0.72 |
| 2 | 28153488 | rs6709355 | 0.217 | 0.591 | 0.519 | -0.314 | -0.019 | -0.047 | -0.047 | 0 | 0.1 | 0.2 | 0.04 | 0 | 0.06 | 0.1 |
| 2 | 28174081 | rs1506535 | 0.676 | 0.67 | 0.469 | -0.257 | 0.014 | 0.041 | 0.041 | 0 | 0.1 | 0.12 | 0.38 | 0.61 | 0.18 | 0.18 |
| 2 | 28203294 | rs898031 | 1.136 | 0.498 | -0.645 | -0.778 | -0.001 | 0.025 | 0.025 | 0 | 0.3 | 0.2 | 0.18 | 0.22 | 0.29 | 0.55 |
| 2 | 28261674 | rs10200337 | 1.578 | 0.545 | -0.8 | -1.738 | 0 | 0.002 | 0.002 | 0 | 0.23 | 0.2 | 0.16 | 0.21 | 0.23 | 0.49 |
| 2 | 28288392 | rs10173528 | 0.066 | 0.587 | -0.76 | -1.69 | 0.012 | 0.055 | 0.055 | 0.053 | 0.53 | 0.38 | 0.37 | 0.22 | 0.55 | 0.61 |
| 2 | 28290347 | rs13414801 | 0.642 | 0.707 | -0.23 | -1.178 | -0.002 | 0 | 0 | 0.541 | 0.7 | 0.35 | 0.5 | 0.4 | 0.63 | 0.71 |
| 2 | 28293299 | rs13409819 | 0.132 | 0.61 | -0.04 | -0.93 | 0.037 | -0.022 | -0.022 | 0 | 0.66 | 0.55 | 0.42 | 0.38 | 0.55 | 0.77 |
| 2 | 28300314 | rs1506534 | -0.491 | 0.326 | 0.065 | -1.121 | -0.025 | -0.113 | -0.113 | 0.618 | 0.17 | 0.36 | 0.09 | 0 | 0.1 | 0.13 |
| 2 | 28301540 | rs937813 | 0.718 | -0.034 | -0.51 | -1.183 | 0.005 | -0.003 | -0.003 | 1.999 | 0.15 | 0.05 | 0.11 | 0.15 | 0.11 | 0.09 |
| 2 | 28305663 | rs13415164 | 1.175 | -0.058 | -0.473 | -1.415 | 0.008 | 0.056 | 0.056 | 0 | 0.38 | 0.25 | 0.2 | 0.29 | 0.35 | 0.56 |
| 2 | 28318008 | rs2063018 | 1.267 | 0.297 | -0.695 | -1.571 | -0.067 | 0.149 | 0.149 | 0 | 0.25 | 0.05 | 0.2 | 0 | 0.28 | 0.05 |
| 2 | 28318879 | rs7578052 | 0.527 | 0.323 | -0.266 | -1.484 | 0.01 | 0.009 | 0.009 | 0 | 0.17 | 0.18 | 0.48 | 0.7 | 0.27 | 0.26 |
| 2 | 28319242 | rs7568903 | 1.878 | 0.296 | -0.163 | -1.435 | -0.003 | 0.006 | 0.006 | 0 | 0.07 | 0.09 | 0.06 | 0.15 | 0.05 | 0.08 |
| 2 | 28320033 | rs7572644 | 1.346 | 0.339 | -0.213 | -1.434 | 0.037 | 0.107 | 0.107 | 0 | 0.1 | 0.13 | 0.42 | 0.56 | 0.22 | 0.18 |
| 2 | 28320181 | rs7558946 | 0.576 | 0.178 | -0.052 | -1.63 | 0.018 | 0.03 | 0.03 | 0 | 0.48 | 0.65 | 0.29 | 0.44 | 0.41 | 0.64 |
| 2 | 28324130 | rs10187665 | 0.891 | 0.254 | 0.071 | -1.528 | 0 | 0.013 | 0.013 | 0.04 | 0.64 | 0.36 | 0.41 | 0.31 | 0.63 | 0.6 |
| 2 | 28342153 | rs1948922 | 1.07 | 0.116 | -0.308 | -1.764 | -0.01 | -0.013 | -0.013 | 0 | 0.19 | 0.12 | 0.44 | 0.55 | 0.29 | 0.24 |
| 2 | 28344285 | rs4401177 | 2.628 | 0.369 | -0.441 | -1.877 | -0.015 | 0.011 | 0.011 | 0 | 0.06 | 0.02 | 0.07 | 0 | 0.11 | 0.01 |
| 2 | 28345789 | rs11889865 | 0.969 | 0.123 | -0.417 | -1.821 | -0.022 | 0.002 | 0.002 | 0 | 0.12 | 0.04 | 0.1 | 0.13 | 0.11 | 0.07 |
| 2 | 28350220 | rs6707054 | 1.895 | -0.049 | -0.153 | -1.897 | 0.012 | 0.004 | 0.004 | 0 | 0.28 | 0.16 | 0.15 | 0.31 | 0.25 | 0.49 |
| 2 | 28350611 | rs12053063 | -1.228 | 0.071 | 0.206 | -1.84 | 0.017 | 0.051 | 0.051 | 0 | 0.81 | 0.78 | 0.56 | 0.47 | 0.73 | 0.76 |
| 2 | 28357270 | rs7581912 | 1.73 | -0.035 | 0.199 | -1.808 | 0.011 | 0.009 | 0.009 | 0 | 0.3 | 0.2 | 0.15 | 0.32 | 0.27 | 0.5 |
| 2 | 28358479 | rs11127133 | -1.389 | 0.081 | 0.202 | -1.517 | -0.001 | 0.007 | 0.007 | 0 | 0.83 | 0.87 | 0.56 | 0.47 | 0.73 | 0.77 |
| 2 | 28371194 | rs1387573 | 0.078 | 0.023 | 0.076 | -1.531 | -0.013 | -0.059 | -0.059 | 10.915 | 0.16 | 0.24 | 0.07 | 0 | 0.08 | 0.12 |
| 2 | 28371475 | rs6707039 | 1.365 | 0.156 | -0.176 | -1.372 | -0.002 | -0.01 | -0.01 | 2.455 | 0.08 | 0.08 | 0.09 | 0.16 | 0.07 | 0.06 |
| 2 | 28379976 | rs11685351 | 0.822 | 0.092 | -0.199 | -1.711 | -0.001 | -0.005 | -0.005 | 0.587 | 0.53 | 0.44 | 0.37 | 0.32 | 0.55 | 0.51 |
| 2 | 28387200 | rs17759655 | 1.522 | 0.281 | -0.716 | -1.594 | -0.002 | 0.001 | 0.001 | 0 | 0.07 | 0.09 | 0.08 | 0.04 | 0.06 | 0.03 |
| 2 | 28389846 | rs17006590 | 1.667 | 0.172 | -0.812 | -1.828 | 0.009 | 0.005 | 0.005 | 0 | 0.27 | 0.16 | 0.14 | 0.31 | 0.24 | 0.44 |
| 2 | 28392794 | rs17759709 | 0.885 | 0.118 | -0.532 | -1.59 | 0.001 | 0.046 | 0.046 | 0.179 | 0.55 | 0.34 | 0.38 | 0.31 | 0.56 | 0.54 |
| 2 | 28394293 | rs17759746 | 0.658 | 0.233 | -0.565 | -1.559 | -0.003 | 0.013 | 0.013 | 0 | 0.12 | 0.11 | 0.1 | 0.04 | 0.11 | 0.04 |
| 2 | 28398324 | rs11127134 | -0.586 | 0.238 | -0.432 | -1.323 | 0 | -0.01 | -0.01 | 0.315 | 0.35 | 0.5 | 0.53 | 0.64 | 0.37 | 0.43 |
| 2 | 28421381 | rs11891642 | 0.003 | 0.144 | -0.528 | -1.7 | -0.002 | -0.075 | -0.075 | 0 | 0.35 | 0.36 | 0.27 | 0 | 0.27 | 0.16 |
| 2 | 28421461 | rs4666042 | -0.446 | 0.445 | -0.207 | -1.108 | 0.018 | 0.062 | 0.062 | 0 | 0.79 | 0.66 | 0.8 | 0.4 | 0.74 | 0.7 |
| 2 | 28424472 | rs17709034 | 1.882 | 0.956 | -0.338 | -1.99 | -0.014 | 0.012 | 0.012 | 0 | 0.06 | 0.02 | 0.08 | 0 | 0.13 | 0.01 |
| 2 | 28425481 | rs6720083 | 0.502 | -0.01 | -0.749 | -2.49 | -0.001 | 0.152 | 0.152 | 0 | 0.41 | 0.22 | 0.55 | 0.99 | 0.41 | 0.7 |
| 2 | 28426725 | rs9808353 | -0.503 | -0.144 | -0.293 | -2.513 | 0.006 | -0.109 | -0.109 | 0 | 0.8 | 0.42 | 0.86 | 0.99 | 0.8 | 0.85 |
| 2 | 28435264 | rs12478271 | 0.314 | -0.36 | -0.442 | -2.606 | -0.003 | -0.071 | -0.071 | 0 | 0.68 | 0.41 | 0.81 | 0.99 | 0.75 | 0.75 |
| 2 | 28436308 | rs4637064 | 0.545 | -0.375 | -0.576 | -2.653 | 0.009 | -0.03 | -0.03 | 2.606 | 0.64 | 0.39 | 0.78 | 0.99 | 0.71 | 0.75 |
| 2 | 28443050 | rs7349418 | 0.366 | -0.368 | -0.374 | -2.728 | -0.001 | 0.073 | 0.073 | 0 | 0.48 | 0.36 | 0.61 | 0.96 | 0.53 | 0.78 |
| 2 | 28448282 | rs7576048 | 0.825 | -0.643 | -0.309 | -2.676 | 0.007 | 0.06 | 0.06 | 0 | 0.37 | 0.26 | 0.52 | 0.99 | 0.37 | 0.68 |
| 2 | 28452956 | rs1968000 | -0.122 | -0.762 | 0.634 | -2.441 | -0.05 | -0.112 | -0.112 | 0.18 | 0.82 | 0.69 | 0.89 | 1 | 0.91 | 0.89 |
| 2 | 28469456 | rs11127135 | 0.228 | -0.766 | 0.577 | -2.443 | -0.074 | -0.15 | -0.15 | 0 | 0.8 | 0.54 | 0.87 | 1 | 0.9 | 0.89 |
| 2 | 28479472 | rs6716971 | 0.205 | -0.77 | 0.629 | -2.433 | -0.047 | -0.075 | -0.075 | 0.002 | 0.15 | 0.24 | 0.08 | 0 | 0.05 | 0.1 |
| 2 | 28485444 | rs4666048 | 0.77 | -0.753 | 0.354 | -2.533 | 0.032 | 0.065 | 0.065 | 0 | 0.63 | 0.28 | 0.77 | 0.99 | 0.71 | 0.74 |
| 2 | 28486416 | rs11684560 | 1.106 | -0.611 | 0.363 | -2.41 | 0.008 | -0.06 | -0.06 | 0.018 | 0.16 | 0.2 | 0.09 | 0.01 | 0.19 | 0.1 |
| 2 | 28493370 | rs7568185 | -0.063 | -0.847 | 0.535 | -2.452 | -0.018 | -0.059 | -0.059 | 0 | 0.18 | 0.21 | 0.1 | 0 | 0.1 | 0.11 |
| 2 | 28493709 | rs4666050 | 1.172 | -0.903 | 0.355 | -2.655 | 0.017 | 0.057 | 0.057 | 0 | 0.33 | 0.19 | 0.47 | 0.95 | 0.31 | 0.67 |
| 2 | 28494000 | rs10203337 | -0.332 | -0.96 | 0.323 | -2.487 | -0.052 | -0.106 | -0.106 | 0 | 0.21 | 0.33 | 0.12 | 0 | 0.11 | 0.15 |
| 2 | 28498321 | rs6547838 | -0.586 | -1.03 | 0.255 | -2.523 | -0.053 | -0.106 | -0.106 | 0 | 0.23 | 0.33 | 0.12 | 0 | 0.11 | 0.15 |
| 2 | 28510347 | rs10495763 | 0.149 | -0.897 | 0.065 | -2.458 | 0.019 | 0.005 | 0.005 | 0 | 0.18 | 0.14 | 0.09 | 0 | 0.1 | 0.11 |
| 2 | 28516271 | rs7567409 | 0.95 | -1.116 | -0.044 | -2.068 | -0.04 | 0.005 | 0.005 | 0.015 | 0.32 | 0.39 | 0.4 | 0.63 | 0.16 | 0.49 |
| 2 | 28520083 | rs10189899 | -0.865 | -0.908 | -0.128 | -2.145 | 0.009 | 0.005 | 0.005 | 0 | 0.77 | 0.77 | 0.78 | 0.65 | 0.61 | 0.67 |
| 2 | 28521057 | rs4665408 | 0.652 | -0.76 | -0.234 | -2.048 | 0 | -0.001 | -0.001 | 0 | 0.54 | 0.45 | 0.65 | 0.64 | 0.5 | 0.51 |
| 2 | 28523995 | rs12621972 | 0.256 | -1.132 | 0.111 | -1.648 | 0.004 | 0 | 0 | 0.002 | 0.34 | 0.32 | 0.43 | 0.65 | 0.21 | 0.47 |
| 2 | 28525459 | rs6721691 | 1.445 | -1.381 | -0.363 | -1.396 | 0.092 | -0.128 | -0.128 | 0.547 | 0.23 | 0.02 | 0.32 | 0.57 | 0.1 | 0.44 |
| 2 | 28542345 | rs6547841 | -0.589 | -1.504 | 0.316 | -1.492 | 0.02 | 0.014 | 0.014 | 0 | 0.54 | 0.56 | 0.47 | 0.63 | 0.26 | 0.66 |
| 2 | 28545109 | rs2279475 | 0.028 | -1.711 | 0.601 | -1.565 | -0.069 | -0.006 | -0.006 | 0.319 | 0.4 | 0.47 | 0.42 | 0.61 | 0.16 | 0.59 |
| 2 | 28546023 | rs12470242 | -0.081 | -1.79 | 0.766 | -1.585 | 0.073 | -0.006 | -0.006 | 0 | 0.4 | 0.21 | 0.42 | 0.58 | 0.2 | 0.56 |
| 2 | 28549350 | rs876251 | 0.049 | -1.753 | 0.743 | -1.566 | 0.11 | 0.04 | 0.04 | 0 | 0.56 | 0.29 | 0.55 | 0.38 | 0.83 | 0.4 |
| 2 | 28549757 | rs2005181 | -1.214 | -1.641 | 0.941 | -1.527 | -0.133 | 0.067 | 0.067 | 0 | 0.76 | 0.56 | 0.63 | 0.39 | 0.93 | 0.52 |
| 2 | 28559717 | rs11127142 | 1.151 | -1.039 | 1.229 | -1.237 | -0.086 | 0.046 | 0.046 | 0.097 | 0.29 | 0.41 | 0.39 | 0.57 | 0.11 | 0.44 |

| | | | | | | | | | | | | | | | | |
|---|---|---|---|---|---|---|---|---|---|---|---|---|---|---|---|---|
| 2 | 28559728 | rs11127143 | 0.807 | 0.296 | 1.627 | -0.668 | -0.041 | -0.052 | -0.052 | 0 | 0.82 | 0.69 | 0.91 | 0.97 | 0.9 | 0.86 |
| 2 | 28566341 | rs7602515 | 1.357 | 0.772 | 1.065 | -0.647 | 0.006 | 0.003 | 0.003 | 0.721 | 0.48 | 0.48 | 0.55 | 0.59 | 0.35 | 0.51 |
| 2 | 28568016 | rs11127145 | -0.144 | 0.512 | 1.298 | -0.382 | 0.016 | 0.035 | 0.035 | 0.032 | 0.17 | 0.21 | 0.19 | 0.36 | 0.23 | 0.25 |
| 2 | 28571021 | rs10174861 | -0.344 | 0.421 | 1.032 | -0.43 | -0.004 | -0.022 | -0.022 | 0 | 0.12 | 0.16 | 0.06 | 0 | 0.05 | 0.1 |
| 2 | 28571241 | rs7568247 | 1.389 | 0.613 | 0.905 | -0.392 | 0 | 0.002 | 0.002 | 0.428 | 0.53 | 0.4 | 0.61 | 0.61 | 0.47 | 0.52 |
| 2 | 28584315 | rs7602568 | -0.193 | 0.832 | 0.581 | -0.564 | 0.002 | 0.007 | 0.007 | 0.041 | 0.84 | 0.83 | 0.82 | 0.66 | 0.78 | 0.77 |
| 2 | 28587020 | rs2175094 | -0.21 | 0.831 | 0.565 | -0.564 | -0.011 | -0.015 | -0.015 | 0.114 | 0.84 | 0.88 | 0.82 | 0.66 | 0.78 | 0.77 |
| 2 | 28587051 | rs7594431 | 0.328 | 0.792 | 0.844 | -0.701 | 0.013 | -0.049 | -0.049 | 0 | 0.17 | 0.23 | 0.15 | 0.02 | 0.25 | 0.12 |
| 2 | 28587069 | rs2175095 | 1.428 | 0.659 | 0.919 | -0.762 | -0.006 | 0.016 | 0.016 | 0 | 0.55 | 0.37 | 0.6 | 0.62 | 0.49 | 0.56 |
| 2 | 28587339 | rs871521 | 0.9 | 0.816 | 0.793 | -0.672 | 0.019 | -0.003 | -0.003 | 2.092 | 0.66 | 0.58 | 0.66 | 0.63 | 0.53 | 0.64 |
| 2 | 28592116 | rs11902329 | -0.245 | 0.885 | -0.055 | -0.53 | -0.03 | -0.039 | -0.039 | 0.364 | 0.19 | 0.09 | 0.18 | 0.35 | 0.24 | 0.23 |
| 2 | 28596064 | rs4665418 | 1.475 | 0.698 | 0.043 | -0.734 | 0.009 | 0.03 | 0.03 | 0 | 0.49 | 0.39 | 0.6 | 0.62 | 0.46 | 0.54 |
| 2 | 28596317 | rs10202096 | -0.933 | 0.855 | 0.021 | -0.628 | -0.001 | -0.135 | -0.135 | 0.073 | 0.3 | 0.49 | 0.23 | 0.04 | 0.3 | 0.22 |
| 2 | 28597305 | rs12714232 | -0.721 | 1.195 | -0.427 | 0.103 | -0.019 | 0 | 0 | 0 | 0.36 | 0.32 | 0.3 | 0.37 | 0.47 | 0.37 |
| 2 | 28599765 | rs13015562 | -1.372 | 1.363 | -0.22 | -0.037 | 0.008 | -0.021 | -0.021 | 0 | 0.12 | 0.18 | 0.13 | 0.02 | 0.17 | 0.21 |
| 2 | 28600239 | rs931948 | 0.642 | 0.983 | 0.011 | -0.002 | 0 | -0.022 | -0.022 | 0 | 0.79 | 0.72 | 0.82 | 0.87 | 0.83 | 0.79 |
| 2 | 28605588 | rs13411730 | -1.688 | 0.752 | 0.942 | -0.037 | 0.061 | 0.021 | 0.021 | 6.264 | 0.09 | 0.45 | 0.09 | 0.05 | 0.06 | 0.16 |
| 2 | 28614794 | rs925255 | -0.266 | 0.847 | 0.169 | -0.306 | -0.022 | 0.014 | 0.014 | 0.853 | 0.33 | 0.25 | 0.28 | 0.24 | 0.46 | 0.5 |
| 2 | 28618161 | rs12473028 | -0.639 | 0.889 | 0.466 | -0.519 | 0.006 | -0.011 | -0.011 | 0 | 0.32 | 0.41 | 0.26 | 0.23 | 0.41 | 0.47 |
| 2 | 28623159 | rs1039823 | -1.374 | -0.085 | 0.278 | -0.918 | -0.003 | 0.107 | 0.107 | 0.013 | 0.47 | 0.76 | 0.33 | 0.22 | 0.48 | 0.55 |
| 2 | 28627629 | rs11127150 | -0.273 | 0.126 | 1.349 | -0.805 | 0.032 | -0.025 | -0.025 | 0 | 0.3 | 0.42 | 0.28 | 0.2 | 0.43 | 0.45 |
| 2 | 28629892 | rs9309662 | -1.037 | 0.298 | 1.051 | -0.814 | -0.014 | 0.042 | 0.042 | 1.616 | 0.37 | 0.75 | 0.29 | 0.2 | 0.43 | 0.52 |
| 2 | 28634790 | rs12624279 | -0.046 | 0.307 | 0.939 | -0.801 | -0.026 | 0.018 | 0.018 | 0.762 | 0.28 | 0.21 | 0.22 | 0.18 | 0.41 | 0.45 |
| 2 | 28635740 | rs7562 | -1.154 | 0.323 | 0.953 | -0.784 | -0.026 | 0.09 | 0.09 | 0.885 | 0.41 | 0.8 | 0.31 | 0.2 | 0.46 | 0.58 |
| 2 | 28636740 | rs2279990 | -0.887 | 0.105 | 0.854 | 0.239 | -0.007 | -0.003 | -0.003 | 1.332 | 0.38 | 0.34 | 0.27 | 0.35 | 0.44 | 0.53 |
| 2 | 28642747 | rs1396733 | 0.483 | 0.19 | 0.895 | 0.725 | -0.004 | -0.005 | -0.005 | 0 | 0.65 | 0.65 | 0.72 | 0.6 | 0.55 | 0.47 |
| 2 | 28645802 | rs7590263 | 0.701 | 0.366 | 0.956 | 0.517 | 0.003 | 0.001 | 0.001 | 0.958 | 0.61 | 0.38 | 0.66 | 0.61 | 0.52 | 0.42 |
| 2 | 28647666 | rs6719666 | -0.42 | -0.235 | -0.768 | -0.538 | 0.036 | 0.026 | 0.026 | 0.13 | 0.6 | 0.34 | 0.68 | 0.3 | 0.7 | 0.68 |
| 2 | 28652529 | rs4666076 | -0.399 | -0.097 | -1.175 | -0.472 | -0.044 | 0.055 | 0.055 | 1.349 | 0.25 | 0.09 | 0.18 | 0.15 | 0.31 | 0.43 |
| 2 | 28655153 | rs10190267 | 0.754 | -0.158 | -1.621 | -0.757 | -0.006 | 0.141 | 0.141 | 0 | 0.51 | 0.23 | 0.57 | 0.17 | 0.52 | 0.42 |
| 2 | 28661201 | rs2940806 | 0.683 | -0.092 | -1.546 | -1.066 | -0.005 | 0.131 | 0.131 | 0 | 0.53 | 0.24 | 0.59 | 0.12 | 0.54 | 0.45 |
| 2 | 28662228 | rs10174692 | 0.039 | -0.373 | -0.99 | -0.77 | 0.008 | 0.008 | 0.008 | 0.817 | 0.27 | 0.11 | 0.23 | 0.73 | 0.24 | 0.26 |
| 2 | 28662264 | rs2338013 | 0.035 | -0.332 | -0.656 | -0.75 | -0.015 | -0.015 | -0.015 | 0.878 | 0.71 | 0.6 | 0.75 | 0.27 | 0.76 | 0.74 |
| 2 | 28662375 | rs10199186 | 1.06 | -0.418 | -0.763 | -0.386 | 0 | -0.043 | -0.043 | 0 | 0.13 | 0 | 0.07 | 0.19 | 0.09 | 0.09 |
| 2 | 28662493 | rs2972052 | 0.836 | -0.079 | -0.816 | -1.351 | -0.007 | 0.006 | 0.006 | 0.163 | 0.4 | 0.34 | 0.57 | 0.16 | 0.51 | 0.51 |
| 2 | 28663333 | rs2940787 | 0.218 | -0.07 | -0.847 | -1.122 | -0.001 | 0.014 | 0.014 | 0.035 | 0.52 | 0.4 | 0.7 | 0.83 | 0.54 | 0.7 |
| 2 | 28667749 | rs13431955 | -0.394 | 0.329 | -0.793 | -1.063 | 0.02 | 0.036 | 0.036 | 0.023 | 0.4 | 0.28 | 0.28 | 0.71 | 0.26 | 0.4 |
| 2 | 28670404 | rs2940797 | 0.463 | 1.168 | -0.988 | -0.739 | -0.016 | 0.032 | 0.032 | 0 | 0.33 | 0.23 | 0.52 | 0.18 | 0.43 | 0.47 |
| 2 | 28671048 | rs2972050 | 0.085 | 1.014 | -0.832 | -0.178 | -0.005 | -0.034 | -0.034 | 0 | 0.55 | 0.48 | 0.65 | 0.21 | 0.57 | 0.67 |
| 2 | 28674572 | rs12622211 | -1.026 | 1.017 | -0.501 | 1.09 | -0.011 | 0.074 | 0.074 | 0 | 0.23 | 0.33 | 0.13 | 0.45 | 0.19 | 0.19 |
| 2 | 28678410 | rs2940803 | -0.854 | 1.716 | -0.935 | 0.527 | -0.033 | 0.013 | 0.013 | 0.752 | 0.25 | 0.16 | 0.4 | 0.18 | 0.3 | 0.41 |
| 2 | 28683174 | rs10186544 | 1.308 | 1.206 | -1.634 | 0.099 | 0.001 | 0.076 | 0.076 | 0 | 0.32 | 0.09 | 0.22 | 0.17 | 0.3 | 0.33 |
| 2 | 28686128 | rs13395604 | 0.518 | 1.336 | -1.448 | 0.064 | 0.006 | 0.011 | 0.011 | 0.304 | 0.09 | 0.05 | 0.14 | 0.04 | 0.04 | 0.08 |
| 2 | 28686332 | rs1581035 | 0.18 | 1.512 | -1.563 | 0.038 | 0.001 | 0.073 | 0.073 | 0 | 0.47 | 0.25 | 0.45 | 0.31 | 0.52 | 0.62 |
| 2 | 28687043 | rs2940790 | 0.172 | 2.035 | -0.095 | 0.636 | 0.019 | 0.185 | 0.185 | 0 | 0.85 | 0.68 | 0.87 | 0.43 | 0.82 | 0.84 |
| 2 | 28690957 | rs12467414 | -0.892 | 1.934 | -0.369 | 0.567 | -0.018 | 0.002 | 0.002 | 0.421 | 0.22 | 0.17 | 0.33 | 0.19 | 0.23 | 0.36 |
| 2 | 28723942 | rs6715256 | -0.532 | 1.835 | -0.341 | 0.545 | -0.008 | 0.087 | 0.087 | 0 | 0.16 | 0.22 | 0.12 | 0.63 | 0.15 | 0.17 |
| 2 | 28731702 | rs12478824 | -0.898 | 2.045 | -0.352 | 0.416 | -0.057 | 0.009 | 0.009 | 0.137 | 0.22 | 0.09 | 0.34 | 0.15 | 0.27 | 0.37 |
| 2 | 28735342 | rs10865499 | -0.16 | 2.113 | -0.086 | 0.318 | -0.024 | -0.002 | -0.002 | 1.206 | 0.14 | 0.09 | 0.22 | 0.11 | 0.22 | 0.29 |
| 2 | 28736582 | rs11885873 | -1.135 | 1.472 | 0.121 | 0.699 | -0.004 | 0.012 | 0.012 | 0 | 0.23 | 0.23 | 0.24 | 0.66 | 0.2 | 0.24 |
| 2 | 28738493 | rs4233726 | -1.591 | 1.748 | 0.371 | 0.688 | 0.024 | 0.002 | 0.002 | 0 | 0.17 | 0.29 | 0.16 | 0.18 | 0.18 | 0.27 |
| 2 | 28747170 | rs4334451 | -0.214 | 1.628 | 0.013 | 1.07 | -0.002 | -0.011 | -0.011 | 0 | 0.22 | 0.22 | 0.15 | 0.39 | 0.21 | 0.19 |
| 2 | 28752555 | rs4473343 | 0.574 | 1.673 | 0.281 | 0.453 | 0.006 | -0.032 | -0.032 | 0 | 0.92 | 0.83 | 0.8 | 1 | 0.93 | 0.96 |
| 2 | 28757368 | rs6547857 | 1.384 | 1.738 | 0.391 | 0.501 | 0.031 | 0.039 | 0.039 | 0 | 0.88 | 0.77 | 0.64 | 0.79 | 0.86 | 0.88 |
| 2 | 28758427 | rs13029005 | 0.866 | 1.694 | -0.19 | -0.01 | -0.064 | 0.118 | 0.118 | 0 | 0.21 | 0.03 | 0.16 | 0.01 | 0.29 | 0.26 |
| 2 | 28760914 | rs7589034 | 1.056 | 1.222 | 0.82 | 0.577 | 0.007 | -0.042 | -0.042 | 0.541 | 0.9 | 0.66 | 0.78 | 0.99 | 0.93 | 0.94 |
| 2 | 28761981 | rs6753929 | 0.136 | 1.101 | 0.472 | 0.471 | -0.002 | 0.021 | 0.021 | 0.459 | 0.27 | 0.09 | 0.3 | 0.22 | 0.24 | 0.2 |
| 2 | 28762315 | rs6744005 | 0.464 | 1.121 | 0.527 | 0.464 | 0.003 | 0.013 | 0.013 | 0 | 0.65 | 0.7 | 0.5 | 0.77 | 0.69 | 0.74 |
| 2 | 28763374 | rs11127162 | 1.096 | 1.164 | 0.113 | 0.567 | -0.019 | 0.108 | 0.108 | 0 | 0.42 | 0.23 | 0.31 | 0.22 | 0.47 | 0.52 |
| 2 | 28765309 | rs10205008 | 1.163 | 1.294 | 0.164 | 0.676 | -0.017 | 0.111 | 0.111 | 0 | 0.42 | 0.16 | 0.31 | 0.21 | 0.48 | 0.52 |
| 2 | 28768251 | rs1881254 | -0.747 | 1.278 | 0.053 | 0.683 | 0 | 0.042 | 0.042 | 0 | 0.49 | 0.41 | 0.47 | 0.78 | 0.45 | 0.42 |
| 2 | 28768427 | rs9309663 | 0.217 | 1.277 | -0.147 | 0.724 | -0.008 | 0.025 | 0.025 | 0.04 | 0.26 | 0.1 | 0.3 | 0.22 | 0.24 | 0.2 |
| 2 | 28770240 | rs11127165 | 1.218 | 1.346 | -0.064 | 0.729 | -0.017 | 0.111 | 0.111 | 0 | 0.42 | 0.22 | 0.31 | 0.21 | 0.47 | 0.52 |
| 2 | 28770802 | rs11127167 | 0.246 | 0.67 | 0.078 | 0.84 | 0.004 | 0.012 | 0.012 | 0.046 | 0.69 | 0.45 | 0.59 | 0.44 | 0.71 | 0.72 |
| 2 | 28772597 | rs13017118 | 0.246 | 0.553 | 0.078 | 0.839 | 0.002 | 0.017 | 0.017 | 0.046 | 0.69 | 0.45 | 0.62 | 0.46 | 0.71 | 0.72 |
| 2 | 28773874 | rs10170463 | 0.246 | 0.553 | 0.078 | 0.836 | 0.002 | 0.018 | 0.018 | 0.046 | 0.69 | 0.45 | 0.62 | 0.47 | 0.71 | 0.72 |
| 2 | 28779228 | rs2141326 | -0.038 | 0.498 | 0.078 | 0.864 | -0.001 | 0.015 | 0.015 | 0 | 0.26 | 0.29 | 0.2 | 0.53 | 0.24 | 0.22 |
| 2 | 28780313 | rs12617735 | 1.012 | 0.538 | -0.127 | 0.858 | -0.012 | 0.075 | 0.075 | 0 | 0.47 | 0.24 | 0.36 | 0.32 | 0.54 | 0.58 |
| 2 | 28780383 | rs17741435 | 0.491 | 0.384 | -0.246 | 1.112 | 0.016 | 0.046 | 0.046 | 0 | 0.21 | 0.12 | 0.26 | 0.15 | 0.18 | 0.15 |

| | | | | | | | | | | | | | | | | |
|---|---|---|---|---|---|---|---|---|---|---|---|---|---|---|---|---|
| 2 | 28782985 | rs6728960 | -0.402 | 0.752 | 0.435 | 1.936 | -0.002 | 0.046 | 0.046 | 0 | 0.23 | 0.32 | 0.16 | 0.53 | 0.2 | 0.21 |
| 2 | 28786640 | rs11679308 | 0.342 | 0.809 | -0.516 | 1.972 | -0.003 | 0.044 | 0.044 | 0.223 | 0.73 | 0.45 | 0.66 | 0.47 | 0.77 | 0.74 |
| 2 | 28800894 | rs877832 | -0.575 | 1.306 | -1.165 | 1.559 | -0.018 | 0.002 | 0.002 | 0 | 0.27 | 0.19 | 0.33 | 0.28 | 0.31 | 0.28 |
| 2 | 28803240 | rs7595633 | -1.144 | 1.068 | -0.553 | 1.329 | 0.001 | 0.006 | 0.006 | 0 | 0.32 | 0.6 | 0.4 | 0.38 | 0.32 | 0.43 |
| 2 | 28807899 | rs4666103 | -0.768 | 1.011 | 0.09 | 1.3 | 0.007 | 0.052 | 0.052 | 0 | 0.25 | 0.42 | 0.32 | 0.37 | 0.21 | 0.34 |
| 2 | 28812285 | rs12468715 | -1.283 | 0.984 | 0.168 | 1.337 | 0.003 | 0.021 | 0.021 | 0 | 0.32 | 0.53 | 0.4 | 0.38 | 0.32 | 0.44 |
| 2 | 28814185 | rs11127175 | -0.517 | 0.815 | 0.454 | 1.172 | 0.008 | 0.022 | 0.022 | 0 | 0.23 | 0.37 | 0.32 | 0.29 | 0.19 | 0.29 |
| 2 | 28814683 | rs11127176 | -0.755 | 0.664 | 0.486 | 1.516 | 0.001 | 0.037 | 0.037 | 0 | 0.25 | 0.42 | 0.36 | 0.38 | 0.2 | 0.34 |
| 2 | 28815110 | rs7559815 | 0.593 | -0.477 | -0.411 | 0.304 | 0.034 | 0.028 | 0.028 | 0.02 | 0.21 | 0.14 | 0.06 | 0 | 0.1 | 0.06 |
| 2 | 28816425 | rs10176136 | 0.549 | -0.117 | -0.841 | -0.142 | 0 | -0.059 | -0.059 | 0 | 0.65 | 0.48 | 0.67 | 0.15 | 0.66 | 0.6 |
| 2 | 28816948 | rs4665429 | 0.589 | -0.023 | -0.775 | -0.239 | -0.004 | -0.036 | -0.036 | 0 | 0.67 | 0.41 | 0.67 | 0.15 | 0.69 | 0.6 |
| 2 | 28817870 | rs10865503 | -0.801 | 0.098 | -0.739 | 0.568 | 0 | -0.004 | -0.004 | 0 | 0.27 | 0.23 | 0.22 | 0.28 | 0.31 | 0.34 |
| 2 | 28818039 | rs12467758 | 0.017 | 0.093 | -0.774 | 0.601 | -0.013 | -0.046 | -0.046 | 0 | 0.2 | 0.09 | 0.15 | 0.27 | 0.2 | 0.26 |
| 2 | 28818931 | rs12993525 | -0.392 | 0.066 | -0.752 | 0.591 | 0.001 | -0.004 | -0.004 | 0 | 0.23 | 0.22 | 0.17 | 0.26 | 0.23 | 0.26 |
| 2 | 28818957 | rs2280733 | 0.547 | -0.076 | -0.291 | 0.736 | 0.018 | 0.282 | 0.282 | 0.035 | 0.85 | 0.62 | 0.82 | 0.46 | 0.83 | 0.78 |
| 2 | 28819398 | rs13026552 | 0.642 | 0.428 | -1.555 | -0.519 | -0.138 | 0.226 | 0.226 | 0 | 0.32 | 0.03 | 0.31 | 0.02 | 0.37 | 0.28 |
| 2 | 28819516 | rs10194430 | 0.749 | 0.829 | -1.837 | -0.719 | -0.007 | 0.101 | 0.101 | 0.618 | 0.52 | 0.33 | 0.43 | 0.18 | 0.49 | 0.51 |
| 2 | 28819603 | rs10171628 | -0.245 | 1.301 | -1.234 | -0.547 | -0.001 | -0.009 | -0.009 | 0.01 | 0.74 | 0.8 | 0.67 | 0.31 | 0.73 | 0.69 |
| 2 | 28820641 | rs1534477 | -1.436 | 1.51 | -1.07 | -0.619 | -0.001 | 0.068 | 0.068 | 0.333 | 0.46 | 0.64 | 0.57 | 0.88 | 0.56 | 0.52 |
| 2 | 28821149 | rs4371315 | 0.303 | 1.538 | -0.782 | -0.063 | -0.017 | -0.031 | -0.031 | 0 | 0.27 | 0.17 | 0.23 | 0.4 | 0.29 | 0.29 |
| 2 | 28833479 | rs1534476 | -0.537 | 1.408 | -0.802 | -0.486 | -0.007 | 0.016 | 0.016 | 0 | 0.37 | 0.3 | 0.47 | 0.87 | 0.45 | 0.46 |
| 2 | 28836285 | rs6750615 | 2.33 | 1.544 | -1.153 | -0.627 | -0.029 | 0.111 | 0.111 | 0 | 0.17 | 0.01 | 0.11 | 0 | 0.14 | 0.06 |
| 2 | 28840691 | rs1534478 | 0.863 | 1.582 | -0.757 | -0.506 | -0.021 | -0.016 | -0.016 | 1.561 | 0.27 | 0.16 | 0.24 | 0.33 | 0.31 | 0.28 |
| 2 | 28851116 | rs11127182 | 1.16 | 1.179 | -0.691 | -0.692 | -0.013 | -0.104 | -0.104 | 0 | 0.28 | 0.47 | 0.19 | 0.06 | 0.22 | 0.11 |
| 2 | 28852926 | rs4530322 | -0.99 | 1.459 | -0.853 | -0.699 | 0.004 | -0.008 | -0.008 | 0 | 0.59 | 0.66 | 0.41 | 0.39 | 0.57 | 0.41 |
| 2 | 28854456 | rs11896668 | 0.663 | 1.743 | -0.235 | -0.395 | -0.009 | -0.012 | -0.012 | 0 | 0.25 | 0.19 | 0.18 | 0.32 | 0.27 | 0.28 |
| 2 | 28854570 | rs6713845 | -0.099 | 1.744 | -0.172 | -0.423 | -0.006 | 0.001 | 0.001 | 0 | 0.33 | 0.28 | 0.23 | 0.32 | 0.36 | 0.3 |
| 2 | 28854958 | rs2199619 | 0.622 | 1.734 | -0.218 | -0.335 | -0.008 | -0.013 | -0.013 | 0.923 | 0.26 | 0.19 | 0.18 | 0.32 | 0.28 | 0.29 |
| 2 | 28861067 | rs11127185 | 1.312 | 1.962 | -0.596 | -0.298 | -0.037 | -0.035 | -0.035 | 0 | 0.11 | 0.05 | 0.11 | 0.16 | 0.19 | 0.17 |
| 2 | 28861205 | rs6738639 | 0.749 | 1.727 | -0.485 | -0.318 | -0.036 | -0.03 | -0.03 | 0.102 | 0.26 | 0.11 | 0.18 | 0.3 | 0.29 | 0.28 |
| 2 | 28866469 | rs2272386 | -0.61 | 1.691 | -0.314 | -0.397 | 0.014 | 0.002 | 0.002 | 0 | 0.37 | 0.5 | 0.28 | 0.36 | 0.42 | 0.33 |
| 2 | 28867201 | rs6741437 | -0.666 | 1.675 | -0.304 | -0.388 | 0.013 | 0.001 | 0.001 | 0 | 0.37 | 0.5 | 0.28 | 0.36 | 0.42 | 0.38 |
| 2 | 28871685 | rs10495766 | 0.488 | 1.694 | -0.057 | -0.298 | 0.041 | 0.004 | 0.004 | 0 | 0.1 | 0.32 | 0.07 | 0.05 | 0.12 | 0.04 |
| 2 | 28876857 | rs11678098 | 0.68 | 1.702 | -0.19 | -0.37 | -0.021 | -0.011 | -0.011 | 1.304 | 0.28 | 0.16 | 0.23 | 0.3 | 0.33 | 0.3 |
| 2 | 28877006 | rs11694841 | 0.511 | 1.691 | -0.192 | -0.376 | 0.002 | 0 | 0 | 0 | 0.29 | 0.31 | 0.24 | 0.3 | 0.32 | 0.29 |
| 2 | 28877585 | rs6547865 | -0.473 | 1.4 | -0.199 | -0.333 | 0.001 | -0.004 | -0.004 | 0 | 0.38 | 0.59 | 0.31 | 0.3 | 0.42 | 0.32 |
| 2 | 28877683 | rs17742402 | 1.598 | 1.179 | -0.678 | -0.727 | -0.008 | 0.108 | 0.108 | 0 | 0.2 | 0.01 | 0.1 | 0.02 | 0.14 | 0.26 |
| 2 | 28892116 | rs10192375 | -1.182 | 1.913 | 0.16 | 0.018 | -0.006 | 0.003 | 0.003 | 0 | 0.44 | 0.61 | 0.46 | 0.39 | 0.51 | 0.36 |
| 2 | 28892960 | rs7586033 | -0.947 | 1.597 | 0.321 | 0.117 | 0.054 | 0.07 | 0.07 | 0.805 | 0.07 | 0.3 | 0.04 | 0.06 | 0.07 | 0.03 |
| 2 | 28897430 | rs13018122 | 1.622 | 1.64 | -0.291 | 0.178 | -0.002 | -0.02 | -0.02 | 0 | 0.06 | 0.1 | 0.02 | 0 | 0.05 | 0.01 |
| 2 | 28902004 | rs4666115 | -0.768 | 1.353 | -0.344 | 0.161 | 0.002 | -0.006 | -0.006 | 0.08 | 0.39 | 0.6 | 0.43 | 0.34 | 0.4 | 0.32 |
| 2 | 28903361 | rs17691678 | -0.279 | 1.085 | -0.053 | 0.232 | -0.006 | -0.007 | -0.007 | 0 | 0.11 | 0.23 | 0.05 | 0.06 | 0.09 | 0.03 |
| 2 | 28905810 | rs17007170 | -1.011 | 1.047 | 0.376 | 0.256 | 0.032 | 0.05 | 0.05 | 0 | 0.07 | 0.28 | 0.04 | 0.06 | 0.05 | 0.01 |
| 2 | 28906644 | rs7586405 | 0.515 | 1.041 | 0.28 | 0.015 | -0.005 | 0.007 | 0.007 | 0 | 0.63 | 0.36 | 0.67 | 0.59 | 0.68 | 0.71 |
| 2 | 28931376 | rs11675572 | -0.533 | 1.169 | -0.128 | -0.247 | -0.008 | 0.031 | 0.031 | 0 | 0.32 | 0.15 | 0.24 | 0.25 | 0.28 | 0.2 |
| 2 | 28951698 | rs4666118 | -0.821 | 1.101 | 0.763 | -0.181 | -0.004 | -0.014 | -0.014 | 0 | 0.36 | 0.42 | 0.28 | 0.28 | 0.29 | 0.2 |
| 2 | 28973883 | rs4372836 | 0.46 | 1.112 | 0.702 | -0.172 | -0.004 | -0.01 | -0.01 | 0 | 0.64 | 0.59 | 0.72 | 0.72 | 0.71 | 0.8 |
| 2 | 28977274 | rs4233728 | 0.609 | 1.771 | 0.808 | -0.226 | 0.002 | -0.005 | -0.005 | 0.502 | 0.62 | 0.39 | 0.61 | 0.71 | 0.62 | 0.78 |
| 2 | 28995226 | rs10183665 | 1.146 | 1.359 | 0.59 | -0.765 | 0.006 | 0.138 | 0.138 | 0 | 0.49 | 0.28 | 0.33 | 0.06 | 0.49 | 0.44 |
| 2 | 29017870 | rs10210252 | 0.627 | 1.238 | 0.768 | -0.581 | 0.022 | 0.122 | 0.122 | 0 | 0.08 | 0.27 | 0.1 | 0.16 | 0.1 | 0.14 |
| 2 | 29022959 | rs11687800 | 0.627 | 1.238 | 0.768 | -0.581 | 0.022 | 0.122 | 0.122 | 0 | 0.08 | 0.27 | 0.1 | 0.16 | 0.1 | 0.14 |
| 2 | 29025479 | rs3190 | 0.596 | 1.242 | 0.801 | -0.63 | 0.001 | 0.008 | 0.008 | 0 | 0.57 | 0.39 | 0.42 | 0.22 | 0.58 | 0.58 |
| 2 | 29030006 | rs12475612 | -0.704 | 1.198 | 0.861 | -0.651 | -0.005 | 0.052 | 0.052 | 0 | 0.4 | 0.19 | 0.55 | 0.78 | 0.41 | 0.42 |
| 2 | 29032673 | rs7601934 | 1.125 | 1.255 | 0.769 | -0.648 | 0.002 | 0 | 0 | 0 | 0.49 | 0.44 | 0.34 | 0.05 | 0.49 | 0.44 |
| 2 | 29032746 | rs6547881 | 1.166 | 1.255 | 0.777 | -0.648 | 0.003 | 0.137 | 0.137 | 0 | 0.49 | 0.28 | 0.33 | 0.06 | 0.49 | 0.44 |
| 2 | 29050318 | rs10187155 | -0.988 | 1.212 | 0.778 | -0.619 | -0.001 | 0.04 | 0.04 | 0 | 0.92 | 0.89 | 0.9 | 0.84 | 0.9 | 0.86 |
| 2 | 29064981 | rs10184527 | -1.042 | 1.21 | 0.789 | -0.62 | 0 | 0.043 | 0.043 | 0 | 0.93 | 0.89 | 0.9 | 0.84 | 0.91 | 0.85 |
| 2 | 29068882 | rs13382966 | 1.155 | 1.2 | 0.821 | -0.641 | 0.006 | 0.121 | 0.121 | 0 | 0.49 | 0.3 | 0.33 | 0.06 | 0.49 | 0.43 |
| 2 | 29070805 | rs6737027 | 1.151 | 1.208 | 0.825 | -0.646 | 0.007 | 0.116 | 0.116 | 0 | 0.49 | 0.3 | 0.33 | 0.06 | 0.49 | 0.43 |
| 2 | 29083850 | rs13025081 | 1.18 | 1.135 | 0.737 | -0.682 | 0.005 | 0.116 | 0.116 | 0 | 0.49 | 0.29 | 0.33 | 0.06 | 0.49 | 0.44 |
| 2 | 29086607 | rs6738736 | -0.839 | 1.134 | 0.741 | -0.678 | -0.002 | 0.035 | 0.035 | 0 | 0.92 | 0.89 | 0.9 | 0.84 | 0.9 | 0.85 |
| 2 | 29087274 | rs6760186 | 1.176 | 1.136 | 0.726 | -0.685 | 0.005 | 0.116 | 0.116 | 0 | 0.49 | 0.29 | 0.33 | 0.06 | 0.49 | 0.44 |
| 2 | 29090582 | rs13001838 | -0.505 | 1.163 | 0.431 | -0.6 | -0.004 | 0.035 | 0.035 | 0 | 0.91 | 0.86 | 0.9 | 0.84 | 0.9 | 0.85 |
| 2 | 29092850 | rs4666122 | -0.711 | 1.155 | 0.598 | -0.636 | 0 | 0.003 | 0.003 | 0 | 0.42 | 0.39 | 0.57 | 0.78 | 0.41 | 0.42 |
| 2 | 29099752 | rs7589735 | -0.704 | 0.978 | 0.89 | -0.538 | 0.012 | 0.111 | 0.111 | 0 | 0.92 | 0.75 | 0.91 | 0.84 | 0.91 | 0.87 |
| 2 | 29100642 | rs4358081 | -0.806 | 1.15 | 0.976 | -0.655 | -0.005 | 0.024 | 0.024 | 0 | 0.44 | 0.38 | 0.62 | 0.78 | 0.46 | 0.43 |
| 2 | 29102888 | rs12465314 | -0.854 | 1.198 | 1.025 | -0.661 | -0.001 | -0.005 | -0.005 | 0 | 0.44 | 0.46 | 0.6 | 0.78 | 0.44 | 0.43 |
| 2 | 29109631 | rs6749462 | 1.267 | 1.013 | 1.185 | -0.776 | -0.025 | 0.205 | 0.205 | 0 | 0.47 | 0.2 | 0.28 | 0.07 | 0.47 | 0.43 |
| 2 | 29120733 | rs4577244 | -0.621 | 1.105 | 1.059 | -0.944 | 0.001 | 0.002 | 0.002 | 0 | 0.2 | 0.2 | 0.45 | 0.79 | 0.23 | 0.3 |

| | | | | | | | | | | | | | | | | |
|---|---|---|---|---|---|---|---|---|---|---|---|---|---|---|---|---|
| 2 | 29128322 | rs6547889 | -1.109 | 1.099 | 1.117 | -0.954 | -0.003 | 0.035 | 0.035 | 0 | 0.93 | 0.86 | 0.92 | 0.86 | 0.92 | 0.87 |
| 2 | 29129418 | rs11688605 | -0.589 | 1.094 | 1.09 | -0.934 | 0.001 | 0.002 | 0.002 | 0 | 0.19 | 0.17 | 0.45 | 0.79 | 0.23 | 0.3 |
| 2 | 29138009 | rs6547892 | 1.44 | 1.089 | 1.103 | -0.949 | -0.004 | 0.087 | 0.087 | 0 | 0.48 | 0.32 | 0.31 | 0.07 | 0.48 | 0.43 |
| 2 | 29145725 | rs9653591 | 0.236 | 1.086 | 1.112 | -0.932 | -0.008 | -0.053 | -0.053 | 0 | 0.67 | 0.52 | 0.76 | 0.86 | 0.7 | 0.73 |
| 2 | 29154418 | rs7561164 | 2.125 | 0.898 | 0.441 | -1.533 | -0.05 | 0.088 | 0.088 | 0 | 0.2 | 0.04 | 0.16 | 0.04 | 0.24 | 0.17 |
| 2 | 29169612 | rs3087649 | -0.276 | 0.706 | 0.531 | -1.325 | 0 | 0.006 | 0.006 | 0 | 0.7 | 0.71 | 0.79 | 0.86 | 0.71 | 0.74 |
| 2 | 29170623 | rs11680458 | -0.351 | 0.734 | 0.252 | -1.249 | 0.001 | 0.001 | 0.001 | 0 | 0.19 | 0.17 | 0.45 | 0.79 | 0.22 | 0.3 |
| 2 | 29180402 | rs13402702 | 2.226 | 0.757 | 0.021 | -1.261 | -0.035 | 0.093 | 0.093 | 0 | 0.15 | 0.01 | 0.11 | 0 | 0.16 | 0.05 |
| 2 | 29181107 | rs10197378 | 0.493 | 0.599 | 0.765 | -0.69 | -0.004 | 0.031 | 0.031 | 0 | 0.23 | 0.15 | 0.12 | 0 | 0.21 | 0.14 |
| 2 | 29184909 | rs6727948 | 1.722 | 0.579 | 0.825 | -0.552 | -0.018 | 0.281 | 0.281 | 0 | 0.5 | 0.18 | 0.47 | 0.08 | 0.55 | 0.52 |
| 2 | 29193017 | rs7595944 | 0.676 | 0.46 | 0.966 | -0.032 | 0 | -0.003 | -0.003 | 0 | 0.67 | 0.56 | 0.63 | 0.66 | 0.67 | 0.7 |
| 2 | 29195674 | rs11127196 | -0.639 | 0.78 | 1.191 | 0.364 | 0.007 | 0.132 | 0.132 | 0 | 0.12 | 0.21 | 0.13 | 0.6 | 0.09 | 0.15 |
| 2 | 29196785 | rs4666157 | 0.036 | 0.813 | 0.859 | 0.438 | 0.018 | 0.081 | 0.081 | 0 | 0.11 | 0.24 | 0.25 | 0.32 | 0.15 | 0.18 |
| 2 | 29202386 | rs6547900 | 0.059 | 0.773 | 0.77 | 0.047 | 0.013 | 0.015 | 0.015 | 0 | 0.85 | 0.64 | 0.82 | 0.83 | 0.84 | 0.73 |
| 2 | 29208656 | rs11676763 | -0.143 | 0.66 | 0.214 | 0.232 | -0.014 | -0.04 | -0.04 | 0.339 | 0.24 | 0.14 | 0.27 | 0.63 | 0.26 | 0.22 |
| 2 | 29213408 | rs7567628 | 1.375 | 0.656 | 0.204 | 0.073 | -0.02 | 0.269 | 0.269 | 0 | 0.49 | 0.16 | 0.45 | 0.05 | 0.51 | 0.44 |
| 2 | 29214293 | rs7567935 | -0.337 | 0.632 | 0.484 | 0.11 | -0.001 | -0.002 | -0.002 | 0 | 0.85 | 0.78 | 0.83 | 0.85 | 0.84 | 0.74 |
| 2 | 29215005 | rs7578007 | -0.849 | 0.63 | 0.439 | 0.113 | -0.005 | 0.024 | 0.024 | 0.236 | 0.36 | 0.7 | 0.38 | 0.79 | 0.33 | 0.29 |
| 2 | 29217896 | rs7563398 | 0.796 | 0.522 | 0.695 | 0.372 | 0.005 | 0.041 | 0.041 | 0 | 0.77 | 0.44 | 0.74 | 0.69 | 0.78 | 0.67 |
| 2 | 29218411 | rs12714251 | 0.847 | 0.533 | 0.149 | 0.375 | 0 | 0.001 | 0.001 | 0 | 0.76 | 0.27 | 0.73 | 0.69 | 0.77 | 0.68 |
| 2 | 29229806 | rs12466400 | 0.423 | 0.588 | 0.471 | 0.308 | 0.004 | 0.133 | 0.133 | 0 | 0.82 | 0.46 | 0.77 | 0.67 | 0.82 | 0.68 |
| 2 | 29231870 | rs11684978 | 0.053 | 0.573 | -0.244 | 0.209 | 0 | 0.018 | 0.018 | 0 | 0.25 | 0.2 | 0.18 | 0.19 | 0.25 | 0.18 |
| 2 | 29233954 | rs6712882 | 1.309 | 0.544 | -0.417 | 0.102 | -0.072 | -0.047 | -0.047 | 0.203 | 0.5 | 0.06 | 0.53 | 0.49 | 0.51 | 0.47 |
| 2 | 29233967 | rs11684304 | -0.834 | 0.514 | 0.037 | 0.171 | -0.002 | -0.003 | -0.003 | 0.834 | 0.36 | 0.63 | 0.29 | 0.35 | 0.34 | 0.25 |
| 2 | 29236919 | rs895588 | 1.232 | 0.552 | -0.686 | 0.092 | -0.074 | -0.071 | -0.071 | 0.289 | 0.49 | 0.06 | 0.53 | 0.48 | 0.51 | 0.47 |
| 2 | 29239078 | rs12987284 | -0.07 | 0.686 | -0.28 | -0.13 | -0.003 | 0.019 | 0.019 | 0 | 0.23 | 0.2 | 0.17 | 0.13 | 0.25 | 0.17 |
| 2 | 29239110 | rs13004554 | -0.025 | 0.607 | -0.264 | -0.113 | 0.005 | 0.009 | 0.009 | 0 | 0.23 | 0.21 | 0.14 | 0.13 | 0.18 | 0.17 |
| 2 | 29246044 | rs6721861 | -0.363 | 0.77 | -0.344 | -0.422 | -0.001 | 0.002 | 0.002 | 0 | 0.35 | 0.62 | 0.4 | 0.7 | 0.32 | 0.25 |
| 2 | 29247894 | rs6739684 | -0.205 | 0.701 | -0.118 | -0.551 | 0.016 | -0.021 | -0.021 | 0 | 0.6 | 0.5 | 0.52 | 0.32 | 0.54 | 0.57 |
| 2 | 29249107 | rs11692447 | -0.008 | 0.32 | -0.487 | -0.569 | 0.02 | 0.032 | 0.032 | 0 | 0.26 | 0.2 | 0.17 | 0.01 | 0.23 | 0.15 |
| 2 | 29251134 | rs17007460 | 0.717 | 0.551 | -0.109 | 0.136 | 0.007 | 0.046 | 0.046 | 0 | 0.07 | 0.08 | 0.07 | 0.17 | 0.07 | 0.09 |
| 2 | 29258193 | rs878292 | -0.489 | 0.494 | -0.307 | -0.193 | -0.016 | 0.009 | 0.009 | 0 | 0.37 | 0.23 | 0.35 | 0.66 | 0.35 | 0.29 |
| 2 | 29260085 | rs6742110 | 0.228 | 0.42 | -0.405 | -0.095 | -0.022 | -0.076 | -0.076 | 0 | 0.26 | 0.15 | 0.28 | 0.51 | 0.25 | 0.19 |
| 2 | 29260106 | rs6756951 | -0.829 | 0.373 | -0.018 | -0.12 | -0.004 | 0.003 | 0.003 | 0 | 0.38 | 0.36 | 0.37 | 0.65 | 0.35 | 0.29 |
| 2 | 29261252 | rs11684613 | 1.638 | 0.267 | -0.32 | -0.389 | 0 | 0.081 | 0.081 | 0 | 0.3 | 0.09 | 0.37 | 0.15 | 0.32 | 0.37 |
| 2 | 29263237 | rs12465721 | 0.954 | 0.279 | -0.305 | -0.362 | 0 | -0.005 | -0.005 | 0 | 0.11 | 0.07 | 0.07 | 0.14 | 0.1 | 0.09 |
| 2 | 29266459 | rs1026044 | 1.297 | 0.433 | -0.579 | -0.576 | -0.078 | 0.211 | 0.211 | 0.016 | 0.35 | 0.01 | 0.4 | 0.03 | 0.38 | 0.29 |
| 2 | 29267713 | rs1992809 | 0.055 | 0.402 | 0.083 | -0.561 | -0.002 | -0.002 | -0.002 | 0 | 0.56 | 0.55 | 0.65 | 0.81 | 0.51 | 0.62 |
| 2 | 29268375 | rs2276551 | 0.715 | 0.447 | -0.225 | -0.565 | -0.001 | 0.098 | 0.098 | 0 | 0.12 | 0.19 | 0.19 | 0.52 | 0.09 | 0.17 |
| 2 | 29270823 | rs921099 | 1 | 0.472 | -0.488 | -0.581 | 0.004 | 0.121 | 0.121 | 0 | 0.41 | 0.16 | 0.45 | 0.18 | 0.41 | 0.38 |
| 2 | 29272472 | rs6547906 | 0.957 | 0.461 | -0.468 | -0.588 | 0.008 | 0.064 | 0.064 | 0 | 0.41 | 0.27 | 0.45 | 0.18 | 0.41 | 0.37 |
| 2 | 29273719 | rs6731719 | -1.127 | 0.53 | -1.093 | -0.638 | 0.035 | 0.084 | 0.084 | 0 | 0.84 | 0.68 | 0.83 | 0.69 | 0.79 | 0.66 |
| 2 | 29274730 | rs7577483 | 0.875 | 0.336 | -1.114 | -0.369 | 0 | -0.004 | -0.004 | 0 | 0.11 | 0.07 | 0.07 | 0.14 | 0.1 | 0.09 |
| 2 | 29277200 | rs1867945 | -1.393 | 0.351 | -0.69 | -0.14 | 0.002 | -0.003 | -0.003 | 0.332 | 0.45 | 0.57 | 0.39 | 0.51 | 0.38 | 0.28 |
| 2 | 29280791 | rs882631 | -0.605 | 0.518 | -1.547 | -0.651 | -0.057 | 0.187 | 0.187 | 0 | 0.28 | 0.02 | 0.16 | 0 | 0.27 | 0.11 |
| 2 | 29285575 | rs1562392 | -1.224 | 0.27 | -0.815 | -0.68 | -0.002 | 0.011 | 0.011 | 0 | 0.56 | 0.58 | 0.49 | 0.83 | 0.53 | 0.52 |
| 2 | 29285631 | rs1562391 | -1.234 | 0.253 | -0.802 | -0.671 | -0.002 | 0.007 | 0.007 | 0 | 0.56 | 0.57 | 0.49 | 0.83 | 0.53 | 0.51 |
| 2 | 29285713 | rs4464231 | -1.222 | 0.283 | -0.784 | -0.67 | -0.002 | 0.009 | 0.009 | 0 | 0.56 | 0.57 | 0.45 | 0.83 | 0.53 | 0.51 |
| 2 | 29286072 | rs958584 | -1.176 | 0.111 | -0.197 | -0.755 | 0 | -0.016 | -0.016 | 0.161 | 0.53 | 0.51 | 0.49 | 0.85 | 0.53 | 0.51 |
| 2 | 29286193 | rs719691 | -1.493 | -0.094 | -0.115 | -0.85 | -0.004 | 0.011 | 0.011 | 0 | 0.59 | 0.66 | 0.51 | 0.85 | 0.53 | 0.52 |
| 2 | 29286670 | rs17007529 | -0.752 | -0.181 | 0.044 | 0.025 | -0.013 | 0.037 | 0.037 | 0 | 0.32 | 0.47 | 0.27 | 0.52 | 0.26 | 0.25 |
| 2 | 29286905 | rs13016696 | -0.606 | -0.187 | 0.219 | 0.022 | -0.003 | 0.011 | 0.011 | 0 | 0.31 | 0.36 | 0.26 | 0.52 | 0.26 | 0.25 |
| 2 | 29287192 | rs10495767 | 0.197 | -0.22 | 0.16 | 0.484 | -0.031 | -0.022 | -0.022 | 0 | 0.21 | 0.04 | 0.15 | 0.18 | 0.21 | 0.1 |
| 2 | 29287227 | rs10196859 | -0.751 | -0.185 | 0.435 | 0.444 | -0.009 | 0.034 | 0.034 | 0 | 0.31 | 0.46 | 0.26 | 0.52 | 0.25 | 0.25 |
| 2 | 29287229 | rs10194257 | -1.064 | -0.146 | 0.362 | 0.464 | -0.018 | 0.033 | 0.033 | 0 | 0.34 | 0.56 | 0.27 | 0.52 | 0.25 | 0.25 |
| 2 | 29289411 | rs17744052 | -0.002 | -0.056 | 0.059 | 0.502 | -0.001 | 0.001 | 0.001 | 0 | 0.23 | 0.16 | 0.17 | 0.17 | 0.22 | 0.1 |
| 2 | 29289501 | rs7569316 | 0.306 | 0.24 | -0.247 | 0.327 | -0.018 | -0.049 | -0.049 | 2.227 | 0.22 | 0.09 | 0.21 | 0.35 | 0.25 | 0.27 |
| 2 | 29293241 | rs6749000 | -1.511 | 0.214 | -0.072 | 0.085 | -0.013 | 0.09 | 0.09 | 0 | 0.57 | 0.72 | 0.53 | 0.86 | 0.53 | 0.47 |
| 2 | 29295676 | rs13385188 | -0.847 | -0.056 | 0.392 | 0.203 | -0.011 | 0.032 | 0.032 | 0 | 0.32 | 0.42 | 0.26 | 0.44 | 0.26 | 0.23 |
| 2 | 29302832 | rs6750016 | -0.448 | 0.145 | -0.084 | -0.663 | 0.003 | 0.11 | 0.11 | 0.552 | 0.46 | 0.31 | 0.31 | 0.12 | 0.46 | 0.33 |
| 2 | 29310048 | rs7568662 | -0.862 | -0.003 | 0.557 | -0.73 | -0.005 | 0.067 | 0.067 | 0 | 0.53 | 0.71 | 0.38 | 0.13 | 0.5 | 0.32 |
| 2 | 29312240 | rs7579035 | -0.459 | -0.119 | 0.616 | -0.951 | -0.007 | -0.155 | -0.155 | 0.768 | 0.28 | 0.52 | 0.21 | 0.02 | 0.28 | 0.09 |
| 2 | 29318077 | rs12714263 | -0.762 | -0.064 | 0.294 | -0.84 | -0.003 | -0.02 | -0.02 | 0 | 0.46 | 0.55 | 0.33 | 0.12 | 0.4 | 0.32 |
| 2 | 29320512 | rs6547910 | 0.005 | 0.212 | -0.096 | -0.508 | 0.011 | 0.045 | 0.045 | 0 | 0.19 | 0.08 | 0.14 | 0.02 | 0.17 | 0.08 |
| 2 | 29335758 | rs4666166 | 0.983 | 0.129 | -0.156 | -0.176 | -0.028 | -0.029 | -0.029 | 0 | 0.34 | 0.21 | 0.55 | 0.62 | 0.42 | 0.49 |
| 2 | 29339777 | rs7604766 | 0.023 | -0.076 | 0.194 | -0.273 | -0.001 | -0.003 | -0.003 | 0 | 0.77 | 0.78 | 0.84 | 0.89 | 0.8 | 0.9 |
| 2 | 29344127 | rs17693414 | 0.003 | 0.057 | 0.074 | -0.482 | 0.006 | -0.026 | -0.026 | 0.951 | 0.07 | 0.11 | 0.06 | 0 | 0.11 | 0.01 |
| 2 | 29352680 | rs1563049 | 0.217 | -0.117 | 0.132 | -0.613 | 0.002 | 0.034 | 0.034 | 0 | 0.21 | 0.1 | 0.15 | 0.02 | 0.2 | 0.08 |
| 2 | 29355303 | rs4666167 | 0.152 | -0.157 | 0.619 | -0.572 | -0.012 | -0.045 | -0.045 | 0 | 0.75 | 0.67 | 0.82 | 0.93 | 0.79 | 0.91 |

| | | | | | | | | | | | | | | | | |
|---|---|---|---|---|---|---|---|---|---|---|---|---|---|---|---|---|
| 2 | 29356728 | rs3768674 | -0.474 | -0.29 | 0.64 | -0.493 | -0.01 | 0.008 | 0.008 | 0 | 0.21 | 0.37 | 0.13 | 0.25 | 0.19 | 0.15 |
| 2 | 29359416 | rs9967678 | 0.243 | -0.032 | 0.086 | -0.505 | 0.001 | -0.014 | -0.014 | 0.345 | 0.06 | 0.08 | 0.05 | 0 | 0.11 | 0.01 |
| 2 | 29359786 | rs17750032 | 0.112 | -0.031 | 0.154 | -0.505 | 0.013 | -0.031 | -0.031 | 0 | 0.07 | 0.11 | 0.05 | 0 | 0.11 | 0.01 |
| 2 | 29365583 | rs3100226 | -1.119 | -0.113 | 0.002 | -0.293 | 0.019 | 0.001 | 0.001 | 0 | 0.46 | 0.67 | 0.32 | 0.44 | 0.4 | 0.24 |
| 2 | 29366830 | rs3213947 | 0.676 | 0.062 | -0.214 | -0.911 | -0.012 | 0.028 | 0.028 | 0.155 | 0.38 | 0.19 | 0.54 | 0.69 | 0.37 | 0.55 |
| 2 | 29369622 | rs13003156 | 0.3 | -0.167 | -0.024 | -1.669 | -0.006 | -0.001 | -0.001 | 0 | 0.17 | 0.16 | 0.08 | 0.2 | 0.09 | 0.16 |
| 2 | 29372029 | rs3100228 | -0.028 | -0.285 | 1.017 | -1.739 | -0.004 | -0.02 | -0.02 | 0 | 0.46 | 0.43 | 0.35 | 0.05 | 0.53 | 0.3 |
| 2 | 29372369 | rs3112916 | -0.518 | -0.257 | 0.727 | -1.698 | -0.018 | -0.002 | -0.002 | 0.473 | 0.83 | 0.81 | 0.92 | 0.8 | 0.91 | 0.84 |
| 2 | 29375902 | rs3100239 | -0.05 | -0.31 | 0.809 | -1.747 | -0.022 | 0.103 | 0.103 | 0.087 | 0.44 | 0.32 | 0.36 | 0.1 | 0.53 | 0.32 |
| 2 | 29376110 | rs3100241 | 0.065 | -0.311 | 0.369 | -1.753 | -0.056 | 0.208 | 0.208 | 0.2 | 0.43 | 0.18 | 0.34 | 0.05 | 0.53 | 0.31 |
| 2 | 29378902 | rs3100244 | 0.601 | -0.149 | 0.039 | -1.879 | -0.017 | 0.114 | 0.114 | 0 | 0.47 | 0.14 | 0.64 | 0.69 | 0.52 | 0.56 |
| 2 | 29382038 | rs2123443 | 1.002 | -0.263 | 0.281 | -1.806 | -0.03 | 0.075 | 0.075 | 0.19 | 0.37 | 0.13 | 0.54 | 0.77 | 0.38 | 0.53 |
| 2 | 29387572 | rs6736999 | 0.289 | -0.282 | 0.46 | -1.744 | -0.009 | -0.009 | -0.009 | 0.608 | 0.17 | 0.15 | 0.08 | 0.13 | 0.09 | 0.15 |
| 2 | 29392680 | rs12620500 | 1.18 | -0.225 | 0.282 | -1.694 | 0.014 | -0.016 | -0.016 | 0 | 0.1 | 0 | 0.03 | 0.1 | 0.05 | 0.16 |
| 2 | 29396270 | rs3100219 | -0.722 | 0.064 | 0.622 | -1.722 | 0.022 | -0.027 | -0.027 | 0.748 | 0.95 | 0.86 | 0.95 | 1 | 0.92 | 0.99 |
| 2 | 29401499 | rs2879454 | -0.421 | 0.226 | 0.492 | -1.818 | -0.004 | 0.066 | 0.066 | 0.005 | 0.46 | 0.71 | 0.39 | 0.1 | 0.53 | 0.31 |
| 2 | 29401544 | rs2879455 | 0.06 | 0.148 | 0.055 | -1.714 | -0.031 | 0.105 | 0.105 | 0.511 | 0.38 | 0.24 | 0.3 | 0.1 | 0.44 | 0.3 |
| 2 | 29411696 | rs1670282 | -0.441 | 0.252 | 0.172 | -1.527 | -0.003 | -0.038 | -0.038 | 0 | 0.88 | 0.87 | 0.94 | 1 | 0.88 | 0.98 |
| 2 | 29412123 | rs1869264 | 0.646 | 0.589 | 0.038 | -2.331 | 0 | 0.002 | 0.002 | 0 | 0.36 | 0.34 | 0.39 | 0.83 | 0.33 | 0.56 |
| 2 | 29413311 | rs17007625 | 0.202 | 0.693 | 0.604 | -2.093 | -0.003 | -0.064 | -0.064 | 0.523 | 0.13 | 0.23 | 0.08 | 0 | 0.12 | 0.03 |
| 2 | 29414339 | rs13025743 | 0.141 | 0.728 | 0.614 | -2.095 | 0 | -0.067 | -0.067 | 0.161 | 0.13 | 0.25 | 0.08 | 0 | 0.12 | 0.03 |
| 2 | 29415014 | rs2278880 | -0.366 | 0.573 | 0.659 | -2.164 | -0.001 | -0.009 | -0.009 | 0 | 0.35 | 0.41 | 0.53 | 0.73 | 0.36 | 0.52 |
| 2 | 29420315 | rs3738870 | 1.056 | 0.614 | 0.581 | -2.075 | -0.005 | -0.011 | -0.011 | 0 | 0.21 | 0.16 | 0.45 | 0.73 | 0.24 | 0.49 |
| 2 | 29431344 | rs3820711 | -0.375 | 0.753 | 0.502 | -2.299 | -0.002 | 0.033 | 0.033 | 0 | 0.71 | 0.8 | 0.81 | 0.85 | 0.74 | 0.84 |
| 2 | 29435447 | rs1615183 | -0.12 | 0.82 | 0.384 | -2.374 | -0.002 | 0.028 | 0.028 | 0 | 0.46 | 0.45 | 0.61 | 0.88 | 0.45 | 0.63 |
| 2 | 29435897 | rs1728827 | -0.901 | 0.827 | -0.071 | -2.447 | 0 | -0.031 | -0.031 | 3.987 | 0.59 | 0.59 | 0.69 | 0.88 | 0.57 | 0.66 |
| 2 | 29437842 | rs12987571 | 0.922 | 0.754 | -0.698 | -2.395 | -0.001 | 0.037 | 0.037 | 0 | 0.12 | 0.04 | 0.06 | 0 | 0.12 | 0.03 |
| 2 | 29438655 | rs11885445 | 0.94 | 0.697 | -0.72 | -2.338 | -0.055 | 0.212 | 0.212 | 0 | 0.35 | 0.02 | 0.22 | 0.05 | 0.35 | 0.32 |
| 2 | 29442426 | rs11681922 | 1.642 | 0.623 | -0.835 | -2.433 | -0.07 | 0.173 | 0.173 | 0 | 0.27 | 0.01 | 0.18 | 0.01 | 0.31 | 0.2 |
| 2 | 29446202 | rs4622670 | 1.057 | 0.691 | -0.598 | -2.245 | -0.067 | 0.204 | 0.204 | 0 | 0.34 | 0.01 | 0.22 | 0.05 | 0.35 | 0.32 |
| 2 | 29446701 | rs12619049 | -1.088 | 0.826 | 0.177 | -1.989 | -0.001 | -0.003 | -0.003 | 0 | 0.44 | 0.43 | 0.3 | 0.16 | 0.42 | 0.19 |
| 2 | 29447108 | rs4387740 | 0.065 | 0.726 | -0.245 | -1.75 | 0 | -0.008 | -0.008 | 0 | 0.69 | 0.61 | 0.77 | 0.76 | 0.73 | 0.83 |
| 2 | 29449005 | rs1317902 | 0.379 | 0.405 | -0.313 | -1.389 | 0 | 0 | 0 | 0 | 0.14 | 0.17 | 0.09 | 0.12 | 0.13 | 0.12 |
| 2 | 29449423 | rs6744522 | 0.426 | 0.576 | -0.358 | -1.521 | -0.017 | 0.17 | 0.17 | 0.407 | 0.47 | 0.26 | 0.35 | 0.05 | 0.55 | 0.34 |
| 2 | 29449819 | rs2293563 | -0.144 | 0.076 | -0.046 | -0.74 | 0.002 | 0.005 | 0.005 | 0 | 0.24 | 0.23 | 0.15 | 0.12 | 0.19 | 0.14 |
| 2 | 29454701 | rs1104869 | 0.783 | 0.454 | 0.696 | -0.011 | 0.008 | 0.089 | 0.089 | 0.022 | 0.89 | 0.57 | 0.9 | 0.86 | 0.92 | 0.86 |
| 2 | 29455738 | rs4666183 | -0.532 | 0.945 | -0.144 | -0.057 | -0.018 | 0.014 | 0.014 | 0.956 | 0.38 | 0.27 | 0.54 | 0.71 | 0.49 | 0.63 |
| 2 | 29459309 | rs13423991 | -1.679 | 0.781 | 0.708 | -0.16 | 0.041 | -0.043 | -0.043 | 0.232 | 0.08 | 0.27 | 0.07 | 0 | 0.09 | 0.02 |
| 2 | 29465429 | rs13391633 | -1.371 | 0.724 | 1.162 | -0.167 | 0.043 | -0.023 | -0.023 | 0 | 0.06 | 0.2 | 0.04 | 0 | 0.08 | 0.02 |
| 2 | 29467231 | rs13382755 | -2.129 | 0.634 | 1.402 | -0.209 | 0.049 | -0.034 | -0.034 | 0 | 0.09 | 0.44 | 0.06 | 0 | 0.07 | 0.02 |
| 2 | 29472936 | rs10197179 | -2.44 | 0.568 | 1.387 | -0.296 | 0.019 | -0.073 | -0.073 | 3.333 | 0.12 | 0.38 | 0.08 | 0 | 0.1 | 0.03 |
| 2 | 29477830 | rs1528424 | 0.543 | 0.453 | 1.346 | -0.368 | -0.005 | -0.015 | -0.015 | 0 | 0.71 | 0.63 | 0.73 | 0.77 | 0.72 | 0.75 |
| 2 | 29482318 | rs11127209 | 0.878 | 0.441 | 1.384 | -0.408 | -0.001 | 0.008 | 0.008 | 0 | 0.65 | 0.35 | 0.69 | 0.78 | 0.66 | 0.75 |
| 2 | 29484101 | rs7561367 | 0.365 | 0.259 | 1.24 | -0.34 | -0.011 | -0.006 | -0.006 | 0 | 0.75 | 0.65 | 0.78 | 0.78 | 0.79 | 0.79 |
| 2 | 29487293 | rs1358513 | 1.132 | 0.093 | 1.536 | -0.409 | -0.026 | -0.016 | -0.016 | 1.251 | 0.73 | 0.5 | 0.77 | 0.79 | 0.79 | 0.79 |
| 2 | 29487405 | rs7572957 | -1.888 | 0.092 | 1.832 | -0.477 | 0.001 | 0.089 | 0.089 | 0.07 | 0.14 | 0.35 | 0.12 | 0.22 | 0.1 | 0.16 |
| 2 | 29491807 | rs2049707 | 2.027 | -0.788 | 1.429 | -1.021 | 0.02 | 0.032 | 0.032 | 1.522 | 0.56 | 0.35 | 0.69 | 0.78 | 0.68 | 0.69 |
| 2 | 29494740 | rs6732925 | 0.431 | -0.682 | 1.535 | -0.8 | 0.022 | 0.23 | 0.23 | 1.351 | 0.93 | 0.66 | 0.92 | 0.8 | 0.98 | 0.96 |
| 2 | 29499956 | rs4666188 | 1.861 | -0.675 | 1.769 | -0.855 | -0.003 | 0.063 | 0.063 | 1.371 | 0.85 | 0.48 | 0.89 | 0.84 | 0.89 | 0.87 |
| 2 | 29502690 | rs4666189 | 0.321 | -0.669 | 1.848 | -0.809 | 0.021 | 0.19 | 0.19 | 1.038 | 0.93 | 0.71 | 0.95 | 0.83 | 0.98 | 0.97 |
| 2 | 29504792 | rs13422564 | -1.093 | -0.686 | 1.662 | -0.869 | -0.001 | -0.001 | -0.001 | 0 | 0.11 | 0.1 | 0.06 | 0 | 0.11 | 0.13 |
| 2 | 29506599 | rs13394509 | -0.102 | -0.695 | 1.66 | -0.871 | -0.003 | -0.019 | -0.019 | 0 | 0.07 | 0.09 | 0.03 | 0 | 0.05 | 0.1 |
| 2 | 29509715 | rs4666191 | 1.913 | -0.707 | 1.77 | -0.873 | -0.022 | 0.004 | 0.004 | 0.727 | 0.81 | 0.47 | 0.88 | 0.84 | 0.87 | 0.85 |
| 2 | 29511480 | rs1881417 | 2.093 | -0.887 | 1.747 | -0.943 | -0.029 | -0.017 | -0.017 | 0 | 0.74 | 0.37 | 0.85 | 0.83 | 0.84 | 0.81 |
| 2 | 29512675 | rs876748 | 2.103 | -0.319 | 1.756 | -1.038 | 0.01 | 0.042 | 0.042 | 2.58 | 0.61 | 0.29 | 0.74 | 0.83 | 0.69 | 0.71 |
| 2 | 29517823 | rs12328429 | -1.144 | -0.344 | 1.712 | -1.03 | 0 | -0.022 | -0.022 | 0.102 | 0.13 | 0.15 | 0.07 | 0 | 0.12 | 0.11 |
| 2 | 29519475 | rs13406263 | -1.222 | -0.344 | 1.705 | -1.029 | -0.003 | -0.047 | -0.047 | 0 | 0.13 | 0.2 | 0.08 | 0 | 0.12 | 0.13 |
| 2 | 29520455 | rs4666195 | 0.454 | -0.34 | 1.641 | -0.924 | 0.023 | 0.198 | 0.198 | 1.827 | 0.92 | 0.66 | 0.93 | 0.83 | 0.98 | 0.96 |
| 2 | 29524432 | rs17716681 | -0.866 | -0.243 | 1.061 | -0.971 | -0.001 | -0.004 | -0.004 | 3.045 | 0.18 | 0.16 | 0.12 | 0.01 | 0.17 | 0.13 |
| 2 | 29533345 | rs6750752 | 1.485 | -0.557 | 1.102 | -0.889 | -0.037 | -0.023 | -0.023 | 0.468 | 0.72 | 0.48 | 0.82 | 0.79 | 0.81 | 0.77 |
| 2 | 29533782 | rs7599783 | 1.16 | -0.618 | 0.155 | -1.571 | 0.058 | 0.114 | 0.114 | 0.501 | 0.51 | 0.29 | 0.71 | 0.78 | 0.61 | 0.57 |
| 2 | 29534329 | rs10432708 | 1.47 | -0.624 | 0.077 | -1.609 | -0.005 | 0.081 | 0.081 | 0.55 | 0.45 | 0.27 | 0.69 | 0.78 | 0.52 | 0.56 |
| 2 | 29535197 | rs4666199 | 1.496 | -0.615 | 0.051 | -1.605 | -0.01 | 0.089 | 0.089 | 0.906 | 0.45 | 0.27 | 0.67 | 0.78 | 0.52 | 0.56 |
| 2 | 29536039 | rs6708752 | -0.584 | -0.37 | -0.177 | -1.359 | -0.005 | -0.008 | -0.008 | 0 | 0.16 | 0.14 | 0.12 | 0 | 0.19 | 0.08 |
| 2 | 29538411 | rs4666200 | 1.34 | -0.506 | -0.41 | -1.255 | 0.048 | 0.058 | 0.058 | 1.877 | 0.64 | 0.12 | 0.74 | 0.79 | 0.72 | 0.75 |
| 2 | 29538860 | rs2339469 | 1.325 | -0.229 | -1.289 | -1.016 | -0.107 | -0.102 | -0.102 | 0 | 0.27 | 0.06 | 0.5 | 0.61 | 0.34 | 0.42 |
| 2 | 29544776 | rs7561975 | 1.214 | -0.175 | -0.378 | -1.003 | 0.015 | 0.037 | 0.037 | 0.634 | 0.65 | 0.24 | 0.75 | 0.85 | 0.71 | 0.78 |
| 2 | 29544920 | rs17727792 | -0.147 | -0.248 | -0.181 | -1.039 | -0.001 | 0 | 0 | 0 | 0.11 | 0.1 | 0.08 | 0.08 | 0.14 | 0.08 |
| 2 | 29545054 | rs13404555 | -0.985 | -0.3 | -0.073 | -1.075 | 0.073 | 0.002 | 0.002 | 0.187 | 0.09 | 0.19 | 0.13 | 0.08 | 0.22 | 0.1 |

| | | | | | | | | | | | | | | | | |
|---|---|---|---|---|---|---|---|---|---|---|---|---|---|---|---|---|
| 2 | 29545378 | rs4666204 | 0.996 | -0.424 | -0.007 | -1.098 | -0.03 | -0.062 | -0.062 | 2.566 | 0.85 | 0.35 | 0.89 | 0.96 | 0.92 | 0.97 |
| 2 | 29546758 | rs6731724 | 1.667 | -0.262 | -0.391 | -0.451 | 0.008 | -0.005 | -0.005 | 0 | 0.69 | 0.25 | 0.7 | 0.69 | 0.75 | 0.63 |
| 2 | 29547526 | rs6722414 | 1.455 | -0.162 | -0.144 | -0.44 | -0.033 | -0.045 | -0.045 | 0 | 0.73 | 0.36 | 0.85 | 0.87 | 0.83 | 0.84 |
| 2 | 29552712 | rs6742761 | -1.576 | 0.224 | -0.343 | -0.064 | 0.003 | 0.005 | 0.005 | 0 | 0.25 | 0.24 | 0.27 | 0.33 | 0.31 | 0.36 |
| 2 | 29554495 | rs17007904 | -0.388 | 0.07 | -0.016 | 0.044 | 0.052 | 0.072 | 0.072 | 0 | 0.08 | 0.23 | 0.11 | 0.16 | 0.12 | 0.12 |
| 2 | 29559519 | rs12465220 | -0.053 | 0.138 | -0.091 | -0.411 | 0 | 0 | 0 | 0 | 0.53 | 0.53 | 0.44 | 0.21 | 0.51 | 0.53 |
| 2 | 29565278 | rs4666205 | 1.086 | 0.257 | -0.433 | -0.237 | -0.132 | -0.037 | -0.037 | 0.888 | 0.3 | 0.07 | 0.5 | 0.74 | 0.37 | 0.52 |
| 2 | 29569562 | rs13000043 | 1.882 | 0.133 | -0.559 | -0.509 | 0.015 | 0.193 | 0.193 | 0 | 0.3 | 0.05 | 0.13 | 0.02 | 0.25 | 0.22 |
| 2 | 29570906 | rs17007920 | 0.484 | 0.143 | 0.007 | -0.502 | 0.008 | -0.005 | -0.005 | 0 | 0.1 | 0.12 | 0.17 | 0.06 | 0.12 | 0.09 |
| 2 | 29572299 | rs12996631 | 0.397 | 0.125 | 0.183 | -0.45 | 0.013 | 0.068 | 0.068 | 0 | 0.44 | 0.28 | 0.38 | 0.22 | 0.42 | 0.53 |
| 2 | 29573090 | rs6547920 | -1.027 | 0.233 | 0.298 | -0.395 | 0 | -0.018 | -0.018 | 0 | 0.9 | 0.91 | 0.95 | 1 | 0.89 | 0.99 |
| 2 | 29575197 | rs6716914 | -1.061 | 0.236 | 0.254 | -0.394 | -0.001 | 0.003 | 0.003 | 0.129 | 0.9 | 0.93 | 0.95 | 1 | 0.89 | 0.99 |
| 2 | 29577405 | rs3927337 | 1.606 | 0.243 | 0.139 | -0.354 | -0.101 | -0.128 | -0.128 | 0 | 0.28 | 0.06 | 0.39 | 0.57 | 0.32 | 0.31 |
| 2 | 29578068 | rs17007931 | 0.071 | 0.574 | 0.364 | -0.471 | -0.008 | -0.011 | -0.011 | 0 | 0.11 | 0.06 | 0.24 | 0.19 | 0.16 | 0.3 |
| 2 | 29580466 | rs10204572 | 0.592 | 0.613 | 0.874 | -0.726 | -0.006 | -0.007 | -0.007 | 0.514 | 0.4 | 0.39 | 0.54 | 0.92 | 0.42 | 0.62 |
| 2 | 29582154 | rs4146674 | 0.713 | 0.63 | 1.169 | -0.66 | 0.03 | -0.04 | -0.04 | 0 | 0.84 | 0.67 | 0.74 | 0.94 | 0.78 | 0.89 |
| 2 | 29582433 | rs4146673 | 1.67 | 0.601 | 1.011 | -0.686 | 0.007 | -0.077 | -0.077 | 0 | 0.73 | 0.44 | 0.67 | 0.94 | 0.68 | 0.87 |
| 2 | 29583501 | rs10199963 | 0.334 | 0.439 | 0.771 | -0.488 | -0.001 | -0.036 | -0.036 | 0.954 | 0.07 | 0.16 | 0.1 | 0 | 0.1 | 0.03 |
| 2 | 29586686 | rs12714277 | 0.382 | 0.409 | 0.698 | -0.481 | 0.014 | -0.02 | -0.02 | 0 | 0.42 | 0.34 | 0.21 | 0.02 | 0.36 | 0.26 |
| 2 | 29586747 | rs12986732 | 0.391 | 0.408 | 0.697 | -0.482 | 0.014 | -0.021 | -0.021 | 0 | 0.42 | 0.34 | 0.21 | 0.02 | 0.36 | 0.26 |
| 2 | 29587488 | rs1528423 | -0.442 | 0.373 | 0.743 | -0.33 | -0.007 | -0.035 | -0.035 | 0.02 | 0.56 | 0.54 | 0.78 | 0.85 | 0.63 | 0.65 |
| 2 | 29589910 | rs11893367 | -1.542 | 0.904 | 1.256 | 0.217 | 0.038 | 0.05 | 0.05 | 0 | 0.11 | 0.21 | 0.09 | 0.15 | 0.13 | 0.13 |
| 2 | 29592200 | rs7602265 | 0.058 | 0.84 | 1.366 | 0.104 | -0.001 | -0.031 | -0.031 | 0.26 | 0.1 | 0.14 | 0.06 | 0 | 0.1 | 0.01 |
| 2 | 29592460 | rs11903143 | -1.947 | 0.812 | 1.304 | 0.027 | 0.032 | -0.013 | -0.013 | 0 | 0.2 | 0.35 | 0.17 | 0.16 | 0.26 | 0.16 |
| 2 | 29595021 | rs17007998 | 0.278 | 0.845 | 1.239 | 0.035 | 0 | -0.013 | -0.013 | 0 | 0.1 | 0.13 | 0.05 | 0 | 0.11 | 0.01 |
| 2 | 29596488 | rs1534544 | 1.731 | 0.856 | 1.32 | 0.022 | 0.024 | -0.038 | -0.038 | 0 | 0.77 | 0.55 | 0.82 | 0.85 | 0.72 | 0.84 |
| 2 | 29599382 | rs6751302 | -0.47 | 0.925 | 0.834 | 0.315 | 0.014 | 0.056 | 0.056 | 0 | 0.06 | 0.09 | 0.05 | 0.19 | 0.1 | 0.12 |
| 2 | 29599866 | rs12621597 | -1.121 | 0.896 | 0.583 | 0.569 | -0.003 | 0.004 | 0.004 | 0 | 0.1 | 0.06 | 0.08 | 0.25 | 0.11 | 0.14 |
| 2 | 29607480 | rs10186332 | 0.295 | 1.304 | 0.818 | -0.029 | 0.004 | -0.005 | -0.005 | 0 | 0.15 | 0.12 | 0.2 | 0.07 | 0.19 | 0.14 |
| 2 | 29609205 | rs12469277 | -0.134 | 1.348 | 0.788 | 0.138 | 0.004 | 0.002 | 0.002 | 0.342 | 0.18 | 0.14 | 0.3 | 0.18 | 0.24 | 0.3 |
| 2 | 29610088 | rs4074943 | -1.312 | 1.297 | 0.945 | 0.033 | 0.024 | -0.023 | -0.023 | 0 | 0.3 | 0.42 | 0.39 | 0.18 | 0.36 | 0.32 |
| 2 | 29612926 | rs10195189 | 1.499 | 1.125 | 0.101 | -0.305 | 0.025 | 0.102 | 0.102 | 0 | 0.32 | 0.16 | 0.15 | 0.01 | 0.29 | 0.24 |
| 2 | 29613214 | rs4666212 | -0.586 | 1.221 | -0.054 | -0.254 | 0.009 | 0.065 | 0.065 | 0 | 0.51 | 0.43 | 0.61 | 0.93 | 0.52 | 0.62 |
| 2 | 29615433 | rs13415294 | 0.019 | 1.323 | -0.404 | 0.104 | -0.006 | -0.001 | -0.001 | 0 | 0.21 | 0.12 | 0.29 | 0.15 | 0.27 | 0.29 |
| 2 | 29618678 | rs11127220 | 0.569 | 1.039 | -0.882 | 0.323 | -0.03 | -0.075 | -0.075 | 0.074 | 0.29 | 0.17 | 0.42 | 0.53 | 0.29 | 0.32 |
| 2 | 29619971 | rs7591913 | 0.049 | 0.933 | -0.623 | 0.111 | -0.018 | 0.052 | 0.052 | 0 | 0.58 | 0.62 | 0.47 | 0.13 | 0.56 | 0.53 |
| 2 | 29624193 | rs6740703 | 0.414 | 0.908 | -0.201 | 0.978 | 0.024 | 0.245 | 0.245 | 0 | 0.93 | 0.78 | 0.92 | 0.65 | 0.94 | 0.87 |
| 2 | 29624787 | rs7590571 | 0.794 | 0.761 | -1.175 | 0.332 | 0 | 0.025 | 0.025 | 0 | 0.55 | 0.13 | 0.54 | 0.45 | 0.57 | 0.56 |
| 2 | 29627861 | rs6547924 | 0.188 | 0.466 | -0.703 | -0.4 | 0 | 0.05 | 0.05 | 0.165 | 0.58 | 0.3 | 0.61 | 0.96 | 0.55 | 0.71 |
| 2 | 29637040 | rs4233734 | 0.966 | 0.414 | -1.079 | -0.093 | -0.047 | 0.068 | 0.068 | 0.995 | 0.46 | 0.11 | 0.51 | 0.69 | 0.44 | 0.51 |
| 2 | 29638906 | rs11674942 | -0.094 | 0.589 | -0.163 | -0.118 | 0.015 | 0.072 | 0.072 | 0 | 0.1 | 0.25 | 0.1 | 0.25 | 0.11 | 0.14 |
| 2 | 29639679 | rs4555301 | -0.696 | 0.371 | 0.24 | -0.613 | 0.003 | -0.037 | -0.037 | 0 | 0.76 | 0.73 | 0.73 | 0.96 | 0.71 | 0.78 |
| 2 | 29643071 | rs11897469 | 0.372 | 0.376 | 0.535 | -0.615 | -0.002 | -0.01 | -0.01 | 0.039 | 0.26 | 0.25 | 0.28 | 0.02 | 0.3 | 0.21 |
| 2 | 29643332 | rs11689426 | 0.326 | 0.186 | 0.696 | 0.001 | -0.018 | 0.025 | 0.025 | 0 | 0.1 | 0.07 | 0.06 | 0 | 0.15 | 0.01 |
| 2 | 29645870 | rs4315478 | -0.212 | 0.15 | 0.981 | 0.013 | 0.018 | 0.05 | 0.05 | 0 | 0.94 | 0.8 | 0.97 | 0.98 | 0.99 | 0.98 |
| 2 | 29646424 | rs11898966 | 0.104 | 0.454 | 0.23 | 0.099 | -0.014 | 0.012 | 0.012 | 0 | 0.1 | 0.07 | 0.07 | 0 | 0.16 | 0.01 |
| 2 | 29654953 | rs7590597 | -0.546 | 0.185 | 0.219 | 0.754 | 0 | 0.02 | 0.02 | 0 | 0.09 | 0.14 | 0.23 | 0.32 | 0.07 | 0.14 |
| 2 | 29666976 | rs4322797 | 0.552 | -0.099 | 0.898 | 0.641 | -0.037 | 0.185 | 0.185 | 0.427 | 0.88 | 0.6 | 0.76 | 0.71 | 0.93 | 0.83 |
| 2 | 29667972 | rs10178439 | 0.399 | -0.094 | 1.011 | 0.651 | -0.034 | 0.155 | 0.155 | 0.08 | 0.88 | 0.63 | 0.76 | 0.72 | 0.93 | 0.83 |
| 2 | 29670834 | rs4456659 | 0.289 | -0.025 | 0.51 | -0.23 | 0 | -0.014 | -0.014 | 0 | 0.57 | 0.48 | 0.44 | 0.67 | 0.56 | 0.51 |
| 2 | 29679463 | rs4346337 | 0.33 | 0.036 | -0.155 | -0.266 | -0.006 | 0.034 | 0.034 | 1.287 | 0.48 | 0.33 | 0.48 | 0.38 | 0.48 | 0.59 |
| 2 | 29683558 | rs7580476 | 1.334 | -0.064 | 0.04 | -0.298 | -0.002 | 0.011 | 0.011 | 0.833 | 0.26 | 0.26 | 0.19 | 0.34 | 0.26 | 0.18 |
| 2 | 29685596 | rs3923028 | 1.621 | -0.098 | -0.289 | -0.295 | 0.001 | -0.03 | -0.03 | 0 | 0.16 | 0.12 | 0.17 | 0.34 | 0.14 | 0.3 |
| 2 | 29691960 | rs12714281 | -0.983 | -0.169 | -0.043 | -0.294 | -0.017 | 0.05 | 0.05 | 0.15 | 0.88 | 0.78 | 0.76 | 0.72 | 0.9 | 0.84 |
| 2 | 29698025 | rs4233739 | 0.136 | -0.197 | 0.027 | -0.387 | 0.038 | 0.036 | 0.036 | 0.44 | 0.54 | 0.31 | 0.51 | 0.37 | 0.59 | 0.6 |
| 2 | 29698164 | rs4493213 | -0.842 | -0.232 | 0.107 | -0.319 | -0.021 | 0.075 | 0.075 | 0.367 | 0.88 | 0.75 | 0.75 | 0.72 | 0.9 | 0.83 |
| 2 | 29698383 | rs4503951 | 1.808 | -0.206 | 0.068 | -0.296 | 0 | -0.033 | -0.033 | 0.032 | 0.16 | 0.1 | 0.16 | 0.31 | 0.14 | 0.29 |
| 2 | 29700165 | rs6547933 | -0.955 | -0.166 | 0.141 | -0.327 | -0.024 | 0.091 | 0.091 | 0.179 | 0.88 | 0.73 | 0.75 | 0.72 | 0.9 | 0.84 |
| 2 | 29702682 | rs7426089 | 1.726 | -0.185 | 0.013 | -0.338 | -0.001 | -0.027 | -0.027 | 0.743 | 0.15 | 0.11 | 0.16 | 0.32 | 0.14 | 0.29 |
| 2 | 29702883 | rs11888731 | 0.59 | -0.252 | 0.096 | -0.439 | -0.012 | -0.001 | -0.001 | 0.143 | 0.33 | 0.5 | 0.24 | 0.34 | 0.28 | 0.2 |
| 2 | 29704715 | rs4528720 | 0.307 | -0.169 | 0.089 | -0.438 | 0.001 | -0.003 | -0.003 | 0 | 0.63 | 0.47 | 0.52 | 0.37 | 0.61 | 0.49 |
| 2 | 29705020 | rs4299297 | -0.1 | -0.188 | 0.101 | -0.427 | 0.008 | 0.032 | 0.032 | 0 | 0.8 | 0.64 | 0.68 | 0.71 | 0.76 | 0.81 |
| 2 | 29706271 | rs7588341 | -0.274 | -0.189 | 0.076 | -0.434 | -0.011 | 0.039 | 0.039 | 0 | 0.73 | 0.52 | 0.58 | 0.39 | 0.73 | 0.54 |
| 2 | 29708619 | rs13383564 | 0.335 | -0.203 | 0.49 | -0.61 | -0.002 | -0.058 | -0.058 | 1.243 | 0.42 | 0.36 | 0.4 | 0.03 | 0.48 | 0.34 |
| 2 | 29709401 | rs3884503 | -0.717 | -0.194 | 0.538 | -0.521 | -0.006 | 0.107 | 0.107 | 0.515 | 0.87 | 0.65 | 0.73 | 0.71 | 0.87 | 0.83 |
| 2 | 29710663 | rs12714287 | 0.551 | -0.275 | 0.416 | -0.596 | 0.002 | 0.014 | 0.014 | 0.059 | 0.39 | 0.28 | 0.33 | 0.68 | 0.37 | 0.47 |
| 2 | 29715388 | rs13000666 | 0.262 | -0.204 | 0.538 | -0.587 | 0.004 | 0.047 | 0.047 | 0.069 | 0.77 | 0.44 | 0.67 | 0.71 | 0.76 | 0.78 |
| 2 | 29715992 | rs6547935 | -0.059 | -0.291 | 0.453 | -0.571 | 0.013 | 0.019 | 0.019 | 0.966 | 0.48 | 0.53 | 0.37 | 0.68 | 0.39 | 0.5 |
| 2 | 29716266 | rs6547936 | -0.851 | -0.27 | 0.365 | -0.428 | -0.006 | 0.077 | 0.077 | 0.12 | 0.88 | 0.73 | 0.75 | 0.71 | 0.87 | 0.82 |

| | | | | | | | | | | | | | | | | |
|---|---|---|---|---|---|---|---|---|---|---|---|---|---|---|---|---|
| 2 | 29719357 | rs4666230 | 0.735 | -0.309 | 0.191 | -0.368 | 0 | 0.013 | 0.013 | 2.015 | 0.31 | 0.18 | 0.2 | 0.29 | 0.31 | 0.21 |
| 2 | 29720219 | rs13407210 | 0.038 | -0.403 | 0.347 | -0.568 | 0.005 | -0.016 | -0.016 | 0.958 | 0.48 | 0.41 | 0.44 | 0.03 | 0.51 | 0.35 |
| 2 | 29721334 | rs9808313 | 0.223 | -0.414 | 0.455 | -0.576 | 0.003 | 0.043 | 0.043 | 0 | 0.46 | 0.35 | 0.4 | 0.03 | 0.51 | 0.35 |
| 2 | 29724183 | rs4261679 | 1.233 | -0.286 | 0.669 | -0.392 | 0 | 0.002 | 0.002 | 0 | 0.24 | 0.23 | 0.18 | 0.32 | 0.25 | 0.2 |
| 2 | 29726078 | rs6734413 | -0.654 | -0.317 | 0.786 | -0.28 | -0.008 | 0.065 | 0.065 | 0.003 | 0.87 | 0.75 | 0.75 | 0.68 | 0.86 | 0.81 |
| 2 | 29727204 | rs10445906 | 1.056 | -0.364 | 0.841 | -0.267 | 0.006 | 0.014 | 0.014 | 0 | 0.26 | 0.3 | 0.2 | 0.36 | 0.28 | 0.2 |
| 2 | 29727528 | rs10445967 | 0.195 | -0.353 | 0.756 | -0.361 | -0.004 | 0.001 | 0.001 | 1.297 | 0.35 | 0.62 | 0.29 | 0.36 | 0.4 | 0.24 |
| 2 | 29728909 | rs12620310 | 1.215 | -0.343 | 0.672 | -0.322 | 0.005 | 0.01 | 0.01 | 0.007 | 0.25 | 0.26 | 0.18 | 0.35 | 0.27 | 0.2 |
| 2 | 29729331 | rs4665462 | 0.367 | -0.411 | 0.518 | -0.353 | -0.045 | 0.034 | 0.034 | 1.142 | 0.53 | 0.06 | 0.47 | 0.32 | 0.51 | 0.59 |
| 2 | 29730492 | rs4433956 | 0.079 | -0.348 | 0.899 | -0.189 | -0.009 | 0.078 | 0.078 | 0 | 0.78 | 0.45 | 0.67 | 0.68 | 0.77 | 0.78 |
| 2 | 29731788 | rs6740714 | 1.004 | -0.138 | 0.739 | -0.408 | -0.002 | 0.001 | 0.001 | 0.013 | 0.32 | 0.25 | 0.21 | 0.32 | 0.33 | 0.23 |
| 2 | 29733601 | rs4665463 | 0.148 | -0.202 | 0.53 | -0.356 | -0.019 | 0.045 | 0.045 | 0.101 | 0.46 | 0.16 | 0.45 | 0.33 | 0.44 | 0.55 |
| 2 | 29733801 | rs6719507 | 0.378 | -0.2 | 0.59 | -0.384 | -0.012 | 0.017 | 0.017 | 1.759 | 0.42 | 0.62 | 0.31 | 0.36 | 0.45 | 0.27 |
| 2 | 29739177 | rs4567889 | 0.103 | -0.197 | 0.298 | -0.299 | -0.019 | -0.01 | -0.01 | 0.951 | 0.46 | 0.15 | 0.44 | 0.4 | 0.44 | 0.57 |
| 2 | 29741526 | rs10187810 | 0.074 | -0.756 | 0.785 | -0.203 | -0.034 | -0.016 | -0.016 | 0 | 0.1 | 0.21 | 0.04 | 0.05 | 0.02 | 0.1 |
| 2 | 29743581 | rs4468777 | -0.102 | -0.767 | 0.669 | -0.303 | 0.026 | 0.025 | 0.025 | 1.901 | 0.55 | 0.63 | 0.35 | 0.38 | 0.59 | 0.27 |
| 2 | 29743733 | rs7578465 | 0.71 | -0.821 | 0.729 | -0.331 | 0.036 | 0.095 | 0.095 | 0.516 | 0.44 | 0.14 | 0.3 | 0.32 | 0.57 | 0.17 |
| 2 | 29744336 | rs4365418 | -0.243 | -0.812 | 0.857 | -0.292 | -0.034 | -0.007 | -0.007 | 0 | 0.11 | 0.49 | 0.06 | 0.05 | 0.03 | 0.1 |
| 2 | 29745284 | rs10171094 | 0.432 | -0.8 | 0.516 | -0.286 | -0.032 | -0.015 | -0.015 | 0 | 0.09 | 0.17 | 0.02 | 0.05 | 0.02 | 0.1 |
| 2 | 29745985 | rs4536609 | 1.107 | -0.912 | 0.254 | -0.352 | 0.028 | -0.098 | -0.098 | 0.494 | 0.16 | 0.01 | 0.17 | 0.3 | 0.1 | 0.34 |
| 2 | 29747174 | rs12467802 | 1.419 | -0.926 | 0.231 | -0.32 | 0.024 | -0.049 | -0.049 | 0.053 | 0.11 | 0.04 | 0.22 | 0.32 | 0.11 | 0.19 |
| 2 | 29748228 | rs7586042 | 1.513 | -0.74 | 0.124 | -0.506 | -0.046 | 0.023 | 0.023 | 0 | 0.31 | 0.18 | 0.22 | 0.3 | 0.38 | 0.14 |
| 2 | 29750729 | rs7594020 | 1.638 | -0.698 | -0.015 | -0.501 | -0.056 | 0.043 | 0.043 | 0.159 | 0.3 | 0.11 | 0.2 | 0.27 | 0.35 | 0.13 |
| 2 | 29750828 | rs12714289 | 0.409 | -0.608 | 0.098 | -0.483 | -0.001 | 0.024 | 0.024 | 0 | 0.5 | 0.33 | 0.46 | 0.63 | 0.51 | 0.41 |
| 2 | 29752367 | rs4666234 | 0.583 | -0.537 | 0.022 | -0.236 | -0.02 | 0.027 | 0.027 | 0 | 0.18 | 0.23 | 0.2 | 0.31 | 0.1 | 0.34 |
| 2 | 29755934 | rs6716559 | -0.161 | -0.251 | 0.091 | -0.312 | 0.01 | 0.025 | 0.025 | 0 | 0.74 | 0.66 | 0.67 | 0.66 | 0.68 | 0.73 |
| 2 | 29758149 | rs12105492 | 1.629 | -0.28 | -0.264 | -0.43 | -0.021 | 0.059 | 0.059 | 0.418 | 0.17 | 0.05 | 0.22 | 0.06 | 0.2 | 0.2 |
| 2 | 29759962 | rs10779970 | 0.187 | -0.54 | 0.103 | -0.549 | -0.038 | 0.046 | 0.046 | 0 | 0.78 | 0.41 | 0.7 | 0.67 | 0.85 | 0.75 |
| 2 | 29763321 | rs7569431 | 1.586 | -0.455 | -0.01 | -0.457 | -0.004 | 0.01 | 0.01 | 0.095 | 0.17 | 0.12 | 0.23 | 0.13 | 0.21 | 0.22 |
| 2 | 29763753 | rs4666243 | -0.597 | -0.526 | -0.105 | -0.421 | 0.015 | 0.041 | 0.041 | 0 | 0.41 | 0.25 | 0.49 | 0.64 | 0.4 | 0.59 |
| 2 | 29764460 | rs13010035 | 0.109 | -0.513 | -0.136 | -0.404 | -0.035 | -0.066 | -0.066 | 0.6 | 0.29 | 0.09 | 0.26 | 0.32 | 0.3 | 0.36 |
| 2 | 29767147 | rs10176482 | -0.73 | -0.598 | -0.028 | -0.445 | 0.027 | 0.011 | 0.011 | 0 | 0.34 | 0.26 | 0.43 | 0.65 | 0.26 | 0.59 |
| 2 | 29767668 | rs4473345 | 1.109 | -0.628 | -0.014 | -0.424 | 0.009 | -0.012 | -0.012 | 1.527 | 0.16 | 0.15 | 0.18 | 0.33 | 0.11 | 0.35 |
| 2 | 29767894 | rs7592571 | 0.357 | -0.643 | -0.046 | -0.407 | 0.014 | -0.019 | -0.019 | 0 | 0.18 | 0.11 | 0.24 | 0.32 | 0.12 | 0.24 |
| 2 | 29769160 | rs4666246 | -0.622 | -0.47 | -0.059 | -0.512 | 0.011 | 0.046 | 0.046 | 0 | 0.41 | 0.27 | 0.5 | 0.64 | 0.41 | 0.6 |
| 2 | 29771510 | rs7567732 | -1.183 | -0.288 | 0.097 | -0.393 | -0.025 | -0.017 | -0.017 | 0 | 0.92 | 0.82 | 0.97 | 0.97 | 0.98 | 0.94 |
| 2 | 29776488 | rs7605970 | 1.641 | -0.22 | -0.049 | -0.425 | -0.052 | 0.062 | 0.062 | 0 | 0.13 | 0.02 | 0.09 | 0 | 0.18 | 0.03 |
| 2 | 29780501 | rs6547945 | -1.03 | -0.287 | 0.236 | -0.346 | -0.025 | -0.02 | -0.02 | 0 | 0.92 | 0.79 | 0.97 | 0.98 | 0.98 | 0.94 |
| 2 | 29783021 | rs4280415 | 0.232 | -0.14 | 0.016 | -0.448 | -0.005 | 0.002 | 0.002 | 0 | 0.43 | 0.48 | 0.43 | 0.54 | 0.41 | 0.34 |
| 2 | 29785679 | rs10193521 | 0.886 | -0.128 | 0.09 | -0.339 | -0.033 | 0.137 | 0.137 | 0.08 | 0.13 | 0.32 | 0.25 | 0.28 | 0.11 | 0.18 |
| 2 | 29785781 | rs12473275 | 1.246 | -0.11 | 0.003 | -0.33 | -0.023 | 0.087 | 0.087 | 0 | 0.12 | 0.2 | 0.23 | 0.28 | 0.11 | 0.18 |
| 2 | 29790622 | rs6547948 | -1.171 | 0.007 | -0.083 | -0.439 | -0.016 | 0.017 | 0.017 | 0 | 0.68 | 0.76 | 0.8 | 0.77 | 0.64 | 0.81 |
| 2 | 29791841 | rs6547949 | -0.147 | -0.123 | -0.117 | -0.523 | 0.012 | 0.019 | 0.019 | 2.074 | 0.46 | 0.6 | 0.68 | 0.77 | 0.44 | 0.74 |
| 2 | 29793174 | rs6742561 | 0.8 | -0.104 | -0.256 | -0.501 | -0.028 | 0.059 | 0.059 | 0 | 0.15 | 0.25 | 0.25 | 0.28 | 0.13 | 0.19 |
| 2 | 29795988 | rs13014477 | -0.645 | -0.058 | -0.402 | -0.569 | -0.004 | 0.003 | 0.003 | 0 | 0.29 | 0.34 | 0.35 | 0.3 | 0.26 | 0.28 |
| 2 | 29796951 | rs5003770 | 0.922 | -0.269 | -0.639 | -0.589 | -0.081 | -0.041 | -0.041 | 0 | 0.4 | 0.11 | 0.32 | 0.59 | 0.43 | 0.46 |
| 2 | 29797217 | rs6547951 | 0.056 | -0.483 | -0.191 | -0.169 | -0.028 | 0.027 | 0.027 | 0 | 0.2 | 0.26 | 0.26 | 0.29 | 0.14 | 0.19 |
| 2 | 29798400 | rs11688563 | 1.21 | -0.112 | -0.081 | -0.221 | 0.019 | -0.041 | -0.041 | 0.033 | 0.14 | 0.19 | 0.1 | 0.04 | 0.18 | 0.04 |
| 2 | 29800482 | rs4128814 | 0.811 | -0.281 | -0.208 | 0.103 | -0.029 | 0.105 | 0.105 | 0 | 0.12 | 0.21 | 0.25 | 0.27 | 0.1 | 0.17 |
| 2 | 29801450 | rs6547955 | 0.616 | -0.369 | -0.494 | 0.075 | 0.009 | -0.015 | -0.015 | 0 | 0.34 | 0.46 | 0.27 | 0.26 | 0.36 | 0.22 |
| 2 | 29802178 | rs6713380 | 0.337 | -0.648 | -0.667 | 0.369 | -0.023 | 0.023 | 0.023 | 0 | 0.17 | 0.23 | 0.23 | 0.28 | 0.1 | 0.18 |
| 2 | 29802440 | rs6757945 | -0.907 | -0.112 | -1.036 | 0.33 | -0.042 | 0.092 | 0.092 | 0 | 0.85 | 0.72 | 0.82 | 0.72 | 0.91 | 0.66 |
| 2 | 29805263 | rs4555303 | -0.529 | 0.139 | -1.025 | 0.321 | -0.001 | 0 | 0 | 0 | 0.67 | 0.69 | 0.59 | 0.62 | 0.66 | 0.47 |
| 2 | 29807890 | rs4381747 | -0.055 | -0.141 | -0.734 | 0.452 | -0.009 | 0.003 | 0.003 | 0 | 0.41 | 0.51 | 0.43 | 0.54 | 0.35 | 0.32 |
| 2 | 29812429 | rs4630725 | 0.029 | 0.1 | -1.088 | 0.325 | 0.022 | 0.01 | 0.01 | 0 | 0.54 | 0.24 | 0.48 | 0.61 | 0.58 | 0.52 |
| 2 | 29814009 | rs6709634 | 0.369 | 0.074 | -1.033 | 0.096 | -0.032 | -0.023 | -0.023 | 0 | 0.4 | 0.28 | 0.43 | 0.54 | 0.47 | 0.4 |
| 2 | 29816800 | rs6547957 | -0.22 | -0.115 | -0.29 | 0.123 | -0.002 | -0.002 | -0.002 | 0 | 0.72 | 0.77 | 0.64 | 0.72 | 0.69 | 0.77 |
| 2 | 29829760 | rs7594717 | 0.402 | -0.211 | -0.522 | 0.386 | -0.034 | -0.122 | -0.122 | 0 | 0.32 | 0.05 | 0.26 | 0.41 | 0.31 | 0.5 |
| 2 | 29835429 | rs7599913 | 0.277 | 0.319 | 0.681 | 0.392 | -0.02 | -0.015 | -0.015 | 0 | 0.84 | 0.9 | 0.7 | 0.76 | 0.73 | 0.8 |
| 2 | 29837275 | rs10208167 | 1.803 | 0.002 | 1.596 | 0.227 | 0.004 | -0.006 | -0.006 | 0.138 | 0.69 | 0.38 | 0.62 | 0.74 | 0.68 | 0.67 |
| 2 | 29838529 | rs9309671 | -0.739 | 0.016 | 1.981 | 0.234 | -0.023 | -0.034 | -0.034 | 0.023 | 0.08 | 0.23 | 0.03 | 0.01 | 0.04 | 0.06 |
| 2 | 29838964 | rs6547962 | 1.762 | 0.005 | 2.068 | 0.24 | 0.026 | 0.001 | 0.001 | 0 | 0.72 | 0.43 | 0.64 | 0.74 | 0.68 | 0.69 |
| 2 | 29843424 | rs6736621 | -0.352 | 0.031 | 2.058 | 0.228 | -0.023 | -0.032 | -0.032 | 0 | 0.07 | 0.13 | 0.02 | 0.01 | 0.04 | 0.06 |
| 2 | 29850379 | rs4233746 | 1.352 | 0.039 | 2.11 | 0.223 | 0.037 | 0.006 | 0.006 | 0 | 0.75 | 0.42 | 0.6 | 0.74 | 0.68 | 0.74 |
| 2 | 29856155 | rs2339554 | 1.374 | 0.062 | 2.142 | 0.235 | 0.057 | 0.013 | 0.013 | 0 | 0.75 | 0.48 | 0.62 | 0.74 | 0.68 | 0.73 |
| 2 | 29860061 | rs2339547 | 0.145 | 0.091 | 2.07 | 0.239 | -0.025 | -0.02 | -0.02 | 0 | 0.85 | 0.91 | 0.68 | 0.76 | 0.72 | 0.79 |
| 2 | 29862915 | rs7606115 | -1.381 | -0.148 | 2.165 | 0.218 | -0.014 | -0.03 | -0.03 | 0 | 0.09 | 0.45 | 0.05 | 0.01 | 0.04 | 0.06 |
| 2 | 29866533 | rs2339540 | 1.857 | -0.113 | 2.076 | 0.215 | 0.009 | -0.009 | -0.009 | 0.102 | 0.7 | 0.43 | 0.61 | 0.74 | 0.67 | 0.67 |
| 2 | 29867599 | rs17008384 | -2.328 | -0.126 | 2.259 | 0.196 | -0.088 | -0.108 | -0.108 | 5.465 | 0.15 | 0.41 | 0.05 | 0.01 | 0.05 | 0.12 |

| | | | | | | | | | | | | | | | | |
|---|---|---|---|---|---|---|---|---|---|---|---|---|---|---|---|---|
| 2 | 29868259 | rs7608034 | -1.038 | -0.121 | 2.358 | 0.198 | -0.026 | -0.042 | -0.042 | 0 | 0.08 | 0.3 | 0.04 | 0.01 | 0.04 | 0.06 |
| 2 | 29870205 | rs10182925 | -1.603 | -0.12 | 2.302 | 0.199 | -0.052 | -0.07 | -0.07 | 0 | 0.11 | 0.31 | 0.04 | 0.01 | 0.04 | 0.11 |
| 2 | 29871991 | rs4666265 | 0.165 | -0.105 | 1.978 | 0.219 | -0.086 | -0.064 | -0.064 | 0 | 0.84 | 0.96 | 0.68 | 0.77 | 0.72 | 0.8 |
| 2 | 29881235 | rs7583871 | -2.301 | -0.52 | 2.398 | 0.314 | -0.086 | -0.104 | -0.104 | 0 | 0.15 | 0.57 | 0.08 | 0.02 | 0.05 | 0.12 |
| 2 | 29889019 | rs2254066 | 0.543 | -0.14 | 1.962 | 0.43 | -0.057 | -0.051 | -0.051 | 0 | 0.89 | 0.96 | 0.75 | 0.78 | 0.81 | 0.87 |
| 2 | 29905337 | rs10200505 | -1.845 | -0.262 | 2.615 | 0.372 | -0.047 | -0.067 | -0.067 | 0 | 0.11 | 0.53 | 0.06 | 0.01 | 0.04 | 0.07 |
| 2 | 29907865 | rs2246276 | 0.351 | -0.143 | 2.468 | 0.4 | -0.053 | -0.048 | -0.048 | 0.01 | 0.89 | 0.97 | 0.76 | 0.79 | 0.83 | 0.87 |
| 2 | 29908726 | rs952455 | 1.241 | -0.16 | 2.351 | 0.392 | 0.021 | 0.014 | 0.014 | 0 | 0.86 | 0.8 | 0.73 | 0.79 | 0.82 | 0.83 |
| 2 | 29913145 | rs6547972 | -0.744 | -0.178 | 2.257 | 0.383 | -0.024 | -0.038 | -0.038 | 0 | 0.07 | 0.14 | 0.02 | 0.01 | 0.03 | 0.06 |
| 2 | 29916548 | rs2541169 | 2.396 | -0.205 | 1.956 | 0.369 | -0.003 | 0.013 | 0.013 | 0 | 0.73 | 0.16 | 0.66 | 0.77 | 0.78 | 0.76 |
| 2 | 29922977 | rs1009024 | 2.424 | -0.232 | 2.098 | 0.385 | 0.002 | -0.003 | -0.003 | 0 | 0.75 | 0.28 | 0.66 | 0.77 | 0.76 | 0.75 |
| 2 | 29924736 | rs2631964 | 1.054 | -0.186 | 2.036 | 0.425 | -0.029 | -0.026 | -0.026 | 0.323 | 0.86 | 0.89 | 0.76 | 0.79 | 0.83 | 0.86 |
| 2 | 29947235 | rs2541194 | 2.282 | -0.226 | 1.957 | 0.386 | -0.001 | 0.002 | 0.002 | 0 | 0.74 | 0.24 | 0.69 | 0.78 | 0.79 | 0.7 |
| 2 | 29947687 | rs17008532 | -1.143 | 0.011 | 2.209 | 0.315 | -0.019 | -0.035 | -0.035 | 0.013 | 0.08 | 0.24 | 0.03 | 0 | 0.04 | 0.05 |
| 2 | 29954652 | rs12998454 | 0.347 | 0.064 | 1.313 | 0.403 | -0.024 | -0.028 | -0.028 | 0 | 0.08 | 0.01 | 0.07 | 0.11 | 0.11 | 0.02 |
| 2 | 29966870 | rs2541191 | 2.231 | 0.127 | 0.811 | 0.495 | -0.002 | 0 | 0 | 0.844 | 0.81 | 0.2 | 0.67 | 0.84 | 0.78 | 0.75 |
| 2 | 29973281 | rs13411840 | -1.823 | -0.882 | 0.467 | 0.489 | -0.052 | -0.067 | -0.067 | 0.981 | 0.11 | 0.72 | 0.09 | 0 | 0.05 | 0.1 |
| 2 | 29981286 | rs2631941 | 1.674 | -0.89 | -0.37 | 0.648 | 0.069 | 0.002 | 0.002 | 0.716 | 0.75 | 0.12 | 0.67 | 0.74 | 0.88 | 0.77 |
| 2 | 29986574 | rs2541180 | 1.512 | -0.848 | -0.227 | 0.655 | 0.028 | 0.005 | 0.005 | 0.02 | 0.76 | 0.19 | 0.69 | 0.75 | 0.88 | 0.75 |
| 2 | 29989203 | rs4666270 | -1.277 | -0.878 | -0.196 | 1.018 | 0.006 | -0.01 | -0.01 | 0 | 0.24 | 0.2 | 0.29 | 0.35 | 0.17 | 0.13 |
| 2 | 29989331 | rs3845689 | -1.786 | -0.775 | 1.057 | 1.104 | -0.014 | 0.08 | 0.08 | 0.487 | 0.15 | 0.58 | 0.26 | 0.28 | 0.12 | 0.14 |
| 2 | 29999161 | rs10495771 | 0.757 | -0.981 | 1.835 | 0.86 | 0 | -0.003 | -0.003 | 0 | 0.94 | 0.67 | 0.95 | 0.98 | 0.99 | 0.94 |
| 2 | 29999860 | rs9973323 | -0.619 | -0.919 | 1.919 | 0.95 | 0.001 | 0.016 | 0.016 | 0 | 0.07 | 0.11 | 0.03 | 0.11 | 0.06 | 0.11 |
| 2 | 30000281 | rs9973358 | -0.683 | -0.916 | 1.94 | 0.963 | -0.024 | -0.021 | -0.021 | 0 | 0.06 | 0.16 | 0.02 | 0.02 | 0.01 | 0.06 |
| 2 | 30002326 | rs7606108 | -1.265 | -0.769 | 1.678 | 1.255 | -0.007 | -0.022 | -0.022 | 0 | 0.18 | 0.16 | 0.31 | 0.39 | 0.19 | 0.14 |
| 2 | 30002690 | rs12466048 | 2.157 | -0.734 | 1.646 | 1.272 | -0.001 | -0.102 | -0.102 | 0 | 0.7 | 0.11 | 0.6 | 0.52 | 0.76 | 0.69 |
| 2 | 30010894 | rs6709517 | -0.947 | -0.73 | 2.195 | 1.036 | -0.017 | -0.015 | -0.015 | 0.034 | 0.06 | 0.21 | 0.02 | 0.02 | 0.01 | 0.06 |
| 2 | 30012952 | rs7571379 | 1.965 | -0.76 | 1.435 | 1.032 | -0.017 | -0.009 | -0.009 | 0 | 0.85 | 0.24 | 0.9 | 0.91 | 0.95 | 0.83 |
| 2 | 30016786 | rs4666275 | 1.749 | -0.729 | 1.449 | 1.036 | -0.018 | -0.015 | -0.015 | 0.131 | 0.85 | 0.29 | 0.89 | 0.9 | 0.92 | 0.82 |
| 2 | 30023373 | rs4395214 | 1.296 | -0.039 | 1.402 | 1.113 | 0.018 | 0.019 | 0.019 | 0.488 | 0.76 | 0.63 | 0.59 | 0.64 | 0.7 | 0.86 |
| 2 | 30026965 | rs6744445 | 1.174 | 0.307 | 1.523 | 1.393 | 0.066 | 0.108 | 0.108 | 0 | 0.8 | 0.63 | 0.61 | 0.61 | 0.72 | 0.76 |
| 2 | 30028900 | rs13030354 | -0.816 | -0.322 | 1.311 | 1.307 | -0.032 | -0.009 | -0.009 | 0 | 0.21 | 0.47 | 0.11 | 0.2 | 0.18 | 0.23 |
| 2 | 30029139 | rs12714291 | -0.179 | -0.169 | 0.994 | 1.433 | 0.021 | 0.064 | 0.064 | 0 | 0.89 | 0.86 | 0.75 | 0.69 | 0.86 | 0.84 |
| 2 | 30029875 | rs12105526 | 0.603 | -0.174 | 0.667 | 1.29 | -0.004 | 0.002 | 0.002 | 0 | 0.06 | 0.04 | 0.09 | 0.04 | 0.11 | 0.06 |
| 2 | 30035602 | rs11690312 | 1.749 | -0.31 | 0.839 | 1.311 | -0.027 | -0.024 | -0.024 | 0 | 0.61 | 0.24 | 0.54 | 0.41 | 0.57 | 0.56 |
| 2 | 30041065 | rs12997218 | -0.566 | -0.173 | 0.557 | 1.196 | -0.008 | 0.024 | 0.024 | 0 | 0.17 | 0.03 | 0.06 | 0.09 | 0.18 | 0.21 |
| 2 | 30041666 | rs4605324 | 1.019 | -0.682 | 0.595 | 0.957 | 0 | 0 | 0 | 0 | 0.71 | 0.73 | 0.56 | 0.57 | 0.74 | 0.56 |
| 2 | 30042217 | rs10187657 | 1.072 | -0.854 | 0.936 | 0.642 | 0.005 | 0.001 | 0.001 | 0 | 0.57 | 0.52 | 0.52 | 0.57 | 0.52 | 0.54 |
| 2 | 30043619 | rs4073186 | -0.254 | -0.84 | 0.209 | 0.555 | 0.005 | -0.014 | -0.014 | 0 | 0.37 | 0.4 | 0.26 | 0.3 | 0.4 | 0.32 |
| 2 | 30045059 | rs4629120 | 1.791 | -0.507 | -0.193 | 0.6 | 0.004 | -0.019 | -0.019 | 0 | 0.33 | 0.24 | 0.28 | 0.38 | 0.34 | 0.31 |
| 2 | 30049010 | rs4265970 | 0.897 | -0.391 | -0.182 | 0.073 | -0.019 | 0.053 | 0.053 | 0 | 0.44 | 0.31 | 0.61 | 0.7 | 0.47 | 0.56 |
| 2 | 30053342 | rs12623676 | 2.17 | -0.389 | -0.395 | 0.39 | -0.036 | -0.109 | -0.109 | 0 | 0.29 | 0.08 | 0.37 | 0.47 | 0.33 | 0.3 |
| 2 | 30057570 | rs6747958 | 0.614 | -0.405 | -0.328 | 0.255 | -0.021 | -0.027 | -0.027 | 0.2 | 0.14 | 0.04 | 0.23 | 0.15 | 0.14 | 0.24 |
| 2 | 30065435 | rs12479392 | 0.045 | -0.311 | 0.04 | 0.057 | -0.012 | -0.048 | -0.048 | 0 | 0.78 | 0.62 | 0.84 | 0.87 | 0.81 | 0.82 |
| 2 | 30066829 | rs4666281 | -1.086 | -0.362 | 0.011 | 0.005 | -0.021 | -0.047 | -0.047 | 0.015 | 0.88 | 0.83 | 0.92 | 1 | 0.93 | 0.93 |
| 2 | 30068116 | rs13012046 | 0.382 | -0.356 | -0.151 | -0.011 | -0.005 | 0.012 | 0.012 | 0 | 0.3 | 0.25 | 0.22 | 0.19 | 0.34 | 0.26 |
| 2 | 30070044 | rs4461225 | -0.35 | -0.381 | -0.098 | 0.029 | -0.011 | 0.022 | 0.022 | 0 | 0.45 | 0.33 | 0.44 | 0.33 | 0.47 | 0.49 |
| 2 | 30071411 | rs4074252 | -0.502 | -0.353 | -0.155 | 0.059 | -0.005 | 0.02 | 0.02 | 0 | 0.47 | 0.37 | 0.45 | 0.34 | 0.49 | 0.51 |
| 2 | 30072993 | rs4665483 | -0.962 | -0.367 | -0.621 | 0.071 | -0.014 | -0.035 | -0.035 | 0 | 0.88 | 0.85 | 0.92 | 1 | 0.93 | 0.93 |
| 2 | 30074066 | rs7575137 | 1.626 | -0.083 | -1.218 | 0.339 | -0.012 | -0.081 | -0.081 | 0.327 | 0.3 | 0.1 | 0.37 | 0.53 | 0.31 | 0.32 |
| 2 | 30074153 | rs7567152 | 1.918 | -0.096 | -1.217 | 0.36 | -0.015 | -0.087 | -0.087 | 0.467 | 0.28 | 0.09 | 0.36 | 0.53 | 0.29 | 0.31 |
| 2 | 30075237 | rs4075919 | 0.09 | -0.043 | -1.254 | -0.331 | 0.001 | 0.09 | 0.09 | 0 | 0.59 | 0.25 | 0.7 | 0.82 | 0.6 | 0.75 |
| 2 | 30075404 | rs6707371 | 0.463 | -0.141 | -1.474 | -0.228 | 0.009 | 0.19 | 0.19 | 0 | 0.49 | 0.15 | 0.62 | 0.77 | 0.45 | 0.62 |
| 2 | 30078957 | rs4581855 | 0.46 | -0.31 | -0.833 | -0.037 | -0.024 | 0.007 | 0.007 | 0 | 0.7 | 0.58 | 0.68 | 0.66 | 0.77 | 0.55 |
| 2 | 30081261 | rs13035485 | -0.267 | -0.271 | -0.82 | -0.007 | 0.001 | -0.02 | -0.02 | 0 | 0.44 | 0.57 | 0.34 | 0.34 | 0.52 | 0.39 |
| 2 | 30081970 | rs7568708 | -0.174 | -0.306 | -0.804 | -0.004 | -0.018 | -0.012 | -0.012 | 0.242 | 0.74 | 0.68 | 0.71 | 0.83 | 0.8 | 0.7 |
| 2 | 30082600 | rs4597472 | -0.211 | -0.273 | -0.744 | 0.017 | -0.011 | -0.007 | -0.007 | 0.578 | 0.74 | 0.7 | 0.71 | 0.83 | 0.8 | 0.66 |
| 2 | 30086291 | rs7564775 | -0.226 | -0.26 | -0.742 | 0.023 | -0.011 | -0.006 | -0.006 | 0.019 | 0.75 | 0.71 | 0.71 | 0.83 | 0.81 | 0.7 |
| 2 | 30087465 | rs12613938 | 2.093 | -0.237 | -0.84 | 0.018 | -0.003 | -0.108 | -0.108 | 0 | 0.28 | 0.07 | 0.37 | 0.47 | 0.28 | 0.27 |
| 2 | 30090341 | rs12620548 | 1.862 | -0.269 | -0.736 | 0.023 | 0.008 | -0.044 | -0.044 | 0 | 0.3 | 0.15 | 0.39 | 0.57 | 0.3 | 0.32 |
| 2 | 30091595 | rs3936219 | 0.078 | -0.119 | -0.55 | 0.095 | -0.001 | -0.028 | -0.028 | 0.157 | 0.4 | 0.42 | 0.3 | 0.09 | 0.49 | 0.23 |
| 2 | 30094209 | rs7574859 | 1.277 | 0.076 | -0.53 | -0.033 | -0.02 | 0.019 | 0.019 | 0 | 0.13 | 0.08 | 0.13 | 0.01 | 0.17 | 0.14 |
| 2 | 30095846 | rs4665485 | 2.677 | 0.162 | -0.71 | -0.126 | 0.004 | -0.058 | -0.058 | 0 | 0.21 | 0.08 | 0.32 | 0.47 | 0.2 | 0.21 |
| 2 | 30097479 | rs13020484 | 0.2 | 0.179 | -0.691 | -0.372 | 0.007 | -0.06 | -0.06 | 0 | 0.4 | 0.44 | 0.3 | 0.09 | 0.49 | 0.24 |
| 2 | 30100133 | rs11127243 | -0.213 | -0.034 | -0.693 | -0.308 | -0.005 | -0.006 | -0.006 | 0 | 0.7 | 0.74 | 0.69 | 0.81 | 0.75 | 0.65 |
| 2 | 30102650 | rs4233750 | -0.266 | 0.034 | -0.615 | -0.251 | -0.006 | -0.008 | -0.008 | 0 | 0.7 | 0.7 | 0.69 | 0.8 | 0.75 | 0.62 |
| 2 | 30110249 | rs12993382 | 1.401 | 0.125 | -0.57 | -0.318 | -0.002 | -0.009 | -0.009 | 0 | 0.27 | 0.25 | 0.17 | 0.06 | 0.32 | 0.11 |
| 2 | 30111840 | rs4545964 | 2.002 | 0.032 | -0.53 | -0.291 | -0.003 | 0.011 | 0.011 | 0 | 0.24 | 0.25 | 0.35 | 0.58 | 0.23 | 0.23 |
| 2 | 30111881 | rs4567890 | -0.218 | 0.093 | -0.554 | -0.341 | 0.003 | 0.063 | 0.063 | 0 | 0.47 | 0.37 | 0.33 | 0.22 | 0.52 | 0.39 |

| | | | | | | | | | | | | | | | | |
|---|---|---|---|---|---|---|---|---|---|---|---|---|---|---|---|---|
| 2 | 30117102 | rs6547982 | 0.231 | -0.152 | 0.149 | 0.053 | -0.009 | -0.019 | -0.019 | 9.371 | 0.67 | 0.6 | 0.66 | 0.79 | 0.71 | 0.61 |
| 2 | 30117954 | rs5018732 | 0.167 | -0.129 | 0.043 | 0.056 | -0.026 | 0.02 | 0.02 | 0 | 0.71 | 0.51 | 0.7 | 0.7 | 0.77 | 0.62 |
| 2 | 30118211 | rs5018731 | 1.561 | -0.253 | -0.108 | 0.268 | -0.007 | -0.019 | -0.019 | 0.975 | 0.47 | 0.24 | 0.47 | 0.48 | 0.54 | 0.34 |
| 2 | 30118386 | rs4520995 | -0.06 | -0.229 | -0.073 | 0.216 | -0.001 | -0.001 | -0.001 | 0 | 0.69 | 0.75 | 0.68 | 0.79 | 0.71 | 0.62 |
| 2 | 30123810 | rs4450565 | -0.068 | -0.185 | -0.125 | 0.247 | 0.001 | 0.001 | 0.001 | 0 | 0.69 | 0.74 | 0.67 | 0.79 | 0.72 | 0.61 |
| 2 | 30125593 | rs4666289 | 0.487 | -0.268 | -0.094 | 0.256 | 0.002 | 0.012 | 0.012 | 0 | 0.63 | 0.7 | 0.59 | 0.78 | 0.65 | 0.57 |
| 2 | 30125934 | rs4233751 | 0.788 | -0.235 | -0.269 | 0.264 | -0.006 | -0.022 | -0.022 | 2.972 | 0.61 | 0.51 | 0.57 | 0.78 | 0.64 | 0.57 |
| 2 | 30132986 | rs7589120 | 0.511 | -0.239 | -0.314 | 0.529 | -0.016 | -0.003 | -0.003 | 0 | 0.42 | 0.25 | 0.47 | 0.57 | 0.43 | 0.46 |
| 2 | 30133304 | rs13031061 | 1.358 | -0.305 | -1.015 | 0.385 | -0.081 | -0.057 | -0.057 | 0 | 0.17 | 0.01 | 0.27 | 0.17 | 0.21 | 0.2 |
| 2 | 30134463 | rs11688109 | 0.588 | -0.932 | -0.555 | 0.569 | -0.113 | 0.021 | 0.021 | 0.07 | 0.38 | 0.06 | 0.39 | 0.24 | 0.43 | 0.28 |
| 2 | 30135236 | rs7605273 | -0.273 | -1.012 | 0.538 | 0.673 | -0.031 | 0.01 | 0.01 | 0 | 0.09 | 0.13 | 0.04 | 0.18 | 0.02 | 0.14 |
| 2 | 30136530 | rs4349303 | -0.164 | -0.56 | 1.077 | 0.593 | 0.001 | -0.055 | -0.055 | 0 | 0.9 | 0.79 | 0.94 | 1 | 0.88 | 0.89 |
| 2 | 30148034 | rs12620211 | 0.327 | -0.564 | 1.44 | 1.099 | -0.012 | 0.129 | 0.129 | 0 | 0.06 | 0.17 | 0.08 | 0.25 | 0.04 | 0.13 |
| 2 | 30149187 | rs13414785 | -0.095 | -0.582 | 1.733 | 0.996 | 0.003 | 0.224 | 0.224 | 0 | 0.06 | 0.31 | 0.09 | 0.27 | 0.04 | 0.14 |
| 2 | 30151617 | rs6547983 | -0.273 | -0.381 | 1.952 | 0.355 | 0 | 0 | 0 | 0 | 0.43 | 0.58 | 0.46 | 0.62 | 0.38 | 0.67 |
| 2 | 30151753 | rs4600601 | -0.314 | -0.379 | 1.959 | 0.356 | 0.003 | 0.001 | 0.001 | 0 | 0.43 | 0.59 | 0.46 | 0.62 | 0.38 | 0.67 |
| 2 | 30156089 | rs10469979 | -0.262 | -0.371 | 1.896 | 0.362 | -0.008 | -0.003 | -0.003 | 0 | 0.43 | 0.48 | 0.45 | 0.62 | 0.38 | 0.67 |
| 2 | 30157386 | rs4274563 | -0.278 | -0.366 | 1.873 | 0.364 | -0.001 | 0 | 0 | 0 | 0.43 | 0.57 | 0.45 | 0.62 | 0.38 | 0.67 |
| 2 | 30157544 | rs10201541 | -0.266 | -0.509 | 1.856 | 0.344 | -0.035 | 0.008 | 0.008 | 0 | 0.09 | 0.29 | 0.06 | 0.08 | 0.02 | 0.14 |
| 2 | 30158494 | rs13412653 | 0.294 | -0.492 | 1.284 | 0.352 | -0.001 | -0.014 | -0.014 | 0.203 | 0.34 | 0.31 | 0.39 | 0.54 | 0.35 | 0.53 |
| 2 | 30160391 | rs10175057 | -0.544 | -0.478 | 1.283 | 0.31 | 0.024 | 0.007 | 0.007 | 0.138 | 0.44 | 0.77 | 0.48 | 0.62 | 0.38 | 0.67 |
| 2 | 30165935 | rs10174538 | -0.512 | -0.488 | 1.03 | 0.299 | -0.024 | 0.032 | 0.032 | 0.572 | 0.1 | 0.47 | 0.09 | 0.08 | 0.02 | 0.14 |
| 2 | 30169886 | rs10187936 | -0.414 | -0.068 | -0.106 | 0.413 | -0.014 | -0.065 | -0.065 | 0 | 0.67 | 0.83 | 0.61 | 0.49 | 0.64 | 0.47 |
| 2 | 30176084 | rs13412672 | 0.383 | -0.157 | 0.21 | 0.254 | -0.033 | -0.045 | -0.045 | 0.011 | 0.09 | 0.15 | 0.02 | 0.01 | 0.02 | 0.09 |
| 2 | 30180657 | rs4510267 | 0.246 | -0.363 | 0.059 | 0.238 | -0.031 | -0.096 | -0.096 | 0.006 | 0.34 | 0.1 | 0.38 | 0.51 | 0.36 | 0.53 |
| 2 | 30184503 | rs13029270 | 0.339 | -0.489 | 0.821 | 0.302 | 0.028 | 0.005 | 0.005 | 0.494 | 0.53 | 0.39 | 0.53 | 0.41 | 0.63 | 0.33 |
| 2 | 30185186 | rs4365502 | 0.143 | -0.506 | 0.855 | 0.283 | 0.019 | -0.014 | -0.014 | 0 | 0.56 | 0.71 | 0.6 | 0.49 | 0.63 | 0.38 |
| 2 | 30185394 | rs6718327 | 0.35 | -0.542 | 1.03 | 0.245 | 0.012 | 0.004 | 0.004 | 0.047 | 0.53 | 0.47 | 0.54 | 0.41 | 0.6 | 0.3 |
| 2 | 30187923 | rs11127248 | 0.004 | -0.066 | 0.903 | 0.542 | -0.036 | -0.086 | -0.086 | 0.128 | 0.36 | 0.11 | 0.39 | 0.48 | 0.38 | 0.56 |
| 2 | 30196778 | rs7590877 | -0.26 | -0.104 | 1.127 | 0.515 | 0.003 | 0.009 | 0.009 | 0 | 0.31 | 0.62 | 0.41 | 0.31 | 0.33 | 0.23 |
| 2 | 30200798 | rs4359705 | -0.362 | -0.107 | 1.23 | 0.528 | 0.001 | 0.011 | 0.011 | 0 | 0.34 | 0.57 | 0.42 | 0.31 | 0.34 | 0.23 |
| 2 | 30210861 | rs4952125 | -0.462 | -0.035 | 0.489 | 0.51 | 0 | -0.013 | -0.013 | 0 | 0.36 | 0.7 | 0.46 | 0.41 | 0.37 | 0.31 |
| 2 | 30210991 | rs11691638 | 0.182 | 0.111 | 0.257 | 0.437 | -0.01 | -0.005 | -0.005 | 0 | 0.25 | 0.16 | 0.33 | 0.2 | 0.29 | 0.18 |
| 2 | 30214473 | rs11127252 | 0.262 | 0.351 | 0.093 | 0.695 | -0.007 | -0.086 | -0.086 | 0 | 0.25 | 0.1 | 0.34 | 0.51 | 0.22 | 0.59 |
| 2 | 30228640 | rs4952167 | -0.775 | 0.639 | -0.038 | 0.582 | -0.009 | 0.089 | 0.089 | 0 | 0.59 | 0.74 | 0.69 | 0.82 | 0.56 | 0.83 |
| 2 | 30237620 | rs7575487 | 0.292 | 0.548 | -0.082 | 0.638 | 0.002 | -0.016 | -0.016 | 0 | 0.27 | 0.23 | 0.35 | 0.51 | 0.23 | 0.59 |
| 2 | 30240720 | rs2361743 | -0.014 | 0.717 | -0.177 | 0.294 | -0.003 | 0.059 | 0.059 | 0 | 0.6 | 0.72 | 0.7 | 0.82 | 0.58 | 0.83 |
| 2 | 30256303 | rs11127254 | 1.172 | 0.625 | 0.499 | 1.56 | 0.051 | 0.307 | 0.307 | 0.071 | 0.94 | 0.68 | 0.85 | 0.7 | 0.97 | 0.76 |
| 2 | 30273535 | rs2290052 | -1.119 | 0.788 | -0.135 | 1.211 | -0.008 | -0.017 | -0.017 | 0 | 0.1 | 0.07 | 0.23 | 0.19 | 0.1 | 0.15 |
| 2 | 30273791 | rs12475473 | -1.119 | 0.788 | 0.128 | 1.211 | 0 | 0.002 | 0.002 | 0 | 0.1 | 0.1 | 0.24 | 0.19 | 0.1 | 0.15 |
| 2 | 30275291 | rs10203090 | -0.948 | 0.579 | -0.896 | 0.328 | -0.02 | 0.117 | 0.117 | 0.655 | 0.46 | 0.89 | 0.5 | 0.31 | 0.46 | 0.56 |
| 2 | 30279480 | rs1562450 | -1.257 | 0.378 | -0.842 | 0.273 | 0.016 | 0.135 | 0.135 | 0.002 | 0.5 | 0.87 | 0.5 | 0.32 | 0.46 | 0.55 |
| 2 | 30282981 | rs1447524 | 0.437 | 0.314 | -0.566 | 0.078 | 0.001 | 0.002 | 0.002 | 0 | 0.1 | 0.08 | 0.04 | 0.08 | 0.1 | 0.09 |
| 2 | 30292478 | rs1447522 | 0.379 | 0.212 | 0.342 | 0.007 | 0.001 | 0.019 | 0.019 | 0 | 0.16 | 0.11 | 0.12 | 0.04 | 0.18 | 0.19 |
| 2 | 30292775 | rs1900589 | -0.421 | 0.228 | 0.685 | 0.099 | 0 | 0.006 | 0.006 | 0 | 0.11 | 0.15 | 0.25 | 0.2 | 0.1 | 0.15 |
| 2 | 30300036 | rs2028983 | -0.527 | -0.706 | 0.096 | 0.199 | 0.032 | -0.083 | -0.083 | 0.049 | 0.24 | 0.07 | 0.25 | 0.55 | 0.18 | 0.32 |
| 2 | 30307179 | rs4952091 | -0.294 | -0.484 | 0.084 | -0.553 | 0.002 | 0.049 | 0.049 | 0.12 | 0.73 | 0.87 | 0.81 | 0.96 | 0.74 | 0.79 |
| 2 | 30310799 | rs2078044 | 0.231 | -0.655 | 0.561 | -0.23 | -0.041 | 0.18 | 0.18 | 6.422 | 0.09 | 0.14 | 0.17 | 0.55 | 0.02 | 0.27 |
| 2 | 30321444 | rs11903016 | 1.869 | -0.092 | 0.409 | -0.635 | -0.041 | 0.064 | 0.064 | 0 | 0.32 | 0.16 | 0.25 | 0.17 | 0.43 | 0.24 |
| 2 | 30325461 | rs10179096 | -0.732 | -0.153 | 0.252 | -0.695 | -0.015 | 0.122 | 0.122 | 0.291 | 0.74 | 0.95 | 0.82 | 0.96 | 0.74 | 0.8 |
| 2 | 30332590 | rs6547992 | -0.006 | -0.097 | 0.579 | -0.738 | 0 | 0.184 | 0.184 | 4.932 | 0.58 | 0.85 | 0.77 | 0.95 | 0.6 | 0.66 |
| 2 | 30338534 | rs4952098 | 0.835 | -0.077 | 0.526 | -0.694 | -0.049 | -0.07 | -0.07 | 0 | 0.11 | 0.01 | 0.3 | 0.22 | 0.13 | 0.15 |
| 2 | 30341725 | rs935486 | -0.313 | -0.126 | 0.722 | -0.669 | -0.03 | 0.121 | 0.121 | 9.216 | 0.13 | 0.16 | 0.18 | 0.55 | 0.04 | 0.27 |
| 2 | 30342750 | rs4952099 | -0.006 | -0.108 | 0.755 | -0.707 | 0.004 | 0.136 | 0.136 | 4.849 | 0.58 | 0.78 | 0.77 | 0.95 | 0.6 | 0.66 |
| 2 | 30346287 | rs4286323 | 0.002 | -0.109 | 0.754 | -0.708 | 0.002 | 0.137 | 0.137 | 3.664 | 0.58 | 0.78 | 0.77 | 0.95 | 0.6 | 0.66 |
| 2 | 30359532 | rs6547994 | 0.343 | -0.1 | 0.874 | -0.577 | -0.034 | 0.246 | 0.246 | 5.698 | 0.1 | 0.2 | 0.17 | 0.54 | 0.02 | 0.26 |
| 2 | 30367224 | rs1509578 | 0.135 | -0.079 | 0.782 | -0.624 | 0.005 | 0.129 | 0.129 | 4.879 | 0.56 | 0.76 | 0.76 | 0.95 | 0.58 | 0.65 |
| 2 | 30372327 | rs3768667 | 1.954 | -0.007 | 0.477 | -0.63 | -0.072 | 0.085 | 0.085 | 0 | 0.32 | 0.1 | 0.25 | 0.18 | 0.43 | 0.23 |
| 2 | 30375186 | rs1345151 | -0.576 | -0.065 | 0.436 | -0.659 | -0.018 | 0.021 | 0.021 | 0 | 0.3 | 0.33 | 0.26 | 0.56 | 0.19 | 0.42 |
| 2 | 30380923 | rs7600457 | 0.94 | -0.06 | 0.393 | -0.59 | -0.004 | 0.039 | 0.039 | 4.058 | 0.42 | 0.28 | 0.42 | 0.74 | 0.45 | 0.51 |
| 2 | 30381505 | rs1137288 | 0.921 | -0.06 | 0.393 | -0.589 | -0.004 | 0.04 | 0.04 | 3.888 | 0.43 | 0.28 | 0.41 | 0.74 | 0.45 | 0.51 |
| 2 | 30381901 | rs4951983 | 0.644 | 0.049 | -0.087 | -0.412 | -0.047 | -0.069 | -0.069 | 1.377 | 0.11 | 0.01 | 0.31 | 0.22 | 0.13 | 0.15 |
| 2 | 30385901 | rs7567202 | -0.106 | 0.033 | 0.067 | -0.419 | 0.044 | 0.123 | 0.123 | 0 | 0.14 | 0.46 | 0.34 | 0.22 | 0.13 | 0.15 |
| 2 | 30386174 | rs6547997 | 0.727 | -0.149 | -0.086 | -0.474 | 0 | 0 | 0 | 0.366 | 0.46 | 0.57 | 0.58 | 0.39 | 0.56 | 0.38 |
| 2 | 30403762 | rs1868073 | 0.312 | 0.472 | -1.664 | -0.788 | 0.01 | 0.099 | 0.099 | 3.986 | 0.54 | 0.71 | 0.75 | 0.93 | 0.59 | 0.6 |
| 2 | 30406902 | rs4264597 | 1.142 | 0.883 | -1.937 | -0.778 | 0.011 | 0.024 | 0.024 | 0 | 0.39 | 0.71 | 0.43 | 0.71 | 0.4 | 0.47 |
| 2 | 30407566 | rs1355207 | -0.277 | 0.994 | -1.756 | -0.422 | 0.002 | 0.029 | 0.029 | 0.037 | 0.74 | 0.82 | 0.83 | 0.93 | 0.77 | 0.78 |
| 2 | 30411085 | rs966498 | -0.156 | 1.181 | -1.703 | 0.247 | -0.065 | -0.066 | -0.066 | 0 | 0.15 | 0.01 | 0.24 | 0.18 | 0.16 | 0.14 |
| 2 | 30418294 | rs12714299 | -0.682 | 0.752 | -2.297 | -0.161 | -0.044 | 0.059 | 0.059 | 0.013 | 0.44 | 0.88 | 0.44 | 0.71 | 0.38 | 0.52 |

| | | | | | | | | | | | | | | | | |
|---|---|---|---|---|---|---|---|---|---|---|---|---|---|---|---|---|
| 2 | 30421533 | rs17009207 | -0.513 | 0.909 | -0.974 | 0.108 | 0.029 | 0.028 | 0.028 | 0 | 0.28 | 0.35 | 0.35 | 0.63 | 0.31 | 0.41 |
| 2 | 30423512 | rs4334534 | -0.466 | 0.908 | -0.922 | 0.108 | 0.027 | 0.025 | 0.025 | 0 | 0.28 | 0.32 | 0.35 | 0.63 | 0.31 | 0.41 |
| 2 | 30426171 | rs4952107 | 0.417 | 0.262 | -1.224 | -0.682 | 0.02 | 0.075 | 0.075 | 0.144 | 0.69 | 0.86 | 0.86 | 0.92 | 0.76 | 0.82 |
| 2 | 30426194 | rs10205388 | 0.526 | -0.097 | -0.36 | -0.903 | -0.027 | -0.047 | -0.047 | 1.333 | 0.09 | 0.29 | 0.04 | 0 | 0.02 | 0.05 |
| 2 | 30426437 | rs11681599 | -0.49 | -0.189 | -0.791 | -1.002 | -0.001 | 0 | 0 | 0 | 0.89 | 0.92 | 0.92 | 1 | 0.91 | 0.94 |
| 2 | 30426554 | rs17736058 | -0.426 | -0.198 | -0.635 | -1.269 | -0.002 | -0.005 | -0.005 | 0 | 0.17 | 0.12 | 0.11 | 0 | 0.14 | 0.09 |
| 2 | 30426674 | rs11693328 | 2.062 | -0.023 | -0.835 | -1.211 | -0.033 | 0.076 | 0.076 | 0 | 0.36 | 0.14 | 0.5 | 0.24 | 0.46 | 0.34 |
| 2 | 30429330 | rs6707974 | -0.39 | -0.047 | -0.553 | -0.948 | -0.051 | -0.06 | -0.06 | 0 | 0.27 | 0.45 | 0.23 | 0.13 | 0.16 | 0.19 |
| 2 | 30429735 | rs4952108 | 0.608 | 0.157 | -1.025 | -0.89 | -0.052 | -0.094 | -0.094 | 0.361 | 0.15 | 0.05 | 0.22 | 0.63 | 0.19 | 0.36 |
| 2 | 30430416 | rs13032691 | 0.444 | 0.146 | -0.847 | -1.091 | 0.007 | 0.006 | 0.006 | 0.097 | 0.18 | 0.14 | 0.17 | 0.13 | 0.14 | 0.14 |
| 2 | 30436249 | rs6718106 | -0.675 | 0.437 | -0.72 | -0.739 | 0 | 0.003 | 0.003 | 0 | 0.46 | 0.58 | 0.42 | 0.65 | 0.46 | 0.62 |
| 2 | 30436467 | rs11887956 | 0.788 | 0.459 | -0.844 | -0.638 | -0.061 | -0.129 | -0.129 | 0.008 | 0.17 | 0.01 | 0.25 | 0.54 | 0.23 | 0.42 |
| 2 | 30441004 | rs7568254 | 1.296 | 0.743 | -0.681 | -0.508 | -0.024 | 0.028 | 0.028 | 0 | 0.39 | 0.11 | 0.49 | 0.24 | 0.44 | 0.35 |
| 2 | 30442158 | rs2055782 | -0.15 | 0.833 | 0.178 | -0.014 | -0.001 | 0.112 | 0.112 | 0.468 | 0.25 | 0.45 | 0.31 | 0.54 | 0.25 | 0.46 |
| 2 | 30445026 | rs7579944 | 0.753 | 0.881 | -0.205 | -0.001 | -0.018 | -0.029 | -0.029 | 0.003 | 0.64 | 0.23 | 0.62 | 0.36 | 0.64 | 0.5 |
| 2 | 30446083 | rs1396836 | 0.192 | 0.867 | -0.104 | -0.2 | -0.009 | 0.088 | 0.088 | 0.085 | 0.74 | 0.5 | 0.69 | 0.46 | 0.75 | 0.54 |
| 2 | 30452959 | rs2362540 | 0.51 | 0.493 | -0.094 | 0.037 | 0.001 | 0 | 0 | 0 | 0.57 | 0.48 | 0.63 | 0.46 | 0.64 | 0.48 |
| 2 | 30456082 | rs6548003 | -0.597 | 0.925 | -1.058 | 0.325 | -0.002 | -0.001 | -0.001 | 0 | 0.37 | 0.39 | 0.43 | 0.53 | 0.37 | 0.5 |
| 2 | 30462920 | rs2102472 | -0.551 | 0.588 | -1.554 | -0.195 | 0.01 | 0.038 | 0.038 | 0 | 0.28 | 0.17 | 0.18 | 0.04 | 0.28 | 0.22 |
| 2 | 30462979 | rs1473591 | -0.052 | 0.894 | -2.248 | 0.178 | -0.062 | -0.132 | -0.132 | 0.319 | 0.23 | 0.03 | 0.36 | 0.55 | 0.28 | 0.42 |
| 2 | 30470178 | rs2089657 | -0.507 | 1.743 | -2.17 | 0.408 | -0.002 | -0.001 | -0.001 | 1.806 | 0.08 | 0.1 | 0.3 | 0.16 | 0.16 | 0.03 |
| 2 | 30473522 | rs3813658 | -0.426 | 1.023 | -2.23 | -0.93 | -0.014 | 0.089 | 0.089 | 0 | 0.54 | 0.37 | 0.63 | 0.16 | 0.52 | 0.3 |
| 2 | 30478386 | rs4952115 | -0.232 | 1.084 | -2.248 | -0.932 | 0.028 | 0.028 | 0.028 | 0.404 | 0.09 | 0.17 | 0.31 | 0.16 | 0.16 | 0.03 |
| 2 | 30478453 | rs2362541 | -0.537 | 0.97 | -2.279 | -1.003 | -0.006 | 0.035 | 0.035 | 0.004 | 0.55 | 0.44 | 0.63 | 0.17 | 0.51 | 0.32 |
| 2 | 30478628 | rs906867 | -0.616 | 1.093 | -2.243 | -0.925 | 0.066 | 0.067 | 0.067 | 1.368 | 0.11 | 0.3 | 0.33 | 0.16 | 0.17 | 0.03 |
| 2 | 30479992 | rs11679132 | 0.654 | 0.224 | -2.717 | -1.392 | -0.009 | 0.055 | 0.055 | 0 | 0.3 | 0.19 | 0.3 | 0.01 | 0.32 | 0.21 |
| 2 | 30480027 | rs11679137 | 0.772 | 0.228 | -2.728 | -1.39 | -0.062 | 0.179 | 0.179 | 0 | 0.29 | 0.06 | 0.28 | 0.01 | 0.32 | 0.21 |
| 2 | 30480865 | rs13386455 | -0.602 | 0.28 | -2.608 | -1.269 | 0.061 | 0.063 | 0.063 | 1.497 | 0.11 | 0.34 | 0.33 | 0.16 | 0.17 | 0.05 |
| 2 | 30481917 | rs9558 | 0.101 | 0.327 | -2.645 | -1.416 | 0.025 | -0.02 | -0.02 | 0 | 0.08 | 0.14 | 0.18 | 0.01 | 0.16 | 0.05 |
| 2 | 30483397 | rs12468820 | 1.392 | 0.359 | -3.255 | -1.004 | -0.049 | -0.095 | -0.095 | 0.084 | 0.13 | 0.01 | 0.15 | 0.37 | 0.16 | 0.31 |
| 2 | 30491556 | rs7604484 | -0.307 | 0.4 | -2.774 | -1.387 | 0.061 | -0.04 | -0.04 | 0.024 | 0.1 | 0.28 | 0.21 | 0.01 | 0.16 | 0.02 |
| 2 | 30494172 | rs17322090 | 0.557 | 0.432 | -2.752 | -1.333 | 0.031 | -0.019 | -0.019 | 0 | 0.06 | 0.14 | 0.16 | 0.01 | 0.11 | 0.02 |
| 2 | 30502177 | rs7569754 | 0.178 | 0.424 | -2.417 | -1.337 | 0.067 | -0.018 | -0.018 | 0 | 0.06 | 0.22 | 0.18 | 0.01 | 0.11 | 0.02 |
| 2 | 30504247 | rs12988733 | -0.858 | 0.186 | -2.497 | -1.645 | -0.007 | 0.05 | 0.05 | 0 | 0.48 | 0.38 | 0.49 | 0.02 | 0.49 | 0.32 |
| 2 | 30507449 | rs11127264 | -0.487 | 0.318 | -2.6 | -1.731 | -0.037 | 0.137 | 0.137 | 0 | 0.42 | 0.22 | 0.36 | 0.02 | 0.48 | 0.28 |
| 2 | 30510147 | rs17394102 | 0.831 | 0.913 | -2.322 | -1.348 | -0.016 | 0.016 | 0.016 | 0 | 0.06 | 0.04 | 0.15 | 0.01 | 0.16 | 0.03 |
| 2 | 30510444 | rs6733372 | -0.351 | 0.697 | -2.186 | -1.389 | 0.006 | -0.086 | -0.086 | 0.008 | 0.38 | 0.45 | 0.42 | 0.02 | 0.5 | 0.24 |
| 2 | 30518711 | rs10210471 | -0.879 | 0.158 | -1.593 | -0.553 | -0.021 | 0.001 | 0.001 | 1.67 | 0.63 | 0.53 | 0.66 | 0.36 | 0.71 | 0.61 |
| 2 | 30522830 | rs11678328 | 0.138 | 0.433 | -1.71 | -0.429 | -0.09 | 0.194 | 0.194 | 0 | 0.27 | 0.02 | 0.2 | 0.01 | 0.33 | 0.23 |
| 2 | 30528297 | rs6760047 | -0.274 | 0.394 | -1.126 | 0.053 | 0 | -0.005 | -0.005 | 0 | 0.37 | 0.43 | 0.42 | 0.27 | 0.43 | 0.37 |
| 2 | 30531189 | rs11683782 | -0.094 | -0.542 | -0.453 | 0.513 | -0.001 | 0.01 | 0.01 | 0 | 0.39 | 0.61 | 0.28 | 0.29 | 0.38 | 0.25 |
| 2 | 30531821 | rs6718743 | -0.215 | -0.052 | -0.229 | 0.806 | 0.005 | 0.013 | 0.013 | 0.421 | 0.38 | 0.36 | 0.47 | 0.29 | 0.34 | 0.43 |
| 2 | 30531922 | rs6758245 | -0.86 | -0.283 | -0.167 | 0.746 | -0.01 | 0.003 | 0.003 | 0 | 0.63 | 0.65 | 0.71 | 0.59 | 0.69 | 0.58 |
| 2 | 30540833 | rs13014895 | 0.085 | 0.155 | -0.951 | 0.52 | -0.003 | 0.019 | 0.019 | 0 | 0.09 | 0.15 | 0.22 | 0.2 | 0.07 | 0.29 |
| 2 | 30549428 | rs4563275 | -0.746 | 0.257 | -1.136 | 0.299 | -0.005 | -0.004 | -0.004 | 0 | 0.75 | 0.77 | 0.75 | 0.59 | 0.61 | 0.83 |
| 2 | 30551366 | rs4434038 | -0.107 | -0.048 | -1.034 | 0.085 | -0.042 | 0.097 | 0.097 | 0.233 | 0.25 | 0.06 | 0.23 | 0.07 | 0.26 | 0.14 |
| 2 | 30556564 | rs7586804 | -0.862 | -0.226 | 0.068 | 0.856 | -0.002 | 0.036 | 0.036 | 0 | 0.24 | 0.3 | 0.35 | 0.44 | 0.21 | 0.53 |
| 2 | 30557749 | rs7604316 | -0.043 | -0.066 | -0.58 | 0.722 | -0.005 | 0.02 | 0.02 | 0 | 0.83 | 0.96 | 0.81 | 0.85 | 0.83 | 0.89 |
| 2 | 30559816 | rs11680937 | 0.418 | -0.079 | -0.301 | 0.661 | 0.009 | -0.032 | -0.032 | 0.681 | 0.62 | 0.26 | 0.45 | 0.48 | 0.67 | 0.37 |
| 2 | 30565269 | rs4321410 | -1.511 | -0.264 | 0.658 | 0.376 | -0.023 | -0.027 | -0.027 | 0 | 0.09 | 0.52 | 0.07 | 0 | 0.01 | 0.02 |
| 2 | 30569156 | rs9967751 | -1.303 | -0.222 | 0.655 | 0.397 | -0.001 | -0.005 | -0.005 | 0 | 0.08 | 0.47 | 0.05 | 0 | 0.01 | 0.02 |
| 2 | 30575898 | rs6729421 | -0.962 | -0.108 | 0.183 | 0.623 | -0.006 | 0.008 | 0.008 | 0 | 0.05 | 0.09 | 0.12 | 0.09 | 0.03 | 0.06 |
| 2 | 30575970 | rs6742446 | -1.134 | -0.129 | 0.329 | 0.612 | -0.009 | 0.015 | 0.015 | 0 | 0.06 | 0.12 | 0.12 | 0.09 | 0.03 | 0.06 |
| 2 | 30577757 | rs6740257 | -1.572 | -0.107 | 0.333 | 0.574 | -0.006 | 0.004 | 0.004 | 3.388 | 0.16 | 0.24 | 0.16 | 0.22 | 0.11 | 0.07 |
| 2 | 30578421 | rs10207504 | -1.742 | -0.598 | 0.426 | 0.009 | -0.027 | -0.033 | -0.033 | 0.417 | 0.37 | 0.48 | 0.29 | 0.31 | 0.26 | 0.24 |
| 2 | 30581537 | rs13004173 | -0.76 | -0.517 | 0.422 | -0.046 | -0.029 | -0.072 | -0.072 | 0.469 | 0.23 | 0.33 | 0.14 | 0.09 | 0.16 | 0.17 |
| 2 | 30581981 | rs6753748 | -1.127 | -0.523 | -0.891 | 0.305 | -0.033 | -0.131 | -0.131 | 2.432 | 0.78 | 0.97 | 0.82 | 0.64 | 0.79 | 0.9 |
| 2 | 30588810 | rs7576009 | -0.407 | -0.644 | 0.105 | -0.044 | -0.007 | -0.039 | -0.039 | 4.515 | 0.23 | 0.22 | 0.22 | 0.1 | 0.21 | 0.2 |
| 2 | 30594615 | rs7578449 | -0.917 | -0.727 | 0.503 | 0.055 | -0.023 | -0.056 | -0.056 | 0.012 | 0.27 | 0.39 | 0.25 | 0.14 | 0.23 | 0.21 |
| 2 | 30598654 | rs4572642 | 1.017 | -0.163 | -0.239 | -0.121 | -0.01 | -0.011 | -0.011 | 0 | 0.18 | 0.11 | 0.16 | 0.26 | 0.21 | 0.36 |
| 2 | 30599878 | rs6719566 | 0.935 | -0.179 | -0.174 | -0.129 | -0.005 | -0.006 | -0.006 | 0.878 | 0.18 | 0.16 | 0.16 | 0.26 | 0.21 | 0.36 |
| 2 | 30600320 | rs11885821 | 0.574 | -0.18 | -0.044 | -0.119 | 0 | 0 | 0 | 0 | 0.21 | 0.24 | 0.2 | 0.29 | 0.23 | 0.38 |
| 2 | 30606663 | rs13033921 | -1.131 | -0.284 | 0.254 | -0.077 | -0.001 | 0.007 | 0.007 | 0 | 0.51 | 0.5 | 0.53 | 0.44 | 0.49 | 0.58 |
| 2 | 30606890 | rs4637169 | 0.362 | -0.31 | 0.37 | -0.093 | -0.002 | -0.003 | -0.003 | 0 | 0.22 | 0.23 | 0.21 | 0.29 | 0.25 | 0.4 |
| 2 | 30607362 | rs6548014 | 0.714 | -0.308 | 0.36 | -0.091 | -0.006 | -0.013 | -0.013 | 0 | 0.2 | 0.14 | 0.19 | 0.29 | 0.23 | 0.38 |
| 2 | 30608160 | rs7583194 | -1.39 | -0.38 | 0.659 | -0.227 | -0.005 | 0.032 | 0.032 | 0 | 0.53 | 0.6 | 0.5 | 0.31 | 0.49 | 0.58 |
| 2 | 30608352 | rs7557751 | 0.314 | -0.396 | 0.811 | -0.245 | -0.022 | -0.047 | -0.047 | 1.558 | 0.12 | 0.31 | 0.06 | 0.03 | 0.04 | 0 |
| 2 | 30608719 | rs7583844 | -1.204 | -0.404 | 0.848 | -0.251 | 0 | 0.004 | 0.004 | 0.258 | 0.49 | 0.41 | 0.46 | 0.28 | 0.47 | 0.56 |
| 2 | 30609786 | rs6725328 | 0.769 | -0.51 | 0.789 | -0.294 | -0.007 | 0.02 | 0.02 | 0 | 0.16 | 0.07 | 0.09 | 0.07 | 0.2 | 0.14 |

| | | | | | | | | | | | | | | | | |
|---|---|---|---|---|---|---|---|---|---|---|---|---|---|---|---|---|
| 2 | 30613090 | rs1357851 | 1.077 | -0.504 | 0.722 | -0.291 | -0.046 | 0.017 | 0.017 | 0 | 0.18 | 0.02 | 0.17 | 0.13 | 0.21 | 0.39 |
| 2 | 30614883 | rs10495775 | -0.696 | -0.501 | 0.801 | -0.292 | -0.014 | -0.058 | -0.058 | 0 | 0.29 | 0.37 | 0.29 | 0.15 | 0.24 | 0.17 |
| 2 | 30623062 | rs829590 | -0.99 | -0.537 | 0.757 | -0.315 | 0.002 | 0.044 | 0.044 | 0.205 | 0.46 | 0.29 | 0.37 | 0.28 | 0.46 | 0.56 |
| 2 | 30624016 | rs12471227 | 0.792 | -0.517 | 0.677 | -0.296 | -0.003 | 0.006 | 0.006 | 0 | 0.21 | 0.16 | 0.19 | 0.16 | 0.23 | 0.41 |
| 2 | 30625264 | rs11899606 | 0.631 | -0.524 | 0.749 | -0.198 | -0.002 | -0.001 | -0.001 | 0 | 0.11 | 0.07 | 0.1 | 0.08 | 0.04 | 0.03 |
| 2 | 30632216 | rs829609 | -0.888 | -0.461 | 0.576 | -0.184 | -0.019 | -0.06 | -0.06 | 0.179 | 0.31 | 0.41 | 0.28 | 0.19 | 0.25 | 0.26 |
| 2 | 30632520 | rs829608 | -0.66 | -0.382 | 0.679 | -0.041 | 0.009 | -0.026 | -0.026 | 0 | 0.2 | 0.39 | 0.26 | 0.16 | 0.21 | 0.24 |
| 2 | 30634928 | rs829605 | -0.931 | -0.431 | 0.65 | -0.134 | -0.005 | -0.052 | -0.052 | 0.152 | 0.24 | 0.47 | 0.15 | 0.16 | 0.24 | 0.19 |
| 2 | 30636263 | rs829602 | -0.28 | -0.906 | -0.247 | -0.909 | 0.027 | 0.037 | 0.037 | 0.857 | 0.65 | 0.73 | 0.85 | 0.93 | 0.77 | 0.66 |
| 2 | 30636916 | rs2076932 | 0.435 | -0.51 | -0.763 | -0.055 | -0.002 | -0.001 | -0.001 | 0 | 0.46 | 0.29 | 0.51 | 0.53 | 0.56 | 0.34 |
| 2 | 30638474 | rs829597 | 0.125 | -0.542 | -0.225 | 0.072 | -0.006 | 0.013 | 0.013 | 0.947 | 0.15 | 0.65 | 0.13 | 0.21 | 0.15 | 0.1 |
| 2 | 30643811 | rs829699 | -0.404 | -0.016 | -0.388 | -0.42 | -0.005 | 0.118 | 0.118 | 0 | 0.56 | 0.72 | 0.72 | 0.94 | 0.55 | 0.64 |
| 2 | 30649220 | rs711237 | -0.62 | -0.069 | 0.152 | -0.108 | 0.017 | 0.01 | 0.01 | 0 | 0.59 | 0.65 | 0.59 | 0.66 | 0.7 | 0.46 |
| 2 | 30652954 | rs13391574 | -0.933 | -0.154 | 0.277 | 0.059 | -0.002 | 0 | 0 | 0 | 0.43 | 0.5 | 0.55 | 0.41 | 0.51 | 0.41 |
| 2 | 30670973 | rs1662955 | 0.609 | -0.008 | -0.13 | 0.513 | 0.003 | 0.021 | 0.021 | 0 | 0.2 | 0.19 | 0.27 | 0.36 | 0.23 | 0.31 |
| 2 | 30677275 | rs829584 | 0.558 | -0.016 | -0.135 | 0.54 | 0.004 | 0.024 | 0.024 | 0 | 0.2 | 0.19 | 0.26 | 0.35 | 0.23 | 0.31 |
| 2 | 30681357 | rs17394450 | 1.424 | -0.101 | -0.179 | 0.27 | -0.026 | 0.017 | 0.017 | 0 | 0.22 | 0.17 | 0.2 | 0.01 | 0.31 | 0.1 |
| 2 | 30682630 | rs829573 | 0.613 | -0.175 | -0.051 | 0.403 | 0.004 | 0.027 | 0.027 | 0 | 0.2 | 0.19 | 0.26 | 0.37 | 0.23 | 0.31 |
| 2 | 30686039 | rs9636423 | -0.791 | -0.391 | -0.197 | 0.319 | 0.03 | 0.045 | 0.045 | 1.016 | 0.62 | 0.32 | 0.8 | 0.73 | 0.71 | 0.58 |
| 2 | 30695786 | rs1357850 | 0.641 | -0.373 | -0.257 | 0.341 | 0.028 | -0.011 | -0.011 | 0 | 0.48 | 0.23 | 0.57 | 0.5 | 0.58 | 0.39 |
| 2 | 30701722 | rs711241 | 1.789 | -0.397 | -0.314 | 0.204 | 0.008 | 0 | 0 | 0 | 0.1 | 0.1 | 0.04 | 0.14 | 0.11 | 0.08 |
| 2 | 30703971 | rs860487 | -2.032 | -0.526 | -0.269 | 0.131 | 0.034 | 0.062 | 0.062 | 0 | 0.75 | 0.81 | 0.9 | 0.94 | 0.84 | 0.71 |
| 2 | 30712016 | rs829672 | -1.008 | -0.508 | -0.153 | 0.152 | -0.044 | -0.071 | -0.071 | 0 | 0.67 | 0.45 | 0.84 | 0.86 | 0.8 | 0.68 |
| 2 | 30714832 | rs10180482 | 0.665 | -0.443 | -0.295 | 0.252 | 0.028 | -0.007 | -0.007 | 0 | 0.47 | 0.26 | 0.58 | 0.5 | 0.57 | 0.37 |
| 2 | 30716307 | rs13033111 | 1.659 | -0.466 | -0.552 | 0.234 | -0.005 | -0.043 | -0.043 | 0 | 0.11 | 0.04 | 0.22 | 0.28 | 0.12 | 0.23 |
| 2 | 30721503 | rs1880540 | 1.01 | -0.46 | -0.55 | 0.17 | -0.073 | 0.058 | 0.058 | 0.357 | 0.36 | 0.19 | 0.36 | 0.21 | 0.46 | 0.14 |
| 2 | 30735763 | rs829634 | -1.407 | -0.544 | -0.28 | -0.075 | -0.047 | -0.073 | -0.073 | 0.039 | 0.69 | 0.45 | 0.84 | 0.87 | 0.81 | 0.7 |
| 2 | 30737707 | rs17009601 | 1.937 | -0.502 | -0.161 | 0.03 | 0.004 | 0.046 | 0.046 | 0.054 | 0.11 | 0.19 | 0.23 | 0.21 | 0.12 | 0.23 |
| 2 | 30740488 | rs17038632 | 1.178 | -0.415 | -0.28 | -0.056 | -0.04 | -0.134 | -0.134 | 0.073 | 0.26 | 0.06 | 0.38 | 0.5 | 0.28 | 0.29 |
| 2 | 30767529 | rs10495776 | -1.161 | -0.51 | -0.048 | 0.025 | 0.028 | 0.008 | 0.008 | 0 | 0.64 | 0.75 | 0.69 | 0.71 | 0.72 | 0.48 |
| 2 | 30770734 | rs6712951 | -0.644 | -0.506 | 0.032 | 0.03 | 0.016 | 0.005 | 0.005 | 0 | 0.58 | 0.32 | 0.62 | 0.64 | 0.69 | 0.47 |
| 2 | 30777269 | rs7586675 | 1.376 | -0.431 | 0.121 | 0.022 | 0.029 | 0.04 | 0.04 | 0.079 | 0.05 | 0.23 | 0.05 | 0.07 | 0.04 | 0.03 |
| 2 | 30792898 | rs11676852 | -0.752 | -0.486 | 0.07 | 0.066 | -0.021 | -0.01 | -0.01 | 0 | 0.59 | 0.49 | 0.64 | 0.66 | 0.69 | 0.47 |
| 2 | 30796361 | rs7583211 | -1.98 | -0.508 | -0.076 | 0.011 | 0.033 | 0.057 | 0.057 | 0 | 0.75 | 0.86 | 0.91 | 0.94 | 0.84 | 0.71 |
| 2 | 30796668 | rs4952151 | 1.376 | -0.491 | -0.104 | 0.031 | -0.003 | -0.032 | -0.032 | 0.674 | 0.15 | 0.09 | 0.23 | 0.25 | 0.14 | 0.24 |
| 2 | 30834985 | rs1869424 | -1.948 | -0.564 | -0.113 | -0.038 | 0.036 | 0.068 | 0.068 | 0 | 0.75 | 0.89 | 0.91 | 0.94 | 0.84 | 0.71 |
| 2 | 30835910 | rs11895131 | 2.164 | -0.473 | -0.047 | 0 | 0.001 | -0.001 | -0.001 | 0 | 0.09 | 0.06 | 0.04 | 0.13 | 0.11 | 0.08 |
| 2 | 30838344 | rs4952000 | 1.993 | -0.496 | -0.152 | 0.089 | -0.002 | 0.003 | 0.003 | 0.042 | 0.1 | 0.11 | 0.22 | 0.21 | 0.12 | 0.23 |
| 2 | 30846262 | rs2593442 | -1.747 | -0.543 | -0.196 | -0.007 | 0.046 | 0.115 | 0.115 | 0 | 0.74 | 0.96 | 0.92 | 0.94 | 0.84 | 0.71 |
| 2 | 30854230 | rs1807334 | 0.539 | -0.488 | -0.126 | -0.142 | 0.005 | -0.025 | -0.025 | 0 | 0.31 | 0.28 | 0.23 | 0.13 | 0.41 | 0.18 |
| 2 | 30856738 | rs12052260 | 0.031 | -0.673 | -0.318 | -0.164 | -0.012 | 0.012 | 0.012 | 0 | 0.36 | 0.56 | 0.47 | 0.59 | 0.32 | 0.32 |
| 2 | 30857157 | rs2609917 | -1.881 | -0.676 | -0.341 | -0.182 | 0.044 | 0.1 | 0.1 | 0 | 0.75 | 0.93 | 0.92 | 0.94 | 0.84 | 0.71 |
| 2 | 30858736 | rs12613783 | 0.756 | -0.625 | -0.128 | -0.096 | 0 | -0.004 | -0.004 | 0 | 0.31 | 0.27 | 0.43 | 0.58 | 0.31 | 0.3 |
| 2 | 30860007 | rs4952002 | -0.481 | -0.62 | -0.147 | -0.17 | 0.002 | 0.012 | 0.012 | 0 | 0.44 | 0.64 | 0.69 | 0.8 | 0.43 | 0.53 |
| 2 | 30861836 | rs7565755 | -0.178 | -0.599 | -0.056 | -0.132 | 0.003 | 0.03 | 0.03 | 0.002 | 0.41 | 0.43 | 0.64 | 0.73 | 0.39 | 0.52 |
| 2 | 30863108 | rs12471868 | 1.932 | -0.557 | -0.395 | -0.031 | -0.004 | -0.005 | -0.005 | 0 | 0.1 | 0.09 | 0.22 | 0.22 | 0.12 | 0.23 |
| 2 | 30863708 | rs10200589 | 1.932 | -0.557 | -0.395 | -0.031 | -0.004 | -0.005 | -0.005 | 1.306 | 0.1 | 0.09 | 0.22 | 0.22 | 0.12 | 0.23 |
| 2 | 30864006 | rs6723023 | 1.932 | -0.558 | -0.395 | -0.031 | -0.004 | -0.005 | -0.005 | 0 | 0.1 | 0.09 | 0.22 | 0.23 | 0.12 | 0.24 |
| 2 | 30864326 | rs2609945 | -2.138 | -0.579 | -0.407 | -0.226 | 0.026 | 0.048 | 0.048 | 0 | 0.77 | 0.86 | 0.92 | 0.94 | 0.84 | 0.72 |
| 2 | 30864789 | rs1060737 | 0.459 | -0.555 | -0.552 | -0.218 | 0.035 | -0.092 | -0.092 | 0 | 0.33 | 0.5 | 0.26 | 0.13 | 0.41 | 0.18 |
| 2 | 30865048 | rs1060734 | -0.142 | -0.573 | -0.53 | -0.18 | 0.006 | 0.087 | 0.087 | 0.003 | 0.41 | 0.27 | 0.62 | 0.73 | 0.39 | 0.52 |
| 2 | 30867573 | rs6548023 | 0.466 | -0.41 | -0.474 | -0.086 | -0.022 | 0.062 | 0.062 | 0 | 0.36 | 0.2 | 0.61 | 0.73 | 0.38 | 0.5 |
| 2 | 30871808 | rs2609954 | -1.238 | -0.518 | -0.52 | -0.131 | 0.008 | 0.005 | 0.005 | 0.017 | 0.69 | 0.77 | 0.87 | 0.87 | 0.79 | 0.68 |
| 2 | 30872064 | rs7583600 | 0.55 | -0.507 | -0.91 | -0.067 | 0 | -0.042 | -0.042 | 0.605 | 0.31 | 0.21 | 0.41 | 0.59 | 0.31 | 0.3 |
| 2 | 30872606 | rs2593463 | -0.273 | -0.472 | -1.121 | 0.009 | 0.015 | 0.019 | 0.019 | 0 | 0.47 | 0.8 | 0.53 | 0.43 | 0.57 | 0.44 |
| 2 | 30873111 | rs11127284 | 0.638 | -0.108 | -1.28 | -0.397 | 0.018 | -0.061 | -0.061 | 0 | 0.26 | 0.34 | 0.24 | 0.08 | 0.34 | 0.13 |
| 2 | 30874229 | rs895662 | -1.355 | -0.462 | -1.946 | -0.057 | 0.018 | 0.101 | 0.101 | 0 | 0.75 | 0.99 | 0.93 | 0.94 | 0.84 | 0.71 |
| 2 | 30878949 | rs2246050 | 0.889 | 0.162 | -1.296 | 0.593 | -0.016 | -0.028 | -0.028 | 0 | 0.19 | 0.05 | 0.18 | 0.24 | 0.19 | 0.1 |
| 2 | 30883211 | rs1447191 | -0.631 | 0.082 | -1.313 | 0.355 | -0.019 | -0.044 | -0.044 | 1.117 | 0.61 | 0.84 | 0.54 | 0.5 | 0.57 | 0.49 |
| 2 | 30884576 | rs1901906 | -0.215 | -0.436 | -1.367 | 0.236 | 0.076 | -0.039 | -0.039 | 0.462 | 0.37 | 0.79 | 0.32 | 0.48 | 0.21 | 0.31 |
| 2 | 30885263 | rs2602772 | -0.062 | -0.413 | -0.992 | 0.245 | -0.004 | -0.001 | -0.001 | 0 | 0.35 | 0.62 | 0.3 | 0.48 | 0.21 | 0.3 |
| 2 | 30888286 | rs4952172 | 0.045 | -0.452 | -0.77 | 0.227 | -0.048 | 0.032 | 0.032 | 0 | 0.34 | 0.52 | 0.3 | 0.48 | 0.21 | 0.31 |
| 2 | 30889161 | rs1901909 | -0.458 | -0.264 | -1.409 | 0.213 | 0.045 | -0.019 | -0.019 | 0.118 | 0.39 | 0.72 | 0.34 | 0.5 | 0.24 | 0.34 |
| 2 | 30900567 | rs6719376 | 0.425 | -0.167 | -1.258 | 0.172 | 0.023 | 0.014 | 0.014 | 0 | 0.28 | 0.18 | 0.27 | 0.27 | 0.22 | 0.23 |
| 2 | 30902939 | rs2593477 | -0.145 | 0.299 | -1.313 | 0.492 | -0.002 | 0.017 | 0.017 | 0.056 | 0.58 | 0.7 | 0.49 | 0.38 | 0.55 | 0.44 |
| 2 | 30905936 | rs2123752 | -0.302 | 0.372 | -1.542 | 0.61 | -0.008 | 0.003 | 0.003 | 0.103 | 0.62 | 0.76 | 0.53 | 0.42 | 0.58 | 0.47 |
| 2 | 30910849 | rs1562417 | -0.081 | 0.321 | -1.531 | 0.748 | 0.04 | -0.007 | -0.007 | 0 | 0.48 | 0.7 | 0.66 | 0.45 | 0.4 | 0.45 |
| 2 | 30914381 | rs547773 | 0.247 | 0.197 | -1.739 | 1.08 | 0 | -0.076 | -0.076 | 0 | 0.16 | 0.03 | 0.09 | 0.32 | 0.15 | 0.26 |
| 2 | 30915386 | rs530076 | -0.431 | 0.151 | -1.182 | 0.458 | -0.003 | -0.001 | -0.001 | 0.13 | 0.54 | 0.49 | 0.36 | 0.48 | 0.58 | 0.44 |

| | | | | | | | | | | | | | | | | |
|---|---|---|---|---|---|---|---|---|---|---|---|---|---|---|---|---|
| 2 | 30917735 | rs543134 | 0.237 | 0.165 | -0.549 | -0.041 | -0.001 | -0.04 | -0.04 | 0 | 0.35 | 0.35 | 0.26 | 0.06 | 0.42 | 0.17 |
| 2 | 30918839 | rs514940 | 0.243 | 0.13 | -0.257 | -0.063 | 0.002 | -0.044 | -0.044 | 1.27 | 0.34 | 0.32 | 0.25 | 0.06 | 0.42 | 0.17 |
| 2 | 30918938 | rs519747 | -0.198 | 0.1 | -0.017 | -0.055 | 0.008 | -0.047 | -0.047 | 0.052 | 0.39 | 0.51 | 0.28 | 0.16 | 0.45 | 0.19 |
| 2 | 30924850 | rs2609942 | 0.728 | -0.23 | -0.581 | 0.225 | -0.004 | -0.002 | -0.002 | 0 | 0.33 | 0.38 | 0.32 | 0.33 | 0.31 | 0.36 |
| 2 | 30927012 | rs7599134 | 1.677 | -0.468 | -0.815 | 0.101 | -0.031 | -0.03 | -0.03 | 0.476 | 0.22 | 0.03 | 0.22 | 0.22 | 0.24 | 0.22 |
| 2 | 30928787 | rs528264 | 0.89 | -0.393 | -0.595 | -0.052 | -0.007 | -0.016 | -0.016 | 0 | 0.34 | 0.27 | 0.25 | 0.06 | 0.42 | 0.17 |
| 2 | 30930165 | rs472161 | 0.121 | -0.477 | -0.685 | 0.091 | 0.005 | 0.062 | 0.062 | 0 | 0.5 | 0.66 | 0.34 | 0.34 | 0.52 | 0.36 |
| 2 | 30938938 | rs17010046 | 0.973 | -0.455 | -0.864 | 0.251 | 0.001 | 0 | 0 | 0 | 0.14 | 0.14 | 0.07 | 0.3 | 0.09 | 0.16 |
| 2 | 30940397 | rs486803 | -1.065 | -0.353 | -0.933 | 0.25 | -0.019 | -0.076 | -0.076 | 0 | 0.83 | 0.96 | 0.67 | 0.7 | 0.84 | 0.71 |
| 2 | 30945896 | rs2280404 | -0.087 | -0.449 | -0.575 | 0.183 | 0.018 | -0.009 | -0.009 | 2.308 | 0.45 | 0.37 | 0.38 | 0.5 | 0.39 | 0.5 |
| 2 | 30949592 | rs2681683 | -0.303 | -0.517 | -0.341 | 0.15 | 0.022 | 0.02 | 0.02 | 0.852 | 0.48 | 0.65 | 0.41 | 0.63 | 0.41 | 0.53 |
| 2 | 30949634 | rs2681682 | -0.303 | -0.517 | -0.341 | 0.173 | 0.022 | 0.021 | 0.021 | 0.852 | 0.48 | 0.65 | 0.41 | 0.63 | 0.41 | 0.53 |
| 2 | 30951486 | rs751062 | -0.337 | -0.512 | -0.307 | 0.147 | 0.008 | 0.008 | 0.008 | 0.142 | 0.48 | 0.47 | 0.4 | 0.63 | 0.42 | 0.53 |
| 2 | 30951919 | rs6748488 | -0.02 | -0.516 | -0.315 | 0.155 | 0.013 | -0.005 | -0.005 | 0 | 0.45 | 0.43 | 0.39 | 0.5 | 0.39 | 0.51 |
| 2 | 30951984 | rs2593430 | -0.444 | -0.5 | -0.39 | 0.167 | 0.029 | 0.029 | 0.029 | 0.845 | 0.49 | 0.66 | 0.41 | 0.64 | 0.42 | 0.53 |
| 2 | 30952987 | rs9679386 | 0.501 | -0.523 | -0.709 | 0.178 | -0.006 | -0.146 | -0.146 | 0 | 0.35 | 0.07 | 0.3 | 0.47 | 0.32 | 0.41 |
| 2 | 30953300 | rs6711743 | 0.501 | -0.523 | -0.709 | 0.178 | -0.006 | -0.146 | -0.146 | 0.254 | 0.35 | 0.07 | 0.3 | 0.47 | 0.32 | 0.41 |
| 2 | 30956090 | rs12052219 | 1.4 | -0.57 | -0.745 | 0.179 | -0.027 | -0.066 | -0.066 | 0 | 0.24 | 0.06 | 0.27 | 0.3 | 0.25 | 0.28 |
| 2 | 30956344 | rs2593434 | 0.128 | -0.483 | -0.744 | 0.03 | -0.026 | 0.166 | 0.166 | 0 | 0.47 | 0.8 | 0.35 | 0.24 | 0.51 | 0.29 |
| 2 | 30957326 | rs2276568 | 2.036 | -0.479 | -0.818 | 0.086 | 0.012 | -0.057 | -0.057 | 0 | 0.11 | 0.01 | 0.04 | 0.17 | 0.07 | 0.13 |
| 2 | 30958137 | rs477229 | -0.379 | -0.445 | -0.768 | 0.097 | -0.036 | 0.018 | 0.018 | 0.03 | 0.58 | 0.87 | 0.39 | 0.41 | 0.58 | 0.42 |
| 2 | 30959262 | rs509380 | -0.403 | -0.44 | -0.512 | 0.234 | -0.003 | 0.051 | 0.051 | 0.12 | 0.79 | 0.66 | 0.62 | 0.58 | 0.81 | 0.69 |
| 2 | 30959827 | rs1879276 | 0.437 | -0.501 | -0.412 | 0.166 | -0.017 | 0.06 | 0.06 | 0 | 0.7 | 0.45 | 0.54 | 0.54 | 0.74 | 0.59 |
| 2 | 30959882 | rs7566361 | 1.391 | -0.369 | -0.675 | 0.06 | -0.001 | -0.034 | -0.034 | 0 | 0.08 | 0.21 | 0.08 | 0.03 | 0.07 | 0.1 |
| 2 | 30960671 | rs2609913 | -0.4 | -0.428 | -0.978 | 0.112 | 0.005 | -0.114 | -0.114 | 0 | 0.72 | 0.94 | 0.62 | 0.54 | 0.76 | 0.57 |
| 2 | 30961646 | rs2593439 | -0.292 | -0.334 | -0.927 | 0.051 | 0.02 | 0.172 | 0.172 | 0 | 0.49 | 0.81 | 0.51 | 0.28 | 0.53 | 0.3 |
| 2 | 30961997 | rs2609914 | 0.096 | 0.291 | -1.318 | -0.361 | 0.031 | -0.087 | -0.087 | 0 | 0.23 | 0.28 | 0.2 | 0.03 | 0.35 | 0.12 |
| 2 | 30962675 | rs10153751 | 1.458 | 0.61 | -1.166 | -0.166 | -0.028 | 0.023 | 0.023 | 0 | 0.1 | 0.05 | 0.09 | 0 | 0.19 | 0.04 |
| 2 | 30963364 | rs1715819 | 0.406 | 0.453 | -1.273 | -0.337 | -0.013 | 0.126 | 0.126 | 0.131 | 0.37 | 0.18 | 0.4 | 0.14 | 0.38 | 0.25 |
| 2 | 30965992 | rs11127287 | 1.259 | 0.221 | -0.929 | -0.022 | -0.001 | 0.046 | 0.046 | 0 | 0.26 | 0.15 | 0.33 | 0.15 | 0.25 | 0.21 |
| 2 | 30966687 | rs2253464 | 1.052 | 0.226 | -0.938 | 0.177 | -0.023 | -0.009 | -0.009 | 0 | 0.31 | 0.11 | 0.31 | 0.27 | 0.32 | 0.29 |
| 2 | 30967074 | rs2280403 | -0.437 | 0.337 | -0.855 | 0.373 | -0.049 | 0.001 | 0.001 | 0 | 0.57 | 0.84 | 0.51 | 0.47 | 0.54 | 0.45 |
| 2 | 30969468 | rs479103 | -0.926 | 0.396 | -0.366 | 0.463 | 0.032 | 0.149 | 0.149 | 0 | 0.89 | 0.79 | 0.8 | 0.61 | 0.85 | 0.74 |
| 2 | 30973825 | rs1879283 | 0.234 | 0.314 | -0.387 | 0.336 | 0.01 | -0.05 | -0.05 | 0 | 0.39 | 0.52 | 0.44 | 0.27 | 0.46 | 0.27 |
| 2 | 30974575 | rs17010141 | 1.435 | 0.149 | -0.605 | 0.342 | 0.021 | -0.077 | -0.077 | 0 | 0.16 | 0.01 | 0.19 | 0.24 | 0.09 | 0.17 |
| 2 | 30976058 | rs2926304 | 0.19 | 0.273 | -0.52 | 0.335 | 0.041 | -0.027 | -0.027 | 1.197 | 0.49 | 0.15 | 0.49 | 0.51 | 0.41 | 0.47 |
| 2 | 30977073 | rs485100 | -0.381 | 0.457 | -0.46 | 0.42 | -0.02 | -0.037 | -0.037 | 0.172 | 0.88 | 0.95 | 0.82 | 0.74 | 0.85 | 0.75 |
| 2 | 30979487 | rs2681676 | -0.87 | 0.419 | -0.816 | 0.554 | -0.063 | -0.054 | -0.054 | 0 | 0.83 | 0.94 | 0.74 | 0.76 | 0.69 | 0.72 |
| 2 | 30979720 | rs17324501 | 1.003 | 0.259 | -0.879 | 0.376 | 0.006 | 0.026 | 0.026 | 0 | 0.27 | 0.22 | 0.17 | 0.1 | 0.26 | 0.25 |
| 2 | 30979871 | rs11688018 | 0.712 | 0.952 | -0.516 | 0.881 | 0.031 | 0.033 | 0.033 | 0.098 | 0.44 | 0.34 | 0.37 | 0.33 | 0.34 | 0.4 |
| 2 | 30980940 | rs508405 | -0.329 | 1.636 | -0.697 | 1.018 | -0.085 | -0.072 | -0.072 | 1.4 | 0.84 | 0.96 | 0.75 | 0.76 | 0.69 | 0.72 |
| 2 | 30982795 | rs536137 | -0.252 | 0.884 | 0.26 | 0.672 | 0.026 | -0.025 | -0.025 | 0.047 | 0.28 | 0.35 | 0.22 | 0.19 | 0.37 | 0.24 |
| 2 | 30983434 | rs2922063 | 1.301 | -0.066 | 1.185 | 0.986 | 0.052 | 0.115 | 0.115 | 0 | 0.9 | 0.64 | 0.92 | 0.83 | 0.91 | 0.86 |
| 2 | 30984850 | rs551356 | 1.205 | -0.296 | 0.504 | 0.587 | -0.002 | -0.003 | -0.003 | 0 | 0.82 | 0.27 | 0.86 | 0.88 | 0.85 | 0.9 |
| 2 | 30986989 | rs579016 | 0.08 | -0.324 | 0.776 | 0.642 | 0.023 | 0.035 | 0.035 | 2.904 | 0.89 | 0.78 | 0.94 | 0.86 | 0.89 | 0.94 |
| 2 | 30988148 | rs6747327 | 0.303 | -0.84 | 0.499 | 0.196 | -0.001 | 0.003 | 0.003 | 0 | 0.6 | 0.63 | 0.45 | 0.27 | 0.61 | 0.63 |
| 2 | 30988375 | rs6548032 | 0.132 | -0.833 | 0.116 | 0.343 | -0.035 | 0.023 | 0.023 | 0 | 0.71 | 0.89 | 0.8 | 0.74 | 0.69 | 0.72 |
| 2 | 30989030 | rs2176840 | -0.277 | -1.182 | 0.658 | 0.319 | -0.015 | -0.02 | -0.02 | 1.157 | 0.12 | 0.4 | 0.06 | 0.09 | 0.05 | 0.06 |
| 2 | 30992877 | rs17324843 | 0.497 | -1.203 | -0.129 | 0.32 | -0.008 | 0.002 | 0.002 | 0.076 | 0.7 | 0.73 | 0.8 | 0.68 | 0.65 | 0.78 |
| 2 | 30993818 | rs12613578 | 0.885 | -1.242 | -0.242 | 0.421 | 0.014 | -0.016 | -0.016 | 0 | 0.09 | 0.03 | 0.31 | 0.43 | 0.07 | 0.09 |
| 2 | 30995027 | rs4952179 | -0.385 | -1.256 | -0.24 | 0.398 | -0.065 | -0.009 | -0.009 | 2.252 | 0.78 | 0.92 | 0.84 | 0.72 | 0.7 | 0.79 |
| 2 | 30995579 | rs17396800 | 0.404 | -1.69 | -0.568 | -0.058 | 0.003 | 0.018 | 0.018 | 0 | 0.47 | 0.61 | 0.65 | 0.6 | 0.45 | 0.42 |
| 2 | 30997898 | rs1464332 | 1.095 | -1.573 | -0.592 | 0.03 | 0.017 | -0.026 | -0.026 | 0 | 0.1 | 0.03 | 0.31 | 0.42 | 0.07 | 0.11 |
| 2 | 30999833 | rs6714182 | 1.213 | -1.64 | -0.604 | -0.19 | 0.019 | -0.05 | -0.05 | 0.151 | 0.33 | 0.46 | 0.32 | 0.19 | 0.36 | 0.3 |
| 2 | 31001167 | rs1568403 | 0.17 | -1.683 | -0.835 | -0.477 | 0.008 | -0.034 | -0.034 | 0.607 | 0.65 | 0.49 | 0.78 | 0.72 | 0.64 | 0.8 |
| 2 | 31001405 | rs17010258 | 0.87 | -1.699 | -1.042 | -0.498 | 0.013 | 0.024 | 0.024 | 0 | 0.25 | 0.22 | 0.16 | 0.07 | 0.21 | 0.37 |
| 2 | 31004365 | rs1520321 | 0.949 | -1.487 | -0.943 | -0.315 | 0.007 | 0.054 | 0.054 | 0.108 | 0.54 | 0.37 | 0.45 | 0.25 | 0.57 | 0.68 |
| 2 | 31005129 | rs2030385 | -0.043 | -1.495 | -0.917 | -0.307 | -0.053 | 0.03 | 0.03 | 0.02 | 0.69 | 0.87 | 0.81 | 0.72 | 0.64 | 0.78 |
| 2 | 31005176 | rs4597573 | 0.271 | -1.252 | -0.841 | -0.477 | -0.007 | 0.037 | 0.037 | 0 | 0.59 | 0.71 | 0.49 | 0.25 | 0.57 | 0.68 |
| 2 | 31005297 | rs4597574 | 0.643 | -1.222 | -0.946 | -0.462 | 0.006 | 0.052 | 0.052 | 0.093 | 0.55 | 0.37 | 0.45 | 0.25 | 0.57 | 0.68 |
| 2 | 31007962 | rs10186446 | 0.538 | -0.492 | -0.605 | -0.415 | 0.006 | -0.003 | -0.003 | 0.24 | 0.14 | 0.13 | 0.13 | 0.17 | 0.21 | 0.15 |
| 2 | 31008142 | rs10495780 | -0.31 | -1.01 | -0.147 | -0.274 | 0.065 | 0.061 | 0.061 | 1.528 | 0.05 | 0.42 | 0.04 | 0 | 0 | 0 |
| 2 | 31009479 | rs6732918 | -0.677 | -0.949 | -1.352 | -0.261 | 0.007 | 0.005 | 0.005 | 0.156 | 0.31 | 0.71 | 0.19 | 0.15 | 0.2 | 0.37 |
| 2 | 31009876 | rs1915126 | -0.054 | -0.252 | -1.724 | 0.243 | -0.041 | -0.001 | -0.001 | 0 | 0.9 | 0.99 | 0.94 | 0.94 | 0.89 | 0.95 |
| 2 | 31010212 | rs13021213 | -0.543 | -0.677 | -1.312 | 0.279 | 0.012 | 0.017 | 0.017 | 0 | 0.16 | 0.13 | 0.11 | 0.08 | 0.13 | 0.33 |
| 2 | 31010924 | rs6709502 | -0.808 | -0.789 | -1.121 | 0.921 | -0.018 | 0.012 | 0.012 | 0.021 | 0.24 | 0.69 | 0.18 | 0.39 | 0.13 | 0.16 |
| 2 | 31014166 | rs1520319 | 0.143 | -0.055 | -1.967 | 0.144 | -0.036 | 0.118 | 0.118 | 0.149 | 0.33 | 0.07 | 0.45 | 0.13 | 0.31 | 0.47 |
| 2 | 31016084 | rs1520317 | -0.547 | 0.509 | -2.324 | 0.637 | -0.034 | -0.075 | -0.075 | 1.778 | 0.78 | 0.93 | 0.74 | 0.48 | 0.73 | 0.72 |
| 2 | 31016239 | rs7576623 | 0.375 | 0.585 | -1.161 | 0.431 | 0.029 | -0.028 | -0.028 | 0 | 0.66 | 0.54 | 0.66 | 0.25 | 0.59 | 0.59 |

| | | | | | | | | | | | | | | | | |
|---|---|---|---|---|---|---|---|---|---|---|---|---|---|---|---|---|
| 2 | 31022484 | rs10203688 | 0.95 | 0.299 | -1.43 | 0.05 | -0.007 | 0.201 | 0.201 | 0.612 | 0.52 | 0.16 | 0.55 | 0.2 | 0.49 | 0.46 |
| 2 | 31022513 | rs17010322 | -0.452 | 0.544 | -0.324 | 0.148 | 0.043 | 0.01 | 0.01 | 0 | 0.14 | 0.31 | 0.11 | 0.16 | 0.16 | 0.12 |
| 2 | 31026737 | rs7597567 | -0.639 | 0.336 | -0.368 | 0.429 | -0.04 | -0.047 | -0.047 | 2.515 | 0.61 | 0.84 | 0.44 | 0.52 | 0.59 | 0.48 |
| 2 | 31027510 | rs13385578 | 0.285 | 0.293 | -0.058 | 0.346 | -0.003 | -0.041 | -0.041 | 0.015 | 0.37 | 0.43 | 0.23 | 0.13 | 0.35 | 0.22 |
| 2 | 31028945 | rs10173959 | -1.314 | 0.224 | -0.137 | 0.417 | 0.024 | 0.067 | 0.067 | 0 | 0.69 | 0.88 | 0.49 | 0.79 | 0.74 | 0.63 |
| 2 | 31029835 | rs1879282 | -1.314 | 0.224 | -0.137 | 0.417 | 0.024 | 0.067 | 0.067 | 0 | 0.69 | 0.88 | 0.49 | 0.79 | 0.74 | 0.63 |
| 2 | 31030713 | rs10177929 | -1.342 | 0.213 | -0.154 | 0.412 | 0.017 | 0.072 | 0.072 | 0 | 0.7 | 0.92 | 0.5 | 0.79 | 0.74 | 0.63 |
| 2 | 31034257 | rs6712121 | -0.073 | 0.205 | 0.375 | 0.571 | -0.003 | 0.001 | 0.001 | 0 | 0.55 | 0.5 | 0.37 | 0.51 | 0.58 | 0.53 |
| 2 | 31037485 | rs6548038 | -1.203 | 0.245 | 0.123 | 0.588 | 0.029 | 0.02 | 0.02 | 0 | 0.69 | 0.83 | 0.49 | 0.7 | 0.74 | 0.63 |
| 2 | 31048617 | rs939371 | 1.066 | 0.186 | 0.022 | 0.576 | -0.009 | 0.075 | 0.075 | 0 | 0.31 | 0.08 | 0.52 | 0.21 | 0.28 | 0.38 |
| 2 | 31061392 | rs7572039 | 0.574 | 0.275 | 0.015 | 0.645 | -0.054 | -0.053 | -0.053 | 0 | 0.08 | 0.01 | 0.08 | 0.2 | 0.16 | 0.1 |
| 2 | 31061536 | rs7560096 | -0.831 | 0.195 | 0.227 | 0.593 | -0.003 | -0.029 | -0.029 | 0 | 0.67 | 0.61 | 0.82 | 0.87 | 0.7 | 0.74 |
| 2 | 31062348 | rs4410329 | 0.894 | 0.149 | -0.246 | 0.598 | -0.023 | -0.078 | -0.078 | 1.415 | 0.22 | 0.06 | 0.17 | 0.37 | 0.25 | 0.28 |
| 2 | 31066592 | rs12993482 | 1.016 | 0.159 | -0.205 | 0.567 | 0.014 | 0.02 | 0.02 | 0 | 0.32 | 0.13 | 0.53 | 0.3 | 0.29 | 0.35 |
| 2 | 31077996 | rs2364430 | 0.144 | 0.231 | 0.1 | 0.621 | -0.008 | 0.02 | 0.02 | 0 | 0.43 | 0.31 | 0.62 | 0.4 | 0.44 | 0.45 |
| 2 | 31079020 | rs6738257 | -0.898 | 0.316 | 0.793 | 0.512 | 0.007 | 0.044 | 0.044 | 0.962 | 0.71 | 0.82 | 0.85 | 0.88 | 0.72 | 0.8 |
| 2 | 31085727 | rs4952188 | 0.611 | 0.357 | -0.064 | 0.601 | -0.045 | 0.014 | 0.014 | 2.362 | 0.51 | 0.1 | 0.68 | 0.59 | 0.52 | 0.64 |
| 2 | 31086044 | rs10469990 | -1.256 | 0.736 | 0.441 | 1.13 | 0.036 | 0.08 | 0.08 | 0 | 0.15 | 0.28 | 0.11 | 0.28 | 0.19 | 0.11 |
| 2 | 31093502 | rs2194452 | -1.297 | 0.489 | 0.294 | 0.9 | -0.017 | 0.042 | 0.042 | 3.351 | 0.15 | 0.35 | 0.07 | 0.18 | 0.09 | 0.07 |
| 2 | 31101646 | rs10167990 | -0.609 | 0.517 | 0.223 | 0.771 | -0.003 | -0.003 | -0.003 | 0.087 | 0.11 | 0.11 | 0.06 | 0.09 | 0.07 | 0.07 |
| 2 | 31107154 | rs3755527 | -1.06 | 0.422 | 0.102 | 0.892 | 0.003 | 0.001 | 0.001 | 0 | 0.41 | 0.59 | 0.36 | 0.57 | 0.36 | 0.4 |
| 2 | 31107181 | rs10184608 | -0.671 | 0.301 | 0.334 | 0.948 | -0.002 | -0.014 | -0.014 | 1.137 | 0.13 | 0.27 | 0.07 | 0.09 | 0.08 | 0.07 |
| 2 | 31109504 | rs11127290 | -0.965 | 0.304 | -0.424 | 0.75 | 0 | 0.019 | 0.019 | 0 | 0.47 | 0.6 | 0.42 | 0.59 | 0.46 | 0.48 |
| 2 | 31112402 | rs11127291 | -0.424 | 0.001 | -1.486 | 0.82 | 0.032 | -0.042 | -0.042 | 0 | 0.38 | 0.8 | 0.37 | 0.42 | 0.29 | 0.33 |
| 2 | 31112433 | rs12623781 | 0.162 | -0.155 | -1.092 | 0.834 | -0.008 | 0.051 | 0.051 | 0 | 0.11 | 0.17 | 0.05 | 0.34 | 0.04 | 0.12 |
| 2 | 31112471 | rs6733757 | 0.139 | 0.121 | -0.972 | 0.72 | 0 | -0.003 | -0.003 | 0 | 0.27 | 0.65 | 0.32 | 0.09 | 0.28 | 0.21 |
| 2 | 31118058 | rs1559497 | -0.565 | 0.206 | -1.017 | 0.647 | 0 | 0 | 0 | 0.008 | 0.56 | 0.51 | 0.48 | 0.55 | 0.48 | 0.55 |
| 2 | 31122598 | rs4952192 | -0.009 | 0.221 | -1.643 | 0.218 | -0.001 | -0.007 | -0.007 | 0.067 | 0.2 | 0.09 | 0.22 | 0.19 | 0.18 | 0.23 |
| 2 | 31125049 | rs7572773 | 1.2 | 0.363 | -0.98 | 0.068 | 0.005 | -0.022 | -0.022 | 1.411 | 0.22 | 0.21 | 0.29 | 0.08 | 0.28 | 0.17 |
| 2 | 31135184 | rs2288101 | -0.275 | -0.069 | -1.13 | 0.527 | -0.003 | 0.017 | 0.017 | 0 | 0.24 | 0.14 | 0.18 | 0.2 | 0.26 | 0.22 |
| 2 | 31146497 | rs10205350 | -0.899 | -0.122 | -0.874 | 0.724 | 0.008 | -0.002 | -0.002 | 0.011 | 0.32 | 0.3 | 0.2 | 0.38 | 0.22 | 0.37 |
| 2 | 31166795 | rs10179187 | -0.899 | 0.23 | -0.683 | 0.898 | 0 | 0 | 0 | 0 | 0.29 | 0.3 | 0.26 | 0.33 | 0.34 | 0.27 |
| 2 | 31171410 | rs889897 | -1.138 | -0.12 | -0.697 | 1.202 | -0.002 | 0.01 | 0.01 | 0 | 0.32 | 0.35 | 0.25 | 0.51 | 0.32 | 0.32 |
| 2 | 31173824 | rs971709 | -1.403 | 0.107 | -0.994 | 1.003 | -0.015 | 0.018 | 0.018 | 0 | 0.66 | 0.82 | 0.71 | 0.73 | 0.68 | 0.68 |
| 2 | 31178818 | rs2288100 | -0.363 | 0.076 | -0.709 | 1.503 | 0.043 | -0.004 | -0.004 | 0 | 0.21 | 0.06 | 0.11 | 0.2 | 0.15 | 0.14 |
| 2 | 31180362 | rs6744118 | -0.479 | 0.042 | 0.076 | 1.651 | 0.065 | 0.265 | 0.265 | 0.3 | 0.12 | 0.42 | 0.17 | 0.39 | 0.16 | 0.17 |
| 2 | 31180935 | rs6741827 | -1.943 | 0.052 | 0.096 | 1.628 | -0.01 | -0.004 | -0.004 | 0 | 0.36 | 0.59 | 0.39 | 0.66 | 0.34 | 0.43 |
| 2 | 31181139 | rs2303325 | 0.282 | 0.055 | -0.258 | 1.476 | -0.01 | 0.054 | 0.054 | 0 | 0.15 | 0.01 | 0.09 | 0.04 | 0.14 | 0.1 |
| 2 | 31182162 | rs11884629 | -0.622 | -0.114 | -0.189 | 1.503 | 0.032 | -0.014 | -0.014 | 0 | 0.22 | 0.15 | 0.18 | 0.26 | 0.15 | 0.19 |
| 2 | 31189304 | rs2303323 | -0.091 | 0.467 | 0.405 | 1.892 | 0.047 | 0.144 | 0.144 | 0 | 0.08 | 0.18 | 0.13 | 0.3 | 0.14 | 0.16 |
| 2 | 31202312 | rs1862972 | -1.657 | 0.348 | 0.503 | 1.996 | -0.009 | 0.13 | 0.13 | 0 | 0.21 | 0.57 | 0.29 | 0.54 | 0.17 | 0.31 |
| 2 | 31204981 | rs4952017 | -0.18 | 0.334 | 0.026 | 1.19 | -0.003 | 0.004 | 0.004 | 0.087 | 0.72 | 0.72 | 0.77 | 0.77 | 0.66 | 0.85 |
| 2 | 31208530 | rs6737493 | 0.065 | 0.411 | 0.064 | 1.1 | 0 | -0.003 | -0.003 | 0 | 0.67 | 0.66 | 0.71 | 0.75 | 0.62 | 0.77 |
| 2 | 31222346 | rs12465448 | -1.047 | 0.641 | 0.229 | 1.585 | -0.002 | 0.032 | 0.032 | 0 | 0.23 | 0.29 | 0.4 | 0.41 | 0.25 | 0.3 |
| 2 | 31230833 | rs867496 | -1.01 | 0.521 | 0.597 | 1.634 | 0 | 0.074 | 0.074 | 0 | 0.22 | 0.36 | 0.38 | 0.45 | 0.25 | 0.29 |
| 2 | 31231371 | rs2365190 | -0.456 | 0.805 | 0.549 | 1.614 | -0.014 | -0.032 | -0.032 | 0.102 | 0.2 | 0.15 | 0.37 | 0.45 | 0.29 | 0.31 |
| 2 | 31232116 | rs8179779 | -0.693 | 0.614 | 1.205 | 1.624 | 0.009 | 0.128 | 0.128 | 0 | 0.21 | 0.4 | 0.37 | 0.45 | 0.25 | 0.3 |
| 2 | 31232512 | rs9636424 | -0.051 | 0.625 | 0.695 | 1.765 | -0.014 | -0.034 | -0.034 | 0 | 0.08 | 0.01 | 0.05 | 0.3 | 0.08 | 0.11 |
| 2 | 31236916 | rs1862969 | -0.59 | 0.807 | 0.726 | 1.454 | -0.007 | 0.006 | 0.006 | 0.508 | 0.73 | 0.73 | 0.73 | 0.85 | 0.63 | 0.78 |
| 2 | 31238524 | rs12987759 | -0.597 | 0.482 | 1.237 | 1.744 | -0.001 | 0.11 | 0.11 | 0 | 0.16 | 0.38 | 0.35 | 0.31 | 0.15 | 0.22 |
| 2 | 31240246 | rs6543592 | -0.687 | 0.259 | 1.009 | 1.272 | -0.002 | 0.047 | 0.047 | 0 | 0.1 | 0.27 | 0.07 | 0.15 | 0.08 | 0.11 |
| 2 | 31241126 | rs4952026 | 0.488 | 0.344 | 0.864 | 1.176 | -0.001 | -0.006 | -0.006 | 3.594 | 0.54 | 0.48 | 0.35 | 0.65 | 0.43 | 0.58 |
| 2 | 31241920 | rs13009779 | -0.651 | 0.467 | 1.312 | 1.138 | -0.001 | 0.001 | 0.001 | 0.048 | 0.34 | 0.6 | 0.5 | 0.38 | 0.29 | 0.34 |
| 2 | 31244635 | rs6543594 | -0.978 | 0.416 | 1.428 | 1.032 | -0.003 | 0 | 0 | 0 | 0.42 | 0.42 | 0.55 | 0.55 | 0.38 | 0.36 |
| 2 | 31246546 | rs10197865 | -1.158 | 0.404 | 1.295 | 1 | 0.016 | -0.002 | -0.002 | 0 | 0.44 | 0.66 | 0.56 | 0.55 | 0.37 | 0.38 |
| 2 | 31248062 | rs6708989 | -0.84 | 0.469 | 1.185 | 1.032 | -0.005 | 0.004 | 0.004 | 0.32 | 0.41 | 0.48 | 0.54 | 0.55 | 0.37 | 0.34 |
| 2 | 31251499 | rs4951960 | -0.284 | 0.514 | 1.095 | 1.057 | -0.001 | -0.004 | -0.004 | 1.47 | 0.32 | 0.28 | 0.47 | 0.41 | 0.3 | 0.31 |
| 2 | 31254348 | rs1862968 | -0.781 | 0.463 | 1.066 | 0.964 | 0.002 | -0.005 | -0.005 | 0 | 0.4 | 0.32 | 0.52 | 0.54 | 0.37 | 0.35 |
| 2 | 31267391 | rs12712304 | -0.154 | 0.592 | 0.101 | 0.996 | -0.01 | -0.054 | -0.054 | 0 | 0.31 | 0.16 | 0.46 | 0.43 | 0.29 | 0.32 |
| 2 | 31271882 | rs1862985 | -0.329 | 0.326 | -0.103 | 0.156 | 0 | -0.048 | -0.048 | 0.001 | 0.74 | 0.74 | 0.84 | 1 | 0.7 | 0.82 |
| 2 | 31271942 | rs1862984 | -0.329 | 0.326 | -0.103 | 0.156 | 0 | -0.048 | -0.048 | 0.085 | 0.74 | 0.74 | 0.84 | 1 | 0.7 | 0.82 |
| 2 | 31273087 | rs4952039 | -0.329 | 0.326 | -0.103 | 0.156 | 0 | -0.048 | -0.048 | 0 | 0.74 | 0.74 | 0.84 | 1 | 0.7 | 0.82 |
| 2 | 31278254 | rs11681174 | 0.379 | -0.103 | -0.298 | 0.489 | -0.002 | -0.001 | -0.001 | 0.192 | 0.1 | 0.1 | 0.06 | 0.13 | 0.07 | 0.05 |
| 2 | 31280917 | rs10211370 | 0.092 | 0.288 | -0.251 | 0.678 | -0.053 | 0.015 | 0.015 | 1.087 | 0.36 | 0.15 | 0.46 | 0.28 | 0.4 | 0.4 |
| 2 | 31283345 | rs8179695 | -0.637 | 0.446 | 0.685 | 0.657 | 0.004 | 0.008 | 0.008 | 0 | 0.61 | 0.54 | 0.63 | 0.45 | 0.6 | 0.49 |
| 2 | 31286910 | rs2161834 | -0.721 | 0.475 | 0.333 | 0.578 | -0.005 | -0.006 | -0.006 | 1.814 | 0.62 | 0.67 | 0.64 | 0.43 | 0.6 | 0.49 |
| 2 | 31291718 | rs11674713 | -1.033 | 0.379 | 0.128 | 0.477 | -0.068 | -0.057 | -0.057 | 0 | 0.64 | 0.81 | 0.62 | 0.46 | 0.53 | 0.5 |
| 2 | 31292787 | rs4362605 | -0.426 | 0.22 | 0.382 | 0.578 | -0.008 | 0.099 | 0.099 | 0 | 0.05 | 0.07 | 0.09 | 0.25 | 0.02 | 0.06 |
| 2 | 31292817 | rs17010651 | -0.198 | 0.001 | 0.608 | 0.336 | -0.006 | -0.004 | -0.004 | 0 | 0.12 | 0.15 | 0.06 | 0.05 | 0.06 | 0.08 |

| | | | | | | | | | | | | | | | | |
|---|---|---|---|---|---|---|---|---|---|---|---|---|---|---|---|---|
| 2 | 31297357 | rs13386542 | -0.055 | -0.014 | 0.625 | 0.234 | 0.009 | 0.008 | 0.008 | 0 | 0.12 | 0.08 | 0.05 | 0.05 | 0.05 | 0.08 |
| 2 | 31297427 | rs13021484 | -0.269 | -0.1 | 0.882 | 0.149 | 0.013 | 0.032 | 0.032 | 0 | 0.48 | 0.69 | 0.56 | 0.4 | 0.46 | 0.38 |
| 2 | 31297524 | rs1862981 | -1.533 | -0.116 | 0.802 | 0.142 | -0.09 | -0.077 | -0.077 | 1.847 | 0.65 | 0.86 | 0.64 | 0.45 | 0.53 | 0.5 |
| 2 | 31300544 | rs4951964 | -0.861 | -0.186 | 1.419 | 0.116 | 0.023 | 0.27 | 0.27 | 0 | 0.07 | 0.27 | 0.11 | 0.25 | 0.02 | 0.06 |
| 2 | 31300892 | rs4951965 | -1.525 | -0.165 | 1.226 | 0.121 | -0.077 | -0.065 | -0.065 | 0 | 0.64 | 0.83 | 0.63 | 0.45 | 0.52 | 0.5 |
| 2 | 31301634 | rs6714508 | 0.088 | -0.232 | 1.224 | 0.132 | 0.012 | 0.01 | 0.01 | 0 | 0.11 | 0.07 | 0.06 | 0.04 | 0.05 | 0.08 |
| 2 | 31304147 | rs2113490 | -0.174 | -0.206 | 1.282 | 0.144 | 0.007 | 0.019 | 0.019 | 0 | 0.47 | 0.62 | 0.54 | 0.4 | 0.46 | 0.38 |
| 2 | 31306055 | rs1862978 | -1.075 | -0.203 | 1.236 | 0.111 | -0.041 | -0.036 | -0.036 | 0.394 | 0.6 | 0.76 | 0.61 | 0.45 | 0.52 | 0.46 |
| 2 | 31308019 | rs13404171 | 0.01 | -0.149 | 1.128 | -1.005 | 0.014 | 0.011 | 0.011 | 0 | 0.12 | 0.07 | 0.05 | 0.05 | 0.05 | 0.08 |
| 2 | 31311548 | rs3919588 | 0.152 | -0.104 | 1.414 | -1.023 | 0.001 | -0.058 | -0.058 | 0 | 0.42 | 0.49 | 0.45 | 0.15 | 0.45 | 0.32 |
| 2 | 31312461 | rs2365195 | 0.188 | -0.1 | 1.427 | -1.022 | -0.001 | -0.046 | -0.046 | 0.204 | 0.42 | 0.46 | 0.44 | 0.15 | 0.44 | 0.32 |
| 2 | 31316509 | rs4561706 | -0.753 | -0.073 | 1.371 | -0.992 | -0.021 | -0.026 | -0.026 | 0.836 | 0.08 | 0.14 | 0.04 | 0.04 | 0.03 | 0.09 |
| 2 | 31320834 | rs11888670 | -0.618 | -0.084 | 1.23 | -1 | -0.011 | 0.04 | 0.04 | 0 | 0.55 | 0.67 | 0.51 | 0.2 | 0.5 | 0.4 |
| 2 | 31320949 | rs6748774 | 0.251 | -0.145 | 0.739 | -0.976 | -0.003 | 0.07 | 0.07 | 0 | 0.41 | 0.3 | 0.43 | 0.15 | 0.44 | 0.32 |
| 2 | 31327850 | rs12996218 | -1.206 | 0.032 | 0.09 | -0.825 | -0.085 | 0.068 | 0.068 | 0 | 0.63 | 0.86 | 0.57 | 0.24 | 0.52 | 0.49 |
| 2 | 31331470 | rs2194463 | -0.673 | -0.365 | 0.119 | -0.783 | -0.036 | -0.021 | -0.021 | 0.106 | 0.13 | 0.19 | 0.06 | 0.05 | 0.04 | 0.1 |
| 2 | 31340646 | rs11687856 | -1.123 | 0.077 | 0.055 | -0.697 | 0.021 | -0.075 | -0.075 | 1.613 | 0.14 | 0.46 | 0.07 | 0.04 | 0.11 | 0.13 |
| 2 | 31344440 | rs10190183 | -1.095 | 0.331 | -0.783 | -0.421 | 0.043 | 0.04 | 0.04 | 0.375 | 0.08 | 0.32 | 0.04 | 0.1 | 0.09 | 0.02 |
| 2 | 31346200 | rs6731612 | 0.552 | 0.519 | -2.29 | -0.627 | -0.096 | 0.042 | 0.042 | 0 | 0.36 | 0.07 | 0.33 | 0.74 | 0.38 | 0.51 |
| 2 | 31348390 | rs2216832 | 0.53 | 0.458 | -2.298 | -1.065 | -0.086 | 0.241 | 0.241 | 5.22 | 0.55 | 0.1 | 0.4 | 0.85 | 0.5 | 0.65 |
| 2 | 31351249 | rs6543604 | 0.147 | 0.832 | -1.999 | -1.142 | -0.044 | 0.075 | 0.075 | 3.133 | 0.76 | 0.87 | 0.63 | 0.98 | 0.64 | 0.82 |
| 2 | 31356428 | rs7560152 | -0.911 | 0.032 | -2.019 | -0.935 | 0.044 | 0.02 | 0.02 | 0 | 0.38 | 0.76 | 0.34 | 0.35 | 0.32 | 0.44 |
| 2 | 31363732 | rs1530394 | 0.56 | 0.478 | -2.818 | -1.569 | -0.097 | 0.121 | 0.121 | 0 | 0.22 | 0.05 | 0.38 | 0 | 0.3 | 0.14 |
| 2 | 31368997 | rs12105231 | 0.353 | 0.024 | -1.983 | -1.283 | 0.013 | 0.009 | 0.009 | 0 | 0.62 | 0.3 | 0.66 | 0.65 | 0.67 | 0.56 |
| 2 | 31372429 | rs6543609 | 0.478 | 0.022 | -2.024 | -1.293 | 0.039 | 0.023 | 0.023 | 0 | 0.61 | 0.18 | 0.66 | 0.65 | 0.67 | 0.57 |
| 2 | 31374733 | rs956747 | -0.828 | 0.057 | -2.011 | -1.278 | 0.013 | 0.009 | 0.009 | 0 | 0.39 | 0.66 | 0.34 | 0.35 | 0.33 | 0.44 |
| 2 | 31380955 | rs6706156 | 0.481 | 0.343 | -2.014 | -1.163 | 0.035 | -0.015 | -0.015 | 0.549 | 0.67 | 0.13 | 0.69 | 0.66 | 0.72 | 0.64 |
| 2 | 31387957 | rs1947027 | -0.611 | 0.162 | -0.581 | -1.537 | -0.03 | -0.051 | -0.051 | 0 | 0.25 | 0.52 | 0.39 | 0.13 | 0.18 | 0.24 |
| 2 | 31392316 | rs1443712 | -0.185 | 0.164 | 0.111 | -1.473 | -0.029 | -0.022 | -0.022 | 0 | 0.21 | 0.39 | 0.35 | 0.16 | 0.13 | 0.19 |
| 2 | 31396186 | rs4952051 | -1.251 | 0.029 | -0.031 | -1.609 | 0.038 | 0.047 | 0.047 | 0.147 | 0.33 | 0.75 | 0.46 | 0.19 | 0.22 | 0.35 |
| 2 | 31396729 | rs17010826 | -0.77 | 0 | 0.076 | -1.584 | -0.021 | -0.07 | -0.07 | 0.765 | 0.14 | 0.26 | 0.1 | 0.02 | 0.09 | 0.13 |
| 2 | 31397332 | rs12992405 | -0.973 | 0.445 | -0.439 | -1.855 | -0.008 | 0.093 | 0.093 | 0 | 0.56 | 0.71 | 0.63 | 0.19 | 0.5 | 0.4 |
| 2 | 31398412 | rs7579175 | -1.411 | 0.614 | 0.076 | -1.24 | 0.008 | 0.013 | 0.013 | 0.076 | 0.35 | 0.7 | 0.47 | 0.18 | 0.26 | 0.34 |
| 2 | 31399988 | rs4592896 | -1.334 | 0.609 | 0.072 | -1.243 | -0.005 | -0.012 | -0.012 | 0 | 0.34 | 0.63 | 0.45 | 0.17 | 0.23 | 0.29 |
| 2 | 31400502 | rs13421721 | -1.485 | 0.592 | 0.067 | -1.237 | 0.018 | 0.032 | 0.032 | 0.04 | 0.36 | 0.73 | 0.47 | 0.18 | 0.25 | 0.33 |
| 2 | 31401763 | rs7564478 | -1.494 | 0.592 | -0.027 | -1.236 | 0.007 | 0.01 | 0.01 | 0.051 | 0.36 | 0.72 | 0.47 | 0.18 | 0.25 | 0.33 |
| 2 | 31407458 | rs4952053 | -1.343 | 0.553 | -0.107 | -1.213 | -0.018 | -0.031 | -0.031 | 0.105 | 0.34 | 0.61 | 0.45 | 0.17 | 0.22 | 0.29 |
| 2 | 31410728 | rs764072 | -0.755 | 0.664 | -0.088 | -1.143 | -0.021 | -0.04 | -0.04 | 0 | 0.27 | 0.59 | 0.39 | 0.16 | 0.19 | 0.26 |
| 2 | 31415263 | rs10204608 | -1.185 | 0.603 | 0.174 | -1.266 | -0.037 | -0.09 | -0.09 | 6.452 | 0.17 | 0.52 | 0.13 | 0.05 | 0.1 | 0.12 |
| 2 | 31415588 | rs4952055 | 0.832 | 1.446 | -0.029 | -0.884 | 0.001 | -0.002 | -0.002 | 0 | 0.56 | 0.48 | 0.5 | 0.62 | 0.59 | 0.65 |
| 2 | 31416213 | rs11124248 | 1.376 | 1.455 | 0.254 | -0.845 | -0.021 | 0.016 | 0.016 | 0.01 | 0.48 | 0.2 | 0.43 | 0.57 | 0.55 | 0.61 |
| 2 | 31417863 | rs4952056 | 0.615 | 1.372 | 0.314 | -0.91 | 0.001 | 0.019 | 0.019 | 0 | 0.16 | 0.24 | 0.28 | 0.37 | 0.16 | 0.17 |
| 2 | 31421582 | rs10188925 | 0.601 | 1.541 | 0.357 | -0.99 | 0.004 | -0.068 | -0.068 | 0 | 0.68 | 0.67 | 0.72 | 0.95 | 0.69 | 0.79 |
| 2 | 31427082 | rs4952060 | -0.691 | 1.576 | 0.272 | -0.978 | 0.027 | -0.077 | -0.077 | 1.506 | 0.82 | 0.79 | 0.81 | 1 | 0.76 | 0.89 |
| 2 | 31431725 | rs11681503 | 1.488 | 1.588 | -0.286 | -0.919 | -0.013 | 0.025 | 0.025 | 0.148 | 0.47 | 0.2 | 0.42 | 0.57 | 0.53 | 0.56 |
| 2 | 31432367 | rs13403357 | 1.303 | 1.823 | -0.486 | -1.132 | -0.043 | 0.036 | 0.036 | 0 | 0.08 | 0 | 0.05 | 0 | 0.12 | 0.04 |
| 2 | 31436358 | rs4952064 | 1.012 | 1.692 | 0.085 | -0.727 | 0 | -0.014 | -0.014 | 0.3 | 0.63 | 0.38 | 0.68 | 0.92 | 0.65 | 0.74 |
| 2 | 31438548 | rs10210053 | 1.161 | 1.713 | 0.064 | -0.722 | -0.002 | 0 | 0 | 0.006 | 0.61 | 0.38 | 0.68 | 0.94 | 0.62 | 0.74 |
| 2 | 31441193 | rs6731524 | 0.113 | 2.123 | 0.251 | -0.123 | -0.001 | -0.025 | -0.025 | 1.365 | 0.15 | 0.12 | 0.25 | 0.34 | 0.12 | 0.19 |
| 2 | 31442747 | rs4952069 | -1.405 | 2.281 | 0.454 | -0.186 | 0 | 0 | 0 | 2.648 | 0.33 | 0.39 | 0.43 | 0.39 | 0.33 | 0.3 |
| 2 | 31444826 | rs13408922 | -1.744 | 2.25 | 0.255 | -0.437 | -0.007 | -0.045 | -0.045 | 0.005 | 0.14 | 0.36 | 0.12 | 0.08 | 0.12 | 0.12 |
| 2 | 31445272 | rs10204574 | 1.844 | 2.355 | -0.009 | -0.287 | 0.077 | -0.009 | -0.009 | 0 | 0.5 | 0.15 | 0.37 | 0.45 | 0.46 | 0.55 |
| 2 | 31445691 | rs13386850 | -1.613 | 2.281 | 0.248 | -0.249 | -0.013 | -0.068 | -0.068 | 0 | 0.16 | 0.54 | 0.13 | 0.08 | 0.12 | 0.13 |
| 2 | 31445795 | rs13386968 | -0.762 | 2.318 | 0.096 | -0.21 | -0.008 | -0.018 | -0.018 | 0 | 0.08 | 0.26 | 0.05 | 0.04 | 0.05 | 0.08 |
| 2 | 31448475 | rs6753831 | -1.39 | 2.416 | -0.222 | -0.192 | 0.001 | -0.004 | -0.004 | 0.024 | 0.36 | 0.68 | 0.46 | 0.53 | 0.29 | 0.33 |
| 2 | 31450522 | rs2344 | -0.356 | 2.626 | -0.25 | 0.059 | 0.003 | 0.017 | 0.017 | 0 | 0.22 | 0.23 | 0.37 | 0.47 | 0.23 | 0.22 |
| 2 | 31451885 | rs6722609 | 0.987 | 2.071 | -0.943 | 0.079 | 0.015 | -0.014 | -0.014 | 0.72 | 0.55 | 0.31 | 0.39 | 0.46 | 0.5 | 0.56 |
| 2 | 31452231 | rs601255 | -0.775 | 2.34 | -0.918 | -0.122 | 0.053 | -0.068 | -0.068 | 0 | 0.15 | 0.2 | 0.18 | 0.01 | 0.26 | 0.13 |
| 2 | 31454545 | rs605832 | 0.047 | 1.394 | -1.98 | 0.308 | -0.026 | -0.042 | -0.042 | 0 | 0.57 | 0.89 | 0.59 | 0.59 | 0.44 | 0.54 |
| 2 | 31454609 | rs13425772 | -0.315 | 1.316 | -1.614 | -0.094 | 0 | -0.006 | -0.006 | 0 | 0.08 | 0.09 | 0.05 | 0.03 | 0.08 | 0.11 |
| 2 | 31459744 | rs1540926 | 0.93 | 1.362 | -2.122 | -0.385 | -0.019 | 0.064 | 0.064 | 0.955 | 0.6 | 0.2 | 0.71 | 0.73 | 0.55 | 0.76 |
| 2 | 31464829 | rs647316 | -0.042 | 1.271 | -1.829 | -0.494 | -0.028 | 0.032 | 0.032 | 0 | 0.83 | 0.92 | 0.85 | 0.93 | 0.71 | 0.92 |
| 2 | 31465278 | rs649509 | 0.033 | 1.188 | -1.465 | -0.465 | 0 | 0 | 0 | 0 | 0.1 | 0.12 | 0.07 | 0.1 | 0.12 | 0.12 |
| 2 | 31467079 | rs2295471 | 1.459 | 1.322 | -1.481 | -0.473 | -0.038 | 0.078 | 0.078 | 0.788 | 0.6 | 0.16 | 0.69 | 0.72 | 0.53 | 0.74 |
| 2 | 31479234 | rs671403 | -0.107 | 1.411 | -1.39 | -0.401 | -0.002 | 0.002 | 0.002 | 0 | 0.1 | 0.12 | 0.07 | 0.13 | 0.12 | 0.13 |
| 2 | 31480495 | rs597644 | 0.51 | 1.447 | -1.403 | -0.684 | -0.045 | 0.057 | 0.057 | 0 | 0.78 | 0.9 | 0.84 | 0.94 | 0.66 | 0.81 |
| 2 | 31482300 | rs625132 | 0.605 | 1.515 | -0.465 | -0.555 | 0.026 | -0.017 | -0.017 | 0 | 0.89 | 0.76 | 0.9 | 0.94 | 0.83 | 0.95 |
| 2 | 31483890 | rs13028669 | 1.646 | 1.239 | 0.326 | -0.638 | 0.011 | -0.042 | -0.042 | 0.544 | 0.69 | 0.5 | 0.79 | 0.77 | 0.67 | 0.71 |
| 2 | 31485125 | rs659396 | -1.224 | 1.474 | -0.112 | -0.505 | 0 | -0.003 | -0.003 | 1.798 | 0.15 | 0.27 | 0.09 | 0.17 | 0.13 | 0.14 |

| | | | | | | | | | | | | | | | | |
|---|---|---|---|---|---|---|---|---|---|---|---|---|---|---|---|---|
| 2 | 31488257 | rs605588 | -0.195 | 1.589 | -0.666 | -0.531 | -0.063 | 0.02 | 0.02 | 0 | 0.87 | 0.98 | 0.9 | 0.94 | 0.79 | 0.87 |
| 2 | 31488554 | rs617163 | -0.053 | 1.607 | -0.625 | -0.524 | 0.001 | -0.001 | -0.001 | 0 | 0.1 | 0.1 | 0.07 | 0.13 | 0.12 | 0.12 |
| 2 | 31489727 | rs644926 | -0.167 | 1.533 | -0.768 | -0.439 | -0.062 | 0.018 | 0.018 | 0 | 0.88 | 0.98 | 0.9 | 0.94 | 0.79 | 0.87 |
| 2 | 31491075 | rs13377 | -0.334 | 1.412 | -0.868 | -0.572 | -0.03 | 0.039 | 0.039 | 0 | 0.07 | 0.01 | 0.07 | 0 | 0.1 | 0.04 |
| 2 | 31491690 | rs678829 | -0.279 | 1.463 | -0.838 | -0.752 | -0.005 | 0.006 | 0.006 | 0.149 | 0.12 | 0.1 | 0.08 | 0.11 | 0.14 | 0.06 |
| 2 | 31493985 | rs7606498 | 1 | 1.088 | -1.932 | -1.199 | -0.016 | 0.051 | 0.051 | 0.249 | 0.3 | 0.09 | 0.2 | 0.26 | 0.24 | 0.52 |
| 2 | 31495278 | rs7593192 | 0.941 | 1.233 | -1.28 | -1.268 | -0.001 | -0.004 | -0.004 | 0 | 0.61 | 0.37 | 0.69 | 0.85 | 0.61 | 0.75 |
| 2 | 31497047 | rs646200 | -0.543 | 0.961 | -0.295 | -1.273 | 0.008 | -0.074 | -0.074 | 0 | 0.8 | 0.66 | 0.83 | 0.97 | 0.79 | 0.83 |
| 2 | 31505021 | rs11124256 | 0.483 | 0.778 | 0.297 | -1.472 | 0.001 | -0.084 | -0.084 | 0.107 | 0.7 | 0.39 | 0.75 | 0.97 | 0.7 | 0.78 |
| 2 | 31509369 | rs12988379 | 0.211 | 0.753 | 0.27 | -1.488 | 0.011 | -0.108 | -0.108 | 0.086 | 0.73 | 0.56 | 0.78 | 0.97 | 0.71 | 0.79 |
| 2 | 31511022 | rs11896460 | -0.028 | 0 | -0.421 | -1.229 | -0.032 | -0.039 | -0.039 | 0.234 | 0.23 | 0.51 | 0.35 | 0.17 | 0.17 | 0.15 |
| 2 | 31512482 | rs590009 | -1.609 | 0.107 | -0.424 | -1.335 | -0.055 | 0.058 | 0.058 | 0 | 0.85 | 0.97 | 0.86 | 0.97 | 0.81 | 0.85 |
| 2 | 31513450 | rs4619655 | 1.155 | -0.036 | -0.803 | -1.409 | 0.002 | 0.054 | 0.054 | 0.045 | 0.56 | 0.33 | 0.47 | 0.85 | 0.57 | 0.67 |
| 2 | 31519001 | rs10178210 | -0.403 | -0.296 | -0.67 | -1.333 | -0.007 | -0.055 | -0.055 | 0 | 0.69 | 0.54 | 0.75 | 0.83 | 0.72 | 0.68 |
| 2 | 31519054 | rs13422633 | 0.638 | -0.362 | -0.644 | -1.361 | -0.004 | -0.063 | -0.063 | 0 | 0.11 | 0.3 | 0.08 | 0 | 0.1 | 0.04 |
| 2 | 31520023 | rs961316 | -0.613 | -0.445 | -0.645 | -1.352 | -0.007 | -0.042 | -0.042 | 0.104 | 0.19 | 0.45 | 0.14 | 0.12 | 0.15 | 0.18 |
| 2 | 31521236 | rs17011242 | 2.339 | -0.481 | -1.049 | -1.158 | -0.111 | -0.166 | -0.166 | 0 | 0.22 | 0.02 | 0.18 | 0.56 | 0.28 | 0.44 |
| 2 | 31521289 | rs604127 | -2.124 | -0.009 | 0.03 | -1.702 | -0.001 | 0.012 | 0.012 | 0 | 0.84 | 0.89 | 0.86 | 0.97 | 0.82 | 0.86 |
| 2 | 31523749 | rs606089 | 0.451 | 0.037 | -1.035 | -1.695 | -0.003 | 0.015 | 0.015 | 0.016 | 0.14 | 0.11 | 0.1 | 0.03 | 0.15 | 0.09 |
| 2 | 31524174 | rs4276068 | 0.515 | 0.084 | -1.355 | -1.637 | -0.005 | 0.099 | 0.099 | 0.234 | 0.54 | 0.32 | 0.4 | 0.83 | 0.52 | 0.67 |
| 2 | 31524264 | rs10179712 | 1.721 | 0.191 | -1.412 | -1.294 | -0.008 | -0.015 | -0.015 | 0 | 0.08 | 0.01 | 0.05 | 0.1 | 0.09 | 0.06 |
| 2 | 31527853 | rs10865088 | 0.496 | 0.313 | -0.818 | -1.536 | 0.006 | 0.089 | 0.089 | 0 | 0.64 | 0.22 | 0.72 | 0.94 | 0.68 | 0.77 |
| 2 | 31537515 | rs207407 | 0.753 | 1.104 | -0.306 | -1.46 | 0.002 | -0.035 | -0.035 | 0 | 0.67 | 0.68 | 0.77 | 0.9 | 0.66 | 0.76 |
| 2 | 31537818 | rs207408 | 0.23 | 1.125 | -0.54 | -1.441 | -0.004 | 0.037 | 0.037 | 0 | 0.75 | 0.84 | 0.82 | 0.9 | 0.76 | 0.81 |
| 2 | 31537931 | rs1560268 | -0.391 | 1.139 | -0.831 | -1.428 | -0.002 | 0.009 | 0.009 | 0 | 0.08 | 0.05 | 0.07 | 0.06 | 0.05 | 0.05 |
| 2 | 31538639 | rs207411 | -0.755 | 1.162 | -0.842 | -1.484 | -0.01 | 0.049 | 0.049 | 0 | 0.82 | 0.93 | 0.9 | 0.96 | 0.81 | 0.85 |
| 2 | 31538823 | rs207412 | 0.993 | 1.115 | -0.792 | -1.572 | -0.032 | 0.142 | 0.142 | 0 | 0.6 | 0.22 | 0.69 | 0.9 | 0.61 | 0.72 |
| 2 | 31540926 | rs207417 | 1.114 | 1.06 | -0.871 | -1.58 | -0.026 | 0.183 | 0.183 | 0 | 0.58 | 0.2 | 0.69 | 0.92 | 0.6 | 0.72 |
| 2 | 31542918 | rs207419 | 0.908 | 1.157 | -0.943 | -1.554 | -0.063 | 0.194 | 0.194 | 0 | 0.6 | 0.17 | 0.68 | 0.91 | 0.59 | 0.72 |
| 2 | 31543124 | rs10153659 | -0.91 | 0.924 | -0.22 | -1.305 | 0.02 | -0.087 | -0.087 | 0 | 0.16 | 0.38 | 0.11 | 0.04 | 0.15 | 0.1 |
| 2 | 31547084 | rs207423 | -1.147 | 0.975 | -1.3 | -1.25 | -0.07 | 0.191 | 0.191 | 0 | 0.54 | 0.88 | 0.33 | 0.23 | 0.47 | 0.49 |
| 2 | 31547773 | rs13392979 | -0.125 | 0.799 | -0.622 | -1.225 | -0.006 | -0.062 | -0.062 | 0 | 0.32 | 0.37 | 0.17 | 0.07 | 0.25 | 0.25 |
| 2 | 31549662 | rs207425 | -0.518 | 0.812 | -0.494 | -1.324 | 0.006 | 0.007 | 0.007 | 0 | 0.4 | 0.61 | 0.21 | 0.13 | 0.32 | 0.4 |
| 2 | 31549757 | rs6756897 | 0.081 | 0.894 | -0.335 | -1.389 | -0.004 | -0.107 | -0.107 | 0 | 0.31 | 0.42 | 0.2 | 0.04 | 0.28 | 0.25 |
| 2 | 31549887 | rs6742051 | -0.402 | 0.904 | -0.253 | -1.361 | -0.001 | -0.055 | -0.055 | 0 | 0.36 | 0.53 | 0.3 | 0.14 | 0.33 | 0.36 |
| 2 | 31549959 | rs207426 | 0.472 | 0.813 | -0.925 | -1.329 | -0.01 | 0.016 | 0.016 | 0 | 0.66 | 0.67 | 0.72 | 0.76 | 0.64 | 0.76 |
| 2 | 31550180 | rs12478111 | -0.907 | 0.847 | -0.953 | -0.825 | -0.013 | -0.002 | -0.002 | 0 | 0.22 | 0.17 | 0.19 | 0.27 | 0.24 | 0.3 |
| 2 | 31552034 | rs10164462 | 0.958 | 0.274 | -0.91 | -0.928 | -0.039 | -0.09 | -0.09 | 0 | 0.23 | 0.36 | 0.09 | 0.04 | 0.14 | 0.17 |
| 2 | 31552902 | rs932559 | 0.774 | 0.718 | -0.998 | -0.487 | -0.046 | -0.028 | -0.028 | 0 | 0.38 | 0.12 | 0.6 | 0.51 | 0.46 | 0.48 |
| 2 | 31553394 | rs207431 | 0.009 | 0.767 | -0.907 | -0.479 | 0.002 | -0.018 | -0.018 | 0 | 0.17 | 0.32 | 0.11 | 0.18 | 0.16 | 0.08 |
| 2 | 31553987 | rs207432 | -0.254 | 0.559 | -1.197 | -0.518 | -0.001 | -0.006 | -0.006 | 0 | 0.56 | 0.55 | 0.71 | 0.69 | 0.63 | 0.55 |
| 2 | 31556986 | rs6543626 | 0.359 | 0.869 | -2.371 | -0.371 | -0.014 | 0.008 | 0.008 | 0 | 0.15 | 0.05 | 0.09 | 0.16 | 0.15 | 0.07 |
| 2 | 31557308 | rs1054889 | 0.746 | 0.86 | -2.389 | -0.408 | -0.056 | 0.001 | 0.001 | 0 | 0.37 | 0.13 | 0.62 | 0.57 | 0.48 | 0.47 |
| 2 | 31557347 | rs6710015 | 0.212 | 0.878 | -2.366 | -0.436 | 0.031 | 0.073 | 0.073 | 0 | 0.45 | 0.76 | 0.27 | 0.3 | 0.33 | 0.46 |
| 2 | 31558306 | rs1042039 | 0.098 | 1.098 | -1.897 | -0.145 | -0.05 | 0.017 | 0.017 | 0 | 0.38 | 0.14 | 0.63 | 0.58 | 0.48 | 0.48 |
| 2 | 31562412 | rs207440 | -0.808 | 0.913 | -1.064 | 0.23 | -0.002 | -0.018 | -0.018 | 0 | 0.06 | 0.05 | 0.03 | 0.01 | 0.05 | 0.06 |
| 2 | 31567929 | rs207455 | 0.84 | 0.987 | -1.283 | 0.664 | -0.023 | -0.037 | -0.037 | 0 | 0.92 | 0.99 | 0.95 | 0.87 | 0.89 | 0.97 |
| 2 | 31568726 | rs9308919 | 0.631 | 1.15 | -0.767 | 0.143 | 0.007 | -0.036 | -0.036 | 2.852 | 0.79 | 0.71 | 0.75 | 0.84 | 0.78 | 0.82 |
| 2 | 31570689 | rs4952085 | 0.672 | 1.067 | -0.434 | 0.123 | 0 | -0.011 | -0.011 | 3.902 | 0.8 | 0.76 | 0.76 | 0.85 | 0.79 | 0.83 |
| 2 | 31571786 | rs1884725 | 0.657 | 1.067 | -0.434 | 0.146 | 0 | -0.01 | -0.01 | 0 | 0.8 | 0.77 | 0.76 | 0.85 | 0.79 | 0.84 |
| 2 | 31587235 | rs1018505 | 0.879 | 1.263 | -0.134 | 0.326 | -0.005 | 0.003 | 0.003 | 0 | 0.78 | 0.78 | 0.73 | 0.84 | 0.76 | 0.73 |
| 2 | 31588561 | rs1429376 | 0.787 | 1.223 | -0.153 | 0.338 | -0.005 | 0.002 | 0.002 | 0 | 0.79 | 0.79 | 0.74 | 0.83 | 0.76 | 0.74 |
| 2 | 31588577 | rs17011356 | 1.154 | 1.024 | -0.388 | 0.412 | -0.013 | 0.015 | 0.015 | 0 | 0.35 | 0.21 | 0.27 | 0.26 | 0.34 | 0.21 |
| 2 | 31588664 | rs12621192 | 0.74 | 0.424 | -0.654 | 0.535 | -0.032 | -0.08 | -0.08 | 0.011 | 0.26 | 0.07 | 0.38 | 0.41 | 0.32 | 0.38 |
| 2 | 31589414 | rs743163 | 1.726 | 0.428 | -0.646 | 0.52 | -0.013 | 0.015 | 0.015 | 0 | 0.35 | 0.21 | 0.27 | 0.26 | 0.34 | 0.21 |
| 2 | 31589654 | rs13387204 | 1.734 | 0.375 | 0.017 | 0.494 | -0.003 | 0.006 | 0.006 | 0 | 0.36 | 0.33 | 0.28 | 0.26 | 0.34 | 0.21 |
| 2 | 31589847 | rs2295475 | 1.101 | 0.227 | -0.132 | 0.557 | -0.043 | -0.068 | -0.068 | 0.025 | 0.21 | 0.07 | 0.37 | 0.33 | 0.3 | 0.37 |
| 2 | 31590156 | rs732436 | 0.259 | 0.077 | 0.371 | 0.443 | 0 | -0.047 | -0.047 | 1.041 | 0.72 | 0.57 | 0.7 | 0.85 | 0.75 | 0.71 |
| 2 | 31590545 | rs2281548 | 2.074 | 0.074 | 0.485 | 0.432 | -0.005 | 0.01 | 0.01 | 0 | 0.35 | 0.28 | 0.27 | 0.26 | 0.34 | 0.21 |
| 2 | 31590759 | rs2281550 | 2.069 | 0.056 | 0.514 | 0.422 | -0.012 | 0.014 | 0.014 | 0 | 0.34 | 0.21 | 0.26 | 0.26 | 0.34 | 0.21 |
| 2 | 31590785 | rs761926 | 1.12 | 0.087 | 0.501 | 0.44 | -0.04 | -0.068 | -0.068 | 0 | 0.21 | 0.07 | 0.36 | 0.34 | 0.3 | 0.38 |
| 2 | 31590917 | rs17011368 | 0.003 | 0.146 | 0.547 | 0.17 | -0.004 | -0.029 | -0.029 | 0 | 0.06 | 0.12 | 0.03 | 0 | 0.05 | 0.03 |
| 2 | 31592323 | rs2043013 | -1.242 | -0.007 | 0.56 | 0.15 | 0.016 | 0.003 | 0.003 | 0 | 0.46 | 0.63 | 0.49 | 0.6 | 0.43 | 0.54 |
| 2 | 31592403 | rs1366817 | -0.711 | 0.056 | 0.524 | 0.19 | -0.006 | 0.004 | 0.004 | 0 | 0.39 | 0.41 | 0.46 | 0.6 | 0.39 | 0.51 |
| 2 | 31598823 | rs2281547 | 1.154 | -0.005 | 0.669 | 0.178 | -0.001 | 0.011 | 0.011 | 0 | 0.39 | 0.3 | 0.29 | 0.28 | 0.36 | 0.24 |
| 2 | 31603134 | rs2163059 | 0.596 | 0.524 | 0.441 | 0.167 | 0.022 | 0.019 | 0.019 | 0.074 | 0.47 | 0.62 | 0.3 | 0.34 | 0.36 | 0.33 |
| 2 | 31605411 | rs6720163 | 0.437 | 0.321 | 0.474 | 0.138 | 0.007 | 0.032 | 0.032 | 0 | 0.5 | 0.64 | 0.35 | 0.34 | 0.44 | 0.34 |
| 2 | 31605447 | rs3769618 | 1.24 | 0.322 | 0.561 | 0.136 | -0.001 | 0.001 | 0.001 | 0.099 | 0.38 | 0.34 | 0.29 | 0.28 | 0.36 | 0.24 |
| 2 | 31605615 | rs1864280 | 0.398 | 0.413 | 0.04 | 0.22 | 0.007 | 0.045 | 0.045 | 0 | 0.51 | 0.71 | 0.35 | 0.34 | 0.44 | 0.35 |

| | | | | | | | | | | | | | | | | |
|---|---|---|---|---|---|---|---|---|---|---|---|---|---|---|---|---|
| 2 | 31606670 | rs4407290 | -0.001 | 0.345 | 0.007 | 0.389 | 0.014 | -0.003 | -0.003 | 0.123 | 0.06 | 0.01 | 0.03 | 0.05 | 0.05 | 0.09 |
| 2 | 31609993 | rs185925 | 0.079 | 0.653 | 0.621 | 0.39 | 0 | 0.001 | 0.001 | 0.274 | 0.75 | 0.78 | 0.59 | 0.41 | 0.69 | 0.58 |
| 2 | 31611029 | rs2073316 | 1.784 | 0.219 | 0.526 | -0.119 | -0.01 | 0.033 | 0.033 | 0 | 0.44 | 0.34 | 0.44 | 0.28 | 0.46 | 0.3 |
| 2 | 31611367 | rs206847 | 0.033 | 0.258 | 0.428 | -0.111 | -0.003 | -0.01 | -0.01 | 1.991 | 0.73 | 0.79 | 0.57 | 0.36 | 0.67 | 0.51 |
| 2 | 31612781 | rs13418515 | -0.375 | 0.276 | 0.496 | -0.143 | -0.005 | -0.056 | -0.056 | 0.87 | 0.18 | 0.18 | 0.1 | 0 | 0.16 | 0.14 |
| 2 | 31613249 | rs206849 | -0.576 | 0.709 | 0.464 | -0.147 | 0.006 | 0.008 | 0.008 | 0.379 | 0.78 | 0.8 | 0.57 | 0.42 | 0.63 | 0.57 |
| 2 | 31614225 | rs13431382 | -0.668 | 0.112 | 0.414 | -0.142 | -0.015 | 0.11 | 0.11 | 0.238 | 0.2 | 0.02 | 0.09 | 0 | 0.18 | 0.15 |
| 2 | 31614900 | rs11904439 | -0.029 | 0.094 | 0.373 | -0.158 | -0.005 | 0.066 | 0.066 | 0.421 | 0.11 | 0.01 | 0.02 | 0 | 0.05 | 0.04 |
| 2 | 31616749 | rs206854 | -0.661 | 0.094 | 0.404 | -0.141 | -0.004 | -0.007 | -0.007 | 0 | 0.79 | 0.82 | 0.6 | 0.42 | 0.71 | 0.59 |
| 2 | 31619101 | rs206857 | -1.405 | -0.016 | 0.409 | -0.201 | -0.02 | -0.074 | -0.074 | 0.371 | 0.25 | 0.29 | 0.12 | 0.07 | 0.19 | 0.15 |
| 2 | 31621598 | rs10175754 | -0.793 | -0.031 | 0.392 | -0.196 | -0.003 | -0.055 | -0.055 | 0.15 | 0.18 | 0.18 | 0.1 | 0 | 0.17 | 0.14 |
| 2 | 31622804 | rs206860 | -0.726 | 0.085 | 0.192 | 0.168 | -0.005 | -0.006 | -0.006 | 0 | 0.81 | 0.87 | 0.65 | 0.55 | 0.73 | 0.61 |
| 2 | 31623810 | rs6714794 | -0.804 | -0.207 | 0.147 | -0.021 | 0.005 | 0.004 | 0.004 | 0 | 0.08 | 0.11 | 0.02 | 0.07 | 0.05 | 0.11 |
| 2 | 31624836 | rs494852 | -2.061 | -0.322 | 0.225 | -0.091 | -0.013 | -0.021 | -0.021 | 0 | 0.25 | 0.29 | 0.13 | 0.12 | 0.2 | 0.25 |
| 2 | 31625610 | rs513311 | 0.025 | -0.314 | 0.242 | -0.083 | -0.026 | -0.015 | -0.015 | 0.072 | 0.08 | 0.15 | 0.01 | 0.06 | 0.02 | 0.01 |
| 2 | 31626636 | rs206798 | -2.17 | -0.32 | 0.239 | -0.09 | -0.014 | -0.021 | -0.021 | 0.146 | 0.25 | 0.28 | 0.13 | 0.12 | 0.2 | 0.25 |
| 2 | 31628705 | rs1366814 | -0.188 | -0.262 | 0.247 | -0.09 | 0.001 | -0.011 | -0.011 | 0 | 0.11 | 0.1 | 0.09 | 0 | 0.14 | 0.12 |
| 2 | 31629030 | rs206803 | -1.202 | -0.271 | 0.252 | -0.092 | -0.015 | -0.039 | -0.039 | 0.187 | 0.18 | 0.25 | 0.1 | 0.07 | 0.17 | 0.13 |
| 2 | 31630382 | rs3769616 | 0.727 | -0.358 | 0.18 | -0.125 | 0.029 | 0.026 | 0.026 | 3.928 | 0.09 | 0.02 | 0.03 | 0.05 | 0.05 | 0.1 |
| 2 | 31630846 | rs206804 | -2.095 | -0.36 | 0.193 | -0.143 | 0 | -0.012 | -0.012 | 0 | 0.29 | 0.29 | 0.14 | 0.11 | 0.22 | 0.28 |
| 2 | 31630867 | rs206805 | -2.07 | -0.359 | 0.194 | -0.143 | 0 | -0.013 | -0.013 | 0 | 0.29 | 0.28 | 0.14 | 0.11 | 0.22 | 0.28 |
| 2 | 31636915 | rs206811 | 0.434 | -0.265 | 0.278 | 0.043 | 0.006 | -0.014 | -0.014 | 0 | 0.67 | 0.76 | 0.57 | 0.36 | 0.66 | 0.49 |
| 2 | 31640455 | rs7575607 | 0.986 | -0.29 | 0.285 | 0.037 | 0.01 | 0.004 | 0.004 | 0 | 0.6 | 0.65 | 0.54 | 0.29 | 0.63 | 0.49 |
| 2 | 31644983 | rs206816 | 0.562 | -0.265 | 0.345 | 0.036 | 0.013 | -0.012 | -0.012 | 0 | 0.65 | 0.74 | 0.55 | 0.35 | 0.64 | 0.46 |
| 2 | 31646126 | rs206818 | 0.179 | -0.299 | 0.391 | 0.073 | -0.021 | -0.006 | -0.006 | 0 | 0.07 | 0.13 | 0.02 | 0.06 | 0.02 | 0.01 |
| 2 | 31656380 | rs7595578 | 1.896 | -0.272 | 0.462 | 0.023 | 0.002 | 0.003 | 0.003 | 0 | 0.47 | 0.53 | 0.45 | 0.29 | 0.48 | 0.33 |
| 2 | 31671769 | rs7579660 | 0.333 | -0.176 | 0.453 | 0.059 | 0 | -0.011 | -0.011 | 0 | 0.09 | 0.07 | 0.07 | 0 | 0.12 | 0.11 |
| 2 | 31683940 | rs206826 | -0.539 | -0.365 | 0.344 | 0.153 | -0.015 | -0.057 | -0.057 | 3.699 | 0.79 | 0.91 | 0.66 | 0.56 | 0.74 | 0.61 |
| 2 | 31686309 | rs206827 | 0.366 | -0.333 | 0.336 | -0.018 | 0.012 | -0.02 | -0.02 | 1.897 | 0.68 | 0.79 | 0.59 | 0.36 | 0.69 | 0.51 |
| 2 | 31693451 | rs761925 | 2.23 | -0.302 | 0.569 | -0.025 | -0.001 | -0.003 | -0.003 | 0 | 0.44 | 0.46 | 0.45 | 0.28 | 0.48 | 0.32 |
| 2 | 31700798 | rs11888215 | 0.129 | -0.3 | 0.479 | -0.03 | 0 | -0.029 | -0.029 | 0 | 0.1 | 0.09 | 0.08 | 0 | 0.13 | 0.11 |
| 2 | 31709417 | rs206834 | -1.392 | -0.357 | 0.706 | -0.085 | -0.017 | -0.045 | -0.045 | 4.179 | 0.28 | 0.32 | 0.17 | 0.12 | 0.24 | 0.22 |
| 2 | 31729141 | rs541263 | -1.106 | -0.377 | 0.936 | -0.167 | -0.076 | -0.094 | -0.094 | 0 | 0.23 | 0.43 | 0.13 | 0.11 | 0.14 | 0.11 |
| 2 | 31738219 | rs2365778 | 1.846 | -0.301 | 0.719 | -0.09 | 0.008 | 0.011 | 0.011 | 0 | 0.48 | 0.48 | 0.5 | 0.29 | 0.5 | 0.34 |
| 2 | 31742467 | rs2224310 | -0.241 | -0.372 | 0.545 | -0.034 | 0 | -0.015 | -0.015 | 0 | 0.77 | 0.82 | 0.67 | 0.4 | 0.76 | 0.57 |
| 2 | 31748207 | rs1884722 | -0.341 | -0.482 | 0.58 | -0.168 | -0.001 | 0.001 | 0.001 | 0 | 0.18 | 0.16 | 0.08 | 0.12 | 0.11 | 0.11 |
| 2 | 31749714 | rs1042578 | -1.015 | -0.527 | 0.709 | -0.165 | -0.037 | -0.028 | -0.028 | 0 | 0.22 | 0.26 | 0.1 | 0.13 | 0.14 | 0.17 |
| 2 | 31750417 | rs9332975 | -0.442 | -0.536 | 0.69 | -0.169 | -0.018 | -0.011 | -0.011 | 0 | 0.19 | 0.2 | 0.08 | 0.12 | 0.12 | 0.11 |
| 2 | 31755485 | rs17011453 | -0.857 | -0.543 | 0.721 | -0.173 | -0.007 | -0.004 | -0.004 | 0 | 0.22 | 0.22 | 0.1 | 0.12 | 0.13 | 0.16 |
| 2 | 31757024 | rs2281546 | -1.176 | -0.549 | 0.676 | -0.175 | -0.026 | -0.019 | -0.019 | 0 | 0.23 | 0.26 | 0.1 | 0.13 | 0.14 | 0.17 |
| 2 | 31763558 | rs12470143 | 2.524 | -0.414 | 0.756 | 0.031 | -0.014 | 0.009 | 0.009 | 0 | 0.41 | 0.3 | 0.47 | 0.34 | 0.46 | 0.31 |
| 2 | 31765556 | rs4952220 | 1.534 | -0.424 | 0.838 | 0.046 | 0.014 | 0.014 | 0.014 | 0 | 0.53 | 0.43 | 0.55 | 0.35 | 0.57 | 0.44 |
| 2 | 31773795 | rs11690596 | -0.941 | -0.441 | 0.901 | 0.047 | -0.075 | -0.057 | -0.057 | 0 | 0.23 | 0.38 | 0.1 | 0.12 | 0.14 | 0.15 |
| 2 | 31782280 | rs2268796 | 1.593 | -0.344 | 0.821 | 0.086 | 0.025 | 0.013 | 0.013 | 0 | 0.53 | 0.38 | 0.56 | 0.36 | 0.59 | 0.43 |
| 2 | 31782791 | rs12467911 | -0.175 | -0.246 | 0.573 | 0.141 | -0.006 | -0.021 | -0.021 | 0 | 0.76 | 0.82 | 0.66 | 0.49 | 0.72 | 0.6 |
| 2 | 31786637 | rs2300697 | 0.69 | -0.255 | 0.557 | 0.1 | -0.009 | -0.027 | -0.027 | 0 | 0.64 | 0.76 | 0.54 | 0.47 | 0.59 | 0.5 |
| 2 | 31786967 | rs2300699 | 0.239 | -0.23 | 0.553 | 0.214 | -0.005 | -0.023 | -0.023 | 0 | 0.73 | 0.8 | 0.62 | 0.47 | 0.7 | 0.58 |
| 2 | 31791793 | rs7594951 | -0.133 | -0.298 | 0.618 | 0.227 | -0.002 | 0.001 | 0.001 | 0 | 0.17 | 0.13 | 0.07 | 0.12 | 0.11 | 0.11 |
| 2 | 31803403 | rs2754530 | 0.471 | -0.221 | 0.744 | 0.273 | -0.004 | 0.069 | 0.069 | 0 | 0.7 | 0.57 | 0.64 | 0.49 | 0.7 | 0.55 |
| 2 | 31805675 | rs522638 | 0.095 | -0.194 | 0.717 | 0.286 | 0.001 | 0.011 | 0.011 | 0 | 0.74 | 0.74 | 0.66 | 0.48 | 0.71 | 0.6 |
| 2 | 31807940 | rs806645 | 0.543 | 0.154 | 0.72 | 0.261 | -0.002 | 0.066 | 0.066 | 0 | 0.71 | 0.59 | 0.64 | 0.49 | 0.7 | 0.55 |
| 2 | 31808300 | rs3754838 | -0.153 | -0.019 | 0.726 | 0.158 | 0.009 | 0.008 | 0.008 | 0 | 0.18 | 0.14 | 0.07 | 0.12 | 0.12 | 0.11 |
| 2 | 31811268 | rs502139 | 0.567 | 0.054 | 0.529 | 0.2 | -0.008 | 0.064 | 0.064 | 0 | 0.69 | 0.56 | 0.64 | 0.49 | 0.69 | 0.53 |
| 2 | 31811870 | rs612224 | -0.173 | 0.063 | 0.399 | 0.383 | -0.009 | -0.028 | -0.028 | 0 | 0.77 | 0.86 | 0.67 | 0.55 | 0.71 | 0.61 |
| 2 | 31834078 | rs623419 | 0.212 | 0.138 | 0.424 | 0.288 | 0.001 | 0.014 | 0.014 | 0 | 0.73 | 0.71 | 0.59 | 0.49 | 0.7 | 0.59 |
| 2 | 31871583 | rs682895 | 0.068 | 0.062 | 0.344 | 0.243 | -0.001 | 0.003 | 0.003 | 0 | 0.75 | 0.75 | 0.65 | 0.49 | 0.72 | 0.6 |
| 2 | 31873476 | rs553464 | 0.913 | 0.171 | 0.367 | 0.196 | -0.001 | 0.005 | 0.005 | 0.052 | 0.66 | 0.65 | 0.56 | 0.47 | 0.67 | 0.51 |
| 2 | 31891571 | rs13027103 | -0.057 | 0.043 | 0.252 | 0.112 | 0.029 | 0.014 | 0.014 | 0 | 0.16 | 0.11 | 0.07 | 0.12 | 0.11 | 0.09 |
| 2 | 31912164 | rs806584 | 2.085 | 0.215 | 0.487 | 0.059 | 0 | 0.004 | 0.004 | 0 | 0.43 | 0.38 | 0.49 | 0.34 | 0.49 | 0.34 |
| 2 | 31933642 | rs806591 | 0.727 | 0.067 | 0.377 | -0.008 | -0.019 | -0.014 | -0.014 | 0 | 0.78 | 0.77 | 0.9 | 0.9 | 0.87 | 0.86 |
| 2 | 31933889 | rs806592 | 0.974 | -0.233 | 0.127 | -0.34 | 0.001 | -0.002 | -0.002 | 3.355 | 0.74 | 0.76 | 0.88 | 0.9 | 0.82 | 0.84 |
| 2 | 31933896 | rs6543636 | 0.365 | -0.053 | 0.052 | -0.302 | 0.001 | -0.006 | -0.006 | 0 | 0.1 | 0.08 | 0.09 | 0 | 0.12 | 0.08 |
| 2 | 31973127 | rs11695295 | -0.277 | -0.026 | 0.176 | -0.224 | 0.001 | 0.008 | 0.008 | 0 | 0.27 | 0.28 | 0.41 | 0.6 | 0.29 | 0.44 |
| 2 | 31986063 | rs223631 | 2.243 | -0.147 | 0.215 | -0.369 | 0 | -0.002 | -0.002 | 0.075 | 0.42 | 0.43 | 0.5 | 0.34 | 0.48 | 0.31 |
| 2 | 32014315 | rs223657 | 0.59 | -0.299 | 0.054 | -0.431 | -0.029 | -0.02 | -0.02 | 0 | 0.78 | 0.73 | 0.89 | 0.88 | 0.86 | 0.83 |
| 2 | 32020049 | rs6735704 | 1.426 | -0.26 | -0.283 | -0.724 | -0.011 | 0.024 | 0.024 | 0 | 0.06 | 0.03 | 0.06 | 0 | 0.1 | 0.04 |
| 2 | 32057759 | rs160804 | 2.558 | -0.181 | -0.209 | -0.599 | 0.01 | 0.002 | 0.002 | 0 | 0.34 | 0.35 | 0.36 | 0.37 | 0.38 | 0.26 |
| 2 | 32070489 | rs11904162 | -0.489 | -0.223 | -0.675 | -0.51 | 0 | -0.001 | -0.001 | 0 | 0.29 | 0.27 | 0.41 | 0.87 | 0.29 | 0.47 |
| 2 | 32071502 | rs6739055 | -0.061 | -0.274 | -0.667 | -0.43 | -0.002 | 0.004 | 0.004 | 0 | 0.24 | 0.18 | 0.33 | 0.79 | 0.27 | 0.44 |

| | | | | | | | | | | | | | | | | |
|---|---|---|---|---|---|---|---|---|---|---|---|---|---|---|---|---|
| 2 | 32103689 | rs4952203 | 0.697 | -0.169 | -0.783 | -0.328 | -0.026 | -0.093 | -0.093 | 0 | 0.18 | 0.04 | 0.23 | 0.43 | 0.22 | 0.39 |
| 2 | 32112652 | rs7591270 | 0.157 | -0.29 | -0.804 | -0.464 | 0.046 | 0.028 | 0.028 | 0 | 0.18 | 0.07 | 0.07 | 0.11 | 0.11 | 0.1 |
| 2 | 32128601 | rs3769611 | -0.27 | 0.138 | -1.397 | -0.772 | -0.001 | -0.011 | -0.011 | 0 | 0.28 | 0.24 | 0.29 | 0.52 | 0.21 | 0.41 |
| 2 | 32183611 | rs7559329 | -0.559 | -0.281 | -0.969 | -0.675 | -0.002 | -0.006 | -0.006 | 0 | 0.24 | 0.23 | 0.11 | 0.12 | 0.16 | 0.17 |
| 2 | 32183767 | rs10439444 | 2.231 | -0.239 | -0.883 | -0.727 | 0.002 | -0.002 | -0.002 | 0 | 0.4 | 0.39 | 0.48 | 0.34 | 0.45 | 0.3 |
| 2 | 32206029 | rs3754835 | 0.772 | -1.128 | -1.088 | -0.648 | -0.001 | -0.009 | -0.009 | 0 | 0.66 | 0.66 | 0.59 | 0.47 | 0.61 | 0.52 |
| 2 | 32212935 | rs2064813 | 1.029 | -1.011 | -1.338 | -0.324 | -0.018 | -0.02 | -0.02 | 5.724 | 0.1 | 0.03 | 0.18 | 0.35 | 0.16 | 0.35 |
| 2 | 32223456 | rs6734180 | 0.863 | -0.973 | -1.222 | -0.603 | 0.003 | -0.011 | -0.011 | 0.713 | 0.1 | 0 | 0.04 | 0.08 | 0.09 | 0.06 |
| 2 | 32247942 | rs6752375 | 2.466 | -1.047 | -1.42 | -0.687 | -0.161 | -0.116 | -0.116 | 0.355 | 0.25 | 0.03 | 0.3 | 0.35 | 0.34 | 0.2 |
| 2 | 32265125 | rs12991444 | 0.091 | -1.298 | -0.457 | 0.026 | -0.002 | -0.024 | -0.024 | 0 | 0.25 | 0.23 | 0.33 | 0.53 | 0.27 | 0.44 |
| 2 | 32278782 | rs7577696 | -1.543 | -1.451 | -0.288 | -0.14 | 0.021 | 0.021 | 0.021 | 0 | 0.48 | 0.4 | 0.42 | 0.65 | 0.4 | 0.61 |
| 2 | 32280699 | rs2365555 | -1.293 | -1.472 | -0.272 | -0.151 | 0.026 | 0.024 | 0.024 | 0 | 0.46 | 0.36 | 0.41 | 0.65 | 0.39 | 0.57 |
| 2 | 32301631 | rs467523 | -1.942 | -1.415 | 0.042 | -0.16 | -0.003 | 0.012 | 0.012 | 0 | 0.57 | 0.65 | 0.51 | 0.65 | 0.53 | 0.65 |
| 2 | 32322495 | rs3769604 | -1.461 | -1.402 | -0.274 | -0.121 | 0.017 | 0.018 | 0.018 | 0 | 0.48 | 0.4 | 0.42 | 0.65 | 0.4 | 0.6 |
| 2 | 32337391 | rs7581340 | -1.428 | -1.387 | -0.273 | -0.111 | 0.039 | 0.037 | 0.037 | 0 | 0.47 | 0.34 | 0.41 | 0.65 | 0.4 | 0.6 |
| 2 | 32339561 | rs11691939 | 0.217 | -1.504 | -0.306 | -0.123 | -0.004 | -0.035 | -0.035 | 0.122 | 0.25 | 0.19 | 0.33 | 0.53 | 0.27 | 0.43 |
| 2 | 32358864 | rs13428933 | 0.111 | -1.599 | -0.308 | -0.171 | 0.01 | 0.059 | 0.059 | 0 | 0.16 | 0.07 | 0.11 | 0 | 0.18 | 0.05 |
| 2 | 32363171 | rs12614686 | -1.113 | -1.6 | -0.28 | -0.168 | 0.01 | 0.004 | 0.004 | 0 | 0.4 | 0.36 | 0.38 | 0.65 | 0.33 | 0.6 |
| 2 | 32381829 | rs3769602 | -0.958 | -1.608 | -0.263 | -0.118 | -0.002 | -0.001 | -0.001 | 0 | 0.4 | 0.42 | 0.38 | 0.66 | 0.33 | 0.6 |
| 2 | 32386659 | rs10208246 | 1.964 | -1.398 | 0.006 | -0.082 | -0.009 | 0.011 | 0.011 | 0 | 0.41 | 0.29 | 0.49 | 0.35 | 0.47 | 0.35 |
| 2 | 32389646 | rs34197800 | 0.072 | -1.233 | -1.035 | 0.048 | -0.004 | -0.038 | -0.038 | 0 | 0.25 | 0.18 | 0.32 | 0.53 | 0.27 | 0.44 |
| 2 | 32391231 | rs13013695 | 0.125 | -1.198 | -1.041 | 0.045 | 0.004 | -0.032 | -0.032 | 0 | 0.25 | 0.18 | 0.32 | 0.53 | 0.25 | 0.43 |
| 2 | 32395315 | rs7581432 | -1.846 | -1.233 | -0.994 | 0.076 | 0.005 | 0.019 | 0.019 | 0 | 0.54 | 0.41 | 0.49 | 0.64 | 0.5 | 0.59 |
| 2 | 32437338 | rs212689 | -0.436 | -1.246 | -1.399 | 0.314 | -0.001 | -0.01 | -0.01 | 1.216 | 0.76 | 0.82 | 0.74 | 0.61 | 0.74 | 0.58 |
| 2 | 32444054 | rs212701 | 0.73 | -1.345 | -1.458 | 0.049 | 0.031 | -0.011 | -0.011 | 0 | 0.57 | 0.72 | 0.66 | 0.46 | 0.62 | 0.42 |
| 2 | 32447718 | rs177083 | 1.398 | -1.347 | -1.447 | 0.043 | 0.013 | 0.004 | 0.004 | 0 | 0.49 | 0.6 | 0.58 | 0.46 | 0.52 | 0.37 |
| 2 | 32449261 | rs6757121 | 0.791 | -1.324 | -1.676 | 0.063 | 0.036 | 0.021 | 0.021 | 0 | 0.17 | 0.06 | 0.07 | 0.11 | 0.11 | 0.11 |
| 2 | 32450348 | rs212704 | 1.342 | -1.265 | -1.627 | 0.096 | 0.019 | 0.003 | 0.003 | 0 | 0.47 | 0.63 | 0.58 | 0.46 | 0.52 | 0.38 |
| 2 | 32473126 | rs212736 | -0.076 | -0.879 | -1.608 | 0.661 | 0.001 | -0.011 | -0.011 | 0 | 0.75 | 0.79 | 0.74 | 0.59 | 0.76 | 0.59 |
| 2 | 32475109 | rs455060 | 0.938 | -0.917 | -1.599 | 0.683 | 0.038 | -0.005 | -0.005 | 0.166 | 0.56 | 0.72 | 0.66 | 0.46 | 0.62 | 0.48 |
| 2 | 32476992 | rs410469 | 0.398 | 0.226 | -1.066 | 0.997 | 0 | -0.016 | -0.016 | 0 | 0.74 | 0.78 | 0.68 | 0.49 | 0.73 | 0.58 |
| 2 | 32478629 | rs7562653 | 0.417 | 0.312 | -1.251 | 1.04 | -0.003 | -0.034 | -0.034 | 0 | 0.75 | 0.82 | 0.68 | 0.49 | 0.73 | 0.58 |
| 2 | 32495395 | rs438408 | 0.506 | 0.41 | -1.104 | 0.868 | -0.001 | -0.002 | -0.002 | 0 | 0.7 | 0.73 | 0.64 | 0.38 | 0.67 | 0.51 |
| 2 | 32495471 | rs409188 | -0.594 | -0.494 | -1.543 | -0.22 | 0.04 | 0.062 | 0.062 | 1.072 | 0.37 | 0.71 | 0.22 | 0.12 | 0.25 | 0.16 |
| 2 | 32502495 | rs387780 | 0.214 | -0.274 | -1.642 | -0.015 | -0.007 | -0.018 | -0.018 | 1.702 | 0.7 | 0.86 | 0.69 | 0.36 | 0.69 | 0.46 |
| 2 | 32503526 | rs398290 | 0.776 | -0.374 | -1.401 | -0.208 | 0.004 | 0.003 | 0.003 | 0 | 0.65 | 0.74 | 0.66 | 0.36 | 0.66 | 0.45 |
| 2 | 32509330 | rs4952258 | 0.812 | -0.349 | -0.907 | -0.643 | 0.011 | -0.045 | -0.045 | 0.571 | 0.09 | 0.19 | 0.19 | 0 | 0.09 | 0.1 |
| 2 | 32510072 | rs390480 | 0.491 | -0.381 | -0.772 | -0.644 | 0.006 | 0.008 | 0.008 | 0 | 0.65 | 0.77 | 0.66 | 0.33 | 0.66 | 0.45 |
| 2 | 32535568 | rs385760 | 2.326 | 0.702 | -1.098 | -0.98 | -0.04 | 0.037 | 0.037 | 0 | 0.07 | 0 | 0.04 | 0 | 0.07 | 0.05 |
| 2 | 32545880 | rs2038537 | -1.376 | 0.063 | -1.111 | -1.282 | 0.009 | 0.033 | 0.033 | 0 | 0.44 | 0.32 | 0.54 | 0.68 | 0.44 | 0.7 |
| 2 | 32550485 | rs6718214 | -2.204 | 0.123 | -0.736 | -1.843 | -0.009 | 0.015 | 0.015 | 0 | 0.63 | 0.69 | 0.64 | 0.69 | 0.58 | 0.74 |
| 2 | 32579759 | rs12612824 | 0.349 | 0.044 | -0.784 | -1.758 | 0 | -0.009 | -0.009 | 0 | 0.3 | 0.2 | 0.32 | 0.63 | 0.31 | 0.53 |
| 2 | 32597234 | rs4952262 | -2.175 | -0.254 | -0.689 | -1.945 | -0.056 | 0.036 | 0.036 | 0.251 | 0.63 | 0.79 | 0.63 | 0.7 | 0.51 | 0.73 |
| 2 | 32611512 | rs4952211 | -1.064 | -0.25 | -0.651 | -2.169 | 0.002 | 0.009 | 0.009 | 0 | 0.45 | 0.55 | 0.54 | 0.65 | 0.44 | 0.59 |
| 2 | 32633569 | rs176400 | -3.829 | -0.166 | -0.948 | -1.935 | -0.122 | -0.079 | -0.079 | 0.016 | 0.77 | 0.98 | 0.73 | 0.71 | 0.68 | 0.76 |
| 2 | 32654636 | rs12712328 | 1.158 | -0.533 | -0.155 | -1.773 | -0.07 | -0.119 | -0.119 | 0.004 | 0.78 | 0.48 | 0.88 | 0.95 | 0.89 | 0.88 |
| 2 | 32661543 | rs12992095 | 2.058 | -0.405 | -0.53 | -1.001 | -0.016 | -0.118 | -0.118 | 0 | 0.19 | 0.03 | 0.2 | 0.53 | 0.2 | 0.45 |
| 2 | 32666137 | rs13409142 | 0.569 | -0.485 | -0.453 | -1.082 | 0.012 | 0.011 | 0.011 | 0 | 0.63 | 0.72 | 0.66 | 0.34 | 0.66 | 0.45 |
| 2 | 32667562 | rs2276592 | 1.746 | -0.391 | -0.556 | -0.97 | 0 | -0.046 | -0.046 | 0.617 | 0.21 | 0.1 | 0.24 | 0.56 | 0.23 | 0.46 |
| 2 | 32679732 | rs3769598 | 0.577 | -0.349 | -0.408 | -1.221 | -0.006 | 0.057 | 0.057 | 0 | 0.16 | 0.05 | 0.1 | 0.05 | 0.15 | 0.08 |
| 2 | 32686582 | rs10181198 | -0.68 | -0.404 | -0.326 | -1.271 | -0.001 | -0.003 | -0.003 | 0 | 0.7 | 0.8 | 0.68 | 0.35 | 0.69 | 0.48 |
| 2 | 32705085 | rs6543657 | 0.482 | -0.292 | -0.391 | -1.235 | 0 | -0.001 | -0.001 | 0 | 0.61 | 0.6 | 0.49 | 0.35 | 0.61 | 0.38 |
| 2 | 32735881 | rs3731579 | -0.963 | -0.207 | -0.508 | -1.187 | 0.001 | -0.009 | -0.009 | 0 | 0.24 | 0.21 | 0.28 | 0.05 | 0.23 | 0.2 |
| 2 | 32751182 | rs11683905 | -0.288 | -0.163 | -0.627 | -1.143 | -0.004 | 0.046 | 0.046 | 0 | 0.2 | 0.11 | 0.26 | 0.05 | 0.21 | 0.18 |
| 2 | 32751688 | rs12466713 | 0.943 | -0.083 | -0.659 | -1.086 | -0.007 | 0.026 | 0.026 | 0 | 0.15 | 0.09 | 0.23 | 0.05 | 0.16 | 0.13 |
| 2 | 32758353 | rs2710628 | 1.939 | -0.017 | -0.609 | -0.942 | -0.02 | 0.017 | 0.017 | 0.541 | 0.37 | 0.26 | 0.36 | 0.32 | 0.44 | 0.29 |
| 2 | 32775984 | rs2243844 | 1.929 | -0.22 | -0.596 | -0.753 | -0.008 | 0.018 | 0.018 | 0 | 0.37 | 0.33 | 0.37 | 0.24 | 0.42 | 0.26 |
| 2 | 32806385 | rs6721331 | 2.14 | -0.366 | -0.533 | -0.891 | -0.006 | -0.027 | -0.027 | 0.251 | 0.05 | 0.1 | 0.02 | 0 | 0.03 | 0.02 |
| 2 | 32808804 | rs2710634 | 1.253 | -0.271 | -0.54 | -0.797 | -0.009 | 0.015 | 0.015 | 0 | 0.42 | 0.38 | 0.54 | 0.3 | 0.51 | 0.39 |
| 2 | 32822957 | rs2710625 | 1.681 | -0.239 | -0.649 | -0.745 | -0.01 | 0.014 | 0.014 | 0 | 0.38 | 0.32 | 0.37 | 0.3 | 0.43 | 0.26 |
| 2 | 32849207 | rs6723226 | -0.627 | -0.115 | -0.776 | -0.247 | 0.001 | 0.012 | 0.012 | 0 | 0.35 | 0.21 | 0.33 | 0.68 | 0.34 | 0.56 |
| 2 | 32861950 | rs2568465 | 0.845 | -0.895 | -0.614 | -0.101 | -0.007 | 0.012 | 0.012 | 0 | 0.63 | 0.62 | 0.55 | 0.55 | 0.67 | 0.43 |
| 2 | 32868255 | rs2710624 | 1.126 | -0.969 | -0.433 | -0.025 | -0.017 | -0.057 | -0.057 | 0 | 0.75 | 0.59 | 0.72 | 0.9 | 0.83 | 0.81 |
| 2 | 32881819 | rs17012046 | 0.899 | -0.924 | -0.345 | 0.145 | -0.001 | 0.002 | 0.002 | 0.008 | 0.1 | 0.09 | 0.12 | 0.13 | 0.08 | 0.05 |
| 2 | 32897055 | rs2295728 | 1.341 | -0.902 | -0.408 | 0.157 | -0.002 | -0.009 | -0.009 | 0 | 0.08 | 0.05 | 0.11 | 0.12 | 0.07 | 0.04 |
| 2 | 32897147 | rs2295730 | 1.235 | -1.095 | -0.467 | 0.421 | 0.009 | -0.027 | -0.027 | 0 | 0.12 | 0.08 | 0.1 | 0.27 | 0.11 | 0.31 |
| 2 | 32900330 | rs7594173 | -0.545 | -1.601 | -0.525 | -0.431 | -0.012 | -0.029 | -0.029 | 0.198 | 0.35 | 0.39 | 0.2 | 0.13 | 0.23 | 0.16 |
| 2 | 32903053 | rs12994364 | 2.162 | -1.129 | -0.767 | 0.03 | -0.009 | -0.003 | -0.003 | 0.206 | 0.05 | 0.03 | 0.03 | 0.21 | 0.06 | 0.02 |
| 2 | 32904596 | rs7563448 | -0.451 | -1.065 | -0.665 | -0.121 | 0.004 | -0.027 | -0.027 | 0.446 | 0.92 | 0.89 | 0.8 | 1 | 0.91 | 0.89 |

| | | | | | | | | | | | | | | | | |
|---|---|---|---|---|---|---|---|---|---|---|---|---|---|---|---|---|
| 2 | 32918112 | rs4952283 | 0.781 | -1.546 | -0.579 | -0.32 | -0.009 | -0.005 | -0.005 | 0 | 0.62 | 0.54 | 0.62 | 0.67 | 0.71 | 0.72 |
| 2 | 32918720 | rs3769589 | 3.104 | -1.484 | -1.015 | -0.422 | -0.036 | -0.023 | -0.023 | 0 | 0.24 | 0.14 | 0.35 | 0.35 | 0.36 | 0.31 |
| 2 | 32922558 | rs958243 | 0.213 | -1.744 | -0.87 | -0.594 | -0.007 | -0.022 | -0.022 | 0 | 0.67 | 0.41 | 0.69 | 0.79 | 0.75 | 0.82 |
| 2 | 32924202 | rs17012106 | 1.821 | -1.873 | -0.815 | -0.625 | -0.03 | -0.044 | -0.044 | 0 | 0.09 | 0.2 | 0.03 | 0 | 0.02 | 0.05 |
| 2 | 32926031 | rs12612455 | -0.125 | -1.964 | -1.093 | -0.642 | 0.018 | 0 | 0 | 0 | 0.31 | 0.22 | 0.2 | 0.29 | 0.26 | 0.47 |
| 2 | 32927576 | rs11893716 | 1.182 | -1.978 | -0.846 | -0.674 | -0.024 | -0.078 | -0.078 | 1.064 | 0.16 | 0.26 | 0.05 | 0 | 0.08 | 0.08 |
| 2 | 32927863 | rs2273659 | 0.934 | -1.792 | -0.865 | -0.521 | -0.001 | -0.066 | -0.066 | 0 | 0.19 | 0.04 | 0.15 | 0.29 | 0.19 | 0.4 |
| 2 | 32928119 | rs7586406 | -0.184 | -1.713 | -0.855 | -0.517 | -0.003 | -0.002 | -0.002 | 0.458 | 0.19 | 0.2 | 0.15 | 0.12 | 0.12 | 0.14 |
| 2 | 32933095 | rs6743661 | 0.907 | -1.512 | -1.134 | -0.295 | -0.029 | -0.102 | -0.102 | 0 | 0.19 | 0.03 | 0.19 | 0.34 | 0.23 | 0.44 |
| 2 | 32940834 | rs3754834 | -1.33 | -1.604 | -1.02 | -0.399 | 0.024 | -0.009 | -0.009 | 0 | 0.52 | 0.33 | 0.36 | 0.46 | 0.45 | 0.58 |
| 2 | 32942791 | rs12712329 | 3.127 | -1.655 | -0.984 | -0.166 | -0.018 | -0.013 | -0.013 | 0 | 0.22 | 0.16 | 0.34 | 0.34 | 0.36 | 0.22 |
| 2 | 32948356 | rs2052998 | 0.647 | -1.528 | -0.934 | 0.112 | -0.016 | -0.078 | -0.078 | 0 | 0.18 | 0.03 | 0.15 | 0.3 | 0.2 | 0.39 |
| 2 | 32958187 | rs1028600 | 1.252 | -1.698 | -0.542 | -0.272 | -0.003 | 0.008 | 0.008 | 0.003 | 0.47 | 0.39 | 0.43 | 0.37 | 0.54 | 0.38 |
| 2 | 32960091 | rs762027 | -0.387 | -1.702 | -0.562 | -0.249 | -0.001 | -0.001 | -0.001 | 0 | 0.8 | 0.72 | 0.72 | 0.76 | 0.84 | 0.89 |
| 2 | 32968121 | rs3769583 | 1.174 | -1.706 | -0.687 | -0.216 | -0.021 | -0.089 | -0.089 | 0.052 | 0.16 | 0.03 | 0.16 | 0.32 | 0.18 | 0.4 |
| 2 | 32969116 | rs3769581 | -0.575 | -1.751 | -0.413 | -0.291 | -0.002 | 0 | 0 | 0 | 0.29 | 0.29 | 0.24 | 0.32 | 0.27 | 0.46 |
| 2 | 32976087 | rs3769579 | 0.189 | -1.938 | -0.292 | -0.287 | -0.016 | -0.005 | -0.005 | 0.462 | 0.14 | 0.22 | 0.09 | 0.1 | 0.06 | 0.05 |
| 2 | 32977741 | rs2116588 | -0.473 | -1.81 | 0.289 | -0.217 | -0.002 | -0.002 | -0.002 | 0 | 0.81 | 0.8 | 0.73 | 0.79 | 0.84 | 0.84 |
| 2 | 32981379 | rs7606648 | 0.916 | -1.869 | 0.432 | -0.315 | -0.011 | -0.059 | -0.059 | 0.053 | 0.18 | 0.21 | 0.05 | 0 | 0.12 | 0.16 |
| 2 | 32983480 | rs2273664 | 2.322 | -1.796 | 0.193 | -0.243 | -0.027 | -0.049 | -0.049 | 0 | 0.08 | 0 | 0.04 | 0.2 | 0.06 | 0.04 |
| 2 | 32985666 | rs13008476 | -0.12 | -1.68 | 0.656 | -0.118 | 0 | -0.006 | -0.006 | 0 | 0.72 | 0.76 | 0.66 | 0.69 | 0.78 | 0.79 |
| 2 | 32986777 | rs4952291 | -0.201 | -1.63 | 0.696 | -0.11 | 0.005 | -0.012 | -0.012 | 0 | 0.24 | 0.21 | 0.2 | 0.35 | 0.22 | 0.44 |
| 2 | 32988331 | rs17012217 | 0.149 | -1.475 | 0.662 | -0.101 | -0.014 | -0.017 | -0.017 | 0 | 0.19 | 0.26 | 0.11 | 0.1 | 0.11 | 0.08 |
| 2 | 32995556 | rs6724076 | 0.086 | -1.477 | 0.676 | -0.102 | -0.016 | -0.019 | -0.019 | 0 | 0.19 | 0.27 | 0.12 | 0.1 | 0.11 | 0.09 |
| 2 | 32998606 | rs2116585 | -1.278 | -1.329 | 0.803 | -0.032 | -0.002 | -0.002 | -0.002 | 0 | 0.54 | 0.54 | 0.34 | 0.41 | 0.43 | 0.58 |
| 2 | 33005238 | rs6729463 | 0.825 | -1.316 | 0.516 | -0.069 | 0.016 | 0.009 | 0.009 | 0 | 0.17 | 0.1 | 0.05 | 0 | 0.1 | 0.16 |
| 2 | 33009932 | rs10490456 | 0.515 | -1.326 | 0.592 | -0.075 | -0.002 | -0.037 | -0.037 | 0.039 | 0.19 | 0.16 | 0.05 | 0 | 0.12 | 0.16 |
| 2 | 33013756 | rs220655 | 0.999 | -1.136 | 0.591 | 0.123 | -0.004 | -0.008 | -0.008 | 2.35 | 0.17 | 0.16 | 0.26 | 0.22 | 0.16 | 0.13 |
| 2 | 33013822 | rs3769570 | -1.086 | -1.184 | 0.733 | 0.088 | 0.003 | 0 | 0 | 0.029 | 0.56 | 0.64 | 0.36 | 0.41 | 0.43 | 0.62 |
| 2 | 33030709 | rs17012300 | 0.225 | -1.163 | 0.87 | -0.104 | -0.016 | -0.036 | -0.036 | 0 | 0.21 | 0.43 | 0.13 | 0.11 | 0.13 | 0.08 |
| 2 | 33031358 | rs9308929 | -0.516 | -1.094 | 0.388 | 0.011 | 0.015 | 0.005 | 0.005 | 0.003 | 0.5 | 0.27 | 0.48 | 0.48 | 0.46 | 0.66 |
| 2 | 33034057 | rs3769567 | -1.629 | -0.63 | 0.463 | 0.26 | -0.025 | -0.023 | -0.023 | 0 | 0.72 | 0.8 | 0.6 | 0.59 | 0.58 | 0.74 |
| 2 | 33037159 | rs927087 | -0.618 | -0.689 | 0.555 | 0.191 | -0.003 | -0.002 | -0.002 | 0 | 0.53 | 0.53 | 0.33 | 0.35 | 0.4 | 0.62 |
| 2 | 33039690 | rs1037626 | 0.777 | -0.553 | 0.187 | 0.218 | 0.006 | 0.004 | 0.004 | 0 | 0.17 | 0.24 | 0.28 | 0.24 | 0.18 | 0.12 |
| 2 | 33051529 | rs4952302 | -1.75 | -0.295 | -0.008 | 0.202 | -0.027 | -0.043 | -0.043 | 0 | 0.74 | 0.83 | 0.6 | 0.56 | 0.59 | 0.73 |
| 2 | 33052763 | rs7570685 | -0.615 | -0.461 | -0.162 | 0.047 | -0.021 | 0.07 | 0.07 | 2.487 | 0.57 | 0.77 | 0.45 | 0.33 | 0.39 | 0.35 |
| 2 | 33057389 | rs10173352 | -1.806 | -0.32 | -0.411 | 0.142 | -0.064 | -0.095 | -0.095 | 1.674 | 0.74 | 0.88 | 0.62 | 0.56 | 0.59 | 0.73 |
| 2 | 33057541 | rs4952272 | -1.582 | -0.343 | -0.445 | 0.096 | -0.072 | -0.103 | -0.103 | 0 | 0.73 | 0.88 | 0.61 | 0.56 | 0.58 | 0.73 |
| 2 | 33062281 | rs6543678 | -1.517 | -0.612 | 0.568 | 0.002 | 0 | 0.008 | 0.008 | 0.015 | 0.57 | 0.66 | 0.37 | 0.34 | 0.44 | 0.64 |
| 2 | 33062481 | rs6543679 | -1.482 | -0.615 | 0.568 | 0 | 0 | 0.008 | 0.008 | 0 | 0.57 | 0.66 | 0.37 | 0.34 | 0.44 | 0.64 |
| 2 | 33080659 | rs17012350 | -0.569 | -0.688 | 0.826 | -0.353 | 0.025 | 0.035 | 0.035 | 0 | 0.37 | 0.24 | 0.18 | 0.12 | 0.24 | 0.27 |
| 2 | 33080673 | rs1455974 | -1.857 | -0.593 | 0.859 | -0.165 | -0.03 | -0.066 | -0.066 | 0 | 0.6 | 0.83 | 0.45 | 0.46 | 0.46 | 0.66 |
| 2 | 33088544 | rs6759807 | 0.176 | -1.135 | 1.014 | -0.314 | 0.001 | -0.029 | -0.029 | 0 | 0.2 | 0.16 | 0.08 | 0 | 0.15 | 0.18 |
| 2 | 33088870 | rs6760225 | -0.909 | -1.108 | 1.221 | -0.387 | -0.007 | -0.074 | -0.074 | 0 | 0.29 | 0.68 | 0.17 | 0 | 0.19 | 0.22 |
| 2 | 33090667 | rs6710845 | -0.208 | -1.559 | 1.339 | -0.195 | -0.039 | -0.121 | -0.121 | 0 | 0.25 | 0.37 | 0.13 | 0 | 0.16 | 0.21 |
| 2 | 33099380 | rs10202440 | -1.284 | -1.175 | 0.975 | 0.066 | -0.002 | -0.019 | -0.019 | 0.314 | 0.43 | 0.51 | 0.25 | 0.11 | 0.3 | 0.26 |
| 2 | 33102494 | rs35930275 | 0.444 | -0.978 | 0.431 | 0.319 | 0.001 | 0.014 | 0.014 | 0 | 0.11 | 0.09 | 0.18 | 0.34 | 0.11 | 0.34 |
| 2 | 33108465 | rs4952315 | 0.236 | -0.998 | 0.427 | 0.203 | -0.015 | 0.164 | 0.164 | 0 | 0.12 | 0.26 | 0.2 | 0.32 | 0.1 | 0.31 |
| 2 | 33114118 | rs4952316 | -0.451 | -0.98 | 0.394 | 0.199 | -0.005 | 0.111 | 0.111 | 0 | 0.14 | 0.27 | 0.23 | 0.34 | 0.17 | 0.37 |
| 2 | 33119383 | rs13033107 | -1.563 | -0.608 | 0.068 | -0.697 | 0 | -0.011 | -0.011 | 0.596 | 0.41 | 0.37 | 0.22 | 0.12 | 0.32 | 0.26 |
| 2 | 33121494 | rs17012363 | -1.19 | -0.521 | -0.222 | -0.657 | 0.002 | -0.001 | -0.001 | 0 | 0.34 | 0.32 | 0.17 | 0.12 | 0.21 | 0.24 |
| 2 | 33121504 | rs6728021 | -0.779 | 0.007 | -1.016 | -0.535 | -0.004 | 0 | 0 | 0 | 0.05 | 0.03 | 0.04 | 0 | 0.11 | 0.02 |
| 2 | 33122143 | rs7607940 | -0.406 | -0.352 | -1.002 | -0.349 | 0.023 | 0.051 | 0.051 | 0 | 0.19 | 0.09 | 0.09 | 0 | 0.14 | 0.14 |
| 2 | 33127080 | rs6716986 | -0.687 | -0.065 | -0.337 | 0.156 | -0.014 | -0.009 | -0.009 | 0 | 0.17 | 0.26 | 0.11 | 0.13 | 0.12 | 0.11 |
| 2 | 33138551 | rs2219232 | -0.503 | -0.591 | -0.272 | 0.181 | -0.03 | 0.019 | 0.019 | 0 | 0.33 | 0.42 | 0.26 | 0.6 | 0.2 | 0.39 |
| 2 | 33144085 | rs7578053 | -0.997 | 0.105 | -1.209 | -0.95 | -0.014 | 0.179 | 0.179 | 3.45 | 0.65 | 0.9 | 0.71 | 0.98 | 0.64 | 0.61 |
| 2 | 33150932 | rs897509 | -0.201 | 0.474 | -0.428 | -1.357 | 0.003 | 0.031 | 0.031 | 1.158 | 0.51 | 0.62 | 0.62 | 0.98 | 0.46 | 0.57 |
| 2 | 33152042 | rs979050 | -0.329 | 0.524 | -0.422 | -1.37 | 0.016 | 0.059 | 0.059 | 0.852 | 0.52 | 0.66 | 0.64 | 0.98 | 0.5 | 0.58 |
| 2 | 33163650 | rs12618959 | -0.021 | 0.414 | -0.241 | -1.666 | 0.027 | 0.142 | 0.142 | 0 | 0.34 | 0.12 | 0.45 | 0.96 | 0.28 | 0.53 |
| 2 | 33169809 | rs150719 | 0.836 | 0.336 | -0.422 | -2.03 | -0.001 | 0.184 | 0.184 | 0 | 0.43 | 0.18 | 0.29 | 0.02 | 0.39 | 0.38 |
| 2 | 33173465 | rs7594155 | 0.163 | 0.105 | -0.246 | -1.851 | -0.011 | -0.046 | -0.046 | 1.815 | 0.13 | 0.17 | 0.1 | 0 | 0.15 | 0.07 |
| 2 | 33173552 | rs6745339 | 0.854 | 0.095 | -0.675 | -1.11 | 0.009 | -0.092 | -0.092 | 0.214 | 0.26 | 0.08 | 0.22 | 0.52 | 0.2 | 0.36 |
| 2 | 33179432 | rs150656 | 0.329 | -0.057 | -0.465 | -1.353 | 0.001 | 0.017 | 0.017 | 0 | 0.52 | 0.42 | 0.56 | 0.23 | 0.47 | 0.51 |
| 2 | 33181959 | rs150660 | 0.34 | -0.121 | -0.428 | -1.371 | 0.005 | 0.044 | 0.044 | 0 | 0.51 | 0.39 | 0.56 | 0.25 | 0.47 | 0.51 |
| 2 | 33183497 | rs219435 | 0.965 | -0.274 | -0.707 | -1.728 | -0.005 | -0.018 | -0.018 | 0 | 0.34 | 0.26 | 0.32 | 0.02 | 0.39 | 0.36 |
| 2 | 33183606 | rs219436 | -0.66 | -0.118 | -0.511 | -1.681 | -0.029 | -0.001 | -0.001 | 0 | 0.67 | 0.6 | 0.75 | 0.29 | 0.77 | 0.61 |
| 2 | 33185302 | rs4952324 | -0.033 | 0.47 | -0.711 | -1.598 | -0.015 | -0.029 | -0.029 | 0 | 0.16 | 0.1 | 0.29 | 0.26 | 0.2 | 0.16 |
| 2 | 33187557 | rs17396246 | 0.907 | 0.48 | -0.716 | -1.643 | -0.03 | 0.021 | 0.021 | 0 | 0.11 | 0.04 | 0.07 | 0.05 | 0.17 | 0.07 |
| 2 | 33187739 | rs150689 | -0.384 | 0.396 | -0.613 | -1.615 | -0.002 | -0.005 | -0.005 | 0 | 0.63 | 0.43 | 0.63 | 0.3 | 0.64 | 0.58 |

| | | | | | | | | | | | | | | | | |
|---|---|---|---|---|---|---|---|---|---|---|---|---|---|---|---|---|
| 2 | 33189370 | rs17012409 | 0.518 | 0.689 | -0.775 | -1.142 | -0.002 | 0.058 | 0.058 | 0 | 0.05 | 0.07 | 0.22 | 0.24 | 0.03 | 0.1 |
| 2 | 33202098 | rs107195 | 0.983 | 0.716 | -1.321 | -1.484 | -0.081 | -0.028 | -0.028 | 0 | 0.64 | 0.17 | 0.43 | 0.68 | 0.57 | 0.58 |
| 2 | 33206458 | rs6714901 | 0.55 | 1.095 | -0.991 | -1.418 | -0.024 | 0.02 | 0.02 | 0 | 0.09 | 0.07 | 0.09 | 0 | 0.19 | 0.09 |
| 2 | 33215833 | rs219152 | -0.548 | 0.877 | -0.267 | -1.404 | 0.072 | 0.062 | 0.062 | 1.772 | 0.95 | 0.66 | 0.97 | 1 | 1 | 1 |
| 2 | 33216626 | rs219154 | 1.628 | 0.91 | -0.955 | -1.719 | -0.052 | 0.085 | 0.085 | 0.071 | 0.19 | 0.07 | 0.19 | 0 | 0.3 | 0.18 |
| 2 | 33216805 | rs10198807 | -0.161 | 0.533 | -1.104 | -1.67 | -0.001 | 0.001 | 0.001 | 0 | 0.29 | 0.31 | 0.33 | 0.27 | 0.25 | 0.23 |
| 2 | 33218037 | rs219158 | 1.777 | 0.625 | -1.204 | -1.702 | -0.063 | 0.078 | 0.078 | 0.252 | 0.2 | 0.08 | 0.21 | 0 | 0.33 | 0.18 |
| 2 | 33219147 | rs13417287 | 0.291 | 0.517 | -1.093 | -1.672 | 0 | -0.037 | -0.037 | 1.184 | 0.25 | 0.22 | 0.11 | 0.01 | 0.23 | 0.16 |
| 2 | 33220291 | rs743237 | 1.51 | 0.47 | -1.589 | -1.627 | 0.023 | 0.018 | 0.018 | 0.341 | 0.15 | 0.07 | 0.03 | 0 | 0.07 | 0.07 |
| 2 | 33228292 | rs219174 | 1.275 | 0.567 | -1.735 | -1.585 | -0.075 | -0.072 | -0.072 | 0.008 | 0.23 | 0.1 | 0.42 | 0.27 | 0.35 | 0.26 |
| 2 | 33230086 | rs219176 | -0.477 | 0.43 | -1.637 | -1.633 | 0.036 | 0.066 | 0.066 | 0 | 0.49 | 0.27 | 0.53 | 0.28 | 0.58 | 0.42 |
| 2 | 33233715 | rs4952279 | 0.827 | 0.47 | -1.596 | -1.531 | 0.011 | -0.06 | -0.06 | 0 | 0.19 | 0.09 | 0.27 | 0.27 | 0.13 | 0.18 |
| 2 | 33233902 | rs4952280 | -0.053 | 0.34 | -1.687 | -1.925 | -0.004 | 0.046 | 0.046 | 1.069 | 0.56 | 0.37 | 0.59 | 0.97 | 0.45 | 0.54 |
| 2 | 33234117 | rs219180 | -0.401 | 0.357 | -1.771 | -1.906 | 0.024 | 0.202 | 0.202 | 0.215 | 0.54 | 0.24 | 0.4 | 0.03 | 0.62 | 0.53 |
| 2 | 33235028 | rs436882 | 0.909 | 0.502 | -1.668 | -1.802 | -0.034 | 0.098 | 0.098 | 1.122 | 0.3 | 0.17 | 0.31 | 0.03 | 0.39 | 0.38 |
| 2 | 33235324 | rs219183 | 0.949 | 0.502 | -1.707 | -1.794 | -0.043 | 0.119 | 0.119 | 1.464 | 0.3 | 0.14 | 0.3 | 0.03 | 0.39 | 0.38 |
| 2 | 33246491 | rs219195 | 0.587 | 0.78 | -1.043 | -1.325 | 0 | 0.001 | 0.001 | 0 | 0.65 | 0.36 | 0.62 | 0.73 | 0.7 | 0.75 |
| 2 | 33248325 | rs2168064 | 0.055 | 0.821 | -0.822 | -1.497 | 0.044 | -0.057 | -0.057 | 0 | 0.11 | 0.21 | 0.15 | 0 | 0.15 | 0.04 |
| 2 | 33249542 | rs219205 | 1.327 | 0.778 | -0.95 | -1.495 | -0.058 | 0.053 | 0.053 | 0.736 | 0.55 | 0.17 | 0.49 | 0.73 | 0.59 | 0.72 |
| 2 | 33250571 | rs4952336 | 0.146 | 0.809 | -0.853 | -1.602 | 0.025 | -0.021 | -0.021 | 0 | 0.28 | 0.19 | 0.36 | 0.28 | 0.19 | 0.3 |
| 2 | 33250843 | rs219206 | 1.808 | 0.828 | -0.878 | -1.58 | -0.006 | 0.047 | 0.047 | 0.073 | 0.46 | 0.14 | 0.39 | 0.72 | 0.5 | 0.57 |
| 2 | 33252278 | rs219208 | 0.893 | 0.807 | -0.768 | -1.646 | -0.003 | -0.036 | -0.036 | 0 | 0.61 | 0.36 | 0.74 | 0.99 | 0.66 | 0.8 |
| 2 | 33260019 | rs1028598 | 1.39 | 0.95 | -0.927 | -1.64 | -0.049 | 0.046 | 0.046 | 0 | 0.08 | 0.01 | 0.06 | 0 | 0.15 | 0.09 |
| 2 | 33263671 | rs10490453 | 1.852 | 1.287 | -1.08 | -1.764 | -0.035 | 0.028 | 0.028 | 0 | 0.06 | 0.01 | 0.05 | 0 | 0.1 | 0.07 |
| 2 | 33263907 | rs2008845 | -0.266 | 0.757 | -0.866 | -1.665 | -0.014 | 0.132 | 0.132 | 0.096 | 0.33 | 0.14 | 0.2 | 0.02 | 0.31 | 0.3 |
| 2 | 33273297 | rs57568 | -0.756 | 0.658 | -0.514 | -1.491 | 0.076 | 0.048 | 0.048 | 1.089 | 0.94 | 0.53 | 0.95 | 1 | 0.99 | 0.99 |
| 2 | 33284382 | rs6753484 | 1.672 | 0.633 | -1.007 | -1.527 | 0.03 | 0.049 | 0.049 | 0 | 0.15 | 0.05 | 0.03 | 0 | 0.07 | 0.06 |
| 2 | 33289577 | rs219082 | 0.558 | 1.001 | -0.472 | -1.575 | 0.092 | -0.125 | -0.125 | 0 | 0.79 | 0.4 | 0.76 | 0.99 | 0.69 | 0.85 |
| 2 | 33295631 | rs439416 | 1.643 | 0.716 | -1.051 | -1.245 | 0.007 | -0.047 | -0.047 | 0.85 | 0.44 | 0.12 | 0.37 | 0.6 | 0.47 | 0.48 |
| 2 | 33296774 | rs368858 | 0.739 | 0.674 | -0.625 | -1.286 | 0.001 | 0.006 | 0.006 | 0 | 0.56 | 0.34 | 0.51 | 0.66 | 0.61 | 0.58 |
| 2 | 33297583 | rs438695 | -0.095 | 0.713 | -0.498 | -1.291 | -0.012 | -0.111 | -0.111 | 0 | 0.8 | 0.4 | 0.82 | 0.99 | 0.85 | 0.79 |
| 2 | 33297929 | rs11896749 | 0.965 | 0.826 | -0.675 | -1.244 | 0.02 | -0.024 | -0.024 | 0.114 | 0.06 | 0.11 | 0.12 | 0 | 0.11 | 0.03 |
| 2 | 33300735 | rs402402 | 1.148 | 0.53 | -0.91 | -1.288 | 0.016 | 0.035 | 0.035 | 0.266 | 0.51 | 0.13 | 0.45 | 0.66 | 0.65 | 0.59 |
| 2 | 33302342 | rs377892 | 1.133 | 0.538 | -0.915 | -1.293 | 0.022 | 0.047 | 0.047 | 0.239 | 0.51 | 0.15 | 0.45 | 0.67 | 0.65 | 0.59 |
| 2 | 33304365 | rs496600 | -0.406 | 0.682 | -0.324 | -1.443 | -0.017 | -0.103 | -0.103 | 5.707 | 0.84 | 0.42 | 0.84 | 0.99 | 0.89 | 0.77 |
| 2 | 33307784 | rs219099 | 0.279 | 0.729 | -0.341 | -1.453 | 0.025 | -0.11 | -0.11 | 0 | 0.75 | 0.39 | 0.76 | 0.99 | 0.71 | 0.67 |
| 2 | 33310894 | rs219104 | -0.439 | 0.381 | -0.049 | -1.168 | 0.069 | 0.052 | 0.052 | 1.562 | 0.94 | 0.5 | 0.95 | 1 | 0.99 | 0.99 |
| 2 | 33319539 | rs13027016 | 1.603 | 0.432 | -0.736 | -1.221 | -0.004 | 0.039 | 0.039 | 0.045 | 0.09 | 0.02 | 0.11 | 0.01 | 0.09 | 0.22 |
| 2 | 33320957 | rs218209 | 0.684 | 0.385 | -0.632 | -1.717 | 0.002 | -0.064 | -0.064 | 0 | 0.56 | 0.43 | 0.75 | 0.98 | 0.64 | 0.64 |
| 2 | 33322077 | rs10169790 | 0.623 | 0.408 | -0.739 | -1.751 | -0.065 | 0.089 | 0.089 | 0.502 | 0.17 | 0.03 | 0.16 | 0.01 | 0.24 | 0.3 |
| 2 | 33322587 | rs218211 | 0.101 | 0.417 | -0.625 | -1.705 | -0.008 | -0.152 | -0.152 | 1.309 | 0.66 | 0.48 | 0.86 | 0.99 | 0.73 | 0.86 |
| 2 | 33324783 | rs10195741 | 1.695 | 0.399 | -0.691 | -1.679 | 0.003 | 0.024 | 0.024 | 0.018 | 0.09 | 0.03 | 0.11 | 0.01 | 0.09 | 0.22 |
| 2 | 33324978 | rs218214 | 0.223 | 0.447 | -0.368 | -1.482 | -0.01 | -0.167 | -0.167 | 3.06 | 0.68 | 0.5 | 0.86 | 1 | 0.74 | 0.86 |
| 2 | 33326484 | rs17012565 | -0.064 | 0.863 | -0.632 | -1.339 | 0.022 | -0.032 | -0.032 | 0 | 0.06 | 0.15 | 0.09 | 0 | 0.1 | 0.04 |
| 2 | 33329957 | rs11124305 | 0.235 | 0.563 | -0.434 | -1.587 | 0 | -0.051 | -0.051 | 0 | 0.26 | 0.29 | 0.14 | 0 | 0.19 | 0.09 |
| 2 | 33331219 | rs218221 | 0.595 | 0.405 | -1.189 | -1.034 | 0.054 | -0.032 | -0.032 | 0 | 0.54 | 0.21 | 0.57 | 0.55 | 0.68 | 0.74 |
| 2 | 33332938 | rs10490450 | 0.819 | 1.049 | -1.157 | -1 | -0.058 | 0.051 | 0.051 | 0 | 0.09 | 0.01 | 0.08 | 0 | 0.17 | 0.07 |
| 2 | 33334699 | rs11904654 | -0.35 | 0.443 | -0.746 | -0.779 | -0.001 | 0.046 | 0.046 | 0.036 | 0.17 | 0.25 | 0.31 | 0.44 | 0.14 | 0.22 |
| 2 | 33334879 | rs218224 | 0.066 | 0.165 | -1.248 | -1.425 | 0.079 | 0.165 | 0.165 | 1.335 | 0.54 | 0.2 | 0.77 | 1 | 0.72 | 0.84 |
| 2 | 33336309 | rs11124307 | 0.706 | 0.87 | -1.491 | -0.92 | -0.007 | -0.001 | -0.001 | 0.36 | 0.31 | 0.25 | 0.37 | 0.37 | 0.38 | 0.44 |
| 2 | 33338110 | rs11124308 | 0.452 | 1.203 | -1.279 | -0.958 | 0.027 | -0.038 | -0.038 | 1.633 | 0.09 | 0.13 | 0.12 | 0 | 0.18 | 0.05 |
| 2 | 33338736 | rs17325551 | 0.875 | 0.787 | -1.477 | -0.981 | 0 | 0.005 | 0.005 | 2.25 | 0.23 | 0.16 | 0.14 | 0.07 | 0.21 | 0.06 |
| 2 | 33341172 | rs7583121 | 0.221 | 1.057 | -1.963 | -0.666 | 0.008 | 0.012 | 0.012 | 0 | 0.47 | 0.34 | 0.37 | 0.48 | 0.43 | 0.5 |
| 2 | 33344570 | rs218193 | 1.331 | 1.33 | -1.569 | -0.452 | -0.014 | 0.029 | 0.029 | 0.044 | 0.06 | 0.02 | 0.04 | 0 | 0.06 | 0.06 |
| 2 | 33348581 | rs218197 | -0.436 | 1.447 | -1.605 | -0.314 | -0.021 | 0.002 | 0.002 | 0.142 | 0.64 | 0.72 | 0.51 | 0.61 | 0.48 | 0.54 |
| 2 | 33358079 | rs897090 | -0.298 | 1.452 | -1.647 | -0.34 | -0.008 | -0.004 | -0.004 | 0 | 0.86 | 0.88 | 0.87 | 0.99 | 0.79 | 0.9 |
| 2 | 33359457 | rs12620839 | 0.28 | 1.511 | -1.744 | -0.144 | -0.023 | -0.04 | -0.04 | 0 | 0.11 | 0.05 | 0.24 | 0.51 | 0.12 | 0.3 |
| 2 | 33360338 | rs1545552 | 0.435 | 1.075 | -1.455 | -1.293 | -0.014 | 0.02 | 0.02 | 0 | 0.77 | 0.83 | 0.83 | 0.99 | 0.71 | 0.86 |
| 2 | 33362643 | rs6705263 | -0.112 | 0.824 | -1.036 | -1.057 | 0 | -0.028 | -0.028 | 0.172 | 0.51 | 0.49 | 0.67 | 0.94 | 0.55 | 0.77 |
| 2 | 33369582 | rs1902049 | 0.418 | 0.44 | -0.694 | -0.182 | -0.014 | 0.064 | 0.064 | 0 | 0.26 | 0.39 | 0.35 | 0.55 | 0.21 | 0.35 |
| 2 | 33370516 | rs17012602 | -0.407 | 0.886 | -0.499 | -0.462 | -0.003 | 0.001 | 0.001 | 0 | 0.39 | 0.33 | 0.24 | 0.16 | 0.4 | 0.24 |
| 2 | 33375133 | rs10432657 | -0.777 | 0.863 | -0.412 | -0.471 | 0 | 0.01 | 0.01 | 0 | 0.47 | 0.57 | 0.52 | 0.36 | 0.55 | 0.38 |
| 2 | 33391901 | rs1947444 | -0.644 | 0.627 | -0.392 | -0.722 | -0.02 | -0.005 | -0.005 | 5.143 | 0.3 | 0.21 | 0.44 | 0.34 | 0.49 | 0.31 |
| 2 | 33392710 | rs17012632 | -0.335 | 0.888 | -0.413 | -0.79 | -0.001 | -0.031 | -0.031 | 0 | 0.35 | 0.26 | 0.19 | 0.14 | 0.34 | 0.21 |
| 2 | 33392911 | rs6728094 | -1.093 | 0.404 | -0.351 | -0.608 | 0.008 | 0.016 | 0.016 | 0 | 0.49 | 0.57 | 0.51 | 0.4 | 0.55 | 0.38 |
| 2 | 33412077 | rs1065324 | -0.901 | 1.192 | 0.306 | -0.193 | 0.007 | 0.003 | 0.003 | 0 | 0.32 | 0.27 | 0.45 | 0.37 | 0.48 | 0.32 |
| 2 | 33412535 | rs6707962 | -1.27 | 1.201 | 0.215 | -0.185 | 0.017 | 0.008 | 0.008 | 0 | 0.36 | 0.52 | 0.49 | 0.39 | 0.5 | 0.32 |
| 2 | 33413640 | rs6712473 | -1.27 | 1.201 | 0.215 | -0.204 | 0.017 | 0.007 | 0.007 | 0 | 0.36 | 0.52 | 0.49 | 0.39 | 0.5 | 0.32 |
| 2 | 33447202 | rs2290448 | -0.854 | 1.404 | -0.732 | -0.142 | -0.038 | -0.014 | -0.014 | 0 | 0.3 | 0.19 | 0.43 | 0.33 | 0.49 | 0.29 |

| | | | | | | | | | | | | | | | | |
|---|---|---|---|---|---|---|---|---|---|---|---|---|---|---|---|---|
| 2 | 33470656 | rs817535 | 0.574 | 1.203 | -0.526 | -0.721 | 0.009 | -0.054 | -0.054 | 0.988 | 0.59 | 0.48 | 0.5 | 0.89 | 0.54 | 0.6 |
| 2 | 33474645 | rs921705 | -1.218 | 1.162 | -0.658 | -0.635 | 0.001 | -0.003 | -0.003 | 5.758 | 0.32 | 0.36 | 0.25 | 0.25 | 0.37 | 0.24 |
| 2 | 33477964 | rs3731575 | 0.353 | 1.026 | -0.772 | -0.122 | -0.029 | -0.019 | -0.019 | 0.152 | 0.17 | 0.1 | 0.17 | 0.21 | 0.25 | 0.16 |
| 2 | 33484210 | rs3769531 | 0.202 | 1.046 | -0.678 | -0.058 | -0.064 | -0.041 | -0.041 | 0 | 0.18 | 0.05 | 0.18 | 0.23 | 0.29 | 0.18 |
| 2 | 33486902 | rs3769536 | -1.188 | 0.924 | -0.36 | -0.083 | 0.005 | -0.012 | -0.012 | 0 | 0.32 | 0.5 | 0.26 | 0.27 | 0.33 | 0.24 |
| 2 | 33488419 | rs12468099 | -0.195 | 0.906 | -0.609 | -0.185 | -0.098 | -0.063 | -0.063 | 0.001 | 0.2 | 0.02 | 0.18 | 0.24 | 0.29 | 0.17 |
| 2 | 33491608 | rs11883651 | -0.935 | 0.881 | -0.622 | -0.066 | -0.118 | -0.164 | -0.164 | 0.093 | 0.25 | 0.02 | 0.22 | 0.41 | 0.34 | 0.27 |
| 2 | 33497335 | rs817529 | 0.284 | 0.86 | -0.602 | -0.154 | -0.022 | 0.024 | 0.024 | 0.427 | 0.27 | 0.21 | 0.46 | 0.09 | 0.39 | 0.35 |
| 2 | 33497656 | rs817530 | -2.191 | 0.878 | -0.83 | -0.116 | 0.042 | -0.059 | -0.059 | 0 | 0.63 | 0.87 | 0.75 | 0.47 | 0.73 | 0.67 |
| 2 | 33499107 | rs3754829 | -0.938 | 0.869 | -0.785 | -0.146 | -0.03 | 0.095 | 0.095 | 0 | 0.43 | 0.71 | 0.54 | 0.11 | 0.48 | 0.42 |
| 2 | 33502505 | rs3769541 | -2.23 | 0.75 | -0.828 | -0.084 | 0.074 | -0.043 | -0.043 | 0 | 0.66 | 0.88 | 0.75 | 0.51 | 0.77 | 0.7 |
| 2 | 33509219 | rs6722904 | 1.271 | 0.706 | -0.819 | -0.149 | 0.018 | 0.037 | 0.037 | 0 | 0.1 | 0.01 | 0.03 | 0.02 | 0.05 | 0.06 |
| 2 | 33514753 | rs11685108 | 0.35 | 0.7 | -0.771 | -0.111 | 0.007 | -0.009 | -0.009 | 0 | 0.14 | 0.11 | 0.06 | 0.16 | 0.07 | 0.16 |
| 2 | 33517458 | rs2123770 | -1.937 | 1.179 | -0.921 | -0.047 | -0.006 | -0.058 | -0.058 | 0.451 | 0.61 | 0.89 | 0.71 | 0.47 | 0.64 | 0.66 |
| 2 | 33525382 | rs935996 | -1.323 | 0.9 | -0.757 | -0.02 | -0.018 | 0.056 | 0.056 | 0 | 0.47 | 0.77 | 0.57 | 0.27 | 0.5 | 0.53 |
| 2 | 33532820 | rs1463313 | -2.356 | 0.881 | -0.714 | 0.066 | 0.065 | -0.054 | -0.054 | 0.129 | 0.68 | 0.88 | 0.76 | 0.5 | 0.78 | 0.69 |
| 2 | 33536151 | rs3769545 | 0.571 | 1.286 | -0.912 | -0.07 | -0.073 | -0.019 | -0.019 | 0.139 | 0.12 | 0.02 | 0.36 | 0.09 | 0.32 | 0.19 |
| 2 | 33540138 | rs2276591 | -1.303 | 0.755 | -0.312 | -0.031 | 0.045 | 0.017 | 0.017 | 1.715 | 0.48 | 0.37 | 0.69 | 0.5 | 0.68 | 0.62 |
| 2 | 33540248 | rs1058840 | -1.075 | 0.765 | -0.326 | -0.015 | 0.031 | 0.028 | 0.028 | 2.561 | 0.45 | 0.32 | 0.63 | 0.31 | 0.65 | 0.5 |
| 2 | 33541702 | rs723042 | -1.549 | 0.401 | -0.278 | 0.027 | 0.057 | 0.016 | 0.016 | 0.514 | 0.54 | 0.37 | 0.71 | 0.52 | 0.76 | 0.64 |
| 2 | 33555177 | rs2167973 | -1.076 | 0.949 | 0.326 | -0.087 | -0.042 | 0.026 | 0.026 | 0.807 | 0.21 | 0.59 | 0.14 | 0.21 | 0.1 | 0.16 |
| 2 | 33555854 | rs9308934 | -0.284 | 1.121 | 0.144 | -0.161 | -0.042 | 0.035 | 0.035 | 0.114 | 0.25 | 0.16 | 0.28 | 0.09 | 0.39 | 0.33 |
| 2 | 33556839 | rs2167972 | -1.253 | 1.092 | 0.267 | -0.197 | 0.045 | 0.022 | 0.022 | 0.003 | 0.3 | 0.52 | 0.36 | 0.28 | 0.43 | 0.41 |
| 2 | 33563575 | rs9678322 | -1.924 | 1.198 | -0.517 | -0.179 | -0.02 | 0.033 | 0.033 | 0 | 0.46 | 0.79 | 0.42 | 0.3 | 0.49 | 0.48 |
| 2 | 33564438 | rs7581438 | -0.825 | 2.404 | -0.484 | -0.151 | -0.044 | 0.035 | 0.035 | 0.19 | 0.25 | 0.16 | 0.28 | 0.09 | 0.4 | 0.33 |
| 2 | 33572555 | rs6706620 | 0.054 | 1.851 | 0.244 | -0.003 | 0.051 | -0.027 | -0.027 | 0 | 0.05 | 0.18 | 0.09 | 0 | 0.1 | 0.05 |
| 2 | 33573090 | rs11895736 | -0.542 | 2.103 | -0.041 | -0.138 | -0.023 | 0.03 | 0.03 | 0.69 | 0.11 | 0.04 | 0.09 | 0.01 | 0.16 | 0.08 |
| 2 | 33574166 | rs17644939 | -0.312 | 1.94 | -0.037 | -0.121 | -0.006 | 0.024 | 0.024 | 0.725 | 0.09 | 0.04 | 0.07 | 0.01 | 0.11 | 0.07 |
| 2 | 33586776 | rs2305502 | -1.705 | 1.561 | 0.778 | 0.136 | -0.001 | 0.045 | 0.045 | 0.079 | 0.19 | 0.52 | 0.19 | 0.21 | 0.19 | 0.26 |
| 2 | 33595197 | rs7566302 | -1.759 | 1.35 | 0.96 | 0.007 | -0.038 | 0.03 | 0.03 | 0.769 | 0.23 | 0.54 | 0.16 | 0.24 | 0.18 | 0.26 |
| 2 | 33596090 | rs11899010 | -1.492 | 1.29 | 1.21 | -0.018 | -0.068 | 0.107 | 0.107 | 0.713 | 0.16 | 0.4 | 0.08 | 0.23 | 0.06 | 0.21 |
| 2 | 33600280 | rs17012892 | 0.761 | 1.301 | 1.085 | -0.02 | 0.004 | 0.007 | 0.007 | 0.006 | 0.05 | 0.06 | 0.03 | 0 | 0.04 | 0.04 |
| 2 | 33602571 | rs3845775 | 0.088 | 1.254 | 1.393 | -0.073 | 0.029 | -0.037 | -0.037 | 0 | 0.07 | 0.18 | 0.08 | 0 | 0.1 | 0.05 |
| 2 | 33604048 | rs6719824 | -1.632 | 1.23 | 1.412 | -0.067 | 0.057 | 0.294 | 0.294 | 0.611 | 0.09 | 0.4 | 0.12 | 0.24 | 0.09 | 0.18 |
| 2 | 33607071 | rs6742198 | -0.186 | 2.208 | 0.448 | -0.07 | -0.076 | -0.022 | -0.022 | 1.359 | 0.88 | 0.99 | 0.91 | 0.91 | 0.8 | 0.86 |
| 2 | 33608957 | rs13404913 | -1.188 | 1.985 | 0.518 | -0.106 | -0.002 | -0.064 | -0.064 | 0 | 0.18 | 0.3 | 0.14 | 0 | 0.19 | 0.11 |
| 2 | 33610073 | rs17012922 | -2.011 | 2.032 | -0.179 | -0.026 | 0.001 | 0.001 | 0.001 | 2.466 | 0.3 | 0.72 | 0.25 | 0.22 | 0.28 | 0.3 |
| 2 | 33613324 | rs13015373 | 0.792 | 1.754 | -0.828 | -0.099 | -0.113 | 0.01 | 0.01 | 0 | 0.22 | 0.02 | 0.26 | 0.14 | 0.33 | 0.32 |
| 2 | 33613354 | rs1869450 | -0.992 | 2.159 | -0.18 | 0.164 | 0.002 | -0.012 | -0.012 | 2.076 | 0.2 | 0.18 | 0.13 | 0.01 | 0.19 | 0.16 |
| 2 | 33613402 | rs1869451 | 0.913 | 2.257 | -0.462 | 0.23 | 0.001 | -0.014 | -0.014 | 0 | 0.08 | 0.06 | 0.06 | 0.01 | 0.1 | 0.07 |
| 2 | 33613593 | rs1017677 | -1.433 | 2.058 | -0.908 | 0.335 | 0.01 | 0.011 | 0.011 | 0.551 | 0.34 | 0.73 | 0.23 | 0.21 | 0.26 | 0.34 |
| 2 | 33614187 | rs642472 | 1.351 | 1.5 | -1.237 | 0.312 | -0.024 | 0.013 | 0.013 | 0.953 | 0.57 | 0.22 | 0.67 | 0.68 | 0.52 | 0.56 |
| 2 | 33616240 | rs6757228 | -1.185 | 1.707 | -0.727 | 0.581 | 0.019 | 0.008 | 0.008 | 0 | 0.29 | 0.62 | 0.25 | 0.31 | 0.37 | 0.34 |
| 2 | 33623306 | rs2034693 | 0.261 | 0.921 | -1.196 | 0.601 | 0.004 | -0.001 | -0.001 | 0 | 0.57 | 0.43 | 0.43 | 0.62 | 0.51 | 0.42 |
| 2 | 33623720 | rs8970 | 0.494 | 0.899 | -2.419 | 0.773 | -0.074 | -0.11 | -0.11 | 0 | 0.44 | 0.05 | 0.36 | 0.52 | 0.42 | 0.28 |
| 2 | 33631442 | rs588252 | 0.376 | 0.608 | -1.41 | 0.616 | 0.004 | -0.004 | -0.004 | 5.169 | 0.73 | 0.83 | 0.78 | 0.73 | 0.74 | 0.58 |
| 2 | 33641932 | rs7607123 | -0.458 | 1.092 | -1.435 | 0.464 | -0.068 | -0.018 | -0.018 | 0.033 | 0.22 | 0.07 | 0.19 | 0.16 | 0.33 | 0.23 |
| 2 | 33645650 | rs7575204 | -1.087 | 0.443 | -1.217 | 0.129 | -0.012 | 0.017 | 0.017 | 0 | 0.37 | 0.28 | 0.29 | 0.24 | 0.4 | 0.34 |
| 2 | 33652677 | rs17646725 | -0.295 | -0.806 | -1.7 | -0.183 | 0.081 | 0.06 | 0.06 | 0 | 0.31 | 0.13 | 0.11 | 0.15 | 0.11 | 0.23 |
| 2 | 33653978 | rs4092337 | -0.445 | -0.609 | -1.124 | -0.524 | -0.006 | 0.052 | 0.052 | 0 | 0.42 | 0.25 | 0.31 | 0.24 | 0.46 | 0.47 |
| 2 | 33657837 | rs604394 | -0.021 | -0.436 | 0.034 | -0.464 | -0.014 | -0.048 | -0.048 | 0 | 0.1 | 0.36 | 0.04 | 0 | 0.03 | 0.01 |
| 2 | 33660964 | rs583942 | 0.042 | 0.401 | -1.08 | -0.408 | -0.097 | -0.064 | -0.064 | 1.041 | 0.84 | 0.99 | 0.76 | 0.79 | 0.74 | 0.83 |
| 2 | 33664350 | rs604788 | -0.661 | -0.34 | -0.337 | -0.446 | -0.039 | -0.105 | -0.105 | 0.14 | 0.21 | 0.41 | 0.24 | 0.05 | 0.14 | 0.12 |
| 2 | 33668178 | rs654291 | -0.798 | 0.289 | -0.83 | -0.017 | -0.002 | -0.004 | -0.004 | 0.602 | 0.17 | 0.16 | 0.24 | 0.21 | 0.19 | 0.19 |
| 2 | 33674384 | rs11695726 | -0.297 | 0.031 | -1.181 | -0.02 | -0.003 | 0.006 | 0.006 | 0.239 | 0.09 | 0 | 0.2 | 0.09 | 0.11 | 0.06 |
| 2 | 33676440 | rs12999228 | -0.232 | 0.06 | -1.362 | 0.231 | 0 | -0.001 | -0.001 | 0 | 0.78 | 0.82 | 0.78 | 0.38 | 0.82 | 0.67 |
| 2 | 33678924 | rs13014054 | 1.445 | -0.091 | -2.061 | -0.535 | -0.093 | 0.068 | 0.068 | 0 | 0.27 | 0.08 | 0.24 | 0.1 | 0.48 | 0.25 |
| 2 | 33679231 | rs2008255 | -0.635 | -0.481 | -1.755 | 0.371 | 0.001 | 0.003 | 0.003 | 0 | 0.82 | 0.8 | 0.73 | 0.68 | 0.86 | 0.79 |
| 2 | 33681479 | rs1979242 | -0.964 | -0.465 | -1.565 | 0.478 | -0.001 | -0.007 | -0.007 | 0.088 | 0.84 | 0.86 | 0.76 | 0.74 | 0.85 | 0.8 |
| 2 | 33682603 | rs6543714 | 0.319 | -0.534 | -0.655 | 0.275 | 0.011 | 0.161 | 0.161 | 0 | 0.23 | 0.44 | 0.2 | 0.54 | 0.17 | 0.27 |
| 2 | 33683727 | rs6728969 | -1.007 | -0.531 | -0.857 | 0.345 | 0 | -0.027 | -0.027 | 0 | 0.84 | 0.88 | 0.75 | 0.72 | 0.86 | 0.79 |
| 2 | 33686059 | rs11124317 | -0.521 | -0.547 | -0.901 | 0.255 | -0.011 | -0.018 | -0.018 | 0 | 0.82 | 0.91 | 0.74 | 0.82 | 0.82 | 0.86 |
| 2 | 33686786 | rs6756985 | -0.505 | -0.866 | -1.045 | 0.085 | -0.008 | -0.017 | -0.017 | 1.152 | 0.79 | 0.93 | 0.74 | 0.82 | 0.82 | 0.87 |
| 2 | 33686901 | rs13388421 | 1.384 | -1.2 | -0.999 | -0.237 | 0.012 | -0.006 | -0.006 | 0.193 | 0.18 | 0.12 | 0.05 | 0.01 | 0.09 | 0.27 |
| 2 | 33688129 | rs17013034 | 1.124 | -0.848 | -0.246 | 0.474 | 0.012 | 0.018 | 0.018 | 0 | 0.15 | 0.2 | 0.12 | 0.2 | 0.13 | 0.18 |
| 2 | 33689158 | rs13398130 | 0.99 | -1.031 | -0.789 | 0.48 | 0.008 | -0.003 | -0.003 | 0 | 0.2 | 0.15 | 0.13 | 0.28 | 0.16 | 0.33 |
| 2 | 33691716 | rs13406014 | 0.415 | -1.037 | -0.792 | 0.48 | 0.038 | 0.021 | 0.021 | 0 | 0.26 | 0.17 | 0.14 | 0.14 | 0.16 | 0.3 |
| 2 | 33691809 | rs2054133 | 1.077 | -0.791 | -1.189 | 0.389 | -0.09 | -0.043 | -0.043 | 0 | 0.43 | 0.07 | 0.43 | 0.34 | 0.53 | 0.42 |
| 2 | 33695073 | rs2887087 | 0.392 | -1.308 | -0.77 | 0.205 | 0.014 | 0.063 | 0.063 | 0 | 0.54 | 0.67 | 0.53 | 0.25 | 0.62 | 0.44 |

| | | | | | | | | | | | | | | | | |
|---|---|---|---|---|---|---|---|---|---|---|---|---|---|---|---|---|
| 2 | 33695424 | rs12712340 | 1.505 | -1.075 | -1.293 | 0.099 | -0.112 | 0.038 | 0.038 | 0 | 0.33 | 0.1 | 0.26 | 0.17 | 0.49 | 0.37 |
| 2 | 33696602 | rs10185666 | -0.59 | -1.017 | -1.313 | -0.17 | 0.002 | 0.002 | 0.002 | 0.061 | 0.79 | 0.77 | 0.73 | 0.84 | 0.84 | 0.87 |
| 2 | 33698944 | rs7576999 | -0.439 | -0.708 | -1.294 | -0.1 | -0.011 | -0.011 | -0.011 | 0 | 0.37 | 0.41 | 0.39 | 0.36 | 0.33 | 0.41 |
| 2 | 33700305 | rs4145831 | -0.921 | 0.014 | -1.704 | 0.377 | -0.002 | 0.003 | 0.003 | 0.258 | 0.85 | 0.88 | 0.74 | 0.86 | 0.85 | 0.87 |
| 2 | 33701632 | rs17594260 | 0.818 | 0.067 | -1.739 | 0.61 | -0.04 | 0.064 | 0.064 | 0.152 | 0.12 | 0.02 | 0.07 | 0.04 | 0.1 | 0.28 |
| 2 | 33703003 | rs11693332 | -0.597 | -0.129 | -1.718 | 0.612 | 0.007 | -0.026 | -0.026 | 0.008 | 0.28 | 0.14 | 0.31 | 0.33 | 0.23 | 0.35 |
| 2 | 33703389 | rs17013095 | -0.429 | 0.035 | -0.916 | 0.714 | -0.026 | 0.018 | 0.018 | 0 | 0.15 | 0.3 | 0.22 | 0.2 | 0.13 | 0.1 |
| 2 | 33703930 | rs13031142 | -0.369 | 0.535 | -1.629 | 0.868 | 0.003 | -0.006 | -0.006 | 0 | 0.51 | 0.25 | 0.5 | 0.48 | 0.56 | 0.45 |
| 2 | 33704123 | rs4670574 | -0.523 | 0.244 | -1.39 | 0.759 | 0.01 | -0.066 | -0.066 | 0.045 | 0.16 | 0.01 | 0.25 | 0.25 | 0.14 | 0.12 |
| 2 | 33704785 | rs6543716 | -1.043 | 0.705 | -0.581 | 0.29 | -0.004 | -0.056 | -0.056 | 0 | 0.24 | 0.65 | 0.25 | 0.06 | 0.22 | 0.1 |
| 2 | 33704969 | rs6758325 | -1.017 | 0.922 | -0.682 | 0.704 | -0.001 | -0.001 | -0.001 | 0.539 | 0.15 | 0.16 | 0.09 | 0.07 | 0.14 | 0.08 |
| 2 | 33709961 | rs17013126 | -0.984 | 0.671 | -0.181 | 0.58 | -0.001 | -0.023 | -0.023 | 0 | 0.15 | 0.25 | 0.1 | 0.08 | 0.13 | 0.09 |
| 2 | 33710289 | rs17013129 | -0.017 | 0.687 | -0.521 | 1.107 | -0.01 | -0.053 | -0.053 | 0 | 0.11 | 0.02 | 0.04 | 0.21 | 0.11 | 0.09 |
| 2 | 33710577 | rs6753711 | -1.736 | 0.635 | -0.032 | 1.14 | -0.006 | 0 | 0 | 0 | 0.27 | 0.42 | 0.28 | 0.29 | 0.22 | 0.25 |
| 2 | 33715742 | rs7561821 | -2.266 | 0.834 | 0.12 | 1.214 | 0.006 | 0.001 | 0.001 | 0 | 0.36 | 0.59 | 0.5 | 0.39 | 0.38 | 0.43 |
| 2 | 33717683 | rs13391694 | -2.21 | 0.766 | -0.002 | 1.178 | 0.008 | -0.001 | -0.001 | 0.836 | 0.36 | 0.55 | 0.48 | 0.37 | 0.37 | 0.39 |
| 2 | 33718255 | rs13408063 | -1.877 | 0.737 | 0.031 | 1.132 | 0.007 | 0.024 | 0.024 | 0 | 0.27 | 0.41 | 0.44 | 0.37 | 0.29 | 0.35 |
| 2 | 33719473 | rs10495790 | -0.128 | 1.006 | 0.1 | 0.614 | -0.008 | 0.02 | 0.02 | 0.11 | 0.09 | 0.02 | 0.04 | 0.04 | 0.08 | 0.1 |
| 2 | 33721496 | rs1531124 | -0.169 | 1.095 | 0.112 | 0.651 | -0.013 | -0.005 | -0.005 | 1.025 | 0.07 | 0.04 | 0.2 | 0.1 | 0.17 | 0.16 |
| 2 | 33722854 | rs10166200 | -1.445 | 0.981 | 0.064 | 0.604 | 0.01 | -0.019 | -0.019 | 0 | 0.23 | 0.34 | 0.28 | 0.16 | 0.26 | 0.27 |
| 2 | 33723184 | rs10495791 | -2.082 | 1.004 | -0.35 | 0.618 | 0.004 | -0.02 | -0.02 | 0 | 0.37 | 0.57 | 0.37 | 0.25 | 0.35 | 0.35 |
| 2 | 33723510 | rs719546 | -0.16 | 0.196 | -0.135 | 0.586 | -0.004 | -0.001 | -0.001 | 0.106 | 0.11 | 0.08 | 0.07 | 0.09 | 0.09 | 0.05 |
| 2 | 33726610 | rs6543719 | -0.952 | 0.31 | -0.193 | 0.677 | -0.028 | -0.055 | -0.055 | 0 | 0.27 | 0.48 | 0.15 | 0.13 | 0.18 | 0.17 |
| 2 | 33733554 | rs6761582 | -0.927 | 0.141 | 0.184 | 1.288 | -0.029 | 0.148 | 0.148 | 0.046 | 0.15 | 0.44 | 0.16 | 0.34 | 0.1 | 0.08 |
| 2 | 33734126 | rs6743600 | -0.273 | 0.477 | -0.491 | 0.356 | -0.019 | 0.041 | 0.041 | 0 | 0.18 | 0.1 | 0.11 | 0.01 | 0.21 | 0.16 |
| 2 | 33736281 | rs10200743 | 0.138 | 1.218 | -1.018 | 0.614 | -0.024 | -0.016 | -0.016 | 0 | 0.8 | 0.89 | 0.86 | 0.78 | 0.73 | 0.73 |
| 2 | 33738613 | rs3754823 | 1.053 | 0.937 | -2.014 | 0.258 | 0.004 | 0.074 | 0.074 | 1.273 | 0.42 | 0.07 | 0.38 | 0.26 | 0.36 | 0.26 |
| 2 | 33739297 | rs2168395 | -0.962 | 1.196 | -1.464 | 0.673 | 0.068 | 0.245 | 0.245 | 0 | 0.14 | 0.41 | 0.15 | 0.41 | 0.21 | 0.14 |
| 2 | 33740855 | rs3845781 | 1.487 | 1.144 | -1.702 | 0.475 | -0.044 | 0.051 | 0.051 | 0 | 0.09 | 0.02 | 0.2 | 0.01 | 0.17 | 0.09 |
| 2 | 33742284 | rs4670648 | -0.319 | 1.025 | -1.613 | 0.569 | -0.034 | -0.031 | -0.031 | 0 | 0.81 | 0.91 | 0.88 | 0.78 | 0.76 | 0.73 |
| 2 | 33744958 | rs7597095 | 1.124 | 0.813 | -1.202 | 0.424 | -0.001 | -0.001 | -0.001 | 0 | 0.66 | 0.32 | 0.69 | 0.36 | 0.64 | 0.54 |
| 2 | 33745208 | rs2305576 | 1.007 | 0.716 | -0.937 | 0.274 | 0.001 | 0.018 | 0.018 | 0 | 0.58 | 0.35 | 0.64 | 0.3 | 0.59 | 0.41 |
| 2 | 33746333 | rs17013215 | -0.404 | 0.654 | -0.719 | 0.524 | -0.02 | 0.059 | 0.059 | 0.608 | 0.23 | 0.65 | 0.25 | 0.48 | 0.18 | 0.34 |
| 2 | 33748838 | rs2305571 | 1.593 | 0.809 | -1.014 | 0.278 | 0.024 | 0.1 | 0.1 | 0.788 | 0.46 | 0.13 | 0.4 | 0.29 | 0.39 | 0.29 |
| 2 | 33756488 | rs17650408 | 1.714 | 0.758 | -0.9 | 0.19 | 0.001 | 0.005 | 0.005 | 0 | 0.08 | 0.05 | 0.04 | 0.06 | 0.07 | 0.12 |
| 2 | 33756520 | rs13404493 | -0.116 | 0.716 | 0.051 | 0.196 | 0.001 | -0.015 | -0.015 | 0.871 | 0.12 | 0.36 | 0.18 | 0.08 | 0.11 | 0.11 |
| 2 | 33757554 | rs11685011 | 0.938 | 0.735 | 0.055 | 0.188 | 0.005 | -0.017 | -0.017 | 2.892 | 0.58 | 0.5 | 0.55 | 0.34 | 0.49 | 0.4 |
| 2 | 33757785 | rs11685205 | 0.409 | 0.783 | 0.091 | 0.203 | 0.035 | 0.01 | 0.01 | 0 | 0.68 | 0.59 | 0.62 | 0.4 | 0.57 | 0.52 |
| 2 | 33762963 | rs17013237 | 0.144 | 0.671 | 0.666 | 0.303 | -0.002 | -0.005 | -0.005 | 0 | 0.1 | 0.19 | 0.16 | 0.11 | 0.11 | 0.11 |
| 2 | 33772525 | rs4258776 | -1.439 | 0.551 | 0.575 | 0.306 | 0 | 0.004 | 0.004 | 0 | 0.34 | 0.58 | 0.47 | 0.46 | 0.36 | 0.48 |
| 2 | 33773536 | rs17013312 | -0.231 | 0.508 | 0.474 | 0.288 | -0.007 | -0.017 | -0.017 | 4.233 | 0.12 | 0.2 | 0.16 | 0.1 | 0.11 | 0.13 |
| 2 | 33775665 | rs17597696 | 1.414 | 0.66 | 0.493 | -0.114 | -0.054 | 0.031 | 0.031 | 0 | 0.11 | 0.02 | 0.21 | 0.02 | 0.18 | 0.13 |
| 2 | 33776743 | rs4491711 | -0.157 | 0.511 | 0.637 | -0.075 | -0.006 | -0.021 | -0.021 | 2.908 | 0.11 | 0.2 | 0.16 | 0.08 | 0.11 | 0.13 |
| 2 | 33780231 | rs10182608 | -0.729 | 0.463 | 0.816 | -0.125 | -0.016 | -0.065 | -0.065 | 0.438 | 0.2 | 0.48 | 0.22 | 0.13 | 0.18 | 0.23 |
| 2 | 33782911 | rs7574183 | -0.737 | 0.455 | 0.568 | -0.26 | -0.011 | -0.027 | -0.027 | 0.163 | 0.14 | 0.38 | 0.18 | 0.1 | 0.11 | 0.12 |
| 2 | 33784161 | rs2278955 | 1.477 | 0.347 | -0.763 | -0.505 | -0.022 | 0.08 | 0.08 | 1.379 | 0.42 | 0.09 | 0.36 | 0.27 | 0.34 | 0.27 |
| 2 | 33784696 | rs17598067 | 1.924 | 0.42 | -0.312 | -0.309 | -0.018 | -0.001 | -0.001 | 0 | 0.07 | 0 | 0.04 | 0.06 | 0.06 | 0.09 |
| 2 | 33788105 | rs17013362 | 1.243 | 0.298 | -0.103 | -0.477 | 0.011 | -0.039 | -0.039 | 0 | 0.09 | 0.13 | 0.06 | 0 | 0.13 | 0.03 |
| 2 | 33788156 | rs3087731 | -0.942 | 0.022 | -0.361 | -0.819 | 0.005 | 0.005 | 0.005 | 0 | 0.3 | 0.3 | 0.25 | 0.09 | 0.27 | 0.16 |
| 2 | 33789032 | rs10514775 | 0.486 | 0.03 | -0.212 | -0.661 | 0.003 | -0.006 | -0.006 | 0.811 | 0.1 | 0.13 | 0.15 | 0.09 | 0.1 | 0.12 |
| 2 | 33789534 | rs6708 | -2.033 | -0.493 | -0.547 | -1.092 | 0.049 | 0.107 | 0.107 | 0.151 | 0.38 | 0.76 | 0.32 | 0.15 | 0.29 | 0.3 |
| 2 | 33790330 | rs6742257 | -1.43 | -0.456 | -0.347 | -1.064 | -0.002 | -0.018 | -0.018 | 0.159 | 0.2 | 0.72 | 0.23 | 0.1 | 0.17 | 0.16 |
| 2 | 33798742 | rs17013391 | -0.773 | -0.591 | 0.031 | -1.133 | 0.004 | -0.033 | -0.033 | 0.525 | 0.06 | 0.15 | 0.04 | 0 | 0.06 | 0.03 |
| 2 | 33801017 | rs10198451 | 0.611 | -0.509 | -0.405 | -0.948 | -0.004 | -0.001 | -0.001 | 0 | 0.64 | 0.41 | 0.65 | 0.37 | 0.61 | 0.52 |
| 2 | 33802902 | rs13388224 | 2.079 | -0.595 | -0.462 | -0.963 | -0.047 | 0.05 | 0.05 | 0.086 | 0.45 | 0.17 | 0.55 | 0.31 | 0.5 | 0.38 |
| 2 | 33802956 | rs4670209 | 0.595 | -0.631 | -0.439 | -0.928 | -0.003 | -0.001 | -0.001 | 0 | 0.64 | 0.41 | 0.65 | 0.37 | 0.61 | 0.52 |
| 2 | 33809294 | rs11066 | -0.579 | -0.493 | -0.222 | -0.752 | -0.011 | -0.099 | -0.099 | 0 | 0.74 | 0.94 | 0.83 | 0.45 | 0.75 | 0.68 |
| 2 | 33814590 | rs17657783 | 1.634 | -0.439 | -0.223 | -0.819 | -0.008 | 0.056 | 0.056 | 0 | 0.46 | 0.25 | 0.56 | 0.3 | 0.5 | 0.39 |
| 2 | 33815948 | rs4627543 | -1.133 | -0.475 | -0.211 | -0.844 | -0.004 | -0.065 | -0.065 | 0 | 0.22 | 0.59 | 0.25 | 0.09 | 0.22 | 0.18 |
| 2 | 33817559 | rs2044644 | 1.702 | -0.523 | -0.637 | -0.85 | -0.06 | 0.041 | 0.041 | 0.141 | 0.45 | 0.13 | 0.56 | 0.3 | 0.5 | 0.39 |
| 2 | 33817881 | rs3023 | -0.204 | -0.648 | -0.652 | -0.775 | 0.014 | -0.065 | -0.065 | 2.242 | 0.68 | 0.9 | 0.81 | 0.39 | 0.73 | 0.57 |
| 2 | 33826877 | rs1375194 | 1.558 | -0.689 | -0.809 | -0.788 | -0.044 | 0.006 | 0.006 | 0.209 | 0.43 | 0.07 | 0.57 | 0.3 | 0.55 | 0.39 |
| 2 | 33827428 | rs2367811 | 0.687 | -0.815 | -0.456 | -0.79 | 0.089 | 0.019 | 0.019 | 0 | 0.54 | 0.25 | 0.72 | 0.38 | 0.66 | 0.53 |
| 2 | 33830319 | rs10173763 | -0.11 | -0.889 | -0.534 | -0.736 | 0.04 | -0.064 | -0.064 | 0.839 | 0.66 | 0.89 | 0.8 | 0.41 | 0.74 | 0.57 |
| 2 | 33832815 | rs6543723 | 1.956 | -0.309 | -0.946 | -0.935 | -0.123 | -0.042 | -0.042 | 1.116 | 0.39 | 0.06 | 0.49 | 0.32 | 0.44 | 0.36 |
| 2 | 33834496 | rs11687252 | 0.838 | -0.239 | -0.557 | -0.996 | 0.007 | 0.008 | 0.008 | 0 | 0.53 | 0.61 | 0.68 | 0.41 | 0.55 | 0.5 |
| 2 | 33836922 | rs7559158 | -0.071 | -0.118 | -0.309 | -1.013 | 0.084 | -0.046 | -0.046 | 0 | 0.09 | 0.31 | 0.1 | 0 | 0.13 | 0.04 |
| 2 | 33845642 | rs6543726 | -0.222 | -0.159 | -0.294 | -1.005 | -0.016 | -0.055 | -0.055 | 0.048 | 0.18 | 0.52 | 0.2 | 0.12 | 0.16 | 0.14 |
| 2 | 33847659 | rs6717058 | -1.695 | -0.087 | -0.259 | -0.962 | -0.002 | 0.006 | 0.006 | 0.467 | 0.28 | 0.68 | 0.3 | 0.14 | 0.27 | 0.31 |

| | | | | | | | | | | | | | | | | |
|---|---|---|---|---|---|---|---|---|---|---|---|---|---|---|---|---|
| 2 | 33850382 | rs6713874 | -0.957 | -0.109 | -0.263 | -0.974 | 0.044 | -0.059 | -0.059 | 0 | 0.2 | 0.55 | 0.25 | 0.12 | 0.22 | 0.17 |
| 2 | 33852489 | rs4670815 | -1.034 | -0.351 | -0.628 | -1.105 | 0.022 | -0.123 | -0.123 | 5.271 | 0.64 | 0.93 | 0.81 | 0.47 | 0.7 | 0.67 |
| 2 | 33854344 | rs1448561 | 1.448 | -0.363 | -0.566 | -1.089 | -0.007 | 0.006 | 0.006 | 0 | 0.41 | 0.37 | 0.53 | 0.32 | 0.45 | 0.37 |
| 2 | 33857236 | rs1448560 | 1.706 | -0.194 | -0.632 | -1.095 | -0.023 | -0.002 | -0.002 | 0 | 0.37 | 0.23 | 0.51 | 0.32 | 0.43 | 0.34 |
| 2 | 33862493 | rs2012014 | 0.389 | 0.01 | 0.08 | -0.856 | -0.018 | -0.036 | -0.036 | 0 | 0.18 | 0.49 | 0.2 | 0.14 | 0.13 | 0.26 |
| 2 | 33863202 | rs11124328 | -2.373 | 0.235 | -0.199 | -0.855 | -0.02 | 0.02 | 0.02 | 0.314 | 0.95 | 0.98 | 0.94 | 1 | 0.89 | 0.99 |
| 2 | 33875088 | rs10188211 | -0.117 | 0.2 | 0.252 | -0.79 | 0.016 | -0.021 | -0.021 | 9.753 | 0.17 | 0.25 | 0.23 | 0.08 | 0.21 | 0.17 |
| 2 | 33877471 | rs7580073 | 0.749 | 0.076 | -0.022 | -0.704 | -0.041 | -0.046 | -0.046 | 0 | 0.12 | 0.39 | 0.07 | 0.06 | 0.05 | 0.15 |
| 2 | 33880853 | rs4670866 | 1.943 | 0.184 | -0.874 | -0.514 | -0.103 | -0.06 | -0.06 | 0 | 0.41 | 0.09 | 0.5 | 0.37 | 0.47 | 0.42 |
| 2 | 33894494 | rs1562614 | 0.959 | 0.554 | -0.621 | -0.45 | 0.027 | 0.02 | 0.02 | 0 | 0.13 | 0.35 | 0.32 | 0.14 | 0.14 | 0.21 |
| 2 | 33895267 | rs6711516 | 1.907 | 0.431 | -0.682 | -0.508 | -0.003 | 0.021 | 0.021 | 0 | 0.41 | 0.29 | 0.37 | 0.31 | 0.43 | 0.31 |
| 2 | 33896461 | rs6543733 | -0.795 | 0.417 | -1.309 | -0.259 | 0.016 | -0.133 | -0.133 | 0 | 0.71 | 0.92 | 0.82 | 0.51 | 0.74 | 0.7 |
| 2 | 33897454 | rs11904039 | 0.323 | 0.608 | -0.634 | -0.017 | 0.019 | 0.016 | 0.016 | 0 | 0.13 | 0.34 | 0.32 | 0.14 | 0.14 | 0.21 |
| 2 | 33903260 | rs1463143 | 1.578 | 0.476 | -1.63 | -0.067 | -0.069 | -0.063 | -0.063 | 0.097 | 0.35 | 0.06 | 0.38 | 0.35 | 0.41 | 0.23 |
| 2 | 33922554 | rs7585514 | 0.987 | 0.164 | -0.635 | -0.186 | -0.021 | -0.027 | -0.027 | 1.633 | 0.06 | 0.2 | 0.01 | 0 | 0.01 | 0 |
| 2 | 33925066 | rs12712350 | -1.199 | 0.357 | -0.867 | -0.273 | 0.012 | 0.014 | 0.014 | 0 | 0.37 | 0.66 | 0.26 | 0.17 | 0.26 | 0.34 |
| 2 | 33925829 | rs1868348 | -0.582 | 0.582 | -1.09 | -0.072 | -0.053 | -0.059 | -0.059 | 1.636 | 0.87 | 0.95 | 0.84 | 0.61 | 0.73 | 0.85 |
| 2 | 33928605 | rs7574123 | -0.297 | 0.676 | -1.382 | -0.242 | -0.086 | -0.075 | -0.075 | 0 | 0.77 | 0.93 | 0.78 | 0.62 | 0.63 | 0.81 |
| 2 | 33928721 | rs7588291 | 2.215 | 0.602 | 0.551 | 0.264 | -0.031 | -0.003 | -0.003 | 0 | 0.49 | 0.19 | 0.55 | 0.44 | 0.49 | 0.5 |
| 2 | 33929419 | rs1375196 | -0.146 | 0.23 | 0.543 | 0.253 | -0.014 | -0.063 | -0.063 | 0 | 0.2 | 0.27 | 0.09 | 0 | 0.13 | 0.08 |
| 2 | 33931617 | rs7585657 | 0.931 | 0.775 | 0.179 | 0.716 | -0.032 | -0.046 | -0.046 | 0 | 0.81 | 0.88 | 0.7 | 0.55 | 0.7 | 0.71 |
| 2 | 33939941 | rs2367955 | -0.74 | 0.9 | 0.403 | 0.125 | 0.031 | 0.029 | 0.029 | 0 | 0.91 | 0.9 | 0.9 | 0.72 | 0.81 | 0.9 |
| 2 | 33940043 | rs1448553 | -0.169 | 0.838 | 0.546 | 0.004 | 0 | -0.002 | -0.002 | 0 | 0.1 | 0.09 | 0.08 | 0.05 | 0.1 | 0.16 |
| 2 | 33940918 | rs12988770 | 1.85 | 0.486 | 0.917 | -0.103 | 0.001 | -0.002 | -0.002 | 0 | 0.44 | 0.39 | 0.51 | 0.43 | 0.44 | 0.46 |
| 2 | 33941926 | rs6755276 | -0.547 | 0.574 | 0.86 | -0.164 | -0.005 | -0.01 | -0.01 | 0 | 0.89 | 0.94 | 0.84 | 0.71 | 0.74 | 0.89 |
| 2 | 33944002 | rs6706223 | 1.966 | 0.468 | 1.171 | -0.204 | -0.006 | -0.02 | -0.02 | 0.109 | 0.43 | 0.29 | 0.57 | 0.43 | 0.44 | 0.45 |
| 2 | 33944025 | rs10469891 | 0.403 | 0.391 | 1.323 | -0.222 | 0.031 | 0.038 | 0.038 | 0 | 0.22 | 0.13 | 0.07 | 0 | 0.16 | 0.1 |
| 2 | 33945714 | rs12471341 | 1.991 | 0.485 | 1.268 | -0.164 | -0.024 | -0.02 | -0.02 | 0 | 0.45 | 0.3 | 0.62 | 0.44 | 0.51 | 0.45 |
| 2 | 33946764 | rs10469892 | -0.991 | 0.476 | 1.346 | -0.166 | -0.048 | -0.023 | -0.023 | 0 | 0.35 | 0.42 | 0.2 | 0.25 | 0.2 | 0.26 |
| 2 | 33947264 | rs2367957 | 1.112 | 0.507 | 1.303 | -0.178 | 0 | 0 | 0 | 0.016 | 0.56 | 0.6 | 0.7 | 0.47 | 0.6 | 0.63 |
| 2 | 33948748 | rs935622 | 1.66 | 0.454 | 1.292 | -0.172 | -0.004 | 0.001 | 0.001 | 0.011 | 0.46 | 0.43 | 0.61 | 0.43 | 0.5 | 0.45 |
| 2 | 33951513 | rs10196645 | 0.332 | 0.464 | 1.173 | -0.301 | -0.026 | -0.003 | -0.003 | 0 | 0.75 | 0.85 | 0.79 | 0.71 | 0.69 | 0.69 |
| 2 | 33952413 | rs4477896 | 0.427 | 0.441 | 1.271 | -0.331 | 0.055 | 0.031 | 0.031 | 0.001 | 0.8 | 0.63 | 0.85 | 0.74 | 0.78 | 0.86 |
| 2 | 33952621 | rs11684598 | 1.818 | 0.315 | 1.177 | -0.478 | 0 | -0.013 | -0.013 | 2.475 | 0.45 | 0.34 | 0.6 | 0.44 | 0.48 | 0.46 |
| 2 | 33955044 | rs6707925 | 0.618 | 0.383 | 1.468 | -0.747 | -0.012 | -0.066 | -0.066 | 0.248 | 0.16 | 0.22 | 0.08 | 0 | 0.14 | 0.1 |
| 2 | 33959349 | rs13387317 | -1.221 | 0.375 | 1.797 | -0.354 | 0.004 | -0.04 | -0.04 | 0 | 0.09 | 0.4 | 0.04 | 0 | 0.02 | 0.01 |
| 2 | 33960341 | rs7581397 | 0.414 | 0.407 | 1.234 | -0.33 | -0.011 | -0.03 | -0.03 | 0 | 0.06 | 0.12 | 0.02 | 0 | 0.01 | 0.01 |
| 2 | 33965251 | rs4257379 | 1.906 | 0.484 | 0.774 | -0.373 | 0.011 | -0.061 | -0.061 | 1.042 | 0.33 | 0.16 | 0.37 | 0.44 | 0.27 | 0.26 |
| 2 | 33969387 | rs4287749 | 0.088 | 0.628 | 0.859 | -0.252 | 0.005 | -0.025 | -0.025 | 0 | 0.13 | 0.12 | 0.22 | 0.01 | 0.19 | 0.19 |
| 2 | 33971813 | rs12995445 | -0.008 | 0.613 | 0.478 | -0.192 | -0.061 | -0.134 | -0.134 | 0 | 0.65 | 0.92 | 0.75 | 0.48 | 0.6 | 0.64 |
| 2 | 33977878 | rs6758248 | 1.446 | 0.058 | 0.832 | -0.044 | 0.015 | -0.029 | -0.029 | 0 | 0.35 | 0.21 | 0.38 | 0.44 | 0.28 | 0.26 |
| 2 | 33978851 | rs6543760 | -1.307 | 0.299 | 1.152 | -0.126 | 0.001 | -0.006 | -0.006 | 0.508 | 0.57 | 0.6 | 0.52 | 0.49 | 0.6 | 0.69 |
| 2 | 33988173 | rs17338700 | 0.437 | 0.574 | 1.431 | -0.195 | 0.002 | -0.011 | -0.011 | 0 | 0.09 | 0.09 | 0.16 | 0.01 | 0.12 | 0.17 |
| 2 | 33989670 | rs5025563 | 0.375 | 0.533 | 1.034 | -0.204 | -0.056 | 0.027 | 0.027 | 0 | 0.27 | 0.1 | 0.23 | 0.24 | 0.41 | 0.21 |
| 2 | 33990715 | rs17247554 | 0.4 | 0.542 | 0.892 | -0.205 | -0.091 | 0.017 | 0.017 | 0 | 0.27 | 0.07 | 0.23 | 0.24 | 0.42 | 0.21 |
| 2 | 33990760 | rs17247561 | 0.327 | 0.537 | 0.928 | -0.206 | -0.049 | 0.029 | 0.029 | 0 | 0.28 | 0.11 | 0.23 | 0.24 | 0.41 | 0.21 |
| 2 | 33993069 | rs11691402 | 0.403 | 0.539 | 0.9 | -0.205 | -0.09 | 0.013 | 0.013 | 0 | 0.27 | 0.06 | 0.22 | 0.24 | 0.41 | 0.21 |
| 2 | 33993207 | rs1543976 | -1.608 | 0.462 | 1.178 | -0.173 | 0.006 | -0.008 | -0.008 | 0 | 0.58 | 0.66 | 0.58 | 0.57 | 0.64 | 0.73 |
| 2 | 33993379 | rs17338771 | 0.298 | 0.38 | 0.565 | -0.11 | -0.07 | 0.029 | 0.029 | 0 | 0.28 | 0.09 | 0.23 | 0.24 | 0.42 | 0.21 |
| 2 | 33994141 | rs7583931 | 0.2 | 0.38 | 0.571 | -0.109 | -0.038 | 0.032 | 0.032 | 0 | 0.29 | 0.15 | 0.23 | 0.24 | 0.42 | 0.21 |
| 2 | 33995410 | rs1533840 | -1.356 | 0.178 | 0.737 | 0.121 | -0.008 | 0.005 | 0.005 | 0.015 | 0.31 | 0.58 | 0.35 | 0.33 | 0.23 | 0.52 |
| 2 | 33995612 | rs12712361 | -1.358 | 0.18 | 0.738 | 0.122 | -0.008 | 0.005 | 0.005 | 0.043 | 0.31 | 0.58 | 0.35 | 0.33 | 0.23 | 0.52 |
| 2 | 33996983 | rs6714970 | -0.889 | -0.571 | 0.69 | 0.362 | -0.048 | 0.054 | 0.054 | 1.355 | 0.22 | 0.54 | 0.2 | 0.31 | 0.1 | 0.33 |
| 2 | 34001967 | rs6735386 | -0.781 | 0.13 | -0.127 | -0.07 | -0.001 | -0.001 | -0.001 | 0 | 0.51 | 0.48 | 0.59 | 0.6 | 0.64 | 0.66 |
| 2 | 34002421 | rs6738691 | -0.846 | 0.148 | -0.421 | -0.061 | 0.011 | 0.004 | 0.004 | 0.042 | 0.52 | 0.56 | 0.59 | 0.6 | 0.64 | 0.66 |
| 2 | 34004920 | rs7576239 | -0.493 | 1.005 | -1.701 | 0.46 | -0.111 | -0.126 | -0.126 | 0.032 | 0.34 | 0.08 | 0.41 | 0.53 | 0.55 | 0.53 |
| 2 | 34019376 | rs6735662 | 0.519 | 0.669 | -0.639 | -0.302 | 0.01 | -0.155 | -0.155 | 0.001 | 0.76 | 0.58 | 0.86 | 1 | 0.77 | 0.82 |
| 2 | 34019809 | rs4290635 | 0.671 | 0.367 | -0.575 | -0.451 | -0.004 | -0.167 | -0.167 | 0.002 | 0.73 | 0.55 | 0.86 | 1 | 0.77 | 0.83 |
| 2 | 34021004 | rs1533845 | 0.519 | 0.551 | -0.655 | -0.527 | 0.019 | -0.166 | -0.166 | 0.027 | 0.74 | 0.54 | 0.84 | 1 | 0.73 | 0.82 |
| 2 | 34022970 | rs10187143 | -0.563 | 0.165 | -1.439 | -0.299 | 0.012 | -0.042 | -0.042 | 0 | 0.25 | 0.12 | 0.36 | 0.71 | 0.26 | 0.25 |
| 2 | 34023020 | rs4313958 | -0.186 | -0.065 | -0.908 | -0.596 | -0.003 | 0.009 | 0.009 | 0.137 | 0.79 | 0.86 | 0.89 | 1 | 0.81 | 0.92 |
| 2 | 34025075 | rs6738183 | -1.842 | -0.066 | -0.321 | -0.114 | -0.002 | -0.051 | -0.051 | 0.465 | 0.09 | 0.23 | 0.04 | 0 | 0.08 | 0.09 |
| 2 | 34028090 | rs6736937 | -0.91 | -0.098 | -0.638 | -0.127 | 0 | -0.006 | -0.006 | 0 | 0.07 | 0.06 | 0.03 | 0 | 0.06 | 0.07 |
| 2 | 34028455 | rs7572718 | -1.331 | -0.482 | -0.726 | 0.452 | 0.022 | -0.025 | -0.025 | 0 | 0.32 | 0.77 | 0.27 | 0.49 | 0.25 | 0.39 |
| 2 | 34030594 | rs4324308 | -1.358 | -0.458 | -0.757 | 0.669 | 0.025 | -0.031 | -0.031 | 0 | 0.33 | 0.78 | 0.37 | 0.59 | 0.25 | 0.4 |
| 2 | 34032345 | rs12621616 | -0.376 | -0.09 | -0.437 | 0.968 | -0.001 | 0.062 | 0.062 | 0 | 0.14 | 0.21 | 0.21 | 0.52 | 0.16 | 0.2 |
| 2 | 34036427 | rs1522581 | -0.039 | -0.48 | -0.615 | 0.343 | 0.007 | 0.011 | 0.011 | 0.053 | 0.61 | 0.68 | 0.77 | 0.79 | 0.65 | 0.75 |
| 2 | 34037163 | rs7559601 | 0.351 | -1.069 | -1.244 | -0.27 | 0.07 | 0.141 | 0.141 | 0.931 | 0.52 | 0.15 | 0.65 | 0.74 | 0.64 | 0.64 |
| 2 | 34038728 | rs6741553 | 0.639 | -1.052 | -1.615 | 0.382 | -0.057 | 0.103 | 0.103 | 0.858 | 0.42 | 0.17 | 0.5 | 0.21 | 0.5 | 0.48 |

| | | | | | | | | | | | | | | | | |
|---|---|---|---|---|---|---|---|---|---|---|---|---|---|---|---|---|
| 2 | 34040204 | rs1522578 | -0.781 | -1.211 | -0.52 | 0.6 | -0.007 | 0.174 | 0.174 | 0.494 | 0.22 | 0.46 | 0.24 | 0.48 | 0.21 | 0.2 |
| 2 | 34043081 | rs13005973 | -0.534 | -1.444 | -0.095 | 1.04 | -0.019 | 0.101 | 0.101 | 0.615 | 0.11 | 0.29 | 0.09 | 0.25 | 0.07 | 0.09 |
| 2 | 34043426 | rs13417671 | -0.56 | -1.479 | -0.093 | 0.949 | -0.001 | -0.029 | -0.029 | 0 | 0.19 | 0.14 | 0.17 | 0.24 | 0.19 | 0.17 |
| 2 | 34044433 | rs1996834 | -0.003 | -1.551 | 0.091 | 0.929 | 0.014 | -0.027 | -0.027 | 0 | 0.62 | 0.32 | 0.67 | 0.43 | 0.69 | 0.65 |
| 2 | 34047785 | rs1543978 | 0.09 | -1.572 | 0.08 | 0.919 | 0.022 | -0.038 | -0.038 | 0.157 | 0.62 | 0.3 | 0.67 | 0.43 | 0.69 | 0.64 |
| 2 | 34048462 | rs7608031 | 1.094 | -1.484 | 0.042 | 0.751 | 0.035 | 0.125 | 0.125 | 1.174 | 0.5 | 0.32 | 0.59 | 0.2 | 0.59 | 0.49 |
| 2 | 34055724 | rs17014142 | -1.416 | -1.531 | 0.099 | 0.917 | -0.018 | 0.063 | 0.063 | 0.485 | 0.23 | 0.3 | 0.17 | 0.45 | 0.17 | 0.23 |
| 2 | 34055950 | rs10179135 | 0.538 | -1.484 | 0.072 | 0.964 | -0.024 | -0.06 | -0.06 | 0 | 0.11 | 0 | 0.09 | 0.2 | 0.1 | 0.13 |
| 2 | 34059750 | rs10204763 | -0.467 | -1.482 | 0.943 | 0.667 | -0.003 | 0.174 | 0.174 | 0 | 0.14 | 0.31 | 0.13 | 0.45 | 0.11 | 0.16 |
| 2 | 34060104 | rs1522591 | 0.023 | -1.406 | 0.754 | 0.378 | -0.005 | -0.004 | -0.004 | 0 | 0.65 | 0.48 | 0.64 | 0.65 | 0.66 | 0.68 |
| 2 | 34061865 | rs869964 | 0.494 | -1.3 | 0.551 | 0.447 | 0.018 | 0.03 | 0.03 | 0.4 | 0.56 | 0.33 | 0.61 | 0.65 | 0.6 | 0.61 |
| 2 | 34067478 | rs4671004 | -0.217 | -0.789 | 0.528 | 0.488 | 0.006 | -0.039 | -0.039 | 0 | 0.92 | 0.83 | 0.89 | 1 | 0.9 | 0.96 |
| 2 | 34074065 | rs1705119 | 1.362 | -0.878 | 0.779 | 0.415 | 0.018 | -0.043 | -0.043 | 0 | 0.92 | 0.74 | 0.92 | 1 | 0.94 | 0.99 |
| 2 | 34074656 | rs1718728 | 1.826 | -1.051 | 1.475 | 1.097 | 0.011 | 0.13 | 0.13 | 0.029 | 0.8 | 0.61 | 0.81 | 0.37 | 0.82 | 0.8 |
| 2 | 34075614 | rs1718730 | 1.579 | -1.065 | 1.773 | 1.07 | 0.078 | -0.01 | -0.01 | 0 | 0.92 | 0.6 | 0.93 | 1 | 0.93 | 0.99 |
| 2 | 34075756 | rs4670303 | 0.477 | -1.607 | 1.207 | 0.491 | -0.003 | -0.008 | -0.008 | 0 | 0.29 | 0.35 | 0.33 | 0.24 | 0.26 | 0.26 |
| 2 | 34083346 | rs1718739 | 0.541 | -1.9 | 0.852 | -0.02 | -0.03 | -0.106 | -0.106 | 1.77 | 0.78 | 0.64 | 0.88 | 0.99 | 0.85 | 0.88 |
| 2 | 34083671 | rs1705118 | 0.871 | -1.864 | 0.883 | 0.002 | 0.056 | -0.021 | -0.021 | 0.012 | 0.92 | 0.66 | 0.93 | 1 | 0.93 | 0.99 |
| 2 | 34085969 | rs773814 | 1.653 | -1.859 | 0.631 | 0 | -0.061 | 0.099 | 0.099 | 0.68 | 0.38 | 0.18 | 0.43 | 0.12 | 0.48 | 0.46 |
| 2 | 34091442 | rs7579347 | 0.939 | -1.768 | 0.852 | 0.267 | 0.003 | -0.036 | -0.036 | 0 | 0.29 | 0.14 | 0.32 | 0.34 | 0.27 | 0.3 |
| 2 | 34094236 | rs810148 | 0.126 | -1.581 | 1.042 | 0.242 | 0.044 | -0.047 | -0.047 | 0 | 0.92 | 0.81 | 0.92 | 1 | 0.87 | 0.97 |
| 2 | 34098507 | rs1718712 | 1.185 | -1.688 | 0.685 | 0.242 | -0.022 | 0.04 | 0.04 | 0.441 | 0.4 | 0.32 | 0.46 | 0.18 | 0.47 | 0.4 |
| 2 | 34103321 | rs1510876 | 1.036 | -1.734 | 0.353 | 0.304 | -0.052 | -0.057 | -0.057 | 0.371 | 0.3 | 0.08 | 0.34 | 0.34 | 0.32 | 0.26 |
| 2 | 34103546 | rs773828 | 0.491 | -1.809 | 0.464 | 0.281 | 0.012 | 0.018 | 0.018 | 0.628 | 0.5 | 0.47 | 0.56 | 0.66 | 0.58 | 0.64 |
| 2 | 34103843 | rs13400136 | -0.457 | -1.737 | 0.66 | 0.45 | 0.003 | 0.107 | 0.107 | 0.173 | 0.12 | 0.16 | 0.11 | 0.49 | 0.12 | 0.24 |
| 2 | 34106664 | rs773834 | 1.42 | -1.775 | 0.025 | 0.359 | 0.009 | 0.094 | 0.094 | 1.156 | 0.81 | 0.4 | 0.8 | 0.52 | 0.82 | 0.75 |
| 2 | 34115198 | rs17014268 | -0.14 | -1.801 | -0.181 | 0.331 | -0.022 | -0.072 | -0.072 | 0 | 0.12 | 0.01 | 0.12 | 0.51 | 0.11 | 0.23 |
| 2 | 34117511 | rs10495797 | 1.264 | -1.779 | -0.256 | 0.301 | -0.029 | -0.086 | -0.086 | 0 | 0.27 | 0.05 | 0.27 | 0.3 | 0.25 | 0.26 |
| 2 | 34119158 | rs1510875 | 1.464 | -1.595 | 0.036 | 0.415 | -0.001 | 0.041 | 0.041 | 1.906 | 0.8 | 0.35 | 0.78 | 0.49 | 0.81 | 0.75 |
| 2 | 34125660 | rs1705152 | 1.366 | -1.369 | -0.35 | 0.046 | -0.081 | 0.072 | 0.072 | 0.637 | 0.37 | 0.17 | 0.44 | 0.18 | 0.48 | 0.38 |
| 2 | 34127078 | rs1705148 | -0.36 | -1.582 | -0.12 | -0.087 | 0.001 | -0.017 | -0.017 | 0 | 0.83 | 0.85 | 0.91 | 1 | 0.91 | 0.89 |
| 2 | 34134854 | rs1705131 | 1.512 | -1.577 | -0.215 | -0.146 | -0.074 | 0.078 | 0.078 | 0.388 | 0.36 | 0.17 | 0.44 | 0.15 | 0.49 | 0.39 |
| 2 | 34135144 | rs13413184 | 0.005 | -1.718 | -0.213 | 0.228 | -0.025 | -0.058 | -0.058 | 4.755 | 0.38 | 0.08 | 0.33 | 0.55 | 0.36 | 0.48 |
| 2 | 34137903 | rs1718718 | 0.402 | -1.741 | 0.424 | 0.191 | 0.004 | 0.047 | 0.047 | 0 | 0.54 | 0.4 | 0.52 | 0.16 | 0.57 | 0.49 |
| 2 | 34144622 | rs7606055 | -1.931 | -1.76 | 0.293 | 0.177 | -0.092 | 0.085 | 0.085 | 0.526 | 0.64 | 0.85 | 0.56 | 0.85 | 0.51 | 0.61 |
| 2 | 34145755 | rs7569283 | -1.782 | -1.672 | 0.837 | 0.302 | 0.089 | 0.254 | 0.254 | 2.309 | 0.08 | 0.45 | 0.15 | 0.3 | 0.07 | 0.04 |
| 2 | 34151860 | rs13414817 | -0.453 | -1.664 | 1.086 | 0.244 | 0 | 0.007 | 0.007 | 0 | 0.44 | 0.28 | 0.4 | 0.54 | 0.43 | 0.47 |
| 2 | 34156809 | rs17014378 | -2.052 | -1.687 | 0.559 | 0.618 | 0.048 | -0.029 | -0.029 | 0 | 0.38 | 0.73 | 0.28 | 0.56 | 0.23 | 0.35 |
| 2 | 34156867 | rs6741754 | -1.812 | -1.815 | 0.142 | 0.312 | -0.063 | 0.049 | 0.049 | 0 | 0.7 | 0.81 | 0.55 | 0.87 | 0.53 | 0.61 |
| 2 | 34163414 | rs13387662 | 1.088 | -1.759 | 0.058 | 0.368 | -0.013 | -0.04 | -0.04 | 0 | 0.35 | 0.07 | 0.29 | 0.32 | 0.33 | 0.31 |
| 2 | 34175977 | rs4309564 | -0.347 | -1.405 | 0.487 | 0.621 | -0.007 | 0.005 | 0.005 | 0.031 | 0.57 | 0.79 | 0.5 | 0.62 | 0.43 | 0.42 |
| 2 | 34185335 | rs4496320 | -0.686 | -1.346 | 0.424 | 0.526 | -0.018 | 0.037 | 0.037 | 0 | 0.61 | 0.68 | 0.58 | 0.87 | 0.53 | 0.55 |
| 2 | 34187256 | rs13410163 | -2.076 | -1.382 | 0.727 | 0.63 | 0.043 | 0.176 | 0.176 | 5.701 | 0.1 | 0.51 | 0.13 | 0.29 | 0.06 | 0.04 |
| 2 | 34195228 | rs1559644 | 1.126 | 0.111 | 0.421 | 0.51 | 0.023 | 0.036 | 0.036 | 0 | 0.71 | 0.63 | 0.69 | 0.6 | 0.61 | 0.73 |
| 2 | 34199734 | rs10189501 | 1.782 | 0.332 | 0.074 | 0.457 | -0.037 | 0.031 | 0.031 | 0.058 | 0.41 | 0.08 | 0.47 | 0.29 | 0.41 | 0.43 |
| 2 | 34199849 | rs17247784 | 1.86 | 0.337 | 0.088 | 0.47 | -0.03 | 0.016 | 0.016 | 0.037 | 0.39 | 0.08 | 0.46 | 0.29 | 0.37 | 0.41 |
| 2 | 34200693 | rs9751085 | -0.807 | 0.505 | 0.534 | 0.126 | -0.011 | -0.001 | -0.001 | 2.492 | 0.85 | 0.91 | 0.8 | 0.87 | 0.76 | 0.88 |
| 2 | 34203838 | rs12162330 | 0.531 | 0.478 | 0.348 | 0.095 | 0.018 | -0.065 | -0.065 | 0.763 | 0.12 | 0.01 | 0.09 | 0.29 | 0.06 | 0.14 |
| 2 | 34204243 | rs13395553 | 0.79 | 0.489 | 0.587 | 0.119 | 0.009 | 0.016 | 0.016 | 0.23 | 0.68 | 0.6 | 0.67 | 0.56 | 0.65 | 0.74 |
| 2 | 34204267 | rs17247791 | 1.561 | 0.496 | 0.49 | 0.132 | -0.036 | 0.004 | 0.004 | 0 | 0.47 | 0.1 | 0.52 | 0.55 | 0.47 | 0.57 |
| 2 | 34207819 | rs12474926 | 1.225 | 0.658 | 0.446 | 0.29 | 0.011 | 0.023 | 0.023 | 0.125 | 0.67 | 0.52 | 0.65 | 0.56 | 0.65 | 0.74 |
| 2 | 34211405 | rs1510868 | -0.005 | 0.677 | 0.407 | 0.282 | 0.095 | 0.009 | 0.009 | 0 | 0.81 | 0.58 | 0.75 | 0.85 | 0.71 | 0.88 |
| 2 | 34214944 | rs13399795 | 1.358 | 0.643 | 0.484 | 0.158 | 0 | 0 | 0 | 0.694 | 0.65 | 0.37 | 0.63 | 0.56 | 0.65 | 0.74 |
| 2 | 34217579 | rs2048406 | 0.933 | 0.643 | 0.369 | 0.158 | 0.008 | 0.018 | 0.018 | 0 | 0.68 | 0.57 | 0.65 | 0.56 | 0.64 | 0.74 |
| 2 | 34222921 | rs12105087 | 0.256 | 0.736 | 0.28 | 0.21 | 0.068 | 0.115 | 0.115 | 0 | 0.78 | 0.51 | 0.68 | 0.57 | 0.7 | 0.84 |
| 2 | 34227952 | rs6715109 | 0.726 | 0.623 | -0.071 | 0.11 | -0.001 | 0.002 | 0.002 | 0.241 | 0.64 | 0.43 | 0.63 | 0.59 | 0.64 | 0.71 |
| 2 | 34234342 | rs6543799 | 1.421 | 0.623 | -0.854 | -0.006 | -0.068 | 0.01 | 0.01 | 0.429 | 0.47 | 0.09 | 0.52 | 0.59 | 0.49 | 0.52 |
| 2 | 34236374 | rs17247805 | 1.166 | 0.452 | -0.143 | -0.485 | -0.003 | -0.005 | -0.005 | 0 | 0.16 | 0.12 | 0.1 | 0.01 | 0.16 | 0.16 |
| 2 | 34237001 | rs9284729 | -0.727 | 0.383 | 0.452 | -0.051 | -0.037 | -0.084 | -0.084 | 0 | 0.15 | 0.34 | 0.07 | 0.01 | 0.08 | 0.1 |
| 2 | 34237472 | rs1965391 | 0.403 | 1.143 | 0.033 | 0.321 | 0.018 | -0.008 | -0.008 | 0.444 | 0.81 | 0.81 | 0.78 | 0.88 | 0.7 | 0.82 |
| 2 | 34245505 | rs11892281 | -0.696 | 0.808 | 0.27 | 0.528 | 0.02 | -0.01 | -0.01 | 1.334 | 0.25 | 0.43 | 0.29 | 0.22 | 0.3 | 0.27 |
| 2 | 34247570 | rs1425012 | 0.205 | 0.798 | 0.425 | 0.549 | 0.036 | 0.015 | 0.015 | 0.023 | 0.11 | 0.16 | 0.24 | 0.21 | 0.27 | 0.19 |
| 2 | 34253356 | rs6748035 | 0.667 | 0.205 | -0.138 | 0.587 | -0.009 | -0.027 | -0.027 | 0 | 0.38 | 0.18 | 0.43 | 0.39 | 0.39 | 0.46 |
| 2 | 34253584 | rs13033324 | -0.356 | 0.157 | 0.241 | 0.561 | 0.053 | 0.04 | 0.04 | 2.373 | 0.26 | 0.58 | 0.41 | 0.38 | 0.38 | 0.26 |
| 2 | 34256774 | rs7588920 | 1.502 | -0.053 | -0.208 | 0.526 | -0.022 | -0.073 | -0.073 | 0 | 0.18 | 0.04 | 0.34 | 0.31 | 0.19 | 0.23 |
| 2 | 34257574 | rs4366890 | 1.116 | 0.23 | 0.502 | 0.554 | 0.007 | -0.002 | -0.002 | 0.2 | 0.11 | 0.1 | 0.23 | 0.21 | 0.27 | 0.19 |
| 2 | 34257592 | rs735379 | 0.952 | 0.227 | 0.57 | 0.549 | 0.021 | 0.046 | 0.046 | 0 | 0.12 | 0.22 | 0.14 | 0.18 | 0.11 | 0.07 |
| 2 | 34258477 | rs6704936 | -0.46 | 0.16 | 0.606 | 0.514 | 0.007 | -0.005 | -0.005 | 0 | 0.27 | 0.39 | 0.3 | 0.24 | 0.3 | 0.28 |
| 2 | 34259736 | rs2216963 | -1.518 | 0.111 | 0.362 | 0.527 | 0.006 | 0.012 | 0.012 | 0.987 | 0.78 | 0.88 | 0.89 | 0.82 | 0.8 | 0.82 |

| | | | | | | | | | | | | | | | | |
|---|---|---|---|---|---|---|---|---|---|---|---|---|---|---|---|---|
| 2 | 34264495 | rs10495805 | 1.235 | 0.121 | 0.203 | 0.537 | -0.014 | -0.046 | -0.046 | 0.12 | 0.38 | 0.12 | 0.43 | 0.4 | 0.39 | 0.47 |
| 2 | 34265211 | rs4671029 | 1.012 | 0.233 | 0.268 | 0.484 | 0.027 | 0.046 | 0.046 | 0 | 0.11 | 0.24 | 0.15 | 0.16 | 0.11 | 0.07 |
| 2 | 34265621 | rs10495806 | -0.368 | 0.267 | -0.066 | 0.203 | -0.008 | 0.001 | 0.001 | 0 | 0.26 | 0.27 | 0.29 | 0.24 | 0.3 | 0.28 |
| 2 | 34267179 | rs2042583 | -0.848 | 1.303 | 0.013 | 0.585 | 0.01 | 0.011 | 0.011 | 0.774 | 0.77 | 0.88 | 0.88 | 0.79 | 0.8 | 0.81 |
| 2 | 34269189 | rs6712647 | 0.626 | 1.454 | -0.103 | 0.516 | -0.026 | -0.018 | -0.018 | 0 | 0.11 | 0.05 | 0.22 | 0.18 | 0.27 | 0.19 |
| 2 | 34272097 | rs745007 | 0.681 | 0.193 | 0.353 | 0.366 | 0.014 | 0.024 | 0.024 | 0.088 | 0.63 | 0.29 | 0.7 | 0.69 | 0.69 | 0.74 |
| 2 | 34273293 | rs6543803 | -0.461 | 0.465 | -0.339 | 0.493 | 0.009 | 0.017 | 0.017 | 1.143 | 0.78 | 0.91 | 0.88 | 0.81 | 0.8 | 0.8 |
| 2 | 34275263 | rs4670314 | -0.419 | 0.49 | -0.361 | 0.477 | 0.005 | 0.009 | 0.009 | 0.933 | 0.78 | 0.88 | 0.88 | 0.81 | 0.8 | 0.81 |
| 2 | 34275757 | rs4670315 | -0.673 | 0.818 | -0.073 | 0.765 | 0.046 | 0.026 | 0.026 | 0 | 0.12 | 0.27 | 0.18 | 0.12 | 0.12 | 0.07 |
| 2 | 34281766 | rs1834282 | 0.18 | 0.19 | -0.648 | 0.639 | 0.001 | 0 | 0 | 0.186 | 0.63 | 0.39 | 0.66 | 0.39 | 0.73 | 0.58 |
| 2 | 34282078 | rs13427419 | -0.436 | -0.651 | -0.52 | -0.29 | 0.021 | 0.023 | 0.023 | 0 | 0.39 | 0.38 | 0.45 | 0.91 | 0.3 | 0.51 |
| 2 | 34282796 | rs1425007 | 0.107 | -1.133 | -0.631 | 0.225 | 0.053 | -0.051 | -0.051 | 0 | 0.25 | 0.12 | 0.19 | 0.59 | 0.18 | 0.27 |
| 2 | 34282953 | rs17014564 | -0.8 | -0.989 | -0.531 | 0.184 | -0.001 | 0.054 | 0.054 | 0 | 0.06 | 0.05 | 0.21 | 0.34 | 0.06 | 0.24 |
| 2 | 34283170 | rs1425005 | 0.918 | -0.628 | -0.808 | -0.699 | -0.005 | 0.026 | 0.026 | 1.841 | 0.19 | 0.12 | 0.12 | 0.06 | 0.18 | 0.15 |
| 2 | 34286221 | rs12617275 | 0.644 | -0.154 | -0.681 | -0.738 | 0.014 | -0.024 | -0.024 | 0 | 0.31 | 0.16 | 0.28 | 0.6 | 0.31 | 0.31 |
| 2 | 34290662 | rs6750420 | -0.913 | -0.481 | -0.01 | -0.984 | 0.017 | 0.148 | 0.148 | 0 | 0.54 | 0.77 | 0.6 | 0.93 | 0.46 | 0.62 |
| 2 | 34291570 | rs17014613 | 0.796 | -0.469 | -0.097 | -0.93 | 0.014 | -0.021 | -0.021 | 0.579 | 0.31 | 0.15 | 0.28 | 0.6 | 0.31 | 0.3 |
| 2 | 34291679 | rs1816151 | -0.24 | -0.528 | 0.331 | -0.975 | 0.006 | -0.012 | -0.012 | 0 | 0.41 | 0.62 | 0.54 | 0.93 | 0.37 | 0.55 |
| 2 | 34293505 | rs17014642 | -0.174 | -0.474 | 0.333 | -0.938 | 0.021 | 0.031 | 0.031 | 0.026 | 0.43 | 0.24 | 0.33 | 0.6 | 0.39 | 0.37 |
| 2 | 34301138 | rs17014644 | -0.724 | -0.484 | 0.476 | -0.903 | -0.029 | 0.041 | 0.041 | 0 | 0.22 | 0.48 | 0.32 | 0.34 | 0.14 | 0.3 |
| 2 | 34301669 | rs17247889 | 1.086 | -0.507 | 0.428 | -0.901 | -0.007 | 0.036 | 0.036 | 0.017 | 0.11 | 0.06 | 0.05 | 0 | 0.08 | 0.07 |
| 2 | 34302743 | rs1425025 | -0.252 | -0.547 | 0.471 | -0.486 | -0.009 | -0.019 | -0.019 | 0 | 0.39 | 0.58 | 0.48 | 0.83 | 0.32 | 0.47 |
| 2 | 34305257 | rs10495809 | -0.314 | -0.521 | 0.223 | -0.43 | 0.03 | -0.017 | -0.017 | 0 | 0.43 | 0.26 | 0.29 | 0.5 | 0.37 | 0.31 |
| 2 | 34305358 | rs13398077 | -0.34 | -0.5 | 0.228 | -0.436 | 0.02 | 0.031 | 0.031 | 0.147 | 0.43 | 0.28 | 0.34 | 0.61 | 0.4 | 0.37 |
| 2 | 34305469 | rs10179470 | 0.519 | -0.507 | 0.225 | -0.441 | 0.017 | -0.017 | -0.017 | 0.964 | 0.32 | 0.16 | 0.28 | 0.6 | 0.31 | 0.3 |
| 2 | 34305524 | rs17339077 | 0.959 | -0.503 | 0.168 | -0.415 | -0.001 | 0.005 | 0.005 | 0 | 0.11 | 0.11 | 0.06 | 0 | 0.09 | 0.07 |
| 2 | 34311016 | rs10184147 | 0.54 | -0.407 | -0.302 | -0.419 | 0.026 | -0.063 | -0.063 | 0 | 0.31 | 0.14 | 0.23 | 0.5 | 0.28 | 0.24 |
| 2 | 34313003 | rs1425021 | 0.383 | -0.082 | -0.229 | -0.573 | 0 | 0.012 | 0.012 | 0 | 0.12 | 0.11 | 0.06 | 0 | 0.09 | 0.07 |
| 2 | 34319511 | rs11688637 | -0.305 | 0.009 | -0.639 | -0.427 | 0.002 | -0.001 | -0.001 | 0 | 0.57 | 0.6 | 0.68 | 0.47 | 0.62 | 0.59 |
| 2 | 34319972 | rs7595485 | -0.331 | 0.054 | -1.093 | -0.467 | -0.017 | 0.143 | 0.143 | 0 | 0.55 | 0.85 | 0.6 | 0.21 | 0.56 | 0.58 |
| 2 | 34324360 | rs2059442 | 1.071 | 0.966 | -0.732 | -0.687 | 0.027 | -0.045 | -0.045 | 0 | 0.11 | 0.19 | 0.21 | 0 | 0.15 | 0.11 |
| 2 | 34326938 | rs17014711 | 1.245 | 0.88 | -0.462 | -0.749 | -0.005 | -0.064 | -0.064 | 0.142 | 0.12 | 0.23 | 0.06 | 0 | 0.08 | 0.06 |
| 2 | 34328691 | rs17014720 | 1.025 | 0.955 | -0.55 | -0.726 | 0.026 | -0.045 | -0.045 | 0 | 0.11 | 0.19 | 0.21 | 0 | 0.15 | 0.12 |
| 2 | 34329248 | rs10495810 | -0.353 | 0.698 | -0.817 | -0.679 | 0.01 | 0.011 | 0.011 | 0 | 0.48 | 0.64 | 0.48 | 0.47 | 0.46 | 0.57 |
| 2 | 34331115 | rs6757934 | -0.716 | 0.605 | -0.69 | -0.626 | -0.001 | 0 | 0 | 0 | 0.57 | 0.59 | 0.67 | 0.46 | 0.63 | 0.63 |
| 2 | 34332491 | rs6704996 | 0.659 | 0.751 | -1.116 | -0.587 | -0.015 | 0.04 | 0.04 | 0 | 0.29 | 0.16 | 0.29 | 0.17 | 0.32 | 0.28 |
| 2 | 34336592 | rs6757137 | -0.958 | 0.211 | -0.873 | -0.594 | -0.016 | 0.046 | 0.046 | 0.825 | 0.37 | 0.8 | 0.48 | 0.28 | 0.35 | 0.4 |
| 2 | 34338541 | rs6707616 | 0.69 | 0.247 | -0.917 | -0.317 | -0.001 | -0.002 | -0.002 | 2.21 | 0.34 | 0.26 | 0.28 | 0.29 | 0.35 | 0.4 |
| 2 | 34339587 | rs6712013 | -0.554 | 0.326 | -0.825 | -0.348 | 0.012 | 0.005 | 0.005 | 0.433 | 0.23 | 0.67 | 0.4 | 0.26 | 0.24 | 0.26 |
| 2 | 34340969 | rs17014748 | 0.183 | 0.251 | -0.28 | -0.344 | -0.011 | -0.009 | -0.009 | 0.918 | 0.18 | 0.32 | 0.12 | 0.11 | 0.12 | 0.15 |
| 2 | 34341570 | rs10187022 | 0.675 | 0.159 | -0.474 | -0.446 | -0.001 | -0.015 | -0.015 | 0 | 0.37 | 0.36 | 0.27 | 0.18 | 0.37 | 0.35 |
| 2 | 34343342 | rs1477507 | -0.791 | 0.176 | -1.045 | -0.424 | 0 | 0.006 | 0.006 | 0.875 | 0.25 | 0.73 | 0.42 | 0.38 | 0.24 | 0.33 |
| 2 | 34346220 | rs17014757 | 1.084 | 0.476 | -1.256 | -0.457 | -0.017 | 0.015 | 0.015 | 0 | 0.08 | 0.04 | 0.19 | 0 | 0.12 | 0.11 |
| 2 | 34346937 | rs1559646 | -1.223 | 0.451 | -1.365 | -0.42 | -0.044 | -0.062 | -0.062 | 0 | 0.6 | 0.93 | 0.68 | 0.56 | 0.6 | 0.69 |
| 2 | 34351759 | rs1364690 | -1.425 | 0.65 | -1.036 | -0.307 | -0.011 | -0.009 | -0.009 | 0.293 | 0.36 | 0.74 | 0.48 | 0.38 | 0.35 | 0.41 |
| 2 | 34355651 | rs4670072 | -1.181 | 0.664 | -1.617 | -0.11 | -0.094 | -0.112 | -0.112 | 0 | 0.65 | 0.97 | 0.69 | 0.57 | 0.61 | 0.69 |
| 2 | 34379845 | rs17470197 | 1.372 | 1.033 | -0.981 | 0.196 | -0.049 | -0.051 | -0.051 | 0 | 0.14 | 0.01 | 0.16 | 0.11 | 0.17 | 0.06 |
| 2 | 34383404 | rs6746903 | 0.857 | 0.802 | -0.742 | -0.218 | -0.011 | 0.033 | 0.033 | 0 | 0.11 | 0.06 | 0.07 | 0 | 0.11 | 0.07 |
| 2 | 34386193 | rs4670075 | 0.202 | 0.819 | -0.498 | 0.094 | -0.01 | -0.009 | -0.009 | 0.152 | 0.08 | 0.06 | 0.16 | 0.27 | 0.12 | 0.14 |
| 2 | 34387353 | rs979019 | 0.599 | 0.112 | 0.042 | -0.248 | -0.01 | 0.002 | 0.002 | 0 | 0.62 | 0.49 | 0.64 | 0.61 | 0.68 | 0.66 |
| 2 | 34395176 | rs4670328 | 0.399 | 0.471 | -0.033 | -0.375 | 0.001 | -0.014 | -0.014 | 0.09 | 0.08 | 0.08 | 0.2 | 0 | 0.11 | 0.12 |
| 2 | 34396126 | rs12614680 | 1.409 | 0.319 | -0.06 | -0.396 | -0.013 | 0.02 | 0.02 | 0 | 0.44 | 0.25 | 0.4 | 0.62 | 0.49 | 0.45 |
| 2 | 34403344 | rs6707582 | -0.983 | 0.144 | 0.317 | -0.464 | 0 | 0.005 | 0.005 | 0 | 0.78 | 0.83 | 0.83 | 0.96 | 0.76 | 0.9 |
| 2 | 34403856 | rs2371930 | 0.605 | 0.046 | 0.777 | -0.428 | -0.001 | -0.001 | -0.001 | 0.039 | 0.61 | 0.62 | 0.64 | 0.58 | 0.64 | 0.67 |
| 2 | 34404072 | rs10174698 | 1.669 | 0.048 | 0.826 | -0.472 | -0.018 | 0.028 | 0.028 | 0 | 0.46 | 0.23 | 0.39 | 0.58 | 0.51 | 0.55 |
| 2 | 34406531 | rs11896442 | 1.204 | 0.053 | 0.794 | -0.458 | -0.001 | -0.012 | -0.012 | 0 | 0.08 | 0.08 | 0.05 | 0 | 0.07 | 0.06 |
| 2 | 34410240 | rs4670331 | 0.441 | 0.004 | 1.014 | -0.457 | -0.02 | 0.008 | 0.008 | 0 | 0.66 | 0.58 | 0.65 | 0.6 | 0.71 | 0.73 |
| 2 | 34410890 | rs12469719 | 1.14 | 0.037 | 0.804 | -0.319 | -0.006 | -0.015 | -0.015 | 0.148 | 0.06 | 0.01 | 0.11 | 0.27 | 0.05 | 0.12 |
| 2 | 34413387 | rs11896394 | -1.656 | -0.039 | 0.931 | -0.369 | 0.001 | 0.008 | 0.008 | 0.171 | 0.3 | 0.6 | 0.43 | 0.37 | 0.26 | 0.33 |
| 2 | 34417339 | rs4670333 | 1.212 | -0.069 | 0.802 | -0.387 | 0.061 | 0.03 | 0.03 | 0 | 0.53 | 0.22 | 0.59 | 0.59 | 0.64 | 0.65 |
| 2 | 34419367 | rs6729698 | -0.858 | -0.047 | 1.198 | -0.332 | 0.039 | 0.169 | 0.169 | 4.478 | 0.05 | 0.31 | 0.06 | 0.09 | 0.01 | 0.04 |
| 2 | 34423760 | rs1075018 | 1.707 | -0.049 | 0.982 | -0.41 | -0.028 | 0.029 | 0.029 | 0 | 0.45 | 0.18 | 0.39 | 0.6 | 0.49 | 0.53 |
| 2 | 34424794 | rs7579205 | 0.058 | -0.062 | 1.073 | -0.456 | 0.009 | 0.013 | 0.013 | 0 | 0.69 | 0.8 | 0.71 | 0.69 | 0.7 | 0.76 |
| 2 | 34426991 | rs11124364 | -0.355 | -0.068 | 1.178 | -0.459 | -0.03 | 0.006 | 0.006 | 1.029 | 0.16 | 0.57 | 0.12 | 0.1 | 0.08 | 0.11 |
| 2 | 34427633 | rs1541636 | -1.103 | -0.063 | 0.871 | -0.472 | 0.001 | -0.002 | -0.002 | 0 | 0.78 | 0.84 | 0.89 | 0.97 | 0.78 | 0.91 |
| 2 | 34430476 | rs4670335 | 1.785 | -0.059 | 0.826 | -0.43 | -0.026 | 0.028 | 0.028 | 0 | 0.45 | 0.22 | 0.4 | 0.58 | 0.5 | 0.53 |
| 2 | 34430530 | rs12472207 | 0.175 | -0.014 | 0.507 | -0.305 | 0.004 | -0.041 | -0.041 | 0 | 0.08 | 0.01 | 0.17 | 0.28 | 0.06 | 0.15 |
| 2 | 34440088 | rs11891597 | 0.646 | 0.006 | 0.724 | -0.37 | 0.001 | 0.015 | 0.015 | 0 | 0.1 | 0.24 | 0.08 | 0.09 | 0.07 | 0.1 |
| 2 | 34442074 | rs6543816 | -0.598 | -0.005 | 0.355 | -0.369 | -0.004 | -0.013 | -0.013 | 0.822 | 0.76 | 0.74 | 0.86 | 0.87 | 0.76 | 0.85 |

| | | | | | | | | | | | | | | | | |
|---|---|---|---|---|---|---|---|---|---|---|---|---|---|---|---|---|
| 2 | 34453722 | rs2161025 | -0.649 | 0.044 | 0.299 | -0.392 | -0.004 | -0.054 | -0.054 | 7.311 | 0.76 | 0.74 | 0.86 | 0.95 | 0.76 | 0.9 |
| 2 | 34455103 | rs2371924 | 1.321 | 0.036 | 0.15 | -0.366 | 0 | 0.002 | 0.002 | 0.559 | 0.51 | 0.56 | 0.44 | 0.58 | 0.5 | 0.53 |
| 2 | 34460541 | rs7574426 | -0.626 | 0.012 | -0.608 | -0.423 | 0.003 | 0.018 | 0.018 | 0 | 0.76 | 0.85 | 0.87 | 0.95 | 0.75 | 0.88 |
| 2 | 34466486 | rs10490441 | 0.008 | -0.166 | -0.451 | -0.126 | 0 | -0.02 | -0.02 | 0 | 0.14 | 0.09 | 0.22 | 0.3 | 0.13 | 0.19 |
| 2 | 34466523 | rs2216491 | -0.073 | -0.042 | -0.626 | -0.313 | -0.02 | 0.063 | 0.063 | 0.499 | 0.68 | 0.84 | 0.69 | 0.95 | 0.63 | 0.76 |
| 2 | 34474473 | rs2371923 | 0.684 | 0.208 | -1.049 | -0.064 | -0.054 | -0.005 | -0.005 | 0 | 0.45 | 0.14 | 0.39 | 0.56 | 0.49 | 0.52 |
| 2 | 34491664 | rs10177156 | 0.329 | 0.147 | -1.332 | 0.123 | 0.009 | 0.008 | 0.008 | 0 | 0.55 | 0.22 | 0.5 | 0.57 | 0.59 | 0.6 |
| 2 | 34499299 | rs10174826 | -0.274 | 0.982 | -1.347 | 0.625 | -0.07 | -0.098 | -0.098 | 0.393 | 0.24 | 0.06 | 0.36 | 0.44 | 0.34 | 0.24 |
| 2 | 34499749 | rs6719309 | 1.092 | 0.09 | -1.235 | -0.045 | -0.007 | 0.006 | 0.006 | 0.007 | 0.46 | 0.34 | 0.5 | 0.55 | 0.5 | 0.5 |
| 2 | 34505480 | rs1978414 | -0.27 | 0.163 | -0.178 | -0.075 | 0.025 | -0.009 | -0.009 | 0.049 | 0.95 | 0.75 | 0.86 | 0.99 | 0.96 | 0.92 |
| 2 | 34509330 | rs2042116 | 0.216 | 0.356 | 0.036 | 0.137 | 0.053 | 0.255 | 0.255 | 0 | 0.09 | 0.33 | 0.25 | 0.33 | 0.11 | 0.19 |
| 2 | 34519344 | rs12465251 | 0.096 | -0.173 | -0.021 | -0.538 | 0.005 | -0.022 | -0.022 | 0.448 | 0.23 | 0.17 | 0.11 | 0 | 0.16 | 0.18 |
| 2 | 34519779 | rs12613086 | 0.207 | -0.429 | -0.424 | -0.629 | -0.052 | -0.071 | -0.071 | 0 | 0.29 | 0.09 | 0.38 | 0.47 | 0.35 | 0.23 |
| 2 | 34520276 | rs249093 | 2.385 | -0.667 | -0.613 | -0.815 | -0.062 | 0.119 | 0.119 | 0 | 0.33 | 0.06 | 0.23 | 0.16 | 0.38 | 0.3 |
| 2 | 34522051 | rs249095 | 2.405 | -0.66 | -0.611 | -0.861 | -0.066 | 0.136 | 0.136 | 0 | 0.33 | 0.05 | 0.21 | 0.12 | 0.37 | 0.29 |
| 2 | 34523233 | rs249099 | 1.411 | -0.492 | -0.235 | -0.626 | 0.024 | -0.042 | -0.042 | 0 | 0.64 | 0.21 | 0.61 | 0.56 | 0.73 | 0.56 |
| 2 | 34529017 | rs4327215 | 0.673 | -0.268 | -0.082 | -0.494 | 0.003 | 0.088 | 0.088 | 0 | 0.09 | 0.13 | 0.26 | 0.37 | 0.11 | 0.2 |
| 2 | 34531245 | rs249113 | 0.38 | -0.321 | 0.025 | -0.68 | -0.003 | -0.046 | -0.046 | 0 | 0.85 | 0.78 | 0.92 | 0.98 | 0.9 | 0.86 |
| 2 | 34537639 | rs13001514 | 0.716 | -0.43 | -0.203 | -0.695 | -0.061 | -0.082 | -0.082 | 0 | 0.23 | 0.07 | 0.35 | 0.45 | 0.32 | 0.22 |
| 2 | 34538100 | rs249106 | 0.509 | -0.701 | 0.064 | -0.89 | -0.086 | -0.081 | -0.081 | 1.832 | 0.82 | 0.58 | 0.93 | 0.95 | 0.92 | 0.81 |
| 2 | 34541186 | rs3910275 | -1.816 | -0.741 | 0.034 | -1.004 | -0.007 | 0.014 | 0.014 | 0 | 0.37 | 0.47 | 0.4 | 0.4 | 0.33 | 0.47 |
| 2 | 34541228 | rs17486979 | 1.349 | -0.667 | -0.073 | -1.048 | -0.024 | 0.043 | 0.043 | 0 | 0.08 | 0.01 | 0.05 | 0 | 0.08 | 0.05 |
| 2 | 34542730 | rs6721933 | 0.774 | -0.532 | 0.226 | -1.11 | -0.004 | -0.003 | -0.003 | 0 | 0.24 | 0.19 | 0.37 | 0.48 | 0.34 | 0.22 |
| 2 | 34542861 | rs12475811 | 0.715 | -0.539 | 0.269 | -1.121 | -0.007 | 0.012 | 0.012 | 0 | 0.12 | 0.11 | 0.27 | 0.38 | 0.14 | 0.23 |
| 2 | 34543150 | rs3845767 | -1.684 | -0.474 | 0.239 | -1.089 | -0.002 | 0.009 | 0.009 | 0 | 0.36 | 0.59 | 0.41 | 0.4 | 0.31 | 0.46 |
| 2 | 34544898 | rs12712378 | 0.696 | -0.435 | 0.234 | -1.061 | -0.004 | -0.004 | -0.004 | 2.615 | 0.24 | 0.19 | 0.37 | 0.46 | 0.34 | 0.21 |
| 2 | 34551566 | rs6543823 | -2.302 | -0.508 | 0.224 | -1.019 | 0.025 | 0.105 | 0.105 | 0 | 0.62 | 0.77 | 0.78 | 0.9 | 0.67 | 0.75 |
| 2 | 34552765 | rs1368934 | -0.991 | -0.438 | 0.302 | -0.897 | -0.011 | 0.08 | 0.08 | 0 | 0.26 | 0.39 | 0.34 | 0.44 | 0.22 | 0.44 |
| 2 | 34554042 | rs10183942 | -1.325 | -0.496 | 0.3 | -0.891 | -0.015 | 0.068 | 0.068 | 0 | 0.28 | 0.51 | 0.36 | 0.4 | 0.23 | 0.45 |
| 2 | 34573155 | rs1434145 | 1.658 | -0.508 | 0.116 | -0.912 | -0.012 | -0.034 | -0.034 | 0 | 0.1 | 0.1 | 0.04 | 0 | 0.07 | 0.08 |
| 2 | 34574054 | rs13411686 | -0.874 | -0.396 | 0.186 | -0.836 | 0.005 | 0.086 | 0.086 | 0 | 0.18 | 0.28 | 0.3 | 0.44 | 0.17 | 0.4 |
| 2 | 34575895 | rs7577875 | -0.663 | -0.407 | 0.047 | -1.123 | 0.017 | 0.01 | 0.01 | 0.018 | 0.37 | 0.52 | 0.44 | 0.46 | 0.43 | 0.29 |
| 2 | 34578706 | rs10166784 | -1.277 | -0.324 | -0.425 | -1.273 | 0.02 | 0.023 | 0.023 | 0.003 | 0.51 | 0.34 | 0.7 | 0.94 | 0.56 | 0.61 |
| 2 | 34580853 | rs13005810 | -1.416 | -0.48 | -0.39 | -1.268 | -0.064 | -0.087 | -0.087 | 0 | 0.2 | 0.56 | 0.09 | 0.05 | 0.1 | 0.17 |
| 2 | 34583568 | rs13026280 | 0.292 | -0.26 | -0.346 | -1.18 | -0.007 | -0.038 | -0.038 | 0 | 0.13 | 0.13 | 0.05 | 0 | 0.09 | 0.09 |
| 2 | 34587546 | rs4670352 | -0.982 | 0.028 | -0.757 | -1.136 | 0 | 0.013 | 0.013 | 0 | 0.39 | 0.41 | 0.42 | 0.47 | 0.4 | 0.39 |
| 2 | 34589009 | rs7600640 | 0.666 | 0.027 | -1.218 | -1.115 | -0.082 | -0.127 | -0.127 | 0 | 0.23 | 0.02 | 0.35 | 0.46 | 0.31 | 0.23 |
| 2 | 34589999 | rs10495818 | -0.801 | 0.183 | -1.067 | -0.953 | -0.047 | -0.093 | -0.093 | 0.024 | 0.21 | 0.1 | 0.33 | 0.47 | 0.25 | 0.3 |
| 2 | 34594367 | rs11124371 | 0.098 | 0.255 | -0.992 | -0.776 | -0.04 | -0.053 | -0.053 | 0.107 | 0.12 | 0.04 | 0.27 | 0.47 | 0.17 | 0.25 |
| 2 | 34594442 | rs10204349 | -1.202 | 0.273 | -0.678 | -0.997 | -0.004 | 0.006 | 0.006 | 0.061 | 0.48 | 0.52 | 0.47 | 0.47 | 0.49 | 0.44 |
| 2 | 34601890 | rs1583365 | -0.136 | 0.49 | -0.661 | -1.272 | 0.006 | 0.005 | 0.005 | 0 | 0.74 | 0.71 | 0.63 | 0.64 | 0.65 | 0.79 |
| 2 | 34601927 | rs1583366 | 2.752 | 0.4 | -1.463 | -1.634 | -0.035 | 0.169 | 0.169 | 0 | 0.28 | 0.05 | 0.18 | 0.05 | 0.27 | 0.15 |
| 2 | 34608896 | rs10495819 | 1.052 | 0.349 | -1.524 | -1.534 | -0.047 | -0.066 | -0.066 | 0.066 | 0.1 | 0.01 | 0.25 | 0.52 | 0.12 | 0.17 |
| 2 | 34611748 | rs6716549 | -0.665 | 0.291 | -1.102 | -1.744 | 0 | 0 | 0 | 0 | 0.32 | 0.33 | 0.36 | 0.58 | 0.26 | 0.51 |
| 2 | 34616306 | rs919933 | -1.406 | 0.338 | -1.702 | -1.764 | -0.042 | 0.052 | 0.052 | 0.684 | 0.85 | 0.99 | 0.91 | 0.92 | 0.83 | 0.95 |
| 2 | 34620100 | rs17339245 | 1.605 | 0.094 | -1.449 | -1.721 | 0.002 | 0.171 | 0.171 | 0 | 0.38 | 0.12 | 0.22 | 0.04 | 0.36 | 0.24 |
| 2 | 34627691 | rs1368930 | -0.152 | 0.15 | -1.364 | -1.631 | -0.016 | -0.001 | -0.001 | 0.165 | 0.7 | 0.77 | 0.6 | 0.64 | 0.57 | 0.72 |
| 2 | 34634302 | rs1368932 | 0.685 | 0.087 | -1.178 | -1.888 | 0.002 | 0 | 0 | 0 | 0.1 | 0.04 | 0.05 | 0 | 0.1 | 0.09 |
| 2 | 34645519 | rs1589267 | 1.153 | 0.128 | -1.149 | -1.857 | -0.013 | 0.207 | 0.207 | 0 | 0.38 | 0.1 | 0.23 | 0.05 | 0.36 | 0.24 |
| 2 | 34650565 | rs13003534 | 0.209 | -0.026 | -0.747 | -1.717 | -0.013 | 0.024 | 0.024 | 0 | 0.58 | 0.68 | 0.34 | 0.08 | 0.49 | 0.6 |
| 2 | 34665412 | rs1560881 | -0.109 | -0.269 | -0.281 | -0.979 | -0.005 | 0.026 | 0.026 | 0 | 0.12 | 0.12 | 0.15 | 0.29 | 0.11 | 0.15 |
| 2 | 34667749 | rs892458 | -1.295 | -0.453 | 0.126 | -1.365 | 0.006 | 0.042 | 0.042 | 0 | 0.45 | 0.63 | 0.54 | 0.95 | 0.43 | 0.5 |
| 2 | 34670602 | rs6712234 | -0.589 | -0.519 | 0.104 | -1.525 | -0.015 | 0.041 | 0.041 | 0.146 | 0.81 | 0.89 | 0.9 | 0.98 | 0.81 | 0.67 |
| 2 | 34671890 | rs10207517 | -1.183 | -0.541 | -0.12 | -1.538 | 0.018 | 0.134 | 0.134 | 0 | 0.46 | 0.71 | 0.55 | 0.95 | 0.42 | 0.5 |
| 2 | 34672921 | rs10495822 | 2.045 | -0.501 | -0.679 | -1.514 | -0.071 | 0.217 | 0.217 | 0 | 0.34 | 0.06 | 0.34 | 0.03 | 0.38 | 0.18 |
| 2 | 34673447 | rs17260930 | 0.848 | -0.541 | -0.499 | -1.429 | -0.013 | -0.016 | -0.016 | 0 | 0.23 | 0.15 | 0.34 | 0.67 | 0.31 | 0.24 |
| 2 | 34678045 | rs1850907 | -0.363 | -0.616 | -0.524 | -1.733 | -0.001 | -0.018 | -0.018 | 0 | 0.35 | 0.29 | 0.47 | 0.94 | 0.38 | 0.36 |
| 2 | 34691083 | rs7583467 | -0.961 | -0.547 | -1.262 | -1.697 | 0.017 | 0.254 | 0.254 | 0 | 0.45 | 0.8 | 0.53 | 0.95 | 0.4 | 0.45 |
| 2 | 34738616 | rs7581230 | -0.699 | -0.229 | -1.374 | -1.544 | 0 | 0 | 0 | 0 | 0.78 | 0.79 | 0.89 | 0.99 | 0.81 | 0.67 |
| 2 | 34739570 | rs10176097 | -0.316 | 0.046 | -1.347 | -1.286 | 0 | -0.031 | -0.031 | 0 | 0.32 | 0.28 | 0.47 | 0.94 | 0.39 | 0.37 |
| 2 | 34741854 | rs6715436 | 0.747 | 0.089 | -1.881 | -0.76 | -0.083 | 0.231 | 0.231 | 0 | 0.34 | 0.06 | 0.35 | 0.03 | 0.39 | 0.19 |
| 2 | 34753390 | rs7340483 | -0.567 | -0.01 | -1.273 | 0.642 | 0.013 | -0.018 | -0.018 | 0 | 0.27 | 0.17 | 0.28 | 0.66 | 0.26 | 0.24 |
| 2 | 34760320 | rs1519916 | -0.958 | -0.036 | -1.604 | 0.768 | 0.034 | -0.017 | -0.017 | 0 | 0.39 | 0.84 | 0.28 | 0.39 | 0.31 | 0.28 |
| 2 | 34760573 | rs11124380 | -0.554 | -0.069 | -0.644 | 0.685 | 0 | -0.002 | -0.002 | 0 | 0.28 | 0.3 | 0.17 | 0.23 | 0.31 | 0.21 |
| 2 | 34776610 | rs11891404 | -0.066 | -0.596 | -0.999 | 0.753 | 0.03 | -0.061 | -0.061 | 0.4 | 0.71 | 0.87 | 0.72 | 0.55 | 0.79 | 0.69 |
| 2 | 34778477 | rs1519918 | -0.378 | -0.388 | 0.103 | 0.971 | 0.021 | 0.159 | 0.159 | 0 | 0.08 | 0.31 | 0.06 | 0.17 | 0.05 | 0.06 |
| 2 | 34790535 | rs10865103 | -0.19 | -0.238 | 0.159 | 0.877 | 0.011 | -0.024 | -0.024 | 5.37 | 0.7 | 0.8 | 0.71 | 0.53 | 0.77 | 0.66 |
| 2 | 34796055 | rs3845770 | 0.393 | -0.259 | 0.27 | 0.222 | 0.054 | 0.021 | 0.021 | 0.049 | 0.6 | 0.18 | 0.61 | 0.36 | 0.72 | 0.57 |
| 2 | 34796176 | rs3901291 | 0.45 | -0.302 | 0.175 | 0.192 | 0.077 | 0.025 | 0.025 | 0 | 0.58 | 0.14 | 0.6 | 0.37 | 0.72 | 0.58 |

| | | | | | | | | | | | | | | | | |
|---|---|---|---|---|---|---|---|---|---|---|---|---|---|---|---|---|
| 2 | 34809821 | rs17015376 | -0.665 | -0.251 | 0.565 | -0.259 | 0.017 | -0.02 | -0.02 | 0 | 0.06 | 0.21 | 0.07 | 0 | 0.06 | 0.03 |
| 2 | 34813997 | rs10165088 | 0.926 | -0.029 | -0.559 | -0.755 | -0.037 | 0.189 | 0.189 | 0.368 | 0.3 | 0.02 | 0.15 | 0.03 | 0.28 | 0.19 |
| 2 | 34814518 | rs1220211 | 0.227 | -0.113 | -0.087 | -0.663 | 0.008 | -0.021 | -0.021 | 0.009 | 0.4 | 0.58 | 0.26 | 0.06 | 0.33 | 0.34 |
| 2 | 34827388 | rs12151473 | 0.644 | 0.328 | -0.765 | -0.103 | -0.093 | 0.043 | 0.043 | 0.392 | 0.42 | 0.03 | 0.36 | 0.28 | 0.46 | 0.3 |
| 2 | 34843169 | rs1527600 | 0.502 | -0.21 | 0.133 | 0.295 | 0.003 | -0.004 | -0.004 | 0.83 | 0.95 | 0.75 | 0.97 | 0.99 | 0.99 | 0.93 |
| 2 | 34868092 | rs11897262 | -0.404 | -0.08 | -0.25 | 0.426 | 0.012 | -0.021 | -0.021 | 0 | 0.35 | 0.24 | 0.29 | 0.55 | 0.27 | 0.32 |
| 2 | 34889395 | rs9917264 | -0.784 | 0.067 | -0.785 | 0.526 | 0.003 | -0.002 | -0.002 | 0 | 0.28 | 0.24 | 0.4 | 0.64 | 0.23 | 0.48 |
| 2 | 34891689 | rs1358358 | 0.073 | 0.317 | -2.043 | -0.316 | -0.064 | 0.108 | 0.108 | 0 | 0.31 | 0.03 | 0.21 | 0.11 | 0.32 | 0.26 |
| 2 | 34892317 | rs1527597 | 0.508 | 0.032 | -2.228 | -0.141 | 0.029 | -0.045 | -0.045 | 0 | 0.34 | 0.19 | 0.24 | 0.54 | 0.25 | 0.42 |
| 2 | 34892832 | rs2178046 | 0.537 | 0.466 | -1.297 | -0.327 | 0.05 | 0.068 | 0.068 | 0.033 | 0.18 | 0.47 | 0.29 | 0.22 | 0.2 | 0.13 |
| 2 | 34900399 | rs2371412 | 0.063 | 0.653 | -1.401 | -0.438 | -0.061 | -0.084 | -0.084 | 0.769 | 0.19 | 0.09 | 0.42 | 0.41 | 0.29 | 0.27 |
| 2 | 34902177 | rs7575839 | 1.198 | 0.38 | -1.643 | -0.528 | -0.08 | 0.144 | 0.144 | 0 | 0.3 | 0.03 | 0.2 | 0.07 | 0.34 | 0.27 |
| 2 | 34905328 | rs17015496 | 0.962 | 0.485 | -1.633 | -0.501 | -0.044 | -0.043 | -0.043 | 0.306 | 0.14 | 0.07 | 0.26 | 0.21 | 0.2 | 0.12 |
| 2 | 34909001 | rs1527616 | -0.353 | 0.197 | -1.643 | -0.774 | -0.017 | 0.089 | 0.089 | 0 | 0.55 | 0.7 | 0.69 | 0.89 | 0.5 | 0.66 |
| 2 | 34909636 | rs12470906 | 1.275 | 0.19 | -1.512 | -0.774 | 0.029 | -0.032 | -0.032 | 0 | 0.31 | 0.21 | 0.23 | 0.53 | 0.19 | 0.39 |
| 2 | 34911127 | rs6740364 | 0.874 | 0.172 | -1.61 | -0.819 | -0.094 | 0.137 | 0.137 | 0 | 0.36 | 0.03 | 0.21 | 0.11 | 0.38 | 0.3 |
| 2 | 34911909 | rs7564039 | -0.744 | 0.217 | -1.48 | -0.797 | 0.031 | -0.053 | -0.053 | 0 | 0.91 | 0.76 | 0.9 | 1 | 0.88 | 0.96 |
| 2 | 34912753 | rs13028105 | 1.654 | 0.029 | -1.848 | -1.092 | -0.084 | 0.161 | 0.161 | 0 | 0.25 | 0.04 | 0.17 | 0.01 | 0.3 | 0.17 |
| 2 | 34912850 | rs11683103 | 1.103 | -0.141 | -1.592 | -0.538 | 0.001 | -0.002 | -0.002 | 0 | 0.72 | 0.35 | 0.68 | 0.73 | 0.68 | 0.72 |
| 2 | 34913212 | rs10210431 | 0.799 | -0.158 | -1.597 | -0.508 | 0.027 | -0.026 | -0.026 | 0 | 0.36 | 0.22 | 0.38 | 0.61 | 0.29 | 0.49 |
| 2 | 34914569 | rs13404871 | -0.349 | -0.289 | -1.382 | -0.411 | 0.005 | 0.053 | 0.053 | 0 | 0.15 | 0.38 | 0.07 | 0.15 | 0.11 | 0.13 |
| 2 | 34915238 | rs7572455 | 0.581 | -0.256 | -1.505 | -0.442 | -0.007 | -0.023 | -0.023 | 0.002 | 0.77 | 0.38 | 0.83 | 0.88 | 0.79 | 0.84 |
| 2 | 34919515 | rs12989663 | 0.587 | -0.187 | -1.42 | -0.124 | 0.045 | -0.051 | -0.051 | 0 | 0.35 | 0.2 | 0.24 | 0.5 | 0.23 | 0.42 |
| 2 | 34923268 | rs6739366 | -0.148 | 0.239 | -0.511 | 0.011 | 0.013 | 0.017 | 0.017 | 0 | 0.2 | 0.34 | 0.27 | 0.21 | 0.22 | 0.19 |
| 2 | 34928080 | rs10201571 | -0.724 | 0.687 | -0.42 | -0.071 | -0.001 | -0.003 | -0.003 | 0 | 0.72 | 0.71 | 0.72 | 0.88 | 0.67 | 0.77 |
| 2 | 34931915 | rs2888082 | 0.519 | 0.546 | -0.06 | -0.04 | 0.064 | 0.059 | 0.059 | 0.001 | 0.08 | 0.29 | 0.17 | 0.12 | 0.11 | 0.11 |
| 2 | 34933685 | rs7580063 | 0.854 | 0.544 | -0.083 | -0.073 | -0.058 | 0.045 | 0.045 | 0 | 0.26 | 0.07 | 0.19 | 0.12 | 0.29 | 0.22 |
| 2 | 34934503 | rs4670097 | 0.184 | 0.543 | 0.036 | -0.045 | -0.002 | -0.002 | -0.002 | 0 | 0.24 | 0.23 | 0.41 | 0.33 | 0.33 | 0.32 |
| 2 | 34942918 | rs6543865 | -0.816 | 0.236 | -0.263 | -0.016 | 0.03 | -0.004 | -0.004 | 0 | 0.54 | 0.64 | 0.64 | 0.46 | 0.63 | 0.53 |
| 2 | 34947118 | rs3731951 | 0.667 | 0.021 | -0.773 | -0.4 | -0.003 | 0.028 | 0.028 | 0 | 0.27 | 0.18 | 0.21 | 0.13 | 0.29 | 0.16 |
| 2 | 34948430 | rs3115248 | 0.597 | -0.212 | -1.242 | -0.501 | -0.044 | -0.006 | -0.006 | 0 | 0.55 | 0.16 | 0.56 | 0.62 | 0.53 | 0.5 |
| 2 | 34952517 | rs3115238 | 0.773 | -0.299 | -0.865 | -0.515 | 0.028 | -0.015 | -0.015 | 0 | 0.39 | 0.27 | 0.4 | 0.59 | 0.29 | 0.37 |
| 2 | 34956399 | rs11897499 | 0.141 | -0.227 | -0.798 | -0.614 | -0.002 | 0.001 | 0.001 | 0 | 0.65 | 0.37 | 0.6 | 0.73 | 0.58 | 0.54 |
| 2 | 34956494 | rs11897621 | -0.475 | -0.229 | -0.841 | -0.65 | 0.02 | -0.003 | -0.003 | 0 | 0.71 | 0.39 | 0.61 | 0.73 | 0.61 | 0.61 |
| 2 | 34962533 | rs1917410 | -0.439 | -0.444 | -0.848 | -0.758 | -0.026 | -0.092 | -0.092 | 0.499 | 0.84 | 0.43 | 0.9 | 0.98 | 0.9 | 0.78 |
| 2 | 34968286 | rs3115263 | -0.203 | -0.412 | -0.962 | -0.76 | 0.012 | 0.028 | 0.028 | 0 | 0.58 | 0.3 | 0.7 | 0.94 | 0.63 | 0.64 |
| 2 | 34970079 | rs3115260 | 0.806 | -0.415 | -0.938 | -0.694 | 0.031 | 0.021 | 0.021 | 0 | 0.39 | 0.27 | 0.36 | 0.7 | 0.26 | 0.41 |
| 2 | 34972791 | rs3108093 | -1.004 | -0.168 | -0.838 | -0.532 | 0.035 | -0.002 | -0.002 | 0 | 0.91 | 0.54 | 0.92 | 0.97 | 0.94 | 0.85 |
| 2 | 34974141 | rs11683299 | 0.974 | 0.111 | -1.837 | -0.412 | -0.1 | -0.048 | -0.048 | 0 | 0.19 | 0.04 | 0.34 | 0.24 | 0.37 | 0.24 |
| 2 | 34976869 | rs3108126 | 1.302 | -0.158 | -1.882 | -0.337 | 0.064 | -0.16 | -0.16 | 0.132 | 0.3 | 0.05 | 0.31 | 0.54 | 0.19 | 0.38 |
| 2 | 34977195 | rs11902219 | -1.636 | -0.173 | -1.626 | -0.604 | -0.003 | -0.015 | -0.015 | 0 | 0.76 | 0.79 | 0.8 | 0.99 | 0.73 | 0.87 |
| 2 | 34977477 | rs1917418 | 1.385 | -0.752 | -1.728 | -0.238 | 0.046 | -0.156 | -0.156 | 0 | 0.26 | 0.04 | 0.32 | 0.54 | 0.17 | 0.35 |
| 2 | 34977980 | rs3108124 | 0.006 | -0.416 | -1.145 | -0.247 | -0.002 | -0.026 | -0.026 | 0 | 0.11 | 0.1 | 0.03 | 0 | 0.07 | 0.02 |
| 2 | 34983529 | rs3115243 | -0.951 | -1.086 | -0.886 | -0.133 | -0.042 | 0.132 | 0.132 | 0 | 0.22 | 0.55 | 0.33 | 0.53 | 0.16 | 0.45 |
| 2 | 34986833 | rs6543870 | -1.62 | -0.815 | -1.597 | -0.484 | -0.126 | 0.012 | 0.012 | 0.013 | 0.7 | 0.97 | 0.62 | 0.81 | 0.61 | 0.77 |
| 2 | 34987206 | rs7591253 | -0.554 | -1.238 | -1.683 | -0.727 | 0.149 | -0.055 | -0.055 | 0.548 | 0.46 | 0.83 | 0.44 | 0.54 | 0.32 | 0.57 |
| 2 | 34988219 | rs4531910 | -0.156 | -1.24 | -1.665 | -0.539 | -0.043 | -0.039 | -0.039 | 0 | 0.15 | 0.57 | 0.09 | 0.08 | 0.1 | 0.23 |
| 2 | 34988474 | rs4439939 | -1.077 | -1.13 | -1.837 | -0.733 | 0.062 | -0.054 | -0.054 | 0 | 0.58 | 0.83 | 0.45 | 0.53 | 0.36 | 0.63 |
| 2 | 34990937 | rs4471866 | 0.647 | -1.066 | -0.621 | -0.758 | -0.016 | -0.04 | -0.04 | 0 | 0.11 | 0.28 | 0.05 | 0.04 | 0.1 | 0.16 |
| 2 | 34995934 | rs4128274 | -0.137 | -0.856 | -0.339 | -0.704 | 0.017 | 0.003 | 0.003 | 0 | 0.27 | 0.39 | 0.22 | 0.29 | 0.29 | 0.22 |
| 2 | 34997372 | rs6747984 | -0.306 | -0.926 | -0.33 | -0.705 | 0 | 0.018 | 0.018 | 0 | 0.48 | 0.39 | 0.29 | 0.29 | 0.46 | 0.32 |
| 2 | 35006395 | rs1438625 | -0.875 | -1.387 | -0.664 | -0.948 | 0.001 | 0.001 | 0.001 | 0 | 0.5 | 0.54 | 0.56 | 0.8 | 0.49 | 0.67 |
| 2 | 35007976 | rs17015616 | 1.887 | -1.575 | -0.639 | -1.091 | -0.006 | -0.038 | -0.038 | 0 | 0.19 | 0.19 | 0.1 | 0 | 0.13 | 0.11 |
| 2 | 35008853 | rs6756796 | 0.268 | -1.714 | -0.478 | -1.026 | -0.005 | 0.056 | 0.056 | 0 | 0.17 | 0.28 | 0.33 | 0.54 | 0.16 | 0.44 |
| 2 | 35009651 | rs7569098 | -2.119 | -1.724 | -0.692 | -1.115 | -0.095 | -0.01 | -0.01 | 0 | 0.73 | 0.97 | 0.67 | 0.8 | 0.63 | 0.78 |
| 2 | 35013851 | rs17015647 | -0.274 | -1.654 | -0.417 | -0.956 | 0.004 | 0.001 | 0.001 | 0 | 0.38 | 0.64 | 0.45 | 0.66 | 0.31 | 0.57 |
| 2 | 35019687 | rs12712404 | -0.091 | -1.476 | -0.118 | -1.366 | 0 | -0.005 | -0.005 | 0 | 0.39 | 0.35 | 0.29 | 0.27 | 0.42 | 0.26 |
| 2 | 35030699 | rs11694339 | 1.604 | -1.459 | -0.255 | -1.348 | -0.01 | -0.039 | -0.039 | 0.255 | 0.11 | 0.12 | 0.03 | 0 | 0.03 | 0.06 |
| 2 | 35032088 | rs2456800 | -0.765 | -1.452 | -0.384 | -1.358 | -0.009 | 0.066 | 0.066 | 0 | 0.53 | 0.62 | 0.33 | 0.27 | 0.46 | 0.32 |
| 2 | 35036487 | rs2118672 | 0.934 | -1.305 | -0.582 | -1.355 | -0.07 | 0.013 | 0.013 | 0 | 0.18 | 0.06 | 0.18 | 0.15 | 0.28 | 0.13 |
| 2 | 35043996 | rs380322 | -0.53 | -1.214 | -0.222 | -1.359 | 0.009 | 0.098 | 0.098 | 0 | 0.49 | 0.6 | 0.31 | 0.15 | 0.42 | 0.31 |
| 2 | 35046665 | rs17261035 | 1.505 | -1.156 | -0.441 | -1.197 | 0.023 | 0.048 | 0.048 | 0 | 0.18 | 0.11 | 0.11 | 0 | 0.14 | 0.1 |
| 2 | 35049995 | rs399420 | -2.27 | -1.061 | -0.726 | -1.333 | -0.054 | 0.142 | 0.142 | 0 | 0.76 | 0.98 | 0.85 | 0.96 | 0.71 | 0.88 |
| 2 | 35052108 | rs6733633 | -0.007 | -1.176 | -0.094 | -0.984 | -0.025 | 0.023 | 0.023 | 0 | 0.33 | 0.19 | 0.2 | 0.27 | 0.34 | 0.28 |
| 2 | 35058421 | rs7559552 | 0.944 | -1.08 | -0.197 | -1.026 | 0.003 | 0.01 | 0.01 | 0 | 0.21 | 0.14 | 0.09 | 0.12 | 0.13 | 0.14 |
| 2 | 35061211 | rs280715 | -1.001 | -1.125 | -0.004 | -1.048 | -0.001 | 0.003 | 0.003 | 0 | 0.57 | 0.54 | 0.33 | 0.27 | 0.48 | 0.38 |
| 2 | 35065246 | rs188113 | 0.967 | -1.091 | 0.132 | -0.636 | -0.013 | -0.015 | -0.015 | 0 | 0.21 | 0.26 | 0.12 | 0.09 | 0.15 | 0.17 |
| 2 | 35068284 | rs10169121 | 1.279 | -0.726 | 0.145 | -0.518 | -0.056 | -0.018 | -0.018 | 4.912 | 0.12 | 0.03 | 0.11 | 0.15 | 0.21 | 0.14 |
| 2 | 35075107 | rs10495829 | -0.403 | -1.096 | 0.386 | -0.456 | 0.013 | 0 | 0 | 0 | 0.07 | 0.34 | 0.04 | 0.02 | 0.04 | 0.16 |

| | | | | | | | | | | | | | | | | |
|---|---|---|---|---|---|---|---|---|---|---|---|---|---|---|---|---|
| 2 | 35085350 | rs280678 | -1.157 | -1.263 | -0.032 | -0.575 | -0.033 | 0.068 | 0.068 | 0 | 0.59 | 0.84 | 0.36 | 0.29 | 0.5 | 0.54 |
| 2 | 35088235 | rs2438427 | -1.028 | -1.287 | 0.029 | -0.56 | -0.02 | 0.061 | 0.061 | 0 | 0.57 | 0.74 | 0.36 | 0.29 | 0.49 | 0.53 |
| 2 | 35089425 | rs280758 | 1.101 | -1.239 | 0.027 | -0.573 | -0.023 | -0.061 | -0.061 | 0 | 0.18 | 0.22 | 0.11 | 0 | 0.11 | 0.09 |
| 2 | 35091308 | rs280757 | 0.657 | -1.038 | 0.219 | -0.222 | 0 | -0.001 | -0.001 | 0 | 0.31 | 0.29 | 0.22 | 0.24 | 0.35 | 0.29 |
| 2 | 35092762 | rs280754 | -0.625 | -1.087 | -0.132 | -0.247 | -0.032 | 0.089 | 0.089 | 0 | 0.55 | 0.82 | 0.34 | 0.29 | 0.47 | 0.55 |
| 2 | 35093293 | rs7600259 | 0.69 | -0.692 | 0.309 | 0.03 | 0.001 | 0.001 | 0.001 | 0 | 0.16 | 0.13 | 0.08 | 0.13 | 0.1 | 0.11 |
| 2 | 35095121 | rs280750 | 0.087 | -0.722 | 0.496 | 0.221 | -0.001 | 0.011 | 0.011 | 0 | 0.47 | 0.44 | 0.24 | 0.34 | 0.43 | 0.53 |
| 2 | 35095373 | rs280749 | 0.485 | -0.807 | 0.408 | 0.192 | -0.04 | 0.04 | 0.04 | 0 | 0.42 | 0.15 | 0.22 | 0.3 | 0.41 | 0.44 |
| 2 | 35095936 | rs280746 | -1.097 | -0.904 | 0.268 | 0.15 | -0.099 | -0.102 | -0.102 | 0 | 0.66 | 0.95 | 0.37 | 0.41 | 0.54 | 0.64 |
| 2 | 35096136 | rs280745 | 0.647 | -0.803 | 0.3 | 0.198 | -0.065 | 0.056 | 0.056 | 0 | 0.4 | 0.11 | 0.21 | 0.25 | 0.41 | 0.44 |
| 2 | 35098351 | rs280743 | 0.222 | -0.715 | 0.415 | 0.205 | 0 | 0.003 | 0.003 | 0 | 0.45 | 0.59 | 0.24 | 0.21 | 0.42 | 0.43 |
| 2 | 35104389 | rs280734 | -0.888 | -0.65 | 0.117 | 0.264 | -0.03 | -0.032 | -0.032 | 0 | 0.63 | 0.8 | 0.35 | 0.41 | 0.54 | 0.62 |
| 2 | 35115848 | rs280780 | -0.254 | -0.321 | 0.602 | 0.125 | 0 | -0.027 | -0.027 | 0.037 | 0.44 | 0.56 | 0.25 | 0.22 | 0.4 | 0.35 |
| 2 | 35115979 | rs280779 | 0.092 | -0.297 | 0.539 | 0.137 | -0.004 | 0.011 | 0.011 | 0 | 0.4 | 0.42 | 0.22 | 0.22 | 0.4 | 0.35 |
| 2 | 35122915 | rs280770 | -0.273 | -0.274 | 0.407 | 0.172 | 0 | -0.01 | -0.01 | 0 | 0.44 | 0.51 | 0.25 | 0.27 | 0.42 | 0.39 |
| 2 | 35123423 | rs280768 | -0.989 | -0.263 | 0.02 | 0.4 | -0.013 | 0.02 | 0.02 | 0 | 0.32 | 0.6 | 0.55 | 0.45 | 0.27 | 0.47 |
| 2 | 35127305 | rs17326572 | 0.658 | -0.11 | -0.432 | 0.084 | -0.066 | 0.009 | 0.009 | 0 | 0.29 | 0.1 | 0.18 | 0.23 | 0.33 | 0.25 |
| 2 | 35127980 | rs13391731 | 0.395 | 0.067 | -0.412 | 0.181 | 0.006 | -0.006 | -0.006 | 0 | 0.34 | 0.4 | 0.2 | 0.27 | 0.34 | 0.33 |
| 2 | 35128057 | rs1553617 | 0.829 | -0.158 | -1.12 | -0.377 | -0.008 | -0.031 | -0.031 | 0.473 | 0.12 | 0.13 | 0.06 | 0 | 0.08 | 0.08 |
| 2 | 35129304 | rs280762 | -0.29 | 0.314 | -1.096 | 0.054 | 0.004 | 0.098 | 0.098 | 0 | 0.6 | 0.83 | 0.74 | 0.79 | 0.6 | 0.73 |
| 2 | 35130378 | rs6543879 | 0.488 | 0.932 | -0.664 | 0.857 | -0.018 | -0.025 | -0.025 | 0 | 0.84 | 0.91 | 0.67 | 0.42 | 0.79 | 0.72 |
| 2 | 35138450 | rs13386292 | 0.147 | 0.377 | 0.305 | 1.039 | -0.003 | 0.293 | 0.293 | 0 | 0.08 | 0.28 | 0.05 | 0.36 | 0.03 | 0.14 |
| 2 | 35143238 | rs2055694 | -0.134 | 0.624 | 0.169 | 1.017 | -0.012 | -0.001 | -0.001 | 0.609 | 0.87 | 0.9 | 0.7 | 0.51 | 0.86 | 0.75 |
| 2 | 35156295 | rs6543881 | 1.062 | 0.328 | 0.579 | 0.763 | 0.031 | 0.206 | 0.206 | 0.007 | 0.79 | 0.51 | 0.51 | 0.51 | 0.83 | 0.72 |
| 2 | 35158357 | rs11883629 | -0.802 | 0.204 | 0.666 | 0.689 | 0.005 | -0.05 | -0.05 | 0 | 0.1 | 0.45 | 0.06 | 0 | 0.04 | 0.01 |
| 2 | 35161672 | rs10175357 | -1.648 | 0.109 | 0.655 | 0.702 | 0.007 | -0.01 | -0.01 | 0 | 0.19 | 0.79 | 0.12 | 0.37 | 0.06 | 0.16 |
| 2 | 35161954 | rs10175571 | -0.422 | 0.116 | 1.107 | 0.713 | -0.028 | 0.241 | 0.241 | 0.733 | 0.11 | 0.56 | 0.07 | 0.37 | 0.03 | 0.15 |
| 2 | 35166346 | rs7609077 | -0.438 | 0.139 | 0.805 | 0.732 | -0.014 | -0.006 | -0.006 | 0 | 0.88 | 0.91 | 0.59 | 0.52 | 0.87 | 0.76 |
| 2 | 35170234 | rs13393137 | -0.014 | 0.113 | 1.207 | 0.912 | -0.002 | 0.326 | 0.326 | 0 | 0.09 | 0.35 | 0.05 | 0.36 | 0.03 | 0.15 |
| 2 | 35190105 | rs11124406 | -1.85 | 0.088 | 1.072 | 0.846 | 0.01 | -0.013 | -0.013 | 0 | 0.19 | 0.79 | 0.12 | 0.34 | 0.06 | 0.16 |
| 2 | 35191207 | rs13386676 | 0.033 | 0.1 | 1.133 | 0.852 | -0.006 | 0.291 | 0.291 | 0 | 0.09 | 0.35 | 0.05 | 0.34 | 0.03 | 0.15 |
| 2 | 35191585 | rs11124407 | 0.495 | 0.723 | 0.571 | 0.233 | -0.053 | -0.009 | -0.009 | 0.503 | 0.08 | 0.01 | 0.1 | 0.09 | 0.21 | 0.11 |
| 2 | 35194150 | rs6760654 | -0.82 | 0.655 | 0.727 | 0.257 | 0.002 | -0.045 | -0.045 | 0 | 0.09 | 0.37 | 0.06 | 0 | 0.04 | 0.01 |
| 2 | 35195262 | rs1507247 | -0.112 | 0.662 | -0.121 | 0.285 | -0.061 | -0.013 | -0.013 | 1.069 | 0.11 | 0.02 | 0.12 | 0.12 | 0.23 | 0.22 |
| 2 | 35204168 | rs10176749 | -1.29 | -0.047 | -0.109 | 0.111 | 0.034 | 0.001 | 0.001 | 0 | 0.41 | 0.74 | 0.29 | 0.41 | 0.38 | 0.43 |
| 2 | 35205952 | rs10204114 | 0.344 | -0.052 | 0.661 | 0.291 | -0.02 | 0.258 | 0.258 | 0 | 0.11 | 0.4 | 0.07 | 0.29 | 0.03 | 0.19 |
| 2 | 35224451 | rs17387182 | -0.468 | -0.165 | 0.611 | 0.046 | -0.002 | -0.056 | -0.056 | 1.966 | 0.26 | 0.34 | 0.17 | 0.12 | 0.29 | 0.22 |
| 2 | 35225041 | rs2290383 | -0.836 | 0.066 | 0.51 | 0.219 | 0.006 | 0.091 | 0.091 | 0 | 0.91 | 0.89 | 0.77 | 0.55 | 0.89 | 0.78 |
| 2 | 35227498 | rs10495831 | 0.093 | 0.348 | 0.606 | 0.1 | 0.005 | -0.017 | -0.017 | 0.294 | 0.2 | 0.2 | 0.17 | 0.12 | 0.32 | 0.23 |
| 2 | 35229511 | rs13396601 | 0.174 | -0.084 | 0.627 | 0.156 | 0.011 | 0.081 | 0.081 | 0 | 0.84 | 0.74 | 0.73 | 0.55 | 0.87 | 0.77 |
| 2 | 35235120 | rs17398711 | 0.526 | -0.084 | 0.648 | 0.136 | 0 | -0.005 | -0.005 | 0.011 | 0.17 | 0.16 | 0.12 | 0.12 | 0.26 | 0.21 |
| 2 | 35244058 | rs2888114 | -1.09 | -0.256 | 0.83 | 0.057 | -0.025 | 0.027 | 0.027 | 0 | 0.33 | 0.54 | 0.19 | 0.41 | 0.3 | 0.42 |
| 2 | 35244395 | rs1009327 | 0.856 | -0.228 | 0.734 | 0.288 | -0.008 | -0.035 | -0.035 | 0 | 0.07 | 0.15 | 0.05 | 0 | 0.03 | 0.02 |
| 2 | 35250422 | rs1453266 | -1.057 | -0.103 | 0.508 | 0.387 | -0.003 | 0.07 | 0.07 | 0 | 0.92 | 0.93 | 0.78 | 0.55 | 0.89 | 0.79 |
| 2 | 35253975 | rs1453268 | -0.598 | -0.2 | 0.457 | 0.338 | -0.011 | 0.033 | 0.033 | 0 | 0.9 | 0.92 | 0.78 | 0.55 | 0.89 | 0.78 |
| 2 | 35261253 | rs1169252 | -1.736 | -0.374 | 0.574 | -0.134 | 0 | -0.062 | -0.062 | 0 | 0.43 | 0.52 | 0.47 | 0.87 | 0.4 | 0.65 |
| 2 | 35262834 | rs1169255 | 0.751 | -0.242 | 0.703 | 0.189 | 0.039 | 0.196 | 0.196 | 0 | 0.09 | 0.18 | 0.28 | 0.51 | 0.11 | 0.22 |
| 2 | 35266175 | rs1993218 | 0.473 | -0.48 | 0.858 | 0.192 | -0.001 | 0.012 | 0.012 | 0 | 0.2 | 0.28 | 0.15 | 0.27 | 0.24 | 0.29 |
| 2 | 35277976 | rs1305581 | -2.201 | -0.527 | 0.169 | -0.001 | 0.018 | 0.119 | 0.119 | 0 | 0.49 | 0.76 | 0.51 | 0.88 | 0.41 | 0.68 |
| 2 | 35293343 | rs12712411 | 1.867 | -0.545 | 0.109 | -0.081 | -0.016 | 0.267 | 0.267 | 0 | 0.48 | 0.1 | 0.48 | 0.12 | 0.57 | 0.34 |
| 2 | 35295078 | rs1955381 | -2.052 | -0.513 | 0.103 | -0.066 | 0.01 | 0.122 | 0.122 | 0 | 0.51 | 0.76 | 0.51 | 0.88 | 0.43 | 0.68 |
| 2 | 35298411 | rs13005421 | -2.127 | -0.534 | 0.102 | -0.062 | 0.016 | 0.128 | 0.128 | 0 | 0.5 | 0.75 | 0.5 | 0.88 | 0.42 | 0.66 |
| 2 | 35300334 | rs1922351 | -1.213 | -0.556 | 0.19 | -0.003 | -0.003 | 0.015 | 0.015 | 0 | 0.41 | 0.63 | 0.48 | 0.8 | 0.42 | 0.55 |
| 2 | 35303167 | rs1317240 | -0.144 | -0.508 | 0.143 | 0.084 | -0.004 | -0.005 | -0.005 | 0 | 0.16 | 0.23 | 0.31 | 0.71 | 0.16 | 0.33 |
| 2 | 35303985 | rs1922355 | 0.008 | -0.493 | 0.105 | 0.069 | -0.008 | -0.01 | -0.01 | 0 | 0.15 | 0.17 | 0.32 | 0.72 | 0.16 | 0.33 |
| 2 | 35305420 | rs13397975 | 0.427 | -0.533 | 0.261 | 0.021 | -0.004 | -0.074 | -0.074 | 0.59 | 0.23 | 0.35 | 0.14 | 0.08 | 0.25 | 0.19 |
| 2 | 35309259 | rs4670434 | 0.561 | -0.531 | -0.12 | 0.039 | 0.052 | 0.081 | 0.081 | 0 | 0.63 | 0.25 | 0.79 | 0.84 | 0.72 | 0.66 |
| 2 | 35309975 | rs2371504 | 1.201 | 0.243 | -0.163 | 0.578 | -0.038 | 0.018 | 0.018 | 0 | 0.07 | 0.01 | 0.26 | 0.52 | 0.11 | 0.22 |
| 2 | 35316932 | rs1997506 | 0.143 | 0.005 | -0.174 | 0.303 | -0.008 | -0.049 | -0.049 | 0 | 0.72 | 0.7 | 0.81 | 0.92 | 0.7 | 0.78 |
| 2 | 35319012 | rs1971830 | 0.039 | 0.043 | -0.658 | 0.326 | 0.009 | 0.024 | 0.024 | 0.095 | 0.72 | 0.83 | 0.83 | 0.92 | 0.7 | 0.78 |
| 2 | 35325586 | rs10495836 | 0.908 | 0.195 | -1.107 | 0.598 | -0.042 | -0.003 | -0.003 | 0 | 0.08 | 0.01 | 0.13 | 0.52 | 0.12 | 0.22 |
| 2 | 35326427 | rs17016153 | 0.434 | 0.111 | -0.949 | 0.559 | -0.006 | 0.099 | 0.099 | 0 | 0.08 | 0.2 | 0.06 | 0.18 | 0.04 | 0.11 |
| 2 | 35339940 | rs2165188 | -0.146 | 0.282 | -1.562 | 0.353 | -0.013 | 0.112 | 0.112 | 0.632 | 0.26 | 0.03 | 0.29 | 0.08 | 0.31 | 0.19 |
| 2 | 35342430 | rs1922366 | -0.124 | 0.529 | -0.937 | 0.252 | -0.048 | -0.021 | -0.021 | 0 | 0.08 | 0.01 | 0.12 | 0.53 | 0.11 | 0.2 |
| 2 | 35355977 | rs12712421 | 0.371 | 0.026 | -1.448 | -1.501 | -0.005 | 0.172 | 0.172 | 0.165 | 0.34 | 0.06 | 0.16 | 0.07 | 0.32 | 0.38 |
| 2 | 35388124 | rs1371501 | -0.143 | 0.504 | -0.533 | -1.108 | 0.031 | 0.096 | 0.096 | 0.014 | 0.15 | 0.22 | 0.25 | 0.67 | 0.18 | 0.36 |
| 2 | 35404011 | rs968059 | -0.831 | 0.198 | -0.775 | -1.71 | -0.06 | 0.1 | 0.1 | 0 | 0.71 | 0.82 | 0.75 | 0.92 | 0.61 | 0.87 |
| 2 | 35406626 | rs10204984 | 0.616 | 0.008 | -0.571 | -1.872 | 0.002 | -0.065 | -0.065 | 0.16 | 0.4 | 0.34 | 0.22 | 0.02 | 0.38 | 0.35 |
| 2 | 35416075 | rs11124419 | 0.771 | 1.008 | -0.036 | -1.037 | -0.036 | -0.004 | -0.004 | 0.037 | 0.08 | 0.02 | 0.22 | 0.6 | 0.15 | 0.28 |

| | | | | | | | | | | | | | | | | |
|---|---|---|---|---|---|---|---|---|---|---|---|---|---|---|---|---|
| 2 | 35428465 | rs13403909 | -0.225 | 0.011 | 0.509 | -0.825 | -0.002 | 0.006 | 0.006 | 0 | 0.58 | 0.57 | 0.36 | 0.36 | 0.46 | 0.5 |
| 2 | 35438101 | rs7597757 | -0.734 | 0.054 | 0.669 | -0.871 | 0.027 | -0.085 | -0.085 | 0 | 0.65 | 0.39 | 0.67 | 0.92 | 0.55 | 0.87 |
| 2 | 35443726 | rs10204994 | 1.103 | -0.057 | 0.517 | -0.989 | 0.033 | 0.037 | 0.037 | 0 | 0.33 | 0.25 | 0.21 | 0.19 | 0.21 | 0.26 |
| 2 | 35454001 | rs17038724 | 1.171 | 0.047 | 0.639 | -0.917 | -0.002 | 0.116 | 0.116 | 0 | 0.06 | 0.14 | 0.03 | 0.15 | 0.04 | 0.1 |
| 2 | 35470727 | rs12613236 | 1.442 | 0.093 | 0.188 | -0.899 | -0.034 | 0.041 | 0.041 | 0.087 | 0.06 | 0.01 | 0.18 | 0.52 | 0.09 | 0.27 |
| 2 | 35470856 | rs1837450 | 1.451 | 0.071 | 0.186 | -0.979 | 0.045 | 0.047 | 0.047 | 0.049 | 0.3 | 0.1 | 0.19 | 0.19 | 0.21 | 0.27 |
| 2 | 35480663 | rs1371505 | -0.951 | -0.563 | 0.687 | -1.439 | -0.093 | 0.126 | 0.126 | 0 | 0.69 | 0.84 | 0.72 | 0.96 | 0.56 | 0.87 |
| 2 | 35482278 | rs1025775 | -0.007 | -0.608 | 0.743 | -1.575 | -0.003 | 0.073 | 0.073 | 0 | 0.2 | 0.09 | 0.12 | 0.03 | 0.21 | 0.13 |
| 2 | 35509352 | rs17327412 | -0.041 | -0.587 | 0.204 | -1.564 | -0.001 | 0.072 | 0.072 | 0 | 0.2 | 0.07 | 0.11 | 0.08 | 0.21 | 0.13 |
| 2 | 35526918 | rs6750486 | 0.02 | -0.521 | 0.338 | -1.517 | 0.004 | 0.098 | 0.098 | 1.352 | 0.09 | 0.34 | 0.05 | 0.11 | 0.05 | 0.1 |
| 2 | 35532579 | rs11689082 | -0.011 | -0.483 | 0.265 | -1.522 | -0.002 | 0.076 | 0.076 | 0 | 0.2 | 0.07 | 0.12 | 0.06 | 0.21 | 0.14 |
| 2 | 35533411 | rs6736209 | 0.298 | -0.344 | 0.526 | -1.331 | -0.068 | 0.19 | 0.19 | 0.22 | 0.4 | 0.19 | 0.34 | 0.03 | 0.53 | 0.14 |
| 2 | 35551496 | rs4366887 | 0.439 | -0.314 | 0.529 | -1.269 | 0.014 | -0.058 | -0.058 | 0 | 0.15 | 0.32 | 0.11 | 0.06 | 0.16 | 0.12 |
| 2 | 35562791 | rs4371344 | -1.288 | -0.332 | 0.34 | -1.284 | 0.005 | 0.001 | 0.001 | 0.372 | 0.64 | 0.95 | 0.64 | 0.28 | 0.72 | 0.47 |
| 2 | 35575340 | rs7590763 | 0.404 | -0.18 | 0.558 | -1.145 | -0.05 | 0.161 | 0.161 | 0 | 0.41 | 0.23 | 0.33 | 0.05 | 0.51 | 0.28 |
| 2 | 35575973 | rs6727683 | -0.404 | -0.179 | 0.553 | -1.139 | -0.032 | 0.158 | 0.158 | 0.243 | 0.52 | 0.87 | 0.4 | 0.22 | 0.55 | 0.39 |
| 2 | 35577771 | rs7584802 | 0.433 | -0.165 | 0.386 | -1.118 | -0.075 | 0.19 | 0.19 | 0 | 0.39 | 0.17 | 0.31 | 0.03 | 0.48 | 0.18 |
| 2 | 35578987 | rs17016449 | -0.724 | -0.157 | 0.482 | -1.091 | -0.011 | -0.015 | -0.015 | 0 | 0.24 | 0.47 | 0.15 | 0.17 | 0.23 | 0.24 |
| 2 | 35584461 | rs7572208 | -0.682 | 0.011 | 0.137 | -1.409 | -0.01 | -0.018 | -0.018 | 0 | 0.24 | 0.44 | 0.14 | 0.17 | 0.23 | 0.24 |
| 2 | 35591234 | rs7598399 | 0.271 | 0.246 | -0.804 | -1.355 | -0.067 | 0.18 | 0.18 | 0 | 0.39 | 0.2 | 0.31 | 0.03 | 0.48 | 0.18 |
| 2 | 35593804 | rs4549067 | 0.448 | 0.292 | -0.781 | -1.306 | 0.008 | 0.106 | 0.106 | 0 | 0.07 | 0.2 | 0.04 | 0.12 | 0.05 | 0.1 |
| 2 | 35597319 | rs4344916 | -0.972 | 0.454 | -1.542 | -1.244 | -0.007 | -0.017 | -0.017 | 0 | 0.62 | 0.47 | 0.44 | 0.22 | 0.71 | 0.51 |
| 2 | 35600357 | rs6722845 | -0.462 | 1.153 | -2.197 | -1.286 | -0.085 | 0.198 | 0.198 | 0 | 0.38 | 0.17 | 0.31 | 0.02 | 0.48 | 0.17 |
| 2 | 35610115 | rs7590663 | -0.726 | 1.02 | -1.694 | -0.915 | -0.003 | 0.02 | 0.02 | 0.953 | 0.26 | 0.25 | 0.16 | 0.04 | 0.28 | 0.19 |
| 2 | 35612621 | rs4670463 | 0.286 | 1.346 | -2.402 | -1.834 | -0.052 | 0.112 | 0.112 | 0.955 | 0.67 | 0.86 | 0.78 | 0.96 | 0.61 | 0.79 |
| 2 | 35613178 | rs6709851 | -0.011 | 1.248 | -1.641 | -0.863 | 0.029 | 0.211 | 0.211 | 0 | 0.14 | 0.35 | 0.28 | 0.45 | 0.16 | 0.19 |
| 2 | 35613198 | rs4346361 | 0.73 | 0.296 | -2.536 | -1.522 | -0.006 | 0.098 | 0.098 | 0.013 | 0.27 | 0.06 | 0.14 | 0.07 | 0.23 | 0.3 |
| 2 | 35613676 | rs6739406 | 0.639 | 0.359 | -2.513 | -1.489 | 0.011 | 0.228 | 0.228 | 0 | 0.13 | 0.34 | 0.27 | 0.45 | 0.13 | 0.18 |
| 2 | 35614344 | rs4381761 | -1.372 | 0.622 | -2.862 | -1.751 | -0.029 | 0.043 | 0.043 | 0.119 | 0.76 | 0.86 | 0.82 | 0.97 | 0.68 | 0.81 |
| 2 | 35614444 | rs17016558 | 1.285 | 0.191 | -2.331 | -1.989 | 0.001 | 0.004 | 0.004 | 0 | 0.16 | 0.17 | 0.26 | 0.45 | 0.1 | 0.29 |
| 2 | 35618483 | rs12476042 | 0.822 | -0.079 | -2.448 | -1.946 | 0.025 | -0.041 | -0.041 | 0 | 0.2 | 0.14 | 0.29 | 0.47 | 0.11 | 0.3 |
| 2 | 35623559 | rs4624361 | -1.18 | -0.195 | -2.578 | -2.109 | -0.068 | 0.108 | 0.108 | 0.097 | 0.69 | 0.86 | 0.75 | 0.96 | 0.54 | 0.8 |
| 2 | 35624746 | rs6756157 | 0.71 | -0.14 | -2.518 | -1.947 | -0.053 | 0.104 | 0.104 | 0 | 0.3 | 0.06 | 0.15 | 0.07 | 0.29 | 0.3 |
| 2 | 35627198 | rs13384335 | 0.765 | -0.276 | -2.256 | -1.747 | 0.001 | 0.182 | 0.182 | 0 | 0.15 | 0.36 | 0.28 | 0.41 | 0.14 | 0.19 |
| 2 | 35627227 | rs7600751 | 0.553 | -0.352 | -2.284 | -1.785 | 0.001 | 0.252 | 0.252 | 0 | 0.16 | 0.5 | 0.29 | 0.41 | 0.14 | 0.19 |
| 2 | 35634511 | rs7580957 | 0.505 | -0.4 | -2.144 | -1.305 | 0.003 | 0.174 | 0.174 | 0 | 0.16 | 0.49 | 0.3 | 0.34 | 0.14 | 0.17 |
| 2 | 35644367 | rs13012972 | 0.299 | -0.138 | -2.5 | -1.348 | -0.103 | 0.06 | 0.06 | 0 | 0.34 | 0.07 | 0.27 | 0.16 | 0.41 | 0.33 |
| 2 | 35647806 | rs17016602 | 0.125 | -0.59 | -2.023 | -0.999 | -0.014 | 0.03 | 0.03 | 0.009 | 0.21 | 0.22 | 0.36 | 0.52 | 0.1 | 0.31 |
| 2 | 35651028 | rs11694690 | -0.847 | -0.669 | -1.909 | -1.061 | -0.028 | -0.039 | -0.039 | 0.066 | 0.39 | 0.51 | 0.53 | 0.8 | 0.27 | 0.49 |
| 2 | 35652873 | rs11890863 | 1.011 | -0.703 | -2.168 | -1.09 | -0.039 | 0.007 | 0.007 | 0 | 0.28 | 0.06 | 0.19 | 0.17 | 0.26 | 0.3 |
| 2 | 35652916 | rs17016629 | 0.273 | -0.738 | -2.067 | -0.98 | 0.023 | -0.15 | -0.15 | 0.529 | 0.18 | 0.01 | 0.31 | 0.51 | 0.1 | 0.31 |
| 2 | 35653722 | rs4670470 | 0.718 | -0.64 | -1.98 | -0.918 | -0.004 | 0.023 | 0.023 | 0 | 0.18 | 0.29 | 0.18 | 0.28 | 0.17 | 0.17 |
| 2 | 35655270 | rs13026558 | 0.377 | -0.542 | -2.145 | -1.113 | -0.082 | 0.06 | 0.06 | 0.017 | 0.35 | 0.07 | 0.26 | 0.17 | 0.39 | 0.33 |
| 2 | 35657891 | rs4280439 | -1.392 | -0.813 | -2.155 | -1.325 | -0.049 | 0.087 | 0.087 | 0.097 | 0.71 | 0.87 | 0.8 | 0.96 | 0.62 | 0.81 |
| 2 | 35659858 | rs4513272 | -1.768 | -0.827 | -1.842 | -1.276 | -0.028 | 0.018 | 0.018 | 0.064 | 0.75 | 0.83 | 0.8 | 0.99 | 0.64 | 0.81 |
| 2 | 35666876 | rs10179616 | -0.581 | -1.065 | -0.873 | -1.343 | -0.013 | -0.063 | -0.063 | 0 | 0.46 | 0.51 | 0.58 | 0.95 | 0.32 | 0.63 |
| 2 | 35670363 | rs4439937 | 0.172 | -1.245 | -0.438 | -0.431 | -0.029 | 0.083 | 0.083 | 0 | 0.21 | 0.28 | 0.32 | 0.48 | 0.13 | 0.3 |
| 2 | 35670651 | rs12619050 | -0.348 | -1.353 | -0.639 | -0.471 | -0.066 | 0.036 | 0.036 | 1.439 | 0.26 | 0.4 | 0.43 | 0.7 | 0.16 | 0.49 |
| 2 | 35672564 | rs6735007 | 0.854 | -1.073 | -0.514 | -0.662 | -0.02 | -0.077 | -0.077 | 0 | 0.16 | 0.21 | 0.06 | 0 | 0.09 | 0.09 |
| 2 | 35673080 | rs13397054 | 0.691 | -0.938 | -0.99 | -0.585 | 0 | 0.002 | 0.002 | 0 | 0.17 | 0.24 | 0.17 | 0.18 | 0.18 | 0.17 |
| 2 | 35675611 | rs4297857 | 0.614 | -1.191 | -1.035 | -0.79 | 0.009 | 0.024 | 0.024 | 0 | 0.51 | 0.46 | 0.65 | 0.73 | 0.6 | 0.58 |
| 2 | 35676282 | rs17469573 | 0.227 | -0.675 | -1.227 | -0.391 | -0.078 | 0.084 | 0.084 | 0.708 | 0.26 | 0.04 | 0.18 | 0.06 | 0.29 | 0.16 |
| 2 | 35688188 | rs10200472 | 0.861 | -0.78 | -0.591 | 0.165 | -0.009 | 0.001 | 0.001 | 0 | 0.14 | 0.17 | 0.12 | 0.15 | 0.08 | 0.14 |
| 2 | 35693719 | rs2719096 | 0.041 | -0.81 | -0.515 | 0.226 | -0.046 | 0.042 | 0.042 | 0 | 0.8 | 0.68 | 0.78 | 0.68 | 0.89 | 0.72 |
| 2 | 35700888 | rs4560089 | -0.768 | -0.587 | -0.584 | -0.16 | -0.007 | -0.034 | -0.034 | 0 | 0.33 | 0.51 | 0.54 | 0.79 | 0.34 | 0.44 |
| 2 | 35709163 | rs17016728 | -0.44 | -0.366 | -0.637 | 0.006 | 0 | 0.008 | 0.008 | 0 | 0.23 | 0.26 | 0.42 | 0.66 | 0.27 | 0.41 |
| 2 | 35713309 | rs2719121 | -0.72 | -0.428 | -1.003 | -0.364 | -0.01 | 0.037 | 0.037 | 0 | 0.82 | 0.9 | 0.88 | 0.95 | 0.82 | 0.86 |
| 2 | 35717726 | rs4380218 | 0.69 | -0.399 | -1.152 | -0.355 | -0.045 | 0.042 | 0.042 | 0.375 | 0.18 | 0.04 | 0.12 | 0.05 | 0.18 | 0.14 |
| 2 | 35729556 | rs7607623 | 0.386 | -0.787 | -0.956 | -0.978 | 0.019 | 0.027 | 0.027 | 0.182 | 0.53 | 0.45 | 0.71 | 0.77 | 0.67 | 0.67 |
| 2 | 35730260 | rs10188049 | -0.542 | -1.024 | -0.989 | -0.847 | -0.016 | 0.001 | 0.001 | 0 | 0.42 | 0.45 | 0.37 | 0.41 | 0.32 | 0.44 |
| 2 | 35730792 | rs11679130 | 1.061 | -0.993 | -1.045 | -1.037 | -0.015 | -0.007 | -0.007 | 0 | 0.42 | 0.27 | 0.54 | 0.58 | 0.56 | 0.53 |
| 2 | 35732012 | rs12712441 | -1.11 | -1.095 | -0.81 | -1.003 | -0.002 | -0.001 | -0.001 | 0.11 | 0.55 | 0.48 | 0.42 | 0.43 | 0.42 | 0.47 |
| 2 | 35732518 | rs17016818 | 2.014 | -1.107 | -0.983 | -1.021 | 0.002 | 0 | 0 | 0 | 0.07 | 0.01 | 0.16 | 0.51 | 0.07 | 0.31 |
| 2 | 35732687 | rs4312465 | 0.687 | -1.195 | -0.981 | -1.135 | 0.003 | -0.004 | -0.004 | 0 | 0.2 | 0.18 | 0.18 | 0.22 | 0.14 | 0.18 |
| 2 | 35732726 | rs11682305 | 0.465 | -1.253 | -0.961 | -1.639 | 0.005 | 0.006 | 0.006 | 0 | 0.35 | 0.31 | 0.25 | 0.21 | 0.27 | 0.29 |
| 2 | 35760604 | rs1371428 | -2.34 | -1.029 | -0.983 | -1.516 | 0.001 | -0.007 | -0.007 | 0 | 0.95 | 0.94 | 0.99 | 1 | 0.98 | 0.99 |
| 2 | 35767002 | rs17400304 | 2.065 | -1.084 | -1.008 | -1.715 | -0.01 | 0.036 | 0.036 | 0 | 0.11 | 0.05 | 0.07 | 0 | 0.1 | 0.05 |
| 2 | 35772759 | rs1371422 | 0.567 | -1.297 | -0.955 | -1.783 | -0.019 | -0.016 | -0.016 | 0.012 | 0.57 | 0.49 | 0.64 | 0.68 | 0.72 | 0.64 |
| 2 | 35773926 | rs2372006 | 1.123 | -1.268 | -1.059 | -1.772 | 0.013 | 0.043 | 0.043 | 0 | 0.47 | 0.32 | 0.56 | 0.68 | 0.59 | 0.61 |

| | | | | | | | | | | | | | | | | |
|---|---|---|---|---|---|---|---|---|---|---|---|---|---|---|---|---|
| 2 | 35777771 | rs1561524 | 1.529 | -1.209 | -1.049 | -1.712 | -0.022 | -0.006 | -0.006 | 1.43 | 0.23 | 0.28 | 0.25 | 0.22 | 0.14 | 0.21 |
| 2 | 35804150 | rs2258 | -0.303 | -1.123 | -1.126 | -1.662 | 0.003 | 0.004 | 0.004 | 0 | 0.45 | 0.58 | 0.42 | 0.22 | 0.38 | 0.28 |
| 2 | 35809575 | rs12471144 | 1.962 | -1.079 | -1.219 | -1.59 | -0.003 | -0.033 | -0.033 | 0.007 | 0.08 | 0.01 | 0.13 | 0.63 | 0.07 | 0.3 |
| 2 | 35819686 | rs2719120 | -0.523 | -1.034 | -0.919 | -1.463 | -0.003 | -0.006 | -0.006 | 0.033 | 0.8 | 0.78 | 0.91 | 0.89 | 0.87 | 0.84 |
| 2 | 35821710 | rs4670477 | 1.074 | -1.002 | -0.904 | -1.502 | 0.021 | 0.069 | 0.069 | 0 | 0.48 | 0.3 | 0.57 | 0.68 | 0.6 | 0.61 |
| 2 | 35823374 | rs1561522 | 1.274 | -0.95 | -1.028 | -1.475 | 0.012 | 0.077 | 0.077 | 0 | 0.46 | 0.21 | 0.55 | 0.68 | 0.59 | 0.62 |
| 2 | 35830334 | rs1371418 | -0.944 | -0.916 | -0.877 | -1.068 | -0.021 | 0.02 | 0.02 | 0 | 0.91 | 0.77 | 0.97 | 0.9 | 0.97 | 0.89 |
| 2 | 35831008 | rs2043914 | 0.514 | -0.993 | -0.835 | -1.15 | -0.006 | -0.006 | -0.006 | 0 | 0.59 | 0.45 | 0.64 | 0.68 | 0.7 | 0.66 |
| 2 | 35831794 | rs1347942 | -0.33 | -0.593 | -1.183 | -1.004 | 0.001 | 0.001 | 0.001 | 0 | 0.33 | 0.28 | 0.33 | 0.22 | 0.29 | 0.23 |
| 2 | 35843803 | rs6736879 | -0.441 | -0.606 | -1.19 | -1.004 | 0.003 | 0.005 | 0.005 | 0 | 0.34 | 0.3 | 0.33 | 0.24 | 0.29 | 0.24 |
| 2 | 35846522 | rs8179828 | 0.504 | -0.562 | -1.246 | -1.055 | -0.01 | -0.005 | -0.005 | 0 | 0.27 | 0.27 | 0.27 | 0.24 | 0.18 | 0.21 |
| 2 | 35848903 | rs13011599 | -0.232 | -0.377 | -1.049 | -1.085 | 0.009 | 0.021 | 0.021 | 0 | 0.38 | 0.3 | 0.35 | 0.24 | 0.34 | 0.25 |
| 2 | 35849684 | rs7562096 | -0.294 | -0.346 | -1.038 | -1.069 | 0.005 | 0.011 | 0.011 | 0 | 0.39 | 0.33 | 0.35 | 0.24 | 0.34 | 0.25 |
| 2 | 35853705 | rs6713533 | 0.143 | 0.281 | -0.558 | -0.523 | 0.004 | 0.017 | 0.017 | 0 | 0.21 | 0.29 | 0.27 | 0.25 | 0.22 | 0.19 |
| 2 | 35859342 | rs10495843 | 0.173 | 0.21 | 0.088 | -0.782 | 0.01 | -0.069 | -0.069 | 5.411 | 0.22 | 0.31 | 0.16 | 0.06 | 0.23 | 0.14 |
| 2 | 35860852 | rs7571190 | 0.378 | -0.007 | 0.013 | -0.842 | -0.024 | 0.034 | 0.034 | 0 | 0.17 | 0.27 | 0.2 | 0.23 | 0.09 | 0.2 |
| 2 | 35868794 | rs1518651 | 0.434 | 0.009 | -0.015 | -0.849 | -0.001 | -0.007 | -0.007 | 0.238 | 0.21 | 0.2 | 0.15 | 0.11 | 0.23 | 0.15 |
| 2 | 35880236 | rs4473362 | -1.241 | 0.216 | -0.031 | -0.867 | 0 | -0.008 | -0.008 | 0 | 0.41 | 0.53 | 0.41 | 0.34 | 0.39 | 0.35 |
| 2 | 35885530 | rs12463508 | -1.43 | 0.215 | -0.082 | -0.867 | -0.001 | 0.042 | 0.042 | 0 | 0.43 | 0.69 | 0.43 | 0.34 | 0.39 | 0.35 |
| 2 | 35896017 | rs6756825 | -1.429 | 0.2 | -0.066 | -0.87 | 0 | 0.04 | 0.04 | 0 | 0.43 | 0.68 | 0.44 | 0.34 | 0.4 | 0.35 |
| 2 | 35899155 | rs7578143 | -0.106 | -0.138 | 0.267 | -0.73 | -0.032 | 0.069 | 0.069 | 0 | 0.18 | 0.4 | 0.22 | 0.25 | 0.1 | 0.19 |
| 2 | 35903080 | rs6543970 | -0.325 | -0.103 | 0.132 | -0.943 | 0.014 | -0.056 | -0.056 | 0 | 0.13 | 0.48 | 0.12 | 0.06 | 0.09 | 0.05 |
| 2 | 35907182 | rs2165167 | 0.59 | -0.242 | -0.02 | -0.87 | -0.025 | 0.037 | 0.037 | 0 | 0.14 | 0.24 | 0.19 | 0.24 | 0.06 | 0.2 |
| 2 | 35907195 | rs6739017 | 0.097 | -0.16 | 0.185 | -1.104 | 0.013 | -0.042 | -0.042 | 0 | 0.11 | 0.29 | 0.11 | 0.06 | 0.09 | 0.05 |
| 2 | 35907337 | rs6710661 | 0.188 | -0.292 | 0.216 | -1.106 | 0.003 | -0.029 | -0.029 | 0 | 0.1 | 0.29 | 0.1 | 0.06 | 0.07 | 0.04 |
| 2 | 35908737 | rs13025239 | 0.643 | -0.301 | 0.113 | -1.156 | 0.003 | -0.035 | -0.035 | 0 | 0.09 | 0.3 | 0.08 | 0.05 | 0.08 | 0.03 |
| 2 | 35909000 | rs925910 | 0.203 | -0.381 | -1.112 | -1.83 | -0.043 | 0.31 | 0.31 | 4.47 | 0.48 | 0.13 | 0.45 | 0.06 | 0.56 | 0.3 |
| 2 | 35910255 | rs1401228 | -0.428 | -0.451 | -1.064 | -1.465 | 0.028 | -0.024 | -0.024 | 0 | 0.27 | 0.76 | 0.31 | 0.42 | 0.14 | 0.24 |
| 2 | 35923095 | rs7602005 | -1.64 | -0.355 | -1.093 | -1.41 | -0.053 | -0.078 | -0.078 | 0 | 0.75 | 0.89 | 0.75 | 0.49 | 0.69 | 0.52 |
| 2 | 35929940 | rs13414263 | -0.982 | -0.377 | -1.42 | -1.678 | -0.09 | -0.054 | -0.054 | 0.001 | 0.61 | 0.91 | 0.67 | 0.45 | 0.54 | 0.4 |
| 2 | 35930877 | rs4670125 | -0.088 | -0.454 | -1.178 | -1.805 | 0.002 | -0.03 | -0.03 | 0.184 | 0.44 | 0.57 | 0.47 | 0.09 | 0.46 | 0.22 |
| 2 | 35933011 | rs11124469 | -0.779 | -0.366 | -1.064 | -1.73 | -0.003 | 0.024 | 0.024 | 1.895 | 0.6 | 0.67 | 0.57 | 0.15 | 0.63 | 0.34 |
| 2 | 35934991 | rs12465166 | 0.621 | -0.21 | -0.857 | -1.754 | -0.024 | 0.077 | 0.077 | 0.225 | 0.25 | 0.14 | 0.33 | 0.04 | 0.35 | 0.18 |
| 2 | 35935526 | rs11683407 | -0.366 | -0.237 | -0.846 | -1.765 | 0 | -0.027 | -0.027 | 0 | 0.36 | 0.63 | 0.43 | 0.09 | 0.43 | 0.2 |
| 2 | 35940628 | rs12105269 | -0.712 | -0.155 | -0.809 | -1.7 | 0.01 | 0.092 | 0.092 | 0.035 | 0.51 | 0.65 | 0.52 | 0.14 | 0.59 | 0.32 |
| 2 | 35947071 | rs10170933 | 0.342 | 0.165 | -0.076 | -1.071 | -0.033 | 0.083 | 0.083 | 0 | 0.15 | 0.27 | 0.19 | 0.36 | 0.07 | 0.18 |
| 2 | 35949516 | rs1401218 | -0.041 | 0.25 | -0.108 | -1.102 | 0 | -0.052 | -0.052 | 0 | 0.87 | 0.77 | 0.9 | 0.94 | 0.87 | 0.96 |
| 2 | 35951818 | rs17471749 | 0.639 | 0.13 | -0.144 | -1.042 | -0.022 | 0.054 | 0.054 | 0 | 0.16 | 0.05 | 0.1 | 0.05 | 0.16 | 0.12 |
| 2 | 35955286 | rs17017365 | -0.492 | -0.334 | 0.406 | -0.725 | -0.005 | -0.053 | -0.053 | 0.879 | 0.12 | 0.43 | 0.11 | 0.05 | 0.07 | 0.04 |
| 2 | 35955381 | rs6755827 | -0.514 | -0.177 | 0.245 | -0.475 | -0.056 | 0.141 | 0.141 | 0 | 0.17 | 0.53 | 0.23 | 0.36 | 0.07 | 0.18 |
| 2 | 35957851 | rs6749172 | -0.2 | -0.048 | -0.255 | -0.412 | -0.032 | 0.099 | 0.099 | 0 | 0.15 | 0.27 | 0.2 | 0.36 | 0.07 | 0.18 |
| 2 | 35967350 | rs3900084 | -1.058 | 0.533 | -0.895 | -0.755 | 0.012 | 0.002 | 0.002 | 0 | 0.54 | 0.85 | 0.57 | 0.5 | 0.55 | 0.42 |
| 2 | 35973963 | rs3914801 | -0.452 | 0.387 | -0.749 | -0.82 | -0.076 | 0.053 | 0.053 | 0.028 | 0.33 | 0.18 | 0.3 | 0.27 | 0.43 | 0.28 |
| 2 | 36001197 | rs17017391 | 0.88 | 0.767 | -0.804 | -0.87 | -0.03 | 0.03 | 0.03 | 0 | 0.08 | 0.03 | 0.11 | 0.01 | 0.17 | 0.06 |
| 2 | 36003061 | rs17017395 | 0.383 | 0.758 | -0.188 | -0.881 | -0.023 | 0.02 | 0.02 | 0 | 0.17 | 0.26 | 0.22 | 0.24 | 0.11 | 0.13 |
| 2 | 36004210 | rs17329328 | -1.06 | 0.746 | -0.253 | -1.015 | -0.011 | 0.015 | 0.015 | 0 | 0.39 | 0.64 | 0.39 | 0.27 | 0.46 | 0.31 |
| 2 | 36004453 | rs1356731 | -2.149 | 0.874 | -0.554 | -1.224 | -0.046 | 0.066 | 0.066 | 0 | 0.89 | 0.99 | 0.95 | 1 | 0.84 | 0.95 |
| 2 | 36004618 | rs7603026 | 0.389 | 0.829 | -0.427 | -1.128 | -0.061 | 0.016 | 0.016 | 0 | 0.22 | 0.09 | 0.19 | 0.23 | 0.25 | 0.21 |
| 2 | 36007771 | rs1160728 | -0.615 | 0.763 | -0.382 | -1.086 | 0.026 | -0.037 | -0.037 | 0 | 0.36 | 0.57 | 0.35 | 0.27 | 0.43 | 0.3 |
| 2 | 36010265 | rs1518670 | -1.248 | 0.653 | -0.48 | -0.75 | 0.012 | -0.015 | -0.015 | 0.297 | 0.53 | 0.82 | 0.58 | 0.55 | 0.54 | 0.44 |
| 2 | 36017688 | rs4670499 | 0.34 | 0.534 | -0.445 | -0.728 | -0.029 | 0.016 | 0.016 | 0 | 0.18 | 0.25 | 0.22 | 0.24 | 0.09 | 0.13 |
| 2 | 36019937 | rs10177485 | -0.597 | 0.723 | -0.521 | -1.071 | -0.006 | -0.005 | -0.005 | 0 | 0.37 | 0.32 | 0.39 | 0.46 | 0.37 | 0.35 |
| 2 | 36020401 | rs10169238 | 0.487 | 0.771 | -0.538 | -1.065 | -0.067 | -0.026 | -0.026 | 0 | 0.21 | 0.09 | 0.19 | 0.28 | 0.28 | 0.22 |
| 2 | 36021687 | rs6745167 | -0.065 | 0.807 | -0.337 | -1.087 | 0.012 | -0.06 | -0.06 | 0.005 | 0.15 | 0.29 | 0.17 | 0.02 | 0.2 | 0.09 |
| 2 | 36021984 | rs1401239 | -1.1 | 0.849 | -0.839 | -0.946 | 0.017 | 0.005 | 0.005 | 0 | 0.52 | 0.61 | 0.56 | 0.48 | 0.58 | 0.44 |
| 2 | 36026347 | rs1518659 | -1.433 | 0.914 | -0.411 | -0.656 | -0.045 | 0.07 | 0.07 | 0 | 0.88 | 0.99 | 0.94 | 1 | 0.84 | 0.95 |
| 2 | 36037158 | rs12475952 | -1.046 | 0.567 | 0.362 | 0.002 | -0.023 | -0.031 | -0.031 | 0 | 0.3 | 0.52 | 0.36 | 0.21 | 0.28 | 0.21 |
| 2 | 36044351 | rs13428332 | -0.925 | 0.795 | 0.565 | 0.192 | 0.01 | 0.025 | 0.025 | 0 | 0.13 | 0.28 | 0.17 | 0.18 | 0.14 | 0.12 |
| 2 | 36044575 | rs17329797 | 0.024 | 0.782 | 0.518 | 0.168 | -0.013 | 0.022 | 0.022 | 0.151 | 0.09 | 0.03 | 0.08 | 0.02 | 0.12 | 0.09 |
| 2 | 36045227 | rs13419901 | -1.565 | 0.603 | 0.854 | 0.057 | 0.021 | -0.001 | -0.001 | 0.45 | 0.32 | 0.75 | 0.4 | 0.35 | 0.25 | 0.26 |
| 2 | 36048221 | rs6747236 | -1.511 | 0.602 | 0.925 | 0.078 | -0.001 | -0.001 | -0.001 | 0 | 0.32 | 0.67 | 0.4 | 0.34 | 0.25 | 0.27 |
| 2 | 36049974 | rs7586691 | 0.381 | 1.282 | 0.862 | -0.032 | -0.081 | -0.01 | -0.01 | 0 | 0.14 | 0.02 | 0.16 | 0.09 | 0.27 | 0.1 |
| 2 | 36057798 | rs13431175 | -0.487 | 1.158 | 0.905 | -0.248 | -0.004 | -0.01 | -0.01 | 0 | 0.12 | 0.08 | 0.15 | 0.17 | 0.14 | 0.11 |
| 2 | 36062875 | rs10203034 | -1.435 | 1.046 | 1.01 | -0.414 | -0.009 | -0.002 | -0.002 | 0 | 0.34 | 0.41 | 0.4 | 0.35 | 0.28 | 0.28 |
| 2 | 36069858 | rs4670517 | 0.211 | 1.08 | 0.959 | -0.368 | -0.015 | 0.001 | 0.001 | 0 | 0.17 | 0.22 | 0.21 | 0.18 | 0.11 | 0.15 |
| 2 | 36070033 | rs4670519 | -1.563 | 1.066 | 1.007 | -0.408 | 0.009 | 0.003 | 0.003 | 0 | 0.36 | 0.66 | 0.43 | 0.34 | 0.28 | 0.28 |
| 2 | 36072931 | rs9678802 | -1.327 | 1.124 | 1.161 | -0.3 | -0.015 | -0.015 | -0.015 | 0 | 0.28 | 0.63 | 0.21 | 0.17 | 0.2 | 0.23 |
| 2 | 36077092 | rs6742996 | -2.104 | 1.284 | 1.078 | -0.344 | -0.038 | 0.068 | 0.068 | 0.016 | 0.46 | 0.84 | 0.45 | 0.33 | 0.5 | 0.37 |
| 2 | 36077848 | rs6543981 | -1.725 | 1.336 | 1.215 | -0.349 | -0.007 | 0.025 | 0.025 | 0.138 | 0.41 | 0.67 | 0.36 | 0.17 | 0.47 | 0.34 |

| | | | | | | | | | | | | | | | |
|---|---|---|---|---|---|---|---|---|---|---|---|---|---|---|---|
| 2 | 36078157 | rs6732309 | 0.322 | 1.377 | 0.723 | -0.372 | -0.036 | 0.035 | 0.035 | 0 | 0.14 | 0.08 | 0.17 | 0.05 | 0.27 | 0.11 |
| 2 | 36078497 | rs13417436 | -1.381 | 1.359 | 0.832 | -0.353 | 0.036 | -0.102 | -0.102 | 0.169 | 0.3 | 0.53 | 0.33 | 0.07 | 0.42 | 0.22 |
| 2 | 36082275 | rs940386 | -0.534 | 1.049 | 0.868 | -0.349 | 0.012 | 0.073 | 0.073 | 0 | 0.05 | 0.18 | 0.1 | 0.15 | 0.03 | 0.03 |
| 2 | 36083263 | rs1879131 | -1.797 | 1.111 | 0.894 | -0.328 | -0.015 | 0.063 | 0.063 | 0.025 | 0.41 | 0.69 | 0.39 | 0.18 | 0.48 | 0.34 |
| 2 | 36083564 | rs2968636 | -0.84 | 1.073 | 0.905 | -0.319 | 0 | -0.064 | -0.064 | 0.016 | 0.15 | 0.3 | 0.15 | 0.02 | 0.15 | 0.1 |
| 2 | 36087642 | rs11885738 | -1.09 | 1.08 | 0.896 | -0.31 | -0.017 | -0.021 | -0.021 | 0 | 0.26 | 0.33 | 0.18 | 0.15 | 0.21 | 0.22 |
| 2 | 36097070 | rs10173700 | -1.903 | 1.12 | 0.901 | -0.291 | -0.013 | 0.054 | 0.054 | 0.027 | 0.42 | 0.68 | 0.39 | 0.21 | 0.48 | 0.33 |
| 2 | 36102001 | rs7584203 | 0.315 | 1.056 | 0.62 | -0.471 | -0.026 | 0.039 | 0.039 | 0 | 0.13 | 0.32 | 0.07 | 0.13 | 0.06 | 0.12 |
| 2 | 36102489 | rs13015448 | -0.427 | 1.155 | 0.612 | -0.479 | -0.014 | 0.05 | 0.05 | 0.34 | 0.14 | 0.06 | 0.13 | 0.01 | 0.16 | 0.1 |
| 2 | 36103785 | rs7591464 | -1.959 | 1.165 | 0.576 | -0.494 | -0.007 | 0.033 | 0.033 | 0.027 | 0.43 | 0.66 | 0.39 | 0.18 | 0.49 | 0.34 |
| 2 | 36104351 | rs17017733 | 0.007 | 1.448 | 0.808 | -0.267 | -0.031 | 0.034 | 0.034 | 0 | 0.15 | 0.08 | 0.16 | 0.03 | 0.26 | 0.11 |
| 2 | 36104933 | rs7575563 | -1.009 | 1.448 | 0.81 | -0.266 | 0.015 | -0.037 | -0.037 | 0 | 0.27 | 0.39 | 0.23 | 0.16 | 0.32 | 0.24 |
| 2 | 36108484 | rs10193157 | 0.099 | 1.478 | 0.838 | -0.234 | -0.027 | 0.037 | 0.037 | 0 | 0.13 | 0.32 | 0.07 | 0.13 | 0.06 | 0.13 |
| 2 | 36109679 | rs1554364 | -2.047 | 1.487 | 0.772 | -0.221 | -0.101 | 0.033 | 0.033 | 0.298 | 0.42 | 0.93 | 0.45 | 0.33 | 0.47 | 0.38 |
| 2 | 36110210 | rs13431783 | -1.1 | 1.482 | 0.802 | -0.299 | 0.009 | -0.042 | -0.042 | 0 | 0.25 | 0.31 | 0.27 | 0.04 | 0.38 | 0.22 |
| 2 | 36110858 | rs6735990 | -1.78 | 1.326 | 0.699 | -0.278 | 0.003 | -0.027 | -0.027 | 0 | 0.37 | 0.58 | 0.33 | 0.16 | 0.43 | 0.34 |
| 2 | 36113122 | rs1401777 | -1.272 | 1.356 | 0.756 | 0.042 | 0.023 | -0.047 | -0.047 | 0 | 0.27 | 0.46 | 0.24 | 0.16 | 0.33 | 0.24 |
| 2 | 36120430 | rs4510197 | -0.274 | 1.279 | 0.661 | 0.079 | -0.014 | 0.035 | 0.035 | 0 | 0.13 | 0.38 | 0.08 | 0.13 | 0.07 | 0.13 |
| 2 | 36120658 | rs2372417 | -0.274 | 1.287 | 0.665 | 0.083 | -0.014 | 0.035 | 0.035 | 0 | 0.13 | 0.38 | 0.08 | 0.13 | 0.07 | 0.13 |
| 2 | 36120755 | rs2177483 | -1.233 | 1.245 | 0.262 | 0.08 | 0 | -0.005 | -0.005 | 0 | 0.29 | 0.67 | 0.28 | 0.17 | 0.34 | 0.24 |
| 2 | 36121342 | rs10171463 | -0.84 | 1.092 | 0.224 | 0.497 | 0.014 | 0.066 | 0.066 | 0 | 0.17 | 0.32 | 0.17 | 0.31 | 0.18 | 0.17 |
| 2 | 36122170 | rs11124487 | -0.912 | 0.54 | 0.068 | -0.07 | -0.022 | -0.071 | -0.071 | 0 | 0.29 | 0.48 | 0.27 | 0.11 | 0.26 | 0.18 |
| 2 | 36122388 | rs1524145 | 0.45 | 0.985 | -0.694 | 0.374 | -0.039 | 0.004 | 0.004 | 0.177 | 0.54 | 0.81 | 0.63 | 0.64 | 0.47 | 0.59 |
| 2 | 36123261 | rs2968576 | -0.54 | 1.037 | -0.782 | 0.303 | -0.011 | 0.014 | 0.014 | 0 | 0.74 | 0.84 | 0.77 | 0.78 | 0.71 | 0.58 |
| 2 | 36123717 | rs2968577 | -0.575 | 1.072 | -0.792 | 0.316 | -0.01 | 0.014 | 0.014 | 0 | 0.74 | 0.84 | 0.77 | 0.79 | 0.71 | 0.64 |
| 2 | 36123824 | rs2949052 | -0.295 | 0.288 | -0.461 | 0.449 | -0.037 | 0.038 | 0.038 | 0.279 | 0.22 | 0.54 | 0.17 | 0.25 | 0.15 | 0.16 |
| 2 | 36124132 | rs2968578 | -0.516 | 0.257 | -0.447 | 0.708 | -0.029 | 0.054 | 0.054 | 0.274 | 0.23 | 0.57 | 0.18 | 0.3 | 0.17 | 0.18 |
| 2 | 36125308 | rs1533969 | -0.453 | 0.216 | -0.456 | 0.682 | -0.036 | 0.023 | 0.023 | 0.384 | 0.23 | 0.61 | 0.17 | 0.25 | 0.15 | 0.17 |
| 2 | 36125664 | rs2949057 | -0.547 | 0.253 | -0.48 | 0.661 | -0.033 | 0.019 | 0.019 | 0.134 | 0.24 | 0.61 | 0.17 | 0.25 | 0.16 | 0.17 |
| 2 | 36132138 | rs935920 | -0.807 | 0.567 | -0.382 | 0.624 | -0.004 | 0.002 | 0.002 | 0 | 0.7 | 0.76 | 0.7 | 0.72 | 0.64 | 0.53 |
| 2 | 36132943 | rs2949065 | -0.693 | 0.563 | -0.388 | 0.622 | -0.006 | 0.003 | 0.003 | 0 | 0.7 | 0.77 | 0.69 | 0.72 | 0.64 | 0.53 |
| 2 | 36135009 | rs11124488 | 0.799 | 0.439 | -0.451 | 0.66 | -0.009 | -0.018 | -0.018 | 0 | 0.47 | 0.18 | 0.53 | 0.49 | 0.49 | 0.37 |
| 2 | 36135389 | rs2968599 | -0.872 | 0.475 | -0.373 | 0.66 | -0.012 | 0.013 | 0.013 | 0 | 0.72 | 0.8 | 0.72 | 0.77 | 0.67 | 0.54 |
| 2 | 36137996 | rs6543987 | 0.088 | 0.325 | 0.43 | 0.79 | 0.01 | 0.063 | 0.063 | 0 | 0.08 | 0.32 | 0.11 | 0.1 | 0.06 | 0.08 |
| 2 | 36139951 | rs2968602 | -0.288 | 0.232 | 0.493 | 0.762 | 0.004 | 0.103 | 0.103 | 0.13 | 0.09 | 0.21 | 0.07 | 0.19 | 0.07 | 0.08 |
| 2 | 36140364 | rs7355533 | 0.391 | 0.208 | 0.402 | 0.723 | 0.009 | 0.011 | 0.011 | 0 | 0.07 | 0.31 | 0.11 | 0.05 | 0.05 | 0.07 |
| 2 | 36147480 | rs1357662 | 0.877 | 0.168 | 0.191 | 0.638 | -0.001 | -0.003 | -0.003 | 0.283 | 0.47 | 0.31 | 0.54 | 0.49 | 0.49 | 0.33 |
| 2 | 36149165 | rs13433044 | 0.859 | 0.319 | 0.309 | 0.542 | -0.011 | -0.006 | -0.006 | 0.198 | 0.06 | 0.14 | 0.07 | 0.05 | 0.04 | 0.09 |
| 2 | 36153965 | rs13397366 | 0.768 | 0.34 | 0.315 | 0.582 | -0.006 | 0.032 | 0.032 | 0 | 0.06 | 0.15 | 0.09 | 0.1 | 0.04 | 0.06 |
| 2 | 36154313 | rs2949030 | -0.155 | 0.137 | 0.402 | 0.315 | 0.006 | -0.022 | -0.022 | 0 | 0.62 | 0.64 | 0.65 | 0.77 | 0.58 | 0.47 |
| 2 | 36162477 | rs2968624 | -0.576 | 0.053 | 0.031 | 0.877 | 0.024 | 0.003 | 0.003 | 0 | 0.68 | 0.6 | 0.61 | 0.37 | 0.62 | 0.5 |
| 2 | 36175047 | rs1510737 | -0.619 | 0.369 | -0.032 | 1.156 | 0.008 | -0.004 | -0.004 | 0 | 0.68 | 0.64 | 0.71 | 0.72 | 0.66 | 0.53 |
| 2 | 36175700 | rs17017895 | -0.218 | 0.271 | -0.197 | 0.733 | -0.001 | -0.004 | -0.004 | 0 | 0.15 | 0.16 | 0.25 | 0.08 | 0.11 | 0.08 |
| 2 | 36176752 | rs13409385 | 0.459 | 0.445 | -0.461 | 0.643 | -0.006 | -0.034 | -0.034 | 0 | 0.11 | 0.17 | 0.09 | 0.05 | 0.08 | 0.07 |
| 2 | 36178802 | rs1721777 | 0.6 | 0.524 | -0.723 | 0.847 | 0.019 | 0.004 | 0.004 | 2.263 | 0.47 | 0.35 | 0.61 | 0.46 | 0.46 | 0.38 |
| 2 | 36178894 | rs1721779 | 1.142 | 0.553 | -0.789 | 0.736 | 0.014 | 0.006 | 0.006 | 4.708 | 0.39 | 0.25 | 0.39 | 0.36 | 0.4 | 0.31 |
| 2 | 36179344 | rs1721781 | 1.194 | 0.569 | -0.827 | 0.684 | 0.014 | 0.001 | 0.001 | 4.367 | 0.38 | 0.22 | 0.38 | 0.36 | 0.4 | 0.31 |
| 2 | 36179888 | rs1620100 | 0.241 | 0.723 | -0.956 | 0.718 | 0.004 | 0.02 | 0.02 | 1.731 | 0.58 | 0.38 | 0.65 | 0.7 | 0.61 | 0.5 |
| 2 | 36180242 | rs13017326 | -0.271 | 0.681 | -0.959 | 0.765 | 0.001 | -0.005 | -0.005 | 1.314 | 0.2 | 0.19 | 0.27 | 0.34 | 0.14 | 0.19 |
| 2 | 36186152 | rs833760 | 0.263 | 0.686 | -0.791 | 0.495 | 0.002 | 0 | 0 | 0 | 0.61 | 0.46 | 0.5 | 0.4 | 0.62 | 0.44 |
| 2 | 36186184 | rs10211245 | -0.117 | 0.661 | -0.828 | 0.346 | -0.011 | -0.054 | -0.054 | 0 | 0.21 | 0.25 | 0.13 | 0.05 | 0.15 | 0.12 |
| 2 | 36188807 | rs13402895 | -0.576 | 0.438 | -0.837 | 0.335 | -0.02 | -0.056 | -0.056 | 0 | 0.21 | 0.25 | 0.11 | 0.03 | 0.12 | 0.21 |
| 2 | 36190919 | rs1912529 | 1.369 | 0.56 | -1.015 | 0.597 | -0.029 | -0.075 | -0.075 | 0 | 0.11 | 0.01 | 0.06 | 0.32 | 0.08 | 0.12 |
| 2 | 36194617 | rs1606033 | 0.854 | 0.77 | -0.859 | 0.362 | 0.027 | -0.03 | -0.03 | 0 | 0.07 | 0.12 | 0.09 | 0 | 0.12 | 0.03 |
| 2 | 36194653 | rs833765 | 1.158 | 0.694 | -1.072 | 0.414 | -0.021 | 0.129 | 0.129 | 1.629 | 0.51 | 0.25 | 0.58 | 0.76 | 0.52 | 0.4 |
| 2 | 36206662 | rs13390347 | -0.558 | 0.593 | -0.812 | 0.364 | -0.021 | -0.088 | -0.088 | 1.588 | 0.17 | 0.34 | 0.12 | 0.04 | 0.11 | 0.16 |
| 2 | 36208336 | rs1533972 | 0.295 | 0.585 | -0.982 | 0.476 | -0.001 | 0 | 0 | 0 | 0.58 | 0.57 | 0.68 | 0.44 | 0.62 | 0.43 |
| 2 | 36209776 | rs2968634 | -1.449 | 0.596 | -0.863 | 0.542 | 0.02 | 0.01 | 0.01 | 0 | 0.8 | 0.68 | 0.79 | 0.79 | 0.76 | 0.64 |
| 2 | 36211712 | rs1705046 | 1.977 | 0.674 | -0.906 | 0.568 | -0.002 | 0.001 | 0.001 | 3.398 | 0.39 | 0.19 | 0.36 | 0.35 | 0.44 | 0.25 |
| 2 | 36216027 | rs833783 | -1.004 | 0.862 | -0.873 | 0.802 | 0.024 | 0.02 | 0.02 | 0.193 | 0.78 | 0.56 | 0.76 | 0.77 | 0.75 | 0.65 |
| 2 | 36220820 | rs833785 | 1.353 | 0.854 | -1.038 | 0.795 | -0.036 | 0.044 | 0.044 | 1.209 | 0.49 | 0.09 | 0.38 | 0.36 | 0.51 | 0.31 |
| 2 | 36234822 | rs6722998 | -0.736 | 0.519 | -0.998 | 0.774 | -0.017 | 0.011 | 0.011 | 0 | 0.28 | 0.35 | 0.15 | 0.32 | 0.17 | 0.23 |
| 2 | 36234933 | rs6738060 | -1.045 | 0.523 | -1.005 | 0.73 | -0.025 | 0.042 | 0.042 | 0 | 0.3 | 0.44 | 0.35 | 0.44 | 0.18 | 0.34 |
| 2 | 36240074 | rs6543990 | 0.067 | 0.748 | -0.148 | 0.975 | 0.04 | 0.025 | 0.025 | 0.75 | 0.26 | 0.57 | 0.46 | 0.3 | 0.32 | 0.46 |
| 2 | 36245072 | rs2634806 | -0.129 | 0.446 | -0.388 | 0.827 | -0.012 | -0.002 | -0.002 | 2.458 | 0.69 | 0.58 | 0.7 | 0.3 | 0.73 | 0.66 |
| 2 | 36248316 | rs10170731 | 1.302 | 0.632 | -0.698 | 1.453 | -0.028 | -0.069 | -0.069 | 0 | 0.11 | 0.01 | 0.12 | 0.21 | 0.13 | 0.17 |
| 2 | 36252307 | rs962378 | 0.199 | 0.629 | 0.223 | 1.585 | 0.006 | -0.003 | -0.003 | 0 | 0.2 | 0.34 | 0.17 | 0.2 | 0.19 | 0.4 |
| 2 | 36252362 | rs305190 | -0.73 | 0.759 | 0.552 | 1.404 | 0.032 | -0.039 | -0.039 | 0 | 0.3 | 0.46 | 0.26 | 0.21 | 0.36 | 0.5 |
| 2 | 36252564 | rs305191 | 0.187 | 0.807 | 0.564 | 1.404 | 0.035 | -0.003 | -0.003 | 0 | 0.21 | 0.37 | 0.2 | 0.21 | 0.28 | 0.44 |

| | | | | | | | | | | | | | | | | |
|---|---|---|---|---|---|---|---|---|---|---|---|---|---|---|---|---|
| 2 | 36254823 | rs10495861 | 0.531 | 0.78 | 0.645 | 1.44 | 0.005 | 0.005 | 0.005 | 0.013 | 0.19 | 0.27 | 0.16 | 0.21 | 0.19 | 0.34 |
| 2 | 36255030 | rs1397443 | -0.427 | 0.781 | 0.616 | 1.404 | 0.024 | -0.043 | -0.043 | 1.372 | 0.1 | 0.12 | 0.1 | 0 | 0.17 | 0.05 |
| 2 | 36255746 | rs1352513 | -0.953 | 0.67 | 0.957 | 1.819 | -0.006 | -0.016 | -0.016 | 0.043 | 0.33 | 0.68 | 0.28 | 0.54 | 0.35 | 0.56 |
| 2 | 36257825 | rs7566742 | -0.676 | 0.794 | 0.934 | 1.295 | 0.034 | -0.03 | -0.03 | 0 | 0.3 | 0.47 | 0.27 | 0.21 | 0.34 | 0.49 |
| 2 | 36259341 | rs12989336 | -0.719 | 1.145 | 1.068 | 1.444 | -0.001 | 0.001 | 0.001 | 0.207 | 0.29 | 0.3 | 0.24 | 0.21 | 0.31 | 0.47 |
| 2 | 36260710 | rs2019606 | 1.957 | 1.002 | 0.918 | 1.331 | 0.013 | 0.191 | 0.191 | 1.093 | 0.5 | 0.14 | 0.36 | 0.21 | 0.5 | 0.39 |
| 2 | 36261234 | rs1906738 | 0.093 | 1.008 | 0.959 | 1.442 | 0.016 | -0.02 | -0.02 | 0 | 0.81 | 0.7 | 0.64 | 0.85 | 0.8 | 0.85 |
| 2 | 36261471 | rs2251598 | 0.154 | 1.016 | 0.997 | 1.44 | -0.009 | -0.026 | -0.026 | 0 | 0.78 | 0.72 | 0.64 | 0.86 | 0.84 | 0.9 |
| 2 | 36262110 | rs10199383 | 1.853 | 1.028 | 1.079 | 1.461 | 0.027 | 0.152 | 0.152 | 0.901 | 0.49 | 0.18 | 0.37 | 0.2 | 0.49 | 0.38 |
| 2 | 36262214 | rs1510728 | 0.551 | 1.155 | 0.736 | 0.994 | 0.009 | -0.026 | -0.026 | 0 | 0.77 | 0.69 | 0.6 | 0.85 | 0.75 | 0.84 |
| 2 | 36264999 | rs12988911 | -0.431 | 1.386 | 0.553 | 1.068 | 0.022 | -0.005 | -0.005 | 1.888 | 0.25 | 0.34 | 0.22 | 0.22 | 0.33 | 0.45 |
| 2 | 36269486 | rs2019397 | -0.963 | 1.313 | 0.502 | 1.04 | -0.001 | -0.021 | -0.021 | 0 | 0.86 | 0.82 | 0.75 | 0.97 | 0.86 | 0.95 |
| 2 | 36270756 | rs4670135 | -0.371 | 1.38 | 0.848 | 1.696 | 0.001 | 0 | 0 | 0.086 | 0.22 | 0.25 | 0.18 | 0.22 | 0.22 | 0.38 |
| 2 | 36273831 | rs2020037 | -0.732 | 1.377 | 0.824 | 1.688 | 0.006 | -0.002 | -0.002 | 0 | 0.24 | 0.29 | 0.19 | 0.23 | 0.25 | 0.41 |
| 2 | 36284939 | rs931401 | -1.091 | 1.083 | 0.536 | 1.977 | -0.005 | -0.006 | -0.006 | 0 | 0.19 | 0.13 | 0.12 | 0.22 | 0.18 | 0.33 |
| 2 | 36284976 | rs6740927 | 1.782 | 1.153 | 1.065 | 1.797 | 0.153 | 0.102 | 0.102 | 0.49 | 0.94 | 0.72 | 0.7 | 0.89 | 0.88 | 0.97 |
| 2 | 36331157 | rs2666138 | -0.34 | 0.94 | 0.925 | 2.408 | -0.008 | -0.008 | -0.008 | 0 | 0.88 | 0.87 | 0.91 | 0.7 | 0.87 | 0.72 |
| 2 | 36331798 | rs2666136 | -0.466 | 0.945 | 0.927 | 2.385 | -0.008 | -0.005 | -0.005 | 0 | 0.89 | 0.87 | 0.91 | 0.7 | 0.87 | 0.73 |
| 2 | 36334070 | rs10199869 | 0.652 | 0.732 | 0.753 | 2.218 | 0.002 | -0.02 | -0.02 | 0.135 | 0.75 | 0.77 | 0.84 | 0.62 | 0.8 | 0.65 |
| 2 | 36353996 | rs7596950 | 0.935 | 0.832 | -0.342 | 1.152 | -0.006 | 0 | 0 | 0.02 | 0.65 | 0.35 | 0.75 | 0.38 | 0.76 | 0.55 |
| 2 | 36355292 | rs1489153 | 0.959 | 0.334 | -0.539 | 0.923 | 0.006 | 0.004 | 0.004 | 0.006 | 0.62 | 0.29 | 0.73 | 0.35 | 0.73 | 0.49 |
| 2 | 36357979 | rs10195029 | 0.898 | 0.286 | -1.444 | 0.648 | 0.006 | 0.036 | 0.036 | 0.561 | 0.57 | 0.13 | 0.61 | 0.36 | 0.65 | 0.41 |
| 2 | 36364164 | rs305175 | 0.504 | 0.182 | -1.128 | 0.792 | 0.009 | 0.006 | 0.006 | 0.061 | 0.64 | 0.31 | 0.67 | 0.35 | 0.73 | 0.41 |
| 2 | 36371417 | rs1167445 | 0.378 | 0.249 | -1.357 | 0.421 | -0.042 | 0.048 | 0.048 | 0.4 | 0.56 | 0.16 | 0.52 | 0.7 | 0.53 | 0.57 |
| 2 | 36374519 | rs1167450 | 0.231 | 0.082 | -0.195 | 0.305 | 0.006 | -0.008 | -0.008 | 0 | 0.6 | 0.48 | 0.55 | 0.71 | 0.54 | 0.57 |
| 2 | 36376699 | rs11904235 | -0.048 | 0.122 | 0.411 | 0.147 | 0.009 | -0.049 | -0.049 | 0 | 0.09 | 0.18 | 0.06 | 0 | 0.08 | 0.09 |
| 2 | 36379720 | rs1167454 | 1.241 | -0.146 | 0.051 | 0.525 | -0.114 | -0.049 | -0.049 | 0 | 0.47 | 0.02 | 0.46 | 0.51 | 0.48 | 0.45 |
| 2 | 36381404 | rs2372441 | -0.734 | 0.011 | 0.179 | -0.177 | 0.025 | -0.025 | -0.025 | 0 | 0.09 | 0.31 | 0.06 | 0.01 | 0.05 | 0.03 |
| 2 | 36386902 | rs12614272 | 0.155 | 0.153 | -0.103 | -0.2 | -0.042 | -0.05 | -0.05 | 0 | 0.1 | 0.04 | 0.12 | 0.26 | 0.15 | 0.19 |
| 2 | 36391086 | rs7561060 | -0.664 | -0.01 | 0.235 | -0.232 | -0.005 | 0.008 | 0.008 | 0.062 | 0.4 | 0.36 | 0.44 | 0.29 | 0.46 | 0.46 |
| 2 | 36392700 | rs1153127 | 1.214 | 0.013 | -0.461 | -0.179 | -0.026 | 0.08 | 0.08 | 0.631 | 0.49 | 0.15 | 0.5 | 0.69 | 0.49 | 0.51 |
| 2 | 36393247 | rs7582800 | -0.194 | 0.086 | -0.353 | -0.083 | -0.018 | -0.006 | -0.006 | 0.013 | 0.32 | 0.18 | 0.38 | 0.3 | 0.37 | 0.38 |
| 2 | 36395705 | rs1167463 | 1.079 | -0.514 | -0.742 | -0.655 | -0.071 | 0.109 | 0.109 | 0 | 0.41 | 0.15 | 0.53 | 0.81 | 0.49 | 0.5 |
| 2 | 36398904 | rs6717847 | 0.473 | -0.631 | -0.361 | -0.812 | -0.001 | 0 | 0 | 0.073 | 0.17 | 0.12 | 0.1 | 0.14 | 0.15 | 0.2 |
| 2 | 36399318 | rs1167459 | 0.866 | -0.745 | -0.485 | -0.936 | 0.003 | 0.158 | 0.158 | 0 | 0.47 | 0.16 | 0.58 | 0.81 | 0.57 | 0.6 |
| 2 | 36408593 | rs11124500 | 0.101 | -0.772 | -0.229 | -0.952 | -0.001 | 0.003 | 0.003 | 0 | 0.2 | 0.15 | 0.1 | 0.15 | 0.18 | 0.21 |
| 2 | 36416792 | rs13405170 | 0.062 | -0.699 | -0.185 | -0.953 | -0.002 | 0.004 | 0.004 | 0 | 0.2 | 0.13 | 0.1 | 0.15 | 0.18 | 0.22 |
| 2 | 36418581 | rs1167441 | 1.504 | -0.717 | -0.201 | -1.03 | -0.071 | 0.104 | 0.104 | 0 | 0.4 | 0.13 | 0.53 | 0.82 | 0.49 | 0.5 |
| 2 | 36420568 | rs1183799 | 0.373 | -0.798 | 0.397 | -1.16 | 0.025 | 0.064 | 0.064 | 0.618 | 0.6 | 0.28 | 0.63 | 0.97 | 0.67 | 0.73 |
| 2 | 36422604 | rs1167436 | 0.418 | -0.724 | 0.798 | -1.022 | 0.074 | -0.001 | -0.001 | 0.007 | 0.92 | 0.55 | 0.95 | 1 | 0.96 | 0.99 |
| 2 | 36423276 | rs1167434 | -0.156 | -0.736 | 0.688 | -1.025 | -0.026 | -0.125 | -0.125 | 0 | 0.69 | 0.48 | 0.7 | 0.97 | 0.74 | 0.82 |
| 2 | 36425140 | rs1167432 | 1.425 | -0.578 | -0.068 | -0.724 | -0.112 | -0.071 | -0.071 | 0 | 0.31 | 0.05 | 0.48 | 0.7 | 0.47 | 0.28 |
| 2 | 36425498 | rs6704612 | 0.027 | -0.893 | 0.702 | -1.323 | 0.005 | -0.034 | -0.034 | 0.168 | 0.26 | 0.36 | 0.29 | 0.1 | 0.24 | 0.34 |
| 2 | 36432391 | rs11124501 | 0.747 | -1.073 | 0.802 | -1.383 | 0.007 | 0.019 | 0.019 | 0 | 0.63 | 0.34 | 0.7 | 0.88 | 0.75 | 0.55 |
| 2 | 36432419 | rs10197906 | 0.378 | -1.068 | 0.892 | -1.342 | -0.011 | -0.016 | -0.016 | 0.824 | 0.08 | 0.09 | 0.01 | 0.02 | 0.03 | 0.06 |
| 2 | 36437449 | rs2204773 | 0.737 | -1.058 | 1.233 | -1.34 | 0.056 | 0.01 | 0.01 | 3.756 | 0.93 | 0.48 | 0.94 | 1 | 0.97 | 0.99 |
| 2 | 36439777 | rs12468624 | 0.455 | -1.042 | 1.03 | -1.367 | -0.013 | -0.062 | -0.062 | 3.508 | 0.67 | 0.53 | 0.73 | 0.88 | 0.78 | 0.55 |
| 2 | 36448149 | rs1881782 | 0.538 | -0.947 | 0.344 | -1.303 | -0.018 | 0.029 | 0.029 | 0 | 0.22 | 0.11 | 0.23 | 0.11 | 0.19 | 0.39 |
| 2 | 36460918 | rs2141537 | -0.73 | -0.773 | -0.415 | -1.099 | 0.02 | 0.11 | 0.11 | 0.62 | 0.38 | 0.83 | 0.32 | 0.12 | 0.26 | 0.43 |
| 2 | 36461251 | rs6743680 | 0.617 | -0.746 | -0.526 | -1.327 | 0.015 | 0.005 | 0.005 | 0 | 0.27 | 0.26 | 0.15 | 0.02 | 0.2 | 0.18 |
| 2 | 36462390 | rs2699150 | -0.513 | 0.033 | -0.82 | -0.833 | -0.078 | 0.114 | 0.114 | 0 | 0.63 | 0.87 | 0.46 | 0.14 | 0.46 | 0.6 |
| 2 | 36464050 | rs1529267 | -0.492 | 0.139 | -0.773 | -0.852 | -0.025 | 0.064 | 0.064 | 0 | 0.64 | 0.81 | 0.44 | 0.15 | 0.41 | 0.62 |
| 2 | 36465942 | rs2161901 | -0.494 | 0.186 | -1.015 | -0.69 | -0.081 | 0.086 | 0.086 | 0.353 | 0.66 | 0.86 | 0.45 | 0.14 | 0.46 | 0.62 |
| 2 | 36477600 | rs2052626 | -0.061 | 0.574 | -0.781 | -0.068 | -0.044 | -0.04 | -0.04 | 0 | 0.09 | 0.01 | 0.25 | 0.63 | 0.14 | 0.17 |
| 2 | 36479648 | rs2699160 | -0.152 | 0.144 | -0.126 | -0.655 | -0.048 | -0.085 | -0.085 | 0 | 0.26 | 0.49 | 0.26 | 0.09 | 0.11 | 0.35 |
| 2 | 36480049 | rs2552007 | 0.555 | 0.112 | -0.39 | -1.219 | 0.03 | -0.012 | -0.012 | 0 | 0.29 | 0.17 | 0.16 | 0.26 | 0.23 | 0.32 |
| 2 | 36482202 | rs2699164 | 0.183 | 0.069 | 0.069 | -1.28 | -0.032 | -0.059 | -0.059 | 0 | 0.26 | 0.53 | 0.26 | 0.09 | 0.11 | 0.35 |
| 2 | 36483942 | rs2246009 | -0.975 | 0.802 | -0.263 | -1.534 | -0.044 | 0.038 | 0.038 | 0 | 0.87 | 0.95 | 0.81 | 0.98 | 0.7 | 0.91 |
| 2 | 36486776 | rs918901 | 0.037 | 0.702 | -0.032 | -1.542 | -0.025 | 0.05 | 0.05 | 0 | 0.66 | 0.8 | 0.6 | 0.92 | 0.61 | 0.59 |
| 2 | 36490476 | rs2699175 | 1.184 | 0.497 | -0.246 | -1.38 | 0.028 | -0.039 | -0.039 | 0 | 0.25 | 0.15 | 0.14 | 0.29 | 0.21 | 0.31 |
| 2 | 36494176 | rs1881778 | -0.912 | 0.743 | 0.213 | -1.461 | -0.028 | 0.024 | 0.024 | 0 | 0.87 | 0.93 | 0.81 | 0.98 | 0.7 | 0.9 |
| 2 | 36494239 | rs1881779 | 0.469 | 0.69 | 0.302 | -1.617 | 0.008 | -0.07 | -0.07 | 0 | 0.59 | 0.57 | 0.53 | 0.89 | 0.51 | 0.53 |
| 2 | 36497171 | rs726946 | 0.302 | 0.656 | 0.454 | -1.606 | 0.105 | -0.113 | -0.113 | 0 | 0.78 | 0.59 | 0.75 | 0.95 | 0.62 | 0.85 |
| 2 | 36497312 | rs726947 | 0.245 | 0.658 | 0.442 | -1.605 | 0.089 | -0.098 | -0.098 | 0 | 0.78 | 0.63 | 0.75 | 0.95 | 0.62 | 0.85 |
| 2 | 36500649 | rs6723999 | 1.2 | 0.58 | 0.226 | -1.606 | -0.001 | -0.015 | -0.015 | 0 | 0.31 | 0.3 | 0.38 | 0.6 | 0.32 | 0.22 |
| 2 | 36506690 | rs2888336 | -0.327 | 0.57 | 0.009 | -1.675 | -0.079 | 0.119 | 0.119 | 0 | 0.68 | 0.88 | 0.59 | 0.93 | 0.59 | 0.57 |
| 2 | 36507357 | rs2552034 | -0.198 | 0.464 | 0.049 | -1.654 | -0.031 | 0.013 | 0.013 | 0.402 | 0.57 | 0.8 | 0.63 | 0.68 | 0.42 | 0.6 |
| 2 | 36508889 | rs7598895 | 0.804 | 0.72 | 0.421 | -1.42 | 0.011 | 0.019 | 0.019 | 0 | 0.22 | 0.14 | 0.22 | 0.05 | 0.11 | 0.33 |
| 2 | 36516173 | rs2161905 | -0.128 | 1.187 | 0.072 | -1.33 | -0.026 | 0.016 | 0.016 | 0 | 0.83 | 0.88 | 0.78 | 0.98 | 0.63 | 0.89 |

| | | | | | | | | | | | | | | | | |
|---|---|---|---|---|---|---|---|---|---|---|---|---|---|---|---|---|
| 2 | 36520350 | rs6733100 | -0.098 | 1.215 | 0.16 | -1.316 | 0.007 | -0.018 | -0.018 | 0 | 0.83 | 0.83 | 0.77 | 0.98 | 0.63 | 0.89 |
| 2 | 36521759 | rs2699187 | -0.496 | 1.091 | 0.334 | -1.149 | -0.003 | 0.016 | 0.016 | 0 | 0.56 | 0.7 | 0.65 | 0.73 | 0.5 | 0.64 |
| 2 | 36521977 | rs2699188 | -0.838 | 1.068 | 0 | -1.123 | 0.061 | 0.027 | 0.027 | 0.418 | 0.6 | 0.91 | 0.78 | 0.74 | 0.67 | 0.68 |
| 2 | 36523782 | rs12712488 | -1.597 | 2.074 | 1.477 | -0.463 | 0.207 | -0.069 | -0.069 | 0 | 0.14 | 0.48 | 0.21 | 0.05 | 0.31 | 0.14 |
| 2 | 36527733 | rs12472996 | -1.58 | 1.989 | 1.19 | -0.404 | 0.063 | 0.017 | 0.017 | 0.001 | 0.3 | 0.46 | 0.42 | 0.31 | 0.45 | 0.52 |
| 2 | 36529151 | rs1030295 | -1.22 | 2.343 | 0.834 | -0.289 | -0.023 | 0.02 | 0.02 | 0 | 0.13 | 0.09 | 0.2 | 0.02 | 0.36 | 0.11 |
| 2 | 36532633 | rs7595790 | -1.058 | 2.122 | 1.086 | -0.283 | -0.01 | -0.071 | -0.071 | 0 | 0.3 | 0.47 | 0.3 | 0.12 | 0.19 | 0.39 |
| 2 | 36533630 | rs4670541 | -1.879 | 2.183 | 1.087 | -0.253 | -0.019 | -0.003 | -0.003 | 0 | 0.41 | 0.81 | 0.47 | 0.36 | 0.48 | 0.55 |
| 2 | 36542575 | rs7560345 | -1.125 | 2.233 | 1.012 | -0.349 | 0.192 | 0.185 | 0.185 | 0.995 | 0.12 | 0.37 | 0.17 | 0.26 | 0.29 | 0.16 |
| 2 | 36545169 | rs6544006 | -0.629 | 2.228 | 1.068 | -0.38 | 0.081 | -0.017 | -0.017 | 1.301 | 0.09 | 0.46 | 0.09 | 0.03 | 0.09 | 0.05 |
| 2 | 36551827 | rs10190416 | -1.125 | 2.275 | 0.943 | -0.353 | 0.193 | 0.182 | 0.182 | 0.937 | 0.12 | 0.37 | 0.17 | 0.26 | 0.29 | 0.17 |
| 2 | 36556928 | rs17407386 | -0.015 | 2.066 | -0.314 | -1.146 | 0.012 | 0.063 | 0.063 | 0 | 0.21 | 0.04 | 0.21 | 0.06 | 0.1 | 0.31 |
| 2 | 36560011 | rs12329047 | -1.126 | 2.35 | -0.002 | -0.684 | 0.165 | -0.086 | -0.086 | 0.047 | 0.17 | 0.44 | 0.23 | 0.05 | 0.37 | 0.15 |
| 2 | 36562973 | rs7608543 | -0.636 | 1.523 | -0.876 | -0.886 | 0.004 | -0.053 | -0.053 | 0 | 0.36 | 0.52 | 0.27 | 0.05 | 0.35 | 0.31 |
| 2 | 36563968 | rs10200367 | -0.385 | 0.736 | -2.178 | -0.07 | -0.02 | -0.112 | -0.112 | 0 | 0.55 | 0.91 | 0.5 | 0.51 | 0.36 | 0.55 |
| 2 | 36569014 | rs7584784 | 0.255 | 0.669 | -2.252 | -0.332 | -0.071 | 0.221 | 0.221 | 0 | 0.39 | 0.12 | 0.45 | 0.06 | 0.6 | 0.33 |
| 2 | 36571060 | rs9308990 | 0.056 | 0.14 | -1.158 | 0.301 | -0.013 | 0.078 | 0.078 | 0 | 0.18 | 0.34 | 0.19 | 0.34 | 0.15 | 0.32 |
| 2 | 36573454 | rs982353 | 0.045 | -0.08 | -1.282 | 0.069 | -0.003 | -0.001 | -0.001 | 0 | 0.74 | 0.73 | 0.77 | 0.73 | 0.79 | 0.67 |
| 2 | 36575659 | rs717805 | -0.41 | -0.028 | -0.826 | -0.361 | 0 | 0.008 | 0.008 | 0.457 | 0.47 | 0.53 | 0.42 | 0.23 | 0.55 | 0.57 |
| 2 | 36578070 | rs10167098 | -1.013 | -0.333 | -0.365 | 0.019 | 0.008 | -0.014 | -0.014 | 0.104 | 0.69 | 0.6 | 0.64 | 0.72 | 0.65 | 0.74 |
| 2 | 36580631 | rs4670544 | 0.154 | -0.523 | -0.516 | -0.613 | -0.016 | 0.047 | 0.047 | 0 | 0.4 | 0.25 | 0.32 | 0.08 | 0.43 | 0.42 |
| 2 | 36588291 | rs3770962 | 0.8 | -0.455 | -0.708 | -0.513 | -0.003 | -0.01 | -0.01 | 0.015 | 0.12 | 0.09 | 0.1 | 0.15 | 0.15 | 0.14 |
| 2 | 36590010 | rs6728742 | -1.014 | -0.654 | -0.601 | -0.054 | -0.001 | 0.017 | 0.017 | 0.246 | 0.64 | 0.44 | 0.6 | 0.45 | 0.64 | 0.7 |
| 2 | 36590023 | rs6757374 | 0.34 | -0.878 | -0.567 | -0.063 | -0.045 | 0.117 | 0.117 | 0 | 0.17 | 0.38 | 0.22 | 0.27 | 0.1 | 0.23 |
| 2 | 36597085 | rs10178804 | 0.685 | -0.365 | -0.786 | -0.315 | -0.047 | 0.197 | 0.197 | 0 | 0.28 | 0.02 | 0.2 | 0.01 | 0.24 | 0.26 |
| 2 | 36601151 | rs3770951 | 1.124 | -0.171 | -0.509 | -0.24 | -0.001 | 0.003 | 0.003 | 0 | 0.2 | 0.22 | 0.25 | 0.19 | 0.18 | 0.13 |
| 2 | 36603763 | rs3770949 | 0.334 | -0.096 | -0.58 | -0.294 | -0.009 | 0.003 | 0.003 | 0 | 0.15 | 0.14 | 0.13 | 0.14 | 0.2 | 0.16 |
| 2 | 36605304 | rs4670546 | -0.628 | -0.229 | -0.351 | -0.226 | 0 | -0.006 | -0.006 | 0 | 0.53 | 0.5 | 0.54 | 0.31 | 0.47 | 0.55 |
| 2 | 36605417 | rs4670547 | 0.607 | -0.186 | -0.772 | -0.438 | -0.043 | 0.194 | 0.194 | 0 | 0.28 | 0.03 | 0.21 | 0.01 | 0.25 | 0.26 |
| 2 | 36610069 | rs3821165 | -0.336 | -0.01 | -0.178 | -0.271 | -0.004 | -0.007 | -0.007 | 0.038 | 0.47 | 0.51 | 0.48 | 0.31 | 0.38 | 0.52 |
| 2 | 36612480 | rs3770945 | -0.171 | -0.067 | -0.066 | -0.092 | 0.001 | 0.015 | 0.015 | 0 | 0.38 | 0.36 | 0.28 | 0.21 | 0.36 | 0.49 |
| 2 | 36614307 | rs4670551 | -0.886 | -0.206 | -0.002 | -0.118 | 0.008 | 0.06 | 0.06 | 0 | 0.53 | 0.71 | 0.53 | 0.7 | 0.47 | 0.67 |
| 2 | 36617528 | rs3770939 | 0.897 | -0.193 | -0.019 | -0.184 | 0.001 | 0.055 | 0.055 | 0 | 0.29 | 0.14 | 0.2 | 0.13 | 0.26 | 0.34 |
| 2 | 36618667 | rs11124513 | 0.007 | -0.23 | 0.109 | -0.268 | -0.007 | 0.01 | 0.01 | 0 | 0.32 | 0.61 | 0.39 | 0.36 | 0.28 | 0.33 |
| 2 | 36619064 | rs4670552 | -0.802 | -0.184 | 0.097 | -0.206 | -0.015 | -0.023 | -0.023 | 0 | 0.6 | 0.71 | 0.59 | 0.45 | 0.53 | 0.65 |
| 2 | 36619764 | rs12465282 | -2.058 | -0.19 | -0.147 | -0.319 | -0.085 | 0.135 | 0.135 | 0.094 | 0.75 | 0.95 | 0.82 | 0.99 | 0.65 | 0.86 |
| 2 | 36621057 | rs17018690 | -0.439 | -0.238 | 0.218 | -0.152 | -0.024 | 0.118 | 0.118 | 0 | 0.23 | 0.47 | 0.32 | 0.57 | 0.21 | 0.37 |
| 2 | 36624137 | rs3821164 | 0.896 | -0.231 | -0.113 | -0.12 | 0.007 | 0.005 | 0.005 | 0 | 0.13 | 0.12 | 0.23 | 0.51 | 0.12 | 0.18 |
| 2 | 36624368 | rs11891181 | 1.184 | -0.209 | -0.228 | -0.309 | -0.019 | 0.06 | 0.06 | 0 | 0.27 | 0.08 | 0.17 | 0.13 | 0.24 | 0.35 |
| 2 | 36625388 | rs3770935 | 1.185 | -0.212 | -0.243 | -0.311 | -0.023 | 0.067 | 0.067 | 0 | 0.27 | 0.08 | 0.16 | 0.13 | 0.23 | 0.34 |
| 2 | 36626078 | rs3755214 | -1.944 | -0.261 | -0.277 | -0.416 | -0.059 | 0.104 | 0.104 | 0.123 | 0.73 | 0.9 | 0.81 | 0.99 | 0.64 | 0.85 |
| 2 | 36627725 | rs4670145 | 1.229 | -0.261 | -0.274 | -0.391 | -0.023 | 0.061 | 0.061 | 0 | 0.27 | 0.07 | 0.17 | 0.13 | 0.23 | 0.34 |
| 2 | 36628642 | rs3770933 | 1.542 | -0.292 | -0.176 | -0.315 | 0 | -0.012 | -0.012 | 0 | 0.2 | 0.15 | 0.25 | 0.21 | 0.17 | 0.11 |
| 2 | 36629212 | rs3770929 | -0.071 | -0.276 | 0.187 | -0.259 | 0.002 | 0.004 | 0.004 | 0 | 0.49 | 0.43 | 0.47 | 0.42 | 0.41 | 0.51 |
| 2 | 36629247 | rs7595245 | -0.672 | -0.217 | 0.267 | -0.289 | 0.004 | -0.025 | -0.025 | 0.222 | 0.61 | 0.48 | 0.64 | 0.84 | 0.53 | 0.64 |
| 2 | 36630268 | rs3821161 | 0.165 | -0.203 | 0.227 | -0.243 | 0.006 | -0.001 | -0.001 | 0 | 0.1 | 0.3 | 0.1 | 0.07 | 0.11 | 0.15 |
| 2 | 36631779 | rs3770924 | -1.717 | -0.176 | 0.142 | -0.276 | -0.011 | 0.011 | 0.011 | 0.05 | 0.72 | 0.81 | 0.75 | 0.91 | 0.64 | 0.8 |
| 2 | 36632040 | rs3770922 | 0.718 | -0.313 | 0.376 | 0.125 | 0.009 | -0.01 | -0.01 | 0 | 0.13 | 0.12 | 0.22 | 0.49 | 0.12 | 0.19 |
| 2 | 36632175 | rs3770920 | -0.748 | -0.36 | 0.391 | 0.019 | 0.01 | 0.03 | 0.03 | 0 | 0.39 | 0.29 | 0.52 | 0.71 | 0.37 | 0.35 |
| 2 | 36634617 | rs3755210 | -0.003 | -0.386 | 0.355 | 0.454 | 0.007 | 0.007 | 0.007 | 0 | 0.1 | 0.3 | 0.1 | 0.07 | 0.11 | 0.16 |
| 2 | 36639259 | rs3770910 | 0.122 | -0.385 | -0.077 | 0.697 | 0.006 | 0.001 | 0.001 | 0 | 0.13 | 0.12 | 0.23 | 0.48 | 0.12 | 0.17 |
| 2 | 36650725 | rs2110928 | -0.153 | -0.427 | 0.058 | 0.664 | -0.01 | 0.076 | 0.076 | 0 | 0.14 | 0.21 | 0.27 | 0.56 | 0.12 | 0.24 |
| 2 | 36653937 | rs11682688 | -0.068 | -0.443 | 0.081 | 0.644 | -0.013 | 0.077 | 0.077 | 0 | 0.14 | 0.2 | 0.26 | 0.56 | 0.11 | 0.24 |
| 2 | 36654772 | rs3770899 | 0.005 | -0.446 | 0.025 | 0.674 | -0.002 | 0.036 | 0.036 | 0 | 0.14 | 0.16 | 0.22 | 0.49 | 0.13 | 0.19 |
| 2 | 36655394 | rs848504 | -0.224 | -0.414 | -0.067 | 0.029 | 0 | -0.005 | -0.005 | 0 | 0.45 | 0.48 | 0.56 | 0.79 | 0.43 | 0.33 |
| 2 | 36657664 | rs848503 | -0.188 | -0.329 | -0.047 | -0.156 | 0.004 | 0.021 | 0.021 | 0 | 0.65 | 0.57 | 0.56 | 0.42 | 0.6 | 0.62 |
| 2 | 36660989 | rs3770888 | 0.55 | -0.319 | -0.409 | -0.021 | 0.002 | 0.017 | 0.017 | 0 | 0.14 | 0.15 | 0.24 | 0.5 | 0.15 | 0.2 |
| 2 | 36668219 | rs2049367 | 0.947 | -0.456 | -0.557 | -0.257 | 0.001 | 0.004 | 0.004 | 0 | 0.29 | 0.28 | 0.23 | 0.12 | 0.29 | 0.4 |
| 2 | 36670011 | rs3732073 | 0.217 | -0.367 | -0.578 | -0.255 | 0.002 | 0.008 | 0.008 | 0 | 0.36 | 0.32 | 0.25 | 0.13 | 0.35 | 0.47 |
| 2 | 36674827 | rs848514 | -0.683 | 0.015 | -0.778 | -0.208 | 0.012 | -0.09 | -0.09 | 0.01 | 0.74 | 0.7 | 0.82 | 0.99 | 0.72 | 0.85 |
| 2 | 36679028 | rs3770876 | -0.046 | 0.067 | -1.822 | -0.074 | 0.063 | 0.095 | 0.095 | 0 | 0.49 | 0.8 | 0.53 | 0.68 | 0.42 | 0.69 |
| 2 | 36680848 | rs711254 | -0.618 | 0.368 | -1.458 | 0.04 | -0.057 | 0.091 | 0.091 | 0 | 0.76 | 0.92 | 0.8 | 0.9 | 0.71 | 0.77 |
| 2 | 36683997 | rs3770874 | -0.191 | 0.017 | -1.157 | 0.568 | 0.037 | -0.022 | -0.022 | 0 | 0.46 | 0.74 | 0.43 | 0.5 | 0.43 | 0.67 |
| 2 | 36687111 | rs888077 | 0.058 | -0.242 | -0.276 | 0.309 | -0.018 | -0.029 | -0.029 | 0 | 0.75 | 0.46 | 0.62 | 0.82 | 0.81 | 0.84 |
| 2 | 36696521 | rs3770858 | 0.891 | -0.48 | -0.346 | 0.511 | 0.006 | 0.037 | 0.037 | 0 | 0.19 | 0.23 | 0.24 | 0.37 | 0.2 | 0.33 |
| 2 | 36696545 | rs3770857 | 0.57 | -1.038 | -1.081 | 0.004 | 0.04 | 0.104 | 0.104 | 0 | 0.33 | 0.09 | 0.2 | 0.11 | 0.26 | 0.34 |
| 2 | 36700639 | rs848551 | 0.684 | -0.971 | -0.598 | 0.138 | -0.007 | 0.018 | 0.018 | 0 | 0.09 | 0.12 | 0.27 | 0.2 | 0.04 | 0.06 |
| 2 | 36704144 | rs848547 | -0.993 | -1.055 | -0.405 | 0.023 | 0.014 | -0.017 | -0.017 | 0 | 0.6 | 0.5 | 0.74 | 0.68 | 0.5 | 0.73 |
| 2 | 36704998 | rs3770852 | 1.901 | -0.741 | -0.359 | 0.252 | -0.076 | -0.045 | -0.045 | 0 | 0.22 | 0.06 | 0.16 | 0.3 | 0.32 | 0.15 |

| | | | | | | | | | | | | | | | | |
|---|---|---|---|---|---|---|---|---|---|---|---|---|---|---|---|---|
| 2 | 36705040 | rs848544 | -0.475 | -0.773 | -0.17 | 0.22 | -0.008 | 0.01 | 0.01 | 0 | 0.34 | 0.59 | 0.53 | 0.59 | 0.27 | 0.43 |
| 2 | 36705747 | rs6720433 | -0.347 | -0.444 | -0.552 | -0.289 | 0.044 | 0.064 | 0.064 | 0 | 0.53 | 0.79 | 0.47 | 0.68 | 0.6 | 0.59 |
| 2 | 36711872 | rs3770844 | 0.524 | -0.563 | 0.046 | -0.544 | 0.007 | -0.054 | -0.054 | 0 | 0.26 | 0.36 | 0.23 | 0.09 | 0.25 | 0.41 |
| 2 | 36712514 | rs1403858 | 0.606 | -0.565 | -0.058 | -0.544 | 0 | -0.009 | -0.009 | 0 | 0.25 | 0.27 | 0.22 | 0.08 | 0.25 | 0.42 |
| 2 | 36712597 | rs4141686 | -0.479 | -0.629 | -0.105 | -0.479 | 0.031 | 0.037 | 0.037 | 0 | 0.57 | 0.74 | 0.51 | 0.7 | 0.67 | 0.7 |
| 2 | 36712905 | rs11893748 | 0.451 | -0.335 | -0.534 | -0.52 | 0 | -0.016 | -0.016 | 0 | 0.26 | 0.28 | 0.22 | 0.09 | 0.25 | 0.43 |
| 2 | 36717819 | rs3899034 | 0.795 | -0.268 | -0.919 | -0.441 | -0.005 | 0.014 | 0.014 | 0 | 0.23 | 0.19 | 0.19 | 0.09 | 0.23 | 0.4 |
| 2 | 36728019 | rs848532 | -0.947 | -0.426 | -1.561 | -0.683 | -0.035 | 0.173 | 0.173 | 0 | 0.69 | 0.94 | 0.79 | 0.98 | 0.69 | 0.81 |
| 2 | 36731860 | rs711249 | -0.534 | -0.369 | -0.892 | -0.097 | 0.021 | -0.026 | -0.026 | 0 | 0.34 | 0.79 | 0.48 | 0.37 | 0.24 | 0.53 |
| 2 | 36735426 | rs3770834 | -0.159 | -0.383 | -0.548 | -0.3 | 0.007 | -0.001 | -0.001 | 0.097 | 0.7 | 0.75 | 0.6 | 0.7 | 0.79 | 0.71 |
| 2 | 36738630 | rs3770832 | 1.025 | -0.072 | -0.387 | -0.345 | -0.001 | 0.026 | 0.026 | 0.18 | 0.34 | 0.28 | 0.25 | 0.15 | 0.38 | 0.22 |
| 2 | 36742955 | rs884215 | -0.002 | 0.025 | 0.063 | -0.219 | -0.014 | -0.021 | -0.021 | 0 | 0.34 | 0.23 | 0.35 | 0.56 | 0.4 | 0.49 |
| 2 | 36747912 | rs3770827 | -0.597 | -0.094 | 0.53 | -0.19 | -0.003 | 0.014 | 0.014 | 0.25 | 0.75 | 0.64 | 0.63 | 0.7 | 0.8 | 0.76 |
| 2 | 36762126 | rs3770821 | 0.798 | -0.308 | 0.541 | -0.276 | 0.006 | 0.052 | 0.052 | 0.331 | 0.36 | 0.24 | 0.25 | 0.13 | 0.37 | 0.22 |
| 2 | 36763205 | rs848518 | 0.322 | -0.336 | 0.893 | -0.2 | 0.003 | 0.002 | 0.002 | 0.242 | 0.43 | 0.42 | 0.52 | 0.39 | 0.4 | 0.28 |
| 2 | 36764402 | rs848516 | -0.437 | -0.548 | 1.074 | 0.031 | -0.04 | 0.113 | 0.113 | 0 | 0.09 | 0.21 | 0.29 | 0.24 | 0.03 | 0.13 |
| 2 | 36767257 | rs848515 | -0.976 | -0.39 | 0.753 | -0.314 | -0.03 | 0.053 | 0.053 | 0 | 0.75 | 0.85 | 0.8 | 0.99 | 0.68 | 0.81 |
| 2 | 36768875 | rs711245 | -0.612 | -0.346 | 0.907 | -0.182 | -0.002 | -0.016 | -0.016 | 0 | 0.39 | 0.58 | 0.54 | 0.83 | 0.3 | 0.53 |
| 2 | 36770192 | rs848607 | -0.326 | -0.376 | 1.016 | 0.157 | -0.001 | -0.001 | -0.001 | 4.953 | 0.35 | 0.4 | 0.48 | 0.37 | 0.28 | 0.49 |
| 2 | 36771309 | rs12712508 | 0.671 | -0.344 | 0.965 | 0.106 | -0.003 | 0 | 0 | 0 | 0.42 | 0.49 | 0.34 | 0.61 | 0.41 | 0.32 |
| 2 | 36771701 | rs2276671 | 0.81 | -0.417 | 0.962 | 0.126 | -0.036 | 0.037 | 0.037 | 0.195 | 0.25 | 0.14 | 0.21 | 0.12 | 0.26 | 0.39 |
| 2 | 36773071 | rs3755196 | 0.957 | -0.335 | 0.961 | 0.131 | 0.013 | 0.047 | 0.047 | 0.274 | 0.37 | 0.26 | 0.26 | 0.16 | 0.37 | 0.27 |
| 2 | 36773260 | rs848605 | -0.309 | -0.154 | 1.038 | 0.209 | -0.001 | -0.004 | -0.004 | 0 | 0.34 | 0.36 | 0.48 | 0.36 | 0.28 | 0.49 |
| 2 | 36778372 | rs848602 | -0.698 | -0.212 | 0.979 | 0.145 | -0.001 | -0.012 | -0.012 | 0 | 0.38 | 0.57 | 0.54 | 0.82 | 0.3 | 0.55 |
| 2 | 36780549 | rs10172196 | 1.323 | -0.274 | 0.853 | 0.125 | 0.006 | 0.045 | 0.045 | 0.054 | 0.32 | 0.2 | 0.21 | 0.12 | 0.34 | 0.15 |
| 2 | 36780952 | rs3770809 | 0.697 | -0.106 | 1.416 | 0.29 | -0.011 | -0.001 | -0.001 | 0 | 0.42 | 0.44 | 0.33 | 0.62 | 0.39 | 0.31 |
| 2 | 36782886 | rs848642 | -0.45 | -0.077 | 1.2 | 0.305 | 0 | -0.016 | -0.016 | 0 | 0.35 | 0.59 | 0.52 | 0.78 | 0.29 | 0.52 |
| 2 | 36791226 | rs13406184 | 0.841 | -0.115 | 0.859 | 0.134 | 0.022 | 0.059 | 0.059 | 1.362 | 0.41 | 0.26 | 0.29 | 0.21 | 0.38 | 0.3 |
| 2 | 36795233 | rs3821150 | 0.397 | 0.108 | 0.367 | 0.006 | -0.054 | 0.051 | 0.051 | 0.209 | 0.26 | 0.09 | 0.22 | 0.12 | 0.28 | 0.41 |
| 2 | 36801999 | rs848622 | 0.073 | 0.042 | 0.495 | -0.1 | -0.01 | 0.064 | 0.064 | 0.981 | 0.56 | 0.62 | 0.66 | 0.87 | 0.54 | 0.55 |
| 2 | 36803863 | rs848596 | -0.028 | 0.021 | 0.383 | -0.11 | -0.028 | 0.128 | 0.128 | 1.17 | 0.57 | 0.71 | 0.67 | 0.87 | 0.54 | 0.56 |
| 2 | 36808732 | rs848637 | -1.107 | -0.282 | 0.473 | -0.045 | -0.014 | 0.051 | 0.051 | 0.994 | 0.76 | 0.81 | 0.81 | 0.99 | 0.71 | 0.82 |
| 2 | 36810133 | rs11691600 | 0.43 | -0.48 | 1.189 | 0.437 | 0.002 | 0.003 | 0.003 | 0.829 | 0.41 | 0.36 | 0.26 | 0.19 | 0.39 | 0.3 |
| 2 | 36810586 | rs14291 | 0.078 | -0.468 | 1.179 | 0.606 | 0.004 | -0.001 | -0.001 | 0 | 0.46 | 0.45 | 0.51 | 0.41 | 0.41 | 0.37 |
| 2 | 36817152 | rs2072535 | -0.008 | -0.235 | 0.49 | 0.569 | 0.028 | 0.012 | 0.012 | 0.743 | 0.45 | 0.64 | 0.35 | 0.65 | 0.39 | 0.33 |
| 2 | 36834707 | rs918067 | 0.521 | -0.099 | -0.082 | 0.643 | -0.016 | 0.043 | 0.043 | 0.051 | 0.34 | 0.16 | 0.33 | 0.23 | 0.39 | 0.2 |
| 2 | 36835678 | rs744917 | 0.14 | -0.031 | 0.159 | 0.46 | -0.001 | -0.006 | -0.006 | 3.95 | 0.42 | 0.43 | 0.58 | 0.69 | 0.45 | 0.24 |
| 2 | 36838600 | rs918066 | 0.096 | 0.042 | -0.108 | 0.478 | 0 | 0.005 | 0.005 | 0 | 0.42 | 0.57 | 0.6 | 0.7 | 0.45 | 0.24 |
| 2 | 36847770 | rs17489237 | 0.885 | 0.465 | -0.828 | 0.13 | -0.038 | 0.035 | 0.035 | 0 | 0.13 | 0.03 | 0.12 | 0.04 | 0.16 | 0.22 |
| 2 | 36848297 | rs4670152 | 1.122 | 0.388 | -0.991 | 0.169 | -0.018 | 0.004 | 0.004 | 2.793 | 0.26 | 0.16 | 0.27 | 0.23 | 0.33 | 0.17 |
| 2 | 36848969 | rs13407899 | -0.809 | 0.094 | -0.42 | 0.34 | -0.011 | -0.006 | -0.006 | 1.122 | 0.09 | 0.13 | 0.05 | 0.08 | 0.06 | 0.14 |
| 2 | 36849220 | rs12612163 | -0.091 | -0.079 | -0.645 | 0.513 | -0.002 | -0.004 | -0.004 | 0 | 0.72 | 0.75 | 0.61 | 0.53 | 0.7 | 0.81 |
| 2 | 36851901 | rs2372692 | 0.342 | 0.021 | -0.817 | -0.184 | -0.004 | -0.027 | -0.027 | 0.727 | 0.55 | 0.52 | 0.71 | 0.79 | 0.6 | 0.47 |
| 2 | 36854548 | rs7600979 | -0.379 | 0.563 | -0.753 | -0.505 | 0.001 | -0.063 | -0.063 | 0 | 0.38 | 0.45 | 0.28 | 0.08 | 0.39 | 0.38 |
| 2 | 36874250 | rs1012668 | -0.744 | 0.684 | -0.948 | -0.035 | -0.004 | 0.005 | 0.005 | 0 | 0.08 | 0.06 | 0.27 | 0.01 | 0.13 | 0.02 |
| 2 | 36877659 | rs10173568 | -0.566 | 0.605 | -0.888 | -0.013 | -0.001 | -0.002 | -0.002 | 0 | 0.46 | 0.47 | 0.54 | 0.4 | 0.45 | 0.58 |
| 2 | 36879521 | rs10167726 | 0.392 | 0.153 | -0.881 | 0.027 | 0.019 | 0.053 | 0.053 | 0 | 0.27 | 0.36 | 0.25 | 0.43 | 0.3 | 0.45 |
| 2 | 36880831 | rs6758291 | -1.017 | 0.264 | -0.817 | -0.19 | -0.002 | -0.001 | -0.001 | 0 | 0.54 | 0.55 | 0.61 | 0.5 | 0.54 | 0.65 |
| 2 | 36883218 | rs759432 | 0.824 | -0.093 | -0.884 | -0.444 | -0.034 | 0.211 | 0.211 | 0.582 | 0.4 | 0.15 | 0.31 | 0.1 | 0.43 | 0.25 |
| 2 | 36884084 | rs17489732 | 0.958 | -0.05 | -1.019 | -0.511 | -0.041 | 0.17 | 0.17 | 0.491 | 0.37 | 0.14 | 0.31 | 0.1 | 0.41 | 0.23 |
| 2 | 36884473 | rs12052493 | -0.666 | 0.217 | -0.576 | -0.071 | -0.007 | -0.022 | -0.022 | 0 | 0.13 | 0.05 | 0.31 | 0.46 | 0.18 | 0.12 |
| 2 | 36884839 | rs6733328 | 0.304 | -0.157 | -0.337 | -0.793 | -0.001 | -0.008 | -0.008 | 0 | 0.27 | 0.31 | 0.19 | 0.11 | 0.3 | 0.19 |
| 2 | 36887718 | rs9308993 | 0.431 | -0.271 | 0.221 | -0.912 | 0.002 | -0.039 | -0.039 | 0.422 | 0.57 | 0.4 | 0.67 | 0.83 | 0.58 | 0.57 |
| 2 | 36890869 | rs1861396 | 0.478 | -0.278 | 0.477 | -0.967 | -0.003 | 0.017 | 0.017 | 0 | 0.57 | 0.32 | 0.67 | 0.83 | 0.58 | 0.6 |
| 2 | 36895344 | rs6726841 | -1.176 | 0.294 | 0.938 | -0.663 | 0.006 | 0.008 | 0.008 | 0 | 0.49 | 0.5 | 0.64 | 0.62 | 0.57 | 0.46 |
| 2 | 36895728 | rs4670582 | 1.254 | 0.193 | 1.008 | -0.769 | 0.018 | 0.026 | 0.026 | 0 | 0.45 | 0.22 | 0.31 | 0.38 | 0.4 | 0.49 |
| 2 | 36900275 | rs2372707 | 0.677 | 0.114 | 0.976 | -0.783 | -0.008 | 0.058 | 0.058 | 0 | 0.73 | 0.39 | 0.54 | 0.5 | 0.71 | 0.75 |
| 2 | 36902056 | rs7587877 | -1.358 | 0.241 | 0.726 | -0.94 | 0.01 | 0.014 | 0.014 | 0.268 | 0.5 | 0.47 | 0.62 | 0.62 | 0.55 | 0.5 |
| 2 | 36902070 | rs7602835 | -1.772 | 0.178 | 0.591 | -0.95 | -0.004 | -0.006 | -0.006 | 0 | 0.55 | 0.53 | 0.64 | 0.62 | 0.57 | 0.53 |
| 2 | 36905013 | rs6714112 | 0.012 | 0.202 | 0.529 | -0.58 | -0.001 | -0.001 | -0.001 | 0 | 0.15 | 0.17 | 0.13 | 0.18 | 0.12 | 0.24 |
| 2 | 36919337 | rs6723801 | 1.103 | 0.239 | 0.514 | -0.585 | -0.003 | -0.008 | -0.008 | 0.377 | 0.69 | 0.33 | 0.49 | 0.37 | 0.67 | 0.66 |
| 2 | 36920273 | rs7604443 | -0.254 | 0.456 | 0.345 | -0.379 | -0.011 | 0.045 | 0.045 | 4.031 | 0.24 | 0.14 | 0.16 | 0.02 | 0.25 | 0.19 |
| 2 | 36924663 | rs12989101 | -1.101 | 0.409 | 0.32 | -0.351 | -0.006 | 0.036 | 0.036 | 1.321 | 0.36 | 0.32 | 0.26 | 0.18 | 0.38 | 0.34 |
| 2 | 36924893 | rs17019460 | 0.277 | 0.467 | 0.108 | -0.181 | -0.009 | -0.01 | -0.01 | 0 | 0.07 | 0.07 | 0.04 | 0.02 | 0.02 | 0.11 |
| 2 | 36926371 | rs17490768 | 0.789 | 0.648 | 0.244 | -0.164 | 0.019 | -0.046 | -0.046 | 0 | 0.65 | 0.44 | 0.72 | 0.82 | 0.59 | 0.69 |
| 2 | 36927283 | rs17019472 | 0.132 | 0.626 | -0.206 | 0.063 | -0.003 | 0.053 | 0.053 | 0 | 0.14 | 0.13 | 0.35 | 0.45 | 0.14 | 0.22 |
| 2 | 36928030 | rs6544028 | -0.844 | 0.741 | -0.474 | 0.18 | 0.058 | 0.049 | 0.049 | 0 | 0.89 | 0.6 | 0.89 | 0.84 | 0.89 | 0.85 |
| 2 | 36928948 | rs12472756 | -1.293 | 0.725 | -0.54 | 0.075 | -0.001 | 0.004 | 0.004 | 0.311 | 0.43 | 0.46 | 0.56 | 0.47 | 0.45 | 0.4 |
| 2 | 36929420 | rs4670591 | 1.673 | 0.733 | -0.649 | 0.08 | 0.005 | 0.036 | 0.036 | 0.174 | 0.47 | 0.19 | 0.34 | 0.37 | 0.44 | 0.46 |

| | | | | | | | | | | | | | | | | |
|---|---|---|---|---|---|---|---|---|---|---|---|---|---|---|---|---|
| 2 | 36930423 | rs6544029 | -1.165 | 1.03 | -0.681 | 0.195 | 0.043 | 0.033 | 0.033 | 0 | 0.9 | 0.66 | 0.89 | 0.84 | 0.89 | 0.85 |
| 2 | 36932843 | rs10460526 | 1.548 | 0.971 | -1.766 | 0.342 | -0.048 | 0.066 | 0.066 | 0 | 0.41 | 0.1 | 0.31 | 0.25 | 0.41 | 0.44 |
| 2 | 36934513 | rs6713326 | -0.21 | 0.923 | -1.397 | 0.577 | -0.053 | -0.004 | -0.004 | 0.076 | 0.18 | 0.04 | 0.13 | 0.15 | 0.24 | 0.15 |
| 2 | 36934887 | rs10182004 | 0.692 | 0.95 | -0.889 | 0.582 | 0.005 | -0.003 | -0.003 | 0 | 0.53 | 0.43 | 0.41 | 0.5 | 0.54 | 0.53 |
| 2 | 36935291 | rs13403956 | -0.847 | 1.242 | -0.674 | 1.173 | 0.007 | 0.06 | 0.06 | 0 | 0.14 | 0.22 | 0.42 | 0.34 | 0.18 | 0.14 |
| 2 | 36935406 | rs12712516 | 0.507 | 1.336 | -0.803 | 0.394 | 0.008 | -0.013 | -0.013 | 0 | 0.75 | 0.68 | 0.78 | 0.82 | 0.7 | 0.62 |
| 2 | 36935570 | rs13029062 | 0.591 | 0.935 | -0.973 | 0.145 | -0.002 | 0.053 | 0.053 | 0 | 0.58 | 0.33 | 0.49 | 0.22 | 0.62 | 0.39 |
| 2 | 36936830 | rs1990640 | -0.983 | 0.49 | -1.136 | 0.216 | -0.036 | 0.098 | 0.098 | 0 | 0.55 | 0.83 | 0.65 | 0.73 | 0.53 | 0.67 |
| 2 | 36937778 | rs7355582 | -1.895 | -0.534 | 0.102 | 1.067 | -0.028 | 0.108 | 0.108 | 0 | 0.13 | 0.39 | 0.13 | 0.26 | 0.08 | 0.3 |
| 2 | 36938368 | rs7593468 | -1.467 | -0.494 | 0.124 | 1.11 | -0.02 | 0.051 | 0.051 | 0 | 0.11 | 0.25 | 0.11 | 0.26 | 0.07 | 0.28 |
| 2 | 36939159 | rs10197807 | -1.555 | -0.493 | 0.143 | 1.11 | -0.024 | 0.064 | 0.064 | 0 | 0.11 | 0.29 | 0.11 | 0.26 | 0.07 | 0.29 |
| 2 | 36940596 | rs11691187 | 1.581 | -0.545 | -0.766 | 0.339 | -0.071 | 0.088 | 0.088 | 0 | 0.43 | 0.06 | 0.25 | 0.28 | 0.42 | 0.34 |
| 2 | 36942266 | rs4670165 | 1.257 | -0.761 | 1.119 | 1.331 | 0.001 | 0.001 | 0.001 | 0 | 0.65 | 0.39 | 0.6 | 0.38 | 0.64 | 0.57 |
| 2 | 36942730 | rs6746047 | 1.895 | -0.74 | 0.609 | 1.267 | 0.026 | 0.11 | 0.11 | 0 | 0.54 | 0.1 | 0.54 | 0.28 | 0.61 | 0.45 |
| 2 | 36946351 | rs7603396 | -0.89 | -0.619 | 1.261 | 1.26 | -0.009 | -0.029 | -0.029 | 0 | 0.07 | 0.27 | 0.05 | 0 | 0.04 | 0.03 |
| 2 | 36946433 | rs721348 | 2.137 | -0.654 | 1.507 | 1.294 | 0.005 | 0.027 | 0.027 | 0 | 0.54 | 0.32 | 0.56 | 0.28 | 0.56 | 0.43 |
| 2 | 36947218 | rs6744274 | 1.219 | -0.477 | 1.637 | 1.44 | -0.021 | -0.031 | -0.031 | 0 | 0.85 | 0.59 | 0.86 | 0.91 | 0.88 | 0.83 |
| 2 | 36948760 | rs10176047 | 1.224 | -0.377 | 0.902 | 1.376 | 0 | -0.006 | -0.006 | 0 | 0.63 | 0.57 | 0.63 | 0.29 | 0.62 | 0.47 |
| 2 | 36955005 | rs4670166 | -1.131 | -0.849 | 1.113 | 1.983 | -0.025 | 0.097 | 0.097 | 0.039 | 0.2 | 0.38 | 0.26 | 0.4 | 0.18 | 0.27 |
| 2 | 36959486 | rs17019636 | -1.049 | -0.853 | 0.642 | 2.006 | 0.008 | -0.014 | -0.014 | 0 | 0.18 | 0.18 | 0.17 | 0.32 | 0.17 | 0.25 |
| 2 | 36959860 | rs1015696 | -0.59 | -0.871 | 0.632 | 1.998 | -0.004 | 0.025 | 0.025 | 0 | 0.13 | 0.18 | 0.12 | 0.32 | 0.11 | 0.22 |
| 2 | 36959885 | rs1015695 | 2.27 | -1.086 | 0.025 | 1.65 | -0.118 | 0.036 | 0.036 | 0.005 | 0.43 | 0.03 | 0.36 | 0.27 | 0.53 | 0.31 |
| 2 | 36961691 | rs10189296 | -0.498 | -0.914 | 0.769 | 1.631 | 0.001 | -0.001 | -0.001 | 0 | 0.15 | 0.09 | 0.12 | 0.19 | 0.16 | 0.16 |
| 2 | 36961723 | rs10201689 | -1.18 | -0.911 | 0.761 | 1.617 | -0.023 | 0.049 | 0.049 | 0 | 0.19 | 0.25 | 0.17 | 0.31 | 0.17 | 0.26 |
| 2 | 36963787 | rs10490664 | -0.611 | -0.892 | 0.289 | 1.18 | -0.08 | 0.109 | 0.109 | 0 | 0.17 | 0.04 | 0.12 | 0.03 | 0.23 | 0.19 |
| 2 | 36963955 | rs4670599 | 2.582 | -1.146 | 0.26 | 1.242 | -0.035 | 0.009 | 0.009 | 0 | 0.44 | 0.19 | 0.38 | 0.4 | 0.5 | 0.38 |
| 2 | 36964751 | rs2160337 | -0.773 | -0.846 | 0.327 | 1.047 | -0.084 | 0.107 | 0.107 | 0 | 0.18 | 0.04 | 0.12 | 0.03 | 0.24 | 0.22 |
| 2 | 36966684 | rs17019662 | -1.313 | -1.002 | 1.352 | 1.231 | -0.071 | 0.123 | 0.123 | 0.541 | 0.19 | 0.41 | 0.34 | 0.38 | 0.09 | 0.24 |
| 2 | 36966828 | rs17019664 | -1.292 | -0.916 | 1.543 | 1.069 | 0.008 | -0.019 | -0.019 | 0 | 0.2 | 0.28 | 0.17 | 0.11 | 0.24 | 0.28 |
| 2 | 36966900 | rs12712518 | 1.958 | -1.164 | 1.703 | 1.127 | 0.025 | 0.024 | 0.024 | 0.432 | 0.66 | 0.33 | 0.77 | 0.81 | 0.83 | 0.64 |
| 2 | 36967402 | rs13411045 | -1.601 | -0.86 | 2.006 | 1.336 | 0.004 | 0.003 | 0.003 | 0.907 | 0.23 | 0.2 | 0.15 | 0.1 | 0.13 | 0.3 |
| 2 | 36967445 | rs13411239 | -1.736 | -0.859 | 2.059 | 1.35 | -0.013 | -0.019 | -0.019 | 0 | 0.24 | 0.23 | 0.15 | 0.1 | 0.13 | 0.29 |
| 2 | 36974472 | rs1072218 | 1.46 | -0.764 | 1.407 | 1.512 | 0.035 | -0.014 | -0.014 | 0.063 | 0.58 | 0.17 | 0.46 | 0.47 | 0.66 | 0.5 |
| 2 | 36977116 | rs12469652 | -0.373 | -0.512 | 1.404 | 1.568 | -0.063 | -0.031 | -0.031 | 0 | 0.23 | 0.11 | 0.4 | 0.34 | 0.31 | 0.3 |
| 2 | 36977577 | rs10207138 | -1.011 | -0.228 | 2.138 | 1.687 | 0.005 | 0.012 | 0.012 | 0 | 0.42 | 0.36 | 0.51 | 0.44 | 0.39 | 0.53 |
| 2 | 36978749 | rs10865116 | -0.896 | -0.452 | 2.089 | 1.705 | 0.007 | 0.009 | 0.009 | 0 | 0.41 | 0.37 | 0.51 | 0.44 | 0.38 | 0.53 |
| 2 | 36979512 | rs4670600 | -0.005 | -0.417 | 1.507 | 1.868 | 0.001 | 0.036 | 0.036 | 0 | 0.89 | 0.84 | 0.93 | 0.86 | 0.89 | 0.86 |
| 2 | 36980698 | rs12997998 | 1.5 | -0.364 | 1.07 | 1.607 | -0.031 | 0.022 | 0.022 | 0.048 | 0.43 | 0.18 | 0.34 | 0.33 | 0.5 | 0.29 |
| 2 | 36985579 | rs2072528 | 0.584 | -0.348 | 1.388 | 1.216 | 0.001 | 0.003 | 0.003 | 0 | 0.62 | 0.69 | 0.49 | 0.61 | 0.64 | 0.52 |
| 2 | 36994439 | rs11124542 | -1.051 | -0.145 | 1.117 | 1.087 | -0.001 | -0.001 | -0.001 | 0.302 | 0.43 | 0.55 | 0.53 | 0.37 | 0.39 | 0.51 |
| 2 | 36996398 | rs4670605 | 1.472 | -0.139 | 1.483 | 1.312 | -0.005 | 0.067 | 0.067 | 5.579 | 0.92 | 0.4 | 0.94 | 0.94 | 0.99 | 0.99 |
| 2 | 37009282 | rs888055 | 0.735 | 0.313 | 0.807 | 1.416 | 0.002 | -0.018 | -0.018 | 0 | 0.73 | 0.65 | 0.78 | 0.81 | 0.77 | 0.56 |
| 2 | 37010574 | rs3821147 | 1.348 | 0.03 | -0.032 | 1.017 | -0.025 | 0.079 | 0.079 | 0 | 0.59 | 0.09 | 0.71 | 0.78 | 0.67 | 0.42 |
| 2 | 37016156 | rs11680457 | -0.284 | -0.035 | 1.557 | 0.818 | -0.001 | -0.014 | -0.014 | 0 | 0.08 | 0.09 | 0.03 | 0 | 0.07 | 0.06 |
| 2 | 37023688 | rs2372713 | 0.709 | 0.034 | 2.098 | 0.668 | -0.002 | -0.042 | -0.042 | 0 | 0.76 | 0.75 | 0.83 | 0.99 | 0.75 | 0.69 |
| 2 | 37024713 | rs12987048 | 1.581 | 0.07 | 2.411 | 0.644 | -0.002 | -0.133 | -0.133 | 0.71 | 0.66 | 0.54 | 0.79 | 0.99 | 0.67 | 0.64 |
| 2 | 37026421 | rs7606098 | 1.469 | 0.063 | 2.422 | 0.851 | -0.013 | -0.046 | -0.046 | 0.013 | 0.71 | 0.47 | 0.78 | 0.82 | 0.75 | 0.51 |
| 2 | 37035934 | rs2072526 | -2.744 | 0.011 | 2.239 | 1.175 | 0.036 | 0.276 | 0.276 | 0.321 | 0.06 | 0.3 | 0.07 | 0.25 | 0.05 | 0.17 |
| 2 | 37039362 | rs13417089 | -1.096 | -0.051 | 1.718 | 0.887 | 0.004 | 0.001 | 0.001 | 0.012 | 0.21 | 0.16 | 0.12 | 0.19 | 0.17 | 0.36 |
| 2 | 37044191 | rs13002466 | 1.414 | -0.135 | 1.913 | 0.642 | -0.03 | -0.117 | -0.117 | 0 | 0.82 | 0.44 | 0.87 | 0.99 | 0.86 | 0.83 |
| 2 | 37046119 | rs4670624 | 1.644 | -0.119 | 1.707 | 0.614 | -0.007 | -0.111 | -0.111 | 0.002 | 0.63 | 0.42 | 0.76 | 0.98 | 0.67 | 0.62 |
| 2 | 37046711 | rs4670627 | 1.999 | 0.044 | 0.779 | 0.508 | 0.031 | 0.168 | 0.168 | 0.119 | 0.53 | 0.14 | 0.68 | 0.79 | 0.58 | 0.41 |
| 2 | 37047353 | rs2160334 | 1.712 | 0.062 | 1.396 | 1.605 | -0.025 | 0.086 | 0.086 | 0 | 0.76 | 0.44 | 0.81 | 0.55 | 0.83 | 0.64 |
| 2 | 37048522 | rs974966 | -1.73 | 0.213 | 1.117 | 1.116 | -0.004 | -0.038 | -0.038 | 0 | 0.39 | 0.51 | 0.5 | 0.23 | 0.35 | 0.5 |
| 2 | 37048724 | rs6747593 | -1.072 | 0.133 | 0.506 | 1.05 | -0.042 | 0.044 | 0.044 | 1.204 | 0.29 | 0.1 | 0.43 | 0.23 | 0.32 | 0.45 |
| 2 | 37048883 | rs888054 | 1.638 | 0.195 | 0.454 | 1.059 | -0.023 | 0.137 | 0.137 | 0.884 | 0.57 | 0.11 | 0.46 | 0.75 | 0.61 | 0.47 |
| 2 | 37051428 | rs13002772 | 2.432 | -0.017 | -0.092 | 1.016 | -0.047 | 0.029 | 0.029 | 0 | 0.47 | 0.09 | 0.39 | 0.33 | 0.55 | 0.38 |
| 2 | 37052585 | rs2075206 | -1.357 | 0.004 | 0.68 | 0.94 | -0.001 | -0.009 | -0.009 | 0 | 0.35 | 0.32 | 0.46 | 0.49 | 0.36 | 0.46 |
| 2 | 37059108 | rs13020894 | 1.726 | 0.062 | 1.204 | 0.553 | -0.007 | 0.024 | 0.024 | 0.018 | 0.88 | 0.53 | 0.93 | 0.85 | 0.95 | 0.87 |
| 2 | 37065320 | rs2540939 | -0.807 | -0.023 | 1.313 | 0.563 | -0.015 | 0.005 | 0.005 | 0 | 0.06 | 0.1 | 0.01 | 0.15 | 0.02 | 0.12 |
| 2 | 37074951 | rs1861436 | 1.928 | -0.002 | 1.749 | 0.568 | 0.003 | 0.054 | 0.054 | 0 | 0.9 | 0.52 | 0.93 | 0.85 | 0.96 | 0.88 |
| 2 | 37117073 | rs888083 | -1.557 | 0.061 | 1.142 | 0.435 | -0.003 | 0 | 0 | 0 | 0.42 | 0.3 | 0.4 | 0.54 | 0.43 | 0.47 |
| 2 | 37121894 | rs11682966 | 0.364 | 0.054 | 1.072 | 0.445 | 0.008 | -0.008 | -0.008 | 0 | 0.05 | 0.04 | 0.01 | 0.15 | 0.02 | 0.1 |
| 2 | 37122552 | rs2717493 | 2.4 | 0.03 | 1.069 | 0.431 | -0.063 | 0.052 | 0.052 | 0 | 0.47 | 0.1 | 0.51 | 0.32 | 0.52 | 0.41 |
| 2 | 37130658 | rs6744560 | -1.17 | 0.014 | 0.968 | 0.321 | -0.001 | 0.012 | 0.012 | 0 | 0.37 | 0.28 | 0.26 | 0.03 | 0.37 | 0.32 |
| 2 | 37136398 | rs10490658 | -0.125 | 0.049 | 0.727 | 0.389 | -0.015 | 0.011 | 0.011 | 0 | 0.07 | 0.01 | 0.16 | 0.51 | 0.09 | 0.17 |
| 2 | 37136638 | rs13418962 | -2.633 | 0.083 | 0.88 | 0.193 | -0.009 | 0.048 | 0.048 | 0 | 0.54 | 0.75 | 0.48 | 0.69 | 0.49 | 0.61 |
| 2 | 37162975 | rs17497197 | -0.224 | 0.208 | 0.111 | 0.065 | -0.027 | 0.072 | 0.072 | 0 | 0.2 | 0.06 | 0.13 | 0.03 | 0.25 | 0.14 |
| 2 | 37178255 | rs10490657 | -1.568 | 0.217 | 0.408 | 0.224 | -0.013 | -0.012 | -0.012 | 0.336 | 0.41 | 0.5 | 0.37 | 0.68 | 0.37 | 0.44 |

| | | | | | | | | | | | | | | | | |
|---|---|---|---|---|---|---|---|---|---|---|---|---|---|---|---|---|
| 2 | 37189123 | rs3845779 | -1.462 | 0.122 | 0.323 | 0.128 | -0.022 | -0.047 | -0.047 | 0 | 0.19 | 0.37 | 0.15 | 0.15 | 0.13 | 0.27 |
| 2 | 37200033 | rs6739171 | -2.075 | 0.416 | -0.032 | -0.391 | -0.008 | 0.013 | 0.013 | 0 | 0.58 | 0.58 | 0.5 | 0.69 | 0.53 | 0.63 |
| 2 | 37204840 | rs12464180 | 1.568 | 0.407 | -0.081 | -0.458 | -0.001 | -0.014 | -0.014 | 0 | 0.05 | 0.04 | 0.03 | 0 | 0.03 | 0.03 |
| 2 | 37206683 | rs6716891 | -0.886 | 0.452 | 0.023 | -0.596 | -0.003 | -0.163 | -0.163 | 0.139 | 0.34 | 0.42 | 0.21 | 0.03 | 0.32 | 0.2 |
| 2 | 37213186 | rs11682782 | -1.398 | 0.4 | -0.182 | -0.393 | -0.005 | -0.075 | -0.075 | 0 | 0.23 | 0.06 | 0.28 | 0.66 | 0.22 | 0.42 |
| 2 | 37230728 | rs1007823 | -0.037 | 0.47 | -0.167 | -0.326 | -0.017 | 0.006 | 0.006 | 0 | 0.07 | 0.01 | 0.16 | 0.53 | 0.09 | 0.17 |
| 2 | 37236821 | rs11694342 | -1.022 | 0.47 | 0.022 | -0.342 | -0.034 | -0.11 | -0.11 | 0 | 0.18 | 0.03 | 0.26 | 0.52 | 0.2 | 0.33 |
| 2 | 37237450 | rs10176270 | -2.23 | 0.417 | -0.026 | -0.499 | -0.003 | 0.012 | 0.012 | 0 | 0.56 | 0.53 | 0.5 | 0.7 | 0.53 | 0.63 |
| 2 | 37241050 | rs17020125 | -1.016 | 0.442 | -0.097 | -0.41 | -0.027 | -0.108 | -0.108 | 0 | 0.18 | 0.05 | 0.27 | 0.53 | 0.19 | 0.33 |
| 2 | 37252235 | rs1008838 | -0.596 | 0.388 | 0.211 | -1.017 | -0.008 | -0.043 | -0.043 | 0 | 0.21 | 0.25 | 0.11 | 0.14 | 0.18 | 0.22 |
| 2 | 37260463 | rs17020147 | 0.219 | 0.411 | 0.146 | -0.996 | 0.002 | -0.075 | -0.075 | 0.054 | 0.17 | 0.26 | 0.13 | 0.02 | 0.15 | 0.08 |
| 2 | 37276651 | rs10495870 | -1.48 | 0.469 | 0.043 | -0.929 | -0.011 | -0.08 | -0.08 | 1.949 | 0.39 | 0.44 | 0.25 | 0.18 | 0.33 | 0.3 |
| 2 | 37277811 | rs13423928 | -1.765 | 0.349 | -0.219 | -0.94 | 0.01 | 0.109 | 0.109 | 0 | 0.51 | 0.59 | 0.34 | 0.17 | 0.44 | 0.46 |
| 2 | 37285412 | rs2540992 | -2.465 | 0.638 | -0.945 | -1.268 | -0.058 | 0.049 | 0.049 | 0.302 | 0.72 | 0.9 | 0.73 | 0.83 | 0.63 | 0.81 |
| 2 | 37290659 | rs2541009 | -2.458 | 0.71 | -0.439 | -1.361 | -0.058 | 0.053 | 0.053 | 0.399 | 0.72 | 0.91 | 0.75 | 0.85 | 0.64 | 0.81 |
| 2 | 37291320 | rs17498534 | 0.13 | 0.553 | -0.224 | -1.574 | -0.002 | 0.026 | 0.026 | 0.716 | 0.23 | 0.16 | 0.12 | 0.05 | 0.22 | 0.23 |
| 2 | 37292890 | rs11903252 | 0.156 | 0.676 | -0.182 | -0.968 | -0.011 | -0.024 | -0.024 | 5.971 | 0.26 | 0.13 | 0.4 | 0.33 | 0.24 | 0.39 |
| 2 | 37306151 | rs17020183 | 1.223 | 0.641 | -0.133 | -1.11 | 0.004 | -0.016 | -0.016 | 0.274 | 0.18 | 0.13 | 0.3 | 0.24 | 0.16 | 0.27 |
| 2 | 37310852 | rs6718647 | 1.006 | 0.659 | -0.09 | -1.187 | 0.001 | -0.021 | -0.021 | 0 | 0.08 | 0.07 | 0.04 | 0 | 0.08 | 0.08 |
| 2 | 37311339 | rs10188586 | 1.006 | 0.659 | -0.09 | -1.187 | 0.001 | -0.021 | -0.021 | 0.159 | 0.08 | 0.07 | 0.04 | 0 | 0.08 | 0.08 |
| 2 | 37317457 | rs6738282 | -0.364 | 1.18 | -0.315 | -1.263 | -0.056 | 0.012 | 0.012 | 0.047 | 0.3 | 0.14 | 0.45 | 0.26 | 0.36 | 0.43 |
| 2 | 37327244 | rs1127906 | -0.803 | 0.697 | 0.142 | -0.998 | 0.004 | 0.013 | 0.013 | 0 | 0.79 | 0.62 | 0.66 | 0.77 | 0.79 | 0.7 |
| 2 | 37336720 | rs4233920 | 0.619 | 0.488 | 0.34 | -0.886 | 0.047 | -0.007 | -0.007 | 0.04 | 0.94 | 0.76 | 0.95 | 1 | 0.95 | 0.93 |
| 2 | 37346151 | rs3770768 | 0.873 | 0.474 | 0.048 | -0.93 | -0.001 | 0.028 | 0.028 | 0.427 | 0.11 | 0.05 | 0.05 | 0.01 | 0.13 | 0.12 |
| 2 | 37354149 | rs4233921 | -1.287 | 0.428 | -0.126 | -0.902 | 0.006 | -0.012 | -0.012 | 0 | 0.37 | 0.29 | 0.3 | 0.59 | 0.34 | 0.39 |
| 2 | 37363107 | rs4648202 | 0.49 | 0.595 | -0.463 | -1.064 | 0.006 | -0.046 | -0.046 | 0.274 | 0.09 | 0.2 | 0.05 | 0 | 0.07 | 0.09 |
| 2 | 37367107 | rs4648186 | 0.633 | 0.659 | -0.861 | -1.045 | 0 | 0.033 | 0.033 | 0.251 | 0.13 | 0.06 | 0.06 | 0.01 | 0.13 | 0.12 |
| 2 | 37367701 | rs4648182 | 1.665 | 0.842 | -0.533 | -0.599 | 0.054 | 0.003 | 0.003 | 0 | 0.05 | 0.23 | 0.04 | 0 | 0.03 | 0.03 |
| 2 | 37368452 | rs2287350 | 1.312 | 1.051 | -1.583 | -0.468 | -0.063 | 0.059 | 0.059 | 0 | 0.33 | 0.08 | 0.32 | 0.19 | 0.41 | 0.24 |
| 2 | 37376314 | rs2307466 | 0.045 | 1.099 | -1.416 | -0.463 | -0.03 | 0.025 | 0.025 | 0 | 0.07 | 0 | 0.07 | 0.01 | 0.07 | 0.1 |
| 2 | 37383315 | rs12712526 | 0.821 | 1.037 | -0.969 | -0.349 | -0.002 | 0.017 | 0.017 | 0 | 0.44 | 0.35 | 0.43 | 0.72 | 0.49 | 0.4 |
| 2 | 37402955 | rs3845785 | -0.041 | 0.579 | -0.701 | -0.87 | 0.005 | -0.004 | -0.004 | 0 | 0.68 | 0.7 | 0.83 | 0.95 | 0.72 | 0.71 |
| 2 | 37407178 | rs4374352 | -0.266 | 0.681 | -0.382 | -0.725 | 0 | -0.001 | -0.001 | 0 | 0.26 | 0.23 | 0.42 | 0.8 | 0.26 | 0.48 |
| 2 | 37413145 | rs13022403 | -0.47 | 0.47 | -0.372 | -0.841 | -0.001 | -0.024 | -0.024 | 0 | 0.72 | 0.72 | 0.85 | 0.95 | 0.73 | 0.71 |
| 2 | 37414227 | rs6544062 | 1.18 | 0.458 | -0.372 | -0.787 | 0 | -0.011 | -0.011 | 0.238 | 0.46 | 0.45 | 0.41 | 0.15 | 0.48 | 0.24 |
| 2 | 37414431 | rs6724820 | 1.944 | 0.46 | -0.486 | -0.866 | -0.012 | 0.089 | 0.089 | 0.203 | 0.35 | 0.12 | 0.33 | 0.15 | 0.39 | 0.2 |
| 2 | 37417052 | rs7583355 | -0.45 | 0.297 | 0.003 | -0.853 | -0.001 | -0.025 | -0.025 | 0 | 0.73 | 0.73 | 0.86 | 0.96 | 0.74 | 0.74 |
| 2 | 37419985 | rs2540975 | -0.768 | 0.141 | -0.092 | -0.925 | -0.011 | 0.079 | 0.079 | 1.383 | 0.78 | 0.92 | 0.9 | 0.96 | 0.77 | 0.76 |
| 2 | 37428743 | rs2540973 | -0.365 | 0.419 | 0.104 | -0.773 | -0.003 | 0.12 | 0.12 | 0 | 0.68 | 0.86 | 0.86 | 0.96 | 0.7 | 0.72 |
| 2 | 37438764 | rs2239650 | 0.157 | 0.579 | 0.142 | 0.085 | 0.003 | 0.085 | 0.085 | 0 | 0.07 | 0.09 | 0.25 | 0.39 | 0.09 | 0.12 |
| 2 | 37438993 | rs1137437 | 1.71 | 0.462 | 0.406 | 0.248 | -0.006 | -0.002 | -0.002 | 0 | 0.37 | 0.26 | 0.37 | 0.35 | 0.43 | 0.23 |
| 2 | 37446288 | rs11689186 | 0.846 | 0.463 | 0.449 | 0.117 | -0.002 | 0.004 | 0.004 | 0 | 0.16 | 0.16 | 0.18 | 0.22 | 0.13 | 0.32 |
| 2 | 37454286 | rs10206380 | 0.672 | 0.599 | 0.733 | 0.168 | -0.005 | 0.015 | 0.015 | 0 | 0.55 | 0.38 | 0.54 | 0.61 | 0.56 | 0.55 |
| 2 | 37458711 | rs3213745 | 1.563 | 0.439 | 0.787 | 0.254 | -0.012 | -0.027 | -0.027 | 0 | 0.37 | 0.16 | 0.33 | 0.37 | 0.42 | 0.23 |
| 2 | 37460234 | rs2372888 | -0.598 | 0.442 | 0.977 | 0.105 | -0.002 | -0.099 | -0.099 | 0 | 0.67 | 0.63 | 0.82 | 0.96 | 0.68 | 0.71 |
| 2 | 37461139 | rs12712528 | 0.552 | 0.41 | 0.947 | 0.108 | 0.011 | 0.069 | 0.069 | 0.215 | 0.51 | 0.38 | 0.6 | 0.73 | 0.55 | 0.39 |
| 2 | 37461786 | rs6743961 | 0.552 | 0.41 | 0.947 | 0.108 | 0.011 | 0.069 | 0.069 | 0.215 | 0.51 | 0.38 | 0.6 | 0.73 | 0.55 | 0.39 |
| 2 | 37463620 | rs2430494 | -0.777 | 0.412 | 1.002 | 0.068 | -0.029 | 0.129 | 0.129 | 0 | 0.68 | 0.89 | 0.85 | 0.96 | 0.68 | 0.71 |
| 2 | 37467264 | rs2041840 | -0.704 | 0.36 | 0.873 | 0.083 | -0.005 | 0.097 | 0.097 | 0 | 0.67 | 0.85 | 0.86 | 0.96 | 0.69 | 0.72 |
| 2 | 37468564 | rs2041837 | -0.448 | 0.404 | 0.828 | 0.104 | -0.008 | -0.15 | -0.15 | 0 | 0.65 | 0.53 | 0.81 | 0.96 | 0.68 | 0.71 |
| 2 | 37473585 | rs11897101 | -1.858 | 0.378 | 0.634 | 0.253 | 0.01 | 0.114 | 0.114 | 0.82 | 0.18 | 0.38 | 0.34 | 0.61 | 0.15 | 0.24 |
| 2 | 37478477 | rs11124570 | -0.551 | 0.41 | 0.45 | 0.32 | 0.006 | 0.065 | 0.065 | 0 | 0.09 | 0.14 | 0.25 | 0.34 | 0.09 | 0.13 |
| 2 | 37479756 | rs10184179 | 2.056 | 0.415 | 0.33 | 0.243 | -0.101 | 0.096 | 0.096 | 0.101 | 0.32 | 0.02 | 0.29 | 0.12 | 0.41 | 0.15 |
| 2 | 37482525 | rs2300892 | 0.931 | 0.676 | 0.622 | 0.307 | -0.006 | -0.036 | -0.036 | 0 | 0.15 | 0.07 | 0.18 | 0.23 | 0.14 | 0.31 |
| 2 | 37489877 | rs3770764 | 1.294 | 0.771 | 0.732 | 0.298 | 0.008 | 0.025 | 0.025 | 0.45 | 0.41 | 0.33 | 0.34 | 0.25 | 0.4 | 0.24 |
| 2 | 37505105 | rs10460527 | -1.596 | 0.669 | 0.428 | 0.157 | 0.017 | 0.082 | 0.082 | 0 | 0.12 | 0.23 | 0.28 | 0.34 | 0.13 | 0.16 |
| 2 | 37506939 | rs3770761 | -0.061 | 0.675 | 0.344 | 0.163 | -0.003 | 0.042 | 0.042 | 0 | 0.07 | 0.08 | 0.24 | 0.34 | 0.08 | 0.12 |
| 2 | 37514564 | rs10178975 | -0.641 | 0.597 | 0.373 | -0.081 | 0.001 | 0.028 | 0.028 | 0 | 0.67 | 0.78 | 0.84 | 0.96 | 0.69 | 0.72 |
| 2 | 37517266 | rs3770759 | 1.664 | 0.619 | 0.321 | -0.013 | -0.01 | -0.048 | -0.048 | 0 | 0.37 | 0.17 | 0.33 | 0.41 | 0.43 | 0.24 |
| 2 | 37527941 | rs6726921 | -0.216 | 0.821 | 0.373 | -0.005 | -0.015 | 0.016 | 0.016 | 0 | 0.07 | 0.05 | 0.04 | 0 | 0.09 | 0.07 |
| 2 | 37536436 | rs4670686 | -0.136 | 0.719 | 0.374 | -0.134 | -0.005 | 0.06 | 0.06 | 0 | 0.59 | 0.71 | 0.81 | 0.96 | 0.62 | 0.69 |
| 2 | 37541060 | rs4670688 | -0.108 | 1.157 | 0.309 | 0.312 | 0.001 | 0.03 | 0.03 | 0 | 0.18 | 0.19 | 0.44 | 0.56 | 0.2 | 0.42 |
| 2 | 37543371 | rs6544066 | -1.366 | 1.158 | 0.279 | 0.304 | 0.023 | 0.053 | 0.053 | 0 | 0.27 | 0.38 | 0.51 | 0.56 | 0.3 | 0.5 |
| 2 | 37561357 | rs4670195 | 0.027 | 0.563 | 0.302 | 0.508 | -0.001 | 0.073 | 0.073 | 0 | 0.17 | 0.2 | 0.42 | 0.56 | 0.16 | 0.41 |
| 2 | 37575287 | rs3821142 | -0.084 | 0.555 | -0.017 | 0.352 | 0.002 | -0.043 | -0.043 | 0.015 | 0.61 | 0.62 | 0.75 | 0.9 | 0.58 | 0.66 |
| 2 | 37583621 | rs2058719 | -0.455 | -0.039 | 0.172 | 1.098 | 0.036 | 0.014 | 0.014 | 1.342 | 0.07 | 0.47 | 0.07 | 0 | 0.01 | 0.03 |
| 2 | 37586246 | rs7606760 | -0.463 | -0.011 | -0.319 | 1.103 | 0.037 | 0.009 | 0.009 | 1.748 | 0.07 | 0.49 | 0.08 | 0 | 0.01 | 0.03 |
| 2 | 37590284 | rs4384764 | -0.32 | 0.215 | -1.521 | 1.165 | 0.026 | -0.004 | -0.004 | 0 | 0.33 | 0.76 | 0.42 | 0.38 | 0.23 | 0.33 |
| 2 | 37592628 | rs2373000 | -0.739 | 0.495 | -1.396 | 1.029 | 0.013 | 0.085 | 0.085 | 0.005 | 0.5 | 0.84 | 0.65 | 0.65 | 0.4 | 0.66 |

| | | | | | | | | | | | | | | | | |
|---|---|---|---|---|---|---|---|---|---|---|---|---|---|---|---|---|
| 2 | 37595525 | rs3770748 | -0.526 | 0.745 | -0.223 | 1.355 | 0.029 | 0.181 | 0.181 | 0 | 0.11 | 0.23 | 0.21 | 0.53 | 0.12 | 0.3 |
| 2 | 37598421 | rs2373001 | -1.068 | 0.43 | 0.034 | 1.321 | 0.008 | 0.26 | 0.26 | 0 | 0.11 | 0.36 | 0.28 | 0.44 | 0.06 | 0.2 |
| 2 | 37599988 | rs4670697 | -0.991 | 0.446 | -0.01 | 1.336 | 0.016 | 0.256 | 0.256 | 0.882 | 0.07 | 0.2 | 0.26 | 0.42 | 0.05 | 0.16 |
| 2 | 37600427 | rs6708310 | -1.805 | 0.514 | -0.075 | 1.274 | 0.013 | 0.189 | 0.189 | 0.184 | 0.09 | 0.21 | 0.27 | 0.42 | 0.08 | 0.17 |
| 2 | 37601208 | rs1024816 | -1.393 | 0.314 | -0.231 | 0.966 | -0.01 | 0.093 | 0.093 | 1.653 | 0.17 | 0.27 | 0.31 | 0.42 | 0.1 | 0.2 |
| 2 | 37602230 | rs7593887 | -0.196 | 0.366 | -0.673 | 0.411 | -0.004 | -0.034 | -0.034 | 0 | 0.1 | 0.11 | 0.05 | 0 | 0.06 | 0.05 |
| 2 | 37603801 | rs888096 | -0.691 | 0.387 | -1.726 | 0.326 | -0.035 | 0.03 | 0.03 | 0 | 0.49 | 0.96 | 0.63 | 0.7 | 0.38 | 0.61 |
| 2 | 37604136 | rs10084460 | 0.425 | 0.935 | -1.79 | -0.245 | -0.053 | 0.189 | 0.189 | 0.176 | 0.27 | 0.01 | 0.2 | 0 | 0.29 | 0.1 |
| 2 | 37617484 | rs10490445 | -0.488 | 1.252 | -0.52 | 1.124 | 0.001 | 0.073 | 0.073 | 0 | 0.11 | 0.1 | 0.18 | 0.55 | 0.13 | 0.19 |
| 2 | 37619469 | rs6544073 | -1.378 | 1.35 | -0.672 | 1.111 | -0.009 | -0.026 | -0.026 | 0 | 0.18 | 0.11 | 0.27 | 0.59 | 0.22 | 0.35 |
| 2 | 37621836 | rs4670200 | -1.86 | 1.229 | 0.178 | 0.947 | -0.001 | 0.008 | 0.008 | 0.471 | 0.26 | 0.66 | 0.29 | 0.64 | 0.23 | 0.34 |
| 2 | 37629222 | rs10173794 | -1.39 | 1.019 | 1.207 | 1.07 | -0.007 | 0.077 | 0.077 | 0 | 0.18 | 0.24 | 0.17 | 0.56 | 0.17 | 0.27 |
| 2 | 37630993 | rs6708931 | -0.658 | 1.132 | 1.271 | 0.714 | 0.041 | -0.007 | -0.007 | 0 | 0.07 | 0.28 | 0.07 | 0.02 | 0.05 | 0.08 |
| 2 | 37646345 | rs6544078 | -0.984 | 1.164 | 0.662 | 0.738 | 0.082 | -0.01 | -0.01 | 0 | 0.09 | 0.46 | 0.14 | 0.03 | 0.07 | 0.09 |
| 2 | 37670926 | rs2373106 | -0.957 | 1.259 | -0.417 | 0.99 | -0.006 | -0.004 | -0.004 | 0.044 | 0.11 | 0.1 | 0.29 | 0.25 | 0.14 | 0.23 |
| 2 | 37676671 | rs7568035 | -1.328 | 1.144 | -0.177 | 0.831 | 0.006 | 0.009 | 0.009 | 0 | 0.21 | 0.68 | 0.37 | 0.26 | 0.2 | 0.31 |
| 2 | 37685248 | rs10460529 | -0.662 | 0.695 | -0.127 | 1.244 | -0.001 | 0.018 | 0.018 | 0 | 0.18 | 0.18 | 0.19 | 0.53 | 0.18 | 0.26 |
| 2 | 37687204 | rs4670720 | -1.182 | 0.243 | -1.274 | 0.366 | -0.089 | -0.057 | -0.057 | 0 | 0.57 | 0.94 | 0.49 | 0.62 | 0.45 | 0.6 |
| 2 | 37687520 | rs6733460 | 0.611 | -0.148 | -1.167 | 0.048 | -0.013 | 0.066 | 0.066 | 0 | 0.25 | 0.06 | 0.12 | 0.09 | 0.2 | 0.19 |
| 2 | 37687770 | rs1024810 | -0.465 | -0.036 | 0.185 | 0.018 | -0.01 | -0.054 | -0.054 | 0 | 0.36 | 0.41 | 0.25 | 0.1 | 0.27 | 0.31 |
| 2 | 37691130 | rs6748955 | -1.325 | -0.16 | 0.166 | -0.038 | 0 | -0.001 | -0.001 | 0 | 0.49 | 0.47 | 0.43 | 0.63 | 0.44 | 0.58 |
| 2 | 37694090 | rs10192261 | -0.61 | -0.168 | 0.641 | -0.045 | -0.002 | -0.005 | -0.005 | 0 | 0.4 | 0.28 | 0.3 | 0.63 | 0.38 | 0.46 |
| 2 | 37695748 | rs2216115 | -1.696 | -0.189 | 0.653 | -0.055 | -0.045 | -0.002 | -0.002 | 0.1 | 0.55 | 0.84 | 0.48 | 0.63 | 0.45 | 0.61 |
| 2 | 37700786 | rs6544081 | 0.244 | -0.015 | 0.478 | 0.086 | -0.002 | 0.054 | 0.054 | 0 | 0.25 | 0.09 | 0.12 | 0.1 | 0.2 | 0.19 |
| 2 | 37703223 | rs6705512 | 0.168 | -0.017 | 0.71 | 0.173 | 0.005 | 0.149 | 0.149 | 0 | 0.19 | 0.41 | 0.23 | 0.52 | 0.18 | 0.29 |
| 2 | 37704766 | rs960139 | -0.515 | 0.42 | -0.215 | -0.023 | -0.023 | 0.044 | 0.044 | 0 | 0.77 | 0.92 | 0.88 | 0.9 | 0.78 | 0.79 |
| 2 | 37713203 | rs12992054 | 0.363 | -0.116 | 0.118 | -0.186 | -0.016 | -0.016 | -0.016 | 0 | 0.43 | 0.31 | 0.49 | 0.46 | 0.48 | 0.39 |
| 2 | 37718181 | rs6708411 | -0.136 | -0.28 | 0.952 | -0.475 | -0.001 | 0.001 | 0.001 | 0 | 0.48 | 0.56 | 0.55 | 0.4 | 0.51 | 0.48 |
| 2 | 37725712 | rs7593333 | 0.446 | -0.319 | 1.031 | -0.492 | -0.032 | -0.021 | -0.021 | 0 | 0.39 | 0.22 | 0.45 | 0.39 | 0.44 | 0.33 |
| 2 | 37728467 | rs11124586 | 0.134 | -0.299 | 1.25 | -0.532 | 0.019 | 0.011 | 0.011 | 0 | 0.48 | 0.27 | 0.43 | 0.6 | 0.45 | 0.46 |
| 2 | 37729133 | rs6544086 | 0.335 | -0.281 | 1.562 | -0.53 | 0.011 | -0.034 | -0.034 | 0.014 | 0.92 | 0.84 | 0.9 | 0.99 | 0.9 | 0.81 |
| 2 | 37731460 | rs2192943 | -0.374 | -0.202 | 1.569 | -0.505 | 0.002 | -0.002 | -0.002 | 0 | 0.55 | 0.48 | 0.54 | 0.6 | 0.53 | 0.52 |
| 2 | 37731665 | rs960902 | 0.085 | -0.203 | 1.531 | -0.505 | -0.002 | 0 | 0 | 1.534 | 0.46 | 0.57 | 0.53 | 0.4 | 0.51 | 0.47 |
| 2 | 37735381 | rs10199137 | 0.126 | -0.237 | 1.255 | -0.515 | 0.001 | 0.009 | 0.009 | 0 | 0.5 | 0.51 | 0.5 | 0.62 | 0.51 | 0.51 |
| 2 | 37739512 | rs4353628 | -0.696 | -0.213 | 0.966 | -0.436 | -0.012 | -0.008 | -0.008 | 0.786 | 0.6 | 0.69 | 0.56 | 0.62 | 0.57 | 0.57 |
| 2 | 37742142 | rs10198396 | -0.119 | -0.009 | 0.128 | -0.335 | 0.005 | 0.018 | 0.018 | 0 | 0.52 | 0.32 | 0.5 | 0.62 | 0.55 | 0.55 |
| 2 | 37748084 | rs7559825 | 0.221 | 0.2 | -1.055 | -0.307 | -0.027 | -0.01 | -0.01 | 0 | 0.3 | 0.19 | 0.39 | 0.27 | 0.36 | 0.3 |
| 2 | 37748367 | rs7559976 | 0.166 | -0.09 | -1.33 | -0.753 | -0.024 | 0.012 | 0.012 | 0 | 0.56 | 0.77 | 0.6 | 0.66 | 0.52 | 0.55 |
| 2 | 37754020 | rs9309009 | -0.049 | -0.539 | -1.416 | -1.406 | -0.007 | -0.004 | -0.004 | 0 | 0.55 | 0.53 | 0.63 | 0.35 | 0.64 | 0.47 |
| 2 | 37768967 | rs6747182 | 1.336 | -0.437 | -1.166 | -1.395 | 0.007 | -0.124 | -0.124 | 0 | 0.23 | 0.36 | 0.23 | 0.03 | 0.25 | 0.15 |
| 2 | 37777752 | rs11124590 | 1.148 | -0.523 | -1.574 | -1.26 | -0.024 | -0.096 | -0.096 | 0 | 0.25 | 0.09 | 0.24 | 0.56 | 0.25 | 0.26 |
| 2 | 37779622 | rs4524108 | 0.514 | -0.233 | -1.59 | -1.142 | -0.094 | -0.022 | -0.022 | 0 | 0.36 | 0.11 | 0.46 | 0.3 | 0.44 | 0.35 |
| 2 | 37787271 | rs4670208 | 0.944 | -0.286 | -1.517 | -1.151 | -0.022 | -0.109 | -0.109 | 0 | 0.3 | 0.08 | 0.26 | 0.57 | 0.28 | 0.39 |
| 2 | 37791321 | rs4633908 | 0.283 | -0.318 | -1.499 | -1.23 | -0.094 | -0.018 | -0.018 | 0 | 0.39 | 0.12 | 0.46 | 0.32 | 0.46 | 0.36 |
| 2 | 37792066 | rs4670733 | 0.863 | -0.313 | -1.242 | -1.294 | -0.016 | -0.099 | -0.099 | 0 | 0.3 | 0.43 | 0.24 | 0.1 | 0.3 | 0.25 |
| 2 | 37797178 | rs11124592 | -0.62 | -0.217 | -1.181 | -1.133 | 0.003 | -0.005 | -0.005 | 0 | 0.63 | 0.67 | 0.73 | 0.41 | 0.71 | 0.59 |
| 2 | 37804408 | rs12712538 | 1.261 | -0.074 | -1.191 | -1.368 | -0.018 | -0.078 | -0.078 | 0.75 | 0.23 | 0.41 | 0.21 | 0.1 | 0.25 | 0.23 |
| 2 | 37804964 | rs6730320 | -0.435 | 0.189 | -0.958 | -0.963 | 0.005 | -0.003 | -0.003 | 0 | 0.63 | 0.69 | 0.71 | 0.37 | 0.71 | 0.61 |
| 2 | 37807010 | rs7560890 | -0.784 | 0.214 | -1.014 | -0.93 | -0.014 | -0.059 | -0.059 | 0.082 | 0.67 | 0.89 | 0.74 | 0.38 | 0.72 | 0.61 |
| 2 | 37807372 | rs6544093 | -0.378 | 0.366 | -1.113 | -0.879 | -0.013 | 0.001 | 0.001 | 0.019 | 0.62 | 0.52 | 0.7 | 0.37 | 0.71 | 0.6 |
| 2 | 37808868 | rs10173816 | -0.709 | 0.447 | -1.409 | -1.035 | 0.003 | 0.002 | 0.002 | 0 | 0.64 | 0.67 | 0.57 | 0.3 | 0.73 | 0.57 |
| 2 | 37811749 | rs2191847 | -0.181 | 1.215 | -1.474 | -0.493 | -0.035 | 0.015 | 0.015 | 0 | 0.27 | 0.13 | 0.24 | 0.2 | 0.33 | 0.22 |
| 2 | 37814827 | rs10490284 | -0.261 | 1.207 | -1.641 | -0.486 | -0.069 | 0.018 | 0.018 | 0 | 0.28 | 0.1 | 0.26 | 0.2 | 0.35 | 0.2 |
| 2 | 37815891 | rs17507744 | 0.05 | 0.848 | -1.605 | -0.628 | -0.045 | 0.034 | 0.034 | 0.854 | 0.21 | 0.04 | 0.17 | 0.09 | 0.22 | 0.11 |
| 2 | 37827519 | rs874427 | -0.265 | 0.724 | -0.708 | -0.84 | 0.004 | -0.078 | -0.078 | 0.131 | 0.59 | 0.53 | 0.69 | 0.94 | 0.59 | 0.6 |
| 2 | 37829977 | rs12712542 | -0.001 | 0.639 | -0.194 | -0.936 | -0.009 | 0.092 | 0.092 | 3.044 | 0.54 | 0.3 | 0.65 | 0.92 | 0.55 | 0.52 |
| 2 | 37831162 | rs4140806 | 0.065 | 0.574 | 0.213 | -0.987 | -0.004 | 0.077 | 0.077 | 6.035 | 0.53 | 0.3 | 0.64 | 0.92 | 0.55 | 0.53 |
| 2 | 37832330 | rs7605481 | -0.418 | 0.557 | 0.3 | -0.997 | 0.003 | -0.099 | -0.099 | 1.232 | 0.59 | 0.46 | 0.69 | 0.94 | 0.6 | 0.6 |
| 2 | 37834333 | rs2058403 | -0.063 | 0.536 | 0.368 | -0.994 | -0.001 | -0.04 | -0.04 | 0 | 0.55 | 0.4 | 0.65 | 0.92 | 0.55 | 0.53 |
| 2 | 37843019 | rs17021120 | -0.567 | 0.463 | 0.411 | -0.977 | 0.04 | 0.008 | 0.008 | 1.396 | 0.07 | 0.25 | 0.05 | 0.02 | 0.06 | 0.08 |
| 2 | 37844646 | rs2011966 | -0.996 | 0.456 | 0.42 | -0.981 | 0.036 | 0.002 | 0.002 | 1.515 | 0.07 | 0.33 | 0.05 | 0.02 | 0.06 | 0.08 |
| 2 | 37848136 | rs2041509 | -0.439 | 0.311 | 0.724 | -1.305 | -0.003 | 0.045 | 0.045 | 0.002 | 0.58 | 0.68 | 0.74 | 0.94 | 0.62 | 0.62 |
| 2 | 37851667 | rs13406277 | -0.892 | 0.37 | 0.422 | -1.237 | 0.002 | -0.023 | -0.023 | 0 | 0.08 | 0.17 | 0.1 | 0.02 | 0.1 | 0.13 |
| 2 | 37855222 | rs6729121 | -0.828 | 0.119 | 0.443 | -1.357 | -0.075 | 0.182 | 0.182 | 0.006 | 0.65 | 0.86 | 0.78 | 0.95 | 0.65 | 0.69 |
| 2 | 37855437 | rs13401336 | -1.261 | -0.073 | 0.863 | -0.854 | 0.012 | -0.021 | -0.021 | 1.026 | 0.08 | 0.24 | 0.1 | 0.02 | 0.1 | 0.12 |
| 2 | 37862717 | rs727805 | -0.596 | -0.323 | 0.069 | -0.077 | -0.009 | -0.137 | -0.137 | 0.011 | 0.26 | 0.05 | 0.42 | 0.54 | 0.23 | 0.41 |
| 2 | 37864069 | rs17021160 | -0.033 | -0.444 | 0.194 | 0.09 | 0.01 | 0.009 | 0.009 | 2.529 | 0.17 | 0.14 | 0.1 | 0.09 | 0.11 | 0.15 |
| 2 | 37865620 | rs2373292 | -1.351 | -0.545 | 0.889 | 0.064 | -0.028 | 0.067 | 0.067 | 0 | 0.28 | 0.52 | 0.46 | 0.55 | 0.23 | 0.41 |
| 2 | 37869494 | rs9742 | -1.741 | -0.66 | 0.883 | -0.145 | -0.013 | 0.027 | 0.027 | 0 | 0.33 | 0.57 | 0.49 | 0.59 | 0.29 | 0.44 |

| | | | | | | | | | | | | | | | | |
|---|---|---|---|---|---|---|---|---|---|---|---|---|---|---|---|---|
| 2 | 37872139 | rs10694 | -1.46 | -0.674 | 0.867 | -0.144 | -0.035 | 0.087 | 0.087 | 0 | 0.29 | 0.47 | 0.45 | 0.6 | 0.24 | 0.41 |
| 2 | 37872508 | rs7735 | -1.623 | -0.676 | 0.718 | -0.145 | -0.033 | 0.047 | 0.047 | 0 | 0.32 | 0.53 | 0.47 | 0.6 | 0.25 | 0.43 |
| 2 | 37872871 | rs15628 | 0.78 | -0.48 | 0.042 | -0.015 | 0.013 | -0.002 | -0.002 | 0 | 0.12 | 0.02 | 0.08 | 0.1 | 0.09 | 0.14 |
| 2 | 37877664 | rs12474331 | -0.072 | -0.453 | 0.061 | 0.078 | -0.014 | 0.014 | 0.014 | 0 | 0.06 | 0 | 0.3 | 0.49 | 0.07 | 0.24 |
| 2 | 37880588 | rs12622958 | -1.109 | -0.546 | 0.384 | -0.094 | -0.016 | 0.049 | 0.049 | 0 | 0.31 | 0.41 | 0.45 | 0.6 | 0.27 | 0.44 |
| 2 | 37886901 | rs4670210 | -0.575 | -0.202 | -0.076 | 0.023 | -0.003 | -0.034 | -0.034 | 0.627 | 0.26 | 0.25 | 0.44 | 0.61 | 0.27 | 0.37 |
| 2 | 37889438 | rs6710421 | 0.75 | -0.116 | -0.354 | -0.244 | -0.008 | -0.008 | -0.008 | 0 | 0.31 | 0.31 | 0.25 | 0.32 | 0.39 | 0.21 |
| 2 | 37893397 | rs11692666 | -0.119 | -0.17 | -0.762 | -0.241 | 0.007 | -0.091 | -0.091 | 0.211 | 0.34 | 0.09 | 0.54 | 0.55 | 0.32 | 0.52 |
| 2 | 37895663 | rs11124596 | -0.033 | -0.092 | -0.771 | -0.23 | -0.009 | -0.097 | -0.097 | 0 | 0.33 | 0.09 | 0.54 | 0.53 | 0.33 | 0.51 |
| 2 | 37897230 | rs4670757 | 0.338 | -0.198 | -0.367 | -0.684 | 0.006 | 0.016 | 0.016 | 0.777 | 0.28 | 0.15 | 0.21 | 0.07 | 0.25 | 0.33 |
| 2 | 37903390 | rs7608326 | -0.36 | -0.084 | 0.147 | -0.648 | 0.022 | -0.05 | -0.05 | 0.198 | 0.09 | 0.25 | 0.06 | 0 | 0.08 | 0.05 |
| 2 | 37908135 | rs2888542 | -0.358 | -0.401 | -0.474 | -1.204 | -0.001 | 0.002 | 0.002 | 0.017 | 0.71 | 0.79 | 0.79 | 0.9 | 0.68 | 0.88 |
| 2 | 37909108 | rs11683907 | -0.221 | -1.48 | 0.04 | -1.277 | -0.024 | -0.085 | -0.085 | 0 | 0.34 | 0.57 | 0.28 | 0.09 | 0.24 | 0.4 |
| 2 | 37911571 | rs11885952 | 0.09 | -1.289 | 0.128 | -1.141 | -0.002 | -0.091 | -0.091 | 0 | 0.16 | 0.32 | 0.11 | 0.01 | 0.12 | 0.13 |
| 2 | 37916060 | rs3845787 | -0.293 | -1.389 | -0.663 | -0.808 | 0.033 | -0.023 | -0.023 | 0 | 0.39 | 0.38 | 0.53 | 0.53 | 0.27 | 0.66 |
| 2 | 37917363 | rs10195538 | -0.82 | -1.27 | -0.779 | -0.796 | 0.005 | -0.004 | -0.004 | 1.443 | 0.57 | 0.61 | 0.65 | 0.54 | 0.39 | 0.78 |
| 2 | 37917773 | rs10172526 | -0.759 | -0.735 | -0.348 | -0.246 | -0.019 | 0.054 | 0.054 | 0 | 0.32 | 0.46 | 0.46 | 0.49 | 0.25 | 0.5 |
| 2 | 37926630 | rs12052821 | -0.143 | -0.581 | -0.334 | -0.211 | -0.006 | 0.15 | 0.15 | 0 | 0.11 | 0.22 | 0.34 | 0.59 | 0.12 | 0.25 |
| 2 | 37926844 | rs7568405 | -0.682 | -0.481 | -0.805 | -1.042 | -0.001 | -0.001 | -0.001 | 0 | 0.61 | 0.45 | 0.76 | 0.65 | 0.62 | 0.71 |
| 2 | 37927161 | rs12471294 | -0.463 | -0.537 | -1.233 | -1.414 | -0.001 | -0.012 | -0.012 | 0.002 | 0.38 | 0.34 | 0.52 | 0.62 | 0.4 | 0.34 |
| 2 | 37932820 | rs2041502 | -0.413 | -0.696 | -1.295 | -1.481 | 0.036 | 0.008 | 0.008 | 0 | 0.44 | 0.25 | 0.59 | 0.62 | 0.36 | 0.66 |
| 2 | 37934470 | rs17463789 | 2.121 | -0.439 | -0.788 | -2.052 | -0.007 | -0.025 | -0.025 | 0.341 | 0.06 | 0.08 | 0.04 | 0.01 | 0.05 | 0.01 |
| 2 | 37936118 | rs12185536 | 0.326 | -0.358 | -0.877 | -2.113 | 0.004 | -0.032 | -0.032 | 2.95 | 0.19 | 0.22 | 0.18 | 0.04 | 0.27 | 0.06 |
| 2 | 37938985 | rs13391045 | 0.443 | -0.305 | -0.911 | -2.106 | -0.007 | -0.008 | -0.008 | 0.769 | 0.18 | 0.18 | 0.17 | 0.04 | 0.27 | 0.05 |
| 2 | 37940542 | rs4670766 | 1.147 | -0.435 | -0.617 | -2.122 | 0.001 | -0.033 | -0.033 | 0 | 0.19 | 0.23 | 0.21 | 0.03 | 0.17 | 0.36 |
| 2 | 37945865 | rs10189364 | 0.517 | -0.457 | -0.635 | -2.137 | -0.001 | -0.025 | -0.025 | 0 | 0.25 | 0.26 | 0.26 | 0.03 | 0.22 | 0.37 |
| 2 | 37949251 | rs1476873 | -0.182 | -0.24 | -0.832 | -1.945 | -0.013 | 0.022 | 0.022 | 0 | 0.24 | 0.22 | 0.22 | 0.04 | 0.33 | 0.07 |
| 2 | 37958369 | rs6725186 | -0.249 | -1.131 | -0.311 | -1.755 | -0.062 | -0.127 | -0.127 | 0 | 0.25 | 0.31 | 0.16 | 0.01 | 0.13 | 0.07 |
| 2 | 37960725 | rs6748492 | -0.684 | -0.827 | -0.842 | -1.839 | 0.04 | 0.102 | 0.102 | 0 | 0.55 | 0.74 | 0.4 | 0.03 | 0.36 | 0.47 |
| 2 | 37963500 | rs539892 | 0.06 | -1.013 | -0.549 | -2.014 | -0.008 | -0.094 | -0.094 | 0 | 0.36 | 0.56 | 0.34 | 0.03 | 0.31 | 0.44 |
| 2 | 37963797 | rs537116 | -0.793 | -0.889 | -0.741 | -1.819 | 0.026 | 0.11 | 0.11 | 0 | 0.55 | 0.72 | 0.43 | 0.07 | 0.4 | 0.49 |
| 2 | 37970848 | rs7600866 | 1.591 | -0.461 | -0.717 | -1.391 | -0.003 | -0.027 | -0.027 | 1.313 | 0.07 | 0.09 | 0.06 | 0 | 0.08 | 0.01 |
| 2 | 37982662 | rs600019 | -0.844 | 0.233 | -0.685 | -0.74 | 0.063 | 0.087 | 0.087 | 0 | 0.76 | 0.59 | 0.67 | 0.61 | 0.67 | 0.77 |
| 2 | 37986275 | rs10167624 | 0.608 | -0.305 | -1.357 | -1.242 | 0.005 | 0.05 | 0.05 | 0 | 0.1 | 0.02 | 0.05 | 0 | 0.09 | 0.03 |
| 2 | 37987124 | rs7566658 | 0.642 | -0.511 | -1.256 | -1.262 | 0.009 | 0.01 | 0.01 | 0 | 0.14 | 0.11 | 0.08 | 0.04 | 0.08 | 0.05 |
| 2 | 37992910 | rs12471513 | -0.116 | -0.154 | -1.651 | -1.504 | -0.01 | 0.12 | 0.12 | 0.005 | 0.36 | 0.16 | 0.31 | 0.02 | 0.37 | 0.43 |
| 2 | 37995727 | rs12615742 | 0.308 | -0.102 | -1.296 | -0.616 | -0.003 | -0.004 | -0.004 | 0 | 0.4 | 0.4 | 0.3 | 0.47 | 0.46 | 0.45 |
| 2 | 37997327 | rs1897094 | 0.377 | 0.38 | -1.695 | -0.75 | -0.007 | 0.013 | 0.013 | 0 | 0.13 | 0.06 | 0.09 | 0 | 0.16 | 0.04 |
| 2 | 37999909 | rs9973704 | -0.004 | 0.121 | -1.822 | -0.946 | 0.013 | 0.07 | 0.07 | 0 | 0.73 | 0.83 | 0.78 | 0.98 | 0.74 | 0.65 |
| 2 | 38002507 | rs7562836 | -0.814 | 0.276 | -1.818 | -0.574 | -0.006 | 0.008 | 0.008 | 0.052 | 0.29 | 0.24 | 0.22 | 0.22 | 0.28 | 0.13 |
| 2 | 38005048 | rs4670214 | -0.221 | -0.13 | -1.4 | 0.041 | -0.001 | -0.051 | -0.051 | 0 | 0.16 | 0.09 | 0.29 | 0.31 | 0.13 | 0.1 |
| 2 | 38008339 | rs4670775 | -0.397 | -0.407 | -1.164 | -0.317 | -0.009 | -0.047 | -0.047 | 0 | 0.34 | 0.42 | 0.34 | 0.09 | 0.25 | 0.43 |
| 2 | 38009280 | rs13404698 | 0.159 | 0.387 | -0.516 | 0.248 | -0.001 | -0.016 | -0.016 | 0 | 0.15 | 0.1 | 0.14 | 0.04 | 0.12 | 0.09 |
| 2 | 38009492 | rs6718541 | 0.056 | 0.374 | -0.523 | 0.242 | 0.002 | -0.009 | -0.009 | 0 | 0.16 | 0.11 | 0.14 | 0.04 | 0.12 | 0.09 |
| 2 | 38010266 | rs10165224 | -0.949 | 0.592 | -0.656 | 0.478 | -0.003 | -0.036 | -0.036 | 0 | 0.25 | 0.41 | 0.2 | 0.12 | 0.19 | 0.21 |
| 2 | 38010296 | rs10165323 | -0.672 | 0.464 | -0.485 | 0.642 | -0.009 | -0.032 | -0.032 | 0 | 0.22 | 0.41 | 0.19 | 0.12 | 0.17 | 0.21 |
| 2 | 38012224 | rs7567760 | -0.327 | 0.646 | -0.402 | 0.721 | 0.022 | 0.04 | 0.04 | 0.199 | 0.05 | 0.19 | 0.05 | 0.09 | 0.06 | 0.12 |
| 2 | 38014191 | rs1367148 | -1.678 | 0.581 | -0.579 | 1.056 | -0.013 | 0.011 | 0.011 | 0 | 0.39 | 0.52 | 0.47 | 0.55 | 0.3 | 0.39 |
| 2 | 38015298 | rs7571582 | -1.512 | 0.633 | -0.583 | 0.606 | -0.009 | 0 | 0 | 0.032 | 0.38 | 0.48 | 0.31 | 0.36 | 0.3 | 0.37 |
| 2 | 38020832 | rs744713 | -0.576 | 0.862 | -0.844 | 0.67 | 0.013 | 0.004 | 0.004 | 0 | 0.09 | 0.16 | 0.07 | 0.08 | 0.12 | 0.19 |
| 2 | 38021498 | rs867574 | 0.826 | 0.867 | -1.067 | 0.8 | -0.003 | -0.001 | -0.001 | 0 | 0.58 | 0.48 | 0.51 | 0.6 | 0.63 | 0.58 |
| 2 | 38021775 | rs10495873 | -0.499 | 0.756 | -1.36 | 1.154 | -0.011 | -0.04 | -0.04 | 0 | 0.12 | 0.06 | 0.12 | 0.24 | 0.12 | 0.15 |
| 2 | 38025151 | rs1430027 | 0.35 | 0.79 | -1.15 | 0.975 | 0.008 | 0.006 | 0.006 | 0 | 0.5 | 0.34 | 0.43 | 0.51 | 0.57 | 0.54 |
| 2 | 38025672 | rs6727303 | -0.998 | 1.054 | -0.526 | 1.673 | 0.077 | 0.212 | 0.212 | 0.074 | 0.06 | 0.24 | 0.15 | 0.19 | 0.08 | 0.11 |
| 2 | 38026026 | rs6544109 | 0.69 | 0.364 | -0.756 | 1.221 | -0.032 | -0.011 | -0.011 | 0 | 0.84 | 0.29 | 0.84 | 0.87 | 0.91 | 0.96 |
| 2 | 38026176 | rs6544110 | 0.236 | 0.357 | -1.987 | 0.94 | 0.069 | 0.018 | 0.018 | 0 | 0.52 | 0.21 | 0.49 | 0.41 | 0.63 | 0.64 |
| 2 | 38027626 | rs1430034 | -0.048 | 0.656 | -2.678 | 0.622 | -0.083 | -0.083 | -0.083 | 2.871 | 0.4 | 0.09 | 0.38 | 0.5 | 0.5 | 0.48 |
| 2 | 38029531 | rs2373305 | -0.317 | 1 | -2.501 | -0.147 | -0.039 | -0.017 | -0.017 | 0.52 | 0.33 | 0.21 | 0.51 | 0.65 | 0.46 | 0.38 |
| 2 | 38030131 | rs1529555 | 0.731 | 0.837 | -2.788 | -0.676 | 0.008 | 0.051 | 0.051 | 0 | 0.19 | 0.07 | 0.07 | 0 | 0.13 | 0.17 |
| 2 | 38035290 | rs1367150 | -1.57 | 0.474 | -2.047 | -0.898 | -0.011 | -0.049 | -0.049 | 0 | 0.84 | 0.73 | 0.9 | 0.97 | 0.85 | 0.84 |
| 2 | 38036428 | rs10865127 | 0.549 | 0.609 | -2.089 | -0.903 | -0.014 | -0.021 | -0.021 | 0 | 0.25 | 0.16 | 0.45 | 0.65 | 0.34 | 0.35 |
| 2 | 38037298 | rs6738882 | 0.646 | 0.535 | -1.754 | -0.704 | 0.008 | 0.039 | 0.039 | 0 | 0.2 | 0.1 | 0.08 | 0 | 0.13 | 0.18 |
| 2 | 38044344 | rs4670779 | 0.336 | 0.591 | -1.304 | -0.078 | -0.064 | -0.03 | -0.03 | 0 | 0.29 | 0.02 | 0.31 | 0.23 | 0.28 | 0.28 |
| 2 | 38047340 | rs11124614 | 0.29 | 0.628 | -0.508 | -0.064 | 0 | -0.001 | -0.001 | 0.634 | 0.31 | 0.3 | 0.31 | 0.31 | 0.28 | 0.29 |
| 2 | 38048671 | rs4670780 | 0.195 | 0.706 | -0.674 | -0.026 | -0.008 | 0 | 0 | 0 | 0.32 | 0.38 | 0.32 | 0.31 | 0.28 | 0.29 |
| 2 | 38048886 | rs7583451 | 0.027 | 0.699 | -0.743 | 0.048 | -0.006 | -0.01 | -0.01 | 0.414 | 0.36 | 0.46 | 0.34 | 0.31 | 0.34 | 0.3 |
| 2 | 38053004 | rs12617882 | -0.581 | 0.806 | -0.382 | 0.867 | 0.001 | 0.034 | 0.034 | 0.147 | 0.18 | 0.2 | 0.35 | 0.62 | 0.21 | 0.31 |
| 2 | 38054087 | rs7558738 | 0.991 | 0.68 | -0.063 | 0.316 | 0.011 | 0.019 | 0.019 | 0 | 0.57 | 0.38 | 0.63 | 0.31 | 0.59 | 0.48 |
| 2 | 38054623 | rs4670222 | 1.091 | 0.986 | 0.627 | 0.248 | 0 | 0.001 | 0.001 | 0 | 0.56 | 0.44 | 0.62 | 0.31 | 0.53 | 0.5 |

| | | | | | | | | | | | | | | | | |
|---|---|---|---|---|---|---|---|---|---|---|---|---|---|---|---|---|
| 2 | 38055097 | rs10495875 | -1.019 | 0.851 | 1.151 | 0.246 | 0.008 | 0.006 | 0.006 | 0.503 | 0.1 | 0.24 | 0.06 | 0.08 | 0.08 | 0.08 |
| 2 | 38055544 | rs6711697 | 0.696 | 0.873 | 1.326 | 0.398 | 0.012 | -0.012 | -0.012 | 0 | 0.67 | 0.75 | 0.7 | 0.39 | 0.7 | 0.58 |
| 2 | 38056403 | rs10206788 | 1.185 | 0.807 | 1.369 | 0.409 | -0.005 | -0.004 | -0.004 | 0.038 | 0.59 | 0.47 | 0.65 | 0.29 | 0.65 | 0.5 |
| 2 | 38056951 | rs4143265 | 0.64 | 0.946 | 1.523 | 0.101 | 0.045 | -0.031 | -0.031 | 0 | 0.85 | 0.52 | 0.86 | 0.93 | 0.85 | 0.96 |
| 2 | 38057628 | rs12712554 | 0.677 | 1.034 | 1.517 | 0.095 | 0.054 | -0.033 | -0.033 | 0.548 | 0.85 | 0.52 | 0.85 | 0.93 | 0.84 | 0.95 |
| 2 | 38060179 | rs12613672 | -0.97 | 0.954 | 1.504 | 0.102 | 0.014 | -0.007 | -0.007 | 0 | 0.37 | 0.26 | 0.31 | 0.64 | 0.33 | 0.48 |
| 2 | 38062895 | rs13019317 | 1.696 | 0.967 | 1.437 | 0.056 | 0.008 | 0.068 | 0.068 | 3.736 | 0.51 | 0.26 | 0.56 | 0.28 | 0.52 | 0.48 |
| 2 | 38066683 | rs17411119 | -0.396 | 0.833 | 1.303 | 0.208 | 0.001 | 0.055 | 0.055 | 0.079 | 0.26 | 0.1 | 0.16 | 0.13 | 0.26 | 0.34 |
| 2 | 38066716 | rs7598641 | -0.758 | 0.829 | 1.405 | 0.214 | 0.003 | 0.015 | 0.015 | 0.825 | 0.28 | 0.21 | 0.18 | 0.2 | 0.26 | 0.35 |
| 2 | 38067593 | rs766231 | 1.718 | 0.921 | 1.419 | 0.178 | -0.002 | 0.019 | 0.019 | 0.244 | 0.53 | 0.36 | 0.56 | 0.35 | 0.52 | 0.49 |
| 2 | 38067671 | rs766232 | 2.01 | 0.987 | 1.408 | 0.134 | 0.012 | 0.028 | 0.028 | 0.218 | 0.51 | 0.35 | 0.54 | 0.35 | 0.43 | 0.49 |
| 2 | 38068167 | rs1346746 | 2.038 | 0.973 | 1.356 | 0.123 | 0.016 | 0.04 | 0.04 | 0.23 | 0.5 | 0.3 | 0.55 | 0.35 | 0.44 | 0.49 |
| 2 | 38069289 | rs1897090 | 0.18 | 0.918 | 1.523 | -0.159 | 0.022 | -0.075 | -0.075 | 1.653 | 0.87 | 0.72 | 0.88 | 1 | 0.85 | 0.97 |
| 2 | 38069839 | rs2115846 | -0.425 | 0.923 | 1.53 | -0.155 | 0.019 | -0.052 | -0.052 | 0.928 | 0.89 | 0.78 | 0.89 | 1 | 0.86 | 0.98 |
| 2 | 38072657 | rs11886923 | -0.557 | 0.907 | 1.411 | -0.111 | 0.011 | 0.156 | 0.156 | 0 | 0.09 | 0.16 | 0.17 | 0.54 | 0.16 | 0.13 |
| 2 | 38077208 | rs748169 | 2.322 | 0.894 | 1.266 | -0.127 | 0.011 | 0.069 | 0.069 | 0.665 | 0.46 | 0.2 | 0.53 | 0.27 | 0.41 | 0.48 |
| 2 | 38080454 | rs6741132 | -0.67 | 0.878 | 1.274 | -0.196 | 0 | -0.003 | -0.003 | 0 | 0.28 | 0.24 | 0.16 | 0.13 | 0.26 | 0.34 |
| 2 | 38084560 | rs2707249 | 1.455 | 0.777 | 1.316 | -0.176 | 0.019 | 0.049 | 0.049 | 0.082 | 0.74 | 0.56 | 0.71 | 0.39 | 0.69 | 0.83 |
| 2 | 38090387 | rs870904 | 2.251 | 0.614 | 1.183 | -0.268 | 0.003 | 0.044 | 0.044 | 1.442 | 0.44 | 0.25 | 0.54 | 0.28 | 0.41 | 0.47 |
| 2 | 38092624 | rs9967771 | -0.08 | 0.43 | 1.205 | -0.247 | 0.014 | 0.03 | 0.03 | 0.053 | 0.23 | 0.14 | 0.14 | 0.1 | 0.19 | 0.27 |
| 2 | 38092846 | rs1009671 | 1.294 | 0.533 | 1.372 | 0.091 | 0.002 | 0.062 | 0.062 | 0.286 | 0.77 | 0.59 | 0.74 | 0.44 | 0.79 | 0.83 |
| 2 | 38099981 | rs2565639 | 1.904 | 0.381 | 1.488 | 0.016 | 0.024 | 0.076 | 0.076 | 0 | 0.81 | 0.59 | 0.77 | 0.31 | 0.79 | 0.86 |
| 2 | 38101439 | rs2565640 | 1.3 | 0.315 | 1.552 | 0.016 | 0.025 | 0.168 | 0.168 | 0 | 0.85 | 0.53 | 0.78 | 0.35 | 0.86 | 0.84 |
| 2 | 38103789 | rs2247278 | 1.356 | 0.314 | 1.587 | 0.015 | 0.021 | 0.147 | 0.147 | 0 | 0.85 | 0.59 | 0.79 | 0.35 | 0.86 | 0.87 |
| 2 | 38104724 | rs7576160 | -0.807 | 0.219 | 1.628 | -0.119 | -0.004 | -0.045 | -0.045 | 0.519 | 0.25 | 0.34 | 0.15 | 0.16 | 0.25 | 0.32 |
| 2 | 38104813 | rs2565646 | 2.101 | 0.15 | 1.317 | -0.16 | 0.008 | 0.103 | 0.103 | 0.073 | 0.56 | 0.2 | 0.62 | 0.19 | 0.61 | 0.53 |
| 2 | 38105208 | rs17021562 | -0.114 | 0.154 | 1.386 | -0.145 | 0 | 0.091 | 0.091 | 0 | 0.07 | 0.1 | 0.14 | 0.63 | 0.07 | 0.11 |
| 2 | 38105877 | rs11886737 | -1.151 | 0.152 | 1.403 | -0.149 | 0 | -0.001 | -0.001 | 0.263 | 0.28 | 0.24 | 0.15 | 0.16 | 0.26 | 0.32 |
| 2 | 38106046 | rs2565649 | 2.326 | 0.162 | 1.389 | -0.155 | -0.01 | 0.122 | 0.122 | 0.11 | 0.54 | 0.19 | 0.61 | 0.19 | 0.54 | 0.52 |
| 2 | 38106831 | rs2707225 | 1.899 | 0.159 | 1.398 | -0.15 | 0.009 | 0.135 | 0.135 | 0.086 | 0.81 | 0.48 | 0.77 | 0.35 | 0.86 | 0.85 |
| 2 | 38107130 | rs6737255 | -0.588 | 0.18 | 1.41 | -0.109 | 0 | 0.02 | 0.02 | 0 | 0.26 | 0.18 | 0.14 | 0.11 | 0.25 | 0.3 |
| 2 | 38110530 | rs2707223 | 1.927 | 0.161 | 1.429 | -0.117 | 0.008 | 0.086 | 0.086 | 0.038 | 0.57 | 0.25 | 0.63 | 0.19 | 0.61 | 0.55 |
| 2 | 38110970 | rs2707221 | 0.624 | 0.202 | 1.422 | -0.092 | -0.005 | -0.085 | -0.085 | 1.653 | 0.73 | 0.63 | 0.84 | 0.88 | 0.75 | 0.68 |
| 2 | 38112226 | rs4670788 | -0.024 | 0.273 | 1.386 | -0.05 | -0.002 | 0.092 | 0.092 | 0 | 0.07 | 0.09 | 0.14 | 0.64 | 0.07 | 0.11 |
| 2 | 38112660 | rs934215 | -1.521 | 0.242 | 1.399 | -0.041 | 0.016 | 0.063 | 0.063 | 0 | 0.17 | 0.35 | 0.22 | 0.65 | 0.2 | 0.13 |
| 2 | 38114189 | rs1835317 | 0.797 | 0.233 | 1.405 | -0.049 | -0.011 | -0.063 | -0.063 | 0 | 0.71 | 0.64 | 0.83 | 0.84 | 0.74 | 0.7 |
| 2 | 38115194 | rs6706045 | -2.029 | 0.324 | 1.483 | 0.082 | 0.007 | -0.033 | -0.033 | 0 | 0.39 | 0.52 | 0.34 | 0.81 | 0.41 | 0.37 |
| 2 | 38115234 | rs17021600 | -0.42 | 0.272 | 1.393 | 0.09 | 0.003 | -0.024 | -0.024 | 0 | 0.07 | 0.16 | 0.05 | 0.01 | 0.06 | 0.02 |
| 2 | 38118608 | rs7582060 | -0.937 | 0.214 | 1.246 | 0.142 | -0.007 | -0.024 | -0.024 | 0 | 0.26 | 0.32 | 0.33 | 0.16 | 0.21 | 0.33 |
| 2 | 38119317 | rs2707273 | 0.405 | 0.406 | 1.268 | 0.139 | -0.017 | -0.076 | -0.076 | 0 | 0.75 | 0.54 | 0.65 | 0.88 | 0.79 | 0.7 |
| 2 | 38120846 | rs2707269 | 0.347 | 0.434 | 1.195 | 0.153 | -0.014 | -0.069 | -0.069 | 0.071 | 0.75 | 0.61 | 0.66 | 0.89 | 0.8 | 0.71 |
| 2 | 38121447 | rs17411686 | -1.386 | 0.518 | 1.232 | 0.939 | 0.009 | 0.033 | 0.033 | 0 | 0.16 | 0.26 | 0.2 | 0.7 | 0.2 | 0.14 |
| 2 | 38139081 | rs4670789 | -1.097 | -0.427 | 0.429 | 0.378 | -0.068 | -0.039 | -0.039 | 1.239 | 0.29 | 0.48 | 0.47 | 0.81 | 0.18 | 0.2 |
| 2 | 38140126 | rs6707023 | -0.953 | -0.494 | 0.991 | 0.578 | -0.006 | -0.001 | -0.001 | 0 | 0.27 | 0.25 | 0.45 | 0.75 | 0.18 | 0.2 |
| 2 | 38140888 | rs17021695 | -1.04 | -0.494 | 1.127 | 0.583 | -0.021 | -0.01 | -0.01 | 0 | 0.27 | 0.32 | 0.45 | 0.81 | 0.18 | 0.2 |
| 2 | 38146418 | rs4670791 | -1.137 | -0.559 | 1.133 | 0.569 | -0.05 | -0.024 | -0.024 | 0 | 0.25 | 0.44 | 0.46 | 0.84 | 0.17 | 0.2 |
| 2 | 38146519 | rs9309016 | -0.292 | -0.413 | 1.28 | 0.259 | 0.01 | -0.033 | -0.033 | 0 | 0.07 | 0.13 | 0.05 | 0 | 0.09 | 0.07 |
| 2 | 38146714 | rs17021734 | -0.83 | -0.496 | 1.37 | 0.243 | 0.036 | -0.019 | -0.019 | 1.666 | 0.22 | 0.09 | 0.43 | 0.75 | 0.17 | 0.19 |
| 2 | 38152707 | rs7584680 | -1.377 | -0.841 | 1.543 | -0.22 | -0.007 | -0.085 | -0.085 | 0 | 0.16 | 0.28 | 0.11 | 0.01 | 0.14 | 0.15 |
| 2 | 38152819 | rs6544115 | -1.926 | -0.768 | 1.036 | -0.057 | -0.011 | -0.013 | -0.013 | 0.365 | 0.3 | 0.51 | 0.21 | 0.23 | 0.3 | 0.26 |
| 2 | 38171937 | rs6544117 | -0.131 | -0.408 | 0.991 | -0.111 | -0.027 | 0.021 | 0.021 | 0 | 0.88 | 0.95 | 0.94 | 0.92 | 0.76 | 0.9 |
| 2 | 38173070 | rs4670798 | -0.115 | -0.396 | 1.057 | -0.113 | -0.018 | 0.016 | 0.016 | 0 | 0.88 | 0.93 | 0.94 | 0.92 | 0.77 | 0.9 |
| 2 | 38174964 | rs6716296 | 1.396 | -0.453 | 1.341 | -0.179 | 0.027 | -0.102 | -0.102 | 0 | 0.73 | 0.5 | 0.82 | 0.89 | 0.64 | 0.75 |
| 2 | 38175574 | rs1900234 | -0.004 | -0.424 | 1.206 | -0.161 | -0.025 | 0.021 | 0.021 | 0 | 0.88 | 0.95 | 0.95 | 0.92 | 0.77 | 0.9 |
| 2 | 38177525 | rs11687301 | 2.328 | -0.533 | 1.425 | 0.089 | 0.011 | 0.151 | 0.151 | 0 | 0.49 | 0.26 | 0.39 | 0.09 | 0.47 | 0.55 |
| 2 | 38178362 | rs2197561 | -1.541 | -0.684 | 1.468 | 0.234 | 0 | -0.001 | -0.001 | 0.086 | 0.31 | 0.34 | 0.47 | 0.66 | 0.25 | 0.28 |
| 2 | 38189887 | rs6544120 | -2.139 | -0.64 | 1.292 | 0.211 | -0.013 | -0.03 | -0.03 | 0 | 0.4 | 0.57 | 0.55 | 0.82 | 0.31 | 0.36 |
| 2 | 38207823 | rs6707190 | -2.51 | -0.656 | 1.042 | 0.195 | 0.001 | 0.283 | 0.283 | 0 | 0.51 | 0.9 | 0.64 | 0.91 | 0.49 | 0.46 |
| 2 | 38208233 | rs1446297 | -1.99 | -0.696 | 1.109 | 0.206 | 0.004 | 0.002 | 0.002 | 0 | 0.4 | 0.62 | 0.55 | 0.82 | 0.31 | 0.37 |
| 2 | 38211778 | rs6738545 | -1.991 | -0.724 | 1.09 | 0.191 | 0.01 | 0.011 | 0.011 | 0 | 0.41 | 0.64 | 0.56 | 0.82 | 0.31 | 0.35 |
| 2 | 38212781 | rs10193252 | -1.635 | -0.775 | 1.048 | 0.272 | -0.02 | -0.007 | -0.007 | 0.17 | 0.26 | 0.63 | 0.2 | 0.21 | 0.2 | 0.23 |
| 2 | 38216416 | rs1446296 | -2.868 | -0.778 | 0.822 | 0.191 | -0.085 | 0.256 | 0.256 | 0 | 0.54 | 0.96 | 0.65 | 0.91 | 0.5 | 0.43 |
| 2 | 38218281 | rs17021813 | -0.043 | -0.72 | 0.883 | 0.241 | 0.016 | -0.112 | -0.112 | 0 | 0.15 | 0.01 | 0.36 | 0.62 | 0.11 | 0.14 |
| 2 | 38219642 | rs6544127 | 0.391 | -0.639 | 1.081 | 0.2 | 0.007 | -0.01 | -0.01 | 0.261 | 0.1 | 0.13 | 0.05 | 0.07 | 0.18 | 0.09 |
| 2 | 38222086 | rs2373327 | -1.838 | -0.653 | 1.204 | 0.227 | -0.004 | -0.012 | -0.012 | 4.876 | 0.4 | 0.6 | 0.55 | 0.81 | 0.31 | 0.33 |
| 2 | 38225019 | rs17412043 | 0.369 | -0.654 | 1.226 | 0.229 | 0.015 | -0.011 | -0.011 | 0 | 0.1 | 0.18 | 0.06 | 0.09 | 0.18 | 0.1 |
| 2 | 38226830 | rs1562285 | -1.156 | -0.703 | 1.362 | 0.251 | 0.01 | -0.013 | -0.013 | 0 | 0.29 | 0.27 | 0.46 | 0.68 | 0.24 | 0.27 |
| 2 | 38227561 | rs2373329 | -1.779 | -0.583 | 1.15 | 0.159 | -0.003 | -0.01 | -0.01 | 0 | 0.4 | 0.63 | 0.56 | 0.82 | 0.31 | 0.34 |
| 2 | 38239356 | rs6544134 | -2.143 | -0.261 | 1.15 | 0.112 | 0.01 | 0.069 | 0.069 | 0 | 0.5 | 0.66 | 0.62 | 0.91 | 0.51 | 0.42 |

| | | | | | | | | | | | | | | | | |
|---|---|---|---|---|---|---|---|---|---|---|---|---|---|---|---|---|
| 2 | 38241710 | rs17412098 | 0.166 | -0.24 | 1.107 | 0.227 | 0.014 | -0.007 | -0.007 | 3.273 | 0.11 | 0.18 | 0.06 | 0.1 | 0.18 | 0.1 |
| 2 | 38244824 | rs11891536 | -1.352 | -0.278 | 1.291 | 0.787 | -0.026 | 0.07 | 0.07 | 0 | 0.3 | 0.49 | 0.47 | 0.55 | 0.23 | 0.26 |
| 2 | 38245716 | rs4670230 | -0.969 | -0.254 | 1.192 | 0.839 | 0.001 | -0.001 | -0.001 | 0 | 0.26 | 0.28 | 0.45 | 0.54 | 0.23 | 0.26 |
| 2 | 38251872 | rs12712574 | -1.323 | -0.204 | 1.199 | 0.506 | -0.014 | 0.013 | 0.013 | 0 | 0.31 | 0.42 | 0.47 | 0.7 | 0.25 | 0.28 |
| 2 | 38253013 | rs7595482 | -1.472 | -0.231 | 1.184 | 0.491 | -0.03 | 0.021 | 0.021 | 0 | 0.32 | 0.55 | 0.48 | 0.7 | 0.24 | 0.28 |
| 2 | 38255921 | rs336046 | -1.969 | 0.262 | 0.089 | -0.319 | -0.027 | 0.195 | 0.195 | 0 | 0.55 | 0.84 | 0.67 | 0.9 | 0.57 | 0.44 |
| 2 | 38261124 | rs336037 | -2.067 | 0.614 | -0.293 | -0.555 | -0.114 | 0.214 | 0.214 | 0 | 0.59 | 0.98 | 0.68 | 0.91 | 0.61 | 0.48 |
| 2 | 38261143 | rs336036 | -1.836 | 0.555 | 0.241 | -0.583 | 0.001 | 0.093 | 0.093 | 0 | 0.57 | 0.71 | 0.66 | 0.91 | 0.61 | 0.46 |
| 2 | 38261467 | rs336035 | -0.811 | 0.526 | 0.508 | -0.74 | 0.019 | -0.07 | -0.07 | 0 | 0.3 | 0.38 | 0.19 | 0.09 | 0.38 | 0.17 |
| 2 | 38265130 | rs336031 | -0.534 | 0.629 | 0.318 | -0.796 | -0.005 | -0.013 | -0.013 | 0.043 | 0.27 | 0.23 | 0.18 | 0.06 | 0.37 | 0.16 |
| 2 | 38266794 | rs17021886 | 0.617 | 0.494 | 0.416 | -0.851 | 0 | -0.019 | -0.019 | 0.889 | 0.07 | 0.08 | 0.04 | 0 | 0.07 | 0.01 |
| 2 | 38266996 | rs336030 | -2.13 | 0.491 | 0.657 | -0.883 | -0.001 | 0.101 | 0.101 | 0 | 0.49 | 0.67 | 0.6 | 0.89 | 0.54 | 0.47 |
| 2 | 38268718 | rs2116232 | 0.844 | 0.471 | 0.439 | -0.843 | 0.005 | -0.051 | -0.051 | 0 | 0.66 | 0.66 | 0.77 | 0.93 | 0.55 | 0.84 |
| 2 | 38272672 | rs10192788 | 2.437 | 0.472 | 0.232 | -0.826 | -0.091 | 0.211 | 0.211 | 0 | 0.41 | 0.03 | 0.31 | 0.09 | 0.38 | 0.52 |
| 2 | 38274908 | rs11124631 | 0.857 | 0.416 | 0.332 | -0.922 | -0.026 | 0.046 | 0.046 | 0 | 0.64 | 0.76 | 0.77 | 0.94 | 0.54 | 0.83 |
| 2 | 38277832 | rs6741148 | -1.452 | 0.168 | 0.634 | -0.48 | -0.032 | 0.058 | 0.058 | 1.268 | 0.14 | 0.7 | 0.13 | 0.16 | 0.08 | 0.14 |
| 2 | 38290916 | rs163084 | -0.011 | 0.549 | 0.548 | -0.925 | 0.002 | -0.034 | -0.034 | 3.718 | 0.18 | 0.19 | 0.15 | 0.02 | 0.18 | 0.1 |
| 2 | 38291704 | rs2256327 | -0.07 | 0.542 | 0.576 | -0.929 | 0.003 | -0.041 | -0.041 | 0 | 0.18 | 0.21 | 0.15 | 0.02 | 0.18 | 0.1 |
| 2 | 38295994 | rs9341266 | 0.23 | 0.386 | 0.924 | -0.868 | 0.005 | -0.031 | -0.031 | 1.402 | 0.07 | 0 | 0.05 | 0.08 | 0.06 | 0.11 |
| 2 | 38296890 | rs2855658 | 1.436 | 0.708 | 0.933 | -0.958 | -0.057 | 0.12 | 0.12 | 0.001 | 0.66 | 0.18 | 0.72 | 0.91 | 0.6 | 0.84 |
| 2 | 38297170 | rs10916 | 0.341 | 0.704 | 1.473 | -0.961 | 0.057 | -0.034 | -0.034 | 0 | 0.87 | 0.69 | 0.89 | 0.94 | 0.78 | 0.95 |
| 2 | 38297515 | rs162562 | 0.918 | 0.696 | 1.565 | -0.965 | 0.026 | -0.037 | -0.037 | 0.267 | 0.84 | 0.33 | 0.87 | 0.92 | 0.78 | 0.93 |
| 2 | 38298139 | rs1800440 | 1.809 | 0.393 | 0.518 | -2.128 | -0.056 | 0.11 | 0.11 | 0.032 | 0.18 | 0.01 | 0.13 | 0.01 | 0.2 | 0.2 |
| 2 | 38298150 | rs1056837 | 0.949 | 0.12 | 0.955 | -1.899 | -0.043 | 0.094 | 0.094 | 0.001 | 0.66 | 0.2 | 0.73 | 0.91 | 0.6 | 0.83 |
| 2 | 38298203 | rs1056836 | 1.012 | 0.116 | 0.9 | -1.901 | -0.056 | 0.123 | 0.123 | 0.002 | 0.66 | 0.18 | 0.72 | 0.91 | 0.6 | 0.83 |
| 2 | 38298877 | rs162561 | -1.178 | 0.211 | 1.006 | -1.82 | 0.029 | -0.005 | -0.005 | 2.1 | 0.92 | 0.87 | 0.93 | 0.94 | 0.83 | 0.95 |
| 2 | 38302794 | rs2551188 | 1.203 | -0.223 | 0.844 | -2.052 | -0.02 | -0.029 | -0.029 | 0 | 0.36 | 0.52 | 0.3 | 0.18 | 0.29 | 0.44 |
| 2 | 38306454 | rs162556 | -0.428 | -0.177 | 0.295 | -2.074 | -0.12 | 0.072 | 0.072 | 0.77 | 0.58 | 0.97 | 0.5 | 0.2 | 0.5 | 0.55 |
| 2 | 38306935 | rs4646429 | 1.536 | -0.15 | 0.44 | -1.979 | 0.014 | 0.04 | 0.04 | 0 | 0.33 | 0.17 | 0.28 | 0.17 | 0.29 | 0.43 |
| 2 | 38307509 | rs162555 | 0.66 | -0.131 | 0.11 | -1.997 | -0.002 | -0.015 | -0.015 | 0.075 | 0.16 | 0.17 | 0.14 | 0.02 | 0.18 | 0.1 |
| 2 | 38307643 | rs10175338 | 1.48 | -0.018 | 0.008 | -1.891 | 0.015 | 0.041 | 0.041 | 0 | 0.33 | 0.16 | 0.28 | 0.17 | 0.29 | 0.44 |
| 2 | 38307861 | rs10175368 | 1.578 | -0.005 | -0.232 | -1.885 | -0.017 | 0.024 | 0.024 | 0 | 0.32 | 0.06 | 0.27 | 0.17 | 0.29 | 0.43 |
| 2 | 38313940 | rs2447753 | 0.331 | 0.483 | 0.991 | -1.641 | 0.009 | -0.047 | -0.047 | 1.204 | 0.16 | 0.18 | 0.14 | 0.02 | 0.17 | 0.09 |
| 2 | 38315020 | rs193103 | 0.629 | 0.655 | 0.862 | -1.623 | 0.077 | -0.037 | -0.037 | 0 | 0.83 | 0.59 | 0.82 | 0.92 | 0.73 | 0.92 |
| 2 | 38315682 | rs149425 | 0.884 | 0.677 | 0.276 | -1.614 | 0.065 | -0.054 | -0.054 | 0 | 0.81 | 0.45 | 0.82 | 0.92 | 0.72 | 0.92 |
| 2 | 38316571 | rs151096 | 1.214 | 0.604 | -0.781 | -1.571 | -0.14 | 0.175 | 0.175 | 0.532 | 0.63 | 0.1 | 0.66 | 0.89 | 0.55 | 0.83 |
| 2 | 38322556 | rs151256 | -0.574 | -0.258 | -0.005 | -1.649 | 0.008 | -0.121 | -0.121 | 0 | 0.24 | 0.5 | 0.19 | 0.03 | 0.22 | 0.11 |
| 2 | 38326907 | rs232557 | -0.99 | -0.629 | -1.044 | -1.84 | -0.155 | 0.065 | 0.065 | 0.437 | 0.64 | 0.96 | 0.42 | 0.19 | 0.46 | 0.5 |
| 2 | 38326921 | rs10865134 | 1.246 | -0.815 | -0.995 | -1.821 | 0.001 | 0.002 | 0.002 | 0 | 0.16 | 0.05 | 0.13 | 0.11 | 0.13 | 0.23 |
| 2 | 38328073 | rs17021971 | 0.508 | -0.943 | -0.807 | -1.965 | -0.001 | -0.015 | -0.015 | 2.13 | 0.08 | 0.08 | 0.04 | 0 | 0.07 | 0.07 |
| 2 | 38328178 | rs232542 | -0.374 | -1.179 | -0.226 | -2.117 | 0.002 | -0.023 | -0.023 | 0 | 0.4 | 0.55 | 0.29 | 0.08 | 0.31 | 0.34 |
| 2 | 38328205 | rs9309022 | 1.459 | -1.099 | -0.268 | -2.03 | -0.001 | 0.05 | 0.05 | 0 | 0.16 | 0.05 | 0.13 | 0.05 | 0.13 | 0.24 |
| 2 | 38331037 | rs17022000 | 1.696 | -1.08 | -0.05 | -1.95 | 0.03 | 0.013 | 0.013 | 1.527 | 0.05 | 0.29 | 0.04 | 0.01 | 0.01 | 0 |
| 2 | 38336269 | rs11886702 | -0.766 | -1.131 | -0.167 | -2.022 | 0.008 | 0.091 | 0.091 | 0.236 | 0.32 | 0.75 | 0.2 | 0.02 | 0.24 | 0.19 |
| 2 | 38336730 | rs10490621 | 0.587 | -1.129 | -0.003 | -2.024 | 0.018 | -0.043 | -0.043 | 0 | 0.08 | 0.17 | 0.04 | 0 | 0.06 | 0.07 |
| 2 | 38338031 | rs11124634 | -1.066 | -1.187 | -0.268 | -2.063 | 0.008 | 0.186 | 0.186 | 0.672 | 0.34 | 0.81 | 0.21 | 0.03 | 0.25 | 0.23 |
| 2 | 38344244 | rs232543 | -0.23 | -1.111 | 0.316 | -1.971 | -0.021 | -0.066 | -0.066 | 0.001 | 0.26 | 0.51 | 0.14 | 0.12 | 0.19 | 0.15 |
| 2 | 38345128 | rs232544 | -0.329 | -1.016 | 0.334 | -1.925 | -0.008 | -0.038 | -0.038 | 0 | 0.25 | 0.63 | 0.15 | 0.13 | 0.19 | 0.15 |
| 2 | 38345461 | rs232545 | -1.744 | -1.104 | 0.075 | -2.076 | -0.064 | 0.213 | 0.213 | 0.268 | 0.53 | 0.94 | 0.35 | 0.18 | 0.39 | 0.45 |
| 2 | 38345836 | rs232547 | 0.353 | -0.842 | 0.524 | -1.901 | 0.004 | -0.124 | -0.124 | 0.532 | 0.21 | 0.38 | 0.12 | 0.04 | 0.18 | 0.11 |
| 2 | 38348653 | rs232553 | -0.444 | -0.871 | 0.097 | -1.889 | -0.018 | -0.051 | -0.051 | 0 | 0.26 | 0.61 | 0.16 | 0.1 | 0.18 | 0.15 |
| 2 | 38350814 | rs3795980 | 1.143 | -0.427 | -0.185 | -1.931 | -0.009 | 0.023 | 0.023 | 0 | 0.15 | 0.03 | 0.13 | 0.08 | 0.13 | 0.21 |
| 2 | 38352639 | rs2432661 | -0.677 | -0.488 | -0.023 | -1.979 | -0.008 | 0.037 | 0.037 | 0 | 0.36 | 0.25 | 0.27 | 0.09 | 0.34 | 0.38 |
| 2 | 38357806 | rs232622 | 0.103 | -0.47 | -0.268 | -2.016 | -0.008 | 0.075 | 0.075 | 0.705 | 0.22 | 0.07 | 0.13 | 0.01 | 0.19 | 0.1 |
| 2 | 38358664 | rs232620 | -1.167 | -0.552 | -0.174 | -2.064 | 0.009 | 0.167 | 0.167 | 1.091 | 0.39 | 0.17 | 0.27 | 0.05 | 0.33 | 0.34 |
| 2 | 38366453 | rs10171555 | -0.695 | -0.59 | 0.259 | -2.101 | -0.012 | -0.092 | -0.092 | 0 | 0.28 | 0.43 | 0.21 | 0.09 | 0.21 | 0.3 |
| 2 | 38367734 | rs2113389 | 0.301 | -0.58 | 0.059 | -2.079 | 0.014 | 0.044 | 0.044 | 0.595 | 0.19 | 0.06 | 0.16 | 0.08 | 0.15 | 0.23 |
| 2 | 38368231 | rs232610 | -1.63 | -0.148 | 0.321 | -2.095 | -0.006 | 0.073 | 0.073 | 0.116 | 0.47 | 0.65 | 0.34 | 0.12 | 0.44 | 0.42 |
| 2 | 38373552 | rs7593657 | -0.261 | -0.596 | 0.391 | -1.957 | -0.004 | -0.015 | -0.015 | 0 | 0.23 | 0.28 | 0.17 | 0.08 | 0.17 | 0.25 |
| 2 | 38375771 | rs232606 | -1.022 | -0.557 | 0.507 | -2.158 | -0.008 | -0.134 | -0.134 | 2.56 | 0.32 | 0.44 | 0.21 | 0.05 | 0.26 | 0.18 |
| 2 | 38376154 | rs232602 | -1.173 | -0.573 | 0.573 | -2.166 | -0.008 | -0.126 | -0.126 | 0.121 | 0.33 | 0.54 | 0.21 | 0.05 | 0.26 | 0.19 |
| 2 | 38377405 | rs184577 | -0.123 | -0.544 | -0.014 | -2.13 | 0 | 0.063 | 0.063 | 0.593 | 0.23 | 0.14 | 0.16 | 0.01 | 0.21 | 0.1 |
| 2 | 38379110 | rs10196846 | -0.528 | -0.504 | 0.158 | -1.795 | -0.014 | -0.052 | -0.052 | 0 | 0.23 | 0.24 | 0.17 | 0.04 | 0.15 | 0.23 |
| 2 | 38379307 | rs11677850 | -0.847 | -0.497 | 0.12 | -1.791 | -0.003 | -0.034 | -0.034 | 0 | 0.28 | 0.29 | 0.19 | 0.1 | 0.21 | 0.3 |
| 2 | 38380958 | rs6737183 | -0.855 | -0.524 | 0.571 | -1.494 | -0.026 | -0.055 | -0.055 | 0 | 0.11 | 0.42 | 0.06 | 0.02 | 0.04 | 0.03 |
| 2 | 38385164 | rs6730133 | -0.783 | -0.644 | 0.572 | -1.472 | 0.019 | 0.046 | 0.046 | 0.17 | 0.08 | 0.39 | 0.05 | 0.04 | 0.03 | 0.1 |
| 2 | 38385902 | rs232578 | -1.247 | -0.722 | 0.152 | -2.149 | 0.005 | 0.116 | 0.116 | 3.293 | 0.45 | 0.73 | 0.36 | 0.14 | 0.39 | 0.39 |
| 2 | 38387227 | rs232574 | 0.473 | -0.705 | -0.121 | -2.206 | -0.096 | 0.146 | 0.146 | 0.053 | 0.25 | 0.04 | 0.25 | 0.04 | 0.32 | 0.22 |
| 2 | 38390311 | rs232572 | -0.934 | -0.824 | -0.554 | -2.749 | 0 | -0.023 | -0.023 | 2.421 | 0.52 | 0.5 | 0.47 | 0.11 | 0.52 | 0.53 |

| | | | | | | | | | | | | | | | | |
|---|---|---|---|---|---|---|---|---|---|---|---|---|---|---|---|---|
| 2 | 38392313 | rs232571 | -1.332 | -1.031 | -0.708 | -2.799 | 0 | -0.057 | -0.057 | 0.586 | 0.58 | 0.55 | 0.49 | 0.12 | 0.56 | 0.54 |
| 2 | 38392952 | rs232570 | -1.717 | -0.582 | -0.8 | -2.796 | 0.001 | 0.078 | 0.078 | 0.092 | 0.64 | 0.8 | 0.54 | 0.17 | 0.6 | 0.55 |
| 2 | 38394363 | rs232569 | 0.922 | -0.618 | -1.009 | -2.769 | -0.063 | 0.117 | 0.117 | 2.011 | 0.25 | 0.09 | 0.26 | 0.04 | 0.32 | 0.21 |
| 2 | 38394739 | rs232567 | -1.055 | -0.593 | -0.911 | -2.719 | 0.001 | -0.058 | -0.058 | 0 | 0.55 | 0.54 | 0.48 | 0.11 | 0.52 | 0.53 |
| 2 | 38398122 | rs2432678 | -0.402 | -0.443 | -1.048 | -2.669 | -0.036 | 0.153 | 0.153 | 2.16 | 0.38 | 0.21 | 0.32 | 0.05 | 0.4 | 0.3 |
| 2 | 38399977 | rs447905 | -0.034 | -0.58 | -1.15 | -2.734 | -0.05 | 0.121 | 0.121 | 2.219 | 0.35 | 0.19 | 0.31 | 0.08 | 0.39 | 0.27 |
| 2 | 38400597 | rs416497 | -0.87 | -0.518 | -0.991 | -2.67 | -0.003 | -0.004 | -0.004 | 0 | 0.55 | 0.56 | 0.49 | 0.15 | 0.51 | 0.47 |
| 2 | 38404386 | rs400681 | -1.116 | -0.385 | -1.002 | -2.626 | -0.003 | 0.088 | 0.088 | 0 | 0.52 | 0.67 | 0.46 | 0.16 | 0.48 | 0.58 |
| 2 | 38405869 | rs13385748 | -0.132 | -0.37 | -0.816 | -2.616 | -0.018 | -0.118 | -0.118 | 0 | 0.25 | 0.5 | 0.2 | 0.07 | 0.19 | 0.33 |
| 2 | 38407966 | rs68352 | -0.445 | -0.51 | -0.981 | -2.57 | 0 | -0.019 | -0.019 | 0 | 0.42 | 0.38 | 0.4 | 0.08 | 0.43 | 0.46 |
| 2 | 38408537 | rs10490623 | 0.592 | -0.502 | -0.979 | -2.56 | 0.008 | 0.001 | 0.001 | 0.12 | 0.09 | 0.2 | 0.07 | 0.08 | 0.06 | 0.06 |
| 2 | 38410270 | rs2447739 | -0.995 | -0.539 | -1.007 | -2.582 | 0.01 | 0.041 | 0.041 | 0 | 0.49 | 0.59 | 0.44 | 0.13 | 0.45 | 0.46 |
| 2 | 38410895 | rs2432671 | -0.731 | -0.538 | -0.944 | -2.49 | -0.002 | 0.039 | 0.039 | 0.006 | 0.36 | 0.28 | 0.31 | 0.17 | 0.33 | 0.37 |
| 2 | 38415333 | rs170815 | -1.177 | -0.323 | -1.177 | -1.697 | 0 | 0 | 0 | 0 | 0.61 | 0.6 | 0.74 | 0.83 | 0.6 | 0.66 |
| 2 | 38420153 | rs232589 | -0.795 | -0.214 | -1.072 | -1.607 | 0.002 | 0.032 | 0.032 | 0 | 0.56 | 0.6 | 0.73 | 0.88 | 0.56 | 0.61 |
| 2 | 38420673 | rs232588 | -0.42 | -0.285 | -1.026 | -1.6 | 0 | 0 | 0 | 2.568 | 0.5 | 0.48 | 0.68 | 0.75 | 0.54 | 0.59 |
| 2 | 38425107 | rs232585 | 0.009 | -0.457 | -1.02 | -1.503 | -0.001 | 0.021 | 0.021 | 0.135 | 0.28 | 0.21 | 0.28 | 0.13 | 0.29 | 0.26 |
| 2 | 38427647 | rs4670823 | -0.122 | -0.407 | -0.891 | -1.274 | -0.005 | 0.009 | 0.009 | 0.841 | 0.3 | 0.49 | 0.46 | 0.71 | 0.28 | 0.4 |
| 2 | 38429763 | rs2261 | -0.793 | -0.497 | -0.892 | -1.457 | -0.007 | 0.041 | 0.041 | 0 | 0.53 | 0.36 | 0.68 | 0.75 | 0.52 | 0.62 |
| 2 | 38431610 | rs7606492 | -0.959 | -1.335 | -0.742 | -1.303 | -0.014 | -0.028 | -0.028 | 0 | 0.4 | 0.53 | 0.54 | 0.82 | 0.3 | 0.56 |
| 2 | 38432325 | rs10174646 | -0.914 | -1.503 | -0.709 | -1.39 | 0.005 | 0.006 | 0.006 | 0 | 0.46 | 0.47 | 0.55 | 0.73 | 0.34 | 0.58 |
| 2 | 38435241 | rs1030278 | -1.453 | -1.566 | -0.626 | -1.359 | 0.015 | 0.02 | 0.02 | 0 | 0.49 | 0.57 | 0.59 | 0.84 | 0.36 | 0.61 |
| 2 | 38436969 | rs4670824 | 0.554 | -1.738 | -0.723 | -1.399 | -0.004 | 0.013 | 0.013 | 0.07 | 0.25 | 0.31 | 0.43 | 0.65 | 0.25 | 0.34 |
| 2 | 38438575 | rs4670826 | 0.35 | -1.763 | -0.756 | -1.435 | -0.007 | 0.009 | 0.009 | 0.014 | 0.27 | 0.36 | 0.43 | 0.69 | 0.26 | 0.38 |
| 2 | 38441983 | rs1871163 | -1.725 | -1.812 | -0.559 | -1.711 | 0.011 | 0.014 | 0.014 | 0.054 | 0.49 | 0.57 | 0.59 | 0.86 | 0.39 | 0.65 |
| 2 | 38443883 | rs4142656 | -1.703 | -1.817 | -0.54 | -1.713 | 0.02 | 0.042 | 0.042 | 0.199 | 0.49 | 0.58 | 0.59 | 0.87 | 0.38 | 0.65 |
| 2 | 38444225 | rs13398184 | -1.857 | -1.836 | -0.423 | -1.725 | 0.023 | 0.08 | 0.08 | 0 | 0.53 | 0.65 | 0.6 | 0.87 | 0.39 | 0.65 |
| 2 | 38445779 | rs10183112 | 0.821 | -1.866 | -0.326 | -1.704 | -0.019 | -0.028 | -0.028 | 0 | 0.15 | 0.18 | 0.11 | 0.05 | 0.08 | 0.2 |
| 2 | 38447523 | rs6723404 | -0.659 | -1.802 | -0.186 | -1.661 | 0.027 | 0.042 | 0.042 | 0.489 | 0.42 | 0.3 | 0.53 | 0.78 | 0.33 | 0.6 |
| 2 | 38450009 | rs4670244 | -1.697 | -1.492 | -0.488 | -1.695 | 0.012 | 0.079 | 0.079 | 0 | 0.53 | 0.66 | 0.61 | 0.88 | 0.42 | 0.67 |
| 2 | 38459930 | rs3856501 | -0.724 | -1.404 | -0.443 | -1.278 | -0.008 | -0.018 | -0.018 | 1.608 | 0.41 | 0.5 | 0.56 | 0.76 | 0.38 | 0.61 |
| 2 | 38463790 | rs2373369 | -0.803 | -1.924 | -1.11 | -1.535 | 0.123 | 0.068 | 0.068 | 0 | 0.44 | 0.87 | 0.5 | 0.73 | 0.25 | 0.46 |
| 2 | 38466625 | rs11124635 | -0.095 | -1.898 | -0.511 | -1.519 | 0.003 | 0.001 | 0.001 | 0 | 0.34 | 0.64 | 0.46 | 0.69 | 0.27 | 0.38 |
| 2 | 38467524 | rs7608905 | 1.269 | -2.066 | -0.605 | -1.575 | -0.015 | 0.005 | 0.005 | 0 | 0.07 | 0.11 | 0.04 | 0.11 | 0.02 | 0.11 |
| 2 | 38469908 | rs12712581 | -1.501 | -2.193 | -0.85 | -1.753 | -0.038 | 0.177 | 0.177 | 0 | 0.56 | 0.96 | 0.63 | 0.85 | 0.43 | 0.7 |
| 2 | 38470492 | rs4101522 | 0.493 | -2.219 | -0.089 | -1.733 | -0.022 | 0.007 | 0.007 | 0 | 0.29 | 0.39 | 0.44 | 0.69 | 0.25 | 0.34 |
| 2 | 38471780 | rs10445918 | 1.298 | -1.766 | -0.22 | -1.576 | -0.015 | 0.005 | 0.005 | 0 | 0.07 | 0.11 | 0.03 | 0.11 | 0.02 | 0.11 |
| 2 | 38473627 | rs7604860 | -0.484 | -1.79 | 0.004 | -1.608 | -0.024 | -0.007 | -0.007 | 0 | 0.13 | 0.36 | 0.05 | 0.12 | 0.05 | 0.16 |
| 2 | 38478237 | rs3911040 | -1.773 | -1.891 | -0.462 | -1.73 | -0.121 | 0.104 | 0.104 | 0 | 0.61 | 0.98 | 0.65 | 0.85 | 0.45 | 0.71 |
| 2 | 38478503 | rs3845790 | -0.443 | -1.572 | 0.24 | -1.196 | -0.075 | 0.055 | 0.055 | 0 | 0.2 | 0.48 | 0.3 | 0.75 | 0.08 | 0.26 |
| 2 | 38482511 | rs7565694 | -0.218 | -1.64 | 0.201 | -1.224 | -0.038 | 0.041 | 0.041 | 0 | 0.19 | 0.27 | 0.34 | 0.69 | 0.07 | 0.26 |
| 2 | 38483796 | rs17022391 | 0.154 | -1.405 | 0.281 | -1.119 | -0.029 | 0.034 | 0.034 | 0 | 0.09 | 0.28 | 0.04 | 0.11 | 0.02 | 0.11 |
| 2 | 38487153 | rs12617934 | 0.544 | -1.294 | 0.087 | -1.162 | -0.037 | 0.072 | 0.072 | 0.014 | 0.19 | 0.37 | 0.31 | 0.65 | 0.1 | 0.16 |
| 2 | 38489285 | rs4670248 | 0.739 | -1.289 | -0.178 | -1.229 | 0.007 | -0.024 | -0.024 | 0.418 | 0.18 | 0.14 | 0.29 | 0.66 | 0.14 | 0.21 |
| 2 | 38490324 | rs13432312 | 0.586 | -1.354 | -0.356 | -1.357 | -0.034 | 0.041 | 0.041 | 0 | 0.12 | 0.01 | 0.11 | 0.05 | 0.13 | 0.23 |
| 2 | 38491227 | rs20505 | 1.092 | -1.277 | 0.083 | -1.225 | -0.002 | -0.034 | -0.034 | 0 | 0.17 | 0.19 | 0.16 | 0.04 | 0.19 | 0.19 |
| 2 | 38491993 | rs4670846 | -0.238 | -1.346 | 0.112 | -1.253 | -0.019 | -0.035 | -0.035 | 0 | 0.28 | 0.54 | 0.22 | 0.14 | 0.22 | 0.3 |
| 2 | 38492587 | rs1972974 | -0.468 | -1.202 | 0.11 | -1.151 | 0.028 | 0.033 | 0.033 | 0.18 | 0.39 | 0.73 | 0.48 | 0.7 | 0.33 | 0.38 |
| 2 | 38502247 | rs13391941 | -0.953 | -0.827 | 0.127 | -0.856 | 0.083 | 0.177 | 0.177 | 6.571 | 0.05 | 0.4 | 0.04 | 0.08 | 0.01 | 0.01 |
| 2 | 38504680 | rs4670855 | 1.076 | -0.731 | -0.49 | -0.511 | 0.008 | -0.053 | -0.053 | 0 | 0.22 | 0.14 | 0.33 | 0.54 | 0.17 | 0.22 |
| 2 | 38509319 | rs12468437 | -0.168 | -0.679 | -0.459 | -1.073 | 0.006 | 0.008 | 0.008 | 0 | 0.27 | 0.7 | 0.23 | 0.18 | 0.21 | 0.32 |
| 2 | 38510608 | rs6743640 | -0.289 | -0.747 | -0.401 | -1.058 | 0.016 | 0.004 | 0.004 | 0 | 0.36 | 0.66 | 0.4 | 0.7 | 0.22 | 0.38 |
| 2 | 38511405 | rs4670858 | -0.523 | -0.125 | -0.423 | -1.047 | 0.017 | 0.03 | 0.03 | 0 | 0.42 | 0.67 | 0.54 | 0.73 | 0.36 | 0.56 |
| 2 | 38513085 | rs2121897 | -0.911 | 0.391 | 0.179 | -0.77 | 0.001 | 0.026 | 0.026 | 0 | 0.35 | 0.68 | 0.36 | 0.12 | 0.37 | 0.49 |
| 2 | 38517515 | rs11894455 | -1.336 | 0.205 | 0.016 | -0.814 | 0.014 | 0.14 | 0.14 | 0 | 0.42 | 0.83 | 0.39 | 0.18 | 0.4 | 0.5 |
| 2 | 38520856 | rs4670861 | -1.297 | 0.297 | 0.108 | -0.799 | 0.001 | 0.105 | 0.105 | 0.018 | 0.4 | 0.8 | 0.39 | 0.19 | 0.4 | 0.5 |
| 2 | 38547749 | rs12712584 | -0.513 | -0.062 | 0.265 | -0.744 | -0.001 | 0.003 | 0.003 | 0.755 | 0.33 | 0.27 | 0.33 | 0.19 | 0.37 | 0.49 |
| 2 | 38571255 | rs10198323 | -0.129 | -0.052 | 0.109 | -0.436 | 0.007 | 0.023 | 0.023 | 0 | 0.23 | 0.35 | 0.21 | 0.25 | 0.28 | 0.39 |
| 2 | 38611011 | rs10182455 | -0.498 | 0.071 | -0.421 | -1.126 | 0.004 | 0.094 | 0.094 | 0 | 0.4 | 0.75 | 0.38 | 0.21 | 0.39 | 0.51 |
| 2 | 38611930 | rs10198160 | -0.29 | 0.139 | -0.353 | -1.079 | 0 | -0.016 | -0.016 | 0 | 0.38 | 0.36 | 0.38 | 0.13 | 0.4 | 0.51 |
| 2 | 38621456 | rs11674467 | -1.134 | 0.142 | -1.797 | -1.145 | -0.09 | 0.07 | 0.07 | 0 | 0.61 | 0.95 | 0.55 | 0.27 | 0.59 | 0.58 |
| 2 | 38623289 | rs11902844 | -0.767 | 0.425 | -0.89 | -0.57 | 0.01 | 0.083 | 0.083 | 0 | 0.42 | 0.74 | 0.4 | 0.25 | 0.4 | 0.53 |
| 2 | 38646846 | rs7568053 | -0.95 | -0.05 | -0.275 | -0.352 | -0.015 | -0.031 | -0.031 | 0 | 0.32 | 0.62 | 0.3 | 0.13 | 0.22 | 0.31 |
| 2 | 38652572 | rs6741107 | -1.422 | -0.235 | 0.343 | -0.447 | -0.05 | -0.096 | -0.096 | 0 | 0.16 | 0.67 | 0.1 | 0 | 0.07 | 0.01 |
| 2 | 38656881 | rs12712595 | -1.053 | 0.49 | 0.062 | -0.066 | -0.009 | 0 | 0 | 0.033 | 0.24 | 0.22 | 0.28 | 0.24 | 0.34 | 0.2 |
| 2 | 38657924 | rs7589540 | -0.875 | 0.61 | 0.208 | -0.154 | -0.008 | -0.041 | -0.041 | 0.359 | 0.09 | 0.19 | 0.07 | 0 | 0.07 | 0.01 |
| 2 | 38665338 | rs11679742 | -1.136 | 0.919 | 1.034 | 0.016 | 0.044 | 0.025 | 0.025 | 0 | 0.23 | 0.31 | 0.27 | 0.27 | 0.39 | 0.26 |
| 2 | 38666024 | rs17603354 | -0.035 | 0.978 | 0.996 | 0.028 | -0.058 | -0.06 | -0.06 | 0 | 0.15 | 0.03 | 0.25 | 0.26 | 0.3 | 0.13 |
| 2 | 38668336 | rs10187298 | -0.403 | 0.933 | 1.192 | 0 | -0.001 | -0.037 | -0.037 | 0.857 | 0.07 | 0.18 | 0.06 | 0 | 0.07 | 0.01 |

| | | | | | | | | | | | | | | | | |
|---|---|---|---|---|---|---|---|---|---|---|---|---|---|---|---|---|
| 2 | 38670529 | rs10205121 | -1.477 | 0.718 | 1.068 | -0.111 | 0.007 | 0.003 | 0.003 | 0 | 0.32 | 0.31 | 0.32 | 0.33 | 0.45 | 0.42 |
| 2 | 38672996 | rs7599196 | -0.339 | 0.776 | 1.706 | -0.035 | 0.049 | 0.031 | 0.031 | 0 | 0.08 | 0.27 | 0.03 | 0.02 | 0.07 | 0.13 |
| 2 | 38673538 | rs17022684 | -0.328 | 0.777 | 1.707 | -0.034 | 0.087 | 0.029 | 0.029 | 0 | 0.08 | 0.27 | 0.04 | 0.02 | 0.09 | 0.13 |
| 2 | 38674466 | rs1368112 | -2.237 | 0.739 | 1.64 | -0.045 | -0.012 | 0 | 0 | 0 | 0.38 | 0.64 | 0.39 | 0.33 | 0.51 | 0.41 |
| 2 | 38675963 | rs12997977 | -0.097 | 0.708 | 1.619 | -0.069 | -0.001 | -0.022 | -0.022 | 0 | 0.07 | 0.13 | 0.06 | 0 | 0.07 | 0.01 |
| 2 | 38677471 | rs6735620 | -2.301 | 0.7 | 1.753 | -0.07 | -0.019 | 0.002 | 0.002 | 0 | 0.39 | 0.64 | 0.39 | 0.33 | 0.51 | 0.4 |
| 2 | 38679551 | rs1396738 | -0.664 | 0.715 | 1.64 | -0.05 | -0.008 | -0.008 | -0.008 | 0 | 0.23 | 0.17 | 0.29 | 0.31 | 0.35 | 0.28 |
| 2 | 38681622 | rs12712605 | -0.591 | 0.715 | 1.651 | -0.05 | -0.023 | -0.024 | -0.024 | 0 | 0.22 | 0.12 | 0.29 | 0.3 | 0.35 | 0.27 |
| 2 | 38687351 | rs367917 | 2.487 | 0.492 | 1.333 | -0.299 | -0.03 | 0.034 | 0.034 | 0.346 | 0.5 | 0.07 | 0.54 | 0.63 | 0.42 | 0.51 |
| 2 | 38687498 | rs11695317 | -2.117 | 0.54 | 1.551 | -0.209 | -0.012 | 0.019 | 0.019 | 0 | 0.41 | 0.67 | 0.41 | 0.37 | 0.5 | 0.38 |
| 2 | 38692619 | rs10168607 | -1.49 | 0.555 | 1.675 | -0.176 | 0.009 | 0.003 | 0.003 | 0 | 0.32 | 0.58 | 0.36 | 0.34 | 0.42 | 0.28 |
| 2 | 38692766 | rs17022754 | -0.303 | 0.548 | 1.738 | -0.209 | 0.037 | -0.031 | -0.031 | 0 | 0.08 | 0.28 | 0.06 | 0 | 0.07 | 0.01 |
| 2 | 38697934 | rs7588370 | -2.013 | 0.549 | 1.572 | -0.135 | -0.01 | 0.033 | 0.033 | 0 | 0.4 | 0.66 | 0.41 | 0.28 | 0.49 | 0.36 |
| 2 | 38701231 | rs6725717 | -1.628 | 0.369 | 1.666 | -0.132 | -0.014 | -0.115 | -0.115 | 0.567 | 0.18 | 0.5 | 0.12 | 0.04 | 0.14 | 0.09 |
| 2 | 38702122 | rs17022787 | -0.207 | 0.371 | 1.69 | -0.131 | -0.005 | -0.03 | -0.03 | 0.011 | 0.09 | 0.15 | 0.05 | 0.04 | 0.07 | 0.09 |
| 2 | 38702337 | rs17022789 | -0.386 | 0.372 | 1.706 | -0.125 | 0.026 | -0.038 | -0.038 | 0 | 0.09 | 0.31 | 0.06 | 0.01 | 0.07 | 0.01 |
| 2 | 38703619 | rs6720959 | -2.334 | 0.472 | 0.974 | -0.066 | 0.013 | 0.029 | 0.029 | 0 | 0.48 | 0.69 | 0.44 | 0.39 | 0.59 | 0.49 |
| 2 | 38704658 | rs7608506 | -1.25 | 0.321 | 0.77 | -0.051 | -0.006 | -0.055 | -0.055 | 2.421 | 0.17 | 0.26 | 0.11 | 0.03 | 0.14 | 0.11 |
| 2 | 38723049 | rs13424546 | -0.305 | 0.697 | 0.19 | 0.319 | -0.078 | -0.047 | -0.047 | 3.556 | 0.19 | 0.03 | 0.23 | 0.22 | 0.31 | 0.19 |
| 2 | 38728911 | rs4490160 | -1.701 | 0.117 | 1.207 | 0.174 | 0.014 | 0.016 | 0.016 | 0 | 0.14 | 0.32 | 0.12 | 0.11 | 0.16 | 0.13 |
| 2 | 38729008 | rs4405737 | -2.509 | -0.008 | 1.44 | 0.157 | -0.012 | -0.003 | -0.003 | 0 | 0.23 | 0.65 | 0.17 | 0.22 | 0.2 | 0.28 |
| 2 | 38730273 | rs7598981 | -1.37 | 0.193 | 1.087 | 0.123 | -0.007 | 0.023 | 0.023 | 0 | 0.11 | 0.3 | 0.08 | 0.1 | 0.09 | 0.13 |
| 2 | 38732293 | rs1125864 | 0.971 | 0.286 | 1.072 | -0.114 | 0.042 | -0.08 | -0.08 | 0.114 | 0.88 | 0.59 | 0.87 | 1 | 0.87 | 0.99 |
| 2 | 38736090 | rs4670876 | -0.33 | 0.354 | 0.604 | 0.061 | 0 | 0 | 0 | 1.543 | 0.73 | 0.72 | 0.7 | 0.75 | 0.69 | 0.62 |
| 2 | 38737000 | rs17421154 | 1.234 | -0.191 | 0.238 | -0.299 | -0.074 | -0.054 | -0.054 | 0.141 | 0.36 | 0.12 | 0.49 | 0.56 | 0.47 | 0.34 |
| 2 | 38740703 | rs7563859 | -0.489 | -0.16 | 0.444 | -0.483 | -0.016 | -0.07 | -0.07 | 2.073 | 0.29 | 0.35 | 0.18 | 0.11 | 0.21 | 0.14 |
| 2 | 38743881 | rs12476717 | 0.779 | -0.123 | -0.33 | -0.334 | -0.084 | 0.052 | 0.052 | 1.282 | 0.43 | 0.11 | 0.54 | 0.65 | 0.49 | 0.48 |
| 2 | 38745624 | rs13423976 | -0.621 | -0.09 | 0.187 | 0.093 | -0.001 | 0 | 0 | 0 | 0.73 | 0.73 | 0.61 | 0.74 | 0.7 | 0.63 |
| 2 | 38746456 | rs7566216 | 0.016 | -0.775 | 0.367 | 0.025 | -0.009 | 0 | 0 | 0 | 0.92 | 0.93 | 0.98 | 0.89 | 0.99 | 0.9 |
| 2 | 38749920 | rs3731843 | -0.805 | -0.376 | 0.66 | 0.127 | 0.022 | -0.007 | -0.007 | 0 | 0.17 | 0.23 | 0.18 | 0.12 | 0.23 | 0.05 |
| 2 | 38757841 | rs7601994 | -0.705 | -0.627 | 0.917 | 0.165 | 0.003 | 0.006 | 0.006 | 0 | 0.52 | 0.51 | 0.62 | 0.75 | 0.41 | 0.52 |
| 2 | 38759645 | rs13392580 | -0.24 | -0.797 | 1.362 | 0.109 | 0.016 | 0.032 | 0.032 | 0 | 0.42 | 0.62 | 0.59 | 0.65 | 0.35 | 0.48 |
| 2 | 38759797 | rs13392800 | -0.134 | -0.808 | 1.261 | 0.129 | -0.003 | -0.006 | -0.006 | 0.022 | 0.4 | 0.39 | 0.56 | 0.64 | 0.35 | 0.39 |
| 2 | 38760294 | rs10164547 | -1.341 | -0.421 | 1.607 | 0.215 | 0.03 | 0.025 | 0.025 | 0 | 0.17 | 0.39 | 0.19 | 0.22 | 0.18 | 0.13 |
| 2 | 38767505 | rs17421378 | 0.126 | -0.303 | 1.456 | 0.321 | 0.076 | 0.047 | 0.047 | 0.002 | 0.06 | 0.15 | 0.11 | 0.11 | 0.11 | 0.02 |
| 2 | 38772507 | rs17493321 | 0.125 | -0.271 | 1.456 | 0.333 | 0.096 | 0.062 | 0.062 | 0.009 | 0.06 | 0.15 | 0.12 | 0.12 | 0.14 | 0.02 |
| 2 | 38772915 | rs7571463 | -1.752 | -0.398 | 1.747 | 0.329 | 0.017 | 0.075 | 0.075 | 0 | 0.19 | 0.47 | 0.16 | 0.29 | 0.16 | 0.17 |
| 2 | 38781589 | rs17022904 | -2.255 | -0.447 | 1.694 | 0.27 | -0.006 | 0.002 | 0.002 | 0 | 0.25 | 0.48 | 0.21 | 0.28 | 0.21 | 0.18 |
| 2 | 38789682 | rs12328034 | -0.485 | -0.286 | 1.893 | 0.296 | 0.002 | 0.002 | 0.002 | 0 | 0.11 | 0.1 | 0.04 | 0.1 | 0.07 | 0.09 |
| 2 | 38799820 | rs1368114 | -1.902 | -0.308 | 1.943 | 0.311 | -0.008 | 0.031 | 0.031 | 0 | 0.2 | 0.47 | 0.15 | 0.27 | 0.14 | 0.18 |
| 2 | 38814517 | rs6544169 | -1.816 | -0.308 | 1.949 | 0.312 | -0.004 | 0.011 | 0.011 | 0 | 0.19 | 0.44 | 0.14 | 0.25 | 0.14 | 0.18 |
| 2 | 38819906 | rs12996029 | 0.21 | -0.292 | 1.891 | 0.314 | 0.063 | 0.023 | 0.023 | 0 | 0.06 | 0.14 | 0.1 | 0.1 | 0.11 | 0.02 |
| 2 | 38820522 | rs6544170 | -0.373 | -0.284 | 1.733 | 0.317 | 0.138 | 0.075 | 0.075 | 0.018 | 0.08 | 0.29 | 0.14 | 0.12 | 0.14 | 0.02 |
| 2 | 38828837 | rs438564 | 0.888 | -0.608 | 1.354 | 0.346 | 0.048 | -0.119 | -0.119 | 0 | 0.31 | 0.05 | 0.44 | 0.44 | 0.23 | 0.31 |
| 2 | 38839323 | rs372139 | -0.787 | -0.507 | 1.462 | 0.162 | 0.036 | 0.071 | 0.071 | 0 | 0.52 | 0.69 | 0.62 | 0.72 | 0.38 | 0.45 |
| 2 | 38845701 | rs17022996 | -1.717 | -0.243 | 1.42 | 0.263 | 0.002 | -0.004 | -0.004 | 0 | 0.21 | 0.6 | 0.17 | 0.25 | 0.17 | 0.18 |
| 2 | 38847132 | rs453182 | -0.857 | -0.176 | 1.413 | 0.286 | -0.025 | 0.074 | 0.074 | 0 | 0.56 | 0.84 | 0.66 | 0.73 | 0.43 | 0.6 |
| 2 | 38853095 | rs17023003 | -2.561 | 0.59 | 2.216 | 0.504 | 0.019 | -0.002 | -0.002 | 0 | 0.22 | 0.51 | 0.29 | 0.26 | 0.19 | 0.22 |
| 2 | 38857632 | rs2163551 | -1.658 | 0.652 | 1.88 | 0.536 | 0.089 | 0.068 | 0.068 | 0 | 0.1 | 0.27 | 0.12 | 0.16 | 0.14 | 0.09 |
| 2 | 38877636 | rs17023025 | -1.167 | 0.68 | 1.167 | 0.539 | 0 | 0 | 0 | 0 | 0.11 | 0.12 | 0.17 | 0.09 | 0.05 | 0.13 |
| 2 | 38881106 | rs6733848 | -0.818 | 0.668 | 1.147 | 0.614 | 0.037 | 0.004 | 0.004 | 0.139 | 0.1 | 0.19 | 0.11 | 0.1 | 0.11 | 0.02 |
| 2 | 38881480 | rs891706 | -1.138 | 0.648 | 0.324 | 0.64 | 0.003 | -0.005 | -0.005 | 0 | 0.23 | 0.63 | 0.19 | 0.25 | 0.2 | 0.18 |
| 2 | 38883974 | rs2163553 | -1.065 | 0.661 | 0.466 | 0.64 | 0.014 | -0.006 | -0.006 | 0.238 | 0.22 | 0.56 | 0.2 | 0.24 | 0.21 | 0.16 |
| 2 | 38893660 | rs3821023 | -1.032 | 0.366 | 1.179 | 1.544 | -0.018 | 0.194 | 0.194 | 0 | 0.15 | 0.57 | 0.15 | 0.41 | 0.12 | 0.26 |
| 2 | 38897810 | rs6751481 | -0.88 | 0.68 | 0.992 | 1.543 | 0 | -0.002 | -0.002 | 0 | 0.46 | 0.37 | 0.54 | 0.54 | 0.5 | 0.64 |
| 2 | 38907605 | rs12712612 | -0.228 | 0.725 | 1.556 | 1.629 | 0.004 | -0.005 | -0.005 | 0 | 0.13 | 0.1 | 0.09 | 0.22 | 0.07 | 0.25 |
| 2 | 38910402 | rs4670885 | -0.232 | 0.724 | 1.67 | 1.629 | -0.005 | 0.006 | 0.006 | 0 | 0.13 | 0.13 | 0.09 | 0.22 | 0.07 | 0.25 |
| 2 | 38913884 | rs4670264 | -1.245 | 0.717 | 1.632 | 1.4 | -0.024 | 0.051 | 0.051 | 0.076 | 0.42 | 0.68 | 0.51 | 0.75 | 0.41 | 0.61 |
| 2 | 38915780 | rs13014787 | -1.501 | 0.715 | 1.598 | 1.579 | -0.034 | 0.057 | 0.057 | 0 | 0.24 | 0.62 | 0.23 | 0.55 | 0.12 | 0.37 |
| 2 | 38920318 | rs12619390 | -0.482 | 0.797 | 1.386 | 1.57 | -0.004 | 0.02 | 0.02 | 0 | 0.09 | 0.07 | 0.02 | 0.23 | 0.05 | 0.09 |
| 2 | 38928981 | rs2043188 | 0.447 | 0.909 | 1.373 | 1.571 | -0.084 | -0.029 | -0.029 | 0.906 | 0.17 | 0.04 | 0.28 | 0.2 | 0.28 | 0.23 |
| 2 | 38932777 | rs4670265 | -1.837 | 0.547 | 1.223 | 1.189 | -0.009 | 0.047 | 0.047 | 0 | 0.44 | 0.67 | 0.58 | 0.73 | 0.49 | 0.53 |
| 2 | 38936539 | rs12987870 | 0.102 | 0.333 | 1.118 | 1.004 | 0.02 | -0.057 | -0.057 | 0 | 0.11 | 0.02 | 0.16 | 0.29 | 0.07 | 0.21 |
| 2 | 38940316 | rs13015612 | 1.382 | 0.273 | 1.309 | 0.467 | -0.01 | -0.075 | -0.075 | 0 | 0.85 | 0.7 | 0.91 | 0.99 | 0.88 | 0.94 |
| 2 | 38943609 | rs1965752 | -0.169 | 0.239 | 1.017 | 0.571 | 0.012 | -0.021 | -0.021 | 0 | 0.94 | 0.99 | 0.98 | 0.92 | 0.97 | 0.89 |
| 2 | 38951666 | rs1997377 | 1.334 | 1.043 | 1.298 | 0.537 | 0.005 | -0.029 | -0.029 | 0.084 | 0.65 | 0.46 | 0.64 | 0.8 | 0.68 | 0.71 |
| 2 | 38955129 | rs7603676 | 1.527 | 1.248 | 1.141 | 0.382 | 0.007 | -0.02 | -0.02 | 0.015 | 0.6 | 0.45 | 0.61 | 0.8 | 0.56 | 0.65 |
| 2 | 38956947 | rs2272445 | 1.437 | 1.339 | 1.158 | 0.399 | 0.009 | -0.022 | -0.022 | 0 | 0.61 | 0.46 | 0.62 | 0.8 | 0.58 | 0.65 |
| 2 | 38960491 | rs3097714 | 0.252 | 1.409 | 1.22 | 0.398 | -0.031 | 0.021 | 0.021 | 0 | 0.19 | 0.12 | 0.27 | 0.1 | 0.25 | 0.18 |

| | | | | | | | | | | | | | | | | |
|---|---|---|---|---|---|---|---|---|---|---|---|---|---|---|---|---|
| 2 | 38960927 | rs3731838 | 1.516 | 1.339 | 1.624 | 0.319 | 0.002 | -0.018 | -0.018 | 0.065 | 0.61 | 0.43 | 0.61 | 0.87 | 0.59 | 0.75 |
| 2 | 38963211 | rs4670891 | 1.699 | 1.354 | 1.6 | 0.333 | -0.032 | 0.087 | 0.087 | 0.09 | 0.58 | 0.29 | 0.6 | 0.8 | 0.59 | 0.64 |
| 2 | 38964531 | rs3097713 | -0.069 | 1.343 | 1.728 | 0.312 | -0.003 | 0.002 | 0.002 | 0 | 0.22 | 0.19 | 0.28 | 0.11 | 0.26 | 0.21 |
| 2 | 38964561 | rs1541877 | 1.403 | 1.288 | 1.787 | 0.291 | 0.006 | -0.04 | -0.04 | 0 | 0.86 | 0.87 | 0.9 | 0.99 | 0.83 | 0.97 |
| 2 | 38980231 | rs13024811 | 0.633 | 1.083 | 1.927 | 0.362 | 0.008 | -0.016 | -0.016 | 0.027 | 0.72 | 0.72 | 0.68 | 0.88 | 0.71 | 0.77 |
| 2 | 38980922 | rs2037875 | 1.691 | 1.049 | 1.872 | 0.324 | -0.025 | 0.107 | 0.107 | 0 | 0.59 | 0.3 | 0.61 | 0.87 | 0.63 | 0.73 |
| 2 | 38991193 | rs10196452 | -1.088 | 1.335 | 1.978 | 0.442 | 0 | -0.016 | -0.016 | 0 | 0.11 | 0.09 | 0.08 | 0.01 | 0.12 | 0.04 |
| 2 | 38995250 | rs3112188 | -0.136 | 1.331 | 1.96 | 0.44 | 0.003 | -0.005 | -0.005 | 0.018 | 0.22 | 0.21 | 0.27 | 0.12 | 0.24 | 0.19 |
| 2 | 38996561 | rs3112190 | 1.446 | 1.325 | 2.024 | 0.45 | 0.03 | -0.073 | -0.073 | 1.235 | 0.87 | 0.57 | 0.89 | 0.99 | 0.88 | 0.96 |
| 2 | 39005205 | rs10173118 | 1.413 | 1.33 | 1.983 | 0.462 | -0.019 | 0.047 | 0.047 | 0.01 | 0.63 | 0.32 | 0.61 | 0.79 | 0.64 | 0.71 |
| 2 | 39008949 | rs1056104 | 0.283 | 1.213 | 1.761 | 0.886 | -0.008 | -0.049 | -0.049 | 0 | 0.08 | 0 | 0.11 | 0.25 | 0.08 | 0.17 |
| 2 | 39030074 | rs13029406 | 1.63 | 1.119 | 1.481 | 0.233 | -0.02 | 0.067 | 0.067 | 0 | 0.61 | 0.3 | 0.57 | 0.79 | 0.64 | 0.7 |
| 2 | 39033590 | rs3099968 | -0.121 | 1.144 | 1.487 | 0.237 | 0.005 | -0.006 | -0.006 | 0 | 0.22 | 0.2 | 0.29 | 0.12 | 0.25 | 0.2 |
| 2 | 39045885 | rs10176932 | 1.752 | 1.285 | 1.165 | 0.162 | -0.066 | 0.097 | 0.097 | 0 | 0.58 | 0.21 | 0.48 | 0.78 | 0.59 | 0.65 |
| 2 | 39050141 | rs3770681 | 1.631 | 1.118 | 1.233 | 0.12 | -0.017 | 0.037 | 0.037 | 0 | 0.64 | 0.33 | 0.58 | 0.8 | 0.66 | 0.7 |
| 2 | 39060756 | rs6746931 | -1.468 | 0.873 | 1.236 | 0.155 | -0.005 | -0.008 | -0.008 | 0 | 0.12 | 0.13 | 0.11 | 0.09 | 0.1 | 0.08 |
| 2 | 39077320 | rs4670266 | 1.256 | 0.894 | 0.838 | 0.124 | 0 | -0.005 | -0.005 | 0.989 | 0.69 | 0.4 | 0.59 | 0.88 | 0.65 | 0.75 |
| 2 | 39082344 | rs7598922 | 1.822 | 0.864 | 0.29 | 0.115 | -0.032 | 0.064 | 0.064 | 0 | 0.62 | 0.3 | 0.57 | 0.8 | 0.63 | 0.72 |
| 2 | 39084757 | rs3112163 | -1.414 | 0.866 | 0.449 | 0.077 | -0.03 | 0.034 | 0.034 | 0 | 0.29 | 0.16 | 0.36 | 0.12 | 0.35 | 0.25 |
| 2 | 39090189 | rs3112165 | 1.666 | 0.853 | 0.483 | 0.072 | -0.018 | -0.067 | -0.067 | 0 | 0.87 | 0.43 | 0.87 | 0.98 | 0.92 | 0.97 |
| 2 | 39098764 | rs3099943 | -0.996 | 0.914 | 0.353 | 0.063 | -0.131 | -0.007 | -0.007 | 0.737 | 0.25 | 0.02 | 0.32 | 0.17 | 0.3 | 0.24 |
| 2 | 39102528 | rs1010859 | -0.862 | 0.88 | 0.48 | 0.087 | -0.017 | 0.02 | 0.02 | 0 | 0.27 | 0.16 | 0.32 | 0.12 | 0.29 | 0.23 |
| 2 | 39103991 | rs3923144 | 1.188 | 0.897 | 0.556 | 0.086 | -0.002 | -0.003 | -0.003 | 0 | 0.68 | 0.44 | 0.61 | 0.87 | 0.68 | 0.76 |
| 2 | 39105781 | rs4670899 | 1.209 | 0.898 | 0.558 | 0.092 | -0.012 | 0.022 | 0.022 | 0 | 0.68 | 0.35 | 0.61 | 0.8 | 0.71 | 0.74 |
| 2 | 39128610 | rs3099959 | -1.029 | 0.915 | 0.562 | 0.175 | 0.005 | -0.003 | -0.003 | 0 | 0.27 | 0.26 | 0.32 | 0.18 | 0.28 | 0.24 |
| 2 | 39129522 | rs10779923 | 2.291 | 0.808 | -0.016 | 0.107 | -0.043 | 0.093 | 0.093 | 0 | 0.57 | 0.27 | 0.54 | 0.78 | 0.6 | 0.71 |
| 2 | 39134127 | rs6754902 | 0.276 | 0.892 | 0.251 | 0.178 | 0.024 | -0.022 | -0.022 | 0.897 | 0.77 | 0.73 | 0.69 | 0.9 | 0.73 | 0.79 |
| 2 | 39135927 | rs6544180 | 0.943 | 0.915 | -0.552 | 0.183 | -0.014 | 0.013 | 0.013 | 0 | 0.71 | 0.35 | 0.63 | 0.81 | 0.72 | 0.76 |
| 2 | 39143803 | rs4670267 | 1.538 | 0.855 | -0.752 | 0.125 | -0.046 | 0.021 | 0.021 | 0.008 | 0.69 | 0.27 | 0.6 | 0.79 | 0.67 | 0.79 |
| 2 | 39154463 | rs373300 | -1.508 | 0.933 | -0.505 | 0.297 | -0.026 | 0.031 | 0.031 | 0 | 0.34 | 0.71 | 0.4 | 0.22 | 0.31 | 0.28 |
| 2 | 39165054 | rs10203995 | 1.863 | -0.227 | -0.552 | 0.791 | 0.05 | -0.05 | -0.05 | 0.012 | 0.77 | 0.18 | 0.79 | 0.71 | 0.9 | 0.73 |
| 2 | 39166576 | rs368451 | 0.285 | -0.078 | -0.214 | 0.619 | 0 | 0.004 | 0.004 | 0 | 0.92 | 0.94 | 0.95 | 0.84 | 0.95 | 0.83 |
| 2 | 39171781 | rs2599074 | 1.086 | -0.101 | -0.107 | 0.581 | 0.002 | -0.003 | -0.003 | 0 | 0.88 | 0.88 | 0.93 | 0.84 | 0.94 | 0.84 |
| 2 | 39176695 | rs413431 | 2.205 | -0.159 | 0.7 | 0.594 | -0.031 | 0.05 | 0.05 | 0 | 0.84 | 0.47 | 0.85 | 0.83 | 0.92 | 0.83 |
| 2 | 39177320 | rs408686 | 0.871 | -0.176 | 0.637 | 0.573 | -0.001 | 0.004 | 0.004 | 0 | 0.89 | 0.85 | 0.92 | 0.85 | 0.93 | 0.84 |
| 2 | 39186339 | rs6544183 | 0.764 | -0.013 | 0.6 | 0.549 | -0.001 | -0.039 | -0.039 | 1.743 | 0.89 | 0.93 | 0.94 | 0.82 | 0.93 | 0.82 |
| 2 | 39204040 | rs12473092 | 2.316 | -0.07 | 0.699 | 0.383 | 0.003 | -0.003 | -0.003 | 0 | 0.76 | 0.28 | 0.8 | 0.78 | 0.84 | 0.79 |
| 2 | 39209047 | rs1043793 | 1.928 | -0.081 | 0.881 | 0.308 | -0.015 | 0.076 | 0.076 | 0 | 0.86 | 0.44 | 0.86 | 0.82 | 0.92 | 0.82 |
| 2 | 39212637 | rs1059310 | 2.021 | 0.23 | 0.324 | 0.127 | -0.021 | 0.009 | 0.009 | 0.115 | 0.55 | 0.16 | 0.65 | 0.72 | 0.63 | 0.61 |
| 2 | 39216873 | rs963731 | 0.701 | 0.158 | 0.614 | 0.215 | -0.013 | 0.036 | 0.036 | 0 | 0.9 | 0.79 | 0.93 | 0.84 | 0.94 | 0.83 |
| 2 | 39242703 | rs10204958 | 2.223 | 0.219 | 0.705 | 0.215 | 0.006 | -0.001 | -0.001 | 0 | 0.76 | 0.28 | 0.77 | 0.79 | 0.82 | 0.79 |
| 2 | 39243129 | rs10184015 | 1.707 | 0.223 | 0.977 | 0.345 | -0.018 | 0.074 | 0.074 | 0 | 0.86 | 0.43 | 0.83 | 0.81 | 0.91 | 0.81 |
| 2 | 39271729 | rs11674846 | 1.47 | 0.254 | 1.073 | 0.346 | 0.011 | 0.134 | 0.134 | 0 | 0.86 | 0.54 | 0.84 | 0.81 | 0.89 | 0.8 |
| 2 | 39276957 | rs1454219 | 1.026 | 0.252 | 0.959 | 0.346 | -0.001 | 0.006 | 0.006 | 0 | 0.88 | 0.81 | 0.85 | 0.81 | 0.89 | 0.8 |
| 2 | 39285372 | rs7607509 | 1.599 | 0.259 | 0.807 | 0.348 | 0.001 | 0.114 | 0.114 | 0 | 0.86 | 0.5 | 0.83 | 0.81 | 0.89 | 0.8 |
| 2 | 39294016 | rs1377862 | 1.727 | 0.209 | 0.434 | 0.238 | -0.03 | 0.016 | 0.016 | 0 | 0.52 | 0.15 | 0.59 | 0.71 | 0.56 | 0.58 |
| 2 | 39305489 | rs297136 | 1.674 | 0.282 | 0.916 | 0.437 | 0.003 | 0.118 | 0.118 | 0 | 0.87 | 0.49 | 0.84 | 0.81 | 0.91 | 0.82 |
| 2 | 39326511 | rs297151 | 0.668 | 0.301 | 0.813 | 0.446 | 0 | 0.004 | 0.004 | 0 | 0.89 | 0.83 | 0.86 | 0.81 | 0.91 | 0.81 |
| 2 | 39328640 | rs297158 | 0.46 | 0.39 | 0.683 | 0.421 | 0.044 | 0.143 | 0.143 | 0 | 0.89 | 0.64 | 0.89 | 0.84 | 0.92 | 0.83 |
| 2 | 39370035 | rs6729036 | -1.652 | 0.343 | 0.355 | 0.103 | -0.018 | -0.02 | -0.02 | 0 | 0.26 | 0.41 | 0.18 | 0.18 | 0.2 | 0.26 |
| 2 | 39373207 | rs10171977 | -1.358 | 0.275 | -0.37 | -0.268 | 0.002 | -0.009 | -0.009 | 0 | 0.55 | 0.52 | 0.42 | 0.29 | 0.55 | 0.42 |
| 2 | 39374982 | rs17023351 | -1.136 | 0.205 | -0.432 | -0.32 | -0.009 | 0.007 | 0.007 | 0.086 | 0.22 | 0.03 | 0.13 | 0.13 | 0.18 | 0.19 |
| 2 | 39378320 | rs1510112 | 1.741 | 0.112 | -0.278 | -0.47 | 0.022 | 0.03 | 0.03 | 0 | 0.69 | 0.24 | 0.78 | 0.82 | 0.77 | 0.72 |
| 2 | 39380978 | rs6544203 | -0.639 | 0.204 | -0.085 | -0.401 | 0 | 0.003 | 0.003 | 0 | 0.45 | 0.46 | 0.38 | 0.28 | 0.47 | 0.4 |
| 2 | 39401658 | rs11124661 | 1.379 | -0.14 | 0.071 | -0.926 | -0.013 | 0 | 0 | 0 | 0.84 | 0.24 | 0.74 | 0.85 | 0.9 | 0.79 |
| 2 | 39404010 | rs7575216 | 0.246 | -0.15 | 0.868 | -0.931 | -0.004 | 0.015 | 0.015 | 0 | 0.89 | 0.78 | 0.8 | 0.88 | 0.93 | 0.83 |
| 2 | 39405570 | rs17665998 | 0.999 | 0.09 | 0.564 | -1.159 | -0.059 | 0.174 | 0.174 | 0 | 0.26 | 0.02 | 0.21 | 0.01 | 0.27 | 0.19 |
| 2 | 39408329 | rs12165133 | 1.56 | -0.177 | 1.024 | -0.828 | -0.013 | -0.001 | -0.001 | 0 | 0.84 | 0.25 | 0.74 | 0.85 | 0.9 | 0.79 |
| 2 | 39424633 | rs2708222 | -1.138 | -0.13 | 0.724 | -0.807 | -0.02 | -0.031 | -0.031 | 0 | 0.95 | 1 | 0.82 | 0.88 | 0.94 | 0.83 |
| 2 | 39443569 | rs6743345 | 2.005 | -0.15 | 0.808 | -0.823 | -0.007 | -0.002 | -0.002 | 0.675 | 0.79 | 0.29 | 0.72 | 0.79 | 0.86 | 0.71 |
| 2 | 39453469 | rs7583170 | 1.025 | -0.12 | 0.794 | -0.844 | 0.002 | 0.005 | 0.005 | 0 | 0.87 | 0.72 | 0.79 | 0.87 | 0.9 | 0.79 |
| 2 | 39463446 | rs11124665 | 1.931 | -0.155 | 0.43 | -0.861 | 0.005 | 0.002 | 0.002 | 0.434 | 0.78 | 0.2 | 0.73 | 0.8 | 0.87 | 0.72 |
| 2 | 39474679 | rs11124667 | 0.739 | 0.355 | 0.086 | -1.067 | -0.038 | 0.188 | 0.188 | 0 | 0.48 | 0.12 | 0.48 | 0.78 | 0.52 | 0.54 |
| 2 | 39477170 | rs6544214 | 1.298 | 0.339 | 0.919 | -0.812 | -0.002 | 0.025 | 0.025 | 0 | 0.81 | 0.54 | 0.76 | 0.8 | 0.86 | 0.72 |
| 2 | 39497758 | rs6756183 | -0.032 | 0.344 | 1.041 | -0.828 | -0.006 | 0.001 | 0.001 | 4.828 | 0.89 | 0.8 | 0.85 | 0.89 | 0.92 | 0.82 |
| 2 | 39500067 | rs10153665 | 0.847 | 0.294 | 1.288 | -0.849 | -0.015 | 0.005 | 0.005 | 3.402 | 0.86 | 0.46 | 0.78 | 0.89 | 0.92 | 0.81 |
| 2 | 39505890 | rs3755177 | -0.083 | 0.301 | 1.106 | -0.846 | -0.005 | 0.001 | 0.001 | 4.001 | 0.89 | 0.82 | 0.85 | 0.91 | 0.92 | 0.82 |
| 2 | 39512437 | rs12151392 | 1.569 | 0.297 | 0.807 | -0.863 | 0.011 | 0.003 | 0.003 | 0.02 | 0.79 | 0.21 | 0.7 | 0.8 | 0.85 | 0.72 |
| 2 | 39518512 | rs10169329 | 1.183 | 0.279 | 0.87 | -0.858 | 0 | 0 | 0 | 0 | 0.83 | 0.21 | 0.76 | 0.85 | 0.9 | 0.79 |

| | | | | | | | | | | | | | | | | |
|---|---|---|---|---|---|---|---|---|---|---|---|---|---|---|---|---|
| 2 | 39522448 | rs10193636 | 1.508 | 0.029 | 0.616 | -1.046 | 0.016 | 0.01 | 0.01 | 1.579 | 0.77 | 0.23 | 0.7 | 0.84 | 0.86 | 0.74 |
| 2 | 39539427 | rs6725784 | -0.032 | 0.031 | 0.984 | -1.037 | -0.009 | -0.027 | -0.027 | 0 | 0.07 | 0.14 | 0.06 | 0.02 | 0.05 | 0.07 |
| 2 | 39543666 | rs3770662 | 0.649 | 0.005 | 0.579 | -1.292 | -0.077 | 0.141 | 0.141 | 0 | 0.44 | 0.12 | 0.47 | 0.78 | 0.49 | 0.52 |
| 2 | 39554376 | rs2024514 | 1.618 | 0.123 | 1.564 | -0.72 | -0.006 | -0.002 | -0.002 | 0 | 0.79 | 0.28 | 0.72 | 0.8 | 0.86 | 0.72 |
| 2 | 39555944 | rs9309036 | 1.779 | 0.119 | 1.395 | -0.707 | 0.011 | 0 | 0 | 0.408 | 0.78 | 0.21 | 0.7 | 0.79 | 0.85 | 0.72 |
| 2 | 39570427 | rs3732053 | 1.194 | 0.086 | 1.536 | -0.723 | -0.022 | 0.003 | 0.003 | 1.482 | 0.84 | 0.33 | 0.74 | 0.84 | 0.9 | 0.78 |
| 2 | 39574984 | rs6753448 | 0.904 | 0.098 | 1.548 | -0.717 | 0.005 | 0.052 | 0.052 | 2.175 | 0.86 | 0.59 | 0.77 | 0.84 | 0.9 | 0.79 |
| 2 | 39588797 | rs12712636 | 0.886 | 0.123 | 1.458 | -0.703 | -0.022 | 0.032 | 0.032 | 2.439 | 0.86 | 0.43 | 0.77 | 0.87 | 0.93 | 0.82 |
| 2 | 39602950 | rs11898704 | -0.489 | 0.109 | 1.383 | -0.727 | -0.013 | -0.03 | -0.03 | 0 | 0.08 | 0.14 | 0.06 | 0.02 | 0.06 | 0.09 |
| 2 | 39604876 | rs12712640 | 1.839 | 0.119 | 1.33 | -0.735 | 0.007 | 0.004 | 0.004 | 0.114 | 0.74 | 0.24 | 0.7 | 0.8 | 0.84 | 0.7 |
| 2 | 39605917 | rs2373530 | 1.662 | 0.145 | 1.344 | -0.71 | -0.02 | -0.006 | -0.006 | 0 | 0.88 | 0.34 | 0.89 | 0.91 | 0.93 | 0.9 |
| 2 | 39620989 | rs6544224 | 0.158 | 0.237 | 1.186 | -0.654 | -0.008 | 0.026 | 0.026 | 4.632 | 0.89 | 0.77 | 0.82 | 0.87 | 0.93 | 0.82 |
| 2 | 39626388 | rs12712643 | 1.754 | 0.207 | 1.248 | -0.676 | -0.031 | -0.01 | -0.01 | 1.28 | 0.76 | 0.52 | 0.73 | 0.8 | 0.83 | 0.69 |
| 2 | 39636469 | rs6712399 | -0.289 | 0.461 | 1.218 | -0.677 | -0.002 | -0.029 | -0.029 | 0 | 0.08 | 0.14 | 0.06 | 0.02 | 0.07 | 0.08 |
| 2 | 39639572 | rs1558854 | 1.342 | 0.606 | 1.556 | -0.509 | -0.011 | 0.028 | 0.028 | 0.234 | 0.83 | 0.47 | 0.76 | 0.81 | 0.88 | 0.77 |
| 2 | 39640893 | rs10170617 | 0.458 | 0.571 | 1.228 | -0.503 | -0.002 | 0.051 | 0.051 | 5.825 | 0.88 | 0.7 | 0.81 | 0.86 | 0.93 | 0.84 |
| 2 | 39641556 | rs6732611 | 0.263 | 0.604 | 1.195 | -0.496 | -0.006 | 0.01 | 0.01 | 4.91 | 0.89 | 0.79 | 0.81 | 0.88 | 0.92 | 0.83 |
| 2 | 39675339 | rs7577097 | 0.321 | 0.616 | 1.187 | -0.47 | 0.048 | 0.039 | 0.039 | 0.483 | 0.92 | 0.62 | 0.95 | 0.95 | 0.96 | 0.94 |
| 2 | 39689746 | rs10865153 | 0.536 | 0.798 | 0.104 | -0.856 | -0.128 | 0.092 | 0.092 | 0 | 0.41 | 0.06 | 0.44 | 0.8 | 0.47 | 0.53 |
| 2 | 39692393 | rs9677244 | -0.116 | 0.741 | 0.11 | -1.009 | 0.004 | 0.236 | 0.236 | 0 | 0.49 | 0.14 | 0.51 | 0.87 | 0.55 | 0.58 |
| 2 | 39712161 | rs2287089 | 0.12 | 0.77 | 0.59 | -1.078 | -0.001 | -0.004 | -0.004 | 0.137 | 0.45 | 0.39 | 0.45 | 0.83 | 0.46 | 0.53 |
| 2 | 39712376 | rs1060691 | 0.163 | 0.81 | 0.331 | -1.056 | -0.064 | 0.201 | 0.201 | 0 | 0.45 | 0.1 | 0.45 | 0.84 | 0.47 | 0.56 |
| 2 | 39728431 | rs11891871 | 0.623 | 0.861 | 0.097 | -0.963 | -0.087 | 0.035 | 0.035 | 0 | 0.38 | 0.03 | 0.28 | 0.77 | 0.39 | 0.48 |
| 2 | 39733850 | rs10210842 | -0.033 | 0.665 | 0.301 | -1.155 | -0.003 | 0.034 | 0.034 | 0 | 0.52 | 0.58 | 0.53 | 0.83 | 0.48 | 0.6 |
| 2 | 39733965 | rs13420542 | 1.331 | 0.551 | -0.146 | -1.288 | -0.068 | 0.145 | 0.145 | 0 | 0.28 | 0.08 | 0.21 | 0.02 | 0.36 | 0.18 |
| 2 | 39735813 | rs13420425 | -1.101 | 0.534 | 0.095 | -1.19 | 0 | -0.039 | -0.039 | 0 | 0.12 | 0.25 | 0.11 | 0.04 | 0.1 | 0.05 |
| 2 | 39740478 | rs7591326 | 0.443 | 0.591 | -0.049 | -1.238 | -0.011 | 0.105 | 0.105 | 0 | 0.45 | 0.29 | 0.46 | 0.86 | 0.44 | 0.56 |
| 2 | 39741928 | rs11124685 | 0.516 | 0.615 | -0.171 | -1.209 | -0.039 | 0.169 | 0.169 | 0 | 0.44 | 0.11 | 0.44 | 0.84 | 0.43 | 0.55 |
| 2 | 39743876 | rs734663 | 0.188 | 0.718 | -0.187 | -1.113 | -0.135 | 0.181 | 0.181 | 0.176 | 0.42 | 0.05 | 0.45 | 0.13 | 0.51 | 0.41 |
| 2 | 39748858 | rs12621670 | 1.284 | 0.773 | -0.443 | -0.502 | -0.007 | -0.065 | -0.065 | 1.101 | 0.1 | 0 | 0.1 | 0.24 | 0.08 | 0.09 |
| 2 | 39752158 | rs6753798 | 0.224 | 1.033 | 0.45 | -0.897 | -0.003 | 0.136 | 0.136 | 0.342 | 0.52 | 0.27 | 0.46 | 0.85 | 0.46 | 0.56 |
| 2 | 39752628 | rs17473802 | 0.905 | 1.081 | 0.227 | -0.905 | -0.071 | 0.212 | 0.212 | 0 | 0.32 | 0.06 | 0.25 | 0.01 | 0.37 | 0.22 |
| 2 | 39752949 | rs10191970 | -0.603 | 1.097 | 0.342 | -0.881 | -0.001 | -0.01 | -0.01 | 0.057 | 0.63 | 0.64 | 0.62 | 0.88 | 0.56 | 0.65 |
| 2 | 39754150 | rs12476783 | 1.256 | 1.058 | -0.244 | -0.867 | -0.087 | 0.172 | 0.172 | 0 | 0.28 | 0.06 | 0.2 | 0.01 | 0.36 | 0.18 |
| 2 | 39758174 | rs4670285 | 0.165 | 1.068 | 0.3 | -0.381 | -0.001 | -0.002 | -0.002 | 0 | 0.08 | 0.07 | 0.18 | 0.11 | 0.11 | 0.17 |
| 2 | 39762982 | rs6745068 | -0.803 | 0.938 | 0.366 | -0.338 | 0.047 | 0.048 | 0.048 | 3.341 | 0.06 | 0.4 | 0.07 | 0.03 | 0.02 | 0.03 |
| 2 | 39766614 | rs7588713 | -0.207 | 1.087 | -0.157 | -0.301 | -0.001 | -0.002 | -0.002 | 0 | 0.08 | 0.07 | 0.18 | 0.12 | 0.12 | 0.17 |
| 2 | 39772004 | rs10196602 | 0.515 | 1.285 | -0.235 | -0.022 | -0.061 | -0.003 | -0.003 | 0 | 0.37 | 0.03 | 0.25 | 0.73 | 0.35 | 0.48 |
| 2 | 39776534 | rs2241162 | -0.421 | 1.472 | 0.009 | -0.195 | 0.002 | 0.167 | 0.167 | 0.752 | 0.53 | 0.19 | 0.48 | 0.85 | 0.5 | 0.57 |
| 2 | 39778109 | rs4670944 | 0.822 | 1.432 | -0.253 | -0.073 | -0.138 | 0.165 | 0.165 | 0.026 | 0.42 | 0.03 | 0.44 | 0.15 | 0.51 | 0.41 |
| 2 | 39789288 | rs6740771 | -0.255 | 1.121 | -0.701 | 0.034 | 0.049 | 0.184 | 0.184 | 0.139 | 0.5 | 0.17 | 0.48 | 0.82 | 0.52 | 0.57 |
| 2 | 39791925 | rs13035388 | 0.546 | 1.141 | -0.928 | -0.001 | -0.02 | 0.183 | 0.183 | 0 | 0.44 | 0.14 | 0.46 | 0.15 | 0.48 | 0.4 |
| 2 | 39809660 | rs11690963 | 0.155 | 1.004 | -1.123 | 0.438 | -0.138 | -0.091 | -0.091 | 0.169 | 0.38 | 0.03 | 0.34 | 0.58 | 0.46 | 0.49 |
| 2 | 39814690 | rs7597939 | -0.147 | 1.018 | -1.174 | 0.638 | 0.03 | 0.084 | 0.084 | 0.044 | 0.52 | 0.21 | 0.5 | 0.67 | 0.55 | 0.59 |
| 2 | 39819339 | rs4670289 | 0.311 | 0.371 | -1.195 | 1.045 | 0.066 | -0.001 | -0.001 | 0 | 0.51 | 0.15 | 0.5 | 0.53 | 0.59 | 0.5 |
| 2 | 39819461 | rs4670947 | 0.57 | 0.214 | 0.048 | 1.15 | 0.022 | 0.113 | 0.113 | 0.026 | 0.93 | 0.71 | 0.84 | 0.88 | 0.96 | 0.89 |
| 2 | 39822983 | rs4670949 | 1.403 | 0.144 | 0.765 | 1.112 | 0.013 | 0.092 | 0.092 | 0.173 | 0.91 | 0.53 | 0.93 | 0.9 | 0.94 | 0.91 |
| 2 | 39837257 | rs2110952 | -0.269 | 0.142 | 0.982 | 1.305 | 0.024 | 0.002 | 0.002 | 0 | 0.43 | 0.35 | 0.33 | 0.44 | 0.37 | 0.49 |
| 2 | 39837382 | rs6544238 | -0.717 | 0.136 | 0.992 | 1.294 | 0.078 | -0.014 | -0.014 | 0 | 0.48 | 0.82 | 0.39 | 0.49 | 0.39 | 0.51 |
| 2 | 39846823 | rs10194567 | 0.067 | 0.144 | 1.103 | 1.203 | 0.027 | 0.018 | 0.018 | 0 | 0.38 | 0.27 | 0.3 | 0.33 | 0.33 | 0.39 |
| 2 | 39847517 | rs2716705 | -0.651 | 0.119 | 1.003 | 1.248 | 0.071 | 0.014 | 0.014 | 0 | 0.47 | 0.87 | 0.39 | 0.42 | 0.39 | 0.49 |
| 2 | 39851299 | rs887969 | -0.422 | 0.045 | 1.104 | 1.235 | -0.014 | -0.009 | -0.009 | 0 | 0.43 | 0.54 | 0.34 | 0.39 | 0.38 | 0.48 |
| 2 | 39851411 | rs887968 | -0.222 | -0.025 | 1.1 | 1.136 | 0.022 | 0.014 | 0.014 | 0 | 0.39 | 0.33 | 0.29 | 0.33 | 0.33 | 0.41 |
| 2 | 39868986 | rs10208929 | -0.01 | 0.167 | 0.416 | 0.29 | 0.009 | 0.069 | 0.069 | 0 | 0.4 | 0.23 | 0.59 | 0.7 | 0.41 | 0.56 |
| 2 | 39874270 | rs6748628 | -0.035 | 0.166 | 0.471 | 0.29 | 0.011 | 0.057 | 0.057 | 0 | 0.4 | 0.27 | 0.59 | 0.7 | 0.41 | 0.56 |
| 2 | 39896800 | rs2716692 | -0.164 | -0.027 | 1.015 | -0.271 | 0.007 | -0.011 | -0.011 | 0 | 0.43 | 0.62 | 0.63 | 0.87 | 0.41 | 0.61 |
| 2 | 39899435 | rs917964 | 0.153 | -0.083 | 1.266 | -0.03 | -0.002 | -0.009 | -0.009 | 0 | 0.38 | 0.39 | 0.58 | 0.74 | 0.4 | 0.56 |
| 2 | 39900326 | rs2540255 | 0.253 | -0.085 | 1.312 | -0.031 | 0.001 | 0.001 | 0.001 | 0 | 0.37 | 0.34 | 0.57 | 0.74 | 0.4 | 0.56 |
| 2 | 39911538 | rs10170462 | 0.412 | 0.015 | 1.4 | -0.111 | 0.003 | 0.059 | 0.059 | 0 | 0.36 | 0.2 | 0.56 | 0.74 | 0.42 | 0.57 |
| 2 | 39953746 | rs2716727 | 0.015 | 0.305 | 1.684 | -0.298 | 0.014 | 0.081 | 0.081 | 0 | 0.44 | 0.74 | 0.6 | 0.76 | 0.42 | 0.57 |
| 2 | 39966594 | rs917962 | -0.049 | 0.255 | 1.779 | -0.278 | -0.001 | 0.075 | 0.075 | 0 | 0.45 | 0.71 | 0.64 | 0.74 | 0.47 | 0.58 |
| 2 | 39969682 | rs6749057 | 0.679 | 0.071 | 1.566 | -0.684 | -0.01 | -0.067 | -0.067 | 2.629 | 0.13 | 0.26 | 0.09 | 0 | 0.1 | 0.05 |
| 2 | 39970438 | rs12991701 | 0.728 | 0.072 | 1.568 | -0.684 | -0.001 | -0.027 | -0.027 | 0 | 0.13 | 0.16 | 0.08 | 0 | 0.1 | 0.05 |
| 2 | 39972721 | rs962262 | 0.93 | -0.053 | 1.355 | -0.761 | -0.002 | 0.072 | 0.072 | 0.124 | 0.41 | 0.2 | 0.58 | 0.73 | 0.45 | 0.54 |
| 2 | 39979125 | rs3815510 | -0.405 | 0.033 | 1.362 | -0.908 | -0.021 | 0.094 | 0.094 | 0 | 0.62 | 0.76 | 0.76 | 0.83 | 0.62 | 0.66 |
| 2 | 39985742 | rs6722610 | 0.07 | -0.14 | 1.943 | -0.53 | 0.004 | 0.039 | 0.039 | 0.175 | 0.46 | 0.65 | 0.64 | 0.81 | 0.47 | 0.6 |
| 2 | 39988703 | rs3732122 | 0.4 | 0.019 | 1.831 | -0.476 | 0 | -0.025 | -0.025 | 0 | 0.42 | 0.48 | 0.62 | 0.74 | 0.48 | 0.58 |
| 2 | 39997694 | rs12624217 | 0.657 | 0.024 | 1.288 | -0.392 | -0.016 | 0.076 | 0.076 | 0 | 0.4 | 0.15 | 0.59 | 0.72 | 0.46 | 0.54 |
| 2 | 40002032 | rs7560642 | 0.399 | 0.045 | 1.28 | -0.42 | -0.008 | 0.089 | 0.089 | 0 | 0.43 | 0.21 | 0.62 | 0.74 | 0.49 | 0.59 |

| | | | | | | | | | | | | | | | | |
|---|---|---|---|---|---|---|---|---|---|---|---|---|---|---|---|---|
| 2 | 40005290 | rs12470473 | -0.079 | 0.099 | 0.765 | -0.35 | 0.014 | 0.179 | 0.179 | 0.082 | 0.5 | 0.18 | 0.33 | 0.27 | 0.51 | 0.41 |
| 2 | 40021197 | rs11678063 | -0.443 | -0.065 | 0.775 | -0.155 | 0.043 | 0.061 | 0.061 | 0 | 0.57 | 0.24 | 0.51 | 0.33 | 0.6 | 0.52 |
| 2 | 40024218 | rs7586524 | 0.367 | -0.066 | 1.281 | -0.084 | 0.002 | 0.002 | 0.002 | 0 | 0.43 | 0.53 | 0.47 | 0.72 | 0.38 | 0.5 |
| 2 | 40027569 | rs9309043 | 0.977 | -0.064 | 1.362 | -0.1 | 0.008 | 0.003 | 0.003 | 0 | 0.36 | 0.24 | 0.42 | 0.65 | 0.36 | 0.48 |
| 2 | 40033717 | rs2192708 | 0.089 | -0.178 | 0.455 | -0.36 | 0.009 | 0.149 | 0.149 | 0 | 0.46 | 0.1 | 0.44 | 0.25 | 0.55 | 0.44 |
| 2 | 40044135 | rs6544257 | 0.422 | -0.088 | 0.821 | -0.111 | 0.02 | 0.03 | 0.03 | 0 | 0.42 | 0.61 | 0.47 | 0.72 | 0.38 | 0.46 |
| 2 | 40046658 | rs11124697 | 0.438 | -0.07 | 0.425 | -0.088 | 0.042 | 0.064 | 0.064 | 0 | 0.42 | 0.73 | 0.48 | 0.72 | 0.38 | 0.47 |
| 2 | 40051216 | rs6717356 | 0.184 | 0.061 | 0.163 | -0.04 | 0.052 | 0.098 | 0.098 | 0.116 | 0.45 | 0.84 | 0.52 | 0.71 | 0.41 | 0.47 |
| 2 | 40069664 | rs4670978 | -0.534 | 1.186 | 0.583 | 0.301 | -0.038 | 0.042 | 0.042 | 0 | 0.44 | 0.16 | 0.43 | 0.35 | 0.44 | 0.46 |
| 2 | 40074903 | rs13430309 | -0.571 | 1.244 | 0.289 | 0.271 | -0.016 | 0.043 | 0.043 | 0 | 0.45 | 0.26 | 0.44 | 0.35 | 0.43 | 0.48 |
| 2 | 40077971 | rs9679549 | 0.53 | 1.138 | -0.438 | 0.246 | 0.031 | 0.018 | 0.018 | 0.318 | 0.51 | 0.2 | 0.47 | 0.56 | 0.5 | 0.51 |
| 2 | 40080343 | rs11692913 | 0.642 | 1.081 | -0.474 | 0.353 | 0.029 | 0.009 | 0.009 | 0 | 0.48 | 0.22 | 0.47 | 0.56 | 0.48 | 0.51 |
| 2 | 40082054 | rs1541564 | -0.449 | 1.51 | -0.606 | 0.302 | -0.076 | -0.017 | -0.017 | 0 | 0.34 | 0.12 | 0.3 | 0.34 | 0.44 | 0.3 |
| 2 | 40087025 | rs2192703 | -0.161 | 1.42 | -0.534 | 0.358 | -0.062 | -0.013 | -0.013 | 0.024 | 0.3 | 0.11 | 0.28 | 0.3 | 0.41 | 0.27 |
| 2 | 40096398 | rs7566309 | 0.384 | 1.423 | -0.425 | 0.37 | 0.001 | 0 | 0 | 0 | 0.86 | 0.81 | 0.83 | 0.87 | 0.87 | 0.82 |
| 2 | 40102764 | rs7562741 | -0.629 | 1.29 | 0.235 | 0.034 | 0 | 0 | 0 | 0 | 0.37 | 0.6 | 0.32 | 0.3 | 0.35 | 0.31 |
| 2 | 40111379 | rs11901530 | -0.877 | 0.965 | -0.34 | 0.114 | -0.038 | 0.03 | 0.03 | 0.504 | 0.41 | 0.76 | 0.43 | 0.31 | 0.45 | 0.35 |
| 2 | 40115378 | rs7597754 | -1.619 | 0.702 | 0.887 | -0.044 | -0.019 | -0.037 | -0.037 | 0 | 0.08 | 0.62 | 0.07 | 0 | 0.01 | 0.05 |
| 2 | 40115727 | rs7571703 | -1.542 | 0.729 | 0.9 | -0.031 | -0.014 | -0.032 | -0.032 | 0 | 0.08 | 0.62 | 0.07 | 0 | 0.01 | 0.05 |
| 2 | 40119060 | rs4670981 | 1.79 | 0.978 | 0.858 | 0.2 | -0.002 | 0.023 | 0.023 | 0.991 | 0.84 | 0.25 | 0.79 | 0.72 | 0.85 | 0.83 |
| 2 | 40119894 | rs6544266 | -1.032 | 0.952 | 1.722 | 0.141 | 0.067 | 0.046 | 0.046 | 0.957 | 0.06 | 0.38 | 0.03 | 0 | 0.01 | 0.05 |
| 2 | 40123679 | rs17024533 | -1.537 | 0.976 | 2.137 | 0.206 | 0.025 | 0.114 | 0.114 | 0 | 0.12 | 0.3 | 0.16 | 0.29 | 0.15 | 0.11 |
| 2 | 40123904 | rs13385459 | -2.352 | 0.931 | 2.411 | 0.185 | -0.016 | -0.024 | -0.024 | 1.197 | 0.09 | 0.59 | 0.1 | 0.03 | 0.01 | 0.06 |
| 2 | 40126281 | rs731353 | 1.242 | 0.535 | 1.766 | -0.334 | 0.003 | 0 | 0 | 0 | 0.69 | 0.31 | 0.69 | 0.71 | 0.77 | 0.74 |
| 2 | 40135076 | rs13415008 | -0.956 | 0.669 | 1.377 | -0.59 | 0.013 | -0.041 | -0.041 | 0.295 | 0.3 | 0.37 | 0.23 | 0.1 | 0.32 | 0.24 |
| 2 | 40136231 | rs2888638 | -0.132 | 0.494 | 1.138 | -0.62 | -0.044 | -0.083 | -0.083 | 0 | 0.13 | 0.29 | 0.1 | 0 | 0.04 | 0.09 |
| 2 | 40137942 | rs2215972 | -1.012 | 0.81 | 1.233 | -0.325 | -0.042 | 0.036 | 0.036 | 0 | 0.29 | 0.16 | 0.22 | 0.19 | 0.35 | 0.28 |
| 2 | 40151882 | rs7567740 | -0.655 | 0.658 | 0.719 | -0.431 | -0.019 | 0.02 | 0.02 | 0 | 0.26 | 0.13 | 0.18 | 0.18 | 0.29 | 0.24 |
| 2 | 40152867 | rs17024612 | 0.701 | 0.894 | 0.49 | -0.024 | -0.028 | 0.009 | 0.009 | 0 | 0.06 | 0.02 | 0.1 | 0.34 | 0.13 | 0.11 |
| 2 | 40152918 | rs17024614 | 0.377 | 0.894 | 0.517 | 0.003 | -0.002 | 0.057 | 0.057 | 0 | 0.07 | 0.05 | 0.13 | 0.38 | 0.13 | 0.11 |
| 2 | 40154844 | rs7559023 | -0.187 | 0.614 | 0.355 | -0.54 | 0.004 | -0.015 | -0.015 | 0 | 0.14 | 0.15 | 0.12 | 0 | 0.08 | 0.09 |
| 2 | 40158363 | rs2058626 | 1.203 | 0.811 | 0.166 | -0.371 | -0.015 | -0.014 | -0.014 | 0 | 0.68 | 0.15 | 0.72 | 0.67 | 0.67 | 0.81 |
| 2 | 40159265 | rs10181603 | 0.748 | 0.766 | 0.472 | -0.382 | -0.013 | 0.05 | 0.05 | 1.323 | 0.84 | 0.25 | 0.86 | 0.67 | 0.88 | 0.91 |
| 2 | 40167555 | rs725375 | -0.059 | 0.823 | 0.406 | -0.597 | -0.003 | -0.021 | -0.021 | 0 | 0.16 | 0.1 | 0.13 | 0 | 0.21 | 0.09 |
| 2 | 40168578 | rs11124710 | 2.261 | 0.956 | -0.006 | -0.68 | -0.062 | -0.074 | -0.074 | 1.909 | 0.5 | 0.04 | 0.55 | 0.49 | 0.51 | 0.54 |
| 2 | 40173905 | rs7584288 | 1.283 | 0.993 | 0.347 | -0.595 | -0.01 | -0.053 | -0.053 | 0 | 0.71 | 0.18 | 0.73 | 0.58 | 0.68 | 0.81 |
| 2 | 40181783 | rs10490338 | 1.164 | 1.004 | 0.78 | -0.595 | 0 | 0.007 | 0.007 | 0.408 | 0.75 | 0.33 | 0.78 | 0.58 | 0.72 | 0.82 |
| 2 | 40182392 | rs720796 | 0.679 | 1.046 | -0.012 | -1.01 | -0.056 | -0.048 | -0.048 | 0 | 0.6 | 0.18 | 0.67 | 0.52 | 0.5 | 0.58 |
| 2 | 40194701 | rs768271 | 0.78 | 0.959 | 0.179 | -0.801 | -0.014 | -0.041 | -0.041 | 0 | 0.69 | 0.19 | 0.73 | 0.59 | 0.68 | 0.8 |
| 2 | 40195548 | rs13015010 | 1.197 | 0.986 | 0.091 | -0.859 | -0.027 | -0.019 | -0.019 | 0 | 0.6 | 0.13 | 0.66 | 0.58 | 0.61 | 0.71 |
| 2 | 40198780 | rs7572370 | 0.562 | 1.021 | 0.26 | -0.827 | -0.002 | -0.001 | -0.001 | 0 | 0.06 | 0.06 | 0.06 | 0.09 | 0.07 | 0.08 |
| 2 | 40202505 | rs17024711 | 0.057 | 0.984 | 0.401 | -0.86 | -0.034 | -0.052 | -0.052 | 0.012 | 0.21 | 0.12 | 0.19 | 0.4 | 0.31 | 0.19 |
| 2 | 40207338 | rs917969 | 1.589 | 0.535 | 0.511 | -0.733 | -0.023 | -0.163 | -0.163 | 0.037 | 0.69 | 0.05 | 0.71 | 0.49 | 0.76 | 0.69 |
| 2 | 40208916 | rs1013954 | 0.968 | 0.647 | 1.085 | -0.667 | -0.016 | 0.044 | 0.044 | 0 | 0.77 | 0.36 | 0.8 | 0.58 | 0.83 | 0.82 |
| 2 | 40215894 | rs10186088 | -0.28 | 0.731 | 1.345 | -0.658 | 0.045 | 0.016 | 0.016 | 0.736 | 0.06 | 0.34 | 0.03 | 0 | 0.02 | 0.01 |
| 2 | 40216202 | rs7599493 | -1.066 | 0.64 | 1.338 | -0.656 | -0.018 | 0.077 | 0.077 | 0 | 0.22 | 0.57 | 0.19 | 0.41 | 0.17 | 0.18 |
| 2 | 40217666 | rs1023584 | 0.132 | 0.629 | 1.367 | -0.704 | -0.028 | -0.095 | -0.095 | 0 | 0.86 | 0.64 | 0.9 | 1 | 0.93 | 0.92 |
| 2 | 40218136 | rs13390026 | -1.09 | 0.636 | 1.338 | -0.697 | -0.017 | 0.053 | 0.053 | 0 | 0.22 | 0.61 | 0.2 | 0.4 | 0.17 | 0.18 |
| 2 | 40221686 | rs6742932 | 0.32 | 0.654 | 1.269 | -0.689 | 0.007 | 0.009 | 0.009 | 0 | 0.06 | 0.1 | 0.07 | 0.11 | 0.07 | 0.09 |
| 2 | 40222206 | rs10865162 | 1.037 | 0.519 | 1.253 | -0.708 | -0.009 | 0.023 | 0.023 | 0 | 0.77 | 0.31 | 0.8 | 0.57 | 0.83 | 0.82 |
| 2 | 40224494 | rs17483267 | 1.653 | 0.542 | 0.656 | -0.794 | -0.048 | 0.036 | 0.036 | 0 | 0.13 | 0.02 | 0.09 | 0.05 | 0.21 | 0.17 |
| 2 | 40225253 | rs10177610 | -0.731 | 0.439 | 1.45 | -0.288 | -0.024 | -0.082 | -0.082 | 0.329 | 0.14 | 0.31 | 0.1 | 0.01 | 0.07 | 0.08 |
| 2 | 40226489 | rs17024784 | 0.009 | 0.495 | 0.899 | -0.251 | 0.017 | 0.207 | 0.207 | 0 | 0.08 | 0.2 | 0.09 | 0.43 | 0.11 | 0.1 |
| 2 | 40241868 | rs17024837 | -0.61 | 0.49 | 0.872 | -0.266 | -0.002 | 0.044 | 0.044 | 0 | 0.16 | 0.21 | 0.17 | 0.42 | 0.16 | 0.17 |
| 2 | 40264540 | rs12990700 | 1.694 | 0.169 | 0.261 | -0.368 | -0.061 | -0.086 | -0.086 | 0 | 0.47 | 0.04 | 0.54 | 0.47 | 0.48 | 0.51 |
| 2 | 40267991 | rs7586183 | -1.453 | -0.377 | 1.481 | -0.679 | -0.003 | 0.005 | 0.005 | 0 | 0.16 | 0.16 | 0.14 | 0.09 | 0.15 | 0.25 |
| 2 | 40270322 | rs6705665 | -1.692 | -0.307 | 1.594 | -0.62 | 0.028 | 0.023 | 0.023 | 0 | 0.09 | 0.22 | 0.08 | 0.1 | 0.1 | 0.19 |
| 2 | 40270847 | rs6719947 | -2.222 | -0.336 | 1.355 | -0.582 | -0.009 | -0.02 | -0.02 | 0 | 0.18 | 0.65 | 0.14 | 0.1 | 0.11 | 0.24 |
| 2 | 40273670 | rs17024951 | -1.758 | -0.262 | 1.242 | -0.558 | -0.064 | -0.146 | -0.146 | 0 | 0.2 | 0.44 | 0.12 | 0 | 0.08 | 0.16 |
| 2 | 40273917 | rs7608512 | -0.647 | -0.186 | 1.19 | -0.519 | 0.005 | -0.025 | -0.025 | 0 | 0.08 | 0.38 | 0.04 | 0 | 0.02 | 0.09 |
| 2 | 40275533 | rs2373788 | 2.15 | -0.188 | 0.844 | -0.529 | 0.026 | 0.063 | 0.063 | 0.932 | 0.7 | 0.21 | 0.79 | 0.9 | 0.83 | 0.7 |
| 2 | 40276550 | rs6751189 | -0.895 | -0.366 | 0.791 | -0.537 | 0.004 | -0.028 | -0.028 | 0 | 0.09 | 0.43 | 0.05 | 0 | 0.02 | 0.08 |
| 2 | 40278909 | rs6759630 | -1.082 | -0.263 | 0.284 | -0.482 | -0.069 | -0.15 | -0.15 | 0.018 | 0.21 | 0.5 | 0.15 | 0 | 0.08 | 0.16 |
| 2 | 40281964 | rs11891674 | -1.331 | 0.394 | -0.111 | -0.136 | 0.009 | -0.03 | -0.03 | 0 | 0.07 | 0.13 | 0.08 | 0.01 | 0.08 | 0.04 |
| 2 | 40282217 | rs11898574 | -1.578 | 0.755 | -0.229 | -0.325 | -0.01 | 0.004 | 0.004 | 0 | 0.08 | 0.07 | 0.08 | 0 | 0.1 | 0.07 |
| 2 | 40283459 | rs1861266 | -1.104 | -0.003 | -0.022 | -0.141 | -0.005 | 0.004 | 0.004 | 0 | 0.4 | 0.45 | 0.35 | 0.5 | 0.31 | 0.44 |
| 2 | 40284963 | rs985059 | -1.305 | -0.08 | -0.298 | -0.59 | -0.016 | 0.015 | 0.015 | 0 | 0.69 | 0.77 | 0.66 | 0.88 | 0.46 | 0.79 |
| 2 | 40288568 | rs6716299 | -0.57 | -0.864 | 0.132 | -0.899 | 0 | -0.002 | -0.002 | 0.171 | 0.28 | 0.29 | 0.29 | 0.43 | 0.22 | 0.41 |
| 2 | 40289267 | rs759361 | -1.687 | -0.868 | 0.086 | -0.973 | -0.006 | 0.001 | 0.001 | 0 | 0.69 | 0.73 | 0.67 | 0.88 | 0.49 | 0.81 |

| | | | | | | | | | | | | | | | | |
|---|---|---|---|---|---|---|---|---|---|---|---|---|---|---|---|---|
| 2 | 40291055 | rs12615150 | 0.308 | -1.055 | 0.392 | -0.878 | 0.017 | -0.021 | -0.021 | 0 | 0.35 | 0.24 | 0.34 | 0.36 | 0.24 | 0.41 |
| 2 | 40291868 | rs17025029 | 0.878 | -1.117 | 0.367 | -0.799 | 0.027 | -0.044 | -0.044 | 1.956 | 0.16 | 0.13 | 0.19 | 0.47 | 0.11 | 0.28 |
| 2 | 40297282 | rs2192723 | 0.462 | -1.307 | 0.51 | -0.914 | 0.007 | -0.01 | -0.01 | 1.9 | 0.2 | 0.2 | 0.22 | 0.47 | 0.14 | 0.32 |
| 2 | 40297772 | rs11898088 | 0.263 | -1.877 | 0.276 | -1.284 | -0.011 | 0.026 | 0.026 | 0.028 | 0.23 | 0.27 | 0.27 | 0.36 | 0.14 | 0.32 |
| 2 | 40306740 | rs11688696 | 0.974 | -1.832 | 0.125 | -1.188 | 0.043 | -0.078 | -0.078 | 0 | 0.18 | 0.08 | 0.21 | 0.44 | 0.13 | 0.29 |
| 2 | 40315899 | rs17025044 | 0.18 | -1.832 | 0.281 | -1.199 | -0.012 | 0.027 | 0.027 | 0 | 0.24 | 0.29 | 0.26 | 0.36 | 0.13 | 0.32 |
| 2 | 40317698 | rs12994093 | 1.001 | -1.636 | -0.071 | -1.265 | -0.033 | 0.031 | 0.031 | 0 | 0.07 | 0.01 | 0.07 | 0 | 0.1 | 0.06 |
| 2 | 40321189 | rs917977 | -1.797 | -1.813 | -0.225 | -1.356 | -0.03 | 0.112 | 0.112 | 0 | 0.59 | 0.81 | 0.6 | 0.84 | 0.41 | 0.71 |
| 2 | 40322900 | rs7422736 | -0.332 | -1.679 | -0.207 | -1.091 | -0.013 | -0.087 | -0.087 | 0.287 | 0.28 | 0.12 | 0.29 | 0.48 | 0.24 | 0.39 |
| 2 | 40323864 | rs13432990 | 1.518 | -1.381 | 0.07 | -1.077 | 0.001 | -0.004 | -0.004 | 0 | 0.14 | 0.09 | 0.1 | 0.01 | 0.11 | 0.11 |
| 2 | 40324846 | rs917976 | -0.516 | -1.045 | 0.454 | -0.656 | -0.043 | 0.071 | 0.071 | 0.099 | 0.26 | 0.5 | 0.28 | 0.32 | 0.13 | 0.32 |
| 2 | 40326142 | rs6726508 | -1.166 | -0.978 | 0.45 | -0.614 | 0.003 | 0.013 | 0.013 | 0 | 0.33 | 0.56 | 0.34 | 0.33 | 0.24 | 0.38 |
| 2 | 40330275 | rs4952441 | 0.553 | -0.076 | 0.608 | -0.841 | 0.055 | -0.109 | -0.109 | 0 | 0.82 | 0.48 | 0.86 | 0.96 | 0.8 | 0.85 |
| 2 | 40331135 | rs3922283 | 1.137 | -0.308 | 0.051 | -0.607 | 0.023 | -0.014 | -0.014 | 4.616 | 0.6 | 0.22 | 0.61 | 0.41 | 0.67 | 0.63 |
| 2 | 40331163 | rs17025097 | 0.28 | -0.39 | -0.139 | -0.591 | 0.039 | -0.069 | -0.069 | 0 | 0.21 | 0.14 | 0.25 | 0.57 | 0.17 | 0.26 |
| 2 | 40334899 | rs4952447 | 1.3 | -0.269 | -0.008 | -0.58 | 0.002 | -0.004 | -0.004 | 0 | 0.57 | 0.28 | 0.57 | 0.41 | 0.56 | 0.59 |
| 2 | 40335459 | rs6726967 | 1.127 | 0.354 | 0.8 | -0.347 | 0.071 | -0.029 | -0.029 | 0.045 | 0.89 | 0.49 | 0.88 | 0.97 | 0.86 | 0.93 |
| 2 | 40335876 | rs6727351 | 0.853 | 0.413 | 0.964 | -0.053 | 0.036 | 0.051 | 0.051 | 0 | 0.92 | 0.64 | 0.92 | 0.91 | 0.95 | 0.96 |
| 2 | 40339986 | rs5551 | -1.444 | 1.261 | 0.409 | 0.197 | -0.001 | 0.064 | 0.064 | 0 | 0.56 | 0.7 | 0.61 | 0.82 | 0.5 | 0.63 |
| 2 | 40341388 | rs5546 | 0.226 | 1.453 | 0.081 | 0.428 | 0.007 | -0.13 | -0.13 | 0 | 0.22 | 0.04 | 0.23 | 0.55 | 0.18 | 0.35 |
| 2 | 40341605 | rs5543 | -0.49 | 1.464 | 0.511 | 0.488 | 0.015 | -0.018 | -0.018 | 0 | 0.27 | 0.18 | 0.29 | 0.64 | 0.24 | 0.39 |
| 2 | 40341660 | rs5542 | -2.1 | 1.405 | 0.728 | 0.434 | 0.006 | 0.075 | 0.075 | 0.009 | 0.53 | 0.69 | 0.61 | 0.81 | 0.5 | 0.61 |
| 2 | 40341847 | rs5541 | 0.299 | 1.406 | 0.548 | 0.48 | 0.006 | -0.134 | -0.134 | 0 | 0.22 | 0.04 | 0.23 | 0.54 | 0.18 | 0.35 |
| 2 | 40345471 | rs13005439 | -1.154 | 1.519 | 0.892 | 0.318 | 0.013 | 0.106 | 0.106 | 0.702 | 0.45 | 0.3 | 0.51 | 0.81 | 0.4 | 0.55 |
| 2 | 40347244 | rs7601672 | 0.284 | 1.522 | 1.5 | 0.3 | 0.121 | -0.017 | -0.017 | 0.436 | 0.93 | 0.61 | 0.91 | 0.99 | 0.9 | 0.93 |
| 2 | 40349094 | rs9309051 | 0.154 | 1.512 | 1.485 | 0.306 | 0 | -0.022 | -0.022 | 0.609 | 0.19 | 0.26 | 0.22 | 0.16 | 0.15 | 0.16 |
| 2 | 40349162 | rs13405865 | -0.39 | 1.546 | 1.255 | 0.384 | 0.008 | -0.01 | -0.01 | 0 | 0.29 | 0.19 | 0.29 | 0.66 | 0.26 | 0.41 |
| 2 | 40350271 | rs2373792 | 1.934 | 1.523 | 1.216 | 0.355 | -0.003 | 0.144 | 0.144 | 0 | 0.45 | 0.19 | 0.4 | 0.18 | 0.5 | 0.37 |
| 2 | 40354688 | rs2110927 | 0.564 | 1.648 | 1.357 | 0.54 | 0.002 | -0.014 | -0.014 | 0 | 0.73 | 0.34 | 0.68 | 0.82 | 0.74 | 0.78 |
| 2 | 40357323 | rs2160364 | 0.336 | 1.614 | 1.467 | 0.55 | -0.001 | -0.005 | -0.005 | 0 | 0.19 | 0.18 | 0.23 | 0.17 | 0.16 | 0.15 |
| 2 | 40358175 | rs765554 | 0.87 | 1.629 | 1.459 | 0.57 | 0.005 | -0.008 | -0.008 | 0 | 0.64 | 0.46 | 0.64 | 0.36 | 0.65 | 0.52 |
| 2 | 40360449 | rs929642 | -2.063 | 1.578 | 1.592 | 0.561 | 0.004 | 0.079 | 0.079 | 0.02 | 0.53 | 0.73 | 0.6 | 0.8 | 0.5 | 0.62 |
| 2 | 40361707 | rs741320 | 0.215 | 1.598 | 1.452 | 0.572 | 0.059 | -0.031 | -0.031 | 0.015 | 0.06 | 0.17 | 0.08 | 0.01 | 0.1 | 0.07 |
| 2 | 40362956 | rs4140823 | -0.202 | 1.646 | 1.412 | 0.588 | 0.009 | -0.034 | -0.034 | 0 | 0.27 | 0.14 | 0.26 | 0.62 | 0.24 | 0.4 |
| 2 | 40363644 | rs2110922 | -1.067 | 1.637 | 1.452 | 0.995 | 0.007 | 0.027 | 0.027 | 0 | 0.34 | 0.45 | 0.35 | 0.52 | 0.34 | 0.46 |
| 2 | 40364895 | rs7573537 | -0.741 | 1.654 | 1.439 | 0.992 | 0.004 | 0.005 | 0.005 | 0 | 0.32 | 0.38 | 0.35 | 0.34 | 0.34 | 0.44 |
| 2 | 40366301 | rs2072532 | -1.958 | 1.635 | 1.161 | -0.058 | 0.008 | 0.105 | 0.105 | 0 | 0.53 | 0.73 | 0.61 | 0.8 | 0.5 | 0.62 |
| 2 | 40372704 | rs4952490 | -0.529 | 1.73 | 1.153 | -0.045 | 0.042 | -0.069 | -0.069 | 0.329 | 0.26 | 0.47 | 0.33 | 0.17 | 0.26 | 0.23 |
| 2 | 40373196 | rs17501526 | 1.917 | 1.717 | 1.006 | -0.059 | -0.019 | 0.17 | 0.17 | 0 | 0.46 | 0.14 | 0.39 | 0.2 | 0.51 | 0.38 |
| 2 | 40373872 | rs444999 | -0.614 | 1.698 | 1.027 | -0.048 | 0.029 | -0.054 | -0.054 | 1.784 | 0.26 | 0.51 | 0.34 | 0.18 | 0.26 | 0.23 |
| 2 | 40376453 | rs2058693 | -0.47 | 1.59 | 0.968 | 0 | 0.006 | -0.006 | -0.006 | 0 | 0.29 | 0.24 | 0.27 | 0.63 | 0.25 | 0.41 |
| 2 | 40377680 | rs387103 | -2.44 | 1.28 | 0.645 | -0.504 | 0.038 | 0.046 | 0.046 | 0 | 0.71 | 0.83 | 0.77 | 0.88 | 0.8 | 0.74 |
| 2 | 40377770 | rs17025271 | 0.032 | 1.534 | 1.154 | -0.139 | 0.04 | -0.033 | -0.033 | 0 | 0.06 | 0.14 | 0.08 | 0.01 | 0.1 | 0.07 |
| 2 | 40378186 | rs432188 | 0.543 | 1.345 | 1.168 | -0.107 | -0.03 | -0.015 | -0.015 | 0 | 0.16 | 0.29 | 0.23 | 0.16 | 0.1 | 0.13 |
| 2 | 40381369 | rs13425134 | -0.386 | 1.376 | 1.15 | -0.085 | -0.027 | -0.007 | -0.007 | 0 | 0.82 | 0.54 | 0.74 | 0.82 | 0.89 | 0.86 |
| 2 | 40384733 | rs452369 | 0.533 | 1.38 | 0.959 | -0.083 | 0.002 | 0.001 | 0.001 | 1.825 | 0.16 | 0.16 | 0.23 | 0.18 | 0.11 | 0.13 |
| 2 | 40386331 | rs7570522 | -0.924 | 1.652 | 0.96 | -0.145 | 0.145 | -0.049 | -0.049 | 0 | 0.1 | 0.32 | 0.12 | 0.01 | 0.14 | 0.07 |
| 2 | 40387489 | rs1861427 | -0.207 | 1.681 | 0.052 | -0.084 | 0.011 | -0.041 | -0.041 | 2.767 | 0.28 | 0.16 | 0.25 | 0.6 | 0.23 | 0.43 |
| 2 | 40388357 | rs2072531 | 1.757 | 1.664 | -0.056 | -0.085 | -0.071 | 0.127 | 0.127 | 0.01 | 0.44 | 0.09 | 0.38 | 0.21 | 0.52 | 0.35 |
| 2 | 40390470 | rs1428571 | -0.391 | 1.398 | 0.286 | -0.021 | 0.004 | -0.013 | -0.013 | 0 | 0.25 | 0.23 | 0.25 | 0.63 | 0.21 | 0.4 |
| 2 | 40391516 | rs2072530 | -0.193 | 1.571 | 0.438 | 0.159 | 0.005 | -0.014 | -0.014 | 0 | 0.18 | 0.17 | 0.22 | 0.64 | 0.15 | 0.46 |
| 2 | 40391789 | rs1005213 | 1.227 | 1.514 | 0.834 | -0.043 | -0.003 | -0.009 | -0.009 | 0 | 0.6 | 0.4 | 0.5 | 0.2 | 0.71 | 0.46 |
| 2 | 40394140 | rs2192919 | 1.924 | 1.849 | 0.736 | -0.139 | -0.021 | 0.167 | 0.167 | 1.714 | 0.51 | 0.22 | 0.4 | 0.17 | 0.56 | 0.4 |
| 2 | 40395937 | rs17025372 | -0.06 | 1.602 | 1.05 | -0.08 | 0.003 | -0.006 | -0.006 | 0.016 | 0.18 | 0.19 | 0.23 | 0.65 | 0.15 | 0.45 |
| 2 | 40396078 | rs404005 | 1.195 | 1.603 | 1.289 | -0.109 | 0.005 | 0.002 | 0.002 | 0 | 0.7 | 0.39 | 0.65 | 0.35 | 0.71 | 0.47 |
| 2 | 40397165 | rs2072529 | 0.072 | 1.572 | 1.187 | -0.056 | 0.002 | -0.003 | -0.003 | 0.265 | 0.18 | 0.19 | 0.22 | 0.65 | 0.12 | 0.43 |
| 2 | 40400312 | rs417591 | -1.066 | 1.562 | 1.21 | -0.099 | 0.003 | -0.056 | -0.056 | 0 | 0.25 | 0.51 | 0.33 | 0.17 | 0.21 | 0.12 |
| 2 | 40400672 | rs408009 | 0.431 | 1.489 | 1.196 | -0.083 | -0.011 | -0.005 | -0.005 | 0 | 0.16 | 0.16 | 0.23 | 0.16 | 0.08 | 0.05 |
| 2 | 40401500 | rs3828295 | 1.094 | 1.521 | 1.213 | -0.105 | -0.015 | -0.024 | -0.024 | 0 | 0.63 | 0.57 | 0.52 | 0.2 | 0.76 | 0.49 |
| 2 | 40403104 | rs106706 | 0.681 | 1.561 | 1.247 | -0.075 | -0.025 | 0.046 | 0.046 | 0 | 0.76 | 0.55 | 0.74 | 0.37 | 0.85 | 0.53 |
| 2 | 40406518 | rs688673 | 0.004 | 1.546 | 1.21 | -0.082 | -0.003 | -0.001 | -0.001 | 0.69 | 0.18 | 0.16 | 0.23 | 0.16 | 0.09 | 0.06 |
| 2 | 40406967 | rs385665 | 0.175 | 1.73 | 1.262 | 0.032 | -0.006 | 0.024 | 0.024 | 0 | 0.81 | 0.72 | 0.76 | 0.35 | 0.86 | 0.54 |
| 2 | 40407867 | rs372072 | 0.345 | 1.729 | 1.233 | 0.016 | -0.002 | 0.006 | 0.006 | 0 | 0.79 | 0.73 | 0.75 | 0.35 | 0.84 | 0.54 |
| 2 | 40411717 | rs376535 | -0.605 | 1.78 | 1.261 | 0.015 | -0.004 | 0.01 | 0.01 | 0.161 | 0.84 | 0.8 | 0.78 | 0.35 | 0.9 | 0.58 |
| 2 | 40413542 | rs426295 | -0.674 | 1.811 | 1.286 | -0.002 | 0.03 | -0.032 | -0.032 | 0 | 0.19 | 0.32 | 0.31 | 0.17 | 0.2 | 0.1 |
| 2 | 40420129 | rs12104840 | 1.972 | 1.803 | 1.215 | -0.053 | -0.03 | 0.201 | 0.201 | 1.144 | 0.52 | 0.11 | 0.4 | 0.19 | 0.63 | 0.43 |
| 2 | 40420497 | rs373857 | -0.722 | 2.04 | 0.947 | -0.323 | -0.001 | -0.001 | -0.001 | 0 | 0.4 | 0.34 | 0.57 | 0.26 | 0.57 | 0.22 |
| 2 | 40420794 | rs369140 | -0.917 | 1.853 | 2.062 | 0.694 | 0.029 | -0.027 | -0.027 | 0.126 | 0.19 | 0.28 | 0.31 | 0.17 | 0.2 | 0.1 |
| 2 | 40423454 | rs381610 | 0.743 | 1.942 | 2.041 | 0.8 | -0.001 | 0.021 | 0.021 | 0 | 0.78 | 0.68 | 0.76 | 0.38 | 0.83 | 0.56 |

| | | | | | | | | | | | | | | | | |
|---|---|---|---|---|---|---|---|---|---|---|---|---|---|---|---|---|
| 2 | 40424444 | rs449383 | 1.881 | 1.985 | 2.083 | 0.784 | 0.005 | 0.002 | 0.002 | 0 | 0.67 | 0.41 | 0.66 | 0.37 | 0.72 | 0.49 |
| 2 | 40429180 | rs13408531 | -1.355 | 1.847 | 2.035 | 0.766 | 0.04 | -0.044 | -0.044 | 0.086 | 0.08 | 0.15 | 0.07 | 0.01 | 0.11 | 0.04 |
| 2 | 40429442 | rs11674102 | 1.419 | 1.815 | 2.046 | 0.787 | -0.019 | -0.027 | -0.027 | 0 | 0.64 | 0.44 | 0.52 | 0.23 | 0.76 | 0.49 |
| 2 | 40435115 | rs408263 | 0.298 | 1.81 | 2.014 | 0.677 | 0.005 | -0.008 | -0.008 | 0.224 | 0.81 | 0.8 | 0.8 | 0.36 | 0.86 | 0.56 |
| 2 | 40438489 | rs399061 | -0.977 | 1.877 | 2.028 | 0.707 | -0.025 | -0.038 | -0.038 | 0 | 0.19 | 0.32 | 0.26 | 0.15 | 0.1 | 0.06 |
| 2 | 40447587 | rs2540904 | 1.493 | 1.864 | 2.061 | 0.703 | 0.02 | 0.004 | 0.004 | 0 | 0.69 | 0.48 | 0.69 | 0.35 | 0.7 | 0.48 |
| 2 | 40451209 | rs4952594 | 2.261 | 1.858 | 2.055 | 0.696 | -0.018 | 0.136 | 0.136 | 0 | 0.54 | 0.21 | 0.43 | 0.21 | 0.61 | 0.44 |
| 2 | 40451724 | rs384994 | -0.255 | 1.936 | 2.073 | 0.73 | -0.02 | -0.023 | -0.023 | 0 | 0.16 | 0.27 | 0.26 | 0.14 | 0.1 | 0.06 |
| 2 | 40453794 | rs10490208 | 2.631 | 1.868 | 1.968 | 0.732 | -0.058 | 0.195 | 0.195 | 0.279 | 0.51 | 0.08 | 0.41 | 0.21 | 0.6 | 0.42 |
| 2 | 40454348 | rs404214 | 0.815 | 0.997 | 2.097 | 0.803 | -0.003 | 0.027 | 0.027 | 0 | 0.77 | 0.64 | 0.76 | 0.36 | 0.82 | 0.55 |
| 2 | 40468418 | rs4952599 | 1.712 | 0.945 | 1.964 | 0.83 | 0.057 | 0.059 | 0.059 | 1.431 | 0.61 | 0.2 | 0.61 | 0.23 | 0.76 | 0.49 |
| 2 | 40469480 | rs615449 | -1.308 | 0.93 | 2.057 | 0.823 | -0.037 | -0.048 | -0.048 | 0.161 | 0.21 | 0.6 | 0.3 | 0.14 | 0.11 | 0.07 |
| 2 | 40469705 | rs371909 | -1.308 | 0.93 | 2.057 | 0.823 | -0.037 | -0.048 | -0.048 | 0 | 0.21 | 0.6 | 0.3 | 0.14 | 0.11 | 0.07 |
| 2 | 40471407 | rs11124737 | 1.393 | 1.106 | 2.093 | 0.908 | 0.015 | 0.017 | 0.017 | 0.461 | 0.65 | 0.3 | 0.62 | 0.22 | 0.76 | 0.5 |
| 2 | 40472864 | rs7602176 | 2.077 | 1.163 | 2.001 | 0.9 | 0.045 | 0.092 | 0.092 | 0 | 0.57 | 0.24 | 0.56 | 0.21 | 0.68 | 0.44 |
| 2 | 40473347 | rs380590 | 0.386 | 1.15 | 2.049 | 0.998 | -0.005 | 0.014 | 0.014 | 0 | 0.81 | 0.74 | 0.89 | 0.38 | 0.87 | 0.55 |
| 2 | 40473425 | rs505473 | -1.028 | 1.248 | 1.836 | 0.944 | 0.013 | -0.031 | -0.031 | 0 | 0.19 | 0.39 | 0.3 | 0.16 | 0.14 | 0.08 |
| 2 | 40476402 | rs7570300 | -1.352 | 0.789 | 1.717 | 1.124 | 0.023 | -0.049 | -0.049 | 4.612 | 0.09 | 0.22 | 0.07 | 0.01 | 0.09 | 0.05 |
| 2 | 40476866 | rs10490209 | -0.378 | 0.805 | 1.237 | 1.604 | -0.036 | -0.129 | -0.129 | 0 | 0.17 | 0.01 | 0.1 | 0.56 | 0.14 | 0.45 |
| 2 | 40477658 | rs380082 | -0.759 | 0.526 | 1.623 | 1.142 | -0.013 | -0.001 | -0.001 | 0 | 0.15 | 0.25 | 0.26 | 0.16 | 0.1 | 0.06 |
| 2 | 40485074 | rs434082 | -0.818 | 0.553 | 1.309 | 1.153 | -0.006 | -0.004 | -0.004 | 0 | 0.14 | 0.22 | 0.26 | 0.16 | 0.1 | 0.06 |
| 2 | 40486939 | rs391378 | -0.93 | 0.698 | 1.239 | 1.357 | 0.002 | -0.01 | -0.01 | 0 | 0.14 | 0.12 | 0.34 | 0.22 | 0.13 | 0.08 |
| 2 | 40489408 | rs7607333 | -0.381 | 0.467 | 1.452 | 1.456 | -0.011 | 0.051 | 0.051 | 0 | 0.17 | 0.22 | 0.11 | 0.56 | 0.11 | 0.41 |
| 2 | 40490009 | rs421576 | -1.606 | 0.337 | 1.816 | 1.275 | -0.003 | 0.009 | 0.009 | 1.379 | 0.2 | 0.38 | 0.4 | 0.24 | 0.2 | 0.12 |
| 2 | 40490288 | rs2540903 | -0.567 | 0.427 | 1.337 | 1.338 | -0.001 | 0.006 | 0.006 | 1.873 | 0.13 | 0.19 | 0.34 | 0.22 | 0.12 | 0.08 |
| 2 | 40493990 | rs3862986 | -1.595 | 0.103 | 1.296 | 1.449 | -0.016 | 0.024 | 0.024 | 0 | 0.27 | 0.62 | 0.21 | 0.6 | 0.18 | 0.46 |
| 2 | 40497091 | rs17025712 | -0.768 | 0.254 | 0.523 | 1.622 | -0.004 | 0.039 | 0.039 | 0 | 0.19 | 0.24 | 0.12 | 0.61 | 0.13 | 0.42 |
| 2 | 40500993 | rs9808490 | -1.521 | 0.338 | -0.141 | 1.323 | 0.008 | 0.022 | 0.022 | 0.131 | 0.4 | 0.62 | 0.51 | 0.72 | 0.34 | 0.51 |
| 2 | 40503246 | rs17025732 | -0.472 | 0.202 | -0.262 | 1.952 | -0.015 | -0.013 | -0.013 | 0 | 0.05 | 0 | 0.02 | 0.2 | 0.04 | 0.16 |
| 2 | 40504788 | rs7571216 | -0.936 | -0.056 | 0.378 | 1.822 | -0.035 | 0.076 | 0.076 | 1.451 | 0.2 | 0.55 | 0.39 | 0.29 | 0.16 | 0.19 |
| 2 | 40509154 | rs2192771 | -0.46 | -0.381 | 0.343 | 1.493 | -0.032 | 0.008 | 0.008 | 0 | 0.34 | 0.47 | 0.29 | 0.37 | 0.25 | 0.35 |
| 2 | 40532749 | rs10210665 | -1.103 | -0.389 | 0.994 | 1.633 | -0.002 | 0.069 | 0.069 | 0.015 | 0.07 | 0.22 | 0.1 | 0.11 | 0.04 | 0.09 |
| 2 | 40538968 | rs2110743 | -1.22 | -0.49 | 1.446 | 1.931 | 0.073 | 0.204 | 0.204 | 0 | 0.1 | 0.45 | 0.22 | 0.25 | 0.1 | 0.11 |
| 2 | 40540620 | rs10207312 | -0.921 | -0.497 | 1.323 | 1.918 | -0.004 | 0.065 | 0.065 | 0.129 | 0.06 | 0.21 | 0.09 | 0.11 | 0.04 | 0.08 |
| 2 | 40546386 | rs13031392 | 0.292 | -0.852 | 0.575 | 1.847 | 0.046 | -0.098 | -0.098 | 0.741 | 0.27 | 0.04 | 0.22 | 0.36 | 0.19 | 0.36 |
| 2 | 40547186 | rs2160237 | -1.488 | -0.824 | 0.575 | 1.785 | 0.086 | 0.125 | 0.125 | 0 | 0.47 | 0.87 | 0.57 | 0.73 | 0.36 | 0.51 |
| 2 | 40555733 | rs4952609 | -0.571 | -0.837 | 1.062 | 1.775 | -0.034 | 0.036 | 0.036 | 0 | 0.34 | 0.53 | 0.33 | 0.45 | 0.26 | 0.36 |
| 2 | 40558573 | rs13005619 | -0.984 | -0.839 | 0.364 | 1.669 | -0.009 | 0.03 | 0.03 | 0 | 0.12 | 0.19 | 0.18 | 0.2 | 0.08 | 0.12 |
| 2 | 40560630 | rs4952404 | 0.223 | -0.872 | -0.222 | 1.457 | 0.006 | -0.098 | -0.098 | 0 | 0.27 | 0.02 | 0.23 | 0.34 | 0.22 | 0.41 |
| 2 | 40564500 | rs13406526 | -0.862 | -0.533 | 0.459 | 1.333 | 0.003 | 0.033 | 0.033 | 0.048 | 0.1 | 0.34 | 0.09 | 0.1 | 0.08 | 0.06 |
| 2 | 40566235 | rs768710 | 0.237 | -0.228 | -0.741 | 1.201 | -0.067 | -0.069 | -0.069 | 3.035 | 0.38 | 0.01 | 0.32 | 0.37 | 0.36 | 0.46 |
| 2 | 40566271 | rs13413412 | -0.819 | 0.339 | -0.008 | 1.099 | 0.046 | 0.04 | 0.04 | 0.259 | 0.11 | 0.46 | 0.13 | 0.1 | 0.12 | 0.06 |
| 2 | 40567743 | rs4952611 | -1.017 | 0.417 | -0.486 | 1.253 | -0.025 | 0 | 0 | 0.018 | 0.63 | 0.74 | 0.65 | 0.66 | 0.55 | 0.66 |
| 2 | 40569746 | rs4521092 | -0.387 | 0.427 | -0.974 | 1.145 | 0.012 | -0.042 | -0.042 | 0 | 0.56 | 0.86 | 0.52 | 0.51 | 0.36 | 0.57 |
| 2 | 40574207 | rs7561383 | -1.057 | 0.794 | -1.026 | 0.53 | -0.061 | 0.07 | 0.07 | 0 | 0.69 | 0.88 | 0.68 | 0.88 | 0.56 | 0.68 |
| 2 | 40574269 | rs11890040 | -0.584 | 0.798 | -0.556 | 0.672 | 0.039 | 0.072 | 0.072 | 0 | 0.08 | 0.17 | 0.19 | 0.2 | 0.09 | 0.08 |
| 2 | 40575809 | rs12712691 | -0.678 | 0.723 | -0.599 | 0.597 | -0.082 | 0.095 | 0.095 | 0 | 0.64 | 0.86 | 0.63 | 0.88 | 0.47 | 0.65 |
| 2 | 40580065 | rs12712692 | -0.412 | 0.758 | -0.579 | 0.651 | -0.016 | 0.005 | 0.005 | 0.128 | 0.62 | 0.73 | 0.47 | 0.69 | 0.44 | 0.61 |
| 2 | 40592311 | rs34408472 | -1.083 | 0.643 | -0.553 | 0.808 | 0.001 | -0.003 | -0.003 | 0 | 0.11 | 0.08 | 0.19 | 0.2 | 0.08 | 0.11 |
| 2 | 40596524 | rs6544334 | -0.677 | 0.327 | -0.798 | 0.483 | 0.004 | -0.023 | -0.023 | 0.238 | 0.6 | 0.87 | 0.64 | 0.36 | 0.65 | 0.55 |
| 2 | 40596632 | rs984707 | -0.705 | 0.326 | -0.799 | 0.478 | 0.003 | -0.03 | -0.03 | 0 | 0.6 | 0.87 | 0.63 | 0.37 | 0.65 | 0.55 |
| 2 | 40598884 | rs7581611 | -0.325 | 0.615 | -0.542 | 0.44 | -0.001 | 0.064 | 0.064 | 0 | 0.54 | 0.67 | 0.46 | 0.17 | 0.59 | 0.47 |
| 2 | 40601179 | rs981739 | -0.915 | 0.638 | -0.579 | 0.448 | -0.04 | -0.045 | -0.045 | 0.109 | 0.62 | 0.92 | 0.64 | 0.37 | 0.64 | 0.54 |
| 2 | 40607504 | rs918001 | -0.027 | 0.782 | -0.701 | 0.402 | -0.01 | 0.152 | 0.152 | 0.658 | 0.51 | 0.77 | 0.43 | 0.16 | 0.56 | 0.43 |
| 2 | 40608306 | rs4952410 | -0.069 | 0.784 | -0.714 | 0.405 | 0.005 | 0.155 | 0.155 | 0.355 | 0.5 | 0.75 | 0.42 | 0.16 | 0.54 | 0.42 |
| 2 | 40613207 | rs4446101 | -0.852 | 0.839 | -0.916 | 0.426 | -0.036 | -0.066 | -0.066 | 0 | 0.61 | 0.98 | 0.67 | 0.36 | 0.69 | 0.55 |
| 2 | 40615520 | rs6742210 | 0.667 | 0.665 | -0.651 | 0.266 | -0.04 | 0.119 | 0.119 | 0 | 0.4 | 0.16 | 0.36 | 0.13 | 0.49 | 0.39 |
| 2 | 40617820 | rs2373859 | -0.636 | 0.562 | -0.754 | 0.391 | -0.06 | -0.077 | -0.077 | 0.003 | 0.6 | 0.98 | 0.67 | 0.36 | 0.66 | 0.53 |
| 2 | 40621060 | rs7557545 | -1.431 | 0.793 | -0.061 | 0.809 | -0.007 | 0.039 | 0.039 | 0 | 0.16 | 0.32 | 0.23 | 0.23 | 0.14 | 0.13 |
| 2 | 40621387 | rs12615002 | -1.425 | 0.731 | -0.01 | 0.853 | 0.014 | 0.05 | 0.05 | 0 | 0.15 | 0.32 | 0.26 | 0.23 | 0.15 | 0.12 |
| 2 | 40626336 | rs10490047 | -0.025 | 0.931 | -0.461 | 0.796 | 0.001 | 0 | 0 | 0 | 0.43 | 0.51 | 0.54 | 0.48 | 0.46 | 0.45 |
| 2 | 40627712 | rs6544336 | 0.129 | 0.307 | -0.649 | 0.613 | 0.007 | 0.01 | 0.01 | 0.392 | 0.58 | 0.37 | 0.58 | 0.33 | 0.71 | 0.35 |
| 2 | 40628222 | rs6730958 | -0.186 | -0.074 | -0.419 | 1.154 | -0.011 | -0.009 | -0.009 | 0.275 | 0.67 | 0.46 | 0.77 | 0.72 | 0.77 | 0.61 |
| 2 | 40628394 | rs6544337 | 0.663 | 0.184 | -0.61 | 1.03 | 0.003 | 0.014 | 0.014 | 2.338 | 0.53 | 0.42 | 0.56 | 0.32 | 0.65 | 0.28 |
| 2 | 40630678 | rs10490046 | 0.243 | -0.088 | -1.107 | 0.485 | -0.003 | 0.001 | 0.001 | 0 | 0.27 | 0.3 | 0.37 | 0.31 | 0.26 | 0.22 |
| 2 | 40632095 | rs2216007 | -0.079 | -0.315 | -0.612 | 0.417 | 0.012 | -0.003 | -0.003 | 0 | 0.37 | 0.23 | 0.31 | 0.37 | 0.25 | 0.5 |
| 2 | 40632431 | rs6732205 | 0.321 | -0.284 | -0.522 | 0.524 | 0.002 | -0.003 | -0.003 | 0 | 0.31 | 0.27 | 0.28 | 0.36 | 0.19 | 0.43 |
| 2 | 40638024 | rs2888677 | -0.129 | -0.359 | -0.553 | 0.151 | 0.025 | 0.063 | 0.063 | 0 | 0.48 | 0.63 | 0.41 | 0.26 | 0.37 | 0.42 |
| 2 | 40638449 | rs4366950 | -0.312 | -0.406 | -0.772 | 0.112 | 0.042 | 0.143 | 0.143 | 0.201 | 0.5 | 0.8 | 0.43 | 0.26 | 0.38 | 0.42 |

| | | | | | | | | | | | | | | | | |
|---|---|---|---|---|---|---|---|---|---|---|---|---|---|---|---|---|
| 2 | 40641410 | rs17466980 | 1.075 | -0.553 | -0.786 | -0.13 | -0.007 | 0.059 | 0.059 | 0.895 | 0.23 | 0.13 | 0.13 | 0 | 0.18 | 0.18 |
| 2 | 40642423 | rs4952621 | 0.105 | -0.627 | -0.743 | -0.122 | 0.001 | 0 | 0 | 1.337 | 0.32 | 0.34 | 0.41 | 0.67 | 0.24 | 0.53 |
| 2 | 40649084 | rs13414307 | -0.774 | -0.556 | -1.022 | -0.24 | -0.038 | 0.051 | 0.051 | 1.486 | 0.6 | 0.77 | 0.48 | 0.26 | 0.48 | 0.55 |
| 2 | 40661305 | rs4952414 | -1.284 | -0.746 | -1.235 | 0.141 | -0.079 | 0.194 | 0.194 | 2.481 | 0.64 | 0.91 | 0.75 | 0.94 | 0.53 | 0.85 |
| 2 | 40662546 | rs4629203 | 0.341 | -0.86 | -0.553 | 0.318 | -0.003 | 0.006 | 0.006 | 0.674 | 0.3 | 0.4 | 0.4 | 0.65 | 0.23 | 0.48 |
| 2 | 40674308 | rs6737745 | -0.414 | -0.585 | -0.799 | -0.253 | 0.006 | 0.232 | 0.232 | 1.22 | 0.54 | 0.81 | 0.67 | 0.94 | 0.41 | 0.75 |
| 2 | 40680149 | rs2301343 | -0.261 | -0.428 | -0.528 | 0.084 | -0.001 | 0.058 | 0.058 | 0.032 | 0.26 | 0.09 | 0.19 | 0.11 | 0.24 | 0.24 |
| 2 | 40681590 | rs2111588 | 0.495 | -0.526 | -0.013 | 0.315 | -0.003 | 0.005 | 0.005 | 0 | 0.29 | 0.34 | 0.35 | 0.35 | 0.21 | 0.32 |
| 2 | 40687437 | rs7597373 | 0.072 | -0.653 | 0.208 | 0.184 | 0.011 | 0.077 | 0.077 | 1.005 | 0.42 | 0.34 | 0.28 | 0.11 | 0.33 | 0.34 |
| 2 | 40688262 | rs2373864 | -0.711 | -0.276 | 0.175 | 0.086 | 0.008 | 0.07 | 0.07 | 1.102 | 0.56 | 0.39 | 0.4 | 0.12 | 0.55 | 0.49 |
| 2 | 40691468 | rs11679585 | 1.448 | -0.296 | 0.119 | 0.131 | -0.035 | 0.11 | 0.11 | 0.087 | 0.22 | 0.07 | 0.11 | 0 | 0.18 | 0.17 |
| 2 | 40710953 | rs10205402 | 1.002 | -0.37 | 0.373 | -0.114 | -0.016 | 0.115 | 0.115 | 1.675 | 0.35 | 0.15 | 0.21 | 0.08 | 0.39 | 0.11 |
| 2 | 40715880 | rs7590554 | -0.087 | -0.104 | 0.892 | 0.581 | -0.029 | 0.078 | 0.078 | 0.014 | 0.74 | 0.44 | 0.7 | 0.6 | 0.81 | 0.63 |
| 2 | 40717245 | rs4952630 | 0.118 | 0.173 | 0.659 | 0.594 | 0.001 | -0.002 | -0.002 | 0 | 0.29 | 0.28 | 0.21 | 0.13 | 0.2 | 0.22 |
| 2 | 40725181 | rs11887097 | -0.755 | 0.231 | 0.636 | 0.807 | 0.001 | -0.003 | -0.003 | 0.453 | 0.42 | 0.61 | 0.55 | 0.56 | 0.4 | 0.43 |
| 2 | 40725202 | rs11897805 | -0.266 | 0.332 | 0.145 | 0.827 | 0.001 | 0.02 | 0.02 | 0 | 0.36 | 0.28 | 0.31 | 0.12 | 0.35 | 0.3 |
| 2 | 40725639 | rs11124743 | -0.297 | 0.338 | 0.156 | 0.854 | 0 | 0.017 | 0.017 | 0 | 0.36 | 0.31 | 0.31 | 0.13 | 0.36 | 0.3 |
| 2 | 40743183 | rs17026114 | -0.854 | 0.081 | 0.409 | 1.6 | 0.018 | 0.349 | 0.349 | 0 | 0.06 | 0.17 | 0.23 | 0.48 | 0.04 | 0.13 |
| 2 | 40761794 | rs3796080 | -0.43 | 0.123 | -0.04 | 1.585 | -0.001 | 0.157 | 0.157 | 0 | 0.05 | 0.06 | 0.22 | 0.45 | 0.04 | 0.13 |
| 2 | 40775133 | rs10179753 | 0.359 | 0.085 | -0.408 | 0.995 | -0.012 | 0.052 | 0.052 | 0.503 | 0.34 | 0.25 | 0.24 | 0.1 | 0.4 | 0.15 |
| 2 | 40778289 | rs7561779 | -1.104 | -0.098 | -0.056 | 1.93 | -0.004 | 0.206 | 0.206 | 0 | 0.06 | 0.14 | 0.07 | 0.39 | 0.05 | 0.1 |
| 2 | 40778325 | rs11674742 | -0.391 | -0.027 | -0.829 | 1.549 | 0.007 | 0.009 | 0.009 | 0 | 0.23 | 0.21 | 0.17 | 0.11 | 0.18 | 0.19 |
| 2 | 40782071 | rs7576549 | -1.384 | 0.689 | -0.689 | 2.328 | 0 | 0.102 | 0.102 | 0 | 0.06 | 0.07 | 0.06 | 0.43 | 0.07 | 0.2 |
| 2 | 40783406 | rs9309056 | -1.458 | 0.969 | -0.523 | 2.304 | 0.011 | 0.123 | 0.123 | 0 | 0.07 | 0.13 | 0.18 | 0.47 | 0.13 | 0.12 |
| 2 | 40783676 | rs7593331 | 0.831 | 0.851 | -0.443 | 2.524 | -0.001 | 0.022 | 0.022 | 0 | 0.86 | 0.84 | 0.67 | 0.71 | 0.82 | 0.76 |
| 2 | 40786281 | rs2373889 | 0.991 | 0.814 | -0.285 | 2.625 | 0.011 | 0.136 | 0.136 | 0 | 0.93 | 0.81 | 0.79 | 0.74 | 0.88 | 0.85 |
| 2 | 40806338 | rs4952421 | 1.333 | 0.85 | 0.158 | 2.566 | 0.033 | 0.15 | 0.15 | 0 | 0.92 | 0.77 | 0.9 | 0.74 | 0.85 | 0.89 |
| 2 | 40818629 | rs2193560 | 2.129 | 0.546 | 0.131 | 2.362 | 0.011 | 0.123 | 0.123 | 0 | 0.87 | 0.63 | 0.86 | 0.73 | 0.82 | 0.88 |
| 2 | 40825531 | rs42815 | 0.561 | 0.167 | 0.094 | 1.097 | 0.002 | 0.006 | 0.006 | 0 | 0.51 | 0.41 | 0.64 | 0.68 | 0.51 | 0.49 |
| 2 | 40825962 | rs6717053 | 0.002 | 0.091 | 0.708 | 1.054 | 0.006 | -0.032 | -0.032 | 0.02 | 0.39 | 0.22 | 0.51 | 0.53 | 0.34 | 0.52 |
| 2 | 40826397 | rs42813 | 1.35 | 0.183 | 0.985 | 1.215 | 0.002 | 0.035 | 0.035 | 0 | 0.87 | 0.75 | 0.85 | 0.72 | 0.81 | 0.86 |
| 2 | 40832632 | rs10195113 | 1.328 | 0.019 | 0.657 | 0.797 | -0.012 | 0.031 | 0.031 | 0 | 0.06 | 0 | 0.03 | 0 | 0.06 | 0.05 |
| 2 | 40837953 | rs42802 | 0.883 | -0.096 | 0.756 | 0.754 | 0 | -0.019 | -0.019 | 0 | 0.49 | 0.49 | 0.34 | 0.17 | 0.48 | 0.34 |
| 2 | 40839463 | rs2216041 | 0.028 | -0.144 | 0.568 | 1.004 | 0.01 | 0.013 | 0.013 | 0 | 0.36 | 0.16 | 0.18 | 0.28 | 0.31 | 0.43 |
| 2 | 40839928 | rs42799 | 1.261 | 0.081 | 0.369 | 0.533 | 0.004 | -0.012 | -0.012 | 0 | 0.28 | 0.3 | 0.17 | 0.16 | 0.27 | 0.14 |
| 2 | 40839969 | rs2192812 | -0.677 | 0.072 | 0.418 | 0.596 | 0 | 0 | 0 | 0 | 0.66 | 0.66 | 0.44 | 0.33 | 0.67 | 0.69 |
| 2 | 40840450 | rs17026346 | -0.41 | 0.125 | 0.462 | 0.607 | 0.006 | -0.006 | -0.006 | 0 | 0.08 | 0.2 | 0.07 | 0.05 | 0.14 | 0.05 |
| 2 | 40843577 | rs42793 | -0.014 | 0.021 | 0.703 | 0.5 | 0.001 | -0.008 | -0.008 | 0 | 0.4 | 0.56 | 0.58 | 0.68 | 0.43 | 0.32 |
| 2 | 40846875 | rs42788 | 0.839 | -0.044 | 0.698 | 0.486 | 0.005 | 0.017 | 0.017 | 0 | 0.32 | 0.39 | 0.53 | 0.65 | 0.3 | 0.27 |
| 2 | 40848550 | rs7560878 | -1.492 | 0.084 | 0.779 | 0.482 | 0.043 | -0.028 | -0.028 | 0 | 0.13 | 0.44 | 0.14 | 0.05 | 0.19 | 0.06 |
| 2 | 40850219 | rs11887013 | -1.545 | -0.054 | 0.745 | 0.527 | 0.027 | 0.495 | 0.495 | 0.004 | 0.06 | 0.31 | 0.35 | 0.49 | 0.03 | 0.13 |
| 2 | 40857899 | rs2373901 | 1.44 | -0.019 | 0.872 | 0.42 | 0.018 | -0.054 | -0.054 | 0 | 0.84 | 0.46 | 0.8 | 0.93 | 0.8 | 0.92 |
| 2 | 40868787 | rs10197644 | 1.011 | -0.568 | 0.497 | 0.116 | 0.053 | 0.065 | 0.065 | 0.002 | 0.22 | 0.07 | 0.07 | 0 | 0.1 | 0.11 |
| 2 | 40875698 | rs12712700 | -0.06 | -0.291 | 0.49 | 0.378 | -0.017 | 0.08 | 0.08 | 0 | 0.54 | 0.18 | 0.31 | 0.29 | 0.53 | 0.66 |
| 2 | 40882419 | rs12468195 | -0.239 | -0.13 | 0.511 | 0.37 | 0.001 | -0.01 | -0.01 | 0 | 0.56 | 0.41 | 0.62 | 0.77 | 0.52 | 0.75 |
| 2 | 40884332 | rs13008680 | 0.519 | -0.159 | 0.365 | 0.514 | -0.001 | 0.072 | 0.072 | 0 | 0.94 | 0.96 | 0.61 | 0.49 | 0.93 | 0.84 |
| 2 | 40893490 | rs2110786 | -0.271 | -0.204 | 0.347 | 0.474 | -0.024 | 0.036 | 0.036 | 0 | 0.58 | 0.18 | 0.34 | 0.31 | 0.57 | 0.68 |
| 2 | 40896385 | rs1861303 | 0.588 | -0.198 | 0.461 | 0.778 | -0.005 | -0.007 | -0.007 | 0 | 0.35 | 0.69 | 0.42 | 0.64 | 0.37 | 0.19 |
| 2 | 40916149 | rs6713004 | -0.013 | -0.233 | 0.268 | 0.716 | -0.021 | 0.081 | 0.081 | 0 | 0.54 | 0.15 | 0.31 | 0.29 | 0.53 | 0.65 |
| 2 | 40931058 | rs6749321 | 0.766 | -0.225 | 0.398 | 0.768 | -0.066 | 0.041 | 0.041 | 0 | 0.34 | 0.15 | 0.24 | 0.21 | 0.41 | 0.19 |
| 2 | 40940311 | rs2888698 | 0.14 | -0.235 | 0.378 | 0.903 | -0.027 | 0.032 | 0.032 | 0 | 0.41 | 0.81 | 0.65 | 0.66 | 0.43 | 0.31 |
| 2 | 40948151 | rs10211572 | -0.025 | -0.241 | 0.37 | 0.922 | -0.036 | 0.063 | 0.063 | 0 | 0.55 | 0.18 | 0.3 | 0.3 | 0.52 | 0.66 |
| 2 | 40949168 | rs1107932 | 1.034 | -0.247 | 0.558 | 0.922 | 0.002 | -0.002 | -0.002 | 0.296 | 0.31 | 0.35 | 0.25 | 0.21 | 0.37 | 0.17 |
| 2 | 40949606 | rs4952645 | -0.89 | -0.246 | 0.537 | 0.92 | -0.011 | 0.007 | 0.007 | 0 | 0.66 | 0.66 | 0.7 | 0.76 | 0.59 | 0.8 |
| 2 | 40958699 | rs12615804 | -0.024 | -0.201 | 0.371 | 0.85 | -0.054 | 0.075 | 0.075 | 0 | 0.54 | 0.12 | 0.3 | 0.29 | 0.52 | 0.66 |
| 2 | 40969323 | rs3851317 | -0.049 | -0.214 | 0.421 | 0.902 | -0.025 | 0.064 | 0.064 | 0 | 0.43 | 0.84 | 0.64 | 0.7 | 0.43 | 0.31 |
| 2 | 40987687 | rs4603810 | 0.549 | -0.141 | 0.313 | 0.913 | -0.01 | 0.004 | 0.004 | 0 | 0.35 | 0.32 | 0.41 | 0.26 | 0.42 | 0.2 |
| 2 | 40990662 | rs7566371 | -0.439 | -0.423 | 0.082 | 0.802 | 0.011 | 0.006 | 0.006 | 0.058 | 0.35 | 0.64 | 0.35 | 0.75 | 0.24 | 0.45 |
| 2 | 41024301 | rs2045342 | 0.089 | 0.248 | 0.485 | 1.14 | -0.014 | 0.007 | 0.007 | 0 | 0.36 | 0.33 | 0.58 | 0.7 | 0.43 | 0.32 |
| 2 | 41036518 | rs6722048 | -0.925 | 0.23 | 0.57 | 1.225 | 0.032 | -0.036 | -0.036 | 1.604 | 0.69 | 0.51 | 0.76 | 0.81 | 0.69 | 0.86 |
| 2 | 41051123 | rs764464 | -0.292 | 0.18 | 0.566 | 1.171 | -0.013 | 0.001 | 0.001 | 0 | 0.63 | 0.23 | 0.4 | 0.33 | 0.63 | 0.71 |
| 2 | 41052544 | rs2168546 | -0.863 | 0.213 | 0.495 | 1.018 | 0.026 | -0.043 | -0.043 | 2.029 | 0.7 | 0.55 | 0.76 | 0.84 | 0.69 | 0.86 |
| 2 | 41057779 | rs17026712 | -1.168 | 0.067 | 0.302 | 0.986 | -0.011 | 0.236 | 0.236 | 0 | 0.14 | 0.38 | 0.4 | 0.53 | 0.15 | 0.2 |
| 2 | 41058665 | rs12712717 | -0.201 | 0.185 | -0.139 | 0.82 | -0.037 | 0.089 | 0.089 | 0 | 0.57 | 0.15 | 0.36 | 0.26 | 0.55 | 0.67 |
| 2 | 41069098 | rs7597016 | -0.18 | 0.28 | -0.997 | 0.83 | -0.03 | 0.058 | 0.058 | 0 | 0.57 | 0.17 | 0.36 | 0.29 | 0.55 | 0.68 |
| 2 | 41073997 | rs12987383 | 0.006 | 0.19 | -1.058 | 0.91 | -0.024 | 0.103 | 0.103 | 0 | 0.52 | 0.14 | 0.31 | 0.29 | 0.5 | 0.62 |
| 2 | 41075429 | rs4952430 | -1.39 | 0.052 | 0.054 | 1.655 | 0.001 | 0.147 | 0.147 | 0.017 | 0.16 | 0.53 | 0.34 | 0.33 | 0.12 | 0.18 |
| 2 | 41079833 | rs6724492 | -0.134 | 0.047 | -0.294 | 1.612 | 0.005 | -0.003 | -0.003 | 0.011 | 0.34 | 0.33 | 0.29 | 0.26 | 0.42 | 0.44 |
| 2 | 41092349 | rs17026752 | -0.142 | -0.4 | 0.387 | 1.71 | -0.016 | 0.152 | 0.152 | 0.121 | 0.1 | 0.31 | 0.34 | 0.31 | 0.06 | 0.06 |

| | | | | | | | | | | | | | | | | |
|---|---|---|---|---|---|---|---|---|---|---|---|---|---|---|---|---|
| 2 | 41096993 | rs9309062 | -0.021 | -0.192 | 0.306 | 1.74 | 0.003 | -0.024 | -0.024 | 0.737 | 0.33 | 0.33 | 0.27 | 0.13 | 0.36 | 0.41 |
| 2 | 41100542 | rs11898602 | -0.859 | -0.206 | 0.702 | 1.808 | -0.001 | -0.001 | -0.001 | 0 | 0.41 | 0.56 | 0.63 | 0.63 | 0.43 | 0.56 |
| 2 | 41101776 | rs17026778 | -0.406 | -0.109 | 0.682 | 1.826 | 0.002 | 0.115 | 0.115 | 0 | 0.12 | 0.27 | 0.36 | 0.31 | 0.08 | 0.1 |
| 2 | 41103223 | rs1379022 | -0.077 | -0.279 | 0.398 | 1.481 | -0.002 | -0.002 | -0.002 | 0 | 0.33 | 0.29 | 0.29 | 0.34 | 0.36 | 0.5 |
| 2 | 41107517 | rs12328759 | 0.417 | -1.046 | 0.008 | 0.722 | 0.058 | 0.077 | 0.077 | 1.398 | 0.3 | 0.14 | 0.14 | 0.05 | 0.18 | 0.18 |
| 2 | 41123765 | rs10203017 | -0.624 | -1.006 | 0.906 | 0.861 | -0.004 | -0.001 | -0.001 | 1.324 | 0.47 | 0.48 | 0.56 | 0.56 | 0.4 | 0.56 |
| 2 | 41123971 | rs7576767 | -0.723 | -1.017 | 0.812 | 0.855 | 0.021 | 0.005 | 0.005 | 0 | 0.48 | 0.61 | 0.58 | 0.56 | 0.4 | 0.56 |
| 2 | 41124963 | rs207603 | 0.061 | -1.027 | 0.893 | 0.829 | 0.005 | 0.007 | 0.007 | 0 | 0.57 | 0.35 | 0.67 | 0.7 | 0.63 | 0.52 |
| 2 | 41132143 | rs10203786 | 0.407 | -0.663 | 1.073 | 1.017 | -0.002 | -0.001 | -0.001 | 0.094 | 0.29 | 0.27 | 0.29 | 0.31 | 0.36 | 0.48 |
| 2 | 41155423 | rs6725737 | 1.051 | -0.668 | 0.923 | 1.098 | 0.02 | 0.233 | 0.233 | 0.064 | 0.91 | 0.59 | 0.71 | 0.74 | 0.96 | 0.94 |
| 2 | 41161464 | rs9309064 | 0.366 | -0.628 | 0.77 | 1.109 | 0.007 | 0.002 | 0.002 | 0.199 | 0.31 | 0.38 | 0.3 | 0.3 | 0.36 | 0.49 |
| 2 | 41180319 | rs11885442 | 0.366 | -0.56 | 0.268 | 0.925 | -0.001 | -0.001 | -0.001 | 0 | 0.31 | 0.28 | 0.29 | 0.31 | 0.37 | 0.48 |
| 2 | 41197708 | rs10205399 | -0.105 | -0.518 | -0.299 | 0.93 | 0 | 0.008 | 0.008 | 0.831 | 0.65 | 0.19 | 0.68 | 0.7 | 0.64 | 0.49 |
| 2 | 41198011 | rs13023575 | 0.183 | -0.5 | -0.298 | 1.035 | -0.002 | -0.03 | -0.03 | 0 | 0.6 | 0.2 | 0.43 | 0.43 | 0.61 | 0.46 |
| 2 | 41198183 | rs7607054 | 0.496 | -0.257 | 0.013 | 1.175 | 0.032 | 0.241 | 0.241 | 0 | 0.93 | 0.7 | 0.72 | 0.74 | 0.97 | 0.94 |
| 2 | 41199040 | rs1981804 | 0.203 | -0.261 | -1.065 | 0.659 | 0.002 | 0.008 | 0.008 | 0.002 | 0.46 | 0.17 | 0.39 | 0.41 | 0.56 | 0.45 |
| 2 | 41201040 | rs17027078 | -1.079 | -0.621 | -0.176 | 0.573 | -0.028 | -0.018 | -0.018 | 0.176 | 0.14 | 0.31 | 0.09 | 0.04 | 0.11 | 0.12 |
| 2 | 41212010 | rs12614285 | -0.137 | -0.468 | -0.773 | 0.52 | 0.005 | -0.002 | -0.002 | 0.001 | 0.35 | 0.38 | 0.35 | 0.64 | 0.43 | 0.3 |
| 2 | 41213624 | rs13022283 | -0.154 | -0.542 | -0.443 | 0.482 | 0.012 | -0.002 | -0.002 | 0 | 0.35 | 0.39 | 0.36 | 0.64 | 0.42 | 0.3 |
| 2 | 41220869 | rs13034585 | 1.197 | -0.915 | -0.892 | -0.169 | -0.085 | 0.107 | 0.107 | 0.064 | 0.26 | 0.09 | 0.38 | 0.02 | 0.36 | 0.26 |
| 2 | 41221458 | rs12990636 | -0.396 | -0.961 | -0.843 | -0.047 | -0.053 | -0.059 | -0.059 | 0.04 | 0.62 | 0.53 | 0.75 | 0.85 | 0.78 | 0.57 |
| 2 | 41225026 | rs6715452 | -1.019 | -0.998 | -0.818 | -0.051 | -0.055 | -0.056 | -0.056 | 0 | 0.69 | 0.59 | 0.77 | 0.85 | 0.82 | 0.64 |
| 2 | 41226427 | rs12988568 | 0.111 | -0.896 | -0.792 | -0.037 | 0.009 | -0.053 | -0.053 | 0 | 0.37 | 0.52 | 0.38 | 0.82 | 0.42 | 0.3 |
| 2 | 41227539 | rs4952434 | 1.188 | -0.681 | -0.886 | 0.387 | -0.021 | -0.001 | -0.001 | 0 | 0.05 | 0 | 0.09 | 0.32 | 0.06 | 0.08 |
| 2 | 41228588 | rs12474136 | -0.209 | -0.732 | -0.815 | 0.092 | -0.01 | -0.016 | -0.016 | 0 | 0.59 | 0.44 | 0.73 | 0.84 | 0.77 | 0.56 |
| 2 | 41230862 | rs12476568 | 0.12 | -0.705 | -0.816 | 0.131 | -0.011 | -0.011 | -0.011 | 0 | 0.56 | 0.45 | 0.69 | 0.8 | 0.7 | 0.52 |
| 2 | 41237919 | rs1862096 | -0.884 | -0.857 | -0.677 | 0.038 | -0.076 | -0.061 | -0.061 | 1.142 | 0.7 | 0.55 | 0.78 | 0.85 | 0.84 | 0.67 |
| 2 | 41238321 | rs7571620 | 1.223 | -0.717 | -0.953 | -0.003 | -0.125 | 0.148 | 0.148 | 0.224 | 0.26 | 0.06 | 0.38 | 0.02 | 0.37 | 0.26 |
| 2 | 41263841 | rs12989241 | -0.081 | -0.359 | -0.887 | -0.023 | -0.003 | 0.018 | 0.018 | 0 | 0.42 | 0.37 | 0.4 | 0.84 | 0.54 | 0.39 |
| 2 | 41267941 | rs6746423 | 0.746 | -0.749 | -0.562 | -0.116 | -0.118 | -0.103 | -0.103 | 0 | 0.19 | 0.47 | 0.11 | 0.06 | 0.06 | 0.15 |
| 2 | 41273631 | rs17027130 | 1.052 | -0.657 | -0.823 | 0.182 | -0.007 | -0.057 | -0.057 | 0.054 | 0.09 | 0.01 | 0.09 | 0.37 | 0.06 | 0.14 |
| 2 | 41275829 | rs7578600 | 2.381 | -0.604 | -0.35 | -0.411 | -0.048 | -0.025 | -0.025 | 0 | 0.12 | 0.16 | 0.05 | 0.04 | 0.02 | 0.07 |
| 2 | 41276504 | rs6544381 | -0.349 | -0.534 | -0.973 | -0.435 | 0.004 | -0.001 | -0.001 | 0.045 | 0.39 | 0.61 | 0.27 | 0.15 | 0.22 | 0.42 |
| 2 | 41279791 | rs11688656 | 1.498 | -0.423 | -1.088 | -0.366 | -0.016 | 0.026 | 0.026 | 0.021 | 0.17 | 0.02 | 0.14 | 0.08 | 0.16 | 0.26 |
| 2 | 41279956 | rs17683610 | 1.445 | -0.305 | -0.927 | -0.38 | -0.049 | 0.094 | 0.094 | 0.005 | 0.22 | 0.07 | 0.35 | 0.01 | 0.27 | 0.24 |
| 2 | 41293934 | rs10196627 | -0.012 | -0.203 | -0.784 | -0.302 | 0.014 | 0 | 0 | 3.21 | 0.43 | 0.6 | 0.47 | 0.08 | 0.31 | 0.4 |
| 2 | 41305118 | rs17684108 | -0.088 | -0.851 | -0.676 | -0.006 | -0.001 | 0.105 | 0.105 | 1.884 | 0.48 | 0.73 | 0.54 | 0.24 | 0.45 | 0.47 |
| 2 | 41305293 | rs17633025 | -0.078 | -0.851 | -0.677 | -0.007 | -0.002 | 0.104 | 0.104 | 1.808 | 0.48 | 0.72 | 0.54 | 0.24 | 0.45 | 0.45 |
| 2 | 41327766 | rs13411627 | 1.558 | -1.039 | -0.272 | -0.032 | -0.104 | 0.048 | 0.048 | 6.424 | 0.14 | 0.47 | 0.1 | 0.16 | 0.03 | 0.1 |
| 2 | 41330027 | rs6705468 | 0.235 | -1.082 | -0.137 | -0.092 | -0.019 | -0.037 | -0.037 | 1.444 | 0.07 | 0.17 | 0.05 | 0 | 0.04 | 0.06 |
| 2 | 41334045 | rs13407981 | 1.455 | -1.06 | -0.234 | -0.003 | -0.11 | 0.056 | 0.056 | 6.711 | 0.15 | 0.47 | 0.09 | 0.18 | 0.03 | 0.12 |
| 2 | 41335222 | rs7585987 | -1.387 | -1.157 | -0.561 | 0.226 | -0.061 | -0.1 | -0.1 | 0 | 0.74 | 0.93 | 0.8 | 0.6 | 0.68 | 0.82 |
| 2 | 41341270 | rs4372953 | 0.497 | -1.276 | -0.252 | -0.506 | -0.029 | 0.021 | 0.021 | 0 | 0.34 | 0.24 | 0.46 | 0.02 | 0.46 | 0.34 |
| 2 | 41341684 | rs13417378 | 1.176 | -1.316 | -0.356 | -0.397 | -0.021 | 0.011 | 0.011 | 0.024 | 0.17 | 0.01 | 0.14 | 0.09 | 0.16 | 0.26 |
| 2 | 41345151 | rs4952352 | 1.075 | -1.371 | -0.258 | -0.401 | -0.001 | -0.001 | -0.001 | 0 | 0.17 | 0.14 | 0.16 | 0.11 | 0.16 | 0.26 |
| 2 | 41345909 | rs10169519 | 0.867 | -1.163 | -0.167 | -1.042 | -0.105 | 0.049 | 0.049 | 8.376 | 0.17 | 0.48 | 0.1 | 0.21 | 0.05 | 0.11 |
| 2 | 41351491 | rs2194436 | 1.144 | -1.157 | -0.36 | -1.052 | -0.091 | 0.066 | 0.066 | 6.995 | 0.16 | 0.42 | 0.09 | 0.24 | 0.05 | 0.11 |
| 2 | 41352620 | rs6544388 | 0.221 | -1.19 | -0.494 | -1.08 | -0.106 | 0.019 | 0.019 | 7.356 | 0.22 | 0.48 | 0.13 | 0.26 | 0.1 | 0.19 |
| 2 | 41358705 | rs17634234 | 1.117 | -1.081 | -0.618 | -1.201 | -0.106 | 0.12 | 0.12 | 0.419 | 0.28 | 0.1 | 0.42 | 0.02 | 0.43 | 0.3 |
| 2 | 41362047 | rs7605382 | 0.279 | -0.927 | -0.536 | -1.122 | 0.019 | -0.13 | -0.13 | 0 | 0.36 | 0.48 | 0.5 | 0.02 | 0.49 | 0.37 |
| 2 | 41366151 | rs7562658 | 0.413 | -0.852 | -0.758 | -1.058 | -0.001 | -0.048 | -0.048 | 2.348 | 0.35 | 0.38 | 0.49 | 0.02 | 0.49 | 0.36 |
| 2 | 41370817 | rs1866904 | -1.149 | -0.977 | -1.273 | -0.998 | 0.006 | 0.012 | 0.012 | 0 | 0.68 | 0.75 | 0.65 | 0.27 | 0.69 | 0.67 |
| 2 | 41373988 | rs7577606 | -0.159 | -0.721 | -0.542 | -0.449 | -0.107 | 0.026 | 0.026 | 7.407 | 0.22 | 0.47 | 0.12 | 0.26 | 0.1 | 0.19 |
| 2 | 41377737 | rs2042461 | 0.251 | -0.371 | -0.705 | -0.42 | 0.003 | -0.026 | -0.026 | 0.414 | 0.32 | 0.33 | 0.45 | 0.09 | 0.45 | 0.3 |
| 2 | 41381344 | rs13421868 | -0.605 | -0.947 | -0.448 | 0.415 | -0.112 | 0.004 | 0.004 | 6.085 | 0.19 | 0.46 | 0.1 | 0.2 | 0.07 | 0.19 |
| 2 | 41381376 | rs10490376 | 0.01 | -0.716 | -0.309 | 0.591 | -0.096 | 0.051 | 0.051 | 7.643 | 0.14 | 0.44 | 0.08 | 0.2 | 0.02 | 0.12 |
| 2 | 41385771 | rs4952440 | -0.203 | -0.237 | -1.734 | 0.393 | 0.018 | 0.045 | 0.045 | 1.783 | 0.43 | 0.17 | 0.57 | 0.72 | 0.37 | 0.53 |
| 2 | 41386562 | rs12613369 | 0.099 | -0.432 | -2.05 | 1.202 | -0.025 | -0.06 | -0.06 | 0 | 0.1 | 0.01 | 0.18 | 0.42 | 0.11 | 0.19 |
| 2 | 41387093 | rs7583193 | 0.744 | 0.176 | -1.936 | 0.372 | -0.002 | 0.003 | 0.003 | 0.271 | 0.33 | 0.62 | 0.24 | 0.22 | 0.36 | 0.36 |
| 2 | 41388121 | rs9309065 | -0.021 | 0.054 | -1.501 | 0.316 | 0.041 | 0.043 | 0.043 | 1.443 | 0.44 | 0.31 | 0.58 | 0.72 | 0.34 | 0.52 |
| 2 | 41391252 | rs17027254 | 0.582 | 0.212 | -1.143 | 0.458 | 0.001 | -0.001 | -0.001 | 0 | 0.32 | 0.62 | 0.25 | 0.22 | 0.38 | 0.36 |
| 2 | 41399187 | rs6732320 | 0.55 | 0.172 | -0.714 | 0.442 | 0.012 | -0.004 | -0.004 | 0 | 0.33 | 0.54 | 0.23 | 0.23 | 0.38 | 0.35 |
| 2 | 41413579 | rs4952443 | -0.052 | 0.077 | -1.065 | 0.381 | 0.04 | 0.005 | 0.005 | 0 | 0.45 | 0.26 | 0.54 | 0.68 | 0.35 | 0.5 |
| 2 | 41414128 | rs12463409 | 1.105 | -0.239 | -1.409 | 0.321 | 0.018 | -0.028 | -0.028 | 0 | 0.28 | 0.02 | 0.3 | 0.26 | 0.19 | 0.2 |
| 2 | 41415129 | rs1899286 | 0.946 | -0.022 | -1.067 | 0.457 | 0.006 | -0.008 | -0.008 | 0 | 0.32 | 0.56 | 0.23 | 0.18 | 0.38 | 0.35 |
| 2 | 41420957 | rs1372165 | -1.283 | 0.012 | -1.151 | 0.372 | -0.015 | 0.02 | 0.02 | 0 | 0.81 | 0.89 | 0.8 | 0.94 | 0.73 | 0.87 |
| 2 | 41424692 | rs11687208 | 0.737 | -0.112 | -0.915 | 0.344 | 0.011 | -0.007 | -0.007 | 0 | 0.34 | 0.51 | 0.24 | 0.24 | 0.38 | 0.36 |
| 2 | 41431085 | rs1372166 | 1.293 | 0.099 | -0.327 | 0.007 | 0.017 | -0.054 | -0.054 | 0 | 0.23 | 0.3 | 0.16 | 0.04 | 0.3 | 0.3 |
| 2 | 41434073 | rs11903277 | 0.252 | -0.035 | -0.239 | 0.09 | 0 | -0.042 | -0.042 | 0 | 0.38 | 0.54 | 0.26 | 0.12 | 0.4 | 0.4 |

| | | | | | | | | | | | | | | | | |
|---|---|---|---|---|---|---|---|---|---|---|---|---|---|---|---|---|
| 2 | 41439870 | rs6544397 | 0.335 | -0.341 | -0.268 | 0.174 | 0.042 | -0.004 | -0.004 | 4.454 | 0.37 | 0.24 | 0.38 | 0.42 | 0.22 | 0.34 |
| 2 | 41444315 | rs1441221 | -0.852 | 0.019 | -0.283 | 0.308 | 0.024 | 0.05 | 0.05 | 0.602 | 0.79 | 0.67 | 0.66 | 0.55 | 0.69 | 0.77 |
| 2 | 41446609 | rs7567386 | -0.859 | 0.025 | -0.364 | 0.332 | 0.005 | 0.012 | 0.012 | 0.499 | 0.78 | 0.79 | 0.67 | 0.55 | 0.68 | 0.77 |
| 2 | 41474388 | rs11894180 | -0.008 | -0.334 | -0.363 | 0.024 | 0 | 0 | 0 | 0 | 0.46 | 0.57 | 0.31 | 0.13 | 0.48 | 0.44 |
| 2 | 41476122 | rs11886398 | 0.764 | -0.437 | -0.914 | -0.151 | -0.041 | 0.138 | 0.138 | 0 | 0.32 | 0.12 | 0.2 | 0.04 | 0.38 | 0.26 |
| 2 | 41486554 | rs17027360 | 1.252 | -0.306 | -0.464 | 0.211 | -0.006 | 0.032 | 0.032 | 0.007 | 0.11 | 0.19 | 0.09 | 0.21 | 0.09 | 0.07 |
| 2 | 41488391 | rs1441234 | 0.012 | -0.513 | -0.43 | 0.159 | 0.003 | 0.003 | 0.003 | 0 | 0.43 | 0.48 | 0.4 | 0.5 | 0.49 | 0.6 |
| 2 | 41489307 | rs1992404 | -0.484 | -0.508 | -0.421 | 0.163 | -0.004 | 0.099 | 0.099 | 0.583 | 0.53 | 0.79 | 0.51 | 0.71 | 0.57 | 0.66 |
| 2 | 41502545 | rs6544404 | -0.41 | -0.511 | 0.003 | 0.135 | 0.029 | -0.051 | -0.051 | 0 | 0.1 | 0.35 | 0.06 | 0 | 0.06 | 0.24 |
| 2 | 41505473 | rs1441231 | 1.505 | -0.479 | 0.049 | 0.257 | 0 | 0.058 | 0.058 | 0 | 0.1 | 0.18 | 0.08 | 0.22 | 0.09 | 0.06 |
| 2 | 41506889 | rs10202299 | -0.651 | -0.657 | -0.15 | 0.204 | 0.004 | 0.005 | 0.005 | 1.114 | 0.55 | 0.74 | 0.52 | 0.54 | 0.58 | 0.67 |
| 2 | 41507059 | rs17027376 | 0.085 | -0.625 | 0.14 | 0.14 | -0.017 | -0.031 | -0.031 | 0.702 | 0.17 | 0.57 | 0.13 | 0.07 | 0.12 | 0.16 |
| 2 | 41509362 | rs1441230 | -0.589 | -0.619 | -0.057 | 0.094 | -0.007 | 0.091 | 0.091 | 0.747 | 0.55 | 0.82 | 0.53 | 0.73 | 0.58 | 0.69 |
| 2 | 41513126 | rs12472156 | -1.254 | -0.748 | -0.236 | -0.212 | -0.02 | 0.044 | 0.044 | 1.077 | 0.78 | 0.89 | 0.81 | 0.95 | 0.72 | 0.82 |
| 2 | 41513438 | rs995345 | 0.562 | -0.781 | -0.008 | -0.146 | 0.043 | 0.002 | 0.002 | 0 | 0.07 | 0.31 | 0.04 | 0 | 0.03 | 0.1 |
| 2 | 41516596 | rs2197036 | -0.377 | -0.753 | -0.633 | -0.032 | -0.01 | 0.127 | 0.127 | 0.825 | 0.56 | 0.83 | 0.53 | 0.75 | 0.58 | 0.67 |
| 2 | 41519239 | rs6727039 | 0.552 | -0.563 | -0.491 | 0.179 | -0.055 | -0.06 | -0.06 | 0 | 0.37 | 0.14 | 0.38 | 0.53 | 0.46 | 0.51 |
| 2 | 41525877 | rs984172 | -1.208 | -0.458 | -0.452 | -0.176 | -0.013 | 0.033 | 0.033 | 0.724 | 0.8 | 0.9 | 0.84 | 0.92 | 0.75 | 0.83 |
| 2 | 41525941 | rs1508115 | 0.035 | -0.278 | -0.225 | 0.13 | -0.055 | 0.033 | 0.033 | 0.007 | 0.39 | 0.16 | 0.41 | 0.65 | 0.49 | 0.53 |
| 2 | 41531398 | rs13385952 | -0.934 | -0.52 | 0.528 | 0.1 | 0.03 | -0.001 | -0.001 | 0 | 0.24 | 0.87 | 0.19 | 0.23 | 0.15 | 0.2 |
| 2 | 41532477 | rs10490381 | -0.378 | -0.495 | 1.23 | -0.044 | -0.032 | -0.046 | -0.046 | 4.327 | 0.19 | 0.53 | 0.14 | 0.1 | 0.13 | 0.13 |
| 2 | 41534961 | rs17027420 | 0.054 | -0.46 | 1.199 | -0.071 | -0.017 | -0.055 | -0.055 | 1.649 | 0.17 | 0.41 | 0.13 | 0.05 | 0.12 | 0.12 |
| 2 | 41540474 | rs12712747 | 0.278 | -0.144 | 0.365 | -0.108 | -0.003 | 0.138 | 0.138 | 0 | 0.54 | 0.2 | 0.62 | 0.91 | 0.52 | 0.65 |
| 2 | 41540966 | rs7602964 | -0.964 | -0.102 | -0.008 | -0.175 | -0.022 | 0.046 | 0.046 | 0 | 0.73 | 0.87 | 0.79 | 0.96 | 0.66 | 0.77 |
| 2 | 41542137 | rs12712748 | 0.389 | 0.04 | -0.505 | 0.075 | -0.037 | 0.1 | 0.1 | 0 | 0.51 | 0.12 | 0.59 | 0.69 | 0.49 | 0.58 |
| 2 | 41556080 | rs1508110 | -0.298 | 0.34 | 0.501 | -0.215 | 0.02 | -0.093 | -0.093 | 1.065 | 0.69 | 0.43 | 0.74 | 0.95 | 0.65 | 0.71 |
| 2 | 41556924 | rs1559468 | -0.801 | 0.521 | 0.416 | -0.213 | -0.044 | 0.091 | 0.091 | 0 | 0.74 | 0.94 | 0.8 | 0.96 | 0.69 | 0.77 |
| 2 | 41559590 | rs1567236 | -0.701 | 0.547 | 0.483 | -0.215 | -0.013 | 0.059 | 0.059 | 0 | 0.74 | 0.84 | 0.79 | 0.96 | 0.7 | 0.77 |
| 2 | 41623189 | rs10192605 | -0.506 | 0.305 | 0.409 | -0.095 | -0.045 | 0.066 | 0.066 | 0 | 0.72 | 0.85 | 0.8 | 0.92 | 0.65 | 0.72 |
| 2 | 41630684 | rs1508118 | -0.335 | 0.257 | 0.632 | -0.118 | -0.01 | 0.015 | 0.015 | 0 | 0.71 | 0.68 | 0.77 | 0.92 | 0.65 | 0.73 |
| 2 | 41633767 | rs6544415 | -0.138 | 0.177 | 1.295 | -0.158 | 0.008 | -0.041 | -0.041 | 0.071 | 0.69 | 0.36 | 0.74 | 0.92 | 0.65 | 0.73 |
| 2 | 41633989 | rs6746520 | -0.329 | 0.188 | 0.53 | -0.152 | 0.001 | -0.012 | -0.012 | 0 | 0.71 | 0.63 | 0.76 | 0.92 | 0.65 | 0.73 |
| 2 | 41636627 | rs4952361 | -0.259 | 0.19 | 0.553 | -0.131 | 0.009 | -0.033 | -0.033 | 0.848 | 0.7 | 0.56 | 0.76 | 0.92 | 0.65 | 0.73 |
| 2 | 41641357 | rs872037 | -0.31 | -0.068 | 0.247 | -0.286 | -0.011 | 0.053 | 0.053 | 0 | 0.68 | 0.78 | 0.79 | 0.96 | 0.66 | 0.76 |
| 2 | 41643650 | rs1508120 | -0.354 | -0.064 | 0.274 | -0.284 | -0.033 | 0.11 | 0.11 | 0 | 0.69 | 0.84 | 0.79 | 0.96 | 0.66 | 0.76 |
| 2 | 41644827 | rs10210181 | -0.004 | -0.082 | 0.332 | 0.321 | 0.003 | 0.003 | 0.003 | 0 | 0.65 | 0.52 | 0.56 | 0.68 | 0.65 | 0.71 |
| 2 | 41653824 | rs906549 | -0.334 | -0.019 | 0.256 | -0.572 | -0.015 | 0.054 | 0.054 | 0 | 0.69 | 0.8 | 0.79 | 0.96 | 0.66 | 0.76 |
| 2 | 41655276 | rs2174883 | 0.085 | -0.016 | 0.208 | -0.567 | 0.007 | -0.068 | -0.068 | 0 | 0.66 | 0.48 | 0.74 | 0.92 | 0.65 | 0.72 |
| 2 | 41656025 | rs1508124 | -0.409 | -0.027 | 0.118 | -0.565 | -0.016 | 0.04 | 0.04 | 0 | 0.69 | 0.73 | 0.78 | 0.92 | 0.65 | 0.71 |
| 2 | 41670280 | rs7574300 | -0.183 | 0.022 | -0.021 | -0.557 | 0.001 | -0.031 | -0.031 | 0 | 0.67 | 0.6 | 0.76 | 0.96 | 0.64 | 0.76 |
| 2 | 41694434 | rs10495889 | 0.52 | 0.016 | -0.83 | -0.497 | -0.018 | -0.088 | -0.088 | 0 | 0.21 | 0.01 | 0.42 | 0.76 | 0.15 | 0.25 |
| 2 | 41701800 | rs8179813 | 0.317 | 0.037 | -0.221 | -0.742 | -0.028 | -0.04 | -0.04 | 0 | 0.35 | 0.42 | 0.52 | 0.84 | 0.25 | 0.36 |
| 2 | 41704150 | rs12996069 | 0.128 | 0.106 | -0.349 | -0.894 | -0.005 | 0.036 | 0.036 | 0.754 | 0.37 | 0.33 | 0.24 | 0.12 | 0.39 | 0.41 |
| 2 | 41704580 | rs12997558 | 0.271 | 0.114 | -0.349 | -0.889 | -0.003 | 0.044 | 0.044 | 0.378 | 0.34 | 0.28 | 0.22 | 0.08 | 0.36 | 0.38 |
| 2 | 41718641 | rs10175292 | 0.371 | 0.088 | -0.469 | -0.893 | -0.013 | 0.083 | 0.083 | 0.585 | 0.33 | 0.22 | 0.21 | 0.1 | 0.36 | 0.37 |
| 2 | 41721609 | rs12474614 | -0.963 | 0.037 | -0.315 | -0.88 | 0.017 | -0.045 | -0.045 | 0 | 0.69 | 0.59 | 0.74 | 0.92 | 0.62 | 0.73 |
| 2 | 41722554 | rs12471569 | 0.26 | 0.008 | -0.338 | -0.777 | -0.02 | -0.022 | -0.022 | 0 | 0.35 | 0.37 | 0.53 | 0.81 | 0.25 | 0.35 |
| 2 | 41737267 | rs13423625 | 0.902 | 0.188 | -0.796 | -0.831 | -0.037 | 0.07 | 0.07 | 0 | 0.25 | 0.08 | 0.18 | 0.08 | 0.32 | 0.21 |
| 2 | 41745916 | rs12463636 | 0.277 | -0.036 | -0.487 | -0.732 | -0.018 | -0.016 | -0.016 | 0 | 0.36 | 0.49 | 0.54 | 0.79 | 0.25 | 0.36 |
| 2 | 41750080 | rs7578278 | -1.078 | -0.043 | -0.283 | -0.963 | 0.013 | -0.038 | -0.038 | 0.864 | 0.7 | 0.63 | 0.75 | 0.92 | 0.63 | 0.74 |
| 2 | 41751055 | rs10205537 | 0.084 | -0.042 | -0.31 | -0.941 | -0.003 | 0.012 | 0.012 | 0.795 | 0.36 | 0.34 | 0.23 | 0.11 | 0.38 | 0.38 |
| 2 | 41755591 | rs4245775 | -1.112 | -0.075 | -0.287 | -0.962 | -0.026 | 0.053 | 0.053 | 0 | 0.7 | 0.79 | 0.76 | 0.91 | 0.62 | 0.73 |
| 2 | 41758888 | rs4468861 | 0.093 | -0.06 | -0.344 | -0.957 | -0.01 | 0.049 | 0.049 | 0.48 | 0.34 | 0.28 | 0.21 | 0.11 | 0.37 | 0.36 |
| 2 | 41765955 | rs12620138 | 0.305 | -0.065 | -0.264 | -0.929 | -0.015 | -0.022 | -0.022 | 0.224 | 0.36 | 0.51 | 0.54 | 0.85 | 0.26 | 0.4 |
| 2 | 41773324 | rs12620745 | 0.787 | -0.202 | -0.279 | -0.886 | -0.018 | -0.001 | -0.001 | 5.098 | 0.31 | 0.36 | 0.52 | 0.73 | 0.24 | 0.35 |
| 2 | 41773756 | rs17027951 | 1.286 | -0.203 | -0.389 | -0.869 | -0.018 | 0.021 | 0.021 | 0 | 0.06 | 0.11 | 0.01 | 0.12 | 0.02 | 0.05 |
| 2 | 41782340 | rs10188010 | 1.512 | 0.006 | -0.605 | -1.159 | -0.026 | 0.054 | 0.054 | 0 | 0.22 | 0.07 | 0.16 | 0.08 | 0.26 | 0.2 |
| 2 | 41784905 | rs10208424 | -1.088 | -0.136 | -0.58 | -1.187 | -0.013 | 0.026 | 0.026 | 0 | 0.69 | 0.73 | 0.76 | 0.95 | 0.59 | 0.76 |
| 2 | 41787981 | rs13026723 | 0.167 | 0.056 | -0.608 | -1.185 | -0.06 | 0.105 | 0.105 | 0.209 | 0.33 | 0.13 | 0.22 | 0.11 | 0.4 | 0.37 |
| 2 | 41790278 | rs6740655 | -0.443 | 0.02 | -0.581 | -1.179 | 0.069 | 0.154 | 0.154 | 0 | 0.47 | 0.75 | 0.6 | 0.88 | 0.31 | 0.56 |
| 2 | 41790301 | rs17027968 | 0.37 | 0.041 | -0.516 | -1.06 | -0.002 | -0.012 | -0.012 | 0 | 0.09 | 0.11 | 0.05 | 0.03 | 0.05 | 0.16 |
| 2 | 41791788 | rs2195496 | -1.239 | 0.149 | -0.691 | -1.052 | -0.047 | 0.105 | 0.105 | 0 | 0.72 | 0.89 | 0.79 | 0.96 | 0.66 | 0.76 |
| 2 | 41795417 | rs1864276 | 0.436 | 0.116 | -0.08 | -0.59 | -0.021 | 0.001 | 0.001 | 0 | 0.06 | 0.24 | 0.03 | 0.04 | 0.01 | 0.04 |
| 2 | 41796861 | rs17027998 | -0.447 | 0.158 | -0.243 | -0.568 | -0.012 | -0.024 | -0.024 | 2.791 | 0.14 | 0.27 | 0.05 | 0.07 | 0.07 | 0.2 |
| 2 | 41802979 | rs6544449 | -0.337 | 0.267 | -0.348 | -0.516 | -0.001 | -0.025 | -0.025 | 0 | 0.39 | 0.62 | 0.27 | 0.16 | 0.35 | 0.4 |
| 2 | 41806652 | rs12712787 | -0.212 | 0.711 | -0.376 | -0.183 | 0.035 | 0.015 | 0.015 | 0 | 0.66 | 0.53 | 0.37 | 0.29 | 0.53 | 0.69 |
| 2 | 41808308 | rs1560262 | -0.009 | 0.328 | -1.039 | -0.171 | 0.001 | 0.019 | 0.019 | 0 | 0.56 | 0.89 | 0.35 | 0.3 | 0.39 | 0.54 |
| 2 | 41814174 | rs1920480 | 0.459 | 0.038 | -0.994 | -0.437 | -0.006 | 0.078 | 0.078 | 0 | 0.41 | 0.17 | 0.28 | 0.17 | 0.42 | 0.44 |
| 2 | 41818847 | rs7556765 | 0.302 | -0.05 | -0.252 | -0.097 | -0.048 | -0.001 | -0.001 | 1.902 | 0.27 | 0.54 | 0.12 | 0.23 | 0.15 | 0.28 |

| | | | | | | | | | | | | | | | | |
|---|---|---|---|---|---|---|---|---|---|---|---|---|---|---|---|---|
| 2 | 41822439 | rs12471180 | -0.889 | 0.28 | -0.557 | 0.062 | -0.099 | -0.108 | -0.108 | 0 | 0.71 | 0.9 | 0.43 | 0.4 | 0.57 | 0.72 |
| 2 | 41826738 | rs2163050 | 0.684 | 0.337 | -0.555 | 0.038 | -0.015 | -0.006 | -0.006 | 0 | 0.15 | 0.04 | 0.11 | 0.06 | 0.14 | 0.18 |
| 2 | 41829089 | rs10865177 | 1.095 | 0.901 | -0.693 | 0.726 | 0.007 | 0.017 | 0.017 | 0.117 | 0.06 | 0.05 | 0.3 | 0.26 | 0.09 | 0.15 |
| 2 | 41832878 | rs7603075 | 0.146 | 0.93 | -0.317 | 0.629 | 0.003 | -0.006 | -0.006 | 1.873 | 0.2 | 0.22 | 0.16 | 0.09 | 0.24 | 0.21 |
| 2 | 41835526 | rs1560259 | -0.319 | 0.701 | -0.055 | 0.518 | 0.01 | 0.003 | 0.003 | 0 | 0.53 | 0.58 | 0.62 | 0.46 | 0.54 | 0.56 |
| 2 | 41836313 | rs7597606 | -0.629 | 0.661 | -0.098 | 0.533 | 0.001 | -0.065 | -0.065 | 0 | 0.58 | 0.87 | 0.68 | 0.53 | 0.59 | 0.6 |
| 2 | 41839007 | rs10205791 | -0.031 | 0.685 | 0.325 | 0.462 | 0.047 | 0.009 | 0.009 | 0.104 | 0.14 | 0.36 | 0.13 | 0.08 | 0.2 | 0.19 |
| 2 | 41839186 | rs7568164 | -0.461 | 0.69 | 0.406 | 0.584 | 0.011 | 0.019 | 0.019 | 0 | 0.49 | 0.66 | 0.62 | 0.45 | 0.53 | 0.57 |
| 2 | 41839915 | rs1920502 | 0.615 | 0.688 | 0.356 | 0.556 | -0.007 | 0.004 | 0.004 | 0 | 0.34 | 0.29 | 0.49 | 0.37 | 0.37 | 0.38 |
| 2 | 41840666 | rs17028106 | -0.383 | 1.071 | 0.702 | 0.652 | 0.022 | 0.006 | 0.006 | 0 | 0.15 | 0.36 | 0.13 | 0.08 | 0.16 | 0.19 |
| 2 | 41848835 | rs13006557 | 0.944 | 0.994 | 0.538 | 0.828 | 0.024 | 0.01 | 0.01 | 0 | 0.45 | 0.35 | 0.27 | 0.33 | 0.38 | 0.36 |
| 2 | 41849795 | rs2374292 | 0.161 | 1.008 | 0.651 | 0.863 | 0.001 | 0.001 | 0.001 | 0 | 0.57 | 0.49 | 0.38 | 0.39 | 0.54 | 0.54 |
| 2 | 41856087 | rs13023970 | 0.545 | 0.978 | 0.754 | 0.879 | 0.046 | 0.024 | 0.024 | 0 | 0.51 | 0.62 | 0.33 | 0.37 | 0.37 | 0.47 |
| 2 | 41857620 | rs17028152 | -0.183 | 1.489 | 1.382 | 1.355 | 0.133 | 0.176 | 0.176 | 2.477 | 0.08 | 0.35 | 0.25 | 0.23 | 0.11 | 0.14 |
| 2 | 41858990 | rs4952494 | -0.168 | 1.529 | 1.146 | 1.329 | -0.104 | -0.073 | -0.073 | 0.689 | 0.68 | 0.92 | 0.65 | 0.6 | 0.63 | 0.69 |
| 2 | 41860854 | rs2059917 | -0.707 | 1.494 | 1.624 | 1.433 | 0.126 | 0.215 | 0.215 | 2.776 | 0.1 | 0.49 | 0.32 | 0.3 | 0.15 | 0.17 |
| 2 | 41867887 | rs13016499 | 0.426 | 1.324 | 1.357 | 1.359 | -0.015 | -0.016 | -0.016 | 0 | 0.6 | 0.69 | 0.41 | 0.38 | 0.51 | 0.54 |
| 2 | 41872719 | rs1920509 | 0.636 | 1.304 | 1.356 | 1.352 | 0.002 | 0.007 | 0.007 | 0 | 0.57 | 0.34 | 0.33 | 0.3 | 0.48 | 0.52 |
| 2 | 41877732 | rs10183045 | 0.044 | 1.655 | 0.858 | 1.289 | 0.02 | 0.011 | 0.011 | 0 | 0.14 | 0.14 | 0.28 | 0.23 | 0.19 | 0.18 |
| 2 | 41885773 | rs6748566 | -0.088 | 1.504 | 0.656 | 1.295 | -0.062 | -0.059 | -0.059 | 0 | 0.73 | 0.93 | 0.67 | 0.61 | 0.69 | 0.72 |
| 2 | 41892171 | rs10198541 | -0.372 | 1.571 | 0.663 | 1.243 | -0.083 | -0.084 | -0.084 | 0 | 0.75 | 0.92 | 0.65 | 0.6 | 0.68 | 0.72 |
| 2 | 41898188 | rs9789345 | -0.711 | 1.737 | 1.078 | 1.21 | 0.079 | 0.097 | 0.097 | 0.463 | 0.14 | 0.32 | 0.31 | 0.3 | 0.19 | 0.21 |
| 2 | 41903118 | rs2043008 | -0.567 | 1.671 | 0.915 | 1.285 | -0.005 | -0.005 | -0.005 | 0 | 0.78 | 0.77 | 0.64 | 0.61 | 0.7 | 0.79 |
| 2 | 41909765 | rs6745319 | 0.738 | 1.309 | 1.006 | 1.239 | 0.001 | 0.002 | 0.002 | 0 | 0.57 | 0.47 | 0.37 | 0.43 | 0.52 | 0.58 |
| 2 | 41925436 | rs4952370 | -0.383 | 1.453 | 0.686 | 1.259 | -0.024 | -0.015 | -0.015 | 0 | 0.14 | 0.06 | 0.28 | 0.23 | 0.19 | 0.21 |
| 2 | 41934802 | rs17028447 | 0.596 | 1.427 | 0.997 | 1.156 | -0.004 | -0.004 | -0.004 | 0 | 0.6 | 0.62 | 0.38 | 0.44 | 0.53 | 0.56 |
| 2 | 41937140 | rs961279 | -0.508 | 1.525 | 0.538 | 1.166 | -0.073 | -0.058 | -0.058 | 0 | 0.74 | 0.94 | 0.67 | 0.68 | 0.69 | 0.77 |
| 2 | 41939950 | rs4952505 | 1.052 | 1.465 | 0.731 | 1.188 | 0.012 | 0.011 | 0.011 | 0 | 0.53 | 0.28 | 0.3 | 0.36 | 0.48 | 0.55 |
| 2 | 41947069 | rs11896329 | 0.444 | 1.442 | 0.68 | 1.145 | -0.006 | -0.003 | -0.003 | 0.849 | 0.59 | 0.38 | 0.52 | 0.56 | 0.46 | 0.61 |
| 2 | 41953286 | rs17611588 | 0.236 | 1.354 | 0.676 | 0.872 | -0.008 | 0.002 | 0.002 | 0.018 | 0.14 | 0.16 | 0.13 | 0.07 | 0.22 | 0.2 |
| 2 | 41954085 | rs6704987 | 1.4 | 1.217 | 0.553 | 0.798 | 0.066 | 0.043 | 0.043 | 0.173 | 0.42 | 0.21 | 0.19 | 0.28 | 0.28 | 0.39 |
| 2 | 41954472 | rs6720861 | 1.161 | 1.267 | 0.526 | 0.816 | -0.018 | -0.011 | -0.011 | 0.072 | 0.45 | 0.49 | 0.21 | 0.28 | 0.28 | 0.39 |
| 2 | 41958105 | rs1997416 | -0.246 | 1.708 | 0.751 | 0.974 | 0.015 | 0.008 | 0.008 | 0.018 | 0.15 | 0.21 | 0.18 | 0.15 | 0.27 | 0.21 |
| 2 | 41962945 | rs749651 | 0.418 | 1.509 | 0.437 | 0.897 | -0.006 | 0.006 | 0.006 | 0 | 0.63 | 0.67 | 0.47 | 0.28 | 0.59 | 0.51 |
| 2 | 41966314 | rs10432635 | -0.204 | 1.542 | 0.395 | 1.053 | 0.002 | -0.004 | -0.004 | 2.392 | 0.17 | 0.15 | 0.36 | 0.24 | 0.13 | 0.27 |
| 2 | 41968222 | rs10180152 | 0.515 | 1.564 | 0.433 | 0.937 | -0.005 | 0.001 | 0.001 | 0 | 0.63 | 0.66 | 0.47 | 0.29 | 0.59 | 0.51 |
| 2 | 41969241 | rs10490641 | 0.951 | 1.634 | 0.283 | 0.805 | -0.001 | -0.008 | -0.008 | 0 | 0.11 | 0.13 | 0.14 | 0.01 | 0.11 | 0.17 |
| 2 | 41979425 | rs7572321 | 1.114 | 1.738 | -0.028 | 0.791 | 0.001 | 0.007 | 0.007 | 0 | 0.54 | 0.53 | 0.39 | 0.27 | 0.44 | 0.43 |
| 2 | 41993806 | rs882585 | -0.901 | 2.007 | -0.445 | 0.986 | -0.009 | -0.013 | -0.013 | 0 | 0.8 | 0.84 | 0.75 | 0.52 | 0.72 | 0.77 |
| 2 | 41996154 | rs17464732 | 0.249 | 2.06 | -0.584 | 0.77 | 0.002 | -0.014 | -0.014 | 1.935 | 0.09 | 0.15 | 0.08 | 0.03 | 0.16 | 0.09 |
| 2 | 41996612 | rs4381822 | 0.129 | 1.798 | -0.55 | 0.876 | -0.001 | -0.001 | -0.001 | 2.822 | 0.6 | 0.46 | 0.59 | 0.39 | 0.59 | 0.45 |
| 2 | 41998764 | rs2005174 | -0.699 | 2.091 | -0.779 | 1.244 | -0.008 | 0.003 | 0.003 | 0 | 0.17 | 0.18 | 0.32 | 0.19 | 0.24 | 0.19 |
| 2 | 42004502 | rs4453726 | 0.068 | 2.146 | -0.675 | 1.659 | 0.001 | 0.006 | 0.006 | 0 | 0.79 | 0.79 | 0.75 | 0.57 | 0.7 | 0.74 |
| 2 | 42004851 | rs6724031 | 0.786 | 2.218 | -0.871 | 1.525 | -0.042 | -0.041 | -0.041 | 0.304 | 0.69 | 0.72 | 0.64 | 0.44 | 0.5 | 0.64 |
| 2 | 42004878 | rs719595 | 0.133 | 1.949 | -0.644 | 1.762 | -0.012 | -0.013 | -0.013 | 0.594 | 0.79 | 0.8 | 0.75 | 0.6 | 0.7 | 0.8 |
| 2 | 42005428 | rs4952511 | -1.03 | 0.787 | 0.155 | 1.297 | 0.014 | 0.08 | 0.08 | 0 | 0.1 | 0.35 | 0.25 | 0.12 | 0.06 | 0.21 |
| 2 | 42008674 | rs13398657 | -0.555 | 0.848 | -0.465 | 1.416 | -0.001 | -0.001 | -0.001 | 0 | 0.26 | 0.25 | 0.28 | 0.4 | 0.31 | 0.26 |
| 2 | 42012806 | rs4633963 | 0.448 | 0.608 | -0.815 | 0.535 | 0.002 | 0.004 | 0.004 | 0 | 0.17 | 0.14 | 0.31 | 0.01 | 0.13 | 0.23 |
| 2 | 42015775 | rs6704827 | -1.114 | 0.03 | -1.677 | 0.579 | -0.147 | -0.072 | -0.072 | 0 | 0.67 | 0.98 | 0.81 | 0.67 | 0.59 | 0.75 |
| 2 | 42018400 | rs4414724 | -0.516 | -0.283 | -0.927 | 0.453 | 0.003 | 0.025 | 0.025 | 0 | 0.44 | 0.62 | 0.61 | 0.2 | 0.42 | 0.51 |
| 2 | 42027041 | rs10204821 | -0.976 | -0.048 | -0.67 | 0.588 | -0.074 | 0.002 | 0.002 | 0.151 | 0.61 | 0.9 | 0.79 | 0.66 | 0.58 | 0.74 |
| 2 | 42038874 | rs4952515 | -0.393 | 0.021 | -0.471 | 0.435 | 0.001 | -0.01 | -0.01 | 0 | 0.38 | 0.6 | 0.58 | 0.2 | 0.4 | 0.46 |
| 2 | 42041338 | rs7599972 | 0.523 | 0.05 | -0.281 | 0.465 | 0.032 | 0.032 | 0.032 | 0 | 0.22 | 0.53 | 0.29 | 0.22 | 0.28 | 0.24 |
| 2 | 42046343 | rs11124848 | 0.299 | 0.039 | -0.271 | 0.455 | 0.014 | 0.011 | 0.011 | 0.011 | 0.23 | 0.6 | 0.3 | 0.22 | 0.28 | 0.24 |
| 2 | 42047032 | rs7599670 | 0.509 | 0.043 | -0.267 | 0.46 | 0.027 | 0.027 | 0.027 | 0 | 0.22 | 0.52 | 0.29 | 0.22 | 0.28 | 0.24 |
| 2 | 42047856 | rs13386861 | 0.508 | 0.042 | -0.265 | 0.468 | 0.025 | 0.028 | 0.028 | 0 | 0.22 | 0.54 | 0.29 | 0.22 | 0.28 | 0.24 |
| 2 | 42049084 | rs11689557 | 0.615 | 0.032 | -0.345 | 0.439 | 0.013 | 0.012 | 0.012 | 0 | 0.21 | 0.31 | 0.26 | 0.22 | 0.28 | 0.24 |
| 2 | 42054278 | rs13392392 | 1.104 | -0.008 | -0.452 | 0.334 | 0.002 | 0.046 | 0.046 | 0 | 0.16 | 0.08 | 0.28 | 0.01 | 0.12 | 0.24 |
| 2 | 42057211 | rs17028820 | 0.662 | -0.096 | -0.468 | 0.464 | -0.015 | 0.009 | 0.009 | 0 | 0.25 | 0.3 | 0.12 | 0.36 | 0.18 | 0.28 |
| 2 | 42062039 | rs4608574 | -0.37 | 0.214 | -1.236 | 0.494 | 0.006 | -0.015 | -0.015 | 1.097 | 0.39 | 0.5 | 0.57 | 0.25 | 0.43 | 0.48 |
| 2 | 42075529 | rs6746569 | -0.699 | 0.316 | -1.294 | 0.583 | -0.02 | -0.013 | -0.013 | 0.209 | 0.61 | 0.76 | 0.66 | 0.6 | 0.56 | 0.74 |
| 2 | 42080624 | rs4305317 | -0.23 | 0.277 | -1.398 | 0.581 | 0.002 | -0.001 | -0.001 | 0 | 0.46 | 0.53 | 0.38 | 0.58 | 0.45 | 0.54 |
| 2 | 42081106 | rs751489 | 0.628 | 0.196 | -1.026 | 0.804 | -0.031 | 0.042 | 0.042 | 0 | 0.21 | 0.38 | 0.11 | 0.34 | 0.15 | 0.25 |
| 2 | 42081856 | rs4331558 | -0.522 | 1.324 | -1.796 | 0.63 | -0.028 | -0.052 | -0.052 | 0.373 | 0.85 | 0.94 | 0.91 | 0.76 | 0.82 | 0.94 |
| 2 | 42085362 | rs11694333 | -0.306 | 1.514 | -1.403 | 0.225 | -0.146 | -0.015 | -0.015 | 0 | 0.26 | 0.02 | 0.53 | 0.17 | 0.38 | 0.23 |
| 2 | 42087252 | rs10167265 | -0.531 | 1.233 | -1.055 | 0.252 | 0.002 | -0.004 | -0.004 | 0 | 0.29 | 0.3 | 0.36 | 0.17 | 0.38 | 0.21 |
| 2 | 42090039 | rs4952521 | -0.017 | 1.033 | -1.307 | -0.063 | 0.019 | 0.096 | 0.096 | 0 | 0.47 | 0.32 | 0.53 | 0.18 | 0.41 | 0.65 |
| 2 | 42090563 | rs6723745 | -0.348 | 1.963 | -0.98 | 0.353 | 0.009 | 0 | 0 | 0.005 | 0.83 | 0.84 | 0.76 | 0.84 | 0.76 | 0.85 |
| 2 | 42095441 | rs4278972 | 0.403 | 0.851 | -1.422 | 0.735 | 0.033 | -0.093 | -0.093 | 0 | 0.26 | 0.07 | 0.27 | 0.44 | 0.18 | 0.42 |

| | | | | | | | | | | | | | | | | |
|---|---|---|---|---|---|---|---|---|---|---|---|---|---|---|---|---|
| 2 | 42098236 | rs4632397 | -0.052 | 1.08 | -1.072 | 0.658 | 0.002 | -0.002 | -0.002 | 0 | 0.62 | 0.53 | 0.44 | 0.64 | 0.45 | 0.71 |
| 2 | 42098839 | rs11695456 | 1.161 | 0.639 | -0.904 | 0.528 | 0.003 | 0.009 | 0.009 | 0.092 | 0.27 | 0.18 | 0.13 | 0.17 | 0.24 | 0.27 |
| 2 | 42100538 | rs11676056 | -0.253 | 1.463 | -0.866 | 0.64 | -0.009 | 0.004 | 0.004 | 0 | 0.78 | 0.78 | 0.56 | 0.85 | 0.63 | 0.75 |
| 2 | 42101473 | rs6731940 | 0.909 | 1.126 | -0.984 | 0.854 | 0 | -0.005 | -0.005 | 2.366 | 0.17 | 0.19 | 0.11 | 0.21 | 0.18 | 0.04 |
| 2 | 42104558 | rs9309077 | -0.259 | 1.842 | -0.823 | 1.048 | -0.064 | 0.004 | 0.004 | 0 | 0.2 | 0.07 | 0.37 | 0.14 | 0.32 | 0.24 |
| 2 | 42109471 | rs13426621 | -0.388 | 1.582 | -0.751 | 1.073 | 0 | -0.003 | -0.003 | 0 | 0.06 | 0.06 | 0.04 | 0.12 | 0.07 | 0.05 |
| 2 | 42111482 | rs4500987 | -0.215 | 1.85 | -0.441 | 1.261 | -0.001 | -0.002 | -0.002 | 0.229 | 0.45 | 0.4 | 0.54 | 0.39 | 0.4 | 0.58 |
| 2 | 42113729 | rs17028942 | -0.441 | 1.936 | -0.311 | 1.211 | 0.028 | -0.009 | -0.009 | 0 | 0.06 | 0.15 | 0.07 | 0.04 | 0.1 | 0.1 |
| 2 | 42114002 | rs13028888 | 0.298 | 1.87 | -0.229 | 1.423 | -0.022 | -0.009 | -0.009 | 0 | 0.13 | 0.07 | 0.17 | 0.14 | 0.18 | 0.14 |
| 2 | 42117362 | rs7601644 | -0.078 | 1.922 | -0.18 | 1.457 | -0.004 | -0.007 | -0.007 | 0 | 0.45 | 0.43 | 0.53 | 0.4 | 0.39 | 0.56 |
| 2 | 42120198 | rs4633961 | -0.066 | 1.927 | -0.457 | 1.49 | 0.004 | 0.013 | 0.013 | 0 | 0.44 | 0.31 | 0.5 | 0.39 | 0.39 | 0.57 |
| 2 | 42120306 | rs2374377 | -0.066 | 1.927 | -0.457 | 1.519 | 0.004 | 0.007 | 0.007 | 0 | 0.44 | 0.31 | 0.5 | 0.41 | 0.39 | 0.57 |
| 2 | 42120443 | rs2374379 | -0.063 | 1.925 | -0.496 | 1.449 | 0.005 | 0.016 | 0.016 | 0 | 0.44 | 0.3 | 0.49 | 0.38 | 0.39 | 0.56 |
| 2 | 42125126 | rs11491283 | 0.453 | 1.979 | -0.661 | 1.269 | -0.032 | -0.002 | -0.002 | 0 | 0.12 | 0.05 | 0.15 | 0.12 | 0.19 | 0.14 |
| 2 | 42129595 | rs4952534 | -0.455 | 1.708 | -0.492 | 1.326 | 0.01 | 0 | 0 | 0 | 0.5 | 0.35 | 0.55 | 0.54 | 0.46 | 0.62 |
| 2 | 42130740 | rs11687338 | -0.727 | 1.552 | -0.383 | 1.437 | -0.009 | 0.005 | 0.005 | 0 | 0.19 | 0.14 | 0.2 | 0.15 | 0.26 | 0.21 |
| 2 | 42131523 | rs4952535 | -0.085 | 1.412 | -0.346 | 1.576 | -0.002 | 0.005 | 0.005 | 0 | 0.37 | 0.14 | 0.32 | 0.38 | 0.36 | 0.51 |
| 2 | 42133049 | rs2374340 | 0.766 | 0.96 | -0.638 | 1.362 | -0.078 | -0.085 | -0.085 | 0 | 0.22 | 0.02 | 0.14 | 0.31 | 0.27 | 0.3 |
| 2 | 42133432 | rs2374341 | -0.538 | 1.026 | -0.103 | 1.437 | 0.006 | 0.009 | 0.009 | 0 | 0.37 | 0.21 | 0.31 | 0.38 | 0.35 | 0.5 |
| 2 | 42133651 | rs10179528 | -1.464 | 1.067 | 0.507 | 1.577 | -0.002 | -0.002 | -0.002 | 0 | 0.54 | 0.47 | 0.71 | 0.58 | 0.58 | 0.64 |
| 2 | 42135100 | rs17772527 | -1.009 | 1.054 | 0.993 | 1.539 | -0.008 | -0.011 | -0.011 | 0 | 0.45 | 0.66 | 0.57 | 0.51 | 0.5 | 0.51 |
| 2 | 42137215 | rs6734260 | -0.745 | 1.154 | 1.065 | 1.502 | 0.001 | 0.001 | 0.001 | 0.077 | 0.41 | 0.56 | 0.47 | 0.5 | 0.45 | 0.52 |
| 2 | 42139094 | rs6741942 | -0.23 | 1.234 | 1.063 | 1.527 | 0.007 | -0.003 | -0.003 | 0 | 0.33 | 0.55 | 0.26 | 0.35 | 0.37 | 0.4 |
| 2 | 42141078 | rs4952537 | -0.786 | 1.24 | 0.937 | 1.52 | -0.003 | -0.003 | -0.003 | 0 | 0.15 | 0.13 | 0.31 | 0.19 | 0.18 | 0.16 |
| 2 | 42141473 | rs2374344 | -0.519 | 1.169 | 0.91 | 1.455 | 0.04 | -0.016 | -0.016 | 0 | 0.11 | 0.27 | 0.15 | 0.07 | 0.16 | 0.09 |
| 2 | 42141552 | rs4952539 | 0.999 | 1.016 | 0.402 | 1.454 | 0.02 | 0.021 | 0.021 | 0 | 0.11 | 0.04 | 0.18 | 0.08 | 0.08 | 0.13 |
| 2 | 42142031 | rs6751325 | -1.887 | 1.154 | 0.407 | 1.46 | 0.012 | -0.038 | -0.038 | 0 | 0.6 | 0.76 | 0.62 | 0.53 | 0.62 | 0.64 |
| 2 | 42148519 | rs4952542 | -1.429 | 1.445 | 0.742 | 1.804 | 0.014 | 0.006 | 0.006 | 0 | 0.47 | 0.68 | 0.73 | 0.57 | 0.56 | 0.51 |
| 2 | 42150310 | rs6723268 | 0.37 | 1.559 | 1.073 | 1.837 | 0.005 | 0.014 | 0.014 | 0 | 0.25 | 0.35 | 0.29 | 0.34 | 0.27 | 0.31 |
| 2 | 42153211 | rs13419919 | -0.191 | 1.561 | 1.122 | 1.624 | 0.01 | -0.013 | -0.013 | 0 | 0.08 | 0.14 | 0.12 | 0.05 | 0.11 | 0.03 |
| 2 | 42155823 | rs17772955 | 0.233 | 1.563 | 1.081 | 1.624 | 0.002 | -0.009 | -0.009 | 0 | 0.07 | 0.08 | 0.11 | 0.05 | 0.11 | 0.03 |
| 2 | 42157310 | rs4952547 | 0.289 | 1.629 | 1.042 | 1.677 | -0.004 | -0.004 | -0.004 | 0 | 0.1 | 0.07 | 0.24 | 0.13 | 0.12 | 0.1 |
| 2 | 42159588 | rs2374390 | -1.581 | 1.562 | 0.991 | 1.932 | 0.016 | 0.041 | 0.041 | 0 | 0.47 | 0.69 | 0.68 | 0.67 | 0.56 | 0.5 |
| 2 | 42160639 | rs2276699 | 0.165 | 1.601 | 0.881 | 1.797 | 0 | -0.003 | -0.003 | 0 | 0.07 | 0.06 | 0.1 | 0.05 | 0.11 | 0.03 |
| 2 | 42163072 | rs11885000 | 0.255 | 1.58 | 1.025 | 2.035 | 0.003 | 0.021 | 0.021 | 0 | 0.26 | 0.35 | 0.29 | 0.45 | 0.29 | 0.35 |
| 2 | 42164215 | rs4952548 | -1.982 | 1.58 | 0.913 | 2.035 | 0.027 | 0.048 | 0.048 | 0 | 0.49 | 0.68 | 0.68 | 0.7 | 0.56 | 0.5 |
| 2 | 42165462 | rs7602635 | -1.909 | 1.508 | 1.079 | 1.993 | 0.04 | 0.006 | 0.006 | 0 | 0.24 | 0.39 | 0.39 | 0.24 | 0.28 | 0.15 |
| 2 | 42171309 | rs6750133 | -1.765 | 1.473 | 0.756 | 1.887 | 0.011 | 0.015 | 0.015 | 0 | 0.48 | 0.67 | 0.66 | 0.66 | 0.55 | 0.5 |
| 2 | 42171566 | rs11673981 | 0.025 | 1.498 | 0.677 | 1.855 | 0.011 | 0.027 | 0.027 | 0 | 0.27 | 0.37 | 0.3 | 0.45 | 0.3 | 0.29 |
| 2 | 42173006 | rs7576650 | -0.604 | 1.342 | 0.44 | 1.488 | 0 | -0.001 | -0.001 | 0 | 0.08 | 0.12 | 0.24 | 0.09 | 0.11 | 0.04 |
| 2 | 42176266 | rs6706641 | -1.117 | 1.247 | 0.478 | 1.601 | 0.006 | 0.004 | 0.004 | 0.691 | 0.13 | 0.22 | 0.11 | 0.14 | 0.14 | 0.13 |
| 2 | 42176797 | rs7598275 | -0.74 | 1.131 | 0.655 | 1.702 | 0.013 | 0.033 | 0.033 | 0 | 0.25 | 0.51 | 0.33 | 0.31 | 0.29 | 0.29 |
| 2 | 42178305 | rs4368377 | -2.159 | 0.819 | 1.142 | 1.614 | 0.013 | 0.136 | 0.136 | 2.862 | 0.08 | 0.55 | 0.12 | 0.1 | 0.03 | 0.08 |
| 2 | 42180912 | rs6737194 | -2.329 | 0.776 | 1.236 | 1.556 | -0.004 | 0.078 | 0.078 | 1.684 | 0.08 | 0.59 | 0.12 | 0.08 | 0.03 | 0.08 |
| 2 | 42182063 | rs6728179 | -2.597 | 1.38 | 0.969 | 2.213 | 0.043 | 0.161 | 0.161 | 0 | 0.19 | 0.61 | 0.23 | 0.47 | 0.22 | 0.21 |
| 2 | 42190583 | rs4952549 | -0.81 | 1.464 | 0.81 | 2.219 | -0.021 | -0.012 | -0.012 | 0 | 0.11 | 0.05 | 0.11 | 0.4 | 0.16 | 0.12 |
| 2 | 42194039 | rs2119172 | -0.645 | 1.516 | 0.698 | 2.186 | -0.05 | -0.065 | -0.065 | 0.17 | 0.1 | 0.01 | 0.1 | 0.39 | 0.13 | 0.1 |
| 2 | 42209530 | rs4952561 | -2.203 | 1.411 | 1.134 | 2.096 | 0.036 | 0.177 | 0.177 | 0.101 | 0.18 | 0.6 | 0.21 | 0.47 | 0.19 | 0.2 |
| 2 | 42211659 | rs12617564 | -0.286 | 1.438 | 0.426 | 1.855 | -0.077 | -0.136 | -0.136 | 0.559 | 0.2 | 0.01 | 0.32 | 0.4 | 0.27 | 0.18 |
| 2 | 42214495 | rs13017167 | -1.95 | 1.264 | 0.937 | 1.895 | 0.066 | 0.197 | 0.197 | 0 | 0.17 | 0.46 | 0.2 | 0.4 | 0.21 | 0.2 |
| 2 | 42227952 | rs1550430 | -1.169 | 0.985 | 1.045 | 1.822 | 0.011 | 0.068 | 0.068 | 0 | 0.09 | 0.15 | 0.36 | 0.33 | 0.1 | 0.17 |
| 2 | 42232439 | rs7570390 | -1.054 | 0.408 | 0.532 | 0.78 | 0.027 | 0.016 | 0.016 | 0.1 | 0.48 | 0.77 | 0.55 | 0.41 | 0.37 | 0.49 |
| 2 | 42236063 | rs6544497 | -0.837 | 0.425 | 0.547 | 0.663 | 0.024 | 0.008 | 0.008 | 0.22 | 0.45 | 0.79 | 0.54 | 0.42 | 0.33 | 0.54 |
| 2 | 42248713 | rs10205005 | 0.49 | 0.296 | 0.682 | 0.348 | 0.013 | 0.053 | 0.053 | 0 | 0.29 | 0.17 | 0.13 | 0 | 0.24 | 0.25 |
| 2 | 42249643 | rs10195970 | 0.156 | 0.295 | 1.067 | 0.351 | -0.026 | -0.157 | -0.157 | 0 | 0.33 | 0.49 | 0.14 | 0 | 0.21 | 0.23 |
| 2 | 42261813 | rs6716080 | -1.085 | 0.328 | 1.16 | 0.656 | 0.01 | 0.006 | 0.006 | 0.076 | 0.45 | 0.74 | 0.57 | 0.41 | 0.35 | 0.41 |
| 2 | 42262132 | rs10173801 | -1.323 | 0.309 | 1.412 | 0.638 | 0.042 | 0.251 | 0.251 | 0.515 | 0.12 | 0.42 | 0.42 | 0.4 | 0.1 | 0.17 |
| 2 | 42263580 | rs10177811 | 0.625 | 0.254 | 1.329 | 0.584 | 0.012 | 0.095 | 0.095 | 0.159 | 0.28 | 0.13 | 0.11 | 0 | 0.21 | 0.23 |
| 2 | 42267428 | rs11898187 | -1.006 | 0.231 | 1.381 | 0.684 | 0.012 | -0.006 | -0.006 | 0 | 0.45 | 0.72 | 0.59 | 0.49 | 0.37 | 0.51 |
| 2 | 42267462 | rs11887431 | 0.1 | 0.247 | 1.484 | 0.694 | -0.011 | -0.045 | -0.045 | 0 | 0.34 | 0.39 | 0.18 | 0.09 | 0.27 | 0.32 |
| 2 | 42268672 | rs4952582 | -0.804 | 0.286 | 1.364 | 0.933 | 0.028 | 0.167 | 0.167 | 0 | 0.1 | 0.22 | 0.4 | 0.4 | 0.1 | 0.17 |
| 2 | 42276921 | rs4952590 | -1.402 | 0.139 | 1.322 | 0.912 | 0.025 | 0.217 | 0.217 | 0 | 0.12 | 0.39 | 0.43 | 0.41 | 0.1 | 0.18 |
| 2 | 42279967 | rs12623480 | -1.87 | 0.136 | 0.433 | 0.232 | -0.066 | 0.148 | 0.148 | 1.101 | 0.72 | 0.96 | 0.81 | 0.98 | 0.69 | 0.71 |
| 2 | 42284884 | rs1044305 | -0.959 | 0.008 | 1.256 | 0.932 | 0.025 | 0.22 | 0.22 | 0 | 0.11 | 0.28 | 0.39 | 0.44 | 0.09 | 0.18 |
| 2 | 42285306 | rs7439 | -1.245 | -0.075 | 1.173 | 0.794 | 0.008 | 0.004 | 0.004 | 0 | 0.46 | 0.63 | 0.55 | 0.52 | 0.38 | 0.49 |
| 2 | 42285642 | rs2425 | -0.115 | -0.117 | 1.071 | 0.606 | 0.019 | 0.044 | 0.044 | 0.139 | 0.34 | 0.22 | 0.15 | 0.08 | 0.29 | 0.3 |
| 2 | 42287306 | rs988958 | -0.163 | -0.09 | 1.074 | 0.598 | 0.009 | 0.013 | 0.013 | 0.046 | 0.34 | 0.27 | 0.16 | 0.08 | 0.28 | 0.3 |
| 2 | 42289666 | rs1440095 | -1.198 | -0.066 | 0.916 | 0.759 | 0.007 | -0.002 | -0.002 | 0 | 0.45 | 0.63 | 0.55 | 0.47 | 0.37 | 0.47 |
| 2 | 42291485 | rs6737623 | -1.451 | -0.496 | 1.274 | 0.077 | 0.052 | 0.131 | 0.131 | 1.243 | 0.06 | 0.36 | 0.06 | 0.09 | 0.05 | 0.1 |

| | | | | | | | | | | | | | | | | |
|---|---|---|---|---|---|---|---|---|---|---|---|---|---|---|---|---|
| 2 | 42296955 | rs13421437 | -1.945 | -0.505 | 1.267 | 0.073 | 0.04 | 0.126 | 0.126 | 2.297 | 0.08 | 0.4 | 0.06 | 0.09 | 0.05 | 0.1 |
| 2 | 42298742 | rs13428594 | -0.731 | -0.426 | 1.136 | 0.04 | -0.012 | -0.054 | -0.054 | 0.591 | 0.39 | 0.56 | 0.18 | 0.09 | 0.29 | 0.31 |
| 2 | 42299028 | rs7608070 | -0.731 | -0.426 | 1.136 | 0.054 | -0.012 | -0.052 | -0.052 | 0.204 | 0.39 | 0.56 | 0.18 | 0.09 | 0.29 | 0.31 |
| 2 | 42299552 | rs6748687 | 1.254 | -0.602 | 0.808 | -0.093 | -0.032 | -0.033 | -0.033 | 0 | 0.06 | 0.15 | 0.02 | 0 | 0.02 | 0.07 |
| 2 | 42300838 | rs10514784 | 0.217 | -0.591 | 0.534 | -0.088 | -0.043 | -0.052 | -0.052 | 0 | 0.1 | 0.35 | 0.04 | 0 | 0.02 | 0.07 |
| 2 | 42301771 | rs893811 | -0.666 | -0.295 | -0.208 | 0.136 | -0.009 | -0.026 | -0.026 | 0 | 0.39 | 0.6 | 0.18 | 0.14 | 0.29 | 0.31 |
| 2 | 42306071 | rs10153685 | -0.627 | -0.127 | -0.608 | 0.229 | 0.012 | 0.022 | 0.022 | 0.341 | 0.39 | 0.71 | 0.19 | 0.12 | 0.29 | 0.33 |
| 2 | 42306258 | rs10153686 | -0.528 | -0.153 | -0.922 | 0.256 | 0.016 | 0.029 | 0.029 | 0.233 | 0.37 | 0.74 | 0.2 | 0.14 | 0.29 | 0.31 |
| 2 | 42306517 | rs6713393 | 0.62 | 0.527 | -1.157 | 1.026 | -0.012 | -0.024 | -0.024 | 0 | 0.81 | 0.88 | 0.7 | 0.54 | 0.74 | 0.82 |
| 2 | 42306722 | rs2581124 | 0.097 | 0.139 | -1.159 | 0.416 | 0.022 | 0.07 | 0.07 | 2.126 | 0.64 | 0.25 | 0.8 | 0.83 | 0.72 | 0.66 |
| 2 | 42306770 | rs6727235 | 0.908 | 0.303 | -0.756 | 1.389 | 0.01 | 0.04 | 0.04 | 0 | 0.84 | 0.83 | 0.69 | 0.59 | 0.78 | 0.86 |
| 2 | 42308573 | rs2466001 | -1.108 | -0.39 | -0.576 | 1.108 | -0.018 | 0.029 | 0.029 | 0 | 0.31 | 0.58 | 0.19 | 0.43 | 0.26 | 0.29 |
| 2 | 42309114 | rs6544499 | -1.25 | -0.697 | -0.66 | 0.861 | 0.028 | -0.011 | -0.011 | 0.003 | 0.41 | 0.71 | 0.26 | 0.58 | 0.3 | 0.43 |
| 2 | 42310322 | rs1866670 | 0.714 | -0.242 | -0.781 | 0.559 | 0.002 | 0.014 | 0.014 | 0.118 | 0.36 | 0.29 | 0.27 | 0.21 | 0.4 | 0.35 |
| 2 | 42310505 | rs2719149 | -0.627 | -0.267 | -1.172 | 0.628 | -0.029 | 0.008 | 0.008 | 0.058 | 0.6 | 0.85 | 0.48 | 0.67 | 0.61 | 0.61 |
| 2 | 42311264 | rs2719150 | -1.207 | -0.142 | -1.407 | 0.609 | -0.124 | -0.07 | -0.07 | 0 | 0.69 | 0.98 | 0.5 | 0.78 | 0.64 | 0.7 |
| 2 | 42312094 | rs7586671 | -0.682 | -0.766 | -0.469 | -0.062 | -0.041 | -0.063 | -0.063 | 0 | 0.16 | 0.56 | 0.08 | 0.12 | 0.07 | 0.12 |
| 2 | 42314135 | rs6737207 | 0.188 | -0.653 | -0.618 | 0.025 | -0.021 | -0.02 | -0.02 | 0 | 0.1 | 0.22 | 0.02 | 0.12 | 0.03 | 0.1 |
| 2 | 42324510 | rs11902318 | -0.28 | -0.396 | -1.652 | 0.155 | 0.026 | 0.077 | 0.077 | 0 | 0.51 | 0.7 | 0.37 | 0.4 | 0.47 | 0.52 |
| 2 | 42329329 | rs7570219 | -0.407 | -0.3 | -1.827 | 0.21 | 0.008 | 0.057 | 0.057 | 0 | 0.54 | 0.71 | 0.39 | 0.41 | 0.49 | 0.53 |
| 2 | 42333345 | rs4953236 | 0.198 | -0.037 | -2.03 | 0.299 | 0.01 | 0.027 | 0.027 | 0.555 | 0.35 | 0.23 | 0.28 | 0.29 | 0.36 | 0.35 |
| 2 | 42335185 | rs2083418 | -0.44 | 0.325 | -2.524 | 0.602 | -0.15 | -0.103 | -0.103 | 0 | 0.64 | 0.97 | 0.41 | 0.46 | 0.5 | 0.63 |
| 2 | 42340127 | rs11687715 | -0.094 | 0.195 | -2.712 | 0.654 | 0.028 | -0.012 | -0.012 | 0 | 0.52 | 0.19 | 0.35 | 0.4 | 0.45 | 0.39 |
| 2 | 42341945 | rs6544504 | -0.482 | -0.961 | -1.968 | 0.667 | -0.071 | 0.059 | 0.059 | 0.077 | 0.27 | 0.52 | 0.2 | 0.32 | 0.17 | 0.24 |
| 2 | 42350487 | rs6733455 | 0.043 | -0.912 | -1.548 | 0.126 | -0.016 | -0.009 | -0.009 | 0.005 | 0.38 | 0.42 | 0.22 | 0.21 | 0.3 | 0.38 |
| 2 | 42353031 | rs7591141 | 1.062 | 0.007 | -1.793 | 0.398 | -0.075 | -0.02 | -0.02 | 0.165 | 0.18 | 0.06 | 0.38 | 0.18 | 0.23 | 0.26 |
| 2 | 42354415 | rs6544505 | 0.333 | 0.013 | -1.871 | 0.203 | -0.006 | -0.009 | -0.009 | 0 | 0.47 | 0.44 | 0.26 | 0.21 | 0.35 | 0.49 |
| 2 | 42354846 | rs2581135 | -1.669 | 0.038 | -2.215 | 0.42 | -0.028 | -0.084 | -0.084 | 0.05 | 0.82 | 0.99 | 0.88 | 0.72 | 0.82 | 0.93 |
| 2 | 42355689 | rs963405 | -0.285 | 0.226 | -1.37 | 0.045 | 0.033 | -0.007 | -0.007 | 0.046 | 0.33 | 0.6 | 0.57 | 0.38 | 0.44 | 0.41 |
| 2 | 42360204 | rs4531983 | 0.4 | -0.18 | -1.14 | -0.178 | -0.001 | 0.045 | 0.045 | 0 | 0.51 | 0.41 | 0.52 | 0.31 | 0.41 | 0.62 |
| 2 | 42360660 | rs4453730 | -0.293 | -0.231 | -1.465 | -0.201 | -0.066 | 0.023 | 0.023 | 0 | 0.57 | 0.85 | 0.44 | 0.3 | 0.55 | 0.54 |
| 2 | 42361047 | rs4953571 | 0.762 | 0.384 | -0.951 | 0.219 | -0.011 | 0.074 | 0.074 | 0 | 0.46 | 0.3 | 0.51 | 0.24 | 0.4 | 0.54 |
| 2 | 42364439 | rs13392988 | 0.039 | 1.603 | -0.512 | 0.717 | 0 | 0 | 0 | 0 | 0.21 | 0.19 | 0.18 | 0.17 | 0.22 | 0.22 |
| 2 | 42370455 | rs4952936 | -0.429 | 1.009 | -0.352 | 0.234 | 0.007 | 0.013 | 0.013 | 0 | 0.45 | 0.61 | 0.7 | 0.41 | 0.58 | 0.54 |
| 2 | 42372986 | rs6544507 | 0.59 | 1.176 | -0.267 | 0.038 | 0.02 | -0.006 | -0.006 | 0 | 0.62 | 0.5 | 0.58 | 0.34 | 0.45 | 0.61 |
| 2 | 42373747 | rs13412252 | 0.241 | 1.231 | -0.759 | 0.264 | -0.045 | -0.065 | -0.065 | 0 | 0.73 | 0.85 | 0.72 | 0.43 | 0.57 | 0.74 |
| 2 | 42398473 | rs11902328 | -0.346 | 0.958 | -0.273 | 0.072 | 0.009 | 0.002 | 0.002 | 0 | 0.48 | 0.44 | 0.73 | 0.42 | 0.52 | 0.66 |
| 2 | 42398918 | rs12622910 | 1.036 | 1.192 | -0.273 | 0.28 | 0.004 | 0.026 | 0.026 | 0.116 | 0.1 | 0.16 | 0.12 | 0.16 | 0.13 | 0.15 |
| 2 | 42405743 | rs10164865 | -0.8 | 1.136 | 0.056 | 0.142 | 0.029 | 0.078 | 0.078 | 0 | 0.83 | 0.75 | 0.82 | 0.42 | 0.67 | 0.87 |
| 2 | 42419829 | rs10193935 | 1.138 | 1.132 | -0.027 | 0.127 | 0.008 | 0.047 | 0.047 | 0.093 | 0.1 | 0.17 | 0.13 | 0.16 | 0.13 | 0.15 |
| 2 | 42420599 | rs2719135 | 0.686 | 1.177 | 0.129 | 0.203 | 0.006 | 0.03 | 0.03 | 0 | 0.17 | 0.16 | 0.17 | 0.58 | 0.33 | 0.13 |
| 2 | 42423645 | rs10183344 | -0.476 | 0.999 | -0.214 | 0.067 | -0.038 | 0.068 | 0.068 | 0 | 0.63 | 0.79 | 0.41 | 0.74 | 0.61 | 0.49 |
| 2 | 42443032 | rs1470052 | 0.484 | 1.111 | -0.028 | 0.049 | 0.02 | 0.042 | 0.042 | 0 | 0.43 | 0.3 | 0.22 | 0.16 | 0.33 | 0.36 |
| 2 | 42451554 | rs2118888 | 0.775 | 0.931 | -0.089 | -0.003 | 0.06 | 0.06 | 0.06 | 0 | 0.41 | 0.24 | 0.17 | 0.15 | 0.24 | 0.34 |
| 2 | 42465484 | rs6544523 | -0.823 | 1.519 | -0.518 | -0.243 | -0.075 | 0.028 | 0.028 | 0 | 0.69 | 0.9 | 0.46 | 0.76 | 0.66 | 0.62 |
| 2 | 42468471 | rs7583046 | -0.291 | 1.472 | 0.051 | -0.351 | 0.073 | -0.024 | -0.024 | 0 | 0.29 | 0.51 | 0.32 | 0.73 | 0.47 | 0.27 |
| 2 | 42470794 | rs6713242 | 1.575 | 1.357 | 0.008 | -0.233 | -0.014 | -0.022 | -0.022 | 0 | 0.09 | 0.06 | 0.11 | 0.12 | 0.13 | 0.13 |
| 2 | 42476676 | rs17029395 | 1.148 | 1.054 | 0.167 | -0.518 | -0.005 | -0.059 | -0.059 | 0.029 | 0.35 | 0.26 | 0.08 | 0.03 | 0.12 | 0.32 |
| 2 | 42479050 | rs17029408 | 1.093 | 1.041 | 0.329 | -0.539 | -0.056 | -0.153 | -0.153 | 1.876 | 0.36 | 0.46 | 0.1 | 0.03 | 0.15 | 0.33 |
| 2 | 42488045 | rs2302346 | -0.728 | 1.439 | 0.252 | -0.319 | 0.021 | -0.019 | -0.019 | 0 | 0.33 | 0.61 | 0.36 | 0.73 | 0.5 | 0.28 |
| 2 | 42494273 | rs6544527 | -0.406 | 1.522 | 0.285 | -0.253 | 0.021 | -0.013 | -0.013 | 0.006 | 0.29 | 0.6 | 0.31 | 0.73 | 0.42 | 0.23 |
| 2 | 42497913 | rs11691536 | 0.967 | 1.584 | 0.237 | -0.072 | -0.038 | -0.061 | -0.061 | 0.172 | 0.15 | 0.06 | 0.15 | 0.58 | 0.28 | 0.11 |
| 2 | 42503030 | rs10490557 | 0.486 | 1.775 | -0.125 | -0.494 | -0.075 | 0.135 | 0.135 | 0 | 0.23 | 0.03 | 0.2 | 0.04 | 0.27 | 0.16 |
| 2 | 42509829 | rs6723361 | 0.131 | 1.496 | 0.163 | -0.464 | 0.071 | 0.193 | 0.193 | 0 | 0.19 | 0.43 | 0.2 | 0.58 | 0.31 | 0.13 |
| 2 | 42514649 | rs6544532 | 0.259 | 1.434 | 0.113 | -0.586 | -0.019 | 0.049 | 0.049 | 0 | 0.58 | 0.68 | 0.37 | 0.73 | 0.52 | 0.45 |
| 2 | 42515388 | rs10202624 | -1.445 | 1.548 | 0.026 | -0.591 | -0.021 | -0.042 | -0.042 | 3.267 | 0.92 | 0.97 | 0.64 | 0.8 | 0.9 | 0.85 |
| 2 | 42515501 | rs10179237 | 0.797 | 1.258 | 0.238 | -0.574 | 0.03 | 0.128 | 0.128 | 0.644 | 0.13 | 0.41 | 0.15 | 0.15 | 0.13 | 0.12 |
| 2 | 42516205 | rs2053349 | 0.709 | 1.263 | 0.243 | -0.532 | 0.027 | 0.125 | 0.125 | 1.51 | 0.13 | 0.42 | 0.15 | 0.15 | 0.13 | 0.12 |
| 2 | 42518025 | rs6544534 | 0.396 | 1.502 | 0.002 | 0.194 | -0.033 | -0.03 | -0.03 | 0.96 | 0.57 | 0.84 | 0.36 | 0.54 | 0.5 | 0.52 |
| 2 | 42518107 | rs6544535 | 1.689 | 1.529 | -0.039 | 0.263 | 0.018 | 0.048 | 0.048 | 0 | 0.1 | 0.16 | 0.13 | 0.35 | 0.24 | 0.07 |
| 2 | 42525308 | rs6544538 | 0.641 | 1.128 | -0.175 | 0.203 | 0.089 | 0.014 | 0.014 | 0 | 0.5 | 0.7 | 0.26 | 0.38 | 0.29 | 0.37 |
| 2 | 42526512 | rs12466769 | -0.291 | 1.139 | -0.446 | -0.179 | -0.107 | 0.024 | 0.024 | 0 | 0.68 | 0.91 | 0.42 | 0.76 | 0.6 | 0.58 |
| 2 | 42532912 | rs10171643 | 1.829 | 1.1 | -0.222 | 0.035 | -0.005 | 0 | 0 | 0 | 0.09 | 0.1 | 0.12 | 0.15 | 0.13 | 0.11 |
| 2 | 42537995 | rs6728830 | -1.997 | 1.086 | -0.282 | 0.112 | -0.009 | -0.03 | -0.03 | 6.137 | 0.95 | 1 | 0.99 | 0.89 | 0.97 | 0.89 |
| 2 | 42543577 | rs6752455 | 1.349 | 1.019 | 0.017 | 0.173 | 0.006 | 0.036 | 0.036 | 1.115 | 0.11 | 0.19 | 0.12 | 0.15 | 0.14 | 0.14 |
| 2 | 42553578 | rs11682029 | -0.306 | 1.08 | -0.063 | 0.082 | -0.028 | 0.053 | 0.053 | 0 | 0.62 | 0.75 | 0.39 | 0.73 | 0.57 | 0.47 |
| 2 | 42559256 | rs6716767 | 1.402 | 0.809 | 0.188 | -0.06 | 0.025 | -0.012 | -0.012 | 0 | 0.36 | 0.26 | 0.08 | 0.04 | 0.13 | 0.32 |
| 2 | 42559288 | rs13375 | 0.657 | 1.008 | 0.258 | 0.176 | 0.031 | 0.13 | 0.13 | 0.877 | 0.13 | 0.37 | 0.15 | 0.17 | 0.14 | 0.14 |
| 2 | 42577170 | rs1981664 | -0.453 | 1.103 | -0.03 | 0.068 | -0.026 | 0.04 | 0.04 | 0 | 0.63 | 0.76 | 0.39 | 0.73 | 0.59 | 0.51 |

| | | | | | | | | | | | | | | | | |
|---|---|---|---|---|---|---|---|---|---|---|---|---|---|---|---|---|
| 2 | 42578149 | rs8162 | -0.384 | 1.028 | 0.454 | -0.024 | 0.06 | -0.007 | -0.007 | 0 | 0.27 | 0.44 | 0.3 | 0.73 | 0.45 | 0.26 |
| 2 | 42583667 | rs10206058 | -0.609 | 1.085 | 0.155 | -0.111 | 0.002 | 0.033 | 0.033 | 0 | 0.64 | 0.76 | 0.41 | 0.72 | 0.65 | 0.54 |
| 2 | 42599917 | rs1992258 | 0.256 | 0.959 | 0.254 | -0.071 | 0.016 | 0.001 | 0.001 | 0 | 0.52 | 0.6 | 0.27 | 0.56 | 0.44 | 0.32 |
| 2 | 42615496 | rs7560797 | 0.546 | 0.437 | 0.303 | -0.235 | -0.014 | -0.017 | -0.017 | 0 | 0.44 | 0.52 | 0.22 | 0.16 | 0.27 | 0.37 |
| 2 | 42617549 | rs13417597 | 0.547 | 0.447 | 0.42 | -0.203 | 0.016 | 0.017 | 0.017 | 0 | 0.47 | 0.37 | 0.21 | 0.16 | 0.28 | 0.39 |
| 2 | 42637749 | rs10169411 | 1.4 | 0.672 | 0.462 | 0.003 | 0.029 | 0.09 | 0.09 | 0.025 | 0.1 | 0.22 | 0.13 | 0.15 | 0.14 | 0.13 |
| 2 | 42638057 | rs222471 | 1.582 | 0.674 | 0.458 | 0.006 | 0.018 | 0.065 | 0.065 | 0 | 0.09 | 0.16 | 0.13 | 0.16 | 0.14 | 0.13 |
| 2 | 42642531 | rs222468 | 1.131 | 0.658 | 0.497 | 0.021 | 0.046 | 0.12 | 0.12 | 0.231 | 0.11 | 0.33 | 0.14 | 0.14 | 0.14 | 0.13 |
| 2 | 42649819 | rs222485 | -0.433 | 0.827 | 0.243 | 0.018 | -0.013 | 0.03 | 0.03 | 0 | 0.62 | 0.74 | 0.36 | 0.73 | 0.59 | 0.49 |
| 2 | 42652658 | rs13015280 | 0.4 | 1.026 | 0.354 | 0.126 | 0.023 | 0.059 | 0.059 | 0 | 0.16 | 0.19 | 0.17 | 0.58 | 0.32 | 0.11 |
| 2 | 42652855 | rs17389988 | 0.815 | 1.057 | 0.012 | -0.005 | -0.044 | 0.051 | 0.051 | 0 | 0.11 | 0.03 | 0.12 | 0.03 | 0.14 | 0.18 |
| 2 | 42656337 | rs92303 | 1.149 | 0.953 | 0.352 | 0.23 | 0.038 | 0.072 | 0.072 | 0.296 | 0.11 | 0.3 | 0.08 | 0.13 | 0.14 | 0.13 |
| 2 | 42660297 | rs11124877 | 1.124 | 1.073 | -0.187 | 0.582 | -0.07 | -0.07 | -0.07 | 0.077 | 0.11 | 0.01 | 0.14 | 0.37 | 0.29 | 0.07 |
| 2 | 42669620 | rs11684852 | 0.383 | 0.978 | -0.083 | 0.151 | -0.018 | -0.014 | -0.014 | 0 | 0.16 | 0.13 | 0.17 | 0.6 | 0.32 | 0.11 |
| 2 | 42677549 | rs17029732 | 0.423 | 1.004 | -0.074 | 0.178 | -0.011 | 0 | 0 | 0 | 0.15 | 0.13 | 0.17 | 0.6 | 0.32 | 0.11 |
| 2 | 42677587 | rs1496328 | 0.575 | 0.769 | -0.468 | -0.086 | -0.015 | 0.03 | 0.03 | 0 | 0.31 | 0.22 | 0.5 | 0.17 | 0.34 | 0.32 |
| 2 | 42688797 | rs1496329 | -1.399 | 1.097 | -0.172 | 0.396 | 0.019 | 0.038 | 0.038 | 0.002 | 0.9 | 0.82 | 0.87 | 0.9 | 0.86 | 0.88 |
| 2 | 42690878 | rs6729011 | 0.494 | 0.136 | -1.109 | -0.303 | 0.015 | 0.078 | 0.078 | 0 | 0.51 | 0.32 | 0.63 | 0.27 | 0.44 | 0.54 |
| 2 | 42690881 | rs10495897 | 0.71 | -0.179 | -1.404 | -0.254 | 0.024 | 0.044 | 0.044 | 0 | 0.22 | 0.13 | 0.06 | 0.04 | 0.1 | 0.22 |
| 2 | 42707354 | rs17029748 | 0.246 | 0.377 | -1.387 | 0.083 | -0.011 | 0.057 | 0.057 | 0 | 0.15 | 0.02 | 0.11 | 0.04 | 0.14 | 0.18 |
| 2 | 42708267 | rs6544550 | 0.193 | 0.959 | -0.909 | 0.571 | -0.011 | -0.003 | -0.003 | 0 | 0.63 | 0.67 | 0.67 | 0.6 | 0.49 | 0.64 |
| 2 | 42712932 | rs13030217 | 0.768 | 0.566 | -1.156 | 0.482 | -0.001 | -0.012 | -0.012 | 0 | 0.26 | 0.21 | 0.2 | 0.28 | 0.28 | 0.34 |
| 2 | 42716984 | rs713611 | 0.769 | 0.476 | -1.235 | 0.335 | 0.039 | 0.132 | 0.132 | 0 | 0.25 | 0.04 | 0.13 | 0.04 | 0.19 | 0.21 |
| 2 | 42717015 | rs1390610 | 0.486 | 1.274 | -0.402 | 0.968 | -0.002 | 0.013 | 0.013 | 0.089 | 0.23 | 0.26 | 0.17 | 0.29 | 0.22 | 0.32 |
| 2 | 42728604 | rs17029761 | 0.438 | 1.333 | -0.157 | 0.974 | 0.001 | -0.002 | -0.002 | 0 | 0.64 | 0.56 | 0.58 | 0.3 | 0.38 | 0.49 |
| 2 | 42735172 | rs13028484 | 0.577 | 1.584 | -0.254 | 1.207 | 0.007 | 0 | 0 | 0.074 | 0.09 | 0.16 | 0.16 | 0.05 | 0.16 | 0.14 |
| 2 | 42739333 | rs924114 | 0.427 | 1.515 | -0.219 | 1.09 | -0.001 | 0.001 | 0.001 | 0 | 0.63 | 0.6 | 0.56 | 0.3 | 0.38 | 0.48 |
| 2 | 42743543 | rs13012795 | -0.623 | 1.043 | 0.051 | 1.046 | -0.002 | 0.02 | 0.02 | 0.002 | 0.35 | 0.5 | 0.33 | 0.36 | 0.38 | 0.45 |
| 2 | 42748348 | rs13033481 | -0.185 | 1.047 | -0.116 | 1.023 | -0.002 | -0.01 | -0.01 | 0 | 0.31 | 0.32 | 0.31 | 0.35 | 0.37 | 0.45 |
| 2 | 42773875 | rs12986795 | -0.392 | 1.113 | -0.201 | 1.107 | -0.001 | 0.001 | 0.001 | 0 | 0.33 | 0.38 | 0.32 | 0.33 | 0.36 | 0.45 |
| 2 | 42781775 | rs4953435 | 0.756 | 1.245 | -0.021 | 1.114 | -0.003 | -0.021 | -0.021 | 1.439 | 0.57 | 0.48 | 0.57 | 0.3 | 0.32 | 0.47 |
| 2 | 42784539 | rs7607881 | -0.741 | 1.361 | -0.19 | 1.201 | 0.009 | 0.02 | 0.02 | 0 | 0.91 | 0.94 | 0.86 | 0.65 | 0.7 | 0.92 |
| 2 | 42803365 | rs2132534 | 0.28 | 1.294 | -0.146 | 1.3 | 0.043 | 0.05 | 0.05 | 4.073 | 0.87 | 0.79 | 0.86 | 0.67 | 0.7 | 0.92 |
| 2 | 42806738 | rs10187433 | 0.366 | 1.287 | 0.02 | 1.227 | 0.086 | 0.118 | 0.118 | 3.954 | 0.87 | 0.72 | 0.86 | 0.65 | 0.71 | 0.92 |
| 2 | 42816929 | rs6544569 | 0.068 | 1.289 | -0.039 | 1.228 | 0.037 | 0.054 | 0.054 | 2.804 | 0.88 | 0.84 | 0.86 | 0.65 | 0.7 | 0.92 |
| 2 | 42856208 | rs6757976 | 0.65 | 1.252 | -0.207 | 1.277 | -0.001 | -0.045 | -0.045 | 0.125 | 0.22 | 0.14 | 0.15 | 0.36 | 0.22 | 0.39 |
| 2 | 42856780 | rs6758688 | -0.324 | 1.353 | -0.143 | 1.312 | 0.017 | 0.029 | 0.029 | 2.139 | 0.89 | 0.88 | 0.87 | 0.66 | 0.71 | 0.92 |
| 2 | 42865441 | rs13027020 | -0.5 | 1.275 | -0.152 | 1.273 | 0.031 | 0.047 | 0.047 | 0 | 0.89 | 0.87 | 0.87 | 0.65 | 0.7 | 0.92 |
| 2 | 42870125 | rs4953527 | 1.152 | 1.209 | -0.113 | 1.094 | -0.007 | -0.022 | -0.022 | 0 | 0.53 | 0.48 | 0.54 | 0.23 | 0.28 | 0.42 |
| 2 | 42876940 | rs11900994 | 0.765 | 1.03 | -0.131 | 1.365 | -0.009 | -0.02 | -0.02 | 0 | 0.22 | 0.14 | 0.15 | 0.28 | 0.22 | 0.31 |
| 2 | 42880771 | rs12996750 | -0.766 | 0.99 | -0.136 | 1.097 | 0.022 | 0.037 | 0.037 | 0 | 0.91 | 0.9 | 0.87 | 0.65 | 0.7 | 0.92 |
| 2 | 42881492 | rs10208367 | -0.723 | 0.988 | -0.142 | 1.258 | -0.054 | -0.044 | -0.044 | 0 | 0.9 | 0.99 | 0.88 | 0.85 | 0.7 | 0.92 |
| 2 | 42881534 | rs10195899 | 0.009 | 0.984 | -0.114 | 1.256 | 0.051 | 0.022 | 0.022 | 4.906 | 0.88 | 0.79 | 0.86 | 0.85 | 0.7 | 0.92 |
| 2 | 42895487 | rs6724638 | 0.092 | 0.95 | -0.171 | 1.266 | 0.036 | 0.014 | 0.014 | 4.53 | 0.88 | 0.81 | 0.86 | 0.84 | 0.7 | 0.92 |
| 2 | 42895816 | rs10865184 | 0.92 | 1.012 | -0.233 | 1.283 | 0.003 | -0.017 | -0.017 | 0 | 0.18 | 0.17 | 0.15 | 0.51 | 0.19 | 0.36 |
| 2 | 42900991 | rs6544586 | -0.132 | 0.213 | -0.365 | 0.922 | -0.001 | -0.008 | -0.008 | 0 | 0.66 | 0.75 | 0.56 | 0.6 | 0.66 | 0.76 |
| 2 | 42903494 | rs8179838 | -0.198 | 0.232 | -0.229 | 0.955 | -0.002 | -0.005 | -0.005 | 0 | 0.67 | 0.74 | 0.54 | 0.6 | 0.64 | 0.74 |
| 2 | 42912035 | rs4953562 | -0.306 | 0.87 | -0.604 | 1.005 | 0.04 | 0.053 | 0.053 | 2.725 | 0.89 | 0.85 | 0.86 | 0.65 | 0.7 | 0.92 |
| 2 | 42912517 | rs4952910 | -0.673 | 0.877 | -0.617 | 1.009 | 0.009 | 0.019 | 0.019 | 0 | 0.91 | 0.91 | 0.87 | 0.64 | 0.7 | 0.92 |
| 2 | 42921899 | rs731205 | 0.623 | 0.669 | -0.561 | 0.905 | 0.068 | 0.072 | 0.072 | 0.918 | 0.8 | 0.66 | 0.72 | 0.59 | 0.53 | 0.84 |
| 2 | 42925459 | rs12472339 | 0.665 | 0.702 | -0.905 | 0.868 | 0.038 | 0.046 | 0.046 | 0.293 | 0.79 | 0.72 | 0.72 | 0.56 | 0.53 | 0.82 |
| 2 | 42935961 | rs4953577 | 0.287 | 0.816 | -1.269 | 0.895 | -0.002 | -0.005 | -0.005 | 0.138 | 0.82 | 0.83 | 0.73 | 0.76 | 0.53 | 0.83 |
| 2 | 42936547 | rs3816186 | 0.01 | 0.871 | -1.44 | 0.895 | -0.023 | -0.019 | -0.019 | 0 | 0.83 | 0.89 | 0.74 | 0.78 | 0.53 | 0.84 |
| 2 | 42942177 | rs10194712 | -0.164 | 0.518 | -1.868 | 0.51 | -0.132 | -0.146 | -0.146 | 0 | 0.69 | 0.96 | 0.5 | 0.56 | 0.51 | 0.81 |
| 2 | 42950753 | rs2119014 | 0.708 | 0.63 | -0.983 | 0.682 | 0.014 | 0.02 | 0.02 | 0.105 | 0.71 | 0.69 | 0.59 | 0.52 | 0.48 | 0.67 |
| 2 | 42971687 | rs1016607 | 0.297 | 1.114 | -0.484 | 0.255 | 0.168 | 0.023 | 0.023 | 1.237 | 0.79 | 0.55 | 0.72 | 0.82 | 0.65 | 0.78 |
| 2 | 42977503 | rs6544593 | -0.159 | 0.638 | -0.354 | 0.093 | 0.002 | 0.01 | 0.01 | 2.197 | 0.28 | 0.37 | 0.3 | 0.63 | 0.35 | 0.43 |
| 2 | 42978232 | rs6717144 | -0.06 | 0.577 | -0.429 | 0.131 | 0 | -0.001 | -0.001 | 0 | 0.27 | 0.3 | 0.28 | 0.62 | 0.33 | 0.43 |
| 2 | 42978436 | rs930420 | 1.043 | 0.486 | -0.474 | 0.021 | 0.026 | 0.038 | 0.038 | 0 | 0.52 | 0.41 | 0.44 | 0.2 | 0.31 | 0.33 |
| 2 | 42985803 | rs6719977 | 0.549 | 0.764 | -0.848 | -0.017 | -0.04 | 0.012 | 0.012 | 0 | 0.74 | 0.86 | 0.74 | 0.84 | 0.55 | 0.77 |
| 2 | 42989592 | rs2278583 | 0.229 | 0.651 | -0.779 | 0.302 | -0.001 | 0.001 | 0.001 | 0.029 | 0.18 | 0.21 | 0.23 | 0.6 | 0.24 | 0.41 |
| 2 | 42989994 | rs2278585 | 0.229 | 0.651 | -0.779 | 0.304 | -0.001 | 0.002 | 0.002 | 0.029 | 0.18 | 0.2 | 0.24 | 0.59 | 0.24 | 0.41 |
| 2 | 42990101 | rs2278586 | 0.229 | 0.651 | -0.779 | 0.305 | -0.001 | 0.001 | 0.001 | 0.029 | 0.18 | 0.2 | 0.24 | 0.6 | 0.24 | 0.41 |
| 2 | 42990522 | rs12712859 | 0.255 | 0.656 | -0.749 | 0.304 | 0 | 0.019 | 0.019 | 0.046 | 0.17 | 0.24 | 0.24 | 0.59 | 0.24 | 0.41 |
| 2 | 42993166 | rs965374 | 0.016 | 1.024 | -0.853 | 0.209 | -0.089 | -0.041 | -0.041 | 0 | 0.84 | 0.95 | 0.79 | 0.82 | 0.62 | 0.78 |
| 2 | 42994484 | rs2304661 | -1.443 | 0.921 | -0.074 | 0.215 | 0.046 | 0.007 | 0.007 | 0 | 0.08 | 0.26 | 0.12 | 0.05 | 0.07 | 0.06 |
| 2 | 42995840 | rs3755540 | -0.183 | 0.985 | -0.719 | 0.133 | 0.001 | 0.001 | 0.001 | 0 | 0.86 | 0.86 | 0.86 | 0.69 | 0.72 | 0.89 |
| 2 | 42995848 | rs3755541 | -0.046 | 1.016 | -0.594 | 0.132 | 0.038 | 0.044 | 0.044 | 0 | 0.85 | 0.79 | 0.86 | 0.66 | 0.71 | 0.89 |
| 2 | 42997614 | rs2241850 | -0.642 | 0.626 | -1.165 | -0.261 | -0.026 | -0.006 | -0.006 | 0 | 0.36 | 0.23 | 0.36 | 0.63 | 0.42 | 0.45 |

| | | | | | | | | | | | | | | | |
|---|---|---|---|---|---|---|---|---|---|---|---|---|---|---|---|
| 2 | 43010739 | rs3821349 | 1.294 | 0.179 | -1.019 | -0.56 | -0.016 | -0.085 | -0.085 | 0 | 0.45 | 0.49 | 0.5 | 0.06 | 0.29 | 0.43 |
| 2 | 43012632 | rs11891403 | -0.054 | -0.274 | -2.041 | -0.605 | -0.071 | 0.062 | 0.062 | 0.801 | 0.58 | 0.18 | 0.42 | 0.72 | 0.56 | 0.58 |
| 2 | 43014289 | rs12613026 | -0.062 | -0.209 | -1.999 | -0.572 | -0.069 | 0.062 | 0.062 | 0 | 0.6 | 0.2 | 0.43 | 0.73 | 0.56 | 0.59 |
| 2 | 43015119 | rs737148 | -0.381 | 0.725 | -1.155 | 0.321 | 0.001 | -0.057 | -0.057 | 0 | 0.28 | 0.14 | 0.25 | 0.58 | 0.28 | 0.29 |
| 2 | 43015719 | rs3816183 | -0.918 | 1.255 | -0.75 | 0.703 | 0.124 | 0.092 | 0.092 | 0 | 0.95 | 0.89 | 0.89 | 0.8 | 0.76 | 0.95 |
| 2 | 43018458 | rs13027051 | -0.438 | 1.161 | -0.474 | 1.006 | -0.008 | 0.038 | 0.038 | 0.628 | 0.17 | 0.19 | 0.15 | 0.53 | 0.09 | 0.2 |
| 2 | 43028132 | rs9309089 | -0.753 | 0.753 | -1.335 | 0.934 | -0.024 | -0.105 | -0.105 | 0 | 0.24 | 0.06 | 0.23 | 0.58 | 0.21 | 0.35 |
| 2 | 43040577 | rs1371404 | 0.302 | 0.768 | -0.911 | 0.975 | -0.008 | -0.017 | -0.017 | 0.034 | 0.63 | 0.65 | 0.61 | 0.53 | 0.46 | 0.64 |
| 2 | 43043116 | rs6717237 | 1.269 | 0.48 | -0.711 | 0.677 | -0.013 | -0.09 | -0.09 | 0 | 0.46 | 0.48 | 0.5 | 0.02 | 0.29 | 0.3 |
| 2 | 43043308 | rs6544601 | -0.358 | 0.873 | -0.964 | 0.887 | -0.075 | -0.072 | -0.072 | 0 | 0.77 | 0.89 | 0.76 | 0.61 | 0.59 | 0.81 |
| 2 | 43050311 | rs1347929 | 0.15 | 0.87 | -0.883 | 1.064 | -0.001 | -0.001 | -0.001 | 0 | 0.2 | 0.2 | 0.18 | 0.18 | 0.21 | 0.37 |
| 2 | 43053772 | rs13013413 | 0.741 | 0.941 | -1.005 | 1.334 | -0.018 | -0.016 | -0.016 | 0 | 0.16 | 0.05 | 0.12 | 0.18 | 0.17 | 0.1 |
| 2 | 43057206 | rs6544602 | -1.348 | 0.758 | -0.926 | 1.123 | -0.001 | 0.021 | 0.021 | 0 | 0.52 | 0.46 | 0.37 | 0.76 | 0.46 | 0.64 |
| 2 | 43058086 | rs11899294 | 0.256 | 0.652 | -1.228 | 1.522 | 0.004 | -0.061 | -0.061 | 0 | 0.1 | 0.02 | 0.04 | 0.35 | 0.07 | 0.14 |
| 2 | 43074225 | rs7355325 | -0.685 | 0.855 | -1.373 | 1.295 | -0.032 | -0.083 | -0.083 | 0.47 | 0.35 | 0.11 | 0.24 | 0.54 | 0.34 | 0.57 |
| 2 | 43076226 | rs13395387 | 0.079 | 0.844 | -1.424 | 1.21 | -0.052 | 0.002 | 0.002 | 0 | 0.24 | 0.06 | 0.17 | 0.17 | 0.26 | 0.39 |
| 2 | 43077509 | rs11896488 | -1.28 | 0.941 | -0.668 | 0.692 | -0.022 | 0.06 | 0.06 | 0 | 0.6 | 0.73 | 0.45 | 0.83 | 0.51 | 0.67 |
| 2 | 43081439 | rs10169957 | -0.473 | 1.29 | 0.667 | 1.004 | 0.345 | 0.204 | 0.204 | 0.016 | 0.93 | 0.52 | 0.85 | 0.86 | 0.76 | 0.92 |
| 2 | 43083070 | rs7585445 | -0.257 | 1.277 | 0.65 | 0.976 | 0.31 | 0.165 | 0.165 | 0.675 | 0.92 | 0.49 | 0.85 | 0.87 | 0.75 | 0.92 |
| 2 | 43084026 | rs13013681 | 0.332 | 0.66 | -0.036 | 0.883 | 0.015 | -0.018 | -0.018 | 0.508 | 0.23 | 0.15 | 0.15 | 0.29 | 0.2 | 0.24 |
| 2 | 43086466 | rs7558605 | 1.284 | 0.214 | -0.128 | 0.653 | 0.022 | 0.028 | 0.028 | 0 | 0.37 | 0.28 | 0.36 | 0.11 | 0.24 | 0.25 |
| 2 | 43091100 | rs6727142 | 0.292 | 0.237 | 0.066 | 0.846 | -0.01 | 0.006 | 0.006 | 0 | 0.25 | 0.31 | 0.17 | 0.29 | 0.22 | 0.26 |
| 2 | 43091550 | rs11687176 | 2.241 | 0.509 | -0.164 | 0.886 | -0.01 | -0.003 | -0.003 | 0 | 0.1 | 0.02 | 0.06 | 0.06 | 0.08 | 0.14 |
| 2 | 43093767 | rs11888901 | 0.18 | 0.869 | -0.009 | 0.89 | 0.003 | 0.002 | 0.002 | 0 | 0.26 | 0.32 | 0.22 | 0.28 | 0.31 | 0.29 |
| 2 | 43095786 | rs4408769 | 1.066 | 0.531 | 0.307 | 0.484 | 0.001 | -0.011 | -0.011 | 1.962 | 0.48 | 0.5 | 0.43 | 0.16 | 0.32 | 0.39 |
| 2 | 43100438 | rs4953672 | -0.162 | 0.814 | 0.499 | 1.185 | 0.038 | 0.049 | 0.049 | 0 | 0.72 | 0.58 | 0.71 | 0.43 | 0.54 | 0.67 |
| 2 | 43102327 | rs7598637 | 0.505 | 0.324 | 0.398 | 0.936 | -0.001 | 0 | 0 | 0 | 0.6 | 0.61 | 0.55 | 0.38 | 0.43 | 0.63 |
| 2 | 43103614 | rs4555392 | 0.572 | -0.13 | 0.373 | 0.692 | 0.031 | 0.032 | 0.032 | 0 | 0.53 | 0.25 | 0.54 | 0.35 | 0.38 | 0.54 |
| 2 | 43105219 | rs12615717 | 0.446 | -0.068 | 0.421 | 0.914 | -0.011 | 0.02 | 0.02 | 1.951 | 0.23 | 0.28 | 0.19 | 0.33 | 0.22 | 0.23 |
| 2 | 43106956 | rs10208080 | 0.03 | -0.135 | 0.224 | 0.894 | -0.033 | 0.055 | 0.055 | 0 | 0.37 | 0.12 | 0.25 | 0.27 | 0.39 | 0.34 |
| 2 | 43107039 | rs3936042 | 0.697 | -0.185 | 0.401 | 0.809 | 0.061 | 0.072 | 0.072 | 0.092 | 0.5 | 0.15 | 0.53 | 0.35 | 0.37 | 0.55 |
| 2 | 43109042 | rs4494803 | 0.101 | -0.122 | 0.244 | 0.879 | -0.035 | 0.075 | 0.075 | 0 | 0.37 | 0.12 | 0.26 | 0.23 | 0.39 | 0.34 |
| 2 | 43109583 | rs11897055 | 0.517 | -0.31 | 0.083 | 0.731 | 0 | -0.001 | -0.001 | 0 | 0.19 | 0.22 | 0.23 | 0.23 | 0.14 | 0.28 |
| 2 | 43111944 | rs13396426 | -0.348 | -0.157 | 0.587 | 0.796 | -0.002 | -0.002 | -0.002 | 0.566 | 0.57 | 0.53 | 0.55 | 0.5 | 0.53 | 0.62 |
| 2 | 43113206 | rs4350800 | -0.54 | 0.156 | 0.358 | 0.968 | 0.12 | 0.196 | 0.196 | 0 | 0.93 | 0.85 | 0.87 | 0.62 | 0.77 | 0.91 |
| 2 | 43117026 | rs6707553 | -0.655 | 0.022 | -0.044 | 0.913 | -0.019 | -0.014 | -0.014 | 0 | 0.61 | 0.72 | 0.55 | 0.51 | 0.54 | 0.64 |
| 2 | 43127816 | rs4434040 | 0.125 | -0.19 | -0.751 | 0.664 | 0.019 | -0.019 | -0.019 | 0 | 0.25 | 0.19 | 0.25 | 0.33 | 0.2 | 0.28 |
| 2 | 43132109 | rs11124899 | 0.354 | -0.148 | -0.507 | 0.772 | 0.004 | -0.005 | -0.005 | 0 | 0.22 | 0.23 | 0.33 | 0.29 | 0.16 | 0.17 |
| 2 | 43132146 | rs11904826 | 0.789 | -0.167 | -0.652 | 0.432 | 0.02 | 0.002 | 0.002 | 0 | 0.3 | 0.11 | 0.29 | 0.27 | 0.31 | 0.48 |
| 2 | 43132547 | rs12986863 | 0.869 | -0.143 | -0.656 | 0.439 | 0.014 | -0.005 | -0.005 | 0 | 0.29 | 0.11 | 0.3 | 0.26 | 0.31 | 0.47 |
| 2 | 43135213 | rs4404328 | 0.162 | 0.31 | -0.029 | 0.587 | 0 | 0 | 0 | 0 | 0.72 | 0.69 | 0.55 | 0.33 | 0.6 | 0.76 |
| 2 | 43135815 | rs12712867 | -0.239 | 0.41 | 0.419 | 0.755 | -0.001 | 0 | 0 | 0 | 0.55 | 0.54 | 0.43 | 0.37 | 0.5 | 0.66 |
| 2 | 43140653 | rs6719452 | 0.523 | 0.494 | 0.05 | 0.544 | 0.024 | 0.072 | 0.072 | 0.068 | 0.43 | 0.11 | 0.33 | 0.28 | 0.41 | 0.62 |
| 2 | 43142947 | rs13020622 | -0.275 | 0.279 | 0.629 | 0.673 | -0.001 | 0 | 0 | 0 | 0.56 | 0.51 | 0.42 | 0.33 | 0.5 | 0.66 |
| 2 | 43143587 | rs3842996 | -0.462 | 0.274 | 0.531 | 0.592 | -0.024 | 0.02 | 0.02 | 0 | 0.61 | 0.73 | 0.57 | 0.32 | 0.45 | 0.41 |
| 2 | 43144956 | rs11682091 | 1.286 | 0.425 | 0.752 | 0.501 | 0.022 | -0.077 | -0.077 | 0.586 | 0.18 | 0.23 | 0.11 | 0.02 | 0.18 | 0.14 |
| 2 | 43148487 | rs4952948 | 0.131 | 0.395 | 0.615 | 0.83 | 0.069 | 0.125 | 0.125 | 1.362 | 0.76 | 0.49 | 0.73 | 0.49 | 0.65 | 0.8 |
| 2 | 43148735 | rs4953685 | 1.341 | 0.196 | 0.607 | 0.816 | 0.011 | 0.002 | 0.002 | 0 | 0.26 | 0.2 | 0.33 | 0.22 | 0.15 | 0.18 |
| 2 | 43149097 | rs4953686 | -0.13 | 0.246 | 0.761 | 0.799 | 0.009 | 0.006 | 0.006 | 0 | 0.5 | 0.53 | 0.44 | 0.42 | 0.43 | 0.6 |
| 2 | 43151040 | rs12990300 | 0.779 | 0.196 | 0.213 | 0.684 | 0.036 | 0.035 | 0.035 | 0 | 0.33 | 0.21 | 0.3 | 0.25 | 0.31 | 0.47 |
| 2 | 43151319 | rs11686965 | 0.063 | 0.355 | 0.418 | 0.639 | 0.003 | -0.029 | -0.029 | 0 | 0.34 | 0.44 | 0.22 | 0.14 | 0.29 | 0.25 |
| 2 | 43151394 | rs10168954 | -0.416 | 0.724 | 0.585 | 0.965 | 0.041 | 0.015 | 0.015 | 0 | 0.75 | 0.72 | 0.76 | 0.64 | 0.56 | 0.78 |
| 2 | 43152320 | rs4953692 | -0.252 | 0.473 | 0.265 | 0.811 | 0.015 | 0.009 | 0.009 | 0 | 0.66 | 0.55 | 0.6 | 0.39 | 0.6 | 0.72 |
| 2 | 43154210 | rs13028927 | 0.369 | 0.018 | 0.155 | 0.724 | 0.015 | 0.007 | 0.007 | 0 | 0.31 | 0.24 | 0.35 | 0.27 | 0.27 | 0.49 |
| 2 | 43156905 | rs2888844 | -0.296 | 0.205 | 0.623 | 0.806 | 0.037 | -0.04 | -0.04 | 0 | 0.77 | 0.52 | 0.78 | 0.85 | 0.73 | 0.83 |
| 2 | 43158832 | rs1550861 | -0.615 | 0.218 | 0.639 | 0.801 | -0.009 | 0.006 | 0.006 | 0.947 | 0.79 | 0.78 | 0.79 | 0.84 | 0.71 | 0.84 |
| 2 | 43159466 | rs11900368 | 0.186 | -0.19 | 1.43 | 1.132 | 0.003 | 0.002 | 0.002 | 0.022 | 0.33 | 0.29 | 0.42 | 0.3 | 0.33 | 0.48 |
| 2 | 43160027 | rs1078579 | -0.714 | -0.233 | 1.375 | 1.067 | 0.023 | 0.057 | 0.057 | 0 | 0.44 | 0.65 | 0.58 | 0.85 | 0.42 | 0.6 |
| 2 | 43160844 | rs895508 | 0.06 | -0.255 | 1.219 | 1.294 | -0.014 | 0.001 | 0.001 | 0 | 0.34 | 0.37 | 0.4 | 0.34 | 0.32 | 0.49 |
| 2 | 43167622 | rs766887 | -0.288 | -0.264 | 0.622 | 1.378 | -0.022 | -0.014 | -0.014 | 0 | 0.38 | 0.49 | 0.44 | 0.33 | 0.34 | 0.5 |
| 2 | 43169272 | rs2166500 | -0.506 | -0.161 | -0.168 | 1.19 | 0.016 | 0.029 | 0.029 | 0 | 0.42 | 0.64 | 0.6 | 0.81 | 0.41 | 0.56 |
| 2 | 43169426 | rs997818 | 0.197 | 0.138 | -1.253 | 1.164 | 0.005 | -0.046 | -0.046 | 0 | 0.32 | 0.08 | 0.37 | 0.34 | 0.33 | 0.48 |
| 2 | 43184504 | rs12476790 | 0.851 | -0.027 | -1.403 | 0.095 | -0.002 | 0.06 | 0.06 | 0.386 | 0.26 | 0.13 | 0.16 | 0 | 0.2 | 0.14 |
| 2 | 43185532 | rs754868 | -0.111 | 0.451 | -1.219 | 0.555 | -0.037 | 0.029 | 0.029 | 0 | 0.6 | 0.79 | 0.5 | 0.68 | 0.45 | 0.49 |
| 2 | 43188499 | rs7590773 | -0.073 | 0.547 | -1.578 | 0.51 | -0.073 | 0.05 | 0.05 | 0 | 0.6 | 0.86 | 0.53 | 0.66 | 0.44 | 0.54 |
| 2 | 43199311 | rs10176316 | -0.295 | 0.868 | -1.045 | 0.697 | 0.021 | 0.036 | 0.036 | 0 | 0.29 | 0.41 | 0.21 | 0.37 | 0.27 | 0.29 |
| 2 | 43200148 | rs7576359 | -0.266 | 0.66 | -1.595 | 0.418 | -0.033 | -0.068 | -0.068 | 0.173 | 0.76 | 0.88 | 0.51 | 0.62 | 0.62 | 0.7 |
| 2 | 43202587 | rs11124908 | 0.295 | 0.425 | -1.697 | 0.219 | -0.035 | 0.021 | 0.021 | 0 | 0.27 | 0.44 | 0.17 | 0.36 | 0.2 | 0.25 |
| 2 | 43203581 | rs3749084 | 1.454 | 0.213 | -1.526 | -0.226 | 0.003 | 0.103 | 0.103 | 0 | 0.3 | 0.1 | 0.18 | 0.09 | 0.25 | 0.23 |

| | | | | | | | | | | | | | | | | |
|---|---|---|---|---|---|---|---|---|---|---|---|---|---|---|---|---|
| 2 | 43215855 | rs7593685 | -0.201 | 0.458 | -1.082 | 0.094 | 0.029 | 0.093 | 0.093 | 0 | 0.25 | 0.43 | 0.52 | 0.39 | 0.37 | 0.32 |
| 2 | 43229909 | rs4953706 | 0.985 | 0.594 | -1.065 | -0.411 | 0.001 | -0.051 | -0.051 | 0.347 | 0.31 | 0.29 | 0.22 | 0.03 | 0.24 | 0.22 |
| 2 | 43232117 | rs17030257 | 0.852 | 0.443 | -1.34 | -0.061 | 0.048 | -0.007 | -0.007 | 0.017 | 0.34 | 0.2 | 0.27 | 0.3 | 0.18 | 0.28 |
| 2 | 43235451 | rs9309095 | 0.212 | 0.448 | -1.464 | -0.231 | -0.005 | 0.133 | 0.133 | 0.926 | 0.55 | 0.69 | 0.36 | 0.13 | 0.44 | 0.37 |
| 2 | 43239425 | rs11686640 | -0.101 | 0.484 | -1.455 | -0.103 | -0.005 | 0.009 | 0.009 | 0.076 | 0.67 | 0.38 | 0.69 | 0.58 | 0.76 | 0.76 |
| 2 | 43241234 | rs10174815 | 1.258 | 1.036 | -1.663 | 0.037 | -0.112 | -0.135 | -0.135 | 0 | 0.22 | 0.02 | 0.19 | 0.4 | 0.31 | 0.32 |
| 2 | 43244845 | rs2166503 | 0.59 | 0.987 | -1.238 | -0.211 | 0.011 | 0.015 | 0.015 | 0 | 0.72 | 0.66 | 0.57 | 0.62 | 0.64 | 0.67 |
| 2 | 43247073 | rs11884157 | 0.707 | 0.641 | -1.177 | -0.443 | -0.011 | -0.048 | -0.048 | 0 | 0.25 | 0.36 | 0.16 | 0.17 | 0.21 | 0.19 |
| 2 | 43247215 | rs4953713 | 1.842 | 0.492 | -1.784 | -0.635 | 0.037 | 0.041 | 0.041 | 0 | 0.26 | 0.15 | 0.17 | 0.03 | 0.16 | 0.13 |
| 2 | 43247227 | rs13421358 | 0.298 | 0.944 | -2.025 | -0.29 | -0.176 | -0.133 | -0.133 | 0 | 0.37 | 0.04 | 0.45 | 0.57 | 0.51 | 0.59 |
| 2 | 43247619 | rs7560093 | -1.165 | 0.543 | -1.878 | -0.175 | -0.008 | -0.005 | -0.005 | 0.518 | 0.94 | 0.99 | 0.85 | 0.78 | 0.96 | 0.94 |
| 2 | 43247753 | rs7560194 | -0.176 | 0.68 | -1.717 | -0.328 | -0.019 | -0.05 | -0.05 | 0.01 | 0.41 | 0.16 | 0.52 | 0.58 | 0.61 | 0.62 |
| 2 | 43249254 | rs10172861 | -0.005 | 0.448 | -0.848 | -0.953 | 0.011 | -0.005 | -0.005 | 0.069 | 0.5 | 0.43 | 0.41 | 0.14 | 0.47 | 0.45 |
| 2 | 43252691 | rs4485603 | -0.276 | 0.955 | -0.668 | -0.595 | 0.014 | 0.012 | 0.012 | 0 | 0.28 | 0.58 | 0.23 | 0.28 | 0.3 | 0.21 |
| 2 | 43253827 | rs2034274 | -0.483 | 0.379 | -0.559 | -0.64 | 0.018 | -0.004 | -0.004 | 0.043 | 0.38 | 0.51 | 0.31 | 0.31 | 0.47 | 0.39 |
| 2 | 43259164 | rs10175942 | -0.523 | 0.061 | -0.545 | -0.586 | 0.019 | 0.021 | 0.021 | 0 | 0.51 | 0.57 | 0.48 | 0.38 | 0.58 | 0.58 |
| 2 | 43263805 | rs7588142 | -0.722 | 0.223 | -0.24 | -0.615 | 0 | -0.016 | -0.016 | 0 | 0.75 | 0.84 | 0.6 | 0.37 | 0.76 | 0.69 |
| 2 | 43270895 | rs4953715 | -0.041 | 0.224 | 0.322 | -0.864 | 0.003 | 0.005 | 0.005 | 0 | 0.63 | 0.48 | 0.69 | 0.42 | 0.64 | 0.55 |
| 2 | 43274679 | rs4952961 | 0.677 | 0.171 | 0.292 | -1.075 | -0.004 | 0.002 | 0.002 | 0.236 | 0.23 | 0.23 | 0.47 | 0.29 | 0.37 | 0.36 |
| 2 | 43278048 | rs9973820 | 0.099 | 0.517 | 0.83 | -0.46 | 0.005 | 0.011 | 0.011 | 0 | 0.64 | 0.53 | 0.71 | 0.42 | 0.64 | 0.56 |
| 2 | 43278660 | rs6756476 | -0.767 | 0.37 | 0.894 | -0.571 | -0.011 | -0.005 | -0.005 | 0 | 0.22 | 0.3 | 0.12 | 0.11 | 0.17 | 0.22 |
| 2 | 43279815 | rs17030315 | 0.07 | 0.356 | 0.601 | -0.522 | -0.018 | 0.007 | 0.007 | 0 | 0.14 | 0.21 | 0.06 | 0.11 | 0.07 | 0.1 |
| 2 | 43281032 | rs7559315 | -0.394 | 0.467 | 0.404 | -0.449 | -0.001 | -0.015 | -0.015 | 0 | 0.72 | 0.79 | 0.76 | 0.42 | 0.72 | 0.67 |
| 2 | 43281211 | rs7601314 | -0.477 | 0.472 | 0.307 | -0.447 | -0.005 | -0.033 | -0.033 | 0 | 0.73 | 0.86 | 0.76 | 0.42 | 0.72 | 0.67 |
| 2 | 43283474 | rs895516 | -0.407 | 0.519 | 0.06 | -0.43 | 0 | 0.017 | 0.017 | 0 | 0.72 | 0.73 | 0.75 | 0.42 | 0.73 | 0.67 |
| 2 | 43284517 | rs11685806 | 0.836 | 0.597 | -0.723 | -0.508 | 0.001 | 0.12 | 0.12 | 0 | 0.5 | 0.33 | 0.61 | 0.31 | 0.55 | 0.45 |
| 2 | 43286610 | rs7582239 | -0.302 | 0.655 | -0.752 | -0.144 | -0.004 | 0.003 | 0.003 | 0 | 0.19 | 0.24 | 0.1 | 0.12 | 0.15 | 0.22 |
| 2 | 43292832 | rs2374459 | 0.414 | 0.827 | -1.53 | -0.686 | 0.019 | 0.077 | 0.077 | 0 | 0.53 | 0.28 | 0.63 | 0.41 | 0.63 | 0.48 |
| 2 | 43299383 | rs17030342 | -0.2 | 0.5 | -1.701 | -1.048 | 0.031 | 0.092 | 0.092 | 0 | 0.22 | 0.08 | 0.06 | 0 | 0.1 | 0.19 |
| 2 | 43301761 | rs7591954 | -0.702 | 0.955 | -2.122 | -1.097 | -0.076 | 0.039 | 0.039 | 0 | 0.76 | 0.96 | 0.55 | 0.18 | 0.67 | 0.66 |
| 2 | 43305926 | rs2374482 | 0.056 | 0.438 | -1.167 | -0.9 | -0.008 | -0.057 | -0.057 | 0 | 0.44 | 0.44 | 0.28 | 0.13 | 0.38 | 0.26 |
| 2 | 43312809 | rs7587206 | -0.236 | -0.01 | -1.189 | -0.435 | 0.04 | 0.061 | 0.061 | 0 | 0.49 | 0.66 | 0.28 | 0.26 | 0.32 | 0.37 |
| 2 | 43314116 | rs6723438 | -0.626 | -0.539 | -1.099 | -0.694 | 0.002 | -0.006 | -0.006 | 0.424 | 0.11 | 0.1 | 0.06 | 0.01 | 0.09 | 0.08 |
| 2 | 43314379 | rs11124915 | 0.528 | -0.77 | -0.774 | -0.55 | 0 | 0.019 | 0.019 | 0.237 | 0.14 | 0.11 | 0.25 | 0.26 | 0.14 | 0.12 |
| 2 | 43314843 | rs1002796 | 0.763 | -0.772 | -0.78 | -0.521 | -0.003 | -0.01 | -0.01 | 0.303 | 0.13 | 0.07 | 0.2 | 0.26 | 0.13 | 0.12 |
| 2 | 43315848 | rs7578467 | -0.609 | -1.775 | -0.78 | -0.641 | 0.035 | -0.004 | -0.004 | 0 | 0.55 | 0.7 | 0.5 | 0.6 | 0.34 | 0.48 |
| 2 | 43317253 | rs6706778 | 0.032 | -1.91 | -0.801 | -0.693 | 0.015 | -0.018 | -0.018 | 0.299 | 0.32 | 0.28 | 0.1 | 0.02 | 0.16 | 0.17 |
| 2 | 43321225 | rs1446571 | -1.103 | -1.045 | -1.167 | -0.215 | -0.094 | -0.069 | -0.069 | 0.486 | 0.69 | 0.89 | 0.67 | 0.62 | 0.5 | 0.6 |
| 2 | 43321742 | rs6752660 | -0.032 | -0.967 | -0.668 | -0.275 | -0.006 | -0.017 | -0.017 | 1.398 | 0.08 | 0.1 | 0.05 | 0.01 | 0.06 | 0.05 |
| 2 | 43330109 | rs4567998 | -0.55 | -1.073 | -0.603 | -0.154 | 0.054 | 0.046 | 0.046 | 0 | 0.48 | 0.64 | 0.38 | 0.39 | 0.28 | 0.42 |
| 2 | 43336638 | rs4952967 | 0.57 | -1.329 | -0.829 | -0.18 | 0.051 | 0.052 | 0.052 | 0.542 | 0.26 | 0.1 | 0.26 | 0.12 | 0.15 | 0.29 |
| 2 | 43338650 | rs11894071 | -0.154 | -1.232 | -0.098 | -0.328 | -0.013 | -0.039 | -0.039 | 0 | 0.09 | 0.26 | 0.05 | 0.01 | 0.04 | 0.04 |
| 2 | 43343806 | rs6741704 | -0.602 | -1.053 | -0.157 | -0.22 | -0.002 | -0.006 | -0.006 | 0 | 0.29 | 0.43 | 0.29 | 0.24 | 0.27 | 0.15 |
| 2 | 43349680 | rs766829 | 0.248 | -1.157 | -0.346 | -0.353 | -0.001 | -0.047 | -0.047 | 0 | 0.3 | 0.25 | 0.09 | 0.01 | 0.15 | 0.21 |
| 2 | 43351007 | rs17030368 | 0.207 | -1.179 | -0.353 | -0.364 | -0.002 | -0.048 | -0.048 | 0.015 | 0.31 | 0.26 | 0.09 | 0.01 | 0.16 | 0.21 |
| 2 | 43353389 | rs747709 | -0.665 | -1.225 | -0.287 | -0.367 | 0.001 | -0.002 | -0.002 | 0 | 0.51 | 0.5 | 0.35 | 0.24 | 0.37 | 0.31 |
| 2 | 43355763 | rs17405635 | -0.653 | -0.691 | -0.237 | 0.283 | -0.001 | -0.001 | -0.001 | 0.485 | 0.28 | 0.32 | 0.28 | 0.26 | 0.25 | 0.14 |
| 2 | 43355955 | rs17405663 | 0.464 | -0.686 | -0.383 | 0.28 | -0.016 | -0.026 | -0.026 | 0 | 0.19 | 0.17 | 0.24 | 0.24 | 0.22 | 0.11 |
| 2 | 43357022 | rs1946539 | -1.108 | 0.401 | -0.566 | 0.563 | -0.051 | -0.065 | -0.065 | 0 | 0.73 | 0.93 | 0.7 | 0.6 | 0.6 | 0.59 |
| 2 | 43359061 | rs12466022 | -0.698 | 0.322 | 0.092 | 0.679 | -0.006 | 0.001 | 0.001 | 0.018 | 0.29 | 0.41 | 0.29 | 0.26 | 0.25 | 0.13 |
| 2 | 43361256 | rs2163226 | -0.689 | 0.487 | -0.137 | 0.794 | 0.002 | 0.019 | 0.019 | 0 | 0.29 | 0.45 | 0.3 | 0.36 | 0.28 | 0.19 |
| 2 | 43362380 | rs17030415 | -0.898 | -0.017 | 0.105 | 1.2 | -0.032 | 0.093 | 0.093 | 0 | 0.17 | 0.3 | 0.24 | 0.37 | 0.1 | 0.16 |
| 2 | 43363752 | rs882502 | -0.132 | 0.457 | -0.544 | 0.401 | -0.017 | 0.03 | 0.03 | 0 | 0.68 | 0.75 | 0.62 | 0.77 | 0.61 | 0.63 |
| 2 | 43371542 | rs1978648 | 0.114 | 0.864 | -0.724 | 0.478 | -0.014 | -0.01 | -0.01 | 0 | 0.87 | 0.91 | 0.69 | 0.76 | 0.74 | 0.86 |
| 2 | 43386134 | rs2374525 | 0.57 | 0.975 | -1.216 | 0.201 | 0.015 | 0.035 | 0.035 | 0 | 0.46 | 0.31 | 0.46 | 0.64 | 0.4 | 0.51 |
| 2 | 43397176 | rs10167465 | 1.026 | 1.454 | 0.024 | 0.902 | 0.059 | 0.032 | 0.032 | 0 | 0.76 | 0.4 | 0.56 | 0.65 | 0.64 | 0.76 |
| 2 | 43397614 | rs12464602 | 1.123 | 1.435 | 0.075 | 1.095 | 0.035 | 0.067 | 0.067 | 0 | 0.73 | 0.39 | 0.55 | 0.52 | 0.64 | 0.72 |
| 2 | 43411001 | rs10193902 | 1.142 | 1.523 | -0.426 | 0.533 | 0.01 | 0 | 0 | 0 | 0.79 | 0.27 | 0.6 | 0.75 | 0.73 | 0.81 |
| 2 | 43417083 | rs7557417 | 0.3 | 1.918 | -0.801 | 0.342 | -0.079 | 0.02 | 0.02 | 0.876 | 0.87 | 0.98 | 0.84 | 0.91 | 0.71 | 0.8 |
| 2 | 43419493 | rs934232 | 1.161 | 1.611 | -0.606 | 0.499 | -0.015 | -0.003 | -0.003 | 2.788 | 0.6 | 0.31 | 0.68 | 0.57 | 0.51 | 0.62 |
| 2 | 43419886 | rs2011896 | 0.811 | 1.356 | -0.476 | 0.018 | 0.061 | 0.033 | 0.033 | 0 | 0.5 | 0.37 | 0.48 | 0.43 | 0.39 | 0.44 |
| 2 | 43423870 | rs4952673 | 1.41 | 2.097 | 0.477 | -0.054 | -0.004 | 0.031 | 0.031 | 0 | 0.6 | 0.33 | 0.66 | 0.73 | 0.47 | 0.66 |
| 2 | 43423915 | rs11897836 | -0.903 | 2.225 | 0.819 | -0.047 | 0.017 | -0.03 | -0.03 | 0.112 | 0.23 | 0.41 | 0.15 | 0.21 | 0.22 | 0.16 |
| 2 | 43428651 | rs4952981 | 1.374 | 2.04 | 0.371 | 0.017 | 0.001 | -0.009 | -0.009 | 0.096 | 0.59 | 0.42 | 0.68 | 0.66 | 0.5 | 0.63 |
| 2 | 43436411 | rs6544633 | -0.041 | 2.048 | -0.602 | -0.092 | -0.11 | 0.028 | 0.028 | 0 | 0.86 | 0.98 | 0.77 | 0.94 | 0.68 | 0.83 |
| 2 | 43436754 | rs2052927 | 0.4 | 1.262 | -0.207 | 0.039 | -0.05 | 0.042 | 0.042 | 0.052 | 0.68 | 0.82 | 0.69 | 0.74 | 0.51 | 0.66 |
| 2 | 43446616 | rs13002812 | 0.57 | 1.554 | -1.677 | 0.293 | -0.055 | 0.041 | 0.041 | 0.014 | 0.57 | 0.1 | 0.57 | 0.63 | 0.5 | 0.44 |
| 2 | 43452183 | rs7933 | -0.493 | 1.663 | -1.628 | -0.219 | 0.013 | 0.077 | 0.077 | 0.033 | 0.46 | 0.82 | 0.38 | 0.24 | 0.37 | 0.55 |
| 2 | 43456734 | rs13407351 | -0.486 | 1.468 | -0.247 | -0.405 | 0.153 | -0.019 | -0.019 | 0 | 0.07 | 0.3 | 0.17 | 0 | 0.11 | 0.04 |

| | | | | | | | | | | | | | | | | |
|---|---|---|---|---|---|---|---|---|---|---|---|---|---|---|---|---|
| 2 | 43460514 | rs1549723 | -1.244 | 1.461 | 0.425 | -0.667 | 0.028 | -0.111 | -0.111 | 0.068 | 0.19 | 0.59 | 0.24 | 0 | 0.17 | 0.23 |
| 2 | 43483044 | rs6755601 | -1.26 | 1.379 | 0.611 | -0.707 | 0.01 | -0.086 | -0.086 | 0.05 | 0.2 | 0.67 | 0.24 | 0.01 | 0.17 | 0.22 |
| 2 | 43496986 | rs6749108 | -0.865 | 1.538 | 0.498 | -1.153 | 0.018 | 0.028 | 0.028 | 0 | 0.44 | 0.38 | 0.22 | 0.21 | 0.36 | 0.54 |
| 2 | 43501686 | rs13399243 | 0.014 | 1.533 | 0.877 | -1.076 | 0.153 | -0.014 | -0.014 | 0 | 0.06 | 0.2 | 0.15 | 0 | 0.11 | 0.04 |
| 2 | 43502477 | rs13026309 | -0.024 | 1.522 | 0.154 | -1.869 | 0.021 | -0.016 | -0.016 | 0 | 0.63 | 0.49 | 0.34 | 0.23 | 0.57 | 0.57 |
| 2 | 43506259 | rs7561670 | 1.14 | 1.437 | 0.404 | -1.917 | 0.013 | -0.002 | -0.002 | 0.796 | 0.27 | 0.15 | 0.15 | 0.23 | 0.22 | 0.33 |
| 2 | 43509617 | rs7599564 | -0.69 | 1.802 | 0.342 | -1.77 | 0.059 | 0.036 | 0.036 | 0 | 0.73 | 0.57 | 0.39 | 0.26 | 0.65 | 0.75 |
| 2 | 43510097 | rs6544639 | -0.446 | 1.73 | 0.507 | -1.717 | 0.173 | -0.024 | -0.024 | 0.471 | 0.08 | 0.37 | 0.16 | 0 | 0.11 | 0.05 |
| 2 | 43514036 | rs11900952 | -0.471 | 1.641 | 0.089 | -1.699 | 0.014 | 0.096 | 0.096 | 0 | 0.46 | 0.81 | 0.39 | 0.23 | 0.36 | 0.51 |
| 2 | 43537711 | rs6714624 | -1.513 | 1.594 | 0.263 | -1.504 | -0.002 | -0.01 | -0.01 | 0 | 0.79 | 0.84 | 0.55 | 0.27 | 0.77 | 0.78 |
| 2 | 43541412 | rs17030668 | 0.783 | 1.669 | 0.362 | -1.57 | -0.007 | 0.033 | 0.033 | 0 | 0.08 | 0.04 | 0.03 | 0 | 0.09 | 0.18 |
| 2 | 43541598 | rs6730757 | -0.176 | 1.737 | 0.665 | -1.582 | 0.031 | 0.013 | 0.013 | 0 | 0.71 | 0.51 | 0.38 | 0.26 | 0.68 | 0.72 |
| 2 | 43551416 | rs10211126 | -1.22 | 1.693 | 0.818 | -1.636 | 0.036 | -0.126 | -0.126 | 0 | 0.19 | 0.42 | 0.22 | 0.01 | 0.18 | 0.23 |
| 2 | 43555506 | rs4340576 | -2.125 | 1.756 | 0.613 | -1.506 | -0.017 | -0.048 | -0.048 | 0.22 | 0.84 | 0.93 | 0.59 | 0.27 | 0.81 | 0.81 |
| 2 | 43566146 | rs4372955 | -1.029 | 1.787 | 0.948 | -1.594 | 0.073 | -0.063 | -0.063 | 2.144 | 0.11 | 0.33 | 0.19 | 0 | 0.11 | 0.06 |
| 2 | 43576997 | rs6544646 | -0.412 | 1.692 | 0.702 | -1.938 | 0 | -0.043 | -0.043 | 0 | 0.48 | 0.58 | 0.34 | 0.03 | 0.4 | 0.41 |
| 2 | 43578419 | rs6544647 | -0.864 | 1.778 | 0.773 | -1.987 | 0.006 | 0.067 | 0.067 | 0 | 0.55 | 0.71 | 0.41 | 0.03 | 0.53 | 0.45 |
| 2 | 43587451 | rs3901678 | -0.817 | 1.869 | 0.843 | -2.036 | -0.005 | 0.095 | 0.095 | 0 | 0.55 | 0.77 | 0.35 | 0.03 | 0.54 | 0.43 |
| 2 | 43590329 | rs11693801 | -1.837 | 1.923 | 0.455 | -1.872 | -0.031 | -0.055 | -0.055 | 6.36 | 0.8 | 0.93 | 0.5 | 0.28 | 0.77 | 0.74 |
| 2 | 43590899 | rs11694173 | 0.362 | 1.889 | 0.498 | -1.92 | -0.034 | 0.036 | 0.036 | 0 | 0.11 | 0.06 | 0.09 | 0.01 | 0.17 | 0.19 |
| 2 | 43592756 | rs7562101 | -1.139 | 1.859 | 0.593 | -1.934 | 0.026 | -0.121 | -0.121 | 0 | 0.18 | 0.44 | 0.14 | 0.01 | 0.17 | 0.2 |
| 2 | 43595460 | rs17406278 | -0.005 | 1.849 | 0.466 | -1.868 | 0 | -0.015 | -0.015 | 0.409 | 0.08 | 0.1 | 0.09 | 0 | 0.13 | 0.03 |
| 2 | 43610027 | rs13408002 | -0.26 | 1.743 | 0.526 | -1.797 | 0.074 | -0.018 | -0.018 | 1.363 | 0.05 | 0.18 | 0.07 | 0 | 0.08 | 0.01 |
| 2 | 43616008 | rs6739828 | -0.508 | 1.838 | 0.469 | -1.753 | 0.001 | -0.009 | -0.009 | 0.532 | 0.48 | 0.57 | 0.44 | 0.06 | 0.44 | 0.39 |
| 2 | 43617726 | rs10495901 | 1.164 | 1.623 | 0.388 | -1.827 | -0.002 | 0.126 | 0.126 | 0 | 0.28 | 0.08 | 0.11 | 0.03 | 0.26 | 0.19 |
| 2 | 43630657 | rs7557989 | -0.077 | 1.7 | 0.437 | -1.939 | -0.01 | -0.074 | -0.074 | 1.475 | 0.41 | 0.51 | 0.23 | 0.03 | 0.33 | 0.31 |
| 2 | 43634135 | rs9677398 | 0.41 | 1.768 | 0.482 | -1.895 | 0.019 | 0.045 | 0.045 | 0 | 0.35 | 0.27 | 0.15 | 0.03 | 0.28 | 0.31 |
| 2 | 43635143 | rs6544656 | 0.445 | 1.759 | 0.477 | -1.901 | 0.027 | 0.043 | 0.043 | 0 | 0.35 | 0.27 | 0.14 | 0.03 | 0.26 | 0.31 |
| 2 | 43638712 | rs13029250 | -0.534 | 1.529 | 0.204 | -2.238 | -0.004 | 0.04 | 0.04 | 0 | 0.57 | 0.68 | 0.34 | 0.1 | 0.55 | 0.51 |
| 2 | 43638838 | rs13429458 | 1.511 | 1.529 | 0.149 | -2.193 | -0.008 | 0.01 | 0.01 | 0 | 0.16 | 0.2 | 0.09 | 0.2 | 0.13 | 0.19 |
| 2 | 43638855 | rs7568365 | -1.352 | 1.644 | 0.31 | -2.081 | -0.021 | -0.036 | -0.036 | 0 | 0.74 | 0.87 | 0.44 | 0.29 | 0.7 | 0.71 |
| 2 | 43641627 | rs7605725 | -0.037 | 1.732 | 0.228 | -2.125 | -0.002 | -0.008 | -0.008 | 0 | 0.35 | 0.38 | 0.23 | 0.26 | 0.37 | 0.4 |
| 2 | 43645675 | rs6739313 | 0.56 | 1.734 | 0.163 | -2.09 | 0.003 | 0.044 | 0.044 | 0 | 0.37 | 0.29 | 0.19 | 0.03 | 0.34 | 0.31 |
| 2 | 43648176 | rs10176241 | -1.491 | 1.697 | 0.04 | -2.015 | -0.07 | -0.057 | -0.057 | 0 | 0.75 | 0.91 | 0.42 | 0.3 | 0.65 | 0.72 |
| 2 | 43656924 | rs4608575 | -0.402 | 1.593 | 0.153 | -2.078 | 0.001 | -0.02 | -0.02 | 0 | 0.49 | 0.57 | 0.3 | 0.03 | 0.48 | 0.37 |
| 2 | 43670183 | rs17030798 | 1.047 | 1.41 | 0.196 | -1.98 | 0.003 | -0.015 | -0.015 | 0 | 0.07 | 0.07 | 0.06 | 0.01 | 0.09 | 0.14 |
| 2 | 43671904 | rs6744642 | -0.871 | 1.535 | 0.215 | -2.16 | -0.001 | -0.002 | -0.002 | 0 | 0.73 | 0.81 | 0.8 | 0.75 | 0.7 | 0.7 |
| 2 | 43675898 | rs6724325 | -0.431 | 0.977 | 0.475 | -2.06 | -0.003 | -0.005 | -0.005 | 0 | 0.05 | 0.22 | 0.04 | 0 | 0.01 | 0.01 |
| 2 | 43686207 | rs2046916 | 1.363 | 0.974 | 0.318 | -2.145 | 0.023 | 0.092 | 0.092 | 0 | 0.28 | 0.15 | 0.13 | 0.03 | 0.24 | 0.18 |
| 2 | 43687879 | rs17030845 | 0.851 | 0.884 | 0.458 | -2.138 | 0.028 | -0.04 | -0.04 | 0.023 | 0.08 | 0.18 | 0.07 | 0.01 | 0.09 | 0.14 |
| 2 | 43695148 | rs6741434 | 1.305 | 0.875 | 0.335 | -2.131 | 0.01 | 0.096 | 0.096 | 0 | 0.29 | 0.14 | 0.13 | 0.03 | 0.26 | 0.19 |
| 2 | 43697975 | rs11124938 | 1.308 | 0.888 | 0.342 | -2.125 | 0.01 | 0.096 | 0.096 | 0 | 0.29 | 0.15 | 0.13 | 0.03 | 0.26 | 0.19 |
| 2 | 43721508 | rs12478601 | -0.897 | 0.929 | 0.37 | -2.077 | -0.046 | -0.017 | -0.017 | 0 | 0.68 | 0.82 | 0.4 | 0.29 | 0.59 | 0.66 |
| 2 | 43732823 | rs7578597 | -0.021 | 0.934 | 0.669 | -1.978 | 0.058 | -0.046 | -0.046 | 0 | 0.1 | 0.31 | 0.07 | 0.01 | 0.09 | 0.13 |
| 2 | 43751480 | rs17030980 | 0.036 | 0.956 | 0.525 | -1.969 | 0.04 | -0.051 | -0.051 | 0 | 0.09 | 0.24 | 0.07 | 0.01 | 0.09 | 0.16 |
| 2 | 43752382 | rs6708660 | -0.227 | 0.877 | 0.343 | -2.126 | 0.016 | 0.055 | 0.055 | 0 | 0.46 | 0.67 | 0.25 | 0.04 | 0.36 | 0.36 |
| 2 | 43779665 | rs1873555 | -0.287 | 0.889 | 0.371 | -2.012 | -0.014 | -0.015 | -0.015 | 0 | 0.37 | 0.41 | 0.23 | 0.26 | 0.34 | 0.47 |
| 2 | 43783141 | rs6725776 | -0.386 | 0.883 | 0.382 | -2.019 | -0.005 | -0.007 | -0.007 | 0 | 0.38 | 0.51 | 0.24 | 0.26 | 0.34 | 0.49 |
| 2 | 43797619 | rs17031056 | 0.658 | 0.816 | 0.278 | -2.039 | 0.019 | -0.005 | -0.005 | 0.069 | 0.27 | 0.15 | 0.15 | 0.25 | 0.23 | 0.32 |
| 2 | 43800174 | rs11898207 | -0.309 | 0.851 | 0.512 | -2.05 | -0.006 | -0.009 | -0.009 | 0.28 | 0.37 | 0.4 | 0.23 | 0.26 | 0.34 | 0.47 |
| 2 | 43801712 | rs11899823 | -0.264 | 0.828 | 0.504 | -2.061 | -0.005 | -0.008 | -0.008 | 0 | 0.36 | 0.4 | 0.23 | 0.26 | 0.34 | 0.44 |
| 2 | 43808065 | rs10179648 | -1.648 | 1.073 | 0.192 | -2.09 | -0.044 | -0.066 | -0.066 | 0.061 | 0.75 | 0.92 | 0.46 | 0.29 | 0.7 | 0.67 |
| 2 | 43809248 | rs17031092 | -0.014 | 0.739 | 0.622 | -1.983 | 0.068 | -0.052 | -0.052 | 0 | 0.11 | 0.36 | 0.1 | 0.01 | 0.11 | 0.16 |
| 2 | 43817505 | rs11897432 | 1.305 | 0.605 | 0.387 | -2.132 | 0.007 | 0.09 | 0.09 | 0 | 0.27 | 0.14 | 0.12 | 0.03 | 0.23 | 0.18 |
| 2 | 43822006 | rs6717791 | 0.003 | 0.903 | 0.789 | -1.766 | 0.056 | -0.047 | -0.047 | 0 | 0.1 | 0.26 | 0.08 | 0.01 | 0.11 | 0.1 |
| 2 | 43834118 | rs6712944 | 0.066 | 0.91 | 0.819 | -1.763 | 0.097 | -0.034 | -0.034 | 0 | 0.09 | 0.34 | 0.09 | 0.01 | 0.11 | 0.1 |
| 2 | 43836687 | rs7558302 | -0.871 | 0.889 | 0.302 | -1.685 | -0.061 | -0.026 | -0.026 | 0 | 0.68 | 0.78 | 0.38 | 0.3 | 0.56 | 0.62 |
| 2 | 43841269 | rs7576643 | -0.218 | 0.883 | 0.472 | -1.802 | 0.059 | -0.048 | -0.048 | 0 | 0.1 | 0.31 | 0.09 | 0.01 | 0.11 | 0.09 |
| 2 | 43850590 | rs11886047 | -0.841 | 0.538 | -0.071 | -2.036 | -0.001 | -0.023 | -0.023 | 0 | 0.19 | 0.21 | 0.13 | 0.01 | 0.19 | 0.16 |
| 2 | 43854142 | rs6736105 | -0.758 | 0.484 | 0.208 | -1.766 | 0.006 | -0.022 | -0.022 | 0 | 0.65 | 0.55 | 0.39 | 0.12 | 0.59 | 0.58 |
| 2 | 43855643 | rs6740739 | 0.851 | 0.066 | 0.264 | -2.154 | 0.002 | 0.057 | 0.057 | 0 | 0.33 | 0.16 | 0.15 | 0.01 | 0.3 | 0.18 |
| 2 | 43856927 | rs6716337 | 0.507 | 0.093 | 0.501 | -1.965 | -0.001 | 0.064 | 0.064 | 0 | 0.41 | 0.23 | 0.23 | 0.03 | 0.38 | 0.28 |
| 2 | 43858579 | rs6710610 | -0.705 | -0.004 | 0.424 | -2.019 | -0.001 | -0.011 | -0.011 | 0.829 | 0.61 | 0.59 | 0.36 | 0.09 | 0.59 | 0.48 |
| 2 | 43859090 | rs13412979 | 0.496 | -0.048 | 0.483 | -2.049 | 0 | -0.018 | -0.018 | 0 | 0.42 | 0.33 | 0.24 | 0.03 | 0.38 | 0.26 |
| 2 | 43862888 | rs12473033 | -1.28 | 0.159 | 0.045 | -1.556 | -0.037 | 0.022 | 0.022 | 0 | 0.68 | 0.82 | 0.51 | 0.21 | 0.61 | 0.57 |
| 2 | 43863866 | rs9309103 | -1.28 | 0.159 | 0.039 | -1.556 | -0.036 | 0.022 | 0.022 | 0 | 0.68 | 0.82 | 0.51 | 0.21 | 0.61 | 0.57 |
| 2 | 43870490 | rs6750909 | -1.27 | 0.123 | 0.032 | -1.258 | -0.029 | -0.009 | -0.009 | 0 | 0.68 | 0.81 | 0.55 | 0.28 | 0.63 | 0.57 |
| 2 | 43875289 | rs7571775 | -0.485 | 0.17 | 0.242 | -1.831 | -0.001 | -0.006 | -0.006 | 3.987 | 0.61 | 0.62 | 0.46 | 0.15 | 0.57 | 0.48 |
| 2 | 43877074 | rs2888878 | -1.385 | 0.33 | 0.2 | -2.261 | -0.032 | 0.018 | 0.018 | 0.006 | 0.7 | 0.8 | 0.5 | 0.2 | 0.63 | 0.56 |

| | | | | | | | | | | | | | | | | |
|---|---|---|---|---|---|---|---|---|---|---|---|---|---|---|---|---|
| 2 | 43879382 | rs6544683 | 0.669 | 0.082 | 0.412 | -2.243 | -0.026 | -0.031 | -0.031 | 0.283 | 0.2 | 0.33 | 0.23 | 0.12 | 0.19 | 0.24 |
| 2 | 43882627 | rs9309105 | -0.715 | 0.281 | 0.191 | -2.357 | 0.011 | -0.029 | -0.029 | 0.135 | 0.64 | 0.6 | 0.46 | 0.15 | 0.57 | 0.49 |
| 2 | 43883711 | rs17335701 | 0.885 | 0.216 | 0.186 | -2.615 | 0.007 | 0.01 | 0.01 | 0 | 0.19 | 0.15 | 0.21 | 0.12 | 0.19 | 0.23 |
| 2 | 43884760 | rs919688 | 0 | 0.2 | 0.428 | -2.616 | 0.003 | 0.032 | 0.032 | 0 | 0.48 | 0.42 | 0.27 | 0.1 | 0.42 | 0.33 |
| 2 | 43886460 | rs13423605 | 0.787 | 0.266 | 0.561 | -2.399 | 0.01 | 0.016 | 0.016 | 0 | 0.05 | 0.14 | 0.04 | 0.05 | 0.03 | 0.07 |
| 2 | 43903066 | rs3828297 | -1.002 | 0.521 | 0.241 | -2.437 | 0.012 | -0.017 | -0.017 | 0 | 0.72 | 0.61 | 0.54 | 0.19 | 0.72 | 0.63 |
| 2 | 43916955 | rs4952996 | 0.692 | 0.017 | 0.28 | -2.362 | -0.005 | -0.017 | -0.017 | 0.002 | 0.15 | 0.17 | 0.2 | 0.08 | 0.15 | 0.16 |
| 2 | 43922928 | rs17031289 | -0.184 | -0.161 | 0.542 | -2.579 | -0.017 | -0.065 | -0.065 | 0.277 | 0.25 | 0.38 | 0.26 | 0.08 | 0.21 | 0.25 |
| 2 | 43929983 | rs4952676 | -1.14 | -0.151 | 0.016 | -2.929 | -0.061 | 0.098 | 0.098 | 0.781 | 0.61 | 0.81 | 0.48 | 0.17 | 0.55 | 0.53 |
| 2 | 43930709 | rs10514787 | 0.845 | -0.128 | -0.241 | -2.971 | -0.007 | 0.072 | 0.072 | 0.028 | 0.29 | 0.16 | 0.17 | 0.05 | 0.32 | 0.22 |
| 2 | 43931176 | rs4953002 | -0.457 | 0.149 | -0.221 | -2.681 | -0.003 | 0.004 | 0.004 | 0.184 | 0.56 | 0.6 | 0.43 | 0.13 | 0.54 | 0.48 |
| 2 | 43935446 | rs12465669 | 0.019 | 0.208 | -0.078 | -2.344 | 0.001 | -0.01 | -0.01 | 0.348 | 0.12 | 0.13 | 0.07 | 0 | 0.1 | 0.1 |
| 2 | 43935725 | rs6716551 | -0.654 | 0.094 | -0.094 | -2.716 | -0.016 | 0.058 | 0.058 | 0.018 | 0.61 | 0.69 | 0.45 | 0.19 | 0.49 | 0.55 |
| 2 | 43936230 | rs6544691 | -0.371 | 0.054 | 0.504 | -2.694 | 0.012 | -0.005 | -0.005 | 0 | 0.07 | 0.3 | 0.03 | 0 | 0.03 | 0 |
| 2 | 43939079 | rs2888881 | -1.947 | 0.804 | -0.121 | -2.85 | -0.064 | -0.052 | -0.052 | 0 | 0.81 | 0.95 | 0.62 | 0.22 | 0.74 | 0.68 |
| 2 | 43939525 | rs13028613 | 1.323 | 0.196 | 0.226 | -2.856 | -0.022 | 0.095 | 0.095 | 0.021 | 0.23 | 0.06 | 0.14 | 0.04 | 0.25 | 0.18 |
| 2 | 43941944 | rs976555 | -1.594 | 0.5 | 0.21 | -2.847 | -0.105 | 0.061 | 0.061 | 0 | 0.68 | 0.87 | 0.51 | 0.17 | 0.56 | 0.63 |
| 2 | 43943116 | rs7582880 | 0.608 | 0.345 | 0.674 | -2.842 | -0.013 | -0.089 | -0.089 | 0 | 0.28 | 0.34 | 0.15 | 0.03 | 0.24 | 0.18 |
| 2 | 43943525 | rs1991739 | -0.867 | 0.508 | 0.874 | -2.781 | 0.008 | 0.015 | 0.015 | 0 | 0.41 | 0.6 | 0.44 | 0.14 | 0.33 | 0.44 |
| 2 | 43946046 | rs1560599 | 0.573 | 0.405 | 0.913 | -2.778 | -0.022 | 0.047 | 0.047 | 0 | 0.2 | 0.08 | 0.31 | 0.1 | 0.21 | 0.23 |
| 2 | 43947318 | rs11885893 | 1.107 | 0.387 | 0.9 | -2.791 | -0.027 | 0.093 | 0.093 | 0.228 | 0.24 | 0.08 | 0.15 | 0.02 | 0.28 | 0.2 |
| 2 | 43948153 | rs1036150 | -0.358 | 0.272 | 1.116 | -2.711 | -0.028 | -0.067 | -0.067 | 0 | 0.2 | 0.4 | 0.11 | 0.05 | 0.15 | 0.2 |
| 2 | 43950929 | rs6740998 | -0.348 | 0.275 | 1.175 | -2.709 | -0.023 | -0.056 | -0.056 | 0 | 0.19 | 0.34 | 0.11 | 0.05 | 0.15 | 0.2 |
| 2 | 43952970 | rs930194 | -1.623 | 0.376 | 0.532 | -2.709 | -0.05 | 0.075 | 0.075 | 0 | 0.66 | 0.82 | 0.57 | 0.17 | 0.57 | 0.63 |
| 2 | 43953752 | rs2374573 | -1.567 | 0.406 | 0.555 | -2.703 | -0.046 | 0.07 | 0.07 | 0 | 0.65 | 0.8 | 0.57 | 0.16 | 0.56 | 0.61 |
| 2 | 43959547 | rs7603323 | -2.016 | 0.148 | 1.707 | -1.692 | 0.053 | 0.091 | 0.091 | 0 | 0.43 | 0.69 | 0.44 | 0.17 | 0.28 | 0.46 |
| 2 | 43964837 | rs1368087 | -1.005 | 0.379 | 1.925 | -1.59 | 0.009 | -0.066 | -0.066 | 0 | 0.25 | 0.4 | 0.4 | 0.1 | 0.28 | 0.28 |
| 2 | 43965007 | rs1368086 | -0.247 | -0.111 | 1.347 | -1.511 | -0.003 | 0.022 | 0.022 | 0 | 0.19 | 0.1 | 0.32 | 0.11 | 0.18 | 0.22 |
| 2 | 43972708 | rs7605900 | -0.383 | -0.132 | 1.365 | -1.685 | -0.001 | 0.013 | 0.013 | 0 | 0.2 | 0.17 | 0.32 | 0.11 | 0.19 | 0.23 |
| 2 | 43995491 | rs7601418 | 0.144 | -0.132 | 1.364 | -1.777 | 0.001 | 0.012 | 0.012 | 0 | 0.14 | 0.1 | 0.12 | 0.02 | 0.14 | 0.17 |
| 2 | 43999182 | rs725238 | -1.979 | -0.305 | 1.652 | -1.884 | 0.013 | 0.019 | 0.019 | 0 | 0.41 | 0.64 | 0.44 | 0.16 | 0.29 | 0.46 |
| 2 | 44000495 | rs6544711 | -2.337 | -0.292 | 1.084 | -1.902 | 0.049 | 0.163 | 0.163 | 0 | 0.44 | 0.89 | 0.48 | 0.16 | 0.33 | 0.51 |
| 2 | 44004010 | rs2288709 | -2.597 | -0.304 | 1.054 | -1.931 | 0.059 | 0.203 | 0.203 | 0 | 0.48 | 0.89 | 0.49 | 0.17 | 0.35 | 0.54 |
| 2 | 44004860 | rs3792026 | -0.061 | -0.086 | 1.353 | -1.744 | 0.005 | 0.011 | 0.011 | 0 | 0.07 | 0.09 | 0.23 | 0.1 | 0.07 | 0.09 |
| 2 | 44010341 | rs4953015 | -0.518 | -0.086 | 1.32 | -1.743 | 0.005 | 0.019 | 0.019 | 0 | 0.08 | 0.18 | 0.24 | 0.1 | 0.07 | 0.09 |
| 2 | 44015409 | rs11695056 | -0.467 | -0.137 | 1.214 | -1.796 | 0.012 | 0.035 | 0.035 | 0 | 0.17 | 0.09 | 0.11 | 0.02 | 0.14 | 0.2 |
| 2 | 44017411 | rs3792017 | -0.894 | -0.107 | 1.35 | -1.755 | 0.029 | 0.109 | 0.109 | 0.156 | 0.09 | 0.3 | 0.27 | 0.14 | 0.08 | 0.11 |
| 2 | 44019799 | rs13420319 | -1.19 | -0.14 | 1.394 | -1.771 | 0.03 | 0.119 | 0.119 | 0.011 | 0.1 | 0.37 | 0.28 | 0.14 | 0.08 | 0.13 |
| 2 | 44022970 | rs1025447 | -0.77 | -0.065 | 1.502 | -1.752 | -0.003 | -0.036 | -0.036 | 0.134 | 0.19 | 0.23 | 0.15 | 0.02 | 0.17 | 0.2 |
| 2 | 44027302 | rs7594731 | -1.656 | -0.088 | 1.427 | -1.783 | 0.002 | -0.04 | -0.04 | 0 | 0.33 | 0.34 | 0.18 | 0.02 | 0.24 | 0.34 |
| 2 | 44030649 | rs1835815 | -2.283 | -0.075 | 1.042 | -1.772 | 0.04 | 0.084 | 0.084 | 0 | 0.42 | 0.72 | 0.46 | 0.18 | 0.32 | 0.47 |
| 2 | 44036438 | rs10173687 | -1.708 | 0.169 | 1.106 | -1.687 | -0.02 | -0.032 | -0.032 | 0.009 | 0.25 | 0.43 | 0.32 | 0.15 | 0.18 | 0.29 |
| 2 | 44036922 | rs10186552 | -1.742 | 0.167 | 1.148 | -1.68 | -0.011 | -0.041 | -0.041 | 0 | 0.24 | 0.36 | 0.31 | 0.12 | 0.18 | 0.28 |
| 2 | 44037333 | rs3792009 | -1.616 | -0.172 | 1.188 | -1.666 | -0.021 | -0.043 | -0.043 | 0.35 | 0.24 | 0.36 | 0.31 | 0.12 | 0.16 | 0.28 |
| 2 | 44039633 | rs4148195 | -1.766 | -0.172 | 1.194 | -1.665 | -0.035 | -0.031 | -0.031 | 0.021 | 0.24 | 0.43 | 0.32 | 0.15 | 0.16 | 0.29 |
| 2 | 44039839 | rs2278357 | -1.59 | -0.167 | 0.99 | -1.662 | -0.02 | -0.042 | -0.042 | 0 | 0.23 | 0.36 | 0.31 | 0.12 | 0.16 | 0.28 |
| 2 | 44039875 | rs2278356 | -2.422 | -0.19 | 0.847 | -1.697 | 0.049 | 0.105 | 0.105 | 0 | 0.45 | 0.72 | 0.47 | 0.18 | 0.33 | 0.51 |
| 2 | 44040401 | rs6720173 | -1.657 | -0.018 | 0.47 | -1.783 | -0.014 | -0.032 | -0.032 | 0 | 0.24 | 0.33 | 0.3 | 0.12 | 0.16 | 0.28 |
| 2 | 44042896 | rs17031664 | -0.633 | -0.002 | -0.401 | -1.904 | 0.016 | 0.018 | 0.018 | 0 | 0.15 | 0.1 | 0.04 | 0 | 0.08 | 0.18 |
| 2 | 44044255 | rs10205816 | -1.802 | 0.221 | -0.407 | -1.565 | 0.007 | 0.009 | 0.009 | 0.041 | 0.34 | 0.69 | 0.43 | 0.09 | 0.27 | 0.39 |
| 2 | 44044680 | rs11891859 | -1.722 | 0.41 | 0.383 | -1.111 | -0.027 | -0.039 | -0.039 | 0 | 0.18 | 0.67 | 0.32 | 0.06 | 0.13 | 0.21 |
| 2 | 44047530 | rs4148189 | -1.444 | 0.508 | 0.325 | -1.056 | -0.024 | -0.042 | -0.042 | 0.055 | 0.16 | 0.6 | 0.31 | 0.05 | 0.12 | 0.2 |
| 2 | 44049120 | rs4245786 | 1.011 | 1.175 | -0.367 | -1.044 | -0.001 | -0.006 | -0.006 | 0 | 0.78 | 0.78 | 0.78 | 0.93 | 0.76 | 0.73 |
| 2 | 44050687 | rs4148188 | -0.494 | 1.008 | -0.578 | -1.342 | 0.005 | 0.031 | 0.031 | 0 | 0.41 | 0.34 | 0.35 | 0.8 | 0.34 | 0.59 |
| 2 | 44050888 | rs4148187 | -0.614 | 0.996 | -0.525 | -1.259 | 0.006 | 0.037 | 0.037 | 0 | 0.4 | 0.29 | 0.35 | 0.78 | 0.33 | 0.58 |
| 2 | 44054123 | rs4289236 | -0.228 | 0.985 | -0.705 | -1.603 | 0.03 | 0.026 | 0.026 | 0 | 0.27 | 0.11 | 0.27 | 0.8 | 0.18 | 0.56 |
| 2 | 44056856 | rs4148179 | -0.925 | 0.9 | 0.471 | -1.607 | 0.005 | 0.028 | 0.028 | 0 | 0.42 | 0.56 | 0.39 | 0.82 | 0.35 | 0.61 |
| 2 | 44065090 | rs6756629 | -0.294 | 0.951 | 0.664 | -1.77 | 0.003 | -0.011 | -0.011 | 0 | 0.06 | 0.1 | 0.1 | 0.01 | 0.08 | 0.04 |
| 2 | 44070780 | rs6720518 | -0.22 | 0.867 | 0.622 | -2.187 | 0.009 | -0.062 | -0.062 | 2.242 | 0.33 | 0.42 | 0.39 | 0.83 | 0.37 | 0.55 |
| 2 | 44070935 | rs6748810 | -0.13 | 0.85 | 0.686 | -2.196 | 0.004 | -0.028 | -0.028 | 0 | 0.32 | 0.31 | 0.38 | 0.84 | 0.37 | 0.55 |
| 2 | 44071743 | rs4148211 | 0.02 | 0.881 | 0.52 | -2.211 | -0.03 | 0.074 | 0.074 | 0 | 0.3 | 0.13 | 0.35 | 0.85 | 0.37 | 0.54 |
| 2 | 44074000 | rs4953023 | -0.045 | 0.798 | 0.288 | -2.108 | 0.005 | -0.012 | -0.012 | 0 | 0.06 | 0.1 | 0.09 | 0.01 | 0.07 | 0.04 |
| 2 | 44080324 | rs6709904 | -0.632 | 0.703 | 0.084 | -2.254 | -0.032 | -0.09 | -0.09 | 2.742 | 0.22 | 0.33 | 0.14 | 0.03 | 0.11 | 0.12 |
| 2 | 44098855 | rs3795860 | -0.188 | 1.019 | 0.106 | -2.245 | 0.003 | -0.026 | -0.026 | 0 | 0.35 | 0.32 | 0.39 | 0.86 | 0.43 | 0.61 |
| 2 | 44099582 | rs4148218 | 0.201 | 0.863 | -0.425 | -2.002 | 0.021 | -0.021 | -0.021 | 0 | 0.16 | 0.29 | 0.32 | 0.12 | 0.2 | 0.16 |
| 2 | 44100849 | rs4245794 | -0.699 | 0.786 | -0.841 | -1.873 | -0.001 | -0.026 | -0.026 | 0 | 0.19 | 0.2 | 0.11 | 0.03 | 0.11 | 0.14 |
| 2 | 44101827 | rs4953028 | -0.396 | 1.05 | -1.381 | -2.117 | -0.003 | 0.078 | 0.078 | 0 | 0.59 | 0.72 | 0.6 | 0.96 | 0.46 | 0.79 |
| 2 | 44102491 | rs4148221 | -0.037 | 1.521 | -1.271 | -2.025 | 0.008 | -0.012 | -0.012 | 0 | 0.11 | 0.17 | 0.16 | 0.02 | 0.18 | 0.11 |
| 2 | 44103017 | rs11685669 | -0.416 | 1.429 | -1.878 | -2.524 | -0.024 | 0.093 | 0.093 | 0 | 0.83 | 0.96 | 0.85 | 1 | 0.81 | 0.92 |

| | | | | | | | | | | | | | | | | |
|---|---|---|---|---|---|---|---|---|---|---|---|---|---|---|---|---|
| 2 | 44107701 | rs4521094 | 0.917 | 0.752 | -1.64 | -1.071 | -0.01 | 0.009 | 0.009 | 0 | 0.31 | 0.31 | 0.26 | 0.6 | 0.22 | 0.56 |
| 2 | 44107742 | rs2954804 | 0.986 | 0.958 | -1.776 | -1.308 | 0.014 | -0.081 | -0.081 | 0.046 | 0.28 | 0.56 | 0.23 | 0.07 | 0.3 | 0.14 |
| 2 | 44109281 | rs2954806 | -0.214 | -0.223 | -1.728 | -1.132 | 0.004 | 0.017 | 0.017 | 0 | 0.36 | 0.33 | 0.48 | 0.9 | 0.31 | 0.58 |
| 2 | 44112052 | rs17578296 | -0.151 | -0.294 | -1.546 | -0.615 | 0.014 | -0.007 | -0.007 | 0 | 0.12 | 0.09 | 0.19 | 0.41 | 0.06 | 0.13 |
| 2 | 44112075 | rs4953030 | 0.835 | -0.302 | -1.442 | -0.349 | 0.007 | 0.058 | 0.058 | 0 | 0.24 | 0.37 | 0.3 | 0.49 | 0.24 | 0.46 |
| 2 | 44112422 | rs2955279 | -0.251 | -0.27 | -1.365 | -0.617 | -0.001 | -0.007 | -0.007 | 0 | 0.35 | 0.42 | 0.48 | 0.8 | 0.3 | 0.57 |
| 2 | 44116100 | rs10445885 | -0.73 | -1.137 | -1.876 | -1.098 | -0.085 | 0.209 | 0.209 | 0.017 | 0.64 | 0.89 | 0.59 | 0.96 | 0.49 | 0.63 |
| 2 | 44116836 | rs2955280 | -0.787 | -1.163 | -1.873 | -1.069 | -0.089 | 0.2 | 0.2 | 0 | 0.64 | 0.9 | 0.64 | 0.96 | 0.5 | 0.64 |
| 2 | 44118189 | rs12465555 | -0.587 | -1.345 | -1.171 | -0.716 | -0.034 | 0.191 | 0.191 | 0 | 0.14 | 0.32 | 0.21 | 0.53 | 0.06 | 0.14 |
| 2 | 44120846 | rs7560946 | -0.63 | -1.321 | -1.361 | -0.734 | 0.002 | 0.001 | 0.001 | 0 | 0.15 | 0.17 | 0.22 | 0.55 | 0.1 | 0.17 |
| 2 | 44120947 | rs4563284 | -0.879 | -1.25 | -1.418 | -0.897 | -0.1 | 0.21 | 0.21 | 0.152 | 0.64 | 0.91 | 0.64 | 0.96 | 0.49 | 0.64 |
| 2 | 44123759 | rs3795859 | 2.066 | -1.469 | -1.319 | -0.682 | -0.007 | -0.092 | -0.092 | 0 | 0.13 | 0.01 | 0.19 | 0.46 | 0.11 | 0.32 |
| 2 | 44135314 | rs13387221 | 0.815 | -1.532 | -0.808 | -0.755 | -0.003 | 0.026 | 0.026 | 0 | 0.2 | 0.26 | 0.25 | 0.47 | 0.18 | 0.34 |
| 2 | 44139197 | rs4953033 | -0.807 | -1.452 | -1.036 | -1.076 | -0.099 | 0.205 | 0.205 | 0.013 | 0.65 | 0.9 | 0.65 | 0.97 | 0.5 | 0.63 |
| 2 | 44145374 | rs7568481 | -0.453 | -1.718 | -1.161 | -1.245 | 0.062 | 0.233 | 0.233 | 0.548 | 0.52 | 0.76 | 0.59 | 0.96 | 0.4 | 0.5 |
| 2 | 44149442 | rs4953037 | -0.63 | -1.746 | -1.203 | -1.262 | 0.037 | 0.281 | 0.281 | 0.111 | 0.53 | 0.84 | 0.6 | 0.97 | 0.42 | 0.54 |
| 2 | 44156440 | rs12712900 | -1.022 | -1.718 | -1.175 | -1.254 | -0.06 | 0.169 | 0.169 | 0.001 | 0.64 | 0.81 | 0.64 | 0.96 | 0.5 | 0.65 |
| 2 | 44162320 | rs17424646 | 1.77 | -1.663 | -1.122 | -1.16 | 0.011 | 0.006 | 0.006 | 0.017 | 0.13 | 0.07 | 0.04 | 0 | 0.1 | 0.14 |
| 2 | 44169889 | rs9309112 | 0.637 | -1.748 | -1.011 | -1.256 | 0.01 | 0.036 | 0.036 | 0 | 0.26 | 0.21 | 0.42 | 0.96 | 0.22 | 0.47 |
| 2 | 44181434 | rs6741740 | 0.782 | -1.817 | -0.841 | -1.293 | -0.042 | -0.16 | -0.16 | 0.088 | 0.31 | 0.38 | 0.18 | 0 | 0.25 | 0.15 |
| 2 | 44185129 | rs4347883 | 0.988 | -1.822 | -0.937 | -1.454 | -0.1 | 0.135 | 0.135 | 0 | 0.32 | 0.17 | 0.29 | 0.02 | 0.43 | 0.22 |
| 2 | 44185306 | rs11886849 | 1.788 | -1.814 | -0.887 | -1.409 | -0.022 | -0.14 | -0.14 | 0 | 0.19 | 0.37 | 0.16 | 0 | 0.17 | 0.03 |
| 2 | 44185345 | rs10206724 | -0.719 | -1.836 | -0.943 | -1.378 | 0.011 | 0.043 | 0.043 | 0 | 0.71 | 0.81 | 0.58 | 0.04 | 0.76 | 0.46 |
| 2 | 44187384 | rs7573769 | 2.044 | -1.861 | -1.007 | -1.076 | -0.001 | -0.11 | -0.11 | 0.144 | 0.17 | 0.03 | 0.23 | 0.51 | 0.17 | 0.39 |
| 2 | 44187829 | rs10190161 | -0.548 | -1.874 | -0.973 | -1.265 | 0.007 | 0.002 | 0.002 | 0 | 0.71 | 0.75 | 0.57 | 0.04 | 0.75 | 0.49 |
| 2 | 44189630 | rs10865196 | 0.122 | -1.838 | -0.958 | -1.261 | 0.03 | 0.051 | 0.051 | 0.023 | 0.57 | 0.68 | 0.51 | 0.04 | 0.64 | 0.33 |
| 2 | 44201259 | rs7593842 | 1.801 | -1.829 | -0.905 | -1.565 | -0.028 | -0.138 | -0.138 | 0 | 0.19 | 0.37 | 0.15 | 0 | 0.17 | 0.02 |
| 2 | 44201376 | rs4953042 | 2.176 | -1.814 | -0.982 | -1.509 | 0 | -0.11 | -0.11 | 0 | 0.16 | 0.03 | 0.27 | 0.52 | 0.15 | 0.39 |
| 2 | 44204717 | rs6736282 | 0.582 | -1.617 | -0.818 | -2.266 | -0.031 | 0.071 | 0.071 | 0 | 0.37 | 0.25 | 0.37 | 0.04 | 0.48 | 0.32 |
| 2 | 44209477 | rs6741066 | -0.454 | -1.538 | -0.906 | -2.146 | 0.013 | 0.023 | 0.023 | 0 | 0.67 | 0.68 | 0.55 | 0.04 | 0.74 | 0.47 |
| 2 | 44211546 | rs6544724 | 0.311 | -1.384 | -0.925 | -2.204 | 0.032 | 0.092 | 0.092 | 0 | 0.51 | 0.62 | 0.47 | 0.04 | 0.58 | 0.32 |
| 2 | 44218637 | rs6756455 | 0.007 | -1.306 | -1.06 | -2.099 | 0.023 | 0.07 | 0.07 | 0 | 0.6 | 0.65 | 0.51 | 0.04 | 0.68 | 0.4 |
| 2 | 44232411 | rs10084473 | 0.674 | -1.052 | -0.914 | -1.851 | -0.078 | 0.089 | 0.089 | 1.665 | 0.3 | 0.17 | 0.32 | 0.04 | 0.44 | 0.31 |
| 2 | 44232704 | rs10084400 | 0.703 | -1.045 | -0.933 | -1.848 | -0.083 | 0.095 | 0.095 | 1.025 | 0.3 | 0.16 | 0.31 | 0.04 | 0.43 | 0.31 |
| 2 | 44232849 | rs4521095 | -0.669 | -1.305 | -0.965 | -2.015 | 0.009 | 0.007 | 0.007 | 0 | 0.63 | 0.67 | 0.49 | 0.04 | 0.66 | 0.47 |
| 2 | 44235543 | rs6544734 | -0.216 | -1.573 | -0.986 | -1.985 | 0.04 | 0.046 | 0.046 | 0 | 0.52 | 0.57 | 0.48 | 0.04 | 0.63 | 0.35 |
| 2 | 44236915 | rs11693923 | 0.767 | -1.073 | -1.441 | -1.766 | -0.138 | 0.175 | 0.175 | 0 | 0.29 | 0.06 | 0.29 | 0.02 | 0.45 | 0.23 |
| 2 | 44237787 | rs4131366 | 0.948 | -0.913 | -1.107 | -1.535 | -0.009 | -0.036 | -0.036 | 0.089 | 0.14 | 0.14 | 0.05 | 0.01 | 0.1 | 0.14 |
| 2 | 44258530 | rs7582693 | -0.099 | -0.781 | -1.341 | -1.254 | 0.068 | 0.217 | 0.217 | 0.277 | 0.47 | 0.16 | 0.41 | 0.13 | 0.61 | 0.49 |
| 2 | 44260347 | rs17496908 | 1.341 | -0.494 | -1.419 | -1.212 | -0.039 | 0.013 | 0.013 | 0 | 0.17 | 0 | 0.09 | 0.08 | 0.17 | 0.15 |
| 2 | 44260752 | rs7590420 | -0.699 | -0.624 | -1.081 | -0.838 | 0 | -0.026 | -0.026 | 0 | 0.89 | 0.94 | 0.79 | 0.58 | 0.93 | 0.84 |
| 2 | 44260914 | rs7421942 | 1.544 | -0.667 | -0.667 | -0.978 | -0.012 | -0.06 | -0.06 | 0 | 0.16 | 0.22 | 0.09 | 0 | 0.11 | 0.01 |
| 2 | 44261892 | rs3853705 | 0.267 | -0.481 | -1.456 | -0.858 | 0.016 | 0.001 | 0.001 | 0.014 | 0.41 | 0.66 | 0.35 | 0.45 | 0.32 | 0.35 |
| 2 | 44265330 | rs6724437 | 1.048 | 0.21 | -1.926 | -1.024 | -0.068 | 0.069 | 0.069 | 0 | 0.21 | 0.1 | 0.23 | 0.02 | 0.33 | 0.13 |
| 2 | 44267800 | rs6761677 | 0.138 | -0.083 | -1.302 | -0.6 | -0.02 | -0.082 | -0.082 | 0 | 0.26 | 0.62 | 0.23 | 0.03 | 0.21 | 0.14 |
| 2 | 44268800 | rs4953045 | 0.321 | 0.308 | -1.475 | 0.36 | -0.002 | -0.101 | -0.101 | 0.055 | 0.72 | 0.92 | 0.71 | 0.56 | 0.77 | 0.66 |
| 2 | 44279622 | rs12613714 | -0.472 | 0.889 | -0.593 | -0.155 | 0.027 | -0.045 | -0.045 | 0 | 0.22 | 0.31 | 0.31 | 0.08 | 0.27 | 0.29 |
| 2 | 44282325 | rs7565864 | -1.16 | 0.337 | -0.496 | -0.818 | 0 | 0.024 | 0.024 | 0.029 | 0.49 | 0.53 | 0.45 | 0.1 | 0.51 | 0.49 |
| 2 | 44283172 | rs17031893 | 0.255 | 0.059 | 0.166 | -0.865 | 0.005 | -0.008 | -0.008 | 0.019 | 0.25 | 0.23 | 0.15 | 0.02 | 0.24 | 0.2 |
| 2 | 44290437 | rs7583854 | 0.029 | 0.151 | 0.013 | -0.842 | -0.008 | -0.045 | -0.045 | 0 | 0.27 | 0.26 | 0.15 | 0.02 | 0.24 | 0.19 |
| 2 | 44291333 | rs9284745 | -0.612 | 0.049 | -0.237 | -0.296 | 0.004 | -0.007 | -0.007 | 0 | 0.9 | 0.89 | 0.81 | 0.64 | 0.89 | 0.87 |
| 2 | 44304047 | rs4953058 | 0.008 | -0.022 | -0.22 | -0.593 | -0.023 | -0.054 | -0.054 | 0 | 0.29 | 0.31 | 0.29 | 0.04 | 0.23 | 0.22 |
| 2 | 44305240 | rs6544738 | 0.228 | 0.407 | -0.566 | -0.268 | 0.003 | -0.022 | -0.022 | 0 | 0.2 | 0.23 | 0.16 | 0.06 | 0.25 | 0.26 |
| 2 | 44306681 | rs10169761 | 0.016 | 0.426 | -0.574 | -0.256 | 0.008 | -0.026 | -0.026 | 0 | 0.22 | 0.26 | 0.29 | 0.08 | 0.25 | 0.29 |
| 2 | 44307450 | rs7607908 | -0.395 | 0.087 | -0.693 | 0.332 | -0.026 | -0.041 | -0.041 | 0.711 | 0.93 | 0.99 | 0.81 | 0.8 | 0.88 | 0.89 |
| 2 | 44312697 | rs10166377 | -0.366 | -0.518 | -0.418 | -0.348 | -0.049 | -0.07 | -0.07 | 0.015 | 0.28 | 0.39 | 0.26 | 0.04 | 0.17 | 0.22 |
| 2 | 44315927 | rs7606187 | -0.287 | -0.435 | -0.356 | -0.646 | -0.001 | -0.025 | -0.025 | 0 | 0.26 | 0.24 | 0.14 | 0.05 | 0.15 | 0.25 |
| 2 | 44316913 | rs11683623 | -0.73 | 1.422 | 0.352 | 0.343 | 0.126 | 0.359 | 0.359 | 1.156 | 0.12 | 0.45 | 0.17 | 0.47 | 0.19 | 0.16 |
| 2 | 44317749 | rs7599435 | -0.69 | 1.419 | 0.521 | 0.366 | 0.117 | 0.34 | 0.34 | 0.802 | 0.12 | 0.4 | 0.16 | 0.47 | 0.19 | 0.16 |
| 2 | 44318510 | rs12612469 | 1.402 | 1.326 | -0.103 | 0.082 | 0.006 | 0.036 | 0.036 | 0.817 | 0.39 | 0.05 | 0.47 | 0.3 | 0.35 | 0.37 |
| 2 | 44319306 | rs7579921 | -0.016 | 1.662 | -0.07 | 0.147 | 0.14 | 0.533 | 0.533 | 1.822 | 0.09 | 0.45 | 0.14 | 0.5 | 0.12 | 0.11 |
| 2 | 44319733 | rs4953063 | 0.811 | 1.652 | -0.108 | 0.128 | -0.001 | -0.022 | -0.022 | 0.031 | 0.48 | 0.48 | 0.61 | 0.81 | 0.47 | 0.48 |
| 2 | 44320255 | rs6705336 | -1.278 | 1.658 | -0.314 | 0.117 | -0.021 | -0.031 | -0.031 | 0 | 0.92 | 0.99 | 0.92 | 0.9 | 0.88 | 0.92 |
| 2 | 44321352 | rs9808443 | 0.123 | 1.632 | 0.206 | 0.443 | -0.002 | 0.017 | 0.017 | 0 | 0.32 | 0.39 | 0.26 | 0.48 | 0.26 | 0.36 |
| 2 | 44322128 | rs7564640 | -0.857 | 1.642 | 0.422 | 0.221 | 0.003 | 0.035 | 0.035 | 0 | 0.47 | 0.62 | 0.33 | 0.08 | 0.41 | 0.43 |
| 2 | 44323135 | rs921680 | 0.878 | 1.597 | 0.537 | 0.301 | -0.006 | 0.044 | 0.044 | 0 | 0.48 | 0.28 | 0.58 | 0.35 | 0.47 | 0.47 |
| 2 | 44324287 | rs17425341 | 1.492 | 1.567 | 0.393 | 0.33 | 0.011 | 0.033 | 0.033 | 0.846 | 0.39 | 0.08 | 0.48 | 0.32 | 0.35 | 0.37 |
| 2 | 44326923 | rs2333825 | 0.511 | 1.556 | 0.744 | 0.763 | 0.015 | 0.035 | 0.035 | 0 | 0.53 | 0.6 | 0.69 | 0.7 | 0.58 | 0.57 |
| 2 | 44327038 | rs7587071 | 0.422 | 1.675 | 0.652 | 0.981 | 0.056 | 0.219 | 0.219 | 0 | 0.06 | 0.23 | 0.1 | 0.36 | 0.1 | 0.1 |

| | | | | | | | | | | | | | | | | |
|---|---|---|---|---|---|---|---|---|---|---|---|---|---|---|---|---|
| 2 | 44328998 | rs17425397 | 1.08 | 1.329 | 0.336 | 0.742 | 0.009 | 0.036 | 0.036 | 0 | 0.08 | 0.03 | 0.04 | 0.01 | 0.06 | 0.06 |
| 2 | 44339612 | rs12622038 | 0.282 | 1.44 | 0.518 | 0.575 | 0.001 | 0.016 | 0.016 | 0 | 0.28 | 0.21 | 0.18 | 0.15 | 0.24 | 0.27 |
| 2 | 44339820 | rs3886714 | -0.629 | 1.452 | 0.386 | 0.813 | 0.004 | 0.03 | 0.03 | 0 | 0.89 | 0.79 | 0.89 | 0.84 | 0.86 | 0.93 |
| 2 | 44341976 | rs1377687 | 1.929 | 1.276 | -0.53 | 0.591 | 0.005 | -0.021 | -0.021 | 0 | 0.33 | 0.04 | 0.46 | 0.31 | 0.33 | 0.31 |
| 2 | 44342583 | rs6752099 | 0.611 | 1.253 | -0.316 | 0.607 | -0.024 | -0.034 | -0.034 | 0 | 0.77 | 0.79 | 0.74 | 0.51 | 0.66 | 0.72 |
| 2 | 44347384 | rs7599136 | -2.152 | 1.632 | -0.612 | 0.713 | -0.007 | -0.026 | -0.026 | 0.801 | 0.68 | 0.96 | 0.52 | 0.73 | 0.65 | 0.71 |
| 2 | 44351536 | rs11124972 | 0.436 | 2.164 | -0.39 | 0.663 | -0.044 | 0.002 | 0.002 | 0 | 0.2 | 0.03 | 0.12 | 0.17 | 0.17 | 0.15 |
| 2 | 44354495 | rs896986 | -1.816 | 2.201 | -0.048 | 0.701 | 0.014 | 0.005 | 0.005 | 0 | 0.49 | 0.6 | 0.38 | 0.51 | 0.51 | 0.47 |
| 2 | 44357936 | rs6712908 | -1.455 | 1.778 | 0.22 | 0.738 | 0.039 | 0.014 | 0.014 | 0.792 | 0.3 | 0.55 | 0.27 | 0.35 | 0.36 | 0.33 |
| 2 | 44358504 | rs7593878 | -0.297 | 1.654 | -0.063 | 0.867 | 0.023 | 0.047 | 0.047 | 0 | 0.13 | 0.23 | 0.13 | 0.31 | 0.13 | 0.17 |
| 2 | 44360400 | rs11124975 | -0.132 | 0.905 | -0.095 | 0.289 | -0.01 | 0.045 | 0.045 | 0 | 0.19 | 0.06 | 0.1 | 0.11 | 0.13 | 0.16 |
| 2 | 44366284 | rs1551319 | -1.198 | 1.016 | 0.184 | 0.389 | -0.02 | 0.04 | 0.04 | 0 | 0.31 | 0.2 | 0.21 | 0.22 | 0.32 | 0.29 |
| 2 | 44369192 | rs17031997 | -1.374 | 0.997 | 0.439 | 0.484 | 0.004 | -0.015 | -0.015 | 0 | 0.33 | 0.4 | 0.22 | 0.26 | 0.31 | 0.3 |
| 2 | 44370198 | rs12465678 | -1.454 | 0.989 | 0.435 | 0.48 | 0.009 | -0.035 | -0.035 | 2.86 | 0.34 | 0.48 | 0.23 | 0.27 | 0.31 | 0.3 |
| 2 | 44371721 | rs2085021 | -0.349 | 0.987 | -0.022 | 0.741 | 0.001 | -0.006 | -0.006 | 0 | 0.12 | 0.09 | 0.09 | 0.22 | 0.13 | 0.12 |
| 2 | 44372909 | rs10166397 | 1.468 | 0.831 | 0.093 | 0.613 | -0.001 | 0.003 | 0.003 | 0 | 0.65 | 0.74 | 0.77 | 0.68 | 0.69 | 0.66 |
| 2 | 44376526 | rs2333821 | 0.488 | 0.863 | 0.488 | 0.613 | 0.035 | 0.001 | 0.001 | 0.044 | 0.85 | 0.68 | 0.79 | 0.85 | 0.79 | 0.87 |
| 2 | 44384185 | rs4146528 | 1.039 | 0.81 | 0.31 | 0.698 | 0.002 | -0.003 | -0.003 | 0 | 0.72 | 0.62 | 0.75 | 0.71 | 0.71 | 0.7 |
| 2 | 44392395 | rs10180864 | 1.621 | 0.501 | -0.569 | -0.116 | -0.001 | 0.166 | 0.166 | 0.699 | 0.24 | 0.01 | 0.13 | 0 | 0.2 | 0.04 |
| 2 | 44395302 | rs1453863 | 1.258 | 1.012 | 1.018 | 1.025 | -0.001 | -0.002 | -0.002 | 2.885 | 0.63 | 0.33 | 0.68 | 0.55 | 0.6 | 0.65 |
| 2 | 44398026 | rs17032022 | -0.257 | 1.139 | 1.519 | 1.045 | 0.001 | -0.009 | -0.009 | 0 | 0.18 | 0.25 | 0.1 | 0.16 | 0.18 | 0.21 |
| 2 | 44404902 | rs13008237 | 0.951 | 1.126 | 1.665 | 1.073 | 0.021 | 0.036 | 0.036 | 0.083 | 0.86 | 0.76 | 0.81 | 0.72 | 0.79 | 0.9 |
| 2 | 44410540 | rs10495915 | -0.292 | 1.15 | 1.802 | 1.205 | -0.005 | -0.002 | -0.002 | 0 | 0.18 | 0.1 | 0.09 | 0.2 | 0.18 | 0.24 |
| 2 | 44437217 | rs6730278 | -0.835 | 1.11 | 2.043 | 1.142 | -0.004 | -0.038 | -0.038 | 0 | 0.22 | 0.49 | 0.13 | 0.17 | 0.18 | 0.24 |
| 2 | 44448632 | rs2053456 | 0.917 | 1.082 | 1.901 | 1.295 | 0.028 | 0.064 | 0.064 | 0 | 0.87 | 0.82 | 0.82 | 0.66 | 0.78 | 0.9 |
| 2 | 44464629 | rs4953074 | 1.597 | 1.545 | 2.039 | 1.415 | 0.121 | 0.168 | 0.168 | 0.34 | 0.87 | 0.54 | 0.79 | 0.71 | 0.79 | 0.9 |
| 2 | 44477588 | rs7598019 | 1.864 | 1.451 | 1.76 | 1.247 | -0.008 | 0.004 | 0.004 | 0.111 | 0.63 | 0.28 | 0.51 | 0.36 | 0.57 | 0.61 |
| 2 | 44482099 | rs12712916 | 1.212 | 1.461 | 1.737 | 1.387 | 0.035 | 0.072 | 0.072 | 0.095 | 0.84 | 0.75 | 0.66 | 0.38 | 0.75 | 0.85 |
| 2 | 44490660 | rs12613771 | 1.565 | 1.472 | 1.764 | 1.369 | 0.047 | 0.096 | 0.096 | 0.728 | 0.82 | 0.61 | 0.64 | 0.37 | 0.75 | 0.84 |
| 2 | 44502788 | rs3738985 | 1.052 | 1.378 | 1.587 | 1.354 | 0.028 | 0.063 | 0.063 | 0.01 | 0.86 | 0.8 | 0.67 | 0.38 | 0.77 | 0.86 |
| 2 | 44504857 | rs713446 | 0.22 | 1.497 | 1.182 | 1.255 | -0.012 | -0.014 | -0.014 | 0.511 | 0.06 | 0.03 | 0.06 | 0.12 | 0.08 | 0.06 |
| 2 | 44507213 | rs10194161 | 1.795 | 1.119 | 1.455 | 1.133 | -0.003 | 0.005 | 0.005 | 0 | 0.61 | 0.29 | 0.52 | 0.35 | 0.59 | 0.61 |
| 2 | 44519142 | rs4952709 | 1.751 | 0.875 | 1.618 | 1.523 | -0.016 | 0.015 | 0.015 | 0 | 0.73 | 0.62 | 0.65 | 0.34 | 0.78 | 0.67 |
| 2 | 44521246 | rs4953081 | 0.785 | 0.932 | 1.7 | 1.594 | -0.001 | -0.004 | -0.004 | 0 | 0.87 | 0.88 | 0.7 | 0.35 | 0.85 | 0.85 |
| 2 | 44524575 | rs4952710 | 2.107 | 0.952 | 1.802 | 1.549 | 0.007 | 0.001 | 0.001 | 0 | 0.68 | 0.52 | 0.57 | 0.33 | 0.65 | 0.65 |
| 2 | 44536249 | rs2340809 | 2.263 | 0.989 | 1.522 | 1.482 | 0 | 0.001 | 0.001 | 0 | 0.66 | 0.26 | 0.54 | 0.33 | 0.66 | 0.65 |
| 2 | 44548282 | rs11124988 | 0.941 | 1.375 | 1.408 | 1.728 | -0.005 | -0.028 | -0.028 | 0.039 | 0.86 | 0.93 | 0.72 | 0.35 | 0.84 | 0.85 |
| 2 | 44554724 | rs4953085 | 1.482 | 1.352 | 1.433 | 1.711 | 0 | -0.036 | -0.036 | 0 | 0.78 | 0.89 | 0.68 | 0.33 | 0.79 | 0.7 |
| 2 | 44557919 | rs9309116 | 1.769 | 1.509 | 1.477 | 1.666 | -0.023 | -0.014 | -0.014 | 0 | 0.74 | 0.84 | 0.61 | 0.31 | 0.64 | 0.66 |
| 2 | 44561532 | rs4953086 | 2.243 | 1.238 | 2.016 | 1.669 | 0.024 | 0.018 | 0.018 | 0 | 0.72 | 0.52 | 0.59 | 0.32 | 0.66 | 0.68 |
| 2 | 44565024 | rs4953089 | 1.893 | 1.247 | 1.788 | 1.688 | -0.004 | -0.003 | -0.003 | 0.278 | 0.73 | 0.76 | 0.62 | 0.31 | 0.66 | 0.68 |
| 2 | 44568694 | rs7596887 | -0.757 | 1.146 | 1.575 | 1.617 | 0.003 | 0.003 | 0.003 | 0 | 0.11 | 0.12 | 0.05 | 0.02 | 0.05 | 0.15 |
| 2 | 44571109 | rs10514790 | 0.038 | 1.162 | 1.475 | 1.625 | 0.005 | 0.015 | 0.015 | 0 | 0.08 | 0.03 | 0.03 | 0.02 | 0.05 | 0.11 |
| 2 | 44584314 | rs698774 | 1.789 | 1.205 | 1.699 | 1.681 | 0 | 0 | 0 | 0 | 0.74 | 0.75 | 0.62 | 0.32 | 0.66 | 0.67 |
| 2 | 44588941 | rs698775 | 1.783 | 1.283 | 1.707 | 1.683 | -0.002 | -0.001 | -0.001 | 0 | 0.74 | 0.75 | 0.61 | 0.32 | 0.65 | 0.69 |
| 2 | 44596768 | rs13406580 | -1.826 | 1.141 | 2.103 | 1.581 | -0.041 | -0.075 | -0.075 | 0 | 0.15 | 0.58 | 0.11 | 0.02 | 0.07 | 0.12 |
| 2 | 44597096 | rs2616450 | 0.428 | 1.16 | 2.073 | 1.985 | -0.015 | -0.013 | -0.013 | 0 | 0.91 | 0.95 | 0.79 | 0.67 | 0.86 | 0.91 |
| 2 | 44598825 | rs6713213 | -2.582 | 1.006 | 2.309 | 1.786 | 0.036 | 0.033 | 0.033 | 3.814 | 0.06 | 0.39 | 0.06 | 0 | 0 | 0 |
| 2 | 44599253 | rs7593926 | -1.932 | 1.013 | 2.306 | 1.786 | -0.037 | -0.066 | -0.066 | 0.33 | 0.16 | 0.65 | 0.12 | 0.02 | 0.07 | 0.12 |
| 2 | 44605518 | rs2616444 | 1.601 | 1.102 | 2.091 | 1.869 | 0.041 | 0.068 | 0.068 | 0 | 0.81 | 0.7 | 0.64 | 0.34 | 0.71 | 0.8 |
| 2 | 44621042 | rs1067343 | 1.392 | 1.028 | 2.002 | 1.879 | 0.006 | 0.017 | 0.017 | 0 | 0.82 | 0.79 | 0.64 | 0.34 | 0.72 | 0.82 |
| 2 | 44624448 | rs1067339 | 2.055 | 0.99 | 1.996 | 1.825 | 0.009 | 0.003 | 0.003 | 0 | 0.7 | 0.58 | 0.59 | 0.32 | 0.65 | 0.69 |
| 2 | 44643077 | rs17032193 | -0.124 | 0.998 | 1.851 | 1.786 | 0.003 | 0.016 | 0.016 | 0 | 0.09 | 0.04 | 0.04 | 0.02 | 0.06 | 0.14 |
| 2 | 44644257 | rs1067386 | 2.365 | 0.943 | 2.258 | 1.804 | 0.002 | -0.001 | -0.001 | 0 | 0.68 | 0.36 | 0.57 | 0.31 | 0.65 | 0.69 |
| 2 | 44652343 | rs1067370 | 2.469 | 0.965 | 2.191 | 1.78 | -0.003 | -0.001 | -0.001 | 0.358 | 0.67 | 0.26 | 0.55 | 0.32 | 0.65 | 0.7 |
| 2 | 44660699 | rs1065785 | 1.764 | 1.019 | 2.075 | 1.897 | 0.029 | 0.029 | 0.029 | 0 | 0.76 | 0.62 | 0.67 | 0.67 | 0.66 | 0.78 |
| 2 | 44669724 | rs1067352 | 1.491 | 1.041 | 1.864 | 1.944 | -0.001 | -0.002 | -0.002 | 0 | 0.78 | 0.76 | 0.81 | 0.67 | 0.68 | 0.78 |
| 2 | 44669946 | rs1067350 | 0.664 | 1.056 | 1.869 | 1.924 | 0.011 | 0.017 | 0.017 | 0 | 0.87 | 0.85 | 0.85 | 0.7 | 0.74 | 0.91 |
| 2 | 44674450 | rs1067406 | 2.218 | 1.028 | 2.048 | 1.965 | 0.01 | 0.001 | 0.001 | 0.023 | 0.69 | 0.51 | 0.71 | 0.29 | 0.66 | 0.68 |
| 2 | 44674861 | rs1067405 | 1.492 | 1.091 | 1.64 | 2.076 | -0.021 | -0.032 | -0.032 | 0 | 0.82 | 0.85 | 0.77 | 0.33 | 0.72 | 0.83 |
| 2 | 44683018 | rs698790 | 0.525 | 1.121 | 1.577 | 1.992 | -0.035 | -0.037 | -0.037 | 0 | 0.88 | 0.93 | 0.86 | 0.69 | 0.74 | 0.91 |
| 2 | 44685685 | rs698793 | 1.52 | 1.109 | 1.634 | 1.925 | -0.021 | -0.031 | -0.031 | 0.085 | 0.81 | 0.84 | 0.77 | 0.31 | 0.72 | 0.83 |
| 2 | 44690514 | rs698798 | -1.359 | 1.355 | 1.612 | 1.875 | 0.027 | 0.037 | 0.037 | 0 | 0.07 | 0.21 | 0.17 | 0.1 | 0.08 | 0.08 |
| 2 | 44697156 | rs1551882 | -0.161 | 1.311 | 1.241 | 1.827 | 0 | 0 | 0 | 0 | 0.09 | 0.06 | 0.05 | 0.02 | 0.08 | 0.12 |
| 2 | 44705615 | rs698813 | 0.469 | 1.347 | 1.037 | 1.94 | -0.034 | -0.036 | -0.036 | 0 | 0.88 | 0.92 | 0.86 | 0.69 | 0.75 | 0.91 |
| 2 | 44706974 | rs698815 | 1.343 | 1.324 | 1.086 | 1.877 | -0.015 | -0.026 | -0.026 | 0 | 0.82 | 0.85 | 0.77 | 0.35 | 0.73 | 0.82 |
| 2 | 44712929 | rs698819 | -1.198 | 1.219 | 0.947 | 1.957 | -0.018 | -0.016 | -0.016 | 0 | 0.07 | 0.03 | 0.16 | 0.26 | 0.09 | 0.08 |
| 2 | 44726639 | rs786412 | -1.833 | 0.901 | 1.394 | 1.671 | -0.022 | -0.018 | -0.018 | 0.416 | 0.21 | 0.49 | 0.25 | 0.15 | 0.15 | 0.33 |
| 2 | 44727772 | rs17581138 | -1.21 | 0.902 | 1.294 | 1.899 | -0.033 | 0.172 | 0.172 | 0.308 | 0.08 | 0.18 | 0.1 | 0.34 | 0.01 | 0.08 |

| | | | | | | | | | | | | | | | | |
|---|---|---|---|---|---|---|---|---|---|---|---|---|---|---|---|---|
| 2 | 44735438 | rs698833 | 1.167 | 1.231 | 1.128 | 1.731 | 0.021 | 0.018 | 0.018 | 0 | 0.78 | 0.75 | 0.81 | 0.67 | 0.68 | 0.78 |
| 2 | 44738763 | rs698838 | 1.643 | 1.485 | 0.686 | 1.069 | -0.023 | 0.004 | 0.004 | 0 | 0.66 | 0.27 | 0.49 | 0.34 | 0.62 | 0.65 |
| 2 | 44740552 | rs13016493 | 1.937 | 1.332 | 0.949 | 0.786 | -0.016 | 0.081 | 0.081 | 0 | 0.57 | 0.31 | 0.38 | 0.2 | 0.53 | 0.57 |
| 2 | 44742517 | rs698779 | -1.534 | 1.335 | 1.652 | 0.787 | 0.073 | -0.032 | -0.032 | 1.266 | 0.08 | 0.33 | 0.09 | 0 | 0.09 | 0.05 |
| 2 | 44744897 | rs698786 | 1.475 | 1.408 | 1.544 | 0.773 | 0.009 | -0.066 | -0.066 | 0.027 | 0.68 | 0.62 | 0.56 | 0.09 | 0.65 | 0.67 |
| 2 | 44746859 | rs12185730 | 1.981 | 1.4 | 1.795 | 0.742 | 0.017 | -0.045 | -0.045 | 0 | 0.61 | 0.5 | 0.52 | 0.2 | 0.57 | 0.63 |
| 2 | 44756361 | rs1223222 | 1.501 | 1.374 | 1.826 | 0.821 | 0.004 | -0.009 | -0.009 | 0 | 0.68 | 0.63 | 0.72 | 0.23 | 0.66 | 0.68 |
| 2 | 44757217 | rs1011797 | 2.007 | 1.394 | 2.002 | 0.884 | 0.023 | 0.001 | 0.001 | 0.022 | 0.71 | 0.44 | 0.57 | 0.29 | 0.64 | 0.73 |
| 2 | 44759714 | rs936039 | 1.181 | 1.387 | 1.983 | 0.88 | 0.039 | 0.056 | 0.056 | 0 | 0.79 | 0.63 | 0.77 | 0.33 | 0.72 | 0.78 |
| 2 | 44759941 | rs2341456 | 1.486 | 1.383 | 1.992 | 0.878 | 0.048 | 0.071 | 0.071 | 0 | 0.77 | 0.53 | 0.75 | 0.32 | 0.72 | 0.78 |
| 2 | 44768202 | rs2341459 | 1.129 | 1.4 | 1.958 | 0.877 | 0.033 | 0.048 | 0.048 | 0 | 0.79 | 0.66 | 0.78 | 0.33 | 0.72 | 0.77 |
| 2 | 44771535 | rs10170837 | 2.162 | 1.387 | 1.915 | 0.802 | -0.016 | 0.031 | 0.031 | 0 | 0.61 | 0.36 | 0.51 | 0.21 | 0.57 | 0.64 |
| 2 | 44774792 | rs755667 | -0.707 | 1.405 | 1.669 | 0.805 | 0.032 | -0.034 | -0.034 | 0 | 0.06 | 0.09 | 0.06 | 0 | 0.09 | 0.05 |
| 2 | 44775683 | rs12712919 | 2 | 1.363 | 1.684 | 0.844 | -0.003 | 0.028 | 0.028 | 0.001 | 0.65 | 0.3 | 0.54 | 0.2 | 0.65 | 0.68 |
| 2 | 44787133 | rs13402122 | -0.485 | 1.611 | 1.679 | 1.369 | 0.043 | 0.063 | 0.063 | 0 | 0.09 | 0.19 | 0.06 | 0.17 | 0.14 | 0.06 |
| 2 | 44795774 | rs17427915 | 0.034 | 1.466 | 1.415 | 1.204 | 0.013 | -0.017 | -0.017 | 0 | 0.11 | 0.01 | 0.06 | 0.09 | 0.06 | 0.09 |
| 2 | 44810215 | rs6752168 | 1.887 | 1.379 | 1.554 | 1.124 | 0.025 | 0.009 | 0.009 | 0 | 0.69 | 0.6 | 0.7 | 0.32 | 0.64 | 0.59 |
| 2 | 44810480 | rs10201401 | 0.435 | 1.432 | 1.559 | 1.197 | 0.001 | 0.088 | 0.088 | 0 | 0.89 | 0.87 | 0.86 | 0.45 | 0.88 | 0.87 |
| 2 | 44825713 | rs17032420 | 1.973 | 1.358 | 1.504 | 1.099 | 0.024 | 0.011 | 0.011 | 0.043 | 0.7 | 0.63 | 0.73 | 0.32 | 0.64 | 0.59 |
| 2 | 44826068 | rs13389981 | 1.973 | 1.358 | 1.486 | 1.099 | 0.023 | 0.01 | 0.01 | 0.043 | 0.7 | 0.63 | 0.73 | 0.32 | 0.64 | 0.59 |
| 2 | 44826173 | rs13403533 | 1.973 | 1.358 | 1.486 | 1.099 | 0.023 | 0.01 | 0.01 | 0.043 | 0.7 | 0.63 | 0.73 | 0.32 | 0.64 | 0.59 |
| 2 | 44831953 | rs17390029 | 0.311 | 1.945 | 1.163 | 1.02 | -0.001 | 0 | 0 | 0.808 | 0.11 | 0.1 | 0.21 | 0.12 | 0.13 | 0.1 |
| 2 | 44845258 | rs934777 | 2.092 | 1.81 | 1.3 | 0.949 | 0.005 | 0 | 0 | 0.237 | 0.67 | 0.4 | 0.67 | 0.32 | 0.64 | 0.59 |
| 2 | 44861910 | rs4953123 | 0.408 | 1.852 | 0.885 | 1.001 | -0.004 | -0.006 | -0.006 | 0 | 0.11 | 0.07 | 0.21 | 0.12 | 0.13 | 0.11 |
| 2 | 44865678 | rs1437747 | 2.377 | 1.773 | 0.886 | 0.918 | -0.006 | 0.063 | 0.063 | 0 | 0.56 | 0.38 | 0.48 | 0.19 | 0.51 | 0.48 |
| 2 | 44871874 | rs13400118 | 1.947 | 1.817 | 0.383 | 1.018 | 0.043 | 0.019 | 0.019 | 0 | 0.7 | 0.58 | 0.72 | 0.32 | 0.64 | 0.59 |
| 2 | 44877010 | rs551660 | -1.541 | 1.754 | 0.385 | 0.932 | 0.034 | -0.018 | -0.018 | 0.223 | 0.19 | 0.28 | 0.13 | 0.13 | 0.24 | 0.28 |
| 2 | 44883762 | rs6711405 | 1.031 | 1.062 | -0.716 | 0.288 | -0.002 | 0.001 | 0.001 | 1.721 | 0.34 | 0.31 | 0.22 | 0.45 | 0.3 | 0.2 |
| 2 | 44885193 | rs473449 | -0.101 | 1.414 | -0.913 | -0.171 | -0.017 | 0.026 | 0.026 | 1.238 | 0.11 | 0.07 | 0.07 | 0 | 0.14 | 0.2 |
| 2 | 44886000 | rs17390344 | 0.075 | 1.285 | -0.646 | 0.153 | 0 | -0.002 | -0.002 | 0 | 0.11 | 0.07 | 0.19 | 0.08 | 0.1 | 0.1 |
| 2 | 44893134 | rs579119 | 0.677 | 1.453 | -0.652 | 0.176 | 0.048 | 0.018 | 0.018 | 0 | 0.72 | 0.58 | 0.73 | 0.29 | 0.66 | 0.7 |
| 2 | 44899976 | rs4953131 | 0.385 | 0.55 | -1.282 | -0.469 | 0.017 | 0.066 | 0.066 | 0.086 | 0.71 | 0.88 | 0.69 | 0.9 | 0.73 | 0.63 |
| 2 | 44905265 | rs1370546 | 1.163 | 0.753 | -0.976 | -1.201 | 0.005 | -0.066 | -0.066 | 6.897 | 0.3 | 0.29 | 0.22 | 0 | 0.41 | 0.09 |
| 2 | 44910413 | rs343974 | 0.011 | 0.415 | -0.824 | -1.302 | 0.009 | 0.032 | 0.032 | 0 | 0.53 | 0.66 | 0.39 | 0.1 | 0.58 | 0.34 |
| 2 | 44932854 | rs4145981 | 0.483 | 0.078 | -0.561 | -0.521 | -0.035 | 0.101 | 0.101 | 0.089 | 0.2 | 0.29 | 0.26 | 0.58 | 0.12 | 0.26 |
| 2 | 44943441 | rs343956 | -1.729 | 0.447 | -1.182 | -0.648 | -0.122 | 0.01 | 0.01 | 4.651 | 0.67 | 0.98 | 0.64 | 0.85 | 0.61 | 0.81 |
| 2 | 44944877 | rs343955 | -1.724 | 0.383 | -1.128 | -0.723 | -0.08 | 0.032 | 0.032 | 4.427 | 0.66 | 0.95 | 0.67 | 0.81 | 0.63 | 0.79 |
| 2 | 44948308 | rs343938 | 0.338 | 0.249 | -0.512 | -0.751 | -0.009 | -0.077 | -0.077 | 0 | 0.19 | 0.3 | 0.16 | 0 | 0.17 | 0.23 |
| 2 | 44985325 | rs12620511 | 0.821 | 0.162 | -0.298 | -0.63 | -0.011 | 0.042 | 0.042 | 0 | 0.19 | 0.19 | 0.28 | 0.55 | 0.12 | 0.2 |
| 2 | 44991100 | rs343935 | 0.591 | 0.473 | -0.842 | -0.795 | -0.015 | 0.075 | 0.075 | 1.363 | 0.17 | 0.06 | 0.11 | 0 | 0.18 | 0.24 |
| 2 | 44992550 | rs343936 | 0.093 | 0.462 | -0.692 | -0.811 | -0.013 | -0.096 | -0.096 | 0 | 0.2 | 0.34 | 0.17 | 0.01 | 0.19 | 0.28 |
| 2 | 44993487 | rs11679997 | 1.037 | 0.62 | -1.119 | -0.791 | -0.028 | -0.031 | -0.031 | 0 | 0.17 | 0.05 | 0.11 | 0.23 | 0.18 | 0.27 |
| 2 | 44999709 | rs343954 | 0.159 | 0.549 | -0.993 | -0.823 | 0 | -0.033 | -0.033 | 0.019 | 0.2 | 0.19 | 0.15 | 0 | 0.19 | 0.25 |
| 2 | 45003555 | rs10495921 | 0.563 | 0.753 | -0.555 | -0.179 | -0.031 | 0.027 | 0.027 | 0 | 0.05 | 0.01 | 0.08 | 0 | 0.15 | 0.02 |
| 2 | 45013538 | rs343939 | -0.55 | 0.229 | -0.311 | 0.014 | -0.026 | 0.034 | 0.034 | 0 | 0.29 | 0.59 | 0.35 | 0.52 | 0.17 | 0.26 |
| 2 | 45013725 | rs17032588 | 0.412 | 0.623 | -0.869 | 0.206 | -0.014 | -0.024 | -0.024 | 1.019 | 0.17 | 0.08 | 0.1 | 0.22 | 0.18 | 0.29 |
| 2 | 45024199 | rs4245799 | -0.09 | 0.46 | -0.655 | 0.363 | -0.011 | 0.067 | 0.067 | 0 | 0.23 | 0.29 | 0.32 | 0.52 | 0.18 | 0.24 |
| 2 | 45029525 | rs4953133 | -0.893 | 0.496 | -1.396 | -0.057 | -0.047 | 0.113 | 0.113 | 0.557 | 0.55 | 0.82 | 0.57 | 0.73 | 0.52 | 0.61 |
| 2 | 45037498 | rs6544770 | -0.823 | 0.502 | -1.448 | -0.22 | -0.083 | 0.205 | 0.205 | 0.019 | 0.56 | 0.88 | 0.73 | 0.87 | 0.5 | 0.66 |
| 2 | 45052611 | rs785679 | -0.805 | 0.366 | -0.947 | 0.018 | -0.007 | 0.069 | 0.069 | 0 | 0.08 | 0.2 | 0.21 | 0.2 | 0.05 | 0.09 |
| 2 | 45059246 | rs341577 | -0.97 | 0.289 | -0.915 | -0.202 | -0.043 | 0.133 | 0.133 | 0 | 0.57 | 0.79 | 0.72 | 0.86 | 0.5 | 0.65 |
| 2 | 45063613 | rs341606 | -0.929 | 0.29 | -0.881 | -0.326 | -0.035 | 0.135 | 0.135 | 0.102 | 0.57 | 0.77 | 0.73 | 0.9 | 0.49 | 0.76 |
| 2 | 45064187 | rs341605 | -0.146 | 0.279 | -1.148 | -0.094 | -0.005 | 0.036 | 0.036 | 0 | 0.06 | 0.12 | 0.21 | 0.21 | 0.05 | 0.17 |
| 2 | 45065588 | rs7565620 | 0.367 | 0.254 | -1.68 | -0.251 | 0.001 | -0.001 | -0.001 | 0 | 0.18 | 0.11 | 0.1 | 0.16 | 0.16 | 0.3 |
| 2 | 45072029 | rs572753 | -1.096 | 0.434 | -1.86 | -0.403 | -0.13 | 0.218 | 0.218 | 0.349 | 0.6 | 0.93 | 0.75 | 0.9 | 0.5 | 0.75 |
| 2 | 45082995 | rs340505 | -0.914 | 0.249 | -2.009 | -0.425 | -0.144 | 0.193 | 0.193 | 0.877 | 0.59 | 0.95 | 0.76 | 0.88 | 0.5 | 0.75 |
| 2 | 45089817 | rs340495 | -1.181 | 0.413 | -2.232 | -0.563 | -0.12 | 0.148 | 0.148 | 1.406 | 0.67 | 0.98 | 0.81 | 0.9 | 0.58 | 0.82 |
| 2 | 45091208 | rs554177 | -0.244 | 0.205 | -1.354 | -0.306 | 0 | -0.004 | -0.004 | 0 | 0.43 | 0.53 | 0.54 | 0.4 | 0.4 | 0.59 |
| 2 | 45095097 | rs11674185 | 0.043 | 0.235 | -1.086 | -0.367 | -0.001 | 0.061 | 0.061 | 0 | 0.38 | 0.24 | 0.23 | 0.18 | 0.34 | 0.44 |
| 2 | 45099750 | rs559396 | -1.673 | 0.31 | -1.155 | -0.416 | -0.1 | 0.153 | 0.153 | 0.571 | 0.68 | 0.97 | 0.81 | 0.91 | 0.58 | 0.82 |
| 2 | 45100208 | rs529177 | -1.277 | 0.291 | -0.888 | -0.451 | -0.059 | 0.143 | 0.143 | 0.008 | 0.64 | 0.84 | 0.77 | 0.91 | 0.53 | 0.76 |
| 2 | 45105675 | rs737448 | -0.582 | 0.441 | -0.211 | 0.116 | 0.025 | -0.027 | -0.027 | 0 | 0.05 | 0.16 | 0.04 | 0 | 0.06 | 0.07 |
| 2 | 45106017 | rs737447 | 0.047 | 0.234 | -0.2 | 0.476 | -0.036 | 0.217 | 0.217 | 0.151 | 0.19 | 0.42 | 0.24 | 0.5 | 0.13 | 0.16 |
| 2 | 45109063 | rs1975288 | -0.952 | 0.264 | -0.397 | 0.022 | 0 | 0 | 0 | 0 | 0.39 | 0.56 | 0.64 | 0.73 | 0.38 | 0.54 |
| 2 | 45111054 | rs508544 | -0.61 | -0.168 | -0.7 | -0.103 | 0.001 | 0.076 | 0.076 | 0 | 0.51 | 0.66 | 0.69 | 0.77 | 0.45 | 0.64 |
| 2 | 45116937 | rs6544776 | -0.978 | -0.319 | -0.073 | 0.238 | -0.033 | 0.081 | 0.081 | 0.635 | 0.17 | 0.47 | 0.35 | 0.25 | 0.08 | 0.26 |
| 2 | 45122342 | rs17041592 | -0.481 | -0.285 | -0.378 | -0.393 | -0.003 | -0.038 | -0.038 | 0.126 | 0.34 | 0.38 | 0.26 | 0.06 | 0.31 | 0.29 |
| 2 | 45123765 | rs6707760 | -1.33 | -0.942 | -0.886 | -0.44 | 0.019 | 0.006 | 0.006 | 3.921 | 0.26 | 0.79 | 0.41 | 0.24 | 0.13 | 0.29 |
| 2 | 45126586 | rs6713476 | -0.223 | -0.408 | -1.182 | 0.151 | -0.002 | -0.014 | -0.014 | 0 | 0.16 | 0.12 | 0.11 | 0.25 | 0.16 | 0.13 |

| | | | | | | | | | | | | | | | | |
|---|---|---|---|---|---|---|---|---|---|---|---|---|---|---|---|---|
| 2 | 45131643 | rs163521 | -1.61 | 0.217 | -1.274 | 0.005 | -0.109 | 0.022 | 0.022 | 0.026 | 0.8 | 0.95 | 0.64 | 0.78 | 0.6 | 0.85 |
| 2 | 45132047 | rs163522 | -1.61 | 0.218 | -1.299 | -0.071 | -0.127 | 0.128 | 0.128 | 0.12 | 0.8 | 0.98 | 0.82 | 0.98 | 0.61 | 0.89 |
| 2 | 45140616 | rs6704663 | 0.109 | -0.418 | -1.233 | 0.111 | 0.004 | -0.013 | -0.013 | 0.243 | 0.19 | 0.19 | 0.29 | 0.5 | 0.17 | 0.2 |
| 2 | 45141180 | rs494904 | 0.076 | -0.616 | -0.517 | -0.145 | 0.002 | -0.012 | -0.012 | 0 | 0.57 | 0.49 | 0.56 | 0.67 | 0.41 | 0.67 |
| 2 | 45142078 | rs1026062 | 0.099 | -0.535 | -0.652 | 0.223 | 0.004 | -0.012 | -0.012 | 0.088 | 0.18 | 0.14 | 0.19 | 0.31 | 0.16 | 0.17 |
| 2 | 45143894 | rs492780 | 0.006 | -0.686 | -0.891 | 0.149 | 0.003 | -0.018 | -0.018 | 2.601 | 0.12 | 0.03 | 0.11 | 0.06 | 0.07 | 0.14 |
| 2 | 45144752 | rs1657855 | -1.201 | -0.751 | -0.373 | 0.107 | -0.036 | 0.064 | 0.064 | 0.312 | 0.17 | 0.43 | 0.16 | 0.22 | 0.07 | 0.21 |
| 2 | 45147724 | rs480385 | -1.476 | -0.781 | -0.513 | 0.069 | -0.01 | 0.01 | 0.01 | 0 | 0.35 | 0.57 | 0.34 | 0.53 | 0.23 | 0.38 |
| 2 | 45150181 | rs519799 | -0.933 | -0.762 | -0.423 | 0.045 | 0.02 | 0.071 | 0.071 | 1.691 | 0.12 | 0.43 | 0.07 | 0.18 | 0.05 | 0.16 |
| 2 | 45151055 | rs511704 | -1.598 | -0.837 | -0.455 | 0.158 | 0.016 | -0.018 | -0.018 | 0.543 | 0.36 | 0.71 | 0.37 | 0.58 | 0.24 | 0.37 |
| 2 | 45152748 | rs2342406 | -0.129 | -0.821 | -1.207 | 0.009 | 0.011 | -0.013 | -0.013 | 0 | 0.23 | 0.18 | 0.22 | 0.37 | 0.19 | 0.18 |
| 2 | 45154908 | rs528301 | -1.822 | -1.271 | -1.726 | -0.514 | -0.137 | 0.16 | 0.16 | 1.687 | 0.76 | 0.97 | 0.79 | 0.98 | 0.57 | 0.87 |
| 2 | 45159091 | rs1004787 | -1.442 | -1.326 | -1.655 | -0.531 | -0.117 | 0.133 | 0.133 | 0.266 | 0.72 | 0.88 | 0.78 | 0.96 | 0.54 | 0.83 |
| 2 | 45159934 | rs4143308 | 0.651 | -1.685 | -1.565 | -0.291 | 0.006 | -0.006 | -0.006 | 0 | 0.3 | 0.22 | 0.35 | 0.33 | 0.2 | 0.39 |
| 2 | 45162618 | rs2271623 | 0.518 | -1.155 | -2.115 | -0.475 | 0 | -0.004 | -0.004 | 0.007 | 0.17 | 0.16 | 0.21 | 0.35 | 0.18 | 0.23 |
| 2 | 45165389 | rs1561227 | -0.63 | -0.99 | -2.067 | -0.563 | -0.053 | 0.11 | 0.11 | 0.042 | 0.65 | 0.78 | 0.72 | 0.92 | 0.54 | 0.79 |
| 2 | 45167570 | rs83995 | -1.465 | -0.753 | -2.088 | -0.638 | -0.126 | 0.158 | 0.158 | 2.11 | 0.76 | 0.97 | 0.81 | 0.97 | 0.59 | 0.88 |
| 2 | 45170515 | rs2673270 | -0.276 | -0.865 | -1.871 | -0.417 | 0.002 | -0.003 | -0.003 | 0 | 0.35 | 0.4 | 0.31 | 0.39 | 0.27 | 0.34 |
| 2 | 45175585 | rs173076 | -0.954 | -0.293 | -1.863 | -0.655 | -0.029 | 0.018 | 0.018 | 0.73 | 0.75 | 0.87 | 0.8 | 0.97 | 0.59 | 0.88 |
| 2 | 45180150 | rs2048774 | -0.418 | -0.199 | -2.017 | 0.058 | 0.003 | -0.069 | -0.069 | 0 | 0.3 | 0.16 | 0.38 | 0.54 | 0.25 | 0.48 |
| 2 | 45185224 | rs163513 | -0.538 | -0.291 | -2.091 | 0.11 | -0.035 | 0.041 | 0.041 | 0 | 0.6 | 0.82 | 0.55 | 0.67 | 0.5 | 0.71 |
| 2 | 45187127 | rs1983312 | -0.605 | 0.414 | -1.208 | 0.919 | -0.006 | 0.014 | 0.014 | 0 | 0.57 | 0.72 | 0.65 | 0.74 | 0.47 | 0.71 |
| 2 | 45189763 | rs4953156 | -0.544 | 0.57 | -0.983 | 1.215 | 0 | -0.006 | -0.006 | 0 | 0.17 | 0.13 | 0.24 | 0.36 | 0.15 | 0.29 |
| 2 | 45200009 | rs908279 | 0.046 | 1.079 | -0.594 | 0.661 | 0.02 | -0.021 | -0.021 | 0 | 0.64 | 0.58 | 0.69 | 0.68 | 0.56 | 0.61 |
| 2 | 45205547 | rs4953161 | 0.806 | 0.562 | -0.693 | 1.31 | 0.018 | -0.019 | -0.019 | 0.02 | 0.43 | 0.16 | 0.45 | 0.44 | 0.37 | 0.33 |
| 2 | 45205666 | rs561078 | -0.671 | 0.488 | 0.107 | 1.4 | 0.002 | 0.303 | 0.303 | 0 | 0.11 | 0.35 | 0.17 | 0.3 | 0.07 | 0.23 |
| 2 | 45207521 | rs4953162 | 0.081 | 0.416 | 0.156 | 1.282 | 0.01 | 0.021 | 0.021 | 0.132 | 0.05 | 0.26 | 0.06 | 0.06 | 0.03 | 0.1 |
| 2 | 45208020 | rs535137 | 0.058 | 0.412 | 0.306 | 1.284 | 0.027 | 0.112 | 0.112 | 0.412 | 0.08 | 0.33 | 0.1 | 0.07 | 0.05 | 0.06 |
| 2 | 45210169 | rs486930 | -0.151 | 0.417 | 0.014 | 1.198 | -0.091 | 0.095 | 0.095 | 0 | 0.66 | 0.8 | 0.74 | 0.8 | 0.54 | 0.67 |
| 2 | 45212258 | rs2917765 | 0.647 | 0.403 | 0.669 | 1.238 | -0.001 | 0 | 0 | 0 | 0.56 | 0.36 | 0.6 | 0.52 | 0.48 | 0.4 |
| 2 | 45213477 | rs1545371 | -0.216 | 0.595 | 0.563 | 1.205 | -0.081 | 0.12 | 0.12 | 0 | 0.68 | 0.81 | 0.77 | 0.91 | 0.56 | 0.74 |
| 2 | 45213745 | rs6743447 | 0.283 | 0.496 | 0.615 | 1.246 | -0.003 | 0.057 | 0.057 | 0.016 | 0.07 | 0.2 | 0.09 | 0.08 | 0.04 | 0.06 |
| 2 | 45217280 | rs2917768 | 0.101 | 0.499 | 0.506 | 1.286 | -0.073 | 0.001 | 0.001 | 0 | 0.63 | 0.78 | 0.73 | 0.65 | 0.51 | 0.51 |
| 2 | 45220348 | rs2917769 | 1.125 | 0.484 | 0.894 | 1.25 | -0.001 | 0 | 0 | 0 | 0.51 | 0.48 | 0.59 | 0.52 | 0.43 | 0.36 |
| 2 | 45221079 | rs767492 | 0.398 | 0.549 | 0.916 | 1.173 | -0.003 | 0.005 | 0.005 | 0 | 0.59 | 0.58 | 0.64 | 0.73 | 0.48 | 0.6 |
| 2 | 45229872 | rs2959253 | -0.098 | 0.514 | 0.911 | 1.087 | -0.054 | 0.068 | 0.068 | 0 | 0.67 | 0.76 | 0.74 | 0.81 | 0.54 | 0.68 |
| 2 | 45232686 | rs2290030 | -1.067 | 0.864 | 0.837 | 1.066 | -0.032 | -0.007 | -0.007 | 0 | 0.95 | 1 | 0.93 | 0.91 | 0.87 | 0.92 |
| 2 | 45234113 | rs2959230 | -0.125 | 0.693 | 0.887 | 1.067 | -0.076 | 0.089 | 0.089 | 0 | 0.68 | 0.78 | 0.74 | 0.81 | 0.54 | 0.69 |
| 2 | 45234363 | rs2959229 | 0.773 | 0.636 | 1.091 | 1.111 | 0.001 | -0.022 | -0.022 | 0 | 0.55 | 0.46 | 0.6 | 0.7 | 0.47 | 0.53 |
| 2 | 45238710 | rs964992 | 0.844 | 0.623 | 1.011 | 1.12 | -0.003 | 0.001 | 0.001 | 0.333 | 0.55 | 0.43 | 0.58 | 0.53 | 0.45 | 0.38 |
| 2 | 45238721 | rs964991 | 1.16 | 0.595 | 0.823 | 1.119 | 0.021 | -0.016 | -0.016 | 0 | 0.52 | 0.1 | 0.48 | 0.46 | 0.44 | 0.35 |
| 2 | 45242008 | rs12712931 | 0.862 | 0.637 | 1.13 | 1.151 | -0.002 | 0.001 | 0.001 | 0 | 0.54 | 0.48 | 0.59 | 0.53 | 0.45 | 0.38 |
| 2 | 45249426 | rs2959265 | 0.2 | 0.715 | 0.947 | 0.952 | 0.016 | -0.026 | -0.026 | 0 | 0.63 | 0.36 | 0.63 | 0.74 | 0.52 | 0.7 |
| 2 | 45250399 | rs4953173 | -0.565 | 0.468 | 0.458 | 0.998 | -0.002 | 0.001 | 0.001 | 0 | 0.06 | 0.06 | 0.05 | 0.1 | 0.03 | 0.14 |
| 2 | 45253117 | rs11898689 | -0.498 | 0.38 | 0.77 | 1.073 | -0.029 | 0.097 | 0.097 | 0.339 | 0.09 | 0.32 | 0.18 | 0.16 | 0.04 | 0.09 |
| 2 | 45253128 | rs2959262 | 0.09 | 0.682 | 0.728 | 1.216 | -0.001 | 0.013 | 0.013 | 0 | 0.9 | 0.91 | 0.84 | 0.69 | 0.82 | 0.74 |
| 2 | 45253578 | rs17032936 | -0.244 | 0.596 | 0.74 | 0.755 | 0.017 | -0.108 | -0.108 | 0.289 | 0.32 | 0.48 | 0.22 | 0.06 | 0.35 | 0.28 |
| 2 | 45254148 | rs17032940 | -0.685 | 0.44 | 0.657 | 1.046 | -0.026 | 0.13 | 0.13 | 0.445 | 0.09 | 0.35 | 0.19 | 0.17 | 0.04 | 0.09 |
| 2 | 45258427 | rs13005516 | 0.21 | 0.403 | -0.437 | 0.217 | -0.001 | 0.001 | 0.001 | 0 | 0.52 | 0.45 | 0.49 | 0.48 | 0.54 | 0.37 |
| 2 | 45260493 | rs4952732 | 0.22 | -0.145 | -0.526 | 0.291 | -0.034 | 0.009 | 0.009 | 0.109 | 0.12 | 0.33 | 0.11 | 0.07 | 0.08 | 0.1 |
| 2 | 45260750 | rs1868022 | 1.671 | 0.291 | -1.426 | 0.009 | -0.01 | 0.034 | 0.034 | 0 | 0.12 | 0.05 | 0.09 | 0 | 0.12 | 0.04 |
| 2 | 45260859 | rs4953174 | -0.911 | 0.357 | -1.544 | -0.03 | -0.077 | 0.098 | 0.098 | 0 | 0.78 | 0.94 | 0.81 | 0.9 | 0.69 | 0.87 |
| 2 | 45261200 | rs13025847 | 1.117 | 0.406 | -1.717 | -0.131 | 0.037 | -0.119 | -0.119 | 0.032 | 0.28 | 0.06 | 0.31 | 0.42 | 0.23 | 0.21 |
| 2 | 45272197 | rs1447107 | -0.394 | 0.173 | -2.217 | -0.268 | -0.033 | -0.055 | -0.055 | 0 | 0.55 | 0.84 | 0.52 | 0.45 | 0.5 | 0.7 |
| 2 | 45273041 | rs921120 | -0.965 | 0.388 | -2.084 | -0.34 | -0.07 | 0.096 | 0.096 | 0 | 0.78 | 0.94 | 0.81 | 0.9 | 0.7 | 0.87 |
| 2 | 45277600 | rs4952738 | -0.333 | 0.15 | -1.853 | -0.162 | -0.015 | 0.017 | 0.017 | 0 | 0.65 | 0.77 | 0.69 | 0.68 | 0.62 | 0.6 |
| 2 | 45280108 | rs13407328 | 0.046 | 0.014 | -1.338 | 0.329 | -0.009 | 0.023 | 0.023 | 0 | 0.12 | 0.17 | 0.11 | 0.23 | 0.06 | 0.27 |
| 2 | 45280704 | rs17032957 | -0.196 | -0.14 | -1.592 | 0.449 | 0.037 | -0.011 | -0.011 | 0.009 | 0.36 | 0.25 | 0.41 | 0.58 | 0.27 | 0.46 |
| 2 | 45281143 | rs13410828 | -0.207 | -0.101 | -0.785 | 0.59 | -0.02 | 0.058 | 0.058 | 0 | 0.13 | 0.31 | 0.12 | 0.21 | 0.06 | 0.28 |
| 2 | 45281360 | rs10191480 | -0.554 | 0.127 | -1.198 | 0.214 | 0 | 0.083 | 0.083 | 0 | 0.52 | 0.37 | 0.56 | 0.75 | 0.51 | 0.65 |
| 2 | 45281541 | rs10191680 | -0.518 | -0.072 | -1.021 | 0.265 | -0.001 | -0.001 | -0.001 | 0 | 0.38 | 0.38 | 0.27 | 0.39 | 0.35 | 0.47 |
| 2 | 45283359 | rs6733636 | -0.561 | -0.413 | -0.837 | 0.234 | -0.004 | -0.012 | -0.012 | 0.014 | 0.43 | 0.47 | 0.44 | 0.7 | 0.34 | 0.45 |
| 2 | 45283491 | rs6748611 | 0.067 | -0.534 | -0.807 | 0.325 | 0 | 0.003 | 0.003 | 0 | 0.2 | 0.17 | 0.16 | 0.23 | 0.18 | 0.19 |
| 2 | 45283518 | rs6748631 | 0.951 | -0.494 | -0.784 | 0.173 | 0 | -0.028 | -0.028 | 0 | 0.13 | 0.12 | 0.09 | 0 | 0.11 | 0.02 |
| 2 | 45284165 | rs2959259 | 0.117 | -0.517 | -0.868 | -0.05 | -0.008 | -0.047 | -0.047 | 0.007 | 0.63 | 0.54 | 0.65 | 0.77 | 0.66 | 0.64 |
| 2 | 45285952 | rs4355165 | 1.294 | -0.36 | -1.276 | -0.162 | -0.059 | 0.082 | 0.082 | 0 | 0.34 | 0.15 | 0.31 | 0.14 | 0.46 | 0.28 |
| 2 | 45293080 | rs2959254 | 0.44 | -0.175 | -0.972 | 0.319 | -0.001 | 0.002 | 0.002 | 0 | 0.56 | 0.47 | 0.61 | 0.49 | 0.61 | 0.5 |
| 2 | 45294327 | rs13024962 | 1.672 | -0.232 | -1.143 | 0.665 | -0.004 | -0.072 | -0.072 | 0 | 0.13 | 0.05 | 0.24 | 0.29 | 0.13 | 0.17 |
| 2 | 45294486 | rs2944665 | 0.426 | -0.085 | -0.973 | 0.255 | 0.003 | 0.019 | 0.019 | 0.025 | 0.56 | 0.58 | 0.46 | 0.18 | 0.58 | 0.37 |

| | | | | | | | | | | | | | | | | |
|---|---|---|---|---|---|---|---|---|---|---|---|---|---|---|---|---|
| 2 | 45296525 | rs868542 | -0.066 | 0.128 | -1.222 | 0.46 | -0.006 | -0.011 | -0.011 | 0 | 0.77 | 0.84 | 0.76 | 0.67 | 0.75 | 0.71 |
| 2 | 45296965 | rs12618367 | -0.087 | 0.02 | -1.206 | 0.642 | 0.002 | -0.01 | -0.01 | 0 | 0.13 | 0.1 | 0.06 | 0.16 | 0.08 | 0.17 |
| 2 | 45297576 | rs12997380 | 1.45 | -0.145 | -1.505 | 0.941 | 0 | -0.085 | -0.085 | 0 | 0.15 | 0.03 | 0.29 | 0.29 | 0.13 | 0.18 |
| 2 | 45297687 | rs6742724 | 0.55 | -0.133 | -1.036 | 1.009 | 0 | 0.125 | 0.125 | 0.254 | 0.06 | 0.19 | 0.04 | 0.15 | 0.01 | 0.11 |
| 2 | 45298085 | rs17319875 | 0.532 | 0.079 | -1.272 | 0.812 | -0.009 | 0.033 | 0.033 | 0 | 0.08 | 0.03 | 0.04 | 0.01 | 0.08 | 0.08 |
| 2 | 45298420 | rs2138648 | 0.511 | 0.122 | -0.9 | 0.685 | 0.004 | -0.002 | -0.002 | 0 | 0.41 | 0.34 | 0.35 | 0.43 | 0.46 | 0.39 |
| 2 | 45311319 | rs2959232 | 0.6 | 0.437 | -0.802 | 0.755 | -0.001 | -0.006 | -0.006 | 0.839 | 0.46 | 0.28 | 0.55 | 0.46 | 0.47 | 0.39 |
| 2 | 45312196 | rs11680125 | 0.887 | 0.771 | -1.079 | 0.571 | 0.047 | 0.036 | 0.036 | 0 | 0.4 | 0.16 | 0.47 | 0.28 | 0.35 | 0.3 |
| 2 | 45312582 | rs11691323 | 0.765 | 0.977 | -1.113 | 0.568 | 0.007 | 0.025 | 0.025 | 0 | 0.4 | 0.1 | 0.47 | 0.29 | 0.37 | 0.31 |
| 2 | 45312967 | rs2959236 | -0.935 | 1.255 | -0.567 | 0.642 | 0.007 | 0.013 | 0.013 | 0 | 0.3 | 0.61 | 0.33 | 0.53 | 0.39 | 0.46 |
| 2 | 45315575 | rs7578888 | 0.541 | 0.976 | -0.082 | 0.842 | -0.004 | 0.061 | 0.061 | 0 | 0.09 | 0.24 | 0.08 | 0.16 | 0.06 | 0.13 |
| 2 | 45316732 | rs908572 | -0.44 | 0.977 | -0.71 | 0.476 | 0.001 | 0.016 | 0.016 | 0 | 0.51 | 0.55 | 0.64 | 0.76 | 0.5 | 0.66 |
| 2 | 45317268 | rs876960 | 0.073 | 0.92 | -1.002 | 0.779 | -0.02 | -0.004 | -0.004 | 0 | 0.72 | 0.8 | 0.69 | 0.67 | 0.66 | 0.71 |
| 2 | 45318465 | rs908571 | -0.018 | 0.791 | -0.334 | 1.133 | 0.008 | -0.007 | -0.007 | 0 | 0.37 | 0.34 | 0.52 | 0.54 | 0.28 | 0.44 |
| 2 | 45319604 | rs10183968 | -0.318 | 1.749 | -0.01 | 1.188 | 0.083 | 0.136 | 0.136 | 0.02 | 0.07 | 0.17 | 0.08 | 0.21 | 0.12 | 0.11 |
| 2 | 45321308 | rs7423619 | 0.203 | 1.067 | -0.501 | 0.63 | 0.02 | 0.032 | 0.032 | 0 | 0.42 | 0.32 | 0.27 | 0.36 | 0.37 | 0.27 |
| 2 | 45328406 | rs1607133 | 0.422 | 1.214 | -0.904 | 0.787 | 0.021 | -0.032 | -0.032 | 0 | 0.37 | 0.2 | 0.41 | 0.46 | 0.31 | 0.35 |
| 2 | 45332395 | rs1517027 | -0.075 | 2.512 | 0.188 | 1.092 | 0.152 | 0.148 | 0.148 | 0 | 0.83 | 0.62 | 0.69 | 0.65 | 0.63 | 0.72 |
| 2 | 45334074 | rs4393800 | 0.787 | 2.466 | 0.493 | 1.002 | 0.091 | 0.104 | 0.104 | 0.637 | 0.74 | 0.49 | 0.55 | 0.54 | 0.56 | 0.58 |
| 2 | 45335413 | rs11125003 | -0.956 | 1.99 | 0.006 | 0.998 | -0.032 | -0.03 | -0.03 | 0 | 0.89 | 0.96 | 0.84 | 0.82 | 0.79 | 0.82 |
| 2 | 45336028 | rs908570 | -0.559 | 1.981 | 0.135 | 0.995 | -0.005 | -0.005 | -0.005 | 0 | 0.88 | 0.91 | 0.83 | 0.82 | 0.79 | 0.83 |
| 2 | 45336859 | rs868714 | 0.177 | 0.986 | 0.051 | 0.689 | 0.01 | 0.02 | 0.02 | 0 | 0.41 | 0.33 | 0.29 | 0.19 | 0.33 | 0.28 |
| 2 | 45339468 | rs12993810 | 1.102 | 0.887 | -0.428 | 0.537 | 0.013 | 0.136 | 0.136 | 0 | 0.28 | 0.04 | 0.11 | 0.07 | 0.21 | 0.13 |
| 2 | 45339491 | rs12464646 | 0.4 | 0.921 | -0.269 | 0.698 | 0.005 | 0.051 | 0.051 | 0 | 0.05 | 0.16 | 0.11 | 0.12 | 0.06 | 0.04 |
| 2 | 45341876 | rs11899053 | 0.699 | 0.908 | -0.34 | 0.455 | 0.021 | 0.053 | 0.053 | 0 | 0.31 | 0.18 | 0.15 | 0.08 | 0.26 | 0.14 |
| 2 | 45342920 | rs7561428 | 0.934 | 1.345 | -0.369 | 0.931 | -0.005 | 0.003 | 0.003 | 0 | 0.07 | 0.04 | 0.11 | 0.18 | 0.11 | 0.17 |
| 2 | 45343740 | rs17033049 | 0.149 | 1.13 | -0.25 | 0.653 | 0 | 0.001 | 0.001 | 0 | 0.17 | 0.14 | 0.16 | 0.09 | 0.14 | 0.18 |
| 2 | 45346666 | rs1517024 | 0.509 | 1.304 | -0.087 | 0.844 | 0.004 | -0.001 | -0.001 | 0 | 0.7 | 0.71 | 0.65 | 0.69 | 0.5 | 0.68 |
| 2 | 45347144 | rs6707629 | -0.347 | 0.508 | 0.275 | 0.511 | -0.025 | -0.04 | -0.04 | 0 | 0.16 | 0.44 | 0.07 | 0.09 | 0.1 | 0.13 |
| 2 | 45348125 | rs6726955 | 1.073 | 0.078 | -0.14 | 0.521 | 0.105 | -0.104 | -0.104 | 0 | 0.29 | 0.09 | 0.36 | 0.41 | 0.14 | 0.37 |
| 2 | 45348277 | rs17033059 | 0.98 | 0.077 | 0.104 | 0.633 | 0.048 | -0.011 | -0.011 | 1.459 | 0.29 | 0.16 | 0.31 | 0.27 | 0.17 | 0.25 |
| 2 | 45349861 | rs6731889 | 1.117 | 0.216 | 0.16 | 0.581 | -0.003 | -0.003 | -0.003 | 0.014 | 0.07 | 0.11 | 0.04 | 0.08 | 0.06 | 0.04 |
| 2 | 45351867 | rs7575297 | -0.095 | 0.143 | 0.297 | 0.563 | -0.025 | -0.012 | -0.012 | 0 | 0.12 | 0.24 | 0.06 | 0.09 | 0.06 | 0.14 |
| 2 | 45352356 | rs6728256 | -1.591 | 1.024 | -0.028 | 0.909 | -0.013 | -0.021 | -0.021 | 0.063 | 0.93 | 0.98 | 0.89 | 0.9 | 0.74 | 0.9 |
| 2 | 45352632 | rs6757058 | -0.274 | 0.684 | 0.093 | 0.826 | 0.042 | 0.017 | 0.017 | 0 | 0.48 | 0.75 | 0.38 | 0.43 | 0.38 | 0.44 |
| 2 | 45354105 | rs17415156 | 1.586 | 0.905 | 0.473 | 0.923 | 0.001 | 0.006 | 0.006 | 0 | 0.28 | 0.22 | 0.2 | 0.19 | 0.26 | 0.27 |
| 2 | 45358521 | rs4952740 | 0.393 | 0.899 | 0.035 | 0.88 | -0.004 | -0.002 | -0.002 | 0 | 0.66 | 0.69 | 0.64 | 0.6 | 0.5 | 0.58 |
| 2 | 45358577 | rs1517013 | 1.273 | 0.87 | -0.076 | 0.9 | 0.006 | -0.004 | -0.004 | 0 | 0.3 | 0.26 | 0.29 | 0.31 | 0.27 | 0.3 |
| 2 | 45359671 | rs973392 | -0.057 | 1.08 | -0.339 | 0.94 | -0.018 | -0.001 | -0.001 | 0.074 | 0.73 | 0.78 | 0.76 | 0.66 | 0.57 | 0.74 |
| 2 | 45363662 | rs9309123 | 1.306 | 0.91 | -0.578 | 1.133 | 0.001 | -0.044 | -0.044 | 0 | 0.28 | 0.11 | 0.27 | 0.34 | 0.28 | 0.3 |
| 2 | 45365176 | rs2102797 | -0.463 | 1.287 | -0.536 | 1.158 | 0.004 | 0.003 | 0.003 | 0 | 0.88 | 0.9 | 0.86 | 0.66 | 0.73 | 0.8 |
| 2 | 45365594 | rs12476150 | 0.869 | 0.321 | -0.557 | 1.059 | 0.008 | -0.028 | -0.028 | 0 | 0.29 | 0.21 | 0.28 | 0.51 | 0.28 | 0.34 |
| 2 | 45372968 | rs4952741 | 0.256 | -0.271 | -0.596 | 0.839 | 0.04 | 0.001 | 0.001 | 0 | 0.3 | 0.11 | 0.34 | 0.26 | 0.2 | 0.35 |
| 2 | 45377907 | rs11125004 | -1.846 | 0.076 | 0.404 | 0.951 | -0.016 | 0.181 | 0.181 | 0 | 0.08 | 0.32 | 0.09 | 0.33 | 0.03 | 0.18 |
| 2 | 45380320 | rs908576 | -1.375 | 0.168 | -0.076 | 0.591 | -0.01 | 0.023 | 0.023 | 0.058 | 0.11 | 0.32 | 0.12 | 0.19 | 0.05 | 0.12 |
| 2 | 45383266 | rs1517025 | 0.434 | -0.209 | -0.934 | 0.123 | 0.015 | 0.029 | 0.029 | 0 | 0.23 | 0.11 | 0.26 | 0.11 | 0.17 | 0.19 |
| 2 | 45383771 | rs11695220 | 0.308 | -0.305 | -1.163 | 0.148 | 0.019 | 0.009 | 0.009 | 0 | 0.12 | 0.03 | 0.07 | 0.1 | 0.1 | 0.09 |
| 2 | 45385062 | rs11685825 | 0.249 | -0.299 | -1.175 | 0.137 | 0.013 | 0.01 | 0.01 | 0 | 0.12 | 0.04 | 0.07 | 0.1 | 0.1 | 0.09 |
| 2 | 45387774 | rs992618 | -1.592 | -0.3 | -1.549 | 0.23 | 0.001 | -0.052 | -0.052 | 0 | 0.81 | 0.92 | 0.87 | 0.66 | 0.8 | 0.78 |
| 2 | 45388312 | rs10179400 | -1.815 | -0.303 | -1.557 | 0.264 | -0.01 | -0.064 | -0.064 | 0 | 0.82 | 0.96 | 0.87 | 0.75 | 0.81 | 0.84 |
| 2 | 45389301 | rs10206235 | -0.266 | -0.35 | -0.979 | -0.132 | -0.033 | -0.049 | -0.049 | 0.7 | 0.22 | 0.5 | 0.24 | 0.14 | 0.16 | 0.21 |
| 2 | 45389983 | rs1356339 | 1.152 | -0.188 | -0.656 | -0.061 | 0.028 | -0.021 | -0.021 | 0 | 0.08 | 0.27 | 0.15 | 0.02 | 0.07 | 0.13 |
| 2 | 45390197 | rs10186146 | 0.474 | 0.204 | -0.843 | -0.293 | 0.055 | 0.092 | 0.092 | 0 | 0.16 | 0.32 | 0.28 | 0.44 | 0.29 | 0.31 |
| 2 | 45390835 | rs12712936 | -0.494 | 0.31 | -1.04 | -0.325 | -0.02 | 0.04 | 0.04 | 0 | 0.68 | 0.47 | 0.63 | 0.59 | 0.74 | 0.58 |
| 2 | 45391048 | rs12712937 | -0.981 | 0.441 | -0.996 | -0.087 | -0.034 | -0.003 | -0.003 | 0.025 | 0.75 | 0.82 | 0.72 | 0.27 | 0.6 | 0.6 |
| 2 | 45391271 | rs994293 | -2.108 | 0.045 | -1.15 | -0.119 | 0.013 | -0.058 | -0.058 | 5.404 | 0.84 | 0.95 | 0.89 | 0.67 | 0.85 | 0.83 |
| 2 | 45402400 | rs7595106 | 1.49 | 0.145 | -0.997 | -0.444 | -0.009 | -0.061 | -0.061 | 0.043 | 0.19 | 0.21 | 0.1 | 0 | 0.15 | 0.09 |
| 2 | 45402571 | rs11125005 | 0.117 | 0.28 | -1.305 | -0.356 | 0.002 | -0.02 | -0.02 | 0 | 0.36 | 0.31 | 0.21 | 0.06 | 0.29 | 0.18 |
| 2 | 45404483 | rs3914642 | 0.775 | 0.46 | -1.197 | -0.058 | -0.001 | 0 | 0 | 0 | 0.08 | 0.2 | 0.13 | 0.05 | 0.03 | 0.15 |
| 2 | 45405049 | rs13418468 | -0.143 | 0.719 | -1.616 | -0.007 | 0.012 | 0.004 | 0.004 | 0 | 0.37 | 0.43 | 0.41 | 0.42 | 0.48 | 0.42 |
| 2 | 45405066 | rs3914643 | -0.354 | 1.189 | -2.079 | 0.583 | -0.06 | -0.061 | -0.061 | 0.084 | 0.33 | 0.21 | 0.42 | 0.55 | 0.46 | 0.53 |
| 2 | 45405160 | rs11674390 | -0.265 | 1.343 | -1.684 | 1.048 | 0.022 | 0.021 | 0.021 | 0 | 0.13 | 0.04 | 0.07 | 0.06 | 0.09 | 0.08 |
| 2 | 45405989 | rs13422236 | -0.197 | 1.848 | -1.393 | 1.365 | -0.013 | 0.006 | 0.006 | 0 | 0.08 | 0.06 | 0.16 | 0.38 | 0.25 | 0.2 |
| 2 | 45408269 | rs6720296 | -0.449 | 0.74 | -1.322 | 0.849 | 0.006 | 0.013 | 0.013 | 0 | 0.47 | 0.37 | 0.59 | 0.46 | 0.56 | 0.48 |
| 2 | 45409988 | rs17033132 | -0.109 | 0.019 | -1.048 | 0.952 | -0.035 | 0.012 | 0.012 | 0 | 0.22 | 0.34 | 0.28 | 0.24 | 0.13 | 0.26 |
| 2 | 45410658 | rs4953204 | -0.738 | 0.472 | -0.726 | 0.807 | 0.022 | -0.034 | -0.034 | 0.372 | 0.07 | 0.2 | 0.11 | 0 | 0.06 | 0.07 |
| 2 | 45411188 | rs2342702 | -0.55 | 0.603 | -1.272 | 0.992 | -0.001 | -0.002 | -0.002 | 0 | 0.36 | 0.47 | 0.5 | 0.41 | 0.35 | 0.42 |
| 2 | 45413618 | rs3851320 | -0.432 | 0.393 | -0.818 | 1.134 | 0.068 | 0.069 | 0.069 | 0.061 | 0.07 | 0.34 | 0.18 | 0.1 | 0.06 | 0.14 |
| 2 | 45413788 | rs3851322 | 0.454 | 0.248 | -0.867 | 1.027 | 0.015 | 0.071 | 0.071 | 0.476 | 0.21 | 0.37 | 0.17 | 0.32 | 0.22 | 0.16 |

| | | | | | | | | | | | | | | | | |
|---|---|---|---|---|---|---|---|---|---|---|---|---|---|---|---|---|
| 2 | 45414728 | rs13384377 | 0.687 | 0.199 | -1.404 | 0.562 | 0.018 | -0.016 | -0.016 | 0.552 | 0.13 | 0.05 | 0.05 | 0.16 | 0.09 | 0.1 |
| 2 | 45420082 | rs7569588 | -0.225 | 0.133 | -1.221 | 0.381 | 0.002 | 0 | 0 | 0 | 0.39 | 0.63 | 0.43 | 0.59 | 0.33 | 0.36 |
| 2 | 45423287 | rs10196185 | -0.027 | 0.157 | -1.041 | 0.129 | -0.002 | 0.013 | 0.013 | 0 | 0.56 | 0.38 | 0.53 | 0.74 | 0.56 | 0.56 |
| 2 | 45424099 | rs3908091 | 0.166 | -0.257 | -0.462 | 0.199 | -0.053 | 0.044 | 0.044 | 0.245 | 0.73 | 0.53 | 0.62 | 0.58 | 0.79 | 0.74 |
| 2 | 45426205 | rs3914636 | -0.549 | 0.071 | -0.531 | 0.178 | -0.004 | 0.005 | 0.005 | 1.839 | 0.81 | 0.81 | 0.84 | 0.73 | 0.81 | 0.83 |
| 2 | 45427012 | rs13383079 | 0.816 | -0.028 | -1.4 | -0.116 | -0.002 | 0.075 | 0.075 | 0 | 0.27 | 0.1 | 0.17 | 0.09 | 0.23 | 0.26 |
| 2 | 45430053 | rs2343408 | 0.378 | 0.486 | -0.788 | 0.358 | 0.003 | 0.007 | 0.007 | 0.099 | 0.48 | 0.49 | 0.41 | 0.4 | 0.39 | 0.61 |
| 2 | 45432763 | rs17033218 | 0.39 | 0.721 | -1.17 | 1.101 | -0.003 | 0.062 | 0.062 | 0 | 0.06 | 0.09 | 0.03 | 0.28 | 0.03 | 0.13 |
| 2 | 45435952 | rs7586888 | -0.362 | 0.836 | -1.235 | 0.498 | -0.001 | 0.002 | 0.002 | 0 | 0.92 | 0.86 | 0.81 | 0.9 | 0.89 | 0.97 |
| 2 | 45437978 | rs6760793 | 0.952 | 1.09 | -1.213 | 0.295 | 0.008 | -0.016 | -0.016 | 0.319 | 0.11 | 0.2 | 0.07 | 0.09 | 0.1 | 0.1 |
| 2 | 45439913 | rs4952752 | -0.025 | 1.317 | -1.98 | 0.159 | -0.095 | -0.01 | -0.01 | 0 | 0.26 | 0.07 | 0.34 | 0.19 | 0.39 | 0.21 |
| 2 | 45447713 | rs11679331 | 0.004 | 1.164 | -1.907 | -0.214 | -0.033 | 0.062 | 0.062 | 0 | 0.22 | 0.1 | 0.12 | 0 | 0.26 | 0.17 |
| 2 | 45448525 | rs12994487 | -0.02 | 1.153 | -1.912 | -0.222 | -0.026 | 0.058 | 0.058 | 0 | 0.23 | 0.12 | 0.12 | 0 | 0.25 | 0.17 |
| 2 | 45449950 | rs7580559 | -0.553 | 1.156 | -2.127 | 0.129 | -0.053 | 0.068 | 0.068 | 0 | 0.43 | 0.16 | 0.51 | 0.2 | 0.52 | 0.37 |
| 2 | 45450533 | rs3908093 | -0.521 | 0.714 | -1.595 | 0.247 | 0.022 | 0.004 | 0.004 | 0 | 0.54 | 0.61 | 0.61 | 0.41 | 0.66 | 0.62 |
| 2 | 45450988 | rs4953210 | -0.122 | 0.519 | -1.603 | 0.412 | 0.001 | -0.002 | -0.002 | 0 | 0.58 | 0.44 | 0.64 | 0.45 | 0.68 | 0.59 |
| 2 | 45452426 | rs9636390 | -0.911 | 0.615 | -1.997 | 0.458 | 0.026 | 0.046 | 0.046 | 0 | 0.82 | 0.93 | 0.91 | 0.92 | 0.84 | 0.85 |
| 2 | 45452838 | rs13413031 | 1.439 | 1.036 | -2.206 | 0.634 | -0.066 | -0.063 | -0.063 | 0 | 0.11 | 0.02 | 0.09 | 0.17 | 0.18 | 0.11 |
| 2 | 45454554 | rs13400787 | -0.03 | 0.987 | -2.232 | 0.64 | -0.041 | -0.038 | -0.038 | 0 | 0.28 | 0.09 | 0.17 | 0.26 | 0.32 | 0.24 |
| 2 | 45454801 | rs13426486 | 1.412 | 1.021 | -2.223 | 0.731 | -0.061 | -0.063 | -0.063 | 1.408 | 0.11 | 0.02 | 0.09 | 0.19 | 0.18 | 0.11 |
| 2 | 45455897 | rs6544792 | -0.87 | 0.725 | -1.91 | 0.689 | 0.016 | 0.019 | 0.019 | 0 | 0.82 | 0.9 | 0.9 | 0.9 | 0.84 | 0.85 |
| 2 | 45457111 | rs883834 | 0.89 | 1.19 | -1.578 | 1.393 | -0.005 | 0.01 | 0.01 | 0 | 0.05 | 0.05 | 0.15 | 0.14 | 0.08 | 0.11 |
| 2 | 45464079 | rs13391740 | 0 | 0.723 | -1.185 | 0.967 | -0.004 | 0.035 | 0.035 | 0.099 | 0.35 | 0.46 | 0.52 | 0.57 | 0.36 | 0.47 |
| 2 | 45465799 | rs869183 | 0.748 | 0.928 | -1.863 | 1.111 | -0.036 | -0.105 | -0.105 | 6.883 | 0.19 | 0.03 | 0.34 | 0.48 | 0.25 | 0.42 |
| 2 | 45466067 | rs3937505 | 0.526 | 1.231 | -1.697 | 0.997 | -0.047 | -0.031 | -0.031 | 0 | 0.13 | 0.04 | 0.14 | 0.17 | 0.22 | 0.13 |
| 2 | 45480142 | rs7569382 | 0.324 | 0.353 | -1.225 | 0.869 | -0.015 | -0.004 | -0.004 | 0 | 0.16 | 0.19 | 0.16 | 0.15 | 0.12 | 0.06 |
| 2 | 45482500 | rs12472971 | -0.148 | 0.113 | -1.428 | 1.262 | -0.01 | 0.022 | 0.022 | 0 | 0.2 | 0.2 | 0.26 | 0.32 | 0.16 | 0.12 |
| 2 | 45482812 | rs7577545 | 0.177 | 0.155 | -1.088 | 0.93 | 0.013 | 0.009 | 0.009 | 0 | 0.2 | 0.36 | 0.19 | 0.17 | 0.19 | 0.16 |
| 2 | 45483869 | rs7595329 | -0.916 | -0.265 | -1.737 | 0.72 | 0.064 | 0.175 | 0.175 | 0 | 0.48 | 0.8 | 0.51 | 0.75 | 0.37 | 0.41 |
| 2 | 45484777 | rs10178040 | -0.716 | -0.196 | -2.364 | 0.284 | -0.112 | -0.069 | -0.069 | 0 | 0.72 | 0.97 | 0.59 | 0.73 | 0.61 | 0.46 |
| 2 | 45485296 | rs17392668 | 1.297 | -0.463 | -2.534 | 0.019 | -0.018 | 0.062 | 0.062 | 0 | 0.27 | 0.16 | 0.18 | 0.09 | 0.28 | 0.13 |
| 2 | 45485938 | rs929901 | -1.83 | -0.386 | -2.542 | 0.096 | -0.103 | 0.015 | 0.015 | 0.12 | 0.81 | 0.97 | 0.69 | 0.85 | 0.66 | 0.66 |
| 2 | 45485957 | rs929902 | 0.043 | -0.421 | -2.009 | 0.116 | 0.016 | 0.003 | 0.003 | 0 | 0.39 | 0.71 | 0.38 | 0.62 | 0.32 | 0.32 |
| 2 | 45486331 | rs7567591 | 0.847 | -0.181 | -1.957 | 0.412 | -0.009 | 0.034 | 0.034 | 0 | 0.25 | 0.47 | 0.26 | 0.28 | 0.19 | 0.12 |
| 2 | 45486867 | rs4953216 | -0.943 | 0.06 | -2.273 | 0.409 | -0.09 | -0.069 | -0.069 | 0.314 | 0.73 | 0.97 | 0.61 | 0.73 | 0.66 | 0.48 |
| 2 | 45487466 | rs7583873 | 0.696 | 0.066 | -2.055 | 0.36 | -0.014 | 0.021 | 0.021 | 0 | 0.26 | 0.52 | 0.28 | 0.29 | 0.2 | 0.16 |
| 2 | 45488003 | rs13394266 | 1.273 | 0.247 | -1.761 | 0.052 | -0.066 | 0.038 | 0.038 | 0 | 0.16 | 0.02 | 0.11 | 0.08 | 0.21 | 0.1 |
| 2 | 45488414 | rs17320808 | -0.354 | 0.142 | -1.263 | 0.327 | 0 | 0 | 0 | 0 | 0.41 | 0.39 | 0.29 | 0.43 | 0.39 | 0.31 |
| 2 | 45492041 | rs7573076 | -1.169 | 0.671 | -1.395 | 0.497 | -0.106 | -0.01 | -0.01 | 0 | 0.78 | 0.97 | 0.7 | 0.82 | 0.71 | 0.72 |
| 2 | 45492077 | rs12993476 | 0.018 | 0.219 | -0.346 | 0.641 | 0.001 | -0.026 | -0.026 | 0.2 | 0.07 | 0.1 | 0.04 | 0.01 | 0.1 | 0.19 |
| 2 | 45492633 | rs10204473 | -2.084 | -0.142 | -0.487 | 0.82 | 0.028 | -0.07 | -0.07 | 0 | 0.43 | 0.9 | 0.41 | 0.51 | 0.35 | 0.46 |
| 2 | 45493703 | rs7594273 | -0.84 | -0.17 | 0.568 | 0.859 | -0.053 | 0.043 | 0.043 | 0 | 0.23 | 0.46 | 0.18 | 0.35 | 0.1 | 0.18 |
| 2 | 45495143 | rs17033293 | 0.966 | -0.142 | 0.517 | 0.816 | 0.008 | -0.001 | -0.001 | 0 | 0.05 | 0.09 | 0.05 | 0.05 | 0.1 | 0.05 |
| 2 | 45496101 | rs17033295 | 1.147 | -0.27 | 0.551 | 0.75 | 0.004 | 0.027 | 0.027 | 0 | 0.1 | 0.04 | 0.06 | 0.01 | 0.06 | 0.04 |
| 2 | 45496813 | rs6721199 | -0.156 | -0.316 | 0.919 | 0.806 | 0.027 | 0.326 | 0.326 | 0.272 | 0.08 | 0.41 | 0.06 | 0.33 | 0.03 | 0.12 |
| 2 | 45498718 | rs7579209 | -1.308 | -0.302 | 0.933 | 0.802 | -0.04 | 0.056 | 0.056 | 0.373 | 0.22 | 0.48 | 0.14 | 0.35 | 0.13 | 0.33 |
| 2 | 45498745 | rs7605521 | -2.538 | -0.248 | 0.865 | 0.949 | 0.022 | -0.055 | -0.055 | 0 | 0.37 | 0.8 | 0.38 | 0.62 | 0.29 | 0.43 |
| 2 | 45500248 | rs11899866 | -1.84 | -0.191 | 0.924 | 0.946 | -0.01 | 0.034 | 0.034 | 0.537 | 0.27 | 0.58 | 0.2 | 0.4 | 0.23 | 0.38 |
| 2 | 45503169 | rs17320892 | 1.29 | -0.15 | 0.847 | 0.951 | 0.002 | 0.024 | 0.024 | 0 | 0.09 | 0.04 | 0.06 | 0.01 | 0.06 | 0.04 |
| 2 | 45503260 | rs870575 | -2.231 | -0.168 | 0.896 | 0.984 | 0.012 | -0.036 | -0.036 | 0 | 0.3 | 0.7 | 0.33 | 0.56 | 0.24 | 0.39 |
| 2 | 45503598 | rs9679711 | 0.881 | -0.158 | 1.064 | 0.963 | -0.001 | -0.025 | -0.025 | 0 | 0.06 | 0.07 | 0.03 | 0 | 0.08 | 0.04 |
| 2 | 45503643 | rs4953219 | -0.792 | -0.161 | 0.995 | 0.984 | 0.035 | 0.092 | 0.092 | 0 | 0.1 | 0.23 | 0.2 | 0.21 | 0.12 | 0.08 |
| 2 | 45505271 | rs6544796 | -1.552 | -0.202 | 1.162 | 0.965 | -0.049 | 0.021 | 0.021 | 0 | 0.24 | 0.42 | 0.14 | 0.34 | 0.12 | 0.29 |
| 2 | 45505432 | rs866922 | 0.536 | 0.011 | 0.903 | 0.912 | -0.003 | -0.027 | -0.027 | 0.048 | 0.12 | 0.13 | 0.08 | 0.02 | 0.09 | 0.18 |
| 2 | 45506771 | rs12476712 | -0.832 | 0.205 | 0.736 | 1.149 | 0.031 | 0.088 | 0.088 | 0 | 0.1 | 0.21 | 0.2 | 0.21 | 0.12 | 0.08 |
| 2 | 45507516 | rs6715182 | 0.057 | 0.301 | -0.05 | 1.049 | 0.001 | -0.001 | -0.001 | 0 | 0.07 | 0.1 | 0.06 | 0.05 | 0.11 | 0.05 |
| 2 | 45510233 | rs12612653 | -0.203 | 0.342 | 0.142 | 1.218 | 0.023 | 0.117 | 0.117 | 0 | 0.11 | 0.27 | 0.09 | 0.38 | 0.13 | 0.15 |
| 2 | 45510971 | rs2042474 | -1.126 | 0.321 | 0.027 | 0.922 | 0.051 | 0.237 | 0.237 | 0.025 | 0.15 | 0.37 | 0.25 | 0.59 | 0.18 | 0.22 |
| 2 | 45512442 | rs2343466 | -1.643 | 0.209 | -0.201 | 0.778 | 0.014 | 0.13 | 0.13 | 0.113 | 0.24 | 0.46 | 0.29 | 0.6 | 0.26 | 0.26 |
| 2 | 45512908 | rs6707333 | -1.307 | 0.609 | -0.512 | 0.98 | 0.003 | 0.022 | 0.022 | 0.88 | 0.3 | 0.62 | 0.36 | 0.61 | 0.31 | 0.3 |
| 2 | 45515092 | rs7602584 | 0.389 | 0.547 | -1.169 | 0.305 | 0.021 | 0.042 | 0.042 | 0 | 0.51 | 0.31 | 0.41 | 0.62 | 0.47 | 0.44 |
| 2 | 45517067 | rs4953222 | 0.099 | 0.551 | -0.98 | -0.217 | -0.023 | 0.087 | 0.087 | 0 | 0.79 | 0.94 | 0.9 | 0.93 | 0.82 | 0.8 |
| 2 | 45521003 | rs2194394 | 0.512 | 0.576 | -0.673 | -0.252 | 0 | -0.004 | -0.004 | 1.218 | 0.87 | 0.87 | 0.93 | 0.93 | 0.89 | 0.83 |
| 2 | 45521185 | rs4953225 | 1.052 | 0.463 | -0.706 | -0.311 | 0.014 | 0.104 | 0.104 | 0.41 | 0.5 | 0.27 | 0.55 | 0.68 | 0.59 | 0.49 |
| 2 | 45525591 | rs17033393 | 0.826 | 0.41 | 0.015 | -0.128 | 0.031 | 0.158 | 0.158 | 0 | 0.05 | 0.18 | 0.14 | 0.19 | 0.06 | 0.04 |
| 2 | 45528422 | rs17033398 | -0.981 | 0.245 | 0.534 | -0.234 | -0.022 | 0.051 | 0.051 | 0 | 0.19 | 0.43 | 0.25 | 0.23 | 0.15 | 0.22 |
| 2 | 45529595 | rs883413 | 1.32 | 0.213 | 0.276 | -0.323 | -0.017 | 0.108 | 0.108 | 2.014 | 0.49 | 0.14 | 0.54 | 0.69 | 0.58 | 0.49 |
| 2 | 45530632 | rs6737172 | -1.413 | 0.557 | 0.595 | -0.218 | 0.053 | 0.096 | 0.096 | 1.872 | 0.43 | 0.73 | 0.4 | 0.23 | 0.31 | 0.35 |
| 2 | 45532011 | rs10169967 | -0.722 | 0.573 | 0.783 | -0.145 | -0.012 | 0.028 | 0.028 | 0 | 0.17 | 0.27 | 0.24 | 0.23 | 0.15 | 0.24 |

| | | | | | | | | | | | | | | | | |
|---|---|---|---|---|---|---|---|---|---|---|---|---|---|---|---|---|
| 2 | 45532366 | rs1862892 | -0.234 | 0.695 | 0.423 | -0.036 | -0.011 | -0.004 | -0.004 | 0 | 0.4 | 0.35 | 0.49 | 0.54 | 0.47 | 0.49 |
| 2 | 45532804 | rs6745203 | 1.406 | 0.415 | 0.748 | 0.057 | -0.025 | -0.026 | -0.026 | 0 | 0.05 | 0.07 | 0.02 | 0 | 0 | 0 |
| 2 | 45533241 | rs13431811 | -0.443 | 0.51 | 0.557 | 0.145 | -0.005 | 0.038 | 0.038 | 0 | 0.15 | 0.26 | 0.24 | 0.25 | 0.15 | 0.25 |
| 2 | 45535161 | rs2216790 | 0.01 | 0.6 | 0.084 | 0.142 | -0.003 | -0.008 | -0.008 | 0 | 0.11 | 0.12 | 0.1 | 0.07 | 0.09 | 0.2 |
| 2 | 45536858 | rs6755279 | -0.799 | 0.724 | 0.36 | 0.361 | 0.01 | 0.02 | 0.02 | 1.708 | 0.36 | 0.67 | 0.39 | 0.25 | 0.29 | 0.34 |
| 2 | 45539196 | rs17321114 | 0.937 | 0.845 | 0.21 | 0.406 | -0.013 | 0.034 | 0.034 | 0 | 0.17 | 0.06 | 0.13 | 0 | 0.18 | 0.1 |
| 2 | 45539725 | rs17033413 | 1.061 | 0.619 | 0.093 | 0.519 | -0.036 | 0.005 | 0.005 | 0.887 | 0.3 | 0.11 | 0.29 | 0.29 | 0.31 | 0.28 |
| 2 | 45543071 | rs7561717 | -0.076 | -0.07 | 0.964 | 1.292 | 0.007 | -0.013 | -0.013 | 0 | 0.65 | 0.68 | 0.67 | 0.53 | 0.63 | 0.63 |
| 2 | 45543090 | rs7575803 | -0.364 | -0.111 | 0.778 | 1.287 | 0.025 | -0.028 | -0.028 | 0 | 0.68 | 0.77 | 0.72 | 0.58 | 0.7 | 0.63 |
| 2 | 45547015 | rs11677320 | -0.506 | -0.034 | 0.812 | 1.12 | -0.005 | 0.016 | 0.016 | 0.978 | 0.16 | 0.27 | 0.17 | 0.18 | 0.11 | 0.14 |
| 2 | 45549305 | rs12473388 | 0.882 | -0.047 | 0.628 | 1.14 | 0.036 | 0.089 | 0.089 | 0.226 | 0.49 | 0.15 | 0.43 | 0.31 | 0.53 | 0.42 |
| 2 | 45549660 | rs11681887 | -1.139 | -0.617 | 0.754 | 0.972 | 0.067 | 0.02 | 0.02 | 0.038 | 0.49 | 0.79 | 0.54 | 0.63 | 0.4 | 0.57 |
| 2 | 45549961 | rs35608719 | 0.413 | -0.476 | 0.298 | 0.864 | 0.023 | -0.025 | -0.025 | 5.633 | 0.28 | 0.24 | 0.28 | 0.42 | 0.27 | 0.3 |
| 2 | 45551530 | rs16909 | -0.747 | -0.294 | -0.207 | 0.992 | 0.046 | 0.011 | 0.011 | 0 | 0.48 | 0.68 | 0.47 | 0.6 | 0.38 | 0.45 |
| 2 | 45554033 | rs6544808 | -0.661 | -0.251 | -0.28 | 0.9 | 0.03 | 0.006 | 0.006 | 0 | 0.47 | 0.66 | 0.48 | 0.6 | 0.39 | 0.36 |
| 2 | 45556388 | rs11904395 | -1.014 | 0.156 | -0.144 | 0.942 | -0.009 | 0.07 | 0.07 | 2.799 | 0.12 | 0.27 | 0.18 | 0.19 | 0.07 | 0.05 |
| 2 | 45560276 | rs2278064 | -1.079 | 0.151 | -0.321 | 0.953 | -0.017 | 0.049 | 0.049 | 0.844 | 0.13 | 0.23 | 0.18 | 0.2 | 0.07 | 0.07 |
| 2 | 45569103 | rs2343468 | -0.021 | -0.033 | -0.877 | 0.113 | 0.009 | 0.028 | 0.028 | 0 | 0.62 | 0.7 | 0.6 | 0.83 | 0.39 | 0.66 |
| 2 | 45571703 | rs6760244 | -0.204 | -0.979 | -0.696 | 0.566 | -0.008 | 0.009 | 0.009 | 0 | 0.25 | 0.31 | 0.24 | 0.31 | 0.16 | 0.23 |
| 2 | 45572181 | rs13036196 | 0.103 | -1.086 | -1.044 | 0.462 | 0.001 | -0.012 | -0.012 | 0.013 | 0.22 | 0.13 | 0.24 | 0.28 | 0.25 | 0.33 |
| 2 | 45573241 | rs2081297 | 0.5 | -1.907 | -1.591 | -0.216 | 0.018 | -0.076 | -0.076 | 0 | 0.33 | 0.13 | 0.32 | 0.46 | 0.31 | 0.37 |
| 2 | 45575110 | rs10460504 | 0.352 | -2.102 | -1.741 | -0.402 | 0.018 | 0.03 | 0.03 | 0.856 | 0.42 | 0.15 | 0.31 | 0.63 | 0.36 | 0.45 |
| 2 | 45575726 | rs3911086 | -1.675 | -2.237 | -1.059 | -0.761 | 0.031 | -0.056 | -0.056 | 0 | 0.78 | 0.71 | 0.75 | 0.97 | 0.61 | 0.83 |
| 2 | 45576017 | rs3915409 | 0.757 | -2.321 | -1.184 | -0.754 | -0.013 | -0.026 | -0.026 | 3.762 | 0.14 | 0.03 | 0.17 | 0.2 | 0.14 | 0.2 |
| 2 | 45577243 | rs11689571 | -0.138 | -2.291 | -1.184 | -0.79 | -0.056 | 0.165 | 0.165 | 0.312 | 0.56 | 0.16 | 0.48 | 0.83 | 0.49 | 0.66 |
| 2 | 45577721 | rs6705684 | 0.853 | -2.432 | -0.51 | -0.818 | -0.062 | -0.114 | -0.114 | 0.058 | 0.17 | 0.36 | 0.12 | 0 | 0.09 | 0.07 |
| 2 | 45579991 | rs13023260 | -0.001 | -2.38 | -0.415 | -0.823 | 0.007 | 0.04 | 0.04 | 0 | 0.55 | 0.34 | 0.42 | 0.64 | 0.44 | 0.53 |
| 2 | 45582319 | rs10167668 | 1.358 | -2.255 | -0.274 | -0.777 | -0.002 | -0.026 | -0.026 | 0.598 | 0.13 | 0.16 | 0.1 | 0 | 0.09 | 0.05 |
| 2 | 45585144 | rs6711122 | -0.191 | -2.468 | 0.036 | -0.821 | -0.064 | -0.028 | -0.028 | 1.946 | 0.23 | 0.59 | 0.28 | 0.15 | 0.12 | 0.2 |
| 2 | 45591275 | rs12328653 | 0.076 | -2.482 | 0.099 | -0.843 | -0.042 | -0.02 | -0.02 | 0 | 0.21 | 0.33 | 0.26 | 0.15 | 0.12 | 0.2 |
| 2 | 45600619 | rs1862070 | 0.35 | -2.465 | 0.108 | -0.856 | -0.063 | -0.115 | -0.115 | 0 | 0.19 | 0.35 | 0.12 | 0 | 0.08 | 0.07 |
| 2 | 45607280 | rs17393522 | 0.937 | -2.298 | -0.159 | -0.933 | -0.018 | 0.011 | 0.011 | 0 | 0.28 | 0.22 | 0.41 | 0.17 | 0.45 | 0.35 |
| 2 | 45617101 | rs6719211 | 0.306 | -2.38 | -0.194 | -0.921 | -0.028 | -0.044 | -0.044 | 0 | 0.25 | 0.35 | 0.18 | 0.08 | 0.18 | 0.2 |
| 2 | 45621627 | rs3770241 | 1.712 | -2.27 | -0.263 | -0.859 | -0.002 | 0 | 0 | 1.051 | 0.1 | 0.08 | 0.08 | 0.08 | 0.13 | 0.14 |
| 2 | 45629700 | rs7581815 | 0.252 | -2.436 | -0.338 | -0.858 | -0.005 | -0.006 | -0.006 | 0.325 | 0.23 | 0.18 | 0.25 | 0.3 | 0.26 | 0.31 |
| 2 | 45635313 | rs7557300 | 0.649 | -2.513 | -0.317 | -0.811 | -0.024 | -0.008 | -0.008 | 0 | 0.18 | 0.27 | 0.25 | 0.15 | 0.1 | 0.16 |
| 2 | 45640334 | rs3755072 | 0.928 | -2.46 | -0.458 | -0.756 | -0.024 | -0.04 | -0.04 | 5.182 | 0.13 | 0.02 | 0.16 | 0.22 | 0.13 | 0.18 |
| 2 | 45661586 | rs3770248 | 1.037 | -2.484 | -0.489 | -0.775 | -0.02 | -0.026 | -0.026 | 0 | 0.12 | 0.04 | 0.16 | 0.21 | 0.13 | 0.13 |
| 2 | 45661587 | rs3770249 | -0.113 | -2.601 | -0.353 | -0.918 | 0.02 | 0.024 | 0.024 | 0 | 0.48 | 0.41 | 0.52 | 0.32 | 0.67 | 0.53 |
| 2 | 45662883 | rs7593154 | -1.497 | -2.734 | -0.43 | -0.973 | -0.011 | -0.115 | -0.115 | 0.056 | 0.69 | 0.95 | 0.78 | 0.47 | 0.77 | 0.7 |
| 2 | 45669349 | rs3755076 | 0.029 | -2.78 | -0.413 | -0.985 | -0.087 | -0.043 | -0.043 | 0.027 | 0.23 | 0.56 | 0.27 | 0.14 | 0.1 | 0.2 |
| 2 | 45673894 | rs6738398 | -1.367 | -2.728 | -0.362 | -0.97 | 0.018 | -0.077 | -0.077 | 0 | 0.68 | 0.92 | 0.79 | 0.47 | 0.77 | 0.69 |
| 2 | 45675684 | rs2193836 | 1.388 | -2.614 | -0.117 | -0.987 | -0.037 | -0.066 | -0.066 | 0 | 0.14 | 0.21 | 0.1 | 0 | 0.06 | 0.05 |
| 2 | 45688221 | rs3770266 | 0.725 | -2.585 | -0.149 | -0.994 | 0.008 | -0.039 | -0.039 | 0 | 0.35 | 0.34 | 0.35 | 0.1 | 0.54 | 0.37 |
| 2 | 45695075 | rs13400867 | 0.571 | -2.662 | -0.088 | -0.94 | -0.056 | -0.015 | -0.015 | 0 | 0.18 | 0.33 | 0.2 | 0.15 | 0.09 | 0.15 |
| 2 | 45700070 | rs17033715 | 1.621 | -2.549 | -0.169 | -0.97 | 0 | -0.023 | -0.023 | 0 | 0.13 | 0.16 | 0.11 | 0 | 0.08 | 0.05 |
| 2 | 45714555 | rs3770272 | 0.81 | -3.033 | -0.611 | -1.287 | 0.001 | -0.028 | -0.028 | 0 | 0.37 | 0.35 | 0.41 | 0.12 | 0.6 | 0.38 |
| 2 | 45717161 | rs3770273 | -1.188 | -3.016 | -0.663 | -1.103 | 0.011 | -0.019 | -0.019 | 0 | 0.71 | 0.76 | 0.77 | 0.47 | 0.81 | 0.7 |
| 2 | 45720658 | rs17033745 | 1.245 | -2.859 | -0.702 | -1.162 | 0 | 0.001 | 0.001 | 0.235 | 0.15 | 0.12 | 0.1 | 0.09 | 0.16 | 0.16 |
| 2 | 45739045 | rs17322370 | 1.287 | -2.961 | -0.899 | -1.247 | -0.095 | 0.107 | 0.107 | 0 | 0.28 | 0.11 | 0.34 | 0.03 | 0.45 | 0.24 |
| 2 | 45740908 | rs12712944 | 0.109 | -2.899 | -0.881 | -1.139 | -0.019 | 0.009 | 0.009 | 0.098 | 0.25 | 0.35 | 0.31 | 0.34 | 0.2 | 0.3 |
| 2 | 45745731 | rs4140775 | 0.389 | -2.856 | -0.802 | -1.099 | -0.06 | -0.018 | -0.018 | 3.68 | 0.19 | 0.35 | 0.19 | 0.14 | 0.09 | 0.15 |
| 2 | 45752140 | rs7573215 | 0.976 | -2.829 | -0.764 | -1.081 | -0.062 | 0 | 0 | 0.011 | 0.15 | 0.35 | 0.16 | 0.14 | 0.07 | 0.13 |
| 2 | 45752214 | rs2112040 | -0.307 | -2.981 | -0.92 | -1.214 | 0.026 | 0.025 | 0.025 | 0 | 0.55 | 0.61 | 0.61 | 0.33 | 0.7 | 0.57 |
| 2 | 45757489 | rs3770283 | -0.897 | -3.004 | -0.765 | -1.199 | 0.006 | 0.006 | 0.006 | 0 | 0.43 | 0.61 | 0.38 | 0.32 | 0.33 | 0.4 |
| 2 | 45765157 | rs13430952 | 1.44 | -2.843 | -0.677 | -1.114 | -0.04 | -0.053 | -0.053 | 0 | 0.12 | 0.19 | 0.06 | 0 | 0.04 | 0.04 |
| 2 | 45769971 | rs755652 | -0.348 | -2.862 | -1.003 | -1.124 | -0.006 | 0.098 | 0.098 | 0 | 0.57 | 0.88 | 0.61 | 0.26 | 0.65 | 0.52 |
| 2 | 45770732 | rs3770292 | -0.067 | -2.823 | -0.824 | -1.133 | -0.027 | 0.173 | 0.173 | 0.051 | 0.47 | 0.77 | 0.52 | 0.17 | 0.52 | 0.39 |
| 2 | 45781771 | rs6734588 | 0.096 | -2.78 | -0.746 | -1.116 | -0.007 | 0.017 | 0.017 | 0.022 | 0.44 | 0.59 | 0.5 | 0.17 | 0.51 | 0.38 |
| 2 | 45796481 | rs2217870 | -0.111 | -2.729 | -0.625 | -0.997 | 0.02 | 0.02 | 0.02 | 0.018 | 0.54 | 0.57 | 0.61 | 0.31 | 0.7 | 0.57 |
| 2 | 45815991 | rs6544836 | -0.034 | -0.489 | -0.549 | -0.912 | -0.017 | -0.114 | -0.114 | 0.718 | 0.24 | 0.6 | 0.21 | 0.01 | 0.2 | 0.07 |
| 2 | 45816569 | rs7576649 | 0 | -0.458 | -0.523 | -0.906 | -0.016 | -0.134 | -0.134 | 1.075 | 0.24 | 0.57 | 0.21 | 0.01 | 0.2 | 0.07 |
| 2 | 45831537 | rs718731 | -0.41 | -0.599 | -0.559 | -0.762 | -0.029 | -0.018 | -0.018 | 0 | 0.29 | 0.59 | 0.22 | 0.2 | 0.21 | 0.28 |
| 2 | 45834076 | rs7591446 | 0.357 | -0.539 | -0.912 | -0.223 | -0.025 | 0.017 | 0.017 | 0.412 | 0.68 | 0.43 | 0.61 | 0.48 | 0.79 | 0.64 |
| 2 | 45836781 | rs7599740 | -1.889 | -0.283 | -1.081 | 0.424 | 0.002 | -0.008 | -0.008 | 0.038 | 0.91 | 0.93 | 0.97 | 0.83 | 0.94 | 0.92 |
| 2 | 45837053 | rs11884064 | 1.357 | -0.525 | -1.51 | 0.199 | 0.009 | -0.009 | -0.009 | 0 | 0.26 | 0.11 | 0.19 | 0.28 | 0.24 | 0.22 |
| 2 | 45839700 | rs7596942 | 1.19 | -0.182 | -0.882 | 0.16 | -0.002 | 0.012 | 0.012 | 0 | 0.15 | 0.18 | 0.3 | 0.22 | 0.14 | 0.14 |
| 2 | 45842064 | rs1522993 | -1.179 | -0.242 | -1.022 | 0.151 | 0.012 | -0.057 | -0.057 | 0 | 0.66 | 0.86 | 0.82 | 0.57 | 0.71 | 0.66 |
| 2 | 45842844 | rs10173144 | 0.179 | -0.001 | -0.456 | -0.026 | -0.053 | 0.029 | 0.029 | 0.515 | 0.19 | 0.42 | 0.21 | 0.18 | 0.11 | 0.14 |

| | | | | | | | | | | | | | | | | |
|---|---|---|---|---|---|---|---|---|---|---|---|---|---|---|---|---|
| 2 | 45843275 | rs1403450 | 1.186 | 0.237 | -0.758 | -0.688 | 0.024 | -0.099 | -0.099 | 0 | 0.22 | 0.37 | 0.23 | 0.02 | 0.29 | 0.22 |
| 2 | 45844714 | rs10209296 | 0.753 | -0.106 | -0.883 | -0.142 | -0.002 | -0.012 | -0.012 | 1.455 | 0.16 | 0.15 | 0.17 | 0 | 0.18 | 0.04 |
| 2 | 45850077 | rs6713298 | -0.399 | -0.485 | -0.77 | 0.439 | 0.023 | 0.008 | 0.008 | 1.353 | 0.4 | 0.67 | 0.26 | 0.37 | 0.27 | 0.41 |
| 2 | 45854864 | rs17323037 | 0.661 | 1.111 | -1.361 | 0.39 | -0.113 | 0.104 | 0.104 | 1.257 | 0.22 | 0.03 | 0.37 | 0.05 | 0.36 | 0.21 |
| 2 | 45855171 | rs4405810 | -0.279 | -0.092 | -0.684 | 1.115 | 0.013 | 0.003 | 0.003 | 0.217 | 0.53 | 0.41 | 0.7 | 0.58 | 0.71 | 0.54 |
| 2 | 45856434 | rs11694157 | -0.704 | -0.443 | -1.244 | 0.882 | 0.028 | 0.015 | 0.015 | 0.535 | 0.6 | 0.6 | 0.78 | 0.68 | 0.77 | 0.66 |
| 2 | 45858240 | rs1589039 | -0.255 | 0.58 | -1.899 | 0.576 | 0.02 | 0.054 | 0.054 | 1.134 | 0.42 | 0.18 | 0.62 | 0.34 | 0.62 | 0.48 |
| 2 | 45864672 | rs11896740 | -0.564 | 0.766 | -1.821 | 0.757 | -0.019 | -0.013 | -0.013 | 0 | 0.16 | 0.07 | 0.15 | 0.11 | 0.17 | 0.13 |
| 2 | 45864710 | rs2344662 | -1.076 | 0.585 | -1.997 | 0.963 | 0 | -0.03 | -0.03 | 0 | 0.44 | 0.74 | 0.38 | 0.44 | 0.43 | 0.45 |
| 2 | 45875839 | rs6722418 | -0.138 | 1.257 | -1.447 | 1.512 | 0.007 | 0.014 | 0.014 | 0.125 | 0.14 | 0.15 | 0.18 | 0.28 | 0.19 | 0.26 |
| 2 | 45884213 | rs666334 | 0.65 | 1.161 | -1.59 | 0.635 | -0.002 | -0.039 | -0.039 | 0 | 0.28 | 0.37 | 0.29 | 0.08 | 0.25 | 0.15 |
| 2 | 45887931 | rs666214 | -0.624 | 1.137 | -1.74 | 0.993 | -0.038 | -0.027 | -0.027 | 0.377 | 0.86 | 0.99 | 0.86 | 0.89 | 0.84 | 0.87 |
| 2 | 45898530 | rs17033965 | 0.074 | 0.994 | -1.511 | 1.09 | -0.002 | -0.004 | -0.004 | 4.319 | 0.14 | 0.12 | 0.1 | 0.24 | 0.16 | 0.25 |
| 2 | 45901755 | rs585156 | 0.594 | 1.162 | -0.675 | 1.164 | 0.021 | 0.043 | 0.043 | 0 | 0.69 | 0.55 | 0.67 | 0.49 | 0.66 | 0.59 |
| 2 | 45902454 | rs556650 | -0.057 | 1.151 | -0.819 | 1.374 | -0.002 | -0.011 | -0.011 | 0 | 0.79 | 0.79 | 0.79 | 0.65 | 0.79 | 0.71 |
| 2 | 45905946 | rs2204204 | 0.296 | 0.076 | -1.301 | 0.79 | 0.049 | 0.002 | 0.002 | 0 | 0.32 | 0.17 | 0.25 | 0.29 | 0.24 | 0.4 |
| 2 | 45908637 | rs12185636 | 0.116 | 0.202 | -1.215 | 0.759 | 0.03 | 0.005 | 0.005 | 0 | 0.34 | 0.23 | 0.27 | 0.29 | 0.3 | 0.4 |
| 2 | 45910616 | rs2285024 | 0.381 | 0.308 | -1.403 | 0.594 | 0.036 | 0.026 | 0.026 | 0 | 0.35 | 0.16 | 0.27 | 0.27 | 0.3 | 0.42 |
| 2 | 45910950 | rs506134 | -0.107 | 0.561 | -0.993 | 0.678 | 0.004 | -0.006 | -0.006 | 0.159 | 0.35 | 0.24 | 0.31 | 0.37 | 0.39 | 0.31 |
| 2 | 45922031 | rs542413 | -0.37 | 0.995 | -0.714 | 0.752 | -0.036 | -0.009 | -0.009 | 0 | 0.75 | 0.78 | 0.66 | 0.69 | 0.65 | 0.7 |
| 2 | 45922470 | rs6720782 | 0.493 | 0.803 | -0.852 | 0.776 | 0.009 | -0.013 | -0.013 | 1.273 | 0.13 | 0.06 | 0.07 | 0.12 | 0.12 | 0.08 |
| 2 | 45925771 | rs629772 | 0.651 | 0.615 | -0.62 | 0.759 | -0.016 | 0.014 | 0.014 | 0 | 0.57 | 0.26 | 0.46 | 0.41 | 0.53 | 0.62 |
| 2 | 45927390 | rs656823 | 1.445 | 0.51 | -0.894 | 0.627 | -0.023 | 0.062 | 0.062 | 0 | 0.15 | 0.05 | 0.15 | 0.05 | 0.19 | 0.12 |
| 2 | 45928086 | rs11674329 | 0.887 | 0.371 | -0.233 | 0.791 | -0.021 | -0.009 | -0.009 | 0 | 0.42 | 0.43 | 0.39 | 0.33 | 0.29 | 0.28 |
| 2 | 45929628 | rs483404 | -0.131 | 0.594 | -0.279 | 0.775 | -0.009 | 0.03 | 0.03 | 0 | 0.29 | 0.1 | 0.23 | 0.18 | 0.33 | 0.21 |
| 2 | 45932849 | rs642200 | 0.801 | 1.254 | -0.106 | 0.626 | 0.017 | -0.01 | -0.01 | 0 | 0.22 | 0.3 | 0.3 | 0.19 | 0.29 | 0.22 |
| 2 | 45937413 | rs6709673 | -1.188 | 1.087 | -0.59 | 0.389 | 0 | 0.004 | 0.004 | 0.012 | 0.84 | 0.84 | 0.84 | 0.95 | 0.82 | 0.88 |
| 2 | 45941915 | rs1464573 | 1.048 | 1.153 | -0.761 | 0.507 | 0.025 | 0.016 | 0.016 | 0 | 0.43 | 0.25 | 0.31 | 0.38 | 0.38 | 0.5 |
| 2 | 45942363 | rs7606454 | 0.671 | 0.963 | -0.839 | 0.396 | 0.002 | -0.01 | -0.01 | 0 | 0.12 | 0.03 | 0.06 | 0.13 | 0.12 | 0.11 |
| 2 | 45946800 | rs6725257 | -1.177 | 0.955 | -0.831 | 0.337 | -0.034 | 0.063 | 0.063 | 0 | 0.85 | 0.97 | 0.85 | 0.94 | 0.81 | 0.9 |
| 2 | 45950270 | rs7577664 | -0.092 | 1.233 | -0.654 | 0.373 | 0.003 | 0 | 0 | 0 | 0.14 | 0.14 | 0.1 | 0.13 | 0.18 | 0.11 |
| 2 | 45953910 | rs1464572 | 1.066 | 1.023 | -0.607 | 0.221 | 0.018 | -0.007 | -0.007 | 0 | 0.41 | 0.32 | 0.33 | 0.4 | 0.35 | 0.54 |
| 2 | 45954324 | rs4557033 | -0.806 | 1.188 | -0.336 | 0.228 | 0 | -0.001 | -0.001 | 0 | 0.39 | 0.33 | 0.4 | 0.39 | 0.46 | 0.36 |
| 2 | 45958251 | rs10201978 | -0.866 | 1.119 | -0.357 | -0.463 | -0.004 | -0.015 | -0.015 | 0 | 0.14 | 0.08 | 0.1 | 0 | 0.18 | 0.08 |
| 2 | 45963001 | rs6740453 | -0.771 | 0.767 | 0.263 | -0.431 | 0.02 | -0.057 | -0.057 | 0.004 | 0.1 | 0.33 | 0.09 | 0 | 0.08 | 0.04 |
| 2 | 45966098 | rs952331 | -0.742 | 0.612 | -0.271 | -0.639 | 0.009 | 0.041 | 0.041 | 0 | 0.54 | 0.72 | 0.56 | 0.33 | 0.53 | 0.34 |
| 2 | 45968233 | rs2176347 | -0.789 | 0.697 | -0.189 | -0.624 | -0.031 | 0.051 | 0.051 | 0 | 0.43 | 0.79 | 0.5 | 0.33 | 0.47 | 0.27 |
| 2 | 45968584 | rs1878383 | 0.042 | 0.84 | 0.13 | -0.44 | -0.027 | -0.007 | -0.007 | 0 | 0.3 | 0.18 | 0.39 | 0.33 | 0.38 | 0.21 |
| 2 | 45968698 | rs1878384 | 0.106 | 0.838 | -0.159 | -0.472 | -0.096 | -0.04 | -0.04 | 0 | 0.29 | 0.09 | 0.38 | 0.32 | 0.38 | 0.21 |
| 2 | 45968987 | rs7424090 | -0.113 | 0.908 | 0.053 | -0.442 | 0.042 | 0.001 | 0.001 | 0 | 0.32 | 0.44 | 0.41 | 0.32 | 0.39 | 0.21 |
| 2 | 45972516 | rs17034091 | 0.293 | 0.402 | -0.193 | -0.745 | -0.014 | -0.089 | -0.089 | 0 | 0.19 | 0.37 | 0.13 | 0.03 | 0.13 | 0.2 |
| 2 | 45976420 | rs11125030 | -0.393 | 0.676 | -0.458 | -0.946 | 0.003 | -0.004 | -0.004 | 1.086 | 0.71 | 0.69 | 0.62 | 0.69 | 0.69 | 0.82 |
| 2 | 45976753 | rs4953245 | 0.484 | 0.627 | -0.822 | -1.126 | 0.007 | 0.07 | 0.07 | 0 | 0.46 | 0.25 | 0.44 | 0.65 | 0.5 | 0.57 |
| 2 | 45977472 | rs6757543 | -1.271 | 1.102 | -0.617 | -1.065 | -0.026 | 0.028 | 0.028 | 0 | 0.93 | 0.98 | 0.92 | 1 | 0.87 | 0.99 |
| 2 | 45977576 | rs11125031 | -0.012 | 0.463 | -0.706 | -0.683 | 0.009 | 0.034 | 0.034 | 0 | 0.3 | 0.38 | 0.38 | 0.42 | 0.37 | 0.38 |
| 2 | 45977815 | rs7579246 | 0.785 | 0.42 | -0.656 | -0.654 | 0.001 | -0.032 | -0.032 | 0.07 | 0.08 | 0.15 | 0.05 | 0 | 0.07 | 0.07 |
| 2 | 45981111 | rs1533476 | -1.003 | 0.441 | -0.466 | -0.681 | -0.013 | -0.015 | -0.015 | 0 | 0.69 | 0.71 | 0.53 | 0.49 | 0.58 | 0.69 |
| 2 | 45981142 | rs940052 | 0.125 | 0.253 | -0.343 | -0.822 | -0.003 | -0.002 | -0.002 | 0.522 | 0.55 | 0.43 | 0.48 | 0.44 | 0.52 | 0.63 |
| 2 | 45981426 | rs6716622 | 0.487 | 0.281 | -0.582 | -0.743 | -0.033 | -0.036 | -0.036 | 0 | 0.14 | 0.27 | 0.05 | 0.06 | 0.06 | 0.07 |
| 2 | 45982030 | rs867286 | 0.942 | 0.683 | -1.266 | -0.523 | -0.002 | 0.007 | 0.007 | 0.068 | 0.18 | 0.15 | 0.33 | 0.39 | 0.31 | 0.33 |
| 2 | 45983518 | rs1468042 | -0.322 | 0.809 | -1.312 | -0.13 | -0.042 | -0.047 | -0.047 | 0 | 0.71 | 0.8 | 0.55 | 0.5 | 0.58 | 0.7 |
| 2 | 45989023 | rs7582320 | -0.255 | -0.388 | -0.745 | 0.068 | -0.006 | 0.021 | 0.021 | 0.612 | 0.36 | 0.4 | 0.38 | 0.52 | 0.35 | 0.48 |
| 2 | 45995350 | rs3924523 | -0.954 | -0.861 | -0.297 | -0.231 | -0.005 | -0.089 | -0.089 | 0.035 | 0.13 | 0.56 | 0.1 | 0 | 0.09 | 0.06 |
| 2 | 45996657 | rs13401638 | 0.63 | -1.475 | -1.1 | -0.433 | 0.04 | 0.066 | 0.066 | 0.12 | 0.32 | 0.02 | 0.22 | 0.15 | 0.23 | 0.26 |
| 2 | 45996694 | rs7592092 | -0.11 | -1.463 | -0.853 | -0.438 | 0.038 | 0.029 | 0.029 | 1.217 | 0.06 | 0.57 | 0.06 | 0 | 0 | 0.01 |
| 2 | 45997209 | rs13404751 | -0.129 | -1.497 | -1.729 | -0.441 | 0.059 | 0.154 | 0.154 | 0.249 | 0.4 | 0.07 | 0.27 | 0.15 | 0.32 | 0.31 |
| 2 | 45997411 | rs13404973 | 0.288 | -1.506 | -1.71 | -0.39 | 0.074 | 0.096 | 0.096 | 0.273 | 0.34 | 0.07 | 0.22 | 0.15 | 0.23 | 0.26 |
| 2 | 45997867 | rs4446102 | 0.437 | -0.929 | -1.726 | -0.108 | -0.022 | -0.023 | -0.023 | 0 | 0.24 | 0.18 | 0.29 | 0.32 | 0.3 | 0.31 |
| 2 | 45999116 | rs4952771 | 0.585 | -0.88 | -1.693 | -0.217 | -0.019 | -0.017 | -0.017 | 0 | 0.23 | 0.17 | 0.29 | 0.28 | 0.3 | 0.29 |
| 2 | 46000408 | rs6758953 | 0.704 | -1.929 | -1.206 | -0.607 | 0.092 | 0.159 | 0.159 | 0 | 0.51 | 0.21 | 0.59 | 0.8 | 0.62 | 0.63 |
| 2 | 46000783 | rs13002562 | 0.183 | -1.986 | -0.903 | -0.742 | 0.023 | 0.049 | 0.049 | 0 | 0.57 | 0.74 | 0.68 | 0.84 | 0.63 | 0.72 |
| 2 | 46002657 | rs13016569 | -0.481 | -1.569 | -1.635 | -0.516 | 0.022 | 0.109 | 0.109 | 0 | 0.69 | 0.87 | 0.72 | 0.9 | 0.74 | 0.78 |
| 2 | 46002777 | rs13017423 | 0.704 | -1.42 | -1.917 | -0.378 | 0.109 | 0.167 | 0.167 | 0 | 0.5 | 0.18 | 0.57 | 0.75 | 0.62 | 0.59 |
| 2 | 46009085 | rs4264601 | 0.799 | -1.505 | -1.74 | -0.28 | 0.073 | 0.127 | 0.127 | 0 | 0.46 | 0.19 | 0.57 | 0.76 | 0.61 | 0.56 |
| 2 | 46010306 | rs6707046 | -0.951 | -1.222 | -0.553 | 0.619 | -0.096 | 0.06 | 0.06 | 0.055 | 0.15 | 0.54 | 0.11 | 0.2 | 0.03 | 0.14 |
| 2 | 46011945 | rs10191412 | -0.926 | -1.048 | -1.566 | 0.562 | 0.074 | 0.002 | 0.002 | 0.189 | 0.3 | 0.81 | 0.22 | 0.25 | 0.17 | 0.22 |
| 2 | 46015851 | rs7558342 | -0.407 | -0.377 | -1.176 | 0.746 | 0.002 | 0.001 | 0.001 | 0 | 0.44 | 0.47 | 0.46 | 0.54 | 0.49 | 0.51 |
| 2 | 46017602 | rs10170073 | -0.484 | -1.092 | -1.662 | 0.615 | 0.015 | -0.037 | -0.037 | 0 | 0.56 | 0.86 | 0.51 | 0.56 | 0.43 | 0.59 |
| 2 | 46018533 | rs11898209 | -0.235 | -1.488 | -1.206 | 0.388 | -0.008 | -0.002 | -0.002 | 0 | 0.28 | 0.63 | 0.22 | 0.2 | 0.17 | 0.21 |

| | | | | | | | | | | | | | | | |
|---|---|---|---|---|---|---|---|---|---|---|---|---|---|---|---|
| 2 | 46021381 | rs4952774 | 0.61 | -1.665 | -1.499 | 0.021 | 0.002 | -0.015 | -0.015 | 0 | 0.16 | 0.15 | 0.21 | 0.02 | 0.09 | 0.19 |
| 2 | 46021606 | rs4952775 | 0.061 | -1.605 | -2.011 | 0.122 | 0.017 | 0.127 | 0.127 | 0 | 0.41 | 0.1 | 0.46 | 0.33 | 0.36 | 0.48 |
| 2 | 46023371 | rs6544852 | 0.497 | -1.634 | -1.543 | 0.162 | -0.001 | 0.002 | 0.002 | 0 | 0.12 | 0.15 | 0.12 | 0.16 | 0.14 | 0.16 |
| 2 | 46023916 | rs12472172 | -0.286 | -1.65 | -1.395 | 0.1 | -0.003 | 0.003 | 0.003 | 0 | 0.59 | 0.56 | 0.61 | 0.45 | 0.48 | 0.54 |
| 2 | 46024212 | rs6720975 | -0.965 | -1.176 | -1.296 | 0.698 | -0.149 | -0.095 | -0.095 | 0 | 0.73 | 0.95 | 0.75 | 0.72 | 0.57 | 0.79 |
| 2 | 46026901 | rs6743144 | -0.457 | -1.188 | -1.065 | 0.664 | -0.058 | -0.017 | -0.017 | 0 | 0.61 | 0.87 | 0.66 | 0.68 | 0.5 | 0.74 |
| 2 | 46028107 | rs10164633 | 0.292 | -1.104 | -0.938 | 0.528 | -0.008 | -0.001 | -0.001 | 0 | 0.32 | 0.39 | 0.35 | 0.32 | 0.24 | 0.36 |
| 2 | 46038801 | rs4296473 | -0.463 | -0.178 | -1.051 | 0.952 | -0.012 | 0.005 | 0.005 | 0 | 0.87 | 0.93 | 0.94 | 0.89 | 0.89 | 0.91 |
| 2 | 46039147 | rs7593800 | -0.17 | -0.121 | -0.766 | 1.098 | -0.014 | 0.012 | 0.012 | 0 | 0.69 | 0.66 | 0.81 | 0.69 | 0.81 | 0.83 |
| 2 | 46040793 | rs4953262 | 0.596 | 0.087 | -1.501 | 0.979 | 0.003 | 0.01 | 0.01 | 0 | 0.5 | 0.46 | 0.57 | 0.41 | 0.57 | 0.56 |
| 2 | 46042478 | rs3886870 | -0.122 | 0.807 | -1.845 | 0.983 | -0.086 | -0.045 | -0.045 | 0.016 | 0.18 | 0.02 | 0.22 | 0.27 | 0.24 | 0.24 |
| 2 | 46045959 | rs935672 | -0.079 | 0.669 | -2.085 | 0.903 | 0.017 | -0.037 | -0.037 | 0.001 | 0.53 | 0.84 | 0.58 | 0.6 | 0.4 | 0.57 |
| 2 | 46049445 | rs11125034 | -1.091 | 0.161 | -0.851 | 0.93 | -0.036 | 0.029 | 0.029 | 0 | 0.17 | 0.5 | 0.24 | 0.19 | 0.11 | 0.15 |
| 2 | 46050036 | rs7559522 | -0.118 | 0.497 | -1.34 | 0.715 | -0.037 | -0.007 | -0.007 | 0 | 0.55 | 0.81 | 0.64 | 0.61 | 0.46 | 0.59 |
| 2 | 46052497 | rs6747706 | -0.934 | 0.223 | 0.04 | 0.86 | 0.077 | 0.132 | 0.132 | 0 | 0.06 | 0.36 | 0.18 | 0.08 | 0.04 | 0.06 |
| 2 | 46054114 | rs10175198 | -0.04 | 0.733 | -0.786 | 0.877 | -0.05 | -0.062 | -0.062 | 0.342 | 0.65 | 0.84 | 0.67 | 0.48 | 0.52 | 0.58 |
| 2 | 46060375 | rs4952779 | 0.11 | 0.755 | -0.846 | 0.897 | -0.06 | -0.092 | -0.092 | 0 | 0.65 | 0.88 | 0.67 | 0.48 | 0.53 | 0.58 |
| 2 | 46062550 | rs6738409 | 0.372 | 0.687 | -0.186 | 0.828 | 0.009 | 0 | 0 | 0 | 0.6 | 0.59 | 0.51 | 0.39 | 0.49 | 0.52 |
| 2 | 46063406 | rs7591351 | 0.495 | 0.713 | -0.12 | 0.852 | -0.002 | 0.002 | 0.002 | 0 | 0.58 | 0.44 | 0.46 | 0.3 | 0.46 | 0.45 |
| 2 | 46066236 | rs6732900 | 0.513 | 0.736 | -0.233 | 0.861 | -0.001 | -0.003 | -0.003 | 0 | 0.55 | 0.47 | 0.46 | 0.37 | 0.4 | 0.49 |
| 2 | 46066261 | rs6761327 | 0.228 | 0.626 | -0.216 | 0.79 | 0.004 | -0.008 | -0.008 | 0 | 0.59 | 0.55 | 0.49 | 0.37 | 0.42 | 0.49 |
| 2 | 46068907 | rs3896679 | -0.676 | 0.637 | -0.221 | 1.02 | 0.004 | -0.006 | -0.006 | 0 | 0.09 | 0.08 | 0.13 | 0.14 | 0.05 | 0.08 |
| 2 | 46073717 | rs6719779 | -0.705 | 1.43 | -0.323 | 1.246 | 0.005 | 0.008 | 0.008 | 1.282 | 0.24 | 0.27 | 0.38 | 0.36 | 0.31 | 0.31 |
| 2 | 46073997 | rs4952781 | 1.132 | 1.448 | -0.343 | 1.166 | 0.003 | 0.011 | 0.011 | 0.271 | 0.53 | 0.47 | 0.44 | 0.25 | 0.39 | 0.43 |
| 2 | 46077038 | rs10166692 | -0.548 | 1.228 | -0.269 | 1.045 | 0.001 | 0.002 | 0.002 | 0 | 0.8 | 0.81 | 0.85 | 0.72 | 0.72 | 0.8 |
| 2 | 46077503 | rs6756452 | 0.8 | 0.199 | -0.785 | 0.351 | -0.054 | -0.078 | -0.078 | 0 | 0.23 | 0.44 | 0.12 | 0.07 | 0.09 | 0.1 |
| 2 | 46078964 | rs7581914 | -0.221 | 1.429 | -1.463 | 0.904 | -0.067 | -0.078 | -0.078 | 0 | 0.22 | 0.12 | 0.41 | 0.52 | 0.4 | 0.37 |
| 2 | 46079213 | rs7558378 | -0.788 | 0.758 | -1.562 | 0.394 | 0.019 | 0.027 | 0.027 | 1.236 | 0.5 | 0.35 | 0.65 | 0.64 | 0.55 | 0.65 |
| 2 | 46082995 | rs4953274 | 0.769 | 1.614 | -1.912 | 0.99 | -0.035 | -0.034 | -0.034 | 0 | 0.1 | 0.04 | 0.17 | 0.31 | 0.21 | 0.17 |
| 2 | 46083842 | rs4953276 | 0.357 | 0.687 | -2.346 | 0.324 | 0.016 | 0.02 | 0.02 | 0 | 0.18 | 0.01 | 0.17 | 0.11 | 0.14 | 0.28 |
| 2 | 46085325 | rs1947195 | 0.173 | 0.138 | -2.096 | -0.088 | -0.021 | 0.001 | 0.001 | 0 | 0.28 | 0.3 | 0.14 | 0.27 | 0.14 | 0.25 |
| 2 | 46087256 | rs4953277 | 1.111 | 0.183 | -2.128 | -0.065 | -0.007 | -0.019 | -0.019 | 0.372 | 0.08 | 0.02 | 0.13 | 0.1 | 0.08 | 0.25 |
| 2 | 46087905 | rs12465045 | 0.458 | 0.02 | -2.255 | -0.183 | -0.024 | 0.031 | 0.031 | 0.28 | 0.18 | 0.02 | 0.18 | 0.1 | 0.15 | 0.29 |
| 2 | 46088544 | rs935653 | -0.139 | -0.198 | -2.236 | -0.215 | 0.01 | 0.01 | 0.01 | 0 | 0.23 | 0.14 | 0.2 | 0.17 | 0.16 | 0.31 |
| 2 | 46089447 | rs3911797 | -0.08 | 0.014 | -2.123 | -0.169 | 0.005 | 0.004 | 0.004 | 0 | 0.35 | 0.34 | 0.43 | 0.7 | 0.29 | 0.43 |
| 2 | 46091060 | rs10179520 | 0.179 | 0.385 | -2.227 | -0.25 | -0.08 | -0.086 | -0.086 | 0 | 0.33 | 0.13 | 0.45 | 0.52 | 0.4 | 0.42 |
| 2 | 46091279 | rs10181963 | 1.75 | 0.256 | -2.111 | -0.162 | -0.049 | -0.089 | -0.089 | 0 | 0.13 | 0.03 | 0.29 | 0.43 | 0.19 | 0.23 |
| 2 | 46093760 | rs10189339 | -1.995 | 0.1 | -1.958 | -0.529 | -0.014 | -0.085 | -0.085 | 0.082 | 0.67 | 0.49 | 0.8 | 0.91 | 0.66 | 0.89 |
| 2 | 46094473 | rs4952787 | 0.167 | -0.095 | -1.851 | -0.409 | 0.022 | 0.037 | 0.037 | 0 | 0.26 | 0.17 | 0.2 | 0.11 | 0.16 | 0.41 |
| 2 | 46096456 | rs11125038 | -0.656 | 0.218 | -1.776 | -0.004 | -0.001 | 0.025 | 0.025 | 3.49 | 0.3 | 0.39 | 0.51 | 0.52 | 0.32 | 0.57 |
| 2 | 46096684 | rs11125039 | -0.852 | 0.11 | -1.816 | -0.383 | 0.029 | 0.007 | 0.007 | 0 | 0.44 | 0.36 | 0.33 | 0.38 | 0.25 | 0.51 |
| 2 | 46097897 | rs1966813 | -0.895 | 0.519 | -1.726 | -0.794 | -0.007 | 0.066 | 0.066 | 4.986 | 0.46 | 0.37 | 0.7 | 0.89 | 0.51 | 0.75 |
| 2 | 46098608 | rs7585985 | -0.664 | 0.583 | -1.456 | -0.496 | -0.011 | -0.024 | -0.024 | 0 | 0.38 | 0.44 | 0.52 | 0.76 | 0.37 | 0.67 |
| 2 | 46100254 | rs2122110 | -0.066 | 1.162 | -1.415 | -0.526 | -0.042 | 0.021 | 0.021 | 0 | 0.15 | 0.06 | 0.2 | 0.09 | 0.21 | 0.18 |
| 2 | 46102346 | rs4245804 | -0.771 | 0.739 | -0.847 | -0.872 | 0.007 | -0.123 | -0.123 | 0 | 0.66 | 0.58 | 0.78 | 0.97 | 0.6 | 0.76 |
| 2 | 46103877 | rs13400225 | -0.334 | 0.771 | -1.674 | -0.704 | -0.018 | 0.215 | 0.215 | 0.369 | 0.56 | 0.15 | 0.69 | 0.97 | 0.57 | 0.77 |
| 2 | 46115025 | rs872288 | 0.307 | 0.655 | -1.837 | -0.31 | -0.013 | -0.038 | -0.038 | 0 | 0.39 | 0.13 | 0.38 | 0.53 | 0.34 | 0.52 |
| 2 | 46122164 | rs10185201 | 0.511 | 0.674 | -1.778 | -0.246 | -0.002 | -0.029 | -0.029 | 0 | 0.17 | 0.11 | 0.3 | 0.42 | 0.21 | 0.24 |
| 2 | 46125865 | rs10201568 | 0.031 | 0.727 | -1.666 | 0.095 | 0.003 | -0.01 | -0.01 | 8.826 | 0.41 | 0.24 | 0.51 | 0.52 | 0.37 | 0.56 |
| 2 | 46129267 | rs1375341 | -0.153 | 0.537 | -1.44 | -0.503 | -0.003 | -0.009 | -0.009 | 0 | 0.76 | 0.87 | 0.83 | 0.99 | 0.66 | 0.88 |
| 2 | 46137869 | rs1868389 | -0.349 | 0.297 | -1.365 | -0.215 | -0.001 | -0.001 | -0.001 | 0 | 0.57 | 0.54 | 0.5 | 0.55 | 0.49 | 0.62 |
| 2 | 46140804 | rs13010241 | 0.464 | 0.216 | -1.961 | -0.486 | -0.107 | 0.094 | 0.094 | 0 | 0.32 | 0.04 | 0.24 | 0.11 | 0.32 | 0.4 |
| 2 | 46158345 | rs13011605 | 0.929 | -0.042 | -2.096 | -0.653 | 0.024 | 0.084 | 0.084 | 0 | 0.15 | 0.01 | 0.05 | 0.01 | 0.08 | 0.06 |
| 2 | 46158523 | rs6720958 | 0.492 | -0.127 | -1.844 | -0.604 | 0.004 | -0.013 | -0.013 | 0 | 0.12 | 0.09 | 0.07 | 0 | 0.08 | 0.11 |
| 2 | 46158690 | rs13036100 | 0.331 | 0.271 | -1.781 | -0.141 | -0.122 | 0.021 | 0.021 | 0 | 0.32 | 0.03 | 0.24 | 0.16 | 0.33 | 0.39 |
| 2 | 46160405 | rs12999695 | 0.098 | 0.266 | -1.382 | -0.143 | -0.017 | 0.033 | 0.033 | 0 | 0.35 | 0.23 | 0.27 | 0.16 | 0.34 | 0.4 |
| 2 | 46165972 | rs10206343 | -0.233 | -0.334 | -0.47 | 0.712 | -0.012 | 0.03 | 0.03 | 0 | 0.19 | 0.22 | 0.35 | 0.67 | 0.14 | 0.35 |
| 2 | 46166063 | rs7578578 | 0.626 | -0.082 | -0.55 | 0.353 | 0.001 | -0.009 | -0.009 | 0 | 0.67 | 0.67 | 0.51 | 0.15 | 0.61 | 0.66 |
| 2 | 46166321 | rs12712955 | 1.325 | -0.457 | -0.479 | -0.081 | 0.004 | 0.072 | 0.072 | 0 | 0.48 | 0.3 | 0.41 | 0.15 | 0.5 | 0.39 |
| 2 | 46167368 | rs2595192 | 0.443 | -0.28 | -0.29 | 0.087 | -0.009 | 0.099 | 0.099 | 0 | 0.14 | 0.18 | 0.33 | 0.59 | 0.13 | 0.19 |
| 2 | 46168929 | rs2711292 | -1.442 | -0.162 | -0.798 | -0.205 | -0.006 | 0.004 | 0.004 | 0.366 | 0.88 | 0.92 | 0.92 | 0.96 | 0.8 | 0.97 |
| 2 | 46171117 | rs2711299 | 0.443 | -0.227 | -0.788 | -0.237 | 0.001 | -0.101 | -0.101 | 2.933 | 0.69 | 0.52 | 0.82 | 0.94 | 0.69 | 0.72 |
| 2 | 46172236 | rs2711297 | 0.729 | -0.18 | -0.976 | 0.555 | 0.006 | -0.022 | -0.022 | 0 | 0.1 | 0.05 | 0.3 | 0.26 | 0.1 | 0.12 |
| 2 | 46174037 | rs12466402 | 0.874 | -0.367 | -0.837 | 0.441 | 0 | 0.006 | 0.006 | 0 | 0.59 | 0.57 | 0.48 | 0.68 | 0.56 | 0.58 |
| 2 | 46176403 | rs6750700 | -0.307 | -0.284 | -1.077 | 0.468 | -0.019 | -0.019 | -0.019 | 0 | 0.78 | 0.86 | 0.59 | 0.7 | 0.68 | 0.85 |
| 2 | 46176962 | rs6751349 | -0.656 | -0.495 | -0.924 | 0.212 | -0.035 | -0.101 | -0.101 | 0 | 0.23 | 0.38 | 0.12 | 0.02 | 0.12 | 0.27 |
| 2 | 46180993 | rs2711293 | 0.132 | 0.677 | -1.805 | 0.911 | -0.018 | -0.014 | -0.014 | 0 | 0.79 | 0.87 | 0.88 | 0.74 | 0.73 | 0.9 |
| 2 | 46181323 | rs1463162 | -0.558 | 0.58 | -1.753 | 0.46 | -0.01 | 0.019 | 0.019 | 0.116 | 0.84 | 0.91 | 0.88 | 0.96 | 0.75 | 0.96 |
| 2 | 46183424 | rs2711305 | 0.532 | 0.71 | -2.116 | 0.5 | -0.045 | 0.058 | 0.058 | 0 | 0.54 | 0.85 | 0.5 | 0.6 | 0.5 | 0.54 |

| | | | | | | | | | | | | | | | | |
|---|---|---|---|---|---|---|---|---|---|---|---|---|---|---|---|---|
| 2 | 46191983 | rs4953292 | 0.464 | 0.462 | -1.644 | 0.54 | 0 | 0 | 0 | 0 | 0.56 | 0.54 | 0.4 | 0.41 | 0.49 | 0.61 |
| 2 | 46192017 | rs2595202 | -0.013 | 0.654 | -1.739 | 0.706 | -0.031 | -0.053 | -0.053 | 0.714 | 0.14 | 0.06 | 0.42 | 0.46 | 0.2 | 0.19 |
| 2 | 46192191 | rs2595203 | 0.719 | 0.753 | -1.786 | 0.776 | -0.053 | -0.067 | -0.067 | 0 | 0.1 | 0.01 | 0.38 | 0.42 | 0.16 | 0.16 |
| 2 | 46192231 | rs6715568 | -0.398 | 0.543 | -1.542 | 0.404 | -0.007 | -0.051 | -0.051 | 1.43 | 0.09 | 0.2 | 0.05 | 0 | 0.06 | 0.09 |
| 2 | 46193347 | rs6706140 | 0.546 | 0.209 | -2.035 | 0.152 | 0 | 0.001 | 0.001 | 0 | 0.44 | 0.54 | 0.42 | 0.34 | 0.43 | 0.54 |
| 2 | 46194334 | rs7607360 | -0.595 | -0.158 | -2.662 | -0.28 | 0.066 | 0.077 | 0.077 | 0.018 | 0.48 | 0.83 | 0.32 | 0.34 | 0.36 | 0.43 |
| 2 | 46195874 | rs2711303 | -1.611 | 0 | -2.401 | -0.316 | -0.027 | 0.063 | 0.063 | 0 | 0.81 | 0.91 | 0.87 | 0.97 | 0.74 | 0.93 |
| 2 | 46196138 | rs2711301 | -1.724 | -0.012 | -2.4 | -0.31 | -0.028 | 0.066 | 0.066 | 0 | 0.81 | 0.93 | 0.88 | 0.97 | 0.76 | 0.93 |
| 2 | 46196932 | rs1562653 | 0.415 | -0.248 | -1.716 | -0.413 | -0.003 | -0.027 | -0.027 | 0 | 0.42 | 0.4 | 0.35 | 0.21 | 0.41 | 0.49 |
| 2 | 46199886 | rs6751933 | 0.824 | -0.346 | -1.604 | -0.339 | -0.005 | -0.012 | -0.012 | 0 | 0.18 | 0.33 | 0.1 | 0.14 | 0.14 | 0.12 |
| 2 | 46202338 | rs753572 | -0.368 | -0.265 | -1.673 | -0.299 | -0.001 | -0.006 | -0.006 | 0.019 | 0.37 | 0.46 | 0.51 | 0.72 | 0.33 | 0.33 |
| 2 | 46203231 | rs751237 | 0.715 | -0.25 | -1.921 | -0.357 | -0.018 | 0.077 | 0.077 | 0 | 0.37 | 0.17 | 0.28 | 0.16 | 0.4 | 0.48 |
| 2 | 46205039 | rs13396424 | 1.295 | -0.112 | -1.831 | -0.025 | -0.029 | -0.02 | -0.02 | 0.581 | 0.1 | 0.05 | 0.4 | 0.58 | 0.17 | 0.16 |
| 2 | 46205320 | rs12464563 | -0.751 | -0.332 | -1.822 | -0.16 | -0.061 | -0.053 | -0.053 | 0 | 0.69 | 0.84 | 0.46 | 0.38 | 0.57 | 0.75 |
| 2 | 46210763 | rs7594827 | 0.545 | -0.517 | -1.777 | -0.341 | 0 | 0.013 | 0.013 | 0 | 0.35 | 0.24 | 0.28 | 0.27 | 0.33 | 0.55 |
| 2 | 46211658 | rs4953294 | 1.865 | -0.334 | -1.866 | -0.365 | -0.047 | 0.027 | 0.027 | 0 | 0.17 | 0.07 | 0.18 | 0.09 | 0.27 | 0.34 |
| 2 | 46217331 | rs2245633 | -0.45 | -0.559 | -1.57 | -0.377 | 0 | -0.001 | -0.001 | 0 | 0.39 | 0.41 | 0.63 | 0.67 | 0.49 | 0.58 |
| 2 | 46217670 | rs2595214 | 0.471 | -0.485 | -1.796 | -0.239 | -0.063 | -0.096 | -0.096 | 0.107 | 0.14 | 0.04 | 0.39 | 0.58 | 0.21 | 0.19 |
| 2 | 46220888 | rs2711302 | 1.622 | -0.553 | -1.802 | 0.004 | -0.053 | -0.058 | -0.058 | 0.048 | 0.1 | 0.02 | 0.36 | 0.56 | 0.16 | 0.16 |
| 2 | 46221755 | rs12622193 | 1.23 | -0.675 | -1.584 | -0.25 | 0.027 | -0.024 | -0.024 | 0 | 0.2 | 0.3 | 0.21 | 0.11 | 0.25 | 0.33 |
| 2 | 46225778 | rs2122113 | -0.302 | -1.215 | -1.812 | -0.444 | -0.007 | 0.096 | 0.096 | 0 | 0.51 | 0.81 | 0.41 | 0.27 | 0.47 | 0.59 |
| 2 | 46225965 | rs2595218 | -0.274 | -1.168 | -1.693 | -0.395 | -0.007 | -0.013 | -0.013 | 0 | 0.38 | 0.32 | 0.5 | 0.56 | 0.39 | 0.3 |
| 2 | 46227378 | rs4953296 | 1.629 | -1.052 | -1.556 | -0.071 | 0.04 | 0.037 | 0.037 | 0 | 0.19 | 0.29 | 0.23 | 0.28 | 0.26 | 0.33 |
| 2 | 46230267 | rs6755308 | 1.012 | -1.094 | -1.685 | -0.119 | -0.005 | -0.044 | -0.044 | 3.996 | 0.23 | 0.28 | 0.12 | 0.04 | 0.19 | 0.12 |
| 2 | 46230383 | rs2595221 | 1.627 | -1.427 | -1.744 | -0.027 | -0.055 | -0.055 | -0.055 | 0 | 0.1 | 0.01 | 0.35 | 0.35 | 0.16 | 0.15 |
| 2 | 46232042 | rs2255094 | -1.213 | -1.558 | -1.745 | -0.258 | 0.027 | 0.036 | 0.036 | 0 | 0.62 | 0.83 | 0.76 | 0.69 | 0.67 | 0.69 |
| 2 | 46235359 | rs7602129 | -0.655 | -1.191 | -1.585 | -0.021 | -0.029 | -0.052 | -0.052 | 0 | 0.85 | 0.9 | 0.61 | 0.65 | 0.8 | 0.79 |
| 2 | 46237041 | rs12465869 | 1.223 | -1.23 | -1.356 | -0.002 | 0.025 | 0.04 | 0.04 | 0 | 0.19 | 0.34 | 0.23 | 0.28 | 0.24 | 0.33 |
| 2 | 46237446 | rs1987070 | 1.6 | -1.417 | -1.818 | -0.27 | -0.002 | 0.006 | 0.006 | 0.149 | 0.24 | 0.15 | 0.14 | 0.22 | 0.27 | 0.2 |
| 2 | 46240240 | rs3738901 | 0.419 | -1.107 | -1.382 | -0.576 | 0.001 | 0.085 | 0.085 | 0 | 0.25 | 0.49 | 0.28 | 0.4 | 0.26 | 0.49 |
| 2 | 46252561 | rs13402503 | 0.951 | -1.586 | -1.412 | -1.52 | -0.018 | -0.074 | -0.074 | 0 | 0.13 | 0.25 | 0.06 | 0 | 0.08 | 0.04 |
| 2 | 46256399 | rs10167883 | -0.203 | -1.697 | -1.631 | -1.669 | -0.007 | -0.024 | -0.024 | 0 | 0.27 | 0.33 | 0.15 | 0.16 | 0.23 | 0.12 |
| 2 | 46256666 | rs6544868 | -1.677 | -1.594 | -2.372 | -1.69 | -0.143 | -0.131 | -0.131 | 0 | 0.8 | 0.98 | 0.49 | 0.39 | 0.62 | 0.59 |
| 2 | 46257880 | rs13024806 | 0.708 | -1.784 | -2.379 | -1.839 | 0.009 | 0.024 | 0.024 | 0 | 0.21 | 0.13 | 0.12 | 0.16 | 0.18 | 0.16 |
| 2 | 46267270 | rs1009399 | -0.261 | -1.753 | -2.347 | -1.828 | -0.06 | 0.085 | 0.085 | 0 | 0.61 | 0.82 | 0.38 | 0.22 | 0.46 | 0.42 |
| 2 | 46269567 | rs1470598 | 0.838 | -1.842 | -2.278 | -1.971 | -0.001 | 0.094 | 0.094 | 0 | 0.36 | 0.16 | 0.23 | 0.12 | 0.35 | 0.26 |
| 2 | 46271484 | rs13420874 | 0.023 | -1.346 | -1.868 | -1.854 | 0.002 | 0.04 | 0.04 | 0 | 0.56 | 0.68 | 0.37 | 0.35 | 0.45 | 0.47 |
| 2 | 46285524 | rs13024907 | 0.048 | -0.676 | -1.572 | -1.35 | 0.003 | -0.003 | -0.003 | 0.211 | 0.58 | 0.59 | 0.39 | 0.4 | 0.51 | 0.46 |
| 2 | 46287743 | rs6751805 | 0.668 | -0.862 | -1.512 | -1.557 | -0.028 | -0.066 | -0.066 | 0.032 | 0.35 | 0.43 | 0.19 | 0.13 | 0.22 | 0.23 |
| 2 | 46290416 | rs921183 | -0.161 | -0.655 | -0.97 | -1.233 | -0.034 | -0.05 | -0.05 | 0 | 0.23 | 0.43 | 0.17 | 0.13 | 0.16 | 0.25 |
| 2 | 46291906 | rs12994911 | 0.847 | -0.235 | -1.9 | -1.414 | -0.079 | 0.062 | 0.062 | 0 | 0.21 | 0.03 | 0.18 | 0.12 | 0.29 | 0.14 |
| 2 | 46293445 | rs13405039 | 0.434 | -0.455 | -1.523 | -1.365 | -0.007 | -0.051 | -0.051 | 0.015 | 0.18 | 0.21 | 0.08 | 0 | 0.12 | 0.06 |
| 2 | 46294811 | rs12712962 | -0.69 | -0.381 | -1.401 | -1.157 | -0.003 | 0.089 | 0.089 | 0 | 0.54 | 0.74 | 0.4 | 0.3 | 0.49 | 0.5 |
| 2 | 46294981 | rs12712963 | -0.201 | 0.105 | -1.601 | -1.304 | -0.004 | 0.071 | 0.071 | 0 | 0.38 | 0.26 | 0.26 | 0.16 | 0.4 | 0.2 |
| 2 | 46297842 | rs6742625 | 0.059 | 0.214 | -1.474 | -1.285 | 0.026 | 0.05 | 0.05 | 0 | 0.44 | 0.63 | 0.32 | 0.04 | 0.38 | 0.21 |
| 2 | 46297914 | rs6742737 | -0.117 | 0.583 | -1.397 | -1.243 | -0.016 | 0.004 | 0.004 | 0 | 0.35 | 0.33 | 0.57 | 0.94 | 0.4 | 0.55 |
| 2 | 46301429 | rs3814371 | 0.114 | 0.3 | -0.639 | 0.108 | 0.047 | 0.354 | 0.354 | 0 | 0.08 | 0.28 | 0.17 | 0.45 | 0.09 | 0.2 |
| 2 | 46308729 | rs12467673 | 0.668 | 0.286 | -1.778 | -0.489 | -0.11 | -0.092 | -0.092 | 3.12 | 0.42 | 0.07 | 0.53 | 0.42 | 0.52 | 0.47 |
| 2 | 46309825 | rs6712557 | -0.017 | 0.215 | -0.933 | -0.9 | -0.001 | -0.076 | -0.076 | 0 | 0.23 | 0.32 | 0.18 | 0.04 | 0.23 | 0.15 |
| 2 | 46311301 | rs11125048 | -0.48 | 0.241 | -1.009 | -0.9 | -0.004 | -0.078 | -0.078 | 0.424 | 0.38 | 0.54 | 0.28 | 0.09 | 0.35 | 0.23 |
| 2 | 46314935 | rs1124787 | -0.134 | -0.154 | -1.178 | -0.974 | 0.031 | 0.071 | 0.071 | 0 | 0.49 | 0.4 | 0.68 | 0.89 | 0.62 | 0.56 |
| 2 | 46319124 | rs3768760 | 0.084 | -0.224 | -1.082 | -0.621 | 0.022 | 0.07 | 0.07 | 0 | 0.47 | 0.35 | 0.66 | 0.84 | 0.61 | 0.56 |
| 2 | 46320880 | rs2345955 | -0.465 | -0.657 | -1.094 | -0.716 | -0.016 | -0.017 | -0.017 | 0 | 0.78 | 0.47 | 0.75 | 0.85 | 0.81 | 0.72 |
| 2 | 46321106 | rs10167555 | -0.24 | -0.806 | -1.221 | -0.9 | -0.012 | -0.027 | -0.027 | 0 | 0.77 | 0.33 | 0.78 | 0.9 | 0.82 | 0.73 |
| 2 | 46321319 | rs12993373 | 1.117 | -0.374 | -2.041 | -0.124 | -0.173 | -0.079 | -0.079 | 0 | 0.36 | 0.05 | 0.52 | 0.4 | 0.5 | 0.31 |
| 2 | 46322951 | rs13024886 | 0.523 | -0.395 | -1.976 | -0.848 | -0.046 | 0.134 | 0.134 | 0 | 0.44 | 0.08 | 0.65 | 0.8 | 0.59 | 0.56 |
| 2 | 46326743 | rs3754573 | 0.004 | -0.153 | -1.639 | -0.649 | -0.007 | -0.035 | -0.035 | 0 | 0.34 | 0.19 | 0.28 | 0.4 | 0.29 | 0.4 |
| 2 | 46333694 | rs12712970 | -0.233 | -0.15 | -1.413 | -0.771 | -0.011 | -0.066 | -0.066 | 4.555 | 0.74 | 0.44 | 0.78 | 0.95 | 0.78 | 0.84 |
| 2 | 46336974 | rs10865212 | -0.09 | 0.097 | -1.388 | -0.795 | -0.005 | -0.067 | -0.067 | 0.156 | 0.73 | 0.45 | 0.77 | 0.95 | 0.74 | 0.83 |
| 2 | 46338536 | rs1597158 | -0.095 | -0.082 | -1.48 | -0.789 | 0.001 | -0.001 | -0.001 | 0 | 0.37 | 0.36 | 0.3 | 0.4 | 0.29 | 0.4 |
| 2 | 46340378 | rs1375055 | -0.366 | -0.015 | -1.508 | -0.92 | -0.01 | -0.087 | -0.087 | 0 | 0.75 | 0.5 | 0.78 | 0.95 | 0.78 | 0.84 |
| 2 | 46342779 | rs3768753 | 0.047 | 0.035 | -1.713 | -0.829 | -0.008 | -0.009 | -0.009 | 0 | 0.33 | 0.14 | 0.26 | 0.32 | 0.26 | 0.38 |
| 2 | 46345345 | rs12616328 | 0.128 | 0.003 | -0.625 | -1.625 | -0.009 | -0.078 | -0.078 | 0 | 0.77 | 0.68 | 0.82 | 0.97 | 0.79 | 0.81 |
| 2 | 46346716 | rs3768751 | -0.532 | -0.12 | -0.88 | -1.902 | 0.011 | 0.007 | 0.007 | 0.509 | 0.4 | 0.64 | 0.19 | 0.15 | 0.27 | 0.3 |
| 2 | 46352453 | rs3754569 | 0.925 | 0.351 | -1.242 | -1.91 | -0.053 | 0.262 | 0.262 | 0.016 | 0.49 | 0.06 | 0.63 | 0.81 | 0.55 | 0.61 |
| 2 | 46352499 | rs10495927 | -0.83 | 0.216 | -0.4 | -2.004 | -0.024 | -0.105 | -0.105 | 0.54 | 0.39 | 0.51 | 0.25 | 0.09 | 0.31 | 0.35 |
| 2 | 46353166 | rs10495928 | -0.91 | 0.109 | -0.024 | -2.117 | -0.016 | -0.062 | -0.062 | 0.369 | 0.38 | 0.47 | 0.28 | 0.16 | 0.32 | 0.33 |
| 2 | 46355051 | rs4953318 | -1.265 | 0.096 | 0.008 | -2.093 | 0.014 | 0.025 | 0.025 | 0.109 | 0.44 | 0.64 | 0.32 | 0.21 | 0.37 | 0.36 |
| 2 | 46362868 | rs13406773 | -0.669 | 0.101 | -0.019 | -2.092 | 0.007 | -0.035 | -0.035 | 0.515 | 0.35 | 0.33 | 0.21 | 0.05 | 0.26 | 0.31 |

| | | | | | | | | | | | | | | | | |
|---|---|---|---|---|---|---|---|---|---|---|---|---|---|---|---|---|
| 2 | 46363699 | rs6724315 | -0.553 | 0.365 | -0.233 | -2.114 | 0.057 | -0.101 | -0.101 | 0.106 | 0.14 | 0.45 | 0.12 | 0 | 0.15 | 0.05 |
| 2 | 46370892 | rs2053797 | 0.201 | 0.476 | -0.391 | -2.08 | 0.007 | 0.007 | 0.007 | 0.014 | 0.21 | 0.18 | 0.15 | 0.05 | 0.18 | 0.24 |
| 2 | 46371214 | rs3754568 | -1.727 | 0.353 | -0.303 | -2.075 | 0.004 | 0.128 | 0.128 | 0 | 0.52 | 0.74 | 0.38 | 0.2 | 0.5 | 0.41 |
| 2 | 46372675 | rs10495930 | -0.184 | 0.388 | -0.112 | -1.712 | 0 | -0.018 | -0.018 | 0.369 | 0.19 | 0.21 | 0.13 | 0.06 | 0.15 | 0.21 |
| 2 | 46373529 | rs7604415 | -0.207 | 0.58 | 0.135 | -1.487 | -0.014 | -0.082 | -0.082 | 0.153 | 0.13 | 0.26 | 0.06 | 0 | 0.1 | 0.07 |
| 2 | 46374529 | rs4953320 | -0.442 | 0.941 | -0.492 | -1.135 | 0.01 | -0.03 | -0.03 | 0 | 0.88 | 0.81 | 0.91 | 0.96 | 0.85 | 0.83 |
| 2 | 46385339 | rs10205024 | 0.831 | 0.802 | -0.567 | -1.586 | 0 | 0.017 | 0.017 | 0 | 0.6 | 0.38 | 0.76 | 0.88 | 0.64 | 0.64 |
| 2 | 46386298 | rs2034360 | 0.113 | 0.846 | -0.215 | -1.839 | -0.001 | -0.02 | -0.02 | 0 | 0.82 | 0.8 | 0.89 | 0.97 | 0.83 | 0.84 |
| 2 | 46387621 | rs2278775 | 1.724 | 0.782 | -0.819 | -2.381 | -0.027 | 0.23 | 0.23 | 0.044 | 0.44 | 0.12 | 0.65 | 0.86 | 0.47 | 0.54 |
| 2 | 46392775 | rs3820729 | 1.029 | 0.835 | -0.387 | -2.294 | 0.001 | 0.002 | 0.002 | 0.01 | 0.13 | 0.09 | 0.09 | 0.1 | 0.1 | 0.16 |
| 2 | 46393815 | rs11125055 | -0.12 | 0.734 | 0.195 | -2.476 | 0.003 | -0.089 | -0.089 | 0 | 0.81 | 0.71 | 0.88 | 0.97 | 0.83 | 0.85 |
| 2 | 46397631 | rs951012 | 1.196 | 0.736 | 0.244 | -2.516 | 0 | -0.051 | -0.051 | 0 | 0.61 | 0.44 | 0.78 | 0.95 | 0.66 | 0.65 |
| 2 | 46400477 | rs281477 | -0.196 | 0.781 | 0.234 | -2.46 | -0.002 | -0.086 | -0.086 | 0 | 0.81 | 0.7 | 0.88 | 0.97 | 0.84 | 0.84 |
| 2 | 46400604 | rs281476 | 0.326 | 0.794 | 0.045 | -2.416 | 0.013 | -0.128 | -0.128 | 0.02 | 0.74 | 0.58 | 0.85 | 0.97 | 0.75 | 0.78 |
| 2 | 46400936 | rs7601785 | 0.719 | 0.854 | 0.031 | -2.037 | 0.007 | -0.068 | -0.068 | 0.322 | 0.15 | 0.24 | 0.1 | 0.01 | 0.17 | 0.09 |
| 2 | 46403837 | rs281472 | 0.467 | 0.613 | -0.501 | -1.67 | -0.01 | -0.038 | -0.038 | 0 | 0.13 | 0.16 | 0.07 | 0.02 | 0.07 | 0.15 |
| 2 | 46407694 | rs281467 | -0.146 | 0.747 | -0.804 | -1.838 | 0.001 | -0.031 | -0.031 | 0 | 0.83 | 0.88 | 0.86 | 0.99 | 0.79 | 0.88 |
| 2 | 46408071 | rs2594489 | -0.458 | 0.18 | -1.777 | -1.169 | 0.058 | -0.024 | -0.024 | 0 | 0.47 | 0.81 | 0.36 | 0.44 | 0.34 | 0.56 |
| 2 | 46409412 | rs2594494 | -0.491 | -0.36 | -1.832 | -1.154 | 0.06 | -0.016 | -0.016 | 0 | 0.41 | 0.76 | 0.31 | 0.41 | 0.23 | 0.44 |
| 2 | 46410143 | rs2592156 | -0.895 | -0.002 | -2.413 | -1.46 | -0.014 | 0.026 | 0.026 | 0 | 0.8 | 0.9 | 0.85 | 0.97 | 0.74 | 0.91 |
| 2 | 46410313 | rs11693321 | 1.14 | -0.047 | -2.95 | -1.8 | -0.03 | 0.112 | 0.112 | 0 | 0.21 | 0.07 | 0.12 | 0.03 | 0.22 | 0.13 |
| 2 | 46411207 | rs281508 | 1.168 | -0.1 | -2.472 | -1.199 | 0.005 | -0.006 | -0.006 | 0.279 | 0.25 | 0.32 | 0.37 | 0.26 | 0.3 | 0.36 |
| 2 | 46411420 | rs281507 | -0.036 | -0.239 | -2.318 | -1.117 | 0.002 | 0.001 | 0.001 | 0 | 0.44 | 0.66 | 0.59 | 0.59 | 0.42 | 0.76 |
| 2 | 46412288 | rs2278774 | 1.347 | 0.209 | -1.658 | -0.427 | -0.009 | 0.036 | 0.036 | 1.421 | 0.09 | 0.12 | 0.08 | 0.3 | 0.03 | 0.01 |
| 2 | 46414434 | rs3738893 | 1.753 | 0.148 | -1.652 | -0.483 | -0.013 | 0.027 | 0.027 | 1.451 | 0.08 | 0.08 | 0.05 | 0.25 | 0.01 | 0.01 |
| 2 | 46414446 | rs14138 | 1.816 | 0.189 | -1.779 | -0.557 | -0.052 | -0.003 | -0.003 | 0 | 0.17 | 0.01 | 0.31 | 0.17 | 0.19 | 0.21 |
| 2 | 46415226 | rs281506 | 1.362 | 0.182 | -1.553 | -0.523 | 0.01 | -0.041 | -0.041 | 0.084 | 0.13 | 0.36 | 0.11 | 0.06 | 0.13 | 0.17 |
| 2 | 46415545 | rs281505 | -0.937 | 0.157 | -1.737 | -0.514 | -0.053 | -0.055 | -0.055 | 0.002 | 0.66 | 0.87 | 0.65 | 0.6 | 0.58 | 0.55 |
| 2 | 46418059 | rs7572993 | 1.434 | 0.048 | -1.482 | -0.248 | -0.009 | 0.019 | 0.019 | 2.076 | 0.08 | 0.07 | 0.03 | 0.27 | 0.01 | 0.01 |
| 2 | 46418606 | rs281499 | 0.309 | 0.094 | -1.372 | -0.299 | -0.005 | 0.055 | 0.055 | 0 | 0.4 | 0.31 | 0.48 | 0.27 | 0.43 | 0.47 |
| 2 | 46419958 | rs281498 | -1.304 | 0.38 | -1.466 | -0.36 | -0.019 | -0.029 | -0.029 | 0 | 0.72 | 0.81 | 0.65 | 0.62 | 0.63 | 0.61 |
| 2 | 46423054 | rs281493 | -0.602 | 0.296 | -1.41 | -0.547 | -0.011 | -0.012 | -0.012 | 0 | 0.53 | 0.74 | 0.56 | 0.57 | 0.5 | 0.5 |
| 2 | 46423756 | rs281491 | -1.495 | 0.374 | -1.421 | -0.538 | -0.043 | -0.082 | -0.082 | 0 | 0.74 | 0.91 | 0.68 | 0.6 | 0.66 | 0.61 |
| 2 | 46423982 | rs281489 | 0.115 | 0.407 | -1.577 | -0.537 | 0 | -0.006 | -0.006 | 0.113 | 0.42 | 0.33 | 0.48 | 0.48 | 0.42 | 0.45 |
| 2 | 46424346 | rs281488 | -1.933 | 0.617 | -1.594 | -0.512 | -0.091 | 0.028 | 0.028 | 0 | 0.8 | 0.99 | 0.67 | 0.84 | 0.74 | 0.75 |
| 2 | 46424401 | rs882586 | -0.484 | 0.578 | -1.466 | -0.479 | 0.004 | 0.007 | 0.007 | 0 | 0.59 | 0.45 | 0.57 | 0.56 | 0.54 | 0.44 |
| 2 | 46425647 | rs183984 | 1.641 | 0.487 | -1.496 | -0.4 | 0.006 | -0.037 | -0.037 | 0 | 0.11 | 0.2 | 0.08 | 0.04 | 0.13 | 0.17 |
| 2 | 46426788 | rs12621370 | 1.347 | 0.463 | -1.971 | -0.333 | 0.037 | -0.04 | -0.04 | 0.365 | 0.15 | 0.07 | 0.09 | 0.28 | 0.04 | 0.06 |
| 2 | 46427455 | rs895761 | 1.299 | 1.622 | -1.556 | 0.081 | 0.002 | 0.032 | 0.032 | 0 | 0.23 | 0.24 | 0.32 | 0.41 | 0.27 | 0.35 |
| 2 | 46427875 | rs17034764 | 1.304 | 1.35 | -1.495 | 0.14 | 0.005 | 0.036 | 0.036 | 0 | 0.23 | 0.24 | 0.3 | 0.41 | 0.27 | 0.34 |
| 2 | 46428362 | rs281518 | 1.142 | 0.535 | -1.435 | 0.231 | -0.031 | 0.006 | 0.006 | 1.276 | 0.26 | 0.42 | 0.19 | 0.29 | 0.18 | 0.23 |
| 2 | 46428466 | rs281517 | 1.022 | 1.206 | -0.679 | 0.85 | 0.003 | 0.003 | 0.003 | 0.005 | 0.42 | 0.43 | 0.27 | 0.32 | 0.33 | 0.31 |
| 2 | 46431304 | rs885181 | 0.569 | 1.456 | -1.726 | 0.879 | -0.118 | 0.033 | 0.033 | 0 | 0.24 | 0.02 | 0.39 | 0.19 | 0.33 | 0.28 |
| 2 | 46452277 | rs4952809 | 0.944 | 0.799 | -1.577 | 1.055 | -0.001 | 0.002 | 0.002 | 0 | 0.21 | 0.19 | 0.26 | 0.36 | 0.19 | 0.33 |
| 2 | 46452317 | rs4952810 | -0.504 | 0.72 | -1.587 | 0.583 | 0.059 | 0.045 | 0.045 | 1.537 | 0.47 | 0.23 | 0.69 | 0.64 | 0.62 | 0.63 |
| 2 | 46452850 | rs1446317 | -1.153 | 0.804 | -1.448 | 0.666 | 0.037 | 0.025 | 0.025 | 0.054 | 0.59 | 0.79 | 0.75 | 0.68 | 0.66 | 0.67 |
| 2 | 46454212 | rs10193766 | 0.111 | 1.657 | -1.984 | 0.938 | -0.149 | 0.081 | 0.081 | 0.001 | 0.28 | 0.03 | 0.41 | 0.18 | 0.4 | 0.3 |
| 2 | 46455316 | rs8179696 | -0.388 | 1.499 | -0.644 | 1.89 | -0.001 | 0.008 | 0.008 | 0 | 0.21 | 0.2 | 0.28 | 0.43 | 0.22 | 0.33 |
| 2 | 46463976 | rs4953334 | -1.398 | 1.591 | -0.429 | 1.688 | 0.021 | 0.047 | 0.047 | 0 | 0.15 | 0.2 | 0.2 | 0.31 | 0.21 | 0.32 |
| 2 | 46468276 | rs11888552 | -1.18 | 1.445 | -0.697 | 1.958 | -0.004 | -0.034 | -0.034 | 0.948 | 0.1 | 0.03 | 0.13 | 0.28 | 0.09 | 0.27 |
| 2 | 46469828 | rs4337520 | -0.427 | 1.374 | -0.477 | 1.571 | -0.005 | -0.009 | -0.009 | 0.187 | 0.22 | 0.27 | 0.15 | 0.15 | 0.18 | 0.11 |
| 2 | 46470109 | rs6742517 | -1.175 | 1.875 | -0.067 | 1.327 | 0.009 | -0.005 | -0.005 | 0 | 0.33 | 0.56 | 0.38 | 0.28 | 0.4 | 0.41 |
| 2 | 46475797 | rs11681778 | 0.108 | 1.61 | -0.215 | 1.352 | 0.016 | 0.02 | 0.02 | 0 | 0.7 | 0.54 | 0.68 | 0.63 | 0.68 | 0.68 |
| 2 | 46475880 | rs7564967 | -0.353 | 1.857 | -0.366 | 1.534 | 0.001 | 0.032 | 0.032 | 0 | 0.9 | 0.88 | 0.85 | 0.75 | 0.91 | 0.81 |
| 2 | 46479107 | rs881547 | 0.62 | 1.454 | -1.089 | 0.552 | 0.01 | 0.002 | 0.002 | 0 | 0.36 | 0.36 | 0.4 | 0.08 | 0.28 | 0.19 |
| 2 | 46479733 | rs11688133 | 0.104 | 1.633 | -1.669 | 0.924 | -0.024 | -0.023 | -0.023 | 0.142 | 0.07 | 0 | 0.12 | 0.34 | 0.08 | 0.29 |
| 2 | 46479838 | rs13027413 | 0.147 | 1.434 | -1.338 | -0.076 | 0.004 | 0.081 | 0.081 | 0 | 0.41 | 0.28 | 0.44 | 0.09 | 0.39 | 0.26 |
| 2 | 46484129 | rs12162357 | -0.17 | 1.249 | -0.917 | 1.204 | -0.002 | 0.048 | 0.048 | 0 | 0.08 | 0.12 | 0.14 | 0.34 | 0.09 | 0.24 |
| 2 | 46484310 | rs17034876 | -0.093 | 0.671 | -1.005 | 0.198 | -0.003 | -0.011 | -0.011 | 0 | 0.65 | 0.64 | 0.59 | 0.52 | 0.72 | 0.55 |
| 2 | 46485130 | rs6727211 | 0.579 | 0.471 | -0.794 | -0.068 | 0.001 | 0.003 | 0.003 | 0 | 0.38 | 0.31 | 0.26 | 0.35 | 0.38 | 0.46 |
| 2 | 46485481 | rs6727698 | 0.018 | 0.519 | -0.983 | -0.096 | -0.007 | 0.007 | 0.007 | 0.308 | 0.24 | 0.4 | 0.37 | 0.33 | 0.27 | 0.37 |
| 2 | 46488081 | rs17745066 | 1.081 | 0.577 | -1.197 | 0.104 | -0.006 | 0.008 | 0.008 | 0 | 0.29 | 0.19 | 0.13 | 0.18 | 0.32 | 0.23 |
| 2 | 46490132 | rs2881321 | -0.546 | 0.005 | -0.788 | -0.001 | -0.019 | -0.019 | -0.019 | 0.231 | 0.31 | 0.48 | 0.43 | 0.33 | 0.34 | 0.35 |
| 2 | 46498675 | rs7592781 | -0.32 | -0.001 | -1.203 | -0.192 | 0.003 | 0.067 | 0.067 | 0 | 0.8 | 0.96 | 0.79 | 1 | 0.87 | 0.92 |
| 2 | 46502593 | rs7581090 | -0.924 | 0.163 | -0.344 | 0.573 | 0.032 | 0.052 | 0.052 | 3.206 | 0.33 | 0.56 | 0.48 | 0.53 | 0.46 | 0.49 |
| 2 | 46504962 | rs12617123 | -0.622 | 0.266 | -0.363 | 0.652 | -0.008 | -0.01 | -0.01 | 0 | 0.25 | 0.2 | 0.35 | 0.39 | 0.38 | 0.42 |
| 2 | 46509657 | rs17802111 | -1.118 | -0.059 | -0.496 | 0.329 | -0.001 | 0.004 | 0.004 | 0 | 0.42 | 0.32 | 0.38 | 0.37 | 0.46 | 0.42 |
| 2 | 46510146 | rs12615992 | -0.341 | 0.178 | -0.291 | 0.521 | 0.027 | 0.014 | 0.014 | 0.772 | 0.23 | 0.33 | 0.31 | 0.29 | 0.4 | 0.31 |
| 2 | 46511939 | rs1000531 | -0.731 | -0.413 | -0.531 | 0.316 | -0.018 | -0.015 | -0.015 | 0 | 0.2 | 0.21 | 0.1 | 0.11 | 0.11 | 0.15 |

| | | | | | | | | | | | | | | | | |
|---|---|---|---|---|---|---|---|---|---|---|---|---|---|---|---|---|
| 2 | 46534338 | rs1867785 | 0.631 | 0.149 | -0.654 | 0.083 | 0.005 | 0.03 | 0.03 | 0 | 0.62 | 0.34 | 0.63 | 0.19 | 0.62 | 0.44 |
| 2 | 46535924 | rs2121266 | 0.564 | 0.031 | 0.097 | 0.627 | 0.001 | -0.003 | -0.003 | 0 | 0.66 | 0.66 | 0.66 | 0.35 | 0.62 | 0.63 |
| 2 | 46542408 | rs7571879 | -1.044 | 0.407 | 0.348 | 0.565 | 0.124 | 0.028 | 0.028 | 0.481 | 0.07 | 0.44 | 0.1 | 0.03 | 0.07 | 0.08 |
| 2 | 46547494 | rs6720535 | 0.855 | 0.075 | -0.319 | 0.842 | -0.01 | 0.009 | 0.009 | 0 | 0.83 | 0.81 | 0.9 | 0.74 | 0.91 | 0.7 |
| 2 | 46552047 | rs4953342 | -1.14 | 0.84 | -0.748 | 1.096 | -0.032 | -0.033 | -0.033 | 0 | 0.14 | 0.09 | 0.31 | 0.65 | 0.29 | 0.25 |
| 2 | 46553966 | rs4952820 | -1.473 | 0.599 | -0.847 | 0.539 | -0.037 | 0.005 | 0.005 | 0.003 | 0.35 | 0.23 | 0.6 | 0.67 | 0.44 | 0.44 |
| 2 | 46558541 | rs6758592 | -1.456 | 0.692 | -0.675 | 0.486 | -0.02 | 0.004 | 0.004 | 0 | 0.36 | 0.27 | 0.62 | 0.66 | 0.46 | 0.45 |
| 2 | 46567276 | rs4953353 | -1.362 | 0.093 | -1.182 | 0.491 | -0.031 | -0.037 | -0.037 | 0 | 0.36 | 0.1 | 0.58 | 0.59 | 0.37 | 0.48 |
| 2 | 46573210 | rs10191091 | -1.286 | -0.617 | -1.184 | -0.613 | -0.011 | 0.054 | 0.054 | 0 | 0.59 | 0.76 | 0.73 | 0.86 | 0.5 | 0.61 |
| 2 | 46577212 | rs6544887 | -1.285 | -0.765 | -0.517 | -0.657 | -0.018 | 0.056 | 0.056 | 0 | 0.6 | 0.78 | 0.73 | 0.85 | 0.5 | 0.63 |
| 2 | 46579675 | rs1868086 | -0.38 | -1.37 | -0.734 | -0.824 | 0.047 | 0.082 | 0.082 | 0 | 0.28 | 0.14 | 0.35 | 0.09 | 0.22 | 0.29 |
| 2 | 46579881 | rs1868088 | -0.449 | -1.558 | -0.818 | -0.919 | 0.065 | 0.123 | 0.123 | 0 | 0.3 | 0.11 | 0.35 | 0.06 | 0.2 | 0.28 |
| 2 | 46581643 | rs4953359 | -2.034 | -1.456 | -0.24 | -0.861 | -0.016 | 0.041 | 0.041 | 0 | 0.6 | 0.7 | 0.76 | 0.86 | 0.53 | 0.64 |
| 2 | 46583235 | rs6743991 | -2.116 | -1.541 | -0.314 | -0.866 | -0.043 | 0.092 | 0.092 | 0 | 0.61 | 0.84 | 0.77 | 0.87 | 0.53 | 0.66 |
| 2 | 46584573 | rs10176396 | -0.692 | -1.451 | -0.288 | -0.866 | -0.005 | -0.024 | -0.024 | 0 | 0.36 | 0.48 | 0.41 | 0.77 | 0.36 | 0.36 |
| 2 | 46584852 | rs6707241 | -2.146 | -1.456 | -0.537 | -0.86 | -0.07 | 0.198 | 0.198 | 0.828 | 0.57 | 0.87 | 0.76 | 0.87 | 0.52 | 0.66 |
| 2 | 46587034 | rs1374748 | -0.651 | -1.399 | -0.212 | -0.854 | -0.024 | -0.03 | -0.03 | 0 | 0.2 | 0.29 | 0.31 | 0.09 | 0.13 | 0.23 |
| 2 | 46590456 | rs1530633 | -2.173 | -1.468 | -0.745 | -0.914 | -0.078 | 0.173 | 0.173 | 0.592 | 0.59 | 0.92 | 0.77 | 0.86 | 0.5 | 0.66 |
| 2 | 46590791 | rs3768728 | -0.486 | -1.551 | -0.638 | -0.862 | 0.018 | 0.019 | 0.019 | 0 | 0.19 | 0.16 | 0.32 | 0.09 | 0.13 | 0.25 |
| 2 | 46593299 | rs2346176 | -1.502 | -1.3 | -0.792 | -0.619 | 0.047 | 0.183 | 0.183 | 0.618 | 0.52 | 0.91 | 0.68 | 0.76 | 0.4 | 0.57 |
| 2 | 46595363 | rs4145836 | -0.445 | -1.123 | -0.739 | -0.613 | -0.029 | 0.001 | 0.001 | 0 | 0.1 | 0 | 0.1 | 0.1 | 0.12 | 0.11 |
| 2 | 46596433 | rs1374749 | -0.967 | -1.126 | 0.004 | -0.67 | -0.002 | -0.018 | -0.018 | 0.548 | 0.47 | 0.5 | 0.71 | 0.8 | 0.44 | 0.56 |
| 2 | 46597827 | rs10178633 | 0.86 | -0.973 | -0.148 | -0.582 | 0.019 | 0.148 | 0.148 | 0.313 | 0.51 | 0.25 | 0.27 | 0.22 | 0.54 | 0.41 |
| 2 | 46599373 | rs7594912 | 1.309 | -0.937 | -0.185 | -0.589 | -0.034 | 0.156 | 0.156 | 1.41 | 0.43 | 0.16 | 0.27 | 0.19 | 0.53 | 0.36 |
| 2 | 46603260 | rs11678465 | 1.338 | -0.622 | -0.37 | -0.615 | -0.105 | 0.169 | 0.169 | 0.091 | 0.41 | 0.06 | 0.23 | 0.2 | 0.49 | 0.37 |
| 2 | 46610605 | rs7577700 | -1.307 | -0.576 | 0.841 | 0.022 | 0.057 | 0.17 | 0.17 | 4.178 | 0.06 | 0.6 | 0.1 | 0.12 | 0.05 | 0.01 |
| 2 | 46614202 | rs1868092 | 1.368 | -0.422 | 0.251 | -0.185 | -0.06 | 0.134 | 0.134 | 0.207 | 0.39 | 0.11 | 0.22 | 0.09 | 0.47 | 0.35 |
| 2 | 46617182 | rs11689694 | 0.83 | -0.489 | 0.257 | -0.153 | 0.003 | 0.005 | 0.005 | 0 | 0.47 | 0.43 | 0.31 | 0.22 | 0.53 | 0.37 |
| 2 | 46632083 | rs9677412 | 1.064 | -0.424 | -0.728 | -0.069 | -0.003 | 0.016 | 0.016 | 0 | 0.43 | 0.32 | 0.28 | 0.32 | 0.53 | 0.37 |
| 2 | 46633667 | rs11689675 | 0.02 | -0.507 | -1.826 | -0.357 | 0.017 | 0.072 | 0.072 | 0.025 | 0.7 | 0.86 | 0.57 | 0.84 | 0.72 | 0.64 |
| 2 | 46633930 | rs1109286 | 0.91 | 0.142 | -2.057 | -0.295 | -0.104 | 0.162 | 0.162 | 1.72 | 0.41 | 0.04 | 0.23 | 0.19 | 0.5 | 0.34 |
| 2 | 46635648 | rs13011481 | 0.906 | 0.029 | -1.997 | -0.289 | -0.098 | 0.197 | 0.197 | 1.585 | 0.41 | 0.05 | 0.46 | 0.18 | 0.56 | 0.37 |
| 2 | 46636967 | rs1447562 | 0.265 | 0.256 | -1.775 | -0.062 | -0.034 | 0 | 0 | 0 | 0.9 | 0.99 | 0.9 | 0.93 | 0.88 | 0.9 |
| 2 | 46641955 | rs2881327 | 0.468 | 1.209 | -0.681 | 0.244 | -0.001 | 0.008 | 0.008 | 1.264 | 0.49 | 0.57 | 0.61 | 0.59 | 0.55 | 0.51 |
| 2 | 46642249 | rs2346177 | 0.821 | 1.121 | -0.63 | 0.205 | 0 | -0.002 | -0.002 | 0 | 0.46 | 0.47 | 0.58 | 0.58 | 0.51 | 0.5 |
| 2 | 46645480 | rs6726089 | 0.466 | 0.968 | -0.451 | 0.177 | 0 | 0.001 | 0.001 | 0.036 | 0.49 | 0.54 | 0.62 | 0.6 | 0.57 | 0.51 |
| 2 | 46657089 | rs7587138 | 0.512 | 1.315 | -0.258 | 0.224 | 0 | 0.013 | 0.013 | 1.168 | 0.49 | 0.56 | 0.6 | 0.59 | 0.52 | 0.5 |
| 2 | 46663172 | rs7556828 | 0.853 | 1.525 | 0.015 | 0.193 | 0.011 | 0.012 | 0.012 | 1.918 | 0.45 | 0.24 | 0.54 | 0.59 | 0.46 | 0.49 |
| 2 | 46675665 | rs2121700 | 0.965 | 1.586 | 0.11 | 0.075 | 0.014 | 0.166 | 0.166 | 1.856 | 0.43 | 0.15 | 0.37 | 0.09 | 0.44 | 0.33 |
| 2 | 46688688 | rs1530621 | 1.06 | 1.487 | 0.025 | 0.05 | -0.012 | 0.262 | 0.262 | 2.515 | 0.42 | 0.05 | 0.34 | 0.09 | 0.42 | 0.31 |
| 2 | 46706468 | rs4953385 | 0.426 | 1.479 | 0.286 | 0.279 | 0.017 | 0.015 | 0.015 | 1.146 | 0.48 | 0.61 | 0.59 | 0.58 | 0.49 | 0.48 |
| 2 | 46709571 | rs2121698 | 1.189 | 1.499 | 0.119 | 0.167 | -0.022 | 0.261 | 0.261 | 2.742 | 0.41 | 0.05 | 0.38 | 0.09 | 0.42 | 0.31 |
| 2 | 46709607 | rs2121697 | 0.427 | 1.492 | 0.19 | 0.171 | 0.018 | 0.022 | 0.022 | 1.235 | 0.48 | 0.6 | 0.59 | 0.62 | 0.49 | 0.48 |
| 2 | 46710405 | rs7562332 | 0.536 | 1.467 | 0.212 | 0.158 | 0.005 | 0.006 | 0.006 | 0.984 | 0.47 | 0.52 | 0.59 | 0.62 | 0.49 | 0.49 |
| 2 | 46712759 | rs2276555 | 0.547 | 1.468 | 0.046 | 0.15 | 0.019 | 0.002 | 0.002 | 1.019 | 0.46 | 0.57 | 0.56 | 0.59 | 0.44 | 0.34 |
| 2 | 46716828 | rs12473626 | 0.449 | 1.451 | 0.052 | 0.072 | 0.023 | 0.011 | 0.011 | 0.837 | 0.47 | 0.6 | 0.56 | 0.61 | 0.44 | 0.42 |
| 2 | 46718090 | rs12986653 | 1.087 | 1.458 | -0.347 | -0.138 | 0.008 | 0.232 | 0.232 | 2.871 | 0.39 | 0.05 | 0.35 | 0.1 | 0.37 | 0.19 |
| 2 | 46718989 | rs2346412 | -0.053 | 1.951 | -0.279 | -0.016 | -0.024 | -0.004 | -0.004 | 0 | 0.9 | 0.94 | 0.86 | 0.91 | 0.86 | 0.89 |
| 2 | 46724016 | rs6713990 | -0.208 | 1.94 | -0.3 | -0.038 | -0.027 | -0.006 | -0.006 | 0 | 0.9 | 0.94 | 0.87 | 0.92 | 0.87 | 0.89 |
| 2 | 46728137 | rs10203773 | -0.097 | 1.778 | 0.233 | -0.38 | 0.016 | -0.097 | -0.097 | 2.204 | 0.23 | 0.46 | 0.2 | 0.04 | 0.2 | 0.31 |
| 2 | 46731454 | rs2053901 | 0.246 | 1.717 | 0.098 | -0.207 | -0.014 | 0.001 | 0.001 | 0 | 0.23 | 0.25 | 0.2 | 0.22 | 0.28 | 0.36 |
| 2 | 46732405 | rs11125080 | 0.583 | 1.758 | 0.353 | -0.07 | 0.021 | 0.084 | 0.084 | 0 | 0.87 | 0.78 | 0.7 | 0.39 | 0.81 | 0.77 |
| 2 | 46733131 | rs11677822 | 0.227 | 1.687 | 0.204 | -0.094 | -0.018 | 0 | 0 | 0.001 | 0.26 | 0.25 | 0.22 | 0.23 | 0.3 | 0.39 |
| 2 | 46738891 | rs12474287 | 0.766 | 1.615 | 0.151 | -0.081 | 0.005 | -0.001 | -0.001 | 0.22 | 0.43 | 0.38 | 0.52 | 0.61 | 0.41 | 0.37 |
| 2 | 46739157 | rs3814045 | 0.793 | 1.61 | 0.148 | -0.142 | 0.004 | 0 | 0 | 0 | 0.43 | 0.37 | 0.52 | 0.62 | 0.41 | 0.37 |
| 2 | 46746745 | rs896210 | -0.585 | 1.614 | 0.103 | -0.961 | 0 | -0.014 | -0.014 | 0.684 | 0.52 | 0.49 | 0.49 | 0.82 | 0.48 | 0.67 |
| 2 | 46746963 | rs4952830 | 1.062 | 1.4 | -0.699 | -0.959 | 0.03 | 0.147 | 0.147 | 0.995 | 0.38 | 0.18 | 0.37 | 0.08 | 0.36 | 0.17 |
| 2 | 46751545 | rs7583756 | 0.621 | 1.703 | -0.345 | -0.728 | 0.046 | 0.021 | 0.021 | 0 | 0.9 | 0.77 | 0.86 | 0.89 | 0.84 | 0.79 |
| 2 | 46751577 | rs7557190 | 0.977 | 1.427 | -0.772 | -1.027 | 0 | 0.212 | 0.212 | 2.517 | 0.39 | 0.05 | 0.36 | 0.1 | 0.38 | 0.18 |
| 2 | 46758746 | rs12997164 | 0.93 | 1.396 | -0.593 | -1.281 | 0 | 0.018 | 0.018 | 0 | 0.24 | 0.19 | 0.32 | 0.1 | 0.25 | 0.16 |
| 2 | 46762596 | rs11676473 | 0.628 | 1.367 | -0.302 | -1.414 | -0.004 | 0.002 | 0.002 | 0 | 0.26 | 0.32 | 0.33 | 0.71 | 0.21 | 0.31 |
| 2 | 46763587 | rs3814043 | -0.365 | 1.783 | -0.389 | -1.592 | -0.027 | 0.027 | 0.027 | 0 | 0.66 | 0.74 | 0.54 | 0.81 | 0.52 | 0.72 |
| 2 | 46768543 | rs1376405 | -1.121 | 1.839 | -0.394 | -1.526 | -0.017 | -0.003 | -0.003 | 0 | 0.9 | 0.94 | 0.87 | 0.92 | 0.78 | 0.88 |
| 2 | 46768559 | rs1868844 | 0.499 | 1.689 | -0.258 | -1.591 | -0.013 | -0.072 | -0.072 | 0.148 | 0.39 | 0.41 | 0.2 | 0.08 | 0.32 | 0.36 |
| 2 | 46774599 | rs1001746 | 0.696 | 1.387 | -0.283 | -1.552 | -0.004 | 0.002 | 0.002 | 0 | 0.27 | 0.33 | 0.34 | 0.71 | 0.23 | 0.3 |
| 2 | 46775059 | rs1001745 | 1.028 | 1.835 | -0.021 | -1.245 | 0.003 | 0.016 | 0.016 | 0 | 0.24 | 0.2 | 0.32 | 0.1 | 0.24 | 0.16 |
| 2 | 46777942 | rs7568909 | -0.621 | 1.905 | -0.156 | -1.149 | -0.031 | -0.003 | -0.003 | 1.095 | 0.89 | 0.94 | 0.84 | 0.91 | 0.71 | 0.87 |
| 2 | 46783890 | rs4142759 | 0.703 | 1.088 | 0.275 | -1.293 | -0.007 | 0.046 | 0.046 | 0 | 0.26 | 0.14 | 0.33 | 0.1 | 0.28 | 0.17 |
| 2 | 46793943 | rs9808496 | 0.722 | 0.701 | 0.523 | -1.418 | -0.066 | -0.065 | -0.065 | 0 | 0.19 | 0.28 | 0.08 | 0.04 | 0.08 | 0.21 |

| | | | | | | | | | | | | | | | | |
|---|---|---|---|---|---|---|---|---|---|---|---|---|---|---|---|---|
| 2 | 46799852 | rs1451153 | -0.833 | 0.789 | 0.58 | -1.447 | 0.092 | 0.193 | 0.193 | 0 | 0.51 | 0.76 | 0.47 | 0.21 | 0.39 | 0.45 |
| 2 | 46800150 | rs1451152 | -0.494 | 0.814 | 0.901 | -1.55 | 0.002 | -0.008 | -0.008 | 0.183 | 0.46 | 0.5 | 0.41 | 0.19 | 0.36 | 0.38 |
| 2 | 46805373 | rs13404342 | 0.607 | 0.763 | 1.14 | -1.408 | -0.066 | -0.072 | -0.072 | 0.701 | 0.19 | 0.32 | 0.08 | 0.04 | 0.09 | 0.2 |
| 2 | 46809621 | rs1057899 | 0.414 | 0.591 | 0.823 | -1.343 | -0.087 | -0.099 | -0.099 | 0.05 | 0.21 | 0.41 | 0.09 | 0.05 | 0.09 | 0.17 |
| 2 | 46812338 | rs1376406 | 0.175 | 0.89 | 0.886 | -1.358 | -0.022 | -0.117 | -0.117 | 0.452 | 0.22 | 0.47 | 0.14 | 0.05 | 0.19 | 0.21 |
| 2 | 46815961 | rs13410076 | 0.553 | 0.789 | 0.771 | -1.346 | -0.076 | -0.083 | -0.083 | 0 | 0.19 | 0.37 | 0.09 | 0.05 | 0.09 | 0.17 |
| 2 | 46819815 | rs2305350 | -0.864 | 0.891 | 0.138 | -1.426 | 0.012 | 0.121 | 0.121 | 0 | 0.49 | 0.63 | 0.48 | 0.19 | 0.48 | 0.44 |
| 2 | 46843631 | rs1531133 | -0.805 | 0.383 | -0.055 | -1.545 | 0.082 | 0.229 | 0.229 | 0 | 0.5 | 0.77 | 0.48 | 0.18 | 0.41 | 0.46 |
| 2 | 46845849 | rs2122715 | 0.265 | 0.285 | -0.062 | -1.598 | -0.007 | 0.05 | 0.05 | 0 | 0.28 | 0.17 | 0.35 | 0.12 | 0.31 | 0.26 |
| 2 | 46848446 | rs6735530 | 0.031 | 0.292 | -0.421 | -1.535 | -0.011 | -0.008 | -0.008 | 0.109 | 0.25 | 0.66 | 0.15 | 0.06 | 0.12 | 0.25 |
| 2 | 46850571 | rs17819022 | 0.62 | 0.28 | -0.571 | -1.618 | -0.065 | 0.114 | 0.114 | 0 | 0.27 | 0.05 | 0.33 | 0.12 | 0.3 | 0.24 |
| 2 | 46852033 | rs3087822 | -0.7 | 0.638 | -0.309 | -1.341 | 0.041 | 0.172 | 0.172 | 0 | 0.53 | 0.76 | 0.48 | 0.18 | 0.42 | 0.49 |
| 2 | 46863872 | rs6544906 | -0.718 | 0.721 | -0.599 | -1.307 | 0.03 | 0.2 | 0.2 | 0 | 0.53 | 0.78 | 0.49 | 0.18 | 0.43 | 0.49 |
| 2 | 46871726 | rs17768226 | 0.168 | 1.531 | -0.201 | -0.758 | -0.064 | 0.068 | 0.068 | 1.649 | 0.19 | 0.03 | 0.26 | 0.1 | 0.22 | 0.21 |
| 2 | 46874746 | rs10495934 | -0.128 | 1.463 | 0.888 | -0.786 | 0.01 | -0.037 | -0.037 | 0 | 0.22 | 0.29 | 0.31 | 0.12 | 0.24 | 0.26 |
| 2 | 46876958 | rs6712910 | -0.731 | 1.364 | 1.149 | -0.891 | -0.011 | -0.089 | -0.089 | 0.175 | 0.27 | 0.47 | 0.34 | 0.13 | 0.24 | 0.32 |
| 2 | 46879069 | rs2881504 | -1.054 | 1.773 | 0.788 | -0.95 | -0.001 | -0.012 | -0.012 | 0.001 | 0.3 | 0.67 | 0.38 | 0.13 | 0.34 | 0.33 |
| 2 | 46881613 | rs13417558 | 0.283 | 1.747 | 0.61 | -0.905 | -0.035 | 0.055 | 0.055 | 2.513 | 0.19 | 0.08 | 0.27 | 0.11 | 0.23 | 0.21 |
| 2 | 46882393 | rs1869836 | -0.781 | 1.657 | 0.761 | -0.909 | 0.006 | -0.024 | -0.024 | 0.034 | 0.27 | 0.69 | 0.37 | 0.13 | 0.31 | 0.31 |
| 2 | 46883739 | rs10204096 | 0.78 | 1.276 | 0.203 | -0.644 | 0.015 | 0.017 | 0.017 | 0.795 | 0.5 | 0.4 | 0.43 | 0.68 | 0.41 | 0.42 |
| 2 | 46883839 | rs6741114 | -0.624 | 1.517 | 0.219 | -0.555 | -0.036 | 0.012 | 0.012 | 0 | 0.74 | 0.83 | 0.78 | 0.83 | 0.62 | 0.72 |
| 2 | 46886360 | rs4953406 | -0.199 | 1.445 | 0.459 | -0.578 | -0.002 | 0.002 | 0.002 | 0 | 0.71 | 0.71 | 0.79 | 0.84 | 0.69 | 0.69 |
| 2 | 46886565 | rs4953408 | 0.276 | 1.495 | 0.618 | -0.633 | -0.009 | 0.012 | 0.012 | 0 | 0.57 | 0.67 | 0.5 | 0.68 | 0.52 | 0.51 |
| 2 | 46888969 | rs17769101 | 0.486 | 1.389 | 0.529 | -0.644 | 0.003 | 0.016 | 0.016 | 0.383 | 0.19 | 0.15 | 0.32 | 0.15 | 0.19 | 0.25 |
| 2 | 46889746 | rs12328738 | 0.901 | 1.44 | 0.725 | -0.667 | 0.022 | 0.029 | 0.029 | 0.524 | 0.51 | 0.34 | 0.43 | 0.68 | 0.43 | 0.44 |
| 2 | 46890538 | rs6742593 | 0.1 | 0.939 | 0.669 | -0.654 | 0.008 | -0.009 | -0.009 | 0 | 0.22 | 0.3 | 0.36 | 0.16 | 0.27 | 0.33 |
| 2 | 46891964 | rs10209278 | 0.564 | 0.92 | 0.264 | -0.164 | 0.056 | 0 | 0 | 0.338 | 0.49 | 0.32 | 0.35 | 0.56 | 0.32 | 0.38 |
| 2 | 46895097 | rs1378763 | 0.549 | 0.914 | 0.271 | 0.072 | 0.053 | 0.004 | 0.004 | 0.416 | 0.49 | 0.31 | 0.36 | 0.48 | 0.33 | 0.38 |
| 2 | 46895324 | rs17035488 | 1.293 | 0.968 | 0.373 | 0.05 | 0 | -0.003 | -0.003 | 0 | 0.15 | 0.21 | 0.32 | 0.15 | 0.18 | 0.15 |
| 2 | 46895485 | rs17035489 | 1.293 | 0.956 | 0.373 | 0.05 | 0.001 | -0.003 | -0.003 | 0 | 0.15 | 0.21 | 0.32 | 0.16 | 0.18 | 0.15 |
| 2 | 46897570 | rs6734725 | -0.608 | 1.012 | 0.261 | 0.016 | -0.049 | -0.037 | -0.037 | 0 | 0.7 | 0.87 | 0.73 | 0.65 | 0.59 | 0.63 |
| 2 | 46899525 | rs6713768 | 0.005 | 0.877 | 0.117 | 0.044 | 0.027 | 0.011 | 0.011 | 0 | 0.65 | 0.52 | 0.68 | 0.65 | 0.52 | 0.59 |
| 2 | 46902667 | rs1466268 | -0.034 | 0.906 | -0.149 | 0.024 | 0.013 | -0.004 | -0.004 | 0.35 | 0.64 | 0.4 | 0.66 | 0.73 | 0.51 | 0.6 |
| 2 | 46913726 | rs1456012 | -0.221 | 0.299 | -1.473 | -0.053 | -0.017 | 0.007 | 0.007 | 3.442 | 0.44 | 0.8 | 0.67 | 0.62 | 0.44 | 0.57 |
| 2 | 46919037 | rs4953413 | -0.239 | -0.069 | -1.695 | 0.051 | -0.007 | 0.019 | 0.019 | 3.261 | 0.53 | 0.85 | 0.74 | 0.62 | 0.57 | 0.63 |
| 2 | 46919332 | rs970711 | -0.925 | -0.031 | -1.872 | 0.148 | -0.045 | -0.082 | -0.082 | 0 | 0.72 | 0.93 | 0.84 | 0.63 | 0.7 | 0.69 |
| 2 | 46925730 | rs4952842 | 1.214 | -0.043 | -1.949 | -0.181 | -0.067 | 0.065 | 0.065 | 0.755 | 0.22 | 0.07 | 0.37 | 0.07 | 0.32 | 0.22 |
| 2 | 46925789 | rs3829835 | 1.135 | -0.076 | -1.585 | -0.197 | -0.03 | 0.042 | 0.042 | 0.816 | 0.22 | 0.12 | 0.37 | 0.07 | 0.32 | 0.22 |
| 2 | 46930776 | rs973491 | 0.84 | -0.821 | -1.298 | 0.059 | -0.033 | 0.047 | 0.047 | 0 | 0.15 | 0.2 | 0.19 | 0.42 | 0.07 | 0.35 |
| 2 | 46932523 | rs10168342 | -1.715 | -1.058 | -1.415 | -0.211 | -0.09 | -0.087 | -0.087 | 0 | 0.72 | 0.89 | 0.8 | 0.54 | 0.6 | 0.71 |
| 2 | 46932666 | rs13411633 | -0.292 | -1.341 | -1.196 | -0.335 | 0.012 | 0.031 | 0.031 | 0 | 0.49 | 0.61 | 0.53 | 0.11 | 0.45 | 0.36 |
| 2 | 46938789 | rs13025984 | 1.638 | -1.18 | -1.467 | -0.209 | -0.029 | 0.041 | 0.041 | 1.291 | 0.23 | 0.12 | 0.38 | 0.07 | 0.33 | 0.22 |
| 2 | 46948175 | rs897529 | -0.004 | -1.267 | -1.028 | -0.252 | 0.002 | 0.001 | 0.001 | 1.364 | 0.42 | 0.6 | 0.61 | 0.49 | 0.44 | 0.57 |
| 2 | 46968051 | rs10167561 | -1.604 | -1.072 | -1.442 | -0.263 | -0.04 | -0.074 | -0.074 | 0 | 0.71 | 0.83 | 0.79 | 0.56 | 0.63 | 0.69 |
| 2 | 46976834 | rs4575769 | 1.378 | -0.926 | -1.534 | -0.404 | -0.004 | 0.004 | 0.004 | 0 | 0.09 | 0.08 | 0.13 | 0.03 | 0.15 | 0.04 |
| 2 | 46987139 | rs17771838 | 1.095 | -0.974 | -1.709 | -0.449 | 0.064 | 0.08 | 0.08 | 0.355 | 0.19 | 0.03 | 0.09 | 0.04 | 0.09 | 0.11 |
| 2 | 46987767 | rs10221923 | 0.833 | -0.984 | -1.727 | -0.466 | 0.063 | 0.095 | 0.095 | 0.134 | 0.2 | 0.03 | 0.1 | 0.04 | 0.11 | 0.12 |
| 2 | 46989869 | rs4953419 | 1.453 | -1.28 | -1.7 | -0.476 | -0.039 | 0.058 | 0.058 | 1.552 | 0.25 | 0.12 | 0.39 | 0.08 | 0.35 | 0.25 |
| 2 | 46995893 | rs13424717 | 0.763 | -1.359 | -1.632 | -0.325 | 0.063 | 0.095 | 0.095 | 0.193 | 0.2 | 0.02 | 0.1 | 0.04 | 0.11 | 0.12 |
| 2 | 47004788 | rs6740539 | -1.329 | -1.319 | -1.585 | -0.063 | 0.098 | 0.007 | 0.007 | 4.577 | 0.46 | 0.87 | 0.43 | 0.49 | 0.31 | 0.38 |
| 2 | 47008964 | rs6756738 | -1.345 | -1.426 | -1.701 | 0.132 | 0.107 | 0.083 | 0.083 | 3.334 | 0.5 | 0.86 | 0.45 | 0.76 | 0.34 | 0.49 |
| 2 | 47009718 | rs11125090 | 1.488 | -1.384 | -1.466 | 0.351 | -0.002 | 0.06 | 0.06 | 0.068 | 0.05 | 0.07 | 0.02 | 0.29 | 0.03 | 0.08 |
| 2 | 47009746 | rs11125091 | -0.102 | -1.623 | -1.014 | 0.202 | 0.004 | 0.027 | 0.027 | 0 | 0.18 | 0.23 | 0.28 | 0.75 | 0.2 | 0.31 |
| 2 | 47011834 | rs9309137 | -0.793 | -1.856 | -0.669 | 0.042 | -0.009 | 0.009 | 0.009 | 0.13 | 0.27 | 0.42 | 0.17 | 0.3 | 0.2 | 0.19 |
| 2 | 47012594 | rs4953420 | -0.762 | -1.936 | -0.926 | -0.365 | -0.001 | 0.006 | 0.006 | 1.614 | 0.56 | 0.72 | 0.69 | 0.5 | 0.48 | 0.63 |
| 2 | 47012616 | rs6544923 | 1.089 | -1.865 | -1.513 | -0.391 | 0.04 | 0.049 | 0.049 | 0 | 0.15 | 0.04 | 0.08 | 0.01 | 0.08 | 0.1 |
| 2 | 47020925 | rs7577004 | -0.865 | -1.823 | -1.509 | -0.347 | -0.016 | 0.002 | 0.002 | 2.326 | 0.57 | 0.83 | 0.69 | 0.49 | 0.46 | 0.62 |
| 2 | 47021070 | rs13414893 | 0.694 | -1.521 | -1.625 | 0.065 | 0.037 | 0.045 | 0.045 | 0 | 0.15 | 0.04 | 0.08 | 0.01 | 0.08 | 0.1 |
| 2 | 47025479 | rs9967800 | 0.006 | -0.873 | -1.315 | 0.53 | -0.014 | 0.03 | 0.03 | 0 | 0.13 | 0.16 | 0.16 | 0.43 | 0.06 | 0.28 |
| 2 | 47028590 | rs17824181 | 0.326 | -0.445 | -1.798 | -0.249 | -0.029 | 0.062 | 0.062 | 0.485 | 0.3 | 0.17 | 0.46 | 0.09 | 0.39 | 0.43 |
| 2 | 47030724 | rs1867822 | -0.507 | -0.463 | -2.083 | 0.137 | -0.023 | -0.06 | -0.06 | 0.084 | 0.6 | 0.87 | 0.67 | 0.64 | 0.58 | 0.66 |
| 2 | 47032546 | rs11884764 | -0.024 | -1.009 | -1.122 | 0.053 | -0.019 | -0.067 | -0.067 | 0 | 0.11 | 0.39 | 0.08 | 0 | 0.03 | 0.02 |
| 2 | 47033078 | rs7565800 | -0.445 | -0.96 | -1.472 | 0.295 | -0.053 | 0.078 | 0.078 | 1.553 | 0.22 | 0.48 | 0.2 | 0.4 | 0.12 | 0.14 |
| 2 | 47039991 | rs935376 | -0.172 | -0.134 | -2.6 | -0.051 | -0.051 | -0.06 | -0.06 | 0 | 0.62 | 0.83 | 0.65 | 0.57 | 0.48 | 0.63 |
| 2 | 47040659 | rs935375 | -0.169 | -0.112 | -2.553 | -0.094 | -0.036 | -0.045 | -0.045 | 0.32 | 0.62 | 0.81 | 0.64 | 0.57 | 0.47 | 0.63 |
| 2 | 47045321 | rs17773492 | 0.923 | 0.233 | -2.512 | 0.201 | -0.067 | 0.065 | 0.065 | 0.539 | 0.27 | 0.06 | 0.36 | 0.15 | 0.32 | 0.36 |
| 2 | 47049409 | rs2278715 | 0.613 | 1.119 | -1.595 | 0.613 | -0.005 | -0.001 | -0.001 | 2.225 | 0.57 | 0.64 | 0.62 | 0.63 | 0.46 | 0.66 |
| 2 | 47049453 | rs1053952 | 0.223 | 1.569 | -1.796 | 1.295 | -0.055 | -0.059 | -0.059 | 0 | 0.77 | 0.91 | 0.74 | 0.68 | 0.62 | 0.74 |
| 2 | 47051551 | rs1374273 | 0.238 | 1.532 | -1.795 | 1.146 | -0.054 | -0.073 | -0.073 | 0 | 0.77 | 0.89 | 0.74 | 0.65 | 0.62 | 0.73 |

| | | | | | | | | | | | | | | | | |
|---|---|---|---|---|---|---|---|---|---|---|---|---|---|---|---|---|
| 2 | 47054981 | rs11125093 | 0.224 | 1.213 | -1.535 | 0.581 | -0.009 | 0.025 | 0.025 | 0 | 0.58 | 0.64 | 0.4 | 0.28 | 0.5 | 0.42 |
| 2 | 47055169 | rs1374274 | -0.165 | 0.192 | -1.441 | 0.089 | -0.006 | -0.011 | -0.011 | 0 | 0.38 | 0.55 | 0.41 | 0.26 | 0.25 | 0.28 |
| 2 | 47057231 | rs17035715 | -0.324 | -0.291 | -1.41 | -0.166 | 0.022 | 0.022 | 0.022 | 0 | 0.22 | 0.13 | 0.16 | 0.1 | 0.13 | 0.14 |
| 2 | 47057325 | rs10173681 | 0.796 | -0.106 | -2.054 | 0.088 | -0.01 | -0.004 | -0.004 | 0.296 | 0.31 | 0.04 | 0.24 | 0.26 | 0.23 | 0.2 |
| 2 | 47058526 | rs7586612 | 0.29 | 0.35 | -1.451 | 0.4 | 0 | 0.001 | 0.001 | 0.014 | 0.25 | 0.27 | 0.42 | 0.48 | 0.25 | 0.33 |
| 2 | 47065699 | rs2121322 | -0.563 | 0.496 | -0.84 | 0.219 | 0 | 0 | 0 | 0 | 0.43 | 0.4 | 0.61 | 0.63 | 0.38 | 0.44 |
| 2 | 47067214 | rs17774013 | 1.18 | 1.047 | -0.548 | 0.139 | 0.001 | 0 | 0 | 0.653 | 0.2 | 0.22 | 0.18 | 0.17 | 0.31 | 0.33 |
| 2 | 47069769 | rs935377 | 0.885 | 0.832 | -0.662 | 0.67 | 0 | -0.023 | -0.023 | 0.017 | 0.2 | 0.1 | 0.3 | 0.19 | 0.21 | 0.23 |
| 2 | 47070827 | rs1579862 | 0.063 | 0.959 | 0.221 | 0.058 | 0.058 | 0.06 | 0.06 | 0 | 0.28 | 0.53 | 0.29 | 0.38 | 0.37 | 0.47 |
| 2 | 47072226 | rs17035766 | -0.259 | 0.917 | -0.005 | -0.009 | 0.039 | 0.001 | 0.001 | 0 | 0.44 | 0.28 | 0.54 | 0.57 | 0.36 | 0.44 |
| 2 | 47073260 | rs17774133 | 0.923 | 1.207 | -0.038 | -0.159 | -0.101 | 0.008 | 0.008 | 0.284 | 0.19 | 0.05 | 0.22 | 0.18 | 0.31 | 0.31 |
| 2 | 47073302 | rs6719049 | 0.686 | 1.213 | 0.062 | -0.15 | -0.006 | 0.001 | 0.001 | 0.119 | 0.22 | 0.22 | 0.24 | 0.23 | 0.35 | 0.33 |
| 2 | 47075393 | rs9789429 | -0.456 | 0.996 | -0.017 | -0.462 | 0.002 | -0.028 | -0.028 | 0 | 0.53 | 0.6 | 0.69 | 0.93 | 0.55 | 0.78 |
| 2 | 47075697 | rs12614366 | -0.614 | -0.021 | 0.202 | -0.073 | 0.029 | 0.007 | 0.007 | 0.017 | 0.46 | 0.65 | 0.57 | 0.56 | 0.36 | 0.45 |
| 2 | 47076673 | rs935374 | 0.369 | 0.142 | -0.029 | 0.127 | 0.003 | -0.017 | -0.017 | 0 | 0.28 | 0.23 | 0.42 | 0.53 | 0.26 | 0.33 |
| 2 | 47077502 | rs6713911 | -0.001 | 0.43 | 0.245 | -0.004 | -0.016 | 0.153 | 0.153 | 0.017 | 0.15 | 0.44 | 0.09 | 0.21 | 0.11 | 0.15 |
| 2 | 47083252 | rs11691225 | 1.272 | 0.565 | -0.487 | -0.345 | -0.036 | 0.01 | 0.01 | 0 | 0.27 | 0.1 | 0.18 | 0.24 | 0.29 | 0.13 |
| 2 | 47084204 | rs2166478 | -0.756 | 0.509 | -0.639 | -0.423 | 0.014 | -0.076 | -0.076 | 0 | 0.63 | 0.85 | 0.47 | 0.52 | 0.63 | 0.55 |
| 2 | 47086006 | rs8622 | 1.442 | 0.578 | -0.532 | -0.446 | -0.018 | 0.01 | 0.01 | 0 | 0.27 | 0.15 | 0.19 | 0.24 | 0.29 | 0.13 |
| 2 | 47086112 | rs9225 | 0.02 | 0.546 | -0.238 | -0.484 | 0.014 | 0.003 | 0.003 | 0 | 0.51 | 0.54 | 0.43 | 0.49 | 0.59 | 0.45 |
| 2 | 47089862 | rs6749716 | 1 | 0.579 | -0.705 | -0.591 | -0.012 | 0.025 | 0.025 | 0 | 0.31 | 0.19 | 0.26 | 0.1 | 0.35 | 0.15 |
| 2 | 47095007 | rs11125098 | 0.579 | 0.571 | -0.812 | -0.48 | -0.039 | -0.002 | -0.002 | 0 | 0.22 | 0.14 | 0.22 | 0.23 | 0.3 | 0.32 |
| 2 | 47095489 | rs4953434 | 0.863 | 0.574 | -0.942 | -0.517 | -0.07 | 0.01 | 0.01 | 0 | 0.2 | 0.05 | 0.19 | 0.15 | 0.26 | 0.27 |
| 2 | 47095560 | rs17035808 | 0.736 | 0.266 | -0.542 | -0.376 | -0.045 | 0.083 | 0.083 | 0 | 0.14 | 0.41 | 0.08 | 0.16 | 0.05 | 0.17 |
| 2 | 47097257 | rs13003608 | 0.752 | 0.825 | -1.282 | -0.2 | -0.017 | 0.014 | 0.014 | 0 | 0.28 | 0.17 | 0.2 | 0.25 | 0.29 | 0.17 |
| 2 | 47098250 | rs13420349 | -1.191 | 0.915 | -0.641 | 0.376 | 0.041 | 0.226 | 0.226 | 0.307 | 0.05 | 0.21 | 0.07 | 0.22 | 0.07 | 0.13 |
| 2 | 47098968 | rs7586009 | -0.52 | 0.941 | -1.25 | -0.085 | 0.002 | -0.015 | -0.015 | 0 | 0.39 | 0.59 | 0.38 | 0.24 | 0.43 | 0.47 |
| 2 | 47111175 | rs6544933 | -0.467 | 1.025 | -1.828 | 0.294 | -0.079 | -0.127 | -0.127 | 0 | 0.76 | 0.92 | 0.5 | 0.49 | 0.68 | 0.53 |
| 2 | 47112985 | rs4334538 | 0.128 | 0.709 | -1.417 | -0.462 | 0.001 | 0 | 0 | 0 | 0.19 | 0.62 | 0.16 | 0.22 | 0.18 | 0.27 |
| 2 | 47113952 | rs1374275 | -0.524 | 0.969 | -2.138 | -0.35 | -0.099 | -0.038 | -0.038 | 0 | 0.72 | 0.89 | 0.53 | 0.35 | 0.61 | 0.55 |
| 2 | 47119469 | rs6749194 | -0.453 | 0.83 | -2.177 | -0.402 | -0.105 | -0.007 | -0.007 | 0 | 0.65 | 0.89 | 0.52 | 0.36 | 0.57 | 0.5 |
| 2 | 47121192 | rs7578781 | -0.005 | 0.403 | -1.742 | -0.615 | 0.01 | 0.082 | 0.082 | 0.085 | 0.34 | 0.17 | 0.21 | 0.12 | 0.32 | 0.2 |
| 2 | 47128954 | rs7586004 | -0.766 | 0.52 | -0.946 | -0.26 | 0.027 | 0.021 | 0.021 | 0 | 0.06 | 0.18 | 0.15 | 0.08 | 0.07 | 0.06 |
| 2 | 47133055 | rs11890177 | -1.227 | -0.073 | -0.34 | -0.952 | -0.012 | -0.058 | -0.058 | 0.041 | 0.24 | 0.62 | 0.26 | 0.08 | 0.19 | 0.13 |
| 2 | 47134904 | rs2289928 | -0.573 | -0.009 | 0.087 | -0.927 | 0.065 | 0.053 | 0.053 | 0 | 0.06 | 0.29 | 0.15 | 0.08 | 0.08 | 0.06 |
| 2 | 47145029 | rs6729621 | -1.376 | 0.149 | -0.921 | -1.153 | -0.063 | -0.072 | -0.072 | 0.207 | 0.66 | 0.95 | 0.55 | 0.38 | 0.65 | 0.51 |
| 2 | 47146046 | rs3087857 | -0.663 | 0.072 | -0.794 | -1.192 | -0.004 | -0.012 | -0.012 | 0 | 0.13 | 0.59 | 0.19 | 0.08 | 0.08 | 0.11 |
| 2 | 47149658 | rs12621059 | 0.986 | 0.204 | -0.721 | -1.403 | -0.026 | 0.001 | 0.001 | 0 | 0.17 | 0.11 | 0.15 | 0.18 | 0.27 | 0.25 |
| 2 | 47153762 | rs11690921 | 1.448 | 0.14 | -0.67 | -1.351 | -0.029 | -0.038 | -0.038 | 0 | 0.34 | 0.1 | 0.46 | 0.62 | 0.35 | 0.49 |
| 2 | 47162739 | rs12712979 | 1.04 | 0.147 | -0.472 | -1.22 | -0.001 | -0.03 | -0.03 | 0.616 | 0.12 | 0.12 | 0.07 | 0 | 0.1 | 0.02 |
| 2 | 47169003 | rs6755303 | 0.306 | 0.139 | -0.42 | -1.209 | -0.007 | -0.028 | -0.028 | 0.048 | 0.15 | 0.31 | 0.19 | 0.08 | 0.12 | 0.1 |
| 2 | 47173359 | rs13404033 | 0.72 | 0.06 | -0.688 | -1.138 | 0.025 | 0.032 | 0.032 | 0 | 0.24 | 0.15 | 0.14 | 0.1 | 0.19 | 0.14 |
| 2 | 47178571 | rs7593939 | 0.092 | 0.101 | -0.728 | -0.971 | 0 | -0.001 | -0.001 | 1.079 | 0.58 | 0.6 | 0.55 | 0.63 | 0.52 | 0.59 |
| 2 | 47181106 | rs2881586 | 0.33 | 0.086 | -0.826 | -1.06 | -0.005 | 0.019 | 0.019 | 0 | 0.54 | 0.62 | 0.54 | 0.68 | 0.5 | 0.56 |
| 2 | 47182032 | rs17035931 | 0.198 | 0.004 | -1.07 | -1.212 | 0.034 | 0.068 | 0.068 | 0.191 | 0.31 | 0.18 | 0.16 | 0.12 | 0.23 | 0.18 |
| 2 | 47182698 | rs1036294 | -1.125 | -0.059 | -1.442 | -0.63 | -0.022 | 0.059 | 0.059 | 0 | 0.62 | 0.84 | 0.64 | 0.82 | 0.57 | 0.71 |
| 2 | 47183650 | rs2278450 | -1.105 | 0.446 | -0.694 | -0.759 | 0.032 | 0.009 | 0.009 | 0.01 | 0.84 | 0.78 | 0.71 | 0.8 | 0.74 | 0.72 |
| 2 | 47184447 | rs6726941 | 1.827 | 0.109 | -1.063 | -0.93 | 0.011 | 0.036 | 0.036 | 0 | 0.11 | 0.06 | 0.04 | 0.01 | 0.07 | 0.05 |
| 2 | 47187393 | rs2241736 | 2.303 | -0.045 | -1.253 | -0.8 | -0.026 | -0.134 | -0.134 | 0.202 | 0.29 | 0.05 | 0.35 | 0.5 | 0.3 | 0.27 |
| 2 | 47189381 | rs10495941 | -0.606 | -0.289 | -0.443 | -1.007 | 0.088 | 0.022 | 0.022 | 0 | 0.79 | 0.49 | 0.68 | 0.78 | 0.73 | 0.64 |
| 2 | 47190873 | rs11674443 | 1.016 | -0.49 | -0.513 | -1.109 | -0.012 | 0.005 | 0.005 | 0 | 0.65 | 0.25 | 0.58 | 0.68 | 0.63 | 0.57 |
| 2 | 47198639 | rs7557193 | -0.786 | -0.45 | -0.088 | -1.127 | 0.082 | 0.024 | 0.024 | 0 | 0.8 | 0.6 | 0.69 | 0.78 | 0.73 | 0.63 |
| 2 | 47198924 | rs4542901 | 0.334 | -0.474 | -0.322 | -1.1 | -0.032 | -0.018 | -0.018 | 0 | 0.13 | 0.21 | 0.1 | 0.1 | 0.11 | 0.08 |
| 2 | 47200025 | rs735888 | 0.383 | -0.55 | -0.383 | -1.177 | 0.01 | -0.016 | -0.016 | 0 | 0.66 | 0.4 | 0.64 | 0.77 | 0.66 | 0.57 |
| 2 | 47201099 | rs6731862 | 0.886 | -0.392 | -0.933 | -1.089 | -0.011 | 0.066 | 0.066 | 0 | 0.54 | 0.21 | 0.54 | 0.67 | 0.56 | 0.49 |
| 2 | 47205483 | rs17036004 | 0.332 | -0.489 | -0.846 | -1.179 | 0.003 | -0.02 | -0.02 | 1.186 | 0.2 | 0.22 | 0.16 | 0 | 0.19 | 0.11 |
| 2 | 47207665 | rs1865260 | 0.422 | -0.478 | -0.928 | -1.174 | 0.004 | -0.012 | -0.012 | 0 | 0.2 | 0.19 | 0.15 | 0 | 0.19 | 0.11 |
| 2 | 47210747 | rs736683 | 0.273 | -0.374 | -0.994 | -1.112 | -0.004 | -0.001 | -0.001 | 0 | 0.7 | 0.36 | 0.6 | 0.7 | 0.66 | 0.55 |
| 2 | 47211273 | rs892381 | 1.743 | -0.293 | -1.137 | -0.889 | -0.033 | -0.009 | -0.009 | 0 | 0.34 | 0.11 | 0.39 | 0.68 | 0.36 | 0.37 |
| 2 | 47217094 | rs6753628 | -1.144 | 0.059 | -0.791 | -0.856 | -0.025 | 0.012 | 0.012 | 3.672 | 0.82 | 0.89 | 0.85 | 0.88 | 0.74 | 0.76 |
| 2 | 47220285 | rs6708635 | -0.633 | -0.293 | -0.338 | -1.093 | -0.01 | -0.081 | -0.081 | 0.646 | 0.31 | 0.43 | 0.22 | 0.1 | 0.27 | 0.22 |
| 2 | 47220840 | rs6751621 | 1.439 | -0.446 | -0.455 | -1.172 | 0.013 | 0.032 | 0.032 | 0 | 0.12 | 0.06 | 0.04 | 0.01 | 0.08 | 0.05 |
| 2 | 47221693 | rs7576915 | 0.155 | -0.411 | -0.311 | -1.183 | 0 | 0.011 | 0.011 | 0 | 0.21 | 0.17 | 0.15 | 0.01 | 0.2 | 0.15 |
| 2 | 47223543 | rs13394109 | -0.822 | -0.43 | -0.231 | -1.188 | -0.021 | -0.084 | -0.084 | 0.013 | 0.3 | 0.42 | 0.21 | 0.1 | 0.25 | 0.22 |
| 2 | 47224927 | rs17481321 | 1.09 | -0.415 | -0.247 | -1.178 | -0.004 | -0.006 | -0.006 | 0 | 0.1 | 0.09 | 0.11 | 0 | 0.12 | 0.09 |
| 2 | 47225087 | rs13432506 | -0.48 | -0.42 | -0.196 | -1.171 | -0.013 | -0.055 | -0.055 | 0.003 | 0.19 | 0.35 | 0.17 | 0.1 | 0.18 | 0.17 |
| 2 | 47227146 | rs13402320 | -0.843 | -0.381 | -0.222 | -1.177 | -0.021 | -0.083 | -0.083 | 0 | 0.31 | 0.43 | 0.21 | 0.11 | 0.26 | 0.22 |
| 2 | 47227689 | rs11125110 | -1.332 | 0.001 | -0.244 | -0.997 | -0.045 | 0.015 | 0.015 | 3.924 | 0.82 | 0.95 | 0.85 | 0.88 | 0.74 | 0.76 |
| 2 | 47231215 | rs13428696 | -0.171 | -0.215 | 0.136 | -0.948 | 0.006 | 0 | 0 | 2.422 | 0.11 | 0.25 | 0.09 | 0.1 | 0.1 | 0.09 |

| | | | | | | | | | | | | | | | | |
|---|---|---|---|---|---|---|---|---|---|---|---|---|---|---|---|---|
| 2 | 47232072 | rs11899526 | -1.53 | -0.431 | 0.213 | -1.193 | -0.003 | -0.059 | -0.059 | 0 | 0.29 | 0.59 | 0.24 | 0.11 | 0.27 | 0.2 |
| 2 | 47232495 | rs1865258 | -0.138 | -0.485 | 0.162 | -1.32 | -0.008 | 0.02 | 0.02 | 0 | 0.64 | 0.71 | 0.62 | 0.79 | 0.63 | 0.59 |
| 2 | 47233048 | rs13407983 | -0.089 | -0.29 | 0.759 | -1.081 | -0.01 | -0.095 | -0.095 | 0 | 0.16 | 0.31 | 0.13 | 0 | 0.14 | 0.09 |
| 2 | 47233749 | rs13424540 | -0.241 | -0.299 | 0.76 | -1.086 | -0.013 | -0.107 | -0.107 | 0 | 0.17 | 0.34 | 0.13 | 0 | 0.14 | 0.08 |
| 2 | 47240012 | rs10194115 | 1.078 | -0.292 | 0.694 | -1.064 | -0.01 | -0.041 | -0.041 | 0 | 0.08 | 0.25 | 0.05 | 0 | 0.05 | 0.01 |
| 2 | 47240149 | rs7581528 | 1.29 | -0.142 | 0.547 | -1.053 | -0.008 | 0.015 | 0.015 | 0 | 0.08 | 0.03 | 0.08 | 0 | 0.09 | 0.07 |
| 2 | 47241075 | rs7558419 | -0.165 | -0.305 | 0.547 | -1.247 | -0.004 | 0.012 | 0.012 | 2.192 | 0.66 | 0.68 | 0.66 | 0.8 | 0.66 | 0.56 |
| 2 | 47241345 | rs7585571 | 0.634 | -0.413 | 0.456 | -1.228 | -0.001 | 0.042 | 0.042 | 2.491 | 0.55 | 0.61 | 0.63 | 0.81 | 0.57 | 0.58 |
| 2 | 47241388 | rs6544951 | 0.462 | -0.467 | 0.472 | -1.34 | -0.022 | 0.023 | 0.023 | 1.704 | 0.62 | 0.7 | 0.69 | 0.94 | 0.53 | 0.59 |
| 2 | 47242340 | rs6739975 | 1.111 | -0.386 | 0.528 | -1.031 | 0.014 | 0.051 | 0.051 | 0.762 | 0.46 | 0.33 | 0.5 | 0.79 | 0.44 | 0.45 |
| 2 | 47243124 | rs1036292 | -0.452 | -0.351 | 0.535 | -1.046 | -0.01 | 0.014 | 0.014 | 1.507 | 0.64 | 0.67 | 0.6 | 0.82 | 0.58 | 0.58 |
| 2 | 47243312 | rs2163982 | 0.885 | -0.325 | 0.046 | -1.025 | 0.005 | 0.063 | 0.063 | 0.3 | 0.47 | 0.35 | 0.52 | 0.79 | 0.49 | 0.47 |
| 2 | 47245022 | rs4953442 | 0.29 | -0.36 | 0.029 | -1.011 | 0 | -0.029 | -0.029 | 0 | 0.54 | 0.5 | 0.55 | 0.79 | 0.52 | 0.48 |
| 2 | 47246899 | rs10164750 | -0.403 | -0.204 | -0.294 | -0.895 | -0.004 | -0.074 | -0.074 | 3.923 | 0.17 | 0.29 | 0.09 | 0.02 | 0.14 | 0.13 |
| 2 | 47248111 | rs919883 | 0.342 | -0.303 | -0.424 | -0.455 | 0.001 | 0.009 | 0.009 | 0.248 | 0.52 | 0.38 | 0.54 | 0.58 | 0.53 | 0.46 |
| 2 | 47248196 | rs919884 | 0.153 | -0.31 | -0.409 | -0.469 | 0.02 | -0.067 | -0.067 | 0 | 0.12 | 0.31 | 0.07 | 0.02 | 0.1 | 0.11 |
| 2 | 47249239 | rs3814036 | 0.746 | -0.149 | -0.918 | -0.274 | -0.028 | -0.016 | -0.016 | 0.099 | 0.43 | 0.2 | 0.51 | 0.56 | 0.49 | 0.37 |
| 2 | 47249816 | rs11695048 | 1.052 | -0.54 | -1.17 | -0.165 | -0.017 | -0.075 | -0.075 | 0 | 0.38 | 0.12 | 0.46 | 0.54 | 0.39 | 0.31 |
| 2 | 47250819 | rs12988535 | -0.323 | -0.438 | -0.844 | -0.464 | -0.018 | 0.134 | 0.134 | 2.112 | 0.45 | 0.32 | 0.25 | 0.13 | 0.41 | 0.53 |
| 2 | 47251125 | rs919880 | -1.396 | -1.474 | 0.242 | 0.272 | -0.003 | -0.018 | -0.018 | 0 | 0.42 | 0.6 | 0.5 | 0.83 | 0.41 | 0.6 |
| 2 | 47251634 | rs6713052 | -2.049 | -1.485 | 0.152 | 0.295 | 0.002 | 0.195 | 0.195 | 0.011 | 0.52 | 0.9 | 0.57 | 0.85 | 0.45 | 0.72 |
| 2 | 47255516 | rs12373629 | 0.314 | -1.372 | 0.471 | 0.432 | 0.002 | -0.001 | -0.001 | 0.736 | 0.07 | 0.09 | 0.1 | 0.05 | 0.12 | 0.06 |
| 2 | 47255723 | rs4952862 | -0.218 | -1.331 | 0.82 | 0.801 | 0.023 | 0.162 | 0.162 | 0 | 0.1 | 0.21 | 0.19 | 0.63 | 0.13 | 0.19 |
| 2 | 47256058 | rs4332965 | -1.42 | -1.442 | 0.973 | 0.844 | -0.018 | -0.016 | -0.016 | 0 | 0.39 | 0.47 | 0.41 | 0.73 | 0.31 | 0.64 |
| 2 | 47256618 | rs7575628 | -1.424 | -0.851 | 1.601 | 1.413 | 0.003 | 0.086 | 0.086 | 0.111 | 0.23 | 0.48 | 0.31 | 0.64 | 0.23 | 0.22 |
| 2 | 47257977 | rs4952863 | -2.166 | -0.863 | 1.609 | 1.382 | 0 | 0 | 0 | 0 | 0.31 | 0.65 | 0.41 | 0.7 | 0.35 | 0.28 |
| 2 | 47257989 | rs6729896 | -0.186 | -1.122 | 1.572 | 1.377 | -0.001 | 0.001 | 0.001 | 0.064 | 0.08 | 0.07 | 0.1 | 0.06 | 0.12 | 0.06 |
| 2 | 47259467 | rs1865257 | -2.222 | -1.136 | 1.617 | 1.448 | -0.001 | 0.008 | 0.008 | 0 | 0.3 | 0.61 | 0.39 | 0.6 | 0.33 | 0.27 |
| 2 | 47260839 | rs7578472 | -1.167 | -1.152 | 1.54 | 1.376 | -0.001 | 0.008 | 0.008 | 0 | 0.17 | 0.13 | 0.14 | 0.09 | 0.2 | 0.16 |
| 2 | 47261639 | rs6717063 | -1.187 | -1.307 | 1.683 | 1.486 | 0.016 | 0.218 | 0.218 | 0 | 0.2 | 0.42 | 0.29 | 0.57 | 0.22 | 0.22 |
| 2 | 47262137 | rs10495942 | 0.76 | -1.262 | 1.214 | 1.402 | 0.006 | 0.023 | 0.023 | 0 | 0.08 | 0.02 | 0.02 | 0.03 | 0.04 | 0.11 |
| 2 | 47262695 | rs11695952 | -2.567 | -1.281 | 1.193 | 1.411 | 0.004 | -0.025 | -0.025 | 0 | 0.39 | 0.81 | 0.42 | 0.65 | 0.38 | 0.38 |
| 2 | 47272245 | rs13423726 | 0.479 | -1.101 | 1.387 | 1.276 | -0.01 | -0.046 | -0.046 | 0 | 0.08 | 0.14 | 0.03 | 0 | 0.04 | 0.07 |
| 2 | 47273668 | rs12712986 | -1.409 | -1.128 | 1.34 | 1.641 | 0.004 | -0.024 | -0.024 | 0 | 0.25 | 0.3 | 0.22 | 0.16 | 0.26 | 0.17 |
| 2 | 47273732 | rs7593339 | -1.309 | -1.332 | 0.897 | 1.874 | -0.045 | 0.049 | 0.049 | 0.145 | 0.27 | 0.37 | 0.31 | 0.51 | 0.14 | 0.3 |
| 2 | 47278610 | rs11693262 | -0.781 | -1.26 | 0.738 | 1.271 | 0 | -0.012 | -0.012 | 0.064 | 0.15 | 0.14 | 0.14 | 0.03 | 0.2 | 0.09 |
| 2 | 47279605 | rs17036168 | -0.376 | -1.22 | 0.938 | 0.82 | -0.009 | 0.013 | 0.013 | 0.679 | 0.17 | 0.2 | 0.27 | 0.6 | 0.07 | 0.28 |
| 2 | 47283557 | rs2436772 | -0.828 | -0.788 | 0.933 | 0.604 | 0 | -0.016 | -0.016 | 0.052 | 0.16 | 0.21 | 0.16 | 0.09 | 0.19 | 0.11 |
| 2 | 47284168 | rs2462590 | -0.819 | -0.77 | 0.945 | 0.614 | 0.001 | -0.018 | -0.018 | 0.865 | 0.16 | 0.21 | 0.16 | 0.09 | 0.19 | 0.1 |
| 2 | 47284506 | rs2692215 | -2.516 | -1.06 | 0.684 | 0.523 | 0.09 | 0.086 | 0.086 | 0 | 0.44 | 0.92 | 0.44 | 0.7 | 0.28 | 0.44 |
| 2 | 47289279 | rs11125114 | 1.378 | -1.02 | 1.201 | 0.429 | -0.014 | -0.045 | -0.045 | 3.988 | 0.74 | 0.41 | 0.81 | 0.93 | 0.8 | 0.86 |
| 2 | 47290537 | rs2692219 | 0.675 | -0.791 | 1.243 | 0.438 | -0.011 | 0.011 | 0.011 | 0 | 0.83 | 0.9 | 0.79 | 0.93 | 0.74 | 0.88 |
| 2 | 47290570 | rs4953456 | -0.669 | -0.752 | 1.499 | 0.484 | -0.04 | 0.075 | 0.075 | 0 | 0.19 | 0.38 | 0.26 | 0.67 | 0.09 | 0.31 |
| 2 | 47295288 | rs919885 | 2.352 | -0.674 | 1.271 | 0.433 | 0.034 | 0.126 | 0.126 | 0 | 0.54 | 0.23 | 0.5 | 0.2 | 0.62 | 0.51 |
| 2 | 47297082 | rs2570515 | 1.749 | 0.114 | 1.491 | 0.638 | -0.001 | -0.059 | -0.059 | 0 | 0.74 | 0.56 | 0.77 | 0.87 | 0.73 | 0.83 |
| 2 | 47297973 | rs9973700 | 1.383 | 0.151 | 1.264 | 0.568 | 0.002 | 0 | 0 | 0 | 0.78 | 0.84 | 0.88 | 0.89 | 0.82 | 0.86 |
| 2 | 47298430 | rs7569495 | 0.694 | 0.435 | 1.33 | 0.524 | -0.001 | -0.013 | -0.013 | 0 | 0.86 | 0.88 | 0.89 | 0.95 | 0.85 | 0.91 |
| 2 | 47302095 | rs1046267 | 1.098 | 0.44 | 0.965 | 0.533 | 0.028 | -0.077 | -0.077 | 0.265 | 0.82 | 0.64 | 0.79 | 0.95 | 0.77 | 0.9 |
| 2 | 47314638 | rs9917276 | -1.102 | 0.061 | 0.598 | 1.245 | -0.051 | 0.147 | 0.147 | 0 | 0.17 | 0.39 | 0.11 | 0.65 | 0.08 | 0.3 |
| 2 | 47315956 | rs6758667 | 0.553 | 0.352 | -0.175 | 0.545 | -0.002 | 0.002 | 0.002 | 0 | 0.88 | 0.89 | 0.92 | 0.96 | 0.88 | 0.92 |
| 2 | 47317631 | rs3739098 | -1.216 | 0.215 | -0.287 | 0.457 | 0.019 | 0.055 | 0.055 | 0 | 0.21 | 0.11 | 0.13 | 0.07 | 0.17 | 0.23 |
| 2 | 47317952 | rs11900463 | -1.595 | 0.318 | 0.29 | 0.295 | -0.016 | -0.051 | -0.051 | 0 | 0.27 | 0.37 | 0.2 | 0.05 | 0.21 | 0.2 |
| 2 | 47318990 | rs10181620 | 1.152 | 0.134 | 0.463 | 1.071 | -0.015 | 0.039 | 0.039 | 0 | 0.85 | 0.79 | 0.9 | 0.67 | 0.87 | 0.7 |
| 2 | 47320846 | rs13382303 | -0.812 | 0.573 | 0.667 | 1.251 | 0 | 0.001 | 0.001 | 0 | 0.23 | 0.25 | 0.32 | 0.25 | 0.24 | 0.2 |
| 2 | 47322222 | rs4952868 | 2.171 | 0.967 | 0.012 | 1.235 | -0.059 | 0.002 | 0.002 | 0 | 0.53 | 0.11 | 0.44 | 0.38 | 0.49 | 0.45 |
| 2 | 47322330 | rs4953463 | 0.535 | 1.047 | 0.107 | 1.414 | 0.003 | 0.08 | 0.08 | 0 | 0.9 | 0.82 | 0.89 | 0.71 | 0.87 | 0.81 |
| 2 | 47323153 | rs2347605 | 0.676 | 1.105 | 0.142 | 1.266 | -0.018 | 0.037 | 0.037 | 0 | 0.84 | 0.75 | 0.89 | 0.7 | 0.85 | 0.68 |
| 2 | 47323432 | rs12470974 | 1.418 | 1.144 | -0.533 | 0.93 | -0.051 | -0.036 | -0.036 | 0 | 0.55 | 0.13 | 0.45 | 0.42 | 0.5 | 0.45 |
| 2 | 47325973 | rs2347611 | 1.384 | 1.388 | -0.192 | 0.282 | -0.022 | 0.109 | 0.109 | 0.014 | 0.55 | 0.27 | 0.39 | 0.78 | 0.52 | 0.58 |
| 2 | 47338623 | rs12988606 | -1.106 | 1.583 | 0.956 | 0.093 | 0.002 | 0.003 | 0.003 | 0 | 0.26 | 0.23 | 0.26 | 0.13 | 0.21 | 0.3 |
| 2 | 47338962 | rs11685529 | 0.767 | 1.709 | 1.261 | 0.108 | 0.138 | 0.085 | 0.085 | 0 | 0.92 | 0.62 | 0.87 | 0.93 | 0.89 | 0.95 |
| 2 | 47339899 | rs949724 | -1.126 | 1.724 | 1.217 | 0.416 | 0.005 | -0.004 | -0.004 | 0 | 0.25 | 0.21 | 0.25 | 0.29 | 0.19 | 0.3 |
| 2 | 47340890 | rs12478385 | 2.179 | 1.799 | 1.154 | 0.349 | -0.104 | 0.028 | 0.028 | 0 | 0.58 | 0.12 | 0.49 | 0.67 | 0.55 | 0.58 |
| 2 | 47342343 | rs12612191 | -0.275 | 1.709 | 1.164 | 0.358 | 0.058 | -0.056 | -0.056 | 0 | 0.14 | 0.02 | 0.05 | 0.23 | 0.05 | 0.05 |
| 2 | 47346185 | rs17036272 | -0.655 | 1.919 | 1.518 | 0.058 | 0.071 | -0.036 | -0.036 | 0 | 0.07 | 0.2 | 0.11 | 0 | 0.13 | 0.06 |
| 2 | 47346329 | rs10210315 | 0.663 | 1.914 | 1.571 | 0.048 | 0.183 | 0.085 | 0.085 | 0.013 | 0.93 | 0.59 | 0.87 | 0.95 | 0.89 | 0.96 |
| 2 | 47348347 | rs1880583 | 2.132 | 1.892 | 1.352 | 0.041 | 0.061 | -0.056 | -0.056 | 0.282 | 0.84 | 0.35 | 0.75 | 0.94 | 0.76 | 0.89 |
| 2 | 47355143 | rs7596776 | 2.501 | 1.718 | 1.159 | 0.033 | -0.114 | 0.02 | 0.02 | 0.845 | 0.62 | 0.09 | 0.49 | 0.73 | 0.56 | 0.6 |
| 2 | 47362229 | rs7596354 | 2.215 | 1.996 | 1.249 | 0.021 | -0.05 | 0.028 | 0.028 | 0.045 | 0.63 | 0.24 | 0.49 | 0.74 | 0.52 | 0.57 |

| | | | | | | | | | | | | | | | | |
|---|---|---|---|---|---|---|---|---|---|---|---|---|---|---|---|---|
| 2 | 47362374 | rs7584243 | 0.708 | 2.055 | 1.453 | 0.038 | 0.133 | 0.109 | 0.109 | 0 | 0.93 | 0.58 | 0.88 | 0.93 | 0.91 | 0.96 |
| 2 | 47368754 | rs1534110 | 1.547 | 2.018 | 1.307 | 0.066 | 0.118 | -0.025 | -0.025 | 0 | 0.86 | 0.45 | 0.76 | 0.93 | 0.76 | 0.89 |
| 2 | 47372425 | rs726117 | 2.354 | 1.715 | 0.9 | -0.115 | -0.037 | 0.039 | 0.039 | 0 | 0.6 | 0.27 | 0.48 | 0.69 | 0.51 | 0.55 |
| 2 | 47374397 | rs1693892 | -1.679 | 1.083 | 1.381 | 0.091 | -0.005 | -0.007 | -0.007 | 0 | 0.26 | 0.37 | 0.29 | 0.24 | 0.25 | 0.34 |
| 2 | 47380145 | rs17036300 | -1.348 | 1.136 | 1.442 | -0.038 | 0.019 | -0.036 | -0.036 | 0 | 0.09 | 0.28 | 0.16 | 0.03 | 0.07 | 0.24 |
| 2 | 47381097 | rs815808 | 2.134 | 1.64 | 1.905 | -0.093 | 0.082 | -0.061 | -0.061 | 0 | 0.84 | 0.59 | 0.72 | 0.9 | 0.8 | 0.69 |
| 2 | 47382300 | rs815805 | -0.393 | 1.684 | 1.931 | -0.028 | -0.003 | -0.036 | -0.036 | 0 | 0.11 | 0.02 | 0.09 | 0.15 | 0.09 | 0.07 |
| 2 | 47382342 | rs815804 | -0.956 | 1.733 | 2.031 | -0.059 | -0.007 | -0.001 | -0.001 | 0.081 | 0.16 | 0.1 | 0.13 | 0.15 | 0.18 | 0.08 |
| 2 | 47385182 | rs10865222 | 2.719 | 1.701 | 1.99 | -0.033 | -0.095 | 0.007 | 0.007 | 0 | 0.64 | 0.13 | 0.54 | 0.67 | 0.59 | 0.6 |
| 2 | 47387187 | rs17036320 | -1.928 | 1.617 | 2.197 | -0.079 | -0.001 | -0.019 | -0.019 | 0 | 0.11 | 0.3 | 0.16 | 0.06 | 0.08 | 0.24 |
| 2 | 47390058 | rs17036325 | -1.507 | 1.716 | 2.251 | -0.028 | 0.015 | 0.001 | 0.001 | 0 | 0.1 | 0.28 | 0.16 | 0.06 | 0.08 | 0.24 |
| 2 | 47392049 | rs815802 | 0.462 | 1.772 | 2.125 | 0.022 | 0.031 | 0.031 | 0.031 | 0 | 0.95 | 0.89 | 0.91 | 0.89 | 0.89 | 0.94 |
| 2 | 47398394 | rs815816 | 1.335 | 1.796 | 2.216 | 0.023 | 0.088 | 0.037 | 0.037 | 0 | 0.92 | 0.83 | 0.87 | 0.89 | 0.86 | 0.93 |
| 2 | 47406539 | rs1723482 | 1.782 | 1.656 | 2.081 | -0.067 | 0.061 | -0.029 | -0.029 | 0 | 0.83 | 0.53 | 0.82 | 0.9 | 0.84 | 0.92 |
| 2 | 47418041 | rs13013607 | -2.446 | 1.694 | 1.961 | -0.206 | 0.104 | -0.039 | -0.039 | 1.997 | 0.09 | 0.36 | 0.1 | 0 | 0.09 | 0.03 |
| 2 | 47423575 | rs7581908 | 2.397 | 1.384 | 1.692 | -0.398 | 0.003 | 0.038 | 0.038 | 0 | 0.73 | 0.24 | 0.67 | 0.91 | 0.79 | 0.71 |
| 2 | 47423591 | rs9808216 | -2.31 | 1.162 | 2.142 | -0.308 | 0.023 | -0.071 | -0.071 | 0.167 | 0.12 | 0.37 | 0.19 | 0.03 | 0.09 | 0.17 |
| 2 | 47426010 | rs6747816 | -2.083 | 1.297 | 1.706 | -0.161 | 0.03 | -0.03 | -0.03 | 1.471 | 0.24 | 0.42 | 0.21 | 0.14 | 0.24 | 0.11 |
| 2 | 47433968 | rs17557422 | -2.207 | 1.017 | 1.341 | -0.03 | 0.033 | 0.02 | 0.02 | 1.53 | 0.31 | 0.84 | 0.25 | 0.27 | 0.2 | 0.28 |
| 2 | 47435823 | rs1021609 | -2.347 | 0.93 | 1.649 | -0.077 | 0 | 0.001 | 0.001 | 0.055 | 0.33 | 0.68 | 0.3 | 0.28 | 0.26 | 0.36 |
| 2 | 47438851 | rs4953477 | -2.102 | 0.956 | 1.764 | -0.08 | -0.001 | 0.003 | 0.003 | 0 | 0.29 | 0.64 | 0.3 | 0.24 | 0.24 | 0.22 |
| 2 | 47448865 | rs904166 | -2.042 | 0.939 | 1.9 | -0.143 | 0.002 | -0.015 | -0.015 | 0 | 0.28 | 0.57 | 0.29 | 0.22 | 0.24 | 0.22 |
| 2 | 47449545 | rs904167 | -2.439 | 0.823 | 1.44 | -0.291 | -0.012 | 0.056 | 0.056 | 0.08 | 0.33 | 0.85 | 0.33 | 0.22 | 0.26 | 0.22 |
| 2 | 47449602 | rs6744385 | -0.962 | 0.877 | 1.642 | -0.222 | -0.004 | 0.006 | 0.006 | 0.011 | 0.14 | 0.32 | 0.12 | 0.13 | 0.1 | 0.07 |
| 2 | 47452710 | rs998587 | -1.234 | 0.909 | 1.089 | -0.215 | 0.028 | -0.026 | -0.026 | 0 | 0.14 | 0.32 | 0.18 | 0.09 | 0.14 | 0.14 |
| 2 | 47454527 | rs6544972 | -2.611 | 0.915 | 1 | -0.166 | -0.02 | 0.032 | 0.032 | 0.686 | 0.35 | 0.9 | 0.33 | 0.25 | 0.27 | 0.35 |
| 2 | 47455238 | rs13430277 | -1.308 | 1.177 | 1.283 | 0.015 | 0.036 | -0.027 | -0.027 | 0.072 | 0.15 | 0.34 | 0.19 | 0.09 | 0.16 | 0.14 |
| 2 | 47461253 | rs17036385 | -1.338 | 0.778 | 0.901 | -0.02 | -0.01 | -0.007 | -0.007 | 0 | 0.13 | 0.22 | 0.08 | 0.13 | 0.09 | 0.06 |
| 2 | 47461518 | rs1880585 | -1.263 | 1.134 | 0.583 | -0.068 | -0.004 | 0.009 | 0.009 | 0.01 | 0.2 | 0.15 | 0.14 | 0.2 | 0.19 | 0.08 |
| 2 | 47462074 | rs7595027 | -1.216 | 1.786 | 1.092 | -0.156 | 0.05 | -0.035 | -0.035 | 0 | 0.2 | 0.37 | 0.16 | 0.18 | 0.23 | 0.08 |
| 2 | 47462144 | rs13429790 | -1.448 | 2.119 | 1.089 | 0.12 | 0.046 | 0.08 | 0.08 | 0 | 0.09 | 0.33 | 0.15 | 0.11 | 0.09 | 0.09 |
| 2 | 47462311 | rs13404793 | -1.468 | 2.115 | 1.065 | 0.118 | 0.041 | 0.074 | 0.074 | 0 | 0.09 | 0.32 | 0.15 | 0.11 | 0.09 | 0.09 |
| 2 | 47462849 | rs6733907 | -1.936 | 1.399 | -0.147 | 0.22 | -0.023 | -0.011 | -0.011 | 0.324 | 0.38 | 0.82 | 0.33 | 0.37 | 0.34 | 0.35 |
| 2 | 47468331 | rs11899692 | -0.937 | 1.003 | -0.392 | 0.182 | -0.001 | -0.002 | -0.002 | 0 | 0.43 | 0.55 | 0.34 | 0.64 | 0.4 | 0.6 |
| 2 | 47468955 | rs7598661 | -1.308 | 0.588 | -1.546 | -0.144 | -0.086 | 0.037 | 0.037 | 0 | 0.62 | 0.94 | 0.59 | 0.78 | 0.54 | 0.76 |
| 2 | 47469711 | rs10191479 | 0.286 | 0.464 | -1.47 | -0.137 | -0.003 | 0.053 | 0.053 | 0 | 0.19 | 0.09 | 0.12 | 0.08 | 0.17 | 0.33 |
| 2 | 47476780 | rs12105518 | 0.168 | 0.443 | -0.857 | -0.145 | -0.002 | -0.008 | -0.008 | 0.393 | 0.25 | 0.25 | 0.28 | 0.14 | 0.22 | 0.19 |
| 2 | 47476816 | rs10209637 | -0.344 | 0.55 | -0.778 | -0.116 | 0.004 | 0.013 | 0.013 | 0.226 | 0.35 | 0.45 | 0.31 | 0.62 | 0.35 | 0.57 |
| 2 | 47477584 | rs904160 | -1.492 | 0.533 | -1.282 | -0.075 | -0.052 | 0.045 | 0.045 | 0 | 0.62 | 0.92 | 0.6 | 0.77 | 0.57 | 0.76 |
| 2 | 47479676 | rs4953485 | -1.617 | 0.65 | -1.427 | -0.244 | -0.015 | -0.011 | -0.011 | 0 | 0.79 | 0.81 | 0.74 | 0.76 | 0.68 | 0.79 |
| 2 | 47480107 | rs10186190 | -0.402 | 0.169 | -1.36 | -0.375 | -0.001 | -0.023 | -0.023 | 0 | 0.31 | 0.65 | 0.28 | 0.57 | 0.29 | 0.4 |
| 2 | 47483458 | rs931640 | 0.437 | 0.162 | -1.291 | -0.407 | 0.001 | 0.006 | 0.006 | 0 | 0.11 | 0.08 | 0.05 | 0.04 | 0.11 | 0.18 |
| 2 | 47483905 | rs11125128 | 0.596 | 0.136 | -0.672 | -0.397 | 0.036 | 0.217 | 0.217 | 1.507 | 0.19 | 0.37 | 0.22 | 0.55 | 0.19 | 0.23 |
| 2 | 47484256 | rs6751203 | -1.03 | -0.003 | -1.512 | -0.435 | 0.007 | -0.015 | -0.015 | 0 | 0.42 | 0.79 | 0.47 | 0.63 | 0.37 | 0.55 |
| 2 | 47486300 | rs13002829 | 1.472 | -0.104 | -1.33 | -0.319 | -0.002 | 0.019 | 0.019 | 0 | 0.1 | 0.03 | 0.17 | 0.04 | 0.08 | 0.14 |
| 2 | 47489587 | rs2055226 | -1.032 | -0.563 | -1.532 | -0.456 | -0.022 | 0.031 | 0.031 | 0 | 0.54 | 0.8 | 0.52 | 0.68 | 0.53 | 0.59 |
| 2 | 47494799 | rs11125132 | 0.78 | -0.529 | -0.805 | 0.241 | -0.002 | -0.001 | -0.001 | 0 | 0.38 | 0.39 | 0.47 | 0.56 | 0.42 | 0.49 |
| 2 | 47502757 | rs7568590 | -0.789 | -0.589 | -0.599 | 0.149 | -0.001 | -0.003 | -0.003 | 0 | 0.65 | 0.73 | 0.61 | 0.58 | 0.66 | 0.72 |
| 2 | 47510324 | rs904162 | 0.326 | -0.623 | -0.501 | -0.112 | -0.004 | 0.015 | 0.015 | 0 | 0.47 | 0.45 | 0.49 | 0.33 | 0.49 | 0.53 |
| 2 | 47511181 | rs11898296 | 0.992 | -0.559 | -0.615 | -0.299 | -0.002 | 0.013 | 0.013 | 0 | 0.37 | 0.32 | 0.42 | 0.24 | 0.4 | 0.34 |
| 2 | 47514011 | rs10204202 | 0.672 | -0.417 | -0.807 | -0.231 | -0.001 | -0.017 | -0.017 | 0 | 0.11 | 0.1 | 0.05 | 0.03 | 0.09 | 0.16 |
| 2 | 47514399 | rs11894871 | -0.607 | -0.577 | -0.695 | -0.053 | 0.001 | 0.001 | 0.001 | 1.358 | 0.69 | 0.71 | 0.68 | 0.42 | 0.68 | 0.72 |
| 2 | 47519899 | rs965721 | -0.525 | -0.856 | -1.839 | -0.079 | -0.087 | -0.087 | -0.087 | 0 | 0.67 | 0.95 | 0.58 | 0.43 | 0.67 | 0.69 |
| 2 | 47520035 | rs10206550 | -0.219 | -0.761 | -0.95 | -0.251 | 0.017 | -0.078 | -0.078 | 2.917 | 0.14 | 0.27 | 0.08 | 0.02 | 0.17 | 0.08 |
| 2 | 47522368 | rs10192326 | -0.217 | -1.214 | -1.695 | 0.101 | 0.073 | 0.039 | 0.039 | 0 | 0.18 | 0.02 | 0.16 | 0.14 | 0.1 | 0.17 |
| 2 | 47522487 | rs10168995 | -0.201 | -1.199 | -1.684 | 0.109 | 0.075 | 0.043 | 0.043 | 0 | 0.19 | 0.04 | 0.17 | 0.14 | 0.1 | 0.17 |
| 2 | 47538477 | rs4341988 | -1.063 | -0.783 | -1.614 | 0.17 | -0.048 | 0.045 | 0.045 | 0 | 0.67 | 0.89 | 0.72 | 0.81 | 0.61 | 0.75 |
| 2 | 47538683 | rs4245814 | -0.349 | -0.202 | -0.723 | 0.531 | 0.013 | 0.009 | 0.009 | 0.147 | 0.13 | 0.13 | 0.23 | 0.09 | 0.07 | 0.16 |
| 2 | 47540532 | rs7604511 | -0.406 | 0.098 | -0.679 | 0.722 | -0.016 | -0.001 | -0.001 | 0.212 | 0.41 | 0.65 | 0.45 | 0.41 | 0.41 | 0.31 |
| 2 | 47552277 | rs6712843 | -0.528 | -0.518 | -0.884 | 0.624 | 0.055 | 0.061 | 0.061 | 3.661 | 0.49 | 0.34 | 0.51 | 0.69 | 0.34 | 0.37 |
| 2 | 47563573 | rs10514804 | 0.023 | -0.722 | -1.286 | 0.979 | 0.015 | -0.018 | -0.018 | 0 | 0.08 | 0.04 | 0.29 | 0.14 | 0.04 | 0.12 |
| 2 | 47564034 | rs4422207 | -0.396 | -0.73 | -0.843 | 0.741 | 0.047 | 0.04 | 0.04 | 3.048 | 0.47 | 0.64 | 0.52 | 0.69 | 0.32 | 0.35 |
| 2 | 47565795 | rs10865224 | -0.677 | -0.213 | -0.81 | 0.548 | 0 | 0.027 | 0.027 | 3.457 | 0.59 | 0.67 | 0.59 | 0.82 | 0.45 | 0.63 |
| 2 | 47570839 | rs13388259 | -0.314 | -0.68 | -0.632 | 1.296 | -0.03 | 0.028 | 0.028 | 0.422 | 0.16 | 0.18 | 0.08 | 0.23 | 0.09 | 0.22 |
| 2 | 47571306 | rs11688926 | -0.747 | -1.19 | -0.701 | 1.305 | 0.004 | -0.001 | -0.001 | 0 | 0.29 | 0.28 | 0.18 | 0.28 | 0.21 | 0.22 |
| 2 | 47599283 | rs10185866 | -1 | -0.091 | -0.641 | 2.197 | -0.001 | 0.002 | 0.002 | 0 | 0.24 | 0.21 | 0.29 | 0.57 | 0.32 | 0.3 |
| 2 | 47599454 | rs17036544 | -0.602 | 0.178 | -0.595 | 2.411 | -0.001 | 0.043 | 0.043 | 0 | 0.05 | 0.07 | 0.14 | 0.26 | 0.09 | 0.13 |
| 2 | 47606683 | rs3923560 | -1.291 | 0.502 | 0.04 | 2.351 | 0.025 | 0.137 | 0.137 | 0 | 0.07 | 0.14 | 0.13 | 0.26 | 0.08 | 0.13 |
| 2 | 47612156 | rs4952886 | -1.271 | 0.508 | 0.063 | 2.418 | 0.03 | 0.11 | 0.11 | 0 | 0.06 | 0.15 | 0.13 | 0.17 | 0.08 | 0.12 |

| | | | | | | | | | | | | | | | | |
|---|---|---|---|---|---|---|---|---|---|---|---|---|---|---|---|---|
| 2 | 47616087 | rs6744343 | -0.578 | 0.328 | 0.567 | 2.68 | 0.001 | 0.06 | 0.06 | 0.521 | 0.11 | 0.26 | 0.08 | 0.25 | 0.09 | 0.14 |
| 2 | 47619408 | rs11692854 | -0.296 | 0.332 | 0.335 | 2.783 | -0.015 | -0.037 | -0.037 | 0 | 0.1 | 0.04 | 0.17 | 0.41 | 0.1 | 0.14 |
| 2 | 47629898 | rs1863332 | -1.2 | -0.089 | 0.452 | 2.264 | -0.006 | 0.03 | 0.03 | 0 | 0.09 | 0.17 | 0.19 | 0.15 | 0.1 | 0.14 |
| 2 | 47646968 | rs4952887 | -1.144 | -0.277 | 0.416 | 2.187 | 0 | 0.01 | 0.01 | 0 | 0.09 | 0.11 | 0.15 | 0.15 | 0.1 | 0.13 |
| 2 | 47657684 | rs6753135 | -0.566 | -0.979 | -0.016 | 1.62 | 0.032 | 0.035 | 0.035 | 0 | 0.19 | 0.13 | 0.09 | 0.01 | 0.12 | 0.12 |
| 2 | 47671565 | rs7607076 | -1.281 | -1.087 | 0.086 | 1.61 | 0.003 | 0.011 | 0.011 | 2.018 | 0.29 | 0.59 | 0.32 | 0.67 | 0.3 | 0.36 |
| 2 | 47672569 | rs1981929 | -2.743 | -0.904 | -0.832 | 1.071 | -0.075 | 0.095 | 0.095 | 0.16 | 0.68 | 0.95 | 0.77 | 0.87 | 0.61 | 0.72 |
| 2 | 47679978 | rs10514805 | 0.478 | -1.076 | -0.506 | 1.027 | 0.068 | 0.09 | 0.09 | 0.004 | 0.16 | 0.02 | 0.04 | 0.01 | 0.08 | 0.11 |
| 2 | 47698702 | rs17036614 | 0.761 | -1.015 | -0.495 | 1.206 | -0.017 | -0.049 | -0.049 | 0 | 0.1 | 0.01 | 0.17 | 0.52 | 0.09 | 0.14 |
| 2 | 47700957 | rs3821227 | -1.749 | -1.037 | -0.295 | 1.147 | 0.008 | 0.011 | 0.011 | 2.888 | 0.42 | 0.71 | 0.38 | 0.68 | 0.39 | 0.44 |
| 2 | 47703984 | rs4608577 | 0.532 | -0.924 | -0.013 | 1.215 | 0.02 | 0.002 | 0.002 | 0 | 0.15 | 0.22 | 0.11 | 0.14 | 0.21 | 0.18 |
| 2 | 47712780 | rs6544991 | 0.341 | -0.953 | 0.035 | 1.28 | 0.025 | 0.078 | 0.078 | 0 | 0.16 | 0.24 | 0.22 | 0.34 | 0.22 | 0.21 |
| 2 | 47714419 | rs6720549 | -1.311 | -0.961 | -0.072 | 0.965 | -0.004 | -0.012 | -0.012 | 0 | 0.31 | 0.71 | 0.34 | 0.69 | 0.31 | 0.35 |
| 2 | 47715944 | rs6544992 | -1.076 | -0.95 | -0.029 | 0.975 | 0.009 | 0.028 | 0.028 | 0.879 | 0.28 | 0.52 | 0.33 | 0.69 | 0.3 | 0.35 |
| 2 | 47740566 | rs876937 | -1.834 | -0.889 | -0.216 | 0.924 | 0.032 | 0.034 | 0.034 | 0.961 | 0.47 | 0.66 | 0.38 | 0.68 | 0.42 | 0.47 |
| 2 | 47750227 | rs6744097 | -2.357 | -1.029 | -0.69 | 0.619 | -0.062 | -0.028 | -0.028 | 0 | 0.67 | 0.87 | 0.51 | 0.7 | 0.56 | 0.58 |
| 2 | 47751239 | rs10495945 | -2.3 | -0.684 | 0.189 | 1.286 | -0.004 | 0.053 | 0.053 | 1.129 | 0.5 | 0.89 | 0.41 | 0.67 | 0.44 | 0.49 |
| 2 | 47754445 | rs11675417 | -1.963 | -0.774 | 0.384 | 1.326 | -0.001 | 0.007 | 0.007 | 0.05 | 0.37 | 0.66 | 0.33 | 0.53 | 0.34 | 0.34 |
| 2 | 47757577 | rs6751501 | -1.166 | -0.8 | -0.021 | 1.237 | 0.045 | 0.103 | 0.103 | 0.546 | 0.25 | 0.46 | 0.29 | 0.53 | 0.29 | 0.27 |
| 2 | 47757623 | rs6722911 | -1.745 | -0.737 | -0.769 | 1.065 | 0.004 | 0.022 | 0.022 | 0.811 | 0.41 | 0.82 | 0.37 | 0.71 | 0.37 | 0.44 |
| 2 | 47763019 | rs7580217 | 0.336 | -0.531 | -0.689 | 1.199 | -0.018 | 0.162 | 0.162 | 2.239 | 0.12 | 0.22 | 0.2 | 0.51 | 0.08 | 0.18 |
| 2 | 47763895 | rs748780 | -1.702 | 0.049 | -1.614 | 0.74 | 0.005 | 0.072 | 0.072 | 0.253 | 0.53 | 0.82 | 0.42 | 0.72 | 0.45 | 0.5 |
| 2 | 47771677 | rs13000537 | -0.84 | 0.575 | -1.027 | 0.908 | 0.126 | 0.123 | 0.123 | 0 | 0.07 | 0.17 | 0.15 | 0.19 | 0.16 | 0.06 |
| 2 | 47779201 | rs12476727 | -0.939 | -0.661 | -1.857 | 0.578 | 0.091 | -0.015 | -0.015 | 0 | 0.44 | 0.84 | 0.54 | 0.59 | 0.31 | 0.53 |
| 2 | 47782555 | rs6752504 | -0.515 | -0.684 | -0.879 | 0.008 | -0.04 | -0.12 | -0.12 | 0 | 0.21 | 0.55 | 0.16 | 0.02 | 0.14 | 0.19 |
| 2 | 47782756 | rs11674800 | -1.086 | -0.281 | -0.841 | 0.675 | 0.003 | 0.052 | 0.052 | 0 | 0.09 | 0.21 | 0.19 | 0.14 | 0.11 | 0.08 |
| 2 | 47784992 | rs1882449 | -0.73 | -0.071 | -0.793 | 0.316 | 0.011 | -0.026 | -0.026 | 0 | 0.12 | 0.3 | 0.09 | 0.05 | 0.11 | 0.1 |
| 2 | 47785013 | rs17568679 | -1.066 | -0.215 | -1.388 | 0.401 | -0.003 | 0.002 | 0.002 | 0.312 | 0.23 | 0.22 | 0.28 | 0.14 | 0.21 | 0.22 |
| 2 | 47787257 | rs11903353 | -0.256 | -0.216 | -1.212 | 0.534 | -0.001 | 0.002 | 0.002 | 0 | 0.14 | 0.12 | 0.21 | 0.1 | 0.12 | 0.12 |
| 2 | 47789775 | rs11125144 | 0.29 | -0.42 | -1.504 | 0.989 | 0.008 | -0.038 | -0.038 | 0 | 0.1 | 0.03 | 0.16 | 0.34 | 0.09 | 0.21 |
| 2 | 47790611 | rs1863334 | -1.321 | 0.195 | -2.011 | -0.068 | -0.062 | 0.048 | 0.048 | 0 | 0.7 | 0.92 | 0.67 | 0.8 | 0.58 | 0.77 |
| 2 | 47791096 | rs2348094 | -0.182 | 0.445 | -1.554 | -0.128 | -0.004 | 0.064 | 0.064 | 0 | 0.21 | 0.12 | 0.12 | 0.06 | 0.19 | 0.23 |
| 2 | 47794426 | rs4953503 | -0.525 | 0.591 | -1.267 | 0.011 | -0.001 | 0.007 | 0.007 | 0 | 0.34 | 0.31 | 0.32 | 0.16 | 0.32 | 0.35 |
| 2 | 47808152 | rs17503541 | -1.143 | 0.986 | -1.066 | 0.086 | 0.001 | -0.015 | -0.015 | 0 | 0.07 | 0.08 | 0.07 | 0 | 0.11 | 0.02 |
| 2 | 47814721 | rs13032306 | -0.843 | 0.885 | -0.94 | -0.112 | -0.002 | -0.009 | -0.009 | 0 | 0.09 | 0.1 | 0.06 | 0 | 0.09 | 0.01 |
| 2 | 47819339 | rs13397771 | -0.462 | 1.021 | -0.671 | 0.11 | 0.004 | 0.012 | 0.012 | 0 | 0.07 | 0.11 | 0.15 | 0.12 | 0.08 | 0.06 |
| 2 | 47821281 | rs4953509 | -0.308 | 1.063 | -0.866 | -0.121 | 0.004 | 0.009 | 0.009 | 0 | 0.64 | 0.35 | 0.71 | 0.76 | 0.62 | 0.59 |
| 2 | 47822310 | rs4521097 | -0.915 | 0.763 | -1.007 | -0.388 | 0 | 0.011 | 0.011 | 0 | 0.36 | 0.47 | 0.39 | 0.39 | 0.43 | 0.37 |
| 2 | 47826681 | rs11694783 | -0.613 | 1.276 | -1.215 | -0.311 | 0.015 | 0.04 | 0.04 | 0 | 0.3 | 0.46 | 0.49 | 0.42 | 0.34 | 0.4 |
| 2 | 47826830 | rs17036703 | -0.613 | 1.276 | -1.215 | -0.304 | 0.015 | 0.044 | 0.044 | 0 | 0.3 | 0.45 | 0.49 | 0.43 | 0.34 | 0.4 |
| 2 | 47829230 | rs4953514 | -1.395 | 1.157 | -1.364 | -0.443 | 0.002 | 0.001 | 0.001 | 0 | 0.43 | 0.47 | 0.59 | 0.44 | 0.5 | 0.55 |
| 2 | 47832420 | rs4953516 | -1.399 | 1.49 | -1.714 | -0.876 | -0.001 | -0.041 | -0.041 | 0 | 0.51 | 0.93 | 0.69 | 0.44 | 0.56 | 0.56 |
| 2 | 47832612 | rs11687554 | 1.498 | 1.494 | -1.678 | -0.878 | -0.032 | 0.035 | 0.035 | 0 | 0.08 | 0.02 | 0.09 | 0 | 0.12 | 0.09 |
| 2 | 47833916 | rs11694647 | 0.454 | 1.486 | -1.297 | -0.873 | 0.022 | -0.047 | -0.047 | 0 | 0.12 | 0.22 | 0.14 | 0 | 0.17 | 0.11 |
| 2 | 47834313 | rs10182660 | -1.026 | 1.362 | -1.341 | -0.918 | -0.009 | -0.003 | -0.003 | 3.329 | 0.45 | 0.64 | 0.63 | 0.44 | 0.51 | 0.56 |
| 2 | 47836070 | rs13408772 | -1.9 | 1.396 | -1.487 | -0.849 | -0.011 | -0.138 | -0.138 | 0.009 | 0.59 | 0.98 | 0.77 | 0.44 | 0.68 | 0.67 |
| 2 | 47837986 | rs11681403 | -0.759 | 1.582 | -1.246 | -0.516 | 0.021 | 0.006 | 0.006 | 0 | 0.37 | 0.75 | 0.51 | 0.34 | 0.28 | 0.37 |
| 2 | 47838874 | rs6716883 | -0.844 | 1.427 | -1.226 | -0.617 | -0.005 | -0.02 | -0.02 | 0.11 | 0.38 | 0.69 | 0.55 | 0.43 | 0.38 | 0.45 |
| 2 | 47844505 | rs6544997 | -0.837 | 1.374 | -1.283 | -0.464 | -0.008 | 0.035 | 0.035 | 0.012 | 0.5 | 0.78 | 0.62 | 0.38 | 0.5 | 0.58 |
| 2 | 47845523 | rs6723080 | -0.776 | 1.684 | -0.312 | -0.275 | 0.064 | -0.086 | -0.086 | 0 | 0.14 | 0.33 | 0.18 | 0 | 0.19 | 0.15 |
| 2 | 47851906 | rs6747656 | -0.782 | 1.334 | -0.311 | 0.674 | 0.018 | 0.121 | 0.121 | 0 | 0.21 | 0.39 | 0.42 | 0.44 | 0.19 | 0.16 |
| 2 | 47854692 | rs1317855 | -1.436 | 1.023 | -0.711 | 0.436 | 0.031 | 0.042 | 0.042 | 1.724 | 0.52 | 0.67 | 0.7 | 0.74 | 0.63 | 0.6 |
| 2 | 47856328 | rs11887626 | -0.395 | 1.333 | -0.649 | 0.332 | 0.009 | -0.019 | -0.019 | 0 | 0.19 | 0.22 | 0.2 | 0.08 | 0.27 | 0.26 |
| 2 | 47857655 | rs7559733 | -1.224 | 0.858 | -0.919 | 0.546 | -0.012 | 0.026 | 0.026 | 0.024 | 0.47 | 0.76 | 0.41 | 0.6 | 0.5 | 0.51 |
| 2 | 47857914 | rs4952894 | -0.663 | 0.965 | -0.553 | 0.708 | -0.006 | 0.015 | 0.015 | 0 | 0.21 | 0.36 | 0.4 | 0.21 | 0.18 | 0.15 |
| 2 | 47866656 | rs4266057 | -1.113 | 0.851 | -1.647 | 0.656 | -0.053 | -0.007 | -0.007 | 0.006 | 0.56 | 0.84 | 0.5 | 0.58 | 0.51 | 0.49 |
| 2 | 47868693 | rs11902856 | 0.368 | 0.956 | -1.852 | 0.469 | -0.111 | 0.119 | 0.119 | 1.029 | 0.21 | 0.03 | 0.24 | 0.04 | 0.32 | 0.31 |
| 2 | 47874713 | rs1574380 | -0.567 | 0.964 | -1.392 | 1.176 | -0.018 | -0.004 | -0.004 | 1.605 | 0.29 | 0.24 | 0.36 | 0.3 | 0.4 | 0.35 |
| 2 | 47877388 | rs17570243 | -0.604 | 0.929 | -1.423 | 1.255 | -0.031 | -0.015 | -0.015 | 0 | 0.3 | 0.21 | 0.37 | 0.34 | 0.4 | 0.36 |
| 2 | 47879202 | rs11125149 | -0.485 | 0.81 | -1.119 | 1.309 | 0.005 | -0.001 | -0.001 | 0.493 | 0.36 | 0.39 | 0.44 | 0.33 | 0.41 | 0.43 |
| 2 | 47880934 | rs1214776 | 0.086 | 0.595 | -1.117 | 1.147 | -0.014 | 0.01 | 0.01 | 0 | 0.22 | 0.33 | 0.12 | 0.27 | 0.19 | 0.19 |
| 2 | 47881550 | rs2969777 | -2.097 | 0.725 | -1.615 | 1.003 | 0.016 | 0.129 | 0.129 | 0 | 0.67 | 0.99 | 0.84 | 0.91 | 0.79 | 0.69 |
| 2 | 47883357 | rs6712934 | 0.134 | 0.642 | -1.373 | 0.949 | 0.024 | -0.047 | -0.047 | 0 | 0.33 | 0.39 | 0.41 | 0.13 | 0.45 | 0.43 |
| 2 | 47886543 | rs2260388 | 0.061 | 0.497 | -1.316 | 1.046 | -0.007 | 0.001 | 0.001 | 0 | 0.23 | 0.26 | 0.12 | 0.28 | 0.19 | 0.2 |
| 2 | 47887090 | rs6729891 | 0.4 | 0.965 | -1.254 | 1.009 | 0.028 | -0.074 | -0.074 | 0.15 | 0.29 | 0.39 | 0.4 | 0.08 | 0.4 | 0.42 |
| 2 | 47890331 | rs2705766 | -1.511 | 0.944 | -1.372 | 1.38 | -0.089 | -0.084 | -0.084 | 0.135 | 0.71 | 0.98 | 0.64 | 0.67 | 0.71 | 0.7 |
| 2 | 47897866 | rs2705775 | -0.021 | 0.205 | -0.997 | 1.388 | 0.004 | 0.009 | 0.009 | 0 | 0.17 | 0.25 | 0.12 | 0.24 | 0.17 | 0.11 |
| 2 | 47901721 | rs1214779 | -1.81 | 0.239 | -1.174 | 1.573 | -0.015 | 0.013 | 0.013 | 0.269 | 0.73 | 0.98 | 0.82 | 0.85 | 0.77 | 0.75 |
| 2 | 47902093 | rs9309149 | 0.152 | 0.196 | -1.018 | 1.607 | 0.001 | 0.002 | 0.002 | 3.211 | 0.48 | 0.45 | 0.62 | 0.33 | 0.55 | 0.58 |

| | | | | | | | | | | | | | | | |
|---|---|---|---|---|---|---|---|---|---|---|---|---|---|---|---|
| 2 | 47902785 | rs1214780 | -0.549 | -0.167 | -1.599 | 1.389 | 0.001 | -0.135 | -0.135 | 0 | 0.67 | 0.95 | 0.77 | 0.5 | 0.7 | 0.69 |
| 2 | 47904096 | rs7603494 | 0.287 | -0.11 | -1.226 | 1.259 | 0.008 | 0.004 | 0.004 | 0.277 | 0.53 | 0.54 | 0.57 | 0.51 | 0.58 | 0.59 |
| 2 | 47906192 | rs2969778 | -0.272 | -0.256 | -1.772 | 0.769 | 0.004 | 0.005 | 0.005 | 0.191 | 0.7 | 0.99 | 0.82 | 0.84 | 0.77 | 0.74 |
| 2 | 47910657 | rs13409431 | -1.01 | -0.082 | -0.378 | 1.301 | -0.035 | 0.108 | 0.108 | 0.285 | 0.17 | 0.37 | 0.2 | 0.35 | 0.14 | 0.13 |
| 2 | 47913358 | rs9789343 | -1.271 | 0.099 | -0.003 | 0.615 | -0.01 | -0.001 | -0.001 | 0 | 0.25 | 0.57 | 0.18 | 0.29 | 0.21 | 0.1 |
| 2 | 47914049 | rs6750911 | -1.141 | -0.005 | 0.104 | 0.172 | -0.032 | -0.125 | -0.125 | 0.839 | 0.23 | 0.55 | 0.14 | 0.01 | 0.15 | 0.07 |
| 2 | 47920541 | rs6713362 | -0.954 | -0.013 | 0.64 | 0.306 | -0.029 | -0.092 | -0.092 | 0.12 | 0.14 | 0.42 | 0.08 | 0.01 | 0.07 | 0.05 |
| 2 | 47933563 | rs17036890 | -0.578 | 0.045 | 0.426 | 0.289 | -0.014 | -0.074 | -0.074 | 0 | 0.12 | 0.26 | 0.05 | 0 | 0.07 | 0.04 |
| 2 | 47934826 | rs2018541 | -0.843 | 0.04 | 0.535 | 0.287 | -0.014 | -0.088 | -0.088 | 0.035 | 0.13 | 0.37 | 0.07 | 0 | 0.07 | 0.04 |
| 2 | 47941304 | rs2348238 | 0.058 | 0.132 | -0.217 | 0.084 | 0.006 | -0.027 | -0.027 | 6.105 | 0.36 | 0.16 | 0.31 | 0.38 | 0.3 | 0.32 |
| 2 | 47942825 | rs4325805 | -0.736 | 0.197 | -0.165 | 0.068 | -0.001 | -0.001 | -0.001 | 0 | 0.5 | 0.44 | 0.38 | 0.39 | 0.42 | 0.38 |
| 2 | 47942902 | rs4588237 | -0.181 | 0.382 | -1.04 | 0.256 | -0.007 | -0.024 | -0.024 | 1.149 | 0.38 | 0.16 | 0.33 | 0.39 | 0.35 | 0.34 |
| 2 | 47947791 | rs797703 | 0.867 | 0.586 | -0.834 | 0.618 | 0.004 | -0.057 | -0.057 | 0 | 0.66 | 0.17 | 0.69 | 0.53 | 0.7 | 0.7 |
| 2 | 47948101 | rs797700 | 0.225 | 0.59 | -0.8 | 0.603 | -0.001 | -0.011 | -0.011 | 0 | 0.74 | 0.26 | 0.73 | 0.53 | 0.76 | 0.73 |
| 2 | 47952231 | rs797689 | 0.212 | 0.599 | -0.788 | 0.594 | -0.001 | 0.03 | 0.03 | 0.546 | 0.74 | 0.34 | 0.73 | 0.53 | 0.76 | 0.73 |
| 2 | 47952777 | rs797685 | 0.262 | 0.586 | -0.791 | 0.577 | 0 | -0.02 | -0.02 | 0 | 0.74 | 0.23 | 0.72 | 0.53 | 0.76 | 0.73 |
| 2 | 47956844 | rs2098241 | -0.544 | 0.56 | -0.531 | 0.166 | -0.006 | -0.041 | -0.041 | 0.059 | 0.08 | 0.16 | 0.05 | 0 | 0.06 | 0.03 |
| 2 | 47982572 | rs441327 | 0.078 | 0.681 | -0.708 | -0.061 | 0 | -0.005 | -0.005 | 1.82 | 0.49 | 0.47 | 0.57 | 0.68 | 0.4 | 0.67 |
| 2 | 47996596 | rs759050 | -0.456 | 1.436 | -0.217 | -0.941 | 0.039 | -0.109 | -0.109 | 0 | 0.25 | 0.32 | 0.22 | 0 | 0.35 | 0.14 |
| 2 | 47999480 | rs17036952 | -0.154 | 1.698 | -0.388 | -1.045 | -0.001 | 0.01 | 0.01 | 0.377 | 0.08 | 0.05 | 0.02 | 0 | 0.07 | 0.07 |
| 2 | 48008574 | rs330792 | 0.113 | 1.308 | -1.222 | -1.724 | 0.008 | 0.094 | 0.094 | 0 | 0.19 | 0.01 | 0.09 | 0.05 | 0.15 | 0.11 |
| 2 | 48009816 | rs3136228 | -0.32 | 1.388 | -0.955 | -1.636 | 0.008 | -0.04 | -0.04 | 0 | 0.19 | 0.21 | 0.2 | 0 | 0.35 | 0.13 |
| 2 | 48013099 | rs3136247 | 0.685 | 2.003 | -0.962 | -1.619 | -0.01 | -0.004 | -0.004 | 0 | 0.13 | 0.08 | 0.12 | 0 | 0.18 | 0.06 |
| 2 | 48035673 | rs2710163 | 0.283 | 2.112 | -0.799 | -1.846 | -0.007 | 0.059 | 0.059 | 0 | 0.41 | 0.16 | 0.49 | 0.67 | 0.38 | 0.54 |
| 2 | 48039744 | rs10490302 | -0.16 | 2.542 | -0.799 | -1.94 | 0.086 | -0.056 | -0.056 | 0 | 0.12 | 0.2 | 0.19 | 0 | 0.3 | 0.1 |
| 2 | 48040285 | rs3732191 | 1.319 | 2.246 | -1.124 | -1.804 | -0.019 | -0.017 | -0.017 | 0.077 | 0.05 | 0.01 | 0.07 | 0.03 | 0.07 | 0.04 |
| 2 | 48041377 | rs330787 | -0.424 | 2.384 | -1.259 | -1.471 | -0.021 | -0.046 | -0.046 | 0.04 | 0.67 | 0.87 | 0.62 | 0.49 | 0.66 | 0.56 |
| 2 | 48044772 | rs2651766 | -0.001 | 2.294 | -1.287 | -1.549 | -0.02 | 0.018 | 0.018 | 0 | 0.61 | 0.85 | 0.53 | 0.4 | 0.59 | 0.45 |
| 2 | 48078838 | rs960106 | -0.067 | 1.954 | -1.263 | -1.883 | 0.001 | -0.072 | -0.072 | 0.011 | 0.6 | 0.53 | 0.61 | 0.88 | 0.58 | 0.75 |
| 2 | 48098708 | rs12463595 | 0.797 | 1.81 | -1.706 | -1.859 | -0.007 | 0.084 | 0.084 | 0 | 0.27 | 0.04 | 0.26 | 0.09 | 0.28 | 0.21 |
| 2 | 48116126 | rs4952896 | -0.905 | 1.8 | -1.743 | -1.863 | -0.042 | 0.077 | 0.077 | 0 | 0.73 | 0.97 | 0.74 | 0.88 | 0.71 | 0.79 |
| 2 | 48124504 | rs12620679 | 0.658 | 1.853 | -1.725 | -1.864 | -0.028 | -0.022 | -0.022 | 0 | 0.32 | 0.1 | 0.44 | 0.6 | 0.34 | 0.44 |
| 2 | 48131936 | rs13035558 | 0.286 | 2.075 | -1.497 | -1.656 | -0.007 | 0.068 | 0.068 | 0 | 0.26 | 0.12 | 0.32 | 0.11 | 0.28 | 0.22 |
| 2 | 48146122 | rs7587289 | -0.453 | 2.076 | -1.41 | -1.531 | 0.049 | -0.029 | -0.029 | 0 | 0.07 | 0.14 | 0.09 | 0 | 0.18 | 0.07 |
| 2 | 48164334 | rs6710889 | -0.456 | 1.949 | -1.577 | -1.565 | -0.04 | 0.128 | 0.128 | 0 | 0.72 | 0.96 | 0.76 | 0.92 | 0.7 | 0.77 |
| 2 | 48167631 | rs4552231 | 0.009 | 1.935 | -1.495 | -1.541 | -0.086 | 0.134 | 0.134 | 0 | 0.66 | 0.84 | 0.67 | 0.92 | 0.52 | 0.71 |
| 2 | 48174199 | rs6729916 | -0.228 | 1.922 | -1.495 | -1.53 | -0.055 | 0.157 | 0.157 | 0 | 0.68 | 0.93 | 0.73 | 0.92 | 0.63 | 0.71 |
| 2 | 48177487 | rs4583515 | 0.616 | 1.468 | -1.841 | -1.402 | 0.064 | 0.083 | 0.083 | 0.283 | 0.51 | 0.81 | 0.35 | 0.41 | 0.37 | 0.45 |
| 2 | 48179497 | rs13382470 | 0.557 | 1.478 | -1.443 | -0.292 | 0.043 | 0.114 | 0.114 | 0 | 0.49 | 0.66 | 0.33 | 0.14 | 0.37 | 0.43 |
| 2 | 48185303 | rs12105860 | -0.628 | 1.349 | -1.824 | -0.245 | -0.035 | 0.135 | 0.135 | 0 | 0.69 | 0.93 | 0.74 | 0.92 | 0.66 | 0.71 |
| 2 | 48196989 | rs6545008 | -1.286 | 0.718 | -1.106 | 0.007 | 0.042 | 0.015 | 0.015 | 0 | 0.6 | 0.73 | 0.77 | 0.58 | 0.73 | 0.56 |
| 2 | 48211719 | rs4953534 | -0.455 | 1.382 | -1.369 | -0.679 | -0.014 | 0.19 | 0.19 | 0 | 0.64 | 0.94 | 0.7 | 0.89 | 0.66 | 0.71 |
| 2 | 48212055 | rs7600005 | 1.04 | 1.25 | -1.161 | -0.81 | 0.007 | 0.029 | 0.029 | 0 | 0.45 | 0.58 | 0.32 | 0.09 | 0.39 | 0.43 |
| 2 | 48217552 | rs999036 | -0.645 | 1.226 | -1.215 | -0.781 | -0.008 | 0.181 | 0.181 | 0 | 0.64 | 0.93 | 0.7 | 0.89 | 0.66 | 0.71 |
| 2 | 48233949 | rs7562885 | -0.014 | 1.163 | -1.055 | -0.812 | -0.012 | 0.016 | 0.016 | 0 | 0.59 | 0.63 | 0.6 | 0.89 | 0.53 | 0.7 |
| 2 | 48242946 | rs12612568 | 1.073 | 1.178 | -1.136 | 0.408 | -0.03 | -0.055 | -0.055 | 0 | 0.09 | 0.01 | 0.25 | 0.35 | 0.11 | 0.13 |
| 2 | 48257961 | rs7580982 | -0.886 | 1.091 | -1.283 | -0.326 | -0.002 | 0.117 | 0.117 | 0 | 0.69 | 0.88 | 0.69 | 0.9 | 0.68 | 0.75 |
| 2 | 48287260 | rs1606930 | -1.705 | 0.777 | -0.506 | -0.383 | -0.032 | -0.034 | -0.034 | 0.999 | 0.23 | 0.44 | 0.29 | 0.81 | 0.17 | 0.25 |
| 2 | 48298383 | rs1878079 | -0.988 | 0.764 | -0.67 | -0.39 | -0.018 | 0.14 | 0.14 | 0 | 0.7 | 0.93 | 0.69 | 0.9 | 0.68 | 0.75 |
| 2 | 48300241 | rs6726144 | 1.52 | 1.005 | -0.321 | -0.589 | 0.003 | -0.013 | -0.013 | 0.003 | 0.39 | 0.32 | 0.25 | 0.09 | 0.28 | 0.45 |
| 2 | 48303182 | rs900511 | -1.032 | 1.005 | -0.419 | -0.333 | 0.001 | 0.1 | 0.1 | 0 | 0.69 | 0.9 | 0.68 | 0.85 | 0.67 | 0.74 |
| 2 | 48304381 | rs6545018 | -0.81 | 0.925 | -0.391 | -0.381 | -0.019 | 0.036 | 0.036 | 0 | 0.65 | 0.7 | 0.6 | 0.88 | 0.55 | 0.73 |
| 2 | 48314490 | rs6740430 | -0.984 | 0.963 | -0.207 | -0.328 | -0.025 | 0.035 | 0.035 | 0 | 0.23 | 0.11 | 0.33 | 0.83 | 0.27 | 0.31 |
| 2 | 48318158 | rs1878078 | -1.414 | 0.655 | -0.483 | -0.723 | -0.045 | 0.098 | 0.098 | 0 | 0.35 | 0.13 | 0.42 | 0.84 | 0.39 | 0.37 |
| 2 | 48344097 | rs6739811 | 1.898 | 0.211 | 0.137 | -0.764 | 0.009 | 0.005 | 0.005 | 0 | 0.27 | 0.16 | 0.14 | 0.06 | 0.16 | 0.28 |
| 2 | 48352405 | rs7566406 | 1.486 | 0.169 | -0.027 | -0.772 | -0.052 | -0.07 | -0.07 | 0.004 | 0.3 | 0.42 | 0.17 | 0.09 | 0.17 | 0.35 |
| 2 | 48361852 | rs2170678 | 0.504 | 0.367 | -0.282 | -0.144 | -0.008 | -0.071 | -0.071 | 0 | 0.13 | 0.02 | 0.26 | 0.52 | 0.13 | 0.24 |
| 2 | 48367696 | rs1842941 | 0.889 | -0.153 | -0.397 | -0.51 | -0.043 | -0.037 | -0.037 | 1.808 | 0.24 | 0.34 | 0.16 | 0.1 | 0.15 | 0.34 |
| 2 | 48371745 | rs4953551 | 0.937 | 0.24 | -0.762 | -0.407 | 0.041 | -0.138 | -0.138 | 0 | 0.25 | 0.03 | 0.28 | 0.52 | 0.16 | 0.27 |
| 2 | 48383746 | rs7570354 | -1.059 | 0.404 | -0.544 | -0.791 | -0.014 | 0.085 | 0.085 | 0 | 0.59 | 0.69 | 0.54 | 0.95 | 0.43 | 0.72 |
| 2 | 48398366 | rs6732149 | -1.476 | 0.192 | -0.563 | -0.728 | -0.024 | 0.038 | 0.038 | 0 | 0.66 | 0.72 | 0.61 | 0.95 | 0.56 | 0.75 |
| 2 | 48408487 | rs2348719 | -0.998 | -0.088 | -0.576 | -0.849 | 0.005 | 0.005 | 0.005 | 0.084 | 0.61 | 0.65 | 0.59 | 0.89 | 0.62 | 0.67 |
| 2 | 48423898 | rs7569620 | -0.523 | -0.08 | -0.46 | -0.723 | 0.002 | -0.029 | -0.029 | 0 | 0.54 | 0.42 | 0.35 | 0.83 | 0.48 | 0.57 |
| 2 | 48426035 | rs7583634 | -1.185 | -0.079 | -0.517 | -0.775 | 0.003 | 0.135 | 0.135 | 0.757 | 0.62 | 0.9 | 0.51 | 0.86 | 0.61 | 0.66 |
| 2 | 48441358 | rs17037195 | 1.593 | -0.301 | -0.424 | -0.956 | -0.049 | -0.073 | -0.073 | 0 | 0.14 | 0.29 | 0.06 | 0.03 | 0.07 | 0.19 |
| 2 | 48443109 | rs6755968 | -0.04 | -0.29 | -0.548 | -1.05 | -0.02 | 0.118 | 0.118 | 1.019 | 0.42 | 0.2 | 0.48 | 0.86 | 0.41 | 0.46 |
| 2 | 48443320 | rs2175138 | -1.047 | -0.013 | -0.824 | -0.864 | 0.014 | 0.149 | 0.149 | 0.396 | 0.61 | 0.83 | 0.63 | 0.9 | 0.61 | 0.65 |
| 2 | 48445680 | rs4375916 | -0.659 | -0.099 | -0.875 | -0.85 | -0.001 | -0.019 | -0.019 | 0 | 0.57 | 0.53 | 0.55 | 0.9 | 0.5 | 0.65 |
| 2 | 48461993 | rs3750272 | 1.212 | -0.564 | -1.294 | -0.458 | 0.005 | -0.16 | -0.16 | 0.032 | 0.23 | 0.02 | 0.29 | 0.56 | 0.16 | 0.3 |

| | | | | | | | | | | | | | | | | |
|---|---|---|---|---|---|---|---|---|---|---|---|---|---|---|---|---|
| 2 | 48463834 | rs1396825 | -1.536 | -0.417 | -1.049 | -0.635 | -0.02 | 0.171 | 0.171 | 0.048 | 0.72 | 0.97 | 0.71 | 0.95 | 0.72 | 0.72 |
| 2 | 48487724 | rs6750123 | -0.426 | -0.075 | -1.529 | -0.487 | 0.007 | 0.165 | 0.165 | 0.274 | 0.61 | 0.88 | 0.63 | 0.9 | 0.59 | 0.65 |
| 2 | 48489733 | rs17397205 | 0.094 | -0.171 | -1.755 | -0.457 | 0.012 | 0.068 | 0.068 | 0.014 | 0.25 | 0.09 | 0.13 | 0.06 | 0.22 | 0.26 |
| 2 | 48490508 | rs2471403 | 0.751 | 0.317 | -1.852 | -0.241 | 0.003 | 0.022 | 0.022 | 0 | 0.51 | 0.63 | 0.62 | 0.51 | 0.48 | 0.47 |
| 2 | 48498115 | rs1509561 | -2.496 | 0.192 | -1.894 | -0.483 | 0.033 | 0.067 | 0.067 | 0 | 0.87 | 0.98 | 0.94 | 0.99 | 0.92 | 0.88 |
| 2 | 48505401 | rs959141 | 0.048 | 0.181 | -1.826 | -0.55 | 0.034 | 0.046 | 0.046 | 0 | 0.59 | 0.76 | 0.8 | 0.93 | 0.68 | 0.61 |
| 2 | 48520539 | rs6545026 | -0.235 | 0.683 | -1.302 | -0.24 | 0.005 | -0.043 | -0.043 | 0 | 0.3 | 0.32 | 0.2 | 0.07 | 0.33 | 0.37 |
| 2 | 48520762 | rs11893741 | 1.938 | 0.646 | -1.188 | -0.424 | 0.002 | 0.02 | 0.02 | 0 | 0.14 | 0.1 | 0.21 | 0.02 | 0.16 | 0.21 |
| 2 | 48528035 | rs10193072 | -1.008 | 0.699 | -1.255 | -0.146 | 0.001 | -0.002 | -0.002 | 0 | 0.4 | 0.38 | 0.28 | 0.53 | 0.38 | 0.57 |
| 2 | 48554191 | rs11676168 | -0.754 | 0.702 | -1.121 | -0.121 | 0.001 | 0.002 | 0.002 | 0.134 | 0.38 | 0.37 | 0.31 | 0.53 | 0.45 | 0.51 |
| 2 | 48558260 | rs6545031 | 0.215 | 0.781 | -1.091 | -0.136 | -0.004 | -0.061 | -0.061 | 0.342 | 0.1 | 0.21 | 0.05 | 0.01 | 0.09 | 0.22 |
| 2 | 48566031 | rs10153773 | -1.544 | 0.639 | -0.64 | -0.224 | 0.031 | -0.014 | -0.014 | 0 | 0.53 | 0.78 | 0.55 | 0.55 | 0.62 | 0.69 |
| 2 | 48579778 | rs7606104 | -1.21 | 0.916 | -0.72 | 0.045 | 0.002 | 0.001 | 0.001 | 0 | 0.36 | 0.35 | 0.31 | 0.47 | 0.44 | 0.46 |
| 2 | 48584474 | rs11679523 | 0.801 | 0.861 | -0.72 | 0.031 | -0.003 | 0.045 | 0.045 | 0 | 0.09 | 0.11 | 0.13 | 0.46 | 0.12 | 0.16 |
| 2 | 48585963 | rs7563625 | -0.025 | 0.883 | -0.749 | 0.016 | -0.005 | -0.073 | -0.073 | 0.565 | 0.25 | 0.21 | 0.13 | 0.01 | 0.21 | 0.27 |
| 2 | 48587741 | rs10495952 | 0.992 | 0.927 | -0.815 | 0.104 | -0.025 | -0.023 | -0.023 | 0 | 0.09 | 0.03 | 0.11 | 0.46 | 0.12 | 0.17 |
| 2 | 48594355 | rs17325488 | 0.017 | 0.854 | -0.741 | -0.071 | 0.011 | 0.017 | 0.017 | 0.061 | 0.25 | 0.12 | 0.19 | 0.02 | 0.22 | 0.27 |
| 2 | 48597559 | rs7562069 | -1.144 | 0.798 | -0.731 | -0.035 | -0.038 | -0.027 | -0.027 | 0.011 | 0.35 | 0.23 | 0.3 | 0.47 | 0.45 | 0.42 |
| 2 | 48603483 | rs6705802 | -1.355 | 0.995 | -0.391 | 0.118 | 0.065 | 0.095 | 0.095 | 0 | 0.22 | 0.37 | 0.24 | 0.46 | 0.33 | 0.32 |
| 2 | 48604978 | rs6718068 | -0.918 | 0.986 | -0.54 | 0.162 | 0.021 | 0.036 | 0.036 | 0 | 0.19 | 0.23 | 0.22 | 0.46 | 0.32 | 0.3 |
| 2 | 48610367 | rs6756596 | -0.818 | 1.031 | -0.578 | 0.196 | 0.002 | 0.037 | 0.037 | 0 | 0.18 | 0.24 | 0.17 | 0.47 | 0.21 | 0.3 |
| 2 | 48625044 | rs10169629 | -0.866 | 1.11 | -0.581 | 0.209 | 0.025 | 0.041 | 0.041 | 0 | 0.18 | 0.25 | 0.22 | 0.47 | 0.32 | 0.3 |
| 2 | 48630053 | rs6747441 | 0.13 | 1.04 | -0.457 | 0.384 | -0.017 | 0.007 | 0.007 | 0 | 0.1 | 0.05 | 0.13 | 0.5 | 0.17 | 0.16 |
| 2 | 48643436 | rs4606978 | -0.298 | 0.932 | -0.595 | 0.247 | 0.001 | 0.012 | 0.012 | 0.083 | 0.27 | 0.24 | 0.23 | 0.5 | 0.31 | 0.35 |
| 2 | 48646399 | rs10454142 | 0.199 | 1.039 | -0.82 | 0.307 | -0.057 | -0.042 | -0.042 | 0 | 0.1 | 0.03 | 0.19 | 0.47 | 0.28 | 0.15 |
| 2 | 48649705 | rs7565792 | -0.509 | 0.852 | -0.672 | -0.319 | -0.002 | 0 | 0 | 1.325 | 0.52 | 0.52 | 0.47 | 0.52 | 0.61 | 0.42 |
| 2 | 48650308 | rs4316980 | 0.198 | 1.042 | -0.789 | -0.183 | -0.002 | -0.006 | -0.006 | 0 | 0.27 | 0.24 | 0.24 | 0.5 | 0.3 | 0.34 |
| 2 | 48653703 | rs4267555 | -0.314 | 0.989 | -0.797 | -0.108 | -0.001 | -0.001 | -0.001 | 0 | 0.3 | 0.32 | 0.32 | 0.57 | 0.42 | 0.44 |
| 2 | 48666736 | rs4434042 | 0.937 | 0.758 | -0.813 | -0.098 | -0.035 | -0.042 | -0.042 | 0 | 0.33 | 0.42 | 0.44 | 0.18 | 0.21 | 0.47 |
| 2 | 48683746 | rs11902595 | 1.77 | 0.711 | -1.031 | -0.105 | -0.001 | 0.008 | 0.008 | 0 | 0.17 | 0.14 | 0.23 | 0.09 | 0.17 | 0.34 |
| 2 | 48686310 | rs4953579 | 0.579 | 0.93 | -1.252 | -0.036 | 0.006 | -0.018 | -0.018 | 0 | 0.58 | 0.46 | 0.54 | 0.23 | 0.48 | 0.65 |
| 2 | 48714938 | rs4561709 | 0.839 | 1.373 | -1.142 | 0.634 | -0.03 | 0.056 | 0.056 | 0.003 | 0.18 | 0.03 | 0.21 | 0.06 | 0.19 | 0.19 |
| 2 | 48715483 | rs11125173 | 0.552 | 1.481 | -0.999 | 1.106 | -0.019 | 0.031 | 0.031 | 0 | 0.08 | 0.05 | 0.07 | 0.46 | 0.15 | 0.11 |
| 2 | 48722755 | rs4443044 | 0.658 | 1.513 | -0.898 | 1.054 | 0.011 | -0.006 | -0.006 | 0 | 0.66 | 0.6 | 0.64 | 0.28 | 0.54 | 0.6 |
| 2 | 48725407 | rs7566996 | 0.926 | 1.435 | -1.004 | 0.402 | -0.002 | -0.033 | -0.033 | 0 | 0.29 | 0.25 | 0.23 | 0.02 | 0.33 | 0.1 |
| 2 | 48732944 | rs4293602 | 0.018 | 2.021 | -0.907 | 1.609 | 0.01 | 0.008 | 0.008 | 0 | 0.9 | 0.88 | 0.8 | 0.26 | 0.72 | 0.81 |
| 2 | 48740008 | rs11674469 | 0.356 | 1.707 | -0.955 | 1.164 | -0.042 | 0.054 | 0.054 | 0.004 | 0.21 | 0.08 | 0.19 | 0.06 | 0.32 | 0.22 |
| 2 | 48740195 | rs4953589 | 0.346 | 1.466 | -0.839 | 1.356 | -0.006 | 0.005 | 0.005 | 0 | 0.63 | 0.61 | 0.64 | 0.19 | 0.47 | 0.57 |
| 2 | 48749047 | rs12615167 | 0.285 | 1.462 | -0.856 | 1.308 | 0.009 | -0.014 | -0.014 | 0 | 0.63 | 0.57 | 0.63 | 0.26 | 0.5 | 0.57 |
| 2 | 48769377 | rs4952916 | -0.002 | 1.991 | -0.611 | 2.378 | 0.013 | 0.135 | 0.135 | 0 | 0.05 | 0.07 | 0.28 | 0.44 | 0.16 | 0.12 |
| 2 | 48776217 | rs4953592 | -1.307 | 2.411 | 0.144 | 2.425 | 0.049 | 0.288 | 0.288 | 0 | 0.08 | 0.22 | 0.28 | 0.44 | 0.14 | 0.12 |
| 2 | 48779064 | rs876970 | 0.204 | 1.914 | -0.789 | 1.556 | 0.008 | 0.006 | 0.006 | 0 | 0.31 | 0.19 | 0.3 | 0.32 | 0.29 | 0.34 |
| 2 | 48781222 | rs6705602 | 0.428 | 2.576 | -0.665 | 1.095 | -0.041 | 0.077 | 0.077 | 0 | 0.13 | 0.01 | 0.07 | 0 | 0.12 | 0.07 |
| 2 | 48787134 | rs2293274 | -0.059 | 2.612 | -0.442 | 1.216 | 0.039 | 0.052 | 0.052 | 0.034 | 0.21 | 0.29 | 0.31 | 0.37 | 0.32 | 0.39 |
| 2 | 48800050 | rs6716878 | 0.38 | 2.374 | -0.595 | 1.059 | -0.035 | 0.105 | 0.105 | 0 | 0.17 | 0.01 | 0.1 | 0 | 0.15 | 0.07 |
| 2 | 48807976 | rs3828340 | -0.012 | 2.402 | -0.26 | 1.117 | 0.035 | 0.056 | 0.056 | 0.048 | 0.21 | 0.28 | 0.32 | 0.39 | 0.33 | 0.4 |
| 2 | 48808594 | rs3749142 | -1.464 | 2.391 | -0.279 | 1.104 | -0.02 | -0.008 | -0.008 | 0.2 | 0.38 | 0.29 | 0.41 | 0.39 | 0.47 | 0.45 |
| 2 | 48808898 | rs3749145 | 0.012 | 2.448 | -0.25 | 1.132 | 0.035 | 0.057 | 0.057 | 0.03 | 0.21 | 0.28 | 0.31 | 0.39 | 0.32 | 0.39 |
| 2 | 48809593 | rs3792234 | -1.224 | 2.139 | -0.16 | 1.228 | -0.018 | -0.002 | -0.002 | 0 | 0.37 | 0.29 | 0.41 | 0.37 | 0.48 | 0.46 |
| 2 | 48810067 | rs13011288 | -1.329 | 1.644 | -0.187 | 1.17 | -0.001 | -0.001 | -0.001 | 0.041 | 0.41 | 0.57 | 0.5 | 0.39 | 0.62 | 0.48 |
| 2 | 48814317 | rs13007591 | -0.22 | 1.7 | -0.232 | 1.143 | 0.014 | 0.013 | 0.013 | 0 | 0.2 | 0.23 | 0.33 | 0.35 | 0.31 | 0.39 |
| 2 | 48821941 | rs10865231 | -0.239 | 1.598 | 0.085 | 1.234 | 0.043 | 0.126 | 0.126 | 0 | 0.21 | 0.43 | 0.34 | 0.46 | 0.27 | 0.4 |
| 2 | 48822540 | rs7574549 | -0.551 | 1.57 | 0.123 | 1.298 | 0.077 | 0.476 | 0.476 | 0 | 0.07 | 0.42 | 0.14 | 0.36 | 0.07 | 0.14 |
| 2 | 48824698 | rs13018198 | -0.276 | 1.589 | -0.429 | 0.505 | -0.037 | 0.075 | 0.075 | 0 | 0.25 | 0.09 | 0.17 | 0 | 0.29 | 0.15 |
| 2 | 48827497 | rs1554612 | -1.422 | 1.322 | -0.009 | 0.652 | -0.001 | -0.004 | -0.004 | 1.488 | 0.45 | 0.57 | 0.51 | 0.43 | 0.56 | 0.54 |
| 2 | 48829030 | rs11901451 | -0.455 | 1.071 | 0.39 | 1.253 | 0.071 | 0.404 | 0.404 | 0 | 0.06 | 0.41 | 0.15 | 0.33 | 0.07 | 0.12 |
| 2 | 48831931 | rs6752945 | 1.003 | 0.758 | 0.202 | 1.061 | -0.036 | 0.153 | 0.153 | 0.006 | 0.13 | 0.39 | 0.16 | 0.32 | 0.08 | 0.16 |
| 2 | 48838955 | rs10208754 | 0.955 | 0.846 | 0.564 | 0.427 | 0.019 | -0.089 | -0.089 | 0 | 0.69 | 0.43 | 0.7 | 0.94 | 0.65 | 0.62 |
| 2 | 48843520 | rs1464727 | 1.126 | 0.81 | 0.526 | 0.923 | 0.005 | -0.001 | -0.001 | 0 | 0.67 | 0.37 | 0.69 | 0.7 | 0.64 | 0.6 |
| 2 | 48845706 | rs6545053 | -0.645 | 0.886 | 0.711 | 0.13 | 0.046 | -0.055 | -0.055 | 1.402 | 0.1 | 0.32 | 0.07 | 0 | 0.1 | 0.07 |
| 2 | 48848295 | rs13035544 | 0.612 | 0.935 | 0.62 | 0.076 | -0.033 | 0.024 | 0.024 | 0.123 | 0.13 | 0.09 | 0.21 | 0.07 | 0.19 | 0.26 |
| 2 | 48850230 | rs10495955 | -0.895 | 0.941 | 0.659 | 0.067 | 0.031 | -0.061 | -0.061 | 0 | 0.23 | 0.35 | 0.28 | 0.07 | 0.29 | 0.32 |
| 2 | 48862902 | rs2349099 | 0.529 | 0.944 | 0.563 | 0.096 | -0.042 | 0.029 | 0.029 | 0.194 | 0.14 | 0.06 | 0.2 | 0.07 | 0.19 | 0.27 |
| 2 | 48873582 | rs12988176 | -0.905 | 0.894 | 0.65 | 0.072 | 0.016 | -0.065 | -0.065 | 0 | 0.11 | 0.26 | 0.07 | 0 | 0.1 | 0.07 |
| 2 | 48874821 | rs10198501 | 1.786 | 0.656 | 0.588 | -0.102 | -0.007 | 0.056 | 0.056 | 0 | 0.61 | 0.24 | 0.63 | 0.93 | 0.61 | 0.61 |
| 2 | 48876377 | rs13021383 | -0.445 | 0.944 | 0.703 | 0.064 | -0.003 | -0.008 | -0.008 | 0 | 0.08 | 0.07 | 0.05 | 0 | 0.1 | 0.07 |
| 2 | 48891287 | rs11677854 | 0.462 | 1.036 | 0.723 | 0.17 | -0.042 | 0.029 | 0.029 | 0.303 | 0.14 | 0.06 | 0.2 | 0.07 | 0.19 | 0.26 |
| 2 | 48899150 | rs3792246 | 1.708 | 0.942 | 0.791 | 0.116 | -0.024 | 0.079 | 0.079 | 0.118 | 0.65 | 0.2 | 0.65 | 0.91 | 0.63 | 0.62 |
| 2 | 48904550 | rs12713012 | -0.352 | 0.771 | 1.028 | -0.067 | 0.066 | -0.054 | -0.054 | 0.623 | 0.18 | 0.35 | 0.25 | 0.08 | 0.25 | 0.28 |

| | | | | | | | | | | | | | | | | |
|---|---|---|---|---|---|---|---|---|---|---|---|---|---|---|---|---|
| 2 | 48907086 | rs6725150 | 1.217 | 0.629 | 1.073 | -0.086 | -0.018 | -0.081 | -0.081 | 0.209 | 0.88 | 0.54 | 0.93 | 1 | 0.92 | 0.95 |
| 2 | 48910133 | rs10865234 | 1.221 | 0.637 | 0.969 | -0.083 | -0.02 | -0.081 | -0.081 | 0.618 | 0.88 | 0.55 | 0.94 | 0.99 | 0.92 | 0.94 |
| 2 | 48912416 | rs11892581 | -0.78 | 0.667 | 0.95 | -0.132 | 0.071 | -0.098 | -0.098 | 0.2 | 0.23 | 0.53 | 0.29 | 0.07 | 0.29 | 0.33 |
| 2 | 48915871 | rs11125179 | 1.325 | 0.739 | 0.583 | -0.089 | -0.008 | 0.012 | 0.012 | 0.085 | 0.66 | 0.25 | 0.66 | 0.92 | 0.64 | 0.61 |
| 2 | 48921375 | rs2293275 | 1.285 | 0.766 | 0.575 | -0.077 | -0.01 | 0.017 | 0.017 | 0 | 0.67 | 0.24 | 0.67 | 0.93 | 0.64 | 0.62 |
| 2 | 48924091 | rs1404057 | 1.156 | 0.121 | 0.19 | 0.261 | -0.015 | -0.077 | -0.077 | 0.116 | 0.86 | 0.75 | 0.93 | 1 | 0.89 | 0.91 |
| 2 | 48925746 | rs6755901 | 0.235 | -0.249 | -0.414 | 0.726 | 0.019 | -0.046 | -0.046 | 1.545 | 0.77 | 0.89 | 0.65 | 0.47 | 0.82 | 0.79 |
| 2 | 48935392 | rs4374421 | -0.062 | -0.499 | -0.462 | -0.394 | 0.031 | 0.09 | 0.09 | 0 | 0.65 | 0.81 | 0.77 | 0.9 | 0.72 | 0.64 |
| 2 | 48937154 | rs12713013 | 0.802 | -0.52 | -0.379 | -0.055 | 0 | 0.067 | 0.067 | 0 | 0.08 | 0.17 | 0.04 | 0.2 | 0.08 | 0.16 |
| 2 | 48937349 | rs12618729 | 0.605 | -0.703 | -0.326 | 0.11 | -0.001 | 0.036 | 0.036 | 0 | 0.14 | 0.23 | 0.3 | 0.25 | 0.14 | 0.06 |
| 2 | 48937576 | rs17326321 | 1.118 | -0.755 | -0.726 | -0.083 | -0.01 | -0.035 | -0.035 | 0 | 0.08 | 0.12 | 0.05 | 0 | 0.07 | 0.06 |
| 2 | 48937614 | rs17326328 | 1.433 | -0.773 | -0.88 | -0.14 | -0.011 | -0.029 | -0.029 | 0 | 0.44 | 0.17 | 0.41 | 0.42 | 0.44 | 0.4 |
| 2 | 48938341 | rs7605712 | -0.619 | -0.923 | -0.847 | -0.01 | -0.004 | -0.002 | -0.002 | 0 | 0.46 | 0.44 | 0.56 | 0.48 | 0.35 | 0.48 |
| 2 | 48938356 | rs7579411 | -1.537 | -0.634 | -0.864 | 0.073 | -0.006 | 0 | 0 | 0 | 0.63 | 0.73 | 0.43 | 0.41 | 0.41 | 0.66 |
| 2 | 48938662 | rs6729809 | 0.062 | -0.522 | -1.576 | 0.039 | 0.031 | 0.057 | 0.057 | 0 | 0.58 | 0.73 | 0.69 | 0.76 | 0.71 | 0.52 |
| 2 | 48939953 | rs7371084 | 0.483 | -0.663 | -1.86 | 0.461 | -0.031 | 0.003 | 0.003 | 0 | 0.05 | 0.01 | 0.24 | 0.28 | 0.15 | 0.06 |
| 2 | 48941428 | rs4953616 | 0.051 | -0.815 | -1.741 | 0.35 | 0.036 | 0.051 | 0.051 | 0 | 0.59 | 0.64 | 0.71 | 0.77 | 0.75 | 0.53 |
| 2 | 48941564 | rs17326446 | 1.447 | -0.843 | -1.968 | 0.378 | -0.106 | -0.031 | -0.031 | 0 | 0.35 | 0.14 | 0.34 | 0.33 | 0.5 | 0.31 |
| 2 | 48944441 | rs4352270 | 1.06 | -0.726 | -1.794 | 0.33 | 0.002 | 0.005 | 0.005 | 0 | 0.2 | 0.18 | 0.15 | 0.12 | 0.08 | 0.19 |
| 2 | 48946167 | rs6711321 | 0.059 | -0.123 | -1.609 | 0.39 | 0 | 0.041 | 0.041 | 0.792 | 0.21 | 0.47 | 0.33 | 0.28 | 0.21 | 0.08 |
| 2 | 48948802 | rs4490239 | 0.492 | 0.367 | -2.165 | 0.276 | -0.038 | 0.035 | 0.035 | 0.019 | 0.15 | 0.03 | 0.13 | 0.1 | 0.15 | 0.27 |
| 2 | 48952574 | rs6545061 | -0.478 | 0.39 | -2.185 | 0.425 | -0.076 | 0.002 | 0.002 | 0 | 0.61 | 0.84 | 0.62 | 0.66 | 0.47 | 0.68 |
| 2 | 48953349 | rs7562693 | -0.344 | 0.689 | -1.972 | 0.556 | 0.003 | 0.016 | 0.016 | 0 | 0.48 | 0.4 | 0.54 | 0.45 | 0.52 | 0.34 |
| 2 | 48958595 | rs4597581 | 0.38 | 1.072 | -2.419 | 0.248 | -0.057 | 0.059 | 0.059 | 0.104 | 0.17 | 0.02 | 0.13 | 0.07 | 0.19 | 0.33 |
| 2 | 48959603 | rs4319975 | -1.274 | 1.002 | -3.223 | 0.366 | 0 | -0.111 | -0.111 | 0.019 | 0.69 | 0.95 | 0.8 | 0.61 | 0.74 | 0.69 |
| 2 | 48960032 | rs10495960 | 0.97 | 1.373 | -3.435 | 0.646 | -0.042 | -0.018 | -0.018 | 0 | 0.07 | 0.01 | 0.27 | 0.33 | 0.18 | 0.06 |
| 2 | 48960317 | rs4637174 | -0.021 | 1.318 | -3.093 | 0.514 | 0.05 | 0.129 | 0.129 | 0.115 | 0.44 | 0.8 | 0.38 | 0.21 | 0.35 | 0.29 |
| 2 | 48962291 | rs17326656 | 0.947 | 1.16 | -2.55 | 0.176 | -0.001 | -0.053 | -0.053 | 0 | 0.24 | 0.26 | 0.15 | 0.01 | 0.22 | 0.08 |
| 2 | 48964420 | rs12998409 | -1.488 | 1.228 | -2.358 | 0.583 | 0.023 | 0.009 | 0.009 | 0.025 | 0.67 | 0.93 | 0.78 | 0.77 | 0.73 | 0.68 |
| 2 | 48965022 | rs4131886 | 0.791 | 1.166 | -2.299 | 0.145 | -0.002 | 0.016 | 0.016 | 0.316 | 0.28 | 0.21 | 0.15 | 0.01 | 0.26 | 0.09 |
| 2 | 48973162 | rs11679263 | 1.009 | 1.235 | -2.297 | 0.302 | -0.039 | -0.017 | -0.017 | 0 | 0.07 | 0.03 | 0.28 | 0.35 | 0.18 | 0.06 |
| 2 | 48976210 | rs4074648 | -1.492 | 1.223 | -2.321 | 0.281 | 0.007 | -0.071 | -0.071 | 0 | 0.67 | 0.94 | 0.78 | 0.69 | 0.72 | 0.66 |
| 2 | 48976886 | rs13020003 | 0.57 | 1.288 | -2.128 | 0.148 | -0.001 | -0.055 | -0.055 | 0 | 0.29 | 0.28 | 0.18 | 0.01 | 0.31 | 0.1 |
| 2 | 48977120 | rs4555391 | -0.157 | 1.275 | -2.105 | 0.201 | -0.018 | 0.084 | 0.084 | 0 | 0.45 | 0.29 | 0.28 | 0.08 | 0.45 | 0.4 |
| 2 | 48983395 | rs7562879 | -0.011 | 1.102 | -1.758 | 0.366 | 0.008 | 0.107 | 0.107 | 0.3 | 0.14 | 0.56 | 0.24 | 0.27 | 0.12 | 0.2 |
| 2 | 48983633 | rs7563126 | 0.238 | 0.918 | -1.729 | 0.385 | -0.029 | 0.138 | 0.138 | 0.439 | 0.13 | 0.53 | 0.21 | 0.27 | 0.07 | 0.19 |
| 2 | 48983789 | rs7563144 | -1.32 | 0.956 | -1.502 | 0.469 | 0.026 | -0.027 | -0.027 | 0 | 0.65 | 0.84 | 0.77 | 0.68 | 0.69 | 0.65 |
| 2 | 48984391 | rs2301267 | -0.101 | 1.07 | -1.716 | 0.399 | -0.004 | 0.001 | 0.001 | 0 | 0.36 | 0.33 | 0.46 | 0.36 | 0.48 | 0.15 |
| 2 | 48996116 | rs4594496 | 0.425 | 1.559 | -1.106 | 1.269 | 0.018 | 0.07 | 0.07 | 0.252 | 0.07 | 0.05 | 0.27 | 0.34 | 0.18 | 0.06 |
| 2 | 48997123 | rs7583082 | -0.497 | 0.899 | -1.006 | 0.951 | -0.022 | 0.004 | 0.004 | 1.795 | 0.28 | 0.55 | 0.32 | 0.32 | 0.22 | 0.49 |
| 2 | 49000874 | rs10185213 | -0.268 | 0.938 | -1.043 | 0.925 | 0.003 | 0.013 | 0.013 | 0.456 | 0.72 | 0.76 | 0.86 | 0.88 | 0.74 | 0.81 |
| 2 | 49003566 | rs10194838 | -1.07 | 1.475 | -0.201 | 1.89 | 0.022 | 0.126 | 0.126 | 0 | 0.14 | 0.49 | 0.23 | 0.24 | 0.12 | 0.18 |
| 2 | 49003623 | rs4453732 | -0.917 | 1.581 | -1.092 | 2.256 | -0.069 | -0.08 | -0.08 | 0 | 0.13 | 0.05 | 0.34 | 0.43 | 0.22 | 0.15 |
| 2 | 49003800 | rs10195116 | -0.407 | 0.376 | -0.355 | 1.728 | -0.031 | 0.099 | 0.099 | 0 | 0.07 | 0.15 | 0.03 | 0.18 | 0.01 | 0.14 |
| 2 | 49006825 | rs4245823 | 1.064 | 0.849 | -0.468 | 1.295 | 0.014 | 0.026 | 0.026 | 0.172 | 0.66 | 0.48 | 0.56 | 0.55 | 0.68 | 0.64 |
| 2 | 49007238 | rs7574612 | 0.989 | 0.848 | -0.528 | 1.127 | 0.003 | 0.011 | 0.011 | 0.134 | 0.64 | 0.4 | 0.44 | 0.45 | 0.67 | 0.6 |
| 2 | 49009112 | rs7595773 | -0.797 | 1.019 | -0.65 | 0.296 | 0.03 | -0.099 | -0.099 | 0.058 | 0.17 | 0.34 | 0.29 | 0.04 | 0.17 | 0.23 |
| 2 | 49011435 | rs4256003 | 0.417 | 0.474 | -1.977 | 0.72 | -0.005 | -0.097 | -0.097 | 0.956 | 0.63 | 0.96 | 0.59 | 0.65 | 0.73 | 0.6 |
| 2 | 49013655 | rs11125183 | -0.712 | 0.72 | -1.385 | 0.096 | 0.004 | 0.018 | 0.018 | 1.954 | 0.55 | 0.55 | 0.46 | 0.8 | 0.62 | 0.68 |
| 2 | 49014171 | rs13429511 | -0.137 | 0.943 | -1.687 | 0.004 | 0.016 | 0.017 | 0.017 | 0.016 | 0.32 | 0.51 | 0.29 | 0.63 | 0.39 | 0.45 |
| 2 | 49015548 | rs12620872 | 0.098 | 1.043 | -1.108 | -0.009 | 0.019 | 0.04 | 0.04 | 0 | 0.28 | 0.36 | 0.25 | 0.62 | 0.34 | 0.42 |
| 2 | 49016904 | rs7589251 | -1.165 | 0.791 | -0.846 | -0.131 | 0.014 | 0.049 | 0.049 | 4.966 | 0.53 | 0.65 | 0.46 | 0.78 | 0.61 | 0.67 |
| 2 | 49026034 | rs4602262 | -0.99 | 1.022 | -0.975 | -0.239 | 0.003 | -0.005 | -0.005 | 0 | 0.22 | 0.7 | 0.2 | 0.17 | 0.23 | 0.26 |
| 2 | 49030757 | rs10185108 | -0.942 | 0.914 | -1.132 | -0.24 | -0.05 | 0.085 | 0.085 | 0.223 | 0.44 | 0.12 | 0.34 | 0.75 | 0.46 | 0.54 |
| 2 | 49030976 | rs10208693 | -1.554 | 1.004 | -0.442 | -0.259 | -0.003 | -0.012 | -0.012 | 0 | 0.56 | 0.45 | 0.46 | 0.78 | 0.63 | 0.72 |
| 2 | 49034015 | rs4538253 | -0.641 | 1.28 | -0.185 | -0.029 | -0.018 | 0.012 | 0.012 | 0.347 | 0.33 | 0.23 | 0.27 | 0.76 | 0.36 | 0.48 |
| 2 | 49035123 | rs9309155 | 0.312 | 1.325 | -0.293 | 0.051 | -0.058 | -0.14 | -0.14 | 0 | 0.23 | 0.03 | 0.16 | 0.59 | 0.26 | 0.34 |
| 2 | 49035714 | rs10183901 | 0.11 | 1.334 | -0.287 | -0.004 | -0.032 | -0.027 | -0.027 | 0 | 0.25 | 0.11 | 0.21 | 0.75 | 0.28 | 0.43 |
| 2 | 49041322 | rs7603311 | -1.682 | 1.509 | -0.103 | 0.031 | 0.004 | 0.128 | 0.128 | 1.06 | 0.48 | 0.75 | 0.41 | 0.78 | 0.54 | 0.65 |
| 2 | 49044584 | rs13003416 | 1.127 | 1.611 | -0.06 | 0.164 | -0.019 | 0.04 | 0.04 | 0 | 0.07 | 0 | 0.04 | 0 | 0.07 | 0.03 |
| 2 | 49051382 | rs4533511 | -0.481 | 1.632 | 0.121 | 0.152 | 0.039 | -0.062 | -0.062 | 0 | 0.14 | 0.26 | 0.11 | 0.02 | 0.2 | 0.18 |
| 2 | 49052255 | rs4482537 | -1.3 | 1.662 | 0.116 | 0.143 | -0.031 | 0.063 | 0.063 | 0.135 | 0.41 | 0.72 | 0.35 | 0.77 | 0.46 | 0.58 |
| 2 | 49052293 | rs6734547 | -0.46 | 1.645 | 0.225 | 0.19 | 0.052 | -0.074 | -0.074 | 0 | 0.15 | 0.33 | 0.11 | 0.02 | 0.2 | 0.18 |
| 2 | 49056717 | rs13001380 | -0.184 | 1.393 | 0.366 | 0.266 | 0.052 | 0.073 | 0.073 | 0 | 0.11 | 0.29 | 0.12 | 0.16 | 0.1 | 0.13 |
| 2 | 49069969 | rs6545073 | -0.943 | 1.799 | 0.183 | 0.705 | 0.084 | 0.135 | 0.135 | 0 | 0.23 | 0.47 | 0.24 | 0.53 | 0.37 | 0.44 |
| 2 | 49070405 | rs6545074 | -1.612 | 1.698 | 0.098 | 0.633 | 0.004 | 0.028 | 0.028 | 1.301 | 0.45 | 0.6 | 0.38 | 0.78 | 0.58 | 0.65 |
| 2 | 49072952 | rs13027348 | -0.787 | 1.76 | 0.113 | 0.709 | -0.003 | 0.001 | 0.001 | 0 | 0.29 | 0.27 | 0.23 | 0.21 | 0.4 | 0.34 |
| 2 | 49073208 | rs3934824 | -0.755 | 1.76 | 0.181 | 0.709 | 0.021 | -0.024 | -0.024 | 0 | 0.3 | 0.41 | 0.24 | 0.21 | 0.4 | 0.34 |
| 2 | 49073556 | rs3935550 | 0.623 | 1.773 | 0.173 | 0.696 | 0.012 | -0.047 | -0.047 | 0 | 0.09 | 0.14 | 0.06 | 0 | 0.09 | 0.04 |

| | | | | | | | | | | | | | | | | |
|---|---|---|---|---|---|---|---|---|---|---|---|---|---|---|---|---|
| 2 | 49073574 | rs10210521 | 0.424 | 1.791 | 0.09 | 0.817 | -0.008 | -0.003 | -0.003 | 1.368 | 0.05 | 0.07 | 0.09 | 0.17 | 0.07 | 0.14 |
| 2 | 49076821 | rs13017739 | -1.231 | 1.725 | 0.141 | 0.845 | 0.002 | 0.001 | 0.001 | 0.603 | 0.36 | 0.39 | 0.31 | 0.44 | 0.49 | 0.5 |
| 2 | 49078132 | rs13030632 | -0.008 | 1.719 | 0.128 | 0.664 | -0.012 | 0.001 | 0.001 | 0.037 | 0.21 | 0.16 | 0.17 | 0.21 | 0.33 | 0.33 |
| 2 | 49082000 | rs12613652 | -0.038 | 1.738 | -0.031 | 0.654 | 0.002 | -0.001 | -0.001 | 5.192 | 0.22 | 0.23 | 0.17 | 0.21 | 0.33 | 0.33 |
| 2 | 49083188 | rs11680420 | 0.926 | 1.776 | 0.174 | 0.727 | 0.013 | 0.101 | 0.101 | 0 | 0.1 | 0.15 | 0.08 | 0.39 | 0.11 | 0.16 |
| 2 | 49086399 | rs17483876 | 0.413 | 1.8 | 0.048 | 0.671 | 0.05 | -0.028 | -0.028 | 0 | 0.09 | 0.19 | 0.08 | 0.02 | 0.2 | 0.17 |
| 2 | 49092438 | rs10187173 | -1.34 | 1.837 | 0.08 | 0.607 | 0.008 | 0.005 | 0.005 | 0 | 0.32 | 0.56 | 0.3 | 0.59 | 0.46 | 0.49 |
| 2 | 49092782 | rs12614310 | 0.877 | 1.787 | -0.035 | 0.599 | 0.006 | -0.015 | -0.015 | 0 | 0.14 | 0.09 | 0.1 | 0.18 | 0.13 | 0.16 |
| 2 | 49093258 | rs10490128 | -0.161 | 1.436 | 0.086 | 0.703 | 0.037 | 0.184 | 0.184 | 0 | 0.14 | 0.25 | 0.16 | 0.57 | 0.18 | 0.28 |
| 2 | 49093975 | rs17556008 | 0.869 | 1.323 | -0.469 | 0.915 | -0.002 | -0.004 | -0.004 | 0 | 0.07 | 0.02 | 0.04 | 0 | 0.07 | 0.03 |
| 2 | 49096214 | rs1558603 | -0.114 | 1.339 | -0.636 | 0.949 | 0.016 | 0.027 | 0.027 | 1.384 | 0.26 | 0.31 | 0.27 | 0.4 | 0.4 | 0.44 |
| 2 | 49096835 | rs733726 | -0.549 | 0.797 | -1.043 | 1.084 | 0.05 | 0.153 | 0.153 | 0 | 0.07 | 0.27 | 0.12 | 0.18 | 0.08 | 0.12 |
| 2 | 49099508 | rs10203937 | -0.452 | 0.775 | -1.486 | 1.098 | 0.006 | 0.011 | 0.011 | 0 | 0.14 | 0.23 | 0.15 | 0.18 | 0.15 | 0.17 |
| 2 | 49110303 | rs17037739 | -0.242 | 0.413 | -2.024 | 0.906 | 0.013 | -0.047 | -0.047 | 0.074 | 0.23 | 0.04 | 0.39 | 0.22 | 0.2 | 0.28 |
| 2 | 49114148 | rs10490126 | -0.577 | 0.774 | -1.52 | 1.095 | -0.002 | -0.006 | -0.006 | 0.981 | 0.16 | 0.18 | 0.16 | 0.2 | 0.15 | 0.21 |
| 2 | 49114231 | rs9789744 | -1.113 | 1.513 | -1.097 | 1.439 | 0.018 | 0.016 | 0.016 | 0.041 | 0.28 | 0.38 | 0.24 | 0.63 | 0.34 | 0.39 |
| 2 | 49114584 | rs9789406 | -0.931 | 0.898 | -2.23 | 0.362 | -0.008 | 0.12 | 0.12 | 3.011 | 0.63 | 0.88 | 0.54 | 0.81 | 0.7 | 0.78 |
| 2 | 49116885 | rs10202351 | -0.509 | 0.682 | -1.562 | 1.102 | 0 | 0.075 | 0.075 | 0 | 0.07 | 0.16 | 0.12 | 0.26 | 0.07 | 0.04 |
| 2 | 49121258 | rs7579295 | 0.439 | 0.195 | -1.947 | 0.768 | 0.012 | 0.014 | 0.014 | 0 | 0.12 | 0.13 | 0.05 | 0.06 | 0.06 | 0.06 |
| 2 | 49122333 | rs7570675 | -0.715 | 0.132 | -1.769 | 0.534 | 0.001 | -0.012 | -0.012 | 0 | 0.4 | 0.58 | 0.62 | 0.73 | 0.46 | 0.49 |
| 2 | 49122750 | rs4953638 | -0.604 | 0.124 | -1.672 | 0.453 | -0.005 | 0.01 | 0.01 | 0 | 0.36 | 0.34 | 0.57 | 0.72 | 0.4 | 0.48 |
| 2 | 49124040 | rs741230 | -1.673 | 0.164 | -2.136 | 0.187 | -0.146 | 0.184 | 0.184 | 0 | 0.62 | 0.97 | 0.77 | 0.91 | 0.58 | 0.71 |
| 2 | 49126802 | rs4953640 | 0.664 | 0.134 | -1.784 | 0.489 | -0.01 | -0.12 | -0.12 | 0 | 0.18 | 0.04 | 0.34 | 0.44 | 0.16 | 0.33 |
| 2 | 49141838 | rs17038295 | 0.025 | -0.035 | -0.501 | 0.455 | -0.028 | -0.083 | -0.083 | 2.566 | 0.19 | 0.45 | 0.14 | 0.06 | 0.11 | 0.15 |
| 2 | 49143829 | rs10490120 | 0.289 | 0.107 | -1.34 | 0.921 | -0.006 | -0.003 | -0.003 | 0.047 | 0.07 | 0.02 | 0.31 | 0.47 | 0.07 | 0.31 |
| 2 | 49153675 | rs6545082 | -0.103 | -0.168 | -1.571 | 0.165 | -0.001 | -0.06 | -0.06 | 0 | 0.23 | 0.23 | 0.15 | 0.01 | 0.24 | 0.12 |
| 2 | 49156580 | rs11891861 | -1.135 | 0.027 | -2.329 | 0.525 | -0.081 | -0.007 | -0.007 | 0 | 0.56 | 0.97 | 0.65 | 0.66 | 0.44 | 0.66 |
| 2 | 49156953 | rs17037869 | 0.297 | 0.389 | -1.214 | 1.379 | 0.011 | 0.007 | 0.007 | 0 | 0.07 | 0.01 | 0.28 | 0.44 | 0.05 | 0.29 |
| 2 | 49158325 | rs989373 | 0.735 | 0.944 | -0.64 | 0.585 | 0.021 | -0.025 | -0.025 | 0.957 | 0.69 | 0.48 | 0.62 | 0.28 | 0.64 | 0.66 |
| 2 | 49161779 | rs988921 | 0.414 | 0.872 | -0.671 | 0.513 | -0.001 | -0.009 | -0.009 | 0.307 | 0.64 | 0.66 | 0.68 | 0.7 | 0.62 | 0.64 |
| 2 | 49161923 | rs1950145 | 1.211 | 0.893 | -0.549 | 1.01 | 0.005 | 0.04 | 0.04 | 0 | 0.52 | 0.24 | 0.53 | 0.41 | 0.52 | 0.55 |
| 2 | 49165092 | rs7565565 | 1.358 | 1.251 | -0.604 | 0.907 | 0.001 | 0.038 | 0.038 | 0.007 | 0.5 | 0.24 | 0.44 | 0.4 | 0.47 | 0.52 |
| 2 | 49169599 | rs1019277 | 0.258 | 1.216 | -0.414 | 0.757 | 0.002 | -0.006 | -0.006 | 0 | 0.72 | 0.7 | 0.75 | 0.69 | 0.74 | 0.74 |
| 2 | 49170035 | rs1024776 | -0.479 | 1.063 | -0.287 | 0.768 | 0 | 0.01 | 0.01 | 0 | 0.19 | 0.31 | 0.19 | 0.2 | 0.2 | 0.16 |
| 2 | 49171837 | rs4952931 | 0.64 | 1.388 | -0.236 | 0.86 | 0.001 | -0.009 | -0.009 | 0 | 0.59 | 0.64 | 0.6 | 0.69 | 0.54 | 0.64 |
| 2 | 49172049 | rs6739570 | 0.113 | 1.262 | -0.086 | 0.96 | 0.024 | 0.187 | 0.187 | 0 | 0.06 | 0.25 | 0.11 | 0.25 | 0.06 | 0.08 |
| 2 | 49175240 | rs17500266 | 1.139 | 0.954 | -0.284 | 0.781 | 0.002 | -0.004 | -0.004 | 0 | 0.52 | 0.36 | 0.5 | 0.48 | 0.52 | 0.55 |
| 2 | 49178436 | rs11125188 | 1.39 | 0.732 | -0.555 | 0.543 | -0.003 | 0.003 | 0.003 | 0 | 0.45 | 0.43 | 0.44 | 0.4 | 0.46 | 0.47 |
| 2 | 49181007 | rs12468889 | 0.82 | 0.726 | -0.635 | 0.613 | 0.028 | -0.015 | -0.015 | 0 | 0.54 | 0.26 | 0.55 | 0.5 | 0.57 | 0.52 |
| 2 | 49183409 | rs4140979 | 0.247 | 0.688 | -0.443 | 0.498 | 0.024 | 0.036 | 0.036 | 0 | 0.62 | 0.69 | 0.73 | 0.74 | 0.66 | 0.61 |
| 2 | 49184327 | rs1922476 | 0.612 | 0.732 | -0.621 | 0.561 | 0.011 | 0.041 | 0.041 | 0 | 0.58 | 0.36 | 0.65 | 0.69 | 0.59 | 0.59 |
| 2 | 49184338 | rs1922475 | -0.148 | 0.75 | -0.465 | 0.575 | 0.046 | 0.069 | 0.069 | 0 | 0.16 | 0.32 | 0.19 | 0.23 | 0.19 | 0.17 |
| 2 | 49184668 | rs12465216 | 1.127 | 0.741 | -0.623 | 0.721 | -0.014 | 0.022 | 0.022 | 0 | 0.05 | 0.04 | 0.13 | 0.24 | 0.09 | 0.07 |
| 2 | 49185716 | rs6545087 | -0.544 | 0.728 | -0.579 | 0.689 | 0.003 | 0.016 | 0.016 | 0 | 0.17 | 0.61 | 0.25 | 0.24 | 0.19 | 0.19 |
| 2 | 49186702 | rs7559066 | 1.517 | 0.421 | -0.761 | 0.525 | -0.046 | -0.026 | -0.026 | 0 | 0.43 | 0.22 | 0.47 | 0.48 | 0.47 | 0.53 |
| 2 | 49187452 | rs6711198 | 0.536 | 0.443 | -0.739 | 0.578 | 0.009 | 0.075 | 0.075 | 0 | 0.07 | 0.17 | 0.14 | 0.23 | 0.09 | 0.07 |
| 2 | 49189921 | rs6166 | 0.848 | 0.364 | -0.685 | 0.494 | 0 | -0.001 | -0.001 | 0.637 | 0.51 | 0.6 | 0.58 | 0.68 | 0.55 | 0.56 |
| 2 | 49191041 | rs6165 | 1.305 | 0.376 | -0.753 | 0.507 | 0.016 | 0.058 | 0.058 | 0 | 0.46 | 0.24 | 0.58 | 0.66 | 0.55 | 0.54 |
| 2 | 49191665 | rs12997223 | 1.702 | 0.351 | -0.859 | 0.468 | 0.001 | 0 | 0 | 0 | 0.07 | 0.09 | 0.18 | 0.12 | 0.06 | 0.2 |
| 2 | 49194702 | rs2300439 | 1.233 | 0.093 | -0.712 | 0.457 | 0.042 | 0.019 | 0.019 | 0 | 0.51 | 0.26 | 0.56 | 0.48 | 0.57 | 0.53 |
| 2 | 49197461 | rs13415618 | 0.898 | 0.121 | -0.679 | 0.477 | 0.002 | 0.055 | 0.055 | 0 | 0.1 | 0.24 | 0.15 | 0.22 | 0.09 | 0.08 |
| 2 | 49197684 | rs759287 | -0.011 | 0.084 | -0.697 | 0.421 | 0.007 | 0.002 | 0.002 | 0 | 0.68 | 0.74 | 0.72 | 0.73 | 0.68 | 0.71 |
| 2 | 49198180 | rs6705106 | -0.046 | 0.082 | -0.729 | 0.41 | 0.001 | -0.008 | -0.008 | 0.654 | 0.69 | 0.82 | 0.72 | 0.73 | 0.68 | 0.71 |
| 2 | 49199448 | rs12713027 | -0.526 | 0.039 | -0.529 | 0.474 | 0.006 | 0.011 | 0.011 | 0 | 0.23 | 0.36 | 0.26 | 0.26 | 0.24 | 0.2 |
| 2 | 49201328 | rs2268363 | -0.172 | -0.139 | -0.506 | 0.726 | -0.001 | 0.07 | 0.07 | 0 | 0.15 | 0.47 | 0.14 | 0.22 | 0.14 | 0.14 |
| 2 | 49201453 | rs6545091 | 0.046 | -0.28 | -0.691 | 0.633 | 0.012 | -0.013 | -0.013 | 0.564 | 0.75 | 0.74 | 0.76 | 0.71 | 0.79 | 0.69 |
| 2 | 49201612 | rs2268361 | 1.472 | -0.31 | -0.757 | 0.627 | 0.065 | -0.011 | -0.011 | 0 | 0.55 | 0.25 | 0.63 | 0.5 | 0.64 | 0.54 |
| 2 | 49202485 | rs989359 | -0.118 | -0.269 | -0.774 | 0.678 | 0.019 | -0.075 | -0.075 | 0.331 | 0.76 | 0.92 | 0.8 | 0.69 | 0.79 | 0.69 |
| 2 | 49205110 | rs2268359 | 0.043 | -0.23 | -0.725 | 0.66 | 0.015 | 0.099 | 0.099 | 0 | 0.07 | 0.21 | 0.18 | 0.27 | 0.11 | 0.09 |
| 2 | 49206571 | rs1558607 | 0.449 | -0.332 | -0.765 | 0.631 | 0.017 | -0.026 | -0.026 | 0 | 0.72 | 0.75 | 0.75 | 0.64 | 0.76 | 0.68 |
| 2 | 49207493 | rs2300437 | 0.256 | -0.343 | -0.645 | 0.51 | -0.031 | 0.033 | 0.033 | 0 | 0.16 | 0.52 | 0.14 | 0.18 | 0.12 | 0.12 |
| 2 | 49209034 | rs1007541 | -0.624 | -0.046 | -0.614 | 0.671 | 0.009 | 0.068 | 0.068 | 0 | 0.09 | 0.21 | 0.21 | 0.27 | 0.11 | 0.08 |
| 2 | 49217916 | rs2072486 | 1.276 | 0.212 | -0.842 | 0.722 | -0.001 | 0.029 | 0.029 | 0 | 0.48 | 0.14 | 0.44 | 0.42 | 0.5 | 0.51 |
| 2 | 49219124 | rs10186089 | 0.255 | -0.424 | -0.468 | 0.771 | 0.001 | 0.028 | 0.028 | 0 | 0.11 | 0.19 | 0.17 | 0.26 | 0.11 | 0.1 |
| 2 | 49220080 | rs1922466 | 0.726 | -0.406 | -0.46 | 0.676 | 0.046 | -0.03 | -0.03 | 0 | 0.58 | 0.27 | 0.62 | 0.52 | 0.65 | 0.55 |
| 2 | 49224580 | rs1922463 | 0.75 | -0.352 | -0.599 | 0.665 | 0.051 | -0.022 | -0.022 | 0 | 0.58 | 0.28 | 0.62 | 0.51 | 0.65 | 0.55 |
| 2 | 49228368 | rs12713029 | 0.039 | -0.635 | -0.493 | 0.219 | 0.025 | 0.013 | 0.013 | 0 | 0.69 | 0.7 | 0.81 | 0.77 | 0.76 | 0.63 |
| 2 | 49230159 | rs917944 | 0.421 | -0.681 | -0.218 | 0.202 | -0.064 | -0.035 | -0.035 | 2.165 | 0.64 | 0.39 | 0.78 | 0.76 | 0.76 | 0.63 |
| 2 | 49230413 | rs1922469 | 0.1 | -0.431 | -0.771 | 0.34 | 0.044 | 0.02 | 0.02 | 0 | 0.69 | 0.75 | 0.81 | 0.76 | 0.76 | 0.63 |

| | | | | | | | | | | | | | | | | |
|---|---|---|---|---|---|---|---|---|---|---|---|---|---|---|---|---|
| 2 | 49244559 | rs2091787 | -0.026 | -0.201 | -0.95 | -0.071 | -0.003 | -0.02 | -0.02 | 0 | 0.53 | 0.55 | 0.65 | 0.78 | 0.53 | 0.56 |
| 2 | 49247832 | rs2349415 | -0.686 | -0.266 | -0.992 | -0.234 | -0.006 | -0.028 | -0.028 | 0 | 0.69 | 0.52 | 0.7 | 0.76 | 0.68 | 0.61 |
| 2 | 49250971 | rs12713034 | 0.024 | -0.041 | -0.82 | 0.674 | -0.049 | 0.012 | 0.012 | 1.385 | 0.36 | 0.23 | 0.27 | 0.29 | 0.43 | 0.34 |
| 2 | 49251260 | rs17038059 | 0.896 | -0.299 | -0.827 | 0.597 | -0.009 | 0.054 | 0.054 | 0.757 | 0.25 | 0.35 | 0.43 | 0.47 | 0.21 | 0.24 |
| 2 | 49251847 | rs9808506 | 0.123 | -0.227 | -0.895 | 0.608 | -0.049 | 0.009 | 0.009 | 1.393 | 0.36 | 0.23 | 0.28 | 0.29 | 0.43 | 0.35 |
| 2 | 49252066 | rs9808508 | 0.116 | -0.072 | -0.997 | 0.596 | -0.122 | -0.029 | -0.029 | 0 | 0.35 | 0.07 | 0.27 | 0.29 | 0.43 | 0.35 |
| 2 | 49257758 | rs17038087 | 1.098 | -0.116 | -0.824 | 0.751 | -0.009 | 0.06 | 0.06 | 1.632 | 0.23 | 0.35 | 0.42 | 0.41 | 0.2 | 0.23 |
| 2 | 49258829 | rs13009588 | 0.938 | 0.137 | -1.286 | 0.661 | -0.089 | -0.062 | -0.062 | 0 | 0.25 | 0.06 | 0.23 | 0.28 | 0.35 | 0.3 |
| 2 | 49259390 | rs17038094 | 0.572 | 0.17 | -0.75 | 0.706 | -0.001 | 0.012 | 0.012 | 0.695 | 0.27 | 0.28 | 0.4 | 0.49 | 0.23 | 0.26 |
| 2 | 49260314 | rs1553476 | 1.451 | 0.235 | -0.61 | 0.494 | -0.011 | -0.019 | -0.019 | 0 | 0.1 | 0.18 | 0.04 | 0.05 | 0.05 | 0.03 |
| 2 | 49265885 | rs6545092 | -0.187 | 0.253 | -0.248 | 0.684 | 0.026 | 0.031 | 0.031 | 1.208 | 0.35 | 0.48 | 0.52 | 0.45 | 0.43 | 0.38 |
| 2 | 49266314 | rs6545094 | -0.076 | 0.259 | -0.237 | 0.687 | 0.01 | 0.012 | 0.012 | 1.777 | 0.34 | 0.38 | 0.48 | 0.45 | 0.43 | 0.36 |
| 2 | 49267193 | rs1277459 | 2.332 | 0.573 | -1.021 | 0.522 | -0.042 | -0.033 | -0.033 | 0 | 0.07 | 0.01 | 0.14 | 0.15 | 0.17 | 0.1 |
| 2 | 49268163 | rs17038110 | 1.412 | 0.323 | -0.764 | 0.591 | 0.005 | 0.006 | 0.006 | 0 | 0.1 | 0.08 | 0.03 | 0.06 | 0.05 | 0.03 |
| 2 | 49268787 | rs10865237 | 1.669 | 0.206 | -0.864 | 0.554 | 0.001 | -0.008 | -0.008 | 0 | 0.18 | 0.12 | 0.17 | 0.2 | 0.16 | 0.19 |
| 2 | 49268804 | rs10865238 | -1.305 | 0.105 | -0.626 | 0.486 | 0 | -0.002 | -0.002 | 0.045 | 0.62 | 0.67 | 0.71 | 0.76 | 0.63 | 0.61 |
| 2 | 49275862 | rs13004879 | 0.241 | -0.024 | -0.868 | 0.679 | -0.047 | -0.04 | -0.04 | 4.595 | 0.42 | 0.18 | 0.46 | 0.48 | 0.5 | 0.49 |
| 2 | 49276242 | rs3913665 | -1.341 | -0.139 | -0.808 | 0.623 | 0.004 | 0.031 | 0.031 | 0 | 0.62 | 0.36 | 0.69 | 0.74 | 0.62 | 0.57 |
| 2 | 49276771 | rs17038169 | 1.912 | -0.008 | -0.832 | 0.673 | -0.005 | -0.005 | -0.005 | 0.329 | 0.18 | 0.09 | 0.16 | 0.17 | 0.16 | 0.17 |
| 2 | 49277823 | rs1504175 | -0.849 | -0.04 | -0.856 | 0.675 | 0.005 | 0.005 | 0.005 | 0 | 0.57 | 0.33 | 0.5 | 0.53 | 0.56 | 0.5 |
| 2 | 49278098 | rs1504178 | 0.934 | 0.005 | -0.904 | 0.699 | -0.06 | -0.061 | -0.061 | 0.297 | 0.25 | 0.09 | 0.3 | 0.3 | 0.35 | 0.31 |
| 2 | 49279644 | rs1504186 | -0.396 | -0.009 | -0.573 | 0.752 | -0.032 | 0.056 | 0.056 | 0 | 0.19 | 0.47 | 0.22 | 0.26 | 0.11 | 0.08 |
| 2 | 49292227 | rs11676909 | 0.64 | 0.112 | -0.657 | 0.704 | -0.054 | -0.052 | -0.052 | 0 | 0.37 | 0.17 | 0.41 | 0.48 | 0.42 | 0.46 |
| 2 | 49293231 | rs11689714 | -0.233 | 0.12 | -0.604 | 0.661 | -0.003 | 0.026 | 0.026 | 0 | 0.46 | 0.63 | 0.61 | 0.68 | 0.48 | 0.52 |
| 2 | 49298586 | rs13007274 | 0.884 | 0.241 | -0.639 | 0.784 | 0.001 | 0.007 | 0.007 | 2.457 | 0.23 | 0.22 | 0.25 | 0.36 | 0.27 | 0.3 |
| 2 | 49300234 | rs12622212 | 1.221 | 0.28 | -0.628 | 0.801 | 0.001 | 0.004 | 0.004 | 2.842 | 0.2 | 0.21 | 0.25 | 0.29 | 0.26 | 0.3 |
| 2 | 49305933 | rs905668 | 1.501 | 0.53 | -0.705 | 0.769 | -0.004 | -0.002 | -0.002 | 2.662 | 0.17 | 0.13 | 0.24 | 0.29 | 0.21 | 0.27 |
| 2 | 49317148 | rs12052611 | -0.901 | 0.467 | -0.681 | 0.459 | 0.003 | 0.023 | 0.023 | 0 | 0.53 | 0.61 | 0.63 | 0.74 | 0.49 | 0.52 |
| 2 | 49330545 | rs7590213 | 0.556 | 0.396 | -0.581 | 0.582 | 0.007 | 0.003 | 0.003 | 1.089 | 0.31 | 0.26 | 0.22 | 0.26 | 0.22 | 0.2 |
| 2 | 49332370 | rs971831 | -0.375 | 0.402 | -0.616 | 0.579 | 0.004 | 0.031 | 0.031 | 0 | 0.35 | 0.5 | 0.46 | 0.53 | 0.32 | 0.34 |
| 2 | 49332877 | rs4510264 | -1.245 | 0.641 | -0.581 | 0.522 | 0 | -0.007 | -0.007 | 0 | 0.6 | 0.6 | 0.69 | 0.76 | 0.58 | 0.54 |
| 2 | 49342052 | rs9309159 | 0.242 | 0.576 | -0.578 | 0.82 | -0.043 | -0.052 | -0.052 | 0 | 0.33 | 0.11 | 0.32 | 0.36 | 0.35 | 0.36 |
| 2 | 49342921 | rs10495966 | 0.736 | 0.955 | -0.225 | 0.953 | 0.014 | 0.093 | 0.093 | 0.296 | 0.21 | 0.32 | 0.41 | 0.48 | 0.23 | 0.3 |
| 2 | 49353428 | rs1394203 | 0.834 | 0.818 | -0.221 | 1.003 | 0.006 | 0.05 | 0.05 | 0 | 0.2 | 0.24 | 0.4 | 0.48 | 0.23 | 0.31 |
| 2 | 49356217 | rs17038310 | 1.293 | 0.8 | -0.2 | 0.917 | 0.016 | -0.009 | -0.009 | 0 | 0.22 | 0.12 | 0.17 | 0.2 | 0.17 | 0.19 |
| 2 | 49356721 | rs10191244 | -0.213 | 0.808 | -0.156 | 0.912 | 0.001 | 0.004 | 0.004 | 0 | 0.42 | 0.44 | 0.56 | 0.68 | 0.39 | 0.49 |
| 2 | 49359676 | rs1553474 | 1.84 | 0.82 | -0.184 | 0.956 | -0.022 | -0.043 | -0.043 | 6.795 | 0.15 | 0.06 | 0.2 | 0.25 | 0.18 | 0.25 |
| 2 | 49362676 | rs10432668 | 0.76 | 0.619 | -0.115 | 0.847 | -0.022 | -0.005 | -0.005 | 0 | 0.25 | 0.42 | 0.19 | 0.2 | 0.17 | 0.19 |
| 2 | 49366047 | rs17038320 | 1.557 | 0.582 | -0.19 | 0.847 | 0.003 | -0.019 | -0.019 | 0.065 | 0.19 | 0.09 | 0.17 | 0.2 | 0.17 | 0.19 |
| 2 | 49371129 | rs1504179 | -0.621 | 0.469 | -0.252 | 0.777 | 0 | -0.004 | -0.004 | 0 | 0.5 | 0.5 | 0.6 | 0.68 | 0.49 | 0.53 |
| 2 | 49375023 | rs13022022 | 0.929 | 0.443 | -0.297 | 0.826 | 0.006 | 0.021 | 0.021 | 0.409 | 0.24 | 0.26 | 0.43 | 0.48 | 0.31 | 0.34 |
| 2 | 49379733 | rs988103 | -0.421 | 0.413 | -0.278 | 0.834 | 0 | 0.047 | 0.047 | 0 | 0.47 | 0.66 | 0.6 | 0.67 | 0.45 | 0.52 |
| 2 | 49381585 | rs1394205 | 0.845 | 0.697 | -0.716 | 0.913 | 0.006 | 0.028 | 0.028 | 0.373 | 0.23 | 0.26 | 0.42 | 0.47 | 0.29 | 0.34 |
| 2 | 49391013 | rs10495963 | 1.762 | 1.033 | -0.968 | 1.005 | -0.013 | -0.013 | -0.013 | 0 | 0.17 | 0.11 | 0.29 | 0.26 | 0.27 | 0.28 |
| 2 | 49394735 | rs13428062 | 0.556 | 0.504 | -1.104 | 0.622 | 0.002 | 0.028 | 0.028 | 0.401 | 0.19 | 0.09 | 0.11 | 0.1 | 0.17 | 0.1 |
| 2 | 49400622 | rs13019040 | 0.026 | -0.102 | -1.363 | 0.68 | -0.014 | 0.043 | 0.043 | 0 | 0.47 | 0.24 | 0.62 | 0.61 | 0.54 | 0.57 |
| 2 | 49415714 | rs4971697 | 1.336 | -0.096 | -1.354 | 0.81 | -0.005 | -0.003 | -0.003 | 0 | 0.23 | 0.21 | 0.37 | 0.45 | 0.34 | 0.33 |
| 2 | 49416106 | rs1027715 | 2.416 | -0.344 | -1.506 | 0.707 | 0.023 | 0.008 | 0.008 | 0 | 0.19 | 0.04 | 0.28 | 0.17 | 0.15 | 0.19 |
| 2 | 49418031 | rs9309160 | -1.597 | -0.42 | -1.202 | 0.812 | 0.004 | 0.001 | 0.001 | 0 | 0.69 | 0.69 | 0.54 | 0.65 | 0.66 | 0.64 |
| 2 | 49418070 | rs10180596 | -0.365 | -0.576 | -1.386 | 0.681 | -0.006 | 0.043 | 0.043 | 0 | 0.53 | 0.36 | 0.49 | 0.68 | 0.55 | 0.53 |
| 2 | 49420410 | rs6720857 | 1.224 | -0.562 | -1.305 | 0.697 | 0.003 | -0.021 | -0.021 | 0 | 0.35 | 0.48 | 0.41 | 0.3 | 0.34 | 0.35 |
| 2 | 49421571 | rs12713042 | -0.018 | -0.496 | -1.269 | 0.767 | 0.005 | -0.001 | -0.001 | 0 | 0.39 | 0.42 | 0.46 | 0.58 | 0.46 | 0.42 |
| 2 | 49426468 | rs13016730 | -0.582 | 0.088 | -0.816 | 1.187 | 0 | -0.004 | -0.004 | 0 | 0.42 | 0.49 | 0.44 | 0.6 | 0.45 | 0.49 |
| 2 | 49427138 | rs11125224 | 0.936 | 0.05 | -0.74 | 1.132 | 0.047 | 0.1 | 0.1 | 1.106 | 0.22 | 0.38 | 0.33 | 0.43 | 0.31 | 0.32 |
| 2 | 49428225 | rs4337506 | 0.648 | 0.073 | -0.728 | 1.122 | -0.007 | 0.031 | 0.031 | 0 | 0.52 | 0.68 | 0.56 | 0.44 | 0.48 | 0.42 |
| 2 | 49431384 | rs13025387 | 0.062 | 0.077 | -0.752 | 1.06 | -0.003 | -0.006 | -0.006 | 0 | 0.33 | 0.32 | 0.35 | 0.54 | 0.36 | 0.44 |
| 2 | 49435907 | rs6739228 | -0.297 | 0.199 | -0.638 | 1.165 | 0.036 | 0.061 | 0.061 | 1.468 | 0.73 | 0.89 | 0.87 | 0.86 | 0.79 | 0.73 |
| 2 | 49436287 | rs12613133 | -0.144 | 0.256 | -0.558 | 1.189 | 0 | 0 | 0 | 0 | 0.4 | 0.47 | 0.38 | 0.55 | 0.43 | 0.49 |
| 2 | 49446588 | rs7609395 | -0.185 | 0.292 | -0.67 | 1.164 | -0.003 | -0.006 | -0.006 | 0.045 | 0.39 | 0.32 | 0.36 | 0.54 | 0.42 | 0.48 |
| 2 | 49452151 | rs7578571 | -0.716 | 0.307 | -0.671 | 1.178 | 0 | 0.019 | 0.019 | 0 | 0.51 | 0.38 | 0.44 | 0.55 | 0.52 | 0.57 |
| 2 | 49463788 | rs9677134 | 1.839 | 0.315 | -0.797 | 1.035 | -0.021 | -0.03 | -0.03 | 4.406 | 0.06 | 0.14 | 0.01 | 0 | 0 | 0 |
| 2 | 49465171 | rs17038496 | -0.124 | 0.329 | -0.777 | 1.02 | -0.012 | -0.076 | -0.076 | 0 | 0.14 | 0.3 | 0.1 | 0 | 0.1 | 0.1 |
| 2 | 49470272 | rs13000772 | 0.924 | 0.403 | -0.972 | 1.19 | -0.027 | -0.019 | -0.019 | 0 | 0.19 | 0.09 | 0.28 | 0.21 | 0.28 | 0.27 |
| 2 | 49480056 | rs12713044 | -0.208 | 0.407 | -0.937 | 1.247 | -0.012 | 0.008 | 0.008 | 0 | 0.32 | 0.25 | 0.35 | 0.22 | 0.37 | 0.37 |
| 2 | 49480306 | rs1427446 | -0.665 | 0.478 | -0.943 | 1.339 | 0 | -0.002 | -0.002 | 0 | 0.52 | 0.52 | 0.43 | 0.36 | 0.49 | 0.56 |
| 2 | 49493299 | rs2350021 | 0.769 | 0.8 | -1.225 | 1.586 | -0.009 | -0.004 | -0.004 | 0.029 | 0.21 | 0.23 | 0.28 | 0.21 | 0.28 | 0.32 |
| 2 | 49498006 | rs9636436 | 0.008 | 0.587 | -1.21 | 1.484 | -0.001 | 0 | 0 | 0 | 0.29 | 0.33 | 0.18 | 0.34 | 0.26 | 0.24 |
| 2 | 49506503 | rs7590557 | 0.11 | 0.624 | -1.338 | 1.186 | 0.033 | 0.053 | 0.053 | 0 | 0.27 | 0.46 | 0.33 | 0.34 | 0.32 | 0.37 |
| 2 | 49507819 | rs10208431 | 0.831 | 0.606 | -1.367 | 1.141 | -0.001 | 0.011 | 0.011 | 0.164 | 0.11 | 0.1 | 0.08 | 0.05 | 0.1 | 0.04 |

| | | | | | | | | | | | | | | | | |
|---|---|---|---|---|---|---|---|---|---|---|---|---|---|---|---|---|
| 2 | 49509374 | rs12992623 | 1.403 | 0.658 | -1.392 | 1.15 | -0.05 | -0.022 | -0.022 | 0 | 0.17 | 0.06 | 0.27 | 0.17 | 0.27 | 0.35 |
| 2 | 49509777 | rs1365862 | 0.699 | 0.593 | -1.394 | 1.122 | 0 | 0 | 0 | 0 | 0.11 | 0.15 | 0.08 | 0.05 | 0.1 | 0.04 |
| 2 | 49510173 | rs7578654 | -0.723 | 0.687 | -1.395 | 1.113 | 0.017 | 0.016 | 0.016 | 0 | 0.38 | 0.51 | 0.43 | 0.41 | 0.45 | 0.43 |
| 2 | 49515633 | rs1344647 | -0.979 | 0.671 | -1.398 | 1.17 | 0 | -0.006 | -0.006 | 0 | 0.54 | 0.59 | 0.43 | 0.6 | 0.54 | 0.58 |
| 2 | 49516618 | rs1365860 | -0.343 | 0.464 | -1.377 | 1.198 | -0.001 | 0.001 | 0.001 | 0 | 0.43 | 0.45 | 0.34 | 0.55 | 0.44 | 0.55 |
| 2 | 49519654 | rs12622975 | 0.287 | 0.292 | -1.431 | 1.055 | 0.033 | -0.008 | -0.008 | 0.041 | 0.27 | 0.13 | 0.14 | 0.27 | 0.22 | 0.21 |
| 2 | 49522163 | rs13412860 | 1.137 | 0.451 | -1.528 | 0.689 | -0.032 | -0.062 | -0.062 | 0 | 0.37 | 0.37 | 0.3 | 0.17 | 0.27 | 0.22 |
| 2 | 49531903 | rs13000527 | 1.091 | 1.027 | -2.292 | 0.959 | -0.054 | -0.016 | -0.016 | 0.574 | 0.18 | 0.07 | 0.28 | 0.17 | 0.31 | 0.37 |
| 2 | 49533498 | rs2350017 | -1.527 | 0.833 | -2.28 | 0.793 | -0.016 | -0.017 | -0.017 | 0 | 0.92 | 0.94 | 0.85 | 0.74 | 0.81 | 0.82 |
| 2 | 49535856 | rs11680024 | 0.388 | 0.682 | -2.037 | 0.35 | -0.001 | -0.003 | -0.003 | 0 | 0.12 | 0.16 | 0.09 | 0.06 | 0.1 | 0.05 |
| 2 | 49537384 | rs7424350 | 0.62 | 0.641 | -2.041 | 0.532 | -0.046 | -0.036 | -0.036 | 0 | 0.18 | 0.07 | 0.25 | 0.21 | 0.3 | 0.36 |
| 2 | 49537795 | rs1991387 | -0.27 | 0.665 | -1.85 | 0.763 | -0.038 | -0.054 | -0.054 | 0 | 0.63 | 0.81 | 0.63 | 0.42 | 0.57 | 0.6 |
| 2 | 49557587 | rs10192212 | -0.725 | 0.614 | -1.196 | 0.449 | -0.001 | -0.002 | -0.002 | 0 | 0.08 | 0.07 | 0.05 | 0.08 | 0.09 | 0.05 |
| 2 | 49559547 | rs7557485 | 0.296 | 0.716 | -1.184 | 0.14 | -0.002 | -0.029 | -0.029 | 0 | 0.46 | 0.44 | 0.31 | 0.2 | 0.45 | 0.32 |
| 2 | 49563508 | rs34058885 | -0.542 | 0.031 | -0.744 | 0.838 | -0.047 | 0.002 | 0.002 | 0 | 0.33 | 0.56 | 0.36 | 0.67 | 0.22 | 0.39 |
| 2 | 49565575 | rs6545107 | -0.338 | 0.113 | -0.694 | 1.169 | -0.019 | 0.04 | 0.04 | 0 | 0.26 | 0.35 | 0.28 | 0.48 | 0.18 | 0.32 |
| 2 | 49571948 | rs817095 | 0.287 | 1.012 | -1.009 | 0.617 | -0.017 | 0.015 | 0.015 | 0 | 0.85 | 0.93 | 0.85 | 0.89 | 0.8 | 0.78 |
| 2 | 49576624 | rs755788 | 1.328 | 1.334 | -0.215 | 0.046 | 0.013 | -0.083 | -0.083 | 0 | 0.71 | 0.55 | 0.75 | 0.85 | 0.75 | 0.74 |
| 2 | 49580566 | rs860994 | -0.412 | 1.368 | 0.023 | 0.031 | -0.006 | -0.004 | -0.004 | 0 | 0.09 | 0.05 | 0.04 | 0 | 0.09 | 0.02 |
| 2 | 49587134 | rs2556407 | -1.964 | 1.308 | 0.133 | 0.314 | 0.012 | -0.017 | -0.017 | 0 | 0.22 | 0.48 | 0.3 | 0.2 | 0.19 | 0.19 |
| 2 | 49588657 | rs817081 | -1.765 | 1.242 | 0.227 | 0.289 | 0.015 | -0.097 | -0.097 | 0 | 0.17 | 0.37 | 0.12 | 0.04 | 0.12 | 0.06 |
| 2 | 49594303 | rs17038608 | -1.162 | 1.21 | 0.719 | 0.444 | 0.079 | 0.189 | 0.189 | 0 | 0.07 | 0.29 | 0.24 | 0.22 | 0.09 | 0.16 |
| 2 | 49596283 | rs817074 | -0.523 | 1.205 | 0.811 | 0.418 | -0.003 | -0.01 | -0.01 | 0 | 0.1 | 0.09 | 0.04 | 0 | 0.09 | 0.02 |
| 2 | 49599540 | rs7583216 | -1.335 | 1.19 | 0.839 | 0.47 | 0.054 | 0.136 | 0.136 | 0 | 0.08 | 0.26 | 0.23 | 0.2 | 0.09 | 0.16 |
| 2 | 49604798 | rs817018 | -0.749 | 1.189 | 0.875 | 0.44 | -0.002 | -0.016 | -0.016 | 0 | 0.11 | 0.1 | 0.05 | 0 | 0.1 | 0.02 |
| 2 | 49605424 | rs817020 | -2.101 | 1.158 | 0.875 | 0.512 | 0.025 | -0.012 | -0.012 | 0 | 0.2 | 0.42 | 0.3 | 0.19 | 0.19 | 0.19 |
| 2 | 49608454 | rs17838375 | -0.452 | 1.145 | 0.828 | 0.511 | 0 | 0.019 | 0.019 | 0 | 0.06 | 0.09 | 0.2 | 0.15 | 0.07 | 0.14 |
| 2 | 49610186 | rs817027 | -1.821 | 1.144 | 0.831 | 0.506 | -0.002 | 0.001 | 0.001 | 0 | 0.17 | 0.17 | 0.3 | 0.16 | 0.17 | 0.17 |
| 2 | 49616270 | rs10469872 | 1.175 | 1.37 | -0.41 | 0.007 | -0.076 | 0.005 | 0.005 | 0 | 0.24 | 0.01 | 0.25 | 0.17 | 0.3 | 0.27 |
| 2 | 49620211 | rs2882306 | 0.904 | 1.488 | 0.143 | 0.083 | 0.079 | -0.091 | -0.091 | 0 | 0.84 | 0.64 | 0.88 | 0.96 | 0.8 | 0.94 |
| 2 | 49623006 | rs817035 | -1.138 | 1.472 | 0.127 | 0.091 | 0.004 | -0.004 | -0.004 | 1.772 | 0.17 | 0.29 | 0.27 | 0.16 | 0.17 | 0.16 |
| 2 | 49630281 | rs1882345 | 1.579 | 1.459 | 0.159 | 0.08 | 0.072 | -0.017 | -0.017 | 1.09 | 0.78 | 0.56 | 0.66 | 0.8 | 0.73 | 0.8 |
| 2 | 49630633 | rs817043 | -0.3 | 1.449 | 0.142 | 0.069 | 0 | -0.033 | -0.033 | 0 | 0.11 | 0.13 | 0.05 | 0 | 0.1 | 0.02 |
| 2 | 49634490 | rs10196852 | 1.121 | 1.453 | 0.149 | 0.068 | 0.084 | -0.106 | -0.106 | 0.1 | 0.83 | 0.61 | 0.87 | 0.96 | 0.8 | 0.94 |
| 2 | 49635681 | rs698840 | 1.016 | 1.376 | 0.11 | 0.116 | 0.108 | 0.097 | 0.097 | 0.016 | 0.89 | 0.66 | 0.72 | 0.8 | 0.83 | 0.81 |
| 2 | 49640979 | rs10170288 | 1.368 | 1.298 | 0.582 | 0.082 | 0.006 | -0.004 | -0.004 | 0.306 | 0.66 | 0.39 | 0.53 | 0.66 | 0.64 | 0.58 |
| 2 | 49641262 | rs1527898 | -0.145 | 1.287 | 0.519 | 0.003 | 0.006 | -0.033 | -0.033 | 0.408 | 0.2 | 0.31 | 0.19 | 0.08 | 0.19 | 0.22 |
| 2 | 49641556 | rs17835319 | 1.412 | 0.632 | -0.405 | 0.238 | 0.005 | -0.038 | -0.038 | 0 | 0.2 | 0.02 | 0.14 | 0.18 | 0.15 | 0.08 |
| 2 | 49642833 | rs17778937 | 1.091 | 0.798 | 0.212 | 0.201 | 0.011 | 0.051 | 0.051 | 0 | 0.45 | 0.16 | 0.39 | 0.33 | 0.42 | 0.3 |
| 2 | 49642896 | rs17835468 | 1.115 | 0.789 | 0.208 | 0.196 | 0.009 | 0.049 | 0.049 | 0 | 0.44 | 0.17 | 0.39 | 0.33 | 0.42 | 0.3 |
| 2 | 49643063 | rs11125238 | 0.171 | 0.504 | 0.49 | 0.287 | -0.002 | 0.005 | 0.005 | 0 | 0.29 | 0.35 | 0.37 | 0.57 | 0.28 | 0.44 |
| 2 | 49644800 | rs17038751 | 1.426 | 0.517 | 0.593 | 0.238 | 0 | -0.022 | -0.022 | 0 | 0.12 | 0.24 | 0.13 | 0.06 | 0.11 | 0.2 |
| 2 | 49646785 | rs10172488 | 1.225 | 0.406 | 0.586 | 0.305 | 0.001 | -0.008 | -0.008 | 0.155 | 0.22 | 0.21 | 0.13 | 0.39 | 0.22 | 0.28 |
| 2 | 49647212 | rs1405959 | -1.36 | 0.445 | 0.633 | 0.257 | -0.011 | 0.049 | 0.049 | 0 | 0.53 | 0.74 | 0.56 | 0.63 | 0.5 | 0.68 |
| 2 | 49649517 | rs817005 | -0.004 | 0.488 | 0.65 | 0.24 | 0.025 | -0.063 | -0.063 | 0 | 0.11 | 0.17 | 0.07 | 0 | 0.12 | 0.05 |
| 2 | 49650145 | rs843840 | -0.397 | 0.49 | 0.615 | 0.241 | 0.024 | -0.075 | -0.075 | 0.403 | 0.13 | 0.23 | 0.07 | 0 | 0.11 | 0.05 |
| 2 | 49651232 | rs7598701 | 1.476 | 0.508 | 0.473 | 0.322 | -0.001 | -0.013 | -0.013 | 0 | 0.12 | 0.18 | 0.13 | 0.06 | 0.11 | 0.2 |
| 2 | 49653672 | rs12105450 | 0.024 | 0.52 | 0.348 | 0.337 | -0.004 | 0.02 | 0.02 | 0 | 0.34 | 0.46 | 0.27 | 0.45 | 0.33 | 0.49 |
| 2 | 49676328 | rs1405969 | 0.45 | 0.567 | 0.491 | 0.314 | 0.002 | 0.009 | 0.009 | 0 | 0.6 | 0.55 | 0.57 | 0.48 | 0.61 | 0.53 |
| 2 | 49678339 | rs10495974 | -0.571 | 0.702 | 0.673 | 0.245 | 0.001 | -0.005 | -0.005 | 0 | 0.25 | 0.31 | 0.4 | 0.24 | 0.24 | 0.37 |
| 2 | 49680236 | rs1921609 | 0.467 | 0.698 | 0.571 | 0.361 | -0.003 | 0.011 | 0.011 | 0.071 | 0.17 | 0.24 | 0.35 | 0.24 | 0.16 | 0.35 |
| 2 | 49680697 | rs4971853 | 0.958 | 0.742 | 0.12 | 0.47 | -0.012 | -0.024 | -0.024 | 0 | 0.05 | 0.01 | 0.21 | 0.21 | 0.05 | 0.16 |
| 2 | 49684567 | rs1405955 | -1.214 | 0.641 | 0.245 | 0.366 | 0.002 | -0.007 | -0.007 | 0 | 0.94 | 0.9 | 0.97 | 0.98 | 0.97 | 0.97 |
| 2 | 49697619 | rs10865241 | 0.984 | 0.408 | -0.274 | 0.154 | -0.021 | 0.015 | 0.015 | 0 | 0.4 | 0.26 | 0.37 | 0.36 | 0.47 | 0.31 |
| 2 | 49700419 | rs10165268 | 1.039 | 0.256 | -0.356 | 0.037 | -0.037 | 0.004 | 0.004 | 0 | 0.37 | 0.21 | 0.37 | 0.35 | 0.47 | 0.32 |
| 2 | 49707932 | rs13427322 | 0.879 | 0.17 | -0.226 | -0.073 | 0 | -0.022 | -0.022 | 2.277 | 0.07 | 0.07 | 0.03 | 0 | 0.06 | 0.01 |
| 2 | 49714129 | rs11888995 | 0.123 | -0.025 | 0.198 | -0.194 | 0.006 | -0.035 | -0.035 | 0 | 0.72 | 0.7 | 0.76 | 0.9 | 0.73 | 0.74 |
| 2 | 49714846 | rs13382313 | 1.927 | -0.062 | -0.363 | -0.392 | -0.028 | -0.016 | -0.016 | 0 | 0.31 | 0.12 | 0.32 | 0.33 | 0.37 | 0.31 |
| 2 | 49715021 | rs12987465 | -0.088 | -0.072 | -0.29 | -0.358 | -0.008 | 0.004 | 0.004 | 0.336 | 0.41 | 0.58 | 0.44 | 0.55 | 0.36 | 0.43 |
| 2 | 49716202 | rs977134 | 0.923 | -0.495 | -0.202 | -0.495 | -0.026 | -0.009 | -0.009 | 0 | 0.18 | 0.21 | 0.22 | 0.18 | 0.08 | 0.19 |
| 2 | 49716853 | rs1405966 | -1.151 | -0.431 | -0.245 | -0.484 | 0.002 | -0.012 | -0.012 | 0.014 | 0.79 | 0.77 | 0.8 | 0.9 | 0.79 | 0.75 |
| 2 | 49722210 | rs4971860 | -0.269 | -0.424 | -0.25 | -0.473 | 0.003 | -0.025 | -0.025 | 0.019 | 0.72 | 0.7 | 0.76 | 0.9 | 0.73 | 0.74 |
| 2 | 49724966 | rs12996690 | 1.029 | -0.316 | -0.312 | -0.447 | -0.01 | -0.022 | -0.022 | 0.052 | 0.21 | 0.19 | 0.21 | 0.39 | 0.28 | 0.24 |
| 2 | 49732935 | rs1553127 | 1.003 | -0.316 | -0.539 | -0.448 | 0.006 | 0.012 | 0.012 | 0 | 0.53 | 0.48 | 0.54 | 0.74 | 0.66 | 0.55 |
| 2 | 49737819 | rs1405965 | -0.107 | -0.297 | -0.664 | -0.446 | 0.001 | 0.021 | 0.021 | 0 | 0.67 | 0.79 | 0.61 | 0.72 | 0.72 | 0.54 |
| 2 | 49739361 | rs6545119 | -0.144 | -0.201 | -0.849 | -0.58 | 0.001 | 0.002 | 0.002 | 0 | 0.77 | 0.77 | 0.71 | 0.79 | 0.83 | 0.73 |
| 2 | 49739812 | rs1527904 | 1.19 | 0.06 | -1.115 | -0.413 | 0.027 | 0.074 | 0.074 | 0 | 0.19 | 0.36 | 0.21 | 0.38 | 0.28 | 0.23 |
| 2 | 49743324 | rs13422448 | 2.439 | 0.141 | -1.513 | -0.462 | -0.068 | 0.057 | 0.057 | 0 | 0.28 | 0.01 | 0.24 | 0.13 | 0.32 | 0.25 |
| 2 | 49744060 | rs17038916 | -0.58 | 0.062 | -1.168 | -0.447 | -0.02 | -0.054 | -0.054 | 0.55 | 0.3 | 0.47 | 0.33 | 0.19 | 0.21 | 0.25 |

| | | | | | | | | | | | | | | | | |
|---|---|---|---|---|---|---|---|---|---|---|---|---|---|---|---|---|
| 2 | 49749038 | rs6709183 | -0.359 | -0.019 | -1.125 | -0.386 | -0.015 | -0.021 | -0.021 | 0 | 0.36 | 0.5 | 0.4 | 0.26 | 0.24 | 0.44 |
| 2 | 49752589 | rs4971621 | 0.91 | 0.158 | -1.553 | -0.199 | -0.025 | 0.014 | 0.014 | 0.172 | 0.48 | 0.15 | 0.48 | 0.43 | 0.53 | 0.53 |
| 2 | 49764298 | rs1116294 | 0.857 | -0.044 | -1.328 | -0.22 | 0.017 | -0.005 | -0.005 | 0 | 0.51 | 0.26 | 0.53 | 0.49 | 0.56 | 0.55 |
| 2 | 49778877 | rs7560656 | 1.08 | -0.38 | -0.66 | -0.129 | -0.009 | -0.036 | -0.036 | 0 | 0.09 | 0.1 | 0.03 | 0 | 0.05 | 0.01 |
| 2 | 49781113 | rs1427414 | 0.489 | -0.406 | -0.528 | -0.142 | -0.02 | -0.06 | -0.06 | 0 | 0.11 | 0.2 | 0.04 | 0 | 0.05 | 0.01 |
| 2 | 49785558 | rs1874912 | 0.105 | -0.404 | -0.501 | -0.153 | -0.024 | -0.069 | -0.069 | 0.001 | 0.12 | 0.2 | 0.04 | 0 | 0.05 | 0.01 |
| 2 | 49788135 | rs17038952 | 1.71 | -0.365 | -0.494 | -0.144 | -0.002 | -0.036 | -0.036 | 0.258 | 0.08 | 0.09 | 0.03 | 0 | 0.05 | 0.01 |
| 2 | 49790063 | rs1995172 | 1.707 | -0.302 | -1.067 | -0.302 | 0.02 | 0.148 | 0.148 | 0 | 0.49 | 0.19 | 0.46 | 0.22 | 0.57 | 0.48 |
| 2 | 49796689 | rs7595953 | 0.444 | -0.379 | -1.122 | -0.268 | -0.022 | -0.066 | -0.066 | 0 | 0.12 | 0.19 | 0.04 | 0 | 0.05 | 0.01 |
| 2 | 49808562 | rs13429217 | 0.491 | -0.244 | -1.121 | -0.042 | -0.02 | -0.007 | -0.007 | 2.623 | 0.21 | 0.21 | 0.21 | 0.19 | 0.09 | 0.19 |
| 2 | 49816534 | rs1593705 | 0.78 | -0.173 | -1.032 | 0.089 | 0.024 | -0.012 | -0.012 | 0 | 0.61 | 0.29 | 0.61 | 0.44 | 0.73 | 0.61 |
| 2 | 49825768 | rs13401858 | 2.097 | -0.14 | -2.197 | -0.465 | -0.043 | 0.006 | 0.006 | 0 | 0.23 | 0.01 | 0.21 | 0.14 | 0.24 | 0.21 |
| 2 | 49829630 | rs1498799 | 1.15 | 0.05 | -1.852 | -0.409 | -0.105 | 0.116 | 0.116 | 0.077 | 0.29 | 0.03 | 0.24 | 0.09 | 0.34 | 0.27 |
| 2 | 49830962 | rs17039054 | -0.111 | -0.212 | -0.941 | 0.13 | -0.014 | -0.01 | -0.01 | 0.307 | 0.22 | 0.21 | 0.23 | 0.18 | 0.1 | 0.18 |
| 2 | 49849268 | rs10179773 | 1.198 | 0.099 | -1.368 | 0.037 | 0.002 | 0.042 | 0.042 | 0 | 0.23 | 0.09 | 0.24 | 0.13 | 0.22 | 0.2 |
| 2 | 49849708 | rs7577053 | 0.104 | -0.088 | -0.937 | 0.347 | -0.001 | 0.049 | 0.049 | 0 | 0.74 | 0.62 | 0.74 | 0.51 | 0.74 | 0.73 |
| 2 | 49850450 | rs1391748 | 0.44 | -0.039 | -1.093 | 0.024 | 0.031 | 0.046 | 0.046 | 0.144 | 0.39 | 0.15 | 0.37 | 0.31 | 0.32 | 0.42 |
| 2 | 49851124 | rs12470758 | 0.563 | -0.11 | -1.065 | 0.549 | 0.001 | -0.007 | -0.007 | 0 | 0.15 | 0.12 | 0.14 | 0.27 | 0.16 | 0.17 |
| 2 | 49852392 | rs1391749 | 1.422 | -0.31 | -1.257 | 0.478 | 0.017 | -0.057 | -0.057 | 0 | 0.09 | 0.01 | 0.17 | 0.22 | 0.05 | 0.16 |
| 2 | 49854117 | rs11895566 | 0.999 | -0.297 | -1.171 | 0.388 | 0.017 | -0.039 | -0.039 | 0.892 | 0.36 | 0.5 | 0.38 | 0.22 | 0.43 | 0.32 |
| 2 | 49855331 | rs4527244 | -0.498 | -0.377 | -1.337 | 0.398 | 0.001 | -0.041 | -0.041 | 0 | 0.82 | 0.92 | 0.86 | 0.72 | 0.83 | 0.82 |
| 2 | 49856202 | rs2114657 | 1.493 | -0.396 | -1.373 | 0.329 | 0.005 | 0.087 | 0.087 | 0.03 | 0.28 | 0.12 | 0.18 | 0.08 | 0.26 | 0.24 |
| 2 | 49856326 | rs976704 | 0.499 | -0.335 | -1.202 | 0.466 | -0.022 | 0.016 | 0.016 | 0.357 | 0.43 | 0.76 | 0.5 | 0.38 | 0.5 | 0.4 |
| 2 | 49857266 | rs1498807 | 0.354 | -0.726 | -1.057 | 0.558 | -0.024 | 0.091 | 0.091 | 0 | 0.77 | 0.62 | 0.76 | 0.55 | 0.82 | 0.76 |
| 2 | 49857536 | rs12713067 | 0.535 | -0.702 | -1.464 | 0.522 | 0.018 | 0.021 | 0.021 | 0 | 0.45 | 0.74 | 0.53 | 0.38 | 0.58 | 0.42 |
| 2 | 49857938 | rs10189404 | -0.905 | -0.704 | -1.708 | 0.54 | 0.022 | -0.03 | -0.03 | 0 | 0.85 | 0.93 | 0.89 | 0.73 | 0.9 | 0.83 |
| 2 | 49858261 | rs13425236 | 0.473 | -0.694 | -1.7 | 0.511 | 0.005 | 0 | 0 | 0.064 | 0.38 | 0.08 | 0.35 | 0.34 | 0.31 | 0.4 |
| 2 | 49861418 | rs1391752 | -0.783 | -0.695 | -1.643 | 0.58 | 0.026 | -0.04 | -0.04 | 0 | 0.85 | 0.93 | 0.89 | 0.7 | 0.9 | 0.83 |
| 2 | 49861898 | rs11125265 | 0.172 | -0.752 | -1.708 | 0.515 | 0.088 | 0.045 | 0.045 | 0.034 | 0.51 | 0.82 | 0.54 | 0.39 | 0.62 | 0.44 |
| 2 | 49862134 | rs7564547 | -0.869 | -0.711 | -1.663 | 0.546 | 0.023 | -0.034 | -0.034 | 0.601 | 0.85 | 0.92 | 0.89 | 0.7 | 0.9 | 0.83 |
| 2 | 49865140 | rs1834782 | 0.212 | -0.733 | -1.689 | 0.539 | 0.082 | 0.03 | 0.03 | 0.209 | 0.48 | 0.82 | 0.54 | 0.39 | 0.62 | 0.44 |
| 2 | 49871330 | rs2350699 | 1.32 | -0.773 | -1.712 | 0.557 | 0.011 | -0.01 | -0.01 | 0 | 0.11 | 0.07 | 0.1 | 0.3 | 0.09 | 0.15 |
| 2 | 49872334 | rs13405761 | 1.757 | -0.635 | -1.602 | 0.597 | 0.004 | 0.031 | 0.031 | 0 | 0.06 | 0.11 | 0.12 | 0.19 | 0.11 | 0.1 |
| 2 | 49872424 | rs1427412 | 1.261 | -0.776 | -1.201 | 0.459 | 0.009 | -0.015 | -0.015 | 0 | 0.35 | 0.34 | 0.38 | 0.2 | 0.48 | 0.32 |
| 2 | 49876591 | rs17794707 | 1.964 | -0.951 | -2.08 | 0.381 | -0.064 | 0.077 | 0.077 | 0.002 | 0.22 | 0.03 | 0.16 | 0.09 | 0.25 | 0.25 |
| 2 | 49878657 | rs17039154 | 1.138 | -1.205 | -1.769 | 0.699 | 0.012 | -0.06 | -0.06 | 6.472 | 0.09 | 0.01 | 0.18 | 0.27 | 0.05 | 0.17 |
| 2 | 49882063 | rs1025248 | -0.342 | -1.296 | -1.795 | 0.698 | 0.03 | -0.047 | -0.047 | 0 | 0.84 | 0.93 | 0.89 | 0.69 | 0.9 | 0.83 |
| 2 | 49883809 | rs10495981 | 1.143 | -1.214 | -1.787 | 0.607 | 0.013 | -0.06 | -0.06 | 0 | 0.09 | 0.01 | 0.18 | 0.26 | 0.05 | 0.18 |
| 2 | 49887106 | rs12713068 | 0.202 | -1.016 | -1.835 | 0.452 | -0.015 | 0.049 | 0.049 | 0 | 0.43 | 0.17 | 0.44 | 0.36 | 0.4 | 0.44 |
| 2 | 49888607 | rs2059713 | 0.398 | -0.99 | -2.012 | 0.441 | -0.063 | 0.048 | 0.048 | 0.021 | 0.41 | 0.08 | 0.42 | 0.32 | 0.4 | 0.44 |
| 2 | 49888676 | rs6545125 | 0.561 | -1.044 | -1.717 | 0.479 | -0.023 | 0.004 | 0.004 | 0.066 | 0.42 | 0.74 | 0.47 | 0.38 | 0.51 | 0.4 |
| 2 | 49894154 | rs10190188 | 0.867 | -0.941 | -1.497 | 0.481 | -0.037 | 0.135 | 0.135 | 0 | 0.33 | 0.11 | 0.24 | 0.11 | 0.34 | 0.25 |
| 2 | 49894534 | rs2350700 | 0.995 | -0.698 | -1.243 | 0.727 | -0.005 | 0.017 | 0.017 | 0 | 0.09 | 0.09 | 0.18 | 0.25 | 0.05 | 0.19 |
| 2 | 49899260 | rs12713070 | -0.086 | -0.789 | -1.34 | 0.656 | 0.025 | 0.029 | 0.029 | 0.34 | 0.49 | 0.24 | 0.51 | 0.59 | 0.45 | 0.57 |
| 2 | 49904589 | rs2882478 | -0.061 | -0.409 | -1.018 | 0.574 | 0.022 | 0.041 | 0.041 | 0 | 0.51 | 0.32 | 0.37 | 0.38 | 0.46 | 0.45 |
| 2 | 49909547 | rs1568287 | -1.124 | -0.229 | -1.012 | 0.897 | 0.019 | -0.04 | -0.04 | 0.183 | 0.25 | 0.75 | 0.31 | 0.46 | 0.18 | 0.31 |
| 2 | 49913955 | rs2350701 | -0.484 | -0.548 | -0.482 | 0.725 | 0 | -0.009 | -0.009 | 0 | 0.55 | 0.58 | 0.52 | 0.69 | 0.46 | 0.58 |
| 2 | 49917644 | rs1518825 | 0.193 | -0.574 | -0.074 | 0.721 | -0.03 | 0.046 | 0.046 | 0.109 | 0.13 | 0.23 | 0.08 | 0.25 | 0.06 | 0.15 |
| 2 | 49926570 | rs1518826 | 1.093 | -0.735 | -0.375 | 0.521 | -0.135 | 0.096 | 0.096 | 0.012 | 0.33 | 0.03 | 0.24 | 0.13 | 0.38 | 0.27 |
| 2 | 49930153 | rs10197605 | -0.248 | -0.47 | -0.121 | 0.522 | 0.01 | 0.028 | 0.028 | 0 | 0.5 | 0.55 | 0.51 | 0.68 | 0.45 | 0.58 |
| 2 | 49941022 | rs17039278 | 0.825 | -0.52 | -0.095 | 0.535 | -0.011 | 0.124 | 0.124 | 0 | 0.09 | 0.25 | 0.08 | 0.26 | 0.06 | 0.15 |
| 2 | 49944137 | rs7595347 | -1.4 | -0.558 | -0.523 | 0.386 | -0.03 | 0.133 | 0.133 | 0 | 0.64 | 0.85 | 0.74 | 0.9 | 0.6 | 0.76 |
| 2 | 49944565 | rs10185615 | 0.971 | -0.545 | -0.538 | 0.445 | -0.076 | 0.093 | 0.093 | 0.064 | 0.35 | 0.08 | 0.26 | 0.16 | 0.4 | 0.29 |
| 2 | 49946543 | rs1159705 | -1.958 | -0.386 | -0.582 | 0.51 | -0.029 | 0.094 | 0.094 | 0 | 0.71 | 0.88 | 0.77 | 0.9 | 0.65 | 0.77 |
| 2 | 49947538 | rs17039201 | 1.368 | -0.416 | -0.509 | 0.705 | -0.001 | -0.035 | -0.035 | 0.577 | 0.08 | 0.01 | 0.18 | 0.3 | 0.06 | 0.16 |
| 2 | 49955497 | rs17039212 | 0.815 | -0.377 | -0.296 | 0.596 | 0.017 | 0.081 | 0.081 | 0 | 0.09 | 0.24 | 0.21 | 0.19 | 0.11 | 0.15 |
| 2 | 49962090 | rs10204080 | 1.477 | -0.254 | -0.382 | 0.472 | -0.026 | 0.097 | 0.097 | 0 | 0.32 | 0.14 | 0.23 | 0.09 | 0.37 | 0.25 |
| 2 | 49963021 | rs7564375 | -0.75 | -0.293 | -0.342 | 0.532 | 0.005 | 0.029 | 0.029 | 0 | 0.87 | 0.76 | 0.78 | 0.82 | 0.87 | 0.84 |
| 2 | 49963077 | rs7593733 | 1.039 | -0.304 | -0.56 | 0.397 | -0.025 | -0.02 | -0.02 | 0 | 0.65 | 0.41 | 0.5 | 0.25 | 0.75 | 0.52 |
| 2 | 49964349 | rs17489439 | 1.61 | 0.035 | -0.577 | 0.518 | -0.076 | 0.042 | 0.042 | 0.858 | 0.27 | 0.04 | 0.21 | 0.15 | 0.34 | 0.26 |
| 2 | 49967393 | rs7562595 | 1.453 | -0.05 | -0.573 | 0.521 | -0.096 | 0.045 | 0.045 | 0.998 | 0.28 | 0.05 | 0.24 | 0.16 | 0.36 | 0.27 |
| 2 | 49972394 | rs2139156 | 0.817 | -0.106 | -0.8 | 0.512 | -0.177 | 0.081 | 0.081 | 0.577 | 0.34 | 0.04 | 0.27 | 0.15 | 0.44 | 0.3 |
| 2 | 49977093 | rs10171772 | 0.268 | -0.284 | -0.49 | 0.684 | -0.003 | -0.004 | -0.004 | 1.253 | 0.42 | 0.37 | 0.45 | 0.45 | 0.44 | 0.45 |
| 2 | 49984263 | rs7594662 | -0.554 | -0.65 | 0.031 | 0.846 | -0.1 | 0.213 | 0.213 | 0 | 0.2 | 0.46 | 0.27 | 0.54 | 0.08 | 0.31 |
| 2 | 49985340 | rs13012739 | 0.655 | -0.544 | -0.34 | 0.451 | -0.027 | 0.144 | 0.144 | 0 | 0.34 | 0.11 | 0.23 | 0.1 | 0.37 | 0.23 |
| 2 | 49990017 | rs7349353 | 0.61 | -1.109 | -0.073 | 0.608 | 0.013 | 0.089 | 0.089 | 0 | 0.34 | 0.12 | 0.21 | 0.19 | 0.32 | 0.29 |
| 2 | 49990368 | rs1518823 | -1.494 | -1.059 | -0.064 | 0.804 | -0.014 | -0.022 | -0.022 | 0 | 0.92 | 0.96 | 0.81 | 0.76 | 0.89 | 0.9 |
| 2 | 49991428 | rs7567970 | 1.062 | -1.364 | 0.07 | 0.859 | -0.025 | 0.026 | 0.026 | 0 | 0.08 | 0.14 | 0.05 | 0.16 | 0.03 | 0.15 |
| 2 | 49992053 | rs7595014 | -0.618 | -1.452 | 0.203 | 0.818 | 0.05 | 0.014 | 0.014 | 0.147 | 0.53 | 0.62 | 0.47 | 0.61 | 0.36 | 0.57 |

| | | | | | | | | | | | | | | | | |
|---|---|---|---|---|---|---|---|---|---|---|---|---|---|---|---|---|
| 2 | 49992135 | rs13406948 | 0.764 | -1.408 | 0.213 | 0.737 | -0.001 | -0.006 | -0.006 | 0.002 | 0.33 | 0.28 | 0.2 | 0.18 | 0.28 | 0.26 |
| 2 | 49997218 | rs870168 | -1.244 | -1.212 | 0.195 | 0.714 | -0.029 | 0.121 | 0.121 | 1.245 | 0.62 | 0.88 | 0.66 | 0.83 | 0.45 | 0.66 |
| 2 | 50004148 | rs7602969 | -1.128 | -1.414 | 0.276 | 0.571 | 0.016 | 0.002 | 0.002 | 0 | 0.57 | 0.63 | 0.48 | 0.62 | 0.39 | 0.57 |
| 2 | 50005959 | rs11903512 | 0.247 | -1.245 | 0.226 | 0.292 | -0.033 | 0.057 | 0.057 | 0 | 0.08 | 0.19 | 0.05 | 0.18 | 0.03 | 0.15 |
| 2 | 50016579 | rs2139162 | 0.489 | -0.708 | -0.022 | 0.428 | 0.005 | 0.009 | 0.009 | 0 | 0.29 | 0.21 | 0.19 | 0.17 | 0.26 | 0.26 |
| 2 | 50022031 | rs17039328 | -0.349 | -0.458 | -0.41 | 0.553 | -0.017 | -0.009 | -0.009 | 0 | 0.43 | 0.51 | 0.26 | 0.36 | 0.31 | 0.44 |
| 2 | 50022788 | rs6736617 | -0.004 | 0.134 | -0.899 | 0.368 | 0.002 | -0.002 | -0.002 | 0 | 0.22 | 0.27 | 0.35 | 0.2 | 0.24 | 0.25 |
| 2 | 50024202 | rs1878856 | -0.093 | -0.205 | -1.441 | -0.109 | 0.058 | 0.108 | 0.108 | 0 | 0.33 | 0.13 | 0.16 | 0.11 | 0.27 | 0.29 |
| 2 | 50028214 | rs7600386 | -0.393 | -0.199 | -0.846 | 0.001 | -0.029 | -0.043 | -0.043 | 0.019 | 0.37 | 0.45 | 0.2 | 0.23 | 0.29 | 0.39 |
| 2 | 50055938 | rs885560 | 1.074 | -0.124 | -1.779 | -0.064 | 0 | 0.05 | 0.05 | 0 | 0.13 | 0.02 | 0.09 | 0 | 0.16 | 0.06 |
| 2 | 50057312 | rs17795521 | -0.431 | -0.184 | -1.917 | -0.157 | 0.058 | 0.133 | 0.133 | 0 | 0.35 | 0.12 | 0.18 | 0.12 | 0.3 | 0.29 |
| 2 | 50057351 | rs13405956 | -0.262 | 1.203 | -1.627 | 0.615 | -0.068 | -0.078 | -0.078 | 0 | 0.87 | 0.97 | 0.8 | 0.68 | 0.77 | 0.84 |
| 2 | 50059808 | rs925931 | 0.641 | 0.649 | -1.269 | 0.118 | 0.004 | 0 | 0 | 0 | 0.13 | 0.07 | 0.1 | 0 | 0.16 | 0.07 |
| 2 | 50060991 | rs6761048 | -0.535 | 0.811 | -1.139 | 0.07 | 0.001 | 0 | 0 | 0 | 0.57 | 0.46 | 0.54 | 0.44 | 0.57 | 0.62 |
| 2 | 50072626 | rs4531970 | -0.263 | 0.82 | -1.484 | 0.007 | -0.051 | 0.061 | 0.061 | 0 | 0.52 | 0.13 | 0.51 | 0.34 | 0.53 | 0.49 |
| 2 | 50074442 | rs13028383 | 1.481 | 0.637 | -1.564 | 0.088 | -0.055 | -0.078 | -0.078 | 4.601 | 0.12 | 0.02 | 0.28 | 0.22 | 0.18 | 0.14 |
| 2 | 50085965 | rs1363047 | 0.45 | 0.242 | -1.742 | -0.219 | 0.04 | 0.059 | 0.059 | 0.015 | 0.24 | 0.08 | 0.12 | 0.12 | 0.18 | 0.27 |
| 2 | 50100865 | rs1592729 | -0.991 | 0.021 | -0.883 | -0.471 | 0.003 | -0.001 | -0.001 | 0 | 0.57 | 0.57 | 0.54 | 0.44 | 0.48 | 0.61 |
| 2 | 50125840 | rs1156742 | -1.224 | 0.078 | -0.843 | -0.594 | -0.023 | -0.006 | -0.006 | 0.183 | 0.6 | 0.62 | 0.56 | 0.42 | 0.51 | 0.65 |
| 2 | 50131767 | rs7558063 | 0.72 | -0.249 | 0.046 | -0.533 | -0.042 | -0.072 | -0.072 | 0.337 | 0.19 | 0.45 | 0.14 | 0.09 | 0.12 | 0.19 |
| 2 | 50142117 | rs971732 | -0.624 | -0.185 | 0.333 | -0.324 | -0.012 | -0.001 | -0.001 | 0 | 0.42 | 0.41 | 0.42 | 0.42 | 0.37 | 0.52 |
| 2 | 50148972 | rs1045881 | 0.594 | -0.338 | 0.181 | -0.405 | 0.027 | 0.032 | 0.032 | 0 | 0.23 | 0.12 | 0.1 | 0.11 | 0.16 | 0.34 |
| 2 | 50149510 | rs17039448 | 0.578 | -0.331 | 0.115 | -0.339 | 0.006 | 0.045 | 0.045 | 0 | 0.18 | 0.22 | 0.3 | 0.31 | 0.19 | 0.17 |
| 2 | 50153280 | rs10495987 | -0.895 | -0.23 | 0.147 | -0.35 | -0.002 | 0 | 0 | 0 | 0.43 | 0.5 | 0.43 | 0.43 | 0.36 | 0.52 |
| 2 | 50157772 | rs12617277 | 0.18 | 0.104 | 0.129 | -0.028 | 0.001 | 0.028 | 0.028 | 0 | 0.19 | 0.25 | 0.29 | 0.3 | 0.19 | 0.17 |
| 2 | 50168239 | rs920035 | 0.765 | 0.203 | -0.199 | 0.03 | -0.003 | -0.006 | -0.006 | 2.662 | 0.16 | 0.12 | 0.28 | 0.3 | 0.19 | 0.17 |
| 2 | 50176965 | rs1421594 | -0.46 | -0.277 | -0.739 | -0.37 | -0.001 | 0 | 0 | 0 | 0.46 | 0.48 | 0.43 | 0.43 | 0.37 | 0.54 |
| 2 | 50179975 | rs2193870 | 1.703 | 0.225 | -1.349 | 0.003 | -0.082 | -0.07 | -0.07 | 0 | 0.15 | 0.01 | 0.2 | 0.24 | 0.24 | 0.2 |
| 2 | 50184372 | rs6713940 | -0.435 | -0.315 | -0.709 | 0.081 | -0.004 | -0.001 | -0.001 | 0 | 0.46 | 0.43 | 0.44 | 0.43 | 0.38 | 0.54 |
| 2 | 50188135 | rs17491881 | 0.878 | -0.413 | -0.912 | -0.048 | 0.036 | 0.068 | 0.068 | 0.092 | 0.26 | 0.1 | 0.12 | 0.1 | 0.18 | 0.29 |
| 2 | 50189738 | rs17039577 | -0.476 | -0.271 | -1.154 | 0.108 | -0.007 | 0 | 0 | 0 | 0.47 | 0.45 | 0.43 | 0.46 | 0.38 | 0.55 |
| 2 | 50193483 | rs17039581 | 1.227 | 0.338 | -1.067 | 0.792 | -0.039 | -0.081 | -0.081 | 0.051 | 0.13 | 0.03 | 0.27 | 0.24 | 0.15 | 0.15 |
| 2 | 50197075 | rs11125280 | -0.194 | 0.022 | -0.775 | 0.672 | -0.011 | 0.007 | 0.007 | 0 | 0.37 | 0.52 | 0.4 | 0.4 | 0.28 | 0.35 |
| 2 | 50201383 | rs4971883 | 0.089 | 0.745 | -0.757 | 0.871 | -0.001 | 0.003 | 0.003 | 0 | 0.26 | 0.29 | 0.47 | 0.46 | 0.3 | 0.33 |
| 2 | 50202021 | rs17439468 | 0.701 | 0.708 | -0.598 | 0.458 | -0.006 | 0.049 | 0.049 | 1.729 | 0.41 | 0.24 | 0.3 | 0.24 | 0.42 | 0.45 |
| 2 | 50202608 | rs10495989 | 1.945 | 0.459 | -0.253 | 0.406 | 0.024 | 0.025 | 0.025 | 0.539 | 0.19 | 0.09 | 0.1 | 0.09 | 0.11 | 0.08 |
| 2 | 50202953 | rs11889551 | 0.059 | 0.454 | -0.03 | 0.424 | 0.006 | -0.006 | -0.006 | 0 | 0.6 | 0.41 | 0.4 | 0.33 | 0.53 | 0.54 |
| 2 | 50209558 | rs2112073 | 0.063 | 0.462 | -0.042 | 0.283 | 0.004 | -0.008 | -0.008 | 1.081 | 0.6 | 0.41 | 0.41 | 0.29 | 0.52 | 0.53 |
| 2 | 50210702 | rs12478797 | -1.368 | 0.475 | -0.044 | 0.296 | 0.015 | 0.034 | 0.034 | 0.018 | 0.86 | 0.71 | 0.87 | 0.79 | 0.84 | 0.87 |
| 2 | 50212116 | rs17492567 | 1.784 | 0.684 | -0.15 | 0.219 | 0.016 | 0.034 | 0.034 | 0.269 | 0.19 | 0.09 | 0.12 | 0.05 | 0.12 | 0.09 |
| 2 | 50214739 | rs1435090 | 0.783 | 0.65 | -0.121 | 0.23 | -0.033 | -0.05 | -0.05 | 0 | 0.26 | 0.21 | 0.47 | 0.52 | 0.33 | 0.34 |
| 2 | 50216143 | rs17439837 | 2.141 | 0.511 | -0.704 | -0.165 | -0.061 | 0.085 | 0.085 | 0.534 | 0.22 | 0.01 | 0.18 | 0.06 | 0.22 | 0.23 |
| 2 | 50217995 | rs1421567 | -1.629 | 0.496 | -0.469 | 0.02 | 0.017 | 0.038 | 0.038 | 0.005 | 0.86 | 0.67 | 0.87 | 0.79 | 0.83 | 0.87 |
| 2 | 50228332 | rs6706472 | 0.647 | 0.544 | -0.088 | 0.046 | 0.013 | 0.041 | 0.041 | 0.045 | 0.29 | 0.46 | 0.49 | 0.51 | 0.32 | 0.33 |
| 2 | 50230334 | rs7604754 | 2.307 | 0.482 | -0.096 | -0.015 | 0.008 | 0.004 | 0.004 | 0.224 | 0.19 | 0.12 | 0.1 | 0.05 | 0.1 | 0.09 |
| 2 | 50231111 | rs1421577 | -1.47 | 0.564 | -0.079 | 0.094 | 0 | 0.004 | 0.004 | 0 | 0.86 | 0.8 | 0.89 | 0.78 | 0.83 | 0.84 |
| 2 | 50232145 | rs1421579 | 0.755 | 0.56 | -0.17 | 0.111 | -0.006 | -0.017 | -0.017 | 0 | 0.28 | 0.28 | 0.47 | 0.49 | 0.31 | 0.33 |
| 2 | 50233352 | rs930295 | -2.052 | 0.563 | -0.367 | 0.141 | -0.043 | -0.055 | -0.055 | 0.008 | 0.88 | 0.99 | 0.9 | 0.79 | 0.84 | 0.84 |
| 2 | 50240644 | rs3796036 | 0.231 | 0.66 | -0.228 | 0.042 | -0.004 | 0.007 | 0.007 | 0 | 0.56 | 0.34 | 0.39 | 0.3 | 0.51 | 0.51 |
| 2 | 50243386 | rs13003111 | 1.923 | 0.841 | -0.458 | 0.348 | -0.022 | 0.002 | 0.002 | 0 | 0.09 | 0.09 | 0.2 | 0.36 | 0.2 | 0.16 |
| 2 | 50243402 | rs6731061 | 0.257 | 0.429 | -0.322 | -0.304 | 0.001 | 0.007 | 0.007 | 0 | 0.56 | 0.51 | 0.31 | 0.21 | 0.44 | 0.45 |
| 2 | 50246249 | rs7586095 | 1.457 | 0.065 | -0.003 | -0.282 | -0.037 | -0.099 | -0.099 | 0 | 0.21 | 0.33 | 0.14 | 0.05 | 0.12 | 0.09 |
| 2 | 50247149 | rs6753833 | 0.468 | -0.151 | -0.103 | -0.389 | 0.017 | 0.023 | 0.023 | 0 | 0.49 | 0.45 | 0.52 | 0.61 | 0.58 | 0.56 |
| 2 | 50247620 | rs10192012 | -0.067 | -0.131 | -0.348 | -0.364 | 0.01 | -0.006 | -0.006 | 0.746 | 0.7 | 0.79 | 0.66 | 0.65 | 0.7 | 0.65 |
| 2 | 50251100 | rs9677968 | 0.459 | -0.145 | -0.145 | -0.4 | 0.009 | 0.014 | 0.014 | 0 | 0.48 | 0.54 | 0.54 | 0.61 | 0.58 | 0.57 |
| 2 | 50252452 | rs1035342 | 0.024 | -0.094 | -0.429 | -0.368 | 0.004 | 0 | 0 | 0 | 0.69 | 0.66 | 0.65 | 0.66 | 0.7 | 0.66 |
| 2 | 50256939 | rs17039676 | 1.325 | -0.109 | -0.185 | -0.384 | -0.044 | -0.114 | -0.114 | 0 | 0.23 | 0.48 | 0.15 | 0.05 | 0.12 | 0.1 |
| 2 | 50257479 | rs12469906 | -0.806 | -0.077 | -0.216 | -0.477 | -0.034 | -0.131 | -0.131 | 0.823 | 0.7 | 0.55 | 0.8 | 0.95 | 0.79 | 0.84 |
| 2 | 50259734 | rs1895132 | -1.793 | 0.422 | -0.221 | 0.13 | -0.007 | -0.029 | -0.029 | 0 | 0.92 | 0.94 | 0.94 | 0.8 | 0.9 | 0.87 |
| 2 | 50259939 | rs1895130 | -0.107 | 0.491 | -0.146 | 0.066 | -0.006 | 0.014 | 0.014 | 1.539 | 0.75 | 0.63 | 0.69 | 0.66 | 0.74 | 0.66 |
| 2 | 50262230 | rs2351152 | 0.398 | 0.542 | -0.568 | 0.101 | 0.004 | -0.012 | -0.012 | 0 | 0.69 | 0.79 | 0.65 | 0.65 | 0.68 | 0.65 |
| 2 | 50268228 | rs1819972 | -0.296 | 0.448 | -0.412 | 0.203 | 0.001 | -0.003 | -0.003 | 0 | 0.77 | 0.78 | 0.72 | 0.66 | 0.77 | 0.67 |
| 2 | 50268555 | rs10191808 | -0.343 | 0.977 | -0.289 | 0.438 | 0.019 | 0.014 | 0.014 | 0 | 0.34 | 0.46 | 0.45 | 0.55 | 0.39 | 0.36 |
| 2 | 50275078 | rs3861561 | 0.176 | 0.741 | -0.405 | 0.401 | 0 | 0 | 0 | 0 | 0.63 | 0.61 | 0.65 | 0.64 | 0.61 | 0.56 |
| 2 | 50276061 | rs12470518 | 0.829 | 0.134 | -0.793 | 0.498 | 0.028 | -0.041 | -0.041 | 0 | 0.17 | 0.11 | 0.3 | 0.41 | 0.1 | 0.23 |
| 2 | 50281638 | rs4971644 | -0.697 | 0.238 | -0.59 | 0.407 | -0.024 | -0.037 | -0.037 | 0 | 0.72 | 0.8 | 0.69 | 0.64 | 0.68 | 0.64 |
| 2 | 50296401 | rs1563021 | 0.114 | 0.092 | 0.538 | 0.314 | -0.002 | 0 | 0 | 0 | 0.63 | 0.6 | 0.64 | 0.63 | 0.61 | 0.56 |
| 2 | 50298963 | rs6748208 | 0.071 | 0.135 | 0.49 | 0.321 | -0.008 | 0.001 | 0.001 | 0 | 0.64 | 0.69 | 0.65 | 0.64 | 0.61 | 0.56 |
| 2 | 50302978 | rs4971552 | 0.565 | 0.329 | 0.668 | 0.201 | -0.002 | -0.001 | -0.001 | 0.242 | 0.56 | 0.61 | 0.54 | 0.46 | 0.49 | 0.47 |

| | | | | | | | | | | | | | | | | |
|---|---|---|---|---|---|---|---|---|---|---|---|---|---|---|---|---|
| 2 | 50303022 | rs4971646 | 0.534 | 0.112 | 0.352 | -0.113 | 0.014 | -0.006 | -0.006 | 0.009 | 0.34 | 0.28 | 0.4 | 0.39 | 0.24 | 0.28 |
| 2 | 50306377 | rs12613363 | 0.447 | 0.112 | 0.188 | -0.151 | -0.035 | 0.006 | 0.006 | 0.348 | 0.35 | 0.47 | 0.42 | 0.4 | 0.24 | 0.28 |
| 2 | 50307245 | rs2351155 | 0.528 | 0.249 | -0.019 | -0.195 | -0.009 | 0.014 | 0.014 | 0 | 0.37 | 0.34 | 0.27 | 0.23 | 0.42 | 0.35 |
| 2 | 50321293 | rs9677302 | 0.681 | 0.187 | -0.478 | -0.161 | -0.006 | -0.016 | -0.016 | 0 | 0.46 | 0.71 | 0.39 | 0.43 | 0.52 | 0.49 |
| 2 | 50322880 | rs1961358 | 0.748 | 0.155 | -0.591 | -0.034 | 0.04 | -0.01 | -0.01 | 0 | 0.33 | 0.19 | 0.39 | 0.37 | 0.23 | 0.28 |
| 2 | 50332400 | rs9309176 | 0.158 | 0.182 | -0.565 | -0.063 | 0.036 | -0.014 | -0.014 | 0.336 | 0.52 | 0.77 | 0.44 | 0.42 | 0.57 | 0.51 |
| 2 | 50342832 | rs990024 | -0.865 | 0.115 | -0.193 | 0.077 | -0.045 | -0.023 | -0.023 | 0 | 0.79 | 0.96 | 0.78 | 0.8 | 0.75 | 0.78 |
| 2 | 50342983 | rs7590544 | 0.176 | 0.068 | 0.215 | 0.022 | -0.007 | -0.002 | -0.002 | 0 | 0.37 | 0.46 | 0.42 | 0.38 | 0.26 | 0.29 |
| 2 | 50358840 | rs17039932 | 0.973 | 0.023 | 0.393 | -0.023 | 0.011 | -0.028 | -0.028 | 0 | 0.19 | 0.14 | 0.33 | 0.38 | 0.13 | 0.24 |
| 2 | 50361388 | rs6545153 | 1.643 | 0.177 | 0.369 | -0.082 | -0.013 | 0.003 | 0.003 | 0 | 0.28 | 0.21 | 0.23 | 0.23 | 0.35 | 0.28 |
| 2 | 50361463 | rs10490240 | 0.523 | 0.151 | 0.487 | -0.095 | 0.001 | -0.003 | -0.003 | 0 | 0.23 | 0.2 | 0.36 | 0.39 | 0.15 | 0.26 |
| 2 | 50366097 | rs17505257 | 2.743 | 0.164 | 0.4 | -0.187 | -0.022 | 0.071 | 0.071 | 0 | 0.2 | 0.05 | 0.12 | 0.05 | 0.24 | 0.19 |
| 2 | 50366158 | rs17039979 | 0.943 | 0.125 | 0.37 | -0.171 | -0.003 | 0.006 | 0.006 | 0 | 0.2 | 0.26 | 0.34 | 0.38 | 0.13 | 0.24 |
| 2 | 50366181 | rs12713088 | -0.844 | 0.081 | 0.074 | -0.208 | -0.042 | -0.075 | -0.075 | 0.208 | 0.69 | 0.88 | 0.7 | 0.61 | 0.64 | 0.59 |
| 2 | 50368229 | rs12467877 | 2.296 | -0.049 | 0.16 | -0.195 | 0.007 | 0.012 | 0.012 | 0 | 0.16 | 0.14 | 0.09 | 0 | 0.13 | 0.05 |
| 2 | 50369150 | rs1584763 | 0.91 | -0.053 | 0.231 | -0.171 | -0.03 | 0.059 | 0.059 | 0 | 0.21 | 0.32 | 0.35 | 0.38 | 0.13 | 0.24 |
| 2 | 50373957 | rs17505495 | 2.32 | -0.025 | 0.05 | -0.197 | 0.006 | 0.06 | 0.06 | 0 | 0.16 | 0.06 | 0.09 | 0 | 0.13 | 0.05 |
| 2 | 50376687 | rs2678228 | 0.215 | 0.037 | 0.118 | -0.175 | 0.002 | 0.023 | 0.023 | 0 | 0.49 | 0.59 | 0.59 | 0.61 | 0.5 | 0.53 |
| 2 | 50386627 | rs970784 | 0.371 | 0.062 | 0.208 | -0.297 | -0.008 | 0.006 | 0.006 | 0 | 0.58 | 0.55 | 0.43 | 0.42 | 0.63 | 0.53 |
| 2 | 50387896 | rs973354 | 0.55 | 0.055 | 0.172 | -0.301 | 0.001 | -0.001 | -0.001 | 0 | 0.56 | 0.47 | 0.42 | 0.42 | 0.63 | 0.53 |
| 2 | 50388064 | rs13410042 | 1.31 | 0.057 | 0.204 | -0.323 | 0.002 | 0.009 | 0.009 | 0 | 0.28 | 0.26 | 0.19 | 0.19 | 0.25 | 0.25 |
| 2 | 50388284 | rs2602003 | -1.151 | 0.066 | 0.057 | -0.315 | -0.026 | 0.009 | 0.009 | 0 | 0.81 | 0.95 | 0.8 | 0.83 | 0.79 | 0.8 |
| 2 | 50388317 | rs1377234 | -1.134 | 0.066 | 0.078 | -0.315 | -0.004 | 0.017 | 0.017 | 0 | 0.81 | 0.91 | 0.79 | 0.83 | 0.79 | 0.8 |
| 2 | 50388475 | rs2688890 | 1.669 | 0.074 | 0.083 | -0.309 | -0.017 | 0.003 | 0.003 | 0 | 0.28 | 0.21 | 0.23 | 0.23 | 0.37 | 0.29 |
| 2 | 50388887 | rs2678229 | -1.119 | 0.082 | 0.04 | -0.296 | -0.004 | 0.016 | 0.016 | 0 | 0.81 | 0.91 | 0.79 | 0.83 | 0.79 | 0.8 |
| 2 | 50390615 | rs7599263 | -1.001 | 0.011 | -0.141 | -0.383 | -0.021 | -0.006 | -0.006 | 0 | 0.43 | 0.86 | 0.62 | 0.6 | 0.36 | 0.4 |
| 2 | 50393271 | rs2678223 | 1.751 | 0.079 | -0.045 | -0.279 | -0.017 | 0.002 | 0.002 | 0 | 0.28 | 0.21 | 0.31 | 0.24 | 0.37 | 0.29 |
| 2 | 50396135 | rs1452769 | 1.001 | 0.04 | 0.047 | -0.287 | -0.033 | 0.114 | 0.114 | 0.005 | 0.21 | 0.47 | 0.35 | 0.42 | 0.14 | 0.24 |
| 2 | 50398712 | rs1403654 | 2.293 | 0.065 | -0.066 | -0.324 | -0.041 | -0.038 | -0.038 | 0 | 0.07 | 0.02 | 0.17 | 0.17 | 0.12 | 0.08 |
| 2 | 50402321 | rs2678219 | 2.601 | 0.021 | 0.115 | -0.247 | -0.025 | 0.074 | 0.074 | 0 | 0.21 | 0.12 | 0.13 | 0.05 | 0.25 | 0.2 |
| 2 | 50409803 | rs17040117 | 0.812 | -0.074 | 0.4 | -0.07 | -0.033 | 0.104 | 0.104 | 0 | 0.2 | 0.43 | 0.34 | 0.4 | 0.13 | 0.23 |
| 2 | 50416835 | rs1452786 | -0.84 | -0.027 | 0.043 | -0.025 | -0.016 | -0.024 | -0.024 | 0 | 0.77 | 0.94 | 0.84 | 0.77 | 0.76 | 0.75 |
| 2 | 50423607 | rs539260 | -0.219 | -0.02 | 0.189 | -0.04 | -0.002 | -0.008 | -0.008 | 0 | 0.65 | 0.78 | 0.76 | 0.63 | 0.64 | 0.57 |
| 2 | 50424350 | rs17040141 | 0.811 | -0.03 | 0.319 | 0.011 | -0.032 | 0.087 | 0.087 | 0 | 0.21 | 0.37 | 0.35 | 0.42 | 0.14 | 0.23 |
| 2 | 50431752 | rs13001872 | 2.647 | 0.166 | 0.318 | -0.013 | -0.006 | -0.017 | -0.017 | 0.01 | 0.13 | 0.08 | 0.12 | 0.17 | 0.14 | 0.11 |
| 2 | 50432463 | rs10187296 | 2.333 | -0.052 | 0.796 | 0.549 | -0.008 | -0.012 | -0.012 | 0 | 0.12 | 0.06 | 0.07 | 0.14 | 0.13 | 0.17 |
| 2 | 50432664 | rs17040199 | 1.519 | -0.042 | 0.911 | 0.513 | -0.002 | -0.041 | -0.041 | 0 | 0.09 | 0.13 | 0.06 | 0 | 0.07 | 0.03 |
| 2 | 50433751 | rs613166 | 2.033 | -0.052 | 0.965 | 0.49 | -0.016 | 0.017 | 0.017 | 0 | 0.35 | 0.32 | 0.28 | 0.22 | 0.43 | 0.3 |
| 2 | 50434741 | rs17040210 | -0.438 | -0.117 | 0.931 | 0.503 | -0.027 | 0.026 | 0.026 | 0 | 0.29 | 0.41 | 0.4 | 0.4 | 0.2 | 0.27 |
| 2 | 50435324 | rs17040212 | -0.478 | -0.141 | 0.811 | 0.485 | -0.007 | 0.016 | 0.016 | 0 | 0.27 | 0.3 | 0.39 | 0.45 | 0.21 | 0.34 |
| 2 | 50437804 | rs1402166 | -0.879 | -0.106 | 0.833 | 0.488 | -0.004 | 0.004 | 0.004 | 0 | 0.33 | 0.34 | 0.43 | 0.44 | 0.25 | 0.34 |
| 2 | 50437852 | rs17508170 | 2.093 | -0.071 | 0.824 | 0.473 | 0.004 | 0.005 | 0.005 | 0 | 0.12 | 0.17 | 0.08 | 0.14 | 0.13 | 0.17 |
| 2 | 50438701 | rs1915222 | 0.182 | -0.263 | 0.496 | 0.512 | -0.007 | -0.097 | -0.097 | 0 | 0.67 | 0.88 | 0.49 | 0.42 | 0.66 | 0.62 |
| 2 | 50439075 | rs1915221 | -0.11 | -0.23 | 0.543 | 0.552 | 0.003 | -0.088 | -0.088 | 0.351 | 0.71 | 0.87 | 0.51 | 0.42 | 0.7 | 0.67 |
| 2 | 50439522 | rs17508435 | 1.879 | -0.219 | 0.428 | 0.517 | -0.009 | 0.045 | 0.045 | 0 | 0.08 | 0.01 | 0.05 | 0 | 0.07 | 0.03 |
| 2 | 50439540 | rs12619144 | 0.357 | -0.224 | 0.507 | 0.522 | -0.021 | -0.017 | -0.017 | 0 | 0.15 | 0.21 | 0.09 | 0.06 | 0.08 | 0.17 |
| 2 | 50440043 | rs17040234 | 0.438 | -0.197 | 0.498 | 0.526 | -0.006 | -0.009 | -0.009 | 0 | 0.15 | 0.16 | 0.1 | 0.06 | 0.1 | 0.18 |
| 2 | 50440176 | rs1715983 | 2.025 | -0.137 | 0.426 | 0.596 | -0.03 | 0.024 | 0.024 | 0 | 0.34 | 0.21 | 0.28 | 0.23 | 0.43 | 0.3 |
| 2 | 50441527 | rs17040235 | 0.31 | -0.575 | 0.208 | 0.691 | 0.015 | -0.041 | -0.041 | 0 | 0.15 | 0.09 | 0.3 | 0.39 | 0.12 | 0.21 |
| 2 | 50442276 | rs6724128 | -0.73 | -0.353 | 0.324 | 0.521 | -0.022 | -0.045 | -0.045 | 0.723 | 0.24 | 0.36 | 0.16 | 0.06 | 0.16 | 0.18 |
| 2 | 50443600 | rs1915220 | 1.322 | 0.174 | 0.088 | 1.121 | 0.004 | 0 | 0 | 0 | 0.08 | 0.03 | 0.12 | 0.37 | 0.09 | 0.17 |
| 2 | 50448539 | rs587928 | 1.05 | 0.253 | 0.614 | 0.521 | 0 | -0.012 | -0.012 | 0 | 0.41 | 0.52 | 0.43 | 0.25 | 0.4 | 0.38 |
| 2 | 50450879 | rs523003 | 0.965 | 0.243 | 0.71 | 0.52 | -0.002 | 0.017 | 0.017 | 0 | 0.4 | 0.42 | 0.42 | 0.25 | 0.4 | 0.39 |
| 2 | 50458008 | rs11681950 | 1.015 | 0.267 | 0.804 | 0.527 | 0.007 | 0.007 | 0.007 | 0 | 0.14 | 0.31 | 0.11 | 0.13 | 0.14 | 0.14 |
| 2 | 50463216 | rs11900179 | 1.203 | 0.272 | 0.543 | 0.529 | 0 | 0 | 0 | 0 | 0.13 | 0.21 | 0.09 | 0.13 | 0.13 | 0.15 |
| 2 | 50464483 | rs1715970 | 0.278 | 0.247 | 0.59 | 0.501 | -0.001 | 0.02 | 0.02 | 0.671 | 0.54 | 0.71 | 0.52 | 0.38 | 0.53 | 0.53 |
| 2 | 50470001 | rs548271 | 1.064 | 0.258 | 0.606 | 0.549 | -0.002 | 0.013 | 0.013 | 0 | 0.4 | 0.4 | 0.42 | 0.27 | 0.4 | 0.38 |
| 2 | 50482571 | rs2678216 | 1.177 | 0.271 | 0.615 | 0.529 | 0 | -0.008 | -0.008 | 0 | 0.4 | 0.47 | 0.43 | 0.27 | 0.4 | 0.39 |
| 2 | 50484360 | rs1915239 | 0.252 | 0.303 | 0.183 | 0.507 | -0.004 | 0.02 | 0.02 | 0.7 | 0.54 | 0.77 | 0.52 | 0.38 | 0.53 | 0.53 |
| 2 | 50489929 | rs13415902 | 0.997 | 0.439 | -0.407 | 0.698 | 0.009 | 0.026 | 0.026 | 0 | 0.14 | 0.27 | 0.09 | 0.16 | 0.13 | 0.14 |
| 2 | 50498605 | rs563148 | 0.049 | 0.499 | -1.298 | 0.704 | -0.034 | -0.061 | -0.061 | 0 | 0.63 | 0.85 | 0.59 | 0.4 | 0.61 | 0.56 |
| 2 | 50505658 | rs4032051 | 0.67 | 0.474 | -0.7 | 0.335 | -0.001 | -0.006 | -0.006 | 0.714 | 0.43 | 0.43 | 0.44 | 0.24 | 0.43 | 0.39 |
| 2 | 50509971 | rs1915234 | -0.928 | 0.573 | -0.722 | 0.462 | -0.003 | 0 | 0 | 0.207 | 0.86 | 0.93 | 0.81 | 0.83 | 0.81 | 0.85 |
| 2 | 50514746 | rs1520526 | -0.896 | 0.517 | -0.729 | 0.48 | -0.003 | 0.001 | 0.001 | 0.314 | 0.86 | 0.92 | 0.81 | 0.84 | 0.82 | 0.85 |
| 2 | 50522118 | rs1715997 | 0.323 | 0.086 | -0.269 | 0.525 | -0.004 | 0.003 | 0.003 | 0 | 0.71 | 0.6 | 0.71 | 0.7 | 0.7 | 0.68 |
| 2 | 50522963 | rs7605803 | 0.167 | 0.012 | 0.091 | 0.424 | -0.018 | 0.069 | 0.069 | 0.35 | 0.22 | 0.32 | 0.21 | 0.41 | 0.17 | 0.3 |
| 2 | 50524684 | rs1879340 | -1.27 | 0.071 | 0.06 | 0.396 | 0 | 0.01 | 0.01 | 0.175 | 0.86 | 0.93 | 0.83 | 0.87 | 0.84 | 0.85 |
| 2 | 50524783 | rs1104979 | -0.612 | 0.016 | 0.19 | 0.371 | 0.008 | -0.015 | -0.015 | 0.202 | 0.32 | 0.3 | 0.27 | 0.39 | 0.25 | 0.33 |
| 2 | 50526915 | rs560734 | -1.256 | 0.063 | -0.246 | 0.346 | 0 | -0.001 | -0.001 | 0 | 0.86 | 0.9 | 0.83 | 0.87 | 0.84 | 0.85 |

| | | | | | | | | | | | | | | | | |
|---|---|---|---|---|---|---|---|---|---|---|---|---|---|---|---|---|
| 2 | 50528952 | rs499598 | 0.123 | 0.062 | -0.359 | 0.315 | 0.002 | 0.027 | 0.027 | 0 | 0.75 | 0.82 | 0.77 | 0.87 | 0.75 | 0.81 |
| 2 | 50529764 | rs17040458 | 0.155 | -0.096 | -0.11 | 0.688 | 0.009 | -0.037 | -0.037 | 5.084 | 0.2 | 0.15 | 0.2 | 0.4 | 0.16 | 0.29 |
| 2 | 50530180 | rs12464203 | 0.252 | 0.235 | -0.015 | 0.909 | -0.001 | 0.003 | 0.003 | 0 | 0.11 | 0.09 | 0.07 | 0 | 0.09 | 0.04 |
| 2 | 50532337 | rs7569775 | 0.283 | 0.318 | -0.482 | 0.84 | -0.055 | -0.129 | -0.129 | 0 | 0.26 | 0.04 | 0.35 | 0.37 | 0.27 | 0.22 |
| 2 | 50540027 | rs10203511 | -1.394 | 0.135 | 0.813 | 1.18 | -0.025 | 0.154 | 0.154 | 0.372 | 0.21 | 0.52 | 0.28 | 0.54 | 0.15 | 0.29 |
| 2 | 50540740 | rs939425 | -2.54 | 0.12 | -0.082 | 0.925 | -0.055 | 0.161 | 0.161 | 1.083 | 0.49 | 0.93 | 0.47 | 0.7 | 0.42 | 0.48 |
| 2 | 50551568 | rs6545163 | -2.248 | -0.213 | 0.36 | 0.822 | 0.034 | -0.082 | -0.082 | 0 | 0.33 | 0.84 | 0.36 | 0.5 | 0.25 | 0.34 |
| 2 | 50563905 | rs9309181 | -2.078 | -0.098 | 0.772 | 0.902 | 0.021 | -0.053 | -0.053 | 0.143 | 0.27 | 0.79 | 0.33 | 0.49 | 0.2 | 0.31 |
| 2 | 50569117 | rs1543973 | -0.322 | 0.054 | 0.975 | 1.024 | -0.001 | 0.057 | 0.057 | 0 | 0.08 | 0.11 | 0.18 | 0.36 | 0.07 | 0.12 |
| 2 | 50594162 | rs4377361 | -1.345 | 0.118 | 1.556 | 0.819 | 0.004 | 0.009 | 0.009 | 0 | 0.37 | 0.46 | 0.42 | 0.57 | 0.4 | 0.32 |
| 2 | 50604829 | rs1520446 | -0.166 | 0.179 | 1.903 | 0.887 | -0.002 | 0.036 | 0.036 | 0 | 0.26 | 0.29 | 0.3 | 0.43 | 0.26 | 0.22 |
| 2 | 50605728 | rs12463670 | -0.623 | 0.18 | 1.844 | 0.88 | 0.008 | 0.009 | 0.009 | 0 | 0.1 | 0.17 | 0.1 | 0.12 | 0.12 | 0.09 |
| 2 | 50606572 | rs4971669 | -2.079 | 0.169 | 2.057 | 0.814 | 0.015 | 0.191 | 0.191 | 0 | 0.46 | 0.77 | 0.61 | 0.93 | 0.44 | 0.49 |
| 2 | 50606642 | rs7564130 | -1.152 | 0.133 | 2.035 | 0.841 | 0 | 0 | 0 | 0 | 0.35 | 0.33 | 0.39 | 0.56 | 0.38 | 0.31 |
| 2 | 50607875 | rs12474009 | -1.838 | 0.128 | 1.959 | 0.876 | -0.011 | 0.096 | 0.096 | 0 | 0.2 | 0.37 | 0.31 | 0.38 | 0.17 | 0.23 |
| 2 | 50607989 | rs10184843 | -0.134 | 0.111 | 1.957 | 0.88 | -0.022 | -0.087 | -0.087 | 0.409 | 0.25 | 0.11 | 0.3 | 0.45 | 0.28 | 0.22 |
| 2 | 50608717 | rs12467043 | -0.118 | 0.116 | 2.032 | 0.882 | -0.008 | -0.049 | -0.049 | 0 | 0.25 | 0.17 | 0.3 | 0.45 | 0.27 | 0.22 |
| 2 | 50611076 | rs1829534 | -2.076 | 0.108 | 2.119 | 0.803 | 0.011 | 0.075 | 0.075 | 0 | 0.45 | 0.65 | 0.61 | 0.81 | 0.44 | 0.45 |
| 2 | 50612316 | rs1851015 | -0.012 | 0.131 | 2.018 | 0.829 | -0.019 | -0.094 | -0.094 | 1.899 | 0.24 | 0.1 | 0.29 | 0.44 | 0.26 | 0.22 |
| 2 | 50614081 | rs4318425 | -1.805 | 0.138 | 2.045 | 0.831 | -0.002 | -0.035 | -0.035 | 0 | 0.43 | 0.56 | 0.59 | 0.81 | 0.44 | 0.45 |
| 2 | 50618708 | rs10191261 | -0.076 | 0.151 | 1.755 | 0.911 | -0.003 | -0.049 | -0.049 | 0 | 0.25 | 0.16 | 0.29 | 0.44 | 0.26 | 0.21 |
| 2 | 50622702 | rs12713097 | -0.008 | 0.144 | 1.726 | 0.908 | -0.01 | -0.07 | -0.07 | 0 | 0.24 | 0.11 | 0.28 | 0.43 | 0.26 | 0.21 |
| 2 | 50624071 | rs9678669 | -2.149 | 0.165 | 1.907 | 0.866 | -0.004 | 0.142 | 0.142 | 0 | 0.47 | 0.74 | 0.63 | 0.82 | 0.47 | 0.47 |
| 2 | 50624617 | rs1520451 | -0.34 | 0.193 | 1.655 | 0.856 | -0.006 | 0 | 0 | 0 | 0.09 | 0.06 | 0.1 | 0 | 0.12 | 0.05 |
| 2 | 50625105 | rs17040569 | -0.285 | 0.197 | 1.755 | 0.882 | 0.001 | 0.006 | 0.006 | 0 | 0.26 | 0.24 | 0.31 | 0.43 | 0.29 | 0.21 |
| 2 | 50633190 | rs13407780 | -1.823 | -0.091 | 1.955 | 0.957 | -0.011 | 0.122 | 0.122 | 0 | 0.2 | 0.56 | 0.31 | 0.38 | 0.18 | 0.19 |
| 2 | 50635445 | rs7599052 | -0.887 | -0.104 | 2.351 | 1.071 | -0.003 | 0.219 | 0.219 | 0 | 0.1 | 0.28 | 0.2 | 0.38 | 0.06 | 0.13 |
| 2 | 50640323 | rs10181522 | -0.136 | -0.163 | 2.063 | 1.025 | -0.005 | -0.064 | -0.064 | 0.124 | 0.25 | 0.16 | 0.29 | 0.44 | 0.26 | 0.27 |
| 2 | 50642287 | rs10188271 | -0.23 | -0.166 | 2.042 | 1.024 | 0.001 | -0.033 | -0.033 | 0 | 0.26 | 0.21 | 0.29 | 0.44 | 0.26 | 0.24 |
| 2 | 50643419 | rs1915170 | -1.553 | -0.012 | 2.313 | 1.118 | -0.007 | 0.169 | 0.169 | 0 | 0.17 | 0.43 | 0.27 | 0.37 | 0.13 | 0.18 |
| 2 | 50644449 | rs12053271 | -1.934 | -0.016 | 1.787 | 1.083 | -0.015 | 0.07 | 0.07 | 0.001 | 0.19 | 0.64 | 0.3 | 0.38 | 0.13 | 0.21 |
| 2 | 50650102 | rs1520439 | -0.544 | -0.119 | 1.375 | 1.053 | 0.008 | -0.028 | -0.028 | 0 | 0.3 | 0.27 | 0.32 | 0.4 | 0.28 | 0.31 |
| 2 | 50653025 | rs7423850 | -0.649 | -0.128 | 1.675 | 1.049 | -0.013 | 0.025 | 0.025 | 0 | 0.31 | 0.45 | 0.34 | 0.4 | 0.29 | 0.31 |
| 2 | 50659515 | rs10490227 | -1.231 | -0.153 | 1.712 | 1.045 | -0.009 | 0.17 | 0.17 | 0 | 0.13 | 0.27 | 0.22 | 0.52 | 0.1 | 0.17 |
| 2 | 50660985 | rs4971675 | -1.925 | -0.198 | 1.486 | 1 | 0.045 | 0.22 | 0.22 | 0 | 0.46 | 0.83 | 0.59 | 0.93 | 0.39 | 0.5 |
| 2 | 50661643 | rs6756219 | -0.513 | -0.093 | 1.17 | 1.091 | 0.007 | -0.04 | -0.04 | 0 | 0.29 | 0.25 | 0.33 | 0.4 | 0.28 | 0.31 |
| 2 | 50668760 | rs10048731 | -0.842 | -0.12 | 1.165 | 1.164 | -0.004 | 0.009 | 0.009 | 0 | 0.33 | 0.39 | 0.34 | 0.5 | 0.29 | 0.37 |
| 2 | 50669861 | rs6709865 | -0.421 | -0.069 | 0.945 | 0.863 | 0.003 | -0.048 | -0.048 | 0 | 0.28 | 0.16 | 0.32 | 0.38 | 0.28 | 0.31 |
| 2 | 50682462 | rs1005431 | -0.535 | 0.013 | 0.676 | 0.829 | 0.004 | -0.01 | -0.01 | 0 | 0.3 | 0.28 | 0.32 | 0.39 | 0.27 | 0.3 |
| 2 | 50682914 | rs9917153 | -1.218 | 0.222 | 1.352 | 1.562 | 0.001 | 0.393 | 0.393 | 0 | 0.1 | 0.34 | 0.21 | 0.54 | 0.07 | 0.16 |
| 2 | 50682959 | rs7590358 | -1.309 | 0.394 | 0.388 | 0.834 | 0.007 | -0.039 | -0.039 | 0 | 0.07 | 0.19 | 0.03 | 0 | 0.05 | 0.04 |
| 2 | 50689260 | rs12997031 | -1.167 | 0.691 | -0.528 | 0.99 | -0.014 | -0.014 | -0.014 | 0 | 0.36 | 0.7 | 0.49 | 0.61 | 0.4 | 0.49 |
| 2 | 50691431 | rs11690362 | -1.167 | 0.704 | -0.544 | 1.003 | -0.017 | -0.019 | -0.019 | 0.053 | 0.36 | 0.7 | 0.49 | 0.59 | 0.4 | 0.49 |
| 2 | 50692168 | rs1520455 | 0.903 | 0.839 | -0.722 | 1.409 | -0.012 | -0.017 | -0.017 | 0.138 | 0.68 | 0.69 | 0.43 | 0.44 | 0.62 | 0.49 |
| 2 | 50692358 | rs1879318 | 0.645 | 0.673 | -0.564 | 1.514 | -0.011 | 0.141 | 0.141 | 0.023 | 0.91 | 0.78 | 0.82 | 0.67 | 0.93 | 0.89 |
| 2 | 50692560 | rs3213756 | -0.661 | 0.806 | -0.767 | 1.537 | -0.036 | -0.063 | -0.063 | 0 | 0.24 | 0.13 | 0.4 | 0.45 | 0.3 | 0.5 |
| 2 | 50698154 | rs2111066 | 0.666 | 0.305 | -0.734 | 1.291 | -0.004 | -0.071 | -0.071 | 1.145 | 0.88 | 0.95 | 0.8 | 0.7 | 0.89 | 0.87 |
| 2 | 50702945 | rs11677389 | -0.235 | 0.232 | -0.659 | 1.249 | -0.061 | -0.001 | -0.001 | 5.448 | 0.24 | 0.1 | 0.39 | 0.21 | 0.34 | 0.43 |
| 2 | 50705043 | rs7595040 | 0.615 | 0.221 | -0.668 | 1.26 | 0 | -0.072 | -0.072 | 0 | 0.87 | 0.96 | 0.8 | 0.7 | 0.89 | 0.89 |
| 2 | 50705098 | rs10176224 | 0.915 | 0.202 | -0.516 | 1.222 | -0.059 | -0.06 | -0.06 | 0 | 0.64 | 0.82 | 0.4 | 0.49 | 0.56 | 0.5 |
| 2 | 50707394 | rs7578902 | 1.124 | 0.154 | 0.327 | 1.188 | 0.007 | 0.007 | 0.007 | 0 | 0.62 | 0.52 | 0.38 | 0.49 | 0.54 | 0.47 |
| 2 | 50707858 | rs11892200 | 0.973 | 0.159 | 0.401 | 1.191 | -0.01 | -0.014 | -0.014 | 0 | 0.63 | 0.63 | 0.39 | 0.49 | 0.57 | 0.49 |
| 2 | 50713012 | rs6721498 | 1.212 | 0.15 | 0.479 | 1.195 | 0.01 | 0.008 | 0.008 | 0 | 0.61 | 0.52 | 0.36 | 0.47 | 0.52 | 0.47 |
| 2 | 50714013 | rs6725307 | -0.629 | 0.182 | 0.49 | 1.049 | 0.063 | 0.342 | 0.342 | 0.268 | 0.11 | 0.43 | 0.08 | 0.4 | 0.08 | 0.09 |
| 2 | 50714226 | rs13011150 | -0.117 | 0.261 | 0.478 | 1.098 | -0.062 | 0 | 0 | 0.218 | 0.23 | 0.11 | 0.4 | 0.2 | 0.35 | 0.4 |
| 2 | 50716768 | rs10183541 | 0.431 | 0.46 | 0.329 | 1.072 | 0 | -0.084 | -0.084 | 0.798 | 0.85 | 0.95 | 0.78 | 0.68 | 0.86 | 0.88 |
| 2 | 50721511 | rs6706713 | 0.803 | 0.477 | 0.376 | 1.008 | -0.015 | -0.012 | -0.012 | 0 | 0.62 | 0.68 | 0.35 | 0.48 | 0.52 | 0.49 |
| 2 | 50733545 | rs13008919 | -1.369 | 0.302 | 0.49 | 1.6 | 0.019 | 0.204 | 0.204 | 0.091 | 0.13 | 0.35 | 0.07 | 0.38 | 0.08 | 0.1 |
| 2 | 50744820 | rs6707531 | 0.83 | 0.464 | 0.3 | 1.723 | -0.03 | -0.101 | -0.101 | 0 | 0.86 | 0.99 | 0.76 | 0.68 | 0.84 | 0.88 |
| 2 | 50750428 | rs7581256 | -1.263 | 0.511 | 0.855 | 1.603 | 0.086 | 0.285 | 0.285 | 0.777 | 0.1 | 0.36 | 0.09 | 0.39 | 0.11 | 0.1 |
| 2 | 50763645 | rs10175222 | -0.803 | 0.111 | 0.464 | 1.3 | -0.006 | 0.061 | 0.061 | 0 | 0.25 | 0.36 | 0.34 | 0.53 | 0.26 | 0.21 |
| 2 | 50766464 | rs7601431 | -1.749 | -0.414 | 0.453 | 0.969 | 0.052 | 0.014 | 0.014 | 1.524 | 0.41 | 0.75 | 0.4 | 0.59 | 0.31 | 0.27 |
| 2 | 50767091 | rs13413672 | -0.985 | -0.376 | 0.853 | 0.981 | 0.009 | 0.038 | 0.038 | 0.334 | 0.05 | 0.29 | 0.04 | 0.07 | 0.02 | 0.03 |
| 2 | 50770218 | rs11125317 | -0.571 | -0.378 | 0.838 | 0.96 | -0.006 | 0.065 | 0.065 | 0 | 0.25 | 0.35 | 0.33 | 0.53 | 0.26 | 0.21 |
| 2 | 50771738 | rs6716892 | -0.77 | -0.402 | 0.853 | 0.948 | -0.009 | 0.061 | 0.061 | 0.81 | 0.27 | 0.36 | 0.34 | 0.52 | 0.26 | 0.21 |
| 2 | 50779296 | rs13412851 | -1.1 | -0.453 | 1.114 | 1.269 | 0.006 | -0.02 | -0.02 | 0 | 0.25 | 0.19 | 0.28 | 0.45 | 0.21 | 0.21 |
| 2 | 50780219 | rs13429627 | -1.302 | -0.442 | 1.319 | 1.277 | -0.021 | 0.076 | 0.076 | 0 | 0.26 | 0.36 | 0.29 | 0.45 | 0.19 | 0.23 |
| 2 | 50780375 | rs13029649 | -0.277 | -0.469 | 1.25 | 1.257 | -0.008 | 0.057 | 0.057 | 0 | 0.11 | 0.22 | 0.09 | 0.15 | 0.1 | 0.07 |
| 2 | 50782599 | rs11675550 | -0.278 | -0.461 | 1.179 | 1.252 | -0.015 | 0.045 | 0.045 | 0 | 0.11 | 0.21 | 0.09 | 0.15 | 0.1 | 0.08 |

| | | | | | | | | | | | | | | | | |
|---|---|---|---|---|---|---|---|---|---|---|---|---|---|---|---|---|
| 2 | 50786770 | rs12713108 | -2.204 | -0.394 | 0.978 | 0.976 | 0.043 | 0.014 | 0.014 | 0.46 | 0.39 | 0.71 | 0.38 | 0.61 | 0.29 | 0.3 |
| 2 | 50790963 | rs10196148 | -1.428 | -0.368 | 1.043 | 1.004 | -0.029 | 0.04 | 0.04 | 0 | 0.3 | 0.43 | 0.33 | 0.54 | 0.25 | 0.23 |
| 2 | 50793781 | rs2351765 | -2.091 | -0.262 | 0.92 | 1.008 | 0.021 | 0.009 | 0.009 | 0.018 | 0.38 | 0.62 | 0.34 | 0.61 | 0.26 | 0.3 |
| 2 | 50798079 | rs10490222 | -1.513 | -0.274 | 0.556 | 0.97 | -0.033 | -0.057 | -0.057 | 0 | 0.16 | 0.41 | 0.07 | 0.07 | 0.07 | 0.08 |
| 2 | 50804469 | rs12613518 | -0.798 | -0.426 | 0.483 | 1.05 | -0.002 | 0.093 | 0.093 | 0 | 0.24 | 0.38 | 0.32 | 0.54 | 0.24 | 0.22 |
| 2 | 50804723 | rs17040856 | 0.69 | -0.438 | 0.058 | 1.291 | -0.034 | -0.073 | -0.073 | 0 | 0.12 | 0.02 | 0.21 | 0.28 | 0.13 | 0.12 |
| 2 | 50805475 | rs725288 | -2.116 | -0.324 | -0.749 | 0.573 | -0.104 | 0.183 | 0.183 | 0 | 0.63 | 0.98 | 0.75 | 0.9 | 0.6 | 0.69 |
| 2 | 50809998 | rs10177104 | -1.686 | -0.365 | -0.679 | 0.604 | 0.087 | 0.076 | 0.076 | 0 | 0.47 | 0.75 | 0.49 | 0.63 | 0.34 | 0.48 |
| 2 | 50812359 | rs13419090 | 0.754 | -0.035 | -0.11 | 0.579 | 0.001 | 0.047 | 0.047 | 0 | 0.12 | 0.12 | 0.22 | 0.32 | 0.12 | 0.14 |
| 2 | 50812891 | rs7584604 | -0.943 | -0.159 | -0.395 | -0.312 | -0.012 | 0.082 | 0.082 | 0 | 0.44 | 0.66 | 0.6 | 0.89 | 0.5 | 0.45 |
| 2 | 50814968 | rs2351767 | -0.415 | 0.006 | -0.588 | -0.142 | 0.028 | -0.058 | -0.058 | 0 | 0.37 | 0.45 | 0.56 | 0.82 | 0.47 | 0.38 |
| 2 | 50816879 | rs7579807 | 0.555 | -0.215 | 0.064 | 0.283 | -0.051 | -0.036 | -0.036 | 0.233 | 0.87 | 0.61 | 0.94 | 0.93 | 0.96 | 0.92 |
| 2 | 50817203 | rs7582692 | -0.478 | 0.312 | -0.469 | 0.517 | 0.001 | 0.005 | 0.005 | 0.037 | 0.44 | 0.49 | 0.44 | 0.53 | 0.55 | 0.58 |
| 2 | 50822050 | rs17040885 | -1.056 | 0.147 | -0.153 | 0.126 | -0.004 | -0.064 | -0.064 | 0 | 0.11 | 0.26 | 0.09 | 0 | 0.09 | 0.06 |
| 2 | 50822463 | rs2473 | -0.193 | 1.196 | -0.873 | -0.307 | -0.025 | 0.021 | 0.021 | 0.057 | 0.86 | 0.93 | 0.87 | 0.96 | 0.76 | 0.89 |
| 2 | 50824331 | rs12713117 | 0.605 | 0.986 | -0.881 | -0.084 | -0.001 | 0.001 | 0.001 | 0 | 0.27 | 0.26 | 0.33 | 0.52 | 0.25 | 0.39 |
| 2 | 50826484 | rs4467312 | 0.155 | 0.248 | -1.283 | -0.665 | 0.019 | 0.093 | 0.093 | 3.86 | 0.32 | 0.16 | 0.27 | 0.14 | 0.27 | 0.2 |
| 2 | 50828633 | rs972112 | -1.417 | 0.326 | -1.299 | -0.633 | -0.091 | 0.043 | 0.043 | 0.122 | 0.86 | 0.99 | 0.88 | 0.96 | 0.76 | 0.89 |
| 2 | 50829144 | rs2216784 | 1.374 | 0.08 | -1.204 | -0.039 | -0.046 | -0.123 | -0.123 | 0 | 0.26 | 0.09 | 0.28 | 0.4 | 0.26 | 0.37 |
| 2 | 50831734 | rs4971686 | 1.192 | 0.106 | -0.865 | 0.057 | 0.002 | 0.05 | 0.05 | 0 | 0.27 | 0.36 | 0.34 | 0.51 | 0.25 | 0.37 |
| 2 | 50851835 | rs10865248 | -0.463 | -0.037 | -0.623 | 0.025 | 0.037 | -0.028 | -0.028 | 0 | 0.71 | 0.56 | 0.66 | 0.73 | 0.6 | 0.63 |
| 2 | 50865334 | rs2194385 | -0.003 | -0.289 | -0.58 | -0.009 | -0.002 | 0.001 | 0.001 | 1.35 | 0.6 | 0.42 | 0.61 | 0.65 | 0.52 | 0.59 |
| 2 | 50865495 | rs2194386 | -0.19 | -0.753 | -1.024 | -0.292 | -0.002 | 0.03 | 0.03 | 0.735 | 0.54 | 0.61 | 0.52 | 0.68 | 0.44 | 0.52 |
| 2 | 50866090 | rs1469160 | 0.313 | -0.86 | -1.203 | -0.362 | -0.046 | 0.13 | 0.13 | 0.256 | 0.25 | 0.04 | 0.22 | 0.06 | 0.26 | 0.15 |
| 2 | 50869459 | rs4518000 | 1.895 | -0.701 | -1.056 | -0.055 | -0.078 | -0.158 | -0.158 | 0 | 0.25 | 0.02 | 0.28 | 0.39 | 0.26 | 0.35 |
| 2 | 50871418 | rs10182729 | 0.386 | -0.733 | -0.926 | -0.05 | 0 | 0 | 0 | 0 | 0.41 | 0.44 | 0.46 | 0.6 | 0.36 | 0.46 |
| 2 | 50871821 | rs2194388 | 0.59 | -0.724 | -1.05 | -0.054 | -0.001 | 0 | 0 | 0.108 | 0.38 | 0.28 | 0.45 | 0.61 | 0.37 | 0.46 |
| 2 | 50873666 | rs2018909 | 1.006 | -0.428 | -1.034 | 0.011 | -0.002 | 0.008 | 0.008 | 1.091 | 0.11 | 0.07 | 0.11 | 0.08 | 0.1 | 0.09 |
| 2 | 50874084 | rs17480512 | 0.155 | -0.389 | -1.206 | -0.089 | 0.017 | 0.019 | 0.019 | 0 | 0.3 | 0.15 | 0.33 | 0.22 | 0.33 | 0.25 |
| 2 | 50874309 | rs918807 | -0.2 | -0.325 | -1.214 | -0.159 | -0.024 | 0.044 | 0.044 | 0.021 | 0.66 | 0.8 | 0.75 | 0.88 | 0.61 | 0.82 |
| 2 | 50874681 | rs969719 | 0.408 | -0.286 | -1.312 | -0.197 | -0.01 | 0.026 | 0.026 | 2.596 | 0.19 | 0.13 | 0.19 | 0.13 | 0.18 | 0.19 |
| 2 | 50875879 | rs858952 | -1.749 | -0.032 | -1.431 | -0.229 | -0.021 | 0.017 | 0.017 | 0 | 0.86 | 0.92 | 0.87 | 0.96 | 0.76 | 0.88 |
| 2 | 50876081 | rs2048979 | 0.672 | -0.361 | -1.332 | 0.114 | -0.002 | 0.001 | 0.001 | 0 | 0.41 | 0.45 | 0.47 | 0.61 | 0.41 | 0.61 |
| 2 | 50877396 | rs858955 | -0.124 | -0.173 | -1.605 | 0.198 | 0.022 | 0.035 | 0.035 | 0 | 0.2 | 0.07 | 0.13 | 0.06 | 0.16 | 0.06 |
| 2 | 50879012 | rs858956 | -0.129 | -0.183 | -1.544 | 0.245 | 0.032 | 0.019 | 0.019 | 0 | 0.19 | 0.07 | 0.11 | 0.12 | 0.14 | 0.07 |
| 2 | 50882657 | rs9752732 | -0.484 | 0.539 | -1.238 | 0.404 | -0.009 | 0.005 | 0.005 | 0 | 0.89 | 0.92 | 0.79 | 0.89 | 0.81 | 0.78 |
| 2 | 50883300 | rs719645 | 0.127 | 0.552 | -1.165 | 0.639 | 0.002 | 0.004 | 0.004 | 0 | 0.27 | 0.38 | 0.23 | 0.31 | 0.23 | 0.18 |
| 2 | 50885859 | rs990590 | -0.263 | 0.305 | -1.061 | 0.945 | -0.008 | 0.011 | 0.011 | 0 | 0.33 | 0.42 | 0.4 | 0.5 | 0.23 | 0.39 |
| 2 | 50889615 | rs858927 | 0.728 | 0.68 | -0.889 | 0.431 | 0.016 | -0.05 | -0.05 | 0.581 | 0.14 | 0.23 | 0.17 | 0.04 | 0.18 | 0.17 |
| 2 | 50889959 | rs991566 | 0.111 | 0.807 | -1.279 | 0.086 | 0.014 | -0.057 | -0.057 | 0 | 0.6 | 0.6 | 0.62 | 0.84 | 0.48 | 0.63 |
| 2 | 50895613 | rs858930 | 0.137 | 0.79 | -1.317 | 0.164 | 0.001 | 0 | 0 | 0 | 0.41 | 0.42 | 0.39 | 0.38 | 0.42 | 0.39 |
| 2 | 50896413 | rs17040937 | 1.432 | 0.867 | -1.294 | 0.335 | -0.003 | -0.02 | -0.02 | 0.18 | 0.26 | 0.14 | 0.2 | 0.29 | 0.24 | 0.21 |
| 2 | 50897456 | rs4246580 | 0.171 | 0.484 | -1.633 | 0.7 | -0.001 | 0.004 | 0.004 | 0 | 0.36 | 0.32 | 0.4 | 0.29 | 0.38 | 0.44 |
| 2 | 50902931 | rs2194390 | -1.87 | 0.808 | -1.84 | 0.583 | -0.032 | 0.014 | 0.014 | 0 | 0.93 | 1 | 0.92 | 0.97 | 0.88 | 0.94 |
| 2 | 50906725 | rs2216789 | 1.469 | 0.798 | -1.285 | 0.633 | -0.018 | 0.037 | 0.037 | 0.383 | 0.12 | 0.21 | 0.18 | 0.3 | 0.04 | 0.13 |
| 2 | 50914485 | rs918808 | -0.716 | 0.788 | -1.101 | 0.458 | -0.037 | 0.069 | 0.069 | 0 | 0.73 | 0.89 | 0.8 | 0.88 | 0.66 | 0.81 |
| 2 | 50916673 | rs9309189 | 0.086 | 0.933 | -1.191 | 0.386 | 0.003 | 0.007 | 0.007 | 0 | 0.49 | 0.57 | 0.46 | 0.65 | 0.39 | 0.48 |
| 2 | 50918967 | rs9309190 | 0.366 | 0.8 | -1.459 | 0.24 | -0.004 | 0.007 | 0.007 | 0 | 0.66 | 0.39 | 0.65 | 0.88 | 0.59 | 0.64 |
| 2 | 50920698 | rs9752126 | 0.091 | 0.606 | -1.308 | 0.408 | -0.015 | 0.013 | 0.013 | 0 | 0.25 | 0.31 | 0.26 | 0.36 | 0.16 | 0.24 |
| 2 | 50923733 | rs17501068 | 2.112 | 0.655 | -1.391 | 0.527 | -0.035 | 0.015 | 0.015 | 0 | 0.11 | 0 | 0.06 | 0.05 | 0.12 | 0.03 |
| 2 | 50923891 | rs994991 | 0.575 | 0.553 | -1.257 | 0.493 | -0.018 | 0.016 | 0.016 | 0.84 | 0.34 | 0.49 | 0.42 | 0.57 | 0.23 | 0.44 |
| 2 | 50924358 | rs17560229 | 1.066 | 0.646 | -1.231 | 0.474 | -0.002 | -0.016 | -0.016 | 0.011 | 0.13 | 0.11 | 0.07 | 0.05 | 0.11 | 0.11 |
| 2 | 50930181 | rs858933 | 0.771 | 0.858 | -1.146 | 0.668 | 0.009 | -0.024 | -0.024 | 0.039 | 0.35 | 0.35 | 0.39 | 0.22 | 0.41 | 0.34 |
| 2 | 50937036 | rs12618157 | 0.303 | 0.992 | -1.203 | 0.518 | 0 | 0 | 0 | 0.602 | 0.56 | 0.55 | 0.64 | 0.75 | 0.49 | 0.64 |
| 2 | 50938133 | rs17040972 | -0.258 | 0.799 | -1.193 | 0.622 | 0.023 | -0.022 | -0.022 | 0 | 0.36 | 0.31 | 0.3 | 0.5 | 0.25 | 0.3 |
| 2 | 50943528 | rs1518553 | 1.517 | 0.72 | -1.037 | 0.7 | -0.011 | 0.025 | 0.025 | 0.408 | 0.11 | 0.19 | 0.18 | 0.3 | 0.04 | 0.14 |
| 2 | 50945471 | rs4971565 | 1.093 | 0.711 | -1.012 | 0.706 | -0.036 | 0.121 | 0.121 | 0.475 | 0.13 | 0.38 | 0.2 | 0.31 | 0.04 | 0.18 |
| 2 | 50947044 | rs17501747 | 2.702 | 1.037 | -1.341 | 0.611 | -0.028 | -0.015 | -0.015 | 0 | 0.17 | 0.01 | 0.17 | 0.13 | 0.14 | 0.13 |
| 2 | 50959683 | rs858951 | 0.437 | 0.962 | -0.845 | 0.696 | 0.003 | -0.003 | -0.003 | 0 | 0.56 | 0.43 | 0.63 | 0.49 | 0.61 | 0.67 |
| 2 | 50968808 | rs10184594 | 0.383 | 0.967 | -0.831 | 0.621 | -0.012 | -0.003 | -0.003 | 0 | 0.39 | 0.57 | 0.42 | 0.39 | 0.3 | 0.37 |
| 2 | 50970292 | rs858938 | 1.082 | 1.324 | -0.745 | 0.473 | 0.033 | 0.032 | 0.032 | 0 | 0.13 | 0.21 | 0.24 | 0.18 | 0.22 | 0.22 |
| 2 | 50970602 | rs10202039 | 0.318 | 1.146 | -0.927 | 0.454 | -0.006 | 0.025 | 0.025 | 0 | 0.39 | 0.63 | 0.44 | 0.24 | 0.42 | 0.39 |
| 2 | 50971951 | rs858939 | -1.841 | 1.035 | -0.883 | 0.568 | -0.038 | -0.03 | -0.03 | 0.298 | 0.93 | 1 | 0.95 | 0.9 | 0.87 | 0.95 |
| 2 | 50972117 | rs17502407 | 1.057 | 0.919 | -0.831 | 0.456 | -0.002 | 0.003 | 0.003 | 0 | 0.1 | 0.04 | 0.06 | 0.04 | 0.11 | 0.11 |
| 2 | 50983186 | rs17041014 | 0.917 | 0.904 | -0.672 | 0.531 | 0.05 | 0.026 | 0.026 | 0 | 0.12 | 0.38 | 0.2 | 0.11 | 0.16 | 0.19 |
| 2 | 50991039 | rs3914722 | 2.098 | 0.742 | -1.095 | 0.468 | -0.013 | 0.018 | 0.018 | 0 | 0.13 | 0 | 0.06 | 0.06 | 0.11 | 0.04 |
| 2 | 50992027 | rs9309195 | -1.806 | 0.815 | -1.035 | 0.566 | -0.022 | -0.041 | -0.041 | 0 | 0.91 | 0.99 | 0.88 | 0.87 | 0.9 | 0.83 |
| 2 | 50992109 | rs9309196 | 0.185 | 0.908 | -1.089 | 0.474 | -0.016 | 0.002 | 0.002 | 0.007 | 0.37 | 0.54 | 0.43 | 0.62 | 0.25 | 0.45 |
| 2 | 50993048 | rs3914724 | 1.436 | 1.055 | -1.159 | 0.434 | -0.002 | 0.004 | 0.004 | 0 | 0.1 | 0.05 | 0.07 | 0.05 | 0.12 | 0.11 |

| | | | | | | | | | | | | | | | |
|---|---|---|---|---|---|---|---|---|---|---|---|---|---|---|---|
| 2 | 50994236 | rs13014442 | 1.71 | 1.091 | -1.18 | 0.414 | -0.036 | 0.084 | 0.084 | 0 | 0.28 | 0.17 | 0.23 | 0.06 | 0.33 | 0.19 |
| 2 | 50994255 | rs17502934 | 1.866 | 1.219 | -1.077 | 0.569 | -0.032 | -0.033 | -0.033 | 0.181 | 0.14 | 0 | 0.08 | 0.15 | 0.15 | 0.07 |
| 2 | 51002606 | rs3850332 | 0.195 | 1.238 | -0.949 | 0.607 | 0.009 | -0.018 | -0.018 | 0 | 0.64 | 0.62 | 0.64 | 0.69 | 0.57 | 0.63 |
| 2 | 51009232 | rs2352078 | 1.625 | 1.215 | -1.04 | 0.589 | -0.035 | -0.02 | -0.02 | 0 | 0.07 | 0.01 | 0.05 | 0.09 | 0.12 | 0.04 |
| 2 | 51013647 | rs9309199 | 0.554 | 1.179 | -0.961 | 0.557 | 0.01 | -0.022 | -0.022 | 0 | 0.57 | 0.6 | 0.62 | 0.7 | 0.45 | 0.61 |
| 2 | 51017111 | rs9309200 | -0.603 | 1.191 | -0.749 | 0.532 | 0.058 | -0.004 | -0.004 | 0 | 0.86 | 0.52 | 0.74 | 0.88 | 0.79 | 0.79 |
| 2 | 51019232 | rs9751737 | -0.782 | 1.174 | -0.692 | 0.759 | 0.022 | 0.031 | 0.031 | 0 | 0.87 | 0.79 | 0.76 | 0.8 | 0.78 | 0.78 |
| 2 | 51021488 | rs17568791 | 0.812 | 1.1 | -0.924 | 0.6 | -0.063 | 0 | 0 | 0.35 | 0.21 | 0.01 | 0.11 | 0.16 | 0.23 | 0.09 |
| 2 | 51025041 | rs3850336 | 1.209 | 0.772 | -0.933 | 0.874 | 0.026 | -0.056 | -0.056 | 0 | 0.23 | 0.07 | 0.3 | 0.3 | 0.14 | 0.27 |
| 2 | 51028662 | rs2352077 | -0.497 | 0.935 | -0.676 | 0.954 | -0.007 | 0.027 | 0.027 | 0 | 0.69 | 0.45 | 0.67 | 0.6 | 0.68 | 0.53 |
| 2 | 51031742 | rs17041042 | -0.34 | 1.049 | -0.742 | 0.796 | 0.023 | -0.053 | -0.053 | 0.225 | 0.34 | 0.43 | 0.37 | 0.2 | 0.42 | 0.23 |
| 2 | 51033083 | rs9284757 | -1.471 | 0.652 | -0.493 | 1.234 | 0.012 | 0.032 | 0.032 | 0 | 0.89 | 0.8 | 0.84 | 0.83 | 0.86 | 0.79 |
| 2 | 51033295 | rs17569028 | 0.937 | 0.648 | -0.685 | 1.071 | 0.001 | -0.006 | -0.006 | 0 | 0.17 | 0.19 | 0.16 | 0.15 | 0.21 | 0.09 |
| 2 | 51041472 | rs3914731 | 1.74 | 0.405 | -1.045 | 1.164 | 0.01 | -0.055 | -0.055 | 0 | 0.16 | 0.04 | 0.08 | 0.2 | 0.11 | 0.22 |
| 2 | 51041603 | rs17041064 | 1.262 | 0.366 | -0.952 | 0.755 | 0.016 | 0.004 | 0.004 | 2.257 | 0.08 | 0.04 | 0.13 | 0.1 | 0.03 | 0.05 |
| 2 | 51044633 | rs10185764 | -1.315 | 0.517 | -0.834 | 0.631 | 0.002 | 0.002 | 0.002 | 0 | 0.9 | 0.86 | 0.83 | 0.88 | 0.85 | 0.8 |
| 2 | 51047313 | rs11125326 | 1.156 | 0.733 | -0.928 | 0.383 | -0.011 | 0.01 | 0.01 | 0 | 0.17 | 0.07 | 0.15 | 0.14 | 0.22 | 0.1 |
| 2 | 51047358 | rs9309203 | -0.193 | 0.609 | -0.811 | 0.37 | 0.028 | 0.033 | 0.033 | 0.561 | 0.6 | 0.25 | 0.45 | 0.27 | 0.69 | 0.42 |
| 2 | 51058745 | rs10167695 | 0.844 | 0.727 | -1.459 | 0.571 | -0.001 | 0.004 | 0.004 | 0 | 0.27 | 0.3 | 0.36 | 0.42 | 0.27 | 0.46 |
| 2 | 51059836 | rs17569702 | 1.418 | 1.275 | -2.273 | 0.018 | -0.023 | 0.036 | 0.036 | 0 | 0.14 | 0 | 0.06 | 0.06 | 0.12 | 0.06 |
| 2 | 51065644 | rs9679539 | 0.599 | 1.152 | -2.359 | -0.124 | -0.086 | 0.156 | 0.156 | 0 | 0.38 | 0.13 | 0.28 | 0.1 | 0.46 | 0.3 |
| 2 | 51068308 | rs17569799 | 0.331 | 1.103 | -2.238 | -0.037 | 0.001 | 0.011 | 0.011 | 0 | 0.2 | 0.13 | 0.16 | 0.14 | 0.22 | 0.11 |
| 2 | 51073449 | rs17504439 | 0.893 | 1.228 | -2.249 | 0.052 | -0.003 | 0.004 | 0.004 | 0 | 0.17 | 0.07 | 0.14 | 0.14 | 0.2 | 0.09 |
| 2 | 51077569 | rs17569963 | 1.285 | 1.44 | -2.287 | 0.027 | -0.029 | 0.045 | 0.045 | 0 | 0.15 | 0.02 | 0.06 | 0.06 | 0.13 | 0.06 |
| 2 | 51079482 | rs2193225 | 0.03 | 1.488 | -2.373 | -0.038 | 0.016 | 0.198 | 0.198 | 0 | 0.46 | 0.16 | 0.33 | 0.17 | 0.53 | 0.35 |
| 2 | 51082754 | rs10490175 | -0.421 | 1.517 | -2.306 | 0.015 | 0.017 | 0.003 | 0.003 | 0 | 0.07 | 0.05 | 0.13 | 0.14 | 0.2 | 0.11 |
| 2 | 51088030 | rs6743452 | 0.482 | 1.207 | -2.405 | -0.067 | -0.033 | 0.063 | 0.063 | 0 | 0.38 | 0.21 | 0.31 | 0.19 | 0.47 | 0.3 |
| 2 | 51093536 | rs11884918 | 0.759 | 1.161 | -2.434 | -0.133 | -0.068 | 0.153 | 0.153 | 0 | 0.35 | 0.13 | 0.23 | 0.07 | 0.37 | 0.22 |
| 2 | 51101583 | rs17570579 | 0.269 | 0.903 | -2.222 | 0.106 | -0.032 | 0.004 | 0.004 | 0 | 0.12 | 0.05 | 0.1 | 0.1 | 0.18 | 0.13 |
| 2 | 51106092 | rs17570759 | 0.162 | 0.855 | -2.298 | 0.084 | -0.042 | 0.055 | 0.055 | 0 | 0.22 | 0.02 | 0.12 | 0.1 | 0.21 | 0.14 |
| 2 | 51108133 | rs983936 | -0.613 | 0.836 | -2.306 | 0.063 | 0.039 | 0.055 | 0.055 | 0.407 | 0.58 | 0.23 | 0.44 | 0.27 | 0.68 | 0.43 |
| 2 | 51108406 | rs6727808 | -0.583 | 0.827 | -1.925 | 0.891 | -0.041 | -0.007 | -0.007 | 0 | 0.23 | 0.05 | 0.14 | 0.21 | 0.25 | 0.21 |
| 2 | 51120335 | rs17041090 | 0.861 | 0.442 | -1.784 | 1.151 | 0.016 | -0.046 | -0.046 | 0 | 0.12 | 0.02 | 0.05 | 0.17 | 0.08 | 0.08 |
| 2 | 51123048 | rs17571325 | -0.209 | 0.493 | -1.879 | 1.111 | -0.003 | -0.017 | -0.017 | 0.959 | 0.2 | 0.03 | 0.11 | 0.19 | 0.19 | 0.16 |
| 2 | 51126078 | rs17512199 | -0.167 | 0.699 | -1.691 | 1.6 | -0.012 | -0.024 | -0.024 | 0 | 0.1 | 0.03 | 0.07 | 0.12 | 0.11 | 0.13 |
| 2 | 51126441 | rs17041094 | -0.126 | 0.71 | -1.687 | 1.619 | -0.025 | -0.041 | -0.041 | 0 | 0.1 | 0.03 | 0.09 | 0.21 | 0.16 | 0.15 |
| 2 | 51127789 | rs17041095 | 0.144 | -0.282 | -1.339 | 1.75 | -0.046 | 0.045 | 0.045 | 0 | 0.25 | 0.62 | 0.34 | 0.49 | 0.11 | 0.31 |
| 2 | 51129285 | rs6741326 | 0.8 | -0.126 | -1.304 | 1.564 | 0 | -0.007 | -0.007 | 0 | 0.07 | 0.04 | 0.08 | 0.01 | 0.09 | 0.11 |
| 2 | 51132555 | rs9677208 | 0.614 | -0.109 | -1.27 | 1.555 | 0.001 | -0.015 | -0.015 | 0.028 | 0.08 | 0.09 | 0.08 | 0.01 | 0.09 | 0.11 |
| 2 | 51135341 | rs1160595 | 0.735 | -0.452 | -1.167 | 1.987 | 0.008 | -0.014 | -0.014 | 0 | 0.18 | 0.14 | 0.19 | 0.36 | 0.11 | 0.25 |
| 2 | 51135474 | rs10490168 | 1.629 | -0.003 | -2.006 | 1.437 | -0.036 | 0.043 | 0.043 | 2.048 | 0.28 | 0.11 | 0.22 | 0.14 | 0.3 | 0.21 |
| 2 | 51141342 | rs1003017 | 1.228 | 0.025 | -1.938 | 1.522 | 0.009 | -0.017 | -0.017 | 0.048 | 0.18 | 0.14 | 0.27 | 0.42 | 0.12 | 0.25 |
| 2 | 51160560 | rs13028294 | 1.563 | 0.09 | -2.021 | 1.264 | -0.036 | 0.028 | 0.028 | 1.941 | 0.28 | 0.11 | 0.22 | 0.16 | 0.31 | 0.21 |
| 2 | 51163990 | rs2193414 | 0.173 | -0.096 | -1 | 1.466 | -0.002 | 0.042 | 0.042 | 0.896 | 0.52 | 0.61 | 0.55 | 0.7 | 0.43 | 0.54 |
| 2 | 51171458 | rs4971709 | 0.39 | -0.157 | -0.77 | 1.536 | -0.054 | 0.121 | 0.121 | 0 | 0.25 | 0.45 | 0.33 | 0.51 | 0.13 | 0.34 |
| 2 | 51181831 | rs6730977 | -0.759 | 0.004 | -0.721 | 1.319 | -0.008 | 0.025 | 0.025 | 0.311 | 0.66 | 0.75 | 0.72 | 0.78 | 0.61 | 0.72 |
| 2 | 51183559 | rs2193417 | -0.462 | 0.101 | -0.807 | 1.261 | -0.002 | 0.019 | 0.019 | 1.402 | 0.61 | 0.64 | 0.69 | 0.76 | 0.59 | 0.68 |
| 2 | 51184223 | rs13409387 | 1.329 | 0.041 | -0.627 | 1.491 | 0.009 | 0.005 | 0.005 | 0 | 0.19 | 0.16 | 0.19 | 0.22 | 0.1 | 0.24 |
| 2 | 51184730 | rs4971711 | -0.384 | 0.074 | -0.845 | 1.456 | -0.002 | 0.002 | 0.002 | 0 | 0.35 | 0.61 | 0.48 | 0.58 | 0.3 | 0.49 |
| 2 | 51186984 | rs10197386 | 0.406 | 0.093 | -0.778 | 1.461 | 0.004 | 0.027 | 0.027 | 1.153 | 0.51 | 0.57 | 0.54 | 0.68 | 0.42 | 0.54 |
| 2 | 51191432 | rs12463983 | 1.477 | 0.365 | -1.037 | 1.089 | -0.088 | 0.04 | 0.04 | 1.295 | 0.3 | 0.08 | 0.25 | 0.19 | 0.36 | 0.22 |
| 2 | 51191693 | rs4971712 | -0.675 | -0.201 | -1.234 | 0.875 | -0.002 | 0.105 | 0.105 | 0 | 0.75 | 0.95 | 0.78 | 0.99 | 0.74 | 0.83 |
| 2 | 51191994 | rs13017431 | 3.078 | -0.216 | -1.567 | 0.687 | -0.07 | -0.007 | -0.007 | 0 | 0.17 | 0.01 | 0.16 | 0.12 | 0.19 | 0.16 |
| 2 | 51193994 | rs7608611 | 0.593 | -0.284 | -1.102 | 0.832 | 0.038 | 0.01 | 0.01 | 0 | 0.5 | 0.39 | 0.45 | 0.43 | 0.61 | 0.48 |
| 2 | 51209909 | rs10203784 | 1.442 | -0.266 | -0.719 | 0.855 | 0.013 | 0.013 | 0.013 | 0 | 0.16 | 0.11 | 0.05 | 0.12 | 0.09 | 0.21 |
| 2 | 51212752 | rs7559936 | 0.213 | -0.093 | -0.944 | 0.877 | -0.003 | 0.021 | 0.021 | 0.19 | 0.43 | 0.29 | 0.51 | 0.38 | 0.44 | 0.41 |
| 2 | 51224091 | rs1558850 | 0.718 | -0.155 | -0.909 | 1.02 | -0.057 | 0.079 | 0.079 | 0 | 0.23 | 0.42 | 0.21 | 0.39 | 0.11 | 0.3 |
| 2 | 51226834 | rs1995583 | -0.985 | -0.145 | -0.943 | 0.934 | -0.017 | 0.005 | 0.005 | 0.595 | 0.66 | 0.71 | 0.71 | 0.74 | 0.55 | 0.71 |
| 2 | 51227254 | rs2193410 | -0.905 | -0.106 | -0.937 | 0.953 | -0.041 | 0.014 | 0.014 | 0.689 | 0.66 | 0.79 | 0.71 | 0.74 | 0.55 | 0.71 |
| 2 | 51237684 | rs12616790 | -0.032 | -0.188 | -0.9 | 1.033 | 0.005 | 0.035 | 0.035 | 0 | 0.09 | 0.1 | 0.29 | 0.45 | 0.04 | 0.18 |
| 2 | 51241789 | rs13418185 | 0.694 | -0.203 | -0.915 | 0.957 | -0.036 | -0.069 | -0.069 | 0 | 0.23 | 0.43 | 0.16 | 0.12 | 0.17 | 0.23 |
| 2 | 51242270 | rs12467557 | -0.141 | -0.117 | -0.954 | 1.026 | 0.006 | 0.03 | 0.03 | 0 | 0.09 | 0.1 | 0.29 | 0.45 | 0.04 | 0.18 |
| 2 | 51247294 | rs13382584 | 1.434 | -0.173 | -1.073 | 0.875 | -0.006 | 0.022 | 0.022 | 1.471 | 0.3 | 0.18 | 0.2 | 0.16 | 0.3 | 0.22 |
| 2 | 51247657 | rs10490162 | -0.464 | -0.011 | -1.368 | 0.992 | 0.004 | -0.023 | -0.023 | 0 | 0.12 | 0.1 | 0.37 | 0.45 | 0.12 | 0.2 |
| 2 | 51250505 | rs7557525 | 0.99 | -0.005 | -1.354 | 0.937 | -0.006 | 0.038 | 0.038 | 1.306 | 0.33 | 0.15 | 0.25 | 0.17 | 0.35 | 0.31 |
| 2 | 51252137 | rs1558852 | -0.725 | -0.049 | -1.314 | 1.011 | 0 | -0.024 | -0.024 | 2.793 | 0.19 | 0.17 | 0.45 | 0.47 | 0.21 | 0.31 |
| 2 | 51258416 | rs2287232 | 1.021 | -0.168 | -1.279 | 0.926 | -0.066 | -0.058 | -0.058 | 0 | 0.19 | 0.49 | 0.07 | 0.12 | 0.07 | 0.2 |
| 2 | 51258887 | rs2287234 | -0.883 | -0.168 | -1.301 | 0.985 | -0.023 | -0.012 | -0.012 | 0 | 0.64 | 0.68 | 0.55 | 0.53 | 0.54 | 0.65 |
| 2 | 51259148 | rs2287235 | -0.322 | -0.16 | -1.31 | 1.001 | -0.001 | -0.035 | -0.035 | 0.608 | 0.17 | 0.07 | 0.29 | 0.26 | 0.2 | 0.26 |

| | | | | | | | | | | | | | | | | |
|---|---|---|---|---|---|---|---|---|---|---|---|---|---|---|---|---|
| 2 | 51272375 | rs11889192 | -0.521 | -0.183 | -1.299 | 0.98 | -0.047 | 0.024 | 0.024 | 0 | 0.3 | 0.52 | 0.29 | 0.36 | 0.18 | 0.35 |
| 2 | 51279394 | rs4971724 | 0.168 | -0.178 | -1.361 | 0.968 | -0.002 | -0.059 | -0.059 | 0.262 | 0.1 | 0.02 | 0.22 | 0.24 | 0.1 | 0.13 |
| 2 | 51279710 | rs12622930 | 1.848 | -0.151 | -1.337 | 0.975 | -0.012 | 0.027 | 0.027 | 2.397 | 0.27 | 0.09 | 0.19 | 0.16 | 0.27 | 0.19 |
| 2 | 51282609 | rs6545190 | -0.951 | -0.113 | -1.344 | 0.968 | -0.041 | 0.015 | 0.015 | 0.63 | 0.66 | 0.8 | 0.71 | 0.75 | 0.55 | 0.71 |
| 2 | 51286169 | rs7606493 | 0.851 | -0.105 | -1.213 | 0.995 | -0.067 | -0.059 | -0.059 | 0.033 | 0.19 | 0.49 | 0.08 | 0.12 | 0.07 | 0.2 |
| 2 | 51292433 | rs7576224 | -0.168 | -0.073 | -1.24 | 1.067 | -0.003 | -0.03 | -0.03 | 0.279 | 0.17 | 0.08 | 0.3 | 0.27 | 0.21 | 0.26 |
| 2 | 51294096 | rs988982 | 0.303 | -0.106 | -1.029 | 1.101 | -0.063 | 0.068 | 0.068 | 0 | 0.23 | 0.53 | 0.2 | 0.36 | 0.1 | 0.3 |
| 2 | 51304238 | rs2163018 | 1.415 | -0.033 | -1.198 | 0.963 | -0.012 | 0.026 | 0.026 | 2.32 | 0.27 | 0.1 | 0.19 | 0.16 | 0.27 | 0.2 |
| 2 | 51306133 | rs1995582 | 0.212 | 0.495 | -1.082 | 0.801 | 0.016 | 0.006 | 0.006 | 1.786 | 0.51 | 0.59 | 0.4 | 0.52 | 0.39 | 0.5 |
| 2 | 51308075 | rs7569904 | -0.569 | 0.743 | -1.258 | 0.664 | -0.026 | -0.018 | -0.018 | 0.2 | 0.74 | 0.87 | 0.81 | 0.74 | 0.7 | 0.74 |
| 2 | 51308895 | rs6720020 | 0.275 | 0.089 | -0.372 | 0.349 | -0.041 | -0.052 | -0.052 | 0 | 0.2 | 0.31 | 0.27 | 0.15 | 0.11 | 0.22 |
| 2 | 51311532 | rs17575302 | 0.662 | 0.535 | -0.161 | 0.708 | 0.031 | 0.033 | 0.033 | 0 | 0.06 | 0.09 | 0.09 | 0.14 | 0.11 | 0.1 |
| 2 | 51316018 | rs6742075 | 0.876 | 0.607 | -0.228 | 0.573 | -0.021 | 0.187 | 0.187 | 0 | 0.43 | 0.13 | 0.25 | 0.14 | 0.41 | 0.28 |
| 2 | 51316822 | rs10208300 | 1.421 | 0.482 | 0.021 | 0.517 | 0.007 | 0.007 | 0.007 | 0 | 0.14 | 0.11 | 0.16 | 0.16 | 0.05 | 0.18 |
| 2 | 51317471 | rs6734633 | -1.436 | 0.602 | -0.009 | 0.673 | -0.015 | -0.006 | -0.006 | 0.244 | 0.73 | 0.86 | 0.8 | 0.75 | 0.7 | 0.69 |
| 2 | 51318000 | rs6545193 | 0.367 | 0.486 | 0.388 | 0.895 | -0.021 | 0 | 0 | 0 | 0.2 | 0.39 | 0.32 | 0.26 | 0.18 | 0.25 |
| 2 | 51324337 | rs1157606 | -1.392 | 0.618 | 0.156 | 0.84 | -0.013 | -0.001 | -0.001 | 1.521 | 0.73 | 0.86 | 0.8 | 0.75 | 0.7 | 0.69 |
| 2 | 51325532 | rs6745471 | 0.435 | 0.466 | 0.331 | 0.75 | -0.026 | -0.014 | -0.014 | 0.09 | 0.2 | 0.36 | 0.32 | 0.2 | 0.17 | 0.24 |
| 2 | 51326497 | rs2195476 | 1.109 | 0.48 | -0.053 | 0.69 | -0.05 | 0.138 | 0.138 | 0.585 | 0.4 | 0.07 | 0.24 | 0.2 | 0.39 | 0.28 |
| 2 | 51331157 | rs735315 | 0.491 | 0.301 | 0.143 | 0.917 | -0.012 | 0.039 | 0.039 | 0.251 | 0.47 | 0.27 | 0.34 | 0.36 | 0.5 | 0.38 |
| 2 | 51339237 | rs12989159 | -0.992 | 0.316 | 0.379 | 0.937 | -0.004 | 0.008 | 0.008 | 0 | 0.7 | 0.8 | 0.8 | 0.78 | 0.71 | 0.74 |
| 2 | 51345474 | rs4971569 | -1.073 | 0.293 | 0.379 | 1.033 | -0.004 | 0.013 | 0.013 | 0 | 0.71 | 0.8 | 0.8 | 0.82 | 0.71 | 0.74 |
| 2 | 51354717 | rs10204611 | 0.856 | 0.106 | 0.577 | 0.627 | -0.006 | -0.031 | -0.031 | 0.129 | 0.31 | 0.37 | 0.35 | 0.17 | 0.29 | 0.31 |
| 2 | 51355852 | rs1013163 | 0.879 | 0.107 | 0.64 | 0.63 | -0.004 | -0.027 | -0.027 | 0 | 0.3 | 0.37 | 0.34 | 0.17 | 0.29 | 0.3 |
| 2 | 51355911 | rs1013164 | -1.387 | 0.092 | 0.584 | 0.63 | -0.004 | 0.007 | 0.007 | 0 | 0.71 | 0.79 | 0.79 | 0.78 | 0.71 | 0.74 |
| 2 | 51357902 | rs7602156 | 0.926 | 0.051 | 0.619 | 0.859 | 0.018 | 0.026 | 0.026 | 0 | 0.12 | 0.15 | 0.16 | 0.19 | 0.16 | 0.14 |
| 2 | 51358887 | rs7579236 | -0.342 | 0.036 | 0.638 | 0.917 | -0.004 | 0.001 | 0.001 | 0 | 0.38 | 0.32 | 0.31 | 0.39 | 0.4 | 0.36 |
| 2 | 51360667 | rs11125335 | -0.615 | 0.042 | 0.627 | 0.84 | 0.002 | -0.003 | -0.003 | 0 | 0.4 | 0.51 | 0.46 | 0.62 | 0.42 | 0.44 |
| 2 | 51367130 | rs10182663 | 0.882 | 0.088 | 0.348 | 0.824 | 0.022 | 0.034 | 0.034 | 0 | 0.12 | 0.17 | 0.16 | 0.19 | 0.16 | 0.15 |
| 2 | 51370461 | rs6727517 | -0.487 | 0.088 | 0.311 | 0.824 | -0.005 | -0.007 | -0.007 | 0.072 | 0.39 | 0.34 | 0.43 | 0.56 | 0.41 | 0.43 |
| 2 | 51386738 | rs13428293 | 0.859 | 0.119 | 0.276 | 0.879 | -0.001 | 0 | 0 | 0 | 0.12 | 0.08 | 0.16 | 0.2 | 0.16 | 0.15 |
| 2 | 51395987 | rs3948613 | -0.506 | 0.038 | 0.319 | 0.775 | -0.001 | -0.002 | -0.002 | 0 | 0.4 | 0.4 | 0.45 | 0.55 | 0.41 | 0.43 |
| 2 | 51398178 | rs2193399 | -0.724 | -0.002 | 0.384 | 0.754 | 0 | 0 | 0 | 0 | 0.42 | 0.6 | 0.47 | 0.55 | 0.41 | 0.44 |
| 2 | 51401196 | rs4408757 | 1.055 | -0.002 | 0.103 | 0.776 | -0.015 | 0.052 | 0.052 | 0 | 0.25 | 0.09 | 0.15 | 0.14 | 0.25 | 0.22 |
| 2 | 51411126 | rs4254538 | -0.662 | 0.04 | 0.196 | 0.779 | -0.001 | 0 | 0 | 0 | 0.42 | 0.48 | 0.45 | 0.55 | 0.41 | 0.44 |
| 2 | 51423122 | rs2193404 | 0.815 | 0.239 | -0.156 | 0.744 | -0.013 | -0.019 | -0.019 | 0 | 0.12 | 0.07 | 0.15 | 0.19 | 0.15 | 0.15 |
| 2 | 51429880 | rs11125339 | 1.389 | 0.007 | -0.22 | 0.606 | 0.008 | 0.003 | 0.003 | 0 | 0.11 | 0.09 | 0.14 | 0.15 | 0.04 | 0.19 |
| 2 | 51434161 | rs1017176 | 0.114 | -0.022 | -0.218 | 0.545 | -0.014 | 0.009 | 0.009 | 0 | 0.36 | 0.19 | 0.3 | 0.33 | 0.4 | 0.35 |
| 2 | 51437416 | rs2058854 | 0.789 | 0.252 | 0.117 | 0.794 | 0.001 | 0.004 | 0.004 | 0 | 0.12 | 0.11 | 0.16 | 0.19 | 0.16 | 0.14 |
| 2 | 51440854 | rs17519847 | 1.018 | 0.257 | 0.086 | 0.786 | -0.007 | -0.009 | -0.009 | 0 | 0.11 | 0.06 | 0.16 | 0.19 | 0.16 | 0.13 |
| 2 | 51445932 | rs10490153 | -0.76 | 0.191 | 0.346 | 0.766 | -0.015 | -0.012 | -0.012 | 0 | 0.53 | 0.69 | 0.52 | 0.57 | 0.52 | 0.57 |
| 2 | 51450802 | rs6716885 | -1.053 | 0.357 | 0.44 | 0.581 | -0.004 | 0.039 | 0.039 | 0 | 0.69 | 0.57 | 0.68 | 0.57 | 0.72 | 0.65 |
| 2 | 51453079 | rs2241225 | -0.682 | 0.43 | 0.479 | 0.571 | 0.003 | 0.014 | 0.014 | 0 | 0.64 | 0.47 | 0.62 | 0.47 | 0.64 | 0.61 |
| 2 | 51455704 | rs13025769 | 2.203 | 0.496 | -0.19 | 0.285 | -0.01 | 0.015 | 0.015 | 0.689 | 0.16 | 0.1 | 0.18 | 0 | 0.21 | 0.06 |
| 2 | 51458570 | rs4141448 | -1.201 | 0.365 | -0.298 | 0.692 | -0.007 | 0.013 | 0.013 | 0 | 0.72 | 0.87 | 0.82 | 0.78 | 0.73 | 0.72 |
| 2 | 51471321 | rs2193392 | -1.239 | 0.432 | -0.249 | 0.759 | -0.019 | -0.005 | -0.005 | 0.486 | 0.78 | 0.87 | 0.83 | 0.78 | 0.72 | 0.73 |
| 2 | 51483528 | rs2193394 | 0.944 | 0.297 | -0.019 | 0.789 | 0.015 | 0.098 | 0.098 | 0 | 0.09 | 0.27 | 0.18 | 0.24 | 0.09 | 0.21 |
| 2 | 51488425 | rs13420545 | -0.133 | 0.257 | -0.359 | 0.689 | 0.013 | 0.015 | 0.015 | 0.051 | 0.38 | 0.25 | 0.34 | 0.38 | 0.34 | 0.45 |
| 2 | 51492867 | rs2193396 | 0.7 | 0.211 | -0.901 | 0.556 | -0.004 | 0.002 | 0.002 | 1.171 | 0.41 | 0.56 | 0.48 | 0.4 | 0.38 | 0.29 |
| 2 | 51492973 | rs1115763 | -0.266 | 0.121 | -1.657 | 0.366 | 0.013 | -0.029 | -0.029 | 0.446 | 0.36 | 0.18 | 0.3 | 0.36 | 0.3 | 0.32 |
| 2 | 51512492 | rs10490146 | 1.586 | 0.064 | -1.678 | -0.207 | -0.041 | 0.151 | 0.151 | 1.154 | 0.26 | 0.03 | 0.2 | 0 | 0.26 | 0.11 |
| 2 | 51513533 | rs12713137 | -0.85 | 0.077 | -1.322 | -0.042 | 0.017 | 0.005 | 0.005 | 1.129 | 0.76 | 0.7 | 0.71 | 0.68 | 0.74 | 0.74 |
| 2 | 51513683 | rs10490145 | 0.971 | -0.15 | -0.935 | 0.364 | 0.005 | 0.022 | 0.022 | 2.86 | 0.37 | 0.23 | 0.34 | 0.24 | 0.35 | 0.36 |
| 2 | 51514659 | rs2216133 | -1.344 | -0.117 | -1.09 | 0.384 | -0.018 | 0.021 | 0.021 | 1.087 | 0.8 | 0.85 | 0.82 | 0.9 | 0.76 | 0.76 |
| 2 | 51516700 | rs7578198 | -0.545 | -0.205 | -1.267 | 0.418 | 0.011 | -0.001 | -0.001 | 0 | 0.74 | 0.67 | 0.69 | 0.68 | 0.72 | 0.72 |
| 2 | 51519639 | rs36016587 | 0.311 | -0.271 | -1.732 | 0.149 | -0.072 | -0.094 | -0.094 | 0 | 0.3 | 0.02 | 0.21 | 0.33 | 0.36 | 0.2 |
| 2 | 51520327 | rs10490142 | 2.152 | -0.185 | -1.307 | 0.081 | -0.012 | 0.097 | 0.097 | 0.805 | 0.27 | 0.1 | 0.2 | 0 | 0.26 | 0.11 |
| 2 | 51528298 | rs7602908 | 2.208 | -0.173 | -1.314 | 0.087 | -0.015 | 0.098 | 0.098 | 1.167 | 0.26 | 0.08 | 0.2 | 0.01 | 0.26 | 0.11 |
| 2 | 51532494 | rs7565012 | 0.667 | -0.227 | -0.764 | 0.243 | -0.007 | 0.002 | 0.002 | 2.911 | 0.42 | 0.49 | 0.46 | 0.4 | 0.38 | 0.36 |
| 2 | 51533191 | rs7568446 | -1.069 | -0.249 | -0.612 | 0.099 | 0.012 | -0.025 | -0.025 | 0 | 0.77 | 0.74 | 0.78 | 0.84 | 0.75 | 0.74 |
| 2 | 51534306 | rs12233091 | -1.455 | -0.197 | -0.769 | 0.071 | -0.023 | 0.032 | 0.032 | 1.173 | 0.81 | 0.9 | 0.82 | 0.91 | 0.77 | 0.76 |
| 2 | 51536930 | rs6709031 | 2.337 | -0.329 | -0.809 | 0.17 | -0.015 | 0.098 | 0.098 | 1.241 | 0.26 | 0.08 | 0.2 | 0 | 0.26 | 0.11 |
| 2 | 51538031 | rs4971739 | 0.043 | -0.341 | -0.73 | 0.218 | -0.013 | 0.134 | 0.134 | 0 | 0.14 | 0.3 | 0.24 | 0.39 | 0.12 | 0.25 |
| 2 | 51544679 | rs7570510 | 0.757 | -0.333 | -0.817 | 0.202 | 0.002 | -0.003 | -0.003 | 0 | 0.41 | 0.39 | 0.45 | 0.4 | 0.38 | 0.36 |
| 2 | 51547046 | rs2160408 | -0.007 | -0.324 | -0.79 | 0.217 | -0.014 | 0.172 | 0.172 | 0 | 0.14 | 0.35 | 0.25 | 0.39 | 0.11 | 0.25 |
| 2 | 51548039 | rs17588882 | 2.244 | -0.251 | -1.03 | 0.042 | -0.025 | 0.126 | 0.126 | 1.306 | 0.26 | 0.05 | 0.2 | 0 | 0.26 | 0.11 |
| 2 | 51550798 | rs17041579 | 0.571 | 0.069 | -0.169 | 0.491 | -0.004 | 0.02 | 0.02 | 0 | 0.7 | 0.59 | 0.69 | 0.58 | 0.73 | 0.66 |
| 2 | 51554047 | rs12713138 | 0.188 | 0.036 | -0.054 | 0.553 | -0.005 | -0.011 | -0.011 | 0 | 0.17 | 0.13 | 0.12 | 0.27 | 0.21 | 0.15 |
| 2 | 51554079 | rs7587510 | -0.135 | 0.076 | 0.04 | 0.481 | 0.001 | -0.013 | -0.013 | 0 | 0.7 | 0.63 | 0.63 | 0.74 | 0.71 | 0.55 |

| | | | | | | | | | | | | | | | | |
|---|---|---|---|---|---|---|---|---|---|---|---|---|---|---|---|---|
| 2 | 51554862 | rs1548898 | -0.032 | -0.009 | -0.104 | 0.554 | -0.014 | 0.18 | 0.18 | 0 | 0.14 | 0.34 | 0.21 | 0.39 | 0.1 | 0.25 |
| 2 | 51556145 | rs1319130 | 0.986 | -0.04 | -0.142 | 0.504 | 0 | -0.001 | -0.001 | 0 | 0.47 | 0.42 | 0.48 | 0.46 | 0.49 | 0.38 |
| 2 | 51559091 | rs1548899 | -0.404 | 0.058 | -0.116 | 0.619 | 0.005 | 0.027 | 0.027 | 0.372 | 0.34 | 0.49 | 0.41 | 0.54 | 0.33 | 0.53 |
| 2 | 51566220 | rs1008618 | 0.368 | -0.073 | -0.446 | 0.665 | -0.006 | 0.014 | 0.014 | 0 | 0.73 | 0.85 | 0.84 | 0.76 | 0.72 | 0.81 |
| 2 | 51566341 | rs17589784 | 1.965 | -0.213 | -0.621 | 0.491 | -0.048 | 0.139 | 0.139 | 0.474 | 0.34 | 0.07 | 0.29 | 0.08 | 0.39 | 0.14 |
| 2 | 51567535 | rs17041599 | 1.851 | -0.18 | -0.043 | 0.538 | 0.026 | -0.027 | -0.027 | 0 | 0.16 | 0.26 | 0.18 | 0.09 | 0.19 | 0.23 |
| 2 | 51569042 | rs6717556 | 0.617 | -0.28 | -0.02 | 0.507 | 0.003 | 0.006 | 0.006 | 0.762 | 0.57 | 0.5 | 0.56 | 0.52 | 0.58 | 0.44 |
| 2 | 51572556 | rs7582583 | 0.667 | -0.282 | -0.013 | 0.522 | 0.001 | 0 | 0 | 0 | 0.57 | 0.53 | 0.54 | 0.46 | 0.55 | 0.43 |
| 2 | 51575056 | rs2160410 | 0.195 | -0.335 | -0.071 | 0.547 | -0.053 | -0.048 | -0.048 | 0.013 | 0.79 | 0.99 | 0.89 | 0.76 | 0.79 | 0.83 |
| 2 | 51576612 | rs2883286 | 0.4 | -0.363 | 0.071 | 0.634 | 0.008 | 0.011 | 0.011 | 1.563 | 0.61 | 0.5 | 0.57 | 0.46 | 0.61 | 0.43 |
| 2 | 51581853 | rs17041624 | 0.828 | -0.39 | 0.065 | 0.681 | 0.052 | 0.153 | 0.153 | 0.165 | 0.22 | 0.5 | 0.36 | 0.42 | 0.25 | 0.46 |
| 2 | 51584418 | rs6545212 | -0.097 | -0.22 | 0.391 | 0.85 | 0.005 | 0.008 | 0.008 | 1.415 | 0.62 | 0.55 | 0.56 | 0.51 | 0.59 | 0.44 |
| 2 | 51593059 | rs6722881 | -0.843 | -0.485 | 0.525 | 0.92 | -0.005 | 0.009 | 0.009 | 0 | 0.27 | 0.32 | 0.27 | 0.44 | 0.23 | 0.3 |
| 2 | 51594480 | rs2110989 | 1.294 | -0.493 | 0.531 | 1.03 | 0.024 | -0.023 | -0.023 | 0.049 | 0.07 | 0.04 | 0.13 | 0.18 | 0.01 | 0.16 |
| 2 | 51612454 | rs5016462 | 0.128 | -0.3 | 0.526 | 0.635 | -0.001 | 0.002 | 0.002 | 0.582 | 0.59 | 0.54 | 0.57 | 0.52 | 0.61 | 0.44 |
| 2 | 51618513 | rs7557697 | 0.093 | -0.283 | 0.449 | 0.603 | -0.002 | -0.003 | -0.003 | 0 | 0.2 | 0.17 | 0.13 | 0.26 | 0.21 | 0.14 |
| 2 | 51626179 | rs10207678 | 0.117 | -0.259 | 0.454 | 0.614 | 0 | 0.001 | 0.001 | 0 | 0.19 | 0.18 | 0.13 | 0.26 | 0.21 | 0.14 |
| 2 | 51633618 | rs1919430 | 0.513 | -0.373 | 0.5 | 0.606 | -0.003 | -0.004 | -0.004 | 0.038 | 0.54 | 0.45 | 0.44 | 0.34 | 0.61 | 0.28 |
| 2 | 51634931 | rs1919429 | 0.172 | -0.131 | 0.325 | 0.742 | -0.007 | -0.003 | -0.003 | 0.168 | 0.21 | 0.16 | 0.14 | 0.25 | 0.23 | 0.15 |
| 2 | 51640946 | rs9309212 | 0.21 | -0.406 | 0.33 | 0.813 | -0.022 | -0.005 | -0.005 | 0 | 0.2 | 0.07 | 0.12 | 0.2 | 0.21 | 0.15 |
| 2 | 51643185 | rs1206401 | -0.813 | -0.473 | 0.352 | 0.803 | -0.012 | 0.046 | 0.046 | 0 | 0.21 | 0.23 | 0.19 | 0.39 | 0.16 | 0.34 |
| 2 | 51643322 | rs13420278 | 0.26 | -0.22 | 0.213 | 0.572 | -0.001 | -0.001 | -0.001 | 0 | 0.2 | 0.18 | 0.14 | 0.25 | 0.21 | 0.15 |
| 2 | 51648736 | rs17525600 | -0.052 | -0.135 | 0.296 | 0.697 | -0.018 | -0.015 | -0.015 | 0.959 | 0.08 | 0.01 | 0.04 | 0.05 | 0.08 | 0.07 |
| 2 | 51651243 | rs2098389 | 0.481 | -0.202 | 0.51 | 0.339 | -0.003 | -0.007 | -0.007 | 0 | 0.58 | 0.41 | 0.41 | 0.27 | 0.62 | 0.28 |
| 2 | 51654878 | rs1015192 | 0.236 | 0.199 | 0.536 | 0.593 | -0.002 | -0.014 | -0.014 | 0 | 0.62 | 0.67 | 0.55 | 0.5 | 0.6 | 0.44 |
| 2 | 51655345 | rs13387812 | 0.141 | 0.227 | 0.404 | 0.606 | -0.005 | -0.02 | -0.02 | 0 | 0.63 | 0.69 | 0.55 | 0.5 | 0.6 | 0.44 |
| 2 | 51663763 | rs10184263 | 0.421 | 0.421 | -0.998 | 0.7 | -0.005 | -0.008 | -0.008 | 0 | 0.17 | 0.15 | 0.11 | 0.27 | 0.18 | 0.13 |
| 2 | 51664304 | rs1206415 | -0.564 | 0.137 | -0.457 | 1.053 | -0.012 | 0.039 | 0.039 | 1.529 | 0.81 | 0.89 | 0.78 | 0.93 | 0.76 | 0.76 |
| 2 | 51664721 | rs6545221 | 0.145 | 0.245 | -0.53 | 1.125 | 0.001 | -0.005 | -0.005 | 0 | 0.59 | 0.38 | 0.52 | 0.5 | 0.59 | 0.42 |
| 2 | 51669730 | rs11563100 | -1.088 | 0.138 | -0.208 | 1.509 | -0.009 | 0.067 | 0.067 | 0 | 0.15 | 0.29 | 0.27 | 0.34 | 0.16 | 0.2 |
| 2 | 51675468 | rs6737381 | -0.814 | 0.089 | -0.111 | 1.619 | 0.003 | 0.007 | 0.007 | 0 | 0.12 | 0.28 | 0.08 | 0.1 | 0.15 | 0.08 |
| 2 | 51680489 | rs11125351 | 0.624 | 0.114 | 0.116 | 0.588 | 0.002 | -0.072 | -0.072 | 0 | 0.28 | 0.23 | 0.24 | 0 | 0.25 | 0.17 |
| 2 | 51685872 | rs7572317 | -0.583 | -0.363 | 0.474 | 1.542 | -0.012 | 0.065 | 0.065 | 0 | 0.11 | 0.21 | 0.08 | 0.29 | 0.08 | 0.26 |
| 2 | 51688180 | rs984795 | -1.231 | -0.397 | 0.52 | 1.605 | -0.039 | 0.22 | 0.22 | 0 | 0.13 | 0.33 | 0.27 | 0.53 | 0.05 | 0.39 |
| 2 | 51689306 | rs991900 | -0.824 | -0.393 | 0.534 | 1.584 | 0.002 | 0.006 | 0.006 | 0 | 0.1 | 0.09 | 0.07 | 0.29 | 0.08 | 0.27 |
| 2 | 51689579 | rs991901 | -0.857 | -0.392 | 0.524 | 1.585 | -0.006 | 0.036 | 0.036 | 0 | 0.1 | 0.13 | 0.08 | 0.28 | 0.08 | 0.26 |
| 2 | 51692449 | rs2353829 | -1.418 | -0.396 | 0.658 | 1.567 | -0.012 | 0.08 | 0.08 | 0.081 | 0.13 | 0.17 | 0.27 | 0.53 | 0.08 | 0.39 |
| 2 | 51692541 | rs11563048 | 0.051 | -0.454 | 1.292 | 1.54 | -0.001 | 0.004 | 0.004 | 0 | 0.11 | 0.13 | 0.06 | 0.1 | 0.14 | 0.07 |
| 2 | 51694815 | rs11563178 | -1.682 | -0.44 | 1.087 | 1.396 | -0.029 | 0.101 | 0.101 | 0 | 0.23 | 0.41 | 0.33 | 0.54 | 0.19 | 0.45 |
| 2 | 51699054 | rs17864503 | -1.265 | -0.223 | 0.868 | 1.343 | -0.001 | -0.012 | -0.012 | 0 | 0.2 | 0.15 | 0.13 | 0.3 | 0.24 | 0.34 |
| 2 | 51702486 | rs6739884 | -1.594 | -0.266 | 1.277 | 1.277 | 0.004 | 0.061 | 0.061 | 0 | 0.22 | 0.52 | 0.15 | 0.3 | 0.24 | 0.34 |
| 2 | 51704524 | rs12996042 | 0.115 | -0.251 | 1.149 | 1.172 | -0.007 | 0.016 | 0.016 | 0 | 0.09 | 0.05 | 0.06 | 0.01 | 0.14 | 0.05 |
| 2 | 51706314 | rs6705941 | -0.536 | -0.898 | 0.63 | 0.943 | 0.023 | -0.015 | -0.015 | 2.212 | 0.79 | 0.88 | 0.88 | 0.8 | 0.85 | 0.88 |
| 2 | 51710533 | rs1919419 | -1.512 | -0.894 | 1.143 | 1.106 | -0.022 | 0.097 | 0.097 | 0.018 | 0.26 | 0.49 | 0.37 | 0.6 | 0.24 | 0.48 |
| 2 | 51711095 | rs17868144 | -0.968 | -0.875 | 1.131 | 1.155 | -0.013 | 0.136 | 0.136 | 0 | 0.14 | 0.22 | 0.3 | 0.52 | 0.1 | 0.41 |
| 2 | 51711823 | rs12995616 | 0.362 | -0.895 | 1.138 | 1.104 | 0.002 | -0.007 | -0.007 | 0.774 | 0.11 | 0.14 | 0.06 | 0.07 | 0.14 | 0.06 |
| 2 | 51716954 | rs6725779 | 0.149 | -0.926 | 0.727 | 1.511 | -0.023 | 0.189 | 0.189 | 0 | 0.08 | 0.26 | 0.24 | 0.33 | 0.02 | 0.33 |
| 2 | 51717718 | rs17862679 | -1.335 | -0.934 | 0.747 | 1.587 | -0.006 | 0.027 | 0.027 | 0.152 | 0.19 | 0.2 | 0.29 | 0.58 | 0.15 | 0.4 |
| 2 | 51718448 | rs13006788 | 0.967 | -0.92 | 0.687 | 1.533 | -0.027 | 0.044 | 0.044 | 0 | 0.08 | 0.02 | 0.05 | 0.01 | 0.11 | 0.05 |
| 2 | 51718820 | rs11563175 | -0.387 | -0.94 | 0.739 | 1.567 | -0.045 | 0.148 | 0.148 | 0.275 | 0.1 | 0.2 | 0.23 | 0.39 | 0.02 | 0.34 |
| 2 | 51720700 | rs1569193 | -1.034 | -0.92 | 0.74 | 1.565 | -0.022 | 0.098 | 0.098 | 0.034 | 0.21 | 0.36 | 0.31 | 0.58 | 0.18 | 0.4 |
| 2 | 51723107 | rs9789477 | 0.837 | -0.967 | -0.186 | 1.059 | -0.151 | -0.119 | -0.119 | 0 | 0.37 | 0.03 | 0.36 | 0.39 | 0.49 | 0.33 |
| 2 | 51729681 | rs10170127 | 1.212 | -1.142 | 0.074 | 0.945 | 0.05 | 0.078 | 0.078 | 1.06 | 0.55 | 0.26 | 0.51 | 0.21 | 0.68 | 0.48 |
| 2 | 51732873 | rs2883423 | 0.966 | -1.01 | 0.072 | 1.077 | 0.018 | -0.006 | -0.006 | 0 | 0.61 | 0.28 | 0.58 | 0.4 | 0.7 | 0.49 |
| 2 | 51736994 | rs1406428 | -0.697 | -0.735 | -0.715 | 1.333 | -0.007 | 0.02 | 0.02 | 0 | 0.19 | 0.21 | 0.29 | 0.41 | 0.16 | 0.39 |
| 2 | 51738393 | rs1528805 | 1.243 | -0.738 | -0.899 | 1.135 | -0.124 | 0.017 | 0.017 | 0.556 | 0.28 | 0.03 | 0.29 | 0.19 | 0.42 | 0.23 |
| 2 | 51742119 | rs2715051 | -1.542 | -0.753 | -0.136 | 1.4 | -0.002 | 0.026 | 0.026 | 0 | 0.95 | 0.99 | 0.95 | 1 | 0.95 | 0.98 |
| 2 | 51753361 | rs12612083 | -0.837 | -0.453 | -0.037 | 1.741 | 0.005 | 0.12 | 0.12 | 0 | 0.06 | 0.1 | 0.09 | 0.28 | 0.05 | 0.28 |
| 2 | 51757112 | rs17862700 | -0.587 | -0.407 | -0.305 | 1.266 | 0.003 | -0.008 | -0.008 | 0 | 0.14 | 0.12 | 0.25 | 0.3 | 0.14 | 0.15 |
| 2 | 51761876 | rs2715091 | -0.065 | -0.492 | -0.068 | 0.72 | -0.019 | -0.008 | -0.008 | 0 | 0.74 | 0.55 | 0.79 | 0.76 | 0.78 | 0.78 |
| 2 | 51770480 | rs1528820 | -0.217 | 0.333 | 0.458 | 0.424 | -0.012 | 0.04 | 0.04 | 0 | 0.39 | 0.32 | 0.34 | 0.18 | 0.42 | 0.36 |
| 2 | 51773393 | rs1358578 | -0.091 | 0.331 | 0.521 | 0.431 | -0.05 | 0.108 | 0.108 | 0 | 0.37 | 0.13 | 0.32 | 0.18 | 0.4 | 0.35 |
| 2 | 51781853 | rs2698028 | -0.277 | 0.358 | 0.641 | 0.217 | -0.011 | 0.076 | 0.076 | 0 | 0.43 | 0.31 | 0.36 | 0.18 | 0.43 | 0.38 |
| 2 | 51786142 | rs2715062 | -0.249 | 0.346 | 0.63 | 0.216 | -0.024 | 0.129 | 0.129 | 0 | 0.43 | 0.21 | 0.34 | 0.18 | 0.43 | 0.38 |
| 2 | 51786520 | rs2698024 | -0.417 | 0.327 | 0.809 | 0.507 | 0 | 0.005 | 0.005 | 0 | 0.46 | 0.58 | 0.4 | 0.32 | 0.43 | 0.39 |
| 2 | 51792223 | rs1609361 | -0.211 | 0.335 | 0.683 | 0.498 | -0.023 | 0.127 | 0.127 | 0 | 0.43 | 0.22 | 0.34 | 0.19 | 0.43 | 0.38 |
| 2 | 51799271 | rs2715076 | -0.354 | 0.363 | 0.535 | 0.475 | -0.001 | 0.003 | 0.003 | 0 | 0.46 | 0.44 | 0.39 | 0.32 | 0.43 | 0.39 |
| 2 | 51804668 | rs4971755 | -0.25 | 0.426 | 0.567 | 0.415 | 0.004 | -0.014 | -0.014 | 0.833 | 0.41 | 0.53 | 0.42 | 0.33 | 0.42 | 0.44 |
| 2 | 51807660 | rs6737616 | 1.165 | -0.08 | 0.24 | 0.537 | 0.044 | -0.019 | -0.019 | 0.111 | 0.22 | 0.08 | 0.27 | 0.24 | 0.1 | 0.26 |

| | | | | | | | | | | | | | | | | |
|---|---|---|---|---|---|---|---|---|---|---|---|---|---|---|---|---|
| 2 | 51810439 | rs6721228 | 0.343 | 0.033 | 0.104 | 0.422 | -0.009 | 0.048 | 0.048 | 0 | 0.38 | 0.29 | 0.34 | 0.19 | 0.39 | 0.4 |
| 2 | 51810484 | rs17868182 | 1.264 | 0.216 | -0.39 | 0.235 | -0.08 | 0.011 | 0.011 | 0 | 0.24 | 0.02 | 0.14 | 0.17 | 0.25 | 0.19 |
| 2 | 51814215 | rs11562998 | 1.352 | 0.065 | -0.02 | 0.569 | 0.03 | -0.037 | -0.037 | 0 | 0.2 | 0.09 | 0.3 | 0.37 | 0.1 | 0.26 |
| 2 | 51815287 | rs12470929 | -0.572 | 0.264 | 0.251 | 0.512 | -0.006 | -0.006 | -0.006 | 0 | 0.58 | 0.61 | 0.63 | 0.56 | 0.44 | 0.66 |
| 2 | 51815380 | rs12478948 | -0.702 | 0.226 | 0.292 | 0.553 | 0.004 | 0.003 | 0.003 | 0.057 | 0.6 | 0.58 | 0.66 | 0.56 | 0.5 | 0.68 |
| 2 | 51820543 | rs13386874 | -0.147 | 0 | -0.23 | 0.75 | 0.008 | -0.001 | -0.001 | 0 | 0.32 | 0.23 | 0.42 | 0.63 | 0.25 | 0.45 |
| 2 | 51821908 | rs12616608 | 0.296 | 0.054 | -0.238 | 0.812 | -0.002 | 0.021 | 0.021 | 0 | 0.12 | 0.09 | 0.12 | 0.26 | 0.15 | 0.17 |
| 2 | 51823943 | rs10190317 | -0.09 | 0.023 | -0.184 | 0.718 | -0.004 | 0 | 0 | 0 | 0.32 | 0.32 | 0.39 | 0.37 | 0.24 | 0.37 |
| 2 | 51824160 | rs10199009 | -0.181 | 0.009 | -0.114 | 0.721 | -0.032 | 0.01 | 0.01 | 0 | 0.33 | 0.4 | 0.4 | 0.37 | 0.23 | 0.38 |
| 2 | 51825042 | rs2883426 | 0.103 | -0.01 | -0.134 | 0.728 | 0.001 | -0.002 | -0.002 | 0 | 0.3 | 0.26 | 0.38 | 0.37 | 0.22 | 0.35 |
| 2 | 51826920 | rs2354382 | 1.127 | -0.029 | -0.234 | 0.727 | -0.026 | 0.022 | 0.022 | 0.174 | 0.21 | 0.21 | 0.3 | 0.36 | 0.1 | 0.3 |
| 2 | 51833271 | rs17864557 | 0.318 | 0.157 | -0.435 | 0.789 | 0.003 | -0.001 | -0.001 | 0 | 0.11 | 0.07 | 0.12 | 0.26 | 0.15 | 0.16 |
| 2 | 51847852 | rs10495999 | 0.684 | 0.274 | -0.808 | 0.471 | -0.11 | 0.092 | 0.092 | 0 | 0.36 | 0.04 | 0.33 | 0.18 | 0.39 | 0.3 |
| 2 | 51849371 | rs2465526 | -0.516 | 0.318 | -0.443 | 0.654 | -0.002 | 0.035 | 0.035 | 0 | 0.14 | 0.21 | 0.15 | 0.26 | 0.18 | 0.19 |
| 2 | 51851017 | rs11563273 | 1.409 | 0.585 | -0.757 | 0.464 | -0.114 | 0.03 | 0.03 | 0 | 0.25 | 0.01 | 0.21 | 0.18 | 0.32 | 0.14 |
| 2 | 51852093 | rs2953299 | -0.086 | 0.063 | -0.308 | 0.833 | -0.001 | 0.03 | 0.03 | 0 | 0.12 | 0.17 | 0.14 | 0.27 | 0.17 | 0.18 |
| 2 | 51852373 | rs2953300 | -0.132 | 0.063 | -0.318 | 0.838 | -0.001 | 0.043 | 0.043 | 0 | 0.13 | 0.19 | 0.14 | 0.25 | 0.17 | 0.18 |
| 2 | 51852755 | rs2675025 | 0.98 | 0.071 | -0.489 | 0.737 | -0.017 | 0.046 | 0.046 | 0 | 0.08 | 0.01 | 0.07 | 0 | 0.12 | 0.02 |
| 2 | 51862691 | rs10496000 | 1.208 | -0.023 | -0.322 | 0.686 | -0.001 | -0.001 | -0.001 | 0.01 | 0.2 | 0.15 | 0.29 | 0.36 | 0.1 | 0.27 |
| 2 | 51864372 | rs11563025 | 1.161 | -0.022 | -0.35 | 0.666 | 0.031 | -0.039 | -0.039 | 0 | 0.2 | 0.08 | 0.29 | 0.36 | 0.1 | 0.27 |
| 2 | 51871946 | rs17864568 | 0.174 | -0.03 | -0.169 | 0.674 | -0.006 | 0.088 | 0.088 | 0 | 0.86 | 0.67 | 0.85 | 0.74 | 0.83 | 0.82 |
| 2 | 51873354 | rs10439476 | 0.324 | -0.055 | -0.209 | 0.649 | -0.008 | -0.001 | -0.001 | 0 | 0.65 | 0.65 | 0.57 | 0.38 | 0.73 | 0.54 |
| 2 | 51876227 | rs952893 | -0.168 | 0.207 | -0.087 | 0.806 | -0.001 | 0.031 | 0.031 | 0 | 0.88 | 0.87 | 0.87 | 0.73 | 0.84 | 0.85 |
| 2 | 51881427 | rs10865256 | 0.617 | 0.397 | -0.258 | 0.487 | -0.039 | 0.077 | 0.077 | 0 | 0.34 | 0.1 | 0.32 | 0.19 | 0.37 | 0.37 |
| 2 | 51890276 | rs966733 | 1.03 | 0.217 | 1.052 | 0.964 | -0.018 | 0.146 | 0.146 | 0.009 | 0.82 | 0.44 | 0.79 | 0.61 | 0.82 | 0.77 |
| 2 | 51890442 | rs966734 | -0.501 | 0.211 | 1.235 | 1.004 | -0.006 | 0.06 | 0.06 | 0 | 0.89 | 0.85 | 0.88 | 0.76 | 0.86 | 0.83 |
| 2 | 51894219 | rs1516178 | 1.117 | 0.218 | 1.247 | 1.003 | -0.018 | 0.152 | 0.152 | 0.004 | 0.82 | 0.46 | 0.8 | 0.62 | 0.82 | 0.77 |
| 2 | 51897741 | rs11563268 | -0.334 | 0.246 | 1.164 | 1.029 | 0.001 | 0.076 | 0.076 | 0 | 0.89 | 0.84 | 0.86 | 0.75 | 0.85 | 0.83 |
| 2 | 51901085 | rs1400104 | 0.682 | 0.239 | 1.208 | 1.007 | 0.008 | 0.085 | 0.085 | 0 | 0.85 | 0.71 | 0.84 | 0.75 | 0.82 | 0.78 |
| 2 | 51908399 | rs959661 | -0.415 | 0.218 | 1.128 | 0.994 | -0.003 | 0.043 | 0.043 | 0 | 0.89 | 0.87 | 0.88 | 0.76 | 0.86 | 0.83 |
| 2 | 51909753 | rs4146703 | 1.342 | 0.167 | 1.531 | 0.977 | -0.013 | 0.066 | 0.066 | 0.854 | 0.81 | 0.32 | 0.79 | 0.61 | 0.82 | 0.77 |
| 2 | 51911211 | rs1400112 | 0.828 | 0.181 | 1.301 | 0.945 | -0.01 | 0.091 | 0.091 | 0.021 | 0.84 | 0.64 | 0.85 | 0.75 | 0.83 | 0.78 |
| 2 | 51920657 | rs6739137 | -0.443 | 0.204 | 1.213 | 0.967 | -0.006 | 0.109 | 0.109 | 0 | 0.89 | 0.79 | 0.87 | 0.74 | 0.86 | 0.84 |
| 2 | 51921531 | rs2354979 | 0.342 | 0.149 | 0.564 | 0.421 | -0.046 | 0.09 | 0.09 | 0 | 0.39 | 0.11 | 0.35 | 0.22 | 0.41 | 0.44 |
| 2 | 51928042 | rs1028145 | -0.709 | 0.285 | 0.763 | 0.529 | -0.001 | -0.026 | -0.026 | 0 | 0.9 | 0.88 | 0.91 | 0.98 | 0.87 | 0.93 |
| 2 | 51928206 | rs1028146 | 0.75 | 0.246 | 1.002 | 0.568 | -0.013 | 0 | 0 | 0 | 0.84 | 0.46 | 0.84 | 0.83 | 0.85 | 0.87 |
| 2 | 51931529 | rs7609540 | -0.001 | 0.258 | 1.018 | 0.549 | 0.007 | -0.076 | -0.076 | 0 | 0.87 | 0.7 | 0.91 | 0.99 | 0.86 | 0.91 |
| 2 | 51931744 | rs17862754 | 1.349 | 0.238 | 0.51 | 0.434 | -0.057 | 0.008 | 0.008 | 0.03 | 0.23 | 0.05 | 0.16 | 0.18 | 0.25 | 0.12 |
| 2 | 51933909 | rs1516184 | 0.735 | 0.312 | 0.951 | 0.487 | -0.021 | -0.01 | -0.01 | 0 | 0.84 | 0.39 | 0.85 | 0.85 | 0.86 | 0.88 |
| 2 | 51934748 | rs4971757 | -0.689 | 0.337 | 0.728 | 0.47 | -0.004 | -0.05 | -0.05 | 0 | 0.9 | 0.82 | 0.92 | 0.98 | 0.88 | 0.93 |
| 2 | 51935903 | rs17874928 | 1.096 | 0.403 | -0.079 | 0.317 | -0.088 | -0.004 | -0.004 | 0.034 | 0.25 | 0.02 | 0.15 | 0.19 | 0.27 | 0.21 |
| 2 | 51940564 | rs12475198 | -0.276 | 0.54 | 0.695 | 0.447 | 0.018 | 0.028 | 0.028 | 0 | 0.88 | 0.68 | 0.84 | 0.84 | 0.85 | 0.9 |
| 2 | 51940718 | rs2354387 | 0.136 | 0.532 | 0.836 | 0.438 | 0.035 | 0.039 | 0.039 | 0 | 0.87 | 0.54 | 0.82 | 0.83 | 0.84 | 0.89 |
| 2 | 51946163 | rs11125373 | 0.64 | 0.539 | 0.749 | 0.404 | 0.014 | -0.015 | -0.015 | 0.025 | 0.83 | 0.42 | 0.79 | 0.85 | 0.81 | 0.87 |
| 2 | 51951101 | rs11693054 | 0.825 | 0.307 | 0.187 | 0.475 | -0.008 | 0.009 | 0.009 | 0 | 0.65 | 0.41 | 0.6 | 0.57 | 0.7 | 0.65 |
| 2 | 51965716 | rs13413603 | 0.877 | 0.347 | -0.001 | 0.429 | 0 | -0.02 | -0.02 | 1.417 | 0.24 | 0.19 | 0.33 | 0.34 | 0.21 | 0.36 |
| 2 | 51967244 | rs17863688 | 1.655 | 0.457 | -0.021 | 0.417 | -0.008 | 0 | 0 | 0 | 0.23 | 0.15 | 0.17 | 0.24 | 0.26 | 0.12 |
| 2 | 51970966 | rs1516191 | 0.281 | 0.306 | 0.328 | 0.426 | 0.007 | -0.005 | -0.005 | 0 | 0.73 | 0.81 | 0.69 | 0.73 | 0.74 | 0.7 |
| 2 | 51975932 | rs10198564 | 1.144 | 0.352 | 0.291 | 0.493 | -0.001 | -0.003 | -0.003 | 0.597 | 0.18 | 0.19 | 0.31 | 0.29 | 0.18 | 0.32 |
| 2 | 51976598 | rs6722793 | 1.451 | 0.349 | 0.277 | 0.486 | -0.009 | -0.001 | -0.001 | 0 | 0.17 | 0.12 | 0.27 | 0.15 | 0.18 | 0.31 |
| 2 | 51979541 | rs952167 | 1.102 | 0.348 | 0.393 | 0.471 | 0.001 | 0.017 | 0.017 | 0 | 0.19 | 0.25 | 0.31 | 0.29 | 0.18 | 0.32 |
| 2 | 51982297 | rs13404286 | 0.036 | 0.307 | 0.458 | 0.319 | 0.005 | -0.004 | -0.004 | 0.249 | 0.68 | 0.62 | 0.57 | 0.72 | 0.64 | 0.63 |
| 2 | 51984344 | rs11562965 | -0.188 | 0.212 | 0.352 | 0.354 | 0.005 | -0.005 | -0.005 | 0 | 0.2 | 0.18 | 0.2 | 0.28 | 0.11 | 0.28 |
| 2 | 51986339 | rs7585968 | -0.989 | 0.218 | 0.476 | 0.324 | 0.023 | -0.001 | -0.001 | 0.743 | 0.43 | 0.73 | 0.56 | 0.56 | 0.32 | 0.63 |
| 2 | 51990959 | rs6709347 | -0.888 | 0.288 | 0.277 | 0.228 | -0.032 | 0.059 | 0.059 | 0 | 0.9 | 0.99 | 0.89 | 1 | 0.84 | 0.97 |
| 2 | 52004166 | rs6708818 | 0.607 | 0.269 | 0.411 | 0.315 | 0.007 | 0 | 0 | 0 | 0.72 | 0.8 | 0.68 | 0.73 | 0.74 | 0.71 |
| 2 | 52010446 | rs13405097 | 1.033 | 0.331 | 0.789 | 0.346 | 0 | 0.006 | 0.006 | 0 | 0.5 | 0.43 | 0.36 | 0.61 | 0.51 | 0.37 |
| 2 | 52012278 | rs1516199 | -0.33 | 0.316 | 0.781 | 0.226 | 0 | -0.015 | -0.015 | 1.816 | 0.91 | 0.95 | 0.95 | 1 | 0.92 | 0.95 |
| 2 | 52017565 | rs6725659 | 0.407 | 0.368 | 0.576 | 0.347 | 0.014 | 0.037 | 0.037 | 0 | 0.23 | 0.53 | 0.38 | 0.27 | 0.23 | 0.37 |
| 2 | 52022010 | rs7564650 | -0.887 | 0.401 | -0.089 | 0.02 | 0.01 | 0.054 | 0.054 | 0 | 0.46 | 0.66 | 0.47 | 0.29 | 0.4 | 0.53 |
| 2 | 52024350 | rs7608332 | 0.361 | 0.347 | 0.091 | 0.023 | -0.001 | 0.004 | 0.004 | 0 | 0.28 | 0.25 | 0.34 | 0.11 | 0.28 | 0.35 |
| 2 | 52025123 | rs11690538 | 1.256 | 0.546 | -0.45 | 0.315 | -0.016 | -0.036 | -0.036 | 0 | 0.41 | 0.23 | 0.36 | 0.47 | 0.44 | 0.33 |
| 2 | 52030134 | rs17863708 | -0.328 | 0.901 | -0.246 | 0.641 | 0.011 | 0.044 | 0.044 | 0 | 0.1 | 0.15 | 0.12 | 0.26 | 0.12 | 0.14 |
| 2 | 52030757 | rs7565870 | 0.237 | 0.805 | -0.811 | -0.149 | -0.038 | 0.082 | 0.082 | 0 | 0.82 | 0.96 | 0.83 | 0.98 | 0.76 | 0.86 |
| 2 | 52033702 | rs1211181 | -0.713 | 0.849 | -0.447 | 0.327 | -0.007 | 0.01 | 0.01 | 0 | 0.78 | 0.82 | 0.74 | 0.9 | 0.78 | 0.68 |
| 2 | 52034035 | rs1543136 | -0.223 | 1.091 | -0.73 | 0.791 | -0.023 | -0.037 | -0.037 | 0 | 0.1 | 0.05 | 0.19 | 0.23 | 0.17 | 0.13 |
| 2 | 52038547 | rs7561923 | -0.785 | 0.587 | -0.695 | -0.203 | -0.001 | -0.041 | -0.041 | 0 | 0.76 | 0.81 | 0.86 | 0.98 | 0.78 | 0.81 |
| 2 | 52038675 | rs11563112 | 0.18 | 0.457 | -0.34 | -0.115 | -0.008 | 0.016 | 0.016 | 0 | 0.63 | 0.3 | 0.58 | 0.74 | 0.6 | 0.56 |
| 2 | 52040842 | rs7595930 | 0.044 | 0.443 | -0.528 | -0.127 | 0.016 | -0.022 | -0.022 | 0 | 0.65 | 0.45 | 0.6 | 0.76 | 0.61 | 0.61 |

| | | | | | | | | | | | | | | | | |
|---|---|---|---|---|---|---|---|---|---|---|---|---|---|---|---|---|
| 2 | 52042513 | rs7603003 | -0.645 | 0.54 | -0.259 | -0.01 | 0.1 | -0.02 | -0.02 | 0 | 0.19 | 0.43 | 0.33 | 0.1 | 0.31 | 0.28 |
| 2 | 52043114 | rs6745103 | -0.557 | 0.492 | -0.172 | 0.095 | 0.08 | 0.115 | 0.115 | 0.158 | 0.16 | 0.45 | 0.29 | 0.25 | 0.2 | 0.28 |
| 2 | 52045220 | rs10184299 | 0.116 | 0.049 | -0.136 | 0.316 | -0.014 | 0.018 | 0.018 | 0.423 | 0.62 | 0.59 | 0.63 | 0.48 | 0.62 | 0.71 |
| 2 | 52045962 | rs13429683 | 0.465 | 0.359 | -0.137 | 0.547 | 0.086 | 0.099 | 0.099 | 0.003 | 0.15 | 0.34 | 0.36 | 0.29 | 0.22 | 0.35 |
| 2 | 52046419 | rs17868260 | 0.841 | 0.255 | -0.418 | 0.583 | -0.035 | -0.029 | -0.029 | 0 | 0.08 | 0.02 | 0.19 | 0.34 | 0.16 | 0.09 |
| 2 | 52047505 | rs2033633 | -1.142 | 0.132 | -0.18 | 0.406 | 0.028 | 0.039 | 0.039 | 1.374 | 0.75 | 0.86 | 0.84 | 0.86 | 0.79 | 0.83 |
| 2 | 52048275 | rs1017419 | -1.167 | 0.104 | -0.245 | 0.409 | 0.035 | 0.044 | 0.044 | 1.496 | 0.75 | 0.91 | 0.85 | 0.86 | 0.8 | 0.84 |
| 2 | 52051069 | rs7590117 | 0.433 | 0.252 | -0.042 | 0.492 | 0.122 | 0.131 | 0.131 | 0 | 0.15 | 0.44 | 0.36 | 0.29 | 0.21 | 0.35 |
| 2 | 52053334 | rs11563133 | 1.003 | -0.133 | -0.335 | 0.411 | -0.009 | -0.104 | -0.104 | 2.199 | 0.23 | 0.05 | 0.24 | 0.37 | 0.22 | 0.17 |
| 2 | 52053430 | rs6545238 | -0.603 | -0.139 | -0.297 | 0.255 | 0.013 | -0.01 | -0.01 | 0.142 | 0.63 | 0.52 | 0.53 | 0.71 | 0.59 | 0.52 |
| 2 | 52055363 | rs10865258 | 0.885 | -0.065 | -0.429 | 0.189 | -0.005 | 0.069 | 0.069 | 0 | 0.36 | 0.18 | 0.23 | 0.18 | 0.36 | 0.29 |
| 2 | 52056672 | rs13387224 | 0.747 | -0.05 | -0.4 | 0.197 | 0.001 | 0.015 | 0.015 | 0 | 0.37 | 0.33 | 0.25 | 0.19 | 0.36 | 0.29 |
| 2 | 52061087 | rs4513332 | 0.432 | 0.174 | -0.389 | 0.354 | 0.001 | 0.004 | 0.004 | 0.037 | 0.51 | 0.46 | 0.43 | 0.53 | 0.47 | 0.47 |
| 2 | 52065162 | rs3861566 | -0.076 | -0.092 | -0.705 | 0.193 | -0.05 | -0.037 | -0.037 | 0.063 | 0.62 | 0.48 | 0.77 | 0.76 | 0.73 | 0.71 |
| 2 | 52065417 | rs3850342 | 0.225 | 0.287 | 0.116 | 0.3 | 0 | -0.001 | -0.001 | 0.569 | 0.57 | 0.4 | 0.48 | 0.45 | 0.6 | 0.53 |
| 2 | 52066351 | rs2033635 | 0.056 | 0.662 | 0.15 | 0.463 | 0.032 | 0.022 | 0.022 | 0 | 0.2 | 0.24 | 0.41 | 0.32 | 0.32 | 0.35 |
| 2 | 52071440 | rs10201204 | 0.411 | 0.681 | -0.015 | 0.508 | 0.117 | 0.174 | 0.174 | 0 | 0.16 | 0.46 | 0.38 | 0.34 | 0.22 | 0.37 |
| 2 | 52072246 | rs12994401 | -0.53 | 0.924 | -0.496 | 0.459 | -0.042 | -0.076 | -0.076 | 3.878 | 0.14 | 0.06 | 0.22 | 0.33 | 0.21 | 0.28 |
| 2 | 52072658 | rs4971764 | -1.111 | 0.708 | -0.209 | 0.259 | -0.018 | -0.028 | -0.028 | 0.247 | 0.76 | 0.65 | 0.87 | 0.91 | 0.81 | 0.89 |
| 2 | 52073146 | rs10207610 | -0.881 | 0.833 | -0.044 | 0.29 | 0.007 | 0 | 0 | 0 | 0.3 | 0.27 | 0.61 | 0.68 | 0.49 | 0.53 |
| 2 | 52073761 | rs10208223 | 0.944 | 0.703 | -0.938 | 0.363 | 0.032 | 0.1 | 0.1 | 0 | 0.46 | 0.18 | 0.26 | 0.25 | 0.35 | 0.37 |
| 2 | 52074452 | rs4971766 | -0.453 | 0.978 | -0.651 | 0.815 | -0.066 | -0.077 | -0.077 | 0 | 0.13 | 0.01 | 0.21 | 0.27 | 0.23 | 0.14 |
| 2 | 52084316 | rs6712068 | -0.196 | 0.512 | -0.183 | 0.531 | 0.002 | 0.006 | 0.006 | 0 | 0.75 | 0.76 | 0.82 | 0.78 | 0.78 | 0.86 |
| 2 | 52084733 | rs17863726 | 0.014 | 1.016 | -0.053 | 0.974 | 0.001 | 0.03 | 0.03 | 0 | 0.07 | 0.08 | 0.12 | 0.25 | 0.11 | 0.12 |
| 2 | 52094467 | rs11562958 | -0.264 | 0.669 | 0.038 | 0.878 | 0.005 | -0.01 | -0.01 | 0 | 0.12 | 0.11 | 0.13 | 0.23 | 0.12 | 0.08 |
| 2 | 52098998 | rs1435802 | 0.231 | 1.109 | -0.149 | 0.604 | -0.04 | -0.027 | -0.027 | 0 | 0.27 | 0.16 | 0.47 | 0.33 | 0.38 | 0.26 |
| 2 | 52100297 | rs12713160 | -0.737 | 1.104 | 0.181 | 0.413 | 0.005 | -0.017 | -0.017 | 0.136 | 0.75 | 0.66 | 0.8 | 0.79 | 0.74 | 0.86 |
| 2 | 52102327 | rs17862811 | 1.744 | 0.833 | 0.347 | 0.426 | 0.033 | 0.048 | 0.048 | 0 | 0.34 | 0.15 | 0.18 | 0.15 | 0.25 | 0.38 |
| 2 | 52102629 | rs6706200 | 1.049 | 1 | 0.372 | 0.256 | 0.003 | 0.002 | 0.002 | 0 | 0.21 | 0.21 | 0.16 | 0.21 | 0.15 | 0.11 |
| 2 | 52114176 | rs984093 | 0.083 | 0.769 | 0.405 | 0.444 | 0.026 | -0.003 | -0.003 | 0 | 0.51 | 0.64 | 0.35 | 0.48 | 0.42 | 0.63 |
| 2 | 52115043 | rs6741100 | 0.358 | 0.709 | 0.548 | 0.448 | 0.038 | -0.005 | -0.005 | 0 | 0.47 | 0.34 | 0.31 | 0.48 | 0.37 | 0.6 |
| 2 | 52126166 | rs17864651 | 1.418 | 0.695 | 0.479 | 0.428 | 0.04 | 0.084 | 0.084 | 0 | 0.33 | 0.1 | 0.18 | 0.12 | 0.23 | 0.37 |
| 2 | 52130099 | rs12994529 | -0.007 | 0.688 | 0.554 | 0.552 | -0.005 | 0.034 | 0.034 | 0.341 | 0.12 | 0.16 | 0.14 | 0.25 | 0.12 | 0.09 |
| 2 | 52137385 | rs6733466 | 0.179 | 0.783 | 0.325 | 0.405 | 0.001 | 0.002 | 0.002 | 0.516 | 0.11 | 0.11 | 0.14 | 0.26 | 0.12 | 0.08 |
| 2 | 52140713 | rs6745758 | 1.353 | 0.77 | 0.229 | 0.07 | 0.036 | 0.09 | 0.09 | 0 | 0.33 | 0.11 | 0.18 | 0.12 | 0.24 | 0.37 |
| 2 | 52145001 | rs6712451 | 0.085 | 0.849 | 0.345 | 0.078 | 0.025 | 0.021 | 0.021 | 0 | 0.48 | 0.62 | 0.35 | 0.36 | 0.4 | 0.46 |
| 2 | 52146364 | rs11125395 | 1.387 | 0.787 | 0.349 | -0.022 | -0.032 | -0.052 | -0.052 | 0.702 | 0.3 | 0.42 | 0.17 | 0.12 | 0.2 | 0.34 |
| 2 | 52153900 | rs737023 | 0.619 | 1.424 | -0.029 | 0.397 | -0.019 | -0.034 | -0.034 | 0.544 | 0.18 | 0.15 | 0.4 | 0.44 | 0.26 | 0.39 |
| 2 | 52156264 | rs1435813 | 1.213 | 1.236 | 0.106 | 0.172 | -0.008 | -0.038 | -0.038 | 0 | 0.35 | 0.51 | 0.21 | 0.12 | 0.28 | 0.29 |
| 2 | 52157626 | rs8179760 | 0.361 | 1.309 | 0.185 | 0.273 | 0.005 | 0.011 | 0.011 | 0 | 0.53 | 0.63 | 0.6 | 0.54 | 0.53 | 0.65 |
| 2 | 52169692 | rs11681667 | 1.744 | 1.039 | -0.324 | 0.102 | 0.019 | 0.062 | 0.062 | 0.176 | 0.11 | 0.01 | 0.04 | 0 | 0.05 | 0.02 |
| 2 | 52171034 | rs1303768 | 0.497 | 1.369 | 0.072 | 0.383 | -0.02 | -0.035 | -0.035 | 2.848 | 0.2 | 0.2 | 0.41 | 0.44 | 0.28 | 0.39 |
| 2 | 52178661 | rs6743164 | 1.609 | 1.29 | 0.104 | 0.153 | 0.027 | 0.059 | 0.059 | 0 | 0.29 | 0.14 | 0.16 | 0.11 | 0.22 | 0.25 |
| 2 | 52182477 | rs1550100 | -0.785 | 1.392 | -0.074 | 0.079 | -0.084 | 0.228 | 0.228 | 0 | 0.69 | 0.98 | 0.81 | 1 | 0.67 | 0.88 |
| 2 | 52198271 | rs13403084 | 0.543 | 1.52 | 0.621 | 0.559 | -0.005 | -0.002 | -0.002 | 1.128 | 0.19 | 0.19 | 0.41 | 0.42 | 0.28 | 0.34 |
| 2 | 52200476 | rs2355173 | 1.66 | 1.189 | 0.283 | 0.383 | 0.03 | 0.078 | 0.078 | 0 | 0.29 | 0.07 | 0.16 | 0.1 | 0.2 | 0.25 |
| 2 | 52202951 | rs2355174 | -0.212 | 1.253 | 0.598 | 0.455 | -0.005 | 0.032 | 0.032 | 0 | 0.66 | 0.76 | 0.76 | 0.76 | 0.68 | 0.76 |
| 2 | 52215641 | rs13022813 | -0.485 | 1.037 | 0.894 | 0.834 | -0.01 | 0.053 | 0.053 | 0 | 0.12 | 0.25 | 0.15 | 0.22 | 0.12 | 0.1 |
| 2 | 52217219 | rs13036114 | -1.483 | 1.065 | 0.923 | 0.802 | 0.008 | 0.086 | 0.086 | 0 | 0.17 | 0.53 | 0.18 | 0.27 | 0.17 | 0.23 |
| 2 | 52219475 | rs6758579 | -0.001 | 0.943 | 0.582 | 0.831 | 0.021 | -0.038 | -0.038 | 0 | 0.1 | 0.04 | 0.11 | 0.2 | 0.1 | 0.1 |
| 2 | 52219924 | rs11677270 | -1.128 | 0.961 | 0.817 | 0.823 | -0.012 | 0.072 | 0.072 | 0 | 0.16 | 0.37 | 0.15 | 0.25 | 0.14 | 0.2 |
| 2 | 52221753 | rs6545247 | -0.52 | 0.917 | 0.668 | 0.85 | -0.023 | 0.082 | 0.082 | 0 | 0.12 | 0.29 | 0.17 | 0.25 | 0.1 | 0.11 |
| 2 | 52222395 | rs2355181 | 0.301 | 1.435 | 0.331 | 0.603 | -0.003 | 0.013 | 0.013 | 0 | 0.45 | 0.38 | 0.41 | 0.35 | 0.44 | 0.4 |
| 2 | 52223396 | rs17042362 | -0.536 | 1.291 | 0.246 | 0.635 | 0 | 0.008 | 0.008 | 0 | 0.11 | 0.13 | 0.19 | 0.25 | 0.18 | 0.14 |
| 2 | 52223651 | rs11892803 | 0.237 | 1.258 | 0.716 | 0.679 | 0.021 | -0.002 | -0.002 | 0 | 0.13 | 0.35 | 0.07 | 0.11 | 0.1 | 0.23 |
| 2 | 52224747 | rs13025813 | -0.437 | 1.26 | 0.349 | 0.694 | -0.014 | -0.059 | -0.059 | 0.81 | 0.11 | 0.07 | 0.18 | 0.23 | 0.18 | 0.14 |
| 2 | 52227074 | rs12468645 | 1.611 | 1.488 | 0.322 | 0.526 | 0.026 | 0.117 | 0.117 | 0 | 0.32 | 0.08 | 0.2 | 0.08 | 0.25 | 0.27 |
| 2 | 52227824 | rs11125398 | 0.953 | 1.832 | 0.288 | 0.547 | 0.005 | 0.005 | 0.005 | 0 | 0.5 | 0.28 | 0.62 | 0.45 | 0.52 | 0.4 |
| 2 | 52229603 | rs13034869 | 0.104 | 1.799 | 0.242 | 0.596 | -0.011 | -0.025 | -0.025 | 0 | 0.11 | 0.08 | 0.18 | 0.23 | 0.16 | 0.13 |
| 2 | 52233463 | rs6736403 | 0.855 | 1.781 | 0.395 | 0.6 | 0.012 | 0.017 | 0.017 | 0 | 0.45 | 0.31 | 0.4 | 0.35 | 0.43 | 0.41 |
| 2 | 52236955 | rs12477454 | -1.045 | 1.794 | 0.733 | 0.867 | 0.003 | 0.017 | 0.017 | 0 | 0.24 | 0.33 | 0.25 | 0.38 | 0.27 | 0.38 |
| 2 | 52243162 | rs6545251 | 0.514 | 1.763 | 0.667 | 0.873 | 0.005 | 0.017 | 0.017 | 0.252 | 0.12 | 0.23 | 0.07 | 0.14 | 0.1 | 0.23 |
| 2 | 52245082 | rs6756931 | -0.261 | 1.976 | 0.719 | 0.759 | -0.006 | -0.031 | -0.031 | 0 | 0.68 | 0.62 | 0.81 | 0.79 | 0.7 | 0.76 |
| 2 | 52245302 | rs6742134 | 0.308 | 1.967 | 0.398 | 0.682 | 0 | -0.01 | -0.01 | 0.003 | 0.13 | 0.23 | 0.06 | 0.09 | 0.1 | 0.24 |
| 2 | 52249401 | rs13389547 | 0.92 | 1.915 | 0.149 | 0.598 | 0.012 | 0.068 | 0.068 | 0.053 | 0.51 | 0.75 | 0.6 | 0.66 | 0.47 | 0.52 |
| 2 | 52255597 | rs4246581 | -0.752 | 1.859 | -0.14 | 0.401 | -0.063 | 0.182 | 0.182 | 0 | 0.74 | 0.98 | 0.85 | 1 | 0.73 | 0.9 |
| 2 | 52260752 | rs10203140 | -0.525 | 2.028 | -0.078 | 0.401 | -0.067 | 0.151 | 0.151 | 0 | 0.72 | 0.94 | 0.82 | 0.97 | 0.68 | 0.89 |
| 2 | 52261308 | rs12473317 | -0.568 | 1.884 | 0.388 | 0.54 | 0.001 | 0.107 | 0.107 | 0 | 0.24 | 0.5 | 0.25 | 0.38 | 0.23 | 0.37 |
| 2 | 52262932 | rs7582793 | -1.327 | 1.874 | -0.038 | 0.305 | -0.03 | 0.127 | 0.127 | 0 | 0.81 | 0.98 | 0.91 | 1 | 0.82 | 0.91 |

| | | | | | | | | | | | | | | | | |
|---|---|---|---|---|---|---|---|---|---|---|---|---|---|---|---|---|
| 2 | 52264332 | rs7595947 | 0.053 | 1.818 | 0.365 | 0.516 | 0 | -0.024 | -0.024 | 0.852 | 0.65 | 0.64 | 0.79 | 0.77 | 0.67 | 0.74 |
| 2 | 52286399 | rs10207605 | 1.046 | 1.804 | 0.4 | 0.709 | -0.01 | -0.019 | -0.019 | 0 | 0.41 | 0.46 | 0.26 | 0.33 | 0.33 | 0.41 |
| 2 | 52287618 | rs11889021 | 0.885 | 1.83 | 0.47 | 0.696 | 0.015 | 0.032 | 0.032 | 0 | 0.42 | 0.71 | 0.28 | 0.34 | 0.34 | 0.41 |
| 2 | 52297892 | rs2355195 | 0.808 | 1.973 | 0.203 | 1.045 | -0.039 | -0.042 | -0.042 | 0.558 | 0.09 | 0.06 | 0.35 | 0.26 | 0.18 | 0.12 |
| 2 | 52298060 | rs10180224 | 0.644 | 1.608 | 0.495 | 0.819 | 0.003 | 0.119 | 0.119 | 0 | 0.05 | 0.17 | 0.03 | 0.2 | 0.03 | 0.14 |
| 2 | 52299636 | rs11899118 | 0.975 | 1.641 | 0.368 | 0.832 | -0.009 | -0.013 | -0.013 | 0 | 0.41 | 0.44 | 0.26 | 0.35 | 0.32 | 0.41 |
| 2 | 52316298 | rs10496005 | 0.455 | 1.676 | -0.183 | 1.15 | -0.025 | -0.027 | -0.027 | 0 | 0.07 | 0.01 | 0.15 | 0.24 | 0.1 | 0.13 |
| 2 | 52318474 | rs7601533 | 0.523 | 1.75 | -0.018 | 0.932 | 0.036 | 0.015 | 0.015 | 0 | 0.51 | 0.7 | 0.42 | 0.54 | 0.43 | 0.54 |
| 2 | 52319233 | rs11886119 | 0.435 | 1.907 | -0.056 | 0.924 | 0.008 | 0.018 | 0.018 | 0 | 0.52 | 0.71 | 0.44 | 0.57 | 0.47 | 0.55 |
| 2 | 52319558 | rs12469644 | -0.867 | 2.015 | -0.098 | 0.731 | -0.007 | 0.099 | 0.099 | 0 | 0.73 | 0.93 | 0.85 | 1 | 0.74 | 0.9 |
| 2 | 52319702 | rs13388814 | 0.109 | 1.532 | -0.021 | 1.155 | 0.025 | 0.007 | 0.007 | 0 | 0.52 | 0.7 | 0.36 | 0.48 | 0.42 | 0.52 |
| 2 | 52324160 | rs13412496 | 1.579 | 1.389 | 0.012 | 0.876 | 0.027 | 0.097 | 0.097 | 0 | 0.31 | 0.13 | 0.18 | 0.11 | 0.25 | 0.26 |
| 2 | 52324258 | rs10181636 | 0.042 | 1.316 | 0.397 | 1.336 | 0 | 0.007 | 0.007 | 0 | 0.53 | 0.61 | 0.45 | 0.56 | 0.51 | 0.54 |
| 2 | 52326591 | rs10199714 | -0.031 | 0.803 | 0.362 | 1.049 | 0.001 | 0.005 | 0.005 | 0 | 0.57 | 0.55 | 0.45 | 0.5 | 0.53 | 0.57 |
| 2 | 52328224 | rs4625950 | 0.121 | 0.885 | 0.181 | 1.087 | 0 | 0 | 0 | 0 | 0.54 | 0.47 | 0.45 | 0.5 | 0.54 | 0.57 |
| 2 | 52331049 | rs13016977 | -1.448 | 0.368 | 0.458 | 1.155 | 0.011 | 0.088 | 0.088 | 0.25 | 0.09 | 0.35 | 0.05 | 0.14 | 0.04 | 0.17 |
| 2 | 52335664 | rs2355803 | -1.035 | 0.163 | -0.049 | 0.964 | 0.001 | -0.001 | -0.001 | 0 | 0.58 | 0.57 | 0.63 | 0.37 | 0.51 | 0.43 |
| 2 | 52338106 | rs11678412 | 0.573 | -0.264 | -0.436 | 0.438 | 0.007 | -0.004 | -0.004 | 0 | 0.2 | 0.17 | 0.1 | 0 | 0.14 | 0.06 |
| 2 | 52338125 | rs2355805 | 0.955 | -0.197 | -0.665 | 0.542 | -0.041 | -0.031 | -0.031 | 0.19 | 0.1 | 0.01 | 0.31 | 0.08 | 0.12 | 0.11 |
| 2 | 52339683 | rs4971790 | -0.594 | -0.319 | -0.587 | 0.477 | -0.004 | -0.023 | -0.023 | 0 | 0.21 | 0.08 | 0.37 | 0.21 | 0.19 | 0.23 |
| 2 | 52342990 | rs12611635 | 0.415 | -0.279 | -0.47 | 0.498 | 0 | -0.008 | -0.008 | 0 | 0.11 | 0.09 | 0.14 | 0.18 | 0.1 | 0.1 |
| 2 | 52343686 | rs7563687 | -0.995 | -0.373 | -0.308 | 0.375 | 0.005 | 0.021 | 0.021 | 0 | 0.39 | 0.31 | 0.3 | 0.21 | 0.33 | 0.21 |
| 2 | 52345248 | rs4971791 | -0.801 | -0.635 | -0.241 | 1.201 | -0.003 | -0.053 | -0.053 | 0 | 0.19 | 0.07 | 0.36 | 0.33 | 0.19 | 0.23 |
| 2 | 52346715 | rs2355807 | -1.707 | -0.737 | 0.086 | 1.091 | 0.019 | 0.057 | 0.057 | 0.559 | 0.48 | 0.38 | 0.55 | 0.73 | 0.4 | 0.56 |
| 2 | 52351522 | rs4356688 | -1.084 | -0.735 | -0.12 | 1.354 | 0.003 | -0.022 | -0.022 | 0 | 0.22 | 0.16 | 0.41 | 0.68 | 0.2 | 0.45 |
| 2 | 52369737 | rs13015643 | -1.085 | -0.92 | -0.011 | 1.651 | 0.002 | 0.005 | 0.005 | 0 | 0.17 | 0.14 | 0.12 | 0.45 | 0.09 | 0.29 |
| 2 | 52371363 | rs7594573 | 0.342 | -1.443 | 0.154 | 1.313 | 0.003 | -0.014 | -0.014 | 0.443 | 0.61 | 0.5 | 0.51 | 0.71 | 0.62 | 0.7 |
| 2 | 52372389 | rs12986646 | 0.128 | -1.331 | -0.35 | 1.357 | 0.041 | -0.01 | -0.01 | 0 | 0.13 | 0.01 | 0.05 | 0.1 | 0.05 | 0.11 |
| 2 | 52382382 | rs10180287 | -1.62 | -1.427 | -0.294 | 1.217 | 0.021 | 0.128 | 0.128 | 1.174 | 0.51 | 0.88 | 0.59 | 0.74 | 0.4 | 0.53 |
| 2 | 52384668 | rs12621097 | 0.425 | -1.147 | -0.109 | 1.206 | -0.04 | 0.01 | 0.01 | 0 | 0.25 | 0.13 | 0.43 | 0.22 | 0.31 | 0.25 |
| 2 | 52395370 | rs7606800 | 0.272 | -1.357 | -0.182 | 1.272 | -0.072 | 0.042 | 0.042 | 0.447 | 0.57 | 0.95 | 0.47 | 0.74 | 0.57 | 0.6 |
| 2 | 52397517 | rs12713179 | -1.046 | -1.031 | 0.911 | 1.851 | 0.013 | 0.027 | 0.027 | 2.775 | 0.51 | 0.45 | 0.59 | 0.73 | 0.41 | 0.54 |
| 2 | 52407060 | rs2103389 | 1.361 | -1.09 | 0.713 | 1.828 | -0.005 | 0.143 | 0.143 | 1.594 | 0.43 | 0.13 | 0.37 | 0.2 | 0.57 | 0.41 |
| 2 | 52430934 | rs2141847 | -1.143 | -0.962 | 0.755 | 2.131 | 0.072 | -0.003 | -0.003 | 0.506 | 0.5 | 0.86 | 0.55 | 0.48 | 0.38 | 0.49 |
| 2 | 52434608 | rs6731958 | 0.951 | -0.994 | 1.146 | 2.216 | -0.02 | 0.056 | 0.056 | 0 | 0.05 | 0.11 | 0.02 | 0.14 | 0.02 | 0.11 |
| 2 | 52437929 | rs3950253 | -0.283 | -1.04 | 1.296 | 2.224 | 0.014 | 0.011 | 0.011 | 2.884 | 0.42 | 0.23 | 0.52 | 0.33 | 0.38 | 0.4 |
| 2 | 52438863 | rs1882475 | -1.471 | -1.002 | 1.171 | 2.197 | 0.004 | 0.161 | 0.161 | 2.552 | 0.55 | 0.87 | 0.63 | 0.8 | 0.43 | 0.59 |
| 2 | 52448661 | rs4971796 | -0.362 | -1.061 | 1.65 | 2.138 | -0.006 | -0.003 | -0.003 | 2.273 | 0.4 | 0.61 | 0.49 | 0.31 | 0.32 | 0.36 |
| 2 | 52457693 | rs17042635 | 0.893 | -1.031 | 1.706 | 2.176 | 0.015 | 0.016 | 0.016 | 1.15 | 0.51 | 0.37 | 0.41 | 0.38 | 0.59 | 0.55 |
| 2 | 52458725 | rs17042637 | 0.109 | -1.018 | 1.714 | 2.157 | -0.023 | 0.123 | 0.123 | 0 | 0.07 | 0.19 | 0.02 | 0.18 | 0.01 | 0.13 |
| 2 | 52461739 | rs6726230 | -0.689 | -1.015 | 1.7 | 2.13 | -0.004 | -0.013 | -0.013 | 0 | 0.45 | 0.49 | 0.53 | 0.19 | 0.39 | 0.37 |
| 2 | 52461984 | rs12995337 | -0.333 | -1.034 | 1.358 | 1.978 | 0.009 | 0.123 | 0.123 | 3.612 | 0.41 | 0.2 | 0.5 | 0.13 | 0.37 | 0.36 |
| 2 | 52465031 | rs2141843 | -1.382 | -0.994 | 1.251 | 1.912 | 0.041 | 0.049 | 0.049 | 3.758 | 0.53 | 0.84 | 0.59 | 0.63 | 0.39 | 0.55 |
| 2 | 52466342 | rs11125414 | -1.376 | -0.993 | 1.237 | 1.887 | 0.043 | 0.105 | 0.105 | 3.394 | 0.53 | 0.79 | 0.59 | 0.74 | 0.4 | 0.55 |
| 2 | 52469618 | rs7586327 | -0.675 | -0.965 | 1.429 | 1.86 | -0.003 | 0.001 | 0.001 | 1.279 | 0.44 | 0.49 | 0.56 | 0.51 | 0.38 | 0.45 |
| 2 | 52473759 | rs12615485 | 0.478 | -0.912 | 0.875 | 1.449 | 0.037 | 0.097 | 0.097 | 1.718 | 0.55 | 0.24 | 0.47 | 0.81 | 0.64 | 0.63 |
| 2 | 52476027 | rs12622126 | 0.31 | -0.37 | 0.333 | 1.131 | -0.028 | 0.06 | 0.06 | 2.204 | 0.55 | 0.77 | 0.44 | 0.72 | 0.55 | 0.58 |
| 2 | 52479251 | rs12472885 | -1.142 | 0.02 | 0.483 | 1.465 | -0.027 | 0.432 | 0.432 | 0 | 0.1 | 0.38 | 0.08 | 0.52 | 0.03 | 0.19 |
| 2 | 52493926 | rs2141844 | -0.61 | 0.632 | -0.717 | 0.127 | -0.004 | 0.053 | 0.053 | 0 | 0.52 | 0.64 | 0.51 | 0.28 | 0.52 | 0.43 |
| 2 | 52505765 | rs13406269 | -0.493 | 0.628 | -0.929 | -0.45 | -0.004 | -0.027 | -0.027 | 0 | 0.3 | 0.58 | 0.34 | 0.2 | 0.26 | 0.19 |
| 2 | 52506774 | rs1997492 | 1.403 | 0.336 | -1.022 | -0.684 | -0.002 | 0.004 | 0.004 | 0.438 | 0.42 | 0.38 | 0.49 | 0.72 | 0.45 | 0.54 |
| 2 | 52507412 | rs17042675 | 0.202 | 0.234 | -0.496 | -0.61 | 0.018 | 0.081 | 0.081 | 0 | 0.05 | 0.19 | 0.11 | 0.13 | 0.03 | 0.06 |
| 2 | 52509039 | rs4971800 | 0.621 | 0.216 | -0.721 | -0.757 | -0.037 | 0.017 | 0.017 | 0.003 | 0.15 | 0.01 | 0.11 | 0.06 | 0.19 | 0.2 |
| 2 | 52511386 | rs13389926 | 0.176 | 0.178 | -0.553 | -0.78 | -0.003 | -0.003 | -0.003 | 0 | 0.12 | 0.11 | 0.15 | 0.01 | 0.13 | 0.06 |
| 2 | 52511933 | rs2464650 | -1.684 | 0.584 | -0.551 | -0.737 | -0.024 | 0.004 | 0.004 | 0 | 0.92 | 1 | 0.87 | 0.96 | 0.93 | 0.93 |
| 2 | 52513418 | rs6723027 | -1.593 | 0.447 | -0.561 | -0.785 | -0.013 | -0.026 | -0.026 | 0 | 0.29 | 0.62 | 0.32 | 0.22 | 0.23 | 0.18 |
| 2 | 52514493 | rs10199014 | -1.091 | 0.5 | -0.57 | -0.771 | -0.006 | -0.013 | -0.013 | 0 | 0.21 | 0.58 | 0.28 | 0.2 | 0.17 | 0.16 |
| 2 | 52520921 | rs7607212 | -1.003 | 0.508 | -0.436 | -0.766 | -0.017 | -0.015 | -0.015 | 0 | 0.21 | 0.54 | 0.27 | 0.2 | 0.16 | 0.13 |
| 2 | 52521126 | rs1922210 | 2.558 | 0.534 | -0.972 | -0.377 | -0.08 | -0.092 | -0.092 | 0.025 | 0.3 | 0.02 | 0.3 | 0.34 | 0.32 | 0.3 |
| 2 | 52523725 | rs10205333 | -0.497 | 0.504 | -0.584 | -0.341 | 0.008 | -0.011 | -0.011 | 0 | 0.17 | 0.36 | 0.25 | 0.13 | 0.15 | 0.12 |
| 2 | 52529983 | rs1424788 | -1.483 | 0.471 | -0.743 | -0.302 | 0.005 | 0.025 | 0.025 | 0 | 0.42 | 0.65 | 0.43 | 0.27 | 0.42 | 0.35 |
| 2 | 52534578 | rs6715421 | -1.369 | 0.449 | -0.483 | -0.17 | -0.002 | -0.015 | -0.015 | 0 | 0.22 | 0.62 | 0.29 | 0.2 | 0.18 | 0.16 |
| 2 | 52535407 | rs12475021 | 0.071 | 0.292 | -0.406 | 0.063 | 0.009 | 0.076 | 0.076 | 0 | 0.05 | 0.19 | 0.1 | 0.13 | 0.02 | 0.06 |
| 2 | 52540372 | rs3109947 | 0.994 | 0.427 | -0.65 | 0.064 | -0.028 | -0.056 | -0.056 | 0 | 0.14 | 0.04 | 0.1 | 0.33 | 0.19 | 0.26 |
| 2 | 52540726 | rs6739969 | 1.272 | 0.334 | -0.482 | -0.131 | 0.011 | 0.024 | 0.024 | 0 | 0.46 | 0.35 | 0.46 | 0.37 | 0.4 | 0.42 |
| 2 | 52547000 | rs2727892 | -0.637 | 0.488 | -0.232 | -0.13 | 0.02 | 0.032 | 0.032 | 0 | 0.32 | 0.39 | 0.37 | 0.53 | 0.36 | 0.39 |
| 2 | 52549472 | rs12464727 | 0.384 | 0.436 | -0.322 | -0.132 | -0.002 | 0.001 | 0.001 | 0 | 0.07 | 0.08 | 0.1 | 0.11 | 0.05 | 0.07 |
| 2 | 52549542 | rs1424784 | 0.8 | 0.401 | -0.427 | -0.165 | -0.033 | -0.081 | -0.081 | 0.46 | 0.19 | 0.04 | 0.12 | 0.33 | 0.22 | 0.28 |
| 2 | 52550366 | rs2552317 | -1.851 | 0.328 | 0.075 | -0.195 | -0.001 | 0.018 | 0.018 | 0 | 0.59 | 0.78 | 0.56 | 0.64 | 0.63 | 0.59 |

| | | | | | | | | | | | | | | | | |
|---|---|---|---|---|---|---|---|---|---|---|---|---|---|---|---|---|
| 2 | 52552297 | rs2244201 | 0.746 | 0.421 | 0.15 | -0.105 | -0.003 | -0.005 | -0.005 | 4.666 | 0.19 | 0.12 | 0.12 | 0.33 | 0.21 | 0.27 |
| 2 | 52562876 | rs17730749 | 0.309 | 0.429 | 0.37 | -0.202 | 0.004 | 0.015 | 0.015 | 0 | 0.22 | 0.19 | 0.17 | 0.12 | 0.25 | 0.2 |
| 2 | 52563643 | rs2552323 | 1.465 | 0.344 | 0.439 | -0.295 | 0.004 | 0.051 | 0.051 | 0 | 0.54 | 0.27 | 0.45 | 0.71 | 0.56 | 0.63 |
| 2 | 52564610 | rs3111858 | 0.595 | 0.337 | 0.517 | -0.3 | -0.016 | -0.049 | -0.049 | 0 | 0.66 | 0.47 | 0.67 | 0.83 | 0.68 | 0.7 |
| 2 | 52566298 | rs2969312 | -0.932 | 0.356 | 0.464 | -0.298 | -0.009 | -0.031 | -0.031 | 0.61 | 0.88 | 0.74 | 0.85 | 0.95 | 0.93 | 0.91 |
| 2 | 52570207 | rs1946312 | 1.288 | 0.362 | 0.233 | -0.296 | 0.006 | 0.01 | 0.01 | 0 | 0.5 | 0.31 | 0.52 | 0.53 | 0.51 | 0.53 |
| 2 | 52570526 | rs13018183 | 0.686 | 0.424 | 0.331 | -0.294 | -0.007 | 0.001 | 0.001 | 0 | 0.1 | 0.08 | 0.15 | 0 | 0.13 | 0.04 |
| 2 | 52572167 | rs2081375 | -0.641 | 0.348 | 0.303 | -0.321 | 0.003 | -0.004 | -0.004 | 0 | 0.3 | 0.36 | 0.38 | 0.23 | 0.35 | 0.28 |
| 2 | 52578154 | rs11904782 | -0.349 | 0.292 | 0.505 | -0.297 | -0.021 | -0.003 | -0.003 | 0 | 0.24 | 0.47 | 0.26 | 0.23 | 0.24 | 0.24 |
| 2 | 52582658 | rs3109948 | 1.212 | 0.411 | 0.636 | -0.227 | -0.016 | 0.105 | 0.105 | 0 | 0.53 | 0.14 | 0.45 | 0.71 | 0.55 | 0.62 |
| 2 | 52586559 | rs7590476 | 0.292 | 0.409 | 0.859 | -0.21 | -0.008 | -0.003 | -0.003 | 1.619 | 0.25 | 0.4 | 0.28 | 0.23 | 0.28 | 0.25 |
| 2 | 52588775 | rs13012563 | 2.023 | 0.426 | 0.786 | -0.17 | 0.003 | 0.001 | 0.001 | 0 | 0.32 | 0.2 | 0.32 | 0.31 | 0.3 | 0.32 |
| 2 | 52594480 | rs11890083 | -0.232 | 0.448 | 0.927 | -0.116 | -0.018 | -0.017 | -0.017 | 0 | 0.3 | 0.59 | 0.31 | 0.24 | 0.3 | 0.27 |
| 2 | 52595591 | rs1424792 | 0.604 | 0.583 | 0.856 | -0.028 | -0.042 | -0.096 | -0.096 | 0.036 | 0.21 | 0.04 | 0.14 | 0.4 | 0.26 | 0.31 |
| 2 | 52596531 | rs17042835 | 0.407 | 0.518 | 0.942 | -0.048 | -0.002 | 0 | 0 | 0.209 | 0.24 | 0.33 | 0.27 | 0.24 | 0.28 | 0.23 |
| 2 | 52599360 | rs17788159 | 0.881 | 0.549 | 0.828 | -0.094 | 0.003 | 0.012 | 0.012 | 0 | 0.21 | 0.17 | 0.16 | 0.02 | 0.25 | 0.17 |
| 2 | 52608484 | rs2727868 | -1.516 | 0.62 | 0.691 | -0.061 | 0.003 | 0.015 | 0.015 | 0 | 0.58 | 0.82 | 0.57 | 0.64 | 0.64 | 0.59 |
| 2 | 52616656 | rs10169856 | -0.009 | 0.584 | 0.792 | -0.03 | -0.02 | -0.015 | -0.015 | 0 | 0.27 | 0.47 | 0.29 | 0.24 | 0.28 | 0.26 |
| 2 | 52623156 | rs1345965 | -0.712 | 0.595 | 0.694 | -0.025 | 0 | 0.011 | 0.011 | 0 | 0.35 | 0.64 | 0.41 | 0.24 | 0.38 | 0.32 |
| 2 | 52624780 | rs4971815 | -0.336 | 0.608 | 0.75 | -0.025 | 0.011 | -0.02 | -0.02 | 0 | 0.31 | 0.55 | 0.4 | 0.24 | 0.36 | 0.29 |
| 2 | 52628235 | rs2552346 | -1.523 | 0.551 | 0.638 | -0.34 | 0.009 | 0.009 | 0.009 | 0 | 0.6 | 0.76 | 0.57 | 0.64 | 0.65 | 0.6 |
| 2 | 52633563 | rs2727865 | -0.838 | 0.66 | 0.627 | -0.25 | 0 | 0 | 0 | 0 | 0.48 | 0.52 | 0.51 | 0.57 | 0.58 | 0.51 |
| 2 | 52635129 | rs2694135 | 0.71 | 0.655 | 0.604 | -0.214 | 0.009 | 0.008 | 0.008 | 0 | 0.49 | 0.43 | 0.48 | 0.42 | 0.4 | 0.44 |
| 2 | 52643615 | rs11673995 | 1.138 | 0.631 | 0.624 | -0.336 | -0.017 | -0.008 | -0.008 | 0 | 0.42 | 0.49 | 0.37 | 0.4 | 0.33 | 0.41 |
| 2 | 52645704 | rs7586951 | 0.441 | 0.752 | 0.661 | -0.304 | -0.002 | 0 | 0 | 0 | 0.52 | 0.68 | 0.59 | 0.52 | 0.46 | 0.51 |
| 2 | 52646336 | rs13399394 | 1.462 | 0.751 | 0.573 | -0.22 | 0.001 | 0.001 | 0.001 | 0 | 0.36 | 0.34 | 0.33 | 0.33 | 0.3 | 0.35 |
| 2 | 52653026 | rs2552347 | 0.603 | 1.291 | 0.632 | 0.009 | 0.005 | -0.004 | -0.004 | 0 | 0.1 | 0.11 | 0.23 | 0.02 | 0.14 | 0.07 |
| 2 | 52653886 | rs1364587 | -0.791 | 1.073 | 0.574 | 0.285 | -0.042 | -0.096 | -0.096 | 0 | 0.83 | 0.97 | 0.86 | 0.58 | 0.79 | 0.74 |
| 2 | 52654662 | rs2694138 | 0.619 | 0.993 | 0.765 | 0.151 | -0.001 | 0.001 | 0.001 | 0.227 | 0.55 | 0.58 | 0.6 | 0.44 | 0.51 | 0.5 |
| 2 | 52655861 | rs12619211 | 0.001 | 0.776 | 0.831 | 0.137 | -0.01 | 0.04 | 0.04 | 0 | 0.24 | 0.26 | 0.16 | 0.51 | 0.2 | 0.32 |
| 2 | 52659874 | rs6730865 | -1.132 | 0.772 | 1.09 | 0.008 | 0.013 | 0.013 | 0.013 | 0 | 0.48 | 0.41 | 0.42 | 0.57 | 0.46 | 0.52 |
| 2 | 52668651 | rs4971598 | 0.528 | 0.805 | 1.013 | 0.049 | 0 | -0.058 | -0.058 | 2.685 | 0.21 | 0.09 | 0.16 | 0.44 | 0.21 | 0.28 |
| 2 | 52690141 | rs10192786 | -1.907 | 1.062 | 1.054 | -0.097 | -0.004 | 0.027 | 0.027 | 0 | 0.61 | 0.8 | 0.67 | 0.69 | 0.62 | 0.62 |
| 2 | 52690413 | rs10193067 | 1.38 | 1.078 | 1.158 | -0.08 | 0 | 0.017 | 0.017 | 0 | 0.07 | 0.18 | 0.13 | 0.09 | 0.05 | 0.04 |
| 2 | 52692596 | rs1424812 | -0.202 | 1.18 | 0.963 | -0.24 | 0.018 | -0.043 | -0.043 | 0.083 | 0.27 | 0.4 | 0.35 | 0.13 | 0.33 | 0.22 |
| 2 | 52694638 | rs12611666 | -1.223 | 1.374 | 0.765 | -0.108 | 0.01 | -0.015 | -0.015 | 0 | 0.59 | 0.52 | 0.56 | 0.69 | 0.54 | 0.57 |
| 2 | 52696419 | rs12052444 | -1.318 | 1.042 | 0.075 | -0.086 | 0 | 0.021 | 0.021 | 0 | 0.6 | 0.75 | 0.66 | 0.69 | 0.62 | 0.59 |
| 2 | 52698940 | rs7588277 | -1.099 | 1.215 | 0.136 | -0.025 | -0.005 | 0.035 | 0.035 | 0 | 0.56 | 0.73 | 0.62 | 0.67 | 0.53 | 0.58 |
| 2 | 52699308 | rs12477282 | -0.166 | 1.081 | 0.367 | -0.134 | -0.001 | -0.008 | -0.008 | 0.418 | 0.24 | 0.3 | 0.25 | 0.11 | 0.24 | 0.18 |
| 2 | 52699456 | rs12477329 | -0.156 | 1.211 | 0.413 | -0.109 | -0.003 | -0.013 | -0.013 | 0 | 0.22 | 0.3 | 0.22 | 0.11 | 0.23 | 0.19 |
| 2 | 52700283 | rs12105524 | 0.309 | 1.212 | 0.348 | -0.146 | -0.002 | -0.031 | -0.031 | 0 | 0.16 | 0.23 | 0.11 | 0.01 | 0.19 | 0.16 |
| 2 | 52703416 | rs1997160 | 0.462 | 1.163 | 0.088 | 0.011 | -0.011 | -0.103 | -0.103 | 0.025 | 0.2 | 0.04 | 0.15 | 0.51 | 0.19 | 0.29 |
| 2 | 52703590 | rs12619788 | -0.599 | 1.01 | 0.812 | -0.067 | 0.005 | 0.001 | 0.001 | 0 | 0.36 | 0.36 | 0.39 | 0.53 | 0.49 | 0.49 |
| 2 | 52704475 | rs2030871 | -0.007 | 1.062 | 0.794 | -0.028 | -0.005 | -0.016 | -0.016 | 0 | 0.27 | 0.17 | 0.22 | 0.52 | 0.3 | 0.34 |
| 2 | 52704836 | rs12469086 | -0.67 | 0.968 | 1.162 | -0.124 | 0.012 | -0.063 | -0.063 | 0.397 | 0.27 | 0.58 | 0.4 | 0.14 | 0.26 | 0.21 |
| 2 | 52705571 | rs13005050 | 0.485 | 1.264 | 1.042 | -0.158 | -0.001 | 0.003 | 0.003 | 0 | 0.11 | 0.16 | 0.22 | 0.15 | 0.11 | 0.09 |
| 2 | 52705959 | rs6545277 | -0.047 | 1.29 | 0.999 | -0.149 | -0.047 | -0.064 | -0.064 | 0.524 | 0.27 | 0.09 | 0.24 | 0.53 | 0.37 | 0.35 |
| 2 | 52706489 | rs13407104 | 0.393 | 0.883 | 0.903 | -0.143 | -0.044 | 0.063 | 0.063 | 0 | 0.16 | 0.05 | 0.18 | 0 | 0.23 | 0.14 |
| 2 | 52707774 | rs1526683 | -0.341 | 0.853 | 1.056 | -0.103 | 0 | 0.004 | 0.004 | 0 | 0.29 | 0.66 | 0.27 | 0.52 | 0.3 | 0.35 |
| 2 | 52709799 | rs17789255 | -0.462 | 0.865 | 0.917 | -0.134 | -0.006 | 0.015 | 0.015 | 0 | 0.25 | 0.22 | 0.37 | 0.14 | 0.26 | 0.2 |
| 2 | 52709926 | rs17789333 | -1.638 | 0.857 | 1.164 | -0.141 | 0.011 | 0.038 | 0.038 | 0 | 0.53 | 0.66 | 0.62 | 0.66 | 0.56 | 0.55 |
| 2 | 52709965 | rs17043016 | -0.133 | 0.895 | 1.125 | -0.102 | 0.008 | 0.042 | 0.042 | 0 | 0.28 | 0.45 | 0.25 | 0.52 | 0.3 | 0.35 |
| 2 | 52716027 | rs10084177 | 0.077 | 0.951 | 0.763 | -0.157 | -0.025 | 0.042 | 0.042 | 0 | 0.17 | 0.08 | 0.19 | 0.01 | 0.23 | 0.16 |
| 2 | 52721889 | rs6760658 | -1.578 | 0.85 | 0.932 | -0.154 | 0.003 | 0.011 | 0.011 | 0 | 0.54 | 0.63 | 0.62 | 0.67 | 0.55 | 0.58 |
| 2 | 52722668 | rs4971604 | -0.081 | 0.903 | 0.948 | -0.209 | 0.03 | -0.097 | -0.097 | 0.57 | 0.19 | 0.41 | 0.21 | 0.01 | 0.23 | 0.18 |
| 2 | 52730021 | rs11891667 | -0.809 | 0.779 | 0.771 | 0.231 | -0.002 | -0.002 | -0.002 | 0 | 0.36 | 0.34 | 0.43 | 0.63 | 0.34 | 0.46 |
| 2 | 52735067 | rs11896983 | -0.065 | 0.886 | 0.061 | 0.312 | -0.037 | -0.074 | -0.074 | 0 | 0.25 | 0.07 | 0.22 | 0.54 | 0.3 | 0.36 |
| 2 | 52740395 | rs6737033 | -0.19 | 0.524 | 1.169 | -0.378 | -0.001 | -0.035 | -0.035 | 4.168 | 0.22 | 0.22 | 0.2 | 0 | 0.19 | 0.13 |
| 2 | 52745557 | rs11125431 | 0.779 | 0.452 | 1.185 | -0.364 | -0.028 | 0.024 | 0.024 | 0 | 0.1 | 0.29 | 0.22 | 0.15 | 0.04 | 0.1 |
| 2 | 52786331 | rs9808457 | -0.031 | 0.439 | 0.974 | -0.494 | -0.024 | 0.081 | 0.081 | 0 | 0.2 | 0.06 | 0.13 | 0 | 0.2 | 0.14 |
| 2 | 52786342 | rs1526677 | -1.812 | 0.464 | 0.831 | -0.257 | -0.013 | 0.03 | 0.03 | 0 | 0.6 | 0.78 | 0.62 | 0.68 | 0.57 | 0.67 |
| 2 | 52790493 | rs1526676 | -1.757 | 0.459 | 0.855 | -0.248 | -0.014 | 0.034 | 0.034 | 0 | 0.59 | 0.78 | 0.61 | 0.68 | 0.55 | 0.65 |
| 2 | 52794107 | rs10865263 | -1.765 | 0.454 | 1.119 | -0.282 | -0.001 | 0.006 | 0.006 | 0 | 0.6 | 0.66 | 0.61 | 0.72 | 0.58 | 0.64 |
| 2 | 52795849 | rs12993516 | -1.796 | 0.446 | 0.978 | -0.279 | -0.02 | 0.032 | 0.032 | 0 | 0.61 | 0.82 | 0.61 | 0.71 | 0.58 | 0.68 |
| 2 | 52797840 | rs10174561 | 0.946 | 0.374 | 1.151 | -0.246 | -0.022 | 0.005 | 0.005 | 0 | 0.1 | 0.17 | 0.2 | 0.15 | 0.04 | 0.08 |
| 2 | 52801074 | rs6720390 | -0.886 | 0.445 | 1.091 | -0.2 | -0.007 | -0.004 | -0.004 | 0 | 0.47 | 0.15 | 0.4 | 0.57 | 0.54 | 0.56 |
| 2 | 52808175 | rs1609012 | 1.037 | 0.455 | 1.078 | -0.299 | -0.015 | 0.003 | 0.003 | 0 | 0.1 | 0.14 | 0.2 | 0.15 | 0.04 | 0.08 |
| 2 | 52810903 | rs7371202 | 0.273 | 0.491 | 1.286 | -0.259 | -0.001 | -0.027 | -0.027 | 0 | 0.21 | 0.2 | 0.21 | 0.05 | 0.21 | 0.22 |
| 2 | 52812248 | rs4625951 | -2.001 | 0.387 | 1.223 | -0.01 | -0.022 | -0.004 | -0.004 | 0 | 0.62 | 0.82 | 0.68 | 0.7 | 0.59 | 0.68 |

| | | | | | | | | | | | | | | | | |
|---|---|---|---|---|---|---|---|---|---|---|---|---|---|---|---|---|
| 2 | 52812988 | rs1384925 | -0.76 | 0.367 | 1.3 | -0.025 | -0.021 | -0.059 | -0.059 | 0.015 | 0.32 | 0.41 | 0.42 | 0.19 | 0.25 | 0.3 |
| 2 | 52814504 | rs2170794 | -0.791 | 0.368 | 1.3 | -0.012 | -0.021 | -0.047 | -0.047 | 0.027 | 0.32 | 0.41 | 0.42 | 0.21 | 0.25 | 0.3 |
| 2 | 52820363 | rs11895971 | -0.135 | 0.562 | 1.176 | 0.049 | 0.013 | 0.045 | 0.045 | 0 | 0.3 | 0.44 | 0.27 | 0.49 | 0.34 | 0.39 |
| 2 | 52820859 | rs11681573 | -0.087 | 0.614 | 0.666 | 0.035 | -0.067 | -0.106 | -0.106 | 9.19 | 0.29 | 0.07 | 0.21 | 0.49 | 0.36 | 0.39 |
| 2 | 52822501 | rs952263 | 0.335 | 0.561 | 0.747 | -0.045 | -0.005 | 0.029 | 0.029 | 0 | 0.21 | 0.14 | 0.21 | 0.05 | 0.21 | 0.22 |
| 2 | 52825943 | rs12464582 | -0.709 | 0.4 | 0.847 | -0.096 | -0.01 | -0.027 | -0.027 | 0 | 0.31 | 0.35 | 0.42 | 0.2 | 0.25 | 0.26 |
| 2 | 52826175 | rs17734324 | 0.417 | 0.463 | 0.577 | -0.1 | -0.021 | 0.059 | 0.059 | 0.002 | 0.19 | 0.09 | 0.22 | 0.05 | 0.23 | 0.17 |
| 2 | 52826401 | rs17043120 | -0.181 | 0.555 | 0.768 | 0.029 | -0.005 | -0.013 | -0.013 | 0 | 0.29 | 0.29 | 0.24 | 0.49 | 0.34 | 0.39 |
| 2 | 52843343 | rs7569652 | 0.09 | 0.531 | 0.571 | 0.148 | 0.005 | 0.096 | 0.096 | 0.129 | 0.19 | 0.34 | 0.18 | 0.44 | 0.2 | 0.29 |
| 2 | 52850368 | rs4672135 | -0.851 | 0.442 | 0.247 | -0.074 | 0 | -0.003 | -0.003 | 0 | 0.4 | 0.65 | 0.53 | 0.26 | 0.37 | 0.36 |
| 2 | 52853564 | rs6545295 | -1.566 | 0.522 | -0.169 | -0.374 | 0.001 | -0.009 | -0.009 | 6.298 | 0.71 | 0.66 | 0.83 | 0.81 | 0.74 | 0.83 |
| 2 | 52853806 | rs1483865 | 0.633 | 0.54 | -0.733 | -0.502 | -0.024 | -0.13 | -0.13 | 0 | 0.23 | 0.06 | 0.25 | 0.48 | 0.25 | 0.35 |
| 2 | 52855689 | rs900660 | 0.301 | -0.101 | -0.497 | -0.816 | -0.023 | -0.012 | -0.012 | 0 | 0.17 | 0.25 | 0.24 | 0.14 | 0.05 | 0.27 |
| 2 | 52856914 | rs871696 | 0.781 | -0.2 | -0.184 | -0.826 | 0.001 | -0.004 | -0.004 | 0 | 0.23 | 0.2 | 0.25 | 0.47 | 0.23 | 0.34 |
| 2 | 52859829 | rs4672216 | -0.992 | -0.401 | -0.634 | -1.163 | -0.016 | 0.091 | 0.091 | 0.196 | 0.45 | 0.29 | 0.55 | 0.31 | 0.5 | 0.46 |
| 2 | 52859840 | rs900663 | -2.53 | -0.55 | -0.518 | -1.291 | 0 | 0.004 | 0.004 | 0.12 | 0.72 | 0.82 | 0.84 | 0.79 | 0.74 | 0.84 |
| 2 | 52868656 | rs350747 | -0.728 | 0.134 | -0.03 | -1.27 | 0.005 | -0.034 | -0.034 | 1.185 | 0.78 | 0.8 | 0.71 | 0.95 | 0.68 | 0.86 |
| 2 | 52871941 | rs350752 | -1.046 | 0.267 | -0.19 | -1.307 | -0.034 | 0.028 | 0.028 | 0 | 0.91 | 0.97 | 0.9 | 0.94 | 0.86 | 0.92 |
| 2 | 52873788 | rs350753 | 0.993 | -0.092 | -0.12 | -1.336 | 0.004 | 0.061 | 0.061 | 2.117 | 0.6 | 0.29 | 0.55 | 0.79 | 0.62 | 0.58 |
| 2 | 52883672 | rs10178513 | -0.595 | -0.089 | 0.187 | -1.098 | -0.079 | -0.012 | -0.012 | 0 | 0.15 | 0.49 | 0.12 | 0.14 | 0.04 | 0.25 |
| 2 | 52885422 | rs10198004 | -0.608 | -0.086 | 0.378 | -1.097 | -0.076 | -0.014 | -0.014 | 0 | 0.15 | 0.41 | 0.12 | 0.14 | 0.04 | 0.27 |
| 2 | 52886040 | rs13418868 | -0.591 | 0.071 | 0.055 | -1.159 | -0.007 | -0.011 | -0.011 | 1.283 | 0.4 | 0.58 | 0.4 | 0.69 | 0.41 | 0.63 |
| 2 | 52887354 | rs6545298 | -0.754 | 0.04 | -0.041 | -1.179 | 0.016 | 0.025 | 0.025 | 0 | 0.42 | 0.71 | 0.41 | 0.69 | 0.41 | 0.67 |
| 2 | 52891171 | rs7578734 | -0.75 | 0.433 | -0.004 | -1.255 | 0 | -0.016 | -0.016 | 0 | 0.79 | 0.82 | 0.69 | 0.94 | 0.72 | 0.86 |
| 2 | 52893165 | rs1377953 | -0.549 | 0.669 | 0.235 | -1.215 | 0.018 | -0.052 | -0.052 | 0 | 0.78 | 0.76 | 0.69 | 0.94 | 0.72 | 0.86 |
| 2 | 52896902 | rs17043185 | 0.916 | 0.466 | 0.513 | -1.171 | -0.04 | 0.004 | 0.004 | 1.655 | 0.1 | 0.33 | 0.1 | 0.12 | 0.03 | 0.2 |
| 2 | 52897938 | rs10173792 | 0.614 | 0.584 | 0.181 | -1.218 | 0.011 | -0.082 | -0.082 | 0 | 0.66 | 0.5 | 0.59 | 0.82 | 0.68 | 0.64 |
| 2 | 52899006 | rs1563200 | 0.82 | 0.642 | -0.048 | -0.965 | -0.013 | -0.028 | -0.028 | 1.224 | 0.26 | 0.19 | 0.39 | 0.57 | 0.37 | 0.39 |
| 2 | 52904760 | rs1157620 | -0.313 | 0.622 | 0.215 | -1.152 | 0.007 | -0.015 | -0.015 | 0.519 | 0.75 | 0.71 | 0.61 | 0.83 | 0.7 | 0.82 |
| 2 | 52905592 | rs4672476 | 0.723 | 0.678 | -0.013 | -1.107 | -0.016 | -0.044 | -0.044 | 0 | 0.27 | 0.16 | 0.38 | 0.56 | 0.36 | 0.5 |
| 2 | 52908627 | rs350792 | -1.35 | 0.859 | -0.475 | -1.304 | -0.004 | 0.07 | 0.07 | 0.369 | 0.53 | 0.78 | 0.69 | 0.69 | 0.55 | 0.72 |
| 2 | 52909087 | rs350793 | 1.742 | 1.037 | -0.848 | -1.169 | -0.053 | 0.086 | 0.086 | 2.129 | 0.14 | 0.01 | 0.18 | 0 | 0.16 | 0.07 |
| 2 | 52910687 | rs350794 | -1.073 | 0.949 | -0.638 | -1.001 | 0.023 | -0.008 | -0.008 | 0.995 | 0.93 | 0.92 | 0.89 | 0.95 | 0.86 | 0.94 |
| 2 | 52915347 | rs4369899 | 0.981 | 0.398 | -0.532 | -0.989 | 0.057 | 0.082 | 0.082 | 0 | 0.47 | 0.24 | 0.21 | 0.27 | 0.34 | 0.34 |
| 2 | 52917512 | rs12477044 | 1.62 | 0.331 | -0.619 | -1.046 | 0.059 | 0.025 | 0.025 | 0 | 0.36 | 0.19 | 0.14 | 0.25 | 0.18 | 0.2 |
| 2 | 52918929 | rs1454382 | 0.454 | 0.258 | -0.578 | -0.693 | -0.06 | -0.071 | -0.071 | 1.227 | 0.59 | 0.88 | 0.37 | 0.48 | 0.46 | 0.44 |
| 2 | 52919630 | rs1454383 | 0.469 | 0.317 | -0.516 | -0.66 | -0.054 | -0.028 | -0.028 | 0.026 | 0.59 | 0.88 | 0.31 | 0.37 | 0.47 | 0.42 |
| 2 | 52922423 | rs2062273 | 1.183 | 0.403 | -0.417 | -0.704 | -0.002 | -0.094 | -0.094 | 0.008 | 0.21 | 0.06 | 0.36 | 0.47 | 0.25 | 0.41 |
| 2 | 52932882 | rs6723934 | 0.912 | 0.436 | 0.09 | -0.703 | 0.001 | -0.003 | -0.003 | 0 | 0.55 | 0.56 | 0.28 | 0.35 | 0.47 | 0.39 |
| 2 | 52933307 | rs1384910 | 0.853 | 0.441 | 0.082 | -0.7 | -0.004 | 0.004 | 0.004 | 0 | 0.56 | 0.66 | 0.29 | 0.35 | 0.47 | 0.38 |
| 2 | 52934145 | rs350784 | -0.998 | 0.793 | -0.125 | -0.592 | -0.002 | 0.006 | 0.006 | 0 | 0.92 | 0.95 | 0.91 | 0.94 | 0.89 | 0.89 |
| 2 | 52935402 | rs2356473 | 0.37 | 0.8 | 0.01 | -0.577 | 0.062 | 0.087 | 0.087 | 0 | 0.07 | 0.32 | 0.05 | 0.1 | 0.11 | 0.05 |
| 2 | 52935725 | rs6545301 | 0.045 | 0.777 | -0.715 | -0.593 | -0.044 | 0.052 | 0.052 | 0.131 | 0.77 | 0.91 | 0.65 | 0.83 | 0.72 | 0.78 |
| 2 | 52938523 | rs350788 | -0.271 | 0.536 | -0.558 | -0.297 | 0.001 | 0.002 | 0.002 | 0.021 | 0.87 | 0.92 | 0.83 | 0.85 | 0.87 | 0.84 |
| 2 | 52938976 | rs10168918 | 2.503 | 0.306 | -1.142 | -0.478 | -0.068 | -0.034 | -0.034 | 0.018 | 0.23 | 0.04 | 0.17 | 0.24 | 0.28 | 0.11 |
| 2 | 52939115 | rs746098 | 0.177 | -0.437 | -0.425 | -0.728 | -0.001 | -0.008 | -0.008 | 0 | 0.18 | 0.31 | 0.08 | 0.13 | 0.16 | 0.23 |
| 2 | 52943646 | rs11892909 | -0.772 | -0.546 | -0.093 | -0.542 | -0.017 | -0.021 | -0.021 | 0.009 | 0.2 | 0.48 | 0.09 | 0.12 | 0.14 | 0.24 |
| 2 | 52944722 | rs350779 | 0.769 | -0.593 | -0.42 | -0.831 | -0.017 | 0.034 | 0.034 | 0 | 0.18 | 0.11 | 0.2 | 0.01 | 0.18 | 0.13 |
| 2 | 52944875 | rs11897448 | 0.227 | -0.684 | -0.258 | -0.806 | -0.016 | 0 | 0 | 0.343 | 0.17 | 0.41 | 0.08 | 0.11 | 0.11 | 0.08 |
| 2 | 52946337 | rs2626385 | 0.829 | -0.657 | -0.356 | -0.811 | -0.013 | -0.043 | -0.043 | 0 | 0.74 | 0.53 | 0.84 | 0.86 | 0.79 | 0.7 |
| 2 | 52947356 | rs1483867 | -0.119 | -0.625 | -0.357 | -0.847 | -0.009 | 0.017 | 0.017 | 0 | 0.26 | 0.19 | 0.27 | 0.03 | 0.26 | 0.22 |
| 2 | 52948828 | rs10173479 | 2.832 | -0.674 | -0.459 | -0.497 | -0.026 | -0.048 | -0.048 | 0 | 0.25 | 0.09 | 0.24 | 0.38 | 0.29 | 0.14 |
| 2 | 52949318 | rs1349882 | 1.07 | -0.651 | -0.342 | -0.498 | 0.001 | 0.007 | 0.007 | 0.041 | 0.43 | 0.48 | 0.33 | 0.5 | 0.42 | 0.37 |
| 2 | 52955892 | rs1305531 | -1.028 | -0.534 | -0.358 | -0.436 | -0.002 | 0 | 0 | 2.997 | 0.91 | 0.89 | 0.86 | 0.87 | 0.91 | 0.85 |
| 2 | 52957024 | rs7605026 | -0.558 | -0.565 | -0.222 | -0.418 | -0.016 | 0.035 | 0.035 | 0 | 0.2 | 0.53 | 0.17 | 0.25 | 0.16 | 0.26 |
| 2 | 52958528 | rs12713213 | 2.75 | -0.391 | -0.545 | -0.254 | -0.086 | -0.071 | -0.071 | 0 | 0.23 | 0.02 | 0.17 | 0.23 | 0.27 | 0.12 |
| 2 | 52962055 | rs1823887 | 1.077 | -0.069 | 0.596 | -0.185 | 0.021 | 0.017 | 0.017 | 2.834 | 0.64 | 0.51 | 0.47 | 0.43 | 0.5 | 0.41 |
| 2 | 52964900 | rs2128960 | 0.135 | 0.1 | 0.847 | -0.188 | 0.009 | 0.038 | 0.038 | 5.589 | 0.78 | 0.71 | 0.59 | 0.46 | 0.68 | 0.56 |
| 2 | 52967969 | rs1454391 | 0.137 | 0.101 | 0.926 | -0.208 | -0.006 | -0.008 | -0.008 | 0 | 0.78 | 0.8 | 0.59 | 0.44 | 0.68 | 0.56 |
| 2 | 52973062 | rs1580092 | 2.067 | -0.133 | 1.298 | -0.317 | 0 | 0.012 | 0.012 | 0 | 0.52 | 0.42 | 0.32 | 0.33 | 0.47 | 0.38 |
| 2 | 52974210 | rs2603375 | 0.159 | -0.14 | 1.184 | -0.357 | 0.01 | 0.002 | 0.002 | 0 | 0.16 | 0.11 | 0.07 | 0 | 0.11 | 0.1 |
| 2 | 52980071 | rs350720 | 1.431 | -0.075 | 1.143 | -0.317 | 0.009 | 0.008 | 0.008 | 3.341 | 0.65 | 0.57 | 0.47 | 0.43 | 0.5 | 0.43 |
| 2 | 52980534 | rs350723 | 0.083 | -0.046 | 1.028 | -0.284 | 0.008 | 0.034 | 0.034 | 5.739 | 0.78 | 0.71 | 0.58 | 0.45 | 0.68 | 0.58 |
| 2 | 52989700 | rs1843042 | 0.569 | -0.13 | 1.02 | -0.268 | 0.03 | 0.038 | 0.038 | 3.824 | 0.74 | 0.64 | 0.55 | 0.44 | 0.58 | 0.54 |
| 2 | 52992061 | rs2045131 | 2.668 | -0.211 | 1.137 | -0.349 | 0.019 | 0.012 | 0.012 | 0 | 0.43 | 0.16 | 0.22 | 0.32 | 0.37 | 0.32 |
| 2 | 52994927 | rs6747507 | -0.469 | -0.159 | 1.116 | 0.01 | 0.007 | 0.047 | 0.047 | 4.19 | 0.83 | 0.77 | 0.65 | 0.58 | 0.75 | 0.66 |
| 2 | 52999305 | rs12615207 | 1.741 | -0.205 | 1.073 | 0.091 | -0.003 | 0.004 | 0.004 | 0 | 0.54 | 0.37 | 0.47 | 0.53 | 0.47 | 0.45 |
| 2 | 53005483 | rs2603392 | -1.016 | -0.233 | 1.024 | 0.021 | -0.005 | -0.066 | -0.066 | 0.207 | 0.24 | 0.33 | 0.17 | 0.03 | 0.2 | 0.18 |
| 2 | 53006142 | rs1451465 | -1.745 | -0.273 | 1.149 | 0.045 | -0.036 | -0.043 | -0.043 | 0 | 0.3 | 0.5 | 0.35 | 0.2 | 0.22 | 0.23 |

| | | | | | | | | | | | | | | | | |
|---|---|---|---|---|---|---|---|---|---|---|---|---|---|---|---|---|
| 2 | 53006402 | rs993563 | -0.529 | -0.183 | 0.932 | 0.116 | 0 | 0.014 | 0.014 | 4.123 | 0.84 | 0.79 | 0.66 | 0.57 | 0.75 | 0.66 |
| 2 | 53010288 | rs2060774 | -0.055 | -0.275 | 0.768 | 0.054 | 0.019 | 0.059 | 0.059 | 0 | 0.16 | 0.07 | 0.08 | 0 | 0.12 | 0.1 |
| 2 | 53012074 | rs785282 | 1.222 | -0.186 | 0.882 | 0.066 | 0.024 | 0.021 | 0.021 | 0 | 0.68 | 0.54 | 0.39 | 0.35 | 0.58 | 0.48 |
| 2 | 53015422 | rs12623673 | 2.061 | -0.185 | 0.516 | 0.066 | 0.042 | 0.005 | 0.005 | 0 | 0.47 | 0.18 | 0.41 | 0.5 | 0.4 | 0.33 |
| 2 | 53018366 | rs12469661 | -0.998 | 0.118 | 0.701 | -0.399 | -0.001 | 0.042 | 0.042 | 0 | 0.12 | 0.29 | 0.26 | 0.2 | 0.09 | 0.13 |
| 2 | 53023825 | rs1454405 | 2.585 | -0.055 | 0 | -0.396 | 0.008 | -0.092 | -0.092 | 0 | 0.4 | 0.06 | 0.22 | 0.46 | 0.34 | 0.17 |
| 2 | 53024050 | rs7559322 | 1.678 | 0.074 | 0.249 | -0.51 | 0 | 0.044 | 0.044 | 0.423 | 0.53 | 0.35 | 0.49 | 0.63 | 0.46 | 0.3 |
| 2 | 53028126 | rs1085209 | -0.32 | -0.034 | 0.563 | -0.108 | 0 | 0.003 | 0.003 | 4.168 | 0.85 | 0.84 | 0.69 | 0.72 | 0.77 | 0.66 |
| 2 | 53030005 | rs785311 | -1.36 | -0.152 | 0.766 | -0.315 | -0.023 | -0.116 | -0.116 | 0 | 0.23 | 0.46 | 0.15 | 0.06 | 0.18 | 0.31 |
| 2 | 53034853 | rs785303 | -1.083 | -0.453 | 1.235 | -0.143 | -0.027 | -0.106 | -0.106 | 0 | 0.18 | 0.37 | 0.12 | 0.03 | 0.13 | 0.28 |
| 2 | 53035849 | rs785302 | -2.557 | -0.33 | 1.156 | -0.272 | -0.005 | 0.035 | 0.035 | 0 | 0.36 | 0.7 | 0.43 | 0.27 | 0.35 | 0.46 |
| 2 | 53037048 | rs785301 | 1.415 | -0.35 | 1.225 | -0.264 | -0.051 | -0.002 | -0.002 | 0 | 0.77 | 0.52 | 0.7 | 0.76 | 0.85 | 0.69 |
| 2 | 53037805 | rs725865 | 0.355 | -0.279 | 1.136 | -0.354 | -0.004 | 0.002 | 0.002 | 0 | 0.07 | 0.07 | 0.09 | 0.03 | 0.1 | 0.08 |
| 2 | 53039770 | rs10865265 | 2.207 | -0.296 | 1.155 | -0.365 | -0.005 | 0.037 | 0.037 | 0 | 0.64 | 0.3 | 0.57 | 0.73 | 0.65 | 0.55 |
| 2 | 53042122 | rs10221577 | -0.797 | -0.282 | 1.189 | -0.409 | 0.007 | -0.013 | -0.013 | 0.554 | 0.12 | 0.14 | 0.13 | 0.04 | 0.2 | 0.1 |
| 2 | 53051473 | rs10490584 | 2.004 | -0.227 | 1.262 | -0.293 | -0.006 | -0.015 | -0.015 | 0 | 0.72 | 0.43 | 0.66 | 0.77 | 0.75 | 0.64 |
| 2 | 53051989 | rs12465316 | 0.261 | -0.388 | 1.614 | -0.274 | -0.042 | -0.086 | -0.086 | 0.025 | 0.87 | 0.7 | 0.93 | 0.99 | 0.93 | 0.8 |
| 2 | 53052480 | rs1549749 | 0.489 | -0.389 | 1.637 | -0.275 | -0.049 | -0.1 | -0.1 | 0 | 0.86 | 0.61 | 0.92 | 0.99 | 0.93 | 0.8 |
| 2 | 53053929 | rs4364055 | 1.939 | -0.385 | 1.549 | -0.271 | -0.036 | -0.011 | -0.011 | 2.13 | 0.76 | 0.43 | 0.69 | 0.77 | 0.85 | 0.69 |
| 2 | 53054366 | rs17043530 | 0.184 | -0.432 | 1.46 | -0.191 | -0.005 | -0.009 | -0.009 | 0 | 0.15 | 0.14 | 0.32 | 0.31 | 0.23 | 0.36 |
| 2 | 53055973 | rs11125449 | 1.901 | -0.324 | 1.509 | -0.234 | -0.036 | -0.011 | -0.011 | 0.486 | 0.76 | 0.43 | 0.69 | 0.77 | 0.85 | 0.69 |
| 2 | 53061834 | rs1542282 | 0.903 | -0.319 | 1.042 | -0.205 | -0.018 | -0.008 | -0.008 | 0.377 | 0.71 | 0.58 | 0.62 | 0.71 | 0.7 | 0.45 |
| 2 | 53072134 | rs12713224 | 1.581 | -0.323 | 0.846 | -0.195 | 0.019 | -0.018 | -0.018 | 0.139 | 0.6 | 0.35 | 0.39 | 0.47 | 0.63 | 0.34 |
| 2 | 53088117 | rs17043603 | 0.002 | -0.297 | 0.694 | -0.185 | -0.029 | 0.101 | 0.101 | 7.577 | 0.06 | 0.14 | 0.17 | 0.19 | 0.01 | 0.05 |
| 2 | 53088412 | rs1451459 | -0.24 | -0.288 | 0.897 | -0.235 | 0.001 | 0.009 | 0.009 | 2.958 | 0.18 | 0.24 | 0.33 | 0.3 | 0.24 | 0.48 |
| 2 | 53095923 | rs10164868 | 2.269 | -0.311 | 0.301 | -0.292 | -0.006 | -0.049 | -0.049 | 1.141 | 0.47 | 0.15 | 0.26 | 0.44 | 0.41 | 0.19 |
| 2 | 53104091 | rs1839770 | 2.064 | 0.049 | 1.239 | -0.151 | 0.003 | 0.012 | 0.012 | 2.294 | 0.52 | 0.41 | 0.43 | 0.62 | 0.42 | 0.25 |
| 2 | 53109005 | rs1864534 | 2.169 | 0.061 | 0.706 | -0.145 | 0.027 | 0.099 | 0.099 | 4.028 | 0.51 | 0.26 | 0.43 | 0.63 | 0.42 | 0.25 |
| 2 | 53123426 | rs1451463 | -1.993 | 0.831 | 1.116 | -0.206 | -0.011 | 0.012 | 0.012 | 3.403 | 0.34 | 0.7 | 0.39 | 0.27 | 0.46 | 0.62 |
| 2 | 53124588 | rs4672065 | -1.121 | 0.908 | 1.078 | -0.164 | 0.022 | -0.011 | -0.011 | 0 | 0.27 | 0.6 | 0.32 | 0.3 | 0.37 | 0.51 |
| 2 | 53132960 | rs17043770 | -0.667 | 0.885 | 0.62 | -0.182 | 0.005 | 0.057 | 0.057 | 0 | 0.06 | 0.11 | 0.2 | 0.22 | 0.07 | 0.13 |
| 2 | 53143884 | rs17039478 | -0.647 | 0.954 | 0.457 | -0.61 | -0.001 | -0.007 | -0.007 | 0 | 0.08 | 0.1 | 0.1 | 0.01 | 0.12 | 0.1 |
| 2 | 53152519 | rs17043834 | -1.035 | 0.917 | 0.919 | -0.621 | 0.006 | 0.031 | 0.031 | 2.992 | 0.23 | 0.31 | 0.43 | 0.42 | 0.28 | 0.49 |
| 2 | 53154899 | rs4672085 | -2.747 | 0.712 | 0.281 | -0.98 | -0.049 | -0.055 | -0.055 | 0.266 | 0.59 | 0.97 | 0.67 | 0.44 | 0.63 | 0.83 |
| 2 | 53164354 | rs2195658 | 0.468 | 0.675 | 0.471 | -0.69 | -0.033 | -0.021 | -0.021 | 0 | 0.05 | 0 | 0.16 | 0.12 | 0.16 | 0.22 |
| 2 | 53174956 | rs7577197 | -1.62 | 0.566 | 0.879 | -0.931 | 0.011 | 0.008 | 0.008 | 4.313 | 0.32 | 0.58 | 0.55 | 0.41 | 0.39 | 0.58 |
| 2 | 53183877 | rs6705923 | -0.362 | 0.574 | 0.796 | -0.901 | 0.006 | -0.032 | -0.032 | 0.442 | 0.1 | 0.18 | 0.12 | 0 | 0.12 | 0.1 |
| 2 | 53188267 | rs2288639 | -0.293 | 0.591 | 0.256 | -0.891 | -0.051 | -0.105 | -0.105 | 0.047 | 0.17 | 0.02 | 0.32 | 0.39 | 0.23 | 0.37 |
| 2 | 53190362 | rs7596997 | -1.978 | 0.491 | 0.34 | -0.968 | -0.003 | 0.004 | 0.004 | 2.227 | 0.39 | 0.79 | 0.51 | 0.41 | 0.41 | 0.63 |
| 2 | 53210079 | rs1533076 | -0.999 | 0.431 | 0.708 | -0.932 | -0.002 | -0.125 | -0.125 | 0.777 | 0.19 | 0.59 | 0.17 | 0.02 | 0.16 | 0.23 |
| 2 | 53212803 | rs17043912 | -0.438 | 0.41 | 0.602 | -0.933 | -0.017 | -0.059 | -0.059 | 1.399 | 0.18 | 0.06 | 0.33 | 0.37 | 0.23 | 0.36 |
| 2 | 53216917 | rs10490577 | -1.552 | 0.443 | 0.705 | -0.95 | 0 | -0.02 | -0.02 | 0.711 | 0.24 | 0.79 | 0.25 | 0.05 | 0.21 | 0.35 |
| 2 | 53222962 | rs6545315 | -1.084 | 0.413 | 0.924 | -0.945 | -0.004 | -0.107 | -0.107 | 0 | 0.19 | 0.49 | 0.16 | 0.02 | 0.17 | 0.24 |
| 2 | 53223478 | rs10197026 | -0.819 | 0.658 | 0.875 | -0.914 | 0.061 | 0.064 | 0.064 | 1.082 | 0.22 | 0.34 | 0.43 | 0.4 | 0.33 | 0.49 |
| 2 | 53224976 | rs10183101 | -1.357 | 0.441 | 0.682 | -1.076 | 0.001 | -0.024 | -0.024 | 0 | 0.36 | 0.67 | 0.22 | 0.1 | 0.28 | 0.32 |
| 2 | 53225133 | rs10185735 | -1.357 | 0.44 | 0.715 | -1.077 | 0.001 | -0.024 | -0.024 | 0 | 0.36 | 0.67 | 0.22 | 0.1 | 0.28 | 0.31 |
| 2 | 53225842 | rs2048310 | -2.678 | 0.313 | 0.472 | -1.099 | -0.074 | -0.12 | -0.12 | 0.264 | 0.61 | 0.97 | 0.65 | 0.51 | 0.62 | 0.82 |
| 2 | 53227130 | rs4635564 | -0.476 | 0.34 | 0.629 | -1.113 | -0.012 | -0.087 | -0.087 | 0 | 0.23 | 0.37 | 0.17 | 0.11 | 0.19 | 0.23 |
| 2 | 53228506 | rs12052449 | -0.467 | 0.372 | 0.56 | -1.069 | -0.067 | -0.086 | -0.086 | 0 | 0.2 | 0.05 | 0.33 | 0.4 | 0.29 | 0.38 |
| 2 | 53231161 | rs6742807 | -0.742 | 0.24 | 0.791 | -1.086 | 0 | -0.099 | -0.099 | 0.415 | 0.18 | 0.53 | 0.15 | 0.02 | 0.15 | 0.19 |
| 2 | 53234750 | rs10198783 | -1.35 | 0.307 | 0.66 | -1.045 | 0.005 | 0 | 0 | 0.509 | 0.4 | 0.47 | 0.5 | 0.49 | 0.47 | 0.6 |
| 2 | 53235554 | rs1446435 | -1.088 | 0.314 | 0.637 | -1.041 | 0.006 | -0.003 | -0.003 | 0 | 0.36 | 0.35 | 0.42 | 0.41 | 0.42 | 0.49 |
| 2 | 53235595 | rs1446436 | 0.214 | 0.347 | 0.576 | -1.074 | -0.014 | -0.035 | -0.035 | 1.21 | 0.16 | 0.07 | 0.32 | 0.39 | 0.22 | 0.36 |
| 2 | 53236165 | rs1446437 | 0.461 | 0.341 | 0.452 | -1.137 | -0.023 | 0.014 | 0.014 | 0 | 0.06 | 0.02 | 0.08 | 0.01 | 0.1 | 0.05 |
| 2 | 53237548 | rs17574300 | 0.022 | 0.292 | 0.472 | -1.119 | -0.013 | 0.052 | 0.052 | 0 | 0.19 | 0.08 | 0.08 | 0.02 | 0.19 | 0.13 |
| 2 | 53246064 | rs1446442 | -1.457 | 0.176 | 0.443 | -1.152 | -0.008 | 0.016 | 0.016 | 0 | 0.39 | 0.19 | 0.33 | 0.42 | 0.42 | 0.5 |
| 2 | 53248330 | rs2121303 | -1.328 | 0.122 | 0.471 | -1.159 | -0.001 | 0.001 | 0.001 | 0 | 0.38 | 0.32 | 0.33 | 0.41 | 0.43 | 0.49 |
| 2 | 53251060 | rs895475 | 0.039 | -0.128 | 0.438 | -1.391 | 0.003 | -0.066 | -0.066 | 0 | 0.19 | 0.22 | 0.08 | 0.02 | 0.2 | 0.12 |
| 2 | 53260274 | rs1446450 | 0.181 | -0.016 | 0.309 | -1.192 | -0.029 | -0.095 | -0.095 | 0.453 | 0.16 | 0.03 | 0.24 | 0.45 | 0.22 | 0.35 |
| 2 | 53261375 | rs17044059 | -0.833 | 0.012 | 0.533 | -1.232 | -0.013 | -0.079 | -0.079 | 0 | 0.18 | 0.47 | 0.16 | 0.03 | 0.14 | 0.24 |
| 2 | 53263116 | rs2121304 | -1.591 | 0.014 | 0.436 | -1.261 | 0 | -0.001 | -0.001 | 0.021 | 0.46 | 0.39 | 0.41 | 0.57 | 0.48 | 0.61 |
| 2 | 53263553 | rs10496011 | -0.483 | -0.074 | 1.052 | -1.92 | -0.013 | 0.082 | 0.082 | 0 | 0.21 | 0.06 | 0.08 | 0.02 | 0.2 | 0.11 |
| 2 | 53263644 | rs1900249 | -1.848 | -0.057 | 1.085 | -1.914 | -0.001 | -0.006 | -0.006 | 0 | 0.23 | 0.77 | 0.25 | 0.12 | 0.19 | 0.36 |
| 2 | 53263871 | rs10496012 | -0.281 | -0.174 | 1.289 | -1.873 | -0.038 | -0.04 | -0.04 | 0 | 0.1 | 0.31 | 0.06 | 0.02 | 0.04 | 0.19 |
| 2 | 53277205 | rs2357016 | -1.231 | 0.126 | 0.696 | -1.707 | 0.009 | -0.072 | -0.072 | 0.685 | 0.25 | 0.27 | 0.17 | 0.05 | 0.26 | 0.26 |
| 2 | 53283334 | rs17189119 | -0.222 | -0.065 | 0.562 | -1.656 | -0.004 | 0.015 | 0.015 | 0 | 0.17 | 0.08 | 0.06 | 0.02 | 0.17 | 0.12 |
| 2 | 53288066 | rs10445935 | -0.613 | -0.026 | 0.594 | -1.637 | -0.002 | -0.07 | -0.07 | 0.742 | 0.2 | 0.21 | 0.07 | 0.02 | 0.17 | 0.12 |
| 2 | 53295106 | rs6717300 | 0.243 | 0.018 | 0.235 | -1.612 | 0.012 | -0.003 | -0.003 | 0 | 0.15 | 0.09 | 0.04 | 0 | 0.08 | 0.08 |
| 2 | 53310440 | rs7604973 | -1.245 | 0.082 | 0.088 | -1.674 | 0.005 | -0.123 | -0.123 | 0.833 | 0.27 | 0.45 | 0.12 | 0.1 | 0.24 | 0.24 |

| | | | | | | | | | | | | | | | | |
|---|---|---|---|---|---|---|---|---|---|---|---|---|---|---|---|---|
| 2 | 53314170 | rs3861569 | -0.71 | 0.165 | -0.084 | -1.591 | 0.01 | -0.017 | -0.017 | 0 | 0.1 | 0.18 | 0.05 | 0.05 | 0.12 | 0.12 |
| 2 | 53321966 | rs12476215 | 0.341 | 0.201 | -0.621 | -1.552 | -0.035 | 0.013 | 0.013 | 0.167 | 0.07 | 0 | 0.16 | 0.06 | 0.11 | 0.07 |
| 2 | 53322243 | rs12477118 | -0.979 | 0.235 | -0.453 | -1.437 | -0.012 | -0.087 | -0.087 | 0 | 0.34 | 0.45 | 0.28 | 0.16 | 0.28 | 0.3 |
| 2 | 53327092 | rs1424968 | -1.378 | 0.443 | -2 | -1.444 | -0.064 | 0.282 | 0.282 | 0 | 0.52 | 0.93 | 0.34 | 0.16 | 0.43 | 0.55 |
| 2 | 53327137 | rs1424969 | 0.264 | 0.764 | -1.599 | -1.163 | -0.026 | 0.017 | 0.017 | 0 | 0.06 | 0 | 0.03 | 0.06 | 0.07 | 0.07 |
| 2 | 53342655 | rs17044122 | -0.145 | 0.52 | -1.321 | -1.024 | -0.002 | -0.06 | -0.06 | 0 | 0.25 | 0.27 | 0.1 | 0.02 | 0.21 | 0.17 |
| 2 | 53352630 | rs9653536 | -0.508 | 0.759 | -1.965 | -0.591 | -0.062 | 0.209 | 0.209 | 0 | 0.54 | 0.84 | 0.45 | 0.19 | 0.47 | 0.56 |
| 2 | 53355627 | rs2113820 | 0 | 0.764 | -1.485 | -0.551 | 0.001 | 0 | 0 | 0 | 0.51 | 0.56 | 0.5 | 0.21 | 0.56 | 0.42 |
| 2 | 53355763 | rs7594954 | -0.86 | 1.743 | -1.286 | -0.198 | 0.005 | 0.002 | 0.002 | 0 | 0.18 | 0.19 | 0.44 | 0.64 | 0.35 | 0.34 |
| 2 | 53358680 | rs17044134 | 0.205 | 1.28 | -1.384 | -0.6 | 0.005 | 0.011 | 0.011 | 0 | 0.11 | 0.1 | 0.15 | 0.09 | 0.08 | 0.08 |
| 2 | 53358953 | rs2034254 | -0.509 | 0.99 | -1.003 | -0.763 | -0.007 | 0.005 | 0.005 | 0.019 | 0.24 | 0.26 | 0.29 | 0.66 | 0.22 | 0.3 |
| 2 | 53359345 | rs1900247 | 0.13 | 0.835 | -0.864 | -0.653 | -0.006 | 0.069 | 0.069 | 0 | 0.17 | 0.32 | 0.27 | 0.63 | 0.17 | 0.24 |
| 2 | 53361731 | rs17189210 | -0.196 | 0.749 | -0.672 | -0.645 | 0.005 | 0.017 | 0.017 | 0 | 0.14 | 0.11 | 0.16 | 0.07 | 0.1 | 0.08 |
| 2 | 53370010 | rs6753476 | -1.139 | 0.511 | -0.287 | -0.909 | 0.011 | -0.035 | -0.035 | 0.26 | 0.26 | 0.32 | 0.3 | 0.11 | 0.31 | 0.26 |
| 2 | 53370427 | rs1374252 | -1.611 | 0.491 | -0.091 | -1.022 | 0.016 | -0.096 | -0.096 | 0 | 0.32 | 0.47 | 0.34 | 0.12 | 0.36 | 0.3 |
| 2 | 53379489 | rs1364676 | -0.795 | 0.629 | 0.33 | -0.605 | 0.109 | 0.014 | 0.014 | 0.07 | 0.13 | 0.28 | 0.19 | 0.13 | 0.25 | 0.23 |
| 2 | 53380974 | rs10186405 | 0.091 | 0.157 | 0.226 | -0.731 | 0.009 | 0.001 | 0.001 | 0 | 0.86 | 0.85 | 0.82 | 0.88 | 0.9 | 0.87 |
| 2 | 53384500 | rs4233969 | 1.825 | 0.059 | 0.216 | -0.826 | 0.001 | -0.018 | -0.018 | 0 | 0.62 | 0.51 | 0.53 | 0.77 | 0.59 | 0.64 |
| 2 | 53385857 | rs10168550 | 0.973 | 0.057 | 0.19 | -0.83 | 0.003 | -0.006 | -0.006 | 0 | 0.75 | 0.7 | 0.67 | 0.83 | 0.67 | 0.71 |
| 2 | 53386014 | rs13021753 | 2.043 | 0.011 | 0.661 | -0.847 | -0.007 | 0.01 | 0.01 | 0 | 0.6 | 0.3 | 0.49 | 0.69 | 0.57 | 0.58 |
| 2 | 53388690 | rs1895697 | 1.785 | 0.02 | 0.616 | -0.87 | -0.002 | 0.002 | 0.002 | 0.047 | 0.63 | 0.59 | 0.53 | 0.76 | 0.59 | 0.66 |
| 2 | 53389260 | rs10181311 | 0.036 | 0.033 | 0.502 | -0.858 | 0.004 | 0.004 | 0.004 | 6.764 | 0.12 | 0.13 | 0.14 | 0.05 | 0.08 | 0.06 |
| 2 | 53389666 | rs11687654 | 1.884 | 0.06 | 0.607 | -0.856 | 0.011 | -0.012 | -0.012 | 0.097 | 0.62 | 0.49 | 0.5 | 0.67 | 0.57 | 0.6 |
| 2 | 53395092 | rs13428097 | -0.036 | 0.244 | 0.261 | -0.752 | 0.004 | 0.004 | 0.004 | 6.7 | 0.12 | 0.13 | 0.13 | 0.07 | 0.08 | 0.06 |
| 2 | 53395407 | rs11892385 | -1.821 | 0.272 | 0.038 | -0.766 | 0.022 | -0.021 | -0.021 | 0.635 | 0.27 | 0.59 | 0.34 | 0.19 | 0.33 | 0.29 |
| 2 | 53398241 | rs4671327 | -1.912 | 0.306 | -0.143 | -0.784 | 0.025 | -0.02 | -0.02 | 0 | 0.3 | 0.57 | 0.35 | 0.23 | 0.35 | 0.31 |
| 2 | 53403042 | rs10496019 | -0.436 | 0.512 | -0.288 | -0.709 | -0.001 | 0 | 0 | 0 | 0.13 | 0.12 | 0.16 | 0.04 | 0.23 | 0.17 |
| 2 | 53407107 | rs1424974 | 0.123 | 0.081 | -0.765 | -0.135 | 0.025 | 0.016 | 0.016 | 0 | 0.84 | 0.91 | 0.82 | 0.89 | 0.88 | 0.86 |
| 2 | 53410123 | rs7603187 | -0.648 | 0.844 | -0.094 | 0.323 | 0.095 | 0.06 | 0.06 | 0.648 | 0.16 | 0.55 | 0.21 | 0.23 | 0.25 | 0.19 |
| 2 | 53410785 | rs6545322 | 0.815 | -0.063 | -0.659 | -0.674 | 0.054 | 0.07 | 0.07 | 1.708 | 0.56 | 0.73 | 0.61 | 0.74 | 0.7 | 0.73 |
| 2 | 53416255 | rs1559631 | -0.166 | -0.618 | -0.231 | -1.179 | 0.006 | 0.002 | 0.002 | 0 | 0.88 | 0.76 | 0.78 | 0.88 | 0.85 | 0.92 |
| 2 | 53417366 | rs2357180 | 0.782 | -1.007 | -0.237 | -1.33 | 0.01 | 0.006 | 0.006 | 3.159 | 0.69 | 0.67 | 0.63 | 0.72 | 0.77 | 0.74 |
| 2 | 53421015 | rs1344595 | 2.023 | -1.24 | -0.118 | -1.558 | 0.075 | 0.088 | 0.088 | 1.268 | 0.49 | 0.2 | 0.51 | 0.67 | 0.64 | 0.66 |
| 2 | 53427383 | rs1424989 | 0.982 | -0.887 | 0.55 | -1.4 | -0.015 | -0.026 | -0.026 | 0 | 0.67 | 0.44 | 0.59 | 0.71 | 0.72 | 0.75 |
| 2 | 53428270 | rs11885974 | 0.189 | -0.962 | 0.476 | -1.438 | -0.041 | -0.03 | -0.03 | 0 | 0.15 | 0.23 | 0.07 | 0.05 | 0.08 | 0.07 |
| 2 | 53428421 | rs11896988 | 0.562 | -0.828 | 0.171 | -1.386 | -0.003 | -0.002 | -0.002 | 2.728 | 0.72 | 0.66 | 0.62 | 0.72 | 0.76 | 0.75 |
| 2 | 53439629 | rs4672264 | -0.138 | -0.73 | -0.036 | -1.332 | 0.048 | 0.021 | 0.021 | 0.5 | 0.19 | 0.08 | 0.17 | 0.17 | 0.1 | 0.18 |
| 2 | 53444766 | rs10165023 | 0.537 | -0.776 | -0.047 | -1.359 | 0.031 | 0.016 | 0.016 | 1.192 | 0.69 | 0.86 | 0.63 | 0.71 | 0.75 | 0.74 |
| 2 | 53450309 | rs10496021 | 0.475 | -0.392 | -0.105 | -1.308 | -0.033 | 0.023 | 0.023 | 0 | 0.05 | 0 | 0.1 | 0.02 | 0.14 | 0.11 |
| 2 | 53453844 | rs6758455 | 0.562 | -1.383 | 0.264 | -1.186 | -0.003 | -0.004 | -0.004 | 0 | 0.13 | 0.11 | 0.06 | 0.04 | 0.1 | 0.05 |
| 2 | 53459016 | rs1364666 | 1.502 | -1.404 | 0.141 | -1.197 | 0.091 | 0.111 | 0.111 | 0.002 | 0.5 | 0.16 | 0.52 | 0.67 | 0.64 | 0.65 |
| 2 | 53462395 | rs1424956 | -1.415 | -0.666 | 0.873 | -0.666 | -0.037 | -0.025 | -0.025 | 0.163 | 0.32 | 0.52 | 0.39 | 0.29 | 0.24 | 0.28 |
| 2 | 53468832 | rs6705480 | 0.126 | -0.278 | -0.825 | -0.727 | 0.012 | -0.027 | -0.027 | 0.402 | 0.1 | 0.09 | 0.07 | 0 | 0.15 | 0.14 |
| 2 | 53469810 | rs4672278 | -0.666 | -1.129 | -0.911 | -0.217 | 0.045 | 0.03 | 0.03 | 0.55 | 0.17 | 0.06 | 0.16 | 0.15 | 0.09 | 0.13 |
| 2 | 53479450 | rs195583 | -1.542 | -0.997 | -0.774 | -0.174 | -0.011 | -0.014 | -0.014 | 0 | 0.3 | 0.48 | 0.39 | 0.29 | 0.26 | 0.25 |
| 2 | 53488034 | rs195592 | -0.907 | -0.464 | -1.461 | 0.129 | -0.005 | 0.001 | 0.001 | 0 | 0.14 | 0.14 | 0.22 | 0.12 | 0.17 | 0.1 |
| 2 | 53493132 | rs195610 | 0.192 | -0.957 | -1.79 | 0.203 | 0.024 | 0.036 | 0.036 | 4.456 | 0.69 | 0.96 | 0.66 | 0.8 | 0.76 | 0.77 |
| 2 | 53495204 | rs1895699 | -0.217 | -0.579 | -1.435 | 0.594 | -0.026 | -0.025 | -0.025 | 0 | 0.23 | 0.03 | 0.3 | 0.23 | 0.21 | 0.21 |
| 2 | 53499062 | rs6756435 | -0.597 | -1.099 | -0.729 | 0.219 | 0.002 | -0.01 | -0.01 | 0 | 0.08 | 0.35 | 0.04 | 0.05 | 0.03 | 0.04 |
| 2 | 53500604 | rs11125480 | -0.647 | -0.963 | -1.452 | 0.781 | 0.048 | -0.015 | -0.015 | 0.015 | 0.29 | 0.19 | 0.44 | 0.71 | 0.17 | 0.34 |
| 2 | 53507604 | rs4672305 | -0.618 | -0.934 | -0.992 | 0.64 | -0.005 | -0.001 | -0.001 | 0.536 | 0.37 | 0.58 | 0.26 | 0.75 | 0.23 | 0.54 |
| 2 | 53510546 | rs6754620 | -0.723 | -1.099 | -0.268 | 0.685 | -0.019 | -0.005 | -0.005 | 0.57 | 0.13 | 0.25 | 0.07 | 0.12 | 0.1 | 0.27 |
| 2 | 53514022 | rs10202079 | -1.057 | -0.704 | -1.499 | -0.123 | -0.092 | 0.137 | 0.137 | 5.234 | 0.65 | 0.95 | 0.68 | 0.88 | 0.65 | 0.91 |
| 2 | 53516993 | rs12473577 | -0.77 | -0.781 | -1.335 | 0.538 | -0.053 | -0.034 | -0.034 | 0 | 0.38 | 0.82 | 0.4 | 0.51 | 0.46 | 0.6 |
| 2 | 53522077 | rs7574199 | -1.041 | -0.938 | -0.158 | 0.65 | 0 | 0.001 | 0.001 | 0.899 | 0.52 | 0.5 | 0.57 | 0.71 | 0.55 | 0.61 |
| 2 | 53524646 | rs6545328 | -1.934 | -0.93 | 0.048 | 0.649 | -0.01 | 0.085 | 0.085 | 4.578 | 0.65 | 0.86 | 0.66 | 0.86 | 0.65 | 0.9 |
| 2 | 53529557 | rs6731364 | -1.144 | -0.94 | 0.285 | 0.652 | 0.001 | 0.005 | 0.005 | 0.767 | 0.53 | 0.57 | 0.6 | 0.76 | 0.55 | 0.66 |
| 2 | 53530783 | rs10164993 | -1.188 | -0.939 | 0.284 | 0.652 | 0.003 | 0.027 | 0.027 | 0.954 | 0.54 | 0.6 | 0.6 | 0.76 | 0.55 | 0.66 |
| 2 | 53531184 | rs12621817 | 0.72 | -0.431 | 0.566 | 1.202 | 0.02 | 0.006 | 0.006 | 0 | 0.1 | 0.11 | 0.26 | 0.16 | 0.27 | 0.22 |
| 2 | 53531829 | rs7588904 | 0.792 | -0.431 | 0.679 | 1.19 | -0.008 | -0.003 | -0.003 | 0 | 0.11 | 0.22 | 0.06 | 0.11 | 0.1 | 0.25 |
| 2 | 53532456 | rs992874 | 0.172 | -0.404 | 0.717 | 1.219 | 0.032 | -0.021 | -0.021 | 0 | 0.26 | 0.13 | 0.28 | 0.36 | 0.19 | 0.32 |
| 2 | 53536813 | rs2161783 | -2.197 | -0.367 | 0.661 | 1.198 | -0.001 | 0.037 | 0.037 | 3.265 | 0.65 | 0.79 | 0.66 | 0.86 | 0.65 | 0.91 |
| 2 | 53537829 | rs6757726 | 0.303 | -0.375 | 0.676 | 1.212 | -0.008 | 0.05 | 0.05 | 0.4 | 0.16 | 0.32 | 0.07 | 0.24 | 0.1 | 0.12 |
| 2 | 53538984 | rs6746636 | -0.623 | -0.43 | 0.624 | 1.181 | 0.013 | -0.002 | -0.002 | 0.9 | 0.38 | 0.35 | 0.33 | 0.46 | 0.29 | 0.56 |
| 2 | 53540121 | rs2216804 | -2.155 | -0.373 | 0.611 | 1.169 | 0 | 0.063 | 0.063 | 3.962 | 0.64 | 0.82 | 0.66 | 0.88 | 0.65 | 0.91 |
| 2 | 53540600 | rs11890869 | -0.86 | -0.065 | 0.948 | 1.44 | 0.047 | 0.065 | 0.065 | 1.903 | 0.27 | 0.43 | 0.33 | 0.4 | 0.36 | 0.35 |
| 2 | 53543619 | rs2194408 | -1.071 | -0.104 | 1.026 | 1.473 | 0.022 | 0.015 | 0.015 | 0 | 0.28 | 0.39 | 0.44 | 0.4 | 0.39 | 0.36 |
| 2 | 53545265 | rs11125482 | -0.464 | 0.207 | 0.955 | 1.484 | 0.004 | 0.001 | 0.001 | 0 | 0.21 | 0.36 | 0.22 | 0.26 | 0.17 | 0.21 |
| 2 | 53547825 | rs7609365 | -2.218 | 0.002 | 0.874 | 1.429 | -0.004 | 0.06 | 0.06 | 0.415 | 0.58 | 0.82 | 0.62 | 0.73 | 0.57 | 0.69 |

| | | | | | | | | | | | | | | | | |
|---|---|---|---|---|---|---|---|---|---|---|---|---|---|---|---|---|
| 2 | 53549075 | rs7576932 | -0.738 | 0.572 | 1.311 | 1.599 | 0.005 | -0.004 | -0.004 | 0 | 0.21 | 0.36 | 0.22 | 0.24 | 0.17 | 0.21 |
| 2 | 53551600 | rs1011572 | -1.469 | 0.558 | 1.259 | 1.623 | 0.042 | 0.025 | 0.025 | 0 | 0.3 | 0.5 | 0.45 | 0.39 | 0.4 | 0.42 |
| 2 | 53552553 | rs13410317 | -0.756 | 0.532 | 1.102 | 1.593 | 0.001 | -0.002 | -0.002 | 0 | 0.21 | 0.33 | 0.21 | 0.24 | 0.17 | 0.21 |
| 2 | 53554943 | rs4672343 | -1.107 | 0.497 | 0.539 | 1.553 | 0.017 | 0.014 | 0.014 | 0.039 | 0.3 | 0.62 | 0.47 | 0.41 | 0.4 | 0.42 |
| 2 | 53561575 | rs12328023 | -1.53 | 0.085 | -0.037 | 1.196 | -0.03 | 0.166 | 0.166 | 1.015 | 0.51 | 0.85 | 0.58 | 0.73 | 0.52 | 0.69 |
| 2 | 53576149 | rs6727931 | -1.651 | 0.029 | -0.092 | 1.17 | -0.081 | 0.151 | 0.151 | 1.1 | 0.52 | 0.93 | 0.59 | 0.72 | 0.52 | 0.69 |
| 2 | 53576769 | rs2194403 | -1.248 | -0.085 | 0.745 | 1.745 | 0 | -0.001 | -0.001 | 0 | 0.42 | 0.6 | 0.53 | 0.44 | 0.45 | 0.59 |
| 2 | 53587240 | rs12466120 | -1.162 | -0.043 | 1.544 | 2.054 | 0.034 | 0.161 | 0.161 | 0.196 | 0.17 | 0.34 | 0.26 | 0.47 | 0.25 | 0.38 |
| 2 | 53593857 | rs10205628 | 0.21 | -0.076 | 1.824 | 1.993 | -0.04 | -0.042 | -0.042 | 1.483 | 0.1 | 0.04 | 0.22 | 0.23 | 0.23 | 0.28 |
| 2 | 53595148 | rs6728653 | -1.052 | -0.138 | 2.081 | 1.951 | 0.025 | 0.315 | 0.315 | 2.799 | 0.06 | 0.27 | 0.04 | 0.27 | 0.02 | 0.1 |
| 2 | 53595922 | rs2059316 | -0.793 | -0.182 | 2.426 | 1.861 | -0.018 | -0.021 | -0.021 | 0.413 | 0.3 | 0.34 | 0.29 | 0.22 | 0.22 | 0.3 |
| 2 | 53597486 | rs7572447 | -0.481 | -0.176 | 2.329 | 1.864 | 0.02 | 0.024 | 0.024 | 0 | 0.27 | 0.11 | 0.27 | 0.22 | 0.2 | 0.3 |
| 2 | 53602776 | rs2042478 | -1.477 | -0.164 | 2.336 | 1.937 | 0.03 | 0.384 | 0.384 | 0.86 | 0.08 | 0.46 | 0.07 | 0.27 | 0.03 | 0.11 |
| 2 | 53609363 | rs1357204 | -2.303 | -0.296 | 2.211 | 1.928 | 0.029 | -0.047 | -0.047 | 0.034 | 0.24 | 0.79 | 0.26 | 0.46 | 0.14 | 0.24 |
| 2 | 53613744 | rs7559945 | -1.69 | -0.276 | 2.313 | 1.889 | 0 | 0.263 | 0.263 | 5.867 | 0.08 | 0.32 | 0.04 | 0.27 | 0.03 | 0.1 |
| 2 | 53615444 | rs11125490 | -1.438 | -0.262 | 2.181 | 1.863 | -0.02 | 0.029 | 0.029 | 0.358 | 0.29 | 0.57 | 0.19 | 0.32 | 0.23 | 0.28 |
| 2 | 53617260 | rs4671394 | -2.528 | -0.163 | 1.566 | 1.898 | -0.03 | 0.177 | 0.177 | 0.942 | 0.5 | 0.87 | 0.58 | 0.74 | 0.52 | 0.69 |
| 2 | 53628263 | rs6725739 | -1.313 | -0.217 | 1.639 | 1.951 | -0.01 | 0.227 | 0.227 | 5.622 | 0.07 | 0.22 | 0.02 | 0.26 | 0.02 | 0.1 |
| 2 | 53629321 | rs4671396 | -2.506 | -0.243 | 1.24 | 1.7 | -0.055 | 0.146 | 0.146 | 1.119 | 0.52 | 0.91 | 0.59 | 0.74 | 0.51 | 0.68 |
| 2 | 53633289 | rs1157474 | -0.098 | 0.026 | 1.076 | 1.92 | -0.003 | 0.027 | 0.027 | 0 | 0.07 | 0.04 | 0.22 | 0.35 | 0.23 | 0.24 |
| 2 | 53633492 | rs6761256 | -0.758 | -0.547 | 1.229 | 1.302 | -0.013 | 0.008 | 0.008 | 0.007 | 0.24 | 0.29 | 0.14 | 0.21 | 0.17 | 0.21 |
| 2 | 53635017 | rs13428153 | -1.26 | -0.385 | 0.913 | 1.399 | -0.023 | -0.052 | -0.052 | 0 | 0.12 | 0.24 | 0.05 | 0 | 0.06 | 0.13 |
| 2 | 53635266 | rs7582207 | -1.398 | -0.235 | 0.738 | 1.65 | -0.071 | 0.096 | 0.096 | 0.282 | 0.23 | 0.47 | 0.22 | 0.33 | 0.13 | 0.27 |
| 2 | 53636324 | rs12475695 | -1.045 | 1.098 | -0.076 | 1.182 | 0.033 | 0.001 | 0.001 | 0 | 0.22 | 0.31 | 0.23 | 0.24 | 0.38 | 0.2 |
| 2 | 53637182 | rs12713247 | -0.943 | 1.197 | -0.626 | 1.005 | -0.016 | 0.013 | 0.013 | 0 | 0.41 | 0.7 | 0.59 | 0.58 | 0.55 | 0.36 |
| 2 | 53644605 | rs11904043 | -0.421 | 2.338 | -0.176 | 0.66 | -0.003 | 0.002 | 0.002 | 0.02 | 0.14 | 0.09 | 0.23 | 0.07 | 0.22 | 0.09 |
| 2 | 53647729 | rs2172645 | 1.556 | 2.466 | 0.476 | 1.304 | 0.02 | 0.035 | 0.035 | 1.089 | 0.68 | 0.6 | 0.43 | 0.52 | 0.44 | 0.52 |
| 2 | 53649620 | rs12994450 | -0.641 | 2.545 | 0.433 | 1.47 | -0.037 | -0.027 | -0.027 | 0 | 0.11 | 0.05 | 0.15 | 0.19 | 0.27 | 0.3 |
| 2 | 53650817 | rs1494018 | -0.226 | 2.698 | 0.433 | 1.466 | -0.016 | -0.014 | -0.014 | 0 | 0.08 | 0.05 | 0.12 | 0.1 | 0.24 | 0.06 |
| 2 | 53650968 | rs10183072 | 0.274 | 2.651 | 0.442 | 1.467 | 0.001 | -0.001 | -0.001 | 0 | 0.13 | 0.09 | 0.22 | 0.08 | 0.22 | 0.08 |
| 2 | 53651911 | rs2131786 | 1.308 | 2.537 | 0.538 | 1.445 | 0.063 | 0 | 0 | 0.693 | 0.86 | 0.76 | 0.8 | 0.89 | 0.68 | 0.85 |
| 2 | 53660931 | rs1521939 | 0.145 | 0.784 | 1.367 | 0.876 | -0.023 | -0.011 | -0.011 | 1.112 | 0.75 | 0.36 | 0.73 | 0.73 | 0.85 | 0.64 |
| 2 | 53667654 | rs4672420 | 1.639 | 0.683 | 0.778 | 0.841 | 0.066 | 0.012 | 0.012 | 0.46 | 0.46 | 0.28 | 0.35 | 0.42 | 0.31 | 0.42 |
| 2 | 53669133 | rs17044715 | -0.853 | 0.536 | 0.578 | 0.908 | 0.011 | -0.012 | -0.012 | 0 | 0.05 | 0.02 | 0.02 | 0.21 | 0.03 | 0.24 |
| 2 | 53670488 | rs1521947 | -1.93 | 0.658 | 0.538 | 0.918 | 0.028 | 0.002 | 0.002 | 0.068 | 0.54 | 0.89 | 0.66 | 0.6 | 0.66 | 0.58 |
| 2 | 53672814 | rs4330141 | 1.67 | 0.404 | 0.673 | 0.891 | 0.018 | 0.016 | 0.016 | 0 | 0.45 | 0.16 | 0.35 | 0.39 | 0.36 | 0.47 |
| 2 | 53674840 | rs6705735 | 0.1 | 0.425 | 0.547 | 0.93 | -0.123 | -0.079 | -0.079 | 0 | 0.76 | 0.92 | 0.61 | 0.69 | 0.51 | 0.77 |
| 2 | 53679392 | rs882250 | 0.584 | 0.352 | 1.046 | 0.851 | 0.001 | 0 | 0 | 0.331 | 0.61 | 0.62 | 0.45 | 0.62 | 0.47 | 0.75 |
| 2 | 53682501 | rs4672433 | 1.897 | 0.297 | 1.027 | 0.833 | 0.021 | 0.016 | 0.016 | 0 | 0.45 | 0.29 | 0.35 | 0.38 | 0.36 | 0.46 |
| 2 | 53687730 | rs1402901 | 0.368 | 0.277 | 0.943 | 0.882 | 0.046 | -0.006 | -0.006 | 0 | 0.22 | 0.11 | 0.18 | 0.3 | 0.09 | 0.27 |
| 2 | 53691289 | rs745685 | 0.523 | 0.362 | 1.114 | 0.796 | -0.029 | -0.02 | -0.02 | 0 | 0.72 | 0.79 | 0.56 | 0.68 | 0.47 | 0.75 |
| 2 | 53693619 | rs10204358 | 1.817 | 0.293 | 1.228 | 0.609 | -0.026 | -0.04 | -0.04 | 0 | 0.13 | 0.11 | 0.03 | 0 | 0.04 | 0.03 |
| 2 | 53695084 | rs903229 | -0.027 | 0.39 | 1.213 | 0.668 | -0.048 | -0.043 | -0.043 | 1.635 | 0.76 | 0.85 | 0.61 | 0.57 | 0.52 | 0.78 |
| 2 | 53696453 | rs7567601 | -1.166 | 0.325 | 1.297 | 0.618 | 0.003 | 0.008 | 0.008 | 0.477 | 0.1 | 0.35 | 0.17 | 0.07 | 0.05 | 0.05 |
| 2 | 53703155 | rs1318578 | -0.599 | 0.637 | 1.022 | 0.645 | -0.152 | -0.15 | -0.15 | 0.027 | 0.79 | 0.97 | 0.65 | 0.56 | 0.56 | 0.84 |
| 2 | 53704641 | rs6727792 | -1.028 | 0.564 | 1.239 | 0.532 | -0.004 | 0.005 | 0.005 | 0.334 | 0.09 | 0.34 | 0.17 | 0.08 | 0.05 | 0.06 |
| 2 | 53706224 | rs10164725 | -0.031 | 0.53 | 0.977 | 0.495 | -0.058 | -0.1 | -0.1 | 0.179 | 0.23 | 0.38 | 0.22 | 0.07 | 0.13 | 0.07 |
| 2 | 53706903 | rs926014 | 1.611 | 0.543 | 0.43 | 0.626 | 0.035 | 0.033 | 0.033 | 0 | 0.47 | 0.25 | 0.35 | 0.36 | 0.37 | 0.46 |
| 2 | 53711697 | rs7604061 | 0.821 | 0.307 | 0.862 | 0.627 | -0.013 | -0.022 | -0.022 | 0 | 0.16 | 0.13 | 0.17 | 0.05 | 0.08 | 0.05 |
| 2 | 53715181 | rs10177230 | -0.707 | 0.431 | 0.096 | 0.524 | -0.021 | -0.03 | -0.03 | 0 | 0.59 | 0.75 | 0.59 | 0.53 | 0.51 | 0.49 |
| 2 | 53718104 | rs6720273 | -0.583 | 0.468 | 0.222 | 0.513 | -0.01 | -0.022 | -0.022 | 0.937 | 0.08 | 0.18 | 0.05 | 0.02 | 0.06 | 0.06 |
| 2 | 53721582 | rs13404343 | 0.721 | 0.47 | 0.357 | 0.55 | -0.002 | 0.002 | 0.002 | 0.088 | 0.33 | 0.24 | 0.32 | 0.3 | 0.37 | 0.16 |
| 2 | 53722595 | rs11900008 | -1.004 | 0.044 | 0.444 | 0.434 | -0.022 | 0.008 | 0.008 | 0.328 | 0.19 | 0.22 | 0.21 | 0.25 | 0.11 | 0.32 |
| 2 | 53723509 | rs10204173 | 0.855 | 0.225 | 0.39 | 0.434 | -0.001 | 0.002 | 0.002 | 0.155 | 0.34 | 0.27 | 0.35 | 0.3 | 0.36 | 0.16 |
| 2 | 53731906 | rs7559951 | -0.641 | 0.243 | 0.384 | 0.398 | -0.006 | -0.022 | -0.022 | 0.175 | 0.55 | 0.71 | 0.58 | 0.53 | 0.47 | 0.45 |
| 2 | 53735050 | rs17268019 | 1.624 | 0.667 | -0.154 | 0.308 | -0.06 | -0.018 | -0.018 | 0 | 0.14 | 0.01 | 0.1 | 0.12 | 0.21 | 0.12 |
| 2 | 53739376 | rs17189470 | 1.026 | 0.506 | 0.335 | 0.349 | -0.005 | -0.033 | -0.033 | 0 | 0.3 | 0.07 | 0.28 | 0.31 | 0.29 | 0.1 |
| 2 | 53743131 | rs13394011 | 0.526 | 0.489 | 0.754 | 0.31 | -0.001 | -0.001 | -0.001 | 0 | 0.34 | 0.31 | 0.36 | 0.3 | 0.35 | 0.15 |
| 2 | 53744110 | rs11887670 | -0.043 | 0.407 | 0.993 | 0.256 | -0.006 | -0.008 | -0.008 | 3.029 | 0.4 | 0.47 | 0.38 | 0.32 | 0.39 | 0.21 |
| 2 | 53745379 | rs7572194 | -1.049 | 0.415 | 0.815 | 0.245 | -0.033 | -0.061 | -0.061 | 0.004 | 0.55 | 0.82 | 0.6 | 0.52 | 0.48 | 0.44 |
| 2 | 53752843 | rs12463516 | 0.991 | 0.68 | 0.593 | 0.28 | -0.034 | -0.046 | -0.046 | 0 | 0.3 | 0.07 | 0.31 | 0.34 | 0.34 | 0.1 |
| 2 | 53757983 | rs6749477 | -1.488 | 0.2 | 0.96 | 0.169 | 0.043 | -0.025 | -0.025 | 0 | 0.07 | 0.29 | 0.06 | 0 | 0.05 | 0.02 |
| 2 | 53764592 | rs11902012 | -1.646 | 0.185 | 0.952 | 0.178 | -0.025 | -0.017 | -0.017 | 0.002 | 0.2 | 0.29 | 0.23 | 0.17 | 0.11 | 0.27 |
| 2 | 53767913 | rs11901158 | -0.466 | 0.252 | 0.794 | 0.184 | -0.004 | -0.029 | -0.029 | 0 | 0.07 | 0.09 | 0.05 | 0 | 0.05 | 0.05 |
| 2 | 53770692 | rs7600140 | -0.931 | 0.251 | 0.729 | 0.187 | 0.004 | -0.051 | -0.051 | 0 | 0.09 | 0.2 | 0.07 | 0 | 0.08 | 0.05 |
| 2 | 53772561 | rs2287511 | -1.02 | 0.241 | 0.229 | 0.222 | -0.042 | -0.031 | -0.031 | 0 | 0.22 | 0.36 | 0.23 | 0.16 | 0.12 | 0.28 |
| 2 | 53773801 | rs4672481 | 0.604 | 0.628 | -1.135 | 0.539 | -0.003 | -0.039 | -0.039 | 0 | 0.29 | 0.08 | 0.29 | 0.32 | 0.29 | 0.1 |
| 2 | 53780790 | rs11681872 | 1.17 | 1.185 | -0.599 | 0.296 | -0.007 | -0.015 | -0.015 | 0 | 0.65 | 0.26 | 0.48 | 0.45 | 0.62 | 0.6 |
| 2 | 53784430 | rs1494020 | -0.303 | 1.486 | -0.316 | 0.33 | 0.001 | 0.077 | 0.077 | 0.095 | 0.08 | 0.1 | 0.12 | 0.39 | 0.13 | 0.11 |

| | | | | | | | | | | | | | | | | |
|---|---|---|---|---|---|---|---|---|---|---|---|---|---|---|---|---|
| 2 | 53785037 | rs6746107 | -0.124 | 1.417 | -0.066 | 0.076 | 0.005 | -0.021 | -0.021 | 0.623 | 0.89 | 0.86 | 0.83 | 1 | 0.84 | 0.93 |
| 2 | 53796167 | rs7566120 | -1.589 | 1.164 | 0.434 | 0.113 | -0.018 | 0.089 | 0.089 | 0.777 | 0.24 | 0.54 | 0.23 | 0.46 | 0.21 | 0.36 |
| 2 | 53800465 | rs6759929 | -0.373 | 1.442 | 0.245 | 0.005 | -0.014 | 0.012 | 0.012 | 1.833 | 0.91 | 0.92 | 0.84 | 1 | 0.86 | 0.91 |
| 2 | 53802786 | rs6739882 | -0.088 | 1.444 | 0.596 | -0.006 | 0.017 | -0.034 | -0.034 | 0.1 | 0.9 | 0.84 | 0.82 | 1 | 0.85 | 0.91 |
| 2 | 53804261 | rs10192011 | 0.077 | 1.277 | 0.376 | -0.36 | -0.005 | 0.003 | 0.003 | 0 | 0.16 | 0.15 | 0.13 | 0.02 | 0.16 | 0.11 |
| 2 | 53808311 | rs17044935 | 1.738 | 1.292 | 0.098 | -0.369 | 0.082 | 0.041 | 0.041 | 4.731 | 0.5 | 0.15 | 0.44 | 0.46 | 0.38 | 0.55 |
| 2 | 53809057 | rs12622794 | 0.35 | 1.366 | 0.287 | -0.39 | 0.036 | -0.092 | -0.092 | 1.624 | 0.74 | 0.57 | 0.67 | 0.98 | 0.67 | 0.79 |
| 2 | 53809294 | rs12623478 | 1.572 | 0.952 | -0.324 | -0.305 | 0.079 | 0.04 | 0.04 | 4.415 | 0.5 | 0.16 | 0.44 | 0.46 | 0.38 | 0.54 |
| 2 | 53810978 | rs888518 | 0.261 | 1.261 | -0.764 | -0.254 | -0.023 | 0.035 | 0.035 | 0 | 0.15 | 0.1 | 0.14 | 0.02 | 0.18 | 0.12 |
| 2 | 53812676 | rs4993917 | 0.497 | 0.43 | -0.81 | -0.081 | -0.011 | 0.079 | 0.079 | 0.011 | 0.58 | 0.75 | 0.56 | 0.83 | 0.47 | 0.61 |
| 2 | 53813500 | rs759718 | 0.153 | 0.516 | -1.169 | -0.089 | -0.062 | 0.07 | 0.07 | 0.123 | 0.63 | 0.83 | 0.73 | 0.83 | 0.56 | 0.74 |
| 2 | 53813848 | rs888520 | -0.002 | 0.528 | -1.163 | -0.082 | -0.062 | 0.06 | 0.06 | 0 | 0.64 | 0.83 | 0.73 | 0.83 | 0.55 | 0.74 |
| 2 | 53820452 | rs17044943 | 1.618 | 0.051 | -1.518 | -0.329 | 0.042 | -0.025 | -0.025 | 0 | 0.32 | 0.1 | 0.2 | 0.35 | 0.25 | 0.16 |
| 2 | 53821511 | rs11682841 | 0.746 | 0.302 | -1.655 | -0.083 | -0.018 | -0.043 | -0.043 | 0 | 0.18 | 0.08 | 0.31 | 0.37 | 0.21 | 0.26 |
| 2 | 53828410 | rs734295 | -0.042 | 0.2 | -1.675 | -0.227 | -0.065 | 0.039 | 0.039 | 0 | 0.66 | 0.82 | 0.76 | 0.8 | 0.58 | 0.75 |
| 2 | 53831467 | rs13420159 | 0.205 | -0.322 | -1.498 | -0.342 | -0.012 | 0.005 | 0.005 | 0 | 0.05 | 0.16 | 0.03 | 0.07 | 0.02 | 0.17 |
| 2 | 53834921 | rs736084 | 1.136 | -0.493 | -2.063 | -0.331 | 0.03 | -0.17 | -0.17 | 0.321 | 0.28 | 0.02 | 0.17 | 0.43 | 0.19 | 0.16 |
| 2 | 53835083 | rs736085 | 0.878 | -0.346 | -1.94 | -0.294 | 0.068 | -0.129 | -0.129 | 1.736 | 0.32 | 0.06 | 0.3 | 0.49 | 0.2 | 0.32 |
| 2 | 53837360 | rs1023600 | 0.303 | 0.319 | -0.999 | -1.037 | 0.012 | -0.095 | -0.095 | 0 | 0.69 | 0.42 | 0.78 | 0.95 | 0.67 | 0.75 |
| 2 | 53837432 | rs10490460 | -0.415 | 0.295 | -1.857 | -0.75 | -0.004 | -0.011 | -0.011 | 0.991 | 0.09 | 0.1 | 0.1 | 0 | 0.06 | 0.13 |
| 2 | 53838838 | rs1011275 | 0.359 | 0.824 | -1.852 | -0.567 | -0.046 | 0.14 | 0.14 | 0 | 0.73 | 0.9 | 0.86 | 0.97 | 0.74 | 0.82 |
| 2 | 53845919 | rs6749860 | -0.915 | 0.51 | -1.335 | -0.067 | 0 | -0.001 | -0.001 | 0 | 0.13 | 0.19 | 0.1 | 0.08 | 0.09 | 0.1 |
| 2 | 53846965 | rs6545359 | 0.126 | 0.936 | -1.482 | 0.192 | -0.011 | -0.003 | -0.003 | 0 | 0.72 | 0.78 | 0.69 | 0.73 | 0.68 | 0.61 |
| 2 | 53851844 | rs17044989 | -0.901 | 0.872 | -0.935 | 0.9 | 0 | -0.001 | -0.001 | 0 | 0.15 | 0.13 | 0.16 | 0.39 | 0.15 | 0.27 |
| 2 | 53852071 | rs17577903 | 1.153 | 0.238 | -1.236 | 0.326 | 0.049 | 0.018 | 0.018 | 0.155 | 0.31 | 0.14 | 0.16 | 0.22 | 0.19 | 0.18 |
| 2 | 53852811 | rs7571459 | -0.748 | 0.853 | -0.978 | 0.1 | -0.028 | 0.013 | 0.013 | 0.501 | 0.63 | 0.83 | 0.54 | 0.69 | 0.56 | 0.55 |
| 2 | 53853291 | rs17578049 | 0.212 | 0.901 | -0.273 | 0.087 | 0.001 | -0.003 | -0.003 | 2.054 | 0.15 | 0.18 | 0.18 | 0.1 | 0.24 | 0.17 |
| 2 | 53854488 | rs11125513 | 1.28 | 0.529 | 0.384 | 0.083 | -0.001 | 0 | 0 | 1.544 | 0.31 | 0.32 | 0.19 | 0.23 | 0.18 | 0.17 |
| 2 | 53856928 | rs741531 | 1.259 | 0.635 | 0.782 | 0.124 | -0.025 | -0.011 | -0.011 | 0 | 0.31 | 0.31 | 0.2 | 0.22 | 0.22 | 0.19 |
| 2 | 53859017 | rs6752239 | -0.927 | 0.522 | 0.484 | 0.079 | -0.044 | 0.013 | 0.013 | 0.353 | 0.64 | 0.89 | 0.56 | 0.7 | 0.58 | 0.54 |
| 2 | 53860007 | rs1468985 | 1.35 | 0.604 | 0.532 | 0.25 | 0.005 | 0.002 | 0.002 | 0 | 0.31 | 0.25 | 0.2 | 0.23 | 0.23 | 0.19 |
| 2 | 53860097 | rs10195829 | -1.145 | 0.855 | 0.861 | 0.56 | 0.023 | 0.031 | 0.031 | 0 | 0.3 | 0.51 | 0.34 | 0.46 | 0.35 | 0.35 |
| 2 | 53861427 | rs13419842 | -1.075 | 0.703 | 1.037 | 1.034 | 0.027 | 0.239 | 0.239 | 0.43 | 0.16 | 0.47 | 0.13 | 0.48 | 0.16 | 0.37 |
| 2 | 53861561 | rs6735360 | 0.326 | 0.056 | 0.196 | 0.385 | 0.027 | 0.033 | 0.033 | 0 | 0.5 | 0.39 | 0.59 | 0.19 | 0.6 | 0.38 |
| 2 | 53864008 | rs6715313 | -1.081 | 1.28 | 0.265 | 0.881 | 0.023 | -0.025 | -0.025 | 0 | 0.07 | 0.13 | 0.07 | 0 | 0.15 | 0.11 |
| 2 | 53864413 | rs12618833 | -0.12 | 0.939 | 0.534 | 1.04 | -0.02 | 0.045 | 0.045 | 0 | 0.11 | 0.22 | 0.13 | 0.16 | 0.07 | 0.09 |
| 2 | 53867187 | rs997644 | 0.126 | 0.94 | 0.46 | 1.068 | 0.001 | -0.001 | -0.001 | 0 | 0.1 | 0.06 | 0.12 | 0.16 | 0.07 | 0.09 |
| 2 | 53872721 | rs10490464 | -0.952 | 1.005 | 0.484 | 1.026 | 0.018 | -0.022 | -0.022 | 0 | 0.07 | 0.11 | 0.07 | 0 | 0.15 | 0.11 |
| 2 | 53872803 | rs10490465 | -0.265 | 0.531 | -0.211 | 1.205 | 0.021 | -0.086 | -0.086 | 0 | 0.27 | 0.14 | 0.14 | 0.45 | 0.2 | 0.41 |
| 2 | 53877608 | rs17045069 | 0.812 | 0.466 | -0.275 | 1.576 | 0.013 | -0.001 | -0.001 | 0.007 | 0.07 | 0.03 | 0.02 | 0.37 | 0.04 | 0.2 |
| 2 | 53882527 | rs2160930 | -2.153 | 1.026 | 0.336 | 1.148 | 0 | 0.003 | 0.003 | 0 | 0.51 | 0.48 | 0.36 | 0.72 | 0.51 | 0.62 |
| 2 | 53886799 | rs11125515 | -1.51 | 0.817 | 0.358 | 1.077 | -0.021 | -0.003 | -0.003 | 0 | 0.39 | 0.44 | 0.28 | 0.63 | 0.28 | 0.51 |
| 2 | 53895843 | rs17045081 | -0.332 | 0.808 | 0.172 | 1.079 | 0.026 | -0.085 | -0.085 | 0 | 0.28 | 0.15 | 0.15 | 0.45 | 0.2 | 0.42 |
| 2 | 53898032 | rs2287512 | 0.448 | 0.901 | 0.215 | 1.083 | -0.002 | -0.002 | -0.002 | 0 | 0.12 | 0.08 | 0.15 | 0.19 | 0.16 | 0.1 |
| 2 | 53903161 | rs6713107 | -1.992 | 0.684 | 0.513 | 0.967 | 0.011 | 0.02 | 0.02 | 0 | 0.5 | 0.58 | 0.34 | 0.63 | 0.42 | 0.61 |
| 2 | 53912128 | rs10490466 | -0.015 | 0.876 | 0.233 | 0.902 | -0.051 | 0.051 | 0.051 | 0 | 0.09 | 0.01 | 0.05 | 0 | 0.15 | 0.1 |
| 2 | 53929855 | rs3755120 | -1.283 | 0.755 | 0.855 | 0.95 | 0.033 | -0.028 | -0.028 | 0 | 0.24 | 0.44 | 0.24 | 0.18 | 0.28 | 0.19 |
| 2 | 53931190 | rs7583622 | -0.294 | 0.616 | 0.813 | 0.994 | 0.014 | -0.034 | -0.034 | 0 | 0.29 | 0.23 | 0.23 | 0.46 | 0.22 | 0.44 |
| 2 | 53932669 | rs7601692 | -2.11 | 0.639 | 0.759 | 0.985 | -0.016 | 0.036 | 0.036 | 0 | 0.53 | 0.75 | 0.48 | 0.64 | 0.5 | 0.64 |
| 2 | 53942367 | rs17045139 | 0.742 | 0.566 | 0.683 | 1.17 | 0.005 | 0.004 | 0.004 | 0 | 0.09 | 0.07 | 0.04 | 0.3 | 0.05 | 0.21 |
| 2 | 53945340 | rs6747749 | 1.222 | 0.315 | 0.12 | 0.607 | 0.005 | 0.007 | 0.007 | 0.191 | 0.3 | 0.25 | 0.24 | 0.26 | 0.23 | 0.28 |
| 2 | 53954123 | rs10198317 | 0.098 | 0.449 | 0.391 | 0.691 | 0.017 | -0.027 | -0.027 | 0 | 0.11 | 0.17 | 0.1 | 0.01 | 0.2 | 0.11 |
| 2 | 53955850 | rs17521008 | 0.491 | 0.326 | 0.25 | 0.75 | 0 | 0.001 | 0.001 | 0 | 0.21 | 0.11 | 0.2 | 0.16 | 0.18 | 0.24 |
| 2 | 53965524 | rs13414183 | 0.607 | 0.427 | 0.514 | 0.734 | 0.005 | 0.014 | 0.014 | 0 | 0.14 | 0.29 | 0.17 | 0.17 | 0.15 | 0.09 |
| 2 | 53968774 | rs918207 | -1.549 | 0.427 | 0.464 | 0.713 | 0.009 | 0.007 | 0.007 | 0 | 0.44 | 0.63 | 0.42 | 0.64 | 0.37 | 0.54 |
| 2 | 53984957 | rs2111623 | -2.069 | 0.447 | 0.466 | 0.68 | -0.003 | 0.03 | 0.03 | 0 | 0.54 | 0.73 | 0.48 | 0.64 | 0.51 | 0.64 |
| 2 | 53989984 | rs10490469 | 0.654 | 0.528 | 0.28 | 0.64 | -0.044 | 0.046 | 0.046 | 0 | 0.08 | 0.02 | 0.05 | 0 | 0.13 | 0.1 |
| 2 | 53991707 | rs10205590 | -1.661 | 0.418 | 0.629 | 0.719 | 0.002 | 0.003 | 0.003 | 0 | 0.41 | 0.54 | 0.33 | 0.59 | 0.39 | 0.5 |
| 2 | 53992859 | rs2287340 | 1.341 | 0.384 | 0.637 | 0.546 | -0.019 | 0.017 | 0.017 | 0.117 | 0.12 | 0.15 | 0.14 | 0.17 | 0.07 | 0.15 |
| 2 | 53993487 | rs17045199 | -1.615 | 0.347 | 0.793 | 0.577 | 0 | -0.001 | -0.001 | 0.524 | 0.38 | 0.35 | 0.3 | 0.46 | 0.35 | 0.53 |
| 2 | 53994820 | rs17045203 | 1.362 | 0.292 | 0.69 | 0.6 | -0.016 | 0.013 | 0.013 | 0.163 | 0.12 | 0.15 | 0.14 | 0.15 | 0.07 | 0.15 |
| 2 | 53995680 | rs4671583 | -1.534 | 0.361 | 0.545 | 0.679 | 0.015 | 0.007 | 0.007 | 0 | 0.44 | 0.64 | 0.42 | 0.64 | 0.34 | 0.54 |
| 2 | 54013527 | rs6545366 | -0.749 | 0.309 | 0.718 | 0.751 | -0.051 | 0.117 | 0.117 | 0 | 0.24 | 0.49 | 0.22 | 0.46 | 0.17 | 0.31 |
| 2 | 54013544 | rs2542585 | -2.018 | 0.391 | 0.496 | 0.705 | -0.011 | 0.029 | 0.029 | 0 | 0.53 | 0.7 | 0.48 | 0.63 | 0.48 | 0.64 |
| 2 | 54014558 | rs2287342 | 1.11 | 0.4 | 0.384 | 0.751 | 0.007 | -0.001 | -0.001 | 0 | 0.09 | 0.06 | 0.03 | 0.29 | 0.05 | 0.18 |
| 2 | 54024626 | rs2692514 | -1.356 | 0.427 | 0.544 | 0.649 | 0.001 | 0.001 | 0.001 | 0 | 0.41 | 0.58 | 0.35 | 0.47 | 0.4 | 0.54 |
| 2 | 54027997 | rs17522496 | 2.814 | 0.431 | 0.192 | 0.4 | -0.04 | 0.1 | 0.1 | 0.431 | 0.17 | 0.01 | 0.09 | 0 | 0.16 | 0.03 |
| 2 | 54039813 | rs2287348 | -0.82 | 0.355 | 0.718 | 0.816 | -0.031 | 0.121 | 0.121 | 0 | 0.24 | 0.49 | 0.22 | 0.46 | 0.19 | 0.29 |
| 2 | 54041983 | rs2542571 | -1.438 | 0.397 | 0.551 | 0.79 | 0.002 | 0.001 | 0.001 | 0 | 0.41 | 0.57 | 0.36 | 0.47 | 0.43 | 0.54 |

| | | | | | | | | | | | | | | | | |
|---|---|---|---|---|---|---|---|---|---|---|---|---|---|---|---|---|
| 2 | 54046646 | rs2542573 | -0.583 | 0.41 | 0.204 | 0.756 | -0.042 | 0.045 | 0.045 | 0 | 0.29 | 0.14 | 0.26 | 0.18 | 0.31 | 0.35 |
| 2 | 54050281 | rs17045292 | 1.381 | 0.393 | 0.256 | 0.845 | 0.015 | -0.043 | -0.043 | 1.111 | 0.11 | 0.01 | 0.12 | 0.18 | 0.07 | 0.1 |
| 2 | 54060151 | rs2357697 | -1.406 | 0.46 | 0.558 | 0.803 | 0.005 | 0.035 | 0.035 | 0.008 | 0.32 | 0.57 | 0.28 | 0.49 | 0.32 | 0.4 |
| 2 | 54075416 | rs805430 | -2.041 | 0.38 | 0.553 | 0.753 | -0.006 | 0.07 | 0.07 | 0 | 0.53 | 0.73 | 0.49 | 0.69 | 0.51 | 0.64 |
| 2 | 54082405 | rs1232 | 0.033 | 0.584 | 0.564 | 0.654 | -0.009 | 0.023 | 0.023 | 0.083 | 0.25 | 0.16 | 0.22 | 0.16 | 0.25 | 0.33 |
| 2 | 54086595 | rs805373 | -1.962 | 0.599 | 0.646 | 0.728 | 0.02 | 0.095 | 0.095 | 0 | 0.5 | 0.68 | 0.48 | 0.73 | 0.44 | 0.63 |
| 2 | 54088024 | rs3755113 | 1.185 | 0.542 | 0.678 | 0.779 | 0.017 | -0.016 | -0.016 | 0.097 | 0.09 | 0.05 | 0.03 | 0.32 | 0.05 | 0.17 |
| 2 | 54089857 | rs805374 | -2.192 | 0.573 | 0.772 | 0.715 | -0.005 | 0.09 | 0.09 | 0 | 0.54 | 0.74 | 0.49 | 0.73 | 0.51 | 0.64 |
| 2 | 54091688 | rs698855 | -0.763 | 0.577 | 0.799 | 0.721 | -0.001 | -0.055 | -0.055 | 0.147 | 0.34 | 0.23 | 0.26 | 0.48 | 0.31 | 0.5 |
| 2 | 54092984 | rs805379 | -2.083 | 0.573 | 0.883 | 0.701 | -0.002 | 0.028 | 0.028 | 0 | 0.53 | 0.63 | 0.48 | 0.73 | 0.5 | 0.64 |
| 2 | 54109239 | rs6713523 | -0.799 | 0.541 | 0.938 | 0.782 | 0.011 | 0.107 | 0.107 | 0 | 0.19 | 0.41 | 0.22 | 0.26 | 0.18 | 0.14 |
| 2 | 54114864 | rs2293361 | 1.255 | 0.571 | 0.673 | 0.774 | 0.021 | -0.067 | -0.067 | 0 | 0.11 | 0.01 | 0.14 | 0.25 | 0.06 | 0.11 |
| 2 | 54120497 | rs805410 | -2.018 | 0.596 | 0.77 | 0.497 | -0.007 | 0.069 | 0.069 | 0 | 0.53 | 0.72 | 0.48 | 0.73 | 0.48 | 0.64 |
| 2 | 54127041 | rs805423 | -1.986 | 0.612 | 0.734 | 0.507 | -0.005 | 0.046 | 0.046 | 0 | 0.52 | 0.64 | 0.47 | 0.73 | 0.49 | 0.65 |
| 2 | 54133744 | rs805316 | -0.039 | 0.501 | 0.571 | 0.55 | -0.03 | 0.033 | 0.033 | 0.003 | 0.26 | 0.11 | 0.22 | 0.16 | 0.27 | 0.36 |
| 2 | 54135052 | rs805318 | -1.629 | 0.596 | 0.888 | 0.276 | 0.001 | 0.02 | 0.02 | 0 | 0.51 | 0.59 | 0.47 | 0.73 | 0.46 | 0.65 |
| 2 | 54164848 | rs805360 | -1.172 | 0.45 | 0.794 | 0.284 | 0.027 | 0.127 | 0.127 | 0 | 0.46 | 0.26 | 0.4 | 0.73 | 0.38 | 0.61 |
| 2 | 54196014 | rs805309 | -1.833 | 0.605 | 0.89 | 0.261 | 0 | 0.044 | 0.044 | 0 | 0.53 | 0.64 | 0.47 | 0.74 | 0.51 | 0.65 |
| 2 | 54210781 | rs10183655 | -0.938 | 0.474 | 1.195 | 0.463 | 0.021 | 0.023 | 0.023 | 0 | 0.37 | 0.5 | 0.31 | 0.47 | 0.43 | 0.53 |
| 2 | 54211184 | rs17045452 | -0.703 | 0.524 | 1.153 | 0.453 | -0.013 | -0.039 | -0.039 | 0 | 0.35 | 0.25 | 0.26 | 0.46 | 0.35 | 0.53 |
| 2 | 54216979 | rs7569954 | -1.238 | 0.504 | 0.935 | 0.136 | -0.002 | -0.022 | -0.022 | 0 | 0.63 | 0.84 | 0.53 | 0.67 | 0.59 | 0.69 |
| 2 | 54220898 | rs6545379 | -2.421 | 0.513 | 0.448 | 0.062 | -0.081 | 0.101 | 0.101 | 4.401 | 0.75 | 0.99 | 0.69 | 0.94 | 0.68 | 0.79 |
| 2 | 54260132 | rs1421617 | -2.179 | 0.477 | 1.033 | 0.556 | 0 | -0.002 | -0.002 | 0 | 0.53 | 0.56 | 0.46 | 0.73 | 0.51 | 0.66 |
| 2 | 54268750 | rs7559384 | -0.356 | 0.458 | 0.695 | 0.69 | -0.009 | -0.078 | -0.078 | 0.165 | 0.3 | 0.16 | 0.25 | 0.46 | 0.29 | 0.5 |
| 2 | 54276502 | rs11683654 | -1.569 | 0.322 | 0.807 | 0.555 | 0.019 | 0.039 | 0.039 | 0.024 | 0.4 | 0.32 | 0.32 | 0.71 | 0.31 | 0.57 |
| 2 | 54278604 | rs7590819 | -0.472 | 0.389 | 0.841 | 0.577 | 0 | -0.008 | -0.008 | 0 | 0.27 | 0.27 | 0.25 | 0.44 | 0.26 | 0.5 |
| 2 | 54279323 | rs1833497 | -1.492 | 0.467 | 0.735 | 0.567 | 0.003 | 0.053 | 0.053 | 0 | 0.4 | 0.29 | 0.36 | 0.74 | 0.39 | 0.58 |
| 2 | 54280129 | rs1862122 | -1.61 | 0.429 | 0.863 | 0.548 | 0.006 | 0.045 | 0.045 | 0 | 0.42 | 0.33 | 0.36 | 0.74 | 0.39 | 0.59 |
| 2 | 54281762 | rs1363062 | -2.601 | 0.535 | 0.802 | 0.543 | -0.004 | 0.054 | 0.054 | 0 | 0.54 | 0.66 | 0.47 | 0.74 | 0.52 | 0.66 |
| 2 | 54282887 | rs3929001 | 0.127 | 0.548 | 0.554 | 0.627 | -0.004 | -0.059 | -0.059 | 0.67 | 0.13 | 0.03 | 0.11 | 0.27 | 0.13 | 0.08 |
| 2 | 54284441 | rs1363061 | 0.515 | 0.502 | 0.485 | 0.484 | -0.022 | 0.03 | 0.03 | 0 | 0.09 | 0.01 | 0.14 | 0.03 | 0.1 | 0.22 |
| 2 | 54285239 | rs1862121 | -1.325 | 0.504 | 0.618 | 0.524 | 0.002 | 0.058 | 0.058 | 0 | 0.4 | 0.27 | 0.36 | 0.74 | 0.39 | 0.59 |
| 2 | 54302039 | rs10865278 | -1.325 | 0.615 | 0.399 | 0.518 | -0.008 | 0.066 | 0.066 | 0 | 0.41 | 0.24 | 0.36 | 0.7 | 0.4 | 0.58 |
| 2 | 54324701 | rs10865279 | -1.297 | 0.606 | 0.355 | 0.465 | -0.013 | 0.083 | 0.083 | 0 | 0.4 | 0.2 | 0.36 | 0.75 | 0.4 | 0.58 |
| 2 | 54328837 | rs10207537 | -2.431 | 0.613 | 0.525 | 0.414 | 0.002 | 0.026 | 0.026 | 0 | 0.54 | 0.61 | 0.49 | 0.75 | 0.53 | 0.64 |
| 2 | 54333078 | rs10176342 | -0.484 | 0.628 | 0.551 | 0.411 | 0.045 | -0.099 | -0.099 | 0 | 0.14 | 0.4 | 0.12 | 0 | 0.14 | 0.06 |
| 2 | 54335433 | rs2287645 | -1.69 | 0.568 | 0.486 | 0.501 | 0.025 | 0.04 | 0.04 | 0 | 0.25 | 0.49 | 0.26 | 0.3 | 0.27 | 0.12 |
| 2 | 54335506 | rs2287643 | -0.483 | 0.525 | 0.327 | 0.547 | -0.002 | -0.077 | -0.077 | 0 | 0.3 | 0.13 | 0.23 | 0.43 | 0.25 | 0.52 |
| 2 | 54341098 | rs12617094 | 0.448 | 0.69 | 0.579 | 0.713 | 0.024 | -0.029 | -0.029 | 9.328 | 0.12 | 0.06 | 0.09 | 0.29 | 0.06 | 0.05 |
| 2 | 54341106 | rs12620359 | -1.096 | 0.692 | 0.779 | 0.696 | 0.006 | -0.01 | -0.01 | 1.879 | 0.24 | 0.18 | 0.2 | 0.29 | 0.19 | 0.17 |
| 2 | 54342955 | rs2287641 | 0.316 | 0.662 | 0.77 | 0.778 | -0.014 | 0.023 | 0.023 | 0.173 | 0.13 | 0.11 | 0.1 | 0.29 | 0.06 | 0.11 |
| 2 | 54358517 | rs10206954 | -1.246 | 0.85 | 0.865 | 0.77 | 0.002 | 0.023 | 0.023 | 0 | 0.25 | 0.36 | 0.23 | 0.31 | 0.22 | 0.17 |
| 2 | 54362975 | rs10186140 | -0.144 | 0.831 | 0.917 | 0.811 | -0.024 | 0.157 | 0.157 | 0 | 0.15 | 0.41 | 0.14 | 0.31 | 0.1 | 0.13 |
| 2 | 54366449 | rs1421619 | 0.608 | 0.784 | 0.786 | 0.818 | 0.034 | -0.042 | -0.042 | 0.059 | 0.12 | 0.05 | 0.09 | 0.29 | 0.06 | 0.12 |
| 2 | 54372118 | rs7572559 | -0.705 | 0.854 | 0.883 | 0.79 | 0.002 | -0.005 | -0.005 | 3.118 | 0.24 | 0.2 | 0.19 | 0.29 | 0.19 | 0.17 |
| 2 | 54372931 | rs10201710 | -1.15 | 0.841 | 0.877 | 0.789 | -0.006 | 0.025 | 0.025 | 0 | 0.27 | 0.51 | 0.25 | 0.31 | 0.23 | 0.18 |
| 2 | 54382335 | rs13399590 | -0.892 | 0.848 | 0.65 | 0.79 | -0.001 | 0.008 | 0.008 | 0 | 0.25 | 0.29 | 0.22 | 0.31 | 0.23 | 0.18 |
| 2 | 54384927 | rs6545388 | -1.953 | 0.801 | 0.531 | 0.603 | 0.008 | 0.056 | 0.056 | 0 | 0.53 | 0.65 | 0.49 | 0.73 | 0.53 | 0.64 |
| 2 | 54393570 | rs6744662 | 0.012 | 0.824 | 0.269 | 1.164 | 0.003 | 0.044 | 0.044 | 0 | 0.16 | 0.18 | 0.16 | 0.41 | 0.19 | 0.15 |
| 2 | 54402934 | rs1215153 | -0.692 | 0.792 | -0.35 | 1.041 | -0.039 | -0.066 | -0.066 | 0 | 0.29 | 0.06 | 0.25 | 0.32 | 0.25 | 0.45 |
| 2 | 54409855 | rs843677 | -2.077 | 0.593 | -0.227 | 0.911 | -0.002 | 0.074 | 0.074 | 0 | 0.49 | 0.67 | 0.48 | 0.72 | 0.46 | 0.58 |
| 2 | 54412539 | rs3850348 | 0.808 | 0.674 | -0.446 | 1.282 | -0.012 | -0.064 | -0.064 | 0.183 | 0.1 | 0.01 | 0.09 | 0.25 | 0.09 | 0.12 |
| 2 | 54420617 | rs13417493 | 0.084 | 0.637 | -0.243 | 1.088 | -0.02 | -0.035 | -0.035 | 0 | 0.21 | 0.13 | 0.22 | 0.38 | 0.27 | 0.19 |
| 2 | 54420839 | rs843671 | -1.839 | 0.495 | -0.053 | 0.883 | 0.007 | 0.04 | 0.04 | 0 | 0.49 | 0.62 | 0.41 | 0.7 | 0.5 | 0.6 |
| 2 | 54424537 | rs1385274 | -0.703 | 0.437 | 0.004 | 0.623 | 0.009 | 0.016 | 0.016 | 0 | 0.31 | 0.24 | 0.2 | 0.3 | 0.27 | 0.46 |
| 2 | 54442697 | rs11894205 | 0.318 | 0.542 | -0.016 | 0.776 | -0.024 | -0.049 | -0.049 | 0.103 | 0.22 | 0.14 | 0.22 | 0.39 | 0.27 | 0.24 |
| 2 | 54448555 | rs843755 | -0.565 | 0.487 | 0.208 | 0.658 | 0 | 0.007 | 0.007 | 0 | 0.28 | 0.21 | 0.19 | 0.3 | 0.26 | 0.39 |
| 2 | 54469283 | rs843653 | -1.672 | 0.561 | 0.545 | 0.809 | -0.003 | -0.006 | -0.006 | 0.001 | 0.39 | 0.67 | 0.32 | 0.46 | 0.38 | 0.52 |
| 2 | 54476678 | rs17045711 | 0.772 | 0.627 | 0.29 | 0.872 | 0.038 | 0.065 | 0.065 | 0.171 | 0.08 | 0.18 | 0.1 | 0.17 | 0.11 | 0.09 |
| 2 | 54479117 | rs843711 | -1.506 | 0.572 | 0.379 | 0.851 | -0.006 | -0.011 | -0.011 | 0 | 0.38 | 0.4 | 0.3 | 0.48 | 0.38 | 0.59 |
| 2 | 54480369 | rs843706 | -1.502 | 0.58 | 0.466 | 0.854 | 0 | 0.001 | 0.001 | 0 | 0.38 | 0.43 | 0.3 | 0.48 | 0.37 | 0.59 |
| 2 | 54481636 | rs10165485 | 0.772 | 0.642 | 0.271 | 0.851 | 0.036 | 0.058 | 0.058 | 0.171 | 0.08 | 0.17 | 0.1 | 0.16 | 0.11 | 0.09 |
| 2 | 54482964 | rs843704 | -0.585 | 0.624 | 0.076 | 0.837 | 0 | 0.008 | 0.008 | 0 | 0.28 | 0.22 | 0.19 | 0.3 | 0.26 | 0.38 |
| 2 | 54490026 | rs843688 | -1.459 | 0.614 | 0.081 | 0.848 | 0.005 | 0.008 | 0.008 | 0 | 0.38 | 0.47 | 0.31 | 0.46 | 0.38 | 0.59 |
| 2 | 54493408 | rs10171249 | 0.552 | 0.687 | 0.072 | 1.004 | 0.06 | 0.105 | 0.105 | 0.003 | 0.09 | 0.19 | 0.12 | 0.17 | 0.11 | 0.13 |
| 2 | 54498212 | rs12465259 | 0.778 | 0.206 | -0.499 | 0.991 | -0.001 | -0.06 | -0.06 | 0.015 | 0.1 | 0.01 | 0.06 | 0.25 | 0.08 | 0.22 |
| 2 | 54501025 | rs843743 | -0.034 | 0.081 | -0.574 | 0.814 | 0.01 | -0.025 | -0.025 | 0 | 0.73 | 0.89 | 0.86 | 0.67 | 0.79 | 0.55 |
| 2 | 54502912 | rs843748 | 0.961 | 0.162 | -0.849 | 0.212 | -0.11 | 0.011 | 0.011 | 0.535 | 0.29 | 0.07 | 0.39 | 0.2 | 0.45 | 0.29 |
| 2 | 54506070 | rs843713 | -0.525 | -0.116 | -0.589 | 0.35 | 0.016 | -0.034 | -0.034 | 0 | 0.62 | 0.73 | 0.74 | 0.48 | 0.65 | 0.52 |

| | | | | | | | | | | | | | | | | |
|---|---|---|---|---|---|---|---|---|---|---|---|---|---|---|---|---|
| 2 | 54508110 | rs843718 | 0.516 | -0.316 | 0.118 | -0.089 | 0 | -0.001 | -0.001 | 0 | 0.37 | 0.39 | 0.53 | 0.32 | 0.38 | 0.36 |
| 2 | 54509549 | rs843719 | -0.271 | -0.327 | -0.252 | -0.13 | -0.007 | 0.061 | 0.061 | 0 | 0.52 | 0.81 | 0.61 | 0.31 | 0.47 | 0.39 |
| 2 | 54513742 | rs843726 | 1.737 | 0.079 | 0.031 | -0.479 | -0.015 | 0.021 | 0.021 | 0.574 | 0.22 | 0.16 | 0.28 | 0.03 | 0.31 | 0.11 |
| 2 | 54527866 | rs964092 | 0.865 | -0.02 | 0.655 | -0.254 | -0.026 | -0.081 | -0.081 | 0.09 | 0.15 | 0.29 | 0.09 | 0 | 0.08 | 0.02 |
| 2 | 54530966 | rs13407896 | 0.839 | -0.013 | 0.711 | -0.251 | -0.029 | -0.098 | -0.098 | 0 | 0.15 | 0.34 | 0.09 | 0 | 0.09 | 0.02 |
| 2 | 54536882 | rs6724542 | 0.592 | 0.132 | 0.494 | -0.18 | -0.005 | -0.027 | -0.027 | 0 | 0.16 | 0.22 | 0.1 | 0.07 | 0.12 | 0.14 |
| 2 | 54546625 | rs985773 | 0.566 | 0.678 | 0.871 | -0.011 | 0.092 | 0.042 | 0.042 | 0 | 0.11 | 0.28 | 0.15 | 0.14 | 0.21 | 0.15 |
| 2 | 54548482 | rs843622 | 0.284 | 0.246 | 0.681 | -0.254 | -0.001 | -0.001 | -0.001 | 0.663 | 0.39 | 0.62 | 0.47 | 0.19 | 0.53 | 0.37 |
| 2 | 54556550 | rs1682139 | 0.503 | 0.136 | 0.926 | -0.099 | 0.005 | 0.021 | 0.021 | 0.164 | 0.45 | 0.58 | 0.47 | 0.19 | 0.57 | 0.37 |
| 2 | 54557481 | rs1727529 | 1.555 | 0.253 | 0.756 | -0.159 | -0.056 | 0.154 | 0.154 | 0.817 | 0.32 | 0.06 | 0.3 | 0.05 | 0.36 | 0.21 |
| 2 | 54559793 | rs843669 | 0.99 | 0.087 | 0.876 | 0.137 | 0.001 | 0.033 | 0.033 | 0.16 | 0.53 | 0.39 | 0.49 | 0.26 | 0.54 | 0.31 |
| 2 | 54567590 | rs12713256 | 0.344 | -0.02 | 0.692 | 0.302 | 0.019 | -0.003 | -0.003 | 0 | 0.63 | 0.68 | 0.66 | 0.38 | 0.75 | 0.46 |
| 2 | 54573489 | rs941351 | 0.062 | 0.009 | 0.579 | 0.138 | 0.019 | 0.006 | 0.006 | 0 | 0.68 | 0.73 | 0.75 | 0.65 | 0.79 | 0.59 |
| 2 | 54574825 | rs11690242 | 0.802 | 0.078 | 0.56 | 0.131 | -0.002 | 0 | 0 | 0 | 0.58 | 0.46 | 0.59 | 0.52 | 0.58 | 0.45 |
| 2 | 54582446 | rs13385514 | 0.756 | 0.096 | 0.008 | 0.091 | 0 | -0.004 | -0.004 | 0.019 | 0.25 | 0.22 | 0.27 | 0.48 | 0.22 | 0.25 |
| 2 | 54583417 | rs13023335 | 1.582 | 0.214 | -0.872 | 0.131 | -0.039 | -0.04 | -0.04 | 0.656 | 0.07 | 0.01 | 0.14 | 0.13 | 0.2 | 0.1 |
| 2 | 54583542 | rs17189869 | 1.621 | 0.056 | -0.814 | 0.016 | -0.027 | 0.123 | 0.123 | 0.998 | 0.33 | 0.09 | 0.31 | 0.05 | 0.37 | 0.2 |
| 2 | 54583590 | rs17189876 | 0.665 | -0.043 | -0.856 | -0.024 | 0.002 | 0.162 | 0.162 | 0 | 0.47 | 0.2 | 0.47 | 0.11 | 0.45 | 0.32 |
| 2 | 54587596 | rs2280718 | 0.66 | -0.221 | -0.489 | 0.419 | -0.009 | -0.002 | -0.002 | 0 | 0.68 | 0.64 | 0.75 | 0.64 | 0.79 | 0.58 |
| 2 | 54599272 | rs4488702 | -0.183 | 0.594 | -0.929 | 0.382 | 0.01 | 0.015 | 0.015 | 0 | 0.14 | 0.1 | 0.16 | 0.07 | 0.09 | 0.11 |
| 2 | 54600556 | rs11904342 | -0.091 | 1.238 | -0.893 | 0.647 | 0.012 | 0 | 0 | 0 | 0.09 | 0.1 | 0.16 | 0.13 | 0.21 | 0.14 |
| 2 | 54608005 | rs6756317 | -0.614 | 0.211 | -1.618 | 0.457 | 0.009 | -0.074 | -0.074 | 0 | 0.53 | 0.96 | 0.51 | 0.57 | 0.4 | 0.53 |
| 2 | 54609774 | rs13408182 | -0.051 | -0.23 | -0.767 | 0.278 | -0.01 | -0.077 | -0.077 | 0 | 0.13 | 0.28 | 0.09 | 0 | 0.1 | 0.05 |
| 2 | 54610789 | rs4131262 | 0.05 | -0.271 | -0.408 | 0.252 | -0.038 | 0.019 | 0.019 | 0 | 0.36 | 0.52 | 0.31 | 0.57 | 0.24 | 0.45 |
| 2 | 54614309 | rs10198244 | -0.019 | -0.263 | -0.548 | 0.268 | -0.017 | 0.006 | 0.006 | 0 | 0.36 | 0.61 | 0.32 | 0.57 | 0.24 | 0.44 |
| 2 | 54615526 | rs6545397 | -0.675 | -0.108 | -1.004 | 0.331 | 0.095 | -0.048 | -0.048 | 0 | 0.5 | 0.96 | 0.41 | 0.57 | 0.35 | 0.5 |
| 2 | 54618361 | rs17045857 | -0.404 | -0.072 | -0.057 | 0.08 | -0.007 | -0.085 | -0.085 | 0 | 0.14 | 0.42 | 0.1 | 0 | 0.11 | 0.06 |
| 2 | 54622582 | rs7601198 | -1.343 | 0.251 | -0.509 | -0.115 | -0.027 | 0.114 | 0.114 | 0 | 0.68 | 0.88 | 0.67 | 0.92 | 0.67 | 0.84 |
| 2 | 54629081 | rs4671925 | -0.334 | 0.187 | -0.152 | -0.202 | 0.001 | -0.051 | -0.051 | 0 | 0.54 | 0.44 | 0.57 | 0.97 | 0.57 | 0.78 |
| 2 | 54630902 | rs4671212 | -0.562 | 0.21 | 0.345 | 0.44 | 0 | -0.003 | -0.003 | 0 | 0.47 | 0.51 | 0.43 | 0.59 | 0.49 | 0.61 |
| 2 | 54631603 | rs6752816 | 0.334 | 0.054 | -0.003 | 0.428 | -0.014 | 0.052 | 0.052 | 0 | 0.09 | 0.25 | 0.12 | 0.16 | 0.08 | 0.25 |
| 2 | 54634288 | rs7592359 | -1.323 | 0.009 | -0.131 | 0.337 | -0.014 | 0.134 | 0.134 | 0 | 0.66 | 0.85 | 0.68 | 0.97 | 0.67 | 0.83 |
| 2 | 54644991 | rs13393949 | 0.329 | 0.252 | 0.009 | 0.934 | 0.001 | -0.002 | -0.002 | 0 | 0.13 | 0.11 | 0.23 | 0.19 | 0.26 | 0.14 |
| 2 | 54645072 | rs7560205 | -0.374 | 0.161 | 0.258 | 0.904 | 0.026 | 0.009 | 0.009 | 0 | 0.2 | 0.34 | 0.27 | 0.2 | 0.3 | 0.23 |
| 2 | 54646852 | rs7565034 | 0.887 | 0.028 | 0.167 | 1.003 | 0.001 | 0 | 0 | 0 | 0.2 | 0.16 | 0.36 | 0.41 | 0.18 | 0.3 |
| 2 | 54647455 | rs17045879 | 1.274 | 0.019 | 0.242 | 0.847 | -0.024 | -0.026 | -0.026 | 0 | 0.06 | 0.17 | 0.03 | 0.01 | 0.04 | 0.1 |
| 2 | 54651933 | rs6734097 | -1.711 | -0.049 | 0.042 | 0.728 | -0.046 | 0.157 | 0.157 | 0 | 0.68 | 0.9 | 0.8 | 0.97 | 0.65 | 0.83 |
| 2 | 54653111 | rs11901520 | 0.865 | -0.067 | -0.001 | 0.762 | -0.002 | 0 | 0 | 0 | 0.26 | 0.26 | 0.16 | 0.3 | 0.17 | 0.32 |
| 2 | 54653561 | rs4393791 | -0.096 | -0.058 | -0.018 | 0.842 | -0.006 | -0.007 | -0.007 | 3.083 | 0.33 | 0.27 | 0.6 | 0.51 | 0.44 | 0.29 |
| 2 | 54660456 | rs4627617 | 1.018 | -0.119 | -0.004 | 0.688 | 0.002 | -0.006 | -0.006 | 0 | 0.2 | 0.15 | 0.36 | 0.37 | 0.18 | 0.19 |
| 2 | 54660542 | rs7597093 | -0.193 | 0.001 | -0.24 | 0.447 | 0.012 | 0.005 | 0.005 | 0 | 0.35 | 0.39 | 0.61 | 0.61 | 0.44 | 0.41 |
| 2 | 54660795 | rs17045898 | 0.824 | -0.045 | -0.338 | 0.442 | -0.003 | 0.001 | 0.001 | 0 | 0.26 | 0.28 | 0.16 | 0.32 | 0.17 | 0.32 |
| 2 | 54661569 | rs6752877 | -1.486 | 0.346 | -1.06 | 0.571 | -0.082 | 0.188 | 0.188 | 0 | 0.68 | 0.95 | 0.8 | 0.97 | 0.64 | 0.83 |
| 2 | 54663715 | rs4671931 | 0.332 | 0.698 | -0.512 | 1.209 | -0.013 | 0.057 | 0.057 | 0.051 | 0.23 | 0.3 | 0.36 | 0.47 | 0.18 | 0.33 |
| 2 | 54668200 | rs4505569 | 0.485 | 0.812 | -0.851 | 1.196 | -0.074 | -0.031 | -0.031 | 0 | 0.12 | 0.01 | 0.22 | 0.13 | 0.24 | 0.09 |
| 2 | 54668247 | rs4306748 | -0.211 | 0.591 | -0.578 | 1.064 | 0.005 | 0.001 | 0.001 | 0 | 0.36 | 0.42 | 0.6 | 0.61 | 0.43 | 0.41 |
| 2 | 54684557 | rs11898505 | -0.706 | 0.746 | -1.69 | 0.387 | -0.132 | 0.116 | 0.116 | 0 | 0.69 | 0.98 | 0.79 | 0.87 | 0.65 | 0.84 |
| 2 | 54690542 | rs7592594 | -0.621 | 0.836 | -1.048 | 0.417 | 0.022 | -0.006 | -0.006 | 0 | 0.36 | 0.54 | 0.44 | 0.64 | 0.45 | 0.49 |
| 2 | 54693490 | rs6755780 | -0.448 | 0.633 | -0.489 | 0.262 | -0.011 | -0.043 | -0.043 | 0.1 | 0.23 | 0.34 | 0.27 | 0.11 | 0.2 | 0.16 |
| 2 | 54703979 | rs6718276 | 0.018 | 0.505 | -0.404 | 0.46 | -0.019 | -0.032 | -0.032 | 0 | 0.2 | 0.11 | 0.3 | 0.39 | 0.29 | 0.29 |
| 2 | 54706092 | rs4129261 | -0.522 | 0.454 | 0.209 | 0.432 | -0.02 | -0.011 | -0.011 | 0.199 | 0.24 | 0.6 | 0.29 | 0.23 | 0.19 | 0.24 |
| 2 | 54708408 | rs10171331 | 0.228 | 0.517 | 0.155 | 0.46 | -0.032 | -0.057 | -0.057 | 0.079 | 0.19 | 0.09 | 0.31 | 0.4 | 0.29 | 0.29 |
| 2 | 54714119 | rs17343815 | 0.904 | 0.58 | -0.073 | 0.445 | -0.028 | 0.018 | 0.018 | 0 | 0.13 | 0.05 | 0.09 | 0.11 | 0.17 | 0.12 |
| 2 | 54720903 | rs7605221 | -1.301 | 0.38 | -0.218 | 0.419 | -0.051 | 0.047 | 0.047 | 0.361 | 0.46 | 0.86 | 0.64 | 0.62 | 0.5 | 0.6 |
| 2 | 54723946 | rs17045932 | -1.288 | 0.195 | 0.312 | 0.456 | -0.001 | 0.065 | 0.065 | 0.891 | 0.11 | 0.5 | 0.05 | 0.13 | 0.03 | 0.17 |
| 2 | 54728082 | rs6545415 | -0.693 | 0.002 | -1.06 | 0.095 | -0.023 | -0.021 | -0.021 | 1.051 | 0.66 | 0.91 | 0.77 | 0.69 | 0.74 | 0.63 |
| 2 | 54738233 | rs17045939 | 1.116 | -0.008 | -0.611 | -0.054 | -0.009 | 0.017 | 0.017 | 0 | 0.15 | 0.1 | 0.1 | 0.12 | 0.17 | 0.17 |
| 2 | 54744093 | rs6545420 | -0.951 | 0.092 | -0.803 | 0.008 | -0.097 | -0.104 | -0.104 | 0.02 | 0.69 | 0.96 | 0.67 | 0.63 | 0.71 | 0.68 |
| 2 | 54746709 | rs6545423 | -0.214 | 0.126 | -1.01 | -0.274 | 0.056 | 0.246 | 0.246 | 0.16 | 0.5 | 0.9 | 0.4 | 0.22 | 0.39 | 0.4 |
| 2 | 54753991 | rs4671224 | 1.013 | 0.209 | -0.947 | -0.15 | -0.068 | -0.11 | -0.11 | 0.872 | 0.2 | 0.07 | 0.38 | 0.45 | 0.36 | 0.31 |
| 2 | 54755229 | rs3796022 | 1.177 | 0.146 | -0.914 | -0.214 | 0.004 | 0.032 | 0.032 | 0.442 | 0.19 | 0.1 | 0.09 | 0.11 | 0.17 | 0.17 |
| 2 | 54760038 | rs4671952 | 0.664 | 0.139 | -0.928 | -0.211 | 0.011 | 0.036 | 0.036 | 0 | 0.22 | 0.25 | 0.48 | 0.45 | 0.37 | 0.31 |
| 2 | 54766158 | rs11904679 | 1.358 | 0.201 | -1.264 | -0.138 | -0.019 | 0.052 | 0.052 | 0.904 | 0.17 | 0.04 | 0.08 | 0.09 | 0.17 | 0.08 |
| 2 | 54771441 | rs4671225 | -1.097 | 0.265 | -1.221 | -0.055 | -0.087 | -0.095 | -0.095 | 0 | 0.7 | 0.97 | 0.77 | 0.67 | 0.74 | 0.7 |
| 2 | 54798596 | rs4671961 | -0.945 | 0.186 | -1.007 | 0.096 | -0.046 | -0.095 | -0.095 | 0.035 | 0.68 | 0.91 | 0.64 | 0.45 | 0.71 | 0.61 |
| 2 | 54801856 | rs7599241 | 0.166 | 0.046 | -0.311 | -0.15 | -0.007 | -0.059 | -0.059 | 0 | 0.26 | 0.62 | 0.26 | 0.11 | 0.21 | 0.24 |
| 2 | 54817683 | rs6545432 | 0.354 | -0.04 | -0.213 | -0.084 | -0.033 | -0.031 | -0.031 | 0.951 | 0.41 | 0.19 | 0.5 | 0.42 | 0.53 | 0.45 |
| 2 | 54823310 | rs6739099 | 1.027 | -0.036 | -0.386 | -0.162 | -0.087 | -0.095 | -0.095 | 0.003 | 0.2 | 0.03 | 0.36 | 0.3 | 0.36 | 0.32 |
| 2 | 54824146 | rs6749802 | 0.165 | -0.08 | -0.202 | -0.176 | 0.002 | -0.004 | -0.004 | 0 | 0.27 | 0.72 | 0.27 | 0.11 | 0.22 | 0.24 |

| | | | | | | | | | | | | | | | | |
|---|---|---|---|---|---|---|---|---|---|---|---|---|---|---|---|---|
| 2 | 54830671 | rs4358162 | 0.918 | -0.106 | 0.133 | -0.19 | -0.01 | -0.081 | -0.081 | 0 | 0.22 | 0.43 | 0.12 | 0.09 | 0.17 | 0.08 |
| 2 | 54831343 | rs7598568 | -1.59 | -0.24 | -0.579 | -0.31 | -0.067 | -0.155 | -0.155 | 0 | 0.75 | 0.97 | 0.78 | 0.46 | 0.79 | 0.68 |
| 2 | 54846400 | rs2941592 | 1.702 | -0.325 | -0.315 | -0.314 | 0.021 | 0.033 | 0.033 | 0 | 0.22 | 0.13 | 0.11 | 0.12 | 0.17 | 0.15 |
| 2 | 54847934 | rs6715538 | -0.422 | -0.259 | -0.232 | -0.35 | -0.004 | 0.002 | 0.002 | 0.141 | 0.44 | 0.6 | 0.62 | 0.4 | 0.56 | 0.46 |
| 2 | 54851300 | rs2971886 | 0.081 | -0.268 | -0.104 | -0.326 | 0.001 | -0.004 | -0.004 | 0.077 | 0.45 | 0.51 | 0.6 | 0.35 | 0.56 | 0.51 |
| 2 | 54858664 | rs2229503 | 1.371 | -0.275 | -0.197 | -0.371 | 0 | 0.038 | 0.038 | 1.109 | 0.2 | 0.13 | 0.1 | 0.08 | 0.17 | 0.08 |
| 2 | 54861825 | rs2941579 | 1.537 | -0.599 | -0.305 | -0.601 | 0.008 | -0.06 | -0.06 | 0.113 | 0.12 | 0.24 | 0.18 | 0.01 | 0.13 | 0.05 |
| 2 | 54863861 | rs2941581 | 0.83 | -0.62 | -0.352 | -0.607 | -0.003 | -0.012 | -0.012 | 0 | 0.23 | 0.23 | 0.22 | 0.13 | 0.2 | 0.18 |
| 2 | 54865527 | rs1510611 | -0.18 | -0.549 | -0.378 | -0.613 | 0.002 | -0.002 | -0.002 | 0 | 0.53 | 0.43 | 0.51 | 0.5 | 0.59 | 0.63 |
| 2 | 54868483 | rs2941583 | -1.425 | -0.554 | -0.536 | -0.565 | -0.082 | -0.178 | -0.178 | 0.012 | 0.69 | 0.97 | 0.72 | 0.52 | 0.73 | 0.7 |
| 2 | 54875799 | rs2941594 | 2.02 | -0.593 | -0.379 | -0.568 | 0.001 | -0.007 | -0.007 | 0.285 | 0.1 | 0.09 | 0.07 | 0 | 0.1 | 0.03 |
| 2 | 54877365 | rs13401431 | 2.089 | -0.616 | -0.302 | -0.555 | -0.016 | -0.025 | -0.025 | 0 | 0.13 | 0.16 | 0.06 | 0.06 | 0.07 | 0.12 |
| 2 | 54881621 | rs2941584 | -0.631 | -0.619 | -0.362 | -0.556 | -0.006 | -0.004 | -0.004 | 0 | 0.54 | 0.52 | 0.68 | 0.4 | 0.68 | 0.58 |
| 2 | 54884870 | rs966003 | -0.493 | -0.571 | -0.641 | -0.58 | 0.043 | 0.199 | 0.199 | 0.013 | 0.48 | 0.82 | 0.38 | 0.22 | 0.39 | 0.34 |
| 2 | 54886347 | rs1052820 | 1.354 | -0.45 | -0.616 | -0.539 | -0.017 | -0.021 | -0.021 | 0.099 | 0.2 | 0.15 | 0.36 | 0.29 | 0.35 | 0.31 |
| 2 | 54888778 | rs2971878 | 1.052 | -0.481 | -0.624 | -0.516 | 0.01 | 0.028 | 0.028 | 0 | 0.23 | 0.15 | 0.21 | 0.15 | 0.2 | 0.17 |
| 2 | 54889015 | rs1047465 | 1.227 | -0.519 | -0.634 | -0.505 | 0.019 | 0.033 | 0.033 | 0 | 0.22 | 0.12 | 0.11 | 0.13 | 0.17 | 0.15 |
| 2 | 54889149 | rs10528 | 1.544 | -0.522 | -0.643 | -0.468 | -0.029 | 0.08 | 0.08 | 1.051 | 0.2 | 0.03 | 0.09 | 0.08 | 0.17 | 0.08 |
| 2 | 54889390 | rs1047499 | -1.391 | -0.52 | -0.574 | -0.361 | -0.075 | -0.181 | -0.181 | 0.31 | 0.7 | 0.97 | 0.74 | 0.52 | 0.75 | 0.71 |
| 2 | 54894043 | rs907135 | -0.87 | -0.461 | 0.085 | -0.416 | -0.004 | 0.046 | 0.046 | 0 | 0.66 | 0.52 | 0.71 | 0.53 | 0.74 | 0.72 |
| 2 | 54898633 | rs3850350 | 1.422 | -0.466 | 0.068 | -0.575 | 0.002 | -0.067 | -0.067 | 0.33 | 0.2 | 0.3 | 0.12 | 0.09 | 0.19 | 0.17 |
| 2 | 54903139 | rs2941588 | -0.223 | -0.433 | 0.12 | -0.501 | 0.002 | 0 | 0 | 0 | 0.44 | 0.4 | 0.58 | 0.41 | 0.49 | 0.42 |
| 2 | 54904463 | rs2971884 | 2.269 | -0.436 | 0.068 | -0.537 | -0.003 | -0.022 | -0.022 | 0.115 | 0.09 | 0.1 | 0.04 | 0 | 0.06 | 0.02 |
| 2 | 54904592 | rs2007020 | -0.292 | -0.388 | -0.073 | -0.564 | 0 | -0.001 | -0.001 | 0 | 0.34 | 0.58 | 0.57 | 0.28 | 0.41 | 0.35 |
| 2 | 54905778 | rs2941589 | -2.456 | -0.144 | -1.035 | -0.532 | -0.037 | 0.052 | 0.052 | 0 | 0.82 | 1 | 0.9 | 0.94 | 0.86 | 0.88 |
| 2 | 54913312 | rs17046153 | 1.255 | -0.101 | -0.716 | -0.386 | -0.065 | -0.101 | -0.101 | 0 | 0.19 | 0.03 | 0.38 | 0.3 | 0.32 | 0.25 |
| 2 | 54914459 | rs2941577 | 2.159 | -0.109 | -0.581 | -0.476 | -0.001 | -0.02 | -0.02 | 0 | 0.1 | 0.11 | 0.07 | 0 | 0.1 | 0.03 |
| 2 | 54915044 | rs6717369 | -0.057 | -0.199 | -0.41 | -0.481 | -0.001 | -0.003 | -0.003 | 0.069 | 0.36 | 0.36 | 0.56 | 0.42 | 0.43 | 0.47 |
| 2 | 54916332 | rs7560775 | 1.114 | -0.161 | -0.3 | -0.738 | -0.032 | -0.102 | -0.102 | 0 | 0.18 | 0.4 | 0.19 | 0.05 | 0.11 | 0.21 |
| 2 | 54916910 | rs354232 | 0.486 | -0.195 | -0.45 | -0.82 | -0.002 | -0.12 | -0.12 | 0 | 0.3 | 0.49 | 0.19 | 0.08 | 0.3 | 0.18 |
| 2 | 54924896 | rs171350 | -0.952 | -0.245 | -0.938 | -0.717 | 0.002 | -0.031 | -0.031 | 0 | 0.64 | 0.8 | 0.74 | 0.42 | 0.73 | 0.62 |
| 2 | 54932516 | rs354211 | -0.364 | -0.398 | -1.015 | -0.588 | 0.037 | 0.185 | 0.185 | 0 | 0.46 | 0.91 | 0.3 | 0.22 | 0.37 | 0.35 |
| 2 | 54939411 | rs13404168 | 1.778 | -0.393 | -1.109 | -0.513 | 0.005 | -0.001 | -0.001 | 0 | 0.14 | 0.16 | 0.07 | 0.05 | 0.08 | 0.13 |
| 2 | 54941951 | rs2567842 | 0.046 | -0.467 | -1.305 | -0.659 | -0.02 | 0.052 | 0.052 | 0.058 | 0.39 | 0.21 | 0.38 | 0.23 | 0.49 | 0.37 |
| 2 | 54942538 | rs2666036 | 0.192 | -0.483 | -1.453 | -0.637 | -0.044 | -0.006 | -0.006 | 0 | 0.38 | 0.13 | 0.37 | 0.31 | 0.48 | 0.37 |
| 2 | 54943102 | rs2631834 | -1.239 | -0.381 | -1.542 | -0.606 | -0.098 | -0.133 | -0.133 | 0.007 | 0.66 | 0.97 | 0.67 | 0.39 | 0.7 | 0.6 |
| 2 | 54954824 | rs354239 | 0.564 | 0.143 | -1.114 | -0.283 | -0.048 | -0.002 | -0.002 | 0 | 0.24 | 0.12 | 0.31 | 0.18 | 0.4 | 0.24 |
| 2 | 54960680 | rs13428703 | 1.024 | 0.101 | -0.51 | -0.194 | -0.001 | 0 | 0 | 0 | 0.19 | 0.19 | 0.11 | 0.17 | 0.19 | 0.14 |
| 2 | 54962352 | rs6752720 | 1.021 | 0.11 | -0.382 | -0.189 | 0.003 | -0.004 | -0.004 | 0.88 | 0.19 | 0.23 | 0.12 | 0.17 | 0.19 | 0.14 |
| 2 | 54971385 | rs354204 | -0.473 | 0.624 | -0.318 | 0.033 | 0.016 | 0.007 | 0.007 | 0 | 0.89 | 0.83 | 0.68 | 0.91 | 0.85 | 0.83 |
| 2 | 54971630 | rs1510613 | -0.411 | 0.554 | -0.59 | -0.243 | 0.016 | 0.023 | 0.023 | 0 | 0.52 | 0.75 | 0.61 | 0.39 | 0.49 | 0.52 |
| 2 | 54972161 | rs7565441 | 0.744 | 0.338 | -0.028 | -0.08 | -0.01 | 0.014 | 0.014 | 0 | 0.17 | 0.3 | 0.19 | 0.19 | 0.19 | 0.21 |
| 2 | 54972476 | rs354205 | 1.761 | 0.554 | -0.666 | -0.295 | -0.036 | 0.068 | 0.068 | 0.287 | 0.17 | 0.04 | 0.19 | 0.04 | 0.18 | 0.14 |
| 2 | 54972648 | rs13401661 | 0.282 | 0.249 | -0.175 | -0.19 | -0.03 | 0.006 | 0.006 | 0 | 0.21 | 0.6 | 0.21 | 0.18 | 0.19 | 0.31 |
| 2 | 54979666 | rs4671972 | 0.36 | 0.429 | -0.319 | -0.239 | 0.006 | 0.014 | 0.014 | 0 | 0.49 | 0.42 | 0.39 | 0.32 | 0.45 | 0.43 |
| 2 | 54987940 | rs11891165 | 1.892 | 0.338 | -0.26 | 0.454 | -0.031 | 0.132 | 0.132 | 0.098 | 0.11 | 0.27 | 0.06 | 0.33 | 0.05 | 0.19 |
| 2 | 54991712 | rs4671975 | -0.122 | 0.243 | -0.307 | 0.405 | -0.005 | 0.004 | 0.004 | 0 | 0.31 | 0.65 | 0.25 | 0.51 | 0.23 | 0.37 |
| 2 | 54995746 | rs2116441 | 0.744 | 0.253 | -0.382 | 0.283 | 0.01 | 0.024 | 0.024 | 0 | 0.46 | 0.38 | 0.33 | 0.32 | 0.44 | 0.39 |
| 2 | 55001405 | rs2116439 | 1.364 | 0.192 | -0.687 | 0.181 | -0.034 | 0.164 | 0.164 | 0 | 0.36 | 0.06 | 0.26 | 0.13 | 0.34 | 0.23 |
| 2 | 55003244 | rs974916 | -0.32 | 0.728 | -0.52 | 0.287 | -0.037 | 0.011 | 0.011 | 0 | 0.62 | 0.82 | 0.48 | 0.64 | 0.54 | 0.6 |
| 2 | 55005649 | rs2033415 | 2.3 | 0.646 | -0.52 | 0.289 | -0.006 | -0.02 | -0.02 | 0.256 | 0.08 | 0.05 | 0.1 | 0.17 | 0.11 | 0.17 |
| 2 | 55007205 | rs2116436 | 0.284 | 0.907 | -0.711 | 0.149 | -0.004 | 0.065 | 0.065 | 0.139 | 0.35 | 0.21 | 0.46 | 0.11 | 0.36 | 0.25 |
| 2 | 55010115 | rs13033894 | 2.078 | 0.731 | -0.515 | 0.236 | 0.002 | 0.089 | 0.089 | 0 | 0.29 | 0.12 | 0.2 | 0.13 | 0.26 | 0.22 |
| 2 | 55010193 | rs6705079 | -0.081 | 0.788 | -0.534 | 0.273 | -0.032 | 0.015 | 0.015 | 0 | 0.61 | 0.79 | 0.46 | 0.63 | 0.5 | 0.56 |
| 2 | 55012202 | rs6713115 | 2.035 | 0.724 | -0.201 | 0.394 | -0.029 | 0.215 | 0.215 | 0.22 | 0.12 | 0.37 | 0.06 | 0.33 | 0.05 | 0.16 |
| 2 | 55016061 | rs11898747 | 1.027 | 0.765 | -0.401 | 0.355 | 0.003 | -0.006 | -0.006 | 0 | 0.41 | 0.4 | 0.3 | 0.49 | 0.35 | 0.38 |
| 2 | 55017837 | rs7564091 | 1.834 | 0.83 | -0.44 | 0.462 | 0.005 | 0.145 | 0.145 | 0.486 | 0.12 | 0.34 | 0.07 | 0.34 | 0.09 | 0.15 |
| 2 | 55019651 | rs164855 | -1.597 | 1.044 | -0.88 | 0.358 | -0.03 | -0.064 | -0.064 | 0 | 0.89 | 0.99 | 0.91 | 0.73 | 0.86 | 0.81 |
| 2 | 55021891 | rs165053 | -0.713 | 1.058 | -0.519 | 0.215 | -0.002 | -0.01 | -0.01 | 0.014 | 0.55 | 0.54 | 0.66 | 0.61 | 0.57 | 0.6 |
| 2 | 55022953 | rs354944 | 0.732 | 1.202 | -0.715 | 0.16 | -0.005 | 0.008 | 0.008 | 0.098 | 0.27 | 0.22 | 0.41 | 0.1 | 0.29 | 0.23 |
| 2 | 55028892 | rs2578716 | 1.424 | 1.263 | -0.957 | 0.005 | -0.07 | 0.11 | 0.11 | 0.256 | 0.19 | 0.02 | 0.19 | 0.02 | 0.2 | 0.15 |
| 2 | 55031023 | rs17345167 | 1.032 | 1.283 | -0.77 | 0.07 | 0.015 | -0.006 | -0.006 | 0.002 | 0.11 | 0.17 | 0.31 | 0.07 | 0.15 | 0.11 |
| 2 | 55031674 | rs2567845 | -0.973 | 1.272 | -0.719 | 0.21 | 0.01 | 0.028 | 0.028 | 0 | 0.87 | 0.81 | 0.89 | 0.75 | 0.83 | 0.82 |
| 2 | 55031983 | rs6705883 | 2.148 | 1.051 | -0.97 | -0.057 | 0.006 | 0.11 | 0.11 | 0 | 0.3 | 0.13 | 0.16 | 0.12 | 0.26 | 0.23 |
| 2 | 55035282 | rs6545443 | 1.24 | 1.011 | -0.877 | 0.237 | 0.006 | -0.022 | -0.022 | 0 | 0.15 | 0.08 | 0.16 | 0.2 | 0.17 | 0.21 |
| 2 | 55039211 | rs2631838 | -0.06 | 1.041 | -0.812 | -0.014 | -0.008 | 0.058 | 0.058 | 0.373 | 0.37 | 0.23 | 0.52 | 0.11 | 0.37 | 0.26 |
| 2 | 55040204 | rs17345285 | 0.715 | 0.997 | -0.643 | -0.024 | -0.002 | 0.019 | 0.019 | 0 | 0.18 | 0.09 | 0.33 | 0.09 | 0.2 | 0.13 |
| 2 | 55040426 | rs2631840 | -1.135 | 0.988 | -0.412 | 0.136 | 0.012 | 0.057 | 0.057 | 0 | 0.88 | 0.72 | 0.89 | 0.76 | 0.86 | 0.75 |
| 2 | 55041322 | rs17046380 | 2.059 | 0.964 | -0.316 | 0.03 | 0.004 | -0.006 | -0.006 | 0 | 0.32 | 0.35 | 0.18 | 0.09 | 0.27 | 0.22 |

| | | | | | | | | | | | | | | | |
|---|---|---|---|---|---|---|---|---|---|---|---|---|---|---|---|
| 2 | 55042463 | rs2567839 | 0.299 | 1.02 | -0.252 | 0.173 | 0.016 | 0.069 | 0.069 | 0 | 0.77 | 0.56 | 0.67 | 0.66 | 0.74 | 0.6 |
| 2 | 55042477 | rs2567969 | -1.979 | 1.026 | -0.526 | 0.207 | -0.006 | -0.013 | -0.013 | 0 | 0.9 | 0.93 | 0.91 | 0.73 | 0.86 | 0.84 |
| 2 | 55044487 | rs2631841 | -1.802 | 1.018 | -0.502 | 0.165 | 0.007 | 0.023 | 0.023 | 0 | 0.89 | 0.82 | 0.9 | 0.77 | 0.86 | 0.75 |
| 2 | 55045017 | rs17417259 | 2.224 | 0.794 | -0.921 | -0.09 | 0.008 | 0.113 | 0.113 | 0 | 0.3 | 0.12 | 0.16 | 0.11 | 0.26 | 0.23 |
| 2 | 55049337 | rs2631842 | 0.627 | 0.91 | -0.96 | -0.096 | -0.01 | 0.028 | 0.028 | 0.161 | 0.27 | 0.18 | 0.44 | 0.1 | 0.28 | 0.28 |
| 2 | 55051827 | rs12999197 | -0.463 | 0.51 | -0.853 | 0.236 | 0.011 | -0.044 | -0.044 | 1.611 | 0.27 | 0.76 | 0.37 | 0.43 | 0.21 | 0.35 |
| 2 | 55052493 | rs10197243 | 1.092 | 0.618 | -0.799 | 0.258 | 0.008 | 0.007 | 0.007 | 0 | 0.11 | 0.14 | 0.28 | 0.11 | 0.13 | 0.15 |
| 2 | 55054378 | rs7569768 | 0.475 | 0.595 | -0.78 | 0.149 | 0.007 | 0.004 | 0.004 | 0 | 0.51 | 0.57 | 0.49 | 0.52 | 0.45 | 0.55 |
| 2 | 55055307 | rs1971105 | 0.466 | 0.59 | -0.744 | 0.207 | 0.022 | 0.037 | 0.037 | 0.007 | 0.51 | 0.74 | 0.39 | 0.59 | 0.44 | 0.56 |
| 2 | 55055628 | rs6723508 | 0.497 | 0.731 | -0.907 | 0.266 | 0.005 | 0.086 | 0.086 | 0 | 0.12 | 0.38 | 0.17 | 0.19 | 0.12 | 0.15 |
| 2 | 55057961 | rs930185 | -1.238 | 0.76 | -1.259 | 0.228 | -0.103 | 0.161 | 0.161 | 0.326 | 0.76 | 0.97 | 0.75 | 1 | 0.68 | 0.82 |
| 2 | 55060556 | rs7565802 | -0.125 | 0.723 | -1.076 | 0.397 | 0.045 | 0.229 | 0.229 | 0.415 | 0.14 | 0.39 | 0.18 | 0.38 | 0.13 | 0.15 |
| 2 | 55062132 | rs11674173 | 2.662 | 0.795 | -1.406 | 0.178 | -0.027 | -0.052 | -0.052 | 0 | 0.09 | 0 | 0.05 | 0.33 | 0.08 | 0.05 |
| 2 | 55063113 | rs1549840 | 1.267 | 0.834 | -1.878 | 0.121 | -0.025 | -0.097 | -0.097 | 0.019 | 0.4 | 0.1 | 0.31 | 0.55 | 0.39 | 0.49 |
| 2 | 55063552 | rs6755779 | -0.786 | 0.609 | -1.681 | 0.277 | 0.008 | -0.096 | -0.096 | 0.607 | 0.36 | 0.87 | 0.44 | 0.43 | 0.3 | 0.32 |
| 2 | 55068451 | rs6754447 | -0.309 | 0.741 | -0.879 | 0.086 | 0.012 | -0.074 | -0.074 | 0.007 | 0.36 | 0.51 | 0.46 | 0.1 | 0.4 | 0.25 |
| 2 | 55071861 | rs17046413 | 1.122 | 0.487 | -1.376 | 0.54 | 0.017 | -0.038 | -0.038 | 0 | 0.13 | 0.04 | 0.2 | 0.28 | 0.08 | 0.22 |
| 2 | 55072474 | rs2116435 | -0.324 | 0.521 | -1.993 | 0.505 | -0.063 | -0.068 | -0.068 | 0 | 0.7 | 0.9 | 0.66 | 0.66 | 0.64 | 0.72 |
| 2 | 55073428 | rs13005511 | 2.571 | 0.514 | -1.907 | 0.364 | 0 | -0.002 | -0.002 | 0 | 0.09 | 0.06 | 0.07 | 0.26 | 0.07 | 0.03 |
| 2 | 55074429 | rs12465327 | -0.278 | 0.421 | -1.258 | 0.25 | 0 | -0.001 | -0.001 | 0 | 0.41 | 0.39 | 0.53 | 0.54 | 0.41 | 0.56 |
| 2 | 55079388 | rs13033818 | 0.797 | 0.41 | -0.9 | 0.146 | 0.003 | 0.035 | 0.035 | 0 | 0.51 | 0.39 | 0.36 | 0.33 | 0.49 | 0.34 |
| 2 | 55080623 | rs12991420 | 0.546 | 0.384 | -1.335 | -0.223 | -0.003 | -0.002 | -0.002 | 0 | 0.3 | 0.33 | 0.19 | 0.27 | 0.23 | 0.22 |
| 2 | 55081376 | rs12713280 | 2.373 | 0.486 | -1.274 | -0.23 | -0.015 | 0.04 | 0.04 | 0 | 0.22 | 0.16 | 0.19 | 0.07 | 0.26 | 0.12 |
| 2 | 55081863 | rs12713282 | -0.005 | 0.357 | -1.361 | -0.181 | -0.018 | 0.019 | 0.019 | 0 | 0.6 | 0.68 | 0.56 | 0.4 | 0.56 | 0.71 |
| 2 | 55082530 | rs1371021 | -1.208 | 0.334 | -1.355 | -0.121 | -0.025 | -0.033 | -0.033 | 0.513 | 0.62 | 0.73 | 0.65 | 0.55 | 0.57 | 0.74 |
| 2 | 55085271 | rs2033413 | -0.62 | 0.312 | -1.386 | -0.114 | 0.055 | -0.008 | -0.008 | 0 | 0.49 | 0.75 | 0.47 | 0.58 | 0.38 | 0.61 |
| 2 | 55087887 | rs1371020 | 1.333 | 0.424 | -1.146 | -0.083 | -0.018 | -0.023 | -0.023 | 0 | 0.22 | 0.12 | 0.25 | 0.22 | 0.26 | 0.14 |
| 2 | 55091223 | rs6545450 | 0.147 | 0.164 | -0.835 | -0.173 | -0.018 | -0.011 | -0.011 | 0 | 0.32 | 0.45 | 0.21 | 0.27 | 0.25 | 0.23 |
| 2 | 55091579 | rs6727776 | -1.182 | 0.245 | -1.124 | -0.406 | 0.01 | -0.116 | -0.116 | 0 | 0.74 | 0.92 | 0.68 | 0.42 | 0.77 | 0.56 |
| 2 | 55093153 | rs6545452 | 0.917 | 0.497 | -0.541 | -0.254 | -0.005 | -0.068 | -0.068 | 0.17 | 0.2 | 0.25 | 0.11 | 0.01 | 0.16 | 0.19 |
| 2 | 55097549 | rs11677058 | 0.462 | 0.61 | -0.763 | 0.009 | 0.004 | 0.02 | 0.02 | 0 | 0.51 | 0.69 | 0.57 | 0.6 | 0.5 | 0.56 |
| 2 | 55102617 | rs10164992 | 0.564 | 0.655 | -0.738 | -0.856 | 0.003 | -0.102 | -0.102 | 2.143 | 0.2 | 0.3 | 0.14 | 0.01 | 0.15 | 0.19 |
| 2 | 55103265 | rs13385982 | 1.951 | 0.867 | -0.958 | -1.052 | -0.015 | 0.102 | 0.102 | 4.557 | 0.28 | 0.16 | 0.19 | 0.06 | 0.27 | 0.1 |
| 2 | 55108504 | rs13410462 | 0.422 | 0.758 | -0.874 | -0.947 | -0.002 | 0.046 | 0.046 | 1.635 | 0.49 | 0.33 | 0.32 | 0.07 | 0.42 | 0.31 |
| 2 | 55115152 | rs9309259 | 0.034 | 0.704 | -0.757 | -0.595 | -0.001 | 0 | 0 | 0.982 | 0.73 | 0.69 | 0.62 | 0.63 | 0.73 | 0.48 |
| 2 | 55117853 | rs4671988 | 0.88 | 0.653 | -0.575 | -0.594 | -0.016 | 0.01 | 0.01 | 0.28 | 0.2 | 0.28 | 0.29 | 0.28 | 0.18 | 0.38 |
| 2 | 55119047 | rs2163617 | -1.386 | 0.723 | -0.88 | -0.809 | -0.044 | 0.097 | 0.097 | 1.757 | 0.63 | 0.83 | 0.72 | 0.82 | 0.64 | 0.76 |
| 2 | 55120179 | rs13422312 | 0.886 | 0.726 | -0.849 | -0.978 | 0.011 | -0.089 | -0.089 | 0 | 0.14 | 0.31 | 0.11 | 0.01 | 0.1 | 0.18 |
| 2 | 55120659 | rs1438818 | -0.055 | 0.741 | -0.826 | -0.95 | -0.006 | 0.017 | 0.017 | 0 | 0.62 | 0.65 | 0.57 | 0.35 | 0.55 | 0.67 |
| 2 | 55121439 | rs6545453 | 0.056 | 0.797 | -0.821 | -0.965 | 0 | -0.009 | -0.009 | 0 | 0.55 | 0.47 | 0.39 | 0.18 | 0.47 | 0.42 |
| 2 | 55122589 | rs6545454 | 0.494 | 0.742 | -0.844 | -1.015 | 0 | -0.046 | -0.046 | 0 | 0.49 | 0.45 | 0.3 | 0.08 | 0.39 | 0.3 |
| 2 | 55124412 | rs891778 | -0.716 | 0.755 | -0.715 | -0.79 | 0.007 | 0.023 | 0.023 | 0 | 0.91 | 0.85 | 0.89 | 0.85 | 0.9 | 0.77 |
| 2 | 55125123 | rs891777 | -0.581 | 0.767 | -0.711 | -0.805 | 0.02 | 0.018 | 0.018 | 0 | 0.9 | 0.75 | 0.89 | 0.9 | 0.9 | 0.87 |
| 2 | 55129171 | rs10201734 | -0.709 | 0.678 | -1.107 | -0.904 | -0.006 | -0.016 | -0.016 | 0 | 0.39 | 0.49 | 0.39 | 0.29 | 0.3 | 0.58 |
| 2 | 55134309 | rs4671992 | -0.081 | 0.664 | -1.272 | -1.034 | -0.001 | -0.035 | -0.035 | 0 | 0.42 | 0.51 | 0.59 | 0.82 | 0.5 | 0.58 |
| 2 | 55146720 | rs17346218 | 1.934 | 0.861 | -1.485 | -1.085 | -0.014 | 0.14 | 0.14 | 4.394 | 0.29 | 0.1 | 0.19 | 0.07 | 0.27 | 0.11 |
| 2 | 55146868 | rs13011849 | -0.349 | 0.944 | -1.401 | -1.06 | 0.002 | -0.02 | -0.02 | 0.148 | 0.4 | 0.45 | 0.43 | 0.68 | 0.44 | 0.31 |
| 2 | 55150611 | rs6545459 | 0.166 | 0.778 | -1.396 | -0.977 | -0.004 | 0.012 | 0.012 | 0 | 0.34 | 0.44 | 0.35 | 0.58 | 0.37 | 0.22 |
| 2 | 55150883 | rs4671995 | 2.837 | 0.68 | -1.623 | -1.068 | 0.009 | -0.041 | -0.041 | 0 | 0.1 | 0.01 | 0.2 | 0.2 | 0.07 | 0.2 |
| 2 | 55151440 | rs4671233 | -1.651 | 0.756 | -1.616 | -1.091 | -0.03 | -0.016 | -0.016 | 0 | 0.94 | 1 | 0.94 | 0.95 | 0.9 | 0.82 |
| 2 | 55152587 | rs13390825 | 0.726 | 0.703 | -1.535 | -1.22 | -0.003 | 0.031 | 0.031 | 1.556 | 0.43 | 0.36 | 0.3 | 0.07 | 0.39 | 0.31 |
| 2 | 55152830 | rs13430119 | 0.155 | 0.808 | -1.388 | -1.168 | -0.004 | -0.109 | -0.109 | 0 | 0.22 | 0.36 | 0.14 | 0 | 0.15 | 0.21 |
| 2 | 55155551 | rs733916 | 2.746 | 0.8 | -1.484 | -1.095 | 0.008 | -0.041 | -0.041 | 0.017 | 0.1 | 0.01 | 0.2 | 0.19 | 0.07 | 0.21 |
| 2 | 55155957 | rs17418194 | 2.438 | 0.78 | -1.566 | -1.319 | -0.03 | 0.11 | 0.11 | 4.25 | 0.23 | 0.08 | 0.14 | 0.03 | 0.22 | 0.09 |
| 2 | 55156296 | rs755229 | -0.546 | 0.785 | -1.517 | -1.321 | 0.001 | -0.024 | -0.024 | 0 | 0.61 | 0.43 | 0.52 | 0.26 | 0.53 | 0.67 |
| 2 | 55157492 | rs4399760 | -0.376 | 0.733 | -1.154 | -0.995 | -0.006 | -0.033 | -0.033 | 0 | 0.32 | 0.4 | 0.37 | 0.23 | 0.27 | 0.56 |
| 2 | 55158082 | rs11689620 | 1.217 | 0.745 | -1.413 | -0.857 | -0.063 | -0.139 | -0.139 | 0 | 0.31 | 0.02 | 0.31 | 0.56 | 0.34 | 0.19 |
| 2 | 55159284 | rs13422342 | 0.336 | 0.653 | -0.919 | -0.812 | 0.023 | -0.095 | -0.095 | 2.498 | 0.15 | 0.34 | 0.11 | 0 | 0.12 | 0.19 |
| 2 | 55167820 | rs6749322 | 0.696 | 0.728 | -0.902 | -0.736 | 0.032 | -0.081 | -0.081 | 7.163 | 0.13 | 0.33 | 0.11 | 0 | 0.11 | 0.16 |
| 2 | 55168119 | rs10166518 | -0.882 | 0.742 | -1.647 | -0.557 | -0.045 | 0.081 | 0.081 | 0.986 | 0.68 | 0.87 | 0.76 | 0.84 | 0.67 | 0.8 |
| 2 | 55179512 | rs11683784 | 0.983 | 0.377 | -1.162 | -0.283 | -0.001 | 0 | 0 | 0.004 | 0.15 | 0.14 | 0.22 | 0.26 | 0.1 | 0.2 |
| 2 | 55180264 | rs3796008 | 0.444 | 0.383 | -1.582 | -0.552 | -0.045 | -0.013 | -0.013 | 0 | 0.4 | 0.12 | 0.41 | 0.59 | 0.44 | 0.32 |
| 2 | 55181382 | rs4672001 | 0.52 | 0.403 | -1.161 | -0.457 | 0.013 | 0.013 | 0.013 | 0.196 | 0.68 | 0.18 | 0.61 | 0.69 | 0.73 | 0.45 |
| 2 | 55182032 | rs11125550 | -0.366 | 0.404 | -0.643 | -0.473 | -0.02 | -0.012 | -0.012 | 0 | 0.27 | 0.61 | 0.33 | 0.26 | 0.18 | 0.37 |
| 2 | 55183627 | rs7584992 | 0.757 | 0.447 | -0.457 | -0.42 | 0.02 | -0.05 | -0.05 | 0.099 | 0.09 | 0.23 | 0.08 | 0 | 0.08 | 0.16 |
| 2 | 55185862 | rs17046514 | 2.191 | 0.391 | -1.04 | -0.394 | 0.017 | -0.049 | -0.049 | 0.092 | 0.12 | 0.01 | 0.2 | 0.2 | 0.07 | 0.19 |
| 2 | 55186394 | rs17346629 | 0.709 | 0.442 | -1.016 | -0.421 | 0.016 | 0.024 | 0.024 | 0.312 | 0.65 | 0.18 | 0.59 | 0.69 | 0.69 | 0.44 |
| 2 | 55192082 | rs2588512 | 1.83 | 0.534 | -1.75 | -0.266 | -0.051 | -0.148 | -0.148 | 0 | 0.19 | 0.01 | 0.17 | 0.5 | 0.21 | 0.15 |
| 2 | 55197967 | rs17046518 | 2.131 | 0.229 | -1.514 | -0.275 | 0.003 | -0.045 | -0.045 | 0 | 0.12 | 0 | 0.16 | 0.17 | 0.08 | 0.19 |

| | | | | | | | | | | | | | | | | |
|---|---|---|---|---|---|---|---|---|---|---|---|---|---|---|---|---|
| 2 | 55199248 | rs2588510 | 1.05 | 0.151 | -1.752 | -0.545 | -0.051 | 0.023 | 0.023 | 0.6 | 0.41 | 0.08 | 0.39 | 0.64 | 0.43 | 0.23 |
| 2 | 55204028 | rs2580765 | 0.703 | 0.1 | -1.504 | -0.545 | -0.035 | 0.066 | 0.066 | 0.779 | 0.44 | 0.1 | 0.4 | 0.66 | 0.44 | 0.24 |
| 2 | 55205124 | rs2588519 | -0.15 | 0.079 | -1.351 | -0.731 | -0.007 | -0.001 | -0.001 | 0 | 0.71 | 0.66 | 0.65 | 0.71 | 0.72 | 0.51 |
| 2 | 55207693 | rs3198123 | 1.115 | 0.001 | -0.969 | -0.802 | 0.006 | -0.064 | -0.064 | 0 | 0.25 | 0.28 | 0.22 | 0.03 | 0.27 | 0.28 |
| 2 | 55214618 | rs2580769 | 0.022 | -0.063 | -1.151 | -0.874 | -0.021 | -0.024 | -0.024 | 1.836 | 0.69 | 0.45 | 0.67 | 0.76 | 0.76 | 0.47 |
| 2 | 55216545 | rs2919129 | 1.402 | 0.031 | -1.182 | -0.865 | -0.021 | 0.03 | 0.03 | 0.692 | 0.41 | 0.11 | 0.39 | 0.63 | 0.42 | 0.23 |
| 2 | 55220584 | rs17046570 | 0.932 | 0.081 | -1.113 | -0.853 | -0.029 | 0.041 | 0.041 | 0 | 0.25 | 0.16 | 0.24 | 0.11 | 0.3 | 0.18 |
| 2 | 55220960 | rs2588517 | -0.054 | 0.031 | -1.12 | -0.826 | -0.004 | 0.003 | 0.003 | 1.613 | 0.71 | 0.65 | 0.64 | 0.67 | 0.7 | 0.51 |
| 2 | 55228392 | rs6545466 | -1.506 | -0.009 | -1.23 | -0.818 | -0.012 | 0.036 | 0.036 | 0.928 | 0.58 | 0.86 | 0.6 | 0.35 | 0.58 | 0.77 |
| 2 | 55231841 | rs6715980 | 0.227 | 0.046 | -1.097 | -0.776 | -0.015 | -0.104 | -0.104 | 0 | 0.14 | 0.52 | 0.12 | 0 | 0.08 | 0.2 |
| 2 | 55237049 | rs17046589 | 1.384 | 0.177 | -1.305 | -0.672 | -0.008 | -0.04 | -0.04 | 0 | 0.18 | 0.04 | 0.23 | 0.25 | 0.19 | 0.35 |
| 2 | 55245220 | rs2864052 | -2.562 | 0.059 | -1.604 | -0.272 | -0.073 | -0.134 | -0.134 | 0.196 | 0.79 | 0.99 | 0.78 | 0.62 | 0.76 | 0.87 |
| 2 | 55251374 | rs7601625 | -0.374 | 0.15 | -1.062 | -0.509 | 0.001 | -0.03 | -0.03 | 0 | 0.37 | 0.62 | 0.36 | 0.12 | 0.4 | 0.42 |
| 2 | 55257677 | rs10496037 | 0.795 | 0.203 | -1.254 | -0.393 | 0.033 | 0.014 | 0.014 | 0 | 0.1 | 0.29 | 0.12 | 0.08 | 0.1 | 0.1 |
| 2 | 55265785 | rs2968787 | 0.035 | 0.121 | -1.343 | -0.404 | -0.009 | 0.015 | 0.015 | 1.191 | 0.56 | 0.6 | 0.5 | 0.64 | 0.5 | 0.43 |
| 2 | 55271875 | rs7584354 | -0.407 | 0.154 | -1.304 | -0.538 | -0.01 | 0.012 | 0.012 | 0 | 0.37 | 0.67 | 0.37 | 0.13 | 0.4 | 0.42 |
| 2 | 55273287 | rs2920898 | -0.881 | 0.213 | -1.299 | -0.372 | -0.004 | 0.001 | 0.001 | 0.137 | 0.92 | 0.97 | 0.92 | 0.94 | 0.87 | 0.76 |
| 2 | 55277858 | rs6545469 | 1.097 | 0.224 | -1.208 | -0.402 | 0.025 | -0.062 | -0.062 | 2.47 | 0.1 | 0.32 | 0.1 | 0 | 0.08 | 0.2 |
| 2 | 55278168 | rs1348528 | 1.147 | 0.304 | -1.625 | -0.327 | 0 | 0.001 | 0.001 | 0 | 0.08 | 0.09 | 0.1 | 0.09 | 0.1 | 0.11 |
| 2 | 55286146 | rs11887323 | 2.019 | 0.149 | -1.66 | -0.361 | -0.007 | -0.006 | -0.006 | 0 | 0.14 | 0.13 | 0.2 | 0.1 | 0.1 | 0.22 |
| 2 | 55295370 | rs2972072 | 1.338 | 0.064 | -1.896 | -0.344 | -0.037 | -0.028 | -0.028 | 0 | 0.41 | 0.07 | 0.34 | 0.59 | 0.42 | 0.22 |
| 2 | 55296816 | rs17046647 | 1.203 | 0.122 | -1.731 | -0.276 | -0.001 | -0.002 | -0.002 | 0 | 0.09 | 0.08 | 0.1 | 0.18 | 0.1 | 0.11 |
| 2 | 55298517 | rs2920881 | 0.475 | 0.046 | -1.589 | -0.327 | -0.004 | -0.054 | -0.054 | 0 | 0.77 | 0.34 | 0.8 | 0.93 | 0.78 | 0.59 |
| 2 | 55301201 | rs2920883 | -2.252 | 0.141 | -1.676 | -0.23 | -0.027 | -0.017 | -0.017 | 0.051 | 0.95 | 1 | 0.94 | 0.94 | 0.9 | 0.82 |
| 2 | 55304458 | rs2920889 | -1.501 | 0.129 | -1.624 | -0.236 | -0.005 | 0.001 | 0.001 | 0 | 0.93 | 0.96 | 0.94 | 0.94 | 0.9 | 0.82 |
| 2 | 55305986 | rs7575107 | 1.844 | -0.157 | -1.828 | -0.114 | 0.008 | -0.061 | -0.061 | 0 | 0.19 | 0.01 | 0.11 | 0.18 | 0.15 | 0.08 |
| 2 | 55307865 | rs2968821 | -0.359 | 0.538 | -1.465 | -0.553 | -0.06 | 0 | 0 | 0 | 0.67 | 0.86 | 0.68 | 0.7 | 0.61 | 0.64 |
| 2 | 55310942 | rs6758258 | 0.349 | 0.346 | -0.984 | -0.446 | 0.016 | -0.092 | -0.092 | 0 | 0.13 | 0.51 | 0.11 | 0 | 0.08 | 0.21 |
| 2 | 55314518 | rs6724138 | 0.186 | 0.404 | -1.3 | -0.362 | -0.016 | -0.038 | -0.038 | 0 | 0.27 | 0.64 | 0.32 | 0.1 | 0.18 | 0.42 |
| 2 | 55316938 | rs2920862 | 0.839 | 0.392 | -1.604 | -0.265 | -0.011 | -0.03 | -0.03 | 2.285 | 0.34 | 0.15 | 0.31 | 0.6 | 0.36 | 0.22 |
| 2 | 55321474 | rs2968808 | 0.954 | 0.41 | -1.754 | -0.1 | -0.027 | -0.153 | -0.153 | 0.147 | 0.36 | 0.04 | 0.3 | 0.45 | 0.33 | 0.22 |
| 2 | 55323183 | rs7607923 | 0.672 | 0.429 | -1.253 | -0.521 | 0.016 | -0.074 | -0.074 | 0 | 0.32 | 0.43 | 0.33 | 0.06 | 0.35 | 0.26 |
| 2 | 55324960 | rs4672016 | -0.585 | 0.354 | -1.167 | -0.534 | 0.004 | 0.01 | 0.01 | 0 | 0.52 | 0.57 | 0.57 | 0.25 | 0.56 | 0.55 |
| 2 | 55325385 | rs2920875 | 0.859 | 0.374 | -1.387 | -0.453 | 0.017 | 0.04 | 0.04 | 0.028 | 0.43 | 0.27 | 0.37 | 0.68 | 0.38 | 0.31 |
| 2 | 55327407 | rs12464796 | 0.885 | 0.441 | -1.527 | -0.199 | -0.005 | 0.01 | 0.01 | 0.211 | 0.18 | 0.1 | 0.24 | 0.19 | 0.21 | 0.28 |
| 2 | 55327550 | rs12464877 | 1.4 | 0.381 | -1.648 | -0.261 | -0.02 | 0.041 | 0.041 | 0.181 | 0.16 | 0.01 | 0.22 | 0.07 | 0.16 | 0.26 |
| 2 | 55338420 | rs2968778 | -0.924 | 0.318 | -1.644 | -0.268 | 0 | 0.013 | 0.013 | 0.842 | 0.81 | 0.97 | 0.77 | 0.86 | 0.82 | 0.63 |
| 2 | 55338488 | rs2920842 | 0.474 | 0.411 | -1.636 | -0.427 | 0.001 | 0.063 | 0.063 | 0 | 0.53 | 0.22 | 0.53 | 0.63 | 0.51 | 0.51 |
| 2 | 55345758 | rs7578257 | 1.308 | 0.711 | -1.137 | -0.93 | 0.063 | -0.131 | -0.131 | 0.053 | 0.21 | 0.41 | 0.21 | 0.03 | 0.24 | 0.22 |
| 2 | 55346488 | rs2920851 | 0.517 | 0.576 | -1.395 | -0.924 | 0 | 0.006 | 0.006 | 0.525 | 0.56 | 0.63 | 0.55 | 0.5 | 0.53 | 0.47 |
| 2 | 55346785 | rs10496039 | 0.652 | 0.545 | -1.468 | -0.998 | -0.002 | 0.016 | 0.016 | 0.428 | 0.28 | 0.24 | 0.32 | 0.23 | 0.32 | 0.28 |
| 2 | 55349726 | rs11892071 | 0.352 | 0.505 | -1.459 | -0.981 | 0.021 | 0.08 | 0.08 | 0.03 | 0.09 | 0.28 | 0.11 | 0.21 | 0.08 | 0.1 |
| 2 | 55351010 | rs2920854 | 0.466 | 0.522 | -1.631 | -1.016 | 0.012 | 0.022 | 0.022 | 0.12 | 0.58 | 0.67 | 0.58 | 0.59 | 0.58 | 0.49 |
| 2 | 55354466 | rs1444261 | -1.634 | 0.492 | -1.616 | -1.001 | 0.002 | 0.004 | 0.004 | 0 | 0.93 | 0.98 | 0.96 | 0.95 | 0.93 | 0.8 |
| 2 | 55354512 | rs1444262 | 0.444 | 0.497 | -1.606 | -0.965 | 0.012 | 0.021 | 0.021 | 0.252 | 0.58 | 0.69 | 0.53 | 0.6 | 0.58 | 0.44 |
| 2 | 55359155 | rs2968800 | 1.528 | 0.338 | -1.822 | -0.856 | 0.016 | -0.101 | -0.101 | 0 | 0.35 | 0.07 | 0.28 | 0.47 | 0.29 | 0.25 |
| 2 | 55369684 | rs6725139 | 0.473 | 0.317 | -1.255 | -0.735 | 0.007 | -0.012 | -0.012 | 2.203 | 0.64 | 0.44 | 0.6 | 0.69 | 0.61 | 0.52 |
| 2 | 55369847 | rs7575639 | -0.222 | 0.521 | -1.553 | -0.656 | -0.014 | 0.013 | 0.013 | 0.012 | 0.3 | 0.68 | 0.38 | 0.26 | 0.32 | 0.28 |
| 2 | 55383624 | rs12470507 | 2.13 | -0.507 | -1.78 | -0.778 | 0.003 | 0.037 | 0.037 | 0.125 | 0.11 | 0.03 | 0.16 | 0.05 | 0.09 | 0.16 |
| 2 | 55384062 | rs12470651 | -0.458 | -0.598 | -1.666 | -0.753 | -0.001 | 0.001 | 0.001 | 0.078 | 0.26 | 0.28 | 0.37 | 0.46 | 0.26 | 0.34 |
| 2 | 55389938 | rs6714830 | -1.187 | -1.329 | -1.633 | -1.216 | -0.007 | 0.014 | 0.014 | 0 | 0.53 | 0.64 | 0.54 | 0.49 | 0.48 | 0.5 |
| 2 | 55391294 | rs7349405 | -0.083 | -1.315 | -1.698 | -1.187 | -0.011 | -0.056 | -0.056 | 0 | 0.13 | 0.04 | 0.17 | 0.28 | 0.11 | 0.15 |
| 2 | 55395732 | rs6717482 | 1.265 | -1.376 | -1.611 | -1.301 | -0.007 | -0.096 | -0.096 | 0.145 | 0.17 | 0.34 | 0.11 | 0.04 | 0.12 | 0.14 |
| 2 | 55404794 | rs14026 | 1.017 | -1.393 | -1.693 | -1.318 | 0.002 | -0.003 | -0.003 | 0 | 0.45 | 0.48 | 0.47 | 0.49 | 0.52 | 0.39 |
| 2 | 55406602 | rs11692206 | -0.314 | -1.49 | -1.716 | -1.329 | -0.003 | -0.02 | -0.02 | 0 | 0.17 | 0.14 | 0.2 | 0.41 | 0.17 | 0.24 |
| 2 | 55420825 | rs6747364 | 0.115 | -1.211 | -1.592 | -1.303 | -0.003 | -0.064 | -0.064 | 0 | 0.35 | 0.37 | 0.34 | 0.08 | 0.31 | 0.26 |
| 2 | 55435164 | rs2118284 | 0.61 | -1.221 | -1.581 | -1.3 | -0.01 | -0.08 | -0.08 | 0 | 0.26 | 0.37 | 0.17 | 0.09 | 0.22 | 0.16 |
| 2 | 55436934 | rs9677948 | 0.676 | -1.215 | -1.58 | -1.296 | -0.008 | -0.078 | -0.078 | 0.741 | 0.25 | 0.37 | 0.17 | 0.09 | 0.21 | 0.15 |
| 2 | 55446356 | rs1437391 | -0.855 | -1.477 | -1.95 | -1.463 | 0.021 | 0.005 | 0.005 | 0 | 0.62 | 0.53 | 0.56 | 0.51 | 0.52 | 0.54 |
| 2 | 55452971 | rs11686867 | 1.075 | -1.456 | -2.241 | -1.576 | -0.164 | -0.092 | -0.092 | 0 | 0.33 | 0.04 | 0.39 | 0.52 | 0.45 | 0.35 |
| 2 | 55460751 | rs6545480 | 0.477 | -1.927 | -1.358 | -1.247 | 0.022 | 0.001 | 0.001 | 0.037 | 0.68 | 0.77 | 0.75 | 0.62 | 0.75 | 0.63 |
| 2 | 55476937 | rs2576711 | 1.468 | -0.698 | -0.384 | -0.374 | -0.01 | -0.005 | -0.005 | 0.877 | 0.65 | 0.42 | 0.58 | 0.58 | 0.69 | 0.51 |
| 2 | 55490819 | rs1056445 | 1.868 | -0.261 | 0.316 | 0.21 | -0.012 | 0.032 | 0.032 | 0 | 0.76 | 0.44 | 0.75 | 0.63 | 0.79 | 0.64 |
| 2 | 55492076 | rs6545482 | -0.08 | -0.317 | -0.115 | 0.302 | -0.006 | -0.015 | -0.015 | 0 | 0.93 | 0.91 | 0.98 | 0.98 | 0.95 | 0.89 |
| 2 | 55505681 | rs2576696 | 1.632 | -0.433 | -0.07 | 0.409 | 0.028 | -0.009 | -0.009 | 0 | 0.75 | 0.22 | 0.84 | 0.64 | 0.85 | 0.54 |
| 2 | 55507386 | rs7579539 | 1.498 | -0.488 | 0.464 | 0.446 | -0.057 | 0.036 | 0.036 | 0 | 0.78 | 0.52 | 0.89 | 0.67 | 0.88 | 0.61 |
| 2 | 55508526 | rs13031346 | 1.704 | -0.523 | 0.628 | 0.316 | -0.011 | -0.062 | -0.062 | 0 | 0.88 | 0.46 | 0.93 | 0.98 | 0.94 | 0.78 |
| 2 | 55508618 | rs6718747 | 0.322 | -0.512 | 0.635 | 0.402 | -0.009 | -0.026 | -0.026 | 1.873 | 0.93 | 0.82 | 0.98 | 0.98 | 0.95 | 0.77 |
| 2 | 55516090 | rs1045613 | -0.379 | -0.517 | 0.517 | 0.601 | 0 | 0.016 | 0.016 | 0 | 0.23 | 0.24 | 0.26 | 0.44 | 0.21 | 0.17 |

| | | | | | | | | | | | | | | | | |
|---|---|---|---|---|---|---|---|---|---|---|---|---|---|---|---|---|
| 2 | 55516323 | rs2589113 | 1.107 | -0.546 | 0.593 | 0.504 | -0.015 | -0.047 | -0.047 | 0.956 | 0.89 | 0.43 | 0.94 | 0.98 | 0.95 | 0.8 |
| 2 | 55516723 | rs2576692 | -0.033 | -0.545 | 0.591 | 0.519 | -0.007 | -0.021 | -0.021 | 0 | 0.93 | 0.86 | 0.98 | 0.98 | 0.95 | 0.8 |
| 2 | 55517491 | rs1047948 | 2.265 | -0.49 | 0.418 | 0.43 | -0.003 | 0.295 | 0.295 | 0 | 0.5 | 0.05 | 0.59 | 0.15 | 0.6 | 0.34 |
| 2 | 55527108 | rs1545121 | 1.734 | -0.512 | 0.625 | 0.551 | -0.038 | -0.072 | -0.072 | 0.022 | 0.84 | 0.37 | 0.91 | 0.94 | 0.91 | 0.78 |
| 2 | 55534186 | rs4484068 | 1.571 | -0.489 | 0.576 | 0.603 | -0.023 | 0.08 | 0.08 | 0 | 0.82 | 0.54 | 0.89 | 0.66 | 0.85 | 0.73 |
| 2 | 55536808 | rs10191785 | -0.203 | -0.484 | 0.57 | 0.562 | -0.007 | -0.024 | -0.024 | 0.702 | 0.07 | 0.3 | 0.04 | 0 | 0.03 | 0.01 |
| 2 | 55539905 | rs10189439 | 1.546 | -0.542 | 0.503 | 0.581 | 0.004 | 0.008 | 0.008 | 0 | 0.61 | 0.64 | 0.66 | 0.14 | 0.65 | 0.58 |
| 2 | 55543840 | rs11125563 | 1.554 | -0.525 | 0.535 | 0.779 | -0.026 | 0.023 | 0.023 | 0 | 0.77 | 0.39 | 0.87 | 0.64 | 0.83 | 0.54 |
| 2 | 55571085 | rs2589077 | -1.801 | -0.239 | 0.706 | 0.997 | 0 | -0.004 | -0.004 | 0 | 0.37 | 0.65 | 0.35 | 0.82 | 0.31 | 0.44 |
| 2 | 55585276 | rs3099084 | -2.074 | -0.125 | 0.637 | 0.868 | 0.003 | 0.003 | 0.003 | 0 | 0.42 | 0.63 | 0.39 | 0.85 | 0.36 | 0.45 |
| 2 | 55585915 | rs12991861 | -0.668 | -0.124 | 0.758 | 0.964 | 0 | 0.011 | 0.011 | 0 | 0.24 | 0.25 | 0.28 | 0.45 | 0.24 | 0.17 |
| 2 | 55587727 | rs2966466 | -0.764 | -0.323 | 0.513 | 0.707 | -0.004 | 0.093 | 0.093 | 0 | 0.11 | 0.27 | 0.07 | 0.31 | 0.09 | 0.25 |
| 2 | 55589364 | rs13019339 | -0.794 | -0.317 | 0.352 | 0.706 | -0.004 | 0.017 | 0.017 | 0 | 0.26 | 0.35 | 0.28 | 0.54 | 0.22 | 0.19 |
| 2 | 55599810 | rs10496042 | -0.782 | -0.313 | 0.28 | 0.706 | -0.004 | 0.021 | 0.021 | 0 | 0.26 | 0.36 | 0.28 | 0.54 | 0.22 | 0.18 |
| 2 | 55619610 | rs11692239 | 1.759 | -0.387 | 0.144 | 0.536 | 0.03 | 0.113 | 0.113 | 0 | 0.58 | 0.22 | 0.62 | 0.15 | 0.67 | 0.54 |
| 2 | 55627639 | rs6709204 | 0.774 | -0.409 | 0.099 | 0.589 | 0.003 | 0.121 | 0.121 | 0 | 0.87 | 0.58 | 0.92 | 0.72 | 0.91 | 0.74 |
| 2 | 55631013 | rs6721972 | 0.015 | -0.345 | -0.01 | 0.659 | -0.034 | 0.057 | 0.057 | 0 | 0.17 | 0.02 | 0.1 | 0.05 | 0.14 | 0.1 |
| 2 | 55639980 | rs6545492 | 0.627 | -0.254 | 0.292 | 0.668 | -0.013 | 0.036 | 0.036 | 0 | 0.85 | 0.6 | 0.9 | 0.8 | 0.88 | 0.73 |
| 2 | 55651886 | rs1902061 | 0.301 | -0.243 | 0.234 | 0.681 | -0.013 | 0.039 | 0.039 | 0 | 0.86 | 0.72 | 0.91 | 0.79 | 0.89 | 0.74 |
| 2 | 55658147 | rs4672045 | 0.449 | -0.251 | -0.335 | 0.59 | -0.036 | 0.066 | 0.066 | 0 | 0.83 | 0.6 | 0.9 | 0.71 | 0.89 | 0.7 |
| 2 | 55659155 | rs11125573 | -0.824 | 0.197 | -0.783 | 0.924 | -0.005 | 0.012 | 0.012 | 0 | 0.27 | 0.35 | 0.29 | 0.57 | 0.24 | 0.17 |
| 2 | 55680496 | rs7593229 | 0.215 | 0.269 | -1.778 | 0.709 | 0.008 | -0.013 | -0.013 | 0.886 | 0.62 | 0.85 | 0.78 | 0.59 | 0.72 | 0.62 |
| 2 | 55682348 | rs7600456 | 0.912 | -0.186 | -1.291 | 0.92 | 0.055 | 0.011 | 0.011 | 0 | 0.5 | 0.28 | 0.7 | 0.41 | 0.67 | 0.42 |
| 2 | 55687618 | rs11694644 | 0.879 | -0.091 | -1.04 | 0.927 | 0.05 | 0.007 | 0.007 | 0 | 0.5 | 0.32 | 0.71 | 0.41 | 0.68 | 0.43 |
| 2 | 55688713 | rs4671246 | -0.279 | -0.349 | -1.376 | 0.663 | -0.005 | -0.084 | -0.084 | 0 | 0.86 | 0.96 | 0.92 | 0.6 | 0.9 | 0.78 |
| 2 | 55696124 | rs7567744 | 0.123 | -0.495 | -1.476 | 0.51 | 0.026 | -0.102 | -0.102 | 0 | 0.75 | 0.94 | 0.88 | 0.59 | 0.85 | 0.72 |
| 2 | 55704333 | rs12612931 | 1.702 | 0.374 | -1.594 | 0.256 | -0.002 | 0.005 | 0.005 | 0 | 0.47 | 0.27 | 0.59 | 0.39 | 0.59 | 0.32 |
| 2 | 55709416 | rs957267 | 1.837 | 0.284 | -1.693 | 0.144 | 0.012 | -0.006 | -0.006 | 0 | 0.46 | 0.28 | 0.68 | 0.39 | 0.62 | 0.34 |
| 2 | 55713102 | rs13011640 | 1.225 | 0.219 | -1.688 | 0.121 | 0.01 | -0.009 | -0.009 | 0.073 | 0.6 | 0.35 | 0.76 | 0.42 | 0.73 | 0.48 |
| 2 | 55723038 | rs10165864 | 1.409 | 0.52 | -2.035 | 0.099 | -0.013 | 0.188 | 0.188 | 0 | 0.48 | 0.73 | 0.54 | 0.15 | 0.52 | 0.36 |
| 2 | 55726222 | rs10184287 | -0.836 | 0.575 | -2.063 | 0.22 | -0.017 | -0.097 | -0.097 | 0.391 | 0.87 | 0.96 | 0.93 | 0.52 | 0.9 | 0.8 |
| 2 | 55727476 | rs12986587 | -0.546 | 0.485 | -2.041 | 0.129 | -0.031 | -0.064 | -0.064 | 0.217 | 0.24 | 0.15 | 0.25 | 0.34 | 0.28 | 0.28 |
| 2 | 55749085 | rs12992408 | -1.293 | 0.468 | -1.553 | 0.042 | 0.015 | 0.08 | 0.08 | 0 | 0.15 | 0.26 | 0.26 | 0.35 | 0.2 | 0.27 |
| 2 | 55757885 | rs17369191 | 2.021 | 0.312 | -1.515 | -0.082 | -0.019 | 0.065 | 0.065 | 0 | 0.43 | 0.31 | 0.45 | 0.09 | 0.52 | 0.21 |
| 2 | 55757939 | rs12987272 | 1.037 | 0.358 | -1.491 | 0.174 | -0.017 | 0 | 0 | 0 | 0.06 | 0.05 | 0.21 | 0.25 | 0.11 | 0.06 |
| 2 | 55764906 | rs17792798 | 1.167 | 0.401 | -1.74 | 0.026 | -0.043 | 0.078 | 0.078 | 0 | 0.13 | 0.01 | 0.08 | 0.01 | 0.12 | 0.14 |
| 2 | 55771161 | rs1045910 | -0.994 | 0.387 | -1.751 | 0.141 | 0.017 | -0.076 | -0.076 | 4.147 | 0.88 | 0.98 | 0.9 | 0.48 | 0.94 | 0.8 |
| 2 | 55775724 | rs8080 | -1.098 | 0.358 | -1.677 | 0.122 | 0.021 | 0.105 | 0.105 | 0 | 0.15 | 0.27 | 0.26 | 0.34 | 0.2 | 0.27 |
| 2 | 55776793 | rs3748945 | 0.606 | 0.366 | -1.678 | 0.127 | -0.002 | 0.041 | 0.041 | 0 | 0.68 | 0.57 | 0.73 | 0.44 | 0.73 | 0.55 |
| 2 | 55784608 | rs1013138 | -1.132 | 0.085 | -1.633 | 0.147 | 0.019 | 0.075 | 0.075 | 0 | 0.15 | 0.26 | 0.26 | 0.29 | 0.2 | 0.27 |
| 2 | 55793476 | rs12993624 | -0.389 | 0.057 | -1.701 | 0.11 | 0 | -0.001 | -0.001 | 1.148 | 0.26 | 0.26 | 0.3 | 0.3 | 0.26 | 0.31 |
| 2 | 55798108 | rs6545506 | 1.924 | 0.053 | -1.726 | 0.187 | -0.02 | 0.07 | 0.07 | 0 | 0.43 | 0.32 | 0.45 | 0.17 | 0.52 | 0.24 |
| 2 | 55812061 | rs4312532 | -1.48 | 0.065 | -1.706 | 0.068 | 0.003 | 0.008 | 0.008 | 0 | 0.45 | 0.68 | 0.45 | 0.4 | 0.42 | 0.59 |
| 2 | 55815997 | rs6545508 | 0.787 | 0.043 | -1.641 | 0.042 | -0.037 | -0.095 | -0.095 | 0 | 0.17 | 0.39 | 0.1 | 0.04 | 0.1 | 0.16 |
| 2 | 55819752 | rs6737995 | -0.158 | 0.047 | -1.671 | 0.064 | -0.004 | 0.005 | 0.005 | 0 | 0.24 | 0.26 | 0.27 | 0.29 | 0.21 | 0.26 |
| 2 | 55829585 | rs12998664 | -0.08 | 0.039 | -1.66 | 0.051 | -0.007 | 0.001 | 0.001 | 0 | 0.1 | 0.1 | 0.22 | 0.3 | 0.14 | 0.18 |
| 2 | 55829967 | rs2586970 | 1.894 | 0.021 | -1.741 | 0.029 | 0 | 0.091 | 0.091 | 0 | 0.46 | 0.32 | 0.48 | 0.16 | 0.58 | 0.31 |
| 2 | 55831415 | rs2627763 | -1.077 | 0.02 | -1.746 | 0.026 | 0.009 | -0.086 | -0.086 | 0 | 0.88 | 0.99 | 0.9 | 0.49 | 0.94 | 0.8 |
| 2 | 55845351 | rs2291336 | -0.461 | 0.029 | -1.73 | 0.031 | 0.011 | 0.095 | 0.095 | 0 | 0.12 | 0.23 | 0.23 | 0.3 | 0.15 | 0.18 |
| 2 | 55846146 | rs736345 | 0.148 | 0.024 | -1.721 | 0.008 | 0.021 | -0.095 | -0.095 | 0.196 | 0.18 | 0.4 | 0.15 | 0.04 | 0.16 | 0.23 |
| 2 | 55849309 | rs782602 | 2.024 | -0.19 | -1.921 | -0.082 | -0.019 | 0.054 | 0.054 | 0 | 0.41 | 0.32 | 0.44 | 0.1 | 0.51 | 0.22 |
| 2 | 55855611 | rs2586949 | -0.585 | -0.233 | -1.919 | 0.082 | 0.015 | -0.092 | -0.092 | 0 | 0.87 | 0.99 | 0.89 | 0.51 | 0.93 | 0.78 |
| 2 | 55860160 | rs782648 | -1.127 | -0.249 | -1.865 | 0.129 | 0.005 | -0.074 | -0.074 | 0 | 0.89 | 0.99 | 0.9 | 0.5 | 0.94 | 0.8 |
| 2 | 55872538 | rs7594497 | 0.303 | -0.254 | -1.757 | 0.097 | 0.084 | 0.062 | 0.062 | 0.253 | 0.06 | 0.41 | 0.06 | 0.04 | 0.04 | 0.08 |
| 2 | 55877113 | rs2627775 | 1.985 | -0.488 | -2.399 | -0.523 | -0.011 | 0.03 | 0.03 | 0 | 0.37 | 0.31 | 0.42 | 0.1 | 0.46 | 0.22 |
| 2 | 55883317 | rs2627765 | 1.454 | -0.546 | -2.213 | -0.401 | -0.01 | 0.079 | 0.079 | 0.431 | 0.56 | 0.32 | 0.49 | 0.14 | 0.58 | 0.31 |
| 2 | 55886602 | rs782606 | 1.633 | -0.5 | -2.186 | -0.384 | -0.019 | 0.089 | 0.089 | 0.016 | 0.45 | 0.31 | 0.46 | 0.09 | 0.53 | 0.23 |
| 2 | 55886780 | rs7575170 | -0.575 | -0.313 | -2.076 | -0.243 | -0.004 | -0.006 | -0.006 | 0.246 | 0.27 | 0.25 | 0.35 | 0.37 | 0.32 | 0.41 |
| 2 | 55894428 | rs12329050 | -1.246 | -0.531 | -2.237 | -0.415 | -0.007 | -0.001 | -0.001 | 0 | 0.37 | 0.66 | 0.41 | 0.35 | 0.36 | 0.49 |
| 2 | 55901317 | rs12617829 | -1.188 | -0.667 | -2.405 | -0.462 | 0.05 | -0.143 | -0.143 | 0 | 0.79 | 0.97 | 0.87 | 0.5 | 0.9 | 0.72 |
| 2 | 55903620 | rs706551 | -1.522 | -0.615 | -2.401 | -0.433 | 0.002 | -0.031 | -0.031 | 0 | 0.91 | 0.98 | 0.9 | 0.5 | 0.94 | 0.79 |
| 2 | 55903711 | rs10496043 | 1.75 | -0.435 | -2.36 | -0.44 | -0.042 | 0.073 | 0.073 | 0 | 0.12 | 0.01 | 0.08 | 0 | 0.11 | 0.14 |
| 2 | 55918189 | rs1443746 | -0.269 | -0.47 | -2.305 | -0.455 | -0.008 | -0.014 | -0.014 | 0 | 0.26 | 0.2 | 0.32 | 0.31 | 0.28 | 0.33 |
| 2 | 55929430 | rs1084525 | 1.532 | -0.389 | -2.53 | -0.575 | -0.018 | 0.037 | 0.037 | 0 | 0.37 | 0.29 | 0.43 | 0.1 | 0.48 | 0.23 |
| 2 | 55941440 | rs11125589 | 0.291 | -0.529 | -2.456 | -0.023 | 0.065 | -0.134 | -0.134 | 0 | 0.25 | 0.04 | 0.14 | 0.55 | 0.13 | 0.4 |
| 2 | 55943068 | rs12623877 | 0.232 | -0.521 | -2.418 | -0.004 | 0.068 | -0.12 | -0.12 | 0 | 0.26 | 0.06 | 0.14 | 0.55 | 0.14 | 0.41 |
| 2 | 55944357 | rs17369212 | 0.289 | 0.106 | -1.966 | -0.402 | -0.023 | 0.037 | 0.037 | 0 | 0.16 | 0.06 | 0.11 | 0.07 | 0.16 | 0.24 |
| 2 | 55958891 | rs10178492 | -0.483 | 0.174 | -1.653 | 0.056 | 0.004 | 0.023 | 0.023 | 0 | 0.1 | 0.13 | 0.22 | 0.26 | 0.16 | 0.11 |
| 2 | 55963418 | rs6729407 | 0.49 | -0.144 | -1.331 | -0.392 | 0.001 | -0.003 | -0.003 | 0 | 0.24 | 0.24 | 0.17 | 0.58 | 0.17 | 0.23 |

| | | | | | | | | | | | | | | | | |
|---|---|---|---|---|---|---|---|---|---|---|---|---|---|---|---|---|
| 2 | 55969580 | rs7568378 | 1.449 | -0.096 | -1.882 | -0.622 | -0.011 | 0.055 | 0.055 | 0 | 0.12 | 0.01 | 0.08 | 0 | 0.09 | 0.13 |
| 2 | 55970281 | rs1346793 | -1.385 | -0.529 | -1.631 | -1.003 | -0.009 | -0.006 | -0.006 | 0 | 0.63 | 0.72 | 0.54 | 0.91 | 0.51 | 0.79 |
| 2 | 55973732 | rs2163237 | -0.201 | 0.136 | -2.053 | -0.435 | -0.034 | 0.025 | 0.025 | 0 | 0.3 | 0.13 | 0.37 | 0.74 | 0.33 | 0.32 |
| 2 | 55983929 | rs1583116 | -0.232 | 0.38 | -1.416 | -0.46 | -0.019 | -0.021 | -0.021 | 0 | 0.21 | 0.12 | 0.3 | 0.26 | 0.26 | 0.26 |
| 2 | 55987068 | rs6755214 | -0.031 | 0.261 | -1.223 | -1.157 | 0.001 | 0.069 | 0.069 | 0 | 0.34 | 0.14 | 0.32 | 0.82 | 0.3 | 0.44 |
| 2 | 55991651 | rs1465624 | -0.889 | 0.22 | -1.066 | -1.792 | -0.043 | 0.182 | 0.182 | 0 | 0.6 | 0.84 | 0.48 | 0.88 | 0.49 | 0.68 |
| 2 | 56004592 | rs7601401 | 0.548 | 0.04 | -0.409 | -1.78 | 0.005 | 0.009 | 0.009 | 0 | 0.32 | 0.26 | 0.3 | 0.83 | 0.22 | 0.29 |
| 2 | 56008904 | rs7577894 | -0.673 | 0.206 | -0.082 | -1.84 | -0.011 | 0.104 | 0.104 | 0 | 0.57 | 0.77 | 0.42 | 0.89 | 0.42 | 0.6 |
| 2 | 56010255 | rs10496053 | 0.286 | 0.207 | 0.193 | -1.811 | 0.013 | -0.034 | -0.034 | 0 | 0.07 | 0.13 | 0.05 | 0.01 | 0.09 | 0.03 |
| 2 | 56010803 | rs4355154 | -0.025 | -0.037 | 0.827 | -2.003 | -0.003 | -0.059 | -0.059 | 1.203 | 0.46 | 0.49 | 0.35 | 0.9 | 0.35 | 0.59 |
| 2 | 56011881 | rs732132 | 0.465 | -0.385 | 1.029 | -2.057 | 0.008 | 0.026 | 0.026 | 0 | 0.33 | 0.25 | 0.26 | 0.89 | 0.28 | 0.49 |
| 2 | 56029444 | rs6712017 | -0.233 | -0.315 | 1.465 | -2.157 | -0.001 | -0.046 | -0.046 | 0 | 0.4 | 0.61 | 0.36 | 0.9 | 0.35 | 0.56 |
| 2 | 56031285 | rs1430189 | -0.117 | -0.322 | 1.451 | -2.161 | 0.003 | 0.018 | 0.018 | 0 | 0.4 | 0.66 | 0.36 | 0.9 | 0.35 | 0.56 |
| 2 | 56032652 | rs7560657 | 1.127 | -0.402 | 1.493 | -2.05 | 0.016 | 0.03 | 0.03 | 0 | 0.28 | 0.16 | 0.24 | 0.81 | 0.21 | 0.27 |
| 2 | 56033427 | rs1430191 | -1.349 | -0.315 | 1.409 | -2.179 | -0.065 | 0.201 | 0.201 | 0 | 0.61 | 0.88 | 0.46 | 0.91 | 0.51 | 0.66 |
| 2 | 56036226 | rs10196888 | -0.982 | -0.29 | 1.473 | -2.099 | 0.053 | 0.101 | 0.101 | 0 | 0.08 | 0.38 | 0.05 | 0.09 | 0.07 | 0.22 |
| 2 | 56036796 | rs6545522 | 0.637 | -0.306 | 1.344 | -2.133 | 0 | 0.011 | 0.011 | 0 | 0.33 | 0.29 | 0.32 | 0.81 | 0.29 | 0.34 |
| 2 | 56039510 | rs4672074 | -0.727 | -0.462 | 0.996 | -2.452 | -0.041 | 0.178 | 0.178 | 0 | 0.58 | 0.82 | 0.45 | 0.9 | 0.45 | 0.63 |
| 2 | 56039930 | rs11125603 | 0.186 | -0.419 | 0.887 | -2.412 | 0.001 | 0.041 | 0.041 | 0 | 0.4 | 0.31 | 0.37 | 0.81 | 0.36 | 0.34 |
| 2 | 56048237 | rs6730976 | 1.066 | -0.299 | 0.565 | -2.247 | 0.001 | 0.028 | 0.028 | 0 | 0.08 | 0.05 | 0.06 | 0 | 0.07 | 0.01 |
| 2 | 56067567 | rs6723018 | 0.017 | -0.282 | 0.752 | -2.262 | 0.002 | -0.027 | -0.027 | 0 | 0.12 | 0.23 | 0.1 | 0.02 | 0.13 | 0.06 |
| 2 | 56068378 | rs10180043 | -0.252 | -0.343 | 0.656 | -2.278 | -0.032 | -0.085 | -0.085 | 0 | 0.29 | 0.45 | 0.16 | 0.09 | 0.22 | 0.32 |
| 2 | 56070352 | rs934278 | -1.586 | -0.509 | -0.238 | -2.326 | -0.127 | 0.107 | 0.107 | 0 | 0.74 | 0.97 | 0.52 | 0.91 | 0.58 | 0.75 |
| 2 | 56082610 | rs6545528 | 0.076 | -0.351 | 0.489 | -2.022 | -0.019 | -0.049 | -0.049 | 0 | 0.26 | 0.4 | 0.17 | 0.06 | 0.18 | 0.18 |
| 2 | 56085261 | rs7582381 | -0.251 | -0.248 | 0.47 | -1.936 | 0.059 | 0.068 | 0.068 | 0 | 0.07 | 0.25 | 0.07 | 0.07 | 0.09 | 0.09 |
| 2 | 56092612 | rs1346787 | -1.293 | -0.273 | -0.102 | -2.127 | -0.053 | 0.141 | 0.141 | 0 | 0.68 | 0.89 | 0.51 | 0.9 | 0.57 | 0.66 |
| 2 | 56096892 | rs3791679 | 1.319 | -0.543 | -0.093 | -1.917 | -0.013 | 0.005 | 0.005 | 0 | 0.27 | 0.03 | 0.24 | 0.77 | 0.23 | 0.25 |
| 2 | 56105934 | rs17047290 | 0.402 | -0.649 | 0.367 | -1.987 | -0.041 | -0.042 | -0.042 | 0.295 | 0.23 | 0.51 | 0.13 | 0.13 | 0.13 | 0.39 |
| 2 | 56108333 | rs1346786 | -0.246 | -0.36 | 0.549 | -1.508 | -0.02 | -0.071 | -0.071 | 0.171 | 0.4 | 0.44 | 0.33 | 0.83 | 0.31 | 0.34 |
| 2 | 56111309 | rs3791675 | 0.568 | -0.2 | -0.462 | -1.329 | 0.003 | 0.011 | 0.011 | 0 | 0.29 | 0.04 | 0.24 | 0.76 | 0.23 | 0.24 |
| 2 | 56118046 | rs10865291 | 0.012 | -0.006 | 0.076 | -1.336 | 0.002 | 0.001 | 0.001 | 0 | 0.42 | 0.37 | 0.31 | 0.77 | 0.35 | 0.43 |
| 2 | 56119657 | rs727878 | -0.008 | -0.008 | 0.14 | -1.351 | -0.002 | -0.012 | -0.012 | 0.129 | 0.43 | 0.4 | 0.32 | 0.77 | 0.35 | 0.42 |
| 2 | 56128027 | rs1344733 | -0.791 | 0.061 | -0.465 | -0.935 | -0.001 | -0.005 | -0.005 | 0.457 | 0.34 | 0.75 | 0.33 | 0.34 | 0.3 | 0.3 |
| 2 | 56129870 | rs3791661 | -0.681 | 0.052 | -0.121 | -0.932 | 0.003 | 0.014 | 0.014 | 0 | 0.33 | 0.58 | 0.31 | 0.35 | 0.3 | 0.3 |
| 2 | 56129880 | rs3791660 | -1.164 | 0.069 | -0.71 | -1.044 | -0.078 | 0.26 | 0.26 | 0 | 0.58 | 0.91 | 0.51 | 0.89 | 0.53 | 0.62 |
| 2 | 56130640 | rs1430195 | -0.701 | 0.074 | -0.587 | -0.93 | 0 | 0.003 | 0.003 | 0 | 0.33 | 0.68 | 0.31 | 0.35 | 0.3 | 0.3 |
| 2 | 56133040 | rs7559906 | -0.491 | 0.33 | -0.901 | -1.487 | -0.033 | 0.204 | 0.204 | 0 | 0.57 | 0.84 | 0.5 | 0.88 | 0.51 | 0.63 |
| 2 | 56135497 | rs4233964 | -0.336 | 0.382 | -0.105 | -0.997 | 0.027 | -0.016 | -0.016 | 0 | 0.06 | 0.08 | 0.11 | 0.03 | 0.11 | 0.12 |
| 2 | 56145270 | rs10496056 | 0.253 | 0.112 | 0.303 | -1.114 | -0.004 | -0.05 | -0.05 | 0 | 0.09 | 0.22 | 0.07 | 0 | 0.07 | 0.02 |
| 2 | 56154316 | rs10469935 | 1.246 | -0.278 | -0.421 | -1.301 | -0.011 | -0.028 | -0.028 | 0 | 0.12 | 0.12 | 0.03 | 0 | 0.02 | 0.06 |
| 2 | 56161538 | rs7608756 | -0.984 | -0.4 | -0.69 | -1.611 | 0.042 | 0.087 | 0.087 | 0 | 0.37 | 0.75 | 0.25 | 0.04 | 0.23 | 0.33 |
| 2 | 56164916 | rs6752225 | -0.696 | 0.537 | -1.187 | -1.645 | -0.054 | 0.057 | 0.057 | 0 | 0.77 | 0.87 | 0.51 | 0.86 | 0.61 | 0.88 |
| 2 | 56170392 | rs6718634 | -0.582 | 0.49 | -0.727 | -1.656 | -0.012 | 0.015 | 0.015 | 0 | 0.76 | 0.79 | 0.5 | 0.86 | 0.61 | 0.89 |
| 2 | 56177013 | rs11685113 | 0.4 | 0.684 | -0.188 | -2.046 | 0.006 | -0.026 | -0.026 | 1.252 | 0.63 | 0.63 | 0.35 | 0.81 | 0.47 | 0.55 |
| 2 | 56184498 | rs4672078 | -0.275 | 0.697 | -0.157 | -2.043 | 0.038 | -0.067 | -0.067 | 1.443 | 0.14 | 0.23 | 0.16 | 0.05 | 0.15 | 0.32 |
| 2 | 56186525 | rs1465684 | 0.981 | 0.695 | -0.288 | -2.043 | -0.054 | 0.055 | 0.055 | 0 | 0.22 | 0.11 | 0.49 | 0.14 | 0.37 | 0.11 |
| 2 | 56189618 | rs1432559 | -0.946 | 0.699 | -0.03 | -2.246 | 0.018 | -0.111 | -0.111 | 0 | 0.22 | 0.5 | 0.21 | 0.05 | 0.2 | 0.22 |
| 2 | 56190273 | rs6735393 | -0.632 | 0.68 | 0.067 | -2.264 | 0.054 | -0.077 | -0.077 | 0.036 | 0.16 | 0.4 | 0.17 | 0.05 | 0.17 | 0.21 |
| 2 | 56191470 | rs2868985 | -2.147 | 0.369 | -0.77 | -1.869 | 0.067 | -0.077 | -0.077 | 1.735 | 0.7 | 0.97 | 0.87 | 0.4 | 0.81 | 0.71 |
| 2 | 56191794 | rs724540 | -1.359 | 0.789 | -0.384 | -1.95 | 0.03 | 0.24 | 0.24 | 0.9 | 0.49 | 0.86 | 0.76 | 0.22 | 0.61 | 0.54 |
| 2 | 56193418 | rs6747490 | -0.819 | 0.81 | -0.375 | -1.918 | -0.007 | 0.069 | 0.069 | 0 | 0.43 | 0.7 | 0.71 | 0.22 | 0.59 | 0.52 |
| 2 | 56198401 | rs17047369 | 0.607 | 0.836 | -0.19 | -1.817 | 0.018 | -0.046 | -0.046 | 0.1 | 0.1 | 0.15 | 0.1 | 0.02 | 0.11 | 0.11 |
| 2 | 56199062 | rs6733889 | -1.026 | 0.656 | -0.601 | -1.727 | 0.043 | 0.183 | 0.183 | 0.617 | 0.49 | 0.8 | 0.76 | 0.22 | 0.62 | 0.53 |
| 2 | 56203798 | rs6545534 | -0.582 | 1.028 | -0.309 | -1.547 | 0.037 | -0.057 | -0.057 | 0 | 0.34 | 0.55 | 0.57 | 0.17 | 0.51 | 0.36 |
| 2 | 56214516 | rs6545535 | -0.487 | 1.004 | -0.554 | -1.701 | 0 | 0.006 | 0.006 | 1.029 | 0.31 | 0.7 | 0.26 | 0.07 | 0.28 | 0.42 |
| 2 | 56216904 | rs7558150 | -1.066 | 0.781 | -0.971 | -1.55 | 0.022 | -0.027 | -0.027 | 0 | 0.89 | 0.96 | 0.91 | 0.87 | 0.92 | 0.92 |
| 2 | 56236854 | rs1368242 | 0.322 | 1.361 | -0.802 | -1.55 | -0.077 | 0.022 | 0.022 | 0 | 0.21 | 0.09 | 0.42 | 0.15 | 0.37 | 0.16 |
| 2 | 56244096 | rs4553860 | -0.387 | 1.118 | -0.028 | -1.501 | -0.004 | -0.08 | -0.08 | 0.654 | 0.27 | 0.53 | 0.21 | 0.06 | 0.23 | 0.36 |
| 2 | 56249038 | rs11886020 | -0.204 | 1.073 | 0.15 | -1.455 | -0.002 | -0.103 | -0.103 | 0.015 | 0.24 | 0.47 | 0.18 | 0.07 | 0.2 | 0.27 |
| 2 | 56251912 | rs11901093 | 0.121 | 1.126 | -0.406 | -1.446 | 0.008 | -0.036 | -0.036 | 0 | 0.16 | 0.22 | 0.15 | 0.05 | 0.18 | 0.22 |
| 2 | 56253505 | rs13418952 | 1.296 | 1.143 | -0.272 | -1.473 | -0.001 | -0.068 | -0.068 | 0 | 0.11 | 0.32 | 0.09 | 0 | 0.08 | 0.09 |
| 2 | 56255832 | rs3850353 | 0.066 | 1.61 | -0.507 | -1.395 | -0.156 | 0.003 | 0.003 | 0 | 0.23 | 0.03 | 0.41 | 0.17 | 0.39 | 0.17 |
| 2 | 56257941 | rs13023665 | -0.13 | 1.709 | 0.13 | -1.198 | 0.008 | -0.03 | -0.03 | 0 | 0.16 | 0.22 | 0.15 | 0.07 | 0.18 | 0.22 |
| 2 | 56265789 | rs17279532 | 0.413 | 1.946 | -0.944 | -0.959 | -0.076 | 0.023 | 0.023 | 0 | 0.16 | 0.05 | 0.39 | 0.13 | 0.3 | 0.18 |
| 2 | 56265813 | rs6753577 | 0.683 | 0.588 | -0.347 | -0.744 | 0.017 | 0.013 | 0.013 | 0 | 0.45 | 0.58 | 0.25 | 0.61 | 0.28 | 0.34 |
| 2 | 56267182 | rs13417586 | 0.162 | 0.638 | -0.229 | -0.564 | -0.001 | -0.013 | -0.013 | 0 | 0.12 | 0.17 | 0.12 | 0.07 | 0.09 | 0.32 |
| 2 | 56269883 | rs7590475 | 0.654 | 0.224 | -0.278 | -0.568 | 0.099 | -0.008 | -0.008 | 2.348 | 0.42 | 0.2 | 0.18 | 0.58 | 0.27 | 0.3 |
| 2 | 56273704 | rs1368243 | -0.102 | 0.801 | 0.081 | -0.647 | 0.018 | 0.001 | 0.001 | 0 | 0.1 | 0.14 | 0.14 | 0.08 | 0.18 | 0.18 |
| 2 | 56274120 | rs2868995 | 0.651 | 0.664 | 0.493 | -0.693 | 0.021 | 0.002 | 0.002 | 0 | 0.43 | 0.39 | 0.24 | 0.58 | 0.28 | 0.44 |

| | | | | | | | | | | | | | | | | |
|---|---|---|---|---|---|---|---|---|---|---|---|---|---|---|---|---|
| 2 | 56280428 | rs891881 | 0.201 | 1.101 | 0.546 | -0.831 | -0.041 | 0.043 | 0.043 | 0 | 0.31 | 0.24 | 0.54 | 0.19 | 0.45 | 0.31 |
| 2 | 56292883 | rs13401550 | 1.945 | 0.607 | 0.523 | -0.743 | 0.004 | -0.008 | -0.008 | 0 | 0.27 | 0.26 | 0.12 | 0.5 | 0.2 | 0.11 |
| 2 | 56297858 | rs7564258 | 1.686 | 0.66 | 0.481 | -0.758 | -0.023 | 0.024 | 0.024 | 0 | 0.29 | 0.36 | 0.16 | 0.51 | 0.21 | 0.25 |
| 2 | 56300279 | rs1432547 | -0.147 | 0.676 | 0.355 | -0.804 | 0.005 | 0.011 | 0.011 | 0 | 0.78 | 0.76 | 0.75 | 0.76 | 0.73 | 0.74 |
| 2 | 56309560 | rs7603021 | -1.871 | 0.689 | 0.166 | -0.747 | 0.006 | -0.014 | -0.014 | 0 | 0.88 | 0.95 | 0.89 | 0.84 | 0.89 | 0.92 |
| 2 | 56315073 | rs973238 | -0.645 | 0.689 | 0.219 | -0.865 | 0.04 | 0.093 | 0.093 | 0.07 | 0.51 | 0.33 | 0.65 | 0.26 | 0.57 | 0.58 |
| 2 | 56316188 | rs17047488 | 1.297 | 0.769 | -0.156 | -0.808 | -0.026 | -0.008 | -0.008 | 0 | 0.1 | 0 | 0.07 | 0.09 | 0.09 | 0.19 |
| 2 | 56326296 | rs3850354 | -1.956 | 0.74 | -0.14 | -0.772 | 0.005 | -0.021 | -0.021 | 0 | 0.88 | 0.95 | 0.9 | 0.82 | 0.88 | 0.93 |
| 2 | 56330148 | rs10166672 | 0.735 | 0.749 | -0.149 | -0.841 | 0.005 | 0.048 | 0.048 | 0.134 | 0.18 | 0.03 | 0.09 | 0.09 | 0.12 | 0.25 |
| 2 | 56331228 | rs1432567 | 0.835 | 0.985 | 0.044 | -0.799 | -0.007 | 0.005 | 0.005 | 0 | 0.28 | 0.26 | 0.53 | 0.17 | 0.37 | 0.38 |
| 2 | 56331375 | rs6545542 | 0.83 | 0.797 | 0.056 | -0.726 | 0.008 | 0.002 | 0.002 | 0 | 0.1 | 0.12 | 0.1 | 0.1 | 0.18 | 0.17 |
| 2 | 56332596 | rs13431320 | 2.112 | 0.712 | 0.176 | -0.732 | 0.022 | -0.032 | -0.032 | 2.201 | 0.27 | 0.21 | 0.12 | 0.46 | 0.19 | 0.1 |
| 2 | 56333730 | rs4347872 | 0.493 | 0.869 | -0.223 | -0.69 | -0.021 | 0.018 | 0.018 | 0 | 0.32 | 0.27 | 0.56 | 0.2 | 0.44 | 0.47 |
| 2 | 56336173 | rs2868994 | 0.289 | 0.888 | -0.18 | -0.68 | -0.024 | 0.021 | 0.021 | 0 | 0.32 | 0.28 | 0.55 | 0.2 | 0.44 | 0.32 |
| 2 | 56338393 | rs7585128 | 1.899 | 0.684 | -0.09 | -0.527 | -0.05 | 0.062 | 0.062 | 0 | 0.28 | 0.45 | 0.16 | 0.42 | 0.2 | 0.13 |
| 2 | 56338546 | rs6545543 | 0.965 | 0.984 | -0.724 | -0.53 | -0.001 | 0.017 | 0.017 | 0 | 0.57 | 0.73 | 0.68 | 0.63 | 0.56 | 0.5 |
| 2 | 56341068 | rs1036165 | -0.471 | 1.106 | -0.661 | -0.423 | -0.002 | -0.007 | -0.007 | 0 | 0.8 | 0.83 | 0.82 | 0.76 | 0.76 | 0.84 |
| 2 | 56347309 | rs2099305 | -0.025 | 1.279 | -0.467 | -0.483 | -0.001 | -0.001 | -0.001 | 0.274 | 0.17 | 0.16 | 0.13 | 0.16 | 0.18 | 0.26 |
| 2 | 56349939 | rs2023514 | -0.478 | 1.284 | -0.537 | -0.488 | -0.016 | 0.018 | 0.018 | 0 | 0.27 | 0.16 | 0.18 | 0.18 | 0.29 | 0.42 |
| 2 | 56354019 | rs1368245 | -0.755 | 1.166 | -0.738 | -0.496 | -0.001 | 0 | 0 | 0.684 | 0.3 | 0.66 | 0.22 | 0.17 | 0.23 | 0.37 |
| 2 | 56354692 | rs4672087 | -0.08 | 1.784 | -0.846 | -0.588 | 0.009 | -0.004 | -0.004 | 0 | 0.1 | 0.13 | 0.19 | 0.06 | 0.22 | 0.29 |
| 2 | 56355315 | rs2021796 | 0.851 | 1.802 | -0.517 | -0.604 | 0.076 | 0.063 | 0.063 | 0.252 | 0.09 | 0.36 | 0.1 | 0.07 | 0.11 | 0.09 |
| 2 | 56357582 | rs929689 | -0.713 | 1.416 | -0.897 | -0.633 | 0.013 | 0.065 | 0.065 | 0 | 0.49 | 0.66 | 0.58 | 0.3 | 0.47 | 0.52 |
| 2 | 56357846 | rs1811272 | -1.414 | 1.353 | -0.907 | -0.617 | 0.07 | 0.01 | 0.01 | 0.022 | 0.59 | 0.78 | 0.77 | 0.36 | 0.68 | 0.81 |
| 2 | 56363377 | rs6751715 | -0.278 | 1.487 | -0.809 | -0.549 | -0.008 | 0.005 | 0.005 | 0 | 0.31 | 0.25 | 0.56 | 0.21 | 0.44 | 0.48 |
| 2 | 56369213 | rs6739384 | -0.105 | 1.533 | -0.806 | -0.508 | 0.007 | -0.002 | -0.002 | 0.043 | 0.1 | 0.12 | 0.19 | 0.07 | 0.22 | 0.28 |
| 2 | 56370879 | rs3850355 | -0.127 | 1.514 | -0.759 | -0.529 | -0.02 | 0.014 | 0.014 | 0 | 0.29 | 0.24 | 0.56 | 0.21 | 0.43 | 0.45 |
| 2 | 56374910 | rs2216322 | 1.01 | 0.834 | -1.009 | -0.375 | -0.002 | 0.002 | 0.002 | 0 | 0.22 | 0.16 | 0.38 | 0.15 | 0.23 | 0.19 |
| 2 | 56376308 | rs2111469 | 0.731 | 0.657 | -0.989 | -0.293 | -0.002 | 0.001 | 0.001 | 0 | 0.26 | 0.22 | 0.44 | 0.18 | 0.32 | 0.4 |
| 2 | 56376564 | rs7609306 | 0.52 | 0.602 | -0.98 | -0.304 | 0.066 | -0.005 | -0.005 | 0.495 | 0.23 | 0.45 | 0.29 | 0.19 | 0.31 | 0.35 |
| 2 | 56376801 | rs17047515 | 1.6 | 0.344 | -1.402 | -0.243 | 0.038 | -0.051 | -0.051 | 0 | 0.27 | 0.2 | 0.13 | 0.48 | 0.18 | 0.1 |
| 2 | 56381698 | rs1835893 | -0.715 | 0.495 | -1.766 | -0.246 | -0.002 | 0.047 | 0.047 | 0 | 0.77 | 0.95 | 0.86 | 0.85 | 0.81 | 0.85 |
| 2 | 56381739 | rs17268598 | 0.621 | 0.536 | -1.451 | -0.002 | 0.006 | -0.014 | -0.014 | 0 | 0.27 | 0.31 | 0.45 | 0.18 | 0.32 | 0.41 |
| 2 | 56383762 | rs6545554 | 0.738 | 1.285 | -1.155 | 0.097 | -0.001 | -0.005 | -0.005 | 1.039 | 0.54 | 0.47 | 0.57 | 0.66 | 0.5 | 0.5 |
| 2 | 56393918 | rs4335989 | 0.196 | 1.111 | -1.442 | 0.462 | -0.002 | 0.005 | 0.005 | 0 | 0.84 | 0.94 | 0.88 | 0.85 | 0.85 | 0.91 |
| 2 | 56396433 | rs10176451 | -0.049 | 1.012 | -1.236 | 0.602 | 0.005 | 0.005 | 0.005 | 0 | 0.4 | 0.69 | 0.45 | 0.3 | 0.37 | 0.27 |
| 2 | 56397566 | rs7589176 | -0.059 | 1.174 | -0.828 | 0.393 | -0.002 | -0.015 | -0.015 | 0 | 0.4 | 0.51 | 0.47 | 0.28 | 0.42 | 0.25 |
| 2 | 56399334 | rs4672095 | -0.165 | 1.123 | -0.754 | 0.352 | -0.001 | -0.01 | -0.01 | 0.065 | 0.4 | 0.56 | 0.47 | 0.27 | 0.42 | 0.25 |
| 2 | 56402808 | rs1861774 | -0.123 | 1.097 | -0.906 | 0.074 | 0.01 | 0.115 | 0.115 | 0 | 0.46 | 0.25 | 0.44 | 0.2 | 0.43 | 0.29 |
| 2 | 56402848 | rs1861773 | 0.163 | 1.797 | -0.741 | -0.117 | 0.038 | -0.038 | -0.038 | 0 | 0.78 | 0.69 | 0.79 | 0.89 | 0.71 | 0.75 |
| 2 | 56403345 | rs7419516 | -0.369 | 1.777 | -0.551 | -0.057 | 0.077 | -0.021 | -0.021 | 1.947 | 0.83 | 0.71 | 0.79 | 0.89 | 0.68 | 0.77 |
| 2 | 56408030 | rs1814277 | 1.178 | 0.949 | -0.929 | 0.162 | 0.065 | -0.022 | -0.022 | 0 | 0.32 | 0.12 | 0.33 | 0.27 | 0.26 | 0.18 |
| 2 | 56411157 | rs960490 | -0.606 | 1.064 | -0.117 | 0.073 | 0.018 | 0.011 | 0.011 | 0 | 0.89 | 0.73 | 0.89 | 0.89 | 0.9 | 0.85 |
| 2 | 56411363 | rs960489 | -0.65 | 1.098 | -0.074 | 0.091 | 0.019 | 0.012 | 0.012 | 0 | 0.89 | 0.73 | 0.89 | 0.89 | 0.9 | 0.83 |
| 2 | 56421972 | rs1016339 | -0.358 | 0.641 | -0.02 | -0.115 | -0.012 | -0.043 | -0.043 | 0.892 | 0.24 | 0.35 | 0.25 | 0.13 | 0.18 | 0.2 |
| 2 | 56422980 | rs6728383 | 0.742 | 0.591 | -0.071 | 0.206 | 0.002 | -0.002 | -0.002 | 0 | 0.09 | 0.09 | 0.06 | 0.17 | 0.08 | 0.11 |
| 2 | 56423598 | rs7594008 | -0.209 | 0.506 | 0.275 | 0.127 | 0.005 | -0.001 | -0.001 | 0 | 0.14 | 0.18 | 0.22 | 0.14 | 0.14 | 0.15 |
| 2 | 56428584 | rs9967877 | -0.241 | 0.493 | 0.119 | -0.053 | 0.006 | -0.013 | -0.013 | 0.228 | 0.13 | 0.19 | 0.1 | 0.1 | 0.13 | 0.13 |
| 2 | 56431594 | rs6756156 | -0.513 | 0.588 | 0.091 | 0.137 | 0.012 | 0.01 | 0.01 | 0 | 0.85 | 0.78 | 0.86 | 0.84 | 0.84 | 0.69 |
| 2 | 56435494 | rs2869526 | 0.183 | 0.249 | -0.657 | -0.024 | 0.029 | 0.035 | 0.035 | 0 | 0.58 | 0.79 | 0.57 | 0.33 | 0.63 | 0.46 |
| 2 | 56438189 | rs6749100 | -0.644 | 0.213 | -0.612 | -0.154 | -0.006 | -0.003 | -0.003 | 0 | 0.38 | 0.62 | 0.49 | 0.42 | 0.49 | 0.54 |
| 2 | 56440166 | rs2216325 | 0.388 | 0.188 | -0.402 | -0.152 | 0.059 | 0.04 | 0.04 | 0 | 0.25 | 0.37 | 0.39 | 0.36 | 0.4 | 0.31 |
| 2 | 56443708 | rs17047623 | 0.814 | 0.349 | -0.307 | -0.152 | 0.28 | -0.018 | -0.018 | 0 | 0.08 | 0.31 | 0.13 | 0 | 0.22 | 0.05 |
| 2 | 56444887 | rs2869527 | -0.197 | 0.157 | -0.441 | -0.189 | 0 | 0 | 0 | 3.773 | 0.8 | 0.84 | 0.84 | 0.65 | 0.83 | 0.74 |
| 2 | 56450038 | rs7579447 | 1.479 | 0.04 | -0.577 | -0.326 | 0.08 | 0.038 | 0.038 | 1.368 | 0.51 | 0.11 | 0.61 | 0.6 | 0.43 | 0.4 |
| 2 | 56450431 | rs6545559 | 0.749 | 0.057 | -0.493 | -0.328 | -0.003 | -0.001 | -0.001 | 0 | 0.61 | 0.55 | 0.75 | 0.62 | 0.68 | 0.57 |
| 2 | 56450856 | rs6733298 | -1.601 | 0.173 | -0.409 | -0.229 | -0.005 | -0.006 | -0.006 | 0 | 0.88 | 0.91 | 0.88 | 0.85 | 0.87 | 0.81 |
| 2 | 56452097 | rs6545560 | 0.741 | 0.182 | -0.502 | -0.237 | 0.286 | -0.02 | -0.02 | 0.003 | 0.08 | 0.33 | 0.13 | 0 | 0.22 | 0.05 |
| 2 | 56456736 | rs6545561 | -1.539 | 0.026 | -0.743 | -0.208 | -0.046 | -0.059 | -0.059 | 0 | 0.89 | 1 | 0.89 | 0.84 | 0.87 | 0.81 |
| 2 | 56457608 | rs17047642 | -0.352 | 0.034 | -0.667 | -0.481 | -0.006 | -0.061 | -0.061 | 0 | 0.15 | 0.32 | 0.1 | 0.03 | 0.12 | 0.14 |
| 2 | 56458345 | rs4672099 | 1.927 | -0.083 | -1.009 | -0.478 | 0.042 | 0.024 | 0.024 | 0.467 | 0.33 | 0.1 | 0.35 | 0.24 | 0.26 | 0.14 |
| 2 | 56459427 | rs2869528 | -0.593 | 0.13 | -0.741 | -0.484 | -0.001 | 0.01 | 0.01 | 0 | 0.81 | 0.82 | 0.84 | 0.65 | 0.83 | 0.76 |
| 2 | 56460013 | rs888279 | 0.28 | 0.153 | -0.671 | -0.464 | -0.003 | 0.022 | 0.022 | 0 | 0.7 | 0.57 | 0.74 | 0.61 | 0.72 | 0.62 |
| 2 | 56461082 | rs12612590 | 0.731 | -0.066 | -1.084 | -0.409 | -0.001 | -0.006 | -0.006 | 0 | 0.6 | 0.31 | 0.62 | 0.59 | 0.46 | 0.45 |
| 2 | 56462994 | rs17047655 | -0.185 | -0.044 | -0.82 | -0.417 | -0.033 | -0.069 | -0.069 | 1.144 | 0.14 | 0.32 | 0.05 | 0.02 | 0.06 | 0.16 |
| 2 | 56464386 | rs7568979 | 0.658 | -0.07 | -1.631 | -0.39 | 0.045 | 0.104 | 0.104 | 0.02 | 0.5 | 0.15 | 0.45 | 0.35 | 0.44 | 0.41 |
| 2 | 56466257 | rs12993884 | 1.33 | 0.006 | -1.507 | -0.262 | 0.014 | -0.07 | -0.07 | 0 | 0.17 | 0.05 | 0.27 | 0.3 | 0.17 | 0.25 |
| 2 | 56466868 | rs969302 | 0.884 | -0.011 | -1.193 | -0.27 | -0.009 | 0.011 | 0.011 | 0 | 0.2 | 0.26 | 0.28 | 0.3 | 0.17 | 0.25 |
| 2 | 56467339 | rs6730327 | -0.975 | 0.167 | -0.974 | -0.393 | 0.002 | 0.021 | 0.021 | 0.352 | 0.44 | 0.7 | 0.49 | 0.33 | 0.54 | 0.48 |

| | | | | | | | | | | | | | | | | |
|---|---|---|---|---|---|---|---|---|---|---|---|---|---|---|---|---|
| 2 | 56472159 | rs2869529 | -0.495 | 0.245 | -1.007 | -0.4 | 0.148 | -0.117 | -0.117 | 0.003 | 0.2 | 0.46 | 0.2 | 0.03 | 0.36 | 0.18 |
| 2 | 56476401 | rs12613716 | 0.341 | -0.046 | -1.138 | -0.276 | 0.027 | -0.017 | -0.017 | 0 | 0.26 | 0.18 | 0.29 | 0.3 | 0.2 | 0.3 |
| 2 | 56480571 | rs759615 | -0.665 | 0.033 | -1.126 | -0.271 | 0.027 | 0.012 | 0.012 | 0 | 0.41 | 0.34 | 0.37 | 0.35 | 0.35 | 0.54 |
| 2 | 56489455 | rs4672103 | 1.903 | 0.333 | -1.157 | 0.011 | 0.02 | -0.032 | -0.032 | 0 | 0.28 | 0.07 | 0.29 | 0.28 | 0.22 | 0.17 |
| 2 | 56490680 | rs214047 | 0.536 | 0.368 | -0.993 | 0.012 | -0.009 | 0.043 | 0.043 | 0 | 0.21 | 0.3 | 0.17 | 0.33 | 0.17 | 0.28 |
| 2 | 56492112 | rs2193485 | -0.057 | 0.539 | -0.918 | 0.031 | 0.015 | -0.007 | -0.007 | 0.067 | 0.36 | 0.29 | 0.38 | 0.36 | 0.33 | 0.38 |
| 2 | 56496589 | rs214045 | -0.587 | 0.691 | -0.881 | -0.051 | 0.171 | -0.099 | -0.099 | 0 | 0.85 | 0.58 | 0.84 | 1 | 0.74 | 0.93 |
| 2 | 56498664 | rs1001471 | 0.878 | 0.256 | -0.857 | 0.521 | 0.011 | -0.136 | -0.136 | 0.771 | 0.24 | 0.03 | 0.29 | 0.35 | 0.21 | 0.26 |
| 2 | 56499879 | rs2111475 | -0.307 | 0.321 | -0.765 | 0.534 | 0.027 | -0.045 | -0.045 | 0.162 | 0.36 | 0.26 | 0.38 | 0.46 | 0.33 | 0.38 |
| 2 | 56501688 | rs214044 | -0.679 | 0.077 | -0.6 | 0.463 | -0.006 | -0.008 | -0.008 | 0.438 | 0.45 | 0.44 | 0.46 | 0.7 | 0.38 | 0.54 |
| 2 | 56501821 | rs7576924 | -0.325 | 0.091 | -0.746 | 0.401 | 0.019 | -0.014 | -0.014 | 2.907 | 0.3 | 0.23 | 0.31 | 0.4 | 0.24 | 0.38 |
| 2 | 56503029 | rs214041 | 0.597 | 1.349 | -0.179 | 0.175 | 0.024 | -0.026 | -0.026 | 0 | 0.75 | 0.68 | 0.8 | 0.84 | 0.65 | 0.74 |
| 2 | 56507299 | rs888281 | -0.655 | 0.807 | 0.345 | 0.412 | 0.015 | 0.06 | 0.06 | 0 | 0.3 | 0.51 | 0.36 | 0.54 | 0.31 | 0.39 |
| 2 | 56507552 | rs1549030 | -0.609 | 0.753 | 0.689 | 0.515 | 0.017 | 0.148 | 0.148 | 0 | 0.23 | 0.47 | 0.29 | 0.51 | 0.23 | 0.3 |
| 2 | 56513819 | rs13410910 | 0.058 | 0.883 | 0.171 | 0.792 | -0.026 | -0.071 | -0.071 | 0 | 0.14 | 0.03 | 0.07 | 0.35 | 0.12 | 0.21 |
| 2 | 56524334 | rs214038 | -0.231 | 1.571 | 0.298 | 0.438 | 0.025 | -0.025 | -0.025 | 0 | 0.92 | 0.89 | 0.87 | 1 | 0.79 | 0.95 |
| 2 | 56527602 | rs7561764 | -1.005 | 0.045 | 0.579 | 0.341 | -0.01 | 0.016 | 0.016 | 0.483 | 0.26 | 0.68 | 0.14 | 0.37 | 0.13 | 0.27 |
| 2 | 56528075 | rs12469239 | -1.372 | 0.065 | 0.535 | 0.497 | -0.003 | 0.005 | 0.005 | 0.195 | 0.3 | 0.67 | 0.38 | 0.54 | 0.23 | 0.37 |
| 2 | 56528373 | rs4671278 | -0.459 | 0.028 | 0.574 | 0.422 | 0.002 | -0.003 | -0.003 | 0 | 0.12 | 0.1 | 0.28 | 0.2 | 0.12 | 0.17 |
| 2 | 56552703 | rs10203280 | -0.88 | -0.3 | 0.744 | 0.311 | -0.008 | 0 | 0 | 0 | 0.18 | 0.19 | 0.3 | 0.22 | 0.13 | 0.21 |
| 2 | 56553865 | rs12616518 | 1.219 | -0.746 | -0.045 | -0.037 | -0.007 | -0.023 | -0.023 | 0 | 0.15 | 0.01 | 0.08 | 0.15 | 0.16 | 0.12 |
| 2 | 56555581 | rs10172020 | -0.455 | -0.616 | 0.515 | -0.002 | 0.01 | 0.005 | 0.005 | 0 | 0.16 | 0.14 | 0.28 | 0.17 | 0.13 | 0.15 |
| 2 | 56560602 | rs6722198 | -1.288 | -0.582 | 0.799 | -0.026 | 0 | -0.001 | -0.001 | 0.162 | 0.34 | 0.68 | 0.38 | 0.53 | 0.25 | 0.36 |
| 2 | 56563822 | rs9808416 | -0.125 | -0.244 | 1.034 | 0.126 | 0.018 | 0.153 | 0.153 | 0 | 0.17 | 0.44 | 0.09 | 0.32 | 0.12 | 0.21 |
| 2 | 56575020 | rs10200604 | -0.63 | -0.301 | 0.834 | -0.357 | -0.025 | -0.03 | -0.03 | 0 | 0.24 | 0.59 | 0.36 | 0.16 | 0.18 | 0.13 |
| 2 | 56580537 | rs17047819 | 0.185 | -0.393 | 0.381 | -0.16 | -0.001 | -0.009 | -0.009 | 0 | 0.18 | 0.18 | 0.07 | 0.27 | 0.13 | 0.2 |
| 2 | 56580933 | rs6760801 | 0.442 | -0.35 | 0.057 | -0.158 | -0.019 | -0.059 | -0.059 | 0 | 0.17 | 0.05 | 0.07 | 0.27 | 0.13 | 0.2 |
| 2 | 56581199 | rs2193481 | -1.016 | -0.439 | 0.284 | -0.275 | 0.01 | -0.008 | -0.008 | 7.072 | 0.41 | 0.68 | 0.41 | 0.42 | 0.31 | 0.32 |
| 2 | 56583628 | rs6545573 | -0.379 | -0.735 | 0.18 | -1.171 | -0.005 | -0.028 | -0.028 | 0 | 0.83 | 0.84 | 0.8 | 0.97 | 0.88 | 0.86 |
| 2 | 56585774 | rs7561980 | -0.64 | -0.74 | 0.303 | -1.173 | -0.012 | -0.069 | -0.069 | 0.175 | 0.85 | 0.73 | 0.79 | 0.97 | 0.88 | 0.86 |
| 2 | 56592639 | rs11691570 | -0.038 | -0.784 | -0.163 | -1.418 | 0.012 | 0.039 | 0.039 | 0.002 | 0.16 | 0.07 | 0.08 | 0.03 | 0.12 | 0.08 |
| 2 | 56593901 | rs6714635 | 1.363 | -0.712 | -0.19 | -1.427 | 0.036 | 0.157 | 0.157 | 0.092 | 0.49 | 0.23 | 0.45 | 0.78 | 0.58 | 0.64 |
| 2 | 56594487 | rs10180182 | -0.57 | -0.481 | -0.238 | -1.146 | 0.017 | 0.03 | 0.03 | 0 | 0.84 | 0.9 | 0.8 | 0.97 | 0.89 | 0.85 |
| 2 | 56598309 | rs17047854 | -0.218 | 0.062 | 0.178 | -0.604 | -0.003 | -0.013 | -0.013 | 0.426 | 0.18 | 0.41 | 0.24 | 0.15 | 0.17 | 0.12 |
| 2 | 56599354 | rs11694209 | -0.427 | 0.323 | -0.515 | -0.396 | 0 | -0.029 | -0.029 | 0 | 0.11 | 0.12 | 0.08 | 0.03 | 0.12 | 0.08 |
| 2 | 56607046 | rs13423780 | -0.439 | 0.106 | -0.573 | -0.558 | -0.007 | -0.043 | -0.043 | 0 | 0.22 | 0.21 | 0.13 | 0.04 | 0.18 | 0.18 |
| 2 | 56607843 | rs17268820 | -0.542 | 0.341 | -0.831 | -0.682 | -0.012 | 0.044 | 0.044 | 0 | 0.09 | 0.01 | 0.07 | 0 | 0.08 | 0.06 |
| 2 | 56608816 | rs2193488 | -0.835 | 0.101 | -0.224 | -0.705 | -0.004 | 0.007 | 0.007 | 0.66 | 0.08 | 0.21 | 0.05 | 0.09 | 0.07 | 0.11 |
| 2 | 56609328 | rs17190254 | 0.228 | -0.027 | -0.154 | -1.103 | -0.002 | -0.015 | -0.015 | 0 | 0.13 | 0.11 | 0.15 | 0.05 | 0.1 | 0.12 |
| 2 | 56611725 | rs11902680 | -1.425 | -0.125 | 0.567 | -1.354 | -0.005 | -0.044 | -0.044 | 0 | 0.34 | 0.58 | 0.29 | 0.13 | 0.29 | 0.29 |
| 2 | 56622471 | rs10191130 | -0.685 | -0.133 | 0.832 | -1.14 | -0.004 | 0.003 | 0.003 | 0.368 | 0.08 | 0.15 | 0.04 | 0.04 | 0.07 | 0.11 |
| 2 | 56634019 | rs4672116 | -0.83 | 0.335 | 1.009 | -0.344 | -0.07 | -0.023 | -0.023 | 0.054 | 0.36 | 0.17 | 0.41 | 0.59 | 0.44 | 0.46 |
| 2 | 56643955 | rs7600814 | -1.276 | 0.283 | 1.503 | 0.216 | -0.024 | 0.013 | 0.013 | 1.128 | 0.42 | 0.75 | 0.5 | 0.52 | 0.49 | 0.5 |
| 2 | 56656894 | rs1861776 | -1.095 | 0.331 | 2.282 | -0.105 | -0.007 | 0.005 | 0.005 | 0.279 | 0.39 | 0.33 | 0.44 | 0.63 | 0.47 | 0.5 |
| 2 | 56666949 | rs10202807 | -1.181 | 0.331 | 2.51 | -0.107 | -0.004 | 0.005 | 0.005 | 0.777 | 0.42 | 0.62 | 0.54 | 0.66 | 0.48 | 0.5 |
| 2 | 56672309 | rs11903884 | -0.789 | 0.341 | 2.492 | -0.096 | -0.014 | -0.004 | -0.004 | 0 | 0.36 | 0.3 | 0.41 | 0.59 | 0.43 | 0.45 |
| 2 | 56682350 | rs17047952 | -0.801 | 0.342 | 2.43 | -0.095 | -0.008 | -0.002 | -0.002 | 0 | 0.36 | 0.34 | 0.41 | 0.59 | 0.43 | 0.45 |
| 2 | 56682547 | rs13397156 | -1.121 | 0.334 | 2.421 | -0.104 | 0.011 | -0.012 | -0.012 | 0 | 0.39 | 0.46 | 0.44 | 0.63 | 0.48 | 0.5 |
| 2 | 56708945 | rs879807 | -1.254 | 0.441 | 2.028 | -0.261 | -0.032 | 0.075 | 0.075 | 0.596 | 0.42 | 0.74 | 0.56 | 0.69 | 0.49 | 0.51 |
| 2 | 56712426 | rs2194803 | 0.207 | 0.509 | 1.91 | -0.217 | -0.005 | -0.013 | -0.013 | 0 | 0.2 | 0.15 | 0.32 | 0.59 | 0.26 | 0.27 |
| 2 | 56730155 | rs17048024 | -0.726 | -0.155 | 1.779 | -0.281 | -0.045 | 0.026 | 0.026 | 0.19 | 0.4 | 0.21 | 0.43 | 0.62 | 0.48 | 0.46 |
| 2 | 56731479 | rs1077270 | -1.465 | -0.829 | 1.474 | -0.559 | -0.063 | 0.25 | 0.25 | 0 | 0.55 | 0.92 | 0.69 | 0.83 | 0.54 | 0.63 |
| 2 | 56739650 | rs982116 | -0.315 | -0.767 | 1.461 | -0.538 | -0.049 | 0.032 | 0.032 | 0.324 | 0.4 | 0.19 | 0.44 | 0.63 | 0.48 | 0.47 |
| 2 | 56751574 | rs987577 | -1.154 | -0.642 | 1.439 | -0.64 | 0.005 | 0.095 | 0.095 | 0 | 0.53 | 0.7 | 0.68 | 0.69 | 0.56 | 0.52 |
| 2 | 56752025 | rs987576 | -0.481 | -0.579 | 1.486 | -0.61 | -0.003 | 0.005 | 0.005 | 0.196 | 0.43 | 0.54 | 0.48 | 0.63 | 0.51 | 0.48 |
| 2 | 56756857 | rs6718973 | -0.871 | -0.573 | 1.507 | -0.622 | -0.002 | 0.08 | 0.08 | 0 | 0.48 | 0.7 | 0.58 | 0.66 | 0.5 | 0.51 |
| 2 | 56757257 | rs6735514 | -1.076 | -0.715 | 1.52 | -0.613 | 0.007 | 0.049 | 0.049 | 0 | 0.53 | 0.67 | 0.61 | 0.66 | 0.56 | 0.53 |
| 2 | 56760592 | rs2870064 | -0.637 | -1.03 | 1.336 | -0.096 | 0.013 | 0.021 | 0.021 | 0 | 0.5 | 0.38 | 0.47 | 0.51 | 0.53 | 0.54 |
| 2 | 56761626 | rs4671282 | -0.39 | -1.026 | 1.436 | -0.543 | -0.014 | 0.062 | 0.062 | 0 | 0.48 | 0.2 | 0.44 | 0.59 | 0.52 | 0.47 |
| 2 | 56762887 | rs7578177 | 0.838 | -0.985 | 0.851 | -0.828 | 0.002 | 0.007 | 0.007 | 0 | 0.1 | 0.1 | 0.05 | 0.06 | 0.09 | 0.03 |
| 2 | 56764646 | rs996846 | -0.17 | -1.04 | 1.126 | -0.856 | 0.001 | 0.002 | 0.002 | 0.557 | 0.48 | 0.47 | 0.54 | 0.63 | 0.56 | 0.52 |
| 2 | 56765038 | rs1160086 | 0.156 | -0.985 | 1.157 | -0.836 | 0.001 | -0.006 | -0.006 | 0.018 | 0.43 | 0.46 | 0.49 | 0.62 | 0.52 | 0.5 |
| 2 | 56766125 | rs7586505 | -1.323 | -0.926 | 0.969 | -0.842 | 0.004 | 0.034 | 0.034 | 0 | 0.63 | 0.84 | 0.59 | 0.69 | 0.66 | 0.55 |
| 2 | 56771307 | rs17039629 | 0.8 | -0.945 | 0.815 | -0.84 | -0.084 | -0.015 | -0.015 | 0.023 | 0.36 | 0.15 | 0.35 | 0.59 | 0.47 | 0.46 |
| 2 | 56771330 | rs17190394 | 0.609 | -0.98 | 0.721 | -0.863 | -0.008 | 0.051 | 0.051 | 0 | 0.14 | 0.02 | 0.06 | 0.06 | 0.14 | 0.04 |
| 2 | 56772337 | rs10490411 | 0.604 | -0.922 | 1.111 | -0.814 | 0.018 | -0.003 | -0.003 | 0.003 | 0.37 | 0.4 | 0.37 | 0.59 | 0.47 | 0.46 |
| 2 | 56784574 | rs6760940 | -0.302 | -0.926 | 1.094 | -0.806 | 0.01 | 0.045 | 0.045 | 0 | 0.49 | 0.66 | 0.51 | 0.62 | 0.53 | 0.51 |
| 2 | 56792506 | rs17048168 | 0.715 | -0.821 | 0.795 | -0.753 | -0.065 | -0.012 | -0.012 | 0 | 0.36 | 0.17 | 0.35 | 0.59 | 0.47 | 0.46 |
| 2 | 56799685 | rs7565891 | -1.545 | -1.237 | 0.39 | -1.022 | -0.038 | 0.096 | 0.096 | 0 | 0.77 | 0.96 | 0.83 | 0.9 | 0.78 | 0.83 |

| | | | | | | | | | | | | | | | | |
|---|---|---|---|---|---|---|---|---|---|---|---|---|---|---|---|---|
| 2 | 56802340 | rs2033016 | 0.722 | -1.083 | 0.637 | -0.884 | -0.024 | 0.022 | 0.022 | 1.193 | 0.4 | 0.27 | 0.46 | 0.62 | 0.52 | 0.5 |
| 2 | 56814463 | rs11125642 | 0.77 | -1.086 | 0.634 | -0.885 | -0.033 | 0.022 | 0.022 | 1.415 | 0.39 | 0.22 | 0.45 | 0.62 | 0.52 | 0.5 |
| 2 | 56817791 | rs12477994 | 0.266 | -0.759 | 0.901 | -0.959 | 0.014 | 0.027 | 0.027 | 0 | 0.38 | 0.21 | 0.41 | 0.31 | 0.32 | 0.46 |
| 2 | 56820468 | rs13401378 | 0.811 | -0.751 | 0.893 | -0.946 | -0.012 | 0.033 | 0.033 | 1.711 | 0.43 | 0.3 | 0.47 | 0.62 | 0.52 | 0.5 |
| 2 | 56822414 | rs2042123 | -0.831 | -0.757 | 1.021 | -0.954 | 0.008 | 0.013 | 0.013 | 0.05 | 0.83 | 0.86 | 0.92 | 0.94 | 0.85 | 0.96 |
| 2 | 56822558 | rs2870066 | 0.594 | -0.894 | 0.9 | -0.944 | -0.009 | 0.137 | 0.137 | 0.281 | 0.36 | 0.07 | 0.32 | 0.15 | 0.32 | 0.32 |
| 2 | 56836468 | rs3948576 | 0.896 | -0.812 | 1.014 | -0.964 | -0.011 | 0.031 | 0.031 | 2.029 | 0.42 | 0.33 | 0.47 | 0.64 | 0.53 | 0.48 |
| 2 | 56837602 | rs13411277 | 0.396 | -0.816 | 1.035 | -0.956 | -0.016 | 0.108 | 0.108 | 3.252 | 0.46 | 0.86 | 0.61 | 0.67 | 0.53 | 0.48 |
| 2 | 56839139 | rs6545583 | -0.632 | -0.821 | 1.08 | -0.968 | 0.01 | 0.01 | 0.01 | 0 | 0.82 | 0.83 | 0.91 | 0.94 | 0.86 | 0.96 |
| 2 | 56840079 | rs2870070 | -0.516 | -0.826 | 1.314 | -0.981 | -0.034 | -0.075 | -0.075 | 0 | 0.81 | 0.7 | 0.91 | 0.94 | 0.86 | 0.96 |
| 2 | 56841117 | rs2112025 | 0.844 | -0.689 | 1.069 | -0.877 | -0.019 | 0.035 | 0.035 | 2.084 | 0.42 | 0.28 | 0.47 | 0.64 | 0.53 | 0.48 |
| 2 | 56841968 | rs12998772 | 1.17 | -0.745 | 0.886 | -0.976 | -0.01 | 0.051 | 0.051 | 0.009 | 0.19 | 0.01 | 0.17 | 0.05 | 0.16 | 0.2 |
| 2 | 56842783 | rs13403703 | 0.795 | -0.635 | 1.067 | -0.887 | -0.001 | 0 | 0 | 2.183 | 0.43 | 0.48 | 0.49 | 0.64 | 0.53 | 0.48 |
| 2 | 56866018 | rs9678179 | 0.451 | -0.74 | 0.405 | -0.881 | -0.021 | 0.071 | 0.071 | 3.935 | 0.44 | 0.84 | 0.53 | 0.65 | 0.53 | 0.48 |
| 2 | 56904198 | rs11890549 | -1.095 | -0.862 | 0.689 | -0.834 | 0.017 | 0.055 | 0.055 | 8.681 | 0.06 | 0.65 | 0.13 | 0.05 | 0.01 | 0.02 |
| 2 | 56906260 | rs17269015 | 1.135 | -0.961 | 0.353 | -0.936 | 0.009 | 0.054 | 0.054 | 0.033 | 0.19 | 0.01 | 0.16 | 0.04 | 0.14 | 0.19 |
| 2 | 56907253 | rs7574093 | -1.264 | -0.553 | 0.788 | -0.685 | -0.001 | 0.034 | 0.034 | 9.081 | 0.06 | 0.68 | 0.13 | 0.04 | 0.01 | 0.02 |
| 2 | 56924658 | rs2042119 | 0.721 | -0.397 | -0.063 | -0.69 | -0.029 | 0.008 | 0.008 | 3.203 | 0.39 | 0.18 | 0.41 | 0.6 | 0.53 | 0.47 |
| 2 | 56928639 | rs4672150 | -1.228 | -0.31 | -0.098 | -0.709 | -0.016 | 0.06 | 0.06 | 0 | 0.81 | 0.93 | 0.76 | 0.91 | 0.83 | 0.84 |
| 2 | 56942624 | rs10207082 | -1.571 | -0.478 | 1.079 | -0.595 | 0.001 | 0.077 | 0.077 | 9.43 | 0.06 | 0.68 | 0.21 | 0.08 | 0.01 | 0.02 |
| 2 | 56964639 | rs7563088 | -0.794 | -0.425 | 1.204 | -0.712 | -0.007 | -0.063 | -0.063 | 0.722 | 0.78 | 0.63 | 0.66 | 0.86 | 0.82 | 0.83 |
| 2 | 56970732 | rs2682837 | -0.456 | -0.693 | 0.887 | -0.752 | 0.013 | 0.009 | 0.009 | 0.251 | 0.39 | 0.41 | 0.36 | 0.3 | 0.29 | 0.4 |
| 2 | 56988751 | rs2544313 | -0.105 | -0.643 | 1.012 | -0.721 | -0.005 | -0.002 | -0.002 | 0 | 0.37 | 0.42 | 0.35 | 0.31 | 0.28 | 0.39 |
| 2 | 56993390 | rs4140686 | 0.189 | -0.311 | 1.138 | -0.632 | 0.006 | 0.025 | 0.025 | 1.262 | 0.45 | 0.62 | 0.42 | 0.6 | 0.54 | 0.45 |
| 2 | 57010309 | rs11125655 | 0.316 | -0.228 | 1.174 | -0.644 | -0.001 | -0.009 | -0.009 | 0 | 0.44 | 0.45 | 0.4 | 0.6 | 0.53 | 0.45 |
| 2 | 57011171 | rs17696579 | 0.623 | -0.235 | 0.723 | -0.606 | -0.013 | -0.001 | -0.001 | 0 | 0.42 | 0.18 | 0.31 | 0.55 | 0.52 | 0.43 |
| 2 | 57017445 | rs976365 | -0.471 | -0.159 | 0.311 | -0.761 | 0.028 | 0.042 | 0.042 | 0 | 0.58 | 0.89 | 0.48 | 0.68 | 0.64 | 0.48 |
| 2 | 57032081 | rs7587789 | 0.203 | 0.075 | 0.605 | -0.595 | 0.002 | 0.011 | 0.011 | 1.086 | 0.45 | 0.58 | 0.41 | 0.6 | 0.53 | 0.45 |
| 2 | 57039250 | rs17048423 | 0.542 | 0.095 | -0.03 | -0.578 | -0.028 | -0.012 | -0.012 | 0.052 | 0.41 | 0.14 | 0.31 | 0.55 | 0.52 | 0.43 |
| 2 | 57040068 | rs17048424 | 1.038 | 0.67 | -0.226 | -0.697 | -0.025 | -0.065 | -0.065 | 0 | 0.26 | 0.1 | 0.2 | 0.53 | 0.32 | 0.27 |
| 2 | 57050007 | rs12987302 | 0.022 | 0.419 | 0.478 | -0.448 | 0.003 | -0.006 | -0.006 | 0 | 0.15 | 0.25 | 0.18 | 0.1 | 0.16 | 0.19 |
| 2 | 57050778 | rs12993505 | 0.431 | 0.408 | 0.329 | -0.472 | -0.001 | -0.004 | -0.004 | 0 | 0.13 | 0.13 | 0.16 | 0.06 | 0.14 | 0.15 |
| 2 | 57051996 | rs1023598 | -0.964 | 0.365 | 0.063 | -0.419 | 0.002 | -0.019 | -0.019 | 6.056 | 0.94 | 0.98 | 0.96 | 0.86 | 0.91 | 0.85 |
| 2 | 57053945 | rs7560622 | 0.181 | 0.439 | 0.109 | -0.702 | 0.014 | 0.076 | 0.076 | 0.022 | 0.52 | 0.75 | 0.37 | 0.68 | 0.54 | 0.52 |
| 2 | 57060131 | rs6715352 | -0.005 | 0.332 | 0.785 | -0.629 | 0.02 | -0.011 | -0.011 | 0 | 0.17 | 0.56 | 0.37 | 0.1 | 0.18 | 0.19 |
| 2 | 57062513 | rs7600489 | 0.529 | 0.363 | 0.67 | -0.379 | 0.015 | 0.061 | 0.061 | 3.247 | 0.47 | 0.37 | 0.34 | 0.23 | 0.55 | 0.45 |
| 2 | 57062894 | rs2870502 | -0.847 | 0.338 | 0.524 | -0.083 | -0.013 | 0.001 | 0.001 | 0 | 0.93 | 0.99 | 0.94 | 0.44 | 0.91 | 0.82 |
| 2 | 57068218 | rs2216413 | -0.558 | 0.402 | 0.612 | -0.231 | 0.026 | 0.068 | 0.068 | 0.559 | 0.48 | 0.73 | 0.64 | 0.28 | 0.41 | 0.41 |
| 2 | 57069147 | rs6545600 | -0.772 | 0.404 | 0.724 | -0.245 | 0.017 | 0.047 | 0.047 | 0 | 0.5 | 0.64 | 0.62 | 0.29 | 0.44 | 0.43 |
| 2 | 57074475 | rs6744438 | -0.343 | 0.364 | 0.813 | -0.231 | 0.026 | 0.015 | 0.015 | 0.353 | 0.2 | 0.55 | 0.37 | 0.17 | 0.22 | 0.26 |
| 2 | 57077745 | rs6741957 | -0.347 | 0.025 | 0.713 | -0.171 | 0.012 | 0.02 | 0.02 | 3.148 | 0.44 | 0.63 | 0.59 | 0.22 | 0.34 | 0.34 |
| 2 | 57101805 | rs2904426 | -0.057 | 0.16 | 0.769 | -0.085 | -0.005 | -0.008 | -0.008 | 0 | 0.59 | 0.59 | 0.65 | 0.3 | 0.65 | 0.62 |
| 2 | 57109560 | rs7422350 | -0.118 | 0.183 | 0.728 | -0.072 | 0.01 | 0.013 | 0.013 | 7.493 | 0.61 | 0.65 | 0.65 | 0.3 | 0.66 | 0.62 |
| 2 | 57118147 | rs10206726 | -0.592 | -0.218 | 0.364 | 0.113 | -0.012 | -0.102 | -0.102 | 0.062 | 0.86 | 0.99 | 0.92 | 0.45 | 0.87 | 0.8 |
| 2 | 57119328 | rs13033180 | 1.078 | -0.197 | 0.566 | -0.122 | -0.006 | 0.162 | 0.162 | 1.761 | 0.46 | 0.21 | 0.32 | 0.24 | 0.52 | 0.45 |
| 2 | 57122342 | rs4672172 | 0.671 | -0.231 | 0.997 | -0.135 | -0.019 | -0.019 | -0.019 | 0 | 0.7 | 0.5 | 0.58 | 0.34 | 0.71 | 0.59 |
| 2 | 57122686 | rs4671301 | 1.151 | -0.177 | 0.892 | -0.098 | 0.003 | 0.007 | 0.007 | 0.465 | 0.23 | 0.23 | 0.25 | 0.11 | 0.19 | 0.15 |
| 2 | 57135651 | rs13400448 | 0.099 | -0.148 | 1.131 | 0.564 | 0.007 | 0.209 | 0.209 | 0.65 | 0.17 | 0.46 | 0.35 | 0.51 | 0.16 | 0.22 |
| 2 | 57137622 | rs7598437 | 0.535 | -0.164 | 1.499 | 0.616 | 0.007 | 0.019 | 0.019 | 0.019 | 0.15 | 0.24 | 0.32 | 0.13 | 0.16 | 0.2 |
| 2 | 57138726 | rs17698880 | 1.04 | -0.265 | 1.438 | 0.611 | 0.013 | 0.166 | 0.166 | 1.951 | 0.47 | 0.17 | 0.32 | 0.3 | 0.54 | 0.48 |
| 2 | 57147590 | rs7589657 | 0.249 | -0.236 | 1.465 | 0.711 | 0 | 0.181 | 0.181 | 0.373 | 0.17 | 0.31 | 0.17 | 0.44 | 0.16 | 0.21 |
| 2 | 57148163 | rs10182943 | 0.127 | -0.235 | 1.472 | 0.699 | 0.004 | 0.191 | 0.191 | 0.581 | 0.17 | 0.41 | 0.34 | 0.5 | 0.16 | 0.24 |
| 2 | 57153537 | rs7585535 | -0.19 | -0.308 | 1.515 | 0.735 | -0.003 | -0.004 | -0.004 | 0 | 0.64 | 0.59 | 0.68 | 0.79 | 0.7 | 0.72 |
| 2 | 57159462 | rs2865258 | -0.753 | -0.495 | 1.164 | 0.749 | -0.017 | -0.029 | -0.029 | 1.645 | 0.87 | 0.98 | 0.92 | 0.89 | 0.87 | 0.83 |
| 2 | 57174489 | rs7578236 | -0.379 | -0.571 | 1.357 | 0.797 | 0.035 | 0.549 | 0.549 | 0.503 | 0.06 | 0.33 | 0.03 | 0.38 | 0.01 | 0.03 |
| 2 | 57177642 | rs4671302 | 1.677 | -0.581 | 1.216 | 0.593 | 0.01 | 0.012 | 0.012 | 2.424 | 0.18 | 0.09 | 0.2 | 0.11 | 0.14 | 0.13 |
| 2 | 57179566 | rs13033220 | -0.182 | -0.17 | 1.124 | 0.467 | 0.011 | 0.022 | 0.022 | 6.59 | 0.61 | 0.7 | 0.51 | 0.73 | 0.64 | 0.65 |
| 2 | 57207748 | rs4672179 | 0.826 | -0.196 | 0.721 | 0.456 | -0.001 | 0 | 0 | 0.055 | 0.14 | 0.16 | 0.14 | 0.07 | 0.11 | 0.17 |
| 2 | 57217196 | rs1946500 | -0.4 | -0.09 | 0.847 | 0.413 | -0.023 | -0.008 | -0.008 | 0 | 0.65 | 0.51 | 0.5 | 0.72 | 0.71 | 0.7 |
| 2 | 57218736 | rs1366797 | -0.652 | 0.214 | 0.672 | 0.47 | 0.013 | 0.016 | 0.016 | 3.696 | 0.61 | 0.77 | 0.45 | 0.67 | 0.63 | 0.67 |
| 2 | 57250415 | rs996839 | -0.477 | 0.334 | 0.539 | 0.51 | -0.072 | -0.061 | -0.061 | 0 | 0.86 | 0.98 | 0.71 | 0.83 | 0.78 | 0.8 |
| 2 | 57254728 | rs2195490 | -0.238 | 0.212 | 0.764 | 0.479 | -0.002 | 0.033 | 0.033 | 3.71 | 0.55 | 0.69 | 0.41 | 0.67 | 0.54 | 0.59 |
| 2 | 57256060 | rs1594143 | 0.861 | 0.142 | 0.466 | 0.426 | 0.014 | 0.022 | 0.022 | 0.626 | 0.17 | 0.07 | 0.15 | 0.07 | 0.14 | 0.19 |
| 2 | 57263345 | rs4099901 | 0.135 | 0.19 | 0.777 | 0.446 | 0.022 | 0.038 | 0.038 | 0 | 0.5 | 0.38 | 0.46 | 0.25 | 0.4 | 0.39 |
| 2 | 57267524 | rs11903925 | 0.871 | 0.236 | 0.912 | 0.484 | -0.005 | -0.006 | -0.006 | 0.119 | 0.34 | 0.33 | 0.24 | 0.58 | 0.39 | 0.4 |
| 2 | 57272711 | rs1429310 | 0.189 | 0.218 | 0.684 | 0.421 | 0.036 | 0.066 | 0.066 | 0 | 0.5 | 0.32 | 0.44 | 0.25 | 0.4 | 0.4 |
| 2 | 57276155 | rs12614157 | 0.81 | 0.301 | 0.776 | 0.45 | 0.023 | -0.073 | -0.073 | 0 | 0.34 | 0.5 | 0.26 | 0.21 | 0.38 | 0.37 |
| 2 | 57280699 | rs2163045 | 0.017 | 0.345 | 0.722 | 0.564 | 0.005 | 0.005 | 0.005 | 0 | 0.52 | 0.45 | 0.46 | 0.63 | 0.39 | 0.44 |
| 2 | 57287411 | rs1116357 | 0.011 | 0.267 | 0.652 | 0.507 | 0.006 | 0.022 | 0.022 | 3.116 | 0.51 | 0.57 | 0.4 | 0.28 | 0.53 | 0.57 |

| | | | | | | | | | | | | | | | | |
|---|---|---|---|---|---|---|---|---|---|---|---|---|---|---|---|---|
| 2 | 57296558 | rs1424642 | -0.312 | 0.322 | 0.571 | 0.626 | -0.005 | -0.009 | -0.009 | 0 | 0.85 | 0.86 | 0.71 | 0.47 | 0.8 | 0.78 |
| 2 | 57300389 | rs10496062 | 0.836 | 0.302 | 0.875 | 0.571 | 0.01 | 0.012 | 0.012 | 0.162 | 0.34 | 0.45 | 0.25 | 0.59 | 0.39 | 0.4 |
| 2 | 57306208 | rs11901902 | 0.05 | 0.3 | 0.891 | 0.543 | 0.001 | 0.013 | 0.013 | 2.392 | 0.5 | 0.47 | 0.39 | 0.66 | 0.52 | 0.6 |
| 2 | 57309754 | rs7563939 | 0.908 | 0.286 | 0.982 | 0.526 | 0.002 | 0.003 | 0.003 | 1.666 | 0.34 | 0.3 | 0.3 | 0.18 | 0.29 | 0.21 |
| 2 | 57332673 | rs1559557 | 0.692 | 0.115 | 1.04 | 0.492 | -0.013 | -0.013 | -0.013 | 0 | 0.64 | 0.5 | 0.51 | 0.75 | 0.66 | 0.61 |
| 2 | 57335044 | rs1345942 | 0.726 | 0.332 | 1.028 | 0.637 | 0.023 | 0.019 | 0.019 | 0.187 | 0.35 | 0.46 | 0.24 | 0.6 | 0.4 | 0.4 |
| 2 | 57344635 | rs12464407 | 1.14 | 0.282 | 0.943 | 0.735 | 0.015 | 0.027 | 0.027 | 1.847 | 0.34 | 0.24 | 0.29 | 0.17 | 0.26 | 0.21 |
| 2 | 57352937 | rs12622970 | 0.924 | 0.299 | 0.976 | 0.744 | 0.009 | 0.017 | 0.017 | 0.248 | 0.16 | 0.06 | 0.16 | 0.07 | 0.14 | 0.2 |
| 2 | 57353334 | rs4672197 | 0.268 | 0.337 | 1.092 | 0.756 | -0.027 | 0.142 | 0.142 | 0.509 | 0.2 | 0.36 | 0.19 | 0.45 | 0.15 | 0.23 |
| 2 | 57353920 | rs4672198 | 1.139 | 0.317 | 1.07 | 0.704 | 0.012 | 0.023 | 0.023 | 2.331 | 0.34 | 0.27 | 0.3 | 0.18 | 0.27 | 0.21 |
| 2 | 57354454 | rs17188812 | 0.943 | 0.342 | 1.008 | 0.698 | -0.01 | 0.012 | 0.012 | 0.07 | 0.33 | 0.3 | 0.23 | 0.22 | 0.39 | 0.37 |
| 2 | 57355662 | rs1593235 | 0.951 | 0.343 | 1.009 | 0.675 | -0.01 | 0.011 | 0.011 | 0.063 | 0.33 | 0.3 | 0.23 | 0.21 | 0.39 | 0.37 |
| 2 | 57365335 | rs17048681 | 0.757 | 0.308 | 1.061 | 0.672 | -0.004 | -0.02 | -0.02 | 0 | 0.66 | 0.54 | 0.52 | 0.38 | 0.65 | 0.58 |
| 2 | 57365499 | rs2161889 | 0.421 | 0.319 | 1.1 | 0.751 | -0.02 | 0.084 | 0.084 | 0.136 | 0.18 | 0.26 | 0.18 | 0.47 | 0.14 | 0.23 |
| 2 | 57366620 | rs17189361 | 0.946 | 0.367 | 0.852 | 0.654 | -0.009 | 0.008 | 0.008 | 0.071 | 0.33 | 0.3 | 0.23 | 0.21 | 0.39 | 0.37 |
| 2 | 57372731 | rs6709593 | 0.603 | 0.037 | 0.608 | 0.599 | 0.011 | 0.014 | 0.014 | 0 | 0.18 | 0.15 | 0.17 | 0.07 | 0.14 | 0.2 |
| 2 | 57374265 | rs1582644 | 1.171 | -0.008 | 0.675 | 0.532 | -0.028 | 0.022 | 0.022 | 0.124 | 0.28 | 0.24 | 0.24 | 0.26 | 0.39 | 0.37 |
| 2 | 57374374 | rs1582645 | -0.207 | -0.141 | 0.828 | 0.628 | 0.006 | 0.01 | 0.01 | 0 | 0.84 | 0.79 | 0.72 | 0.85 | 0.8 | 0.81 |
| 2 | 57388379 | rs7562542 | 0.13 | -0.054 | 0.542 | 0.581 | 0.013 | 0.029 | 0.029 | 0.1 | 0.54 | 0.57 | 0.48 | 0.25 | 0.41 | 0.42 |
| 2 | 57393245 | rs6545625 | -0.09 | -0.045 | 0.451 | 0.434 | 0 | -0.001 | -0.001 | 0.318 | 0.47 | 0.49 | 0.41 | 0.27 | 0.52 | 0.57 |
| 2 | 57394031 | rs6760508 | -0.164 | -0.029 | 0.215 | 0.441 | 0.004 | 0.019 | 0.019 | 2.837 | 0.48 | 0.58 | 0.42 | 0.28 | 0.52 | 0.57 |
| 2 | 57398024 | rs727282 | 0.345 | 0.491 | -0.081 | 0.778 | 0 | 0.002 | 0.002 | 0 | 0.83 | 0.83 | 0.71 | 0.45 | 0.79 | 0.78 |
| 2 | 57399830 | rs6545632 | 0.415 | 0.473 | -0.091 | 0.769 | 0 | 0 | 0 | 0 | 0.82 | 0.83 | 0.71 | 0.45 | 0.79 | 0.78 |
| 2 | 57402640 | rs877993 | 0.099 | 0.518 | -0.089 | 0.754 | -0.001 | 0.021 | 0.021 | 0 | 0.85 | 0.83 | 0.72 | 0.45 | 0.83 | 0.79 |
| 2 | 57404022 | rs12713336 | 0.35 | 0.504 | -0.076 | 0.757 | 0 | 0.003 | 0.003 | 0 | 0.83 | 0.82 | 0.72 | 0.45 | 0.8 | 0.77 |
| 2 | 57405885 | rs1424635 | -0.263 | 0.597 | -0.386 | 0.603 | -0.051 | -0.036 | -0.036 | 0 | 0.86 | 0.98 | 0.74 | 0.86 | 0.81 | 0.81 |
| 2 | 57416841 | rs13030956 | 0.659 | 0.654 | -0.082 | 0.312 | -0.009 | 0.001 | 0.001 | 0.776 | 0.28 | 0.25 | 0.23 | 0.16 | 0.39 | 0.36 |
| 2 | 57427953 | rs6755060 | 0.153 | 0.626 | -0.324 | 0.459 | 0.008 | 0.024 | 0.024 | 0 | 0.54 | 0.58 | 0.5 | 0.3 | 0.46 | 0.43 |
| 2 | 57429923 | rs7423025 | -0.759 | 0.759 | -1.031 | 0.548 | 0.005 | 0.037 | 0.037 | 6.175 | 0.57 | 0.89 | 0.61 | 0.64 | 0.62 | 0.56 |
| 2 | 57430709 | rs1981900 | 0.696 | 0.841 | -1.243 | 0.682 | -0.005 | -0.014 | -0.014 | 0 | 0.06 | 0.03 | 0.06 | 0.11 | 0.09 | 0.08 |
| 2 | 57431622 | rs1582642 | -0.795 | 0.907 | -1.417 | 0.497 | -0.056 | 0.078 | 0.078 | 8.663 | 0.55 | 0.93 | 0.6 | 0.66 | 0.59 | 0.58 |
| 2 | 57435578 | rs1345941 | 1.644 | 0.999 | -1.683 | 0.451 | -0.036 | -0.023 | -0.023 | 0.452 | 0.19 | 0.01 | 0.17 | 0.09 | 0.18 | 0.24 |
| 2 | 57436631 | rs7604249 | -0.163 | 1.108 | -1.064 | 0.55 | 0.02 | -0.016 | -0.016 | 0.161 | 0.36 | 0.7 | 0.31 | 0.53 | 0.31 | 0.4 |
| 2 | 57441643 | rs12052504 | -0.024 | 1.279 | -0.896 | 0.562 | -0.018 | 0.007 | 0.007 | 5.096 | 0.2 | 0.21 | 0.29 | 0.17 | 0.3 | 0.17 |
| 2 | 57441710 | rs4390805 | -0.053 | 1.278 | -0.868 | 0.561 | -0.01 | 0.004 | 0.004 | 4.822 | 0.2 | 0.24 | 0.3 | 0.17 | 0.3 | 0.17 |
| 2 | 57445471 | rs4290697 | -1.19 | 0.954 | -0.882 | 0.492 | -0.011 | 0.03 | 0.03 | 8.236 | 0.58 | 0.81 | 0.58 | 0.61 | 0.6 | 0.52 |
| 2 | 57450236 | rs4608569 | -1.068 | 1.117 | -1.005 | 0.478 | -0.006 | 0.092 | 0.092 | 5.465 | 0.54 | 0.77 | 0.59 | 0.64 | 0.58 | 0.56 |
| 2 | 57464756 | rs13417963 | -1.507 | 1.151 | -0.159 | 0.433 | 0 | 0.003 | 0.003 | 9.732 | 0.61 | 0.65 | 0.63 | 0.72 | 0.68 | 0.6 |
| 2 | 57482289 | rs12622457 | 0.552 | 1.144 | -0.254 | 0.569 | 0.01 | 0.019 | 0.019 | 0 | 0.3 | 0.17 | 0.25 | 0.16 | 0.29 | 0.29 |
| 2 | 57483669 | rs4271799 | 0.587 | 1.105 | -0.103 | 0.564 | 0.017 | 0.02 | 0.02 | 0 | 0.29 | 0.19 | 0.24 | 0.14 | 0.28 | 0.28 |
| 2 | 57488302 | rs4619596 | 0.397 | 1.143 | 0.013 | 0.706 | 0 | 0.018 | 0.018 | 0.054 | 0.09 | 0.09 | 0.06 | 0.35 | 0.08 | 0.11 |
| 2 | 57507510 | rs1589603 | 0.885 | 1.338 | -0.14 | 0.622 | -0.066 | -0.026 | -0.026 | 0.415 | 0.11 | 0.01 | 0.22 | 0.11 | 0.19 | 0.12 |
| 2 | 57514841 | rs1916141 | 0.365 | 1.259 | 0.485 | 0.684 | 0.019 | 0.201 | 0.201 | 0.394 | 0.09 | 0.24 | 0.07 | 0.35 | 0.08 | 0.11 |
| 2 | 57516048 | rs7603447 | -0.432 | 1.289 | 0.436 | 0.663 | 0.004 | 0.136 | 0.136 | 0 | 0.14 | 0.25 | 0.14 | 0.45 | 0.16 | 0.19 |
| 2 | 57518011 | rs1403335 | 0.69 | 1.241 | 0.171 | 0.575 | 0.013 | 0.02 | 0.02 | 0 | 0.3 | 0.17 | 0.25 | 0.14 | 0.29 | 0.29 |
| 2 | 57532112 | rs17048747 | -0.033 | 1.333 | 0.24 | 0.642 | -0.061 | -0.04 | -0.04 | 2.289 | 0.16 | 0.04 | 0.29 | 0.24 | 0.27 | 0.2 |
| 2 | 57542882 | rs13408500 | -1.03 | 1.284 | 0.579 | 0.631 | -0.001 | 0.006 | 0.006 | 0.668 | 0.21 | 0.25 | 0.17 | 0.46 | 0.23 | 0.21 |
| 2 | 57547114 | rs1357377 | -2.141 | 1.313 | 0.63 | 0.58 | -0.007 | 0.052 | 0.052 | 10.474 | 0.63 | 0.85 | 0.67 | 0.71 | 0.7 | 0.62 |
| 2 | 57549451 | rs11125710 | -2.174 | 1.303 | 0.631 | 0.575 | -0.024 | 0.055 | 0.055 | 10.515 | 0.64 | 0.9 | 0.67 | 0.71 | 0.7 | 0.62 |
| 2 | 57549639 | rs11125711 | -1.027 | 1.283 | 0.661 | 0.595 | 0.005 | 0.012 | 0.012 | 0 | 0.51 | 0.5 | 0.41 | 0.6 | 0.52 | 0.49 |
| 2 | 57563358 | rs4671306 | 1.758 | 1.22 | 0.312 | 0.517 | -0.028 | -0.041 | -0.041 | 0 | 0.21 | 0.01 | 0.17 | 0.09 | 0.18 | 0.24 |
| 2 | 57571600 | rs17193105 | -2.021 | 1.299 | 0.675 | 0.549 | -0.003 | -0.002 | -0.002 | 0 | 0.41 | 0.6 | 0.47 | 0.63 | 0.52 | 0.37 |
| 2 | 57574704 | rs13396309 | 0.035 | 1.336 | 0.819 | 0.556 | 0.001 | -0.007 | -0.007 | 0 | 0.12 | 0.18 | 0.12 | 0.12 | 0.16 | 0.1 |
| 2 | 57577034 | rs13390366 | -2.112 | 1.372 | 0.872 | 0.551 | 0.008 | 0.004 | 0.004 | 8.49 | 0.41 | 0.54 | 0.46 | 0.63 | 0.52 | 0.37 |
| 2 | 57579667 | rs10181692 | 0.922 | 1.032 | 0.724 | 0.621 | 0 | -0.014 | -0.014 | 0 | 0.07 | 0.09 | 0.04 | 0 | 0.07 | 0.03 |
| 2 | 57582921 | rs7602729 | 0.271 | 1.044 | 0.772 | 0.679 | -0.002 | 0.064 | 0.064 | 0.199 | 0.09 | 0.15 | 0.06 | 0.35 | 0.08 | 0.11 |
| 2 | 57585692 | rs12464561 | 0.742 | 1.044 | 0.73 | 0.48 | -0.005 | -0.018 | -0.018 | 0 | 0.09 | 0.17 | 0.07 | 0.04 | 0.11 | 0.04 |
| 2 | 57588405 | rs13409605 | 0.616 | 1.058 | 0.562 | 0.533 | 0 | -0.003 | -0.003 | 0.321 | 0.07 | 0.07 | 0.04 | 0 | 0.07 | 0.03 |
| 2 | 57589477 | rs10204326 | 1.386 | 0.979 | 0.606 | 0.526 | 0.003 | 0.001 | 0.001 | 0 | 0.22 | 0.21 | 0.2 | 0.09 | 0.18 | 0.25 |
| 2 | 57595408 | rs6743780 | -0.409 | 0.929 | 0.762 | 0.553 | 0.005 | -0.007 | -0.007 | 0 | 0.12 | 0.25 | 0.12 | 0.12 | 0.16 | 0.1 |
| 2 | 57607449 | rs1518681 | -0.778 | 0.983 | 0.496 | 0.395 | -0.001 | 0.001 | 0.001 | 0.103 | 0.48 | 0.46 | 0.55 | 0.71 | 0.49 | 0.62 |
| 2 | 57614310 | rs10210959 | -0.422 | 0.997 | 0.488 | 0.555 | 0 | 0.006 | 0.006 | 0 | 0.44 | 0.62 | 0.42 | 0.56 | 0.41 | 0.53 |
| 2 | 57618446 | rs2203805 | -0.932 | 1.015 | 0.505 | 0.556 | -0.006 | 0.091 | 0.091 | 0 | 0.49 | 0.68 | 0.56 | 0.71 | 0.5 | 0.62 |
| 2 | 57633070 | rs1850672 | 0.236 | 1.078 | 0.514 | 0.631 | -0.001 | 0.006 | 0.006 | 0.161 | 0.34 | 0.38 | 0.21 | 0.45 | 0.33 | 0.43 |
| 2 | 57634432 | rs13428949 | -1.994 | 1.186 | 0.228 | 0.454 | -0.085 | 0.051 | 0.051 | 2.586 | 0.63 | 0.95 | 0.66 | 0.72 | 0.62 | 0.68 |
| 2 | 57644319 | rs17048806 | 0.169 | 1.314 | 0.465 | 0.669 | 0.007 | 0.106 | 0.106 | 0.646 | 0.1 | 0.17 | 0.09 | 0.39 | 0.13 | 0.15 |
| 2 | 57645506 | rs7601574 | 1.513 | 1.278 | 0.243 | 0.565 | -0.023 | 0.057 | 0.057 | 0 | 0.23 | 0.04 | 0.1 | 0.06 | 0.19 | 0.28 |
| 2 | 57645820 | rs2871608 | 1.009 | 1.291 | 0.276 | 0.609 | -0.024 | -0.031 | -0.031 | 0 | 0.09 | 0 | 0.19 | 0.11 | 0.09 | 0.1 |
| 2 | 57658970 | rs2310825 | 0.379 | 1.244 | 0.544 | 0.569 | -0.001 | -0.001 | -0.001 | 0.097 | 0.31 | 0.31 | 0.28 | 0.19 | 0.29 | 0.38 |

| | | | | | | | | | | | | | | | | |
|---|---|---|---|---|---|---|---|---|---|---|---|---|---|---|---|---|
| 2 | 57660908 | rs2310826 | 1.189 | 1.272 | 0.506 | 0.575 | -0.004 | -0.034 | -0.034 | 0.764 | 0.24 | 0.29 | 0.13 | 0.06 | 0.19 | 0.28 |
| 2 | 57669304 | rs17048826 | -0.021 | 1.311 | 0.403 | 0.658 | 0.003 | 0.009 | 0.009 | 0 | 0.11 | 0.16 | 0.2 | 0.17 | 0.17 | 0.12 |
| 2 | 57673796 | rs1028189 | -1.586 | 1.606 | 0.061 | 0.714 | -0.032 | -0.035 | -0.035 | 0 | 0.94 | 0.99 | 0.89 | 0.87 | 0.89 | 0.96 |
| 2 | 57676255 | rs17048831 | 1.301 | 1.596 | 0.547 | 0.748 | 0 | 0 | 0 | 6.692 | 0.23 | 0.18 | 0.11 | 0.05 | 0.19 | 0.28 |
| 2 | 57677416 | rs2139123 | -1.239 | 1.638 | 0.451 | 0.697 | -0.007 | 0.099 | 0.099 | 0.012 | 0.52 | 0.74 | 0.58 | 0.7 | 0.51 | 0.63 |
| 2 | 57689942 | rs17048834 | 1.463 | 1.698 | -0.122 | 0.553 | -0.038 | 0.013 | 0.013 | 0.097 | 0.09 | 0.01 | 0.06 | 0.13 | 0.11 | 0.18 |
| 2 | 57690033 | rs7589636 | -0.659 | 1.687 | 0.113 | 0.683 | 0.008 | 0.11 | 0.11 | 0.09 | 0.23 | 0.42 | 0.32 | 0.5 | 0.23 | 0.27 |
| 2 | 57694030 | rs13024352 | 0.959 | 1.442 | -0.208 | 0.741 | 0.001 | -0.019 | -0.019 | 0 | 0.07 | 0.1 | 0.06 | 0.02 | 0.09 | 0.04 |
| 2 | 57696957 | rs17048844 | 0.344 | 1.451 | -0.307 | 1.008 | -0.003 | 0.18 | 0.18 | 0.944 | 0.07 | 0.14 | 0.05 | 0.38 | 0.07 | 0.12 |
| 2 | 57708917 | rs952397 | -1.454 | 1.456 | -0.887 | 0.766 | -0.004 | -0.101 | -0.101 | 0 | 0.63 | 0.89 | 0.67 | 0.52 | 0.64 | 0.81 |
| 2 | 57714717 | rs10518819 | -1.178 | 1.411 | -0.474 | 1.008 | -0.003 | -0.005 | -0.005 | 0 | 0.27 | 0.28 | 0.3 | 0.63 | 0.32 | 0.43 |
| 2 | 57727701 | rs10165125 | 0.862 | 1.151 | -0.345 | 0.813 | -0.008 | -0.053 | -0.053 | 0.021 | 0.29 | 0.34 | 0.13 | 0.07 | 0.2 | 0.27 |
| 2 | 57728868 | rs2243931 | -0.857 | 1.219 | -0.248 | 0.912 | 0.017 | 0.037 | 0.037 | 0 | 0.48 | 0.68 | 0.51 | 0.32 | 0.42 | 0.48 |
| 2 | 57730190 | rs17048858 | 0.218 | 1.387 | -0.102 | 1.165 | -0.005 | 0.091 | 0.091 | 0 | 0.08 | 0.14 | 0.05 | 0.29 | 0.08 | 0.05 |
| 2 | 57731262 | rs2246820 | -0.034 | 1.38 | 0.01 | 1.103 | -0.007 | -0.003 | -0.003 | 0 | 0.15 | 0.19 | 0.23 | 0.13 | 0.14 | 0.13 |
| 2 | 57731944 | rs2264692 | 1.116 | 1.399 | -0.048 | 1.11 | -0.012 | -0.012 | -0.012 | 0 | 0.06 | 0.03 | 0.19 | 0.09 | 0.09 | 0.1 |
| 2 | 57732408 | rs6545653 | -2.281 | 1.28 | -0.045 | 1.041 | 0.003 | 0.039 | 0.039 | 2.091 | 0.69 | 0.9 | 0.67 | 0.81 | 0.68 | 0.84 |
| 2 | 57739441 | rs1881020 | -0.244 | 1.231 | -0.137 | 0.999 | 0.004 | 0.003 | 0.003 | 0 | 0.41 | 0.63 | 0.35 | 0.32 | 0.36 | 0.44 |
| 2 | 57742834 | rs820968 | -1.319 | 1.264 | -0.165 | 1.022 | 0 | -0.006 | -0.006 | 3.998 | 0.58 | 0.65 | 0.59 | 0.51 | 0.55 | 0.75 |
| 2 | 57743543 | rs11125715 | -0.127 | 0.878 | 0.094 | 1.177 | -0.026 | -0.028 | -0.028 | 0 | 0.11 | 0.18 | 0.06 | 0.03 | 0.07 | 0.03 |
| 2 | 57755496 | rs2264599 | -1.268 | 1.057 | -0.183 | 1.557 | -0.003 | -0.007 | -0.007 | 0 | 0.51 | 0.85 | 0.41 | 0.51 | 0.47 | 0.49 |
| 2 | 57755753 | rs6742652 | -0.342 | 1.384 | 0.288 | 1.797 | -0.003 | -0.004 | -0.004 | 1.987 | 0.36 | 0.3 | 0.18 | 0.37 | 0.35 | 0.35 |
| 2 | 57757289 | rs820983 | -0.199 | 1.366 | 0.327 | 1.787 | 0.092 | 0.269 | 0.269 | 0 | 0.07 | 0.34 | 0.2 | 0.23 | 0.1 | 0.14 |
| 2 | 57757646 | rs820984 | -0.872 | 1.266 | -0.349 | 1.445 | -0.012 | 0.018 | 0.018 | 1.455 | 0.44 | 0.7 | 0.39 | 0.6 | 0.46 | 0.49 |
| 2 | 57758608 | rs10460564 | -0.328 | 0.938 | -0.233 | 1.439 | 0.007 | -0.001 | -0.001 | 0.376 | 0.3 | 0.29 | 0.21 | 0.25 | 0.25 | 0.3 |
| 2 | 57763213 | rs13392012 | -1.45 | 0.45 | 0.985 | 2.154 | -0.015 | 0.354 | 0.354 | 0.996 | 0.1 | 0.48 | 0.16 | 0.33 | 0.03 | 0.11 |
| 2 | 57766378 | rs4671309 | 0.465 | 0.937 | 0.772 | 2.228 | -0.018 | -0.006 | -0.006 | 0.739 | 0.86 | 0.87 | 0.88 | 0.78 | 0.84 | 0.73 |
| 2 | 57767972 | rs13021425 | 0.72 | 0.938 | 0.882 | 2.361 | 0.027 | 0.059 | 0.059 | 0 | 0.84 | 0.68 | 0.76 | 0.66 | 0.84 | 0.7 |
| 2 | 57768677 | rs1527624 | 0.947 | 0.914 | 0.722 | 2.259 | -0.029 | -0.005 | -0.005 | 0 | 0.81 | 0.85 | 0.73 | 0.76 | 0.79 | 0.67 |
| 2 | 57772923 | rs9309298 | -0.411 | 0.884 | 0.815 | 2.007 | -0.003 | 0.069 | 0.069 | 0.539 | 0.08 | 0.28 | 0.03 | 0.08 | 0.03 | 0.06 |
| 2 | 57773233 | rs6722978 | 1.925 | 0.885 | 0.837 | 2.07 | 0.001 | 0.03 | 0.03 | 0.225 | 0.71 | 0.36 | 0.59 | 0.44 | 0.76 | 0.55 |
| 2 | 57774450 | rs1527627 | 1 | 0.96 | 0.817 | 2.071 | -0.029 | -0.036 | -0.036 | 0.528 | 0.8 | 0.86 | 0.73 | 0.65 | 0.79 | 0.66 |
| 2 | 57775544 | rs9309300 | 1.019 | 0.982 | 0.755 | 1.986 | 0.015 | 0.053 | 0.053 | 0 | 0.79 | 0.65 | 0.62 | 0.53 | 0.79 | 0.62 |
| 2 | 57780007 | rs17048911 | -0.21 | 1.129 | 0.863 | 1.714 | 0.021 | -0.022 | -0.022 | 0.024 | 0.08 | 0.16 | 0.12 | 0.03 | 0.15 | 0.06 |
| 2 | 57781712 | rs6747684 | -0.553 | 1.186 | 0.603 | 1.863 | 0.075 | 0.079 | 0.079 | 0 | 0.09 | 0.29 | 0.16 | 0.13 | 0.18 | 0.1 |
| 2 | 57793672 | rs4144515 | -0.252 | 1.059 | 0.679 | 1.782 | 0.002 | 0.08 | 0.08 | 0 | 0.07 | 0.25 | 0.03 | 0.09 | 0.02 | 0.06 |
| 2 | 57794837 | rs12713341 | 0.083 | 1.323 | 0.521 | 2.073 | 0.007 | 0.067 | 0.067 | 0 | 0.92 | 0.9 | 0.88 | 0.58 | 0.88 | 0.71 |
| 2 | 57799040 | rs10204095 | 0.31 | 1.315 | 0.556 | 2.071 | 0.011 | 0.045 | 0.045 | 0 | 0.07 | 0.2 | 0.15 | 0.14 | 0.07 | 0.06 |
| 2 | 57800605 | rs11125718 | 1.575 | 1.213 | -0.087 | 1.737 | -0.075 | 0.161 | 0.161 | 1.814 | 0.47 | 0.05 | 0.5 | 0.17 | 0.45 | 0.31 |
| 2 | 57801443 | rs13384136 | -1.924 | 1.299 | 0.008 | 1.711 | -0.006 | -0.013 | -0.013 | 0 | 0.28 | 0.76 | 0.46 | 0.27 | 0.3 | 0.24 |
| 2 | 57805439 | rs10196737 | -1.52 | 1.296 | 0.023 | 1.682 | -0.002 | -0.008 | -0.008 | 0 | 0.23 | 0.75 | 0.31 | 0.25 | 0.25 | 0.18 |
| 2 | 57812484 | rs10185414 | 0.089 | 1.28 | 0.224 | 1.662 | 0.008 | 0.092 | 0.092 | 0.302 | 0.07 | 0.29 | 0.04 | 0.09 | 0.02 | 0.06 |
| 2 | 57820945 | rs1918454 | 0.4 | 1.32 | 0.151 | 1.633 | 0.065 | -0.011 | -0.011 | 0 | 0.07 | 0.22 | 0.12 | 0.03 | 0.15 | 0.06 |
| 2 | 57821416 | rs11125720 | 0.728 | 1.101 | -0.341 | 1.521 | 0.028 | 0.057 | 0.057 | 0 | 0.25 | 0.11 | 0.08 | 0.15 | 0.19 | 0.19 |
| 2 | 57826028 | rs11125721 | 2.049 | 1.03 | -0.387 | 1.569 | -0.08 | 0.121 | 0.121 | 3.636 | 0.39 | 0.03 | 0.34 | 0.15 | 0.39 | 0.25 |
| 2 | 57835525 | rs17048978 | 1.22 | 0.925 | -0.303 | 1.591 | -0.007 | 0.003 | 0.003 | 0 | 0.05 | 0.02 | 0.15 | 0.02 | 0.05 | 0.06 |
| 2 | 57835603 | rs17048981 | 0.522 | 0.888 | -0.067 | 1.589 | 0.01 | 0.098 | 0.098 | 0.019 | 0.06 | 0.25 | 0.04 | 0.09 | 0.02 | 0.06 |
| 2 | 57842453 | rs17810870 | 0.66 | 0.982 | -0.664 | 1.555 | 0.045 | 0.062 | 0.062 | 0 | 0.25 | 0.11 | 0.08 | 0.15 | 0.17 | 0.19 |
| 2 | 57845016 | rs1405826 | 0.213 | 1.029 | -0.619 | 1.599 | -0.011 | -0.054 | -0.054 | 0 | 0.62 | 0.81 | 0.66 | 0.41 | 0.66 | 0.45 |
| 2 | 57854484 | rs2695595 | -0.64 | 1.151 | -0.587 | 1.629 | -0.011 | 0.013 | 0.013 | 0 | 0.93 | 0.95 | 0.92 | 0.69 | 0.91 | 0.75 |
| 2 | 57857099 | rs6748662 | 0.62 | 0.921 | -0.493 | 1.195 | 0.014 | -0.028 | -0.028 | 0 | 0.07 | 0.2 | 0.06 | 0.02 | 0.08 | 0.03 |
| 2 | 57864042 | rs1380706 | 1.514 | 0.973 | -0.99 | 1.186 | -0.056 | 0.116 | 0.116 | 5.125 | 0.4 | 0.03 | 0.34 | 0.15 | 0.39 | 0.25 |
| 2 | 57877166 | rs820777 | 1.545 | 0.997 | -0.667 | 1.329 | -0.044 | -0.02 | -0.02 | 0 | 0.6 | 0.26 | 0.52 | 0.44 | 0.55 | 0.48 |
| 2 | 57882889 | rs820772 | 0.54 | 1.189 | -0.346 | 1.539 | 0.026 | 0.053 | 0.053 | 0 | 0.83 | 0.71 | 0.82 | 0.58 | 0.79 | 0.67 |
| 2 | 57883073 | rs2030944 | -0.967 | 1.051 | -0.215 | 1.38 | 0.003 | -0.012 | -0.012 | 1.81 | 0.23 | 0.47 | 0.31 | 0.15 | 0.23 | 0.19 |
| 2 | 57883201 | rs2030945 | -0.985 | 0.956 | -0.418 | 1.302 | 0.009 | 0.026 | 0.026 | 1.837 | 0.42 | 0.61 | 0.38 | 0.3 | 0.4 | 0.36 |
| 2 | 57884390 | rs820769 | -0.153 | 1.094 | -0.404 | 1.454 | -0.036 | -0.04 | -0.04 | 0.066 | 0.91 | 0.97 | 0.91 | 0.71 | 0.89 | 0.73 |
| 2 | 57884675 | rs2311732 | -0.921 | 0.666 | -0.223 | 1.052 | -0.001 | -0.011 | -0.011 | 0 | 0.22 | 0.5 | 0.31 | 0.15 | 0.23 | 0.18 |
| 2 | 57888034 | rs10496071 | 0.072 | 0.598 | 0.17 | 1.057 | -0.022 | 0.068 | 0.068 | 0.091 | 0.07 | 0.21 | 0.03 | 0.1 | 0.02 | 0.06 |
| 2 | 57888655 | rs10496072 | 0.87 | 0.567 | 0.073 | 1.017 | 0.017 | 0.051 | 0.051 | 0 | 0.22 | 0.08 | 0.07 | 0.15 | 0.18 | 0.19 |
| 2 | 57889352 | rs7586538 | -0.689 | 0.567 | 0.145 | 1.011 | -0.014 | 0.04 | 0.04 | 0.417 | 0.13 | 0.3 | 0.2 | 0.12 | 0.08 | 0.14 |
| 2 | 57896756 | rs1460256 | 0.038 | 0.569 | 0.124 | 1.09 | 0.029 | 0.049 | 0.049 | 0 | 0.11 | 0.24 | 0.19 | 0.15 | 0.16 | 0.13 |
| 2 | 57899062 | rs2199271 | -0.546 | 0.55 | 0.142 | 1.093 | -0.004 | -0.006 | -0.006 | 0 | 0.36 | 0.37 | 0.28 | 0.3 | 0.35 | 0.33 |
| 2 | 57901997 | rs17049036 | -0.378 | 0.68 | 0.272 | 1.216 | -0.01 | 0.032 | 0.032 | 1.557 | 0.1 | 0.2 | 0.18 | 0.11 | 0.06 | 0.1 |
| 2 | 57905695 | rs1380697 | -1.577 | 0.713 | 0.14 | 1.228 | -0.005 | 0.033 | 0.033 | 3.242 | 0.24 | 0.55 | 0.41 | 0.3 | 0.23 | 0.24 |
| 2 | 57905815 | rs17812960 | 0.714 | -0.095 | -0.808 | 0.652 | 0.06 | 0.116 | 0.116 | 0 | 0.31 | 0.08 | 0.09 | 0.15 | 0.19 | 0.2 |
| 2 | 57906216 | rs1380699 | -0.73 | 0.159 | -0.384 | 0.937 | 0.001 | 0.021 | 0.021 | 0 | 0.16 | 0.27 | 0.23 | 0.14 | 0.13 | 0.15 |
| 2 | 57909360 | rs10496075 | 0.187 | 0.021 | -0.919 | 0.84 | 0.055 | 0.07 | 0.07 | 0 | 0.41 | 0.17 | 0.17 | 0.27 | 0.33 | 0.29 |
| 2 | 57911867 | rs17049058 | -0.666 | 0.778 | 0.124 | 1.538 | 0.035 | 0.059 | 0.059 | 1.102 | 0.1 | 0.53 | 0.08 | 0.12 | 0.1 | 0.07 |

| | | | | | | | | | | | | | | | | |
|---|---|---|---|---|---|---|---|---|---|---|---|---|---|---|---|---|
| 2 | 57921232 | rs10196493 | -0.633 | 0.491 | -0.827 | 1.22 | -0.021 | 0.066 | 0.066 | 0 | 0.65 | 0.97 | 0.83 | 0.75 | 0.67 | 0.7 |
| 2 | 57922033 | rs6732639 | -1.308 | 0.274 | 0.06 | 1.098 | 0.013 | 0.177 | 0.177 | 1.306 | 0.1 | 0.65 | 0.17 | 0.25 | 0.08 | 0.12 |
| 2 | 57923006 | rs13012916 | 1.098 | 0.728 | -0.314 | 1.237 | -0.064 | 0.013 | 0.013 | 0.177 | 0.35 | 0.04 | 0.33 | 0.18 | 0.37 | 0.25 |
| 2 | 57929951 | rs13427078 | 0.664 | 0.806 | -0.135 | 1.266 | 0.002 | 0.061 | 0.061 | 0 | 0.05 | 0.11 | 0.11 | 0.16 | 0.05 | 0.08 |
| 2 | 57930015 | rs10193760 | -0.408 | 0.757 | -0.444 | 1.245 | -0.058 | -0.023 | -0.023 | 0.931 | 0.55 | 0.91 | 0.72 | 0.5 | 0.56 | 0.44 |
| 2 | 57933881 | rs3732135 | -0.532 | 0.985 | -0.218 | 0.663 | 0.037 | 0.046 | 0.046 | 0.098 | 0.1 | 0.34 | 0.07 | 0.11 | 0.09 | 0.06 |
| 2 | 57934120 | rs2871780 | 0.461 | 0.959 | -1.253 | 0.526 | -0.023 | 0.049 | 0.049 | 0 | 0.4 | 0.17 | 0.51 | 0.21 | 0.42 | 0.31 |
| 2 | 57937373 | rs17227657 | 0.119 | 0.939 | -1.304 | 0.342 | -0.013 | 0.01 | 0.01 | 0 | 0.09 | 0.03 | 0.18 | 0.06 | 0.09 | 0.07 |
| 2 | 57937744 | rs11125728 | 0.23 | 0.91 | -1.064 | 0.56 | -0.001 | 0.069 | 0.069 | 0 | 0.47 | 0.27 | 0.55 | 0.22 | 0.47 | 0.33 |
| 2 | 57941416 | rs6712575 | -0.966 | 1.565 | -1.022 | 0.531 | -0.027 | 0.012 | 0.012 | 3.496 | 0.28 | 0.76 | 0.39 | 0.22 | 0.41 | 0.49 |
| 2 | 57941597 | rs12612495 | -1.499 | 1.988 | -0.418 | 0.919 | 0.041 | 0.036 | 0.036 | 0 | 0.09 | 0.15 | 0.11 | 0.27 | 0.17 | 0.19 |
| 2 | 57943044 | rs10496077 | -0.542 | 1.565 | -0.624 | 0.994 | 0 | -0.021 | -0.021 | 0 | 0.08 | 0.09 | 0.1 | 0.19 | 0.1 | 0.1 |
| 2 | 57948003 | rs17049085 | -1.101 | 1.751 | -0.745 | 0.837 | -0.011 | -0.023 | -0.023 | 0 | 0.09 | 0.11 | 0.13 | 0.19 | 0.12 | 0.12 |
| 2 | 57950104 | rs10180540 | -1.589 | 1.642 | -0.441 | 0.817 | 0 | -0.001 | -0.001 | 0.173 | 0.06 | 0.08 | 0.05 | 0.07 | 0.06 | 0.1 |
| 2 | 57970943 | rs6722461 | -0.39 | 1.716 | 0.317 | 0.835 | 0.003 | -0.003 | -0.003 | 0 | 0.64 | 0.64 | 0.59 | 0.62 | 0.59 | 0.58 |
| 2 | 57999642 | rs10204882 | -1.223 | 1.623 | 0.347 | 0.676 | -0.001 | 0.004 | 0.004 | 0.092 | 0.22 | 0.22 | 0.2 | 0.15 | 0.19 | 0.15 |
| 2 | 58012833 | rs1568452 | 1.27 | 1.808 | 0.672 | 0.623 | 0.026 | -0.031 | -0.031 | 0.067 | 0.4 | 0.13 | 0.38 | 0.34 | 0.37 | 0.35 |
| 2 | 58026655 | rs10496080 | -0.478 | 1.791 | 0.649 | 0.606 | 0.009 | -0.015 | -0.015 | 0 | 0.16 | 0.21 | 0.14 | 0.1 | 0.17 | 0.13 |
| 2 | 58041482 | rs2953441 | -0.781 | 1.729 | 0.615 | 0.751 | -0.039 | 0.006 | 0.006 | 3.35 | 0.67 | 0.87 | 0.64 | 0.63 | 0.62 | 0.58 |
| 2 | 58041936 | rs2717046 | 1.315 | 1.727 | 0.548 | 0.755 | 0.022 | -0.041 | -0.041 | 0 | 0.4 | 0.1 | 0.37 | 0.34 | 0.37 | 0.36 |
| 2 | 58046683 | rs2717054 | -0.539 | 1.736 | 1.018 | 0.887 | 0.009 | -0.007 | -0.007 | 0 | 0.66 | 0.63 | 0.62 | 0.63 | 0.62 | 0.57 |
| 2 | 58061127 | rs2717076 | -0.69 | 1.713 | 0.862 | 0.806 | -0.122 | -0.045 | -0.045 | 0 | 0.67 | 0.97 | 0.65 | 0.63 | 0.63 | 0.58 |
| 2 | 58068741 | rs1106090 | 0.156 | 1.916 | 0.837 | 0.688 | -0.005 | 0.01 | 0.01 | 0 | 0.61 | 0.49 | 0.54 | 0.47 | 0.61 | 0.55 |
| 2 | 58070482 | rs1851140 | 1.14 | 1.884 | 0.237 | 0.608 | -0.079 | -0.059 | -0.059 | 0 | 0.47 | 0.04 | 0.39 | 0.36 | 0.47 | 0.42 |
| 2 | 58073897 | rs2176931 | 0.737 | 1.844 | 0.419 | 0.521 | -0.079 | -0.018 | -0.018 | 0 | 0.27 | 0.01 | 0.3 | 0.25 | 0.27 | 0.33 |
| 2 | 58092754 | rs13405071 | -0.796 | 1.501 | 1.035 | 0.637 | 0.014 | 0.003 | 0.003 | 0 | 0.06 | 0.13 | 0.05 | 0.08 | 0.06 | 0.1 |
| 2 | 58110969 | rs2717063 | -0.561 | 1.529 | 0.369 | 0.663 | -0.154 | -0.052 | -0.052 | 0 | 0.65 | 0.97 | 0.62 | 0.62 | 0.58 | 0.58 |
| 2 | 58124359 | rs2678919 | 0.956 | 1.485 | 1.023 | 0.582 | -0.004 | 0 | 0 | 1.244 | 0.4 | 0.36 | 0.34 | 0.38 | 0.31 | 0.35 |
| 2 | 58131532 | rs17049243 | 1.055 | 1.418 | 1.531 | 0.707 | 0.015 | -0.017 | -0.017 | 0 | 0.07 | 0.16 | 0.09 | 0.02 | 0.08 | 0.02 |
| 2 | 58133522 | rs2678907 | 1.043 | 1.42 | 1.626 | 0.694 | 0.034 | -0.025 | -0.025 | 0.014 | 0.42 | 0.16 | 0.36 | 0.39 | 0.38 | 0.36 |
| 2 | 58135872 | rs2678905 | -0.704 | 1.416 | 1.651 | 0.697 | -0.027 | 0.023 | 0.023 | 1.39 | 0.64 | 0.78 | 0.63 | 0.67 | 0.59 | 0.58 |
| 2 | 58138581 | rs2678902 | 1.014 | 1.381 | 1.546 | 0.726 | 0.034 | -0.02 | -0.02 | 0 | 0.42 | 0.16 | 0.36 | 0.39 | 0.38 | 0.36 |
| 2 | 58143267 | rs11887238 | -0.818 | 1.362 | 1.501 | 0.748 | 0.024 | 0.022 | 0.022 | 0 | 0.05 | 0.17 | 0.06 | 0.08 | 0.04 | 0.12 |
| 2 | 58153676 | rs10189138 | -0.899 | 1.364 | 1.297 | 0.849 | -0.015 | 0.014 | 0.014 | 0 | 0.17 | 0.28 | 0.19 | 0.16 | 0.13 | 0.08 |
| 2 | 58158283 | rs2678880 | -0.607 | 1.349 | 1.008 | 0.648 | -0.019 | 0.019 | 0.019 | 1.242 | 0.63 | 0.76 | 0.63 | 0.67 | 0.59 | 0.58 |
| 2 | 58161520 | rs7596775 | -1.601 | 1.393 | 0.604 | 0.705 | -0.001 | -0.002 | -0.002 | 0 | 0.23 | 0.77 | 0.28 | 0.28 | 0.22 | 0.21 |
| 2 | 58169494 | rs2717040 | -0.686 | 1.384 | 0.501 | 0.644 | -0.079 | 0.029 | 0.029 | 1.154 | 0.64 | 0.9 | 0.64 | 0.67 | 0.59 | 0.58 |
| 2 | 58192905 | rs17049270 | 1.075 | 1.379 | 0.797 | 0.631 | 0.004 | -0.018 | -0.018 | 0 | 0.07 | 0.1 | 0.09 | 0.01 | 0.08 | 0.02 |
| 2 | 58193774 | rs17049272 | -0.774 | 1.354 | 0.794 | 0.739 | -0.003 | 0.01 | 0.01 | 0.013 | 0.12 | 0.23 | 0.15 | 0.2 | 0.12 | 0.1 |
| 2 | 58203057 | rs4672226 | 1.011 | 0.997 | 0.359 | 0.684 | -0.084 | -0.148 | -0.148 | 0.574 | 0.4 | 0.05 | 0.38 | 0.47 | 0.41 | 0.46 |
| 2 | 58210239 | rs6732310 | -0.766 | 1.058 | 0.476 | 0.715 | -0.091 | 0.014 | 0.014 | 0.765 | 0.65 | 0.94 | 0.66 | 0.68 | 0.6 | 0.58 |
| 2 | 58212803 | rs1518396 | -0.989 | 1.167 | 0.947 | 0.775 | -0.011 | 0.068 | 0.068 | 0.35 | 0.13 | 0.39 | 0.17 | 0.2 | 0.11 | 0.1 |
| 2 | 58212839 | rs1518397 | -0.453 | 1.179 | 0.645 | 0.78 | 0.003 | -0.009 | -0.009 | 0 | 0.12 | 0.16 | 0.14 | 0.2 | 0.11 | 0.09 |
| 2 | 58222928 | rs2312147 | -0.911 | 1.137 | 0.569 | 0.733 | -0.108 | -0.005 | -0.005 | 0 | 0.67 | 0.93 | 0.66 | 0.67 | 0.61 | 0.6 |
| 2 | 58230233 | rs6755324 | -1.076 | 1.188 | 1.203 | 0.85 | 0 | 0.077 | 0.077 | 2.072 | 0.12 | 0.57 | 0.15 | 0.11 | 0.1 | 0.07 |
| 2 | 58233464 | rs7561688 | 0.7 | 1.115 | 0.732 | 0.858 | -0.024 | -0.067 | -0.067 | 0 | 0.47 | 0.06 | 0.42 | 0.53 | 0.42 | 0.49 |
| 2 | 58235386 | rs1518400 | -2 | 0.936 | 0.361 | 0.782 | 0.004 | -0.007 | -0.007 | 0 | 0.25 | 0.85 | 0.28 | 0.22 | 0.22 | 0.21 |
| 2 | 58237076 | rs6719889 | -1.586 | 1.08 | 0.341 | 0.87 | 0.01 | -0.005 | -0.005 | 0 | 0.2 | 0.85 | 0.24 | 0.14 | 0.18 | 0.1 |
| 2 | 58257450 | rs17830749 | 0.687 | 1.119 | 0.31 | 0.918 | -0.012 | -0.034 | -0.034 | 0 | 0.07 | 0.01 | 0.08 | 0.09 | 0.09 | 0.06 |
| 2 | 58258689 | rs12478718 | 0.022 | 1.114 | 0.543 | 0.907 | -0.004 | 0.011 | 0.011 | 0.955 | 0.09 | 0.17 | 0.11 | 0.11 | 0.09 | 0.07 |
| 2 | 58268266 | rs1518403 | 0.098 | 1.098 | 0.485 | 0.902 | 0.005 | -0.018 | -0.018 | 0.714 | 0.09 | 0.06 | 0.1 | 0.11 | 0.09 | 0.1 |
| 2 | 58275666 | rs986512 | -1.086 | 1.584 | 0.21 | 0.839 | -0.096 | -0.067 | -0.067 | 0 | 0.7 | 0.91 | 0.64 | 0.59 | 0.64 | 0.58 |
| 2 | 58304878 | rs1439269 | -2.099 | 1.579 | 0.241 | 1.185 | -0.012 | -0.023 | -0.023 | 0.257 | 0.25 | 0.82 | 0.29 | 0.25 | 0.27 | 0.24 |
| 2 | 58309374 | rs7577299 | 0.221 | 1.543 | 0.784 | 1.052 | 0.078 | -0.005 | -0.005 | 0.82 | 0.09 | 0.42 | 0.11 | 0.02 | 0.09 | 0.03 |
| 2 | 58316814 | rs1051061 | 0.828 | 1.48 | 0.577 | 1.102 | -0.001 | -0.021 | -0.021 | 0 | 0.45 | 0.07 | 0.37 | 0.41 | 0.4 | 0.36 |
| 2 | 58318398 | rs13386998 | -0.626 | 1.547 | 0.581 | 1.13 | -0.001 | -0.001 | -0.001 | 1.682 | 0.07 | 0.07 | 0.06 | 0.1 | 0.08 | 0.15 |
| 2 | 58322052 | rs13424099 | -0.222 | 1.514 | 0.584 | 1.085 | 0.001 | -0.001 | -0.001 | 0.085 | 0.06 | 0.06 | 0.05 | 0.08 | 0.07 | 0.15 |
| 2 | 58332757 | rs10198382 | -0.609 | 1.526 | 0.678 | 1.085 | -0.001 | 0 | 0 | 0 | 0.07 | 0.06 | 0.06 | 0.09 | 0.08 | 0.15 |
| 2 | 58334239 | rs13033987 | -0.773 | 1.495 | 0.692 | 1.048 | 0.005 | -0.003 | -0.003 | 0 | 0.07 | 0.08 | 0.05 | 0.08 | 0.08 | 0.15 |
| 2 | 58338086 | rs13011711 | 0.813 | 1.47 | 0.481 | 1.032 | 0.016 | -0.038 | -0.038 | 1.155 | 0.45 | 0.11 | 0.38 | 0.42 | 0.4 | 0.36 |
| 2 | 58354422 | rs3771211 | 1.448 | 1.436 | 0.702 | 1.053 | -0.003 | 0.001 | 0.001 | 0 | 0.06 | 0.04 | 0.07 | 0.02 | 0.09 | 0.02 |
| 2 | 58377014 | rs12713372 | -0.266 | 1.482 | 0.912 | 1.092 | 0.009 | 0.011 | 0.011 | 2.86 | 0.6 | 0.52 | 0.55 | 0.48 | 0.56 | 0.44 |
| 2 | 58382490 | rs848293 | -0.273 | 1.429 | 0.648 | 1.073 | -0.042 | -0.042 | -0.042 | 1.714 | 0.43 | 0.79 | 0.47 | 0.5 | 0.44 | 0.56 |
| 2 | 58382691 | rs17049378 | -0.371 | 1.456 | 0.66 | 1.064 | 0 | -0.001 | -0.001 | 0.012 | 0.15 | 0.21 | 0.18 | 0.08 | 0.18 | 0.07 |
| 2 | 58385353 | rs848292 | -1.437 | 1.524 | 0.425 | 1.086 | -0.004 | 0.005 | 0.005 | 2.207 | 0.6 | 0.96 | 0.68 | 0.69 | 0.65 | 0.7 |
| 2 | 58387000 | rs3732136 | -2.19 | 1.58 | 0.434 | 1.251 | -0.028 | -0.059 | -0.059 | 0 | 0.27 | 0.92 | 0.31 | 0.28 | 0.26 | 0.26 |
| 2 | 58388696 | rs848291 | -1.328 | 1.58 | 0.428 | 1.253 | -0.025 | -0.063 | -0.063 | 0 | 0.58 | 0.95 | 0.64 | 0.59 | 0.62 | 0.64 |
| 2 | 58390547 | rs10445896 | 1.028 | 1.598 | 0.837 | 1.22 | 0.081 | 0.012 | 0.012 | 0 | 0.07 | 0.3 | 0.09 | 0.02 | 0.09 | 0.03 |
| 2 | 58392789 | rs848288 | 0.974 | 1.574 | 0.666 | 1.236 | -0.051 | 0.019 | 0.019 | 0.437 | 0.3 | 0.11 | 0.33 | 0.29 | 0.35 | 0.38 |

| | | | | | | | | | | | | | | | | |
|---|---|---|---|---|---|---|---|---|---|---|---|---|---|---|---|---|
| 2 | 58394543 | rs848286 | 0.619 | 1.625 | 0.727 | 1.257 | -0.03 | -0.009 | -0.009 | 0 | 0.33 | 0.21 | 0.37 | 0.4 | 0.38 | 0.44 |
| 2 | 58410972 | rs768298 | -2.271 | 1.6 | 0.565 | 1.284 | -0.02 | -0.067 | -0.067 | 0 | 0.28 | 0.85 | 0.33 | 0.39 | 0.29 | 0.31 |
| 2 | 58412472 | rs848280 | -1.407 | 1.576 | 0.516 | 1.409 | 0.01 | 0.05 | 0.05 | 2.422 | 0.59 | 0.94 | 0.67 | 0.72 | 0.65 | 0.69 |
| 2 | 58417817 | rs17049394 | 0.129 | 1.317 | 1.197 | 1.568 | 0 | -0.001 | -0.001 | 0.298 | 0.09 | 0.12 | 0.1 | 0.09 | 0.09 | 0.05 |
| 2 | 58423312 | rs11903212 | -1.016 | 1.34 | 1.364 | 1.456 | 0.021 | 0.12 | 0.12 | 2.055 | 0.1 | 0.25 | 0.1 | 0.22 | 0.08 | 0.18 |
| 2 | 58423643 | rs13006005 | -0.317 | 1.348 | 1.1 | 1.462 | -0.029 | -0.041 | -0.041 | 0 | 0.07 | 0 | 0.08 | 0.22 | 0.08 | 0.18 |
| 2 | 58441542 | rs17049417 | 0.097 | 1.34 | 1.234 | 1.398 | 0.001 | 0.001 | 0.001 | 0.226 | 0.09 | 0.11 | 0.1 | 0.05 | 0.09 | 0.05 |
| 2 | 58450569 | rs3771220 | -1.685 | 1.33 | 0.724 | 1.401 | 0.036 | 0.176 | 0.176 | 1.461 | 0.13 | 0.56 | 0.16 | 0.32 | 0.11 | 0.25 |
| 2 | 58455672 | rs6761301 | -2.68 | 1.355 | 0.396 | 1.283 | -0.055 | -0.117 | -0.117 | 0 | 0.32 | 0.93 | 0.37 | 0.39 | 0.33 | 0.32 |
| 2 | 58463541 | rs11889211 | -0.775 | 1.369 | 0.579 | 1.246 | 0.006 | -0.025 | -0.025 | 0 | 0.16 | 0.36 | 0.19 | 0.07 | 0.18 | 0.07 |
| 2 | 58470914 | rs17049425 | -2.318 | 1.365 | 0.071 | 1.444 | -0.02 | -0.078 | -0.078 | 0 | 0.3 | 0.85 | 0.35 | 0.41 | 0.31 | 0.36 |
| 2 | 58475916 | rs2192689 | -0.087 | 1.227 | 0.103 | 1.13 | -0.009 | 0.002 | 0.002 | 0.807 | 0.1 | 0.02 | 0.09 | 0.05 | 0.1 | 0.07 |
| 2 | 58478179 | rs13411119 | -2.171 | 1.247 | 0.128 | 1.19 | -0.004 | -0.031 | -0.031 | 0 | 0.27 | 0.83 | 0.3 | 0.29 | 0.27 | 0.3 |
| 2 | 58481589 | rs17398871 | -0.27 | 1.12 | 0.216 | 1.061 | 0.003 | 0.001 | 0.001 | 0.719 | 0.11 | 0.09 | 0.11 | 0.06 | 0.11 | 0.11 |
| 2 | 58488026 | rs11894186 | -0.562 | 1.368 | 0.308 | 1.031 | 0 | 0 | 0 | 0.119 | 0.47 | 0.51 | 0.39 | 0.52 | 0.47 | 0.55 |
| 2 | 58492345 | rs9789483 | -0.075 | 1.388 | 0.584 | 1.001 | 0.008 | -0.018 | -0.018 | 0.092 | 0.39 | 0.27 | 0.31 | 0.51 | 0.38 | 0.51 |
| 2 | 58492441 | rs9789714 | -0.083 | 1.382 | 0.672 | 1.005 | 0.004 | -0.006 | -0.006 | 0 | 0.41 | 0.37 | 0.31 | 0.51 | 0.38 | 0.52 |
| 2 | 58493260 | rs7593369 | -0.058 | 1.418 | 0.558 | 0.993 | 0.013 | -0.021 | -0.021 | 0 | 0.39 | 0.26 | 0.3 | 0.51 | 0.38 | 0.51 |
| 2 | 58493541 | rs9973912 | -0.137 | 1.345 | 0.533 | 1.077 | -0.005 | 0.062 | 0.062 | 0.093 | 0.08 | 0.14 | 0.06 | 0.19 | 0.06 | 0.15 |
| 2 | 58497451 | rs6739403 | 0.534 | 1.633 | 0.094 | 0.778 | -0.002 | 0 | 0 | 3.014 | 0.47 | 0.48 | 0.54 | 0.48 | 0.42 | 0.43 |
| 2 | 58498481 | rs10490251 | 0.97 | 1.477 | -0.577 | 0.679 | -0.052 | -0.034 | -0.034 | 0.236 | 0.3 | 0.02 | 0.23 | 0.31 | 0.32 | 0.36 |
| 2 | 58503341 | rs6706369 | -0.644 | 1.357 | -0.521 | 0.739 | -0.117 | 0.012 | 0.012 | 0.791 | 0.63 | 0.98 | 0.71 | 0.69 | 0.57 | 0.63 |
| 2 | 58509628 | rs10490250 | -0.196 | 1.112 | 0.296 | 0.923 | 0.017 | 0.019 | 0.019 | 0 | 0.16 | 0.26 | 0.13 | 0.2 | 0.18 | 0.19 |
| 2 | 58513505 | rs7355437 | -1.186 | 1.033 | 0.807 | 0.878 | 0.002 | 0.003 | 0.003 | 0.587 | 0.2 | 0.7 | 0.17 | 0.19 | 0.18 | 0.19 |
| 2 | 58514666 | rs2215960 | -0.735 | 1.039 | 0.917 | 0.881 | 0.014 | 0.015 | 0.015 | 0.234 | 0.19 | 0.63 | 0.16 | 0.19 | 0.18 | 0.19 |
| 2 | 58525659 | rs9646985 | 0.169 | 1.153 | 0.608 | 0.98 | 0.005 | 0.006 | 0.006 | 0 | 0.13 | 0.17 | 0.1 | 0.18 | 0.16 | 0.18 |
| 2 | 58525741 | rs7595474 | -0.791 | 1.149 | 0.575 | 0.973 | 0.021 | 0.027 | 0.027 | 0 | 0.18 | 0.38 | 0.14 | 0.2 | 0.18 | 0.21 |
| 2 | 58559427 | rs11903838 | 0.191 | 0.696 | -0.353 | 0.823 | 0.011 | -0.032 | -0.032 | 0 | 0.22 | 0.12 | 0.2 | 0.35 | 0.19 | 0.26 |
| 2 | 58565395 | rs17049463 | 0.457 | 0.684 | -0.473 | 0.77 | 0.004 | -0.059 | -0.059 | 0 | 0.21 | 0.05 | 0.19 | 0.33 | 0.18 | 0.26 |
| 2 | 58565501 | rs6708046 | -1.064 | 0.594 | -0.245 | 0.738 | 0.01 | -0.037 | -0.037 | 0 | 0.39 | 0.73 | 0.32 | 0.51 | 0.36 | 0.46 |
| 2 | 58567163 | rs12619492 | 0.421 | 0.488 | 0.021 | 0.68 | 0.009 | -0.021 | -0.021 | 0 | 0.22 | 0.14 | 0.2 | 0.33 | 0.18 | 0.28 |
| 2 | 58569066 | rs6734368 | 0.407 | 0.499 | 0.355 | 0.686 | 0.009 | -0.021 | -0.021 | 0 | 0.22 | 0.14 | 0.2 | 0.33 | 0.18 | 0.28 |
| 2 | 58569214 | rs6708936 | 0.149 | 1.258 | 0.18 | 0.495 | 0.026 | -0.013 | -0.013 | 0 | 0.07 | 0.12 | 0.25 | 0.02 | 0.15 | 0.13 |
| 2 | 58580782 | rs17049476 | 0.816 | 1.078 | 0.057 | 0.64 | 0.008 | -0.029 | -0.029 | 0 | 0.19 | 0.07 | 0.19 | 0.29 | 0.16 | 0.24 |
| 2 | 58581018 | rs6761469 | -1.004 | 0.175 | -0.548 | 0.245 | -0.043 | -0.053 | -0.053 | 0 | 0.62 | 0.82 | 0.65 | 0.54 | 0.52 | 0.65 |
| 2 | 58584218 | rs6545681 | -0.158 | 0.213 | -0.505 | 0.22 | 0.008 | 0.06 | 0.06 | 0.402 | 0.42 | 0.75 | 0.48 | 0.25 | 0.38 | 0.39 |
| 2 | 58592685 | rs7588962 | -0.149 | 0.218 | -0.377 | 0.226 | 0.004 | 0.023 | 0.023 | 0.108 | 0.42 | 0.66 | 0.48 | 0.25 | 0.38 | 0.39 |
| 2 | 58597957 | rs887956 | 1.384 | 0.181 | -0.465 | 0.24 | 0.011 | -0.05 | -0.05 | 0.329 | 0.2 | 0.07 | 0.18 | 0.34 | 0.16 | 0.26 |
| 2 | 58602481 | rs722764 | 1.325 | 0.171 | -0.47 | 0.235 | 0.012 | -0.05 | -0.05 | 0.944 | 0.2 | 0.07 | 0.19 | 0.34 | 0.16 | 0.26 |
| 2 | 58607850 | rs6757964 | -0.096 | 0.177 | -0.151 | 0.1 | 0.002 | 0.017 | 0.017 | 0 | 0.41 | 0.64 | 0.48 | 0.2 | 0.38 | 0.38 |
| 2 | 58615998 | rs10192578 | 0.123 | -0.051 | -0.254 | -0.102 | 0.011 | -0.053 | -0.053 | 0.516 | 0.27 | 0.48 | 0.34 | 0.15 | 0.28 | 0.29 |
| 2 | 58623301 | rs12713384 | -2.033 | -0.006 | -0.428 | 0.101 | -0.052 | -0.062 | -0.062 | 1.282 | 0.79 | 0.96 | 0.73 | 0.74 | 0.72 | 0.85 |
| 2 | 58628304 | rs4672239 | -0.247 | -0.148 | 0.078 | 0.12 | 0 | -0.026 | -0.026 | 2.049 | 0.33 | 0.61 | 0.4 | 0.15 | 0.32 | 0.34 |
| 2 | 58629752 | rs4444510 | 0.844 | -0.21 | -0.134 | 0.164 | 0.004 | -0.04 | -0.04 | 0 | 0.19 | 0.11 | 0.18 | 0.38 | 0.16 | 0.29 |
| 2 | 58649144 | rs7577048 | -0.277 | -0.223 | -0.039 | 0.212 | 0 | -0.033 | -0.033 | 0.215 | 0.33 | 0.52 | 0.48 | 0.18 | 0.31 | 0.35 |
| 2 | 58650355 | rs4672240 | -1.073 | -0.08 | -0.104 | 0.198 | -0.031 | 0.001 | 0.001 | 0.394 | 0.53 | 0.82 | 0.67 | 0.56 | 0.48 | 0.65 |
| 2 | 58656792 | rs6734591 | 0.065 | 0.049 | 0.155 | 0.206 | 0.137 | 0.036 | 0.036 | 0 | 0.11 | 0.38 | 0.32 | 0.06 | 0.17 | 0.16 |
| 2 | 58662069 | rs12613228 | 1.926 | 0.01 | 0.039 | 0.356 | -0.007 | -0.021 | -0.021 | 0.17 | 0.14 | 0.03 | 0.15 | 0.17 | 0.12 | 0.19 |
| 2 | 58664988 | rs2192661 | 0.014 | 0.219 | 0.394 | 0.499 | 0.149 | 0.043 | 0.043 | 0 | 0.11 | 0.38 | 0.32 | 0.06 | 0.17 | 0.17 |
| 2 | 58666521 | rs6753024 | -0.01 | -0.081 | -0.013 | 0.501 | -0.01 | 0.001 | 0.001 | 0 | 0.27 | 0.33 | 0.2 | 0.32 | 0.19 | 0.28 |
| 2 | 58685318 | rs3732138 | 1.511 | 0.023 | -0.213 | 0.572 | 0.002 | 0 | 0 | 0 | 0.16 | 0.13 | 0.15 | 0.17 | 0.12 | 0.18 |
| 2 | 58690295 | rs17399334 | 1.536 | 0.035 | -0.207 | 0.578 | 0.002 | 0 | 0 | 0.064 | 0.16 | 0.13 | 0.15 | 0.17 | 0.12 | 0.18 |
| 2 | 58704449 | rs6743916 | -0.451 | -0.232 | 0.438 | 0.578 | -0.016 | 0.029 | 0.029 | 1.994 | 0.35 | 0.49 | 0.32 | 0.48 | 0.28 | 0.48 |
| 2 | 58705069 | rs10174892 | -0.955 | -0.286 | 0.546 | 0.566 | 0.017 | -0.016 | -0.016 | 0 | 0.4 | 0.69 | 0.36 | 0.51 | 0.3 | 0.5 |
| 2 | 58713430 | rs11894830 | 0.574 | -0.244 | 0.742 | 0.606 | -0.001 | 0.002 | 0.002 | 1.251 | 0.18 | 0.17 | 0.15 | 0.3 | 0.16 | 0.28 |
| 2 | 58715610 | rs6738399 | -0.014 | -0.256 | 0.93 | 0.594 | -0.013 | 0.041 | 0.041 | 0.62 | 0.22 | 0.41 | 0.2 | 0.32 | 0.18 | 0.3 |
| 2 | 58740603 | rs6545684 | -0.526 | -0.336 | 1.008 | 0.552 | -0.016 | 0.013 | 0.013 | 0.864 | 0.25 | 0.66 | 0.22 | 0.34 | 0.18 | 0.3 |
| 2 | 58744670 | rs7557639 | -0.719 | -0.42 | 0.992 | 0.499 | 0.016 | -0.012 | -0.012 | 3.376 | 0.41 | 0.69 | 0.35 | 0.5 | 0.3 | 0.5 |
| 2 | 58751249 | rs13007356 | 1.815 | -0.066 | 0.19 | 0.453 | -0.057 | -0.062 | -0.062 | 0.008 | 0.19 | 0.05 | 0.18 | 0.26 | 0.26 | 0.14 |
| 2 | 58760640 | rs10490086 | 1.858 | -0.027 | 0.03 | 0.445 | -0.048 | -0.06 | -0.06 | 0 | 0.19 | 0.05 | 0.18 | 0.26 | 0.25 | 0.14 |
| 2 | 58765789 | rs17269091 | 1.889 | -0.072 | 0.033 | 0.371 | 0.012 | 0 | 0 | 0 | 0.17 | 0.05 | 0.11 | 0.17 | 0.12 | 0.18 |
| 2 | 58767826 | rs11896571 | -0.475 | -0.034 | 0.164 | 0.411 | 0 | 0.009 | 0.009 | 0 | 0.45 | 0.76 | 0.45 | 0.59 | 0.44 | 0.45 |
| 2 | 58777169 | rs6732504 | -0.505 | -0.032 | 0.259 | 0.396 | 0.001 | 0.006 | 0.006 | 0 | 0.45 | 0.69 | 0.44 | 0.59 | 0.45 | 0.46 |
| 2 | 58785543 | rs11125757 | -0.426 | -0.026 | 0.19 | 0.4 | 0 | 0 | 0 | 0 | 0.44 | 0.58 | 0.39 | 0.57 | 0.44 | 0.46 |
| 2 | 58792232 | rs6736526 | -0.406 | 0.011 | -0.09 | 0.426 | -0.004 | -0.002 | -0.002 | 0 | 0.42 | 0.71 | 0.4 | 0.59 | 0.44 | 0.46 |
| 2 | 58794524 | rs994836 | 0.844 | 0.044 | -0.159 | 0.467 | -0.006 | -0.015 | -0.015 | 0.407 | 0.17 | 0.36 | 0.16 | 0.13 | 0.13 | 0.21 |
| 2 | 58794665 | rs994834 | -0.946 | 0.022 | -0.351 | 0.438 | -0.012 | 0.053 | 0.053 | 2.989 | 0.57 | 0.77 | 0.54 | 0.75 | 0.55 | 0.63 |
| 2 | 58795436 | rs933525 | -1.72 | 0.546 | -0.474 | 0.113 | -0.007 | 0.005 | 0.005 | 0 | 0.89 | 0.92 | 0.7 | 0.95 | 0.8 | 0.85 |
| 2 | 58802370 | rs17039689 | 1.764 | 0.052 | -0.35 | 0.337 | -0.071 | -0.051 | -0.051 | 0.223 | 0.17 | 0.03 | 0.18 | 0.24 | 0.26 | 0.14 |

| | | | | | | | | | | | | | | | | |
|---|---|---|---|---|---|---|---|---|---|---|---|---|---|---|---|---|
| 2 | 58807544 | rs6712660 | 0.57 | 0.006 | 0.1 | 0.356 | -0.002 | -0.009 | -0.009 | 2.529 | 0.17 | 0.36 | 0.16 | 0.13 | 0.13 | 0.21 |
| 2 | 58820417 | rs2902140 | 0.653 | 0.187 | -0.471 | 0.441 | -0.008 | -0.05 | -0.05 | 0 | 0.18 | 0.09 | 0.15 | 0.34 | 0.21 | 0.2 |
| 2 | 58823770 | rs7604277 | 0.104 | 0.192 | -0.626 | 0.134 | 0.002 | -0.002 | -0.002 | 1.168 | 0.4 | 0.37 | 0.33 | 0.41 | 0.38 | 0.37 |
| 2 | 58824738 | rs6723710 | -0.336 | 0.208 | -1.079 | 0.095 | 0 | -0.012 | -0.012 | 0 | 0.53 | 0.5 | 0.46 | 0.74 | 0.57 | 0.54 |
| 2 | 58825311 | rs7577494 | -1.46 | 0.267 | -1.218 | -0.086 | -0.015 | 0.032 | 0.032 | 0.066 | 0.76 | 0.85 | 0.66 | 0.91 | 0.73 | 0.79 |
| 2 | 58826638 | rs7607950 | 0.153 | 0.217 | -1.335 | -0.058 | -0.004 | 0.05 | 0.05 | 0 | 0.33 | 0.45 | 0.3 | 0.49 | 0.33 | 0.39 |
| 2 | 58828431 | rs12612428 | 2.273 | 0.207 | -1.32 | -0.058 | -0.002 | 0.008 | 0.008 | 0 | 0.16 | 0.04 | 0.15 | 0.16 | 0.14 | 0.2 |
| 2 | 58828603 | rs17049586 | 1.356 | 0.213 | -0.828 | 0.406 | -0.01 | -0.023 | -0.023 | 0 | 0.22 | 0.11 | 0.17 | 0.24 | 0.24 | 0.14 |
| 2 | 58830445 | rs7569678 | -1.326 | 0.264 | -0.783 | 0.398 | -0.006 | 0.023 | 0.023 | 0 | 0.59 | 0.66 | 0.5 | 0.76 | 0.59 | 0.59 |
| 2 | 58832407 | rs17269126 | 1.979 | 0.237 | -1.058 | 0.408 | -0.035 | -0.038 | -0.038 | 0.991 | 0.11 | 0.01 | 0.11 | 0.13 | 0.11 | 0.19 |
| 2 | 58832708 | rs6727462 | -1.113 | 0.28 | -0.806 | 0.178 | -0.02 | 0.083 | 0.083 | 0 | 0.59 | 0.79 | 0.5 | 0.76 | 0.59 | 0.58 |
| 2 | 58833651 | rs11887255 | 0.534 | 0.323 | -0.838 | 0.31 | 0.006 | -0.009 | -0.009 | 0 | 0.26 | 0.18 | 0.2 | 0.27 | 0.26 | 0.19 |
| 2 | 58834185 | rs6545689 | 0.879 | 0.551 | -0.913 | 0.443 | -0.001 | -0.018 | -0.018 | 0.015 | 0.23 | 0.12 | 0.18 | 0.26 | 0.25 | 0.17 |
| 2 | 58842418 | rs13029039 | -0.388 | 0.727 | -0.505 | 0.781 | 0.008 | 0.042 | 0.042 | 0 | 0.33 | 0.48 | 0.37 | 0.49 | 0.37 | 0.41 |
| 2 | 58847953 | rs6545694 | 0.164 | 0.602 | -0.344 | 0.687 | 0.004 | 0.037 | 0.037 | 0 | 0.36 | 0.53 | 0.37 | 0.47 | 0.41 | 0.35 |
| 2 | 58854574 | rs2041813 | -0.855 | 0.503 | -0.676 | 0.284 | 0.006 | -0.003 | -0.003 | 0.389 | 0.68 | 0.75 | 0.69 | 0.65 | 0.71 | 0.72 |
| 2 | 58857542 | rs4672251 | 0.115 | 0.536 | -0.481 | 0.222 | -0.01 | 0.018 | 0.018 | 0 | 0.28 | 0.19 | 0.31 | 0.2 | 0.3 | 0.33 |
| 2 | 58866584 | rs7572387 | 0.493 | 0.543 | -0.192 | 0.322 | -0.009 | -0.019 | -0.019 | 0 | 0.37 | 0.3 | 0.38 | 0.46 | 0.44 | 0.4 |
| 2 | 58877236 | rs1158041 | -0.182 | 0.624 | -0.097 | 0.197 | 0.01 | 0.015 | 0.015 | 0 | 0.52 | 0.57 | 0.6 | 0.64 | 0.57 | 0.56 |
| 2 | 58892049 | rs10490092 | 1.317 | 0.7 | -0.164 | 0.115 | 0.001 | -0.009 | -0.009 | 0 | 0.13 | 0.16 | 0.09 | 0.08 | 0.12 | 0.11 |
| 2 | 58898820 | rs7561534 | 0.421 | 0.683 | -0.236 | 0.103 | 0.005 | 0.005 | 0.005 | 0 | 0.39 | 0.52 | 0.5 | 0.54 | 0.44 | 0.44 |
| 2 | 58899650 | rs1990642 | -0.218 | 0.669 | -0.261 | 0.11 | 0.005 | 0.01 | 0.01 | 0 | 0.51 | 0.54 | 0.58 | 0.62 | 0.56 | 0.55 |
| 2 | 58913139 | rs10865306 | -0.782 | 0.837 | -0.628 | 0.139 | -0.048 | -0.05 | -0.05 | 0 | 0.92 | 0.99 | 0.82 | 0.82 | 0.85 | 0.96 |
| 2 | 58915608 | rs17614895 | 1.678 | 0.82 | -0.659 | 0.049 | -0.035 | 0.035 | 0.035 | 0 | 0.13 | 0.02 | 0.08 | 0.08 | 0.12 | 0.11 |
| 2 | 58922570 | rs17549476 | 1.698 | 0.82 | -0.668 | 0.02 | -0.039 | 0.055 | 0.055 | 0 | 0.12 | 0 | 0.08 | 0.03 | 0.12 | 0.11 |
| 2 | 58923091 | rs1981731 | -0.766 | 0.845 | -0.664 | 0.091 | -0.049 | -0.053 | -0.053 | 1.751 | 0.92 | 0.99 | 0.83 | 0.78 | 0.86 | 0.96 |
| 2 | 58935023 | rs17615036 | 1.396 | 0.772 | -0.674 | -0.002 | -0.038 | 0.059 | 0.059 | 0 | 0.13 | 0.01 | 0.09 | 0.03 | 0.13 | 0.12 |
| 2 | 58935282 | rs4671328 | 0.032 | 0.584 | -0.627 | -0.092 | 0.002 | 0.004 | 0.004 | 0 | 0.52 | 0.54 | 0.6 | 0.57 | 0.57 | 0.63 |
| 2 | 58937463 | rs11884767 | 1.628 | 0.363 | -0.704 | -0.272 | -0.002 | -0.011 | -0.011 | 0 | 0.13 | 0.09 | 0.08 | 0 | 0.14 | 0.2 |
| 2 | 58944760 | rs13416859 | 1.676 | 0.247 | -0.937 | -0.29 | 0.059 | 0.025 | 0.025 | 0.35 | 0.24 | 0.05 | 0.09 | 0.19 | 0.14 | 0.22 |
| 2 | 58945843 | rs17615381 | 1.661 | 0.435 | -0.886 | -0.272 | -0.04 | 0.055 | 0.055 | 0 | 0.12 | 0.01 | 0.08 | 0.02 | 0.12 | 0.1 |
| 2 | 58946567 | rs17550035 | 1.352 | 0.409 | -0.803 | -0.222 | -0.021 | 0.058 | 0.058 | 0 | 0.14 | 0.03 | 0.1 | 0 | 0.15 | 0.2 |
| 2 | 58949839 | rs1796434 | 1.031 | 0.394 | -0.519 | -0.234 | 0.002 | -0.048 | -0.048 | 0.211 | 0.15 | 0.15 | 0.12 | 0 | 0.17 | 0.2 |
| 2 | 58950863 | rs993532 | 1.632 | 0.449 | -0.654 | -0.286 | -0.042 | 0.056 | 0.056 | 0 | 0.13 | 0.02 | 0.08 | 0.02 | 0.14 | 0.11 |
| 2 | 58955953 | rs2708146 | -0.432 | 0.289 | -0.44 | -0.276 | 0 | 0.005 | 0.005 | 0 | 0.55 | 0.59 | 0.34 | 0.22 | 0.44 | 0.53 |
| 2 | 58965211 | rs1641155 | 0.382 | 0.549 | -0.074 | -0.108 | -0.002 | -0.007 | -0.007 | 0 | 0.37 | 0.43 | 0.22 | 0.22 | 0.3 | 0.34 |
| 2 | 58965713 | rs2540325 | 1.803 | 0.301 | -0.032 | -0.2 | 0.048 | 0.018 | 0.018 | 0.856 | 0.23 | 0.1 | 0.11 | 0.19 | 0.15 | 0.21 |
| 2 | 58966082 | rs1796435 | 1.306 | 0.496 | 0.07 | -0.104 | 0.042 | 0.028 | 0.028 | 0.611 | 0.27 | 0.2 | 0.12 | 0.19 | 0.15 | 0.22 |
| 2 | 58971625 | rs17049716 | 0.02 | 0.95 | -0.004 | -0.105 | 0.076 | -0.01 | -0.01 | 2.124 | 0.06 | 0.14 | 0.08 | 0.01 | 0.1 | 0.04 |
| 2 | 58975143 | rs12986742 | -0.736 | 0.649 | -0.275 | -0.046 | -0.001 | 0.003 | 0.003 | 0 | 0.54 | 0.59 | 0.42 | 0.44 | 0.46 | 0.42 |
| 2 | 58978523 | rs17550821 | 1.207 | 1.242 | -0.034 | 0.216 | -0.048 | 0.032 | 0.032 | 0 | 0.09 | 0.01 | 0.09 | 0.02 | 0.16 | 0.11 |
| 2 | 58981325 | rs6759519 | -1.237 | 1.176 | 0.004 | 0.196 | 0 | 0.004 | 0.004 | 0.152 | 0.63 | 0.59 | 0.54 | 0.45 | 0.6 | 0.56 |
| 2 | 58987726 | rs10490095 | 0.293 | 1.264 | -0.094 | 0.238 | 0.056 | -0.009 | -0.009 | 0 | 0.05 | 0.12 | 0.06 | 0.01 | 0.09 | 0.04 |
| 2 | 58990485 | rs2862874 | -1.146 | 1.157 | -0.487 | 0.296 | 0.001 | 0.007 | 0.007 | 0.108 | 0.63 | 0.58 | 0.54 | 0.44 | 0.6 | 0.59 |
| 2 | 58997712 | rs10210385 | 0.007 | 0.54 | -0.791 | 0.325 | 0 | -0.001 | -0.001 | 0 | 0.43 | 0.58 | 0.29 | 0.27 | 0.36 | 0.42 |
| 2 | 58999505 | rs10204566 | -0.814 | 0.824 | -0.646 | 0.761 | -0.006 | -0.004 | -0.004 | 1.375 | 0.63 | 0.64 | 0.54 | 0.44 | 0.56 | 0.58 |
| 2 | 59037042 | rs7597339 | -0.378 | 0.886 | -0.948 | 0.624 | -0.015 | -0.022 | -0.022 | 0.412 | 0.18 | 0.07 | 0.23 | 0.22 | 0.21 | 0.15 |
| 2 | 59039998 | rs12617233 | -0.515 | 1.237 | -0.389 | 0.728 | -0.004 | -0.006 | -0.006 | 0 | 0.66 | 0.69 | 0.56 | 0.5 | 0.59 | 0.65 |
| 2 | 59041432 | rs2862733 | 0.466 | 1.197 | 0.118 | 0.573 | 0.005 | 0.006 | 0.006 | 0 | 0.33 | 0.33 | 0.5 | 0.56 | 0.44 | 0.38 |
| 2 | 59042847 | rs4671330 | -0.219 | 1.019 | 0.111 | 0.211 | -0.001 | 0.031 | 0.031 | 0 | 0.66 | 0.78 | 0.65 | 0.76 | 0.68 | 0.65 |
| 2 | 59047919 | rs2160206 | -0.308 | 1.088 | 0.111 | 0.207 | -0.002 | 0.047 | 0.047 | 0 | 0.67 | 0.86 | 0.67 | 0.76 | 0.71 | 0.68 |
| 2 | 59047973 | rs10182324 | -0.301 | 1.109 | 0.709 | 0.202 | 0.071 | -0.042 | -0.042 | 0.523 | 0.11 | 0.35 | 0.1 | 0.01 | 0.13 | 0.03 |
| 2 | 59065808 | rs7593324 | 1.202 | 0.92 | 0.567 | 0.167 | 0.008 | 0.003 | 0.003 | 0 | 0.23 | 0.21 | 0.08 | 0.19 | 0.11 | 0.2 |
| 2 | 59072516 | rs7592415 | -0.166 | 1.013 | 0.642 | 0.17 | -0.016 | -0.016 | -0.016 | 0 | 0.35 | 0.59 | 0.18 | 0.24 | 0.24 | 0.31 |
| 2 | 59072969 | rs17552189 | 1.38 | 1.065 | 0.557 | 0.257 | -0.035 | 0.036 | 0.036 | 0.575 | 0.26 | 0.13 | 0.26 | 0.14 | 0.39 | 0.21 |
| 2 | 59074334 | rs17049775 | 0.865 | 0.57 | 0.729 | 0.687 | 0.035 | -0.004 | -0.004 | 0 | 0.22 | 0.12 | 0.09 | 0.22 | 0.11 | 0.27 |
| 2 | 59082438 | rs10198586 | -0.517 | 0.712 | 0.821 | 0.671 | 0.023 | -0.04 | -0.04 | 0.823 | 0.1 | 0.23 | 0.07 | 0.01 | 0.12 | 0.05 |
| 2 | 59083361 | rs17049782 | 0.193 | 0.753 | 0.81 | 0.696 | -0.003 | 0.011 | 0.011 | 0 | 0.05 | 0.05 | 0.11 | 0.2 | 0.06 | 0.07 |
| 2 | 59089629 | rs17049785 | -1.421 | 0.495 | 0.833 | 0.697 | 0.02 | 0.023 | 0.023 | 0 | 0.15 | 0.28 | 0.17 | 0.22 | 0.16 | 0.11 |
| 2 | 59090390 | rs2215964 | -0.878 | 0.445 | 0.724 | 0.675 | -0.018 | 0.008 | 0.008 | 0 | 0.39 | 0.55 | 0.28 | 0.44 | 0.27 | 0.38 |
| 2 | 59091613 | rs11125762 | -0.345 | 0.582 | 0.75 | 0.558 | 0.037 | -0.038 | -0.038 | 0.54 | 0.09 | 0.26 | 0.08 | 0.01 | 0.12 | 0.05 |
| 2 | 59094190 | rs6719683 | -0.707 | 0.563 | 0.272 | 0.602 | -0.02 | 0.003 | 0.003 | 0 | 0.37 | 0.54 | 0.26 | 0.41 | 0.26 | 0.32 |
| 2 | 59095800 | rs12618463 | 0.642 | 0.208 | -0.413 | 0.584 | 0.027 | 0.005 | 0.005 | 0 | 0.22 | 0.14 | 0.07 | 0.19 | 0.11 | 0.19 |
| 2 | 59099013 | rs17049794 | -0.426 | 0.982 | 0.519 | 1.26 | 0.038 | 0.196 | 0.196 | 0 | 0.05 | 0.21 | 0.13 | 0.24 | 0.04 | 0.05 |
| 2 | 59100543 | rs734926 | 0.905 | 1.534 | -0.075 | 0.561 | -0.002 | 0.01 | 0.01 | 0 | 0.53 | 0.38 | 0.34 | 0.32 | 0.43 | 0.5 |
| 2 | 59100938 | rs17049796 | -0.251 | 1.741 | -1.204 | 0.844 | -0.044 | -0.156 | -0.156 | 0 | 0.2 | 0.02 | 0.38 | 0.59 | 0.22 | 0.37 |
| 2 | 59104524 | rs2192692 | 0.912 | 2.137 | -1.108 | 0.675 | -0.054 | -0.029 | -0.029 | 0 | 0.66 | 0.87 | 0.47 | 0.33 | 0.57 | 0.56 |
| 2 | 59104941 | rs12621847 | -0.969 | 1.881 | -0.812 | 0.88 | -0.004 | -0.033 | -0.033 | 0 | 0.23 | 0.16 | 0.44 | 0.64 | 0.27 | 0.35 |
| 2 | 59106418 | rs17617873 | 0.76 | 2.249 | -0.499 | 1.101 | -0.01 | 0.013 | 0.013 | 0.813 | 0.05 | 0.02 | 0.05 | 0.04 | 0.09 | 0.05 |

| | | | | | | | | | | | | | | | | |
|---|---|---|---|---|---|---|---|---|---|---|---|---|---|---|---|---|
| 2 | 59106786 | rs42848 | -2.141 | 2.247 | -0.298 | 1.102 | 0.004 | 0.013 | 0.013 | 0 | 0.24 | 0.31 | 0.46 | 0.68 | 0.32 | 0.36 |
| 2 | 59109155 | rs42844 | 0.255 | 3.098 | -0.328 | 1.164 | -0.042 | -0.041 | -0.041 | 0 | 0.93 | 0.99 | 0.81 | 0.82 | 0.86 | 0.89 |
| 2 | 59111389 | rs42842 | -0.906 | 3.086 | -0.2 | 1.135 | 0.023 | -0.042 | -0.042 | 0.248 | 0.12 | 0.3 | 0.12 | 0.05 | 0.15 | 0.05 |
| 2 | 59113045 | rs42837 | 0.536 | 3.094 | -0.259 | 1.142 | -0.048 | -0.045 | -0.045 | 0 | 0.92 | 0.99 | 0.79 | 0.82 | 0.84 | 0.87 |
| 2 | 59113179 | rs42836 | 0.301 | 2.913 | -0.249 | 1.151 | -0.044 | -0.042 | -0.042 | 0.306 | 0.93 | 0.99 | 0.8 | 0.82 | 0.86 | 0.87 |
| 2 | 59114739 | rs6758100 | 1.823 | 2.898 | -0.193 | 1.178 | 0.073 | 0.026 | 0.026 | 0 | 0.81 | 0.61 | 0.67 | 0.73 | 0.7 | 0.79 |
| 2 | 59119018 | rs2058563 | 0.787 | 2.737 | -1.434 | 0.706 | -0.075 | 0.058 | 0.058 | 0.042 | 0.23 | 0.04 | 0.19 | 0.09 | 0.31 | 0.15 |
| 2 | 59119064 | rs42829 | -1.015 | 2.47 | -0.549 | 1.277 | -0.002 | -0.001 | -0.001 | 0 | 0.1 | 0.11 | 0.11 | 0.05 | 0.16 | 0.08 |
| 2 | 59129941 | rs10490098 | -0.591 | 2.28 | -0.43 | 1.651 | -0.002 | 0.093 | 0.093 | 0 | 0.08 | 0.1 | 0.21 | 0.46 | 0.1 | 0.16 |
| 2 | 59133513 | rs1861100 | 1.528 | 2.515 | 0.002 | 1.157 | 0.055 | 0.034 | 0.034 | 0.018 | 0.81 | 0.58 | 0.69 | 0.72 | 0.75 | 0.84 |
| 2 | 59138000 | rs10490099 | -1.265 | 1.723 | 0.103 | 1.269 | 0.001 | 0.07 | 0.07 | 0.009 | 0.16 | 0.4 | 0.18 | 0.27 | 0.15 | 0.12 |
| 2 | 59145840 | rs2160091 | -1.396 | 2.176 | -0.35 | 1.345 | -0.006 | -0.007 | -0.007 | 0 | 0.16 | 0.15 | 0.17 | 0.28 | 0.2 | 0.21 |
| 2 | 59147007 | rs12621461 | 0.2 | 2.205 | -0.548 | 1.961 | -0.025 | 0.045 | 0.045 | 0 | 0.05 | 0 | 0.17 | 0.44 | 0.08 | 0.15 |
| 2 | 59147994 | rs10490100 | -1.006 | 2.718 | -0.298 | 1.639 | -0.005 | -0.005 | -0.005 | 0.768 | 0.17 | 0.17 | 0.21 | 0.27 | 0.25 | 0.2 |
| 2 | 59150956 | rs12328778 | -0.434 | 2.274 | 0.61 | 1.167 | 0.029 | -0.014 | -0.014 | 1.219 | 0.08 | 0.46 | 0.09 | 0 | 0.08 | 0.03 |
| 2 | 59155105 | rs7573672 | -1.825 | 1.742 | 0.524 | 1.118 | -0.01 | 0.047 | 0.047 | 0.367 | 0.29 | 0.59 | 0.44 | 0.63 | 0.31 | 0.33 |
| 2 | 59156648 | rs4672270 | -2.281 | 1.7 | 0.351 | 0.94 | -0.003 | -0.004 | -0.004 | 0 | 0.4 | 0.68 | 0.54 | 0.65 | 0.44 | 0.38 |
| 2 | 59162140 | rs2215849 | -0.365 | 1.289 | 0.088 | 0.393 | -0.009 | -0.036 | -0.036 | 0.344 | 0.75 | 0.66 | 0.83 | 0.95 | 0.74 | 0.66 |
| 2 | 59162425 | rs1073975 | -2.115 | 1.185 | 0.009 | 0.33 | -0.03 | 0.064 | 0.064 | 1.029 | 0.56 | 0.84 | 0.61 | 0.69 | 0.58 | 0.63 |
| 2 | 59166461 | rs9973935 | 1.584 | 1.108 | 0.011 | 0.533 | 0.001 | 0 | 0 | 0 | 0.52 | 0.47 | 0.47 | 0.47 | 0.5 | 0.38 |
| 2 | 59167905 | rs1079815 | -0.377 | 1.096 | -0.215 | 0.525 | 0.005 | 0.024 | 0.024 | 0 | 0.26 | 0.33 | 0.39 | 0.43 | 0.31 | 0.26 |
| 2 | 59169043 | rs11675817 | 2.462 | 1.112 | -0.241 | 0.531 | -0.018 | 0.054 | 0.054 | 0.408 | 0.39 | 0.16 | 0.35 | 0.26 | 0.38 | 0.29 |
| 2 | 59170650 | rs2080217 | 1.486 | 1.114 | -0.24 | 0.53 | 0 | -0.001 | -0.001 | 0 | 0.53 | 0.53 | 0.48 | 0.48 | 0.5 | 0.39 |
| 2 | 59171779 | rs12615223 | 1.572 | 1.959 | -0.184 | 0.586 | -0.001 | -0.001 | -0.001 | 0 | 0.54 | 0.53 | 0.4 | 0.47 | 0.44 | 0.38 |
| 2 | 59172847 | rs2080216 | -2.294 | 1.835 | -0.315 | 0.666 | -0.017 | 0.001 | 0.001 | 1.056 | 0.43 | 0.76 | 0.64 | 0.65 | 0.51 | 0.41 |
| 2 | 59175260 | rs17554668 | 1.016 | 1.597 | -0.354 | 0.7 | 0.004 | -0.01 | -0.01 | 0.314 | 0.1 | 0.1 | 0.1 | 0.02 | 0.13 | 0.06 |
| 2 | 59178456 | rs13027265 | 2.457 | 1.505 | -0.719 | 0.441 | 0.031 | 0.025 | 0.025 | 0.99 | 0.34 | 0.15 | 0.24 | 0.26 | 0.27 | 0.18 |
| 2 | 59178687 | rs713457 | 1.23 | 1.58 | -0.678 | 0.357 | 0.001 | -0.006 | -0.006 | 1.782 | 0.1 | 0.09 | 0.1 | 0.02 | 0.13 | 0.04 |
| 2 | 59183439 | rs7586333 | 0.494 | 1.617 | -0.809 | 0.518 | 0.004 | 0.02 | 0.02 | 0 | 0.1 | 0.16 | 0.12 | 0.22 | 0.13 | 0.12 |
| 2 | 59184123 | rs17049848 | 0.606 | 1.396 | -1.158 | 0.709 | 0.001 | -0.076 | -0.076 | 0 | 0.11 | 0 | 0.22 | 0.41 | 0.09 | 0.17 |
| 2 | 59187207 | rs12615247 | 1.944 | 1.539 | -1.014 | 0.618 | 0.033 | 0.005 | 0.005 | 0.985 | 0.35 | 0.15 | 0.25 | 0.31 | 0.27 | 0.25 |
| 2 | 59188638 | rs2863300 | 0.736 | 1.973 | -0.605 | 0.728 | 0.027 | -0.011 | -0.011 | 0.653 | 0.68 | 0.47 | 0.65 | 0.74 | 0.64 | 0.65 |
| 2 | 59193664 | rs17049851 | 0.257 | 1.623 | -1.001 | 1.198 | -0.013 | -0.01 | -0.01 | 0.044 | 0.05 | 0.01 | 0.13 | 0.23 | 0.09 | 0.11 |
| 2 | 59194640 | rs4383319 | -0.987 | 1.711 | -0.674 | 0.453 | 0.002 | -0.008 | -0.008 | 0 | 0.34 | 0.59 | 0.29 | 0.23 | 0.38 | 0.35 |
| 2 | 59199847 | rs2863301 | -0.802 | 1.923 | -0.93 | 0.8 | -0.02 | 0 | 0 | 0 | 0.26 | 0.19 | 0.26 | 0.25 | 0.32 | 0.23 |
| 2 | 59204906 | rs2192463 | 0.251 | 2.024 | -1.045 | 0.752 | -0.037 | 0.012 | 0.012 | 0 | 0.91 | 0.98 | 0.87 | 0.93 | 0.83 | 0.91 |
| 2 | 59205954 | rs7577730 | -1.08 | 1.946 | -1.27 | 0.915 | -0.026 | -0.02 | -0.02 | 0.151 | 0.57 | 0.85 | 0.53 | 0.53 | 0.55 | 0.73 |
| 2 | 59207605 | rs7570921 | -0.262 | 1.665 | -0.575 | 0.704 | -0.027 | 0.008 | 0.008 | 1.78 | 0.26 | 0.64 | 0.1 | 0.24 | 0.13 | 0.32 |
| 2 | 59213427 | rs763694 | -0.348 | 1.934 | -0.154 | 0.883 | 0 | 0 | 0 | 0 | 0.36 | 0.57 | 0.26 | 0.27 | 0.35 | 0.41 |
| 2 | 59219263 | rs6746590 | -0.51 | 1.95 | -0.199 | 0.876 | 0.001 | 0.007 | 0.007 | 0.057 | 0.37 | 0.68 | 0.27 | 0.27 | 0.35 | 0.41 |
| 2 | 59221939 | rs10169123 | -0.878 | 1.875 | -0.35 | 0.879 | -0.02 | 0.065 | 0.065 | 0.968 | 0.5 | 0.79 | 0.36 | 0.33 | 0.45 | 0.56 |
| 2 | 59226413 | rs12993019 | 0.521 | 1.672 | 0.267 | 0.3 | -0.023 | 0.011 | 0.011 | 3.853 | 0.26 | 0.19 | 0.26 | 0.2 | 0.36 | 0.26 |
| 2 | 59227300 | rs11125766 | -0.764 | 1.609 | 0.559 | 0.252 | 0.001 | -0.017 | -0.017 | 1.052 | 0.44 | 0.51 | 0.42 | 0.29 | 0.53 | 0.48 |
| 2 | 59227529 | rs6758107 | -0.55 | 1.596 | 0.582 | 0.246 | -0.02 | 0.059 | 0.059 | 2.649 | 0.42 | 0.24 | 0.39 | 0.28 | 0.53 | 0.48 |
| 2 | 59228502 | rs6704661 | 0.395 | 1.535 | 1.065 | 0.251 | -0.026 | -0.022 | -0.022 | 0 | 0.11 | 0.23 | 0.04 | 0.04 | 0.06 | 0.12 |
| 2 | 59232709 | rs7585990 | -0.665 | 1.617 | 0.567 | 0.283 | -0.009 | 0.054 | 0.054 | 2.765 | 0.43 | 0.31 | 0.4 | 0.29 | 0.53 | 0.49 |
| 2 | 59234869 | rs17620941 | 0.335 | 1.701 | 0.259 | 0.226 | -0.007 | 0.026 | 0.026 | 0 | 0.16 | 0.1 | 0.13 | 0.06 | 0.16 | 0.22 |
| 2 | 59236502 | rs4672274 | -1.358 | 1.697 | 0.201 | 0.349 | -0.033 | -0.054 | -0.054 | 0 | 0.57 | 0.93 | 0.56 | 0.44 | 0.59 | 0.68 |
| 2 | 59236891 | rs7567417 | -1.125 | 2.488 | 0.629 | 0.572 | 0 | -0.002 | -0.002 | 0 | 0.44 | 0.52 | 0.5 | 0.38 | 0.52 | 0.55 |
| 2 | 59239368 | rs11687358 | -0.886 | 2.41 | -0.339 | 0.535 | -0.008 | 0.008 | 0.008 | 1.201 | 0.41 | 0.37 | 0.45 | 0.35 | 0.47 | 0.53 |
| 2 | 59240153 | rs6545712 | 0.636 | 1.725 | -0.348 | 0.532 | 0.005 | 0.029 | 0.029 | 1.581 | 0.59 | 0.8 | 0.49 | 0.64 | 0.44 | 0.44 |
| 2 | 59240476 | rs7581819 | -0.365 | 0.859 | -0.071 | 0.421 | 0.005 | 0.027 | 0.027 | 0 | 0.14 | 0.07 | 0.08 | 0.06 | 0.1 | 0.22 |
| 2 | 59260307 | rs9309307 | 0.109 | 1.338 | -1.004 | 0.649 | -0.028 | 0.04 | 0.04 | 7.077 | 0.58 | 0.76 | 0.56 | 0.68 | 0.51 | 0.52 |
| 2 | 59266958 | rs978624 | -1.219 | 1.495 | -0.304 | 0.968 | 0.048 | 0.037 | 0.037 | 0 | 0.07 | 0.2 | 0.16 | 0.1 | 0.13 | 0.08 |
| 2 | 59271782 | rs10176695 | 0.651 | 2.018 | -0.916 | 1.481 | -0.048 | -0.048 | -0.048 | 0.897 | 0.92 | 0.99 | 0.72 | 0.75 | 0.85 | 0.79 |
| 2 | 59273136 | rs2540049 | 0.579 | 2.032 | -0.911 | 1.49 | -0.044 | -0.042 | -0.042 | 0 | 0.93 | 0.99 | 0.72 | 0.75 | 0.85 | 0.82 |
| 2 | 59276451 | rs1544626 | -0.214 | 1.432 | -1.776 | 1.017 | 0.045 | 0.089 | 0.089 | 1.228 | 0.42 | 0.14 | 0.2 | 0.29 | 0.3 | 0.4 |
| 2 | 59276560 | rs1544625 | -0.447 | 1.169 | -1.98 | 0.663 | 0.053 | 0.136 | 0.136 | 0 | 0.48 | 0.17 | 0.28 | 0.29 | 0.36 | 0.43 |
| 2 | 59282442 | rs991965 | -0.49 | 0.771 | -0.908 | 1.392 | 0.006 | -0.001 | -0.001 | 0 | 0.53 | 0.46 | 0.4 | 0.56 | 0.36 | 0.55 |
| 2 | 59282591 | rs991964 | -0.795 | 0.852 | -0.773 | 1.518 | -0.058 | -0.003 | -0.003 | 0 | 0.61 | 0.88 | 0.51 | 0.66 | 0.46 | 0.58 |
| 2 | 59297140 | rs10169897 | 1.17 | 0.579 | -1.064 | 1.588 | 0.015 | 0.01 | 0.01 | 1.782 | 0.29 | 0.12 | 0.13 | 0.28 | 0.21 | 0.31 |
| 2 | 59301572 | rs17049905 | 0.017 | 0.606 | 0.321 | 1.851 | 0.062 | 0.177 | 0.177 | 0 | 0.08 | 0.42 | 0.11 | 0.15 | 0.1 | 0.04 |
| 2 | 59301917 | rs13027140 | 0.324 | 1.217 | -0.142 | 1.167 | -0.001 | -0.001 | -0.001 | 0.337 | 0.27 | 0.23 | 0.43 | 0.33 | 0.36 | 0.36 |
| 2 | 59302938 | rs17049906 | 0.783 | 1.187 | 0.066 | 1.142 | 0.003 | 0.04 | 0.04 | 0 | 0.07 | 0.12 | 0.09 | 0.15 | 0.1 | 0.04 |
| 2 | 59305625 | rs1016287 | -0.558 | 1.105 | 0.199 | 1.072 | 0.003 | 0.018 | 0.018 | 0.105 | 0.71 | 0.8 | 0.72 | 0.76 | 0.72 | 0.77 |
| 2 | 59308101 | rs7599151 | -0.224 | 1.07 | 0.026 | 1.133 | 0.001 | -0.017 | -0.017 | 0.474 | 0.64 | 0.8 | 0.63 | 0.63 | 0.62 | 0.77 |
| 2 | 59310698 | rs11125769 | 0.477 | 1.092 | -0.09 | 1.063 | -0.048 | 0.054 | 0.054 | 0 | 0.24 | 0.09 | 0.33 | 0.1 | 0.36 | 0.38 |
| 2 | 59311536 | rs980183 | -0.355 | 1.075 | -0.125 | 1.001 | -0.005 | 0.066 | 0.066 | 1.626 | 0.64 | 0.83 | 0.75 | 0.85 | 0.62 | 0.81 |
| 2 | 59312140 | rs10490105 | 1.241 | 1.07 | -0.229 | 0.95 | 0.023 | 0.037 | 0.037 | 2.427 | 0.34 | 0.13 | 0.17 | 0.29 | 0.25 | 0.35 |

| | | | | | | | | | | | | | | | | |
|---|---|---|---|---|---|---|---|---|---|---|---|---|---|---|---|---|
| 2 | 59331066 | rs12615810 | 0.121 | 1.202 | 0.058 | 1.001 | -0.035 | -0.029 | -0.029 | 0 | 0.26 | 0.12 | 0.43 | 0.33 | 0.35 | 0.39 |
| 2 | 59337008 | rs13025124 | -0.342 | 1.149 | 0.201 | 0.976 | 0.004 | 0.003 | 0.003 | 0 | 0.28 | 0.68 | 0.46 | 0.33 | 0.36 | 0.36 |
| 2 | 59339111 | rs11686642 | -0.395 | 0.97 | 0.336 | 1.143 | 0.003 | 0.007 | 0.007 | 0 | 0.16 | 0.21 | 0.15 | 0.18 | 0.25 | 0.24 |
| 2 | 59343477 | rs17049931 | -0.101 | 0.951 | 0.709 | 0.967 | -0.017 | -0.037 | -0.037 | 0.047 | 0.15 | 0.28 | 0.11 | 0.07 | 0.09 | 0.03 |
| 2 | 59344469 | rs7602061 | -0.401 | 0.941 | 0.692 | 0.967 | 0.002 | 0.005 | 0.005 | 0 | 0.17 | 0.22 | 0.15 | 0.19 | 0.25 | 0.31 |
| 2 | 59346419 | rs11887991 | 1.641 | 1.064 | 0.347 | 0.913 | 0.012 | 0.092 | 0.092 | 0.987 | 0.36 | 0.14 | 0.18 | 0.24 | 0.28 | 0.32 |
| 2 | 59346956 | rs7597683 | -0.688 | 1.173 | 0.391 | 1.173 | 0.002 | -0.002 | -0.002 | 0.199 | 0.69 | 0.67 | 0.55 | 0.72 | 0.66 | 0.65 |
| 2 | 59356681 | rs2902572 | -0.833 | 1.142 | 0.436 | 1.059 | -0.019 | 0.005 | 0.005 | 0 | 0.7 | 0.86 | 0.57 | 0.72 | 0.66 | 0.65 |
| 2 | 59373344 | rs12713392 | -0.644 | 1.149 | 0.423 | 1.119 | 0.001 | -0.003 | -0.003 | 0 | 0.68 | 0.68 | 0.55 | 0.72 | 0.65 | 0.65 |
| 2 | 59388342 | rs12052355 | -1.141 | 1.19 | 0.633 | 1.114 | 0.007 | 0.026 | 0.026 | 0.017 | 0.25 | 0.56 | 0.26 | 0.27 | 0.31 | 0.29 |
| 2 | 59394998 | rs6744164 | -0.366 | 1.137 | 0.622 | 1.129 | 0 | -0.008 | -0.008 | 0 | 0.59 | 0.65 | 0.44 | 0.51 | 0.63 | 0.6 |
| 2 | 59401058 | rs7588875 | -0.187 | 1.156 | 0.696 | 1.176 | 0.005 | 0.007 | 0.007 | 0.009 | 0.16 | 0.23 | 0.18 | 0.17 | 0.27 | 0.24 |
| 2 | 59403078 | rs12611535 | 0.875 | 1.156 | 0.571 | 1.121 | 0.002 | 0.013 | 0.013 | 1.367 | 0.41 | 0.38 | 0.27 | 0.32 | 0.37 | 0.35 |
| 2 | 59404651 | rs6720604 | -1.158 | 1.072 | 0.309 | 1.127 | 0.001 | 0.003 | 0.003 | 0.577 | 0.28 | 0.7 | 0.35 | 0.26 | 0.33 | 0.31 |
| 2 | 59409979 | rs6741197 | -0.347 | 1.039 | 0.016 | 1.094 | -0.01 | -0.037 | -0.037 | 0 | 0.61 | 0.84 | 0.52 | 0.55 | 0.62 | 0.61 |
| 2 | 59421047 | rs1861108 | 0.736 | 0.911 | 0.157 | 1.216 | -0.004 | 0.021 | 0.021 | 0 | 0.37 | 0.59 | 0.25 | 0.48 | 0.32 | 0.41 |
| 2 | 59423256 | rs10196564 | -0.204 | 1.002 | 0.099 | 1.285 | -0.014 | -0.02 | -0.02 | 0.105 | 0.17 | 0.12 | 0.26 | 0.24 | 0.27 | 0.2 |
| 2 | 59429192 | rs7584083 | 0.812 | 1.333 | 0.055 | 1.196 | 0.002 | -0.009 | -0.009 | 0 | 0.35 | 0.41 | 0.38 | 0.23 | 0.4 | 0.47 |
| 2 | 59429807 | rs10175975 | 2.232 | 1.114 | -0.238 | 1.124 | -0.046 | 0.058 | 0.058 | 0 | 0.23 | 0.04 | 0.1 | 0.14 | 0.21 | 0.23 |
| 2 | 59435583 | rs13015251 | 0.147 | 0.439 | -0.073 | 1.203 | 0.002 | 0.035 | 0.035 | 0 | 0.12 | 0.18 | 0.12 | 0.26 | 0.12 | 0.08 |
| 2 | 59439450 | rs12993357 | 0.109 | 0.443 | -0.058 | 1.206 | 0.002 | 0.038 | 0.038 | 0 | 0.12 | 0.19 | 0.12 | 0.26 | 0.12 | 0.09 |
| 2 | 59444675 | rs4672279 | -1.081 | 0.15 | 0.046 | 1.068 | 0.028 | 0.023 | 0.023 | 1.28 | 0.65 | 0.78 | 0.8 | 0.75 | 0.79 | 0.74 |
| 2 | 59453954 | rs4671340 | -0.127 | 0.438 | 0.128 | 1.343 | 0.006 | 0.04 | 0.04 | 0 | 0.15 | 0.19 | 0.28 | 0.38 | 0.24 | 0.13 |
| 2 | 59456143 | rs7599431 | -0.482 | 0.604 | 0.347 | 1.307 | 0.046 | 0.012 | 0.012 | 0 | 0.08 | 0.27 | 0.23 | 0.06 | 0.12 | 0.2 |
| 2 | 59456623 | rs2192567 | -1.506 | 0.628 | 0.309 | 1.318 | 0.021 | 0.035 | 0.035 | 0 | 0.31 | 0.46 | 0.38 | 0.46 | 0.4 | 0.27 |
| 2 | 59457661 | rs10192541 | 0.063 | 0.639 | 0.223 | 1.119 | -0.003 | -0.001 | -0.001 | 0 | 0.14 | 0.12 | 0.1 | 0.11 | 0.17 | 0.13 |
| 2 | 59465518 | rs1861143 | 0.955 | 0.586 | 0.122 | 1.092 | -0.009 | 0.014 | 0.014 | 0 | 0.36 | 0.31 | 0.42 | 0.28 | 0.42 | 0.47 |
| 2 | 59466891 | rs10490111 | 0.337 | 0.553 | 0.246 | 1.009 | -0.001 | -0.007 | -0.007 | 0 | 0.12 | 0.12 | 0.08 | 0.03 | 0.15 | 0.12 |
| 2 | 59467016 | rs7423968 | 0.932 | 0.163 | 0.02 | 0.915 | -0.015 | 0.012 | 0.012 | 0 | 0.42 | 0.3 | 0.46 | 0.41 | 0.44 | 0.51 |
| 2 | 59467329 | rs10490112 | 1.078 | 0.376 | -0.006 | 0.942 | -0.035 | 0.028 | 0.028 | 0 | 0.11 | 0.01 | 0.07 | 0.03 | 0.15 | 0.12 |
| 2 | 59467945 | rs2215894 | 0.546 | 0.401 | 0.115 | 1.055 | 0.01 | 0.015 | 0.015 | 0 | 0.13 | 0.16 | 0.18 | 0.2 | 0.21 | 0.1 |
| 2 | 59471298 | rs10210132 | -1.113 | 0.32 | 0.149 | 1.064 | 0.009 | -0.005 | -0.005 | 1.737 | 0.67 | 0.72 | 0.82 | 0.9 | 0.81 | 0.75 |
| 2 | 59474051 | rs1990558 | -0.192 | 0.37 | 0.025 | 1.164 | -0.007 | -0.004 | -0.004 | 0 | 0.54 | 0.51 | 0.64 | 0.62 | 0.66 | 0.61 |
| 2 | 59474387 | rs12990714 | -0.851 | 0.653 | -0.3 | 1.09 | 0.007 | 0.003 | 0.003 | 0 | 0.24 | 0.31 | 0.26 | 0.23 | 0.36 | 0.22 |
| 2 | 59478292 | rs2192562 | 0.646 | 0.642 | -0.531 | 0.87 | -0.027 | 0.104 | 0.104 | 0 | 0.42 | 0.23 | 0.3 | 0.26 | 0.47 | 0.43 |
| 2 | 59478988 | rs5008666 | 0.788 | 0.7 | 0.152 | 1.161 | 0.031 | 0.074 | 0.074 | 0 | 0.74 | 0.52 | 0.56 | 0.62 | 0.68 | 0.69 |
| 2 | 59484780 | rs17050030 | 1.42 | 0.499 | -0.19 | 1.148 | 0.003 | -0.02 | -0.02 | 0 | 0.06 | 0.02 | 0.03 | 0.12 | 0.03 | 0.03 |
| 2 | 59494725 | rs13023187 | 1.117 | 0.435 | -0.18 | 1.134 | -0.029 | -0.009 | -0.009 | 1.336 | 0.34 | 0.17 | 0.29 | 0.37 | 0.34 | 0.31 |
| 2 | 59496548 | rs12986770 | 1.166 | 0.575 | -0.303 | 1.285 | -0.035 | -0.031 | -0.031 | 0 | 0.08 | 0.02 | 0.14 | 0.21 | 0.18 | 0.09 |
| 2 | 59497734 | rs12994597 | -0.766 | 0.451 | 0.478 | 1.134 | -0.004 | 0.014 | 0.014 | 0 | 0.57 | 0.61 | 0.64 | 0.74 | 0.52 | 0.58 |
| 2 | 59502592 | rs4325734 | 0.028 | 0.363 | 0.603 | 1.118 | 0 | -0.001 | -0.001 | 0.165 | 0.46 | 0.45 | 0.55 | 0.62 | 0.47 | 0.54 |
| 2 | 59503431 | rs4672283 | -0.792 | 0.337 | 0.669 | 1.07 | 0 | -0.055 | -0.055 | 0.903 | 0.86 | 0.79 | 0.91 | 0.99 | 0.84 | 0.88 |
| 2 | 59508624 | rs1344456 | 0.783 | 0.075 | 0.416 | 0.938 | -0.003 | -0.008 | -0.008 | 11.817 | 0.72 | 0.66 | 0.84 | 0.87 | 0.78 | 0.83 |
| 2 | 59509803 | rs17050074 | 0.505 | 0.172 | 0.092 | 1.03 | 0.017 | -0.003 | -0.003 | 0 | 0.14 | 0.09 | 0.07 | 0.12 | 0.06 | 0.05 |
| 2 | 59511950 | rs7608940 | 0.032 | 0.274 | 0.304 | 1.205 | 0.041 | 0.176 | 0.176 | 0 | 0.07 | 0.2 | 0.28 | 0.29 | 0.1 | 0.18 |
| 2 | 59515129 | rs12464814 | 0.842 | 0.261 | -0.008 | 0.696 | -0.039 | -0.012 | -0.012 | 0 | 0.42 | 0.18 | 0.43 | 0.6 | 0.4 | 0.35 |
| 2 | 59516617 | rs17050091 | 1.603 | 0.263 | 0.132 | 0.753 | -0.001 | 0.009 | 0.009 | 0 | 0.05 | 0.08 | 0.05 | 0.05 | 0.04 | 0.03 |
| 2 | 59521784 | rs11125784 | 1.215 | 0.174 | 0.088 | 0.689 | -0.019 | -0.044 | -0.044 | 0 | 0.35 | 0.25 | 0.4 | 0.56 | 0.35 | 0.31 |
| 2 | 59523041 | rs6713914 | 0.016 | 0.062 | 0.189 | 0.565 | 0.001 | 0.005 | 0.005 | 0 | 0.44 | 0.35 | 0.44 | 0.35 | 0.44 | 0.46 |
| 2 | 59524395 | rs12622464 | -0.363 | 0.139 | 0.127 | 0.582 | -0.006 | 0.002 | 0.002 | 0.13 | 0.19 | 0.21 | 0.13 | 0.19 | 0.1 | 0.1 |
| 2 | 59530400 | rs4672284 | 0.929 | 0.385 | 0.394 | 0.738 | 0.006 | 0.026 | 0.026 | 0 | 0.36 | 0.47 | 0.45 | 0.55 | 0.38 | 0.39 |
| 2 | 59540020 | rs11901069 | 0.979 | 0.39 | 0.416 | 0.698 | 0 | 0.001 | 0.001 | 0 | 0.35 | 0.37 | 0.44 | 0.53 | 0.39 | 0.34 |
| 2 | 59542854 | rs13421114 | 0.387 | 0.386 | 0.198 | 0.698 | 0.01 | 0.054 | 0.054 | 0.02 | 0.08 | 0.32 | 0.08 | 0.08 | 0.04 | 0.1 |
| 2 | 59544127 | rs11125786 | 1.721 | 0.4 | 0.16 | 0.709 | -0.057 | -0.088 | -0.088 | 0 | 0.28 | 0.11 | 0.36 | 0.48 | 0.34 | 0.3 |
| 2 | 59544419 | rs1344461 | 1.028 | 0.407 | 0.153 | 0.661 | 0 | 0.002 | 0.002 | 0 | 0.35 | 0.41 | 0.45 | 0.56 | 0.37 | 0.4 |
| 2 | 59545416 | rs17643867 | 0.53 | 0.32 | 0.028 | 0.608 | 0.023 | 0.057 | 0.057 | 1.047 | 0.39 | 0.11 | 0.21 | 0.22 | 0.35 | 0.26 |
| 2 | 59553286 | rs2540023 | 0.243 | 0.313 | 0.029 | 0.637 | -0.004 | -0.006 | -0.006 | 0 | 0.74 | 0.68 | 0.81 | 0.83 | 0.76 | 0.8 |
| 2 | 59563744 | rs2561883 | 1.424 | 0.338 | -0.328 | 0.549 | -0.02 | -0.033 | -0.033 | 1.091 | 0.3 | 0.2 | 0.42 | 0.49 | 0.35 | 0.35 |
| 2 | 59565198 | rs2540015 | 0.077 | 0.031 | -0.038 | 0.382 | -0.019 | -0.052 | -0.052 | 0 | 0.78 | 0.54 | 0.84 | 0.92 | 0.8 | 0.77 |
| 2 | 59580736 | rs17644631 | 1.021 | 0.185 | -1.135 | -0.059 | 0.024 | 0.128 | 0.128 | 0 | 0.23 | 0.03 | 0.07 | 0 | 0.16 | 0.15 |
| 2 | 59582383 | rs17050137 | 0.283 | 0.41 | -0.753 | 0.604 | -0.012 | -0.039 | -0.039 | 0.323 | 0.17 | 0.11 | 0.14 | 0.29 | 0.18 | 0.13 |
| 2 | 59585303 | rs12612292 | -0.089 | 0.019 | -0.491 | 0.1 | 0 | -0.027 | -0.027 | 0.271 | 0.81 | 0.84 | 0.86 | 0.99 | 0.79 | 0.84 |
| 2 | 59586372 | rs7606152 | 0.279 | -0.083 | 0.149 | 0.503 | -0.025 | 0.046 | 0.046 | 0.216 | 0.16 | 0.28 | 0.06 | 0.18 | 0.07 | 0.04 |
| 2 | 59593107 | rs2058586 | -0.677 | -0.337 | -0.147 | 0.255 | 0.011 | -0.006 | -0.006 | 0.693 | 0.42 | 0.64 | 0.44 | 0.37 | 0.37 | 0.35 |
| 2 | 59601449 | rs2418495 | 1.219 | -0.509 | -0.489 | 0.033 | -0.012 | -0.026 | -0.026 | 0 | 0.43 | 0.19 | 0.48 | 0.62 | 0.5 | 0.53 |
| 2 | 59603319 | rs2900638 | -0.793 | -0.399 | 0.108 | 0.211 | -0.003 | 0 | 0 | 0.17 | 0.44 | 0.47 | 0.44 | 0.37 | 0.38 | 0.36 |
| 2 | 59605576 | rs4432437 | 0.134 | -0.047 | 0.136 | 0.11 | -0.007 | -0.008 | -0.008 | 0 | 0.79 | 0.68 | 0.84 | 0.84 | 0.82 | 0.78 |
| 2 | 59605724 | rs4413146 | 0.645 | 0.127 | 0.597 | 0.113 | 0.146 | 0.017 | 0.017 | 0 | 0.94 | 0.68 | 0.88 | 1 | 0.9 | 0.92 |
| 2 | 59610535 | rs7603446 | -1.627 | -0.457 | -0.261 | -0.052 | -0.044 | -0.037 | -0.037 | 0 | 0.6 | 0.84 | 0.47 | 0.39 | 0.54 | 0.47 |

| | | | | | | | | | | | | | | | | |
|---|---|---|---|---|---|---|---|---|---|---|---|---|---|---|---|---|
| 2 | 59610711 | rs7577424 | 1.051 | -0.741 | -0.037 | -0.109 | -0.057 | 0.019 | 0.019 | 3.317 | 0.14 | 0.33 | 0.13 | 0.11 | 0.03 | 0.03 |
| 2 | 59611195 | rs11694432 | 0.165 | -0.783 | 0.006 | -0.254 | -0.003 | 0.014 | 0.014 | 0 | 0.19 | 0.11 | 0.19 | 0.12 | 0.26 | 0.19 |
| 2 | 59612358 | rs6734833 | -2.073 | -0.767 | -0.107 | -0.24 | -0.064 | -0.046 | -0.046 | 0 | 0.58 | 0.9 | 0.46 | 0.37 | 0.53 | 0.46 |
| 2 | 59612917 | rs6750614 | -0.84 | -0.686 | 0.049 | -0.224 | -0.031 | -0.019 | -0.019 | 0 | 0.33 | 0.46 | 0.32 | 0.22 | 0.29 | 0.21 |
| 2 | 59644487 | rs4672289 | 0.892 | -0.537 | -0.422 | 0.077 | -0.026 | -0.064 | -0.064 | 0 | 0.48 | 0.09 | 0.53 | 0.53 | 0.53 | 0.52 |
| 2 | 59651584 | rs6749972 | 0.921 | -0.55 | 0.124 | 0.097 | 0.005 | -0.023 | -0.023 | 0 | 0.48 | 0.21 | 0.52 | 0.52 | 0.5 | 0.49 |
| 2 | 59652486 | rs6725245 | -1.425 | -0.424 | 0.179 | 0.084 | 0.023 | -0.008 | -0.008 | 0 | 0.91 | 0.84 | 0.89 | 0.88 | 0.85 | 0.83 |
| 2 | 59655381 | rs11693937 | 1.274 | -0.058 | -0.669 | 0.403 | -0.071 | -0.069 | -0.069 | 0 | 0.16 | 0.01 | 0.19 | 0.24 | 0.2 | 0.19 |
| 2 | 59661521 | rs17584600 | 1.108 | -0.159 | -0.376 | 0.003 | 0.003 | -0.019 | -0.019 | 0 | 0.47 | 0.26 | 0.53 | 0.52 | 0.51 | 0.51 |
| 2 | 59661788 | rs17645334 | 0.855 | -0.137 | -0.827 | 0.081 | -0.049 | -0.05 | -0.05 | 0.283 | 0.21 | 0.06 | 0.2 | 0.24 | 0.27 | 0.28 |
| 2 | 59668401 | rs17050170 | 0.696 | -0.058 | -0.774 | -0.163 | 0.016 | -0.004 | -0.004 | 0 | 0.14 | 0.08 | 0.13 | 0.11 | 0.04 | 0.04 |
| 2 | 59669528 | rs6737829 | 0.701 | 0.021 | -0.798 | -0.144 | -0.016 | -0.007 | -0.007 | 0 | 0.22 | 0.12 | 0.21 | 0.24 | 0.28 | 0.29 |
| 2 | 59673530 | rs12464531 | 1.059 | -0.032 | -0.85 | -0.167 | -0.04 | -0.032 | -0.032 | 0.562 | 0.47 | 0.13 | 0.52 | 0.54 | 0.52 | 0.51 |
| 2 | 59675929 | rs6741153 | -0.411 | -0.028 | -0.197 | -0.098 | -0.003 | 0.003 | 0.003 | 0 | 0.78 | 0.75 | 0.76 | 0.77 | 0.8 | 0.78 |
| 2 | 59694749 | rs13018470 | 1.224 | 0.104 | -0.309 | -0.706 | -0.043 | -0.056 | -0.056 | 0 | 0.41 | 0.14 | 0.48 | 0.54 | 0.41 | 0.38 |
| 2 | 59699929 | rs2160142 | -0.061 | 0.37 | -0.052 | -0.588 | 0.004 | -0.01 | -0.01 | 0.525 | 0.73 | 0.7 | 0.8 | 0.79 | 0.73 | 0.69 |
| 2 | 59738664 | rs12151705 | -0.215 | 0.296 | 0.23 | -0.313 | -0.003 | -0.003 | -0.003 | 0 | 0.26 | 0.61 | 0.32 | 0.24 | 0.28 | 0.27 |
| 2 | 59742478 | rs741229 | -0.084 | 0.368 | -0.031 | -0.33 | 0.006 | -0.007 | -0.007 | 0 | 0.25 | 0.45 | 0.3 | 0.23 | 0.28 | 0.26 |
| 2 | 59744766 | rs6748990 | -0.201 | 0.335 | -0.732 | -0.239 | 0.009 | -0.002 | -0.002 | 0.258 | 0.91 | 0.96 | 0.9 | 0.88 | 0.93 | 0.93 |
| 2 | 59745743 | rs2041719 | -0.093 | 0.482 | -0.769 | -0.143 | -0.013 | -0.047 | -0.047 | 1.151 | 0.23 | 0.13 | 0.27 | 0.32 | 0.28 | 0.28 |
| 2 | 59747044 | rs7593123 | -0.858 | 0.417 | -0.273 | -0.433 | 0.001 | 0.007 | 0.007 | 0.466 | 0.42 | 0.44 | 0.42 | 0.47 | 0.46 | 0.52 |
| 2 | 59747106 | rs2540102 | 1.239 | 0.32 | -0.425 | -0.843 | 0.012 | -0.024 | -0.024 | 0 | 0.43 | 0.35 | 0.5 | 0.53 | 0.38 | 0.32 |
| 2 | 59747932 | rs2418862 | -1.282 | 0.421 | -0.505 | -0.804 | -0.003 | -0.011 | -0.011 | 0 | 0.89 | 0.94 | 0.94 | 1 | 0.86 | 0.86 |
| 2 | 59749401 | rs2110565 | -0.462 | 0.361 | -0.297 | -0.759 | -0.008 | -0.021 | -0.021 | 0 | 0.4 | 0.29 | 0.4 | 0.47 | 0.45 | 0.51 |
| 2 | 59750886 | rs17050204 | 0.799 | 0.497 | -0.381 | -0.664 | -0.001 | -0.002 | -0.002 | 0.409 | 0.18 | 0.17 | 0.13 | 0.15 | 0.16 | 0.23 |
| 2 | 59753331 | rs2674061 | 1.153 | 0.304 | -0.647 | -0.861 | -0.001 | -0.015 | -0.015 | 0 | 0.31 | 0.35 | 0.43 | 0.51 | 0.29 | 0.29 |
| 2 | 59761338 | rs2540094 | -1.316 | 0.419 | -0.53 | -0.82 | -0.008 | -0.005 | -0.005 | 0 | 0.89 | 0.95 | 0.94 | 1 | 0.86 | 0.86 |
| 2 | 59762594 | rs2540087 | 1.106 | 0.331 | -0.613 | -0.882 | -0.001 | -0.016 | -0.016 | 0 | 0.33 | 0.35 | 0.4 | 0.52 | 0.31 | 0.33 |
| 2 | 59764478 | rs2215907 | -1.263 | 0.33 | -0.678 | -0.866 | 0.012 | -0.043 | -0.043 | 0 | 0.89 | 0.91 | 0.94 | 1 | 0.86 | 0.87 |
| 2 | 59770353 | rs2674052 | 0.836 | 0.196 | -0.726 | -0.886 | -0.003 | -0.048 | -0.048 | 0 | 0.34 | 0.26 | 0.41 | 0.53 | 0.32 | 0.34 |
| 2 | 59779478 | rs6740378 | 0.102 | 0.113 | -0.468 | -0.897 | -0.026 | -0.056 | -0.056 | 0.557 | 0.2 | 0.36 | 0.17 | 0.05 | 0.13 | 0.1 |
| 2 | 59780472 | rs13026344 | 0.91 | 0.622 | -1.125 | -0.577 | 0.015 | -0.036 | -0.036 | 0 | 0.08 | 0.07 | 0.05 | 0 | 0.1 | 0.13 |
| 2 | 59788722 | rs13420520 | -0.062 | 0.13 | -1.241 | -0.305 | 0 | 0.005 | 0.005 | 0 | 0.47 | 0.53 | 0.4 | 0.32 | 0.49 | 0.41 |
| 2 | 59789443 | rs10202884 | -0.244 | 0.25 | -1.147 | -0.422 | -0.001 | -0.002 | -0.002 | 4.091 | 0.52 | 0.57 | 0.55 | 0.47 | 0.52 | 0.5 |
| 2 | 59790316 | rs1035102 | -0.037 | 0.665 | -1.55 | 0.016 | -0.024 | -0.027 | -0.027 | 1.497 | 0.6 | 0.79 | 0.59 | 0.44 | 0.57 | 0.61 |
| 2 | 59791233 | rs10209275 | -0.016 | 0.744 | -1.252 | 0.337 | -0.003 | -0.003 | -0.003 | 0 | 0.15 | 0.11 | 0.23 | 0.19 | 0.2 | 0.26 |
| 2 | 59792964 | rs11125800 | -0.722 | 0.47 | -1.414 | 0.513 | 0.021 | 0.018 | 0.018 | 0.178 | 0.39 | 0.22 | 0.39 | 0.27 | 0.36 | 0.54 |
| 2 | 59793184 | rs12052704 | -0.052 | 0.341 | -1.437 | 0.518 | 0.006 | -0.015 | -0.015 | 0.088 | 0.25 | 0.13 | 0.32 | 0.3 | 0.25 | 0.36 |
| 2 | 59793800 | rs2058514 | -0.702 | 0.759 | -1.29 | 0.919 | 0.004 | 0.019 | 0.019 | 0.154 | 0.22 | 0.27 | 0.32 | 0.33 | 0.28 | 0.4 |
| 2 | 59794128 | rs17647479 | -0.31 | 0.727 | -1.309 | 0.896 | 0.011 | 0.004 | 0.004 | 0.03 | 0.19 | 0.11 | 0.23 | 0.14 | 0.16 | 0.27 |
| 2 | 59794609 | rs2539669 | -1.623 | 0.613 | -1.097 | 0.205 | -0.002 | -0.011 | -0.011 | 0 | 0.62 | 0.68 | 0.63 | 0.5 | 0.65 | 0.74 |
| 2 | 59794882 | rs2539668 | -0.598 | 0.781 | -1.167 | 0.77 | 0.003 | 0.021 | 0.021 | 0.089 | 0.23 | 0.26 | 0.34 | 0.37 | 0.29 | 0.4 |
| 2 | 59796020 | rs2674075 | -0.299 | 0.774 | -1.104 | 0.676 | -0.001 | 0.001 | 0.001 | 0.117 | 0.21 | 0.2 | 0.3 | 0.32 | 0.27 | 0.38 |
| 2 | 59800893 | rs13035467 | 0.099 | 0.393 | -1.326 | 0.537 | -0.045 | 0.019 | 0.019 | 0 | 0.19 | 0.02 | 0.15 | 0.08 | 0.22 | 0.23 |
| 2 | 59803687 | rs6545742 | -0.345 | -0.033 | -0.812 | 0.391 | -0.046 | -0.032 | -0.032 | 0.06 | 0.29 | 0.43 | 0.16 | 0.23 | 0.18 | 0.17 |
| 2 | 59803819 | rs17050260 | 0.265 | 0.077 | -1.409 | 0.732 | -0.051 | -0.081 | -0.081 | 0 | 0.21 | 0.01 | 0.16 | 0.25 | 0.22 | 0.26 |
| 2 | 59812464 | rs6760859 | -0.775 | 0.42 | -0.5 | 0.707 | 0.006 | 0.045 | 0.045 | 2.393 | 0.52 | 0.62 | 0.4 | 0.76 | 0.45 | 0.45 |
| 2 | 59813297 | rs2539709 | 0.144 | 0.409 | -0.082 | 1.003 | 0.001 | 0.009 | 0.009 | 2.634 | 0.58 | 0.7 | 0.7 | 0.64 | 0.59 | 0.72 |
| 2 | 59818647 | rs1861031 | 0.411 | 0.382 | 0.048 | 1.233 | 0.001 | 0.006 | 0.006 | 1.083 | 0.54 | 0.61 | 0.62 | 0.27 | 0.56 | 0.61 |
| 2 | 59822418 | rs1861030 | 0.826 | 0.28 | 0.478 | 1.187 | 0.024 | 0.088 | 0.088 | 2.353 | 0.5 | 0.28 | 0.59 | 0.27 | 0.56 | 0.61 |
| 2 | 59824104 | rs6545745 | -1.462 | 0.247 | 0.556 | 1.228 | 0.042 | 0.437 | 0.437 | 1.661 | 0.06 | 0.41 | 0.11 | 0.38 | 0.03 | 0.1 |
| 2 | 59824460 | rs17050288 | 0.243 | 0.13 | 0.347 | 1.004 | -0.014 | -0.086 | -0.086 | 0 | 0.23 | 0.26 | 0.12 | 0 | 0.18 | 0.06 |
| 2 | 59825380 | rs2674034 | 0.785 | 0.123 | 0.375 | 1.015 | 0.005 | 0.03 | 0.03 | 2.637 | 0.53 | 0.67 | 0.62 | 0.27 | 0.56 | 0.62 |
| 2 | 59851653 | rs1861029 | -0.771 | 0.15 | 0.22 | 1.002 | -0.054 | -0.12 | -0.12 | 1.702 | 0.79 | 0.99 | 0.84 | 0.63 | 0.81 | 0.77 |
| 2 | 59852603 | rs2539683 | -0.388 | -0.074 | 0.256 | 0.883 | -0.006 | 0.011 | 0.011 | 0 | 0.74 | 0.75 | 0.81 | 0.63 | 0.79 | 0.77 |
| 2 | 59858364 | rs13019632 | 0.421 | 0.092 | 0.027 | 0.791 | -0.055 | 0.104 | 0.104 | 1.193 | 0.16 | 0.01 | 0.12 | 0 | 0.19 | 0.07 |
| 2 | 59873126 | rs2080167 | 0.184 | -0.13 | 0.34 | 0.94 | -0.013 | 0.003 | 0.003 | 0 | 0.69 | 0.44 | 0.72 | 0.28 | 0.76 | 0.66 |
| 2 | 59879776 | rs9917142 | 1.411 | -0.078 | -0.301 | 0.315 | 0.007 | 0.018 | 0.018 | 0.147 | 0.3 | 0.24 | 0.26 | 0.17 | 0.24 | 0.24 |
| 2 | 59880310 | rs268066 | 0.308 | -0.044 | -0.047 | 0.577 | 0.007 | 0.001 | 0.001 | 0 | 0.66 | 0.33 | 0.72 | 0.28 | 0.76 | 0.66 |
| 2 | 59882782 | rs268070 | -1.432 | 0.002 | -0.092 | 0.608 | -0.063 | -0.121 | -0.121 | 0 | 0.81 | 0.99 | 0.85 | 0.63 | 0.8 | 0.79 |
| 2 | 59886357 | rs2244983 | -0.187 | 0.038 | -0.001 | 0.423 | -0.012 | 0.004 | 0.004 | 0 | 0.7 | 0.43 | 0.73 | 0.28 | 0.76 | 0.69 |
| 2 | 59893561 | rs268062 | -0.104 | 0.035 | 0.227 | 0.412 | -0.016 | 0.007 | 0.007 | 0 | 0.7 | 0.62 | 0.74 | 0.28 | 0.76 | 0.66 |
| 2 | 59898341 | rs268065 | 0.079 | 0.068 | 0.168 | 0.404 | -0.019 | 0 | 0 | 0 | 0.68 | 0.48 | 0.73 | 0.28 | 0.76 | 0.65 |
| 2 | 59903052 | rs2110330 | 0.489 | 0.175 | 0.049 | 0.327 | -0.002 | -0.006 | -0.006 | 0.251 | 0.55 | 0.53 | 0.63 | 0.28 | 0.59 | 0.6 |
| 2 | 59911126 | rs17650159 | 0.283 | 0.544 | -0.295 | 0.333 | -0.022 | 0.067 | 0.067 | 2.025 | 0.18 | 0.06 | 0.13 | 0 | 0.2 | 0.07 |
| 2 | 59919277 | rs155568 | -1.416 | 0.42 | -0.612 | 0.288 | -0.055 | 0.04 | 0.04 | 2.784 | 0.79 | 0.93 | 0.66 | 0.89 | 0.64 | 0.68 |
| 2 | 59941799 | rs197085 | -1.089 | 0.024 | 0.322 | 0.723 | 0.003 | 0.022 | 0.022 | 0 | 0.48 | 0.59 | 0.41 | 0.75 | 0.41 | 0.44 |
| 2 | 59947184 | rs2072471 | -0.12 | 0.384 | 0.223 | 1.143 | 0 | 0.039 | 0.039 | 0.981 | 0.17 | 0.21 | 0.18 | 0.36 | 0.16 | 0.15 |
| 2 | 59949495 | rs11691352 | -0.566 | 0.32 | 0.339 | 0.775 | -0.004 | -0.011 | -0.011 | 0 | 0.36 | 0.41 | 0.34 | 0.72 | 0.34 | 0.36 |

| | | | | | | | | | | | | | | | | |
|---|---|---|---|---|---|---|---|---|---|---|---|---|---|---|---|---|
| 2 | 59951465 | rs13028903 | -0.914 | 0.061 | 0.81 | 0.794 | -0.001 | -0.004 | -0.004 | 0.599 | 0.39 | 0.39 | 0.36 | 0.19 | 0.43 | 0.29 |
| 2 | 59952274 | rs10176878 | 0.307 | 0.019 | 1.084 | 0.754 | 0.014 | -0.08 | -0.08 | 0.136 | 0.15 | 0.23 | 0.11 | 0 | 0.16 | 0.06 |
| 2 | 59952846 | rs6545747 | -1.524 | 0.017 | 0.839 | 1.152 | -0.01 | -0.008 | -0.008 | 0.059 | 0.52 | 0.66 | 0.45 | 0.52 | 0.49 | 0.42 |
| 2 | 59953475 | rs17050389 | 0.225 | 0.126 | 0.685 | 1.275 | -0.038 | -0.001 | -0.001 | 0 | 0.25 | 0.03 | 0.22 | 0.18 | 0.23 | 0.24 |
| 2 | 59955616 | rs9309316 | 0.262 | -0.03 | 0.813 | 1.358 | -0.048 | 0.108 | 0.108 | 1.15 | 0.12 | 0.26 | 0.1 | 0.34 | 0.06 | 0.15 |
| 2 | 59959062 | rs10167515 | -0.771 | 0.066 | 0.456 | 1.255 | -0.006 | 0.005 | 0.005 | 0 | 0.37 | 0.33 | 0.3 | 0.5 | 0.26 | 0.32 |
| 2 | 59970212 | rs12471799 | 0.347 | 0.24 | -0.275 | 1.183 | -0.06 | -0.035 | -0.035 | 1.076 | 0.23 | 0.01 | 0.21 | 0.18 | 0.21 | 0.22 |
| 2 | 59970477 | rs17656897 | 0.339 | 0.4 | -0.027 | 0.992 | -0.024 | 0.037 | 0.037 | 0.04 | 0.13 | 0.07 | 0.11 | 0 | 0.16 | 0.06 |
| 2 | 59973741 | rs11679028 | -0.013 | 0.387 | 0.218 | 1.197 | 0.025 | 0.036 | 0.036 | 0.124 | 0.15 | 0.3 | 0.13 | 0.21 | 0.17 | 0.07 |
| 2 | 59985441 | rs10187463 | -0.693 | 0.277 | 0.037 | 1.208 | -0.025 | -0.006 | -0.006 | 0.446 | 0.38 | 0.42 | 0.25 | 0.68 | 0.27 | 0.29 |
| 2 | 59986066 | rs6545750 | 0.132 | 0.427 | -0.259 | 1.213 | -0.055 | 0.035 | 0.035 | 0 | 0.26 | 0.41 | 0.17 | 0.36 | 0.19 | 0.22 |
| 2 | 59987410 | rs2165763 | -1.693 | 0.806 | -0.468 | 1.125 | -0.003 | 0.024 | 0.024 | 0.319 | 0.58 | 0.73 | 0.5 | 0.68 | 0.57 | 0.45 |
| 2 | 59987774 | rs11125811 | 0.012 | 0.898 | -0.632 | 1.178 | -0.008 | 0 | 0 | 0 | 0.14 | 0.15 | 0.18 | 0.13 | 0.17 | 0.14 |
| 2 | 59989621 | rs11125812 | -1.475 | 1.096 | -0.841 | 1.166 | -0.004 | 0.023 | 0.023 | 0.389 | 0.59 | 0.76 | 0.5 | 0.68 | 0.57 | 0.47 |
| 2 | 59991047 | rs11125813 | 0.133 | 1.452 | -0.595 | 1.233 | 0.011 | 0.01 | 0.01 | 0 | 0.15 | 0.18 | 0.13 | 0.19 | 0.17 | 0.07 |
| 2 | 59998513 | rs12471948 | -0.33 | 1.597 | -0.937 | 1.133 | -0.044 | -0.037 | -0.037 | 0 | 0.08 | 0.02 | 0.14 | 0.12 | 0.12 | 0.14 |
| 2 | 60000136 | rs4672315 | 0.55 | 1.575 | -0.807 | 1.207 | -0.012 | -0.005 | -0.005 | 0 | 0.57 | 0.66 | 0.63 | 0.56 | 0.53 | 0.63 |
| 2 | 60012090 | rs1025876 | 0.538 | 1.402 | -0.184 | 1.372 | 0.002 | 0.004 | 0.004 | 0 | 0.57 | 0.49 | 0.65 | 0.47 | 0.57 | 0.64 |
| 2 | 60012802 | rs1019264 | 0.828 | 2.01 | -0.344 | 1.111 | 0.013 | 0.001 | 0.001 | 0 | 0.51 | 0.42 | 0.55 | 0.47 | 0.41 | 0.59 |
| 2 | 60014129 | rs722889 | 0.359 | 1.651 | -0.37 | 1.229 | -0.025 | -0.036 | -0.036 | 2.432 | 0.07 | 0.02 | 0.06 | 0.13 | 0.08 | 0.12 |
| 2 | 60016819 | rs2419061 | -0.348 | 1.581 | -0.1 | 1.183 | -0.003 | -0.003 | -0.003 | 3.319 | 0.66 | 0.75 | 0.72 | 0.6 | 0.65 | 0.76 |
| 2 | 60021962 | rs1584003 | 0.63 | 1.404 | -0.116 | 1.067 | 0.001 | 0 | 0 | 0.031 | 0.53 | 0.51 | 0.61 | 0.47 | 0.53 | 0.62 |
| 2 | 60024272 | rs6735549 | 1.348 | 0.311 | -0.802 | 0.767 | 0.045 | -0.011 | -0.011 | 0.012 | 0.19 | 0.05 | 0.18 | 0.21 | 0.11 | 0.24 |
| 2 | 60025963 | rs1372876 | -0.7 | 0.404 | -0.502 | 0.749 | -0.002 | 0.003 | 0.003 | 0 | 0.64 | 0.69 | 0.72 | 0.67 | 0.62 | 0.76 |
| 2 | 60027032 | rs1442873 | -0.315 | 0.357 | -0.491 | 0.742 | -0.003 | -0.005 | -0.005 | 0 | 0.6 | 0.58 | 0.68 | 0.59 | 0.62 | 0.72 |
| 2 | 60029072 | rs4671355 | -0.337 | 0.345 | -0.494 | 0.642 | -0.006 | -0.033 | -0.033 | 0 | 0.6 | 0.51 | 0.73 | 0.81 | 0.65 | 0.77 |
| 2 | 60029857 | rs6545754 | -0.58 | 0.338 | -0.484 | 0.639 | -0.001 | -0.022 | -0.022 | 0 | 0.63 | 0.58 | 0.74 | 0.81 | 0.65 | 0.78 |
| 2 | 60030987 | rs1442881 | -0.787 | 0.325 | -0.519 | 0.643 | 0 | -0.003 | -0.003 | 0 | 0.65 | 0.69 | 0.75 | 0.81 | 0.64 | 0.79 |
| 2 | 60033484 | rs7565849 | -0.617 | 0.281 | -0.605 | 0.659 | 0.002 | 0.013 | 0.013 | 0 | 0.63 | 0.71 | 0.71 | 0.67 | 0.65 | 0.74 |
| 2 | 60034937 | rs4671356 | -0.351 | 0.434 | -0.839 | 0.558 | -0.01 | -0.035 | -0.035 | 0 | 0.59 | 0.53 | 0.74 | 0.81 | 0.65 | 0.77 |
| 2 | 60040489 | rs7572396 | 1.274 | 0.259 | -1.527 | 0.499 | 0.041 | -0.022 | -0.022 | 0.121 | 0.16 | 0.05 | 0.15 | 0.26 | 0.08 | 0.26 |
| 2 | 60051642 | rs17399704 | 0.974 | 0.864 | -1.404 | 0.023 | 0.003 | 0.047 | 0.047 | 0 | 0.17 | 0.08 | 0.06 | 0 | 0.12 | 0.04 |
| 2 | 60052940 | rs17399724 | 0.799 | 1.27 | -1.267 | 0.476 | -0.024 | -0.02 | -0.02 | 0 | 0.18 | 0.08 | 0.26 | 0.21 | 0.27 | 0.24 |
| 2 | 60055010 | rs1013344 | -0.902 | 1.024 | -1.023 | 0.357 | 0.011 | -0.002 | -0.002 | 1.775 | 0.44 | 0.64 | 0.58 | 0.38 | 0.59 | 0.47 |
| 2 | 60055238 | rs1035092 | 0.721 | 1.256 | -0.17 | 0.4 | 0.051 | 0.098 | 0.098 | 0 | 0.11 | 0.31 | 0.16 | 0.18 | 0.12 | 0.16 |
| 2 | 60056624 | rs13025401 | 0.296 | 1.36 | -0.234 | 0.274 | -0.019 | 0.015 | 0.015 | 0.004 | 0.09 | 0.05 | 0.11 | 0 | 0.17 | 0.06 |
| 2 | 60056802 | rs11891920 | 0.52 | 1.198 | 0.033 | 0.211 | 0.014 | 0.081 | 0.081 | 0 | 0.16 | 0.46 | 0.18 | 0.18 | 0.12 | 0.18 |
| 2 | 60060788 | rs6760129 | -1.149 | 1.56 | -0.122 | 0.176 | -0.025 | -0.008 | -0.008 | 0.081 | 0.43 | 0.68 | 0.49 | 0.41 | 0.45 | 0.41 |
| 2 | 60061191 | rs6545755 | 0.638 | 1.403 | -0.144 | 0.272 | 0.012 | 0.092 | 0.092 | 0 | 0.16 | 0.45 | 0.17 | 0.21 | 0.12 | 0.17 |
| 2 | 60064055 | rs7580417 | -0.396 | 1.4 | -0.165 | 0.255 | -0.001 | 0.03 | 0.03 | 0 | 0.33 | 0.48 | 0.39 | 0.41 | 0.31 | 0.37 |
| 2 | 60064501 | rs7602575 | 0.705 | 1.377 | 0.055 | 0.297 | 0.009 | 0.068 | 0.068 | 0 | 0.15 | 0.34 | 0.17 | 0.21 | 0.12 | 0.18 |
| 2 | 60068526 | rs6713648 | 0.738 | 1.344 | 0.098 | 0.326 | -0.001 | 0.041 | 0.041 | 0 | 0.14 | 0.27 | 0.16 | 0.2 | 0.12 | 0.17 |
| 2 | 60074136 | rs11682656 | -0.08 | 1.567 | 0.084 | 0.167 | 0.037 | -0.056 | -0.056 | 0 | 0.12 | 0.17 | 0.13 | 0 | 0.19 | 0.06 |
| 2 | 60074591 | rs929616 | -1.119 | 1.439 | 0.015 | 0.163 | -0.008 | 0 | 0 | 0 | 0.45 | 0.69 | 0.53 | 0.41 | 0.5 | 0.43 |
| 2 | 60079250 | rs2419402 | -1.289 | 1.29 | -0.033 | 0.068 | -0.001 | 0.007 | 0.007 | 0 | 0.47 | 0.69 | 0.54 | 0.41 | 0.5 | 0.46 |
| 2 | 60082872 | rs11888922 | -0.46 | 1.321 | 0.175 | 0.028 | 0.057 | -0.086 | -0.086 | 0.017 | 0.14 | 0.32 | 0.15 | 0 | 0.2 | 0.07 |
| 2 | 60086515 | rs7600300 | -0.013 | 1.289 | 0.164 | 0.052 | 0 | 0.007 | 0.007 | 0 | 0.32 | 0.32 | 0.38 | 0.4 | 0.3 | 0.38 |
| 2 | 60094706 | rs730402 | -1.036 | 1.294 | 0.047 | 0.043 | -0.008 | -0.003 | -0.003 | 0.005 | 0.44 | 0.64 | 0.53 | 0.41 | 0.49 | 0.42 |
| 2 | 60097447 | rs763297 | -1.214 | 1.011 | -0.622 | -0.241 | -0.016 | -0.099 | -0.099 | 0.556 | 0.65 | 0.92 | 0.67 | 0.51 | 0.62 | 0.55 |
| 2 | 60098316 | rs9653546 | 0.073 | 1.133 | -0.337 | -0.258 | 0.033 | -0.055 | -0.055 | 0 | 0.12 | 0.18 | 0.14 | 0 | 0.19 | 0.06 |
| 2 | 60100727 | rs13403546 | 1.131 | 0.953 | -0.365 | -0.077 | 0.001 | 0.053 | 0.053 | 0 | 0.15 | 0.3 | 0.17 | 0.21 | 0.12 | 0.18 |
| 2 | 60101212 | rs6545760 | -0.532 | 0.455 | -1.095 | -0.532 | -0.005 | 0.03 | 0.03 | 0 | 0.52 | 0.81 | 0.55 | 0.38 | 0.55 | 0.44 |
| 2 | 60103775 | rs17400093 | 0.027 | 0.459 | -0.969 | -0.561 | 0.001 | -0.007 | -0.007 | 0 | 0.37 | 0.25 | 0.44 | 0.37 | 0.37 | 0.51 |
| 2 | 60105275 | rs17328175 | 0.557 | 0.671 | -0.73 | -0.65 | -0.002 | 0.002 | 0.002 | 1.658 | 0.27 | 0.25 | 0.29 | 0.2 | 0.3 | 0.22 |
| 2 | 60105682 | rs1005038 | -0.873 | 0.549 | -0.761 | -0.642 | 0.024 | 0.048 | 0.048 | 0 | 0.55 | 0.74 | 0.59 | 0.35 | 0.59 | 0.41 |
| 2 | 60107055 | rs7557059 | -0.961 | 0.487 | -0.74 | -0.588 | 0.028 | 0.04 | 0.04 | 0 | 0.57 | 0.77 | 0.6 | 0.35 | 0.61 | 0.45 |
| 2 | 60107632 | rs17400114 | 1.968 | 0.664 | -0.543 | -0.257 | -0.023 | -0.021 | -0.021 | 0 | 0.08 | 0.01 | 0.15 | 0.15 | 0.09 | 0.13 |
| 2 | 60111222 | rs12991146 | 1.776 | 0.582 | -0.074 | -0.312 | -0.022 | -0.056 | -0.056 | 0 | 0.1 | 0.21 | 0.06 | 0 | 0.06 | 0.02 |
| 2 | 60112579 | rs6728812 | 1.75 | 0.581 | -0.086 | -0.309 | -0.018 | -0.054 | -0.054 | 0.488 | 0.1 | 0.27 | 0.06 | 0 | 0.06 | 0.02 |
| 2 | 60112742 | rs17400149 | 0.336 | 0.974 | -0.417 | -0.109 | -0.003 | 0.001 | 0.001 | 1.332 | 0.26 | 0.25 | 0.28 | 0.2 | 0.3 | 0.25 |
| 2 | 60114261 | rs11682272 | -0.583 | 1.042 | 0.344 | -0.034 | 0.095 | -0.012 | -0.012 | 0.28 | 0.12 | 0.47 | 0.11 | 0.05 | 0.15 | 0.03 |
| 2 | 60114444 | rs4672326 | -0.759 | 0.149 | -0.293 | 0.514 | -0.029 | -0.06 | -0.06 | 0.251 | 0.9 | 0.99 | 0.9 | 0.57 | 0.89 | 0.77 |
| 2 | 60115422 | rs6719345 | 0.143 | -0.211 | -0.084 | -0.099 | 0.001 | -0.005 | -0.005 | 0 | 0.45 | 0.39 | 0.68 | 0.5 | 0.56 | 0.55 |
| 2 | 60117297 | rs10865313 | 0.404 | -0.52 | -0.225 | -0.187 | -0.017 | -0.044 | -0.044 | 0 | 0.42 | 0.18 | 0.66 | 0.49 | 0.57 | 0.6 |
| 2 | 60117491 | rs11675976 | -0.687 | -0.677 | -0.053 | -0.272 | 0.015 | -0.047 | -0.047 | 0.049 | 0.6 | 0.79 | 0.77 | 0.5 | 0.65 | 0.64 |
| 2 | 60125363 | rs11674994 | 1.958 | -0.628 | 0.409 | -0.281 | -0.016 | -0.057 | -0.057 | 3.382 | 0.1 | 0.22 | 0.05 | 0 | 0.05 | 0.01 |
| 2 | 60131753 | rs11888341 | -0.458 | -0.665 | 0.373 | -0.275 | 0.022 | -0.02 | -0.02 | 1.091 | 0.57 | 0.71 | 0.76 | 0.5 | 0.64 | 0.56 |
| 2 | 60133158 | rs1344293 | 0.149 | -0.571 | 0.412 | -0.264 | 0.006 | 0.032 | 0.032 | 0 | 0.49 | 0.37 | 0.6 | 0.34 | 0.55 | 0.52 |
| 2 | 60133565 | rs17050485 | 0.01 | -0.624 | 0.276 | -0.041 | -0.008 | 0.05 | 0.05 | 0 | 0.14 | 0.17 | 0.11 | 0.39 | 0.12 | 0.23 |

| | | | | | | | | | | | | | | | | |
|---|---|---|---|---|---|---|---|---|---|---|---|---|---|---|---|---|
| 2 | 60135348 | rs1863162 | -0.551 | -0.61 | 0.251 | -0.07 | -0.001 | 0.014 | 0.014 | 0 | 0.19 | 0.18 | 0.23 | 0.53 | 0.2 | 0.34 |
| 2 | 60135937 | rs1030321 | -0.683 | -0.625 | 0.224 | -0.153 | -0.011 | -0.041 | -0.041 | 0 | 0.63 | 0.55 | 0.8 | 0.87 | 0.71 | 0.79 |
| 2 | 60140608 | rs1559621 | -0.831 | -0.609 | 0.254 | -0.615 | -0.011 | -0.098 | -0.098 | 0 | 0.64 | 0.55 | 0.82 | 0.95 | 0.71 | 0.81 |
| 2 | 60145579 | rs6545767 | -1.443 | -1.088 | 0.151 | -0.858 | 0.009 | 0.033 | 0.033 | 0 | 0.76 | 0.89 | 0.88 | 1 | 0.78 | 0.87 |
| 2 | 60155276 | rs17050495 | 0.868 | -0.854 | 0.466 | -0.609 | -0.047 | -0.1 | -0.1 | 0 | 0.15 | 0.36 | 0.1 | 0.01 | 0.08 | 0.02 |
| 2 | 60155693 | rs17050497 | -0.793 | -0.853 | -0.211 | -0.538 | 0.019 | -0.042 | -0.042 | 0 | 0.63 | 0.77 | 0.8 | 0.49 | 0.69 | 0.66 |
| 2 | 60157097 | rs13005495 | 0.351 | -0.581 | -0.412 | -0.49 | 0.009 | -0.041 | -0.041 | 0 | 0.44 | 0.2 | 0.67 | 0.47 | 0.59 | 0.59 |
| 2 | 60160790 | rs12713413 | 0.084 | -0.261 | -0.628 | -0.209 | 0.01 | -0.001 | -0.001 | 0.225 | 0.52 | 0.6 | 0.7 | 0.48 | 0.6 | 0.6 |
| 2 | 60178342 | rs890067 | -0.208 | -0.291 | -0.812 | -0.235 | 0.009 | -0.004 | -0.004 | 0 | 0.54 | 0.66 | 0.7 | 0.5 | 0.59 | 0.59 |
| 2 | 60179313 | rs6735059 | 0.941 | 0.046 | -0.845 | -0.414 | 0.001 | -0.001 | -0.001 | 0 | 0.27 | 0.29 | 0.27 | 0.22 | 0.34 | 0.33 |
| 2 | 60179482 | rs6545769 | -1.514 | -0.18 | -1.784 | -0.842 | -0.054 | 0.174 | 0.174 | 0 | 0.75 | 0.99 | 0.85 | 1 | 0.76 | 0.86 |
| 2 | 60181642 | rs10490065 | 1.646 | -0.001 | -1.581 | -0.533 | -0.017 | -0.025 | -0.025 | 0 | 0.06 | 0.05 | 0.12 | 0.16 | 0.06 | 0.14 |
| 2 | 60182071 | rs7556782 | -0.849 | -0.279 | -1.469 | -0.699 | 0.005 | -0.028 | -0.028 | 0 | 0.62 | 0.75 | 0.73 | 0.49 | 0.63 | 0.6 |
| 2 | 60183216 | rs12471714 | 1.608 | 0.263 | -1.265 | -0.417 | -0.023 | -0.031 | -0.031 | 0 | 0.06 | 0.03 | 0.12 | 0.16 | 0.06 | 0.14 |
| 2 | 60203917 | rs12476772 | 0.4 | 0.341 | -1.276 | -0.693 | 0.051 | -0.064 | -0.064 | 0.058 | 0.26 | 0.52 | 0.29 | 0.15 | 0.33 | 0.21 |
| 2 | 60209439 | rs4671360 | 1.652 | 0.345 | -1.935 | -0.436 | -0.028 | -0.025 | -0.025 | 2.1 | 0.1 | 0.02 | 0.23 | 0.11 | 0.14 | 0.09 |
| 2 | 60217093 | rs4672337 | 0.379 | 0.07 | -1.826 | -0.437 | -0.03 | -0.034 | -0.034 | 0 | 0.11 | 0.04 | 0.15 | 0.15 | 0.1 | 0.14 |
| 2 | 60225873 | rs6545772 | 0.185 | -0.158 | -1.457 | -0.472 | 0.026 | -0.009 | -0.009 | 0 | 0.35 | 0.47 | 0.47 | 0.24 | 0.45 | 0.35 |
| 2 | 60230714 | rs17050525 | 1.378 | -0.227 | -1.525 | -0.533 | 0 | -0.002 | -0.002 | 0 | 0.05 | 0.09 | 0.04 | 0.01 | 0.03 | 0.01 |
| 2 | 60234327 | rs11125823 | -0.84 | -0.384 | -2.023 | -0.664 | 0.031 | -0.004 | -0.004 | 0 | 0.55 | 0.74 | 0.71 | 0.52 | 0.62 | 0.64 |
| 2 | 60235568 | rs11896330 | -0.92 | -0.673 | -2.362 | -1.088 | -0.037 | -0.096 | -0.096 | 0 | 0.64 | 0.91 | 0.74 | 0.54 | 0.62 | 0.61 |
| 2 | 60237213 | rs12373776 | -0.279 | -0.389 | -2.325 | -0.877 | 0.003 | -0.003 | -0.003 | 0.061 | 0.46 | 0.66 | 0.67 | 0.51 | 0.55 | 0.55 |
| 2 | 60255313 | rs11888990 | 0.696 | -0.293 | -2.268 | -0.932 | 0.001 | 0.053 | 0.053 | 0.159 | 0.12 | 0.05 | 0.04 | 0.01 | 0.08 | 0.04 |
| 2 | 60255546 | rs953755 | 0.973 | -0.248 | -2.019 | -1.12 | 0.031 | -0.078 | -0.078 | 0.739 | 0.24 | 0.27 | 0.24 | 0.04 | 0.36 | 0.23 |
| 2 | 60255567 | rs953754 | 0.305 | -0.293 | -2.502 | -0.935 | 0.018 | 0.005 | 0.005 | 0 | 0.34 | 0.49 | 0.49 | 0.3 | 0.46 | 0.33 |
| 2 | 60257677 | rs12477781 | -0.519 | 0.004 | -2.846 | -0.396 | -0.02 | -0.001 | -0.001 | 0 | 0.2 | 0.06 | 0.2 | 0.22 | 0.15 | 0.27 |
| 2 | 60260639 | rs2215897 | -0.925 | -0.038 | -2.842 | -0.386 | -0.009 | 0.032 | 0.032 | 0 | 0.24 | 0.13 | 0.23 | 0.22 | 0.18 | 0.3 |
| 2 | 60268260 | rs13016030 | -0.003 | 0.101 | -2.562 | -0.295 | 0.009 | 0.03 | 0.03 | 0.192 | 0.12 | 0.09 | 0.04 | 0.01 | 0.08 | 0.04 |
| 2 | 60273795 | rs7607085 | -0.471 | 0.178 | -2.858 | -0.203 | -0.109 | -0.128 | -0.128 | 0 | 0.55 | 0.96 | 0.44 | 0.51 | 0.55 | 0.51 |
| 2 | 60279418 | rs994972 | -0.159 | 0.304 | -2.767 | -0.172 | -0.087 | -0.07 | -0.07 | 0 | 0.5 | 0.93 | 0.41 | 0.51 | 0.51 | 0.51 |
| 2 | 60283805 | rs10187705 | -0.779 | 0.968 | -1.664 | 0.422 | -0.002 | -0.001 | -0.001 | 0 | 0.48 | 0.67 | 0.66 | 0.57 | 0.53 | 0.51 |
| 2 | 60284614 | rs1861151 | -0.188 | 1.128 | -1.164 | 0.672 | 0.003 | -0.002 | -0.002 | 0.416 | 0.25 | 0.33 | 0.29 | 0.29 | 0.22 | 0.34 |
| 2 | 60284697 | rs7603779 | -0.249 | 0.983 | -1.144 | 0.691 | 0.005 | -0.005 | -0.005 | 0 | 0.26 | 0.35 | 0.28 | 0.27 | 0.26 | 0.32 |
| 2 | 60285100 | rs980329 | -0.301 | 1.037 | -1.166 | 0.711 | 0.002 | -0.003 | -0.003 | 0.195 | 0.27 | 0.33 | 0.31 | 0.29 | 0.26 | 0.34 |
| 2 | 60289424 | rs10185622 | -0.756 | 0.555 | -1.129 | 0.734 | 0.006 | 0.003 | 0.003 | 0.142 | 0.52 | 0.65 | 0.69 | 0.6 | 0.61 | 0.61 |
| 2 | 60290360 | rs2192580 | -0.202 | 0.331 | -0.629 | 1.288 | 0.009 | 0.046 | 0.046 | 0 | 0.17 | 0.3 | 0.36 | 0.33 | 0.26 | 0.19 |
| 2 | 60291204 | rs11890420 | -0.625 | -0.528 | -0.095 | 1.271 | -0.018 | 0.014 | 0.014 | 0 | 0.37 | 0.51 | 0.29 | 0.5 | 0.25 | 0.3 |
| 2 | 60295277 | rs17030159 | -0.406 | -0.961 | 0.879 | 1.785 | -0.018 | 0.02 | 0.02 | 0 | 0.07 | 0.03 | 0.17 | 0.51 | 0.06 | 0.29 |
| 2 | 60296223 | rs10178652 | -1.976 | -1.054 | 1.154 | 1.398 | 0.001 | 0.005 | 0.005 | 0 | 0.34 | 0.58 | 0.36 | 0.62 | 0.35 | 0.39 |
| 2 | 60299010 | rs882924 | -2.027 | -1.054 | 1.179 | 1.417 | -0.002 | 0.008 | 0.008 | 0 | 0.34 | 0.59 | 0.36 | 0.6 | 0.34 | 0.38 |
| 2 | 60299441 | rs4671369 | -1.966 | -1.045 | 1.17 | 1.4 | 0.002 | 0.009 | 0.009 | 0 | 0.34 | 0.56 | 0.36 | 0.62 | 0.35 | 0.39 |
| 2 | 60299921 | rs17329014 | -0.511 | -0.833 | 0.911 | 0.529 | -0.014 | -0.001 | -0.001 | 0 | 0.15 | 0.1 | 0.16 | 0 | 0.26 | 0.08 |
| 2 | 60302746 | rs13394010 | -1.899 | -0.988 | 1.133 | 0.464 | -0.002 | -0.072 | -0.072 | 0.886 | 0.33 | 0.51 | 0.27 | 0.1 | 0.36 | 0.17 |
| 2 | 60305110 | rs10203396 | -1.945 | -1.001 | 1.252 | 0.455 | -0.002 | -0.04 | -0.04 | 0 | 0.34 | 0.55 | 0.27 | 0.1 | 0.35 | 0.17 |
| 2 | 60321554 | rs17027670 | -0.162 | -0.915 | 1.249 | 0.5 | -0.006 | -0.024 | -0.024 | 0 | 0.05 | 0.1 | 0.02 | 0 | 0.03 | 0.02 |
| 2 | 60323975 | rs917929 | -1.442 | -1.014 | 1.054 | 0.383 | -0.002 | -0.051 | -0.051 | 0 | 0.3 | 0.35 | 0.21 | 0.09 | 0.32 | 0.14 |
| 2 | 60324127 | rs13415334 | -1.838 | -0.938 | 0.441 | 0.47 | 0.009 | 0.041 | 0.041 | 1.196 | 0.4 | 0.67 | 0.28 | 0.12 | 0.4 | 0.19 |
| 2 | 60331164 | rs17329188 | -0.858 | -0.267 | 0.504 | 1.292 | -0.005 | -0.004 | -0.004 | 0 | 0.16 | 0.18 | 0.24 | 0.23 | 0.27 | 0.2 |
| 2 | 60331399 | rs972337 | -2.224 | -0.558 | 0.48 | 1.464 | 0.003 | -0.012 | -0.012 | 0 | 0.41 | 0.8 | 0.43 | 0.57 | 0.41 | 0.37 |
| 2 | 60333732 | rs887930 | 0.105 | -0.453 | 0.341 | 1.458 | -0.001 | 0.008 | 0.008 | 0 | 0.07 | 0.03 | 0.14 | 0.2 | 0.1 | 0.13 |
| 2 | 60333953 | rs7588487 | -0.813 | -0.504 | 0.532 | 1.418 | -0.011 | 0 | 0 | 0 | 0.23 | 0.22 | 0.24 | 0.24 | 0.29 | 0.19 |
| 2 | 60338227 | rs6737141 | -1.914 | -0.721 | 1.167 | 1.531 | -0.042 | 0.11 | 0.11 | 0.361 | 0.17 | 0.53 | 0.21 | 0.31 | 0.14 | 0.17 |
| 2 | 60338651 | rs872277 | -0.716 | -0.71 | 1.287 | 1.545 | 0.01 | 0.082 | 0.082 | 0 | 0.07 | 0.28 | 0.06 | 0.09 | 0.03 | 0.04 |
| 2 | 60339170 | rs17029624 | -1.657 | -0.684 | 1.036 | 1.517 | -0.038 | 0.141 | 0.141 | 0.093 | 0.17 | 0.48 | 0.2 | 0.31 | 0.14 | 0.15 |
| 2 | 60356628 | rs11691580 | -0.882 | -0.969 | 0.694 | 0.815 | -0.05 | -0.046 | -0.046 | 0.021 | 0.27 | 0.43 | 0.11 | 0.08 | 0.13 | 0.11 |
| 2 | 60375583 | rs9808521 | -0.901 | -0.982 | 0.722 | 0.797 | -0.066 | -0.064 | -0.064 | 1.986 | 0.27 | 0.47 | 0.12 | 0.08 | 0.13 | 0.12 |
| 2 | 60376614 | rs10186608 | -0.306 | -1.081 | 0.82 | 1.137 | 0.006 | 0.088 | 0.088 | 0 | 0.08 | 0.21 | 0.08 | 0.2 | 0.05 | 0.11 |
| 2 | 60376723 | rs10175707 | -1.386 | -1.018 | 0.665 | 1.137 | 0 | 0 | 0 | 0.066 | 0.35 | 0.7 | 0.2 | 0.28 | 0.18 | 0.24 |
| 2 | 60385252 | rs7591835 | -0.297 | -1.019 | 0.215 | 0.989 | -0.04 | -0.038 | -0.038 | 0.026 | 0.2 | 0.23 | 0.06 | 0.05 | 0.09 | 0.09 |
| 2 | 60389854 | rs6759758 | -0.234 | -1.013 | 0.233 | 0.98 | -0.045 | -0.047 | -0.047 | 0 | 0.19 | 0.24 | 0.06 | 0.05 | 0.09 | 0.09 |
| 2 | 60398416 | rs4672367 | -0.054 | -0.274 | -0.539 | 1.587 | -0.008 | 0.026 | 0.026 | 0 | 0.08 | 0.03 | 0.34 | 0.42 | 0.2 | 0.19 |
| 2 | 60403018 | rs6545786 | -0.849 | -0.773 | -0.339 | 1.163 | -0.05 | -0.013 | -0.013 | 0 | 0.32 | 0.61 | 0.18 | 0.31 | 0.16 | 0.25 |
| 2 | 60408305 | rs7585504 | -1.17 | 0.045 | -1.7 | 0.675 | -0.128 | -0.043 | -0.043 | 0 | 0.73 | 0.98 | 0.76 | 0.82 | 0.68 | 0.72 |
| 2 | 60409379 | rs10490066 | 0.21 | -0.536 | -1.611 | 0.176 | 0.019 | 0.115 | 0.115 | 1.57 | 0.23 | 0.05 | 0.13 | 0 | 0.19 | 0.14 |
| 2 | 60425653 | rs13406467 | -0.078 | -0.482 | -0.689 | 0.647 | -0.011 | 0.027 | 0.027 | 0.693 | 0.07 | 0.25 | 0.04 | 0.14 | 0.02 | 0.01 |
| 2 | 60441184 | rs9309322 | -0.256 | -0.662 | -0.648 | 1.037 | 0.004 | -0.007 | -0.007 | 0 | 0.43 | 0.45 | 0.54 | 0.6 | 0.45 | 0.36 |
| 2 | 60449069 | rs11892403 | -0.492 | -0.339 | 0.135 | 1.181 | -0.011 | 0.068 | 0.068 | 0 | 0.12 | 0.24 | 0.08 | 0.16 | 0.05 | 0.16 |
| 2 | 60461578 | rs191692 | -0.932 | -0.054 | -0.479 | 0.717 | -0.041 | 0.033 | 0.033 | 0 | 0.7 | 0.86 | 0.69 | 0.84 | 0.59 | 0.55 |
| 2 | 60461671 | rs7579491 | -0.645 | -0.374 | 0.386 | 1.177 | -0.048 | 0.092 | 0.092 | 0.006 | 0.2 | 0.5 | 0.15 | 0.32 | 0.1 | 0.17 |

| | | | | | | | | | | | | | | | | |
|---|---|---|---|---|---|---|---|---|---|---|---|---|---|---|---|---|
| 2 | 60475257 | rs359242 | 0.474 | -0.393 | 0.146 | 1.123 | 0.015 | 0.042 | 0.042 | 0 | 0.48 | 0.24 | 0.5 | 0.38 | 0.48 | 0.36 |
| 2 | 60480973 | rs359250 | -1.556 | -0.392 | 0.023 | 1.115 | -0.141 | -0.014 | -0.014 | 0 | 0.75 | 0.98 | 0.7 | 0.85 | 0.61 | 0.55 |
| 2 | 60484507 | rs359255 | -1.395 | -0.349 | 0.274 | 1.122 | -0.068 | 0.025 | 0.025 | 0 | 0.73 | 0.9 | 0.7 | 0.85 | 0.61 | 0.56 |
| 2 | 60485161 | rs359257 | 0.548 | -0.383 | 0.32 | 1.129 | 0.002 | 0.048 | 0.048 | 0 | 0.5 | 0.26 | 0.53 | 0.38 | 0.51 | 0.37 |
| 2 | 60486154 | rs1553935 | -0.181 | -0.893 | 0.947 | 1.194 | -0.021 | 0.074 | 0.074 | 0 | 0.13 | 0.22 | 0.1 | 0.3 | 0.05 | 0.16 |
| 2 | 60487902 | rs17017706 | -1.071 | -0.906 | 1.211 | 1.185 | -0.048 | 0.096 | 0.096 | 0 | 0.2 | 0.45 | 0.14 | 0.33 | 0.09 | 0.16 |
| 2 | 60494345 | rs4672376 | -0.607 | -0.673 | 1.569 | 1.152 | 0.02 | -0.009 | -0.009 | 0 | 0.64 | 0.55 | 0.64 | 0.75 | 0.54 | 0.54 |
| 2 | 60495038 | rs4671379 | -1.404 | -0.616 | 1.318 | 1.131 | -0.078 | 0.026 | 0.026 | 0 | 0.73 | 0.89 | 0.69 | 0.85 | 0.58 | 0.57 |
| 2 | 60497135 | rs4671382 | -1.091 | -0.648 | 1.565 | 1.169 | 0.026 | -0.002 | -0.002 | 0.373 | 0.7 | 0.66 | 0.66 | 0.72 | 0.56 | 0.55 |
| 2 | 60498211 | rs17019696 | 0.074 | -0.752 | 1.594 | 1.209 | 0.002 | -0.003 | -0.003 | 0 | 0.12 | 0.09 | 0.08 | 0.24 | 0.05 | 0.15 |
| 2 | 60500158 | rs7587924 | -0.145 | -0.764 | 1.651 | 1.202 | -0.009 | 0.013 | 0.013 | 0 | 0.13 | 0.2 | 0.1 | 0.24 | 0.05 | 0.16 |
| 2 | 60500484 | rs2110398 | -1.176 | -0.694 | 1.611 | 1.196 | -0.008 | 0 | 0 | 0 | 0.7 | 0.75 | 0.67 | 0.72 | 0.56 | 0.55 |
| 2 | 60501421 | rs955658 | -1.477 | -0.675 | 1.484 | 1.184 | -0.077 | 0.026 | 0.026 | 0 | 0.73 | 0.89 | 0.7 | 0.85 | 0.58 | 0.57 |
| 2 | 60503230 | rs907574 | -1.27 | -0.772 | 1.837 | 1.183 | -0.055 | 0.079 | 0.079 | 0 | 0.21 | 0.49 | 0.14 | 0.33 | 0.09 | 0.18 |
| 2 | 60503453 | rs2901001 | -0.285 | -0.75 | 1.735 | 1.176 | -0.02 | 0.036 | 0.036 | 0.014 | 0.09 | 0.27 | 0.05 | 0.1 | 0.04 | 0.02 |
| 2 | 60509450 | rs17021086 | -1.064 | -0.742 | 1.512 | 1.238 | -0.037 | 0.041 | 0.041 | 0.014 | 0.19 | 0.4 | 0.13 | 0.27 | 0.09 | 0.16 |
| 2 | 60514993 | rs4671386 | -0.976 | -0.627 | 1.559 | 1.209 | -0.017 | -0.005 | -0.005 | 0 | 0.69 | 0.76 | 0.67 | 0.64 | 0.57 | 0.53 |
| 2 | 60516388 | rs2110399 | -1.621 | -0.601 | 1.219 | 1.184 | -0.091 | 0.03 | 0.03 | 0 | 0.75 | 0.88 | 0.7 | 0.87 | 0.6 | 0.55 |
| 2 | 60517086 | rs7597821 | -0.03 | -1.334 | 0.691 | 0.827 | -0.001 | 0 | 0 | 0.863 | 0.36 | 0.63 | 0.3 | 0.32 | 0.37 | 0.28 |
| 2 | 60517108 | rs10211066 | -0.522 | -1.45 | 0.598 | 0.915 | 0.002 | 0.036 | 0.036 | 0 | 0.44 | 0.66 | 0.31 | 0.32 | 0.4 | 0.3 |
| 2 | 60517299 | rs7598252 | -0.873 | -1.478 | 0.781 | 1.233 | -0.016 | -0.027 | -0.027 | 0 | 0.63 | 0.71 | 0.58 | 0.56 | 0.52 | 0.41 |
| 2 | 60518350 | rs1468739 | -1.966 | -1.393 | 0.285 | 1.2 | -0.176 | -0.024 | -0.024 | 0 | 0.74 | 0.98 | 0.69 | 0.85 | 0.59 | 0.57 |
| 2 | 60519588 | rs2058550 | -1.512 | -1.416 | 0.512 | 1.243 | -0.056 | -0.016 | -0.016 | 0 | 0.71 | 0.82 | 0.66 | 0.69 | 0.56 | 0.54 |
| 2 | 60522691 | rs11687445 | -0.019 | -1.479 | 0.304 | 1.049 | 0.024 | 0.058 | 0.058 | 0 | 0.53 | 0.23 | 0.52 | 0.36 | 0.48 | 0.4 |
| 2 | 60523559 | rs907573 | -1.604 | -1.139 | 0.062 | 0.931 | -0.029 | 0.01 | 0.01 | 0 | 0.73 | 0.78 | 0.68 | 0.85 | 0.58 | 0.58 |
| 2 | 60528759 | rs7581043 | -0.317 | -1.659 | -0.009 | 1.107 | -0.079 | 0.002 | 0.002 | 4.145 | 0.22 | 0.59 | 0.17 | 0.24 | 0.08 | 0.19 |
| 2 | 60529026 | rs4671387 | 0.073 | -1.512 | -0.684 | 0.837 | -0.028 | 0.088 | 0.088 | 0.189 | 0.4 | 0.15 | 0.41 | 0.18 | 0.46 | 0.24 |
| 2 | 60529903 | rs7597268 | -0.28 | -1.625 | -0.594 | 1.082 | 0.023 | 0.012 | 0.012 | 0.035 | 0.5 | 0.59 | 0.48 | 0.47 | 0.51 | 0.4 |
| 2 | 60530702 | rs876207 | 0.722 | -1.206 | -0.884 | 0.908 | -0.021 | 0.003 | 0.003 | 0 | 0.19 | 0.14 | 0.34 | 0.13 | 0.32 | 0.1 |
| 2 | 60531740 | rs7592241 | -1.051 | -0.84 | -1.008 | 0.94 | -0.027 | -0.001 | -0.001 | 0 | 0.68 | 0.72 | 0.62 | 0.73 | 0.58 | 0.52 |
| 2 | 60532247 | rs17039727 | -0.172 | -1.021 | -0.722 | 1.364 | -0.004 | -0.066 | -0.066 | 0 | 0.12 | 0.01 | 0.14 | 0.24 | 0.08 | 0.18 |
| 2 | 60533240 | rs12611562 | -0.791 | -0.942 | 0.037 | 1.347 | -0.007 | 0.035 | 0.035 | 0 | 0.11 | 0.22 | 0.12 | 0.18 | 0.08 | 0.17 |
| 2 | 60533913 | rs7586170 | -0.544 | -0.42 | 0.067 | 1.311 | 0.095 | 0.025 | 0.025 | 0 | 0.09 | 0.16 | 0.14 | 0.13 | 0.23 | 0.1 |
| 2 | 60534199 | rs6545803 | -0.029 | -1.105 | -0.043 | 1.222 | -0.013 | 0.006 | 0.006 | 0 | 0.31 | 0.38 | 0.34 | 0.39 | 0.25 | 0.31 |
| 2 | 60534338 | rs17024964 | 0.036 | -1.072 | -0.923 | 1.239 | 0.01 | -0.089 | -0.089 | 0 | 0.3 | 0.06 | 0.32 | 0.39 | 0.25 | 0.31 |
| 2 | 60537741 | rs13416098 | -0.704 | -0.97 | -0.394 | 0.9 | 0.1 | 0.058 | 0.058 | 0.047 | 0.5 | 0.3 | 0.48 | 0.65 | 0.34 | 0.54 |
| 2 | 60538609 | rs6545804 | -1.545 | -1.262 | -0.057 | 0.773 | 0.019 | -0.014 | -0.014 | 1.018 | 0.63 | 0.5 | 0.52 | 0.73 | 0.58 | 0.58 |
| 2 | 60538727 | rs6750077 | -0.673 | -0.615 | 0.4 | 0.796 | 0.111 | -0.005 | -0.005 | 0 | 0.16 | 0.39 | 0.2 | 0.13 | 0.28 | 0.1 |
| 2 | 60541853 | rs7595905 | -1.281 | -1.377 | -0.02 | 0.634 | -0.01 | 0.014 | 0.014 | 0.038 | 0.6 | 0.66 | 0.57 | 0.89 | 0.58 | 0.58 |
| 2 | 60542657 | rs13383059 | -0.809 | -1.379 | -0.202 | 1.199 | -0.04 | 0.052 | 0.052 | 0 | 0.17 | 0.31 | 0.18 | 0.29 | 0.08 | 0.19 |
| 2 | 60547567 | rs11125833 | -1.219 | -1.516 | -1.125 | 0.745 | -0.012 | 0.111 | 0.111 | 0 | 0.51 | 0.78 | 0.56 | 0.69 | 0.52 | 0.39 |
| 2 | 60553143 | rs7567242 | -0.494 | -1.557 | -0.891 | 0.734 | -0.055 | 0.022 | 0.022 | 2.234 | 0.26 | 0.67 | 0.16 | 0.36 | 0.12 | 0.24 |
| 2 | 60554216 | rs10193182 | -0.886 | -0.905 | -0.908 | 0.658 | -0.013 | 0.017 | 0.017 | 0.965 | 0.52 | 0.77 | 0.55 | 0.55 | 0.54 | 0.35 |
| 2 | 60560560 | rs13423091 | -0.258 | -0.941 | -0.433 | 1.011 | -0.037 | 0.09 | 0.09 | 0 | 0.18 | 0.24 | 0.23 | 0.42 | 0.09 | 0.22 |
| 2 | 60566266 | rs48314 | 0.996 | -1.037 | -1.383 | 0.508 | -0.086 | 0.098 | 0.098 | 0 | 0.39 | 0.06 | 0.33 | 0.22 | 0.35 | 0.26 |
| 2 | 60569348 | rs243087 | 1.001 | -1.042 | -1.386 | 0.505 | -0.086 | 0.097 | 0.097 | 0 | 0.39 | 0.06 | 0.3 | 0.22 | 0.34 | 0.26 |
| 2 | 60572982 | rs13408761 | 0.033 | -0.969 | -0.839 | 0.671 | -0.04 | 0.086 | 0.086 | 0 | 0.19 | 0.26 | 0.24 | 0.42 | 0.11 | 0.25 |
| 2 | 60573870 | rs243083 | -0.132 | -1.259 | -0.13 | 0.51 | -0.02 | 0.003 | 0.003 | 0 | 0.63 | 0.65 | 0.57 | 0.65 | 0.45 | 0.51 |
| 2 | 60575895 | rs6757208 | 0.414 | -1.106 | -0.029 | 0.693 | -0.028 | 0.079 | 0.079 | 0 | 0.17 | 0.21 | 0.22 | 0.43 | 0.09 | 0.25 |
| 2 | 60578843 | rs10180282 | -0.955 | -1.457 | -0.537 | 0.119 | 0.002 | 0.059 | 0.059 | 0 | 0.5 | 0.61 | 0.62 | 0.79 | 0.57 | 0.72 |
| 2 | 60584819 | rs243021 | 0.712 | -0.973 | -0.165 | 0.207 | 0.005 | -0.004 | -0.004 | 0 | 0.59 | 0.39 | 0.54 | 0.65 | 0.45 | 0.49 |
| 2 | 60586661 | rs10207503 | -0.458 | -0.42 | -0.135 | 0.426 | -0.022 | 0.017 | 0.017 | 0 | 0.19 | 0.35 | 0.13 | 0.24 | 0.13 | 0.38 |
| 2 | 60590257 | rs243052 | -0.4 | 1.121 | -0.646 | 0.41 | -0.012 | -0.014 | -0.014 | 0 | 0.09 | 0.04 | 0.22 | 0.1 | 0.33 | 0.09 |
| 2 | 60592027 | rs243050 | -0.514 | 1.194 | -0.644 | 0.4 | -0.013 | -0.012 | -0.012 | 1.922 | 0.1 | 0.05 | 0.23 | 0.1 | 0.33 | 0.09 |
| 2 | 60595793 | rs243044 | -0.505 | 1.188 | -0.579 | 0.397 | 0.001 | -0.008 | -0.008 | 0.179 | 0.1 | 0.07 | 0.23 | 0.1 | 0.33 | 0.09 |
| 2 | 60597565 | rs243039 | -0.492 | 1.187 | -0.317 | 0.405 | 0.073 | 0.012 | 0.012 | 0 | 0.1 | 0.14 | 0.23 | 0.1 | 0.33 | 0.1 |
| 2 | 60598672 | rs4672388 | -0.593 | 0.322 | -0.192 | 0.409 | -0.022 | 0.041 | 0.041 | 0 | 0.2 | 0.3 | 0.28 | 0.71 | 0.12 | 0.58 |
| 2 | 60601376 | rs243035 | -1.149 | 0.377 | -0.394 | 0.687 | 0 | -0.001 | -0.001 | 0 | 0.3 | 0.27 | 0.5 | 0.56 | 0.48 | 0.47 |
| 2 | 60602892 | rs243034 | -0.884 | -0.08 | -0.472 | -0.032 | 0.017 | 0.025 | 0.025 | 0 | 0.5 | 0.57 | 0.44 | 0.28 | 0.62 | 0.46 |
| 2 | 60605030 | rs4671389 | -0.516 | -0.248 | 0.113 | -0.076 | 0.03 | 0.321 | 0.321 | 0 | 0.07 | 0.24 | 0.19 | 0.45 | 0.06 | 0.27 |
| 2 | 60612457 | rs2137283 | -0.366 | 0.67 | -0.218 | -0.709 | -0.03 | 0.016 | 0.016 | 0.053 | 0.21 | 0.17 | 0.22 | 0.01 | 0.42 | 0.14 |
| 2 | 60613776 | rs243081 | -0.982 | 0.495 | -0.099 | -0.687 | -0.001 | -0.002 | -0.002 | 0 | 0.26 | 0.26 | 0.51 | 0.61 | 0.48 | 0.47 |
| 2 | 60624939 | rs9309326 | -0.039 | 0.037 | -0.09 | -0.985 | -0.051 | 0.023 | 0.023 | 0 | 0.3 | 0.44 | 0.26 | 0.39 | 0.16 | 0.3 |
| 2 | 60626246 | rs3948623 | 0.067 | 0.215 | -0.065 | -0.774 | -0.005 | 0.105 | 0.105 | 0.252 | 0.09 | 0.15 | 0.3 | 0.61 | 0.07 | 0.36 |
| 2 | 60633364 | rs13009393 | -0.495 | 0.391 | -0.172 | -1.283 | 0.031 | -0.072 | -0.072 | 0 | 0.23 | 0.3 | 0.23 | 0.01 | 0.42 | 0.14 |
| 2 | 60641329 | rs13028240 | 0.093 | 0.663 | -0.141 | -1.239 | 0.148 | -0.038 | -0.038 | 0.076 | 0.09 | 0.29 | 0.18 | 0.01 | 0.29 | 0.11 |
| 2 | 60676363 | rs17028222 | -0.757 | 0.025 | -0.387 | -1.026 | -0.006 | 0.01 | 0.01 | 0 | 0.22 | 0.23 | 0.36 | 0.72 | 0.19 | 0.4 |
| 2 | 60679942 | rs17331129 | 0.776 | -0.235 | -0.788 | -1.233 | 0.019 | -0.013 | -0.013 | 0 | 0.19 | 0.11 | 0.24 | 0.24 | 0.14 | 0.1 |
| 2 | 60698397 | rs12477097 | -1.107 | 0.212 | -0.571 | -1.034 | -0.009 | 0.003 | 0.003 | 0 | 0.25 | 0.38 | 0.39 | 0.75 | 0.24 | 0.44 |

| | | | | | | | | | | | | | | | | |
|---|---|---|---|---|---|---|---|---|---|---|---|---|---|---|---|---|
| 2 | 60707588 | rs1123573 | 0.18 | 0.469 | -1.094 | -1.627 | 0.014 | -0.108 | -0.108 | 0.089 | 0.31 | 0.36 | 0.24 | 0.02 | 0.39 | 0.27 |
| 2 | 60707894 | rs7579014 | -0.566 | 0.824 | -1.14 | -1.794 | -0.007 | -0.118 | -0.118 | 2.513 | 0.4 | 0.49 | 0.39 | 0.03 | 0.64 | 0.33 |
| 2 | 60713235 | rs10189857 | 0.401 | 0.328 | -1.796 | -1.718 | -0.005 | 0.013 | 0.013 | 0 | 0.39 | 0.28 | 0.49 | 0.73 | 0.42 | 0.71 |
| 2 | 60714861 | rs6545816 | 0.037 | 0.35 | -1.735 | -1.685 | 0.002 | 0.043 | 0.043 | 0 | 0.5 | 0.34 | 0.55 | 0.73 | 0.54 | 0.74 |
| 2 | 60715179 | rs6545817 | -0.084 | 0.371 | -1.87 | -1.675 | -0.001 | 0.061 | 0.061 | 0 | 0.49 | 0.29 | 0.54 | 0.73 | 0.53 | 0.73 |
| 2 | 60717399 | rs13031396 | 0.881 | 0.529 | -1.912 | -1.612 | -0.007 | 0.013 | 0.013 | 0 | 0.09 | 0.05 | 0.16 | 0 | 0.26 | 0.07 |
| 2 | 60718043 | rs1427407 | -1.242 | 0.026 | -1.904 | -1.643 | 0.039 | 0.006 | 0.006 | 0 | 0.69 | 0.76 | 0.77 | 0.74 | 0.85 | 0.88 |
| 2 | 60719074 | rs1896294 | -0.534 | -0.015 | -2.056 | -1.746 | 0.1 | 0.21 | 0.21 | 0 | 0.5 | 0.73 | 0.46 | 0.03 | 0.69 | 0.39 |
| 2 | 60719970 | rs766432 | -1.279 | -0.048 | -2.055 | -1.708 | 0.027 | 0.002 | 0.002 | 0 | 0.69 | 0.73 | 0.77 | 0.74 | 0.85 | 0.87 |
| 2 | 60720246 | rs11886868 | -1.002 | 0.719 | -1.426 | -1.923 | 0.101 | 0.214 | 0.214 | 0 | 0.5 | 0.74 | 0.46 | 0.03 | 0.69 | 0.41 |
| 2 | 60720589 | rs10195871 | -0.82 | 0.721 | -1.443 | -1.916 | 0.099 | 0.195 | 0.195 | 0 | 0.5 | 0.72 | 0.46 | 0.03 | 0.69 | 0.41 |
| 2 | 60722040 | rs6706648 | -0.379 | 1.368 | -0.954 | -1.557 | 0.086 | -0.112 | -0.112 | 0.059 | 0.2 | 0.44 | 0.23 | 0.01 | 0.32 | 0.14 |
| 2 | 60723266 | rs7565301 | 0.836 | 0.821 | -0.904 | -0.919 | 0.04 | -0.079 | -0.079 | 2.784 | 0.18 | 0.26 | 0.17 | 0.02 | 0.25 | 0.22 |
| 2 | 60729294 | rs10184550 | -0.088 | 0.471 | -1.309 | -0.69 | -0.004 | 0.106 | 0.106 | 0 | 0.55 | 0.72 | 0.36 | 0.19 | 0.56 | 0.35 |
| 2 | 60736837 | rs6724431 | 0.398 | 0.489 | -1.137 | -1.187 | -0.005 | -0.027 | -0.027 | 0 | 0.38 | 0.62 | 0.27 | 0.03 | 0.39 | 0.26 |
| 2 | 60738227 | rs1896297 | 0.512 | -0.065 | -0.548 | -0.497 | 0.011 | 0.011 | 0.011 | 0 | 0.55 | 0.43 | 0.65 | 0.75 | 0.63 | 0.79 |
| 2 | 60744331 | rs168565 | 0.291 | -0.15 | -0.421 | -0.512 | -0.022 | -0.018 | -0.018 | 0.012 | 0.57 | 0.46 | 0.66 | 0.75 | 0.68 | 0.79 |
| 2 | 60744872 | rs356991 | 0.232 | -0.166 | -0.44 | -0.52 | -0.009 | -0.007 | -0.007 | 0 | 0.58 | 0.51 | 0.67 | 0.75 | 0.68 | 0.79 |
| 2 | 60747192 | rs168562 | 0.157 | -0.142 | -0.465 | -0.517 | -0.002 | -0.001 | -0.001 | 0 | 0.59 | 0.56 | 0.68 | 0.75 | 0.68 | 0.79 |
| 2 | 60757419 | rs2457660 | 0.509 | 0.171 | -0.735 | -0.455 | 0.024 | 0.035 | 0.035 | 0 | 0.54 | 0.35 | 0.64 | 0.74 | 0.61 | 0.78 |
| 2 | 60760032 | rs2556376 | 0.252 | 0.273 | -0.796 | 0.042 | -0.005 | 0.016 | 0.016 | 0 | 0.79 | 0.73 | 0.77 | 0.73 | 0.82 | 0.84 |
| 2 | 60760637 | rs2556377 | 0.338 | 0.265 | -0.803 | 0.055 | -0.004 | 0.014 | 0.014 | 0 | 0.78 | 0.73 | 0.77 | 0.73 | 0.81 | 0.83 |
| 2 | 60764941 | rs10427250 | 0.334 | 0.244 | -0.898 | -0.054 | -0.002 | -0.018 | -0.018 | 0.014 | 0.08 | 0.06 | 0.03 | 0 | 0.07 | 0.03 |
| 2 | 60774773 | rs971563 | 0.626 | 0.193 | -0.857 | 0.008 | -0.003 | 0.012 | 0.012 | 0 | 0.79 | 0.74 | 0.77 | 0.74 | 0.82 | 0.84 |
| 2 | 60783572 | rs3765154 | 0.167 | 0.584 | -1.09 | 0.175 | 0.007 | 0.004 | 0.004 | 0.712 | 0.76 | 0.69 | 0.62 | 0.28 | 0.7 | 0.56 |
| 2 | 60784033 | rs17029711 | 0.415 | -0.491 | -0.934 | -0.504 | -0.064 | -0.047 | -0.047 | 0.692 | 0.25 | 0.32 | 0.26 | 0.11 | 0.12 | 0.1 |
| 2 | 60785937 | rs12328348 | 1.172 | 0.026 | -1.343 | -0.595 | -0.056 | 0.086 | 0.086 | 0 | 0.37 | 0.18 | 0.29 | 0.16 | 0.46 | 0.44 |
| 2 | 60789053 | rs13382983 | -0.468 | -0.026 | -1.187 | -0.573 | -0.032 | -0.058 | -0.058 | 0 | 0.37 | 0.5 | 0.32 | 0.1 | 0.24 | 0.12 |
| 2 | 60790985 | rs1510478 | 0.703 | 0.922 | -1.274 | 0.516 | -0.052 | -0.037 | -0.037 | 0.511 | 0.08 | 0 | 0.14 | 0.34 | 0.17 | 0.27 |
| 2 | 60797896 | rs2901066 | -0.144 | 0.32 | -0.918 | -0.595 | -0.029 | -0.047 | -0.047 | 0.037 | 0.21 | 0.47 | 0.3 | 0.1 | 0.17 | 0.11 |
| 2 | 60799121 | rs356982 | 0.715 | 0.831 | -1.564 | -0.929 | 0.009 | 0.037 | 0.037 | 0 | 0.45 | 0.39 | 0.45 | 0.76 | 0.4 | 0.41 |
| 2 | 60800641 | rs971742 | -0.757 | 1.074 | -1.405 | -0.841 | -0.002 | 0.029 | 0.029 | 0 | 0.44 | 0.6 | 0.46 | 0.24 | 0.45 | 0.51 |
| 2 | 60805397 | rs357004 | 0.168 | 1.27 | -0.88 | -0.967 | 0.003 | -0.022 | -0.022 | 0 | 0.6 | 0.42 | 0.45 | 0.88 | 0.46 | 0.77 |
| 2 | 60811584 | rs356999 | -0.532 | 1.469 | -0.765 | -0.865 | 0.01 | -0.02 | -0.02 | 0 | 0.72 | 0.66 | 0.67 | 0.89 | 0.63 | 0.85 |
| 2 | 60817254 | rs2098985 | 0.053 | 1.743 | -0.638 | -0.623 | 0.06 | 0.024 | 0.024 | 0 | 0.07 | 0.16 | 0.21 | 0.11 | 0.18 | 0.09 |
| 2 | 60822323 | rs356986 | 0.367 | 1.315 | -0.655 | -0.872 | -0.023 | 0.085 | 0.085 | 0 | 0.54 | 0.28 | 0.44 | 0.73 | 0.45 | 0.63 |
| 2 | 60832662 | rs2042799 | -0.08 | 0.838 | -0.346 | -0.953 | -0.008 | -0.029 | -0.029 | 0 | 0.61 | 0.51 | 0.65 | 0.83 | 0.63 | 0.71 |
| 2 | 60835999 | rs842944 | -0.064 | 0.378 | -0.043 | -1.742 | 0.004 | -0.043 | -0.043 | 0 | 0.59 | 0.39 | 0.75 | 0.98 | 0.62 | 0.84 |
| 2 | 60838156 | rs2059710 | -0.273 | 0.422 | 0.28 | -1.819 | -0.016 | -0.13 | -0.13 | 0 | 0.2 | 0.5 | 0.12 | 0.02 | 0.18 | 0.1 |
| 2 | 60842571 | rs735133 | -0.182 | 0.365 | 0.262 | -1.824 | -0.019 | -0.118 | -0.118 | 2.404 | 0.2 | 0.44 | 0.12 | 0.02 | 0.18 | 0.1 |
| 2 | 60845560 | rs842777 | -1.402 | 0.338 | 0.315 | -1.847 | -0.014 | -0.108 | -0.108 | 0 | 0.77 | 0.64 | 0.85 | 1 | 0.8 | 0.93 |
| 2 | 60846858 | rs4672396 | 0.976 | 0.492 | -0.759 | -1.889 | -0.005 | 0.013 | 0.013 | 0 | 0.28 | 0.18 | 0.32 | 0.17 | 0.27 | 0.17 |
| 2 | 60862874 | rs1467068 | -0.422 | 0.606 | -0.484 | -1.832 | 0.017 | -0.059 | -0.059 | 0 | 0.25 | 0.55 | 0.31 | 0.11 | 0.31 | 0.16 |
| 2 | 60863833 | rs6732653 | -0.445 | 0.597 | -0.489 | -1.836 | 0.016 | -0.06 | -0.06 | 1.167 | 0.25 | 0.55 | 0.31 | 0.11 | 0.31 | 0.17 |
| 2 | 60868614 | rs842767 | -0.128 | 0.578 | -0.559 | -1.882 | -0.002 | 0.053 | 0.053 | 0 | 0.54 | 0.32 | 0.56 | 0.89 | 0.48 | 0.77 |
| 2 | 60872851 | rs842764 | -0.214 | 0.315 | -0.634 | -1.819 | 0.008 | 0.091 | 0.091 | 1.141 | 0.58 | 0.28 | 0.6 | 0.9 | 0.59 | 0.78 |
| 2 | 60876626 | rs842762 | -2 | 0.346 | -0.654 | -1.786 | -0.001 | 0.065 | 0.065 | 0.417 | 0.79 | 0.92 | 0.87 | 1 | 0.79 | 0.94 |
| 2 | 60878133 | rs842760 | 0.572 | 0.223 | -0.74 | -1.491 | 0.006 | -0.007 | -0.007 | 2.028 | 0.28 | 0.21 | 0.25 | 0.69 | 0.24 | 0.6 |
| 2 | 60881975 | rs6705125 | 1.041 | 0.478 | -0.506 | -1.393 | 0.029 | 0.025 | 0.025 | 0 | 0.05 | 0.13 | 0.2 | 0.1 | 0.14 | 0.07 |
| 2 | 60888202 | rs881952 | 0.089 | 0.223 | -0.617 | -1.672 | -0.006 | 0.018 | 0.018 | 0 | 0.32 | 0.29 | 0.44 | 0.81 | 0.37 | 0.63 |
| 2 | 60893016 | rs17031423 | 0.675 | 0.072 | -0.546 | -1.911 | -0.022 | -0.123 | -0.123 | 0 | 0.31 | 0.39 | 0.18 | 0 | 0.24 | 0.16 |
| 2 | 60899462 | rs17537498 | 1.184 | 0.528 | -0.935 | -1.767 | -0.032 | 0.032 | 0.032 | 0.154 | 0.06 | 0.01 | 0.07 | 0 | 0.14 | 0.05 |
| 2 | 60902430 | rs934796 | 0.112 | 0.246 | -0.652 | -1.79 | 0 | 0.021 | 0.021 | 0 | 0.51 | 0.57 | 0.55 | 0.8 | 0.54 | 0.67 |
| 2 | 60905416 | rs4672400 | -0.511 | 0.431 | -0.795 | -1.718 | -0.052 | 0.115 | 0.115 | 0 | 0.65 | 0.88 | 0.62 | 0.85 | 0.62 | 0.78 |
| 2 | 60910033 | rs10198826 | -0.183 | 0.387 | -0.837 | -1.692 | -0.018 | 0.044 | 0.044 | 0 | 0.59 | 0.72 | 0.5 | 0.75 | 0.49 | 0.7 |
| 2 | 60914577 | rs6732794 | 0.001 | 0.41 | -0.975 | -1.655 | -0.006 | 0.041 | 0.041 | 0 | 0.56 | 0.67 | 0.44 | 0.75 | 0.48 | 0.63 |
| 2 | 60914648 | rs13006409 | 0.594 | 0.605 | -1.1 | -1.531 | -0.028 | -0.024 | -0.024 | 0 | 0.07 | 0.02 | 0.11 | 0.09 | 0.13 | 0.09 |
| 2 | 60914729 | rs17031887 | 0.485 | -0.037 | -0.871 | -1.824 | 0.017 | 0.004 | 0.004 | 0.35 | 0.28 | 0.16 | 0.13 | 0 | 0.21 | 0.06 |
| 2 | 60928367 | rs17537701 | 0.273 | 0.708 | -0.853 | -1.517 | -0.013 | 0.012 | 0.012 | 0.951 | 0.12 | 0.06 | 0.11 | 0 | 0.21 | 0.08 |
| 2 | 60931870 | rs7573815 | 0.056 | 0.241 | -0.942 | -1.49 | 0.017 | 0.028 | 0.028 | 0 | 0.62 | 0.75 | 0.79 | 0.88 | 0.68 | 0.7 |
| 2 | 60957929 | rs13420053 | 0.225 | 0.771 | -1.231 | -1.325 | -0.004 | 0.06 | 0.06 | 0 | 0.37 | 0.25 | 0.25 | 0.1 | 0.33 | 0.11 |
| 2 | 60957993 | rs13011212 | 0.9 | 0.68 | -1.4 | -1.568 | -0.001 | 0.008 | 0.008 | 0 | 0.21 | 0.17 | 0.15 | 0.04 | 0.22 | 0.12 |
| 2 | 60958989 | rs7562375 | 0.842 | 0.659 | -1.406 | -1.574 | 0.001 | 0.009 | 0.009 | 0 | 0.21 | 0.17 | 0.15 | 0.04 | 0.22 | 0.19 |
| 2 | 60968070 | rs7579240 | 0.636 | 0.672 | -1.434 | -1.574 | -0.018 | 0.097 | 0.097 | 0 | 0.36 | 0.15 | 0.23 | 0.1 | 0.33 | 0.11 |
| 2 | 60972786 | rs1866207 | -0.357 | 0.634 | -1.265 | -1.798 | -0.01 | -0.036 | -0.036 | 0 | 0.4 | 0.53 | 0.3 | 0.15 | 0.34 | 0.26 |
| 2 | 60975288 | rs13395150 | 1.156 | 0.603 | -1.283 | -1.781 | 0.003 | -0.007 | -0.007 | 0 | 0.15 | 0.1 | 0.12 | 0.1 | 0.12 | 0.07 |
| 2 | 60982796 | rs12471897 | -0.691 | 0.563 | -1.305 | -1.839 | 0.003 | -0.039 | -0.039 | 0 | 0.58 | 0.56 | 0.41 | 0.14 | 0.54 | 0.3 |
| 2 | 60984923 | rs10196947 | 0.934 | 0.56 | -1.228 | -1.855 | -0.005 | -0.061 | -0.061 | 12.601 | 0.24 | 0.33 | 0.17 | 0.04 | 0.22 | 0.13 |
| 2 | 60994945 | rs7599341 | -0.69 | 0.55 | -1.264 | -1.844 | 0.003 | -0.034 | -0.034 | 1.522 | 0.58 | 0.53 | 0.4 | 0.15 | 0.54 | 0.3 |

| | | | | | | | | | | | | | | | | |
|---|---|---|---|---|---|---|---|---|---|---|---|---|---|---|---|---|
| 2 | 61003193 | rs11687809 | 1.179 | 0.576 | -1.451 | -1.807 | 0.002 | -0.01 | -0.01 | 0.814 | 0.15 | 0.09 | 0.12 | 0.1 | 0.12 | 0.07 |
| 2 | 61007782 | rs10496089 | -1.38 | 0.654 | -1.386 | -1.286 | -0.003 | 0.023 | 0.023 | 0 | 0.73 | 0.72 | 0.68 | 0.57 | 0.71 | 0.8 |
| 2 | 61019699 | rs7600065 | 1.722 | 0.686 | -2.103 | -1.95 | -0.029 | 0.086 | 0.086 | 0.258 | 0.19 | 0.06 | 0.11 | 0 | 0.2 | 0.04 |
| 2 | 61027921 | rs11678813 | 0.714 | 0.579 | -1.941 | -1.66 | 0.006 | -0.007 | -0.007 | 0 | 0.16 | 0.11 | 0.12 | 0.1 | 0.11 | 0.14 |
| 2 | 61029363 | rs17479462 | 0.265 | 0.837 | -1.835 | -1.641 | 0 | -0.017 | -0.017 | 0 | 0.23 | 0.23 | 0.17 | 0.04 | 0.23 | 0.15 |
| 2 | 61041155 | rs17033754 | 0.446 | 0.639 | -1.803 | -1.563 | -0.005 | -0.037 | -0.037 | 0 | 0.21 | 0.29 | 0.11 | 0.04 | 0.18 | 0.1 |
| 2 | 61043834 | rs13034020 | 0.896 | 1.059 | -2.241 | -1.303 | -0.01 | 0.022 | 0.022 | 0.159 | 0.17 | 0.09 | 0.11 | 0 | 0.18 | 0.04 |
| 2 | 61059383 | rs6545832 | -0.062 | 0.912 | -2.565 | -1.575 | -0.075 | 0.159 | 0.159 | 0 | 0.62 | 0.85 | 0.68 | 0.98 | 0.47 | 0.86 |
| 2 | 61066383 | rs6735168 | 0.13 | 0.842 | -2.397 | -1.176 | 0.013 | 0.003 | 0.003 | 0 | 0.18 | 0.12 | 0.14 | 0.1 | 0.13 | 0.13 |
| 2 | 61066666 | rs1432295 | -0.307 | 0.816 | -2.695 | -1.492 | -0.069 | 0.084 | 0.084 | 0.263 | 0.73 | 0.87 | 0.74 | 0.99 | 0.57 | 0.9 |
| 2 | 61068451 | rs11125858 | -0.496 | 0.675 | -2.406 | -1.36 | -0.003 | 0 | 0 | 0.181 | 0.83 | 0.86 | 0.86 | 0.9 | 0.89 | 0.88 |
| 2 | 61071656 | rs4672410 | 0.631 | 1.05 | -2.571 | -1.518 | -0.028 | 0.098 | 0.098 | 0.035 | 0.61 | 0.84 | 0.66 | 0.88 | 0.55 | 0.81 |
| 2 | 61075189 | rs1177206 | -1.08 | 0.963 | -2.265 | -1.46 | -0.005 | 0.036 | 0.036 | 0 | 0.4 | 0.29 | 0.36 | 0.11 | 0.45 | 0.21 |
| 2 | 61077693 | rs1177210 | 0.597 | 0.947 | -2.386 | -1.494 | -0.015 | 0 | 0 | 0 | 0.15 | 0.13 | 0.11 | 0.1 | 0.09 | 0.11 |
| 2 | 61078350 | rs1177211 | -1.057 | 0.971 | -2.375 | -1.481 | -0.015 | 0.079 | 0.079 | 0.041 | 0.39 | 0.19 | 0.35 | 0.12 | 0.45 | 0.19 |
| 2 | 61087829 | rs2901182 | 0.365 | 1.468 | -2.589 | -1.525 | -0.055 | 0.042 | 0.042 | 0 | 0.1 | 0.01 | 0.13 | 0.01 | 0.18 | 0.04 |
| 2 | 61091222 | rs842634 | -1.207 | 1.451 | -3.194 | -0.99 | -0.095 | -0.137 | -0.137 | 0 | 0.82 | 0.98 | 0.72 | 0.55 | 0.75 | 0.37 |
| 2 | 61099420 | rs13019590 | 0.234 | 1.542 | -3.089 | -1.369 | -0.114 | 0.135 | 0.135 | 0.399 | 0.23 | 0.03 | 0.22 | 0.01 | 0.36 | 0.08 |
| 2 | 61118113 | rs12713428 | 0.934 | 0.878 | -3.21 | -1.663 | 0.043 | 0.088 | 0.088 | 0.051 | 0.35 | 0.2 | 0.17 | 0.01 | 0.25 | 0.11 |
| 2 | 61122317 | rs34023434 | 2.075 | 1.072 | -3.09 | -1.467 | -0.043 | 0.064 | 0.064 | 0 | 0.12 | 0.02 | 0.1 | 0 | 0.18 | 0.05 |
| 2 | 61179160 | rs10176601 | 0.78 | 0.643 | -2.916 | -1.387 | 0.034 | 0.033 | 0.033 | 0.203 | 0.36 | 0.24 | 0.18 | 0.01 | 0.25 | 0.12 |
| 2 | 61183657 | rs11691111 | -0.154 | 1.27 | -3.075 | -0.941 | -0.016 | -0.055 | -0.055 | 0.265 | 0.27 | 0.14 | 0.4 | 0.5 | 0.29 | 0.2 |
| 2 | 61186829 | rs13003464 | 0.071 | 0.932 | -3.412 | -1.612 | 0.057 | 0.359 | 0.359 | 1.524 | 0.51 | 0.91 | 0.31 | 0.04 | 0.36 | 0.17 |
| 2 | 61209875 | rs7562347 | -0.029 | 1.338 | -3.447 | -1.649 | -0.005 | 0.344 | 0.344 | 0.434 | 0.52 | 0.91 | 0.32 | 0.04 | 0.43 | 0.18 |
| 2 | 61217542 | rs10188217 | -0.283 | 1.423 | -3.3 | -1.565 | -0.101 | 0.288 | 0.288 | 0.125 | 0.57 | 0.93 | 0.38 | 0.05 | 0.5 | 0.19 |
| 2 | 61220015 | rs10182774 | 1.403 | 1.028 | -3.411 | -1.64 | 0.036 | 0.138 | 0.138 | 0.293 | 0.31 | 0.1 | 0.15 | 0.03 | 0.24 | 0.1 |
| 2 | 61223290 | rs717304 | 0.263 | 0.98 | -3.279 | -1.711 | -0.008 | 0.043 | 0.043 | 0 | 0.18 | 0.08 | 0.14 | 0.01 | 0.19 | 0.07 |
| 2 | 61223666 | rs10195800 | 1.218 | 0.909 | -3.139 | -1.505 | 0.036 | 0.138 | 0.138 | 0.325 | 0.31 | 0.1 | 0.15 | 0.02 | 0.23 | 0.1 |
| 2 | 61232705 | rs9309333 | 1.096 | 1.21 | -2.086 | -0.883 | 0.008 | 0.01 | 0.01 | 0 | 0.31 | 0.26 | 0.17 | 0.05 | 0.24 | 0.11 |
| 2 | 61233289 | rs2564118 | -0.786 | 1.466 | -2.329 | -0.37 | -0.077 | -0.101 | -0.101 | 0.134 | 0.62 | 0.93 | 0.62 | 0.49 | 0.55 | 0.29 |
| 2 | 61235855 | rs1182688 | -0.099 | 1.44 | -1.537 | -0.365 | -0.001 | 0 | 0 | 0 | 0.29 | 0.29 | 0.42 | 0.43 | 0.31 | 0.19 |
| 2 | 61240882 | rs12998880 | 1.953 | 1.668 | -1.293 | -0.525 | -0.019 | 0.014 | 0.014 | 0 | 0.12 | 0.05 | 0.12 | 0.03 | 0.2 | 0.04 |
| 2 | 61244663 | rs12988616 | 1.188 | 1.658 | -1.278 | -0.533 | 0.075 | -0.044 | -0.044 | 0.136 | 0.15 | 0.43 | 0.15 | 0.03 | 0.2 | 0.04 |
| 2 | 61249676 | rs12713431 | 1.645 | 1.597 | -1.421 | -0.59 | 0.017 | 0.112 | 0.112 | 0 | 0.3 | 0.1 | 0.16 | 0.06 | 0.24 | 0.11 |
| 2 | 61254033 | rs10496090 | 2.161 | 1.683 | -1.472 | -0.552 | -0.036 | 0.057 | 0.057 | 0 | 0.11 | 0.02 | 0.09 | 0 | 0.17 | 0.03 |
| 2 | 61254954 | rs6728872 | 0.293 | 1.641 | -1.36 | -0.561 | 0.006 | 0.156 | 0.156 | 1.213 | 0.46 | 0.76 | 0.33 | 0.08 | 0.44 | 0.15 |
| 2 | 61260063 | rs1177234 | -0.731 | 1.476 | -1.369 | -0.295 | -0.08 | -0.11 | -0.11 | 6.025 | 0.62 | 0.92 | 0.61 | 0.49 | 0.55 | 0.29 |
| 2 | 61268292 | rs2698189 | 1.05 | 1.492 | -1.142 | -0.311 | -0.033 | 0.055 | 0.055 | 0 | 0.1 | 0.01 | 0.08 | 0 | 0.1 | 0.05 |
| 2 | 61291174 | rs2698195 | 2.188 | 1.267 | -1.089 | -0.225 | -0.005 | 0.032 | 0.032 | 0 | 0.06 | 0 | 0.03 | 0 | 0.04 | 0.04 |
| 2 | 61304271 | rs1729659 | -0.28 | 1.458 | -1.01 | -0.202 | 0.002 | 0.012 | 0.012 | 0 | 0.3 | 0.39 | 0.43 | 0.42 | 0.32 | 0.18 |
| 2 | 61306754 | rs1177298 | -0.687 | 1.511 | -1.66 | -0.354 | -0.087 | -0.111 | -0.111 | 0 | 0.62 | 0.95 | 0.64 | 0.5 | 0.57 | 0.29 |
| 2 | 61307982 | rs12713433 | 1.26 | 1.419 | -1.443 | -0.376 | 0.024 | 0.046 | 0.046 | 0 | 0.29 | 0.16 | 0.14 | 0.06 | 0.2 | 0.11 |
| 2 | 61323840 | rs1177299 | -0.872 | 2.126 | -1.301 | 0.297 | -0.036 | -0.028 | -0.028 | 0 | 0.82 | 0.97 | 0.76 | 0.77 | 0.81 | 0.53 |
| 2 | 61330217 | rs1729672 | 0.865 | 1.462 | -1.143 | 0.542 | 0.061 | 0.035 | 0.035 | 0.107 | 0.4 | 0.13 | 0.26 | 0.34 | 0.29 | 0.16 |
| 2 | 61338832 | rs1177265 | 0.155 | 1.444 | -0.556 | 0.823 | -0.003 | -0.003 | -0.003 | 0 | 0.6 | 0.41 | 0.58 | 0.5 | 0.57 | 0.34 |
| 2 | 61346133 | rs1177290 | -0.151 | 1.48 | -0.541 | 0.866 | -0.009 | -0.008 | -0.008 | 0 | 0.63 | 0.68 | 0.6 | 0.51 | 0.58 | 0.34 |
| 2 | 61348804 | rs1177283 | -0.315 | 1.579 | -0.761 | 0.83 | -0.103 | -0.083 | -0.083 | 0 | 0.66 | 0.91 | 0.56 | 0.43 | 0.54 | 0.36 |
| 2 | 61370819 | rs11498 | 0.479 | 1.518 | -0.114 | 0.586 | 0.003 | 0.009 | 0.009 | 0.234 | 0.52 | 0.47 | 0.36 | 0.36 | 0.42 | 0.25 |
| 2 | 61376463 | rs720201 | 0.521 | 1.591 | -0.115 | 0.57 | 0.009 | 0.018 | 0.018 | 0.347 | 0.52 | 0.45 | 0.35 | 0.36 | 0.41 | 0.25 |
| 2 | 61379286 | rs1665272 | -0.575 | 1.732 | -0.135 | 0.764 | 0.029 | 0.022 | 0.022 | 0 | 0.72 | 0.65 | 0.62 | 0.64 | 0.58 | 0.61 |
| 2 | 61391305 | rs1729662 | -0.663 | 1.739 | -0.237 | 0.773 | -0.003 | -0.003 | -0.003 | 0.075 | 0.73 | 0.75 | 0.62 | 0.63 | 0.59 | 0.6 |
| 2 | 61405008 | rs746032 | 0.4 | 1.546 | -0.47 | 0.761 | -0.038 | -0.096 | -0.096 | 0 | 0.18 | 0.02 | 0.24 | 0.29 | 0.18 | 0.36 |
| 2 | 61405795 | rs2600665 | -0.828 | 1.868 | -0.392 | 0.833 | -0.002 | -0.002 | -0.002 | 0 | 0.73 | 0.74 | 0.66 | 0.68 | 0.66 | 0.63 |
| 2 | 61406438 | rs17006750 | 1.361 | 1.853 | -0.151 | 0.564 | 0.029 | -0.055 | -0.055 | 0.003 | 0.1 | 0.21 | 0.08 | 0.02 | 0.15 | 0.08 |
| 2 | 61407245 | rs6741734 | 0.118 | 1.788 | -0.106 | 0.616 | -0.004 | 0.009 | 0.009 | 0 | 0.2 | 0.25 | 0.25 | 0.28 | 0.17 | 0.36 |
| 2 | 61410346 | rs777584 | 0.624 | 1.698 | -0.268 | 0.644 | -0.001 | -0.003 | -0.003 | 0 | 0.52 | 0.5 | 0.35 | 0.33 | 0.42 | 0.24 |
| 2 | 61413723 | rs1809028 | -1.722 | 1.903 | -0.671 | 0.729 | -0.061 | -0.107 | -0.107 | 0 | 0.83 | 0.99 | 0.78 | 0.68 | 0.8 | 0.66 |
| 2 | 61435847 | rs4671397 | -1.667 | 1.204 | -0.895 | 0.372 | -0.006 | -0.081 | -0.081 | 0.303 | 0.84 | 0.98 | 0.85 | 0.67 | 0.88 | 0.92 |
| 2 | 61448493 | rs11889205 | 0.561 | 1.614 | -0.031 | 0.57 | 0.045 | -0.075 | -0.075 | 0.15 | 0.12 | 0.36 | 0.09 | 0.02 | 0.14 | 0.08 |
| 2 | 61448776 | rs2290324 | 1.552 | 1.705 | -0.01 | 0.633 | 0 | 0.006 | 0.006 | 0 | 0.18 | 0.16 | 0.23 | 0.33 | 0.2 | 0.35 |
| 2 | 61454010 | rs13388389 | 0.99 | 1.678 | -0.008 | 0.52 | 0.052 | -0.064 | -0.064 | 0.317 | 0.11 | 0.31 | 0.1 | 0.02 | 0.14 | 0.08 |
| 2 | 61461412 | rs7588041 | -1.206 | 1.627 | -0.104 | 0.459 | 0.033 | 0.029 | 0.029 | 0.953 | 0.57 | 0.75 | 0.67 | 0.62 | 0.66 | 0.81 |
| 2 | 61473148 | rs4671401 | 1.266 | 1.721 | 0.012 | 0.618 | 0.004 | 0.019 | 0.019 | 0 | 0.19 | 0.25 | 0.28 | 0.31 | 0.23 | 0.36 |
| 2 | 61482261 | rs10496091 | 0.274 | 1.641 | -0.04 | 0.21 | 0.063 | -0.119 | -0.119 | 0.403 | 0.19 | 0.36 | 0.19 | 0.02 | 0.29 | 0.1 |
| 2 | 61491514 | rs12474871 | 1.76 | 1.563 | -0.157 | 0.319 | 0.004 | -0.022 | -0.022 | 0 | 0.08 | 0.07 | 0.1 | 0 | 0.15 | 0.03 |
| 2 | 61500322 | rs7583383 | -0.45 | 1.446 | -0.221 | 0.37 | -0.002 | -0.003 | -0.003 | 0 | 0.39 | 0.46 | 0.49 | 0.6 | 0.37 | 0.71 |
| 2 | 61519408 | rs2167566 | 0.985 | 0.666 | 0.046 | 0.577 | 0.019 | 0.016 | 0.016 | 0 | 0.27 | 0.39 | 0.34 | 0.32 | 0.34 | 0.37 |
| 2 | 61524382 | rs11889702 | 0.957 | 0.663 | -0.041 | 0.409 | 0.008 | -0.031 | -0.031 | 0 | 0.1 | 0.15 | 0.07 | 0.02 | 0.14 | 0.06 |
| 2 | 61540642 | rs2694633 | -1.166 | 0.54 | -0.329 | 0.401 | 0.036 | 0.02 | 0.02 | 0.906 | 0.59 | 0.85 | 0.7 | 0.63 | 0.68 | 0.81 |

| | | | | | | | | | | | | | | | | |
|---|---|---|---|---|---|---|---|---|---|---|---|---|---|---|---|---|
| 2 | 61548857 | rs17482440 | -0.242 | 0.585 | -0.146 | 0.459 | -0.011 | 0.016 | 0.016 | 0.878 | 0.44 | 0.33 | 0.58 | 0.6 | 0.52 | 0.74 |
| 2 | 61566536 | rs17008405 | 0.985 | 0.603 | -0.047 | 0.431 | 0.023 | -0.052 | -0.052 | 0 | 0.1 | 0.21 | 0.08 | 0.02 | 0.14 | 0.08 |
| 2 | 61594498 | rs10177225 | -0.257 | 0.562 | -0.14 | 0.465 | -0.008 | 0.011 | 0.011 | 0.145 | 0.44 | 0.36 | 0.58 | 0.6 | 0.52 | 0.73 |
| 2 | 61596180 | rs2694642 | 1.276 | 0.589 | -0.156 | 0.576 | -0.005 | -0.003 | -0.003 | 0 | 0.25 | 0.24 | 0.33 | 0.31 | 0.34 | 0.38 |
| 2 | 61604429 | rs17008755 | 0.473 | 0.569 | -0.212 | 0.56 | -0.003 | -0.031 | -0.031 | 0 | 0.16 | 0.09 | 0.23 | 0.29 | 0.16 | 0.35 |
| 2 | 61638952 | rs4672441 | -0.927 | 0.358 | -0.103 | 0.435 | 0.039 | 0.026 | 0.026 | 0.688 | 0.55 | 0.8 | 0.66 | 0.63 | 0.64 | 0.81 |
| 2 | 61647901 | rs778155 | 1.247 | 0.48 | -0.194 | 0.518 | -0.004 | -0.002 | -0.002 | 0 | 0.26 | 0.24 | 0.45 | 0.33 | 0.35 | 0.38 |
| 2 | 61665336 | rs2592352 | -1.411 | 0.471 | -0.007 | 0.485 | 0.015 | 0.01 | 0.01 | 0 | 0.75 | 0.71 | 0.56 | 0.67 | 0.65 | 0.62 |
| 2 | 61669248 | rs2421203 | -1.212 | 0.457 | 0.028 | 0.479 | 0.026 | 0.017 | 0.017 | 0 | 0.73 | 0.64 | 0.56 | 0.67 | 0.65 | 0.62 |
| 2 | 61673394 | rs1186706 | -0.007 | 0.35 | -0.045 | 0.625 | 0.005 | -0.008 | -0.008 | 0 | 0.58 | 0.45 | 0.43 | 0.3 | 0.51 | 0.29 |
| 2 | 61684564 | rs778139 | 0.29 | 0.324 | 0.018 | 0.613 | -0.005 | 0.014 | 0.014 | 0.359 | 0.54 | 0.42 | 0.43 | 0.3 | 0.5 | 0.25 |
| 2 | 61703856 | rs4430924 | -1.537 | 0.429 | -0.376 | 0.866 | -0.018 | -0.018 | -0.018 | 0 | 0.75 | 0.82 | 0.56 | 0.63 | 0.65 | 0.6 |
| 2 | 61710573 | rs11125883 | 0.737 | 0.34 | -0.78 | 0.693 | 0.024 | -0.031 | -0.031 | 0.083 | 0.45 | 0.11 | 0.35 | 0.42 | 0.37 | 0.2 |
| 2 | 61731478 | rs6735330 | 0.726 | 0.564 | -0.362 | 0.755 | 0.038 | -0.071 | -0.071 | 0 | 0.12 | 0.29 | 0.1 | 0.02 | 0.15 | 0.06 |
| 2 | 61749690 | rs3732171 | -1.127 | 0.467 | -0.45 | 0.769 | 0.039 | 0.023 | 0.023 | 0.016 | 0.73 | 0.58 | 0.54 | 0.66 | 0.63 | 0.61 |
| 2 | 61757721 | rs10186325 | -1.108 | 0.483 | -0.441 | 0.803 | 0.037 | 0.046 | 0.046 | 0.041 | 0.72 | 0.57 | 0.54 | 0.57 | 0.63 | 0.6 |
| 2 | 61782047 | rs778757 | 0.242 | -0.082 | -0.782 | 0.735 | -0.025 | -0.015 | -0.015 | 0 | 0.54 | 0.16 | 0.32 | 0.4 | 0.43 | 0.51 |
| 2 | 61798829 | rs2441381 | -0.856 | 0.237 | -0.549 | 0.851 | 0.036 | 0.013 | 0.013 | 0 | 0.71 | 0.54 | 0.54 | 0.66 | 0.62 | 0.59 |
| 2 | 61803309 | rs17010833 | 1.397 | 0.315 | -0.617 | 1.012 | 0.003 | 0.014 | 0.014 | 0 | 0.05 | 0.01 | 0.12 | 0.28 | 0.06 | 0.04 |
| 2 | 61825142 | rs7574631 | -0.967 | 0.341 | -0.488 | 0.636 | 0.027 | 0.011 | 0.011 | 0 | 0.72 | 0.63 | 0.55 | 0.67 | 0.62 | 0.59 |
| 2 | 61839925 | rs13004371 | 0.51 | 0.388 | -0.844 | 0.474 | -0.038 | -0.015 | -0.015 | 0 | 0.14 | 0.02 | 0.09 | 0.08 | 0.18 | 0.13 |
| 2 | 61840018 | rs7594191 | -1.202 | 0.218 | -0.652 | 0.384 | 0.033 | 0.016 | 0.016 | 0 | 0.73 | 0.6 | 0.54 | 0.66 | 0.62 | 0.56 |
| 2 | 61841370 | rs6752673 | 0.711 | -0.255 | -0.709 | -0.495 | -0.032 | -0.137 | -0.137 | 0 | 0.25 | 0.41 | 0.12 | 0.02 | 0.15 | 0.07 |
| 2 | 61852247 | rs2421333 | -0.329 | 0.051 | -0.956 | -0.155 | 0 | 0 | 0 | 0.174 | 0.45 | 0.58 | 0.36 | 0.32 | 0.42 | 0.42 |
| 2 | 61857177 | rs748569 | -0.349 | 0.14 | -1.418 | -0.475 | 0.018 | 0.049 | 0.049 | 0.859 | 0.73 | 0.83 | 0.88 | 0.9 | 0.81 | 0.85 |
| 2 | 61860989 | rs10195376 | 0.394 | 0.656 | -1.232 | -0.551 | -0.005 | 0.002 | 0.002 | 0 | 0.68 | 0.68 | 0.77 | 0.88 | 0.61 | 0.73 |
| 2 | 61863427 | rs12151427 | -0.145 | 0.706 | -1.232 | -0.513 | 0 | -0.002 | -0.002 | 0 | 0.11 | 0.13 | 0.15 | 0.1 | 0.26 | 0.12 |
| 2 | 61864069 | rs4378794 | -0.24 | 0.286 | -1.141 | -0.879 | -0.004 | 0.015 | 0.015 | 1.271 | 0.55 | 0.6 | 0.34 | 0.18 | 0.5 | 0.27 |
| 2 | 61874015 | rs1345207 | 0.12 | -0.217 | -1.141 | -0.871 | 0.005 | 0.001 | 0.001 | 0.869 | 0.43 | 0.57 | 0.27 | 0.23 | 0.24 | 0.3 |
| 2 | 61888702 | rs4671408 | -0.119 | 0.436 | -1.079 | -0.465 | -0.005 | -0.001 | -0.001 | 0.018 | 0.36 | 0.37 | 0.51 | 0.54 | 0.48 | 0.4 |
| 2 | 61895745 | rs1476447 | 0.89 | 0.711 | -1.402 | -0.825 | -0.054 | -0.125 | -0.125 | 0.673 | 0.18 | 0.01 | 0.4 | 0.57 | 0.2 | 0.46 |
| 2 | 61898552 | rs4444511 | 0.633 | -0.296 | -0.571 | -1.864 | -0.025 | -0.16 | -0.16 | 0.023 | 0.33 | 0.57 | 0.15 | 0.01 | 0.2 | 0.11 |
| 2 | 61899141 | rs4233975 | -0.808 | -0.284 | -0.589 | -1.977 | -0.014 | -0.042 | -0.042 | 0.346 | 0.78 | 0.76 | 0.88 | 0.93 | 0.84 | 0.87 |
| 2 | 61919387 | rs13393399 | 0.751 | -0.249 | 0.005 | -1.873 | -0.018 | -0.13 | -0.13 | 0 | 0.32 | 0.46 | 0.14 | 0.01 | 0.2 | 0.12 |
| 2 | 61928100 | rs4672448 | -1.151 | -0.272 | 0.036 | -1.861 | -0.007 | -0.022 | -0.022 | 0 | 0.82 | 0.83 | 0.9 | 0.93 | 0.86 | 0.87 |
| 2 | 61929733 | rs10166986 | -0.172 | -0.175 | -0.001 | -2.007 | 0 | -0.021 | -0.021 | 0.283 | 0.46 | 0.55 | 0.31 | 0.11 | 0.38 | 0.23 |
| 2 | 61938725 | rs12468414 | 0.52 | -0.13 | -0.041 | -2.053 | -0.016 | 0.002 | 0.002 | 0 | 0.14 | 0.05 | 0.19 | 0.1 | 0.18 | 0.13 |
| 2 | 61941368 | rs10165566 | -0.387 | -0.174 | 0.08 | -2.1 | 0.002 | -0.004 | -0.004 | 0 | 0.46 | 0.59 | 0.33 | 0.11 | 0.38 | 0.25 |
| 2 | 61944755 | rs2178295 | -1.296 | 0.022 | 0.133 | -2.02 | -0.007 | -0.026 | -0.026 | 0.337 | 0.82 | 0.83 | 0.9 | 0.95 | 0.85 | 0.9 |
| 2 | 61949300 | rs10174761 | 1.337 | -0.15 | 0.159 | -2.069 | -0.001 | -0.051 | -0.051 | 0 | 0.23 | 0.27 | 0.1 | 0.01 | 0.14 | 0.14 |
| 2 | 61953674 | rs1186889 | -0.692 | -0.107 | -0.066 | -2.037 | 0.001 | -0.01 | -0.01 | 0 | 0.45 | 0.61 | 0.33 | 0.11 | 0.37 | 0.27 |
| 2 | 61962902 | rs1881612 | 0.984 | 0.174 | -0.177 | -1.925 | -0.078 | 0.103 | 0.103 | 0 | 0.36 | 0.11 | 0.58 | 0.84 | 0.51 | 0.61 |
| 2 | 61967109 | rs12479403 | 0.156 | 0.253 | 0.303 | -1.389 | 0.001 | -0.003 | -0.003 | 0 | 0.13 | 0.17 | 0.2 | 0.1 | 0.17 | 0.12 |
| 2 | 61971220 | rs6751214 | 0.429 | 0.248 | 0.438 | -1.434 | -0.002 | -0.057 | -0.057 | 0 | 0.26 | 0.32 | 0.1 | 0.01 | 0.15 | 0.14 |
| 2 | 61971253 | rs1186834 | 1.435 | 0.174 | 0.441 | -1.358 | -0.003 | -0.034 | -0.034 | 3.672 | 0.07 | 0.12 | 0.04 | 0 | 0.06 | 0.01 |
| 2 | 61975658 | rs1186839 | 0.904 | 0.164 | 0.389 | -1.364 | -0.008 | -0.042 | -0.042 | 6.783 | 0.08 | 0.17 | 0.04 | 0 | 0.06 | 0.01 |
| 2 | 61978303 | rs13389550 | -0.738 | 0.179 | 0.313 | -1.352 | 0.002 | 0.011 | 0.011 | 0.066 | 0.4 | 0.4 | 0.28 | 0.11 | 0.32 | 0.26 |
| 2 | 61985827 | rs1186859 | -1.167 | 0.146 | 0.146 | -1.264 | 0.004 | 0.014 | 0.014 | 0.878 | 0.47 | 0.61 | 0.33 | 0.13 | 0.38 | 0.27 |
| 2 | 61988050 | rs11681128 | 0.777 | 0.209 | -0.009 | -1.25 | 0.015 | 0.121 | 0.121 | 0 | 0.23 | 0.04 | 0.06 | 0.01 | 0.12 | 0.13 |
| 2 | 61999572 | rs6738532 | -0.817 | 0.299 | 0.033 | -1.207 | -0.001 | -0.009 | -0.009 | 0 | 0.4 | 0.43 | 0.29 | 0.13 | 0.32 | 0.26 |
| 2 | 62000627 | rs13413178 | 0.083 | 0.113 | -0.065 | -1.154 | -0.021 | -0.097 | -0.097 | 0 | 0.28 | 0.36 | 0.09 | 0.01 | 0.13 | 0.14 |
| 2 | 62014427 | rs13020758 | 0.894 | 0.666 | 0.16 | -0.381 | 0 | -0.034 | -0.034 | 0 | 0.06 | 0.16 | 0.04 | 0 | 0.06 | 0.01 |
| 2 | 62016451 | rs10865326 | -0.894 | 0.93 | 0.145 | -0.269 | 0.031 | 0.006 | 0.006 | 0.149 | 0.17 | 0.39 | 0.22 | 0.15 | 0.21 | 0.15 |
| 2 | 62026934 | rs1534644 | -0.594 | 0.829 | -0.263 | -0.207 | -0.007 | -0.009 | -0.009 | 0 | 0.14 | 0.1 | 0.19 | 0.15 | 0.17 | 0.15 |
| 2 | 62027085 | rs1534645 | -1.35 | 0.713 | 0.271 | -0.394 | -0.004 | -0.022 | -0.022 | 0 | 0.23 | 0.45 | 0.25 | 0.15 | 0.23 | 0.16 |
| 2 | 62036610 | rs11688210 | -0.782 | 0.718 | -0.1 | -0.62 | -0.006 | 0.004 | 0.004 | 0 | 0.21 | 0.17 | 0.25 | 0.16 | 0.25 | 0.2 |
| 2 | 62038088 | rs1534647 | 0.868 | 0.451 | -0.985 | -1.523 | 0.016 | 0.161 | 0.161 | 0 | 0.28 | 0.04 | 0.08 | 0.01 | 0.16 | 0.17 |
| 2 | 62038198 | rs1534648 | -1.235 | 0.192 | -0.229 | -1.629 | -0.029 | 0.065 | 0.065 | 1.59 | 0.6 | 0.74 | 0.4 | 0.17 | 0.47 | 0.38 |
| 2 | 62042152 | rs10191166 | -0.867 | 0.067 | 0.112 | -1.522 | -0.006 | -0.027 | -0.027 | 0 | 0.3 | 0.42 | 0.3 | 0.16 | 0.31 | 0.21 |
| 2 | 62044253 | rs7589969 | -0.715 | 0.287 | -0.061 | -1.459 | 0.001 | -0.003 | -0.003 | 0.078 | 0.5 | 0.54 | 0.3 | 0.16 | 0.39 | 0.37 |
| 2 | 62057317 | rs11125894 | -0.024 | 0.193 | -0.861 | -1.464 | -0.002 | 0 | 0 | 0.038 | 0.2 | 0.19 | 0.19 | 0.15 | 0.2 | 0.22 |
| 2 | 62065759 | rs6545910 | 1.849 | 0.228 | -1.188 | -0.964 | -0.024 | -0.045 | -0.045 | 0 | 0.07 | 0.01 | 0.21 | 0.17 | 0.1 | 0.07 |
| 2 | 62067433 | rs17513722 | 1.762 | 0.405 | -1.424 | -1.472 | -0.024 | 0.054 | 0.054 | 0 | 0.22 | 0.04 | 0.14 | 0.09 | 0.22 | 0.12 |
| 2 | 62069358 | rs11125895 | 0.491 | 0.564 | -1.3 | -1.407 | -0.006 | -0.006 | -0.006 | 1.059 | 0.16 | 0.11 | 0.18 | 0.15 | 0.17 | 0.21 |
| 2 | 62069528 | rs11125896 | 0.112 | 0.566 | -1.318 | -1.386 | -0.002 | 0 | 0 | 0.776 | 0.21 | 0.19 | 0.19 | 0.15 | 0.21 | 0.22 |
| 2 | 62079267 | rs7569467 | 1.191 | 0.511 | -1.258 | -1.388 | 0.006 | -0.026 | -0.026 | 0 | 0.28 | 0.24 | 0.09 | 0.01 | 0.17 | 0.16 |
| 2 | 62079895 | rs11125898 | -0.004 | 0.521 | -1.278 | -1.378 | -0.003 | 0.002 | 0.002 | 0.445 | 0.22 | 0.19 | 0.21 | 0.15 | 0.23 | 0.22 |
| 2 | 62089938 | rs7581802 | 1.515 | 0.683 | -1.207 | -0.782 | -0.069 | -0.084 | -0.084 | 0 | 0.14 | 0.02 | 0.24 | 0.34 | 0.24 | 0.31 |
| 2 | 62101344 | rs2901428 | -0.508 | 0.6 | -0.733 | -0.64 | 0.001 | 0.003 | 0.003 | 0 | 0.49 | 0.47 | 0.29 | 0.16 | 0.4 | 0.37 |

| | | | | | | | | | | | | | | | | |
|---|---|---|---|---|---|---|---|---|---|---|---|---|---|---|---|---|
| 2 | 62120665 | rs7558608 | -0.221 | 0.752 | -0.633 | -0.282 | -0.004 | 0.009 | 0.009 | 0 | 0.34 | 0.25 | 0.21 | 0.21 | 0.33 | 0.36 |
| 2 | 62131773 | rs7606632 | -0.047 | 1.387 | 0.074 | 0.108 | 0.083 | 0.146 | 0.146 | 0.223 | 0.05 | 0.38 | 0.11 | 0.07 | 0.02 | 0.05 |
| 2 | 62178266 | rs6755447 | -0.475 | 1.365 | 0.191 | 0.097 | 0.068 | 0.131 | 0.131 | 1.028 | 0.06 | 0.43 | 0.11 | 0.07 | 0.02 | 0.05 |
| 2 | 62179264 | rs7607266 | 1.258 | 1.435 | -0.004 | 0.13 | -0.006 | -0.062 | -0.062 | 0 | 0.23 | 0.09 | 0.33 | 0.39 | 0.27 | 0.28 |
| 2 | 62193438 | rs11677744 | -1.756 | 1.93 | 1.133 | 0.615 | 0.005 | 0.061 | 0.061 | 0.338 | 0.19 | 0.6 | 0.25 | 0.29 | 0.18 | 0.27 |
| 2 | 62197878 | rs6755858 | -1.234 | 1.823 | 1.131 | 0.568 | -0.002 | 0.011 | 0.011 | 2.741 | 0.93 | 0.97 | 0.95 | 1 | 0.9 | 0.99 |
| 2 | 62220914 | rs11695621 | -0.434 | 1.689 | 1.173 | 0.58 | 0.084 | 0.149 | 0.149 | 0.119 | 0.05 | 0.38 | 0.11 | 0.07 | 0.02 | 0.05 |
| 2 | 62230970 | rs7574498 | 0.376 | 1.643 | 1.17 | 0.626 | 0.009 | -0.045 | -0.045 | 0 | 0.16 | 0.19 | 0.07 | 0.01 | 0.16 | 0.09 |
| 2 | 62247451 | rs7571389 | -0.382 | 1.804 | 1.108 | 1.045 | 0.007 | 0.021 | 0.021 | 0 | 0.11 | 0.13 | 0.21 | 0.21 | 0.15 | 0.21 |
| 2 | 62249619 | rs10084368 | -0.757 | 1.803 | 1.107 | 1.034 | 0.076 | 0.139 | 0.139 | 0.292 | 0.06 | 0.41 | 0.1 | 0.07 | 0.02 | 0.05 |
| 2 | 62276973 | rs10518958 | 0.674 | 1.962 | 0.922 | 1.058 | -0.033 | 0.038 | 0.038 | 0 | 0.1 | 0.02 | 0.04 | 0.01 | 0.12 | 0.08 |
| 2 | 62298745 | rs6743108 | -2.094 | 1.113 | 1.432 | 1.314 | 0.005 | 0.005 | 0.005 | 1.483 | 0.27 | 0.63 | 0.35 | 0.27 | 0.3 | 0.27 |
| 2 | 62317646 | rs4123561 | -0.566 | 0.891 | 1.065 | 1.037 | 0.01 | 0.027 | 0.027 | 0 | 0.11 | 0.12 | 0.24 | 0.23 | 0.16 | 0.22 |
| 2 | 62321492 | rs11674110 | -0.638 | 0.897 | 1.024 | 1.039 | 0.003 | 0.011 | 0.011 | 0 | 0.11 | 0.07 | 0.24 | 0.23 | 0.17 | 0.23 |
| 2 | 62349503 | rs11125908 | -0.558 | 0.683 | 1.015 | 1.069 | -0.073 | -0.046 | -0.046 | 0 | 0.2 | 0.04 | 0.32 | 0.24 | 0.33 | 0.26 |
| 2 | 62366688 | rs733517 | -0.699 | 0.7 | 0.714 | 1.054 | -0.016 | -0.004 | -0.004 | 0 | 0.22 | 0.16 | 0.33 | 0.23 | 0.33 | 0.26 |
| 2 | 62381474 | rs3943540 | -0.942 | 0.739 | 0.645 | 1.102 | 0.002 | -0.003 | -0.003 | 0.001 | 0.28 | 0.65 | 0.38 | 0.23 | 0.33 | 0.29 |
| 2 | 62430208 | rs1008725 | 0.155 | 0.254 | -0.397 | 0.351 | 0 | -0.001 | -0.001 | 0.193 | 0.61 | 0.61 | 0.59 | 0.7 | 0.68 | 0.63 |
| 2 | 62432133 | rs7605321 | 0.346 | -0.083 | -0.596 | 0.187 | 0.003 | 0.002 | 0.002 | 0.408 | 0.57 | 0.58 | 0.59 | 0.64 | 0.66 | 0.7 |
| 2 | 62438681 | rs4671416 | -0.578 | -0.828 | -0.368 | -0.172 | 0.004 | 0.004 | 0.004 | 0 | 0.35 | 0.66 | 0.45 | 0.26 | 0.28 | 0.29 |
| 2 | 62442438 | rs7570305 | -0.808 | -0.971 | -0.346 | -0.253 | -0.008 | 0.03 | 0.03 | 0.593 | 0.18 | 0.57 | 0.29 | 0.2 | 0.14 | 0.19 |
| 2 | 62446914 | rs10182572 | -0.858 | -0.547 | -0.503 | -0.384 | -0.012 | 0.026 | 0.026 | 0.776 | 0.18 | 0.6 | 0.29 | 0.22 | 0.14 | 0.2 |
| 2 | 62448399 | rs4672482 | -1.285 | -0.228 | -1.36 | -0.651 | -0.08 | -0.067 | -0.067 | 0 | 0.62 | 0.88 | 0.66 | 0.44 | 0.51 | 0.64 |
| 2 | 62448855 | rs10189468 | -0.838 | 0.021 | -0.923 | -0.562 | -0.011 | 0.021 | 0.021 | 0.705 | 0.19 | 0.61 | 0.29 | 0.21 | 0.14 | 0.22 |
| 2 | 62450631 | rs3811616 | 0.135 | -0.027 | -0.586 | -0.597 | 0.022 | 0.054 | 0.054 | 0 | 0.09 | 0.13 | 0.25 | 0.24 | 0.13 | 0.22 |
| 2 | 62450743 | rs1136151 | -1.221 | -0.045 | -0.534 | -0.606 | -0.006 | 0.005 | 0.005 | 1.358 | 0.2 | 0.72 | 0.31 | 0.25 | 0.14 | 0.23 |
| 2 | 62451422 | rs13393608 | -0.077 | 0.42 | -0.8 | -0.389 | 0.02 | 0.052 | 0.052 | 0 | 0.09 | 0.13 | 0.25 | 0.24 | 0.13 | 0.22 |
| 2 | 62453884 | rs4319920 | 0.239 | 0.2 | -1.074 | -0.758 | -0.009 | 0.045 | 0.045 | 1.024 | 0.2 | 0.09 | 0.08 | 0.01 | 0.19 | 0.11 |
| 2 | 62474984 | rs7584792 | -0.496 | 0.185 | -0.83 | -0.025 | 0 | 0.001 | 0.001 | 0.074 | 0.43 | 0.54 | 0.45 | 0.55 | 0.41 | 0.49 |
| 2 | 62475676 | rs13397746 | 0.372 | 0.638 | -0.443 | -0.467 | -0.009 | -0.012 | -0.012 | 1.18 | 0.15 | 0.14 | 0.13 | 0 | 0.21 | 0.12 |
| 2 | 62481483 | rs7584033 | -0.142 | 0.56 | -0.341 | -0.498 | 0.048 | -0.133 | -0.133 | 0 | 0.18 | 0.44 | 0.15 | 0 | 0.21 | 0.11 |
| 2 | 62491643 | rs11689568 | -0.05 | -0.022 | -1.189 | -0.346 | 0.014 | 0.068 | 0.068 | 0.083 | 0.36 | 0.17 | 0.3 | 0.18 | 0.31 | 0.31 |
| 2 | 62493960 | rs12713456 | -0.8 | -0.358 | -1.067 | -0.646 | 0.048 | -0.072 | -0.072 | 0 | 0.7 | 0.49 | 0.71 | 0.89 | 0.56 | 0.71 |
| 2 | 62495457 | rs10193629 | 1.395 | -1.007 | -1.289 | -0.915 | 0.033 | 0.024 | 0.024 | 5.758 | 0.15 | 0.1 | 0.05 | 0 | 0.07 | 0.02 |
| 2 | 62496723 | rs12620241 | 0.175 | -1.495 | -0.799 | -0.26 | -0.01 | 0.011 | 0.011 | 0 | 0.17 | 0.22 | 0.16 | 0.19 | 0.14 | 0.11 |
| 2 | 62497170 | rs10200191 | -0.121 | -1.449 | -0.63 | -0.318 | -0.009 | -0.011 | -0.011 | 0 | 0.55 | 0.52 | 0.56 | 0.68 | 0.43 | 0.62 |
| 2 | 62502387 | rs11676235 | 0.445 | -1.557 | -0.755 | -0.354 | 0.008 | 0.01 | 0.01 | 2.774 | 0.39 | 0.32 | 0.51 | 0.69 | 0.35 | 0.6 |
| 2 | 62503017 | rs4563212 | 0.309 | -1.579 | -0.841 | -0.143 | 0.018 | -0.04 | -0.04 | 0 | 0.16 | 0.08 | 0.24 | 0.22 | 0.14 | 0.12 |
| 2 | 62506598 | rs7599801 | 0.179 | -2 | -0.956 | -0.029 | -0.059 | 0.024 | 0.024 | 0.692 | 0.25 | 0.39 | 0.12 | 0.37 | 0.13 | 0.2 |
| 2 | 62516544 | rs6713082 | 1.611 | -1.954 | -1.461 | -0.433 | -0.041 | 0.128 | 0.128 | 0 | 0.21 | 0.01 | 0.23 | 0.02 | 0.22 | 0.12 |
| 2 | 62521244 | rs4672495 | -1.128 | -1.919 | -1.434 | -0.202 | -0.075 | 0.003 | 0.003 | 0.023 | 0.72 | 0.98 | 0.69 | 0.82 | 0.68 | 0.76 |
| 2 | 62523878 | rs4672498 | -1.089 | -1.684 | -1.196 | -0.391 | 0.015 | 0 | 0 | 2.13 | 0.38 | 0.73 | 0.26 | 0.65 | 0.28 | 0.51 |
| 2 | 62525703 | rs7578484 | 1.154 | -1.049 | -0.726 | -0.423 | 0.015 | -0.041 | -0.041 | 0 | 0.24 | 0.24 | 0.2 | 0.04 | 0.33 | 0.28 |
| 2 | 62533045 | rs3924500 | 0.093 | -1.504 | -0.525 | 0.019 | -0.109 | 0.038 | 0.038 | 1.116 | 0.3 | 0.46 | 0.12 | 0.45 | 0.12 | 0.29 |
| 2 | 62541900 | rs10195033 | -0.549 | -0.966 | -0.737 | 0.066 | 0.004 | 0.025 | 0.025 | 0 | 0.65 | 0.82 | 0.48 | 0.76 | 0.63 | 0.63 |
| 2 | 62542667 | rs4672501 | 0.048 | -0.972 | -0.421 | 0.5 | -0.06 | 0.11 | 0.11 | 0.337 | 0.25 | 0.49 | 0.16 | 0.59 | 0.15 | 0.21 |
| 2 | 62549284 | rs4672503 | 0.363 | -0.566 | -0.341 | 0.556 | -0.011 | 0 | 0 | 0 | 0.39 | 0.46 | 0.33 | 0.44 | 0.28 | 0.28 |
| 2 | 62549902 | rs9309343 | -0.132 | -0.596 | -0.292 | 0.532 | 0.015 | 0.006 | 0.006 | 0 | 0.48 | 0.63 | 0.38 | 0.62 | 0.36 | 0.49 |
| 2 | 62551472 | rs10865331 | -0.459 | -0.274 | -0.065 | 0.413 | 0.022 | 0.009 | 0.009 | 0 | 0.51 | 0.46 | 0.61 | 0.54 | 0.62 | 0.56 |
| 2 | 62552779 | rs12475982 | 1.599 | 0.474 | -0.495 | 0.053 | -0.07 | 0.063 | 0.063 | 0.148 | 0.15 | 0.03 | 0.14 | 0 | 0.26 | 0.07 |
| 2 | 62562310 | rs12612759 | -0.471 | -0.328 | -0.248 | 0.332 | -0.018 | 0.026 | 0.026 | 0 | 0.27 | 0.37 | 0.3 | 0.36 | 0.16 | 0.39 |
| 2 | 62571117 | rs11681733 | 1.135 | -0.153 | -0.639 | 0.236 | -0.128 | 0.092 | 0.092 | 0 | 0.21 | 0.03 | 0.24 | 0.03 | 0.41 | 0.2 |
| 2 | 62576006 | rs4470332 | -0.054 | -0.726 | -0.484 | 0.43 | 0.017 | -0.007 | -0.007 | 0 | 0.16 | 0.13 | 0.05 | 0.16 | 0.05 | 0.2 |
| 2 | 62581284 | rs10865332 | -0.93 | -0.788 | -0.259 | 0.287 | -0.002 | -0.001 | -0.001 | 0 | 0.5 | 0.5 | 0.55 | 0.45 | 0.56 | 0.58 |
| 2 | 62593531 | rs11901283 | -0.293 | -1.125 | -0.133 | 0.567 | -0.034 | 0.011 | 0.011 | 0 | 0.15 | 0.27 | 0.07 | 0.12 | 0.07 | 0.25 |
| 2 | 62593632 | rs7603526 | -0.319 | -1.069 | 0.083 | 0.573 | -0.01 | 0.05 | 0.05 | 0 | 0.14 | 0.19 | 0.21 | 0.29 | 0.09 | 0.11 |
| 2 | 62599559 | rs4671421 | -0.747 | -1.111 | 0.107 | 0.578 | -0.005 | 0.004 | 0.004 | 0.145 | 0.21 | 0.2 | 0.22 | 0.29 | 0.11 | 0.28 |
| 2 | 62602378 | rs6755155 | -1.616 | -0.982 | 0.052 | 0.502 | -0.028 | 0.031 | 0.031 | 0 | 0.34 | 0.49 | 0.4 | 0.56 | 0.22 | 0.49 |
| 2 | 62608855 | rs13027346 | 0.006 | -0.881 | -0.226 | 0.43 | 0.009 | 0.015 | 0.015 | 0.188 | 0.15 | 0.04 | 0.13 | 0.15 | 0.11 | 0.3 |
| 2 | 62608898 | rs4328619 | 0.802 | -0.893 | -0.381 | 0.041 | -0.138 | 0.15 | 0.15 | 0 | 0.28 | 0.06 | 0.27 | 0.03 | 0.48 | 0.25 |
| 2 | 62609378 | rs11125916 | -0.187 | -0.886 | -0.338 | 0.233 | 0.02 | -0.037 | -0.037 | 0.015 | 0.27 | 0.17 | 0.33 | 0.51 | 0.18 | 0.43 |
| 2 | 62611707 | rs6545930 | -1.411 | -0.956 | -0.087 | 0.151 | 0.02 | -0.001 | -0.001 | 0 | 0.64 | 0.66 | 0.67 | 0.66 | 0.7 | 0.77 |
| 2 | 62612045 | rs6713620 | -0.982 | -0.742 | -0.117 | 0.537 | -0.033 | 0.016 | 0.016 | 0 | 0.34 | 0.4 | 0.38 | 0.63 | 0.22 | 0.52 |
| 2 | 62617142 | rs6743832 | -0.561 | -0.668 | -0.48 | 0.574 | 0 | -0.001 | -0.001 | 0 | 0.28 | 0.25 | 0.34 | 0.52 | 0.18 | 0.44 |
| 2 | 62629315 | rs6742894 | -0.086 | -0.204 | -0.392 | 0.079 | 0.003 | -0.021 | -0.021 | 1.681 | 0.36 | 0.6 | 0.33 | 0.14 | 0.52 | 0.33 |
| 2 | 62635367 | rs6717002 | 0.257 | -0.315 | -0.558 | -0.291 | -0.001 | -0.029 | -0.029 | 0.301 | 0.53 | 0.52 | 0.5 | 0.13 | 0.41 | 0.28 |
| 2 | 62637252 | rs4672510 | 0.454 | 0.475 | -0.948 | 0.67 | -0.034 | -0.052 | -0.052 | 0 | 0.84 | 0.92 | 0.79 | 0.63 | 0.77 | 0.79 |
| 2 | 62643510 | rs4672511 | 0.32 | -0.105 | -1.034 | 0.582 | 0.001 | 0.009 | 0.009 | 0 | 0.88 | 0.93 | 0.86 | 0.89 | 0.88 | 0.82 |
| 2 | 62644214 | rs6545935 | 0.008 | -0.568 | -0.919 | 0.493 | -0.005 | 0.001 | 0.001 | 0 | 0.42 | 0.33 | 0.54 | 0.63 | 0.53 | 0.46 |

| | | | | | | | | | | | | | | | | |
|---|---|---|---|---|---|---|---|---|---|---|---|---|---|---|---|---|
| 2 | 62662704 | rs13412212 | -0.094 | -0.731 | -0.912 | 0.353 | -0.016 | 0 | 0 | 0 | 0.41 | 0.65 | 0.44 | 0.44 | 0.45 | 0.39 |
| 2 | 62666178 | rs9808571 | 0.962 | -0.558 | -0.727 | 0.203 | -0.024 | 0.019 | 0.019 | 0 | 0.22 | 0.17 | 0.23 | 0.02 | 0.32 | 0.17 |
| 2 | 62666220 | rs4233976 | 0.106 | -0.637 | -0.196 | 0.179 | 0.066 | 0.052 | 0.052 | 0.101 | 0.29 | 0.48 | 0.39 | 0.44 | 0.4 | 0.36 |
| 2 | 62667392 | rs12464504 | 1.375 | -1.433 | -0.942 | 0.21 | 0.036 | -0.058 | -0.058 | 0 | 0.16 | 0.05 | 0.16 | 0.26 | 0.11 | 0.09 |
| 2 | 62673967 | rs12475689 | 1.188 | -1.387 | -1.438 | -0.069 | -0.109 | 0.049 | 0.049 | 0 | 0.22 | 0.05 | 0.24 | 0.06 | 0.36 | 0.32 |
| 2 | 62676906 | rs6727349 | -0.217 | -1.764 | -1.331 | -0.124 | 0.025 | 0.003 | 0.003 | 0.191 | 0.36 | 0.72 | 0.38 | 0.67 | 0.24 | 0.32 |
| 2 | 62679582 | rs6723856 | 0.33 | -1.547 | -1.71 | -0.319 | 0.051 | 0.03 | 0.03 | 0 | 0.52 | 0.31 | 0.45 | 0.37 | 0.42 | 0.27 |
| 2 | 62680153 | rs10205398 | -0.495 | -1.033 | -1.787 | -0.285 | -0.066 | -0.07 | -0.07 | 0.01 | 0.86 | 0.99 | 0.86 | 0.73 | 0.8 | 0.77 |
| 2 | 62680333 | rs10205621 | -0.876 | -1.026 | -1.691 | -0.812 | 0.004 | 0.001 | 0.001 | 0 | 0.71 | 0.71 | 0.78 | 0.93 | 0.72 | 0.81 |
| 2 | 62681169 | rs2067864 | 1.28 | -0.229 | -1.932 | -0.642 | -0.058 | 0.029 | 0.029 | 0 | 0.21 | 0.1 | 0.24 | 0 | 0.38 | 0.17 |
| 2 | 62686776 | rs4672524 | 0.002 | -1.056 | -1.532 | -0.177 | -0.01 | -0.005 | -0.005 | 0 | 0.39 | 0.36 | 0.42 | 0.6 | 0.47 | 0.55 |
| 2 | 62687380 | rs11693346 | -0.061 | -1.216 | -1.446 | -0.297 | -0.009 | -0.007 | -0.007 | 0 | 0.62 | 0.46 | 0.71 | 0.73 | 0.69 | 0.7 |
| 2 | 62693830 | rs4413143 | 0.633 | -0.535 | -1.583 | -0.483 | -0.035 | 0.071 | 0.071 | 0 | 0.41 | 0.21 | 0.49 | 0.68 | 0.45 | 0.46 |
| 2 | 62694693 | rs3924400 | -0.219 | 0.008 | -1.063 | -0.035 | 0 | -0.001 | -0.001 | 0 | 0.84 | 0.8 | 0.84 | 0.76 | 0.78 | 0.78 |
| 2 | 62697989 | rs11689888 | 0.054 | -0.076 | -1.296 | -0.181 | -0.033 | -0.076 | -0.076 | 1.097 | 0.17 | 0.26 | 0.06 | 0 | 0.08 | 0.17 |
| 2 | 62699790 | rs4350731 | 0.582 | 0.081 | -1.421 | -0.071 | -0.004 | -0.017 | -0.017 | 0 | 0.67 | 0.61 | 0.78 | 0.76 | 0.7 | 0.62 |
| 2 | 62703006 | rs11686508 | -0.294 | 0.406 | -2.022 | 0.454 | -0.073 | -0.012 | -0.012 | 0 | 0.39 | 0.1 | 0.33 | 0.36 | 0.41 | 0.5 |
| 2 | 62712386 | rs12616584 | -0.414 | 0.819 | -1.585 | 0.896 | -0.025 | -0.032 | -0.032 | 0 | 0.05 | 0.01 | 0.06 | 0.15 | 0.06 | 0.12 |
| 2 | 62712483 | rs10183370 | -0.821 | 0.844 | -0.992 | 0.57 | 0.012 | -0.035 | -0.035 | 0.002 | 0.07 | 0.13 | 0.04 | 0.01 | 0.07 | 0.09 |
| 2 | 62713533 | rs6545946 | -0.61 | 0.705 | -0.616 | 0.626 | -0.008 | -0.01 | -0.01 | 0 | 0.23 | 0.34 | 0.24 | 0.17 | 0.22 | 0.18 |
| 2 | 62715566 | rs11690138 | 0.995 | 0.813 | -0.732 | 0.69 | -0.006 | -0.024 | -0.024 | 0 | 0.13 | 0.05 | 0.18 | 0.16 | 0.14 | 0.08 |
| 2 | 62719582 | rs12989129 | 1.032 | 0.814 | -0.692 | 0.691 | -0.002 | -0.011 | -0.011 | 0 | 0.13 | 0.08 | 0.18 | 0.16 | 0.14 | 0.08 |
| 2 | 62720684 | rs6727061 | 0.183 | 1.273 | -0.574 | 0.829 | -0.005 | -0.005 | -0.005 | 0 | 0.76 | 0.78 | 0.62 | 0.57 | 0.65 | 0.77 |
| 2 | 62722292 | rs10211381 | 0.66 | 0.968 | -0.534 | 0.684 | -0.006 | -0.006 | -0.006 | 0 | 0.56 | 0.56 | 0.42 | 0.4 | 0.44 | 0.6 |
| 2 | 62724752 | rs10200802 | -0.702 | 0.555 | -0.793 | 0.42 | -0.001 | 0.078 | 0.078 | 0 | 0.49 | 0.4 | 0.54 | 0.72 | 0.44 | 0.51 |
| 2 | 62727154 | rs7582882 | 0.907 | 0.644 | -0.945 | 0.787 | 0.013 | -0.025 | -0.025 | 0 | 0.18 | 0.13 | 0.19 | 0.31 | 0.13 | 0.17 |
| 2 | 62727902 | rs4233979 | 0.378 | 0.822 | -1.011 | 1.017 | 0.013 | -0.047 | -0.047 | 0 | 0.23 | 0.16 | 0.29 | 0.45 | 0.19 | 0.24 |
| 2 | 62728105 | rs13396048 | 0.92 | 0.807 | -0.971 | 0.944 | 0.013 | -0.025 | -0.025 | 0.851 | 0.18 | 0.13 | 0.19 | 0.31 | 0.13 | 0.17 |
| 2 | 62739979 | rs7570024 | 1.072 | 0.9 | -0.533 | 0.508 | 0.002 | 0.02 | 0.02 | 0 | 0.32 | 0.19 | 0.2 | 0.17 | 0.34 | 0.4 |
| 2 | 62751619 | rs6545950 | -0.322 | 0.686 | -0.468 | 0.77 | 0.02 | -0.024 | -0.024 | 0 | 0.66 | 0.71 | 0.64 | 0.58 | 0.73 | 0.75 |
| 2 | 62752921 | rs7424968 | 0.491 | 0.693 | -0.289 | 0.633 | 0.003 | 0.004 | 0.004 | 0 | 0.5 | 0.51 | 0.53 | 0.59 | 0.57 | 0.7 |
| 2 | 62754382 | rs12471388 | -0.97 | 0.244 | -0.469 | 0.822 | 0 | 0 | 0 | 0 | 0.8 | 0.79 | 0.8 | 0.65 | 0.87 | 0.82 |
| 2 | 62756049 | rs6726083 | -0.87 | 0.233 | -0.346 | 0.816 | -0.019 | 0.025 | 0.025 | 0 | 0.79 | 0.73 | 0.75 | 0.65 | 0.88 | 0.82 |
| 2 | 62757393 | rs10496097 | 1.594 | 0.322 | -0.512 | 0.71 | 0.003 | 0.02 | 0.02 | 0.207 | 0.15 | 0.06 | 0.16 | 0.09 | 0.13 | 0.08 |
| 2 | 62758283 | rs17575420 | 0.743 | 0.551 | -0.552 | 0.618 | -0.022 | 0.043 | 0.043 | 0 | 0.12 | 0.04 | 0.08 | 0 | 0.13 | 0.05 |
| 2 | 62758933 | rs6753189 | 0.067 | 0.516 | -0.433 | 0.693 | 0.003 | -0.003 | -0.003 | 0 | 0.68 | 0.68 | 0.71 | 0.66 | 0.75 | 0.78 |
| 2 | 62761855 | rs6750204 | -0.734 | 0.486 | -0.368 | 0.679 | 0.023 | 0.003 | 0.003 | 0 | 0.5 | 0.63 | 0.48 | 0.43 | 0.38 | 0.3 |
| 2 | 62763347 | rs901532 | 0.967 | 0.542 | -0.344 | 0.695 | 0.019 | -0.038 | -0.038 | 0 | 0.19 | 0.14 | 0.2 | 0.33 | 0.13 | 0.19 |
| 2 | 62764247 | rs4608500 | -0.991 | 1.014 | -0.445 | 0.614 | -0.002 | 0.012 | 0.012 | 0 | 0.55 | 0.66 | 0.59 | 0.65 | 0.52 | 0.5 |
| 2 | 62774415 | rs17575679 | 0.234 | 0.821 | -0.561 | 0.369 | -0.024 | 0.045 | 0.045 | 0 | 0.14 | 0.05 | 0.1 | 0 | 0.15 | 0.07 |
| 2 | 62781499 | rs7580494 | -0.755 | 0.61 | -0.358 | 0.589 | 0.006 | 0.003 | 0.003 | 0 | 0.53 | 0.6 | 0.48 | 0.4 | 0.41 | 0.35 |
| 2 | 62782186 | rs2054837 | 1.082 | 0.914 | -0.132 | 0.585 | -0.003 | -0.007 | -0.007 | 0 | 0.17 | 0.18 | 0.15 | 0.08 | 0.13 | 0.09 |
| 2 | 62790805 | rs7576389 | -0.377 | 0.885 | 0.204 | 0.446 | 0.006 | -0.09 | -0.09 | 0 | 0.32 | 0.49 | 0.27 | 0.08 | 0.3 | 0.17 |
| 2 | 62793514 | rs17025805 | -0.399 | 0.557 | 0.292 | 0.806 | -0.006 | 0.166 | 0.166 | 0 | 0.1 | 0.24 | 0.24 | 0.36 | 0.1 | 0.17 |
| 2 | 62793940 | rs6709361 | 0.168 | 0.661 | 0.238 | 0.71 | 0.003 | -0.006 | -0.006 | 0.094 | 0.25 | 0.26 | 0.36 | 0.44 | 0.21 | 0.25 |
| 2 | 62794350 | rs11125928 | -1.381 | 0.531 | -0.099 | 0.547 | -0.029 | -0.075 | -0.075 | 0 | 0.46 | 0.88 | 0.53 | 0.44 | 0.43 | 0.34 |
| 2 | 62795483 | rs7601084 | -0.558 | 0.811 | 0.176 | 0.466 | 0.043 | -0.115 | -0.115 | 0.048 | 0.18 | 0.36 | 0.14 | 0 | 0.21 | 0.09 |
| 2 | 62797517 | rs7596224 | -1.699 | 0.739 | -0.318 | 0.609 | 0.003 | 0.006 | 0.006 | 0 | 0.33 | 0.64 | 0.36 | 0.37 | 0.38 | 0.42 |
| 2 | 62797738 | rs13417654 | -0.718 | 0.587 | -0.374 | 0.878 | -0.002 | 0.116 | 0.116 | 0 | 0.13 | 0.27 | 0.19 | 0.35 | 0.13 | 0.33 |
| 2 | 62805224 | rs17025954 | -0.074 | 0.652 | -1.19 | 0.836 | -0.022 | -0.035 | -0.035 | 0 | 0.09 | 0.01 | 0.19 | 0.15 | 0.14 | 0.07 |
| 2 | 62815730 | rs1527966 | -0.867 | 0.33 | -1.785 | 0.536 | -0.008 | -0.017 | -0.017 | 0.878 | 0.88 | 0.96 | 0.92 | 0.86 | 0.9 | 0.9 |
| 2 | 62816083 | rs11679934 | 1.937 | -0.022 | -2.164 | 0.463 | 0.009 | -0.064 | -0.064 | 0 | 0.17 | 0.02 | 0.2 | 0.18 | 0.13 | 0.11 |
| 2 | 62831112 | rs12617373 | 1.768 | 0.064 | -2.09 | 0.468 | -0.022 | -0.017 | -0.017 | 0 | 0.05 | 0.01 | 0.04 | 0.05 | 0.06 | 0.08 |
| 2 | 62887884 | rs6727888 | -0.708 | -0.242 | -2.454 | -0.27 | 0.057 | 0.09 | 0.09 | 0 | 0.51 | 0.78 | 0.63 | 0.74 | 0.65 | 0.71 |
| 2 | 62890171 | rs1038570 | -0.99 | -0.409 | -2.513 | -0.403 | 0.014 | 0.092 | 0.092 | 0.84 | 0.7 | 0.89 | 0.84 | 0.91 | 0.78 | 0.83 |
| 2 | 62891474 | rs6545969 | -0.75 | -0.404 | -2.506 | -0.341 | 0.051 | 0.08 | 0.08 | 0 | 0.51 | 0.76 | 0.63 | 0.73 | 0.64 | 0.7 |
| 2 | 62914811 | rs17577380 | 0.158 | -0.28 | -2.476 | -0.95 | 0.035 | 0.079 | 0.079 | 3.402 | 0.31 | 0.17 | 0.13 | 0 | 0.21 | 0.12 |
| 2 | 62924309 | rs360805 | 1.143 | -0.214 | -2.357 | -1.001 | -0.001 | -0.008 | -0.008 | 0 | 0.22 | 0.21 | 0.17 | 0.08 | 0.21 | 0.15 |
| 2 | 62939397 | rs360804 | 1.393 | -0.248 | -2.316 | -0.785 | 0.008 | 0.023 | 0.023 | 1.004 | 0.38 | 0.16 | 0.34 | 0.27 | 0.34 | 0.25 |
| 2 | 62946658 | rs1906198 | 0.909 | -0.478 | -2.457 | -0.884 | 0.049 | 0.078 | 0.078 | 0 | 0.48 | 0.27 | 0.29 | 0.25 | 0.37 | 0.26 |
| 2 | 62988169 | rs4671050 | 1.26 | -0.384 | -2.587 | -0.849 | 0.009 | 0.025 | 0.025 | 0.222 | 0.41 | 0.3 | 0.36 | 0.23 | 0.35 | 0.26 |
| 2 | 63073194 | rs11682530 | 1.099 | 0.46 | -2.376 | -0.339 | 0.031 | 0.041 | 0.041 | 0 | 0.47 | 0.32 | 0.37 | 0.27 | 0.37 | 0.26 |
| 2 | 63075033 | rs2421816 | -1.446 | 0.512 | -2.661 | -0.285 | 0.054 | -0.017 | -0.017 | 3.124 | 0.54 | 0.91 | 0.57 | 0.59 | 0.62 | 0.73 |
| 2 | 63126755 | rs11901070 | 0.207 | 0.504 | -2.594 | -0.954 | 0.003 | 0.016 | 0.016 | 1.386 | 0.24 | 0.17 | 0.12 | 0 | 0.19 | 0.11 |
| 2 | 63131731 | rs721048 | 1.665 | 0.703 | -2.693 | -0.821 | -0.038 | 0.022 | 0.022 | 0 | 0.09 | 0.01 | 0.19 | 0.05 | 0.17 | 0.1 |
| 2 | 63134879 | rs2710646 | 1.791 | 0.71 | -2.715 | -0.817 | -0.042 | 0.017 | 0.017 | 0 | 0.09 | 0.01 | 0.19 | 0.05 | 0.17 | 0.1 |
| 2 | 63156832 | rs4671454 | 2.477 | 0.402 | -2.624 | -0.833 | 0.029 | -0.045 | -0.045 | 0.8 | 0.2 | 0.04 | 0.2 | 0.19 | 0.14 | 0.12 |
| 2 | 63176139 | rs17432615 | 2.194 | 0.567 | -2.624 | -0.817 | -0.032 | 0.008 | 0.008 | 0 | 0.06 | 0.02 | 0.08 | 0.03 | 0.11 | 0.08 |
| 2 | 63187114 | rs12987254 | 1.28 | 0.381 | -2.74 | -0.883 | 0.002 | 0.064 | 0.064 | 0.011 | 0.25 | 0.03 | 0.21 | 0.14 | 0.21 | 0.14 |

| | | | | | | | | | | | | | | | | |
|---|---|---|---|---|---|---|---|---|---|---|---|---|---|---|---|---|
| 2 | 63200294 | rs2539979 | -1.408 | 0.443 | -2.682 | -1.119 | -0.009 | 0.12 | 0.12 | 0.011 | 0.74 | 0.96 | 0.86 | 0.92 | 0.78 | 0.86 |
| 2 | 63208460 | rs2539980 | -0.803 | 0.336 | -2.553 | -1.177 | 0.005 | 0.006 | 0.006 | 0 | 0.46 | 0.55 | 0.6 | 0.75 | 0.64 | 0.73 |
| 2 | 63209314 | rs12612046 | 2.386 | 0.452 | -2.613 | -1.169 | -0.031 | 0.007 | 0.007 | 0 | 0.06 | 0.01 | 0.07 | 0.03 | 0.1 | 0.08 |
| 2 | 63213970 | rs2710647 | -0.461 | 0.453 | -2.406 | -1.156 | 0.015 | -0.014 | -0.014 | 0 | 0.4 | 0.48 | 0.52 | 0.71 | 0.53 | 0.65 |
| 2 | 63231534 | rs2553042 | -1.272 | 0.007 | -2.76 | -1.133 | 0.005 | 0.116 | 0.116 | 0 | 0.74 | 0.95 | 0.86 | 0.92 | 0.78 | 0.86 |
| 2 | 63272724 | rs2018650 | 2.364 | 0.006 | -2.649 | -0.973 | 0.002 | -0.076 | -0.076 | 0.312 | 0.2 | 0.01 | 0.2 | 0.19 | 0.14 | 0.12 |
| 2 | 63298150 | rs11125946 | 0.865 | 0.105 | -2.296 | -1.059 | 0.017 | 0.018 | 0.018 | 0 | 0.51 | 0.45 | 0.39 | 0.18 | 0.3 | 0.26 |
| 2 | 63301164 | rs6545977 | 0.197 | 0.48 | -2.266 | -1.011 | 0.012 | -0.018 | -0.018 | 0.269 | 0.6 | 0.49 | 0.49 | 0.23 | 0.5 | 0.36 |
| 2 | 63343399 | rs6754628 | -0.296 | 0.972 | -2.046 | -0.66 | 0.004 | 0.021 | 0.021 | 0 | 0.46 | 0.58 | 0.61 | 0.71 | 0.61 | 0.68 |
| 2 | 63344312 | rs13013948 | 2.194 | 0.757 | -2.177 | -0.746 | 0.017 | 0.031 | 0.031 | 0.432 | 0.21 | 0.01 | 0.11 | 0.09 | 0.11 | 0.15 |
| 2 | 63345507 | rs2162014 | 0.584 | 0.955 | -2.142 | -0.754 | -0.003 | -0.005 | -0.005 | 0 | 0.35 | 0.51 | 0.41 | 0.65 | 0.39 | 0.56 |
| 2 | 63346702 | rs13002109 | -0.435 | 1 | -2.2 | -0.817 | 0 | -0.003 | -0.003 | 0 | 0.61 | 0.42 | 0.53 | 0.26 | 0.52 | 0.41 |
| 2 | 63347964 | rs11696083 | 1.927 | 0.976 | -2.616 | -0.578 | 0.03 | 0.043 | 0.043 | 0.341 | 0.22 | 0.03 | 0.12 | 0.09 | 0.12 | 0.14 |
| 2 | 63363355 | rs13010733 | 1.381 | 1.077 | -2.581 | -0.649 | -0.018 | 0.029 | 0.029 | 0 | 0.19 | 0.06 | 0.2 | 0.13 | 0.19 | 0.13 |
| 2 | 63374063 | rs4671458 | 1.376 | 1.147 | -2.563 | -0.545 | -0.052 | -0.05 | -0.05 | 3.172 | 0.08 | 0.02 | 0.26 | 0.14 | 0.19 | 0.1 |
| 2 | 63374600 | rs11685079 | 2.236 | 0.979 | -2.598 | -0.665 | 0.003 | 0.029 | 0.029 | 0.012 | 0.19 | 0.01 | 0.11 | 0.08 | 0.11 | 0.14 |
| 2 | 63397974 | rs17408841 | 1.449 | 1.197 | -2.514 | -0.616 | 0.012 | 0.126 | 0.126 | 0 | 0.24 | 0.03 | 0.13 | 0.05 | 0.23 | 0.14 |
| 2 | 63401230 | rs13411767 | 0.677 | 1.293 | -2.439 | -0.555 | -0.043 | 0.052 | 0.052 | 0 | 0.31 | 0.11 | 0.32 | 0.15 | 0.41 | 0.24 |
| 2 | 63405218 | rs13009169 | 1.101 | 0.77 | -2.627 | -0.654 | 0.009 | 0.148 | 0.148 | 0.026 | 0.38 | 0.09 | 0.3 | 0.13 | 0.29 | 0.27 |
| 2 | 63430175 | rs12151606 | 1.364 | 1.06 | -2.395 | -0.546 | -0.055 | 0.13 | 0.13 | 0 | 0.23 | 0.03 | 0.16 | 0.06 | 0.3 | 0.15 |
| 2 | 63447682 | rs2421916 | -0.742 | 1.05 | -2.18 | -0.372 | 0.015 | 0.069 | 0.069 | 0 | 0.48 | 0.75 | 0.54 | 0.67 | 0.59 | 0.65 |
| 2 | 63460744 | rs17348207 | 1.527 | 1.407 | -2.08 | -0.215 | -0.04 | -0.038 | -0.038 | 0 | 0.07 | 0.02 | 0.19 | 0.1 | 0.19 | 0.1 |
| 2 | 63472250 | rs12329383 | 0.547 | 1.153 | -2.191 | -0.367 | -0.116 | 0.051 | 0.051 | 0 | 0.32 | 0.07 | 0.35 | 0.15 | 0.49 | 0.24 |
| 2 | 63474793 | rs11125954 | 0.002 | 0.858 | -2.266 | -0.456 | 0.065 | 0.065 | 0.065 | 0 | 0.5 | 0.12 | 0.61 | 0.33 | 0.67 | 0.38 |
| 2 | 63482704 | rs7601558 | -0.861 | 0.858 | -1.994 | -0.438 | -0.001 | -0.011 | -0.011 | 0 | 0.87 | 0.82 | 0.93 | 0.92 | 0.89 | 0.92 |
| 2 | 63511435 | rs2138798 | 0.467 | 0.757 | -1.105 | -0.426 | 0.006 | 0.181 | 0.181 | 0 | 0.13 | 0.28 | 0.19 | 0.6 | 0.13 | 0.41 |
| 2 | 63542732 | rs6545988 | -1.742 | 0.945 | -1.693 | -0.68 | -0.015 | 0.1 | 0.1 | 0.013 | 0.76 | 0.93 | 0.84 | 0.95 | 0.7 | 0.85 |
| 2 | 63548716 | rs7585664 | -1.852 | 0.999 | -1.725 | -0.682 | -0.038 | 0.125 | 0.125 | 0 | 0.77 | 0.98 | 0.85 | 0.95 | 0.72 | 0.85 |
| 2 | 63556849 | rs2421882 | -0.368 | 0.864 | -1.167 | -0.566 | -0.001 | 0.004 | 0.004 | 0 | 0.6 | 0.49 | 0.64 | 0.35 | 0.56 | 0.42 |
| 2 | 63564683 | rs7566860 | 1.489 | 0.795 | -1.463 | -0.65 | -0.005 | 0.007 | 0.007 | 0 | 0.19 | 0.09 | 0.27 | 0.17 | 0.19 | 0.13 |
| 2 | 63571808 | rs1949411 | 0.893 | 0.634 | -1.425 | -0.632 | 0.028 | 0.058 | 0.058 | 0 | 0.3 | 0.19 | 0.16 | 0.08 | 0.18 | 0.21 |
| 2 | 63593212 | rs1922422 | -0.352 | 0.64 | -1.335 | -0.466 | -0.004 | 0.003 | 0.003 | 0 | 0.61 | 0.46 | 0.63 | 0.35 | 0.57 | 0.42 |
| 2 | 63596836 | rs11683229 | 1.844 | 0.58 | -1.587 | -0.609 | 0 | 0.017 | 0.017 | 0.002 | 0.19 | 0.01 | 0.11 | 0.09 | 0.11 | 0.14 |
| 2 | 63652403 | rs10164844 | -0.27 | 0.858 | -0.897 | -0.458 | 0 | 0.028 | 0.028 | 0 | 0.49 | 0.64 | 0.41 | 0.28 | 0.43 | 0.32 |
| 2 | 63692447 | rs2034270 | 1.249 | 1.179 | -1.368 | -0.545 | -0.03 | -0.097 | -0.097 | 0 | 0.19 | 0.04 | 0.36 | 0.65 | 0.22 | 0.38 |
| 2 | 63716223 | rs2028887 | -1.215 | 1.029 | -0.644 | -0.973 | -0.012 | -0.118 | -0.118 | 0.335 | 0.72 | 0.6 | 0.82 | 0.98 | 0.7 | 0.8 |
| 2 | 63744282 | rs33977056 | 0.796 | 1.17 | -0.499 | -1.171 | 0.043 | 0.107 | 0.107 | 0 | 0.37 | 0.21 | 0.18 | 0.07 | 0.21 | 0.22 |
| 2 | 63754275 | rs2421954 | 1.347 | 1.24 | -0.406 | -1.141 | 0.002 | 0.02 | 0.02 | 0 | 0.21 | 0.21 | 0.39 | 0.72 | 0.23 | 0.4 |
| 2 | 63771769 | rs12472865 | 1.532 | 1.331 | -0.626 | -1.112 | -0.045 | -0.04 | -0.04 | 0 | 0.08 | 0.01 | 0.19 | 0.1 | 0.21 | 0.08 |
| 2 | 63780605 | rs2422008 | 0.004 | 1.305 | -0.592 | -1.111 | -0.009 | 0.054 | 0.054 | 0 | 0.47 | 0.32 | 0.4 | 0.26 | 0.44 | 0.31 |
| 2 | 63813458 | rs3754540 | 1.372 | 1.347 | -0.693 | -1.019 | -0.003 | -0.013 | -0.013 | 0 | 0.2 | 0.14 | 0.38 | 0.7 | 0.22 | 0.42 |
| 2 | 63815206 | rs1109027 | 1.245 | 1.352 | -0.885 | -0.98 | -0.048 | -0.042 | -0.042 | 0 | 0.08 | 0.01 | 0.2 | 0.1 | 0.21 | 0.08 |
| 2 | 63815801 | rs2278718 | 0.909 | 1.264 | -0.882 | -1.004 | -0.052 | -0.05 | -0.05 | 1.513 | 0.09 | 0.02 | 0.22 | 0.19 | 0.24 | 0.09 |
| 2 | 63819348 | rs10200146 | -0.022 | 1.104 | -0.844 | -1.017 | -0.021 | 0.065 | 0.065 | 0 | 0.46 | 0.25 | 0.4 | 0.26 | 0.44 | 0.3 |
| 2 | 63909132 | rs6546020 | 0.084 | 0.812 | -0.851 | -1.101 | 0.002 | 0.062 | 0.062 | 0 | 0.46 | 0.65 | 0.47 | 0.23 | 0.58 | 0.33 |
| 2 | 63911761 | rs11125981 | 0.693 | 0.743 | -1.06 | -1.11 | 0.025 | 0.068 | 0.068 | 0.057 | 0.3 | 0.17 | 0.15 | 0.08 | 0.19 | 0.19 |
| 2 | 63917073 | rs17028097 | -0.024 | 0.706 | -0.578 | -1.245 | 0.062 | -0.026 | -0.026 | 0.143 | 0.08 | 0.32 | 0.06 | 0 | 0.06 | 0.06 |
| 2 | 63917221 | rs10168461 | -0.91 | 0.731 | -1.358 | -1.331 | 0.015 | 0.03 | 0.03 | 0.384 | 0.56 | 0.83 | 0.51 | 0.28 | 0.55 | 0.45 |
| 2 | 63921775 | rs6546024 | -0.469 | 0.756 | -1.378 | -1.532 | 0.026 | 0.154 | 0.154 | 0 | 0.51 | 0.75 | 0.48 | 0.16 | 0.59 | 0.43 |
| 2 | 63922369 | rs10209140 | -0.779 | 0.817 | -1.304 | -1.516 | 0.021 | -0.012 | -0.012 | 0.233 | 0.26 | 0.64 | 0.35 | 0.14 | 0.36 | 0.26 |
| 2 | 63923561 | rs6546025 | -1.649 | 0.667 | -1.323 | -1.447 | -0.007 | -0.046 | -0.046 | 0 | 0.81 | 0.89 | 0.64 | 0.24 | 0.79 | 0.61 |
| 2 | 63926197 | rs6741298 | -0.745 | 0.684 | -1.38 | -1.521 | 0.021 | 0.08 | 0.08 | 0.163 | 0.54 | 0.78 | 0.5 | 0.22 | 0.55 | 0.44 |
| 2 | 63939417 | rs6730863 | 1.262 | 0.545 | -1.402 | -1.618 | 0.018 | 0.071 | 0.071 | 0 | 0.28 | 0.11 | 0.15 | 0.02 | 0.26 | 0.16 |
| 2 | 63943257 | rs17619717 | 0.342 | 0.599 | -1.607 | -1.667 | -0.049 | 0.04 | 0.04 | 1.537 | 0.36 | 0.19 | 0.36 | 0.17 | 0.4 | 0.26 |
| 2 | 63962974 | rs6709123 | -0.033 | 0.553 | -1.519 | -1.674 | 0.041 | 0.121 | 0.121 | 1.47 | 0.44 | 0.69 | 0.29 | 0.06 | 0.38 | 0.32 |
| 2 | 64029271 | rs6710230 | -0.444 | 0.581 | -1.721 | -1.743 | -0.005 | 0.001 | 0.001 | 0.837 | 0.83 | 0.82 | 0.73 | 0.91 | 0.72 | 0.88 |
| 2 | 64041944 | rs10201844 | 1.769 | 0.451 | -1.708 | -1.65 | -0.017 | -0.054 | -0.054 | 0.075 | 0.19 | 0.07 | 0.33 | 0.76 | 0.21 | 0.42 |
| 2 | 64058819 | rs10187013 | 1.582 | 0.338 | -1.445 | -1.635 | -0.015 | -0.027 | -0.027 | 0.048 | 0.2 | 0.08 | 0.33 | 0.79 | 0.22 | 0.41 |
| 2 | 64067708 | rs4293553 | 2.091 | 0.296 | -1.531 | -0.837 | -0.032 | -0.047 | -0.047 | 0 | 0.17 | 0.04 | 0.26 | 0.25 | 0.19 | 0.24 |
| 2 | 64068788 | rs6710763 | -0.499 | 0.205 | -1.261 | -1.138 | -0.006 | -0.026 | -0.026 | 0 | 0.68 | 0.65 | 0.82 | 0.92 | 0.78 | 0.75 |
| 2 | 64071232 | rs4671535 | 2.354 | 0.745 | -1.472 | -1.047 | -0.037 | -0.034 | -0.034 | 0 | 0.06 | 0.01 | 0.18 | 0.07 | 0.15 | 0.06 |
| 2 | 64075076 | rs6546032 | -1.48 | 0.296 | -1.421 | -1.128 | 0.006 | 0.051 | 0.051 | 0 | 0.78 | 0.87 | 0.85 | 0.92 | 0.83 | 0.85 |
| 2 | 64089748 | rs6732746 | -1.151 | 0.327 | -1.331 | -1.195 | 0.011 | 0.05 | 0.05 | 0 | 0.74 | 0.84 | 0.84 | 0.92 | 0.81 | 0.83 |
| 2 | 64091533 | rs6755646 | -1.163 | 0.335 | -1.329 | -1.191 | 0.011 | 0.048 | 0.048 | 0 | 0.74 | 0.83 | 0.84 | 0.92 | 0.81 | 0.83 |
| 2 | 64112428 | rs12713498 | -0.366 | 0.296 | -0.427 | -1.324 | -0.013 | -0.094 | -0.094 | 0 | 0.65 | 0.54 | 0.77 | 0.92 | 0.75 | 0.77 |
| 2 | 64116589 | rs6546038 | -1.598 | -0.401 | -1.083 | -1.673 | 0.035 | -0.019 | -0.019 | 0 | 0.71 | 0.92 | 0.66 | 0.23 | 0.77 | 0.54 |
| 2 | 64117944 | rs10166971 | 1.643 | 0.019 | -0.827 | -1.673 | 0.015 | 0 | 0 | 0.103 | 0.09 | 0.09 | 0.22 | 0.07 | 0.23 | 0.08 |
| 2 | 64118877 | rs6546039 | -2.31 | -0.262 | -0.95 | -1.631 | -0.014 | -0.054 | -0.054 | 0 | 0.8 | 0.93 | 0.67 | 0.23 | 0.78 | 0.56 |
| 2 | 64133562 | rs6710437 | 0.747 | -0.136 | -0.676 | -1.571 | 0.033 | -0.041 | -0.041 | 0 | 0.08 | 0.26 | 0.06 | 0 | 0.06 | 0.06 |

| | | | | | | | | | | | | | | | | |
|---|---|---|---|---|---|---|---|---|---|---|---|---|---|---|---|---|
| 2 | 64168149 | rs6723613 | 1.164 | -0.23 | -1.658 | -1.759 | -0.008 | 0.172 | 0.172 | 0 | 0.28 | 0.01 | 0.16 | 0.03 | 0.25 | 0.18 |
| 2 | 64168232 | rs6708611 | 1.629 | -0.22 | -1.647 | -1.751 | -0.009 | 0.146 | 0.146 | 0 | 0.25 | 0.01 | 0.15 | 0.03 | 0.23 | 0.15 |
| 2 | 64168682 | rs12991346 | -0.352 | 0.077 | -1.174 | -1.436 | 0.011 | -0.03 | -0.03 | 0 | 0.41 | 0.51 | 0.42 | 0.17 | 0.45 | 0.34 |
| 2 | 64190732 | rs12466966 | 2.151 | 0.455 | -1.592 | -2.554 | -0.048 | -0.036 | -0.036 | 0 | 0.08 | 0.01 | 0.21 | 0.07 | 0.21 | 0.07 |
| 2 | 64193684 | rs11683408 | 1.5 | 0.414 | -1.7 | -2.721 | 0.02 | 0.031 | 0.031 | 0 | 0.2 | 0.07 | 0.12 | 0.08 | 0.12 | 0.11 |
| 2 | 64200119 | rs17562423 | 1.206 | 0.479 | -1.706 | -2.761 | 0.02 | 0.039 | 0.039 | 0 | 0.22 | 0.07 | 0.14 | 0.08 | 0.14 | 0.14 |
| 2 | 64240609 | rs11682249 | 0.21 | 0.099 | -1.627 | -2.705 | -0.071 | 0.049 | 0.049 | 0 | 0.34 | 0.09 | 0.36 | 0.16 | 0.4 | 0.24 |
| 2 | 64255895 | rs7609385 | 1.417 | 0.468 | -1.082 | -2.44 | 0.039 | -0.029 | -0.029 | 2.699 | 0.06 | 0.13 | 0.04 | 0 | 0.06 | 0.06 |
| 2 | 64257746 | rs329478 | -1.308 | 0.175 | -1.815 | -2.396 | 0.055 | 0.072 | 0.072 | 0 | 0.68 | 0.83 | 0.84 | 0.92 | 0.82 | 0.83 |
| 2 | 64263530 | rs329489 | -0.154 | 0.193 | -1.545 | -2.265 | 0.001 | 0.011 | 0.011 | 0.026 | 0.52 | 0.53 | 0.58 | 0.84 | 0.55 | 0.64 |
| 2 | 64279606 | rs10171434 | -0.226 | 0.361 | -2.143 | -2.155 | 0.001 | -0.016 | -0.016 | 0 | 0.07 | 0.16 | 0.06 | 0.04 | 0.08 | 0.09 |
| 2 | 64281498 | rs7560803 | -0.05 | 0.347 | -2.132 | -2.145 | 0.001 | -0.015 | -0.015 | 0.078 | 0.07 | 0.14 | 0.05 | 0.04 | 0.08 | 0.09 |
| 2 | 64291780 | rs329483 | -0.165 | 0.4 | -2.331 | -2.297 | -0.004 | 0.061 | 0.061 | 0.499 | 0.49 | 0.6 | 0.55 | 0.79 | 0.49 | 0.63 |
| 2 | 64292988 | rs17753048 | 1.405 | 0.171 | -2.637 | -2.482 | -0.034 | 0.092 | 0.092 | 0 | 0.19 | 0.03 | 0.15 | 0.04 | 0.22 | 0.14 |
| 2 | 64293274 | rs2919222 | -0.456 | 0.192 | -2.283 | -2.247 | 0.02 | 0.022 | 0.022 | 0 | 0.58 | 0.63 | 0.77 | 0.87 | 0.72 | 0.72 |
| 2 | 64299693 | rs10198990 | -0.767 | 0.706 | -2.577 | -1.772 | -0.009 | 0.006 | 0.006 | 0.935 | 0.23 | 0.23 | 0.3 | 0.13 | 0.34 | 0.2 |
| 2 | 64307320 | rs2917793 | 0.143 | 0.536 | -3.111 | -1.884 | -0.085 | 0.108 | 0.108 | 0 | 0.37 | 0.17 | 0.48 | 0.87 | 0.49 | 0.47 |
| 2 | 64308048 | rs2581047 | -1.724 | 0.548 | -3.258 | -2.021 | -0.075 | 0.067 | 0.067 | 0 | 0.81 | 0.99 | 0.8 | 0.96 | 0.75 | 0.83 |
| 2 | 64312142 | rs17028375 | -0.862 | 0.685 | -2.966 | -1.955 | 0.012 | -0.025 | -0.025 | 0 | 0.09 | 0.18 | 0.08 | 0.04 | 0.12 | 0.09 |
| 2 | 64316687 | rs329502 | -0.561 | 0.393 | -3.161 | -2.095 | 0.029 | 0.161 | 0.161 | 0 | 0.46 | 0.29 | 0.54 | 0.88 | 0.58 | 0.6 |
| 2 | 64320746 | rs10496105 | 1.704 | 0.166 | -3.292 | -2.182 | 0.003 | 0.003 | 0.003 | 0 | 0.14 | 0.06 | 0.18 | 0.08 | 0.11 | 0.16 |
| 2 | 64321823 | rs3731962 | -0.917 | 0.815 | -3.088 | -1.833 | -0.01 | -0.028 | -0.028 | 0 | 0.08 | 0.19 | 0.06 | 0.03 | 0.05 | 0.04 |
| 2 | 64324076 | rs329499 | -1.772 | 0.723 | -3.238 | -1.963 | -0.067 | 0.066 | 0.066 | 0 | 0.81 | 0.98 | 0.81 | 0.96 | 0.74 | 0.82 |
| 2 | 64327592 | rs329497 | 0.725 | 0.952 | -3.304 | -1.939 | -0.061 | -0.057 | -0.057 | 3.163 | 0.29 | 0.07 | 0.31 | 0.75 | 0.3 | 0.4 |
| 2 | 64334456 | rs6718082 | 1.053 | 0.645 | -3.246 | -1.828 | 0.039 | 0.082 | 0.082 | 0 | 0.37 | 0.69 | 0.29 | 0.07 | 0.23 | 0.3 |
| 2 | 64335650 | rs13408188 | -0.879 | 0.794 | -2.817 | -1.412 | -0.008 | -0.029 | -0.029 | 0 | 0.07 | 0.18 | 0.06 | 0.03 | 0.05 | 0.04 |
| 2 | 64339277 | rs2576837 | 1.273 | 0.8 | -2.871 | -1.414 | -0.028 | -0.094 | -0.094 | 0 | 0.26 | 0.04 | 0.29 | 0.75 | 0.23 | 0.38 |
| 2 | 64342469 | rs7349403 | 1.207 | 0.956 | -2.837 | -1.379 | -0.051 | -0.036 | -0.036 | 0 | 0.08 | 0.01 | 0.16 | 0.09 | 0.21 | 0.1 |
| 2 | 64342626 | rs11679748 | 0.872 | 0.33 | -2.636 | -1.235 | 0 | -0.055 | -0.055 | 0.141 | 0.28 | 0.11 | 0.3 | 0.75 | 0.25 | 0.4 |
| 2 | 64345645 | rs7604199 | 0.414 | 0.416 | -2.562 | -1.182 | 0.019 | 0.002 | 0.002 | 0 | 0.33 | 0.25 | 0.35 | 0.78 | 0.29 | 0.42 |
| 2 | 64369336 | rs10496107 | 0.008 | 0.894 | -2.452 | -1.154 | -0.058 | 0.049 | 0.049 | 0 | 0.68 | 0.8 | 0.61 | 0.87 | 0.51 | 0.71 |
| 2 | 64371569 | rs6717575 | 0.056 | 0.63 | -2.511 | -1.261 | 0.033 | 0.174 | 0.174 | 0 | 0.42 | 0.84 | 0.36 | 0.13 | 0.33 | 0.31 |
| 2 | 64374041 | rs6712700 | -0.081 | 0.479 | -2.486 | -1.205 | -0.055 | 0.19 | 0.19 | 0 | 0.51 | 0.89 | 0.49 | 0.19 | 0.47 | 0.42 |
| 2 | 64379672 | rs10170666 | -0.081 | 0.466 | -2.443 | -1.205 | -0.048 | 0.19 | 0.19 | 0 | 0.51 | 0.87 | 0.5 | 0.19 | 0.46 | 0.42 |
| 2 | 64387765 | rs11125991 | -0.134 | 0.412 | -1.854 | -1.086 | 0.005 | 0.11 | 0.11 | 0 | 0.5 | 0.76 | 0.5 | 0.26 | 0.48 | 0.43 |
| 2 | 64400608 | rs17754672 | 0.111 | 0.307 | -1.437 | -1.122 | -0.002 | 0.006 | 0.006 | 0 | 0.45 | 0.49 | 0.44 | 0.23 | 0.43 | 0.42 |
| 2 | 64401060 | rs4144251 | 0.929 | 0.264 | -1.803 | -1.151 | 0.031 | -0.069 | -0.069 | 0 | 0.28 | 0.12 | 0.28 | 0.67 | 0.24 | 0.42 |
| 2 | 64402689 | rs1881712 | -1.077 | 0.277 | -1.761 | -1.153 | -0.073 | 0.06 | 0.06 | 0 | 0.79 | 0.96 | 0.76 | 0.91 | 0.68 | 0.85 |
| 2 | 64407710 | rs12999721 | -0.191 | -0.04 | -1.222 | -1.033 | -0.003 | 0.037 | 0.037 | 0 | 0.47 | 0.58 | 0.48 | 0.23 | 0.44 | 0.42 |
| 2 | 64413221 | rs6710681 | 0.13 | 0.109 | -1.056 | -0.907 | -0.018 | 0.015 | 0.015 | 0 | 0.62 | 0.35 | 0.54 | 0.75 | 0.6 | 0.64 |
| 2 | 64414077 | rs6546056 | -0.134 | 0.221 | -1.062 | -0.874 | 0.002 | -0.003 | -0.003 | 0.263 | 0.65 | 0.43 | 0.56 | 0.77 | 0.64 | 0.64 |
| 2 | 64420236 | rs2422116 | 0.009 | 0.233 | -1.015 | -0.888 | 0.013 | -0.011 | -0.011 | 0 | 0.65 | 0.53 | 0.56 | 0.75 | 0.63 | 0.65 |
| 2 | 64424920 | rs2118303 | 0.792 | -0.219 | -1.501 | -1.006 | 0.103 | 0.143 | 0.143 | 0 | 0.3 | 0.04 | 0.17 | 0.05 | 0.23 | 0.15 |
| 2 | 64426384 | rs6734073 | 0.151 | 0.441 | -0.558 | -0.707 | -0.001 | -0.003 | -0.003 | 0 | 0.42 | 0.59 | 0.34 | 0.64 | 0.38 | 0.53 |
| 2 | 64432911 | rs11688386 | 0.332 | 0.57 | -0.382 | -0.722 | 0.025 | 0.016 | 0.016 | 0 | 0.51 | 0.31 | 0.56 | 0.36 | 0.46 | 0.38 |
| 2 | 64434512 | rs2164734 | -0.402 | 0.59 | -0.193 | -0.681 | 0.002 | -0.007 | -0.007 | 0 | 0.57 | 0.65 | 0.63 | 0.41 | 0.5 | 0.4 |
| 2 | 64438890 | rs11125993 | 0.152 | 0.583 | -0.028 | -0.721 | 0.013 | 0.007 | 0.007 | 0 | 0.53 | 0.34 | 0.56 | 0.36 | 0.45 | 0.38 |
| 2 | 64441643 | rs1370388 | -0.61 | 0.609 | -0.166 | -0.713 | -0.004 | -0.011 | -0.011 | 0.132 | 0.6 | 0.64 | 0.63 | 0.39 | 0.51 | 0.4 |
| 2 | 64445563 | rs1898445 | -0.935 | 1.104 | -0.351 | -0.874 | 0.022 | -0.03 | -0.03 | 0.054 | 0.86 | 0.8 | 0.87 | 0.96 | 0.78 | 0.85 |
| 2 | 64453486 | rs2555435 | 0.094 | 0.609 | -0.263 | -0.511 | 0.006 | 0.008 | 0.008 | 0.06 | 0.36 | 0.44 | 0.36 | 0.64 | 0.37 | 0.52 |
| 2 | 64455674 | rs871776 | 1.142 | 0.576 | -0.393 | -0.4 | -0.039 | -0.081 | -0.081 | 0 | 0.26 | 0.13 | 0.28 | 0.56 | 0.29 | 0.44 |
| 2 | 64463958 | rs17677161 | 1.883 | 0.152 | -0.673 | -0.91 | 0.068 | 0.144 | 0.144 | 0 | 0.23 | 0.02 | 0.1 | 0.01 | 0.17 | 0.11 |
| 2 | 64471185 | rs11677415 | 1.141 | 0.947 | 0.125 | -0.363 | -0.003 | -0.005 | -0.005 | 0 | 0.07 | 0.1 | 0.04 | 0.03 | 0.05 | 0.07 |
| 2 | 64474069 | rs6750096 | 0.484 | 0.934 | -0.116 | -0.307 | -0.033 | -0.065 | -0.065 | 0 | 0.29 | 0.16 | 0.29 | 0.6 | 0.29 | 0.47 |
| 2 | 64479060 | rs1025209 | 0.355 | 0.944 | -0.096 | -0.277 | -0.019 | -0.047 | -0.047 | 0.04 | 0.3 | 0.2 | 0.3 | 0.6 | 0.29 | 0.5 |
| 2 | 64485465 | rs2251764 | 0.429 | 0.937 | -0.06 | -0.79 | -0.014 | -0.023 | -0.023 | 0 | 0.31 | 0.24 | 0.35 | 0.64 | 0.28 | 0.47 |
| 2 | 64485812 | rs2114432 | -0.564 | 0.95 | 0.213 | -0.965 | -0.001 | 0.004 | 0.004 | 0 | 0.42 | 0.4 | 0.43 | 0.69 | 0.38 | 0.57 |
| 2 | 64486499 | rs1358624 | -0.996 | 1.219 | 0.332 | -0.984 | -0.002 | -0.014 | -0.014 | 0.586 | 0.49 | 0.48 | 0.56 | 0.78 | 0.51 | 0.66 |
| 2 | 64488679 | rs2698541 | 0.349 | 1.386 | -0.24 | -0.886 | -0.011 | -0.015 | -0.015 | 0 | 0.32 | 0.22 | 0.35 | 0.65 | 0.29 | 0.47 |
| 2 | 64494913 | rs17739675 | 0.8 | 1.557 | -0.465 | -0.924 | 0.047 | 0.009 | 0.009 | 0 | 0.06 | 0.09 | 0.13 | 0.06 | 0.13 | 0.1 |
| 2 | 64495627 | rs1529102 | -0.976 | 1.269 | -0.452 | -0.963 | -0.001 | -0.009 | -0.009 | 0.01 | 0.48 | 0.56 | 0.55 | 0.79 | 0.5 | 0.65 |
| 2 | 64499629 | rs2698527 | -0.093 | 1.037 | -0.993 | -0.531 | -0.002 | -0.017 | -0.017 | 0 | 0.31 | 0.23 | 0.35 | 0.65 | 0.27 | 0.46 |
| 2 | 64503895 | rs2698530 | 0.047 | 0.955 | -1.244 | -0.485 | -0.015 | -0.051 | -0.051 | 0 | 0.28 | 0.14 | 0.32 | 0.64 | 0.24 | 0.46 |
| 2 | 64509799 | rs10200259 | -0.283 | 0.971 | -0.667 | -2.219 | -0.003 | -0.073 | -0.073 | 0.008 | 0.33 | 0.45 | 0.26 | 0.12 | 0.28 | 0.36 |
| 2 | 64509954 | rs2698537 | 0.725 | 0.922 | -0.562 | -2.341 | -0.005 | -0.004 | -0.004 | 0 | 0.09 | 0.21 | 0.08 | 0.09 | 0.08 | 0.08 |
| 2 | 64510119 | rs2433384 | -0.261 | 1.091 | -1.14 | -2.782 | -0.016 | 0.068 | 0.068 | 0 | 0.37 | 0.3 | 0.55 | 0.87 | 0.41 | 0.47 |
| 2 | 64511082 | rs2114430 | 0.242 | 1.038 | -0.775 | -2.67 | -0.028 | 0.042 | 0.042 | 0 | 0.29 | 0.19 | 0.48 | 0.77 | 0.33 | 0.39 |
| 2 | 64513207 | rs7600002 | 0.385 | 1.058 | -0.867 | -2.662 | -0.053 | 0.042 | 0.042 | 0 | 0.28 | 0.1 | 0.47 | 0.77 | 0.33 | 0.39 |
| 2 | 64517459 | rs1030406 | -0.143 | 1.04 | -0.756 | -2.752 | 0 | -0.014 | -0.014 | 0 | 0.42 | 0.62 | 0.33 | 0.19 | 0.36 | 0.42 |

| | | | | | | | | | | | | | | | | |
|---|---|---|---|---|---|---|---|---|---|---|---|---|---|---|---|---|
| 2 | 64522307 | rs2555454 | -1.252 | 0.937 | -0.557 | -2.706 | -0.003 | 0.112 | 0.112 | 0 | 0.71 | 0.93 | 0.83 | 0.98 | 0.69 | 0.83 |
| 2 | 64523758 | rs1025212 | -0.252 | 0.84 | -0.1 | -2.935 | 0.004 | -0.049 | -0.049 | 0 | 0.31 | 0.52 | 0.23 | 0.17 | 0.31 | 0.36 |
| 2 | 64533523 | rs1426712 | -0.643 | 0.88 | 0.118 | -2.895 | 0.004 | 0.013 | 0.013 | 0 | 0.51 | 0.62 | 0.48 | 0.22 | 0.52 | 0.53 |
| 2 | 64541233 | rs919126 | 1.908 | 0.855 | 0.151 | -2.909 | -0.005 | -0.036 | -0.036 | 0 | 0.15 | 0.08 | 0.3 | 0.76 | 0.15 | 0.3 |
| 2 | 64544552 | rs883021 | -1.288 | 0.606 | 0.328 | -2.9 | 0.004 | -0.016 | -0.016 | 0 | 0.68 | 0.8 | 0.79 | 0.98 | 0.67 | 0.84 |
| 2 | 64547448 | rs7568533 | 0.668 | 0.651 | 0.182 | -2.902 | 0.002 | 0.01 | 0.01 | 0.037 | 0.25 | 0.21 | 0.26 | 0.06 | 0.21 | 0.17 |
| 2 | 64550332 | rs17740892 | 0.94 | 0.687 | 0.177 | -2.884 | -0.002 | 0.063 | 0.063 | 0.018 | 0.23 | 0.08 | 0.26 | 0.06 | 0.2 | 0.17 |
| 2 | 64551328 | rs10187246 | 0.704 | 0.698 | 0.248 | -2.889 | 0.037 | -0.015 | -0.015 | 0 | 0.2 | 0.34 | 0.19 | 0.16 | 0.29 | 0.3 |
| 2 | 64553440 | rs919123 | -0.412 | 0.289 | 0.201 | -2.676 | 0.001 | 0.016 | 0.016 | 0 | 0.51 | 0.62 | 0.48 | 0.22 | 0.52 | 0.53 |
| 2 | 64553673 | rs7565077 | 1.491 | 0.369 | -0.328 | -2.371 | -0.006 | -0.016 | -0.016 | 2.949 | 0.16 | 0.11 | 0.31 | 0.76 | 0.17 | 0.31 |
| 2 | 64555550 | rs4671561 | 1.466 | 0.723 | -0.302 | -2.289 | -0.001 | -0.022 | -0.022 | 0 | 0.19 | 0.08 | 0.32 | 0.77 | 0.18 | 0.31 |
| 2 | 64559621 | rs1365475 | -0.524 | 0.708 | -0.189 | -2.466 | 0.02 | 0.06 | 0.06 | 0 | 0.5 | 0.64 | 0.49 | 0.19 | 0.54 | 0.51 |
| 2 | 64561680 | rs1426704 | -1.501 | 0.701 | -0.507 | -2.535 | 0.001 | 0.145 | 0.145 | 0 | 0.76 | 0.98 | 0.86 | 0.97 | 0.78 | 0.88 |
| 2 | 64565228 | rs1426701 | -0.64 | 0.873 | -0.106 | -2.292 | 0.037 | 0.135 | 0.135 | 0 | 0.51 | 0.78 | 0.49 | 0.18 | 0.54 | 0.5 |
| 2 | 64567550 | rs7591530 | 0.149 | 1.162 | -0.095 | -2.358 | 0.015 | -0.032 | -0.032 | 0 | 0.25 | 0.34 | 0.26 | 0.13 | 0.28 | 0.3 |
| 2 | 64568628 | rs6731818 | -0.816 | 1.012 | -0.264 | -2.357 | 0.011 | 0.03 | 0.03 | 0 | 0.49 | 0.6 | 0.48 | 0.17 | 0.54 | 0.48 |
| 2 | 64574128 | rs13008221 | 0.073 | 1.096 | -0.521 | -2.283 | 0.006 | -0.028 | -0.028 | 0.077 | 0.25 | 0.29 | 0.24 | 0.13 | 0.26 | 0.29 |
| 2 | 64588987 | rs7559505 | -0.722 | 0.441 | -0.827 | -1.938 | 0.01 | 0.05 | 0.05 | 0 | 0.72 | 0.89 | 0.81 | 0.98 | 0.72 | 0.88 |
| 2 | 64594552 | rs4300812 | -0.737 | 0.464 | -0.833 | -1.916 | 0.01 | 0.055 | 0.055 | 0 | 0.72 | 0.89 | 0.81 | 0.97 | 0.72 | 0.89 |
| 2 | 64595522 | rs7604487 | 0.013 | 0.236 | -0.607 | -2.048 | -0.008 | -0.049 | -0.049 | 1.605 | 0.61 | 0.54 | 0.7 | 0.83 | 0.65 | 0.63 |
| 2 | 64606265 | rs2288298 | -1.293 | 0.447 | -1.069 | -2.06 | -0.068 | 0.011 | 0.011 | 0.145 | 0.85 | 0.99 | 0.89 | 0.93 | 0.79 | 0.83 |
| 2 | 64607964 | rs13412599 | 0.468 | -0.358 | -0.513 | -1.666 | -0.024 | -0.022 | -0.022 | 0 | 0.3 | 0.41 | 0.27 | 0.18 | 0.21 | 0.25 |
| 2 | 64608409 | rs17690928 | 1.222 | -0.184 | -0.751 | -1.674 | -0.001 | 0.034 | 0.034 | 0 | 0.13 | 0.04 | 0.11 | 0 | 0.1 | 0.08 |
| 2 | 64609979 | rs1346236 | -0.158 | -0.057 | -0.652 | -1.595 | 0 | -0.013 | -0.013 | 0.243 | 0.56 | 0.64 | 0.67 | 0.89 | 0.48 | 0.64 |
| 2 | 64612323 | rs11892779 | 0.829 | -0.205 | -1.122 | -1.447 | 0.013 | 0.013 | 0.013 | 0 | 0.23 | 0.21 | 0.17 | 0.07 | 0.14 | 0.07 |
| 2 | 64620637 | rs7578955 | -0.586 | 0.486 | -1.706 | -1.351 | -0.098 | 0.092 | 0.092 | 0.593 | 0.73 | 0.92 | 0.8 | 0.93 | 0.58 | 0.74 |
| 2 | 64622008 | rs1365480 | 0.514 | 0.533 | -1.639 | -1.098 | -0.009 | -0.037 | -0.037 | 0.78 | 0.24 | 0.12 | 0.39 | 0.7 | 0.26 | 0.45 |
| 2 | 64623733 | rs10496112 | -0.067 | 0.125 | -1.147 | -1.483 | 0.033 | 0.08 | 0.08 | 0 | 0.38 | 0.2 | 0.32 | 0.08 | 0.28 | 0.17 |
| 2 | 64630299 | rs872241 | -0.082 | 0.42 | -0.743 | -1.614 | 0.004 | -0.012 | -0.012 | 0.159 | 0.48 | 0.55 | 0.25 | 0.13 | 0.35 | 0.38 |
| 2 | 64631084 | rs869798 | -0.056 | 0.408 | -0.532 | -1.621 | -0.002 | -0.05 | -0.05 | 0 | 0.48 | 0.52 | 0.24 | 0.1 | 0.35 | 0.32 |
| 2 | 64643871 | rs7599508 | -0.27 | 0.936 | -0.282 | -1.733 | 0.008 | -0.023 | -0.023 | 0 | 0.17 | 0.21 | 0.2 | 0.07 | 0.23 | 0.13 |
| 2 | 64644762 | rs11678307 | -0.25 | 1.299 | 0.137 | -2.038 | 0.049 | -0.021 | -0.021 | 0 | 0.27 | 0.42 | 0.37 | 0.76 | 0.36 | 0.34 |
| 2 | 64656409 | rs13391261 | 0.887 | 1.286 | 0.385 | -2.578 | 0.011 | -0.075 | -0.075 | 0 | 0.67 | 0.41 | 0.57 | 0.9 | 0.58 | 0.63 |
| 2 | 64661404 | rs4671582 | -0.138 | 1.326 | 0.692 | -2.619 | 0.011 | -0.068 | -0.068 | 2.615 | 0.8 | 0.68 | 0.68 | 0.94 | 0.74 | 0.74 |
| 2 | 64662732 | rs7591656 | 0.553 | 1.3 | 0.874 | -2.634 | 0.019 | -0.119 | -0.119 | 0.532 | 0.76 | 0.41 | 0.64 | 0.92 | 0.71 | 0.69 |
| 2 | 64663169 | rs9309360 | 0.549 | 0.87 | 0.306 | -2.858 | -0.001 | 0.001 | 0.001 | 0 | 0.26 | 0.22 | 0.33 | 0.79 | 0.28 | 0.35 |
| 2 | 64665661 | rs6733373 | -0.144 | 1.151 | 0.62 | -2.543 | 0.024 | -0.1 | -0.1 | 0.062 | 0.79 | 0.57 | 0.67 | 0.94 | 0.74 | 0.74 |
| 2 | 64668297 | rs10496116 | -0.275 | 1.23 | 0.124 | -2.458 | 0.031 | -0.046 | -0.046 | 0 | 0.12 | 0.26 | 0.11 | 0.04 | 0.17 | 0.11 |
| 2 | 64680967 | rs6729083 | 0.267 | 1.137 | 0.065 | -2.22 | 0.019 | -0.079 | -0.079 | 0.033 | 0.77 | 0.6 | 0.67 | 0.92 | 0.71 | 0.7 |
| 2 | 64687665 | rs10496117 | 0.735 | 0.895 | -0.319 | -2.387 | 0.015 | -0.024 | -0.024 | 0 | 0.23 | 0.33 | 0.33 | 0.79 | 0.27 | 0.34 |
| 2 | 64687777 | rs10496118 | 0.625 | 0.834 | -0.67 | -1.899 | 0.016 | -0.079 | -0.079 | 0.208 | 0.77 | 0.6 | 0.67 | 0.92 | 0.71 | 0.67 |
| 2 | 64690510 | rs7355745 | -0.327 | 0.678 | -1.182 | -1.523 | 0.004 | -0.033 | -0.033 | 0 | 0.89 | 0.85 | 0.9 | 0.98 | 0.88 | 0.87 |
| 2 | 64692041 | rs918292 | -1.018 | 0.89 | -1.7 | -1.398 | -0.044 | 0.037 | 0.037 | 0 | 0.92 | 0.99 | 0.9 | 0.98 | 0.87 | 0.92 |
| 2 | 64695893 | rs12996082 | 0.392 | 0.382 | -2.087 | -1.693 | -0.009 | 0.092 | 0.092 | 0 | 0.24 | 0.03 | 0.15 | 0.08 | 0.15 | 0.15 |
| 2 | 64698397 | rs6546073 | -0.142 | 0.604 | -1.486 | -0.956 | 0.002 | -0.001 | -0.001 | 0 | 0.65 | 0.56 | 0.51 | 0.35 | 0.63 | 0.52 |
| 2 | 64699137 | rs11126005 | -0.279 | 0.957 | -1.66 | -0.553 | -0.004 | 0.002 | 0.002 | 0 | 0.27 | 0.25 | 0.29 | 0.2 | 0.38 | 0.36 |
| 2 | 64700471 | rs12617093 | 0.118 | 0.26 | -1.883 | -0.232 | 0.007 | 0.13 | 0.13 | 0 | 0.52 | 0.8 | 0.55 | 0.73 | 0.41 | 0.55 |
| 2 | 64702082 | rs1421336 | -0.616 | -0.368 | -1.601 | 0.512 | 0.022 | -0.023 | -0.023 | 0 | 0.14 | 0.05 | 0.1 | 0.24 | 0.06 | 0.14 |
| 2 | 64703286 | rs1345516 | -1.19 | -0.516 | -1.21 | 0.156 | -0.007 | -0.014 | -0.014 | 0 | 0.43 | 0.44 | 0.43 | 0.35 | 0.37 | 0.52 |
| 2 | 64707211 | rs870312 | -0.31 | -0.7 | -0.911 | 0.197 | 0.024 | -0.011 | -0.011 | 0 | 0.15 | 0.12 | 0.08 | 0.2 | 0.06 | 0.14 |
| 2 | 64707623 | rs4671593 | -0.986 | -0.791 | -1.041 | 0.298 | 0.032 | -0.051 | -0.051 | 1.258 | 0.21 | 0.11 | 0.17 | 0.34 | 0.13 | 0.31 |
| 2 | 64713532 | rs888527 | 0.476 | -0.693 | -0.864 | -0.069 | -0.011 | -0.057 | -0.057 | 0.178 | 0.61 | 0.57 | 0.6 | 0.84 | 0.62 | 0.48 |
| 2 | 64717119 | rs12477656 | 1.013 | -1.023 | -0.685 | 0.409 | 0.071 | 0.07 | 0.07 | 0.113 | 0.53 | 0.13 | 0.51 | 0.62 | 0.6 | 0.43 |
| 2 | 64719236 | rs13409095 | 0.565 | -1.044 | -0.465 | 0.327 | 0.006 | 0.01 | 0.01 | 0 | 0.57 | 0.41 | 0.54 | 0.8 | 0.6 | 0.43 |
| 2 | 64720254 | rs6756027 | -1.262 | -1.132 | -0.338 | 0.682 | 0.052 | 0.055 | 0.055 | 0 | 0.48 | 0.79 | 0.47 | 0.35 | 0.42 | 0.54 |
| 2 | 64722216 | rs12104828 | -1.3 | -1.058 | -0.204 | 1.118 | 0.05 | -0.025 | -0.025 | 0 | 0.49 | 0.84 | 0.47 | 0.45 | 0.43 | 0.55 |
| 2 | 64725466 | rs6546079 | -1.347 | -1.098 | -0.144 | 1.072 | 0.041 | -0.053 | -0.053 | 0.005 | 0.49 | 0.89 | 0.47 | 0.45 | 0.43 | 0.55 |
| 2 | 64726449 | rs13002043 | -1.348 | -1.1 | -0.152 | 1.071 | 0.037 | -0.058 | -0.058 | 0 | 0.49 | 0.9 | 0.47 | 0.45 | 0.43 | 0.55 |
| 2 | 64737208 | rs11686051 | -0.845 | -1.215 | 0.396 | 0.993 | 0.006 | 0.023 | 0.023 | 0.022 | 0.38 | 0.29 | 0.29 | 0.2 | 0.38 | 0.41 |
| 2 | 64744358 | rs6731663 | -1.351 | -1.258 | 0.379 | 1.033 | 0.073 | -0.033 | -0.033 | 0 | 0.44 | 0.75 | 0.31 | 0.44 | 0.33 | 0.48 |
| 2 | 64758410 | rs12713515 | -0.859 | -1.178 | 0.445 | 1.002 | 0.005 | 0.008 | 0.008 | 0.216 | 0.38 | 0.34 | 0.28 | 0.23 | 0.33 | 0.39 |
| 2 | 64769481 | rs2042091 | -1.034 | -1.204 | 0.421 | 1.006 | -0.018 | -0.03 | -0.03 | 0 | 0.41 | 0.48 | 0.35 | 0.27 | 0.34 | 0.44 |
| 2 | 64773779 | rs1978404 | -1.631 | -1.232 | 0.35 | 1.041 | 0.099 | -0.046 | -0.046 | 0 | 0.48 | 0.91 | 0.38 | 0.5 | 0.35 | 0.53 |
| 2 | 64779509 | rs3770740 | -0.85 | -1.231 | 0.417 | 1.06 | 0.039 | 0.45 | 0.45 | 5.893 | 0.06 | 0.43 | 0.04 | 0.23 | 0.01 | 0.08 |
| 2 | 64781613 | rs3770737 | -1.012 | -1.252 | 0.425 | 1.022 | 0.007 | 0.026 | 0.026 | 0.012 | 0.4 | 0.33 | 0.31 | 0.23 | 0.39 | 0.4 |
| 2 | 64788138 | rs3770734 | -1.57 | -1.23 | 0.43 | 1.05 | -0.006 | 0.395 | 0.395 | 5.223 | 0.08 | 0.43 | 0.09 | 0.29 | 0.01 | 0.14 |
| 2 | 64790171 | rs2287529 | -1.503 | -1.22 | 0.505 | 1.069 | 0.058 | -0.018 | -0.018 | 0 | 0.47 | 0.82 | 0.39 | 0.49 | 0.4 | 0.53 |
| 2 | 64800452 | rs17700924 | 1.645 | -1.354 | 0.004 | 0.656 | -0.005 | 0.042 | 0.042 | 0 | 0.08 | 0 | 0.05 | 0 | 0.08 | 0.02 |
| 2 | 64807294 | rs1001291 | -0.141 | -1.384 | 0.148 | 0.761 | -0.002 | -0.012 | -0.012 | 0 | 0.3 | 0.31 | 0.25 | 0.16 | 0.28 | 0.29 |

| | | | | | | | | | | | | | | | | |
|---|---|---|---|---|---|---|---|---|---|---|---|---|---|---|---|---|
| 2 | 64813124 | rs6747349 | -0.649 | -1.36 | 0.415 | 0.742 | -0.009 | 0.2 | 0.2 | 0 | 0.14 | 0.39 | 0.09 | 0.28 | 0.12 | 0.2 |
| 2 | 64814549 | rs3770728 | -0.127 | -1.385 | 0.337 | 0.727 | 0.001 | 0.003 | 0.003 | 0.015 | 0.3 | 0.28 | 0.24 | 0.17 | 0.27 | 0.29 |
| 2 | 64814765 | rs3770725 | -0.788 | -1.422 | 0.356 | 0.727 | -0.024 | 0.195 | 0.195 | 0 | 0.15 | 0.44 | 0.09 | 0.28 | 0.12 | 0.2 |
| 2 | 64814837 | rs3770724 | -1.425 | -1.379 | 0.045 | 0.623 | 0.053 | -0.029 | -0.029 | 0.057 | 0.46 | 0.77 | 0.34 | 0.45 | 0.4 | 0.49 |
| 2 | 64823848 | rs1206983 | 0.98 | -1.063 | 0.048 | 0.739 | -0.023 | 0.148 | 0.148 | 0 | 0.78 | 0.45 | 0.78 | 0.57 | 0.78 | 0.66 |
| 2 | 64824348 | rs10211119 | -0.014 | -0.763 | 0.073 | 0.333 | -0.009 | -0.036 | -0.036 | 0 | 0.22 | 0.29 | 0.18 | 0.09 | 0.17 | 0.18 |
| 2 | 64826469 | rs11682570 | -0.736 | -0.88 | 0.338 | 0.498 | -0.05 | -0.034 | -0.034 | 0.012 | 0.29 | 0.51 | 0.22 | 0.16 | 0.18 | 0.27 |
| 2 | 64826959 | rs10171427 | -0.282 | -0.725 | 0.373 | 0.46 | -0.02 | -0.039 | -0.039 | 0.496 | 0.25 | 0.32 | 0.14 | 0.08 | 0.16 | 0.2 |
| 2 | 64829151 | rs13020375 | -0.658 | -0.414 | 0.367 | 0.476 | 0.016 | 0.044 | 0.044 | 0 | 0.47 | 0.61 | 0.45 | 0.25 | 0.38 | 0.49 |
| 2 | 64829465 | rs4671599 | -1.393 | 0.173 | -0.402 | 0.842 | -0.064 | -0.059 | -0.059 | 0 | 0.71 | 0.89 | 0.66 | 0.61 | 0.59 | 0.73 |
| 2 | 64833920 | rs1861404 | -0.039 | -0.381 | 0.069 | 0.761 | -0.018 | -0.04 | -0.04 | 0 | 0.4 | 0.51 | 0.37 | 0.19 | 0.29 | 0.35 |
| 2 | 64836095 | rs4671600 | -0.792 | 0.025 | -0.686 | 1.067 | 0.016 | -0.029 | -0.029 | 0.058 | 0.32 | 0.75 | 0.33 | 0.44 | 0.26 | 0.4 |
| 2 | 64839959 | rs11126017 | 1.319 | 0.377 | -0.819 | 1.129 | -0.144 | -0.033 | -0.033 | 0 | 0.25 | 0.02 | 0.31 | 0.24 | 0.39 | 0.16 |
| 2 | 64840180 | rs7585112 | -0.142 | -0.066 | 0.442 | 1.349 | -0.042 | 0.062 | 0.062 | 0 | 0.08 | 0.17 | 0.06 | 0.17 | 0.01 | 0.2 |
| 2 | 64845543 | rs6734765 | -0.613 | -0.08 | 0.306 | 1.3 | -0.054 | 0.215 | 0.215 | 1.646 | 0.11 | 0.4 | 0.17 | 0.28 | 0.02 | 0.24 |
| 2 | 64852557 | rs7557491 | 0.311 | 0.753 | -0.448 | 0.722 | -0.035 | -0.022 | -0.022 | 0 | 0.11 | 0.03 | 0.13 | 0.13 | 0.16 | 0.11 |
| 2 | 64856267 | rs7605342 | 0.613 | 0.065 | -0.188 | 0.053 | -0.012 | -0.067 | -0.067 | 0 | 0.7 | 0.65 | 0.86 | 0.95 | 0.76 | 0.77 |
| 2 | 64857836 | rs10186940 | 0.192 | 0.15 | -0.519 | 0.112 | -0.001 | 0.08 | 0.08 | 0.465 | 0.76 | 0.92 | 0.88 | 0.95 | 0.78 | 0.8 |
| 2 | 64859679 | rs7451 | -0.083 | 0.092 | -0.483 | 0.143 | 0.001 | 0.052 | 0.052 | 0 | 0.8 | 0.91 | 0.92 | 0.97 | 0.82 | 0.81 |
| 2 | 64862055 | rs2070063 | -0.115 | 0.27 | -0.372 | 0.617 | 0.002 | 0.003 | 0.003 | 0 | 0.36 | 0.34 | 0.44 | 0.45 | 0.39 | 0.38 |
| 2 | 64867783 | rs1477035 | -0.255 | 0.295 | -0.287 | 0.559 | -0.003 | -0.003 | -0.003 | 0 | 0.37 | 0.37 | 0.44 | 0.48 | 0.44 | 0.37 |
| 2 | 64872736 | rs4671603 | 0.105 | -0.241 | -0.081 | 0.375 | -0.022 | -0.023 | -0.023 | 0.042 | 0.8 | 0.81 | 0.92 | 0.93 | 0.89 | 0.8 |
| 2 | 64873442 | rs6546094 | -0.222 | -0.171 | -0.517 | 0.41 | 0.036 | 0.038 | 0.038 | 0 | 0.83 | 0.96 | 0.94 | 0.93 | 0.9 | 0.81 |
| 2 | 64895272 | rs10189932 | -0.462 | -0.168 | -0.665 | 0.493 | 0.023 | 0.005 | 0.005 | 0 | 0.86 | 1 | 0.97 | 0.94 | 0.94 | 0.88 |
| 2 | 64905729 | rs4671606 | 0.506 | -0.109 | 0.198 | -0.03 | -0.006 | -0.042 | -0.042 | 2.394 | 0.64 | 0.4 | 0.82 | 0.89 | 0.73 | 0.73 |
| 2 | 64906295 | rs6546096 | 0.106 | -0.122 | 0.121 | -0.037 | -0.022 | 0.086 | 0.086 | 0 | 0.7 | 0.89 | 0.87 | 0.89 | 0.74 | 0.73 |
| 2 | 64918070 | rs3796051 | 1.222 | -0.226 | -0.08 | 0.335 | -0.024 | -0.057 | -0.057 | 0.063 | 0.44 | 0.17 | 0.5 | 0.52 | 0.48 | 0.49 |
| 2 | 64928603 | rs12471768 | 0.639 | -0.34 | 0.377 | 0.152 | -0.007 | -0.062 | -0.062 | 0.055 | 0.65 | 0.41 | 0.81 | 0.92 | 0.71 | 0.73 |
| 2 | 64936472 | rs6733507 | 0.434 | -0.158 | 0.394 | 0.449 | 0.014 | -0.007 | -0.007 | 0 | 0.18 | 0.24 | 0.14 | 0.11 | 0.26 | 0.16 |
| 2 | 64948153 | rs4671609 | 1.101 | -0.51 | -0.076 | 0.753 | -0.004 | 0 | 0 | 0.416 | 0.54 | 0.5 | 0.63 | 0.58 | 0.59 | 0.47 |
| 2 | 64948378 | rs6760401 | -0.622 | -0.579 | 0.025 | 0.736 | -0.038 | -0.027 | -0.027 | 0 | 0.84 | 0.75 | 0.95 | 0.95 | 0.95 | 0.88 |
| 2 | 64961036 | rs6744311 | 0.226 | -0.463 | -0.091 | 0.74 | -0.001 | 0 | 0 | 0 | 0.67 | 0.65 | 0.83 | 0.84 | 0.68 | 0.72 |
| 2 | 64971551 | rs11680278 | 0.494 | -0.124 | -0.457 | 0.959 | 0.008 | -0.012 | -0.012 | 0 | 0.12 | 0.11 | 0.19 | 0.26 | 0.09 | 0.25 |
| 2 | 64983495 | rs6723426 | 0.921 | -0.032 | -0.558 | 0.837 | -0.041 | 0.006 | 0.006 | 0 | 0.15 | 0.09 | 0.13 | 0.11 | 0.27 | 0.15 |
| 2 | 64993045 | rs2136451 | -0.656 | -0.248 | -0.435 | 0.527 | 0.002 | -0.016 | -0.016 | 0 | 0.3 | 0.39 | 0.31 | 0.24 | 0.31 | 0.31 |
| 2 | 64994005 | rs847808 | -0.31 | 0.007 | -1.209 | 0.446 | -0.032 | 0.027 | 0.027 | 0.08 | 0.82 | 1 | 0.92 | 0.95 | 0.87 | 0.81 |
| 2 | 65004346 | rs17608651 | 0.496 | 0.421 | -0.932 | 0.572 | -0.051 | 0.064 | 0.064 | 0 | 0.17 | 0.07 | 0.11 | 0.02 | 0.24 | 0.18 |
| 2 | 65007900 | rs6721629 | 0.32 | -0.291 | -0.662 | 1.124 | 0.011 | -0.017 | -0.017 | 0 | 0.1 | 0.05 | 0.18 | 0.22 | 0.08 | 0.15 |
| 2 | 65008738 | rs929699 | -0.392 | -0.931 | -0.534 | 0.162 | 0.005 | 0.004 | 0.004 | 0.02 | 0.45 | 0.63 | 0.41 | 0.28 | 0.32 | 0.29 |
| 2 | 65008756 | rs929698 | -0.934 | -0.881 | -0.507 | 0.395 | -0.011 | -0.005 | -0.005 | 0.335 | 0.61 | 0.72 | 0.61 | 0.46 | 0.5 | 0.46 |
| 2 | 65008812 | rs929697 | -0.298 | -1.156 | -0.702 | 0.171 | 0.011 | 0.011 | 0.011 | 0 | 0.32 | 0.28 | 0.38 | 0.26 | 0.26 | 0.36 |
| 2 | 65009646 | rs7579416 | 0.789 | -1.291 | -0.815 | 0.055 | 0.001 | 0.02 | 0.02 | 0.142 | 0.2 | 0.16 | 0.13 | 0.05 | 0.19 | 0.31 |
| 2 | 65010120 | rs6732883 | -0.636 | -1.575 | -0.582 | 0.264 | 0.014 | 0.002 | 0.002 | 0 | 0.36 | 0.32 | 0.36 | 0.3 | 0.3 | 0.51 |
| 2 | 65010397 | rs17608884 | 1.664 | -1.457 | -1.252 | 0.084 | -0.03 | 0.068 | 0.068 | 0.297 | 0.16 | 0.03 | 0.1 | 0.05 | 0.15 | 0.04 |
| 2 | 65010556 | rs6546102 | -1.543 | -1.31 | -0.682 | 0.554 | -0.047 | -0.056 | -0.056 | 0 | 0.65 | 0.83 | 0.62 | 0.53 | 0.49 | 0.61 |
| 2 | 65010688 | rs6704589 | -0.122 | -0.892 | -0.136 | 0.88 | -0.01 | 0.014 | 0.014 | 0 | 0.13 | 0.15 | 0.2 | 0.25 | 0.08 | 0.18 |
| 2 | 65023697 | rs17029444 | -0.459 | -0.511 | 0.391 | 0.56 | -0.009 | 0.026 | 0.026 | 0.102 | 0.07 | 0.22 | 0.03 | 0.07 | 0.02 | 0.05 |
| 2 | 65027624 | rs11683197 | 0.149 | -0.71 | 0.094 | 0.336 | 0.011 | 0.059 | 0.059 | 0 | 0.29 | 0.08 | 0.25 | 0.17 | 0.26 | 0.4 |
| 2 | 65029203 | rs6724824 | -1.071 | -0.687 | 0.611 | 0.582 | -0.002 | -0.001 | -0.001 | 0.043 | 0.43 | 0.53 | 0.39 | 0.37 | 0.34 | 0.52 |
| 2 | 65036516 | rs11126021 | 0.862 | 0.012 | 0.348 | 0.725 | -0.003 | -0.008 | -0.008 | 0.042 | 0.55 | 0.67 | 0.58 | 0.54 | 0.55 | 0.46 |
| 2 | 65043456 | rs4671614 | 2.063 | 0.276 | 1.24 | 1.323 | 0.024 | 0.099 | 0.099 | 0 | 0.79 | 0.49 | 0.81 | 0.66 | 0.8 | 0.83 |
| 2 | 65045992 | rs7608749 | 1.454 | 0.252 | 1.166 | 1.16 | 0.046 | 0.011 | 0.011 | 0 | 0.86 | 0.6 | 0.91 | 0.86 | 0.86 | 0.93 |
| 2 | 65053216 | rs6706514 | -0.826 | 0.002 | 0.693 | 1.147 | -0.014 | 0.041 | 0.041 | 0 | 0.12 | 0.16 | 0.11 | 0.22 | 0.07 | 0.13 |
| 2 | 65056358 | rs12373586 | 0.356 | 0.837 | 0.602 | 0.688 | 0.04 | 0 | 0 | 0 | 0.89 | 0.8 | 0.89 | 0.87 | 0.77 | 0.91 |
| 2 | 65057763 | rs7607348 | 0.561 | 0.881 | 0.57 | 0.653 | 0.023 | -0.024 | -0.024 | 0 | 0.06 | 0.1 | 0.05 | 0.01 | 0.11 | 0.05 |
| 2 | 65059603 | rs6738590 | 0.254 | 0.445 | 0.347 | 0.395 | -0.007 | -0.016 | -0.016 | 0 | 0.71 | 0.7 | 0.78 | 0.79 | 0.77 | 0.88 |
| 2 | 65066311 | rs741477 | 0.997 | 0.72 | -0.339 | 0.33 | -0.028 | 0.027 | 0.027 | 0 | 0.05 | 0.01 | 0.04 | 0.03 | 0.12 | 0.05 |
| 2 | 65078728 | rs13026295 | 1.16 | 0.336 | -0.377 | 0.519 | -0.016 | 0.019 | 0.019 | 4.182 | 0.47 | 0.21 | 0.45 | 0.43 | 0.54 | 0.55 |
| 2 | 65080772 | rs7600672 | -0.232 | -0.284 | -0.104 | -0.029 | 0.001 | -0.001 | -0.001 | 0 | 0.28 | 0.27 | 0.19 | 0.06 | 0.23 | 0.13 |
| 2 | 65082367 | rs6546106 | -0.817 | -0.014 | -0.18 | 0.37 | -0.005 | -0.009 | -0.009 | 0 | 0.39 | 0.53 | 0.34 | 0.31 | 0.34 | 0.23 |
| 2 | 65084066 | rs7573247 | -0.414 | -0.124 | 0.162 | 0.226 | -0.026 | -0.071 | -0.071 | 0 | 0.13 | 0.34 | 0.05 | 0.02 | 0.06 | 0.04 |
| 2 | 65084171 | rs7602986 | -0.599 | -0.096 | 0.125 | 0.695 | -0.045 | 0.14 | 0.14 | 0.508 | 0.24 | 0.54 | 0.26 | 0.5 | 0.15 | 0.2 |
| 2 | 65089291 | rs4296412 | 0.212 | -0.135 | -0.975 | 0.37 | 0.008 | 0.008 | 0.008 | 0 | 0.37 | 0.27 | 0.3 | 0.2 | 0.31 | 0.4 |
| 2 | 65091601 | rs2160782 | -0.33 | 0.737 | -1.443 | 0.734 | -0.086 | 0.007 | 0.007 | 0.145 | 0.67 | 0.91 | 0.64 | 0.73 | 0.6 | 0.64 |
| 2 | 65096936 | rs17029539 | -0.348 | 0.091 | -0.793 | 0.595 | 0.003 | 0.002 | 0.002 | 1.125 | 0.38 | 0.7 | 0.39 | 0.6 | 0.34 | 0.41 |
| 2 | 65097201 | rs2063307 | -0.577 | -0.305 | -0.489 | 0.131 | -0.01 | -0.041 | -0.041 | 0.091 | 0.15 | 0.21 | 0.15 | 0.05 | 0.13 | 0.12 |
| 2 | 65097855 | rs9678117 | 0.534 | -0.358 | -0.735 | 0.516 | 0.011 | 0.001 | 0.001 | 2.471 | 0.74 | 0.23 | 0.82 | 0.72 | 0.82 | 0.8 |
| 2 | 65101579 | rs2048262 | 0.218 | -0.311 | -0.569 | 0.382 | -0.003 | -0.008 | -0.008 | 0 | 0.91 | 0.9 | 0.91 | 0.94 | 0.91 | 0.97 |
| 2 | 65102741 | rs6736423 | -0.259 | 0.472 | -0.267 | -0.036 | 0.031 | -0.034 | -0.034 | 0 | 0.27 | 0.32 | 0.24 | 0.14 | 0.38 | 0.16 |

| | | | | | | | | | | | | | | | | |
|---|---|---|---|---|---|---|---|---|---|---|---|---|---|---|---|---|
| 2 | 65103983 | rs6546108 | -0.901 | -0.077 | -0.251 | 0.437 | -0.024 | -0.024 | -0.024 | 0.078 | 0.45 | 0.68 | 0.51 | 0.51 | 0.5 | 0.33 |
| 2 | 65106350 | rs918197 | -1.179 | -0.276 | -0.233 | 0.598 | -0.02 | 0.006 | 0.006 | 2.041 | 0.48 | 0.78 | 0.58 | 0.54 | 0.51 | 0.41 |
| 2 | 65106369 | rs17029550 | -0.369 | -0.314 | -0.014 | 0.501 | -0.013 | -0.028 | -0.028 | 0 | 0.12 | 0.32 | 0.15 | 0.04 | 0.07 | 0.21 |
| 2 | 65109039 | rs1995829 | -1.138 | -0.258 | -0.323 | 0.443 | -0.012 | 0.013 | 0.013 | 1.899 | 0.49 | 0.74 | 0.58 | 0.54 | 0.51 | 0.42 |
| 2 | 65110275 | rs10490211 | 0.585 | -0.234 | -0.644 | 0.367 | 0 | 0 | 0 | 0 | 0.29 | 0.24 | 0.21 | 0.32 | 0.27 | 0.32 |
| 2 | 65110310 | rs6546110 | -0.162 | 0.631 | -0.713 | 0.19 | 0.005 | 0.035 | 0.035 | 0 | 0.5 | 0.56 | 0.47 | 0.29 | 0.5 | 0.58 |
| 2 | 65113981 | rs2058905 | 0.744 | 1.127 | -0.473 | 0.796 | -0.002 | 0.002 | 0.002 | 0 | 0.06 | 0.04 | 0.15 | 0.18 | 0.06 | 0.2 |
| 2 | 65114005 | rs2058904 | -0.719 | 1.115 | -0.519 | 0.81 | 0.018 | 0.05 | 0.05 | 0.776 | 0.24 | 0.34 | 0.42 | 0.41 | 0.29 | 0.3 |
| 2 | 65114067 | rs918195 | -0.339 | 1.294 | -0.632 | 0.899 | 0.009 | 0.027 | 0.027 | 0.328 | 0.21 | 0.2 | 0.41 | 0.41 | 0.29 | 0.3 |
| 2 | 65128290 | rs6761711 | 0.749 | 1.16 | -0.614 | 0.192 | 0.001 | -0.053 | -0.053 | 5.395 | 0.25 | 0.37 | 0.21 | 0.13 | 0.26 | 0.21 |
| 2 | 65130948 | rs13022383 | -0.21 | 1.155 | -0.741 | 0.653 | 0.041 | 0.146 | 0.146 | 0.152 | 0.19 | 0.42 | 0.29 | 0.4 | 0.27 | 0.12 |
| 2 | 65135370 | rs4671628 | -1.336 | 1.462 | -1.709 | 0.288 | -0.121 | 0.049 | 0.049 | 0 | 0.78 | 0.98 | 0.79 | 0.85 | 0.68 | 0.78 |
| 2 | 65141149 | rs17029619 | 0.746 | 1.405 | -1.493 | 0.764 | -0.031 | -0.077 | -0.077 | 0.109 | 0.11 | 0.01 | 0.2 | 0.43 | 0.15 | 0.11 |
| 2 | 65142754 | rs6740379 | -0.791 | 0.806 | -1.604 | 0.445 | 0.001 | 0.006 | 0.006 | 0 | 0.42 | 0.38 | 0.45 | 0.54 | 0.42 | 0.29 |
| 2 | 65144657 | rs4671629 | -0.536 | 0.856 | -1.576 | 0.372 | -0.048 | -0.083 | -0.083 | 0 | 0.21 | 0.1 | 0.31 | 0.43 | 0.32 | 0.12 |
| 2 | 65146446 | rs1974689 | 0.825 | -0.026 | -1.486 | 0.542 | -0.011 | -0.006 | -0.006 | 0 | 0.2 | 0.27 | 0.13 | 0.11 | 0.09 | 0.15 |
| 2 | 65156458 | rs918060 | 0.24 | 0.125 | -1.575 | 0.618 | -0.036 | -0.104 | -0.104 | 0.433 | 0.13 | 0.01 | 0.24 | 0.43 | 0.17 | 0.11 |
| 2 | 65159596 | rs1035165 | -0.241 | 0.218 | -1.357 | 0.408 | 0.026 | -0.019 | -0.019 | 0 | 0.35 | 0.31 | 0.37 | 0.51 | 0.25 | 0.26 |
| 2 | 65162318 | rs7571146 | 0.355 | -0.028 | -0.825 | -0.322 | -0.005 | -0.047 | -0.047 | 0.06 | 0.11 | 0.21 | 0.07 | 0 | 0.08 | 0.1 |
| 2 | 65170092 | rs12466049 | -0.25 | -0.276 | -1.179 | -0.156 | 0.054 | -0.021 | -0.021 | 0 | 0.37 | 0.21 | 0.39 | 0.46 | 0.27 | 0.46 |
| 2 | 65171762 | rs17029659 | 0.87 | 0.174 | -0.646 | -0.302 | 0.091 | -0.028 | -0.028 | 0 | 0.06 | 0.16 | 0.07 | 0 | 0.15 | 0.01 |
| 2 | 65174408 | rs7590631 | 0.15 | 0.088 | -0.139 | -0.344 | 0.143 | -0.021 | -0.021 | 0 | 0.08 | 0.31 | 0.09 | 0 | 0.15 | 0.02 |
| 2 | 65180817 | rs4671635 | -0.247 | -0.33 | -0.683 | -0.464 | 0.058 | -0.045 | -0.045 | 0.105 | 0.37 | 0.26 | 0.37 | 0.45 | 0.23 | 0.34 |
| 2 | 65193391 | rs2017320 | 1.011 | 0.143 | -0.516 | -0.29 | -0.001 | 0.006 | 0.006 | 0 | 0.12 | 0.13 | 0.07 | 0.01 | 0.1 | 0.11 |
| 2 | 65200618 | rs4671636 | -0.254 | 0.489 | -0.542 | -0.068 | 0.006 | 0 | 0 | 2.934 | 0.54 | 0.72 | 0.48 | 0.45 | 0.42 | 0.56 |
| 2 | 65212178 | rs6546118 | 0.564 | 1.186 | -0.316 | 0.01 | 0.025 | 0.067 | 0.067 | 0.209 | 0.11 | 0.16 | 0.18 | 0.27 | 0.17 | 0.16 |
| 2 | 65212520 | rs9677813 | 0.857 | 1.447 | -0.488 | -0.143 | -0.082 | 0.048 | 0.048 | 0.845 | 0.18 | 0.07 | 0.2 | 0.1 | 0.31 | 0.23 |
| 2 | 65213104 | rs6712664 | 0.639 | 1.157 | -0.441 | 0.039 | 0.031 | 0.055 | 0.055 | 0.663 | 0.11 | 0.18 | 0.12 | 0.24 | 0.17 | 0.16 |
| 2 | 65215030 | rs11675218 | 0.422 | -0.115 | -0.992 | 0.1 | -0.001 | -0.001 | -0.001 | 1.234 | 0.25 | 0.3 | 0.37 | 0.54 | 0.13 | 0.19 |
| 2 | 65221184 | rs1541580 | -0.075 | 0.498 | -1.218 | -0.469 | 0.069 | 0.105 | 0.105 | 0.544 | 0.62 | 0.85 | 0.79 | 0.88 | 0.76 | 0.58 |
| 2 | 65231245 | rs11891502 | -0.539 | 1.475 | -0.36 | 0.095 | 0.098 | -0.025 | -0.025 | 0.319 | 0.06 | 0.12 | 0.08 | 0 | 0.14 | 0.01 |
| 2 | 65234410 | rs11689731 | -0.95 | 0.834 | -0.132 | -0.064 | -0.011 | -0.026 | -0.026 | 0 | 0.18 | 0.32 | 0.11 | 0.13 | 0.13 | 0.11 |
| 2 | 65234613 | rs7583682 | 0.64 | 0.11 | -0.852 | -0.904 | 0.03 | 0.068 | 0.068 | 0.005 | 0.81 | 0.93 | 0.89 | 0.97 | 0.86 | 0.97 |
| 2 | 65235333 | rs6546119 | -0.565 | 0.324 | -1.119 | -0.677 | -0.012 | -0.009 | -0.009 | 0 | 0.73 | 0.76 | 0.4 | 0.37 | 0.57 | 0.44 |
| 2 | 65243321 | rs6728107 | -0.56 | -0.51 | -0.957 | -1.236 | 0.007 | -0.004 | -0.004 | 0 | 0.23 | 0.2 | 0.09 | 0.02 | 0.19 | 0.13 |
| 2 | 65245365 | rs759458 | 0.772 | -1.15 | -0.758 | -1.43 | 0.081 | 0.123 | 0.123 | 6.944 | 0.38 | 0.14 | 0.19 | 0.11 | 0.24 | 0.3 |
| 2 | 65246123 | rs2075209 | 0.962 | -1.931 | -0.568 | -1.694 | 0.01 | -0.008 | -0.008 | 0.023 | 0.28 | 0.2 | 0.15 | 0.32 | 0.18 | 0.25 |
| 2 | 65248271 | rs7563131 | 0.354 | -1.829 | -0.459 | -1.718 | -0.033 | -0.091 | -0.091 | 0.017 | 0.16 | 0.27 | 0.1 | 0.01 | 0.12 | 0.09 |
| 2 | 65249922 | rs7850 | 0.573 | -1.772 | -1.037 | -1.69 | -0.001 | -0.024 | -0.024 | 0 | 0.15 | 0.12 | 0.1 | 0.01 | 0.12 | 0.1 |
| 2 | 65255435 | rs6546121 | 0.249 | -1.602 | -1.39 | -1.617 | -0.001 | -0.022 | -0.022 | 0 | 0.13 | 0.14 | 0.07 | 0.01 | 0.14 | 0.11 |
| 2 | 65265995 | rs7601712 | -0.85 | -1.507 | -1.626 | -1.47 | 0.104 | 0.15 | 0.15 | 0 | 0.49 | 0.83 | 0.36 | 0.24 | 0.4 | 0.29 |
| 2 | 65272362 | rs2540953 | -0.884 | -0.747 | -1.312 | -1.041 | -0.1 | -0.061 | -0.061 | 0 | 0.73 | 0.89 | 0.47 | 0.32 | 0.52 | 0.47 |
| 2 | 65276049 | rs1009360 | -0.325 | -0.891 | -1.095 | -1.152 | 0.001 | 0.01 | 0.01 | 3.242 | 0.62 | 0.71 | 0.41 | 0.31 | 0.44 | 0.44 |
| 2 | 65278339 | rs6711537 | 0.49 | -1.482 | -0.716 | -1.484 | -0.024 | -0.06 | -0.06 | 0.613 | 0.11 | 0.21 | 0.03 | 0 | 0.05 | 0.02 |
| 2 | 65279223 | rs2540950 | 0.334 | -1.787 | -0.33 | -1.651 | 0.045 | 0.036 | 0.036 | 2.485 | 0.5 | 0.39 | 0.37 | 0.31 | 0.36 | 0.41 |
| 2 | 65287896 | rs2249105 | 0.168 | -1.846 | -0.253 | -1.523 | 0 | 0 | 0 | 0.974 | 0.51 | 0.49 | 0.37 | 0.33 | 0.38 | 0.41 |
| 2 | 65291546 | rs2241160 | -0.689 | -1.906 | -0.253 | -1.576 | -0.002 | 0.001 | 0.001 | 0.967 | 0.64 | 0.71 | 0.43 | 0.33 | 0.44 | 0.44 |
| 2 | 65296798 | rs7572857 | 1.765 | -2.492 | -0.772 | -1.79 | 0.005 | 0.052 | 0.052 | 0.853 | 0.24 | 0.01 | 0.12 | 0.09 | 0.17 | 0.22 |
| 2 | 65298839 | rs17849707 | 0.577 | -2.683 | -0.775 | -1.869 | 0.065 | -0.003 | -0.003 | 0.299 | 0.36 | 0.15 | 0.27 | 0.33 | 0.23 | 0.3 |
| 2 | 65300995 | rs2723090 | 1.041 | -2.713 | -0.658 | -1.866 | 0 | -0.02 | -0.02 | 0 | 0.11 | 0.11 | 0.08 | 0 | 0.12 | 0.08 |
| 2 | 65309223 | rs6546125 | 1.002 | -2.857 | -0.759 | -1.888 | 0.058 | -0.044 | -0.044 | 0.201 | 0.31 | 0.08 | 0.26 | 0.33 | 0.19 | 0.28 |
| 2 | 65309296 | rs10496123 | 1.961 | -1.462 | -1.143 | -0.686 | -0.099 | -0.095 | -0.095 | 0 | 0.17 | 0.03 | 0.24 | 0.32 | 0.27 | 0.29 |
| 2 | 65312217 | rs4671638 | -0.775 | -1.92 | -0.418 | -1.143 | -0.016 | -0.042 | -0.042 | 1.204 | 0.62 | 0.63 | 0.79 | 0.91 | 0.74 | 0.71 |
| 2 | 65315903 | rs1420185 | 0.184 | -1.899 | -0.422 | -1.13 | 0.081 | 0.053 | 0.053 | 1.942 | 0.48 | 0.32 | 0.62 | 0.67 | 0.66 | 0.63 |
| 2 | 65324089 | rs3771105 | 0.968 | -1.723 | -0.626 | -1.017 | 0.001 | 0.01 | 0.01 | 0 | 0.09 | 0.07 | 0.15 | 0.24 | 0.05 | 0.08 |
| 2 | 65326568 | rs9784068 | 2.221 | -1.785 | -0.559 | -1.063 | -0.021 | -0.029 | -0.029 | 0.989 | 0.24 | 0.27 | 0.13 | 0.09 | 0.14 | 0.21 |
| 2 | 65328694 | rs10195328 | 0.031 | -1.799 | -0.575 | -1.078 | 0.013 | -0.005 | -0.005 | 0.139 | 0.78 | 0.86 | 0.78 | 0.76 | 0.82 | 0.84 |
| 2 | 65331659 | rs2302631 | -0.112 | -1.793 | -0.582 | -1.076 | -0.011 | -0.005 | -0.005 | 0.134 | 0.54 | 0.56 | 0.64 | 0.67 | 0.68 | 0.63 |
| 2 | 65341374 | rs17029863 | 2.068 | -1.878 | -0.542 | -1.058 | -0.012 | -0.017 | -0.017 | 2.597 | 0.25 | 0.26 | 0.13 | 0.09 | 0.14 | 0.21 |
| 2 | 65355744 | rs10185243 | 0.016 | -1.769 | -0.504 | -1.236 | 0.02 | 0.014 | 0.014 | 1.569 | 0.5 | 0.49 | 0.65 | 0.67 | 0.67 | 0.61 |
| 2 | 65364702 | rs12713533 | -0.432 | -1.479 | -0.345 | -1.023 | -0.016 | -0.005 | -0.005 | 0 | 0.59 | 0.49 | 0.67 | 0.67 | 0.73 | 0.7 |
| 2 | 65369964 | rs984748 | -0.729 | -1.516 | -0.415 | -1.045 | 0.073 | 0.003 | 0.003 | 4.133 | 0.63 | 0.86 | 0.7 | 0.67 | 0.73 | 0.7 |
| 2 | 65370766 | rs2422441 | -0.119 | -1.486 | -0.451 | -1.034 | 0.023 | 0.011 | 0.011 | 4.653 | 0.57 | 0.42 | 0.66 | 0.67 | 0.7 | 0.7 |
| 2 | 65371112 | rs4671120 | -0.393 | -1.275 | -0.776 | -0.893 | -0.026 | -0.007 | -0.007 | 0.918 | 0.59 | 0.57 | 0.67 | 0.67 | 0.73 | 0.7 |
| 2 | 65378927 | rs10179474 | -0.385 | -0.388 | -0.679 | -0.783 | 0.006 | -0.006 | -0.006 | 0 | 0.88 | 0.93 | 0.92 | 1 | 0.88 | 0.92 |
| 2 | 65390283 | rs12470270 | 0.19 | 0.123 | -0.474 | -0.713 | 0.005 | 0.009 | 0.009 | 0 | 0.24 | 0.31 | 0.44 | 0.72 | 0.28 | 0.39 |
| 2 | 65392042 | rs6546132 | -0.627 | -0.009 | -0.533 | -0.984 | 0.007 | 0.02 | 0.02 | 0 | 0.42 | 0.36 | 0.53 | 0.72 | 0.38 | 0.51 |
| 2 | 65394714 | rs11681377 | -0.791 | 0.179 | -0.659 | -1.033 | 0.013 | 0.043 | 0.043 | 0 | 0.47 | 0.32 | 0.57 | 0.68 | 0.46 | 0.58 |
| 2 | 65395946 | rs11903118 | -0.655 | 1.212 | -0.097 | -0.815 | 0.065 | 0.016 | 0.016 | 0 | 0.15 | 0.26 | 0.21 | 0.21 | 0.28 | 0.21 |

| | | | | | | | | | | | | | | | | |
|---|---|---|---|---|---|---|---|---|---|---|---|---|---|---|---|---|
| 2 | 65402878 | rs2024438 | -1.331 | 0.859 | -0.188 | -1.316 | 0.016 | 0.022 | 0.022 | 0.644 | 0.63 | 0.72 | 0.78 | 0.89 | 0.73 | 0.83 |
| 2 | 65414296 | rs2110793 | -0.972 | 1.253 | 0.369 | -1.443 | 0.008 | -0.09 | -0.09 | 0 | 0.26 | 0.64 | 0.18 | 0.01 | 0.3 | 0.18 |
| 2 | 65417487 | rs12329001 | -0.534 | 0.962 | 0.536 | -1.413 | -0.073 | -0.141 | -0.141 | 0 | 0.21 | 0.5 | 0.12 | 0 | 0.11 | 0.15 |
| 2 | 65421515 | rs2110795 | 0.011 | 0.912 | 0.723 | -1.44 | -0.007 | -0.037 | -0.037 | 4.157 | 0.16 | 0.13 | 0.09 | 0 | 0.11 | 0.11 |
| 2 | 65434832 | rs13416763 | -0.152 | 1.072 | 0.885 | -1.332 | 0.032 | 0.029 | 0.029 | 3.294 | 0.05 | 0.35 | 0.02 | 0 | 0 | 0.04 |
| 2 | 65443314 | rs6546135 | 0.037 | 1.036 | 0.542 | -1.376 | 0.001 | -0.021 | -0.021 | 0 | 0.15 | 0.11 | 0.08 | 0 | 0.1 | 0.12 |
| 2 | 65448897 | rs9309369 | -1.003 | 1.381 | 0.572 | -1.42 | 0.034 | -0.136 | -0.136 | 1.707 | 0.26 | 0.52 | 0.18 | 0.01 | 0.33 | 0.19 |
| 2 | 65454644 | rs2008312 | -1.016 | 1.38 | 0.571 | -1.42 | 0.033 | -0.134 | -0.134 | 0 | 0.26 | 0.52 | 0.18 | 0.01 | 0.33 | 0.19 |
| 2 | 65461831 | rs7589984 | -1.072 | 1.365 | 0.564 | -1.429 | 0.007 | -0.068 | -0.068 | 0.549 | 0.26 | 0.66 | 0.19 | 0.01 | 0.33 | 0.2 |
| 2 | 65475061 | rs12463661 | 1.196 | 1.31 | 0.281 | -1.302 | -0.005 | 0.003 | 0.003 | 0 | 0.63 | 0.37 | 0.6 | 0.77 | 0.52 | 0.55 |
| 2 | 65480620 | rs17703416 | 0.664 | 1.434 | -0.14 | -1.431 | -0.001 | -0.001 | -0.001 | 0 | 0.33 | 0.32 | 0.41 | 0.66 | 0.27 | 0.38 |
| 2 | 65490659 | rs268866 | 0.205 | 1.464 | -0.099 | -1.472 | 0.002 | -0.023 | -0.023 | 0 | 0.56 | 0.56 | 0.36 | 0.07 | 0.58 | 0.32 |
| 2 | 65492486 | rs4671647 | 0.947 | 1.587 | -0.186 | -1.401 | 0.006 | 0.02 | 0.02 | 0 | 0.31 | 0.14 | 0.4 | 0.7 | 0.27 | 0.39 |
| 2 | 65495283 | rs4671124 | 0.457 | 1.735 | -0.447 | -1.451 | -0.002 | -0.005 | -0.005 | 0 | 0.35 | 0.31 | 0.6 | 0.91 | 0.35 | 0.35 |
| 2 | 65495887 | rs268875 | -1.016 | 1.681 | -0.371 | -0.98 | -0.002 | 0.012 | 0.012 | 0 | 0.91 | 0.9 | 0.79 | 0.71 | 0.87 | 0.73 |
| 2 | 65496251 | rs4513276 | 0.819 | 1.709 | -0.357 | -1.281 | 0.001 | 0.004 | 0.004 | 0 | 0.31 | 0.27 | 0.41 | 0.7 | 0.27 | 0.38 |
| 2 | 65512068 | rs7556894 | 0.848 | 1.484 | -0.321 | -1.334 | -0.02 | -0.065 | -0.065 | 0.444 | 0.37 | 0.45 | 0.22 | 0.11 | 0.26 | 0.17 |
| 2 | 65545090 | rs7601027 | -0.739 | 1.931 | -0.775 | -0.992 | 0.008 | 0.003 | 0.003 | 0 | 0.52 | 0.44 | 0.74 | 0.84 | 0.67 | 0.73 |
| 2 | 65545702 | rs7601514 | 0.247 | 2.056 | -1.11 | -1.065 | -0.001 | 0.026 | 0.026 | 0 | 0.32 | 0.11 | 0.5 | 0.76 | 0.27 | 0.49 |
| 2 | 65546422 | rs7604944 | -0.006 | 1.723 | -0.792 | -1.028 | 0.017 | -0.043 | -0.043 | 0 | 0.17 | 0.28 | 0.18 | 0.08 | 0.23 | 0.21 |
| 2 | 65546879 | rs10170788 | -0.195 | 1.738 | -1.356 | -0.654 | -0.06 | 0.031 | 0.031 | 0 | 0.33 | 0.11 | 0.51 | 0.76 | 0.39 | 0.51 |
| 2 | 65554415 | rs1542994 | 0.005 | 1.057 | -1.3 | -0.581 | 0.006 | 0.015 | 0.015 | 0 | 0.29 | 0.14 | 0.47 | 0.76 | 0.25 | 0.43 |
| 2 | 65555720 | rs6743585 | -0.506 | -0.045 | -1.654 | -1.048 | 0.07 | 0.045 | 0.045 | 1.791 | 0.33 | 0.76 | 0.31 | 0.15 | 0.18 | 0.22 |
| 2 | 65557163 | rs1039767 | 1.679 | -0.112 | -1.944 | -1.3 | -0.005 | 0.014 | 0.014 | 0.005 | 0.25 | 0.01 | 0.15 | 0.17 | 0.19 | 0.08 |
| 2 | 65559027 | rs2271198 | -0.929 | 0.447 | -1.571 | -1.364 | -0.012 | 0.127 | 0.127 | 0.804 | 0.58 | 0.84 | 0.65 | 0.78 | 0.57 | 0.66 |
| 2 | 65561392 | rs4671658 | 0.77 | 0.679 | -1.626 | -1.312 | -0.071 | -0.08 | -0.08 | 0 | 0.27 | 0.06 | 0.39 | 0.68 | 0.3 | 0.47 |
| 2 | 65568634 | rs11694684 | 1.192 | 0.407 | -1.29 | -1.397 | -0.01 | -0.001 | -0.001 | 0 | 0.15 | 0.11 | 0.12 | 0.09 | 0.04 | 0.15 |
| 2 | 65579228 | rs6711791 | -0.92 | 0.484 | -0.52 | -1.488 | 0.005 | 0.091 | 0.091 | 0.014 | 0.54 | 0.69 | 0.62 | 0.78 | 0.52 | 0.66 |
| 2 | 65581200 | rs11695373 | 0.061 | 0.517 | -0.52 | -1.505 | 0.014 | 0.096 | 0.096 | 0.273 | 0.42 | 0.17 | 0.52 | 0.77 | 0.34 | 0.62 |
| 2 | 65592325 | rs6738087 | -1.103 | 0.508 | -0.353 | -1.491 | -0.035 | 0.139 | 0.139 | 0.146 | 0.57 | 0.9 | 0.66 | 0.78 | 0.58 | 0.66 |
| 2 | 65605194 | rs174847 | 0.828 | 0.766 | -0.126 | -1.344 | 0.024 | 0.15 | 0.15 | 0 | 0.46 | 0.18 | 0.36 | 0.23 | 0.45 | 0.36 |
| 2 | 65606653 | rs1194849 | 0.37 | 0.866 | -0.085 | -1.293 | 0.024 | 0.092 | 0.092 | 0.66 | 0.53 | 0.71 | 0.4 | 0.23 | 0.45 | 0.36 |
| 2 | 65607580 | rs268135 | -0.763 | 0.821 | -0.119 | -1.292 | -0.023 | -0.006 | -0.006 | 0.661 | 0.82 | 0.78 | 0.84 | 0.91 | 0.91 | 0.85 |
| 2 | 65608363 | rs268134 | -0.435 | 0.859 | -0.118 | -1.291 | 0.006 | -0.007 | -0.007 | 0 | 0.8 | 0.77 | 0.8 | 0.91 | 0.75 | 0.84 |
| 2 | 65614613 | rs1039764 | 0.609 | 0.862 | -0.126 | -1.294 | -0.001 | -0.012 | -0.012 | 0 | 0.51 | 0.55 | 0.39 | 0.23 | 0.45 | 0.36 |
| 2 | 65623189 | rs964505 | 0.448 | 0.743 | -0.654 | -1.138 | -0.07 | -0.077 | -0.077 | 0.727 | 0.27 | 0.06 | 0.39 | 0.68 | 0.3 | 0.47 |
| 2 | 65653199 | rs6755128 | -0.494 | 0.752 | -0.628 | -1.282 | -0.075 | -0.078 | -0.078 | 0 | 0.71 | 0.93 | 0.59 | 0.35 | 0.68 | 0.51 |
| 2 | 65656969 | rs7569084 | 0.032 | 0.832 | -0.441 | -1.416 | -0.013 | 0.151 | 0.151 | 0 | 0.55 | 0.82 | 0.44 | 0.17 | 0.58 | 0.32 |
| 2 | 65665641 | rs2860773 | 0.418 | 1.04 | 0.106 | -1.385 | 0.001 | -0.013 | -0.013 | 0 | 0.49 | 0.49 | 0.36 | 0.17 | 0.4 | 0.3 |
| 2 | 65677729 | rs1866050 | 1.389 | 0.796 | -0.445 | -1.448 | 0.065 | 0.088 | 0.088 | 0 | 0.33 | 0.12 | 0.25 | 0.17 | 0.24 | 0.11 |
| 2 | 65691760 | rs754110 | -0.957 | 0.867 | 0.237 | -1.006 | -0.001 | -0.02 | -0.02 | 0.031 | 0.51 | 0.49 | 0.61 | 0.82 | 0.54 | 0.7 |
| 2 | 65691896 | rs17476501 | 0.716 | 1.446 | -0.318 | -0.8 | -0.046 | 0.004 | 0.004 | 0 | 0.08 | 0.01 | 0.08 | 0.09 | 0.11 | 0.01 |
| 2 | 65701166 | rs840966 | 0.489 | 1.332 | -0.027 | -0.872 | 0.028 | 0.108 | 0.108 | 0 | 0.46 | 0.7 | 0.47 | 0.17 | 0.43 | 0.31 |
| 2 | 65701757 | rs840967 | -0.278 | 1.354 | -0.907 | -0.798 | -0.128 | -0.137 | -0.137 | 0 | 0.75 | 0.96 | 0.62 | 0.44 | 0.62 | 0.5 |
| 2 | 65702803 | rs840969 | -0.551 | 1.332 | -0.903 | -0.794 | -0.025 | -0.134 | -0.134 | 0 | 0.77 | 0.97 | 0.68 | 0.44 | 0.79 | 0.52 |
| 2 | 65705581 | rs840974 | 0.876 | 0.431 | -0.719 | -0.751 | 0.005 | 0.13 | 0.13 | 0 | 0.41 | 0.18 | 0.43 | 0.17 | 0.43 | 0.31 |
| 2 | 65706325 | rs10519034 | 0.435 | 0.552 | -1.143 | -0.658 | 0.001 | -0.045 | -0.045 | 0 | 0.29 | 0.03 | 0.13 | 0.27 | 0.19 | 0.19 |
| 2 | 65707760 | rs840976 | 0.415 | 0.381 | -0.825 | -0.747 | 0.039 | 0.138 | 0.138 | 0 | 0.46 | 0.74 | 0.45 | 0.16 | 0.41 | 0.28 |
| 2 | 65710328 | rs2244364 | 0.702 | 0.521 | -0.82 | -0.382 | -0.005 | -0.049 | -0.049 | 0 | 0.42 | 0.49 | 0.4 | 0.15 | 0.42 | 0.31 |
| 2 | 65710688 | rs2244259 | -0.464 | 0.997 | -2.091 | 0.021 | -0.035 | -0.04 | -0.04 | 0 | 0.78 | 0.84 | 0.59 | 0.43 | 0.63 | 0.54 |
| 2 | 65715444 | rs10451656 | -0.566 | 0.132 | -1.82 | 0.259 | 0.057 | 0.004 | 0.004 | 0.011 | 0.34 | 0.14 | 0.16 | 0.28 | 0.21 | 0.23 |
| 2 | 65718174 | rs13393157 | -0.587 | 0.731 | -1.719 | -0.096 | -0.01 | 0.101 | 0.101 | 0 | 0.39 | 0.16 | 0.27 | 0.17 | 0.38 | 0.16 |
| 2 | 65719839 | rs17536001 | -0.122 | 0.666 | -1.742 | -0.362 | -0.027 | 0.055 | 0.055 | 0 | 0.28 | 0.1 | 0.2 | 0.17 | 0.27 | 0.15 |
| 2 | 65719941 | rs840952 | 1.063 | 1.171 | -1.517 | -0.395 | 0.018 | 0.083 | 0.083 | 0 | 0.51 | 0.75 | 0.61 | 0.73 | 0.46 | 0.78 |
| 2 | 65723237 | rs10203913 | -0.739 | 1.154 | -1.724 | -0.487 | -0.005 | 0.122 | 0.122 | 0 | 0.42 | 0.17 | 0.3 | 0.17 | 0.44 | 0.16 |
| 2 | 65723956 | rs2917816 | 0.711 | 0.844 | -1.733 | 0.617 | -0.072 | -0.041 | -0.041 | 1.044 | 0.56 | 0.13 | 0.52 | 0.46 | 0.56 | 0.73 |
| 2 | 65724073 | rs840955 | 0.633 | 0.814 | -0.526 | 1.006 | 0.044 | 0.138 | 0.138 | 0 | 0.87 | 0.75 | 0.73 | 0.64 | 0.83 | 0.85 |
| 2 | 65726375 | rs840959 | -0.53 | 0.415 | -1.263 | 0.668 | 0.016 | -0.043 | -0.043 | 0 | 0.21 | 0.06 | 0.17 | 0.28 | 0.17 | 0.11 |
| 2 | 65726755 | rs840961 | -0.344 | 0.534 | -1.108 | 0.584 | -0.001 | -0.012 | -0.012 | 0 | 0.12 | 0.06 | 0.11 | 0.25 | 0.14 | 0.04 |
| 2 | 65726877 | rs6717262 | 1.236 | 0.384 | -1.02 | 0.219 | -0.012 | 0.027 | 0.027 | 0 | 0.36 | 0.3 | 0.43 | 0.25 | 0.41 | 0.49 |
| 2 | 65731846 | rs6708230 | 0.631 | -0.019 | -0.474 | 0.012 | -0.031 | -0.027 | -0.027 | 0.753 | 0.72 | 0.49 | 0.78 | 0.77 | 0.79 | 0.9 |
| 2 | 65732023 | rs840978 | -0.397 | -0.027 | -0.254 | 0.008 | -0.038 | -0.116 | -0.116 | 0.017 | 0.83 | 0.63 | 0.9 | 0.99 | 0.89 | 0.94 |
| 2 | 65734749 | rs702927 | 0.687 | 0.049 | -0.4 | 0.066 | 0.002 | -0.007 | -0.007 | 0 | 0.11 | 0.06 | 0.11 | 0.22 | 0.11 | 0.04 |
| 2 | 65740698 | rs11126040 | 1.019 | 0.402 | -0.55 | 0.005 | -0.036 | 0.028 | 0.028 | 0 | 0.11 | 0.03 | 0.08 | 0.04 | 0.12 | 0.04 |
| 2 | 65756965 | rs702942 | -0.948 | 0.252 | -0.696 | 0.159 | 0 | 0.007 | 0.007 | 0 | 0.41 | 0.34 | 0.42 | 0.38 | 0.4 | 0.25 |
| 2 | 65758355 | rs3845816 | 0.74 | 0.273 | -0.694 | 0.155 | -0.01 | 0.012 | 0.012 | 0 | 0.58 | 0.69 | 0.57 | 0.62 | 0.58 | 0.74 |
| 2 | 65759671 | rs3911862 | 1.523 | 0.205 | -0.616 | 0.107 | -0.018 | 0.008 | 0.008 | 0 | 0.42 | 0.34 | 0.49 | 0.61 | 0.5 | 0.69 |
| 2 | 65765212 | rs702902 | -0.372 | 0.779 | -0.269 | 0.6 | -0.014 | -0.018 | -0.018 | 0.484 | 0.2 | 0.08 | 0.18 | 0.27 | 0.21 | 0.08 |
| 2 | 65772155 | rs9309375 | 0.009 | 0.813 | -0.293 | 0.609 | -0.003 | 0.033 | 0.033 | 0 | 0.21 | 0.13 | 0.22 | 0.12 | 0.18 | 0.18 |

| | | | | | | | | | | | | | | | | |
|---|---|---|---|---|---|---|---|---|---|---|---|---|---|---|---|---|
| 2 | 65772291 | rs702919 | -1.367 | 0.908 | 0.09 | 0.572 | 0 | 0 | 0 | 0 | 0.43 | 0.39 | 0.45 | 0.39 | 0.43 | 0.25 |
| 2 | 65773712 | rs12614525 | 0.334 | 1.023 | 0.13 | 0.625 | 0.009 | -0.014 | -0.014 | 0 | 0.11 | 0.11 | 0.1 | 0.05 | 0.13 | 0.03 |
| 2 | 65783877 | rs1641464 | -1.162 | 0.895 | -0.221 | 0.756 | 0.005 | 0.02 | 0.02 | 0 | 0.51 | 0.45 | 0.41 | 0.69 | 0.54 | 0.4 |
| 2 | 65785369 | rs4671131 | 1.256 | 0.882 | -0.208 | 0.754 | 0.009 | 0.025 | 0.025 | 0 | 0.46 | 0.41 | 0.51 | 0.27 | 0.43 | 0.5 |
| 2 | 65786773 | rs11126046 | 0.955 | 0.874 | -0.195 | 0.721 | 0.012 | 0.031 | 0.031 | 0 | 0.51 | 0.58 | 0.58 | 0.31 | 0.46 | 0.6 |
| 2 | 65789219 | rs1673448 | -0.213 | 0.946 | -0.419 | 0.938 | -0.054 | -0.045 | -0.045 | 0.541 | 0.22 | 0.09 | 0.21 | 0.32 | 0.32 | 0.15 |
| 2 | 65791089 | rs1641474 | -0.903 | 0.953 | -0.371 | 0.931 | 0.035 | 0.043 | 0.043 | 0.576 | 0.28 | 0.43 | 0.25 | 0.4 | 0.34 | 0.15 |
| 2 | 65791581 | rs897876 | -0.415 | 0.953 | -0.351 | 0.937 | 0.03 | 0.038 | 0.038 | 0 | 0.24 | 0.31 | 0.23 | 0.4 | 0.33 | 0.15 |
| 2 | 65800982 | rs11677815 | 0.966 | 0.813 | -0.573 | 1.169 | 0.007 | 0.018 | 0.018 | 0 | 0.47 | 0.41 | 0.56 | 0.32 | 0.44 | 0.56 |
| 2 | 65803661 | rs1194480 | -0.396 | 0.781 | -0.319 | 0.659 | 0.019 | 0.007 | 0.007 | 0 | 0.15 | 0.28 | 0.15 | 0.18 | 0.21 | 0.07 |
| 2 | 65804266 | rs12472695 | -0.571 | 0.465 | -0.906 | 0.65 | -0.002 | 0.012 | 0.012 | 0.066 | 0.32 | 0.28 | 0.23 | 0.26 | 0.3 | 0.28 |
| 2 | 65810588 | rs1673459 | -0.083 | -0.508 | -1.06 | 0.902 | 0.054 | -0.037 | -0.037 | 0.665 | 0.7 | 0.83 | 0.62 | 0.69 | 0.8 | 0.82 |
| 2 | 65816307 | rs6546162 | -0.067 | -0.633 | -0.925 | 0.964 | -0.005 | 0.015 | 0.015 | 0 | 0.15 | 0.16 | 0.27 | 0.31 | 0.06 | 0.14 |
| 2 | 65823020 | rs11695927 | -1.18 | -0.716 | -0.137 | 0.872 | 0.001 | -0.001 | -0.001 | 0 | 0.25 | 0.73 | 0.36 | 0.41 | 0.13 | 0.29 |
| 2 | 65832209 | rs6710386 | -1.189 | -0.778 | -0.081 | 0.864 | 0.002 | -0.003 | -0.003 | 0 | 0.25 | 0.72 | 0.36 | 0.41 | 0.13 | 0.28 |
| 2 | 65836286 | rs17030469 | -0.339 | -0.808 | 0.439 | 0.884 | -0.021 | 0.066 | 0.066 | 0 | 0.18 | 0.28 | 0.31 | 0.31 | 0.11 | 0.15 |
| 2 | 65838800 | rs11902125 | -0.709 | -0.81 | 0.931 | 0.832 | 0.03 | 0.065 | 0.065 | 0 | 0.06 | 0.23 | 0.04 | 0.1 | 0.05 | 0.14 |
| 2 | 65838854 | rs6546167 | -1.298 | -0.82 | 0.667 | 0.822 | -0.006 | 0.008 | 0.008 | 0 | 0.26 | 0.71 | 0.37 | 0.41 | 0.16 | 0.29 |
| 2 | 65844734 | rs6760617 | -0.084 | -0.574 | 0.453 | 1.043 | 0.012 | -0.025 | -0.025 | 0 | 0.15 | 0.09 | 0.29 | 0.24 | 0.11 | 0.11 |
| 2 | 65849494 | rs971410 | -0.489 | 0.264 | 0.042 | 0.56 | -0.114 | 0.049 | 0.049 | 0.067 | 0.67 | 0.98 | 0.73 | 0.83 | 0.57 | 0.79 |
| 2 | 65851312 | rs7593455 | -1.997 | -0.059 | 1.228 | 1.041 | 0.041 | 0.392 | 0.392 | 0 | 0.11 | 0.44 | 0.31 | 0.4 | 0.1 | 0.21 |
| 2 | 65862378 | rs1457451 | -1.549 | -0.029 | 1.505 | 1.128 | 0.03 | 0.042 | 0.042 | 0 | 0.08 | 0.18 | 0.13 | 0.18 | 0.1 | 0.3 |
| 2 | 65877962 | rs7579084 | -1.375 | 0.046 | 1.467 | 1.072 | 0.054 | 0.413 | 0.413 | 0 | 0.08 | 0.43 | 0.24 | 0.29 | 0.06 | 0.09 |
| 2 | 65883314 | rs7574634 | 0.496 | 0.328 | 0.982 | 0.636 | 0.042 | 0.047 | 0.047 | 0 | 0.45 | 0.3 | 0.25 | 0.36 | 0.31 | 0.38 |
| 2 | 65889672 | rs840778 | 1.2 | 0.539 | 0.866 | 0.703 | 0 | 0.05 | 0.05 | 1.271 | 0.89 | 0.85 | 0.67 | 0.59 | 0.85 | 0.76 |
| 2 | 65890333 | rs840776 | 0.924 | 0.276 | 0.617 | 0.511 | -0.139 | 0.047 | 0.047 | 0 | 0.31 | 0.03 | 0.25 | 0.14 | 0.42 | 0.18 |
| 2 | 65892430 | rs1819852 | 0.411 | 0.213 | 1.169 | 0.491 | 0.006 | -0.01 | -0.01 | 0 | 0.35 | 0.34 | 0.31 | 0.19 | 0.48 | 0.2 |
| 2 | 65892607 | rs2860850 | -0.498 | -0.235 | 0.849 | 0.118 | -0.02 | -0.022 | -0.022 | 0 | 0.71 | 0.85 | 0.68 | 0.64 | 0.67 | 0.5 |
| 2 | 65895471 | rs13397571 | 0.807 | -0.127 | 1.046 | 0.03 | 0.022 | -0.008 | -0.008 | 0 | 0.27 | 0.19 | 0.06 | 0.02 | 0.11 | 0.08 |
| 2 | 65899519 | rs6546173 | -0.221 | 0.443 | 1.406 | 0.497 | 0.136 | 0.021 | 0.021 | 0 | 0.05 | 0.28 | 0.07 | 0 | 0.08 | 0.01 |
| 2 | 65904240 | rs4146116 | -0.399 | 0.394 | 1.08 | 0.578 | -0.007 | 0.018 | 0.018 | 0.55 | 0.44 | 0.63 | 0.66 | 0.64 | 0.52 | 0.49 |
| 2 | 65904304 | rs4146117 | 0.952 | 0.469 | 0.643 | 0.677 | -0.096 | 0.002 | 0.002 | 0 | 0.29 | 0.06 | 0.28 | 0.21 | 0.4 | 0.17 |
| 2 | 65904864 | rs1477054 | -1.673 | 0.353 | 1.423 | 1.056 | 0.068 | 0.419 | 0.419 | 0 | 0.08 | 0.45 | 0.31 | 0.33 | 0.07 | 0.13 |
| 2 | 65923957 | rs9678488 | 0 | 0.434 | 0.537 | 0.522 | 0.001 | 0.001 | 0.001 | 0.073 | 0.37 | 0.38 | 0.45 | 0.46 | 0.43 | 0.46 |
| 2 | 65924769 | rs9678380 | -1.864 | 0.344 | 0.875 | 0.571 | -0.008 | 0.01 | 0.01 | 3.183 | 0.12 | 0.6 | 0.12 | 0.1 | 0.06 | 0.16 |
| 2 | 65924908 | rs1019365 | -1.248 | 0.417 | 0.561 | 0.572 | 0.074 | -0.025 | -0.025 | 0 | 0.08 | 0.58 | 0.08 | 0 | 0.08 | 0 |
| 2 | 65925377 | rs4671681 | -0.904 | 0.162 | -0.677 | 0.546 | -0.04 | 0.06 | 0.06 | 3.842 | 0.65 | 0.9 | 0.75 | 0.76 | 0.59 | 0.66 |
| 2 | 65926939 | rs11891634 | -1.504 | 0.123 | -0.208 | 0.554 | 0.021 | 0.04 | 0.04 | 0.157 | 0.13 | 0.39 | 0.16 | 0.19 | 0.12 | 0.27 |
| 2 | 65928971 | rs4671682 | 0.367 | 0.293 | -1.323 | 0.538 | -0.025 | -0.041 | -0.041 | 0.003 | 0.27 | 0.08 | 0.28 | 0.27 | 0.28 | 0.18 |
| 2 | 65929932 | rs17030660 | -0.778 | 0.186 | -0.406 | 0.495 | 0.002 | 0.003 | 0.003 | 0 | 0.05 | 0.09 | 0.06 | 0.08 | 0.07 | 0.11 |
| 2 | 65934798 | rs12622132 | -0.666 | -0.575 | -0.353 | 0.066 | -0.062 | -0.081 | -0.081 | 0 | 0.29 | 0.39 | 0.23 | 0.12 | 0.15 | 0.31 |
| 2 | 65935157 | rs17030670 | -1.208 | -0.438 | -0.12 | 0.696 | -0.011 | 0.023 | 0.023 | 0 | 0.14 | 0.21 | 0.16 | 0.23 | 0.13 | 0.22 |
| 2 | 65935393 | rs6546174 | -0.525 | 0.024 | -0.62 | 0.636 | -0.006 | 0.015 | 0.015 | 0 | 0.45 | 0.35 | 0.4 | 0.64 | 0.49 | 0.6 |
| 2 | 65938598 | rs4671683 | 0.186 | 0.615 | -0.919 | 0.726 | -0.096 | 0.063 | 0.063 | 0 | 0.76 | 0.97 | 0.83 | 0.89 | 0.69 | 0.79 |
| 2 | 65938740 | rs12713544 | 1.316 | 0.09 | -0.274 | 1.106 | 0 | 0.005 | 0.005 | 0 | 0.52 | 0.41 | 0.43 | 0.38 | 0.49 | 0.53 |
| 2 | 65946512 | rs10490617 | -0.547 | -0.416 | 0.181 | 1.903 | 0.004 | 0.01 | 0.01 | 0 | 0.09 | 0.12 | 0.13 | 0.1 | 0.12 | 0.15 |
| 2 | 65948094 | rs7580807 | 0.163 | 0.036 | 0.576 | 1.771 | 0.018 | -0.013 | -0.013 | 0 | 0.71 | 0.64 | 0.74 | 0.84 | 0.59 | 0.77 |
| 2 | 65956150 | rs2111464 | 1.065 | -0.274 | 1.113 | 1.84 | -0.003 | -0.001 | -0.001 | 0 | 0.56 | 0.61 | 0.48 | 0.44 | 0.51 | 0.53 |
| 2 | 65961112 | rs6733631 | -1.982 | -0.318 | 1.256 | 1.851 | 0.09 | 0.12 | 0.12 | 1.871 | 0.07 | 0.4 | 0.07 | 0.06 | 0.06 | 0.03 |
| 2 | 65970775 | rs2287282 | -0.198 | -0.219 | 1.265 | 1.908 | -0.042 | 0.06 | 0.06 | 0 | 0.73 | 0.85 | 0.8 | 0.9 | 0.66 | 0.78 |
| 2 | 65972163 | rs12997745 | 1.579 | -0.18 | 1.09 | 1.981 | 0.011 | 0.057 | 0.057 | 0 | 0.51 | 0.27 | 0.41 | 0.39 | 0.46 | 0.53 |
| 2 | 65975064 | rs17030748 | -0.859 | -0.157 | 1.061 | 2.01 | 0.013 | -0.047 | -0.047 | 0 | 0.15 | 0.07 | 0.24 | 0.4 | 0.14 | 0.22 |
| 2 | 65977863 | rs2009778 | -0.019 | -0.143 | 0.992 | 1.986 | -0.001 | -0.011 | -0.011 | 0 | 0.72 | 0.68 | 0.78 | 0.9 | 0.65 | 0.78 |
| 2 | 65979640 | rs4671140 | -1.759 | -0.518 | 1.111 | 2.542 | -0.007 | 0.068 | 0.068 | 0.066 | 0.2 | 0.24 | 0.38 | 0.53 | 0.19 | 0.25 |
| 2 | 65991365 | rs2193467 | -1.264 | -0.294 | 1.001 | 2.527 | -0.011 | 0.077 | 0.077 | 0.206 | 0.15 | 0.17 | 0.24 | 0.4 | 0.14 | 0.22 |
| 2 | 65993579 | rs4671141 | -1.201 | -0.253 | 0.552 | 2.237 | 0.006 | 0.09 | 0.09 | 0.35 | 0.16 | 0.2 | 0.29 | 0.42 | 0.19 | 0.21 |
| 2 | 66001417 | rs11126058 | -1.479 | -0.657 | 0.298 | 2.268 | -0.029 | 0.041 | 0.041 | 0.796 | 0.23 | 0.63 | 0.3 | 0.58 | 0.13 | 0.22 |
| 2 | 66001846 | rs7597061 | -1.205 | -0.527 | 0.423 | 2.082 | -0.042 | 0.119 | 0.119 | 0.981 | 0.2 | 0.58 | 0.26 | 0.49 | 0.13 | 0.21 |
| 2 | 66006558 | rs6546180 | -0.866 | -0.116 | 0.041 | 1.701 | 0.005 | 0.008 | 0.008 | 0 | 0.35 | 0.56 | 0.47 | 0.38 | 0.4 | 0.19 |
| 2 | 66009594 | rs10172064 | -0.358 | -0.372 | 0.419 | 2.019 | -0.02 | 0.163 | 0.163 | 0 | 0.1 | 0.22 | 0.07 | 0.31 | 0.05 | 0.15 |
| 2 | 66012032 | rs10179158 | -2.057 | -0.248 | 0.574 | 1.701 | -0.006 | 0.008 | 0.008 | 0.093 | 0.24 | 0.65 | 0.2 | 0.35 | 0.18 | 0.28 |
| 2 | 66012093 | rs10181534 | 0.114 | -0.257 | 0.564 | 1.821 | -0.022 | 0.206 | 0.206 | 0 | 0.1 | 0.22 | 0.1 | 0.37 | 0.05 | 0.15 |
| 2 | 66013031 | rs10193514 | 0.032 | -0.256 | 0.633 | 1.886 | -0.022 | 0.099 | 0.099 | 0 | 0.1 | 0.19 | 0.07 | 0.22 | 0.05 | 0.14 |
| 2 | 66014460 | rs13400930 | 0.118 | -0.257 | 0.681 | 1.784 | -0.02 | 0.14 | 0.14 | 0 | 0.1 | 0.22 | 0.07 | 0.29 | 0.05 | 0.14 |
| 2 | 66015141 | rs10173939 | -2.138 | -0.283 | 0.652 | 1.784 | -0.005 | 0.013 | 0.013 | 0.371 | 0.25 | 0.68 | 0.23 | 0.44 | 0.18 | 0.29 |
| 2 | 66021675 | rs11693109 | -0.89 | -0.276 | 0.66 | 1.758 | -0.009 | -0.006 | -0.006 | 0.726 | 0.12 | 0.09 | 0.13 | 0.14 | 0.13 | 0.15 |
| 2 | 66024928 | rs12617585 | -2.213 | -0.467 | 0.749 | 1.735 | -0.005 | 0.006 | 0.006 | 0 | 0.3 | 0.64 | 0.24 | 0.49 | 0.21 | 0.32 |
| 2 | 66030433 | rs13388557 | -2.146 | -0.621 | 0.723 | 1.678 | -0.021 | 0.013 | 0.013 | 0 | 0.28 | 0.38 | 0.19 | 0.42 | 0.2 | 0.31 |
| 2 | 66030980 | rs6758005 | -2.141 | -0.642 | 0.708 | 1.654 | -0.022 | 0.013 | 0.013 | 0 | 0.28 | 0.39 | 0.19 | 0.41 | 0.19 | 0.31 |

| | | | | | | | | | | | | | | | | |
|---|---|---|---|---|---|---|---|---|---|---|---|---|---|---|---|---|
| 2 | 66036540 | rs3919602 | 0.435 | -0.328 | 0.166 | 1.643 | -0.081 | -0.074 | -0.074 | 0.674 | 0.23 | 0.03 | 0.35 | 0.29 | 0.29 | 0.15 |
| 2 | 66040362 | rs7596231 | -2.695 | -0.755 | 0.888 | 1.535 | 0.03 | -0.021 | -0.021 | 0 | 0.32 | 0.75 | 0.25 | 0.48 | 0.19 | 0.31 |
| 2 | 66045064 | rs6743363 | 1.419 | -0.708 | 0.842 | 1.405 | 0 | -0.011 | -0.011 | 0 | 0.86 | 0.82 | 0.9 | 0.94 | 0.87 | 0.87 |
| 2 | 66051513 | rs890092 | -1.928 | -0.693 | 0.927 | 1.531 | -0.037 | 0.036 | 0.036 | 0.532 | 0.23 | 0.49 | 0.17 | 0.35 | 0.16 | 0.25 |
| 2 | 66053473 | rs3845821 | -1.761 | -0.703 | 0.934 | 1.627 | -0.019 | 0.039 | 0.039 | 0 | 0.1 | 0.25 | 0.07 | 0.14 | 0.04 | 0.08 |
| 2 | 66055467 | rs9309379 | -0.884 | -0.504 | 0.813 | 1.696 | -0.036 | 0.141 | 0.141 | 0 | 0.12 | 0.34 | 0.08 | 0.28 | 0.05 | 0.12 |
| 2 | 66058256 | rs7570905 | -1.823 | -0.415 | 0.213 | 1.596 | -0.015 | 0.019 | 0.019 | 0.034 | 0.28 | 0.62 | 0.22 | 0.43 | 0.19 | 0.29 |
| 2 | 66060863 | rs6744162 | -1.767 | -0.251 | 0.078 | 1.747 | 0 | 0.003 | 0.003 | 0 | 0.23 | 0.26 | 0.2 | 0.26 | 0.24 | 0.28 |
| 2 | 66062059 | rs7597074 | -1.12 | -0.282 | 0.13 | 1.785 | 0.007 | 0.007 | 0.007 | 0 | 0.07 | 0.12 | 0.11 | 0.1 | 0.11 | 0.14 |
| 2 | 66062158 | rs6736318 | -1.48 | -0.288 | 0.536 | 1.833 | -0.03 | 0.046 | 0.046 | 0 | 0.17 | 0.49 | 0.08 | 0.3 | 0.09 | 0.15 |
| 2 | 66068976 | rs10179903 | -1.837 | -0.632 | 0.462 | 1.587 | -0.046 | 0.051 | 0.051 | 0 | 0.28 | 0.4 | 0.14 | 0.42 | 0.2 | 0.32 |
| 2 | 66071365 | rs6751687 | -2.625 | -0.636 | 0.515 | 1.67 | 0.023 | -0.052 | -0.052 | 0.217 | 0.37 | 0.71 | 0.26 | 0.53 | 0.32 | 0.44 |
| 2 | 66072634 | rs11681842 | -2.892 | -0.653 | 0.52 | 1.681 | 0.036 | -0.051 | -0.051 | 0 | 0.4 | 0.79 | 0.27 | 0.57 | 0.33 | 0.47 |
| 2 | 66080617 | rs11889877 | -2.204 | -0.687 | 0.716 | 1.804 | -0.003 | 0.013 | 0.013 | 0.245 | 0.19 | 0.25 | 0.16 | 0.22 | 0.17 | 0.24 |
| 2 | 66082204 | rs2194782 | -2.695 | -0.799 | 0.731 | 1.926 | 0.039 | -0.051 | -0.051 | 1.021 | 0.31 | 0.73 | 0.23 | 0.56 | 0.22 | 0.39 |
| 2 | 66082215 | rs2901807 | -2 | -0.8 | 0.86 | 1.96 | 0.024 | 0.034 | 0.034 | 0.607 | 0.1 | 0.27 | 0.12 | 0.12 | 0.11 | 0.13 |
| 2 | 66086769 | rs6750138 | -2.198 | -0.893 | 0.856 | 1.914 | -0.029 | 0.01 | 0.01 | 0 | 0.24 | 0.61 | 0.13 | 0.47 | 0.12 | 0.26 |
| 2 | 66087571 | rs7563053 | -0.742 | -0.92 | 0.904 | 1.909 | -0.055 | 0.174 | 0.174 | 0 | 0.12 | 0.39 | 0.05 | 0.31 | 0.06 | 0.16 |
| 2 | 66089359 | rs6719655 | -0.408 | -0.882 | 1.045 | 1.928 | -0.051 | 0.111 | 0.111 | 0 | 0.11 | 0.25 | 0.04 | 0.31 | 0.06 | 0.16 |
| 2 | 66089760 | rs6720085 | -2.039 | -0.879 | 0.926 | 1.908 | -0.071 | 0.051 | 0.051 | 0 | 0.23 | 0.55 | 0.13 | 0.47 | 0.12 | 0.26 |
| 2 | 66094705 | rs976394 | -2.068 | -0.457 | 0.725 | 1.594 | -0.012 | 0.06 | 0.06 | 1.124 | 0.2 | 0.43 | 0.21 | 0.25 | 0.16 | 0.24 |
| 2 | 66095536 | rs993468 | -2.078 | -0.454 | 0.737 | 1.543 | -0.003 | 0.05 | 0.05 | 0.284 | 0.19 | 0.37 | 0.24 | 0.21 | 0.16 | 0.24 |
| 2 | 66097430 | rs6730703 | -1.731 | -0.496 | 0.706 | 1.399 | -0.039 | 0.007 | 0.007 | 0.054 | 0.13 | 0.31 | 0.11 | 0.13 | 0.07 | 0.1 |
| 2 | 66098825 | rs2553050 | -0.869 | -0.346 | -0.377 | 0.927 | 0.009 | 0.041 | 0.041 | 0 | 0.51 | 0.71 | 0.62 | 0.7 | 0.56 | 0.62 |
| 2 | 66101048 | rs13413944 | -0.919 | -0.337 | -0.195 | 0.937 | -0.005 | 0.037 | 0.037 | 0 | 0.27 | 0.48 | 0.37 | 0.4 | 0.28 | 0.27 |
| 2 | 66104784 | rs2553053 | -1.385 | 0.065 | -0.395 | 0.98 | -0.02 | 0.056 | 0.056 | 0 | 0.47 | 0.76 | 0.61 | 0.72 | 0.53 | 0.61 |
| 2 | 66104881 | rs2631669 | -1.358 | 0.062 | -0.384 | 0.979 | -0.013 | 0.045 | 0.045 | 0 | 0.47 | 0.72 | 0.61 | 0.72 | 0.53 | 0.61 |
| 2 | 66111639 | rs2194788 | -0.979 | -0.176 | -0.025 | 0.99 | 0.003 | 0.023 | 0.023 | 0 | 0.21 | 0.27 | 0.34 | 0.35 | 0.25 | 0.26 |
| 2 | 66112394 | rs990307 | -1.99 | -0.387 | -0.02 | 1 | -0.017 | 0.077 | 0.077 | 0 | 0.29 | 0.49 | 0.46 | 0.46 | 0.28 | 0.32 |
| 2 | 66123593 | rs11684515 | 0.206 | -0.254 | -0.11 | 0.961 | -0.025 | 0.036 | 0.036 | 0.188 | 0.07 | 0 | 0.04 | 0 | 0.1 | 0.1 |
| 2 | 66124371 | rs2194789 | -2.049 | -0.344 | -0.01 | 0.984 | -0.015 | 0.097 | 0.097 | 0 | 0.27 | 0.5 | 0.45 | 0.46 | 0.26 | 0.31 |
| 2 | 66127164 | rs12469561 | -1.245 | -0.259 | -0.202 | 0.884 | 0 | 0.023 | 0.023 | 0 | 0.21 | 0.27 | 0.33 | 0.35 | 0.24 | 0.25 |
| 2 | 66135696 | rs2553049 | -1.215 | -0.001 | -0.215 | 0.881 | 0.01 | 0.028 | 0.028 | 0 | 0.2 | 0.26 | 0.33 | 0.36 | 0.24 | 0.24 |
| 2 | 66142233 | rs11126064 | 0.328 | -0.139 | -0.58 | 0.832 | -0.052 | -0.082 | -0.082 | 0 | 0.12 | 0.01 | 0.27 | 0.37 | 0.15 | 0.19 |
| 2 | 66148466 | rs6728803 | -2.413 | -0.18 | -1.379 | 0.437 | -0.097 | 0.079 | 0.079 | 0.108 | 0.54 | 0.93 | 0.64 | 0.75 | 0.48 | 0.63 |
| 2 | 66149357 | rs6546197 | -1.913 | -0.15 | -0.88 | 0.217 | 0.002 | 0.013 | 0.013 | 0 | 0.4 | 0.47 | 0.54 | 0.51 | 0.41 | 0.43 |
| 2 | 66149484 | rs7582590 | -0.765 | -0.175 | -0.687 | 0.114 | -0.001 | -0.073 | -0.073 | 0.112 | 0.12 | 0.34 | 0.06 | 0 | 0.09 | 0.08 |
| 2 | 66150153 | rs17031137 | -0.267 | -0.139 | -0.751 | 0.155 | 0 | -0.001 | -0.001 | 0 | 0.16 | 0.11 | 0.21 | 0.14 | 0.17 | 0.16 |
| 2 | 66158857 | rs11695386 | -1.419 | -0.12 | -0.691 | 0.162 | -0.009 | -0.043 | -0.043 | 0 | 0.28 | 0.43 | 0.27 | 0.15 | 0.26 | 0.24 |
| 2 | 66160691 | rs9653501 | -1.323 | -0.108 | -0.822 | 0.169 | -0.006 | -0.032 | -0.032 | 0 | 0.27 | 0.37 | 0.26 | 0.16 | 0.26 | 0.24 |
| 2 | 66162120 | rs7598654 | -1.335 | -0.098 | -0.868 | 0.16 | -0.007 | -0.039 | -0.039 | 0 | 0.27 | 0.39 | 0.26 | 0.14 | 0.26 | 0.24 |
| 2 | 66162412 | rs7598978 | -1.125 | -0.085 | -0.823 | 0.167 | -0.001 | -0.005 | -0.005 | 0.03 | 0.25 | 0.23 | 0.25 | 0.14 | 0.26 | 0.24 |
| 2 | 66165446 | rs1825430 | 0.723 | 0.04 | -1.038 | 0.286 | 0.003 | 0.002 | 0.002 | 0 | 0.07 | 0.07 | 0.02 | 0.04 | 0.02 | 0.08 |
| 2 | 66172098 | rs11126068 | -1.211 | 0.193 | -1.254 | 0.309 | 0 | -0.002 | -0.002 | 0.227 | 0.32 | 0.28 | 0.51 | 0.52 | 0.32 | 0.35 |
| 2 | 66172199 | rs17604526 | 0.884 | 0.233 | -1.211 | 0.294 | 0.004 | -0.018 | -0.018 | 0 | 0.05 | 0.1 | 0.03 | 0 | 0.05 | 0.05 |
| 2 | 66173226 | rs1155746 | -1.117 | 0.822 | -0.274 | 1.279 | -0.001 | 0.002 | 0.002 | 0.002 | 0.22 | 0.2 | 0.25 | 0.14 | 0.21 | 0.21 |
| 2 | 66173904 | rs11126069 | -1.639 | 0.768 | -0.367 | 0.947 | 0.001 | 0.004 | 0.004 | 0 | 0.38 | 0.4 | 0.54 | 0.52 | 0.37 | 0.4 |
| 2 | 66196670 | rs987217 | -1.035 | 0.626 | 0.066 | 0.481 | -0.006 | -0.085 | -0.085 | 0 | 0.15 | 0.34 | 0.11 | 0.01 | 0.13 | 0.1 |
| 2 | 66198055 | rs2017570 | -1.246 | 0.693 | 0.51 | 1.033 | 0.01 | 0.081 | 0.081 | 0 | 0.13 | 0.25 | 0.18 | 0.28 | 0.13 | 0.18 |
| 2 | 66200648 | rs1385167 | -1.243 | 0.697 | 0.517 | 1.07 | 0.01 | 0.085 | 0.085 | 0 | 0.13 | 0.24 | 0.18 | 0.29 | 0.13 | 0.19 |
| 2 | 66201731 | rs6716270 | -1.784 | 0.469 | 0.731 | 1.022 | -0.013 | 0.11 | 0.11 | 0.382 | 0.2 | 0.44 | 0.21 | 0.33 | 0.15 | 0.26 |
| 2 | 66204983 | rs6707056 | -1.167 | 0.35 | 1.028 | 0.808 | 0.01 | 0.046 | 0.046 | 0 | 0.13 | 0.26 | 0.18 | 0.21 | 0.13 | 0.18 |
| 2 | 66207377 | rs1030327 | -3.027 | 0.486 | 0.88 | 0.663 | 0.001 | 0.05 | 0.05 | 0 | 0.39 | 0.86 | 0.35 | 0.26 | 0.32 | 0.4 |
| 2 | 66209485 | rs10180226 | -2.398 | 0.495 | 0.956 | 0.623 | -0.003 | 0 | 0 | 0 | 0.28 | 0.56 | 0.25 | 0.25 | 0.22 | 0.31 |
| 2 | 66209626 | rs4398261 | -2.914 | 0.47 | 0.939 | 0.611 | 0 | -0.005 | -0.005 | 0 | 0.35 | 0.62 | 0.32 | 0.26 | 0.32 | 0.38 |
| 2 | 66211610 | rs10187079 | -3.334 | 0.456 | 0.832 | 0.61 | 0.004 | 0.053 | 0.053 | 0 | 0.39 | 0.85 | 0.35 | 0.26 | 0.31 | 0.4 |
| 2 | 66211789 | rs10210513 | -1.524 | 0.461 | 0.919 | 0.642 | 0.003 | 0.064 | 0.064 | 0.261 | 0.16 | 0.51 | 0.2 | 0.21 | 0.13 | 0.18 |
| 2 | 66219395 | rs17606067 | -0.701 | 0.63 | 0.686 | 0.635 | -0.004 | 0.01 | 0.01 | 0 | 0.1 | 0.09 | 0.08 | 0.01 | 0.11 | 0.1 |
| 2 | 66221977 | rs17546835 | -1.571 | 0.633 | 0.764 | 0.617 | 0.006 | -0.044 | -0.044 | 1.119 | 0.14 | 0.22 | 0.1 | 0.01 | 0.16 | 0.13 |
| 2 | 66228539 | rs13429266 | -1.166 | 0.642 | 0.876 | 0.652 | -0.008 | 0.011 | 0.011 | 0 | 0.14 | 0.38 | 0.14 | 0.09 | 0.08 | 0.14 |
| 2 | 66232736 | rs1484483 | -0.59 | 0.644 | 0.85 | 0.643 | -0.007 | 0.02 | 0.02 | 0 | 0.09 | 0.06 | 0.08 | 0.01 | 0.11 | 0.1 |
| 2 | 66238005 | rs6719428 | -2.689 | 0.716 | 0.925 | 0.745 | 0.003 | -0.004 | -0.004 | 0 | 0.3 | 0.48 | 0.3 | 0.26 | 0.28 | 0.35 |
| 2 | 66242561 | rs17481447 | 0.036 | 0.656 | 0.873 | 0.712 | 0.008 | -0.029 | -0.029 | 0 | 0.06 | 0.13 | 0.03 | 0 | 0.05 | 0.04 |
| 2 | 66244404 | rs1531905 | -2.841 | 0.77 | 0.795 | 0.75 | 0 | -0.003 | -0.003 | 1.645 | 0.35 | 0.6 | 0.33 | 0.26 | 0.33 | 0.37 |
| 2 | 66251743 | rs2699780 | 2.135 | 0.7 | 0.744 | 0.759 | -0.01 | 0.012 | 0.012 | 0 | 0.76 | 0.46 | 0.75 | 0.74 | 0.83 | 0.77 |
| 2 | 66258662 | rs12713561 | 1.429 | 0.679 | 0.934 | 0.756 | 0.017 | 0.058 | 0.058 | 0 | 0.85 | 0.66 | 0.79 | 0.79 | 0.87 | 0.84 |
| 2 | 66258701 | rs2699783 | -0.077 | 0.714 | 0.926 | 0.777 | -0.009 | -0.018 | -0.018 | 0.327 | 0.92 | 0.96 | 0.91 | 0.83 | 0.92 | 0.88 |
| 2 | 66265838 | rs17481962 | -0.405 | 0.718 | 0.947 | 0.745 | -0.001 | -0.01 | -0.01 | 0 | 0.09 | 0.09 | 0.07 | 0.01 | 0.1 | 0.09 |
| 2 | 66272243 | rs7600106 | 0.002 | 0.651 | 0.702 | 0.806 | -0.009 | 0.001 | 0.001 | 0.376 | 0.09 | 0.19 | 0.04 | 0.06 | 0.04 | 0.08 |

| | | | | | | | | | | | | | | | | |
|---|---|---|---|---|---|---|---|---|---|---|---|---|---|---|---|---|
| 2 | 66277661 | rs1032110 | 1.94 | 0.693 | 0.239 | 0.799 | -0.004 | 0.007 | 0.007 | 0 | 0.71 | 0.32 | 0.7 | 0.77 | 0.72 | 0.71 |
| 2 | 66285566 | rs900884 | 2.061 | 0.762 | 0.207 | 0.979 | -0.004 | 0.01 | 0.01 | 0.012 | 0.61 | 0.32 | 0.6 | 0.67 | 0.67 | 0.62 |
| 2 | 66285871 | rs4671709 | -1.664 | 1.079 | -0.41 | 1.045 | 0.009 | 0 | 0 | 2.153 | 0.52 | 0.49 | 0.58 | 0.53 | 0.52 | 0.51 |
| 2 | 66286114 | rs6546205 | -1.87 | 1.569 | -0.234 | 0.915 | 0.029 | 0.093 | 0.093 | 0.461 | 0.51 | 0.81 | 0.45 | 0.37 | 0.52 | 0.53 |
| 2 | 66287178 | rs7571668 | -1.128 | 1.469 | -0.226 | 0.902 | 0.013 | 0.017 | 0.017 | 0 | 0.19 | 0.25 | 0.3 | 0.24 | 0.23 | 0.37 |
| 2 | 66290514 | rs10198404 | -0.803 | 1.416 | -0.457 | 0.474 | 0 | -0.032 | -0.032 | 0 | 0.33 | 0.63 | 0.18 | 0.12 | 0.29 | 0.18 |
| 2 | 66293343 | rs6546206 | -2.502 | 1.213 | -0.718 | 0.812 | -0.046 | 0.003 | 0.003 | 0.22 | 0.63 | 0.97 | 0.69 | 0.76 | 0.61 | 0.73 |
| 2 | 66300380 | rs9678491 | 1.042 | 1.317 | -0.267 | 0.697 | 0.002 | 0.011 | 0.011 | 0 | 0.66 | 0.4 | 0.51 | 0.48 | 0.68 | 0.62 |
| 2 | 66302093 | rs2627857 | 1.828 | 1.234 | -0.231 | 0.731 | -0.014 | 0.035 | 0.035 | 0 | 0.55 | 0.34 | 0.4 | 0.32 | 0.55 | 0.35 |
| 2 | 66302873 | rs10496131 | 0.759 | 0.989 | -0.156 | 0.392 | 0.001 | -0.012 | -0.012 | 0 | 0.17 | 0.19 | 0.08 | 0.12 | 0.17 | 0.06 |
| 2 | 66303200 | rs17031450 | -0.56 | 1.083 | 0.387 | 0.694 | 0.003 | 0.037 | 0.037 | 0 | 0.1 | 0.21 | 0.09 | 0.16 | 0.11 | 0.26 |
| 2 | 66305945 | rs13409331 | -0.762 | 1.103 | 0.315 | 0.61 | 0.014 | -0.038 | -0.038 | 1.307 | 0.11 | 0.29 | 0.11 | 0.03 | 0.09 | 0.08 |
| 2 | 66309172 | rs6710236 | 0.769 | 0.864 | -0.111 | 0.723 | 0.018 | 0.079 | 0.079 | 0.075 | 0.73 | 0.49 | 0.59 | 0.51 | 0.73 | 0.68 |
| 2 | 66326853 | rs7586464 | -1.338 | 0.176 | -0.051 | 0.268 | -0.009 | 0.075 | 0.075 | 0 | 0.23 | 0.49 | 0.34 | 0.31 | 0.21 | 0.33 |
| 2 | 66329532 | rs984578 | -1.039 | 0.055 | -0.952 | 0.005 | -0.029 | 0.007 | 0.007 | 0.147 | 0.53 | 0.81 | 0.55 | 0.46 | 0.48 | 0.52 |
| 2 | 66331816 | rs4313962 | -1.191 | 0.006 | -1.144 | -0.019 | -0.063 | -0.019 | -0.019 | 0 | 0.55 | 0.88 | 0.56 | 0.47 | 0.48 | 0.52 |
| 2 | 66332914 | rs902146 | -0.731 | -0.085 | -0.987 | 0.09 | -0.013 | -0.043 | -0.043 | 0.534 | 0.13 | 0.04 | 0.22 | 0.3 | 0.13 | 0.28 |
| 2 | 66334171 | rs1819125 | -1.334 | -0.426 | -1.034 | -0.362 | 0.042 | 0.001 | 0.001 | 0.816 | 0.36 | 0.78 | 0.4 | 0.37 | 0.26 | 0.42 |
| 2 | 66350062 | rs1000087 | -1.703 | -0.332 | -0.302 | -0.455 | -0.023 | -0.003 | -0.003 | 0 | 0.53 | 0.75 | 0.54 | 0.52 | 0.47 | 0.49 |
| 2 | 66350457 | rs1532118 | 0.215 | -0.163 | -0.03 | -0.372 | 0.009 | -0.021 | -0.021 | 0 | 0.21 | 0.29 | 0.16 | 0.16 | 0.23 | 0.11 |
| 2 | 66354489 | rs17556229 | 1.32 | -0.151 | -0.307 | -0.318 | 0 | -0.008 | -0.008 | 0 | 0.05 | 0.03 | 0.1 | 0 | 0.02 | 0.03 |
| 2 | 66355189 | rs7581689 | -1.515 | -0.268 | -0.384 | -0.446 | -0.019 | 0.004 | 0.004 | 0.018 | 0.56 | 0.71 | 0.53 | 0.48 | 0.46 | 0.53 |
| 2 | 66357905 | rs10169352 | -1.59 | 0.56 | -0.099 | -0.111 | 0 | 0 | 0 | 0 | 0.45 | 0.53 | 0.52 | 0.5 | 0.46 | 0.5 |
| 2 | 66358238 | rs4671716 | 0.211 | 0.546 | 0.005 | -0.13 | -0.008 | -0.045 | -0.045 | 0.266 | 0.08 | 0.19 | 0.13 | 0 | 0.04 | 0.09 |
| 2 | 66359326 | rs13033204 | 0.156 | 0.569 | 0.029 | -0.136 | 0.009 | -0.035 | -0.035 | 0 | 0.22 | 0.25 | 0.17 | 0.14 | 0.23 | 0.13 |
| 2 | 66361227 | rs7565180 | 0.079 | 0.611 | -0.067 | -0.103 | -0.01 | -0.022 | -0.022 | 2.333 | 0.15 | 0.11 | 0.22 | 0.35 | 0.19 | 0.29 |
| 2 | 66361926 | rs6728380 | -0.796 | 0.549 | -0.152 | -0.046 | 0.015 | 0.014 | 0.014 | 0.167 | 0.39 | 0.5 | 0.42 | 0.5 | 0.42 | 0.42 |
| 2 | 66364645 | rs9309383 | -0.774 | 0.732 | -0.94 | 0.235 | -0.003 | -0.001 | -0.001 | 1.689 | 0.44 | 0.58 | 0.53 | 0.49 | 0.48 | 0.49 |
| 2 | 66379446 | rs13000105 | -0.058 | 1.214 | -1.462 | 0.006 | -0.09 | 0.062 | 0.062 | 0 | 0.17 | 0.02 | 0.11 | 0.06 | 0.24 | 0.07 |
| 2 | 66382388 | rs6712489 | -0.31 | 1.679 | -0.619 | 0.228 | 0.079 | 0.116 | 0.116 | 0.018 | 0.21 | 0.51 | 0.33 | 0.37 | 0.28 | 0.31 |
| 2 | 66383467 | rs2046795 | -1.512 | 0.506 | -1.034 | -0.54 | 0.011 | -0.001 | -0.001 | 0 | 0.58 | 0.68 | 0.6 | 0.44 | 0.63 | 0.6 |
| 2 | 66387107 | rs6748113 | 0.105 | 0.471 | -0.089 | -0.411 | 0.02 | 0.03 | 0.03 | 0 | 0.17 | 0.18 | 0.29 | 0.37 | 0.27 | 0.28 |
| 2 | 66388139 | rs7600569 | -0.505 | 0.472 | -0.126 | -0.41 | 0.074 | 0.107 | 0.107 | 0 | 0.21 | 0.52 | 0.33 | 0.37 | 0.28 | 0.31 |
| 2 | 66393650 | rs1997383 | -0.244 | 0.483 | 0.09 | -0.379 | 0.041 | 0.051 | 0.051 | 0.145 | 0.18 | 0.3 | 0.3 | 0.36 | 0.28 | 0.3 |
| 2 | 66400834 | rs13383903 | -0.232 | 0.84 | 0.224 | -0.211 | 0.033 | 0.075 | 0.075 | 0 | 0.08 | 0.12 | 0.19 | 0.3 | 0.15 | 0.22 |
| 2 | 66400889 | rs13384003 | -0.878 | 0.832 | 0.629 | -0.217 | 0.1 | 0.185 | 0.185 | 0.02 | 0.1 | 0.35 | 0.21 | 0.3 | 0.15 | 0.25 |
| 2 | 66402505 | rs2971851 | -0.669 | 0.322 | -0.373 | -0.049 | -0.084 | -0.098 | -0.098 | 0.097 | 0.68 | 0.93 | 0.74 | 0.6 | 0.64 | 0.68 |
| 2 | 66410946 | rs2951282 | 0.26 | -0.009 | -0.26 | 0.028 | 0.003 | -0.047 | -0.047 | 0 | 0.41 | 0.76 | 0.53 | 0.51 | 0.4 | 0.44 |
| 2 | 66417209 | rs2971848 | 0.873 | -0.492 | -0.212 | -0.306 | -0.037 | -0.032 | -0.032 | 0.006 | 0.28 | 0.48 | 0.28 | 0.22 | 0.21 | 0.15 |
| 2 | 66423991 | rs2932252 | 1.138 | -0.447 | -0.329 | -0.27 | -0.029 | -0.02 | -0.02 | 0 | 0.27 | 0.36 | 0.27 | 0.22 | 0.21 | 0.12 |
| 2 | 66430063 | rs2951288 | 1.125 | -0.407 | -0.185 | -0.274 | -0.015 | -0.014 | -0.014 | 0 | 0.27 | 0.29 | 0.27 | 0.23 | 0.22 | 0.12 |
| 2 | 66450555 | rs10196975 | -0.745 | -0.805 | -0.017 | -0.4 | -0.044 | -0.059 | -0.059 | 3.702 | 0.28 | 0.57 | 0.13 | 0.08 | 0.13 | 0.25 |
| 2 | 66452546 | rs2951289 | -0.431 | -0.422 | -0.163 | 0.068 | -0.013 | -0.02 | -0.02 | 0.112 | 0.87 | 0.91 | 0.82 | 0.73 | 0.82 | 0.75 |
| 2 | 66453086 | rs1358698 | 1.028 | -0.398 | 0.04 | 0.087 | -0.003 | -0.002 | -0.002 | 0 | 0.6 | 0.39 | 0.68 | 0.65 | 0.67 | 0.47 |
| 2 | 66455372 | rs1489697 | 0.194 | -0.456 | 0.301 | 0.048 | 0.009 | 0.025 | 0.025 | 0.048 | 0.83 | 0.75 | 0.81 | 0.74 | 0.81 | 0.72 |
| 2 | 66455866 | rs975443 | 1.079 | -0.486 | 0.292 | 0.024 | 0.002 | 0.002 | 0.002 | 1.073 | 0.6 | 0.36 | 0.68 | 0.65 | 0.67 | 0.47 |
| 2 | 66456180 | rs6546217 | -0.216 | -0.437 | 0.302 | 0.03 | -0.026 | -0.042 | -0.042 | 0 | 0.23 | 0.37 | 0.12 | 0.09 | 0.14 | 0.25 |
| 2 | 66459248 | rs7609526 | -0.115 | -0.434 | 0.359 | 0.053 | -0.032 | -0.047 | -0.047 | 0 | 0.22 | 0.42 | 0.12 | 0.09 | 0.13 | 0.23 |
| 2 | 66460709 | rs9309384 | 0.173 | -0.423 | 0.29 | 0.077 | 0.002 | 0.002 | 0.002 | 0 | 0.2 | 0.23 | 0.1 | 0.09 | 0.13 | 0.19 |
| 2 | 66464350 | rs1002897 | -0.013 | -0.242 | 0.322 | 0.346 | -0.001 | 0.001 | 0.001 | 0 | 0.12 | 0.1 | 0.15 | 0.24 | 0.18 | 0.25 |
| 2 | 66467515 | rs7566973 | -0.487 | -0.415 | 0.329 | 0.716 | 0 | 0.138 | 0.138 | 0 | 0.9 | 0.76 | 0.75 | 0.69 | 0.89 | 0.79 |
| 2 | 66498775 | rs1565328 | -0.405 | -0.382 | -0.175 | 0.725 | -0.008 | 0.029 | 0.029 | 0 | 0.89 | 0.71 | 0.84 | 0.86 | 0.9 | 0.76 |
| 2 | 66515700 | rs7562304 | -0.621 | -0.111 | -0.738 | 0.835 | 0.026 | 0.045 | 0.045 | 0 | 0.17 | 0.38 | 0.17 | 0.23 | 0.19 | 0.26 |
| 2 | 66522294 | rs4407237 | 0.721 | 0.185 | -1.187 | 0.93 | -0.036 | -0.084 | -0.084 | 0 | 0.29 | 0.02 | 0.31 | 0.31 | 0.26 | 0.25 |
| 2 | 66530723 | rs4075737 | -0.997 | 0.212 | 0.263 | 0.848 | 0.015 | 0.075 | 0.075 | 0.102 | 0.18 | 0.33 | 0.17 | 0.29 | 0.19 | 0.32 |
| 2 | 66537344 | rs4414666 | 0.791 | -0.931 | -0.639 | -0.108 | 0.099 | 0.051 | 0.051 | 0 | 0.49 | 0.07 | 0.55 | 0.59 | 0.65 | 0.45 |
| 2 | 66567399 | rs1519107 | -0.539 | -1.365 | 0.145 | -0.407 | -0.002 | 0.003 | 0.003 | 0.959 | 0.25 | 0.74 | 0.36 | 0.38 | 0.23 | 0.33 |
| 2 | 66567896 | rs2860235 | -1.598 | -1.544 | 0.479 | -0.511 | 0.078 | 0.062 | 0.062 | 0 | 0.46 | 0.83 | 0.43 | 0.38 | 0.35 | 0.39 |
| 2 | 66568115 | rs17501374 | 0.296 | -1.601 | 0.344 | -0.537 | 0.013 | 0.09 | 0.09 | 0 | 0.15 | 0.01 | 0.06 | 0 | 0.11 | 0.05 |
| 2 | 66568270 | rs869578 | -0.396 | -1.595 | 0.628 | -0.527 | -0.004 | 0.043 | 0.043 | 1.656 | 0.23 | 0.44 | 0.33 | 0.38 | 0.23 | 0.33 |
| 2 | 66580863 | rs17031999 | 0.061 | -1.39 | 0.69 | -0.56 | -0.004 | -0.032 | -0.032 | 0 | 0.15 | 0.14 | 0.07 | 0 | 0.11 | 0.05 |
| 2 | 66585152 | rs17032007 | 0.061 | -1.374 | 0.562 | -0.553 | -0.004 | -0.032 | -0.032 | 0 | 0.15 | 0.14 | 0.07 | 0 | 0.11 | 0.05 |
| 2 | 66587063 | rs2946375 | -0.719 | -1.338 | 0.568 | -0.598 | -0.014 | -0.036 | -0.036 | 0 | 0.1 | 0.1 | 0.03 | 0 | 0.03 | 0.04 |
| 2 | 66593246 | rs952267 | -1.684 | -1.295 | 1.084 | -0.685 | 0.03 | 0.048 | 0.048 | 0.322 | 0.47 | 0.63 | 0.34 | 0.37 | 0.41 | 0.45 |
| 2 | 66602521 | rs10182624 | -1.354 | -1.338 | 1.488 | -0.711 | 0.014 | 0.019 | 0.019 | 0 | 0.4 | 0.33 | 0.3 | 0.37 | 0.38 | 0.4 |
| 2 | 66605631 | rs1467229 | -1.828 | -1.346 | 1.34 | -0.747 | 0.044 | 0.082 | 0.082 | 0.048 | 0.48 | 0.75 | 0.37 | 0.33 | 0.41 | 0.49 |
| 2 | 66606448 | rs10496134 | 0.298 | -1.339 | 1.401 | -0.764 | 0.004 | -0.016 | -0.016 | 0 | 0.15 | 0.12 | 0.08 | 0 | 0.12 | 0.05 |
| 2 | 66607003 | rs6752427 | -1.613 | -0.989 | 1.845 | -0.305 | 0.001 | 0.001 | 0.001 | 2.169 | 0.33 | 0.68 | 0.28 | 0.32 | 0.29 | 0.42 |
| 2 | 66608972 | rs6760326 | -2.052 | -1.026 | 1.608 | -0.319 | 0.015 | 0.003 | 0.003 | 0.956 | 0.38 | 0.91 | 0.3 | 0.37 | 0.31 | 0.58 |

| | | | | | | | | | | | | | | | | |
|---|---|---|---|---|---|---|---|---|---|---|---|---|---|---|---|---|
| 2 | 66610296 | rs13397921 | -0.887 | -1 | 1.724 | -0.332 | -0.005 | -0.031 | -0.031 | 0.45 | 0.06 | 0.15 | 0.03 | 0 | 0.03 | 0.06 |
| 2 | 66625375 | rs7604762 | -1.675 | -1.073 | 1.707 | -0.312 | 0.001 | 0.002 | 0.002 | 2.509 | 0.34 | 0.7 | 0.32 | 0.33 | 0.3 | 0.42 |
| 2 | 66627412 | rs13013095 | -0.976 | -1.054 | 1.984 | -0.284 | 0.001 | 0.006 | 0.006 | 0.982 | 0.26 | 0.41 | 0.29 | 0.33 | 0.28 | 0.36 |
| 2 | 66628286 | rs1519118 | -0.765 | -0.979 | 2.105 | -0.337 | -0.007 | 0.046 | 0.046 | 0 | 0.05 | 0.17 | 0.01 | 0.07 | 0 | 0.17 |
| 2 | 66651744 | rs1519116 | -1.909 | -0.884 | 1.466 | -0.321 | -0.01 | -0.01 | -0.01 | 0 | 0.24 | 0.39 | 0.26 | 0.26 | 0.23 | 0.53 |
| 2 | 66652032 | rs1519113 | -1.222 | -0.876 | 1.516 | -0.315 | 0.006 | 0.02 | 0.02 | 0.368 | 0.16 | 0.26 | 0.24 | 0.25 | 0.21 | 0.46 |
| 2 | 66655256 | rs7604427 | -0.385 | -0.682 | 0.782 | -0.43 | -0.006 | 0.007 | 0.007 | 0.683 | 0.66 | 0.57 | 0.53 | 0.39 | 0.71 | 0.83 |
| 2 | 66657939 | rs756292 | -0.36 | -0.716 | 0.737 | -0.424 | -0.007 | 0.01 | 0.01 | 0.638 | 0.66 | 0.53 | 0.52 | 0.39 | 0.7 | 0.83 |
| 2 | 66665146 | rs2280334 | -0.04 | -0.57 | 0.594 | -0.343 | -0.008 | -0.005 | -0.005 | 0 | 0.59 | 0.48 | 0.52 | 0.39 | 0.68 | 0.76 |
| 2 | 66687843 | rs6716792 | 0.414 | -0.693 | 0.108 | -0.381 | -0.002 | 0.056 | 0.056 | 3.167 | 0.51 | 0.62 | 0.4 | 0.26 | 0.48 | 0.47 |
| 2 | 66692508 | rs17032103 | -0.412 | -0.748 | 0.525 | -0.024 | -0.007 | -0.009 | -0.009 | 0 | 0.05 | 0.02 | 0.1 | 0.16 | 0.11 | 0.27 |
| 2 | 66693328 | rs6755563 | -2.469 | -1.036 | 0.815 | -0.163 | -0.04 | -0.031 | -0.031 | 0.331 | 0.25 | 0.68 | 0.24 | 0.26 | 0.16 | 0.39 |
| 2 | 66695321 | rs6727352 | -2.477 | -1.031 | 0.847 | -0.16 | -0.029 | -0.023 | -0.023 | 0 | 0.25 | 0.7 | 0.24 | 0.26 | 0.16 | 0.39 |
| 2 | 66702070 | rs6755135 | -2.477 | -1.041 | 1.044 | -0.164 | -0.029 | -0.024 | -0.024 | 0 | 0.25 | 0.7 | 0.24 | 0.26 | 0.16 | 0.39 |
| 2 | 66703251 | rs6730722 | -1.44 | -1.017 | 0.917 | -0.154 | -0.012 | 0.01 | 0.01 | 0.009 | 0.11 | 0.53 | 0.11 | 0.09 | 0.03 | 0.09 |
| 2 | 66705785 | rs7595378 | -2.047 | -0.996 | 0.592 | -0.144 | -0.068 | -0.058 | -0.058 | 0 | 0.19 | 0.67 | 0.15 | 0.09 | 0.05 | 0.12 |
| 2 | 66714597 | rs17032114 | -1.104 | -0.135 | -0.299 | 0.309 | -0.007 | 0.008 | 0.008 | 0 | 0.08 | 0.28 | 0.09 | 0.09 | 0.03 | 0.09 |
| 2 | 66720099 | rs4671729 | -0.855 | 0.25 | -1.039 | 0.404 | -0.001 | -0.002 | -0.002 | 0 | 0.06 | 0.05 | 0.1 | 0.17 | 0.11 | 0.27 |
| 2 | 66724142 | rs6722685 | -0.006 | 0.307 | -1.595 | 0.335 | 0.058 | 0.09 | 0.09 | 0.127 | 0.52 | 0.2 | 0.62 | 0.59 | 0.63 | 0.43 |
| 2 | 66726453 | rs4671730 | -0.543 | 0.882 | -0.752 | 0.909 | 0.05 | 0.06 | 0.06 | 0.185 | 0.27 | 0.48 | 0.52 | 0.49 | 0.39 | 0.35 |
| 2 | 66726568 | rs4547518 | 0.748 | 0.116 | -1.367 | -0.211 | -0.007 | 0.097 | 0.097 | 2.158 | 0.36 | 0.18 | 0.2 | 0.08 | 0.34 | 0.22 |
| 2 | 66729420 | rs12713567 | -0.486 | -0.557 | -0.728 | -0.097 | -0.006 | 0.012 | 0.012 | 0.657 | 0.57 | 0.63 | 0.64 | 0.68 | 0.56 | 0.68 |
| 2 | 66743591 | rs6721499 | -0.936 | -0.678 | -0.587 | -0.192 | -0.02 | 0.099 | 0.099 | 1.092 | 0.63 | 0.84 | 0.74 | 0.79 | 0.63 | 0.73 |
| 2 | 66744603 | rs3890755 | -0.387 | -0.554 | 0.203 | 0.559 | 0.008 | 0.005 | 0.005 | 4.287 | 0.5 | 0.39 | 0.52 | 0.48 | 0.5 | 0.56 |
| 2 | 66745663 | rs7586211 | -0.381 | -0.489 | 0.559 | 0.592 | 0.177 | 0.251 | 0.251 | 0 | 0.94 | 0.64 | 0.92 | 0.86 | 0.92 | 0.94 |
| 2 | 66749610 | rs4430933 | -0.895 | -0.607 | 0.713 | 0.506 | 0.006 | -0.021 | -0.021 | 0.717 | 0.6 | 0.44 | 0.73 | 0.78 | 0.6 | 0.8 |
| 2 | 66750152 | rs6742861 | 1.549 | -0.618 | 0.449 | 0.416 | -0.078 | 0.17 | 0.17 | 10.706 | 0.31 | 0.08 | 0.17 | 0.07 | 0.32 | 0.13 |
| 2 | 66758422 | rs6710341 | 0.465 | -0.844 | 0.382 | 0.462 | 0.006 | -0.008 | -0.008 | 0 | 0.2 | 0.18 | 0.18 | 0.29 | 0.15 | 0.34 |
| 2 | 66760311 | rs4433986 | -1.279 | -0.838 | 0.406 | 0.517 | 0.012 | -0.04 | -0.04 | 0 | 0.61 | 0.49 | 0.74 | 0.81 | 0.6 | 0.8 |
| 2 | 66760905 | rs4316931 | 1.693 | -0.827 | 0.143 | 0.527 | -0.065 | -0.079 | -0.079 | 0 | 0.18 | 0.02 | 0.28 | 0.22 | 0.27 | 0.19 |
| 2 | 66764308 | rs12469063 | 0.221 | -0.861 | 0.024 | 0.474 | -0.033 | -0.059 | -0.059 | 0 | 0.25 | 0.02 | 0.19 | 0.29 | 0.24 | 0.27 |
| 2 | 66771047 | rs997153 | 1.34 | -0.8 | 0.259 | 0.444 | 0.001 | -0.008 | -0.008 | 0 | 0.37 | 0.37 | 0.25 | 0.19 | 0.4 | 0.19 |
| 2 | 66776729 | rs2300477 | 1.419 | -0.853 | -0.4 | 0.468 | -0.03 | 0.065 | 0.065 | 3.563 | 0.49 | 0.04 | 0.43 | 0.3 | 0.59 | 0.32 |
| 2 | 66777521 | rs2192954 | 0.358 | -0.495 | 0.098 | 0.652 | 0.003 | -0.005 | -0.005 | 0 | 0.6 | 0.71 | 0.45 | 0.48 | 0.55 | 0.55 |
| 2 | 66781453 | rs2300478 | -0.126 | -0.379 | 0.111 | 0.822 | 0.011 | 0 | 0 | 0.002 | 0.26 | 0.11 | 0.22 | 0.29 | 0.25 | 0.27 |
| 2 | 66783940 | rs2284706 | 0.263 | 0.047 | 0.206 | 0.771 | 0.096 | 0.088 | 0.088 | 0 | 0.05 | 0.15 | 0.09 | 0.12 | 0.1 | 0.06 |
| 2 | 66786405 | rs10202912 | 0.679 | -0.162 | -0.02 | 0.656 | 0.002 | -0.013 | -0.013 | 0.382 | 0.58 | 0.56 | 0.48 | 0.3 | 0.55 | 0.44 |
| 2 | 66788314 | rs2300484 | 1.023 | -0.431 | -0.03 | 0.281 | 0.002 | -0.011 | -0.011 | 0 | 0.34 | 0.35 | 0.18 | 0.07 | 0.27 | 0.19 |
| 2 | 66792109 | rs12614369 | -0.314 | -0.617 | 0.253 | 1.268 | 0 | -0.007 | -0.007 | 0.534 | 0.18 | 0.14 | 0.18 | 0.24 | 0.2 | 0.23 |
| 2 | 66792291 | rs6735071 | 0.956 | -0.787 | 0.704 | 1.524 | -0.043 | 0.053 | 0.053 | 0 | 0.8 | 0.65 | 0.84 | 0.71 | 0.85 | 0.74 |
| 2 | 66797138 | rs2216124 | -0.615 | -0.437 | -0.246 | 1.673 | 0.021 | -0.009 | -0.009 | 0.491 | 0.09 | 0.01 | 0.09 | 0.11 | 0.04 | 0.13 |
| 2 | 66803893 | rs10184250 | -0.787 | 0.269 | -0.513 | 1.172 | -0.004 | -0.003 | -0.003 | 0 | 0.25 | 0.23 | 0.18 | 0.24 | 0.24 | 0.32 |
| 2 | 66808036 | rs4671737 | -0.542 | 0.398 | -0.959 | 1.412 | 0.005 | -0.023 | -0.023 | 0 | 0.26 | 0.18 | 0.38 | 0.65 | 0.26 | 0.35 |
| 2 | 66812892 | rs17032269 | -0.748 | 0.506 | -0.437 | 1.617 | -0.01 | 0.021 | 0.021 | 0 | 0.37 | 0.55 | 0.43 | 0.54 | 0.35 | 0.41 |
| 2 | 66815275 | rs6706382 | -0.672 | 0.54 | -0.358 | 1.652 | -0.005 | 0.01 | 0.01 | 0 | 0.35 | 0.62 | 0.43 | 0.56 | 0.32 | 0.38 |
| 2 | 66815660 | rs12987658 | 1.083 | 0.453 | -0.982 | 1.26 | -0.059 | 0.155 | 0.155 | 0 | 0.46 | 0.09 | 0.45 | 0.21 | 0.58 | 0.41 |
| 2 | 66816409 | rs10865356 | -0.486 | 0.498 | -0.545 | 1.436 | -0.009 | 0.015 | 0.015 | 0 | 0.35 | 0.61 | 0.42 | 0.53 | 0.29 | 0.38 |
| 2 | 66817402 | rs984306 | 0.646 | 0.704 | 0.084 | 1.457 | -0.003 | 0.03 | 0.03 | 0 | 0.73 | 0.59 | 0.77 | 0.6 | 0.76 | 0.71 |
| 2 | 66819030 | rs2861111 | 0.72 | 1.015 | 0.289 | 1.498 | 0.005 | -0.012 | -0.012 | 0.106 | 0.63 | 0.37 | 0.63 | 0.49 | 0.66 | 0.67 |
| 2 | 66821247 | rs17032300 | -0.639 | 1.322 | -0.136 | 1.547 | -0.028 | -0.041 | -0.041 | 0.76 | 0.15 | 0.11 | 0.17 | 0.32 | 0.19 | 0.21 |
| 2 | 66821263 | rs4671741 | -0.414 | 1.346 | 0.588 | 1.391 | -0.019 | 0.029 | 0.029 | 0.003 | 0.31 | 0.35 | 0.31 | 0.47 | 0.24 | 0.24 |
| 2 | 66825474 | rs2192963 | 1.292 | 1.465 | 0.914 | 1.552 | 0.062 | 0.009 | 0.009 | 0.026 | 0.8 | 0.44 | 0.74 | 0.82 | 0.73 | 0.73 |
| 2 | 66827052 | rs10195834 | 1.061 | 1.613 | 1.017 | 1.412 | 0.077 | 0.079 | 0.079 | 0 | 0.89 | 0.66 | 0.78 | 0.85 | 0.82 | 0.83 |
| 2 | 66827585 | rs2192964 | 0.874 | 1.596 | 1.005 | 1.413 | 0.069 | 0.089 | 0.089 | 0 | 0.91 | 0.67 | 0.8 | 0.86 | 0.87 | 0.84 |
| 2 | 66831138 | rs6749832 | -0.777 | 1.565 | 0.908 | 1.648 | 0.018 | 0.169 | 0.169 | 0.144 | 0.07 | 0.22 | 0.13 | 0.29 | 0.07 | 0.1 |
| 2 | 66834215 | rs12713569 | 1.27 | 1.416 | 0.549 | 1.407 | 0 | 0.19 | 0.19 | 0 | 0.53 | 0.14 | 0.47 | 0.21 | 0.58 | 0.49 |
| 2 | 66836319 | rs1981736 | -1.797 | 1.394 | 1.231 | 1.456 | 0.031 | 0.189 | 0.189 | 0 | 0.14 | 0.58 | 0.18 | 0.32 | 0.16 | 0.24 |
| 2 | 66876624 | rs6725996 | 0.566 | 1.274 | 1.33 | 1.279 | -0.013 | -0.045 | -0.045 | 0.449 | 0.09 | 0.17 | 0.04 | 0 | 0.05 | 0.09 |
| 2 | 66878488 | rs11695019 | 2.522 | 1.048 | 0.894 | 1.104 | -0.087 | 0.154 | 0.154 | 0.018 | 0.4 | 0.03 | 0.39 | 0.17 | 0.4 | 0.36 |
| 2 | 66881333 | rs2861172 | 0.596 | 0.946 | 0.902 | 1.13 | -0.002 | -0.091 | -0.091 | 0.094 | 0.19 | 0.03 | 0.14 | 0.33 | 0.18 | 0.17 |
| 2 | 66891388 | rs7593846 | -1.624 | 1.035 | 1.082 | 1.206 | 0 | -0.001 | -0.001 | 0.862 | 0.25 | 0.7 | 0.22 | 0.33 | 0.2 | 0.33 |
| 2 | 66896718 | rs9941639 | -0.732 | 1.041 | 1.275 | 1.202 | 0.021 | -0.086 | -0.086 | 0 | 0.17 | 0.4 | 0.11 | 0.02 | 0.17 | 0.25 |
| 2 | 66898267 | rs10490189 | -0.036 | 1.009 | 0.99 | 1.216 | 0.001 | 0.269 | 0.269 | 3.085 | 0.09 | 0.3 | 0.11 | 0.31 | 0.03 | 0.08 |
| 2 | 66900737 | rs6745277 | 0.343 | 1.112 | 0.43 | 0.914 | 0.001 | -0.004 | -0.004 | 0.219 | 0.06 | 0.07 | 0.04 | 0.02 | 0.09 | 0.16 |
| 2 | 66905657 | rs13025385 | 2.001 | 1.057 | 0.17 | 0.759 | -0.078 | 0.152 | 0.152 | 0 | 0.4 | 0.03 | 0.39 | 0.16 | 0.4 | 0.34 |
| 2 | 66906040 | rs10206999 | -0.704 | 1.334 | 0.795 | 0.826 | 0.031 | -0.034 | -0.034 | 0 | 0.16 | 0.26 | 0.13 | 0.05 | 0.23 | 0.25 |
| 2 | 66909353 | rs6546256 | -1.463 | 1.265 | 1.103 | 0.843 | 0.04 | -0.058 | -0.058 | 0.014 | 0.22 | 0.64 | 0.21 | 0.06 | 0.3 | 0.31 |
| 2 | 66909648 | rs7607842 | -0.096 | 1.258 | 1.398 | 0.839 | 0.069 | -0.04 | -0.04 | 0.713 | 0.11 | 0.33 | 0.08 | 0.03 | 0.14 | 0.1 |
| 2 | 66910712 | rs933533 | -1.274 | 1.077 | 1.311 | 1.122 | 0.02 | 0.098 | 0.098 | 0.161 | 0.2 | 0.5 | 0.18 | 0.33 | 0.17 | 0.18 |

| | | | | | | | | | | | | | | | | |
|---|---|---|---|---|---|---|---|---|---|---|---|---|---|---|---|---|
| 2 | 66911701 | rs13018192 | -0.255 | 1.092 | 1.164 | 1.097 | 0.083 | -0.046 | -0.046 | 0 | 0.12 | 0.41 | 0.09 | 0.03 | 0.14 | 0.1 |
| 2 | 66911725 | rs7562380 | 0.488 | 0.869 | 0.288 | 1.208 | 0.012 | -0.055 | -0.055 | 0 | 0.19 | 0.05 | 0.15 | 0.32 | 0.18 | 0.15 |
| 2 | 66911948 | rs13018438 | -1.981 | 0.903 | 0.186 | 1.179 | 0.024 | 0.105 | 0.105 | 0 | 0.5 | 0.79 | 0.46 | 0.69 | 0.55 | 0.55 |
| 2 | 66913252 | rs7583376 | -1.717 | 1.044 | 0.341 | 1.048 | -0.021 | -0.015 | -0.015 | 1.025 | 0.32 | 0.75 | 0.32 | 0.36 | 0.38 | 0.4 |
| 2 | 66917717 | rs7565020 | -0.503 | 1.034 | -0.171 | 1.094 | -0.058 | -0.074 | -0.074 | 0 | 0.31 | 0.14 | 0.33 | 0.44 | 0.44 | 0.41 |
| 2 | 66919316 | rs17679928 | 0.562 | 1.012 | -0.449 | 0.891 | -0.007 | 0.029 | 0.029 | 0 | 0.08 | 0.02 | 0.04 | 0.02 | 0.09 | 0.1 |
| 2 | 66919360 | rs10200612 | -1.433 | 0.915 | -0.046 | 0.947 | 0.006 | 0.046 | 0.046 | 0 | 0.5 | 0.38 | 0.47 | 0.72 | 0.57 | 0.59 |
| 2 | 66927859 | rs17680434 | 0.056 | 1.425 | -0.452 | 0.621 | -0.053 | -0.027 | -0.027 | 0 | 0.09 | 0.01 | 0.1 | 0.11 | 0.21 | 0.23 |
| 2 | 66930461 | rs2192955 | -0.239 | 0.894 | -0.131 | 0.482 | 0.01 | -0.051 | -0.051 | 0.642 | 0.89 | 0.78 | 0.94 | 0.98 | 0.92 | 0.89 |
| 2 | 66936494 | rs6705424 | 1.031 | 1.153 | -0.701 | 0.669 | 0.045 | 0.025 | 0.025 | 0 | 0.51 | 0.75 | 0.47 | 0.46 | 0.39 | 0.4 |
| 2 | 66937329 | rs10490191 | 0.838 | 1.071 | -0.68 | 0.606 | 0.009 | 0.01 | 0.01 | 0 | 0.54 | 0.77 | 0.49 | 0.46 | 0.42 | 0.41 |
| 2 | 66953286 | rs10201517 | -0.164 | 0.983 | -0.765 | 0.672 | 0.002 | -0.008 | -0.008 | 0.172 | 0.77 | 0.78 | 0.69 | 0.57 | 0.81 | 0.76 |
| 2 | 66954363 | rs6748591 | -0.308 | 1.045 | -0.485 | 0.331 | -0.003 | -0.054 | -0.054 | 0 | 0.12 | 0.21 | 0.05 | 0.01 | 0.08 | 0.11 |
| 2 | 66956404 | rs1111234 | 0.035 | 1.108 | -1.398 | 0.326 | 0.011 | -0.051 | -0.051 | 0.41 | 0.32 | 0.2 | 0.36 | 0.43 | 0.28 | 0.26 |
| 2 | 66956820 | rs17680993 | -0.634 | 1.832 | -1.23 | 0.151 | -0.03 | 0.005 | 0.005 | 0 | 0.14 | 0.07 | 0.17 | 0.12 | 0.3 | 0.24 |
| 2 | 66963401 | rs6714705 | -0.308 | 1.395 | -1.348 | -0.095 | -0.003 | 0.001 | 0.001 | 0.208 | 0.44 | 0.59 | 0.48 | 0.45 | 0.5 | 0.47 |
| 2 | 66963775 | rs6757438 | 0.116 | 1.348 | -1.65 | -0.288 | -0.16 | -0.12 | -0.12 | 0.075 | 0.35 | 0.06 | 0.41 | 0.45 | 0.46 | 0.44 |
| 2 | 66972843 | rs7566062 | -0.21 | 1.296 | -1.271 | -0.534 | 0.012 | -0.046 | -0.046 | 0.005 | 0.21 | 0.31 | 0.2 | 0.08 | 0.23 | 0.16 |
| 2 | 66973628 | rs17681226 | 1.138 | 1.707 | -2.041 | -0.503 | -0.04 | 0.022 | 0.022 | 0.019 | 0.23 | 0.03 | 0.25 | 0.14 | 0.21 | 0.31 |
| 2 | 66974287 | rs13000961 | -1.048 | 1.131 | -1.616 | -0.234 | -0.009 | 0.009 | 0.009 | 0 | 0.64 | 0.6 | 0.64 | 0.51 | 0.7 | 0.58 |
| 2 | 66977032 | rs17681403 | 0.426 | 1.232 | -1.702 | -0.491 | -0.035 | 0.041 | 0.041 | 0 | 0.15 | 0.06 | 0.16 | 0.07 | 0.19 | 0.14 |
| 2 | 66992263 | rs2216921 | -0.432 | 0.892 | -1.086 | -0.158 | -0.004 | 0.015 | 0.015 | 0 | 0.6 | 0.66 | 0.48 | 0.36 | 0.58 | 0.43 |
| 2 | 66999666 | rs11126096 | -0.087 | 0.894 | -1.614 | 0.019 | 0.01 | 0.07 | 0.07 | 0 | 0.55 | 0.2 | 0.47 | 0.36 | 0.58 | 0.4 |
| 2 | 67008311 | rs10168064 | -0.862 | 1.009 | -1.473 | 0.345 | -0.042 | 0.005 | 0.005 | 0 | 0.41 | 0.79 | 0.44 | 0.41 | 0.44 | 0.5 |
| 2 | 67010348 | rs17032536 | -0.159 | 1.17 | -0.851 | 0.438 | 0.049 | 0.016 | 0.016 | 0.119 | 0.13 | 0.6 | 0.13 | 0.14 | 0.15 | 0.22 |
| 2 | 67014761 | rs2194659 | -0.984 | 1.166 | -0.919 | 0.381 | 0.001 | 0 | 0 | 0 | 0.35 | 0.64 | 0.4 | 0.41 | 0.41 | 0.48 |
| 2 | 67022351 | rs4671748 | 0.126 | 1.201 | -1.013 | 0.431 | -0.038 | -0.014 | -0.014 | 0 | 0.22 | 0.09 | 0.17 | 0.23 | 0.27 | 0.21 |
| 2 | 67026142 | rs6727188 | -1.277 | 0.95 | -0.598 | 0.269 | -0.009 | 0.004 | 0.004 | 0.049 | 0.44 | 0.75 | 0.34 | 0.45 | 0.45 | 0.56 |
| 2 | 67030447 | rs17631057 | -0.802 | 0.679 | 0.142 | 0.209 | -0.003 | -0.013 | -0.013 | 0 | 0.34 | 0.42 | 0.25 | 0.3 | 0.33 | 0.36 |
| 2 | 67041677 | rs17032567 | -0.87 | 0.78 | -0.259 | 0.658 | 0.007 | -0.019 | -0.019 | 0 | 0.4 | 0.76 | 0.33 | 0.46 | 0.42 | 0.41 |
| 2 | 67045142 | rs11126099 | -0.45 | 0.553 | -0.678 | 0.412 | -0.012 | -0.006 | -0.006 | 0.147 | 0.61 | 0.7 | 0.6 | 0.63 | 0.54 | 0.61 |
| 2 | 67046565 | rs10865358 | -0.903 | 0.619 | -1.243 | 0.192 | -0.002 | 0.113 | 0.113 | 1.085 | 0.54 | 0.84 | 0.67 | 0.75 | 0.6 | 0.63 |
| 2 | 67046865 | rs6546266 | -0.294 | 0.527 | -0.637 | 0.236 | 0 | -0.003 | -0.003 | 0.02 | 0.41 | 0.44 | 0.54 | 0.58 | 0.48 | 0.44 |
| 2 | 67047640 | rs1114117 | 0.906 | 0.755 | -0.352 | 0.477 | 0.013 | 0.039 | 0.039 | 0.033 | 0.13 | 0.2 | 0.35 | 0.26 | 0.2 | 0.19 |
| 2 | 67048161 | rs11682718 | 2.121 | 0.862 | -0.526 | 0.774 | -0.028 | -0.031 | -0.031 | 2.461 | 0.05 | 0 | 0.19 | 0.08 | 0.06 | 0.06 |
| 2 | 67050752 | rs13018425 | 1.551 | 0.332 | -0.762 | 0.299 | 0.008 | 0.008 | 0.008 | 0 | 0.24 | 0.15 | 0.19 | 0.19 | 0.16 | 0.29 |
| 2 | 67055115 | rs17032597 | 0.921 | 0.156 | -1.046 | 0.137 | -0.006 | 0.042 | 0.042 | 0 | 0.28 | 0.17 | 0.19 | 0.09 | 0.28 | 0.15 |
| 2 | 67055919 | rs17032602 | 1.85 | 0.203 | -0.752 | 0.268 | 0.002 | 0.005 | 0.005 | 0 | 0.08 | 0.12 | 0.09 | 0.17 | 0.09 | 0.18 |
| 2 | 67060126 | rs3919596 | 0.919 | 0.259 | -0.774 | 0.281 | -0.009 | -0.007 | -0.007 | 0 | 0.33 | 0.35 | 0.29 | 0.35 | 0.25 | 0.47 |
| 2 | 67060996 | rs7566349 | 0.443 | 0.705 | -0.839 | 0.786 | 0.012 | 0.036 | 0.036 | 0.007 | 0.14 | 0.2 | 0.35 | 0.26 | 0.2 | 0.2 |
| 2 | 67067669 | rs1559603 | -0.919 | 0.429 | -1.351 | 0.788 | -0.004 | -0.028 | -0.028 | 0 | 0.81 | 0.87 | 0.85 | 0.71 | 0.76 | 0.87 |
| 2 | 67072059 | rs2081399 | 0.203 | 0.922 | -1.291 | 0.671 | -0.001 | -0.001 | -0.001 | 0.77 | 0.24 | 0.28 | 0.38 | 0.29 | 0.31 | 0.23 |
| 2 | 67072082 | rs2081398 | -0.956 | 0.477 | -1.818 | 0.385 | 0.012 | -0.056 | -0.056 | 0 | 0.54 | 0.83 | 0.65 | 0.53 | 0.57 | 0.59 |
| 2 | 67072125 | rs2098819 | 0.466 | 0.543 | -1.605 | 0.433 | 0 | -0.001 | -0.001 | 0 | 0.25 | 0.3 | 0.39 | 0.27 | 0.31 | 0.23 |
| 2 | 67075166 | rs6732212 | 1.08 | 0.254 | -1.728 | 0.206 | -0.01 | -0.025 | -0.025 | 1.705 | 0.08 | 0.12 | 0.03 | 0.01 | 0.05 | 0.07 |
| 2 | 67077111 | rs1110581 | -1.937 | 0.2 | -2.143 | 0.303 | -0.032 | -0.096 | -0.096 | 0 | 0.86 | 0.99 | 0.9 | 0.66 | 0.82 | 0.87 |
| 2 | 67077597 | rs2216925 | 0.317 | 0.126 | -2.197 | -0.019 | 0.029 | 0.1 | 0.1 | 0 | 0.44 | 0.26 | 0.31 | 0.19 | 0.37 | 0.37 |
| 2 | 67080778 | rs17032637 | 1.247 | 0.108 | -2.237 | 0.04 | -0.01 | -0.002 | -0.002 | 0 | 0.09 | 0.07 | 0.09 | 0.07 | 0.09 | 0.06 |
| 2 | 67081507 | rs12713575 | 1.409 | 0.064 | -2.323 | -0.042 | 0.024 | 0.066 | 0.066 | 0 | 0.29 | 0.14 | 0.2 | 0.13 | 0.24 | 0.28 |
| 2 | 67081562 | rs6728659 | 0.86 | 0.14 | -2.302 | -0.061 | 0.035 | 0.082 | 0.082 | 0 | 0.36 | 0.26 | 0.23 | 0.14 | 0.28 | 0.35 |
| 2 | 67090833 | rs1126147 | 2.602 | -0.086 | -2.65 | -0.226 | 0.004 | 0.078 | 0.078 | 0 | 0.21 | 0.05 | 0.15 | 0.09 | 0.18 | 0.14 |
| 2 | 67091841 | rs11902129 | 0.839 | 0.078 | -2.057 | -0.099 | 0.057 | 0.062 | 0.062 | 0 | 0.26 | 0.39 | 0.47 | 0.35 | 0.33 | 0.27 |
| 2 | 67094541 | rs1110585 | 0.283 | 0.062 | -2.187 | -0.31 | 0.019 | 0.138 | 0.138 | 0 | 0.44 | 0.22 | 0.31 | 0.18 | 0.38 | 0.38 |
| 2 | 67107863 | rs10176733 | 1.275 | 0.092 | -2.262 | -0.315 | 0.009 | 0.078 | 0.078 | 0 | 0.29 | 0.12 | 0.2 | 0.14 | 0.27 | 0.28 |
| 2 | 67114505 | rs6714117 | 1.131 | 0.046 | -2.196 | -0.352 | -0.019 | -0.033 | -0.033 | 0 | 0.13 | 0.23 | 0.1 | 0.1 | 0.1 | 0.2 |
| 2 | 67115437 | rs11677614 | 0.598 | 0.047 | -2.337 | -0.382 | 0.016 | 0.118 | 0.118 | 0 | 0.38 | 0.21 | 0.25 | 0.15 | 0.35 | 0.35 |
| 2 | 67115938 | rs13031245 | 0.125 | 0.081 | -2.335 | -0.399 | 0.015 | 0.155 | 0.155 | 0 | 0.47 | 0.3 | 0.33 | 0.19 | 0.43 | 0.38 |
| 2 | 67117415 | rs7562476 | 0.376 | 0.23 | -2.296 | 0.003 | -0.011 | 0.011 | 0.011 | 0 | 0.21 | 0.33 | 0.33 | 0.33 | 0.18 | 0.39 |
| 2 | 67121698 | rs10490198 | 1.322 | 0.317 | -2.63 | -0.011 | -0.027 | 0.002 | 0.002 | 0 | 0.09 | 0.05 | 0.09 | 0.04 | 0.11 | 0.02 |
| 2 | 67123397 | rs13010493 | 2.409 | 0.175 | -2.794 | -0.123 | -0.017 | 0.077 | 0.077 | 0 | 0.21 | 0.04 | 0.16 | 0.09 | 0.23 | 0.14 |
| 2 | 67125383 | rs3732047 | 1.496 | 0.559 | -2.58 | 0.187 | -0.123 | 0.025 | 0.025 | 0 | 0.21 | 0.01 | 0.23 | 0.07 | 0.29 | 0.17 |
| 2 | 67129177 | rs7587400 | -1.038 | 0.528 | -2.515 | 0.281 | 0.017 | -0.074 | -0.074 | 0 | 0.78 | 0.87 | 0.85 | 0.6 | 0.79 | 0.8 |
| 2 | 67131631 | rs10519066 | 0.49 | 0.565 | -2.443 | 0.23 | -0.013 | 0.023 | 0.023 | 0 | 0.16 | 0.13 | 0.12 | 0.05 | 0.16 | 0.09 |
| 2 | 67144798 | rs10519071 | 2.099 | 0.574 | -2.439 | 0.242 | -0.026 | -0.013 | -0.013 | 0.581 | 0.07 | 0.01 | 0.08 | 0.04 | 0.09 | 0.02 |
| 2 | 67145433 | rs6546272 | -1.206 | 0.51 | -2.476 | 0.316 | 0.047 | -0.043 | -0.043 | 0 | 0.54 | 0.81 | 0.68 | 0.47 | 0.58 | 0.58 |
| 2 | 67150295 | rs4671759 | 0.375 | 0.59 | -2.224 | 0.447 | -0.001 | 0.001 | 0.001 | 0 | 0.19 | 0.23 | 0.32 | 0.35 | 0.17 | 0.39 |
| 2 | 67150881 | rs11674518 | 1.171 | 0.73 | -2.141 | 0.42 | -0.002 | 0.003 | 0.003 | 0.205 | 0.08 | 0.09 | 0.04 | 0.01 | 0.08 | 0.07 |
| 2 | 67158316 | rs6546273 | -2.037 | 0.691 | -2.186 | 0.583 | -0.001 | -0.078 | -0.078 | 0 | 0.86 | 0.96 | 0.9 | 0.68 | 0.85 | 0.87 |
| 2 | 67159182 | rs17032772 | 1.187 | 0.817 | -2.027 | 0.408 | -0.022 | -0.007 | -0.007 | 0 | 0.08 | 0.03 | 0.09 | 0.05 | 0.09 | 0.02 |

| | | | | | | | | | | | | | | | | |
|---|---|---|---|---|---|---|---|---|---|---|---|---|---|---|---|---|
| 2 | 67163302 | rs6546274 | -0.529 | 1.048 | -1.992 | 0.327 | -0.001 | -0.001 | -0.001 | 0 | 0.31 | 0.64 | 0.33 | 0.13 | 0.34 | 0.22 |
| 2 | 67167431 | rs1362479 | 1.734 | 0.899 | -2.102 | 0.425 | -0.001 | -0.012 | -0.012 | 0 | 0.06 | 0.09 | 0.03 | 0.01 | 0.05 | 0.06 |
| 2 | 67172941 | rs4671761 | 0.781 | 0.792 | -2.182 | 0.587 | -0.013 | -0.054 | -0.054 | 1.354 | 0.27 | 0.1 | 0.22 | 0.37 | 0.28 | 0.21 |
| 2 | 67177439 | rs1362483 | 1.519 | 0.801 | -2.121 | 0.622 | -0.002 | -0.022 | -0.022 | 0.445 | 0.1 | 0.04 | 0.26 | 0.32 | 0.12 | 0.31 |
| 2 | 67182916 | rs12620024 | 0.987 | 0.676 | -2.04 | 0.522 | -0.009 | -0.025 | -0.025 | 0.762 | 0.25 | 0.13 | 0.21 | 0.32 | 0.28 | 0.17 |
| 2 | 67184938 | rs1420370 | -1.548 | 0.701 | -2.273 | 0.267 | 0.021 | 0.073 | 0.073 | 0 | 0.71 | 0.94 | 0.85 | 0.85 | 0.78 | 0.79 |
| 2 | 67191073 | rs1420371 | 1.521 | 1.018 | -2.19 | 0.303 | -0.014 | 0.013 | 0.013 | 0 | 0.13 | 0.1 | 0.14 | 0.05 | 0.19 | 0.09 |
| 2 | 67195459 | rs10210389 | 0.619 | 1.037 | -1.704 | 0.371 | 0.039 | 0.03 | 0.03 | 0 | 0.13 | 0.37 | 0.14 | 0.09 | 0.1 | 0.12 |
| 2 | 67196296 | rs13409540 | 0.857 | 0.977 | -1.677 | 0.402 | 0.028 | -0.045 | -0.045 | 0.008 | 0.24 | 0.39 | 0.2 | 0.09 | 0.22 | 0.16 |
| 2 | 67197076 | rs4671162 | -1.499 | 0.659 | -1.847 | 0.434 | 0.009 | 0.05 | 0.05 | 0 | 0.72 | 0.93 | 0.85 | 0.84 | 0.77 | 0.77 |
| 2 | 67197568 | rs17032810 | 1.774 | 0.713 | -1.737 | 0.654 | -0.001 | 0.01 | 0.01 | 0.002 | 0.13 | 0.11 | 0.15 | 0.38 | 0.16 | 0.14 |
| 2 | 67198954 | rs10197180 | 1.011 | 0.696 | -1.658 | 0.46 | 0.011 | -0.057 | -0.057 | 0.134 | 0.26 | 0.39 | 0.22 | 0.08 | 0.22 | 0.16 |
| 2 | 67204629 | rs1548900 | -0.682 | 0.653 | -2.24 | 0.781 | -0.011 | 0.028 | 0.028 | 0.286 | 0.43 | 0.36 | 0.6 | 0.76 | 0.52 | 0.6 |
| 2 | 67204642 | rs12471464 | 0.929 | 0.7 | -2.236 | 0.743 | -0.003 | -0.004 | -0.004 | 0 | 0.14 | 0.12 | 0.31 | 0.23 | 0.18 | 0.31 |
| 2 | 67217705 | rs981718 | -0.041 | 0.614 | -2.768 | 0.789 | -0.005 | -0.046 | -0.046 | 0 | 0.3 | 0.21 | 0.29 | 0.49 | 0.35 | 0.29 |
| 2 | 67221738 | rs7601922 | -1.524 | 0.629 | -2.887 | 0.607 | 0.027 | 0.018 | 0.018 | 0.06 | 0.77 | 0.88 | 0.88 | 0.84 | 0.82 | 0.88 |
| 2 | 67222446 | rs977253 | 0.334 | 1.081 | -2.361 | 1.127 | 0 | 0.019 | 0.019 | 0 | 0.19 | 0.14 | 0.14 | 0.07 | 0.19 | 0.12 |
| 2 | 67223326 | rs1420376 | -1.358 | 1.183 | -2.07 | 1.77 | -0.024 | -0.039 | -0.039 | 0 | 0.3 | 0.17 | 0.48 | 0.68 | 0.36 | 0.53 |
| 2 | 67224057 | rs1345354 | -0.593 | 0.877 | -1.451 | 1.534 | -0.003 | -0.008 | -0.008 | 0.639 | 0.15 | 0.11 | 0.17 | 0.38 | 0.18 | 0.16 |
| 2 | 67237214 | rs13029838 | -0.109 | 0.53 | -1.747 | 1.114 | 0.015 | -0.088 | -0.088 | 0 | 0.28 | 0.14 | 0.17 | 0.39 | 0.23 | 0.31 |
| 2 | 67240028 | rs6727258 | -0.784 | 0.884 | -0.614 | 0.306 | 0.002 | -0.011 | -0.011 | 0 | 0.46 | 0.33 | 0.31 | 0.4 | 0.42 | 0.38 |
| 2 | 67241174 | rs6734275 | -0.627 | 0.76 | -1.051 | -0.037 | 0.021 | -0.003 | -0.003 | 0 | 0.46 | 0.34 | 0.41 | 0.45 | 0.36 | 0.47 |
| 2 | 67244299 | rs4635520 | 1.703 | 1.034 | -1.258 | -0.197 | -0.004 | -0.019 | -0.019 | 0 | 0.25 | 0.1 | 0.34 | 0.26 | 0.24 | 0.18 |
| 2 | 67247936 | rs12621498 | -0.19 | 1.087 | -0.432 | -0.165 | 0.037 | -0.026 | -0.026 | 3.51 | 0.82 | 0.64 | 0.72 | 0.79 | 0.8 | 0.8 |
| 2 | 67254136 | rs17032888 | 0.301 | 0.824 | -0.553 | -0.088 | 0.009 | -0.015 | -0.015 | 0.012 | 0.38 | 0.33 | 0.22 | 0.41 | 0.3 | 0.46 |
| 2 | 67258675 | rs1362475 | 0.227 | 0.836 | -0.556 | -0.082 | 0.008 | -0.012 | -0.012 | 0 | 0.38 | 0.34 | 0.23 | 0.42 | 0.3 | 0.46 |
| 2 | 67260549 | rs2080400 | 0.287 | 1.032 | -0.372 | -0.063 | -0.004 | 0.015 | 0.015 | 0 | 0.41 | 0.36 | 0.49 | 0.33 | 0.47 | 0.35 |
| 2 | 67260723 | rs10192430 | -0.252 | 0.965 | -0.382 | 0.036 | 0.001 | 0.013 | 0.013 | 0.032 | 0.54 | 0.58 | 0.57 | 0.36 | 0.58 | 0.43 |
| 2 | 67262646 | rs10519086 | 0.789 | 1.023 | -0.6 | 0.106 | -0.006 | -0.002 | -0.002 | 0 | 0.06 | 0.05 | 0.07 | 0.06 | 0.12 | 0.11 |
| 2 | 67263143 | rs10496139 | -0.373 | 0.763 | -0.208 | 0.195 | 0.005 | 0.098 | 0.098 | 0 | 0.2 | 0.37 | 0.16 | 0.32 | 0.19 | 0.25 |
| 2 | 67264563 | rs869354 | -1.005 | 0.745 | -0.212 | 0.179 | 0.011 | 0.078 | 0.078 | 0.044 | 0.26 | 0.42 | 0.23 | 0.37 | 0.31 | 0.37 |
| 2 | 67264964 | rs2098390 | 1.041 | 0.541 | -0.406 | 0.016 | -0.002 | 0.005 | 0.005 | 2.163 | 0.38 | 0.29 | 0.34 | 0.34 | 0.38 | 0.3 |
| 2 | 67265215 | rs1420360 | -0.372 | 0.39 | -0.166 | -0.215 | -0.013 | -0.059 | -0.059 | 0 | 0.19 | 0.33 | 0.21 | 0.05 | 0.15 | 0.25 |
| 2 | 67265783 | rs1420361 | -1.311 | 0.359 | -0.205 | -0.211 | 0 | 0.01 | 0.01 | 0.587 | 0.65 | 0.73 | 0.7 | 0.76 | 0.7 | 0.79 |
| 2 | 67265979 | rs12993985 | 0.121 | 0.378 | -0.118 | -0.018 | 0.005 | 0.198 | 0.198 | 0 | 0.11 | 0.26 | 0.11 | 0.31 | 0.09 | 0.19 |
| 2 | 67266483 | rs17032925 | 0.173 | 0.325 | -0.326 | -0.195 | 0.003 | 0.006 | 0.006 | 0.015 | 0.12 | 0.08 | 0.18 | 0.05 | 0.11 | 0.22 |
| 2 | 67266707 | rs7579521 | 1.184 | 0.233 | -0.517 | -0.185 | -0.021 | -0.044 | -0.044 | 0 | 0.33 | 0.14 | 0.31 | 0.36 | 0.34 | 0.22 |
| 2 | 67269295 | rs12622277 | 1.088 | 0.637 | -0.279 | -0.142 | -0.027 | -0.001 | -0.001 | 3.857 | 0.39 | 0.2 | 0.41 | 0.52 | 0.46 | 0.39 |
| 2 | 67271291 | rs746784 | -0.99 | 0.678 | -0.038 | -0.647 | 0 | 0.017 | 0.017 | 0.001 | 0.66 | 0.76 | 0.7 | 0.79 | 0.7 | 0.77 |
| 2 | 67276947 | rs6546277 | -0.274 | 0.655 | 0.181 | -0.562 | -0.005 | -0.035 | -0.035 | 0 | 0.16 | 0.25 | 0.08 | 0.08 | 0.14 | 0.11 |
| 2 | 67278099 | rs4671770 | -1.337 | 0.632 | 0.219 | -0.583 | -0.016 | -0.061 | -0.061 | 0.057 | 0.24 | 0.42 | 0.23 | 0.12 | 0.18 | 0.28 |
| 2 | 67282576 | rs11676618 | -0.399 | 0.649 | 0.209 | -0.571 | -0.004 | -0.033 | -0.033 | 0 | 0.57 | 0.53 | 0.53 | 0.75 | 0.66 | 0.61 |
| 2 | 67291941 | rs1833218 | 0.953 | 0.607 | 0.079 | -0.618 | -0.003 | 0.053 | 0.053 | 0 | 0.46 | 0.32 | 0.53 | 0.67 | 0.56 | 0.53 |
| 2 | 67294023 | rs1362478 | -0.451 | 0.612 | 0.073 | -0.596 | 0 | 0.003 | 0.003 | 0 | 0.61 | 0.65 | 0.7 | 0.73 | 0.64 | 0.68 |
| 2 | 67294779 | rs1420365 | -0.831 | 0.631 | 0.109 | -0.555 | -0.001 | -0.004 | -0.004 | 3.297 | 0.67 | 0.7 | 0.72 | 0.73 | 0.67 | 0.71 |
| 2 | 67301732 | rs12713590 | -0.742 | 0.971 | -0.341 | -0.665 | 0.003 | -0.026 | -0.026 | 0 | 0.66 | 0.67 | 0.71 | 0.79 | 0.64 | 0.83 |
| 2 | 67301846 | rs1921267 | -0.924 | 0.332 | -0.213 | -0.566 | -0.005 | -0.027 | -0.027 | 0.04 | 0.68 | 0.69 | 0.76 | 0.79 | 0.74 | 0.84 |
| 2 | 67302169 | rs17590703 | 1.516 | 0.377 | -0.496 | -0.544 | -0.043 | -0.016 | -0.016 | 0 | 0.1 | 0.01 | 0.13 | 0.09 | 0.12 | 0.11 |
| 2 | 67302208 | rs13404435 | -0.442 | 0.157 | 0.058 | -0.522 | -0.008 | -0.038 | -0.038 | 0.048 | 0.17 | 0.28 | 0.09 | 0.09 | 0.14 | 0.15 |
| 2 | 67302300 | rs17032976 | -1.122 | 0.175 | 0.101 | -0.516 | 0.002 | -0.029 | -0.029 | 0 | 0.19 | 0.42 | 0.14 | 0.15 | 0.19 | 0.25 |
| 2 | 67302866 | rs13405122 | -0.994 | 0.178 | 0.102 | -0.515 | 0.004 | -0.025 | -0.025 | 0 | 0.19 | 0.39 | 0.13 | 0.15 | 0.19 | 0.25 |
| 2 | 67306158 | rs13399908 | -1.002 | 0.141 | 0.03 | -0.498 | 0 | -0.051 | -0.051 | 0.441 | 0.24 | 0.46 | 0.16 | 0.15 | 0.23 | 0.28 |
| 2 | 67308243 | rs10199998 | -1.132 | 0.142 | 0.029 | -0.498 | 0.002 | -0.046 | -0.046 | 0 | 0.24 | 0.47 | 0.16 | 0.15 | 0.24 | 0.28 |
| 2 | 67311244 | rs752937 | -0.298 | 0.186 | 0.145 | -0.48 | 0.003 | -0.022 | -0.022 | 0 | 0.18 | 0.27 | 0.11 | 0.07 | 0.2 | 0.23 |
| 2 | 67312459 | rs1921275 | 1.336 | 0.13 | 0.049 | -0.542 | -0.051 | 0.074 | 0.074 | 0 | 0.43 | 0.15 | 0.48 | 0.6 | 0.49 | 0.42 |
| 2 | 67317774 | rs6751541 | 0.328 | 0.172 | 0.098 | -0.535 | 0.016 | 0.054 | 0.054 | 0 | 0.06 | 0.23 | 0.05 | 0.09 | 0.03 | 0.05 |
| 2 | 67323400 | rs10196486 | -0.948 | 0.129 | 0.025 | -0.571 | -0.002 | -0.017 | -0.017 | 0.096 | 0.69 | 0.72 | 0.76 | 0.8 | 0.74 | 0.82 |
| 2 | 67327778 | rs7603473 | 0.035 | 0.142 | 0.045 | -0.543 | 0.006 | -0.023 | -0.023 | 0 | 0.17 | 0.26 | 0.12 | 0.07 | 0.19 | 0.22 |
| 2 | 67331198 | rs4671773 | -0.881 | 0.14 | -0.048 | -0.573 | -0.002 | -0.015 | -0.015 | 0.021 | 0.69 | 0.72 | 0.76 | 0.8 | 0.74 | 0.82 |
| 2 | 67333139 | rs17033085 | 0.482 | 0.192 | -0.007 | -0.415 | 0.011 | 0.047 | 0.047 | 0 | 0.05 | 0.2 | 0.05 | 0.09 | 0.03 | 0.05 |
| 2 | 67334338 | rs1921294 | 0.903 | 0.232 | -0.41 | -0.356 | -0.051 | 0.095 | 0.095 | 0 | 0.44 | 0.15 | 0.49 | 0.63 | 0.5 | 0.43 |
| 2 | 67334411 | rs6719951 | -0.492 | 0.138 | -0.577 | -0.489 | -0.017 | -0.079 | -0.079 | 0 | 0.38 | 0.51 | 0.19 | 0.13 | 0.31 | 0.33 |
| 2 | 67341817 | rs1921290 | 0.087 | 0.105 | -0.77 | -0.343 | -0.001 | -0.021 | -0.021 | 0.09 | 0.58 | 0.52 | 0.7 | 0.72 | 0.63 | 0.69 |
| 2 | 67345557 | rs1921287 | 0.138 | 0.087 | -0.8 | -0.387 | 0 | -0.002 | -0.002 | 0.001 | 0.58 | 0.49 | 0.69 | 0.72 | 0.62 | 0.69 |
| 2 | 67353028 | rs2178524 | 0.347 | -0.434 | -0.95 | 0.097 | -0.001 | -0.004 | -0.004 | 0 | 0.48 | 0.49 | 0.61 | 0.59 | 0.59 | 0.59 |
| 2 | 67365786 | rs13014894 | 1.259 | -0.504 | -0.61 | 0.266 | -0.021 | -0.039 | -0.039 | 0 | 0.26 | 0.16 | 0.39 | 0.36 | 0.35 | 0.27 |
| 2 | 67368780 | rs6721279 | 0.044 | -0.679 | -0.54 | 0.239 | -0.002 | -0.003 | -0.003 | 0 | 0.48 | 0.51 | 0.62 | 0.57 | 0.58 | 0.56 |
| 2 | 67370269 | rs6743774 | 1.149 | -0.479 | -0.679 | 0.391 | 0.009 | 0.095 | 0.095 | 0 | 0.05 | 0.13 | 0.14 | 0.21 | 0.07 | 0.24 |
| 2 | 67377875 | rs4671779 | 1.275 | -0.487 | -0.746 | 0.368 | 0 | 0.05 | 0.05 | 0 | 0.05 | 0.06 | 0.13 | 0.21 | 0.07 | 0.23 |

| | | | | | | | | | | | | | | | | |
|---|---|---|---|---|---|---|---|---|---|---|---|---|---|---|---|---|
| 2 | 67381644 | rs7600464 | 0.702 | -0.587 | -0.992 | 0.222 | -0.077 | -0.081 | -0.081 | 0 | 0.13 | 0.02 | 0.19 | 0.34 | 0.23 | 0.12 |
| 2 | 67387721 | rs10199442 | -0.367 | -0.637 | -0.522 | 0.074 | 0.006 | -0.008 | -0.008 | 0.228 | 0.61 | 0.56 | 0.64 | 0.57 | 0.61 | 0.57 |
| 2 | 67388232 | rs4671784 | 0.808 | -0.615 | -0.645 | -0.289 | 0 | 0.057 | 0.057 | 0 | 0.06 | 0.09 | 0.14 | 0.21 | 0.07 | 0.23 |
| 2 | 67388797 | rs6739469 | -0.078 | -0.617 | -0.677 | -0.327 | 0.012 | 0.011 | 0.011 | 3.595 | 0.34 | 0.34 | 0.23 | 0.21 | 0.23 | 0.3 |
| 2 | 67398320 | rs12613227 | 1.194 | 0.111 | -0.965 | 0.027 | -0.048 | -0.045 | -0.045 | 0.128 | 0.07 | 0.01 | 0.16 | 0.22 | 0.17 | 0.07 |
| 2 | 67406029 | rs17592856 | 1.051 | -0.718 | -0.292 | -0.296 | 0.008 | 0.015 | 0.015 | 0 | 0.29 | 0.27 | 0.41 | 0.56 | 0.36 | 0.38 |
| 2 | 67406814 | rs10180549 | 0.077 | -0.932 | -0.6 | -0.621 | -0.017 | 0.042 | 0.042 | 0 | 0.43 | 0.35 | 0.41 | 0.24 | 0.5 | 0.34 |
| 2 | 67407060 | rs7563789 | 0.147 | -0.779 | -1.124 | -0.653 | 0.026 | 0.008 | 0.008 | 1.176 | 0.19 | 0.11 | 0.05 | 0 | 0.07 | 0.05 |
| 2 | 67412681 | rs17527701 | 0.346 | -0.482 | -1.373 | -0.561 | -0.036 | 0.025 | 0.025 | 0.053 | 0.28 | 0.17 | 0.35 | 0.2 | 0.4 | 0.28 |
| 2 | 67413254 | rs17033180 | 0.67 | -0.627 | -1.306 | -0.485 | 0.018 | -0.005 | -0.005 | 0.127 | 0.15 | 0.13 | 0.13 | 0.17 | 0.07 | 0.21 |
| 2 | 67423735 | rs10779952 | -0.89 | -0.343 | -2.164 | -0.301 | 0 | 0.028 | 0.028 | 0 | 0.51 | 0.42 | 0.51 | 0.38 | 0.53 | 0.52 |
| 2 | 67424193 | rs7590628 | -0.817 | -0.518 | -2.33 | -0.511 | 0.028 | 0.108 | 0.108 | 0 | 0.48 | 0.31 | 0.39 | 0.2 | 0.45 | 0.33 |
| 2 | 67429146 | rs11683984 | -0.883 | 0.206 | -2.607 | 0.161 | -0.024 | 0.092 | 0.092 | 0 | 0.54 | 0.23 | 0.43 | 0.28 | 0.52 | 0.38 |
| 2 | 67437597 | rs10206916 | 0.53 | 0.233 | -2.607 | 0.141 | 0.065 | 0.09 | 0.09 | 1.957 | 0.26 | 0.09 | 0.1 | 0.08 | 0.14 | 0.09 |
| 2 | 67438060 | rs11126115 | -0.156 | 0.55 | -2.391 | 0.211 | -0.035 | 0.027 | 0.027 | 0 | 0.27 | 0.13 | 0.31 | 0.19 | 0.37 | 0.28 |
| 2 | 67439004 | rs1921301 | 0.276 | 0.468 | -2.479 | 0.44 | -0.057 | -0.016 | -0.016 | 0.263 | 0.17 | 0.02 | 0.17 | 0.14 | 0.25 | 0.13 |
| 2 | 67444692 | rs1921266 | -0.408 | -0.019 | -1.884 | 0.601 | 0.043 | 0 | 0 | 0 | 0.29 | 0.19 | 0.22 | 0.25 | 0.2 | 0.27 |
| 2 | 67446590 | rs17033208 | -0.455 | 0.648 | -1.693 | 0.709 | -0.015 | -0.015 | -0.015 | 0 | 0.37 | 0.31 | 0.4 | 0.54 | 0.43 | 0.4 |
| 2 | 67449899 | rs6714614 | -0.422 | 0.45 | -1.11 | 0.374 | -0.015 | -0.036 | -0.036 | 0.182 | 0.81 | 0.91 | 0.73 | 0.73 | 0.77 | 0.72 |
| 2 | 67452013 | rs11894761 | 0.007 | 0.133 | -1.024 | -0.388 | 0.001 | -0.003 | -0.003 | 0 | 0.47 | 0.54 | 0.41 | 0.65 | 0.51 | 0.48 |
| 2 | 67453172 | rs1384741 | -0.433 | -0.328 | -0.944 | -0.937 | -0.003 | -0.002 | -0.002 | 0.251 | 0.67 | 0.66 | 0.65 | 0.73 | 0.69 | 0.68 |
| 2 | 67457481 | rs17033231 | 1.381 | 0.02 | -1.432 | -0.58 | 0.004 | -0.003 | -0.003 | 0 | 0.16 | 0.1 | 0.08 | 0.16 | 0.14 | 0.16 |
| 2 | 67459829 | rs1384746 | -0.462 | -0.394 | -1.592 | -0.917 | -0.008 | -0.006 | -0.006 | 0.264 | 0.55 | 0.64 | 0.34 | 0.55 | 0.53 | 0.42 |
| 2 | 67462012 | rs11126116 | 0.955 | -0.374 | -1.753 | -0.859 | 0.02 | 0.004 | 0.004 | 0 | 0.21 | 0.08 | 0.1 | 0.16 | 0.17 | 0.16 |
| 2 | 67464147 | rs1545861 | 1.058 | -0.122 | -1.7 | -0.718 | -0.003 | 0.018 | 0.018 | 0 | 0.2 | 0.15 | 0.26 | 0.08 | 0.19 | 0.2 |
| 2 | 67465753 | rs17033254 | 1.812 | -0.038 | -1.635 | -0.382 | -0.052 | -0.029 | -0.029 | 0 | 0.08 | 0.01 | 0.12 | 0.43 | 0.19 | 0.17 |
| 2 | 67470196 | rs7577524 | 0.217 | -0.122 | -1.871 | -0.793 | 0.01 | 0.058 | 0.058 | 0.013 | 0.34 | 0.2 | 0.15 | 0.17 | 0.25 | 0.27 |
| 2 | 67472630 | rs13012056 | 0.937 | -0.001 | -1.77 | -0.678 | -0.053 | -0.09 | -0.09 | 0.723 | 0.16 | 0.06 | 0.16 | 0.46 | 0.25 | 0.25 |
| 2 | 67473625 | rs1466521 | 1.243 | -0.151 | -1.607 | -0.805 | 0.002 | 0.004 | 0.004 | 0 | 0.17 | 0.16 | 0.1 | 0.08 | 0.11 | 0.16 |
| 2 | 67475306 | rs6754760 | 0.158 | -0.242 | -1.489 | -0.869 | -0.017 | -0.076 | -0.076 | 0 | 0.34 | 0.55 | 0.34 | 0.05 | 0.28 | 0.16 |
| 2 | 67479214 | rs11126117 | -1.84 | 0.108 | -1.707 | -0.448 | -0.028 | -0.042 | -0.042 | 0 | 0.8 | 0.92 | 0.76 | 0.74 | 0.76 | 0.71 |
| 2 | 67482322 | rs6754571 | -0.165 | -0.047 | -0.945 | -0.759 | 0.015 | 0.007 | 0.007 | 0.598 | 0.47 | 0.6 | 0.45 | 0.11 | 0.38 | 0.32 |
| 2 | 67485028 | rs10496144 | 1.737 | 0.19 | -0.983 | -0.451 | -0.034 | -0.013 | -0.013 | 0 | 0.08 | 0.03 | 0.1 | 0.41 | 0.17 | 0.16 |
| 2 | 67490638 | rs6745786 | 1.191 | -0.318 | -0.639 | -0.758 | -0.001 | -0.014 | -0.014 | 1.368 | 0.18 | 0.16 | 0.1 | 0.06 | 0.1 | 0.17 |
| 2 | 67492984 | rs10168151 | 0.007 | -0.228 | -0.585 | -0.722 | -0.01 | -0.061 | -0.061 | 0 | 0.44 | 0.49 | 0.42 | 0.1 | 0.31 | 0.33 |
| 2 | 67495579 | rs2170704 | -0.197 | -0.272 | -0.817 | -0.64 | 0.026 | 0.052 | 0.052 | 0.214 | 0.48 | 0.62 | 0.58 | 0.21 | 0.37 | 0.45 |
| 2 | 67496866 | rs1483344 | -1.468 | -0.15 | -1.15 | -0.216 | -0.007 | -0.026 | -0.026 | 0 | 0.78 | 0.84 | 0.78 | 0.61 | 0.75 | 0.7 |
| 2 | 67501511 | rs966779 | -1.513 | -0.119 | -1.401 | -0.129 | -0.02 | -0.063 | -0.063 | 4.415 | 0.78 | 0.89 | 0.78 | 0.61 | 0.75 | 0.7 |
| 2 | 67506724 | rs17033369 | 1.246 | -0.306 | -1.204 | -0.195 | 0.002 | -0.011 | -0.011 | 0 | 0.05 | 0.01 | 0.18 | 0.08 | 0.08 | 0.11 |
| 2 | 67508174 | rs9941589 | 0.195 | -0.608 | -1.159 | 0.422 | 0.006 | -0.024 | -0.024 | 0 | 0.24 | 0.18 | 0.11 | 0.46 | 0.23 | 0.29 |
| 2 | 67508925 | rs6719169 | 1.056 | -1.113 | -0.512 | -0.499 | -0.051 | -0.079 | -0.079 | 0.092 | 0.17 | 0.32 | 0.09 | 0.05 | 0.07 | 0.18 |
| 2 | 67513152 | rs11900586 | -0.192 | -1.377 | -0.668 | -0.825 | -0.008 | -0.024 | -0.024 | 0 | 0.3 | 0.63 | 0.31 | 0.1 | 0.16 | 0.31 |
| 2 | 67515489 | rs1483349 | 1.291 | -0.966 | -0.758 | -1.035 | -0.002 | 0.02 | 0.02 | 0.001 | 0.17 | 0.13 | 0.06 | 0.01 | 0.16 | 0.07 |
| 2 | 67515512 | rs6740565 | 0.965 | -0.939 | -1.204 | -1.117 | 0.009 | 0.042 | 0.042 | 0 | 0.2 | 0.05 | 0.23 | 0.14 | 0.17 | 0.2 |
| 2 | 67520850 | rs7568231 | -1.606 | -0.309 | -1.074 | -0.761 | -0.038 | -0.046 | -0.046 | 0 | 0.65 | 0.75 | 0.67 | 0.56 | 0.59 | 0.69 |
| 2 | 67521740 | rs7559393 | -1.108 | -0.363 | -0.855 | -0.799 | -0.01 | -0.013 | -0.013 | 0 | 0.63 | 0.61 | 0.66 | 0.56 | 0.58 | 0.67 |
| 2 | 67523685 | rs12713595 | -2.322 | -0.311 | -0.925 | -0.794 | -0.023 | -0.041 | -0.041 | 0.057 | 0.77 | 0.84 | 0.71 | 0.56 | 0.69 | 0.72 |
| 2 | 67525162 | rs10172022 | 1.612 | -0.57 | -0.459 | -0.953 | -0.019 | -0.035 | -0.035 | 0 | 0.16 | 0.19 | 0.08 | 0.05 | 0.08 | 0.19 |
| 2 | 67527590 | rs12991257 | 0.91 | -0.604 | -0.433 | -1.002 | 0.003 | 0.011 | 0.011 | 0 | 0.2 | 0.19 | 0.34 | 0.37 | 0.29 | 0.25 |
| 2 | 67528120 | rs6727523 | 3.112 | -0.634 | -0.379 | -1.05 | 0 | 0.006 | 0.006 | 0 | 0.12 | 0.11 | 0.04 | 0 | 0.11 | 0.04 |
| 2 | 67528927 | rs10170342 | 0.016 | -0.701 | -0.293 | -1.107 | -0.026 | -0.106 | -0.106 | 0 | 0.37 | 0.55 | 0.16 | 0.07 | 0.24 | 0.28 |
| 2 | 67529018 | rs10496146 | 2.419 | -0.683 | -0.22 | -1.066 | -0.003 | -0.024 | -0.024 | 0.463 | 0.14 | 0.19 | 0.07 | 0.05 | 0.12 | 0.11 |
| 2 | 67530125 | rs10176209 | 1.537 | -0.667 | -0.213 | -1.059 | -0.048 | -0.078 | -0.078 | 0 | 0.17 | 0.29 | 0.09 | 0.05 | 0.08 | 0.19 |
| 2 | 67530280 | rs1483333 | -2.414 | -0.434 | -0.636 | -0.858 | -0.034 | -0.059 | -0.059 | 1.391 | 0.77 | 0.87 | 0.72 | 0.56 | 0.69 | 0.72 |
| 2 | 67536442 | rs2062206 | -0.159 | -0.342 | -0.645 | -0.769 | -0.008 | -0.014 | -0.014 | 0 | 0.36 | 0.32 | 0.42 | 0.45 | 0.42 | 0.37 |
| 2 | 67536465 | rs2062205 | 0.721 | -0.304 | -0.654 | -0.755 | 0.006 | 0.021 | 0.021 | 0 | 0.2 | 0.19 | 0.34 | 0.4 | 0.3 | 0.25 |
| 2 | 67537804 | rs874236 | 3.012 | -0.261 | -0.519 | -0.753 | -0.002 | 0.034 | 0.034 | 0 | 0.11 | 0.08 | 0.04 | 0 | 0.1 | 0.04 |
| 2 | 67539226 | rs7572809 | 0.093 | -0.118 | -0.373 | -0.555 | -0.008 | -0.012 | -0.012 | 0 | 0.35 | 0.28 | 0.41 | 0.43 | 0.41 | 0.34 |
| 2 | 67554283 | rs4671792 | -0.289 | -0.336 | -0.434 | -0.879 | 0.001 | -0.005 | -0.005 | 0 | 0.31 | 0.61 | 0.32 | 0.16 | 0.24 | 0.37 |
| 2 | 67555446 | rs961328 | 0.589 | 0.006 | -0.788 | -0.925 | -0.015 | -0.02 | -0.02 | 0 | 0.36 | 0.25 | 0.38 | 0.41 | 0.39 | 0.33 |
| 2 | 67557025 | rs13394751 | 0.062 | -0.025 | -0.36 | -0.94 | -0.03 | -0.082 | -0.082 | 0 | 0.29 | 0.41 | 0.3 | 0.16 | 0.24 | 0.37 |
| 2 | 67566281 | rs979043 | -0.961 | 0.139 | -0.41 | -0.926 | -0.008 | -0.019 | -0.019 | 0 | 0.65 | 0.68 | 0.67 | 0.57 | 0.63 | 0.69 |
| 2 | 67578422 | rs6546296 | -2.11 | 0.199 | -0.63 | -1.002 | -0.038 | -0.075 | -0.075 | 0 | 0.8 | 0.92 | 0.73 | 0.57 | 0.73 | 0.73 |
| 2 | 67593803 | rs962856 | -0.981 | 0.183 | -0.506 | -0.961 | -0.024 | -0.05 | -0.05 | 0 | 0.66 | 0.79 | 0.68 | 0.57 | 0.63 | 0.69 |
| 2 | 67606326 | rs11126123 | -0.029 | 0.104 | -0.222 | -1.068 | -0.001 | -0.001 | -0.001 | 0 | 0.44 | 0.42 | 0.56 | 0.57 | 0.46 | 0.54 |
| 2 | 67622348 | rs4671800 | -1.497 | 0.218 | -0.311 | -0.92 | -0.018 | -0.031 | -0.031 | 0 | 0.76 | 0.81 | 0.71 | 0.58 | 0.7 | 0.74 |
| 2 | 67624302 | rs13016812 | -1.485 | 0.216 | -0.322 | -0.921 | -0.022 | -0.039 | -0.039 | 0 | 0.76 | 0.81 | 0.72 | 0.58 | 0.7 | 0.74 |
| 2 | 67630476 | rs13036061 | 2.147 | 0.1 | 0.034 | -0.993 | -0.007 | -0.046 | -0.046 | 0.072 | 0.16 | 0.2 | 0.05 | 0 | 0.1 | 0.04 |
| 2 | 67630980 | rs3770657 | -2.128 | 0.18 | 0.049 | -1.007 | -0.018 | -0.035 | -0.035 | 0.835 | 0.8 | 0.87 | 0.72 | 0.57 | 0.73 | 0.73 |

| | | | | | | | | | | | | | | | | |
|---|---|---|---|---|---|---|---|---|---|---|---|---|---|---|---|---|
| 2 | 67631395 | rs10496153 | 2.408 | 0.182 | 0.035 | -1.006 | 0.01 | 0.012 | 0.012 | 0 | 0.14 | 0.12 | 0.04 | 0 | 0.1 | 0.04 |
| 2 | 67632125 | rs3770655 | -2.12 | 0.185 | -0.022 | -1.004 | -0.033 | -0.063 | -0.063 | 0 | 0.8 | 0.92 | 0.72 | 0.57 | 0.73 | 0.73 |
| 2 | 67633072 | rs10185288 | -1.935 | 0.227 | 0.071 | -0.96 | 0.001 | 0.011 | 0.011 | 0 | 0.79 | 0.78 | 0.7 | 0.57 | 0.73 | 0.73 |
| 2 | 67639769 | rs1486134 | -2.031 | 0.34 | -0.187 | -0.93 | -0.01 | -0.018 | -0.018 | 0.217 | 0.79 | 0.85 | 0.72 | 0.57 | 0.73 | 0.73 |
| 2 | 67640278 | rs13424211 | 1.074 | 0.299 | -0.086 | -0.888 | -0.005 | -0.049 | -0.049 | 0.045 | 0.21 | 0.25 | 0.13 | 0.07 | 0.16 | 0.21 |
| 2 | 67666133 | rs659116 | -0.268 | 0.273 | -0.58 | -0.658 | 0 | -0.007 | -0.007 | 0 | 0.44 | 0.36 | 0.55 | 0.51 | 0.47 | 0.5 |
| 2 | 67668847 | rs632647 | 0.735 | 0.213 | -0.765 | -0.689 | -0.022 | -0.033 | -0.033 | 0 | 0.36 | 0.22 | 0.47 | 0.45 | 0.4 | 0.37 |
| 2 | 67671433 | rs642915 | -0.268 | 0.295 | -0.756 | -0.739 | 0.001 | -0.001 | -0.001 | 0 | 0.45 | 0.41 | 0.55 | 0.51 | 0.45 | 0.5 |
| 2 | 67673780 | rs600524 | 0.341 | 0.318 | -0.848 | -0.923 | 0 | 0.001 | 0.001 | 0.103 | 0.32 | 0.29 | 0.38 | 0.59 | 0.4 | 0.52 |
| 2 | 67674538 | rs13033289 | 2.214 | 0.447 | -1.055 | -1.078 | -0.006 | 0.054 | 0.054 | 0 | 0.15 | 0.06 | 0.06 | 0 | 0.14 | 0.06 |
| 2 | 67682091 | rs623457 | -0.659 | 0.42 | -0.72 | -1.162 | -0.004 | -0.001 | -0.001 | 0 | 0.67 | 0.67 | 0.54 | 0.65 | 0.6 | 0.77 |
| 2 | 67683703 | rs605980 | -2.091 | 0.451 | -0.708 | -1.149 | -0.025 | -0.026 | -0.026 | 0.169 | 0.85 | 0.91 | 0.61 | 0.66 | 0.74 | 0.83 |
| 2 | 67684209 | rs681825 | 0.573 | 0.321 | -0.6 | -1.239 | 0 | 0.008 | 0.008 | 0 | 0.31 | 0.34 | 0.36 | 0.59 | 0.35 | 0.51 |
| 2 | 67685092 | rs686880 | -1.047 | 0.332 | -0.823 | -1.216 | -0.066 | -0.022 | -0.022 | 0.436 | 0.7 | 0.85 | 0.55 | 0.66 | 0.6 | 0.77 |
| 2 | 67686789 | rs658602 | 0.49 | 0.453 | -0.599 | -0.75 | -0.002 | 0.015 | 0.015 | 1.298 | 0.45 | 0.33 | 0.41 | 0.59 | 0.49 | 0.57 |
| 2 | 67687629 | rs6731529 | -0.076 | -0.36 | -0.758 | -1.168 | 0.007 | -0.021 | -0.021 | 0 | 0.37 | 0.34 | 0.18 | 0.07 | 0.24 | 0.26 |
| 2 | 67702012 | rs17033632 | 1.001 | -0.023 | -0.696 | -0.831 | 0.005 | -0.014 | -0.014 | 0 | 0.18 | 0.22 | 0.11 | 0.01 | 0.15 | 0.06 |
| 2 | 67702146 | rs11126126 | 0.322 | -0.57 | -0.395 | -1.088 | -0.018 | -0.031 | -0.031 | 0 | 0.15 | 0.22 | 0.08 | 0.07 | 0.12 | 0.23 |
| 2 | 67702819 | rs1521041 | -0.353 | -0.303 | -0.513 | -0.568 | 0 | 0 | 0 | 0 | 0.44 | 0.49 | 0.36 | 0.42 | 0.39 | 0.46 |
| 2 | 67705080 | rs6760732 | -0.353 | -0.185 | -0.353 | -0.245 | -0.005 | -0.001 | -0.001 | 0 | 0.44 | 0.53 | 0.37 | 0.42 | 0.38 | 0.46 |
| 2 | 67705898 | rs6719623 | -0.326 | -0.207 | -0.431 | -0.262 | -0.005 | 0 | 0 | 2.356 | 0.44 | 0.48 | 0.36 | 0.42 | 0.38 | 0.47 |
| 2 | 67711185 | rs6419640 | -0.6 | 0.311 | -0.652 | 0.049 | -0.002 | -0.002 | -0.002 | 0 | 0.63 | 0.73 | 0.47 | 0.49 | 0.54 | 0.57 |
| 2 | 67711468 | rs6756612 | -0.726 | 0.873 | -0.718 | 0.239 | 0.02 | 0.037 | 0.037 | 0.375 | 0.8 | 0.82 | 0.57 | 0.6 | 0.72 | 0.65 |
| 2 | 67712269 | rs1552357 | 0.512 | 0.432 | -0.995 | 0.232 | -0.003 | -0.05 | -0.05 | 0 | 0.25 | 0.12 | 0.26 | 0.36 | 0.26 | 0.24 |
| 2 | 67713437 | rs1011090 | 0.439 | 0.201 | -0.549 | 0.215 | -0.014 | 0.031 | 0.031 | 0 | 0.23 | 0.26 | 0.16 | 0.34 | 0.21 | 0.18 |
| 2 | 67716030 | rs1521043 | -0.22 | 0.229 | -0.834 | -0.43 | 0.008 | 0.017 | 0.017 | 0 | 0.67 | 0.65 | 0.46 | 0.51 | 0.62 | 0.42 |
| 2 | 67716539 | rs1568472 | 0.51 | 0.669 | -0.728 | -0.471 | -0.007 | -0.026 | -0.026 | 0 | 0.2 | 0.25 | 0.22 | 0.11 | 0.17 | 0.34 |
| 2 | 67718290 | rs7584345 | -0.185 | 0.91 | -0.417 | -0.307 | 0.073 | 0.104 | 0.104 | 0 | 0.77 | 0.61 | 0.47 | 0.61 | 0.7 | 0.66 |
| 2 | 67722867 | rs7573319 | 1.259 | 0.635 | -0.467 | -0.81 | 0.001 | -0.041 | -0.041 | 0.381 | 0.25 | 0.21 | 0.15 | 0 | 0.26 | 0.09 |
| 2 | 67723513 | rs1521045 | -1.388 | 0.639 | -0.444 | -0.687 | -0.023 | -0.033 | -0.033 | 0 | 0.83 | 0.92 | 0.62 | 0.6 | 0.77 | 0.73 |
| 2 | 67723565 | rs17033681 | 0.445 | 0.569 | -0.267 | -0.653 | 0.019 | 0.002 | 0.002 | 1.744 | 0.52 | 0.39 | 0.3 | 0.56 | 0.45 | 0.55 |
| 2 | 67725604 | rs10170407 | 1.506 | 0.495 | -0.032 | -0.595 | -0.029 | -0.035 | -0.035 | 0 | 0.12 | 0.31 | 0.09 | 0.03 | 0.1 | 0.05 |
| 2 | 67727152 | rs1521049 | -0.525 | 0.534 | -0.095 | -0.508 | 0.036 | 0.106 | 0.106 | 0 | 0.79 | 0.69 | 0.59 | 0.61 | 0.77 | 0.69 |
| 2 | 67737672 | rs1357011 | 1.22 | 0.372 | -0.723 | -0.677 | -0.052 | -0.117 | -0.117 | 0.013 | 0.14 | 0.03 | 0.09 | 0.48 | 0.16 | 0.21 |
| 2 | 67739603 | rs6748393 | 1.935 | 0.308 | -0.679 | -0.798 | 0.004 | -0.04 | -0.04 | 0 | 0.08 | 0.19 | 0.06 | 0 | 0.08 | 0.03 |
| 2 | 67741792 | rs1402440 | 0.526 | 0.431 | -0.648 | -0.733 | -0.03 | -0.034 | -0.034 | 0.474 | 0.21 | 0.12 | 0.56 | 0.41 | 0.33 | 0.32 |
| 2 | 67744538 | rs1568471 | 1.08 | 0.114 | -0.506 | -0.77 | -0.049 | -0.072 | -0.072 | 0 | 0.31 | 0.41 | 0.16 | 0.1 | 0.17 | 0.23 |
| 2 | 67746422 | rs2683684 | -1.302 | 0.148 | -0.665 | -0.778 | 0 | -0.013 | -0.013 | 0 | 0.69 | 0.68 | 0.82 | 0.93 | 0.67 | 0.78 |
| 2 | 67746433 | rs2683683 | -2.334 | 0.146 | -0.685 | -0.781 | -0.015 | 0.064 | 0.064 | 2.183 | 0.77 | 0.87 | 0.88 | 0.93 | 0.75 | 0.81 |
| 2 | 67747535 | rs2683681 | -0.468 | 0.234 | -0.85 | -0.727 | -0.049 | 0.123 | 0.123 | 0 | 0.37 | 0.16 | 0.65 | 0.9 | 0.49 | 0.53 |
| 2 | 67748234 | rs17033716 | 0.946 | 0.015 | -0.901 | -0.625 | -0.041 | -0.121 | -0.121 | 0 | 0.3 | 0.41 | 0.15 | 0.03 | 0.17 | 0.22 |
| 2 | 67751727 | rs17033724 | 2.506 | 0.29 | -1.618 | -0.403 | -0.029 | -0.065 | -0.065 | 0.014 | 0.1 | 0.01 | 0.06 | 0.27 | 0.11 | 0.07 |
| 2 | 67752547 | rs2683690 | -0.378 | 0.338 | -1.557 | -0.641 | 0 | 0.006 | 0.006 | 0 | 0.27 | 0.27 | 0.32 | 0.49 | 0.34 | 0.3 |
| 2 | 67758663 | rs2683687 | -0.038 | 0.262 | -1.844 | -0.677 | 0.048 | -0.035 | -0.035 | 0 | 0.43 | 0.23 | 0.22 | 0.51 | 0.33 | 0.46 |
| 2 | 67760781 | rs2861633 | 0.887 | -0.17 | -1.611 | -1.197 | 0.052 | 0.076 | 0.076 | 3.357 | 0.27 | 0.13 | 0.13 | 0.03 | 0.17 | 0.25 |
| 2 | 67762906 | rs13413974 | 0.221 | 0.078 | -1.28 | -1.06 | 0.014 | -0.077 | -0.077 | 0 | 0.13 | 0.28 | 0.23 | 0 | 0.18 | 0.08 |
| 2 | 67766949 | rs1581370 | -0.375 | 0.038 | -2.18 | -1.072 | -0.009 | 0.168 | 0.168 | 1.478 | 0.58 | 0.15 | 0.6 | 0.9 | 0.57 | 0.67 |
| 2 | 67771586 | rs1568468 | -0.043 | 0.403 | -2.271 | -0.802 | -0.162 | 0.074 | 0.074 | 0 | 0.3 | 0.04 | 0.46 | 0.87 | 0.4 | 0.42 |
| 2 | 67775047 | rs1009280 | 2.342 | 0.212 | -1.985 | -0.028 | -0.029 | -0.015 | -0.015 | 0.122 | 0.07 | 0 | 0.2 | 0.37 | 0.06 | 0.16 |
| 2 | 67775280 | rs744565 | 1.252 | 0.18 | -2.282 | -0.288 | -0.026 | 0.023 | 0.023 | 0 | 0.16 | 0.03 | 0.26 | 0.09 | 0.18 | 0.19 |
| 2 | 67776518 | rs2861638 | -0.252 | 0.134 | -2.442 | -0.438 | -0.044 | -0.092 | -0.092 | 0.356 | 0.43 | 0.07 | 0.47 | 0.5 | 0.52 | 0.53 |
| 2 | 67780749 | rs10182267 | 0.845 | -0.084 | -1.408 | -0.477 | 0.027 | -0.025 | -0.025 | 0 | 0.08 | 0.34 | 0.07 | 0 | 0.08 | 0.02 |
| 2 | 67793236 | rs13398896 | -0.328 | -0.286 | -1.436 | -0.567 | -0.016 | -0.076 | -0.076 | 0 | 0.12 | 0.5 | 0.11 | 0.01 | 0.09 | 0.1 |
| 2 | 67793326 | rs4671187 | 1.24 | -0.18 | -1.901 | -0.505 | -0.097 | -0.171 | -0.171 | 1.638 | 0.23 | 0.02 | 0.46 | 0.46 | 0.25 | 0.37 |
| 2 | 67793687 | rs7602356 | 2.055 | -0.004 | -1.996 | -0.471 | -0.084 | -0.083 | -0.083 | 0.182 | 0.15 | 0.01 | 0.1 | 0.29 | 0.21 | 0.16 |
| 2 | 67798503 | rs10190391 | 1.575 | -0.261 | -1.79 | -0.725 | 0.081 | 0.018 | 0.018 | 0 | 0.07 | 0.42 | 0.07 | 0 | 0.07 | 0.02 |
| 2 | 67800520 | rs6546315 | 0.693 | -0.351 | -1.797 | -0.793 | -0.047 | -0.059 | -0.059 | 0.01 | 0.24 | 0.08 | 0.18 | 0.42 | 0.29 | 0.32 |
| 2 | 67803820 | rs2861647 | 2.138 | -0.911 | -1.923 | -1.139 | -0.037 | -0.03 | -0.03 | 0.452 | 0.19 | 0.28 | 0.14 | 0.11 | 0.13 | 0.05 |
| 2 | 67806472 | rs12620828 | -0.976 | -0.942 | -1.989 | -1.213 | -0.029 | -0.149 | -0.149 | 0 | 0.69 | 0.39 | 0.81 | 0.99 | 0.71 | 0.76 |
| 2 | 67814778 | rs7582772 | 0.336 | -0.963 | -1.872 | -1.116 | -0.016 | -0.073 | -0.073 | 0.411 | 0.12 | 0.56 | 0.11 | 0.01 | 0.1 | 0.1 |
| 2 | 67825685 | rs2902021 | 1.406 | -0.879 | -1.938 | -1.086 | -0.022 | -0.028 | -0.028 | 0 | 0.25 | 0.31 | 0.19 | 0.15 | 0.18 | 0.07 |
| 2 | 67828078 | rs17033861 | 1.418 | -0.863 | -1.929 | -1.077 | -0.022 | -0.028 | -0.028 | 0 | 0.25 | 0.31 | 0.2 | 0.15 | 0.18 | 0.09 |
| 2 | 67833878 | rs6731513 | 0.402 | -1.341 | -1.547 | -1.048 | 0.003 | 0.006 | 0.006 | 0 | 0.33 | 0.39 | 0.22 | 0.44 | 0.34 | 0.41 |
| 2 | 67836507 | rs2861683 | -0.096 | -1.362 | -1.668 | -1.1 | 0.004 | 0.003 | 0.003 | 1.853 | 0.46 | 0.38 | 0.63 | 0.57 | 0.4 | 0.44 |
| 2 | 67838474 | rs2861688 | 1.468 | -1.298 | -1.678 | -1.074 | -0.032 | -0.095 | -0.095 | 2.524 | 0.24 | 0.09 | 0.49 | 0.46 | 0.27 | 0.38 |
| 2 | 67841006 | rs10189870 | 2.051 | -1.266 | -1.574 | -1.064 | 0.071 | 0.015 | 0.015 | 0 | 0.06 | 0.34 | 0.07 | 0 | 0.07 | 0.02 |
| 2 | 67847595 | rs7579880 | -0.177 | -1.327 | -1.642 | -1.099 | -0.032 | -0.095 | -0.095 | 3.436 | 0.14 | 0.61 | 0.11 | 0.01 | 0.1 | 0.11 |
| 2 | 67848751 | rs2861696 | 0.999 | -1.361 | -1.74 | -1.11 | -0.034 | -0.099 | -0.099 | 0.01 | 0.27 | 0.09 | 0.5 | 0.46 | 0.29 | 0.39 |
| 2 | 67852992 | rs6708126 | 2.139 | -1.297 | -2.007 | -0.973 | -0.038 | 0.009 | 0.009 | 3.85 | 0.18 | 0.28 | 0.13 | 0.2 | 0.11 | 0.06 |

| | | | | | | | | | | | | | | | | |
|---|---|---|---|---|---|---|---|---|---|---|---|---|---|---|---|---|
| 2 | 67853013 | rs7598832 | -0.053 | -1.112 | -1.915 | -0.876 | -0.004 | 0.003 | 0.003 | 0 | 0.36 | 0.62 | 0.6 | 0.38 | 0.36 | 0.46 |
| 2 | 67853282 | rs12992403 | 1.433 | -1.001 | -1.974 | -0.904 | -0.092 | -0.094 | -0.094 | 3.399 | 0.27 | 0.06 | 0.19 | 0.42 | 0.32 | 0.34 |
| 2 | 67864090 | rs980228 | -0.255 | -1.418 | -1.326 | -1.578 | -0.033 | -0.086 | -0.086 | 0 | 0.13 | 0.61 | 0.11 | 0.01 | 0.1 | 0.1 |
| 2 | 67864675 | rs4141819 | -0.924 | -1.193 | -1.74 | -1.427 | 0.01 | 0.011 | 0.011 | 0 | 0.64 | 0.72 | 0.79 | 0.88 | 0.69 | 0.82 |
| 2 | 67865595 | rs10490725 | 1.59 | -1.097 | -1.887 | -1.392 | -0.06 | -0.012 | -0.012 | 0.044 | 0.18 | 0.01 | 0.27 | 0.11 | 0.2 | 0.21 |
| 2 | 67866542 | rs10490726 | 1.386 | -1.603 | -0.843 | -1.285 | 0.001 | 0.001 | 0.001 | 4.703 | 0.16 | 0.13 | 0.11 | 0.11 | 0.11 | 0.05 |
| 2 | 67867456 | rs1367476 | 0.395 | -0.879 | -0.38 | -0.736 | -0.029 | -0.031 | -0.031 | 0 | 0.07 | 0.21 | 0.04 | 0.01 | 0.03 | 0.09 |
| 2 | 67868756 | rs1023692 | -0.551 | -0.394 | -0.901 | -0.489 | -0.004 | -0.019 | -0.019 | 0 | 0.73 | 0.75 | 0.86 | 0.88 | 0.76 | 0.83 |
| 2 | 67870225 | rs7572685 | -0.425 | -0.587 | -0.68 | -0.092 | -0.009 | -0.014 | -0.014 | 0 | 0.09 | 0.15 | 0.2 | 0.05 | 0.1 | 0.12 |
| 2 | 67870799 | rs4671194 | -0.103 | -0.684 | -0.526 | 0.077 | 0.005 | -0.005 | -0.005 | 0 | 0.35 | 0.31 | 0.26 | 0.58 | 0.32 | 0.25 |
| 2 | 67872571 | rs2902028 | 0.2 | -0.848 | -0.443 | 0.016 | 0.039 | -0.032 | -0.032 | 0.255 | 0.36 | 0.18 | 0.24 | 0.54 | 0.27 | 0.25 |
| 2 | 67878108 | rs1430779 | -0.112 | -0.459 | 0.357 | 0.096 | 0.054 | -0.029 | -0.029 | 0 | 0.79 | 0.64 | 0.84 | 0.83 | 0.76 | 0.86 |
| 2 | 67878328 | rs1430780 | 0.664 | -0.519 | 0.625 | 0.02 | 0.038 | -0.066 | -0.066 | 0 | 0.72 | 0.52 | 0.8 | 0.82 | 0.7 | 0.83 |
| 2 | 67880148 | rs12476951 | 1.362 | -0.776 | 0.414 | -0.023 | -0.117 | 0.166 | 0.166 | 0 | 0.37 | 0.08 | 0.56 | 0.1 | 0.47 | 0.46 |
| 2 | 67890323 | rs4671821 | 1.109 | -1.019 | 0.798 | 0.047 | -0.032 | 0.041 | 0.041 | 0 | 0.39 | 0.27 | 0.59 | 0.27 | 0.5 | 0.6 |
| 2 | 67892813 | rs971096 | -0.736 | -0.879 | 0.784 | 0.037 | 0.053 | -0.063 | -0.063 | 0 | 0.84 | 0.7 | 0.87 | 0.94 | 0.8 | 0.87 |
| 2 | 67892890 | rs17034087 | 0.291 | -0.904 | 0.47 | 0.122 | 0.032 | -0.148 | -0.148 | 0 | 0.34 | 0.03 | 0.22 | 0.55 | 0.26 | 0.24 |
| 2 | 67897708 | rs12478802 | 0.504 | -0.888 | 0.499 | 0.104 | 0.051 | -0.021 | -0.021 | 0 | 0.32 | 0.12 | 0.23 | 0.64 | 0.24 | 0.26 |
| 2 | 67903473 | rs10205913 | 0.988 | -0.771 | 0.705 | 0.054 | -0.031 | 0.054 | 0.054 | 0 | 0.42 | 0.29 | 0.61 | 0.27 | 0.5 | 0.52 |
| 2 | 67903629 | rs10182504 | -0.005 | -0.84 | 1.003 | 0.004 | -0.04 | -0.013 | -0.013 | 0.106 | 0.38 | 0.47 | 0.23 | 0.65 | 0.24 | 0.25 |
| 2 | 67904632 | rs11681312 | 1.175 | -0.846 | 0.985 | 0.002 | -0.032 | 0.092 | 0.092 | 0 | 0.41 | 0.28 | 0.6 | 0.11 | 0.5 | 0.46 |
| 2 | 67905772 | rs2861713 | 0.466 | -0.848 | 1.22 | -0.004 | 0.039 | -0.111 | -0.111 | 0.144 | 0.76 | 0.56 | 0.83 | 0.92 | 0.74 | 0.77 |
| 2 | 67906813 | rs13418956 | 0.926 | -0.87 | 1.188 | 0.019 | -0.006 | 0.013 | 0.013 | 0 | 0.7 | 0.33 | 0.8 | 0.81 | 0.7 | 0.72 |
| 2 | 67913224 | rs6733362 | -1.006 | -0.819 | 1.178 | 0.031 | 0.043 | -0.04 | -0.04 | 0 | 0.86 | 0.76 | 0.87 | 0.93 | 0.8 | 0.8 |
| 2 | 67917864 | rs4671831 | 1.784 | -0.766 | 1.073 | -0.043 | -0.132 | 0.158 | 0.158 | 0.201 | 0.32 | 0.04 | 0.54 | 0.08 | 0.43 | 0.42 |
| 2 | 67918196 | rs4671836 | 1.48 | -0.8 | 1.099 | -0.037 | -0.023 | 0.021 | 0.021 | 0 | 0.65 | 0.26 | 0.77 | 0.65 | 0.63 | 0.65 |
| 2 | 67919706 | rs7586232 | -0.512 | -0.803 | 1.119 | -0.038 | -0.004 | -0.012 | -0.012 | 0 | 0.45 | 0.51 | 0.26 | 0.81 | 0.3 | 0.35 |
| 2 | 67920009 | rs1978656 | 0.964 | -0.739 | 1.049 | -0.041 | 0.034 | -0.05 | -0.05 | 0 | 0.71 | 0.51 | 0.8 | 0.77 | 0.68 | 0.7 |
| 2 | 67920930 | rs4671839 | 1.246 | -0.874 | 0.949 | -0.057 | -0.019 | 0.018 | 0.018 | 0 | 0.68 | 0.21 | 0.79 | 0.72 | 0.7 | 0.67 |
| 2 | 67921015 | rs12465119 | 0.49 | -0.925 | 0.929 | -0.003 | 0.024 | -0.139 | -0.139 | 0.244 | 0.34 | 0.03 | 0.23 | 0.55 | 0.27 | 0.25 |
| 2 | 67923868 | rs6711052 | -1.154 | -0.838 | 1.131 | -0.045 | -0.014 | 0.001 | 0.001 | 0.97 | 0.15 | 0.33 | 0.08 | 0.14 | 0.1 | 0.08 |
| 2 | 67924370 | rs6735177 | 0.101 | -0.84 | 1.143 | -0.046 | -0.008 | -0.036 | -0.036 | 0 | 0.07 | 0.2 | 0.03 | 0 | 0.05 | 0.02 |
| 2 | 67927293 | rs6747212 | 0.145 | -0.857 | 1.281 | -0.046 | 0.024 | 0.001 | 0.001 | 0 | 0.37 | 0.29 | 0.24 | 0.61 | 0.27 | 0.31 |
| 2 | 67929132 | rs10496149 | 0.944 | -0.871 | 1.242 | -0.053 | 0.041 | -0.114 | -0.114 | 0 | 0.3 | 0.03 | 0.2 | 0.59 | 0.2 | 0.22 |
| 2 | 67929673 | rs17034152 | 0.155 | -0.862 | 1.291 | -0.054 | -0.011 | -0.035 | -0.035 | 0 | 0.07 | 0.17 | 0.03 | 0 | 0.05 | 0.02 |
| 2 | 67931078 | rs2060042 | -0.891 | -0.839 | 1.296 | -0.067 | 0.046 | -0.029 | -0.029 | 0 | 0.86 | 0.78 | 0.88 | 0.92 | 0.79 | 0.83 |
| 2 | 67931187 | rs4233941 | -1.327 | -0.839 | 1.308 | -0.067 | 0.008 | -0.004 | -0.004 | 0 | 0.87 | 0.89 | 0.89 | 0.92 | 0.8 | 0.83 |
| 2 | 67933079 | rs1430770 | 1.37 | -0.839 | 1.184 | -0.063 | 0.021 | -0.036 | -0.036 | 0.541 | 0.66 | 0.47 | 0.77 | 0.77 | 0.62 | 0.72 |
| 2 | 67935402 | rs12465679 | 0.56 | -0.71 | 1.206 | 0.02 | 0.051 | -0.014 | -0.014 | 0 | 0.33 | 0.17 | 0.2 | 0.6 | 0.2 | 0.28 |
| 2 | 67936401 | rs17034184 | 0.202 | -0.689 | 1.151 | -0.017 | -0.002 | 0.012 | 0.012 | 0 | 0.08 | 0.13 | 0.06 | 0.14 | 0.05 | 0.06 |
| 2 | 67937412 | rs6724226 | -0.907 | -0.67 | 1.079 | -0.021 | 0.047 | -0.029 | -0.029 | 0 | 0.86 | 0.78 | 0.88 | 0.92 | 0.79 | 0.83 |
| 2 | 67940962 | rs10188512 | 1.631 | -0.709 | 1.12 | 0.051 | -0.017 | 0.031 | 0.031 | 0.018 | 0.33 | 0.28 | 0.57 | 0.17 | 0.43 | 0.42 |
| 2 | 67942480 | rs724568 | 1.091 | -0.711 | 0.982 | 0.04 | -0.023 | 0.05 | 0.05 | 0.276 | 0.64 | 0.25 | 0.76 | 0.76 | 0.62 | 0.65 |
| 2 | 67944609 | rs17034207 | 0.014 | -0.626 | 0.844 | -0.14 | -0.01 | -0.036 | -0.036 | 0 | 0.07 | 0.19 | 0.03 | 0 | 0.05 | 0.02 |
| 2 | 67945371 | rs4671842 | -0.571 | -0.585 | 0.79 | -0.127 | 0.031 | -0.034 | -0.034 | 1.008 | 0.81 | 0.77 | 0.86 | 0.91 | 0.72 | 0.79 |
| 2 | 67951861 | rs1367457 | 1.354 | -0.501 | 0.303 | -0.128 | -0.045 | 0.083 | 0.083 | 0.019 | 0.34 | 0.18 | 0.56 | 0.17 | 0.43 | 0.44 |
| 2 | 67954356 | rs17034246 | -0.684 | -0.408 | 0.135 | 0.114 | -0.008 | -0.036 | -0.036 | 0 | 0.07 | 0.21 | 0.03 | 0.01 | 0.05 | 0.08 |
| 2 | 67955280 | rs2861708 | 0.763 | -0.559 | -0.04 | 0.544 | 0.016 | -0.012 | -0.012 | 1.038 | 0.2 | 0.18 | 0.14 | 0.26 | 0.12 | 0.18 |
| 2 | 67955919 | rs17034251 | -0.691 | -0.435 | 0.05 | 0.667 | 0.01 | 0.048 | 0.048 | 2.134 | 0.08 | 0.3 | 0.06 | 0.13 | 0.06 | 0.12 |
| 2 | 67956730 | rs6758705 | 0.273 | -0.435 | -0.149 | 0.692 | -0.001 | -0.002 | -0.002 | 2.555 | 0.59 | 0.46 | 0.74 | 0.64 | 0.64 | 0.64 |
| 2 | 67956823 | rs6758822 | 0.163 | -0.369 | -0.2 | 0.696 | -0.001 | -0.001 | -0.001 | 0 | 0.61 | 0.44 | 0.71 | 0.59 | 0.67 | 0.57 |
| 2 | 67957066 | rs12477687 | 1.072 | 0.282 | 0.146 | 0.171 | -0.047 | 0.055 | 0.055 | 0.018 | 0.33 | 0.17 | 0.54 | 0.21 | 0.44 | 0.37 |
| 2 | 67959183 | rs4671198 | -0.98 | 0.281 | 0.204 | 0.166 | -0.004 | 0 | 0 | 0 | 0.84 | 0.89 | 0.87 | 0.85 | 0.77 | 0.8 |
| 2 | 67959729 | rs10865365 | -0.593 | 0.309 | -0.21 | 0.204 | 0.05 | -0.016 | -0.016 | 0 | 0.81 | 0.65 | 0.84 | 0.85 | 0.76 | 0.8 |
| 2 | 67959865 | rs11885596 | -1.221 | 0.276 | -0.204 | 0.205 | 0.013 | 0.099 | 0.099 | 4.023 | 0.12 | 0.51 | 0.08 | 0.17 | 0.08 | 0.16 |
| 2 | 67960538 | rs7607241 | 1.011 | 0.281 | -0.282 | 0.196 | -0.08 | 0.084 | 0.084 | 0.043 | 0.33 | 0.08 | 0.53 | 0.2 | 0.44 | 0.39 |
| 2 | 67962437 | rs17034279 | 0.672 | 0.171 | -0.287 | 0.164 | 0.06 | -0.136 | -0.136 | 0.577 | 0.36 | 0.05 | 0.22 | 0.45 | 0.23 | 0.24 |
| 2 | 67962988 | rs13411734 | 0.004 | 0.211 | -0.241 | 0.156 | 0 | -0.001 | -0.001 | 0 | 0.44 | 0.6 | 0.62 | 0.38 | 0.53 | 0.56 |
| 2 | 67975586 | rs4671848 | 0.417 | 0.705 | -1.721 | -0.243 | -0.034 | 0.027 | 0.027 | 0.254 | 0.69 | 0.25 | 0.74 | 0.8 | 0.63 | 0.69 |
| 2 | 67977997 | rs12713602 | -0.02 | 0.541 | -1.328 | 0.284 | 0.046 | 0.034 | 0.034 | 0 | 0.44 | 0.28 | 0.29 | 0.63 | 0.32 | 0.36 |
| 2 | 67982191 | rs2116051 | 0.384 | 1.251 | -1.134 | 0.138 | -0.017 | 0.024 | 0.024 | 0 | 0.76 | 0.87 | 0.81 | 0.89 | 0.68 | 0.81 |
| 2 | 67983034 | rs11126151 | -0.727 | 0.769 | -0.748 | 0.827 | -0.006 | 0.017 | 0.017 | 0 | 0.25 | 0.28 | 0.37 | 0.51 | 0.25 | 0.38 |
| 2 | 67984498 | rs6730348 | -1.346 | 0.861 | 0.389 | 0.583 | 0.053 | -0.018 | -0.018 | 0 | 0.08 | 0.41 | 0.18 | 0 | 0.04 | 0.06 |
| 2 | 67989075 | rs1346870 | 0.809 | 0.834 | -0.25 | 0.362 | 0 | 0.054 | 0.054 | 0.004 | 0.5 | 0.31 | 0.34 | 0.32 | 0.52 | 0.42 |
| 2 | 67993265 | rs12713604 | 0.814 | 0.957 | -0.124 | 0.78 | 0 | 0.053 | 0.053 | 0 | 0.5 | 0.31 | 0.34 | 0.32 | 0.52 | 0.41 |
| 2 | 67999042 | rs13014491 | 0.265 | 1.051 | -0.539 | 0.857 | -0.009 | 0.05 | 0.05 | 0 | 0.59 | 0.74 | 0.54 | 0.34 | 0.57 | 0.51 |
| 2 | 68003307 | rs17034385 | -0.371 | 0.871 | -0.661 | 1.319 | 0.007 | -0.061 | -0.061 | 0.277 | 0.19 | 0.08 | 0.33 | 0.34 | 0.18 | 0.31 |
| 2 | 68003733 | rs934312 | 0.047 | 1.011 | -0.299 | 1.086 | -0.009 | -0.023 | -0.023 | 0 | 0.74 | 0.83 | 0.87 | 0.65 | 0.74 | 0.77 |
| 2 | 68006463 | rs4671851 | 0.085 | 1.002 | -0.254 | 0.559 | 0 | -0.013 | -0.013 | 0 | 0.75 | 0.71 | 0.86 | 0.91 | 0.75 | 0.85 |

| | | | | | | | | | | | | | | | | |
|---|---|---|---|---|---|---|---|---|---|---|---|---|---|---|---|---|
| 2 | 68009259 | rs1897179 | 0.339 | 0.974 | -0.331 | 0.569 | 0.005 | -0.052 | -0.052 | 0.652 | 0.69 | 0.53 | 0.79 | 0.91 | 0.72 | 0.82 |
| 2 | 68010701 | rs881645 | 0.281 | 1.342 | -0.288 | 0.922 | 0.021 | 0.038 | 0.038 | 0 | 0.71 | 0.48 | 0.68 | 0.47 | 0.61 | 0.8 |
| 2 | 68010800 | rs7571144 | -0.151 | 0.83 | -0.514 | 0.642 | -0.004 | -0.006 | -0.006 | 0 | 0.1 | 0.12 | 0.04 | 0.07 | 0.05 | 0.12 |
| 2 | 68011742 | rs17551494 | 1.189 | 0.636 | -1.31 | 0.161 | -0.017 | 0.033 | 0.033 | 0 | 0.26 | 0.1 | 0.18 | 0.11 | 0.25 | 0.24 |
| 2 | 68012459 | rs1430778 | 0.916 | 0.947 | -0.935 | 0.029 | -0.008 | 0.003 | 0.003 | 0 | 0.27 | 0.21 | 0.27 | 0.19 | 0.32 | 0.38 |
| 2 | 68019342 | rs17034390 | 0.473 | 0.784 | -0.78 | 0.142 | -0.004 | -0.005 | -0.005 | 0 | 0.11 | 0.13 | 0.05 | 0.08 | 0.05 | 0.12 |
| 2 | 68024446 | rs1346866 | -0.797 | 0.852 | -0.287 | 0.171 | 0.001 | 0.032 | 0.032 | 0 | 0.75 | 0.74 | 0.73 | 0.43 | 0.69 | 0.81 |
| 2 | 68025340 | rs11674863 | 0.883 | 0.026 | -0.302 | -0.467 | 0.023 | 0.023 | 0.023 | 0 | 0.26 | 0.15 | 0.12 | 0.02 | 0.18 | 0.22 |
| 2 | 68026807 | rs10469954 | -0.25 | 0.01 | -0.23 | -0.41 | -0.028 | -0.013 | -0.013 | 0 | 0.16 | 0.31 | 0.26 | 0.15 | 0.09 | 0.15 |
| 2 | 68027155 | rs13429160 | 0.886 | 0.159 | -0.482 | -0.345 | 0.029 | 0.045 | 0.045 | 0 | 0.25 | 0.13 | 0.12 | 0.02 | 0.18 | 0.23 |
| 2 | 68028751 | rs4308115 | -0.502 | 0.62 | -0.489 | -0.141 | 0.017 | -0.001 | -0.001 | 0.757 | 0.67 | 0.63 | 0.52 | 0.31 | 0.46 | 0.57 |
| 2 | 68030319 | rs11126157 | 0.32 | 0.587 | -0.643 | -0.224 | 0.029 | 0.04 | 0.04 | 0 | 0.35 | 0.29 | 0.17 | 0.13 | 0.23 | 0.34 |
| 2 | 68031025 | rs11126159 | 0.049 | 1.322 | -0.59 | 0.581 | -0.001 | -0.001 | -0.001 | 0 | 0.67 | 0.69 | 0.53 | 0.36 | 0.46 | 0.61 |
| 2 | 68035962 | rs6715551 | -0.896 | 1.561 | -0.36 | 0.741 | 0.082 | 0.037 | 0.037 | 0 | 0.28 | 0.4 | 0.41 | 0.65 | 0.43 | 0.47 |
| 2 | 68041842 | rs7588898 | -0.392 | 0.707 | -0.867 | 0.732 | 0.005 | -0.004 | -0.004 | 0 | 0.44 | 0.35 | 0.49 | 0.39 | 0.39 | 0.45 |
| 2 | 68043435 | rs1367464 | -0.223 | 0.594 | -0.87 | 0.763 | -0.014 | -0.032 | -0.032 | 2.171 | 0.42 | 0.18 | 0.46 | 0.39 | 0.38 | 0.44 |
| 2 | 68049568 | rs10199432 | -1.502 | 0.21 | -1.383 | 0.311 | -0.004 | 0.166 | 0.166 | 0 | 0.7 | 0.96 | 0.76 | 0.91 | 0.69 | 0.81 |
| 2 | 68050599 | rs7561904 | -0.204 | 0.144 | -0.955 | 0.533 | 0.002 | -0.006 | -0.006 | 0 | 0.44 | 0.39 | 0.5 | 0.4 | 0.39 | 0.44 |
| 2 | 68053984 | rs12989344 | 0.116 | 0.141 | -0.951 | 0.51 | -0.019 | -0.04 | -0.04 | 0.178 | 0.41 | 0.21 | 0.47 | 0.4 | 0.39 | 0.44 |
| 2 | 68059696 | rs13385981 | -0.233 | 0.143 | -0.989 | 0.512 | 0.005 | -0.005 | -0.005 | 0 | 0.45 | 0.4 | 0.5 | 0.4 | 0.38 | 0.44 |
| 2 | 68060244 | rs6729128 | 1.262 | 0.17 | -1.304 | 0.523 | -0.044 | 0.022 | 0.022 | 1.362 | 0.24 | 0.07 | 0.21 | 0.13 | 0.25 | 0.21 |
| 2 | 68060996 | rs12104671 | -0.804 | -0.388 | -0.614 | 0.77 | -0.008 | 0.015 | 0.015 | 0 | 0.2 | 0.27 | 0.29 | 0.27 | 0.14 | 0.23 |
| 2 | 68063154 | rs2654472 | -0.564 | -0.332 | -0.768 | 0.803 | 0 | -0.004 | -0.004 | 0 | 0.18 | 0.18 | 0.26 | 0.27 | 0.14 | 0.22 |
| 2 | 68067589 | rs958979 | -0.304 | -0.275 | -0.922 | 0.831 | -0.001 | -0.028 | -0.028 | 0.266 | 0.17 | 0.1 | 0.26 | 0.26 | 0.14 | 0.24 |
| 2 | 68069867 | rs1820987 | -0.173 | -0.324 | -1.046 | 0.762 | 0.016 | 0.041 | 0.041 | 0 | 0.56 | 0.86 | 0.51 | 0.65 | 0.57 | 0.67 |
| 2 | 68076780 | rs6723375 | -0.175 | 0.024 | -0.85 | 0.823 | -0.019 | -0.013 | -0.013 | 2.154 | 0.9 | 0.95 | 0.76 | 0.88 | 0.9 | 0.85 |
| 2 | 68077707 | rs2861754 | 0.53 | -0.346 | -0.158 | 0.992 | -0.009 | 0.013 | 0.013 | 0.018 | 0.34 | 0.41 | 0.28 | 0.49 | 0.34 | 0.42 |
| 2 | 68080882 | rs905495 | 0.6 | -0.326 | -0.427 | 0.923 | -0.037 | -0.01 | -0.01 | 0 | 0.37 | 0.25 | 0.43 | 0.35 | 0.51 | 0.28 |
| 2 | 68086034 | rs11126162 | -0.946 | -0.153 | 0.3 | 1.699 | -0.014 | 0.053 | 0.053 | 0.432 | 0.16 | 0.29 | 0.06 | 0.41 | 0.09 | 0.29 |
| 2 | 68086045 | rs13030887 | 1.451 | -0.203 | 0.329 | 1.777 | -0.022 | 0.09 | 0.09 | 0.458 | 0.74 | 0.43 | 0.7 | 0.44 | 0.83 | 0.58 |
| 2 | 68086910 | rs1393820 | -0.492 | 0.205 | -0.449 | 1.496 | -0.027 | -0.042 | -0.042 | 0 | 0.93 | 1 | 0.78 | 0.85 | 0.92 | 0.87 |
| 2 | 68089831 | rs7557514 | -0.497 | 0.195 | -0.456 | 1.569 | -0.024 | -0.044 | -0.044 | 0.802 | 0.93 | 1 | 0.71 | 0.82 | 0.92 | 0.85 |
| 2 | 68091229 | rs6741698 | 1.481 | -0.312 | 0.309 | 1.645 | -0.025 | 0.021 | 0.021 | 0 | 0.78 | 0.64 | 0.66 | 0.75 | 0.87 | 0.7 |
| 2 | 68092380 | rs6761173 | 0.512 | -0.814 | -0.058 | 1.558 | -0.002 | 0.005 | 0.005 | 0.756 | 0.33 | 0.45 | 0.21 | 0.42 | 0.34 | 0.35 |
| 2 | 68092404 | rs905493 | 0.835 | -0.813 | -0.09 | 1.583 | -0.032 | 0.033 | 0.033 | 1.453 | 0.44 | 0.17 | 0.45 | 0.32 | 0.54 | 0.35 |
| 2 | 68097169 | rs1503243 | 0.161 | -0.616 | -0.106 | 1.857 | 0 | 0.003 | 0.003 | 0.305 | 0.41 | 0.42 | 0.26 | 0.51 | 0.39 | 0.46 |
| 2 | 68097760 | rs1503242 | -1.035 | -0.547 | 0.001 | 1.825 | -0.031 | -0.051 | -0.051 | 0 | 0.92 | 1 | 0.7 | 0.8 | 0.91 | 0.82 |
| 2 | 68100701 | rs1503241 | 0.247 | -0.54 | 0.134 | 2.225 | -0.002 | 0.005 | 0.005 | 0.047 | 0.41 | 0.42 | 0.26 | 0.51 | 0.39 | 0.47 |
| 2 | 68101715 | rs13032405 | 0.901 | -0.502 | -0.151 | 1.987 | -0.041 | 0.066 | 0.066 | 0.845 | 0.39 | 0.15 | 0.4 | 0.23 | 0.48 | 0.28 |
| 2 | 68103954 | rs7563733 | 0.555 | -0.454 | -0.078 | 1.939 | -0.001 | 0.094 | 0.094 | 0.938 | 0.45 | 0.26 | 0.42 | 0.21 | 0.49 | 0.32 |
| 2 | 68107869 | rs11679294 | 0.282 | -0.52 | 0.483 | 1.938 | -0.002 | -0.002 | -0.002 | 0.001 | 0.36 | 0.41 | 0.24 | 0.36 | 0.34 | 0.3 |
| 2 | 68109308 | rs11688030 | 0.069 | -0.5 | 0.434 | 1.945 | -0.002 | -0.009 | -0.009 | 0 | 0.38 | 0.55 | 0.25 | 0.32 | 0.34 | 0.3 |
| 2 | 68109558 | rs10182649 | -0.705 | -0.012 | 0.184 | 1.988 | -0.008 | -0.008 | -0.008 | 8.168 | 0.89 | 0.93 | 0.6 | 0.83 | 0.85 | 0.81 |
| 2 | 68115179 | rs11126167 | 0.555 | -0.014 | 0.145 | 1.849 | -0.016 | -0.018 | -0.018 | 0 | 0.45 | 0.22 | 0.33 | 0.5 | 0.5 | 0.47 |
| 2 | 68120579 | rs13422556 | 0.037 | 0.021 | 0.528 | 1.889 | -0.015 | 0.031 | 0.031 | 1.247 | 0.07 | 0.14 | 0.02 | 0.07 | 0.02 | 0.07 |
| 2 | 68121443 | rs867022 | -0.585 | 0.005 | 0.394 | 2.188 | -0.051 | -0.063 | -0.063 | 0 | 0.88 | 0.99 | 0.6 | 0.79 | 0.84 | 0.8 |
| 2 | 68122154 | rs13391665 | 0.11 | 0.017 | 0.508 | 2.109 | -0.011 | 0.032 | 0.032 | 1.247 | 0.07 | 0.14 | 0.02 | 0.07 | 0.02 | 0.07 |
| 2 | 68125000 | rs11689443 | 0.199 | -0.07 | 0.905 | 2.165 | -0.003 | -0.007 | -0.007 | 0 | 0.37 | 0.38 | 0.24 | 0.36 | 0.34 | 0.33 |
| 2 | 68128082 | rs7558015 | 0.262 | -0.086 | 1.047 | 2.155 | 0.002 | 0.006 | 0.006 | 0 | 0.37 | 0.3 | 0.23 | 0.35 | 0.34 | 0.33 |
| 2 | 68138786 | rs1503236 | 0.961 | -0.155 | 0.937 | 2.114 | -0.008 | 0.022 | 0.022 | 0 | 0.42 | 0.28 | 0.33 | 0.35 | 0.47 | 0.42 |
| 2 | 68139931 | rs1503239 | 0.313 | -0.163 | 0.847 | 2.183 | 0.003 | 0.016 | 0.016 | 0 | 0.51 | 0.63 | 0.37 | 0.59 | 0.5 | 0.47 |
| 2 | 68141262 | rs6750910 | 0.175 | -0.126 | 0.858 | 2.052 | -0.002 | -0.002 | -0.002 | 0 | 0.38 | 0.37 | 0.24 | 0.36 | 0.34 | 0.33 |
| 2 | 68141355 | rs13014970 | 0.32 | -0.24 | 0.847 | 2.147 | -0.001 | -0.001 | -0.001 | 1.197 | 0.48 | 0.49 | 0.35 | 0.43 | 0.5 | 0.47 |
| 2 | 68143595 | rs872108 | 0.503 | -0.261 | 0.669 | 1.897 | -0.004 | 0 | 0 | 0 | 0.46 | 0.28 | 0.33 | 0.59 | 0.49 | 0.46 |
| 2 | 68148917 | rs9967692 | -0.32 | -0.295 | 0.475 | 1.579 | 0 | 0 | 0 | 0 | 0.46 | 0.48 | 0.63 | 0.49 | 0.47 | 0.43 |
| 2 | 68149455 | rs6704741 | -0.913 | -0.209 | -0.147 | 1.521 | -0.023 | -0.026 | -0.026 | 0 | 0.58 | 0.81 | 0.68 | 0.56 | 0.53 | 0.58 |
| 2 | 68151411 | rs1393819 | -0.737 | 0.199 | -0.622 | 1.07 | -0.029 | -0.025 | -0.025 | 0.234 | 0.62 | 0.75 | 0.74 | 0.54 | 0.56 | 0.76 |
| 2 | 68151619 | rs6741945 | 0.954 | 0.049 | -0.469 | 0.9 | -0.016 | 0.005 | 0.005 | 0 | 0.13 | 0.28 | 0.24 | 0.17 | 0.13 | 0.09 |
| 2 | 68152012 | rs1503237 | -0.895 | 0.238 | -0.896 | 0.939 | -0.021 | -0.018 | -0.018 | 0 | 0.62 | 0.72 | 0.74 | 0.54 | 0.56 | 0.76 |
| 2 | 68157039 | rs17578007 | 1.518 | 0.539 | -1.251 | 0.505 | -0.005 | 0.005 | 0.005 | 0 | 0.2 | 0.11 | 0.25 | 0.16 | 0.23 | 0.33 |
| 2 | 68164638 | rs2089024 | -0.463 | 0.595 | -1.4 | 0.23 | -0.014 | 0.035 | 0.035 | 0 | 0.32 | 0.25 | 0.39 | 0.25 | 0.39 | 0.2 |
| 2 | 68165196 | rs6726612 | 1.066 | 0.882 | -1.253 | -0.017 | -0.015 | 0.006 | 0.006 | 0.178 | 0.16 | 0.13 | 0.15 | 0.04 | 0.25 | 0.09 |
| 2 | 68165892 | rs10496155 | 0.902 | 0.711 | -1.021 | 0.206 | 0.016 | -0.024 | -0.024 | 0 | 0.11 | 0.15 | 0.12 | 0.04 | 0.17 | 0.09 |
| 2 | 68166080 | rs1032807 | 0.391 | -0.08 | -0.973 | 0.437 | -0.002 | -0.006 | -0.006 | 0 | 0.57 | 0.53 | 0.55 | 0.41 | 0.5 | 0.72 |
| 2 | 68174567 | rs11681748 | 0.357 | 0.178 | 0.001 | 0.24 | 0.017 | -0.036 | -0.036 | 0 | 0.15 | 0.24 | 0.15 | 0.1 | 0.17 | 0.15 |
| 2 | 68176433 | rs17034705 | 0.796 | 0.16 | 0.25 | -0.015 | 0.012 | -0.07 | -0.07 | 0 | 0.13 | 0.25 | 0.09 | 0 | 0.1 | 0.12 |
| 2 | 68181448 | rs7564090 | 0.545 | -0.058 | -0.005 | -0.066 | 0.001 | -0.027 | -0.027 | 0 | 0.34 | 0.41 | 0.41 | 0.24 | 0.37 | 0.39 |
| 2 | 68181823 | rs881664 | -0.932 | -0.198 | -0.288 | 0.243 | 0.045 | -0.052 | -0.052 | 0 | 0.66 | 0.91 | 0.61 | 0.6 | 0.69 | 0.63 |
| 2 | 68182025 | rs905488 | -0.942 | -0.196 | -0.287 | 0.244 | 0.044 | -0.053 | -0.053 | 0 | 0.66 | 0.91 | 0.61 | 0.6 | 0.69 | 0.63 |

| | | | | | | | | | | | | | | | | |
|---|---|---|---|---|---|---|---|---|---|---|---|---|---|---|---|---|
| 2 | 68187992 | rs905487 | 0.161 | -0.099 | -0.096 | 0.212 | -0.009 | 0.038 | 0.038 | 0 | 0.45 | 0.58 | 0.47 | 0.19 | 0.5 | 0.47 |
| 2 | 68188887 | rs1393806 | -1.237 | -0.26 | -0.181 | 0.378 | 0.025 | -0.057 | -0.057 | 0 | 0.7 | 0.91 | 0.67 | 0.68 | 0.72 | 0.68 |
| 2 | 68190624 | rs4671863 | 0.667 | -0.246 | 0.037 | 0.256 | 0.007 | 0.008 | 0.008 | 0 | 0.11 | 0.19 | 0.25 | 0.14 | 0.15 | 0.31 |
| 2 | 68192266 | rs12713614 | 0.485 | -0.368 | 0.166 | 0.162 | 0.003 | -0.036 | -0.036 | 0 | 0.21 | 0.26 | 0.28 | 0.11 | 0.25 | 0.18 |
| 2 | 68192880 | rs7566701 | -2.347 | -0.383 | 0.103 | 0.175 | 0.003 | -0.062 | -0.062 | 4.135 | 0.78 | 0.93 | 0.84 | 0.72 | 0.82 | 0.76 |
| 2 | 68195179 | rs969536 | -0.792 | -0.3 | 0.417 | 0.527 | 0.013 | -0.022 | -0.022 | 0 | 0.32 | 0.46 | 0.54 | 0.25 | 0.39 | 0.49 |
| 2 | 68196982 | rs10191711 | -1.53 | -0.346 | 0.563 | 0.538 | -0.015 | 0.017 | 0.017 | 0 | 0.68 | 0.62 | 0.74 | 0.62 | 0.76 | 0.7 |
| 2 | 68198367 | rs12713624 | -1.861 | -0.316 | 0.575 | 0.55 | -0.012 | 0.011 | 0.011 | 0 | 0.72 | 0.63 | 0.8 | 0.72 | 0.78 | 0.75 |
| 2 | 68201544 | rs10171662 | -1.563 | -0.367 | 0.61 | 0.552 | -0.009 | 0.01 | 0.01 | 0 | 0.69 | 0.66 | 0.74 | 0.62 | 0.75 | 0.68 |
| 2 | 68204413 | rs17034763 | -0.641 | -0.404 | 0.951 | 0.807 | -0.004 | -0.002 | -0.002 | 0 | 0.47 | 0.42 | 0.46 | 0.52 | 0.5 | 0.49 |
| 2 | 68206828 | rs12713625 | -0.977 | -0.399 | 0.875 | 0.807 | -0.001 | 0.02 | 0.02 | 0 | 0.52 | 0.67 | 0.53 | 0.61 | 0.53 | 0.55 |
| 2 | 68214370 | rs974366 | 0.216 | -0.521 | 0.298 | 0.766 | -0.019 | -0.026 | -0.026 | 0 | 0.34 | 0.14 | 0.21 | 0.38 | 0.35 | 0.17 |
| 2 | 68214819 | rs11126170 | 1.613 | -0.436 | 0.211 | 0.827 | -0.012 | -0.016 | -0.016 | 0 | 0.19 | 0.06 | 0.12 | 0.17 | 0.24 | 0.05 |
| 2 | 68221677 | rs4671866 | -0.794 | -0.319 | 0.277 | 0.507 | 0 | -0.002 | -0.002 | 0.251 | 0.27 | 0.33 | 0.36 | 0.25 | 0.24 | 0.46 |
| 2 | 68222281 | rs34422503 | 0.816 | -0.283 | -0.052 | 0.206 | -0.015 | -0.019 | -0.019 | 0 | 0.08 | 0.04 | 0.21 | 0.16 | 0.12 | 0.29 |
| 2 | 68222743 | rs10183044 | -0.259 | -0.37 | 0.054 | 0.019 | 0 | -0.042 | -0.042 | 0.247 | 0.45 | 0.47 | 0.31 | 0.17 | 0.5 | 0.24 |
| 2 | 68224745 | rs2221338 | -0.427 | -0.411 | 0.047 | 0.12 | -0.037 | 0.095 | 0.095 | 0 | 0.11 | 0.3 | 0.08 | 0.3 | 0.04 | 0.09 |
| 2 | 68227555 | rs1503248 | 0.41 | -0.382 | 0.041 | 0.107 | 0.005 | -0.019 | -0.019 | 0 | 0.34 | 0.33 | 0.22 | 0.17 | 0.4 | 0.14 |
| 2 | 68229753 | rs4671202 | -0.89 | -0.374 | 0.019 | 0.101 | 0.006 | 0.004 | 0.004 | 0.542 | 0.54 | 0.64 | 0.38 | 0.47 | 0.54 | 0.32 |
| 2 | 68234640 | rs964450 | -0.783 | -0.543 | 0.096 | 0.187 | 0.01 | 0.004 | 0.004 | 0 | 0.51 | 0.7 | 0.35 | 0.53 | 0.51 | 0.29 |
| 2 | 68235796 | rs17034835 | 0.714 | -0.404 | -0.184 | 0.299 | -0.004 | 0.128 | 0.128 | 0.067 | 0.07 | 0.32 | 0.03 | 0.19 | 0.01 | 0.02 |
| 2 | 68235872 | rs17034837 | 1.586 | -0.393 | -0.464 | 0.292 | 0.018 | 0.005 | 0.005 | 0 | 0.06 | 0.02 | 0.04 | 0.06 | 0.06 | 0.06 |
| 2 | 68237578 | rs11678194 | -0.72 | -0.395 | -0.337 | 0.27 | 0.003 | -0.001 | -0.001 | 0 | 0.47 | 0.6 | 0.35 | 0.52 | 0.5 | 0.28 |
| 2 | 68238321 | rs7586459 | 0.671 | -0.289 | -0.44 | 0.159 | -0.018 | 0.025 | 0.025 | 0.27 | 0.3 | 0.19 | 0.22 | 0.16 | 0.39 | 0.13 |
| 2 | 68238580 | rs7586762 | 0.294 | -0.049 | -0.246 | 0.146 | 0.011 | -0.026 | -0.026 | 0 | 0.36 | 0.5 | 0.3 | 0.27 | 0.42 | 0.2 |
| 2 | 68240538 | rs7594119 | -0.576 | 0.148 | -0.5 | 0.16 | -0.003 | 0.001 | 0.001 | 0 | 0.62 | 0.59 | 0.71 | 0.62 | 0.66 | 0.62 |
| 2 | 68250296 | rs7558466 | -0.607 | 0.473 | -0.789 | 0.105 | -0.012 | 0.019 | 0.019 | 0 | 0.56 | 0.71 | 0.58 | 0.43 | 0.55 | 0.49 |
| 2 | 68253616 | rs6546353 | -0.044 | 0.395 | -0.786 | 0.573 | 0.003 | 0.002 | 0.002 | 0 | 0.26 | 0.24 | 0.19 | 0.22 | 0.25 | 0.12 |
| 2 | 68257398 | rs1955369 | -0.626 | 0.697 | -0.379 | 0.734 | 0.009 | 0.013 | 0.013 | 0 | 0.27 | 0.4 | 0.46 | 0.32 | 0.25 | 0.4 |
| 2 | 68260795 | rs7576733 | -1.118 | 0.491 | -0.343 | 0.242 | -0.001 | -0.03 | -0.03 | 0.011 | 0.13 | 0.18 | 0.35 | 0.01 | 0.13 | 0.12 |
| 2 | 68261015 | rs10439493 | -0.329 | -0.531 | -1.15 | -0.002 | 0.02 | 0.01 | 0.01 | 0 | 0.28 | 0.71 | 0.19 | 0.27 | 0.17 | 0.19 |
| 2 | 68262703 | rs17034864 | -0.104 | -0.705 | -0.441 | -0.084 | 0.017 | 0.004 | 0.004 | 0 | 0.23 | 0.19 | 0.14 | 0.25 | 0.16 | 0.18 |
| 2 | 68267191 | rs7582426 | 0.564 | 0.256 | -0.811 | -0.181 | -0.009 | 0.008 | 0.008 | 0 | 0.8 | 0.79 | 0.65 | 0.91 | 0.75 | 0.82 |
| 2 | 68274097 | rs9807970 | -0.153 | -0.09 | 0.304 | 0.111 | -0.046 | 0.018 | 0.018 | 0 | 0.26 | 0.42 | 0.14 | 0.29 | 0.18 | 0.33 |
| 2 | 68281759 | rs10203061 | -0.344 | -0.08 | 0.506 | 0.115 | -0.039 | 0.007 | 0.007 | 0 | 0.27 | 0.54 | 0.15 | 0.31 | 0.18 | 0.34 |
| 2 | 68285641 | rs4671871 | -0.212 | 0.276 | 0.433 | 0.292 | -0.004 | 0.008 | 0.008 | 0 | 0.11 | 0.06 | 0.24 | 0.02 | 0.1 | 0.11 |
| 2 | 68302228 | rs6761122 | -1.333 | 0.001 | 0.343 | 0.278 | -0.015 | -0.003 | -0.003 | 0 | 0.27 | 0.59 | 0.37 | 0.25 | 0.14 | 0.24 |
| 2 | 68306739 | rs17034913 | -0.508 | 0.234 | 0.361 | 0.081 | 0.005 | -0.031 | -0.031 | 0 | 0.12 | 0.16 | 0.26 | 0.02 | 0.12 | 0.16 |
| 2 | 68310101 | rs1868403 | -0.755 | 0.102 | 0.188 | -0.085 | -0.004 | 0.002 | 0.002 | 0 | 0.23 | 0.22 | 0.34 | 0.09 | 0.26 | 0.25 |
| 2 | 68313496 | rs12616172 | -1.221 | -0.254 | -0.688 | -0.003 | -0.019 | 0.13 | 0.13 | 0 | 0.53 | 0.73 | 0.66 | 0.77 | 0.47 | 0.55 |
| 2 | 68322579 | rs11681944 | -0.59 | -0.257 | -1.132 | 0.564 | -0.112 | -0.098 | -0.098 | 0 | 0.36 | 0.06 | 0.52 | 0.41 | 0.43 | 0.46 |
| 2 | 68330834 | rs4671873 | -1.452 | -0.706 | -0.764 | 0.564 | 0.027 | 0.124 | 0.124 | 0.981 | 0.53 | 0.78 | 0.62 | 0.75 | 0.42 | 0.52 |
| 2 | 68335961 | rs2122128 | 0.151 | -0.902 | -0.268 | 0.608 | 0.009 | 0.014 | 0.014 | 0 | 0.09 | 0.05 | 0.07 | 0.35 | 0.06 | 0.08 |
| 2 | 68337745 | rs13391405 | -1.455 | -0.94 | 0.155 | 0.551 | -0.001 | -0.019 | -0.019 | 0 | 0.49 | 0.47 | 0.59 | 0.76 | 0.43 | 0.52 |
| 2 | 68338613 | rs17034987 | -0.486 | -0.579 | -0.169 | 0.804 | 0.028 | -0.076 | -0.076 | 0.824 | 0.11 | 0.01 | 0.08 | 0.34 | 0.07 | 0.08 |
| 2 | 68342443 | rs1075454 | 0.067 | -0.5 | -0.103 | 1.068 | 0.015 | -0.029 | -0.029 | 0 | 0.08 | 0.01 | 0.12 | 0.37 | 0.05 | 0.08 |
| 2 | 68343030 | rs744636 | -0.13 | -0.107 | -0.746 | 0.364 | 0.027 | 0.004 | 0.004 | 0.115 | 0.46 | 0.1 | 0.47 | 0.38 | 0.41 | 0.66 |
| 2 | 68350615 | rs751759 | -0.218 | 0.061 | -0.038 | 0.072 | 0.035 | 0.012 | 0.012 | 0.026 | 0.51 | 0.36 | 0.44 | 0.5 | 0.43 | 0.65 |
| 2 | 68350891 | rs6734622 | -1.369 | 0.674 | -0.743 | -0.274 | -0.138 | -0.083 | -0.083 | 0 | 0.75 | 0.98 | 0.68 | 0.66 | 0.63 | 0.83 |
| 2 | 68356917 | rs34842123 | 0.94 | 0.246 | -0.18 | -0.451 | -0.014 | 0.005 | 0.005 | 0.202 | 0.16 | 0.16 | 0.2 | 0.15 | 0.22 | 0.16 |
| 2 | 68359897 | rs4078978 | 1.053 | -0.154 | -0.451 | -0.72 | 0.022 | 0.063 | 0.063 | 0 | 0.15 | 0.03 | 0.09 | 0 | 0.07 | 0.11 |
| 2 | 68367863 | rs7581776 | 1.147 | -0.205 | 0.127 | -0.732 | -0.018 | -0.02 | -0.02 | 0 | 0.05 | 0.15 | 0.01 | 0 | 0 | 0.01 |
| 2 | 68372154 | rs12463718 | 1.289 | -0.311 | 0.003 | -0.751 | 0.033 | -0.083 | -0.083 | 0 | 0.33 | 0.09 | 0.35 | 0.49 | 0.33 | 0.54 |
| 2 | 68376376 | rs2278789 | 0.345 | 0.131 | 0.832 | -0.434 | 0.056 | 0.025 | 0.025 | 0.101 | 0.51 | 0.37 | 0.45 | 0.49 | 0.4 | 0.65 |
| 2 | 68391087 | rs12713631 | 0.373 | 0.103 | 0.648 | -0.414 | 0.053 | -0.025 | -0.025 | 0 | 0.19 | 0.45 | 0.22 | 0.15 | 0.22 | 0.16 |
| 2 | 68402012 | rs1060842 | -1.516 | 0.191 | 0.106 | -0.408 | -0.149 | -0.08 | -0.08 | 0.131 | 0.75 | 0.98 | 0.69 | 0.65 | 0.64 | 0.82 |
| 2 | 68402360 | rs13003580 | 0.998 | 0.176 | 0.201 | -0.434 | -0.004 | 0.001 | 0.001 | 0.321 | 0.16 | 0.18 | 0.2 | 0.15 | 0.22 | 0.16 |
| 2 | 68413348 | rs12994570 | 0.91 | 0.187 | 0.238 | -0.448 | 0.023 | -0.011 | -0.011 | 0 | 0.17 | 0.27 | 0.2 | 0.15 | 0.22 | 0.16 |
| 2 | 68433831 | rs6736178 | -1.399 | 0.444 | -0.844 | -0.302 | -0.149 | -0.083 | -0.083 | 0 | 0.75 | 0.98 | 0.69 | 0.64 | 0.64 | 0.82 |
| 2 | 68477948 | rs2029091 | -0.339 | -0.097 | -1.077 | 0.046 | 0.062 | 0.03 | 0.03 | 0.166 | 0.51 | 0.33 | 0.46 | 0.5 | 0.42 | 0.66 |
| 2 | 68507833 | rs17035173 | 0.321 | 0.041 | -1.538 | 0.304 | 0.008 | -0.088 | -0.088 | 1.615 | 0.32 | 0.06 | 0.4 | 0.51 | 0.35 | 0.55 |
| 2 | 68514628 | rs11674241 | -0.01 | -0.064 | -0.985 | -0.315 | 0.009 | 0.043 | 0.043 | 0 | 0.19 | 0.13 | 0.11 | 0.01 | 0.17 | 0.17 |
| 2 | 68515530 | rs6740317 | 0.49 | 0.332 | -0.434 | -0.243 | -0.025 | -0.006 | -0.006 | 4.71 | 0.38 | 0.18 | 0.45 | 0.56 | 0.44 | 0.52 |
| 2 | 68518280 | rs7600326 | -0.813 | 0.237 | 0.107 | -0.27 | -0.001 | -0.006 | -0.006 | 0.014 | 0.59 | 0.47 | 0.57 | 0.57 | 0.61 | 0.69 |
| 2 | 68519544 | rs10496161 | 0.094 | 0.341 | -0.022 | -0.254 | 0.003 | 0.029 | 0.029 | 0 | 0.22 | 0.17 | 0.14 | 0.01 | 0.21 | 0.17 |
| 2 | 68519877 | rs3732042 | 0.51 | 0.122 | 0.166 | -0.036 | -0.027 | 0.015 | 0.015 | 0 | 0.12 | 0.19 | 0.03 | 0.17 | 0.04 | 0.14 |
| 2 | 68529682 | rs6746740 | -0.025 | 0.563 | -0.425 | -0.443 | 0.004 | 0.016 | 0.016 | 0 | 0.56 | 0.39 | 0.56 | 0.71 | 0.61 | 0.72 |
| 2 | 68531637 | rs12995380 | -0.457 | 1.687 | -0.669 | -0.191 | 0.005 | -0.003 | -0.003 | 0 | 0.11 | 0.14 | 0.27 | 0.12 | 0.25 | 0.13 |
| 2 | 68537909 | rs6729652 | -1.041 | 1.088 | -1.115 | -0.75 | -0.059 | 0.009 | 0.009 | 1.205 | 0.78 | 0.94 | 0.83 | 0.76 | 0.7 | 0.89 |

| | | | | | | | | | | | | | | | | |
|---|---|---|---|---|---|---|---|---|---|---|---|---|---|---|---|---|
| 2 | 68540378 | rs10169617 | 0.179 | 0.068 | -0.487 | -1.298 | 0.012 | 0.013 | 0.013 | 0 | 0.49 | 0.55 | 0.39 | 0.71 | 0.35 | 0.6 |
| 2 | 68542498 | rs6546371 | -0.876 | 0.7 | -0.672 | -1.523 | -0.035 | 0.006 | 0.006 | 0.007 | 0.79 | 0.88 | 0.71 | 0.79 | 0.69 | 0.79 |
| 2 | 68543294 | rs10179521 | 0.865 | -0.022 | -0.699 | -1.915 | 0.016 | 0.009 | 0.009 | 0 | 0.16 | 0.11 | 0.09 | 0.02 | 0.07 | 0.08 |
| 2 | 68545940 | rs3796056 | -0.815 | 0.114 | -0.415 | -1.863 | 0 | 0 | 0 | 0 | 0.77 | 0.76 | 0.69 | 0.79 | 0.69 | 0.82 |
| 2 | 68546374 | rs4671898 | -0.755 | -0.079 | -0.179 | -2.073 | 0 | -0.014 | -0.014 | 0 | 0.5 | 0.54 | 0.49 | 0.16 | 0.46 | 0.41 |
| 2 | 68546449 | rs13025842 | 0.696 | -0.04 | -0.718 | -2.149 | -0.026 | 0.04 | 0.04 | 0 | 0.28 | 0.16 | 0.3 | 0.08 | 0.33 | 0.22 |
| 2 | 68548160 | rs13383631 | 0.631 | -0.37 | -0.523 | -1.748 | -0.026 | -0.03 | -0.03 | 0.117 | 0.15 | 0.23 | 0.1 | 0.04 | 0.06 | 0.08 |
| 2 | 68553195 | rs4671900 | 1.115 | -0.095 | -0.909 | -1.522 | 0.004 | -0.001 | -0.001 | 0 | 0.27 | 0.27 | 0.27 | 0.68 | 0.22 | 0.42 |
| 2 | 68554641 | rs6546374 | -1.021 | 0.184 | -1.328 | -1.668 | -0.097 | -0.01 | -0.01 | 0.667 | 0.77 | 0.95 | 0.68 | 0.77 | 0.67 | 0.84 |
| 2 | 68555700 | rs6717329 | -0.271 | 0.133 | -1.268 | -1.707 | -0.013 | 0.23 | 0.23 | 0 | 0.51 | 0.81 | 0.42 | 0.1 | 0.45 | 0.42 |
| 2 | 68559450 | rs17035306 | 1.473 | 0.215 | -1.342 | -1.617 | 0.03 | -0.018 | -0.018 | 0.274 | 0.25 | 0.1 | 0.26 | 0.69 | 0.22 | 0.38 |
| 2 | 68561460 | rs4671903 | -0.828 | 0.31 | -1.318 | -1.604 | -0.08 | 0.001 | 0.001 | 0.132 | 0.77 | 0.92 | 0.68 | 0.78 | 0.67 | 0.79 |
| 2 | 68569316 | rs1443651 | -0.026 | 0.316 | -1.102 | -1.78 | 0.01 | 0.025 | 0.025 | 0.01 | 0.46 | 0.64 | 0.41 | 0.08 | 0.41 | 0.39 |
| 2 | 68569763 | rs4255948 | -1.193 | 0.428 | -1.391 | -1.703 | -0.097 | -0.016 | -0.016 | 0 | 0.78 | 0.95 | 0.69 | 0.78 | 0.68 | 0.84 |
| 2 | 68571153 | rs6759864 | 0.041 | 0.468 | -1.322 | -1.694 | 0.014 | 0.074 | 0.074 | 0 | 0.46 | 0.66 | 0.41 | 0.08 | 0.41 | 0.38 |
| 2 | 68573781 | rs4671908 | 0.469 | 0.24 | -1.272 | -1.086 | 0.015 | 0.006 | 0.006 | 0.447 | 0.29 | 0.14 | 0.26 | 0.7 | 0.27 | 0.41 |
| 2 | 68578358 | rs6723149 | 0.909 | 0.207 | -1.289 | -1.215 | 0.032 | -0.037 | -0.037 | 0 | 0.3 | 0.08 | 0.25 | 0.69 | 0.23 | 0.46 |
| 2 | 68578675 | rs6546383 | -0.963 | 0.607 | -1.77 | -1.612 | -0.086 | 0.074 | 0.074 | 1.067 | 0.79 | 0.95 | 0.71 | 0.93 | 0.69 | 0.81 |
| 2 | 68578758 | rs17528215 | 0.867 | 0.192 | -1.798 | -1.895 | -0.038 | 0.025 | 0.025 | 0.231 | 0.26 | 0.1 | 0.22 | 0.17 | 0.33 | 0.19 |
| 2 | 68579931 | rs7578047 | 0.339 | 0.055 | -0.661 | -1.906 | -0.017 | -0.033 | -0.033 | 0 | 0.14 | 0.35 | 0.21 | 0.06 | 0.11 | 0.14 |
| 2 | 68579972 | rs7604812 | -0.299 | 0.031 | -0.165 | -1.918 | -0.04 | -0.066 | -0.066 | 0.282 | 0.18 | 0.53 | 0.23 | 0.07 | 0.13 | 0.17 |
| 2 | 68583528 | rs7592526 | -0.702 | 0.031 | 0.104 | -1.963 | 0.002 | 0.027 | 0.027 | 2.166 | 0.44 | 0.64 | 0.35 | 0.21 | 0.45 | 0.35 |
| 2 | 68585819 | rs10865374 | 0.456 | 0.07 | 0.194 | -1.952 | 0.013 | 0.025 | 0.025 | 0 | 0.46 | 0.63 | 0.49 | 0.75 | 0.39 | 0.59 |
| 2 | 68587477 | rs7595717 | 1.586 | 0.044 | 0.271 | -1.97 | 0.015 | -0.006 | -0.006 | 0 | 0.33 | 0.22 | 0.32 | 0.69 | 0.27 | 0.51 |
| 2 | 68587588 | rs6546387 | -1.031 | 0.04 | 0.217 | -1.98 | -0.067 | 0.081 | 0.081 | 1.373 | 0.77 | 0.95 | 0.72 | 0.93 | 0.71 | 0.82 |
| 2 | 68592078 | rs11889922 | 0.184 | 0.003 | 0.355 | -2.037 | 0.035 | 0.071 | 0.071 | 0 | 0.5 | 0.7 | 0.61 | 0.76 | 0.4 | 0.63 |
| 2 | 68592315 | rs3732046 | 0.184 | 0.003 | 0.355 | -2.037 | 0.035 | 0.071 | 0.071 | 0 | 0.5 | 0.7 | 0.61 | 0.76 | 0.4 | 0.63 |
| 2 | 68595829 | rs1822470 | 0.082 | 0.001 | 0.522 | -2.032 | 0.022 | 0.043 | 0.043 | 0 | 0.5 | 0.67 | 0.6 | 0.76 | 0.4 | 0.63 |
| 2 | 68595939 | rs7567378 | 0.869 | 0.039 | 0.548 | -2.026 | 0.004 | -0.029 | -0.029 | 0 | 0.11 | 0.31 | 0.1 | 0.03 | 0.1 | 0.07 |
| 2 | 68596469 | rs10174380 | 0.428 | 0.026 | 0.549 | -2.036 | -0.002 | 0.012 | 0.012 | 0.345 | 0.29 | 0.23 | 0.22 | 0.19 | 0.32 | 0.23 |
| 2 | 68598955 | rs17035378 | 0.839 | 0.02 | 0.567 | -2.017 | 0.01 | 0.009 | 0.009 | 0 | 0.39 | 0.36 | 0.51 | 0.73 | 0.3 | 0.56 |
| 2 | 68602258 | rs4671912 | 0.731 | 0.029 | 0.511 | -2.013 | -0.014 | -0.013 | -0.013 | 0 | 0.4 | 0.45 | 0.52 | 0.73 | 0.3 | 0.56 |
| 2 | 68602859 | rs7605863 | 0.172 | 0.046 | 0.49 | -2.014 | 0.044 | 0.092 | 0.092 | 0 | 0.5 | 0.74 | 0.61 | 0.76 | 0.4 | 0.63 |
| 2 | 68607947 | rs3816281 | 1.586 | 0.035 | 0.456 | -1.999 | -0.002 | 0 | 0 | 0 | 0.33 | 0.3 | 0.42 | 0.69 | 0.27 | 0.51 |
| 2 | 68613710 | rs2070171 | 0.714 | 0.041 | 0.447 | -2.033 | -0.004 | 0.018 | 0.018 | 0 | 0.28 | 0.18 | 0.21 | 0.17 | 0.31 | 0.21 |
| 2 | 68616085 | rs2120334 | 1.655 | 0.053 | 0.316 | -2.017 | -0.001 | 0 | 0 | 0.428 | 0.32 | 0.28 | 0.41 | 0.69 | 0.27 | 0.5 |
| 2 | 68621338 | rs17530191 | 0.233 | 0.207 | -0.146 | -1.956 | -0.003 | 0.038 | 0.038 | 0 | 0.31 | 0.17 | 0.22 | 0.19 | 0.32 | 0.24 |
| 2 | 68622914 | rs1063479 | -1.125 | 0.201 | -0.169 | -1.97 | -0.081 | 0.088 | 0.088 | 0.659 | 0.78 | 0.96 | 0.82 | 0.93 | 0.71 | 0.83 |
| 2 | 68622990 | rs6725743 | -0.179 | 0.18 | -0.095 | -1.985 | -0.019 | -0.043 | -0.043 | 1.452 | 0.16 | 0.38 | 0.12 | 0.04 | 0.12 | 0.11 |
| 2 | 68623302 | rs6742200 | -0.567 | 0.215 | -0.09 | -1.978 | 0 | 0.003 | 0.003 | 0.83 | 0.42 | 0.5 | 0.33 | 0.22 | 0.43 | 0.31 |
| 2 | 68623652 | rs1867313 | -1.133 | 0.218 | -0.088 | -1.976 | -0.081 | 0.088 | 0.088 | 0.659 | 0.78 | 0.96 | 0.82 | 0.93 | 0.71 | 0.83 |
| 2 | 68624260 | rs8761 | -0.598 | 0.236 | -0.054 | -1.93 | 0 | 0.002 | 0.002 | 0.842 | 0.42 | 0.5 | 0.33 | 0.22 | 0.43 | 0.31 |
| 2 | 68639273 | rs13011577 | 0.574 | 0.327 | -0.319 | -1.818 | -0.018 | 0.03 | 0.03 | 0 | 0.27 | 0.12 | 0.21 | 0.17 | 0.31 | 0.2 |
| 2 | 68647095 | rs7595037 | 0.451 | 0.402 | 0.124 | -1.736 | 0.017 | -0.061 | -0.061 | 0 | 0.64 | 0.51 | 0.69 | 0.9 | 0.57 | 0.74 |
| 2 | 68652580 | rs11126185 | 1.178 | 0.714 | -0.178 | -1.494 | 0.012 | -0.038 | -0.038 | 0.235 | 0.71 | 0.33 | 0.73 | 0.89 | 0.66 | 0.67 |
| 2 | 68658790 | rs6419647 | 0.027 | 0.897 | -0.532 | -1.535 | -0.064 | 0.05 | 0.05 | 0 | 0.87 | 0.96 | 0.86 | 0.95 | 0.77 | 0.86 |
| 2 | 68660126 | rs6720557 | 1.233 | 0.694 | -0.39 | -1.588 | 0.001 | -0.017 | -0.017 | 0 | 0.66 | 0.31 | 0.65 | 0.88 | 0.64 | 0.66 |
| 2 | 68663534 | rs7596633 | 0.139 | 0.939 | -0.603 | -1.239 | -0.039 | 0.007 | 0.007 | 0 | 0.92 | 1 | 0.93 | 0.96 | 0.87 | 0.89 |
| 2 | 68665732 | rs6719948 | 1.761 | 0.853 | 0.316 | -1.478 | 0.028 | -0.075 | -0.075 | 1.418 | 0.77 | 0.47 | 0.77 | 0.94 | 0.76 | 0.81 |
| 2 | 68665945 | rs7421154 | -0.822 | 0.611 | 0.505 | -1.48 | 0.028 | -0.023 | -0.023 | 0.012 | 0.11 | 0.27 | 0.14 | 0.03 | 0.11 | 0.08 |
| 2 | 68668459 | rs10196560 | 1.353 | -0.02 | -0.401 | -2.087 | 0.019 | 0.136 | 0.136 | 0 | 0.49 | 0.27 | 0.56 | 0.87 | 0.45 | 0.57 |
| 2 | 68671494 | rs7424155 | 1.611 | 0.259 | -0.012 | -1.772 | -0.006 | -0.071 | -0.071 | 0 | 0.78 | 0.46 | 0.79 | 0.94 | 0.8 | 0.84 |
| 2 | 68679539 | rs10168414 | 1.547 | 0.257 | 0.022 | -1.817 | 0.015 | -0.074 | -0.074 | 0 | 0.83 | 0.46 | 0.84 | 0.95 | 0.84 | 0.92 |
| 2 | 68719022 | rs7419739 | 0.391 | 0.087 | -0.343 | -1.511 | -0.112 | -0.002 | -0.002 | 0 | 0.41 | 0.06 | 0.31 | 0.37 | 0.48 | 0.45 |
| 2 | 68728483 | rs7563017 | -1.877 | -0.918 | 0.341 | -1.016 | -0.039 | -0.043 | -0.043 | 0 | 0.12 | 0.47 | 0.05 | 0.04 | 0.05 | 0.05 |
| 2 | 68729992 | rs11902811 | -1.996 | -0.925 | 0.275 | -1.02 | -0.043 | -0.047 | -0.047 | 0 | 0.12 | 0.48 | 0.05 | 0.04 | 0.05 | 0.05 |
| 2 | 68733389 | rs10208940 | -2.34 | -0.98 | 0.234 | -1.046 | -0.051 | -0.1 | -0.1 | 0 | 0.26 | 0.6 | 0.12 | 0.04 | 0.13 | 0.15 |
| 2 | 68759471 | rs13415763 | -0.455 | -0.794 | 0.413 | -0.93 | -0.011 | -0.049 | -0.049 | 0.953 | 0.12 | 0.2 | 0.07 | 0.01 | 0.07 | 0.07 |
| 2 | 68763075 | rs10172469 | -0.43 | -0.785 | 0.413 | -0.926 | -0.006 | -0.034 | -0.034 | 0 | 0.13 | 0.15 | 0.07 | 0.01 | 0.07 | 0.07 |
| 2 | 68765205 | rs13404469 | -0.43 | -0.785 | 0.413 | -0.926 | -0.005 | -0.03 | -0.03 | 0.012 | 0.13 | 0.15 | 0.07 | 0.01 | 0.07 | 0.07 |
| 2 | 68770276 | rs7560955 | -0.43 | -0.785 | 0.413 | -0.91 | -0.006 | -0.031 | -0.031 | 0.369 | 0.13 | 0.15 | 0.07 | 0.01 | 0.07 | 0.07 |
| 2 | 68774100 | rs12713646 | -1.04 | -0.822 | 0.375 | -0.938 | -0.009 | -0.039 | -0.039 | 0 | 0.16 | 0.19 | 0.08 | 0.01 | 0.08 | 0.09 |
| 2 | 68799454 | rs6546420 | -2.229 | -0.697 | 0.448 | -0.844 | -0.052 | -0.1 | -0.1 | 2.279 | 0.24 | 0.6 | 0.12 | 0.04 | 0.12 | 0.12 |
| 2 | 68807614 | rs4129711 | -1.534 | -0.655 | 0.384 | -0.822 | -0.03 | -0.033 | -0.033 | 0 | 0.1 | 0.33 | 0.04 | 0.03 | 0.05 | 0.05 |
| 2 | 68811427 | rs10180884 | -0.606 | -0.63 | 0.364 | -0.814 | -0.016 | -0.062 | -0.062 | 0.346 | 0.13 | 0.24 | 0.07 | 0.01 | 0.07 | 0.07 |
| 2 | 68833572 | rs7580719 | -1.519 | -0.567 | 0.062 | -0.719 | -0.052 | -0.09 | -0.09 | 0.515 | 0.23 | 0.58 | 0.11 | 0.05 | 0.12 | 0.12 |
| 2 | 68851403 | rs7608248 | -1.651 | -0.522 | -0.034 | -0.714 | -0.042 | -0.086 | -0.086 | 0 | 0.25 | 0.6 | 0.12 | 0.05 | 0.13 | 0.12 |
| 2 | 68858318 | rs13016618 | -0.809 | -0.509 | 0.018 | -0.604 | -0.013 | -0.028 | -0.028 | 0.872 | 0.18 | 0.24 | 0.09 | 0.05 | 0.11 | 0.11 |
| 2 | 68861119 | rs6736893 | -0.631 | -0.466 | -0.702 | -0.628 | 0.004 | -0.01 | -0.01 | 0 | 0.72 | 0.86 | 0.77 | 0.68 | 0.66 | 0.74 |

| | | | | | | | | | | | | | | | | |
|---|---|---|---|---|---|---|---|---|---|---|---|---|---|---|---|---|
| 2 | 68865293 | rs7567869 | -1.573 | -0.284 | 0.208 | -0.357 | -0.049 | -0.087 | -0.087 | 0 | 0.22 | 0.59 | 0.12 | 0.05 | 0.12 | 0.11 |
| 2 | 68873071 | rs7570797 | -1.389 | -0.105 | 0.463 | -0.07 | -0.011 | 0.005 | 0.005 | 1.449 | 0.1 | 0.47 | 0.05 | 0.04 | 0.05 | 0.05 |
| 2 | 68884537 | rs6731427 | -1.133 | 0.328 | 0.406 | 0.365 | 0.009 | 0.01 | 0.01 | 0 | 0.28 | 0.65 | 0.3 | 0.38 | 0.32 | 0.4 |
| 2 | 68884666 | rs4854436 | -1.717 | 0.154 | -0.323 | 0.682 | -0.052 | 0.061 | 0.061 | 0.114 | 0.51 | 0.91 | 0.47 | 0.64 | 0.49 | 0.69 |
| 2 | 68888708 | rs4854480 | -1.76 | 0.256 | 0.299 | 0.655 | 0.018 | 0.042 | 0.042 | 0.1 | 0.48 | 0.72 | 0.31 | 0.64 | 0.42 | 0.63 |
| 2 | 68890104 | rs6754841 | -1.268 | 0.201 | 0.509 | 0.566 | 0 | 0.001 | 0.001 | 0 | 0.34 | 0.64 | 0.25 | 0.64 | 0.35 | 0.54 |
| 2 | 68895542 | rs6754173 | -1.197 | 0.216 | 0.607 | 0.573 | 0.001 | 0.003 | 0.003 | 0 | 0.33 | 0.61 | 0.25 | 0.64 | 0.35 | 0.54 |
| 2 | 68898708 | rs10178143 | -0.197 | 0.027 | 0.947 | 0.446 | 0.036 | 0 | 0 | 0 | 0.06 | 0.29 | 0.05 | 0 | 0.04 | 0.05 |
| 2 | 68901038 | rs13029650 | 0.603 | 0.012 | 0.194 | 0.528 | 0.008 | 0.003 | 0.003 | 0 | 0.09 | 0.01 | 0.04 | 0.12 | 0.04 | 0.05 |
| 2 | 68901608 | rs3969864 | -0.363 | 0.077 | 0.292 | 0.525 | 0.003 | -0.002 | -0.002 | 0 | 0.16 | 0.09 | 0.21 | 0 | 0.14 | 0.16 |
| 2 | 68902702 | rs10203944 | -0.496 | 0.164 | 0.316 | 0.74 | 0.024 | 0.252 | 0.252 | 0.016 | 0.12 | 0.42 | 0.13 | 0.24 | 0.11 | 0.18 |
| 2 | 68911266 | rs13423988 | -0.798 | 0.224 | 0.239 | 0.823 | 0.034 | 0.247 | 0.247 | 0.434 | 0.16 | 0.42 | 0.29 | 0.37 | 0.19 | 0.28 |
| 2 | 68913500 | rs13033526 | 0.358 | 0.078 | 0.159 | 0.669 | 0.009 | -0.003 | -0.003 | 0 | 0.12 | 0.08 | 0.06 | 0 | 0.07 | 0.1 |
| 2 | 68918043 | rs12477648 | -0.729 | 0.129 | -0.345 | 0.593 | -0.001 | -0.001 | -0.001 | 0.066 | 0.34 | 0.34 | 0.32 | 0.34 | 0.37 | 0.43 |
| 2 | 68918682 | rs4854491 | -1.214 | 0.077 | -0.415 | 0.546 | 0.001 | 0.003 | 0.003 | 0 | 0.43 | 0.34 | 0.35 | 0.46 | 0.41 | 0.48 |
| 2 | 68933876 | rs13000469 | 0.663 | 0.469 | -0.717 | 0.92 | -0.004 | -0.005 | -0.005 | 0 | 0.09 | 0 | 0.03 | 0.12 | 0.04 | 0.05 |
| 2 | 68947023 | rs4854447 | -0.366 | 0.469 | -0.596 | 0.879 | -0.05 | -0.079 | -0.079 | 0 | 0.31 | 0.12 | 0.29 | 0.45 | 0.34 | 0.38 |
| 2 | 68951477 | rs10172004 | -1.355 | 0.354 | 0.261 | 1.332 | 0.013 | 0.17 | 0.17 | 0.805 | 0.17 | 0.52 | 0.3 | 0.37 | 0.2 | 0.28 |
| 2 | 68957910 | rs13393229 | -1.351 | 0.152 | 0.475 | 1.439 | 0.009 | 0.3 | 0.3 | 0.454 | 0.15 | 0.47 | 0.14 | 0.36 | 0.13 | 0.21 |
| 2 | 68960650 | rs34093192 | 0.068 | -0.001 | 0.217 | 0.977 | 0.001 | 0.01 | 0.01 | 0 | 0.1 | 0.06 | 0.09 | 0.07 | 0.07 | 0.06 |
| 2 | 68965078 | rs4854502 | -1.041 | 0.26 | 0.836 | 1.555 | 0.021 | 0.112 | 0.112 | 0.182 | 0.07 | 0.18 | 0.09 | 0.27 | 0.07 | 0.14 |
| 2 | 68969904 | rs7608597 | 0.798 | 0.139 | 1.154 | 1.805 | -0.008 | 0.028 | 0.028 | 0 | 0.76 | 0.71 | 0.76 | 0.65 | 0.79 | 0.81 |
| 2 | 68970181 | rs10176108 | 0.77 | -0.015 | 0.25 | 0.948 | -0.011 | 0.018 | 0.018 | 0 | 0.35 | 0.18 | 0.38 | 0.31 | 0.36 | 0.44 |
| 2 | 68975706 | rs4854506 | 0.45 | -0.321 | 0.345 | 0.444 | -0.02 | 0.008 | 0.008 | 0.424 | 0.65 | 0.5 | 0.71 | 0.65 | 0.72 | 0.79 |
| 2 | 68979302 | rs2280244 | 1.634 | 0.045 | 0.468 | 0.543 | -0.008 | 0.072 | 0.072 | 0 | 0.27 | 0.16 | 0.25 | 0.07 | 0.3 | 0.2 |
| 2 | 68979569 | rs2280242 | -0.157 | 0.034 | 0.882 | 0.937 | 0.007 | -0.037 | -0.037 | 0.564 | 0.79 | 0.88 | 0.8 | 0.66 | 0.84 | 0.82 |
| 2 | 68983217 | rs4309612 | 1.22 | -0.052 | 0.871 | 0.982 | 0.001 | -0.001 | -0.001 | 0 | 0.62 | 0.37 | 0.64 | 0.38 | 0.65 | 0.66 |
| 2 | 68984896 | rs4281935 | -0.125 | -0.043 | 0.702 | 0.999 | 0.001 | -0.001 | -0.001 | 0.051 | 0.79 | 0.81 | 0.8 | 0.66 | 0.84 | 0.84 |
| 2 | 68989691 | rs4241341 | -0.312 | -0.039 | 0.576 | 1.001 | 0.003 | -0.055 | -0.055 | 0 | 0.8 | 0.92 | 0.81 | 0.66 | 0.84 | 0.84 |
| 2 | 68992798 | rs4854451 | -0.503 | -0.043 | 0.713 | 0.988 | 0.038 | 0.082 | 0.082 | 0 | 0.16 | 0.42 | 0.16 | 0.27 | 0.18 | 0.18 |
| 2 | 68994498 | rs6720182 | -0.159 | -0.104 | 0.348 | 1.282 | 0.003 | -0.056 | -0.056 | 0 | 0.8 | 0.92 | 0.81 | 0.66 | 0.84 | 0.84 |
| 2 | 68995389 | rs12232891 | 0.631 | 0.014 | 0.425 | 1.314 | 0.009 | 0.076 | 0.076 | 0 | 0.06 | 0.07 | 0.08 | 0.27 | 0.07 | 0.13 |
| 2 | 68996668 | rs13413887 | 0.761 | -0.06 | 0.554 | 0.951 | -0.004 | 0.033 | 0.033 | 0 | 0.72 | 0.63 | 0.72 | 0.39 | 0.77 | 0.71 |
| 2 | 68998919 | rs4627610 | -0.05 | -0.04 | 0.312 | 1.137 | 0.002 | -0.015 | -0.015 | 0 | 0.84 | 0.89 | 0.84 | 0.5 | 0.86 | 0.75 |
| 2 | 69004457 | rs4854512 | 0.349 | -0.004 | 0.21 | 0.775 | -0.008 | 0.015 | 0.015 | 0 | 0.43 | 0.3 | 0.43 | 0.37 | 0.41 | 0.42 |
| 2 | 69006123 | rs4854515 | 0.756 | 0.003 | 0.052 | 0.759 | -0.011 | 0.016 | 0.016 | 0 | 0.38 | 0.21 | 0.33 | 0.31 | 0.36 | 0.4 |
| 2 | 69009304 | rs2271834 | 0.422 | 0.005 | 0.272 | 0.413 | 0 | -0.007 | -0.007 | 1.007 | 0.39 | 0.33 | 0.4 | 0.11 | 0.44 | 0.32 |
| 2 | 69010557 | rs2311412 | 0.125 | -0.025 | 0.26 | 0.448 | -0.001 | 0.027 | 0.027 | 0 | 0.44 | 0.38 | 0.51 | 0.18 | 0.5 | 0.34 |
| 2 | 69011925 | rs6546435 | 0.787 | -0.007 | -0.2 | 0.772 | -0.023 | -0.011 | -0.011 | 0.012 | 0.38 | 0.18 | 0.33 | 0.34 | 0.37 | 0.39 |
| 2 | 69012522 | rs2311413 | -0.024 | 0.056 | 0.031 | 0.924 | -0.002 | 0.033 | 0.033 | 0 | 0.82 | 0.84 | 0.86 | 0.51 | 0.86 | 0.74 |
| 2 | 69013007 | rs11677065 | 0.347 | -0.13 | -0.203 | 0.732 | 0.004 | 0.032 | 0.032 | 0 | 0.44 | 0.57 | 0.33 | 0.3 | 0.46 | 0.36 |
| 2 | 69017824 | rs4854516 | 0.404 | -0.084 | -0.47 | 0.819 | -0.007 | -0.028 | -0.028 | 0 | 0.07 | 0.02 | 0.06 | 0.15 | 0.05 | 0.14 |
| 2 | 69017904 | rs12477343 | 1.038 | 0.007 | -0.544 | 0.894 | -0.04 | -0.047 | -0.047 | 0 | 0.33 | 0.15 | 0.43 | 0.36 | 0.36 | 0.38 |
| 2 | 69019272 | rs4854517 | -1.341 | -0.088 | -0.547 | 0.912 | 0.006 | -0.015 | -0.015 | 0.764 | 0.9 | 0.92 | 0.94 | 0.89 | 0.93 | 0.91 |
| 2 | 69019549 | rs10211539 | 0.321 | -0.065 | 0.133 | 0.815 | -0.002 | -0.013 | -0.013 | 2.621 | 0.43 | 0.47 | 0.33 | 0.32 | 0.46 | 0.36 |
| 2 | 69023704 | rs12713663 | 0.31 | -0.164 | -0.303 | 0.586 | 0.012 | -0.057 | -0.057 | 0.614 | 0.16 | 0.45 | 0.17 | 0.06 | 0.14 | 0.16 |
| 2 | 69024089 | rs4854519 | 0.399 | -0.088 | -1.01 | 0.76 | -0.039 | -0.071 | -0.071 | 0 | 0.35 | 0.09 | 0.21 | 0.42 | 0.35 | 0.33 |
| 2 | 69029707 | rs4854457 | 0.518 | 0.059 | -1.034 | 0.655 | -0.025 | -0.122 | -0.122 | 0.011 | 0.39 | 0.09 | 0.34 | 0.51 | 0.36 | 0.48 |
| 2 | 69029770 | rs6727322 | 0.023 | 0.101 | -0.879 | 0.228 | -0.009 | 0.059 | 0.059 | 0 | 0.65 | 0.74 | 0.68 | 0.87 | 0.64 | 0.66 |
| 2 | 69030378 | rs17035940 | -0.174 | 0.25 | 0.1 | 0.376 | 0.136 | 0.051 | 0.051 | 0.056 | 0.05 | 0.34 | 0.12 | 0.05 | 0.07 | 0.05 |
| 2 | 69031595 | rs10496167 | 1.144 | 0.074 | -0.247 | 0.419 | -0.001 | -0.004 | -0.004 | 0 | 0.18 | 0.16 | 0.2 | 0.32 | 0.2 | 0.12 |
| 2 | 69034042 | rs17543981 | 0.222 | 0.337 | -0.685 | -0.39 | -0.014 | 0.038 | 0.038 | 2.814 | 0.29 | 0.22 | 0.28 | 0.09 | 0.4 | 0.44 |
| 2 | 69034157 | rs12992898 | -0.777 | 0.198 | -0.635 | -0.045 | -0.012 | -0.004 | -0.004 | 0 | 0.58 | 0.38 | 0.57 | 0.48 | 0.56 | 0.72 |
| 2 | 69036013 | rs11891985 | 0.801 | 0.076 | -0.742 | 0.044 | 0.025 | -0.053 | -0.053 | 0 | 0.27 | 0.14 | 0.28 | 0.4 | 0.16 | 0.28 |
| 2 | 69040500 | rs2280310 | 0.756 | 0.038 | -0.051 | 0.015 | -0.004 | 0.028 | 0.028 | 0 | 0.25 | 0.29 | 0.32 | 0.44 | 0.18 | 0.26 |
| 2 | 69042093 | rs1316845 | 0.212 | -0.032 | 0.359 | -0.267 | -0.004 | 0.09 | 0.09 | 0.411 | 0.57 | 0.63 | 0.64 | 0.85 | 0.52 | 0.42 |
| 2 | 69044629 | rs11897459 | 1.831 | -0.169 | 0.11 | -0.066 | -0.049 | -0.074 | -0.074 | 0 | 0.19 | 0.02 | 0.28 | 0.26 | 0.15 | 0.14 |
| 2 | 69045158 | rs10197806 | 1.851 | -0.087 | 0.285 | -0.113 | -0.006 | -0.018 | -0.018 | 0 | 0.29 | 0.23 | 0.32 | 0.41 | 0.33 | 0.17 |
| 2 | 69045773 | rs12713666 | 0.353 | -0.168 | 0.371 | -0.205 | 0 | -0.001 | -0.001 | 0 | 0.61 | 0.42 | 0.6 | 0.49 | 0.72 | 0.61 |
| 2 | 69045953 | rs7557267 | -1.992 | -0.215 | 0.434 | -0.168 | 0.008 | -0.023 | -0.023 | 0 | 0.93 | 0.78 | 0.96 | 0.99 | 0.95 | 0.93 |
| 2 | 69046164 | rs17545022 | 1.058 | -0.11 | 0.03 | -0.323 | -0.023 | 0.074 | 0.074 | 0 | 0.16 | 0.06 | 0.11 | 0.01 | 0.18 | 0.18 |
| 2 | 69049941 | rs10177248 | 1.452 | -0.297 | 0.395 | 0.044 | 0.007 | 0.039 | 0.039 | 0 | 0.31 | 0.35 | 0.32 | 0.41 | 0.34 | 0.17 |
| 2 | 69051320 | rs7605681 | 0.155 | -0.226 | 0.345 | -0.2 | -0.001 | -0.011 | -0.011 | 0.022 | 0.55 | 0.54 | 0.62 | 0.84 | 0.5 | 0.42 |
| 2 | 69052991 | rs2311421 | 0.254 | -0.3 | 0.362 | -0.139 | -0.001 | -0.008 | -0.008 | 0 | 0.53 | 0.45 | 0.58 | 0.83 | 0.45 | 0.45 |
| 2 | 69053557 | rs9902 | -0.83 | -0.316 | 0.05 | -0.241 | -0.032 | 0.059 | 0.059 | 1.969 | 0.66 | 0.77 | 0.68 | 0.92 | 0.54 | 0.58 |
| 2 | 69054755 | rs3892168 | -0.734 | -0.317 | 0.093 | -0.228 | 0.003 | -0.049 | -0.049 | 0 | 0.65 | 0.66 | 0.67 | 0.92 | 0.54 | 0.58 |
| 2 | 69063909 | rs10496166 | 0.293 | -0.54 | -0.525 | 0.043 | -0.034 | 0.052 | 0.052 | 0.368 | 0.27 | 0.47 | 0.31 | 0.44 | 0.16 | 0.27 |
| 2 | 69064232 | rs8179547 | 0.874 | -0.395 | -0.71 | 0.114 | -0.003 | 0.01 | 0.01 | 0.078 | 0.35 | 0.33 | 0.32 | 0.41 | 0.34 | 0.25 |
| 2 | 69064431 | rs2311422 | 1.857 | -0.405 | -2.057 | 0.246 | -0.026 | -0.063 | -0.063 | 0 | 0.17 | 0.02 | 0.26 | 0.25 | 0.12 | 0.14 |

| | | | | | | | | | | | | | | | | |
|---|---|---|---|---|---|---|---|---|---|---|---|---|---|---|---|---|
| 2 | 69065316 | rs10182268 | -0.323 | -0.113 | -1.869 | 0.328 | 0.059 | -0.002 | -0.002 | 0 | 0.37 | 0.86 | 0.39 | 0.35 | 0.21 | 0.27 |
| 2 | 69066077 | rs12052847 | 1.325 | 0.332 | -1.546 | 0.651 | -0.08 | -0.156 | -0.156 | 0 | 0.25 | 0.02 | 0.28 | 0.39 | 0.3 | 0.18 |
| 2 | 69069905 | rs4854525 | 1.267 | 0.34 | -1.53 | 0.653 | -0.066 | -0.143 | -0.143 | 0.843 | 0.26 | 0.04 | 0.29 | 0.39 | 0.3 | 0.18 |
| 2 | 69072331 | rs7591722 | -0.135 | -0.07 | -0.895 | 0.295 | -0.013 | -0.014 | -0.014 | 0 | 0.33 | 0.48 | 0.3 | 0.26 | 0.21 | 0.2 |
| 2 | 69072557 | rs4854526 | 0.442 | -0.462 | -1.625 | 0.381 | -0.025 | 0.026 | 0.026 | 0 | 0.44 | 0.17 | 0.48 | 0.58 | 0.51 | 0.38 |
| 2 | 69072802 | rs4854527 | 0.788 | -0.33 | -1.966 | 0.261 | 0.026 | -0.005 | -0.005 | 3.642 | 0.42 | 0.21 | 0.39 | 0.37 | 0.38 | 0.3 |
| 2 | 69072857 | rs10182291 | -0.859 | -0.457 | -1.594 | 0.414 | 0.03 | -0.054 | -0.054 | 2.93 | 0.3 | 0.85 | 0.37 | 0.46 | 0.19 | 0.32 |
| 2 | 69073928 | rs11903756 | -0.49 | -0.403 | -1.033 | -0.209 | 0.012 | 0.01 | 0.01 | 0.731 | 0.24 | 0.22 | 0.11 | 0.04 | 0.15 | 0.19 |
| 2 | 69075016 | rs4422194 | -0.228 | 0.221 | -0.557 | 0.078 | 0 | 0.002 | 0.002 | 0 | 0.19 | 0.18 | 0.18 | 0.2 | 0.19 | 0.19 |
| 2 | 69075164 | rs2311424 | 0.615 | 0.174 | -0.279 | 0.168 | 0 | 0.001 | 0.001 | 0 | 0.56 | 0.44 | 0.58 | 0.34 | 0.46 | 0.46 |
| 2 | 69077287 | rs10496169 | -0.177 | 0.324 | 0.024 | -0.049 | 0.018 | -0.01 | -0.01 | 2.65 | 0.72 | 0.42 | 0.73 | 0.76 | 0.65 | 0.7 |
| 2 | 69081746 | rs10171076 | -0.856 | 0.375 | 0.039 | 0.087 | -0.017 | 0.038 | 0.038 | 0 | 0.72 | 0.8 | 0.82 | 0.87 | 0.67 | 0.7 |
| 2 | 69091772 | rs13401986 | 0.229 | -0.016 | 0.2 | 0.209 | -0.012 | -0.017 | -0.017 | 0 | 0.08 | 0.21 | 0.06 | 0.03 | 0.06 | 0.07 |
| 2 | 69095152 | rs2312078 | -0.45 | 0.368 | -0.479 | 0.169 | 0.029 | -0.028 | -0.028 | 0.02 | 0.69 | 0.51 | 0.76 | 0.76 | 0.64 | 0.64 |
| 2 | 69097862 | rs3817368 | -0.83 | 0.433 | -0.615 | -0.016 | 0 | 0 | 0 | 0 | 0.72 | 0.72 | 0.77 | 0.76 | 0.65 | 0.68 |
| 2 | 69098972 | rs3762613 | -0.825 | 0.436 | -0.648 | -0.014 | -0.002 | 0.001 | 0.001 | 0 | 0.72 | 0.72 | 0.77 | 0.76 | 0.65 | 0.69 |
| 2 | 69107710 | rs12987243 | 0.692 | 0.349 | -2.108 | 0.178 | -0.016 | -0.058 | -0.058 | 0 | 0.17 | 0.1 | 0.15 | 0.45 | 0.19 | 0.18 |
| 2 | 69110610 | rs4854528 | -1.159 | 0.35 | -2.336 | -0.106 | -0.003 | 0.007 | 0.007 | 0 | 0.72 | 0.78 | 0.85 | 0.88 | 0.71 | 0.67 |
| 2 | 69111794 | rs2312083 | 1.772 | 0.327 | -3.093 | -0.039 | -0.091 | -0.107 | -0.107 | 0.118 | 0.26 | 0.02 | 0.45 | 0.28 | 0.31 | 0.23 |
| 2 | 69115331 | rs17036196 | 0.348 | -0.211 | -2.485 | -0.874 | -0.003 | -0.026 | -0.026 | 0 | 0.12 | 0.13 | 0.08 | 0 | 0.07 | 0.12 |
| 2 | 69125605 | rs12611735 | 0.651 | 0.268 | -2.811 | -0.685 | -0.116 | 0.058 | 0.058 | 0 | 0.41 | 0.09 | 0.56 | 0.72 | 0.47 | 0.35 |
| 2 | 69126125 | rs10192176 | -0.487 | 0.779 | -2.01 | -0.156 | -0.02 | 0.033 | 0.033 | 0 | 0.75 | 0.85 | 0.85 | 0.87 | 0.71 | 0.69 |
| 2 | 69135362 | rs13021388 | -0.646 | 0.233 | -1.693 | 0.296 | 0.024 | 0.029 | 0.029 | 0.886 | 0.23 | 0.11 | 0.18 | 0.13 | 0.18 | 0.16 |
| 2 | 69139026 | rs10191936 | -0.443 | 0.465 | -1.037 | 0.578 | -0.014 | -0.004 | -0.004 | 0 | 0.3 | 0.55 | 0.3 | 0.32 | 0.23 | 0.19 |
| 2 | 69139137 | rs12999343 | 0.033 | 0.854 | -1.421 | 0.562 | -0.002 | 0.022 | 0.022 | 0 | 0.41 | 0.26 | 0.53 | 0.36 | 0.42 | 0.23 |
| 2 | 69139834 | rs4419260 | -0.621 | 1.07 | -1.509 | 0.805 | -0.041 | -0.022 | -0.022 | 1.681 | 0.11 | 0.02 | 0.1 | 0.18 | 0.17 | 0.08 |
| 2 | 69142158 | rs4629190 | -0.375 | 0.68 | -1.337 | 0.817 | 0.006 | 0.03 | 0.03 | 3.25 | 0.54 | 0.68 | 0.5 | 0.58 | 0.52 | 0.37 |
| 2 | 69142254 | rs2312070 | -0.143 | 0.569 | -1.12 | 1.164 | -0.028 | -0.005 | -0.005 | 0 | 0.28 | 0.57 | 0.2 | 0.27 | 0.24 | 0.18 |
| 2 | 69146026 | rs4450639 | 0.296 | 0.207 | -0.646 | 1.124 | -0.057 | -0.041 | -0.041 | 0 | 0.21 | 0.58 | 0.13 | 0.12 | 0.11 | 0.1 |
| 2 | 69147566 | rs7603541 | -0.556 | 0.158 | -0.781 | 1.169 | -0.001 | -0.011 | -0.011 | 1.672 | 0.45 | 0.66 | 0.44 | 0.45 | 0.4 | 0.28 |
| 2 | 69147960 | rs10169464 | -0.55 | 0.153 | -0.784 | 1.152 | -0.001 | 0.025 | 0.025 | 0.093 | 0.45 | 0.65 | 0.43 | 0.33 | 0.38 | 0.2 |
| 2 | 69148310 | rs10183139 | -0.381 | -0.272 | -0.476 | 1.266 | 0 | 0.015 | 0.015 | 0 | 0.93 | 0.93 | 0.93 | 0.76 | 0.92 | 0.76 |
| 2 | 69150125 | rs10166239 | 0.239 | -0.442 | 0.281 | 0.964 | -0.051 | -0.111 | -0.111 | 0 | 0.21 | 0.6 | 0.12 | 0 | 0.09 | 0.02 |
| 2 | 69151257 | rs6714171 | -0.401 | -0.335 | 0.18 | 1.303 | -0.005 | -0.002 | -0.002 | 0.045 | 0.25 | 0.66 | 0.32 | 0.18 | 0.15 | 0.1 |
| 2 | 69152944 | rs10197208 | -0.348 | -0.32 | 0.18 | 1.36 | -0.001 | -0.001 | -0.001 | 0 | 0.25 | 0.66 | 0.33 | 0.36 | 0.16 | 0.17 |
| 2 | 69154148 | rs11126204 | 0.065 | -0.711 | -0.046 | 1.001 | -0.011 | -0.051 | -0.051 | 0 | 0.9 | 0.98 | 0.92 | 0.75 | 0.91 | 0.73 |
| 2 | 69154483 | rs6546449 | 0.607 | -0.806 | 0.929 | 0.966 | -0.029 | 0.007 | 0.007 | 0.184 | 0.72 | 0.62 | 0.81 | 0.62 | 0.81 | 0.64 |
| 2 | 69155326 | rs4337503 | 0.381 | -0.865 | 1.289 | 0.935 | 0.032 | -0.021 | -0.021 | 0 | 0.74 | 0.85 | 0.82 | 0.62 | 0.82 | 0.64 |
| 2 | 69158120 | rs6705747 | 0.778 | -0.98 | 1.439 | 0.84 | 0.009 | 0 | 0 | 0 | 0.69 | 0.72 | 0.81 | 0.62 | 0.79 | 0.62 |
| 2 | 69170857 | rs12620517 | 0.553 | -0.99 | 1.085 | 0.796 | -0.004 | -0.044 | -0.044 | 0 | 0.88 | 0.98 | 0.91 | 0.75 | 0.89 | 0.72 |
| 2 | 69171896 | rs11126206 | 0.012 | -0.987 | 1.079 | 0.826 | -0.013 | -0.056 | -0.056 | 0.486 | 0.9 | 1 | 0.92 | 0.75 | 0.91 | 0.73 |
| 2 | 69173570 | rs1128272 | -0.63 | -1.031 | 1.421 | 0.839 | -0.093 | 0.016 | 0.016 | 0.995 | 0.27 | 0.5 | 0.16 | 0.38 | 0.12 | 0.31 |
| 2 | 69193133 | rs6546454 | 0.508 | -0.013 | 0.604 | 0.844 | -0.005 | 0.007 | 0.007 | 0 | 0.55 | 0.28 | 0.55 | 0.38 | 0.6 | 0.48 |
| 2 | 69198388 | rs4254535 | 0.422 | 0.08 | 1.065 | 0.942 | 0.023 | 0.011 | 0.011 | 0 | 0.27 | 0.39 | 0.25 | 0.26 | 0.27 | 0.18 |
| 2 | 69198766 | rs7422789 | 1.081 | -0.25 | 1.404 | 1.083 | -0.024 | 0.044 | 0.044 | 0 | 0.81 | 0.64 | 0.79 | 0.64 | 0.86 | 0.66 |
| 2 | 69201894 | rs4417751 | 0.426 | -0.055 | 0.466 | 0.903 | 0.009 | -0.015 | -0.015 | 0 | 0.73 | 0.7 | 0.64 | 0.72 | 0.61 | 0.61 |
| 2 | 69236358 | rs6743488 | -1.84 | -1.29 | 1.549 | 1.943 | -0.068 | 0.207 | 0.207 | 0.6 | 0.16 | 0.57 | 0.25 | 0.52 | 0.07 | 0.24 |
| 2 | 69236515 | rs6743513 | -1.921 | -1.301 | 1.489 | 1.938 | -0.067 | 0.199 | 0.199 | 0.487 | 0.16 | 0.58 | 0.25 | 0.52 | 0.07 | 0.24 |
| 2 | 69263120 | rs4318420 | -2.254 | -1.286 | 1.967 | 2.312 | -0.037 | 0.309 | 0.309 | 1.779 | 0.11 | 0.61 | 0.16 | 0.4 | 0.05 | 0.11 |
| 2 | 69266291 | rs6743357 | -1.484 | -1.311 | 2.179 | 2.403 | -0.027 | 0.24 | 0.24 | 0 | 0.07 | 0.19 | 0.21 | 0.49 | 0.04 | 0.17 |
| 2 | 69267134 | rs4366940 | 0.085 | -1.449 | 1.645 | 2.237 | 0 | 0 | 0 | 0.047 | 0.06 | 0.08 | 0.15 | 0.04 | 0.04 | 0.02 |
| 2 | 69269809 | rs4314026 | -1.225 | -2.04 | 1.315 | 1.846 | 0.016 | -0.014 | -0.014 | 0 | 0.21 | 0.8 | 0.16 | 0.25 | 0.06 | 0.08 |
| 2 | 69270926 | rs11887395 | -1.658 | -1.855 | 1.137 | 1.438 | -0.052 | 0.005 | 0.005 | 0.006 | 0.3 | 0.59 | 0.45 | 0.7 | 0.14 | 0.31 |
| 2 | 69296510 | rs6723425 | -1.352 | -1.912 | 0.012 | 1.414 | 0.034 | -0.027 | -0.027 | 1.256 | 0.27 | 0.79 | 0.22 | 0.38 | 0.14 | 0.22 |
| 2 | 69297283 | rs6727307 | -0.127 | -0.978 | -0.473 | 1.226 | -0.019 | -0.025 | -0.025 | 0 | 0.64 | 0.73 | 0.66 | 0.52 | 0.55 | 0.47 |
| 2 | 69301308 | rs13427290 | -0.265 | -0.736 | -0.629 | 1.293 | -0.001 | 0.034 | 0.034 | 0.015 | 0.1 | 0.17 | 0.08 | 0.23 | 0.1 | 0.15 |
| 2 | 69305747 | rs4854538 | 0.542 | -0.674 | 0.116 | 1.329 | -0.002 | 0.001 | 0.001 | 1.267 | 0.61 | 0.41 | 0.55 | 0.31 | 0.53 | 0.44 |
| 2 | 69312969 | rs11686284 | -0.553 | -0.584 | -0.125 | 0.904 | -0.024 | 0.01 | 0.01 | 0 | 0.75 | 0.87 | 0.83 | 0.78 | 0.67 | 0.66 |
| 2 | 69313884 | rs7591556 | 0.292 | -1.011 | -0.074 | 0.556 | 0.005 | -0.008 | -0.008 | 0 | 0.13 | 0.11 | 0.05 | 0.01 | 0.06 | 0.08 |
| 2 | 69314667 | rs10167468 | -0.297 | -0.655 | 0.269 | 0.864 | 0.007 | 0.045 | 0.045 | 0.009 | 0.11 | 0.18 | 0.11 | 0.25 | 0.12 | 0.17 |
| 2 | 69317925 | rs4854546 | -1.096 | -0.381 | -0.047 | 0.69 | -0.021 | 0.022 | 0.022 | 0 | 0.94 | 0.98 | 0.95 | 0.98 | 0.9 | 0.89 |
| 2 | 69321709 | rs4527239 | 0.834 | -0.616 | 0.294 | 0.723 | -0.005 | -0.004 | -0.004 | 0 | 0.58 | 0.39 | 0.69 | 0.51 | 0.5 | 0.39 |
| 2 | 69328218 | rs1106518 | 0.788 | -0.544 | -0.18 | 0.758 | -0.017 | -0.003 | -0.003 | 0.005 | 0.2 | 0.1 | 0.19 | 0.18 | 0.25 | 0.25 |
| 2 | 69328883 | rs4435491 | 0.527 | -0.502 | 0.053 | 0.815 | 0.011 | 0.049 | 0.049 | 0 | 0.85 | 0.71 | 0.89 | 0.73 | 0.84 | 0.73 |
| 2 | 69329772 | rs4542888 | 1.068 | -0.536 | -0.448 | 0.75 | 0.011 | 0.06 | 0.06 | 0.117 | 0.5 | 0.32 | 0.57 | 0.37 | 0.47 | 0.4 |
| 2 | 69330118 | rs6710260 | -0.264 | -0.257 | -0.978 | 0.615 | 0.044 | 0.028 | 0.028 | 1.904 | 0.31 | 0.83 | 0.26 | 0.22 | 0.17 | 0.14 |
| 2 | 69330284 | rs4241349 | 0.279 | -0.283 | -0.981 | 0.607 | 0.023 | 0.036 | 0.036 | 1.045 | 0.64 | 0.22 | 0.7 | 0.75 | 0.76 | 0.75 |
| 2 | 69330445 | rs10179227 | -0.15 | -0.397 | -0.9 | 0.647 | 0.029 | 0.015 | 0.015 | 2.31 | 0.29 | 0.76 | 0.25 | 0.22 | 0.13 | 0.14 |
| 2 | 69334220 | rs11689690 | 0.455 | 0.523 | -1.457 | 0.704 | -0.086 | -0.033 | -0.033 | 0 | 0.27 | 0.03 | 0.24 | 0.22 | 0.33 | 0.27 |

| | | | | | | | | | | | | | | | | |
|---|---|---|---|---|---|---|---|---|---|---|---|---|---|---|---|---|
| 2 | 69342533 | rs4854547 | -0.182 | 0.761 | -0.496 | 0.875 | -0.001 | -0.001 | -0.001 | 0 | 0.33 | 0.33 | 0.38 | 0.57 | 0.37 | 0.47 |
| 2 | 69342970 | rs6710147 | -0.042 | 1.304 | -0.506 | 0.893 | -0.008 | -0.02 | -0.02 | 0 | 0.15 | 0.09 | 0.25 | 0.3 | 0.19 | 0.19 |
| 2 | 69343149 | rs7564112 | -1.049 | 1.367 | 0.287 | 0.412 | 0.081 | -0.018 | -0.018 | 0 | 0.11 | 0.37 | 0.06 | 0.06 | 0.11 | 0.12 |
| 2 | 69346155 | rs6760880 | -0.016 | 1.215 | 0.058 | 0.243 | 0.016 | -0.049 | -0.049 | 0 | 0.13 | 0.18 | 0.2 | 0.02 | 0.15 | 0.1 |
| 2 | 69350431 | rs7584948 | -1.146 | 0.604 | 0.83 | 0.01 | -0.008 | -0.084 | -0.084 | 0 | 0.24 | 0.56 | 0.28 | 0.11 | 0.22 | 0.14 |
| 2 | 69354700 | rs4358156 | -0.742 | 0.424 | 0.129 | -0.524 | -0.066 | 0.1 | 0.1 | 0 | 0.65 | 0.84 | 0.67 | 0.9 | 0.57 | 0.56 |
| 2 | 69363033 | rs12105936 | 0.506 | 0.596 | 0.501 | -0.098 | 0.007 | -0.037 | -0.037 | 0 | 0.13 | 0.14 | 0.2 | 0.02 | 0.15 | 0.14 |
| 2 | 69368247 | rs11673753 | -0.118 | 0.367 | 0.602 | 0.754 | -0.006 | 0.127 | 0.127 | 0 | 0.06 | 0.11 | 0.09 | 0.3 | 0.05 | 0.06 |
| 2 | 69370771 | rs3890787 | -1.333 | 0.33 | 0.794 | 0.753 | -0.012 | 0.034 | 0.034 | 0 | 0.3 | 0.47 | 0.37 | 0.41 | 0.28 | 0.26 |
| 2 | 69370976 | rs4488695 | -0.429 | 0.64 | 0.687 | 0.935 | 0.005 | 0.046 | 0.046 | 0 | 0.2 | 0.27 | 0.29 | 0.32 | 0.2 | 0.22 |
| 2 | 69377069 | rs9989781 | -1.172 | 0.591 | 1.119 | 1.58 | 0.045 | 0.449 | 0.449 | 0 | 0.08 | 0.43 | 0.11 | 0.32 | 0.04 | 0.08 |
| 2 | 69377605 | rs6708226 | -0.223 | 0.652 | 0.35 | 1.09 | -0.025 | 0.015 | 0.015 | 0 | 0.66 | 0.73 | 0.47 | 0.76 | 0.53 | 0.73 |
| 2 | 69380879 | rs9808255 | -1.007 | 0.957 | 0.875 | 1.409 | 0.024 | 0.192 | 0.192 | 0 | 0.17 | 0.45 | 0.21 | 0.54 | 0.12 | 0.15 |
| 2 | 69384107 | rs12105268 | -0.349 | 0.896 | 0.684 | 0.813 | 0 | -0.012 | -0.012 | 0 | 0.64 | 0.69 | 0.53 | 0.89 | 0.48 | 0.71 |
| 2 | 69391521 | rs13016276 | 0.971 | 0.606 | 0.503 | 0.808 | 0.104 | 0.153 | 0.153 | 0 | 0.46 | 0.16 | 0.33 | 0.3 | 0.37 | 0.49 |
| 2 | 69392619 | rs11126220 | -0.465 | 0.602 | 0.632 | 0.767 | -0.086 | 0.125 | 0.125 | 0 | 0.66 | 0.91 | 0.55 | 0.89 | 0.48 | 0.72 |
| 2 | 69393713 | rs12620093 | -0.801 | 0.525 | 1.038 | 1.127 | 0.008 | 0.232 | 0.232 | 0 | 0.17 | 0.43 | 0.17 | 0.47 | 0.1 | 0.22 |
| 2 | 69394848 | rs7565538 | -0.214 | 0.513 | 1.053 | 0.937 | -0.006 | 0.034 | 0.034 | 0 | 0.62 | 0.75 | 0.49 | 0.77 | 0.42 | 0.71 |
| 2 | 69396387 | rs10188804 | -0.218 | 0.689 | 0.889 | 0.878 | -0.045 | 0.146 | 0.146 | 0 | 0.63 | 0.88 | 0.52 | 0.89 | 0.43 | 0.71 |
| 2 | 69398263 | rs4299364 | 1.28 | 0.53 | 0.789 | 0.92 | 0.096 | 0.106 | 0.106 | 0 | 0.42 | 0.09 | 0.29 | 0.31 | 0.32 | 0.43 |
| 2 | 69402025 | rs6546484 | -0.781 | 1.047 | 0.656 | 0.677 | -0.132 | 0.08 | 0.08 | 0 | 0.74 | 0.98 | 0.59 | 0.92 | 0.6 | 0.84 |
| 2 | 69408045 | rs11126223 | 0.075 | 1.066 | 0.98 | 0.799 | 0.01 | -0.031 | -0.031 | 2.836 | 0.6 | 0.55 | 0.48 | 0.77 | 0.46 | 0.7 |
| 2 | 69410325 | rs10182789 | -1.157 | 0.616 | 0.921 | 0.766 | 0.035 | 0.364 | 0.364 | 0 | 0.13 | 0.38 | 0.18 | 0.44 | 0.11 | 0.26 |
| 2 | 69411517 | rs6732795 | 0.22 | 0.67 | 0.007 | 0.083 | 0.039 | 0.102 | 0.102 | 0.045 | 0.56 | 0.78 | 0.49 | 0.85 | 0.4 | 0.68 |
| 2 | 69422629 | rs7591558 | -0.284 | 0.226 | -0.896 | -0.214 | -0.082 | 0.147 | 0.147 | 0 | 0.67 | 0.95 | 0.58 | 0.92 | 0.61 | 0.73 |
| 2 | 69431269 | rs7564412 | -0.257 | -0.135 | -0.367 | -0.596 | 0.014 | 0.075 | 0.075 | 0 | 0.5 | 0.64 | 0.54 | 0.89 | 0.55 | 0.61 |
| 2 | 69435243 | rs11126226 | 1.328 | -0.092 | -0.771 | -0.747 | -0.006 | 0.004 | 0.004 | 0 | 0.15 | 0.11 | 0.09 | 0.02 | 0.17 | 0.13 |
| 2 | 69437291 | rs4464317 | -0.015 | -0.171 | -0.61 | -0.339 | -0.05 | 0.049 | 0.049 | 0 | 0.32 | 0.11 | 0.44 | 0.77 | 0.41 | 0.48 |
| 2 | 69438988 | rs13035541 | -0.154 | -0.2 | -0.234 | -0.339 | -0.001 | 0 | 0 | 0 | 0.34 | 0.31 | 0.44 | 0.77 | 0.39 | 0.48 |
| 2 | 69443231 | rs4854553 | -0.017 | -0.084 | -0.313 | 0.055 | -0.026 | -0.031 | -0.031 | 0 | 0.3 | 0.13 | 0.45 | 0.67 | 0.36 | 0.47 |
| 2 | 69444581 | rs11680427 | -0.151 | -0.086 | -0.314 | 0.053 | -0.048 | -0.027 | -0.027 | 0 | 0.31 | 0.13 | 0.46 | 0.67 | 0.4 | 0.48 |
| 2 | 69446038 | rs12997939 | 0.339 | -0.096 | -0.514 | 0.91 | -0.072 | -0.066 | -0.066 | 0 | 0.27 | 0.06 | 0.24 | 0.29 | 0.34 | 0.36 |
| 2 | 69451757 | rs6751798 | -0.327 | -0.008 | -0.098 | 0.145 | -0.001 | 0.004 | 0.004 | 0 | 0.42 | 0.38 | 0.52 | 0.7 | 0.44 | 0.53 |
| 2 | 69487977 | rs7558134 | -1.573 | -0.097 | 1.092 | 0.093 | -0.011 | -0.009 | -0.009 | 0 | 0.07 | 0.64 | 0.04 | 0.02 | 0 | 0.09 |
| 2 | 69489816 | rs6546493 | -1.593 | 0.227 | 0.041 | -0.132 | -0.095 | 0.004 | 0.004 | 0 | 0.73 | 0.97 | 0.68 | 0.81 | 0.67 | 0.87 |
| 2 | 69513030 | rs7585658 | -0.847 | 0.281 | -0.007 | -0.216 | -0.018 | 0.028 | 0.028 | 0 | 0.65 | 0.76 | 0.65 | 0.76 | 0.6 | 0.71 |
| 2 | 69518726 | rs6709954 | 0.233 | 0.059 | 0.026 | -0.033 | -0.015 | -0.003 | -0.003 | 0 | 0.4 | 0.52 | 0.53 | 0.63 | 0.37 | 0.42 |
| 2 | 69520581 | rs11683511 | 0.267 | -0.046 | -0.194 | 0.015 | 0.004 | 0.001 | 0.001 | 0 | 0.41 | 0.29 | 0.52 | 0.61 | 0.41 | 0.43 |
| 2 | 69520795 | rs13030314 | -0.171 | 0.064 | -0.715 | -0.266 | 0.02 | 0.066 | 0.066 | 0 | 0.46 | 0.66 | 0.57 | 0.76 | 0.42 | 0.57 |
| 2 | 69521831 | rs12713676 | -1.458 | 0.409 | -1.405 | -0.002 | -0.097 | 0.01 | 0.01 | 0 | 0.75 | 0.97 | 0.68 | 0.82 | 0.67 | 0.82 |
| 2 | 69523838 | rs11126231 | -0.503 | 0.23 | -0.567 | 0.315 | -0.001 | -0.001 | -0.001 | 0 | 0.42 | 0.47 | 0.51 | 0.62 | 0.42 | 0.5 |
| 2 | 69527166 | rs6546497 | -1.42 | 0.343 | -0.454 | 0.023 | -0.013 | 0.037 | 0.037 | 0 | 0.62 | 0.72 | 0.62 | 0.79 | 0.59 | 0.64 |
| 2 | 69528697 | rs7422275 | -0.472 | 0.423 | -0.716 | 0.436 | -0.013 | -0.116 | -0.116 | 0 | 0.38 | 0.12 | 0.46 | 0.49 | 0.37 | 0.36 |
| 2 | 69528858 | rs11687792 | 0.528 | 0.304 | -0.893 | 0.698 | 0.007 | -0.056 | -0.056 | 0 | 0.11 | 0.01 | 0.08 | 0.19 | 0.1 | 0.11 |
| 2 | 69558112 | rs12473304 | 0.788 | 0.399 | -0.484 | 0.168 | -0.009 | 0.012 | 0.012 | 0 | 0.15 | 0.09 | 0.09 | 0.04 | 0.18 | 0.13 |
| 2 | 69560024 | rs12466648 | 0.582 | 0.388 | -0.496 | 0.156 | -0.012 | 0.02 | 0.02 | 0 | 0.16 | 0.09 | 0.1 | 0.04 | 0.2 | 0.14 |
| 2 | 69565345 | rs10164798 | 1.675 | 0.321 | -0.271 | 0.103 | -0.008 | 0.03 | 0.03 | 0 | 0.39 | 0.25 | 0.38 | 0.21 | 0.41 | 0.36 |
| 2 | 69567461 | rs4625963 | -1.37 | 0.26 | -0.557 | -0.042 | -0.046 | 0.047 | 0.047 | 0 | 0.89 | 0.99 | 0.92 | 0.97 | 0.87 | 0.87 |
| 2 | 69569930 | rs4416261 | 1.106 | 0.427 | 0.341 | 0.327 | 0.001 | -0.006 | -0.006 | 3.967 | 0.44 | 0.64 | 0.4 | 0.44 | 0.41 | 0.51 |
| 2 | 69573029 | rs6722492 | 0.111 | 0.439 | 0.318 | 0.333 | -0.041 | -0.107 | -0.107 | 0.583 | 0.6 | 0.89 | 0.52 | 0.48 | 0.59 | 0.63 |
| 2 | 69578292 | rs6546503 | 0.174 | 0.442 | 0.319 | 0.334 | -0.042 | -0.108 | -0.108 | 0.135 | 0.6 | 0.89 | 0.52 | 0.48 | 0.59 | 0.63 |
| 2 | 69606336 | rs13422656 | 1.591 | 0.446 | 0.298 | 0.326 | -0.005 | 0.028 | 0.028 | 0 | 0.4 | 0.29 | 0.38 | 0.21 | 0.41 | 0.35 |
| 2 | 69622299 | rs5017686 | 0.228 | 0.423 | 0.243 | 0.388 | -0.008 | 0.068 | 0.068 | 0 | 0.6 | 0.84 | 0.5 | 0.29 | 0.6 | 0.52 |
| 2 | 69625157 | rs10182682 | 0.074 | 0.428 | 0.118 | 0.396 | -0.07 | -0.146 | -0.146 | 0.713 | 0.61 | 0.95 | 0.53 | 0.48 | 0.6 | 0.63 |
| 2 | 69642970 | rs10175692 | 0.076 | 0.422 | 0.114 | 0.393 | -0.07 | -0.146 | -0.146 | 0.545 | 0.61 | 0.95 | 0.53 | 0.49 | 0.6 | 0.63 |
| 2 | 69646504 | rs7605572 | 0.015 | 0.449 | 0.271 | 0.394 | 0.004 | 0.026 | 0.026 | 0 | 0.19 | 0.38 | 0.13 | 0.23 | 0.18 | 0.23 |
| 2 | 69655790 | rs4581940 | 1.271 | 0.488 | 0.309 | 0.377 | 0.001 | -0.011 | -0.011 | 0 | 0.42 | 0.52 | 0.4 | 0.25 | 0.42 | 0.41 |
| 2 | 69664976 | rs13416004 | 1.517 | 0.494 | 0.029 | 0.376 | -0.007 | 0.027 | 0.027 | 0 | 0.4 | 0.28 | 0.39 | 0.22 | 0.41 | 0.38 |
| 2 | 69667987 | rs6758228 | 0.955 | 0.446 | 0.061 | 0.491 | -0.017 | -0.027 | -0.027 | 0 | 0.46 | 0.81 | 0.45 | 0.45 | 0.43 | 0.54 |
| 2 | 69686286 | rs7577689 | -0.045 | 0.435 | -0.18 | 0.425 | -0.064 | -0.146 | -0.146 | 0 | 0.63 | 0.94 | 0.55 | 0.49 | 0.62 | 0.65 |
| 2 | 69687224 | rs4441525 | 1.375 | 0.348 | -0.03 | 0.39 | -0.01 | 0.046 | 0.046 | 0 | 0.42 | 0.28 | 0.4 | 0.21 | 0.43 | 0.38 |
| 2 | 69693006 | rs9309428 | 1.289 | 0.463 | -0.027 | 0.415 | -0.006 | 0.057 | 0.057 | 0 | 0.43 | 0.26 | 0.26 | 0.22 | 0.44 | 0.39 |
| 2 | 69696001 | rs7420531 | -1.472 | 0.26 | -0.009 | 0.424 | -0.015 | 0.061 | 0.061 | 0 | 0.57 | 0.74 | 0.74 | 0.78 | 0.54 | 0.6 |
| 2 | 69698158 | rs3552 | -1.496 | 0.266 | 0.09 | 0.434 | -0.016 | 0.057 | 0.057 | 0 | 0.57 | 0.74 | 0.74 | 0.79 | 0.54 | 0.6 |
| 2 | 69702628 | rs12151791 | -0.196 | 0.336 | -0.12 | 0.484 | -0.041 | -0.177 | -0.177 | 0 | 0.34 | 0.04 | 0.44 | 0.51 | 0.35 | 0.33 |
| 2 | 69704941 | rs2312205 | -0.015 | 0.372 | 0.399 | 0.532 | 0.004 | 0.024 | 0.024 | 0 | 0.18 | 0.34 | 0.25 | 0.24 | 0.19 | 0.24 |
| 2 | 69715845 | rs2312207 | -1.533 | 0.362 | 0.033 | 0.441 | -0.019 | 0.06 | 0.06 | 0 | 0.57 | 0.73 | 0.73 | 0.79 | 0.54 | 0.62 |
| 2 | 69723710 | rs7577851 | -0.589 | 0.363 | 0.128 | 0.563 | -0.004 | 0.019 | 0.019 | 0 | 0.22 | 0.67 | 0.29 | 0.28 | 0.19 | 0.28 |
| 2 | 69741609 | rs2276689 | -0.763 | 0.683 | 0.109 | 0.753 | -0.05 | -0.137 | -0.137 | 0 | 0.39 | 0.06 | 0.45 | 0.51 | 0.39 | 0.32 |

| | | | | | | | | | | | | | | | | |
|---|---|---|---|---|---|---|---|---|---|---|---|---|---|---|---|---|
| 2 | 69744800 | rs6748361 | -1.472 | 1.139 | 0.832 | 0.752 | 0.003 | 0.085 | 0.085 | 0 | 0.46 | 0.72 | 0.67 | 0.75 | 0.42 | 0.54 |
| 2 | 69746296 | rs7603932 | -1.3 | 1.118 | 1.252 | 0.718 | 0.04 | -0.004 | -0.004 | 0 | 0.08 | 0.15 | 0.05 | 0.05 | 0.11 | 0.11 |
| 2 | 69750113 | rs7565783 | -0.671 | 1.101 | 1.068 | 0.715 | -0.005 | 0.033 | 0.033 | 0 | 0.36 | 0.1 | 0.57 | 0.7 | 0.32 | 0.46 |
| 2 | 69759415 | rs2276687 | -1.616 | 1.046 | 1.05 | 0.677 | 0.004 | 0.109 | 0.109 | 0 | 0.48 | 0.72 | 0.67 | 0.76 | 0.44 | 0.58 |
| 2 | 69788597 | rs12467072 | -0.916 | 1.308 | 1.235 | 1.282 | -0.029 | -0.041 | -0.041 | 0 | 0.36 | 0.2 | 0.45 | 0.55 | 0.39 | 0.37 |
| 2 | 69794350 | rs4852283 | -0.897 | 1.377 | 1.5 | 1.337 | -0.012 | -0.02 | -0.02 | 0.036 | 0.36 | 0.27 | 0.45 | 0.56 | 0.37 | 0.39 |
| 2 | 69796259 | rs11126237 | -1.036 | 1.315 | 1.545 | 1.411 | 0.001 | -0.014 | -0.014 | 0 | 0.36 | 0.39 | 0.59 | 0.76 | 0.35 | 0.55 |
| 2 | 69797008 | rs6546534 | -0.401 | 1.323 | 1.205 | 1.615 | 0 | -0.047 | -0.047 | 1.582 | 0.29 | 0.19 | 0.42 | 0.48 | 0.27 | 0.25 |
| 2 | 69812069 | rs6759804 | -0.98 | 1.673 | 1.824 | 1.712 | -0.031 | -0.016 | -0.016 | 0.004 | 0.37 | 0.27 | 0.45 | 0.57 | 0.39 | 0.36 |
| 2 | 69813458 | rs12987661 | -1.283 | 1.803 | 1.795 | 1.709 | 0.028 | -0.004 | -0.004 | 0 | 0.08 | 0.1 | 0.03 | 0.04 | 0.14 | 0.09 |
| 2 | 69813713 | rs10865378 | -1.839 | 1.691 | 2.2 | 1.678 | -0.007 | 0.09 | 0.09 | 0 | 0.47 | 0.74 | 0.65 | 0.76 | 0.43 | 0.66 |
| 2 | 69824151 | rs10177462 | -1.704 | 1.395 | 2.326 | 1.861 | 0.062 | 0.328 | 0.328 | 0.834 | 0.05 | 0.37 | 0.19 | 0.19 | 0.01 | 0.21 |
| 2 | 69833382 | rs6546535 | -1.07 | 1.461 | 2.095 | 1.879 | -0.012 | -0.003 | -0.003 | 1.874 | 0.37 | 0.61 | 0.61 | 0.69 | 0.26 | 0.55 |
| 2 | 69846510 | rs17036825 | -1.861 | 1.452 | 1.676 | 1.897 | 0.065 | 0.347 | 0.347 | 1.808 | 0.06 | 0.44 | 0.19 | 0.18 | 0.01 | 0.21 |
| 2 | 69864236 | rs6727435 | -0.478 | 1.481 | 0.979 | 1.843 | 0.013 | -0.083 | -0.083 | 0 | 0.31 | 0.13 | 0.42 | 0.5 | 0.25 | 0.34 |
| 2 | 69877490 | rs7567691 | -1.471 | 1.493 | 0.644 | 1.535 | 0.002 | 0.092 | 0.092 | 0 | 0.44 | 0.78 | 0.59 | 0.8 | 0.36 | 0.6 |
| 2 | 69940780 | rs35907069 | -1.658 | 2.237 | 0.974 | 1.852 | 0.084 | 0.035 | 0.035 | 0 | 0.05 | 0.16 | 0.03 | 0.05 | 0.1 | 0.04 |
| 2 | 69941287 | rs12612409 | -1.658 | 2.237 | 0.974 | 1.852 | 0.084 | 0.035 | 0.035 | 0 | 0.05 | 0.16 | 0.03 | 0.05 | 0.1 | 0.04 |
| 2 | 69955760 | rs2312548 | -1.445 | 1.541 | 0.655 | 1.706 | 0.01 | 0.056 | 0.056 | 0.108 | 0.47 | 0.72 | 0.69 | 0.73 | 0.54 | 0.58 |
| 2 | 69979290 | rs10177224 | -1.503 | 1.967 | 1.27 | 2.556 | 0.172 | 0.299 | 0.299 | 0 | 0.08 | 0.3 | 0.36 | 0.36 | 0.2 | 0.12 |
| 2 | 69980193 | rs7563401 | -1.482 | 1.803 | 1.412 | 2.403 | 0.027 | -0.011 | -0.011 | 0 | 0.37 | 0.49 | 0.62 | 0.73 | 0.47 | 0.46 |
| 2 | 69982091 | rs722265 | -0.784 | 1.593 | 0.777 | 2.203 | 0.008 | -0.001 | -0.001 | 0.259 | 0.3 | 0.18 | 0.27 | 0.38 | 0.27 | 0.34 |
| 2 | 69987384 | rs6727897 | -1.399 | 1.461 | 0.866 | 1.771 | 0.011 | -0.02 | -0.02 | 0 | 0.39 | 0.5 | 0.63 | 0.81 | 0.47 | 0.49 |
| 2 | 69995606 | rs12993700 | -0.979 | 1.202 | 1.202 | 2.519 | -0.004 | 0.008 | 0.008 | 0 | 0.28 | 0.32 | 0.49 | 0.55 | 0.24 | 0.31 |
| 2 | 70009035 | rs6546546 | -0.755 | 1.055 | 2.06 | 2.482 | -0.013 | 0.001 | 0.001 | 0 | 0.26 | 0.32 | 0.24 | 0.35 | 0.24 | 0.31 |
| 2 | 70013191 | rs4852334 | -2.908 | 1.132 | 2.16 | 2.655 | 0.016 | 0.121 | 0.121 | 0.023 | 0.07 | 0.28 | 0.28 | 0.19 | 0.02 | 0.11 |
| 2 | 70015181 | rs10496171 | -0.802 | 0.823 | 1.919 | 2.375 | 0.008 | 0.002 | 0.002 | 0 | 0.29 | 0.27 | 0.24 | 0.33 | 0.24 | 0.32 |
| 2 | 70017022 | rs6546548 | -1.689 | 1.233 | 1.733 | 2.595 | -0.01 | -0.019 | -0.019 | 0 | 0.39 | 0.68 | 0.41 | 0.48 | 0.46 | 0.42 |
| 2 | 70023469 | rs7588022 | -0.903 | 1.358 | 1.512 | 2.586 | -0.079 | 0.001 | 0.001 | 0 | 0.31 | 0.12 | 0.28 | 0.3 | 0.39 | 0.3 |
| 2 | 70028904 | rs4853025 | -1.143 | 1.336 | 1.82 | 2.619 | -0.045 | -0.001 | -0.001 | 0 | 0.32 | 0.53 | 0.51 | 0.41 | 0.24 | 0.29 |
| 2 | 70028976 | rs12988897 | -0.421 | 1.335 | 1.636 | 2.62 | 0.01 | -0.024 | -0.024 | 0 | 0.27 | 0.08 | 0.22 | 0.31 | 0.24 | 0.29 |
| 2 | 70029310 | rs3771542 | -0.914 | 1.34 | 1.668 | 2.616 | 0.036 | -0.018 | -0.018 | 0 | 0.3 | 0.23 | 0.5 | 0.41 | 0.24 | 0.29 |
| 2 | 70030827 | rs6730981 | -1.859 | 1.35 | 1.82 | 2.64 | 0.06 | -0.097 | -0.097 | 0 | 0.39 | 0.91 | 0.62 | 0.58 | 0.28 | 0.38 |
| 2 | 70033649 | rs17037076 | -2.213 | 1.257 | 2.141 | 2.554 | 0.077 | 0.078 | 0.078 | 3.322 | 0.11 | 0.63 | 0.18 | 0.17 | 0.2 | 0.1 |
| 2 | 70040144 | rs13432169 | -2.565 | 1.194 | 2.292 | 2.508 | 0.068 | 0.157 | 0.157 | 0.35 | 0.06 | 0.45 | 0.29 | 0.1 | 0 | 0 |
| 2 | 70043387 | rs10173964 | -2.565 | 1.194 | 2.293 | 2.508 | 0.067 | 0.155 | 0.155 | 0.35 | 0.06 | 0.45 | 0.29 | 0.1 | 0 | 0 |
| 2 | 70056365 | rs2278933 | -1.214 | 1.297 | 2.327 | 2.574 | 0.166 | 0.162 | 0.162 | 1.065 | 0.08 | 0.3 | 0.17 | 0.18 | 0.2 | 0.1 |
| 2 | 70059824 | rs4853036 | -1.216 | 1.23 | 2.203 | 2.588 | -0.011 | 0.006 | 0.006 | 0 | 0.34 | 0.59 | 0.51 | 0.49 | 0.26 | 0.37 |
| 2 | 70080397 | rs6711065 | -2.276 | 1.175 | 1.038 | 2.281 | -0.077 | 0.019 | 0.019 | 0.16 | 0.54 | 0.96 | 0.73 | 0.72 | 0.54 | 0.64 |
| 2 | 70084599 | rs11902295 | -0.812 | 1.387 | 1.546 | 2.612 | 0.009 | -0.015 | -0.015 | 0 | 0.29 | 0.13 | 0.22 | 0.37 | 0.26 | 0.37 |
| 2 | 70091246 | rs10865380 | -1.842 | 1.442 | 1.939 | 2.525 | -0.009 | 0.001 | 0.001 | 0 | 0.42 | 0.67 | 0.67 | 0.67 | 0.46 | 0.57 |
| 2 | 70117074 | rs2054046 | -1.441 | 1.499 | 1.94 | 2.504 | 0.159 | 0.077 | 0.077 | 0.153 | 0.08 | 0.29 | 0.12 | 0.13 | 0.2 | 0.1 |
| 2 | 70120608 | rs6738783 | -0.817 | 1.416 | 1.821 | 2.596 | 0.01 | -0.037 | -0.037 | 0 | 0.29 | 0.13 | 0.26 | 0.39 | 0.24 | 0.34 |
| 2 | 70128717 | rs7598283 | -1.822 | 1.497 | 1.43 | 2.601 | -0.047 | -0.058 | -0.058 | 0.105 | 0.41 | 0.82 | 0.4 | 0.54 | 0.44 | 0.54 |
| 2 | 70153848 | rs10496175 | -0.602 | 1.418 | 0.835 | 2.717 | -0.009 | -0.049 | -0.049 | 0.016 | 0.26 | 0.1 | 0.26 | 0.36 | 0.24 | 0.34 |
| 2 | 70157158 | rs10196546 | -1.366 | 1.491 | 0.634 | 2.669 | 0.038 | -0.036 | -0.036 | 0 | 0.35 | 0.81 | 0.32 | 0.42 | 0.25 | 0.46 |
| 2 | 70172116 | rs11686934 | -1.479 | 1.462 | 1.028 | 2.641 | 0 | 0 | 0 | 0 | 0.36 | 0.39 | 0.4 | 0.62 | 0.45 | 0.58 |
| 2 | 70172587 | rs6756513 | -1.529 | 1.62 | 1.158 | 2.765 | 0.071 | 0.048 | 0.048 | 0.078 | 0.12 | 0.25 | 0.14 | 0.29 | 0.27 | 0.35 |
| 2 | 70173629 | rs10496176 | -0.743 | 0.775 | 1.47 | 2.705 | 0.011 | -0.034 | -0.034 | 0 | 0.23 | 0.17 | 0.51 | 0.42 | 0.16 | 0.24 |
| 2 | 70174766 | rs4853177 | -0.918 | 0.751 | 1.761 | 2.704 | -0.006 | 0.018 | 0.018 | 0 | 0.24 | 0.25 | 0.51 | 0.42 | 0.16 | 0.24 |
| 2 | 70182805 | rs7583236 | -0.989 | 0.662 | 2.301 | 2.726 | -0.005 | 0.029 | 0.029 | 0.04 | 0.17 | 0.27 | 0.2 | 0.29 | 0.11 | 0.22 |
| 2 | 70187173 | rs3821276 | -1.549 | 0.594 | 2.161 | 2.723 | -0.02 | 0.047 | 0.047 | 0.617 | 0.23 | 0.66 | 0.37 | 0.38 | 0.11 | 0.34 |
| 2 | 70187538 | rs3087933 | -1.489 | 0.59 | 2.201 | 2.681 | -0.026 | 0.072 | 0.072 | 0.488 | 0.22 | 0.6 | 0.37 | 0.38 | 0.11 | 0.34 |
| 2 | 70196497 | rs13423562 | -0.876 | 0.256 | 1.885 | 2.365 | -0.037 | -0.01 | -0.01 | 0.277 | 0.21 | 0.35 | 0.13 | 0.23 | 0.07 | 0.27 |
| 2 | 70202613 | rs7558246 | -1.089 | 0.601 | 1.946 | 2.678 | 0.068 | 0.096 | 0.096 | 0 | 0.15 | 0.27 | 0.16 | 0.31 | 0.31 | 0.18 |
| 2 | 70225397 | rs6546562 | -0.612 | 0.571 | 1.984 | 2.679 | -0.034 | -0.009 | -0.009 | 0 | 0.2 | 0.3 | 0.12 | 0.24 | 0.07 | 0.27 |
| 2 | 70237360 | rs4853252 | -1.246 | 0.547 | 2.062 | 2.799 | -0.063 | 0.033 | 0.033 | 0.099 | 0.24 | 0.57 | 0.37 | 0.32 | 0.07 | 0.34 |
| 2 | 70238098 | rs12614930 | -1.246 | 0.547 | 2.062 | 2.799 | -0.063 | 0.033 | 0.033 | 0.099 | 0.24 | 0.57 | 0.37 | 0.32 | 0.07 | 0.34 |
| 2 | 70263260 | rs13016716 | -1.021 | 0.563 | 2.074 | 2.815 | 0.067 | 0.105 | 0.105 | 0 | 0.14 | 0.26 | 0.16 | 0.32 | 0.29 | 0.17 |
| 2 | 70283210 | rs4852424 | -0.808 | 0.603 | 2.074 | 2.445 | -0.035 | 0.027 | 0.027 | 0 | 0.21 | 0.3 | 0.36 | 0.31 | 0.07 | 0.34 |
| 2 | 70310878 | rs3755388 | -1.323 | 0.6 | 2.071 | 2.445 | -0.066 | 0.028 | 0.028 | 0.081 | 0.24 | 0.6 | 0.37 | 0.31 | 0.07 | 0.34 |
| 2 | 70316364 | rs3732261 | -0.676 | 0.67 | 1.784 | 2.428 | -0.032 | -0.015 | -0.015 | 0 | 0.2 | 0.3 | 0.12 | 0.22 | 0.08 | 0.27 |
| 2 | 70321659 | rs3732264 | -0.88 | 0.699 | 1.412 | 2.442 | -0.052 | -0.027 | -0.027 | 0 | 0.22 | 0.38 | 0.13 | 0.22 | 0.08 | 0.27 |
| 2 | 70338344 | rs7585767 | -1.741 | -0.021 | -0.118 | 1.834 | -0.122 | -0.088 | -0.088 | 0 | 0.61 | 0.97 | 0.71 | 0.68 | 0.48 | 0.79 |
| 2 | 70360262 | rs12992553 | -1.458 | 0.573 | 0.713 | 1.732 | -0.006 | 0.016 | 0.016 | 0 | 0.28 | 0.75 | 0.25 | 0.3 | 0.3 | 0.28 |
| 2 | 70368923 | rs6708331 | -1.023 | 0.596 | 1.101 | 1.738 | 0.174 | 0.082 | 0.082 | 0 | 0.07 | 0.32 | 0.11 | 0.11 | 0.22 | 0.08 |
| 2 | 70372029 | rs6720799 | -0.846 | 0.582 | 1.141 | 1.754 | 0.109 | 0.046 | 0.046 | 0 | 0.07 | 0.22 | 0.1 | 0.11 | 0.22 | 0.08 |
| 2 | 70372369 | rs728476 | -0.653 | 0.155 | 1.144 | 1.538 | -0.05 | -0.038 | -0.038 | 0 | 0.2 | 0.5 | 0.15 | 0.19 | 0.07 | 0.2 |
| 2 | 70373677 | rs6743318 | -0.769 | 0.154 | 1.026 | 1.649 | -0.043 | 0.023 | 0.023 | 0.084 | 0.21 | 0.56 | 0.39 | 0.29 | 0.07 | 0.28 |

| | | | | | | | | | | | | | | | | |
|---|---|---|---|---|---|---|---|---|---|---|---|---|---|---|---|---|
| 2 | 70389900 | rs11677137 | -0.796 | 0.333 | 0.763 | 1.542 | -0.037 | -0.049 | -0.049 | 0 | 0.21 | 0.45 | 0.16 | 0.19 | 0.11 | 0.21 |
| 2 | 70429052 | rs13403416 | -0.199 | 0.349 | 0.403 | 1.485 | -0.008 | 0.052 | 0.052 | 0 | 0.09 | 0.27 | 0.09 | 0.16 | 0.07 | 0.04 |
| 2 | 70439404 | rs1545357 | 0.25 | 0.367 | 0.124 | 1.494 | 0.004 | -0.035 | -0.035 | 0.733 | 0.08 | 0.01 | 0.08 | 0.16 | 0.07 | 0.04 |
| 2 | 70445020 | rs10519207 | 0.25 | 0.363 | 0.15 | 1.581 | 0.007 | -0.012 | -0.012 | 0.314 | 0.08 | 0.05 | 0.08 | 0.16 | 0.07 | 0.04 |
| 2 | 70446785 | rs2921711 | -0.344 | 0.638 | -0.119 | 1.677 | -0.041 | -0.056 | -0.056 | 0.039 | 0.21 | 0.1 | 0.21 | 0.39 | 0.31 | 0.31 |
| 2 | 70469555 | rs1004831 | 0.377 | 0.628 | -0.134 | 1.685 | 0.004 | -0.036 | -0.036 | 2.905 | 0.08 | 0.01 | 0.08 | 0.16 | 0.07 | 0.04 |
| 2 | 70481867 | rs2592183 | -0.394 | 0.643 | -0.059 | 1.652 | -0.029 | -0.04 | -0.04 | 1.568 | 0.21 | 0.14 | 0.21 | 0.38 | 0.31 | 0.31 |
| 2 | 70508379 | rs4853438 | 0.183 | 0.708 | -0.181 | 1.474 | -0.03 | -0.041 | -0.041 | 0 | 0.79 | 0.86 | 0.79 | 0.61 | 0.68 | 0.63 |
| 2 | 70509689 | rs2862185 | -0.023 | 0.247 | 0.321 | 1.536 | 0.003 | -0.035 | -0.035 | 0 | 0.08 | 0.01 | 0.08 | 0.16 | 0.07 | 0.04 |
| 2 | 70520498 | rs2272532 | -0.868 | 0.191 | 0.614 | 1.508 | -0.019 | 0.018 | 0.018 | 0 | 0.1 | 0.21 | 0.09 | 0.16 | 0.07 | 0.04 |
| 2 | 70523141 | rs7459 | -0.032 | 0.223 | 0.387 | 1.524 | 0.007 | -0.027 | -0.027 | 0 | 0.08 | 0.02 | 0.08 | 0.16 | 0.07 | 0.04 |
| 2 | 70557801 | rs17615710 | -0.559 | 0.251 | -0.373 | 1.486 | -0.008 | -0.032 | -0.032 | 0 | 0.13 | 0.01 | 0.1 | 0.17 | 0.1 | 0.07 |
| 2 | 70560939 | rs6751950 | -0.024 | 0.054 | -0.523 | 1.185 | -0.018 | 0.002 | 0.002 | 0 | 0.31 | 0.53 | 0.2 | 0.3 | 0.19 | 0.38 |
| 2 | 70562650 | rs4852161 | -1.081 | 0.123 | -1.028 | 1.258 | 0.094 | 0.072 | 0.072 | 0 | 0.48 | 0.82 | 0.57 | 0.6 | 0.3 | 0.56 |
| 2 | 70571103 | rs17654377 | -0.257 | 0.385 | -0.831 | 1.248 | -0.005 | 0.034 | 0.034 | 0.621 | 0.21 | 0.11 | 0.14 | 0.18 | 0.2 | 0.06 |
| 2 | 70572693 | rs11126264 | -1.003 | 0.357 | -0.609 | 1.356 | 0.003 | 0.001 | 0.001 | 0 | 0.5 | 0.44 | 0.27 | 0.5 | 0.43 | 0.42 |
| 2 | 70574318 | rs1871238 | -0.381 | 0.136 | -0.939 | 1.285 | 0.008 | -0.012 | -0.012 | 0 | 0.32 | 0.68 | 0.37 | 0.41 | 0.2 | 0.37 |
| 2 | 70579922 | rs1871239 | -0.653 | 0.855 | -1.524 | 1.146 | -0.068 | -0.04 | -0.04 | 0 | 0.42 | 0.16 | 0.26 | 0.45 | 0.47 | 0.36 |
| 2 | 70580161 | rs1871241 | -1.155 | 0.902 | -1.036 | 1.3 | 0.01 | 0.022 | 0.022 | 0.011 | 0.51 | 0.65 | 0.55 | 0.58 | 0.52 | 0.47 |
| 2 | 70587543 | rs7584070 | -1.413 | 1.021 | 0.548 | 1.316 | 0.033 | 0.111 | 0.111 | 0 | 0.13 | 0.55 | 0.34 | 0.22 | 0.13 | 0.2 |
| 2 | 70590129 | rs1351170 | -0.785 | 1.007 | 0.526 | 1.312 | 0.041 | 0.068 | 0.068 | 0 | 0.1 | 0.27 | 0.24 | 0.16 | 0.13 | 0.15 |
| 2 | 70597309 | rs11900489 | -1.504 | 1.575 | 0.127 | 1.21 | 0.026 | 0.011 | 0.011 | 0.202 | 0.29 | 0.54 | 0.47 | 0.38 | 0.37 | 0.27 |
| 2 | 70604385 | rs1871243 | -1.831 | 1.769 | 0.173 | 1.402 | 0.082 | 0.075 | 0.075 | 0 | 0.52 | 0.8 | 0.63 | 0.69 | 0.64 | 0.54 |
| 2 | 70606777 | rs6546594 | -1.652 | 1.787 | 0.097 | 1.376 | 0.028 | 0.015 | 0.015 | 0 | 0.49 | 0.59 | 0.54 | 0.62 | 0.64 | 0.48 |
| 2 | 70607612 | rs2169842 | 0.96 | 1.883 | -0.002 | 1.393 | -0.006 | -0.012 | -0.012 | 1.068 | 0.21 | 0.19 | 0.14 | 0.3 | 0.25 | 0.26 |
| 2 | 70610874 | rs11681491 | 0.351 | 1.943 | 0.056 | 1.347 | -0.055 | 0.026 | 0.026 | 0.662 | 0.16 | 0.05 | 0.14 | 0.16 | 0.24 | 0.07 |
| 2 | 70615184 | rs10496179 | -0.819 | 1.721 | 0.629 | 1.725 | 0.038 | 0.064 | 0.064 | 0 | 0.09 | 0.24 | 0.24 | 0.16 | 0.12 | 0.15 |
| 2 | 70616143 | rs10182620 | 0.469 | 1.732 | 0.406 | 1.819 | -0.001 | 0.001 | 0.001 | 0 | 0.17 | 0.17 | 0.13 | 0.23 | 0.21 | 0.21 |
| 2 | 70625197 | rs1382454 | -0.812 | 1.323 | 0.759 | 1.602 | 0 | 0.09 | 0.09 | 0 | 0.06 | 0.24 | 0.09 | 0.09 | 0.01 | 0.08 |
| 2 | 70628925 | rs2169835 | -1.837 | 1.543 | 0.752 | 1.561 | 0.136 | 0.144 | 0.144 | 0 | 0.14 | 0.5 | 0.29 | 0.32 | 0.25 | 0.21 |
| 2 | 70639877 | rs1904212 | -1.393 | 1.034 | 1.068 | 1.474 | 0.068 | 0.106 | 0.106 | 0.002 | 0.12 | 0.46 | 0.25 | 0.17 | 0.13 | 0.14 |
| 2 | 70647902 | rs4852178 | -1.897 | 1.008 | 0.814 | 1.456 | -0.011 | 0.021 | 0.021 | 0.002 | 0.19 | 0.64 | 0.34 | 0.25 | 0.17 | 0.23 |
| 2 | 70649775 | rs12996793 | -1.793 | 1.024 | 0.873 | 1.46 | -0.012 | 0.044 | 0.044 | 0 | 0.19 | 0.56 | 0.34 | 0.25 | 0.17 | 0.23 |
| 2 | 70659116 | rs494537 | -1.418 | 0.984 | 0.379 | 1.092 | 0.033 | 0.068 | 0.068 | 0 | 0.53 | 0.7 | 0.65 | 0.67 | 0.61 | 0.6 |
| 2 | 70668585 | rs472384 | -1.462 | 0.925 | 0.347 | 1.141 | 0.003 | 0.007 | 0.007 | 0 | 0.93 | 0.9 | 0.89 | 0.94 | 0.94 | 0.97 |
| 2 | 70670283 | rs11890652 | -0.005 | 0.917 | 0.476 | 1.071 | -0.009 | -0.016 | -0.016 | 0 | 0.12 | 0.11 | 0.04 | 0.03 | 0.05 | 0.13 |
| 2 | 70670873 | rs9284774 | 0.127 | 1.014 | 0.232 | 1.196 | 0 | 0.029 | 0.029 | 0 | 0.12 | 0.19 | 0.32 | 0.18 | 0.12 | 0.17 |
| 2 | 70671184 | rs10865381 | 0.875 | 1.165 | -0.242 | 1.271 | -0.037 | -0.046 | -0.046 | 0 | 0.19 | 0.12 | 0.14 | 0.28 | 0.26 | 0.23 |
| 2 | 70676098 | rs3732253 | 0.85 | 0.876 | -0.275 | 1.219 | -0.047 | -0.057 | -0.057 | 0.279 | 0.19 | 0.1 | 0.13 | 0.28 | 0.26 | 0.23 |
| 2 | 70676639 | rs538118 | 1.456 | 0.735 | -0.28 | 1.114 | 0.023 | 0.079 | 0.079 | 0 | 0.45 | 0.17 | 0.34 | 0.33 | 0.39 | 0.39 |
| 2 | 70677994 | rs2166975 | 0.567 | 0.785 | 0.003 | 1.537 | -0.029 | -0.037 | -0.037 | 0 | 0.2 | 0.13 | 0.13 | 0.29 | 0.26 | 0.23 |
| 2 | 70684451 | rs930655 | -1.526 | 0.653 | -0.163 | 1.258 | -0.009 | 0.04 | 0.04 | 0 | 0.5 | 0.9 | 0.57 | 0.6 | 0.55 | 0.59 |
| 2 | 70687052 | rs1880039 | -0.139 | 0.69 | 0.32 | 1.38 | -0.003 | -0.003 | -0.003 | 0.555 | 0.13 | 0.2 | 0.31 | 0.08 | 0.12 | 0.17 |
| 2 | 70692557 | rs3732247 | 0.819 | 0.687 | 0.486 | 1.437 | -0.067 | -0.107 | -0.107 | 0 | 0.19 | 0.06 | 0.14 | 0.38 | 0.26 | 0.22 |
| 2 | 70695562 | rs3771515 | -0.078 | 0.325 | 0.814 | 1.392 | -0.001 | -0.001 | -0.001 | 0 | 0.12 | 0.12 | 0.31 | 0.08 | 0.12 | 0.17 |
| 2 | 70696953 | rs17005644 | -0.825 | 0.272 | 0.89 | 1.417 | 0.002 | 0.019 | 0.019 | 0 | 0.16 | 0.32 | 0.16 | 0.21 | 0.16 | 0.12 |
| 2 | 70697667 | rs3771514 | -0.939 | 0.295 | 1.214 | 1.343 | 0.025 | 0.015 | 0.015 | 0 | 0.17 | 0.37 | 0.33 | 0.17 | 0.23 | 0.21 |
| 2 | 70698531 | rs4852608 | -0.653 | 0.341 | 0.908 | 1.206 | 0.014 | 0.008 | 0.008 | 0 | 0.39 | 0.53 | 0.54 | 0.53 | 0.44 | 0.39 |
| 2 | 70701076 | rs3821264 | -1.347 | 0.264 | 0.847 | 1.136 | 0.002 | 0.02 | 0.02 | 0 | 0.53 | 0.68 | 0.59 | 0.6 | 0.49 | 0.51 |
| 2 | 70705402 | rs17005666 | -0.018 | 0.203 | 0.959 | 1.121 | -0.01 | -0.018 | -0.018 | 0.039 | 0.13 | 0.13 | 0.05 | 0.03 | 0.06 | 0.09 |
| 2 | 70708261 | rs1807968 | -1.196 | 0.165 | 1.143 | 1.118 | -0.001 | -0.002 | -0.002 | 0.625 | 0.52 | 0.56 | 0.59 | 0.6 | 0.51 | 0.52 |
| 2 | 70710420 | rs3849383 | -0.542 | 0.155 | 1.139 | 1.188 | -0.001 | 0 | 0 | 0.101 | 0.39 | 0.46 | 0.55 | 0.66 | 0.45 | 0.44 |
| 2 | 70712005 | rs3771504 | -1.766 | 0.138 | 0.962 | 1.283 | -0.06 | 0.043 | 0.043 | 0 | 0.57 | 0.98 | 0.65 | 0.69 | 0.62 | 0.58 |
| 2 | 70712860 | rs17005692 | 0.068 | 0.226 | 1.039 | 1.314 | -0.002 | -0.006 | -0.006 | 0 | 0.13 | 0.12 | 0.05 | 0.03 | 0.05 | 0.09 |
| 2 | 70717078 | rs3911077 | 1.133 | 0.135 | 0.767 | 1.325 | -0.049 | -0.043 | -0.043 | 0 | 0.23 | 0.14 | 0.19 | 0.38 | 0.33 | 0.21 |
| 2 | 70717507 | rs3771502 | -0.687 | -0.115 | 0.92 | 1.252 | -0.014 | 0.023 | 0.023 | 0 | 0.17 | 0.31 | 0.37 | 0.19 | 0.12 | 0.23 |
| 2 | 70725352 | rs3771494 | -1.53 | -0.499 | 0.203 | 1.012 | -0.023 | -0.014 | -0.014 | 0 | 0.34 | 0.58 | 0.46 | 0.22 | 0.2 | 0.32 |
| 2 | 70728877 | rs3771492 | -0.635 | -0.43 | 0.431 | 1.026 | -0.017 | 0.072 | 0.072 | 1.839 | 0.1 | 0.45 | 0.12 | 0.1 | 0.02 | 0 |
| 2 | 70729253 | rs3821261 | -0.735 | -0.356 | -0.223 | 1.032 | -0.05 | -0.018 | -0.018 | 0 | 0.21 | 0.67 | 0.17 | 0.14 | 0.07 | 0.09 |
| 2 | 70732852 | rs3755377 | -0.446 | 0.038 | -0.494 | 0.958 | -0.007 | -0.005 | -0.005 | 0 | 0.47 | 0.79 | 0.4 | 0.54 | 0.43 | 0.31 |
| 2 | 70733526 | rs404420 | -1.175 | 0.198 | -0.859 | 0.557 | 0.024 | 0.084 | 0.084 | 0 | 0.66 | 0.89 | 0.63 | 0.77 | 0.72 | 0.68 |
| 2 | 70736219 | rs454305 | -1.011 | 0.22 | -0.307 | 0.701 | 0.01 | 0.016 | 0.016 | 0 | 0.37 | 0.71 | 0.36 | 0.26 | 0.35 | 0.38 |
| 2 | 70748334 | rs731461 | 0.977 | -0.046 | -0.814 | 0.163 | -0.079 | -0.008 | -0.008 | 0.545 | 0.25 | 0.03 | 0.16 | 0.21 | 0.3 | 0.18 |
| 2 | 70750531 | rs426081 | -1.28 | -0.806 | 0.395 | 0.171 | -0.011 | -0.034 | -0.034 | 0 | 0.33 | 0.64 | 0.26 | 0.1 | 0.17 | 0.32 |
| 2 | 70752195 | rs6708768 | -0.382 | -0.651 | 0.227 | 0.223 | -0.031 | -0.074 | -0.074 | 0.074 | 0.13 | 0.31 | 0.06 | 0 | 0.06 | 0.1 |
| 2 | 70753882 | rs374640 | -0.961 | -0.499 | -0.476 | 0.403 | 0.01 | 0.01 | 0.01 | 0 | 0.35 | 0.7 | 0.29 | 0.18 | 0.2 | 0.33 |
| 2 | 70756057 | rs446086 | -1.365 | 0.187 | 0.078 | 1.195 | 0.044 | 0.117 | 0.117 | 0 | 0.14 | 0.29 | 0.13 | 0.34 | 0.25 | 0.19 |
| 2 | 70757358 | rs11466212 | -0.216 | 0.098 | -0.41 | 1.114 | -0.018 | -0.026 | -0.026 | 0 | 0.11 | 0.02 | 0.25 | 0.13 | 0.1 | 0.2 |
| 2 | 70758123 | rs7562947 | -0.179 | -0.646 | 0.127 | 0.758 | -0.064 | -0.058 | -0.058 | 0 | 0.2 | 0.59 | 0.11 | 0.09 | 0.09 | 0.18 |

| | | | | | | | | | | | | | | | | |
|---|---|---|---|---|---|---|---|---|---|---|---|---|---|---|---|---|
| 2 | 70759161 | rs428225 | -1.455 | -0.945 | -0.395 | 0.221 | 0.049 | 0.076 | 0.076 | 0 | 0.46 | 0.67 | 0.47 | 0.22 | 0.32 | 0.42 |
| 2 | 70764688 | rs432203 | -1.664 | -0.986 | -0.229 | 0.344 | -0.013 | 0 | 0 | 0 | 0.56 | 0.66 | 0.56 | 0.43 | 0.54 | 0.58 |
| 2 | 70766165 | rs418647 | -1.665 | -0.995 | -0.248 | 0.34 | -0.013 | -0.013 | -0.013 | 0 | 0.56 | 0.66 | 0.57 | 0.49 | 0.55 | 0.58 |
| 2 | 70769365 | rs1523300 | -0.982 | -1.035 | -0.247 | 0.208 | 0.001 | 0.008 | 0.008 | 0 | 0.38 | 0.63 | 0.27 | 0.27 | 0.41 | 0.27 |
| 2 | 70771302 | rs6546610 | -1.213 | -0.824 | -0.101 | 0.468 | -0.009 | -0.015 | -0.015 | 0 | 0.36 | 0.29 | 0.46 | 0.58 | 0.44 | 0.5 |
| 2 | 70772455 | rs10489984 | 0.378 | -0.822 | -0.226 | 0.41 | 0.028 | 0.066 | 0.066 | 0 | 0.16 | 0.08 | 0.07 | 0 | 0.11 | 0.03 |
| 2 | 70776384 | rs12473408 | -1.345 | -0.739 | 0.351 | 0.512 | -0.001 | 0 | 0 | 0 | 0.4 | 0.6 | 0.49 | 0.59 | 0.45 | 0.51 |
| 2 | 70777550 | rs7568707 | -0.843 | -0.919 | -0.28 | 0.403 | 0.038 | 0.033 | 0.033 | 0 | 0.33 | 0.18 | 0.37 | 0.24 | 0.25 | 0.35 |
| 2 | 70783422 | rs540006 | 0.604 | -1.049 | -0.363 | 0.821 | -0.015 | -0.033 | -0.033 | 0 | 0.07 | 0 | 0.11 | 0.14 | 0.07 | 0.14 |
| 2 | 70789532 | rs17040266 | -0.884 | -1.394 | -0.172 | 0.533 | 0.048 | 0.007 | 0.007 | 0 | 0.31 | 0.16 | 0.19 | 0.29 | 0.19 | 0.26 |
| 2 | 70796257 | rs2215021 | 0.342 | -1.307 | -0.208 | 0.415 | -0.039 | -0.047 | -0.047 | 0 | 0.25 | 0.12 | 0.36 | 0.36 | 0.38 | 0.33 |
| 2 | 70796339 | rs713111 | -0.234 | -1.43 | 0.065 | 0.124 | -0.029 | -0.065 | -0.065 | 0.908 | 0.21 | 0.28 | 0.1 | 0 | 0.11 | 0.03 |
| 2 | 70802535 | rs452710 | -1.291 | -1.147 | 0.28 | 0.469 | 0.01 | -0.002 | -0.002 | 0 | 0.57 | 0.52 | 0.58 | 0.47 | 0.57 | 0.58 |
| 2 | 70808147 | rs10489986 | -0.658 | -1.243 | 0.081 | 0.347 | -0.012 | -0.023 | -0.023 | 0 | 0.28 | 0.27 | 0.18 | 0.13 | 0.18 | 0.25 |
| 2 | 70809219 | rs10489987 | -0.518 | -1.251 | -0.327 | 0.342 | 0.029 | 0.039 | 0.039 | 0 | 0.27 | 0.18 | 0.18 | 0.13 | 0.17 | 0.24 |
| 2 | 70810641 | rs428148 | -0.149 | -0.555 | -0.238 | 0.782 | -0.042 | -0.024 | -0.024 | 0 | 0.28 | 0.12 | 0.38 | 0.34 | 0.4 | 0.33 |
| 2 | 70817894 | rs413056 | -0.595 | -0.23 | -0.593 | 0.714 | -0.002 | -0.004 | -0.004 | 0 | 0.46 | 0.49 | 0.51 | 0.63 | 0.49 | 0.64 |
| 2 | 70823386 | rs6546612 | -0.279 | 0.022 | -1.346 | 0.83 | 0.008 | -0.032 | -0.032 | 0 | 0.56 | 0.76 | 0.37 | 0.52 | 0.44 | 0.7 |
| 2 | 70823521 | rs436640 | -0.024 | 0.144 | -1.587 | 0.738 | -0.014 | -0.006 | -0.006 | 0 | 0.62 | 0.78 | 0.55 | 0.56 | 0.55 | 0.61 |
| 2 | 70825231 | rs2108481 | -0.338 | 0.167 | -1.351 | 0.608 | 0.011 | 0.09 | 0.09 | 0 | 0.67 | 0.84 | 0.81 | 0.87 | 0.68 | 0.71 |
| 2 | 70825480 | rs2108483 | 0.101 | 0.262 | -1.117 | 0.868 | 0.048 | 0.025 | 0.025 | 0 | 0.49 | 0.33 | 0.65 | 0.57 | 0.61 | 0.38 |
| 2 | 70826216 | rs11126282 | 0.098 | 0.246 | -1.067 | 0.86 | 0.053 | 0.027 | 0.027 | 0 | 0.49 | 0.3 | 0.65 | 0.57 | 0.61 | 0.38 |
| 2 | 70829851 | rs11677281 | 1.037 | 0.288 | -1.359 | 0.824 | -0.107 | 0.054 | 0.054 | 0 | 0.34 | 0.07 | 0.32 | 0.21 | 0.38 | 0.2 |
| 2 | 70830628 | rs12466990 | 0.566 | 0.457 | -0.74 | 1.231 | -0.002 | 0.054 | 0.054 | 0 | 0.05 | 0.05 | 0.27 | 0.33 | 0.14 | 0.11 |
| 2 | 70831819 | rs10166358 | -0.704 | -0.058 | -0.184 | 1.45 | 0.003 | 0.118 | 0.118 | 1.172 | 0.19 | 0.35 | 0.35 | 0.43 | 0.2 | 0.16 |
| 2 | 70834323 | rs7591505 | -0.923 | -0.422 | 0.265 | 0.957 | -0.016 | -0.058 | -0.058 | 0 | 0.2 | 0.41 | 0.21 | 0.09 | 0.14 | 0.14 |
| 2 | 70835618 | rs17005935 | -0.48 | -0.388 | 0.305 | 0.986 | 0.001 | -0.036 | -0.036 | 0.355 | 0.17 | 0.36 | 0.19 | 0.09 | 0.13 | 0.14 |
| 2 | 70836266 | rs17005940 | 1.057 | -0.371 | 0.236 | 0.973 | -0.004 | -0.004 | -0.004 | 0 | 0.08 | 0.03 | 0.1 | 0.08 | 0.08 | 0.06 |
| 2 | 70838190 | rs13389582 | -0.406 | -0.359 | 0.287 | 0.765 | 0.003 | 0.004 | 0.004 | 0 | 0.07 | 0.05 | 0.08 | 0 | 0.05 | 0.08 |
| 2 | 70840415 | rs13396913 | -0.334 | -0.344 | 0.408 | 0.781 | 0.025 | -0.014 | -0.014 | 0.176 | 0.06 | 0.25 | 0.03 | 0 | 0.03 | 0.05 |
| 2 | 70842774 | rs9309438 | 0.087 | -0.779 | 0.025 | 0.459 | -0.002 | -0.013 | -0.013 | 0 | 0.29 | 0.24 | 0.33 | 0.43 | 0.33 | 0.36 |
| 2 | 70844394 | rs13405786 | 0.365 | -0.576 | 0.061 | -0.052 | -0.002 | 0.01 | 0.01 | 1.087 | 0.13 | 0.1 | 0.13 | 0.07 | 0.12 | 0.1 |
| 2 | 70847531 | rs2079538 | 1.1 | -0.392 | -0.81 | -0.263 | -0.118 | 0.037 | 0.037 | 0.059 | 0.29 | 0.03 | 0.3 | 0.19 | 0.36 | 0.17 |
| 2 | 70853832 | rs13399586 | -0.592 | -0.409 | 0.177 | -0.462 | 0 | 0.042 | 0.042 | 0 | 0.49 | 0.61 | 0.54 | 0.36 | 0.5 | 0.33 |
| 2 | 70857483 | rs6735669 | -0.356 | -0.402 | 0.728 | -0.545 | -0.005 | -0.086 | -0.086 | 0.694 | 0.2 | 0.49 | 0.25 | 0.07 | 0.16 | 0.14 |
| 2 | 70858190 | rs1859917 | 0.483 | -0.071 | 1.124 | -0.536 | 0.004 | -0.025 | -0.025 | 0 | 0.36 | 0.47 | 0.33 | 0.29 | 0.36 | 0.3 |
| 2 | 70858565 | rs7355719 | 0.444 | -0.047 | 1.199 | -0.558 | 0.003 | -0.026 | -0.026 | 0 | 0.35 | 0.47 | 0.33 | 0.28 | 0.35 | 0.21 |
| 2 | 70859248 | rs7606257 | 0.566 | -0.05 | 1.2 | -0.559 | 0.004 | -0.018 | -0.018 | 0 | 0.34 | 0.43 | 0.32 | 0.28 | 0.35 | 0.21 |
| 2 | 70863351 | rs1990209 | -0.24 | -0.002 | 1.009 | -0.599 | 0.003 | -0.014 | -0.014 | 0 | 0.44 | 0.44 | 0.44 | 0.35 | 0.44 | 0.27 |
| 2 | 70865498 | rs11903447 | 1.288 | -0.036 | 1.03 | -0.577 | 0.012 | -0.016 | -0.016 | 0.001 | 0.11 | 0.22 | 0.13 | 0.07 | 0.09 | 0.06 |
| 2 | 70866891 | rs7560329 | -0.609 | -0.04 | 0.907 | -0.582 | 0.006 | 0.059 | 0.059 | 0 | 0.5 | 0.64 | 0.55 | 0.35 | 0.5 | 0.32 |
| 2 | 70867644 | rs12468912 | -0.702 | -0.037 | 0.802 | -0.58 | 0.003 | 0.071 | 0.071 | 0 | 0.51 | 0.7 | 0.56 | 0.35 | 0.5 | 0.32 |
| 2 | 70869473 | rs6733938 | 0.32 | -0.029 | 0.751 | -0.586 | 0.022 | -0.044 | -0.044 | 0.364 | 0.15 | 0.4 | 0.18 | 0.07 | 0.13 | 0.09 |
| 2 | 70872293 | rs880848 | -0.533 | 0.041 | 0.432 | -0.578 | 0.012 | 0.057 | 0.057 | 0 | 0.5 | 0.67 | 0.57 | 0.33 | 0.53 | 0.31 |
| 2 | 70872817 | rs11898642 | -0.69 | 0.043 | 0.006 | -0.569 | 0.001 | 0.091 | 0.091 | 0 | 0.52 | 0.82 | 0.57 | 0.34 | 0.53 | 0.31 |
| 2 | 70874972 | rs2041078 | 1.461 | 0.161 | -0.397 | -0.754 | -0.121 | 0.118 | 0.118 | 0 | 0.27 | 0.02 | 0.3 | 0.11 | 0.35 | 0.19 |
| 2 | 70878684 | rs12617676 | 0.311 | 0.157 | 0.012 | -0.682 | -0.005 | 0.007 | 0.007 | 0 | 0.41 | 0.35 | 0.45 | 0.18 | 0.45 | 0.26 |
| 2 | 70881592 | rs10197417 | 0.321 | 0.271 | -0.011 | -0.684 | -0.001 | -0.01 | -0.01 | 0.092 | 0.38 | 0.35 | 0.42 | 0.18 | 0.4 | 0.25 |
| 2 | 70889498 | rs17698193 | 0.11 | -0.526 | -0.188 | -0.687 | 0.009 | 0.009 | 0.009 | 0 | 0.27 | 0.2 | 0.18 | 0.16 | 0.21 | 0.29 |
| 2 | 70891515 | rs13003996 | 0.85 | -0.604 | -0.348 | -0.722 | 0.031 | 0.049 | 0.049 | 0.058 | 0.22 | 0.05 | 0.12 | 0.16 | 0.16 | 0.24 |
| 2 | 70892697 | rs3771467 | -1.587 | -0.478 | -0.201 | -0.715 | 0.054 | 0.073 | 0.073 | 0 | 0.49 | 0.77 | 0.4 | 0.37 | 0.38 | 0.43 |
| 2 | 70896023 | rs740388 | 0.315 | -0.462 | 0.071 | -0.744 | -0.006 | -0.068 | -0.068 | 0 | 0.12 | 0.24 | 0.12 | 0 | 0.11 | 0.09 |
| 2 | 70896062 | rs758062 | -1.56 | -0.444 | -0.602 | -0.746 | 0.063 | 0.076 | 0.076 | 0 | 0.49 | 0.77 | 0.4 | 0.37 | 0.38 | 0.43 |
| 2 | 70898744 | rs3771466 | -0.377 | -0.5 | -0.163 | -0.759 | -0.02 | -0.006 | -0.006 | 0 | 0.36 | 0.47 | 0.27 | 0.36 | 0.28 | 0.34 |
| 2 | 70898822 | rs3771464 | -0.387 | -0.502 | -0.145 | -0.76 | -0.02 | -0.006 | -0.006 | 0 | 0.36 | 0.48 | 0.27 | 0.36 | 0.28 | 0.33 |
| 2 | 70899002 | rs3771463 | -1.663 | -0.483 | -0.184 | -0.768 | 0.042 | 0.069 | 0.069 | 0 | 0.49 | 0.78 | 0.42 | 0.37 | 0.43 | 0.44 |
| 2 | 70901699 | rs11894520 | 0.307 | -0.409 | -0.041 | -0.81 | 0.002 | -0.053 | -0.053 | 0 | 0.12 | 0.25 | 0.14 | 0 | 0.13 | 0.1 |
| 2 | 70906115 | rs2074815 | -0.33 | -0.523 | -0.132 | -0.792 | -0.023 | 0.005 | 0.005 | 0 | 0.34 | 0.47 | 0.27 | 0.38 | 0.27 | 0.31 |
| 2 | 70909820 | rs11893633 | 0.135 | -0.596 | -0.28 | -0.864 | -0.014 | -0.028 | -0.028 | 1.246 | 0.05 | 0.16 | 0.07 | 0 | 0.04 | 0.04 |
| 2 | 70912868 | rs12470211 | 1.928 | -0.329 | -0.862 | -1.014 | -0.117 | 0.065 | 0.065 | 0 | 0.26 | 0.03 | 0.28 | 0.14 | 0.31 | 0.2 |
| 2 | 70915347 | rs7340146 | -0.19 | -0.647 | -0.697 | -0.874 | -0.029 | 0.018 | 0.018 | 0 | 0.34 | 0.43 | 0.25 | 0.43 | 0.25 | 0.3 |
| 2 | 70917093 | rs13401890 | -0.674 | -0.62 | -0.661 | -0.862 | 0.006 | -0.001 | -0.001 | 0 | 0.4 | 0.6 | 0.33 | 0.44 | 0.31 | 0.34 |
| 2 | 70921539 | rs4852700 | -2.231 | -0.573 | -0.653 | -0.944 | 0.006 | -0.053 | -0.053 | 0 | 0.76 | 0.83 | 0.71 | 0.58 | 0.75 | 0.59 |
| 2 | 70923854 | rs759463 | -0.627 | -0.544 | -0.713 | -0.805 | -0.027 | 0.003 | 0.003 | 0 | 0.38 | 0.45 | 0.31 | 0.44 | 0.31 | 0.32 |
| 2 | 70928436 | rs12476158 | 1.58 | -0.695 | -1.509 | -0.91 | -0.017 | 0.006 | 0.006 | 0 | 0.14 | 0.01 | 0.12 | 0.12 | 0.11 | 0.07 |
| 2 | 70932355 | rs3771456 | 0.732 | -0.639 | -1.36 | -0.83 | 0.002 | -0.014 | -0.014 | 0 | 0.16 | 0.05 | 0.12 | 0.2 | 0.15 | 0.19 |
| 2 | 70936407 | rs7597655 | -0.712 | -0.563 | -1.182 | -0.767 | -0.006 | 0 | 0 | 0 | 0.39 | 0.55 | 0.3 | 0.44 | 0.27 | 0.33 |
| 2 | 70936861 | rs3771452 | -1.378 | -0.525 | -1.657 | -0.856 | -0.009 | -0.041 | -0.041 | 0 | 0.66 | 0.76 | 0.63 | 0.57 | 0.63 | 0.52 |
| 2 | 70938472 | rs6546619 | -0.753 | -0.66 | -1.257 | -0.64 | 0.041 | -0.001 | -0.001 | 0 | 0.4 | 0.68 | 0.31 | 0.44 | 0.27 | 0.33 |

| | | | | | | | | | | | | | | | | |
|---|---|---|---|---|---|---|---|---|---|---|---|---|---|---|---|---|
| 2 | 70942964 | rs3755369 | -0.047 | -1.118 | -0.817 | -0.631 | -0.008 | -0.033 | -0.033 | 0 | 0.27 | 0.37 | 0.19 | 0.21 | 0.25 | 0.21 |
| 2 | 70943138 | rs3771450 | -0.748 | -1.36 | -1.653 | -0.919 | -0.153 | 0.143 | 0.143 | 0 | 0.61 | 0.95 | 0.58 | 0.78 | 0.54 | 0.7 |
| 2 | 70943629 | rs11126290 | 0.46 | -1.255 | -1.045 | -0.723 | -0.029 | 0.004 | 0.004 | 0 | 0.33 | 0.46 | 0.35 | 0.57 | 0.28 | 0.47 |
| 2 | 70944362 | rs3821257 | -0.723 | -0.635 | -1.249 | -0.801 | -0.121 | 0.096 | 0.096 | 0 | 0.77 | 0.97 | 0.67 | 0.9 | 0.65 | 0.78 |
| 2 | 70944880 | rs2052273 | -1.474 | -0.405 | -1.157 | -0.706 | -0.079 | 0.081 | 0.081 | 0.527 | 0.84 | 0.98 | 0.75 | 0.98 | 0.74 | 0.88 |
| 2 | 70944965 | rs3755363 | -0.039 | -1.603 | -0.119 | -0.343 | -0.023 | -0.038 | -0.038 | 0.14 | 0.32 | 0.55 | 0.23 | 0.21 | 0.25 | 0.2 |
| 2 | 70945251 | rs3755361 | -1.421 | -1.524 | 0.011 | -0.554 | -0.112 | 0.119 | 0.119 | 0.813 | 0.72 | 0.95 | 0.64 | 0.9 | 0.59 | 0.72 |
| 2 | 70947575 | rs17040269 | 1.187 | -1.578 | 0.062 | -0.465 | 0.002 | 0 | 0 | 0.288 | 0.11 | 0.09 | 0.14 | 0.11 | 0.09 | 0.06 |
| 2 | 70949978 | rs3771437 | -0.114 | -1.292 | 0.415 | -0.327 | -0.017 | -0.031 | -0.031 | 0 | 0.33 | 0.44 | 0.23 | 0.21 | 0.26 | 0.21 |
| 2 | 70951020 | rs2128 | -0.227 | -1.292 | 0.382 | -0.334 | 0.024 | 0.072 | 0.072 | 0 | 0.51 | 0.75 | 0.46 | 0.62 | 0.45 | 0.59 |
| 2 | 70951723 | rs3771434 | -0.859 | -1.319 | 0.298 | -0.356 | -0.078 | 0.1 | 0.1 | 0 | 0.61 | 0.89 | 0.59 | 0.73 | 0.54 | 0.65 |
| 2 | 70951928 | rs3771433 | -0.301 | -1.164 | 0.479 | -0.211 | -0.003 | -0.039 | -0.039 | 0 | 0.34 | 0.57 | 0.32 | 0.16 | 0.31 | 0.2 |
| 2 | 70952091 | rs7584508 | 0.417 | -1.127 | 0.493 | -0.234 | -0.008 | -0.108 | -0.108 | 0 | 0.25 | 0.5 | 0.18 | 0.05 | 0.21 | 0.14 |
| 2 | 70957503 | rs3771426 | -0.091 | -0.963 | 0.184 | -0.105 | 0.009 | -0.014 | -0.014 | 0 | 0.18 | 0.12 | 0.18 | 0.28 | 0.15 | 0.13 |
| 2 | 70965196 | rs4852706 | -0.563 | -0.914 | 0.443 | -0.092 | 0.005 | 0.007 | 0.007 | 0 | 0.4 | 0.35 | 0.3 | 0.33 | 0.36 | 0.26 |
| 2 | 70973247 | rs1861451 | 0.846 | -1 | 0.524 | -0.119 | -0.017 | 0.026 | 0.026 | 0 | 0.27 | 0.35 | 0.28 | 0.57 | 0.23 | 0.44 |
| 2 | 70974890 | rs3755351 | -0.592 | -0.873 | 0.214 | -0.133 | 0.005 | 0.008 | 0.008 | 0 | 0.4 | 0.34 | 0.3 | 0.33 | 0.36 | 0.26 |
| 2 | 70980886 | rs17006285 | -0.058 | -0.432 | 0.074 | 0.277 | 0.015 | -0.046 | -0.046 | 0 | 0.17 | 0.08 | 0.17 | 0.3 | 0.15 | 0.13 |
| 2 | 70981146 | rs3771408 | -1.017 | 0.017 | 0.365 | -0.215 | -0.117 | 0.146 | 0.146 | 0 | 0.7 | 0.96 | 0.7 | 0.91 | 0.59 | 0.73 |
| 2 | 71003968 | rs11676833 | 0.012 | -0.134 | 0.594 | -0.096 | 0.001 | 0.016 | 0.016 | 0.045 | 0.55 | 0.47 | 0.53 | 0.74 | 0.46 | 0.6 |
| 2 | 71006246 | rs7596564 | 0.137 | -0.135 | 0.342 | -0.097 | 0.008 | 0.089 | 0.089 | 0.12 | 0.54 | 0.27 | 0.52 | 0.74 | 0.46 | 0.63 |
| 2 | 71006389 | rs2024454 | -1.012 | 0.196 | -0.569 | 0.031 | -0.083 | 0.117 | 0.117 | 0 | 0.71 | 0.93 | 0.7 | 0.91 | 0.6 | 0.74 |
| 2 | 71007394 | rs2024457 | 0.841 | 0.179 | -0.374 | 0.189 | 0.002 | 0.039 | 0.039 | 0 | 0.1 | 0.19 | 0.14 | 0.16 | 0.11 | 0.1 |
| 2 | 71009303 | rs2080393 | 0.841 | 0.179 | -0.491 | 0.189 | 0 | 0.022 | 0.022 | 0 | 0.1 | 0.15 | 0.13 | 0.17 | 0.11 | 0.1 |
| 2 | 71012604 | rs7566541 | -1.027 | 0.182 | -0.633 | 0.142 | -0.146 | 0.18 | 0.18 | 0 | 0.68 | 0.97 | 0.7 | 0.91 | 0.59 | 0.73 |
| 2 | 71014322 | rs10184254 | -0.21 | -0.011 | -0.515 | 0.253 | 0.013 | 0.074 | 0.074 | 0 | 0.55 | 0.73 | 0.55 | 0.74 | 0.45 | 0.62 |
| 2 | 71020254 | rs6546638 | -0.334 | -0.031 | -0.109 | 1.19 | -0.039 | 0.033 | 0.033 | 0 | 0.29 | 0.45 | 0.17 | 0.33 | 0.2 | 0.2 |
| 2 | 71024411 | rs6735921 | 0.509 | 0.829 | -0.196 | 1.152 | 0.008 | 0.016 | 0.016 | 0 | 0.88 | 0.8 | 0.85 | 0.85 | 0.86 | 0.89 |
| 2 | 71028889 | rs7560275 | -1.237 | 0.509 | 0.295 | 1.376 | 0.002 | 0.176 | 0.176 | 0 | 0.1 | 0.36 | 0.12 | 0.19 | 0.08 | 0.11 |
| 2 | 71029020 | rs6546640 | 0.317 | 0.534 | -0.288 | 0.685 | -0.011 | 0.053 | 0.053 | 0 | 0.8 | 0.91 | 0.75 | 0.9 | 0.81 | 0.83 |
| 2 | 71029439 | rs6712273 | 1.14 | 1.081 | -0.194 | 0.522 | -0.004 | -0.004 | -0.004 | 0.396 | 0.7 | 0.75 | 0.65 | 0.7 | 0.67 | 0.76 |
| 2 | 71036482 | rs2287101 | -1.755 | 0.879 | 0.684 | 0.793 | 0.009 | 0.219 | 0.219 | 0.623 | 0.11 | 0.36 | 0.09 | 0.24 | 0.06 | 0.08 |
| 2 | 71038565 | rs17656058 | 0.906 | 0.803 | -0.882 | -0.246 | -0.074 | 0.161 | 0.161 | 0.02 | 0.3 | 0.04 | 0.19 | 0.11 | 0.32 | 0.29 |
| 2 | 71043461 | rs722896 | 0.91 | 0.629 | -0.712 | -0.316 | -0.008 | -0.061 | -0.061 | 0 | 0.29 | 0.12 | 0.4 | 0.52 | 0.31 | 0.43 |
| 2 | 71044211 | rs2075221 | 1.01 | 0.375 | -0.952 | -0.393 | -0.116 | -0.139 | -0.139 | 0 | 0.28 | 0.03 | 0.4 | 0.52 | 0.38 | 0.44 |
| 2 | 71049495 | rs1548888 | 0.431 | -0.078 | -0.32 | -0.586 | 0 | -0.002 | -0.002 | 0 | 0.58 | 0.48 | 0.6 | 0.51 | 0.69 | 0.61 |
| 2 | 71049753 | rs1548891 | 0.416 | -0.079 | -0.32 | -0.587 | -0.001 | -0.001 | -0.001 | 0 | 0.58 | 0.48 | 0.6 | 0.51 | 0.69 | 0.61 |
| 2 | 71050401 | rs1005429 | 0.395 | -0.223 | -0.203 | -0.588 | -0.015 | 0.016 | 0.016 | 0 | 0.81 | 0.73 | 0.86 | 0.63 | 0.91 | 0.79 |
| 2 | 71056098 | rs10206319 | 0.089 | -0.313 | -0.113 | -0.638 | 0.003 | -0.011 | -0.011 | 0 | 0.7 | 0.72 | 0.66 | 0.66 | 0.74 | 0.74 |
| 2 | 71058184 | rs13421115 | 1.337 | -0.132 | -0.213 | -1.277 | -0.001 | -0.041 | -0.041 | 0 | 0.26 | 0.2 | 0.17 | 0 | 0.27 | 0.16 |
| 2 | 71058226 | rs2080390 | 0.336 | -0.188 | -0.108 | -0.718 | -0.022 | 0.046 | 0.046 | 0 | 0.21 | 0.04 | 0.24 | 0.08 | 0.22 | 0.17 |
| 2 | 71058795 | rs3213747 | -0.204 | -0.111 | 0.079 | -0.495 | -0.017 | -0.006 | -0.006 | 0 | 0.13 | 0.22 | 0.05 | 0.14 | 0.06 | 0.13 |
| 2 | 71058906 | rs3213749 | -0.215 | -0.11 | 0.08 | -0.494 | -0.016 | -0.006 | -0.006 | 0.259 | 0.13 | 0.22 | 0.05 | 0.14 | 0.06 | 0.13 |
| 2 | 71059763 | rs10204437 | -0.524 | -0.057 | 0.245 | -0.487 | -0.03 | -0.04 | -0.04 | 0.441 | 0.15 | 0.37 | 0.06 | 0.14 | 0.1 | 0.15 |
| 2 | 71060037 | rs6712863 | -0.571 | -0.003 | -0.125 | -0.461 | -0.025 | -0.031 | -0.031 | 0.26 | 0.16 | 0.33 | 0.06 | 0.14 | 0.1 | 0.15 |
| 2 | 71061108 | rs17662453 | 0.233 | 0.135 | -0.405 | -0.474 | -0.004 | 0.006 | 0.006 | 0 | 0.17 | 0.11 | 0.22 | 0.1 | 0.18 | 0.14 |
| 2 | 71061399 | rs17719042 | -0.72 | 0.178 | -0.587 | -0.378 | -0.001 | 0.018 | 0.018 | 0 | 0.31 | 0.29 | 0.28 | 0.26 | 0.29 | 0.33 |
| 2 | 71062648 | rs10489990 | 0.254 | 0.207 | -1.468 | -0.026 | -0.011 | 0.002 | 0.002 | 0 | 0.32 | 0.13 | 0.4 | 0.31 | 0.36 | 0.3 |
| 2 | 71063066 | rs2075225 | -0.306 | 0.357 | -1.578 | 0.58 | 0.083 | 0.032 | 0.032 | 0 | 0.48 | 0.8 | 0.36 | 0.39 | 0.34 | 0.61 |
| 2 | 71063410 | rs3755349 | 0.418 | -0.215 | -1.758 | 0.552 | 0.004 | -0.015 | -0.015 | 2.721 | 0.25 | 0.11 | 0.15 | 0.25 | 0.21 | 0.41 |
| 2 | 71063725 | rs3755348 | 1.217 | -0.372 | -1.403 | 0.494 | -0.011 | -0.001 | -0.001 | 0 | 0.15 | 0.18 | 0.18 | 0.14 | 0.13 | 0.18 |
| 2 | 71063942 | rs4852708 | 1.217 | -0.372 | -1.403 | 0.494 | -0.011 | -0.001 | -0.001 | 0.466 | 0.15 | 0.18 | 0.18 | 0.14 | 0.13 | 0.18 |
| 2 | 71066277 | rs7557453 | 0.35 | -0.474 | -0.687 | 0.427 | -0.034 | -0.019 | -0.019 | 0 | 0.2 | 0.39 | 0.19 | 0.14 | 0.14 | 0.2 |
| 2 | 71066601 | rs17006467 | 0.383 | -0.474 | -0.686 | 0.427 | -0.033 | -0.018 | -0.018 | 0 | 0.2 | 0.39 | 0.19 | 0.14 | 0.14 | 0.2 |
| 2 | 71067853 | rs6723629 | 0.42 | -0.267 | -0.514 | 0.472 | 0.001 | -0.025 | -0.025 | 0 | 0.24 | 0.09 | 0.15 | 0.25 | 0.2 | 0.4 |
| 2 | 71068400 | rs1030046 | -1.471 | -0.343 | -0.613 | 0.483 | -0.012 | -0.01 | -0.01 | 0 | 0.81 | 0.82 | 0.76 | 0.71 | 0.7 | 0.91 |
| 2 | 71068623 | rs17006471 | 0.801 | -0.285 | -0.814 | 0.516 | 0.009 | 0.001 | 0.001 | 0 | 0.48 | 0.21 | 0.58 | 0.46 | 0.49 | 0.49 |
| 2 | 71077902 | rs7565578 | 0.188 | -0.186 | -0.93 | 0.668 | 0.012 | -0.002 | -0.002 | 0 | 0.25 | 0.14 | 0.15 | 0.25 | 0.2 | 0.4 |
| 2 | 71093812 | rs4852716 | 0.519 | -0.02 | -1.092 | 0.573 | -0.002 | 0.005 | 0.005 | 0 | 0.51 | 0.21 | 0.6 | 0.47 | 0.52 | 0.49 |
| 2 | 71101403 | rs6546646 | 0.221 | -0.131 | -1.252 | 0.695 | 0.004 | -0.012 | -0.012 | 0 | 0.22 | 0.11 | 0.09 | 0.3 | 0.16 | 0.27 |
| 2 | 71101426 | rs6546647 | -0.877 | -0.725 | -1.032 | 0.393 | 0.004 | 0.003 | 0.003 | 0 | 0.36 | 0.37 | 0.18 | 0.32 | 0.27 | 0.44 |
| 2 | 71101847 | rs6748323 | -0.283 | -0.268 | -1.417 | 0.604 | -0.01 | 0.002 | 0.002 | 0.465 | 0.54 | 0.73 | 0.56 | 0.61 | 0.52 | 0.52 |
| 2 | 71118757 | rs7594319 | -0.971 | -0.145 | -1.528 | 0.713 | -0.04 | 0.011 | 0.011 | 0.03 | 0.66 | 0.89 | 0.62 | 0.7 | 0.59 | 0.68 |
| 2 | 71130394 | rs3771403 | 1.5 | 0.013 | -1.217 | 0.807 | 0.015 | 0.003 | 0.003 | 0.069 | 0.16 | 0.2 | 0.21 | 0.15 | 0.25 | 0.06 |
| 2 | 71133014 | rs3771395 | 0.894 | -0.662 | -0.942 | 0.864 | -0.008 | -0.001 | -0.001 | 0 | 0.2 | 0.18 | 0.11 | 0.27 | 0.13 | 0.29 |
| 2 | 71136033 | rs6758537 | -0.214 | -0.939 | -0.881 | 0.985 | -0.009 | 0.013 | 0.013 | 0 | 0.31 | 0.33 | 0.14 | 0.52 | 0.22 | 0.46 |
| 2 | 71138276 | rs17006519 | 1.618 | -0.842 | -1.343 | 1.067 | -0.004 | -0.045 | -0.045 | 0 | 0.1 | 0.04 | 0.04 | 0.13 | 0.08 | 0.18 |
| 2 | 71138685 | rs11681519 | 1.307 | -0.866 | -1.032 | 0.699 | 0.006 | 0.001 | 0.001 | 0 | 0.2 | 0.16 | 0.24 | 0.18 | 0.17 | 0.19 |
| 2 | 71140960 | rs7584322 | -0.731 | -0.849 | -1.388 | 0.482 | -0.001 | 0.047 | 0.047 | 0 | 0.55 | 0.84 | 0.45 | 0.66 | 0.54 | 0.57 |

| | | | | | | | | | | | | | | | | |
|---|---|---|---|---|---|---|---|---|---|---|---|---|---|---|---|---|
| 2 | 71145643 | rs3771391 | -1.68 | -0.571 | -1.312 | 0.327 | -0.068 | 0.013 | 0.013 | 0 | 0.75 | 0.98 | 0.69 | 0.84 | 0.72 | 0.8 |
| 2 | 71152850 | rs6748268 | 1.449 | -0.369 | -0.684 | 0.367 | 0.021 | -0.001 | -0.001 | 0 | 0.18 | 0.18 | 0.17 | 0.15 | 0.29 | 0.09 |
| 2 | 71153154 | rs3771389 | 0.005 | -0.892 | -0.487 | 0.335 | -0.02 | 0.005 | 0.005 | 0 | 0.43 | 0.53 | 0.39 | 0.49 | 0.26 | 0.46 |
| 2 | 71162773 | rs17663700 | -0.069 | -0.366 | -0.314 | 0.424 | 0.012 | 0.001 | 0.001 | 0 | 0.34 | 0.38 | 0.26 | 0.36 | 0.42 | 0.32 |
| 2 | 71164634 | rs7588170 | 0.053 | -0.076 | -0.024 | 0.261 | 0.006 | -0.023 | -0.023 | 0 | 0.3 | 0.39 | 0.39 | 0.22 | 0.33 | 0.31 |
| 2 | 71166840 | rs10185490 | -0.65 | -0.377 | -0.043 | 0.211 | -0.011 | 0.023 | 0.023 | 0 | 0.58 | 0.62 | 0.57 | 0.67 | 0.55 | 0.66 |
| 2 | 71167025 | rs11695103 | -0.513 | -0.417 | 0.409 | 0.198 | -0.007 | 0.019 | 0.019 | 0.083 | 0.54 | 0.38 | 0.53 | 0.67 | 0.52 | 0.64 |
| 2 | 71167141 | rs6750310 | 0.845 | -0.496 | 0.167 | 0.149 | -0.009 | 0.016 | 0.016 | 0 | 0.25 | 0.18 | 0.34 | 0.14 | 0.3 | 0.23 |
| 2 | 71167577 | rs10165187 | 0.033 | -0.284 | 0.372 | 0.489 | -0.026 | 0.035 | 0.035 | 0 | 0.3 | 0.38 | 0.19 | 0.53 | 0.21 | 0.42 |
| 2 | 71169705 | rs2266917 | -1.008 | 0.032 | -0.517 | 0.391 | 0.037 | -0.021 | -0.021 | 0.093 | 0.78 | 0.6 | 0.8 | 0.83 | 0.73 | 0.84 |
| 2 | 71170807 | rs2266918 | -0.317 | -0.5 | -0.974 | 0.848 | 0.047 | -0.068 | -0.068 | 0 | 0.31 | 0.1 | 0.17 | 0.53 | 0.21 | 0.41 |
| 2 | 71173408 | rs2239484 | -0.138 | -0.019 | -0.713 | 1.007 | 0.001 | 0.2 | 0.2 | 0 | 0.14 | 0.24 | 0.15 | 0.47 | 0.13 | 0.26 |
| 2 | 71176660 | rs1006049 | -0.458 | 0.245 | -1.208 | 0.24 | 0.001 | 0 | 0 | 0 | 0.31 | 0.28 | 0.4 | 0.38 | 0.39 | 0.32 |
| 2 | 71182900 | rs10210235 | -0.732 | 0.601 | -1.584 | 0.324 | 0.005 | -0.003 | -0.003 | 0 | 0.42 | 0.55 | 0.51 | 0.42 | 0.54 | 0.53 |
| 2 | 71190384 | rs2072462 | 0.561 | 0.513 | -2.261 | 0.169 | -0.023 | 0.061 | 0.061 | 0 | 0.53 | 0.15 | 0.49 | 0.65 | 0.46 | 0.54 |
| 2 | 71190545 | rs2072463 | 0.55 | 0.523 | -2.255 | 0.174 | -0.024 | 0.06 | 0.06 | 0 | 0.53 | 0.15 | 0.49 | 0.66 | 0.46 | 0.54 |
| 2 | 71197623 | rs7562668 | 0.085 | 1.507 | -2.116 | 0.772 | -0.141 | -0.079 | -0.079 | 0.067 | 0.72 | 0.95 | 0.6 | 0.67 | 0.53 | 0.7 |
| 2 | 71202508 | rs17721027 | 0.164 | 1.183 | -1.523 | 0.71 | 0.002 | -0.001 | -0.001 | 0 | 0.46 | 0.43 | 0.34 | 0.43 | 0.47 | 0.41 |
| 2 | 71203004 | rs17664561 | -0.193 | 1.387 | -1.729 | 0.894 | -0.061 | -0.04 | -0.04 | 0 | 0.12 | 0.01 | 0.26 | 0.12 | 0.22 | 0.08 |
| 2 | 71203077 | rs2058552 | 0.581 | 1.065 | -1.678 | 0.904 | 0.027 | 0.022 | 0.022 | 0 | 0.45 | 0.25 | 0.25 | 0.36 | 0.35 | 0.5 |
| 2 | 71211815 | rs3796098 | -0.604 | 0.94 | -1.052 | 0.942 | -0.021 | -0.024 | -0.024 | 0.775 | 0.16 | 0.08 | 0.13 | 0.16 | 0.23 | 0.09 |
| 2 | 71217864 | rs4852737 | -0.532 | 0.551 | -0.96 | 0.598 | 0.019 | 0.083 | 0.083 | 0.107 | 0.14 | 0.01 | 0.24 | 0 | 0.06 | 0.13 |
| 2 | 71220793 | rs2276684 | -0.475 | 0.548 | -0.906 | 0.615 | -0.019 | -0.017 | -0.017 | 0 | 0.16 | 0.11 | 0.13 | 0.16 | 0.23 | 0.1 |
| 2 | 71221822 | rs2302561 | -1.92 | 0.54 | 0.029 | 0.602 | -0.009 | -0.019 | -0.019 | 0 | 0.34 | 0.59 | 0.45 | 0.19 | 0.32 | 0.22 |
| 2 | 71234671 | rs6760496 | -1.663 | 0.166 | 0.419 | 0.569 | -0.056 | -0.138 | -0.138 | 0.101 | 0.2 | 0.43 | 0.27 | 0.01 | 0.11 | 0.13 |
| 2 | 71235034 | rs10196871 | -1.134 | 0.319 | -0.275 | 0.773 | 0.002 | -0.019 | -0.019 | 0.603 | 0.14 | 0.15 | 0.25 | 0 | 0.06 | 0.12 |
| 2 | 71253907 | rs407439 | -0.963 | 0.707 | -0.369 | 1.741 | -0.002 | 0.029 | 0.029 | 0 | 0.08 | 0.06 | 0.09 | 0.28 | 0.08 | 0.22 |
| 2 | 71263768 | rs13420242 | 1.215 | 0.542 | -0.198 | 1.459 | -0.008 | 0.041 | 0.041 | 5.322 | 0.75 | 0.4 | 0.68 | 0.5 | 0.83 | 0.52 |
| 2 | 71274196 | rs2418891 | -0.178 | 0.626 | -0.468 | 0.706 | 0 | 0 | 0 | 0 | 0.41 | 0.55 | 0.43 | 0.72 | 0.43 | 0.54 |
| 2 | 71275700 | rs7576317 | -0.748 | 0.88 | -0.359 | 0.659 | 0 | 0.003 | 0.003 | 0 | 0.26 | 0.24 | 0.31 | 0.37 | 0.25 | 0.43 |
| 2 | 71285678 | rs6725116 | 0.634 | 0.884 | -0.255 | 0.646 | 0.014 | 0.105 | 0.105 | 0 | 0.07 | 0.11 | 0.13 | 0.23 | 0.08 | 0.26 |
| 2 | 71285874 | rs10167593 | 1.474 | 0.673 | -0.232 | 0.58 | -0.021 | 0.057 | 0.057 | 0.265 | 0.22 | 0.1 | 0.16 | 0.12 | 0.24 | 0.13 |
| 2 | 71288817 | rs9636398 | -0.169 | 0.707 | 0.306 | 0.607 | 0 | 0.004 | 0.004 | 0 | 0.22 | 0.21 | 0.27 | 0.61 | 0.17 | 0.25 |
| 2 | 71288912 | rs9636399 | -1.04 | 0.701 | 0.314 | 0.489 | 0 | -0.042 | -0.042 | 0.154 | 0.3 | 0.34 | 0.4 | 0.85 | 0.26 | 0.5 |
| 2 | 71291058 | rs11126315 | -0.791 | 0.709 | 0.307 | 0.493 | 0 | -0.017 | -0.017 | 0 | 0.28 | 0.29 | 0.39 | 0.85 | 0.25 | 0.5 |
| 2 | 71292664 | rs11675048 | -0.756 | 0.569 | 0.835 | 0.487 | -0.002 | 0.054 | 0.054 | 0 | 0.28 | 0.39 | 0.32 | 0.42 | 0.23 | 0.43 |
| 2 | 71294128 | rs11890238 | 0.616 | 0.573 | 0.823 | 0.492 | 0.016 | 0.106 | 0.106 | 0 | 0.08 | 0.13 | 0.13 | 0.24 | 0.09 | 0.26 |
| 2 | 71295054 | rs2287329 | 0.406 | 0.543 | 0.733 | 0.475 | 0.001 | 0.018 | 0.018 | 0 | 0.18 | 0.11 | 0.17 | 0.14 | 0.14 | 0.16 |
| 2 | 71297057 | rs888352 | 1.601 | 0.338 | 0.562 | 0.296 | -0.027 | 0.076 | 0.076 | 1.108 | 0.22 | 0.06 | 0.17 | 0.1 | 0.25 | 0.14 |
| 2 | 71297695 | rs2287328 | 0.481 | 0.266 | 0.735 | 0.332 | 0 | 0.014 | 0.014 | 0 | 0.17 | 0.12 | 0.17 | 0.14 | 0.14 | 0.16 |
| 2 | 71297982 | rs2287327 | -1.331 | 0.009 | 1.305 | 0.226 | 0 | 0.01 | 0.01 | 0 | 0.35 | 0.56 | 0.35 | 0.4 | 0.3 | 0.51 |
| 2 | 71305359 | rs2302794 | 0.075 | 0.019 | 1.291 | -0.055 | 0.021 | 0.045 | 0.045 | 2.628 | 0.41 | 0.32 | 0.3 | 0.25 | 0.32 | 0.32 |
| 2 | 71308656 | rs17429341 | 0.386 | 0.066 | 1.192 | -0.075 | -0.001 | -0.013 | -0.013 | 0.58 | 0.09 | 0.07 | 0.09 | 0 | 0.15 | 0.09 |
| 2 | 71309586 | rs1558899 | 0.865 | -0.048 | 1.265 | 0.187 | 0.023 | 0.086 | 0.086 | 0 | 0.08 | 0.11 | 0.15 | 0.21 | 0.1 | 0.26 |
| 2 | 71311643 | rs4852246 | 0.296 | -0.109 | 1.226 | 0.098 | 0.018 | 0.095 | 0.095 | 0.612 | 0.38 | 0.16 | 0.29 | 0.25 | 0.33 | 0.31 |
| 2 | 71313262 | rs6546667 | -0.826 | 0 | 1.293 | 0.033 | 0.005 | -0.006 | -0.006 | 0 | 0.52 | 0.59 | 0.53 | 0.96 | 0.44 | 0.65 |
| 2 | 71320364 | rs6744843 | 0.485 | 0.159 | 1.383 | 0.606 | 0.014 | 0.066 | 0.066 | 0 | 0.08 | 0.09 | 0.15 | 0.22 | 0.1 | 0.26 |
| 2 | 71327146 | rs6546670 | 0.185 | 0.201 | 1.436 | 0.577 | 0.072 | 0.192 | 0.192 | 0 | 0.08 | 0.25 | 0.16 | 0.22 | 0.1 | 0.26 |
| 2 | 71330331 | rs7558220 | 0.028 | -0.028 | 0.936 | 0.385 | 0.016 | 0.004 | 0.004 | 0 | 0.14 | 0.13 | 0.11 | 0.15 | 0.05 | 0.16 |
| 2 | 71334998 | rs6546674 | -1.008 | 0.023 | 1.096 | 0.171 | -0.048 | 0.051 | 0.051 | 0 | 0.25 | 0.54 | 0.26 | 0.68 | 0.15 | 0.32 |
| 2 | 71335872 | rs10198989 | -0.653 | 0.322 | 1.227 | -0.185 | 0.062 | -0.043 | -0.043 | 0.048 | 0.91 | 0.72 | 0.92 | 1 | 0.86 | 0.93 |
| 2 | 71338585 | rs9653504 | -0.797 | 0.265 | 1.194 | -0.191 | -0.037 | 0.037 | 0.037 | 0.312 | 0.26 | 0.4 | 0.26 | 0.68 | 0.17 | 0.31 |
| 2 | 71342142 | rs17583054 | 0.528 | 0.981 | 1.307 | 0.124 | 0.032 | 0.118 | 0.118 | 0 | 0.07 | 0.13 | 0.12 | 0.27 | 0.11 | 0.23 |
| 2 | 71360625 | rs1813160 | -0.754 | 0.802 | 0.984 | -0.197 | 0.025 | 0.018 | 0.018 | 0 | 0.27 | 0.43 | 0.4 | 0.27 | 0.3 | 0.37 |
| 2 | 71365280 | rs357758 | -0.481 | 0.807 | 1.002 | -0.195 | -0.013 | -0.01 | -0.01 | 0 | 0.25 | 0.18 | 0.39 | 0.26 | 0.29 | 0.36 |
| 2 | 71376201 | rs1458868 | 1.553 | 0.501 | 1.337 | 0.09 | -0.038 | 0.081 | 0.081 | 0 | 0.58 | 0.23 | 0.45 | 0.28 | 0.53 | 0.47 |
| 2 | 71376999 | rs6574 | -0.907 | 0.951 | 1.221 | -0.059 | -0.011 | -0.008 | -0.008 | 0.344 | 0.25 | 0.18 | 0.39 | 0.26 | 0.29 | 0.36 |
| 2 | 71377745 | rs357741 | -1.036 | 0.982 | 0.784 | -0.049 | 0.027 | 0.022 | 0.022 | 0 | 0.27 | 0.44 | 0.41 | 0.28 | 0.3 | 0.37 |
| 2 | 71380052 | rs7604412 | 0.875 | 0.55 | 0.288 | -0.802 | 0.012 | -0.12 | -0.12 | 1.974 | 0.73 | 0.63 | 0.67 | 0.99 | 0.68 | 0.85 |
| 2 | 71381984 | rs357728 | 0.191 | 0.448 | 0.356 | -0.692 | 0.058 | -0.024 | -0.024 | 0.027 | 0.94 | 0.79 | 0.93 | 0.99 | 0.89 | 0.97 |
| 2 | 71382495 | rs11674899 | -0.514 | 0.201 | 0.143 | -0.761 | 0.006 | -0.018 | -0.018 | 0.018 | 0.19 | 0.13 | 0.26 | 0 | 0.18 | 0.11 |
| 2 | 71403284 | rs417263 | 0.738 | 0.284 | 1.132 | -0.929 | 0.033 | -0.137 | -0.137 | 0 | 0.8 | 0.56 | 0.87 | 1 | 0.73 | 0.87 |
| 2 | 71403616 | rs413693 | 0.066 | 0.08 | 0.744 | -1.214 | 0 | 0.007 | 0.007 | 0 | 0.18 | 0.17 | 0.11 | 0.14 | 0.14 | 0.17 |
| 2 | 71408929 | rs12478186 | 2.213 | -0.222 | 0.163 | -1.266 | -0.069 | 0.13 | 0.13 | 0.053 | 0.39 | 0.05 | 0.47 | 0.11 | 0.41 | 0.18 |
| 2 | 71409350 | rs412014 | -0.499 | -0.286 | 0.245 | -0.771 | 0.001 | 0.002 | 0.002 | 0 | 0.28 | 0.33 | 0.22 | 0.33 | 0.23 | 0.53 |
| 2 | 71413984 | rs3732238 | 0.178 | -0.631 | 0.055 | -0.951 | -0.014 | 0.032 | 0.032 | 0 | 0.15 | 0.27 | 0.13 | 0.2 | 0.09 | 0.36 |
| 2 | 71414007 | rs3821248 | 0.178 | -0.631 | 0.076 | -0.951 | -0.014 | 0.032 | 0.032 | 0 | 0.15 | 0.27 | 0.13 | 0.2 | 0.09 | 0.37 |
| 2 | 71414645 | rs3187463 | 0.41 | -0.004 | 0.4 | -0.63 | 0.022 | 0.083 | 0.083 | 0.179 | 0.11 | 0.26 | 0.13 | 0.2 | 0.09 | 0.36 |
| 2 | 71417134 | rs357776 | -0.461 | 0.065 | 0.536 | -0.651 | 0.001 | 0.003 | 0.003 | 0 | 0.28 | 0.33 | 0.22 | 0.33 | 0.23 | 0.53 |

| | | | | | | | | | | | | | | | | |
|---|---|---|---|---|---|---|---|---|---|---|---|---|---|---|---|---|
| 2 | 71428191 | rs357762 | -0.036 | 0.076 | 0.766 | -0.612 | 0.02 | -0.102 | -0.102 | 0 | 0.8 | 0.65 | 0.87 | 1 | 0.73 | 0.87 |
| 2 | 71431885 | rs12615297 | -0.125 | -0.041 | 0.669 | -0.48 | 0.004 | 0.02 | 0.02 | 0 | 0.13 | 0.1 | 0.17 | 0.55 | 0.09 | 0.16 |
| 2 | 71433860 | rs13030137 | 0.61 | 0.037 | 0.701 | -0.548 | -0.002 | -0.02 | -0.02 | 0 | 0.08 | 0.08 | 0.05 | 0.01 | 0.06 | 0.08 |
| 2 | 71436854 | rs741472 | -1.046 | 0.024 | 0.687 | -0.526 | 0.02 | 0.012 | 0.012 | 0.158 | 0.23 | 0.31 | 0.31 | 0.73 | 0.24 | 0.55 |
| 2 | 71438784 | rs4852771 | 1.457 | 0.021 | 0.658 | -0.591 | 0.001 | -0.018 | -0.018 | 2.275 | 0.57 | 0.47 | 0.58 | 0.11 | 0.58 | 0.28 |
| 2 | 71443645 | rs4852772 | -0.038 | -0.058 | 0.719 | -0.443 | -0.024 | 0.028 | 0.028 | 0 | 0.81 | 0.89 | 0.88 | 0.83 | 0.8 | 0.76 |
| 2 | 71448455 | rs763713 | -1.343 | -0.046 | 0.797 | -0.238 | 0.002 | -0.018 | -0.018 | 0.725 | 0.89 | 0.94 | 0.95 | 0.81 | 0.92 | 0.87 |
| 2 | 71461604 | rs13016081 | 0.654 | -0.021 | 0.934 | -0.598 | -0.002 | -0.049 | -0.049 | 0 | 0.13 | 0.17 | 0.09 | 0 | 0.18 | 0.09 |
| 2 | 71464128 | rs918192 | -0.192 | 0.015 | 0.847 | -0.53 | 0.016 | -0.053 | -0.053 | 2.963 | 0.12 | 0.19 | 0.11 | 0.01 | 0.16 | 0.11 |
| 2 | 71467971 | rs6713034 | 0.335 | 0.014 | 0.906 | -0.467 | 0.012 | 0.064 | 0.064 | 0.01 | 0.11 | 0.22 | 0.11 | 0.18 | 0.09 | 0.36 |
| 2 | 71471154 | rs1870384 | 0.475 | 0.013 | 0.911 | -0.467 | 0.013 | 0.058 | 0.058 | 0 | 0.1 | 0.16 | 0.11 | 0.18 | 0.09 | 0.36 |
| 2 | 71486250 | rs12476972 | 1.466 | -0.006 | 0.819 | -0.491 | 0.004 | -0.048 | -0.048 | 0 | 0.62 | 0.5 | 0.62 | 0.11 | 0.64 | 0.32 |
| 2 | 71488944 | rs2609505 | 0.858 | 0.034 | 0.595 | -0.482 | -0.016 | 0.009 | 0.009 | 0 | 0.08 | 0.05 | 0.07 | 0.01 | 0.12 | 0.11 |
| 2 | 71507414 | rs11126328 | 1.278 | 0.009 | 0.667 | -0.461 | 0.016 | -0.013 | -0.013 | 0 | 0.66 | 0.54 | 0.74 | 0.65 | 0.67 | 0.4 |
| 2 | 71509666 | rs2542540 | -0.055 | 0.117 | 1.002 | -0.477 | -0.012 | -0.015 | -0.015 | 0 | 0.85 | 0.78 | 0.92 | 0.85 | 0.89 | 0.87 |
| 2 | 71516191 | rs6725892 | 1.001 | 0.032 | 1.083 | -0.425 | -0.001 | -0.001 | -0.001 | 0.185 | 0.75 | 0.77 | 0.75 | 0.29 | 0.77 | 0.69 |
| 2 | 71522113 | rs7419694 | -0.729 | 0.055 | 1.098 | -0.452 | -0.004 | -0.005 | -0.005 | 0 | 0.88 | 0.86 | 0.94 | 0.84 | 0.92 | 0.87 |
| 2 | 71525478 | rs2216373 | -0.402 | 0.09 | 1.022 | -0.455 | -0.006 | 0.018 | 0.018 | 0 | 0.13 | 0.11 | 0.19 | 0.56 | 0.16 | 0.18 |
| 2 | 71525602 | rs12713744 | 0.007 | -0.055 | 1.115 | -0.555 | -0.017 | -0.1 | -0.1 | 0.227 | 0.16 | 0.35 | 0.1 | 0 | 0.18 | 0.1 |
| 2 | 71525698 | rs7590738 | 2.287 | -0.13 | 0.909 | -0.6 | 0.027 | 0.04 | 0.04 | 0 | 0.53 | 0.19 | 0.62 | 0.3 | 0.56 | 0.56 |
| 2 | 71529775 | rs7579108 | -0.965 | -0.146 | 0.951 | -0.676 | -0.041 | -0.127 | -0.127 | 0 | 0.22 | 0.58 | 0.14 | 0 | 0.21 | 0.13 |
| 2 | 71531413 | rs2542547 | -1.015 | -0.074 | 0.885 | -0.604 | 0.004 | 0.007 | 0.007 | 0 | 0.89 | 0.93 | 0.95 | 0.84 | 0.92 | 0.87 |
| 2 | 71558924 | rs2257136 | -0.862 | -0.069 | 0.762 | -0.602 | -0.001 | -0.002 | -0.002 | 0 | 0.88 | 0.87 | 0.94 | 0.84 | 0.92 | 0.87 |
| 2 | 71583191 | rs2542501 | 0.57 | -0.014 | 0.738 | -0.675 | -0.003 | -0.01 | -0.01 | 0.02 | 0.09 | 0.06 | 0.07 | 0.01 | 0.13 | 0.11 |
| 2 | 71584272 | rs17006902 | 0.616 | -0.06 | 0.932 | -0.728 | -0.007 | -0.064 | -0.064 | 0 | 0.12 | 0.21 | 0.08 | 0 | 0.16 | 0.09 |
| 2 | 71589618 | rs11691076 | -0.379 | -0.056 | 0.651 | -0.623 | -0.007 | 0.015 | 0.015 | 0 | 0.13 | 0.09 | 0.19 | 0.55 | 0.16 | 0.18 |
| 2 | 71589743 | rs2542509 | -1.415 | -0.098 | 0.713 | -0.661 | -0.002 | -0.082 | -0.082 | 0.054 | 0.31 | 0.62 | 0.2 | 0.01 | 0.34 | 0.23 |
| 2 | 71615534 | rs2542528 | 0.675 | -0.058 | 0.659 | -0.659 | 0 | 0 | 0 | 0 | 0.12 | 0.11 | 0.06 | 0.15 | 0.08 | 0.13 |
| 2 | 71631109 | rs3732235 | 0.646 | -0.097 | 0.684 | -0.644 | -0.002 | -0.002 | -0.002 | 0 | 0.12 | 0.13 | 0.06 | 0.16 | 0.08 | 0.13 |
| 2 | 71633275 | rs3732233 | 2.257 | -0.137 | 0.665 | -0.757 | 0.023 | 0.037 | 0.037 | 0 | 0.53 | 0.16 | 0.64 | 0.29 | 0.55 | 0.57 |
| 2 | 71633389 | rs6714975 | 2.231 | -0.121 | 0.671 | -0.749 | 0.024 | 0.036 | 0.036 | 0 | 0.53 | 0.22 | 0.62 | 0.29 | 0.55 | 0.57 |
| 2 | 71654175 | rs1804020 | -1.202 | -0.077 | 0.487 | -0.559 | -0.003 | -0.007 | -0.007 | 0 | 0.25 | 0.23 | 0.24 | 0.7 | 0.24 | 0.31 |
| 2 | 71680632 | rs6546693 | -0.396 | -0.242 | 0.093 | -0.303 | -0.048 | 0.157 | 0.157 | 0 | 0.15 | 0.33 | 0.1 | 0.48 | 0.07 | 0.14 |
| 2 | 71685529 | rs13406110 | 0.226 | 0.001 | -0.329 | -0.579 | -0.008 | 0.079 | 0.079 | 0.067 | 0.2 | 0.06 | 0.12 | 0.13 | 0.22 | 0.21 |
| 2 | 71688249 | rs4268945 | -0.446 | -0.257 | -0.049 | -0.414 | 0.001 | 0.055 | 0.055 | 0 | 0.28 | 0.42 | 0.29 | 0.59 | 0.28 | 0.26 |
| 2 | 71688588 | rs11898330 | -0.39 | -0.073 | -0.269 | -0.548 | -0.021 | -0.094 | -0.094 | 3.3 | 0.3 | 0.44 | 0.13 | 0.13 | 0.24 | 0.26 |
| 2 | 71689050 | rs7582270 | 0.275 | -0.074 | -1.167 | -0.104 | 0.009 | -0.07 | -0.07 | 1.625 | 0.18 | 0.12 | 0.16 | 0.44 | 0.15 | 0.15 |
| 2 | 71695111 | rs6738477 | -0.275 | 0.412 | -1.042 | -0.058 | 0.009 | -0.054 | -0.054 | 0 | 0.32 | 0.42 | 0.23 | 0.26 | 0.35 | 0.33 |
| 2 | 71701352 | rs4428032 | -0.534 | 0.605 | -1.834 | 0.287 | -0.045 | -0.088 | -0.088 | 0.003 | 0.62 | 0.88 | 0.58 | 0.56 | 0.6 | 0.56 |
| 2 | 71703364 | rs12477276 | 0.097 | 0.824 | -1.3 | -0.544 | 0.018 | -0.086 | -0.086 | 0.015 | 0.3 | 0.45 | 0.34 | 0.14 | 0.38 | 0.24 |
| 2 | 71703945 | rs10865385 | -0.347 | 0.489 | -1.233 | -0.029 | -0.011 | -0.008 | -0.008 | 0.523 | 0.35 | 0.49 | 0.27 | 0.63 | 0.26 | 0.31 |
| 2 | 71706697 | rs4852787 | -0.196 | 0.582 | -1.571 | -0.803 | -0.012 | 0.116 | 0.116 | 0 | 0.57 | 0.27 | 0.65 | 0.79 | 0.47 | 0.68 |
| 2 | 71708732 | rs12998057 | -0.322 | -0.025 | -0.665 | -0.29 | -0.024 | 0.024 | 0.024 | 1.046 | 0.18 | 0.26 | 0.07 | 0.21 | 0.11 | 0.19 |
| 2 | 71710068 | rs10187354 | 1.107 | 0.136 | -1.067 | -1.073 | -0.032 | 0.161 | 0.161 | 0 | 0.4 | 0.16 | 0.31 | 0.14 | 0.51 | 0.28 |
| 2 | 71710615 | rs13035702 | -0.26 | 0.137 | -0.319 | -0.625 | -0.037 | 0.06 | 0.06 | 0.579 | 0.18 | 0.34 | 0.07 | 0.26 | 0.1 | 0.19 |
| 2 | 71713738 | rs4309600 | -0.699 | 0.226 | -0.535 | -0.856 | -0.001 | -0.009 | -0.009 | 1.443 | 0.22 | 0.19 | 0.32 | 0.1 | 0.18 | 0.22 |
| 2 | 71714223 | rs4536672 | 0.622 | 0.284 | -0.892 | -1.042 | -0.036 | -0.026 | -0.026 | 0.365 | 0.78 | 0.89 | 0.84 | 0.77 | 0.76 | 0.78 |
| 2 | 71715665 | rs4852260 | 1.6 | -0.083 | -0.982 | -1.473 | -0.029 | 0.129 | 0.129 | 0.228 | 0.39 | 0.17 | 0.29 | 0.13 | 0.51 | 0.26 |
| 2 | 71716494 | rs7574062 | 0.372 | 0.255 | -0.632 | -0.794 | 0.007 | -0.035 | -0.035 | 0 | 0.09 | 0.03 | 0.04 | 0.23 | 0.06 | 0.16 |
| 2 | 71717062 | rs4270372 | 0.495 | 0.28 | -0.606 | -0.789 | -0.04 | -0.049 | -0.049 | 0.545 | 0.78 | 0.91 | 0.84 | 0.75 | 0.77 | 0.78 |
| 2 | 71722096 | rs4852261 | 0.419 | 0.371 | 0.148 | -1.066 | 0.037 | -0.042 | -0.042 | 0.007 | 0.92 | 0.79 | 0.92 | 1 | 0.86 | 0.97 |
| 2 | 71722232 | rs4852792 | 0.589 | 0.082 | 0.289 | -1.156 | -0.001 | -0.009 | -0.009 | 0 | 0.64 | 0.58 | 0.61 | 0.29 | 0.67 | 0.48 |
| 2 | 71724254 | rs4852794 | 0.85 | 0.084 | 0.485 | -1.197 | 0.001 | -0.024 | -0.024 | 0.474 | 0.61 | 0.52 | 0.57 | 0.27 | 0.61 | 0.45 |
| 2 | 71724362 | rs6749480 | 0.081 | -0.204 | 0.67 | -1.181 | -0.014 | -0.03 | -0.03 | 0.719 | 0.09 | 0.13 | 0.02 | 0 | 0.03 | 0.03 |
| 2 | 71724396 | rs11689987 | -0.815 | -0.169 | 0.774 | -1.187 | 0 | 0.026 | 0.026 | 0.671 | 0.08 | 0.12 | 0.05 | 0.15 | 0.06 | 0.13 |
| 2 | 71729786 | rs12713752 | 0.47 | -0.205 | 1.066 | -1.612 | -0.007 | -0.077 | -0.077 | 0 | 0.66 | 0.57 | 0.63 | 0.13 | 0.73 | 0.39 |
| 2 | 71730055 | rs12713753 | 1.186 | -0.327 | 0.634 | -1.734 | 0.02 | 0.065 | 0.065 | 2.854 | 0.53 | 0.34 | 0.57 | 0.11 | 0.64 | 0.35 |
| 2 | 71733582 | rs12469262 | 1.58 | -0.205 | 0.091 | -1.675 | -0.09 | 0.284 | 0.284 | 0.328 | 0.45 | 0.12 | 0.49 | 0.12 | 0.53 | 0.32 |
| 2 | 71733941 | rs9309457 | -0.326 | -0.187 | 0.679 | -0.684 | -0.004 | 0.017 | 0.017 | 0.902 | 0.78 | 0.72 | 0.75 | 0.43 | 0.84 | 0.53 |
| 2 | 71734357 | rs9677918 | 0.305 | -0.216 | 0.894 | -1.496 | -0.015 | -0.059 | -0.059 | 0 | 0.7 | 0.59 | 0.66 | 0.17 | 0.78 | 0.44 |
| 2 | 71736806 | rs4852797 | 1.086 | -0.132 | 0.966 | -1.595 | 0 | -0.016 | -0.016 | 2.819 | 0.57 | 0.42 | 0.58 | 0.15 | 0.66 | 0.37 |
| 2 | 71737334 | rs12467684 | 1.171 | -0.142 | 1.07 | -1.769 | 0.003 | 0.038 | 0.038 | 2.566 | 0.56 | 0.38 | 0.57 | 0.13 | 0.65 | 0.37 |
| 2 | 71737389 | rs11690862 | 0.488 | -0.062 | 1.276 | -1.646 | 0.007 | 0.005 | 0.005 | 0 | 0.09 | 0.12 | 0.07 | 0.11 | 0.11 | 0.1 |
| 2 | 71737519 | rs11679925 | 0.976 | -0.074 | 1.203 | -1.646 | 0.001 | -0.004 | -0.004 | 1 | 0.08 | 0.07 | 0.03 | 0.11 | 0.06 | 0.09 |
| 2 | 71738385 | rs3924297 | 0.074 | 0.018 | 1.157 | -1.781 | -0.003 | -0.041 | -0.041 | 0.085 | 0.1 | 0.2 | 0.09 | 0 | 0.1 | 0.04 |
| 2 | 71739734 | rs7581887 | -1.789 | -0.003 | 1.26 | -1.834 | 0.003 | -0.058 | -0.058 | 0 | 0.31 | 0.53 | 0.21 | 0.13 | 0.29 | 0.2 |
| 2 | 71742978 | rs13428194 | -0.351 | 0.098 | 0.763 | -1.788 | 0.016 | -0.025 | -0.025 | 0 | 0.11 | 0.15 | 0.09 | 0.03 | 0.13 | 0.07 |
| 2 | 71743376 | rs11903960 | -0.242 | 0.168 | 0.81 | -1.724 | 0.049 | -0.005 | -0.005 | 0 | 0.05 | 0.14 | 0.07 | 0.03 | 0.09 | 0.04 |
| 2 | 71747235 | rs11675897 | -0.753 | 0.018 | 0.688 | -1.785 | 0.007 | 0 | 0 | 0 | 0.79 | 0.89 | 0.71 | 0.25 | 0.84 | 0.53 |

| | | | | | | | | | | | | | | | | |
|---|---|---|---|---|---|---|---|---|---|---|---|---|---|---|---|---|
| 2 | 71754430 | rs4350773 | 1.668 | 0.134 | 0.65 | -2.077 | -0.019 | 0.134 | 0.134 | 0.058 | 0.45 | 0.21 | 0.49 | 0.11 | 0.54 | 0.3 |
| 2 | 71756645 | rs6713986 | 0.538 | 0.14 | 1.221 | -1.538 | -0.003 | -0.012 | -0.012 | 0 | 0.69 | 0.65 | 0.62 | 0.23 | 0.72 | 0.49 |
| 2 | 71757432 | rs12329322 | 1.204 | 0.145 | 1.251 | -1.535 | -0.005 | 0.018 | 0.018 | 0.001 | 0.07 | 0.17 | 0.03 | 0.1 | 0.04 | 0.08 |
| 2 | 71758834 | rs13408415 | 1.041 | 0.131 | 1.018 | -1.558 | -0.013 | -0.099 | -0.099 | 2.216 | 0.62 | 0.47 | 0.59 | 0.12 | 0.67 | 0.39 |
| 2 | 71773740 | rs4852805 | 0.932 | 0.37 | 0.217 | -1.257 | -0.008 | -0.067 | -0.067 | 0 | 0.65 | 0.45 | 0.58 | 0.2 | 0.68 | 0.4 |
| 2 | 71774011 | rs4852806 | -0.135 | 0.543 | -0.633 | -0.735 | 0.005 | 0.003 | 0.003 | 0 | 0.81 | 0.86 | 0.72 | 0.22 | 0.86 | 0.49 |
| 2 | 71780215 | rs2303596 | 0.212 | 0.826 | -0.927 | -0.756 | 0.001 | 0.041 | 0.041 | 0.113 | 0.69 | 0.78 | 0.64 | 0.22 | 0.73 | 0.46 |
| 2 | 71782242 | rs734248 | 0.179 | 0.723 | -0.888 | -0.766 | -0.002 | 0.031 | 0.031 | 0.009 | 0.69 | 0.75 | 0.62 | 0.21 | 0.72 | 0.46 |
| 2 | 71791577 | rs17007067 | -0.734 | 0.892 | -0.404 | -0.684 | 0.01 | -0.067 | -0.067 | 0 | 0.13 | 0.24 | 0.09 | 0.01 | 0.17 | 0.07 |
| 2 | 71797176 | rs10183967 | -0.182 | 1.026 | -1.228 | -0.967 | -0.037 | 0.146 | 0.146 | 0 | 0.55 | 0.83 | 0.55 | 0.22 | 0.5 | 0.47 |
| 2 | 71802951 | rs11680124 | 0.126 | 0.399 | -1.582 | -0.04 | -0.037 | -0.097 | -0.097 | 1.251 | 0.19 | 0.06 | 0.2 | 0.35 | 0.25 | 0.07 |
| 2 | 71805522 | rs10169824 | -0.484 | 0.637 | -0.716 | -0.547 | -0.002 | -0.001 | -0.001 | 0.014 | 0.56 | 0.64 | 0.53 | 0.8 | 0.56 | 0.59 |
| 2 | 71806112 | rs7568834 | -0.569 | 0.599 | -0.742 | -0.57 | -0.002 | -0.008 | -0.008 | 0 | 0.57 | 0.65 | 0.53 | 0.82 | 0.57 | 0.59 |
| 2 | 71813055 | rs13413696 | -0.294 | 0.315 | -0.241 | -0.552 | -0.017 | -0.012 | -0.012 | 0 | 0.08 | 0.26 | 0.1 | 0.02 | 0.03 | 0.02 |
| 2 | 71817003 | rs1473228 | -0.868 | 0.095 | -0.525 | -0.702 | -0.028 | 0.057 | 0.057 | 0 | 0.68 | 0.81 | 0.64 | 0.81 | 0.61 | 0.73 |
| 2 | 71818820 | rs9973420 | -0.588 | -0.136 | -0.411 | -0.606 | -0.002 | 0.003 | 0.003 | 1.921 | 0.32 | 0.37 | 0.25 | 0.36 | 0.29 | 0.16 |
| 2 | 71819017 | rs12474121 | 1.497 | -0.084 | -0.988 | -0.373 | -0.049 | -0.102 | -0.102 | 0 | 0.27 | 0.02 | 0.28 | 0.35 | 0.27 | 0.27 |
| 2 | 71819471 | rs11126337 | 0.042 | 0.06 | 0.296 | -0.168 | -0.001 | 0.071 | 0.071 | 0.115 | 0.22 | 0.35 | 0.22 | 0.36 | 0.24 | 0.09 |
| 2 | 71821847 | rs17616999 | 0.618 | -0.159 | -0.509 | -0.071 | -0.002 | -0.054 | -0.054 | 0 | 0.19 | 0.1 | 0.18 | 0.37 | 0.2 | 0.09 |
| 2 | 71823762 | rs1529406 | -0.792 | -0.085 | -0.694 | -0.153 | -0.051 | -0.087 | -0.087 | 0 | 0.66 | 0.9 | 0.68 | 0.56 | 0.62 | 0.42 |
| 2 | 71826077 | rs934062 | 1.292 | -0.094 | -0.875 | -0.107 | -0.083 | -0.126 | -0.126 | 0.57 | 0.28 | 0.02 | 0.29 | 0.36 | 0.31 | 0.39 |
| 2 | 71826102 | rs2303602 | -0.206 | -0.179 | -0.018 | -0.428 | 0.001 | -0.024 | -0.024 | 0 | 0.4 | 0.43 | 0.42 | 0.2 | 0.42 | 0.27 |
| 2 | 71827555 | rs13028056 | 0.683 | -0.144 | -0.743 | -0.3 | -0.079 | 0.075 | 0.075 | 0 | 0.26 | 0.04 | 0.17 | 0.15 | 0.33 | 0.19 |
| 2 | 71830689 | rs1529409 | 0.852 | -0.585 | 0.52 | -0.062 | -0.007 | -0.014 | -0.014 | 0 | 0.89 | 0.8 | 0.94 | 0.91 | 0.93 | 0.79 |
| 2 | 71834676 | rs12472134 | -0.591 | -0.492 | -0.196 | -0.033 | -0.108 | -0.093 | -0.093 | 0 | 0.66 | 0.95 | 0.69 | 0.57 | 0.6 | 0.44 |
| 2 | 71835788 | rs4290680 | -1.542 | -0.543 | 0.824 | 0.049 | 0.001 | 0.041 | 0.041 | 3.238 | 0.1 | 0.51 | 0.1 | 0.05 | 0.04 | 0.04 |
| 2 | 71836525 | rs13030767 | -0.641 | -0.614 | 0.631 | 0 | -0.084 | -0.086 | -0.086 | 0.589 | 0.62 | 0.92 | 0.64 | 0.52 | 0.58 | 0.44 |
| 2 | 71838597 | rs2303606 | 0.155 | -0.488 | 0.919 | 0.1 | -0.004 | 0.001 | 0.001 | 0 | 0.48 | 0.36 | 0.53 | 0.48 | 0.54 | 0.39 |
| 2 | 71838852 | rs2016605 | -1.31 | -0.352 | 1.408 | 0.22 | 0.023 | 0.044 | 0.044 | 0.281 | 0.08 | 0.33 | 0.09 | 0.04 | 0.04 | 0.04 |
| 2 | 71841175 | rs13409105 | -1.804 | -0.296 | 0.768 | 0.226 | 0.066 | 0.074 | 0.074 | 1.054 | 0.07 | 0.47 | 0.09 | 0.03 | 0.04 | 0.01 |
| 2 | 71856229 | rs227772 | 0.306 | -0.285 | -0.425 | 0.125 | -0.012 | -0.007 | -0.007 | 0 | 0.45 | 0.33 | 0.59 | 0.51 | 0.51 | 0.38 |
| 2 | 71856682 | rs227771 | 0.677 | -0.254 | -0.732 | 0.139 | -0.067 | -0.064 | -0.064 | 0.731 | 0.41 | 0.15 | 0.49 | 0.48 | 0.5 | 0.38 |
| 2 | 71860893 | rs847622 | 0.247 | 0.335 | -0.715 | 0.395 | 0 | 0.002 | 0.002 | 0.014 | 0.5 | 0.47 | 0.61 | 0.51 | 0.51 | 0.43 |
| 2 | 71861587 | rs17718067 | -0.001 | 0.733 | -1.933 | 0.684 | -0.061 | -0.08 | -0.08 | 0 | 0.36 | 0.05 | 0.28 | 0.4 | 0.36 | 0.36 |
| 2 | 71864377 | rs227768 | 0.814 | 0.987 | -0.987 | 0.726 | -0.003 | -0.001 | -0.001 | 0 | 0.95 | 0.96 | 0.96 | 0.92 | 0.91 | 0.83 |
| 2 | 71865287 | rs13387541 | -1.511 | 0.466 | -0.025 | 0.412 | -0.033 | -0.054 | -0.054 | 0 | 0.13 | 0.38 | 0.3 | 0.04 | 0.03 | 0.05 |
| 2 | 71866842 | rs227782 | 1.167 | 0.642 | -0.116 | 0.663 | 0.011 | 0.012 | 0.012 | 0.324 | 0.72 | 0.42 | 0.53 | 0.65 | 0.7 | 0.65 |
| 2 | 71871039 | rs84181 | 0.783 | 0.599 | 0.311 | 0.758 | 0.001 | 0.011 | 0.011 | 0 | 0.9 | 0.86 | 0.71 | 0.81 | 0.88 | 0.82 |
| 2 | 71875036 | rs227777 | 0.638 | 0.558 | 0.265 | 0.857 | -0.034 | -0.057 | -0.057 | 0 | 0.91 | 0.99 | 0.71 | 0.79 | 0.88 | 0.83 |
| 2 | 71880211 | rs1736168 | 2.261 | 0.672 | 0.256 | 0.739 | -0.005 | 0.002 | 0.002 | 1.665 | 0.67 | 0.37 | 0.52 | 0.62 | 0.7 | 0.61 |
| 2 | 71891361 | rs2303599 | 1.645 | 0.884 | 0.611 | 0.947 | 0.015 | 0.091 | 0.091 | 0 | 0.87 | 0.67 | 0.68 | 0.69 | 0.87 | 0.81 |
| 2 | 71895157 | rs11674060 | -0.998 | 0.987 | 0.856 | 1.134 | 0.031 | 0 | 0 | 0.27 | 0.13 | 0.29 | 0.07 | 0.1 | 0.16 | 0.21 |
| 2 | 71897075 | rs3791829 | 0.962 | 0.81 | 0.992 | 1.153 | 0.011 | 0.068 | 0.068 | 0 | 0.91 | 0.81 | 0.87 | 0.76 | 0.9 | 0.83 |
| 2 | 71899180 | rs3791833 | -0.773 | 1.86 | 0.978 | 1.356 | 0.001 | -0.001 | -0.001 | 0 | 0.14 | 0.13 | 0.09 | 0.12 | 0.18 | 0.24 |
| 2 | 71900805 | rs13024390 | 2.384 | 2.134 | 1.299 | 1.21 | 0.007 | -0.005 | -0.005 | 0 | 0.64 | 0.43 | 0.64 | 0.64 | 0.62 | 0.56 |
| 2 | 71901286 | rs2559081 | 2.228 | 2.101 | 1.383 | 1.243 | -0.034 | 0.033 | 0.033 | 1.254 | 0.8 | 0.58 | 0.77 | 0.7 | 0.86 | 0.79 |
| 2 | 71906171 | rs1863812 | 0.685 | 2.094 | 1.238 | 1.24 | -0.017 | -0.059 | -0.059 | 0 | 0.9 | 0.99 | 0.89 | 0.76 | 0.9 | 0.83 |
| 2 | 71906278 | rs17718530 | -0.824 | 2.421 | 1.246 | 1.254 | -0.005 | 0.004 | 0.004 | 0.006 | 0.16 | 0.15 | 0.12 | 0.07 | 0.26 | 0.24 |
| 2 | 71906725 | rs4436990 | 2.448 | 2.499 | 1.39 | 1.255 | 0.014 | -0.008 | -0.008 | 0 | 0.64 | 0.47 | 0.65 | 0.65 | 0.61 | 0.57 |
| 2 | 71907149 | rs10187598 | 0.919 | 2.458 | 1.424 | 1.25 | -0.005 | 0.024 | 0.024 | 0 | 0.89 | 0.84 | 0.87 | 0.75 | 0.9 | 0.83 |
| 2 | 71907425 | rs10198903 | 1.704 | 2.489 | 1.408 | 1.248 | -0.015 | -0.002 | -0.002 | 0 | 0.73 | 0.83 | 0.78 | 0.69 | 0.68 | 0.58 |
| 2 | 71907930 | rs7558412 | 0.741 | 2.257 | 1.06 | 1.054 | 0.003 | -0.013 | -0.013 | 0 | 0.84 | 0.91 | 0.86 | 0.73 | 0.84 | 0.77 |
| 2 | 71909822 | rs2559082 | -1.284 | 2.391 | 1.046 | 1.042 | -0.014 | 0.02 | 0.02 | 0.089 | 0.2 | 0.12 | 0.13 | 0.08 | 0.25 | 0.26 |
| 2 | 71914438 | rs1961080 | 1.927 | 1.839 | 0.495 | 0.576 | 0 | 0 | 0 | 0 | 0.47 | 0.52 | 0.51 | 0.56 | 0.48 | 0.36 |
| 2 | 71917617 | rs2162625 | -1.024 | 2.419 | 0.879 | 0.727 | 0.001 | 0 | 0 | 0 | 0.2 | 0.17 | 0.19 | 0.09 | 0.31 | 0.29 |
| 2 | 71919512 | rs12713761 | -0.959 | 2.354 | 0.893 | 0.723 | 0.012 | -0.011 | -0.011 | 0 | 0.19 | 0.22 | 0.17 | 0.09 | 0.27 | 0.29 |
| 2 | 71920075 | rs2900975 | -1.827 | 2.237 | 0.851 | 0.729 | 0.037 | -0.047 | -0.047 | 0 | 0.27 | 0.47 | 0.3 | 0.12 | 0.33 | 0.31 |
| 2 | 71920555 | rs2114809 | -1.576 | 2.246 | 0.836 | 0.72 | 0.033 | -0.05 | -0.05 | 0 | 0.26 | 0.39 | 0.21 | 0.09 | 0.32 | 0.3 |
| 2 | 71929775 | rs750355 | 1.866 | 1.614 | 0.802 | 1.131 | -0.008 | 0.041 | 0.041 | 0 | 0.79 | 0.56 | 0.69 | 0.68 | 0.83 | 0.78 |
| 2 | 71932953 | rs6736164 | 1.282 | 1.625 | 0.371 | 1.204 | 0.001 | -0.01 | -0.01 | 3.318 | 0.84 | 0.9 | 0.85 | 0.71 | 0.85 | 0.78 |
| 2 | 71941150 | rs7598653 | 1.435 | 2.106 | 0.61 | 1.509 | 0.033 | 0.024 | 0.024 | 0 | 0.64 | 0.46 | 0.41 | 0.49 | 0.6 | 0.46 |
| 2 | 71943026 | rs10192575 | -0.779 | 1.828 | 0.479 | 1.266 | 0.043 | -0.025 | -0.025 | 0 | 0.09 | 0.51 | 0.08 | 0.01 | 0.07 | 0.01 |
| 2 | 71943247 | rs2589513 | 0.136 | 2.153 | 0.114 | 1.385 | -0.11 | -0.037 | -0.037 | 0 | 0.85 | 0.98 | 0.63 | 0.9 | 0.69 | 0.66 |
| 2 | 71945432 | rs934069 | -1.397 | 1.612 | 0.391 | 1.47 | -0.006 | 0.049 | 0.049 | 0.196 | 0.22 | 0.64 | 0.26 | 0.4 | 0.15 | 0.17 |
| 2 | 71946135 | rs2004421 | -1.403 | 1.59 | 0.391 | 1.47 | -0.009 | 0.049 | 0.049 | 0.185 | 0.22 | 0.64 | 0.26 | 0.4 | 0.14 | 0.17 |
| 2 | 71950376 | rs4852822 | 2.478 | 1.493 | -0.145 | 1.298 | -0.143 | -0.081 | -0.081 | 0 | 0.55 | 0.03 | 0.51 | 0.48 | 0.57 | 0.47 |
| 2 | 71953686 | rs6749644 | 1.067 | 1.45 | 0.23 | 1.472 | -0.004 | -0.001 | -0.001 | 0 | 0.79 | 0.79 | 0.79 | 0.83 | 0.73 | 0.65 |
| 2 | 71974813 | rs11681627 | -2.232 | 1.322 | 0.454 | 1.62 | -0.034 | 0.043 | 0.043 | 0 | 0.38 | 0.77 | 0.36 | 0.19 | 0.38 | 0.45 |
| 2 | 71975921 | rs4123814 | -0.044 | 1.152 | 0.386 | 1.631 | -0.006 | -0.02 | -0.02 | 0 | 0.11 | 0.03 | 0.15 | 0.45 | 0.08 | 0.18 |

| | | | | | | | | | | | | | | | | |
|---|---|---|---|---|---|---|---|---|---|---|---|---|---|---|---|---|
| 2 | 71986420 | rs7570634 | -1.543 | 1.206 | 1.297 | 1.364 | 0.014 | -0.034 | -0.034 | 0.011 | 0.27 | 0.59 | 0.29 | 0.11 | 0.31 | 0.33 |
| 2 | 71989267 | rs6746698 | -2.405 | 1.229 | 0.659 | 1.478 | -0.076 | -0.091 | -0.091 | 0.017 | 0.42 | 0.91 | 0.47 | 0.53 | 0.39 | 0.52 |
| 2 | 71993330 | rs10175361 | -2.402 | 1.082 | 0.632 | 1.513 | -0.058 | -0.09 | -0.09 | 0 | 0.42 | 0.91 | 0.52 | 0.54 | 0.37 | 0.52 |
| 2 | 71993963 | rs6748598 | -1.031 | 1.077 | 0.777 | 1.515 | 0 | -0.002 | -0.002 | 0.906 | 0.23 | 0.21 | 0.3 | 0.11 | 0.28 | 0.32 |
| 2 | 71997634 | rs4852826 | -2.211 | 1.153 | 0.277 | 1.573 | -0.059 | -0.091 | -0.091 | 0 | 0.42 | 0.91 | 0.52 | 0.54 | 0.36 | 0.52 |
| 2 | 71999008 | rs12623780 | -0.869 | 1.151 | 0.454 | 1.625 | 0.002 | 0.003 | 0.003 | 0 | 0.14 | 0.12 | 0.16 | 0.43 | 0.09 | 0.19 |
| 2 | 72005343 | rs6711634 | -1.506 | 1.177 | 0.829 | 1.603 | -0.001 | -0.001 | -0.001 | 0.117 | 0.4 | 0.61 | 0.5 | 0.59 | 0.37 | 0.55 |
| 2 | 72010747 | rs6546721 | -1.708 | 1.209 | -0.033 | 1.485 | -0.01 | 0.014 | 0.014 | 0 | 0.43 | 0.87 | 0.51 | 0.62 | 0.37 | 0.55 |
| 2 | 72011314 | rs11684597 | -1.67 | 1.214 | 0.087 | 1.347 | 0.002 | 0.038 | 0.038 | 0 | 0.43 | 0.81 | 0.51 | 0.66 | 0.37 | 0.56 |
| 2 | 72013617 | rs1809265 | -0.629 | 0.403 | 0.428 | 0.962 | -0.019 | 0.027 | 0.027 | 0.265 | 0.29 | 0.36 | 0.36 | 0.66 | 0.19 | 0.27 |
| 2 | 72014622 | rs6712030 | -1.429 | 0.43 | 1.091 | 0.687 | 0.062 | 0.02 | 0.02 | 0.653 | 0.06 | 0.5 | 0.05 | 0 | 0.02 | 0.02 |
| 2 | 72022087 | rs2419777 | -1.503 | 0.764 | 0.581 | 0.653 | 0.005 | 0.034 | 0.034 | 0 | 0.49 | 0.62 | 0.54 | 0.72 | 0.4 | 0.44 |
| 2 | 72023218 | rs12713764 | -1.375 | 0.753 | 0.57 | 0.648 | 0.001 | 0.023 | 0.023 | 0.129 | 0.47 | 0.62 | 0.54 | 0.72 | 0.4 | 0.44 |
| 2 | 72027858 | rs10206242 | -1.189 | 0.862 | 0.257 | 1.02 | 0.018 | -0.037 | -0.037 | 0 | 0.39 | 0.75 | 0.46 | 0.57 | 0.28 | 0.29 |
| 2 | 72030760 | rs2419778 | -0.739 | 0.854 | 0.189 | 0.407 | -0.003 | 0.029 | 0.029 | 0.404 | 0.47 | 0.55 | 0.5 | 0.17 | 0.46 | 0.31 |
| 2 | 72031172 | rs898234 | -0.384 | 0.747 | 0.368 | 0.445 | 0.013 | -0.026 | -0.026 | 0 | 0.38 | 0.47 | 0.5 | 0.23 | 0.41 | 0.29 |
| 2 | 72031828 | rs2125637 | -0.916 | 0.724 | 0.199 | 0.616 | -0.005 | 0.057 | 0.057 | 0.054 | 0.49 | 0.61 | 0.55 | 0.23 | 0.46 | 0.32 |
| 2 | 72033801 | rs7588101 | -0.459 | 0.451 | 0.875 | 1.02 | 0.045 | 0.019 | 0.019 | 0.021 | 0.13 | 0.33 | 0.11 | 0.09 | 0.15 | 0.18 |
| 2 | 72035671 | rs17007417 | 0.191 | 0.158 | 0.708 | 1.266 | 0.005 | 0.014 | 0.014 | 0 | 0.89 | 0.85 | 0.68 | 0.84 | 0.9 | 0.79 |
| 2 | 72035866 | rs17485056 | -0.047 | -0.119 | 0.548 | 0.992 | -0.005 | 0.024 | 0.024 | 0 | 0.4 | 0.19 | 0.32 | 0.26 | 0.55 | 0.26 |
| 2 | 72037487 | rs11684170 | 1.41 | -0.302 | 0.638 | 0.908 | 0.034 | -0.107 | -0.107 | 0 | 0.38 | 0.13 | 0.27 | 0.58 | 0.33 | 0.48 |
| 2 | 72038203 | rs11679843 | 1.151 | -0.43 | 1.012 | 0.936 | -0.03 | 0.036 | 0.036 | 0.132 | 0.83 | 0.46 | 0.62 | 0.81 | 0.91 | 0.79 |
| 2 | 72044808 | rs6546723 | 0.488 | -0.379 | 0.676 | 0.944 | -0.008 | 0.008 | 0.008 | 0 | 0.86 | 0.8 | 0.64 | 0.82 | 0.91 | 0.75 |
| 2 | 72050664 | rs1459269 | 0.654 | -0.39 | 0.792 | 0.936 | -0.017 | 0.019 | 0.019 | 0 | 0.85 | 0.71 | 0.64 | 0.81 | 0.91 | 0.73 |
| 2 | 72062531 | rs6705555 | 1.297 | -0.406 | 0.488 | 0.977 | 0.018 | -0.116 | -0.116 | 0 | 0.39 | 0.12 | 0.28 | 0.58 | 0.35 | 0.44 |
| 2 | 72066907 | rs1471219 | -0.273 | -0.382 | 1.191 | 0.99 | 0.006 | 0.013 | 0.013 | 0.199 | 0.44 | 0.33 | 0.35 | 0.27 | 0.56 | 0.31 |
| 2 | 72073715 | rs13427905 | -0.329 | -0.328 | 1.21 | 0.99 | -0.002 | 0.032 | 0.032 | 0 | 0.9 | 0.85 | 0.69 | 0.82 | 0.92 | 0.78 |
| 2 | 72077960 | rs11895277 | 0.46 | -0.348 | 1.492 | 0.972 | 0.003 | 0.113 | 0.113 | 0 | 0.87 | 0.63 | 0.67 | 0.76 | 0.91 | 0.75 |
| 2 | 72083272 | rs2169027 | -0.246 | -0.368 | 1.15 | 0.994 | -0.007 | 0.059 | 0.059 | 0 | 0.9 | 0.82 | 0.7 | 0.78 | 0.94 | 0.8 |
| 2 | 72083780 | rs7596753 | 0.37 | -0.388 | 1.141 | 0.981 | -0.013 | 0.041 | 0.041 | 0 | 0.87 | 0.8 | 0.7 | 0.76 | 0.93 | 0.8 |
| 2 | 72088761 | rs11687017 | 0.915 | -0.498 | 1.152 | 0.969 | 0.028 | -0.015 | -0.015 | 0.251 | 0.43 | 0.33 | 0.33 | 0.58 | 0.37 | 0.46 |
| 2 | 72089937 | rs10208677 | -0.398 | -0.419 | 1.165 | 0.973 | -0.001 | -0.007 | -0.007 | 0 | 0.46 | 0.42 | 0.36 | 0.21 | 0.56 | 0.32 |
| 2 | 72090576 | rs10175577 | -0.223 | -0.048 | 0.843 | 1.142 | 0 | 0.04 | 0.04 | 0 | 0.91 | 0.84 | 0.7 | 0.8 | 0.93 | 0.81 |
| 2 | 72092652 | rs7565433 | -0.173 | -0.044 | 0.845 | 1.088 | 0 | 0.033 | 0.033 | 0 | 0.9 | 0.86 | 0.7 | 0.81 | 0.93 | 0.81 |
| 2 | 72096377 | rs6737513 | -0.132 | -0.776 | 0.969 | 1.111 | 0.01 | 0.012 | 0.012 | 0.106 | 0.58 | 0.77 | 0.43 | 0.46 | 0.45 | 0.48 |
| 2 | 72097591 | rs13425927 | -1.712 | -0.604 | 1.588 | 1.403 | 0.035 | 0.185 | 0.185 | 0.112 | 0.08 | 0.27 | 0.1 | 0.25 | 0.08 | 0.17 |
| 2 | 72099440 | rs1459262 | 0.474 | -0.665 | 1.03 | 1.176 | 0.03 | 0.036 | 0.036 | 0.284 | 0.44 | 0.34 | 0.23 | 0.42 | 0.38 | 0.35 |
| 2 | 72101633 | rs1380263 | -0.29 | -0.706 | 1.036 | 1.077 | 0.001 | 0.012 | 0.012 | 0.06 | 0.54 | 0.7 | 0.35 | 0.67 | 0.48 | 0.55 |
| 2 | 72102665 | rs11678520 | 0.441 | -1.426 | 1.058 | 1.117 | 0.032 | -0.018 | -0.018 | 0 | 0.39 | 0.22 | 0.26 | 0.48 | 0.33 | 0.4 |
| 2 | 72105601 | rs6710683 | -0.612 | -1.386 | 0.952 | 1.098 | 0.032 | 0.02 | 0.02 | 0 | 0.55 | 0.87 | 0.35 | 0.68 | 0.41 | 0.49 |
| 2 | 72107826 | rs881455 | -2.008 | -1.259 | 1.787 | 1.212 | 0.085 | 0.092 | 0.092 | 0 | 0.07 | 0.38 | 0.11 | 0.1 | 0.05 | 0.1 |
| 2 | 72111155 | rs6546730 | -0.237 | -1.382 | 1.647 | 1.185 | 0.092 | 0.011 | 0.011 | 0 | 0.51 | 0.71 | 0.29 | 0.51 | 0.36 | 0.44 |
| 2 | 72111651 | rs7584587 | 0.449 | -1.324 | 1.428 | 1.176 | 0 | 0 | 0 | 0 | 0.41 | 0.38 | 0.21 | 0.41 | 0.29 | 0.34 |
| 2 | 72116933 | rs6546731 | -0.177 | -1.309 | 0.865 | 0.735 | 0.061 | 0.032 | 0.032 | 0.226 | 0.52 | 0.68 | 0.32 | 0.67 | 0.38 | 0.47 |
| 2 | 72126420 | rs744027 | -0.035 | -1.293 | 0.462 | 0.722 | 0.149 | 0.047 | 0.047 | 0.076 | 0.51 | 0.75 | 0.28 | 0.62 | 0.34 | 0.39 |
| 2 | 72131032 | rs6712232 | -0.619 | -0.8 | 0.873 | 0.57 | 0.049 | -0.003 | -0.003 | 2.618 | 0.05 | 0.23 | 0.09 | 0.03 | 0.05 | 0.02 |
| 2 | 72138330 | rs9973794 | 0.053 | -0.917 | 0.949 | 0.516 | 0.02 | 0.008 | 0.008 | 0.668 | 0.06 | 0.22 | 0.02 | 0 | 0.03 | 0.01 |
| 2 | 72145170 | rs1380268 | -0.985 | -0.863 | 0.426 | 0.541 | 0.04 | -0.032 | -0.032 | 0 | 0.1 | 0.52 | 0.11 | 0.04 | 0.08 | 0.03 |
| 2 | 72150898 | rs6735281 | -1.068 | -0.881 | 0.063 | 0.54 | 0.013 | -0.045 | -0.045 | 2.554 | 0.11 | 0.57 | 0.09 | 0.04 | 0.08 | 0.03 |
| 2 | 72155152 | rs908614 | 0.121 | -0.652 | -1.147 | 0.708 | -0.01 | 0.005 | 0.005 | 0 | 0.39 | 0.1 | 0.63 | 0.24 | 0.54 | 0.52 |
| 2 | 72160412 | rs6753449 | 0.255 | -0.81 | -1.296 | 0.455 | 0.022 | 0.048 | 0.048 | 0 | 0.44 | 0.18 | 0.62 | 0.36 | 0.6 | 0.52 |
| 2 | 72161465 | rs4852275 | -0.45 | -1.584 | -1.301 | 0.229 | 0.018 | -0.076 | -0.076 | 0 | 0.67 | 0.85 | 0.78 | 0.55 | 0.7 | 0.62 |
| 2 | 72164293 | rs7570117 | -0.922 | -1.433 | -0.911 | 0.061 | -0.037 | 0.01 | 0.01 | 0.856 | 0.13 | 0.64 | 0.11 | 0.14 | 0.06 | 0.07 |
| 2 | 72167136 | rs12052842 | -1.194 | -1.412 | -0.567 | 0.126 | -0.033 | 0.001 | 0.001 | 0.82 | 0.14 | 0.63 | 0.1 | 0.13 | 0.07 | 0.04 |
| 2 | 72169353 | rs7588891 | 0.337 | -1.611 | -1.214 | 0.11 | -0.025 | -0.109 | -0.109 | 0.452 | 0.67 | 0.95 | 0.63 | 0.47 | 0.69 | 0.73 |
| 2 | 72173427 | rs6546733 | -0.089 | -1.73 | -0.329 | 0.017 | -0.027 | 0.117 | 0.117 | 0 | 0.22 | 0.58 | 0.23 | 0.5 | 0.2 | 0.21 |
| 2 | 72187212 | rs13010378 | 0.98 | -1.67 | -0.357 | -0.4 | 0.031 | 0.086 | 0.086 | 0 | 0.57 | 0.17 | 0.61 | 0.3 | 0.62 | 0.72 |
| 2 | 72189807 | rs954163 | 0.34 | -1.671 | -0.445 | -0.402 | -0.023 | -0.05 | -0.05 | 0.046 | 0.66 | 0.93 | 0.68 | 0.43 | 0.66 | 0.74 |
| 2 | 72191462 | rs6723028 | -1.612 | -1.468 | 0.126 | -0.751 | -0.01 | -0.029 | -0.029 | 0 | 0.14 | 0.59 | 0.16 | 0.08 | 0.12 | 0.04 |
| 2 | 72191662 | rs6736570 | -0.125 | -1.758 | -0.676 | -0.958 | -0.009 | -0.007 | -0.007 | 0 | 0.65 | 0.93 | 0.7 | 0.33 | 0.66 | 0.71 |
| 2 | 72195993 | rs7569239 | 0.393 | -1.516 | -0.054 | -1.076 | 0.01 | 0.031 | 0.031 | 0 | 0.48 | 0.59 | 0.6 | 0.29 | 0.53 | 0.62 |
| 2 | 72203991 | rs7558552 | -1.22 | -0.731 | 0.492 | -0.798 | 0.015 | -0.019 | -0.019 | 0 | 0.12 | 0.27 | 0.08 | 0.08 | 0.09 | 0.04 |
| 2 | 72206006 | rs6722569 | -0.128 | -0.297 | 0.284 | -1.111 | -0.014 | 0.108 | 0.108 | 0 | 0.51 | 0.81 | 0.62 | 0.32 | 0.49 | 0.53 |
| 2 | 72219476 | rs17007663 | 0.333 | -0.32 | 0.618 | -1.031 | -0.017 | -0.03 | -0.03 | 0 | 0.06 | 0.15 | 0.04 | 0 | 0.05 | 0.02 |
| 2 | 72220565 | rs4144821 | -0.866 | -0.26 | 0.265 | -1.189 | -0.068 | -0.081 | -0.081 | 0 | 0.63 | 0.98 | 0.74 | 0.37 | 0.67 | 0.6 |
| 2 | 72221149 | rs11899600 | -1.532 | 0.1 | 0.975 | -1.057 | -0.008 | 0.013 | 0.013 | 3.029 | 0.11 | 0.53 | 0.07 | 0.08 | 0.07 | 0.03 |
| 2 | 72234860 | rs17559295 | 1.432 | 0.29 | 0.388 | -1.054 | -0.04 | 0.013 | 0.013 | 0 | 0.08 | 0 | 0.11 | 0.05 | 0.12 | 0.04 |
| 2 | 72235688 | rs4405788 | -1.046 | 0.202 | 0.978 | -0.953 | -0.003 | 0.052 | 0.052 | 0 | 0.57 | 0.75 | 0.65 | 0.32 | 0.56 | 0.57 |
| 2 | 72236929 | rs4852833 | -0.613 | 0.132 | 1.152 | -0.991 | 0.005 | 0.069 | 0.069 | 0 | 0.5 | 0.66 | 0.6 | 0.31 | 0.51 | 0.51 |

| | | | | | | | | | | | | | | | | |
|---|---|---|---|---|---|---|---|---|---|---|---|---|---|---|---|---|
| 2 | 72239692 | rs883313 | 0.301 | 0.069 | 0.921 | -1.011 | -0.067 | 0.088 | 0.088 | 0 | 0.39 | 0.11 | 0.52 | 0.24 | 0.45 | 0.51 |
| 2 | 72241295 | rs10208040 | -0.315 | 0.084 | 1.157 | -1.024 | -0.004 | 0.015 | 0.015 | 0 | 0.46 | 0.59 | 0.55 | 0.31 | 0.48 | 0.53 |
| 2 | 72241504 | rs7568074 | -1.966 | 0.032 | 1.158 | -1.045 | -0.027 | -0.059 | -0.059 | 0 | 0.18 | 0.41 | 0.11 | 0.08 | 0.12 | 0.09 |
| 2 | 72242013 | rs2176407 | -1.03 | 0.015 | 0.826 | -0.988 | -0.005 | 0.074 | 0.074 | 0.18 | 0.55 | 0.77 | 0.65 | 0.31 | 0.54 | 0.58 |
| 2 | 72251751 | rs1878503 | -0.08 | 0.469 | 0.607 | -0.857 | -0.068 | 0.146 | 0.146 | 0 | 0.44 | 0.11 | 0.55 | 0.22 | 0.45 | 0.53 |
| 2 | 72254126 | rs4852837 | -0.217 | 0.435 | 0.818 | -0.804 | -0.015 | 0.099 | 0.099 | 0 | 0.47 | 0.27 | 0.58 | 0.23 | 0.49 | 0.54 |
| 2 | 72254272 | rs4852838 | -0.877 | 0.519 | 0.418 | -0.765 | -0.01 | 0.06 | 0.06 | 0 | 0.56 | 0.75 | 0.64 | 0.31 | 0.55 | 0.57 |
| 2 | 72264864 | rs4852279 | -0.201 | 0.573 | -0.193 | -0.209 | -0.005 | 0.009 | 0.009 | 0 | 0.41 | 0.4 | 0.53 | 0.34 | 0.44 | 0.54 |
| 2 | 72270951 | rs1400681 | -0.405 | 0.638 | 0.051 | 0.164 | -0.003 | 0.016 | 0.016 | 0 | 0.08 | 0.06 | 0.24 | 0.02 | 0.08 | 0.08 |
| 2 | 72273058 | rs6750558 | -0.902 | 0.458 | 0.573 | -0.011 | -0.017 | -0.024 | -0.024 | 0 | 0.35 | 0.56 | 0.28 | 0.31 | 0.27 | 0.27 |
| 2 | 72305463 | rs13406776 | 0.162 | 0.214 | 0.829 | 0.033 | 0 | -0.014 | -0.014 | 0 | 0.08 | 0.09 | 0.24 | 0.02 | 0.08 | 0.08 |
| 2 | 72306479 | rs7601186 | -1.898 | 0.267 | 0.662 | -0.029 | 0.045 | 0.102 | 0.102 | 0 | 0.47 | 0.91 | 0.55 | 0.37 | 0.35 | 0.36 |
| 2 | 72308674 | rs11126361 | -1.109 | 0.262 | 0.797 | -0.025 | 0.017 | 0.019 | 0.019 | 0 | 0.37 | 0.74 | 0.31 | 0.35 | 0.27 | 0.26 |
| 2 | 72309757 | rs6737062 | -0.797 | 0.348 | 0.797 | -0.114 | -0.026 | -0.053 | -0.053 | 0.222 | 0.33 | 0.54 | 0.22 | 0.25 | 0.25 | 0.24 |
| 2 | 72310763 | rs17007729 | -1.084 | 0.417 | 1.288 | 0.202 | 0.023 | -0.035 | -0.035 | 0 | 0.12 | 0.37 | 0.27 | 0.05 | 0.1 | 0.08 |
| 2 | 72319253 | rs10168803 | -1.27 | 0.434 | 1.617 | 0.587 | -0.003 | -0.009 | -0.009 | 0 | 0.24 | 0.32 | 0.32 | 0.26 | 0.22 | 0.2 |
| 2 | 72320853 | rs7568849 | -2.278 | 0.211 | 1.383 | 0.572 | 0.023 | 0.034 | 0.034 | 0.01 | 0.41 | 0.9 | 0.56 | 0.37 | 0.32 | 0.33 |
| 2 | 72325404 | rs10194900 | -1.437 | 0.51 | 2.228 | 0.84 | 0.023 | 0.1 | 0.1 | 0 | 0.25 | 0.53 | 0.29 | 0.44 | 0.27 | 0.16 |
| 2 | 72334559 | rs6716351 | -2.524 | 0.529 | 2.322 | 1.174 | -0.017 | -0.056 | -0.056 | 0 | 0.32 | 0.89 | 0.42 | 0.43 | 0.29 | 0.24 |
| 2 | 72340493 | rs10203870 | -1.434 | 0.62 | 2.984 | 1.227 | 0.05 | 0.15 | 0.15 | 0 | 0.14 | 0.33 | 0.14 | 0.4 | 0.21 | 0.11 |
| 2 | 72345674 | rs6546738 | -1.437 | 0.743 | 2.937 | 1.125 | 0.053 | 0.209 | 0.209 | 0 | 0.13 | 0.36 | 0.13 | 0.4 | 0.16 | 0.11 |
| 2 | 72345739 | rs7589043 | -1.977 | 0.729 | 2.203 | 1.135 | -0.001 | -0.002 | -0.002 | 0 | 0.27 | 0.76 | 0.32 | 0.41 | 0.24 | 0.22 |
| 2 | 72348725 | rs6546741 | -2.046 | 0.725 | 1.92 | 1.125 | -0.009 | -0.025 | -0.025 | 0 | 0.28 | 0.8 | 0.34 | 0.41 | 0.29 | 0.22 |
| 2 | 72353138 | rs4852856 | -2.156 | 0.633 | 1.812 | 1.104 | -0.001 | -0.047 | -0.047 | 0 | 0.3 | 0.86 | 0.33 | 0.43 | 0.24 | 0.24 |
| 2 | 72355518 | rs7606254 | -1.57 | 0.148 | 1.848 | 1.067 | -0.024 | 0.044 | 0.044 | 0 | 0.23 | 0.43 | 0.25 | 0.34 | 0.17 | 0.22 |
| 2 | 72356195 | rs887844 | -1.889 | 0.249 | 1.042 | 0.494 | 0.024 | 0.058 | 0.058 | 0 | 0.43 | 0.77 | 0.39 | 0.68 | 0.38 | 0.37 |
| 2 | 72356370 | rs2420201 | -1.264 | 0.562 | 1.489 | 0.47 | -0.015 | -0.043 | -0.043 | 0 | 0.1 | 0.13 | 0.14 | 0 | 0.05 | 0.08 |
| 2 | 72356449 | rs707718 | -1.702 | 0.774 | 1.232 | 0.716 | 0.005 | -0.012 | -0.012 | 0.147 | 0.27 | 0.76 | 0.29 | 0.55 | 0.2 | 0.3 |
| 2 | 72361960 | rs2241057 | -1.893 | 1.557 | 1.49 | -0.427 | 0.039 | -0.015 | -0.015 | 0 | 0.12 | 0.29 | 0.21 | 0.08 | 0.17 | 0.1 |
| 2 | 72394893 | rs194242 | -1.63 | 2.011 | 1.414 | -1.074 | 0.031 | -0.118 | -0.118 | 0 | 0.25 | 0.58 | 0.31 | 0.03 | 0.3 | 0.14 |
| 2 | 72405665 | rs17007922 | -1.024 | 2.336 | 2.096 | -1.078 | 0.079 | -0.052 | -0.052 | 0 | 0.1 | 0.38 | 0.1 | 0 | 0.1 | 0.01 |
| 2 | 72405833 | rs41413 | -2.74 | 2.256 | 2.217 | -1.198 | -0.003 | -0.02 | -0.02 | 0 | 0.24 | 0.74 | 0.35 | 0.02 | 0.33 | 0.12 |
| 2 | 72409750 | rs41409 | -0.787 | 2.243 | 2.15 | -1.183 | -0.034 | 0.024 | 0.024 | 0 | 0.08 | 0.02 | 0.2 | 0.02 | 0.18 | 0.11 |
| 2 | 72418689 | rs41403 | -2.406 | 2.186 | 3.148 | -1.237 | 0.095 | -0.119 | -0.119 | 6.747 | 0.18 | 0.54 | 0.29 | 0.02 | 0.27 | 0.11 |
| 2 | 72419055 | rs41402 | -2.801 | 2.202 | 3.511 | -1.244 | 0.051 | -0.1 | -0.1 | 0 | 0.22 | 0.59 | 0.32 | 0.02 | 0.32 | 0.12 |
| 2 | 72425292 | rs41393 | -2.138 | 2.176 | 3.386 | -1.245 | 0.113 | -0.104 | -0.104 | 0 | 0.17 | 0.44 | 0.29 | 0.02 | 0.27 | 0.11 |
| 2 | 72430862 | rs194235 | -3.063 | 2.206 | 3.09 | -1.26 | -0.031 | 0.059 | 0.059 | 5.038 | 0.24 | 0.85 | 0.34 | 0.02 | 0.3 | 0.11 |
| 2 | 72430888 | rs7597679 | -2.427 | 2.225 | 3.139 | -1.262 | 0.004 | -0.059 | -0.059 | 0.469 | 0.16 | 0.73 | 0.15 | 0 | 0.15 | 0.02 |
| 2 | 72433296 | rs13417114 | -2.218 | 2.217 | 3.126 | -1.254 | 0.051 | -0.048 | -0.048 | 0.658 | 0.12 | 0.27 | 0.23 | 0.02 | 0.2 | 0.1 |
| 2 | 72464241 | rs1159766 | -2.584 | 2.331 | 2.845 | -1.3 | 0.001 | -0.032 | -0.032 | 0 | 0.21 | 0.74 | 0.32 | 0.02 | 0.26 | 0.11 |
| 2 | 72483919 | rs10205108 | -2.749 | 2.32 | 2.833 | -1.305 | -0.007 | -0.008 | -0.008 | 2.049 | 0.22 | 0.78 | 0.32 | 0.02 | 0.26 | 0.11 |
| 2 | 72485039 | rs17007984 | -1.404 | 2.286 | 2.992 | -1.309 | 0.076 | -0.067 | -0.067 | 0 | 0.11 | 0.51 | 0.11 | 0 | 0.11 | 0.01 |
| 2 | 72501904 | rs17008015 | -1.897 | 2.309 | 2.856 | -1.316 | 0.084 | -0.084 | -0.084 | 0 | 0.13 | 0.54 | 0.14 | 0 | 0.16 | 0.02 |
| 2 | 72505475 | rs4852288 | -2.742 | 2.104 | 2.721 | -1.385 | -0.002 | -0.033 | -0.033 | 0 | 0.22 | 0.76 | 0.32 | 0.01 | 0.26 | 0.12 |
| 2 | 72507640 | rs10204415 | -1.734 | 2.133 | 2.694 | -1.363 | 0.007 | -0.031 | -0.031 | 0.083 | 0.1 | 0.17 | 0.2 | 0.01 | 0.15 | 0.11 |
| 2 | 72522769 | rs13409513 | -2.165 | 2.096 | 2.661 | -1.322 | 0.028 | -0.033 | -0.033 | 0 | 0.08 | 0.27 | 0.09 | 0.01 | 0.1 | 0.08 |
| 2 | 72533963 | rs9309464 | -2.857 | 2.115 | 2.611 | -1.342 | -0.049 | 0.131 | 0.131 | 3.46 | 0.21 | 0.95 | 0.19 | 0.01 | 0.2 | 0.09 |
| 2 | 72535478 | rs9309465 | -2.16 | 2.113 | 2.666 | -1.341 | -0.002 | -0.04 | -0.04 | 0.545 | 0.14 | 0.75 | 0.1 | 0 | 0.1 | 0.01 |
| 2 | 72544130 | rs10173381 | -3.031 | 2.107 | 2.648 | -1.331 | -0.049 | 0.133 | 0.133 | 6.383 | 0.22 | 0.96 | 0.19 | 0.01 | 0.2 | 0.09 |
| 2 | 72560841 | rs11903912 | -1.638 | 2.107 | 2.713 | -1.333 | 0.016 | -0.076 | -0.076 | 0.284 | 0.12 | 0.63 | 0.1 | 0 | 0.1 | 0.01 |
| 2 | 72584121 | rs13390754 | -2.972 | 2.109 | 2.662 | -1.335 | -0.01 | 0.044 | 0.044 | 3.246 | 0.2 | 0.87 | 0.18 | 0.01 | 0.2 | 0.09 |
| 2 | 72648432 | rs7596126 | -1.785 | 2.108 | 2.732 | -1.34 | 0.003 | -0.066 | -0.066 | 1.015 | 0.13 | 0.68 | 0.11 | 0 | 0.1 | 0.01 |
| 2 | 72650937 | rs640610 | -2.958 | 2.107 | 2.72 | -1.342 | -0.022 | 0.102 | 0.102 | 6.377 | 0.21 | 0.92 | 0.19 | 0.01 | 0.2 | 0.09 |
| 2 | 72684766 | rs674863 | -1.78 | 2.032 | 2.718 | -1.349 | 0.009 | -0.03 | -0.03 | 0 | 0.07 | 0.19 | 0.08 | 0.01 | 0.1 | 0.07 |
| 2 | 72703425 | rs632670 | -2.172 | 2.032 | 2.739 | -1.349 | 0.039 | -0.03 | -0.03 | 0.107 | 0.08 | 0.33 | 0.1 | 0.01 | 0.1 | 0.07 |
| 2 | 72707874 | rs653220 | -3.134 | 2.3 | 2.541 | -1.538 | -0.063 | 0.163 | 0.163 | 4.846 | 0.24 | 0.95 | 0.21 | 0.01 | 0.23 | 0.09 |
| 2 | 72735509 | rs2138683 | 1.735 | 2.306 | 2.61 | -1.52 | 0.033 | -0.029 | -0.029 | 0 | 0.92 | 0.68 | 0.89 | 0.99 | 0.9 | 0.93 |
| 2 | 72749995 | rs10460572 | 1.444 | 2.381 | 2.215 | -1.777 | 0.025 | -0.031 | -0.031 | 0 | 0.92 | 0.74 | 0.9 | 0.99 | 0.9 | 0.93 |
| 2 | 72752939 | rs4852873 | 2.813 | 2.376 | 2.188 | -1.792 | -0.068 | 0.16 | 0.16 | 6.009 | 0.76 | 0.04 | 0.79 | 0.99 | 0.77 | 0.91 |
| 2 | 72758775 | rs4852875 | 1.775 | 2.386 | 2.214 | -1.784 | 0.038 | -0.03 | -0.03 | 0 | 0.91 | 0.63 | 0.89 | 0.99 | 0.9 | 0.93 |
| 2 | 72774203 | rs764766 | -1.478 | 2.388 | 2.372 | -1.784 | 0.059 | -0.094 | -0.094 | 0 | 0.14 | 0.54 | 0.09 | 0 | 0.13 | 0.02 |
| 2 | 72789552 | rs4364045 | 2.748 | 2.381 | 2.153 | -1.789 | -0.071 | 0.154 | 0.154 | 2.9 | 0.77 | 0.04 | 0.79 | 0.99 | 0.77 | 0.91 |
| 2 | 72795730 | rs1446890 | 1.709 | 2.315 | 2.165 | -1.816 | 0.014 | -0.039 | -0.039 | 0.031 | 0.91 | 0.74 | 0.9 | 0.99 | 0.9 | 0.93 |
| 2 | 72834955 | rs17008315 | -1.571 | 2.294 | 2.116 | -1.828 | 0.026 | -0.098 | -0.098 | 0 | 0.15 | 0.6 | 0.11 | 0 | 0.13 | 0.02 |
| 2 | 72882532 | rs4852886 | 2.72 | 2.299 | 2.065 | -1.829 | -0.022 | 0.085 | 0.085 | 2.67 | 0.76 | 0.1 | 0.79 | 0.99 | 0.77 | 0.91 |
| 2 | 72941246 | rs17008360 | -2.142 | 2.242 | 2.083 | -1.816 | 0 | 0.035 | 0.035 | 3.936 | 0.18 | 0.86 | 0.12 | 0 | 0.12 | 0.02 |
| 2 | 72975980 | rs990569 | 0.517 | 2.258 | 2.082 | -1.801 | -0.012 | 0.016 | 0.016 | 0 | 0.95 | 0.97 | 0.92 | 0.99 | 0.89 | 0.93 |
| 2 | 72993493 | rs6720889 | -0.92 | 2.174 | 2.602 | -1.807 | 0.031 | -0.063 | -0.063 | 0 | 0.11 | 0.26 | 0.07 | 0 | 0.12 | 0.01 |
| 2 | 73013836 | rs7593084 | -1.683 | 2.217 | 1.411 | -1.779 | 0.024 | -0.054 | -0.054 | 0.052 | 0.19 | 0.6 | 0.31 | 0.05 | 0.19 | 0.05 |

| | | | | | | | | | | | | | | | | |
|---|---|---|---|---|---|---|---|---|---|---|---|---|---|---|---|---|
| 2 | 73041727 | rs4852893 | 0.632 | 2.164 | 1.145 | -1.752 | -0.002 | 0.003 | 0.003 | 0 | 0.94 | 0.94 | 0.93 | 0.99 | 0.9 | 0.94 |
| 2 | 73075222 | rs6758431 | 1.525 | 2.153 | 1.214 | -1.758 | 0.007 | -0.046 | -0.046 | 6.198 | 0.91 | 0.73 | 0.91 | 0.99 | 0.91 | 0.94 |
| 2 | 73078673 | rs1508060 | 2.443 | 2.175 | 0.808 | -1.723 | -0.073 | 0.105 | 0.105 | 3.075 | 0.71 | 0.04 | 0.59 | 0.93 | 0.72 | 0.88 |
| 2 | 73105933 | rs2048172 | 2.302 | 1.874 | 1.383 | -1.493 | -0.005 | -0.063 | -0.063 | 0.717 | 0.88 | 0.62 | 0.9 | 0.99 | 0.9 | 0.94 |
| 2 | 73109017 | rs12611708 | 1.869 | 1.903 | 1.212 | -1.479 | 0.009 | -0.041 | -0.041 | 0 | 0.91 | 0.71 | 0.92 | 0.99 | 0.91 | 0.94 |
| 2 | 73113524 | rs2421095 | 1.957 | 1.907 | 1.157 | -1.477 | 0.001 | -0.051 | -0.051 | 0 | 0.9 | 0.7 | 0.92 | 0.99 | 0.9 | 0.94 |
| 2 | 73131167 | rs1561245 | -1.885 | 2.222 | -0.368 | -1.784 | -0.062 | 0.13 | 0.13 | 0.479 | 0.35 | 0.86 | 0.45 | 0.12 | 0.38 | 0.38 |
| 2 | 73157395 | rs999494 | -1.118 | 2.096 | -0.638 | -1.905 | 0 | 0.001 | 0.001 | 0 | 0.18 | 0.16 | 0.31 | 0.13 | 0.19 | 0.37 |
| 2 | 73158476 | rs10196456 | -1.713 | 2.03 | -0.613 | -1.975 | 0.001 | -0.029 | -0.029 | 0 | 0.22 | 0.73 | 0.25 | 0.06 | 0.24 | 0.1 |
| 2 | 73161551 | rs2077586 | -1.348 | 2.343 | -0.686 | -1.993 | 0.007 | -0.007 | -0.007 | 0 | 0.22 | 0.24 | 0.4 | 0.15 | 0.27 | 0.31 |
| 2 | 73162541 | rs12470805 | -0.895 | 2.345 | -0.807 | -1.981 | 0.003 | -0.013 | -0.013 | 0.193 | 0.18 | 0.2 | 0.32 | 0.08 | 0.23 | 0.27 |
| 2 | 73167944 | rs11126387 | -1.504 | 1.721 | -0.902 | -1.745 | -0.121 | 0.153 | 0.153 | 0.169 | 0.42 | 0.96 | 0.53 | 0.16 | 0.48 | 0.4 |
| 2 | 73172445 | rs2421097 | -1.173 | 1.441 | -0.733 | -1.761 | -0.012 | 0.027 | 0.027 | 0.006 | 0.42 | 0.71 | 0.5 | 0.16 | 0.47 | 0.39 |
| 2 | 73174848 | rs17504249 | -0.388 | 0.754 | -1.438 | -1.074 | -0.071 | 0.066 | 0.066 | 0 | 0.33 | 0.09 | 0.47 | 0.19 | 0.34 | 0.35 |
| 2 | 73179990 | rs879511 | -1.392 | 0.334 | -0.127 | -0.494 | -0.042 | -0.06 | -0.06 | 0.054 | 0.11 | 0.77 | 0.07 | 0 | 0.02 | 0.01 |
| 2 | 73199619 | rs11898850 | -0.009 | 0.746 | -0.679 | 0.397 | 0.01 | 0.003 | 0.003 | 0 | 0.52 | 0.62 | 0.29 | 0.47 | 0.44 | 0.48 |
| 2 | 73204824 | rs10209859 | -1.152 | 1.229 | -0.439 | 0.848 | 0.01 | 0.087 | 0.087 | 2.593 | 0.22 | 0.65 | 0.3 | 0.47 | 0.22 | 0.21 |
| 2 | 73210411 | rs17008541 | 0.165 | 1.162 | 0.232 | 0.815 | 0.014 | 0.061 | 0.061 | 0.117 | 0.11 | 0.19 | 0.22 | 0.3 | 0.15 | 0.16 |
| 2 | 73219922 | rs893258 | 1.461 | 1.047 | -0.657 | 0.267 | -0.111 | -0.104 | -0.104 | 0.675 | 0.58 | 0.04 | 0.58 | 0.49 | 0.54 | 0.7 |
| 2 | 73226090 | rs12233055 | -1.179 | 1.003 | 0.159 | 0.625 | 0.031 | 0.144 | 0.144 | 0.491 | 0.2 | 0.43 | 0.28 | 0.49 | 0.22 | 0.22 |
| 2 | 73226259 | rs732180 | -1.552 | 0.979 | 0.304 | 0.613 | 0.003 | 0.059 | 0.059 | 3.57 | 0.23 | 0.7 | 0.31 | 0.49 | 0.22 | 0.26 |
| 2 | 73230087 | rs17504810 | 0.538 | 1.026 | -0.108 | 0.608 | -0.002 | -0.007 | -0.007 | 0.798 | 0.07 | 0.06 | 0.03 | 0.19 | 0.06 | 0.1 |
| 2 | 73259564 | rs17008608 | 0.673 | 1.039 | -0.049 | 0.588 | -0.005 | 0.028 | 0.028 | 0 | 0.09 | 0.08 | 0.2 | 0.35 | 0.16 | 0.16 |
| 2 | 73278990 | rs6710739 | -1.136 | 0.946 | -0.191 | 0.537 | 0.005 | 0.114 | 0.114 | 1.113 | 0.22 | 0.56 | 0.28 | 0.56 | 0.21 | 0.28 |
| 2 | 73300661 | rs1046139 | 1.055 | 0.879 | -0.482 | 0.399 | -0.07 | -0.029 | -0.029 | 0 | 0.29 | 0.02 | 0.17 | 0.21 | 0.29 | 0.29 |
| 2 | 73310099 | rs10180995 | -1.615 | 1.13 | 0.59 | 0.842 | 0.012 | 0.089 | 0.089 | 0.937 | 0.23 | 0.6 | 0.29 | 0.56 | 0.22 | 0.28 |
| 2 | 73330954 | rs1430351 | -1.565 | 1.142 | 0.622 | 0.846 | 0.01 | 0.125 | 0.125 | 0.702 | 0.22 | 0.52 | 0.27 | 0.56 | 0.21 | 0.28 |
| 2 | 73333985 | rs1430349 | -1.401 | 1.181 | 0.606 | 0.854 | 0.03 | 0.136 | 0.136 | 0.37 | 0.22 | 0.46 | 0.27 | 0.56 | 0.22 | 0.29 |
| 2 | 73339430 | rs4143146 | 0.084 | 1.063 | 0.358 | 0.873 | -0.002 | 0 | 0 | 0 | 0.89 | 0.91 | 0.93 | 0.83 | 0.87 | 0.87 |
| 2 | 73343973 | rs6710755 | 1.401 | 1.047 | 0.252 | 0.791 | 0.006 | 0.034 | 0.034 | 0.122 | 0.73 | 0.36 | 0.67 | 0.43 | 0.71 | 0.68 |
| 2 | 73352605 | rs2115916 | 1.504 | 1.167 | 0.18 | 0.757 | 0.003 | 0.011 | 0.011 | 0.193 | 0.72 | 0.32 | 0.67 | 0.43 | 0.7 | 0.68 |
| 2 | 73357527 | rs2060001 | 0.226 | 1.12 | -0.13 | 0.504 | -0.014 | 0.009 | 0.009 | 0 | 0.9 | 0.93 | 0.94 | 0.87 | 0.87 | 0.87 |
| 2 | 73358292 | rs10865397 | 1.153 | 1.028 | 0.121 | 0.331 | 0.001 | 0.001 | 0.001 | 0.164 | 0.59 | 0.35 | 0.58 | 0.58 | 0.5 | 0.63 |
| 2 | 73361948 | rs9989755 | -1.548 | 0.537 | -0.023 | 0.21 | -0.073 | -0.082 | -0.082 | 0 | 0.42 | 0.91 | 0.39 | 0.41 | 0.47 | 0.35 |
| 2 | 73368662 | rs10048837 | -0.008 | 0.508 | 0.433 | 0.161 | -0.029 | -0.049 | -0.049 | 0 | 0.12 | 0.22 | 0.04 | 0.02 | 0.07 | 0.1 |
| 2 | 73374096 | rs2043100 | -0.608 | 0.513 | 0.881 | 0.125 | 0.051 | 0.141 | 0.141 | 0 | 0.23 | 0.54 | 0.3 | 0.38 | 0.31 | 0.21 |
| 2 | 73378241 | rs768851 | -1.174 | 0.499 | 0.852 | 0.117 | 0 | 0.001 | 0.001 | 0 | 0.29 | 0.72 | 0.37 | 0.39 | 0.4 | 0.25 |
| 2 | 73392069 | rs13409784 | -0.772 | 0.429 | 0.819 | 0.081 | -0.006 | -0.051 | -0.051 | 0.086 | 0.18 | 0.25 | 0.08 | 0.02 | 0.16 | 0.14 |
| 2 | 73401781 | rs6710726 | -1.437 | 0.413 | 0.548 | 0.124 | -0.004 | -0.03 | -0.03 | 0 | 0.36 | 0.77 | 0.36 | 0.41 | 0.36 | 0.31 |
| 2 | 73407442 | rs2043095 | -1.837 | 0.296 | 0.272 | 0.06 | -0.048 | -0.039 | -0.039 | 0 | 0.44 | 0.85 | 0.43 | 0.4 | 0.47 | 0.34 |
| 2 | 73413182 | rs17348756 | 0.583 | 0.284 | 0.639 | 0.05 | 0.043 | -0.028 | -0.028 | 0.137 | 0.06 | 0.17 | 0.05 | 0.01 | 0.09 | 0.04 |
| 2 | 73416873 | rs11886257 | -2.065 | 0.267 | 0.625 | 0.043 | -0.056 | -0.048 | -0.048 | 0.817 | 0.44 | 0.9 | 0.44 | 0.4 | 0.47 | 0.34 |
| 2 | 73418291 | rs12471813 | -1.503 | 0.275 | 0.617 | 0.047 | 0.002 | -0.016 | -0.016 | 0 | 0.38 | 0.72 | 0.38 | 0.4 | 0.38 | 0.29 |
| 2 | 73421207 | rs13009610 | -0.284 | 0.318 | 0.883 | 0.02 | -0.006 | -0.016 | -0.016 | 0 | 0.22 | 0.21 | 0.3 | 0.38 | 0.29 | 0.2 |
| 2 | 73425019 | rs6711278 | -2.342 | -0.304 | -0.163 | -0.313 | -0.16 | -0.037 | -0.037 | 0 | 0.61 | 0.97 | 0.49 | 0.56 | 0.54 | 0.49 |
| 2 | 73427222 | rs10189022 | -1.064 | 0.373 | 0.734 | -0.036 | 0.016 | -0.126 | -0.126 | 1.057 | 0.19 | 0.49 | 0.13 | 0.02 | 0.18 | 0.15 |
| 2 | 73442516 | rs17008864 | -0.731 | 0.089 | 0.842 | 0.041 | -0.021 | -0.098 | -0.098 | 0 | 0.15 | 0.49 | 0.08 | 0.01 | 0.1 | 0.1 |
| 2 | 73451404 | rs11893547 | 0.107 | 0.096 | 0.953 | 0.045 | -0.027 | -0.051 | -0.051 | 0.583 | 0.12 | 0.2 | 0.03 | 0.01 | 0.07 | 0.1 |
| 2 | 73455747 | rs2288631 | -0.079 | 0.221 | 0.684 | 0.094 | -0.028 | -0.087 | -0.087 | 0 | 0.24 | 0.11 | 0.29 | 0.37 | 0.3 | 0.22 |
| 2 | 73457552 | rs2288630 | -1.903 | 0.248 | 0.693 | 0.079 | -0.036 | -0.052 | -0.052 | 0 | 0.44 | 0.85 | 0.43 | 0.38 | 0.47 | 0.32 |
| 2 | 73461192 | rs2231422 | -1.488 | 0.245 | 0.73 | 0.087 | 0.154 | 0.01 | 0.01 | 0 | 0.08 | 0.44 | 0.09 | 0 | 0.1 | 0.01 |
| 2 | 73461483 | rs1053329 | -0.034 | 0.255 | -0.115 | 0.103 | -0.032 | -0.085 | -0.085 | 0.255 | 0.23 | 0.11 | 0.29 | 0.37 | 0.3 | 0.21 |
| 2 | 73464808 | rs10207625 | -1.794 | 0.308 | -0.129 | 0.067 | -0.05 | -0.056 | -0.056 | 0.863 | 0.44 | 0.85 | 0.43 | 0.38 | 0.47 | 0.31 |
| 2 | 73466592 | rs1474298 | -0.055 | 0.312 | -0.134 | 0.071 | -0.024 | -0.069 | -0.069 | 0 | 0.24 | 0.12 | 0.29 | 0.37 | 0.3 | 0.22 |
| 2 | 73470615 | rs12464589 | -0.329 | 0.28 | -0.12 | 0.075 | 0.096 | 0.009 | 0.009 | 0 | 0.05 | 0.24 | 0.05 | 0 | 0.08 | 0.01 |
| 2 | 73471653 | rs1018342 | -0.133 | 0.219 | -1.474 | 0.132 | -0.025 | -0.071 | -0.071 | 0 | 0.24 | 0.12 | 0.29 | 0.37 | 0.31 | 0.24 |
| 2 | 73472012 | rs12465780 | -0.845 | -0.064 | -1.626 | 0.436 | -0.053 | -0.066 | -0.066 | 0 | 0.44 | 0.88 | 0.43 | 0.38 | 0.48 | 0.33 |
| 2 | 73483057 | rs1045902 | -0.8 | -0.122 | -1.611 | 0.438 | -0.052 | -0.055 | -0.055 | 0 | 0.44 | 0.85 | 0.43 | 0.38 | 0.48 | 0.34 |
| 2 | 73500128 | rs7609407 | -0.85 | -1.299 | -0.817 | 0.513 | 0.025 | -0.016 | -0.016 | 0 | 0.33 | 0.26 | 0.15 | 0.4 | 0.17 | 0.31 |
| 2 | 73521166 | rs6718289 | -1.213 | -0.698 | -0.203 | 1.077 | -0.059 | 0.09 | 0.09 | 3.787 | 0.19 | 0.65 | 0.22 | 0.4 | 0.07 | 0.22 |
| 2 | 73525537 | rs1522928 | -1.18 | -0.681 | -0.203 | 1.03 | -0.057 | 0.081 | 0.081 | 3.789 | 0.19 | 0.65 | 0.22 | 0.37 | 0.07 | 0.22 |
| 2 | 73529077 | rs17349049 | -0.235 | 0.463 | -1.205 | 0.132 | -0.088 | 0.103 | 0.103 | 0 | 0.17 | 0.01 | 0.16 | 0.01 | 0.22 | 0.12 |
| 2 | 73530309 | rs6546818 | 0.605 | 0.349 | -1.231 | 0.271 | -0.001 | -0.006 | -0.006 | 0.491 | 0.83 | 0.9 | 0.92 | 0.8 | 0.84 | 0.83 |
| 2 | 73537537 | rs6711843 | 0.972 | 0.171 | 0.038 | 0.189 | -0.015 | -0.004 | -0.004 | 0 | 0.8 | 0.59 | 0.75 | 0.8 | 0.83 | 0.83 |
| 2 | 73556155 | rs1522927 | -1.368 | -0.089 | 0.451 | 0.196 | 0.008 | -0.009 | -0.009 | 0.834 | 0.29 | 0.62 | 0.26 | 0.21 | 0.29 | 0.26 |
| 2 | 73587331 | rs6546824 | -1.451 | -0.163 | 1.548 | 0.14 | 0.015 | -0.013 | -0.013 | 0 | 0.29 | 0.63 | 0.27 | 0.21 | 0.29 | 0.25 |
| 2 | 73601606 | rs7586361 | -1.505 | -0.159 | 1.446 | 0.141 | -0.001 | 0 | 0 | 0.381 | 0.3 | 0.69 | 0.27 | 0.21 | 0.29 | 0.25 |
| 2 | 73641326 | rs12995433 | 1.58 | -0.464 | 1.706 | -0.087 | -0.007 | 0.029 | 0.029 | 0.068 | 0.63 | 0.29 | 0.71 | 0.79 | 0.66 | 0.72 |
| 2 | 73649928 | rs1881245 | -1.266 | -0.397 | 1.665 | -0.153 | -0.003 | 0.007 | 0.007 | 0.1 | 0.24 | 0.79 | 0.38 | 0.01 | 0.22 | 0.15 |

| | | | | | | | | | | | | | | | | |
|---|---|---|---|---|---|---|---|---|---|---|---|---|---|---|---|---|
| 2 | 73651967 | rs3813227 | -1.443 | -0.406 | 1.64 | -0.157 | -0.004 | 0.083 | 0.083 | 0.416 | 0.26 | 0.88 | 0.39 | 0.01 | 0.23 | 0.15 |
| 2 | 73675669 | rs2037814 | 0.901 | -0.398 | 1.636 | -0.146 | -0.001 | -0.018 | -0.018 | 0 | 0.85 | 0.91 | 0.94 | 0.79 | 0.88 | 0.84 |
| 2 | 73675844 | rs7598901 | 2.127 | -0.413 | 1.641 | -0.153 | -0.033 | 0.099 | 0.099 | 0.502 | 0.6 | 0.1 | 0.56 | 0.78 | 0.65 | 0.69 |
| 2 | 73677833 | rs6546836 | -1.278 | -0.412 | 1.673 | -0.156 | 0 | -0.001 | -0.001 | 0.102 | 0.25 | 0.79 | 0.38 | 0.01 | 0.23 | 0.16 |
| 2 | 73679280 | rs6546838 | -1.44 | -0.412 | 1.649 | -0.156 | -0.003 | 0.074 | 0.074 | 0.426 | 0.26 | 0.86 | 0.39 | 0.01 | 0.23 | 0.16 |
| 2 | 73685852 | rs6749680 | 2.363 | -0.736 | 1.32 | -0.402 | -0.029 | 0.155 | 0.155 | 0.228 | 0.54 | 0.1 | 0.55 | 0.78 | 0.62 | 0.68 |
| 2 | 73694764 | rs780396 | 1.127 | -0.748 | 1.338 | -0.409 | 0.003 | -0.002 | -0.002 | 0 | 0.79 | 0.87 | 0.93 | 0.79 | 0.85 | 0.84 |
| 2 | 73698518 | rs10189574 | -1.425 | -0.546 | 1.49 | -0.251 | -0.003 | 0.085 | 0.085 | 0.33 | 0.26 | 0.88 | 0.39 | 0.01 | 0.23 | 0.16 |
| 2 | 73716810 | rs3820700 | -1.046 | -0.558 | 1.495 | -0.253 | -0.003 | -0.019 | -0.019 | 0 | 0.15 | 0.09 | 0.05 | 0.2 | 0.13 | 0.16 |
| 2 | 73717567 | rs2056486 | -1.364 | -0.553 | 1.503 | -0.264 | -0.003 | 0.074 | 0.074 | 0.426 | 0.26 | 0.86 | 0.39 | 0.01 | 0.23 | 0.16 |
| 2 | 73717656 | rs10193972 | -1.364 | -0.553 | 1.503 | -0.264 | -0.003 | 0.074 | 0.074 | 0.426 | 0.26 | 0.86 | 0.39 | 0.01 | 0.23 | 0.16 |
| 2 | 73746923 | rs11884776 | -0.695 | -0.54 | 1.63 | -0.269 | 0.018 | -0.128 | -0.128 | 0 | 0.21 | 0.57 | 0.23 | 0.01 | 0.21 | 0.15 |
| 2 | 73774019 | rs13008860 | -2.909 | -0.636 | 1.448 | 0.752 | -0.07 | -0.04 | -0.04 | 0.244 | 0.47 | 0.98 | 0.48 | 0.63 | 0.39 | 0.42 |
| 2 | 73785358 | rs6546847 | -1.414 | -0.397 | 1.648 | 0.399 | -0.018 | 0.116 | 0.116 | 0.432 | 0.26 | 0.87 | 0.39 | 0.01 | 0.23 | 0.16 |
| 2 | 73818936 | rs13391552 | -0.981 | -0.4 | 1.836 | 0.398 | 0.008 | -0.114 | -0.114 | 0 | 0.23 | 0.63 | 0.24 | 0.01 | 0.22 | 0.15 |
| 2 | 73825770 | rs2948441 | 0.912 | -0.405 | 1.907 | 0.413 | 0.002 | -0.014 | -0.014 | 3.245 | 0.85 | 0.87 | 0.94 | 0.8 | 0.88 | 0.85 |
| 2 | 73827099 | rs6732440 | 2.063 | -0.408 | 1.91 | 0.41 | -0.026 | 0.09 | 0.09 | 0.383 | 0.61 | 0.13 | 0.7 | 0.79 | 0.66 | 0.7 |
| 2 | 73828538 | rs1052161 | 1.984 | -0.415 | 1.962 | 0.644 | -0.009 | 0.024 | 0.024 | 0.124 | 0.62 | 0.25 | 0.71 | 0.76 | 0.66 | 0.69 |
| 2 | 73837751 | rs6546856 | -1.35 | -0.317 | 1.834 | 0.61 | -0.013 | 0.068 | 0.068 | 0.465 | 0.26 | 0.85 | 0.39 | 0.01 | 0.24 | 0.15 |
| 2 | 73857002 | rs4852940 | 1.131 | -0.505 | 1.904 | 0.895 | 0.003 | -0.019 | -0.019 | 0 | 0.84 | 0.86 | 0.94 | 0.66 | 0.87 | 0.84 |
| 2 | 73859184 | rs12472502 | 2.06 | -0.423 | 1.925 | 0.892 | -0.012 | -0.016 | -0.016 | 0 | 0.62 | 0.16 | 0.67 | 0.66 | 0.67 | 0.69 |
| 2 | 73860348 | rs6546862 | 2.086 | -0.465 | 1.912 | 0.576 | -0.006 | 0.025 | 0.025 | 0.195 | 0.62 | 0.18 | 0.67 | 0.69 | 0.67 | 0.7 |
| 2 | 73869666 | rs2280506 | 1.01 | -0.461 | 1.935 | 0.562 | 0.001 | -0.022 | -0.022 | 0 | 0.84 | 0.88 | 0.94 | 0.7 | 0.87 | 0.84 |
| 2 | 73878352 | rs6759452 | 0.045 | -0.468 | 2.129 | 0.535 | -0.022 | 0.048 | 0.048 | 0 | 0.15 | 0.06 | 0.19 | 0 | 0.15 | 0.12 |
| 2 | 73878827 | rs7607989 | 1.124 | -0.356 | 2.049 | 0.507 | 0 | -0.024 | -0.024 | 0 | 0.84 | 0.87 | 0.93 | 0.7 | 0.86 | 0.84 |
| 2 | 73885096 | rs2421668 | 2.183 | -0.376 | 2.051 | 0.521 | -0.006 | 0.065 | 0.065 | 0.169 | 0.62 | 0.2 | 0.67 | 0.78 | 0.67 | 0.71 |
| 2 | 73899719 | rs2421586 | 2.203 | -0.376 | 2.049 | 0.515 | -0.008 | 0.027 | 0.027 | 0.225 | 0.62 | 0.18 | 0.67 | 0.69 | 0.67 | 0.7 |
| 2 | 73900329 | rs2421587 | 1.133 | -0.449 | 2.023 | 0.583 | -0.004 | 0.031 | 0.031 | 0 | 0.84 | 0.8 | 0.93 | 0.7 | 0.88 | 0.84 |
| 2 | 73927931 | rs4852974 | 1.334 | -0.256 | 1.929 | 0.58 | -0.019 | -0.012 | -0.012 | 0 | 0.7 | 0.86 | 0.77 | 0.71 | 0.74 | 0.73 |
| 2 | 73932044 | rs11126414 | 1.679 | -0.259 | 1.982 | 0.622 | -0.002 | 0.002 | 0.002 | 0.001 | 0.64 | 0.37 | 0.59 | 0.64 | 0.67 | 0.69 |
| 2 | 73937901 | rs7599453 | 1.802 | -0.284 | 1.97 | 0.286 | -0.004 | 0.033 | 0.033 | 0 | 0.63 | 0.29 | 0.59 | 0.71 | 0.69 | 0.69 |
| 2 | 73938295 | rs4852977 | 1.589 | -0.287 | 2.02 | 0.264 | 0 | -0.003 | -0.003 | 0 | 0.65 | 0.41 | 0.59 | 0.71 | 0.69 | 0.69 |
| 2 | 73955571 | rs17009372 | -1.105 | -0.348 | 1.893 | 0.274 | -0.041 | -0.035 | -0.035 | 0 | 0.24 | 0.04 | 0.2 | 0.23 | 0.2 | 0.25 |
| 2 | 73956114 | rs12713793 | 1.941 | -0.277 | 1.89 | 0.294 | 0.003 | 0.024 | 0.024 | 0 | 0.62 | 0.3 | 0.58 | 0.69 | 0.69 | 0.69 |
| 2 | 73957124 | rs7210 | 1.908 | -0.303 | 1.875 | 0.281 | 0.002 | 0.027 | 0.027 | 0 | 0.62 | 0.29 | 0.58 | 0.7 | 0.69 | 0.69 |
| 2 | 73966087 | rs2421676 | 1.768 | -0.314 | 2.266 | 0.355 | 0.005 | -0.016 | -0.016 | 0 | 0.65 | 0.51 | 0.6 | 0.68 | 0.69 | 0.69 |
| 2 | 73973877 | rs11126416 | 1.816 | -0.329 | 2.331 | 0.32 | 0.004 | -0.02 | -0.02 | 0 | 0.65 | 0.5 | 0.6 | 0.68 | 0.69 | 0.69 |
| 2 | 73986421 | rs12713798 | 2.192 | -0.331 | 2.315 | 0.318 | 0.004 | 0.024 | 0.024 | 0 | 0.62 | 0.29 | 0.58 | 0.68 | 0.69 | 0.69 |
| 2 | 73994074 | rs2567599 | -0.039 | -0.338 | 2.384 | 0.316 | 0 | -0.027 | -0.027 | 0 | 0.95 | 0.91 | 0.97 | 0.99 | 0.94 | 0.97 |
| 2 | 73998973 | rs13033744 | 1.807 | -0.319 | 2.343 | 0.324 | -0.001 | -0.014 | -0.014 | 0 | 0.64 | 0.47 | 0.6 | 0.67 | 0.69 | 0.69 |
| 2 | 73999654 | rs2421559 | 2.029 | -0.313 | 2.307 | 0.334 | 0 | 0.001 | 0.001 | 0 | 0.63 | 0.35 | 0.59 | 0.67 | 0.69 | 0.69 |
| 2 | 74007136 | rs2272051 | 1.823 | -0.285 | 2.329 | 0.292 | -0.001 | -0.017 | -0.017 | 0 | 0.64 | 0.43 | 0.6 | 0.7 | 0.69 | 0.69 |
| 2 | 74051612 | rs10193032 | 1.788 | -0.437 | 2.345 | 0.232 | -0.019 | -0.027 | -0.027 | 0.131 | 0.64 | 0.48 | 0.61 | 0.7 | 0.74 | 0.72 |
| 2 | 74054272 | rs17009396 | -2.285 | -0.121 | 2.62 | 0.289 | 0.02 | 0.022 | 0.022 | 0.082 | 0.06 | 0.31 | 0.14 | 0.07 | 0.02 | 0.03 |
| 2 | 74056564 | rs3813229 | -3.04 | -0.065 | 2.372 | 0.393 | -0.001 | -0.003 | -0.003 | 0 | 0.34 | 0.74 | 0.42 | 0.37 | 0.29 | 0.31 |
| 2 | 74066461 | rs1474335 | -1.821 | -0.001 | 2.032 | 0.419 | -0.01 | -0.023 | -0.023 | 0 | 0.23 | 0.19 | 0.22 | 0.32 | 0.22 | 0.27 |
| 2 | 74087290 | rs1025424 | -1.461 | 0.132 | 2.153 | 0.145 | 0.004 | -0.068 | -0.068 | 0.263 | 0.18 | 0.27 | 0.22 | 0.01 | 0.18 | 0.1 |
| 2 | 74087298 | rs2303987 | -0.984 | 0.15 | 2.356 | 0.271 | 0.016 | 0.18 | 0.18 | 0 | 0.06 | 0.19 | 0.03 | 0.27 | 0.05 | 0.21 |
| 2 | 74091725 | rs11126418 | -1.66 | 0.134 | 2.44 | 0.232 | 0 | -0.025 | -0.025 | 2.025 | 0.06 | 0.13 | 0.04 | 0 | 0.05 | 0.01 |
| 2 | 74091805 | rs3738846 | -2.894 | 0.187 | 2.56 | 0.221 | -0.002 | -0.004 | -0.004 | 0 | 0.3 | 0.77 | 0.41 | 0.35 | 0.26 | 0.33 |
| 2 | 74092119 | rs11126419 | -2.159 | 0.213 | 2.513 | 0.203 | 0.042 | -0.038 | -0.038 | 0 | 0.1 | 0.47 | 0.2 | 0.07 | 0.09 | 0.04 |
| 2 | 74093652 | rs3813230 | -2.478 | 0.22 | 2.5 | 0.208 | 0.012 | -0.137 | -0.137 | 6.763 | 0.27 | 0.5 | 0.38 | 0.08 | 0.26 | 0.14 |
| 2 | 74097364 | rs17009429 | -1.019 | 0.318 | 2.18 | 0.375 | 0.015 | 0.179 | 0.179 | 0 | 0.06 | 0.19 | 0.03 | 0.27 | 0.05 | 0.21 |
| 2 | 74110021 | rs10519327 | -2.374 | 0.302 | 1.977 | 0.281 | 0.024 | 0.147 | 0.147 | 0.61 | 0.15 | 0.45 | 0.2 | 0.35 | 0.13 | 0.24 |
| 2 | 74114397 | rs11890369 | -1.235 | 0.176 | 1.685 | 0.27 | -0.019 | -0.026 | -0.026 | 0.319 | 0.08 | 0.2 | 0.13 | 0.07 | 0.04 | 0.03 |
| 2 | 74116536 | rs2163539 | -1.326 | 0.125 | 1.305 | 0.254 | 0.013 | 0.065 | 0.065 | 0 | 0.13 | 0.25 | 0.08 | 0.29 | 0.15 | 0.23 |
| 2 | 74117499 | rs7600452 | -1.555 | -0.009 | 1.288 | 0.242 | -0.033 | -0.046 | -0.046 | 0 | 0.11 | 0.4 | 0.14 | 0.07 | 0.04 | 0.03 |
| 2 | 74120181 | rs2070581 | -1.614 | 0.415 | 0.728 | 0.454 | 0.024 | 0.031 | 0.031 | 0 | 0.27 | 0.46 | 0.26 | 0.3 | 0.27 | 0.32 |
| 2 | 74120785 | rs10176602 | -1.313 | 0.316 | -0.134 | 0.5 | -0.003 | -0.009 | -0.009 | 0 | 0.24 | 0.21 | 0.2 | 0.27 | 0.22 | 0.31 |
| 2 | 74122223 | rs13415017 | -0.469 | 0.462 | -0.72 | -0.791 | 0 | -0.076 | -0.076 | 0 | 0.38 | 0.37 | 0.4 | 0 | 0.31 | 0.15 |
| 2 | 74122345 | rs13024800 | -0.148 | 0.718 | -1.312 | -0.074 | -0.058 | -0.031 | -0.031 | 0 | 0.33 | 0.12 | 0.28 | 0.31 | 0.44 | 0.44 |
| 2 | 74122953 | rs1632296 | -0.159 | -0.238 | -0.99 | 0.231 | 0 | -0.053 | -0.053 | 0 | 0.79 | 0.88 | 0.85 | 0.54 | 0.81 | 0.65 |
| 2 | 74124342 | rs831540 | 0.306 | -0.536 | -0.139 | 0.072 | -0.019 | 0.048 | 0.048 | 0 | 0.75 | 0.61 | 0.83 | 0.61 | 0.81 | 0.67 |
| 2 | 74125550 | rs7557701 | -0.134 | -0.87 | -0.415 | -0.589 | 0.007 | 0.046 | 0.046 | 0 | 0.26 | 0.14 | 0.11 | 0.07 | 0.19 | 0.13 |
| 2 | 74126457 | rs2249441 | -0.283 | -0.879 | 0.14 | -0.418 | -0.007 | 0.025 | 0.025 | 0 | 0.76 | 0.72 | 0.84 | 0.6 | 0.8 | 0.65 |
| 2 | 74127289 | rs1721240 | 0.883 | -0.883 | 0.095 | -0.478 | -0.001 | -0.008 | -0.008 | 0 | 0.45 | 0.36 | 0.69 | 0.5 | 0.59 | 0.5 |
| 2 | 74127845 | rs1618091 | 1.093 | -1.011 | -0.303 | -0.624 | -0.07 | -0.062 | -0.062 | 0.031 | 0.41 | 0.09 | 0.68 | 0.54 | 0.6 | 0.49 |
| 2 | 74128369 | rs1632564 | 0.992 | -1.043 | 0.003 | -0.624 | -0.015 | -0.024 | -0.024 | 0 | 0.42 | 0.18 | 0.68 | 0.52 | 0.61 | 0.51 |
| 2 | 74129389 | rs7586415 | -0.316 | -1.267 | 0.476 | -0.819 | 0.004 | -0.016 | -0.016 | 0 | 0.3 | 0.28 | 0.12 | 0.07 | 0.18 | 0.14 |

| | | | | | | | | | | | | | | | | |
|---|---|---|---|---|---|---|---|---|---|---|---|---|---|---|---|---|
| 2 | 74129895 | rs1721244 | 1.055 | -1.334 | 0.727 | -0.806 | 0 | 0 | 0 | 0 | 0.44 | 0.42 | 0.71 | 0.52 | 0.61 | 0.54 |
| 2 | 74133975 | rs831551 | -0.135 | -1.692 | 0.815 | -0.402 | 0.003 | -0.008 | -0.008 | 0 | 0.75 | 0.75 | 0.84 | 0.59 | 0.8 | 0.68 |
| 2 | 74135732 | rs702460 | 0.58 | -1.547 | 0.505 | -0.35 | 0.015 | 0.002 | 0.002 | 0 | 0.55 | 0.39 | 0.76 | 0.53 | 0.69 | 0.61 |
| 2 | 74135898 | rs756128 | 0.58 | -1.547 | 0.505 | -0.35 | 0.015 | 0.002 | 0.002 | 0 | 0.55 | 0.39 | 0.76 | 0.53 | 0.69 | 0.61 |
| 2 | 74139996 | rs10169974 | 0.562 | -1.372 | 0.31 | -0.266 | -0.019 | -0.002 | -0.002 | 1.364 | 0.56 | 0.48 | 0.76 | 0.53 | 0.7 | 0.62 |
| 2 | 74142783 | rs6711332 | -0.85 | -1.225 | 0.55 | -0.601 | -0.052 | -0.121 | -0.121 | 0 | 0.26 | 0.46 | 0.14 | 0.07 | 0.17 | 0.14 |
| 2 | 74144110 | rs1623605 | 1.206 | -0.926 | 0.363 | -0.21 | 0.065 | 0.019 | 0.019 | 0 | 0.52 | 0.24 | 0.55 | 0.53 | 0.6 | 0.57 |
| 2 | 74146869 | rs1721238 | 0.316 | -1.037 | 0.943 | -0.224 | -0.019 | 0.022 | 0.022 | 0 | 0.67 | 0.59 | 0.6 | 0.51 | 0.69 | 0.64 |
| 2 | 74151700 | rs13411881 | -0.073 | -1.054 | 0.855 | -0.345 | 0.004 | -0.055 | -0.055 | 0.12 | 0.09 | 0.25 | 0.06 | 0 | 0.06 | 0.07 |
| 2 | 74155641 | rs7557705 | 0.438 | -1.01 | 0.659 | -0.323 | -0.001 | -0.018 | -0.018 | 0 | 0.08 | 0.08 | 0.04 | 0 | 0.06 | 0.08 |
| 2 | 74159734 | rs6737156 | 0.4 | -1.019 | 0.611 | -0.324 | -0.009 | -0.042 | -0.042 | 0 | 0.08 | 0.15 | 0.04 | 0 | 0.04 | 0.07 |
| 2 | 74172133 | rs831550 | 0.241 | -1.052 | 0.672 | -0.231 | -0.009 | 0.011 | 0.011 | 0 | 0.68 | 0.65 | 0.61 | 0.52 | 0.69 | 0.64 |
| 2 | 74183673 | rs4852323 | 1.536 | -1.127 | -0.521 | -0.458 | -0.003 | -0.009 | -0.009 | 0 | 0.41 | 0.31 | 0.5 | 0.53 | 0.53 | 0.47 |
| 2 | 74186781 | rs951409 | 0.982 | -1.164 | -0.557 | -0.378 | 0.074 | 0.003 | 0.003 | 0 | 0.54 | 0.26 | 0.56 | 0.53 | 0.64 | 0.5 |
| 2 | 74187061 | rs7568797 | -0.354 | -0.474 | -0.55 | 0.032 | -0.003 | -0.056 | -0.056 | 0 | 0.1 | 0.2 | 0.04 | 0 | 0.05 | 0.08 |
| 2 | 74194587 | rs10169997 | -0.147 | -0.512 | -1.359 | 0.471 | 0.027 | -0.151 | -0.151 | 0.003 | 0.73 | 0.96 | 0.65 | 0.52 | 0.79 | 0.65 |
| 2 | 74197109 | rs870722 | -0.71 | 0.039 | -0.446 | -0.119 | 0.003 | -0.026 | -0.026 | 1.112 | 0.3 | 0.66 | 0.23 | 0.01 | 0.29 | 0.21 |
| 2 | 74200833 | rs940296 | -2.106 | 0.485 | 1.083 | 1.078 | 0.069 | 0.228 | 0.228 | 1.481 | 0.11 | 0.47 | 0.24 | 0.21 | 0.14 | 0.11 |
| 2 | 74202578 | rs4852324 | -1.543 | 0.489 | 1.139 | 1.111 | 0.076 | 0.303 | 0.303 | 1.898 | 0.08 | 0.42 | 0.16 | 0.19 | 0.09 | 0.08 |
| 2 | 74210442 | rs6714060 | -2.42 | 0.609 | 1.681 | 1.153 | 0.048 | 0.143 | 0.143 | 2.432 | 0.1 | 0.61 | 0.25 | 0.16 | 0.14 | 0.07 |
| 2 | 74225434 | rs13416937 | -2.658 | 0.524 | 2.456 | 0.854 | 0.048 | -0.043 | -0.043 | 0 | 0.09 | 0.28 | 0.14 | 0 | 0.12 | 0.04 |
| 2 | 74232251 | rs13404452 | -2.28 | 0.609 | 2.201 | 0.862 | 0.022 | -0.041 | -0.041 | 0.02 | 0.07 | 0.13 | 0.12 | 0 | 0.13 | 0.04 |
| 2 | 74245777 | rs6546886 | -1.699 | 0.455 | 1.82 | 0.6 | 0.001 | 0.016 | 0.016 | 0.679 | 0.26 | 0.8 | 0.3 | 0.11 | 0.32 | 0.16 |
| 2 | 74269279 | rs13016198 | -1.111 | 0.407 | 2.457 | 0.576 | 0.009 | -0.061 | -0.061 | 0.095 | 0.2 | 0.27 | 0.19 | 0.01 | 0.3 | 0.12 |
| 2 | 74272257 | rs4853007 | -1.563 | 0.395 | 2.467 | 0.572 | 0.01 | -0.033 | -0.033 | 0 | 0.23 | 0.43 | 0.28 | 0.11 | 0.32 | 0.16 |
| 2 | 74306056 | rs10164943 | -1.769 | 0.377 | 2.607 | 0.562 | 0.006 | -0.043 | -0.043 | 0 | 0.24 | 0.55 | 0.29 | 0.11 | 0.32 | 0.16 |
| 2 | 74309401 | rs13014576 | -1.588 | 0.393 | 2.343 | 0.569 | 0.006 | -0.028 | -0.028 | 0 | 0.23 | 0.39 | 0.26 | 0.11 | 0.31 | 0.16 |
| 2 | 74313080 | rs828872 | -0.037 | 0.301 | 1.228 | 1.053 | 0.001 | -0.032 | -0.032 | 0 | 0.85 | 0.94 | 0.77 | 0.5 | 0.9 | 0.74 |
| 2 | 74316414 | rs7593050 | -0.769 | 0.405 | 0.973 | 0.73 | 0 | -0.026 | -0.026 | 0 | 0.19 | 0.19 | 0.17 | 0.01 | 0.3 | 0.11 |
| 2 | 74322469 | rs7589197 | -1.625 | 0.307 | 0.362 | 1.037 | 0.002 | 0.005 | 0.005 | 0 | 0.41 | 0.72 | 0.42 | 0.59 | 0.4 | 0.44 |
| 2 | 74323500 | rs828869 | 1.546 | 0.275 | -0.499 | 0.952 | -0.044 | 0.004 | 0.004 | 0.251 | 0.56 | 0.05 | 0.47 | 0.37 | 0.6 | 0.53 |
| 2 | 74327453 | rs6738709 | -0.753 | 0.478 | -0.247 | 0.779 | -0.026 | 0.029 | 0.029 | 0.007 | 0.21 | 0.08 | 0.12 | 0.13 | 0.25 | 0.16 |
| 2 | 74334462 | rs828867 | 1.189 | 0.564 | -0.868 | 0.539 | -0.036 | -0.017 | -0.017 | 0.015 | 0.46 | 0.18 | 0.43 | 0.48 | 0.52 | 0.51 |
| 2 | 74334790 | rs6546891 | -0.823 | 0.789 | 0.162 | 0.729 | 0.018 | 0.089 | 0.089 | 0 | 0.23 | 0.4 | 0.43 | 0.47 | 0.21 | 0.32 |
| 2 | 74337353 | rs13025827 | 0.304 | 0.734 | 0.471 | 0.089 | 0.003 | -0.008 | -0.008 | 0 | 0.32 | 0.28 | 0.44 | 0.51 | 0.27 | 0.48 |
| 2 | 74338942 | rs7570950 | -0.146 | 0.794 | -0.02 | 0.189 | 0.019 | 0.136 | 0.136 | 0 | 0.18 | 0.36 | 0.38 | 0.43 | 0.17 | 0.29 |
| 2 | 74344656 | rs6707619 | -0.36 | 0.84 | -0.281 | 0.002 | 0.005 | 0.069 | 0.069 | 0 | 0.2 | 0.31 | 0.39 | 0.37 | 0.19 | 0.29 |
| 2 | 74345206 | rs4452177 | -0.319 | 0.841 | -0.272 | 0.013 | 0.003 | 0.065 | 0.065 | 0 | 0.2 | 0.29 | 0.39 | 0.37 | 0.18 | 0.28 |
| 2 | 74345780 | rs1026150 | 0.81 | 0.908 | -1.129 | -0.023 | -0.008 | 0.034 | 0.034 | 0 | 0.26 | 0.3 | 0.3 | 0.49 | 0.21 | 0.45 |
| 2 | 74347666 | rs1667618 | -0.348 | 1.021 | -0.997 | 0.012 | 0 | 0.056 | 0.056 | 0.019 | 0.45 | 0.32 | 0.66 | 0.85 | 0.4 | 0.74 |
| 2 | 74358901 | rs2122290 | -0.268 | 0.876 | -1.368 | 0.478 | 0 | -0.004 | -0.004 | 0 | 0.19 | 0.19 | 0.38 | 0.41 | 0.17 | 0.25 |
| 2 | 74361279 | rs702463 | -0.183 | 1.163 | -1.745 | 0.645 | -0.094 | -0.089 | -0.089 | 0 | 0.43 | 0.09 | 0.41 | 0.51 | 0.48 | 0.55 |
| 2 | 74361641 | rs702464 | -0.168 | 1.18 | -1.784 | 0.653 | -0.102 | -0.097 | -0.097 | 0.118 | 0.43 | 0.08 | 0.41 | 0.5 | 0.48 | 0.56 |
| 2 | 74366459 | rs860455 | -0.051 | 1.119 | -1.561 | 0.581 | -0.025 | 0.042 | 0.042 | 0.032 | 0.54 | 0.79 | 0.67 | 0.6 | 0.53 | 0.37 |
| 2 | 74370612 | rs17009668 | -0.005 | 1.003 | -1.648 | 0.961 | -0.001 | -0.039 | -0.039 | 1.221 | 0.13 | 0.08 | 0.32 | 0.36 | 0.1 | 0.19 |
| 2 | 74375804 | rs828882 | 0.416 | 0.958 | -1.208 | 0.881 | -0.003 | 0.026 | 0.026 | 0 | 0.4 | 0.68 | 0.35 | 0.25 | 0.44 | 0.19 |
| 2 | 74392940 | rs11126426 | -0.629 | 0.915 | -0.981 | 0.531 | -0.027 | 0.064 | 0.064 | 0 | 0.57 | 0.8 | 0.61 | 0.75 | 0.4 | 0.77 |
| 2 | 74401657 | rs4083173 | 0.09 | 0.792 | 0.147 | 0.581 | -0.003 | 0.088 | 0.088 | 0 | 0.14 | 0.24 | 0.33 | 0.36 | 0.1 | 0.16 |
| 2 | 74406055 | rs1667601 | 0.814 | 0.909 | 0.495 | 0.557 | -0.001 | 0.001 | 0.001 | 0 | 0.37 | 0.34 | 0.31 | 0.25 | 0.44 | 0.21 |
| 2 | 74411281 | rs828860 | -0.332 | 0.909 | -0.148 | 0.545 | -0.042 | 0.046 | 0.046 | 0.238 | 0.54 | 0.86 | 0.64 | 0.61 | 0.54 | 0.37 |
| 2 | 74426038 | rs1723285 | 0.533 | 1.053 | 0.082 | 0.658 | -0.037 | 0.113 | 0.113 | 0 | 0.36 | 0.14 | 0.22 | 0.14 | 0.44 | 0.16 |
| 2 | 74448655 | rs7587117 | 0.737 | 1.234 | 0.041 | 0.625 | -0.017 | 0.087 | 0.087 | 0.174 | 0.32 | 0.11 | 0.18 | 0.13 | 0.37 | 0.16 |
| 2 | 74451762 | rs12053233 | 0.954 | 1.245 | -0.015 | 0.708 | -0.045 | 0.032 | 0.032 | 0.307 | 0.3 | 0.05 | 0.18 | 0.2 | 0.36 | 0.14 |
| 2 | 74452259 | rs828863 | 1.222 | 0.729 | -0.28 | 0.434 | -0.003 | 0.075 | 0.075 | 0.125 | 0.13 | 0 | 0.03 | 0 | 0.1 | 0.04 |
| 2 | 74461634 | rs895988 | -0.419 | 0.836 | 0.015 | 0.99 | 0.008 | 0.041 | 0.041 | 1.653 | 0.46 | 0.68 | 0.63 | 0.66 | 0.46 | 0.34 |
| 2 | 74463566 | rs13006863 | 1.013 | 0.775 | -0.427 | 0.852 | -0.011 | 0.04 | 0.04 | 0.025 | 0.29 | 0.11 | 0.17 | 0.2 | 0.33 | 0.14 |
| 2 | 74466152 | rs7561242 | -0.526 | 0.607 | -0.414 | 1.056 | 0.005 | 0.116 | 0.116 | 0.528 | 0.51 | 0.81 | 0.71 | 0.7 | 0.56 | 0.37 |
| 2 | 74466594 | rs4853018 | 0.516 | 0.635 | -0.753 | 0.928 | -0.055 | 0.069 | 0.069 | 0 | 0.33 | 0.14 | 0.23 | 0.2 | 0.43 | 0.17 |
| 2 | 74480659 | rs17009774 | -0.453 | 0.124 | -0.249 | 1.982 | 0.005 | -0.054 | -0.054 | 0 | 0.1 | 0.03 | 0.21 | 0.36 | 0.06 | 0.17 |
| 2 | 74482892 | rs7602215 | -1.367 | -0.616 | 0.365 | 1.997 | -0.04 | 0.032 | 0.032 | 0.919 | 0.26 | 0.34 | 0.14 | 0.34 | 0.14 | 0.23 |
| 2 | 74483688 | rs11902148 | -0.982 | -0.756 | 0.35 | 2.153 | -0.054 | 0.017 | 0.017 | 1.036 | 0.19 | 0.31 | 0.1 | 0.18 | 0.1 | 0.15 |
| 2 | 74485697 | rs3771724 | -2.348 | -1.02 | 0.878 | 2.363 | 0.033 | 0.029 | 0.029 | 3.484 | 0.33 | 0.73 | 0.36 | 0.76 | 0.17 | 0.27 |
| 2 | 74486357 | rs1006502 | -2.342 | -1.017 | 0.879 | 2.364 | 0.033 | 0.029 | 0.029 | 3.573 | 0.33 | 0.73 | 0.36 | 0.76 | 0.17 | 0.27 |
| 2 | 74501510 | rs6723819 | -0.768 | -0.983 | 1.19 | 2.315 | -0.005 | 0 | 0 | 1.154 | 0.18 | 0.16 | 0.09 | 0.23 | 0.1 | 0.15 |
| 2 | 74515158 | rs7560244 | -1.762 | -0.531 | 1.617 | 2.515 | -0.029 | 0.137 | 0.137 | 0.557 | 0.17 | 0.34 | 0.11 | 0.35 | 0.12 | 0.22 |
| 2 | 74517251 | rs2191309 | -2.178 | -0.522 | 1.691 | 2.374 | 0 | 0.387 | 0.387 | 4.582 | 0.09 | 0.55 | 0.24 | 0.42 | 0.04 | 0.07 |
| 2 | 74534691 | rs9789404 | -2.236 | -0.406 | 1.983 | 2.324 | -0.04 | 0.199 | 0.199 | 0.056 | 0.19 | 0.49 | 0.17 | 0.44 | 0.12 | 0.23 |
| 2 | 74540094 | rs2421844 | -2.374 | -0.418 | 1.976 | 2.318 | -0.045 | 0.187 | 0.187 | 0 | 0.2 | 0.49 | 0.18 | 0.44 | 0.12 | 0.23 |
| 2 | 74548807 | rs10211612 | -2.217 | -0.409 | 2.014 | 2.338 | -0.036 | 0.163 | 0.163 | 0.043 | 0.2 | 0.5 | 0.19 | 0.44 | 0.13 | 0.24 |

| | | | | | | | | | | | | | | | | |
|---|---|---|---|---|---|---|---|---|---|---|---|---|---|---|---|---|
| 2 | 74569118 | rs3755441 | -1.747 | -0.42 | 2.384 | 2.346 | 0.001 | 0.019 | 0.019 | 0 | 0.16 | 0.16 | 0.14 | 0.44 | 0.12 | 0.22 |
| 2 | 74570654 | rs3771742 | -1.739 | -0.425 | 2.35 | 2.343 | 0.017 | -0.054 | -0.054 | 0 | 0.16 | 0.09 | 0.13 | 0.44 | 0.12 | 0.22 |
| 2 | 74573906 | rs6546903 | -1.597 | -0.385 | 2.35 | 2.343 | 0.012 | -0.047 | -0.047 | 0.317 | 0.15 | 0.09 | 0.13 | 0.44 | 0.12 | 0.22 |
| 2 | 74583478 | rs3771747 | -1.696 | -0.384 | 2.35 | 2.323 | 0.011 | -0.03 | -0.03 | 0.141 | 0.16 | 0.11 | 0.13 | 0.42 | 0.12 | 0.22 |
| 2 | 74586811 | rs9309484 | -2.506 | -0.394 | 2.489 | 2.316 | -0.042 | 0.191 | 0.191 | 0.02 | 0.2 | 0.47 | 0.18 | 0.43 | 0.12 | 0.23 |
| 2 | 74604673 | rs3213672 | -2.254 | -0.246 | 2.528 | 2.401 | -0.038 | 0.231 | 0.231 | 0.022 | 0.18 | 0.47 | 0.15 | 0.43 | 0.09 | 0.22 |
| 2 | 74616870 | rs6740802 | -2.608 | -0.261 | 2.702 | 2.396 | -0.036 | 0.165 | 0.165 | 1.112 | 0.19 | 0.51 | 0.16 | 0.43 | 0.12 | 0.24 |
| 2 | 74618760 | rs7570851 | -1.781 | -0.264 | 2.673 | 2.398 | 0.01 | -0.021 | -0.021 | 0 | 0.15 | 0.11 | 0.12 | 0.43 | 0.12 | 0.23 |
| 2 | 74621660 | rs12328185 | -2.599 | -0.264 | 2.724 | 2.402 | -0.039 | 0.166 | 0.166 | 1.145 | 0.19 | 0.52 | 0.16 | 0.43 | 0.12 | 0.27 |
| 2 | 74639637 | rs10187076 | -2.808 | -0.259 | 2.795 | 2.398 | -0.03 | 0.154 | 0.154 | 1.182 | 0.19 | 0.53 | 0.23 | 0.44 | 0.14 | 0.29 |
| 2 | 74648368 | rs2268424 | -1.834 | -0.255 | 2.794 | 2.4 | 0.008 | -0.036 | -0.036 | 0 | 0.15 | 0.11 | 0.13 | 0.43 | 0.13 | 0.28 |
| 2 | 74649934 | rs3732293 | -1.861 | -0.261 | 2.799 | 2.441 | -0.001 | 0.018 | 0.018 | 0 | 0.11 | 0.11 | 0.1 | 0.28 | 0.09 | 0.22 |
| 2 | 74664467 | rs2268418 | -2.88 | -0.37 | 2.917 | 2.827 | -0.029 | 0.184 | 0.184 | 1.032 | 0.19 | 0.5 | 0.17 | 0.44 | 0.13 | 0.29 |
| 2 | 74665250 | rs11694367 | -2.225 | -0.341 | 2.332 | 2.844 | 0.009 | -0.03 | -0.03 | 0 | 0.16 | 0.12 | 0.13 | 0.44 | 0.13 | 0.28 |
| 2 | 74666075 | rs11695896 | -2.259 | -0.344 | 2.332 | 2.931 | 0.009 | -0.027 | -0.027 | 1.245 | 0.16 | 0.12 | 0.13 | 0.5 | 0.13 | 0.29 |
| 2 | 74669515 | rs7556852 | -2.619 | -0.295 | 2.177 | 2.313 | 0.01 | 0.017 | 0.017 | 0 | 0.21 | 0.14 | 0.29 | 0.82 | 0.18 | 0.39 |
| 2 | 74725178 | rs17009998 | -2.16 | 0.505 | 1.608 | 2.108 | 0.017 | 0.005 | 0.005 | 0 | 0.18 | 0.14 | 0.26 | 0.82 | 0.16 | 0.33 |
| 2 | 74725804 | rs10454145 | -2.17 | 0.506 | 1.608 | 2.117 | 0.018 | 0.004 | 0.004 | 0 | 0.19 | 0.14 | 0.26 | 0.82 | 0.15 | 0.33 |
| 2 | 74745173 | rs3815476 | 0.033 | 0.56 | 1.143 | 2.185 | -0.017 | -0.025 | -0.025 | 0 | 0.05 | 0.01 | 0.05 | 0.15 | 0.07 | 0.08 |
| 2 | 74746080 | rs6709863 | -2.444 | 0.617 | 1.199 | 2.17 | 0.006 | 0.001 | 0.001 | 0.051 | 0.18 | 0.2 | 0.26 | 0.82 | 0.16 | 0.34 |
| 2 | 74758934 | rs2241028 | -0.033 | 0.876 | 0.961 | 1.696 | -0.017 | -0.025 | -0.025 | 0 | 0.05 | 0.01 | 0.05 | 0.15 | 0.07 | 0.08 |
| 2 | 74841569 | rs363691 | -1.351 | 0.738 | 1.769 | 1.707 | -0.014 | -0.004 | -0.004 | 0 | 0.09 | 0.04 | 0.11 | 0.46 | 0.13 | 0.27 |
| 2 | 74845940 | rs10187609 | -0.941 | 0.807 | 1.961 | 0.999 | -0.001 | -0.004 | -0.004 | 0.051 | 0.06 | 0.23 | 0.02 | 0 | 0 | 0.03 |
| 2 | 74849953 | rs13407946 | -1.453 | 0.826 | 1.932 | 1.008 | -0.006 | -0.013 | -0.013 | 0 | 0.07 | 0.29 | 0.03 | 0 | 0 | 0.03 |
| 2 | 74856123 | rs205651 | -0.769 | 0.718 | 0.905 | 0.166 | -0.127 | 0.275 | 0.275 | 0.523 | 0.61 | 0.94 | 0.57 | 0.99 | 0.48 | 0.79 |
| 2 | 74895348 | rs11900406 | -1.168 | 0.498 | 2.364 | 1.458 | 0.036 | 0.033 | 0.033 | 0 | 0.06 | 0.37 | 0.03 | 0 | 0 | 0.03 |
| 2 | 74916183 | rs17010185 | -1.613 | 0.547 | 2.436 | 1.665 | 0.075 | 0.425 | 0.425 | 0.601 | 0.06 | 0.38 | 0.08 | 0.34 | 0.06 | 0.2 |
| 2 | 74960055 | rs6710960 | -1.525 | 0.484 | 1.626 | 1.546 | -0.04 | 0.297 | 0.297 | 0.035 | 0.16 | 0.51 | 0.23 | 0.46 | 0.08 | 0.13 |
| 2 | 74961016 | rs205628 | 0.567 | 0.795 | 1.18 | 1.493 | 0.047 | 0.12 | 0.12 | 0.127 | 0.88 | 0.76 | 0.86 | 0.65 | 0.83 | 0.66 |
| 2 | 74961853 | rs10185931 | -1.526 | 0.646 | 1.107 | 1.476 | -0.046 | 0.228 | 0.228 | 0.346 | 0.16 | 0.58 | 0.23 | 0.46 | 0.07 | 0.12 |
| 2 | 74968240 | rs205621 | 1.115 | 1.106 | 0.659 | 0.991 | 0.021 | 0.102 | 0.102 | 0.882 | 0.82 | 0.46 | 0.85 | 0.66 | 0.83 | 0.61 |
| 2 | 74968610 | rs205618 | 1.092 | 1.114 | 0.636 | 0.993 | 0.025 | 0.108 | 0.108 | 0.838 | 0.82 | 0.46 | 0.85 | 0.66 | 0.83 | 0.61 |
| 2 | 74969763 | rs205612 | -0.385 | 1.306 | -0.473 | -0.436 | 0.001 | -0.06 | -0.06 | 0 | 0.38 | 0.41 | 0.21 | 0.1 | 0.41 | 0.31 |
| 2 | 74971385 | rs6546920 | 0.013 | 1.345 | -1.547 | -0.071 | -0.066 | 0.095 | 0.095 | 0 | 0.58 | 0.88 | 0.73 | 0.73 | 0.54 | 0.64 |
| 2 | 74972081 | rs205611 | -0.46 | 1.333 | -1.182 | -0.202 | -0.015 | 0.032 | 0.032 | 0.034 | 0.17 | 0.08 | 0.08 | 0 | 0.2 | 0.05 |
| 2 | 74973952 | rs205609 | -0.644 | 1.689 | -1.672 | 0.471 | -0.079 | -0.134 | -0.134 | 0 | 0.78 | 0.92 | 0.67 | 0.5 | 0.68 | 0.63 |
| 2 | 74974965 | rs6715368 | 1.076 | 1.244 | -0.638 | 0.367 | 0.007 | 0.008 | 0.008 | 0.011 | 0.5 | 0.47 | 0.54 | 0.36 | 0.41 | 0.48 |
| 2 | 74975361 | rs6718760 | 1.224 | 1.259 | -0.64 | 0.365 | 0.007 | 0.008 | 0.008 | 0.027 | 0.5 | 0.47 | 0.54 | 0.36 | 0.41 | 0.48 |
| 2 | 74989749 | rs13405664 | 1.232 | 1.135 | 0.187 | 0.326 | 0.027 | 0.034 | 0.034 | 0 | 0.48 | 0.36 | 0.52 | 0.36 | 0.4 | 0.47 |
| 2 | 74990885 | rs7574100 | 0.189 | 1.206 | 0.111 | 0.274 | -0.05 | -0.022 | -0.022 | 1.177 | 0.67 | 0.84 | 0.64 | 0.37 | 0.63 | 0.58 |
| 2 | 74991580 | rs3913433 | 0.135 | 1.163 | 0.066 | 0.273 | -0.064 | -0.034 | -0.034 | 0 | 0.68 | 0.86 | 0.64 | 0.37 | 0.63 | 0.58 |
| 2 | 74996512 | rs6740974 | -0.477 | 1.181 | 0.166 | 0.207 | 0.04 | -0.115 | -0.115 | 0.9 | 0.18 | 0.37 | 0.11 | 0.01 | 0.23 | 0.1 |
| 2 | 74999114 | rs11691168 | -0.544 | 1.209 | 0.084 | 0.234 | 0.053 | -0.139 | -0.139 | 0 | 0.19 | 0.47 | 0.11 | 0.01 | 0.22 | 0.1 |
| 2 | 75004753 | rs6737045 | -0.652 | 1.151 | -0.121 | 0.574 | 0.015 | -0.045 | -0.045 | 0 | 0.21 | 0.55 | 0.12 | 0.11 | 0.23 | 0.11 |
| 2 | 75008284 | rs4853048 | 0.046 | 1.214 | -0.522 | 0.718 | -0.1 | -0.133 | -0.133 | 0 | 0.69 | 0.9 | 0.65 | 0.47 | 0.63 | 0.6 |
| 2 | 75011686 | rs13013272 | 0.284 | 1.129 | -0.322 | 0.63 | 0 | -0.029 | -0.029 | 0.718 | 0.16 | 0.16 | 0.08 | 0.01 | 0.18 | 0.08 |
| 2 | 75012327 | rs6546923 | 0.336 | 1.385 | -0.255 | 1.215 | -0.089 | -0.117 | -0.117 | 0 | 0.71 | 0.89 | 0.64 | 0.47 | 0.62 | 0.56 |
| 2 | 75025709 | rs7586593 | -0.523 | 1.454 | -0.228 | 1.159 | -0.084 | -0.124 | -0.124 | 0 | 0.78 | 0.91 | 0.66 | 0.52 | 0.67 | 0.56 |
| 2 | 75032435 | rs624572 | 0.585 | 1.275 | 0.023 | 0.822 | -0.026 | 0.026 | 0.026 | 0 | 0.62 | 0.7 | 0.61 | 0.28 | 0.57 | 0.46 |
| 2 | 75034144 | rs648309 | 0.656 | 1.288 | -0.068 | 0.816 | 0 | -0.003 | -0.003 | 0 | 0.6 | 0.42 | 0.59 | 0.27 | 0.57 | 0.46 |
| 2 | 75037843 | rs682369 | 0.023 | 1.788 | -0.37 | 1.296 | -0.018 | -0.017 | -0.017 | 5.701 | 0.84 | 0.85 | 0.7 | 0.43 | 0.74 | 0.61 |
| 2 | 75039614 | rs581720 | 0.445 | 1.573 | -0.036 | 1.315 | 0.01 | 0.013 | 0.013 | 1.312 | 0.77 | 0.66 | 0.67 | 0.67 | 0.72 | 0.63 |
| 2 | 75041874 | rs642247 | 0.959 | 1.857 | 0.091 | 1.1 | -0.015 | 0.004 | 0.004 | 1.317 | 0.63 | 0.65 | 0.58 | 0.61 | 0.53 | 0.54 |
| 2 | 75042972 | rs627152 | 1.019 | 1.85 | 0.297 | 1.099 | 0.004 | -0.002 | -0.002 | 1.354 | 0.63 | 0.6 | 0.58 | 0.6 | 0.53 | 0.54 |
| 2 | 75050732 | rs17678046 | -0.004 | 1.764 | 0.205 | 1.119 | 0.002 | -0.003 | -0.003 | 1.102 | 0.15 | 0.13 | 0.11 | 0.11 | 0.19 | 0.11 |
| 2 | 75051823 | rs17678088 | 0.067 | 1.772 | 0.209 | 1.111 | 0.003 | -0.004 | -0.004 | 4.259 | 0.14 | 0.14 | 0.1 | 0.11 | 0.19 | 0.09 |
| 2 | 75053174 | rs4853059 | -0.078 | 1.83 | 0.037 | 1.106 | -0.045 | -0.033 | -0.033 | 0 | 0.79 | 0.93 | 0.7 | 0.71 | 0.73 | 0.64 |
| 2 | 75059013 | rs619850 | 0.687 | 1.558 | 0.13 | 1.161 | 0.022 | -0.007 | -0.007 | 0 | 0.63 | 0.52 | 0.57 | 0.6 | 0.53 | 0.55 |
| 2 | 75060099 | rs656489 | 0.64 | 1.572 | -0.101 | 1.171 | 0.005 | -0.002 | -0.002 | 0 | 0.63 | 0.62 | 0.57 | 0.6 | 0.53 | 0.54 |
| 2 | 75065473 | rs602898 | -0.774 | 1.526 | -0.343 | 1.316 | 0.026 | 0.078 | 0.078 | 0.278 | 0.22 | 0.57 | 0.18 | 0.36 | 0.24 | 0.21 |
| 2 | 75065650 | rs603743 | -0.273 | 1.567 | -0.972 | 1.417 | -0.024 | -0.039 | -0.039 | 0 | 0.8 | 0.92 | 0.6 | 0.65 | 0.74 | 0.62 |
| 2 | 75070216 | rs3771791 | 0.288 | 1.082 | -1.695 | 0.688 | -0.036 | 0.098 | 0.098 | 5.376 | 0.29 | 0.1 | 0.16 | 0.12 | 0.28 | 0.07 |
| 2 | 75070263 | rs3821310 | 0.178 | 1.213 | -0.971 | 1.255 | 0.013 | 0.095 | 0.095 | 0.013 | 0.1 | 0.24 | 0.09 | 0.2 | 0.1 | 0.11 |
| 2 | 75071010 | rs11893422 | -0.226 | 1.105 | -1.112 | 1.18 | -0.028 | -0.069 | -0.069 | 0.149 | 0.76 | 0.85 | 0.57 | 0.51 | 0.7 | 0.58 |
| 2 | 75072923 | rs3821309 | 0.003 | 1.167 | -0.821 | 1.167 | 0.033 | 0.084 | 0.084 | 0.045 | 0.1 | 0.24 | 0.09 | 0.19 | 0.11 | 0.12 |
| 2 | 75074055 | rs28362967 | 0.868 | 0.63 | -1.698 | 0.836 | -0.029 | 0.015 | 0.015 | 0 | 0.29 | 0.05 | 0.3 | 0.23 | 0.32 | 0.33 |
| 2 | 75074967 | rs681900 | -0.081 | 0.646 | -1.275 | 0.969 | -0.003 | -0.013 | -0.013 | 0 | 0.26 | 0.31 | 0.13 | 0.17 | 0.22 | 0.18 |
| 2 | 75078201 | rs3755454 | -0.142 | 0.525 | -0.965 | 1.338 | 0 | 0.126 | 0.126 | 1.165 | 0.07 | 0.58 | 0.07 | 0.17 | 0.03 | 0.07 |
| 2 | 75082895 | rs3771783 | -0.038 | 0.785 | -1.353 | 1.716 | -0.01 | -0.042 | -0.042 | 0 | 0.14 | 0.06 | 0.38 | 0.32 | 0.17 | 0.28 |

| | | | | | | | | | | | | | | | | |
|---|---|---|---|---|---|---|---|---|---|---|---|---|---|---|---|---|
| 2 | 75085163 | rs3771781 | 0.038 | 0.632 | -1.799 | 1.344 | -0.014 | -0.023 | -0.023 | 0 | 0.58 | 0.76 | 0.64 | 0.55 | 0.56 | 0.48 |
| 2 | 75086181 | rs651071 | -0.201 | 0.444 | -1.165 | 2.02 | 0.011 | -0.004 | -0.004 | 0 | 0.24 | 0.14 | 0.09 | 0.25 | 0.17 | 0.2 |
| 2 | 75087490 | rs634849 | 0.106 | 0.804 | -0.922 | 2 | -0.003 | -0.033 | -0.033 | 0 | 0.21 | 0.14 | 0.09 | 0.32 | 0.19 | 0.19 |
| 2 | 75090155 | rs4853066 | -1.061 | 0.896 | -0.407 | 2.199 | -0.002 | -0.009 | -0.009 | 0 | 0.14 | 0.12 | 0.39 | 0.32 | 0.17 | 0.24 |
| 2 | 75094125 | rs3771770 | -0.903 | 0.976 | -0.737 | 2.121 | -0.005 | -0.023 | -0.023 | 0 | 0.14 | 0.07 | 0.38 | 0.3 | 0.16 | 0.24 |
| 2 | 75098183 | rs6732614 | 0.23 | 1.705 | -0.799 | 1.737 | -0.074 | -0.098 | -0.098 | 0.788 | 0.81 | 0.93 | 0.53 | 0.58 | 0.68 | 0.65 |
| 2 | 75099872 | rs13384240 | -0.3 | 1.334 | -0.513 | 1.306 | 0.002 | 0.003 | 0.003 | 0.457 | 0.24 | 0.22 | 0.17 | 0.16 | 0.18 | 0.17 |
| 2 | 75106441 | rs1872837 | 1.182 | 1.26 | 0.143 | 1.419 | -0.005 | 0.01 | 0.01 | 0.55 | 0.57 | 0.36 | 0.36 | 0.32 | 0.51 | 0.52 |
| 2 | 75108744 | rs2306805 | -0.201 | 1.087 | -0.063 | 1.329 | 0.006 | 0.005 | 0.005 | 0.91 | 0.22 | 0.19 | 0.15 | 0.16 | 0.17 | 0.16 |
| 2 | 75113657 | rs2229626 | 0.986 | 1.1 | -0.192 | 1.159 | -0.005 | 0.038 | 0.038 | 0 | 0.54 | 0.75 | 0.35 | 0.41 | 0.47 | 0.47 |
| 2 | 75114730 | rs923273 | 1.485 | 0.949 | -0.064 | 1.054 | 0.033 | 0.166 | 0.166 | 0.643 | 0.46 | 0.12 | 0.28 | 0.26 | 0.41 | 0.44 |
| 2 | 75115108 | rs10194657 | 0.567 | 1.478 | 0.284 | 1.339 | 0.018 | 0.046 | 0.046 | 1.132 | 0.71 | 0.42 | 0.47 | 0.48 | 0.63 | 0.64 |
| 2 | 75115687 | rs17010504 | -1.507 | 1.017 | 0.306 | 1.298 | 0.05 | 0.209 | 0.209 | 1.843 | 0.05 | 0.33 | 0.02 | 0.11 | 0.01 | 0.04 |
| 2 | 75116627 | rs4853069 | 0.07 | 1.263 | -0.356 | 1.298 | -0.128 | -0.111 | -0.111 | 0.126 | 0.74 | 0.93 | 0.47 | 0.59 | 0.57 | 0.66 |
| 2 | 75120525 | rs10496196 | 0.551 | 1.134 | -0.456 | 1.293 | -0.008 | -0.046 | -0.046 | 0 | 0.16 | 0.06 | 0.13 | 0.18 | 0.14 | 0.16 |
| 2 | 75120713 | rs10496197 | 0.506 | 1.115 | -0.092 | 1.121 | 0.039 | 0.025 | 0.025 | 0.279 | 0.67 | 0.55 | 0.44 | 0.58 | 0.56 | 0.62 |
| 2 | 75121645 | rs10865404 | 0.7 | 0.965 | -0.599 | 0.632 | 0.034 | 0.087 | 0.087 | 0.318 | 0.47 | 0.26 | 0.31 | 0.28 | 0.41 | 0.39 |
| 2 | 75125907 | rs7597871 | 0.415 | 0.842 | -0.256 | 0.95 | -0.029 | 0.004 | 0.004 | 0.322 | 0.76 | 0.42 | 0.84 | 0.78 | 0.84 | 0.77 |
| 2 | 75128656 | rs7596427 | 0.34 | 0.841 | -0.226 | 0.951 | -0.034 | 0.008 | 0.008 | 0.157 | 0.76 | 0.45 | 0.84 | 0.79 | 0.84 | 0.77 |
| 2 | 75138842 | rs12989728 | 1.026 | 1.097 | -1.706 | 1.023 | -0.051 | -0.05 | -0.05 | 0 | 0.49 | 0.1 | 0.43 | 0.5 | 0.54 | 0.44 |
| 2 | 75139785 | rs1002026 | -0.283 | 1.501 | -1.702 | 1.107 | -0.034 | -0.045 | -0.045 | 0.713 | 0.2 | 0.08 | 0.33 | 0.3 | 0.22 | 0.27 |
| 2 | 75144084 | rs11686274 | 0.201 | 1.352 | -1.446 | 0.904 | -0.029 | -0.023 | -0.023 | 0 | 0.76 | 0.9 | 0.82 | 0.78 | 0.76 | 0.71 |
| 2 | 75146521 | rs4853077 | 0.4 | 1.548 | -2.132 | 0.942 | -0.151 | -0.021 | -0.021 | 0 | 0.66 | 0.94 | 0.67 | 0.69 | 0.57 | 0.61 |
| 2 | 75146534 | rs10187337 | -0.446 | 1.343 | -0.783 | 1.693 | -0.016 | 0.111 | 0.111 | 0.378 | 0.22 | 0.51 | 0.37 | 0.46 | 0.15 | 0.27 |
| 2 | 75146717 | rs13008916 | 0.274 | 1.168 | -0.479 | 1.148 | -0.003 | -0.003 | -0.003 | 0 | 0.67 | 0.39 | 0.78 | 0.72 | 0.69 | 0.65 |
| 2 | 75148429 | rs8179569 | 0.643 | 1.243 | -0.663 | 1.332 | -0.025 | -0.003 | -0.003 | 0 | 0.63 | 0.77 | 0.5 | 0.4 | 0.5 | 0.45 |
| 2 | 75156603 | rs11884580 | -0.53 | 1.328 | -0.556 | 1.245 | 0.022 | 0.011 | 0.011 | 0.194 | 0.29 | 0.53 | 0.19 | 0.32 | 0.36 | 0.31 |
| 2 | 75163550 | rs965078 | -0.956 | 0.999 | -1.268 | 1.159 | -0.017 | 0.003 | 0.003 | 3.663 | 0.47 | 0.8 | 0.54 | 0.57 | 0.52 | 0.6 |
| 2 | 75168664 | rs7573356 | -0.57 | 0.258 | -1.055 | 0.92 | -0.025 | -0.022 | -0.022 | 0 | 0.42 | 0.31 | 0.59 | 0.58 | 0.54 | 0.67 |
| 2 | 75170336 | rs11688381 | -0.403 | 0.101 | -1.021 | 0.818 | -0.047 | -0.045 | -0.045 | 0.073 | 0.37 | 0.23 | 0.57 | 0.58 | 0.53 | 0.66 |
| 2 | 75186337 | rs2287102 | -0.128 | 0.3 | -0.84 | 0.973 | 0.004 | -0.002 | -0.002 | 0 | 0.21 | 0.28 | 0.45 | 0.2 | 0.27 | 0.38 |
| 2 | 75190024 | rs12713820 | 1.202 | -0.008 | 0.113 | 0.859 | -0.002 | -0.001 | -0.001 | 0 | 0.54 | 0.45 | 0.4 | 0.42 | 0.46 | 0.31 |
| 2 | 75202015 | rs13401471 | 1.074 | -0.531 | -0.149 | 0.513 | 0.002 | -0.005 | -0.005 | 0 | 0.21 | 0.19 | 0.24 | 0.25 | 0.17 | 0.13 |
| 2 | 75220996 | rs6546942 | -0.157 | -0.702 | 0.703 | 0.522 | 0.006 | 0.018 | 0.018 | 0 | 0.74 | 0.61 | 0.85 | 0.63 | 0.73 | 0.66 |
| 2 | 75223516 | rs923276 | 0.831 | -0.725 | 0.66 | 0.507 | -0.007 | -0.004 | -0.004 | 0.983 | 0.53 | 0.3 | 0.39 | 0.42 | 0.46 | 0.29 |
| 2 | 75227997 | rs4853094 | -1.574 | -0.678 | 0.577 | 0.509 | -0.047 | -0.056 | -0.056 | 0.363 | 0.84 | 0.93 | 0.87 | 0.63 | 0.75 | 0.72 |
| 2 | 75229665 | rs17563336 | 1.929 | -0.671 | 0.544 | 0.459 | -0.051 | 0.016 | 0.016 | 0 | 0.29 | 0.03 | 0.14 | 0.18 | 0.26 | 0.18 |
| 2 | 75248797 | rs6740220 | -1.219 | -0.39 | 1.171 | 0.659 | -0.049 | -0.059 | -0.059 | 0.564 | 0.83 | 0.93 | 0.87 | 0.62 | 0.75 | 0.72 |
| 2 | 75249187 | rs4853100 | -0.947 | -0.382 | 1.172 | 0.661 | 0.006 | 0.01 | 0.01 | 0 | 0.82 | 0.79 | 0.86 | 0.63 | 0.75 | 0.73 |
| 2 | 75264029 | rs10515918 | 0.378 | -0.464 | 1.05 | 0.685 | -0.074 | -0.022 | -0.022 | 0 | 0.19 | 0.01 | 0.42 | 0.14 | 0.28 | 0.31 |
| 2 | 75268480 | rs2422090 | 0.717 | -0.686 | 1.277 | 0.618 | 0.002 | 0.003 | 0.003 | 2.927 | 0.58 | 0.45 | 0.44 | 0.42 | 0.56 | 0.29 |
| 2 | 75268803 | rs4614953 | -0.953 | -0.632 | 1.171 | 0.737 | -0.015 | -0.008 | -0.008 | 0 | 0.81 | 0.83 | 0.73 | 0.72 | 0.72 | 0.72 |
| 2 | 75269064 | rs13012537 | 0.57 | -0.95 | 1.199 | 0.486 | -0.008 | -0.009 | -0.009 | 0 | 0.58 | 0.59 | 0.41 | 0.48 | 0.46 | 0.41 |
| 2 | 75269221 | rs10202905 | 0.051 | -0.778 | 0.634 | 0.532 | 0.001 | -0.016 | -0.016 | 0.389 | 0.05 | 0.06 | 0.05 | 0 | 0.11 | 0.01 |
| 2 | 75276081 | rs12713828 | 0.889 | -0.893 | 0.51 | 0.552 | -0.005 | -0.005 | -0.005 | 0.062 | 0.52 | 0.28 | 0.38 | 0.43 | 0.45 | 0.28 |
| 2 | 75281054 | rs17010698 | 0.088 | -1.602 | 0.989 | 1.002 | -0.015 | 0.061 | 0.061 | 0 | 0.14 | 0.2 | 0.18 | 0.33 | 0.14 | 0.19 |
| 2 | 75281597 | rs11680315 | 0.238 | -1.52 | 0.766 | 0.956 | -0.001 | 0.022 | 0.022 | 0 | 0.13 | 0.23 | 0.27 | 0.21 | 0.14 | 0.26 |
| 2 | 75283508 | rs6546951 | -0.562 | -1.611 | 0.678 | 0.923 | -0.031 | 0.077 | 0.077 | 0.005 | 0.18 | 0.53 | 0.29 | 0.25 | 0.14 | 0.32 |
| 2 | 75287106 | rs4439987 | -1.444 | -1.806 | -0.429 | 0.65 | -0.168 | 0.038 | 0.038 | 0.009 | 0.67 | 0.96 | 0.62 | 0.78 | 0.54 | 0.67 |
| 2 | 75287819 | rs3755456 | 0.805 | -1.852 | -0.117 | 0.709 | -0.021 | -0.056 | -0.056 | 0.018 | 0.11 | 0.03 | 0.25 | 0.21 | 0.14 | 0.26 |
| 2 | 75290700 | rs13404785 | -0.385 | -1.744 | 0.135 | 0.662 | -0.013 | 0.093 | 0.093 | 0 | 0.17 | 0.45 | 0.42 | 0.31 | 0.15 | 0.33 |
| 2 | 75291510 | rs3755457 | -0.417 | -1.742 | 0.031 | 0.663 | -0.011 | 0.113 | 0.113 | 0 | 0.17 | 0.52 | 0.44 | 0.3 | 0.15 | 0.32 |
| 2 | 75291521 | rs3755458 | 0.425 | -1.8 | -0.004 | 0.703 | -0.031 | 0.057 | 0.057 | 0.024 | 0.14 | 0.17 | 0.14 | 0.31 | 0.06 | 0.16 |
| 2 | 75293157 | rs11688000 | 0.192 | -1.566 | -0.35 | 0.77 | -0.005 | -0.043 | -0.043 | 0 | 0.34 | 0.18 | 0.5 | 0.42 | 0.37 | 0.38 |
| 2 | 75295900 | rs768601 | -0.561 | -1.039 | 0.15 | 0.759 | -0.016 | 0.088 | 0.088 | 0 | 0.16 | 0.43 | 0.42 | 0.3 | 0.14 | 0.33 |
| 2 | 75301763 | rs6546952 | -0.654 | -0.557 | -0.875 | 0.702 | -0.047 | 0.07 | 0.07 | 0 | 0.65 | 0.82 | 0.72 | 0.8 | 0.51 | 0.64 |
| 2 | 75307296 | rs6715420 | -0.866 | -0.479 | -0.285 | 1.285 | 0 | 0.05 | 0.05 | 0 | 0.13 | 0.28 | 0.39 | 0.24 | 0.13 | 0.28 |
| 2 | 75314020 | rs11680998 | -0.889 | -0.322 | -1.329 | 1.363 | -0.014 | -0.065 | -0.065 | 5.117 | 0.14 | 0.03 | 0.38 | 0.24 | 0.13 | 0.29 |
| 2 | 75319659 | rs3771811 | -0.781 | -1.044 | -0.691 | 1.233 | -0.066 | 0.027 | 0.027 | 0 | 0.26 | 0.35 | 0.18 | 0.31 | 0.09 | 0.23 |
| 2 | 75321180 | rs34242711 | -1.198 | -1.149 | -0.609 | 1.202 | -0.023 | -0.001 | -0.001 | 0 | 0.25 | 0.29 | 0.13 | 0.3 | 0.14 | 0.25 |
| 2 | 75321603 | rs3755459 | -1.139 | -1.398 | -0.376 | 1.389 | -0.019 | 0.004 | 0.004 | 0.027 | 0.37 | 0.4 | 0.48 | 0.61 | 0.36 | 0.56 |
| 2 | 75321635 | rs3755460 | -0.871 | -1.097 | -0.421 | 1.251 | 0.006 | 0.002 | 0.002 | 0 | 0.5 | 0.48 | 0.48 | 0.55 | 0.44 | 0.47 |
| 2 | 75323267 | rs1861457 | -0.226 | -1.542 | -0.773 | 1.013 | 0.013 | -0.007 | -0.007 | 0 | 0.69 | 0.84 | 0.67 | 0.67 | 0.68 | 0.57 |
| 2 | 75327682 | rs17010738 | -0.19 | -2.138 | -0.365 | 1.343 | -0.035 | 0.053 | 0.053 | 0 | 0.13 | 0.24 | 0.04 | 0.29 | 0.03 | 0.1 |
| 2 | 75329306 | rs13417600 | -0.599 | -2.752 | -0.407 | 0.818 | 0.046 | -0.04 | -0.04 | 0 | 0.26 | 0.17 | 0.32 | 0.56 | 0.12 | 0.28 |
| 2 | 75338380 | rs3771817 | -0.651 | -2.728 | -0.255 | 0.611 | 0.027 | -0.037 | -0.037 | 0 | 0.25 | 0.19 | 0.32 | 0.48 | 0.1 | 0.27 |
| 2 | 75338619 | rs6747886 | 1.035 | -2.832 | -0.256 | 0.482 | -0.006 | -0.072 | -0.072 | 0.242 | 0.42 | 0.49 | 0.34 | 0.12 | 0.55 | 0.3 |
| 2 | 75339314 | rs7600024 | -0.814 | -2.747 | -0.475 | 0.552 | 0.005 | -0.061 | -0.061 | 0 | 0.71 | 0.91 | 0.66 | 0.61 | 0.66 | 0.57 |
| 2 | 75345195 | rs6741029 | -0.488 | -2.734 | -0.415 | 0.521 | 0.009 | -0.012 | -0.012 | 0 | 0.67 | 0.71 | 0.65 | 0.59 | 0.65 | 0.57 |

| | | | | | | | | | | | | | | | | |
|---|---|---|---|---|---|---|---|---|---|---|---|---|---|---|---|---|
| 2 | 75351100 | rs3771818 | 0.323 | -2.686 | -0.453 | -0.001 | 0.025 | -0.016 | -0.016 | 0 | 0.15 | 0.11 | 0.28 | 0.18 | 0.08 | 0.17 |
| 2 | 75351948 | rs3771823 | 0.775 | -2.317 | -0.253 | -0.111 | -0.006 | -0.082 | -0.082 | 0.075 | 0.41 | 0.45 | 0.32 | 0.12 | 0.53 | 0.3 |
| 2 | 75363044 | rs735668 | 1.132 | -1.96 | -0.395 | -0.313 | -0.012 | 0.034 | 0.034 | 0 | 0.36 | 0.28 | 0.3 | 0.12 | 0.5 | 0.23 |
| 2 | 75363186 | rs741418 | -0.281 | -2.175 | -0.866 | -0.125 | 0.058 | 0.108 | 0.108 | 1.027 | 0.56 | 0.83 | 0.39 | 0.29 | 0.58 | 0.4 |
| 2 | 75366627 | rs3771832 | 0.754 | -1.827 | -0.334 | -0.313 | -0.002 | -0.082 | -0.082 | 0 | 0.4 | 0.45 | 0.33 | 0.14 | 0.52 | 0.28 |
| 2 | 75366937 | rs3771833 | 0.556 | -2.182 | -0.624 | -0.279 | 0.024 | -0.009 | -0.009 | 0 | 0.21 | 0.15 | 0.23 | 0.19 | 0.11 | 0.17 |
| 2 | 75368740 | rs11126453 | 0.613 | -2.203 | -0.684 | -0.261 | 0.058 | -0.024 | -0.024 | 0 | 0.2 | 0.09 | 0.24 | 0.19 | 0.09 | 0.19 |
| 2 | 75368797 | rs975664 | 0.453 | -1.47 | -0.426 | -0.445 | 0 | -0.069 | -0.069 | 0 | 0.4 | 0.63 | 0.33 | 0.03 | 0.5 | 0.25 |
| 2 | 75370716 | rs11902071 | 0.532 | -1.364 | -0.306 | -0.37 | -0.001 | -0.083 | -0.083 | 0 | 0.41 | 0.54 | 0.32 | 0.13 | 0.52 | 0.29 |
| 2 | 75371588 | rs10208860 | 0.325 | -1.396 | -0.452 | -0.366 | 0.002 | -0.007 | -0.007 | 0 | 0.42 | 0.62 | 0.33 | 0.13 | 0.52 | 0.29 |
| 2 | 75374043 | rs3771835 | 0.441 | -1.189 | -0.702 | -0.261 | 0 | -0.074 | -0.074 | 0.513 | 0.39 | 0.43 | 0.33 | 0.13 | 0.52 | 0.29 |
| 2 | 75380952 | rs3771836 | 0.375 | -0.704 | -0.95 | 0.262 | -0.019 | 0.089 | 0.089 | 0 | 0.46 | 0.17 | 0.51 | 0.24 | 0.5 | 0.33 |
| 2 | 75382391 | rs3755468 | -0.225 | -0.817 | -0.545 | 0.523 | 0.001 | -0.005 | -0.005 | 0 | 0.6 | 0.65 | 0.57 | 0.5 | 0.55 | 0.5 |
| 2 | 75382710 | rs10168354 | -0.11 | -0.799 | -0.547 | 0.531 | 0.007 | -0.002 | -0.002 | 0 | 0.58 | 0.6 | 0.57 | 0.51 | 0.54 | 0.5 |
| 2 | 75384549 | rs759588 | 0.776 | -0.454 | -1.005 | 0.543 | -0.049 | -0.032 | -0.032 | 0 | 0.38 | 0.1 | 0.49 | 0.38 | 0.44 | 0.46 |
| 2 | 75386316 | rs3821316 | 0.425 | -0.752 | -0.499 | 0.466 | -0.002 | 0.003 | 0.003 | 0 | 0.51 | 0.56 | 0.58 | 0.47 | 0.56 | 0.52 |
| 2 | 75387146 | rs3771845 | 0.115 | -0.958 | -0.844 | 0.65 | 0 | -0.001 | -0.001 | 0 | 0.16 | 0.15 | 0.04 | 0.15 | 0.06 | 0.26 |
| 2 | 75387834 | rs6733933 | -1.114 | -0.325 | -1.224 | 0.677 | -0.033 | -0.033 | -0.033 | 0 | 0.86 | 0.92 | 0.84 | 0.77 | 0.82 | 0.91 |
| 2 | 75390741 | rs17010840 | 0.429 | -0.412 | -1.056 | 0.617 | 0.013 | -0.004 | -0.004 | 0.083 | 0.13 | 0.06 | 0.18 | 0.13 | 0.09 | 0.12 |
| 2 | 75392831 | rs2216307 | 0.372 | 0.003 | -0.75 | 0.545 | -0.001 | 0.002 | 0.002 | 0 | 0.66 | 0.61 | 0.79 | 0.65 | 0.7 | 0.7 |
| 2 | 75399430 | rs3771848 | -0.612 | 0.376 | -0.811 | 0.349 | 0.004 | -0.023 | -0.023 | 0 | 0.05 | 0.07 | 0.03 | 0.02 | 0.07 | 0.03 |
| 2 | 75401614 | rs3771853 | -0.427 | 0.127 | -1.033 | 0.375 | 0.013 | 0.019 | 0.019 | 0 | 0.57 | 0.73 | 0.57 | 0.34 | 0.66 | 0.54 |
| 2 | 75402065 | rs12477554 | -0.185 | 0.059 | -0.928 | 0.473 | 0.004 | 0.025 | 0.025 | 0 | 0.52 | 0.62 | 0.54 | 0.33 | 0.6 | 0.52 |
| 2 | 75406823 | rs4627599 | -0.083 | 0.266 | -1.133 | 0.432 | -0.005 | 0.005 | 0.005 | 0.753 | 0.39 | 0.39 | 0.48 | 0.3 | 0.51 | 0.46 |
| 2 | 75411057 | rs2160654 | -0.561 | -0.02 | -1.958 | 0.547 | -0.034 | 0.01 | 0.01 | 0 | 0.49 | 0.9 | 0.36 | 0.38 | 0.45 | 0.42 |
| 2 | 75415419 | rs2193409 | 0.003 | 0.008 | -1.16 | 0.491 | -0.006 | 0.002 | 0.002 | 0 | 0.38 | 0.54 | 0.41 | 0.38 | 0.33 | 0.38 |
| 2 | 75416296 | rs4853119 | 0.955 | -0.009 | -1.172 | 0.23 | -0.015 | 0.022 | 0.022 | 0.792 | 0.21 | 0.16 | 0.31 | 0.07 | 0.26 | 0.1 |
| 2 | 75419745 | rs10204684 | 0.644 | -0.003 | -1.243 | 0.271 | -0.019 | 0.021 | 0.021 | 0 | 0.07 | 0.04 | 0.03 | 0.01 | 0.08 | 0.04 |
| 2 | 75427878 | rs2111375 | 1.02 | 0.035 | -1.901 | 0.436 | -0.017 | -0.046 | -0.046 | 0 | 0.34 | 0.01 | 0.3 | 0.33 | 0.3 | 0.32 |
| 2 | 75429228 | rs1477157 | -0.87 | -0.038 | -1.594 | 0.222 | -0.03 | 0.04 | 0.04 | 0 | 0.49 | 0.9 | 0.5 | 0.39 | 0.46 | 0.42 |
| 2 | 75437182 | rs4853122 | 1.015 | -0.148 | -1.171 | 0.176 | 0.002 | -0.027 | -0.027 | 0 | 0.22 | 0.26 | 0.3 | 0.11 | 0.22 | 0.1 |
| 2 | 75440136 | rs6725947 | -0.498 | -0.117 | -0.566 | 0.375 | -0.03 | 0.009 | 0.009 | 0 | 0.26 | 0.55 | 0.17 | 0.28 | 0.17 | 0.31 |
| 2 | 75444073 | rs11684394 | 1.213 | -0.07 | -1.486 | 0.271 | -0.088 | -0.017 | -0.017 | 0.506 | 0.28 | 0.03 | 0.39 | 0.23 | 0.29 | 0.19 |
| 2 | 75449983 | rs1468900 | 0.41 | -0.065 | -1.111 | 0.432 | -0.05 | 0.105 | 0.105 | 0.253 | 0.17 | 0.33 | 0.12 | 0.29 | 0.09 | 0.28 |
| 2 | 75451047 | rs6717216 | -0.475 | 0.149 | -2.301 | 0.44 | -0.123 | -0.063 | -0.063 | 0 | 0.58 | 0.91 | 0.59 | 0.58 | 0.51 | 0.51 |
| 2 | 75457783 | rs4852364 | -0.205 | 0.223 | -2.02 | 0.385 | -0.058 | 0.064 | 0.064 | 1.468 | 0.55 | 0.83 | 0.59 | 0.32 | 0.47 | 0.36 |
| 2 | 75459144 | rs7568535 | -0.013 | 0.243 | -2.224 | 0.382 | -0.033 | 0.005 | 0.005 | 0.172 | 0.69 | 0.81 | 0.74 | 0.73 | 0.65 | 0.57 |
| 2 | 75460060 | rs980545 | -0.142 | 0.213 | -1.901 | -0.134 | -0.018 | -0.016 | -0.016 | 0 | 0.14 | 0.69 | 0.14 | 0.07 | 0.08 | 0.16 |
| 2 | 75460555 | rs12328696 | -0.404 | 0.354 | -1.854 | -0.031 | -0.017 | 0.048 | 0.048 | 1.546 | 0.54 | 0.7 | 0.59 | 0.33 | 0.48 | 0.36 |
| 2 | 75461177 | rs13017967 | 0.55 | 0.526 | -2.342 | 0.299 | -0.058 | -0.003 | -0.003 | 0 | 0.32 | 0.06 | 0.41 | 0.25 | 0.32 | 0.16 |
| 2 | 75465817 | rs4853125 | -0.058 | 0.492 | -1.967 | 0.394 | -0.028 | -0.029 | -0.029 | 0 | 0.32 | 0.05 | 0.39 | 0.7 | 0.32 | 0.51 |
| 2 | 75470713 | rs1861430 | -0.483 | 0.615 | -0.929 | 0.307 | 0.001 | -0.009 | -0.009 | 0.025 | 0.59 | 0.57 | 0.6 | 0.69 | 0.58 | 0.44 |
| 2 | 75471057 | rs7576919 | -0.457 | 0.607 | -0.847 | 0.303 | 0.001 | -0.011 | -0.011 | 0.238 | 0.59 | 0.52 | 0.59 | 0.69 | 0.58 | 0.43 |
| 2 | 75473916 | rs7588010 | 0.226 | 0.565 | -0.981 | 0.39 | -0.005 | 0.02 | 0.02 | 0.988 | 0.37 | 0.29 | 0.39 | 0.27 | 0.36 | 0.22 |
| 2 | 75479258 | rs4508624 | 0.706 | 0.652 | -0.953 | 0.252 | -0.005 | -0.009 | -0.009 | 0.449 | 0.3 | 0.34 | 0.33 | 0.26 | 0.28 | 0.17 |
| 2 | 75479798 | rs12612891 | 1.099 | 0.752 | -1.261 | 0.296 | -0.032 | -0.148 | -0.148 | 0 | 0.2 | 0.02 | 0.19 | 0.42 | 0.21 | 0.21 |
| 2 | 75482223 | rs10490307 | 0.094 | 0.919 | -0.368 | -0.045 | -0.011 | -0.024 | -0.024 | 0 | 0.3 | 0.39 | 0.33 | 0.26 | 0.28 | 0.16 |
| 2 | 75482375 | rs10490308 | -0.654 | 1.202 | -0.504 | 0.305 | 0.001 | -0.027 | -0.027 | 0 | 0.11 | 0.11 | 0.07 | 0.01 | 0.13 | 0.07 |
| 2 | 75498798 | rs12469135 | -0.862 | 1.292 | -0.051 | 0.81 | 0.043 | 0.058 | 0.058 | 0 | 0.1 | 0.25 | 0.3 | 0.23 | 0.19 | 0.15 |
| 2 | 75501720 | rs9309489 | -1.036 | 0.35 | 0.517 | 0.937 | -0.007 | 0.025 | 0.025 | 0.218 | 0.26 | 0.32 | 0.34 | 0.39 | 0.25 | 0.29 |
| 2 | 75502853 | rs10205827 | -1.334 | 0.286 | 0.495 | 0.906 | -0.022 | 0.045 | 0.045 | 0.354 | 0.28 | 0.51 | 0.35 | 0.4 | 0.25 | 0.3 |
| 2 | 75503116 | rs10208470 | -1.192 | 0.256 | 0.554 | 0.891 | -0.015 | 0.041 | 0.041 | 0.361 | 0.27 | 0.43 | 0.34 | 0.39 | 0.25 | 0.3 |
| 2 | 75504977 | rs17010981 | 0.015 | 0.542 | 0.245 | 0.722 | -0.002 | 0 | 0 | 0 | 0.08 | 0.07 | 0.28 | 0.22 | 0.19 | 0.14 |
| 2 | 75505999 | rs2422153 | 0.685 | -0.183 | -1.131 | 0.305 | 0.017 | 0.017 | 0.017 | 0 | 0.28 | 0.02 | 0.25 | 0.17 | 0.2 | 0.32 |
| 2 | 75512548 | rs2192925 | 0.355 | -0.166 | -0.819 | 0.488 | 0.006 | 0.018 | 0.018 | 0 | 0.09 | 0.1 | 0.29 | 0.26 | 0.18 | 0.15 |
| 2 | 75512668 | rs2192926 | -0.347 | -0.242 | -0.339 | 0.443 | 0.023 | 0.024 | 0.024 | 0 | 0.13 | 0.19 | 0.32 | 0.26 | 0.24 | 0.21 |
| 2 | 75515611 | rs6546962 | -1.332 | -0.024 | -0.346 | 0.029 | 0.034 | 0.028 | 0.028 | 0.008 | 0.76 | 0.69 | 0.75 | 0.63 | 0.64 | 0.71 |
| 2 | 75521456 | rs17011008 | 0.97 | -0.441 | 0.147 | -0.032 | -0.024 | 0.001 | 0.001 | 0.288 | 0.19 | 0.24 | 0.06 | 0.14 | 0.07 | 0.15 |
| 2 | 75524061 | rs4853131 | -1.505 | -0.796 | 0.205 | -0.235 | -0.021 | -0.013 | -0.013 | 0 | 0.71 | 0.81 | 0.74 | 0.61 | 0.6 | 0.71 |
| 2 | 75526489 | rs17581492 | 1.361 | -0.832 | 0.367 | -0.352 | 0.005 | 0 | 0 | 0.19 | 0.15 | 0.14 | 0.12 | 0.01 | 0.12 | 0.05 |
| 2 | 75541243 | rs6546964 | -1.589 | -0.757 | 0.316 | -0.274 | -0.02 | -0.014 | -0.014 | 0 | 0.72 | 0.81 | 0.74 | 0.61 | 0.6 | 0.72 |
| 2 | 75541353 | rs7578998 | 1.545 | -0.866 | 0.292 | -0.319 | 0.005 | -0.001 | -0.001 | 0.005 | 0.17 | 0.15 | 0.05 | 0.13 | 0.07 | 0.15 |
| 2 | 75549380 | rs7562368 | -0.311 | -0.622 | 0.556 | -0.328 | -0.016 | 0.054 | 0.054 | 0 | 0.53 | 0.27 | 0.6 | 0.64 | 0.5 | 0.68 |
| 2 | 75551754 | rs759445 | 0.763 | -0.703 | 0.353 | -0.374 | -0.107 | -0.178 | -0.178 | 0.521 | 0.29 | 0.03 | 0.52 | 0.51 | 0.37 | 0.45 |
| 2 | 75554032 | rs7565730 | 0.371 | -0.876 | 0.307 | -0.539 | 0.031 | 0.029 | 0.029 | 0 | 0.27 | 0.16 | 0.08 | 0.13 | 0.16 | 0.28 |
| 2 | 75554692 | rs6739089 | -0.459 | -0.79 | 0.49 | -0.469 | 0.059 | 0.077 | 0.077 | 0.636 | 0.39 | 0.77 | 0.2 | 0.13 | 0.22 | 0.21 |
| 2 | 75555202 | rs3771871 | 1.58 | -0.719 | 0.407 | -0.416 | 0.005 | 0 | 0 | 0 | 0.18 | 0.15 | 0.06 | 0.13 | 0.09 | 0.15 |
| 2 | 75561061 | rs17582810 | 1.281 | -0.691 | 0.358 | -0.379 | 0.012 | 0.005 | 0.005 | 0.37 | 0.21 | 0.15 | 0.07 | 0.13 | 0.13 | 0.21 |
| 2 | 75561691 | rs10191221 | 0.897 | -0.497 | 0.595 | -0.304 | -0.007 | -0.115 | -0.115 | 0.259 | 0.17 | 0.35 | 0.13 | 0.01 | 0.13 | 0.05 |

| | | | | | | | | | | | | | | | | |
|---|---|---|---|---|---|---|---|---|---|---|---|---|---|---|---|---|
| 2 | 75562499 | rs11681817 | 1.585 | -0.493 | 0.456 | -0.301 | -0.025 | 0.083 | 0.083 | 0 | 0.14 | 0.02 | 0.1 | 0.01 | 0.13 | 0.05 |
| 2 | 75562760 | rs741321 | 1.532 | -0.452 | 0.389 | -0.145 | -0.042 | -0.038 | -0.038 | 0 | 0.08 | 0.03 | 0.43 | 0.33 | 0.19 | 0.14 |
| 2 | 75562933 | rs759446 | -0.08 | -0.625 | 0.519 | -0.288 | -0.025 | -0.051 | -0.051 | 0 | 0.4 | 0.45 | 0.19 | 0.14 | 0.27 | 0.28 |
| 2 | 75568216 | rs7567259 | 1.149 | -0.611 | 0.164 | -0.164 | 0.029 | 0.016 | 0.016 | 0.219 | 0.24 | 0.13 | 0.07 | 0.13 | 0.14 | 0.24 |
| 2 | 75572718 | rs2110933 | -0.391 | -0.275 | -0.06 | 0.028 | -0.032 | -0.04 | -0.04 | 0 | 0.43 | 0.23 | 0.63 | 0.53 | 0.49 | 0.48 |
| 2 | 75577646 | rs17011222 | 1.356 | -0.26 | -0.222 | -0.054 | -0.042 | -0.033 | -0.033 | 0 | 0.19 | 0.02 | 0.1 | 0.19 | 0.18 | 0.29 |
| 2 | 75577730 | rs2192921 | -0.564 | -0.164 | 0.326 | 0.033 | 0.012 | -0.014 | -0.014 | 0 | 0.61 | 0.6 | 0.63 | 0.67 | 0.52 | 0.72 |
| 2 | 75578615 | rs4852370 | -1.407 | -0.014 | 0.379 | 0.113 | 0.025 | 0.009 | 0.009 | 0 | 0.74 | 0.71 | 0.73 | 0.68 | 0.63 | 0.76 |
| 2 | 75582351 | rs4852371 | 0.079 | -0.212 | -0.141 | -0.094 | 0 | -0.001 | -0.001 | 0 | 0.29 | 0.27 | 0.19 | 0.19 | 0.27 | 0.29 |
| 2 | 75584273 | rs6718666 | 0.178 | -0.466 | -0.119 | -0.048 | 0.012 | 0.005 | 0.005 | 0 | 0.44 | 0.56 | 0.17 | 0.36 | 0.27 | 0.46 |
| 2 | 75588230 | rs6749524 | -0.445 | -0.4 | -0.11 | -0.02 | 0.004 | 0.015 | 0.015 | 0 | 0.57 | 0.63 | 0.26 | 0.31 | 0.38 | 0.5 |
| 2 | 75594626 | rs715161 | 0.982 | -0.458 | -0.397 | 0.083 | -0.056 | -0.016 | -0.016 | 0.952 | 0.28 | 0.38 | 0.07 | 0.18 | 0.1 | 0.21 |
| 2 | 75604668 | rs6546985 | -1.544 | 0.078 | -0.947 | 0.316 | -0.027 | -0.02 | -0.02 | 0 | 0.77 | 0.8 | 0.74 | 0.64 | 0.66 | 0.73 |
| 2 | 75608486 | rs1861429 | -0.538 | 0.83 | -0.672 | 0.206 | -0.015 | -0.008 | -0.008 | 1.064 | 0.31 | 0.17 | 0.56 | 0.34 | 0.39 | 0.27 |
| 2 | 75617495 | rs13429274 | -0.463 | 1.241 | -0.464 | -0.046 | -0.009 | -0.012 | -0.012 | 0 | 0.41 | 0.35 | 0.6 | 0.49 | 0.47 | 0.46 |
| 2 | 75620383 | rs10181895 | -0.987 | 1.092 | -0.434 | 0.048 | 0.011 | -0.002 | -0.002 | 0 | 0.53 | 0.58 | 0.7 | 0.5 | 0.59 | 0.51 |
| 2 | 75627059 | rs6546992 | 0.62 | 1.092 | -0.504 | 0.003 | 0.002 | 0 | 0 | 0.296 | 0.49 | 0.52 | 0.31 | 0.48 | 0.4 | 0.49 |
| 2 | 75629885 | rs6758300 | 0.609 | 1.257 | -0.606 | 0.4 | 0.002 | 0.006 | 0.006 | 0 | 0.57 | 0.49 | 0.36 | 0.48 | 0.47 | 0.54 |
| 2 | 75629909 | rs6546993 | 0.345 | 1.804 | -0.517 | 0.274 | 0.028 | -0.016 | -0.016 | 0.119 | 0.82 | 0.77 | 0.85 | 0.87 | 0.72 | 0.79 |
| 2 | 75630833 | rs6704951 | -0.253 | 0.934 | -0.813 | 0.484 | 0.005 | -0.021 | -0.021 | 0.044 | 0.2 | 0.1 | 0.32 | 0.29 | 0.16 | 0.22 |
| 2 | 75630998 | rs7570957 | -0.042 | 0.699 | -0.306 | 0.047 | -0.009 | -0.049 | -0.049 | 0 | 0.14 | 0.31 | 0.05 | 0.01 | 0.09 | 0.14 |
| 2 | 75632086 | rs720069 | -0.872 | 1.081 | -1.039 | 0.04 | -0.08 | -0.118 | -0.118 | 0 | 0.75 | 0.96 | 0.64 | 0.59 | 0.6 | 0.7 |
| 2 | 75642987 | rs735881 | -0.687 | 1.115 | 0.088 | -0.32 | 0.084 | -0.087 | -0.087 | 1.93 | 0.13 | 0.38 | 0.21 | 0 | 0.16 | 0.07 |
| 2 | 75644607 | rs2110937 | -1.154 | 1.127 | -1.065 | -0.303 | -0.001 | -0.014 | -0.014 | 2.031 | 0.29 | 0.74 | 0.27 | 0.01 | 0.26 | 0.22 |
| 2 | 75651136 | rs2058702 | -0.496 | 1.405 | -1.482 | 0.185 | -0.021 | 0.014 | 0.014 | 0 | 0.09 | 0.05 | 0.19 | 0 | 0.17 | 0.07 |
| 2 | 75652737 | rs6722299 | -0.316 | 1.021 | -1.578 | 0.361 | 0.014 | -0.012 | -0.012 | 0.051 | 0.34 | 0.7 | 0.27 | 0.37 | 0.19 | 0.31 |
| 2 | 75654059 | rs7564504 | -0.325 | 1.037 | -1.876 | 0.369 | 0.043 | -0.028 | -0.028 | 0 | 0.36 | 0.76 | 0.28 | 0.37 | 0.2 | 0.33 |
| 2 | 75654625 | rs6737535 | -0.899 | 1.113 | -1.901 | 0.37 | -0.004 | -0.012 | -0.012 | 0 | 0.44 | 0.8 | 0.46 | 0.38 | 0.36 | 0.37 |
| 2 | 75656149 | rs6744901 | -0.557 | 0.411 | -1.739 | 0.579 | 0.055 | -0.029 | -0.029 | 0.643 | 0.37 | 0.78 | 0.28 | 0.37 | 0.19 | 0.32 |
| 2 | 75658780 | rs6546997 | 0.469 | 0.673 | -2.134 | 0.333 | -0.017 | 0.021 | 0.021 | 0 | 0.81 | 0.93 | 0.81 | 0.9 | 0.83 | 0.85 |
| 2 | 75666206 | rs6547000 | -0.228 | 2.036 | -2.288 | 1.214 | -0.007 | -0.01 | -0.01 | 0.051 | 0.13 | 0.14 | 0.11 | 0.29 | 0.14 | 0.29 |
| 2 | 75668451 | rs11126470 | -0.084 | 1.574 | -2.165 | 1.076 | 0.011 | 0.019 | 0.019 | 3.346 | 0.76 | 0.36 | 0.82 | 0.7 | 0.76 | 0.72 |
| 2 | 75669070 | rs13031576 | 0.485 | 1.359 | -3.034 | 0.437 | 0.014 | 0.197 | 0.197 | 2.08 | 0.33 | 0.02 | 0.16 | 0.06 | 0.29 | 0.19 |
| 2 | 75672583 | rs11903584 | 0.729 | 1.231 | -1.762 | 0.344 | -0.02 | 0.003 | 0.003 | 0.886 | 0.18 | 0.28 | 0.09 | 0.18 | 0.09 | 0.25 |
| 2 | 75672647 | rs7581555 | 1.119 | 1.382 | -1.641 | 0.526 | -0.031 | -0.06 | -0.06 | 0 | 0.26 | 0.04 | 0.36 | 0.34 | 0.27 | 0.24 |
| 2 | 75673718 | rs12611634 | -0.49 | 0.991 | -1.396 | 0.427 | 0.037 | 0.039 | 0.039 | 0 | 0.66 | 0.56 | 0.54 | 0.52 | 0.57 | 0.73 |
| 2 | 75675860 | rs10201616 | 0.023 | 0.847 | -0.596 | 0.691 | 0.024 | 0.094 | 0.094 | 0 | 0.15 | 0.33 | 0.3 | 0.28 | 0.17 | 0.27 |
| 2 | 75680801 | rs6714189 | -0.326 | 0.078 | -0.883 | 0.21 | -0.031 | -0.034 | -0.034 | 0.038 | 0.32 | 0.38 | 0.37 | 0.16 | 0.21 | 0.35 |
| 2 | 75681330 | rs10208759 | -0.887 | 0.253 | -1.298 | 0.725 | 0.074 | -0.021 | -0.021 | 0 | 0.43 | 0.85 | 0.42 | 0.55 | 0.3 | 0.56 |
| 2 | 75681462 | rs10496198 | -0.367 | 0.745 | -1.466 | 0.353 | 0 | -0.001 | -0.001 | 0 | 0.36 | 0.63 | 0.45 | 0.73 | 0.45 | 0.47 |
| 2 | 75683350 | rs12994030 | 1.46 | 1.055 | -1.889 | 0.577 | -0.083 | -0.093 | -0.093 | 0 | 0.17 | 0.02 | 0.25 | 0.31 | 0.24 | 0.18 |
| 2 | 75683911 | rs7564244 | 1.107 | 0.849 | -1.756 | 0.466 | 0.001 | -0.024 | -0.024 | 0.039 | 0.06 | 0.1 | 0.14 | 0 | 0.08 | 0.07 |
| 2 | 75693318 | rs6711412 | -1.492 | 0.719 | -1.869 | 0.453 | -0.053 | 0.132 | 0.132 | 2.418 | 0.71 | 0.94 | 0.84 | 0.9 | 0.71 | 0.83 |
| 2 | 75693541 | rs12612199 | 0.058 | 0.096 | -1.203 | 0.661 | -0.002 | 0.087 | 0.087 | 0 | 0.19 | 0.31 | 0.17 | 0.38 | 0.19 | 0.29 |
| 2 | 75698352 | rs4852379 | 0.212 | -0.097 | -0.857 | 0.433 | -0.069 | -0.045 | -0.045 | 0 | 0.27 | 0.46 | 0.27 | 0.16 | 0.15 | 0.28 |
| 2 | 75702671 | rs13019578 | -0.43 | 0.924 | -1.211 | 0.833 | -0.017 | -0.018 | -0.018 | 0 | 0.29 | 0.27 | 0.5 | 0.38 | 0.41 | 0.33 |
| 2 | 75705619 | rs2539992 | 0.03 | 1.056 | -1.544 | 0.918 | -0.046 | -0.068 | -0.068 | 0 | 0.22 | 0.08 | 0.36 | 0.38 | 0.32 | 0.31 |
| 2 | 75707925 | rs6739249 | -0.61 | 0.474 | -1.721 | 0.183 | -0.036 | 0.071 | 0.071 | 0.469 | 0.61 | 0.84 | 0.7 | 0.78 | 0.58 | 0.73 |
| 2 | 75708979 | rs2588499 | 0.348 | 0.6 | -1.887 | 0.113 | -0.068 | 0.003 | 0.003 | 0.657 | 0.15 | 0.02 | 0.2 | 0.09 | 0.2 | 0.19 |
| 2 | 75709160 | rs2588498 | -0.359 | 0.063 | -1.902 | -0.222 | 0.011 | 0.076 | 0.076 | 0 | 0.23 | 0.07 | 0.16 | 0.08 | 0.19 | 0.23 |
| 2 | 75710745 | rs1158392 | -0.433 | -0.3 | -0.967 | 0.366 | -0.014 | 0.125 | 0.125 | 0 | 0.18 | 0.37 | 0.27 | 0.51 | 0.14 | 0.25 |
| 2 | 75710803 | rs6723042 | 0.012 | -0.585 | -0.643 | -0.077 | -0.011 | -0.062 | -0.062 | 0 | 0.19 | 0.24 | 0.11 | 0.03 | 0.16 | 0.15 |
| 2 | 75711650 | rs11686817 | 0.218 | -0.577 | -0.496 | -0.021 | 0.001 | 0.023 | 0.023 | 0 | 0.14 | 0.14 | 0.25 | 0.51 | 0.13 | 0.25 |
| 2 | 75714591 | rs6717602 | -1.418 | -0.849 | -0.48 | -0.349 | 0.012 | -0.004 | -0.004 | 0 | 0.46 | 0.56 | 0.41 | 0.54 | 0.34 | 0.46 |
| 2 | 75714764 | rs11126471 | -1.342 | -0.81 | -0.471 | -0.33 | -0.011 | 0.005 | 0.005 | 0 | 0.44 | 0.46 | 0.4 | 0.54 | 0.34 | 0.45 |
| 2 | 75720372 | rs11126472 | 0.313 | -0.683 | -0.271 | -0.508 | -0.006 | -0.027 | -0.027 | 0 | 0.09 | 0.08 | 0.04 | 0 | 0.05 | 0.04 |
| 2 | 75720664 | rs6741778 | -0.959 | -0.763 | -0.08 | -0.614 | -0.046 | -0.158 | -0.158 | 0 | 0.32 | 0.46 | 0.16 | 0.03 | 0.21 | 0.2 |
| 2 | 75728903 | rs2016839 | -0.39 | 0.125 | 0.069 | -0.026 | 0.02 | 0.033 | 0.033 | 0 | 0.25 | 0.32 | 0.38 | 0.38 | 0.33 | 0.32 |
| 2 | 75734347 | rs7564143 | -0.463 | 0.197 | 0.492 | 0.011 | 0.032 | 0.047 | 0.047 | 0 | 0.26 | 0.4 | 0.38 | 0.41 | 0.34 | 0.32 |
| 2 | 75737551 | rs2160102 | -1.149 | -0.198 | 0.959 | -0.192 | -0.003 | -0.009 | -0.009 | 2.815 | 0.33 | 0.72 | 0.4 | 0.41 | 0.34 | 0.34 |
| 2 | 75738954 | rs6714783 | 0.37 | -0.161 | 1.068 | -0.194 | 0.013 | -0.044 | -0.044 | 0.121 | 0.15 | 0.32 | 0.1 | 0.1 | 0.15 | 0.13 |
| 2 | 75744918 | rs2286995 | -1.828 | -0.745 | 0.516 | -0.597 | 0.015 | 0.058 | 0.058 | 0.633 | 0.51 | 0.8 | 0.47 | 0.39 | 0.44 | 0.41 |
| 2 | 75745081 | rs2286994 | -0.951 | -0.563 | 0.977 | -0.417 | 0.002 | 0.005 | 0.005 | 0.278 | 0.31 | 0.61 | 0.39 | 0.39 | 0.34 | 0.34 |
| 2 | 75750667 | rs17011425 | -0.281 | -0.454 | 0.911 | -0.268 | 0.04 | 0.129 | 0.129 | 0.094 | 0.16 | 0.43 | 0.3 | 0.3 | 0.19 | 0.2 |
| 2 | 75752431 | rs6720653 | 0.736 | -0.378 | 0.855 | -0.23 | 0.032 | 0.087 | 0.087 | 0 | 0.12 | 0.2 | 0.29 | 0.3 | 0.19 | 0.2 |
| 2 | 75764175 | rs6754778 | 0.341 | -0.35 | 0.618 | -0.164 | 0.044 | 0.111 | 0.111 | 0 | 0.13 | 0.3 | 0.29 | 0.3 | 0.19 | 0.2 |
| 2 | 75767709 | rs2588495 | 0.718 | -0.483 | -0.14 | -0.222 | -0.015 | -0.026 | -0.026 | 0 | 0.59 | 0.31 | 0.55 | 0.47 | 0.53 | 0.57 |
| 2 | 75768077 | rs6713572 | -1.627 | -0.586 | -0.487 | -0.439 | 0.003 | 0.048 | 0.048 | 1.419 | 0.49 | 0.82 | 0.46 | 0.42 | 0.45 | 0.38 |
| 2 | 75768098 | rs10489977 | 0.529 | -0.235 | -0.357 | -0.123 | 0.045 | 0.106 | 0.106 | 0 | 0.14 | 0.32 | 0.3 | 0.32 | 0.21 | 0.2 |

| | | | | | | | | | | | | | | | | |
|---|---|---|---|---|---|---|---|---|---|---|---|---|---|---|---|---|
| 2 | 75775195 | rs11899583 | 0.846 | -0.483 | -1.1 | -0.52 | 0.015 | 0.076 | 0.076 | 17.442 | 0.14 | 0.02 | 0.05 | 0 | 0.08 | 0.04 |
| 2 | 75775779 | rs13036115 | 0.902 | -0.111 | -0.6 | -0.266 | 0.021 | -0.017 | -0.017 | 0 | 0.12 | 0.17 | 0.09 | 0.1 | 0.15 | 0.11 |
| 2 | 75778286 | rs6732989 | -0.907 | -0.611 | -1.539 | -0.071 | 0.014 | -0.019 | -0.019 | 0 | 0.77 | 0.99 | 0.82 | 0.82 | 0.83 | 0.84 |
| 2 | 75779654 | rs12053281 | 0.772 | -0.292 | -1.58 | 0.025 | -0.092 | 0.14 | 0.14 | 0 | 0.39 | 0.08 | 0.43 | 0.14 | 0.44 | 0.4 |
| 2 | 75788396 | rs759255 | 1.327 | -0.043 | -1.014 | 0.223 | 0.011 | -0.007 | -0.007 | 5.392 | 0.11 | 0.2 | 0.2 | 0.11 | 0.14 | 0.09 |
| 2 | 75788703 | rs1473167 | -0.065 | -0.302 | -0.8 | 0.4 | 0.002 | 0.008 | 0.008 | 0.21 | 0.5 | 0.5 | 0.43 | 0.43 | 0.49 | 0.52 |
| 2 | 75790665 | rs7589605 | -2.256 | -0.467 | -1.211 | 0.453 | 0.008 | 0.01 | 0.01 | 0 | 0.85 | 0.92 | 0.93 | 0.96 | 0.87 | 0.9 |
| 2 | 75792143 | rs17011478 | 2.117 | -0.426 | -1.235 | 0.758 | 0.019 | -0.02 | -0.02 | 0.768 | 0.07 | 0.01 | 0.12 | 0.11 | 0.03 | 0.05 |
| 2 | 75792267 | rs17739099 | 1.521 | -0.282 | -1.21 | 0.77 | -0.037 | -0.051 | -0.051 | 3.619 | 0.14 | 0.06 | 0.17 | 0.31 | 0.22 | 0.22 |
| 2 | 75796950 | rs17677380 | 1.191 | -0.374 | -1.233 | 0.3 | -0.049 | 0.041 | 0.041 | 0.084 | 0.3 | 0.07 | 0.35 | 0.21 | 0.32 | 0.33 |
| 2 | 75797440 | rs7595746 | -2.298 | -0.316 | -0.913 | 0.264 | 0.007 | 0.008 | 0.008 | 0 | 0.85 | 0.92 | 0.94 | 0.96 | 0.87 | 0.91 |
| 2 | 75801693 | rs17011498 | 0.713 | -0.394 | -0.688 | 0.31 | -0.029 | -0.002 | -0.002 | 0 | 0.37 | 0.09 | 0.48 | 0.32 | 0.35 | 0.38 |
| 2 | 75802151 | rs6709627 | 1.766 | -0.131 | -0.272 | 0.214 | 0.001 | -0.002 | -0.002 | 0 | 0.11 | 0.12 | 0.2 | 0.1 | 0.14 | 0.08 |
| 2 | 75803049 | rs10048809 | 1.286 | -0.228 | -0.198 | 0.292 | 0.018 | 0.002 | 0.002 | 0 | 0.11 | 0.19 | 0.05 | 0.15 | 0.14 | 0.13 |
| 2 | 75804641 | rs11675369 | 0.894 | -0.243 | -0.144 | 0.285 | 0.029 | 0.004 | 0.004 | 0 | 0.12 | 0.28 | 0.06 | 0.15 | 0.14 | 0.13 |
| 2 | 75808132 | rs1075054 | 1.275 | -0.184 | -0.448 | 0.315 | 0.018 | 0.003 | 0.003 | 0 | 0.1 | 0.19 | 0.06 | 0.15 | 0.14 | 0.13 |
| 2 | 75810755 | rs2110487 | -1.918 | -0.318 | -0.843 | 0.332 | 0.011 | 0.012 | 0.012 | 0 | 0.85 | 0.93 | 0.94 | 0.95 | 0.87 | 0.91 |
| 2 | 75814908 | rs730048 | 0.249 | -0.036 | -0.865 | 0.403 | 0 | -0.005 | -0.005 | 0 | 0.36 | 0.25 | 0.51 | 0.35 | 0.37 | 0.4 |
| 2 | 75815249 | rs6747987 | -0.801 | -0.1 | -0.627 | 0.149 | 0.005 | 0.012 | 0.012 | 0.009 | 0.71 | 0.8 | 0.89 | 0.82 | 0.73 | 0.76 |
| 2 | 75820691 | rs10203769 | 0.794 | -0.09 | -0.897 | 0.163 | 0 | 0 | 0 | 0 | 0.19 | 0.19 | 0.29 | 0.19 | 0.14 | 0.13 |
| 2 | 75823162 | rs1000585 | -0.041 | 0.019 | -0.624 | 0.245 | 0.011 | 0.015 | 0.015 | 0 | 0.51 | 0.47 | 0.6 | 0.62 | 0.57 | 0.62 |
| 2 | 75823747 | rs7574273 | 0.681 | 0.244 | -0.883 | 0.338 | -0.041 | -0.065 | -0.065 | 0 | 0.31 | 0.14 | 0.38 | 0.46 | 0.37 | 0.35 |
| 2 | 75825819 | rs917235 | -0.236 | 0.245 | -0.826 | 0.06 | 0 | 0.013 | 0.013 | 0 | 0.49 | 0.61 | 0.6 | 0.62 | 0.52 | 0.64 |
| 2 | 75829459 | rs11675652 | -0.393 | 0.229 | -0.752 | 0.153 | -0.002 | -0.004 | -0.004 | 0 | 0.43 | 0.51 | 0.47 | 0.37 | 0.39 | 0.44 |
| 2 | 75830221 | rs11677196 | 0.508 | 0.056 | -0.912 | 0.054 | -0.017 | 0.021 | 0.021 | 0 | 0.3 | 0.2 | 0.3 | 0.19 | 0.35 | 0.34 |
| 2 | 75832122 | rs10178367 | -1.49 | 0.093 | -1.479 | 0.028 | -0.037 | 0.048 | 0.048 | 1.902 | 0.73 | 0.95 | 0.86 | 0.88 | 0.73 | 0.84 |
| 2 | 75834937 | rs714938 | 2.171 | 0.115 | -1.473 | 0.042 | -0.018 | 0.006 | 0.006 | 0 | 0.1 | 0.02 | 0.05 | 0.07 | 0.09 | 0.04 |
| 2 | 75837011 | rs740270 | 2.079 | 0.121 | -1.165 | 0.088 | 0 | -0.007 | -0.007 | 0 | 0.12 | 0.1 | 0.19 | 0.11 | 0.15 | 0.15 |
| 2 | 75839376 | rs17011534 | 0.944 | 0.116 | -0.786 | 0.314 | 0.01 | 0.116 | 0.116 | 0 | 0.18 | 0.37 | 0.21 | 0.4 | 0.19 | 0.24 |
| 2 | 75839733 | rs6732511 | -2.209 | 0.207 | -1.341 | 0.163 | -0.025 | 0.029 | 0.029 | 0.459 | 0.83 | 0.98 | 0.92 | 0.95 | 0.83 | 0.88 |
| 2 | 75844627 | rs875756 | 0.452 | 0.255 | -0.92 | 0.301 | -0.004 | 0.006 | 0.006 | 0 | 0.15 | 0.16 | 0.21 | 0.22 | 0.11 | 0.11 |
| 2 | 75844679 | rs875755 | 0.935 | 0.136 | -1.336 | -0.008 | 0 | 0.052 | 0.052 | 0.454 | 0.21 | 0.07 | 0.22 | 0.12 | 0.19 | 0.22 |
| 2 | 75846545 | rs10166389 | -1.436 | 0.103 | -1.21 | 0.062 | -0.01 | -0.057 | -0.057 | 0.1 | 0.68 | 0.78 | 0.73 | 0.55 | 0.67 | 0.72 |
| 2 | 75846558 | rs7588344 | -1.578 | 0.095 | -1.211 | 0.057 | -0.013 | -0.055 | -0.055 | 0.273 | 0.69 | 0.8 | 0.73 | 0.55 | 0.67 | 0.72 |
| 2 | 75847704 | rs714479 | -0.827 | 0.068 | -1.028 | 0.011 | -0.001 | 0.005 | 0.005 | 1.235 | 0.57 | 0.65 | 0.67 | 0.46 | 0.56 | 0.67 |
| 2 | 75848679 | rs2422229 | -0.948 | 0.081 | -1.036 | -0.019 | -0.009 | -0.013 | -0.013 | 0.009 | 0.58 | 0.73 | 0.68 | 0.48 | 0.56 | 0.68 |
| 2 | 75849673 | rs11690601 | -0.383 | 0.368 | -0.379 | 0.187 | 0.002 | 0 | 0 | 0 | 0.35 | 0.55 | 0.45 | 0.35 | 0.36 | 0.43 |
| 2 | 75853788 | rs6547014 | -0.916 | 0.392 | -0.343 | 0.21 | -0.005 | 0.001 | 0.001 | 0 | 0.45 | 0.57 | 0.5 | 0.43 | 0.48 | 0.5 |
| 2 | 75859919 | rs7577050 | 0.381 | 0.379 | -0.126 | 0.21 | 0.004 | -0.002 | -0.002 | 0.693 | 0.27 | 0.31 | 0.29 | 0.21 | 0.36 | 0.35 |
| 2 | 75862200 | rs13397241 | -1.148 | 0.3 | -0.254 | 0.148 | -0.019 | -0.002 | -0.002 | 0 | 0.47 | 0.79 | 0.53 | 0.43 | 0.48 | 0.5 |
| 2 | 75862932 | rs1859710 | 1.478 | 0.616 | -0.065 | 0.498 | -0.068 | -0.045 | -0.045 | 0.488 | 0.16 | 0.03 | 0.22 | 0.14 | 0.26 | 0.3 |
| 2 | 75865277 | rs1017051 | -0.248 | 0.482 | -0.185 | 0.607 | -0.008 | -0.005 | -0.005 | 0 | 0.46 | 0.39 | 0.49 | 0.51 | 0.5 | 0.56 |
| 2 | 75866236 | rs7570229 | -0.134 | 0.49 | -0.232 | 0.611 | -0.015 | -0.011 | -0.011 | 0 | 0.46 | 0.3 | 0.48 | 0.51 | 0.5 | 0.56 |
| 2 | 75869828 | rs12105510 | 0.677 | 0.45 | -0.329 | 0.697 | 0.014 | -0.008 | -0.008 | 0.006 | 0.28 | 0.18 | 0.24 | 0.35 | 0.23 | 0.22 |
| 2 | 75872246 | rs1001893 | 0.457 | 0.427 | -0.041 | 0.707 | 0.028 | 0.069 | 0.069 | 0 | 0.08 | 0.16 | 0.19 | 0.18 | 0.08 | 0.11 |
| 2 | 75873217 | rs2422230 | -0.389 | 0.361 | -0.002 | 0.707 | 0.001 | 0.004 | 0.004 | 0 | 0.51 | 0.43 | 0.67 | 0.69 | 0.56 | 0.63 |
| 2 | 75877562 | rs3771872 | 1.944 | 0.502 | -0.247 | 0.68 | -0.011 | 0.004 | 0.004 | 0.197 | 0.09 | 0.06 | 0.05 | 0.08 | 0.11 | 0.02 |
| 2 | 75877650 | rs1990145 | 0.317 | 0.479 | -0.362 | 0.722 | 0.005 | -0.014 | -0.014 | 0 | 0.34 | 0.18 | 0.3 | 0.42 | 0.29 | 0.26 |
| 2 | 75877812 | rs3771875 | -1.316 | 0.456 | 0.087 | 0.671 | -0.006 | -0.027 | -0.027 | 0.01 | 0.64 | 0.61 | 0.72 | 0.81 | 0.68 | 0.71 |
| 2 | 75882399 | rs17689863 | 1.816 | 0.48 | -0.171 | 0.721 | -0.067 | -0.068 | -0.068 | 0 | 0.15 | 0.02 | 0.23 | 0.16 | 0.24 | 0.3 |
| 2 | 75882855 | rs3732304 | 1.816 | 0.48 | -0.171 | 0.721 | -0.067 | -0.068 | -0.068 | 0 | 0.15 | 0.02 | 0.23 | 0.16 | 0.24 | 0.3 |
| 2 | 75883276 | rs10165899 | 1.559 | 0.475 | -0.114 | 0.719 | -0.022 | -0.024 | -0.024 | 0.032 | 0.16 | 0.1 | 0.23 | 0.16 | 0.24 | 0.3 |
| 2 | 75883390 | rs917236 | -0.625 | 0.434 | -0.107 | 0.702 | -0.015 | -0.037 | -0.037 | 0 | 0.33 | 0.78 | 0.46 | 0.38 | 0.39 | 0.45 |
| 2 | 75883878 | rs13397865 | -0.427 | 0.433 | -0.082 | 0.701 | -0.005 | -0.03 | -0.03 | 0.517 | 0.32 | 0.75 | 0.46 | 0.38 | 0.37 | 0.42 |
| 2 | 75887579 | rs7577196 | -0.608 | 0.441 | -0.17 | 0.711 | -0.016 | -0.035 | -0.035 | 0 | 0.33 | 0.78 | 0.46 | 0.38 | 0.39 | 0.45 |
| 2 | 75887834 | rs13426287 | -0.608 | 0.441 | -0.17 | 0.711 | -0.016 | -0.037 | -0.037 | 0 | 0.33 | 0.79 | 0.46 | 0.38 | 0.39 | 0.45 |
| 2 | 75888160 | rs6547016 | -0.643 | 0.423 | -0.179 | 0.703 | -0.015 | -0.036 | -0.036 | 0 | 0.34 | 0.78 | 0.46 | 0.38 | 0.39 | 0.45 |
| 2 | 75899732 | rs13385913 | 0.396 | 0.411 | -0.06 | 0.711 | 0.01 | 0.077 | 0.077 | 1.463 | 0.14 | 0.51 | 0.21 | 0.22 | 0.13 | 0.12 |
| 2 | 75900915 | rs718967 | -0.144 | 0.398 | -0.156 | 0.704 | -0.012 | 0.01 | 0.01 | 3.766 | 0.17 | 0.71 | 0.22 | 0.22 | 0.13 | 0.12 |
| 2 | 75907351 | rs6722682 | 0.759 | 0.401 | -0.071 | 0.701 | 0.052 | -0.014 | -0.014 | 0 | 0.13 | 0.53 | 0.08 | 0.08 | 0.13 | 0.06 |
| 2 | 75915035 | rs1803196 | -0.567 | 0.437 | -0.263 | 0.72 | -0.013 | -0.035 | -0.035 | 0 | 0.34 | 0.79 | 0.46 | 0.38 | 0.39 | 0.43 |
| 2 | 75923413 | rs7560262 | -0.86 | 0.434 | -0.546 | 0.607 | -0.05 | -0.036 | -0.036 | 3.396 | 0.4 | 0.8 | 0.52 | 0.41 | 0.47 | 0.47 |
| 2 | 75926565 | rs2298948 | 0.783 | 0.522 | -0.976 | 0.501 | -0.037 | -0.056 | -0.056 | 0 | 0.32 | 0.07 | 0.3 | 0.39 | 0.3 | 0.28 |
| 2 | 75931600 | rs2298945 | -0.435 | 0.516 | -0.707 | 0.595 | -0.013 | -0.035 | -0.035 | 0 | 0.33 | 0.79 | 0.46 | 0.38 | 0.39 | 0.42 |
| 2 | 75938251 | rs10179871 | 0.607 | 0.614 | -0.823 | 0.652 | 0.029 | 0.064 | 0.064 | 1.334 | 0.14 | 0.38 | 0.19 | 0.22 | 0.15 | 0.15 |
| 2 | 75940599 | rs3755476 | -2.249 | 0.474 | -1.091 | 0.516 | -0.035 | 0.044 | 0.044 | 0.124 | 0.78 | 0.99 | 0.89 | 0.92 | 0.79 | 0.92 |
| 2 | 75940864 | rs3755477 | -0.061 | 0.392 | -0.378 | 1.031 | 0.005 | 0.01 | 0.01 | 0 | 0.29 | 0.34 | 0.43 | 0.38 | 0.39 | 0.43 |
| 2 | 75945344 | rs6718646 | -0.419 | 0.408 | -0.224 | 0.887 | -0.002 | 0.025 | 0.025 | 0 | 0.52 | 0.45 | 0.64 | 0.78 | 0.53 | 0.69 |
| 2 | 75953851 | rs2158815 | -1.218 | 0.596 | 0.463 | 1.871 | -0.019 | -0.038 | -0.038 | 0 | 0.41 | 0.88 | 0.49 | 0.66 | 0.36 | 0.37 |

| | | | | | | | | | | | | | | | | |
|---|---|---|---|---|---|---|---|---|---|---|---|---|---|---|---|---|
| 2 | 75954038 | rs1004391 | 0.143 | 0.611 | 0.897 | 1.624 | -0.004 | 0.006 | 0.006 | 0.462 | 0.24 | 0.34 | 0.21 | 0.36 | 0.17 | 0.17 |
| 2 | 75964293 | rs10166482 | -1.581 | 0.668 | 0.838 | 1.575 | -0.005 | 0.06 | 0.06 | 0.186 | 0.45 | 0.72 | 0.5 | 0.81 | 0.4 | 0.52 |
| 2 | 75966134 | rs7607937 | -0.492 | 0.626 | 1.063 | 1.725 | 0.012 | 0.043 | 0.043 | 0 | 0.29 | 0.51 | 0.42 | 0.64 | 0.26 | 0.44 |
| 2 | 75972793 | rs2422237 | -2.363 | 0.723 | 1.103 | 1.754 | 0.019 | 0.052 | 0.052 | 0.466 | 0.17 | 0.61 | 0.29 | 0.33 | 0.17 | 0.33 |
| 2 | 75977535 | rs11675641 | -1.337 | 0.69 | 1.153 | 1.761 | 0.015 | 0.121 | 0.121 | 0 | 0.12 | 0.23 | 0.23 | 0.31 | 0.1 | 0.28 |
| 2 | 75979275 | rs1859707 | -1.58 | 0.636 | 0.911 | 1.709 | -0.008 | 0.049 | 0.049 | 0 | 0.44 | 0.7 | 0.52 | 0.79 | 0.41 | 0.54 |
| 2 | 75991045 | rs7419897 | -2.356 | 0.798 | 1.086 | 1.846 | 0.015 | 0.043 | 0.043 | 0.875 | 0.17 | 0.6 | 0.3 | 0.33 | 0.17 | 0.33 |
| 2 | 75991584 | rs17011730 | 1.396 | 0.828 | 1.005 | 1.896 | -0.012 | -0.025 | -0.025 | 0 | 0.07 | 0.02 | 0.04 | 0.11 | 0.1 | 0.03 |
| 2 | 75991794 | rs11126485 | -1.889 | 0.803 | 1.231 | 1.936 | 0.05 | 0.175 | 0.175 | 0 | 0.13 | 0.35 | 0.26 | 0.33 | 0.13 | 0.32 |
| 2 | 75993443 | rs9789774 | 0.819 | 0.827 | 1.05 | 1.642 | -0.047 | -0.074 | -0.074 | 0 | 0.18 | 0.03 | 0.2 | 0.35 | 0.17 | 0.18 |
| 2 | 75995540 | rs13383643 | -1.124 | 0.88 | 1.444 | 1.167 | 0.136 | 0.002 | 0.002 | 0.28 | 0.1 | 0.38 | 0.09 | 0.03 | 0.11 | 0.02 |
| 2 | 76010976 | rs2190355 | -0.502 | 0.352 | 0.846 | 1.679 | 0.005 | 0.332 | 0.332 | 0 | 0.05 | 0.14 | 0.24 | 0.47 | 0.04 | 0.26 |
| 2 | 76035099 | rs4361150 | 0.381 | 0.406 | 0.138 | 0.895 | -0.02 | -0.002 | -0.002 | 0.533 | 0.22 | 0.09 | 0.21 | 0.27 | 0.2 | 0.23 |
| 2 | 76041848 | rs7573336 | -0.338 | 0.269 | 0.426 | 0.842 | 0 | -0.048 | -0.048 | 0 | 0.29 | 0.43 | 0.25 | 0.25 | 0.23 | 0.2 |
| 2 | 76057275 | rs12616488 | -0.07 | 0.245 | 0.224 | 0.871 | -0.015 | 0.009 | 0.009 | 0 | 0.27 | 0.19 | 0.24 | 0.29 | 0.25 | 0.25 |
| 2 | 76068710 | rs6710469 | -0.513 | 0.199 | 0.352 | 0.774 | 0 | -0.032 | -0.032 | 0 | 0.32 | 0.44 | 0.27 | 0.28 | 0.26 | 0.24 |
| 2 | 76069618 | rs11682079 | -1.498 | -0.051 | 0.014 | 0.646 | -0.04 | 0.315 | 0.315 | 0 | 0.51 | 0.92 | 0.57 | 0.89 | 0.4 | 0.56 |
| 2 | 76079609 | rs4452179 | 0.063 | -0.062 | 0.222 | 0.842 | -0.015 | 0.224 | 0.224 | 0.243 | 0.08 | 0.18 | 0.24 | 0.52 | 0.03 | 0.26 |
| 2 | 76080701 | rs4456732 | -0.428 | -0.008 | -0.237 | 0.416 | -0.002 | -0.032 | -0.032 | 0 | 0.33 | 0.52 | 0.27 | 0.28 | 0.26 | 0.25 |
| 2 | 76081681 | rs3935887 | 0.284 | -0.004 | -0.559 | 0.365 | -0.03 | 0.034 | 0.034 | 0.097 | 0.27 | 0.08 | 0.22 | 0.25 | 0.24 | 0.2 |
| 2 | 76088361 | rs11896839 | -1.246 | 0.16 | -0.446 | 0.143 | 0.04 | 0.036 | 0.036 | 0.122 | 0.08 | 0.38 | 0.05 | 0.03 | 0.03 | 0.06 |
| 2 | 76088880 | rs10194197 | -0.742 | 0.225 | -0.624 | 0.176 | 0.053 | 0.049 | 0.049 | 0 | 0.07 | 0.3 | 0.04 | 0.03 | 0.03 | 0.06 |
| 2 | 76091644 | rs6547035 | -0.833 | 0.296 | -1.738 | 0.267 | -0.027 | 0.236 | 0.236 | 0 | 0.5 | 0.91 | 0.57 | 0.83 | 0.4 | 0.56 |
| 2 | 76109228 | rs6547036 | -0.47 | 0.231 | -1.52 | 0.371 | 0.022 | 0.038 | 0.038 | 0.084 | 0.61 | 0.79 | 0.61 | 0.74 | 0.67 | 0.55 |
| 2 | 76109411 | rs6547037 | -0.13 | 0.218 | -1.456 | 0.296 | 0.024 | 0.052 | 0.052 | 0 | 0.55 | 0.71 | 0.59 | 0.71 | 0.62 | 0.5 |
| 2 | 76112863 | rs12612334 | 0.314 | 0.327 | -0.508 | 0.542 | -0.008 | 0.03 | 0.03 | 0 | 0.45 | 0.37 | 0.42 | 0.26 | 0.49 | 0.33 |
| 2 | 76120604 | rs6751526 | 0.494 | 0.352 | -0.495 | 0.811 | 0.062 | 0.373 | 0.373 | 0 | 0.1 | 0.34 | 0.16 | 0.51 | 0.14 | 0.17 |
| 2 | 76121914 | rs10204120 | -1.169 | 0.177 | -0.339 | 0.719 | -0.006 | 0.104 | 0.104 | 0 | 0.2 | 0.65 | 0.22 | 0.47 | 0.19 | 0.22 |
| 2 | 76124770 | rs17011907 | 0.503 | 0.244 | -0.037 | 0.781 | 0.042 | 0.305 | 0.305 | 0 | 0.1 | 0.25 | 0.16 | 0.45 | 0.14 | 0.17 |
| 2 | 76132622 | rs6706336 | -1.163 | 0.178 | -0.428 | 0.492 | -0.015 | 0.051 | 0.051 | 0.105 | 0.65 | 0.92 | 0.63 | 0.79 | 0.67 | 0.55 |
| 2 | 76139406 | rs4852387 | -1.148 | 0.217 | -0.43 | 0.667 | -0.018 | 0.014 | 0.014 | 0.112 | 0.65 | 0.92 | 0.63 | 0.75 | 0.66 | 0.53 |
| 2 | 76153088 | rs11888390 | -0.515 | 0.031 | 0.026 | 0.353 | 0.004 | -0.034 | -0.034 | 0 | 0.75 | 0.68 | 0.72 | 0.92 | 0.72 | 0.82 |
| 2 | 76153643 | rs7558098 | -1.267 | 0.035 | -0.238 | 0.35 | -0.04 | 0.075 | 0.075 | 0 | 0.8 | 0.94 | 0.77 | 0.95 | 0.76 | 0.86 |
| 2 | 76160203 | rs2422306 | 0.213 | -0.077 | -0.201 | 0.318 | -0.023 | -0.019 | -0.019 | 0.085 | 0.63 | 0.23 | 0.49 | 0.42 | 0.58 | 0.65 |
| 2 | 76161574 | rs2422307 | 0.258 | -0.219 | -0.226 | -0.174 | 0.042 | 0.041 | 0.041 | 0 | 0.33 | 0.17 | 0.23 | 0.16 | 0.24 | 0.36 |
| 2 | 76163817 | rs7588316 | -1.253 | 0.482 | 0.208 | -0.105 | 0.046 | 0.021 | 0.021 | 0 | 0.08 | 0.46 | 0.06 | 0.02 | 0.04 | 0.05 |
| 2 | 76187099 | rs1807143 | 0.452 | 0.799 | -0.429 | -0.104 | 0.001 | -0.001 | -0.001 | 0 | 0.22 | 0.25 | 0.34 | 0.25 | 0.24 | 0.17 |
| 2 | 76189402 | rs7567436 | -1.023 | 1.034 | 0.5 | 0.478 | 0.05 | 0.426 | 0.426 | 0.168 | 0.08 | 0.35 | 0.29 | 0.46 | 0.11 | 0.14 |
| 2 | 76193226 | rs1372303 | 0.125 | 1.196 | 0.564 | 0.198 | 0.023 | -0.046 | -0.046 | 0 | 0.08 | 0.23 | 0.06 | 0.01 | 0.08 | 0.11 |
| 2 | 76206795 | rs4853186 | -0.162 | 1.09 | 0.311 | -0.036 | 0.001 | -0.045 | -0.045 | 0 | 0.18 | 0.23 | 0.1 | 0.03 | 0.17 | 0.21 |
| 2 | 76209834 | rs4853187 | -0.797 | 1.068 | 0.397 | 0.015 | 0.013 | 0.07 | 0.07 | 0 | 0.23 | 0.33 | 0.36 | 0.52 | 0.27 | 0.35 |
| 2 | 76213310 | rs2583537 | -0.966 | 0.892 | -0.078 | -0.49 | 0.009 | -0.101 | -0.101 | 0 | 0.74 | 0.65 | 0.85 | 0.98 | 0.69 | 0.89 |
| 2 | 76226375 | rs6708139 | -0.385 | 0.758 | -0.109 | -0.398 | 0.006 | 0.044 | 0.044 | 0 | 0.22 | 0.28 | 0.36 | 0.51 | 0.27 | 0.36 |
| 2 | 76232882 | rs317229 | -0.49 | 0.873 | -0.341 | -0.382 | 0.005 | 0.036 | 0.036 | 0 | 0.72 | 0.64 | 0.61 | 0.49 | 0.64 | 0.75 |
| 2 | 76236858 | rs17012004 | -0.719 | 0.713 | -0.317 | -0.043 | 0.01 | 0.071 | 0.071 | 0 | 0.23 | 0.37 | 0.37 | 0.52 | 0.27 | 0.36 |
| 2 | 76240361 | rs436000 | -0.096 | 0.57 | -0.262 | -0.388 | 0.002 | 0.01 | 0.01 | 0 | 0.44 | 0.39 | 0.29 | 0.24 | 0.37 | 0.47 |
| 2 | 76242326 | rs3843860 | 0.164 | 0.52 | -0.418 | -0.387 | -0.005 | 0.029 | 0.029 | 0 | 0.33 | 0.57 | 0.45 | 0.54 | 0.35 | 0.23 |
| 2 | 76243116 | rs3843862 | -1.036 | 0.72 | -0.562 | -0.514 | -0.065 | 0.04 | 0.04 | 0.402 | 0.69 | 0.89 | 0.7 | 0.77 | 0.62 | 0.65 |
| 2 | 76244948 | rs427171 | 0.183 | 0.713 | -0.316 | -0.565 | -0.05 | 0.025 | 0.025 | 0.05 | 0.37 | 0.18 | 0.5 | 0.7 | 0.45 | 0.43 |
| 2 | 76250543 | rs12614261 | 0.083 | 0.645 | -0.171 | -0.459 | 0.003 | 0.173 | 0.173 | 0 | 0.08 | 0.14 | 0.28 | 0.5 | 0.12 | 0.12 |
| 2 | 76272837 | rs413828 | -0.775 | 0.554 | -0.163 | -0.613 | 0.001 | 0.002 | 0.002 | 0 | 0.64 | 0.67 | 0.76 | 0.76 | 0.69 | 0.56 |
| 2 | 76281624 | rs7560937 | 0.636 | 0.511 | -0.17 | -0.607 | 0.008 | 0.013 | 0.013 | 0 | 0.35 | 0.28 | 0.17 | 0.21 | 0.27 | 0.42 |
| 2 | 76287089 | rs2028420 | -0.186 | 0.525 | -0.246 | -0.519 | -0.001 | 0.127 | 0.127 | 0 | 0.09 | 0.13 | 0.28 | 0.5 | 0.12 | 0.13 |
| 2 | 76289108 | rs1440887 | 1.355 | 0.344 | -0.378 | -0.724 | 0.029 | 0.03 | 0.03 | 0 | 0.26 | 0.07 | 0.16 | 0.18 | 0.18 | 0.38 |
| 2 | 76302733 | rs451469 | 1.012 | 0.359 | -0.365 | -0.791 | -0.074 | 0.028 | 0.028 | 1.013 | 0.31 | 0.06 | 0.31 | 0.23 | 0.33 | 0.31 |
| 2 | 76303913 | rs404892 | 0.259 | 0.421 | -0.06 | -0.728 | -0.015 | 0.023 | 0.023 | 0.062 | 0.39 | 0.29 | 0.59 | 0.73 | 0.46 | 0.44 |
| 2 | 76312868 | rs6722106 | 1.606 | 0.42 | -0.066 | -0.704 | -0.004 | 0.006 | 0.006 | 0 | 0.08 | 0.05 | 0.05 | 0.05 | 0.09 | 0.05 |
| 2 | 76314728 | rs389616 | -0.353 | 0.308 | 0.146 | -0.608 | 0.029 | 0.056 | 0.056 | 0 | 0.58 | 0.27 | 0.72 | 0.76 | 0.68 | 0.55 |
| 2 | 76333150 | rs1524210 | 0.667 | 0.345 | 0.273 | -0.579 | 0.018 | 0.031 | 0.031 | 0 | 0.34 | 0.2 | 0.17 | 0.21 | 0.27 | 0.42 |
| 2 | 76335351 | rs10188424 | 1.124 | 0.355 | 0.28 | -0.613 | -0.009 | -0.13 | -0.13 | 0.759 | 0.25 | 0.45 | 0.17 | 0.03 | 0.23 | 0.1 |
| 2 | 76337346 | rs17012180 | 0.298 | 0.349 | 0.298 | -0.55 | 0 | 0.116 | 0.116 | 0 | 0.08 | 0.09 | 0.27 | 0.5 | 0.12 | 0.13 |
| 2 | 76339746 | rs6709581 | 0.695 | 0.296 | 0.218 | -0.582 | 0.007 | 0.011 | 0.011 | 0 | 0.35 | 0.3 | 0.17 | 0.22 | 0.28 | 0.42 |
| 2 | 76343190 | rs10496200 | -0.137 | 0.36 | 0.112 | -0.512 | 0.002 | 0.227 | 0.227 | 0 | 0.09 | 0.22 | 0.28 | 0.5 | 0.12 | 0.13 |
| 2 | 76346138 | rs6744439 | 0.92 | 0.443 | -0.46 | -0.536 | -0.047 | 0.034 | 0.034 | 0 | 0.31 | 0.14 | 0.31 | 0.22 | 0.34 | 0.33 |
| 2 | 76361115 | rs2422416 | 1.027 | 0.444 | -0.502 | -0.529 | -0.06 | 0.031 | 0.031 | 0 | 0.3 | 0.09 | 0.31 | 0.22 | 0.33 | 0.33 |
| 2 | 76372316 | rs1595968 | 1.027 | 0.444 | -0.502 | -0.529 | -0.06 | 0.031 | 0.031 | 0 | 0.3 | 0.1 | 0.31 | 0.22 | 0.33 | 0.33 |
| 2 | 76374665 | rs7559751 | 1.508 | 0.457 | -0.455 | -0.486 | -0.013 | -0.001 | -0.001 | 0.21 | 0.24 | 0.1 | 0.25 | 0.22 | 0.24 | 0.3 |
| 2 | 76375711 | rs10204325 | 0.222 | 0.499 | -0.322 | -0.456 | -0.002 | 0.059 | 0.059 | 0 | 0.42 | 0.26 | 0.22 | 0.21 | 0.37 | 0.46 |
| 2 | 76382249 | rs1996974 | -0.107 | -0.438 | 0.118 | -0.026 | 0.004 | 0.061 | 0.061 | 0 | 0.7 | 0.55 | 0.52 | 0.48 | 0.66 | 0.79 |

| | | | | | | | | | | | | | | | | |
|---|---|---|---|---|---|---|---|---|---|---|---|---|---|---|---|---|
| 2 | 76383589 | rs13027159 | -1.023 | -0.491 | 0.321 | -0.055 | -0.003 | -0.005 | -0.005 | 0 | 0.78 | 0.76 | 0.84 | 0.95 | 0.73 | 0.91 |
| 2 | 76385277 | rs1440882 | 0.109 | -0.474 | 0.375 | -0.056 | -0.003 | 0.073 | 0.073 | 0 | 0.27 | 0.41 | 0.46 | 0.5 | 0.3 | 0.18 |
| 2 | 76385773 | rs10179941 | -0.975 | -0.577 | 0.209 | -0.243 | -0.017 | 0.015 | 0.015 | 0 | 0.63 | 0.69 | 0.64 | 0.69 | 0.58 | 0.64 |
| 2 | 76386381 | rs12469625 | 1.566 | -0.579 | 0.175 | -0.236 | -0.085 | 0.01 | 0.01 | 0.047 | 0.29 | 0.05 | 0.31 | 0.23 | 0.34 | 0.31 |
| 2 | 76387187 | rs1440881 | -0.046 | -0.572 | 0.395 | -0.218 | 0.009 | 0.034 | 0.034 | 0 | 0.68 | 0.57 | 0.5 | 0.41 | 0.61 | 0.77 |
| 2 | 76389455 | rs12465096 | -1.168 | -0.521 | 0.002 | -0.249 | 0.015 | 0.012 | 0.012 | 0.526 | 0.06 | 0.25 | 0.04 | 0.06 | 0.05 | 0.02 |
| 2 | 76393883 | rs12622977 | 0.663 | -0.44 | -0.63 | -0.16 | -0.048 | 0.002 | 0.002 | 0.93 | 0.35 | 0.12 | 0.55 | 0.68 | 0.42 | 0.45 |
| 2 | 76404263 | rs11126503 | 0.427 | -0.64 | -0.525 | -0.078 | 0.02 | 0.105 | 0.105 | 0 | 0.51 | 0.23 | 0.55 | 0.74 | 0.56 | 0.55 |
| 2 | 76409872 | rs13421819 | 0.584 | -0.611 | 0.105 | 0.117 | -0.011 | 0.006 | 0.006 | 0 | 0.14 | 0.13 | 0.06 | 0.16 | 0.07 | 0.29 |
| 2 | 76419240 | rs12996043 | 1.268 | -0.758 | 0.043 | -0.06 | -0.011 | 0.014 | 0.014 | 0 | 0.14 | 0.1 | 0.13 | 0.03 | 0.21 | 0.04 |
| 2 | 76419466 | rs11687064 | 0.352 | -0.818 | 0.571 | -0.065 | -0.044 | 0.038 | 0.038 | 0 | 0.16 | 0.42 | 0.08 | 0.16 | 0.07 | 0.29 |
| 2 | 76420089 | rs6759778 | 0.086 | -0.778 | 0.326 | -0.016 | -0.003 | -0.021 | -0.021 | 0 | 0.68 | 0.42 | 0.78 | 0.8 | 0.67 | 0.65 |
| 2 | 76442464 | rs17012308 | 0.376 | -0.656 | 0.163 | 0.592 | 0 | 0.086 | 0.086 | 0 | 0.06 | 0.06 | 0.18 | 0.46 | 0.07 | 0.18 |
| 2 | 76448948 | rs9917265 | 0.577 | -0.976 | 0.631 | 0.383 | 0.002 | 0.007 | 0.007 | 0 | 0.61 | 0.34 | 0.4 | 0.26 | 0.61 | 0.46 |
| 2 | 76453762 | rs11891956 | -0.088 | -0.941 | 0.521 | 0.402 | -0.013 | -0.022 | -0.022 | 0.022 | 0.32 | 0.53 | 0.4 | 0.26 | 0.29 | 0.31 |
| 2 | 76464500 | rs1405343 | 1.143 | -1.012 | 0.288 | 0.293 | -0.015 | 0.164 | 0.164 | 0 | 0.51 | 0.15 | 0.34 | 0.24 | 0.51 | 0.42 |
| 2 | 76468390 | rs9973760 | 0.714 | -1.064 | 0.492 | 0.448 | 0.009 | -0.01 | -0.01 | 3.486 | 0.14 | 0.08 | 0.08 | 0.22 | 0.06 | 0.26 |
| 2 | 76472334 | rs11126510 | 0.518 | -1.049 | 0.477 | 0.64 | 0.011 | -0.015 | -0.015 | 0.853 | 0.14 | 0.08 | 0.08 | 0.28 | 0.06 | 0.3 |
| 2 | 76478835 | rs4852399 | 0.277 | -1.167 | 0.081 | 0.441 | 0.002 | 0 | 0 | 0 | 0.43 | 0.44 | 0.41 | 0.44 | 0.41 | 0.48 |
| 2 | 76482416 | rs12468296 | 0.777 | -1.057 | -0.095 | 0.547 | -0.007 | 0.001 | 0.001 | 0 | 0.37 | 0.32 | 0.37 | 0.39 | 0.39 | 0.41 |
| 2 | 76486806 | rs10865419 | 0.683 | -1.319 | -0.121 | 0.63 | 0.014 | -0.016 | -0.016 | 0 | 0.16 | 0.16 | 0.2 | 0.39 | 0.07 | 0.33 |
| 2 | 76488564 | rs2860732 | 0.065 | -1.353 | 0.182 | 0.411 | -0.003 | 0.071 | 0.071 | 0 | 0.52 | 0.35 | 0.57 | 0.8 | 0.46 | 0.75 |
| 2 | 76490572 | rs1526649 | -0.214 | -1.427 | 0.25 | 0.468 | -0.004 | 0 | 0 | 0 | 0.79 | 0.76 | 0.62 | 0.3 | 0.86 | 0.52 |
| 2 | 76490982 | rs1980322 | -0.032 | -1.442 | 0.343 | 0.459 | -0.023 | 0.013 | 0.013 | 0 | 0.22 | 0.26 | 0.38 | 0.69 | 0.11 | 0.49 |
| 2 | 76491890 | rs13407506 | 1.006 | -1.285 | 0.176 | 0.413 | -0.035 | 0.069 | 0.069 | 0 | 0.36 | 0.23 | 0.23 | 0.11 | 0.4 | 0.27 |
| 2 | 76501978 | rs1000459 | 0.029 | -1.254 | 0.354 | 0.253 | -0.017 | -0.064 | -0.064 | 0.217 | 0.66 | 0.54 | 0.81 | 0.86 | 0.72 | 0.82 |
| 2 | 76503910 | rs1568984 | 0.604 | -1.271 | 0.338 | 0.351 | -0.005 | 0.003 | 0.003 | 0 | 0.6 | 0.49 | 0.63 | 0.53 | 0.66 | 0.63 |
| 2 | 76504256 | rs1004630 | 1.041 | -1.291 | 0.289 | 0.281 | -0.023 | -0.003 | -0.003 | 0 | 0.38 | 0.25 | 0.37 | 0.41 | 0.41 | 0.49 |
| 2 | 76509738 | rs1526624 | 1.205 | -1.368 | 0.236 | 0.311 | 0.015 | -0.011 | -0.011 | 0.157 | 0.13 | 0.09 | 0.2 | 0.38 | 0.06 | 0.27 |
| 2 | 76509864 | rs17754897 | 1.7 | -1.311 | 0.297 | 0.22 | 0.01 | -0.015 | -0.015 | 0 | 0.16 | 0.24 | 0.24 | 0.06 | 0.26 | 0.06 |
| 2 | 76522519 | rs13397093 | 0.32 | -1.406 | 0.336 | 0.185 | -0.049 | -0.086 | -0.086 | 0.179 | 0.29 | 0.43 | 0.16 | 0.13 | 0.2 | 0.15 |
| 2 | 76526320 | rs13006506 | 0.345 | -1.282 | 0.32 | 0.334 | 0 | 0.001 | 0.001 | 0 | 0.29 | 0.33 | 0.45 | 0.44 | 0.32 | 0.32 |
| 2 | 76535474 | rs6747727 | -0.339 | -1.463 | 0.144 | 0.181 | -0.031 | -0.038 | -0.038 | 0 | 0.79 | 0.54 | 0.85 | 0.86 | 0.8 | 0.83 |
| 2 | 76535759 | rs17012485 | 1.436 | -1.31 | 0.192 | 0.283 | 0.017 | -0.013 | -0.013 | 0 | 0.12 | 0.08 | 0.2 | 0.38 | 0.06 | 0.25 |
| 2 | 76537604 | rs885187 | -0.688 | -1.217 | 0.178 | 0.281 | -0.012 | 0.002 | 0.002 | 0 | 0.39 | 0.49 | 0.43 | 0.44 | 0.3 | 0.36 |
| 2 | 76545193 | rs13422380 | 0.394 | -1.226 | 0.172 | 0.476 | 0 | 0.003 | 0.003 | 0 | 0.28 | 0.34 | 0.37 | 0.39 | 0.31 | 0.31 |
| 2 | 76554776 | rs10167686 | 0.671 | -0.837 | 0.716 | 0.647 | 0.017 | 0.169 | 0.169 | 0 | 0.94 | 0.77 | 0.73 | 0.69 | 0.92 | 0.79 |
| 2 | 76556338 | rs925991 | 0.998 | -1.263 | 0.767 | 0.366 | 0.063 | -0.007 | -0.007 | 1.741 | 0.58 | 0.25 | 0.56 | 0.5 | 0.69 | 0.62 |
| 2 | 76564897 | rs1850899 | -1.018 | -1.227 | 0.775 | 0.375 | 0.011 | -0.001 | -0.001 | 1.785 | 0.39 | 0.57 | 0.42 | 0.49 | 0.28 | 0.35 |
| 2 | 76569676 | rs12463599 | 1.182 | -1.348 | 1.005 | 0.162 | -0.01 | 0.005 | 0.005 | 0 | 0.11 | 0.1 | 0.07 | 0.17 | 0.1 | 0.14 |
| 2 | 76571440 | rs6758645 | 0.967 | -1.397 | 1.274 | 0.116 | 0.002 | -0.017 | -0.017 | 0 | 0.18 | 0.13 | 0.05 | 0.01 | 0.09 | 0.02 |
| 2 | 76578575 | rs4853238 | 1.628 | -1.306 | 0.576 | 0.083 | -0.082 | -0.078 | -0.078 | 0.045 | 0.42 | 0.07 | 0.4 | 0.47 | 0.4 | 0.57 |
| 2 | 76579521 | rs4853241 | -0.091 | -1.296 | 0.704 | 0.042 | -0.005 | 0.081 | 0.081 | 0 | 0.08 | 0.19 | 0.29 | 0.3 | 0.09 | 0.19 |
| 2 | 76580070 | rs6758293 | 0.955 | -1.311 | 0.74 | -0.003 | 0.049 | 0.001 | 0.001 | 0.023 | 0.59 | 0.28 | 0.56 | 0.5 | 0.69 | 0.63 |
| 2 | 76580601 | rs6730042 | 1.258 | -1.31 | 0.728 | -0.002 | 0.075 | 0.003 | 0.003 | 0.069 | 0.58 | 0.22 | 0.56 | 0.5 | 0.69 | 0.63 |
| 2 | 76583236 | rs7420310 | 0.533 | -1.169 | 0.771 | 0.08 | -0.027 | 0.084 | 0.084 | 0 | 0.79 | 0.58 | 0.62 | 0.52 | 0.78 | 0.65 |
| 2 | 76585748 | rs4569502 | 0.636 | -0.754 | 0.902 | 0.301 | 0 | 0.017 | 0.017 | 0 | 0.06 | 0.05 | 0.27 | 0.3 | 0.09 | 0.17 |
| 2 | 76586332 | rs17515858 | 1.492 | -0.729 | 0.818 | 0.275 | -0.023 | 0.025 | 0.025 | 0 | 0.16 | 0.13 | 0.16 | 0.04 | 0.29 | 0.06 |
| 2 | 76599289 | rs11891955 | -0.259 | -1.314 | 1.206 | 0.917 | -0.028 | -0.038 | -0.038 | 4.287 | 0.3 | 0.33 | 0.12 | 0.19 | 0.18 | 0.16 |
| 2 | 76602785 | rs17012581 | -1.256 | -0.321 | 1.179 | 1.656 | -0.009 | 0.103 | 0.103 | 0 | 0.08 | 0.2 | 0.3 | 0.34 | 0.09 | 0.18 |
| 2 | 76605834 | rs908957 | 1.546 | -0.307 | 1.119 | 1.441 | 0.005 | 0.132 | 0.132 | 0 | 0.89 | 0.65 | 0.68 | 0.68 | 0.88 | 0.79 |
| 2 | 76606093 | rs908959 | 1.521 | -0.299 | 1.06 | 1.445 | 0.004 | 0.128 | 0.128 | 0 | 0.89 | 0.66 | 0.68 | 0.68 | 0.88 | 0.79 |
| 2 | 76606274 | rs17012588 | -1.136 | -0.297 | 0.992 | 1.446 | -0.005 | 0.044 | 0.044 | 0 | 0.08 | 0.15 | 0.29 | 0.31 | 0.09 | 0.19 |
| 2 | 76613760 | rs950290 | 1.551 | -0.446 | 0.874 | 1.436 | -0.022 | 0.146 | 0.146 | 0 | 0.83 | 0.51 | 0.65 | 0.57 | 0.83 | 0.76 |
| 2 | 76614158 | rs1996634 | -1.33 | -0.229 | -0.188 | 1.399 | -0.007 | 0.081 | 0.081 | 0 | 0.14 | 0.27 | 0.33 | 0.41 | 0.15 | 0.22 |
| 2 | 76618512 | rs908963 | 0.588 | -0.344 | -0.25 | 1.536 | -0.02 | 0.107 | 0.107 | 0 | 0.89 | 0.76 | 0.71 | 0.58 | 0.92 | 0.81 |
| 2 | 76618920 | rs1519898 | 0.256 | -0.702 | -1.071 | 1.762 | 0.002 | -0.107 | -0.107 | 0 | 0.15 | 0 | 0.2 | 0.32 | 0.1 | 0.21 |
| 2 | 76619122 | rs1519899 | 0.529 | -0.571 | -0.778 | 0.894 | 0 | -0.001 | -0.001 | 0 | 0.55 | 0.6 | 0.62 | 0.28 | 0.58 | 0.56 |
| 2 | 76620126 | rs17575434 | 0.46 | -0.078 | -1.429 | 0.295 | -0.026 | 0.084 | 0.084 | 0 | 0.28 | 0.1 | 0.26 | 0.08 | 0.38 | 0.14 |
| 2 | 76625060 | rs13017817 | 1.01 | -0.117 | -1.178 | 0.124 | -0.02 | 0.022 | 0.022 | 1.545 | 0.23 | 0.16 | 0.1 | 0.1 | 0.25 | 0.35 |
| 2 | 76625113 | rs10169401 | -0.35 | -0.111 | -0.854 | 0.64 | 0.007 | 0.201 | 0.201 | 0.287 | 0.88 | 0.6 | 0.78 | 0.7 | 0.92 | 0.81 |
| 2 | 76627129 | rs1879191 | -0.25 | -0.153 | -1.228 | 0.396 | -0.006 | -0.005 | -0.005 | 0.006 | 0.63 | 0.35 | 0.63 | 0.59 | 0.68 | 0.45 |
| 2 | 76627963 | rs1879192 | 0.914 | -0.035 | -1.458 | 0.473 | 0.003 | -0.135 | -0.135 | 0 | 0.25 | 0.01 | 0.36 | 0.34 | 0.22 | 0.16 |
| 2 | 76630436 | rs11683883 | -0.156 | 1.192 | -0.453 | 0.049 | -0.029 | 0.036 | 0.036 | 0 | 0.27 | 0.18 | 0.22 | 0.06 | 0.39 | 0.17 |
| 2 | 76641171 | rs7605051 | -0.696 | 0.538 | -0.36 | 0.448 | -0.002 | -0.033 | -0.033 | 0.02 | 0.31 | 0.56 | 0.37 | 0.17 | 0.34 | 0.25 |
| 2 | 76641791 | rs6742852 | 0.102 | 0.999 | -0.917 | 0.452 | -0.053 | 0.063 | 0.063 | 0 | 0.19 | 0.04 | 0.18 | 0.06 | 0.29 | 0.12 |
| 2 | 76643267 | rs1568378 | 0.482 | 0.87 | -0.159 | 1.004 | 0.016 | 0.048 | 0.048 | 0.06 | 0.88 | 0.74 | 0.94 | 0.8 | 0.89 | 0.77 |
| 2 | 76648993 | rs12614728 | -0.111 | 0.281 | -1.414 | 0.799 | -0.009 | -0.16 | -0.16 | 0 | 0.31 | 0.01 | 0.27 | 0.4 | 0.25 | 0.32 |
| 2 | 76649038 | rs1607639 | -0.395 | 0.391 | -1.761 | 0.371 | 0.019 | 0.098 | 0.098 | 0 | 0.51 | 0.17 | 0.38 | 0.66 | 0.45 | 0.52 |

| | | | | | | | | | | | | | | | | |
|---|---|---|---|---|---|---|---|---|---|---|---|---|---|---|---|---|
| 2 | 76654561 | rs1401841 | 0.147 | 0.214 | -1.267 | -0.309 | 0 | -0.004 | -0.004 | 0 | 0.42 | 0.43 | 0.53 | 0.24 | 0.55 | 0.49 |
| 2 | 76656726 | rs10196661 | -1.051 | 0.053 | -1.617 | 0 | 0.023 | -0.143 | -0.143 | 0 | 0.69 | 0.94 | 0.7 | 0.49 | 0.78 | 0.69 |
| 2 | 76657685 | rs4622759 | 1.207 | 0.687 | -1.578 | -0.374 | -0.051 | 0.07 | 0.07 | 0.305 | 0.23 | 0.08 | 0.35 | 0.09 | 0.35 | 0.19 |
| 2 | 76657950 | rs4481012 | 1.957 | 0.404 | -0.939 | -0.234 | -0.003 | -0.034 | -0.034 | 0 | 0.07 | 0.12 | 0.04 | 0 | 0.04 | 0.01 |
| 2 | 76674191 | rs1316911 | -0.121 | 0.528 | -0.813 | -0.421 | -0.01 | -0.12 | -0.12 | 0 | 0.83 | 0.49 | 0.88 | 1 | 0.86 | 0.97 |
| 2 | 76682789 | rs1921249 | -0.066 | 0.466 | -0.734 | -0.437 | -0.01 | 0.006 | 0.006 | 0.743 | 0.57 | 0.55 | 0.58 | 0.49 | 0.66 | 0.67 |
| 2 | 76682847 | rs1921250 | -1.197 | 0.452 | -0.676 | -0.457 | 0.001 | -0.034 | -0.034 | 0 | 0.88 | 0.86 | 0.89 | 1 | 0.85 | 0.96 |
| 2 | 76683260 | rs7563300 | 0.68 | 0.738 | -1.411 | -0.443 | -0.025 | -0.007 | -0.007 | 0 | 0.42 | 0.25 | 0.43 | 0.45 | 0.49 | 0.5 |
| 2 | 76686174 | rs1921251 | 1.669 | 0.686 | -2.022 | -0.159 | 0.004 | -0.153 | -0.153 | 0 | 0.24 | 0.01 | 0.19 | 0.48 | 0.16 | 0.26 |
| 2 | 76687485 | rs1882237 | 1.189 | 0.725 | -1.893 | -0.387 | -0.005 | 0.007 | 0.007 | 0 | 0.08 | 0.06 | 0.11 | 0.03 | 0.1 | 0.09 |
| 2 | 76688846 | rs13010366 | 0.536 | 0.964 | -2.095 | -0.557 | -0.013 | 0.029 | 0.029 | 0.004 | 0.33 | 0.23 | 0.4 | 0.22 | 0.35 | 0.37 |
| 2 | 76689207 | rs17012723 | 1.681 | 0.987 | -2.091 | -0.542 | -0.049 | 0.003 | 0.003 | 1.745 | 0.23 | 0.04 | 0.28 | 0.18 | 0.24 | 0.28 |
| 2 | 76693358 | rs17012735 | 1.392 | 0.992 | -2.051 | -0.52 | -0.009 | 0.011 | 0.011 | 0.001 | 0.07 | 0.04 | 0.11 | 0.03 | 0.1 | 0.1 |
| 2 | 76696790 | rs11688462 | -0.453 | 1.072 | -2.095 | -0.601 | 0.007 | -0.006 | -0.006 | 0.053 | 0.58 | 0.45 | 0.59 | 0.49 | 0.68 | 0.68 |
| 2 | 76706357 | rs6720979 | 1.77 | 1.191 | -2.197 | -0.557 | -0.003 | 0 | 0 | 0 | 0.16 | 0.17 | 0.11 | 0.19 | 0.18 | 0.15 |
| 2 | 76707046 | rs1921243 | 0.818 | 1.149 | -2.229 | -0.572 | -0.055 | 0.102 | 0.102 | 0.009 | 0.54 | 0.13 | 0.68 | 0.7 | 0.51 | 0.66 |
| 2 | 76708697 | rs13026371 | -0.022 | 1.182 | -2.135 | -0.469 | 0 | -0.001 | -0.001 | 0 | 0.28 | 0.35 | 0.19 | 0.28 | 0.34 | 0.3 |
| 2 | 76716538 | rs7603169 | 1.102 | 1.181 | -2.167 | -0.521 | -0.036 | -0.03 | -0.03 | 0.587 | 0.2 | 0.1 | 0.15 | 0.26 | 0.27 | 0.23 |
| 2 | 76721928 | rs11683701 | 0.976 | 1.19 | -2.154 | -0.523 | -0.016 | -0.011 | -0.011 | 0 | 0.21 | 0.16 | 0.16 | 0.26 | 0.27 | 0.23 |
| 2 | 76722019 | rs12613882 | 1.906 | 1.165 | -2.207 | -0.546 | 0.022 | -0.156 | -0.156 | 0.192 | 0.24 | 0.03 | 0.29 | 0.51 | 0.17 | 0.28 |
| 2 | 76726228 | rs2860780 | 0.097 | 1.216 | -2.158 | -0.539 | 0.009 | 0.096 | 0.096 | 0 | 0.47 | 0.27 | 0.45 | 0.78 | 0.44 | 0.51 |
| 2 | 76726553 | rs1921226 | 0.746 | 1.232 | -2.175 | -0.566 | -0.039 | 0.082 | 0.082 | 0 | 0.56 | 0.19 | 0.68 | 0.73 | 0.5 | 0.65 |
| 2 | 76726686 | rs1997448 | -1.32 | 1.235 | -2.139 | -0.618 | 0.011 | -0.117 | -0.117 | 1.262 | 0.84 | 0.51 | 0.87 | 1 | 0.84 | 0.96 |
| 2 | 76727046 | rs13002864 | 0.976 | 1.345 | -2.381 | -0.539 | -0.024 | 0.035 | 0.035 | 0 | 0.29 | 0.18 | 0.32 | 0.19 | 0.31 | 0.36 |
| 2 | 76741695 | rs12477015 | 0.682 | 1.272 | -2.461 | -0.402 | 0.027 | -0.067 | -0.067 | 0.476 | 0.33 | 0.15 | 0.4 | 0.55 | 0.27 | 0.38 |
| 2 | 76741847 | rs7604513 | 0.409 | 1.271 | -2.478 | -0.424 | -0.047 | 0.09 | 0.09 | 0 | 0.56 | 0.2 | 0.68 | 0.73 | 0.51 | 0.65 |
| 2 | 76744940 | rs11126537 | -0.852 | 1.439 | -1.785 | -0.149 | -0.005 | 0.109 | 0.109 | 0.81 | 0.54 | 0.3 | 0.56 | 0.81 | 0.54 | 0.6 |
| 2 | 76745098 | rs17040477 | 0.884 | 1.395 | -1.824 | -0.149 | -0.005 | 0.007 | 0.007 | 0.119 | 0.07 | 0.04 | 0.1 | 0.03 | 0.1 | 0.1 |
| 2 | 76745159 | rs11126538 | -0.563 | 1.319 | -1.655 | -0.104 | -0.008 | 0.14 | 0.14 | 1.004 | 0.53 | 0.24 | 0.55 | 0.81 | 0.54 | 0.6 |
| 2 | 76746597 | rs11692079 | 1.205 | 1.275 | -1.727 | 0.006 | -0.049 | -0.018 | -0.018 | 0.895 | 0.13 | 0.02 | 0.09 | 0.17 | 0.18 | 0.13 |
| 2 | 76752478 | rs17012839 | -0.182 | 1.104 | -1.248 | -0.098 | 0.001 | 0.059 | 0.059 | 1.207 | 0.55 | 0.36 | 0.57 | 0.81 | 0.58 | 0.63 |
| 2 | 76757244 | rs1921237 | 0.719 | 1.588 | -0.712 | 0.642 | 0.003 | 0.003 | 0.003 | 0.058 | 0.79 | 0.8 | 0.83 | 0.82 | 0.77 | 0.72 |
| 2 | 76760606 | rs12052843 | 0.97 | 1.506 | -0.754 | 0.636 | -0.001 | 0 | 0 | 0 | 0.77 | 0.79 | 0.82 | 0.81 | 0.75 | 0.72 |
| 2 | 76762757 | rs17012903 | 0.505 | 1.709 | -1.204 | 1.096 | -0.015 | -0.077 | -0.077 | 0 | 0.16 | 0.05 | 0.17 | 0.45 | 0.17 | 0.26 |
| 2 | 76764537 | rs7564033 | 0.373 | 1.722 | -1.196 | 1.068 | -0.001 | -0.001 | -0.001 | 0 | 0.82 | 0.85 | 0.83 | 0.82 | 0.78 | 0.78 |
| 2 | 76764790 | rs11126543 | 2.157 | 1.726 | -1.695 | 0.692 | -0.072 | 0.1 | 0.1 | 3.383 | 0.43 | 0.05 | 0.39 | 0.27 | 0.4 | 0.39 |
| 2 | 76768976 | rs12613217 | 2.728 | 2.265 | -2.095 | 0.481 | 0.018 | 0.122 | 0.122 | 8.382 | 0.32 | 0.02 | 0.11 | 0.09 | 0.22 | 0.2 |
| 2 | 76769627 | rs2141732 | 0.295 | 2.058 | -0.617 | 0.754 | 0.076 | 0.137 | 0.137 | 0.037 | 0.06 | 0.28 | 0.15 | 0.09 | 0.08 | 0.04 |
| 2 | 76770013 | rs2141733 | 0.276 | 2.011 | -0.768 | 0.519 | -0.002 | -0.001 | -0.001 | 0 | 0.77 | 0.82 | 0.82 | 0.81 | 0.75 | 0.72 |
| 2 | 76777279 | rs7592517 | -0.554 | 1.966 | -0.461 | 0.501 | 0 | 0.065 | 0.065 | 0.063 | 0.31 | 0.51 | 0.41 | 0.55 | 0.35 | 0.33 |
| 2 | 76780044 | rs1519995 | -0.506 | 2.078 | -0.56 | 0.466 | -0.029 | 0.036 | 0.036 | 0 | 0.91 | 0.98 | 0.95 | 0.97 | 0.89 | 0.9 |
| 2 | 76783281 | rs6725794 | 0.498 | 2.032 | -0.386 | 0.452 | -0.003 | -0.032 | -0.032 | 0 | 0.09 | 0.17 | 0.09 | 0 | 0.1 | 0.03 |
| 2 | 76785084 | rs991725 | 2.218 | 2.083 | -0.373 | 0.499 | 0.001 | 0.109 | 0.109 | 3.541 | 0.46 | 0.22 | 0.4 | 0.26 | 0.4 | 0.4 |
| 2 | 76786845 | rs2139502 | -0.168 | 2.188 | 0.006 | 0.916 | 0.013 | 0.168 | 0.168 | 1.799 | 0.23 | 0.43 | 0.33 | 0.55 | 0.26 | 0.29 |
| 2 | 76796030 | rs12464655 | -0.1 | 2.101 | -0.04 | 0.872 | -0.003 | 0.002 | 0.002 | 0 | 0.91 | 0.93 | 0.96 | 0.96 | 0.9 | 0.89 |
| 2 | 76796561 | rs963552 | -1.211 | 1.575 | 0.47 | 1.095 | -0.007 | 0.016 | 0.016 | 2.131 | 0.3 | 0.53 | 0.41 | 0.66 | 0.3 | 0.41 |
| 2 | 76811778 | rs10203812 | 2.046 | 1.662 | 0.657 | 1.062 | 0.001 | 0.086 | 0.086 | 1.168 | 0.49 | 0.33 | 0.25 | 0.22 | 0.45 | 0.47 |
| 2 | 76814757 | rs7607201 | 0.075 | 1.594 | 1.022 | 1.362 | 0.06 | 0.171 | 0.171 | 0 | 0.08 | 0.26 | 0.12 | 0.16 | 0.1 | 0.08 |
| 2 | 76816071 | rs10201906 | -0.005 | 1.654 | 0.905 | 1.54 | -0.002 | 0.001 | 0.001 | 0 | 0.91 | 0.92 | 0.8 | 0.96 | 0.91 | 0.89 |
| 2 | 76821113 | rs11897408 | -1.415 | 1.633 | 0.919 | 1.537 | -0.001 | -0.025 | -0.025 | 0.355 | 0.3 | 0.27 | 0.43 | 0.56 | 0.35 | 0.29 |
| 2 | 76824150 | rs2139498 | -1.407 | 1.645 | 0.887 | 1.546 | -0.002 | -0.035 | -0.035 | 0.454 | 0.3 | 0.25 | 0.43 | 0.56 | 0.36 | 0.3 |
| 2 | 76826142 | rs10153813 | 1.556 | 1.706 | 0.925 | 1.555 | -0.002 | 0 | 0 | 0.038 | 0.59 | 0.67 | 0.37 | 0.39 | 0.55 | 0.55 |
| 2 | 76836344 | rs1401888 | -0.608 | 1.355 | 1.388 | 1.457 | -0.004 | 0.238 | 0.238 | 0.709 | 0.18 | 0.49 | 0.24 | 0.56 | 0.17 | 0.32 |
| 2 | 76839833 | rs7582080 | 0.971 | 1.42 | 1.296 | 1.367 | -0.008 | 0.018 | 0.018 | 0 | 0.76 | 0.79 | 0.67 | 0.92 | 0.76 | 0.86 |
| 2 | 76844026 | rs1474177 | 2.07 | 1.375 | 1.457 | 1.369 | 0.003 | 0.018 | 0.018 | 0.476 | 0.55 | 0.31 | 0.38 | 0.35 | 0.55 | 0.52 |
| 2 | 76844466 | rs1154641 | 0.942 | 1.349 | 1.461 | 1.343 | -0.001 | -0.003 | -0.003 | 0 | 0.77 | 0.77 | 0.67 | 0.92 | 0.76 | 0.86 |
| 2 | 76860212 | rs6716298 | -0.064 | 1.367 | 1.416 | 1.388 | 0 | 0.101 | 0.101 | 2.236 | 0.16 | 0.25 | 0.23 | 0.55 | 0.17 | 0.32 |
| 2 | 76865585 | rs7566719 | 2.182 | 1.586 | 1.346 | 1.371 | 0.002 | 0.142 | 0.142 | 0.454 | 0.55 | 0.2 | 0.31 | 0.22 | 0.56 | 0.51 |
| 2 | 76870303 | rs7585161 | 0.7 | 1.537 | 0.992 | 1.339 | 0.022 | 0.024 | 0.024 | 0 | 0.09 | 0.03 | 0.04 | 0 | 0.06 | 0.01 |
| 2 | 76870739 | rs9332421 | 1.913 | 1.617 | 1.048 | 1.414 | 0 | 0.002 | 0.002 | 0.334 | 0.56 | 0.38 | 0.39 | 0.37 | 0.55 | 0.52 |
| 2 | 76874141 | rs10195080 | -0.356 | 1.725 | 0.611 | 0.955 | -0.032 | -0.004 | -0.004 | 0 | 0.93 | 0.99 | 0.79 | 0.97 | 0.92 | 0.89 |
| 2 | 76881210 | rs7606395 | 0.031 | 1.747 | 0.744 | 1.033 | 0 | -0.018 | -0.018 | 0.132 | 0.79 | 0.75 | 0.71 | 0.94 | 0.81 | 0.87 |
| 2 | 76889411 | rs2901780 | -1.216 | 1.402 | 0.315 | 1.327 | 0 | -0.005 | -0.005 | 0 | 0.37 | 0.4 | 0.56 | 0.71 | 0.37 | 0.44 |
| 2 | 76890567 | rs1519982 | -1.136 | 1.465 | 0.59 | 2.096 | 0 | 0.037 | 0.037 | 0.047 | 0.2 | 0.29 | 0.33 | 0.68 | 0.25 | 0.31 |
| 2 | 76911782 | rs985343 | -1.384 | 1.431 | 0.498 | 2.017 | 0.036 | 0.18 | 0.18 | 0 | 0.18 | 0.41 | 0.28 | 0.49 | 0.22 | 0.31 |
| 2 | 76912285 | rs11126551 | -1.116 | 1.576 | 0.142 | 1.932 | 0.029 | 0.099 | 0.099 | 0 | 0.16 | 0.28 | 0.27 | 0.44 | 0.23 | 0.3 |
| 2 | 76915107 | rs985812 | -0.998 | 1.459 | -0.117 | 1.633 | 0.026 | 0.052 | 0.052 | 0 | 0.32 | 0.49 | 0.34 | 0.5 | 0.36 | 0.43 |
| 2 | 76915391 | rs11689433 | -0.766 | 1.477 | -0.239 | 1.702 | -0.004 | -0.006 | -0.006 | 0.126 | 0.16 | 0.14 | 0.17 | 0.23 | 0.17 | 0.15 |
| 2 | 76916042 | rs11126552 | 0.02 | 0.907 | -1.146 | 0.944 | 0.038 | -0.059 | -0.059 | 0 | 0.49 | 0.1 | 0.49 | 0.53 | 0.57 | 0.58 |

| | | | | | | | | | | | | | | | | |
|---|---|---|---|---|---|---|---|---|---|---|---|---|---|---|---|---|
| 2 | 76917583 | rs1879219 | -0.137 | 0.504 | -0.757 | 0.627 | -0.005 | 0.027 | 0.027 | 0 | 0.66 | 0.53 | 0.76 | 0.56 | 0.65 | 0.75 |
| 2 | 76918138 | rs6745127 | 0.994 | 0.299 | -0.594 | 0.522 | -0.018 | -0.009 | -0.009 | 0.565 | 0.43 | 0.26 | 0.46 | 0.48 | 0.5 | 0.53 |
| 2 | 76918322 | rs955949 | 0.218 | 0.298 | -0.48 | 0.594 | -0.002 | 0.001 | 0.001 | 0 | 0.59 | 0.42 | 0.56 | 0.56 | 0.58 | 0.7 |
| 2 | 76921182 | rs11676468 | -0.272 | 0.407 | -0.489 | 0.624 | -0.015 | 0.03 | 0.03 | 0 | 0.67 | 0.54 | 0.79 | 0.6 | 0.71 | 0.78 |
| 2 | 76927056 | rs12469595 | 0.387 | 0.673 | -0.718 | 0.729 | 0.039 | 0.018 | 0.018 | 0.088 | 0.5 | 0.25 | 0.5 | 0.49 | 0.56 | 0.57 |
| 2 | 76941049 | rs1519998 | 0.491 | 0.712 | -1.476 | 0.749 | -0.016 | -0.049 | -0.049 | 1.282 | 0.46 | 0.16 | 0.48 | 0.53 | 0.53 | 0.55 |
| 2 | 76943150 | rs7568302 | -0.424 | 0.279 | -0.634 | 0.385 | -0.012 | -0.022 | -0.022 | 0.014 | 0.31 | 0.57 | 0.32 | 0.18 | 0.28 | 0.21 |
| 2 | 76943205 | rs4852410 | -0.999 | 0.179 | -0.169 | 0.625 | 0.001 | 0.007 | 0.007 | 0 | 0.37 | 0.46 | 0.43 | 0.36 | 0.44 | 0.37 |
| 2 | 76943863 | rs10185473 | -1.081 | 0.309 | -0.493 | -0.215 | -0.003 | 0.015 | 0.015 | 0.334 | 0.37 | 0.66 | 0.47 | 0.25 | 0.37 | 0.37 |
| 2 | 76947107 | rs7589991 | -1.439 | 0.236 | -0.806 | -0.056 | -0.015 | -0.004 | -0.004 | 0 | 0.51 | 0.82 | 0.58 | 0.43 | 0.54 | 0.52 |
| 2 | 76947356 | rs1519999 | -1.471 | 0.201 | -0.825 | -0.073 | -0.019 | -0.009 | -0.009 | 0 | 0.51 | 0.82 | 0.58 | 0.43 | 0.54 | 0.52 |
| 2 | 76947750 | rs4618084 | -1.422 | 0.274 | -0.678 | -0.06 | -0.001 | 0.005 | 0.005 | 4.656 | 0.51 | 0.77 | 0.58 | 0.43 | 0.54 | 0.52 |
| 2 | 76947840 | rs11901729 | -0.873 | 0.289 | -0.287 | -0.142 | -0.01 | 0.029 | 0.029 | 0.7 | 0.27 | 0.49 | 0.2 | 0.33 | 0.23 | 0.32 |
| 2 | 76948043 | rs4387821 | -1.326 | 0.383 | -0.886 | -0.001 | -0.012 | -0.003 | -0.003 | 0 | 0.52 | 0.8 | 0.58 | 0.43 | 0.54 | 0.52 |
| 2 | 76948525 | rs12470091 | 0.385 | 0.479 | -0.624 | 0.066 | 0.006 | 0.002 | 0.002 | 0 | 0.85 | 0.87 | 0.94 | 0.88 | 0.9 | 0.77 |
| 2 | 76948994 | rs10208376 | 0.233 | 0.442 | -0.245 | -0.091 | -0.01 | -0.077 | -0.077 | 0 | 0.22 | 0.28 | 0.33 | 0.04 | 0.23 | 0.18 |
| 2 | 76949033 | rs12105118 | 0.533 | 0.472 | -0.517 | -0.079 | -0.001 | -0.022 | -0.022 | 0 | 0.21 | 0.2 | 0.33 | 0.04 | 0.23 | 0.16 |
| 2 | 76949074 | rs17404871 | 0.823 | 0.932 | -1.148 | 0.15 | -0.037 | -0.013 | -0.013 | 0.13 | 0.06 | 0.01 | 0.09 | 0.04 | 0.13 | 0.07 |
| 2 | 76949101 | rs10210863 | -0.873 | 0.693 | -0.938 | 0.255 | -0.013 | 0.05 | 0.05 | 0.237 | 0.46 | 0.7 | 0.5 | 0.26 | 0.48 | 0.45 |
| 2 | 76956302 | rs1969314 | 0.806 | 0.422 | -1.451 | 0.85 | -0.057 | -0.147 | -0.147 | 0 | 0.19 | 0.01 | 0.25 | 0.44 | 0.17 | 0.24 |
| 2 | 76957088 | rs12053161 | -0.564 | -0.523 | -0.838 | 0.541 | -0.03 | 0.016 | 0.016 | 0 | 0.15 | 0.2 | 0.06 | 0.24 | 0.05 | 0.2 |
| 2 | 76957370 | rs10172954 | -0.583 | -0.582 | -0.592 | 0.531 | -0.048 | 0.036 | 0.036 | 0 | 0.16 | 0.27 | 0.07 | 0.26 | 0.05 | 0.21 |
| 2 | 76961427 | rs971121 | -0.842 | -0.485 | -0.477 | 0.152 | 0.037 | 0.039 | 0.039 | 0.779 | 0.44 | 0.74 | 0.46 | 0.36 | 0.35 | 0.38 |
| 2 | 76969846 | rs6730814 | -1.001 | -0.483 | -0.425 | 0.143 | 0.046 | 0.06 | 0.06 | 1.582 | 0.47 | 0.75 | 0.47 | 0.36 | 0.37 | 0.46 |
| 2 | 76970796 | rs13009537 | 0.41 | -0.318 | -0.495 | 0.239 | 0.009 | -0.008 | -0.008 | 0 | 0.08 | 0.06 | 0.03 | 0.13 | 0.02 | 0.13 |
| 2 | 76972429 | rs4853269 | 0.43 | 0.135 | -0.045 | 0.123 | -0.003 | -0.052 | -0.052 | 0.334 | 0.25 | 0.34 | 0.37 | 0.09 | 0.24 | 0.16 |
| 2 | 76975383 | rs7609111 | -0.743 | 0.177 | 0.22 | 0.519 | 0 | 0 | 0 | 0.878 | 0.54 | 0.59 | 0.53 | 0.4 | 0.51 | 0.47 |
| 2 | 76975730 | rs6547099 | -0.123 | 0.065 | 0.343 | 0.391 | -0.011 | 0.049 | 0.049 | 0 | 0.07 | 0.28 | 0.03 | 0.11 | 0.02 | 0.1 |
| 2 | 76976577 | rs13396159 | 0.341 | 0.156 | 0.108 | 0.048 | -0.003 | -0.059 | -0.059 | 0.047 | 0.26 | 0.34 | 0.34 | 0.07 | 0.25 | 0.23 |
| 2 | 76976626 | rs12713881 | -0.968 | 0.081 | 0.067 | 0.444 | 0 | 0 | 0 | 0 | 0.56 | 0.47 | 0.64 | 0.55 | 0.5 | 0.59 |
| 2 | 76981827 | rs4564799 | -0.535 | -0.156 | -0.023 | 0.512 | -0.008 | 0.022 | 0.022 | 0 | 0.35 | 0.43 | 0.33 | 0.5 | 0.25 | 0.44 |
| 2 | 76985361 | rs7608234 | -1.222 | -0.095 | -0.054 | 0.525 | -0.004 | -0.004 | -0.004 | 0 | 0.6 | 0.49 | 0.67 | 0.61 | 0.59 | 0.65 |
| 2 | 76987268 | rs1400848 | -0.205 | -0.143 | -0.064 | 0.805 | -0.001 | 0 | 0 | 0 | 0.3 | 0.3 | 0.31 | 0.47 | 0.24 | 0.35 |
| 2 | 76989568 | rs9917184 | 0.985 | -0.135 | -0.386 | 0.766 | -0.009 | 0.016 | 0.016 | 0.023 | 0.21 | 0.11 | 0.1 | 0.16 | 0.16 | 0.1 |
| 2 | 76989998 | rs1517779 | -0.59 | -0.272 | 0.028 | 0.95 | -0.013 | 0.009 | 0.009 | 0 | 0.33 | 0.5 | 0.33 | 0.37 | 0.24 | 0.34 |
| 2 | 76994984 | rs13008894 | -0.923 | -0.431 | -0.135 | 0.881 | 0.007 | -0.005 | -0.005 | 2.334 | 0.17 | 0.13 | 0.21 | 0.43 | 0.1 | 0.38 |
| 2 | 76997488 | rs2222179 | -1.028 | -0.423 | -0.198 | 0.794 | -0.006 | 0.001 | 0.001 | 0 | 0.4 | 0.53 | 0.34 | 0.55 | 0.27 | 0.45 |
| 2 | 77014110 | rs17404976 | 0.195 | -0.405 | -0.351 | 0.463 | 0.01 | 0.182 | 0.182 | 0 | 0.45 | 0.19 | 0.46 | 0.15 | 0.51 | 0.28 |
| 2 | 77016245 | rs7585584 | -0.206 | -0.247 | -0.017 | 0.665 | 0.007 | 0.002 | 0.002 | 0 | 0.55 | 0.38 | 0.68 | 0.43 | 0.6 | 0.51 |
| 2 | 77016349 | rs12988164 | -0.795 | -0.115 | -0.414 | 0.663 | 0.005 | -0.012 | -0.012 | 0 | 0.64 | 0.65 | 0.71 | 0.54 | 0.63 | 0.65 |
| 2 | 77018329 | rs4853273 | -0.971 | -0.055 | -0.789 | 0.408 | -0.011 | 0.044 | 0.044 | 0.222 | 0.67 | 0.81 | 0.69 | 0.78 | 0.61 | 0.65 |
| 2 | 77020219 | rs10178612 | 0.524 | 0.017 | -0.746 | 0.365 | -0.016 | 0.014 | 0.014 | 0 | 0.09 | 0.06 | 0.08 | 0.03 | 0.15 | 0.07 |
| 2 | 77021928 | rs10170138 | -0.086 | -0.095 | -0.107 | 0.612 | 0.017 | 0.005 | 0.005 | 0 | 0.11 | 0.21 | 0.09 | 0.12 | 0.15 | 0.09 |
| 2 | 77029323 | rs7561233 | -1.413 | -0.241 | -0.188 | 0.329 | -0.019 | 0.033 | 0.033 | 1.858 | 0.64 | 0.73 | 0.66 | 0.77 | 0.54 | 0.59 |
| 2 | 77031247 | rs6718129 | -1.225 | -0.511 | -0.461 | 0.208 | 0.029 | 0.057 | 0.057 | 1.1 | 0.45 | 0.7 | 0.41 | 0.69 | 0.38 | 0.52 |
| 2 | 77041264 | rs2102827 | -1.068 | -0.524 | -0.002 | 0.278 | 0.004 | 0.028 | 0.028 | 0.524 | 0.54 | 0.59 | 0.61 | 0.74 | 0.4 | 0.53 |
| 2 | 77047442 | rs1517772 | -1.035 | -0.967 | -0.17 | 0.201 | -0.012 | -0.012 | -0.012 | 0 | 0.35 | 0.52 | 0.54 | 0.7 | 0.3 | 0.45 |
| 2 | 77048896 | rs7607547 | -1.562 | -0.981 | -0.229 | 0.17 | 0.02 | 0.1 | 0.1 | 0.185 | 0.48 | 0.67 | 0.61 | 0.77 | 0.41 | 0.53 |
| 2 | 77049269 | rs4852419 | -0.829 | -0.941 | -0.131 | 0.219 | -0.006 | 0.009 | 0.009 | 0 | 0.33 | 0.37 | 0.53 | 0.61 | 0.29 | 0.43 |
| 2 | 77049296 | rs4853278 | -0.159 | -0.964 | -0.018 | 0.609 | -0.001 | -0.013 | -0.013 | 0.094 | 0.26 | 0.19 | 0.46 | 0.39 | 0.27 | 0.33 |
| 2 | 77053916 | rs10520165 | 1.078 | -0.91 | -0.379 | 0.681 | 0.017 | -0.086 | -0.086 | 0.013 | 0.12 | 0.01 | 0.21 | 0.29 | 0.1 | 0.26 |
| 2 | 77055883 | rs1517774 | -1.52 | -0.669 | -0.288 | 0.156 | -0.019 | 0.062 | 0.062 | 0.559 | 0.57 | 0.67 | 0.63 | 0.77 | 0.46 | 0.54 |
| 2 | 77059199 | rs1914508 | -2.253 | -1.085 | -0.199 | 0.421 | -0.058 | 0.057 | 0.057 | 2.86 | 0.66 | 0.89 | 0.7 | 0.79 | 0.56 | 0.6 |
| 2 | 77074251 | rs1517776 | -0.576 | -0.6 | 0.757 | 0.907 | -0.003 | 0.007 | 0.007 | 0 | 0.28 | 0.24 | 0.34 | 0.24 | 0.28 | 0.17 |
| 2 | 77078101 | rs4340548 | -1.7 | -0.567 | 1.336 | 1.195 | -0.01 | 0.006 | 0.006 | 0 | 0.32 | 0.51 | 0.53 | 0.63 | 0.28 | 0.38 |
| 2 | 77082852 | rs17013242 | -0.357 | -0.875 | 1.275 | 1.183 | 0.022 | -0.036 | -0.036 | 0 | 0.14 | 0.08 | 0.25 | 0.49 | 0.1 | 0.27 |
| 2 | 77082875 | rs17013248 | -0.176 | -0.881 | 1.431 | 1.21 | 0.009 | 0.005 | 0.005 | 0 | 0.13 | 0.12 | 0.22 | 0.46 | 0.09 | 0.28 |
| 2 | 77084241 | rs13412373 | -2.196 | -0.877 | 1.448 | 1.12 | 0.03 | 0.064 | 0.064 | 3.14 | 0.05 | 0.37 | 0.03 | 0.04 | 0 | 0.07 |
| 2 | 77085351 | rs7607019 | -2.143 | -0.791 | 1.256 | 1.088 | -0.007 | -0.001 | -0.001 | 0 | 0.35 | 0.62 | 0.55 | 0.62 | 0.31 | 0.4 |
| 2 | 77088825 | rs17333276 | 0.737 | -0.734 | 0.577 | 1.133 | -0.059 | 0.015 | 0.015 | 0 | 0.12 | 0.01 | 0.24 | 0.09 | 0.17 | 0.07 |
| 2 | 77094790 | rs721390 | -1.05 | -0.772 | 0.612 | 1.139 | -0.016 | -0.078 | -0.078 | 0 | 0.27 | 0.08 | 0.49 | 0.58 | 0.27 | 0.33 |
| 2 | 77103474 | rs10191141 | -1.992 | -0.875 | 1.126 | 0.998 | -0.026 | -0.028 | -0.028 | 0.01 | 0.35 | 0.5 | 0.53 | 0.71 | 0.28 | 0.43 |
| 2 | 77113949 | rs17405288 | 0.263 | -0.524 | 1.005 | 1.248 | -0.036 | 0.011 | 0.011 | 0 | 0.13 | 0.03 | 0.25 | 0.09 | 0.17 | 0.08 |
| 2 | 77116764 | rs985234 | -0.753 | -0.509 | 1.023 | 1.476 | -0.036 | -0.135 | -0.135 | 0.164 | 0.23 | 0.06 | 0.4 | 0.49 | 0.26 | 0.38 |
| 2 | 77119346 | rs17013296 | -1.095 | -0.496 | 1.489 | 1.49 | -0.001 | -0.015 | -0.015 | 0 | 0.26 | 0.26 | 0.46 | 0.57 | 0.27 | 0.4 |
| 2 | 77122673 | rs914389 | 0.657 | -0.547 | 1.334 | 1.545 | 0.002 | -0.032 | -0.032 | 0 | 0.08 | 0.01 | 0.14 | 0.36 | 0.07 | 0.23 |
| 2 | 77123866 | rs953855 | -0.833 | -0.557 | 1.581 | 1.452 | 0.013 | 0.056 | 0.056 | 0 | 0.21 | 0.41 | 0.39 | 0.31 | 0.21 | 0.13 |
| 2 | 77125218 | rs1929458 | -0.905 | -0.563 | 1.533 | 1.452 | 0.015 | 0.057 | 0.057 | 0 | 0.21 | 0.43 | 0.4 | 0.31 | 0.21 | 0.13 |
| 2 | 77129922 | rs4852420 | 0.07 | -0.402 | 1.61 | 1.205 | 0.011 | -0.037 | -0.037 | 0 | 0.11 | 0.14 | 0.09 | 0.04 | 0.16 | 0.08 |

| | | | | | | | | | | | | | | | | |
|---|---|---|---|---|---|---|---|---|---|---|---|---|---|---|---|---|
| 2 | 77138339 | rs10189940 | -1.091 | -0.442 | 1.339 | 1.285 | -0.032 | 0.158 | 0.158 | 0 | 0.15 | 0.37 | 0.27 | 0.59 | 0.09 | 0.29 |
| 2 | 77143070 | rs7592470 | -1.613 | -0.436 | 1.322 | 1.227 | -0.008 | -0.012 | -0.012 | 0.104 | 0.3 | 0.43 | 0.53 | 0.71 | 0.28 | 0.43 |
| 2 | 77151777 | rs10084200 | 0.159 | -0.405 | 1.161 | 1.344 | 0.007 | 0.027 | 0.027 | 0 | 0.1 | 0.09 | 0.18 | 0.46 | 0.08 | 0.26 |
| 2 | 77161916 | rs13417401 | -1.482 | -0.453 | 1.473 | 1.11 | 0.002 | -0.009 | -0.009 | 3.171 | 0.18 | 0.6 | 0.2 | 0.19 | 0.18 | 0.12 |
| 2 | 77170201 | rs10520170 | -1.024 | -0.44 | 1.658 | 1.096 | 0.058 | -0.103 | -0.103 | 0 | 0.14 | 0.46 | 0.12 | 0.03 | 0.18 | 0.08 |
| 2 | 77172693 | rs1921627 | -0.844 | -0.5 | 1.327 | 1.094 | -0.022 | -0.074 | -0.074 | 1.487 | 0.26 | 0.09 | 0.44 | 0.56 | 0.26 | 0.4 |
| 2 | 77174354 | rs10204605 | -1.555 | -0.503 | 1.339 | 1.089 | -0.003 | -0.01 | -0.01 | 2.291 | 0.18 | 0.64 | 0.2 | 0.19 | 0.18 | 0.11 |
| 2 | 77178484 | rs17405711 | 0.505 | -0.53 | 0.962 | 1.065 | -0.062 | 0.02 | 0.02 | 0 | 0.14 | 0 | 0.26 | 0.07 | 0.18 | 0.06 |
| 2 | 77188061 | rs4853285 | 0.464 | -0.531 | 0.963 | 1.074 | -0.058 | 0.005 | 0.005 | 0 | 0.14 | 0.01 | 0.26 | 0.11 | 0.18 | 0.13 |
| 2 | 77202747 | rs1446710 | -2.423 | -0.714 | 1.071 | 1.076 | 0.002 | 0.125 | 0.125 | 1.225 | 0.45 | 0.84 | 0.63 | 0.75 | 0.43 | 0.51 |
| 2 | 77209245 | rs6547106 | -1.639 | -0.867 | 1.617 | 1.133 | -0.008 | 0.041 | 0.041 | 0.044 | 0.27 | 0.49 | 0.35 | 0.65 | 0.26 | 0.38 |
| 2 | 77210501 | rs13389562 | -0.599 | -0.849 | 1.663 | 1.259 | -0.021 | 0.076 | 0.076 | 0 | 0.18 | 0.25 | 0.19 | 0.41 | 0.12 | 0.28 |
| 2 | 77213066 | rs7561189 | -1.683 | -0.849 | 1.694 | 1.258 | -0.015 | 0.099 | 0.099 | 0 | 0.26 | 0.47 | 0.31 | 0.56 | 0.25 | 0.36 |
| 2 | 77213578 | rs13413951 | -0.3 | -0.821 | 1.539 | 1.125 | 0.004 | -0.033 | -0.033 | 0 | 0.18 | 0.32 | 0.27 | 0.1 | 0.18 | 0.14 |
| 2 | 77217310 | rs10520176 | -2.794 | -0.989 | 0.685 | 0.836 | -0.117 | 0.099 | 0.099 | 0.656 | 0.6 | 0.95 | 0.69 | 0.79 | 0.55 | 0.61 |
| 2 | 77222810 | rs1470504 | -2.226 | -0.749 | 1.441 | 1.351 | 0 | 0.004 | 0.004 | 0.811 | 0.41 | 0.69 | 0.55 | 0.6 | 0.42 | 0.48 |
| 2 | 77226087 | rs12988557 | -0.22 | -0.608 | 1.078 | 1.17 | -0.023 | 0.025 | 0.025 | 0 | 0.09 | 0.04 | 0.09 | 0.03 | 0.16 | 0.08 |
| 2 | 77228622 | rs10520177 | 0.068 | -0.644 | 1.133 | 1.246 | -0.016 | 0.017 | 0.017 | 0 | 0.14 | 0.07 | 0.25 | 0.1 | 0.16 | 0.14 |
| 2 | 77234621 | rs11899047 | -1.18 | -0.724 | 1.514 | 1.304 | 0.019 | 0.19 | 0.19 | 6.261 | 0.07 | 0.47 | 0.1 | 0.15 | 0.01 | 0.03 |
| 2 | 77234767 | rs17013592 | -1.835 | -0.783 | 1.344 | 1.374 | -0.001 | -0.002 | -0.002 | 0 | 0.36 | 0.35 | 0.48 | 0.5 | 0.4 | 0.47 |
| 2 | 77235219 | rs1446704 | -1.517 | -0.787 | 1.157 | 1.372 | -0.051 | -0.114 | -0.114 | 0 | 0.34 | 0.11 | 0.46 | 0.5 | 0.4 | 0.47 |
| 2 | 77246331 | rs6547115 | -2.487 | -0.772 | 0.959 | 1.329 | 0.004 | 0.048 | 0.048 | 2.074 | 0.45 | 0.89 | 0.59 | 0.67 | 0.42 | 0.51 |
| 2 | 77248641 | rs17089 | -0.77 | -0.714 | 0.954 | 1.444 | -0.013 | -0.088 | -0.088 | 0 | 0.2 | 0.09 | 0.22 | 0.41 | 0.24 | 0.33 |
| 2 | 77256291 | rs10201530 | -2.5 | -0.771 | 1.072 | 1.372 | -0.019 | 0.032 | 0.032 | 2.677 | 0.45 | 0.92 | 0.59 | 0.67 | 0.42 | 0.5 |
| 2 | 77264269 | rs17334527 | 0.226 | -0.798 | 1.234 | 1.522 | -0.049 | -0.011 | -0.011 | 0 | 0.14 | 0 | 0.24 | 0.11 | 0.16 | 0.14 |
| 2 | 77264354 | rs13033902 | -0.984 | -0.654 | 1.473 | 1.599 | -0.001 | 0.011 | 0.011 | 0 | 0.21 | 0.24 | 0.22 | 0.41 | 0.24 | 0.32 |
| 2 | 77267654 | rs1348827 | -2.465 | -0.673 | 1.46 | 1.576 | 0.003 | -0.058 | -0.058 | 0.533 | 0.4 | 0.88 | 0.56 | 0.59 | 0.4 | 0.47 |
| 2 | 77270019 | rs17406465 | 0.73 | -0.563 | 1.333 | 1.73 | -0.054 | -0.036 | -0.036 | 0 | 0.11 | 0 | 0.24 | 0.14 | 0.17 | 0.08 |
| 2 | 77274083 | rs6547119 | -1.031 | -0.57 | 1.558 | 1.745 | -0.001 | 0.009 | 0.009 | 0 | 0.19 | 0.22 | 0.2 | 0.28 | 0.22 | 0.29 |
| 2 | 77284280 | rs13397044 | -1.108 | -0.614 | 1.447 | 1.684 | 0 | -0.001 | -0.001 | 0.036 | 0.2 | 0.18 | 0.2 | 0.3 | 0.22 | 0.3 |
| 2 | 77284790 | rs1446689 | -1.755 | -0.571 | 1.56 | 1.692 | -0.02 | 0.04 | 0.04 | 0.789 | 0.27 | 0.62 | 0.3 | 0.44 | 0.23 | 0.33 |
| 2 | 77287190 | rs10175781 | -1.884 | -0.627 | 1.241 | 1.435 | -0.01 | -0.001 | -0.001 | 0.725 | 0.16 | 0.65 | 0.17 | 0.18 | 0.15 | 0.1 |
| 2 | 77288585 | rs7562487 | -1.611 | -0.598 | 1.184 | 1.384 | 0.004 | 0.001 | 0.001 | 2.482 | 0.26 | 0.66 | 0.26 | 0.32 | 0.32 | 0.17 |
| 2 | 77292449 | rs13412552 | -1.832 | -1.034 | 1.168 | 1.399 | -0.002 | 0 | 0 | 0 | 0.34 | 0.67 | 0.37 | 0.66 | 0.26 | 0.42 |
| 2 | 77297536 | rs3910998 | -1.318 | -0.81 | 0.989 | 1.205 | 0.027 | 0.004 | 0.004 | 0.534 | 0.22 | 0.51 | 0.2 | 0.25 | 0.24 | 0.19 |
| 2 | 77326832 | rs3820749 | 0.435 | -0.693 | -0.165 | 1.126 | -0.057 | -0.032 | -0.032 | 0 | 0.16 | 0 | 0.09 | 0.16 | 0.18 | 0.11 |
| 2 | 77333401 | rs6547121 | -1.389 | -0.899 | 0.282 | 1.136 | -0.015 | -0.008 | -0.008 | 0 | 0.38 | 0.6 | 0.39 | 0.68 | 0.29 | 0.44 |
| 2 | 77337464 | rs7593559 | -0.517 | -0.775 | 0.127 | 0.69 | -0.036 | 0.11 | 0.11 | 0 | 0.43 | 0.17 | 0.44 | 0.28 | 0.48 | 0.31 |
| 2 | 77339370 | rs4852429 | -1.399 | -0.924 | 0.777 | 0.887 | -0.011 | 0.007 | 0.007 | 0 | 0.57 | 0.64 | 0.68 | 0.65 | 0.54 | 0.62 |
| 2 | 77339606 | rs13398218 | 0.704 | -1.017 | 0.977 | 1.142 | -0.021 | 0.074 | 0.074 | 0 | 0.08 | 0.1 | 0.14 | 0.22 | 0.04 | 0.19 |
| 2 | 77343756 | rs11901327 | -1.266 | -1.173 | 1.205 | 1.018 | -0.005 | 0.007 | 0.007 | 0 | 0.57 | 0.58 | 0.61 | 0.67 | 0.51 | 0.61 |
| 2 | 77345491 | rs1176782 | -2.216 | -0.715 | 1.585 | 0.666 | 0.005 | 0.113 | 0.113 | 1.6 | 0.09 | 0.42 | 0.17 | 0.2 | 0.04 | 0.12 |
| 2 | 77351013 | rs10186579 | 0.248 | -0.737 | 1.223 | 0.58 | 0 | 0.001 | 0.001 | 0 | 0.23 | 0.19 | 0.31 | 0.18 | 0.25 | 0.21 |
| 2 | 77351450 | rs13383149 | -1.036 | -0.589 | 1.015 | 0.6 | -0.002 | 0.013 | 0.013 | 0 | 0.47 | 0.39 | 0.5 | 0.45 | 0.48 | 0.43 |
| 2 | 77360561 | rs6547123 | -0.779 | -0.723 | 1.226 | 1.231 | -0.008 | 0.001 | 0.001 | 0 | 0.32 | 0.58 | 0.43 | 0.41 | 0.27 | 0.38 |
| 2 | 77377286 | rs4853301 | -0.644 | -0.254 | -0.524 | 1.009 | -0.02 | 0.047 | 0.047 | 0 | 0.3 | 0.43 | 0.44 | 0.49 | 0.27 | 0.4 |
| 2 | 77383715 | rs17013760 | 0.094 | 0.164 | -1.248 | 0.673 | -0.007 | 0.028 | 0.028 | 0 | 0.42 | 0.21 | 0.29 | 0.4 | 0.47 | 0.37 |
| 2 | 77387208 | rs2919058 | 0.378 | -0.444 | -1.146 | 0.382 | 0.043 | 0.07 | 0.07 | 0.942 | 0.63 | 0.17 | 0.71 | 0.68 | 0.73 | 0.63 |
| 2 | 77395150 | rs1837426 | 0.243 | -0.728 | -0.292 | 0.206 | 0 | -0.001 | -0.001 | 0.151 | 0.62 | 0.68 | 0.66 | 0.63 | 0.64 | 0.7 |
| 2 | 77404112 | rs1837425 | -1.475 | -0.957 | 0.734 | 0.231 | 0.02 | 0.056 | 0.056 | 0 | 0.07 | 0.41 | 0.05 | 0.02 | 0.01 | 0.07 |
| 2 | 77411737 | rs1030956 | 0.122 | -1.032 | 1.247 | 0.236 | -0.008 | 0.031 | 0.031 | 0 | 0.66 | 0.82 | 0.68 | 0.64 | 0.65 | 0.69 |
| 2 | 77412859 | rs17013773 | 0.077 | -1.428 | 1.34 | 1.021 | -0.017 | 0.092 | 0.092 | 4.549 | 0.08 | 0.12 | 0.03 | 0.23 | 0.04 | 0.13 |
| 2 | 77414507 | rs4369888 | 1.278 | -1.524 | 1.481 | 0.95 | 0.037 | 0.038 | 0.038 | 0 | 0.48 | 0.16 | 0.6 | 0.38 | 0.59 | 0.48 |
| 2 | 77429297 | rs11126593 | 0.001 | -1.547 | 1.714 | 1.068 | -0.025 | 0.124 | 0.124 | 5.319 | 0.08 | 0.13 | 0.06 | 0.32 | 0.04 | 0.14 |
| 2 | 77429576 | rs10865428 | -1.742 | -1.554 | 1.701 | 1.044 | -0.085 | 0.186 | 0.186 | 0.274 | 0.17 | 0.57 | 0.11 | 0.34 | 0.06 | 0.21 |
| 2 | 77437279 | rs10171377 | 1.356 | -1.525 | 1.48 | 1.007 | -0.011 | 0.059 | 0.059 | 0.128 | 0.47 | 0.07 | 0.56 | 0.31 | 0.58 | 0.47 |
| 2 | 77441127 | rs4852430 | -0.164 | -1.48 | 1.507 | 0.994 | -0.025 | 0.05 | 0.05 | 0 | 0.66 | 0.89 | 0.71 | 0.66 | 0.64 | 0.69 |
| 2 | 77464752 | rs7581365 | -2.269 | -1.269 | 2.376 | 1.189 | -0.026 | 0.325 | 0.325 | 0.447 | 0.1 | 0.38 | 0.09 | 0.33 | 0.06 | 0.22 |
| 2 | 77465662 | rs7558341 | -1.832 | -1.269 | 2.305 | 1.188 | -0.016 | 0.28 | 0.28 | 0.235 | 0.08 | 0.27 | 0.08 | 0.33 | 0.06 | 0.22 |
| 2 | 77468540 | rs6705164 | 0.588 | -1.275 | 2.494 | 1.107 | 0.002 | 0.003 | 0.003 | 0.012 | 0.54 | 0.6 | 0.63 | 0.32 | 0.58 | 0.47 |
| 2 | 77479554 | rs3943682 | -2.473 | -1.169 | 2.457 | 1.034 | -0.004 | 0.317 | 0.317 | 0.086 | 0.12 | 0.42 | 0.1 | 0.34 | 0.09 | 0.25 |
| 2 | 77479587 | rs1470076 | -0.011 | -1.161 | 2.148 | 0.98 | 0.005 | 0.065 | 0.065 | 0.063 | 0.61 | 0.83 | 0.71 | 0.66 | 0.64 | 0.7 |
| 2 | 77483276 | rs13414522 | -2.738 | -1.071 | 3.04 | 1.421 | 0.051 | 0.041 | 0.041 | 3.333 | 0.06 | 0.5 | 0.05 | 0 | 0 | 0 |
| 2 | 77493970 | rs6729161 | -2.856 | -1.028 | 2.742 | 1.185 | -0.053 | -0.033 | -0.033 | 0 | 0.12 | 0.65 | 0.07 | 0.03 | 0.02 | 0.07 |
| 2 | 77497242 | rs12611712 | -0.989 | -1.083 | 1.658 | 1.19 | 0.005 | -0.066 | -0.066 | 0.207 | 0.18 | 0.07 | 0.12 | 0.45 | 0.18 | 0.24 |
| 2 | 77501554 | rs10179301 | -1.259 | -1.601 | 2.084 | 1.131 | -0.074 | 0.057 | 0.057 | 0 | 0.15 | 0.46 | 0.11 | 0.15 | 0.04 | 0.19 |
| 2 | 77515995 | rs10180141 | -1.245 | -1.635 | 1.881 | 1.125 | -0.071 | 0.047 | 0.047 | 0 | 0.14 | 0.31 | 0.1 | 0.14 | 0.03 | 0.18 |
| 2 | 77520451 | rs13416390 | -1.406 | -1.651 | 1.869 | 1.115 | -0.081 | 0.065 | 0.065 | 0 | 0.15 | 0.42 | 0.1 | 0.14 | 0.03 | 0.18 |
| 2 | 77524377 | rs10179549 | -2.268 | -1.964 | 1.727 | 1.145 | 0.061 | -0.007 | -0.007 | 1.151 | 0.35 | 0.78 | 0.23 | 0.59 | 0.17 | 0.37 |

| | | | | | | | | | | | | | | | | |
|---|---|---|---|---|---|---|---|---|---|---|---|---|---|---|---|---|
| 2 | 77529369 | rs290047 | -1.359 | -1.639 | 1.799 | 1.282 | -0.051 | 0.051 | 0.051 | 0 | 0.15 | 0.42 | 0.13 | 0.14 | 0.05 | 0.18 |
| 2 | 77529951 | rs290046 | -1.959 | -1.594 | 1.62 | 1.236 | -0.012 | 0.001 | 0.001 | 0 | 0.32 | 0.67 | 0.25 | 0.6 | 0.22 | 0.38 |
| 2 | 77536249 | rs401204 | -1.917 | -1.564 | 1.533 | 1.267 | -0.06 | 0.026 | 0.026 | 0 | 0.3 | 0.53 | 0.23 | 0.6 | 0.22 | 0.37 |
| 2 | 77538592 | rs11126601 | -1.373 | -1.401 | 1.233 | 1.283 | -0.025 | 0.059 | 0.059 | 0 | 0.21 | 0.36 | 0.17 | 0.49 | 0.17 | 0.28 |
| 2 | 77541883 | rs12328137 | -1.179 | -0.938 | 1.012 | 1.306 | -0.002 | 0.06 | 0.06 | 0 | 0.16 | 0.22 | 0.12 | 0.42 | 0.19 | 0.15 |
| 2 | 77550421 | rs13402802 | -0.705 | -0.923 | 0.738 | 1.336 | -0.007 | -0.033 | -0.033 | 1.143 | 0.13 | 0.05 | 0.1 | 0.42 | 0.19 | 0.16 |
| 2 | 77553930 | rs290039 | -1.912 | -0.985 | 1.067 | 1.044 | -0.001 | -0.001 | -0.001 | 0 | 0.32 | 0.7 | 0.22 | 0.6 | 0.27 | 0.31 |
| 2 | 77572079 | rs1354592 | 1.685 | -1.487 | -0.192 | 0.737 | 0.053 | 0.128 | 0.128 | 0.311 | 0.57 | 0.14 | 0.69 | 0.21 | 0.65 | 0.47 |
| 2 | 77579102 | rs1505226 | 1.466 | -1.616 | 0.268 | 0.805 | 0.01 | 0.001 | 0.001 | 1.777 | 0.73 | 0.34 | 0.83 | 0.7 | 0.88 | 0.69 |
| 2 | 77580531 | rs13026667 | 0.579 | -1.594 | 0.462 | 0.674 | -0.023 | -0.033 | -0.033 | 1.989 | 0.88 | 0.81 | 0.95 | 0.94 | 0.95 | 0.92 |
| 2 | 77581832 | rs12151596 | -1.111 | -1.503 | -0.11 | 0.663 | -0.002 | 0.11 | 0.11 | 0 | 0.06 | 0.18 | 0.03 | 0.18 | 0.04 | 0.12 |
| 2 | 77583963 | rs924901 | 1.683 | -1.55 | -0.855 | 0.217 | 0.062 | 0.199 | 0.199 | 1.238 | 0.51 | 0.12 | 0.67 | 0.19 | 0.63 | 0.46 |
| 2 | 77586295 | rs398543 | 1.341 | -0.853 | -0.85 | 0.278 | 0.015 | 0.12 | 0.12 | 0.757 | 0.54 | 0.15 | 0.65 | 0.21 | 0.6 | 0.48 |
| 2 | 77589901 | rs1505229 | -0.732 | -0.578 | -0.456 | 0.398 | 0.013 | 0.019 | 0.019 | 1.286 | 0.32 | 0.46 | 0.27 | 0.62 | 0.34 | 0.38 |
| 2 | 77591743 | rs1505231 | 1.132 | -0.621 | -0.783 | 0.039 | -0.003 | 0.046 | 0.046 | 0 | 0.48 | 0.42 | 0.55 | 0.16 | 0.53 | 0.43 |
| 2 | 77596340 | rs11126606 | -0.278 | -0.252 | -1.031 | 0.83 | -0.029 | -0.06 | -0.06 | 0 | 0.25 | 0.12 | 0.25 | 0.55 | 0.37 | 0.28 |
| 2 | 77602477 | rs11693018 | -0.274 | -0.242 | -1.025 | 0.816 | -0.029 | -0.059 | -0.059 | 0 | 0.25 | 0.12 | 0.24 | 0.55 | 0.37 | 0.28 |
| 2 | 77603802 | rs2439567 | -1.67 | -0.077 | -0.89 | 0.603 | -0.004 | 0.101 | 0.101 | 0 | 0.48 | 0.3 | 0.43 | 0.85 | 0.5 | 0.53 |
| 2 | 77612121 | rs427294 | -1.52 | 0.257 | 0.092 | 1.199 | 0.007 | 0.007 | 0.007 | 0.012 | 0.35 | 0.38 | 0.36 | 0.61 | 0.43 | 0.46 |
| 2 | 77618401 | rs9309515 | -1.014 | 0.352 | 0.01 | 1.188 | -0.046 | 0.002 | 0.002 | 0.581 | 0.13 | 0.39 | 0.05 | 0.09 | 0.05 | 0.2 |
| 2 | 77627119 | rs433156 | -1.145 | 0.997 | -0.512 | 1.325 | 0.004 | 0.025 | 0.025 | 1.113 | 0.3 | 0.38 | 0.33 | 0.53 | 0.37 | 0.3 |
| 2 | 77636909 | rs393564 | -0.743 | 1.274 | -1.121 | 1.055 | -0.052 | -0.038 | -0.038 | 0 | 0.23 | 0.07 | 0.22 | 0.27 | 0.35 | 0.25 |
| 2 | 77639237 | rs424198 | -1.25 | 1.359 | -0.67 | 0.602 | 0.002 | -0.024 | -0.024 | 1.054 | 0.18 | 0.21 | 0.13 | 0.06 | 0.2 | 0.17 |
| 2 | 77651108 | rs10209045 | -0.366 | 1.014 | -0.068 | 0.361 | 0.001 | -0.016 | -0.016 | 0.74 | 0.35 | 0.4 | 0.23 | 0.18 | 0.35 | 0.4 |
| 2 | 77656160 | rs13403846 | -0.444 | 0.955 | -0.069 | 0.323 | -0.001 | -0.01 | -0.01 | 1.055 | 0.36 | 0.37 | 0.23 | 0.18 | 0.35 | 0.38 |
| 2 | 77659859 | rs10171661 | -0.514 | 0.923 | -0.003 | 0.709 | -0.001 | -0.012 | -0.012 | 1.323 | 0.37 | 0.42 | 0.24 | 0.28 | 0.35 | 0.42 |
| 2 | 77660813 | rs17751162 | -0.359 | 0.984 | -0.072 | 0.725 | -0.002 | 0.006 | 0.006 | 0 | 0.35 | 0.29 | 0.23 | 0.29 | 0.36 | 0.42 |
| 2 | 77663264 | rs1922807 | -0.622 | 1.028 | -0.713 | 0.859 | 0.002 | 0.006 | 0.006 | 0 | 0.52 | 0.71 | 0.48 | 0.42 | 0.53 | 0.55 |
| 2 | 77670072 | rs4145833 | 0.287 | 0.75 | -0.891 | 0.699 | -0.002 | -0.003 | -0.003 | 0 | 0.69 | 0.59 | 0.78 | 0.71 | 0.7 | 0.77 |
| 2 | 77683562 | rs1882681 | 0.373 | 0.523 | -1.911 | 0.674 | 0.005 | 0.004 | 0.004 | 0 | 0.66 | 0.85 | 0.74 | 0.7 | 0.67 | 0.63 |
| 2 | 77687101 | rs1922809 | -0.425 | 0.518 | -1.961 | 0.767 | 0.003 | -0.021 | -0.021 | 0 | 0.38 | 0.25 | 0.42 | 0.51 | 0.4 | 0.49 |
| 2 | 77688109 | rs12614202 | -0.851 | 0.776 | -2.346 | 0.889 | 0.034 | 0.018 | 0.018 | 0 | 0.48 | 0.29 | 0.29 | 0.62 | 0.43 | 0.63 |
| 2 | 77689016 | rs17014128 | 0.092 | 0.712 | -2.343 | 0.791 | 0.004 | -0.025 | -0.025 | 0 | 0.24 | 0.13 | 0.1 | 0.25 | 0.2 | 0.32 |
| 2 | 77689799 | rs2901847 | -1.382 | 0.879 | -1.759 | 0.472 | 0.011 | -0.029 | -0.029 | 0.01 | 0.66 | 0.48 | 0.72 | 0.78 | 0.63 | 0.83 |
| 2 | 77691733 | rs17700606 | -0.198 | 0.879 | -1.891 | 0.28 | -0.005 | 0.003 | 0.003 | 0.451 | 0.21 | 0.18 | 0.25 | 0.15 | 0.28 | 0.36 |
| 2 | 77692584 | rs2861079 | -0.71 | 0.362 | -2.354 | 0.201 | -0.014 | -0.002 | -0.002 | 0 | 0.54 | 0.27 | 0.51 | 0.53 | 0.54 | 0.73 |
| 2 | 77695237 | rs12468802 | 1.129 | 0.243 | -2.172 | 0.267 | -0.004 | -0.002 | -0.002 | 0.041 | 0.06 | 0.08 | 0.22 | 0.03 | 0.04 | 0.03 |
| 2 | 77695496 | rs1922806 | 0.004 | 0.148 | -1.506 | 0.228 | 0.022 | -0.015 | -0.015 | 0.725 | 0.4 | 0.3 | 0.49 | 0.42 | 0.31 | 0.4 |
| 2 | 77697995 | rs2861078 | 0.267 | 0.028 | -1.249 | 0.125 | -0.008 | 0.01 | 0.01 | 0 | 0.62 | 0.66 | 0.49 | 0.75 | 0.57 | 0.65 |
| 2 | 77698670 | rs1922824 | 1.17 | 0.197 | -1.154 | 0.204 | 0.009 | 0.011 | 0.011 | 0 | 0.27 | 0.4 | 0.22 | 0.36 | 0.3 | 0.28 |
| 2 | 77705469 | rs12713914 | -0.309 | 0.155 | -1.35 | -0.126 | 0.008 | -0.004 | -0.004 | 0 | 0.9 | 0.92 | 0.87 | 0.86 | 0.93 | 0.91 |
| 2 | 77711560 | rs12997025 | 0.711 | 0.01 | -1.684 | -0.601 | 0.012 | 0.095 | 0.095 | 0 | 0.27 | 0.13 | 0.22 | 0.05 | 0.27 | 0.29 |
| 2 | 77721499 | rs17648731 | 1.048 | 0.277 | -1.847 | -1.112 | -0.001 | -0.013 | -0.013 | 0 | 0.12 | 0.09 | 0.1 | 0.02 | 0.21 | 0.18 |
| 2 | 77722736 | rs11126614 | 1.05 | 0.261 | -1.716 | -1.148 | 0.013 | 0.06 | 0.06 | 0 | 0.27 | 0.16 | 0.21 | 0.05 | 0.27 | 0.29 |
| 2 | 77727037 | rs9309516 | -1.028 | 0.138 | -1.687 | -1.082 | 0.008 | -0.006 | -0.006 | 0 | 0.91 | 0.93 | 0.88 | 0.86 | 0.93 | 0.91 |
| 2 | 77729748 | rs10206763 | 1.113 | -0.095 | -1.349 | -0.821 | 0.007 | -0.031 | -0.031 | 0 | 0.4 | 0.51 | 0.46 | 0.26 | 0.41 | 0.23 |
| 2 | 77730541 | rs1344255 | 0.977 | 0.053 | -1.441 | -0.746 | -0.001 | 0.011 | 0.011 | 0.193 | 0.65 | 0.74 | 0.49 | 0.28 | 0.66 | 0.51 |
| 2 | 77734812 | rs1922813 | -1.138 | 0.005 | -1.748 | -0.655 | 0.01 | -0.02 | -0.02 | 0 | 0.91 | 0.97 | 0.88 | 0.86 | 0.93 | 0.92 |
| 2 | 77735664 | rs689553 | 0.317 | 0.214 | -1.707 | -0.692 | -0.029 | 0.195 | 0.195 | 0.003 | 0.38 | 0.18 | 0.27 | 0.05 | 0.39 | 0.34 |
| 2 | 77736014 | rs17014206 | -0.087 | 0.419 | -1.334 | -0.126 | 0.004 | 0.037 | 0.037 | 0.071 | 0.09 | 0.07 | 0.1 | 0.53 | 0.05 | 0.2 |
| 2 | 77742249 | rs10445951 | 1.475 | 0.643 | -0.794 | -0.523 | 0.007 | -0.005 | -0.005 | 0 | 0.24 | 0.3 | 0.2 | 0.22 | 0.29 | 0.18 |
| 2 | 77747669 | rs11677111 | 1.143 | 0.317 | -1.105 | -0.672 | -0.012 | 0.042 | 0.042 | 0 | 0.11 | 0.05 | 0.04 | 0 | 0.08 | 0.03 |
| 2 | 77750482 | rs2901848 | 1.386 | 0.712 | -0.855 | -0.597 | -0.005 | 0.002 | 0.002 | 0 | 0.22 | 0.2 | 0.22 | 0.2 | 0.28 | 0.21 |
| 2 | 77754710 | rs2178759 | -0.413 | 0.099 | -0.729 | -1.316 | -0.002 | -0.041 | -0.041 | 0 | 0.62 | 0.68 | 0.73 | 0.93 | 0.68 | 0.75 |
| 2 | 77755115 | rs17014234 | 1.474 | -0.045 | -0.781 | -0.843 | -0.004 | -0.011 | -0.011 | 0 | 0.27 | 0.26 | 0.23 | 0.38 | 0.28 | 0.35 |
| 2 | 77755436 | rs1922818 | 1.419 | -0.065 | -0.524 | -0.852 | 0.002 | 0.011 | 0.011 | 0 | 0.27 | 0.36 | 0.23 | 0.38 | 0.28 | 0.35 |
| 2 | 77757132 | rs2861049 | 1.423 | -0.078 | -0.531 | -0.859 | 0.002 | 0.012 | 0.012 | 0 | 0.27 | 0.36 | 0.23 | 0.38 | 0.28 | 0.35 |
| 2 | 77760819 | rs2141938 | -0.097 | -0.076 | -0.624 | -0.854 | -0.015 | -0.021 | -0.021 | 0 | 0.58 | 0.53 | 0.72 | 0.75 | 0.68 | 0.63 |
| 2 | 77763824 | rs13027448 | 0.812 | 0.161 | -0.996 | -0.697 | -0.109 | -0.151 | -0.151 | 0.037 | 0.22 | 0.03 | 0.23 | 0.39 | 0.28 | 0.31 |
| 2 | 77772143 | rs6717256 | 1.231 | -0.108 | -1.017 | -0.989 | -0.004 | 0 | 0 | 0 | 0.09 | 0.06 | 0.11 | 0.01 | 0.13 | 0.07 |
| 2 | 77774970 | rs9789540 | -0.099 | -0.031 | -0.903 | -0.845 | -0.066 | -0.09 | -0.09 | 0 | 0.32 | 0.16 | 0.49 | 0.48 | 0.4 | 0.38 |
| 2 | 77779249 | rs12713918 | 0.04 | -0.147 | -0.822 | -1.002 | -0.011 | -0.047 | -0.047 | 0 | 0.56 | 0.54 | 0.72 | 0.86 | 0.65 | 0.68 |
| 2 | 77780037 | rs2901849 | -0.469 | -0.025 | -1.075 | -0.929 | -0.013 | 0.008 | 0.008 | 0 | 0.35 | 0.34 | 0.51 | 0.66 | 0.39 | 0.48 |
| 2 | 77780227 | rs13021967 | 2.026 | -0.371 | -1.467 | -1.159 | 0.019 | 0.02 | 0.02 | 0 | 0.16 | 0.08 | 0.12 | 0 | 0.14 | 0.06 |
| 2 | 77780499 | rs6547150 | 0.106 | -0.016 | -1.181 | -0.789 | -0.011 | -0.031 | -0.031 | 0.063 | 0.74 | 0.74 | 0.85 | 0.86 | 0.8 | 0.76 |
| 2 | 77781489 | rs6547152 | 0.396 | 0.371 | -1.432 | -0.68 | -0.001 | 0 | 0 | 0.087 | 0.78 | 0.74 | 0.88 | 0.73 | 0.81 | 0.8 |
| 2 | 77781935 | rs17014308 | -0.344 | 0.315 | -1.407 | -0.681 | -0.016 | -0.043 | -0.043 | 0 | 0.32 | 0.21 | 0.31 | 0.45 | 0.34 | 0.48 |
| 2 | 77783184 | rs11895756 | 0.551 | 0.447 | -1.284 | -0.505 | 0.018 | 0.086 | 0.086 | 0 | 0.07 | 0.21 | 0.04 | 0.21 | 0.06 | 0.11 |
| 2 | 77783335 | rs6740463 | 1.151 | 0.485 | -1.503 | -0.486 | 0.003 | 0.008 | 0.008 | 0 | 0.47 | 0.52 | 0.57 | 0.27 | 0.46 | 0.32 |

| | | | | | | | | | | | | | | | | |
|---|---|---|---|---|---|---|---|---|---|---|---|---|---|---|---|---|
| 2 | 77783575 | rs2901850 | 0.532 | 0.929 | -1.806 | 0.074 | -0.063 | -0.099 | -0.099 | 0 | 0.13 | 0.01 | 0.18 | 0.39 | 0.18 | 0.28 |
| 2 | 77786252 | rs6711151 | -0.876 | 0.886 | -1.783 | -0.182 | -0.045 | -0.013 | -0.013 | 2.324 | 0.37 | 0.23 | 0.53 | 0.58 | 0.46 | 0.42 |
| 2 | 77786509 | rs6724806 | -0.526 | 0.931 | -1.719 | -0.157 | -0.023 | -0.062 | -0.062 | 0 | 0.3 | 0.17 | 0.26 | 0.49 | 0.35 | 0.37 |
| 2 | 77786742 | rs6547155 | -0.736 | 1.088 | -1.623 | -0.462 | -0.008 | 0.023 | 0.023 | 0 | 0.69 | 0.77 | 0.75 | 0.81 | 0.68 | 0.8 |
| 2 | 77787896 | rs4473409 | -0.77 | 0.686 | -1.578 | -0.381 | -0.029 | 0.047 | 0.047 | 0 | 0.64 | 0.77 | 0.71 | 0.8 | 0.61 | 0.79 |
| 2 | 77788869 | rs4853318 | 0.787 | 0.663 | -1.751 | -0.156 | -0.013 | -0.017 | -0.017 | 0 | 0.06 | 0.02 | 0.25 | 0.08 | 0.07 | 0.05 |
| 2 | 77797064 | rs7566976 | -0.917 | 0.493 | -1.537 | -0.34 | -0.008 | 0.022 | 0.022 | 0.003 | 0.7 | 0.78 | 0.75 | 0.84 | 0.71 | 0.82 |
| 2 | 77805397 | rs11677059 | -0.694 | 0.323 | -2.207 | -0.494 | -0.131 | 0.195 | 0.195 | 0.732 | 0.6 | 0.96 | 0.67 | 0.89 | 0.57 | 0.79 |
| 2 | 77826046 | rs13412598 | 0.975 | 0.599 | -2.076 | 0.202 | -0.03 | -0.061 | -0.061 | 0 | 0.11 | 0.03 | 0.16 | 0.39 | 0.14 | 0.3 |
| 2 | 77826374 | rs7592577 | 1.186 | 0.373 | -1.917 | -0.047 | 0 | -0.001 | -0.001 | 1.701 | 0.09 | 0.11 | 0.05 | 0.06 | 0.07 | 0.06 |
| 2 | 77833759 | rs12469613 | 0.228 | 0.491 | -1.698 | 0.069 | -0.007 | -0.011 | -0.011 | 0 | 0.17 | 0.15 | 0.43 | 0.48 | 0.21 | 0.37 |
| 2 | 77836798 | rs13393843 | -0.478 | 0.263 | -1.341 | -0.173 | 0.002 | 0.005 | 0.005 | 0 | 0.31 | 0.34 | 0.5 | 0.53 | 0.37 | 0.41 |
| 2 | 77848881 | rs4490221 | -1.788 | 0.108 | -1.222 | -0.552 | -0.001 | -0.002 | -0.002 | 0 | 0.68 | 0.71 | 0.7 | 0.85 | 0.67 | 0.8 |
| 2 | 77855707 | rs10865430 | -0.3 | 0.209 | -1.7 | -0.259 | 0.037 | 0.01 | 0.01 | 0 | 0.34 | 0.17 | 0.19 | 0.34 | 0.3 | 0.42 |
| 2 | 77856415 | rs12052760 | 0.28 | 0.567 | -1.637 | 0.166 | 0.019 | -0.017 | -0.017 | 0.008 | 0.2 | 0.08 | 0.13 | 0.24 | 0.21 | 0.34 |
| 2 | 77857564 | rs4337498 | 0.119 | 1.081 | -1.082 | 0.41 | -0.004 | 0 | 0 | 0 | 0.16 | 0.1 | 0.19 | 0.42 | 0.19 | 0.31 |
| 2 | 77857764 | rs4485592 | -0.956 | 0.841 | -0.934 | -0.011 | 0.004 | -0.004 | -0.004 | 0 | 0.13 | 0.17 | 0.3 | 0.08 | 0.17 | 0.07 |
| 2 | 77879421 | rs11685470 | -0.851 | 0.572 | -0.859 | -0.115 | 0.005 | -0.044 | -0.044 | 3.02 | 0.12 | 0.18 | 0.1 | 0 | 0.15 | 0.01 |
| 2 | 77889872 | rs6707409 | -0.426 | -0.026 | -1.471 | 0.28 | 0.02 | -0.059 | -0.059 | 0 | 0.27 | 0.08 | 0.33 | 0.35 | 0.22 | 0.37 |
| 2 | 77892612 | rs4513323 | 0.128 | -0.57 | -1.13 | -0.66 | -0.037 | -0.149 | -0.149 | 1.573 | 0.36 | 0.49 | 0.17 | 0.02 | 0.2 | 0.25 |
| 2 | 77893810 | rs10190071 | -0.98 | -0.484 | -0.912 | -0.573 | 0.006 | -0.005 | -0.005 | 0 | 0.61 | 0.51 | 0.49 | 0.36 | 0.42 | 0.61 |
| 2 | 77918355 | rs10185229 | -0.564 | -0.526 | -0.344 | -0.803 | -0.025 | -0.134 | -0.134 | 2.488 | 0.39 | 0.51 | 0.17 | 0.02 | 0.23 | 0.25 |
| 2 | 77923927 | rs4539817 | -0.628 | -0.132 | -0.206 | 0.371 | -0.012 | 0.171 | 0.171 | 0 | 0.12 | 0.22 | 0.27 | 0.46 | 0.08 | 0.17 |
| 2 | 77928314 | rs7598757 | -1.039 | -0.283 | -0.217 | 0.015 | 0 | 0.061 | 0.061 | 0 | 0.31 | 0.47 | 0.39 | 0.52 | 0.27 | 0.42 |
| 2 | 77936605 | rs1487033 | 0.478 | 0.097 | -0.687 | -0.264 | -0.014 | 0.025 | 0.025 | 0.094 | 0.4 | 0.19 | 0.35 | 0.31 | 0.42 | 0.47 |
| 2 | 77938924 | rs17014417 | 0.062 | 0.006 | -0.474 | -0.141 | -0.005 | 0.193 | 0.193 | 0 | 0.11 | 0.21 | 0.26 | 0.46 | 0.08 | 0.16 |
| 2 | 77940273 | rs729756 | -1.454 | 0.022 | -0.418 | -0.387 | 0.01 | -0.039 | -0.039 | 0 | 0.73 | 0.6 | 0.73 | 0.83 | 0.7 | 0.88 |
| 2 | 77943385 | rs1542678 | 0.598 | -0.091 | -0.926 | -0.147 | -0.058 | -0.03 | -0.03 | 0 | 0.4 | 0.09 | 0.36 | 0.4 | 0.44 | 0.5 |
| 2 | 77944581 | rs11126631 | 0.468 | -0.085 | -0.876 | -0.144 | -0.037 | -0.016 | -0.016 | 0 | 0.4 | 0.14 | 0.36 | 0.39 | 0.43 | 0.48 |
| 2 | 77946175 | rs10520195 | 0.646 | -0.187 | -0.877 | -0.293 | 0.003 | -0.009 | -0.009 | 0.012 | 0.12 | 0.12 | 0.06 | 0 | 0.1 | 0.17 |
| 2 | 77947122 | rs10169036 | 0.089 | -0.101 | -0.825 | -0.194 | -0.017 | 0.019 | 0.019 | 0 | 0.47 | 0.24 | 0.42 | 0.45 | 0.5 | 0.56 |
| 2 | 77947988 | rs11676317 | -0.224 | 0.022 | -0.46 | 0.073 | -0.007 | -0.005 | -0.005 | 4.036 | 0.54 | 0.43 | 0.43 | 0.53 | 0.5 | 0.57 |
| 2 | 77955068 | rs2129495 | -1.063 | 0.377 | 0.243 | 0.393 | 0.032 | 0.247 | 0.247 | 1.762 | 0.13 | 0.43 | 0.29 | 0.39 | 0.12 | 0.15 |
| 2 | 77963782 | rs7591531 | -0.518 | 0.376 | 0.228 | 0.421 | 0.024 | 0.176 | 0.176 | 0 | 0.11 | 0.25 | 0.28 | 0.38 | 0.12 | 0.12 |
| 2 | 77965325 | rs6547158 | -0.169 | 0.252 | -0.319 | -0.059 | 0.003 | 0 | 0 | 0.093 | 0.46 | 0.54 | 0.35 | 0.51 | 0.41 | 0.67 |
| 2 | 77965521 | rs6729102 | -0.162 | 0.247 | -0.322 | -0.062 | 0.003 | 0 | 0 | 0 | 0.46 | 0.53 | 0.35 | 0.51 | 0.41 | 0.67 |
| 2 | 77977262 | rs4538235 | -1.308 | 0.287 | -0.289 | 0.022 | 0.002 | 0.116 | 0.116 | 0.189 | 0.21 | 0.5 | 0.34 | 0.35 | 0.18 | 0.2 |
| 2 | 77979650 | rs6722155 | 0.993 | 0.248 | -0.556 | 0.022 | 0.003 | 0.011 | 0.011 | 0 | 0.08 | 0.05 | 0.04 | 0 | 0.07 | 0.08 |
| 2 | 77989607 | rs1487036 | -0.476 | 0.264 | -0.312 | 0.122 | 0 | 0.126 | 0.126 | 0 | 0.09 | 0.18 | 0.26 | 0.3 | 0.1 | 0.1 |
| 2 | 77990389 | rs13410776 | 0.802 | 0.257 | -0.514 | 0.028 | -0.017 | -0.086 | -0.086 | 0 | 0.28 | 0.12 | 0.25 | 0.5 | 0.27 | 0.49 |
| 2 | 77990617 | rs4268948 | 0.634 | 0.3 | -0.425 | 0.049 | 0.005 | -0.029 | -0.029 | 0 | 0.31 | 0.24 | 0.26 | 0.5 | 0.27 | 0.49 |
| 2 | 78001566 | rs992351 | -0.215 | 0.453 | -0.314 | 0.011 | -0.001 | 0.007 | 0.007 | 0 | 0.5 | 0.37 | 0.4 | 0.54 | 0.5 | 0.69 |
| 2 | 78005594 | rs1117351 | -0.611 | 0.423 | -0.291 | -0.024 | 0.001 | -0.038 | -0.038 | 0.091 | 0.55 | 0.5 | 0.62 | 0.86 | 0.53 | 0.8 |
| 2 | 78006805 | rs2087144 | -0.434 | 0.429 | -0.185 | 0.067 | 0 | 0.098 | 0.098 | 0 | 0.1 | 0.16 | 0.26 | 0.31 | 0.11 | 0.09 |
| 2 | 78017869 | rs7564473 | -0.358 | 0.329 | -0.119 | -0.088 | 0 | 0.002 | 0.002 | 0 | 0.51 | 0.58 | 0.6 | 0.86 | 0.48 | 0.76 |
| 2 | 78018076 | rs1386382 | 0.408 | 0.335 | -0.14 | -0.083 | -0.002 | -0.016 | -0.016 | 0 | 0.38 | 0.35 | 0.34 | 0.53 | 0.37 | 0.59 |
| 2 | 78025242 | rs10177440 | 0.103 | 0.439 | 0.076 | 0.034 | -0.002 | 0 | 0 | 0 | 0.48 | 0.46 | 0.38 | 0.53 | 0.46 | 0.66 |
| 2 | 78027063 | rs1586422 | -0.873 | 0.408 | 0.159 | 0.023 | -0.002 | 0.044 | 0.044 | 0 | 0.61 | 0.75 | 0.66 | 0.86 | 0.59 | 0.84 |
| 2 | 78027298 | rs1586421 | -0.84 | 0.426 | 0.043 | 0.031 | 0 | 0.014 | 0.014 | 0 | 0.6 | 0.7 | 0.65 | 0.85 | 0.58 | 0.77 |
| 2 | 78028556 | rs2035620 | -1.509 | 0.471 | -0.043 | 0.554 | -0.009 | 0.09 | 0.09 | 0 | 0.13 | 0.25 | 0.27 | 0.34 | 0.11 | 0.12 |
| 2 | 78034230 | rs12471971 | -0.057 | 0.625 | -0.282 | 0.075 | 0 | -0.004 | -0.004 | 0 | 0.07 | 0.07 | 0.04 | 0.01 | 0.07 | 0.02 |
| 2 | 78039666 | rs4575747 | -0.198 | 0.837 | -0.298 | -0.21 | -0.002 | 0.001 | 0.001 | 0 | 0.68 | 0.7 | 0.66 | 0.88 | 0.64 | 0.88 |
| 2 | 78042083 | rs6726826 | -0.924 | 1.057 | -0.619 | -0.033 | -0.117 | 0.035 | 0.035 | 0 | 0.77 | 0.98 | 0.74 | 0.91 | 0.66 | 0.9 |
| 2 | 78046515 | rs2861216 | 0.413 | 0.828 | -0.251 | 0.114 | 0 | -0.001 | -0.001 | 0 | 0.43 | 0.44 | 0.39 | 0.49 | 0.43 | 0.61 |
| 2 | 78046792 | rs2861215 | 0.444 | 0.828 | -0.263 | 0.103 | 0 | -0.001 | -0.001 | 0.563 | 0.42 | 0.44 | 0.38 | 0.49 | 0.42 | 0.61 |
| 2 | 78048788 | rs12465947 | 0.411 | 0.828 | -0.106 | 0.097 | -0.002 | -0.002 | -0.002 | 0 | 0.43 | 0.47 | 0.39 | 0.49 | 0.41 | 0.61 |
| 2 | 78061074 | rs992870 | 0.305 | 0.665 | -0.393 | 0.183 | -0.007 | -0.005 | -0.005 | 0 | 0.48 | 0.23 | 0.44 | 0.52 | 0.53 | 0.61 |
| 2 | 78062898 | rs2171326 | -0.417 | 1.339 | 0.059 | 0.4 | -0.008 | 0.01 | 0.01 | 0 | 0.75 | 0.79 | 0.76 | 0.92 | 0.7 | 0.88 |
| 2 | 78065404 | rs1487044 | -0.469 | 1.261 | -0.192 | 0.413 | 0 | -0.014 | -0.014 | 0 | 0.76 | 0.76 | 0.75 | 0.94 | 0.71 | 0.89 |
| 2 | 78074314 | rs6742440 | 0.051 | 0.89 | -0.558 | 0.662 | -0.018 | 0.096 | 0.096 | 0 | 0.19 | 0.3 | 0.35 | 0.4 | 0.15 | 0.2 |
| 2 | 78075083 | rs7594767 | 0.114 | 1.09 | -1.383 | 0.684 | -0.01 | 0.001 | 0.001 | 0.017 | 0.52 | 0.23 | 0.39 | 0.51 | 0.49 | 0.63 |
| 2 | 78079485 | rs7557384 | 0.322 | 1.15 | -0.986 | 1.02 | -0.006 | -0.011 | -0.011 | 0.036 | 0.61 | 0.31 | 0.4 | 0.5 | 0.58 | 0.51 |
| 2 | 78079746 | rs11899064 | -0.519 | 1.332 | -0.417 | 1.1 | 0.018 | 0.03 | 0.03 | 0 | 0.33 | 0.56 | 0.43 | 0.44 | 0.39 | 0.38 |
| 2 | 78092512 | rs6716793 | -0.8 | 0.923 | -0.554 | 0.977 | -0.035 | 0.003 | 0.003 | 0 | 0.89 | 0.99 | 0.77 | 0.96 | 0.85 | 0.93 |
| 2 | 78093107 | rs11126639 | -0.533 | 0.84 | 0.336 | 0.929 | 0.038 | 0.002 | 0.002 | 0.583 | 0.06 | 0.27 | 0.04 | 0.01 | 0.04 | 0.06 |
| 2 | 78096219 | rs13398639 | -1.21 | 0.777 | 0.638 | 0.966 | 0.05 | 0.138 | 0.138 | 0.36 | 0.08 | 0.34 | 0.14 | 0.15 | 0.08 | 0.1 |
| 2 | 78100362 | rs13387483 | -1.512 | 0.699 | 0.994 | 1.095 | 0.048 | 0.231 | 0.231 | 1.847 | 0.09 | 0.46 | 0.15 | 0.22 | 0.08 | 0.1 |
| 2 | 78102968 | rs6751378 | -0.631 | 0.617 | 0.429 | 0.656 | 0.011 | 0 | 0 | 6.441 | 0.38 | 0.71 | 0.49 | 0.56 | 0.37 | 0.53 |
| 2 | 78111574 | rs12713927 | 0.326 | 0.7 | 0.593 | 0.306 | 0.004 | 0.026 | 0.026 | 0 | 0.83 | 0.71 | 0.7 | 0.8 | 0.78 | 0.86 |

| | | | | | | | | | | | | | | | |
|---|---|---|---|---|---|---|---|---|---|---|---|---|---|---|---|
| 2 | 78114852 | rs13022252 | 0.897 | 0.491 | -0.3 | -0.773 | 0.015 | 0.058 | 0.058 | 0 | 0.23 | 0.09 | 0.11 | 0 | 0.16 | 0.07 |
| 2 | 78114950 | rs13021944 | 0.266 | 0.47 | 0.228 | -0.732 | -0.005 | -0.051 | -0.051 | 0.167 | 0.29 | 0.36 | 0.16 | 0.04 | 0.24 | 0.15 |
| 2 | 78120305 | rs1897616 | 0.716 | 0.62 | -0.271 | -1.062 | -0.001 | 0.062 | 0.062 | 0 | 0.39 | 0.23 | 0.51 | 0.84 | 0.42 | 0.68 |
| 2 | 78123113 | rs7570807 | 0.794 | 0.718 | -0.993 | -1.042 | 0.006 | 0.021 | 0.021 | 0 | 0.28 | 0.22 | 0.15 | 0.02 | 0.25 | 0.12 |
| 2 | 78124509 | rs1836032 | -1.015 | 1.034 | -0.954 | -0.388 | -0.017 | -0.014 | -0.014 | 0 | 0.88 | 0.98 | 0.82 | 0.92 | 0.85 | 0.92 |
| 2 | 78124975 | rs6719192 | 0.216 | 1.074 | -0.941 | -0.473 | -0.04 | 0.028 | 0.028 | 0 | 0.78 | 0.89 | 0.73 | 0.89 | 0.72 | 0.87 |
| 2 | 78125034 | rs6719220 | -0.848 | 0.665 | -0.827 | -0.238 | -0.028 | -0.024 | -0.024 | 0 | 0.88 | 0.99 | 0.82 | 0.92 | 0.85 | 0.92 |
| 2 | 78126898 | rs10192753 | -0.657 | 0.626 | -0.263 | -0.121 | 0 | -0.034 | -0.034 | 0 | 0.08 | 0.11 | 0.04 | 0.02 | 0.05 | 0.03 |
| 2 | 78132296 | rs17597326 | -0.775 | 0.671 | -0.458 | -0.073 | -0.001 | -0.009 | -0.009 | 0 | 0.07 | 0.09 | 0.04 | 0.02 | 0.04 | 0.03 |
| 2 | 78133560 | rs7602889 | 0.003 | 1.177 | -0.75 | -0.241 | -0.042 | 0.005 | 0.005 | 0 | 0.76 | 0.86 | 0.68 | 0.83 | 0.67 | 0.75 |
| 2 | 78145340 | rs1433197 | 0.062 | 1.245 | -0.887 | -0.355 | -0.066 | 0.024 | 0.024 | 0 | 0.74 | 0.88 | 0.71 | 0.85 | 0.66 | 0.75 |
| 2 | 78147175 | rs1405580 | -1.012 | 0.941 | -0.621 | -0.478 | -0.007 | -0.021 | -0.021 | 0 | 0.09 | 0.17 | 0.05 | 0.03 | 0.04 | 0.05 |
| 2 | 78170534 | rs1368502 | 0.011 | 1.047 | -0.599 | -0.496 | -0.032 | 0.023 | 0.023 | 0 | 0.74 | 0.87 | 0.74 | 0.85 | 0.68 | 0.76 |
| 2 | 78174003 | rs1405584 | 0.752 | 0.947 | 0.045 | -0.538 | 0.004 | -0.012 | -0.012 | 0 | 0.65 | 0.59 | 0.68 | 0.82 | 0.64 | 0.71 |
| 2 | 78178739 | rs6757727 | 0.759 | 0.937 | 0.04 | -0.543 | 0.003 | -0.01 | -0.01 | 0 | 0.65 | 0.59 | 0.68 | 0.82 | 0.64 | 0.71 |
| 2 | 78185777 | rs6715299 | 0.872 | 0.469 | -0.242 | -0.272 | -0.013 | -0.018 | -0.018 | 0 | 0.28 | 0.19 | 0.4 | 0.32 | 0.34 | 0.42 |
| 2 | 78190318 | rs961371 | -0.103 | 0.488 | -0.32 | -0.38 | -0.039 | 0.029 | 0.029 | 0 | 0.7 | 0.8 | 0.71 | 0.83 | 0.65 | 0.74 |
| 2 | 78214766 | rs1609077 | -1.417 | 0.715 | -0.541 | -0.409 | -0.012 | 0.008 | 0.008 | 0.413 | 0.86 | 0.92 | 0.83 | 0.9 | 0.85 | 0.91 |
| 2 | 78234004 | rs13015404 | 1.016 | 0.756 | -0.453 | -0.393 | -0.019 | 0.005 | 0.005 | 0 | 0.12 | 0.06 | 0.1 | 0.06 | 0.16 | 0.15 |
| 2 | 78240557 | rs10195907 | 1.939 | 0.253 | -0.267 | -0.865 | 0.006 | -0.016 | -0.016 | 0 | 0.23 | 0.16 | 0.12 | 0 | 0.18 | 0.11 |
| 2 | 78245876 | rs12328912 | 1.079 | 0.281 | -0.23 | -0.824 | -0.008 | 0.001 | 0.001 | 0 | 0.13 | 0.1 | 0.1 | 0.07 | 0.17 | 0.15 |
| 2 | 78251191 | rs9973347 | 0.824 | 0.265 | -0.133 | -0.831 | 0.01 | -0.005 | -0.005 | 0.369 | 0.14 | 0.21 | 0.12 | 0.07 | 0.17 | 0.15 |
| 2 | 78253852 | rs1918253 | 0.452 | 0.268 | 0.135 | -0.856 | 0.015 | -0.04 | -0.04 | 0 | 0.63 | 0.48 | 0.67 | 0.82 | 0.64 | 0.72 |
| 2 | 78254270 | rs10207002 | -0.388 | 0.289 | 0.231 | -0.839 | 0.037 | -0.054 | -0.054 | 0 | 0.22 | 0.5 | 0.17 | 0.09 | 0.21 | 0.18 |
| 2 | 78256829 | rs10178085 | -0.025 | 0.267 | 0.101 | -0.814 | -0.001 | 0.069 | 0.069 | 0 | 0.46 | 0.66 | 0.37 | 0.09 | 0.4 | 0.29 |
| 2 | 78259923 | rs12463423 | 0.595 | 0.349 | -0.201 | -0.672 | 0.015 | 0.037 | 0.037 | 0 | 0.3 | 0.48 | 0.51 | 0.38 | 0.32 | 0.42 |
| 2 | 78260079 | rs12713938 | 0.07 | 0.298 | -0.262 | -0.721 | 0 | 0.001 | 0.001 | 0 | 0.55 | 0.44 | 0.32 | 0.53 | 0.53 | 0.49 |
| 2 | 78262099 | rs1405588 | 0.169 | 0.257 | -0.281 | -0.753 | 0.012 | 0.012 | 0.012 | 0 | 0.35 | 0.48 | 0.53 | 0.38 | 0.38 | 0.44 |
| 2 | 78262379 | rs1918256 | 1.947 | 0.288 | -0.308 | -0.858 | 0.004 | -0.025 | -0.025 | 3.933 | 0.23 | 0.17 | 0.12 | 0 | 0.18 | 0.11 |
| 2 | 78265453 | rs1543282 | -0.85 | 0.303 | -0.237 | -0.824 | 0.018 | 0.071 | 0.071 | 2.151 | 0.59 | 0.73 | 0.71 | 0.9 | 0.67 | 0.8 |
| 2 | 78265948 | rs1405587 | -0.815 | 0.318 | -0.235 | -0.8 | 0.011 | 0.035 | 0.035 | 1.79 | 0.58 | 0.66 | 0.7 | 0.89 | 0.67 | 0.79 |
| 2 | 78271797 | rs1527186 | -0.051 | 0.491 | -0.231 | -0.759 | 0.012 | -0.056 | -0.056 | 0 | 0.74 | 0.53 | 0.63 | 0.86 | 0.76 | 0.86 |
| 2 | 78280763 | rs6735096 | 2.173 | 0.328 | -0.303 | -0.914 | 0.01 | 0.014 | 0.014 | 0 | 0.23 | 0.13 | 0.12 | 0 | 0.18 | 0.1 |
| 2 | 78280821 | rs7584763 | 0.884 | 0.277 | -0.613 | -0.781 | -0.02 | -0.119 | -0.119 | 1.075 | 0.16 | 0.01 | 0.1 | 0.45 | 0.17 | 0.25 |
| 2 | 78284331 | rs4853367 | 0.852 | 0.36 | -0.577 | -0.826 | -0.047 | -0.057 | -0.057 | 0 | 0.29 | 0.08 | 0.36 | 0.35 | 0.36 | 0.42 |
| 2 | 78287339 | rs6721524 | -0.38 | 0.479 | -0.361 | -0.9 | 0.072 | 0.017 | 0.017 | 0.049 | 0.09 | 0.43 | 0.17 | 0.04 | 0.08 | 0.05 |
| 2 | 78292782 | rs13022668 | 0.554 | 0.743 | -0.289 | -0.666 | 0 | -0.008 | -0.008 | 0 | 0.58 | 0.51 | 0.45 | 0.21 | 0.56 | 0.38 |
| 2 | 78293138 | rs12713943 | 1.435 | 0.446 | -0.423 | -0.777 | 0 | -0.028 | -0.028 | 0 | 0.29 | 0.29 | 0.18 | 0.07 | 0.29 | 0.19 |
| 2 | 78294642 | rs17014915 | -0.464 | 0.531 | -0.294 | -0.471 | -0.009 | 0.077 | 0.077 | 0.164 | 0.25 | 0.39 | 0.27 | 0.55 | 0.25 | 0.33 |
| 2 | 78296074 | rs17014922 | -0.35 | 0.526 | -0.305 | -0.473 | -0.006 | 0.065 | 0.065 | 0.114 | 0.24 | 0.37 | 0.27 | 0.55 | 0.24 | 0.33 |
| 2 | 78296145 | rs4853368 | 0.881 | 0.553 | -0.569 | -0.519 | 0 | -0.013 | -0.013 | 4.005 | 0.54 | 0.46 | 0.42 | 0.21 | 0.51 | 0.37 |
| 2 | 78298989 | rs2033751 | -1.105 | 0.636 | -1.068 | -0.425 | -0.069 | -0.069 | -0.069 | 0 | 0.8 | 0.97 | 0.7 | 0.76 | 0.74 | 0.69 |
| 2 | 78300998 | rs13018490 | 1.403 | 0.588 | -0.893 | -0.443 | 0.024 | 0 | 0 | 0 | 0.08 | 0.16 | 0.06 | 0.06 | 0.1 | 0.11 |
| 2 | 78301575 | rs13388243 | 1.502 | 0.4 | -1.009 | -0.512 | 0.023 | 0.126 | 0.126 | 0 | 0.44 | 0.21 | 0.35 | 0.16 | 0.39 | 0.25 |
| 2 | 78301879 | rs10182886 | 0.6 | 0.209 | -0.882 | -0.289 | 0.019 | -0.071 | -0.071 | 0 | 0.14 | 0.04 | 0.08 | 0.49 | 0.13 | 0.25 |
| 2 | 78304795 | rs11126651 | -0.217 | 0.479 | -0.38 | -0.428 | 0.021 | -0.002 | -0.002 | 0 | 0.13 | 0.18 | 0.15 | 0.12 | 0.19 | 0.21 |
| 2 | 78307412 | rs1370639 | -0.678 | 0.335 | -0.653 | -0.56 | -0.011 | 0.068 | 0.068 | 0.09 | 0.61 | 0.84 | 0.67 | 0.77 | 0.63 | 0.71 |
| 2 | 78308569 | rs7572214 | -0.784 | 0.315 | -0.361 | -0.577 | 0.027 | -0.001 | -0.001 | 0 | 0.18 | 0.63 | 0.19 | 0.16 | 0.19 | 0.25 |
| 2 | 78308594 | rs1370638 | 0.554 | 0.273 | -0.439 | -0.496 | 0.003 | -0.009 | -0.009 | 0.414 | 0.42 | 0.18 | 0.47 | 0.62 | 0.44 | 0.45 |
| 2 | 78311453 | rs7569428 | 0.679 | 0.268 | -0.453 | -0.514 | -0.002 | -0.02 | -0.02 | 0 | 0.41 | 0.16 | 0.47 | 0.61 | 0.43 | 0.44 |
| 2 | 78311502 | rs1437986 | -0.807 | 0.492 | -0.403 | -0.642 | -0.022 | 0.043 | 0.043 | 0 | 0.82 | 0.92 | 0.78 | 0.91 | 0.81 | 0.89 |
| 2 | 78313046 | rs7573663 | -0.686 | 0.361 | 0.152 | -0.518 | 0.087 | 0.115 | 0.115 | 0.485 | 0.05 | 0.47 | 0.05 | 0.04 | 0.02 | 0.04 |
| 2 | 78314245 | rs1437993 | 0.535 | 0.44 | -0.389 | -0.526 | 0.003 | -0.024 | -0.024 | 0 | 0.38 | 0.15 | 0.37 | 0.65 | 0.41 | 0.41 |
| 2 | 78318600 | rs1370641 | -1.06 | 1.173 | 0.35 | -0.771 | 0.008 | -0.004 | -0.004 | 0 | 0.3 | 0.69 | 0.52 | 0.25 | 0.39 | 0.41 |
| 2 | 78321095 | rs9646992 | -0.016 | 1.288 | 0.17 | -0.798 | -0.007 | 0.005 | 0.005 | 0.056 | 0.21 | 0.17 | 0.33 | 0.15 | 0.29 | 0.24 |
| 2 | 78325476 | rs10197148 | -0.003 | 1.417 | -0.277 | -0.769 | -0.055 | 0.019 | 0.019 | 0 | 0.2 | 0.05 | 0.31 | 0.15 | 0.28 | 0.24 |
| 2 | 78330028 | rs7571032 | -1.145 | 1.366 | -0.036 | -0.784 | 0.093 | -0.015 | -0.015 | 0 | 0.11 | 0.53 | 0.08 | 0.04 | 0.11 | 0.18 |
| 2 | 78332806 | rs10173591 | -0.143 | 1.369 | -0.046 | -0.785 | 0.01 | -0.002 | -0.002 | 0 | 0.21 | 0.2 | 0.36 | 0.22 | 0.29 | 0.25 |
| 2 | 78341245 | rs1370637 | -0.817 | 1.406 | -0.17 | -0.782 | 0.003 | -0.003 | -0.003 | 0.155 | 0.26 | 0.69 | 0.4 | 0.22 | 0.31 | 0.28 |
| 2 | 78342416 | rs1347723 | -1.164 | 1.392 | -0.135 | -0.768 | 0.001 | -0.001 | -0.001 | 0 | 0.31 | 0.67 | 0.42 | 0.26 | 0.39 | 0.42 |
| 2 | 78346731 | rs10208377 | -1.212 | 1.63 | 0.278 | -0.443 | 0.009 | 0.006 | 0.006 | 0 | 0.12 | 0.14 | 0.13 | 0.2 | 0.2 | 0.23 |
| 2 | 78347877 | rs9309526 | -0.68 | 1.365 | 0.309 | -0.673 | -0.001 | -0.046 | -0.046 | 0 | 0.21 | 0.57 | 0.14 | 0.09 | 0.19 | 0.14 |
| 2 | 78348702 | rs9309527 | -0.543 | 1.282 | 0.31 | -0.668 | -0.006 | -0.013 | -0.013 | 0 | 0.2 | 0.6 | 0.15 | 0.16 | 0.17 | 0.13 |
| 2 | 78352550 | rs13425498 | -1.555 | 1.295 | 0.323 | -0.802 | 0.006 | -0.028 | -0.028 | 0 | 0.25 | 0.61 | 0.17 | 0.18 | 0.25 | 0.26 |
| 2 | 78353572 | rs723705 | -0.545 | 1.277 | 0.332 | -0.798 | -0.006 | -0.031 | -0.031 | 0 | 0.2 | 0.59 | 0.15 | 0.14 | 0.17 | 0.13 |
| 2 | 78359187 | rs17015011 | 0.163 | 1.232 | 0.585 | -0.884 | -0.04 | -0.089 | -0.089 | 0.215 | 0.14 | 0.39 | 0.06 | 0 | 0.07 | 0.05 |
| 2 | 78374332 | rs12990795 | 1.831 | 1.426 | 0.022 | -1.109 | -0.037 | 0.051 | 0.051 | 0 | 0.1 | 0.02 | 0.09 | 0 | 0.13 | 0.03 |
| 2 | 78374950 | rs13015524 | 0.575 | 1.125 | 0.461 | -0.688 | 0.005 | -0.002 | -0.002 | 0 | 0.07 | 0.06 | 0.07 | 0.06 | 0.12 | 0.11 |
| 2 | 78381263 | rs7586511 | -0.373 | 1.129 | 0.521 | -0.687 | 0 | -0.068 | -0.068 | 0.152 | 0.17 | 0.5 | 0.08 | 0.07 | 0.14 | 0.17 |

| | | | | | | | | | | | | | | | | |
|---|---|---|---|---|---|---|---|---|---|---|---|---|---|---|---|---|
| 2 | 78381862 | rs12991296 | 0.168 | 1.056 | -0.068 | -0.634 | 0.01 | 0.004 | 0.004 | 0 | 0.08 | 0.12 | 0.1 | 0.11 | 0.12 | 0.11 |
| 2 | 78384746 | rs6708183 | -1.449 | 0.791 | -0.119 | -0.762 | -0.003 | 0.017 | 0.017 | 5.214 | 0.34 | 0.68 | 0.2 | 0.19 | 0.35 | 0.34 |
| 2 | 78395467 | rs6547190 | 0.033 | 0.173 | 0.43 | -0.371 | -0.018 | -0.081 | -0.081 | 0 | 0.13 | 0.49 | 0.06 | 0 | 0.06 | 0.03 |
| 2 | 78400037 | rs6748053 | -0.394 | 0.171 | 0.313 | -0.369 | -0.037 | -0.041 | -0.041 | 0 | 0.14 | 0.61 | 0.07 | 0.07 | 0.07 | 0.05 |
| 2 | 78409539 | rs6724320 | 0.04 | 0.205 | 0.373 | -0.404 | -0.016 | -0.08 | -0.08 | 3.396 | 0.13 | 0.48 | 0.05 | 0 | 0.06 | 0.03 |
| 2 | 78416572 | rs17015041 | -1.06 | 0.319 | 0.326 | -0.508 | -0.017 | -0.088 | -0.088 | 1.11 | 0.19 | 0.6 | 0.11 | 0.05 | 0.15 | 0.17 |
| 2 | 78423757 | rs2164850 | -1.368 | 0.196 | 0.329 | -0.5 | -0.021 | -0.041 | -0.041 | 0 | 0.21 | 0.65 | 0.11 | 0.13 | 0.16 | 0.19 |
| 2 | 78428973 | rs2901929 | -1.24 | 0.233 | 0.348 | -0.486 | -0.018 | -0.043 | -0.043 | 0 | 0.2 | 0.64 | 0.11 | 0.13 | 0.16 | 0.19 |
| 2 | 78434830 | rs1595422 | -1.095 | 0.203 | 0.438 | -0.504 | -0.014 | -0.022 | -0.022 | 0.313 | 0.19 | 0.7 | 0.11 | 0.13 | 0.14 | 0.18 |
| 2 | 78451939 | rs1347725 | -0.264 | 0.177 | 0.474 | -0.502 | -0.043 | -0.042 | -0.042 | 0 | 0.14 | 0.6 | 0.06 | 0.07 | 0.06 | 0.04 |
| 2 | 78452934 | rs1526382 | 0.159 | 0.179 | 0.544 | -0.504 | -0.028 | -0.087 | -0.087 | 2.177 | 0.13 | 0.48 | 0.05 | 0 | 0.05 | 0.03 |
| 2 | 78457741 | rs6547194 | 0.16 | 0.18 | 0.544 | -0.504 | -0.028 | -0.087 | -0.087 | 0 | 0.13 | 0.48 | 0.05 | 0 | 0.05 | 0.03 |
| 2 | 78459066 | rs6743609 | -1.448 | 0.018 | 0.304 | -0.633 | -0.01 | -0.021 | -0.021 | 0 | 0.23 | 0.7 | 0.12 | 0.13 | 0.16 | 0.21 |
| 2 | 78465567 | rs11883520 | -0.894 | -0.042 | 0.347 | -0.657 | 0.023 | 0.006 | 0.006 | 0 | 0.05 | 0.1 | 0.05 | 0.07 | 0.09 | 0.14 |
| 2 | 78467412 | rs6715934 | -1.712 | -0.019 | -0.071 | -0.967 | 0.011 | 0.028 | 0.028 | 4.13 | 0.33 | 0.71 | 0.34 | 0.15 | 0.27 | 0.34 |
| 2 | 78471314 | rs2839828 | -1.119 | -0.049 | -0.01 | -0.984 | -0.032 | -0.073 | -0.073 | 0 | 0.27 | 0.58 | 0.28 | 0.09 | 0.17 | 0.21 |
| 2 | 78481166 | rs17015088 | -0.623 | -0.106 | 0.043 | -0.916 | -0.038 | -0.042 | -0.042 | 1.332 | 0.19 | 0.68 | 0.09 | 0.08 | 0.08 | 0.18 |
| 2 | 78485584 | rs17015097 | -0.27 | -0.1 | 0.052 | -0.916 | -0.058 | -0.098 | -0.098 | 0.037 | 0.17 | 0.59 | 0.07 | 0.02 | 0.06 | 0.07 |
| 2 | 78489746 | rs11885621 | -0.679 | -0.041 | -0.151 | -0.893 | 0.004 | -0.004 | -0.004 | 3.832 | 0.06 | 0.07 | 0.05 | 0.05 | 0.09 | 0.14 |
| 2 | 78497142 | rs12464158 | 1.142 | -0.101 | -1.237 | -1.004 | -0.053 | -0.064 | -0.064 | 0.377 | 0.32 | 0.1 | 0.44 | 0.64 | 0.32 | 0.41 |
| 2 | 78511542 | rs1589931 | -1.615 | -0.158 | -1.052 | -0.915 | -0.017 | -0.05 | -0.05 | 0 | 0.83 | 0.96 | 0.9 | 0.78 | 0.83 | 0.84 |
| 2 | 78516288 | rs17566368 | -0.906 | -0.231 | -0.943 | -0.943 | 0.031 | 0.072 | 0.072 | 12.394 | 0.26 | 0.82 | 0.15 | 0.13 | 0.19 | 0.21 |
| 2 | 78532705 | rs11897364 | -0.736 | 0.276 | -0.163 | -0.634 | 0.03 | -0.004 | -0.004 | 0 | 0.06 | 0.14 | 0.05 | 0.06 | 0.09 | 0.13 |
| 2 | 78542946 | rs990276 | -0.223 | 0.001 | 0.209 | -0.653 | -0.019 | -0.014 | -0.014 | 0 | 0.64 | 0.55 | 0.83 | 0.74 | 0.73 | 0.69 |
| 2 | 78580198 | rs6730492 | -0.509 | 0.105 | 0.546 | -0.608 | 0.058 | 0.032 | 0.032 | 0 | 0.06 | 0.25 | 0.05 | 0.06 | 0.06 | 0.12 |
| 2 | 78594007 | rs9309532 | 0.005 | -0.06 | 0.432 | -0.612 | -0.014 | -0.006 | -0.006 | 0 | 0.62 | 0.39 | 0.81 | 0.69 | 0.72 | 0.69 |
| 2 | 78617965 | rs2138490 | 0.317 | -0.116 | 0.403 | -0.613 | 0.038 | 0.034 | 0.034 | 1.736 | 0.58 | 0.27 | 0.78 | 0.69 | 0.68 | 0.57 |
| 2 | 78626927 | rs1960632 | -0.208 | -0.158 | 0.811 | -0.606 | -0.01 | -0.008 | -0.008 | 0 | 0.65 | 0.66 | 0.83 | 0.74 | 0.74 | 0.69 |
| 2 | 78631587 | rs1455380 | -0.218 | -0.159 | 0.874 | -0.607 | -0.01 | -0.008 | -0.008 | 0 | 0.65 | 0.65 | 0.82 | 0.74 | 0.73 | 0.69 |
| 2 | 78633880 | rs2861609 | 0.338 | -0.159 | 0.867 | -0.607 | 0.036 | 0.033 | 0.033 | 1.746 | 0.58 | 0.27 | 0.78 | 0.68 | 0.68 | 0.57 |
| 2 | 78638423 | rs6759570 | -0.222 | -0.163 | 0.872 | -0.583 | -0.027 | -0.012 | -0.012 | 0 | 0.64 | 0.56 | 0.82 | 0.7 | 0.74 | 0.69 |
| 2 | 78644536 | rs11126667 | 0.5 | -0.133 | 0.704 | -0.575 | 0.058 | 0.025 | 0.025 | 2.162 | 0.56 | 0.14 | 0.77 | 0.65 | 0.67 | 0.57 |
| 2 | 78654537 | rs11691877 | 0.343 | -0.136 | 0.716 | -0.576 | 0.03 | 0.027 | 0.027 | 1.462 | 0.58 | 0.29 | 0.78 | 0.68 | 0.67 | 0.57 |
| 2 | 78655712 | rs1455390 | 0.511 | -0.123 | 0.586 | -0.576 | 0.06 | 0.027 | 0.027 | 2.429 | 0.56 | 0.14 | 0.77 | 0.63 | 0.67 | 0.57 |
| 2 | 78668935 | rs2861610 | -0.246 | -0.109 | 0.538 | -0.566 | -0.026 | -0.011 | -0.011 | 0 | 0.64 | 0.53 | 0.83 | 0.7 | 0.75 | 0.7 |
| 2 | 78680056 | rs6726060 | 0.513 | 0.198 | -0.023 | -0.12 | -0.042 | -0.077 | -0.077 | 0 | 0.27 | 0.08 | 0.49 | 0.4 | 0.31 | 0.32 |
| 2 | 78684017 | rs13426270 | -0.326 | 0.218 | 0.214 | -0.134 | 0.044 | 0.033 | 0.033 | 0 | 0.06 | 0.27 | 0.04 | 0.07 | 0.06 | 0.1 |
| 2 | 78693252 | rs6547205 | 0.134 | 0.168 | 0.183 | -0.179 | 0.003 | -0.01 | -0.01 | 0 | 0.39 | 0.58 | 0.35 | 0.33 | 0.45 | 0.38 |
| 2 | 78696239 | rs7580939 | 1.345 | 0.123 | 0.192 | -0.191 | -0.033 | 0.024 | 0.024 | 1.468 | 0.3 | 0.09 | 0.29 | 0.24 | 0.38 | 0.27 |
| 2 | 78699428 | rs7606396 | 0.828 | 0.12 | 0.24 | -0.18 | 0 | -0.004 | -0.004 | 0.311 | 0.32 | 0.31 | 0.3 | 0.24 | 0.38 | 0.27 |
| 2 | 78707767 | rs11681421 | 0.433 | 0.172 | 0.206 | -0.172 | 0.004 | -0.006 | -0.006 | 0 | 0.37 | 0.39 | 0.33 | 0.33 | 0.45 | 0.38 |
| 2 | 78710833 | rs17015382 | 0.565 | 0.116 | 0.122 | -0.14 | -0.035 | -0.077 | -0.077 | 1.297 | 0.27 | 0.09 | 0.49 | 0.4 | 0.29 | 0.31 |
| 2 | 78711422 | rs1516248 | 0.555 | 0.108 | 0.117 | -0.141 | -0.035 | -0.093 | -0.093 | 1.625 | 0.27 | 0.09 | 0.49 | 0.41 | 0.29 | 0.32 |
| 2 | 78714978 | rs6761959 | 0.404 | 0.174 | 0.24 | -0.136 | 0.009 | -0.011 | -0.011 | 0 | 0.37 | 0.41 | 0.33 | 0.33 | 0.45 | 0.38 |
| 2 | 78718429 | rs6735382 | 1.107 | 0.244 | 0.059 | -0.122 | -0.056 | 0.032 | 0.032 | 1.207 | 0.3 | 0.09 | 0.29 | 0.25 | 0.39 | 0.27 |
| 2 | 78737668 | rs6547206 | 1.079 | 0.191 | 0.069 | -0.13 | -0.033 | 0.028 | 0.028 | 1.101 | 0.3 | 0.1 | 0.29 | 0.25 | 0.38 | 0.27 |
| 2 | 78737871 | rs1516255 | -0.448 | 0.208 | 0.104 | -0.101 | 0.017 | 0.036 | 0.036 | 0 | 0.06 | 0.21 | 0.04 | 0.11 | 0.07 | 0.12 |
| 2 | 78740761 | rs17015412 | -0.525 | 0.206 | 0.112 | -0.102 | 0.029 | 0.052 | 0.052 | 0 | 0.06 | 0.24 | 0.04 | 0.11 | 0.07 | 0.12 |
| 2 | 78740927 | rs13422559 | -0.525 | 0.206 | 0.112 | -0.102 | 0.029 | 0.052 | 0.052 | 0 | 0.06 | 0.24 | 0.04 | 0.11 | 0.07 | 0.12 |
| 2 | 78744697 | rs1158752 | 0.273 | 0.204 | 0.151 | -0.129 | 0.011 | -0.01 | -0.01 | 0 | 0.38 | 0.52 | 0.35 | 0.36 | 0.45 | 0.39 |
| 2 | 78754012 | rs6740971 | -0.658 | 0.175 | 0.057 | -0.144 | -0.008 | -0.006 | -0.006 | 0.068 | 0.65 | 0.6 | 0.84 | 0.75 | 0.74 | 0.69 |
| 2 | 78755686 | rs12477341 | -0.104 | 0.191 | 0.032 | -0.145 | -0.006 | -0.012 | -0.012 | 0 | 0.33 | 0.32 | 0.53 | 0.5 | 0.36 | 0.43 |
| 2 | 78756447 | rs17643638 | 2.157 | -0.003 | -0.269 | -0.389 | -0.031 | 0.046 | 0.046 | 0 | 0.11 | 0.02 | 0.07 | 0 | 0.13 | 0.03 |
| 2 | 78757904 | rs12713963 | 0.416 | 0.365 | -0.006 | -0.143 | -0.014 | 0.016 | 0.016 | 0.872 | 0.35 | 0.27 | 0.33 | 0.28 | 0.44 | 0.38 |
| 2 | 78803313 | rs1239108 | 0.38 | 0.622 | -0.211 | -0.157 | -0.031 | 0.05 | 0.05 | 0.411 | 0.37 | 0.11 | 0.32 | 0.26 | 0.41 | 0.34 |
| 2 | 78805831 | rs1239105 | 0.277 | 0.216 | -0.123 | -0.079 | -0.068 | 0.062 | 0.062 | 0.85 | 0.41 | 0.11 | 0.35 | 0.3 | 0.48 | 0.45 |
| 2 | 78805852 | rs1239104 | -1.144 | 0.23 | -0.107 | -0.038 | -0.014 | 0.002 | 0.002 | 0.828 | 0.7 | 0.57 | 0.84 | 0.7 | 0.75 | 0.69 |
| 2 | 78810352 | rs2861618 | -0.111 | 0.172 | -0.141 | 0.038 | -0.015 | 0.045 | 0.045 | 0 | 0.3 | 0.58 | 0.49 | 0.43 | 0.27 | 0.26 |
| 2 | 78817822 | rs2177032 | 0.894 | 0.26 | -0.334 | -0.324 | -0.039 | 0.042 | 0.042 | 0.635 | 0.31 | 0.13 | 0.31 | 0.24 | 0.39 | 0.32 |
| 2 | 78826234 | rs1717554 | -0.667 | 0.14 | -0.189 | -0.32 | 0 | -0.001 | -0.001 | 0 | 0.62 | 0.64 | 0.8 | 0.72 | 0.69 | 0.67 |
| 2 | 78826406 | rs1519624 | -0.036 | 0.184 | -0.134 | -0.29 | -0.001 | 0.029 | 0.029 | 0 | 0.29 | 0.38 | 0.47 | 0.46 | 0.29 | 0.28 |
| 2 | 78830137 | rs10208240 | -0.152 | 0.228 | -0.239 | -0.295 | -0.002 | 0.056 | 0.056 | 0 | 0.31 | 0.53 | 0.48 | 0.52 | 0.3 | 0.36 |
| 2 | 78837866 | rs2139928 | -0.944 | 0.401 | -0.315 | -0.187 | -0.001 | -0.001 | -0.001 | 2.506 | 0.7 | 0.69 | 0.85 | 0.76 | 0.73 | 0.74 |
| 2 | 78841778 | rs17040590 | -0.871 | 0.604 | -0.532 | 0.132 | 0.002 | -0.005 | -0.005 | 0 | 0.42 | 0.43 | 0.54 | 0.5 | 0.37 | 0.43 |
| 2 | 78843902 | rs1402675 | -0.512 | 1.023 | -0.836 | 0.503 | 0.005 | 0.02 | 0.02 | 0 | 0.23 | 0.24 | 0.45 | 0.51 | 0.29 | 0.3 |
| 2 | 78845631 | rs17015524 | -0.763 | 0.859 | -0.727 | 0.48 | 0.01 | 0.016 | 0.016 | 0 | 0.25 | 0.28 | 0.42 | 0.56 | 0.3 | 0.37 |
| 2 | 78857870 | rs2946582 | -0.086 | 0.695 | -0.776 | -0.328 | 0.002 | 0.005 | 0.005 | 0 | 0.46 | 0.39 | 0.52 | 0.39 | 0.57 | 0.43 |
| 2 | 78859874 | rs1401737 | -0.277 | 0.711 | -0.754 | -0.314 | 0.036 | -0.016 | -0.016 | 0 | 0.15 | 0.26 | 0.15 | 0.13 | 0.19 | 0.12 |
| 2 | 78860071 | rs1239084 | 1.099 | 0.649 | -0.769 | -0.319 | -0.081 | 0.052 | 0.052 | 0 | 0.31 | 0.07 | 0.36 | 0.25 | 0.38 | 0.29 |

| | | | | | | | | | | | | | | | | |
|---|---|---|---|---|---|---|---|---|---|---|---|---|---|---|---|---|
| 2 | 78870238 | rs2037451 | -0.25 | 0.643 | -0.715 | -0.25 | 0.057 | 0.009 | 0.009 | 0.083 | 0.15 | 0.32 | 0.15 | 0.16 | 0.19 | 0.12 |
| 2 | 78873621 | rs1239102 | -0.668 | 0.772 | -0.579 | -0.232 | -0.008 | -0.009 | -0.009 | 0 | 0.6 | 0.53 | 0.58 | 0.41 | 0.64 | 0.53 |
| 2 | 78900269 | rs6716040 | 0.195 | 0.472 | -0.755 | -0.152 | 0.041 | -0.013 | -0.013 | 0 | 0.11 | 0.18 | 0.14 | 0.09 | 0.18 | 0.1 |
| 2 | 78914837 | rs17584593 | 0.537 | 0.435 | -0.721 | -0.027 | 0.027 | 0.015 | 0.015 | 0 | 0.14 | 0.05 | 0.06 | 0.06 | 0.07 | 0.19 |
| 2 | 78927328 | rs1669754 | 0.901 | 0.591 | -0.793 | -0.08 | -0.083 | 0.09 | 0.09 | 0 | 0.35 | 0.07 | 0.38 | 0.25 | 0.41 | 0.31 |
| 2 | 78937122 | rs11883911 | 0.419 | 0.781 | -0.626 | 0.04 | 0.034 | -0.012 | -0.012 | 0 | 0.11 | 0.17 | 0.13 | 0.08 | 0.16 | 0.1 |
| 2 | 78945755 | rs1452044 | -0.248 | 0.776 | -0.665 | 0.052 | 0.066 | 0.005 | 0.005 | 0.943 | 0.13 | 0.31 | 0.15 | 0.14 | 0.17 | 0.12 |
| 2 | 78968942 | rs4853406 | 0.496 | 0.769 | -0.932 | 0.062 | 0.03 | 0.04 | 0.04 | 0 | 0.16 | 0.04 | 0.06 | 0.06 | 0.09 | 0.21 |
| 2 | 78978011 | rs11687801 | -0.68 | 0.968 | -0.807 | 0.11 | -0.016 | 0.017 | 0.017 | 0 | 0.65 | 0.52 | 0.59 | 0.47 | 0.66 | 0.64 |
| 2 | 78999216 | rs6739221 | -1.084 | 1.255 | -0.652 | 0.316 | 0.033 | 0.014 | 0.014 | 0.488 | 0.13 | 0.22 | 0.14 | 0.15 | 0.17 | 0.11 |
| 2 | 79001780 | rs2167558 | -0.569 | 1.271 | -0.716 | 0.349 | 0.023 | 0 | 0 | 0 | 0.11 | 0.14 | 0.13 | 0.1 | 0.17 | 0.08 |
| 2 | 79001963 | rs17015634 | -0.711 | 1.262 | -0.741 | 0.269 | 0.024 | 0.016 | 0.016 | 0.421 | 0.12 | 0.2 | 0.2 | 0.17 | 0.18 | 0.11 |
| 2 | 79007948 | rs6751026 | -0.836 | 1.116 | -0.823 | 0.092 | 0.001 | -0.024 | -0.024 | 0.218 | 0.28 | 0.34 | 0.27 | 0.15 | 0.26 | 0.24 |
| 2 | 79011998 | rs10178977 | -1.005 | 1.729 | -1.305 | -0.015 | -0.038 | 0.041 | 0.041 | 0 | 0.15 | 0.04 | 0.11 | 0.06 | 0.21 | 0.27 |
| 2 | 79012163 | rs7578411 | -1.676 | 2.172 | -0.43 | 0.191 | 0.042 | -0.014 | -0.014 | 0 | 0.27 | 0.43 | 0.28 | 0.21 | 0.37 | 0.54 |
| 2 | 79012740 | rs17015646 | -0.121 | 2.21 | -0.869 | 1.149 | -0.013 | 0.022 | 0.022 | 0 | 0.07 | 0.01 | 0.33 | 0.41 | 0.12 | 0.14 |
| 2 | 79018187 | rs1869083 | -0.814 | 2.197 | -1.352 | 0.35 | -0.068 | 0.104 | 0.104 | 0 | 0.24 | 0.06 | 0.18 | 0.06 | 0.3 | 0.35 |
| 2 | 79037108 | rs7598432 | -0.866 | 2.29 | -0.523 | 0.365 | 0.003 | -0.001 | -0.001 | 0 | 0.11 | 0.13 | 0.17 | 0.09 | 0.21 | 0.12 |
| 2 | 79038224 | rs17586318 | 0.496 | 1.726 | -1.285 | 0.102 | 0.01 | 0.098 | 0.098 | 0.006 | 0.5 | 0.12 | 0.38 | 0.33 | 0.5 | 0.64 |
| 2 | 79039355 | rs896530 | -0.644 | 1.619 | -0.703 | 0.082 | -0.037 | 0.021 | 0.021 | 0.018 | 0.64 | 0.58 | 0.59 | 0.44 | 0.72 | 0.77 |
| 2 | 79045079 | rs7569615 | -0.64 | 1.367 | -1.015 | 0.073 | 0.008 | -0.005 | -0.005 | 0 | 0.13 | 0.18 | 0.17 | 0.09 | 0.21 | 0.14 |
| 2 | 79047205 | rs1020567 | 0.008 | 1.465 | -1.036 | 0.041 | 0.01 | -0.002 | -0.002 | 0 | 0.57 | 0.39 | 0.52 | 0.44 | 0.63 | 0.74 |
| 2 | 79055423 | rs10189572 | 0.842 | 1.493 | -1.228 | 0.008 | -0.036 | 0.058 | 0.058 | 0 | 0.43 | 0.14 | 0.34 | 0.32 | 0.4 | 0.59 |
| 2 | 79056092 | rs17586938 | 1.552 | 1.702 | -0.844 | -0.346 | 0.018 | 0.064 | 0.064 | 0 | 0.33 | 0.19 | 0.23 | 0.16 | 0.23 | 0.17 |
| 2 | 79061185 | rs2177098 | 0.379 | 1.671 | -0.376 | -0.243 | 0 | 0.001 | 0.001 | 0 | 0.49 | 0.48 | 0.44 | 0.37 | 0.4 | 0.44 |
| 2 | 79063788 | rs2140039 | 0.031 | 1.699 | -0.373 | -0.318 | -0.001 | 0.001 | 0.001 | 0 | 0.57 | 0.59 | 0.57 | 0.39 | 0.55 | 0.54 |
| 2 | 79064180 | rs17015724 | -0.231 | 1.691 | -0.454 | -0.2 | -0.008 | -0.002 | -0.002 | 0 | 0.61 | 0.47 | 0.58 | 0.4 | 0.63 | 0.59 |
| 2 | 79069569 | rs4853414 | -0.03 | 1.742 | -0.695 | -0.119 | -0.018 | -0.022 | -0.022 | 0 | 0.11 | 0.07 | 0.13 | 0.15 | 0.18 | 0.23 |
| 2 | 79072931 | rs7579074 | 1.476 | 1.701 | -0.558 | -0.235 | -0.002 | -0.01 | -0.01 | 0 | 0.35 | 0.35 | 0.3 | 0.21 | 0.23 | 0.17 |
| 2 | 79085411 | rs17015751 | 0.413 | 1.384 | -0.958 | 0.378 | -0.001 | -0.038 | -0.038 | 0 | 0.09 | 0.01 | 0.22 | 0.24 | 0.07 | 0.15 |
| 2 | 79088516 | rs4853418 | 1.725 | 1.428 | -0.902 | 0.063 | 0.016 | 0.051 | 0.051 | 0 | 0.33 | 0.21 | 0.23 | 0.16 | 0.23 | 0.17 |
| 2 | 79089793 | rs17647989 | -0.264 | 1.666 | -0.981 | 0.274 | -0.008 | -0.01 | -0.01 | 0 | 0.14 | 0.11 | 0.17 | 0.2 | 0.2 | 0.28 |
| 2 | 79090327 | rs4853420 | -0.264 | 1.671 | -0.979 | 0.277 | -0.008 | -0.01 | -0.01 | 0 | 0.14 | 0.11 | 0.17 | 0.2 | 0.2 | 0.26 |
| 2 | 79093607 | rs11883728 | 0.445 | 1.459 | -0.954 | 0.129 | 0.002 | -0.003 | -0.003 | 0 | 0.56 | 0.48 | 0.58 | 0.42 | 0.48 | 0.45 |
| 2 | 79096243 | rs1521973 | -0.19 | 1.181 | -1.102 | 0.018 | 0.001 | -0.001 | -0.001 | 0 | 0.2 | 0.2 | 0.28 | 0.19 | 0.19 | 0.24 |
| 2 | 79100224 | rs1402906 | -0.756 | 1.578 | -1.126 | -0.19 | -0.005 | 0.002 | 0.002 | 0 | 0.9 | 0.9 | 0.84 | 0.98 | 0.79 | 0.89 |
| 2 | 79101215 | rs6761240 | -0.165 | 1.327 | -1.239 | -0.329 | -0.007 | 0.005 | 0.005 | 0 | 0.36 | 0.42 | 0.26 | 0.59 | 0.32 | 0.46 |
| 2 | 79104576 | rs6708376 | -1.389 | 1.494 | -1.156 | -0.316 | -0.003 | 0.017 | 0.017 | 0 | 0.54 | 0.58 | 0.59 | 0.79 | 0.51 | 0.69 |
| 2 | 79110274 | rs10520215 | 2.746 | 1.639 | -1.376 | -0.221 | -0.031 | -0.067 | -0.067 | 0.05 | 0.13 | 0.02 | 0.04 | 0.28 | 0.16 | 0.12 |
| 2 | 79113604 | rs1996768 | 2.273 | 1.696 | -1.182 | -0.644 | 0.016 | 0.061 | 0.061 | 0 | 0.36 | 0.23 | 0.25 | 0.19 | 0.28 | 0.21 |
| 2 | 79113640 | rs1996767 | -0.791 | 1.621 | -1.177 | -0.477 | -0.007 | 0.004 | 0.004 | 0 | 0.9 | 0.91 | 0.84 | 0.98 | 0.79 | 0.89 |
| 2 | 79116722 | rs4853422 | -0.448 | 1.677 | -1.228 | -0.525 | -0.036 | 0.036 | 0.036 | 0 | 0.86 | 0.93 | 0.82 | 0.98 | 0.74 | 0.87 |
| 2 | 79116878 | rs13395260 | 1.62 | 1.674 | -1.166 | -0.484 | -0.016 | 0.018 | 0.018 | 0 | 0.67 | 0.72 | 0.5 | 0.78 | 0.56 | 0.65 |
| 2 | 79121367 | rs4300858 | -0.489 | 1.805 | -0.813 | -0.186 | -0.005 | 0.007 | 0.007 | 0 | 0.91 | 0.93 | 0.82 | 0.97 | 0.78 | 0.88 |
| 2 | 79122483 | rs1915751 | -0.105 | 1.696 | -0.856 | -0.221 | 0 | 0.002 | 0.002 | 0 | 0.15 | 0.18 | 0.18 | 0.19 | 0.16 | 0.2 |
| 2 | 79124814 | rs2139974 | -0.638 | 1.786 | -0.942 | -0.235 | 0.017 | -0.013 | -0.013 | 0 | 0.91 | 0.88 | 0.83 | 0.97 | 0.8 | 0.89 |
| 2 | 79125686 | rs13014483 | 2.185 | 1.655 | -1.242 | -0.384 | -0.054 | 0.15 | 0.15 | 0.046 | 0.36 | 0.02 | 0.26 | 0.16 | 0.29 | 0.2 |
| 2 | 79130310 | rs4145992 | 1.009 | 1.871 | -0.484 | -0.095 | 0.079 | -0.015 | -0.015 | 0 | 0.75 | 0.52 | 0.68 | 0.81 | 0.61 | 0.67 |
| 2 | 79133151 | rs12612243 | -0.175 | 1.776 | -0.36 | -0.041 | 0.083 | 0.048 | 0.048 | 0 | 0.92 | 0.7 | 0.87 | 0.93 | 0.88 | 0.97 |
| 2 | 79135349 | rs17648900 | -0.633 | 1.765 | -0.516 | -0.094 | 0.005 | -0.004 | -0.004 | 0.644 | 0.2 | 0.2 | 0.24 | 0.18 | 0.27 | 0.3 |
| 2 | 79139515 | rs7566244 | 0.527 | 1.729 | -0.467 | -0.081 | 0.001 | -0.014 | -0.014 | 0 | 0.81 | 0.71 | 0.81 | 0.86 | 0.8 | 0.8 |
| 2 | 79140963 | rs4853423 | 0.891 | 1.771 | -0.489 | -0.091 | 0.014 | -0.012 | -0.012 | 1.179 | 0.78 | 0.7 | 0.7 | 0.82 | 0.68 | 0.69 |
| 2 | 79143946 | rs11903030 | -0.482 | 1.846 | -0.308 | -0.037 | 0.194 | 0.112 | 0.112 | 0 | 0.07 | 0.28 | 0.2 | 0.1 | 0.18 | 0.13 |
| 2 | 79144604 | rs1949661 | 1.263 | 1.742 | -0.552 | -0.096 | 0.036 | -0.008 | -0.008 | 0 | 0.72 | 0.48 | 0.63 | 0.75 | 0.62 | 0.66 |
| 2 | 79146708 | rs1915762 | -1.121 | 1.681 | -0.79 | -0.1 | 0.007 | -0.022 | -0.022 | 0 | 0.95 | 0.9 | 0.94 | 1 | 0.91 | 0.99 |
| 2 | 79152829 | rs10208925 | -0.082 | 1.486 | -0.887 | -0.356 | 0.006 | 0.006 | 0.006 | 0 | 0.39 | 0.32 | 0.38 | 0.6 | 0.37 | 0.44 |
| 2 | 79152922 | rs2861680 | -1.172 | 1.565 | -0.855 | -0.409 | 0.001 | -0.039 | -0.039 | 0 | 0.57 | 0.52 | 0.61 | 0.77 | 0.62 | 0.75 |
| 2 | 79155348 | rs10200751 | -0.234 | 1.684 | -0.945 | -0.473 | 0.081 | 0.016 | 0.016 | 0.362 | 0.89 | 0.61 | 0.85 | 0.93 | 0.85 | 0.97 |
| 2 | 79160743 | rs17691986 | 2.327 | 1.522 | -0.998 | -0.474 | -0.019 | 0.129 | 0.129 | 0 | 0.32 | 0.09 | 0.23 | 0.16 | 0.24 | 0.22 |
| 2 | 79161609 | rs6761498 | -0.887 | 1.586 | -1.202 | -0.471 | -0.001 | -0.011 | -0.011 | 2.084 | 0.54 | 0.53 | 0.51 | 0.75 | 0.49 | 0.65 |
| 2 | 79162080 | rs11904716 | 1.45 | 1.049 | -1.158 | -0.445 | -0.004 | -0.027 | -0.027 | 1.603 | 0.35 | 0.4 | 0.32 | 0.23 | 0.26 | 0.23 |
| 2 | 79164165 | rs10165807 | 1.179 | 0.992 | -1.744 | -0.107 | -0.035 | -0.097 | -0.097 | 0 | 0.22 | 0.02 | 0.15 | 0.31 | 0.21 | 0.13 |
| 2 | 79167769 | rs986386 | 0.963 | 1.313 | -0.95 | -0.739 | -0.007 | -0.021 | -0.021 | 0 | 0.34 | 0.38 | 0.3 | 0.24 | 0.24 | 0.24 |
| 2 | 79170700 | rs1992886 | 1.428 | 1.151 | -1.224 | -0.647 | -0.061 | 0.101 | 0.101 | 0.01 | 0.31 | 0.02 | 0.22 | 0.17 | 0.23 | 0.22 |
| 2 | 79175671 | rs17015937 | 0.357 | 1.316 | -0.716 | -0.264 | 0 | -0.001 | -0.001 | 0 | 0.11 | 0.13 | 0.11 | 0.13 | 0.11 | 0.2 |
| 2 | 79176572 | rs1261232 | -0.44 | 1.554 | -0.799 | -0.044 | 0.006 | 0 | 0 | 0 | 0.65 | 0.71 | 0.64 | 0.41 | 0.67 | 0.62 |
| 2 | 79178130 | rs11126689 | -1.331 | 2.002 | -0.65 | 0.176 | -0.007 | 0.002 | 0.002 | 0 | 0.12 | 0.1 | 0.18 | 0.03 | 0.25 | 0.16 |
| 2 | 79182821 | rs1261234 | -0.811 | 1.439 | -0.574 | 0.277 | -0.001 | -0.012 | -0.012 | 0 | 0.09 | 0.08 | 0.06 | 0 | 0.12 | 0.01 |
| 2 | 79185600 | rs17692933 | 0.352 | 1.179 | -0.844 | 0.471 | -0.03 | 0.041 | 0.041 | 0 | 0.26 | 0.12 | 0.26 | 0.16 | 0.29 | 0.24 |

| | | | | | | | | | | | | | | | | |
|---|---|---|---|---|---|---|---|---|---|---|---|---|---|---|---|---|
| 2 | 79189199 | rs13027536 | -0.216 | 1.167 | -0.974 | 0.357 | -0.021 | 0.008 | 0.008 | 0 | 0.06 | 0.01 | 0.05 | 0.02 | 0.08 | 0.06 |
| 2 | 79190408 | rs10520219 | -1.474 | 0.985 | -0.357 | 0.512 | -0.001 | 0.032 | 0.032 | 0 | 0.13 | 0.27 | 0.17 | 0.13 | 0.11 | 0.19 |
| 2 | 79190499 | rs10520221 | -0.43 | 1.023 | -0.329 | 0.552 | 0.002 | -0.014 | -0.014 | 0 | 0.36 | 0.42 | 0.33 | 0.25 | 0.36 | 0.37 |
| 2 | 79191047 | rs17015957 | -1.378 | 0.949 | -0.245 | 0.52 | 0.004 | 0.046 | 0.046 | 0 | 0.15 | 0.46 | 0.18 | 0.13 | 0.11 | 0.19 |
| 2 | 79194418 | rs1261238 | -1.332 | 0.671 | -1.097 | 0.379 | -0.036 | 0.196 | 0.196 | 0.465 | 0.68 | 0.97 | 0.77 | 0.91 | 0.75 | 0.79 |
| 2 | 79196630 | rs13384662 | -0.875 | 0.549 | 0.187 | 1.371 | -0.005 | 0.065 | 0.065 | 0 | 0.07 | 0.17 | 0.02 | 0.1 | 0.02 | 0.12 |
| 2 | 79198168 | rs11683503 | -1.706 | 0.542 | -0.024 | 1.345 | -0.013 | 0.067 | 0.067 | 0 | 0.13 | 0.42 | 0.15 | 0.13 | 0.08 | 0.19 |
| 2 | 79198620 | rs11126690 | -1.197 | 0.552 | -0.167 | 1.348 | -0.005 | -0.012 | -0.012 | 0 | 0.41 | 0.53 | 0.35 | 0.27 | 0.38 | 0.43 |
| 2 | 79205672 | rs12618775 | -0.311 | 0.617 | -1.437 | 1.206 | -0.002 | 0.135 | 0.135 | 0 | 0.05 | 0.05 | 0.24 | 0.51 | 0.02 | 0.14 |
| 2 | 79208014 | rs17749772 | -0.153 | 0.65 | -1.479 | 1.168 | -0.057 | 0.043 | 0.043 | 0 | 0.26 | 0.05 | 0.19 | 0.14 | 0.28 | 0.24 |
| 2 | 79208026 | rs12617352 | -1.087 | 0.384 | -0.578 | 1.754 | -0.019 | 0.203 | 0.203 | 0 | 0.07 | 0.11 | 0.25 | 0.5 | 0.03 | 0.14 |
| 2 | 79223215 | rs1666768 | 0.1 | 0.828 | -0.859 | 0.348 | -0.001 | -0.006 | -0.006 | 0.453 | 0.54 | 0.59 | 0.33 | 0.24 | 0.54 | 0.48 |
| 2 | 79223593 | rs17016000 | -0.914 | 0.666 | -0.314 | 0.898 | 0.002 | 0.067 | 0.067 | 0 | 0.23 | 0.5 | 0.29 | 0.3 | 0.2 | 0.25 |
| 2 | 79223902 | rs4853428 | -1.287 | 0.909 | -0.978 | 1.246 | -0.037 | -0.061 | -0.061 | 0.219 | 0.15 | 0.02 | 0.17 | 0.24 | 0.15 | 0.15 |
| 2 | 79224687 | rs1666769 | -0.349 | 0.045 | -0.821 | 0.994 | -0.01 | -0.006 | -0.006 | 0.227 | 0.59 | 0.55 | 0.78 | 0.7 | 0.66 | 0.73 |
| 2 | 79225690 | rs402221 | 0.159 | -0.291 | -0.882 | 0.971 | 0.034 | 0.014 | 0.014 | 0 | 0.32 | 0.21 | 0.24 | 0.28 | 0.16 | 0.43 |
| 2 | 79225952 | rs384761 | -0.326 | -1.015 | -0.701 | 0.758 | 0.004 | 0.003 | 0.003 | 0 | 0.49 | 0.45 | 0.36 | 0.4 | 0.3 | 0.56 |
| 2 | 79227053 | rs10179468 | 0.529 | -1.294 | -0.823 | 0.765 | 0 | 0 | 0 | 0 | 0.07 | 0.1 | 0.06 | 0.1 | 0.06 | 0.14 |
| 2 | 79227099 | rs811526 | -0.509 | -1.79 | -0.692 | 0.428 | 0.017 | 0.023 | 0.023 | 0.038 | 0.56 | 0.59 | 0.76 | 0.79 | 0.66 | 0.73 |
| 2 | 79227243 | rs283819 | -0.711 | -1.763 | -0.621 | 0.346 | 0.024 | 0.024 | 0.024 | 0.199 | 0.6 | 0.63 | 0.8 | 0.79 | 0.72 | 0.74 |
| 2 | 79227275 | rs7565256 | 0.095 | -1.616 | -0.538 | 0.374 | -0.014 | -0.039 | -0.039 | 0 | 0.23 | 0.35 | 0.16 | 0.11 | 0.15 | 0.15 |
| 2 | 79227583 | rs283822 | -0.281 | -1.292 | -0.23 | 0.373 | -0.01 | 0 | 0 | 0 | 0.4 | 0.54 | 0.39 | 0.41 | 0.28 | 0.54 |
| 2 | 79228257 | rs10520223 | -0.122 | -1.306 | -0.245 | 0.451 | -0.001 | 0 | 0 | 0 | 0.4 | 0.41 | 0.52 | 0.47 | 0.54 | 0.32 |
| 2 | 79230585 | rs10520227 | 2.002 | -1.666 | -0.228 | 0.426 | -0.005 | -0.001 | -0.001 | 0 | 0.15 | 0.07 | 0.09 | 0.13 | 0.05 | 0.15 |
| 2 | 79230697 | rs10179963 | 0.032 | -1.698 | 0.036 | 0.405 | -0.003 | -0.001 | -0.001 | 0 | 0.39 | 0.37 | 0.51 | 0.47 | 0.53 | 0.32 |
| 2 | 79233079 | rs283826 | 0.64 | -1.674 | 0.208 | 0.445 | 0 | -0.001 | -0.001 | 0 | 0.21 | 0.12 | 0.13 | 0.2 | 0.21 | 0.26 |
| 2 | 79236614 | rs10520228 | 1.498 | -1.278 | 0.054 | 0.352 | -0.007 | -0.01 | -0.01 | 1.219 | 0.18 | 0.15 | 0.11 | 0.13 | 0.12 | 0.17 |
| 2 | 79239278 | rs405595 | -0.144 | -1.28 | 0.146 | 0.488 | -0.013 | 0.049 | 0.049 | 0 | 0.24 | 0.43 | 0.3 | 0.32 | 0.22 | 0.39 |
| 2 | 79243597 | rs434922 | 0.205 | -1.304 | 0.142 | 0.509 | -0.002 | -0.001 | -0.001 | 0.001 | 0.22 | 0.25 | 0.14 | 0.2 | 0.22 | 0.26 |
| 2 | 79244207 | rs393149 | -0.038 | -1.285 | 0.082 | 0.464 | -0.009 | 0.04 | 0.04 | 0 | 0.24 | 0.39 | 0.3 | 0.31 | 0.22 | 0.39 |
| 2 | 79246108 | rs1023825 | -0.694 | -1.412 | 0.047 | 0.387 | -0.005 | 0.001 | 0.001 | 0 | 0.54 | 0.41 | 0.6 | 0.59 | 0.59 | 0.48 |
| 2 | 79252761 | rs368506 | 0.025 | -1.356 | 0.084 | 0.423 | -0.01 | 0.046 | 0.046 | 0 | 0.24 | 0.4 | 0.31 | 0.32 | 0.22 | 0.39 |
| 2 | 79253150 | rs369967 | 0.053 | -1.353 | 0.052 | 0.424 | -0.009 | 0.041 | 0.041 | 0 | 0.23 | 0.39 | 0.31 | 0.32 | 0.22 | 0.39 |
| 2 | 79254261 | rs452267 | -2.611 | -1.306 | -0.111 | 0.392 | 0.018 | 0.026 | 0.026 | 0 | 0.78 | 0.84 | 0.9 | 0.91 | 0.8 | 0.86 |
| 2 | 79255588 | rs374577 | -1.019 | -1.362 | -0.074 | 0.383 | 0.005 | -0.004 | -0.004 | 0.479 | 0.42 | 0.56 | 0.42 | 0.45 | 0.34 | 0.56 |
| 2 | 79257948 | rs565716 | -2.565 | -1.282 | -0.071 | 0.421 | 0.022 | 0.034 | 0.034 | 0 | 0.77 | 0.84 | 0.9 | 0.91 | 0.8 | 0.86 |
| 2 | 79258144 | rs372679 | -2.606 | -1.278 | -0.074 | 0.442 | 0.022 | 0.032 | 0.032 | 0 | 0.78 | 0.85 | 0.9 | 0.91 | 0.8 | 0.86 |
| 2 | 79263703 | rs283838 | 0.651 | -1.387 | 0 | 0.421 | -0.002 | -0.001 | -0.001 | 0 | 0.17 | 0.18 | 0.09 | 0.13 | 0.16 | 0.19 |
| 2 | 79265567 | rs13397545 | 0.497 | -1.611 | -0.059 | 0.221 | -0.023 | 0.039 | 0.039 | 0 | 0.23 | 0.37 | 0.32 | 0.31 | 0.17 | 0.36 |
| 2 | 79267708 | rs283842 | -1.117 | -1.626 | -0.085 | 0.186 | -0.002 | -0.004 | -0.004 | 0 | 0.6 | 0.64 | 0.79 | 0.78 | 0.68 | 0.7 |
| 2 | 79277543 | rs446861 | -1.16 | -1.557 | -0.468 | 0.22 | 0.041 | 0.055 | 0.055 | 0 | 0.61 | 0.82 | 0.81 | 0.78 | 0.68 | 0.72 |
| 2 | 79284271 | rs402180 | -1.538 | -1.523 | -0.446 | 0.269 | 0.021 | -0.012 | -0.012 | 0 | 0.71 | 0.72 | 0.71 | 0.73 | 0.76 | 0.65 |
| 2 | 79290106 | rs13405626 | 0.122 | -1.251 | -0.458 | 0.345 | -0.002 | 0.06 | 0.06 | 0 | 0.07 | 0.18 | 0.21 | 0.19 | 0.05 | 0.21 |
| 2 | 79291934 | rs364446 | 0.581 | -1.365 | -0.495 | 0.251 | -0.009 | -0.008 | -0.008 | 0 | 0.31 | 0.26 | 0.44 | 0.49 | 0.48 | 0.37 |
| 2 | 79293342 | rs378492 | -0.964 | -1.371 | -0.709 | 0.205 | 0.027 | 0.002 | 0.002 | 0 | 0.55 | 0.65 | 0.6 | 0.6 | 0.64 | 0.49 |
| 2 | 79293494 | rs13004360 | 1.282 | -1.071 | -0.923 | -0.149 | 0.002 | 0.005 | 0.005 | 0 | 0.17 | 0.08 | 0.11 | 0.13 | 0.14 | 0.18 |
| 2 | 79295653 | rs1448212 | -0.05 | -0.779 | -1.028 | 0.065 | -0.011 | -0.023 | -0.023 | 0.095 | 0.41 | 0.4 | 0.28 | 0.24 | 0.29 | 0.31 |
| 2 | 79297847 | rs283904 | -1.636 | -0.634 | -1.589 | 0.199 | -0.01 | 0.017 | 0.017 | 0 | 0.77 | 0.99 | 0.9 | 0.9 | 0.8 | 0.87 |
| 2 | 79298364 | rs1448213 | -0.024 | -0.461 | -1.35 | 0.101 | 0.006 | 0.033 | 0.033 | 0 | 0.52 | 0.43 | 0.68 | 0.77 | 0.54 | 0.54 |
| 2 | 79298471 | rs1522858 | -0.554 | -0.749 | -1.243 | 0.278 | 0.007 | -0.003 | -0.003 | 2.505 | 0.47 | 0.64 | 0.52 | 0.53 | 0.54 | 0.47 |
| 2 | 79300515 | rs6710164 | 0.735 | -1.35 | -0.804 | 0.369 | 0.041 | 0.043 | 0.043 | 0 | 0.39 | 0.28 | 0.18 | 0.16 | 0.22 | 0.22 |
| 2 | 79301265 | rs1448214 | -0.216 | -1.321 | -0.74 | 0.481 | 0.002 | 0.014 | 0.014 | 0.082 | 0.38 | 0.5 | 0.67 | 0.56 | 0.56 | 0.42 |
| 2 | 79303418 | rs1608234 | 1.256 | -1.009 | -0.941 | 0.691 | -0.024 | -0.02 | -0.02 | 0.161 | 0.21 | 0.14 | 0.27 | 0.42 | 0.38 | 0.34 |
| 2 | 79310943 | rs1522856 | 0.343 | -1.077 | -1.041 | 0.618 | -0.002 | 0.001 | 0.001 | 0 | 0.07 | 0.1 | 0.26 | 0.09 | 0.12 | 0.05 |
| 2 | 79311773 | rs7583314 | -0.402 | -1.415 | -1.036 | 0.715 | -0.001 | -0.002 | -0.002 | 0 | 0.36 | 0.38 | 0.6 | 0.53 | 0.53 | 0.42 |
| 2 | 79313815 | rs2073445 | -0.46 | -1.658 | -0.833 | 0.5 | -0.023 | -0.01 | -0.01 | 0 | 0.67 | 0.61 | 0.78 | 0.7 | 0.72 | 0.57 |
| 2 | 79315285 | rs2073443 | 0.956 | -1.637 | -0.853 | 0.535 | -0.02 | 0.039 | 0.039 | 0 | 0.09 | 0.15 | 0.27 | 0.15 | 0.03 | 0.07 |
| 2 | 79316422 | rs1375050 | 1.272 | -1.631 | -0.931 | 0.453 | 0.022 | 0.064 | 0.064 | 0 | 0.36 | 0.25 | 0.24 | 0.13 | 0.31 | 0.2 |
| 2 | 79317456 | rs2049269 | 1.406 | -1.445 | -1.203 | 0.551 | -0.015 | -0.013 | -0.013 | 0 | 0.21 | 0.16 | 0.26 | 0.42 | 0.38 | 0.32 |
| 2 | 79319089 | rs10206626 | -0.178 | -1.687 | -1.176 | 0.385 | 0.007 | 0.005 | 0.005 | 0.43 | 0.38 | 0.51 | 0.69 | 0.6 | 0.53 | 0.43 |
| 2 | 79323413 | rs11126695 | 1.571 | -1.695 | -1.531 | 0.221 | 0.023 | 0.061 | 0.061 | 0.203 | 0.28 | 0.02 | 0.09 | 0.11 | 0.2 | 0.16 |
| 2 | 79326215 | rs283895 | 0.586 | -1.844 | -1.32 | 0.376 | 0.024 | 0.014 | 0.014 | 0 | 0.38 | 0.19 | 0.25 | 0.32 | 0.35 | 0.46 |
| 2 | 79332083 | rs283851 | -0.811 | -1.734 | -1.025 | 0.389 | -0.011 | -0.052 | -0.052 | 0.113 | 0.88 | 0.71 | 0.83 | 0.97 | 0.91 | 0.98 |
| 2 | 79338929 | rs13407679 | 1.251 | -1.718 | -1.412 | 0.567 | -0.057 | -0.057 | -0.057 | 0.078 | 0.25 | 0.15 | 0.32 | 0.5 | 0.41 | 0.34 |
| 2 | 79345504 | rs2117727 | 1.192 | -1.719 | -1.476 | 0.942 | -0.063 | -0.03 | -0.03 | 0.145 | 0.26 | 0.15 | 0.32 | 0.3 | 0.41 | 0.34 |
| 2 | 79347050 | rs1349077 | 1.253 | -1.47 | -1.519 | 0.371 | -0.049 | -0.056 | -0.056 | 0 | 0.24 | 0.15 | 0.32 | 0.49 | 0.39 | 0.34 |
| 2 | 79347745 | rs283890 | -0.279 | -1.565 | -1.338 | 0.314 | 0.011 | 0.02 | 0.02 | 0.257 | 0.62 | 0.51 | 0.51 | 0.47 | 0.49 | 0.64 |
| 2 | 79348282 | rs2070708 | 0.086 | -1.535 | -1.065 | 0.51 | -0.047 | -0.023 | -0.023 | 0.003 | 0.25 | 0.34 | 0.26 | 0.16 | 0.13 | 0.18 |
| 2 | 79350505 | rs12072 | 0.644 | -0.429 | -1.373 | 0.887 | -0.047 | -0.054 | -0.054 | 0 | 0.24 | 0.15 | 0.31 | 0.5 | 0.39 | 0.34 |

| | | | | | | | | | | | | | | | | |
|---|---|---|---|---|---|---|---|---|---|---|---|---|---|---|---|---|
| 2 | 79352763 | rs4853434 | -1.054 | -0.485 | -1.412 | 0.827 | 0.002 | -0.004 | -0.004 | 0.854 | 0.85 | 0.86 | 0.92 | 0.92 | 0.81 | 0.74 |
| 2 | 79353898 | rs7608639 | -0.283 | 0.123 | -1.446 | 1.259 | -0.047 | -0.053 | -0.053 | 0 | 0.29 | 0.2 | 0.4 | 0.53 | 0.43 | 0.36 |
| 2 | 79359874 | rs17753959 | 0.466 | 0.006 | -1.227 | 1.033 | 0.002 | -0.002 | -0.002 | 0 | 0.05 | 0.05 | 0.11 | 0.07 | 0.04 | 0.03 |
| 2 | 79363251 | rs6721752 | -0.304 | -0.141 | -0.903 | 0.951 | -0.026 | 0.051 | 0.051 | 0.017 | 0.22 | 0.48 | 0.41 | 0.32 | 0.18 | 0.11 |
| 2 | 79363415 | rs3769505 | 0.565 | 0.176 | -1.325 | 1.003 | -0.096 | -0.083 | -0.083 | 0 | 0.22 | 0.1 | 0.34 | 0.41 | 0.39 | 0.31 |
| 2 | 79364601 | rs3819315 | 0.948 | -0.089 | -1.446 | 0.666 | 0.004 | 0.021 | 0.021 | 0 | 0.11 | 0.03 | 0.05 | 0.07 | 0.08 | 0.18 |
| 2 | 79364824 | rs879021 | 0.526 | 0.078 | -1.335 | 0.746 | -0.061 | -0.045 | -0.045 | 0 | 0.26 | 0.17 | 0.36 | 0.39 | 0.41 | 0.33 |
| 2 | 79364979 | rs892867 | -1.004 | -0.213 | -1.327 | 0.597 | 0.002 | 0.001 | 0.001 | 0.103 | 0.59 | 0.66 | 0.8 | 0.74 | 0.66 | 0.69 |
| 2 | 79366600 | rs3806561 | -0.321 | 0.269 | -1.152 | 0.993 | 0.031 | 0.031 | 0.031 | 0 | 0.31 | 0.44 | 0.6 | 0.52 | 0.46 | 0.35 |
| 2 | 79370426 | rs1589614 | -0.597 | -0.355 | -1.458 | 0.779 | 0.001 | -0.001 | -0.001 | 0 | 0.14 | 0.14 | 0.35 | 0.33 | 0.16 | 0.17 |
| 2 | 79371625 | rs1589615 | 0.176 | -0.318 | -1.717 | 0.74 | 0.008 | -0.007 | -0.007 | 0 | 0.1 | 0.03 | 0.18 | 0.12 | 0.1 | 0.09 |
| 2 | 79388587 | rs7588571 | -1.035 | -0.659 | -2.208 | 0.309 | 0.057 | 0.028 | 0.028 | 0 | 0.38 | 0.89 | 0.54 | 0.32 | 0.35 | 0.24 |
| 2 | 79400697 | rs17754357 | 0.502 | -0.376 | -1.813 | 0.249 | -0.01 | 0.045 | 0.045 | 0.288 | 0.09 | 0.01 | 0.06 | 0 | 0.07 | 0.02 |
| 2 | 79404926 | rs17016433 | 0.108 | -0.175 | -1.539 | -0.104 | 0.005 | -0.024 | -0.024 | 0.202 | 0.16 | 0.18 | 0.13 | 0.05 | 0.17 | 0.17 |
| 2 | 79405562 | rs986389 | 0.029 | -1.311 | -1.072 | -0.464 | -0.068 | -0.053 | -0.053 | 0 | 0.2 | 0.44 | 0.12 | 0.1 | 0.14 | 0.2 |
| 2 | 79410158 | rs892865 | -1.7 | -2.12 | -1.687 | -0.153 | -0.051 | -0.117 | -0.117 | 0 | 0.61 | 0.93 | 0.7 | 0.53 | 0.48 | 0.51 |
| 2 | 79415793 | rs9309546 | -0.213 | -2.18 | -1.652 | -0.206 | 0 | -0.035 | -0.035 | 0 | 0.31 | 0.38 | 0.31 | 0.14 | 0.29 | 0.21 |
| 2 | 79417632 | rs17016473 | 0.724 | -2.211 | -1.639 | -0.215 | -0.004 | -0.006 | -0.006 | 0 | 0.14 | 0.17 | 0.18 | 0.09 | 0.11 | 0.03 |
| 2 | 79417943 | rs10496214 | 1.988 | -2.259 | -1.69 | -0.158 | 0.021 | -0.025 | -0.025 | 1.96 | 0.08 | 0 | 0.17 | 0.13 | 0.05 | 0.03 |
| 2 | 79418487 | rs11126702 | -0.3 | -2.247 | -1.682 | -0.106 | -0.001 | -0.009 | -0.009 | 0.34 | 0.31 | 0.38 | 0.39 | 0.25 | 0.29 | 0.26 |
| 2 | 79419891 | rs1991992 | 0.975 | -2.204 | -1.724 | -0.091 | 0.003 | -0.021 | -0.021 | 0 | 0.17 | 0.2 | 0.13 | 0.05 | 0.17 | 0.17 |
| 2 | 79420044 | rs7592506 | -1.695 | -2.113 | -1.923 | -0.063 | -0.057 | -0.081 | -0.081 | 0 | 0.6 | 0.93 | 0.68 | 0.48 | 0.48 | 0.48 |
| 2 | 79432784 | rs17016549 | 1.505 | -2.044 | -1.874 | 0.013 | 0.022 | -0.028 | -0.028 | 2.336 | 0.1 | 0 | 0.18 | 0.14 | 0.06 | 0.03 |
| 2 | 79433627 | rs13407231 | -0.745 | -2.214 | -1.802 | -0.101 | -0.011 | -0.023 | -0.023 | 0 | 0.41 | 0.49 | 0.49 | 0.27 | 0.34 | 0.25 |
| 2 | 79438263 | rs6734557 | -0.479 | -2.261 | -1.723 | -0.142 | -0.044 | 0.003 | 0.003 | 0 | 0.35 | 0.5 | 0.37 | 0.33 | 0.27 | 0.28 |
| 2 | 79445309 | rs12994866 | 1.197 | -2.08 | -1.369 | -0.219 | -0.012 | -0.031 | -0.031 | 0.039 | 0.39 | 0.21 | 0.29 | 0.53 | 0.53 | 0.48 |
| 2 | 79447701 | rs12467106 | -1.365 | -2.05 | -1.315 | -0.269 | 0.02 | -0.095 | -0.095 | 0 | 0.85 | 0.71 | 0.87 | 0.99 | 0.86 | 0.8 |
| 2 | 79447879 | rs1529788 | -1.47 | -2.053 | -1.312 | -0.271 | 0.025 | -0.091 | -0.091 | 1.503 | 0.85 | 0.72 | 0.87 | 0.99 | 0.86 | 0.8 |
| 2 | 79448921 | rs12713982 | 0.804 | -2.317 | -1.484 | -0.388 | 0.102 | 0.028 | 0.028 | 0.192 | 0.52 | 0.15 | 0.46 | 0.6 | 0.64 | 0.5 |
| 2 | 79450227 | rs13404338 | 1.536 | -2.48 | -1.587 | -0.427 | -0.061 | -0.086 | -0.086 | 0.108 | 0.36 | 0.11 | 0.28 | 0.52 | 0.52 | 0.47 |
| 2 | 79450316 | rs4853444 | 0.606 | -2.298 | -1.399 | -0.632 | -0.049 | -0.027 | -0.027 | 0 | 0.19 | 0.44 | 0.14 | 0.12 | 0.16 | 0.2 |
| 2 | 79451065 | rs1434197 | 0.022 | -2.094 | -1.282 | -0.616 | -0.003 | -0.053 | -0.053 | 0 | 0.69 | 0.66 | 0.69 | 0.91 | 0.74 | 0.76 |
| 2 | 79460830 | rs13014809 | 0.812 | -1.828 | -2.071 | -0.299 | 0.017 | -0.014 | -0.014 | 0.121 | 0.25 | 0.04 | 0.16 | 0.22 | 0.18 | 0.26 |
| 2 | 79462603 | rs10779960 | 0.643 | -1.611 | -1.537 | -0.177 | 0.016 | 0.022 | 0.022 | 0 | 0.28 | 0.31 | 0.28 | 0.46 | 0.44 | 0.43 |
| 2 | 79468305 | rs1996867 | -0.062 | -2.148 | -1.175 | -0.249 | 0.004 | 0.01 | 0.01 | 0 | 0.49 | 0.58 | 0.55 | 0.62 | 0.6 | 0.51 |
| 2 | 79472697 | rs735358 | -0.334 | -1.558 | -0.816 | -0.155 | 0.013 | -0.06 | -0.06 | 4.412 | 0.73 | 0.59 | 0.71 | 0.83 | 0.72 | 0.75 |
| 2 | 79476846 | rs7607254 | 0.171 | -1.598 | -1.103 | -0.125 | 0.071 | 0.029 | 0.029 | 0 | 0.38 | 0.19 | 0.35 | 0.32 | 0.23 | 0.31 |
| 2 | 79478859 | rs6710298 | 0.509 | -1.201 | -0.632 | 0.106 | -0.009 | -0.006 | -0.006 | 0 | 0.31 | 0.28 | 0.35 | 0.51 | 0.48 | 0.41 |
| 2 | 79478988 | rs6739387 | 0.116 | -1.275 | -0.444 | 0.181 | 0.012 | 0.007 | 0.007 | 0 | 0.35 | 0.4 | 0.35 | 0.5 | 0.49 | 0.43 |
| 2 | 79481038 | rs1434207 | -0.517 | -1.504 | -0.237 | 0.003 | 0.012 | 0.045 | 0.045 | 0 | 0.47 | 0.65 | 0.52 | 0.66 | 0.61 | 0.53 |
| 2 | 79481297 | rs2861789 | 0.802 | -1.516 | -0.21 | 0.219 | -0.021 | -0.019 | -0.019 | 0 | 0.31 | 0.25 | 0.34 | 0.44 | 0.48 | 0.41 |
| 2 | 79482688 | rs934516 | -0.186 | -1.466 | -0.129 | 0.196 | -0.001 | -0.013 | -0.013 | 0 | 0.43 | 0.46 | 0.51 | 0.66 | 0.6 | 0.51 |
| 2 | 79486250 | rs6757830 | -0.268 | -1.154 | -0.214 | 0.242 | -0.009 | -0.006 | -0.006 | 0 | 0.09 | 0.17 | 0.02 | 0.07 | 0.05 | 0.13 |
| 2 | 79488285 | rs934510 | -0.577 | -1.162 | -0.25 | 0.345 | 0.009 | -0.003 | -0.003 | 0 | 0.48 | 0.56 | 0.51 | 0.55 | 0.59 | 0.52 |
| 2 | 79489923 | rs17705745 | 1.162 | -1.186 | -0.277 | 0.401 | -0.006 | -0.003 | -0.003 | 0 | 0.27 | 0.25 | 0.32 | 0.32 | 0.44 | 0.29 |
| 2 | 79495572 | rs1434189 | -0.67 | -1.181 | -0.319 | 0.41 | 0.021 | 0.031 | 0.031 | 0 | 0.48 | 0.68 | 0.52 | 0.6 | 0.6 | 0.52 |
| 2 | 79497295 | rs11674663 | 1.715 | -1.606 | -0.722 | 0.329 | 0.028 | -0.052 | -0.052 | 0 | 0.18 | 0.02 | 0.05 | 0.18 | 0.1 | 0.06 |
| 2 | 79497558 | rs10190281 | -0.541 | -1.467 | -0.348 | 0.406 | -0.033 | 0.056 | 0.056 | 0.031 | 0.21 | 0.43 | 0.2 | 0.28 | 0.16 | 0.23 |
| 2 | 79497829 | rs17016675 | 0.86 | -1.478 | -0.382 | 0.41 | -0.011 | -0.006 | -0.006 | 0 | 0.11 | 0.16 | 0.16 | 0.13 | 0.1 | 0.1 |
| 2 | 79498507 | rs1368952 | 1.406 | -1.554 | -0.535 | 0.308 | 0.002 | -0.05 | -0.05 | 0.98 | 0.25 | 0.04 | 0.18 | 0.25 | 0.18 | 0.28 |
| 2 | 79499259 | rs1368953 | -0.07 | -0.98 | 0.103 | 0.488 | -0.006 | -0.002 | -0.002 | 0 | 0.4 | 0.35 | 0.49 | 0.45 | 0.55 | 0.39 |
| 2 | 79501362 | rs12622013 | 0.451 | -0.967 | 0.11 | 0.506 | -0.013 | 0.051 | 0.051 | 0 | 0.11 | 0.22 | 0.17 | 0.18 | 0.11 | 0.11 |
| 2 | 79502890 | rs1434191 | -1.025 | -0.963 | 0.062 | 0.493 | 0.022 | 0.031 | 0.031 | 0 | 0.5 | 0.65 | 0.52 | 0.6 | 0.6 | 0.52 |
| 2 | 79504663 | rs1368954 | -0.417 | -0.935 | -0.267 | 0.277 | -0.004 | -0.009 | -0.009 | 0 | 0.45 | 0.61 | 0.51 | 0.55 | 0.56 | 0.5 |
| 2 | 79505248 | rs13009491 | 0.639 | -0.502 | -0.491 | 0.4 | -0.026 | -0.006 | -0.006 | 0.569 | 0.29 | 0.22 | 0.33 | 0.33 | 0.44 | 0.29 |
| 2 | 79507303 | rs7591792 | -0.987 | -0.752 | 0.061 | 0.191 | -0.011 | 0.002 | 0.002 | 0 | 0.09 | 0.2 | 0.02 | 0.09 | 0.05 | 0.13 |
| 2 | 79508503 | rs6547228 | 0.553 | -0.633 | -0.054 | -0.256 | 0 | -0.024 | -0.024 | 0.358 | 0.76 | 0.76 | 0.78 | 0.87 | 0.77 | 0.83 |
| 2 | 79508931 | rs7596189 | 1.319 | -0.967 | -0.032 | -0.265 | 0.04 | 0.043 | 0.043 | 0 | 0.59 | 0.21 | 0.65 | 0.68 | 0.68 | 0.68 |
| 2 | 79509292 | rs11126705 | 1.808 | -1.009 | -0.071 | -0.419 | 0.032 | 0.132 | 0.132 | 0 | 0.51 | 0.28 | 0.59 | 0.74 | 0.59 | 0.64 |
| 2 | 79516139 | rs12623011 | 0.69 | -0.995 | -0.05 | -0.408 | 0.001 | 0.012 | 0.012 | 0 | 0.68 | 0.75 | 0.72 | 0.82 | 0.72 | 0.79 |
| 2 | 79517133 | rs11126706 | 1.72 | -1.184 | 0.251 | -0.418 | -0.001 | -0.008 | -0.008 | 0.107 | 0.52 | 0.49 | 0.61 | 0.74 | 0.6 | 0.64 |
| 2 | 79518889 | rs732260 | -0.559 | -1.198 | 0.356 | -0.468 | -0.012 | -0.082 | -0.082 | 0 | 0.24 | 0.4 | 0.17 | 0.08 | 0.21 | 0.18 |
| 2 | 79520327 | rs1434170 | 0.693 | -1.124 | 0.281 | -0.454 | 0.011 | -0.04 | -0.04 | 0 | 0.09 | 0.1 | 0.07 | 0 | 0.11 | 0.04 |
| 2 | 79521270 | rs11889051 | 0.551 | -1.17 | 0.418 | -0.466 | 0 | -0.026 | -0.026 | 0 | 0.13 | 0.26 | 0.1 | 0.08 | 0.12 | 0.16 |
| 2 | 79521952 | rs12471468 | 2.074 | -1.291 | 0.408 | -0.475 | 0.077 | 0.101 | 0.101 | 0 | 0.52 | 0.17 | 0.61 | 0.67 | 0.64 | 0.65 |
| 2 | 79522589 | rs10167813 | 0.755 | -1.447 | 0.6 | -0.265 | -0.031 | -0.029 | -0.029 | 0 | 0.74 | 0.62 | 0.77 | 0.82 | 0.84 | 0.83 |
| 2 | 79523292 | rs13402353 | 1.037 | -1.426 | 0.529 | -0.254 | -0.017 | -0.013 | -0.013 | 0 | 0.11 | 0.26 | 0.09 | 0.08 | 0.09 | 0.14 |
| 2 | 79524148 | rs934512 | 0.576 | -1.433 | 0.484 | -0.258 | 0.037 | 0.031 | 0.031 | 0 | 0.76 | 0.88 | 0.79 | 0.82 | 0.84 | 0.83 |
| 2 | 79525428 | rs1434205 | -0.748 | -1.032 | 0.844 | -0.059 | -0.006 | -0.034 | -0.034 | 0.137 | 0.22 | 0.69 | 0.17 | 0.08 | 0.18 | 0.16 |

| | | | | | | | | | | | | | | | | |
|---|---|---|---|---|---|---|---|---|---|---|---|---|---|---|---|---|
| 2 | 79526043 | rs11892349 | -0.005 | -0.951 | 0.864 | -0.021 | 0.071 | -0.052 | -0.052 | 0.077 | 0.1 | 0.41 | 0.08 | 0 | 0.09 | 0.02 |
| 2 | 79528694 | rs10520235 | -0.671 | -0.795 | 0.31 | 0.105 | -0.001 | 0.002 | 0.002 | 0.021 | 0.09 | 0.12 | 0.13 | 0.21 | 0.08 | 0.11 |
| 2 | 79529472 | rs6760650 | -1.14 | -0.834 | 0.249 | 0.021 | -0.019 | -0.071 | -0.071 | 0 | 0.3 | 0.48 | 0.23 | 0.11 | 0.23 | 0.2 |
| 2 | 79530813 | rs10200371 | -1.964 | -0.835 | 0.244 | 0.049 | -0.01 | -0.021 | -0.021 | 0.236 | 0.38 | 0.54 | 0.33 | 0.26 | 0.31 | 0.31 |
| 2 | 79536397 | rs13422993 | -0.419 | -0.821 | 0.306 | 0.057 | -0.038 | -0.032 | -0.032 | 0 | 0.22 | 0.33 | 0.17 | 0.11 | 0.14 | 0.18 |
| 2 | 79537278 | rs17016741 | 0.821 | -0.832 | 0.283 | 0.035 | 0.014 | -0.037 | -0.037 | 0 | 0.08 | 0.1 | 0.06 | 0 | 0.09 | 0.02 |
| 2 | 79538168 | rs13418113 | -1.189 | -0.792 | 0.269 | 0.058 | -0.02 | -0.06 | -0.06 | 0.352 | 0.31 | 0.5 | 0.23 | 0.11 | 0.24 | 0.2 |
| 2 | 79539758 | rs17016767 | 0.024 | -0.881 | -0.15 | 0.351 | -0.009 | -0.027 | -0.027 | 0 | 0.07 | 0.01 | 0.1 | 0.15 | 0.08 | 0.11 |
| 2 | 79541025 | rs13412922 | -1.209 | -0.911 | -0.098 | 0.366 | 0.011 | -0.008 | -0.008 | 0 | 0.19 | 0.14 | 0.18 | 0.25 | 0.13 | 0.15 |
| 2 | 79542116 | rs2060787 | -1.042 | -0.912 | -0.102 | 0.364 | 0.012 | 0.004 | 0.004 | 0.022 | 0.19 | 0.13 | 0.18 | 0.18 | 0.13 | 0.14 |
| 2 | 79546297 | rs6757696 | -1.445 | -0.933 | -0.06 | 0.361 | -0.026 | 0.004 | 0.004 | 0 | 0.3 | 0.35 | 0.27 | 0.32 | 0.22 | 0.29 |
| 2 | 79547342 | rs17016820 | 0.874 | -0.814 | -0.037 | 0.203 | -0.009 | 0.003 | 0.003 | 0.228 | 0.1 | 0.21 | 0.09 | 0.08 | 0.09 | 0.14 |
| 2 | 79549374 | rs10520237 | -1.113 | -0.743 | -0.338 | 0.197 | 0.01 | 0.002 | 0.002 | 0.028 | 0.19 | 0.14 | 0.18 | 0.18 | 0.13 | 0.14 |
| 2 | 79550787 | rs13429593 | 0.38 | -0.661 | -0.349 | 0.195 | -0.002 | -0.002 | -0.002 | 0.085 | 0.11 | 0.12 | 0.07 | 0.04 | 0.05 | 0.03 |
| 2 | 79554690 | rs17016879 | -0.499 | -0.427 | -0.255 | 0.322 | -0.015 | 0.005 | 0.005 | 0.043 | 0.18 | 0.3 | 0.2 | 0.22 | 0.17 | 0.26 |
| 2 | 79556909 | rs6706637 | -0.674 | 0.04 | -1.069 | 0.356 | -0.032 | 0.01 | 0.01 | 0 | 0.55 | 0.79 | 0.67 | 0.46 | 0.56 | 0.49 |
| 2 | 79556993 | rs6706744 | -0.057 | -0.078 | -0.477 | -0.191 | -0.002 | -0.079 | -0.079 | 0 | 0.25 | 0.23 | 0.16 | 0 | 0.23 | 0.08 |
| 2 | 79561648 | rs1469763 | 1.278 | 0.027 | -1.092 | -0.035 | -0.029 | -0.02 | -0.02 | 0.701 | 0.33 | 0.17 | 0.23 | 0.27 | 0.3 | 0.31 |
| 2 | 79563202 | rs10186222 | 0.292 | -0.244 | -0.939 | -0.219 | 0.01 | -0.007 | -0.007 | 2.322 | 0.2 | 0.13 | 0.11 | 0 | 0.14 | 0.05 |
| 2 | 79563519 | rs10496221 | 0.724 | 0.299 | -0.538 | 0.645 | -0.004 | -0.015 | -0.015 | 0 | 0.1 | 0.06 | 0.12 | 0.29 | 0.13 | 0.12 |
| 2 | 79563718 | rs17016899 | 0.71 | 0.311 | -0.584 | 0.624 | -0.009 | -0.026 | -0.026 | 0.148 | 0.1 | 0.04 | 0.1 | 0.25 | 0.13 | 0.12 |
| 2 | 79568621 | rs10865442 | -0.249 | -0.385 | -0.678 | 0.173 | -0.006 | 0.102 | 0.102 | 0 | 0.79 | 0.97 | 0.9 | 1 | 0.83 | 0.93 |
| 2 | 79573515 | rs1434208 | 1.162 | -0.327 | -0.566 | 0.303 | 0.013 | 0.059 | 0.059 | 0 | 0.46 | 0.24 | 0.32 | 0.26 | 0.43 | 0.49 |
| 2 | 79575187 | rs726236 | 0.655 | -0.411 | -0.012 | 0.565 | 0.04 | 0.045 | 0.045 | 0 | 0.19 | 0.31 | 0.47 | 0.35 | 0.28 | 0.26 |
| 2 | 79581175 | rs966878 | -0.021 | -0.193 | -0.304 | 0.48 | -0.007 | 0.016 | 0.016 | 0 | 0.74 | 0.81 | 0.83 | 0.96 | 0.73 | 0.83 |
| 2 | 79586600 | rs17016923 | 1.355 | -0.278 | -0.747 | 0.618 | 0.043 | 0.031 | 0.031 | 0 | 0.41 | 0.27 | 0.28 | 0.26 | 0.33 | 0.39 |
| 2 | 79586998 | rs11897194 | 1.371 | -0.281 | -0.744 | 0.617 | 0.049 | 0.033 | 0.033 | 0 | 0.41 | 0.21 | 0.28 | 0.26 | 0.33 | 0.39 |
| 2 | 79588189 | rs13011795 | 0.606 | -0.254 | -0.735 | 0.62 | 0.024 | 0.046 | 0.046 | 0 | 0.53 | 0.36 | 0.35 | 0.6 | 0.43 | 0.57 |
| 2 | 79593298 | rs1434173 | 0.277 | 0.181 | -0.835 | 0.841 | 0.003 | -0.006 | -0.006 | 0 | 0.11 | 0.09 | 0.06 | 0.35 | 0.1 | 0.17 |
| 2 | 79596675 | rs4143799 | 0.616 | 0.109 | -0.621 | 0.797 | -0.006 | 0.004 | 0.004 | 0 | 0.62 | 0.71 | 0.77 | 0.63 | 0.63 | 0.67 |
| 2 | 79599061 | rs6547231 | -0.185 | 0.385 | -0.886 | 0.636 | -0.005 | 0.002 | 0.002 | 0 | 0.13 | 0.1 | 0.09 | 0.04 | 0.17 | 0.14 |
| 2 | 79601323 | rs4852481 | -0.057 | -0.35 | -1.145 | 0.98 | -0.04 | -0.066 | -0.066 | 0 | 0.75 | 0.94 | 0.86 | 0.64 | 0.76 | 0.8 |
| 2 | 79603321 | rs11683598 | 1.141 | -0.643 | -1.103 | 1.029 | 0.047 | -0.02 | -0.02 | 0 | 0.41 | 0.23 | 0.37 | 0.36 | 0.33 | 0.38 |
| 2 | 79605911 | rs17760461 | 0.034 | -0.632 | -0.355 | 0.797 | 0.016 | -0.018 | -0.018 | 0.689 | 0.29 | 0.33 | 0.44 | 0.18 | 0.41 | 0.4 |
| 2 | 79606195 | rs11691007 | -0.3 | -0.531 | -0.279 | 1.039 | -0.063 | 0 | 0 | 0 | 0.75 | 0.98 | 0.86 | 0.77 | 0.77 | 0.86 |
| 2 | 79607267 | rs6741335 | 0.062 | -0.522 | 0.099 | 1.042 | -0.001 | -0.011 | -0.011 | 0.697 | 0.72 | 0.69 | 0.84 | 0.75 | 0.75 | 0.85 |
| 2 | 79608891 | rs6547234 | 0.17 | -0.576 | 0.19 | 1.107 | -0.007 | 0.003 | 0.003 | 0 | 0.71 | 0.54 | 0.81 | 0.65 | 0.75 | 0.85 |
| 2 | 79612064 | rs10171522 | 0.028 | -0.482 | -0.204 | 1.435 | -0.027 | -0.018 | -0.018 | 0.049 | 0.28 | 0.22 | 0.44 | 0.3 | 0.4 | 0.39 |
| 2 | 79618040 | rs995772 | -0.466 | -0.741 | -0.736 | 1.182 | -0.032 | 0.038 | 0.038 | 0 | 0.75 | 0.97 | 0.87 | 0.78 | 0.78 | 0.81 |
| 2 | 79618231 | rs6751052 | 0.812 | -0.885 | -0.626 | 0.914 | 0.043 | -0.014 | -0.014 | 0 | 0.45 | 0.72 | 0.42 | 0.49 | 0.37 | 0.38 |
| 2 | 79623646 | rs1901335 | 0.081 | -0.537 | -0.581 | 0.578 | -0.001 | -0.041 | -0.041 | 0 | 0.75 | 0.8 | 0.85 | 0.96 | 0.79 | 0.84 |
| 2 | 79624698 | rs1451540 | -0.143 | -0.612 | -1.263 | 0.739 | -0.018 | 0.107 | 0.107 | 0 | 0.75 | 0.98 | 0.87 | 0.92 | 0.8 | 0.83 |
| 2 | 79626629 | rs1584666 | -0.293 | -0.479 | -0.719 | 0.657 | -0.003 | -0.039 | -0.039 | 0.166 | 0.75 | 0.75 | 0.84 | 0.93 | 0.77 | 0.85 |
| 2 | 79632061 | rs11691832 | 0.497 | -0.294 | -1.478 | 0.786 | 0.005 | 0.007 | 0.007 | 0 | 0.27 | 0.2 | 0.18 | 0.23 | 0.23 | 0.25 |
| 2 | 79632347 | rs2100290 | -0.136 | 0.339 | -1.387 | 1.068 | -0.067 | -0.032 | -0.032 | 0 | 0.34 | 0.19 | 0.5 | 0.38 | 0.48 | 0.46 |
| 2 | 79641996 | rs6729991 | -0.089 | -0.178 | -0.969 | 0.882 | 0.025 | 0.008 | 0.008 | 0 | 0.39 | 0.33 | 0.28 | 0.34 | 0.27 | 0.46 |
| 2 | 79643276 | rs13384184 | -0.306 | 0.793 | -0.692 | 1.1 | -0.002 | -0.004 | -0.004 | 0 | 0.81 | 0.83 | 0.58 | 0.69 | 0.65 | 0.69 |
| 2 | 79651633 | rs10865443 | 0.484 | 0.638 | -0.601 | 0.964 | 0.046 | 0.02 | 0.02 | 1.309 | 0.68 | 0.45 | 0.48 | 0.38 | 0.53 | 0.56 |
| 2 | 79655517 | rs6714400 | 0.895 | 0.476 | -0.584 | 1.278 | 0.024 | -0.001 | -0.001 | 0.252 | 0.4 | 0.33 | 0.34 | 0.37 | 0.3 | 0.44 |
| 2 | 79657338 | rs6722266 | 0.23 | 0.533 | -0.634 | 1.203 | 0.054 | 0.054 | 0.054 | 0 | 0.5 | 0.69 | 0.43 | 0.63 | 0.41 | 0.55 |
| 2 | 79667252 | rs7564015 | 0.225 | 0.519 | -0.386 | 1.203 | 0.043 | 0.04 | 0.04 | 0 | 0.5 | 0.67 | 0.42 | 0.62 | 0.4 | 0.55 |
| 2 | 79679809 | rs678206 | -0.457 | 0.538 | 0.223 | 1.244 | -0.009 | -0.007 | -0.007 | 0.73 | 0.78 | 0.81 | 0.57 | 0.67 | 0.64 | 0.67 |
| 2 | 79688779 | rs410120 | 0.334 | 0.329 | 0.3 | 1.202 | 0.065 | 0.049 | 0.049 | 0 | 0.48 | 0.68 | 0.38 | 0.63 | 0.33 | 0.52 |
| 2 | 79689572 | rs1864548 | 0.097 | 0.411 | 0.482 | 1.171 | -0.005 | -0.006 | -0.006 | 0 | 0.38 | 0.4 | 0.22 | 0.31 | 0.33 | 0.23 |
| 2 | 79698996 | rs2250249 | 0.755 | 0.347 | 0.484 | 1.187 | -0.02 | 0 | 0 | 0 | 0.42 | 0.48 | 0.36 | 0.53 | 0.31 | 0.49 |
| 2 | 79699411 | rs1864549 | 0.141 | 0.368 | 0.387 | 1.132 | -0.013 | 0.084 | 0.084 | 0.811 | 0.06 | 0.21 | 0.04 | 0.12 | 0.02 | 0.03 |
| 2 | 79702156 | rs386271 | 0.763 | 0.311 | 0.863 | 1.103 | 0.033 | -0.004 | -0.004 | 0 | 0.4 | 0.3 | 0.34 | 0.38 | 0.3 | 0.45 |
| 2 | 79705678 | rs13405677 | -0.625 | 0.254 | 0.812 | 1.086 | 0.012 | 0.001 | 0.001 | 0 | 0.54 | 0.38 | 0.62 | 0.48 | 0.67 | 0.51 |
| 2 | 79709327 | rs6547242 | -1.066 | 0.227 | 0.295 | 1.067 | 0.064 | -0.033 | -0.033 | 0.561 | 0.62 | 0.84 | 0.67 | 0.62 | 0.7 | 0.55 |
| 2 | 79720299 | rs1430637 | -0.076 | 0.909 | 0.539 | 1.319 | 0.004 | -0.001 | -0.001 | 0 | 0.23 | 0.2 | 0.41 | 0.31 | 0.37 | 0.33 |
| 2 | 79725584 | rs1430638 | -1.06 | 0.397 | 0.438 | 1.241 | 0.009 | 0.011 | 0.011 | 0.2 | 0.36 | 0.62 | 0.54 | 0.58 | 0.49 | 0.41 |
| 2 | 79747625 | rs1583148 | -0.417 | 0.287 | -0.123 | 1.193 | -0.001 | -0.023 | -0.023 | 0 | 0.73 | 0.86 | 0.59 | 0.67 | 0.72 | 0.68 |
| 2 | 79756265 | rs6745583 | 1.353 | -0.358 | 0.058 | 0.63 | 0 | -0.028 | -0.028 | 0 | 0.35 | 0.34 | 0.14 | 0.07 | 0.21 | 0.17 |
| 2 | 79760238 | rs6719738 | -0.629 | -0.357 | 0.424 | 0.624 | -0.007 | 0.01 | 0.01 | 0 | 0.73 | 0.69 | 0.6 | 0.68 | 0.73 | 0.69 |
| 2 | 79761016 | rs6547243 | -0.781 | -0.198 | 0.406 | 0.657 | -0.011 | -0.005 | -0.005 | 0 | 0.33 | 0.34 | 0.46 | 0.61 | 0.52 | 0.43 |
| 2 | 79766537 | rs1864551 | 1.618 | -0.61 | 0.107 | 0.316 | -0.003 | -0.019 | -0.019 | 0.311 | 0.32 | 0.31 | 0.11 | 0.07 | 0.18 | 0.22 |
| 2 | 79766703 | rs6707658 | -0.633 | -0.591 | 0.002 | 0.338 | 0.017 | -0.037 | -0.037 | 0 | 0.7 | 0.92 | 0.61 | 0.67 | 0.72 | 0.66 |
| 2 | 79768731 | rs6716273 | -0.573 | -0.462 | 0.086 | 0.421 | -0.007 | -0.005 | -0.005 | 0 | 0.33 | 0.36 | 0.45 | 0.43 | 0.52 | 0.39 |
| 2 | 79772054 | rs17017233 | 0.099 | -0.004 | 0.165 | 0.505 | -0.035 | -0.046 | -0.046 | 0 | 0.25 | 0.19 | 0.37 | 0.51 | 0.39 | 0.37 |

| | | | | | | | | | | | | | | | | |
|---|---|---|---|---|---|---|---|---|---|---|---|---|---|---|---|---|
| 2 | 79773708 | rs6707264 | -0.339 | -0.001 | 0.16 | 0.497 | -0.047 | -0.042 | -0.042 | 0 | 0.28 | 0.22 | 0.41 | 0.62 | 0.46 | 0.41 |
| 2 | 79776291 | rs880813 | -0.105 | -0.155 | 0.312 | 0.44 | -0.015 | 0.013 | 0.013 | 0 | 0.64 | 0.54 | 0.54 | 0.66 | 0.67 | 0.55 |
| 2 | 79779796 | rs7588649 | 2.225 | -0.263 | 0.355 | 0.311 | 0.021 | 0.032 | 0.032 | 0 | 0.26 | 0.12 | 0.1 | 0.01 | 0.18 | 0.12 |
| 2 | 79781587 | rs17017249 | 0.829 | 0.403 | 0.26 | 0.705 | -0.054 | -0.05 | -0.05 | 0 | 0.15 | 0.05 | 0.29 | 0.27 | 0.26 | 0.3 |
| 2 | 79781745 | rs2566539 | -0.016 | -0.501 | 0.722 | 1.105 | -0.011 | 0.003 | 0.003 | 0 | 0.68 | 0.63 | 0.56 | 0.73 | 0.7 | 0.66 |
| 2 | 79783547 | rs2685162 | 0.356 | -0.486 | 0.738 | 1.01 | -0.019 | 0.022 | 0.022 | 0 | 0.64 | 0.41 | 0.56 | 0.56 | 0.69 | 0.52 |
| 2 | 79784187 | rs1972755 | -0.153 | -0.54 | 0.978 | 0.938 | -0.007 | 0.004 | 0.004 | 0 | 0.73 | 0.71 | 0.67 | 0.75 | 0.73 | 0.73 |
| 2 | 79793629 | rs2566541 | 1.132 | -0.496 | 0.827 | 0.351 | 0.001 | -0.001 | -0.001 | 0.156 | 0.57 | 0.39 | 0.49 | 0.59 | 0.6 | 0.61 |
| 2 | 79795361 | rs2566542 | -0.557 | -0.898 | 0.928 | 0.547 | 0.013 | -0.072 | -0.072 | 0.034 | 0.8 | 0.97 | 0.66 | 0.75 | 0.81 | 0.8 |
| 2 | 79801922 | rs2685151 | 1.24 | -0.915 | 0.479 | 0.522 | -0.004 | 0.002 | 0.002 | 0 | 0.63 | 0.38 | 0.55 | 0.62 | 0.7 | 0.68 |
| 2 | 79803287 | rs17017286 | -1.054 | -0.865 | 0.657 | 0.536 | 0.044 | -0.042 | -0.042 | 0 | 0.08 | 0.23 | 0.08 | 0 | 0.09 | 0.01 |
| 2 | 79811317 | rs11126736 | 1.187 | -0.895 | 0.508 | 0.519 | -0.005 | 0.008 | 0.008 | 0.007 | 0.66 | 0.37 | 0.54 | 0.64 | 0.69 | 0.71 |
| 2 | 79813196 | rs1430645 | 0.91 | -1.015 | 0.845 | 0.682 | 0.036 | 0.008 | 0.008 | 0.044 | 0.66 | 0.76 | 0.61 | 0.76 | 0.74 | 0.77 |
| 2 | 79813622 | rs7598670 | 0.737 | -0.912 | 0.974 | 0.744 | -0.002 | 0 | 0 | 0 | 0.71 | 0.67 | 0.63 | 0.64 | 0.77 | 0.71 |
| 2 | 79817143 | rs1549765 | -0.296 | -0.803 | 0.708 | 0.816 | 0.005 | -0.007 | -0.007 | 0 | 0.8 | 0.82 | 0.67 | 0.74 | 0.84 | 0.84 |
| 2 | 79818350 | rs2566550 | -0.462 | -0.798 | 0.555 | 0.81 | 0.027 | -0.065 | -0.065 | 0.547 | 0.81 | 0.95 | 0.68 | 0.76 | 0.84 | 0.84 |
| 2 | 79822618 | rs6750164 | 0.951 | -0.639 | 0.791 | 0.86 | -0.02 | 0.007 | 0.007 | 0 | 0.61 | 0.47 | 0.54 | 0.63 | 0.68 | 0.7 |
| 2 | 79827274 | rs7597183 | -0.939 | -0.56 | 0.637 | 0.973 | 0.066 | 0.11 | 0.11 | 0.424 | 0.07 | 0.36 | 0.08 | 0.14 | 0.07 | 0.1 |
| 2 | 79828929 | rs919537 | 0.37 | -0.593 | 0.275 | 0.979 | 0.054 | -0.006 | -0.006 | 0.092 | 0.69 | 0.88 | 0.63 | 0.75 | 0.79 | 0.79 |
| 2 | 79849617 | rs17017351 | -1.345 | 0.097 | 0.689 | 1.142 | 0.084 | 0.121 | 0.121 | 0.499 | 0.07 | 0.42 | 0.08 | 0.13 | 0.07 | 0.11 |
| 2 | 79855622 | rs6740609 | 0.32 | -0.239 | 0.05 | 0.972 | -0.007 | 0.003 | 0.003 | 0.5 | 0.75 | 0.76 | 0.52 | 0.73 | 0.8 | 0.85 |
| 2 | 79856831 | rs17715064 | -0.345 | 0.353 | -0.344 | 1.01 | -0.064 | -0.046 | -0.046 | 0.34 | 0.24 | 0.07 | 0.44 | 0.23 | 0.34 | 0.12 |
| 2 | 79859101 | rs4852501 | -0.424 | 0.056 | 0.173 | 1.042 | -0.005 | 0.058 | 0.058 | 0 | 0.14 | 0.28 | 0.23 | 0.28 | 0.14 | 0.19 |
| 2 | 79864370 | rs17017378 | 0.705 | 0.02 | 0.13 | 1.071 | -0.003 | 0.034 | 0.034 | 0 | 0.11 | 0.12 | 0.05 | 0.37 | 0.1 | 0.23 |
| 2 | 79864902 | rs17017379 | 0.386 | -0.054 | 0.228 | 1.034 | -0.01 | 0.06 | 0.06 | 0 | 0.12 | 0.14 | 0.05 | 0.37 | 0.1 | 0.23 |
| 2 | 79866608 | rs2033335 | -0.687 | -0.322 | 0.415 | 0.675 | 0.005 | 0 | 0 | 0 | 0.36 | 0.44 | 0.51 | 0.34 | 0.42 | 0.35 |
| 2 | 79867668 | rs1430642 | -1.154 | -0.249 | 0.378 | 0.453 | -0.002 | 0.024 | 0.024 | 0 | 0.48 | 0.41 | 0.55 | 0.71 | 0.51 | 0.58 |
| 2 | 79869978 | rs10209612 | -0.13 | -0.256 | 0.13 | 0.438 | -0.007 | 0.086 | 0.086 | 0.024 | 0.16 | 0.38 | 0.23 | 0.28 | 0.13 | 0.19 |
| 2 | 79873004 | rs3795994 | -1.727 | -0.233 | -0.495 | 0.527 | -0.067 | 0.134 | 0.134 | 0 | 0.6 | 0.96 | 0.76 | 0.79 | 0.59 | 0.71 |
| 2 | 79882260 | rs13395022 | -0.392 | -0.2 | 0.086 | 0.662 | -0.019 | 0.093 | 0.093 | 0 | 0.2 | 0.31 | 0.16 | 0.46 | 0.16 | 0.34 |
| 2 | 79885036 | rs17716058 | 0.898 | -0.211 | -0.047 | 0.654 | -0.005 | -0.021 | -0.021 | 0 | 0.07 | 0.02 | 0.08 | 0.07 | 0.07 | 0.12 |
| 2 | 79885936 | rs11126738 | -0.143 | 0.032 | 0.27 | 0.41 | -0.004 | 0.007 | 0.007 | 0 | 0.23 | 0.31 | 0.23 | 0.7 | 0.24 | 0.41 |
| 2 | 79887830 | rs11687661 | 0.754 | 0.12 | 0.254 | 0.135 | -0.002 | -0.011 | -0.011 | 0 | 0.19 | 0.18 | 0.31 | 0.04 | 0.25 | 0.09 |
| 2 | 79888869 | rs13006976 | 0.644 | 0.225 | 0.273 | 0.08 | -0.006 | -0.001 | -0.001 | 0.025 | 0.05 | 0.03 | 0.05 | 0 | 0.08 | 0.03 |
| 2 | 79890100 | rs2861854 | -1.059 | 0.112 | 0.431 | 0.049 | -0.004 | 0.058 | 0.058 | 0 | 0.49 | 0.63 | 0.56 | 0.74 | 0.51 | 0.5 |
| 2 | 79890172 | rs12464912 | -0.239 | 0.082 | 0.4 | 0.069 | -0.022 | -0.026 | -0.026 | 0 | 0.12 | 0.09 | 0.14 | 0.25 | 0.18 | 0.1 |
| 2 | 79890344 | rs2861855 | -1.601 | 0.083 | 0.41 | 0.053 | 0.006 | 0.043 | 0.043 | 0 | 0.57 | 0.66 | 0.61 | 0.79 | 0.59 | 0.67 |
| 2 | 79890866 | rs7590351 | 0.32 | 0.065 | 0.187 | 0.104 | 0.002 | 0.029 | 0.029 | 0 | 0.15 | 0.26 | 0.23 | 0.25 | 0.14 | 0.27 |
| 2 | 79892257 | rs1567529 | -0.918 | 0.116 | 0.236 | 0.554 | -0.014 | -0.014 | -0.014 | 0 | 0.28 | 0.42 | 0.24 | 0.75 | 0.26 | 0.46 |
| 2 | 79893775 | rs10190617 | -0.381 | 0.002 | 0.105 | 0.363 | -0.034 | 0.134 | 0.134 | 0 | 0.22 | 0.41 | 0.16 | 0.46 | 0.17 | 0.35 |
| 2 | 79894144 | rs10175610 | -0.025 | 0.144 | 0.042 | 0.367 | -0.007 | 0.007 | 0.007 | 0 | 0.21 | 0.17 | 0.32 | 0.04 | 0.25 | 0.09 |
| 2 | 79894172 | rs4852507 | 0.011 | 0.174 | -0.239 | 0.383 | -0.016 | 0.037 | 0.037 | 0 | 0.21 | 0.13 | 0.32 | 0.04 | 0.25 | 0.09 |
| 2 | 79897781 | rs17017476 | -0.827 | 0.132 | -0.971 | 1.24 | -0.006 | 0.032 | 0.032 | 0 | 0.19 | 0.22 | 0.15 | 0.46 | 0.16 | 0.34 |
| 2 | 79902900 | rs17727983 | -0.497 | 0.201 | -1.665 | 0.294 | -0.003 | 0.093 | 0.093 | 0 | 0.27 | 0.07 | 0.16 | 0.14 | 0.25 | 0.31 |
| 2 | 79903266 | rs1355414 | -1.113 | 0.205 | -1.191 | 0.901 | 0.004 | -0.004 | -0.004 | 1.502 | 0.44 | 0.35 | 0.46 | 0.44 | 0.41 | 0.43 |
| 2 | 79904049 | rs17017518 | 0.069 | 0.023 | -0.486 | 0.868 | 0.042 | 0.017 | 0.017 | 1.311 | 0.07 | 0.33 | 0.1 | 0.02 | 0.04 | 0.11 |
| 2 | 79905813 | rs4422182 | 0.291 | -0.052 | -0.449 | 1.008 | -0.015 | -0.013 | -0.013 | 0.543 | 0.14 | 0.06 | 0.2 | 0.13 | 0.16 | 0.07 |
| 2 | 79906106 | rs2861857 | -0.808 | -0.748 | -0.613 | 0.864 | -0.008 | 0.006 | 0.006 | 0 | 0.64 | 0.66 | 0.63 | 0.87 | 0.54 | 0.6 |
| 2 | 79906739 | rs4852508 | -0.73 | -0.889 | -0.517 | 0.742 | 0 | 0.05 | 0.05 | 0.016 | 0.15 | 0.33 | 0.16 | 0.22 | 0.13 | 0.13 |
| 2 | 79907315 | rs4258819 | -0.596 | 0.1 | -0.745 | 0.205 | 0.006 | -0.064 | -0.064 | 0.658 | 0.72 | 0.55 | 0.75 | 0.92 | 0.69 | 0.67 |
| 2 | 79907588 | rs6709350 | -0.284 | 0.265 | -0.823 | 0.102 | 0.01 | -0.04 | -0.04 | 0 | 0.71 | 0.62 | 0.72 | 0.91 | 0.64 | 0.64 |
| 2 | 79908507 | rs17017537 | 0.5 | 0.017 | -0.499 | -0.071 | -0.003 | -0.038 | -0.038 | 0 | 0.1 | 0.14 | 0.05 | 0 | 0.1 | 0.15 |
| 2 | 79909989 | rs13030183 | 0.176 | -0.359 | -0.831 | -0.048 | -0.024 | -0.005 | -0.005 | 0.046 | 0.36 | 0.21 | 0.54 | 0.37 | 0.41 | 0.37 |
| 2 | 79910116 | rs1511179 | -1.817 | -0.756 | -0.528 | -0.283 | 0.013 | -0.055 | -0.055 | 0 | 0.7 | 0.6 | 0.68 | 0.9 | 0.66 | 0.73 |
| 2 | 79910349 | rs12471455 | 1.625 | -0.74 | -0.559 | -0.274 | -0.001 | -0.026 | -0.026 | 1.67 | 0.09 | 0.12 | 0.05 | 0 | 0.09 | 0.14 |
| 2 | 79911834 | rs12472740 | -0.488 | -0.727 | -0.581 | -0.064 | -0.015 | 0.016 | 0.016 | 0 | 0.44 | 0.32 | 0.58 | 0.38 | 0.51 | 0.5 |
| 2 | 79914226 | rs10183482 | 1.964 | -0.818 | -0.73 | -0.001 | 0.021 | -0.114 | -0.114 | 0.284 | 0.16 | 0.02 | 0.06 | 0.42 | 0.11 | 0.12 |
| 2 | 79914884 | rs10184380 | 1.316 | -0.795 | -0.65 | 0.021 | 0.022 | -0.109 | -0.109 | 0.426 | 0.21 | 0.04 | 0.11 | 0.44 | 0.15 | 0.14 |
| 2 | 79915152 | rs6719934 | -0.491 | -0.852 | -0.543 | -0.065 | 0.026 | -0.024 | -0.024 | 0 | 0.35 | 0.26 | 0.21 | 0.55 | 0.27 | 0.42 |
| 2 | 79915275 | rs11684245 | 0.308 | -0.828 | -0.486 | -0.076 | -0.002 | 0 | 0 | 0 | 0.37 | 0.34 | 0.53 | 0.37 | 0.41 | 0.37 |
| 2 | 79917201 | rs1511181 | -0.485 | -0.825 | -0.487 | -0.072 | -0.005 | 0.008 | 0.008 | 0 | 0.45 | 0.4 | 0.59 | 0.37 | 0.5 | 0.5 |
| 2 | 79918572 | rs6732862 | -0.196 | -0.825 | -0.62 | -0.047 | 0.006 | -0.009 | -0.009 | 0.105 | 0.33 | 0.3 | 0.15 | 0.53 | 0.25 | 0.36 |
| 2 | 79921161 | rs11677290 | 0.828 | -0.752 | -0.987 | -0.072 | -0.017 | 0 | 0 | 0 | 0.31 | 0.14 | 0.45 | 0.29 | 0.33 | 0.32 |
| 2 | 79922802 | rs11695685 | 0.958 | -0.729 | -0.972 | -0.061 | -0.019 | -0.005 | -0.005 | 0 | 0.29 | 0.13 | 0.44 | 0.28 | 0.32 | 0.31 |
| 2 | 79925969 | rs6547260 | 1.409 | -0.762 | -1.021 | 0.017 | 0.022 | -0.081 | -0.081 | 0 | 0.2 | 0.08 | 0.11 | 0.45 | 0.15 | 0.14 |
| 2 | 79928312 | rs955749 | 0.16 | -0.564 | -0.908 | -0.143 | -0.005 | -0.001 | -0.001 | 0 | 0.08 | 0.04 | 0.09 | 0.09 | 0.09 | 0.06 |
| 2 | 79930269 | rs13028762 | 0.437 | -0.496 | -0.798 | -0.17 | -0.01 | 0.006 | 0.006 | 0 | 0.32 | 0.18 | 0.46 | 0.29 | 0.34 | 0.34 |
| 2 | 79932294 | rs6738702 | -1.539 | -0.452 | -0.387 | -0.328 | 0.028 | -0.09 | -0.09 | 0 | 0.72 | 0.54 | 0.76 | 0.94 | 0.69 | 0.8 |
| 2 | 79933595 | rs11902209 | -0.912 | -0.29 | -0.504 | -0.277 | -0.001 | -0.009 | -0.009 | 0.372 | 0.55 | 0.53 | 0.65 | 0.81 | 0.51 | 0.72 |

| | | | | | | | | | | | | | | | | |
|---|---|---|---|---|---|---|---|---|---|---|---|---|---|---|---|---|
| 2 | 79933683 | rs11682196 | 0.576 | -1.067 | -0.31 | 0.096 | 0.002 | 0.022 | 0.022 | 0.363 | 0.18 | 0.08 | 0.12 | 0.13 | 0.18 | 0.09 |
| 2 | 79934085 | rs17017634 | 0.564 | -1.716 | 0.1 | 0.425 | -0.04 | 0.025 | 0.025 | 1.448 | 0.23 | 0.3 | 0.11 | 0.4 | 0.14 | 0.23 |
| 2 | 79934288 | rs17017645 | -1.138 | -1.771 | -0.008 | 0.388 | -0.018 | 0.008 | 0.008 | 0.365 | 0.34 | 0.58 | 0.21 | 0.58 | 0.23 | 0.39 |
| 2 | 79934864 | rs7602740 | 1.01 | -1.796 | 0.08 | 0.453 | -0.001 | -0.002 | -0.002 | 0 | 0.22 | 0.21 | 0.1 | 0.39 | 0.13 | 0.23 |
| 2 | 79935003 | rs7576479 | -2.59 | -1.648 | -0.368 | -0.081 | -0.045 | 0.056 | 0.056 | 0 | 0.63 | 0.81 | 0.52 | 0.78 | 0.5 | 0.6 |
| 2 | 79936139 | rs4852513 | -0.951 | -1.598 | -0.365 | -0.036 | -0.005 | -0.001 | -0.001 | 0.207 | 0.36 | 0.59 | 0.23 | 0.6 | 0.22 | 0.4 |
| 2 | 79936545 | rs13007809 | 0.068 | -0.931 | -0.169 | -0.257 | -0.07 | 0.024 | 0.024 | 0 | 0.22 | 0.02 | 0.16 | 0.13 | 0.25 | 0.11 |
| 2 | 79941174 | rs6725030 | -0.201 | -0.77 | 0.491 | -0.393 | -0.008 | -0.058 | -0.058 | 0.766 | 0.1 | 0.25 | 0.07 | 0 | 0.08 | 0.12 |
| 2 | 79941692 | rs11895953 | -1.801 | -0.919 | 0.343 | -0.358 | -0.043 | -0.03 | -0.03 | 0.218 | 0.23 | 0.52 | 0.14 | 0.14 | 0.16 | 0.31 |
| 2 | 79942703 | rs7579935 | -2.58 | -0.809 | -0.309 | -0.381 | 0.018 | 0.077 | 0.077 | 0.054 | 0.45 | 0.81 | 0.42 | 0.28 | 0.42 | 0.47 |
| 2 | 79945441 | rs13034159 | -0.28 | -0.919 | -0.152 | -0.198 | -0.025 | 0.036 | 0.036 | 0 | 0.18 | 0.09 | 0.13 | 0.1 | 0.21 | 0.1 |
| 2 | 79956356 | rs11693318 | -0.488 | -1.003 | -0.09 | -0.173 | -0.005 | 0.022 | 0.022 | 0 | 0.21 | 0.13 | 0.14 | 0.1 | 0.24 | 0.11 |
| 2 | 79963574 | rs10496231 | -0.457 | -0.981 | -0.054 | -0.126 | -0.003 | -0.004 | -0.004 | 0.428 | 0.19 | 0.16 | 0.14 | 0.22 | 0.23 | 0.2 |
| 2 | 79966298 | rs13007685 | -1.796 | -0.997 | -0.227 | -0.126 | 0.026 | 0.044 | 0.044 | 0 | 0.41 | 0.7 | 0.23 | 0.23 | 0.33 | 0.38 |
| 2 | 79968075 | rs17017756 | -0.651 | -2.43 | 0.132 | -0.04 | -0.018 | -0.015 | -0.015 | 0.011 | 0.23 | 0.33 | 0.25 | 0.16 | 0.18 | 0.18 |
| 2 | 79970319 | rs13035866 | -0.476 | -2.486 | 0.834 | -0.01 | 0.022 | -0.011 | -0.011 | 0 | 0.19 | 0.32 | 0.12 | 0.17 | 0.22 | 0.11 |
| 2 | 79971248 | rs6724922 | -1.241 | -2.745 | 0.783 | -0.443 | -0.029 | -0.098 | -0.098 | 0 | 0.19 | 0.33 | 0.07 | 0 | 0.12 | 0.16 |
| 2 | 79973543 | rs13033608 | -1.532 | -2.673 | 0.563 | 0.131 | 0 | -0.022 | -0.022 | 0 | 0.34 | 0.45 | 0.19 | 0.23 | 0.33 | 0.34 |
| 2 | 79979268 | rs4852515 | -0.983 | -3.319 | 0.995 | 0.202 | 0.025 | 0.04 | 0.04 | 0 | 0.46 | 0.37 | 0.38 | 0.78 | 0.35 | 0.54 |
| 2 | 79982689 | rs925404 | -1.647 | -3.883 | 1.005 | 0.237 | 0.072 | 0.126 | 0.126 | 0 | 0.55 | 0.8 | 0.44 | 0.78 | 0.36 | 0.61 |
| 2 | 79983244 | rs9808114 | -1.677 | -3.863 | 1.112 | 0.285 | 0.065 | 0.099 | 0.099 | 0 | 0.54 | 0.73 | 0.42 | 0.78 | 0.36 | 0.6 |
| 2 | 79987825 | rs1022152 | -1.354 | -3.77 | 1.155 | 0.794 | -0.012 | 0.005 | 0.005 | 0 | 0.39 | 0.57 | 0.39 | 0.42 | 0.31 | 0.47 |
| 2 | 79989434 | rs7584972 | -1.605 | -3.15 | 1.066 | 0.983 | -0.002 | 0.009 | 0.009 | 0.767 | 0.24 | 0.67 | 0.33 | 0.27 | 0.21 | 0.2 |
| 2 | 79991610 | rs13001342 | 0.684 | -2.971 | 0.362 | 0.871 | 0.016 | -0.031 | -0.031 | 7.737 | 0.23 | 0.07 | 0.3 | 0.23 | 0.19 | 0.1 |
| 2 | 79992038 | rs7592956 | -0.87 | -2.407 | 0.878 | 1.194 | 0.033 | 0.022 | 0.022 | 0 | 0.14 | 0.24 | 0.28 | 0.17 | 0.19 | 0.1 |
| 2 | 79995558 | rs6709353 | -1.75 | -2.631 | 0.261 | 1.153 | 0.079 | 0.095 | 0.095 | 1.725 | 0.44 | 0.79 | 0.44 | 0.77 | 0.29 | 0.58 |
| 2 | 79995904 | rs17017837 | -0.651 | -2.222 | 0.353 | 1.681 | -0.018 | -0.075 | -0.075 | 0.114 | 0.18 | 0.07 | 0.26 | 0.53 | 0.2 | 0.26 |
| 2 | 79996773 | rs1397696 | -2 | -2.404 | 0.484 | 1.356 | 0.09 | 0.075 | 0.075 | 2.887 | 0.42 | 0.83 | 0.38 | 0.75 | 0.26 | 0.59 |
| 2 | 80002274 | rs1159457 | 0.848 | -2.389 | 0.33 | 1.179 | 0.023 | 0.024 | 0.024 | 0 | 0.1 | 0.06 | 0.02 | 0 | 0.05 | 0.07 |
| 2 | 80002523 | rs11126742 | -0.304 | -2.435 | 0.897 | 1.473 | -0.054 | 0.164 | 0.164 | 0.092 | 0.14 | 0.4 | 0.11 | 0.4 | 0.07 | 0.17 |
| 2 | 80007503 | rs6736573 | -0.453 | -2.423 | 1.028 | 1.537 | -0.001 | -0.004 | -0.004 | 0 | 0.18 | 0.3 | 0.27 | 0.14 | 0.16 | 0.16 |
| 2 | 80007717 | rs2861911 | -1.426 | -2.444 | 1.059 | 1.56 | -0.025 | 0.047 | 0.047 | 0.515 | 0.27 | 0.47 | 0.32 | 0.32 | 0.19 | 0.41 |
| 2 | 80007727 | rs17017866 | -0.373 | -2.435 | 0.994 | 1.557 | -0.014 | 0.044 | 0.044 | 0 | 0.08 | 0.14 | 0.04 | 0.18 | 0.03 | 0.21 |
| 2 | 80010375 | rs994984 | -2.058 | -2.418 | 0.948 | 1.553 | 0.085 | 0.022 | 0.022 | 3.035 | 0.41 | 0.87 | 0.39 | 0.7 | 0.25 | 0.58 |
| 2 | 80011752 | rs2861907 | 1.177 | -2.368 | 0.411 | 1.465 | -0.037 | -0.02 | -0.02 | 0 | 0.19 | 0.05 | 0.24 | 0.19 | 0.26 | 0.1 |
| 2 | 80012056 | rs7575462 | -0.194 | -2.363 | 0.464 | 1.105 | -0.057 | 0.184 | 0.184 | 0.404 | 0.15 | 0.49 | 0.07 | 0.39 | 0.06 | 0.18 |
| 2 | 80013260 | rs975051 | -2.269 | -2.366 | 0.31 | 0.669 | -0.183 | 0.256 | 0.256 | 0.187 | 0.63 | 0.98 | 0.71 | 0.96 | 0.53 | 0.73 |
| 2 | 80015386 | rs1397687 | -2.098 | -1.896 | 0.654 | 1.002 | 0.082 | 0.004 | 0.004 | 3.501 | 0.39 | 0.87 | 0.42 | 0.69 | 0.25 | 0.54 |
| 2 | 80016594 | rs7600023 | -0.342 | -1.817 | 0.339 | 1.057 | -0.066 | 0.145 | 0.145 | 0.755 | 0.16 | 0.46 | 0.11 | 0.39 | 0.06 | 0.18 |
| 2 | 80023852 | rs6751684 | 0.589 | -1.535 | -0.489 | 0.826 | -0.001 | -0.002 | -0.002 | 0 | 0.06 | 0.06 | 0.24 | 0.12 | 0.12 | 0.09 |
| 2 | 80025940 | rs4284854 | -1.421 | -1.454 | -0.729 | 0.78 | 0.058 | -0.026 | -0.026 | 0.028 | 0.39 | 0.89 | 0.43 | 0.67 | 0.25 | 0.54 |
| 2 | 80026432 | rs1160581 | -1.31 | -1.256 | -0.7 | 0.885 | 0.051 | 0.013 | 0.013 | 0.386 | 0.41 | 0.76 | 0.43 | 0.67 | 0.27 | 0.57 |
| 2 | 80035372 | rs7605056 | 0.191 | -0.569 | -1.34 | 0.741 | 0.03 | 0.02 | 0.02 | 0.212 | 0.08 | 0.15 | 0.25 | 0.12 | 0.14 | 0.13 |
| 2 | 80039791 | rs2102548 | -1.236 | -0.496 | -2.089 | 0.508 | -0.189 | 0.236 | 0.236 | 0.096 | 0.64 | 0.97 | 0.71 | 0.95 | 0.53 | 0.74 |
| 2 | 80040994 | rs7585451 | -0.55 | -0.352 | -0.618 | 1.705 | -0.004 | 0.039 | 0.039 | 0.022 | 0.22 | 0.57 | 0.33 | 0.26 | 0.19 | 0.29 |
| 2 | 80045133 | rs1511196 | -0.977 | -0.198 | -1.143 | 1.783 | 0.033 | -0.049 | -0.049 | 0 | 0.14 | 0.07 | 0.18 | 0.38 | 0.06 | 0.29 |
| 2 | 80054826 | rs6734305 | 0.1 | 0.175 | -0.927 | 1.564 | 0.014 | 0.016 | 0.016 | 0 | 0.53 | 0.66 | 0.43 | 0.39 | 0.46 | 0.47 |
| 2 | 80055917 | rs12713993 | 0.528 | -0.565 | -0.07 | 1.256 | 0.005 | 0.025 | 0.025 | 0.21 | 0.41 | 0.33 | 0.35 | 0.13 | 0.4 | 0.42 |
| 2 | 80056097 | rs10180951 | -0.462 | -0.723 | -0.505 | 1.524 | -0.001 | 0 | 0 | 0.127 | 0.89 | 0.9 | 0.86 | 0.82 | 0.92 | 0.77 |
| 2 | 80059495 | rs2861904 | 0.974 | -0.678 | -0.518 | 1.515 | -0.004 | 0.137 | 0.137 | 0.241 | 0.39 | 0.2 | 0.34 | 0.12 | 0.41 | 0.41 |
| 2 | 80064042 | rs6742151 | -0.205 | -0.72 | 0.3 | 1.727 | -0.036 | 0.003 | 0.003 | 0.75 | 0.12 | 0.3 | 0.06 | 0.13 | 0.05 | 0.05 |
| 2 | 80067886 | rs2974180 | -0.623 | -0.627 | 0.063 | 1.825 | -0.138 | -0.09 | -0.09 | 0 | 0.65 | 0.91 | 0.59 | 0.46 | 0.54 | 0.7 |
| 2 | 80077093 | rs12477070 | 0.682 | -0.817 | 0.56 | 1.741 | -0.018 | 0.014 | 0.014 | 0 | 0.83 | 0.81 | 0.84 | 0.71 | 0.9 | 0.75 |
| 2 | 80078285 | rs17339556 | 0.777 | 0.2 | 0.101 | 1.824 | -0.072 | 0.015 | 0.015 | 0 | 0.22 | 0.06 | 0.15 | 0.13 | 0.3 | 0.08 |
| 2 | 80081182 | rs2861914 | 0.602 | 0.218 | 0.587 | 1.812 | 0.003 | -0.003 | -0.003 | 3.593 | 0.23 | 0.2 | 0.17 | 0.12 | 0.3 | 0.08 |
| 2 | 80081856 | rs2974154 | 0.04 | 0.403 | 0.426 | 1.821 | -0.01 | 0.006 | 0.006 | 0 | 0.86 | 0.91 | 0.82 | 0.86 | 0.83 | 0.75 |
| 2 | 80083806 | rs2974157 | 0.102 | 0.396 | 0.548 | 1.811 | -0.005 | 0.004 | 0.004 | 0 | 0.86 | 0.88 | 0.82 | 0.88 | 0.84 | 0.75 |
| 2 | 80084011 | rs7600078 | -0.556 | 0.317 | 0.703 | 1.967 | 0 | -0.001 | -0.001 | 0 | 0.08 | 0.06 | 0.03 | 0.15 | 0.03 | 0.03 |
| 2 | 80084691 | rs1368918 | 0.011 | 0.333 | 0.677 | 1.943 | -0.01 | 0.007 | 0.007 | 0 | 0.86 | 0.91 | 0.82 | 0.86 | 0.83 | 0.74 |
| 2 | 80086434 | rs1368919 | -1.09 | 0.246 | 0.891 | 2.052 | 0.003 | -0.032 | -0.032 | 0 | 0.42 | 0.43 | 0.52 | 0.73 | 0.42 | 0.34 |
| 2 | 80086475 | rs2916492 | -0.459 | 0.337 | 0.772 | 2.057 | 0.004 | 0.002 | 0.002 | 0.014 | 0.34 | 0.32 | 0.48 | 0.58 | 0.39 | 0.3 |
| 2 | 80086888 | rs2916493 | -0.299 | 0.084 | 1.172 | 2.274 | -0.014 | 0.235 | 0.235 | 0 | 0.13 | 0.28 | 0.33 | 0.45 | 0.1 | 0.22 |
| 2 | 80089717 | rs2043453 | 1.273 | -0.267 | 1.416 | 1.806 | -0.07 | -0.067 | -0.067 | 0.409 | 0.78 | 0.63 | 0.8 | 0.91 | 0.92 | 0.78 |
| 2 | 80090527 | rs7560784 | -2.428 | 0.658 | 1.644 | 2.488 | 0.06 | 0.309 | 0.309 | 0.244 | 0.14 | 0.37 | 0.27 | 0.55 | 0.17 | 0.26 |
| 2 | 80092547 | rs6547286 | -2.517 | 0.778 | 1.567 | 2.483 | 0.053 | 0.194 | 0.194 | 0.212 | 0.14 | 0.32 | 0.33 | 0.58 | 0.18 | 0.26 |
| 2 | 80094614 | rs7582888 | -1.365 | 0.825 | 1.414 | 2.902 | 0.052 | 0.05 | 0.05 | 0 | 0.07 | 0.11 | 0.09 | 0.23 | 0.17 | 0.17 |
| 2 | 80095216 | rs6736489 | -1.143 | 0.351 | 1.418 | 2.449 | -0.033 | 0.026 | 0.026 | 0 | 0.13 | 0.3 | 0.21 | 0.11 | 0.07 | 0.12 |
| 2 | 80097279 | rs3755095 | -1.882 | 0.351 | 1.138 | 2.257 | -0.091 | 0.077 | 0.077 | 0.024 | 0.22 | 0.45 | 0.47 | 0.65 | 0.1 | 0.27 |
| 2 | 80102308 | rs6707690 | 0.169 | 1.219 | -0.048 | 1.498 | -0.064 | 0.072 | 0.072 | 0.081 | 0.24 | 0.07 | 0.21 | 0.12 | 0.31 | 0.15 |

| | | | | | | | | | | | | | | | | |
|---|---|---|---|---|---|---|---|---|---|---|---|---|---|---|---|---|
| 2 | 80104027 | rs4852536 | -1.236 | 1.023 | 0.854 | 2.069 | -0.037 | 0.034 | 0.034 | 0 | 0.15 | 0.35 | 0.28 | 0.19 | 0.09 | 0.14 |
| 2 | 80104331 | rs4852537 | -0.72 | 1.308 | 0.387 | 2.522 | -0.003 | 0.131 | 0.131 | 0 | 0.06 | 0.09 | 0.16 | 0.48 | 0.02 | 0.11 |
| 2 | 80104527 | rs2974161 | 0.913 | 1.685 | 0.287 | 2.026 | 0.028 | 0.05 | 0.05 | 0 | 0.79 | 0.71 | 0.73 | 0.43 | 0.7 | 0.64 |
| 2 | 80110642 | rs1434108 | 0.054 | 1.499 | -0.443 | 1.628 | -0.037 | -0.033 | -0.033 | 0 | 0.87 | 0.98 | 0.9 | 0.79 | 0.89 | 0.93 |
| 2 | 80115288 | rs983217 | 0.582 | 1.358 | -1.142 | 1.666 | -0.015 | 0.041 | 0.041 | 0 | 0.19 | 0.08 | 0.15 | 0.08 | 0.23 | 0.11 |
| 2 | 80115384 | rs2916510 | 1.751 | 0.845 | -1.287 | 1.144 | 0.045 | 0.173 | 0.173 | 0 | 0.4 | 0.15 | 0.17 | 0.06 | 0.32 | 0.32 |
| 2 | 80120235 | rs10496234 | -0.653 | 1.871 | 0.486 | 1.68 | -0.005 | -0.001 | -0.001 | 0 | 0.35 | 0.36 | 0.45 | 0.32 | 0.47 | 0.43 |
| 2 | 80124540 | rs1540441 | 0.238 | 2.064 | 0.505 | 1.795 | -0.011 | -0.026 | -0.026 | 0 | 0.24 | 0.16 | 0.33 | 0.29 | 0.29 | 0.24 |
| 2 | 80126127 | rs6740491 | 1.453 | 1.761 | 0.37 | 1.011 | -0.002 | -0.049 | -0.049 | 0 | 0.14 | 0.17 | 0.09 | 0 | 0.12 | 0.1 |
| 2 | 80126192 | rs2117289 | 0.562 | 1.655 | 0.057 | 1.209 | -0.003 | -0.038 | -0.038 | 0 | 0.27 | 0.2 | 0.37 | 0.59 | 0.3 | 0.3 |
| 2 | 80130964 | rs10865447 | -0.516 | 1.793 | 0.199 | 1.233 | 0.073 | 0.053 | 0.053 | 0.062 | 0.16 | 0.42 | 0.27 | 0.13 | 0.22 | 0.28 |
| 2 | 80134915 | rs17261195 | 0.672 | 1.63 | -0.615 | 1.283 | 0.002 | 0.001 | 0.001 | 0 | 0.11 | 0.14 | 0.24 | 0.09 | 0.17 | 0.18 |
| 2 | 80137331 | rs12172883 | 0.851 | 1.169 | -0.863 | 1.24 | -0.034 | 0.066 | 0.066 | 5.257 | 0.15 | 0.04 | 0.11 | 0.03 | 0.18 | 0.12 |
| 2 | 80142822 | rs2916515 | -0.591 | 1.065 | -0.595 | 1.19 | -0.001 | -0.001 | -0.001 | 0 | 0.69 | 0.66 | 0.6 | 0.78 | 0.56 | 0.68 |
| 2 | 80143021 | rs2916516 | 0.586 | 0.883 | -0.614 | 1.266 | 0.004 | -0.017 | -0.017 | 0.307 | 0.56 | 0.41 | 0.3 | 0.35 | 0.48 | 0.52 |
| 2 | 80144546 | rs2974168 | -0.245 | 1.271 | -1.089 | 1.021 | -0.021 | 0.042 | 0.042 | 0 | 0.65 | 0.74 | 0.75 | 0.84 | 0.56 | 0.67 |
| 2 | 80144828 | rs2916518 | -0.398 | 1.3 | -1.081 | 0.873 | -0.02 | 0.009 | 0.009 | 0 | 0.65 | 0.72 | 0.6 | 0.78 | 0.56 | 0.67 |
| 2 | 80144920 | rs2916519 | -0.398 | 1.3 | -1.081 | 0.873 | -0.02 | 0.009 | 0.009 | 0 | 0.65 | 0.72 | 0.6 | 0.78 | 0.55 | 0.67 |
| 2 | 80146561 | rs7572401 | 0.082 | 1.035 | -0.89 | 0.894 | -0.012 | 0.026 | 0.026 | 0.095 | 0.09 | 0.04 | 0.07 | 0.03 | 0.1 | 0.07 |
| 2 | 80151340 | rs10204907 | -1.007 | 0.858 | -0.482 | 1.383 | -0.005 | 0.056 | 0.056 | 0 | 0.08 | 0.16 | 0.28 | 0.24 | 0.08 | 0.13 |
| 2 | 80152516 | rs1816611 | -1.196 | 1.502 | -0.047 | 1.565 | 0.026 | 0.029 | 0.029 | 0 | 0.31 | 0.58 | 0.58 | 0.59 | 0.37 | 0.4 |
| 2 | 80155493 | rs2974149 | -1.017 | 1.701 | -0.135 | 1.533 | 0.001 | 0.021 | 0.021 | 0 | 0.11 | 0.17 | 0.32 | 0.26 | 0.14 | 0.14 |
| 2 | 80156026 | rs2916529 | -0.898 | 1.704 | -0.027 | 1.535 | 0.003 | 0.034 | 0.034 | 0 | 0.11 | 0.18 | 0.33 | 0.26 | 0.14 | 0.15 |
| 2 | 80156246 | rs2916530 | -0.892 | 1.221 | -0.74 | 0.842 | -0.044 | 0.127 | 0.127 | 0.713 | 0.69 | 0.9 | 0.77 | 0.88 | 0.63 | 0.71 |
| 2 | 80165319 | rs2916532 | 1.03 | 1.096 | -0.495 | 0.941 | -0.003 | -0.002 | -0.002 | 0.3 | 0.41 | 0.39 | 0.23 | 0.31 | 0.33 | 0.33 |
| 2 | 80169400 | rs2916480 | 1.318 | 0.94 | -0.394 | 0.971 | 0.073 | 0.093 | 0.093 | 0.579 | 0.35 | 0.19 | 0.13 | 0.07 | 0.23 | 0.21 |
| 2 | 80171873 | rs1030733 | -1.504 | 0.826 | -0.563 | 1.119 | 0.009 | 0.007 | 0.007 | 8.032 | 0.87 | 0.9 | 0.96 | 0.95 | 0.92 | 0.88 |
| 2 | 80171888 | rs1030734 | 0.914 | 0.915 | -0.106 | 1.136 | -0.024 | -0.01 | -0.01 | 0 | 0.17 | 0.1 | 0.13 | 0.25 | 0.23 | 0.26 |
| 2 | 80173213 | rs2974139 | -0.543 | 0.871 | 0.25 | 1.158 | -0.059 | 0.017 | 0.017 | 7.774 | 0.83 | 0.57 | 0.82 | 0.88 | 0.92 | 0.88 |
| 2 | 80173900 | rs10172596 | 0.105 | 1.065 | 0.237 | 1.278 | 0.067 | 0.082 | 0.082 | 0 | 0.27 | 0.46 | 0.42 | 0.36 | 0.39 | 0.28 |
| 2 | 80174923 | rs4143780 | 0.799 | 1.056 | 0.134 | 1.213 | -0.032 | -0.018 | -0.018 | 0 | 0.23 | 0.13 | 0.28 | 0.28 | 0.39 | 0.28 |
| 2 | 80175253 | rs1434104 | 0.846 | 0.997 | 0.473 | 0.841 | 0.006 | -0.044 | -0.044 | 0.137 | 0.58 | 0.56 | 0.35 | 0.14 | 0.54 | 0.38 |
| 2 | 80175510 | rs1434105 | 1.575 | 1.028 | 0.073 | 0.843 | -0.054 | 0.059 | 0.059 | 0 | 0.18 | 0.08 | 0.19 | 0.06 | 0.29 | 0.16 |
| 2 | 80175600 | rs1434106 | -0.025 | 0.965 | 0.07 | 0.871 | -0.02 | 0.014 | 0.014 | 1.019 | 0.71 | 0.81 | 0.84 | 0.7 | 0.72 | 0.63 |
| 2 | 80175705 | rs2916486 | 1.383 | 0.304 | -0.064 | 0.728 | 0.084 | 0.115 | 0.115 | 0.197 | 0.37 | 0.2 | 0.15 | 0.07 | 0.25 | 0.22 |
| 2 | 80177744 | rs12472674 | -0.275 | 0.432 | 0.574 | 1.254 | 0.006 | -0.019 | -0.019 | 0 | 0.41 | 0.32 | 0.61 | 0.61 | 0.57 | 0.53 |
| 2 | 80178514 | rs1836208 | 0.339 | 0.382 | 0.594 | 1.369 | 0.016 | 0.027 | 0.027 | 0 | 0.51 | 0.61 | 0.58 | 0.61 | 0.43 | 0.46 |
| 2 | 80180246 | rs2974134 | 1.373 | -0.041 | 0.196 | 0.947 | 0.052 | 0.131 | 0.131 | 0.058 | 0.35 | 0.18 | 0.15 | 0.01 | 0.25 | 0.17 |
| 2 | 80182551 | rs4591366 | -1.222 | 0.165 | 0.459 | 1.552 | 0 | 0.127 | 0.127 | 0 | 0.14 | 0.23 | 0.28 | 0.52 | 0.14 | 0.28 |
| 2 | 80185073 | rs4852169 | -0.041 | 0.029 | -0.159 | 1.323 | 0.003 | 0.006 | 0.006 | 0.258 | 0.08 | 0.08 | 0.23 | 0.27 | 0.08 | 0.14 |
| 2 | 80185314 | rs892454 | -0.44 | 0.455 | -0.477 | 0.757 | -0.081 | 0.113 | 0.113 | 0.061 | 0.64 | 0.86 | 0.72 | 0.85 | 0.52 | 0.6 |
| 2 | 80185514 | rs2861960 | -0.528 | 0.21 | -0.547 | 0.643 | 0.021 | 0.055 | 0.055 | 0.939 | 0.82 | 0.95 | 0.92 | 0.93 | 0.86 | 0.81 |
| 2 | 80185885 | rs747921 | 0.624 | -0.07 | -0.522 | 0.604 | -0.012 | -0.023 | -0.023 | 0 | 0.45 | 0.24 | 0.28 | 0.52 | 0.4 | 0.44 |
| 2 | 80186107 | rs13032014 | 1.128 | 0.498 | -0.284 | 0.846 | -0.053 | -0.017 | -0.017 | 0.916 | 0.24 | 0.13 | 0.39 | 0.17 | 0.38 | 0.24 |
| 2 | 80187969 | rs6752828 | -0.298 | 0.209 | -0.494 | 0.538 | 0.004 | -0.002 | -0.002 | 1.095 | 0.43 | 0.35 | 0.44 | 0.6 | 0.53 | 0.53 |
| 2 | 80188021 | rs2060396 | 1.142 | 0.103 | -0.386 | 0.474 | -0.017 | 0.002 | 0.002 | 0 | 0.38 | 0.57 | 0.43 | 0.35 | 0.26 | 0.24 |
| 2 | 80188476 | rs12997292 | 1.297 | 0.415 | -0.779 | 0.289 | -0.049 | 0.053 | 0.053 | 0 | 0.24 | 0.13 | 0.28 | 0.08 | 0.38 | 0.24 |
| 2 | 80189383 | rs10173073 | -0.454 | 0.117 | -0.562 | 0.302 | 0.013 | 0.06 | 0.06 | 0 | 0.45 | 0.68 | 0.58 | 0.69 | 0.53 | 0.54 |
| 2 | 80191985 | rs2974132 | -0.073 | 0.057 | -1.147 | 0.116 | -0.002 | 0.01 | 0.01 | 0.043 | 0.58 | 0.69 | 0.68 | 0.65 | 0.56 | 0.5 |
| 2 | 80192352 | rs2916490 | 0.778 | -0.326 | -1.675 | 0.201 | 0.072 | 0.048 | 0.048 | 0.536 | 0.41 | 0.12 | 0.24 | 0.31 | 0.3 | 0.31 |
| 2 | 80192435 | rs2861913 | 0.833 | 0.367 | -1.148 | 0.035 | -0.01 | 0.011 | 0.011 | 0 | 0.23 | 0.19 | 0.26 | 0.09 | 0.32 | 0.24 |
| 2 | 80194080 | rs13001262 | -0.588 | -0.429 | -1.104 | 0.478 | 0.08 | -0.014 | -0.014 | 0 | 0.51 | 0.86 | 0.46 | 0.49 | 0.62 | 0.57 |
| 2 | 80194625 | rs13028697 | 0.722 | -0.232 | -1.153 | 0.355 | -0.051 | 0.071 | 0.071 | 0 | 0.29 | 0.16 | 0.26 | 0.09 | 0.43 | 0.3 |
| 2 | 80196736 | rs542255 | 0.937 | -0.559 | -1.491 | 0.587 | 0.047 | -0.085 | -0.085 | 0.01 | 0.32 | 0.11 | 0.43 | 0.47 | 0.2 | 0.29 |
| 2 | 80196987 | rs17018346 | -0.382 | -0.424 | -1.38 | 0.595 | -0.038 | -0.075 | -0.075 | 0.146 | 0.2 | 0.09 | 0.14 | 0.4 | 0.25 | 0.3 |
| 2 | 80197438 | rs671802 | -0.061 | -0.494 | -0.965 | 0.248 | 0.003 | -0.014 | -0.014 | 0 | 0.79 | 0.81 | 0.86 | 1 | 0.79 | 0.89 |
| 2 | 80197462 | rs671772 | 1.118 | -0.576 | -0.221 | 0.427 | -0.001 | 0.003 | 0.003 | 0.017 | 0.62 | 0.42 | 0.76 | 0.59 | 0.63 | 0.61 |
| 2 | 80197843 | rs17261321 | 0.031 | -0.514 | -0.202 | 0.643 | 0.011 | 0.024 | 0.024 | 0 | 0.48 | 0.41 | 0.6 | 0.65 | 0.39 | 0.49 |
| 2 | 80198597 | rs490805 | 0.297 | -0.27 | -0.217 | 0.816 | -0.001 | 0.018 | 0.018 | 0 | 0.9 | 0.87 | 0.94 | 0.78 | 0.91 | 0.89 |
| 2 | 80199239 | rs518046 | 1.185 | -0.545 | -0.015 | 0.919 | -0.022 | 0.037 | 0.037 | 0 | 0.65 | 0.49 | 0.66 | 0.49 | 0.71 | 0.72 |
| 2 | 80200955 | rs478083 | 0.855 | -0.536 | -0.042 | 0.86 | -0.024 | 0.03 | 0.03 | 0 | 0.7 | 0.54 | 0.75 | 0.57 | 0.76 | 0.74 |
| 2 | 80204329 | rs648361 | 0.292 | -0.765 | -0.772 | 0.528 | 0.024 | 0.167 | 0.167 | 0.271 | 0.51 | 0.17 | 0.36 | 0.23 | 0.63 | 0.41 |
| 2 | 80213680 | rs544483 | 0.667 | -0.569 | 0.027 | 0.484 | 0.022 | 0.025 | 0.025 | 0 | 0.73 | 0.61 | 0.42 | 0.33 | 0.72 | 0.63 |
| 2 | 80214554 | rs598453 | 0.549 | -0.703 | -0.423 | 0.307 | -0.002 | 0.025 | 0.025 | 0 | 0.48 | 0.38 | 0.33 | 0.33 | 0.57 | 0.42 |
| 2 | 80214820 | rs609682 | 1.503 | -0.675 | -0.724 | 0.279 | -0.054 | -0.012 | -0.012 | 1.795 | 0.32 | 0.12 | 0.25 | 0.32 | 0.44 | 0.34 |
| 2 | 80215228 | rs13389996 | 1.651 | -0.678 | -0.795 | 0.271 | 0.02 | 0.102 | 0.102 | 0 | 0.23 | 0.09 | 0.07 | 0 | 0.15 | 0.21 |
| 2 | 80216176 | rs17018359 | 0.564 | -0.483 | -0.871 | 0.684 | -0.026 | -0.057 | -0.057 | 0 | 0.1 | 0.01 | 0.04 | 0.21 | 0.08 | 0.13 |
| 2 | 80216399 | rs556003 | 0.531 | -0.393 | -0.631 | 0.652 | 0.001 | -0.012 | -0.012 | 0 | 0.59 | 0.46 | 0.33 | 0.33 | 0.6 | 0.55 |
| 2 | 80217014 | rs13394481 | 0.599 | -0.415 | -0.534 | 0.561 | -0.017 | -0.041 | -0.041 | 0 | 0.26 | 0.31 | 0.09 | 0.06 | 0.16 | 0.21 |

| | | | | | | | | | | | | | | | | |
|---|---|---|---|---|---|---|---|---|---|---|---|---|---|---|---|---|
| 2 | 80219150 | rs489974 | 0.173 | 0.363 | -0.79 | 1.376 | -0.029 | -0.038 | -0.038 | 0.063 | 0.93 | 0.99 | 0.86 | 0.78 | 0.92 | 0.93 |
| 2 | 80221392 | rs17018376 | -0.658 | 0.08 | -0.38 | 0.984 | -0.007 | 0.071 | 0.071 | 0 | 0.14 | 0.25 | 0.25 | 0.29 | 0.11 | 0.21 |
| 2 | 80224755 | rs4852547 | -0.123 | 0.198 | -0.905 | -0.201 | 0 | -0.01 | -0.01 | 0 | 0.58 | 0.51 | 0.56 | 0.85 | 0.52 | 0.67 |
| 2 | 80226782 | rs563702 | 0.02 | -0.47 | -1.525 | -0.534 | 0.058 | 0.071 | 0.071 | 0 | 0.3 | 0.17 | 0.22 | 0.12 | 0.18 | 0.2 |
| 2 | 80228740 | rs1368909 | 0.254 | -0.333 | -1.423 | -0.901 | -0.038 | 0.009 | 0.009 | 0.145 | 0.35 | 0.18 | 0.47 | 0.73 | 0.42 | 0.57 |
| 2 | 80228863 | rs1897775 | -0.352 | -0.761 | -1.032 | -1.273 | 0.004 | 0.139 | 0.139 | 0 | 0.5 | 0.66 | 0.55 | 0.87 | 0.46 | 0.69 |
| 2 | 80230038 | rs1368910 | -0.217 | -1.504 | -0.444 | -1.434 | -0.075 | -0.078 | -0.078 | 5.427 | 0.71 | 0.57 | 0.62 | 0.92 | 0.85 | 0.88 |
| 2 | 80230127 | rs374511 | 0.851 | -0.561 | -0.973 | -1.793 | -0.024 | 0.031 | 0.031 | 0 | 0.41 | 0.23 | 0.48 | 0.77 | 0.57 | 0.45 |
| 2 | 80230424 | rs6741085 | 0.074 | -0.469 | -1.027 | -1.812 | -0.079 | 0.066 | 0.066 | 0.652 | 0.26 | 0.08 | 0.19 | 0.08 | 0.29 | 0.4 |
| 2 | 80231175 | rs431413 | -1.63 | -0.772 | -0.855 | -1.741 | 0.06 | 0.049 | 0.049 | 0.363 | 0.79 | 0.95 | 0.7 | 0.92 | 0.89 | 0.9 |
| 2 | 80231469 | rs560737 | 0.406 | -0.491 | -0.64 | -1.803 | 0.001 | 0 | 0 | 0 | 0.52 | 0.38 | 0.51 | 0.76 | 0.66 | 0.48 |
| 2 | 80234681 | rs12471345 | 0.284 | -0.868 | -0.786 | -1.687 | 0.005 | 0.024 | 0.024 | 0.033 | 0.19 | 0.09 | 0.12 | 0.08 | 0.14 | 0.34 |
| 2 | 80236251 | rs423412 | 0.592 | -0.99 | -0.879 | -1.764 | 0.063 | 0.053 | 0.053 | 0 | 0.45 | 0.22 | 0.5 | 0.76 | 0.7 | 0.48 |
| 2 | 80238375 | rs53916 | 0.702 | -1.059 | -0.928 | -1.803 | 0.051 | 0.04 | 0.04 | 0 | 0.43 | 0.22 | 0.5 | 0.75 | 0.7 | 0.48 |
| 2 | 80241463 | rs7597545 | -0.396 | -1.119 | -0.678 | -1.665 | -0.021 | -0.008 | -0.008 | 0.751 | 0.66 | 0.66 | 0.65 | 0.82 | 0.85 | 0.82 |
| 2 | 80242693 | rs6711371 | 0.307 | -0.585 | -0.948 | -1.375 | 0.051 | 0.028 | 0.028 | 0 | 0.43 | 0.22 | 0.51 | 0.72 | 0.7 | 0.48 |
| 2 | 80252665 | rs13402286 | -0.431 | -0.182 | -0.376 | -1.175 | -0.006 | 0.041 | 0.041 | 0.396 | 0.05 | 0.22 | 0.01 | 0.04 | 0 | 0.06 |
| 2 | 80259424 | rs921637 | -0.618 | 0.143 | -1.64 | -1.771 | -0.002 | 0.153 | 0.153 | 0 | 0.53 | 0.75 | 0.32 | 0.13 | 0.48 | 0.46 |
| 2 | 80261170 | rs6729700 | 0.536 | 0.224 | -1.668 | -1.798 | -0.052 | 0.215 | 0.215 | 0.236 | 0.45 | 0.17 | 0.65 | 0.86 | 0.49 | 0.53 |
| 2 | 80266700 | rs8179672 | 1.343 | 0.44 | -2.116 | -1.609 | 0.004 | 0.036 | 0.036 | 0.056 | 0.24 | 0.13 | 0.22 | 0.11 | 0.21 | 0.15 |
| 2 | 80266897 | rs10170918 | 1.435 | 0.668 | -2.224 | -1.819 | -0.02 | 0.016 | 0.016 | 0 | 0.15 | 0.12 | 0.16 | 0.1 | 0.28 | 0.13 |
| 2 | 80272344 | rs10865448 | -0.896 | 0.213 | -2.431 | -1.813 | 0.008 | 0.135 | 0.135 | 0 | 0.75 | 0.91 | 0.94 | 0.98 | 0.83 | 0.67 |
| 2 | 80272811 | rs10865451 | 1.057 | 0.077 | -2.092 | -1.339 | -0.073 | -0.04 | -0.04 | 0.005 | 0.22 | 0.03 | 0.44 | 0.76 | 0.29 | 0.37 |
| 2 | 80277135 | rs2861966 | -0.759 | -0.005 | -1.537 | -1.233 | -0.02 | 0.196 | 0.196 | 0.037 | 0.53 | 0.78 | 0.33 | 0.13 | 0.48 | 0.47 |
| 2 | 80283255 | rs9973600 | -0.277 | 0.548 | -1.198 | -0.935 | -0.012 | 0.038 | 0.038 | 0 | 0.41 | 0.34 | 0.24 | 0.1 | 0.45 | 0.45 |
| 2 | 80286981 | rs7596307 | 0.02 | 0.545 | -1.55 | -0.678 | -0.033 | 0.099 | 0.099 | 0 | 0.6 | 0.77 | 0.74 | 0.82 | 0.56 | 0.51 |
| 2 | 80293874 | rs1900270 | 0.437 | 0.154 | -0.921 | 0.219 | -0.009 | 0 | 0 | 0 | 0.61 | 0.39 | 0.59 | 0.39 | 0.7 | 0.4 |
| 2 | 80298707 | rs1484465 | 0.086 | 0.381 | -1.199 | 0.812 | -0.01 | -0.056 | -0.056 | 0.375 | 0.85 | 0.95 | 0.72 | 0.48 | 0.83 | 0.71 |
| 2 | 80299064 | rs1484466 | -0.989 | 0.299 | -1.365 | 0.346 | 0.006 | -0.001 | -0.001 | 0 | 0.3 | 0.11 | 0.42 | 0.7 | 0.29 | 0.65 |
| 2 | 80306270 | rs4852550 | -0.605 | 0.174 | -1.196 | -0.129 | -0.103 | 0.045 | 0.045 | 0 | 0.73 | 0.92 | 0.84 | 0.82 | 0.62 | 0.85 |
| 2 | 80310648 | rs716487 | -0.721 | -0.635 | 0.248 | 0.199 | -0.005 | 0.07 | 0.07 | 0 | 0.24 | 0.33 | 0.52 | 0.57 | 0.22 | 0.34 |
| 2 | 80312122 | rs1843131 | -0.707 | -0.645 | 0.381 | 0.229 | -0.005 | 0.051 | 0.051 | 0 | 0.24 | 0.28 | 0.52 | 0.53 | 0.22 | 0.37 |
| 2 | 80315216 | rs10181663 | -0.375 | -0.911 | 0.479 | -0.005 | -0.035 | 0.036 | 0.036 | 0.75 | 0.17 | 0.59 | 0.11 | 0.14 | 0.1 | 0.35 |
| 2 | 80317685 | rs2902109 | 0.276 | -0.867 | 0.481 | 0.122 | 0 | 0.014 | 0.014 | 0 | 0.17 | 0.14 | 0.46 | 0.49 | 0.16 | 0.28 |
| 2 | 80326330 | rs17018612 | 0.695 | -0.717 | 0.739 | 0.177 | 0.001 | -0.002 | -0.002 | 0 | 0.11 | 0.09 | 0.07 | 0.12 | 0.09 | 0.34 |
| 2 | 80329535 | rs6720989 | 0.354 | -1.112 | 0.54 | 0.021 | 0.002 | 0 | 0 | 0 | 0.21 | 0.16 | 0.5 | 0.5 | 0.19 | 0.27 |
| 2 | 80332151 | rs1031442 | 0.338 | -1.088 | 0.661 | 0.027 | -0.003 | 0.024 | 0.024 | 0 | 0.21 | 0.17 | 0.45 | 0.52 | 0.19 | 0.27 |
| 2 | 80335727 | rs7600625 | -0.731 | -1.145 | 1.123 | 0.011 | -0.015 | -0.042 | -0.042 | 0 | 0.21 | 0.3 | 0.14 | 0.09 | 0.16 | 0.39 |
| 2 | 80340234 | rs1017632 | 0.234 | -1.061 | 1.132 | 0.085 | -0.001 | 0.01 | 0.01 | 0.303 | 0.22 | 0.21 | 0.45 | 0.49 | 0.2 | 0.28 |
| 2 | 80342507 | rs12463927 | -1.675 | -1.056 | 1.118 | 0.052 | 0.055 | 0.095 | 0.095 | 0 | 0.45 | 0.76 | 0.66 | 0.69 | 0.34 | 0.68 |
| 2 | 80343419 | rs1900265 | -1.21 | -1.034 | 1.144 | 0.052 | -0.011 | 0 | 0 | 0 | 0.41 | 0.45 | 0.58 | 0.58 | 0.35 | 0.65 |
| 2 | 80359253 | rs3849389 | 1.39 | -0.452 | 0.696 | -0.087 | -0.067 | 0.089 | 0.089 | 0 | 0.38 | 0.19 | 0.23 | 0.23 | 0.49 | 0.23 |
| 2 | 80365193 | rs11685451 | -0.764 | -0.736 | 1.024 | -0.061 | -0.019 | 0.008 | 0.008 | 0 | 0.21 | 0.43 | 0.24 | 0.18 | 0.15 | 0.43 |
| 2 | 80371312 | rs1867805 | 0.51 | -0.686 | 0.881 | -0.034 | 0.004 | 0.007 | 0.007 | 0 | 0.5 | 0.51 | 0.32 | 0.35 | 0.58 | 0.36 |
| 2 | 80371730 | rs6547294 | 0.51 | -0.686 | 0.881 | -0.026 | 0.004 | 0.007 | 0.007 | 0 | 0.5 | 0.51 | 0.32 | 0.37 | 0.58 | 0.36 |
| 2 | 80372017 | rs7558009 | -0.352 | -0.723 | 0.728 | -0.032 | 0.013 | -0.035 | -0.035 | 0 | 0.63 | 0.83 | 0.53 | 0.47 | 0.67 | 0.69 |
| 2 | 80373393 | rs11686939 | 0.356 | -0.61 | 0.688 | -0.008 | 0.012 | 0.029 | 0.029 | 0.005 | 0.52 | 0.69 | 0.34 | 0.36 | 0.58 | 0.36 |
| 2 | 80373740 | rs17018689 | -1.653 | -0.429 | 1.31 | 0.229 | -0.011 | 0.007 | 0.007 | 2.689 | 0.13 | 0.46 | 0.07 | 0.07 | 0.06 | 0.1 |
| 2 | 80382666 | rs7564961 | -1.772 | -0.482 | 1.3 | 0.371 | -0.018 | 0.059 | 0.059 | 4.17 | 0.1 | 0.48 | 0.07 | 0.07 | 0.03 | 0.06 |
| 2 | 80384006 | rs6746626 | -1.132 | -0.476 | 0.901 | 0.353 | -0.037 | -0.01 | -0.01 | 0.918 | 0.2 | 0.65 | 0.25 | 0.16 | 0.12 | 0.38 |
| 2 | 80392892 | rs1319241 | -0.979 | -0.424 | 0.97 | 0.407 | -0.036 | 0.045 | 0.045 | 0.001 | 0.18 | 0.53 | 0.23 | 0.18 | 0.12 | 0.38 |
| 2 | 80399212 | rs11893350 | -0.625 | -0.521 | 1.348 | 0.571 | -0.021 | 0.026 | 0.026 | 0 | 0.14 | 0.38 | 0.2 | 0.13 | 0.09 | 0.33 |
| 2 | 80399551 | rs1319550 | 0.132 | -0.871 | 0.785 | 0.223 | 0.001 | 0.001 | 0.001 | 0 | 0.54 | 0.59 | 0.55 | 0.38 | 0.62 | 0.61 |
| 2 | 80401497 | rs4852559 | 1.266 | -0.877 | 0.499 | 0.219 | -0.079 | 0.071 | 0.071 | 0 | 0.38 | 0.11 | 0.34 | 0.25 | 0.52 | 0.27 |
| 2 | 80406262 | rs1989614 | 0.008 | -0.633 | 0.849 | 0.187 | 0.026 | 0.011 | 0.011 | 0.007 | 0.57 | 0.81 | 0.57 | 0.38 | 0.63 | 0.62 |
| 2 | 80413336 | rs894829 | -1.078 | -0.965 | 0.34 | 0.071 | 0.006 | 0.031 | 0.031 | 0 | 0.85 | 0.98 | 0.92 | 0.94 | 0.89 | 0.9 |
| 2 | 80418483 | rs7601708 | 1.407 | -0.904 | 0.583 | 0.103 | -0.044 | 0.072 | 0.072 | 0 | 0.4 | 0.2 | 0.35 | 0.28 | 0.52 | 0.28 |
| 2 | 80418601 | rs1446106 | -0.055 | -1.004 | 1.343 | 0.138 | -0.02 | 0.012 | 0.012 | 0 | 0.15 | 0.45 | 0.2 | 0.09 | 0.1 | 0.33 |
| 2 | 80421509 | rs1899705 | -0.135 | -1.07 | 1.585 | 0.113 | -0.025 | 0.013 | 0.013 | 0 | 0.15 | 0.54 | 0.21 | 0.1 | 0.1 | 0.34 |
| 2 | 80424134 | rs7604121 | -0.36 | -1.077 | 1.64 | 0.112 | -0.034 | 0.004 | 0.004 | 0.057 | 0.16 | 0.61 | 0.21 | 0.1 | 0.1 | 0.34 |
| 2 | 80443877 | rs1979024 | 0.92 | -1.228 | 1.363 | 0.523 | -0.017 | -0.06 | -0.06 | 0.64 | 0.14 | 0.03 | 0.28 | 0.38 | 0.19 | 0.14 |
| 2 | 80445512 | rs7590636 | -0.099 | -1.201 | 1.567 | 0.417 | -0.029 | 0.009 | 0.009 | 0 | 0.15 | 0.44 | 0.21 | 0.09 | 0.09 | 0.36 |
| 2 | 80447262 | rs1816909 | 1.67 | -1.262 | 1.499 | 0.499 | -0.03 | 0.079 | 0.079 | 0 | 0.41 | 0.2 | 0.35 | 0.27 | 0.51 | 0.26 |
| 2 | 80448882 | rs1006607 | -0.131 | -1.241 | 1.542 | 0.533 | -0.032 | 0.011 | 0.011 | 0 | 0.16 | 0.5 | 0.21 | 0.1 | 0.1 | 0.36 |
| 2 | 80451184 | rs1965833 | -0.744 | -1.26 | 1.497 | 0.539 | -0.045 | 0.021 | 0.021 | 0 | 0.19 | 0.58 | 0.25 | 0.16 | 0.13 | 0.42 |
| 2 | 80453570 | rs13416818 | -0.089 | -1.223 | 1.57 | 0.528 | -0.028 | 0.012 | 0.012 | 0 | 0.15 | 0.44 | 0.21 | 0.09 | 0.09 | 0.36 |
| 2 | 80456138 | rs2862024 | 0.128 | -1.197 | 1.076 | 0.583 | 0 | -0.044 | -0.044 | 0 | 0.61 | 0.86 | 0.6 | 0.43 | 0.64 | 0.68 |
| 2 | 80469759 | rs6738640 | -1.322 | -1.171 | 1.163 | 0.683 | 0.064 | 0.255 | 0.255 | 3.236 | 0.06 | 0.45 | 0.14 | 0.09 | 0.01 | 0.04 |
| 2 | 80470464 | rs2587149 | 0.21 | -1.252 | 0.84 | 0.594 | 0.008 | -0.02 | -0.02 | 0 | 0.58 | 0.82 | 0.57 | 0.43 | 0.62 | 0.67 |

| | | | | | | | | | | | | | | | | |
|---|---|---|---|---|---|---|---|---|---|---|---|---|---|---|---|---|
| 2 | 80474158 | rs383452 | 0.74 | -1.272 | 0.805 | 0.583 | 0.028 | 0.052 | 0.052 | 0 | 0.52 | 0.34 | 0.43 | 0.34 | 0.61 | 0.63 |
| 2 | 80476142 | rs465119 | 0.746 | -1.228 | 0.818 | 0.945 | 0.033 | -0.013 | -0.013 | 0 | 0.56 | 0.28 | 0.46 | 0.47 | 0.62 | 0.64 |
| 2 | 80478274 | rs160864 | 1.069 | -1.325 | 0.67 | 0.842 | 0.02 | 0.006 | 0.006 | 0 | 0.49 | 0.36 | 0.44 | 0.47 | 0.56 | 0.3 |
| 2 | 80485796 | rs17018914 | 0.625 | -1.158 | -0.027 | 0.774 | 0.002 | -0.001 | -0.001 | 1.357 | 0.09 | 0.08 | 0.04 | 0.01 | 0.07 | 0.33 |
| 2 | 80488774 | rs318369 | 1.574 | -1.224 | -0.175 | 0.776 | -0.042 | -0.029 | -0.029 | 0 | 0.42 | 0.09 | 0.4 | 0.42 | 0.52 | 0.26 |
| 2 | 80492708 | rs318363 | 1.767 | -1.264 | -0.424 | 0.755 | -0.094 | -0.078 | -0.078 | 0.557 | 0.4 | 0.04 | 0.4 | 0.42 | 0.52 | 0.26 |
| 2 | 80495229 | rs735719 | 0.439 | -0.96 | -0.069 | 0.526 | -0.004 | -0.015 | -0.015 | 0.335 | 0.1 | 0.1 | 0.03 | 0.01 | 0.07 | 0.33 |
| 2 | 80501998 | rs310776 | -0.798 | -1.322 | -0.711 | 0.308 | 0.034 | 0.061 | 0.061 | 0 | 0.83 | 0.98 | 0.92 | 0.95 | 0.9 | 0.94 |
| 2 | 80504562 | rs310774 | 0.148 | -0.991 | -0.247 | 0.633 | 0.011 | 0.028 | 0.028 | 0 | 0.57 | 0.69 | 0.59 | 0.59 | 0.62 | 0.66 |
| 2 | 80504787 | rs2587147 | 0.85 | -0.67 | -0.464 | 0.771 | -0.005 | 0.004 | 0.004 | 0 | 0.43 | 0.37 | 0.53 | 0.54 | 0.52 | 0.3 |
| 2 | 80514672 | rs1930 | -1.43 | -0.403 | -0.698 | 0.979 | -0.012 | -0.01 | -0.01 | 0 | 0.46 | 0.81 | 0.49 | 0.4 | 0.47 | 0.58 |
| 2 | 80517753 | rs1446110 | 0.093 | -0.308 | -0.819 | 1.112 | -0.103 | -0.084 | -0.084 | 0 | 0.23 | 0.04 | 0.33 | 0.37 | 0.37 | 0.25 |
| 2 | 80521341 | rs2862286 | -1.558 | -0.249 | -0.678 | 0.902 | -0.008 | -0.017 | -0.017 | 0 | 0.47 | 0.8 | 0.51 | 0.42 | 0.51 | 0.6 |
| 2 | 80524178 | rs6712681 | -0.873 | 0.063 | -0.117 | 1.273 | 0.023 | 0.028 | 0.028 | 0 | 0.33 | 0.39 | 0.44 | 0.47 | 0.42 | 0.31 |
| 2 | 80529956 | rs6733871 | -1.272 | -1.874 | 0.301 | 1.328 | -0.033 | 0.047 | 0.047 | 3.128 | 0.31 | 0.51 | 0.32 | 0.48 | 0.18 | 0.28 |
| 2 | 80538419 | rs1446109 | -0.251 | -1.583 | 0.256 | 1.438 | 0.025 | 0.093 | 0.093 | 0 | 0.06 | 0.1 | 0.15 | 0.28 | 0.09 | 0.1 |
| 2 | 80540600 | rs17018984 | 0.124 | -1.574 | 0.126 | 1.456 | -0.013 | 0.025 | 0.025 | 0 | 0.05 | 0.02 | 0.13 | 0.25 | 0.09 | 0.09 |
| 2 | 80542744 | rs9808084 | -1.841 | -2.116 | 0.088 | 1.189 | 0.05 | -0.002 | -0.002 | 0 | 0.43 | 0.73 | 0.39 | 0.53 | 0.26 | 0.29 |
| 2 | 80551464 | rs1006154 | -1.264 | -2.135 | 0.182 | 1.065 | 0.005 | 0 | 0 | 0 | 0.2 | 0.2 | 0.15 | 0.13 | 0.15 | 0.15 |
| 2 | 80553514 | rs1972956 | -1.823 | -2.112 | 0.399 | 1.016 | -0.009 | 0.001 | 0.001 | 0 | 0.39 | 0.62 | 0.37 | 0.53 | 0.28 | 0.28 |
| 2 | 80560504 | rs6750519 | -0.857 | -1.593 | 0.811 | 1.259 | -0.013 | -0.03 | -0.03 | 0 | 0.07 | 0.15 | 0.04 | 0 | 0.04 | 0.03 |
| 2 | 80563130 | rs1562775 | -0.418 | -1.717 | 0.616 | 1.334 | 0.002 | 0.003 | 0.003 | 0 | 0.09 | 0.05 | 0.06 | 0.06 | 0.08 | 0.08 |
| 2 | 80570625 | rs11885868 | -2.423 | -1.928 | 0.974 | 1.436 | 0.011 | -0.043 | -0.043 | 0 | 0.31 | 0.78 | 0.31 | 0.43 | 0.24 | 0.34 |
| 2 | 80572750 | rs1443896 | -1.933 | -1.987 | 1.78 | 1.163 | -0.024 | -0.055 | -0.055 | 0 | 0.11 | 0.35 | 0.05 | 0 | 0.05 | 0.05 |
| 2 | 80574477 | rs1947033 | -0.321 | -2 | 1.84 | 1.302 | 0.034 | 0.169 | 0.169 | 0.044 | 0.06 | 0.33 | 0.03 | 0.1 | 0.02 | 0.1 |
| 2 | 80574834 | rs1947032 | -1.872 | -1.962 | 1.816 | 1.404 | 0.02 | 0.13 | 0.13 | 0 | 0.19 | 0.39 | 0.31 | 0.51 | 0.19 | 0.3 |
| 2 | 80580875 | rs149248 | 0.42 | -2.632 | 1.202 | 0.975 | 0.031 | 0.038 | 0.038 | 0 | 0.17 | 0.07 | 0.06 | 0.09 | 0.09 | 0.22 |
| 2 | 80581801 | rs723524 | -0.683 | -2.13 | 1.463 | 1.279 | -0.052 | -0.077 | -0.077 | 0 | 0.13 | 0.02 | 0.27 | 0.38 | 0.17 | 0.17 |
| 2 | 80584769 | rs17019074 | -1.648 | -2.167 | 1.728 | 1.233 | 0.021 | 0.105 | 0.105 | 0 | 0.18 | 0.31 | 0.3 | 0.39 | 0.18 | 0.2 |
| 2 | 80587715 | rs11126761 | -1.549 | -2.159 | 1.686 | 1.136 | 0.032 | 0.157 | 0.157 | 0.122 | 0.07 | 0.4 | 0.05 | 0.13 | 0.05 | 0.09 |
| 2 | 80590889 | rs1838432 | -0.833 | -2.172 | 1.865 | 1.138 | -0.002 | 0.108 | 0.108 | 0 | 0.06 | 0.24 | 0.04 | 0.12 | 0.04 | 0.09 |
| 2 | 80600505 | rs13407218 | -0.705 | -2.166 | 1.805 | 1.137 | 0.023 | 0.138 | 0.138 | 0.643 | 0.07 | 0.38 | 0.03 | 0.1 | 0.01 | 0.12 |
| 2 | 80602524 | rs7593219 | -1.516 | -2.162 | 1.805 | 1.224 | 0.006 | 0.039 | 0.039 | 0 | 0.18 | 0.17 | 0.29 | 0.37 | 0.18 | 0.2 |
| 2 | 80607670 | rs10496240 | -2.749 | -2.166 | 1.55 | 1.189 | 0.01 | -0.031 | -0.031 | 2.023 | 0.29 | 0.78 | 0.35 | 0.54 | 0.22 | 0.33 |
| 2 | 80608909 | rs2166129 | -0.798 | -2.09 | 1.278 | 1.204 | -0.05 | -0.075 | -0.075 | 0 | 0.13 | 0.02 | 0.27 | 0.42 | 0.17 | 0.17 |
| 2 | 80613607 | rs6547309 | -1.924 | -2.484 | 1.053 | 0.946 | 0.073 | -0.022 | -0.022 | 0 | 0.5 | 0.93 | 0.43 | 0.58 | 0.34 | 0.51 |
| 2 | 80619803 | rs11896302 | -1.287 | -2.229 | 1.868 | 0.656 | -0.032 | 0.022 | 0.022 | 0 | 0.16 | 0.41 | 0.11 | 0.18 | 0.12 | 0.16 |
| 2 | 80630354 | rs1992632 | -1.863 | -2.15 | 1.197 | 0.675 | -0.09 | 0.097 | 0.097 | 0 | 0.57 | 0.97 | 0.52 | 0.79 | 0.38 | 0.66 |
| 2 | 80636937 | rs10190500 | -1.035 | -1.724 | 2.041 | 0.377 | -0.003 | -0.019 | -0.019 | 0 | 0.09 | 0.45 | 0.03 | 0 | 0.01 | 0.06 |
| 2 | 80641365 | rs11902575 | -1.663 | -1.692 | 1.008 | 0.425 | -0.007 | -0.002 | -0.002 | 0 | 0.23 | 0.74 | 0.13 | 0.14 | 0.12 | 0.21 |
| 2 | 80652386 | rs994593 | -0.731 | -0.895 | 0.992 | 0.565 | -0.001 | -0.021 | -0.021 | 0 | 0.11 | 0.2 | 0.09 | 0.06 | 0.11 | 0.07 |
| 2 | 80655311 | rs7605418 | -0.112 | -0.607 | 0.755 | 0.951 | 0.021 | 0.149 | 0.149 | 0 | 0.05 | 0.08 | 0.3 | 0.51 | 0.1 | 0.1 |
| 2 | 80656878 | rs1444541 | -1.159 | -0.85 | 0.613 | 0.09 | -0.008 | 0.226 | 0.226 | 0.01 | 0.52 | 0.85 | 0.59 | 0.92 | 0.39 | 0.63 |
| 2 | 80657752 | rs10201249 | -1.193 | -0.483 | 1.247 | 0.677 | 0.001 | 0.043 | 0.043 | 0 | 0.13 | 0.4 | 0.09 | 0.15 | 0.08 | 0.07 |
| 2 | 80668722 | rs6747004 | -0.439 | -0.319 | 0.818 | 0.573 | 0.004 | -0.025 | -0.025 | 0 | 0.1 | 0.3 | 0.07 | 0.07 | 0.08 | 0.06 |
| 2 | 80680830 | rs895024 | -0.782 | -0.566 | 0.759 | 0.55 | -0.003 | -0.011 | -0.011 | 0 | 0.42 | 0.55 | 0.54 | 0.72 | 0.33 | 0.55 |
| 2 | 80685229 | rs7600784 | -1.022 | -0.69 | 0.842 | 0.546 | 0.028 | 0.081 | 0.081 | 0.204 | 0.44 | 0.72 | 0.56 | 0.73 | 0.33 | 0.55 |
| 2 | 80690431 | rs6715806 | -0.709 | -0.674 | 1.231 | 0.537 | -0.006 | -0.027 | -0.027 | 0.184 | 0.37 | 0.45 | 0.47 | 0.73 | 0.31 | 0.54 |
| 2 | 80692107 | rs2117277 | -0.536 | -0.751 | 0.546 | 0.567 | -0.02 | -0.013 | -0.013 | 0 | 0.34 | 0.17 | 0.52 | 0.63 | 0.32 | 0.54 |
| 2 | 80702160 | rs4528777 | -1.065 | -0.69 | 0.448 | 0.602 | -0.007 | -0.008 | -0.008 | 0 | 0.4 | 0.52 | 0.54 | 0.62 | 0.32 | 0.53 |
| 2 | 80709918 | rs722559 | -0.762 | -0.425 | -0.602 | 1.036 | -0.033 | 0.022 | 0.022 | 1.021 | 0.06 | 0.01 | 0.35 | 0.5 | 0.1 | 0.11 |
| 2 | 80711589 | rs11902138 | -0.808 | -0.69 | -0.652 | 0.659 | 0.023 | 0.001 | 0.001 | 0 | 0.35 | 0.24 | 0.45 | 0.63 | 0.26 | 0.5 |
| 2 | 80728120 | rs7576475 | -0.694 | 0.169 | -1.045 | 0.529 | 0 | -0.007 | -0.007 | 0 | 0.26 | 0.15 | 0.24 | 0.33 | 0.27 | 0.41 |
| 2 | 80731847 | rs1434076 | -1.199 | -0.017 | -1.238 | -0.379 | 0.029 | 0.212 | 0.212 | 0.119 | 0.48 | 0.79 | 0.55 | 0.94 | 0.4 | 0.52 |
| 2 | 80734826 | rs1368900 | 0.343 | -0.07 | -0.947 | -0.897 | 0.015 | -0.107 | -0.107 | 0 | 0.38 | 0.49 | 0.4 | 0.06 | 0.4 | 0.47 |
| 2 | 80734971 | rs17261683 | 0.258 | 0.019 | -1.692 | -0.857 | -0.012 | 0.009 | 0.009 | 0 | 0.24 | 0.08 | 0.22 | 0.23 | 0.25 | 0.24 |
| 2 | 80735034 | rs1368898 | -0.158 | -0.004 | -1.068 | -0.338 | 0.008 | 0.015 | 0.015 | 0.181 | 0.63 | 0.71 | 0.64 | 0.32 | 0.65 | 0.71 |
| 2 | 80736075 | rs2117276 | -0.319 | 0.112 | -1.18 | -0.08 | 0.005 | -0.027 | -0.027 | 0.642 | 0.7 | 0.82 | 0.69 | 0.39 | 0.71 | 0.78 |
| 2 | 80741406 | rs6742713 | 0.291 | 0.085 | -1.047 | -0.136 | 0.015 | 0.124 | 0.124 | 0.489 | 0.44 | 0.83 | 0.38 | 0.17 | 0.4 | 0.44 |
| 2 | 80743194 | rs1434075 | 0.695 | -0.136 | -0.209 | -0.034 | 0.003 | -0.044 | -0.044 | 1.956 | 0.37 | 0.54 | 0.35 | 0.12 | 0.42 | 0.43 |
| 2 | 80744297 | rs744957 | 0.134 | -0.313 | -0.327 | 0.511 | -0.034 | 0 | 0 | 0.13 | 0.22 | 0.09 | 0.16 | 0.16 | 0.27 | 0.23 |
| 2 | 80748807 | rs6750107 | 1.186 | -0.626 | -0.167 | 0.299 | 0.007 | -0.042 | -0.042 | 0.244 | 0.33 | 0.39 | 0.32 | 0.12 | 0.37 | 0.4 |
| 2 | 80756301 | rs17340143 | 0.788 | -0.572 | 0.451 | 0.732 | 0.008 | -0.006 | -0.006 | 0 | 0.4 | 0.48 | 0.41 | 0.41 | 0.43 | 0.58 |
| 2 | 80757130 | rs7558519 | 1.127 | -0.578 | 0.447 | 0.671 | 0.013 | -0.064 | -0.064 | 2.88 | 0.37 | 0.47 | 0.36 | 0.13 | 0.41 | 0.43 |
| 2 | 80757507 | rs11685198 | -0.156 | -0.452 | 0.168 | 0.333 | 0.004 | 0.095 | 0.095 | 0 | 0.22 | 0.05 | 0.11 | 0 | 0.2 | 0.14 |
| 2 | 80761078 | rs13422536 | 0.625 | -0.442 | 0.214 | 0.37 | -0.003 | 0.066 | 0.066 | 2.376 | 0.45 | 0.74 | 0.43 | 0.35 | 0.43 | 0.59 |
| 2 | 80761192 | rs12611756 | -0.775 | -0.407 | 0.171 | 0.422 | -0.038 | -0.053 | -0.053 | 0.46 | 0.67 | 0.88 | 0.56 | 0.43 | 0.63 | 0.75 |
| 2 | 80761692 | rs1368901 | 0.569 | -0.212 | 0.637 | 0.771 | 0.003 | -0.014 | -0.014 | 0 | 0.41 | 0.54 | 0.41 | 0.34 | 0.43 | 0.58 |
| 2 | 80770808 | rs13423141 | 0.73 | -0.111 | 0.892 | 0.674 | -0.001 | 0.001 | 0.001 | 2.339 | 0.38 | 0.37 | 0.39 | 0.35 | 0.43 | 0.58 |

| | | | | | | | | | | | | | | | | |
|---|---|---|---|---|---|---|---|---|---|---|---|---|---|---|---|---|
| 2 | 80771190 | rs7605261 | 0.572 | -0.014 | 0.39 | 0.752 | 0.008 | -0.022 | -0.022 | 0.576 | 0.4 | 0.47 | 0.39 | 0.35 | 0.43 | 0.58 |
| 2 | 80774809 | rs4852581 | 0.401 | 0.092 | -0.551 | 0.761 | -0.014 | 0.01 | 0.01 | 2.548 | 0.44 | 0.69 | 0.43 | 0.44 | 0.44 | 0.63 |
| 2 | 80786548 | rs3770310 | -0.4 | 0.209 | -0.7 | 0.685 | -0.048 | 0.044 | 0.044 | 2.965 | 0.48 | 0.1 | 0.52 | 0.36 | 0.53 | 0.26 |
| 2 | 80789481 | rs1434061 | -0.333 | 0.279 | -0.817 | 0.476 | 0.002 | 0.024 | 0.024 | 0 | 0.22 | 0.13 | 0.15 | 0.12 | 0.19 | 0.24 |
| 2 | 80790287 | rs3770313 | 1.261 | 0.092 | -0.026 | 0.468 | -0.003 | -0.004 | -0.004 | 0 | 0.39 | 0.34 | 0.35 | 0.37 | 0.42 | 0.58 |
| 2 | 80791001 | rs1434063 | 0.667 | 0.032 | 0.514 | 0.509 | -0.009 | -0.044 | -0.044 | 0.105 | 0.73 | 0.65 | 0.82 | 0.84 | 0.8 | 0.74 |
| 2 | 80791939 | rs3770317 | -0.489 | 0.039 | 0.496 | 0.502 | -0.006 | -0.043 | -0.043 | 0 | 0.23 | 0.31 | 0.17 | 0.13 | 0.19 | 0.24 |
| 2 | 80792792 | rs12714010 | -0.181 | 0.049 | 0.363 | 0.523 | -0.003 | 0.026 | 0.026 | 3.491 | 0.63 | 0.47 | 0.51 | 0.51 | 0.63 | 0.84 |
| 2 | 80795159 | rs3770321 | -0.147 | -0.489 | 0.235 | 0.542 | -0.001 | 0.002 | 0.002 | 0 | 0.18 | 0.14 | 0.14 | 0.14 | 0.17 | 0.23 |
| 2 | 80795887 | rs10192914 | 0.179 | -0.474 | 0.043 | 0.569 | -0.006 | 0.001 | 0.001 | 0 | 0.91 | 0.9 | 0.95 | 0.86 | 0.96 | 0.96 |
| 2 | 80797905 | rs3755105 | -0.384 | -0.27 | 0.264 | 0.384 | 0.003 | -0.057 | -0.057 | 0 | 0.2 | 0.38 | 0.15 | 0.14 | 0.17 | 0.23 |
| 2 | 80802775 | rs1821411 | 0.262 | -0.139 | -0.619 | 0.239 | -0.016 | 0.042 | 0.042 | 0 | 0.15 | 0.05 | 0.1 | 0 | 0.16 | 0.11 |
| 2 | 80804156 | rs216616 | -0.612 | 0.003 | -0.371 | 0.598 | 0.013 | 0.001 | 0.001 | 3.469 | 0.57 | 0.39 | 0.65 | 0.62 | 0.64 | 0.43 |
| 2 | 80805144 | rs216618 | -0.978 | -0.396 | -0.369 | 0.223 | 0.008 | 0.003 | 0.003 | 0 | 0.18 | 0.14 | 0.14 | 0.09 | 0.09 | 0.16 |
| 2 | 80805192 | rs216619 | -0.849 | -0.135 | -0.476 | 0.501 | 0.009 | 0.005 | 0.005 | 0 | 0.48 | 0.29 | 0.56 | 0.56 | 0.44 | 0.33 |
| 2 | 80805522 | rs17019386 | -0.71 | -0.129 | 0.169 | 0.549 | 0.025 | 0.069 | 0.069 | 1.289 | 0.06 | 0.33 | 0.07 | 0.07 | 0.02 | 0.01 |
| 2 | 80809406 | rs17019409 | 0.512 | -0.141 | -0.068 | 0.629 | -0.046 | -0.086 | -0.086 | 0 | 0.31 | 0.13 | 0.42 | 0.51 | 0.36 | 0.17 |
| 2 | 80810423 | rs6547316 | 0.245 | -0.174 | 0.166 | 0.625 | -0.005 | -0.007 | -0.007 | 0 | 0.33 | 0.3 | 0.45 | 0.55 | 0.36 | 0.17 |
| 2 | 80814388 | rs216626 | -1.344 | -0.698 | 0.482 | 0.533 | -0.062 | -0.032 | -0.032 | 0 | 0.28 | 0.57 | 0.23 | 0.2 | 0.11 | 0.19 |
| 2 | 80814794 | rs216627 | -1.194 | -0.256 | 0.374 | 0.687 | -0.009 | 0.008 | 0.008 | 0.895 | 0.57 | 0.64 | 0.61 | 0.65 | 0.46 | 0.35 |
| 2 | 80822772 | rs216640 | -1.266 | -0.332 | 0.497 | 0.646 | -0.075 | -0.042 | -0.042 | 0 | 0.27 | 0.52 | 0.22 | 0.2 | 0.11 | 0.21 |
| 2 | 80831481 | rs2302875 | 0.858 | -0.268 | 0.43 | 0.679 | -0.057 | -0.082 | -0.082 | 1.317 | 0.3 | 0.08 | 0.37 | 0.42 | 0.35 | 0.15 |
| 2 | 80839203 | rs216676 | -0.297 | -0.417 | 0.579 | 0.617 | -0.011 | 0.008 | 0.008 | 0 | 0.12 | 0.19 | 0.11 | 0.1 | 0.08 | 0.06 |
| 2 | 80843935 | rs2058956 | -1.327 | -0.158 | 0.399 | 0.616 | -0.006 | 0.002 | 0.002 | 0.418 | 0.58 | 0.64 | 0.62 | 0.62 | 0.51 | 0.37 |
| 2 | 80847702 | rs3770364 | -1.467 | -0.261 | 0.438 | 0.469 | -0.086 | -0.015 | -0.015 | 0 | 0.22 | 0.52 | 0.19 | 0.19 | 0.07 | 0.17 |
| 2 | 80849020 | rs3770365 | 0.243 | -0.228 | 0.378 | 0.637 | -0.063 | -0.063 | -0.063 | 0 | 0.36 | 0.18 | 0.44 | 0.44 | 0.43 | 0.2 |
| 2 | 80849811 | rs3770369 | -1.261 | -0.196 | 0.449 | 0.618 | -0.014 | 0.014 | 0.014 | 0.042 | 0.57 | 0.69 | 0.61 | 0.63 | 0.47 | 0.36 |
| 2 | 80856465 | rs7580421 | 0.843 | -0.157 | 0.316 | 0.616 | -0.083 | -0.1 | -0.1 | 4.61 | 0.3 | 0.07 | 0.39 | 0.49 | 0.39 | 0.18 |
| 2 | 80864323 | rs10490510 | -1.042 | -0.631 | 0.012 | 0.354 | -0.003 | 0.003 | 0.003 | 0 | 0.59 | 0.62 | 0.6 | 0.67 | 0.55 | 0.53 |
| 2 | 80869195 | rs7582779 | -1.507 | -0.627 | 0.015 | 0.378 | -0.052 | -0.009 | -0.009 | 0 | 0.64 | 0.81 | 0.64 | 0.67 | 0.53 | 0.53 |
| 2 | 80871975 | rs3770381 | 1.113 | -0.585 | -0.289 | 0.488 | -0.081 | -0.114 | -0.114 | 0 | 0.3 | 0.07 | 0.38 | 0.47 | 0.39 | 0.17 |
| 2 | 80875611 | rs11445 | 1.201 | -0.672 | -0.287 | 0.477 | -0.067 | -0.113 | -0.113 | 0 | 0.29 | 0.06 | 0.36 | 0.43 | 0.34 | 0.17 |
| 2 | 80898579 | rs7574764 | -0.159 | -0.973 | 0.074 | 0.327 | -0.003 | 0.047 | 0.047 | 0 | 0.09 | 0.3 | 0.06 | 0.11 | 0.07 | 0.16 |
| 2 | 80900986 | rs6718076 | -1.178 | -0.97 | -0.135 | 0.37 | -0.096 | -0.013 | -0.013 | 0 | 0.21 | 0.51 | 0.16 | 0.24 | 0.08 | 0.3 |
| 2 | 80907225 | rs11691644 | -0.489 | -0.501 | -0.392 | 0.332 | -0.006 | -0.049 | -0.049 | 0 | 0.1 | 0.19 | 0.07 | 0 | 0.07 | 0.01 |
| 2 | 80913850 | rs4852589 | 0.688 | -0.384 | -0.941 | 0.567 | -0.048 | -0.08 | -0.08 | 0 | 0.31 | 0.1 | 0.38 | 0.48 | 0.39 | 0.21 |
| 2 | 80915114 | rs1370339 | -0.83 | -0.495 | -1.374 | 0.154 | -0.144 | 0.157 | 0.157 | 0.601 | 0.57 | 0.97 | 0.56 | 0.82 | 0.48 | 0.53 |
| 2 | 80920342 | rs1437359 | -0.085 | -0.456 | -1.08 | 0.146 | -0.002 | 0.087 | 0.087 | 0.525 | 0.44 | 0.83 | 0.48 | 0.7 | 0.42 | 0.33 |
| 2 | 80929441 | rs996742 | -1.089 | -0.647 | -0.566 | 0.12 | 0.03 | 0.126 | 0.126 | 0 | 0.48 | 0.78 | 0.56 | 0.8 | 0.42 | 0.63 |
| 2 | 80939019 | rs6757169 | -0.651 | -0.73 | -0.232 | 0.152 | 0.001 | -0.025 | -0.025 | 0 | 0.36 | 0.6 | 0.5 | 0.77 | 0.38 | 0.5 |
| 2 | 80941466 | rs1437368 | -1.121 | -0.839 | 0.291 | 0.686 | -0.025 | 0.058 | 0.058 | 0 | 0.13 | 0.3 | 0.11 | 0.23 | 0.09 | 0.29 |
| 2 | 80942951 | rs7607618 | -1.289 | -0.914 | 0.213 | 0.672 | -0.043 | 0.096 | 0.096 | 0.206 | 0.16 | 0.46 | 0.15 | 0.23 | 0.1 | 0.31 |
| 2 | 80945496 | rs13419774 | -1.723 | -1.02 | -0.121 | 0.594 | -0.06 | 0.01 | 0.01 | 0.067 | 0.27 | 0.62 | 0.2 | 0.26 | 0.14 | 0.43 |
| 2 | 80950629 | rs6755493 | 0.901 | -0.705 | -0.065 | 1.136 | 0.024 | 0.068 | 0.068 | 0 | 0.19 | 0.28 | 0.37 | 0.46 | 0.28 | 0.2 |
| 2 | 80951823 | rs1265750 | 1.498 | -1.397 | -0.076 | 0.961 | -0.043 | -0.004 | -0.004 | 0 | 0.71 | 0.34 | 0.8 | 0.77 | 0.85 | 0.56 |
| 2 | 80970449 | rs1370336 | -0.263 | -0.927 | 0.477 | 1.41 | -0.023 | 0.06 | 0.06 | 0 | 0.15 | 0.27 | 0.1 | 0.27 | 0.1 | 0.15 |
| 2 | 80972177 | rs17019687 | 0.965 | -0.919 | 0.787 | 1.315 | -0.011 | -0.02 | -0.02 | 0 | 0.06 | 0.17 | 0.04 | 0.01 | 0.01 | 0.04 |
| 2 | 80974735 | rs6731158 | 2.232 | -0.272 | 0.434 | 1.599 | -0.046 | -0.052 | -0.052 | 0 | 0.16 | 0.08 | 0.35 | 0.31 | 0.25 | 0.24 |
| 2 | 80984824 | rs6547331 | -0.653 | 0.05 | 0.452 | 1.52 | -0.048 | -0.06 | -0.06 | 0 | 0.72 | 0.82 | 0.56 | 0.54 | 0.64 | 0.59 |
| 2 | 80999206 | rs1845017 | -0.599 | 0.056 | 0.224 | 1.348 | -0.013 | -0.015 | -0.015 | 0 | 0.9 | 0.94 | 0.93 | 0.87 | 0.89 | 0.83 |
| 2 | 81000427 | rs11682698 | 2.345 | -0.264 | 0.129 | 1.12 | -0.06 | 0.147 | 0.147 | 0 | 0.46 | 0.07 | 0.38 | 0.25 | 0.52 | 0.34 |
| 2 | 81004626 | rs10179482 | 1.759 | -0.493 | 0.492 | 0.975 | 0 | -0.012 | -0.012 | 0 | 0.52 | 0.49 | 0.44 | 0.27 | 0.51 | 0.42 |
| 2 | 81018549 | rs1826413 | 2.461 | -0.278 | 0.487 | 1.115 | -0.08 | 0.135 | 0.135 | 0 | 0.46 | 0.04 | 0.38 | 0.25 | 0.51 | 0.35 |
| 2 | 81025111 | rs7561634 | -1.053 | -0.307 | 0.741 | 1.298 | -0.059 | -0.074 | -0.074 | 0 | 0.73 | 0.84 | 0.57 | 0.54 | 0.64 | 0.6 |
| 2 | 81061238 | rs1978819 | 2.331 | -0.29 | 0.609 | 1.247 | -0.081 | 0.127 | 0.127 | 0 | 0.46 | 0.05 | 0.39 | 0.25 | 0.52 | 0.35 |
| 2 | 81061501 | rs17019785 | 0.185 | -0.307 | 1.114 | 1.298 | -0.019 | 0.049 | 0.049 | 0 | 0.16 | 0.25 | 0.12 | 0.27 | 0.11 | 0.16 |
| 2 | 81086610 | rs10496244 | 1.258 | -0.178 | 1.115 | 1.219 | 0 | 0.131 | 0.131 | 0 | 0.5 | 0.2 | 0.42 | 0.26 | 0.52 | 0.39 |
| 2 | 81089974 | rs6749488 | -0.847 | -0.288 | 0.966 | 1.224 | -0.023 | 0.055 | 0.055 | 0 | 0.18 | 0.31 | 0.12 | 0.28 | 0.12 | 0.16 |
| 2 | 81094008 | rs12467735 | -0.163 | -0.289 | 0.743 | 1.18 | 0 | 0.036 | 0.036 | 0.114 | 0.14 | 0.23 | 0.14 | 0.29 | 0.13 | 0.1 |
| 2 | 81098202 | rs11678886 | 0.654 | -0.024 | 0.488 | 1.124 | 0.005 | -0.004 | -0.004 | 0.324 | 0.56 | 0.34 | 0.52 | 0.52 | 0.6 | 0.43 |
| 2 | 81102264 | rs2131255 | 0.218 | 0.01 | 0.16 | 1.189 | -0.033 | -0.017 | -0.017 | 0.049 | 0.57 | 0.78 | 0.45 | 0.46 | 0.5 | 0.38 |
| 2 | 81107544 | rs1492549 | -0.324 | 0.021 | 0.093 | 1.088 | -0.071 | -0.064 | -0.064 | 0.097 | 0.65 | 0.87 | 0.52 | 0.47 | 0.55 | 0.43 |
| 2 | 81115259 | rs6732417 | 2.43 | 0.11 | -0.03 | 1.231 | -0.08 | -0.09 | -0.09 | 2.864 | 0.2 | 0.03 | 0.39 | 0.3 | 0.28 | 0.31 |
| 2 | 81120551 | rs6710603 | -0.533 | -0.168 | 0.336 | 1.138 | -0.022 | 0.043 | 0.043 | 4.804 | 0.14 | 0.29 | 0.12 | 0.21 | 0.09 | 0.08 |
| 2 | 81125236 | rs17019859 | -0.704 | -0.178 | 0.29 | 1.064 | -0.025 | 0.007 | 0.007 | 2.597 | 0.2 | 0.34 | 0.15 | 0.24 | 0.12 | 0.09 |
| 2 | 81126233 | rs17019864 | -0.429 | -0.166 | -0.177 | 1.062 | -0.068 | -0.058 | -0.058 | 0 | 0.65 | 0.83 | 0.51 | 0.47 | 0.55 | 0.44 |
| 2 | 81129793 | rs902885 | -1.029 | 0.235 | 0.133 | 1.253 | -0.044 | 0.011 | 0.011 | 2.167 | 0.22 | 0.44 | 0.15 | 0.24 | 0.12 | 0.09 |
| 2 | 81140473 | rs13007790 | 1.184 | 0.607 | -0.594 | 1.353 | -0.125 | -0.123 | -0.123 | 1.706 | 0.23 | 0.04 | 0.4 | 0.37 | 0.34 | 0.47 |
| 2 | 81143279 | rs1845025 | -0.364 | 0.678 | -0.526 | 0.891 | -0.064 | -0.052 | -0.052 | 0.169 | 0.68 | 0.81 | 0.51 | 0.47 | 0.54 | 0.44 |

| | | | | | | | | | | | | | | | | |
|---|---|---|---|---|---|---|---|---|---|---|---|---|---|---|---|---|
| 2 | 81165670 | rs17614582 | 1.552 | 0.604 | -0.585 | 0.835 | 0.001 | 0.107 | 0.107 | 1.299 | 0.43 | 0.19 | 0.34 | 0.23 | 0.44 | 0.3 |
| 2 | 81176175 | rs4432478 | -0.326 | 0.894 | -0.679 | 0.523 | -0.068 | -0.062 | -0.062 | 0 | 0.69 | 0.85 | 0.52 | 0.47 | 0.56 | 0.44 |
| 2 | 81178803 | rs35513010 | 1.64 | 0.442 | 0.436 | 0.181 | -0.056 | -0.075 | -0.075 | 0 | 0.23 | 0.1 | 0.4 | 0.37 | 0.33 | 0.33 |
| 2 | 81179374 | rs10210142 | 0.714 | 0.28 | 0.539 | 0.205 | -0.013 | -0.026 | -0.026 | 0.418 | 0.81 | 0.86 | 0.89 | 0.71 | 0.77 | 0.77 |
| 2 | 81182131 | rs4302235 | 2.122 | 0.168 | 0.854 | 0.084 | -0.002 | -0.008 | -0.008 | 0 | 0.65 | 0.32 | 0.71 | 0.55 | 0.67 | 0.57 |
| 2 | 81191209 | rs4462808 | 0.192 | 0.185 | 1.384 | 0.038 | 0.024 | 0.045 | 0.045 | 0 | 0.51 | 0.36 | 0.4 | 0.27 | 0.41 | 0.36 |
| 2 | 81201683 | rs4389353 | -0.143 | 0.183 | 1.37 | 0.049 | 0.024 | 0.053 | 0.053 | 0.071 | 0.56 | 0.71 | 0.46 | 0.33 | 0.4 | 0.42 |
| 2 | 81207413 | rs10520291 | 0.282 | 0.266 | 1.427 | 0.186 | 0.003 | 0.009 | 0.009 | 0 | 0.09 | 0.09 | 0.04 | 0.33 | 0.06 | 0.16 |
| 2 | 81228591 | rs4852599 | 0.81 | 0.409 | 1.503 | 0.2 | -0.043 | -0.028 | -0.028 | 0 | 0.33 | 0.19 | 0.48 | 0.38 | 0.45 | 0.41 |
| 2 | 81286993 | rs12465404 | -1.009 | 0.664 | 1.848 | 0.135 | 0.013 | 0.111 | 0.111 | 0 | 0.18 | 0.34 | 0.14 | 0.41 | 0.16 | 0.28 |
| 2 | 81287257 | rs6718491 | -1.358 | 0.656 | 2.034 | 0.171 | 0.014 | 0.2 | 0.2 | 0 | 0.2 | 0.49 | 0.17 | 0.47 | 0.16 | 0.34 |
| 2 | 81289914 | rs12476834 | -0.979 | 0.658 | 2.09 | 0.156 | 0.035 | -0.004 | -0.004 | 0 | 0.1 | 0.24 | 0.12 | 0.11 | 0.11 | 0.1 |
| 2 | 81291355 | rs12470945 | -1.577 | 0.671 | 1.823 | 0.158 | 0.062 | 0.051 | 0.051 | 0.372 | 0.12 | 0.42 | 0.15 | 0.17 | 0.11 | 0.16 |
| 2 | 81307430 | rs6712750 | 0.421 | 0.63 | 1.654 | 0.164 | 0.002 | -0.011 | -0.011 | 0 | 0.09 | 0.07 | 0.03 | 0.3 | 0.07 | 0.2 |
| 2 | 81309339 | rs12467439 | 0.825 | 0.539 | 1.603 | 0.046 | -0.011 | 0.002 | 0.002 | 1.468 | 0.37 | 0.33 | 0.49 | 0.33 | 0.46 | 0.4 |
| 2 | 81309453 | rs12467480 | -0.804 | 0.527 | 1.067 | -0.017 | 0.029 | 0.118 | 0.118 | 1.943 | 0.61 | 0.9 | 0.67 | 0.79 | 0.63 | 0.75 |
| 2 | 81313893 | rs4852193 | 0.079 | 0.699 | 1.407 | 0.042 | 0.003 | 0.003 | 0.003 | 1.504 | 0.47 | 0.6 | 0.62 | 0.38 | 0.55 | 0.52 |
| 2 | 81321126 | rs11126794 | 0.015 | 0.735 | 1.469 | 0.064 | 0.01 | 0.005 | 0.005 | 2.04 | 0.47 | 0.66 | 0.63 | 0.43 | 0.55 | 0.52 |
| 2 | 81331311 | rs1870114 | 1.746 | 0.86 | 1.448 | -0.002 | -0.047 | -0.044 | -0.044 | 0 | 0.28 | 0.11 | 0.41 | 0.32 | 0.34 | 0.32 |
| 2 | 81346430 | rs7577855 | 1.002 | 0.555 | 1.351 | 0.048 | -0.016 | -0.012 | -0.012 | 0 | 0.36 | 0.25 | 0.45 | 0.43 | 0.42 | 0.42 |
| 2 | 81358993 | rs984903 | -0.131 | 0.491 | 1.596 | 0.188 | 0.024 | -0.027 | -0.027 | 0 | 0.06 | 0.13 | 0.1 | 0.01 | 0.09 | 0.07 |
| 2 | 81385884 | rs1457338 | -0.65 | 0.419 | 1.425 | 0.11 | -0.005 | 0.09 | 0.09 | 0.105 | 0.65 | 0.95 | 0.7 | 0.82 | 0.68 | 0.79 |
| 2 | 81389365 | rs1457341 | 0.265 | 0.434 | 1.669 | 0.123 | -0.004 | -0.018 | -0.018 | 4.259 | 0.53 | 0.49 | 0.55 | 0.73 | 0.59 | 0.66 |
| 2 | 81392921 | rs6738160 | 1.104 | 0.426 | 1.663 | 0.153 | -0.019 | -0.008 | -0.008 | 0.048 | 0.41 | 0.26 | 0.5 | 0.45 | 0.53 | 0.49 |
| 2 | 81400247 | rs7609197 | 0.7 | 0.234 | 1.699 | 0.16 | 0.003 | 0.002 | 0.002 | 0.692 | 0.45 | 0.46 | 0.53 | 0.51 | 0.52 | 0.54 |
| 2 | 81404222 | rs2198935 | 1.159 | 0.241 | 1.699 | 0.141 | -0.019 | -0.009 | -0.009 | 0 | 0.4 | 0.25 | 0.48 | 0.45 | 0.47 | 0.48 |
| 2 | 81407428 | rs7589441 | -0.582 | 0.223 | 1.708 | 0.092 | 0.026 | 0.074 | 0.074 | 0.234 | 0.65 | 0.86 | 0.69 | 0.79 | 0.7 | 0.77 |
| 2 | 81408898 | rs13402918 | -1.267 | 0.261 | 1.718 | 0.068 | 0.047 | 0.023 | 0.023 | 2.679 | 0.05 | 0.33 | 0.05 | 0 | 0.02 | 0 |
| 2 | 81420899 | rs1968758 | 1.567 | 0.252 | 1.004 | 0.075 | 0.027 | -0.021 | -0.021 | 0.097 | 0.23 | 0.02 | 0.22 | 0.19 | 0.16 | 0.17 |
| 2 | 81421702 | rs1982543 | 1.158 | 0.301 | 1.058 | 0.105 | -0.026 | -0.006 | -0.006 | 0 | 0.39 | 0.22 | 0.47 | 0.43 | 0.46 | 0.52 |
| 2 | 81427187 | rs2010447 | -0.639 | 0.265 | 1.045 | 0.126 | 0.008 | 0.074 | 0.074 | 0.056 | 0.64 | 0.94 | 0.69 | 0.81 | 0.68 | 0.77 |
| 2 | 81430162 | rs754361 | -1.326 | 0.265 | 1.22 | 0.137 | 0.003 | 0.078 | 0.078 | 0 | 0.25 | 0.58 | 0.21 | 0.38 | 0.22 | 0.27 |
| 2 | 81440526 | rs6710284 | -0.696 | 0.245 | 1.21 | 0.135 | 0.03 | 0.05 | 0.05 | 0 | 0.66 | 0.86 | 0.7 | 0.82 | 0.72 | 0.79 |
| 2 | 81444443 | rs6547354 | -1.294 | 0.384 | 1.101 | 0.204 | 0.005 | 0.045 | 0.045 | 0 | 0.26 | 0.64 | 0.23 | 0.38 | 0.24 | 0.26 |
| 2 | 81447075 | rs12619674 | 0.594 | 0.312 | 1.429 | 0.409 | -0.013 | -0.001 | -0.001 | 0 | 0.4 | 0.29 | 0.48 | 0.44 | 0.48 | 0.52 |
| 2 | 81464186 | rs4852612 | 0.741 | 0.235 | 1.265 | 0.403 | -0.013 | 0.001 | 0.001 | 0 | 0.4 | 0.28 | 0.47 | 0.43 | 0.46 | 0.51 |
| 2 | 81469658 | rs7557987 | -0.716 | -0.008 | 0.965 | 0.23 | -0.026 | 0.106 | 0.106 | 0.622 | 0.6 | 0.9 | 0.61 | 0.79 | 0.58 | 0.69 |
| 2 | 81472511 | rs12617125 | 0.953 | -0.046 | 1.065 | 0.164 | -0.001 | 0.01 | 0.01 | 0 | 0.77 | 0.69 | 0.87 | 0.74 | 0.77 | 0.8 |
| 2 | 81475647 | rs4456730 | -0.469 | -0.01 | 0.614 | 0.145 | -0.007 | 0.006 | 0.006 | 0.003 | 0.9 | 0.95 | 0.93 | 1 | 0.87 | 0.96 |
| 2 | 81486134 | rs6711618 | 1.197 | -0.094 | 0.431 | 0.348 | -0.042 | -0.02 | -0.02 | 0.315 | 0.31 | 0.11 | 0.42 | 0.32 | 0.36 | 0.36 |
| 2 | 81505707 | rs7581118 | 0.912 | -0.1 | 0.622 | 0.497 | -0.019 | 0.007 | 0.007 | 0.001 | 0.4 | 0.27 | 0.48 | 0.4 | 0.45 | 0.48 |
| 2 | 81521290 | rs7594123 | -0.605 | -0.36 | 0.532 | 0.528 | -0.008 | 0.028 | 0.028 | 0.008 | 0.2 | 0.27 | 0.12 | 0.32 | 0.18 | 0.22 |
| 2 | 81523033 | rs6547361 | -0.834 | -0.304 | -0.285 | 0.556 | -0.031 | 0.075 | 0.075 | 0 | 0.23 | 0.43 | 0.13 | 0.32 | 0.18 | 0.22 |
| 2 | 81546274 | rs7420125 | -0.856 | -0.221 | -0.455 | 0.595 | -0.038 | 0.007 | 0.007 | 0.333 | 0.67 | 0.95 | 0.66 | 0.75 | 0.68 | 0.7 |
| 2 | 81565786 | rs4430978 | -1.042 | -0.133 | -0.381 | 0.721 | -0.031 | 0.066 | 0.066 | 0 | 0.24 | 0.59 | 0.15 | 0.36 | 0.18 | 0.22 |
| 2 | 81582626 | rs1550121 | -0.294 | 0.291 | -1.027 | 1.004 | 0.003 | 0.04 | 0.04 | 0.005 | 0.64 | 0.83 | 0.64 | 0.75 | 0.65 | 0.65 |
| 2 | 81583557 | rs7589293 | 0.233 | 0.44 | -1.074 | 0.738 | 0.013 | 0.002 | 0.002 | 0.283 | 0.56 | 0.68 | 0.61 | 0.46 | 0.6 | 0.55 |
| 2 | 81591163 | rs6547367 | 0.076 | 0.51 | -0.833 | 0.187 | 0 | -0.003 | -0.003 | 0.132 | 0.61 | 0.59 | 0.61 | 0.74 | 0.65 | 0.7 |
| 2 | 81596818 | rs7580205 | -0.979 | 0.55 | -0.44 | -0.108 | 0.009 | -0.043 | -0.043 | 0 | 0.09 | 0.24 | 0.08 | 0.01 | 0.12 | 0.03 |
| 2 | 81607450 | rs1595071 | 0.127 | 0.722 | -1.298 | -0.51 | -0.029 | 0.067 | 0.067 | 0.68 | 0.65 | 0.85 | 0.69 | 0.82 | 0.61 | 0.69 |
| 2 | 81614001 | rs2862499 | 0.137 | 0.74 | -1.224 | -0.517 | -0.029 | 0.067 | 0.067 | 0.632 | 0.65 | 0.86 | 0.69 | 0.82 | 0.61 | 0.68 |
| 2 | 81623974 | rs6547369 | -0.345 | 0.224 | -1.048 | -1.421 | 0.013 | 0.002 | 0.002 | 0.525 | 0.43 | 0.33 | 0.2 | 0.07 | 0.33 | 0.21 |
| 2 | 81628019 | rs1435887 | -0.388 | -0.091 | -0.658 | -1.617 | -0.002 | -0.003 | -0.003 | 0 | 0.25 | 0.27 | 0.15 | 0.08 | 0.25 | 0.12 |
| 2 | 81633206 | rs966308 | -0.52 | -0.281 | -0.463 | -1.619 | 0 | -0.033 | -0.033 | 3.64 | 0.28 | 0.37 | 0.15 | 0.13 | 0.25 | 0.14 |
| 2 | 81638922 | rs11692929 | 0.276 | 0.57 | -0.577 | -1.324 | 0.029 | -0.011 | -0.011 | 0 | 0.65 | 0.48 | 0.63 | 0.68 | 0.54 | 0.44 |
| 2 | 81639912 | rs11903374 | -0.75 | 0.069 | 0.226 | -1.711 | -0.018 | -0.054 | -0.054 | 0 | 0.15 | 0.42 | 0.09 | 0.09 | 0.07 | 0.04 |
| 2 | 81642505 | rs1435883 | -1.832 | 0.102 | 0.45 | -1.619 | 0.01 | -0.058 | -0.058 | 1.398 | 0.28 | 0.63 | 0.21 | 0.15 | 0.3 | 0.12 |
| 2 | 81665310 | rs1347504 | -0.934 | 0.596 | 0.08 | -1.311 | 0.014 | 0.034 | 0.034 | 0.362 | 0.82 | 0.74 | 0.85 | 0.8 | 0.79 | 0.68 |
| 2 | 81676394 | rs11890114 | -0.832 | 0.607 | 0.517 | -1.398 | -0.031 | -0.053 | -0.053 | 0.637 | 0.19 | 0.46 | 0.13 | 0.15 | 0.14 | 0.07 |
| 2 | 81722849 | rs17659465 | 1.09 | 0.57 | 0.228 | -1.038 | -0.044 | -0.001 | -0.001 | 0 | 0.15 | 0.04 | 0.17 | 0.13 | 0.2 | 0.18 |
| 2 | 81731317 | rs12474704 | -0.034 | 0.312 | 0.693 | -1.013 | -0.019 | -0.043 | -0.043 | 1.202 | 0.13 | 0.31 | 0.07 | 0.06 | 0.07 | 0.03 |
| 2 | 81741750 | rs6752979 | -1.676 | 0.333 | 0.704 | -0.965 | 0.017 | -0.06 | -0.06 | 0 | 0.26 | 0.63 | 0.2 | 0.16 | 0.29 | 0.12 |
| 2 | 81755599 | rs11889709 | 0.556 | 0.097 | 0.324 | -1.048 | 0 | 0.003 | 0.003 | 0 | 0.06 | 0.05 | 0.05 | 0.1 | 0.09 | 0.04 |
| 2 | 81757391 | rs764977 | -1.832 | 0.013 | 0.34 | -1.157 | 0.012 | -0.08 | -0.08 | 0 | 0.28 | 0.56 | 0.19 | 0.16 | 0.31 | 0.12 |
| 2 | 81759688 | rs6725541 | -1.037 | 0.132 | 0.418 | -1.079 | 0.03 | -0.111 | -0.111 | 0 | 0.22 | 0.51 | 0.14 | 0.07 | 0.22 | 0.08 |
| 2 | 81768548 | rs12476748 | 0.782 | 0.131 | 0.359 | -1.082 | -0.003 | 0.016 | 0.016 | 0 | 0.1 | 0.01 | 0.04 | 0.06 | 0.07 | 0.03 |
| 2 | 81808100 | rs6547375 | -1.294 | 0.118 | 0.409 | -1.092 | 0.015 | -0.085 | -0.085 | 0.043 | 0.22 | 0.57 | 0.15 | 0.06 | 0.22 | 0.06 |
| 2 | 81813281 | rs7607015 | -0.646 | 0.056 | 0.576 | -1.094 | -0.042 | -0.068 | -0.068 | 1.8 | 0.15 | 0.5 | 0.08 | 0.06 | 0.07 | 0.03 |
| 2 | 81819551 | rs7570909 | -1.011 | 0.075 | 0.461 | -0.9 | -0.056 | 0.035 | 0.035 | 0.311 | 0.21 | 0.09 | 0.2 | 0.18 | 0.3 | 0.12 |

| | | | | | | | | | | | | | | | | |
|---|---|---|---|---|---|---|---|---|---|---|---|---|---|---|---|---|
| 2 | 81828640 | rs6547376 | -0.67 | 0.08 | 0.437 | -0.94 | 0.076 | -0.078 | -0.078 | 0.021 | 0.12 | 0.58 | 0.12 | 0 | 0.16 | 0.04 |
| 2 | 81829389 | rs11685793 | -0.965 | 0.065 | 0.42 | -0.951 | -0.023 | -0.063 | -0.063 | 0.786 | 0.17 | 0.61 | 0.13 | 0.06 | 0.13 | 0.04 |
| 2 | 81832706 | rs6751593 | -1.761 | 0.07 | 0.408 | -0.95 | 0.01 | -0.056 | -0.056 | 0 | 0.25 | 0.63 | 0.21 | 0.06 | 0.29 | 0.08 |
| 2 | 81837644 | rs17020525 | 0.183 | 0.036 | 0.226 | -0.878 | 0.001 | 0.034 | 0.034 | 0.494 | 0.12 | 0.06 | 0.09 | 0.06 | 0.13 | 0.04 |
| 2 | 81844729 | rs10520321 | 0.161 | 0.022 | 0.225 | -0.878 | 0.004 | 0.035 | 0.035 | 0.478 | 0.12 | 0.06 | 0.07 | 0.06 | 0.12 | 0.04 |
| 2 | 81848922 | rs17020583 | 0.087 | 0.02 | 0.224 | -0.879 | 0.005 | 0.036 | 0.036 | 0.422 | 0.13 | 0.06 | 0.07 | 0.06 | 0.12 | 0.04 |
| 2 | 81852144 | rs12470076 | -0.893 | -0.497 | 0.316 | -1.184 | -0.022 | -0.019 | -0.019 | 0 | 0.21 | 0.27 | 0.11 | 0.19 | 0.15 | 0.09 |
| 2 | 81858897 | rs6547379 | 0.401 | -0.382 | 0.404 | -1.224 | 0.103 | 0.01 | 0.01 | 0 | 0.09 | 0.21 | 0.11 | 0.12 | 0.16 | 0.08 |
| 2 | 81872922 | rs12052359 | 2.506 | -0.684 | -0.269 | -0.752 | -0.006 | -0.074 | -0.074 | 0 | 0.31 | 0.02 | 0.43 | 0.34 | 0.26 | 0.33 |
| 2 | 81876709 | rs11126826 | -2.031 | -0.407 | 0.651 | -0.726 | 0 | 0.02 | 0.02 | 3.628 | 0.32 | 0.69 | 0.23 | 0.19 | 0.31 | 0.13 |
| 2 | 81885942 | rs6547382 | -1.808 | -0.401 | 0.655 | -0.743 | 0.001 | 0.004 | 0.004 | 1.87 | 0.31 | 0.67 | 0.2 | 0.19 | 0.29 | 0.16 |
| 2 | 81911025 | rs10174219 | -1.958 | -0.409 | 0.638 | -0.717 | -0.001 | 0.032 | 0.032 | 2.916 | 0.32 | 0.71 | 0.22 | 0.19 | 0.31 | 0.16 |
| 2 | 81912847 | rs10180671 | -1.09 | -0.402 | 0.751 | -0.719 | -0.067 | -0.105 | -0.105 | 0 | 0.22 | 0.48 | 0.1 | 0.07 | 0.13 | 0.09 |
| 2 | 81918015 | rs12464395 | -2.008 | -0.396 | 0.634 | -0.715 | -0.001 | 0.033 | 0.033 | 3.638 | 0.32 | 0.71 | 0.22 | 0.19 | 0.31 | 0.16 |
| 2 | 81919965 | rs7609125 | 0.701 | -0.415 | 0.638 | -0.724 | 0.006 | 0.006 | 0.006 | 0.335 | 0.09 | 0.08 | 0.04 | 0.01 | 0.07 | 0.04 |
| 2 | 81921062 | rs934892 | -2.028 | -0.401 | 0.597 | -0.717 | -0.001 | 0.033 | 0.033 | 3.457 | 0.32 | 0.71 | 0.22 | 0.19 | 0.31 | 0.16 |
| 2 | 81930807 | rs4555368 | 1.279 | -0.34 | 0.552 | -0.696 | -0.022 | 0.019 | 0.019 | 0 | 0.07 | 0.02 | 0.08 | 0 | 0.16 | 0.04 |
| 2 | 81931197 | rs17020666 | -0.906 | -0.365 | 0.647 | -0.708 | 0.041 | -0.118 | -0.118 | 0.025 | 0.21 | 0.46 | 0.14 | 0.06 | 0.22 | 0.08 |
| 2 | 81933316 | rs11691608 | -0.444 | -0.36 | 0.636 | -0.705 | 0.005 | -0.023 | -0.023 | 0 | 0.17 | 0.15 | 0.11 | 0.06 | 0.21 | 0.08 |
| 2 | 81934345 | rs1922029 | 0.334 | -0.209 | 0.755 | -0.553 | -0.017 | -0.036 | -0.036 | 0.699 | 0.09 | 0.17 | 0.04 | 0.01 | 0.07 | 0.04 |
| 2 | 81944962 | rs17020687 | 0.407 | -0.123 | 0.698 | -0.502 | -0.002 | -0.009 | -0.009 | 0 | 0.09 | 0.11 | 0.04 | 0.01 | 0.07 | 0.04 |
| 2 | 81946409 | rs1432187 | -0.239 | -0.114 | 0.643 | -0.485 | -0.012 | 0.002 | 0.002 | 1.062 | 0.11 | 0.23 | 0.09 | 0.13 | 0.09 | 0.08 |
| 2 | 81946861 | rs1368048 | -1.077 | -0.139 | 0.234 | -0.499 | 0.047 | -0.06 | -0.06 | 0.248 | 0.21 | 0.47 | 0.17 | 0.13 | 0.25 | 0.12 |
| 2 | 81969986 | rs11681750 | 0.418 | 0.193 | -0.317 | -0.502 | 0.005 | 0.022 | 0.022 | 0.1 | 0.06 | 0.01 | 0.04 | 0 | 0.08 | 0.07 |
| 2 | 81978821 | rs1221194 | -1.01 | -0.151 | -0.397 | -0.4 | 0.001 | 0.008 | 0.008 | 0.064 | 0.46 | 0.44 | 0.31 | 0.25 | 0.48 | 0.42 |
| 2 | 81988382 | rs1237758 | -0.287 | -0.278 | -0.548 | -0.516 | -0.009 | 0.008 | 0.008 | 0 | 0.33 | 0.25 | 0.27 | 0.21 | 0.4 | 0.33 |
| 2 | 81991507 | rs1108799 | 0.447 | -0.21 | -0.645 | -0.616 | -0.003 | 0.008 | 0.008 | 0 | 0.2 | 0.23 | 0.16 | 0.21 | 0.22 | 0.3 |
| 2 | 81993943 | rs11684035 | 1.631 | -0.116 | -0.727 | -0.576 | -0.048 | 0.06 | 0.06 | 0 | 0.13 | 0.03 | 0.11 | 0 | 0.18 | 0.05 |
| 2 | 82017373 | rs1432155 | 1.917 | -0.183 | -0.37 | -0.771 | -0.009 | 0.039 | 0.039 | 0.063 | 0.46 | 0.31 | 0.63 | 0.73 | 0.48 | 0.54 |
| 2 | 82019665 | rs6717916 | 0.816 | -0.276 | -0.184 | -0.628 | -0.017 | -0.012 | -0.012 | 0.725 | 0.62 | 0.49 | 0.76 | 0.73 | 0.66 | 0.6 |
| 2 | 82021040 | rs10188944 | 1.074 | 0.232 | -0.228 | -0.788 | -0.001 | -0.019 | -0.019 | 0 | 0.57 | 0.47 | 0.69 | 0.74 | 0.53 | 0.54 |
| 2 | 82024197 | rs10171375 | 1.087 | 0.233 | -0.227 | -0.793 | -0.001 | -0.018 | -0.018 | 0.05 | 0.57 | 0.46 | 0.69 | 0.74 | 0.53 | 0.54 |
| 2 | 82027767 | rs1368038 | -2.253 | 0.155 | -0.12 | -0.729 | 0.002 | 0.041 | 0.041 | 0 | 0.87 | 0.8 | 0.88 | 0.8 | 0.85 | 0.77 |
| 2 | 82028095 | rs1368041 | 0.144 | 0.143 | -0.155 | -0.759 | -0.006 | 0.001 | 0.001 | 0 | 0.74 | 0.63 | 0.81 | 0.78 | 0.73 | 0.67 |
| 2 | 82039382 | rs11126830 | -1.9 | 0.169 | -0.152 | -0.444 | -0.001 | 0.098 | 0.098 | 0 | 0.86 | 0.74 | 0.79 | 0.63 | 0.86 | 0.73 |
| 2 | 82046291 | rs17020914 | 2.545 | 0.197 | -1.269 | -0.415 | -0.047 | -0.075 | -0.075 | 0.032 | 0.28 | 0.01 | 0.39 | 0.35 | 0.26 | 0.35 |
| 2 | 82047642 | rs12151616 | 1.694 | 0.43 | -0.939 | -0.871 | 0.009 | -0.041 | -0.041 | 0 | 0.14 | 0.13 | 0.12 | 0 | 0.2 | 0.05 |
| 2 | 82047715 | rs10166031 | 1.875 | 0.317 | -0.802 | -0.855 | 0.01 | -0.017 | -0.017 | 0.019 | 0.41 | 0.2 | 0.46 | 0.58 | 0.36 | 0.45 |
| 2 | 82052112 | rs1432170 | -0.805 | 0.263 | -0.572 | -0.992 | -0.006 | 0.013 | 0.013 | 0 | 0.76 | 0.71 | 0.74 | 0.63 | 0.78 | 0.67 |
| 2 | 82075453 | rs11126837 | 1.149 | 0.216 | -0.634 | -1.05 | 0.009 | 0.011 | 0.011 | 0.234 | 0.54 | 0.33 | 0.58 | 0.58 | 0.54 | 0.49 |
| 2 | 82077473 | rs11674245 | 0.892 | 0.204 | -0.45 | -1.116 | -0.004 | -0.039 | -0.039 | 0.85 | 0.08 | 0.13 | 0.07 | 0 | 0.09 | 0.06 |
| 2 | 82079857 | rs1432180 | 1.436 | 0.139 | -0.426 | -1.187 | -0.017 | 0.038 | 0.038 | 0 | 0.15 | 0.07 | 0.1 | 0.04 | 0.19 | 0.13 |
| 2 | 82093538 | rs13029269 | 1.949 | 0.189 | -0.429 | -1.195 | 0.005 | -0.037 | -0.037 | 0 | 0.14 | 0.13 | 0.1 | 0 | 0.19 | 0.06 |
| 2 | 82093632 | rs17021075 | 1.88 | 0.173 | -0.524 | -1.128 | -0.004 | 0.024 | 0.024 | 0 | 0.06 | 0.02 | 0.05 | 0 | 0.09 | 0.07 |
| 2 | 82094566 | rs6715448 | -0.816 | 0.017 | -0.553 | -1.179 | 0 | -0.005 | -0.005 | 0.011 | 0.71 | 0.73 | 0.69 | 0.6 | 0.69 | 0.6 |
| 2 | 82106093 | rs10520309 | -0.598 | 0.131 | -0.616 | -1.114 | 0.008 | -0.068 | -0.068 | 0 | 0.12 | 0.33 | 0.1 | 0.01 | 0.13 | 0.1 |
| 2 | 82111016 | rs12469697 | 1.822 | 0.036 | -0.764 | -1.157 | -0.011 | -0.031 | -0.031 | 0 | 0.38 | 0.16 | 0.47 | 0.59 | 0.35 | 0.44 |
| 2 | 82135708 | rs11895085 | 0.461 | 0.051 | -0.848 | -1.171 | 0.002 | 0.001 | 0.001 | 0 | 0.59 | 0.58 | 0.61 | 0.6 | 0.58 | 0.53 |
| 2 | 82166534 | rs2685724 | -2.059 | 0.054 | -1.013 | -1.17 | -0.01 | -0.025 | -0.025 | 0 | 0.62 | 0.84 | 0.53 | 0.4 | 0.65 | 0.57 |
| 2 | 82168465 | rs17640902 | 1.103 | 0.016 | -1.026 | -1.125 | -0.01 | 0.021 | 0.021 | 0 | 0.18 | 0.14 | 0.13 | 0.05 | 0.21 | 0.13 |
| 2 | 82170262 | rs2685710 | -0.846 | 0.009 | -1.093 | -1.1 | 0.01 | -0.016 | -0.016 | 0.28 | 0.41 | 0.82 | 0.37 | 0.4 | 0.35 | 0.43 |
| 2 | 82171067 | rs10209091 | 2.421 | 0.034 | -1.104 | -0.879 | -0.001 | 0 | 0 | 0 | 0.11 | 0.11 | 0.05 | 0.17 | 0.1 | 0.08 |
| 2 | 82171348 | rs17566909 | 1.551 | 0.07 | -1.028 | -1.095 | -0.055 | 0.082 | 0.082 | 0 | 0.15 | 0.02 | 0.1 | 0 | 0.19 | 0.06 |
| 2 | 82171825 | rs2947961 | 1.592 | -0.034 | -0.592 | -0.781 | -0.015 | 0.039 | 0.039 | 0 | 0.17 | 0.21 | 0.19 | 0.34 | 0.1 | 0.26 |
| 2 | 82205627 | rs2685200 | -0.161 | 0.145 | -0.998 | -0.935 | -0.002 | 0.004 | 0.004 | 0 | 0.34 | 0.43 | 0.33 | 0.39 | 0.31 | 0.39 |
| 2 | 82208620 | rs12614042 | 1.158 | 0.116 | -1.711 | -1.003 | 0.018 | -0.057 | -0.057 | 0.005 | 0.33 | 0.15 | 0.41 | 0.55 | 0.26 | 0.45 |
| 2 | 82213961 | rs11686261 | 0.355 | -0.033 | -1.599 | -1.165 | 0.014 | 0.006 | 0.006 | 0 | 0.55 | 0.19 | 0.59 | 0.57 | 0.6 | 0.58 |
| 2 | 82217592 | rs17568010 | 1.958 | 0.296 | -1.64 | -1.189 | -0.029 | 0.028 | 0.028 | 0 | 0.06 | 0.02 | 0.06 | 0 | 0.12 | 0.04 |
| 2 | 82222407 | rs1402778 | 1.144 | 0.163 | -1.576 | -1.221 | -0.004 | 0.017 | 0.017 | 0 | 0.19 | 0.14 | 0.13 | 0.05 | 0.2 | 0.09 |
| 2 | 82225608 | rs1521694 | 1.157 | 0.175 | -2.027 | -1.288 | -0.066 | -0.138 | -0.138 | 0 | 0.33 | 0.07 | 0.46 | 0.44 | 0.35 | 0.46 |
| 2 | 82227252 | rs17021319 | 0.333 | 0.554 | -0.861 | -1.086 | 0 | -0.047 | -0.047 | 0 | 0.08 | 0.17 | 0.05 | 0 | 0.1 | 0.07 |
| 2 | 82230841 | rs909044 | 0.333 | 0.554 | -0.861 | -1.086 | 0 | -0.047 | -0.047 | 0 | 0.08 | 0.17 | 0.05 | 0 | 0.1 | 0.07 |
| 2 | 82237352 | rs7355329 | -0.826 | 0.599 | -1.506 | -0.982 | -0.002 | -0.006 | -0.006 | 0 | 0.78 | 0.76 | 0.75 | 0.63 | 0.78 | 0.71 |
| 2 | 82241600 | rs13027453 | 0.53 | 0.511 | -1.866 | -0.643 | -0.019 | 0.011 | 0.011 | 0 | 0.48 | 0.14 | 0.52 | 0.59 | 0.48 | 0.54 |
| 2 | 82248401 | rs1608009 | 0.647 | 0.301 | -1.685 | -0.666 | -0.003 | 0.014 | 0.014 | 0 | 0.46 | 0.13 | 0.64 | 0.62 | 0.45 | 0.49 |
| 2 | 82248571 | rs1581386 | -0.474 | 0.386 | -1.053 | -0.741 | -0.001 | -0.001 | -0.001 | 0 | 0.34 | 0.32 | 0.35 | 0.2 | 0.46 | 0.31 |
| 2 | 82268707 | rs7609503 | -0.06 | 0.338 | -1.376 | -0.923 | -0.102 | 0.128 | 0.128 | 0 | 0.28 | 0.1 | 0.23 | 0.04 | 0.41 | 0.24 |
| 2 | 82287744 | rs2685211 | -0.836 | 0.2 | -1.287 | -0.74 | -0.001 | 0.034 | 0.034 | 0 | 0.94 | 0.94 | 0.88 | 0.73 | 0.94 | 0.94 |
| 2 | 82292908 | rs2566748 | -0.715 | 0.157 | -1.039 | -0.823 | -0.042 | 0.017 | 0.017 | 0 | 0.35 | 0.24 | 0.36 | 0.29 | 0.47 | 0.3 |

| | | | | | | | | | | | | | | | | |
|---|---|---|---|---|---|---|---|---|---|---|---|---|---|---|---|---|
| 2 | 82299945 | rs1357144 | 0.531 | 0.496 | -1.119 | -0.925 | -0.017 | 0.002 | 0.002 | 0 | 0.14 | 0.12 | 0.12 | 0 | 0.24 | 0.11 |
| 2 | 82315179 | rs2116045 | -0.168 | -0.074 | -0.938 | -1.166 | 0.02 | 0.044 | 0.044 | 0 | 0.68 | 0.76 | 0.83 | 0.84 | 0.77 | 0.62 |
| 2 | 82320474 | rs1344739 | -1.331 | -0.091 | -0.935 | -1.169 | -0.006 | 0 | 0 | 0 | 0.83 | 0.81 | 0.96 | 0.88 | 0.95 | 0.76 |
| 2 | 82320654 | rs1344740 | -0.16 | -0.043 | -0.902 | -1.156 | -0.008 | -0.004 | -0.004 | 0.158 | 0.21 | 0.19 | 0.23 | 0.26 | 0.3 | 0.17 |
| 2 | 82323314 | rs7605223 | 0.52 | -0.057 | -0.917 | -1.163 | -0.024 | 0.149 | 0.149 | 0 | 0.54 | 0.71 | 0.72 | 0.85 | 0.53 | 0.52 |
| 2 | 82328861 | rs2052955 | 1.607 | -0.4 | -1.038 | -1.448 | -0.058 | -0.037 | -0.037 | 0.218 | 0.36 | 0.12 | 0.54 | 0.59 | 0.44 | 0.44 |
| 2 | 82328907 | rs17582064 | 1.134 | 0.062 | -0.722 | -1.105 | -0.06 | 0.068 | 0.068 | 0 | 0.13 | 0.03 | 0.11 | 0.01 | 0.23 | 0.11 |
| 2 | 82331873 | rs10167584 | 1.127 | -0.019 | -0.295 | -1.037 | -0.001 | 0.006 | 0.006 | 0 | 0.46 | 0.38 | 0.58 | 0.6 | 0.47 | 0.46 |
| 2 | 82338371 | rs17021454 | 0.321 | -0.137 | -0.645 | -0.415 | 0.002 | 0.003 | 0.003 | 0 | 0.06 | 0.03 | 0.12 | 0.26 | 0.06 | 0.06 |
| 2 | 82344428 | rs4852213 | 0.831 | -0.082 | -0.63 | -0.386 | -0.001 | 0.007 | 0.007 | 0 | 0.05 | 0.03 | 0.12 | 0.26 | 0.06 | 0.06 |
| 2 | 82351371 | rs4852669 | -0.082 | -0.308 | -0.654 | -0.537 | -0.072 | -0.046 | -0.046 | 0.299 | 0.79 | 0.42 | 0.92 | 0.9 | 0.94 | 0.76 |
| 2 | 82357490 | rs6752176 | 0.783 | 0.412 | -1.691 | -0.365 | -0.052 | -0.015 | -0.015 | 0 | 0.09 | 0 | 0.09 | 0.05 | 0.17 | 0.1 |
| 2 | 82357720 | rs12464109 | 0.293 | 0.463 | -1.853 | -0.41 | -0.039 | -0.06 | -0.06 | 0.856 | 0.19 | 0.06 | 0.23 | 0.29 | 0.23 | 0.2 |
| 2 | 82360446 | rs7558635 | 0.545 | 0.311 | -1.396 | -0.238 | -0.052 | -0.015 | -0.015 | 0.014 | 0.09 | 0 | 0.09 | 0.05 | 0.17 | 0.1 |
| 2 | 82369359 | rs6547406 | -0.585 | -0.287 | -1.057 | -0.27 | -0.009 | -0.002 | -0.002 | 0 | 0.84 | 0.81 | 0.95 | 0.9 | 0.94 | 0.76 |
| 2 | 82375877 | rs13017348 | -0.308 | 0.191 | -1.283 | 0.014 | -0.02 | -0.069 | -0.069 | 0.147 | 0.18 | 0.05 | 0.2 | 0.3 | 0.22 | 0.13 |
| 2 | 82380703 | rs1430713 | -0.282 | 0.189 | -1.264 | 0.029 | -0.019 | -0.064 | -0.064 | 0 | 0.18 | 0.05 | 0.2 | 0.3 | 0.22 | 0.13 |
| 2 | 82380867 | rs1430716 | -0.228 | 0.185 | -1.268 | 0.014 | -0.042 | -0.093 | -0.093 | 0 | 0.18 | 0.02 | 0.2 | 0.3 | 0.22 | 0.13 |
| 2 | 82381293 | rs17021504 | -0.357 | -0.101 | -0.368 | -0.164 | 0.006 | -0.005 | -0.005 | 0 | 0.05 | 0.21 | 0.02 | 0 | 0 | 0 |
| 2 | 82390498 | rs1864564 | -0.197 | -0.151 | 0.27 | -0.181 | -0.007 | -0.035 | -0.035 | 3.622 | 0.18 | 0.08 | 0.2 | 0.3 | 0.22 | 0.13 |
| 2 | 82390641 | rs1864565 | -0.266 | -0.151 | 0.274 | -0.181 | -0.002 | -0.019 | -0.019 | 0.074 | 0.18 | 0.11 | 0.2 | 0.3 | 0.22 | 0.13 |
| 2 | 82396211 | rs934310 | 1.09 | -0.194 | -0.045 | -0.848 | 0.003 | -0.007 | -0.007 | 0 | 0.37 | 0.38 | 0.51 | 0.57 | 0.32 | 0.41 |
| 2 | 82410462 | rs1430726 | 0.545 | -0.031 | 0.085 | -1.172 | 0.01 | 0.011 | 0.011 | 0 | 0.49 | 0.41 | 0.61 | 0.57 | 0.52 | 0.5 |
| 2 | 82416244 | rs17644951 | 1.218 | -0.001 | -0.016 | -1.16 | -0.062 | 0.07 | 0.07 | 0 | 0.13 | 0.03 | 0.11 | 0 | 0.21 | 0.09 |
| 2 | 82427409 | rs17040762 | 0.334 | -0.107 | -0.133 | -1.219 | -0.001 | 0.015 | 0.015 | 0 | 0.21 | 0.21 | 0.23 | 0.3 | 0.23 | 0.12 |
| 2 | 82427826 | rs6736459 | 1.076 | -0.004 | -0.402 | -1.024 | -0.062 | 0.069 | 0.069 | 0 | 0.12 | 0.03 | 0.1 | 0 | 0.21 | 0.09 |
| 2 | 82432988 | rs4852681 | 1.089 | -0.035 | -0.373 | -1.011 | 0.001 | -0.004 | -0.004 | 0 | 0.36 | 0.38 | 0.49 | 0.57 | 0.31 | 0.41 |
| 2 | 82462133 | rs7577033 | 0.398 | 0.041 | -0.193 | -1.13 | -0.003 | -0.02 | -0.02 | 0.296 | 0.13 | 0.13 | 0.11 | 0 | 0.21 | 0.09 |
| 2 | 82476873 | rs11896872 | -0.915 | 0.019 | -0.213 | -1.149 | -0.005 | 0.044 | 0.044 | 1.982 | 0.23 | 0.86 | 0.17 | 0 | 0.21 | 0.09 |
| 2 | 82477944 | rs10208061 | 1.439 | -0.227 | -0.396 | -1.131 | -0.007 | -0.003 | -0.003 | 0.04 | 0.07 | 0.08 | 0.02 | 0.06 | 0.03 | 0.04 |
| 2 | 82482510 | rs6707963 | -0.798 | 0.035 | -0.179 | -1.095 | 0 | -0.049 | -0.049 | 0 | 0.23 | 0.76 | 0.17 | 0 | 0.22 | 0.09 |
| 2 | 82488467 | rs11893649 | -0.148 | -0.135 | 0.279 | -0.923 | 0.004 | -0.08 | -0.08 | 0 | 0.12 | 0.46 | 0.06 | 0.01 | 0.07 | 0.04 |
| 2 | 82493580 | rs4128656 | -0.127 | -0.132 | 0.28 | -0.921 | 0.005 | -0.079 | -0.079 | 0 | 0.12 | 0.46 | 0.06 | 0.01 | 0.07 | 0.04 |
| 2 | 82499876 | rs4393782 | -1.545 | -0.442 | -0.479 | -0.964 | 0.014 | -0.022 | -0.022 | 1.2 | 0.9 | 1 | 0.97 | 0.89 | 0.92 | 0.77 |
| 2 | 82509665 | rs6733403 | -1.013 | -0.321 | -0.64 | -1.505 | 0.016 | 0.075 | 0.075 | 0.673 | 0.31 | 0.81 | 0.19 | 0.02 | 0.26 | 0.11 |
| 2 | 82510902 | rs12714093 | 1.373 | -0.404 | -1.592 | -1.832 | -0.139 | -0.051 | -0.051 | 0.078 | 0.28 | 0.04 | 0.63 | 0.72 | 0.4 | 0.46 |
| 2 | 82512797 | rs13028564 | -0.441 | -0.07 | 0.044 | -1.827 | -0.003 | -0.14 | -0.14 | 0.253 | 0.21 | 0.51 | 0.14 | 0.01 | 0.19 | 0.07 |
| 2 | 82513843 | rs7423219 | -0.475 | -0.084 | 0.441 | -1.833 | -0.001 | -0.108 | -0.108 | 0.955 | 0.2 | 0.37 | 0.11 | 0.01 | 0.19 | 0.07 |
| 2 | 82514844 | rs6547418 | -1.247 | -0.095 | 0.227 | -1.839 | 0.009 | 0.073 | 0.073 | 0.805 | 0.31 | 0.8 | 0.2 | 0.02 | 0.27 | 0.09 |
| 2 | 82522552 | rs13011567 | 0.155 | -0.069 | 0.325 | -1.814 | -0.001 | -0.018 | -0.018 | 0 | 0.16 | 0.14 | 0.1 | 0.01 | 0.19 | 0.07 |
| 2 | 82524725 | rs12151746 | -1.151 | -0.118 | 0.446 | -1.813 | 0.002 | -0.025 | -0.025 | 0.017 | 0.31 | 0.71 | 0.19 | 0.02 | 0.27 | 0.09 |
| 2 | 82527769 | rs6734995 | -0.536 | -0.172 | 0.458 | -1.839 | 0.011 | 0.008 | 0.008 | 0.123 | 0.05 | 0.28 | 0.03 | 0 | 0 | 0 |
| 2 | 82537976 | rs7563467 | -1.118 | -0.104 | 0.072 | -1.807 | 0.002 | -0.025 | -0.025 | 0 | 0.31 | 0.72 | 0.19 | 0.02 | 0.27 | 0.09 |
| 2 | 82540277 | rs11895031 | -1.118 | -0.104 | 0.059 | -1.807 | 0.002 | -0.022 | -0.022 | 0 | 0.31 | 0.72 | 0.19 | 0.02 | 0.27 | 0.09 |
| 2 | 82578928 | rs1427648 | -1.129 | -0.171 | 0.758 | -1.882 | -0.008 | -0.099 | -0.099 | 0 | 0.28 | 0.62 | 0.18 | 0.02 | 0.24 | 0.14 |
| 2 | 82585098 | rs13009422 | 0.566 | -0.211 | 0.833 | -1.908 | 0.003 | -0.021 | -0.021 | 0 | 0.12 | 0.13 | 0.1 | 0.01 | 0.17 | 0.08 |
| 2 | 82607433 | rs11890108 | -1.033 | -0.303 | 0.927 | -1.866 | 0.006 | 0.002 | 0.002 | 4.068 | 0.08 | 0.47 | 0.04 | 0 | 0 | 0 |
| 2 | 82613120 | rs1365953 | -1.405 | -0.328 | 0.827 | -1.906 | -0.012 | -0.111 | -0.111 | 0.244 | 0.3 | 0.61 | 0.19 | 0.02 | 0.25 | 0.14 |
| 2 | 82615016 | rs11887253 | -1.315 | -0.323 | 0.807 | -1.904 | -0.01 | -0.12 | -0.12 | 0 | 0.29 | 0.58 | 0.19 | 0.02 | 0.25 | 0.14 |
| 2 | 82665676 | rs1365951 | -1.97 | -0.729 | -0.216 | -1.588 | -0.02 | -0.008 | -0.008 | 0 | 0.55 | 0.95 | 0.37 | 0.34 | 0.34 | 0.3 |
| 2 | 82668227 | rs6713854 | -1 | -0.433 | 0.268 | -1.872 | -0.008 | -0.166 | -0.166 | 0 | 0.27 | 0.51 | 0.18 | 0.02 | 0.25 | 0.14 |
| 2 | 82679503 | rs2902368 | -1.57 | -0.597 | 0.211 | -1.96 | -0.007 | -0.087 | -0.087 | 0 | 0.37 | 0.62 | 0.22 | 0.03 | 0.29 | 0.2 |
| 2 | 82698316 | rs11126867 | 0.942 | -0.873 | 0.462 | -1.865 | -0.006 | -0.062 | -0.062 | 0.16 | 0.1 | 0.25 | 0.05 | 0 | 0.08 | 0.06 |
| 2 | 82706914 | rs6705383 | -1.005 | -0.877 | 0.402 | -1.869 | -0.008 | -0.164 | -0.164 | 0 | 0.27 | 0.52 | 0.18 | 0.02 | 0.25 | 0.14 |
| 2 | 82717155 | rs11681331 | 0.387 | -0.869 | 0.336 | -1.865 | -0.018 | 0.054 | 0.054 | 0 | 0.15 | 0.09 | 0.11 | 0.02 | 0.17 | 0.08 |
| 2 | 82720679 | rs1549581 | -0.441 | -0.861 | 0.355 | -1.851 | -0.009 | 0.053 | 0.053 | 0 | 0.24 | 0.14 | 0.15 | 0.02 | 0.25 | 0.14 |
| 2 | 82736253 | rs6746730 | -1.422 | -0.794 | 0.217 | -1.809 | -0.008 | -0.093 | -0.093 | 0 | 0.37 | 0.62 | 0.22 | 0.03 | 0.29 | 0.2 |
| 2 | 82744841 | rs7579676 | 0.388 | -0.611 | 0.177 | -1.672 | 0.016 | -0.051 | -0.051 | 0 | 0.13 | 0.22 | 0.1 | 0.02 | 0.15 | 0.07 |
| 2 | 82745838 | rs6733738 | 0.931 | -0.775 | -0.167 | -1.736 | -0.029 | -0.034 | -0.034 | 0 | 0.06 | 0.16 | 0.01 | 0 | 0 | 0 |
| 2 | 82746544 | rs1469370 | -1.374 | -0.867 | -0.22 | -1.775 | -0.022 | -0.117 | -0.117 | 0.675 | 0.36 | 0.54 | 0.19 | 0.04 | 0.26 | 0.18 |
| 2 | 82760396 | rs2862943 | -1.418 | -0.884 | -0.248 | -1.787 | -0.022 | -0.114 | -0.114 | 0.115 | 0.36 | 0.54 | 0.19 | 0.04 | 0.26 | 0.18 |
| 2 | 82761573 | rs2862944 | -2.213 | -0.531 | -0.744 | -1.692 | -0.155 | 0.002 | 0.002 | 2.683 | 0.59 | 0.95 | 0.6 | 0.27 | 0.48 | 0.33 |
| 2 | 82761799 | rs2862945 | -0.698 | -1.186 | -0.438 | -1.569 | -0.031 | -0.121 | -0.121 | 0 | 0.27 | 0.53 | 0.16 | 0.04 | 0.19 | 0.11 |
| 2 | 82771228 | rs6724687 | -1.101 | -1.193 | -0.398 | -1.583 | -0.01 | -0.076 | -0.076 | 0 | 0.34 | 0.44 | 0.18 | 0.04 | 0.26 | 0.18 |
| 2 | 82775177 | rs6761541 | -1.268 | -1.193 | -0.455 | -1.583 | -0.022 | -0.113 | -0.113 | 0.198 | 0.36 | 0.54 | 0.19 | 0.04 | 0.26 | 0.18 |
| 2 | 82781005 | rs2902382 | 1.472 | -1.223 | -0.475 | -1.607 | -0.037 | 0.043 | 0.043 | 0 | 0.11 | 0.01 | 0.09 | 0.02 | 0.15 | 0.07 |
| 2 | 82788806 | rs7581293 | -1.959 | -1.172 | -0.397 | -1.615 | 0.053 | 0.104 | 0.104 | 3.208 | 0.52 | 0.93 | 0.38 | 0.25 | 0.34 | 0.28 |
| 2 | 82799733 | rs6547435 | -1.428 | -0.572 | -0.083 | -1.748 | -0.021 | -0.11 | -0.11 | 0 | 0.36 | 0.56 | 0.2 | 0.04 | 0.26 | 0.18 |
| 2 | 82804806 | rs2862982 | -0.602 | -0.576 | -0.067 | -1.735 | -0.055 | -0.124 | -0.124 | 0 | 0.25 | 0.47 | 0.11 | 0.03 | 0.11 | 0.12 |

| | | | | | | | | | | | | | | | | |
|---|---|---|---|---|---|---|---|---|---|---|---|---|---|---|---|---|
| 2 | 82814223 | rs6756461 | -0.602 | -0.576 | -0.058 | -1.735 | -0.055 | -0.123 | -0.123 | 0.491 | 0.25 | 0.47 | 0.11 | 0.03 | 0.11 | 0.12 |
| 2 | 82814510 | rs6757052 | -1.366 | -0.551 | -0.062 | -1.73 | -0.023 | -0.116 | -0.116 | 0.634 | 0.36 | 0.54 | 0.19 | 0.04 | 0.26 | 0.19 |
| 2 | 82828982 | rs7584969 | 1.3 | -0.581 | 0.004 | -1.712 | -0.012 | -0.025 | -0.025 | 0 | 0.09 | 0.25 | 0.05 | 0.02 | 0.04 | 0.04 |
| 2 | 82839226 | rs11886368 | 1.299 | -0.576 | -0.105 | -1.709 | -0.012 | -0.025 | -0.025 | 0 | 0.09 | 0.25 | 0.05 | 0.02 | 0.04 | 0.04 |
| 2 | 82839774 | rs7567118 | -1.152 | -0.43 | -0.58 | -1.638 | -0.022 | -0.11 | -0.11 | 0.022 | 0.36 | 0.56 | 0.19 | 0.04 | 0.26 | 0.18 |
| 2 | 82857815 | rs1344278 | -0.479 | -0.617 | -0.378 | -1.514 | -0.061 | -0.141 | -0.141 | 0 | 0.26 | 0.55 | 0.12 | 0.03 | 0.13 | 0.12 |
| 2 | 82859580 | rs12104529 | -1.106 | -0.371 | -0.657 | -1.718 | 0.009 | -0.01 | -0.01 | 0 | 0.43 | 0.64 | 0.27 | 0.05 | 0.35 | 0.21 |
| 2 | 82867347 | rs1910233 | -0.54 | -1.028 | -0.119 | -1.606 | -0.034 | 0.044 | 0.044 | 0 | 0.21 | 0.43 | 0.22 | 0.28 | 0.16 | 0.17 |
| 2 | 82883860 | rs1393504 | 0.718 | -1.043 | -0.03 | -1.607 | -0.003 | 0.105 | 0.105 | 0 | 0.11 | 0.25 | 0.18 | 0.27 | 0.08 | 0.07 |
| 2 | 82886562 | rs6706314 | 0.879 | -1.253 | 0.152 | -1.737 | -0.007 | -0.029 | -0.029 | 0.007 | 0.13 | 0.19 | 0.08 | 0.02 | 0.09 | 0.07 |
| 2 | 82886959 | rs17022099 | 0.545 | -1.227 | 0.059 | -1.706 | -0.005 | 0.134 | 0.134 | 0.108 | 0.11 | 0.35 | 0.18 | 0.27 | 0.08 | 0.07 |
| 2 | 82892794 | rs6731289 | -2.306 | -1.46 | -0.167 | -1.81 | 0.094 | 0.023 | 0.023 | 1.515 | 0.43 | 0.95 | 0.32 | 0.31 | 0.25 | 0.24 |
| 2 | 82900043 | rs12052614 | -0.808 | -1.323 | 0.088 | -1.742 | -0.028 | 0.049 | 0.049 | 0 | 0.2 | 0.41 | 0.21 | 0.28 | 0.15 | 0.17 |
| 2 | 82905879 | rs2863020 | -1.647 | -1.257 | -0.798 | -2.27 | -0.123 | 0.195 | 0.195 | 0.012 | 0.68 | 0.99 | 0.83 | 0.96 | 0.68 | 0.75 |
| 2 | 82918725 | rs10496272 | -0.747 | -1.35 | 0.073 | -1.615 | -0.026 | 0.051 | 0.051 | 0 | 0.2 | 0.41 | 0.21 | 0.28 | 0.15 | 0.17 |
| 2 | 82935171 | rs17022214 | 0.786 | -1.216 | -0.482 | -1.614 | -0.004 | 0.037 | 0.037 | 0 | 0.08 | 0 | 0.04 | 0.01 | 0.08 | 0.11 |
| 2 | 82941261 | rs10865463 | 0.637 | -1.034 | -0.054 | -1.318 | -0.003 | 0.106 | 0.106 | 0 | 0.11 | 0.25 | 0.17 | 0.27 | 0.08 | 0.07 |
| 2 | 82941825 | rs7559513 | 0.29 | -0.891 | -0.128 | -1.294 | -0.024 | -0.075 | -0.075 | 0 | 0.15 | 0.39 | 0.11 | 0.03 | 0.09 | 0.07 |
| 2 | 82956517 | rs1532731 | 1.498 | -0.732 | -1.047 | -1.07 | -0.087 | -0.141 | -0.141 | 0 | 0.23 | 0.04 | 0.44 | 0.55 | 0.32 | 0.39 |
| 2 | 82957817 | rs17022288 | -0.116 | -1.745 | -0.051 | -1.425 | -0.012 | -0.053 | -0.053 | 0 | 0.19 | 0.25 | 0.13 | 0.03 | 0.15 | 0.07 |
| 2 | 82991635 | rs17022383 | -0.913 | -1.845 | 0.331 | -1.605 | -0.046 | -0.142 | -0.142 | 0.793 | 0.27 | 0.5 | 0.18 | 0.04 | 0.19 | 0.08 |
| 2 | 82993400 | rs11678338 | -0.892 | -1.851 | 0.348 | -1.608 | -0.044 | -0.138 | -0.138 | 3.345 | 0.27 | 0.47 | 0.17 | 0.04 | 0.19 | 0.08 |
| 2 | 83004733 | rs7558379 | -1.996 | -2.041 | 0.056 | -1.786 | 0.049 | 0.086 | 0.086 | 0.87 | 0.53 | 0.83 | 0.39 | 0.32 | 0.35 | 0.24 |
| 2 | 83013886 | rs13001234 | 0.276 | -1.566 | 0.236 | -1.468 | -0.001 | 0.006 | 0.006 | 0 | 0.2 | 0.21 | 0.21 | 0.28 | 0.16 | 0.15 |
| 2 | 83020550 | rs13012745 | 0.308 | -1.537 | 0.141 | -1.46 | -0.001 | 0.001 | 0.001 | 0 | 0.2 | 0.2 | 0.21 | 0.28 | 0.16 | 0.15 |
| 2 | 83037125 | rs17022447 | -0.108 | -1.595 | 0.279 | -2.428 | -0.02 | -0.083 | -0.083 | 0 | 0.21 | 0.32 | 0.13 | 0.03 | 0.16 | 0.07 |
| 2 | 83040357 | rs972827 | -0.063 | -1.153 | 0.852 | -1.927 | -0.001 | -0.001 | -0.001 | 0.005 | 0.21 | 0.2 | 0.21 | 0.23 | 0.15 | 0.15 |
| 2 | 83041653 | rs974260 | -2.056 | -1.258 | 0.845 | -2 | 0.002 | 0.002 | 0.002 | 0.524 | 0.38 | 0.62 | 0.35 | 0.26 | 0.29 | 0.2 |
| 2 | 83047423 | rs9678502 | -0.539 | -1.218 | 0.918 | -2.148 | -0.01 | -0.052 | -0.052 | 1.052 | 0.12 | 0.28 | 0.09 | 0.03 | 0.1 | 0.05 |
| 2 | 83058160 | rs17022548 | -0.061 | -1.212 | 0.849 | -2.122 | -0.002 | -0.005 | -0.005 | 0 | 0.2 | 0.25 | 0.21 | 0.2 | 0.15 | 0.14 |
| 2 | 83059970 | rs10496276 | -1.255 | -1.112 | 0.667 | -2.217 | -0.011 | -0.054 | -0.054 | 2.073 | 0.16 | 0.25 | 0.12 | 0.03 | 0.15 | 0.06 |
| 2 | 83078896 | rs6547446 | -0.167 | -1.022 | 0.619 | -2.064 | -0.002 | -0.013 | -0.013 | 1.805 | 0.2 | 0.26 | 0.22 | 0.19 | 0.15 | 0.14 |
| 2 | 83082084 | rs12998947 | -0.443 | -1.037 | 0.667 | -2.071 | -0.007 | -0.023 | -0.023 | 0 | 0.22 | 0.34 | 0.23 | 0.19 | 0.15 | 0.15 |
| 2 | 83093602 | rs17022690 | -0.611 | -0.948 | 0.427 | -2.051 | -0.006 | -0.049 | -0.049 | 0 | 0.11 | 0.26 | 0.1 | 0.03 | 0.12 | 0.05 |
| 2 | 83098489 | rs1961190 | -0.066 | -0.923 | 0.367 | -2.045 | -0.004 | 0.007 | 0.007 | 0 | 0.19 | 0.12 | 0.2 | 0.19 | 0.15 | 0.15 |
| 2 | 83107695 | rs10184915 | -0.998 | -0.924 | 0.415 | -2.044 | -0.003 | -0.022 | -0.022 | 0 | 0.3 | 0.4 | 0.3 | 0.22 | 0.27 | 0.2 |
| 2 | 83120727 | rs10469974 | 2.174 | -1.108 | 1.1 | -1.663 | 0.049 | 0.042 | 0.042 | 1.79 | 0.7 | 0.15 | 0.76 | 0.79 | 0.85 | 0.8 |
| 2 | 83129299 | rs17022737 | -0.854 | -0.623 | 1.657 | -1.389 | -0.008 | -0.006 | -0.006 | 0 | 0.05 | 0.14 | 0.03 | 0.02 | 0.02 | 0.03 |
| 2 | 83136642 | rs1420761 | 0.557 | -0.624 | 1.843 | -1.393 | 0.007 | 0.018 | 0.018 | 0 | 0.92 | 0.86 | 0.86 | 0.81 | 0.92 | 0.87 |
| 2 | 83151323 | rs10496278 | 1.046 | -0.633 | 1.964 | -1.397 | 0.012 | 0.031 | 0.031 | 0 | 0.91 | 0.79 | 0.85 | 0.81 | 0.92 | 0.87 |
| 2 | 83153359 | rs897383 | 2.637 | -0.673 | 1.948 | -1.405 | -0.021 | 0.004 | 0.004 | 0.764 | 0.81 | 0.53 | 0.78 | 0.79 | 0.87 | 0.83 |
| 2 | 83157987 | rs7577594 | 2.776 | -0.624 | 1.971 | -1.406 | -0.025 | -0.002 | -0.002 | 0 | 0.81 | 0.47 | 0.78 | 0.79 | 0.87 | 0.83 |
| 2 | 83163347 | rs7340436 | 0.592 | -0.604 | 1.981 | -1.396 | 0.009 | 0.022 | 0.022 | 0 | 0.92 | 0.85 | 0.86 | 0.82 | 0.92 | 0.87 |
| 2 | 83178268 | rs11890594 | -0.727 | -0.575 | 1.707 | -1.382 | -0.002 | -0.001 | -0.001 | 0 | 0.08 | 0.08 | 0.14 | 0.18 | 0.08 | 0.13 |
| 2 | 83183292 | rs10196573 | -2.53 | -0.569 | 1.701 | -1.373 | -0.019 | 0.004 | 0.004 | 1.252 | 0.19 | 0.46 | 0.21 | 0.21 | 0.13 | 0.17 |
| 2 | 83195125 | rs17022826 | -2.172 | -0.531 | 1.396 | -1.355 | -0.009 | 0.003 | 0.003 | 0.098 | 0.17 | 0.28 | 0.2 | 0.21 | 0.13 | 0.17 |
| 2 | 83197589 | rs12615558 | -2.172 | -0.531 | 1.413 | -1.355 | -0.009 | 0.003 | 0.003 | 0.098 | 0.17 | 0.28 | 0.2 | 0.21 | 0.13 | 0.17 |
| 2 | 83205910 | rs7585648 | -2.326 | -0.504 | 1.353 | -1.354 | -0.012 | 0.004 | 0.004 | 0.592 | 0.18 | 0.39 | 0.21 | 0.21 | 0.13 | 0.17 |
| 2 | 83220312 | rs1372083 | -1.869 | -0.522 | 1.246 | -1.358 | -0.002 | 0.003 | 0.003 | 0.184 | 0.14 | 0.28 | 0.18 | 0.17 | 0.13 | 0.13 |
| 2 | 83223524 | rs17022896 | -2.378 | -0.532 | 1.33 | -1.427 | -0.013 | -0.006 | -0.006 | 0 | 0.17 | 0.39 | 0.2 | 0.17 | 0.13 | 0.16 |
| 2 | 83224879 | rs13425508 | -2.472 | -0.572 | 1.317 | -1.409 | -0.017 | 0 | 0 | 0.44 | 0.18 | 0.38 | 0.2 | 0.19 | 0.13 | 0.16 |
| 2 | 83228251 | rs17022928 | -2.295 | -0.564 | 1.322 | -1.407 | -0.001 | 0.011 | 0.011 | 0.703 | 0.15 | 0.38 | 0.2 | 0.17 | 0.13 | 0.13 |
| 2 | 83250940 | rs11887951 | 0.695 | -0.468 | 0.914 | -1.463 | -0.01 | 0.026 | 0.026 | 0 | 0.06 | 0.03 | 0.05 | 0 | 0.08 | 0.03 |
| 2 | 83251041 | rs17022985 | -1.394 | -0.573 | 1.143 | -1.274 | -0.015 | 0.004 | 0.004 | 0.069 | 0.06 | 0.19 | 0.03 | 0.03 | 0 | 0.03 |
| 2 | 83261859 | rs17023054 | -2.35 | -0.637 | 1.338 | -1.323 | -0.028 | 0.005 | 0.005 | 0.105 | 0.17 | 0.41 | 0.2 | 0.19 | 0.12 | 0.14 |
| 2 | 83262540 | rs17023057 | -1.427 | -0.646 | 1.438 | -1.327 | -0.01 | 0.009 | 0.009 | 0.512 | 0.06 | 0.24 | 0.02 | 0.03 | 0 | 0.02 |
| 2 | 83267197 | rs17023068 | -2.526 | -0.652 | 1.473 | -1.33 | -0.029 | 0.005 | 0.005 | 0.197 | 0.18 | 0.42 | 0.2 | 0.19 | 0.12 | 0.15 |
| 2 | 83273436 | rs11890115 | -1.441 | -0.649 | 1.475 | -1.328 | -0.001 | 0.005 | 0.005 | 0 | 0.11 | 0.14 | 0.17 | 0.16 | 0.11 | 0.12 |
| 2 | 83276705 | rs1367260 | -1.441 | -0.649 | 1.475 | -1.328 | -0.001 | 0.005 | 0.005 | 0 | 0.11 | 0.14 | 0.17 | 0.16 | 0.11 | 0.12 |
| 2 | 83278233 | rs17040798 | -2.882 | -0.701 | 1.689 | -1.353 | -0.04 | -0.003 | -0.003 | 0.3 | 0.19 | 0.52 | 0.21 | 0.19 | 0.12 | 0.14 |
| 2 | 83282256 | rs2863064 | -1.645 | -0.629 | 1.285 | -1.321 | -0.003 | 0.003 | 0.003 | 0 | 0.12 | 0.14 | 0.17 | 0.16 | 0.11 | 0.12 |
| 2 | 83291464 | rs17023115 | -1.46 | -0.601 | 1.299 | -1.307 | -0.001 | 0.005 | 0.005 | 0 | 0.11 | 0.14 | 0.17 | 0.16 | 0.11 | 0.12 |
| 2 | 83292565 | rs10496288 | -1.46 | -0.601 | 1.299 | -1.307 | -0.001 | 0.005 | 0.005 | 0 | 0.11 | 0.14 | 0.17 | 0.16 | 0.11 | 0.13 |
| 2 | 83307240 | rs2902404 | -1.113 | -0.61 | 1.296 | -1.312 | 0.001 | 0.017 | 0.017 | 0 | 0.08 | 0.16 | 0.15 | 0.16 | 0.08 | 0.11 |
| 2 | 83332983 | rs17023156 | -2.463 | -0.629 | 1.375 | -1.338 | -0.009 | 0.004 | 0.004 | 0.093 | 0.14 | 0.33 | 0.19 | 0.16 | 0.11 | 0.12 |
| 2 | 83335815 | rs963064 | -1.74 | -0.625 | 1.368 | -1.334 | 0.001 | 0.028 | 0.028 | 0.723 | 0.1 | 0.29 | 0.17 | 0.15 | 0.08 | 0.11 |
| 2 | 83336271 | rs1430275 | -1.74 | -0.625 | 1.368 | -1.334 | 0.001 | 0.028 | 0.028 | 0.723 | 0.1 | 0.29 | 0.17 | 0.15 | 0.08 | 0.11 |
| 2 | 83342154 | rs10496293 | -1.092 | -0.621 | 1.352 | -1.332 | 0.001 | 0.015 | 0.015 | 0 | 0.08 | 0.15 | 0.15 | 0.15 | 0.08 | 0.11 |
| 2 | 83351149 | rs7596981 | -2.43 | -0.641 | 1.363 | -1.363 | -0.016 | -0.006 | -0.006 | 0 | 0.16 | 0.45 | 0.2 | 0.16 | 0.11 | 0.12 |

| | | | | | | | | | | | | | | | | |
|---|---|---|---|---|---|---|---|---|---|---|---|---|---|---|---|---|
| 2 | 83357496 | rs17023252 | -0.889 | -0.585 | 1.163 | -1.33 | 0.007 | 0.023 | 0.023 | 0 | 0.07 | 0.16 | 0.15 | 0.15 | 0.08 | 0.11 |
| 2 | 83383702 | rs12469609 | -0.874 | -0.189 | 0.564 | -1.825 | -0.019 | 0.024 | 0.024 | 0 | 0.39 | 0.37 | 0.52 | 0.89 | 0.54 | 0.5 |
| 2 | 83386258 | rs11126889 | 0.659 | -0.208 | 0.538 | -1.802 | -0.141 | -0.036 | -0.036 | 0 | 0.25 | 0.04 | 0.34 | 0.72 | 0.43 | 0.39 |
| 2 | 83386889 | rs11687402 | -0.658 | -0.121 | 0.582 | -1.868 | -0.021 | 0.025 | 0.025 | 0 | 0.37 | 0.36 | 0.52 | 0.9 | 0.53 | 0.48 |
| 2 | 83390541 | rs6743753 | -0.345 | -0.132 | 0.527 | -1.821 | -0.049 | 0.079 | 0.079 | 0 | 0.34 | 0.22 | 0.49 | 0.88 | 0.52 | 0.49 |
| 2 | 83391938 | rs962033 | -1.68 | -0.411 | 0.468 | -2.135 | -0.027 | 0.168 | 0.168 | 0.324 | 0.59 | 0.85 | 0.58 | 0.91 | 0.58 | 0.59 |
| 2 | 83392185 | rs17314018 | 0.75 | 0.011 | 0.286 | -1.867 | -0.135 | -0.02 | -0.02 | 0 | 0.26 | 0.04 | 0.35 | 0.76 | 0.43 | 0.39 |
| 2 | 83395285 | rs17314045 | 0.18 | 0.013 | 0.348 | -1.858 | -0.04 | 0.027 | 0.027 | 0 | 0.3 | 0.18 | 0.37 | 0.76 | 0.46 | 0.4 |
| 2 | 83397203 | rs6750287 | -0.416 | 0.006 | 0.37 | -1.87 | -0.044 | 0.074 | 0.074 | 0 | 0.36 | 0.22 | 0.51 | 0.88 | 0.54 | 0.5 |
| 2 | 83416545 | rs17023305 | -0.258 | 0.02 | 0.489 | -1.754 | -0.002 | -0.005 | -0.005 | 0 | 0.07 | 0.04 | 0.14 | 0.13 | 0.08 | 0.1 |
| 2 | 83418770 | rs985697 | -0.719 | -0.034 | 0.597 | -1.887 | 0.006 | -0.051 | -0.051 | 0 | 0.41 | 0.49 | 0.53 | 0.88 | 0.54 | 0.5 |
| 2 | 83424531 | rs1430286 | 0.037 | -0.05 | 0.682 | -1.883 | -0.015 | 0.014 | 0.014 | 0 | 0.32 | 0.32 | 0.39 | 0.76 | 0.46 | 0.4 |
| 2 | 83436237 | rs11126894 | 0.707 | -0.074 | 0.639 | -1.904 | -0.143 | -0.021 | -0.021 | 0 | 0.26 | 0.03 | 0.35 | 0.75 | 0.44 | 0.39 |
| 2 | 83438064 | rs6547460 | -0.736 | -0.044 | 0.706 | -1.904 | 0.007 | -0.065 | -0.065 | 0 | 0.4 | 0.48 | 0.54 | 0.9 | 0.54 | 0.5 |
| 2 | 83442194 | rs7596070 | 0.249 | -0.041 | 0.695 | -1.901 | -0.062 | 0.032 | 0.032 | 0 | 0.29 | 0.13 | 0.38 | 0.76 | 0.46 | 0.4 |
| 2 | 83445609 | rs7587978 | -0.736 | -0.043 | 0.682 | -1.904 | 0.008 | -0.068 | -0.068 | 0 | 0.4 | 0.48 | 0.54 | 0.9 | 0.54 | 0.5 |
| 2 | 83445623 | rs7587984 | -0.773 | -0.106 | 0.673 | -1.885 | 0.004 | 0.034 | 0.034 | 0 | 0.08 | 0.16 | 0.15 | 0.14 | 0.08 | 0.1 |
| 2 | 83446208 | rs7591274 | -0.586 | 0.201 | 0.457 | -2.108 | -0.045 | 0.102 | 0.102 | 0 | 0.39 | 0.24 | 0.52 | 0.89 | 0.54 | 0.51 |
| 2 | 83448159 | rs6547465 | -0.818 | 0.333 | 0.632 | -1.923 | 0.007 | -0.073 | -0.073 | 0.179 | 0.39 | 0.48 | 0.53 | 0.9 | 0.51 | 0.49 |
| 2 | 83463172 | rs12470218 | -0.656 | 0.351 | 0.608 | -1.901 | -0.011 | 0.012 | 0.012 | 0 | 0.38 | 0.37 | 0.52 | 0.9 | 0.51 | 0.49 |
| 2 | 83464860 | rs7605657 | 0.856 | 0.352 | 0.601 | -1.885 | -0.119 | -0.034 | -0.034 | 0.155 | 0.24 | 0.03 | 0.34 | 0.75 | 0.4 | 0.38 |
| 2 | 83465188 | rs7422703 | -0.668 | 0.352 | 0.602 | -1.885 | -0.011 | 0.012 | 0.012 | 0 | 0.38 | 0.37 | 0.52 | 0.9 | 0.51 | 0.49 |
| 2 | 83467585 | rs7563837 | 0.425 | 0.37 | 0.614 | -1.875 | -0.06 | 0.02 | 0.02 | 0 | 0.28 | 0.11 | 0.36 | 0.76 | 0.42 | 0.39 |
| 2 | 83472006 | rs7422290 | 0.847 | 0.37 | 0.587 | -1.873 | -0.119 | -0.034 | -0.034 | 0 | 0.24 | 0.03 | 0.34 | 0.75 | 0.4 | 0.38 |
| 2 | 83480213 | rs4129274 | -1.181 | 0.427 | 0.643 | -1.851 | -0.003 | 0.026 | 0.026 | 0 | 0.1 | 0.26 | 0.19 | 0.14 | 0.09 | 0.1 |
| 2 | 83480886 | rs12622959 | -0.623 | 0.438 | 0.582 | -1.862 | 0.007 | -0.069 | -0.069 | 0 | 0.39 | 0.48 | 0.53 | 0.9 | 0.5 | 0.48 |
| 2 | 83489250 | rs12615094 | -1.355 | 0.386 | 0.523 | -1.75 | 0.001 | 0.106 | 0.106 | 0.031 | 0.57 | 0.8 | 0.54 | 0.89 | 0.59 | 0.54 |
| 2 | 83497255 | rs10187267 | -1.275 | 0.458 | 0.397 | -1.71 | 0 | -0.02 | -0.02 | 2.081 | 0.56 | 0.63 | 0.53 | 0.89 | 0.59 | 0.54 |
| 2 | 83500317 | rs11126908 | -0.139 | 0.483 | 0.433 | -1.731 | 0.076 | -0.045 | -0.045 | 0.109 | 0.1 | 0.29 | 0.05 | 0.02 | 0.12 | 0.1 |
| 2 | 83507543 | rs6419673 | -1.228 | 0.397 | 0.404 | -1.737 | -0.002 | -0.055 | -0.055 | 0.012 | 0.54 | 0.52 | 0.52 | 0.89 | 0.59 | 0.54 |
| 2 | 83509062 | rs6731262 | -1.236 | 0.399 | 0.407 | -1.736 | -0.004 | -0.065 | -0.065 | 0 | 0.54 | 0.53 | 0.52 | 0.89 | 0.59 | 0.54 |
| 2 | 83513890 | rs7423195 | -0.731 | 0.399 | 0.415 | -1.687 | -0.004 | 0.113 | 0.113 | 1.083 | 0.46 | 0.28 | 0.38 | 0.76 | 0.52 | 0.5 |
| 2 | 83522999 | rs6547481 | -1.225 | 0.367 | 0.185 | -1.703 | 0.007 | 0.01 | 0.01 | 0 | 0.54 | 0.44 | 0.52 | 0.87 | 0.6 | 0.55 |
| 2 | 83537379 | rs12618878 | -1.211 | 0.329 | 0.179 | -1.7 | 0.016 | 0.041 | 0.041 | 0.017 | 0.53 | 0.41 | 0.52 | 0.87 | 0.61 | 0.55 |
| 2 | 83538976 | rs6547485 | -0.172 | 0.238 | 0.092 | -1.741 | -0.02 | 0.033 | 0.033 | 1.307 | 0.37 | 0.3 | 0.37 | 0.75 | 0.52 | 0.47 |
| 2 | 83541302 | rs6547486 | -0.247 | 0.283 | 0.085 | -1.75 | -0.017 | 0.038 | 0.038 | 0.869 | 0.37 | 0.3 | 0.38 | 0.76 | 0.53 | 0.47 |
| 2 | 83557502 | rs3884626 | -1.014 | 0.764 | -0.305 | -1.52 | 0.001 | -0.041 | -0.041 | 0.343 | 0.56 | 0.61 | 0.55 | 0.91 | 0.62 | 0.58 |
| 2 | 83560840 | rs7422781 | 0.175 | 0.764 | -0.865 | -1.533 | -0.046 | 0.093 | 0.093 | 3.582 | 0.38 | 0.14 | 0.34 | 0.75 | 0.43 | 0.4 |
| 2 | 83570115 | rs11894798 | -0.614 | 0.335 | -0.918 | -1.218 | 0.014 | 0.035 | 0.035 | 0.081 | 0.57 | 0.71 | 0.56 | 0.91 | 0.62 | 0.58 |
| 2 | 83590342 | rs7563569 | -0.669 | 0.343 | -1.003 | -1.224 | 0.014 | 0.033 | 0.033 | 0.056 | 0.57 | 0.7 | 0.57 | 0.91 | 0.62 | 0.58 |
| 2 | 83601285 | rs1991542 | 1.059 | 0.558 | -0.998 | -0.086 | 0.003 | -0.008 | -0.008 | 0.923 | 0.94 | 0.96 | 0.87 | 0.87 | 0.93 | 0.93 |
| 2 | 83605769 | rs1896853 | 1.174 | 0.694 | -1.159 | -0.038 | -0.008 | -0.028 | -0.028 | 0.171 | 0.89 | 0.96 | 0.86 | 0.83 | 0.87 | 0.9 |
| 2 | 83610841 | rs2043014 | -0.275 | 0.441 | -2.261 | -0.936 | 0.013 | 0.233 | 0.233 | 0.376 | 0.45 | 0.15 | 0.58 | 0.85 | 0.41 | 0.39 |
| 2 | 83614035 | rs1035979 | -0.923 | 0.723 | -2.87 | -1.611 | -0.059 | 0.111 | 0.111 | 0.005 | 0.74 | 0.92 | 0.83 | 0.93 | 0.68 | 0.65 |
| 2 | 83614315 | rs7581633 | 0.191 | 0.406 | -1.563 | -1.764 | 0.014 | -0.159 | -0.159 | 2.523 | 0.25 | 0.42 | 0.23 | 0.01 | 0.27 | 0.15 |
| 2 | 83619117 | rs7600094 | 0.258 | 0.485 | -1.592 | -1.931 | 0.116 | -0.044 | -0.044 | 0 | 0.1 | 0.37 | 0.15 | 0.01 | 0.13 | 0.11 |
| 2 | 83620727 | rs1366819 | -0.052 | 0.466 | -1.55 | -1.873 | 0.024 | -0.166 | -0.166 | 0 | 0.24 | 0.43 | 0.23 | 0.01 | 0.27 | 0.15 |
| 2 | 83622161 | rs2043015 | 1.394 | 0.197 | -1.974 | -1.162 | -0.045 | -0.138 | -0.138 | 0.502 | 0.32 | 0.06 | 0.41 | 0.53 | 0.3 | 0.27 |
| 2 | 83632334 | rs10179347 | 0.436 | 0.275 | -1.473 | -1.364 | 0.026 | 0.141 | 0.141 | 0.491 | 0.45 | 0.26 | 0.58 | 0.85 | 0.41 | 0.39 |
| 2 | 83633730 | rs1896848 | -1.016 | 0.307 | -1.383 | -1.261 | -0.007 | 0.008 | 0.008 | 0 | 0.72 | 0.75 | 0.82 | 0.87 | 0.67 | 0.63 |
| 2 | 83637055 | rs17380093 | -0.018 | 0.53 | -1.545 | -1.199 | -0.017 | 0.271 | 0.271 | 0 | 0.49 | 0.2 | 0.65 | 0.87 | 0.45 | 0.52 |
| 2 | 83659667 | rs981525 | 0.197 | 0.49 | -0.85 | -1.06 | 0.009 | 0.038 | 0.038 | 1.858 | 0.47 | 0.4 | 0.61 | 0.84 | 0.44 | 0.42 |
| 2 | 83664988 | rs1366825 | 0.27 | 0.377 | -0.094 | -1.114 | 0.006 | 0.173 | 0.173 | 2.083 | 0.45 | 0.2 | 0.6 | 0.84 | 0.44 | 0.42 |
| 2 | 83671221 | rs17023511 | 0.774 | 0.31 | -0.247 | -0.972 | -0.008 | -0.042 | -0.042 | 0.347 | 0.11 | 0.03 | 0.18 | 0.19 | 0.15 | 0.1 |
| 2 | 83679699 | rs2361508 | -1.595 | 0.076 | -0.346 | -1.233 | -0.096 | 0.084 | 0.084 | 1.541 | 0.79 | 0.97 | 0.79 | 0.93 | 0.69 | 0.69 |
| 2 | 83680196 | rs1429395 | 0.174 | 0.084 | -0.29 | -1.169 | 0.014 | 0.02 | 0.02 | 2.3 | 0.51 | 0.64 | 0.62 | 0.92 | 0.44 | 0.53 |
| 2 | 83687932 | rs919431 | 0.206 | 0.093 | -0.21 | -1.164 | 0.014 | 0.019 | 0.019 | 2.219 | 0.5 | 0.64 | 0.62 | 0.92 | 0.44 | 0.53 |
| 2 | 83700783 | rs7580607 | 0.555 | 0.036 | -0.325 | -1.137 | -0.019 | -0.135 | -0.135 | 0.091 | 0.26 | 0.34 | 0.17 | 0 | 0.24 | 0.1 |
| 2 | 83703964 | rs2115619 | 1.161 | 0.031 | -0.158 | -0.646 | -0.045 | -0.146 | -0.146 | 0.169 | 0.32 | 0.06 | 0.4 | 0.53 | 0.29 | 0.27 |
| 2 | 83718206 | rs11126917 | 0.242 | 0.085 | 0.033 | -0.949 | 0.014 | 0.122 | 0.122 | 2.253 | 0.45 | 0.29 | 0.6 | 0.84 | 0.44 | 0.42 |
| 2 | 83739650 | rs1429377 | -0.293 | 0.018 | -0.186 | -1.016 | 0.009 | -0.011 | -0.011 | 0 | 0.5 | 0.63 | 0.62 | 0.92 | 0.44 | 0.53 |
| 2 | 83741398 | rs9631074 | -0.707 | -0.054 | -0.141 | -0.798 | -0.015 | 0.138 | 0.138 | 0 | 0.06 | 0.23 | 0.04 | 0.2 | 0.02 | 0.15 |
| 2 | 83748050 | rs1429388 | -0.293 | 0.035 | -0.683 | -0.942 | 0.007 | 0.1 | 0.1 | 0.904 | 0.55 | 0.78 | 0.43 | 0.3 | 0.57 | 0.63 |
| 2 | 83753288 | rs1366827 | 0.322 | 0.378 | -0.599 | -1.033 | -0.005 | 0.012 | 0.012 | 0.247 | 0.45 | 0.35 | 0.36 | 0.2 | 0.48 | 0.39 |
| 2 | 83753456 | rs1366828 | -0.922 | 0.476 | -0.794 | -1.068 | -0.052 | -0.027 | -0.027 | 0.276 | 0.67 | 0.8 | 0.5 | 0.4 | 0.59 | 0.72 |
| 2 | 83763938 | rs993736 | 0.834 | 0.399 | -0.094 | -1.15 | 0.028 | 0.07 | 0.07 | 0.634 | 0.38 | 0.18 | 0.29 | 0.2 | 0.29 | 0.35 |
| 2 | 83787422 | rs11901677 | -0.092 | 0.426 | 0.445 | -1.072 | 0.004 | 0.054 | 0.054 | 0.245 | 0.11 | 0.32 | 0.07 | 0.17 | 0.09 | 0.21 |
| 2 | 83790116 | rs10202054 | -0.489 | 0.412 | 0.535 | -1.137 | -0.013 | -0.007 | -0.007 | 0.025 | 0.21 | 0.33 | 0.16 | 0.2 | 0.22 | 0.31 |
| 2 | 83792506 | rs13430367 | -0.489 | 0.412 | 0.526 | -1.137 | -0.014 | -0.007 | -0.007 | 0 | 0.21 | 0.33 | 0.16 | 0.2 | 0.22 | 0.31 |

| | | | | | | | | | | | | | | | | |
|---|---|---|---|---|---|---|---|---|---|---|---|---|---|---|---|---|
| 2 | 83809428 | rs1906127 | -0.936 | 0.47 | 0.383 | -1.093 | -0.083 | -0.043 | -0.043 | 0 | 0.66 | 0.84 | 0.51 | 0.4 | 0.58 | 0.72 |
| 2 | 83829652 | rs17023719 | -0.169 | 0.352 | 0.527 | -1.123 | 0.003 | -0.001 | -0.001 | 0.093 | 0.17 | 0.18 | 0.15 | 0.19 | 0.18 | 0.25 |
| 2 | 83831871 | rs11888141 | -1.038 | 0.422 | 0.329 | -1.176 | -0.095 | -0.055 | -0.055 | 0 | 0.67 | 0.85 | 0.51 | 0.4 | 0.58 | 0.72 |
| 2 | 83831891 | rs12714104 | -0.993 | 0.475 | 0.556 | -1.145 | -0.027 | -0.009 | -0.009 | 0.081 | 0.64 | 0.71 | 0.5 | 0.4 | 0.58 | 0.68 |
| 2 | 83841076 | rs4144323 | 0.51 | 0.59 | 0.704 | -1.269 | 0.002 | 0.051 | 0.051 | 0.16 | 0.45 | 0.29 | 0.35 | 0.21 | 0.39 | 0.39 |
| 2 | 83842902 | rs13413501 | -0.733 | 0.352 | 0.849 | -1.196 | -0.02 | -0.014 | -0.014 | 0 | 0.23 | 0.38 | 0.17 | 0.19 | 0.23 | 0.31 |
| 2 | 83845469 | rs13007209 | 0.734 | 0.364 | 0.783 | -1.194 | 0.007 | 0.032 | 0.032 | 0.009 | 0.42 | 0.29 | 0.33 | 0.21 | 0.35 | 0.37 |
| 2 | 83850912 | rs2362235 | -0.905 | 0.404 | 0.676 | -1.149 | -0.03 | -0.01 | -0.01 | 0.101 | 0.64 | 0.69 | 0.49 | 0.4 | 0.58 | 0.68 |
| 2 | 83857943 | rs10173571 | -0.668 | 0.404 | 1.103 | -0.953 | 0.003 | 0.034 | 0.034 | 0 | 0.11 | 0.28 | 0.07 | 0.16 | 0.09 | 0.14 |
| 2 | 83863422 | rs11681252 | 0.376 | 0.318 | 0.959 | -1.101 | 0.008 | 0.028 | 0.028 | 0.003 | 0.4 | 0.29 | 0.33 | 0.2 | 0.34 | 0.33 |
| 2 | 83869897 | rs7566925 | -0.348 | 0.404 | 0.461 | -0.937 | 0.018 | 0.054 | 0.054 | 0.433 | 0.52 | 0.68 | 0.4 | 0.38 | 0.44 | 0.47 |
| 2 | 83875739 | rs4144322 | 0.29 | 0.409 | 0.455 | -0.96 | 0.02 | 0.011 | 0.011 | 1.044 | 0.42 | 0.31 | 0.34 | 0.38 | 0.32 | 0.38 |
| 2 | 83880421 | rs6738222 | -0.272 | 0.437 | -0.142 | -0.751 | 0.008 | 0.046 | 0.046 | 0.429 | 0.52 | 0.71 | 0.42 | 0.39 | 0.44 | 0.54 |
| 2 | 83886934 | rs6750156 | 0.389 | 0.333 | 0.309 | -0.808 | -0.001 | -0.009 | -0.009 | 1.154 | 0.4 | 0.36 | 0.34 | 0.2 | 0.34 | 0.35 |
| 2 | 83891406 | rs7590611 | 0.618 | 0.249 | 0.201 | -0.47 | -0.015 | 0.101 | 0.101 | 0 | 0.39 | 0.25 | 0.38 | 0.14 | 0.44 | 0.23 |
| 2 | 83891679 | rs7590960 | 0.342 | 0.298 | 0.093 | 0.21 | -0.026 | 0.017 | 0.017 | 0 | 0.44 | 0.25 | 0.61 | 0.61 | 0.47 | 0.32 |
| 2 | 83898441 | rs12328621 | 0.657 | 0.156 | 0.922 | 0.615 | -0.01 | -0.036 | -0.036 | 0 | 0.74 | 0.41 | 0.82 | 0.86 | 0.74 | 0.66 |
| 2 | 83908841 | rs7575993 | -0.388 | 0.112 | 1.3 | 0.646 | -0.009 | -0.076 | -0.076 | 0 | 0.21 | 0.38 | 0.15 | 0.07 | 0.23 | 0.08 |
| 2 | 83912152 | rs1868657 | -0.775 | 0.115 | 1.525 | 0.613 | 0 | 0.046 | 0.046 | 0.033 | 0.25 | 0.29 | 0.38 | 0.52 | 0.27 | 0.17 |
| 2 | 83913808 | rs12619893 | 1.282 | 0.116 | 1.572 | 0.592 | -0.005 | 0.042 | 0.042 | 0 | 0.71 | 0.51 | 0.59 | 0.4 | 0.7 | 0.65 |
| 2 | 83919123 | rs10197667 | 1.36 | 0.106 | 1.553 | 0.582 | 0 | -0.001 | -0.001 | 0 | 0.69 | 0.38 | 0.58 | 0.4 | 0.7 | 0.57 |
| 2 | 83922229 | rs2966376 | 1.28 | 0.108 | 1.549 | 0.583 | -0.005 | 0.042 | 0.042 | 0 | 0.71 | 0.51 | 0.59 | 0.4 | 0.7 | 0.65 |
| 2 | 83928010 | rs12477956 | 1.349 | 0.059 | 1.324 | 0.476 | 0 | -0.001 | -0.001 | 0 | 0.69 | 0.38 | 0.58 | 0.43 | 0.7 | 0.58 |
| 2 | 83932640 | rs2129392 | -0.981 | 0.141 | 0.749 | 0.413 | -0.008 | 0.085 | 0.085 | 0 | 0.26 | 0.39 | 0.39 | 0.49 | 0.27 | 0.16 |
| 2 | 83947833 | rs12476812 | -0.521 | 0.152 | 0.616 | 0.186 | -0.003 | -0.068 | -0.068 | 0 | 0.23 | 0.34 | 0.17 | 0.06 | 0.27 | 0.09 |
| 2 | 83948943 | rs11897350 | -0.17 | 0.126 | 0.749 | 0.173 | -0.002 | 0.01 | 0.01 | 0.262 | 0.21 | 0.17 | 0.16 | 0.06 | 0.27 | 0.09 |
| 2 | 83951665 | rs10084276 | -0.555 | 0.087 | 0.649 | 0.154 | -0.005 | -0.063 | -0.063 | 0 | 0.24 | 0.34 | 0.17 | 0.06 | 0.27 | 0.09 |
| 2 | 83955777 | rs13383442 | 0.292 | 0.037 | 0.627 | 0.105 | -0.023 | 0.044 | 0.044 | 0 | 0.16 | 0.06 | 0.11 | 0 | 0.24 | 0.07 |
| 2 | 83956861 | rs10203758 | -0.778 | 0.087 | 0.563 | 0.14 | -0.005 | -0.065 | -0.065 | 0 | 0.24 | 0.34 | 0.17 | 0.06 | 0.27 | 0.09 |
| 2 | 83964080 | rs7559876 | -0.778 | 0.087 | 0.563 | 0.14 | -0.005 | -0.065 | -0.065 | 0 | 0.24 | 0.34 | 0.17 | 0.06 | 0.27 | 0.09 |
| 2 | 83983335 | rs3100111 | -1.915 | -0.042 | 0.606 | 0.268 | -0.003 | 0.005 | 0.005 | 0.153 | 0.36 | 0.56 | 0.44 | 0.5 | 0.31 | 0.38 |
| 2 | 84001716 | rs2218792 | -0.317 | 0.008 | 0.414 | 0.299 | 0.002 | 0.037 | 0.037 | 0 | 0.1 | 0.07 | 0.26 | 0.44 | 0.05 | 0.29 |
| 2 | 84006399 | rs10183410 | -0.706 | 0.124 | 0.469 | 0.116 | -0.004 | -0.062 | -0.062 | 0 | 0.23 | 0.29 | 0.17 | 0.06 | 0.27 | 0.09 |
| 2 | 84023157 | rs1451777 | -0.823 | 0.152 | 0.451 | 0.131 | -0.004 | -0.059 | -0.059 | 0.367 | 0.24 | 0.29 | 0.17 | 0.06 | 0.27 | 0.16 |
| 2 | 84030512 | rs7603217 | -1.493 | 0.094 | 0.576 | 0.334 | -0.009 | 0.038 | 0.038 | 0 | 0.31 | 0.54 | 0.42 | 0.51 | 0.3 | 0.24 |
| 2 | 84031764 | rs921338 | -1.777 | -0.034 | 0.565 | 0.312 | -0.002 | 0.001 | 0.001 | 0 | 0.33 | 0.61 | 0.42 | 0.52 | 0.27 | 0.38 |
| 2 | 84035363 | rs10167628 | -0.655 | 0.043 | 0.448 | -0.04 | -0.014 | -0.067 | -0.067 | 0.658 | 0.22 | 0.32 | 0.16 | 0.06 | 0.23 | 0.09 |
| 2 | 84062502 | rs7370256 | -0.461 | 0.129 | 0.553 | 0.423 | -0.009 | 0.136 | 0.136 | 0 | 0.12 | 0.23 | 0.3 | 0.57 | 0.07 | 0.11 |
| 2 | 84064355 | rs6547515 | 0.814 | -0.003 | 0.902 | 0.443 | 0.045 | 0.107 | 0.107 | 1.306 | 0.95 | 0.73 | 0.98 | 0.93 | 0.98 | 0.82 |
| 2 | 84074675 | rs7589446 | -0.691 | -0.018 | 0.959 | 0.442 | -0.013 | 0.2 | 0.2 | 0 | 0.13 | 0.31 | 0.3 | 0.57 | 0.07 | 0.11 |
| 2 | 84112346 | rs13409715 | 1.385 | -0.009 | -0.006 | -0.256 | -0.041 | -0.036 | -0.036 | 0 | 0.26 | 0.05 | 0.19 | 0.24 | 0.26 | 0.36 |
| 2 | 84116856 | rs10186197 | -0.922 | 0.156 | 0.386 | -0.378 | -0.025 | 0.282 | 0.282 | 0 | 0.15 | 0.46 | 0.31 | 0.55 | 0.07 | 0.18 |
| 2 | 84117460 | rs7587040 | 0.543 | 0.152 | -0.072 | -0.933 | 0.012 | -0.02 | -0.02 | 2.416 | 0.22 | 0.29 | 0.23 | 0.13 | 0.26 | 0.14 |
| 2 | 84145733 | rs6714853 | 0.228 | 0.096 | -0.145 | -1.004 | -0.012 | 0.023 | 0.023 | 0 | 0.63 | 0.69 | 0.67 | 0.87 | 0.53 | 0.8 |
| 2 | 84147567 | rs11883841 | 0.215 | 0.092 | -0.139 | -0.99 | 0.006 | 0.03 | 0.03 | 0.314 | 0.08 | 0.44 | 0.07 | 0.06 | 0.03 | 0.02 |
| 2 | 84155146 | rs4832353 | -0.779 | 0.09 | -0.195 | -0.956 | -0.032 | 0.254 | 0.254 | 0.026 | 0.16 | 0.52 | 0.31 | 0.55 | 0.07 | 0.2 |
| 2 | 84173414 | rs11687214 | 1.795 | 0.153 | -0.395 | -0.984 | -0.078 | 0.027 | 0.027 | 0.911 | 0.46 | 0.07 | 0.35 | 0.32 | 0.45 | 0.6 |
| 2 | 84186478 | rs4144836 | 0.475 | 0.364 | 0.005 | -0.595 | 0 | 0.17 | 0.17 | 0 | 0.08 | 0.13 | 0.24 | 0.48 | 0.05 | 0.09 |
| 2 | 84189927 | rs17024323 | 0.303 | 0.373 | -0.123 | -0.772 | 0.014 | -0.027 | -0.027 | 0 | 0.2 | 0.3 | 0.2 | 0.12 | 0.23 | 0.13 |
| 2 | 84195984 | rs2885856 | -0.823 | 0.344 | 0.103 | -0.628 | -0.002 | 0.272 | 0.272 | 0 | 0.13 | 0.33 | 0.26 | 0.49 | 0.07 | 0.18 |
| 2 | 84196973 | rs4586641 | -0.68 | 0.385 | 0.063 | -0.773 | 0.023 | -0.025 | -0.025 | 0 | 0.12 | 0.41 | 0.13 | 0.06 | 0.1 | 0.05 |
| 2 | 84198445 | rs7566018 | 1.124 | 0.384 | -0.051 | -0.747 | -0.006 | 0.011 | 0.011 | 0 | 0.06 | 0.2 | 0.07 | 0.06 | 0.03 | 0.02 |
| 2 | 84212663 | rs1584312 | -1.102 | 0.532 | -0.276 | -0.898 | -0.004 | -0.047 | -0.047 | 0 | 0.25 | 0.58 | 0.28 | 0.12 | 0.23 | 0.16 |
| 2 | 84219974 | rs1348954 | -0.281 | 0.479 | -0.158 | -0.887 | -0.022 | -0.002 | -0.002 | 0.053 | 0.09 | 0.34 | 0.07 | 0.07 | 0.03 | 0.09 |
| 2 | 84225912 | rs17024359 | 0.112 | 0.51 | -0.104 | -0.929 | 0.038 | -0.031 | -0.031 | 0 | 0.14 | 0.2 | 0.2 | 0.05 | 0.2 | 0.06 |
| 2 | 84226391 | rs17316184 | 1.44 | 0.507 | -0.24 | -1.084 | -0.029 | 0.025 | 0.025 | 0 | 0.06 | 0.02 | 0.04 | 0.04 | 0.11 | 0.04 |
| 2 | 84228877 | rs4832367 | 0.069 | 0.548 | -0.187 | -1.037 | -0.022 | 0.001 | 0.001 | 0.171 | 0.09 | 0.3 | 0.07 | 0.07 | 0.03 | 0.02 |
| 2 | 84246586 | rs1447537 | -1.529 | 0.565 | -0.268 | -0.942 | -0.001 | -0.006 | -0.006 | 0 | 0.3 | 0.65 | 0.51 | 0.61 | 0.27 | 0.24 |
| 2 | 84248820 | rs1007899 | 0.195 | 0.511 | 0.003 | -0.965 | -0.021 | 0.001 | 0.001 | 0.15 | 0.08 | 0.26 | 0.07 | 0.07 | 0.03 | 0.02 |
| 2 | 84252099 | rs6711260 | 0.414 | 0.548 | -0.062 | -1.326 | 0.012 | -0.044 | -0.044 | 0.331 | 0.14 | 0.28 | 0.11 | 0.06 | 0.19 | 0.13 |
| 2 | 84257316 | rs12621764 | 0.806 | 0.392 | 0.16 | -1.102 | 0.005 | -0.011 | -0.011 | 0 | 0.11 | 0.12 | 0.15 | 0.05 | 0.13 | 0.03 |
| 2 | 84263861 | rs11690211 | 0.931 | 0.404 | -0.189 | -1.277 | -0.001 | -0.032 | -0.032 | 0.485 | 0.72 | 0.37 | 0.73 | 0.95 | 0.71 | 0.85 |
| 2 | 84267429 | rs2238 | -1.21 | 0.384 | -0.265 | -1.235 | -0.038 | 0.062 | 0.062 | 0 | 0.89 | 0.99 | 0.92 | 1 | 0.84 | 0.96 |
| 2 | 84277089 | rs12614368 | 0.165 | 0.167 | -0.235 | -1.159 | 0.063 | -0.03 | -0.03 | 0 | 0.13 | 0.44 | 0.18 | 0.05 | 0.13 | 0.1 |
| 2 | 84280413 | rs7608563 | -1.234 | 0.081 | -0.432 | -1.148 | -0.009 | 0.021 | 0.021 | 0.758 | 0.24 | 0.65 | 0.46 | 0.62 | 0.17 | 0.19 |
| 2 | 84285913 | rs11901149 | -0.18 | 0.088 | -0.209 | -0.991 | 0.037 | -0.042 | -0.042 | 3.577 | 0.14 | 0.53 | 0.18 | 0.08 | 0.13 | 0.03 |
| 2 | 84287417 | rs13431069 | -0.263 | 0.087 | -0.211 | -0.991 | 0.029 | -0.045 | -0.045 | 0.04 | 0.14 | 0.55 | 0.18 | 0.08 | 0.13 | 0.03 |
| 2 | 84291074 | rs7558055 | -1.579 | 0.331 | 0.039 | -0.626 | -0.008 | 0.017 | 0.017 | 0.633 | 0.24 | 0.66 | 0.46 | 0.65 | 0.17 | 0.19 |
| 2 | 84293335 | rs7579710 | -0.687 | 0.543 | 0.28 | -0.755 | 0.011 | -0.052 | -0.052 | 0 | 0.15 | 0.59 | 0.19 | 0.09 | 0.13 | 0.1 |

| | | | | | | | | | | | | | | | | |
|---|---|---|---|---|---|---|---|---|---|---|---|---|---|---|---|---|
| 2 | 84302018 | rs13412176 | -0.455 | 0.604 | 0.293 | -0.725 | 0.05 | -0.035 | -0.035 | 0 | 0.14 | 0.5 | 0.18 | 0.08 | 0.13 | 0.03 |
| 2 | 84305184 | rs11904255 | -0.938 | 0.59 | 0.225 | -0.732 | 0 | -0.061 | -0.061 | 0 | 0.17 | 0.56 | 0.18 | 0.09 | 0.12 | 0.1 |
| 2 | 84307936 | rs2197694 | 0.163 | 0.925 | 0.106 | -0.567 | 0.044 | -0.02 | -0.02 | 0 | 0.11 | 0.3 | 0.17 | 0.08 | 0.12 | 0.03 |
| 2 | 84324021 | rs1597039 | -0.084 | 1.382 | -0.348 | -0.318 | 0.057 | 0.034 | 0.034 | 0 | 0.19 | 0.42 | 0.19 | 0.22 | 0.27 | 0.23 |
| 2 | 84328335 | rs11126949 | -0.464 | 1.273 | -1.001 | -0.35 | 0.031 | 0.009 | 0.009 | 0 | 0.25 | 0.63 | 0.3 | 0.3 | 0.35 | 0.25 |
| 2 | 84348628 | rs11674417 | 0.29 | 1.046 | -1.664 | -0.149 | -0.077 | 0.053 | 0.053 | 1.264 | 0.3 | 0.05 | 0.34 | 0.12 | 0.31 | 0.47 |
| 2 | 84350015 | rs12612847 | 0.478 | 0.847 | -1.662 | -0.222 | 0 | 0.104 | 0.104 | 0.775 | 0.53 | 0.78 | 0.43 | 0.3 | 0.56 | 0.33 |
| 2 | 84356320 | rs17388631 | 0.478 | 0.943 | -1.742 | -0.215 | -0.126 | 0.04 | 0.04 | 1.97 | 0.3 | 0.02 | 0.34 | 0.12 | 0.33 | 0.48 |
| 2 | 84359610 | rs10445960 | -0.138 | 0.53 | -1.575 | -0.029 | 0.008 | -0.078 | -0.078 | 0 | 0.87 | 0.98 | 0.78 | 0.43 | 0.88 | 0.89 |
| 2 | 84366620 | rs4436973 | 0.709 | 0.617 | -1.39 | -0.083 | -0.004 | 0.019 | 0.019 | 0.451 | 0.45 | 0.61 | 0.37 | 0.28 | 0.49 | 0.31 |
| 2 | 84368347 | rs1007874 | 0.149 | 0.782 | -2.029 | -0.121 | -0.017 | 0.063 | 0.063 | 1.782 | 0.37 | 0.2 | 0.39 | 0.13 | 0.37 | 0.57 |
| 2 | 84370890 | rs6547526 | 0.705 | 0.497 | -1.752 | 0.824 | 0.006 | -0.116 | -0.116 | 0.901 | 0.84 | 0.98 | 0.77 | 0.41 | 0.86 | 0.71 |
| 2 | 84371635 | rs1358192 | 0.601 | 1.082 | -1.29 | 0.448 | -0.007 | 0.005 | 0.005 | 0.687 | 0.67 | 0.32 | 0.61 | 0.68 | 0.57 | 0.43 |
| 2 | 84371943 | rs2364236 | 0.991 | 0.691 | -1.253 | 0.572 | -0.001 | -0.004 | -0.004 | 3.492 | 0.89 | 0.96 | 0.92 | 0.9 | 0.89 | 0.91 |
| 2 | 84377246 | rs12466737 | -0.806 | -0.048 | -1.349 | 0.041 | 0.029 | 0.041 | 0.041 | 0 | 0.19 | 0.08 | 0.07 | 0.05 | 0.15 | 0.09 |
| 2 | 84378583 | rs10496307 | -0.867 | -0.254 | -0.633 | 0.522 | -0.032 | 0.007 | 0.007 | 0 | 0.17 | 0.26 | 0.15 | 0.21 | 0.13 | 0.13 |
| 2 | 84381680 | rs6706539 | 0.698 | -0.077 | -1.651 | -0.122 | -0.023 | 0.068 | 0.068 | 0 | 0.44 | 0.2 | 0.45 | 0.33 | 0.48 | 0.26 |
| 2 | 84385203 | rs6737955 | -0.668 | -0.644 | -0.767 | -0.061 | -0.066 | -0.101 | -0.101 | 0.159 | 0.27 | 0.53 | 0.13 | 0.06 | 0.17 | 0.19 |
| 2 | 84385564 | rs4832376 | -1.008 | -0.456 | -1.619 | -0.109 | 0.051 | 0 | 0 | 2.115 | 0.34 | 0.79 | 0.37 | 0.35 | 0.23 | 0.35 |
| 2 | 84407064 | rs17024618 | -0.649 | 0.269 | -2.043 | 0.186 | -0.063 | -0.107 | -0.107 | 0.128 | 0.18 | 0.02 | 0.37 | 0.69 | 0.25 | 0.34 |
| 2 | 84410242 | rs17024623 | 0.008 | 0.343 | -1.949 | 0.607 | -0.049 | -0.057 | -0.057 | 0 | 0.11 | 0.01 | 0.11 | 0.19 | 0.17 | 0.15 |
| 2 | 84419732 | rs10496308 | 0.131 | 0.011 | -2.958 | -0.53 | -0.03 | 0.156 | 0.156 | 0.008 | 0.3 | 0.01 | 0.14 | 0.05 | 0.27 | 0.11 |
| 2 | 84423973 | rs2886028 | -0.328 | 0.277 | -2.504 | -1.054 | -0.003 | -0.001 | -0.001 | 0 | 0.79 | 0.24 | 0.7 | 0.78 | 0.78 | 0.79 |
| 2 | 84430329 | rs2364241 | -0.244 | 0.249 | -2.191 | -1.062 | 0.009 | 0.002 | 0.002 | 0.635 | 0.8 | 0.33 | 0.71 | 0.79 | 0.78 | 0.8 |
| 2 | 84433896 | rs11126954 | -0.184 | 0.214 | -1.765 | -1.109 | 0.002 | 0 | 0 | 0 | 0.79 | 0.27 | 0.7 | 0.78 | 0.79 | 0.79 |
| 2 | 84437352 | rs12714120 | -0.339 | 0.175 | -1.075 | -1.088 | 0.053 | -0.039 | -0.039 | 0.083 | 0.81 | 0.49 | 0.72 | 0.88 | 0.79 | 0.8 |
| 2 | 84443628 | rs10178271 | -0.197 | 0.174 | -1.048 | -1.088 | 0.027 | -0.033 | -0.033 | 0 | 0.81 | 0.41 | 0.72 | 0.88 | 0.79 | 0.79 |
| 2 | 84446193 | rs7590843 | 1.064 | -0.098 | -1.906 | -1.669 | -0.132 | 0.246 | 0.246 | 1.114 | 0.36 | 0.02 | 0.19 | 0.04 | 0.39 | 0.27 |
| 2 | 84460865 | rs4020792 | -0.225 | 0.098 | -1.169 | -1.225 | 0.014 | 0.003 | 0.003 | 0.549 | 0.8 | 0.32 | 0.7 | 0.78 | 0.79 | 0.79 |
| 2 | 84503719 | rs2364271 | -0.256 | 0.043 | -1.177 | -1.479 | 0.032 | -0.04 | -0.04 | 0.471 | 0.8 | 0.41 | 0.72 | 0.88 | 0.79 | 0.78 |
| 2 | 84505969 | rs10451629 | -0.203 | 0.042 | -1.199 | -1.479 | 0.005 | -0.012 | -0.012 | 0 | 0.79 | 0.3 | 0.71 | 0.84 | 0.79 | 0.77 |
| 2 | 84509017 | rs731331 | 0.536 | -0.154 | -1.724 | -1.614 | -0.058 | 0.013 | 0.013 | 0 | 0.27 | 0.03 | 0.28 | 0.17 | 0.28 | 0.26 |
| 2 | 84517162 | rs6547551 | 0.435 | -0.452 | -1.116 | -1.518 | -0.006 | 0.064 | 0.064 | 0.772 | 0.73 | 0.14 | 0.69 | 0.84 | 0.79 | 0.69 |
| 2 | 84518438 | rs1968722 | 0.274 | -0.474 | -0.865 | -1.529 | 0.002 | 0.012 | 0.012 | 0.563 | 0.74 | 0.26 | 0.69 | 0.88 | 0.79 | 0.7 |
| 2 | 84518609 | rs1531047 | 0.21 | -0.494 | -0.875 | -1.538 | 0.001 | 0.009 | 0.009 | 0.394 | 0.75 | 0.26 | 0.69 | 0.88 | 0.79 | 0.7 |
| 2 | 84522961 | rs10177482 | -0.872 | -0.432 | -1.043 | -1.528 | -0.009 | 0.063 | 0.063 | 0.978 | 0.29 | 0.88 | 0.31 | 0.17 | 0.23 | 0.31 |
| 2 | 84528756 | rs11686691 | -0.88 | -0.431 | -1.04 | -1.528 | -0.006 | 0.064 | 0.064 | 1.06 | 0.29 | 0.88 | 0.32 | 0.18 | 0.23 | 0.31 |
| 2 | 84532773 | rs10193228 | -0.679 | -0.398 | -0.908 | -1.569 | 0.002 | 0.012 | 0.012 | 0.162 | 0.27 | 0.74 | 0.3 | 0.14 | 0.23 | 0.31 |
| 2 | 84536960 | rs7572995 | -0.001 | -0.406 | -0.629 | -1.594 | -0.002 | -0.029 | -0.029 | 0 | 0.19 | 0.32 | 0.22 | 0.13 | 0.15 | 0.14 |
| 2 | 84537078 | rs17024868 | -0.606 | -0.218 | -0.347 | -1.72 | 0.052 | -0.033 | -0.033 | 0 | 0.09 | 0.5 | 0.1 | 0 | 0.08 | 0.17 |
| 2 | 84550200 | rs1531045 | -0.002 | -0.294 | -0.55 | -1.948 | -0.001 | 0 | 0 | 0 | 0.36 | 0.31 | 0.28 | 0.16 | 0.37 | 0.38 |
| 2 | 84550988 | rs6715601 | -1.23 | -0.473 | -0.826 | -2.154 | -0.077 | 0.062 | 0.062 | 0.028 | 0.58 | 0.94 | 0.51 | 0.3 | 0.56 | 0.51 |
| 2 | 84551902 | rs6761772 | 0.135 | -0.464 | -0.572 | -2.077 | 0.005 | -0.022 | -0.022 | 0 | 0.18 | 0.54 | 0.2 | 0.14 | 0.14 | 0.11 |
| 2 | 84552604 | rs11690708 | 1.306 | -0.241 | -0.679 | -1.975 | -0.015 | 0.002 | 0.002 | 0 | 0.11 | 0.04 | 0.15 | 0.1 | 0.13 | 0.05 |
| 2 | 84554692 | rs11126959 | 0.107 | -0.261 | -0.53 | -2.048 | -0.001 | 0.025 | 0.025 | 0 | 0.36 | 0.28 | 0.28 | 0.16 | 0.37 | 0.38 |
| 2 | 84559150 | rs13414758 | -0.902 | -0.415 | -0.257 | -1.928 | -0.008 | 0.005 | 0.005 | 0 | 0.7 | 0.75 | 0.75 | 0.8 | 0.61 | 0.56 |
| 2 | 84573946 | rs1840403 | -0.94 | -0.636 | 0.027 | -1.807 | 0.011 | 0.061 | 0.061 | 0 | 0.41 | 0.27 | 0.32 | 0.09 | 0.37 | 0.21 |
| 2 | 84576048 | rs1840404 | -0.323 | -0.571 | 0.629 | -1.71 | 0.041 | -0.002 | -0.002 | 0 | 0.08 | 0.41 | 0.05 | 0.01 | 0.06 | 0.02 |
| 2 | 84603760 | rs4832076 | 0.29 | -0.68 | -0.064 | -1.961 | -0.026 | 0.091 | 0.091 | 0.003 | 0.19 | 0.03 | 0.19 | 0.03 | 0.2 | 0.11 |
| 2 | 84615472 | rs10496309 | -0.469 | -0.695 | 0.488 | -1.427 | -0.005 | -0.02 | -0.02 | 0 | 0.2 | 0.24 | 0.1 | 0.07 | 0.11 | 0.08 |
| 2 | 84618582 | rs11904045 | -0.858 | -0.83 | 0.901 | -1.174 | 0.017 | 0.029 | 0.029 | 1.161 | 0.09 | 0.38 | 0.06 | 0.06 | 0.06 | 0.03 |
| 2 | 84625023 | rs10496310 | 0.185 | -0.847 | 0.55 | -1.196 | -0.017 | -0.021 | -0.021 | 0 | 0.09 | 0.13 | 0.03 | 0.01 | 0.01 | 0.06 |
| 2 | 84657404 | rs4831972 | 0.975 | -0.852 | 0.321 | -1.201 | -0.04 | -0.039 | -0.039 | 0.114 | 0.84 | 0.75 | 0.93 | 0.92 | 0.94 | 0.91 |
| 2 | 84662093 | rs6753640 | 0.114 | -0.796 | 0.249 | -1.134 | -0.031 | -0.036 | -0.036 | 0 | 0.09 | 0.14 | 0.03 | 0.01 | 0.01 | 0.06 |
| 2 | 84673252 | rs10210248 | 1.441 | -0.898 | 0.137 | -1.346 | -0.016 | -0.03 | -0.03 | 0 | 0.72 | 0.37 | 0.76 | 0.9 | 0.83 | 0.82 |
| 2 | 84686999 | rs3795867 | 1.003 | -0.949 | 0.479 | -1.338 | -0.076 | -0.076 | -0.076 | 0.129 | 0.82 | 0.47 | 0.91 | 0.92 | 0.94 | 0.91 |
| 2 | 84696003 | rs7575751 | 1.025 | -0.788 | -0.173 | -1.72 | -0.04 | 0.079 | 0.079 | 0 | 0.4 | 0.24 | 0.32 | 0.21 | 0.47 | 0.47 |
| 2 | 84712523 | rs10182687 | 1.135 | -1.019 | 0.545 | -1.377 | -0.032 | -0.049 | -0.049 | 0 | 0.84 | 0.67 | 0.8 | 0.95 | 0.88 | 0.86 |
| 2 | 84719390 | rs4441484 | 0.137 | -0.919 | 0.451 | -1.345 | 0.025 | -0.013 | -0.013 | 2.293 | 0.93 | 0.68 | 0.95 | 1 | 0.94 | 0.95 |
| 2 | 84756049 | rs4832089 | -0.912 | -0.867 | 0.153 | -1.319 | -0.001 | -0.023 | -0.023 | 0 | 0.95 | 0.89 | 0.97 | 1 | 0.94 | 0.95 |
| 2 | 84756450 | rs4832091 | -0.912 | -0.867 | 0.153 | -1.301 | -0.001 | -0.019 | -0.019 | 1.679 | 0.95 | 0.89 | 0.97 | 0.99 | 0.94 | 0.95 |
| 2 | 84769058 | rs1460698 | 0.483 | -1.179 | 0.183 | -1.477 | -0.031 | -0.057 | -0.057 | 0 | 0.86 | 0.78 | 0.94 | 0.99 | 0.94 | 0.89 |
| 2 | 84782750 | rs13034783 | 0.186 | -1.19 | -0.028 | -1.475 | -0.015 | -0.002 | -0.002 | 0.008 | 0.09 | 0.02 | 0.14 | 0.05 | 0.1 | 0.1 |
| 2 | 84785748 | rs4484047 | 0.697 | -1.211 | 0.358 | -1.517 | -0.072 | -0.116 | -0.116 | 0 | 0.83 | 0.49 | 0.91 | 0.99 | 0.94 | 0.9 |
| 2 | 84786174 | rs7559623 | -0.574 | -1.103 | -0.087 | -1.463 | 0 | -0.027 | -0.027 | 0 | 0.94 | 0.83 | 0.96 | 1 | 0.94 | 0.95 |
| 2 | 84800605 | rs11891970 | -0.632 | -0.991 | -0.239 | -1.55 | 0 | -0.027 | -0.027 | 0 | 0.94 | 0.83 | 0.96 | 1 | 0.95 | 0.95 |
| 2 | 84800898 | rs13002679 | 0.84 | -0.931 | -0.701 | -1.653 | -0.055 | 0.108 | 0.108 | 0.177 | 0.38 | 0.18 | 0.33 | 0.19 | 0.51 | 0.47 |
| 2 | 84808483 | rs4832094 | -0.084 | -1 | -0.525 | -1.527 | -0.001 | -0.016 | -0.016 | 0 | 0.86 | 0.85 | 0.93 | 0.99 | 0.92 | 0.9 |
| 2 | 84822549 | rs12466574 | 1.373 | -0.478 | -0.458 | -1.914 | -0.086 | -0.007 | -0.007 | 0 | 0.15 | 0.03 | 0.21 | 0.12 | 0.29 | 0.21 |

| | | | | | | | | | | | | | | | | |
|---|---|---|---|---|---|---|---|---|---|---|---|---|---|---|---|---|
| 2 | 84823346 | rs13013322 | 0.89 | -0.993 | -0.007 | -1.653 | -0.021 | -0.019 | -0.019 | 0 | 0.73 | 0.38 | 0.76 | 0.86 | 0.82 | 0.79 |
| 2 | 84834592 | rs4832103 | 0.071 | -0.983 | 0.121 | -1.515 | -0.069 | -0.116 | -0.116 | 0 | 0.83 | 0.47 | 0.9 | 0.99 | 0.92 | 0.9 |
| 2 | 84839270 | rs4832104 | -0.125 | -0.986 | 0.132 | -1.517 | -0.065 | -0.116 | -0.116 | 0 | 0.16 | 0.46 | 0.1 | 0.01 | 0.08 | 0.1 |
| 2 | 84846683 | rs11889456 | 0.103 | -0.99 | 0.147 | -1.519 | -0.049 | -0.089 | -0.089 | 0 | 0.15 | 0.32 | 0.08 | 0.01 | 0.08 | 0.1 |
| 2 | 84849386 | rs11888840 | -0.121 | -0.985 | 0.059 | -1.517 | -0.065 | -0.116 | -0.116 | 0.041 | 0.16 | 0.46 | 0.09 | 0.01 | 0.08 | 0.1 |
| 2 | 84855366 | rs4513310 | 0.868 | -1.092 | -0.199 | -1.627 | 0.007 | 0 | 0 | 0.033 | 0.36 | 0.13 | 0.44 | 0.67 | 0.33 | 0.32 |
| 2 | 84862843 | rs4313996 | -0.623 | -0.981 | 0.576 | -1.199 | -0.068 | -0.114 | -0.114 | 0 | 0.17 | 0.49 | 0.09 | 0.01 | 0.08 | 0.1 |
| 2 | 84867252 | rs7572042 | -0.399 | -0.966 | 0.496 | -1.197 | -0.05 | -0.089 | -0.089 | 0.167 | 0.16 | 0.32 | 0.08 | 0.01 | 0.08 | 0.1 |
| 2 | 84868419 | rs7590097 | -0.446 | -0.964 | 0.411 | -1.196 | -0.056 | -0.098 | -0.098 | 0.241 | 0.16 | 0.38 | 0.09 | 0.01 | 0.08 | 0.1 |
| 2 | 84899144 | rs12185616 | 1.09 | -1.154 | 0.089 | -1.576 | -0.058 | -0.095 | -0.095 | 0 | 0.75 | 0.44 | 0.86 | 0.93 | 0.86 | 0.88 |
| 2 | 84905096 | rs11126974 | 0.487 | -1.415 | -0.036 | -1.482 | 0.042 | 0.059 | 0.059 | 0 | 0.54 | 0.32 | 0.57 | 0.8 | 0.66 | 0.53 |
| 2 | 84906723 | rs2178103 | -0.592 | -0.781 | 0.546 | -1.548 | 0.008 | 0.022 | 0.022 | 0 | 0.16 | 0.1 | 0.17 | 0.09 | 0.14 | 0.1 |
| 2 | 84906915 | rs1918690 | 1.427 | -0.885 | 1.098 | -1.601 | -0.025 | -0.064 | -0.064 | 1.448 | 0.76 | 0.62 | 0.78 | 0.91 | 0.8 | 0.84 |
| 2 | 84921433 | rs1192295 | -0.261 | -0.874 | 1.097 | -1.619 | 0.007 | -0.031 | -0.031 | 0 | 0.08 | 0.26 | 0.04 | 0 | 0.06 | 0.06 |
| 2 | 84924378 | rs1192291 | -2.414 | -0.95 | 1.225 | -1.664 | -0.017 | -0.039 | -0.039 | 0.231 | 0.33 | 0.59 | 0.26 | 0.12 | 0.23 | 0.2 |
| 2 | 84936200 | rs1192266 | -0.818 | -0.913 | 1.204 | -1.635 | -0.043 | -0.078 | -0.078 | 0 | 0.15 | 0.34 | 0.08 | 0.03 | 0.09 | 0.1 |
| 2 | 84945679 | rs1192343 | -2.448 | -0.906 | 0.614 | -1.641 | -0.017 | -0.039 | -0.039 | 0 | 0.34 | 0.57 | 0.26 | 0.12 | 0.23 | 0.2 |
| 2 | 84950006 | rs1192336 | -1.937 | -0.772 | 0.384 | -1.495 | -0.056 | -0.087 | -0.087 | 0 | 0.29 | 0.46 | 0.23 | 0.12 | 0.19 | 0.17 |
| 2 | 84967896 | rs13407438 | -0.133 | -0.575 | 0.035 | -1.447 | 0.006 | 0.008 | 0.008 | 0 | 0.12 | 0.08 | 0.15 | 0.09 | 0.1 | 0.07 |
| 2 | 84971398 | rs13383748 | -0.394 | -0.576 | 0.03 | -1.447 | -0.013 | -0.018 | -0.018 | 0 | 0.13 | 0.2 | 0.16 | 0.09 | 0.1 | 0.07 |
| 2 | 84975925 | rs13004624 | 0.706 | -0.176 | -0.181 | -1.31 | -0.075 | -0.031 | -0.031 | 0 | 0.14 | 0.04 | 0.22 | 0.19 | 0.31 | 0.16 |
| 2 | 84982251 | rs1192350 | -2.323 | -1.106 | 0.464 | -1.31 | -0.04 | -0.064 | -0.064 | 1.058 | 0.29 | 0.41 | 0.23 | 0.12 | 0.18 | 0.17 |
| 2 | 85008373 | rs1192375 | -0.47 | -1.022 | 0.42 | -1.264 | 0.001 | 0.009 | 0.009 | 0 | 0.12 | 0.08 | 0.16 | 0.09 | 0.11 | 0.07 |
| 2 | 85024731 | rs1192395 | -0.604 | -1.012 | 0.395 | -1.259 | 0.001 | 0.006 | 0.006 | 0 | 0.13 | 0.1 | 0.16 | 0.09 | 0.11 | 0.07 |
| 2 | 85046918 | rs7584703 | -0.592 | -1.045 | 0.485 | -1.27 | -0.029 | -0.036 | -0.036 | 4.936 | 0.09 | 0.24 | 0.05 | 0.03 | 0.03 | 0.04 |
| 2 | 85050488 | rs11674022 | -1.586 | -1.081 | 0.475 | -1.293 | -0.062 | -0.08 | -0.08 | 0.765 | 0.16 | 0.34 | 0.07 | 0.03 | 0.08 | 0.1 |
| 2 | 85051122 | rs1649292 | -1.357 | -0.993 | 0.317 | -1.253 | -0.012 | -0.022 | -0.022 | 0.046 | 0.2 | 0.21 | 0.2 | 0.11 | 0.14 | 0.11 |
| 2 | 85055822 | rs1192277 | -0.758 | -0.668 | 0.333 | -1.179 | 0.001 | 0.005 | 0.005 | 0 | 0.13 | 0.11 | 0.16 | 0.09 | 0.11 | 0.07 |
| 2 | 85058053 | rs7579460 | -1.063 | -0.632 | 0.318 | -1.165 | -0.031 | -0.043 | -0.043 | 0 | 0.09 | 0.25 | 0.03 | 0 | 0.05 | 0.06 |
| 2 | 85059063 | rs1192273 | -1.611 | -0.534 | -0.038 | -1.1 | -0.033 | -0.047 | -0.047 | 1.817 | 0.22 | 0.35 | 0.2 | 0.1 | 0.15 | 0.14 |
| 2 | 85066245 | rs736711 | -0.754 | -0.152 | -0.519 | -0.93 | -0.001 | -0.002 | -0.002 | 1.411 | 0.12 | 0.12 | 0.17 | 0.1 | 0.1 | 0.08 |
| 2 | 85068632 | rs10514646 | -0.584 | -0.425 | -1.267 | -1.406 | 0.008 | 0.075 | 0.075 | 1.251 | 0.22 | 0.03 | 0.22 | 0.07 | 0.16 | 0.14 |
| 2 | 85073697 | rs1529385 | 0.688 | 0.073 | -1.159 | -1.686 | 0.037 | 0.131 | 0.131 | 0 | 0.38 | 0.18 | 0.25 | 0.05 | 0.29 | 0.29 |
| 2 | 85074092 | rs12993644 | -2.028 | 0.87 | -0.315 | -0.888 | 0.037 | -0.077 | -0.077 | 0 | 0.16 | 0.3 | 0.23 | 0.01 | 0.23 | 0.1 |
| 2 | 85077774 | rs10514645 | -2.013 | 0.346 | -0.236 | -0.842 | -0.005 | 0.005 | 0.005 | 0 | 0.14 | 0.13 | 0.2 | 0.01 | 0.17 | 0.09 |
| 2 | 85080251 | rs1365899 | -2.332 | 0.364 | -0.126 | -0.662 | 0.019 | -0.081 | -0.081 | 0.02 | 0.15 | 0.36 | 0.18 | 0.01 | 0.18 | 0.13 |
| 2 | 85083503 | rs890617 | -2 | 0.404 | -0.022 | -0.521 | 0.009 | -0.052 | -0.052 | 1.073 | 0.11 | 0.37 | 0.16 | 0.01 | 0.1 | 0.11 |
| 2 | 85090036 | rs11683239 | -1.722 | 0.442 | -0.244 | -0.512 | 0.01 | -0.047 | -0.047 | 0 | 0.11 | 0.27 | 0.16 | 0.01 | 0.13 | 0.12 |
| 2 | 85091910 | rs12464608 | -0.939 | 0.33 | -0.29 | -0.515 | 0.059 | 0.006 | 0.006 | 0.001 | 0.06 | 0.26 | 0.14 | 0.01 | 0.07 | 0.07 |
| 2 | 85096254 | rs999401 | -0.458 | 0.368 | -0.616 | -0.083 | -0.03 | -0.139 | -0.139 | 0.029 | 0.8 | 0.99 | 0.85 | 0.37 | 0.81 | 0.67 |
| 2 | 85096481 | rs1000919 | -2.205 | 0.586 | 0.426 | -0.257 | 0.008 | -0.053 | -0.053 | 0 | 0.11 | 0.33 | 0.16 | 0.01 | 0.12 | 0.12 |
| 2 | 85115111 | rs6728804 | -1.61 | 0.919 | 0.235 | -0.035 | 0.016 | -0.008 | -0.008 | 0.196 | 0.13 | 0.26 | 0.31 | 0.08 | 0.16 | 0.09 |
| 2 | 85118358 | rs724012 | -0.958 | 0.919 | 0.202 | -0.06 | 0.027 | 0 | 0 | 0 | 0.1 | 0.2 | 0.3 | 0.08 | 0.16 | 0.09 |
| 2 | 85124209 | rs12469611 | -1.415 | 0.922 | 0.091 | -0.053 | 0.018 | -0.011 | -0.011 | 0.207 | 0.14 | 0.37 | 0.32 | 0.08 | 0.16 | 0.09 |
| 2 | 85140165 | rs2086175 | -0.989 | 0.866 | -0.324 | -0.215 | 0.003 | -0.072 | -0.072 | 0.052 | 0.18 | 0.25 | 0.23 | 0.02 | 0.16 | 0.18 |
| 2 | 85147568 | rs1382506 | 0.378 | 1.216 | -1.112 | -0.451 | -0.077 | 0.094 | 0.094 | 0.116 | 0.72 | 0.93 | 0.7 | 0.91 | 0.65 | 0.76 |
| 2 | 85163536 | rs4832128 | 0.605 | 1.194 | -1.098 | 0.29 | -0.081 | -0.028 | -0.028 | 0.396 | 0.64 | 0.82 | 0.61 | 0.62 | 0.58 | 0.61 |
| 2 | 85168571 | rs1979403 | 0.966 | 0.868 | -0.072 | 0.219 | 0.012 | -0.006 | -0.006 | 0 | 0.6 | 0.46 | 0.59 | 0.61 | 0.57 | 0.61 |
| 2 | 85169420 | rs6726213 | 0.226 | 0.888 | -0.153 | 0.228 | -0.116 | 0.133 | 0.133 | 0.24 | 0.7 | 0.97 | 0.71 | 0.95 | 0.65 | 0.71 |
| 2 | 85173517 | rs7591327 | 0.573 | 0.87 | -0.009 | 0.229 | -0.07 | -0.021 | -0.021 | 0 | 0.64 | 0.8 | 0.61 | 0.61 | 0.57 | 0.6 |
| 2 | 85190325 | rs936540 | 0.893 | 0.846 | 0.048 | 0.214 | 0.011 | -0.007 | -0.007 | 0 | 0.6 | 0.44 | 0.59 | 0.62 | 0.57 | 0.6 |
| 2 | 85190907 | rs7583930 | 0.506 | 0.903 | -0.305 | 0.23 | -0.067 | -0.024 | -0.024 | 1.161 | 0.65 | 0.8 | 0.63 | 0.62 | 0.59 | 0.67 |
| 2 | 85191263 | rs7572990 | 0.607 | 0.899 | -0.325 | 0.228 | -0.068 | -0.019 | -0.019 | 0 | 0.64 | 0.78 | 0.61 | 0.62 | 0.57 | 0.6 |
| 2 | 85204425 | rs17025763 | -2.6 | 0.649 | 0.537 | 0.685 | 0.037 | 0.212 | 0.212 | 4.328 | 0.08 | 0.39 | 0.11 | 0.34 | 0.08 | 0.12 |
| 2 | 85253059 | rs11896509 | -2.26 | 0.656 | 0.064 | 0.688 | 0.014 | 0.127 | 0.127 | 0.409 | 0.08 | 0.23 | 0.1 | 0.34 | 0.08 | 0.12 |
| 2 | 85283713 | rs4832131 | -0.264 | 0.425 | -0.862 | 0.31 | 0.016 | 0.074 | 0.074 | 0.067 | 0.36 | 0.17 | 0.59 | 0.82 | 0.33 | 0.44 |
| 2 | 85286519 | rs2272514 | -1.588 | 0.773 | -0.266 | 1.005 | 0.025 | 0.158 | 0.158 | 2.282 | 0.06 | 0.21 | 0.12 | 0.33 | 0.08 | 0.05 |
| 2 | 85287207 | rs6720075 | 1.228 | 0.61 | 0.372 | 1.045 | 0.03 | 0.086 | 0.086 | 0.051 | 0.93 | 0.79 | 0.94 | 0.9 | 0.94 | 0.87 |
| 2 | 85287288 | rs9808242 | -0.547 | 0.297 | -0.087 | 0.382 | 0.003 | 0.003 | 0.003 | 0 | 0.22 | 0.19 | 0.23 | 0.12 | 0.15 | 0.21 |
| 2 | 85287948 | rs10175174 | -0.111 | 1.381 | -0.581 | 0.099 | -0.073 | 0.072 | 0.072 | 0.782 | 0.19 | 0.04 | 0.24 | 0.04 | 0.3 | 0.15 |
| 2 | 85290623 | rs2583544 | 0.488 | 1.12 | -0.779 | 0.641 | 0.002 | -0.004 | -0.004 | 1.052 | 0.29 | 0.2 | 0.45 | 0.4 | 0.24 | 0.38 |
| 2 | 85291829 | rs11690856 | 0.725 | 1.064 | -0.772 | 0.224 | 0.022 | 0.041 | 0.041 | 0.913 | 0.42 | 0.39 | 0.2 | 0.26 | 0.33 | 0.38 |
| 2 | 85296388 | rs4019832 | 0.356 | 1.129 | -1.433 | 0.338 | -0.099 | 0.074 | 0.074 | 1.552 | 0.23 | 0.03 | 0.25 | 0.07 | 0.36 | 0.2 |
| 2 | 85303481 | rs2702035 | -0.88 | 0.805 | -1.026 | 0.273 | 0.004 | 0.028 | 0.028 | 1.564 | 0.6 | 0.64 | 0.79 | 0.77 | 0.65 | 0.69 |
| 2 | 85309672 | rs2583551 | 0.258 | 0.388 | -1.331 | 0.07 | 0.039 | -0.077 | -0.077 | 0 | 0.27 | 0.06 | 0.43 | 0.43 | 0.22 | 0.39 |
| 2 | 85316885 | rs7567580 | -0.25 | 0.902 | -0.582 | -0.963 | 0.018 | -0.033 | -0.033 | 1.123 | 0.63 | 0.46 | 0.74 | 0.84 | 0.54 | 0.72 |
| 2 | 85317168 | rs13392751 | -0.068 | 0.725 | -0.448 | -1.011 | -0.021 | 0.055 | 0.055 | 2.081 | 0.59 | 0.23 | 0.72 | 0.82 | 0.54 | 0.7 |
| 2 | 85318939 | rs7561211 | -0.015 | 0.671 | -0.426 | -1.435 | -0.001 | -0.001 | -0.001 | 0 | 0.46 | 0.4 | 0.68 | 0.79 | 0.46 | 0.6 |
| 2 | 85321746 | rs2583569 | -0.09 | 0.646 | -0.149 | -1.447 | -0.002 | 0.006 | 0.006 | 0.03 | 0.47 | 0.51 | 0.69 | 0.79 | 0.46 | 0.6 |

| | | | | | | | | | | | | | | | | |
|---|---|---|---|---|---|---|---|---|---|---|---|---|---|---|---|---|
| 2 | 85331425 | rs2583558 | -0.07 | 0.679 | -0.146 | -1.41 | -0.002 | 0.005 | 0.005 | 0.045 | 0.47 | 0.51 | 0.69 | 0.79 | 0.46 | 0.6 |
| 2 | 85331686 | rs2583556 | 0.266 | 0.787 | -0.441 | -1.565 | -0.005 | -0.012 | -0.012 | 0 | 0.35 | 0.35 | 0.59 | 0.76 | 0.3 | 0.56 |
| 2 | 85333976 | rs4832135 | -0.418 | 1.029 | -0.484 | -1.624 | -0.001 | 0.001 | 0.001 | 1.085 | 0.45 | 0.58 | 0.41 | 0.22 | 0.52 | 0.35 |
| 2 | 85334034 | rs4832136 | 0.024 | 1.088 | -0.421 | -1.657 | 0.025 | -0.021 | -0.021 | 0.739 | 0.13 | 0.23 | 0.23 | 0.07 | 0.18 | 0.1 |
| 2 | 85339385 | rs7560476 | 0.024 | 1.088 | -0.421 | -1.657 | 0.025 | -0.021 | -0.021 | 0.596 | 0.13 | 0.23 | 0.23 | 0.07 | 0.18 | 0.1 |
| 2 | 85339718 | rs11685301 | 1.241 | 0.926 | -0.506 | -1.984 | -0.014 | -0.007 | -0.007 | 0 | 0.15 | 0.12 | 0.12 | 0.13 | 0.24 | 0.17 |
| 2 | 85339839 | rs11884056 | 1.468 | 0.909 | -0.461 | -1.995 | 0.007 | 0.005 | 0.005 | 7.535 | 0.15 | 0.09 | 0.05 | 0.02 | 0.1 | 0.08 |
| 2 | 85340366 | rs1432265 | 0.15 | 0.943 | -0.501 | -1.946 | -0.002 | -0.009 | -0.009 | 0.538 | 0.31 | 0.34 | 0.19 | 0.15 | 0.34 | 0.26 |
| 2 | 85343124 | rs6709816 | 0.231 | 1.048 | -0.472 | -1.876 | 0.024 | -0.021 | -0.021 | 0.566 | 0.14 | 0.23 | 0.23 | 0.07 | 0.18 | 0.09 |
| 2 | 85357333 | rs2583554 | 1.163 | 0.484 | -0.769 | -1.935 | -0.019 | -0.044 | -0.044 | 0 | 0.27 | 0.06 | 0.41 | 0.72 | 0.24 | 0.48 |
| 2 | 85358556 | rs6728374 | -0.662 | 0.778 | -0.473 | -1.506 | 0.012 | -0.014 | -0.014 | 0 | 0.35 | 0.55 | 0.36 | 0.2 | 0.43 | 0.34 |
| 2 | 85364093 | rs10195517 | 0.507 | 1.445 | -0.922 | -0.948 | 0.02 | 0.019 | 0.019 | 0 | 0.14 | 0.19 | 0.16 | 0.17 | 0.26 | 0.21 |
| 2 | 85366971 | rs6754757 | -0.512 | 0.875 | -0.99 | -1.445 | 0.007 | -0.007 | -0.007 | 0.915 | 0.36 | 0.39 | 0.36 | 0.2 | 0.45 | 0.32 |
| 2 | 85375132 | rs2568218 | -0.286 | -0.059 | -0.771 | -1.808 | 0.024 | 0.038 | 0.038 | 0 | 0.6 | 0.72 | 0.62 | 0.91 | 0.62 | 0.77 |
| 2 | 85375282 | rs17025940 | 0.665 | -0.178 | -0.771 | -1.885 | 0.009 | 0.007 | 0.007 | 0 | 0.16 | 0.14 | 0.06 | 0.02 | 0.1 | 0.06 |
| 2 | 85377979 | rs2568222 | 0.152 | 0.288 | -0.613 | -1.474 | -0.001 | -0.023 | -0.023 | 0 | 0.43 | 0.56 | 0.52 | 0.88 | 0.47 | 0.7 |
| 2 | 85381416 | rs17762406 | -0.3 | -0.099 | -1.069 | -0.617 | 0.016 | 0.04 | 0.04 | 7.277 | 0.13 | 0.04 | 0.08 | 0.01 | 0.12 | 0.07 |
| 2 | 85389685 | rs6547597 | -0.401 | -0.446 | -0.434 | -0.716 | -0.059 | -0.095 | -0.095 | 2.498 | 0.16 | 0.31 | 0.06 | 0.01 | 0.04 | 0.07 |
| 2 | 85389705 | rs1560585 | 0.008 | -0.388 | -0.504 | -0.739 | 0 | 0.004 | 0.004 | 0 | 0.09 | 0.05 | 0.04 | 0.04 | 0.06 | 0.05 |
| 2 | 85392216 | rs11691660 | -0.742 | -0.102 | -0.985 | -0.578 | -0.001 | -0.002 | -0.002 | 0.984 | 0.38 | 0.34 | 0.26 | 0.26 | 0.37 | 0.35 |
| 2 | 85394611 | rs11691849 | -1.017 | -0.168 | -0.856 | -0.752 | 0.005 | 0.011 | 0.011 | 0 | 0.37 | 0.26 | 0.23 | 0.3 | 0.33 | 0.45 |
| 2 | 85395104 | rs11682047 | -0.729 | -0.167 | -0.947 | -0.637 | -0.004 | 0.044 | 0.044 | 0 | 0.32 | 0.1 | 0.23 | 0.22 | 0.32 | 0.41 |
| 2 | 85401958 | rs2568197 | -1.024 | -0.436 | -1.317 | -1.482 | 0.005 | 0.104 | 0.104 | 0.292 | 0.5 | 0.75 | 0.48 | 0.26 | 0.47 | 0.53 |
| 2 | 85402493 | rs17025989 | 1.349 | -0.536 | -1.358 | -1.534 | 0.01 | 0.005 | 0.005 | 0 | 0.08 | 0.06 | 0.03 | 0.04 | 0.03 | 0.05 |
| 2 | 85403546 | rs2568198 | 0.388 | -0.47 | -1.442 | -1.617 | -0.053 | 0 | 0 | 0 | 0.24 | 0.08 | 0.2 | 0.2 | 0.29 | 0.31 |
| 2 | 85408298 | rs7594695 | 0.321 | 0.017 | -1.463 | -1.862 | 0.008 | 0.01 | 0.01 | 0 | 0.15 | 0.13 | 0.08 | 0.05 | 0.09 | 0.07 |
| 2 | 85410089 | rs2583545 | -0.738 | -0.13 | -0.439 | -1.942 | 0.001 | -0.039 | -0.039 | 0 | 0.35 | 0.52 | 0.26 | 0.2 | 0.33 | 0.37 |
| 2 | 85414140 | rs2583547 | -1.798 | -0.218 | -0.177 | -1.967 | -0.036 | 0.085 | 0.085 | 1.108 | 0.54 | 0.85 | 0.5 | 0.27 | 0.48 | 0.51 |
| 2 | 85414647 | rs13432342 | 1.011 | -0.151 | -0.054 | -1.971 | -0.008 | -0.031 | -0.031 | 0 | 0.15 | 0.23 | 0.22 | 0.07 | 0.13 | 0.15 |
| 2 | 85420124 | rs2568210 | -0.688 | -0.24 | -0.021 | -1.972 | 0.003 | -0.024 | -0.024 | 0 | 0.35 | 0.48 | 0.27 | 0.21 | 0.34 | 0.37 |
| 2 | 85420870 | rs880092 | 0.03 | -0.263 | 0.023 | -1.99 | -0.013 | -0.053 | -0.053 | 0 | 0.21 | 0.27 | 0.27 | 0.07 | 0.19 | 0.16 |
| 2 | 85421691 | rs874838 | 0.971 | -0.248 | 0.011 | -1.964 | -0.008 | -0.031 | -0.031 | 0 | 0.14 | 0.23 | 0.22 | 0.07 | 0.14 | 0.15 |
| 2 | 85422993 | rs747078 | 1.208 | -0.171 | -0.072 | -1.903 | -0.044 | -0.003 | -0.003 | 2.354 | 0.18 | 0.08 | 0.17 | 0.16 | 0.25 | 0.3 |
| 2 | 85426269 | rs6733735 | 1.285 | -0.261 | 0.08 | -1.844 | -0.018 | -0.007 | -0.007 | 0.078 | 0.1 | 0.23 | 0.04 | 0.04 | 0.03 | 0.06 |
| 2 | 85432404 | rs17026015 | 1.298 | -0.261 | 0.041 | -1.813 | -0.017 | -0.003 | -0.003 | 0 | 0.1 | 0.24 | 0.04 | 0.05 | 0.03 | 0.05 |
| 2 | 85434540 | rs4832148 | -2.491 | -0.218 | -0.966 | -1.928 | 0.016 | 0.043 | 0.043 | 0 | 0.77 | 0.92 | 0.85 | 0.89 | 0.78 | 0.92 |
| 2 | 85439553 | rs2043228 | -0.677 | -0.138 | -0.816 | -1.883 | -0.003 | 0.05 | 0.05 | 0 | 0.46 | 0.67 | 0.49 | 0.29 | 0.47 | 0.54 |
| 2 | 85443924 | rs6709476 | -0.501 | -0.308 | -0.748 | -1.863 | 0.001 | 0.001 | 0.001 | 3.161 | 0.22 | 0.72 | 0.26 | 0.08 | 0.09 | 0.11 |
| 2 | 85453593 | rs6743132 | -0.518 | -0.121 | -0.615 | -1.799 | -0.005 | -0.006 | -0.006 | 0 | 0.2 | 0.74 | 0.12 | 0.08 | 0.09 | 0.14 |
| 2 | 85459682 | rs723465 | 1.412 | 0.09 | -0.948 | -1.906 | -0.075 | -0.007 | -0.007 | 1.491 | 0.26 | 0.03 | 0.18 | 0.18 | 0.27 | 0.26 |
| 2 | 85462602 | rs12477490 | 1.326 | 0.245 | -0.945 | -1.811 | -0.065 | -0.104 | -0.104 | 0 | 0.14 | 0.01 | 0.37 | 0.58 | 0.19 | 0.32 |
| 2 | 85468002 | rs10205484 | 0.339 | 0.095 | -0.555 | -1.967 | 0.019 | -0.043 | -0.043 | 0 | 0.08 | 0.27 | 0.08 | 0.01 | 0.08 | 0.01 |
| 2 | 85470087 | rs6547602 | -0.167 | 0.237 | -1.043 | -2.035 | 0.039 | 0.106 | 0.106 | 0.656 | 0.48 | 0.82 | 0.32 | 0.27 | 0.37 | 0.34 |
| 2 | 85470281 | rs11675205 | 1.622 | 0.257 | -1.211 | -2.012 | -0.04 | -0.005 | -0.005 | 0.151 | 0.27 | 0.04 | 0.16 | 0.16 | 0.23 | 0.23 |
| 2 | 85470987 | rs6725799 | -0.618 | 0.24 | -1.112 | -1.97 | -0.095 | 0.032 | 0.032 | 0.031 | 0.58 | 0.92 | 0.39 | 0.32 | 0.5 | 0.49 |
| 2 | 85471459 | rs7566763 | 0.717 | 0.563 | -1.007 | -1.79 | -0.097 | -0.115 | -0.115 | 0 | 0.17 | 0.02 | 0.45 | 0.61 | 0.26 | 0.36 |
| 2 | 85471980 | rs6758778 | 0.245 | -0.174 | -0.091 | -1.571 | -0.005 | -0.022 | -0.022 | 0.108 | 0.12 | 0.36 | 0.07 | 0.06 | 0.06 | 0.08 |
| 2 | 85480107 | rs17763748 | 1.06 | -0.046 | -0.904 | -1.546 | -0.034 | 0 | 0 | 0.044 | 0.32 | 0.18 | 0.22 | 0.26 | 0.32 | 0.26 |
| 2 | 85480422 | rs7580690 | 0.516 | -0.514 | -0.782 | -1.458 | 0.01 | 0.008 | 0.008 | 0.372 | 0.28 | 0.23 | 0.15 | 0.07 | 0.22 | 0.14 |
| 2 | 85480715 | rs17763853 | 1.765 | -0.49 | -0.876 | -1.445 | -0.035 | -0.003 | -0.003 | 0.18 | 0.32 | 0.18 | 0.22 | 0.26 | 0.32 | 0.26 |
| 2 | 85480817 | rs7581133 | 1.003 | -0.444 | -0.893 | -1.46 | -0.052 | 0.027 | 0.027 | 0 | 0.38 | 0.19 | 0.27 | 0.26 | 0.4 | 0.27 |
| 2 | 85483003 | rs2043229 | 0.39 | 0.332 | 0.158 | -0.724 | 0.01 | 0.016 | 0.016 | 0.636 | 0.67 | 0.57 | 0.44 | 0.33 | 0.62 | 0.4 |
| 2 | 85491365 | rs11689667 | 1.713 | 0.274 | -0.317 | -0.717 | -0.054 | 0.071 | 0.071 | 0.111 | 0.46 | 0.15 | 0.28 | 0.27 | 0.43 | 0.3 |
| 2 | 85492141 | rs908301 | 0.916 | 0.301 | -0.185 | -0.649 | -0.003 | 0.025 | 0.025 | 0 | 0.56 | 0.39 | 0.35 | 0.29 | 0.51 | 0.31 |
| 2 | 85502264 | rs12714137 | -0.62 | 0.335 | -0.148 | -0.661 | -0.005 | -0.011 | -0.011 | 0 | 0.88 | 0.89 | 0.92 | 0.95 | 0.87 | 0.9 |
| 2 | 85502419 | rs12714138 | 1.426 | 0.297 | -0.328 | -0.826 | 0.006 | 0.015 | 0.015 | 0.714 | 0.68 | 0.27 | 0.65 | 0.83 | 0.69 | 0.65 |
| 2 | 85503139 | rs6745529 | 1.975 | 0.302 | -0.484 | -0.667 | -0.129 | -0.022 | -0.022 | 0 | 0.45 | 0.05 | 0.28 | 0.28 | 0.43 | 0.3 |
| 2 | 85505602 | rs6732834 | 1.386 | 0.293 | -0.483 | -0.729 | -0.009 | -0.043 | -0.043 | 2.274 | 0.11 | 0.13 | 0.07 | 0 | 0.08 | 0.08 |
| 2 | 85506104 | rs6736348 | 0.504 | 0.292 | -0.141 | -0.553 | -0.023 | -0.059 | -0.059 | 0.138 | 0.77 | 0.43 | 0.72 | 0.85 | 0.78 | 0.66 |
| 2 | 85506160 | rs6736366 | 1.293 | 0.277 | -0.428 | -0.537 | 0.002 | -0.002 | -0.002 | 0 | 0.69 | 0.3 | 0.65 | 0.84 | 0.69 | 0.65 |
| 2 | 85507657 | rs10204252 | 0.648 | 0.277 | -0.374 | -0.452 | -0.001 | 0.003 | 0.003 | 0 | 0.23 | 0.2 | 0.38 | 0.55 | 0.26 | 0.35 |
| 2 | 85508065 | rs10206876 | 0.431 | 0.254 | -0.393 | -0.443 | 0.002 | 0.027 | 0.027 | 0 | 0.24 | 0.26 | 0.38 | 0.56 | 0.26 | 0.35 |
| 2 | 85508850 | rs6547608 | 1.088 | 0.353 | -0.746 | -0.195 | 0.002 | 0.052 | 0.052 | 0 | 0.45 | 0.3 | 0.26 | 0.22 | 0.38 | 0.29 |
| 2 | 85510425 | rs7601117 | 1.243 | 0.661 | -1.755 | -0.276 | -0.086 | -0.023 | -0.023 | 0 | 0.44 | 0.06 | 0.25 | 0.28 | 0.39 | 0.29 |
| 2 | 85512062 | rs17713693 | 0.005 | 1.021 | -1.064 | 0.085 | -0.044 | 0.051 | 0.051 | 0 | 0.1 | 0.01 | 0.09 | 0.01 | 0.15 | 0.04 |
| 2 | 85512961 | rs6719501 | 0.061 | 0.624 | -0.762 | 0.334 | -0.05 | -0.042 | -0.042 | 0 | 0.6 | 0.81 | 0.47 | 0.44 | 0.47 | 0.56 |
| 2 | 85514932 | rs6748174 | -1.14 | 0.442 | 0.116 | 0.232 | -0.012 | 0.004 | 0.004 | 0 | 0.05 | 0.23 | 0.1 | 0.05 | 0 | 0.04 |
| 2 | 85518483 | rs7572226 | -1.332 | 0.764 | -0.023 | 0.21 | 0.004 | -0.008 | -0.008 | 0 | 0.09 | 0.19 | 0.15 | 0.07 | 0.09 | 0.16 |
| 2 | 85523276 | rs4832156 | 0.672 | 1.15 | -0.061 | -0.804 | 0.002 | -0.084 | -0.084 | 0 | 0.59 | 0.53 | 0.68 | 0.9 | 0.5 | 0.67 |

| | | | | | | | | | | | | | | | | |
|---|---|---|---|---|---|---|---|---|---|---|---|---|---|---|---|---|
| 2 | 85523541 | rs4832159 | -0.201 | 1.731 | -0.582 | -0.866 | -0.127 | 0.086 | 0.086 | 0.046 | 0.77 | 0.99 | 0.8 | 0.93 | 0.66 | 0.91 |
| 2 | 85524369 | rs6737773 | -0.035 | 1.714 | -0.489 | -0.486 | -0.001 | -0.013 | -0.013 | 2.485 | 0.05 | 0.02 | 0.03 | 0.09 | 0.04 | 0.11 |
| 2 | 85531807 | rs12714139 | -0.296 | 1.963 | -0.539 | -0.688 | -0.097 | 0.041 | 0.041 | 0 | 0.83 | 0.98 | 0.83 | 0.93 | 0.7 | 0.92 |
| 2 | 85537312 | rs9248 | 1.186 | 2.061 | 0.505 | -0.961 | 0.029 | -0.08 | -0.08 | 0 | 0.66 | 0.53 | 0.72 | 0.87 | 0.52 | 0.67 |
| 2 | 85545490 | rs7428 | 1.803 | 0.921 | -0.129 | -0.576 | 0.021 | -0.1 | -0.1 | 0 | 0.42 | 0.07 | 0.59 | 0.5 | 0.38 | 0.49 |
| 2 | 85546052 | rs1053561 | 0.702 | 0.987 | -0.064 | -0.557 | 0.033 | -0.084 | -0.084 | 0 | 0.66 | 0.52 | 0.72 | 0.87 | 0.52 | 0.67 |
| 2 | 85546192 | rs1053560 | 0.371 | 0.959 | 0.04 | -0.58 | 0.066 | -0.12 | -0.12 | 0 | 0.72 | 0.58 | 0.75 | 0.96 | 0.54 | 0.68 |
| 2 | 85546615 | rs1061782 | 0.371 | 0.959 | 0.04 | -0.58 | 0.066 | -0.12 | -0.12 | 0 | 0.72 | 0.58 | 0.75 | 0.96 | 0.54 | 0.68 |
| 2 | 85546932 | rs4832164 | 0.371 | 0.959 | 0.04 | -0.58 | 0.066 | -0.12 | -0.12 | 0 | 0.72 | 0.58 | 0.75 | 0.96 | 0.54 | 0.67 |
| 2 | 85547429 | rs4459734 | 0.246 | 0.952 | 0.372 | -0.583 | 0.01 | -0.045 | -0.045 | 0 | 0.73 | 0.68 | 0.75 | 0.96 | 0.54 | 0.67 |
| 2 | 85547963 | rs1554110 | 0.246 | 0.952 | 0.372 | -0.583 | 0.01 | -0.045 | -0.045 | 0 | 0.73 | 0.68 | 0.75 | 0.96 | 0.54 | 0.67 |
| 2 | 85548639 | rs6547611 | -0.194 | 0.949 | 0.513 | -0.592 | -0.039 | 0.028 | 0.028 | 0 | 0.77 | 0.83 | 0.8 | 0.96 | 0.6 | 0.72 |
| 2 | 85549527 | rs3184781 | 0.913 | 0.918 | 0.453 | -0.58 | 0.033 | -0.084 | -0.084 | 0 | 0.66 | 0.52 | 0.72 | 0.87 | 0.52 | 0.67 |
| 2 | 85549547 | rs3184780 | 0.913 | 0.918 | 0.453 | -0.58 | 0.033 | -0.084 | -0.084 | 8.245 | 0.66 | 0.52 | 0.72 | 0.87 | 0.52 | 0.67 |
| 2 | 85552261 | rs2292653 | 1.008 | 0.805 | 0.775 | -0.514 | 0.008 | -0.004 | -0.004 | 0 | 0.12 | 0.12 | 0.06 | 0.29 | 0.08 | 0.06 |
| 2 | 85553784 | rs1044973 | 0.559 | 1.028 | 0.807 | -0.616 | 0.067 | -0.075 | -0.075 | 0.055 | 0.71 | 0.55 | 0.72 | 0.88 | 0.53 | 0.66 |
| 2 | 85555262 | rs10460585 | -0.244 | 1.191 | 0.857 | -0.626 | -0.018 | 0.002 | 0.002 | 0 | 0.78 | 0.81 | 0.81 | 0.96 | 0.62 | 0.74 |
| 2 | 85555478 | rs10460587 | 0.401 | 1.187 | 0.799 | -0.606 | 0.074 | -0.061 | -0.061 | 0 | 0.73 | 0.59 | 0.73 | 0.87 | 0.55 | 0.69 |
| 2 | 85562172 | rs3923229 | -1.105 | 1.532 | 0.55 | -0.616 | -0.083 | 0.016 | 0.016 | 0 | 0.86 | 0.98 | 0.85 | 0.92 | 0.74 | 0.93 |
| 2 | 85565634 | rs6747513 | 0.686 | 1.385 | 1.086 | -0.69 | 0.016 | -0.045 | -0.045 | 0 | 0.69 | 0.4 | 0.75 | 0.88 | 0.62 | 0.72 |
| 2 | 85568810 | rs1969260 | 2.086 | 1.026 | 0.935 | -0.685 | 0.019 | -0.097 | -0.097 | 0 | 0.41 | 0.13 | 0.6 | 0.52 | 0.38 | 0.48 |
| 2 | 85569294 | rs999939 | 0.198 | 0.966 | 1.125 | -0.663 | -0.011 | 0.016 | 0.016 | 0.085 | 0.18 | 0.2 | 0.08 | 0.27 | 0.13 | 0.08 |
| 2 | 85573033 | rs908302 | 0.653 | 1.109 | 1.296 | -0.664 | 0.053 | -0.149 | -0.149 | 0 | 0.7 | 0.51 | 0.78 | 0.96 | 0.62 | 0.72 |
| 2 | 85574015 | rs4474882 | 0.614 | 1.132 | 1.208 | -0.65 | 0.045 | -0.13 | -0.13 | 0.454 | 0.7 | 0.57 | 0.78 | 0.96 | 0.62 | 0.72 |
| 2 | 85594106 | rs7340340 | -1.687 | 0.902 | 0.446 | -0.65 | 0.053 | -0.031 | -0.031 | 0.243 | 0.58 | 0.71 | 0.4 | 0.51 | 0.65 | 0.52 |
| 2 | 85596553 | rs6716601 | -0.445 | 1.304 | 0.737 | -0.716 | -0.003 | 0.008 | 0.008 | 0 | 0.28 | 0.31 | 0.16 | 0.35 | 0.25 | 0.25 |
| 2 | 85603064 | rs6741368 | -0.433 | 1.299 | 0.734 | -0.718 | -0.003 | 0.009 | 0.009 | 0 | 0.28 | 0.31 | 0.16 | 0.35 | 0.25 | 0.25 |
| 2 | 85605304 | rs6731223 | -0.344 | 1.526 | 0.48 | -1.044 | 0.001 | -0.01 | -0.01 | 0 | 0.9 | 0.9 | 0.93 | 0.96 | 0.85 | 0.85 |
| 2 | 85606311 | rs7558094 | 0.893 | 1.672 | 0.65 | -0.869 | 0.029 | 0.014 | 0.014 | 0 | 0.06 | 0.11 | 0.04 | 0.13 | 0.09 | 0.16 |
| 2 | 85615012 | rs1135604 | -1.559 | 1.804 | 0.692 | -0.976 | -0.011 | -0.024 | -0.024 | 0 | 0.45 | 0.69 | 0.29 | 0.49 | 0.43 | 0.44 |
| 2 | 85622059 | rs6886 | -1.823 | 1.865 | 0.717 | -1.038 | 0.07 | -0.04 | -0.04 | 0 | 0.58 | 0.75 | 0.4 | 0.5 | 0.66 | 0.51 |
| 2 | 85622317 | rs11690650 | -1.113 | 2.201 | 1.076 | -0.925 | 0.076 | -0.022 | -0.022 | 1.129 | 0.16 | 0.3 | 0.13 | 0.12 | 0.22 | 0.25 |
| 2 | 85627714 | rs11684413 | 1.045 | 2.211 | 0.733 | -0.925 | -0.054 | 0.051 | 0.051 | 0.004 | 0.13 | 0.04 | 0.11 | 0 | 0.22 | 0.06 |
| 2 | 85628556 | rs2002444 | -0.343 | 1.783 | 0.835 | -0.824 | 0.001 | -0.005 | -0.005 | 0 | 0.09 | 0.08 | 0.08 | 0.02 | 0.12 | 0.09 |
| 2 | 85628983 | rs2229668 | -0.315 | 1.807 | 0.846 | -0.808 | 0.003 | 0 | 0 | 0 | 0.08 | 0.08 | 0.08 | 0.08 | 0.12 | 0.1 |
| 2 | 85638323 | rs11682055 | -0.707 | 1.575 | 0.902 | -0.843 | 0.062 | -0.082 | -0.082 | 0 | 0.31 | 0.45 | 0.22 | 0.11 | 0.4 | 0.26 |
| 2 | 85654299 | rs1877954 | -0.53 | 1.117 | 0.751 | -0.592 | -0.021 | 0.057 | 0.057 | 0 | 0.22 | 0.5 | 0.17 | 0.27 | 0.2 | 0.15 |
| 2 | 85658232 | rs11674612 | -0.49 | 1.682 | 0.685 | -0.567 | 0.05 | -0.05 | -0.05 | 2.875 | 0.27 | 0.39 | 0.18 | 0.13 | 0.35 | 0.27 |
| 2 | 85662650 | rs6547618 | -1.284 | 0.881 | -0.193 | -0.233 | 0.097 | 0.052 | 0.052 | 0.25 | 0.52 | 0.73 | 0.47 | 0.58 | 0.63 | 0.52 |
| 2 | 85664399 | rs12473017 | -1.243 | 0.907 | -0.153 | -0.234 | 0.091 | 0.055 | 0.055 | 0.272 | 0.52 | 0.72 | 0.47 | 0.58 | 0.62 | 0.51 |
| 2 | 85665669 | rs2292655 | -0.347 | 1.006 | 0.328 | -0.047 | -0.013 | 0.066 | 0.066 | 0 | 0.19 | 0.25 | 0.21 | 0.36 | 0.14 | 0.2 |
| 2 | 85668988 | rs11682343 | -0.157 | 1.568 | 0.434 | -0.198 | 0.022 | -0.004 | -0.004 | 0 | 0.26 | 0.3 | 0.2 | 0.22 | 0.41 | 0.24 |
| 2 | 85670488 | rs4832175 | -1.164 | 1.428 | 0.397 | -0.139 | 0.003 | -0.013 | -0.013 | 0 | 0.39 | 0.45 | 0.35 | 0.29 | 0.51 | 0.34 |
| 2 | 85672868 | rs4832176 | -1.692 | 0.841 | -0.568 | -0.077 | 0.098 | -0.001 | -0.001 | 0 | 0.6 | 0.84 | 0.52 | 0.6 | 0.7 | 0.62 |
| 2 | 85673991 | rs908307 | -0.948 | 1.128 | -0.244 | -0.161 | -0.004 | 0.006 | 0.006 | 0 | 0.39 | 0.55 | 0.33 | 0.46 | 0.3 | 0.44 |
| 2 | 85674576 | rs7594872 | 0.09 | 1.551 | 0.046 | -0.057 | 0.057 | -0.011 | -0.011 | 0 | 0.21 | 0.29 | 0.2 | 0.14 | 0.4 | 0.17 |
| 2 | 85678425 | rs7583141 | -0.663 | 1.764 | 0.123 | -0.1 | 0.03 | -0.026 | -0.026 | 0 | 0.25 | 0.33 | 0.19 | 0.14 | 0.34 | 0.17 |
| 2 | 85683099 | rs17026352 | -0.23 | 1.942 | 0.138 | 0.016 | 0.053 | -0.008 | -0.008 | 2.831 | 0.18 | 0.3 | 0.17 | 0.14 | 0.32 | 0.21 |
| 2 | 85684210 | rs6716108 | -1.533 | 0.653 | -0.756 | -0.267 | 0.037 | -0.042 | -0.042 | 0.1 | 0.75 | 0.85 | 0.59 | 0.61 | 0.81 | 0.65 |
| 2 | 85692999 | rs11695848 | -1.193 | 1.796 | 0.098 | -0.862 | 0.052 | -0.081 | -0.081 | 0.018 | 0.25 | 0.6 | 0.25 | 0.09 | 0.37 | 0.25 |
| 2 | 85696551 | rs12616455 | -0.194 | 1.827 | 0.004 | -0.904 | 0.013 | -0.032 | -0.032 | 0 | 0.79 | 0.67 | 0.81 | 0.91 | 0.76 | 0.94 |
| 2 | 85699750 | rs7577642 | 0.04 | 1.79 | 0.245 | -0.94 | 0.023 | -0.059 | -0.059 | 0 | 0.78 | 0.61 | 0.8 | 0.89 | 0.72 | 0.88 |
| 2 | 85700924 | rs960066 | 0.475 | 2.294 | -0.306 | -1.113 | -0.025 | 0.019 | 0.019 | 0 | 0.09 | 0.06 | 0.12 | 0.04 | 0.23 | 0.07 |
| 2 | 85703308 | rs6714421 | 1.25 | 1.998 | -0.238 | -1.333 | 0.012 | 0.102 | 0.102 | 0.043 | 0.58 | 0.73 | 0.6 | 0.81 | 0.38 | 0.7 |
| 2 | 85707654 | rs11680227 | -1.296 | 2.24 | 0.258 | -0.937 | 0.018 | -0.024 | -0.024 | 1.157 | 0.2 | 0.24 | 0.22 | 0.08 | 0.33 | 0.17 |
| 2 | 85709069 | rs35700171 | -0.244 | 2.144 | 0.499 | -0.778 | 0.021 | -0.012 | -0.012 | 0 | 0.1 | 0.14 | 0.13 | 0.05 | 0.16 | 0.06 |
| 2 | 85723742 | rs4331503 | -0.101 | 1.551 | 0.644 | -0.601 | -0.009 | -0.002 | -0.002 | 0 | 0.1 | 0.14 | 0.07 | 0.09 | 0.06 | 0.16 |
| 2 | 85731858 | rs6750610 | -0.107 | 1.994 | 0.168 | -0.604 | 0.036 | -0.011 | -0.011 | 0 | 0.1 | 0.13 | 0.15 | 0.07 | 0.26 | 0.06 |
| 2 | 85738374 | rs7591175 | -1.547 | 1.238 | -0.525 | -0.497 | -0.058 | 0.024 | 0.024 | 1.047 | 0.44 | 0.82 | 0.39 | 0.38 | 0.49 | 0.34 |
| 2 | 85738944 | rs2044475 | 0.581 | 1.395 | -0.396 | -0.717 | -0.029 | 0.026 | 0.026 | 0.016 | 0.09 | 0.04 | 0.13 | 0.02 | 0.25 | 0.06 |
| 2 | 85747371 | rs10496316 | -0.008 | 1 | 0.264 | -0.823 | -0.032 | -0.086 | -0.086 | 0 | 0.21 | 0.37 | 0.14 | 0.08 | 0.13 | 0.22 |
| 2 | 85748719 | rs17735885 | 0.762 | 0.959 | 0.359 | -0.843 | 0.002 | 0.002 | 0.002 | 0 | 0.16 | 0.13 | 0.1 | 0.07 | 0.12 | 0.2 |
| 2 | 85755357 | rs6739015 | -1.691 | 0.985 | 0.566 | -0.77 | -0.03 | 0.008 | 0.008 | 0.194 | 0.38 | 0.75 | 0.37 | 0.38 | 0.45 | 0.35 |
| 2 | 85762209 | rs1446669 | -1.013 | 1.001 | 0.624 | -0.761 | 0.076 | 0.064 | 0.064 | 0 | 0.2 | 0.4 | 0.25 | 0.32 | 0.34 | 0.16 |
| 2 | 85767735 | rs2028900 | -1.889 | 0.967 | 0.678 | -0.842 | -0.03 | 0.027 | 0.027 | 0.371 | 0.43 | 0.72 | 0.36 | 0.38 | 0.48 | 0.35 |
| 2 | 85769975 | rs2043675 | -0.472 | 0.976 | 0.682 | -0.838 | 0.105 | 0.097 | 0.097 | 0 | 0.17 | 0.39 | 0.22 | 0.31 | 0.31 | 0.14 |
| 2 | 85780107 | rs10179904 | 1.037 | 0.961 | 0.68 | -0.844 | 0.006 | 0.015 | 0.015 | 0 | 0.16 | 0.12 | 0.1 | 0.07 | 0.11 | 0.2 |
| 2 | 85780536 | rs699664 | -1.528 | 0.943 | 0.698 | -0.86 | 0.007 | 0.002 | 0.002 | 0 | 0.26 | 0.66 | 0.27 | 0.31 | 0.37 | 0.15 |
| 2 | 85794297 | rs10187424 | -1.771 | 0.869 | 0.657 | -0.821 | -0.016 | 0.008 | 0.008 | 0 | 0.39 | 0.69 | 0.36 | 0.39 | 0.44 | 0.35 |

| | | | | | | | | | | | | | | | | |
|---|---|---|---|---|---|---|---|---|---|---|---|---|---|---|---|---|
| 2 | 85803542 | rs6757263 | -1.544 | 0.876 | 0.582 | -0.809 | -0.008 | 0.001 | 0.001 | 0.025 | 0.37 | 0.64 | 0.35 | 0.39 | 0.44 | 0.35 |
| 2 | 85804096 | rs10166612 | 0.993 | 0.889 | 0.475 | -0.808 | 0.006 | 0.016 | 0.016 | 0 | 0.16 | 0.11 | 0.1 | 0.07 | 0.11 | 0.2 |
| 2 | 85806266 | rs3731828 | -0.514 | 0.934 | 0.547 | -0.786 | 0.102 | 0.098 | 0.098 | 0 | 0.17 | 0.4 | 0.22 | 0.31 | 0.3 | 0.14 |
| 2 | 85808737 | rs1009 | -1.169 | 0.893 | 0.587 | -0.801 | -0.001 | 0 | 0 | 0 | 0.36 | 0.6 | 0.35 | 0.39 | 0.43 | 0.35 |
| 2 | 85808871 | rs1058588 | -1.085 | 0.9 | 0.371 | -0.798 | 0.018 | -0.001 | -0.001 | 0 | 0.36 | 0.51 | 0.34 | 0.39 | 0.43 | 0.35 |
| 2 | 85813834 | rs12714147 | 0.332 | 0.576 | -0.489 | -0.731 | 0.017 | 0.046 | 0.046 | 0 | 0.21 | 0.12 | 0.15 | 0.08 | 0.16 | 0.25 |
| 2 | 85814984 | rs10206961 | -1.491 | 0.665 | -0.731 | -0.729 | -0.053 | 0.017 | 0.017 | 1.939 | 0.44 | 0.92 | 0.38 | 0.37 | 0.42 | 0.37 |
| 2 | 85816334 | rs1254900 | -1.358 | 0.246 | -1.2 | -0.75 | 0.073 | -0.179 | -0.179 | 0.265 | 0.67 | 0.97 | 0.58 | 0.51 | 0.8 | 0.64 |
| 2 | 85818488 | rs719023 | 0.891 | 0.343 | -0.124 | -0.595 | -0.001 | -0.022 | -0.022 | 0 | 0.57 | 0.45 | 0.39 | 0.12 | 0.61 | 0.41 |
| 2 | 85818886 | rs14976 | -0.238 | 0.944 | 0.428 | -0.616 | -0.027 | 0.06 | 0.06 | 0 | 0.25 | 0.15 | 0.2 | 0.09 | 0.28 | 0.26 |
| 2 | 85822682 | rs2232739 | 0.758 | 0.821 | 0.667 | -0.523 | -0.001 | -0.012 | -0.012 | 0 | 0.07 | 0.05 | 0.04 | 0 | 0.07 | 0.01 |
| 2 | 85824251 | rs6643 | -0.82 | 0.822 | 0.831 | -0.488 | 0.095 | 0.426 | 0.426 | 3.416 | 0.06 | 0.52 | 0.08 | 0.3 | 0.04 | 0.06 |
| 2 | 85826132 | rs1044158 | -1.648 | 0.826 | 0.593 | -0.55 | -0.008 | 0.026 | 0.026 | 4.556 | 0.41 | 0.77 | 0.32 | 0.38 | 0.37 | 0.32 |
| 2 | 85826721 | rs11891495 | -0.242 | 0.84 | 0.58 | -0.547 | -0.029 | 0.069 | 0.069 | 0 | 0.24 | 0.14 | 0.18 | 0.08 | 0.25 | 0.24 |
| 2 | 85828014 | rs6733550 | -1.584 | 0.847 | 0.605 | -0.53 | -0.015 | 0.03 | 0.03 | 5.112 | 0.41 | 0.84 | 0.32 | 0.38 | 0.37 | 0.31 |
| 2 | 85852143 | rs7569073 | 0.538 | 1.041 | 0.166 | -0.664 | 0.013 | 0.045 | 0.045 | 0 | 0.17 | 0.09 | 0.1 | 0.07 | 0.1 | 0.19 |
| 2 | 85886013 | rs7316 | 0.03 | 0.693 | 0.176 | -0.314 | 0.016 | 0.087 | 0.087 | 1.233 | 0.12 | 0.26 | 0.21 | 0.22 | 0.1 | 0.07 |
| 2 | 85889525 | rs2040349 | -0.517 | 0.742 | -0.344 | -0.132 | -0.019 | 0.037 | 0.037 | 4.968 | 0.41 | 0.84 | 0.44 | 0.36 | 0.37 | 0.31 |
| 2 | 85893741 | rs1130866 | -0.126 | 0.805 | -1.191 | 0.007 | -0.028 | 0.037 | 0.037 | 0.705 | 0.61 | 0.8 | 0.42 | 0.26 | 0.54 | 0.46 |
| 2 | 85895704 | rs3024791 | -0.216 | 0.46 | -0.573 | -0.198 | -0.014 | -0.03 | -0.03 | 0.679 | 0.21 | 0.28 | 0.12 | 0.11 | 0.13 | 0.21 |
| 2 | 85898143 | rs934774 | 0.771 | 0.581 | -1.68 | -0.075 | -0.049 | -0.124 | -0.124 | 0.055 | 0.37 | 0.05 | 0.39 | 0.48 | 0.37 | 0.39 |
| 2 | 85898898 | rs12714149 | 0.749 | 0.556 | -1.694 | -0.044 | -0.052 | -0.137 | -0.137 | 0 | 0.37 | 0.06 | 0.4 | 0.5 | 0.37 | 0.42 |
| 2 | 85902569 | rs1030862 | 0.588 | 0.427 | -0.411 | -0.093 | -0.006 | 0.012 | 0.012 | 0 | 0.42 | 0.45 | 0.45 | 0.53 | 0.38 | 0.47 |
| 2 | 85911339 | rs7355681 | -0.368 | 0.447 | 0.616 | -0.535 | -0.012 | -0.009 | -0.009 | 0 | 0.69 | 0.74 | 0.59 | 0.63 | 0.58 | 0.7 |
| 2 | 85912973 | rs6714324 | 0.076 | 0.28 | 0.373 | -0.659 | -0.013 | -0.062 | -0.062 | 0.158 | 0.17 | 0.32 | 0.11 | 0.04 | 0.16 | 0.2 |
| 2 | 85913162 | rs17026540 | 0.721 | 0.378 | 0.266 | -0.643 | 0.025 | 0.005 | 0.005 | 0 | 0.47 | 0.39 | 0.46 | 0.56 | 0.4 | 0.36 |
| 2 | 85916487 | rs4832183 | -0.12 | 0.483 | 0.055 | -0.762 | -0.025 | -0.002 | -0.002 | 0 | 0.63 | 0.8 | 0.58 | 0.65 | 0.61 | 0.65 |
| 2 | 85921353 | rs1866140 | 1.51 | 0.252 | -0.375 | -0.846 | -0.059 | -0.088 | -0.088 | 0.079 | 0.36 | 0.05 | 0.38 | 0.58 | 0.37 | 0.34 |
| 2 | 85921992 | rs2043760 | -0.034 | 0.225 | 0.517 | -0.794 | 0 | -0.011 | -0.011 | 0 | 0.63 | 0.53 | 0.57 | 0.65 | 0.66 | 0.69 |
| 2 | 85924088 | rs11887600 | -0.194 | 0.237 | 0.505 | -0.82 | -0.013 | -0.038 | -0.038 | 0 | 0.09 | 0.2 | 0.07 | 0 | 0.08 | 0.03 |
| 2 | 85924131 | rs11887686 | -0.313 | 0.233 | 0.502 | -0.822 | -0.014 | -0.04 | -0.04 | 0 | 0.09 | 0.2 | 0.07 | 0 | 0.08 | 0.03 |
| 2 | 85924729 | rs11127 | 1.618 | 0.112 | 0.861 | -0.899 | -0.077 | -0.066 | -0.066 | 0.546 | 0.37 | 0.06 | 0.4 | 0.59 | 0.4 | 0.36 |
| 2 | 85925680 | rs4240202 | 1.236 | 0.138 | 0.875 | -0.887 | -0.001 | -0.002 | -0.002 | 0.065 | 0.42 | 0.48 | 0.43 | 0.59 | 0.4 | 0.36 |
| 2 | 85932210 | rs12714151 | -0.209 | 0.131 | 0.717 | -0.852 | -0.001 | 0 | 0 | 0 | 0.64 | 0.62 | 0.57 | 0.64 | 0.64 | 0.58 |
| 2 | 85933367 | rs6547632 | -0.138 | 0.138 | 0.71 | -0.872 | -0.009 | 0 | 0 | 2.127 | 0.65 | 0.74 | 0.58 | 0.65 | 0.64 | 0.55 |
| 2 | 85934115 | rs17026595 | -0.434 | 0.473 | 1.265 | -0.709 | 0.021 | -0.087 | -0.087 | 0 | 0.18 | 0.44 | 0.11 | 0.05 | 0.16 | 0.17 |
| 2 | 85935282 | rs7577293 | -1.343 | 0.57 | 0.868 | -0.664 | -0.008 | -0.067 | -0.067 | 0.054 | 0.26 | 0.56 | 0.18 | 0.07 | 0.24 | 0.22 |
| 2 | 85938000 | rs4525696 | -0.077 | 0.922 | 0.483 | -0.412 | 0.013 | -0.038 | -0.038 | 0 | 0.14 | 0.2 | 0.09 | 0.04 | 0.16 | 0.17 |
| 2 | 85941074 | rs10188985 | 0.734 | 0.717 | 0.132 | -0.421 | -0.026 | 0.041 | 0.041 | 0.138 | 0.64 | 0.76 | 0.61 | 0.8 | 0.61 | 0.45 |
| 2 | 85946708 | rs751249 | 0.73 | 0.806 | 0.282 | -0.383 | -0.014 | 0.015 | 0.015 | 0.041 | 0.68 | 0.76 | 0.62 | 0.83 | 0.62 | 0.49 |
| 2 | 85948793 | rs6759087 | -0.35 | 1.028 | 0.662 | -0.457 | 0.11 | -0.091 | -0.091 | 0.062 | 0.17 | 0.43 | 0.24 | 0.04 | 0.22 | 0.22 |
| 2 | 85952162 | rs4832005 | -0.165 | 0.813 | -0.278 | -0.383 | -0.023 | 0.001 | 0.001 | 0 | 0.83 | 0.99 | 0.87 | 0.9 | 0.84 | 0.67 |
| 2 | 85954544 | rs2366812 | -0.528 | 0.758 | -0.084 | -0.497 | -0.003 | -0.016 | -0.016 | 0 | 0.23 | 0.23 | 0.21 | 0.05 | 0.26 | 0.2 |
| 2 | 85962551 | rs2164807 | 1.726 | 0.382 | -0.859 | 0.011 | -0.052 | -0.12 | -0.12 | 0.41 | 0.39 | 0.07 | 0.43 | 0.51 | 0.43 | 0.34 |
| 2 | 85963780 | rs6756074 | -0.579 | 0.446 | -0.213 | 0.179 | 0.002 | 0.029 | 0.029 | 0.417 | 0.18 | 0.44 | 0.19 | 0.22 | 0.15 | 0.09 |
| 2 | 85964356 | rs4832006 | -0.059 | 0.08 | 0.228 | 0.248 | 0.003 | -0.008 | -0.008 | 0.253 | 0.81 | 0.84 | 0.84 | 0.74 | 0.83 | 0.66 |
| 2 | 85965769 | rs1561283 | 0.224 | 0.209 | 0.154 | 0.167 | -0.072 | 0.094 | 0.094 | 0.472 | 0.2 | 0.03 | 0.18 | 0.05 | 0.23 | 0.2 |
| 2 | 85969520 | rs6743301 | 0.073 | 0.177 | 0.255 | 0.336 | -0.007 | 0.106 | 0.106 | 4.894 | 0.09 | 0.37 | 0.12 | 0.17 | 0.05 | 0.06 |
| 2 | 85970703 | rs13430978 | 0.703 | 0.16 | -0.237 | 0.303 | -0.038 | 0.025 | 0.025 | 0 | 0.62 | 0.83 | 0.66 | 0.7 | 0.61 | 0.48 |
| 2 | 85975594 | rs17026610 | -0.663 | -0.428 | 0.514 | 0.828 | -0.019 | 0.102 | 0.102 | 6.221 | 0.11 | 0.44 | 0.12 | 0.18 | 0.05 | 0.06 |
| 2 | 85976645 | rs1465823 | 0.604 | 0.085 | -0.483 | 0.687 | 0.003 | -0.006 | -0.006 | 0 | 0.82 | 0.87 | 0.85 | 0.79 | 0.86 | 0.68 |
| 2 | 85986112 | rs7558427 | 0.974 | -0.151 | -0.019 | 0.778 | -0.028 | 0.047 | 0.047 | 0.188 | 0.79 | 0.47 | 0.81 | 0.73 | 0.86 | 0.66 |
| 2 | 85995648 | rs4832198 | 0.143 | 0 | -0.706 | -0.217 | -0.023 | 0.008 | 0.008 | 0 | 0.35 | 0.31 | 0.32 | 0.02 | 0.48 | 0.31 |
| 2 | 86001255 | rs1465821 | -0.184 | -0.349 | 0.083 | 0.44 | -0.008 | 0.016 | 0.016 | 0 | 0.42 | 0.68 | 0.34 | 0.25 | 0.49 | 0.4 |
| 2 | 86001628 | rs6746541 | 0.608 | -0.548 | 0.394 | 0.501 | 0.002 | 0.002 | 0.002 | 0 | 0.46 | 0.47 | 0.52 | 0.71 | 0.38 | 0.32 |
| 2 | 86002303 | rs1048084 | 0.725 | -0.142 | 0.68 | 0.449 | -0.004 | -0.002 | -0.002 | 0.062 | 0.58 | 0.36 | 0.63 | 0.49 | 0.51 | 0.46 |
| 2 | 86004894 | rs934775 | -1.357 | 0.336 | 0.467 | -0.186 | 0.01 | 0.115 | 0.115 | 0 | 0.39 | 0.79 | 0.27 | 0.16 | 0.34 | 0.44 |
| 2 | 86005932 | rs4832199 | 0.332 | 0.304 | 0.885 | -0.255 | 0.007 | -0.081 | -0.081 | 0 | 0.22 | 0.25 | 0.17 | 0.01 | 0.24 | 0.17 |
| 2 | 86010243 | rs869778 | -1.715 | 0.194 | 0.995 | -0.127 | -0.025 | 0 | 0 | 0.527 | 0.16 | 0.48 | 0.09 | 0.14 | 0.1 | 0.27 |
| 2 | 86013029 | rs13386681 | -0.748 | 0.191 | 0.712 | -0.114 | -0.001 | 0.002 | 0.002 | 2.129 | 0.09 | 0.11 | 0.06 | 0.07 | 0.07 | 0.26 |
| 2 | 86015121 | rs13402783 | -1.359 | 0.171 | 0.663 | -0.147 | 0.01 | 0.023 | 0.023 | 0.012 | 0.13 | 0.07 | 0.07 | 0.08 | 0.09 | 0.25 |
| 2 | 86017065 | rs1914821 | -1.389 | 0.325 | -0.038 | -0.153 | -0.041 | 0.115 | 0.115 | 0.014 | 0.4 | 0.78 | 0.31 | 0.15 | 0.46 | 0.44 |
| 2 | 86018569 | rs13388951 | -0.184 | 0.55 | -0.016 | -0.176 | 0.026 | -0.093 | -0.093 | 0 | 0.26 | 0.55 | 0.22 | 0.01 | 0.37 | 0.17 |
| 2 | 86021919 | rs10180924 | -0.927 | 0.454 | -0.847 | 0.04 | -0.07 | 0.138 | 0.138 | 0.374 | 0.4 | 0.85 | 0.3 | 0.17 | 0.42 | 0.44 |
| 2 | 86022788 | rs1356626 | -1.128 | 0.719 | -0.76 | 0.431 | -0.029 | 0.066 | 0.066 | 0 | 0.46 | 0.8 | 0.35 | 0.31 | 0.5 | 0.5 |
| 2 | 86022929 | rs10207756 | -1.038 | 1.03 | 0.026 | 0.6 | 0.161 | 0.321 | 0.321 | 0.457 | 0.05 | 0.34 | 0.06 | 0.23 | 0.1 | 0.03 |
| 2 | 86023022 | rs10184538 | -1.164 | 0.857 | -1.158 | 0.101 | -0.019 | 0.135 | 0.135 | 0 | 0.45 | 0.8 | 0.32 | 0.24 | 0.45 | 0.44 |
| 2 | 86023398 | rs1878901 | -0.797 | 0.635 | -0.999 | 0.279 | 0.009 | 0.011 | 0.011 | 1.074 | 0.36 | 0.63 | 0.23 | 0.13 | 0.28 | 0.5 |
| 2 | 86023425 | rs1878902 | -1.165 | 0.677 | -1.948 | 0.471 | -0.041 | 0.024 | 0.024 | 0 | 0.46 | 0.85 | 0.31 | 0.37 | 0.45 | 0.55 |

| | | | | | | | | | | | | | | | | |
|---|---|---|---|---|---|---|---|---|---|---|---|---|---|---|---|---|
| 2 | 86025331 | rs6706567 | -1.437 | 0.13 | -1.063 | 0.27 | -0.024 | -0.011 | -0.011 | 1.905 | 0.14 | 0.72 | 0.08 | 0.11 | 0.08 | 0.3 |
| 2 | 86028257 | rs13405488 | -0.476 | 0.609 | -1.62 | 0.298 | -0.018 | 0.059 | 0.059 | 0 | 0.28 | 0.1 | 0.19 | 0.14 | 0.27 | 0.16 |
| 2 | 86032343 | rs11690161 | -1.053 | 1.001 | -2.32 | 0.523 | -0.056 | -0.002 | -0.002 | 2.622 | 0.51 | 0.96 | 0.4 | 0.42 | 0.5 | 0.57 |
| 2 | 86033879 | rs10203272 | -0.94 | 1.119 | -0.914 | 0.49 | 0.049 | 0.183 | 0.183 | 0.24 | 0.13 | 0.44 | 0.1 | 0.28 | 0.13 | 0.03 |
| 2 | 86038750 | rs1105882 | -1.38 | 0.766 | -1.641 | 0.772 | -0.085 | -0.134 | -0.134 | 1.42 | 0.56 | 0.96 | 0.46 | 0.54 | 0.53 | 0.59 |
| 2 | 86046509 | rs2139199 | 0.099 | 0.933 | -1.217 | 0.522 | -0.018 | 0.025 | 0.025 | 0.354 | 0.26 | 0.17 | 0.21 | 0.15 | 0.32 | 0.16 |
| 2 | 86047505 | rs6753123 | -0.006 | 0.84 | -0.534 | 0.598 | -0.011 | 0.003 | 0.003 | 0 | 0.18 | 0.25 | 0.14 | 0.26 | 0.14 | 0.42 |
| 2 | 86048593 | rs11676487 | -0.318 | 0.875 | -0.562 | 0.545 | 0 | 0.013 | 0.013 | 0 | 0.22 | 0.25 | 0.16 | 0.4 | 0.2 | 0.43 |
| 2 | 86051235 | rs1518987 | -1.172 | 0.873 | -0.222 | 0.484 | -0.007 | 0.015 | 0.015 | 0.049 | 0.45 | 0.41 | 0.34 | 0.4 | 0.45 | 0.57 |
| 2 | 86051597 | rs2049015 | -1.976 | 0.812 | -1.119 | 0.124 | -0.071 | -0.158 | -0.158 | 0.371 | 0.63 | 0.97 | 0.47 | 0.55 | 0.62 | 0.64 |
| 2 | 86055154 | rs11127000 | -1.59 | 0.771 | -1.071 | 0.081 | -0.056 | 0.013 | 0.013 | 0.152 | 0.59 | 0.91 | 0.42 | 0.4 | 0.55 | 0.64 |
| 2 | 86055946 | rs17431365 | 0.981 | 0.771 | -1.054 | -0.014 | -0.033 | 0.079 | 0.079 | 0.027 | 0.21 | 0.08 | 0.16 | 0.01 | 0.25 | 0.14 |
| 2 | 86061013 | rs2366946 | -1.74 | 0.841 | -1.021 | 0.108 | -0.025 | -0.078 | -0.078 | 1.916 | 0.62 | 0.86 | 0.45 | 0.56 | 0.6 | 0.65 |
| 2 | 86062425 | rs4832202 | -1.762 | 0.834 | -1.038 | 0.123 | -0.041 | -0.102 | -0.102 | 0 | 0.62 | 0.92 | 0.49 | 0.58 | 0.6 | 0.65 |
| 2 | 86062855 | rs2139200 | -1.736 | 0.841 | -0.913 | 0.127 | -0.016 | -0.063 | -0.063 | 0.59 | 0.62 | 0.85 | 0.48 | 0.58 | 0.6 | 0.65 |
| 2 | 86063044 | rs13420465 | -1.289 | 0.802 | -0.704 | 0.149 | 0.011 | -0.021 | -0.021 | 0 | 0.39 | 0.69 | 0.26 | 0.54 | 0.34 | 0.5 |
| 2 | 86063465 | rs11900847 | -1.331 | 0.753 | -0.611 | 0.213 | 0.023 | -0.023 | -0.023 | 0.871 | 0.39 | 0.73 | 0.25 | 0.53 | 0.32 | 0.5 |
| 2 | 86065256 | rs11127001 | -0.337 | 0.548 | 0.045 | 0.005 | -0.014 | -0.084 | -0.084 | 0 | 0.87 | 0.69 | 0.94 | 1 | 0.89 | 0.93 |
| 2 | 86074512 | rs13020072 | -0.646 | 1.036 | -0.049 | 0.333 | -0.001 | 0 | 0 | 0 | 0.28 | 0.24 | 0.23 | 0.54 | 0.3 | 0.47 |
| 2 | 86085145 | rs13025142 | 1.629 | 0.825 | -0.389 | 0.175 | -0.042 | -0.125 | -0.125 | 3.242 | 0.36 | 0.1 | 0.45 | 0.41 | 0.4 | 0.35 |
| 2 | 86086855 | rs17026701 | 0.417 | 0.839 | -0.2 | 0.17 | 0 | 0.044 | 0.044 | 0 | 0.05 | 0.1 | 0.06 | 0.14 | 0.05 | 0.02 |
| 2 | 86092253 | rs1533605 | 1.446 | 0.741 | 0.215 | 0.123 | -0.001 | 0.001 | 0.001 | 1.513 | 0.4 | 0.47 | 0.55 | 0.44 | 0.41 | 0.36 |
| 2 | 86094179 | rs4832211 | -0.629 | 0.763 | 0.279 | 0.075 | 0 | -0.054 | -0.054 | 0.02 | 0.69 | 0.71 | 0.79 | 0.98 | 0.7 | 0.82 |
| 2 | 86095201 | rs4553826 | 1.296 | 0.799 | 0.255 | 0.099 | -0.001 | 0.001 | 0.001 | 1.539 | 0.4 | 0.47 | 0.55 | 0.44 | 0.41 | 0.36 |
| 2 | 86096042 | rs12614498 | 0.74 | 0.798 | 0.181 | 0.113 | -0.016 | -0.03 | -0.03 | 0 | 0.18 | 0.06 | 0.15 | 0.29 | 0.19 | 0.37 |
| 2 | 86100563 | rs6547636 | 0.458 | 0.766 | 0.236 | 0.1 | 0.003 | 0.044 | 0.044 | 1.79 | 0.49 | 0.63 | 0.64 | 0.69 | 0.51 | 0.44 |
| 2 | 86101899 | rs2030275 | -0.367 | 0.773 | 0.245 | 0.107 | -0.008 | -0.019 | -0.019 | 0.056 | 0.26 | 0.17 | 0.23 | 0.54 | 0.3 | 0.46 |
| 2 | 86102178 | rs17026729 | 0.376 | 0.684 | 0.404 | 0.366 | -0.003 | 0.023 | 0.023 | 0 | 0.05 | 0.07 | 0.05 | 0.13 | 0.05 | 0.02 |
| 2 | 86104729 | rs6721125 | 1.042 | 0.677 | 0.443 | 0.354 | -0.001 | 0.001 | 0.001 | 0.515 | 0.39 | 0.46 | 0.56 | 0.44 | 0.41 | 0.36 |
| 2 | 86114252 | rs1878904 | 0.141 | 0.447 | 0.262 | 0.121 | 0 | -0.009 | -0.009 | 0 | 0.4 | 0.59 | 0.61 | 0.73 | 0.43 | 0.43 |
| 2 | 86117958 | rs12714154 | 1.516 | 0.428 | 0.322 | 0.235 | -0.019 | -0.089 | -0.089 | 0 | 0.27 | 0.13 | 0.52 | 0.47 | 0.33 | 0.35 |
| 2 | 86122874 | rs1401383 | 1.047 | 0.437 | 0.375 | 0.24 | -0.003 | 0.019 | 0.019 | 0 | 0.31 | 0.41 | 0.54 | 0.47 | 0.33 | 0.35 |
| 2 | 86125299 | rs2438307 | 0.246 | 0.57 | -0.1 | 0.311 | 0.015 | 0.077 | 0.077 | 0.111 | 0.9 | 0.82 | 0.92 | 0.74 | 0.9 | 0.92 |
| 2 | 86128563 | rs4832219 | 1.258 | 0.538 | -0.315 | 0.34 | -0.021 | -0.104 | -0.104 | 0 | 0.28 | 0.11 | 0.52 | 0.47 | 0.33 | 0.35 |
| 2 | 86139180 | rs4292089 | 0.132 | 0.564 | -0.089 | 0.411 | 0.014 | 0.093 | 0.093 | 0 | 0.06 | 0.15 | 0.06 | 0.15 | 0.05 | 0.02 |
| 2 | 86149532 | rs10185207 | 1.073 | 0.548 | -0.684 | 0.499 | -0.027 | -0.148 | -0.148 | 0.365 | 0.29 | 0.07 | 0.47 | 0.48 | 0.33 | 0.36 |
| 2 | 86155271 | rs6547647 | -0.715 | 0.609 | -0.44 | 0.461 | -0.006 | -0.021 | -0.021 | 0.041 | 0.65 | 0.76 | 0.5 | 0.43 | 0.61 | 0.56 |
| 2 | 86165443 | rs11691934 | 0.487 | 0.36 | -0.238 | 0.167 | 0.01 | -0.016 | -0.016 | 0 | 0.63 | 0.6 | 0.72 | 0.7 | 0.63 | 0.67 |
| 2 | 86165765 | rs10167259 | 0.19 | 0.347 | -0.21 | 0.084 | 0.002 | -0.029 | -0.029 | 0 | 0.25 | 0.22 | 0.2 | 0.04 | 0.26 | 0.16 |
| 2 | 86167439 | rs12714159 | 1.005 | -0.034 | 0.013 | 0.854 | 0 | -0.006 | -0.006 | 0 | 0.63 | 0.35 | 0.73 | 0.62 | 0.68 | 0.67 |
| 2 | 86167602 | rs13431245 | 0.24 | -0.021 | 0.131 | 0.38 | 0.004 | -0.069 | -0.069 | 0 | 0.22 | 0.25 | 0.18 | 0.02 | 0.21 | 0.16 |
| 2 | 86167678 | rs11689401 | 1.631 | 0.003 | -0.395 | 0.141 | 0.013 | -0.075 | -0.075 | 0 | 0.23 | 0.08 | 0.41 | 0.49 | 0.24 | 0.29 |
| 2 | 86167829 | rs2033770 | 0.606 | -0.068 | -0.399 | 0.015 | 0 | -0.004 | -0.004 | 0.018 | 0.67 | 0.38 | 0.75 | 0.72 | 0.74 | 0.71 |
| 2 | 86172274 | rs1370699 | 0.606 | -0.079 | -0.362 | 0.043 | 0.002 | 0.007 | 0.007 | 0.001 | 0.67 | 0.42 | 0.75 | 0.62 | 0.74 | 0.71 |
| 2 | 86177322 | rs735738 | 0.685 | -0.236 | -0.5 | -0.082 | -0.028 | 0.148 | 0.148 | 0 | 0.44 | 0.25 | 0.33 | 0.11 | 0.5 | 0.42 |
| 2 | 86184486 | rs6547651 | -0.681 | -0.199 | -0.318 | 0.07 | -0.001 | 0.053 | 0.053 | 0 | 0.7 | 0.81 | 0.53 | 0.15 | 0.67 | 0.55 |
| 2 | 86185291 | rs6713002 | -1.033 | -0.199 | -0.291 | 0.212 | -0.008 | -0.052 | -0.052 | 0 | 0.73 | 0.88 | 0.57 | 0.39 | 0.7 | 0.57 |
| 2 | 86187595 | rs1866242 | -0.391 | -0.216 | -0.146 | -0.116 | 0 | -0.009 | -0.009 | 0 | 0.68 | 0.7 | 0.52 | 0.15 | 0.67 | 0.55 |
| 2 | 86214360 | rs1866245 | -0.307 | -0.32 | -0.265 | -0.257 | 0.008 | 0.054 | 0.054 | 0.011 | 0.62 | 0.72 | 0.48 | 0.15 | 0.63 | 0.52 |
| 2 | 86226539 | rs1978842 | -0.831 | -0.34 | -0.348 | -0.392 | 0.029 | 0.054 | 0.054 | 0 | 0.29 | 0.79 | 0.2 | 0.12 | 0.14 | 0.21 |
| 2 | 86239772 | rs1469981 | 0.16 | -0.346 | -0.005 | -0.395 | -0.006 | -0.052 | -0.052 | 0 | 0.22 | 0.31 | 0.16 | 0.12 | 0.14 | 0.17 |
| 2 | 86248309 | rs4832233 | -1.178 | -0.346 | -0.197 | -0.275 | -0.049 | -0.096 | -0.096 | 0 | 0.72 | 0.95 | 0.53 | 0.46 | 0.67 | 0.58 |
| 2 | 86253581 | rs10173155 | -1.173 | -0.329 | -0.256 | -0.267 | -0.049 | -0.096 | -0.096 | 0.314 | 0.72 | 0.95 | 0.54 | 0.46 | 0.67 | 0.58 |
| 2 | 86258784 | rs2278536 | 1.098 | -0.513 | -0.196 | -0.208 | -0.166 | 0.175 | 0.175 | 0 | 0.38 | 0.03 | 0.29 | 0.12 | 0.49 | 0.36 |
| 2 | 86263288 | rs3770090 | -1.131 | -0.452 | 0.244 | -0.025 | -0.005 | 0.168 | 0.168 | 0.027 | 0.1 | 0.54 | 0.08 | 0.23 | 0.04 | 0.06 |
| 2 | 86292656 | rs311563 | 0.063 | -0.432 | 0.213 | -0.11 | -0.011 | 0.076 | 0.076 | 0 | 0.84 | 0.71 | 0.87 | 0.66 | 0.88 | 0.86 |
| 2 | 86296566 | rs311572 | 0.302 | -0.436 | 0.048 | -0.112 | -0.016 | 0.085 | 0.085 | 0 | 0.83 | 0.68 | 0.86 | 0.66 | 0.88 | 0.86 |
| 2 | 86299122 | rs311579 | 0.587 | -0.479 | -0.244 | -0.251 | -0.098 | -0.087 | -0.087 | 0 | 0.36 | 0.05 | 0.52 | 0.54 | 0.39 | 0.47 |
| 2 | 86302103 | rs3815840 | 0.002 | -0.409 | 0.017 | -0.122 | 0.003 | 0.063 | 0.063 | 0 | 0.85 | 0.71 | 0.86 | 0.75 | 0.89 | 0.9 |
| 2 | 86307128 | rs12328100 | 1.452 | -0.297 | -0.341 | -0.323 | -0.076 | 0.164 | 0.164 | 0 | 0.35 | 0.05 | 0.35 | 0.11 | 0.43 | 0.32 |
| 2 | 86307624 | rs17584578 | 0.312 | -0.345 | 0.176 | -0.108 | 0.008 | 0.139 | 0.139 | 0 | 0.06 | 0.24 | 0.04 | 0.23 | 0.04 | 0.02 |
| 2 | 86310110 | rs2288118 | 1.085 | -0.375 | 0.216 | -0.162 | -0.024 | 0.021 | 0.021 | 0.052 | 0.77 | 0.41 | 0.83 | 0.66 | 0.84 | 0.82 |
| 2 | 86315741 | rs2288120 | 1.347 | -0.376 | 0.207 | -0.163 | -0.01 | 0.004 | 0.004 | 0.2 | 0.75 | 0.31 | 0.82 | 0.66 | 0.84 | 0.82 |
| 2 | 86330726 | rs10779967 | -0.947 | -0.314 | 0.269 | -0.186 | -0.002 | -0.049 | -0.049 | 0 | 0.88 | 0.78 | 0.89 | 0.98 | 0.91 | 0.92 |
| 2 | 86333193 | rs2303340 | 0.575 | -0.345 | 0.338 | -0.146 | -0.003 | 0.098 | 0.098 | 0 | 0.06 | 0.18 | 0.04 | 0.23 | 0.04 | 0.02 |
| 2 | 86335208 | rs11679343 | 0.854 | -0.365 | 0.401 | -0.148 | -0.021 | 0.054 | 0.054 | 0.018 | 0.79 | 0.52 | 0.83 | 0.66 | 0.84 | 0.86 |
| 2 | 86343316 | rs17618119 | 1.312 | -0.323 | 0.076 | -0.046 | -0.056 | -0.117 | -0.117 | 0 | 0.16 | 0.02 | 0.16 | 0.41 | 0.16 | 0.25 |
| 2 | 86345536 | rs2367202 | 0.538 | -0.534 | 0.322 | -0.451 | -0.01 | 0.081 | 0.081 | 0.089 | 0.8 | 0.54 | 0.84 | 0.66 | 0.84 | 0.86 |
| 2 | 86346180 | rs2303339 | 0.574 | -0.528 | 0.29 | -0.41 | -0.003 | 0.066 | 0.066 | 0 | 0.06 | 0.18 | 0.04 | 0.14 | 0.04 | 0.02 |

| | | | | | | | | | | | | | | | |
|---|---|---|---|---|---|---|---|---|---|---|---|---|---|---|---|
| 2 | 86348960 | rs4832249 | 1.186 | -0.549 | 0.294 | -0.457 | -0.021 | 0.015 | 0.015 | 0.019 | 0.76 | 0.39 | 0.83 | 0.66 | 0.84 | 0.82 |
| 2 | 86362204 | rs7606421 | 1.13 | -0.553 | 0.302 | -0.459 | -0.025 | 0.023 | 0.023 | 0 | 0.77 | 0.39 | 0.83 | 0.66 | 0.84 | 0.82 |
| 2 | 86365607 | rs11395 | 1.173 | -0.612 | 0.107 | -0.577 | -0.092 | 0.172 | 0.172 | 0 | 0.39 | 0.06 | 0.37 | 0.16 | 0.48 | 0.35 |
| 2 | 86365884 | rs7598787 | 1.236 | -0.659 | 0.095 | -0.716 | -0.015 | 0.004 | 0.004 | 0.209 | 0.74 | 0.36 | 0.82 | 0.66 | 0.81 | 0.8 |
| 2 | 86368392 | rs10174095 | 0.317 | -0.64 | 0.114 | -0.711 | -0.018 | 0.08 | 0.08 | 0 | 0.06 | 0.18 | 0.04 | 0.22 | 0.03 | 0.02 |
| 2 | 86369700 | rs4832027 | -0.438 | -0.628 | 0.145 | -0.723 | -0.008 | -0.066 | -0.066 | 0.092 | 0.86 | 0.75 | 0.88 | 0.98 | 0.88 | 0.9 |
| 2 | 86371012 | rs876242 | 0.737 | -0.611 | 0.108 | -0.7 | -0.001 | -0.027 | -0.027 | 0 | 0.44 | 0.5 | 0.41 | 0.15 | 0.48 | 0.39 |
| 2 | 86376089 | rs1124685 | 0.183 | -0.752 | -0.156 | -0.69 | -0.081 | -0.1 | -0.1 | 0.397 | 0.31 | 0.05 | 0.43 | 0.59 | 0.35 | 0.4 |
| 2 | 86376474 | rs1124686 | 0.813 | -0.05 | 0.839 | -0.57 | -0.067 | 0.197 | 0.197 | 0 | 0.41 | 0.09 | 0.39 | 0.14 | 0.5 | 0.41 |
| 2 | 86383171 | rs6547660 | 0.674 | 0.078 | 1.233 | -0.445 | -0.012 | 0.054 | 0.054 | 0 | 0.4 | 0.31 | 0.39 | 0.15 | 0.48 | 0.35 |
| 2 | 86398785 | rs12477274 | 0.844 | 0.071 | 1.221 | -0.448 | -0.083 | 0.175 | 0.175 | 0 | 0.39 | 0.06 | 0.37 | 0.15 | 0.48 | 0.35 |
| 2 | 86400084 | rs6707029 | 0.508 | 0.09 | 1.181 | -0.438 | -0.001 | -0.001 | -0.001 | 0 | 0.43 | 0.44 | 0.4 | 0.15 | 0.48 | 0.39 |
| 2 | 86405804 | rs6749090 | 0.182 | 0.169 | 1.139 | -0.398 | 0 | -0.027 | -0.027 | 0.121 | 0.47 | 0.52 | 0.42 | 0.25 | 0.51 | 0.44 |
| 2 | 86406556 | rs17027011 | -0.229 | 0.159 | 1.183 | -0.39 | -0.009 | 0.077 | 0.077 | 0 | 0.06 | 0.18 | 0.04 | 0.22 | 0.04 | 0.02 |
| 2 | 86422816 | rs9653508 | 0.145 | 0.212 | 1.127 | -0.462 | 0 | -0.027 | -0.027 | 0.232 | 0.47 | 0.52 | 0.42 | 0.25 | 0.51 | 0.44 |
| 2 | 86436067 | rs3770059 | 0.179 | 0.401 | 1.156 | -0.654 | -0.006 | 0.08 | 0.08 | 0 | 0.47 | 0.35 | 0.57 | 0.86 | 0.43 | 0.53 |
| 2 | 86436514 | rs1540391 | 0.821 | 0.446 | 0.611 | -0.7 | -0.055 | 0.159 | 0.159 | 0 | 0.37 | 0.14 | 0.28 | 0.12 | 0.46 | 0.36 |
| 2 | 86437551 | rs7565645 | 0.905 | 0.14 | 0.847 | -0.669 | -0.036 | -0.071 | -0.071 | 0.211 | 0.83 | 0.31 | 0.85 | 0.99 | 0.92 | 0.92 |
| 2 | 86438146 | rs2241432 | -1.159 | 0.279 | 0.429 | -0.527 | -0.114 | -0.068 | -0.068 | 2.4 | 0.63 | 0.98 | 0.47 | 0.45 | 0.63 | 0.54 |
| 2 | 86438600 | rs17438451 | 0.61 | 0.326 | 0.458 | -0.834 | -0.122 | 0.016 | 0.016 | 0 | 0.4 | 0.02 | 0.53 | 0.66 | 0.4 | 0.53 |
| 2 | 86447273 | rs7583021 | -0.358 | 0.262 | 0.942 | -0.783 | -0.003 | -0.049 | -0.049 | 0 | 0.89 | 0.77 | 0.89 | 0.98 | 0.91 | 0.93 |
| 2 | 86448231 | rs749581 | 1.467 | 0.194 | 0.75 | -0.774 | 0.029 | -0.008 | -0.008 | 0 | 0.77 | 0.17 | 0.82 | 0.69 | 0.87 | 0.84 |
| 2 | 86451127 | rs7422930 | 0.863 | 0.24 | 0.622 | -0.883 | -0.054 | 0.156 | 0.156 | 0 | 0.37 | 0.15 | 0.29 | 0.09 | 0.47 | 0.38 |
| 2 | 86451502 | rs1010239 | 0.883 | 0.239 | 0.637 | -0.886 | -0.086 | -0.003 | -0.003 | 0 | 0.38 | 0.06 | 0.53 | 0.6 | 0.37 | 0.46 |
| 2 | 86453919 | rs4832257 | 1.433 | 0.169 | 0.762 | -0.844 | 0.012 | 0.001 | 0.001 | 2.082 | 0.78 | 0.22 | 0.82 | 0.78 | 0.87 | 0.88 |
| 2 | 86454914 | rs11127018 | 1.19 | 0.227 | 1.034 | -1.048 | -0.039 | -0.079 | -0.079 | 0 | 0.81 | 0.43 | 0.85 | 0.91 | 0.88 | 0.85 |
| 2 | 86462747 | rs6758173 | -0.216 | 0.333 | 0.985 | -1.01 | 0 | 0.009 | 0.009 | 0 | 0.5 | 0.58 | 0.34 | 0.35 | 0.54 | 0.46 |
| 2 | 86473740 | rs718292 | 0.705 | 0.125 | 0.761 | -1.002 | -0.133 | 0.007 | 0.007 | 0.175 | 0.39 | 0.04 | 0.59 | 0.59 | 0.46 | 0.43 |
| 2 | 86475443 | rs6547667 | -1.39 | 0.056 | 1.12 | -1.022 | 0.02 | 0.015 | 0.015 | 2.145 | 0.07 | 0.56 | 0.04 | 0 | 0 | 0.03 |
| 2 | 86476588 | rs6714362 | 0.671 | 0.046 | 0.985 | -1.068 | -0.039 | 0.083 | 0.083 | 0 | 0.39 | 0.1 | 0.24 | 0.29 | 0.41 | 0.4 |
| 2 | 86479287 | rs17438687 | 0.475 | 0.158 | 0.96 | -0.958 | -0.085 | 0.065 | 0.065 | 0.078 | 0.42 | 0.08 | 0.61 | 0.66 | 0.5 | 0.49 |
| 2 | 86481835 | rs2276625 | 0.651 | 0.181 | 1.04 | -0.944 | -0.045 | 0.039 | 0.039 | 0.004 | 0.4 | 0.18 | 0.6 | 0.6 | 0.46 | 0.43 |
| 2 | 86488265 | rs4832265 | 0.682 | 0.188 | 0.723 | -0.892 | -0.135 | -0.024 | -0.024 | 0.158 | 0.38 | 0.04 | 0.6 | 0.58 | 0.46 | 0.43 |
| 2 | 86502663 | rs11687392 | 0.584 | 0.198 | 0.73 | -0.886 | -0.085 | 0.011 | 0.011 | 0.209 | 0.39 | 0.11 | 0.6 | 0.59 | 0.46 | 0.43 |
| 2 | 86509388 | rs1863059 | -0.074 | 0.225 | 0.869 | -0.895 | -0.001 | 0.01 | 0.01 | 0 | 0.48 | 0.61 | 0.66 | 0.65 | 0.5 | 0.52 |
| 2 | 86523440 | rs1863065 | 0.63 | 0.125 | 1.025 | -0.772 | -0.023 | -0.053 | -0.053 | 0 | 0.85 | 0.56 | 0.88 | 0.95 | 0.91 | 0.91 |
| 2 | 86526127 | rs6760222 | -0.066 | 0.069 | 1.14 | -0.81 | -0.001 | -0.01 | -0.01 | 0 | 0.05 | 0.2 | 0.03 | 0.06 | 0.04 | 0.09 |
| 2 | 86528938 | rs6547673 | 0.097 | 0.112 | 0.964 | -1.03 | 0 | 0.003 | 0.003 | 0 | 0.46 | 0.46 | 0.28 | 0.36 | 0.45 | 0.49 |
| 2 | 86531748 | rs11127026 | 0.767 | 0.095 | 1.02 | -1.038 | -0.017 | 0.064 | 0.064 | 0 | 0.4 | 0.14 | 0.24 | 0.3 | 0.42 | 0.4 |
| 2 | 86533551 | rs6718596 | 0.175 | 0.205 | 1.211 | -1.083 | 0.002 | -0.041 | -0.041 | 0 | 0.87 | 0.75 | 0.86 | 0.95 | 0.84 | 0.91 |
| 2 | 86534521 | rs13024436 | 0.775 | 0.341 | 1.001 | -0.947 | -0.117 | -0.03 | -0.03 | 0 | 0.36 | 0.03 | 0.55 | 0.59 | 0.39 | 0.42 |
| 2 | 86538070 | rs6716825 | -0.629 | 0.374 | 1.032 | -0.914 | 0.017 | 0.005 | 0.005 | 0 | 0.06 | 0.31 | 0.04 | 0.06 | 0.04 | 0.09 |
| 2 | 86558975 | rs11675985 | -0.69 | 0.459 | 0.732 | -0.865 | 0.013 | -0.025 | -0.025 | 0 | 0.12 | 0.4 | 0.09 | 0.05 | 0.12 | 0.07 |
| 2 | 86559148 | rs1863051 | -0.546 | 0.452 | -0.167 | -0.921 | 0.003 | 0.071 | 0.071 | 0 | 0.55 | 0.74 | 0.68 | 0.7 | 0.54 | 0.58 |
| 2 | 86566701 | rs12471787 | 0.42 | 0.617 | -0.696 | -1.647 | -0.138 | 0.171 | 0.171 | 0 | 0.37 | 0.03 | 0.56 | 0.83 | 0.42 | 0.48 |
| 2 | 86571415 | rs13401223 | -0.902 | 0.216 | -0.52 | -2.165 | 0.039 | 0.052 | 0.052 | 3.121 | 0.29 | 0.83 | 0.15 | 0.14 | 0.17 | 0.17 |
| 2 | 86583240 | rs6756078 | -0.346 | 0.176 | 0.562 | -2.155 | 0.022 | 0.055 | 0.055 | 0.153 | 0.07 | 0.33 | 0.04 | 0.08 | 0.03 | 0.02 |
| 2 | 86583721 | rs12714182 | -0.69 | 0.246 | 0.252 | -2.285 | -0.028 | 0.137 | 0.137 | 0 | 0.62 | 0.86 | 0.68 | 0.91 | 0.59 | 0.58 |
| 2 | 86584250 | rs13405197 | -0.768 | 0.016 | 0.613 | -2.266 | -0.013 | -0.035 | -0.035 | 0 | 0.26 | 0.69 | 0.12 | 0.08 | 0.14 | 0.1 |
| 2 | 86596001 | rs6756146 | -0.627 | 0.181 | 0.51 | -2.276 | -0.005 | 0.121 | 0.121 | 0 | 0.62 | 0.84 | 0.68 | 0.93 | 0.6 | 0.56 |
| 2 | 86599296 | rs2113608 | 0.236 | 0.263 | 1.016 | -2.258 | -0.004 | 0.004 | 0.004 | 0 | 0.46 | 0.64 | 0.62 | 0.93 | 0.49 | 0.53 |
| 2 | 86611284 | rs1033289 | 0.267 | 0.285 | 1.223 | -2.261 | -0.006 | 0.029 | 0.029 | 0 | 0.46 | 0.64 | 0.62 | 0.93 | 0.49 | 0.53 |
| 2 | 86612651 | rs908791 | 0.991 | 0.32 | 1.18 | -2.224 | -0.032 | 0.064 | 0.064 | 0.146 | 0.36 | 0.23 | 0.54 | 0.79 | 0.43 | 0.42 |
| 2 | 86612691 | rs908790 | -0.555 | 0.309 | 1.078 | -2.229 | -0.004 | 0.145 | 0.145 | 0 | 0.61 | 0.85 | 0.68 | 0.93 | 0.6 | 0.56 |
| 2 | 86627316 | rs1105865 | -1.099 | 0.346 | 1.162 | -2.118 | 0.039 | 0.059 | 0.059 | 0 | 0.05 | 0.19 | 0.06 | 0.15 | 0.06 | 0.05 |
| 2 | 86630192 | rs12617075 | 1.043 | 0.376 | 0.696 | -2.215 | -0.065 | 0.121 | 0.121 | 0.353 | 0.36 | 0.17 | 0.54 | 0.78 | 0.43 | 0.42 |
| 2 | 86630752 | rs11903855 | -0.344 | 0.439 | 0.816 | -2.114 | 0.005 | 0.089 | 0.089 | 0 | 0.6 | 0.82 | 0.68 | 0.93 | 0.6 | 0.56 |
| 2 | 86639100 | rs11678201 | 0.007 | 0.466 | 0.862 | -2.09 | 0.006 | 0.004 | 0.004 | 0 | 0.55 | 0.43 | 0.63 | 0.87 | 0.57 | 0.47 |
| 2 | 86643733 | rs11894251 | -1.223 | 0.571 | 0.999 | -2.354 | -0.003 | -0.052 | -0.052 | 0 | 0.33 | 0.68 | 0.21 | 0.15 | 0.3 | 0.11 |
| 2 | 86649747 | rs1033287 | -1.104 | 0.847 | 0.927 | -2.546 | -0.003 | -0.051 | -0.051 | 0.468 | 0.33 | 0.67 | 0.21 | 0.15 | 0.3 | 0.11 |
| 2 | 86653699 | rs17027257 | -1.091 | 0.884 | 1.328 | -2.586 | 0.004 | -0.062 | -0.062 | 1.936 | 0.16 | 0.55 | 0.14 | 0.15 | 0.16 | 0.09 |
| 2 | 86677496 | rs11682587 | -0.143 | 0.845 | 1.79 | -2.601 | 0.008 | -0.033 | -0.033 | 0 | 0.1 | 0.25 | 0.09 | 0.07 | 0.14 | 0.07 |
| 2 | 86693826 | rs34605051 | 0.632 | 1.119 | 1.811 | -2.54 | -0.001 | -0.038 | -0.038 | 0 | 0.16 | 0.12 | 0.07 | 0 | 0.14 | 0.02 |
| 2 | 86715696 | rs1518812 | 0.209 | 1.124 | 1.509 | -2.59 | -0.008 | -0.122 | -0.122 | 0 | 0.19 | 0.33 | 0.09 | 0 | 0.14 | 0.02 |
| 2 | 86732524 | rs3201448 | 1.284 | 1.064 | 1.406 | -2.394 | -0.101 | 0.138 | 0.138 | 1.466 | 0.38 | 0.09 | 0.46 | 0.73 | 0.46 | 0.48 |
| 2 | 86737649 | rs966477 | 0.16 | 1.071 | 1.379 | -2.374 | -0.004 | -0.094 | -0.094 | 0 | 0.54 | 0.57 | 0.68 | 0.94 | 0.63 | 0.57 |
| 2 | 86766153 | rs7557796 | -0.094 | 1.073 | 1.226 | -2.367 | 0.005 | 0.03 | 0.03 | 0 | 0.62 | 0.63 | 0.58 | 0.78 | 0.63 | 0.52 |
| 2 | 86790716 | rs17027296 | 0.154 | 1.076 | 1.139 | -2.329 | -0.007 | -0.142 | -0.142 | 0 | 0.19 | 0.37 | 0.09 | 0 | 0.14 | 0.02 |
| 2 | 86793121 | rs6752371 | 0.069 | 1.083 | 0.64 | -2.327 | 0.005 | 0.015 | 0.015 | 0 | 0.58 | 0.56 | 0.55 | 0.73 | 0.6 | 0.5 |

| | | | | | | | | | | | | | | | | |
|---|---|---|---|---|---|---|---|---|---|---|---|---|---|---|---|---|
| 2 | 86793868 | rs13015620 | 1.083 | 1.147 | 0.091 | -2.222 | -0.101 | 0.135 | 0.135 | 1.356 | 0.38 | 0.09 | 0.46 | 0.73 | 0.46 | 0.48 |
| 2 | 86818630 | rs10184879 | 1.085 | 1.152 | 0.094 | -2.22 | -0.101 | 0.135 | 0.135 | 1.433 | 0.38 | 0.09 | 0.46 | 0.72 | 0.46 | 0.48 |
| 2 | 86819096 | rs11894098 | 0.048 | 1.073 | 0.207 | -2.144 | -0.001 | -0.005 | -0.005 | 0.109 | 0.09 | 0.1 | 0.06 | 0.01 | 0.1 | 0.04 |
| 2 | 86826508 | rs11127047 | -0.085 | 0.995 | 0.4 | -2.006 | 0.005 | 0.039 | 0.039 | 0 | 0.62 | 0.67 | 0.57 | 0.76 | 0.62 | 0.52 |
| 2 | 86841775 | rs308907 | -0.211 | 0.631 | -0.212 | -2.201 | 0.027 | 0.079 | 0.079 | 0 | 0.64 | 0.79 | 0.7 | 0.94 | 0.69 | 0.55 |
| 2 | 86846489 | rs308901 | -0.252 | 0.616 | -0.27 | -2.197 | 0.029 | 0.096 | 0.096 | 0 | 0.65 | 0.82 | 0.72 | 0.92 | 0.7 | 0.57 |
| 2 | 86850022 | rs396354 | -0.252 | 0.616 | -0.271 | -2.2 | 0.029 | 0.088 | 0.088 | 0 | 0.65 | 0.81 | 0.71 | 0.94 | 0.7 | 0.57 |
| 2 | 86853529 | rs433655 | 0.133 | 0.578 | -0.306 | -2.382 | 0.029 | 0.196 | 0.196 | 0 | 0.56 | 0.86 | 0.7 | 0.94 | 0.62 | 0.56 |
| 2 | 86855977 | rs308911 | -0.226 | 0.771 | -0.44 | -2.192 | 0.025 | 0.091 | 0.091 | 0 | 0.65 | 0.8 | 0.71 | 0.94 | 0.69 | 0.56 |
| 2 | 86877360 | rs7564251 | -1.234 | 0.614 | 0.366 | -1.608 | -0.022 | -0.01 | -0.01 | 0 | 0.18 | 0.3 | 0.17 | 0.18 | 0.14 | 0.12 |
| 2 | 86923178 | rs7558827 | -0.398 | -0.147 | 0.307 | -2.765 | 0.005 | 0.015 | 0.015 | 0.925 | 0.16 | 0.33 | 0.19 | 0.18 | 0.15 | 0.12 |
| 2 | 88410731 | rs1441640 | -0.826 | -0.679 | -0.781 | -0.318 | 0.01 | 0.009 | 0.009 | 0.004 | 0.69 | 0.81 | 0.59 | 0.78 | 0.72 | 0.8 |
| 2 | 88411904 | rs2970889 | 0.043 | -0.732 | -0.497 | 0.033 | -0.046 | 0.032 | 0.032 | 1.613 | 0.31 | 0.57 | 0.16 | 0.58 | 0.18 | 0.55 |
| 2 | 88415087 | rs1441642 | 0.844 | 0.428 | -1.672 | -0.563 | -0.079 | 0.018 | 0.018 | 0 | 0.27 | 0.09 | 0.29 | 0.14 | 0.35 | 0.16 |
| 2 | 88415739 | rs2970892 | -0.534 | 0.135 | -1.51 | -0.637 | 0.016 | 0.024 | 0.024 | 0.281 | 0.61 | 0.26 | 0.54 | 0.77 | 0.68 | 0.76 |
| 2 | 88417387 | rs2970895 | -0.899 | 0.157 | -1.397 | -0.447 | -0.001 | 0.006 | 0.006 | 0 | 0.48 | 0.57 | 0.32 | 0.61 | 0.47 | 0.66 |
| 2 | 88423510 | rs1545223 | 0.095 | 0.488 | -2.206 | -0.163 | -0.024 | 0.017 | 0.017 | 0.015 | 0.23 | 0.14 | 0.35 | 0.21 | 0.35 | 0.14 |
| 2 | 88424066 | rs2241883 | -0.524 | 1.046 | -1.466 | 0.013 | -0.025 | 0.019 | 0.019 | 0.011 | 0.24 | 0.14 | 0.35 | 0.21 | 0.35 | 0.14 |
| 2 | 88428571 | rs2919872 | 0.719 | 0.657 | -1.215 | 0.159 | -0.026 | 0.031 | 0.031 | 0 | 0.38 | 0.22 | 0.48 | 0.19 | 0.47 | 0.22 |
| 2 | 88432981 | rs2860399 | 0.283 | 0.809 | -1.147 | 0.247 | -0.001 | 0.001 | 0.001 | 0 | 0.68 | 0.35 | 0.65 | 0.46 | 0.73 | 0.42 |
| 2 | 88433305 | rs2970924 | -0.091 | 0.807 | -0.771 | 0.381 | 0 | 0.009 | 0.009 | 0 | 0.89 | 0.84 | 0.77 | 0.86 | 0.87 | 0.76 |
| 2 | 88438050 | rs4246598 | 1.33 | 0.703 | -0.531 | 0.278 | -0.006 | 0.022 | 0.022 | 0.36 | 0.43 | 0.33 | 0.48 | 0.21 | 0.48 | 0.23 |
| 2 | 88446892 | rs10865489 | 0.91 | 1.19 | -0.338 | 0.287 | 0.065 | 0.034 | 0.034 | 0 | 0.83 | 0.64 | 0.51 | 0.78 | 0.76 | 0.73 |
| 2 | 88460399 | rs13010480 | 1.819 | 1.075 | -0.395 | 0.226 | -0.015 | 0.075 | 0.075 | 0.695 | 0.37 | 0.08 | 0.21 | 0.13 | 0.37 | 0.2 |
| 2 | 88463237 | rs10185660 | 0.07 | 1.043 | -0.057 | 0.442 | -0.012 | 0.132 | 0.132 | 0 | 0.22 | 0.41 | 0.13 | 0.47 | 0.14 | 0.38 |
| 2 | 88464840 | rs4359651 | 1.037 | 1.154 | -0.065 | 0.488 | 0.058 | 0.033 | 0.033 | 0 | 0.83 | 0.64 | 0.51 | 0.78 | 0.76 | 0.73 |
| 2 | 88464926 | rs4246599 | -0.461 | 1.15 | -0.142 | 0.477 | 0.017 | 0.006 | 0.006 | 0 | 0.92 | 0.83 | 0.78 | 0.93 | 0.88 | 0.89 |
| 2 | 88465476 | rs4633926 | -0.71 | 1.167 | -0.328 | 0.486 | -0.005 | -0.002 | -0.002 | 1.43 | 0.93 | 0.94 | 0.79 | 0.93 | 0.88 | 0.89 |
| 2 | 88471796 | rs11683461 | -0.216 | 1.203 | -0.176 | 0.482 | -0.004 | -0.002 | -0.002 | 0.141 | 0.21 | 0.16 | 0.36 | 0.55 | 0.24 | 0.41 |
| 2 | 88472280 | rs7569216 | -0.62 | 0.935 | -0.258 | -0.217 | -0.004 | 0 | 0 | 1.875 | 0.93 | 0.94 | 0.78 | 0.94 | 0.88 | 0.91 |
| 2 | 88472791 | rs4129190 | -0.11 | 0.919 | -0.252 | -0.215 | -0.002 | -0.003 | -0.003 | 0 | 0.92 | 0.91 | 0.78 | 0.86 | 0.88 | 0.87 |
| 2 | 88476288 | rs2176569 | 0.245 | 0.538 | -0.144 | -0.259 | -0.001 | -0.019 | -0.019 | 1.444 | 0.23 | 0.28 | 0.15 | 0.18 | 0.22 | 0.14 |
| 2 | 88476616 | rs1136562 | 0.237 | 0.538 | -0.143 | -0.259 | -0.001 | -0.019 | -0.019 | 0 | 0.24 | 0.28 | 0.15 | 0.19 | 0.22 | 0.14 |
| 2 | 88484808 | rs2292869 | 0.14 | 0.517 | 0.161 | 0.548 | -0.042 | -0.095 | -0.095 | 0 | 0.13 | 0.01 | 0.1 | 0.56 | 0.13 | 0.37 |
| 2 | 88492058 | rs1914733 | 1.282 | 0.511 | 0.378 | 0.463 | 0.004 | 0.004 | 0.004 | 0 | 0.72 | 0.47 | 0.65 | 0.38 | 0.72 | 0.53 |
| 2 | 88494994 | rs12621922 | -0.368 | 0.585 | 0.345 | 0.644 | -0.001 | 0.017 | 0.017 | 0.237 | 0.15 | 0.13 | 0.12 | 0.56 | 0.13 | 0.35 |
| 2 | 88498277 | rs1878813 | -0.364 | 0.568 | 0.362 | 0.566 | -0.001 | 0 | 0 | 0 | 0.92 | 0.91 | 0.79 | 0.94 | 0.87 | 0.9 |
| 2 | 88514990 | rs7571019 | 0.903 | 0.386 | 0.387 | 0.524 | 0 | 0 | 0 | 0.387 | 0.69 | 0.74 | 0.65 | 0.37 | 0.71 | 0.51 |
| 2 | 88521793 | rs2363731 | -0.033 | 0.673 | 0.253 | 0.299 | 0.024 | 0.003 | 0.003 | 0 | 0.08 | 0.1 | 0.28 | 0.07 | 0.12 | 0.06 |
| 2 | 88525965 | rs4260241 | -0.397 | -0.191 | 0.319 | 0.813 | -0.001 | 0.021 | 0.021 | 0.425 | 0.14 | 0.09 | 0.11 | 0.44 | 0.13 | 0.36 |
| 2 | 88531886 | rs11683364 | -0.446 | -0.014 | -0.398 | 0.903 | 0 | 0.048 | 0.048 | 0.42 | 0.14 | 0.12 | 0.12 | 0.44 | 0.13 | 0.36 |
| 2 | 88537343 | rs11898335 | -0.694 | -0.061 | -0.542 | 0.879 | -0.005 | 0.134 | 0.134 | 0.166 | 0.17 | 0.27 | 0.13 | 0.45 | 0.15 | 0.45 |
| 2 | 88539371 | rs11127090 | 0.056 | -0.053 | -0.645 | 0.651 | -0.004 | 0.018 | 0.018 | 0 | 0.85 | 0.93 | 0.78 | 0.94 | 0.84 | 0.87 |
| 2 | 88541064 | rs11689291 | -0.868 | -0.009 | -0.361 | 0.919 | 0.001 | 0.203 | 0.203 | 0.268 | 0.16 | 0.31 | 0.16 | 0.47 | 0.14 | 0.4 |
| 2 | 88541437 | rs4972049 | 0.356 | 0.185 | -1.348 | 0.006 | 0.006 | 0.078 | 0.078 | 0 | 0.69 | 0.83 | 0.71 | 0.93 | 0.7 | 0.81 |
| 2 | 88541669 | rs11127091 | -0.707 | -0.149 | -1.369 | 0.105 | -0.012 | -0.035 | -0.035 | 0 | 0.39 | 0.13 | 0.39 | 0.54 | 0.37 | 0.45 |
| 2 | 88549952 | rs13431169 | -1.201 | -0.889 | -0.471 | -0.62 | -0.01 | -0.036 | -0.036 | 0.455 | 0.25 | 0.69 | 0.27 | 0.04 | 0.15 | 0.22 |
| 2 | 88554351 | rs6707137 | -1.229 | -0.758 | 0.723 | -0.504 | 0.059 | 0.01 | 0.01 | 0.516 | 0.07 | 0.31 | 0.06 | 0.04 | 0.04 | 0.05 |
| 2 | 88557666 | rs2139099 | -2.137 | -0.899 | 1.058 | -0.457 | -0.039 | -0.058 | -0.058 | 0.013 | 0.2 | 0.64 | 0.12 | 0.06 | 0.11 | 0.17 |
| 2 | 88559331 | rs7606234 | -1.061 | -0.95 | 1.328 | -0.459 | -0.051 | -0.043 | -0.043 | 0 | 0.13 | 0.31 | 0.05 | 0.02 | 0.07 | 0.12 |
| 2 | 88574827 | rs6728787 | -1.635 | -0.92 | 1.606 | -0.468 | -0.072 | -0.066 | -0.066 | 0.099 | 0.15 | 0.46 | 0.08 | 0.02 | 0.06 | 0.14 |
| 2 | 88591175 | rs6722369 | -1.419 | -1.041 | 1.172 | -0.528 | -0.017 | -0.01 | -0.01 | 1.706 | 0.15 | 0.21 | 0.09 | 0.08 | 0.11 | 0.14 |
| 2 | 88596890 | rs1271103 | -1.56 | -0.818 | 1.246 | -0.485 | -0.004 | -0.003 | -0.003 | 0 | 0.14 | 0.22 | 0.1 | 0.1 | 0.12 | 0.15 |
| 2 | 88615224 | rs1258391 | -1.504 | -0.771 | 1.266 | -0.675 | -0.004 | -0.004 | -0.004 | 0 | 0.14 | 0.22 | 0.1 | 0.1 | 0.12 | 0.15 |
| 2 | 88625879 | rs1258396 | -2.076 | -0.73 | 1.249 | -0.673 | -0.003 | -0.026 | -0.026 | 0 | 0.2 | 0.34 | 0.13 | 0.1 | 0.17 | 0.15 |
| 2 | 88639638 | rs1258384 | -2.035 | -0.717 | 1.364 | -0.552 | 0.003 | -0.016 | -0.016 | 0 | 0.2 | 0.51 | 0.26 | 0.14 | 0.17 | 0.14 |
| 2 | 88648331 | rs1258429 | -2.021 | -0.681 | 1.266 | -0.607 | 0.002 | -0.026 | -0.026 | 0 | 0.19 | 0.39 | 0.13 | 0.12 | 0.17 | 0.12 |
| 2 | 88709968 | rs13425454 | -1.002 | -0.205 | 0.535 | -0.328 | 0.096 | 0.047 | 0.047 | 1.651 | 0.08 | 0.43 | 0.07 | 0.05 | 0.06 | 0.06 |
| 2 | 88723471 | rs2364560 | -0.836 | -0.23 | 0.901 | -0.324 | 0.05 | 0.043 | 0.043 | 0.008 | 0.05 | 0.23 | 0.05 | 0.05 | 0.05 | 0.04 |
| 2 | 97400324 | rs6727384 | -1.023 | -0.118 | 0.636 | -0.228 | -0.004 | -0.01 | -0.01 | 0.405 | 0.33 | 0.72 | 0.29 | 0.4 | 0.39 | 0.2 |
| 2 | 97410949 | rs6746896 | -0.068 | -0.247 | 0.237 | -0.199 | -0.035 | -0.047 | -0.047 | 0 | 0.19 | 0.1 | 0.19 | 0.33 | 0.31 | 0.16 |
| 2 | 97413488 | rs2314398 | -0.181 | -0.178 | 0.375 | -0.182 | -0.008 | -0.012 | -0.012 | 0 | 0.21 | 0.19 | 0.2 | 0.33 | 0.3 | 0.16 |
| 2 | 97483783 | rs7608621 | -1.839 | -1.222 | 0.729 | 0.257 | 0.018 | 0.005 | 0.005 | 2.682 | 0.2 | 0.86 | 0.13 | 0.19 | 0.1 | 0.08 |
| 2 | 97489870 | rs7594727 | -1.952 | -0.874 | 0.951 | 0.404 | 0.018 | 0.005 | 0.005 | 2.7 | 0.2 | 0.86 | 0.13 | 0.19 | 0.1 | 0.08 |
| 2 | 97525099 | rs6576989 | -1.488 | -0.836 | 0.267 | 0.436 | 0.021 | 0.003 | 0.003 | 2.969 | 0.2 | 0.86 | 0.13 | 0.21 | 0.1 | 0.09 |
| 2 | 97553747 | rs12613171 | 0.339 | -0.945 | 0.311 | 0.326 | 0.046 | -0.006 | -0.006 | 0 | 0.38 | 0.24 | 0.44 | 0.68 | 0.16 | 0.71 |
| 2 | 97563745 | rs4243786 | -0.783 | -1.325 | 0.408 | 0.233 | 0.026 | 0.02 | 0.02 | 0.011 | 0.55 | 0.61 | 0.56 | 0.75 | 0.27 | 0.75 |
| 2 | 97567389 | rs7603256 | -1.905 | -0.782 | 0.954 | 0.411 | 0.016 | 0.004 | 0.004 | 4.476 | 0.23 | 0.82 | 0.16 | 0.22 | 0.14 | 0.08 |
| 2 | 97572798 | rs6576990 | -1.612 | -0.729 | 1.007 | 0.367 | -0.042 | -0.019 | -0.019 | 0 | 0.21 | 0.49 | 0.13 | 0.14 | 0.14 | 0.06 |

| | | | | | | | | | | | | | | | | |
|---|---|---|---|---|---|---|---|---|---|---|---|---|---|---|---|---|
| 2 | 97578329 | rs17119656 | -1.106 | -0.326 | 0.278 | 0.347 | 0.008 | 0.01 | 0.01 | 0 | 0.14 | 0.34 | 0.11 | 0.13 | 0.14 | 0.06 |
| 2 | 97585936 | rs11900466 | -1.121 | -0.328 | 0.301 | 0.414 | 0.007 | 0.04 | 0.04 | 0 | 0.07 | 0.3 | 0.04 | 0.08 | 0.04 | 0.02 |
| 2 | 97595991 | rs10874468 | 0.705 | -0.704 | -0.684 | 0.277 | 0.054 | 0.065 | 0.065 | 1.933 | 0.47 | 0.17 | 0.43 | 0.31 | 0.65 | 0.27 |
| 2 | 97599266 | rs13383143 | -0.826 | -0.595 | -0.351 | 0.306 | -0.012 | -0.003 | -0.003 | 0 | 0.08 | 0.18 | 0.04 | 0.08 | 0.06 | 0.02 |
| 2 | 97611396 | rs10202644 | -0.641 | -0.666 | -0.617 | 0.444 | 0.03 | -0.127 | -0.127 | 5.4 | 0.71 | 0.94 | 0.59 | 0.42 | 0.81 | 0.32 |
| 2 | 97613616 | rs1624844 | -0.656 | -0.661 | -0.629 | 0.441 | 0.025 | -0.129 | -0.129 | 0.292 | 0.71 | 0.96 | 0.59 | 0.5 | 0.81 | 0.33 |
| 2 | 97617140 | rs1257019 | -0.594 | -0.677 | -0.372 | 0.433 | 0.035 | -0.079 | -0.079 | 0 | 0.7 | 0.92 | 0.58 | 0.5 | 0.81 | 0.33 |
| 2 | 97623413 | rs1796043 | 0.716 | -0.88 | -0.691 | 0.276 | 0.032 | 0.008 | 0.008 | 3.66 | 0.47 | 0.03 | 0.43 | 0.25 | 0.69 | 0.26 |
| 2 | 97637905 | rs6715329 | -0.859 | -0.86 | -0.249 | 0.276 | 0.03 | -0.033 | -0.033 | 0 | 0.7 | 0.84 | 0.6 | 0.43 | 0.9 | 0.34 |
| 2 | 97640502 | rs1257016 | -0.869 | -0.874 | 0.298 | 0.281 | -0.009 | 0.005 | 0.005 | 0 | 0.69 | 0.74 | 0.59 | 0.38 | 0.9 | 0.32 |
| 2 | 97643576 | rs1796048 | 0.533 | -0.953 | 0.444 | 0.226 | 0.167 | 0.065 | 0.065 | 2.6 | 0.5 | 0.08 | 0.45 | 0.25 | 0.77 | 0.27 |
| 2 | 97654678 | rs1257029 | -0.824 | -0.96 | 0.432 | 0.224 | -0.012 | 0.011 | 0.011 | 0 | 0.7 | 0.74 | 0.58 | 0.38 | 0.9 | 0.32 |
| 2 | 97655963 | rs1257026 | 0.474 | -0.873 | 0.412 | 0.191 | 0.169 | 0.078 | 0.078 | 2.735 | 0.5 | 0.13 | 0.45 | 0.25 | 0.77 | 0.27 |
| 2 | 97714323 | rs1148591 | 0.343 | -0.673 | 0.733 | 0.174 | 0.107 | -0.005 | -0.005 | 0 | 0.54 | 0.3 | 0.48 | 0.43 | 0.8 | 0.3 |
| 2 | 97757509 | rs1257084 | 0.665 | -0.613 | 1.16 | 0.132 | 0.165 | 0.069 | 0.069 | 1.779 | 0.5 | 0.09 | 0.47 | 0.25 | 0.79 | 0.28 |
| 2 | 97764450 | rs1257097 | 0.448 | -0.565 | 1.185 | 0.166 | 0.139 | 0.055 | 0.055 | 0.909 | 0.53 | 0.2 | 0.48 | 0.26 | 0.79 | 0.28 |
| 2 | 97845100 | rs10179966 | -0.546 | -0.613 | 1.395 | 0.131 | -0.033 | -0.064 | -0.064 | 0.004 | 0.18 | 0.46 | 0.12 | 0.06 | 0.12 | 0.05 |
| 2 | 97846850 | rs11164114 | -1.205 | -0.594 | 0.982 | 0.185 | 0.058 | -0.134 | -0.134 | 1.075 | 0.74 | 0.97 | 0.63 | 0.49 | 0.92 | 0.35 |
| 2 | 97860223 | rs4530406 | -0.975 | -0.58 | 1.031 | 0.202 | 0.009 | -0.014 | -0.014 | 0 | 0.72 | 0.81 | 0.61 | 0.44 | 0.92 | 0.34 |
| 2 | 97870317 | rs3926004 | 0.354 | -0.535 | 0.809 | 0.173 | 0.117 | 0.013 | 0.013 | 0.81 | 0.53 | 0.24 | 0.49 | 0.38 | 0.8 | 0.3 |
| 2 | 97871314 | rs12994584 | 0.354 | -0.535 | 0.809 | 0.192 | 0.117 | 0.054 | 0.054 | 0.81 | 0.53 | 0.24 | 0.49 | 0.26 | 0.79 | 0.28 |
| 2 | 97877591 | rs10171740 | -0.337 | -0.266 | 1.348 | 0.357 | -0.023 | 0.01 | 0.01 | 0.832 | 0.05 | 0.15 | 0.02 | 0.06 | 0.01 | 0.01 |
| 2 | 97881267 | rs6756986 | -1.355 | -0.323 | 1.669 | 0.321 | -0.043 | -0.056 | -0.056 | 0.05 | 0.2 | 0.55 | 0.13 | 0.1 | 0.12 | 0.06 |
| 2 | 98017804 | rs11901495 | -1.135 | -0.334 | 2.263 | 0.289 | -0.038 | -0.008 | -0.008 | 0 | 0.18 | 0.43 | 0.11 | 0.2 | 0.12 | 0.06 |
| 2 | 98019683 | rs12989766 | 0.639 | 0.375 | 1.617 | -0.12 | -0.033 | -0.001 | -0.001 | 0 | 0.36 | 0.24 | 0.43 | 0.24 | 0.62 | 0.2 |
| 2 | 98024837 | rs7598442 | 0.262 | 0.437 | 1.595 | -0.173 | -0.002 | 0.002 | 0.002 | 0 | 0.39 | 0.38 | 0.44 | 0.37 | 0.67 | 0.21 |
| 2 | 98166928 | rs11691779 | 0.046 | 0.346 | 1.569 | -0.137 | 0.003 | -0.003 | -0.003 | 0.579 | 0.43 | 0.34 | 0.44 | 0.3 | 0.65 | 0.2 |
| 2 | 98261802 | rs11904110 | 0.01 | 0.256 | 1.337 | -0.187 | -0.015 | -0.055 | -0.055 | 0 | 0.13 | 0.35 | 0.1 | 0.02 | 0.09 | 0.03 |
| 2 | 98265491 | rs1470625 | -0.547 | 0.289 | 0.943 | -0.184 | 0.064 | 0.012 | 0.012 | 0 | 0.53 | 0.75 | 0.54 | 0.37 | 0.74 | 0.24 |
| 2 | 98271946 | rs1550995 | -0.494 | 0.291 | 0.734 | -0.183 | 0.061 | 0.016 | 0.016 | 0 | 0.53 | 0.71 | 0.54 | 0.37 | 0.74 | 0.24 |
| 2 | 98277804 | rs873886 | -1.015 | 0.951 | 0.375 | -0.189 | 0.007 | -0.125 | -0.125 | 0 | 0.59 | 0.97 | 0.56 | 0.43 | 0.74 | 0.25 |
| 2 | 98286235 | rs11123849 | -0.633 | 1.077 | 0.472 | -0.207 | 0.067 | 0.007 | 0.007 | 0 | 0.54 | 0.78 | 0.54 | 0.37 | 0.74 | 0.24 |
| 2 | 98309882 | rs11123861 | 0.334 | 1.389 | -0.102 | -0.038 | -0.042 | 0.007 | 0.007 | 0 | 0.36 | 0.1 | 0.4 | 0.22 | 0.65 | 0.19 |
| 2 | 98327875 | rs17695937 | 0.797 | 1.881 | -0.645 | -0.328 | 0.003 | 0.035 | 0.035 | 0 | 0.21 | 0.16 | 0.18 | 0.12 | 0.23 | 0.07 |
| 2 | 98329197 | rs13034349 | 0.249 | 2.239 | -0.709 | -0.014 | -0.046 | -0.044 | -0.044 | 0 | 0.17 | 0.07 | 0.22 | 0.23 | 0.39 | 0.12 |
| 2 | 98330052 | rs2276645 | 0.259 | 2.231 | -0.953 | 0.021 | -0.124 | -0.113 | -0.113 | 0 | 0.16 | 0.01 | 0.23 | 0.23 | 0.42 | 0.14 |
| 2 | 98334450 | rs11123869 | -1.446 | 1.232 | -1.666 | 0.119 | 0.006 | -0.098 | -0.098 | 8.276 | 0.64 | 0.97 | 0.53 | 0.41 | 0.76 | 0.31 |
| 2 | 98336352 | rs13420683 | -0.704 | 1.029 | -1.39 | -0.068 | 0.009 | 0 | 0 | 0 | 0.34 | 0.77 | 0.28 | 0.23 | 0.29 | 0.11 |
| 2 | 98339513 | rs13033383 | 0.684 | 1.379 | -1.322 | -0.228 | -0.019 | -0.002 | -0.002 | 0 | 0.17 | 0.09 | 0.2 | 0.01 | 0.36 | 0.11 |
| 2 | 98340364 | rs6712517 | -2.055 | 1.113 | -1.631 | 0.072 | -0.002 | -0.126 | -0.126 | 0 | 0.66 | 0.96 | 0.68 | 0.47 | 0.75 | 0.3 |
| 2 | 98345086 | rs6714710 | -1.246 | 1.128 | -0.75 | 0.038 | 0 | -0.001 | -0.001 | 0.009 | 0.37 | 0.64 | 0.47 | 0.35 | 0.51 | 0.21 |
| 2 | 98353847 | rs11686881 | 0.773 | 1.006 | -0.924 | -0.092 | -0.029 | 0.014 | 0.014 | 0 | 0.17 | 0.09 | 0.33 | 0.1 | 0.36 | 0.11 |
| 2 | 98362907 | rs746004 | -0.163 | 0.73 | -0.524 | -0.359 | -0.011 | 0.015 | 0.015 | 0.122 | 0.16 | 0.25 | 0.12 | 0.14 | 0.12 | 0.08 |
| 2 | 98365250 | rs1466020 | -2.271 | 0.615 | -1.005 | -0.383 | 0.023 | -0.046 | -0.046 | 0 | 0.65 | 0.97 | 0.57 | 0.32 | 0.77 | 0.3 |
| 2 | 98372980 | rs6875 | -0.045 | 0.776 | -0.409 | -0.48 | 0 | -0.007 | -0.007 | 0 | 0.29 | 0.32 | 0.21 | 0.06 | 0.27 | 0.11 |
| 2 | 98373006 | rs5865 | -1.157 | 0.725 | -0.109 | -0.411 | 0.019 | 0.02 | 0.02 | 0 | 0.49 | 0.61 | 0.43 | 0.29 | 0.65 | 0.25 |
| 2 | 98391887 | rs3821053 | -0.688 | 0.587 | -0.243 | -0.67 | 0 | -0.014 | -0.014 | 0.102 | 0.38 | 0.41 | 0.29 | 0.06 | 0.35 | 0.13 |
| 2 | 98393231 | rs17489287 | 0.69 | 0.776 | 0.007 | -0.644 | -0.004 | -0.003 | -0.003 | 0 | 0.18 | 0.16 | 0.18 | 0.05 | 0.31 | 0.12 |
| 2 | 98407168 | rs17022373 | 1.409 | 0.301 | 0.151 | -0.666 | -0.022 | -0.032 | -0.032 | 0 | 0.06 | 0.17 | 0.03 | 0 | 0.02 | 0 |
| 2 | 98424802 | rs4311104 | 0.356 | 0.331 | 0.126 | -0.608 | 0.022 | 0.009 | 0.009 | 0 | 0.21 | 0.24 | 0.22 | 0.19 | 0.39 | 0.15 |
| 2 | 98436208 | rs11899171 | -2.376 | 0.331 | 0.038 | -0.831 | 0.041 | -0.007 | -0.007 | 0 | 0.65 | 0.9 | 0.54 | 0.29 | 0.77 | 0.28 |
| 2 | 98451717 | rs6738401 | 0.376 | 0.339 | 0.074 | -0.836 | 0 | -0.006 | -0.006 | 0.108 | 0.29 | 0.3 | 0.21 | 0.06 | 0.27 | 0.11 |
| 2 | 98458002 | rs6744331 | -2.473 | 0.295 | 0.072 | -0.823 | 0.021 | -0.049 | -0.049 | 0 | 0.66 | 0.97 | 0.56 | 0.29 | 0.78 | 0.3 |
| 2 | 98461802 | rs10208070 | -0.443 | 0.317 | 0.022 | -0.889 | -0.001 | -0.118 | -0.118 | 4.213 | 0.38 | 0.53 | 0.29 | 0.08 | 0.36 | 0.13 |
| 2 | 98462658 | rs12712163 | -0.382 | 0.318 | 0.009 | -0.895 | 0 | -0.013 | -0.013 | 0 | 0.37 | 0.41 | 0.29 | 0.06 | 0.35 | 0.13 |
| 2 | 98471378 | rs10176201 | -2.448 | 0.251 | -0.014 | -0.901 | 0.033 | -0.012 | -0.012 | 0.586 | 0.66 | 0.9 | 0.56 | 0.29 | 0.78 | 0.3 |
| 2 | 98479581 | rs11680018 | 0.438 | 0.794 | -0.077 | -1.463 | -0.003 | -0.003 | -0.003 | 0 | 0.18 | 0.17 | 0.22 | 0.04 | 0.39 | 0.12 |
| 2 | 98562116 | rs7588227 | -0.97 | 0.447 | -0.136 | -1.146 | 0 | -0.015 | -0.015 | 1.378 | 0.37 | 0.41 | 0.28 | 0.07 | 0.35 | 0.13 |
| 2 | 98620765 | rs7600204 | -0.002 | 0.483 | -0.468 | -1.043 | -0.001 | -0.004 | -0.004 | 0 | 0.08 | 0.11 | 0.08 | 0.04 | 0.09 | 0.03 |
| 2 | 98622697 | rs10209970 | -0.541 | 0.505 | -0.405 | -0.935 | -0.004 | -0.083 | -0.083 | 0.644 | 0.3 | 0.43 | 0.21 | 0.07 | 0.27 | 0.11 |
| 2 | 98629966 | rs12989231 | 0.046 | 0.196 | -1.017 | -1.111 | -0.038 | 0.135 | 0.135 | 0.106 | 0.36 | 0.17 | 0.38 | 0.04 | 0.63 | 0.24 |
| 2 | 98636132 | rs17025744 | -0.262 | -0.571 | -0.305 | -0.753 | -0.093 | 0.028 | 0.028 | 3.333 | 0.2 | 0.47 | 0.08 | 0.23 | 0.08 | 0.15 |
| 2 | 98637504 | rs17025778 | 0.304 | -0.625 | 0.05 | -0.956 | -0.027 | -0.03 | -0.03 | 0.437 | 0.19 | 0.25 | 0.08 | 0.08 | 0.09 | 0.11 |
| 2 | 98640981 | rs6543250 | -0.951 | -0.003 | -0.122 | -1.097 | -0.043 | 0.053 | 0.053 | 0.09 | 0.67 | 0.87 | 0.47 | 0.25 | 0.61 | 0.36 |
| 2 | 98653406 | rs11898149 | 0.44 | -0.4 | 0.045 | -0.983 | -0.058 | -0.067 | -0.067 | 0 | 0.2 | 0.31 | 0.09 | 0.08 | 0.1 | 0.05 |
| 2 | 98675606 | rs298904 | -0.41 | -0.368 | -0.108 | -0.743 | 0.028 | 0.045 | 0.045 | 0.107 | 0.56 | 0.72 | 0.57 | 0.93 | 0.32 | 0.62 |
| 2 | 98677164 | rs17026292 | -0.25 | -0.456 | -0.125 | -0.685 | -0.092 | -0.053 | -0.053 | 0 | 0.27 | 0.55 | 0.11 | 0.17 | 0.1 | 0.11 |
| 2 | 98686445 | rs921426 | -0.463 | -0.498 | -0.193 | -0.746 | 0.019 | 0.004 | 0.004 | 0 | 0.55 | 0.69 | 0.58 | 0.92 | 0.32 | 0.71 |
| 2 | 98686948 | rs10186392 | 0.215 | -0.342 | -0.091 | -0.762 | 0.001 | 0.002 | 0.002 | 0 | 0.55 | 0.47 | 0.6 | 0.56 | 0.58 | 0.76 |

| | | | | | | | | | | | | | | | |
|---|---|---|---|---|---|---|---|---|---|---|---|---|---|---|---|
| 2 | 98687752 | rs6729411 | 0.338 | 0.119 | 0.138 | -0.879 | 0.02 | -0.109 | -0.109 | 0.004 | 0.31 | 0.37 | 0.26 | 0.04 | 0.46 | 0.2 |
| 2 | 98696912 | rs11124008 | 0.172 | 0.147 | 0.228 | -0.799 | -0.007 | 0.007 | 0.007 | 0 | 0.11 | 0.28 | 0.06 | 0.1 | 0.07 | 0.08 |
| 2 | 98696966 | rs6753823 | -1.098 | -0.082 | -0.382 | -0.969 | -0.091 | 0.179 | 0.179 | 0.827 | 0.54 | 0.96 | 0.35 | 0.24 | 0.49 | 0.39 |
| 2 | 98701043 | rs6543280 | 0.427 | 0.28 | 0.093 | -0.883 | 0.026 | -0.153 | -0.153 | 9.113 | 0.29 | 0.5 | 0.25 | 0.04 | 0.4 | 0.19 |
| 2 | 98703659 | rs10211067 | -0.485 | 0.32 | -0.21 | -0.817 | -0.011 | 0.074 | 0.074 | 0 | 0.39 | 0.79 | 0.3 | 0.14 | 0.46 | 0.27 |
| 2 | 98710243 | rs298916 | -0.173 | 0.541 | -0.093 | -0.724 | -0.008 | -0.025 | -0.025 | 0.003 | 0.36 | 0.63 | 0.23 | 0.17 | 0.28 | 0.21 |
| 2 | 98714445 | rs6705251 | 1.082 | 0.291 | -0.282 | -0.875 | -0.034 | -0.035 | -0.035 | 6.122 | 0.15 | 0.15 | 0.05 | 0.06 | 0.05 | 0.04 |
| 2 | 98716393 | rs6713479 | -1.048 | 0.495 | -0.39 | -0.672 | -0.127 | 0.098 | 0.098 | 0.715 | 0.53 | 0.96 | 0.36 | 0.31 | 0.54 | 0.29 |
| 2 | 98726903 | rs6733249 | 0.775 | 0.222 | 0.117 | -0.652 | -0.024 | -0.082 | -0.082 | 4.785 | 0.26 | 0.42 | 0.18 | 0.12 | 0.21 | 0.14 |
| 2 | 98736225 | rs2305355 | 0.353 | 0.33 | 0.109 | -0.436 | -0.004 | 0.044 | 0.044 | 0 | 0.09 | 0.25 | 0.05 | 0.13 | 0.07 | 0.08 |
| 2 | 98749890 | rs13008613 | 0.901 | 0.148 | 0.092 | -0.593 | -0.053 | -0.056 | -0.056 | 0 | 0.15 | 0.22 | 0.05 | 0.06 | 0.05 | 0.04 |
| 2 | 98755252 | rs7559678 | -0.923 | 0.665 | -0.075 | -0.275 | 0.023 | 0.014 | 0.014 | 0.177 | 0.36 | 0.47 | 0.25 | 0.35 | 0.46 | 0.21 |
| 2 | 98761589 | rs13422327 | 1.012 | 0.054 | -0.16 | -0.488 | -0.045 | -0.142 | -0.142 | 0 | 0.22 | 0.45 | 0.12 | 0.02 | 0.15 | 0.12 |
| 2 | 98763159 | rs17426403 | 0.584 | 0.501 | -0.135 | 0.041 | -0.041 | -0.056 | -0.056 | 0 | 0.09 | 0.01 | 0.08 | 0.18 | 0.13 | 0.07 |
| 2 | 98764923 | rs6744417 | -0.259 | 0.544 | 0.205 | 0.02 | 0.042 | -0.062 | -0.062 | 0.413 | 0.24 | 0.42 | 0.22 | 0.17 | 0.35 | 0.11 |
| 2 | 98766550 | rs6748458 | -0.268 | 0.295 | -0.321 | -0.146 | -0.031 | -0.038 | -0.038 | 0.018 | 0.28 | 0.51 | 0.13 | 0.19 | 0.16 | 0.16 |
| 2 | 98774780 | rs9973945 | 0.878 | 0.02 | -0.901 | -0.265 | -0.067 | -0.093 | -0.093 | 0 | 0.26 | 0.06 | 0.54 | 0.61 | 0.36 | 0.59 |
| 2 | 98775298 | rs4851839 | 0.866 | 0.017 | -0.903 | -0.266 | -0.067 | -0.093 | -0.093 | 0.032 | 0.26 | 0.06 | 0.54 | 0.61 | 0.36 | 0.59 |
| 2 | 98777992 | rs7605986 | -0.076 | -0.023 | -0.8 | -0.294 | -0.03 | -0.036 | -0.036 | 0 | 0.28 | 0.52 | 0.13 | 0.19 | 0.16 | 0.16 |
| 2 | 98781847 | rs13022257 | 0.902 | 0.036 | -1.422 | -0.032 | -0.095 | -0.16 | -0.16 | 0 | 0.26 | 0.04 | 0.46 | 0.53 | 0.36 | 0.6 |
| 2 | 98784275 | rs2309389 | -0.565 | 0.459 | -1.099 | 0.066 | -0.022 | 0.086 | 0.086 | 0 | 0.56 | 0.7 | 0.68 | 0.8 | 0.52 | 0.75 |
| 2 | 98787951 | rs12712221 | -0.547 | 0.499 | -1.105 | 0.015 | -0.02 | 0.057 | 0.057 | 0 | 0.55 | 0.72 | 0.61 | 0.73 | 0.53 | 0.75 |
| 2 | 98790763 | rs12473335 | -0.309 | 0.984 | -0.862 | 0.544 | -0.002 | 0.001 | 0.001 | 0 | 0.14 | 0.1 | 0.07 | 0.14 | 0.11 | 0.11 |
| 2 | 98798577 | rs13428410 | -0.917 | 1.157 | -0.301 | 1.046 | 0.091 | 0.338 | 0.338 | 0 | 0.06 | 0.32 | 0.15 | 0.24 | 0.07 | 0.04 |
| 2 | 98803025 | rs2122752 | -0.34 | 1.584 | -0.742 | 0.58 | -0.025 | 0.037 | 0.037 | 0 | 0.89 | 0.95 | 0.89 | 0.98 | 0.84 | 0.95 |
| 2 | 98807030 | rs2871213 | 0.92 | 1.28 | -1.053 | 0.367 | -0.001 | -0.001 | -0.001 | 0.115 | 0.53 | 0.51 | 0.31 | 0.32 | 0.4 | 0.34 |
| 2 | 98814054 | rs7574579 | 0.198 | 1.581 | -1.483 | 0.879 | -0.088 | -0.103 | -0.103 | 0.903 | 0.19 | 0.04 | 0.25 | 0.41 | 0.32 | 0.52 |
| 2 | 98824172 | rs17028800 | 0.412 | 1.923 | -1.574 | 0.703 | 0.009 | -0.114 | -0.114 | 0 | 0.18 | 0.04 | 0.21 | 0.47 | 0.13 | 0.48 |
| 2 | 98828807 | rs13431656 | -1.422 | 2.398 | -0.9 | 0.44 | 0.015 | -0.038 | -0.038 | 0 | 0.13 | 0.19 | 0.13 | 0.01 | 0.26 | 0.07 |
| 2 | 98839367 | rs13430738 | -0.663 | 2.356 | -0.848 | 0.459 | 0.025 | -0.031 | -0.031 | 0 | 0.08 | 0.15 | 0.08 | 0 | 0.19 | 0.05 |
| 2 | 98845348 | rs11673807 | 1.355 | 2.421 | -0.757 | 0.397 | 0.038 | -0.007 | -0.007 | 0 | 0.76 | 0.64 | 0.52 | 0.8 | 0.6 | 0.85 |
| 2 | 98848414 | rs4851951 | 0.298 | 2.302 | -0.744 | 0.445 | -0.002 | -0.002 | -0.002 | 0 | 0.85 | 0.85 | 0.62 | 0.81 | 0.8 | 0.9 |
| 2 | 98868978 | rs11901562 | 1.574 | 2.443 | -0.749 | 0.394 | 0.029 | -0.011 | -0.011 | 0 | 0.72 | 0.64 | 0.5 | 0.8 | 0.54 | 0.84 |
| 2 | 98872039 | rs4851122 | 1.563 | 2.442 | -0.734 | 0.393 | 0.029 | -0.011 | -0.011 | 0 | 0.72 | 0.64 | 0.5 | 0.8 | 0.54 | 0.84 |
| 2 | 98873599 | rs7606591 | 1.574 | 2.463 | -0.741 | 0.41 | 0.028 | -0.01 | -0.01 | 0 | 0.72 | 0.63 | 0.5 | 0.8 | 0.54 | 0.84 |
| 2 | 98879818 | rs4851954 | -1.732 | 2.438 | -0.735 | 0.399 | 0.016 | -0.005 | -0.005 | 0 | 0.27 | 0.34 | 0.5 | 0.2 | 0.46 | 0.16 |
| 2 | 98885925 | rs2221416 | -1.838 | 2.438 | -0.735 | 0.399 | 0.029 | -0.011 | -0.011 | 0 | 0.28 | 0.37 | 0.5 | 0.2 | 0.46 | 0.16 |
| 2 | 98892107 | rs10203437 | -1.567 | 2.475 | -0.725 | 0.397 | 0.048 | -0.008 | -0.008 | 0 | 0.24 | 0.36 | 0.49 | 0.2 | 0.42 | 0.16 |
| 2 | 98892800 | rs17427684 | -0.544 | 2.432 | -0.824 | 0.407 | -0.02 | -0.022 | -0.022 | 0 | 0.15 | 0.12 | 0.38 | 0.19 | 0.21 | 0.09 |
| 2 | 98898220 | rs17493655 | -1.678 | 2.419 | -0.832 | 0.394 | -0.048 | 0.012 | 0.012 | 0 | 0.26 | 0.19 | 0.42 | 0.2 | 0.46 | 0.16 |
| 2 | 98898672 | rs17493788 | -0.576 | 2.363 | -0.81 | 0.423 | -0.008 | -0.009 | -0.009 | 0.232 | 0.15 | 0.15 | 0.32 | 0.19 | 0.21 | 0.09 |
| 2 | 98910858 | rs6725543 | -0.442 | 2.377 | -0.818 | 0.423 | -0.001 | -0.001 | -0.001 | 0 | 0.15 | 0.18 | 0.32 | 0.19 | 0.21 | 0.09 |
| 2 | 98912271 | rs6715989 | -1.583 | 2.338 | -0.983 | 0.274 | 0.001 | -0.001 | -0.001 | 0.329 | 0.25 | 0.27 | 0.4 | 0.2 | 0.3 | 0.17 |
| 2 | 98919820 | rs999900 | 1.323 | 1.818 | -1.235 | 0.037 | 0.043 | 0 | 0 | 0.237 | 0.23 | 0.11 | 0.1 | 0.18 | 0.17 | 0.16 |
| 2 | 98928929 | rs1466944 | -1.754 | 1.563 | -1.119 | 0.001 | 0.014 | -0.014 | -0.014 | 0 | 0.32 | 0.39 | 0.48 | 0.2 | 0.5 | 0.22 |
| 2 | 98941836 | rs1504680 | -1.276 | 1.49 | -1.136 | 0.103 | -0.003 | 0 | 0 | 0.004 | 0.56 | 0.58 | 0.59 | 0.41 | 0.67 | 0.41 |
| 2 | 98949400 | rs7571001 | -1.652 | 1.808 | -1.291 | -0.152 | 0.016 | -0.013 | -0.013 | 0 | 0.25 | 0.36 | 0.4 | 0.18 | 0.31 | 0.16 |
| 2 | 98953400 | rs13394706 | -0.504 | 1.941 | -1.629 | -0.068 | -0.02 | -0.013 | -0.013 | 0 | 0.15 | 0.16 | 0.32 | 0.17 | 0.22 | 0.09 |
| 2 | 98955040 | rs2102270 | -1.136 | 1.963 | -1.633 | -0.072 | -0.016 | 0.001 | 0.001 | 0.135 | 0.2 | 0.22 | 0.36 | 0.17 | 0.26 | 0.14 |
| 2 | 98966130 | rs17429299 | -0.003 | 1.863 | -1.66 | -0.217 | -0.038 | 0.048 | 0.048 | 0 | 0.11 | 0.05 | 0.11 | 0.01 | 0.17 | 0.04 |
| 2 | 98967700 | rs2134936 | -1.048 | 1.816 | -1.645 | -0.215 | -0.032 | 0.033 | 0.033 | 0 | 0.2 | 0.15 | 0.18 | 0.01 | 0.27 | 0.11 |
| 2 | 98974389 | rs937729 | -1.207 | 1.876 | -1.569 | 0.018 | -0.029 | -0.006 | -0.006 | 0 | 0.2 | 0.1 | 0.43 | 0.16 | 0.27 | 0.1 |
| 2 | 98975294 | rs1546352 | -0.933 | 1.905 | -1.576 | 0.019 | -0.007 | -0.017 | -0.017 | 0 | 0.12 | 0.08 | 0.34 | 0.15 | 0.14 | 0.08 |
| 2 | 98986510 | rs6727412 | -0.362 | 1.699 | -1.406 | -0.017 | -0.007 | -0.011 | -0.011 | 0.166 | 0.13 | 0.08 | 0.13 | 0.01 | 0.17 | 0.07 |
| 2 | 98989062 | rs11123677 | 0.417 | 0.687 | -1.271 | 0.316 | 0.023 | -0.015 | -0.015 | 0 | 0.26 | 0.19 | 0.38 | 0.33 | 0.21 | 0.17 |
| 2 | 98989700 | rs3769754 | -0.43 | 0.976 | -1.322 | 0.358 | -0.003 | -0.002 | -0.002 | 0.035 | 0.41 | 0.47 | 0.52 | 0.38 | 0.44 | 0.35 |
| 2 | 98989956 | rs11691306 | -0.171 | 1.096 | -1.565 | 0.376 | -0.05 | 0.019 | 0.019 | 0 | 0.08 | 0 | 0.07 | 0.03 | 0.19 | 0.12 |
| 2 | 98992113 | rs2055593 | -0.028 | 0.153 | -1.268 | 0.597 | -0.022 | 0.002 | 0.002 | 1.538 | 0.33 | 0.44 | 0.45 | 0.35 | 0.25 | 0.2 |
| 2 | 98994414 | rs937725 | -0.064 | 0.544 | -1.674 | 0.938 | -0.004 | -0.015 | -0.015 | 0 | 0.63 | 0.81 | 0.73 | 0.41 | 0.7 | 0.46 |
| 2 | 98994641 | rs4146045 | -1.228 | 0.194 | -1.299 | 1.662 | -0.006 | 0.052 | 0.052 | 0 | 0.09 | 0.12 | 0.14 | 0.57 | 0.09 | 0.1 |
| 2 | 98998009 | rs1394431 | -0.045 | 0.092 | -1.491 | 0.774 | 0.027 | 0.055 | 0.055 | 0 | 0.45 | 0.39 | 0.4 | 0.18 | 0.38 | 0.23 |
| 2 | 98998344 | rs1995531 | -0.457 | 0.258 | -1.678 | 1.632 | -0.01 | -0.026 | -0.026 | 1.45 | 0.91 | 0.98 | 0.97 | 0.68 | 0.9 | 0.84 |
| 2 | 98999373 | rs7589617 | 0.836 | -0.116 | -0.931 | 1.338 | -0.01 | -0.016 | -0.016 | 0 | 0.06 | 0.23 | 0.01 | 0.02 | 0.01 | 0 |
| 2 | 99000814 | rs3769747 | 0.255 | -0.108 | -1.005 | 1.524 | -0.013 | 0.047 | 0.047 | 0 | 0.84 | 0.74 | 0.95 | 0.63 | 0.89 | 0.83 |
| 2 | 99001403 | rs3769745 | -0.124 | 0.069 | -1.495 | 1.165 | -0.01 | -0.032 | -0.032 | 0 | 0.21 | 0.13 | 0.37 | 0.32 | 0.23 | 0.35 |
| 2 | 99002448 | rs4851130 | 0.615 | -0.556 | -1.759 | 0.66 | 0.033 | 0.003 | 0.003 | 0.07 | 0.14 | 0.01 | 0.19 | 0.05 | 0.06 | 0.06 |
| 2 | 99016975 | rs1857745 | -0.605 | -0.763 | -1.164 | 0.776 | 0.01 | -0.003 | -0.003 | 0 | 0.24 | 0.27 | 0.26 | 0.3 | 0.2 | 0.28 |
| 2 | 99083764 | rs6542781 | -1.089 | -0.617 | -1.061 | 0.807 | -0.036 | -0.107 | -0.107 | 0 | 0.82 | 0.97 | 0.77 | 0.57 | 0.75 | 0.78 |
| 2 | 99088018 | rs3754891 | -0.594 | -0.637 | -1.288 | 0.544 | 0.001 | 0 | 0 | 0.738 | 0.3 | 0.39 | 0.33 | 0.32 | 0.28 | 0.32 |

| | | | | | | | | | | | | | | | | |
|---|---|---|---|---|---|---|---|---|---|---|---|---|---|---|---|---|
| 2 | 99107677 | rs3754886 | -0.558 | -0.607 | -1.27 | 0.53 | 0.003 | -0.01 | -0.01 | 0 | 0.29 | 0.14 | 0.32 | 0.3 | 0.28 | 0.31 |
| 2 | 99129040 | rs6753380 | -0.744 | -0.607 | -1.265 | 0.53 | 0.011 | 0.003 | 0.003 | 0.864 | 0.3 | 0.28 | 0.33 | 0.32 | 0.28 | 0.32 |
| 2 | 99135738 | rs2278214 | -0.297 | -0.659 | -1.302 | 0.502 | -0.026 | -0.038 | -0.038 | 0 | 0.24 | 0.09 | 0.31 | 0.3 | 0.28 | 0.31 |
| 2 | 99147872 | rs222 | 1.229 | -0.843 | -1.407 | -0.086 | 0.01 | -0.007 | -0.007 | 0 | 0.25 | 0.38 | 0.26 | 0.22 | 0.2 | 0.29 |
| 2 | 99149946 | rs2278211 | 0.378 | -0.857 | -1.769 | -0.123 | -0.038 | -0.052 | -0.052 | 0.382 | 0.23 | 0.07 | 0.3 | 0.3 | 0.27 | 0.28 |
| 2 | 99154229 | rs2278208 | -0.274 | -0.897 | -1.776 | -0.12 | -0.001 | -0.012 | -0.012 | 0.121 | 0.28 | 0.12 | 0.3 | 0.29 | 0.26 | 0.31 |
| 2 | 99156296 | rs6734569 | 1.445 | -0.897 | -1.767 | -0.207 | -0.025 | 0.109 | 0.109 | 0.008 | 0.26 | 0.1 | 0.18 | 0.03 | 0.28 | 0.17 |
| 2 | 99185867 | rs3754876 | 0.775 | -0.793 | -1.516 | 0.022 | -0.061 | 0.034 | 0.034 | 0.172 | 0.12 | 0.01 | 0.1 | 0.05 | 0.17 | 0.1 |
| 2 | 99204428 | rs6736755 | -0.875 | -0.87 | -1.384 | 0.067 | 0.003 | 0.001 | 0.001 | 0.658 | 0.31 | 0.38 | 0.32 | 0.31 | 0.27 | 0.32 |
| 2 | 99208209 | rs6542833 | 1.037 | -0.878 | -1.403 | 0.055 | -0.002 | -0.004 | -0.004 | 0.078 | 0.25 | 0.44 | 0.26 | 0.23 | 0.19 | 0.29 |
| 2 | 99209314 | rs13402379 | -0.887 | -0.448 | -1.815 | -0.153 | 0.002 | 0.001 | 0.001 | 0.762 | 0.3 | 0.39 | 0.31 | 0.31 | 0.27 | 0.32 |
| 2 | 99210559 | rs13016305 | 1.67 | -0.716 | -2.126 | -0.265 | -0.077 | 0.077 | 0.077 | 0 | 0.16 | 0.01 | 0.1 | 0.04 | 0.18 | 0.26 |
| 2 | 99225645 | rs11896729 | -1.127 | -0.602 | -1.395 | 0.194 | 0.01 | 0.002 | 0.002 | 0.787 | 0.3 | 0.23 | 0.3 | 0.31 | 0.27 | 0.32 |
| 2 | 99226021 | rs3203856 | -1.012 | -0.603 | -1.396 | 0.193 | -0.001 | -0.009 | -0.009 | 0.222 | 0.29 | 0.16 | 0.29 | 0.29 | 0.27 | 0.31 |
| 2 | 99232312 | rs1516085 | -1.088 | -0.587 | -1.38 | 0.21 | 0.003 | 0.001 | 0.001 | 0.687 | 0.31 | 0.39 | 0.31 | 0.31 | 0.27 | 0.32 |
| 2 | 99233894 | rs3769698 | -1.016 | -0.575 | -1.539 | 0.198 | 0.012 | 0.004 | 0.004 | 0.82 | 0.3 | 0.24 | 0.3 | 0.31 | 0.27 | 0.32 |
| 2 | 99235169 | rs17513468 | -1.016 | -0.575 | -1.539 | 0.198 | 0.012 | 0.004 | 0.004 | 0.82 | 0.3 | 0.24 | 0.3 | 0.31 | 0.27 | 0.32 |
| 2 | 99237792 | rs11892744 | -0.511 | -0.554 | -1.777 | 0.219 | -0.023 | -0.042 | -0.042 | 0.073 | 0.22 | 0.16 | 0.27 | 0.29 | 0.25 | 0.28 |
| 2 | 99239931 | rs12618769 | -0.437 | -0.356 | -1.315 | -0.059 | 0.004 | 0.011 | 0.011 | 0.022 | 0.2 | 0.26 | 0.11 | 0.07 | 0.17 | 0.14 |
| 2 | 99247820 | rs7560357 | -0.585 | -0.331 | -1.332 | -0.047 | 0.009 | 0.024 | 0.024 | 0 | 0.22 | 0.25 | 0.12 | 0.1 | 0.18 | 0.16 |
| 2 | 99264217 | rs2871234 | 0.062 | 0.215 | -1.134 | 0.428 | -0.002 | 0.001 | 0.001 | 0 | 0.43 | 0.56 | 0.45 | 0.58 | 0.4 | 0.55 |
| 2 | 99272802 | rs11679648 | 1.561 | 0.273 | -1.722 | 0.034 | -0.094 | 0.098 | 0.098 | 0.481 | 0.2 | 0.01 | 0.17 | 0.03 | 0.27 | 0.1 |
| 2 | 99279354 | rs1028144 | -0.493 | 0.286 | -1.581 | 0.343 | -0.076 | 0 | 0 | 0.298 | 0.5 | 0.93 | 0.57 | 0.64 | 0.46 | 0.67 |
| 2 | 99283348 | rs17021869 | 0.42 | 0.494 | -0.418 | 0.41 | 0.005 | 0.058 | 0.058 | 0 | 0.16 | 0.34 | 0.3 | 0.22 | 0.13 | 0.29 |
| 2 | 99301833 | rs10165386 | 0.367 | 0.419 | -0.696 | 0.189 | -0.001 | 0.002 | 0.002 | 0.157 | 0.31 | 0.33 | 0.33 | 0.14 | 0.34 | 0.21 |
| 2 | 99304794 | rs885036 | -0.805 | 0.504 | -0.989 | 0.249 | -0.033 | 0.057 | 0.057 | 0.007 | 0.47 | 0.79 | 0.55 | 0.36 | 0.5 | 0.49 |
| 2 | 99327311 | rs10196218 | -0.821 | 0.518 | -0.283 | 0.423 | -0.001 | 0.005 | 0.005 | 0.336 | 0.22 | 0.66 | 0.31 | 0.27 | 0.22 | 0.36 |
| 2 | 99339122 | rs4851150 | -0.889 | 0.486 | -0.046 | 0.457 | -0.004 | 0.033 | 0.033 | 0.021 | 0.22 | 0.63 | 0.4 | 0.34 | 0.2 | 0.39 |
| 2 | 99350109 | rs1016124 | -0.921 | 0.472 | 0.085 | 0.45 | -0.005 | 0.033 | 0.033 | 0.004 | 0.22 | 0.61 | 0.4 | 0.34 | 0.2 | 0.39 |
| 2 | 99359096 | rs6743770 | -0.753 | 0.976 | -0.248 | 0.123 | 0.006 | 0.033 | 0.033 | 0 | 0.46 | 0.63 | 0.51 | 0.28 | 0.56 | 0.5 |
| 2 | 99359117 | rs6733957 | 0.343 | 0.851 | 0.552 | 0.108 | 0.012 | 0.02 | 0.02 | 0 | 0.15 | 0.25 | 0.23 | 0.21 | 0.17 | 0.25 |
| 2 | 99362181 | rs17021908 | 1.46 | 0.573 | -0.063 | 0.391 | -0.026 | -0.105 | -0.105 | 0 | 0.16 | 0.01 | 0.12 | 0.33 | 0.15 | 0.17 |
| 2 | 99362400 | rs1913621 | -0.653 | 0.646 | 0.034 | 0.158 | -0.004 | 0.025 | 0.025 | 0.024 | 0.22 | 0.64 | 0.4 | 0.34 | 0.2 | 0.39 |
| 2 | 99363790 | rs11899659 | 0.684 | 0.727 | -0.066 | 0.131 | 0.019 | -0.046 | -0.046 | 0.018 | 0.3 | 0.37 | 0.34 | 0.13 | 0.4 | 0.26 |
| 2 | 99363908 | rs4539843 | 1.228 | 0.746 | -0.299 | 0.095 | -0.108 | 0.16 | 0.16 | 0.469 | 0.26 | 0.05 | 0.21 | 0.01 | 0.38 | 0.14 |
| 2 | 99366626 | rs1567951 | -1.478 | 0.669 | -0.402 | 0.215 | -0.039 | -0.099 | -0.099 | 0.975 | 0.67 | 0.99 | 0.74 | 0.68 | 0.72 | 0.73 |
| 2 | 99369757 | rs11123751 | -0.654 | 1.446 | 0.343 | 0.039 | 0.009 | 0.065 | 0.065 | 0 | 0.2 | 0.46 | 0.38 | 0.3 | 0.19 | 0.33 |
| 2 | 99373720 | rs6712597 | -1.041 | 1.509 | 0.231 | 0.013 | 0.005 | 0.048 | 0.048 | 0 | 0.48 | 0.7 | 0.61 | 0.37 | 0.55 | 0.54 |
| 2 | 99378801 | rs6711574 | 0.653 | 1.794 | -0.225 | -0.252 | -0.084 | 0.16 | 0.16 | 0.418 | 0.27 | 0.06 | 0.2 | 0.01 | 0.36 | 0.16 |
| 2 | 99382537 | rs10168498 | 0.877 | 1.697 | 0.256 | -0.012 | 0.059 | -0.012 | -0.012 | 0 | 0.06 | 0.12 | 0.09 | 0.02 | 0.13 | 0.11 |
| 2 | 99383330 | rs13000014 | -0.184 | 1.752 | 0.592 | -0.089 | 0.038 | -0.073 | -0.073 | 0 | 0.34 | 0.51 | 0.3 | 0.13 | 0.43 | 0.27 |
| 2 | 99384299 | rs17021941 | 0.459 | 1.74 | 0.603 | -0.086 | 0.007 | -0.043 | -0.043 | 0 | 0.28 | 0.38 | 0.21 | 0.11 | 0.3 | 0.17 |
| 2 | 99384371 | rs12615689 | -0.584 | 1.607 | 0.459 | -0.054 | 0.014 | -0.04 | -0.04 | 0.126 | 0.39 | 0.54 | 0.44 | 0.19 | 0.48 | 0.5 |
| 2 | 99417433 | rs10182043 | 0.314 | 1.563 | 0.714 | 0.208 | 0.004 | -0.045 | -0.045 | 0 | 0.33 | 0.44 | 0.33 | 0.18 | 0.32 | 0.16 |
| 2 | 99422558 | rs2871244 | -0.229 | 1.585 | 0.625 | 0.28 | -0.002 | -0.007 | -0.007 | 0 | 0.05 | 0.03 | 0.07 | 0.11 | 0.07 | 0.08 |
| 2 | 99425838 | rs1356424 | -0.455 | 1.608 | 0.746 | 0.263 | -0.002 | 0.012 | 0.012 | 0 | 0.42 | 0.68 | 0.5 | 0.33 | 0.4 | 0.44 |
| 2 | 99450831 | rs1517682 | -0.373 | 1.426 | 0.482 | 0.283 | 0.002 | -0.002 | -0.002 | 0 | 0.56 | 0.52 | 0.46 | 0.54 | 0.49 | 0.34 |
| 2 | 99454227 | rs1983352 | -0.399 | 1.583 | 0.689 | 0.361 | -0.002 | 0.013 | 0.013 | 0 | 0.42 | 0.68 | 0.52 | 0.33 | 0.41 | 0.42 |
| 2 | 99463025 | rs12712037 | 0.494 | 1.521 | 0.808 | 0.377 | -0.004 | 0.016 | 0.016 | 0 | 0.32 | 0.24 | 0.34 | 0.17 | 0.32 | 0.15 |
| 2 | 99464530 | rs6542839 | -0.294 | 1.521 | 0.579 | 0.444 | -0.001 | -0.003 | -0.003 | 0 | 0.4 | 0.6 | 0.43 | 0.27 | 0.4 | 0.21 |
| 2 | 99468341 | rs1949424 | 0.078 | 1.542 | 0.626 | 0.441 | 0.001 | -0.013 | -0.013 | 0.47 | 0.36 | 0.6 | 0.47 | 0.23 | 0.35 | 0.38 |
| 2 | 99485421 | rs4851165 | -0.843 | 1.525 | 0.584 | 0.503 | 0.01 | 0.066 | 0.066 | 0 | 0.06 | 0.14 | 0.29 | 0.2 | 0.09 | 0.24 |
| 2 | 99486318 | rs6760065 | 0.189 | 1.495 | 0.646 | 0.484 | 0.005 | -0.048 | -0.048 | 0 | 0.35 | 0.52 | 0.38 | 0.19 | 0.34 | 0.17 |
| 2 | 99491635 | rs883773 | 0.209 | 1.51 | 0.434 | 0.492 | 0 | -0.013 | -0.013 | 2.472 | 0.36 | 0.61 | 0.38 | 0.19 | 0.34 | 0.18 |
| 2 | 99501698 | rs1878585 | -0.191 | 1.5 | 0.396 | 0.486 | -0.006 | 0.028 | 0.028 | 0 | 0.41 | 0.69 | 0.53 | 0.25 | 0.42 | 0.4 |
| 2 | 99509605 | rs10172158 | -0.274 | 1.503 | 0.352 | 0.491 | -0.009 | -0.002 | -0.002 | 0.026 | 0.42 | 0.75 | 0.67 | 0.38 | 0.43 | 0.41 |
| 2 | 99518319 | rs7423519 | 0.364 | 1.5 | 0.64 | 0.34 | 0.004 | -0.024 | -0.024 | 0.215 | 0.33 | 0.39 | 0.38 | 0.2 | 0.34 | 0.18 |
| 2 | 99520608 | rs4851173 | -0.158 | 1.552 | 0.428 | 0.375 | -0.007 | 0.006 | 0.006 | 0 | 0.4 | 0.74 | 0.66 | 0.36 | 0.42 | 0.43 |
| 2 | 99531842 | rs17758170 | 0.308 | 1.43 | 0.493 | 0.407 | 0.01 | -0.053 | -0.053 | 0.646 | 0.34 | 0.46 | 0.52 | 0.21 | 0.34 | 0.18 |
| 2 | 99537569 | rs908672 | -0.124 | 1.431 | 0.316 | 0.357 | -0.004 | 0.013 | 0.013 | 0 | 0.39 | 0.71 | 0.66 | 0.3 | 0.41 | 0.43 |
| 2 | 99543105 | rs1607255 | 1.121 | 1.493 | 0.209 | 0.365 | -0.038 | -0.122 | -0.122 | 0.324 | 0.17 | 0.01 | 0.13 | 0.37 | 0.19 | 0.2 |
| 2 | 99546423 | rs10208260 | -0.594 | 1.921 | 1.011 | -0.161 | -0.005 | -0.019 | -0.019 | 0.164 | 0.41 | 0.54 | 0.4 | 0.27 | 0.33 | 0.19 |
| 2 | 99548333 | rs13385547 | 0.05 | 2.065 | 1.063 | -0.172 | 0.036 | -0.035 | -0.035 | 0 | 0.09 | 0.12 | 0.12 | 0.01 | 0.19 | 0.11 |
| 2 | 99549154 | rs13397765 | -0.575 | 1.274 | 1.314 | 0.194 | -0.004 | -0.006 | -0.006 | 0 | 0.35 | 0.56 | 0.45 | 0.32 | 0.31 | 0.4 |
| 2 | 99558970 | rs10181422 | -1.074 | 0.87 | 1.103 | 0.135 | 0.033 | 0.036 | 0.036 | 0.86 | 0.42 | 0.79 | 0.47 | 0.33 | 0.3 | 0.36 |
| 2 | 99651963 | rs896884 | -0.394 | 0.833 | 1.531 | 0.079 | 0.013 | 0.001 | 0.001 | 2.759 | 0.07 | 0.34 | 0.04 | 0.04 | 0.02 | 0.19 |
| 2 | 99684929 | rs2045057 | -1.073 | 0.766 | 1.547 | 0.074 | 0.001 | -0.014 | -0.014 | 0.041 | 0.38 | 0.62 | 0.35 | 0.12 | 0.26 | 0.33 |
| 2 | 99692829 | rs10181344 | -1.279 | 0.768 | 1.51 | 0.076 | 0.034 | 0.036 | 0.036 | 0.866 | 0.42 | 0.79 | 0.47 | 0.33 | 0.3 | 0.36 |
| 2 | 99699464 | rs12475034 | -0.168 | 0.756 | 1.553 | 0.063 | 0.001 | 0.028 | 0.028 | 0 | 0.29 | 0.17 | 0.3 | 0.08 | 0.24 | 0.14 |

| | | | | | | | | | | | | | | | | |
|---|---|---|---|---|---|---|---|---|---|---|---|---|---|---|---|---|
| 2 | 99701153 | rs1563108 | -0.926 | 1.355 | 1.61 | 0.442 | 0.008 | 0.006 | 0.006 | 0 | 0.36 | 0.73 | 0.45 | 0.33 | 0.28 | 0.35 |
| 2 | 99705903 | rs11123761 | 0.246 | 1.326 | 1.631 | 0.447 | 0.027 | -0.078 | -0.078 | 0 | 0.68 | 0.5 | 0.64 | 0.87 | 0.65 | 0.62 |
| 2 | 99724512 | rs11683188 | 0.703 | 1.367 | 1.569 | 0.473 | -0.019 | 0.018 | 0.018 | 0 | 0.62 | 0.21 | 0.52 | 0.66 | 0.61 | 0.59 |
| 2 | 99732741 | rs12613624 | 0.703 | 1.367 | 1.549 | 0.471 | -0.02 | 0.018 | 0.018 | 0 | 0.62 | 0.21 | 0.52 | 0.66 | 0.61 | 0.59 |
| 2 | 99735052 | rs11688004 | 0.686 | 1.376 | 1.548 | 0.476 | -0.015 | 0.016 | 0.016 | 0 | 0.62 | 0.25 | 0.52 | 0.66 | 0.61 | 0.59 |
| 2 | 99742161 | rs11695379 | 0.504 | 1.38 | 1.548 | 0.443 | 0.002 | -0.03 | -0.03 | 0.296 | 0.65 | 0.37 | 0.63 | 0.87 | 0.65 | 0.62 |
| 2 | 99749350 | rs4588238 | 0.494 | 1.38 | 1.557 | 0.443 | 0.004 | -0.037 | -0.037 | 0 | 0.65 | 0.4 | 0.63 | 0.87 | 0.65 | 0.62 |
| 2 | 99773201 | rs2516829 | 0.294 | 1.39 | 1.528 | 0.441 | 0.024 | -0.077 | -0.077 | 0.045 | 0.67 | 0.49 | 0.65 | 0.87 | 0.65 | 0.62 |
| 2 | 99778985 | rs2632277 | 0.435 | 1.372 | 1.511 | 0.494 | -0.005 | -0.002 | -0.002 | 0 | 0.64 | 0.33 | 0.62 | 0.87 | 0.63 | 0.61 |
| 2 | 99779131 | rs2516835 | 0.648 | 1.372 | 1.511 | 0.494 | -0.02 | 0.018 | 0.018 | 0 | 0.62 | 0.21 | 0.52 | 0.66 | 0.61 | 0.59 |
| 2 | 99795360 | rs2171359 | 0.451 | 1.388 | 1.392 | 0.502 | 0.004 | -0.037 | -0.037 | 0.463 | 0.65 | 0.41 | 0.63 | 0.87 | 0.65 | 0.62 |
| 2 | 99847890 | rs7603462 | 1.439 | 1.678 | 0.748 | 0.958 | 0.154 | 0.162 | 0.162 | 0 | 0.91 | 0.59 | 0.72 | 0.82 | 0.84 | 0.72 |
| 2 | 99857670 | rs7579999 | 0.684 | 1.734 | 0.407 | 0.709 | 0.116 | 0.01 | 0.01 | 0 | 0.95 | 0.77 | 0.95 | 0.95 | 0.88 | 0.75 |
| 2 | 99858556 | rs4851196 | -0.234 | 1.482 | 0.262 | -0.054 | 0.001 | -0.005 | -0.005 | 0 | 0.29 | 0.26 | 0.33 | 0.08 | 0.25 | 0.14 |
| 2 | 99859863 | rs17760953 | -0.179 | 1.4 | 0.284 | -0.095 | 0.004 | 0.029 | 0.029 | 0 | 0.28 | 0.19 | 0.32 | 0.08 | 0.25 | 0.14 |
| 2 | 99871370 | rs11123775 | -0.662 | 1.576 | 0.09 | 0.062 | -0.002 | 0.005 | 0.005 | 0 | 0.42 | 0.44 | 0.62 | 0.31 | 0.45 | 0.26 |
| 2 | 99872861 | rs11676581 | 0.947 | 1.555 | 0.811 | 0.004 | 0.027 | 0.041 | 0.041 | 0.059 | 0.51 | 0.24 | 0.31 | 0.64 | 0.43 | 0.48 |
| 2 | 99875018 | rs12053519 | 0.084 | 1.519 | 0.802 | -0.055 | 0.019 | 0.046 | 0.046 | 0 | 0.09 | 0.14 | 0.3 | 0.25 | 0.2 | 0.12 |
| 2 | 99876244 | rs2200578 | 0.105 | 1.514 | 0.836 | -0.055 | -0.021 | -0.016 | -0.016 | 0 | 0.09 | 0.06 | 0.3 | 0.25 | 0.2 | 0.12 |
| 2 | 99881228 | rs7567462 | 0.103 | 1.513 | 0.863 | -0.059 | 0.115 | 0.013 | 0.013 | 0.221 | 0.93 | 0.69 | 0.94 | 0.95 | 0.88 | 0.74 |
| 2 | 99898998 | rs2309615 | 0.103 | 1.513 | 0.863 | -0.059 | 0.115 | 0.013 | 0.013 | 0.221 | 0.93 | 0.69 | 0.94 | 0.95 | 0.88 | 0.74 |
| 2 | 99899463 | rs13009884 | 1.064 | 1.499 | 0.855 | -0.036 | 0.025 | 0.037 | 0.037 | 0.042 | 0.51 | 0.25 | 0.31 | 0.63 | 0.43 | 0.48 |
| 2 | 99900248 | rs13033740 | 1.064 | 1.499 | 0.876 | -0.036 | 0.027 | 0.039 | 0.039 | 0.042 | 0.51 | 0.24 | 0.31 | 0.63 | 0.43 | 0.48 |
| 2 | 99909958 | rs1922624 | 0.12 | 1.486 | 0.882 | -0.046 | 0.009 | 0.006 | 0.006 | 0 | 0.32 | 0.23 | 0.31 | 0.06 | 0.24 | 0.14 |
| 2 | 99913998 | rs1922625 | 0.116 | 1.371 | 0.89 | -0.027 | 0.115 | 0.013 | 0.013 | 0.221 | 0.93 | 0.69 | 0.94 | 0.95 | 0.87 | 0.74 |
| 2 | 99924918 | rs4851200 | 0.953 | 1.333 | 0.856 | -0.014 | 0.021 | 0.03 | 0.03 | 0 | 0.52 | 0.27 | 0.32 | 0.63 | 0.43 | 0.48 |
| 2 | 99932255 | rs7589893 | 0.338 | 1.429 | 0.86 | -0.047 | 0.006 | -0.075 | -0.075 | 0 | 0.59 | 0.39 | 0.6 | 0.89 | 0.6 | 0.6 |
| 2 | 99938942 | rs12464785 | -0.715 | 1.494 | 0.84 | -0.022 | 0 | 0.004 | 0.004 | 0 | 0.42 | 0.46 | 0.62 | 0.31 | 0.43 | 0.27 |
| 2 | 99946836 | rs13013984 | 1.047 | 1.502 | 0.849 | -0.023 | 0.016 | 0.041 | 0.041 | 0.07 | 0.51 | 0.24 | 0.31 | 0.64 | 0.45 | 0.48 |
| 2 | 99955047 | rs17022584 | 0.841 | 1.247 | 0.93 | -0.149 | -0.024 | -0.024 | -0.024 | 0 | 0.06 | 0.09 | 0.02 | 0 | 0.01 | 0 |
| 2 | 99959845 | rs13018274 | 1.273 | 1.254 | 0.904 | -0.145 | 0.016 | 0.041 | 0.041 | 0.07 | 0.51 | 0.24 | 0.31 | 0.64 | 0.45 | 0.48 |
| 2 | 99975870 | rs12623620 | 1.363 | 0.455 | -0.242 | -0.247 | 0.034 | -0.014 | -0.014 | 0 | 0.24 | 0.01 | 0.09 | 0.25 | 0.14 | 0.16 |
| 2 | 99995517 | rs11896520 | 1.024 | 0.628 | -0.006 | -0.25 | -0.03 | 0.08 | 0.08 | 0 | 0.27 | 0.09 | 0.3 | 0.01 | 0.28 | 0.14 |
| 2 | 100022772 | rs717454 | -0.273 | 0.64 | 0.031 | -0.205 | 0.001 | 0.012 | 0.012 | 0 | 0.42 | 0.41 | 0.62 | 0.31 | 0.43 | 0.27 |
| 2 | 100032449 | rs13402784 | -0.316 | 0.705 | 0.032 | -0.204 | 0.091 | 0.003 | 0.003 | 0 | 0.08 | 0.29 | 0.08 | 0.06 | 0.14 | 0.26 |
| 2 | 100046038 | rs12479064 | 0.456 | 0.626 | -0.374 | -0.545 | 0.001 | -0.024 | -0.024 | 0 | 0.26 | 0.25 | 0.3 | 0.06 | 0.21 | 0.12 |
| 2 | 100055506 | rs3087386 | -1.056 | 0.635 | -0.534 | -0.329 | 0.019 | 0.037 | 0.037 | 0 | 0.5 | 0.73 | 0.7 | 0.36 | 0.57 | 0.53 |
| 2 | 100057489 | rs1451245 | 0.451 | 0.604 | -0.545 | -0.37 | -0.002 | 0.018 | 0.018 | 0 | 0.09 | 0.1 | 0.3 | 0.25 | 0.16 | 0.12 |
| 2 | 100058424 | rs10186223 | 0.259 | 0.611 | -0.539 | -0.378 | 0.123 | 0.02 | 0.02 | 0.417 | 0.07 | 0.33 | 0.06 | 0.05 | 0.12 | 0.25 |
| 2 | 100058870 | rs3087403 | 0.166 | 0.613 | -0.562 | -0.377 | 0.004 | 0.006 | 0.006 | 0 | 0.34 | 0.3 | 0.34 | 0.06 | 0.28 | 0.15 |
| 2 | 100060304 | rs10207435 | 0.39 | 0.621 | -0.548 | -0.363 | 0.104 | 0.011 | 0.011 | 0.189 | 0.06 | 0.29 | 0.06 | 0.05 | 0.12 | 0.25 |
| 2 | 100062163 | rs3792135 | 0.416 | 0.619 | -0.566 | -0.36 | -0.002 | 0.018 | 0.018 | 0 | 0.09 | 0.1 | 0.3 | 0.25 | 0.16 | 0.12 |
| 2 | 100075804 | rs10207981 | -0.504 | 0.662 | -0.506 | -0.351 | 0.076 | 0.113 | 0.113 | 0.013 | 0.14 | 0.35 | 0.33 | 0.29 | 0.27 | 0.37 |
| 2 | 100077966 | rs769105 | -0.352 | 0.568 | -0.623 | -0.303 | 0.001 | 0.017 | 0.017 | 0 | 0.43 | 0.4 | 0.64 | 0.31 | 0.45 | 0.28 |
| 2 | 100113278 | rs7595242 | 1.067 | 0.363 | -0.796 | -0.26 | -0.032 | -0.033 | -0.033 | 0 | 0.06 | 0.15 | 0.02 | 0 | 0.01 | 0 |
| 2 | 100123030 | rs6707827 | -0.778 | 0.277 | -1.406 | -0.347 | -0.101 | -0.161 | -0.161 | 0.97 | 0.75 | 0.99 | 0.81 | 0.61 | 0.71 | 0.74 |
| 2 | 100137660 | rs2309586 | -0.414 | 0.354 | -1.094 | -0.175 | 0.015 | 0.064 | 0.064 | 0 | 0.51 | 0.76 | 0.69 | 0.28 | 0.57 | 0.55 |
| 2 | 100149336 | rs17765112 | -0.096 | 0.373 | -1.441 | -0.267 | 0.006 | 0.039 | 0.039 | 0 | 0.51 | 0.52 | 0.6 | 0.21 | 0.58 | 0.53 |
| 2 | 100152365 | rs4850905 | -1.253 | 0.402 | -2.291 | -0.032 | -0.091 | -0.163 | -0.163 | 0.561 | 0.8 | 0.99 | 0.76 | 0.56 | 0.75 | 0.81 |
| 2 | 100171928 | rs3792118 | -0.982 | -0.558 | -1.6 | 0.388 | 0.063 | -0.058 | -0.058 | 3.084 | 0.44 | 0.89 | 0.4 | 0.48 | 0.32 | 0.4 |
| 2 | 100172081 | rs3792119 | 0.838 | -0.663 | -1.682 | 0.367 | -0.021 | -0.039 | -0.039 | 0 | 0.21 | 0.02 | 0.1 | 0.22 | 0.14 | 0.22 |
| 2 | 100174645 | rs10179116 | -1.098 | -0.628 | -1.65 | 0.539 | 0.064 | -0.084 | -0.084 | 0.092 | 0.44 | 0.89 | 0.4 | 0.51 | 0.31 | 0.41 |
| 2 | 100176775 | rs13008878 | 0.469 | -0.54 | -1.484 | 0.396 | 0.017 | -0.001 | -0.001 | 0.438 | 0.24 | 0.1 | 0.12 | 0.23 | 0.15 | 0.28 |
| 2 | 100184719 | rs3828312 | -0.828 | -0.514 | -1.383 | 0.651 | 0.032 | -0.047 | -0.047 | 0 | 0.22 | 0.8 | 0.32 | 0.35 | 0.19 | 0.18 |
| 2 | 100190049 | rs744145 | -0.878 | -0.55 | -1.482 | 0.573 | 0.04 | -0.055 | -0.055 | 0 | 0.23 | 0.81 | 0.33 | 0.36 | 0.2 | 0.19 |
| 2 | 100193898 | rs3792125 | 0.28 | 0.261 | -1.175 | 0.993 | 0.014 | 0.042 | 0.042 | 0.078 | 0.23 | 0.11 | 0.09 | 0.15 | 0.14 | 0.28 |
| 2 | 100195156 | rs3792126 | -0.837 | 0.724 | 0.116 | 1.482 | 0.031 | 0.297 | 0.297 | 0 | 0.12 | 0.35 | 0.13 | 0.35 | 0.15 | 0.17 |
| 2 | 100205498 | rs11123788 | 0.537 | 1.237 | -0.266 | 0.723 | 0.022 | -0.011 | -0.011 | 0 | 0.72 | 0.62 | 0.64 | 0.77 | 0.67 | 0.81 |
| 2 | 100210642 | rs1047265 | 0.454 | 1.196 | -0.905 | 0.443 | -0.021 | 0.02 | 0.02 | 0.071 | 0.19 | 0.09 | 0.12 | 0.09 | 0.2 | 0.25 |
| 2 | 100217449 | rs3792133 | -0.04 | 1.606 | -0.983 | 0.406 | -0.024 | -0.027 | -0.027 | 0 | 0.09 | 0.03 | 0.16 | 0.12 | 0.12 | 0.1 |
| 2 | 100230334 | rs7567996 | -1.464 | 1.506 | -0.374 | 0.208 | -0.001 | -0.003 | -0.003 | 0 | 0.48 | 0.49 | 0.46 | 0.28 | 0.49 | 0.43 |
| 2 | 100232743 | rs12616127 | -1.701 | 1.453 | -0.084 | 0.184 | 0.028 | 0.05 | 0.05 | 0 | 0.5 | 0.64 | 0.49 | 0.32 | 0.54 | 0.44 |
| 2 | 100248860 | rs11682649 | 0.406 | 1.598 | 0.065 | 0.263 | 0.062 | 0.011 | 0.011 | 1.045 | 0.77 | 0.51 | 0.65 | 0.76 | 0.7 | 0.83 |
| 2 | 100250011 | rs12474603 | -1.461 | 1.701 | 0.111 | 0.214 | -0.001 | 0.005 | 0.005 | 0 | 0.44 | 0.57 | 0.48 | 0.31 | 0.52 | 0.48 |
| 2 | 100252159 | rs1357719 | -0.534 | 1.65 | 0.229 | 0.185 | -0.01 | 0.036 | 0.036 | 0.156 | 0.84 | 0.91 | 0.84 | 0.92 | 0.83 | 0.91 |
| 2 | 100258343 | rs4851218 | 0.826 | 1.653 | 0.253 | 0.208 | 0.005 | -0.053 | -0.053 | 0.12 | 0.61 | 0.5 | 0.69 | 0.84 | 0.61 | 0.6 |
| 2 | 100264145 | rs6745568 | -0.52 | 1.7 | 0.152 | 0.239 | -0.008 | 0.03 | 0.03 | 0.2 | 0.84 | 0.88 | 0.84 | 0.92 | 0.83 | 0.91 |
| 2 | 100272937 | rs1916959 | 1.068 | 1.39 | 0.241 | 0.141 | -0.001 | 0.001 | 0.001 | 0 | 0.57 | 0.41 | 0.52 | 0.66 | 0.49 | 0.55 |
| 2 | 100273893 | rs6728983 | 0.551 | 1.437 | -0.024 | 0.185 | -0.044 | 0.046 | 0.046 | 0 | 0.63 | 0.82 | 0.69 | 0.76 | 0.58 | 0.62 |

| | | | | | | | | | | | | | | | | |
|---|---|---|---|---|---|---|---|---|---|---|---|---|---|---|---|---|
| 2 | 100275925 | rs11676683 | 0.822 | 1.378 | 0.068 | 0.184 | -0.029 | 0.005 | 0.005 | 0 | 0.59 | 0.74 | 0.54 | 0.66 | 0.49 | 0.55 |
| 2 | 100281046 | rs12615018 | 0.602 | 1.497 | -0.142 | 0.202 | -0.061 | -0.039 | -0.039 | 0 | 0.18 | 0.01 | 0.11 | 0.13 | 0.2 | 0.28 |
| 2 | 100291105 | rs6710594 | 0.153 | 1.451 | -0.101 | 0.198 | -0.04 | 0.009 | 0.009 | 0 | 0.82 | 0.9 | 0.7 | 0.83 | 0.71 | 0.85 |
| 2 | 100291413 | rs6542886 | 0.806 | 1.142 | -0.415 | -0.104 | -0.024 | 0.023 | 0.023 | 0 | 0.58 | 0.82 | 0.56 | 0.66 | 0.48 | 0.55 |
| 2 | 100295638 | rs10189976 | 0.806 | 1.07 | -0.026 | -0.139 | 0.001 | -0.003 | -0.003 | 0 | 0.58 | 0.58 | 0.56 | 0.67 | 0.5 | 0.56 |
| 2 | 100296415 | rs10205134 | 0.694 | 1.071 | 0.086 | -0.139 | -0.041 | 0.01 | 0.01 | 0 | 0.6 | 0.81 | 0.58 | 0.67 | 0.51 | 0.56 |
| 2 | 100307874 | rs882855 | 0.396 | 1.139 | 0.328 | -0.142 | -0.005 | 0.022 | 0.022 | 0 | 0.63 | 0.74 | 0.73 | 0.79 | 0.63 | 0.64 |
| 2 | 100324766 | rs12712058 | 0.446 | 1.096 | 0.3 | -0.142 | 0.005 | -0.005 | -0.005 | 0.067 | 0.22 | 0.27 | 0.15 | 0.13 | 0.24 | 0.29 |
| 2 | 100328931 | rs6708267 | -0.127 | 1.089 | 0.285 | -0.155 | 0.001 | -0.004 | -0.004 | 0 | 0.25 | 0.27 | 0.16 | 0.13 | 0.27 | 0.3 |
| 2 | 100335761 | rs6542889 | 0.988 | 1.082 | 0.261 | -0.139 | -0.002 | 0.007 | 0.007 | 0 | 0.55 | 0.61 | 0.57 | 0.67 | 0.5 | 0.59 |
| 2 | 100337252 | rs13431191 | 0.804 | 1.052 | 0.112 | -0.136 | 0 | -0.004 | -0.004 | 0 | 0.58 | 0.46 | 0.55 | 0.67 | 0.49 | 0.58 |
| 2 | 100346964 | rs2309723 | 0.833 | 0.947 | 0.214 | 0.049 | 0.007 | -0.014 | -0.014 | 0 | 0.57 | 0.49 | 0.67 | 0.71 | 0.55 | 0.67 |
| 2 | 100358932 | rs1404177 | 0.148 | 1.005 | 0.25 | 0.017 | 0.037 | -0.086 | -0.086 | 0 | 0.79 | 0.55 | 0.8 | 0.91 | 0.78 | 0.92 |
| 2 | 100360350 | rs7601137 | 0.84 | 0.578 | -0.327 | -0.624 | 0.004 | -0.01 | -0.01 | 0 | 0.16 | 0.16 | 0.25 | 0.01 | 0.12 | 0.15 |
| 2 | 100364339 | rs6744510 | -0.902 | 0.643 | -0.268 | -0.546 | -0.009 | 0.03 | 0.03 | 0 | 0.83 | 0.92 | 0.86 | 0.91 | 0.83 | 0.94 |
| 2 | 100376139 | rs6756821 | -1.686 | 0.547 | -0.289 | -0.541 | -0.045 | 0 | 0 | 2.566 | 0.9 | 1 | 0.86 | 0.91 | 0.86 | 0.94 |
| 2 | 100377321 | rs6721652 | -0.604 | 0.521 | -0.242 | -0.557 | 0 | 0.023 | 0.023 | 0.062 | 0.84 | 0.92 | 0.85 | 0.92 | 0.85 | 0.94 |
| 2 | 100379253 | rs17023082 | 1.565 | -0.008 | -0.389 | -0.234 | 0.014 | -0.042 | -0.042 | 0 | 0.27 | 0.08 | 0.21 | 0.39 | 0.25 | 0.23 |
| 2 | 100379717 | rs7589376 | 0.293 | 0.032 | -0.279 | -0.343 | 0.012 | -0.097 | -0.097 | 0 | 0.76 | 0.53 | 0.66 | 0.9 | 0.78 | 0.81 |
| 2 | 100383580 | rs6746134 | 1.422 | 0.131 | -0.392 | -0.319 | -0.089 | -0.066 | -0.066 | 0.022 | 0.22 | 0.01 | 0.14 | 0.2 | 0.26 | 0.29 |
| 2 | 100405753 | rs10204617 | 0.052 | 0.188 | 0.142 | -0.177 | 0.011 | -0.027 | -0.027 | 2.25 | 0.79 | 0.44 | 0.8 | 0.84 | 0.81 | 0.91 |
| 2 | 100410644 | rs10496342 | 0.72 | 0.258 | -0.007 | -0.13 | -0.001 | -0.001 | -0.001 | 0 | 0.13 | 0.1 | 0.1 | 0.19 | 0.16 | 0.17 |
| 2 | 100421767 | rs1568786 | 0.134 | 0.224 | 0.046 | -0.218 | 0.001 | -0.027 | -0.027 | 0 | 0.78 | 0.41 | 0.8 | 0.83 | 0.8 | 0.91 |
| 2 | 100424310 | rs10185087 | 0.73 | 0.223 | -0.24 | -0.223 | -0.008 | -0.008 | -0.008 | 1.019 | 0.65 | 0.29 | 0.61 | 0.64 | 0.62 | 0.69 |
| 2 | 100441349 | rs1366765 | 0.414 | 0.324 | 0.038 | -0.162 | 0 | -0.002 | -0.002 | 0 | 0.7 | 0.36 | 0.6 | 0.63 | 0.74 | 0.8 |
| 2 | 100445030 | rs7425117 | 0.414 | 0.324 | 0.038 | -0.174 | 0 | -0.001 | -0.001 | 0 | 0.7 | 0.36 | 0.6 | 0.64 | 0.74 | 0.81 |
| 2 | 100457033 | rs12473053 | 0.299 | 0.318 | -0.052 | -0.192 | 0.03 | -0.034 | -0.034 | 0 | 0.18 | 0.05 | 0.21 | 0.18 | 0.08 | 0.15 |
| 2 | 100459435 | rs6542902 | -1.462 | 0.338 | 0.277 | -0.183 | 0.089 | 0.088 | 0.088 | 0 | 0.89 | 0.6 | 0.84 | 0.83 | 0.85 | 0.94 |
| 2 | 100462110 | rs4850915 | -0.086 | 0.336 | 0.285 | -0.185 | 0.007 | -0.003 | -0.003 | 0 | 0.2 | 0.15 | 0.23 | 0.18 | 0.12 | 0.17 |
| 2 | 100464187 | rs6756248 | 1.268 | 0.334 | 0.125 | -0.259 | -0.009 | -0.047 | -0.047 | 0.113 | 0.12 | 0.18 | 0.09 | 0.01 | 0.09 | 0.07 |
| 2 | 100466155 | rs17023216 | 0.793 | 0.544 | 0.024 | -0.04 | -0.004 | -0.002 | -0.002 | 0 | 0.06 | 0.07 | 0.03 | 0 | 0.1 | 0.12 |
| 2 | 100473686 | rs7576327 | 0.754 | 0.278 | 0.227 | 0.214 | -0.004 | 0.001 | 0.001 | 0 | 0.68 | 0.42 | 0.59 | 0.67 | 0.73 | 0.77 |
| 2 | 100474071 | rs17023232 | -0.239 | 0.243 | -0.03 | 0.153 | 0.013 | -0.01 | -0.01 | 0 | 0.2 | 0.11 | 0.25 | 0.18 | 0.13 | 0.17 |
| 2 | 100480281 | rs1346623 | 0.252 | 0.767 | 0.473 | -0.287 | -0.003 | -0.001 | -0.001 | 0 | 0.51 | 0.5 | 0.59 | 0.57 | 0.39 | 0.46 |
| 2 | 100480919 | rs2309687 | 0.375 | 0.737 | 0.537 | -0.302 | 0.051 | -0.012 | -0.012 | 0 | 0.48 | 0.17 | 0.56 | 0.57 | 0.39 | 0.46 |
| 2 | 100480951 | rs6715454 | 0.274 | 0.844 | 0.504 | -0.305 | -0.01 | 0.012 | 0.012 | 0 | 0.37 | 0.3 | 0.25 | 0.25 | 0.42 | 0.46 |
| 2 | 100482934 | rs11691830 | 0.004 | 0.793 | 0.486 | -0.306 | 0.005 | 0.002 | 0.002 | 0 | 0.53 | 0.61 | 0.62 | 0.57 | 0.44 | 0.48 |
| 2 | 100487001 | rs11123802 | -0.007 | 0.79 | 0.338 | -0.283 | 0.014 | -0.003 | -0.003 | 0.195 | 0.53 | 0.68 | 0.62 | 0.55 | 0.43 | 0.48 |
| 2 | 100493439 | rs4850917 | 0.253 | 0.851 | 0.315 | -0.425 | 0.049 | 0.016 | 0.016 | 0 | 0.49 | 0.3 | 0.57 | 0.57 | 0.4 | 0.46 |
| 2 | 100508244 | rs1366762 | -0.992 | 0.689 | -0.093 | -0.425 | -0.048 | -0.025 | -0.025 | 0 | 0.87 | 0.98 | 0.84 | 0.83 | 0.81 | 0.92 |
| 2 | 100516134 | rs6735012 | -0.227 | 0.429 | 0.543 | -0.798 | -0.009 | 0.018 | 0.018 | 0 | 0.43 | 0.37 | 0.27 | 0.21 | 0.47 | 0.48 |
| 2 | 100520274 | rs6733591 | -0.037 | 0.814 | 0.44 | -0.454 | 0.014 | -0.018 | -0.018 | 0 | 0.32 | 0.36 | 0.19 | 0.18 | 0.39 | 0.42 |
| 2 | 100534633 | rs10183501 | -0.447 | 0.964 | -0.453 | -0.126 | -0.006 | 0.01 | 0.01 | 0.013 | 0.35 | 0.66 | 0.22 | 0.18 | 0.4 | 0.43 |
| 2 | 100558451 | rs11897706 | -0.993 | 0.845 | -0.09 | -0.183 | 0.011 | -0.071 | -0.071 | 3.046 | 0.13 | 0.27 | 0.05 | 0 | 0.12 | 0.08 |
| 2 | 100576807 | rs11887597 | -0.624 | 0.654 | 0.481 | -0.132 | -0.004 | 0.024 | 0.024 | 0 | 0.51 | 0.74 | 0.39 | 0.57 | 0.46 | 0.49 |
| 2 | 100584535 | rs7576469 | -0.513 | 0.514 | 0.8 | -0.084 | 0.02 | -0.095 | -0.095 | 0.581 | 0.14 | 0.38 | 0.07 | 0 | 0.12 | 0.08 |
| 2 | 100596865 | rs10202379 | -0.505 | 0.458 | 0.495 | -0.241 | -0.001 | 0.006 | 0.006 | 0 | 0.33 | 0.65 | 0.28 | 0.5 | 0.31 | 0.29 |
| 2 | 100602759 | rs13032879 | 1.232 | 0.462 | 0.326 | -0.229 | -0.014 | -0.112 | -0.112 | 0.119 | 0.17 | 0.02 | 0.18 | 0.5 | 0.17 | 0.2 |
| 2 | 100621265 | rs6736239 | -0.583 | 0.328 | 0.816 | -0.417 | 0.009 | -0.105 | -0.105 | 0.282 | 0.16 | 0.64 | 0.1 | 0 | 0.14 | 0.08 |
| 2 | 100636757 | rs11685341 | -1.006 | 0.356 | 0.529 | -0.047 | -0.013 | 0.028 | 0.028 | 0 | 0.54 | 0.66 | 0.41 | 0.37 | 0.47 | 0.48 |
| 2 | 100637331 | rs11675333 | 0.854 | 0.262 | 0.453 | -0.16 | 0.021 | 0.072 | 0.072 | 0 | 0.22 | 0.05 | 0.12 | 0.07 | 0.17 | 0.21 |
| 2 | 100647872 | rs1115910 | -1.034 | 0.261 | 0.451 | -0.075 | -0.014 | 0.025 | 0.025 | 0 | 0.55 | 0.67 | 0.42 | 0.37 | 0.48 | 0.57 |
| 2 | 100662618 | rs9308833 | -0.394 | 0.261 | 0.579 | -0.098 | 0.018 | 0.019 | 0.019 | 0 | 0.45 | 0.33 | 0.37 | 0.37 | 0.36 | 0.43 |
| 2 | 100665673 | rs2028137 | -1.819 | 0.376 | -0.298 | -0.201 | -0.046 | -0.02 | -0.02 | 0 | 0.88 | 0.99 | 0.91 | 0.89 | 0.85 | 0.91 |
| 2 | 100671449 | rs4851244 | 1.362 | 0.236 | -0.293 | -0.186 | -0.021 | -0.069 | -0.069 | 0.181 | 0.21 | 0.06 | 0.21 | 0.28 | 0.18 | 0.2 |
| 2 | 100678198 | rs11695782 | 1.25 | 0.279 | -0.233 | -0.26 | 0.01 | 0.023 | 0.023 | 0 | 0.22 | 0.12 | 0.13 | 0.09 | 0.17 | 0.21 |
| 2 | 100679024 | rs11681737 | 0.077 | 0.317 | -0.23 | -0.248 | 0.02 | 0.021 | 0.021 | 0 | 0.44 | 0.19 | 0.34 | 0.37 | 0.35 | 0.41 |
| 2 | 100683550 | rs17023459 | 0.356 | 0.522 | -0.094 | -0.102 | 0.052 | -0.037 | -0.037 | 0 | 0.07 | 0.19 | 0.05 | 0 | 0.12 | 0.14 |
| 2 | 100687778 | rs1370355 | -1.142 | 0.65 | -0.464 | 0.125 | -0.051 | -0.084 | -0.084 | 0.912 | 0.84 | 0.96 | 0.88 | 0.6 | 0.76 | 0.82 |
| 2 | 100729293 | rs4851250 | 1.285 | 0.558 | -0.303 | 0.076 | -0.029 | -0.083 | -0.083 | 0 | 0.21 | 0.03 | 0.2 | 0.28 | 0.18 | 0.2 |
| 2 | 100735567 | rs6743817 | 0.043 | 0.455 | 0.623 | -0.465 | 0.102 | -0.024 | -0.024 | 0.358 | 0.07 | 0.29 | 0.06 | 0 | 0.12 | 0.08 |
| 2 | 100735856 | rs10199628 | 0.438 | 0.464 | 0.634 | -0.461 | 0.049 | -0.034 | -0.034 | 0 | 0.06 | 0.14 | 0.05 | 0 | 0.12 | 0.08 |
| 2 | 100739732 | rs17437538 | 1.032 | -0.021 | 0.675 | -0.796 | 0.018 | 0.021 | 0.021 | 0 | 0.21 | 0.17 | 0.09 | 0.09 | 0.11 | 0.18 |
| 2 | 100739970 | rs6706188 | -0.402 | -0.107 | 0.788 | -0.835 | 0.006 | 0.003 | 0.003 | 0 | 0.31 | 0.72 | 0.18 | 0.09 | 0.24 | 0.26 |
| 2 | 100746395 | rs2118280 | -0.023 | -0.108 | 0.468 | -0.752 | 0.003 | 0.006 | 0.006 | 0 | 0.52 | 0.45 | 0.37 | 0.63 | 0.39 | 0.4 |
| 2 | 100746573 | rs11692215 | 0.794 | -0.288 | 0.578 | -0.838 | 0.003 | -0.013 | -0.013 | 0 | 0.31 | 0.22 | 0.28 | 0.55 | 0.27 | 0.24 |
| 2 | 100746935 | rs11885529 | -0.448 | -0.184 | 0.658 | -0.857 | -0.002 | 0.01 | 0.01 | 0 | 0.56 | 0.56 | 0.41 | 0.63 | 0.5 | 0.47 |
| 2 | 100747357 | rs6713524 | 1.295 | -0.282 | 0.128 | -0.791 | -0.045 | -0.089 | -0.089 | 0 | 0.27 | 0.06 | 0.26 | 0.55 | 0.27 | 0.23 |
| 2 | 100753490 | rs4851251 | 1.875 | -0.605 | -0.261 | -1.114 | -0.076 | -0.004 | -0.004 | 0.25 | 0.14 | 0.02 | 0.35 | 0.11 | 0.26 | 0.21 |

| | | | | | | | | | | | | | | | | |
|---|---|---|---|---|---|---|---|---|---|---|---|---|---|---|---|---|
| 2 | 100757308 | rs4850920 | 1.47 | -0.639 | -0.048 | -0.934 | -0.008 | -0.045 | -0.045 | 0.545 | 0.27 | 0.17 | 0.27 | 0.48 | 0.27 | 0.23 |
| 2 | 100759030 | rs17023563 | 0.346 | -0.679 | 0.01 | -0.956 | -0.026 | -0.032 | -0.032 | 0 | 0.06 | 0.17 | 0.02 | 0 | 0.01 | 0.01 |
| 2 | 100759078 | rs13003982 | 0.66 | -0.557 | 0.021 | -0.917 | 0.011 | 0.021 | 0.021 | 0 | 0.44 | 0.24 | 0.33 | 0.55 | 0.38 | 0.38 |
| 2 | 100759810 | rs11682173 | 1.451 | -0.593 | 0.002 | -0.906 | -0.008 | -0.045 | -0.045 | 0 | 0.27 | 0.17 | 0.26 | 0.48 | 0.27 | 0.23 |
| 2 | 100763006 | rs12712066 | -0.415 | -0.735 | -0.308 | -0.984 | 0.003 | -0.009 | -0.009 | 0 | 0.68 | 0.57 | 0.73 | 0.66 | 0.66 | 0.63 |
| 2 | 100765077 | rs2009094 | 0.08 | -0.462 | -0.57 | -0.614 | 0.006 | 0.01 | 0.01 | 0 | 0.45 | 0.39 | 0.35 | 0.55 | 0.38 | 0.38 |
| 2 | 100767082 | rs11123814 | 0.067 | -0.46 | -0.586 | -0.633 | 0.007 | 0.012 | 0.012 | 0 | 0.46 | 0.4 | 0.35 | 0.55 | 0.38 | 0.38 |
| 2 | 100783653 | rs11123816 | 0.866 | -0.055 | -1.521 | -0.273 | -0.057 | -0.14 | -0.14 | 0 | 0.27 | 0.03 | 0.26 | 0.47 | 0.27 | 0.23 |
| 2 | 100801867 | rs4851263 | -1.308 | 0.049 | -1.277 | -0.245 | -0.017 | -0.062 | -0.062 | 0 | 0.89 | 0.99 | 0.94 | 0.73 | 0.88 | 0.97 |
| 2 | 100803778 | rs6722241 | 0.323 | 0.089 | -0.52 | -0.221 | -0.002 | 0.018 | 0.018 | 0 | 0.3 | 0.4 | 0.31 | 0.57 | 0.28 | 0.25 |
| 2 | 100805996 | rs11691869 | 0.889 | 0.077 | -0.764 | -0.14 | -0.112 | -0.154 | -0.154 | 0.122 | 0.25 | 0.02 | 0.29 | 0.54 | 0.35 | 0.24 |
| 2 | 100813499 | rs6740838 | -0.527 | -0.492 | -0.632 | -0.498 | -0.003 | -0.002 | -0.002 | 0 | 0.57 | 0.46 | 0.43 | 0.6 | 0.61 | 0.48 |
| 2 | 100820431 | rs11689201 | 0.794 | -0.024 | -0.914 | -0.092 | -0.14 | -0.15 | -0.15 | 0.012 | 0.26 | 0.02 | 0.29 | 0.56 | 0.38 | 0.25 |
| 2 | 100833898 | rs13432804 | 0.094 | -0.476 | -0.043 | -0.125 | 0.003 | -0.031 | -0.031 | 0 | 0.06 | 0.14 | 0.03 | 0 | 0.07 | 0.06 |
| 2 | 100849928 | rs17023791 | -1.498 | -0.692 | 0.607 | -0.245 | -0.006 | -0.065 | -0.065 | 0 | 0.11 | 0.38 | 0.15 | 0.01 | 0.08 | 0.17 |
| 2 | 100868488 | rs17023825 | -0.41 | -0.937 | 0.692 | -0.656 | -0.02 | -0.088 | -0.088 | 0.443 | 0.37 | 0.46 | 0.27 | 0.08 | 0.28 | 0.27 |
| 2 | 100870862 | rs4851274 | -0.625 | -0.943 | 0.481 | -0.236 | 0.009 | -0.002 | -0.002 | 0 | 0.63 | 0.65 | 0.47 | 0.63 | 0.68 | 0.53 |
| 2 | 100871330 | rs6722320 | 0.122 | -1.586 | 0.808 | -0.427 | -0.035 | -0.097 | -0.097 | 0 | 0.33 | 0.47 | 0.15 | 0.08 | 0.21 | 0.26 |
| 2 | 100873975 | rs13394110 | 0.956 | -1.492 | 0.482 | -0.46 | 0.012 | 0.026 | 0.026 | 0 | 0.26 | 0.15 | 0.34 | 0.03 | 0.18 | 0.19 |
| 2 | 100876789 | rs4438499 | 0.587 | -1.545 | 0.749 | -0.318 | -0.003 | -0.002 | -0.002 | 0.727 | 0.35 | 0.34 | 0.28 | 0.58 | 0.51 | 0.51 |
| 2 | 100878197 | rs4851278 | -0.583 | -1.544 | 0.36 | -0.305 | 0.024 | -0.004 | -0.004 | 0 | 0.63 | 0.72 | 0.64 | 0.61 | 0.69 | 0.72 |
| 2 | 100883151 | rs13397010 | 0.968 | -1.375 | 0.17 | -0.372 | 0.008 | 0.002 | 0.002 | 0 | 0.26 | 0.2 | 0.13 | 0.03 | 0.18 | 0.18 |
| 2 | 100885992 | rs2309814 | -1.255 | -1.538 | -0.204 | -0.28 | -0.001 | 0.001 | 0.001 | 0 | 0.89 | 0.9 | 0.94 | 0.75 | 0.89 | 0.95 |
| 2 | 100887393 | rs4851281 | 0.429 | -1.636 | -0.292 | -0.569 | 0.029 | 0.055 | 0.055 | 0 | 0.31 | 0.24 | 0.35 | 0.03 | 0.2 | 0.2 |
| 2 | 100892018 | rs4622752 | 1.285 | -1.597 | -0.524 | -0.03 | -0.179 | -0.173 | -0.173 | 0 | 0.28 | 0.02 | 0.24 | 0.5 | 0.49 | 0.28 |
| 2 | 100893113 | rs6542924 | -0.853 | -1.758 | -0.568 | -0.307 | -0.055 | 0.218 | 0.218 | 0 | 0.65 | 0.93 | 0.82 | 0.96 | 0.7 | 0.8 |
| 2 | 100898081 | rs6542925 | 0.542 | -1.78 | -0.013 | -0.127 | -0.003 | -0.004 | -0.004 | 1.539 | 0.28 | 0.25 | 0.44 | 0.41 | 0.18 | 0.27 |
| 2 | 100898634 | rs2309818 | -0.325 | -1.797 | 0.53 | -0.271 | -0.001 | -0.055 | -0.055 | 0.008 | 0.58 | 0.58 | 0.7 | 0.94 | 0.65 | 0.56 |
| 2 | 100899665 | rs2309819 | -0.325 | -1.797 | 0.53 | -0.271 | -0.001 | -0.055 | -0.055 | 0 | 0.58 | 0.58 | 0.7 | 0.94 | 0.65 | 0.56 |
| 2 | 100900304 | rs2309821 | -0.677 | -1.811 | 0.597 | -0.291 | -0.013 | 0.111 | 0.111 | 0 | 0.62 | 0.81 | 0.72 | 0.95 | 0.66 | 0.75 |
| 2 | 100900379 | rs3748928 | 0.632 | -1.789 | 0.329 | -0.261 | 0 | -0.005 | -0.005 | 0 | 0.28 | 0.25 | 0.44 | 0.4 | 0.17 | 0.27 |
| 2 | 100900406 | rs2309822 | -0.849 | -1.633 | -0.041 | -0.309 | -0.086 | 0.23 | 0.23 | 0 | 0.65 | 0.96 | 0.73 | 0.96 | 0.66 | 0.76 |
| 2 | 100910578 | rs4850925 | -0.097 | -1.679 | 0.331 | 0.119 | -0.082 | 0.039 | 0.039 | 1.818 | 0.35 | 0.47 | 0.46 | 0.46 | 0.19 | 0.28 |
| 2 | 100915772 | rs4851287 | -0.755 | -1.566 | -0.217 | -0.032 | -0.027 | 0.131 | 0.131 | 0.005 | 0.65 | 0.89 | 0.74 | 0.94 | 0.68 | 0.56 |
| 2 | 100916315 | rs11123823 | 0.922 | -0.853 | 0.028 | 0.452 | -0.186 | -0.172 | -0.172 | 0 | 0.27 | 0.03 | 0.24 | 0.49 | 0.49 | 0.28 |
| 2 | 100936276 | rs11123826 | 0.389 | -0.785 | 0.558 | 0.221 | -0.076 | 0.023 | 0.023 | 0.122 | 0.34 | 0.14 | 0.48 | 0.78 | 0.49 | 0.29 |
| 2 | 100954659 | rs7581011 | -0.295 | -0.95 | 1.231 | 0.115 | -0.013 | 0.025 | 0.025 | 0 | 0.42 | 0.68 | 0.52 | 0.78 | 0.51 | 0.3 |
| 2 | 100956949 | rs2309798 | 0.283 | -0.946 | 1.642 | 0.138 | 0.026 | -0.007 | -0.007 | 0.735 | 0.36 | 0.55 | 0.49 | 0.72 | 0.49 | 0.29 |
| 2 | 100966244 | rs876439 | -1.641 | -1.011 | 1.878 | 0.179 | -0.04 | 0.009 | 0.009 | 0.428 | 0.1 | 0.23 | 0.12 | 0.09 | 0.03 | 0.22 |
| 2 | 100975297 | rs13404965 | -1.374 | -0.98 | 1.523 | 0.162 | -0.027 | -0.005 | -0.005 | 0.025 | 0.1 | 0.55 | 0.13 | 0.03 | 0.02 | 0.21 |
| 2 | 100976685 | rs11685401 | -0.465 | -1.005 | 1.349 | 0.126 | -0.018 | 0.046 | 0.046 | 0 | 0.44 | 0.69 | 0.59 | 0.75 | 0.52 | 0.51 |
| 2 | 100987387 | rs1437972 | -1.478 | -1.021 | 1.42 | 0.163 | -0.041 | 0.03 | 0.03 | 2.532 | 0.11 | 0.56 | 0.14 | 0.09 | 0.02 | 0.21 |
| 2 | 101005145 | rs4850931 | -0.08 | -1.005 | 1.476 | 0.105 | 0.021 | -0.032 | -0.032 | 0 | 0.39 | 0.46 | 0.56 | 0.81 | 0.5 | 0.5 |
| 2 | 101006787 | rs7587333 | -0.635 | -1.009 | 1.494 | 0.107 | 0.059 | 0.101 | 0.101 | 0.384 | 0.05 | 0.35 | 0.11 | 0.03 | 0.01 | 0.21 |
| 2 | 101007039 | rs4149523 | -0.58 | -1.06 | 1.567 | 0.088 | -0.028 | 0.117 | 0.117 | 0 | 0.46 | 0.78 | 0.6 | 0.81 | 0.52 | 0.51 |
| 2 | 101007178 | rs1030902 | 0.506 | -1.044 | 1.586 | 0.107 | -0.026 | 0.003 | 0.003 | 0.1 | 0.34 | 0.27 | 0.47 | 0.72 | 0.49 | 0.29 |
| 2 | 101008233 | rs6755285 | -1.119 | -1.039 | 1.586 | 0.113 | 0.034 | 0.123 | 0.123 | 0.551 | 0.06 | 0.36 | 0.11 | 0.09 | 0.01 | 0.21 |
| 2 | 101009082 | rs1057056 | -1.121 | -1.038 | 1.544 | 0.066 | 0.038 | 0.13 | 0.13 | 0.525 | 0.06 | 0.37 | 0.11 | 0.09 | 0.01 | 0.21 |
| 2 | 101010082 | rs3748930 | -0.165 | -1.018 | 1.448 | 0.107 | -0.015 | 0.021 | 0.021 | 0.878 | 0.6 | 0.63 | 0.58 | 0.81 | 0.5 | 0.5 |
| 2 | 101011618 | rs6754869 | -1.766 | -0.995 | 1.466 | -0.007 | 0.011 | 0.104 | 0.104 | 1.538 | 0.08 | 0.47 | 0.12 | 0.09 | 0.01 | 0.21 |
| 2 | 101011877 | rs2241810 | 0.628 | -1 | 1.342 | -0.018 | -0.104 | -0.015 | -0.015 | 0.345 | 0.33 | 0.1 | 0.45 | 0.72 | 0.49 | 0.29 |
| 2 | 101013649 | rs4149518 | 0.472 | -0.67 | 1.561 | 0.161 | -0.106 | -0.012 | -0.012 | 0 | 0.33 | 0.1 | 0.45 | 0.72 | 0.49 | 0.29 |
| 2 | 101014247 | rs4149517 | -0.724 | -0.675 | 1.635 | 0.167 | -0.025 | 0.109 | 0.109 | 0.01 | 0.46 | 0.78 | 0.6 | 0.8 | 0.52 | 0.51 |
| 2 | 101014363 | rs2241809 | 0.61 | -0.665 | 1.644 | 0.172 | -0.183 | -0.067 | -0.067 | 1.284 | 0.32 | 0.03 | 0.45 | 0.71 | 0.49 | 0.29 |
| 2 | 101018411 | rs4149516 | -2.034 | -0.664 | 1.681 | 0.141 | -0.058 | 0.008 | 0.008 | 1.849 | 0.12 | 0.51 | 0.13 | 0.09 | 0.03 | 0.22 |
| 2 | 101018445 | rs4149515 | -1.407 | -0.661 | 1.677 | 0.143 | 0.029 | 0.13 | 0.13 | 0.39 | 0.07 | 0.44 | 0.11 | 0.09 | 0.01 | 0.21 |
| 2 | 101019663 | rs4850932 | 1.351 | -0.653 | 1.55 | 0.09 | 0.054 | 0.027 | 0.027 | 0 | 0.89 | 0.55 | 0.84 | 0.91 | 0.9 | 0.78 |
| 2 | 101020137 | rs7575422 | -1.788 | -0.66 | 1.466 | 0.105 | 0.019 | 0.112 | 0.112 | 0.536 | 0.07 | 0.44 | 0.11 | 0.09 | 0.01 | 0.21 |
| 2 | 101020624 | rs6725672 | -1.269 | -0.661 | 1.478 | 0.105 | 0.03 | 0.117 | 0.117 | 0.544 | 0.06 | 0.35 | 0.11 | 0.09 | 0.01 | 0.21 |
| 2 | 101020898 | rs4149514 | -1.193 | -0.658 | 1.276 | 0.107 | 0.031 | 0.133 | 0.133 | 0.301 | 0.07 | 0.45 | 0.11 | 0.09 | 0.01 | 0.21 |
| 2 | 101021090 | rs2099611 | -0.836 | -0.657 | 1.273 | 0.11 | -0.025 | 0.109 | 0.109 | 0 | 0.46 | 0.78 | 0.6 | 0.81 | 0.52 | 0.51 |
| 2 | 101021135 | rs6758622 | -1.467 | -0.685 | 1.283 | 0.109 | 0.019 | 0.112 | 0.112 | 0.536 | 0.07 | 0.44 | 0.11 | 0.09 | 0.01 | 0.21 |
| 2 | 101022886 | rs4149512 | 0.378 | -0.671 | 1.244 | 0.106 | -0.101 | -0.015 | -0.015 | 0.065 | 0.33 | 0.1 | 0.45 | 0.72 | 0.49 | 0.29 |
| 2 | 101023635 | rs4149510 | 0.38 | -0.682 | 1.238 | 0.1 | -0.101 | -0.016 | -0.016 | 0 | 0.32 | 0.1 | 0.45 | 0.72 | 0.49 | 0.29 |
| 2 | 101023699 | rs4149509 | -1.463 | -0.679 | 1.275 | -0.043 | 0.02 | 0.114 | 0.114 | 0.536 | 0.07 | 0.45 | 0.12 | 0.09 | 0.01 | 0.21 |
| 2 | 101025811 | rs10172841 | -2.03 | -0.686 | 1.277 | -0.046 | -0.058 | 0.008 | 0.008 | 1.849 | 0.12 | 0.51 | 0.13 | 0.09 | 0.03 | 0.22 |
| 2 | 101027400 | rs7582096 | -1.528 | -0.682 | 1.239 | -0.048 | 0.016 | 0.095 | 0.095 | 0.332 | 0.08 | 0.44 | 0.12 | 0.09 | 0.02 | 0.22 |
| 2 | 101028236 | rs7585713 | -1.569 | -0.682 | 1.222 | -0.048 | 0.014 | 0.093 | 0.093 | 0.434 | 0.08 | 0.45 | 0.12 | 0.09 | 0.02 | 0.22 |
| 2 | 101029002 | rs4851313 | -0.901 | -0.679 | 1.038 | 0.132 | -0.011 | 0.074 | 0.074 | 0 | 0.46 | 0.79 | 0.6 | 0.77 | 0.52 | 0.51 |

| | | | | | | | | | | | | | | | | |
|---|---|---|---|---|---|---|---|---|---|---|---|---|---|---|---|---|
| 2 | 101031561 | rs3828193 | 0.462 | -0.573 | 0.822 | 0.172 | -0.177 | -0.11 | -0.11 | 0 | 0.32 | 0.03 | 0.45 | 0.68 | 0.49 | 0.29 |
| 2 | 101032745 | rs4149507 | 0.338 | -0.536 | 0.886 | 0.191 | -0.101 | -0.051 | -0.051 | 1.107 | 0.33 | 0.1 | 0.45 | 0.68 | 0.49 | 0.29 |
| 2 | 101032816 | rs4149506 | -2.096 | -0.196 | 1.838 | 0.023 | 0.017 | 0.104 | 0.104 | 0.806 | 0.08 | 0.47 | 0.12 | 0.09 | 0.01 | 0.22 |
| 2 | 101038647 | rs13400841 | -2.164 | -0.554 | 2.013 | 0.193 | 0.011 | 0.09 | 0.09 | 0.884 | 0.08 | 0.47 | 0.12 | 0.09 | 0.02 | 0.23 |
| 2 | 101039956 | rs2309801 | -2.164 | -0.554 | 2.013 | 0.193 | 0.011 | 0.09 | 0.09 | 0.884 | 0.08 | 0.47 | 0.12 | 0.09 | 0.02 | 0.22 |
| 2 | 101045508 | rs6542950 | -1.786 | 0.031 | 1.611 | 0.017 | 0.01 | -0.042 | -0.042 | 0.078 | 0.16 | 0.65 | 0.19 | 0.07 | 0.17 | 0.25 |
| 2 | 101048668 | rs12328503 | -1.417 | 0.173 | 0.896 | 0.102 | 0.045 | -0.026 | -0.026 | 0 | 0.13 | 0.31 | 0.16 | 0.07 | 0.16 | 0.24 |
| 2 | 101052766 | rs4851316 | -0.29 | -0.594 | 0.289 | 0.055 | -0.015 | -0.006 | -0.006 | 0 | 0.4 | 0.4 | 0.46 | 0.58 | 0.27 | 0.47 |
| 2 | 101065563 | rs12622229 | 0.807 | -0.857 | -0.17 | 0.262 | 0.021 | -0.095 | -0.095 | 0 | 0.15 | 0.01 | 0.06 | 0.32 | 0.09 | 0.25 |
| 2 | 101066051 | rs6716842 | -1.005 | -0.34 | -0.167 | -0.793 | -0.05 | 0.195 | 0.195 | 0 | 0.6 | 0.87 | 0.68 | 0.92 | 0.48 | 0.78 |
| 2 | 101066296 | rs4850933 | -1 | -0.325 | -0.247 | -0.824 | -0.074 | 0.227 | 0.227 | 0 | 0.61 | 0.91 | 0.68 | 1 | 0.48 | 0.81 |
| 2 | 101067435 | rs17024338 | 0.5 | 0.061 | 0.351 | -0.272 | 0.01 | -0.019 | -0.019 | 0.074 | 0.06 | 0.15 | 0.27 | 0.02 | 0.07 | 0.06 |
| 2 | 101086151 | rs10166865 | 0.534 | 0.028 | 0.493 | -0.14 | 0.011 | 0.043 | 0.043 | 0 | 0.17 | 0.28 | 0.19 | 0.35 | 0.2 | 0.3 |
| 2 | 101086894 | rs13385681 | 0.534 | 0.028 | 0.493 | -0.14 | 0.011 | 0.043 | 0.043 | 0 | 0.17 | 0.28 | 0.19 | 0.35 | 0.2 | 0.3 |
| 2 | 101086966 | rs13411940 | 0.534 | 0.028 | 0.493 | -0.14 | 0.011 | 0.043 | 0.043 | 0 | 0.17 | 0.28 | 0.19 | 0.35 | 0.2 | 0.3 |
| 2 | 101088840 | rs753439 | -0.298 | -0.088 | 0.446 | -0.2 | -0.004 | 0.066 | 0.066 | 0 | 0.25 | 0.55 | 0.24 | 0.43 | 0.23 | 0.37 |
| 2 | 101096960 | rs1519654 | -1.136 | -0.235 | -0.181 | -0.877 | -0.103 | 0.206 | 0.206 | 0 | 0.63 | 0.92 | 0.69 | 1 | 0.5 | 0.84 |
| 2 | 101098494 | rs6735337 | -0.107 | -0.296 | 0.181 | -0.791 | 0 | 0.058 | 0.058 | 0 | 0.25 | 0.54 | 0.24 | 0.43 | 0.24 | 0.36 |
| 2 | 101100231 | rs1589246 | 0.443 | -0.379 | 0.473 | -0.848 | 0.028 | -0.015 | -0.015 | 0 | 0.07 | 0.22 | 0.27 | 0.02 | 0.07 | 0.06 |
| 2 | 101103245 | rs17024409 | 0.516 | -0.374 | 0.635 | -0.835 | 0.002 | 0.016 | 0.016 | 0 | 0.2 | 0.23 | 0.21 | 0.35 | 0.21 | 0.28 |
| 2 | 101103384 | rs10183360 | 0.119 | -0.386 | 0.454 | -0.842 | 0.003 | 0.051 | 0.051 | 0 | 0.23 | 0.37 | 0.22 | 0.43 | 0.22 | 0.36 |
| 2 | 101121719 | rs964559 | -1.391 | -0.46 | 0.195 | -0.955 | -0.106 | 0.196 | 0.196 | 0 | 0.63 | 0.92 | 0.69 | 1 | 0.5 | 0.85 |
| 2 | 101131552 | rs869840 | 0.456 | -0.366 | 0.431 | -0.758 | 0.009 | 0.04 | 0.04 | 0 | 0.21 | 0.33 | 0.22 | 0.35 | 0.22 | 0.3 |
| 2 | 101140614 | rs6542966 | 0.578 | -0.379 | 0.406 | -0.712 | 0.028 | 0.152 | 0.152 | 0 | 0.18 | 0.42 | 0.2 | 0.4 | 0.2 | 0.28 |
| 2 | 101143716 | rs17024489 | 0.579 | -0.57 | -0.089 | -0.886 | 0.053 | -0.077 | -0.077 | 1.973 | 0.31 | 0.11 | 0.17 | 0.51 | 0.21 | 0.44 |
| 2 | 101145430 | rs2176733 | -1.384 | -0.527 | -0.14 | -0.952 | -0.131 | 0.225 | 0.225 | 0 | 0.64 | 0.94 | 0.69 | 1 | 0.5 | 0.85 |
| 2 | 101150154 | rs7596463 | 0.109 | -0.606 | 0.015 | -0.852 | 0.046 | -0.013 | -0.013 | 0 | 0.36 | 0.25 | 0.2 | 0.58 | 0.23 | 0.52 |
| 2 | 101158259 | rs11123835 | -1.497 | 0.215 | -0.468 | -1.25 | -0.027 | 0.076 | 0.076 | 0 | 0.73 | 0.9 | 0.8 | 1 | 0.68 | 0.91 |
| 2 | 101159761 | rs4851329 | -0.466 | 0.152 | -0.144 | -1.199 | 0.005 | -0.026 | -0.026 | 0 | 0.23 | 0.29 | 0.42 | 0.09 | 0.27 | 0.19 |
| 2 | 101161050 | rs9917224 | -0.5 | -0.151 | -0.031 | -1.009 | 0.001 | -0.025 | -0.025 | 0.426 | 0.32 | 0.75 | 0.58 | 0.49 | 0.3 | 0.4 |
| 2 | 101167924 | rs4851333 | 0.088 | -0.137 | 0.256 | -1.115 | -0.006 | -0.059 | -0.059 | 0 | 0.13 | 0.26 | 0.31 | 0.02 | 0.1 | 0.14 |
| 2 | 101173256 | rs3748929 | 0.557 | -0.372 | -0.318 | -1.217 | 0.049 | -0.026 | -0.026 | 0.478 | 0.33 | 0.21 | 0.14 | 0.58 | 0.22 | 0.49 |
| 2 | 101183475 | rs2946589 | -0.898 | 0.14 | -0.76 | -1.412 | -0.101 | 0.165 | 0.165 | 0 | 0.64 | 0.89 | 0.71 | 1 | 0.52 | 0.87 |
| 2 | 101192499 | rs2970997 | 0.387 | 0.22 | -0.634 | -1.702 | 0.007 | -0.104 | -0.104 | 1.109 | 0.62 | 0.6 | 0.51 | 0.97 | 0.59 | 0.76 |
| 2 | 101196111 | rs13005184 | 0.295 | 0.17 | -0.722 | -1.742 | 0.002 | -0.101 | -0.101 | 0.663 | 0.64 | 0.63 | 0.52 | 0.97 | 0.63 | 0.79 |
| 2 | 101201669 | rs7594990 | 0.827 | -0.517 | -0.459 | -1.23 | 0 | 0.099 | 0.099 | 0.102 | 0.21 | 0.38 | 0.26 | 0.46 | 0.22 | 0.29 |
| 2 | 101205999 | rs11678350 | 0.421 | -0.633 | -0.175 | -1.063 | -0.005 | -0.038 | -0.038 | 0.009 | 0.12 | 0.16 | 0.3 | 0.02 | 0.09 | 0.13 |
| 2 | 101206265 | rs13392981 | 0.516 | -0.747 | -0.211 | -1.101 | -0.039 | 0.088 | 0.088 | 0 | 0.24 | 0.4 | 0.13 | 0.51 | 0.17 | 0.42 |
| 2 | 101213038 | rs11123838 | -0.562 | -0.636 | -0.423 | -1.027 | -0.027 | 0.008 | 0.008 | 0 | 0.41 | 0.52 | 0.45 | 0.52 | 0.27 | 0.54 |
| 2 | 101223405 | rs4851346 | -0.136 | -0.324 | -0.83 | -0.427 | -0.003 | 0.022 | 0.022 | 0 | 0.17 | 0.2 | 0.33 | 0.34 | 0.17 | 0.39 |
| 2 | 101227896 | rs2971014 | -0.736 | -0.367 | -1.278 | -1.334 | -0.009 | 0.056 | 0.056 | 0.795 | 0.56 | 0.69 | 0.69 | 1 | 0.49 | 0.85 |
| 2 | 101229192 | rs11123839 | 0.51 | 0.035 | -1.081 | -0.482 | -0.005 | 0 | 0 | 0 | 0.16 | 0.08 | 0.11 | 0.18 | 0.16 | 0.18 |
| 2 | 101236680 | rs1879121 | -0.592 | 0.023 | -0.91 | -0.413 | -0.041 | -0.02 | -0.02 | 0 | 0.42 | 0.24 | 0.59 | 0.56 | 0.51 | 0.7 |
| 2 | 101241404 | rs1533657 | -1.088 | -0.308 | -0.872 | -1.073 | 0.042 | 0.048 | 0.048 | 0.162 | 0.74 | 0.86 | 0.9 | 0.92 | 0.89 | 0.9 |
| 2 | 101242680 | rs2946601 | -0.315 | -0.109 | -0.881 | -0.89 | -0.03 | -0.006 | -0.006 | 0 | 0.43 | 0.22 | 0.59 | 0.5 | 0.52 | 0.66 |
| 2 | 101244758 | rs2222282 | -1.788 | -0.036 | -0.889 | -0.878 | 0.037 | 0.072 | 0.072 | 0.007 | 0.79 | 0.92 | 0.9 | 0.99 | 0.87 | 0.89 |
| 2 | 101247565 | rs7570854 | 1.193 | -0.194 | -0.211 | -0.895 | 0.038 | 0.07 | 0.07 | 0 | 0.27 | 0.42 | 0.27 | 0.45 | 0.33 | 0.21 |
| 2 | 101257623 | rs7605067 | 0.389 | -0.198 | -0.259 | -0.882 | 0.008 | 0.015 | 0.015 | 0.011 | 0.49 | 0.64 | 0.37 | 0.45 | 0.45 | 0.32 |
| 2 | 101269677 | rs4396744 | -0.36 | 0.269 | -0.35 | -0.724 | -0.054 | -0.006 | -0.006 | 0 | 0.67 | 0.85 | 0.71 | 0.59 | 0.61 | 0.39 |
| 2 | 101280697 | rs4513317 | 0.148 | 0.101 | -0.35 | -0.871 | 0.075 | 0.123 | 0.123 | 0.348 | 0.41 | 0.21 | 0.5 | 0.09 | 0.31 | 0.22 |
| 2 | 101286023 | rs10201136 | -0.93 | 0.579 | 0.159 | -0.464 | 0.015 | 0.018 | 0.018 | 0 | 0.87 | 0.76 | 0.89 | 0.76 | 0.85 | 0.65 |
| 2 | 101291137 | rs4851365 | 0.394 | 0.573 | 0.085 | -0.577 | -0.065 | 0.137 | 0.137 | 1.539 | 0.53 | 0.1 | 0.59 | 0.29 | 0.53 | 0.47 |
| 2 | 101294579 | rs4270369 | 0.359 | 0.552 | 0.674 | -0.55 | 0.041 | 0.064 | 0.064 | 0.104 | 0.15 | 0.44 | 0.12 | 0.28 | 0.15 | 0.35 |
| 2 | 101299150 | rs2971017 | 0.801 | 0.558 | 0.695 | -0.547 | 0.011 | 0.014 | 0.014 | 0.141 | 0.13 | 0.2 | 0.09 | 0.24 | 0.15 | 0.34 |
| 2 | 101305327 | rs2175793 | 1.354 | 0.524 | 0.802 | -0.593 | -0.032 | -0.056 | -0.056 | 2.867 | 0.24 | 0.15 | 0.25 | 0.43 | 0.31 | 0.18 |
| 2 | 101308711 | rs2942906 | 0.794 | 0.484 | 0.776 | -0.548 | 0.016 | 0.021 | 0.021 | 0.12 | 0.13 | 0.23 | 0.1 | 0.24 | 0.15 | 0.34 |
| 2 | 101315849 | rs2942878 | 0.644 | 0.483 | 0.516 | -0.55 | 0.022 | 0.035 | 0.035 | 0.012 | 0.13 | 0.23 | 0.1 | 0.28 | 0.15 | 0.36 |
| 2 | 101318308 | rs2942883 | -0.354 | 0.477 | 0.278 | -0.578 | -0.014 | -0.001 | -0.001 | 5.144 | 0.21 | 0.41 | 0.14 | 0.27 | 0.17 | 0.36 |
| 2 | 101319712 | rs10175405 | 0.021 | 0.598 | -0.448 | -0.987 | 0.008 | 0.158 | 0.158 | 0.731 | 0.49 | 0.07 | 0.48 | 0.28 | 0.43 | 0.41 |
| 2 | 101321075 | rs1028045 | -0.059 | 0.303 | -0.409 | -1.117 | -0.058 | 0.22 | 0.22 | 0.503 | 0.5 | 0.09 | 0.5 | 0.26 | 0.5 | 0.45 |
| 2 | 101323275 | rs2175792 | 0.431 | 0.288 | -0.211 | -1.111 | 0.006 | -0.014 | -0.014 | 1.518 | 0.39 | 0.41 | 0.44 | 0.7 | 0.49 | 0.54 |
| 2 | 101323578 | rs2942884 | 0.472 | -0.2 | -0.294 | -1.089 | 0.001 | -0.013 | -0.013 | 0 | 0.4 | 0.57 | 0.42 | 0.7 | 0.46 | 0.54 |
| 2 | 101325207 | rs13384306 | -0.509 | 0.359 | -0.39 | -0.72 | -0.01 | -0.048 | -0.048 | 3.218 | 0.08 | 0.2 | 0.11 | 0 | 0.06 | 0.03 |
| 2 | 101337982 | rs11123842 | -0.107 | 0.103 | -1.185 | -0.714 | -0.004 | 0.137 | 0.137 | 0 | 0.54 | 0.85 | 0.47 | 0.74 | 0.48 | 0.57 |
| 2 | 101338073 | rs12623994 | 0.376 | 0.663 | -1.147 | -0.437 | -0.004 | 0.04 | 0.04 | 0 | 0.43 | 0.31 | 0.41 | 0.7 | 0.47 | 0.56 |
| 2 | 101338509 | rs4851370 | -0.416 | 1.624 | -1.451 | 0.393 | -0.044 | 0.025 | 0.025 | 0 | 0.24 | 0.08 | 0.4 | 0.2 | 0.29 | 0.26 |
| 2 | 101338644 | rs1513628 | -0.467 | 1.327 | -1.436 | 0.405 | -0.068 | 0.038 | 0.038 | 0 | 0.26 | 0.08 | 0.45 | 0.2 | 0.37 | 0.27 |
| 2 | 101339816 | rs12616239 | -1.351 | 0.27 | -0.992 | 1.553 | -0.003 | 0.034 | 0.034 | 0 | 0.14 | 0.18 | 0.2 | 0.47 | 0.09 | 0.2 |
| 2 | 101343140 | rs2137671 | 0.158 | -0.047 | -0.864 | 1.491 | -0.014 | 0.038 | 0.038 | 0 | 0.68 | 0.49 | 0.48 | 0.54 | 0.74 | 0.52 |

| | | | | | | | | | | | | | | | | |
|---|---|---|---|---|---|---|---|---|---|---|---|---|---|---|---|---|
| 2 | 101343387 | rs12621996 | 1.08 | -0.185 | -1.836 | 0.956 | -0.161 | 0.215 | 0.215 | 0.766 | 0.4 | 0.01 | 0.28 | 0.08 | 0.54 | 0.17 |
| 2 | 101346696 | rs7579862 | -0.542 | -0.251 | -1.611 | 1.181 | -0.02 | 0.063 | 0.063 | 0.348 | 0.27 | 0.48 | 0.21 | 0.46 | 0.19 | 0.36 |
| 2 | 101346959 | rs17024686 | -0.406 | -0.333 | -1.636 | 1.388 | 0.004 | -0.015 | -0.015 | 0 | 0.24 | 0.22 | 0.13 | 0.34 | 0.19 | 0.31 |
| 2 | 101349735 | rs12469700 | -0.174 | -0.388 | -1.777 | 1.686 | -0.015 | -0.044 | -0.044 | 0.017 | 0.17 | 0.05 | 0.16 | 0.19 | 0.15 | 0.15 |
| 2 | 101371896 | rs17024700 | -0.218 | -0.445 | -1.743 | 1.675 | 0.006 | -0.023 | -0.023 | 0 | 0.17 | 0.05 | 0.16 | 0.15 | 0.12 | 0.14 |
| 2 | 101372680 | rs12691175 | -1.321 | -0.662 | -1.022 | 1.538 | -0.003 | 0.002 | 0.002 | 0 | 0.79 | 0.83 | 0.63 | 0.9 | 0.76 | 0.64 |
| 2 | 101376496 | rs7582332 | -1.311 | -0.758 | -0.129 | 1.983 | -0.003 | 0.006 | 0.006 | 0 | 0.27 | 0.25 | 0.27 | 0.53 | 0.16 | 0.31 |
| 2 | 101376778 | rs6542985 | -1.726 | -0.8 | 0.299 | 2.012 | -0.041 | 0.017 | 0.017 | 0 | 0.3 | 0.48 | 0.3 | 0.64 | 0.18 | 0.4 |
| 2 | 101378095 | rs6740105 | -0.16 | -0.715 | 0.338 | 2.013 | -0.014 | 0.091 | 0.091 | 0 | 0.08 | 0.1 | 0.1 | 0.37 | 0.04 | 0.17 |
| 2 | 101383053 | rs11897738 | -0.727 | -0.761 | 0.391 | 1.804 | -0.029 | 0.008 | 0.008 | 0 | 0.21 | 0.35 | 0.19 | 0.16 | 0.12 | 0.14 |
| 2 | 101395850 | rs10166048 | -1.191 | -0.11 | 0.204 | 1.823 | -0.045 | -0.106 | -0.106 | 0 | 0.29 | 0.06 | 0.24 | 0.38 | 0.22 | 0.33 |
| 2 | 101408323 | rs6740234 | -1.359 | -0.499 | 0.988 | 1.766 | -0.021 | 0.005 | 0.005 | 0 | 0.32 | 0.57 | 0.31 | 0.63 | 0.17 | 0.4 |
| 2 | 101409601 | rs7577100 | 0.475 | -0.579 | 0.937 | 1.723 | -0.018 | 0.002 | 0.002 | 0 | 0.07 | 0.1 | 0.09 | 0.09 | 0.03 | 0.01 |
| 2 | 101409898 | rs7593910 | -1.569 | -0.552 | 0.766 | 1.654 | -0.007 | 0.001 | 0.001 | 0 | 0.32 | 0.61 | 0.31 | 0.63 | 0.17 | 0.4 |
| 2 | 101413381 | rs11694387 | -1.049 | -0.557 | -0.096 | 1.123 | -0.018 | 0.053 | 0.053 | 0 | 0.71 | 0.87 | 0.59 | 0.81 | 0.69 | 0.68 |
| 2 | 101417047 | rs7419651 | 0.087 | -0.528 | 0.001 | 1.418 | 0.001 | 0.004 | 0.004 | 0 | 0.14 | 0.11 | 0.15 | 0.11 | 0.12 | 0.14 |
| 2 | 101417970 | rs11686712 | 1.244 | -0.474 | -0.531 | 1.153 | -0.177 | 0.199 | 0.199 | 0 | 0.34 | 0.01 | 0.23 | 0.09 | 0.47 | 0.25 |
| 2 | 101418410 | rs10194962 | -0.945 | -0.73 | 0.585 | 1.997 | -0.061 | 0.244 | 0.244 | 0 | 0.14 | 0.32 | 0.13 | 0.41 | 0.06 | 0.13 |
| 2 | 101419887 | rs1435510 | 0.381 | -0.735 | 0.688 | 1.979 | -0.023 | 0.007 | 0.007 | 0 | 0.07 | 0.14 | 0.08 | 0.08 | 0.03 | 0.02 |
| 2 | 101421854 | rs6754281 | -1.685 | -0.717 | 0.468 | 1.959 | 0.034 | -0.016 | -0.016 | 0.624 | 0.31 | 0.72 | 0.31 | 0.6 | 0.18 | 0.27 |
| 2 | 101423777 | rs17024792 | -0.892 | -0.684 | 0.509 | 1.99 | -0.066 | 0.245 | 0.245 | 0 | 0.15 | 0.34 | 0.13 | 0.4 | 0.06 | 0.14 |
| 2 | 101432710 | rs12475500 | -0.26 | -0.575 | 0.42 | 1.837 | -0.009 | 0.045 | 0.045 | 0 | 0.16 | 0.36 | 0.18 | 0.2 | 0.12 | 0.13 |
| 2 | 101434684 | rs3811561 | -1.32 | -0.611 | -0.135 | 1.741 | 0.024 | -0.014 | -0.014 | 0 | 0.31 | 0.69 | 0.3 | 0.59 | 0.18 | 0.26 |
| 2 | 101439596 | rs6542992 | -1.304 | -0.611 | -0.134 | 1.658 | 0.021 | -0.025 | -0.025 | 0 | 0.3 | 0.69 | 0.3 | 0.52 | 0.18 | 0.26 |
| 2 | 101449302 | rs3860455 | -1.336 | -0.822 | -0.13 | 1.62 | -0.005 | 0 | 0 | 0.03 | 0.28 | 0.71 | 0.28 | 0.25 | 0.14 | 0.19 |
| 2 | 101450679 | rs3888170 | -1.745 | -0.822 | -0.183 | 1.786 | 0.02 | -0.011 | -0.011 | 0 | 0.34 | 0.71 | 0.3 | 0.55 | 0.17 | 0.32 |
| 2 | 101451819 | rs3849381 | 0.518 | 0.369 | -0.788 | 0.878 | -0.089 | 0.012 | 0.012 | 0.016 | 0.13 | 0.02 | 0.16 | 0.06 | 0.33 | 0.1 |
| 2 | 101452840 | rs7587573 | -0.509 | -0.301 | -0.028 | 0.983 | -0.013 | 0.034 | 0.034 | 0 | 0.17 | 0.26 | 0.23 | 0.23 | 0.11 | 0.19 |
| 2 | 101456989 | rs10165970 | 0.397 | -0.784 | -0.605 | 0.224 | 0.026 | 0.015 | 0.015 | 0.099 | 0.32 | 0.19 | 0.14 | 0.09 | 0.18 | 0.21 |
| 2 | 101460043 | rs17024869 | 0.14 | -0.346 | -0.72 | 0.485 | 0 | 0.002 | 0.002 | 0.653 | 0.09 | 0.07 | 0.11 | 0.05 | 0.09 | 0.06 |
| 2 | 101465271 | rs1561002 | 0.051 | -0.291 | -1.044 | 0.632 | -0.023 | -0.023 | -0.023 | 0.643 | 0.75 | 0.92 | 0.58 | 0.8 | 0.76 | 0.81 |
| 2 | 101470349 | rs965519 | 1.633 | -0.617 | -1 | 0.318 | 0.023 | 0.053 | 0.053 | 0 | 0.24 | 0.1 | 0.14 | 0.08 | 0.18 | 0.26 |
| 2 | 101473434 | rs7598826 | 0.264 | -0.777 | -0.531 | 0.795 | 0.004 | 0.002 | 0.002 | 0 | 0.42 | 0.66 | 0.34 | 0.41 | 0.57 | 0.42 |
| 2 | 101479014 | rs1811399 | 0.67 | -0.543 | 0.228 | 1.182 | 0.001 | -0.001 | -0.001 | 0 | 0.67 | 0.66 | 0.64 | 0.7 | 0.78 | 0.74 |
| 2 | 101480401 | rs983287 | -0.046 | -0.512 | -0.035 | 1.078 | 0.026 | -0.007 | -0.007 | 0 | 0.73 | 0.83 | 0.67 | 0.78 | 0.81 | 0.81 |
| 2 | 101480885 | rs2043534 | 1.173 | -0.418 | 0.077 | 1.079 | 0.072 | 0.08 | 0.08 | 0 | 0.56 | 0.25 | 0.43 | 0.26 | 0.71 | 0.61 |
| 2 | 101487468 | rs12472321 | 0.997 | -0.398 | 0.047 | 1.006 | -0.011 | -0.001 | -0.001 | 0 | 0.58 | 0.44 | 0.45 | 0.42 | 0.71 | 0.61 |
| 2 | 101503416 | rs1369482 | -0.281 | -0.422 | 0.24 | 1.204 | -0.031 | 0.105 | 0.105 | 0.155 | 0.1 | 0.24 | 0.19 | 0.34 | 0.06 | 0.14 |
| 2 | 101506423 | rs6740935 | 1.101 | -0.064 | -0.098 | 0.805 | 0.013 | 0.026 | 0.026 | 0 | 0.49 | 0.41 | 0.29 | 0.2 | 0.39 | 0.55 |
| 2 | 101508899 | rs12712083 | 0.287 | 0.024 | -0.496 | 0.943 | -0.062 | -0.026 | -0.026 | 0 | 0.62 | 0.87 | 0.51 | 0.7 | 0.45 | 0.69 |
| 2 | 101512543 | rs17654772 | 0.732 | 0.913 | -0.37 | 0.784 | -0.067 | -0.002 | -0.002 | 0 | 0.1 | 0.01 | 0.14 | 0.06 | 0.28 | 0.1 |
| 2 | 101518910 | rs10194413 | 0.31 | 0.452 | -0.029 | 0.751 | -0.015 | 0.024 | 0.024 | 0 | 0.85 | 0.8 | 0.59 | 0.64 | 0.9 | 0.81 |
| 2 | 101521777 | rs2117714 | -0.786 | 0.278 | -0.253 | 0.439 | -0.022 | -0.005 | -0.005 | 0.048 | 0.59 | 0.53 | 0.43 | 0.56 | 0.71 | 0.52 |
| 2 | 101522266 | rs4851377 | -0.142 | 0.245 | -0.402 | 0.185 | 0 | -0.002 | -0.002 | 0 | 0.43 | 0.41 | 0.34 | 0.56 | 0.54 | 0.43 |
| 2 | 101524476 | rs11123853 | 0.492 | 0.817 | -0.613 | 0.36 | 0.001 | -0.006 | -0.006 | 0 | 0.16 | 0.11 | 0.1 | 0.45 | 0.13 | 0.16 |
| 2 | 101528047 | rs7570190 | 2.363 | 0.69 | -0.606 | 0.267 | 0.012 | 0.02 | 0.02 | 0 | 0.24 | 0.2 | 0.14 | 0.1 | 0.17 | 0.24 |
| 2 | 101529415 | rs356655 | 0.721 | 0.752 | -0.697 | 0.298 | -0.027 | -0.009 | -0.009 | 0 | 0.4 | 0.5 | 0.56 | 0.43 | 0.28 | 0.46 |
| 2 | 101529818 | rs11894671 | 0.649 | 0.948 | -0.719 | 0.51 | 0.031 | -0.002 | -0.002 | 0 | 0.1 | 0.12 | 0.13 | 0.07 | 0.23 | 0.09 |
| 2 | 101531943 | rs6729727 | 0.495 | 0.81 | -0.729 | 0.516 | 0.009 | -0.02 | -0.02 | 0 | 0.15 | 0.08 | 0.08 | 0.27 | 0.09 | 0.15 |
| 2 | 101532830 | rs11894322 | 0.591 | 0.842 | -0.846 | 0.522 | 0.036 | 0.022 | 0.022 | 0 | 0.11 | 0.12 | 0.14 | 0.24 | 0.28 | 0.1 |
| 2 | 101536192 | rs356643 | 0.923 | 0.66 | -0.941 | 0.074 | 0 | -0.001 | -0.001 | 0 | 0.58 | 0.59 | 0.68 | 0.48 | 0.47 | 0.66 |
| 2 | 101537438 | rs356642 | -0.824 | 0.346 | -1.322 | 0.417 | -0.007 | 0.007 | 0.007 | 0 | 0.84 | 0.89 | 0.91 | 1 | 0.83 | 0.9 |
| 2 | 101540415 | rs356652 | 0.314 | 0.316 | -1.422 | 0.494 | 0.003 | -0.065 | -0.065 | 0 | 0.1 | 0.01 | 0.4 | 0.33 | 0.09 | 0.19 |
| 2 | 101541798 | rs356651 | -0.611 | 0.455 | -1.229 | 0.114 | -0.011 | 0.031 | 0.031 | 0 | 0.84 | 0.92 | 0.91 | 1 | 0.83 | 0.91 |
| 2 | 101546716 | rs356649 | -0.497 | 0.529 | -1.225 | 0.107 | -0.019 | 0.033 | 0.033 | 0 | 0.82 | 0.88 | 0.87 | 1 | 0.78 | 0.9 |
| 2 | 101546879 | rs13414074 | 0.218 | -0.201 | -0.909 | 0.077 | -0.023 | -0.006 | -0.006 | 0 | 0.08 | 0.31 | 0.02 | 0.02 | 0 | 0.05 |
| 2 | 101547520 | rs356647 | 0.26 | 0.165 | -1.817 | 0.548 | 0.003 | -0.036 | -0.036 | 0 | 0.1 | 0.04 | 0.4 | 0.33 | 0.09 | 0.19 |
| 2 | 101549843 | rs3754675 | 0.26 | 0.165 | -1.817 | 0.561 | 0.003 | -0.036 | -0.036 | 0 | 0.1 | 0.04 | 0.4 | 0.33 | 0.09 | 0.19 |
| 2 | 101550181 | rs13025524 | 0.857 | 0.44 | -1.916 | 0.543 | -0.015 | 0.02 | 0.02 | 0 | 0.31 | 0.24 | 0.24 | 0.23 | 0.33 | 0.29 |
| 2 | 101550987 | rs3768984 | 0.031 | -0.234 | -1.951 | 0.653 | 0.028 | -0.048 | -0.048 | 0 | 0.31 | 0.13 | 0.16 | 0.33 | 0.25 | 0.35 |
| 2 | 101553614 | rs17025005 | -0.185 | -0.41 | -1.819 | 0.373 | 0.061 | 0.011 | 0.011 | 0 | 0.28 | 0.11 | 0.15 | 0.24 | 0.17 | 0.14 |
| 2 | 101557566 | rs11674199 | 0.334 | 0.298 | -1.35 | -0.366 | 0.004 | 0.023 | 0.023 | 0.301 | 0.41 | 0.31 | 0.24 | 0.04 | 0.34 | 0.22 |
| 2 | 101558984 | rs7605434 | -0.266 | 0.321 | -1.335 | -0.328 | 0 | 0 | 0 | 0 | 0.09 | 0.09 | 0.07 | 0.08 | 0.12 | 0.27 |
| 2 | 101559604 | rs895522 | -0.5 | -0.03 | -0.886 | 0.478 | -0.025 | -0.024 | -0.024 | 0.755 | 0.66 | 0.72 | 0.67 | 0.57 | 0.56 | 0.62 |
| 2 | 101560758 | rs4851383 | 0.601 | -0.054 | -0.868 | 0.483 | 0.003 | 0.039 | 0.039 | 0 | 0.51 | 0.4 | 0.31 | 0.13 | 0.47 | 0.49 |
| 2 | 101561638 | rs4851384 | 0.61 | -0.016 | -0.852 | 0.896 | 0.004 | -0.046 | -0.046 | 1.899 | 0.11 | 0.03 | 0.33 | 0.37 | 0.09 | 0.12 |
| 2 | 101565151 | rs12712085 | -0.414 | 0.144 | -0.812 | 0.758 | -0.02 | -0.03 | -0.03 | 0 | 0.64 | 0.76 | 0.68 | 0.57 | 0.58 | 0.71 |
| 2 | 101566070 | rs3739005 | 0.282 | 0.13 | -0.571 | 0.718 | -0.002 | -0.002 | -0.002 | 0.288 | 0.57 | 0.44 | 0.62 | 0.5 | 0.52 | 0.59 |
| 2 | 101566234 | rs3820787 | 0.138 | 0.435 | -1.024 | 0.98 | 0.002 | -0.027 | -0.027 | 0 | 0.65 | 0.76 | 0.41 | 0.49 | 0.63 | 0.74 |

| | | | | | | | | | | | | | | | | |
|---|---|---|---|---|---|---|---|---|---|---|---|---|---|---|---|---|
| 2 | 101570530 | rs12612424 | -0.266 | 0.448 | -0.734 | 1.069 | 0 | 0.006 | 0.006 | 0.164 | 0.15 | 0.16 | 0.37 | 0.39 | 0.15 | 0.25 |
| 2 | 101571705 | rs4851390 | -0.804 | 0.474 | -0.679 | 1.008 | 0 | 0.004 | 0.004 | 1.286 | 0.19 | 0.2 | 0.41 | 0.47 | 0.23 | 0.5 |
| 2 | 101576199 | rs882272 | 1.446 | 0.416 | -0.374 | 0.681 | -0.009 | 0.116 | 0.116 | 0 | 0.42 | 0.21 | 0.28 | 0.07 | 0.41 | 0.29 |
| 2 | 101578458 | rs6747755 | -0.92 | 0.503 | 0.074 | 0.918 | -0.001 | 0.011 | 0.011 | 0.259 | 0.28 | 0.39 | 0.4 | 0.74 | 0.26 | 0.44 |
| 2 | 101578542 | rs6738097 | 0.246 | 0.522 | -0.019 | 0.999 | 0.015 | 0.043 | 0.043 | 0 | 0.15 | 0.3 | 0.07 | 0.28 | 0.14 | 0.15 |
| 2 | 101579454 | rs12622050 | -0.623 | 0.508 | -0.023 | 0.998 | 0.001 | 0.01 | 0.01 | 0.26 | 0.26 | 0.32 | 0.38 | 0.7 | 0.24 | 0.25 |
| 2 | 101581946 | rs3768986 | -0.572 | 0.424 | 0.204 | 1.037 | -0.024 | 0.164 | 0.164 | 0 | 0.17 | 0.31 | 0.35 | 0.51 | 0.11 | 0.3 |
| 2 | 101584895 | rs2289950 | -0.567 | 0.429 | 0.206 | 1.035 | -0.024 | 0.136 | 0.136 | 0 | 0.17 | 0.29 | 0.35 | 0.46 | 0.11 | 0.31 |
| 2 | 101587455 | rs1562313 | -0.006 | 0.714 | -0.118 | 1.065 | 0.026 | 0.024 | 0.024 | 0 | 0.17 | 0.3 | 0.11 | 0.28 | 0.23 | 0.2 |
| 2 | 101591304 | rs2305160 | -0.568 | 0.746 | -1.078 | 1.035 | -0.064 | 0 | 0 | 0 | 0.75 | 0.93 | 0.71 | 0.86 | 0.67 | 0.75 |
| 2 | 101595475 | rs1542178 | -0.578 | 0.757 | -1.076 | 1.022 | -0.061 | -0.033 | -0.033 | 0 | 0.75 | 0.93 | 0.71 | 0.84 | 0.68 | 0.75 |
| 2 | 101596624 | rs3768988 | 0.117 | 0.592 | -0.735 | 1.437 | 0.001 | 0 | 0 | 0 | 0.1 | 0.06 | 0.32 | 0.36 | 0.09 | 0.1 |
| 2 | 101599071 | rs6719437 | 1.082 | 0.301 | -1.142 | 0.944 | -0.005 | 0.049 | 0.049 | 0.627 | 0.36 | 0.11 | 0.22 | 0.26 | 0.32 | 0.37 |
| 2 | 101599087 | rs6719533 | 0.261 | 0.236 | -1.186 | 1.263 | 0.005 | 0.034 | 0.034 | 0.062 | 0.49 | 0.16 | 0.39 | 0.49 | 0.42 | 0.53 |
| 2 | 101599294 | rs3754677 | 0.747 | 0.463 | -0.76 | 1.356 | 0.004 | -0.015 | -0.015 | 0 | 0.61 | 0.2 | 0.49 | 0.36 | 0.63 | 0.62 |
| 2 | 101599375 | rs3754678 | 0.753 | 0.494 | -0.798 | 1.373 | 0.002 | -0.023 | -0.023 | 0 | 0.61 | 0.17 | 0.49 | 0.37 | 0.63 | 0.62 |
| 2 | 101599674 | rs3768990 | 0.945 | 0.936 | -0.913 | 1.08 | -0.041 | -0.015 | -0.015 | 2.28 | 0.56 | 0.17 | 0.42 | 0.38 | 0.51 | 0.54 |
| 2 | 101599767 | rs3768991 | 1.026 | 0.953 | -0.92 | 1.059 | -0.043 | -0.017 | -0.017 | 0 | 0.55 | 0.16 | 0.42 | 0.37 | 0.51 | 0.54 |
| 2 | 101602100 | rs9653466 | 0.247 | 0.887 | -0.008 | 0.905 | 0.004 | -0.027 | -0.027 | 0.054 | 0.07 | 0.23 | 0.07 | 0 | 0.07 | 0.04 |
| 2 | 101603812 | rs11123857 | -0.624 | 1.238 | 0.319 | 0.806 | 0.088 | -0.103 | -0.103 | 0.194 | 0.16 | 0.37 | 0.16 | 0.01 | 0.26 | 0.11 |
| 2 | 101609827 | rs9223 | 1.779 | 0.842 | -0.218 | 0.698 | 0.016 | 0.081 | 0.081 | 1.199 | 0.37 | 0.08 | 0.3 | 0.22 | 0.3 | 0.43 |
| 2 | 101612051 | rs2305158 | 0.579 | 1.025 | -0.365 | 0.751 | 0 | 0 | 0 | 0.019 | 0.21 | 0.19 | 0.17 | 0.13 | 0.26 | 0.13 |
| 2 | 101612568 | rs3739008 | 1.94 | 0.971 | -0.428 | 0.728 | -0.006 | 0.065 | 0.065 | 1.225 | 0.36 | 0.06 | 0.3 | 0.22 | 0.3 | 0.41 |
| 2 | 101612615 | rs1053096 | 0.116 | 0.992 | -0.225 | 0.753 | 0.003 | 0.04 | 0.04 | 0.609 | 0.68 | 0.46 | 0.57 | 0.38 | 0.69 | 0.6 |
| 2 | 101620136 | rs2278722 | 0.691 | 0.991 | 0.112 | 1.007 | -0.004 | 0.079 | 0.079 | 0 | 0.17 | 0.35 | 0.25 | 0.43 | 0.16 | 0.16 |
| 2 | 101622728 | rs10934 | 2.311 | 0.999 | -0.314 | 0.948 | 0.017 | 0.131 | 0.131 | 1.49 | 0.36 | 0.09 | 0.3 | 0.17 | 0.29 | 0.4 |
| 2 | 101624471 | rs746924 | 0.401 | 1.001 | -0.223 | 1.201 | 0.001 | 0.011 | 0.011 | 0.917 | 0.67 | 0.47 | 0.57 | 0.29 | 0.68 | 0.59 |
| 2 | 101628324 | rs4850956 | 0.861 | 1.117 | -0.235 | 1.183 | 0.007 | -0.006 | -0.006 | 0 | 0.59 | 0.41 | 0.5 | 0.29 | 0.55 | 0.53 |
| 2 | 101628998 | rs1302 | -0.993 | 1.127 | -0.325 | 1.216 | -0.043 | -0.045 | -0.045 | 2.712 | 0.86 | 0.98 | 0.83 | 0.71 | 0.84 | 0.75 |
| 2 | 101629210 | rs13017465 | 0.702 | 1.12 | -0.299 | 1.1 | -0.011 | -0.005 | -0.005 | 0 | 0.21 | 0.09 | 0.19 | 0.12 | 0.26 | 0.13 |
| 2 | 101637009 | rs17663443 | 1.071 | 1.117 | -0.281 | 1.086 | 0 | -0.014 | -0.014 | 0 | 0.08 | 0.06 | 0.06 | 0 | 0.13 | 0.05 |
| 2 | 101638888 | rs3739014 | 0.361 | 1.095 | -0.183 | 1.105 | 0.001 | 0.006 | 0.006 | 0.815 | 0.67 | 0.52 | 0.55 | 0.29 | 0.68 | 0.59 |
| 2 | 101642260 | rs10185855 | -0.229 | 1.038 | -0.458 | 1.086 | 0.019 | -0.024 | -0.024 | 0 | 0.31 | 0.37 | 0.25 | 0.13 | 0.39 | 0.19 |
| 2 | 101644299 | rs2289951 | 1.282 | 0.961 | -0.792 | 1.323 | -0.042 | -0.077 | -0.077 | 0.924 | 0.12 | 0.02 | 0.23 | 0.3 | 0.14 | 0.14 |
| 2 | 101644855 | rs11683877 | 2.385 | 0.766 | -0.81 | 0.998 | -0.022 | 0.109 | 0.109 | 1.18 | 0.34 | 0.04 | 0.29 | 0.17 | 0.27 | 0.4 |
| 2 | 101646144 | rs3739015 | 0.82 | 0.861 | -0.469 | 1.241 | -0.011 | -0.013 | -0.013 | 0 | 0.2 | 0.1 | 0.19 | 0.15 | 0.27 | 0.13 |
| 2 | 101649427 | rs10179599 | 0.547 | 0.531 | -0.544 | 1.223 | -0.013 | 0.005 | 0.005 | 0 | 0.21 | 0.31 | 0.19 | 0.13 | 0.21 | 0.12 |
| 2 | 101656726 | rs2289953 | 0.784 | 0.664 | -0.479 | 1.343 | 0.001 | 0.003 | 0.003 | 0.309 | 0.67 | 0.35 | 0.56 | 0.31 | 0.68 | 0.59 |
| 2 | 101661540 | rs4850958 | -0.281 | 0.689 | -0.496 | 1.327 | -0.027 | 0.119 | 0.119 | 0.498 | 0.19 | 0.63 | 0.29 | 0.4 | 0.14 | 0.16 |
| 2 | 101663937 | rs6543009 | -0.625 | 0.897 | -0.673 | 1.284 | -0.053 | -0.061 | -0.061 | 0 | 0.86 | 0.97 | 0.85 | 0.67 | 0.82 | 0.75 |
| 2 | 101665829 | rs11123859 | -0.668 | 0.458 | -0.674 | 1.01 | 0.001 | 0.033 | 0.033 | 0.139 | 0.5 | 0.34 | 0.49 | 0.37 | 0.53 | 0.41 |
| 2 | 101670928 | rs9677214 | -0.38 | 0.302 | -0.661 | 0.974 | 0 | -0.003 | -0.003 | 0 | 0.35 | 0.31 | 0.44 | 0.37 | 0.36 | 0.31 |
| 2 | 101676663 | rs17025231 | 1.318 | 0.377 | -0.934 | 1.106 | -0.039 | -0.066 | -0.066 | 0 | 0.1 | 0.01 | 0.24 | 0.27 | 0.12 | 0.17 |
| 2 | 101678170 | rs7585873 | -0.798 | 0.403 | -0.782 | 1.171 | -0.03 | -0.06 | -0.06 | 0.068 | 0.87 | 0.97 | 0.86 | 0.68 | 0.85 | 0.76 |
| 2 | 101690336 | rs2121370 | 0.359 | 0.427 | -0.635 | 1.218 | 0 | 0.019 | 0.019 | 0 | 0.68 | 0.53 | 0.76 | 0.56 | 0.68 | 0.66 |
| 2 | 101691001 | rs6543012 | -1.243 | 0.622 | -1.385 | 1.274 | -0.052 | -0.064 | -0.064 | 0 | 0.9 | 0.99 | 0.88 | 0.68 | 0.87 | 0.76 |
| 2 | 101693332 | rs4851398 | 1.081 | 0.521 | -1.384 | 1.085 | -0.108 | -0.037 | -0.037 | 0 | 0.44 | 0.05 | 0.56 | 0.43 | 0.45 | 0.57 |
| 2 | 101694009 | rs11676053 | 0.248 | 0.508 | -0.739 | 1.031 | -0.005 | -0.017 | -0.017 | 0 | 0.22 | 0.29 | 0.18 | 0.09 | 0.23 | 0.1 |
| 2 | 101696649 | rs4851400 | 0.626 | 0.677 | -1.211 | 1.35 | -0.027 | -0.076 | -0.076 | 0 | 0.13 | 0.01 | 0.25 | 0.29 | 0.13 | 0.2 |
| 2 | 101698782 | rs6543017 | -0.95 | 0.751 | -1.358 | 1.538 | -0.024 | -0.047 | -0.047 | 0.239 | 0.88 | 0.97 | 0.95 | 0.78 | 0.88 | 0.8 |
| 2 | 101699184 | rs11678957 | 0.557 | 0.56 | -1.208 | 1.455 | -0.001 | -0.004 | -0.004 | 0 | 0.19 | 0.18 | 0.23 | 0.24 | 0.19 | 0.1 |
| 2 | 101699594 | rs2309938 | -1.149 | 0.396 | -0.544 | 1.268 | 0 | 0.001 | 0.001 | 1.588 | 0.49 | 0.53 | 0.57 | 0.63 | 0.5 | 0.4 |
| 2 | 101703771 | rs7584650 | -0.072 | 0.386 | -0.628 | 1.265 | 0 | 0.001 | 0.001 | 0 | 0.29 | 0.31 | 0.34 | 0.39 | 0.3 | 0.3 |
| 2 | 101710796 | rs6755081 | 0.846 | 0.421 | -0.75 | 1.241 | -0.011 | -0.019 | -0.019 | 0 | 0.18 | 0.09 | 0.22 | 0.24 | 0.19 | 0.1 |
| 2 | 101729443 | rs796063 | -1.013 | 0.381 | -0.241 | 1.56 | 0 | -0.002 | -0.002 | 0.019 | 0.48 | 0.54 | 0.38 | 0.63 | 0.5 | 0.4 |
| 2 | 101730288 | rs6543018 | -1.729 | 0.344 | -0.229 | 1.31 | 0 | 0.002 | 0.002 | 1.343 | 0.62 | 0.64 | 0.63 | 0.8 | 0.63 | 0.6 |
| 2 | 101731071 | rs796054 | 0.967 | 0.491 | -0.592 | 1.38 | -0.028 | -0.031 | -0.031 | 0.335 | 0.13 | 0.03 | 0.06 | 0.18 | 0.13 | 0.22 |
| 2 | 101731135 | rs2165427 | 1.066 | 0.501 | -0.489 | 1.376 | -0.004 | -0.036 | -0.036 | 0 | 0.13 | 0.04 | 0.25 | 0.29 | 0.13 | 0.2 |
| 2 | 101732898 | rs796055 | -2.253 | 0.543 | -0.733 | 0.795 | -0.01 | 0.099 | 0.099 | 0.601 | 0.73 | 0.94 | 0.72 | 0.91 | 0.76 | 0.85 |
| 2 | 101737379 | rs2033956 | 1.196 | 0.857 | -0.636 | 1.008 | -0.028 | -0.079 | -0.079 | 0.236 | 0.13 | 0.02 | 0.27 | 0.3 | 0.16 | 0.2 |
| 2 | 101751685 | rs11898628 | 0.6 | 0.811 | -0.183 | 1.009 | 0.007 | -0.009 | -0.009 | 0 | 0.16 | 0.33 | 0.08 | 0.15 | 0.14 | 0.23 |
| 2 | 101753234 | rs2165426 | 1.133 | 0.805 | -0.182 | 1.006 | -0.032 | -0.013 | -0.013 | 0 | 0.14 | 0.03 | 0.05 | 0.15 | 0.14 | 0.23 |
| 2 | 101756929 | rs11694166 | 0.842 | 0.797 | -0.745 | 0.966 | -0.006 | -0.041 | -0.041 | 0 | 0.2 | 0.08 | 0.19 | 0.26 | 0.16 | 0.17 |
| 2 | 101759350 | rs17025300 | 1.402 | 0.911 | -0.643 | 0.497 | 0 | -0.009 | -0.009 | 1.487 | 0.14 | 0.18 | 0.1 | 0.07 | 0.17 | 0.03 |
| 2 | 101761744 | rs6749166 | -0.369 | 0.887 | -0.584 | 0.53 | 0.006 | 0.035 | 0.035 | 0.606 | 0.3 | 0.46 | 0.36 | 0.44 | 0.3 | 0.43 |
| 2 | 101763227 | rs10207371 | -1.57 | 1.108 | -0.936 | 0.728 | -0.005 | -0.007 | -0.007 | 0 | 0.65 | 0.77 | 0.56 | 0.59 | 0.63 | 0.63 |
| 2 | 101789124 | rs1192779 | -1.552 | 0.792 | -0.899 | 0.618 | 0.009 | -0.015 | -0.015 | 0 | 0.69 | 0.77 | 0.62 | 0.57 | 0.73 | 0.73 |
| 2 | 101801651 | rs4850962 | -0.298 | 1.083 | -0.89 | 0.652 | -0.016 | -0.014 | -0.014 | 0 | 0.27 | 0.21 | 0.38 | 0.33 | 0.39 | 0.24 |
| 2 | 101802078 | rs4851411 | 0.269 | 0.982 | -0.555 | 0.716 | 0 | 0.002 | 0.002 | 0.024 | 0.15 | 0.15 | 0.3 | 0.27 | 0.22 | 0.21 |

| | | | | | | | | | | | | | | | | |
|---|---|---|---|---|---|---|---|---|---|---|---|---|---|---|---|---|
| 2 | 101816249 | rs6742826 | -0.314 | 0.71 | -0.604 | 0.601 | 0.004 | -0.004 | -0.004 | 0.049 | 0.41 | 0.49 | 0.25 | 0.35 | 0.41 | 0.4 |
| 2 | 101824546 | rs1192827 | 0.037 | 0.463 | -0.623 | 0.694 | -0.002 | 0.004 | 0.004 | 0 | 0.7 | 0.7 | 0.59 | 0.61 | 0.69 | 0.7 |
| 2 | 101826225 | rs12712090 | 0.459 | 0.576 | -1.105 | 0.144 | -0.037 | 0.134 | 0.134 | 0.045 | 0.46 | 0.23 | 0.26 | 0.28 | 0.46 | 0.33 |
| 2 | 101835501 | rs1660695 | -0.069 | 0.828 | -0.24 | 0.336 | 0.032 | -0.03 | -0.03 | 0 | 0.94 | 0.83 | 0.95 | 1 | 0.93 | 0.92 |
| 2 | 101856926 | rs13025261 | 0.102 | 0.801 | -0.821 | 0.265 | -0.004 | -0.04 | -0.04 | 0 | 0.41 | 0.2 | 0.45 | 0.53 | 0.43 | 0.35 |
| 2 | 101868316 | rs11887349 | 0.915 | 0.568 | -0.094 | -0.031 | -0.003 | -0.001 | -0.001 | 0 | 0.27 | 0.25 | 0.19 | 0.25 | 0.26 | 0.18 |
| 2 | 101870862 | rs935067 | -0.109 | 0.497 | 0.021 | -0.118 | 0.018 | -0.062 | -0.062 | 0 | 0.85 | 0.53 | 0.9 | 0.93 | 0.86 | 0.83 |
| 2 | 101881980 | rs13015194 | 1.516 | 0.313 | -0.482 | -0.327 | -0.038 | 0.06 | 0.06 | 0 | 0.24 | 0.03 | 0.14 | 0.14 | 0.25 | 0.16 |
| 2 | 101887491 | rs1660711 | 0.161 | 0.044 | -0.25 | -0.222 | -0.019 | -0.005 | -0.005 | 0 | 0.43 | 0.26 | 0.44 | 0.4 | 0.43 | 0.48 |
| 2 | 101897871 | rs1192793 | -0.287 | 0.027 | 0.061 | -0.243 | 0.022 | -0.097 | -0.097 | 1.179 | 0.86 | 0.63 | 0.91 | 1 | 0.87 | 0.89 |
| 2 | 101898222 | rs1192794 | 1.481 | 0.031 | -0.118 | -0.238 | -0.044 | -0.035 | -0.035 | 0 | 0.15 | 0.01 | 0.25 | 0.16 | 0.17 | 0.3 |
| 2 | 101900500 | rs1192795 | -1.138 | 0.039 | 0.014 | -0.248 | 0.03 | -0.031 | -0.031 | 1.411 | 0.94 | 0.83 | 0.95 | 1 | 0.92 | 0.92 |
| 2 | 101911385 | rs3828194 | 0.673 | 0.036 | -0.236 | -0.223 | 0.004 | -0.025 | -0.025 | 0 | 0.42 | 0.26 | 0.46 | 0.53 | 0.44 | 0.4 |
| 2 | 101917365 | rs12617343 | -0.4 | 0.075 | -0.308 | -0.224 | -0.014 | 0 | 0 | 0 | 0.57 | 0.7 | 0.56 | 0.62 | 0.57 | 0.52 |
| 2 | 101924004 | rs4851416 | -0.407 | 0.059 | -0.394 | -0.23 | -0.008 | 0.045 | 0.045 | 1.92 | 0.43 | 0.77 | 0.46 | 0.32 | 0.39 | 0.36 |
| 2 | 101926811 | rs7568067 | 0.886 | 0.138 | -0.368 | -0.237 | -0.002 | 0 | 0 | 1.172 | 0.27 | 0.26 | 0.35 | 0.25 | 0.25 | 0.18 |
| 2 | 101926957 | rs6543026 | -1.285 | 0.003 | -0.424 | -0.168 | -0.042 | -0.033 | -0.033 | 0 | 0.84 | 0.99 | 0.91 | 0.86 | 0.83 | 0.7 |
| 2 | 101928997 | rs2056716 | 1.039 | -0.572 | 0.157 | 0.011 | 0.002 | 0.001 | 0.001 | 2.085 | 0.7 | 0.31 | 0.64 | 0.78 | 0.79 | 0.78 |
| 2 | 101949770 | rs1972163 | 0.214 | -0.618 | 0.594 | -0.035 | 0.007 | -0.114 | -0.114 | 0 | 0.83 | 0.62 | 0.9 | 1 | 0.86 | 0.92 |
| 2 | 101949911 | rs2177996 | 0.615 | -0.657 | 0.611 | 0.276 | -0.004 | -0.051 | -0.051 | 0 | 0.81 | 0.5 | 0.73 | 0.93 | 0.84 | 0.85 |
| 2 | 101958174 | rs11684238 | 0.583 | -0.653 | 0.622 | 0.217 | -0.017 | 0.059 | 0.059 | 0 | 0.11 | 0.22 | 0.25 | 0.22 | 0.07 | 0.13 |
| 2 | 101958326 | rs4851423 | 1.128 | -0.632 | 0.471 | 0.221 | 0 | 0 | 0 | 0.797 | 0.7 | 0.31 | 0.64 | 0.77 | 0.79 | 0.79 |
| 2 | 101964726 | rs10166900 | 1.192 | -0.839 | 0.574 | 0.202 | -0.014 | -0.007 | -0.007 | 0 | 0.7 | 0.39 | 0.65 | 0.77 | 0.81 | 0.79 |
| 2 | 101965367 | rs4539813 | -0.041 | -0.847 | 0.775 | 0.172 | -0.006 | -0.059 | -0.059 | 0 | 0.85 | 0.82 | 0.92 | 1 | 0.88 | 0.93 |
| 2 | 101966038 | rs11677690 | 1.165 | -1.035 | 0.147 | 0.008 | -0.057 | 0.043 | 0.043 | 0.831 | 0.26 | 0.05 | 0.17 | 0.17 | 0.32 | 0.36 |
| 2 | 101971618 | rs6753727 | 0.825 | -1.041 | 0.221 | 0.023 | -0.025 | 0.01 | 0.01 | 0 | 0.44 | 0.32 | 0.47 | 0.6 | 0.48 | 0.43 |
| 2 | 101973286 | rs11677377 | 0.938 | -1.06 | 0.288 | 0.038 | -0.011 | -0.006 | -0.006 | 0 | 0.69 | 0.39 | 0.65 | 0.77 | 0.81 | 0.77 |
| 2 | 101981794 | rs4280474 | 0.048 | -1.042 | 0.27 | 0.024 | -0.012 | -0.098 | -0.098 | 0 | 0.84 | 0.69 | 0.91 | 1 | 0.88 | 0.92 |
| 2 | 101982603 | rs6729593 | 0.229 | -1.057 | 0.301 | 0.134 | -0.016 | 0.021 | 0.021 | 0 | 0.13 | 0.18 | 0.25 | 0.23 | 0.07 | 0.14 |
| 2 | 101991907 | rs10186962 | 0.026 | -0.886 | -0.252 | 0.181 | 0.003 | -0.012 | -0.012 | 0 | 0.36 | 0.3 | 0.27 | 0.46 | 0.37 | 0.33 |
| 2 | 101992474 | rs4405786 | -0.926 | -1.108 | 0.202 | 0.765 | -0.058 | -0.087 | -0.087 | 0 | 0.74 | 0.94 | 0.67 | 0.69 | 0.67 | 0.64 |
| 2 | 101994731 | rs7584951 | -0.312 | -0.909 | 0.402 | 0.737 | -0.024 | -0.035 | -0.035 | 0 | 0.18 | 0.29 | 0.23 | 0.07 | 0.12 | 0.15 |
| 2 | 101997738 | rs9308846 | 0.505 | -0.612 | -0.04 | 0.92 | -0.008 | -0.009 | -0.009 | 2.836 | 0.44 | 0.38 | 0.32 | 0.55 | 0.5 | 0.4 |
| 2 | 102006527 | rs11692240 | -0.736 | -0.569 | 0.177 | 1.078 | 0.006 | 0.011 | 0.011 | 0 | 0.51 | 0.57 | 0.54 | 0.39 | 0.46 | 0.6 |
| 2 | 102014754 | rs6716306 | 0.773 | -0.672 | 0.074 | 1.308 | -0.015 | -0.007 | -0.007 | 0 | 0.45 | 0.17 | 0.28 | 0.49 | 0.53 | 0.4 |
| 2 | 102017186 | rs4241207 | 0.261 | -0.771 | 0.357 | 1.255 | -0.028 | 0.02 | 0.02 | 0 | 0.77 | 0.63 | 0.55 | 0.8 | 0.83 | 0.83 |
| 2 | 102019801 | rs10865041 | 0.412 | -0.803 | 0.428 | 1.075 | -0.01 | -0.01 | -0.01 | 0 | 0.27 | 0.13 | 0.23 | 0.32 | 0.29 | 0.43 |
| 2 | 102020770 | rs7577705 | -0.369 | -0.709 | 0.553 | 0.905 | -0.022 | -0.036 | -0.036 | 1.527 | 0.22 | 0.34 | 0.29 | 0.08 | 0.17 | 0.17 |
| 2 | 102029548 | rs1554070 | 0.692 | -0.477 | 0.033 | 1.072 | -0.015 | -0.003 | -0.003 | 0 | 0.23 | 0.11 | 0.21 | 0.21 | 0.28 | 0.33 |
| 2 | 102029739 | rs7603471 | 0.861 | -0.693 | 0.476 | 1.226 | -0.012 | 0.007 | 0.007 | 0 | 0.73 | 0.33 | 0.49 | 0.71 | 0.81 | 0.76 |
| 2 | 102032171 | rs6543042 | 0.997 | -0.706 | 0.389 | 1.245 | 0.015 | -0.01 | -0.01 | 0 | 0.72 | 0.27 | 0.45 | 0.71 | 0.82 | 0.81 |
| 2 | 102032791 | rs908134 | -0.575 | -0.599 | 1.246 | 1.391 | -0.02 | 0.081 | 0.081 | 0.422 | 0.1 | 0.32 | 0.07 | 0.15 | 0.03 | 0.08 |
| 2 | 102036455 | rs7565429 | 0.55 | -0.463 | 0.689 | 1.181 | -0.005 | 0.002 | 0.002 | 0 | 0.25 | 0.18 | 0.17 | 0.21 | 0.3 | 0.38 |
| 2 | 102040283 | rs2310127 | 1.223 | -0.41 | 0.359 | 1.017 | -0.056 | 0 | 0 | 0.015 | 0.47 | 0.15 | 0.27 | 0.51 | 0.49 | 0.42 |
| 2 | 102042298 | rs17025554 | -0.4 | -0.427 | 1.159 | 0.928 | 0.005 | -0.076 | -0.076 | 0 | 0.14 | 0.31 | 0.17 | 0 | 0.13 | 0.07 |
| 2 | 102045328 | rs6543047 | 0.722 | -0.475 | 0.683 | 1.243 | -0.006 | 0.034 | 0.034 | 0 | 0.55 | 0.31 | 0.36 | 0.43 | 0.52 | 0.43 |
| 2 | 102047481 | rs17805225 | 0.251 | -0.458 | 1.112 | 1.469 | -0.003 | 0.014 | 0.014 | 0 | 0.3 | 0.24 | 0.18 | 0.18 | 0.32 | 0.36 |
| 2 | 102050264 | rs4851439 | 1.073 | -0.461 | 1.148 | 1.434 | 0.031 | 0.312 | 0.312 | 0 | 0.85 | 0.55 | 0.54 | 0.62 | 0.85 | 0.79 |
| 2 | 102051912 | rs17025611 | 0.813 | -0.609 | 1.009 | 1.321 | -0.024 | -0.025 | -0.025 | 0 | 0.05 | 0.07 | 0.04 | 0 | 0.01 | 0 |
| 2 | 102053439 | rs6732292 | 0.323 | -0.499 | 0.947 | 1.289 | -0.004 | 0.013 | 0.013 | 0 | 0.3 | 0.24 | 0.18 | 0.19 | 0.33 | 0.36 |
| 2 | 102059797 | rs12712101 | 1.105 | -0.48 | 0.806 | 1.355 | -0.008 | 0.094 | 0.094 | 0.047 | 0.5 | 0.17 | 0.32 | 0.43 | 0.52 | 0.42 |
| 2 | 102059854 | rs12712102 | 1.105 | -0.48 | 0.806 | 1.355 | -0.008 | 0.094 | 0.094 | 0.047 | 0.5 | 0.17 | 0.32 | 0.43 | 0.52 | 0.42 |
| 2 | 102064729 | rs7583866 | -0.168 | -0.516 | 0.837 | 1.402 | 0 | -0.004 | -0.004 | 0 | 0.35 | 0.3 | 0.22 | 0.19 | 0.33 | 0.36 |
| 2 | 102065501 | rs11682765 | 0.353 | -0.483 | 0.747 | 1.538 | 0.001 | 0.177 | 0.177 | 0 | 0.94 | 0.89 | 0.67 | 0.67 | 0.92 | 0.85 |
| 2 | 102078997 | rs2138002 | -0.293 | -0.469 | 1.024 | 1.221 | -0.007 | -0.031 | -0.031 | 0 | 0.36 | 0.46 | 0.22 | 0.24 | 0.34 | 0.36 |
| 2 | 102083183 | rs4851455 | 0.683 | -0.498 | 0.822 | 1.359 | -0.006 | 0.007 | 0.007 | 0 | 0.63 | 0.42 | 0.39 | 0.52 | 0.64 | 0.63 |
| 2 | 102088370 | rs4629180 | 0.112 | -0.488 | 0.399 | 1.576 | -0.002 | -0.033 | -0.033 | 0 | 0.72 | 0.87 | 0.49 | 0.7 | 0.72 | 0.73 |
| 2 | 102095080 | rs10183203 | 0.617 | -0.5 | 0.565 | 1.597 | 0 | 0.001 | 0.001 | 0 | 0.67 | 0.7 | 0.42 | 0.51 | 0.66 | 0.65 |
| 2 | 102104814 | rs11688698 | -1.181 | -0.4 | 0.583 | 1.671 | 0.046 | 0.21 | 0.21 | 0 | 0.05 | 0.21 | 0.08 | 0.19 | 0.07 | 0.07 |
| 2 | 102129420 | rs1995826 | 0.597 | -0.643 | 0.364 | 1.278 | -0.013 | 0.013 | 0.013 | 0 | 0.64 | 0.51 | 0.39 | 0.52 | 0.67 | 0.63 |
| 2 | 102138384 | rs2048252 | 0.627 | -0.726 | 0.079 | 1.237 | 0.001 | -0.001 | -0.001 | 0 | 0.63 | 0.6 | 0.41 | 0.52 | 0.67 | 0.63 |
| 2 | 102145661 | rs6745920 | 0.396 | -0.229 | -0.202 | 1.081 | -0.003 | -0.031 | -0.031 | 0 | 0.65 | 0.72 | 0.61 | 0.62 | 0.61 | 0.69 |
| 2 | 102148022 | rs13426360 | -0.613 | 0.08 | -0.706 | 0.896 | 0.002 | 0.002 | 0.002 | 0 | 0.61 | 0.61 | 0.74 | 0.65 | 0.53 | 0.7 |
| 2 | 102155961 | rs7569002 | 1.28 | 0.234 | -0.895 | 0.803 | -0.049 | 0.085 | 0.085 | 0 | 0.27 | 0.07 | 0.14 | 0.1 | 0.26 | 0.21 |
| 2 | 102157560 | rs11891427 | 0.104 | 0.127 | -0.711 | 0.826 | 0.006 | 0.009 | 0.009 | 0 | 0.39 | 0.47 | 0.23 | 0.35 | 0.44 | 0.32 |
| 2 | 102160250 | rs7561182 | 1.105 | 0.093 | -0.798 | 0.718 | -0.021 | 0.072 | 0.072 | 0 | 0.29 | 0.12 | 0.15 | 0.1 | 0.26 | 0.21 |
| 2 | 102160442 | rs7575166 | 0.175 | 0.138 | -0.666 | 0.825 | 0.006 | 0.01 | 0.01 | 0 | 0.39 | 0.47 | 0.23 | 0.35 | 0.44 | 0.32 |
| 2 | 102166870 | rs4308140 | 0.512 | -0.148 | -0.835 | 0.743 | 0.021 | 0.04 | 0.04 | 0.14 | 0.41 | 0.24 | 0.19 | 0.2 | 0.3 | 0.3 |
| 2 | 102168016 | rs6543074 | 0.319 | -0.179 | -0.683 | 0.76 | 0.006 | 0.009 | 0.009 | 0.146 | 0.43 | 0.4 | 0.21 | 0.23 | 0.3 | 0.36 |

| | | | | | | | | | | | | | | | | |
|---|---|---|---|---|---|---|---|---|---|---|---|---|---|---|---|---|
| 2 | 102176693 | rs7607789 | -0.018 | -0.155 | -0.587 | 0.808 | 0 | 0 | 0 | 0.054 | 0.51 | 0.53 | 0.28 | 0.38 | 0.46 | 0.49 |
| 2 | 102178524 | rs7589339 | 0.257 | -0.279 | -0.297 | 0.707 | 0.01 | 0.013 | 0.013 | 0 | 0.46 | 0.59 | 0.23 | 0.24 | 0.3 | 0.34 |
| 2 | 102179247 | rs17201799 | 1.349 | -0.285 | -0.515 | 0.571 | -0.041 | 0.121 | 0.121 | 0 | 0.3 | 0.08 | 0.16 | 0.09 | 0.27 | 0.23 |
| 2 | 102181444 | rs11123891 | -0.887 | -0.121 | 0.124 | 1.243 | 0.004 | 0.028 | 0.028 | 0 | 0.19 | 0.18 | 0.37 | 0.39 | 0.26 | 0.38 |
| 2 | 102188552 | rs12617031 | -1.025 | -0.057 | 0.149 | 1.222 | 0.061 | 0.132 | 0.132 | 0 | 0.07 | 0.22 | 0.07 | 0.17 | 0.14 | 0.13 |
| 2 | 102194026 | rs3900546 | -0.722 | -0.136 | 0.107 | 1.166 | -0.004 | 0.023 | 0.023 | 0 | 0.28 | 0.59 | 0.3 | 0.41 | 0.29 | 0.22 |
| 2 | 102205282 | rs12712108 | -0.701 | -0.217 | 0.236 | 1.06 | 0 | -0.014 | -0.014 | 0 | 0.35 | 0.3 | 0.37 | 0.53 | 0.36 | 0.35 |
| 2 | 102205736 | rs7603851 | -1.555 | -0.203 | 0.201 | 1.032 | 0.003 | 0.103 | 0.103 | 0 | 0.5 | 0.68 | 0.7 | 0.72 | 0.47 | 0.59 |
| 2 | 102211111 | rs11680510 | -0.649 | -0.211 | 0.058 | 1.046 | 0 | -0.014 | -0.014 | 0.088 | 0.35 | 0.29 | 0.37 | 0.52 | 0.36 | 0.35 |
| 2 | 102215229 | rs11884391 | -0.272 | -0.049 | -0.07 | 1.13 | 0.001 | 0.036 | 0.036 | 0 | 0.14 | 0.22 | 0.31 | 0.19 | 0.11 | 0.24 |
| 2 | 102227623 | rs10186803 | 0.344 | -0.009 | -0.189 | 0.924 | -0.01 | -0.015 | -0.015 | 0.34 | 0.16 | 0.13 | 0.17 | 0.33 | 0.24 | 0.28 |
| 2 | 102233731 | rs6543081 | -0.416 | -0.472 | 0.202 | 0.66 | -0.015 | 0.138 | 0.138 | 0 | 0.15 | 0.3 | 0.3 | 0.33 | 0.1 | 0.25 |
| 2 | 102234065 | rs6720988 | -0.802 | -0.383 | -0.224 | 0.7 | 0.007 | -0.002 | -0.002 | 0 | 0.38 | 0.63 | 0.55 | 0.39 | 0.26 | 0.33 |
| 2 | 102243860 | rs6755294 | -0.694 | -0.287 | -0.365 | 0.142 | 0.006 | 0.03 | 0.03 | 0 | 0.56 | 0.42 | 0.79 | 0.82 | 0.64 | 0.71 |
| 2 | 102247207 | rs2310116 | -0.743 | -0.283 | -0.332 | 0.144 | -0.003 | -0.035 | -0.035 | 0 | 0.56 | 0.59 | 0.8 | 0.82 | 0.64 | 0.71 |
| 2 | 102252504 | rs1876802 | -1.075 | -0.286 | -0.804 | 0.148 | -0.026 | 0.134 | 0.134 | 0.057 | 0.61 | 0.88 | 0.82 | 0.82 | 0.64 | 0.72 |
| 2 | 102255114 | rs10496357 | -0.01 | 0.117 | -0.928 | 0.967 | -0.029 | -0.087 | -0.087 | 0.687 | 0.12 | 0.01 | 0.16 | 0.33 | 0.14 | 0.22 |
| 2 | 102261596 | rs4851475 | -0.339 | -0.242 | -0.788 | 0.708 | 0.023 | 0.049 | 0.049 | 0.687 | 0.51 | 0.27 | 0.6 | 0.55 | 0.6 | 0.62 |
| 2 | 102267257 | rs13407493 | -0.259 | -0.324 | -0.404 | 0.375 | 0.02 | 0.132 | 0.132 | 0.132 | 0.05 | 0.27 | 0.01 | 0.13 | 0.01 | 0.03 |
| 2 | 102274098 | rs4851484 | -0.338 | -0.045 | -1.473 | -0.158 | -0.027 | 0.057 | 0.057 | 0 | 0.6 | 0.8 | 0.57 | 0.77 | 0.56 | 0.59 |
| 2 | 102274546 | rs7598866 | 0.611 | 0.486 | -1.612 | -0.153 | -0.046 | 0.097 | 0.097 | 0.092 | 0.3 | 0.09 | 0.14 | 0.14 | 0.3 | 0.22 |
| 2 | 102288186 | rs4280476 | -0.121 | 0.272 | -1.156 | -0.384 | -0.002 | -0.021 | -0.021 | 0.613 | 0.31 | 0.29 | 0.34 | 0.12 | 0.31 | 0.25 |
| 2 | 102292286 | rs2016677 | 0.336 | 0.093 | -1.459 | -0.3 | -0.026 | 0.154 | 0.154 | 0 | 0.48 | 0.19 | 0.48 | 0.72 | 0.51 | 0.58 |
| 2 | 102294890 | rs6543085 | -0.164 | -0.01 | -1.293 | -0.027 | 0.011 | 0.023 | 0.023 | 0 | 0.5 | 0.61 | 0.33 | 0.66 | 0.41 | 0.63 |
| 2 | 102302978 | rs4851487 | -0.251 | -0.385 | -1.038 | -0.309 | 0.003 | -0.01 | -0.01 | 0 | 0.45 | 0.63 | 0.37 | 0.54 | 0.42 | 0.57 |
| 2 | 102315057 | rs1567385 | 1.377 | -0.477 | -0.361 | -0.629 | -0.063 | -0.046 | -0.046 | 0 | 0.22 | 0.05 | 0.31 | 0.23 | 0.29 | 0.16 |
| 2 | 102320640 | rs17809691 | 1.424 | -0.49 | -0.346 | -0.636 | -0.064 | -0.047 | -0.047 | 0 | 0.22 | 0.05 | 0.31 | 0.23 | 0.29 | 0.16 |
| 2 | 102325324 | rs11674694 | 0.938 | -0.471 | -0.211 | -0.595 | -0.038 | -0.09 | -0.09 | 1.524 | 0.33 | 0.11 | 0.25 | 0.46 | 0.35 | 0.52 |
| 2 | 102336865 | rs4851491 | -0.638 | -0.573 | 0 | -0.644 | -0.001 | 0.029 | 0.029 | 1.624 | 0.69 | 0.91 | 0.7 | 0.76 | 0.71 | 0.73 |
| 2 | 102342511 | rs11894820 | 1.025 | -0.582 | 0.243 | -0.733 | 0.001 | -0.014 | -0.014 | 0 | 0.07 | 0.14 | 0.07 | 0.07 | 0.06 | 0.05 |
| 2 | 102343062 | rs6543092 | -0.596 | -0.604 | 0.161 | -0.713 | -0.001 | 0.05 | 0.05 | 1.603 | 0.69 | 0.91 | 0.71 | 0.79 | 0.71 | 0.73 |
| 2 | 102345345 | rs12476979 | 0.963 | -0.59 | 0.165 | -0.702 | -0.037 | -0.092 | -0.092 | 0 | 0.33 | 0.11 | 0.25 | 0.49 | 0.35 | 0.52 |
| 2 | 102359923 | rs4384813 | -0.487 | -0.591 | 0.457 | -0.708 | -0.021 | -0.046 | -0.046 | 0 | 0.12 | 0.65 | 0.14 | 0.07 | 0.07 | 0.05 |
| 2 | 102362333 | rs7420579 | 1.609 | -0.576 | 0.37 | -0.678 | -0.063 | -0.049 | -0.049 | 0.013 | 0.22 | 0.05 | 0.31 | 0.23 | 0.29 | 0.16 |
| 2 | 102367562 | rs17810475 | 1.609 | -0.576 | 0.37 | -0.678 | -0.06 | -0.046 | -0.046 | 0.333 | 0.22 | 0.05 | 0.31 | 0.23 | 0.29 | 0.16 |
| 2 | 102372131 | rs6543094 | -1.837 | -0.638 | 0.262 | -0.857 | -0.062 | 0.012 | 0.012 | 1.488 | 0.83 | 0.99 | 0.85 | 0.9 | 0.79 | 0.84 |
| 2 | 102374959 | rs11123901 | -0.006 | -0.666 | 0.42 | -0.867 | -0.003 | -0.011 | -0.011 | 0 | 0.67 | 0.67 | 0.66 | 0.79 | 0.7 | 0.72 |
| 2 | 102432368 | rs4851502 | 0.957 | -0.601 | -0.016 | -0.776 | 0.021 | 0.011 | 0.011 | 0.025 | 0.14 | 0.06 | 0.14 | 0.11 | 0.08 | 0.11 |
| 2 | 102440166 | rs2286240 | 0.963 | -0.597 | -0.081 | -0.737 | -0.045 | -0.104 | -0.104 | 9.639 | 0.33 | 0.09 | 0.25 | 0.48 | 0.35 | 0.51 |
| 2 | 102457090 | rs4850990 | -0.04 | -0.606 | -0.503 | -0.979 | -0.032 | 0.009 | 0.009 | 0.078 | 0.58 | 0.9 | 0.53 | 0.41 | 0.53 | 0.4 |
| 2 | 102462574 | rs17802002 | 1.455 | -0.598 | -0.324 | -0.996 | -0.007 | 0.047 | 0.047 | 0 | 0.22 | 0.11 | 0.09 | 0.1 | 0.18 | 0.18 |
| 2 | 102467321 | rs972372 | -0.81 | -0.645 | -0.361 | -1.014 | -0.031 | -0.072 | -0.072 | 0.011 | 0.71 | 0.88 | 0.67 | 0.51 | 0.62 | 0.5 |
| 2 | 102472459 | rs1139583 | 1.438 | -0.546 | -0.299 | -0.99 | -0.007 | 0.047 | 0.047 | 0 | 0.22 | 0.11 | 0.09 | 0.1 | 0.18 | 0.19 |
| 2 | 102478352 | rs13425874 | 0.737 | -0.663 | -0.063 | -0.964 | 0.013 | -0.011 | -0.011 | 0 | 0.09 | 0.3 | 0.08 | 0.05 | 0.06 | 0.05 |
| 2 | 102479029 | rs3771903 | -0.041 | -0.65 | -0.48 | -0.933 | -0.02 | 0.017 | 0.017 | 0.076 | 0.57 | 0.86 | 0.52 | 0.4 | 0.53 | 0.39 |
| 2 | 102508146 | rs6705975 | 0.009 | -0.579 | -0.435 | -0.918 | -0.036 | 0.014 | 0.014 | 0.075 | 0.57 | 0.89 | 0.53 | 0.4 | 0.53 | 0.39 |
| 2 | 102531388 | rs4851508 | 0.484 | -0.505 | -0.356 | -0.903 | -0.001 | 0.006 | 0.006 | 0.382 | 0.36 | 0.76 | 0.44 | 0.3 | 0.36 | 0.21 |
| 2 | 102533882 | rs7603475 | -1.744 | -0.398 | -0.246 | -0.833 | -0.008 | 0.021 | 0.021 | 0 | 0.78 | 0.84 | 0.68 | 0.84 | 0.75 | 0.78 |
| 2 | 102535394 | rs4851509 | 1.002 | -0.775 | -0.107 | -0.445 | 0.017 | 0.027 | 0.027 | 0 | 0.26 | 0.41 | 0.35 | 0.28 | 0.31 | 0.16 |
| 2 | 102535941 | rs13011687 | -0.298 | -0.747 | -1.247 | -0.441 | 0.012 | -0.099 | -0.099 | 0.187 | 0.27 | 0.1 | 0.22 | 0.46 | 0.26 | 0.45 |
| 2 | 102537802 | rs17746734 | 1.368 | -0.682 | -1.267 | -0.394 | -0.065 | -0.081 | -0.081 | 0 | 0.23 | 0.04 | 0.3 | 0.28 | 0.3 | 0.15 |
| 2 | 102546801 | rs2056400 | 0.802 | -0.718 | -0.876 | -0.073 | -0.043 | -0.096 | -0.096 | 0.085 | 0.13 | 0.01 | 0.18 | 0.4 | 0.17 | 0.34 |
| 2 | 102562344 | rs1541435 | -0.542 | -0.779 | -0.869 | -0.08 | 0.004 | -0.106 | -0.106 | 0 | 0.27 | 0.1 | 0.22 | 0.44 | 0.25 | 0.45 |
| 2 | 102569724 | rs12467695 | 0.679 | -0.582 | -0.807 | 0.062 | -0.048 | -0.097 | -0.097 | 0 | 0.13 | 0.01 | 0.18 | 0.39 | 0.18 | 0.34 |
| 2 | 102573478 | rs1859716 | -0.966 | -0.548 | -0.622 | -0.468 | -0.017 | 0.034 | 0.034 | 0 | 0.72 | 0.81 | 0.64 | 0.86 | 0.68 | 0.73 |
| 2 | 102574838 | rs1019031 | -0.966 | -0.548 | -0.622 | -0.468 | -0.017 | 0.034 | 0.034 | 0 | 0.72 | 0.81 | 0.64 | 0.86 | 0.68 | 0.73 |
| 2 | 102576098 | rs2190360 | -0.707 | -0.665 | 0.1 | -0.53 | 0.015 | -0.092 | -0.092 | 0 | 0.69 | 0.53 | 0.61 | 0.86 | 0.67 | 0.73 |
| 2 | 102581224 | rs2190358 | 0.81 | -0.752 | 0.175 | -0.597 | 0.002 | 0.011 | 0.011 | 0 | 0.27 | 0.64 | 0.38 | 0.27 | 0.31 | 0.1 |
| 2 | 102589726 | rs6715873 | 0.79 | -0.986 | 0.204 | -0.591 | 0.009 | 0.011 | 0.011 | 0 | 0.28 | 0.63 | 0.38 | 0.27 | 0.33 | 0.11 |
| 2 | 102598561 | rs7566831 | -1.459 | -1.041 | 0.027 | -0.62 | -0.048 | 0.095 | 0.095 | 0 | 0.74 | 0.91 | 0.77 | 0.95 | 0.68 | 0.79 |
| 2 | 102598940 | rs13006661 | -0.275 | -1.065 | 0.112 | -0.591 | 0.041 | 0.027 | 0.027 | 0 | 0.43 | 0.15 | 0.38 | 0.67 | 0.35 | 0.62 |
| 2 | 102601271 | rs4851518 | 1.162 | -1.144 | 0.132 | -0.364 | 0.005 | 0.004 | 0.004 | 0 | 0.24 | 0.28 | 0.34 | 0.26 | 0.33 | 0.11 |
| 2 | 102601393 | rs4851519 | 0.6 | -1.566 | 0.05 | -0.066 | 0.055 | 0.042 | 0.042 | 0.584 | 0.22 | 0.06 | 0.17 | 0.2 | 0.1 | 0.17 |
| 2 | 102604649 | rs12467316 | -0.478 | -1.356 | 0.013 | 0.028 | 0.045 | -0.015 | -0.015 | 0 | 0.38 | 0.13 | 0.36 | 0.68 | 0.3 | 0.57 |
| 2 | 102610721 | rs11886877 | -0.022 | -1.36 | 1.078 | 0.616 | -0.036 | 0.041 | 0.041 | 0 | 0.18 | 0.42 | 0.18 | 0.23 | 0.15 | 0.33 |
| 2 | 102610828 | rs4851522 | -0.091 | -1.385 | 1.189 | 0.591 | -0.041 | 0.033 | 0.033 | 0 | 0.18 | 0.5 | 0.18 | 0.21 | 0.14 | 0.33 |
| 2 | 102613299 | rs757918 | -1.154 | -1.511 | 0.846 | 0.535 | -0.061 | 0.073 | 0.073 | 0 | 0.59 | 0.86 | 0.67 | 0.8 | 0.55 | 0.66 |
| 2 | 102613581 | rs1108338 | -0.68 | -1.608 | 0.898 | 0.618 | -0.013 | 0.001 | 0.001 | 0 | 0.37 | 0.58 | 0.36 | 0.53 | 0.26 | 0.54 |
| 2 | 102617204 | rs7561460 | -0.152 | -1.549 | 0.923 | 0.626 | 0.01 | 0.022 | 0.022 | 0.019 | 0.37 | 0.48 | 0.47 | 0.48 | 0.43 | 0.46 |

| | | | | | | | | | | | | | | | | |
|---|---|---|---|---|---|---|---|---|---|---|---|---|---|---|---|---|
| 2 | 102617452 | rs11691240 | -0.152 | -1.549 | 0.923 | 0.646 | 0.01 | 0.025 | 0.025 | 0.019 | 0.37 | 0.48 | 0.47 | 0.46 | 0.43 | 0.46 |
| 2 | 102620729 | rs17026386 | 0.877 | -1.575 | 0.627 | 0.648 | 0.024 | -0.015 | -0.015 | 0 | 0.19 | 0.09 | 0.17 | 0.33 | 0.11 | 0.21 |
| 2 | 102622376 | rs4851527 | -1.246 | -1.562 | 0.529 | 0.65 | -0.071 | 0.074 | 0.074 | 0 | 0.59 | 0.89 | 0.67 | 0.79 | 0.55 | 0.67 |
| 2 | 102622780 | rs3755481 | 1.557 | -1.578 | 0.509 | 0.662 | -0.085 | -0.087 | -0.087 | 0.085 | 0.19 | 0.04 | 0.28 | 0.26 | 0.29 | 0.13 |
| 2 | 102623005 | rs13408303 | 0.761 | -1.582 | 0.779 | 0.628 | 0.005 | -0.005 | -0.005 | 0 | 0.14 | 0.12 | 0.15 | 0.2 | 0.14 | 0.28 |
| 2 | 102623875 | rs719248 | -1.191 | -1.547 | 0.834 | 0.514 | -0.079 | 0.073 | 0.073 | 0 | 0.6 | 0.91 | 0.67 | 0.79 | 0.55 | 0.67 |
| 2 | 102628509 | rs3218872 | 0.174 | -1.547 | 1.187 | 0.635 | -0.036 | -0.008 | -0.008 | 0 | 0.22 | 0.33 | 0.19 | 0.3 | 0.12 | 0.21 |
| 2 | 102633347 | rs3218911 | 0.245 | -1.531 | 1.067 | 0.643 | -0.022 | -0.004 | -0.004 | 0.197 | 0.22 | 0.27 | 0.19 | 0.3 | 0.12 | 0.21 |
| 2 | 102639209 | rs2072474 | -0.26 | -1.271 | 0.585 | 0.61 | -0.031 | -0.034 | -0.034 | 0 | 0.28 | 0.51 | 0.23 | 0.28 | 0.16 | 0.29 |
| 2 | 102641201 | rs3218977 | 0.4 | -1.069 | 0.192 | 0.869 | 0.002 | -0.007 | -0.007 | 0 | 0.15 | 0.13 | 0.16 | 0.25 | 0.15 | 0.29 |
| 2 | 102646391 | rs7589525 | 0.967 | -1.025 | 0.454 | 0.839 | -0.03 | -0.036 | -0.036 | 0 | 0.2 | 0.12 | 0.3 | 0.27 | 0.27 | 0.13 |
| 2 | 102648609 | rs4851531 | -0.508 | -1.306 | 0.441 | 0.726 | -0.002 | 0 | 0 | 0 | 0.47 | 0.58 | 0.52 | 0.53 | 0.42 | 0.36 |
| 2 | 102649900 | rs4851532 | 0.497 | -1.3 | 0.263 | 0.722 | 0.032 | 0.021 | 0.021 | 0.37 | 0.23 | 0.11 | 0.19 | 0.27 | 0.15 | 0.23 |
| 2 | 102654228 | rs4850994 | 0.683 | -1.181 | 0.32 | 0.737 | 0 | 0.001 | 0.001 | 0.074 | 0.14 | 0.14 | 0.15 | 0.23 | 0.15 | 0.28 |
| 2 | 102659194 | rs974950 | -0.402 | -1.172 | 0.219 | 0.677 | 0.001 | -0.002 | -0.002 | 0.393 | 0.46 | 0.5 | 0.52 | 0.55 | 0.42 | 0.36 |
| 2 | 102662172 | rs17026510 | 0.426 | -1.217 | -0.129 | 0.531 | 0.036 | 0.031 | 0.031 | 0.401 | 0.23 | 0.1 | 0.2 | 0.27 | 0.15 | 0.23 |
| 2 | 102662888 | rs10185424 | -0.861 | -1.104 | 0.255 | 0.538 | -0.01 | 0.018 | 0.018 | 0 | 0.6 | 0.77 | 0.67 | 0.72 | 0.56 | 0.56 |
| 2 | 102663628 | rs2310173 | -1.214 | -1.234 | 0.023 | 0.472 | -0.061 | -0.017 | -0.017 | 0 | 0.63 | 0.89 | 0.57 | 0.69 | 0.56 | 0.61 |
| 2 | 102665259 | rs11889289 | -1.183 | -1.507 | 0.295 | 0.229 | -0.096 | 0.128 | 0.128 | 0 | 0.62 | 0.92 | 0.6 | 0.84 | 0.5 | 0.71 |
| 2 | 102665552 | rs7607548 | 0.031 | -1.741 | 0.618 | 0.442 | 0 | 0.001 | 0.001 | 0.228 | 0.35 | 0.34 | 0.46 | 0.49 | 0.39 | 0.46 |
| 2 | 102670809 | rs2310175 | 0.618 | -1.746 | 0.586 | 0.433 | 0.046 | -0.046 | -0.046 | 0 | 0.21 | 0.09 | 0.09 | 0.34 | 0.11 | 0.24 |
| 2 | 102671067 | rs11123905 | 1.349 | -1.801 | 0.432 | 0.544 | -0.032 | -0.04 | -0.04 | 0 | 0.18 | 0.1 | 0.3 | 0.27 | 0.24 | 0.26 |
| 2 | 102674501 | rs11689480 | -1.237 | -1.772 | 0.497 | 0.504 | -0.008 | 0.012 | 0.012 | 0 | 0.58 | 0.61 | 0.57 | 0.83 | 0.5 | 0.71 |
| 2 | 102675594 | rs888001 | 0.46 | -1.493 | 0.874 | 0.805 | -0.004 | 0.01 | 0.01 | 1.54 | 0.16 | 0.18 | 0.16 | 0.23 | 0.16 | 0.2 |
| 2 | 102677765 | rs12623383 | 0.657 | -1.612 | 0.735 | 0.742 | 0.022 | -0.037 | -0.037 | 0 | 0.14 | 0.07 | 0.07 | 0.34 | 0.07 | 0.21 |
| 2 | 102680601 | rs7604529 | -0.384 | -1.33 | 0.938 | 0.174 | -0.033 | 0.043 | 0.043 | 2.144 | 0.2 | 0.49 | 0.19 | 0.23 | 0.16 | 0.2 |
| 2 | 102683031 | rs2192751 | 0.065 | -1.177 | 0.333 | 0.236 | -0.008 | 0.009 | 0.009 | 0 | 0.37 | 0.53 | 0.4 | 0.61 | 0.31 | 0.49 |
| 2 | 102683386 | rs868251 | 0.048 | -1.19 | 0.3 | 0.23 | -0.008 | 0.008 | 0.008 | 0 | 0.37 | 0.54 | 0.41 | 0.61 | 0.31 | 0.49 |
| 2 | 102683452 | rs868252 | -0.817 | -0.981 | -0.341 | 0.135 | -0.031 | 0.188 | 0.188 | 1.436 | 0.54 | 0.87 | 0.58 | 0.83 | 0.47 | 0.69 |
| 2 | 102685411 | rs6755229 | -0.758 | -0.936 | -0.473 | 0.137 | -0.006 | 0.108 | 0.108 | 0.482 | 0.54 | 0.74 | 0.56 | 0.85 | 0.47 | 0.72 |
| 2 | 102688158 | rs10203724 | -0.652 | -0.973 | -0.317 | 0.39 | -0.055 | -0.027 | -0.027 | 0 | 0.3 | 0.45 | 0.16 | 0.21 | 0.17 | 0.36 |
| 2 | 102688853 | rs17026582 | -0.504 | -0.504 | -0.623 | 0.327 | -0.003 | -0.01 | -0.01 | 1.057 | 0.3 | 0.61 | 0.48 | 0.23 | 0.27 | 0.23 |
| 2 | 102689020 | rs13401717 | -0.805 | -0.24 | -1.011 | -0.002 | -0.012 | 0.022 | 0.022 | 0.223 | 0.38 | 0.71 | 0.45 | 0.23 | 0.41 | 0.44 |
| 2 | 102689031 | rs13427957 | -0.761 | -0.263 | -0.647 | 0.067 | 0.007 | -0.013 | -0.013 | 0 | 0.37 | 0.58 | 0.44 | 0.23 | 0.41 | 0.43 |
| 2 | 102690578 | rs7569218 | -1.163 | -0.191 | -1.083 | 0.061 | 0.023 | 0.118 | 0.118 | 0 | 0.49 | 0.73 | 0.56 | 0.82 | 0.56 | 0.67 |
| 2 | 102690727 | rs12624116 | 0.507 | 0.049 | -1.426 | 0.767 | -0.002 | -0.019 | -0.019 | 0 | 0.16 | 0.13 | 0.18 | 0.49 | 0.2 | 0.21 |
| 2 | 102691310 | rs1558644 | 0.516 | 0.058 | -1.516 | 0.794 | -0.038 | -0.128 | -0.128 | 0.055 | 0.16 | 0.03 | 0.18 | 0.55 | 0.21 | 0.21 |
| 2 | 102701352 | rs12477295 | 0.734 | 0.102 | -1.558 | 0.718 | -0.072 | -0.022 | -0.022 | 0 | 0.17 | 0.05 | 0.3 | 0.18 | 0.32 | 0.25 |
| 2 | 102707179 | rs10199359 | -0.292 | -0.267 | -1.034 | 0.749 | -0.002 | 0.015 | 0.015 | 0 | 0.22 | 0.26 | 0.21 | 0.55 | 0.22 | 0.39 |
| 2 | 102711009 | rs933494 | 0.706 | 0.395 | -1.341 | 0.577 | -0.097 | -0.032 | -0.032 | 0 | 0.17 | 0.05 | 0.32 | 0.19 | 0.35 | 0.26 |
| 2 | 102711031 | rs933493 | -0.873 | -0.363 | -1.538 | 0.25 | 0.029 | 0.059 | 0.059 | 0.788 | 0.62 | 0.8 | 0.83 | 0.7 | 0.71 | 0.68 |
| 2 | 102717337 | rs10490571 | 0.963 | -0.131 | -1.105 | 0.148 | -0.061 | -0.012 | -0.012 | 0.075 | 0.19 | 0.1 | 0.32 | 0.19 | 0.35 | 0.26 |
| 2 | 102723017 | rs13029804 | 1.025 | -0.512 | -0.849 | 0.247 | -0.03 | -0.04 | -0.04 | 0.78 | 0.23 | 0.45 | 0.29 | 0.17 | 0.2 | 0.13 |
| 2 | 102726661 | rs12712127 | -0.131 | -0.598 | -1.105 | 0.309 | 0.008 | 0.023 | 0.023 | 0.294 | 0.45 | 0.71 | 0.64 | 0.36 | 0.57 | 0.39 |
| 2 | 102730307 | rs887998 | -1.407 | -0.949 | -1.495 | 0.086 | 0.021 | 0.123 | 0.123 | 0 | 0.72 | 0.92 | 0.88 | 0.91 | 0.79 | 0.84 |
| 2 | 102730440 | rs874954 | -0.148 | -0.619 | -1.02 | 0.344 | 0.006 | 0.016 | 0.016 | 0.22 | 0.45 | 0.64 | 0.63 | 0.37 | 0.57 | 0.39 |
| 2 | 102732778 | rs13007967 | 0.497 | -0.735 | -0.836 | 0.239 | -0.075 | -0.06 | -0.06 | 0.713 | 0.25 | 0.56 | 0.24 | 0.16 | 0.12 | 0.21 |
| 2 | 102736588 | rs7557377 | 0.496 | -0.739 | -0.838 | 0.263 | -0.075 | -0.046 | -0.046 | 0 | 0.25 | 0.56 | 0.25 | 0.18 | 0.12 | 0.21 |
| 2 | 102736867 | rs12328681 | 0.782 | -0.859 | -1.025 | 0.276 | -0.001 | -0.01 | -0.01 | 0.512 | 0.16 | 0.21 | 0.13 | 0.19 | 0.19 | 0.22 |
| 2 | 102737937 | rs12712130 | -0.352 | -0.731 | -1.095 | 0.323 | 0.058 | 0.052 | 0.052 | 0 | 0.41 | 0.77 | 0.36 | 0.37 | 0.31 | 0.43 |
| 2 | 102738545 | rs6713468 | -0.352 | -0.731 | -1.095 | 0.323 | 0.058 | 0.052 | 0.052 | 0 | 0.41 | 0.77 | 0.36 | 0.37 | 0.31 | 0.43 |
| 2 | 102740459 | rs4851541 | 0.54 | -0.505 | -1.362 | 0.823 | -0.018 | -0.019 | -0.019 | 0 | 0.25 | 0.2 | 0.45 | 0.44 | 0.44 | 0.32 |
| 2 | 102746083 | rs4851543 | 1.612 | -0.429 | -1.422 | 0.92 | -0.032 | -0.029 | -0.029 | 0.01 | 0.07 | 0 | 0.12 | 0.25 | 0.11 | 0.06 |
| 2 | 102746276 | rs7588201 | -0.015 | -0.384 | -1.284 | 0.941 | 0.003 | 0.002 | 0.002 | 0 | 0.39 | 0.67 | 0.35 | 0.34 | 0.3 | 0.43 |
| 2 | 102747313 | rs1024794 | 1.749 | -0.331 | -1.463 | 0.969 | -0.109 | -0.073 | -0.073 | 0 | 0.16 | 0.02 | 0.31 | 0.22 | 0.33 | 0.26 |
| 2 | 102749240 | rs4851544 | 0.816 | -0.772 | -1.291 | 0.696 | -0.068 | -0.067 | -0.067 | 0 | 0.24 | 0.12 | 0.44 | 0.47 | 0.44 | 0.32 |
| 2 | 102757216 | rs4141631 | -1.087 | -0.752 | -1.313 | 0.487 | 0.013 | 0.048 | 0.048 | 0 | 0.66 | 0.96 | 0.81 | 0.81 | 0.75 | 0.75 |
| 2 | 102761240 | rs11883987 | -0.011 | -0.86 | -0.838 | 0.799 | 0.02 | -0.001 | -0.001 | 2.742 | 0.46 | 0.66 | 0.69 | 0.56 | 0.57 | 0.45 |
| 2 | 102761538 | rs11123911 | 1.201 | -0.776 | -0.675 | 0.801 | -0.037 | -0.024 | -0.024 | 0.037 | 0.19 | 0.4 | 0.2 | 0.17 | 0.11 | 0.12 |
| 2 | 102763837 | rs13035227 | 1.229 | -0.833 | -0.634 | 0.823 | -0.039 | -0.014 | -0.014 | 0 | 0.19 | 0.4 | 0.2 | 0.17 | 0.11 | 0.12 |
| 2 | 102763960 | rs1558642 | -1.132 | -1.005 | -1 | 0.672 | -0.012 | 0.051 | 0.051 | 0 | 0.69 | 0.98 | 0.84 | 0.86 | 0.77 | 0.78 |
| 2 | 102769373 | rs2192752 | -1.132 | -1.005 | -1 | 0.666 | -0.012 | 0.047 | 0.047 | 0 | 0.69 | 0.98 | 0.84 | 0.86 | 0.77 | 0.78 |
| 2 | 102769786 | rs949963 | 0.457 | -1.019 | -0.665 | 0.754 | -0.007 | 0.045 | 0.045 | 0 | 0.19 | 0.32 | 0.14 | 0.26 | 0.19 | 0.32 |
| 2 | 102770738 | rs2287049 | -0.248 | -0.944 | -0.968 | 0.756 | 0.042 | -0.015 | -0.015 | 0.416 | 0.38 | 0.72 | 0.33 | 0.43 | 0.3 | 0.44 |
| 2 | 102774988 | rs3917243 | 0.56 | -0.477 | -0.841 | 0.795 | -0.044 | -0.032 | -0.032 | 0 | 0.31 | 0.15 | 0.5 | 0.43 | 0.44 | 0.34 |
| 2 | 102776202 | rs13019803 | 1.24 | -0.689 | -0.124 | 0.498 | -0.036 | -0.005 | -0.005 | 0 | 0.16 | 0.3 | 0.16 | 0.17 | 0.08 | 0.08 |
| 2 | 102776518 | rs3917254 | 0.424 | -0.895 | -0.086 | 0.49 | -0.044 | -0.007 | -0.007 | 0.614 | 0.22 | 0.32 | 0.18 | 0.22 | 0.1 | 0.21 |
| 2 | 102782166 | rs3917291 | 1.926 | -1.225 | 0.046 | 0.181 | 0.019 | 0.008 | 0.008 | 0.932 | 0.14 | 0.11 | 0.04 | 0 | 0.05 | 0.01 |
| 2 | 102784574 | rs13020778 | -0.632 | -0.907 | -0.625 | -0.092 | -0.028 | -0.057 | -0.057 | 0 | 0.45 | 0.91 | 0.6 | 0.49 | 0.45 | 0.38 |

| | | | | | | | | | | | | | | | | |
|---|---|---|---|---|---|---|---|---|---|---|---|---|---|---|---|---|
| 2 | 102788125 | rs3917304 | 0.853 | -0.861 | -0.217 | -0.14 | 0.026 | 0.085 | 0.085 | 0 | 0.22 | 0.3 | 0.48 | 0.48 | 0.31 | 0.35 |
| 2 | 102788774 | rs3771200 | 1.681 | -0.925 | -0.483 | -0.336 | -0.071 | -0.107 | -0.107 | 0 | 0.29 | 0.09 | 0.26 | 0.43 | 0.36 | 0.37 |
| 2 | 102792875 | rs3917320 | 2.683 | -0.917 | -0.664 | -0.345 | 0.019 | 0.009 | 0.009 | 0.984 | 0.14 | 0.11 | 0.04 | 0 | 0.05 | 0.01 |
| 2 | 102793907 | rs3917325 | 2.683 | -0.917 | -0.728 | -0.345 | 0.03 | 0.032 | 0.032 | 0.984 | 0.14 | 0.08 | 0.04 | 0 | 0.05 | 0.01 |
| 2 | 102794282 | rs2110726 | 1.955 | -0.958 | -0.797 | -0.292 | -0.111 | -0.133 | -0.133 | 0.833 | 0.27 | 0.03 | 0.25 | 0.37 | 0.35 | 0.35 |
| 2 | 102794541 | rs3917328 | 2.622 | -0.784 | -0.633 | -0.374 | 0.03 | 0.031 | 0.031 | 1.136 | 0.14 | 0.08 | 0.04 | 0 | 0.05 | 0.01 |
| 2 | 102794603 | rs3732131 | 0.594 | -0.624 | -0.459 | -0.149 | 0.069 | 0.204 | 0.204 | 0 | 0.05 | 0.2 | 0.36 | 0.23 | 0.08 | 0.09 |
| 2 | 102799152 | rs6744454 | -1.48 | -0.686 | -0.662 | -0.22 | -0.023 | 0.145 | 0.145 | 0 | 0.78 | 0.97 | 0.86 | 0.99 | 0.82 | 0.88 |
| 2 | 102802255 | rs10184597 | 0.737 | -0.41 | -0.146 | 0.083 | 0.018 | 0.061 | 0.061 | 0 | 0.21 | 0.23 | 0.47 | 0.48 | 0.31 | 0.35 |
| 2 | 102804035 | rs2241132 | 0.805 | -0.691 | 0.187 | 0.107 | -0.031 | 0.023 | 0.023 | 0.388 | 0.22 | 0.58 | 0.43 | 0.31 | 0.16 | 0.08 |
| 2 | 102810168 | rs1558648 | 1.224 | -0.453 | -0.108 | -0.056 | -0.001 | -0.007 | -0.007 | 0 | 0.19 | 0.24 | 0.1 | 0.08 | 0.14 | 0.03 |
| 2 | 102822879 | rs1922296 | 1.2 | -0.267 | -0.203 | -0.036 | -0.017 | -0.045 | -0.045 | 0 | 0.28 | 0.12 | 0.25 | 0.39 | 0.33 | 0.38 |
| 2 | 102829013 | rs1922292 | 0.892 | -0.299 | -0.104 | -0.056 | -0.006 | -0.017 | -0.017 | 0 | 0.31 | 0.22 | 0.26 | 0.39 | 0.34 | 0.39 |
| 2 | 102835909 | rs1922294 | -1.6 | -0.421 | -0.102 | -0.264 | 0.011 | 0.003 | 0.003 | 0 | 0.82 | 0.82 | 0.9 | 0.97 | 0.9 | 0.89 |
| 2 | 102840348 | rs3771188 | 0.193 | 0.302 | 0.236 | 0.21 | 0.107 | 0.196 | 0.196 | 0 | 0.06 | 0.22 | 0.38 | 0.21 | 0.12 | 0.07 |
| 2 | 102840507 | rs3771185 | 0.465 | 0.257 | 0.141 | 0.14 | 0.024 | 0.009 | 0.009 | 0 | 0.34 | 0.34 | 0.63 | 0.61 | 0.45 | 0.45 |
| 2 | 102842124 | rs2302621 | -0.509 | 0.166 | 0.067 | 0.09 | 0.011 | 0.054 | 0.054 | 0 | 0.55 | 0.68 | 0.74 | 0.68 | 0.6 | 0.48 |
| 2 | 102842467 | rs2302620 | 0.442 | 0.325 | 0.104 | 0.214 | 0.125 | 0.214 | 0.214 | 0 | 0.05 | 0.21 | 0.38 | 0.2 | 0.12 | 0.07 |
| 2 | 102849081 | rs955754 | 0.227 | 0.27 | -0.091 | 0.099 | -0.025 | 0 | 0 | 0 | 0.26 | 0.53 | 0.45 | 0.28 | 0.19 | 0.09 |
| 2 | 102851708 | rs2302612 | 0.227 | 0.274 | -0.095 | 0.139 | -0.024 | 0.001 | 0.001 | 0.658 | 0.26 | 0.54 | 0.45 | 0.29 | 0.19 | 0.1 |
| 2 | 102852802 | rs10167431 | -0.436 | 0.022 | -0.366 | 0.102 | 0 | 0.013 | 0.013 | 0 | 0.53 | 0.56 | 0.68 | 0.67 | 0.51 | 0.47 |
| 2 | 102852980 | rs2287040 | 1.059 | 0.337 | -0.523 | 0.416 | -0.06 | -0.086 | -0.086 | 0 | 0.27 | 0.08 | 0.23 | 0.36 | 0.32 | 0.38 |
| 2 | 102854054 | rs7583215 | -0.99 | 0.602 | 0.001 | 0.564 | 0.068 | 0.203 | 0.203 | 0 | 0.09 | 0.32 | 0.4 | 0.31 | 0.15 | 0.1 |
| 2 | 102855065 | rs11692230 | 0.367 | 0.444 | -0.717 | 0.09 | 0.011 | 0.024 | 0.024 | 0.109 | 0.47 | 0.21 | 0.3 | 0.37 | 0.39 | 0.42 |
| 2 | 102855241 | rs6752467 | -0.622 | 0.399 | -0.038 | -0.876 | -0.018 | -0.083 | -0.083 | 4.36 | 0.8 | 0.72 | 0.88 | 0.96 | 0.85 | 0.87 |
| 2 | 102855298 | rs6752589 | -0.666 | 0.521 | -0.179 | -0.745 | -0.011 | 0.014 | 0.014 | 0.001 | 0.8 | 0.87 | 0.56 | 0.8 | 0.75 | 0.82 |
| 2 | 102862641 | rs1035131 | 0.639 | 0.564 | -0.417 | -1.154 | -0.034 | -0.005 | -0.005 | 0.083 | 0.31 | 0.18 | 0.34 | 0.67 | 0.36 | 0.62 |
| 2 | 102864310 | rs13018912 | 2.336 | 0.189 | -0.614 | -1.389 | 0.04 | 0.031 | 0.031 | 3.406 | 0.17 | 0.09 | 0.05 | 0 | 0.05 | 0.01 |
| 2 | 102864681 | rs17689452 | 0.815 | 0.297 | -0.539 | -1.359 | -0.038 | -0.005 | -0.005 | 0 | 0.31 | 0.17 | 0.34 | 0.67 | 0.36 | 0.62 |
| 2 | 102866273 | rs12712133 | -0.321 | 0.244 | -0.062 | -1.491 | -0.007 | 0.019 | 0.019 | 0 | 0.63 | 0.62 | 0.52 | 0.78 | 0.56 | 0.7 |
| 2 | 102878891 | rs12475055 | 0.738 | 0.074 | -0.031 | -1.711 | -0.055 | 0.058 | 0.058 | 0.013 | 0.34 | 0.19 | 0.37 | 0.84 | 0.45 | 0.67 |
| 2 | 102879464 | rs10189629 | 2.16 | 0.314 | 0.213 | -1.641 | 0.02 | 0.006 | 0.006 | 0 | 0.11 | 0.25 | 0.13 | 0.07 | 0.16 | 0.08 |
| 2 | 102880823 | rs11123915 | -0.377 | 0.287 | 0.215 | -1.659 | 0 | -0.046 | -0.046 | 1.268 | 0.66 | 0.6 | 0.55 | 0.95 | 0.65 | 0.76 |
| 2 | 102883975 | rs11692065 | 2.097 | 0.285 | 0.006 | -1.534 | 0.02 | 0.009 | 0.009 | 0 | 0.11 | 0.25 | 0.13 | 0.07 | 0.15 | 0.08 |
| 2 | 102887128 | rs11674302 | 2.074 | 0.292 | -0.155 | -1.543 | 0.031 | 0.014 | 0.014 | 0 | 0.11 | 0.3 | 0.13 | 0.07 | 0.15 | 0.08 |
| 2 | 102888565 | rs1922288 | -0.528 | 0.209 | -0.628 | -1.493 | 0.013 | 0.04 | 0.04 | 0 | 0.71 | 0.77 | 0.62 | 0.95 | 0.76 | 0.78 |
| 2 | 102909277 | rs4851003 | -0.126 | 0.481 | -0.42 | -1.136 | -0.017 | -0.159 | -0.159 | 0 | 0.33 | 0.57 | 0.21 | 0 | 0.27 | 0.12 |
| 2 | 102917788 | rs4399750 | 0.221 | 0.824 | -0.586 | -0.56 | 0.004 | 0.004 | 0.004 | 2.291 | 0.58 | 0.43 | 0.36 | 0.57 | 0.49 | 0.44 |
| 2 | 102926511 | rs10178436 | -0.004 | 0.764 | -0.387 | -0.565 | -0.021 | -0.019 | -0.019 | 0 | 0.61 | 0.71 | 0.38 | 0.59 | 0.5 | 0.44 |
| 2 | 102926981 | rs11685424 | 0.037 | 0.761 | -0.331 | -0.594 | -0.013 | -0.012 | -0.012 | 1.309 | 0.6 | 0.68 | 0.37 | 0.56 | 0.49 | 0.44 |
| 2 | 102930948 | rs12712135 | -0.003 | 0.765 | -0.366 | -0.595 | -0.021 | -0.02 | -0.02 | 1.328 | 0.61 | 0.71 | 0.38 | 0.59 | 0.5 | 0.44 |
| 2 | 102938389 | rs1420089 | 0.576 | 1.068 | -0.286 | -0.648 | 0.056 | -0.031 | -0.031 | 2.665 | 0.06 | 0.21 | 0.06 | 0 | 0.12 | 0.01 |
| 2 | 102939036 | rs13001325 | 0.612 | 0.874 | -0.539 | -0.801 | 0.054 | 0.038 | 0.038 | 1.403 | 0.47 | 0.14 | 0.24 | 0.47 | 0.34 | 0.35 |
| 2 | 102953444 | rs17639215 | 1.634 | 1.139 | -0.501 | -0.829 | -0.011 | 0.006 | 0.006 | 0 | 0.07 | 0.03 | 0.08 | 0 | 0.13 | 0.11 |
| 2 | 102955468 | rs1041973 | 0.489 | 0.63 | -0.604 | -1.152 | -0.007 | -0.017 | -0.017 | 0 | 0.43 | 0.57 | 0.2 | 0.15 | 0.24 | 0.14 |
| 2 | 102957716 | rs1420101 | 0.239 | 0.688 | -0.769 | -1.132 | 0.053 | 0.047 | 0.047 | 0.854 | 0.5 | 0.36 | 0.25 | 0.46 | 0.34 | 0.35 |
| 2 | 102959260 | rs12999517 | 1.285 | 0.802 | -0.777 | -1.178 | 0.046 | 0.045 | 0.045 | 1.885 | 0.32 | 0.27 | 0.08 | 0.06 | 0.12 | 0.06 |
| 2 | 102960584 | rs12712142 | 0.437 | 0.878 | -0.7 | -1.087 | 0.029 | 0.041 | 0.041 | 0.505 | 0.49 | 0.36 | 0.26 | 0.47 | 0.37 | 0.35 |
| 2 | 102960824 | rs2160203 | 0.547 | 1.009 | -0.493 | -1.043 | 0.063 | -0.039 | -0.039 | 0 | 0.17 | 0.49 | 0.18 | 0.09 | 0.27 | 0.09 |
| 2 | 102965153 | rs17696376 | 1.47 | 1.003 | -0.892 | -1.022 | 0.012 | -0.019 | -0.019 | 0.165 | 0.05 | 0.08 | 0.06 | 0 | 0.12 | 0.01 |
| 2 | 102966067 | rs1921622 | 0.076 | 0.964 | -1.069 | -1.091 | -0.014 | -0.004 | -0.004 | 0 | 0.56 | 0.18 | 0.33 | 0.49 | 0.51 | 0.46 |
| 2 | 102968007 | rs4988956 | -0.257 | 1.014 | -1.048 | -1.113 | -0.022 | 0.024 | 0.024 | 0.061 | 0.28 | 0.75 | 0.29 | 0.13 | 0.4 | 0.21 |
| 2 | 102968075 | rs4988957 | -0.257 | 1.014 | -1.048 | -1.113 | -0.022 | 0.024 | 0.024 | 0.061 | 0.28 | 0.75 | 0.29 | 0.13 | 0.4 | 0.21 |
| 2 | 102968211 | rs10192036 | -0.257 | 1.014 | -1.048 | -1.113 | -0.022 | 0.024 | 0.024 | 0.061 | 0.28 | 0.74 | 0.29 | 0.13 | 0.4 | 0.2 |
| 2 | 102968285 | rs4988958 | -0.257 | 1.014 | -1.048 | -1.113 | -0.022 | 0.024 | 0.024 | 0.061 | 0.28 | 0.75 | 0.29 | 0.13 | 0.4 | 0.21 |
| 2 | 102968356 | rs10192157 | -0.257 | 1.014 | -1.048 | -1.113 | -0.022 | 0.024 | 0.024 | 0.061 | 0.28 | 0.75 | 0.29 | 0.13 | 0.4 | 0.21 |
| 2 | 102971865 | rs13015714 | -1.632 | 0.977 | -1.091 | -1.116 | 0.011 | -0.088 | -0.088 | 0 | 0.76 | 0.89 | 0.53 | 0.59 | 0.78 | 0.55 |
| 2 | 102977730 | rs10204837 | -0.134 | 1.05 | -0.997 | -1.096 | -0.007 | 0.006 | 0.006 | 0.004 | 0.27 | 0.67 | 0.27 | 0.13 | 0.4 | 0.21 |
| 2 | 102979028 | rs2287037 | 0.417 | 0.997 | -1.013 | -1.096 | 0.037 | 0.058 | 0.058 | 1.452 | 0.49 | 0.22 | 0.26 | 0.46 | 0.38 | 0.35 |
| 2 | 102981018 | rs10515921 | 1.707 | 1.151 | -0.9 | -0.973 | -0.001 | -0.008 | -0.008 | 0.351 | 0.08 | 0.05 | 0.08 | 0 | 0.13 | 0.11 |
| 2 | 102986154 | rs2160202 | 1.367 | 1.012 | -1.083 | -1 | -0.046 | -0.061 | -0.061 | 0 | 0.18 | 0.07 | 0.17 | 0.41 | 0.25 | 0.28 |
| 2 | 102986170 | rs2058623 | -1.594 | 1.004 | -1.123 | -1.052 | 0.009 | -0.103 | -0.103 | 0 | 0.75 | 0.89 | 0.53 | 0.58 | 0.78 | 0.56 |
| 2 | 102986222 | rs3771166 | -0.008 | 0.95 | -1.055 | -1.134 | -0.007 | 0.005 | 0.005 | 0.013 | 0.27 | 0.67 | 0.27 | 0.13 | 0.4 | 0.21 |
| 2 | 102991369 | rs6758936 | -0.239 | 0.944 | -1.056 | -1.058 | -0.027 | 0.112 | 0.112 | 1.797 | 0.57 | 0.74 | 0.34 | 0.17 | 0.53 | 0.24 |
| 2 | 103001402 | rs1035130 | 1.27 | 0.997 | -1.043 | -0.992 | -0.011 | -0.014 | -0.014 | 0.009 | 0.18 | 0.14 | 0.19 | 0.42 | 0.25 | 0.3 |
| 2 | 103003265 | rs2241116 | 1.687 | 1.151 | -1.097 | -0.917 | -0.032 | -0.03 | -0.03 | 0.103 | 0.15 | 0.05 | 0.15 | 0.24 | 0.2 | 0.25 |
| 2 | 103011552 | rs4851005 | 0.971 | 1.248 | -1.023 | -0.975 | -0.042 | -0.037 | -0.037 | 0 | 0.2 | 0.1 | 0.23 | 0.42 | 0.36 | 0.3 |
| 2 | 103013962 | rs3732126 | 1.439 | 1.302 | -1.095 | -0.961 | -0.036 | -0.048 | -0.048 | 0.261 | 0.17 | 0.06 | 0.18 | 0.42 | 0.25 | 0.3 |
| 2 | 103014302 | rs1135354 | 1.216 | 1.291 | -1.09 | -0.966 | -0.013 | -0.017 | -0.017 | 0 | 0.18 | 0.14 | 0.19 | 0.42 | 0.25 | 0.3 |

| | | | | | | | | | | | | | | | | |
|---|---|---|---|---|---|---|---|---|---|---|---|---|---|---|---|---|
| 2 | 103034440 | rs11465670 | 1.794 | 1.295 | -0.514 | -1.019 | 0.03 | -0.027 | -0.027 | 0 | 0.08 | 0.15 | 0.09 | 0.02 | 0.13 | 0.1 |
| 2 | 103035044 | rs1420106 | -1.378 | 0.96 | -0.671 | -0.991 | 0.007 | -0.109 | -0.109 | 0 | 0.75 | 0.89 | 0.53 | 0.58 | 0.78 | 0.55 |
| 2 | 103035468 | rs2293223 | 1.501 | 0.519 | -0.882 | -1.241 | -0.024 | -0.057 | -0.057 | 0 | 0.23 | 0.37 | 0.16 | 0.09 | 0.18 | 0.1 |
| 2 | 103050925 | rs2110735 | -1.545 | 0.543 | -0.989 | -1.197 | 0.003 | -0.111 | -0.111 | 0 | 0.76 | 0.9 | 0.53 | 0.58 | 0.78 | 0.56 |
| 2 | 103052206 | rs2110734 | -1.568 | 0.553 | -0.984 | -1.192 | 0.003 | -0.111 | -0.111 | 0.19 | 0.76 | 0.9 | 0.53 | 0.58 | 0.78 | 0.56 |
| 2 | 103054556 | rs2058659 | -0.432 | 0.604 | -0.9 | -1.269 | -0.013 | 0.061 | 0.061 | 1.824 | 0.57 | 0.67 | 0.34 | 0.17 | 0.53 | 0.26 |
| 2 | 103057043 | rs17027173 | 1.489 | 0.745 | -0.933 | -1.256 | 0.05 | -0.041 | -0.041 | 0 | 0.12 | 0.19 | 0.14 | 0.02 | 0.25 | 0.11 |
| 2 | 103057892 | rs11465702 | 2.439 | 0.715 | -0.828 | -1.016 | -0.01 | 0.017 | 0.017 | 1.027 | 0.1 | 0.06 | 0.1 | 0.01 | 0.15 | 0.07 |
| 2 | 103060851 | rs3771150 | 0.954 | 0.989 | -0.815 | -0.8 | -0.035 | -0.051 | -0.051 | 1.443 | 0.18 | 0.07 | 0.18 | 0.42 | 0.25 | 0.3 |
| 2 | 103062880 | rs6734736 | -1.275 | 0.989 | -0.91 | -0.778 | 0.001 | -0.115 | -0.115 | 0 | 0.76 | 0.9 | 0.53 | 0.58 | 0.78 | 0.56 |
| 2 | 103066858 | rs11465730 | 0.331 | 0.964 | -0.884 | -0.82 | -0.001 | -0.001 | -0.001 | 0 | 0.41 | 0.56 | 0.34 | 0.51 | 0.43 | 0.38 |
| 2 | 103068787 | rs7559479 | -1.286 | 0.934 | -0.917 | -0.797 | 0.001 | -0.115 | -0.115 | 0 | 0.76 | 0.9 | 0.53 | 0.58 | 0.78 | 0.56 |
| 2 | 103070568 | rs917997 | -1.286 | 0.939 | -0.929 | -0.794 | 0.001 | -0.115 | -0.115 | 2.765 | 0.76 | 0.9 | 0.53 | 0.58 | 0.78 | 0.56 |
| 2 | 103072220 | rs11886793 | 0.41 | 0.942 | -0.919 | -0.817 | -0.001 | -0.024 | -0.024 | 0 | 0.35 | 0.34 | 0.19 | 0.07 | 0.34 | 0.18 |
| 2 | 103088777 | rs1403552 | 1.846 | 0.93 | -0.833 | -0.782 | -0.022 | -0.058 | -0.058 | 0.062 | 0.23 | 0.37 | 0.16 | 0.09 | 0.18 | 0.08 |
| 2 | 103090127 | rs17027255 | 0.878 | 1.013 | -0.921 | -0.737 | 0.004 | 0.009 | 0.009 | 0 | 0.18 | 0.17 | 0.18 | 0.41 | 0.26 | 0.3 |
| 2 | 103092270 | rs13018263 | 1.181 | 0.328 | -0.892 | -0.706 | -0.005 | 0.007 | 0.007 | 3.163 | 0.24 | 0.36 | 0.2 | 0.28 | 0.22 | 0.11 |
| 2 | 103092513 | rs1468788 | -0.036 | 0.35 | -0.894 | -0.726 | 0.004 | 0.013 | 0.013 | 0.341 | 0.52 | 0.53 | 0.33 | 0.3 | 0.56 | 0.45 |
| 2 | 103094213 | rs759382 | -0.053 | 0.364 | -0.667 | -0.083 | 0.002 | -0.109 | -0.109 | 0 | 0.75 | 0.88 | 0.52 | 0.57 | 0.79 | 0.58 |
| 2 | 103095404 | rs11692304 | 1.296 | 0.458 | -0.251 | -0.287 | -0.015 | 0.085 | 0.085 | 0 | 0.46 | 0.28 | 0.33 | 0.23 | 0.52 | 0.35 |
| 2 | 103096695 | rs6543146 | 0.471 | 0.084 | 0.304 | -0.348 | 0.001 | 0.001 | 0.001 | 0.669 | 0.67 | 0.71 | 0.66 | 0.38 | 0.64 | 0.52 |
| 2 | 103098793 | rs6737325 | 0.09 | 0.018 | 0.248 | -0.302 | -0.011 | -0.019 | -0.019 | 0 | 0.18 | 0.27 | 0.06 | 0.06 | 0.09 | 0.08 |
| 2 | 103117283 | rs6741235 | 0.066 | 0.225 | 0.141 | -0.225 | -0.001 | -0.001 | -0.001 | 0 | 0.76 | 0.74 | 0.73 | 0.32 | 0.76 | 0.66 |
| 2 | 103120769 | rs4851017 | 0.044 | 0.224 | 0.154 | -0.211 | -0.001 | -0.001 | -0.001 | 0 | 0.76 | 0.74 | 0.73 | 0.32 | 0.76 | 0.66 |
| 2 | 103125736 | rs10195948 | -0.104 | 0.029 | 0.262 | -0.127 | 0.003 | -0.01 | -0.01 | 0 | 0.78 | 0.76 | 0.76 | 0.49 | 0.8 | 0.69 |
| 2 | 103131679 | rs1476999 | 1.311 | -0.042 | 0.395 | -0.248 | 0.033 | 0.053 | 0.053 | 0 | 0.5 | 0.3 | 0.61 | 0.41 | 0.58 | 0.46 |
| 2 | 103148169 | rs741285 | 0.465 | 0.04 | 0.499 | -0.29 | -0.001 | 0.018 | 0.018 | 0.198 | 0.68 | 0.67 | 0.68 | 0.53 | 0.68 | 0.56 |
| 2 | 103149100 | rs1014286 | 0.465 | 0.04 | 0.499 | -0.29 | -0.001 | 0.018 | 0.018 | 0.198 | 0.68 | 0.67 | 0.68 | 0.53 | 0.68 | 0.56 |
| 2 | 103149162 | rs2008159 | 0.465 | 0.04 | 0.499 | -0.29 | -0.001 | 0.018 | 0.018 | 0.198 | 0.68 | 0.67 | 0.68 | 0.53 | 0.68 | 0.56 |
| 2 | 103149182 | rs2008157 | 0.465 | 0.04 | 0.499 | -0.29 | -0.001 | 0.018 | 0.018 | 0.198 | 0.68 | 0.67 | 0.68 | 0.53 | 0.68 | 0.56 |
| 2 | 103155069 | rs6761291 | 1.451 | -0.079 | 0.589 | -0.366 | 0.031 | 0.06 | 0.06 | 0 | 0.5 | 0.3 | 0.61 | 0.39 | 0.58 | 0.47 |
| 2 | 103167757 | rs11123936 | 0.209 | -0.078 | 0.628 | -0.361 | -0.011 | -0.003 | -0.003 | 0 | 0.17 | 0.23 | 0.06 | 0.14 | 0.09 | 0.08 |
| 2 | 103168977 | rs10193009 | 0.64 | 0.125 | 0.727 | -0.142 | -0.001 | 0.029 | 0.029 | 0.655 | 0.68 | 0.64 | 0.69 | 0.53 | 0.67 | 0.55 |
| 2 | 103176411 | rs4851619 | -0.107 | -0.193 | 0.484 | -0.09 | 0.002 | -0.071 | -0.071 | 0 | 0.79 | 0.9 | 0.78 | 0.56 | 0.8 | 0.68 |
| 2 | 103180530 | rs2192756 | 0.936 | 0.018 | 0.856 | -0.179 | 0.005 | -0.025 | -0.025 | 0.116 | 0.58 | 0.46 | 0.57 | 0.27 | 0.49 | 0.37 |
| 2 | 103180864 | rs11681159 | 0.018 | 0.204 | 0.637 | -0.031 | -0.016 | -0.003 | -0.003 | 0 | 0.2 | 0.17 | 0.18 | 0.24 | 0.32 | 0.34 |
| 2 | 103205175 | rs17027517 | -1.097 | 0.149 | 0.472 | 0.052 | -0.024 | 0.009 | 0.009 | 0.009 | 0.34 | 0.77 | 0.52 | 0.35 | 0.39 | 0.55 |
| 2 | 103208610 | rs1861229 | -0.034 | 0.024 | 0.543 | 0.047 | 0.006 | -0.109 | -0.109 | 1.123 | 0.83 | 0.98 | 0.83 | 0.51 | 0.86 | 0.74 |
| 2 | 103210221 | rs7573475 | -1.121 | 0.222 | 0.817 | -0.45 | -0.006 | 0.003 | 0.003 | 0 | 0.33 | 0.68 | 0.52 | 0.33 | 0.4 | 0.55 |
| 2 | 103213050 | rs7593772 | -1.387 | 0.227 | 1.2 | -0.501 | 0.083 | 0.066 | 0.066 | 0 | 0.08 | 0.44 | 0.09 | 0.09 | 0.06 | 0.07 |
| 2 | 103215935 | rs11673795 | 0.253 | 0.253 | 1.073 | -0.492 | -0.015 | 0.003 | 0.003 | 0 | 0.21 | 0.18 | 0.42 | 0.22 | 0.33 | 0.36 |
| 2 | 103225209 | rs6543166 | -0.798 | 0.354 | 1.102 | -0.469 | -0.016 | 0.011 | 0.011 | 0 | 0.31 | 0.75 | 0.52 | 0.3 | 0.39 | 0.43 |
| 2 | 103232093 | rs6708609 | -1.509 | 0.346 | 1.265 | -0.47 | 0.061 | 0.072 | 0.072 | 0 | 0.09 | 0.4 | 0.09 | 0.11 | 0.06 | 0.15 |
| 2 | 103232630 | rs2058612 | -0.66 | -0.075 | 1.075 | -0.436 | -0.016 | 0.011 | 0.011 | 0 | 0.31 | 0.75 | 0.52 | 0.3 | 0.39 | 0.43 |
| 2 | 103234720 | rs2110637 | -0.937 | -0.075 | 1.044 | -0.466 | 0.002 | -0.002 | -0.002 | 0 | 0.34 | 0.68 | 0.54 | 0.32 | 0.39 | 0.55 |
| 2 | 103252467 | rs4851621 | 0.604 | 0.22 | 0.841 | -0.396 | -0.024 | 0.041 | 0.041 | 0 | 0.6 | 0.22 | 0.42 | 0.66 | 0.52 | 0.53 |
| 2 | 103253943 | rs17775237 | 0.658 | 0.354 | -0.039 | -0.373 | -0.011 | 0.017 | 0.017 | 0.018 | 0.1 | 0 | 0.03 | 0.06 | 0.08 | 0.02 |
| 2 | 103256547 | rs10181400 | 0.556 | 0.705 | 0.417 | -0.174 | 0.002 | -0.004 | -0.004 | 2.944 | 0.63 | 0.61 | 0.44 | 0.66 | 0.52 | 0.54 |
| 2 | 103257149 | rs10184340 | 0.431 | 0.763 | 0.34 | -0.143 | -0.073 | 0.061 | 0.061 | 0.225 | 0.66 | 0.81 | 0.47 | 0.75 | 0.52 | 0.54 |
| 2 | 103257560 | rs12998183 | 0.539 | 0.69 | 0.388 | -0.103 | -0.01 | 0.022 | 0.022 | 0 | 0.58 | 0.63 | 0.42 | 0.66 | 0.47 | 0.49 |
| 2 | 103260445 | rs4851622 | 0.408 | 0.93 | -0.693 | -0.206 | -0.027 | 0.063 | 0.063 | 0.017 | 0.6 | 0.65 | 0.43 | 0.75 | 0.48 | 0.5 |
| 2 | 103268511 | rs4140836 | -0.01 | 0.606 | -0.587 | -0.092 | 0 | 0.01 | 0.01 | 0 | 0.82 | 0.88 | 0.92 | 0.93 | 0.8 | 0.9 |
| 2 | 103272957 | rs10178585 | 0.946 | 0.886 | -0.393 | -0.073 | 0.007 | 0.001 | 0.001 | 0 | 0.47 | 0.19 | 0.35 | 0.48 | 0.37 | 0.42 |
| 2 | 103276384 | rs4851629 | 1.087 | 0.678 | -0.46 | -0.215 | -0.017 | -0.016 | -0.016 | 0.103 | 0.42 | 0.21 | 0.35 | 0.48 | 0.4 | 0.41 |
| 2 | 103277619 | rs7561632 | -0.544 | 0.715 | 0.637 | -0.16 | 0.056 | -0.007 | -0.007 | 0 | 0.07 | 0.38 | 0.07 | 0 | 0.05 | 0.07 |
| 2 | 103281417 | rs11123944 | 1.066 | 0.595 | 0.675 | -0.068 | -0.06 | -0.066 | -0.066 | 1.102 | 0.44 | 0.08 | 0.36 | 0.48 | 0.41 | 0.38 |
| 2 | 103292693 | rs6731896 | 0.197 | 0.727 | 0.538 | -0.103 | -0.013 | 0.07 | 0.07 | 0 | 0.69 | 0.79 | 0.77 | 0.85 | 0.59 | 0.78 |
| 2 | 103297515 | rs962207 | 0.24 | 0.69 | 0.483 | -0.089 | -0.004 | 0.037 | 0.037 | 0 | 0.68 | 0.76 | 0.77 | 0.85 | 0.6 | 0.78 |
| 2 | 103297704 | rs7606208 | 0.546 | 0.79 | 0.308 | 0.396 | -0.033 | 0.005 | 0.005 | 0 | 0.45 | 0.14 | 0.36 | 0.44 | 0.41 | 0.38 |
| 2 | 103301198 | rs6716756 | 0.253 | 0.95 | -1.013 | 0.223 | -0.007 | 0.009 | 0.009 | 0 | 0.1 | 0.1 | 0.12 | 0.08 | 0.17 | 0.1 |
| 2 | 103301268 | rs7608617 | 0.26 | 0.714 | -0.986 | 0.481 | -0.017 | -0.024 | -0.024 | 1.352 | 0.45 | 0.21 | 0.36 | 0.48 | 0.41 | 0.39 |
| 2 | 103313018 | rs10173702 | 0.386 | 0.119 | -0.619 | 0.597 | -0.011 | -0.001 | -0.001 | 0.332 | 0.8 | 0.73 | 0.86 | 0.76 | 0.81 | 0.79 |
| 2 | 103313748 | rs878539 | 0.907 | 0.132 | -0.325 | 0.853 | -0.018 | 0.017 | 0.017 | 0.875 | 0.6 | 0.47 | 0.65 | 0.53 | 0.65 | 0.59 |
| 2 | 103315418 | rs7607987 | 0.093 | 0.246 | -0.251 | 0.785 | 0.004 | -0.002 | -0.002 | 0 | 0.78 | 0.78 | 0.82 | 0.75 | 0.76 | 0.75 |
| 2 | 103324575 | rs17833353 | 1.323 | 0.497 | -1.126 | -0.108 | -0.02 | 0.04 | 0.04 | 0 | 0.25 | 0.18 | 0.3 | 0.02 | 0.26 | 0.07 |
| 2 | 103327476 | rs11123946 | -0.56 | 0.445 | -0.536 | -0.147 | -0.005 | -0.002 | -0.002 | 0.006 | 0.78 | 0.74 | 0.82 | 0.78 | 0.76 | 0.76 |
| 2 | 103328105 | rs17776951 | 1.555 | 0.446 | -0.57 | -0.153 | -0.043 | 0.076 | 0.076 | 0.023 | 0.23 | 0.08 | 0.28 | 0 | 0.26 | 0.07 |
| 2 | 103330790 | rs6757508 | 0.051 | 0.312 | 0.086 | -0.203 | -0.001 | 0.004 | 0.004 | 0 | 0.53 | 0.47 | 0.51 | 0.77 | 0.5 | 0.69 |
| 2 | 103334969 | rs1052431 | 0.735 | 0.115 | 0.102 | -0.036 | 0 | -0.008 | -0.008 | 0.2 | 0.34 | 0.3 | 0.33 | 0.57 | 0.35 | 0.52 |

| | | | | | | | | | | | | | | | |
|---|---|---|---|---|---|---|---|---|---|---|---|---|---|---|---|
| 2 | 103340541 | rs6759290 | -0.45 | 0.015 | 0.254 | 0.067 | -0.004 | -0.003 | -0.003 | 0 | 0.77 | 0.74 | 0.75 | 0.78 | 0.74 | 0.75 |
| 2 | 103347857 | rs2732824 | 0.432 | 0.071 | 0.939 | 0.083 | -0.005 | -0.002 | -0.002 | 0 | 0.22 | 0.34 | 0.18 | 0.22 | 0.23 | 0.25 |
| 2 | 103349230 | rs2540319 | -0.539 | 0.156 | 0.86 | 0.108 | 0.005 | -0.012 | -0.012 | 0 | 0.39 | 0.28 | 0.34 | 0.4 | 0.38 | 0.41 |
| 2 | 103351736 | rs955915 | 0.158 | 0.148 | 0.967 | 0.073 | -0.012 | 0.002 | 0.002 | 0.068 | 0.23 | 0.52 | 0.21 | 0.23 | 0.23 | 0.25 |
| 2 | 103376157 | rs2540304 | -0.809 | 0.202 | 1.055 | 0.189 | 0.008 | -0.01 | -0.01 | 5.578 | 0.43 | 0.63 | 0.46 | 0.43 | 0.41 | 0.45 |
| 2 | 103377786 | rs2540299 | -0.029 | 0.161 | 1.154 | 0.171 | -0.011 | -0.003 | -0.003 | 0 | 0.24 | 0.49 | 0.29 | 0.24 | 0.26 | 0.27 |
| 2 | 103382425 | rs2080294 | 0.935 | 0.103 | 1.001 | 0.1 | 0.006 | -0.026 | -0.026 | 0.078 | 0.19 | 0.14 | 0.18 | 0.19 | 0.15 | 0.17 |
| 2 | 103385808 | rs2540290 | 0.404 | 0.152 | 1.165 | 0.074 | -0.001 | -0.002 | -0.002 | 0.276 | 0.22 | 0.21 | 0.24 | 0.22 | 0.26 | 0.27 |
| 2 | 103389792 | rs2732846 | -0.934 | 0.174 | 0.983 | 0.246 | -0.002 | 0.016 | 0.016 | 4.398 | 0.51 | 0.57 | 0.49 | 0.56 | 0.5 | 0.65 |
| 2 | 103398334 | rs2732860 | -1.059 | 0.187 | 1.01 | 0.289 | -0.008 | 0.059 | 0.059 | 4.143 | 0.52 | 0.66 | 0.5 | 0.58 | 0.49 | 0.65 |
| 2 | 103406699 | rs2732811 | -1.367 | 0.224 | 0.841 | -0.035 | 0.011 | -0.05 | -0.05 | 0 | 0.76 | 0.87 | 0.79 | 0.57 | 0.74 | 0.74 |
| 2 | 103409685 | rs7604365 | 1.074 | 0.295 | 0.911 | 0.146 | -0.008 | 0.017 | 0.017 | 1.167 | 0.2 | 0.28 | 0.2 | 0.21 | 0.15 | 0.2 |
| 2 | 103413658 | rs2310358 | 0.986 | 0.576 | 0.985 | 0.069 | -0.031 | -0.038 | -0.038 | 0 | 0.08 | 0 | 0.05 | 0.12 | 0.08 | 0.17 |
| 2 | 103425677 | rs2080292 | -0.452 | 0.514 | 1.152 | -0.048 | -0.001 | 0.007 | 0.007 | 0.063 | 0.33 | 0.51 | 0.31 | 0.35 | 0.34 | 0.45 |
| 2 | 103425860 | rs2080291 | -0.015 | 0.629 | 0.563 | 0.009 | 0.001 | 0.003 | 0.003 | 0 | 0.3 | 0.25 | 0.29 | 0.35 | 0.34 | 0.45 |
| 2 | 103432337 | rs2732822 | -0.025 | 0.595 | 0.61 | 0.053 | 0.001 | 0.003 | 0.003 | 0 | 0.3 | 0.25 | 0.29 | 0.35 | 0.34 | 0.45 |
| 2 | 103445960 | rs7602289 | 0.847 | 0.832 | 0.777 | -0.257 | 0.025 | -0.155 | -0.155 | 0 | 0.25 | 0.39 | 0.31 | 0 | 0.25 | 0.08 |
| 2 | 103460719 | rs13410064 | 0.447 | 0.949 | 0.83 | -0.351 | 0.017 | -0.12 | -0.12 | 0 | 0.3 | 0.55 | 0.35 | 0 | 0.32 | 0.13 |
| 2 | 103475598 | rs870155 | 0.693 | 0.824 | 0.937 | 0.086 | 0 | 0.079 | 0.079 | 0.152 | 0.1 | 0.19 | 0.16 | 0.19 | 0.13 | 0.21 |
| 2 | 103499457 | rs2160205 | 0.781 | 0.818 | 1.014 | 0.096 | 0.008 | 0.09 | 0.09 | 0.006 | 0.1 | 0.22 | 0.1 | 0.19 | 0.12 | 0.21 |
| 2 | 103505482 | rs1473170 | -0.425 | 0.49 | 0.695 | 0.01 | 0.009 | -0.008 | -0.008 | 0 | 0.59 | 0.67 | 0.69 | 0.37 | 0.62 | 0.55 |
| 2 | 103509684 | rs10181783 | 1.257 | 0.595 | 0.412 | 0.184 | -0.018 | -0.033 | -0.033 | 0 | 0.18 | 0.07 | 0.26 | 0.18 | 0.19 | 0.21 |
| 2 | 103539431 | rs11123950 | 0.568 | 0.093 | 0.001 | 0.401 | 0 | -0.007 | -0.007 | 0 | 0.18 | 0.18 | 0.2 | 0.28 | 0.18 | 0.22 |
| 2 | 103546444 | rs13388810 | 0.893 | 0.16 | 0.127 | 0.46 | 0.011 | 0.069 | 0.069 | 0.003 | 0.1 | 0.21 | 0.06 | 0.15 | 0.11 | 0.21 |
| 2 | 103553133 | rs4569500 | -0.194 | 0.12 | -0.195 | 0.435 | 0 | -0.003 | -0.003 | 0 | 0.41 | 0.44 | 0.59 | 0.47 | 0.39 | 0.46 |
| 2 | 103557981 | rs6738764 | 0.362 | 0.099 | -0.312 | 0.535 | 0.003 | -0.006 | -0.006 | 0.058 | 0.19 | 0.23 | 0.2 | 0.29 | 0.16 | 0.22 |
| 2 | 103558556 | rs6726414 | -0.969 | 0.261 | -0.346 | 0.535 | -0.025 | 0.158 | 0.158 | 0 | 0.52 | 0.86 | 0.69 | 0.7 | 0.51 | 0.67 |
| 2 | 103559294 | rs4241213 | -0.151 | 0.263 | -0.034 | 0.522 | 0.008 | 0.028 | 0.028 | 0.41 | 0.32 | 0.54 | 0.49 | 0.41 | 0.35 | 0.44 |
| 2 | 103559677 | rs13004328 | 1.504 | 0.286 | -0.245 | 0.587 | -0.047 | -0.05 | -0.05 | 0 | 0.19 | 0.04 | 0.37 | 0.18 | 0.23 | 0.23 |
| 2 | 103561126 | rs11691730 | 0.4 | 0.26 | -0.178 | 0.447 | -0.024 | 0.017 | 0.017 | 0.921 | 0.22 | 0.4 | 0.21 | 0.29 | 0.16 | 0.24 |
| 2 | 103561609 | rs17028085 | 0.074 | 0.263 | -0.287 | 0.481 | 0.04 | 0.083 | 0.083 | 0 | 0.09 | 0.2 | 0.1 | 0.15 | 0.12 | 0.2 |
| 2 | 103562368 | rs10195542 | -0.881 | 0.154 | -1.608 | -0.045 | 0.002 | 0.002 | 0.002 | 0 | 0.74 | 0.91 | 0.85 | 0.7 | 0.74 | 0.76 |
| 2 | 103568193 | rs11899727 | 0.383 | 0.456 | -1.546 | -0.188 | -0.012 | 0.008 | 0.008 | 0.786 | 0.22 | 0.33 | 0.2 | 0.29 | 0.16 | 0.24 |
| 2 | 103572929 | rs2310351 | 1.763 | 0.506 | -1.477 | -0.07 | -0.034 | 0.111 | 0.111 | 0 | 0.22 | 0.05 | 0.16 | 0 | 0.23 | 0.09 |
| 2 | 103582093 | rs6735868 | -0.111 | 0.42 | -0.925 | 0.592 | -0.002 | 0 | 0 | 0 | 0.24 | 0.29 | 0.25 | 0.28 | 0.2 | 0.29 |
| 2 | 103585422 | rs2242290 | 0.339 | 0.549 | -1.082 | 0.564 | -0.035 | -0.048 | -0.048 | 0 | 0.23 | 0.06 | 0.32 | 0.22 | 0.25 | 0.24 |
| 2 | 103587575 | rs956966 | 0.219 | 0.401 | -0.484 | 0.384 | -0.001 | 0.001 | 0.001 | 0.251 | 0.52 | 0.59 | 0.54 | 0.71 | 0.54 | 0.55 |
| 2 | 103593540 | rs11123953 | -1.37 | 0.259 | 0.681 | 0.327 | 0.008 | 0.023 | 0.023 | 1.295 | 0.09 | 0.62 | 0.06 | 0.06 | 0.03 | 0.01 |
| 2 | 103598208 | rs6730289 | -1.092 | 0.262 | 1.204 | 0.328 | 0.071 | 0.09 | 0.09 | 0 | 0.07 | 0.52 | 0.06 | 0.06 | 0.03 | 0.01 |
| 2 | 103598575 | rs6543194 | -1.192 | 0.778 | 0.745 | 0.29 | 0.021 | -0.012 | -0.012 | 0.013 | 0.21 | 0.67 | 0.18 | 0.21 | 0.22 | 0.32 |
| 2 | 103617979 | rs7587833 | -0.447 | 0.968 | 0.343 | 0.272 | -0.026 | -0.008 | -0.008 | 0.084 | 0.92 | 0.94 | 0.91 | 0.86 | 0.84 | 0.74 |
| 2 | 103624188 | rs1542601 | 0.988 | 0.042 | -0.264 | -0.099 | -0.026 | 0.111 | 0.111 | 0 | 0.39 | 0.13 | 0.31 | 0.14 | 0.4 | 0.25 |
| 2 | 103641828 | rs6759830 | -0.377 | 0.062 | -0.485 | 0.342 | -0.074 | 0.017 | 0.017 | 0 | 0.65 | 0.89 | 0.59 | 0.61 | 0.59 | 0.51 |
| 2 | 103680790 | rs13402350 | -0.875 | -0.396 | 0.271 | 0.633 | -0.033 | 0.098 | 0.098 | 0.559 | 0.19 | 0.68 | 0.21 | 0.44 | 0.12 | 0.16 |
| 2 | 103680914 | rs17028190 | 0.588 | -0.359 | 0.323 | 0.544 | 0.002 | 0.001 | 0.001 | 0 | 0.09 | 0.09 | 0.05 | 0.04 | 0.06 | 0.02 |
| 2 | 103722263 | rs12386183 | -0.965 | -0.497 | 0.489 | 0.601 | -0.03 | 0.094 | 0.094 | 0.38 | 0.19 | 0.68 | 0.21 | 0.44 | 0.13 | 0.16 |
| 2 | 103742570 | rs17028232 | 0.067 | -0.241 | 0.176 | 1.13 | -0.016 | 0.257 | 0.257 | 0 | 0.05 | 0.18 | 0.13 | 0.32 | 0.05 | 0.13 |
| 2 | 103753030 | rs6709515 | -0.108 | 0.127 | -0.616 | 0.423 | -0.114 | 0.033 | 0.033 | 0.587 | 0.6 | 0.94 | 0.59 | 0.6 | 0.54 | 0.44 |
| 2 | 103761689 | rs1881182 | -0.525 | -0.638 | -0.525 | -0.05 | 0.002 | 0.001 | 0.001 | 0 | 0.4 | 0.58 | 0.52 | 0.72 | 0.34 | 0.32 |
| 2 | 103767455 | rs1728260 | -0.265 | -0.454 | -0.329 | -0.692 | 0.001 | -0.033 | -0.033 | 0 | 0.85 | 0.81 | 0.87 | 1 | 0.75 | 0.9 |
| 2 | 103772597 | rs10496364 | 0.601 | -0.653 | -0.62 | -0.649 | -0.027 | -0.027 | -0.027 | 0.009 | 0.3 | 0.22 | 0.42 | 0.64 | 0.38 | 0.48 |
| 2 | 103773825 | rs4319952 | -0.211 | -1.045 | -0.026 | -0.917 | -0.002 | 0.005 | 0.005 | 0 | 0.73 | 0.7 | 0.7 | 0.65 | 0.77 | 0.73 |
| 2 | 103782330 | rs11674158 | 1.012 | -1.106 | 0.403 | -1.27 | -0.02 | -0.058 | -0.058 | 0.309 | 0.14 | 0.18 | 0.07 | 0 | 0.1 | 0.11 |
| 2 | 103791847 | rs11895342 | -0.506 | -1.277 | 0.584 | -1.685 | -0.004 | -0.132 | -0.132 | 0.074 | 0.38 | 0.43 | 0.24 | 0 | 0.37 | 0.26 |
| 2 | 103813251 | rs1590057 | 0.129 | -0.873 | 0.921 | -1.408 | -0.003 | -0.035 | -0.035 | 0 | 0.24 | 0.23 | 0.17 | 0 | 0.27 | 0.15 |
| 2 | 103816804 | rs1430068 | -0.255 | -0.968 | 1.451 | -1.456 | -0.001 | -0.014 | -0.014 | 0 | 0.28 | 0.78 | 0.21 | 0 | 0.27 | 0.15 |
| 2 | 103856812 | rs2043071 | 1.216 | -0.963 | -0.346 | -1.005 | -0.135 | -0.137 | -0.137 | 0 | 0.31 | 0.02 | 0.49 | 0.62 | 0.38 | 0.49 |
| 2 | 103873312 | rs1430079 | 0.066 | -0.254 | 0.186 | -0.985 | 0.013 | -0.067 | -0.067 | 0.196 | 0.19 | 0.26 | 0.16 | 0 | 0.25 | 0.14 |
| 2 | 103900735 | rs17413833 | 0.26 | -0.237 | -0.004 | -0.977 | -0.007 | -0.02 | -0.02 | 0 | 0.18 | 0.16 | 0.15 | 0 | 0.25 | 0.14 |
| 2 | 103910701 | rs1430074 | 0.707 | -1.067 | 0.518 | -0.126 | -0.001 | -0.008 | -0.008 | 0 | 0.85 | 0.94 | 0.81 | 0.84 | 0.83 | 0.87 |
| 2 | 103950575 | rs10191917 | 1.292 | -1.034 | 0.175 | -0.109 | -0.048 | -0.033 | -0.033 | 0 | 0.75 | 0.48 | 0.76 | 0.84 | 0.81 | 0.86 |
| 2 | 103968651 | rs11123959 | 1.099 | -0.828 | 0.398 | 0.092 | -0.043 | -0.026 | -0.026 | 0 | 0.8 | 0.46 | 0.84 | 0.87 | 0.85 | 0.94 |
| 2 | 103981461 | rs10469842 | -1.509 | -0.981 | 0.947 | -0.084 | -0.059 | -0.058 | -0.058 | 2.274 | 0.15 | 0.53 | 0.09 | 0.05 | 0.04 | 0.04 |
| 2 | 103987135 | rs1433312 | -2.134 | -1.2 | 0.768 | -0.098 | -0.039 | -0.018 | -0.018 | 0.324 | 0.18 | 0.31 | 0.12 | 0.12 | 0.08 | 0.07 |
| 2 | 103994511 | rs6707103 | -2.089 | -1.288 | 1.011 | -0.141 | -0.013 | -0.003 | -0.003 | 0 | 0.14 | 0.2 | 0.1 | 0.11 | 0.08 | 0.08 |
| 2 | 103999574 | rs7565678 | -2.489 | -1.59 | 0.697 | -0.562 | -0.052 | -0.071 | -0.071 | 1.054 | 0.29 | 0.52 | 0.17 | 0.14 | 0.17 | 0.16 |
| 2 | 104017337 | rs10496370 | -2.264 | -1.507 | 0.923 | -0.448 | -0.051 | -0.023 | -0.023 | 0 | 0.18 | 0.39 | 0.12 | 0.12 | 0.08 | 0.08 |
| 2 | 104019740 | rs10496372 | -2.078 | -1.489 | 0.806 | -0.439 | -0.037 | -0.015 | -0.015 | 0 | 0.17 | 0.3 | 0.11 | 0.11 | 0.08 | 0.08 |
| 2 | 104036433 | rs13390641 | -1.764 | -1.492 | 0.686 | -0.441 | -0.021 | -0.007 | -0.007 | 0 | 0.16 | 0.23 | 0.1 | 0.11 | 0.08 | 0.08 |

| | | | | | | | | | | | | | | | | |
|---|---|---|---|---|---|---|---|---|---|---|---|---|---|---|---|---|
| 2 | 104052459 | rs13429196 | -1.645 | -1.49 | 0.691 | -0.435 | -0.054 | -0.018 | -0.018 | 1.623 | 0.13 | 0.38 | 0.09 | 0.09 | 0.05 | 0.03 |
| 2 | 104055007 | rs10177199 | -2.025 | -1.487 | 0.664 | -0.436 | -0.052 | -0.018 | -0.018 | 0 | 0.17 | 0.36 | 0.11 | 0.12 | 0.08 | 0.08 |
| 2 | 104060568 | rs7562738 | -0.748 | -1.317 | 0.185 | -1.241 | 0.021 | -0.096 | -0.096 | 0 | 0.62 | 0.56 | 0.69 | 0.91 | 0.52 | 0.81 |
| 2 | 104067175 | rs266064 | 1.016 | -1.735 | -0.303 | -0.663 | -0.024 | -0.123 | -0.123 | 1.301 | 0.3 | 0.02 | 0.47 | 0.58 | 0.27 | 0.42 |
| 2 | 104073034 | rs817225 | -0.107 | -1.639 | 0.014 | -1.056 | 0.015 | 0.204 | 0.204 | 0 | 0.51 | 0.26 | 0.58 | 0.9 | 0.44 | 0.73 |
| 2 | 104076420 | rs266069 | -0.053 | -1.641 | -0.027 | -1.057 | 0.005 | 0.278 | 0.278 | 0 | 0.51 | 0.2 | 0.58 | 0.9 | 0.44 | 0.73 |
| 2 | 104079435 | rs266074 | -0.966 | -1.615 | 0.079 | -1.073 | -0.028 | 0.021 | 0.021 | 0.082 | 0.62 | 0.33 | 0.67 | 0.98 | 0.5 | 0.8 |
| 2 | 104079777 | rs10496374 | -0.418 | -1.485 | 0.159 | -0.856 | -0.012 | -0.002 | -0.002 | 1.947 | 0.11 | 0.13 | 0.09 | 0.08 | 0.06 | 0.07 |
| 2 | 104113458 | rs6761396 | -0.05 | -1.495 | 0.021 | -1.051 | 0.006 | 0.18 | 0.18 | 0 | 0.47 | 0.72 | 0.41 | 0.08 | 0.56 | 0.26 |
| 2 | 104115429 | rs6543213 | 0.891 | -1.576 | 0.218 | -1.118 | 0.029 | -0.148 | -0.148 | 0.133 | 0.36 | 0.44 | 0.3 | 0.02 | 0.5 | 0.19 |
| 2 | 104115536 | rs12712175 | -0.566 | -1.598 | 0.204 | -1.087 | -0.028 | -0.013 | -0.013 | 4.331 | 0.12 | 0.28 | 0.1 | 0.06 | 0.06 | 0.08 |
| 2 | 104142548 | rs1022298 | 0.429 | -1.758 | 0.17 | -1.189 | -0.015 | -0.013 | -0.013 | 0 | 0.4 | 0.66 | 0.33 | 0.02 | 0.51 | 0.23 |
| 2 | 104145165 | rs985864 | 0.196 | -1.702 | 0.255 | -1.043 | -0.01 | -0.001 | -0.001 | 0.002 | 0.08 | 0.08 | 0.08 | 0.06 | 0.05 | 0.04 |
| 2 | 104165823 | rs2167541 | -1.476 | -1.704 | 0.403 | -1.079 | -0.035 | -0.03 | -0.03 | 0 | 0.14 | 0.46 | 0.11 | 0.07 | 0.06 | 0.08 |
| 2 | 104209003 | rs10188273 | -0.65 | -2.006 | 0.011 | -1.414 | 0.022 | 0.166 | 0.166 | 0 | 0.61 | 0.82 | 0.45 | 0.08 | 0.67 | 0.38 |
| 2 | 104210674 | rs11690639 | -0.196 | -2.018 | 0.04 | -1.434 | 0.012 | 0.029 | 0.029 | 0 | 0.17 | 0.08 | 0.11 | 0.07 | 0.14 | 0.13 |
| 2 | 104237468 | rs2167539 | 1.188 | -2.001 | 0.187 | -1.421 | -0.004 | -0.06 | -0.06 | 0.04 | 0.34 | 0.27 | 0.28 | 0.01 | 0.51 | 0.21 |
| 2 | 104240205 | rs11123965 | -0.207 | -1.972 | 0.065 | -1.402 | 0.021 | 0.261 | 0.261 | 0 | 0.52 | 0.82 | 0.42 | 0.08 | 0.58 | 0.3 |
| 2 | 104243342 | rs10182738 | -0.186 | -1.958 | 0.084 | -1.395 | 0.027 | 0.223 | 0.223 | 0 | 0.52 | 0.78 | 0.41 | 0.08 | 0.58 | 0.3 |
| 2 | 104244870 | rs1451985 | 0.317 | -1.931 | 0.067 | -1.379 | 0.002 | 0 | 0 | 0 | 0.08 | 0.06 | 0.07 | 0.07 | 0.05 | 0.05 |
| 2 | 104245978 | rs4851661 | 0.196 | -1.911 | 0.119 | -1.352 | 0.008 | 0.067 | 0.067 | 0 | 0.47 | 0.6 | 0.39 | 0.08 | 0.57 | 0.26 |
| 2 | 104254564 | rs264963 | -0.439 | -1.928 | 0.431 | -1.36 | 0.007 | 0.058 | 0.058 | 0 | 0.53 | 0.4 | 0.61 | 0.92 | 0.43 | 0.74 |
| 2 | 104273554 | rs264921 | -0.464 | -1.929 | 0.388 | -1.361 | 0.007 | 0.06 | 0.06 | 0 | 0.54 | 0.4 | 0.61 | 0.92 | 0.44 | 0.74 |
| 2 | 104276502 | rs171939 | -0.249 | -1.934 | 0.324 | -1.361 | 0.037 | 0.181 | 0.181 | 0 | 0.5 | 0.31 | 0.6 | 0.92 | 0.42 | 0.7 |
| 2 | 104281272 | rs264927 | -0.049 | -2.193 | 0.065 | -0.438 | 0.003 | 0.034 | 0.034 | 0 | 0.44 | 0.2 | 0.38 | 0.61 | 0.41 | 0.64 |
| 2 | 104298541 | rs264943 | -1.125 | -2.263 | 0.546 | -0.626 | -0.011 | -0.017 | -0.017 | 0 | 0.59 | 0.69 | 0.7 | 0.99 | 0.48 | 0.78 |
| 2 | 104303311 | rs264952 | 0.198 | -2.267 | 0.483 | -0.453 | 0.003 | 0.089 | 0.089 | 0 | 0.42 | 0.21 | 0.51 | 0.7 | 0.4 | 0.67 |
| 2 | 104320510 | rs11675245 | 0.185 | -2.259 | 0.566 | -0.5 | 0.006 | 0.048 | 0.048 | 0 | 0.43 | 0.26 | 0.39 | 0.62 | 0.4 | 0.67 |
| 2 | 104335186 | rs1947565 | -0.163 | -2.191 | 0.609 | -0.466 | 0.019 | 0.064 | 0.064 | 0 | 0.48 | 0.3 | 0.4 | 0.63 | 0.41 | 0.71 |
| 2 | 104348213 | rs6543232 | -1.146 | -1.657 | 0.657 | -0.567 | -0.002 | 0.104 | 0.104 | 0 | 0.17 | 0.41 | 0.33 | 0.37 | 0.13 | 0.14 |
| 2 | 104361420 | rs9308868 | 0.027 | -1.715 | 0.297 | -0.559 | -0.007 | 0.102 | 0.102 | 0.308 | 0.5 | 0.24 | 0.41 | 0.62 | 0.46 | 0.75 |
| 2 | 104366056 | rs767048 | -0.271 | -1.439 | 0.565 | -0.012 | 0.01 | -0.008 | -0.008 | 4.605 | 0.09 | 0.05 | 0.27 | 0.36 | 0.06 | 0.07 |
| 2 | 104392496 | rs1457453 | -0.259 | -1.474 | 0.44 | 0.498 | 0 | 0.004 | 0.004 | 0 | 0.19 | 0.2 | 0.19 | 0.31 | 0.21 | 0.34 |
| 2 | 104412205 | rs11123969 | -1.545 | -1.646 | -0.09 | 0.237 | 0.025 | 0.065 | 0.065 | 0 | 0.42 | 0.83 | 0.53 | 0.72 | 0.32 | 0.48 |
| 2 | 104415982 | rs10177144 | -1.524 | -1.635 | 0.025 | 0.246 | 0.015 | 0.037 | 0.037 | 0 | 0.39 | 0.76 | 0.52 | 0.72 | 0.31 | 0.47 |
| 2 | 104421572 | rs12623591 | -0.338 | -1.662 | 0.128 | 0.819 | -0.003 | 0.077 | 0.077 | 0 | 0.07 | 0.08 | 0.13 | 0.37 | 0.04 | 0.07 |
| 2 | 104426491 | rs7605293 | -0.225 | -1.722 | 0.431 | 0.825 | 0.022 | 0.098 | 0.098 | 0.265 | 0.16 | 0.37 | 0.17 | 0.25 | 0.12 | 0.26 |
| 2 | 104430410 | rs4131749 | -0.387 | -2.061 | 0.61 | 0.673 | -0.013 | -0.032 | -0.032 | 0 | 0.15 | 0.37 | 0.21 | 0.08 | 0.12 | 0.11 |
| 2 | 104432433 | rs17029159 | -0.188 | -2.07 | 0.613 | 0.746 | -0.006 | 0.052 | 0.052 | 0 | 0.07 | 0.08 | 0.12 | 0.32 | 0.03 | 0.07 |
| 2 | 104432714 | rs11889924 | -1.435 | -2.079 | 0.589 | 0.73 | -0.048 | 0.093 | 0.093 | 0.518 | 0.23 | 0.48 | 0.34 | 0.41 | 0.16 | 0.18 |
| 2 | 104444273 | rs2678664 | 0.677 | -2.261 | 0.154 | 0.351 | 0.023 | 0.02 | 0.02 | 0.459 | 0.48 | 0.43 | 0.44 | 0.3 | 0.63 | 0.46 |
| 2 | 104449203 | rs17029224 | 0.002 | -2.198 | 0.234 | 0.363 | -0.003 | -0.01 | -0.01 | 2.993 | 0.13 | 0.35 | 0.2 | 0.09 | 0.11 | 0.11 |
| 2 | 104452617 | rs2678659 | 0.781 | -2.221 | -0.135 | 0.389 | 0.03 | -0.022 | -0.022 | 1.553 | 0.29 | 0.06 | 0.22 | 0.29 | 0.21 | 0.38 |
| 2 | 104452843 | rs2678658 | -0.223 | -2.327 | 0.057 | 0.406 | 0.034 | -0.012 | -0.012 | 0 | 0.4 | 0.29 | 0.48 | 0.62 | 0.26 | 0.44 |
| 2 | 104458637 | rs2678657 | 0.815 | -2.392 | -0.322 | 0.484 | 0.038 | -0.055 | -0.055 | 1.42 | 0.24 | 0.06 | 0.2 | 0.32 | 0.17 | 0.3 |
| 2 | 104458763 | rs2582953 | 0.148 | -2.314 | -0.235 | 0.137 | 0.001 | -0.003 | -0.003 | 0 | 0.3 | 0.27 | 0.44 | 0.62 | 0.21 | 0.34 |
| 2 | 104472743 | rs1822144 | -0.329 | -2.046 | -0.573 | 0.16 | -0.021 | -0.032 | -0.032 | 0 | 0.19 | 0.26 | 0.11 | 0.08 | 0.11 | 0.14 |
| 2 | 104481325 | rs2570500 | -0.045 | -1.869 | -1.17 | 0.386 | 0.023 | -0.027 | -0.027 | 0.477 | 0.28 | 0.21 | 0.34 | 0.62 | 0.2 | 0.34 |
| 2 | 104489351 | rs3860446 | -1.349 | -1.515 | -1.982 | -0.471 | 0.09 | 0.379 | 0.379 | 0.063 | 0.54 | 0.96 | 0.63 | 1 | 0.3 | 0.67 |
| 2 | 104498739 | rs6717408 | -1.47 | -1.382 | -2.044 | -0.524 | 0.033 | 0.372 | 0.372 | 0 | 0.55 | 0.96 | 0.64 | 1 | 0.34 | 0.7 |
| 2 | 104502085 | rs10496375 | -0.225 | -1.859 | -1.255 | -0.148 | 0.003 | 0.032 | 0.032 | 0 | 0.21 | 0.3 | 0.29 | 0.28 | 0.19 | 0.3 |
| 2 | 104502188 | rs7582619 | -0.583 | -1.929 | -0.68 | 0.429 | -0.092 | 0.039 | 0.039 | 0.061 | 0.3 | 0.52 | 0.19 | 0.42 | 0.13 | 0.29 |
| 2 | 104506942 | rs17029335 | -0.049 | -1.827 | -0.065 | 0.609 | -0.032 | 0.088 | 0.088 | 0 | 0.08 | 0.3 | 0.03 | 0.17 | 0.01 | 0.07 |
| 2 | 104512226 | rs2570501 | 0.887 | -1.173 | -0.752 | 0.233 | -0.055 | 0.065 | 0.065 | 0.643 | 0.17 | 0.05 | 0.15 | 0 | 0.3 | 0.13 |
| 2 | 104515963 | rs1550533 | 1.118 | -1.439 | -0.913 | 0.256 | 0.05 | -0.047 | -0.047 | 5.899 | 0.18 | 0.07 | 0.15 | 0.24 | 0.11 | 0.21 |
| 2 | 104521151 | rs1441105 | -1.684 | -1.409 | -0.362 | 0.335 | 0.009 | 0.009 | 0.009 | 0 | 0.49 | 0.57 | 0.47 | 0.7 | 0.44 | 0.46 |
| 2 | 104524841 | rs13031410 | 1.912 | -1.508 | -0.498 | 0.344 | 0.018 | -0.065 | -0.065 | 0 | 0.13 | 0 | 0.13 | 0.21 | 0.09 | 0.2 |
| 2 | 104530177 | rs2570479 | -1.171 | -1.125 | -0.083 | 0.488 | 0.035 | 0.031 | 0.031 | 0 | 0.49 | 0.66 | 0.56 | 0.72 | 0.43 | 0.53 |
| 2 | 104534826 | rs2610681 | -0.812 | -1.172 | -0.197 | 0.517 | -0.006 | -0.002 | -0.002 | 0 | 0.45 | 0.5 | 0.33 | 0.43 | 0.44 | 0.42 |
| 2 | 104535716 | rs1441104 | -1.695 | -1.074 | -0.675 | 0.425 | -0.16 | 0.302 | 0.302 | 0 | 0.62 | 0.97 | 0.7 | 1 | 0.51 | 0.74 |
| 2 | 104537499 | rs1992393 | 0.146 | -0.898 | 0.182 | 1.154 | -0.017 | 0.119 | 0.119 | 0 | 0.07 | 0.25 | 0.02 | 0.17 | 0.02 | 0.07 |
| 2 | 104549992 | rs10195724 | 0.02 | -0.897 | 0.171 | 1.181 | -0.029 | 0.111 | 0.111 | 0 | 0.08 | 0.25 | 0.02 | 0.18 | 0.02 | 0.07 |
| 2 | 104563894 | rs12464237 | 0.227 | -0.299 | -0.503 | 0.835 | 0.001 | 0.095 | 0.095 | 0 | 0.11 | 0.19 | 0.12 | 0.28 | 0.06 | 0.21 |
| 2 | 104572318 | rs2570481 | -0.662 | 0.151 | -0.99 | 0.101 | 0.004 | 0.081 | 0.081 | 0.097 | 0.5 | 0.67 | 0.64 | 0.72 | 0.49 | 0.63 |
| 2 | 104573005 | rs2119507 | 0.449 | 0.039 | -1.294 | -0.209 | -0.009 | 0.016 | 0.016 | 0 | 0.14 | 0.15 | 0.06 | 0.23 | 0.1 | 0.12 |
| 2 | 104575444 | rs2570483 | -2.139 | 0.376 | -1.811 | -0.872 | -0.022 | 0.065 | 0.065 | 0 | 0.87 | 0.97 | 0.9 | 1 | 0.87 | 0.96 |
| 2 | 104578923 | rs2582974 | -1.134 | 0.418 | -1.724 | -0.837 | -0.113 | 0.265 | 0.265 | 0.018 | 0.64 | 0.97 | 0.72 | 1 | 0.6 | 0.75 |
| 2 | 104580503 | rs12613775 | 0.125 | 0.4 | -1.382 | -0.854 | -0.014 | 0.042 | 0.042 | 0 | 0.17 | 0.25 | 0.08 | 0.24 | 0.12 | 0.14 |
| 2 | 104580975 | rs2582973 | -0.16 | 0.546 | -1.187 | -0.622 | 0 | 0.007 | 0.007 | 0.259 | 0.45 | 0.58 | 0.61 | 0.71 | 0.45 | 0.54 |

| | | | | | | | | | | | | | | | | |
|---|---|---|---|---|---|---|---|---|---|---|---|---|---|---|---|---|
| 2 | 104582013 | rs2375937 | 0.172 | 0.571 | -1.464 | -0.895 | -0.03 | -0.004 | -0.004 | 0.166 | 0.32 | 0.58 | 0.51 | 0.71 | 0.21 | 0.45 |
| 2 | 104591465 | rs1816754 | 0.09 | -0.365 | -1.454 | -0.179 | -0.005 | 0.118 | 0.118 | 0 | 0.09 | 0.19 | 0.16 | 0.44 | 0.05 | 0.34 |
| 2 | 104592311 | rs1441094 | -0.372 | -0.207 | -1.363 | -0.647 | -0.006 | -0.008 | -0.008 | 0 | 0.25 | 0.3 | 0.26 | 0.19 | 0.21 | 0.14 |
| 2 | 104597455 | rs2610683 | -1.025 | -0.148 | -0.965 | -0.502 | 0.035 | 0.038 | 0.038 | 1.662 | 0.47 | 0.7 | 0.58 | 0.72 | 0.34 | 0.57 |
| 2 | 104599615 | rs1372101 | 0.259 | -0.202 | -1.045 | -0.516 | -0.027 | 0.03 | 0.03 | 4.278 | 0.28 | 0.38 | 0.34 | 0.52 | 0.14 | 0.46 |
| 2 | 104614308 | rs2376056 | 0.175 | -0.258 | -1.053 | -0.524 | -0.018 | 0.025 | 0.025 | 3.448 | 0.29 | 0.38 | 0.34 | 0.52 | 0.16 | 0.47 |
| 2 | 104617456 | rs2889275 | 0.193 | -0.089 | -1.197 | -0.684 | 0.002 | 0.002 | 0.002 | 0 | 0.17 | 0.17 | 0.22 | 0.21 | 0.18 | 0.1 |
| 2 | 104632493 | rs7581688 | -1.404 | -0.675 | -1.314 | -0.88 | -0.047 | 0.296 | 0.296 | 1.936 | 0.6 | 0.95 | 0.63 | 1 | 0.41 | 0.68 |
| 2 | 104634850 | rs12474171 | -1.399 | -0.664 | -1.308 | -0.875 | -0.046 | 0.297 | 0.297 | 1.987 | 0.6 | 0.95 | 0.63 | 1 | 0.41 | 0.68 |
| 2 | 104638469 | rs17417183 | 1.497 | -0.655 | -1.192 | -0.84 | -0.005 | -0.028 | -0.028 | 0 | 0.11 | 0.12 | 0.08 | 0 | 0.11 | 0.04 |
| 2 | 104641510 | rs17345319 | 1.524 | -0.651 | -1.197 | -0.838 | -0.004 | -0.026 | -0.026 | 0 | 0.11 | 0.11 | 0.08 | 0 | 0.11 | 0.04 |
| 2 | 104646414 | rs2376070 | -1.933 | -0.109 | -1.19 | -0.727 | -0.053 | 0.061 | 0.061 | 0.418 | 0.86 | 0.96 | 0.88 | 1 | 0.79 | 0.94 |
| 2 | 104653581 | rs13426125 | -0.197 | -1.211 | -0.321 | 0.2 | 0 | -0.001 | -0.001 | 0.098 | 0.09 | 0.11 | 0.1 | 0.13 | 0.07 | 0.13 |
| 2 | 104689444 | rs2376064 | -1.479 | -0.837 | -1.354 | -0.071 | -0.06 | 0.285 | 0.285 | 1.38 | 0.61 | 0.95 | 0.63 | 0.99 | 0.41 | 0.68 |
| 2 | 104699282 | rs2889279 | -0.289 | -1.123 | -0.614 | 0.615 | -0.038 | 0.056 | 0.056 | 3.166 | 0.29 | 0.42 | 0.36 | 0.47 | 0.17 | 0.45 |
| 2 | 104707682 | rs13392864 | -0.319 | -1.097 | -0.631 | 0.605 | -0.03 | 0.042 | 0.042 | 3.303 | 0.3 | 0.42 | 0.36 | 0.47 | 0.17 | 0.45 |
| 2 | 104713962 | rs2376057 | -0.785 | -0.413 | -1.34 | 0.069 | 0.082 | 0.285 | 0.285 | 4.316 | 0.53 | 0.87 | 0.59 | 1 | 0.35 | 0.6 |
| 2 | 104714131 | rs2889276 | -0.774 | -0.335 | -1.129 | 0.372 | 0.083 | 0.126 | 0.126 | 4.102 | 0.49 | 0.73 | 0.46 | 0.75 | 0.33 | 0.55 |
| 2 | 104719185 | rs4487106 | -1.258 | -0.017 | -0.779 | 0.998 | -0.001 | 0.001 | 0.001 | 0 | 0.09 | 0.07 | 0.09 | 0.06 | 0.06 | 0.14 |
| 2 | 104729566 | rs12987666 | -1.236 | -0.034 | -0.691 | 1.248 | 0.003 | -0.01 | -0.01 | 0 | 0.09 | 0.07 | 0.09 | 0.12 | 0.05 | 0.16 |
| 2 | 104730171 | rs12613102 | -1.236 | -0.034 | -0.691 | 1.248 | 0.003 | -0.01 | -0.01 | 0 | 0.09 | 0.07 | 0.09 | 0.12 | 0.05 | 0.16 |
| 2 | 104733337 | rs12618077 | -1.236 | -0.034 | -0.717 | 1.248 | 0.001 | -0.022 | -0.022 | 0 | 0.09 | 0.05 | 0.08 | 0.11 | 0.05 | 0.16 |
| 2 | 104736734 | rs10496382 | -0.431 | -0.145 | -0.589 | 0.969 | 0 | -0.016 | -0.016 | 0 | 0.09 | 0.09 | 0.06 | 0 | 0.07 | 0.09 |
| 2 | 104742560 | rs13404302 | -0.713 | -0.267 | 0.362 | 2.074 | 0.014 | 0.396 | 0.396 | 0 | 0.06 | 0.28 | 0.26 | 0.45 | 0.04 | 0.23 |
| 2 | 104749844 | rs12994143 | -1.073 | -0.204 | 0.165 | 1.938 | 0.001 | -0.022 | -0.022 | 0.044 | 0.09 | 0.06 | 0.08 | 0.11 | 0.06 | 0.16 |
| 2 | 104798151 | rs4613305 | -1.965 | 0.215 | 1.581 | 2.379 | -0.001 | -0.038 | -0.038 | 0 | 0.15 | 0.17 | 0.13 | 0.02 | 0.13 | 0.13 |
| 2 | 104826331 | rs7597477 | -2.257 | -0.098 | 1.601 | 2.635 | -0.004 | 0.054 | 0.054 | 0 | 0.09 | 0.16 | 0.08 | 0.23 | 0.06 | 0.22 |
| 2 | 104841582 | rs10170023 | -1.226 | -0.343 | 0.896 | 2.373 | 0.006 | -0.058 | -0.058 | 0 | 0.18 | 0.1 | 0.21 | 0.59 | 0.14 | 0.31 |
| 2 | 104856148 | rs4851675 | 0.673 | -0.249 | 0.481 | 1.875 | -0.004 | -0.053 | -0.053 | 0.135 | 0.63 | 0.17 | 0.53 | 0.59 | 0.64 | 0.59 |
| 2 | 104859949 | rs7590841 | 0.069 | -0.334 | 1.053 | 1.381 | -0.005 | -0.061 | -0.061 | 0 | 0.16 | 0.19 | 0.11 | 0.02 | 0.17 | 0.16 |
| 2 | 104881377 | rs7580071 | -0.176 | -0.446 | 0.918 | 1.266 | -0.009 | -0.056 | -0.056 | 0 | 0.17 | 0.21 | 0.11 | 0.02 | 0.17 | 0.17 |
| 2 | 104888087 | rs4539815 | 0.281 | -0.412 | 1.168 | 1.417 | 0.053 | 0.108 | 0.108 | 6.129 | 0.83 | 0.46 | 0.66 | 0.76 | 0.83 | 0.88 |
| 2 | 104888405 | rs12992682 | 1.167 | -0.622 | 0.48 | 1.706 | -0.015 | -0.051 | -0.051 | 0.036 | 0.62 | 0.15 | 0.48 | 0.57 | 0.62 | 0.58 |
| 2 | 104911786 | rs13391945 | -0.445 | -1.246 | 1.221 | 1.99 | -0.039 | 0.071 | 0.071 | 6.01 | 0.07 | 0.14 | 0.03 | 0.2 | 0.01 | 0.13 |
| 2 | 104916482 | rs11892570 | 1.037 | -1.38 | 0.947 | 1.883 | -0.003 | 0.016 | 0.016 | 0 | 0.8 | 0.77 | 0.74 | 0.43 | 0.84 | 0.77 |
| 2 | 104930232 | rs4851683 | 2.101 | -1.838 | 0.587 | 1.457 | 0.142 | 0.093 | 0.093 | 0.289 | 0.57 | 0.13 | 0.54 | 0.23 | 0.76 | 0.55 |
| 2 | 104931550 | rs6713996 | 0.538 | -1.468 | 0.87 | 1.909 | -0.001 | 0.044 | 0.044 | 0 | 0.91 | 0.89 | 0.91 | 0.65 | 0.91 | 0.8 |
| 2 | 104936545 | rs7568278 | -0.41 | -1.483 | 1.508 | 1.837 | 0.003 | -0.012 | -0.012 | 0 | 0.06 | 0.26 | 0.12 | 0.01 | 0.05 | 0.03 |
| 2 | 104942169 | rs4851686 | -1.855 | -1.476 | 1.501 | 1.708 | -0.04 | 0.027 | 0.027 | 0.539 | 0.27 | 0.5 | 0.28 | 0.65 | 0.2 | 0.38 |
| 2 | 104944485 | rs13390052 | -0.183 | -1.226 | 1.603 | 2.232 | -0.031 | 0.093 | 0.093 | 8.411 | 0.07 | 0.11 | 0.03 | 0.27 | 0.01 | 0.11 |
| 2 | 104948911 | rs1519627 | -1.485 | -1.281 | 1.654 | 2.2 | -0.039 | 0.073 | 0.073 | 0 | 0.24 | 0.44 | 0.32 | 0.41 | 0.17 | 0.29 |
| 2 | 104950103 | rs1109458 | -0.096 | -1.261 | 1.458 | 2.201 | -0.034 | 0.134 | 0.134 | 7.596 | 0.07 | 0.15 | 0.04 | 0.31 | 0.01 | 0.1 |
| 2 | 104950170 | rs11895798 | -1.594 | -0.864 | 1.04 | 1.776 | -0.038 | 0.07 | 0.07 | 0 | 0.24 | 0.44 | 0.32 | 0.41 | 0.16 | 0.29 |
| 2 | 104954640 | rs2376158 | 1.179 | -0.551 | 0.91 | 1.19 | -0.018 | -0.031 | -0.031 | 0 | 0.79 | 0.42 | 0.84 | 0.82 | 0.85 | 0.7 |
| 2 | 104957361 | rs4851687 | 1.906 | -0.958 | 0.492 | 1.038 | 0.083 | 0.089 | 0.089 | 1.332 | 0.66 | 0.07 | 0.73 | 0.76 | 0.83 | 0.66 |
| 2 | 104976101 | rs7585377 | -2.491 | -0.139 | 1.515 | 2.077 | 0.046 | 0.198 | 0.198 | 0 | 0.14 | 0.39 | 0.35 | 0.46 | 0.18 | 0.14 |
| 2 | 104983267 | rs7570682 | -2.996 | -0.13 | 2.025 | 2.085 | 0.089 | 0.237 | 0.237 | 0 | 0.14 | 0.52 | 0.3 | 0.4 | 0.22 | 0.14 |
| 2 | 104983639 | rs7598451 | -2.279 | -0.089 | 2.269 | 2.071 | 0.048 | 0.126 | 0.126 | 0 | 0.11 | 0.23 | 0.24 | 0.4 | 0.19 | 0.16 |
| 2 | 104991388 | rs17029778 | -2.609 | -0.104 | 2.248 | 2.195 | 0.034 | 0.118 | 0.118 | 0 | 0.13 | 0.22 | 0.24 | 0.43 | 0.2 | 0.17 |
| 2 | 104998427 | rs12615991 | -1.234 | -0.18 | 2.091 | 2.425 | -0.033 | -0.028 | -0.028 | 0 | 0.06 | 0.02 | 0.12 | 0.21 | 0.13 | 0.11 |
| 2 | 104998581 | rs17029789 | -2.292 | -0.192 | 2.143 | 2.51 | -0.008 | 0.027 | 0.027 | 0 | 0.06 | 0.12 | 0.1 | 0.19 | 0.03 | 0.08 |
| 2 | 105012664 | rs908916 | -1.234 | -0.177 | 2.155 | 2.519 | -0.033 | -0.028 | -0.028 | 0.173 | 0.06 | 0.02 | 0.12 | 0.21 | 0.12 | 0.11 |
| 2 | 105012855 | rs4464293 | -3.221 | -0.223 | 2.272 | 2.468 | 0.013 | 0.116 | 0.116 | 0 | 0.14 | 0.24 | 0.22 | 0.39 | 0.16 | 0.19 |
| 2 | 105019418 | rs7425226 | -2.032 | -0.161 | 1.756 | 2.177 | 0.002 | 0.007 | 0.007 | 0 | 0.31 | 0.64 | 0.29 | 0.38 | 0.32 | 0.32 |
| 2 | 105034927 | rs7572010 | -1.337 | -0.17 | 1.759 | 1.595 | 0.005 | -0.014 | -0.014 | 0 | 0.22 | 0.32 | 0.22 | 0.1 | 0.29 | 0.25 |
| 2 | 105039687 | rs2889323 | -2.649 | -0.226 | 1.422 | 1.846 | -0.011 | -0.082 | -0.082 | 1.534 | 0.38 | 0.88 | 0.58 | 0.48 | 0.36 | 0.42 |
| 2 | 105053265 | rs11888862 | -1.07 | -0.329 | 1.729 | 1.651 | 0.01 | -0.017 | -0.017 | 0 | 0.2 | 0.3 | 0.18 | 0.12 | 0.25 | 0.21 |
| 2 | 105056854 | rs6543255 | -2.109 | -0.381 | 1.913 | 1.802 | 0.005 | 0.004 | 0.004 | 0 | 0.28 | 0.57 | 0.45 | 0.31 | 0.32 | 0.33 |
| 2 | 105061195 | rs6729889 | -1.017 | -0.33 | 1.817 | 1.729 | 0.017 | -0.029 | -0.029 | 0 | 0.21 | 0.36 | 0.18 | 0.12 | 0.26 | 0.23 |
| 2 | 105085140 | rs6749443 | -2.197 | -0.373 | 1.284 | 1.813 | 0.001 | -0.022 | -0.022 | 0.006 | 0.31 | 0.73 | 0.48 | 0.38 | 0.32 | 0.34 |
| 2 | 105093698 | rs1033302 | 2.076 | -0.672 | 0.328 | 1.358 | -0.006 | -0.093 | -0.093 | 0 | 0.5 | 0.03 | 0.4 | 0.53 | 0.59 | 0.56 |
| 2 | 105097129 | rs934956 | -1.681 | 0.049 | 1.381 | 1.442 | 0.011 | 0.014 | 0.014 | 0 | 0.23 | 0.47 | 0.2 | 0.18 | 0.26 | 0.23 |
| 2 | 105097977 | rs3849367 | -1.071 | -0.202 | 0.226 | 0.697 | -0.029 | 0.1 | 0.1 | 0.221 | 0.19 | 0.04 | 0.13 | 0.04 | 0.21 | 0.18 |
| 2 | 105105148 | rs6743263 | -1.401 | 0.193 | 0.812 | 0.816 | -0.014 | -0.107 | -0.107 | 0 | 0.26 | 0.48 | 0.23 | 0.08 | 0.22 | 0.23 |
| 2 | 105110002 | rs6753471 | -2.282 | 0.209 | 0.732 | 1.026 | -0.009 | -0.003 | -0.003 | 0 | 0.35 | 0.57 | 0.45 | 0.36 | 0.31 | 0.37 |
| 2 | 105114879 | rs12620665 | -0.751 | 0.224 | 0.342 | 1.142 | -0.005 | 0.054 | 0.054 | 0.625 | 0.21 | 0.09 | 0.19 | 0.08 | 0.2 | 0.24 |
| 2 | 105115341 | rs12987646 | -0.987 | 0.25 | 0.177 | 1.164 | -0.001 | -0.001 | -0.001 | 0 | 0.09 | 0.07 | 0.22 | 0.27 | 0.09 | 0.13 |
| 2 | 105116184 | rs1439671 | -1.264 | 0.261 | 0.225 | 1.149 | -0.011 | 0.001 | 0.001 | 0 | 0.33 | 0.51 | 0.44 | 0.36 | 0.3 | 0.37 |
| 2 | 105139339 | rs11123995 | -0.307 | 0.679 | -0.608 | 1.069 | -0.007 | -0.023 | -0.023 | 0 | 0.53 | 0.83 | 0.58 | 0.48 | 0.44 | 0.47 |

| | | | | | | | | | | | | | | | | |
|---|---|---|---|---|---|---|---|---|---|---|---|---|---|---|---|---|
| 2 | 105141788 | rs1866555 | 0.38 | 0.484 | -0.561 | 0.666 | 0 | 0.144 | 0.144 | 0 | 0.37 | 0.12 | 0.3 | 0.15 | 0.29 | 0.41 |
| 2 | 105147650 | rs17029922 | -0.751 | 0.146 | 0.471 | 0.595 | -0.031 | -0.038 | -0.038 | 0 | 0.15 | 0.27 | 0.08 | 0.07 | 0.09 | 0.1 |
| 2 | 105153455 | rs1946878 | 0.474 | 0.442 | 0.758 | 0.474 | 0.007 | 0.019 | 0.019 | 0 | 0.34 | 0.28 | 0.25 | 0.06 | 0.26 | 0.36 |
| 2 | 105160414 | rs10205261 | 0.46 | 0.5 | 1.021 | 0.84 | -0.003 | -0.051 | -0.051 | 0 | 0.36 | 0.38 | 0.26 | 0.06 | 0.27 | 0.36 |
| 2 | 105167888 | rs714753 | 0.74 | 0.465 | 1.063 | 0.833 | 0 | 0.119 | 0.119 | 0 | 0.32 | 0.16 | 0.24 | 0.06 | 0.26 | 0.37 |
| 2 | 105168190 | rs1439669 | -0.509 | 0.377 | 1.122 | 1.101 | 0.016 | -0.011 | -0.011 | 0 | 0.5 | 0.91 | 0.49 | 0.53 | 0.4 | 0.43 |
| 2 | 105170353 | rs6733728 | 0.562 | 0.428 | 1.265 | 1.045 | -0.003 | -0.053 | -0.053 | 0 | 0.34 | 0.36 | 0.26 | 0.06 | 0.26 | 0.37 |
| 2 | 105178452 | rs6744638 | 0.503 | 0.414 | 1.45 | 1.077 | -0.008 | -0.066 | -0.066 | 0 | 0.34 | 0.42 | 0.29 | 0.12 | 0.27 | 0.37 |
| 2 | 105183312 | rs17029978 | -1.158 | 0.762 | 1.419 | 1.55 | -0.018 | 0.193 | 0.193 | 1.039 | 0.16 | 0.58 | 0.25 | 0.46 | 0.14 | 0.07 |
| 2 | 105187436 | rs11123997 | -0.41 | 1.234 | 1.288 | 1.452 | 0.031 | 0.114 | 0.114 | 0 | 0.07 | 0.12 | 0.08 | 0.25 | 0.13 | 0.03 |
| 2 | 105188004 | rs6716044 | -0.9 | 1.188 | 1.018 | 0.709 | -0.013 | 0.101 | 0.101 | 0.656 | 0.16 | 0.51 | 0.23 | 0.22 | 0.12 | 0.06 |
| 2 | 105189350 | rs7598356 | -1.328 | 1.598 | 0.882 | 0.578 | 0.052 | 0.014 | 0.014 | 0.01 | 0.23 | 0.51 | 0.41 | 0.22 | 0.29 | 0.16 |
| 2 | 105189937 | rs7571700 | -0.661 | 0.935 | 0.478 | 0.204 | 0 | -0.013 | -0.013 | 0 | 0.07 | 0.08 | 0.07 | 0 | 0.12 | 0.03 |
| 2 | 105193300 | rs1866558 | -0.249 | 0.657 | 0.513 | 0.151 | -0.004 | 0.022 | 0.022 | 0 | 0.53 | 0.67 | 0.55 | 0.37 | 0.49 | 0.52 |
| 2 | 105197447 | rs12479172 | -0.098 | 0.57 | 0.774 | 0.146 | 0.001 | 0.034 | 0.034 | 0 | 0.49 | 0.39 | 0.46 | 0.3 | 0.48 | 0.43 |
| 2 | 105198201 | rs2289357 | -0.511 | 0.464 | 0.653 | 0.312 | -0.006 | -0.002 | -0.002 | 0 | 0.61 | 0.77 | 0.59 | 0.37 | 0.56 | 0.54 |
| 2 | 105219317 | rs11679786 | -1.272 | 0.616 | -0.926 | 0.534 | -0.026 | 0.032 | 0.032 | 0.72 | 0.15 | 0.06 | 0.24 | 0.08 | 0.2 | 0.24 |
| 2 | 105223181 | rs12991560 | -1.072 | 0.622 | -0.923 | 0.243 | -0.025 | 0.046 | 0.046 | 1.002 | 0.15 | 0.06 | 0.2 | 0.02 | 0.2 | 0.24 |
| 2 | 105237863 | rs1866557 | 0.694 | -0.486 | -0.519 | 0.902 | -0.029 | 0.019 | 0.019 | 0.719 | 0.73 | 0.34 | 0.79 | 0.54 | 0.88 | 0.63 |
| 2 | 105238181 | rs1866556 | -1.737 | -0.538 | 0.448 | 0.595 | -0.027 | -0.011 | -0.011 | 0 | 0.14 | 0.34 | 0.05 | 0.12 | 0.08 | 0.03 |
| 2 | 105248454 | rs10204762 | -0.431 | -0.663 | 0.322 | -0.061 | -0.001 | -0.016 | -0.016 | 0 | 0.29 | 0.35 | 0.25 | 0.19 | 0.31 | 0.12 |
| 2 | 105253184 | rs7567189 | -1.2 | -1.219 | -0.553 | 0.011 | -0.099 | -0.131 | -0.131 | 0 | 0.6 | 0.94 | 0.55 | 0.51 | 0.47 | 0.39 |
| 2 | 105253814 | rs6543262 | -0.133 | -1.046 | 0.503 | -0.306 | -0.005 | -0.03 | -0.03 | 0 | 0.32 | 0.59 | 0.27 | 0.19 | 0.32 | 0.12 |
| 2 | 105255306 | rs7595911 | 0.072 | -1.123 | 0.438 | -0.46 | 0.002 | -0.045 | -0.045 | 0 | 0.28 | 0.2 | 0.19 | 0 | 0.29 | 0.09 |
| 2 | 105270919 | rs17030065 | -0.538 | -1.029 | 0.965 | -0.101 | 0.023 | 0.126 | 0.126 | 1.014 | 0.07 | 0.4 | 0.07 | 0.16 | 0.01 | 0.02 |
| 2 | 105278158 | rs6743006 | -0.438 | -1.548 | 1.283 | -0.351 | -0.058 | -0.051 | -0.051 | 0 | 0.29 | 0.53 | 0.19 | 0.23 | 0.15 | 0.13 |
| 2 | 105283526 | rs11899001 | 0.409 | -1.339 | 0.978 | -0.624 | 0.004 | 0.049 | 0.049 | 0 | 0.26 | 0.14 | 0.2 | 0 | 0.29 | 0.11 |
| 2 | 105296110 | rs2222186 | 0.63 | -1.333 | 0.921 | 0.143 | 0.059 | -0.005 | -0.005 | 0 | 0.64 | 0.22 | 0.64 | 0.59 | 0.81 | 0.72 |
| 2 | 105300588 | rs1517862 | 0.545 | -0.992 | 1.326 | 0.387 | 0.03 | 0.169 | 0.169 | 0 | 0.93 | 0.71 | 0.84 | 0.81 | 0.97 | 0.86 |
| 2 | 105304396 | rs6727249 | 0.533 | -1.085 | 1.293 | 0.432 | 0.001 | 0.148 | 0.148 | 0 | 0.93 | 0.74 | 0.85 | 0.79 | 0.98 | 0.86 |
| 2 | 105305426 | rs6713364 | 0.677 | -1.215 | 0.518 | -0.168 | 0.078 | 0.05 | 0.05 | 0.387 | 0.49 | 0.22 | 0.54 | 0.56 | 0.66 | 0.64 |
| 2 | 105307941 | rs10178865 | -0.452 | -1.354 | 0.99 | -0.021 | -0.038 | 0.05 | 0.05 | 0.069 | 0.11 | 0.24 | 0.07 | 0.21 | 0.03 | 0.03 |
| 2 | 105309317 | rs4851704 | 0.077 | -1.353 | 1.053 | -0.032 | 0 | 0.122 | 0.122 | 0 | 0.93 | 0.77 | 0.84 | 0.78 | 0.98 | 0.87 |
| 2 | 105314914 | rs2037262 | 0.783 | -1.643 | 0.727 | -0.286 | 0.027 | 0.008 | 0.008 | 0.661 | 0.53 | 0.33 | 0.63 | 0.54 | 0.64 | 0.67 |
| 2 | 105314970 | rs17627584 | -0.019 | -1.572 | 0.603 | -0.365 | 0.01 | 0.052 | 0.052 | 0.532 | 0.33 | 0.2 | 0.22 | 0.03 | 0.31 | 0.19 |
| 2 | 105335047 | rs17679248 | 0.341 | -1.447 | 0.566 | -0.285 | -0.033 | 0.13 | 0.13 | 0 | 0.25 | 0.06 | 0.19 | 0.04 | 0.26 | 0.18 |
| 2 | 105335971 | rs1517859 | 1.08 | -1.584 | 0.807 | -0.094 | 0.036 | 0.005 | 0.005 | 0.379 | 0.55 | 0.3 | 0.64 | 0.54 | 0.68 | 0.67 |
| 2 | 105340625 | rs1517854 | 1.493 | -0.697 | 0.429 | -0.008 | -0.126 | -0.102 | -0.102 | 1.724 | 0.34 | 0.04 | 0.45 | 0.43 | 0.44 | 0.54 |
| 2 | 105341447 | rs6735422 | 0.615 | -0.694 | 0.612 | 0.001 | 0 | 0.073 | 0.073 | 0 | 0.27 | 0.13 | 0.2 | 0.1 | 0.27 | 0.2 |
| 2 | 105342436 | rs6721646 | 0.997 | -0.475 | 0.519 | 0.155 | -0.003 | 0.011 | 0.011 | 0 | 0.21 | 0.15 | 0.17 | 0.12 | 0.24 | 0.14 |
| 2 | 105343111 | rs4851709 | -0.294 | -0.647 | 0.816 | 0.367 | -0.005 | 0.069 | 0.069 | 0 | 0.95 | 0.87 | 0.93 | 0.79 | 0.98 | 0.91 |
| 2 | 105350891 | rs17020852 | 0.055 | -0.655 | 0.883 | 0.361 | 0.004 | 0.012 | 0.012 | 0 | 0.33 | 0.27 | 0.26 | 0.26 | 0.29 | 0.22 |
| 2 | 105351097 | rs17020855 | 0.723 | -0.641 | 0.871 | 0.37 | 0.04 | -0.001 | -0.001 | 0 | 0.56 | 0.27 | 0.64 | 0.53 | 0.68 | 0.69 |
| 2 | 105354926 | rs4851712 | -0.49 | -0.567 | 0.377 | 0.402 | -0.006 | -0.012 | -0.012 | 0 | 0.38 | 0.4 | 0.33 | 0.27 | 0.3 | 0.23 |
| 2 | 105356894 | rs2037261 | -0.255 | -0.212 | 0.209 | 0.593 | -0.002 | 0.114 | 0.114 | 0 | 0.95 | 0.88 | 0.93 | 0.79 | 0.98 | 0.91 |
| 2 | 105359354 | rs11899487 | -0.213 | -0.153 | -0.323 | 0.603 | -0.002 | -0.007 | -0.007 | 0.019 | 0.33 | 0.34 | 0.28 | 0.26 | 0.29 | 0.22 |
| 2 | 105360898 | rs10203687 | 0.487 | 0.197 | -0.664 | 0.982 | 0.005 | 0.139 | 0.139 | 0 | 0.94 | 0.83 | 0.92 | 0.77 | 0.97 | 0.9 |
| 2 | 105366165 | rs1110143 | 0.149 | 0.499 | -1.669 | 0.63 | -0.028 | -0.115 | -0.115 | 2.045 | 0.34 | 0.07 | 0.39 | 0.51 | 0.3 | 0.52 |
| 2 | 105378957 | rs12615966 | 0.01 | 0.154 | -1.853 | 0.086 | 0.015 | 0.018 | 0.018 | 0.081 | 0.19 | 0.13 | 0.19 | 0.09 | 0.14 | 0.07 |
| 2 | 105385872 | rs17030308 | -0.034 | 0.04 | -1.264 | 0.031 | -0.01 | -0.012 | -0.012 | 0 | 0.2 | 0.3 | 0.21 | 0.09 | 0.14 | 0.07 |
| 2 | 105395302 | rs442203 | -0.613 | 0.215 | -1.347 | -0.935 | 0.026 | 0.055 | 0.055 | 0 | 0.56 | 0.73 | 0.68 | 0.9 | 0.64 | 0.72 |
| 2 | 105401824 | rs376337 | -0.657 | 0.172 | -1.299 | -0.956 | 0.032 | 0.096 | 0.096 | 0 | 0.56 | 0.78 | 0.68 | 0.9 | 0.64 | 0.72 |
| 2 | 105406549 | rs435454 | -0.85 | 0.437 | -1.629 | -1.163 | 0.019 | 0.133 | 0.133 | 1.384 | 0.48 | 0.75 | 0.6 | 0.91 | 0.46 | 0.52 |
| 2 | 105432142 | rs4851716 | -0.021 | 0.114 | -1.279 | -1.262 | 0.004 | -0.025 | -0.025 | 2.902 | 0.3 | 0.4 | 0.4 | 0.76 | 0.34 | 0.44 |
| 2 | 105442170 | rs7596328 | 0.162 | 0.081 | -1.14 | -1.314 | 0.006 | -0.009 | -0.009 | 0 | 0.25 | 0.35 | 0.38 | 0.76 | 0.3 | 0.41 |
| 2 | 105455303 | rs3739154 | 0.905 | -0.045 | -1.348 | -1.382 | 0.008 | 0.042 | 0.042 | 0.58 | 0.21 | 0.11 | 0.2 | 0.09 | 0.16 | 0.1 |
| 2 | 105464087 | rs11124005 | -0.801 | 0.436 | -1.14 | -1.473 | 0.004 | 0.029 | 0.029 | 1.398 | 0.46 | 0.68 | 0.6 | 0.91 | 0.47 | 0.52 |
| 2 | 105466651 | rs6753397 | 0.274 | 0.315 | -0.787 | -0.253 | -0.002 | 0.135 | 0.135 | 0 | 0.22 | 0.4 | 0.32 | 0.56 | 0.26 | 0.4 |
| 2 | 105469864 | rs6723328 | -0.944 | 0.411 | -0.986 | -0.768 | -0.002 | -0.02 | -0.02 | 2.511 | 0.64 | 0.72 | 0.72 | 0.92 | 0.65 | 0.59 |
| 2 | 105473698 | rs732278 | 0.154 | 0.191 | -1.127 | -0.639 | -0.001 | 0.007 | 0.007 | 0 | 0.39 | 0.27 | 0.45 | 0.68 | 0.42 | 0.45 |
| 2 | 105477036 | rs13410373 | 1.156 | 0.183 | -1.087 | -0.642 | -0.001 | -0.004 | -0.004 | 0 | 0.21 | 0.24 | 0.23 | 0.08 | 0.2 | 0.13 |
| 2 | 105478347 | rs11687658 | -0.965 | -0.003 | -1.097 | -0.718 | -0.004 | -0.006 | -0.006 | 0.274 | 0.65 | 0.74 | 0.73 | 0.91 | 0.65 | 0.6 |
| 2 | 105481442 | rs1159964 | -0.153 | 0.071 | -0.924 | -0.39 | -0.006 | -0.007 | -0.007 | 0 | 0.4 | 0.5 | 0.45 | 0.69 | 0.42 | 0.45 |
| 2 | 105486125 | rs1376614 | 0.394 | 0.529 | -0.416 | 0.189 | -0.001 | -0.004 | -0.004 | 0 | 0.3 | 0.27 | 0.36 | 0.63 | 0.36 | 0.39 |
| 2 | 105501615 | rs6743963 | 0.115 | 0.698 | -0.696 | 0.235 | -0.013 | 0.007 | 0.007 | 0.021 | 0.33 | 0.35 | 0.33 | 0.6 | 0.26 | 0.39 |
| 2 | 105504322 | rs1026220 | -0.502 | 0.801 | -0.776 | 0.076 | 0 | -0.021 | -0.021 | 0 | 0.56 | 0.59 | 0.67 | 0.88 | 0.59 | 0.52 |
| 2 | 105507589 | rs6719286 | 0.804 | 0.878 | -1.055 | 0.14 | -0.007 | -0.006 | -0.006 | 3.472 | 0.24 | 0.22 | 0.33 | 0.28 | 0.37 | 0.15 |
| 2 | 105516725 | rs993557 | -0.868 | 0.373 | -1.011 | 0.09 | -0.004 | 0.071 | 0.071 | 1.028 | 0.6 | 0.76 | 0.68 | 0.88 | 0.62 | 0.55 |
| 2 | 105523791 | rs1005999 | 0.546 | 0.548 | -0.693 | 0.518 | -0.003 | -0.006 | -0.006 | 0.017 | 0.24 | 0.22 | 0.29 | 0.28 | 0.24 | 0.14 |

| | | | | | | | | | | | | | | | | |
|---|---|---|---|---|---|---|---|---|---|---|---|---|---|---|---|---|
| 2 | 105532887 | rs896357 | 0.446 | 0.465 | -0.402 | 0.724 | -0.003 | 0.006 | 0.006 | 0 | 0.34 | 0.36 | 0.37 | 0.45 | 0.37 | 0.42 |
| 2 | 105540676 | rs1947307 | -0.944 | 0.481 | -0.434 | 0.739 | -0.015 | 0.038 | 0.038 | 0.434 | 0.65 | 0.8 | 0.69 | 0.83 | 0.63 | 0.56 |
| 2 | 105544888 | rs6734108 | 0.474 | 0.446 | -0.481 | 0.72 | -0.001 | -0.026 | -0.026 | 0 | 0.3 | 0.21 | 0.32 | 0.38 | 0.26 | 0.15 |
| 2 | 105549404 | rs10496388 | -0.692 | 0.577 | -0.453 | 0.506 | 0.004 | -0.024 | -0.024 | 0 | 0.64 | 0.62 | 0.7 | 0.83 | 0.64 | 0.57 |
| 2 | 105565363 | rs1367175 | -0.839 | 0.497 | -0.835 | 0.54 | -0.131 | 0.08 | 0.08 | 1.937 | 0.68 | 0.99 | 0.71 | 0.86 | 0.65 | 0.64 |
| 2 | 105566954 | rs13386928 | 0.486 | 0.438 | -0.522 | 0.681 | -0.004 | 0.016 | 0.016 | 0 | 0.3 | 0.41 | 0.33 | 0.37 | 0.25 | 0.14 |
| 2 | 105578331 | rs1583108 | 0.672 | 0.403 | -0.501 | 0.564 | -0.001 | 0 | 0 | 0.212 | 0.34 | 0.33 | 0.36 | 0.46 | 0.37 | 0.44 |
| 2 | 105589679 | rs12611661 | 0.504 | 0.481 | -0.623 | 0.658 | 0.001 | -0.016 | -0.016 | 0 | 0.3 | 0.25 | 0.31 | 0.35 | 0.25 | 0.14 |
| 2 | 105591354 | rs10204200 | 0.478 | 0.497 | -0.684 | 0.666 | 0 | -0.003 | -0.003 | 0 | 0.3 | 0.33 | 0.32 | 0.35 | 0.25 | 0.14 |
| 2 | 105593028 | rs7608291 | -0.468 | 0.585 | -1.051 | 0.217 | -0.029 | 0.046 | 0.046 | 1.566 | 0.66 | 0.86 | 0.7 | 0.8 | 0.64 | 0.58 |
| 2 | 105613339 | rs9653448 | 0.044 | 0.62 | -0.79 | 0.643 | -0.013 | 0.011 | 0.011 | 0 | 0.35 | 0.46 | 0.36 | 0.42 | 0.28 | 0.24 |
| 2 | 105620400 | rs11884065 | -0.096 | 0.852 | -0.927 | 0.824 | 0.01 | -0.001 | -0.001 | 0 | 0.39 | 0.57 | 0.42 | 0.65 | 0.46 | 0.49 |
| 2 | 105637431 | rs10205766 | -0.108 | 0.857 | -0.931 | 0.851 | 0.009 | -0.001 | -0.001 | 0 | 0.39 | 0.57 | 0.42 | 0.65 | 0.46 | 0.49 |
| 2 | 105648817 | rs2376275 | 0.322 | 0.88 | -0.932 | 0.812 | 0.007 | 0.004 | 0.004 | 0 | 0.34 | 0.43 | 0.37 | 0.39 | 0.41 | 0.39 |
| 2 | 105680566 | rs7604397 | -1.018 | 1.064 | -1.183 | 0.331 | -0.005 | 0.015 | 0.015 | 0 | 0.74 | 0.77 | 0.68 | 0.89 | 0.72 | 0.86 |
| 2 | 105681812 | rs2217420 | -0.753 | 1.041 | -1.228 | 0.336 | -0.032 | 0.053 | 0.053 | 0.567 | 0.69 | 0.9 | 0.72 | 0.83 | 0.7 | 0.63 |
| 2 | 105690136 | rs12712196 | -0.758 | 1.041 | -1.195 | 0.335 | -0.024 | 0.05 | 0.05 | 0.087 | 0.69 | 0.89 | 0.72 | 0.83 | 0.7 | 0.63 |
| 2 | 105690215 | rs12620928 | 0.06 | 0.979 | -1.152 | 0.377 | 0.017 | -0.017 | -0.017 | 0 | 0.34 | 0.29 | 0.34 | 0.37 | 0.28 | 0.24 |
| 2 | 105697068 | rs7562875 | 1.805 | 0.936 | -1.419 | 0.129 | -0.047 | 0.054 | 0.054 | 0.045 | 0.11 | 0 | 0.12 | 0.02 | 0.14 | 0.17 |
| 2 | 105700545 | rs12618715 | 1.075 | 0.919 | -1.178 | 0.351 | 0.006 | 0.016 | 0.016 | 0 | 0.25 | 0.21 | 0.28 | 0.11 | 0.22 | 0.13 |
| 2 | 105701114 | rs12619439 | 1.217 | 0.906 | -1.172 | 0.344 | 0.004 | 0.035 | 0.035 | 0 | 0.24 | 0.12 | 0.28 | 0.11 | 0.22 | 0.13 |
| 2 | 105711904 | rs12617795 | 1.203 | 0.467 | -1.135 | 0.372 | 0.005 | 0.034 | 0.034 | 0 | 0.24 | 0.13 | 0.28 | 0.11 | 0.22 | 0.13 |
| 2 | 105720155 | rs12612044 | 0.06 | 0.464 | -1.013 | 0.386 | 0.015 | -0.017 | -0.017 | 0 | 0.34 | 0.28 | 0.34 | 0.37 | 0.28 | 0.24 |
| 2 | 105729122 | rs10187600 | 0.436 | -0.178 | -1.215 | 0.425 | 0.005 | -0.033 | -0.033 | 0 | 0.09 | 0.05 | 0.06 | 0.25 | 0.06 | 0.1 |
| 2 | 105733847 | rs11687311 | 1.171 | 0.503 | -1.256 | 0.339 | -0.043 | 0.037 | 0.037 | 0 | 0.24 | 0.11 | 0.32 | 0.13 | 0.35 | 0.15 |
| 2 | 105739439 | rs13034052 | 0.689 | 0.543 | -1.21 | 0.393 | -0.014 | 0.015 | 0.015 | 0 | 0.34 | 0.41 | 0.32 | 0.47 | 0.27 | 0.36 |
| 2 | 105742131 | rs7567068 | 0.997 | 0.72 | -1.38 | 0.258 | -0.043 | 0.04 | 0.04 | 0 | 0.25 | 0.11 | 0.32 | 0.13 | 0.35 | 0.15 |
| 2 | 105746447 | rs6729109 | 0.217 | 0.819 | -1.37 | 0.275 | 0.008 | -0.001 | -0.001 | 3.396 | 0.37 | 0.46 | 0.4 | 0.36 | 0.41 | 0.25 |
| 2 | 105747758 | rs6730532 | 0.687 | 0.61 | -1.828 | -0.094 | -0.004 | 0.093 | 0.093 | 0.047 | 0.17 | 0.04 | 0.1 | 0.02 | 0.17 | 0.1 |
| 2 | 105754029 | rs11680324 | -0.05 | 0.801 | -1.598 | 0.37 | -0.042 | 0.008 | 0.008 | 0 | 0.25 | 0.05 | 0.26 | 0.22 | 0.27 | 0.33 |
| 2 | 105757732 | rs11673925 | -0.049 | 0.829 | -1.579 | 0.381 | -0.055 | 0.016 | 0.016 | 0 | 0.25 | 0.04 | 0.27 | 0.19 | 0.27 | 0.33 |
| 2 | 105763818 | rs1125383 | -0.199 | 0.742 | -1.038 | 0.753 | 0.003 | 0.081 | 0.081 | 0 | 0.78 | 0.61 | 0.68 | 0.54 | 0.8 | 0.67 |
| 2 | 105767133 | rs4600676 | 0.07 | 0.668 | -0.373 | 0.859 | 0.005 | 0.05 | 0.05 | 0.144 | 0.76 | 0.42 | 0.66 | 0.36 | 0.79 | 0.64 |
| 2 | 105771650 | rs4605389 | 0.788 | 0.468 | -0.376 | 0.725 | 0.012 | 0.169 | 0.169 | 0 | 0.49 | 0.22 | 0.38 | 0.17 | 0.51 | 0.31 |
| 2 | 105776156 | rs10169497 | 0.067 | 0.556 | -0.382 | 1.02 | -0.009 | 0.083 | 0.083 | 0 | 0.78 | 0.61 | 0.69 | 0.54 | 0.83 | 0.67 |
| 2 | 105784318 | rs12621507 | 0.555 | 0.287 | -1.003 | 0.394 | -0.021 | 0.064 | 0.064 | 0 | 0.45 | 0.17 | 0.24 | 0.33 | 0.48 | 0.33 |
| 2 | 105784872 | rs13422730 | 0.344 | 0.214 | -0.787 | 0.728 | -0.001 | 0.004 | 0.004 | 0.78 | 0.67 | 0.32 | 0.64 | 0.57 | 0.72 | 0.77 |
| 2 | 105786209 | rs13028605 | 1.018 | 0.002 | -0.246 | 0.47 | -0.023 | -0.06 | -0.06 | 0 | 0.11 | 0.31 | 0.07 | 0 | 0.05 | 0.02 |
| 2 | 105787410 | rs1437400 | -0.571 | 0.136 | -0.556 | 0.536 | -0.02 | -0.017 | -0.017 | 0 | 0.79 | 0.94 | 0.6 | 0.76 | 0.76 | 0.54 |
| 2 | 105790639 | rs6761331 | 1.289 | 0.027 | -0.367 | 0.277 | -0.001 | -0.034 | -0.034 | 0 | 0.46 | 0.53 | 0.37 | 0.13 | 0.48 | 0.28 |
| 2 | 105791078 | rs2679882 | -0.73 | 0.11 | -0.952 | 0.493 | -0.064 | -0.035 | -0.035 | 1.126 | 0.77 | 0.99 | 0.6 | 0.75 | 0.74 | 0.56 |
| 2 | 105791914 | rs13388299 | 1.372 | -0.032 | -0.572 | 0.413 | -0.016 | 0.099 | 0.099 | 0 | 0.44 | 0.31 | 0.26 | 0.13 | 0.48 | 0.28 |
| 2 | 105792850 | rs2679870 | 0.214 | 0.16 | -0.476 | 1.355 | -0.008 | 0.146 | 0.146 | 0 | 0.11 | 0.18 | 0.1 | 0.34 | 0.06 | 0.14 |
| 2 | 105795508 | rs6543284 | 1.01 | 0.059 | -1.004 | 0.944 | -0.007 | -0.014 | -0.014 | 0 | 0.6 | 0.81 | 0.41 | 0.39 | 0.57 | 0.36 |
| 2 | 105795581 | rs2576800 | 0.3 | -0.451 | -0.886 | 1.374 | -0.007 | 0.019 | 0.019 | 0 | 0.15 | 0.12 | 0.12 | 0.31 | 0.09 | 0.12 |
| 2 | 105796650 | rs6709948 | 0.433 | -0.247 | -0.854 | 1.045 | 0.024 | -0.028 | -0.028 | 0 | 0.64 | 0.82 | 0.45 | 0.4 | 0.68 | 0.41 |
| 2 | 105797790 | rs2679852 | -0.792 | -0.425 | -0.44 | 0.997 | -0.004 | 0.001 | 0.001 | 0 | 0.36 | 0.31 | 0.55 | 0.6 | 0.28 | 0.61 |
| 2 | 105798292 | rs2679851 | -0.841 | -0.354 | -0.708 | 0.271 | -0.028 | 0.021 | 0.021 | 0 | 0.72 | 0.75 | 0.8 | 0.93 | 0.53 | 0.79 |
| 2 | 105799763 | rs9653475 | 1.593 | -0.449 | -0.558 | 0.419 | 0.016 | 0.038 | 0.038 | 0 | 0.31 | 0.22 | 0.15 | 0.09 | 0.24 | 0.14 |
| 2 | 105800952 | rs6736495 | -0.058 | -0.438 | -0.628 | 0.923 | -0.028 | 0.027 | 0.027 | 0 | 0.2 | 0.24 | 0.29 | 0.31 | 0.13 | 0.24 |
| 2 | 105801309 | rs2576797 | 0.637 | -0.347 | -1.026 | 1.144 | -0.012 | -0.083 | -0.083 | 0 | 0.16 | 0.02 | 0.26 | 0.29 | 0.15 | 0.37 |
| 2 | 105801651 | rs2376400 | 0.741 | -0.179 | -0.519 | 0.896 | -0.003 | 0 | 0 | 0.092 | 0.35 | 0.37 | 0.18 | 0.33 | 0.27 | 0.21 |
| 2 | 105808662 | rs2576791 | 0.312 | 1.089 | 0.046 | 1.205 | 0.003 | -0.003 | -0.003 | 0 | 0.58 | 0.69 | 0.48 | 0.59 | 0.48 | 0.61 |
| 2 | 105811219 | rs2576789 | 0.391 | 1.042 | 0.126 | 1.182 | 0.003 | -0.002 | -0.002 | 0.224 | 0.57 | 0.64 | 0.4 | 0.58 | 0.48 | 0.62 |
| 2 | 105813164 | rs2576783 | 0.674 | 1.038 | -0.271 | 0.49 | 0.027 | 0.066 | 0.066 | 0.227 | 0.54 | 0.23 | 0.38 | 0.38 | 0.47 | 0.61 |
| 2 | 105813839 | rs6754553 | -0.116 | 1.257 | -0.439 | -0.022 | 0.003 | 0.023 | 0.023 | 0 | 0.26 | 0.21 | 0.19 | 0.08 | 0.25 | 0.14 |
| 2 | 105832193 | rs11683715 | 0.372 | 1.668 | -0.301 | 0.508 | 0.063 | 0.135 | 0.135 | 0.001 | 0.76 | 0.57 | 0.54 | 0.51 | 0.65 | 0.71 |
| 2 | 105833093 | rs7584136 | 0.466 | 1.542 | -0.483 | 0.302 | -0.005 | 0.111 | 0.111 | 0.033 | 0.53 | 0.29 | 0.37 | 0.22 | 0.49 | 0.38 |
| 2 | 105835945 | rs6705097 | 0.628 | 1.503 | -0.678 | 0.1 | -0.044 | 0.127 | 0.127 | 0.046 | 0.24 | 0.04 | 0.17 | 0.02 | 0.3 | 0.19 |
| 2 | 105836936 | rs4851739 | 0.208 | 0.925 | 0.013 | 0.625 | 0.032 | 0.148 | 0.148 | 0 | 0.09 | 0.13 | 0.3 | 0.48 | 0.16 | 0.21 |
| 2 | 105837598 | rs6712932 | 0.455 | 0.885 | -0.35 | 0.605 | 0.051 | 0.094 | 0.094 | 0 | 0.45 | 0.29 | 0.32 | 0.22 | 0.33 | 0.27 |
| 2 | 105844490 | rs17030706 | 0.289 | 1.136 | -0.349 | 0.773 | 0.033 | 0.163 | 0.163 | 0 | 0.09 | 0.16 | 0.28 | 0.48 | 0.14 | 0.19 |
| 2 | 105846287 | rs13020904 | -0.185 | 0.612 | -0.404 | 0.635 | 0.033 | -0.009 | -0.009 | 0.862 | 0.85 | 0.64 | 0.84 | 0.92 | 0.8 | 0.9 |
| 2 | 105846891 | rs13022179 | -0.356 | 0.637 | -0.373 | 0.523 | 0.021 | -0.023 | -0.023 | 0.047 | 0.86 | 0.7 | 0.85 | 0.95 | 0.8 | 0.91 |
| 2 | 105852832 | rs1545122 | -0.631 | 0.798 | -1.407 | 0.23 | -0.004 | -0.01 | -0.01 | 0 | 0.87 | 0.85 | 0.85 | 1 | 0.8 | 0.92 |
| 2 | 105855232 | rs4851747 | -0.752 | 0.863 | -2.272 | 0.342 | -0.009 | 0.01 | 0.01 | 0 | 0.88 | 0.89 | 0.85 | 0.92 | 0.81 | 0.91 |
| 2 | 105857819 | rs17636399 | 0.398 | 0.431 | -2.255 | 0.441 | -0.107 | -0.022 | -0.022 | 0 | 0.29 | 0.03 | 0.22 | 0.22 | 0.32 | 0.43 |
| 2 | 105865967 | rs17687727 | -0.342 | -0.254 | -1.3 | 0.612 | 0.003 | 0.001 | 0.001 | 0 | 0.27 | 0.28 | 0.19 | 0.14 | 0.16 | 0.15 |
| 2 | 105875798 | rs12471644 | -0.821 | 0.234 | -1.43 | 1.563 | 0 | -0.028 | -0.028 | 0 | 0.07 | 0.01 | 0.22 | 0.34 | 0.06 | 0.12 |

| | | | | | | | | | | | | | | | | |
|---|---|---|---|---|---|---|---|---|---|---|---|---|---|---|---|---|
| 2 | 105881547 | rs4851752 | -0.549 | -0.001 | -2.324 | 0.554 | -0.019 | 0.125 | 0.125 | 0.132 | 0.22 | 0.01 | 0.14 | 0.02 | 0.18 | 0.06 |
| 2 | 105882061 | rs2460255 | 0.455 | 1.107 | -2.954 | 0.651 | -0.061 | -0.044 | -0.044 | 0 | 0.1 | 0.01 | 0.22 | 0.14 | 0.21 | 0.25 |
| 2 | 105882683 | rs1866039 | 0.021 | 1.34 | -2.379 | 1.429 | 0.016 | 0.184 | 0.184 | 0 | 0.13 | 0.25 | 0.21 | 0.52 | 0.13 | 0.27 |
| 2 | 105882761 | rs1866040 | -0.503 | 0.923 | -2.256 | 0.993 | 0.005 | 0.025 | 0.025 | 0 | 0.29 | 0.5 | 0.32 | 0.69 | 0.28 | 0.35 |
| 2 | 105885961 | rs2241797 | -0.348 | 0.219 | -2.743 | 0.064 | -0.01 | -0.013 | -0.013 | 0 | 0.59 | 0.47 | 0.64 | 0.28 | 0.66 | 0.52 |
| 2 | 105887549 | rs10202544 | 1.299 | 0.286 | -2.552 | 0.729 | 0.001 | 0.036 | 0.036 | 0 | 0.09 | 0.1 | 0.19 | 0.38 | 0.09 | 0.18 |
| 2 | 105889349 | rs11676273 | 0.818 | -0.068 | -2.773 | 0.267 | -0.062 | 0.038 | 0.038 | 0 | 0.19 | 0.01 | 0.22 | 0.08 | 0.22 | 0.12 |
| 2 | 105890971 | rs2576736 | -1.242 | -0.124 | -2.591 | 0.299 | 0.032 | 0.023 | 0.023 | 0 | 0.71 | 0.83 | 0.87 | 0.81 | 0.79 | 0.79 |
| 2 | 105897003 | rs2241799 | 2.034 | -0.035 | -2.527 | 0.348 | -0.061 | -0.044 | -0.044 | 0 | 0.1 | 0.01 | 0.22 | 0.15 | 0.21 | 0.25 |
| 2 | 105901179 | rs3792045 | 0.848 | -0.159 | -0.75 | 0.421 | 0.012 | 0.152 | 0.152 | 0 | 0.12 | 0.22 | 0.21 | 0.53 | 0.13 | 0.27 |
| 2 | 105901255 | rs6725410 | 1.656 | -0.164 | -0.487 | 0.282 | -0.001 | 0.01 | 0.01 | 0 | 0.06 | 0.12 | 0.03 | 0.03 | 0.04 | 0.09 |
| 2 | 105910513 | rs1030877 | -0.472 | -0.196 | -0.629 | 0.251 | 0.008 | 0.011 | 0.011 | 0 | 0.59 | 0.6 | 0.66 | 0.28 | 0.66 | 0.52 |
| 2 | 105914443 | rs2679889 | 0.542 | 0.129 | -0.437 | 0.123 | -0.007 | 0.013 | 0.013 | 0 | 0.27 | 0.19 | 0.3 | 0.19 | 0.31 | 0.35 |
| 2 | 105918939 | rs2576751 | 0.531 | 0.126 | -0.439 | 0.122 | -0.007 | 0.013 | 0.013 | 0 | 0.27 | 0.19 | 0.3 | 0.18 | 0.31 | 0.35 |
| 2 | 105935166 | rs6727677 | 0.745 | 0.153 | -0.713 | 0.001 | -0.077 | 0.037 | 0.037 | 0 | 0.18 | 0.01 | 0.26 | 0.08 | 0.24 | 0.12 |
| 2 | 105938107 | rs2679876 | -1.509 | 0.06 | -0.59 | 0.066 | 0.018 | 0.032 | 0.032 | 0 | 0.64 | 0.69 | 0.83 | 0.83 | 0.72 | 0.78 |
| 2 | 105938378 | rs2164718 | -0.669 | 0.099 | -0.477 | 0.145 | -0.019 | 0.074 | 0.074 | 0 | 0.46 | 0.68 | 0.58 | 0.75 | 0.49 | 0.66 |
| 2 | 105948446 | rs10460493 | 0.723 | 0.112 | -0.458 | 0.226 | 0.024 | 0.263 | 0.263 | 0 | 0.13 | 0.36 | 0.23 | 0.54 | 0.14 | 0.25 |
| 2 | 105950631 | rs1054250 | 0.545 | 0.157 | -0.817 | 0.05 | -0.015 | 0.027 | 0.027 | 0 | 0.26 | 0.15 | 0.3 | 0.18 | 0.31 | 0.34 |
| 2 | 105952417 | rs4851758 | 1.436 | 0.083 | -0.906 | -0.019 | -0.01 | 0.077 | 0.077 | 0 | 0.15 | 0.01 | 0.11 | 0.02 | 0.12 | 0.05 |
| 2 | 105953646 | rs2033724 | -0.131 | 0.136 | -0.833 | 0.007 | -0.006 | 0.016 | 0.016 | 0 | 0.33 | 0.22 | 0.34 | 0.2 | 0.35 | 0.4 |
| 2 | 105960047 | rs2576766 | 2.009 | 0.371 | -0.726 | 0.162 | -0.059 | -0.042 | -0.042 | 0.118 | 0.1 | 0.01 | 0.17 | 0.16 | 0.19 | 0.26 |
| 2 | 105960333 | rs2576767 | 2.009 | 0.371 | -0.726 | 0.186 | -0.059 | -0.044 | -0.044 | 0 | 0.1 | 0.01 | 0.17 | 0.16 | 0.19 | 0.26 |
| 2 | 105970650 | rs13012632 | 0.489 | 0.271 | -0.844 | 0.055 | -0.016 | 0.025 | 0.025 | 0 | 0.17 | 0.08 | 0.16 | 0.03 | 0.2 | 0.12 |
| 2 | 105974978 | rs2576776 | -0.378 | -0.097 | -0.894 | -0.013 | -0.025 | 0.036 | 0.036 | 0 | 0.38 | 0.23 | 0.45 | 0.21 | 0.5 | 0.45 |
| 2 | 105976344 | rs12476328 | -1.355 | -0.226 | -1.073 | -0.04 | 0.027 | 0.046 | 0.046 | 0 | 0.64 | 0.71 | 0.84 | 0.83 | 0.73 | 0.78 |
| 2 | 105977776 | rs3087523 | 1.696 | 0.203 | -1.173 | -0.133 | -0.027 | 0.037 | 0.037 | 0 | 0.1 | 0.01 | 0.11 | 0.02 | 0.11 | 0.04 |
| 2 | 105979506 | rs2278501 | -0.321 | 0.021 | -0.861 | -0.08 | -0.007 | 0.022 | 0.022 | 0 | 0.4 | 0.28 | 0.54 | 0.21 | 0.52 | 0.48 |
| 2 | 105982753 | rs2576778 | 1.295 | 0.378 | -1.231 | 0.24 | -0.038 | -0.03 | -0.03 | 0 | 0.12 | 0.05 | 0.2 | 0.2 | 0.21 | 0.22 |
| 2 | 105985228 | rs880427 | -0.976 | -0.06 | -1.77 | 0.148 | 0.021 | -0.013 | -0.013 | 0 | 0.63 | 0.8 | 0.82 | 0.68 | 0.67 | 0.66 |
| 2 | 105989716 | rs6738207 | -0.431 | 0.703 | -1.468 | 0.225 | -0.008 | -0.018 | -0.018 | 0 | 0.4 | 0.67 | 0.43 | 0.56 | 0.4 | 0.48 |
| 2 | 105992771 | rs10193943 | -0.355 | 0.63 | -1.639 | 0.262 | -0.01 | -0.033 | -0.033 | 0 | 0.41 | 0.74 | 0.43 | 0.56 | 0.39 | 0.51 |
| 2 | 105997385 | rs10190186 | 0.108 | 0.398 | -1.023 | 0.179 | -0.035 | -0.064 | -0.064 | 0.043 | 0.15 | 0.41 | 0.06 | 0.05 | 0.07 | 0.07 |
| 2 | 105998329 | rs6543289 | -1.227 | 0.706 | -2.036 | 0.286 | 0.025 | -0.032 | -0.032 | 0 | 0.67 | 0.78 | 0.82 | 0.66 | 0.68 | 0.7 |
| 2 | 105998627 | rs6753046 | -0.693 | 0.953 | -1.447 | 0.509 | -0.011 | -0.008 | -0.008 | 0 | 0.42 | 0.65 | 0.43 | 0.52 | 0.4 | 0.53 |
| 2 | 106009852 | rs12614625 | 0.796 | 1.216 | -1.683 | 0.728 | -0.028 | -0.009 | -0.009 | 0 | 0.14 | 0.06 | 0.17 | 0.15 | 0.21 | 0.22 |
| 2 | 106020941 | rs11124031 | 0.249 | 1.021 | -1.73 | 0.812 | -0.014 | -0.054 | -0.054 | 0.749 | 0.25 | 0.08 | 0.4 | 0.42 | 0.21 | 0.25 |
| 2 | 106024451 | rs4374396 | -0.15 | 0.854 | -1.106 | 0.466 | 0.005 | -0.005 | -0.005 | 0 | 0.34 | 0.43 | 0.37 | 0.31 | 0.4 | 0.45 |
| 2 | 106032291 | rs2376740 | 0.361 | 0.425 | -1.473 | 0.292 | -0.021 | 0.012 | 0.012 | 0 | 0.38 | 0.21 | 0.44 | 0.37 | 0.4 | 0.3 |
| 2 | 106034305 | rs7572930 | 0.309 | -0.018 | -1.43 | 0.071 | -0.013 | -0.025 | -0.025 | 1.878 | 0.12 | 0.17 | 0.12 | 0.02 | 0.09 | 0.14 |
| 2 | 106035580 | rs1914748 | -0.132 | 0.101 | -0.804 | 0.145 | 0.002 | 0.003 | 0.003 | 0 | 0.51 | 0.54 | 0.42 | 0.53 | 0.46 | 0.49 |
| 2 | 106036457 | rs6543294 | 0.331 | 0.098 | -0.637 | 0.147 | -0.008 | -0.016 | -0.016 | 0.394 | 0.39 | 0.25 | 0.3 | 0.52 | 0.39 | 0.39 |
| 2 | 106037635 | rs11690543 | 0.719 | 0.124 | -0.685 | 0.327 | -0.015 | -0.062 | -0.062 | 0 | 0.35 | 0.16 | 0.28 | 0.47 | 0.35 | 0.34 |
| 2 | 106039695 | rs4381801 | 0.487 | 0.267 | -0.568 | 0.211 | 0.003 | -0.008 | -0.008 | 0.193 | 0.4 | 0.44 | 0.47 | 0.36 | 0.43 | 0.35 |
| 2 | 106042902 | rs10194273 | 0.468 | -0.771 | -0.765 | 0.255 | 0.058 | 0.093 | 0.093 | 0 | 0.65 | 0.83 | 0.79 | 0.87 | 0.76 | 0.7 |
| 2 | 106042958 | rs10169988 | 1.231 | -0.774 | -0.393 | 0.446 | 0.036 | 0.003 | 0.003 | 0 | 0.53 | 0.37 | 0.59 | 0.43 | 0.65 | 0.54 |
| 2 | 106045085 | rs11694681 | -0.192 | -0.598 | -0.469 | 0.684 | 0.044 | 0.063 | 0.063 | 0 | 0.22 | 0.1 | 0.24 | 0.09 | 0.18 | 0.16 |
| 2 | 106045613 | rs7594621 | -0.424 | -0.506 | -0.373 | 0.852 | 0.021 | 0.003 | 0.003 | 0 | 0.27 | 0.16 | 0.29 | 0.22 | 0.25 | 0.28 |
| 2 | 106046662 | rs6735260 | 0.346 | 0.201 | 0.103 | 0.708 | 0.024 | -0.033 | -0.033 | 0 | 0.2 | 0.26 | 0.17 | 0.07 | 0.26 | 0.19 |
| 2 | 106046707 | rs6753085 | -0.304 | 0.247 | -0.195 | 0.801 | 0.023 | 0.009 | 0.009 | 0 | 0.27 | 0.55 | 0.28 | 0.35 | 0.28 | 0.31 |
| 2 | 106046789 | rs6750100 | -0.231 | 0.306 | -0.649 | 0.83 | 0.03 | 0.014 | 0.014 | 0 | 0.26 | 0.46 | 0.27 | 0.35 | 0.28 | 0.3 |
| 2 | 106050658 | rs11680431 | -0.013 | -0.06 | -1.436 | 0.734 | 0 | -0.008 | -0.008 | 0.427 | 0.1 | 0 | 0.12 | 0.1 | 0.07 | 0.17 |
| 2 | 106051956 | rs4851772 | -0.483 | -0.099 | -0.994 | 1.104 | -0.044 | 0.063 | 0.063 | 0.055 | 0.82 | 0.64 | 0.87 | 0.69 | 0.9 | 0.84 |
| 2 | 106051979 | rs11903583 | -0.355 | 0.855 | -1.215 | 1.32 | -0.006 | -0.006 | -0.006 | 0 | 0.4 | 0.4 | 0.57 | 0.48 | 0.45 | 0.47 |
| 2 | 106053343 | rs7583367 | -0.247 | 0.978 | -2.48 | 0.995 | -0.164 | -0.087 | -0.087 | 0.063 | 0.36 | 0.02 | 0.43 | 0.37 | 0.43 | 0.4 |
| 2 | 106059545 | rs11124034 | -0.319 | 0.969 | -1.979 | 1.136 | -0.033 | -0.045 | -0.045 | 0 | 0.38 | 0.16 | 0.42 | 0.45 | 0.41 | 0.43 |
| 2 | 106059781 | rs13411044 | -0.622 | 0.841 | -1.168 | 1.603 | -0.013 | 0.004 | 0.004 | 0.02 | 0.44 | 0.53 | 0.53 | 0.57 | 0.37 | 0.6 |
| 2 | 106061515 | rs6543295 | 0.816 | 1.442 | -0.989 | 1.521 | -0.015 | 0.029 | 0.029 | 0.161 | 0.79 | 0.87 | 0.66 | 0.93 | 0.72 | 0.67 |
| 2 | 106065506 | rs884876 | 0.791 | 1.46 | -1.087 | 0.897 | -0.021 | 0.035 | 0.035 | 0.888 | 0.78 | 0.84 | 0.66 | 0.93 | 0.68 | 0.64 |
| 2 | 106071159 | rs6543297 | 0.618 | 1.271 | -0.746 | 1.366 | 0.01 | 0.003 | 0.003 | 0 | 0.69 | 0.61 | 0.61 | 0.68 | 0.65 | 0.59 |
| 2 | 106076110 | rs10203748 | -1.024 | 0.45 | -0.451 | 1.838 | -0.019 | 0.012 | 0.012 | 0 | 0.17 | 0.3 | 0.18 | 0.16 | 0.13 | 0.19 |
| 2 | 106076400 | rs6752609 | -0.667 | 0.829 | -0.153 | 1.703 | 0.013 | 0.003 | 0.003 | 1.975 | 0.19 | 0.24 | 0.21 | 0.16 | 0.23 | 0.2 |
| 2 | 106078289 | rs7608329 | -1.273 | 0.232 | 0.499 | 1.938 | -0.01 | 0.15 | 0.15 | 0.454 | 0.06 | 0.23 | 0.23 | 0.24 | 0.04 | 0.1 |
| 2 | 106080746 | rs6711988 | -1.789 | 0.015 | 0.678 | 1.787 | -0.029 | 0.086 | 0.086 | 8.429 | 0.14 | 0.31 | 0.28 | 0.32 | 0.07 | 0.18 |
| 2 | 106083368 | rs10177620 | -0.636 | -0.112 | 0.405 | 0.716 | -0.018 | -0.066 | -0.066 | 0 | 0.36 | 0.45 | 0.31 | 0.22 | 0.33 | 0.31 |
| 2 | 106085803 | rs6543300 | -0.57 | -0.494 | 0.39 | 0.501 | -0.019 | -0.008 | -0.008 | 0.583 | 0.16 | 0.22 | 0.08 | 0.16 | 0.06 | 0.14 |
| 2 | 106090331 | rs2048982 | 0.782 | -0.175 | 0.162 | 0.447 | -0.036 | -0.036 | -0.036 | 0 | 0.29 | 0.23 | 0.27 | 0.4 | 0.34 | 0.28 |
| 2 | 106091003 | rs6543302 | -0.142 | -0.191 | 0.386 | 0.523 | -0.023 | 0.01 | 0.01 | 2.387 | 0.18 | 0.22 | 0.21 | 0.27 | 0.09 | 0.23 |
| 2 | 106092054 | rs1401208 | 1.713 | -0.221 | -0.025 | 0.266 | 0.006 | -0.051 | -0.051 | 0 | 0.17 | 0.05 | 0.21 | 0.25 | 0.17 | 0.1 |

| | | | | | | | | | | | | | | | | |
|---|---|---|---|---|---|---|---|---|---|---|---|---|---|---|---|---|
| 2 | 106092217 | rs1401209 | 1.661 | -0.238 | -0.033 | 0.231 | 0.007 | -0.054 | -0.054 | 2.021 | 0.17 | 0.05 | 0.22 | 0.25 | 0.17 | 0.1 |
| 2 | 106093381 | rs17652317 | 1.793 | -0.134 | 0.239 | 0.001 | 0 | 0.013 | 0.013 | 0.157 | 0.23 | 0.15 | 0.25 | 0.07 | 0.29 | 0.22 |
| 2 | 106093436 | rs2203801 | -0.376 | -0.018 | 0.087 | 0.085 | -0.009 | 0.009 | 0.009 | 0.13 | 0.9 | 0.94 | 0.96 | 1 | 0.88 | 0.84 |
| 2 | 106097519 | rs1607420 | 0.624 | -0.028 | 0.147 | 0.272 | -0.013 | -0.014 | -0.014 | 0 | 0.3 | 0.3 | 0.28 | 0.4 | 0.34 | 0.3 |
| 2 | 106101243 | rs7601617 | -0.408 | 0.061 | 0.241 | -0.347 | -0.01 | -0.089 | -0.089 | 0 | 0.72 | 0.6 | 0.81 | 0.93 | 0.7 | 0.76 |
| 2 | 106102328 | rs12997792 | 1.117 | 0.58 | -0.131 | -0.379 | -0.027 | 0.018 | 0.018 | 0 | 0.12 | 0.07 | 0.14 | 0 | 0.19 | 0.07 |
| 2 | 106103606 | rs1518614 | -0.723 | 0.428 | 0.039 | -0.549 | -0.011 | 0.014 | 0.014 | 0 | 0.34 | 0.32 | 0.48 | 0.24 | 0.39 | 0.46 |
| 2 | 106104856 | rs13006682 | 0.552 | 0.565 | 0.18 | -0.48 | 0.104 | 0.013 | 0.013 | 0 | 0.16 | 0.29 | 0.37 | 0.17 | 0.31 | 0.18 |
| 2 | 106106221 | rs11901258 | 0.571 | 0.505 | -0.175 | -0.391 | -0.013 | -0.019 | -0.019 | 0 | 0.16 | 0.06 | 0.31 | 0.23 | 0.19 | 0.3 |
| 2 | 106106996 | rs11896170 | -0.892 | 0.028 | -0.251 | -0.596 | 0.026 | 0.074 | 0.074 | 0.035 | 0.41 | 0.71 | 0.33 | 0.17 | 0.32 | 0.56 |
| 2 | 106110184 | rs1401212 | 0.116 | -0.177 | -0.147 | -0.492 | -0.001 | -0.001 | -0.001 | 0 | 0.23 | 0.22 | 0.34 | 0.24 | 0.23 | 0.35 |
| 2 | 106115555 | rs6707139 | -2.153 | -0.105 | -0.806 | -0.231 | -0.044 | -0.057 | -0.057 | 0.432 | 0.79 | 0.94 | 0.75 | 0.73 | 0.71 | 0.88 |
| 2 | 106116942 | rs13426368 | 1.819 | -0.062 | -0.663 | -0.199 | 0.015 | -0.026 | -0.026 | 0 | 0.23 | 0.18 | 0.16 | 0.32 | 0.23 | 0.14 |
| 2 | 106119891 | rs12994383 | 1.816 | 0.135 | -0.644 | -0.251 | -0.046 | 0.05 | 0.05 | 0 | 0.12 | 0.05 | 0.13 | 0 | 0.15 | 0.07 |
| 2 | 106125913 | rs6543306 | -0.02 | -0.05 | -0.3 | -0.251 | 0.01 | -0.01 | -0.01 | 0 | 0.44 | 0.71 | 0.58 | 0.55 | 0.55 | 0.35 |
| 2 | 106126100 | rs4851783 | 0.258 | -0.162 | 0.086 | -0.326 | -0.003 | -0.092 | -0.092 | 0 | 0.29 | 0.49 | 0.24 | 0.09 | 0.23 | 0.37 |
| 2 | 106128407 | rs13412180 | 0.625 | -0.041 | -0.063 | -0.202 | 0.057 | 0.076 | 0.076 | 0 | 0.07 | 0.13 | 0.27 | 0.23 | 0.17 | 0.14 |
| 2 | 106129071 | rs10865067 | 0.218 | -0.269 | -0.139 | -0.321 | -0.007 | -0.001 | -0.001 | 0 | 0.43 | 0.33 | 0.5 | 0.62 | 0.48 | 0.42 |
| 2 | 106130054 | rs2376739 | -2.39 | -0.28 | -0.219 | -0.331 | -0.047 | -0.07 | -0.07 | 0 | 0.79 | 0.94 | 0.74 | 0.71 | 0.71 | 0.88 |
| 2 | 106130142 | rs17031046 | 1.224 | -0.282 | -0.249 | -0.338 | -0.021 | -0.037 | -0.037 | 0.018 | 0.3 | 0.23 | 0.42 | 0.55 | 0.4 | 0.27 |
| 2 | 106131851 | rs4851787 | -1.876 | -0.146 | 0.093 | -0.282 | -0.002 | -0.003 | -0.003 | 0 | 0.77 | 0.79 | 0.73 | 0.71 | 0.71 | 0.88 |
| 2 | 106131935 | rs17031063 | 0.516 | -0.204 | 0.149 | -0.334 | -0.002 | 0.041 | 0.041 | 0.554 | 0.28 | 0.23 | 0.21 | 0.09 | 0.22 | 0.37 |
| 2 | 106132986 | rs6713907 | -1.589 | -0.164 | 0.611 | 0.004 | 0.008 | 0.022 | 0.022 | 0 | 0.76 | 0.73 | 0.73 | 0.71 | 0.71 | 0.88 |
| 2 | 106133488 | rs939008 | 1.461 | -0.159 | 0.34 | -0.007 | 0.028 | 0.047 | 0.047 | 0 | 0.13 | 0.01 | 0.07 | 0.07 | 0.08 | 0.15 |
| 2 | 106134743 | rs17031089 | 0.954 | -0.216 | 0.328 | -0.059 | 0.016 | 0.016 | 0.016 | 0 | 0.16 | 0.13 | 0.08 | 0.07 | 0.09 | 0.18 |
| 2 | 106134934 | rs12471907 | -0.534 | -0.227 | 0.352 | -0.011 | -0.004 | 0.003 | 0.003 | 0.523 | 0.6 | 0.5 | 0.63 | 0.57 | 0.64 | 0.58 |
| 2 | 106136714 | rs12712211 | 0.034 | -0.127 | 0.133 | -0.332 | -0.005 | 0.037 | 0.037 | 0.137 | 0.54 | 0.38 | 0.37 | 0.4 | 0.45 | 0.54 |
| 2 | 106138304 | rs12468349 | 0.586 | -0.111 | -0.096 | -0.263 | 0.004 | -0.001 | -0.001 | 0.138 | 0.2 | 0.22 | 0.35 | 0.22 | 0.25 | 0.3 |
| 2 | 106139216 | rs13426275 | 1.185 | -0.171 | 0.058 | -0.295 | -0.04 | -0.037 | -0.037 | 0 | 0.17 | 0.28 | 0.09 | 0.08 | 0.07 | 0.3 |
| 2 | 106139293 | rs12473672 | -1.13 | 0.375 | 0.246 | 0.046 | 0.005 | 0.043 | 0.043 | 0 | 0.75 | 0.71 | 0.73 | 0.62 | 0.71 | 0.84 |
| 2 | 106140152 | rs1568261 | 0.06 | 0.323 | -0.085 | -0.071 | -0.001 | 0.004 | 0.004 | 0.004 | 0.54 | 0.49 | 0.38 | 0.4 | 0.45 | 0.54 |
| 2 | 106149270 | rs6543316 | -1.073 | 0.707 | -0.347 | 0.329 | -0.04 | -0.062 | -0.062 | 0 | 0.79 | 0.92 | 0.74 | 0.65 | 0.7 | 0.85 |
| 2 | 106153496 | rs4851793 | -0.02 | 0.859 | 0.475 | 0.327 | 0.045 | 0.04 | 0.04 | 0 | 0.69 | 0.47 | 0.43 | 0.47 | 0.53 | 0.6 |
| 2 | 106154536 | rs12987492 | 0.836 | 0.435 | 0.325 | 0.1 | -0.042 | 0.053 | 0.053 | 0 | 0.54 | 0.18 | 0.34 | 0.39 | 0.48 | 0.38 |
| 2 | 106156342 | rs11675788 | 1.066 | 0.034 | 0.572 | 0.066 | 0.023 | 0.012 | 0.012 | 0 | 0.12 | 0.09 | 0.07 | 0.09 | 0.06 | 0.19 |
| 2 | 106157515 | rs17031192 | 1.066 | 0.034 | 0.538 | 0.068 | 0.023 | 0.012 | 0.012 | 0 | 0.12 | 0.09 | 0.07 | 0.09 | 0.06 | 0.19 |
| 2 | 106159626 | rs7575977 | -0.64 | 0.054 | 0.818 | 0.096 | 0.051 | 0.105 | 0.105 | 0 | 0.73 | 0.52 | 0.61 | 0.57 | 0.64 | 0.77 |
| 2 | 106160334 | rs11682070 | -1.55 | 0.037 | 0.811 | 0.09 | -0.055 | -0.077 | -0.077 | 0 | 0.8 | 0.95 | 0.74 | 0.65 | 0.7 | 0.85 |
| 2 | 106160498 | rs6543317 | -1.423 | 0.023 | 0.815 | 0.095 | -0.041 | -0.059 | -0.059 | 0 | 0.79 | 0.92 | 0.73 | 0.65 | 0.69 | 0.85 |
| 2 | 106160606 | rs6726692 | 0.638 | 0.039 | 0.605 | 0.038 | -0.007 | 0.01 | 0.01 | 0 | 0.57 | 0.36 | 0.35 | 0.38 | 0.46 | 0.41 |
| 2 | 106160899 | rs6727113 | -0.401 | 0.066 | 0.447 | 0.135 | -0.005 | -0.002 | -0.002 | 0 | 0.19 | 0.28 | 0.27 | 0.18 | 0.18 | 0.39 |
| 2 | 106163524 | rs6723920 | 1.193 | 0.045 | -0.322 | -0.121 | 0.073 | 0.019 | 0.019 | 0 | 0.41 | 0.2 | 0.26 | 0.4 | 0.29 | 0.32 |
| 2 | 106164853 | rs2889453 | 0.652 | -0.055 | -0.286 | -0.184 | -0.007 | -0.008 | -0.008 | 0 | 0.17 | 0.19 | 0.12 | 0.07 | 0.1 | 0.19 |
| 2 | 106165605 | rs6738475 | -0.367 | 0.016 | -0.265 | -0.143 | 0.005 | 0.008 | 0.008 | 0 | 0.64 | 0.41 | 0.55 | 0.57 | 0.51 | 0.68 |
| 2 | 106166928 | rs6733046 | 0.826 | 0.049 | -0.934 | -0.126 | 0.053 | -0.008 | -0.008 | 0 | 0.46 | 0.21 | 0.44 | 0.49 | 0.41 | 0.49 |
| 2 | 106170826 | rs17188560 | 0.819 | 0.078 | -0.936 | -0.113 | 0.035 | 0.038 | 0.038 | 0 | 0.17 | 0.07 | 0.11 | 0.07 | 0.1 | 0.19 |
| 2 | 106174649 | rs4561691 | 1.076 | 0.361 | -0.828 | -0.064 | -0.054 | 0.016 | 0.016 | 0 | 0.14 | 0.04 | 0.16 | 0.08 | 0.19 | 0.16 |
| 2 | 106175739 | rs877328 | -0.316 | -0.026 | -0.766 | -0.062 | -0.019 | -0.029 | -0.029 | 0 | 0.63 | 0.29 | 0.54 | 0.56 | 0.51 | 0.68 |
| 2 | 106178712 | rs17031275 | 0.391 | 0.051 | -0.782 | -0.217 | 0.036 | 0 | 0 | 0.055 | 0.17 | 0.07 | 0.11 | 0.12 | 0.1 | 0.19 |
| 2 | 106178815 | rs875429 | 0.334 | 0.144 | -0.195 | -0.245 | 0.057 | 0.009 | 0.009 | 0.266 | 0.06 | 0.18 | 0.19 | 0.05 | 0.12 | 0.17 |
| 2 | 106179399 | rs874773 | -0.273 | 0.118 | -0.25 | -0.25 | -0.004 | -0.004 | -0.004 | 0.365 | 0.22 | 0.25 | 0.29 | 0.17 | 0.22 | 0.36 |
| 2 | 106181868 | rs10170383 | -0.859 | 0.042 | 0.043 | -0.637 | -0.008 | -0.036 | -0.036 | 0 | 0.37 | 0.6 | 0.22 | 0.12 | 0.26 | 0.3 |
| 2 | 106182953 | rs6729061 | -1.06 | 0.078 | 0.032 | -0.642 | 0 | -0.002 | -0.002 | 0 | 0.41 | 0.63 | 0.4 | 0.17 | 0.38 | 0.47 |
| 2 | 106184379 | rs17031301 | -0.286 | 0.02 | 0.207 | -0.665 | -0.014 | -0.016 | -0.016 | 0 | 0.06 | 0.26 | 0.03 | 0 | 0 | 0.02 |
| 2 | 106184438 | rs10496397 | 1.137 | 0.189 | -0.201 | -0.6 | -0.016 | 0.006 | 0.006 | 0 | 0.13 | 0.11 | 0.08 | 0 | 0.15 | 0.09 |
| 2 | 106189955 | rs11891488 | -0.92 | 0.14 | -0.276 | -0.555 | 0.006 | 0.042 | 0.042 | 0 | 0.42 | 0.7 | 0.41 | 0.17 | 0.38 | 0.47 |
| 2 | 106195093 | rs2084779 | -0.695 | -0.102 | -0.346 | 0.068 | 0.011 | 0.005 | 0.005 | 0 | 0.69 | 0.7 | 0.67 | 0.62 | 0.58 | 0.77 |
| 2 | 106197398 | rs1518611 | -0.473 | -0.643 | -0.965 | -0.203 | 0.018 | 0.018 | 0.018 | 0.521 | 0.38 | 0.3 | 0.26 | 0.13 | 0.24 | 0.34 |
| 2 | 106209113 | rs6705579 | -0.192 | -0.238 | -1.317 | 0.518 | -0.009 | -0.049 | -0.049 | 0 | 0.28 | 0.2 | 0.3 | 0.54 | 0.26 | 0.32 |
| 2 | 106210350 | rs10201413 | -0.189 | -1.591 | -0.776 | 0.139 | -0.066 | -0.032 | -0.032 | 0 | 0.34 | 0.4 | 0.19 | 0.22 | 0.16 | 0.19 |
| 2 | 106211726 | rs2167462 | -0.427 | -1.047 | -0.415 | -0.085 | -0.01 | -0.033 | -0.033 | 0.171 | 0.4 | 0.43 | 0.4 | 0.76 | 0.38 | 0.61 |
| 2 | 106214189 | rs6761943 | 0.851 | -1.153 | -0.4 | 0.456 | -0.003 | -0.004 | -0.004 | 1.883 | 0.15 | 0.14 | 0.17 | 0.21 | 0.18 | 0.09 |
| 2 | 106215749 | rs17031337 | 0.419 | -1.269 | -0.323 | 0.87 | 0.002 | -0.008 | -0.008 | 0 | 0.21 | 0.2 | 0.2 | 0.36 | 0.11 | 0.44 |
| 2 | 106217330 | rs6543322 | -1.609 | -0.808 | -0.963 | 0.209 | -0.049 | 0.078 | 0.078 | 0.804 | 0.68 | 0.96 | 0.79 | 0.79 | 0.71 | 0.83 |
| 2 | 106220799 | rs7576878 | -1.225 | -0.943 | -0.297 | 0.124 | 0.077 | 0.182 | 0.182 | 0.039 | 0.49 | 0.77 | 0.53 | 0.79 | 0.42 | 0.67 |
| 2 | 106221754 | rs17196498 | 2.166 | -0.92 | -0.25 | 0.103 | -0.014 | 0.009 | 0.009 | 0 | 0.1 | 0.06 | 0.1 | 0 | 0.15 | 0.03 |
| 2 | 106223220 | rs7590257 | 0.358 | -0.997 | 0.229 | 0.145 | 0.013 | 0.034 | 0.034 | 0.195 | 0.2 | 0.39 | 0.2 | 0.22 | 0.19 | 0.12 |
| 2 | 106225207 | rs2044902 | -0.586 | -1.01 | 0.172 | 0.182 | 0.002 | 0.053 | 0.053 | 0.694 | 0.27 | 0.57 | 0.33 | 0.43 | 0.31 | 0.24 |
| 2 | 106225295 | rs17196637 | 1.824 | -1.054 | 0.152 | 0.095 | -0.06 | 0.056 | 0.056 | 0 | 0.18 | 0.06 | 0.25 | 0 | 0.29 | 0.15 |

| | | | | | | | | | | | | | | | | |
|---|---|---|---|---|---|---|---|---|---|---|---|---|---|---|---|---|
| 2 | 106227721 | rs10496399 | 0.595 | -1.115 | 0.143 | 0.126 | 0.018 | -0.022 | -0.022 | 0 | 0.25 | 0.21 | 0.31 | 0.58 | 0.22 | 0.53 |
| 2 | 106232863 | rs10207508 | -1.738 | -1.053 | 0.104 | 0.116 | -0.01 | 0.093 | 0.093 | 0.611 | 0.68 | 0.88 | 0.77 | 0.79 | 0.7 | 0.83 |
| 2 | 106236355 | rs17031393 | 1.915 | -0.96 | 0.359 | 0.154 | 0.025 | -0.06 | -0.06 | 0 | 0.11 | 0.17 | 0.11 | 0 | 0.15 | 0.03 |
| 2 | 106236536 | rs7564636 | 0.52 | -0.963 | 0.334 | 0.219 | -0.015 | 0.095 | 0.095 | 0 | 0.09 | 0.2 | 0.09 | 0.22 | 0.04 | 0.08 |
| 2 | 106237751 | rs12474808 | -0.726 | -0.739 | 0.171 | 0.015 | 0.017 | 0.033 | 0.033 | 0 | 0.5 | 0.64 | 0.53 | 0.58 | 0.4 | 0.67 |
| 2 | 106238994 | rs13021658 | 1.798 | -0.61 | -0.444 | -0.061 | -0.044 | 0.039 | 0.039 | 0 | 0.17 | 0.06 | 0.24 | 0 | 0.27 | 0.14 |
| 2 | 106241808 | rs896444 | 0.652 | -0.714 | -0.431 | 0.281 | 0.013 | -0.016 | -0.016 | 1.72 | 0.23 | 0.2 | 0.25 | 0.35 | 0.1 | 0.45 |
| 2 | 106243636 | rs6712413 | -0.672 | -0.436 | -0.823 | 0.344 | -0.016 | -0.032 | -0.032 | 1.751 | 0.58 | 0.78 | 0.62 | 0.45 | 0.53 | 0.58 |
| 2 | 106243696 | rs4851816 | 0.557 | -0.515 | -0.871 | 0.44 | 0.043 | -0.061 | -0.061 | 0.061 | 0.37 | 0.17 | 0.32 | 0.54 | 0.33 | 0.35 |
| 2 | 106244673 | rs6543324 | 0.39 | -0.509 | -0.751 | 0.029 | 0.003 | 0.001 | 0.001 | 0.149 | 0.39 | 0.34 | 0.29 | 0.24 | 0.31 | 0.14 |
| 2 | 106245692 | rs4851817 | 1.165 | -0.804 | -1.38 | 0.24 | 0.062 | 0.04 | 0.04 | 2.138 | 0.28 | 0.12 | 0.18 | 0.24 | 0.17 | 0.11 |
| 2 | 106245742 | rs6743358 | -0.171 | -0.277 | -0.838 | 0.333 | -0.011 | 0.013 | 0.013 | 1.558 | 0.38 | 0.64 | 0.39 | 0.22 | 0.45 | 0.26 |
| 2 | 106247640 | rs4851073 | -0.357 | -0.818 | -0.539 | 0.884 | -0.065 | 0.079 | 0.079 | 0 | 0.25 | 0.58 | 0.28 | 0.43 | 0.13 | 0.43 |
| 2 | 106247818 | rs4851076 | -0.205 | 0.919 | -0.717 | 0.11 | 0.034 | -0.056 | -0.056 | 0.999 | 0.8 | 0.74 | 0.7 | 0.98 | 0.63 | 0.85 |
| 2 | 106248663 | rs11124049 | 0.428 | 0.495 | -1.231 | 0.106 | -0.005 | 0.027 | 0.027 | 0 | 0.17 | 0.08 | 0.12 | 0.02 | 0.21 | 0.22 |
| 2 | 106249412 | rs12478680 | -0.236 | 0.333 | -1.293 | 0.899 | 0 | -0.035 | -0.035 | 0 | 0.23 | 0.11 | 0.25 | 0.46 | 0.24 | 0.35 |
| 2 | 106250759 | rs876585 | -0.213 | 0.228 | -1.488 | 0.955 | -0.006 | -0.067 | -0.067 | 0 | 0.24 | 0.11 | 0.27 | 0.35 | 0.26 | 0.39 |
| 2 | 106250893 | rs10201112 | 0.135 | 0.82 | -0.693 | 0.367 | 0.071 | -0.093 | -0.093 | 0 | 0.17 | 0.29 | 0.22 | 0.01 | 0.24 | 0.11 |
| 2 | 106252991 | rs4851819 | -0.4 | 0.827 | -1.167 | 0.77 | -0.09 | -0.028 | -0.028 | 0 | 0.79 | 0.9 | 0.63 | 0.73 | 0.6 | 0.79 |
| 2 | 106256187 | rs11889938 | 0.75 | 0.09 | -1.214 | 0.257 | -0.009 | -0.04 | -0.04 | 0 | 0.15 | 0.13 | 0.06 | 0 | 0.07 | 0.18 |
| 2 | 106258025 | rs4851079 | 0.164 | -0.104 | -0.73 | 0.7 | 0.026 | 0.021 | 0.021 | 0 | 0.67 | 0.45 | 0.53 | 0.59 | 0.53 | 0.59 |
| 2 | 106259024 | rs17031438 | 1.217 | -0.636 | -1.012 | -0.008 | 0.019 | 0.003 | 0.003 | 0 | 0.28 | 0.19 | 0.24 | 0.35 | 0.27 | 0.26 |
| 2 | 106260780 | rs4851821 | 0.327 | -0.83 | -0.74 | -0.086 | 0.011 | 0.011 | 0.011 | 0 | 0.43 | 0.4 | 0.29 | 0.26 | 0.34 | 0.39 |
| 2 | 106261133 | rs4851822 | 0.845 | -0.813 | -0.975 | -0.148 | 0.061 | 0.081 | 0.081 | 0 | 0.31 | 0.16 | 0.23 | 0.14 | 0.2 | 0.22 |
| 2 | 106262719 | rs12467322 | 2.066 | -0.442 | -0.986 | 0.189 | -0.004 | -0.005 | -0.005 | 0 | 0.11 | 0.06 | 0.22 | 0.14 | 0.13 | 0.13 |
| 2 | 106263078 | rs7572653 | 1.263 | -0.792 | -0.283 | 0.035 | -0.025 | -0.018 | -0.018 | 0 | 0.28 | 0.35 | 0.16 | 0.22 | 0.17 | 0.21 |
| 2 | 106266007 | rs10183242 | -0.386 | -0.664 | -0.222 | 0.217 | -0.014 | -0.001 | -0.001 | 0 | 0.56 | 0.69 | 0.59 | 0.59 | 0.53 | 0.63 |
| 2 | 106266677 | rs4851824 | 0.248 | -0.733 | -0.17 | 0.215 | -0.023 | -0.007 | -0.007 | 0 | 0.39 | 0.59 | 0.28 | 0.33 | 0.26 | 0.47 |
| 2 | 106266695 | rs4851825 | 1.133 | -0.699 | -0.189 | 0.254 | -0.018 | 0.003 | 0.003 | 0 | 0.3 | 0.37 | 0.22 | 0.33 | 0.21 | 0.35 |
| 2 | 106269399 | rs17031461 | 1.614 | -0.749 | -0.096 | 0.099 | -0.017 | -0.044 | -0.044 | 0 | 0.08 | 0.13 | 0.04 | 0 | 0.04 | 0.05 |
| 2 | 106279090 | rs1597695 | 0.26 | -0.828 | 0.01 | 0.084 | 0.02 | 0.002 | 0.002 | 0 | 0.42 | 0.71 | 0.35 | 0.45 | 0.36 | 0.48 |
| 2 | 106287156 | rs7563743 | -1.212 | -0.836 | -0.115 | 0.06 | -0.101 | -0.085 | -0.085 | 0 | 0.67 | 0.89 | 0.64 | 0.59 | 0.58 | 0.71 |
| 2 | 106291637 | rs7565674 | 1.367 | -0.799 | 0.127 | 0.115 | 0.019 | 0.045 | 0.045 | 0 | 0.06 | 0.27 | 0.02 | 0.09 | 0.03 | 0.14 |
| 2 | 106292197 | rs10175607 | 0.78 | -0.702 | 0.052 | 0.165 | -0.007 | 0.019 | 0.019 | 0.459 | 0.11 | 0.19 | 0.11 | 0.11 | 0.09 | 0.25 |
| 2 | 106294737 | rs4851831 | 0.388 | -0.733 | -0.113 | 0.099 | 0.001 | 0.001 | 0.001 | 0 | 0.48 | 0.49 | 0.53 | 0.59 | 0.46 | 0.49 |
| 2 | 106295955 | rs4851081 | -0.003 | -1.425 | -0.014 | 0.306 | 0.022 | 0.002 | 0.002 | 0 | 0.42 | 0.71 | 0.35 | 0.45 | 0.36 | 0.48 |
| 2 | 106303174 | rs13025485 | 2.094 | -0.741 | -0.745 | 0.055 | -0.088 | 0.084 | 0.084 | 0 | 0.22 | 0.04 | 0.23 | 0 | 0.34 | 0.15 |
| 2 | 106303243 | rs1037381 | -0.157 | -0.788 | -0.483 | 0.445 | -0.019 | -0.006 | -0.006 | 0 | 0.38 | 0.62 | 0.34 | 0.36 | 0.26 | 0.39 |
| 2 | 106309119 | rs2122967 | -0.677 | -0.662 | -0.824 | 0.339 | -0.011 | -0.125 | -0.125 | 0.057 | 0.72 | 0.92 | 0.73 | 0.59 | 0.74 | 0.76 |
| 2 | 106310212 | rs11884955 | 0.347 | -0.695 | -0.701 | 0.282 | 0.009 | 0.022 | 0.022 | 0 | 0.6 | 0.63 | 0.5 | 0.35 | 0.64 | 0.46 |
| 2 | 106311348 | rs6709683 | 0.179 | -0.306 | -0.037 | 0.75 | 0.001 | -0.054 | -0.054 | 0 | 0.09 | 0.25 | 0.07 | 0 | 0.07 | 0.14 |
| 2 | 106315312 | rs13393585 | 0.207 | -0.187 | -0.259 | 0.585 | -0.036 | -0.005 | -0.005 | 0 | 0.26 | 0.37 | 0.25 | 0.29 | 0.17 | 0.21 |
| 2 | 106324719 | rs336506 | -0.336 | 0.756 | -0.173 | 0.474 | 0.012 | -0.021 | -0.021 | 0 | 0.27 | 0.35 | 0.38 | 0.18 | 0.31 | 0.23 |
| 2 | 106325753 | rs336507 | 0.711 | 0.502 | -0.748 | 0.318 | -0.02 | -0.018 | -0.018 | 0.315 | 0.62 | 0.43 | 0.59 | 0.32 | 0.66 | 0.48 |
| 2 | 106333074 | rs6758695 | 0.601 | 0.747 | -0.744 | 0.343 | -0.014 | -0.031 | -0.031 | 0.557 | 0.15 | 0.06 | 0.22 | 0.15 | 0.17 | 0.1 |
| 2 | 106338654 | rs4851089 | -1.077 | 0.708 | -0.724 | 0.519 | 0.028 | 0.059 | 0.059 | 0 | 0.26 | 0.53 | 0.36 | 0.38 | 0.32 | 0.32 |
| 2 | 106338685 | rs955799 | 2.123 | 0.684 | -0.998 | 0.46 | -0.04 | 0.129 | 0.129 | 0 | 0.35 | 0.08 | 0.21 | 0.14 | 0.34 | 0.25 |
| 2 | 106340164 | rs10496401 | 0.236 | 1.285 | -0.371 | 0.921 | -0.004 | -0.021 | -0.021 | 0.1 | 0.15 | 0.1 | 0.22 | 0.16 | 0.17 | 0.1 |
| 2 | 106340814 | rs10514814 | -0.558 | 1.294 | -0.37 | 0.93 | -0.019 | -0.046 | -0.046 | 0 | 0.19 | 0.11 | 0.27 | 0.27 | 0.22 | 0.2 |
| 2 | 106343836 | rs4556998 | -0.689 | 1.127 | 0.05 | 0.881 | -0.004 | 0.019 | 0.019 | 0.041 | 0.19 | 0.46 | 0.25 | 0.16 | 0.19 | 0.1 |
| 2 | 106345945 | rs17265921 | 0.111 | 1.126 | -0.137 | 0.882 | -0.002 | -0.018 | -0.018 | 0 | 0.16 | 0.11 | 0.22 | 0.16 | 0.17 | 0.1 |
| 2 | 106346234 | rs11899929 | -0.875 | 1.169 | -0.368 | 0.912 | 0.018 | 0.045 | 0.045 | 0 | 0.22 | 0.56 | 0.31 | 0.28 | 0.27 | 0.21 |
| 2 | 106355247 | rs2011352 | 0.321 | 1.363 | -0.613 | 0.741 | 0.024 | -0.004 | -0.004 | 0 | 0.88 | 0.78 | 0.89 | 0.88 | 0.87 | 0.73 |
| 2 | 106358929 | rs4851837 | 1.132 | 1.281 | -0.809 | 0.79 | 0.002 | -0.001 | -0.001 | 1.882 | 0.68 | 0.32 | 0.62 | 0.69 | 0.7 | 0.63 |
| 2 | 106360675 | rs12992944 | 1.964 | 1.29 | -1.156 | 0.733 | -0.045 | 0.143 | 0.143 | 0 | 0.42 | 0.06 | 0.23 | 0.21 | 0.37 | 0.34 |
| 2 | 106366022 | rs997136 | 1.277 | 1.288 | -1.033 | 0.771 | -0.036 | -0.009 | -0.009 | 0.256 | 0.66 | 0.16 | 0.55 | 0.67 | 0.61 | 0.62 |
| 2 | 106372358 | rs13017109 | 1.813 | 1.401 | -1.307 | 0.542 | -0.02 | 0.135 | 0.135 | 0 | 0.41 | 0.05 | 0.2 | 0.2 | 0.34 | 0.32 |
| 2 | 106372671 | rs12712216 | -0.228 | 1.504 | -1.031 | 0.556 | 0.001 | -0.014 | -0.014 | 0 | 0.91 | 0.9 | 0.92 | 0.99 | 0.9 | 0.87 |
| 2 | 106376569 | rs1114882 | -1.229 | 1.367 | -0.75 | 0.647 | -0.003 | 0.019 | 0.019 | 0.009 | 0.46 | 0.61 | 0.67 | 0.68 | 0.52 | 0.42 |
| 2 | 106378341 | rs4851843 | -0.14 | 1.574 | -0.907 | 1.007 | -0.038 | -0.089 | -0.089 | 0.832 | 0.27 | 0.14 | 0.38 | 0.48 | 0.34 | 0.31 |
| 2 | 106380105 | rs3769485 | -0.485 | 1.458 | -1.337 | 0.984 | -0.115 | -0.162 | -0.162 | 0.107 | 0.33 | 0.04 | 0.39 | 0.46 | 0.37 | 0.34 |
| 2 | 106381443 | rs11691815 | -0.632 | 1.578 | -1.294 | 0.949 | -0.095 | -0.115 | -0.115 | 0 | 0.34 | 0.09 | 0.41 | 0.47 | 0.41 | 0.35 |
| 2 | 106386294 | rs10201738 | 1.247 | 1.913 | -1.142 | 0.909 | 0.032 | 0.007 | 0.007 | 0.237 | 0.54 | 0.68 | 0.47 | 0.41 | 0.42 | 0.4 |
| 2 | 106391338 | rs4851864 | 0.467 | 2.032 | -1.071 | 0.504 | 0.061 | 0 | 0 | 0.473 | 0.88 | 0.77 | 0.88 | 0.88 | 0.84 | 0.74 |
| 2 | 106393694 | rs12475901 | 0.889 | 2.062 | -1.328 | 0.641 | -0.05 | -0.035 | -0.035 | 0 | 0.17 | 0.01 | 0.27 | 0.18 | 0.18 | 0.09 |
| 2 | 106397496 | rs10196773 | -0.19 | 2.152 | -1.141 | 0.652 | 0.012 | -0.013 | -0.013 | 0 | 0.91 | 0.91 | 0.91 | 0.99 | 0.86 | 0.85 |
| 2 | 106401975 | rs6543336 | 0.459 | 2.092 | -0.963 | 0.682 | 0.068 | 0.002 | 0.002 | 0.645 | 0.88 | 0.74 | 0.88 | 0.88 | 0.85 | 0.79 |
| 2 | 106402795 | rs7602442 | 1.244 | 1.977 | -1.217 | 0.86 | 0.045 | 0.004 | 0.004 | 0.961 | 0.51 | 0.69 | 0.5 | 0.49 | 0.45 | 0.41 |
| 2 | 106405502 | rs13401952 | -0.709 | 1.744 | -1.022 | 0.84 | -0.017 | 0.142 | 0.142 | 0 | 0.14 | 0.49 | 0.28 | 0.21 | 0.14 | 0.09 |

| | | | | | | | | | | | | | | | | |
|---|---|---|---|---|---|---|---|---|---|---|---|---|---|---|---|---|
| 2 | 106410049 | rs6706946 | -0.616 | 1.873 | -0.954 | 0.859 | -0.009 | 0.148 | 0.148 | 0.021 | 0.13 | 0.53 | 0.28 | 0.22 | 0.14 | 0.09 |
| 2 | 106412191 | rs9646959 | 2.496 | 1.844 | -0.988 | 0.826 | 0.052 | 0.024 | 0.024 | 2.521 | 0.36 | 0.11 | 0.19 | 0.28 | 0.27 | 0.32 |
| 2 | 106413031 | rs17031793 | -0.282 | 1.958 | -1.02 | 0.909 | -0.044 | -0.035 | -0.035 | 0.431 | 0.13 | 0.03 | 0.09 | 0.17 | 0.17 | 0.26 |
| 2 | 106413077 | rs2047699 | -0.973 | 1.864 | -0.891 | 0.625 | 0 | 0.088 | 0.088 | 0.016 | 0.14 | 0.6 | 0.32 | 0.22 | 0.18 | 0.09 |
| 2 | 106415229 | rs17208505 | -0.549 | 1.856 | -0.946 | 0.633 | -0.125 | -0.07 | -0.07 | 1.375 | 0.38 | 0.05 | 0.38 | 0.38 | 0.4 | 0.38 |
| 2 | 106415679 | rs13392889 | -0.608 | 1.621 | -0.866 | 0.651 | -0.007 | 0.152 | 0.152 | 0 | 0.13 | 0.52 | 0.28 | 0.22 | 0.14 | 0.08 |
| 2 | 106419622 | rs10496403 | 1.919 | 1.613 | -1.381 | 0.605 | 0.054 | 0.026 | 0.026 | 2.274 | 0.37 | 0.11 | 0.19 | 0.27 | 0.27 | 0.32 |
| 2 | 106425713 | rs6707820 | -0.691 | 1.367 | -1.044 | 0.546 | -0.012 | 0.08 | 0.08 | 0 | 0.16 | 0.57 | 0.27 | 0.23 | 0.15 | 0.1 |
| 2 | 106428720 | rs12053259 | 1.671 | 1.39 | -1.31 | 0.539 | 0.009 | -0.006 | -0.006 | 0 | 0.39 | 0.08 | 0.34 | 0.33 | 0.36 | 0.37 |
| 2 | 106429319 | rs1549769 | 0.713 | 1.538 | -1.299 | 0.533 | 0.042 | 0.005 | 0.005 | 0.307 | 0.89 | 0.82 | 0.89 | 0.87 | 0.85 | 0.84 |
| 2 | 106433478 | rs7589342 | 1.747 | 1.465 | -1.34 | 0.502 | 0.014 | -0.006 | -0.006 | 0 | 0.39 | 0.08 | 0.34 | 0.33 | 0.36 | 0.37 |
| 2 | 106438936 | rs1465641 | 1.679 | 1.667 | -1.204 | 0.589 | 0.019 | 0.029 | 0.029 | 1.69 | 0.41 | 0.18 | 0.36 | 0.34 | 0.4 | 0.38 |
| 2 | 106439181 | rs1465639 | 1.679 | 1.667 | -1.204 | 0.589 | 0.019 | 0.029 | 0.029 | 1.69 | 0.41 | 0.18 | 0.36 | 0.34 | 0.4 | 0.38 |
| 2 | 106443111 | rs12995849 | 0.505 | 1.816 | -1.248 | 0.569 | -0.057 | -0.027 | -0.027 | 0 | 0.62 | 0.85 | 0.68 | 0.55 | 0.59 | 0.5 |
| 2 | 106444843 | rs6749913 | 0.896 | 1.873 | -1.025 | 0.671 | 0.026 | 0.002 | 0.002 | 0.276 | 0.9 | 0.87 | 0.88 | 0.87 | 0.85 | 0.84 |
| 2 | 106457648 | rs2163349 | -0.639 | 1.776 | -0.865 | 0.697 | -0.011 | 0.023 | 0.023 | 0.002 | 0.21 | 0.58 | 0.33 | 0.2 | 0.23 | 0.13 |
| 2 | 106460814 | rs4627591 | 0.304 | 1.799 | -1.003 | 0.627 | -0.01 | -0.032 | -0.032 | 0 | 0.25 | 0.09 | 0.22 | 0.31 | 0.22 | 0.27 |
| 2 | 106462205 | rs4851878 | 0.346 | 1.89 | -0.956 | 0.671 | -0.013 | -0.034 | -0.034 | 0 | 0.25 | 0.09 | 0.22 | 0.31 | 0.22 | 0.25 |
| 2 | 106462256 | rs6543342 | 0.05 | 1.889 | -0.984 | 0.662 | -0.041 | -0.009 | -0.009 | 0 | 0.71 | 0.82 | 0.73 | 0.67 | 0.67 | 0.68 |
| 2 | 106465351 | rs13396833 | -0.838 | 1.868 | -0.897 | 0.715 | -0.005 | 0.04 | 0.04 | 0 | 0.23 | 0.6 | 0.33 | 0.29 | 0.26 | 0.17 |
| 2 | 106468039 | rs6745887 | 1.765 | 1.704 | -0.993 | 0.667 | 0.047 | 0.029 | 0.029 | 1.504 | 0.42 | 0.18 | 0.34 | 0.36 | 0.36 | 0.42 |
| 2 | 106477092 | rs3769504 | -1.808 | 1.576 | -1.105 | 0.553 | -0.006 | 0.01 | 0.01 | 0 | 0.58 | 0.66 | 0.62 | 0.63 | 0.58 | 0.52 |
| 2 | 106479284 | rs17031883 | -0.419 | 1.434 | -1.064 | 0.538 | 0 | 0.012 | 0.012 | 0 | 0.12 | 0.08 | 0.21 | 0.27 | 0.1 | 0.13 |
| 2 | 106487382 | rs3754806 | 0.557 | 1.473 | -1.36 | 0.593 | -0.078 | -0.095 | -0.095 | 0 | 0.24 | 0.01 | 0.21 | 0.31 | 0.22 | 0.25 |
| 2 | 106490021 | rs3769497 | -0.48 | 1.569 | -1.102 | 0.738 | -0.001 | -0.005 | -0.005 | 0.001 | 0.05 | 0.05 | 0.05 | 0.03 | 0.04 | 0.09 |
| 2 | 106495876 | rs3769496 | 0.497 | 1.565 | -1.05 | 0.793 | 0.007 | 0 | 0 | 0 | 0.07 | 0.04 | 0.17 | 0.16 | 0.05 | 0.08 |
| 2 | 106505478 | rs6740343 | 0.906 | 1.653 | -0.97 | 0.729 | 0.022 | 0.024 | 0.024 | 0.051 | 0.75 | 0.4 | 0.72 | 0.7 | 0.68 | 0.78 |
| 2 | 106509907 | rs10469862 | 1.576 | 1.203 | -1.024 | 0.37 | 0.009 | -0.009 | -0.009 | 0.152 | 0.37 | 0.13 | 0.34 | 0.37 | 0.35 | 0.4 |
| 2 | 106511846 | rs933793 | 1.267 | 1.196 | -0.975 | 0.447 | 0.009 | 0.014 | 0.014 | 0.005 | 0.4 | 0.19 | 0.36 | 0.38 | 0.39 | 0.43 |
| 2 | 106533298 | rs1477516 | -0.909 | 0.971 | -0.6 | 0.662 | 0.009 | -0.005 | -0.005 | 0.168 | 0.24 | 0.47 | 0.26 | 0.2 | 0.3 | 0.21 |
| 2 | 106535038 | rs17031993 | -0.871 | 1.002 | -1.045 | 0.677 | -0.001 | -0.008 | -0.008 | 0 | 0.25 | 0.57 | 0.27 | 0.2 | 0.31 | 0.21 |
| 2 | 106535569 | rs12232973 | 0.069 | 0.85 | -1.576 | 0.852 | 0.01 | -0.011 | -0.011 | 0.98 | 0.08 | 0.04 | 0.07 | 0.11 | 0.04 | 0.06 |
| 2 | 106536134 | rs11895604 | 0.506 | 1.151 | -1.594 | 0.768 | -0.034 | 0.004 | 0.004 | 0.183 | 0.39 | 0.19 | 0.35 | 0.38 | 0.39 | 0.45 |
| 2 | 106539419 | rs6736352 | -1.015 | 1.274 | -0.763 | 0.742 | 0.008 | -0.008 | -0.008 | 0 | 0.18 | 0.24 | 0.14 | 0.11 | 0.2 | 0.11 |
| 2 | 106541229 | rs2098836 | 0.67 | 0.707 | -0.878 | 1.01 | -0.014 | 0.018 | 0.018 | 0.194 | 0.75 | 0.7 | 0.89 | 0.67 | 0.82 | 0.8 |
| 2 | 106550301 | rs17721404 | -0.342 | 1.389 | -1.879 | 0.741 | -0.142 | -0.068 | -0.068 | 0.021 | 0.3 | 0.05 | 0.3 | 0.33 | 0.44 | 0.38 |
| 2 | 106551158 | rs6748289 | -0.681 | 1.073 | -1.946 | 0.768 | 0.008 | -0.042 | -0.042 | 0.173 | 0.42 | 0.19 | 0.56 | 0.44 | 0.56 | 0.47 |
| 2 | 106554990 | rs2889606 | -0.676 | 0.596 | -1.331 | 0.513 | -0.005 | -0.009 | -0.009 | 0 | 0.55 | 0.48 | 0.73 | 0.67 | 0.64 | 0.58 |
| 2 | 106555857 | rs17665024 | 0.316 | 0.91 | -1.547 | 0.474 | -0.039 | -0.015 | -0.015 | 0 | 0.17 | 0.03 | 0.28 | 0.12 | 0.19 | 0.09 |
| 2 | 106557690 | rs4851100 | 0.105 | 0.157 | -1.192 | 0.016 | 0.002 | 0.029 | 0.029 | 0 | 0.6 | 0.68 | 0.59 | 0.78 | 0.63 | 0.69 |
| 2 | 106558545 | rs10490697 | 1.593 | 0.044 | -1.066 | 0.025 | 0.002 | 0.01 | 0.01 | 0 | 0.16 | 0.1 | 0.08 | 0.1 | 0.14 | 0.2 |
| 2 | 106568457 | rs1079039 | 0.136 | -0.19 | -0.734 | -0.032 | -0.014 | -0.002 | -0.002 | 0 | 0.39 | 0.52 | 0.22 | 0.41 | 0.25 | 0.38 |
| 2 | 106569996 | rs4851885 | -0.085 | -0.235 | -0.73 | -0.081 | 0.029 | 0.013 | 0.013 | 0 | 0.55 | 0.41 | 0.75 | 0.59 | 0.7 | 0.59 |
| 2 | 106571717 | rs3849370 | 1.404 | 0.649 | -0.929 | -0.038 | -0.005 | -0.004 | -0.004 | 0 | 0.1 | 0.07 | 0.22 | 0.21 | 0.15 | 0.17 |
| 2 | 106572176 | rs2567195 | 0.237 | 0.51 | -0.448 | -0.051 | -0.001 | 0.007 | 0.007 | 0 | 0.35 | 0.59 | 0.41 | 0.43 | 0.33 | 0.36 |
| 2 | 106574745 | rs17032053 | 1.472 | 0.391 | -0.512 | -0.064 | -0.008 | -0.007 | -0.007 | 0.213 | 0.22 | 0.28 | 0.16 | 0.22 | 0.18 | 0.19 |
| 2 | 106579824 | rs13427564 | -0.057 | 0.636 | -0.826 | -0.216 | -0.055 | 0.002 | 0.002 | 0 | 0.3 | 0.19 | 0.52 | 0.27 | 0.41 | 0.27 |
| 2 | 106579871 | rs17032063 | 1.862 | 0.372 | -0.759 | -0.076 | 0.018 | -0.01 | -0.01 | 0 | 0.15 | 0.09 | 0.06 | 0.19 | 0.08 | 0.19 |
| 2 | 106582663 | rs1863216 | 0.148 | 0.419 | -0.367 | -0.151 | -0.002 | -0.002 | -0.002 | 0 | 0.47 | 0.53 | 0.56 | 0.62 | 0.41 | 0.51 |
| 2 | 106583817 | rs7605126 | 1.047 | 0.573 | -0.342 | -0.207 | 0.002 | -0.024 | -0.024 | 0 | 0.29 | 0.39 | 0.35 | 0.21 | 0.28 | 0.22 |
| 2 | 106584554 | rs12997193 | -0.07 | 0.507 | -0.594 | -0.364 | 0.038 | -0.11 | -0.11 | 0 | 0.82 | 0.56 | 0.86 | 0.98 | 0.78 | 0.9 |
| 2 | 106586411 | rs11675026 | 0.878 | 0.487 | -1.275 | -0.191 | 0.019 | -0.009 | -0.009 | 0 | 0.24 | 0.08 | 0.18 | 0.22 | 0.17 | 0.22 |
| 2 | 106587548 | rs1111513 | -0.719 | 0.686 | -0.989 | 0.073 | -0.027 | -0.046 | -0.046 | 0.011 | 0.7 | 0.91 | 0.78 | 0.67 | 0.68 | 0.66 |
| 2 | 106588730 | rs6726474 | -0.599 | 0.877 | -0.162 | 0.117 | 0.029 | 0.009 | 0.009 | 0 | 0.26 | 0.42 | 0.37 | 0.26 | 0.3 | 0.24 |
| 2 | 106592564 | rs6742317 | -0.03 | 0.015 | 0.102 | 0.311 | -0.021 | 0.109 | 0.109 | 0 | 0.77 | 0.43 | 0.59 | 0.45 | 0.79 | 0.72 |
| 2 | 106594966 | rs6543356 | -0.647 | -0.008 | 0.789 | 0.026 | -0.011 | 0.021 | 0.021 | 0 | 0.83 | 0.74 | 0.73 | 0.74 | 0.84 | 0.85 |
| 2 | 106606105 | rs3860448 | 0.791 | -0.404 | 0.935 | -0.114 | -0.02 | 0.026 | 0.026 | 0 | 0.16 | 0.22 | 0.24 | 0.26 | 0.12 | 0.13 |
| 2 | 106610454 | rs1647646 | -0.192 | -0.62 | 0.19 | -0.724 | -0.02 | -0.024 | -0.024 | 0 | 0.4 | 0.5 | 0.37 | 0.69 | 0.31 | 0.54 |
| 2 | 106615576 | rs1869070 | 0.58 | -0.834 | 0.159 | -0.837 | 0.018 | -0.009 | -0.009 | 0.007 | 0.29 | 0.2 | 0.28 | 0.67 | 0.25 | 0.52 |
| 2 | 106620083 | rs1669518 | -1.202 | -0.538 | -0.12 | -1.18 | 0 | 0.001 | 0.001 | 0 | 0.63 | 0.66 | 0.54 | 0.7 | 0.61 | 0.49 |
| 2 | 106621670 | rs6725326 | 0.815 | -0.535 | -0.151 | -1.171 | 0.015 | -0.004 | -0.004 | 0 | 0.3 | 0.23 | 0.28 | 0.67 | 0.25 | 0.52 |
| 2 | 106626988 | rs2052672 | 0.877 | -0.452 | -0.327 | -1.112 | -0.047 | 0.061 | 0.061 | 0 | 0.22 | 0.09 | 0.18 | 0.01 | 0.3 | 0.11 |
| 2 | 106627281 | rs1669492 | -1.132 | -0.424 | -0.348 | -1.087 | 0 | 0.006 | 0.006 | 0 | 0.63 | 0.69 | 0.55 | 0.7 | 0.61 | 0.66 |
| 2 | 106632393 | rs1647681 | 0.039 | 0.348 | -0.312 | -1.185 | 0.011 | -0.084 | -0.084 | 0.212 | 0.25 | 0.27 | 0.19 | 0.01 | 0.31 | 0.11 |
| 2 | 106636741 | rs10490701 | -0.21 | 0.344 | -0.946 | -1.087 | -0.074 | 0.12 | 0.12 | 0 | 0.28 | 0.09 | 0.23 | 0.08 | 0.37 | 0.11 |
| 2 | 106637411 | rs41386450 | 0.468 | 0.07 | -1.182 | -1.04 | 0.032 | 0.027 | 0.027 | 0 | 0.3 | 0.19 | 0.12 | 0.18 | 0.2 | 0.25 |
| 2 | 106637427 | rs10182846 | -1.172 | 0.05 | -0.768 | -1.047 | 0 | -0.011 | -0.011 | 0 | 0.59 | 0.48 | 0.36 | 0.25 | 0.57 | 0.36 |
| 2 | 106638813 | rs10186668 | -0.385 | 0.53 | -0.303 | -0.668 | 0.01 | -0.06 | -0.06 | 4.184 | 0.3 | 0.32 | 0.24 | 0.08 | 0.37 | 0.11 |
| 2 | 106641803 | rs1647686 | 0.62 | 0.738 | -0.023 | -0.463 | 0.016 | -0.118 | -0.118 | 2.384 | 0.67 | 0.53 | 0.76 | 0.93 | 0.64 | 0.81 |

| | | | | | | | | | | | | | | | | |
|---|---|---|---|---|---|---|---|---|---|---|---|---|---|---|---|---|
| 2 | 106643279 | rs1647689 | 0.255 | 0.5 | -0.018 | -0.367 | 0.006 | -0.148 | -0.148 | 1.481 | 0.74 | 0.56 | 0.79 | 0.97 | 0.73 | 0.84 |
| 2 | 106644662 | rs1647695 | 0.898 | 0.418 | -0.415 | -0.31 | -0.002 | -0.013 | -0.013 | 0 | 0.57 | 0.55 | 0.71 | 0.76 | 0.61 | 0.63 |
| 2 | 106644772 | rs17032192 | 0.927 | 0.318 | -0.506 | -0.311 | 0.015 | -0.061 | -0.061 | 0 | 0.16 | 0.01 | 0.07 | 0.21 | 0.09 | 0.21 |
| 2 | 106649971 | rs2377442 | -1.265 | 0.479 | -0.369 | 0.171 | -0.001 | 0 | 0 | 0 | 0.45 | 0.46 | 0.46 | 0.51 | 0.42 | 0.51 |
| 2 | 106650081 | rs2377443 | 0.021 | 1.113 | -0.272 | 0.566 | 0.02 | 0.015 | 0.015 | 0 | 0.18 | 0.24 | 0.31 | 0.26 | 0.27 | 0.18 |
| 2 | 106653676 | rs1005725 | -0.075 | 0.743 | -0.573 | 0.305 | 0.022 | -0.035 | -0.035 | 6.925 | 0.2 | 0.06 | 0.11 | 0.25 | 0.13 | 0.29 |
| 2 | 106653926 | rs6726744 | 0.303 | 1.172 | 0.039 | 0.064 | 0.034 | -0.057 | -0.057 | 0 | 0.1 | 0.27 | 0.08 | 0.01 | 0.14 | 0.1 |
| 2 | 106654006 | rs6708041 | -0.81 | 1.56 | 0.565 | -0.196 | 0.075 | -0.125 | -0.125 | 0 | 0.2 | 0.46 | 0.16 | 0.04 | 0.27 | 0.15 |
| 2 | 106654062 | rs11901318 | -0.039 | 1.287 | 0.588 | 0.074 | 0.02 | -0.028 | -0.028 | 0 | 0.1 | 0.14 | 0.07 | 0.03 | 0.13 | 0.06 |
| 2 | 106657452 | rs12612811 | 0.137 | 1.668 | 0.29 | 0.494 | -0.022 | -0.044 | -0.044 | 0.155 | 0.13 | 0.07 | 0.34 | 0.46 | 0.19 | 0.22 |
| 2 | 106657799 | rs6543371 | 0.084 | 1.542 | 0.331 | 0.34 | 0.004 | -0.016 | -0.016 | 0 | 0.24 | 0.11 | 0.16 | 0.27 | 0.2 | 0.34 |
| 2 | 106658192 | rs12465304 | -1.634 | 1.346 | 0.578 | 0.102 | 0 | -0.002 | -0.002 | 0 | 0.52 | 0.46 | 0.59 | 0.77 | 0.53 | 0.65 |
| 2 | 106660701 | rs2377445 | 0.812 | 1.808 | 1.175 | 0.447 | 0.021 | 0.064 | 0.064 | 0 | 0.78 | 0.63 | 0.56 | 0.47 | 0.72 | 0.66 |
| 2 | 106665898 | rs11893869 | 0.341 | 1.834 | 1.146 | 0.417 | -0.001 | -0.012 | -0.012 | 0 | 0.13 | 0.14 | 0.08 | 0.03 | 0.13 | 0.06 |
| 2 | 106672133 | rs732081 | 1.336 | 1.828 | 1.146 | 0.479 | 0.016 | 0.008 | 0.008 | 1.412 | 0.64 | 0.41 | 0.47 | 0.43 | 0.58 | 0.56 |
| 2 | 106672429 | rs1869060 | 0.614 | 1.862 | 1.206 | 0.467 | 0.018 | 0.047 | 0.047 | 1.106 | 0.74 | 0.61 | 0.57 | 0.48 | 0.68 | 0.72 |
| 2 | 106673532 | rs17032235 | 0.92 | 1.822 | 1.15 | 0.2 | 0.009 | 0.013 | 0.013 | 0.302 | 0.7 | 0.61 | 0.46 | 0.43 | 0.65 | 0.7 |
| 2 | 106674092 | rs7595084 | 1.425 | 1.881 | 1.211 | 0.216 | 0.006 | -0.001 | -0.001 | 0.045 | 0.62 | 0.4 | 0.36 | 0.41 | 0.55 | 0.55 |
| 2 | 106674925 | rs17032247 | -0.632 | 1.883 | 1.097 | 0.225 | 0.042 | -0.042 | -0.042 | 0.345 | 0.09 | 0.21 | 0.1 | 0.02 | 0.1 | 0.15 |
| 2 | 106676043 | rs17032252 | -0.632 | 1.883 | 1.046 | 0.225 | 0.042 | -0.042 | -0.042 | 0 | 0.09 | 0.22 | 0.1 | 0.02 | 0.1 | 0.15 |
| 2 | 106676525 | rs873766 | 0.343 | 1.981 | 0.618 | 0.26 | 0.018 | -0.021 | -0.021 | 0 | 0.07 | 0.09 | 0.1 | 0.02 | 0.09 | 0.12 |
| 2 | 106678223 | rs4255978 | 1.074 | 1.598 | 0.483 | 0.513 | 0.015 | 0.02 | 0.02 | 0.186 | 0.71 | 0.59 | 0.46 | 0.43 | 0.65 | 0.7 |
| 2 | 106679037 | rs13386274 | 0.272 | 1.424 | 0.132 | 0.029 | 0.003 | -0.04 | -0.04 | 2.386 | 0.84 | 0.75 | 0.88 | 0.97 | 0.83 | 0.9 |
| 2 | 106679941 | rs2377446 | -0.278 | 1.353 | -0.003 | 0.396 | 0.003 | -0.012 | -0.012 | 1.212 | 0.21 | 0.29 | 0.13 | 0.14 | 0.2 | 0.3 |
| 2 | 106682622 | rs4420718 | -0.875 | 1.373 | -0.291 | 0.643 | 0.024 | 0.044 | 0.044 | 0 | 0.34 | 0.56 | 0.51 | 0.5 | 0.4 | 0.52 |
| 2 | 106690137 | rs4851111 | -0.004 | 0.836 | -0.765 | 0.074 | 0 | -0.004 | -0.004 | 0 | 0.44 | 0.4 | 0.39 | 0.62 | 0.44 | 0.62 |
| 2 | 106692818 | rs17271098 | 0.776 | 0.532 | -0.897 | -0.659 | -0.017 | 0.041 | 0.041 | 0 | 0.23 | 0.12 | 0.14 | 0 | 0.22 | 0.14 |
| 2 | 106693142 | rs7584287 | 0.607 | 0.652 | -0.819 | -0.058 | 0.003 | 0.006 | 0.006 | 0 | 0.42 | 0.43 | 0.34 | 0.49 | 0.44 | 0.57 |
| 2 | 106700313 | rs1901460 | 0.043 | 0.101 | -0.781 | -0.087 | -0.001 | 0 | 0 | 0 | 0.69 | 0.65 | 0.61 | 0.6 | 0.72 | 0.77 |
| 2 | 106705507 | rs7587204 | -0.309 | -0.093 | -0.885 | -0.199 | 0 | 0.005 | 0.005 | 0 | 0.79 | 0.78 | 0.64 | 0.6 | 0.77 | 0.78 |
| 2 | 106705747 | rs12621836 | 0.356 | -0.083 | -0.948 | -0.19 | -0.004 | -0.076 | -0.076 | 0 | 0.18 | 0.09 | 0.35 | 0.4 | 0.23 | 0.22 |
| 2 | 106714142 | rs10191679 | -0.238 | -0.076 | -0.948 | -0.189 | 0.005 | -0.022 | -0.022 | 0 | 0.79 | 0.81 | 0.65 | 0.61 | 0.77 | 0.78 |
| 2 | 106714317 | rs17032320 | 0.778 | -0.085 | -0.952 | -0.263 | 0.006 | -0.003 | -0.003 | 0 | 0.09 | 0.06 | 0.03 | 0 | 0.05 | 0.04 |
| 2 | 106721945 | rs725216 | 0.254 | -0.027 | -0.84 | -0.152 | 0 | 0.005 | 0.005 | 0 | 0.76 | 0.72 | 0.63 | 0.55 | 0.74 | 0.73 |
| 2 | 106722953 | rs17032398 | 1.392 | -0.47 | -0.991 | -0.657 | -0.034 | 0.132 | 0.132 | 0 | 0.22 | 0.33 | 0.17 | 0.46 | 0.16 | 0.21 |
| 2 | 106725928 | rs11695995 | -0.049 | -0.348 | -0.57 | -0.434 | 0.003 | 0.206 | 0.206 | 1.194 | 0.92 | 0.74 | 0.74 | 0.64 | 0.91 | 0.8 |
| 2 | 106729992 | rs6543374 | 1.203 | -0.112 | -0.896 | -0.771 | 0 | 0.001 | 0.001 | 0.3 | 0.41 | 0.34 | 0.35 | 0.56 | 0.4 | 0.44 |
| 2 | 106732545 | rs737101 | 0.304 | -0.014 | -0.908 | -0.761 | 0 | 0.001 | 0.001 | 0 | 0.6 | 0.47 | 0.56 | 0.58 | 0.62 | 0.59 |
| 2 | 106736313 | rs11681404 | 0.366 | -0.026 | -0.924 | -0.767 | -0.014 | 0.064 | 0.064 | 0 | 0.3 | 0.19 | 0.17 | 0.06 | 0.29 | 0.21 |
| 2 | 106737335 | rs1869056 | -0.007 | -0.036 | -0.908 | -0.695 | -0.005 | 0.2 | 0.2 | 1.68 | 0.91 | 0.7 | 0.74 | 0.65 | 0.91 | 0.8 |
| 2 | 106742778 | rs17213510 | 1.45 | 0.125 | -1 | -0.737 | -0.028 | 0.068 | 0.068 | 0 | 0.21 | 0.08 | 0.12 | 0 | 0.2 | 0.12 |
| 2 | 106765133 | rs6543378 | 0.571 | -0.009 | -0.938 | -0.679 | 0.001 | -0.003 | -0.003 | 0 | 0.46 | 0.43 | 0.47 | 0.59 | 0.48 | 0.51 |
| 2 | 106766480 | rs11685300 | -0.094 | 0.019 | -0.893 | -0.714 | 0.001 | 0.064 | 0.064 | 0 | 0.43 | 0.32 | 0.28 | 0.02 | 0.41 | 0.23 |
| 2 | 106775223 | rs4012753 | -0.148 | 0.164 | -0.638 | -0.283 | 0.013 | 0.223 | 0.223 | 1.512 | 0.93 | 0.79 | 0.76 | 0.67 | 0.92 | 0.8 |
| 2 | 106780726 | rs7591409 | 0.94 | 0.076 | -0.466 | -0.334 | 0.016 | 0.231 | 0.231 | 1.003 | 0.87 | 0.6 | 0.68 | 0.6 | 0.87 | 0.73 |
| 2 | 106790116 | rs4567955 | 0.028 | 0.007 | -0.603 | -0.646 | -0.005 | 0.063 | 0.063 | 0 | 0.39 | 0.26 | 0.26 | 0.02 | 0.37 | 0.21 |
| 2 | 106823652 | rs13006272 | 0.864 | 0.449 | -0.415 | -0.125 | 0.02 | 0.163 | 0.163 | 0.998 | 0.87 | 0.67 | 0.68 | 0.6 | 0.84 | 0.72 |
| 2 | 106839924 | rs4851946 | 0.561 | 0.153 | -0.45 | -0.422 | 0.011 | 0.003 | 0.003 | 0.224 | 0.49 | 0.31 | 0.39 | 0.51 | 0.45 | 0.53 |
| 2 | 106849382 | rs12712247 | -0.171 | 0.18 | -0.387 | -0.443 | -0.007 | 0.004 | 0.004 | 0 | 0.42 | 0.4 | 0.36 | 0.07 | 0.44 | 0.28 |
| 2 | 106852762 | rs4514893 | 0.491 | 0.224 | -0.483 | -0.383 | 0.014 | 0.004 | 0.004 | 0.165 | 0.49 | 0.32 | 0.39 | 0.51 | 0.44 | 0.53 |
| 2 | 106878354 | rs10187480 | -0.746 | 0.438 | -0.008 | 0.298 | 0.004 | 0.105 | 0.105 | 0 | 0.07 | 0.11 | 0.3 | 0.42 | 0.11 | 0.2 |
| 2 | 106886935 | rs11673736 | -0.342 | 0.469 | 0.059 | 0.094 | 0.017 | 0.03 | 0.03 | 0 | 0.3 | 0.45 | 0.4 | 0.43 | 0.32 | 0.32 |
| 2 | 106887870 | rs7419384 | -0.736 | 0.526 | -0.013 | 0.134 | -0.006 | -0.006 | -0.006 | 0 | 0.35 | 0.35 | 0.4 | 0.42 | 0.35 | 0.36 |
| 2 | 106889381 | rs17032739 | 0.888 | 0.352 | 0.638 | 0.671 | 0.006 | 0.103 | 0.103 | 0.012 | 0.09 | 0.17 | 0.04 | 0.19 | 0.06 | 0.21 |
| 2 | 106891087 | rs7583604 | 0.756 | 0.427 | 0.474 | 0.42 | 0.007 | 0.005 | 0.005 | 0.85 | 0.93 | 0.73 | 0.96 | 0.96 | 0.97 | 0.95 |
| 2 | 106895212 | rs6719028 | -0.65 | 0.474 | 0.287 | 0.297 | -0.001 | -0.082 | -0.082 | 0 | 0.77 | 0.69 | 0.86 | 0.99 | 0.8 | 0.88 |
| 2 | 106897262 | rs11124101 | 0.332 | 0.489 | 0.316 | 0.756 | -0.006 | 0.081 | 0.081 | 0 | 0.05 | 0.06 | 0.19 | 0.43 | 0.1 | 0.2 |
| 2 | 106898105 | rs4449127 | -0.2 | 0.533 | 0.333 | 0.322 | 0.009 | -0.045 | -0.045 | 1.945 | 0.08 | 0.17 | 0.06 | 0 | 0.08 | 0.03 |
| 2 | 106900546 | rs7585986 | -0.244 | 0.832 | 0.336 | 0.246 | 0.003 | -0.114 | -0.114 | 0 | 0.74 | 0.58 | 0.84 | 0.96 | 0.79 | 0.85 |
| 2 | 106901623 | rs11124102 | 0.194 | 0.859 | 0.362 | 0.593 | 0.004 | 0.114 | 0.114 | 0 | 0.06 | 0.11 | 0.16 | 0.47 | 0.1 | 0.2 |
| 2 | 106904475 | rs13405655 | 0.336 | 0.72 | 0.401 | 0.041 | 0.005 | -0.11 | -0.11 | 0 | 0.67 | 0.54 | 0.74 | 0.95 | 0.7 | 0.81 |
| 2 | 106907544 | rs4234111 | 0.365 | 0.714 | 0.377 | 0.036 | 0.001 | -0.113 | -0.113 | 0.006 | 0.65 | 0.41 | 0.73 | 0.95 | 0.7 | 0.81 |
| 2 | 106909658 | rs894391 | -0.108 | 0.723 | 0.388 | 0.213 | 0.004 | 0.089 | 0.089 | 0 | 0.14 | 0.22 | 0.2 | 0.6 | 0.15 | 0.23 |
| 2 | 106912841 | rs13395525 | 0.877 | 0.628 | 0.305 | 0.067 | 0.003 | 0.011 | 0.011 | 0 | 0.51 | 0.4 | 0.49 | 0.35 | 0.49 | 0.56 |
| 2 | 106930993 | rs4643539 | 0.72 | 0.32 | 0.274 | 0.086 | 0.001 | 0.012 | 0.012 | 0 | 0.52 | 0.41 | 0.57 | 0.35 | 0.55 | 0.57 |
| 2 | 106934129 | rs4308118 | 0.922 | 0.325 | 0.246 | 0.12 | 0.006 | 0.044 | 0.044 | 0 | 0.5 | 0.32 | 0.57 | 0.31 | 0.54 | 0.57 |
| 2 | 106938041 | rs13429003 | -0.24 | 0.069 | 0.777 | -0.141 | -0.001 | -0.084 | -0.084 | 0 | 0.79 | 0.65 | 0.85 | 0.95 | 0.82 | 0.87 |
| 2 | 106945278 | rs6543406 | 0.076 | 0.027 | 0.721 | -0.128 | -0.016 | -0.072 | -0.072 | 0 | 0.76 | 0.62 | 0.82 | 0.95 | 0.81 | 0.86 |
| 2 | 106949542 | rs4234115 | 0.073 | 0.026 | 0.515 | -0.111 | -0.008 | -0.026 | -0.026 | 0 | 0.76 | 0.67 | 0.81 | 0.91 | 0.79 | 0.85 |

| | | | | | | | | | | | | | | | | |
|---|---|---|---|---|---|---|---|---|---|---|---|---|---|---|---|---|
| 2 | 106951208 | rs11677201 | 0.384 | 0.048 | 0.06 | -0.101 | -0.021 | -0.046 | -0.046 | 0 | 0.72 | 0.38 | 0.79 | 0.91 | 0.79 | 0.85 |
| 2 | 106954287 | rs4441466 | 0.21 | 0.01 | 0.114 | -0.039 | -0.031 | -0.055 | -0.055 | 0.041 | 0.74 | 0.4 | 0.8 | 0.91 | 0.81 | 0.88 |
| 2 | 106965073 | rs6543408 | -0.382 | -0.037 | -0.478 | 0.056 | 0.014 | 0.049 | 0.049 | 0 | 0.75 | 0.86 | 0.81 | 0.87 | 0.83 | 0.88 |
| 2 | 106974619 | rs4234103 | 0.901 | -0.018 | -0.622 | 0.403 | 0.04 | 0.009 | 0.009 | 0.114 | 0.54 | 0.21 | 0.64 | 0.49 | 0.66 | 0.69 |
| 2 | 106983692 | rs13413684 | -0.535 | 0.257 | -0.087 | 0.596 | -0.002 | 0.232 | 0.232 | 0 | 0.18 | 0.44 | 0.14 | 0.38 | 0.15 | 0.19 |
| 2 | 106990423 | rs13416767 | 0.193 | 0.506 | -0.491 | 0.1 | 0 | -0.001 | -0.001 | 0.419 | 0.61 | 0.45 | 0.68 | 0.59 | 0.68 | 0.71 |
| 2 | 107091864 | rs12471101 | -0.438 | 1.289 | -0.682 | -0.469 | -0.013 | 0.012 | 0.012 | 0.187 | 0.11 | 0.09 | 0.1 | 0.05 | 0.2 | 0.17 |
| 2 | 107102418 | rs4362555 | -0.35 | 0.998 | -0.58 | -1.221 | -0.004 | -0.159 | -0.159 | 0 | 0.62 | 0.5 | 0.73 | 0.99 | 0.65 | 0.89 |
| 2 | 107135170 | rs7575343 | 0.294 | 0.961 | -0.718 | -1.003 | 0.001 | 0.003 | 0.003 | 0 | 0.48 | 0.48 | 0.52 | 0.76 | 0.45 | 0.43 |
| 2 | 107142845 | rs4676222 | -0.4 | 0.701 | -0.215 | -1.304 | -0.036 | -0.117 | -0.117 | 0 | 0.83 | 0.61 | 0.9 | 1 | 0.9 | 0.93 |
| 2 | 107142890 | rs4455150 | -0.076 | 0.674 | -0.077 | -1.321 | 0.034 | -0.02 | -0.02 | 0.807 | 0.07 | 0.26 | 0.05 | 0 | 0.04 | 0.02 |
| 2 | 107151134 | rs4281904 | 0.655 | 1.194 | -0.187 | -1.336 | 0.002 | 0.005 | 0.005 | 0.482 | 0.51 | 0.46 | 0.51 | 0.14 | 0.54 | 0.38 |
| 2 | 107152142 | rs4234108 | 0.509 | 1.432 | -0.404 | -0.855 | -0.006 | 0.012 | 0.012 | 0.298 | 0.12 | 0.1 | 0.23 | 0.58 | 0.17 | 0.32 |
| 2 | 107153049 | rs6733787 | 1.806 | 1.334 | -0.423 | -0.966 | 0.019 | 0.052 | 0.052 | 0 | 0.09 | 0.01 | 0.05 | 0 | 0.06 | 0.04 |
| 2 | 107153121 | rs6746717 | 1.806 | 1.334 | -0.423 | -0.966 | 0.019 | 0.052 | 0.052 | 0 | 0.09 | 0.01 | 0.05 | 0 | 0.06 | 0.04 |
| 2 | 107154331 | rs6729848 | 0.815 | 1.361 | -0.272 | -0.99 | -0.001 | -0.03 | -0.03 | 0 | 0.52 | 0.5 | 0.5 | 0.12 | 0.54 | 0.38 |
| 2 | 107161001 | rs4503985 | 0.261 | 1.426 | -0.21 | -0.972 | 0.002 | -0.033 | -0.033 | 0 | 0.62 | 0.6 | 0.57 | 0.15 | 0.6 | 0.41 |
| 2 | 107166211 | rs6743616 | -1.231 | 1.902 | -0.105 | -1.094 | 0.011 | 0.019 | 0.019 | 0.143 | 0.56 | 0.57 | 0.58 | 0.89 | 0.62 | 0.77 |
| 2 | 107175283 | rs11904823 | -0.41 | 2.106 | -0.234 | -1.32 | -0.024 | 0.041 | 0.041 | 0 | 0.37 | 0.25 | 0.45 | 0.87 | 0.48 | 0.7 |
| 2 | 107176040 | rs4381779 | -0.208 | 2.102 | 0.007 | -0.923 | 0.034 | -0.031 | -0.031 | 0 | 0.91 | 0.83 | 0.89 | 0.96 | 0.84 | 0.84 |
| 2 | 107177898 | rs11890579 | 0.341 | 2.043 | -0.362 | -0.722 | -0.007 | 0.008 | 0.008 | 0.491 | 0.12 | 0.1 | 0.22 | 0.55 | 0.17 | 0.31 |
| 2 | 107180865 | rs6722500 | 0.549 | 1.792 | -0.435 | -0.615 | -0.009 | 0.004 | 0.004 | 0.56 | 0.59 | 0.38 | 0.54 | 0.39 | 0.53 | 0.45 |
| 2 | 107199075 | rs2125904 | -0.535 | 1.581 | -0.774 | -0.763 | -0.001 | 0.012 | 0.012 | 0.16 | 0.48 | 0.43 | 0.52 | 0.7 | 0.44 | 0.61 |
| 2 | 107199518 | rs968914 | -1.256 | 1.703 | -0.633 | -0.846 | 0.014 | -0.027 | -0.027 | 0 | 0.95 | 0.89 | 0.95 | 1 | 0.94 | 0.99 |
| 2 | 107203992 | rs2035030 | 0.62 | 1.672 | -0.561 | -0.834 | 0.003 | 0.015 | 0.015 | 0.218 | 0.46 | 0.41 | 0.43 | 0.3 | 0.49 | 0.37 |
| 2 | 107204110 | rs13028368 | 0.342 | 1.81 | -0.566 | -0.883 | -0.009 | -0.027 | -0.027 | 0 | 0.23 | 0.3 | 0.25 | 0.15 | 0.17 | 0.24 |
| 2 | 107218831 | rs1117160 | 0.484 | 1.883 | -0.591 | -0.936 | -0.018 | -0.028 | -0.028 | 0.362 | 0.22 | 0.3 | 0.2 | 0.15 | 0.15 | 0.24 |
| 2 | 107225761 | rs12998193 | 0.518 | 1.884 | -0.61 | -0.936 | -0.018 | -0.028 | -0.028 | 0 | 0.22 | 0.3 | 0.2 | 0.15 | 0.15 | 0.24 |
| 2 | 107230276 | rs956354 | 1.556 | 1.631 | -0.773 | -1.139 | 0.013 | -0.028 | -0.028 | 0 | 0.15 | 0.07 | 0.1 | 0.25 | 0.12 | 0.2 |
| 2 | 107231903 | rs17217480 | 2.153 | 1.749 | -0.726 | -1.252 | 0.011 | 0.052 | 0.052 | 1.62 | 0.24 | 0.16 | 0.24 | 0.02 | 0.26 | 0.1 |
| 2 | 107234192 | rs2125905 | 0.199 | 1.837 | -0.646 | -1.191 | -0.007 | 0.055 | 0.055 | 0 | 0.53 | 0.64 | 0.53 | 0.21 | 0.53 | 0.41 |
| 2 | 107242240 | rs10203595 | 1.448 | 1.781 | -0.639 | -1.191 | -0.001 | -0.061 | -0.061 | 0.006 | 0.3 | 0.31 | 0.27 | 0.02 | 0.32 | 0.12 |
| 2 | 107248302 | rs6725607 | -0.939 | 1.798 | -0.357 | -0.727 | 0.022 | -0.031 | -0.031 | 0 | 0.94 | 0.82 | 0.94 | 1 | 0.94 | 0.99 |
| 2 | 107255552 | rs6704791 | 0.61 | 1.73 | -0.388 | -0.708 | -0.007 | 0.059 | 0.059 | 0 | 0.53 | 0.64 | 0.53 | 0.2 | 0.52 | 0.41 |
| 2 | 107260916 | rs6543429 | 0.054 | 1.852 | -0.312 | -0.597 | -0.009 | -0.014 | -0.014 | 0 | 0.68 | 0.71 | 0.63 | 0.45 | 0.64 | 0.61 |
| 2 | 107261300 | rs6543431 | 0.219 | 1.852 | -0.322 | -0.597 | 0.015 | 0.027 | 0.027 | 0.526 | 0.67 | 0.59 | 0.62 | 0.45 | 0.64 | 0.61 |
| 2 | 107262432 | rs13432204 | -0.388 | 1.795 | -0.236 | -0.865 | 0.007 | -0.053 | -0.053 | 0 | 0.2 | 0.55 | 0.24 | 0.11 | 0.18 | 0.23 |
| 2 | 107269417 | rs6710590 | 1.841 | 1.531 | -0.649 | -1.058 | -0.012 | 0.144 | 0.144 | 0 | 0.3 | 0.1 | 0.24 | 0.03 | 0.34 | 0.1 |
| 2 | 107272996 | rs17285281 | 0.331 | 1.912 | -0.335 | -0.698 | 0.001 | -0.01 | -0.01 | 0 | 0.08 | 0.07 | 0.08 | 0.04 | 0.08 | 0.08 |
| 2 | 107276339 | rs6746723 | 0.453 | 2.147 | -0.431 | -0.488 | -0.018 | 0.014 | 0.014 | 2.294 | 0.09 | 0.03 | 0.22 | 0.56 | 0.15 | 0.28 |
| 2 | 107292475 | rs11695678 | -0.992 | 2.191 | -0.83 | -0.492 | 0.021 | -0.076 | -0.076 | 0 | 0.23 | 0.57 | 0.21 | 0.08 | 0.24 | 0.21 |
| 2 | 107304977 | rs10171234 | 0.298 | 2.13 | -1.174 | -0.311 | 0.002 | -0.011 | -0.011 | 0 | 0.08 | 0.07 | 0.08 | 0.04 | 0.08 | 0.08 |
| 2 | 107312537 | rs11124128 | 0.099 | 2.234 | -1.469 | -0.302 | 0.053 | 0.135 | 0.135 | 2.238 | 0.1 | 0.17 | 0.25 | 0.56 | 0.21 | 0.31 |
| 2 | 107318188 | rs17033164 | 1.119 | 1.718 | -2.088 | -0.474 | 0.006 | -0.018 | -0.018 | 0.9 | 0.06 | 0 | 0.12 | 0.08 | 0.03 | 0.12 |
| 2 | 107319073 | rs13396045 | 0.034 | 1.979 | -1.788 | -0.367 | 0.003 | -0.01 | -0.01 | 0 | 0.08 | 0.09 | 0.08 | 0.04 | 0.08 | 0.08 |
| 2 | 107324349 | rs11686358 | -1.145 | 2.186 | -1.655 | -0.113 | 0.113 | 0.018 | 0.018 | 0 | 0.05 | 0.13 | 0.11 | 0.05 | 0.14 | 0.09 |
| 2 | 107324499 | rs1606266 | 0.761 | 1.908 | -1.979 | -0.244 | -0.027 | 0.018 | 0.018 | 0.049 | 0.39 | 0.17 | 0.43 | 0.66 | 0.46 | 0.4 |
| 2 | 107329883 | rs7562312 | -0.911 | 2.624 | -0.901 | 0.119 | 0.012 | 0.017 | 0.017 | 2.017 | 0.12 | 0.09 | 0.23 | 0.67 | 0.23 | 0.36 |
| 2 | 107332141 | rs7599975 | -1.011 | 2.637 | -0.886 | 0.126 | 0.034 | 0.043 | 0.043 | 1.199 | 0.13 | 0.13 | 0.24 | 0.67 | 0.23 | 0.36 |
| 2 | 107338258 | rs6543438 | -0.566 | 2.46 | -0.882 | 0.034 | 0 | -0.123 | -0.123 | 0 | 0.23 | 0.61 | 0.16 | 0.01 | 0.23 | 0.1 |
| 2 | 107350916 | rs4487086 | 0.305 | 2.461 | -0.811 | 0.326 | -0.011 | 0.013 | 0.013 | 1.152 | 0.08 | 0.03 | 0.15 | 0.63 | 0.11 | 0.29 |
| 2 | 107385240 | rs6721068 | -0.144 | 2.468 | -0.851 | 0.208 | 0.016 | 0.076 | 0.076 | 1.081 | 0.91 | 0.89 | 0.84 | 0.36 | 0.86 | 0.71 |
| 2 | 107399193 | rs1375000 | -0.456 | 2.428 | -0.901 | 0.111 | 0 | -0.102 | -0.102 | 0 | 0.28 | 0.63 | 0.17 | 0.02 | 0.28 | 0.12 |
| 2 | 107399365 | rs2241992 | 0.266 | 2.436 | -0.91 | 0.122 | -0.007 | 0.017 | 0.017 | 1.038 | 0.08 | 0.04 | 0.15 | 0.64 | 0.11 | 0.29 |
| 2 | 107399759 | rs1868255 | -0.658 | 1.986 | -1.021 | 0.251 | 0.048 | -0.005 | -0.005 | 0 | 0.06 | 0.08 | 0.14 | 0.04 | 0.14 | 0.19 |
| 2 | 107400636 | rs1006413 | -0.833 | 0.969 | -1.495 | 0.204 | -0.011 | 0.07 | 0.07 | 0 | 0.43 | 0.74 | 0.46 | 0.7 | 0.53 | 0.6 |
| 2 | 107411525 | rs10182443 | -0.638 | 0.297 | -0.225 | 0.424 | -0.016 | -0.018 | -0.018 | 0 | 0.17 | 0.27 | 0.08 | 0.13 | 0.15 | 0.2 |
| 2 | 107411613 | rs4577275 | -0.187 | -0.421 | 0.423 | -0.164 | -0.027 | -0.06 | -0.06 | 0 | 0.84 | 0.69 | 0.93 | 0.98 | 0.9 | 0.92 |
| 2 | 107412071 | rs10185327 | 0.455 | -0.455 | 0.364 | -0.187 | -0.001 | -0.004 | -0.004 | 0 | 0.11 | 0.07 | 0.04 | 0.04 | 0.06 | 0.09 |
| 2 | 107412293 | rs11891978 | 0.25 | -0.914 | 0.363 | 0.768 | -0.007 | 0.087 | 0.087 | 0.057 | 0.09 | 0.08 | 0.12 | 0.48 | 0.07 | 0.27 |
| 2 | 107415834 | rs17653405 | 0.397 | -0.951 | 0.729 | 0.645 | 0.006 | 0.01 | 0.01 | 0 | 0.13 | 0.09 | 0.05 | 0.02 | 0.1 | 0.08 |
| 2 | 107420011 | rs10202530 | -0.504 | -0.94 | 0.663 | 0.641 | -0.017 | -0.022 | -0.022 | 0 | 0.17 | 0.26 | 0.08 | 0.13 | 0.15 | 0.2 |
| 2 | 107420143 | rs10496407 | 1.623 | -0.789 | -0.045 | -0.266 | -0.058 | 0.145 | 0.145 | 0.53 | 0.38 | 0.22 | 0.33 | 0.04 | 0.46 | 0.13 |
| 2 | 107420409 | rs9308902 | 0.561 | -0.816 | -0.071 | -0.249 | 0.01 | 0.04 | 0.04 | 0.075 | 0.53 | 0.43 | 0.41 | 0.18 | 0.61 | 0.34 |
| 2 | 107423188 | rs1470591 | 1.753 | -0.624 | 0.006 | -0.081 | -0.079 | 0.173 | 0.173 | 0.364 | 0.37 | 0.18 | 0.33 | 0.04 | 0.45 | 0.13 |
| 2 | 107423395 | rs1470590 | 0.587 | -0.553 | 0.157 | -0.004 | 0.003 | 0.012 | 0.012 | 0.107 | 0.56 | 0.45 | 0.47 | 0.17 | 0.61 | 0.34 |
| 2 | 107424428 | rs10427207 | -0.252 | -0.135 | 0.085 | 0.273 | 0.01 | 0.005 | 0.005 | 0.356 | 0.29 | 0.33 | 0.34 | 0.31 | 0.38 | 0.34 |
| 2 | 107425876 | rs10191793 | -0.671 | -0.241 | 0.404 | 0.316 | 0.003 | 0.149 | 0.149 | 2.149 | 0.16 | 0.28 | 0.17 | 0.51 | 0.17 | 0.35 |
| 2 | 107426060 | rs10182025 | 0.202 | -0.243 | 0.413 | 0.344 | -0.011 | 0.115 | 0.115 | 0.696 | 0.1 | 0.12 | 0.13 | 0.5 | 0.08 | 0.3 |

| | | | | | | | | | | | | | | | | |
|---|---|---|---|---|---|---|---|---|---|---|---|---|---|---|---|---|
| 2 | 107431226 | rs10514795 | -0.508 | -0.23 | 0.42 | 0.352 | 0.005 | 0.169 | 0.169 | 1.584 | 0.14 | 0.27 | 0.17 | 0.5 | 0.15 | 0.34 |
| 2 | 107440303 | rs7607147 | 1.034 | -0.22 | 0.325 | 0.284 | 0.022 | -0.028 | -0.028 | 0 | 0.05 | 0.13 | 0.04 | 0 | 0.09 | 0.05 |
| 2 | 107443369 | rs12622433 | 0.224 | -0.347 | 0.505 | 0.361 | -0.02 | 0.214 | 0.214 | 0.089 | 0.1 | 0.2 | 0.14 | 0.5 | 0.06 | 0.27 |
| 2 | 107444176 | rs6543445 | 1.751 | -0.658 | 0.28 | 0.025 | -0.039 | 0.124 | 0.124 | 0.013 | 0.37 | 0.18 | 0.42 | 0.12 | 0.42 | 0.23 |
| 2 | 107445707 | rs1868256 | -0.516 | -0.805 | 0.482 | 0.131 | -0.013 | -0.023 | -0.023 | 1.276 | 0.76 | 0.76 | 0.9 | 0.92 | 0.86 | 0.85 |
| 2 | 107448206 | rs6712885 | -0.534 | -0.808 | 0.615 | 0.135 | -0.005 | -0.01 | -0.01 | 0 | 0.77 | 0.77 | 0.9 | 0.92 | 0.85 | 0.85 |
| 2 | 107450525 | rs12615112 | 0.116 | -0.707 | 0.767 | 0.299 | -0.011 | 0.246 | 0.246 | 0.003 | 0.1 | 0.24 | 0.15 | 0.49 | 0.08 | 0.28 |
| 2 | 107450589 | rs3796111 | -1.148 | -0.723 | 0.69 | 0.273 | -0.008 | 0.224 | 0.224 | 0.429 | 0.19 | 0.41 | 0.3 | 0.57 | 0.18 | 0.36 |
| 2 | 107456723 | rs1979083 | -0.935 | -0.621 | 0.807 | 0.105 | 0.058 | 0.037 | 0.037 | 0.301 | 0.1 | 0.44 | 0.17 | 0.08 | 0.11 | 0.09 |
| 2 | 107459974 | rs3796110 | -0.072 | -0.562 | 0.584 | 0.135 | 0.013 | 0.019 | 0.019 | 0.803 | 0.08 | 0.15 | 0.15 | 0.14 | 0.1 | 0.16 |
| 2 | 107466891 | rs1375005 | -0.602 | -0.84 | 0.528 | -0.029 | -0.002 | -0.014 | -0.014 | 0 | 0.78 | 0.79 | 0.91 | 0.97 | 0.86 | 0.93 |
| 2 | 107471527 | rs17033373 | 0.923 | -0.787 | 0.953 | 0.258 | 0.012 | 0.034 | 0.034 | 0.206 | 0.06 | 0.11 | 0.14 | 0.12 | 0.09 | 0.08 |
| 2 | 107474733 | rs11124137 | 0.414 | -0.924 | 0.947 | 0.454 | -0.07 | 0.042 | 0.042 | 0.01 | 0.68 | 0.55 | 0.77 | 0.51 | 0.82 | 0.66 |
| 2 | 107474931 | rs6543453 | 0.414 | -0.924 | 0.947 | 0.454 | -0.07 | 0.042 | 0.042 | 0.01 | 0.68 | 0.55 | 0.77 | 0.51 | 0.82 | 0.66 |
| 2 | 107477619 | rs11679344 | 0.376 | -0.824 | 0.653 | 0.304 | -0.025 | -0.021 | -0.021 | 0 | 0.23 | 0.14 | 0.21 | 0.3 | 0.27 | 0.37 |
| 2 | 107482017 | rs10865075 | 0.359 | -0.86 | 0.707 | 0.328 | -0.01 | 0.004 | 0.004 | 0 | 0.65 | 0.64 | 0.65 | 0.38 | 0.73 | 0.57 |
| 2 | 107484185 | rs11681787 | 0.663 | -0.834 | 0.257 | 0.343 | 0.04 | -0.013 | -0.013 | 0 | 0.62 | 0.35 | 0.62 | 0.38 | 0.71 | 0.57 |
| 2 | 107490565 | rs11680654 | 0.61 | -0.832 | 0.252 | 0.374 | 0.015 | -0.007 | -0.007 | 0 | 0.62 | 0.4 | 0.63 | 0.4 | 0.71 | 0.57 |
| 2 | 107493123 | rs4676093 | 1.497 | -0.87 | 0.135 | 0.3 | -0.045 | 0.035 | 0.035 | 0 | 0.4 | 0.25 | 0.56 | 0.27 | 0.51 | 0.29 |
| 2 | 107493639 | rs2108208 | 0.374 | -0.672 | 0.151 | 0.246 | -0.019 | -0.032 | -0.032 | 0 | 0.22 | 0.15 | 0.31 | 0.38 | 0.27 | 0.3 |
| 2 | 107497202 | rs6729112 | 1.338 | -0.744 | 0.15 | 0.207 | -0.045 | -0.051 | -0.051 | 0 | 0.46 | 0.21 | 0.54 | 0.53 | 0.53 | 0.48 |
| 2 | 107499041 | rs4676309 | 0.667 | -0.807 | 0.444 | -0.004 | -0.006 | -0.044 | -0.044 | 0 | 0.66 | 0.43 | 0.83 | 0.97 | 0.75 | 0.83 |
| 2 | 107499573 | rs1448119 | 0.081 | -0.558 | 0.56 | 0.144 | 0.02 | -0.01 | -0.01 | 0 | 0.08 | 0.11 | 0.05 | 0.03 | 0.12 | 0.09 |
| 2 | 107504369 | rs10166025 | -1.137 | -0.902 | 0.431 | -0.057 | 0 | -0.018 | -0.018 | 0 | 0.27 | 0.66 | 0.13 | 0.03 | 0.18 | 0.1 |
| 2 | 107507702 | rs6709151 | 0.173 | -0.972 | 0.762 | -0.092 | 0 | -0.008 | -0.008 | 0 | 0.17 | 0.18 | 0.08 | 0.03 | 0.16 | 0.09 |
| 2 | 107516533 | rs12712262 | -0.955 | -1.023 | 0.135 | -0.13 | 0.017 | -0.063 | -0.063 | 0 | 0.73 | 0.87 | 0.72 | 0.56 | 0.78 | 0.7 |
| 2 | 107523541 | rs1404143 | 2.209 | -0.749 | 0.433 | -0.154 | -0.052 | 0.096 | 0.096 | 0.004 | 0.33 | 0.18 | 0.4 | 0.09 | 0.42 | 0.12 |
| 2 | 107523808 | rs12712263 | -0.909 | -0.652 | 0.312 | -0.031 | 0.008 | -0.052 | -0.052 | 0.031 | 0.73 | 0.84 | 0.72 | 0.56 | 0.76 | 0.67 |
| 2 | 107532159 | rs11889164 | 0.993 | -0.66 | 0.659 | 0.226 | -0.007 | 0.072 | 0.072 | 0 | 0.12 | 0.11 | 0.15 | 0.41 | 0.09 | 0.35 |
| 2 | 107547777 | rs12105611 | -0.212 | -0.665 | 0.946 | -0.105 | -0.012 | -0.047 | -0.047 | 0.02 | 0.19 | 0.24 | 0.09 | 0.03 | 0.16 | 0.1 |
| 2 | 107550844 | rs6730588 | 0.889 | -0.173 | 0.386 | -0.193 | -0.015 | -0.015 | -0.015 | 0 | 0.42 | 0.3 | 0.52 | 0.48 | 0.48 | 0.45 |
| 2 | 107554490 | rs17033552 | 0.374 | -0.175 | 0.37 | -0.194 | -0.003 | -0.018 | -0.018 | 0.01 | 0.17 | 0.16 | 0.07 | 0.03 | 0.16 | 0.1 |
| 2 | 107558907 | rs13033073 | 0.877 | -0.126 | 0.328 | -0.158 | -0.019 | -0.02 | -0.02 | 0 | 0.42 | 0.29 | 0.52 | 0.48 | 0.48 | 0.45 |
| 2 | 107567363 | rs6742683 | 0.177 | -0.297 | 0.31 | -0.235 | 0.012 | -0.001 | -0.001 | 0.525 | 0.51 | 0.41 | 0.57 | 0.53 | 0.59 | 0.55 |
| 2 | 107570090 | rs1034816 | -0.816 | -0.105 | 0.151 | -0.206 | -0.013 | -0.037 | -0.037 | 1.211 | 0.65 | 0.75 | 0.6 | 0.53 | 0.63 | 0.55 |
| 2 | 107574487 | rs12712270 | -0.805 | -0.243 | 0.158 | -0.176 | -0.019 | -0.039 | -0.039 | 0.039 | 0.65 | 0.75 | 0.59 | 0.53 | 0.63 | 0.54 |
| 2 | 107581130 | rs11684876 | 0.069 | -0.294 | 0.329 | -0.277 | -0.002 | -0.001 | -0.001 | 0 | 0.52 | 0.51 | 0.56 | 0.57 | 0.59 | 0.57 |
| 2 | 107584422 | rs12477245 | 1.821 | -0.344 | 0.103 | -0.345 | 0.025 | 0.019 | 0.019 | 1.254 | 0.1 | 0.04 | 0.02 | 0 | 0.04 | 0.01 |
| 2 | 107587099 | rs11884940 | 1.671 | -0.356 | 0.112 | -0.352 | 0.002 | -0.01 | -0.01 | 0.651 | 0.11 | 0.08 | 0.02 | 0 | 0.04 | 0 |
| 2 | 107593110 | rs987987 | -0.873 | -0.244 | -0.158 | -0.319 | -0.002 | -0.029 | -0.029 | 0 | 0.67 | 0.79 | 0.64 | 0.6 | 0.7 | 0.65 |
| 2 | 107601737 | rs7566171 | 0.431 | 0.07 | -0.345 | -0.396 | 0.028 | 0.037 | 0.037 | 0.004 | 0.49 | 0.26 | 0.41 | 0.4 | 0.39 | 0.38 |
| 2 | 107602873 | rs11885179 | 0.205 | 0.078 | -0.333 | -0.409 | -0.001 | 0.003 | 0.003 | 0 | 0.53 | 0.44 | 0.46 | 0.42 | 0.46 | 0.47 |
| 2 | 107603011 | rs4676096 | 1.106 | 0.093 | -0.727 | -0.329 | -0.032 | -0.034 | -0.034 | 2.884 | 0.37 | 0.15 | 0.37 | 0.4 | 0.35 | 0.38 |
| 2 | 107614545 | rs11887254 | 1.539 | -0.018 | -0.424 | -0.338 | -0.038 | -0.058 | -0.058 | 0 | 0.12 | 0.16 | 0.04 | 0 | 0.05 | 0 |
| 2 | 107614969 | rs12469093 | 0.932 | 0.163 | -0.666 | -0.42 | -0.038 | -0.015 | -0.015 | 0.59 | 0.4 | 0.17 | 0.4 | 0.4 | 0.4 | 0.38 |
| 2 | 107621748 | rs2906601 | -0.126 | 0.189 | -0.517 | -0.432 | 0.005 | -0.003 | -0.003 | 0.022 | 0.57 | 0.53 | 0.49 | 0.43 | 0.52 | 0.47 |
| 2 | 107623734 | rs11124147 | 1.209 | 0.149 | -0.897 | -0.499 | -0.018 | 0.076 | 0.076 | 0.275 | 0.46 | 0.28 | 0.32 | 0.26 | 0.44 | 0.34 |
| 2 | 107637902 | rs2906607 | -0.444 | 0.502 | -1.269 | -0.433 | -0.057 | 0.051 | 0.051 | 0 | 0.75 | 0.9 | 0.81 | 0.85 | 0.68 | 0.83 |
| 2 | 107640555 | rs4676099 | 1.517 | 0.334 | -1.3 | -0.443 | -0.023 | 0.051 | 0.051 | 0 | 0.41 | 0.22 | 0.42 | 0.27 | 0.39 | 0.38 |
| 2 | 107641898 | rs13390663 | -0.016 | 0.278 | -1.136 | -0.496 | -0.008 | 0.004 | 0.004 | 0 | 0.33 | 0.58 | 0.39 | 0.57 | 0.29 | 0.46 |
| 2 | 107642482 | rs12614346 | 2.039 | 0.168 | -1.347 | -0.75 | -0.041 | 0.097 | 0.097 | 0 | 0.34 | 0.13 | 0.24 | 0.11 | 0.31 | 0.24 |
| 2 | 107648312 | rs12464185 | 0.97 | -0.011 | 0.071 | -0.632 | -0.019 | -0.036 | -0.036 | 2.867 | 0.11 | 0.11 | 0.03 | 0 | 0.04 | 0.02 |
| 2 | 107673715 | rs2140588 | 0.685 | 1.101 | 0.343 | -0.168 | 0.001 | -0.017 | -0.017 | 0 | 0.7 | 0.68 | 0.77 | 0.86 | 0.69 | 0.84 |
| 2 | 107677369 | rs6760504 | 1.153 | 1.171 | 0.658 | 0.076 | 0.007 | 0.004 | 0.004 | 0 | 0.63 | 0.39 | 0.56 | 0.6 | 0.56 | 0.68 |
| 2 | 107677489 | rs2949222 | -1.088 | 1.235 | 0.53 | -0.383 | -0.01 | 0.02 | 0.02 | 0 | 0.32 | 0.29 | 0.24 | 0.14 | 0.36 | 0.17 |
| 2 | 107677690 | rs2949224 | -1.434 | 1.331 | 1.011 | 0.039 | 0.008 | -0.016 | -0.016 | 0 | 0.37 | 0.41 | 0.36 | 0.22 | 0.44 | 0.28 |
| 2 | 107679474 | rs2030675 | -1.332 | 2.029 | 0.887 | 0.395 | -0.002 | -0.001 | -0.001 | 0 | 0.26 | 0.29 | 0.34 | 0.31 | 0.32 | 0.18 |
| 2 | 107680238 | rs10496410 | 0.119 | 2.115 | 0.889 | 0.405 | 0.034 | 0.054 | 0.054 | 0.257 | 0.07 | 0.11 | 0.2 | 0.18 | 0.12 | 0.05 |
| 2 | 107684694 | rs2222522 | -1.005 | 0.945 | 0.7 | 0.573 | -0.01 | 0.02 | 0.02 | 0 | 0.19 | 0.17 | 0.14 | 0.13 | 0.2 | 0.13 |
| 2 | 107705986 | rs6755974 | 1.139 | 0.876 | 1.084 | 0.907 | 0.05 | 0.059 | 0.059 | 2.855 | 0.8 | 0.6 | 0.72 | 0.68 | 0.75 | 0.87 |
| 2 | 107706123 | rs13427867 | -1.785 | -0.099 | 1.683 | 1.112 | 0.012 | 0.051 | 0.051 | 1.417 | 0.14 | 0.42 | 0.26 | 0.16 | 0.1 | 0.1 |
| 2 | 107709517 | rs4077162 | -0.049 | 0.104 | 1.04 | 1.555 | -0.003 | -0.13 | -0.13 | 1.532 | 0.15 | 0.01 | 0.25 | 0.5 | 0.12 | 0.33 |
| 2 | 107714710 | rs6419584 | 2.442 | 0.538 | 2.031 | 1.182 | 0.027 | 0.125 | 0.125 | 0 | 0.92 | 0.68 | 0.76 | 0.81 | 0.93 | 0.92 |
| 2 | 107728660 | rs4676318 | 2.461 | 0.535 | 1.984 | 1.095 | 0.028 | 0.17 | 0.17 | 0.011 | 0.91 | 0.66 | 0.75 | 0.79 | 0.93 | 0.92 |
| 2 | 107732029 | rs4283419 | -1.195 | 0.495 | 1.844 | 1.063 | 0.019 | 0.269 | 0.269 | 0 | 0.06 | 0.31 | 0.2 | 0.22 | 0.04 | 0.08 |
| 2 | 107736937 | rs7589263 | 2.427 | 0.499 | 1.583 | 1.064 | 0.005 | 0.149 | 0.149 | 0.09 | 0.86 | 0.44 | 0.7 | 0.78 | 0.89 | 0.87 |
| 2 | 107741454 | rs6759688 | 2.398 | 0.517 | 1.546 | 1.073 | 0.009 | 0.157 | 0.157 | 0.031 | 0.87 | 0.44 | 0.7 | 0.78 | 0.89 | 0.87 |
| 2 | 107744643 | rs3923303 | -1.462 | 0.66 | 1.574 | 1.175 | 0.03 | 0.275 | 0.275 | 0 | 0.08 | 0.41 | 0.28 | 0.22 | 0.07 | 0.1 |
| 2 | 107753988 | rs4319926 | 0.913 | 0.837 | 0.797 | 1.037 | 0.02 | -0.023 | -0.023 | 0 | 0.93 | 0.85 | 0.92 | 0.99 | 0.89 | 0.94 |

| | | | | | | | | | | | | | | | | |
|---|---|---|---|---|---|---|---|---|---|---|---|---|---|---|---|---|
| 2 | 107778931 | rs1541496 | -1.302 | 0.784 | 0.384 | 1.093 | 0.013 | 0.078 | 0.078 | 0 | 0.16 | 0.61 | 0.35 | 0.24 | 0.17 | 0.15 |
| 2 | 107781838 | rs2907904 | 0.496 | 0.713 | -1.106 | 0.462 | -0.006 | 0.004 | 0.004 | 0 | 0.89 | 0.97 | 0.77 | 0.91 | 0.91 | 0.93 |
| 2 | 107781903 | rs6732006 | -0.322 | 0.15 | -2.213 | -0.416 | 0.019 | 0.118 | 0.118 | 0.428 | 0.28 | 0.07 | 0.15 | 0.07 | 0.21 | 0.19 |
| 2 | 107782407 | rs2971657 | 0.337 | 0.154 | -2.876 | -0.181 | -0.015 | -0.074 | -0.074 | 0.068 | 0.72 | 0.93 | 0.64 | 0.36 | 0.76 | 0.63 |
| 2 | 107783240 | rs10170038 | 0.18 | 0.69 | -3.21 | -1.045 | -0.018 | -0.01 | -0.01 | 0 | 0.21 | 0.14 | 0.23 | 0.19 | 0.33 | 0.18 |
| 2 | 107784393 | rs1029864 | -0.162 | 0.57 | -3.423 | -1.538 | -0.017 | 0.018 | 0.018 | 0 | 0.63 | 0.86 | 0.52 | 0.31 | 0.64 | 0.47 |
| 2 | 107784665 | rs2971653 | 0.292 | 0.346 | -3.904 | -2 | 0.036 | 0.119 | 0.119 | 3.05 | 0.46 | 0.84 | 0.37 | 0.23 | 0.42 | 0.3 |
| 2 | 107786016 | rs2907907 | 0.065 | 0.214 | -3.886 | -1.982 | -0.015 | 0.124 | 0.124 | 2.92 | 0.54 | 0.86 | 0.38 | 0.23 | 0.47 | 0.32 |
| 2 | 107786443 | rs1990409 | -0.742 | 0.131 | -3.886 | -1.982 | -0.012 | -0.019 | -0.019 | 0 | 0.71 | 0.88 | 0.54 | 0.3 | 0.69 | 0.49 |
| 2 | 107786827 | rs2971652 | 1.587 | -0.55 | -3.364 | -2.325 | -0.024 | -0.08 | -0.08 | 0.047 | 0.13 | 0.53 | 0.1 | 0.02 | 0.07 | 0.04 |
| 2 | 107789903 | rs10196004 | -0.035 | -0.505 | -3.473 | -2.371 | 0.033 | 0.092 | 0.092 | 0.719 | 0.46 | 0.19 | 0.4 | 0.27 | 0.59 | 0.38 |
| 2 | 107790491 | rs2079917 | -0.948 | -0.612 | -3.551 | -2.419 | -0.004 | -0.008 | -0.008 | 0 | 0.68 | 0.87 | 0.52 | 0.3 | 0.68 | 0.48 |
| 2 | 107791876 | rs17718331 | 1.344 | -0.265 | -3.46 | -2.38 | -0.06 | 0.042 | 0.042 | 5.017 | 0.15 | 0.01 | 0.11 | 0.07 | 0.17 | 0.13 |
| 2 | 107794246 | rs2215482 | 0.017 | -0.304 | -3.49 | -2.413 | 0.051 | 0.074 | 0.074 | 2.82 | 0.31 | 0.78 | 0.2 | 0.05 | 0.19 | 0.21 |
| 2 | 107794867 | rs13390271 | 0.749 | -0.057 | -3.334 | -2.259 | -0.028 | 0.041 | 0.041 | 2.796 | 0.18 | 0.09 | 0.16 | 0.08 | 0.22 | 0.18 |
| 2 | 107795527 | rs17718567 | 1.442 | -0.105 | -3.24 | -2.181 | -0.003 | 0.011 | 0.011 | 0 | 0.09 | 0.04 | 0.08 | 0.02 | 0.1 | 0.1 |
| 2 | 107796639 | rs17020992 | 1.245 | -0.396 | -2.273 | -2.333 | -0.01 | -0.065 | -0.065 | 0 | 0.12 | 0.48 | 0.12 | 0.02 | 0.07 | 0.05 |
| 2 | 107796734 | rs2889744 | 1.568 | -0.528 | -2.511 | -2.502 | -0.005 | -0.073 | -0.073 | 0.869 | 0.23 | 0.06 | 0.38 | 0.68 | 0.19 | 0.43 |
| 2 | 107798929 | rs6744712 | -0.431 | -0.557 | -2.342 | -2.606 | 0 | 0.072 | 0.072 | 0 | 0.38 | 0.7 | 0.29 | 0.11 | 0.34 | 0.27 |
| 2 | 107802658 | rs10439406 | 1.337 | -0.518 | -2.287 | -2.545 | -0.004 | 0.003 | 0.003 | 6.764 | 0.15 | 0.13 | 0.1 | 0.07 | 0.17 | 0.12 |
| 2 | 107804217 | rs2079918 | -0.95 | -0.597 | -2.098 | -2.593 | 0.024 | -0.091 | -0.091 | 0 | 0.86 | 0.56 | 0.82 | 0.97 | 0.86 | 0.9 |
| 2 | 107807644 | rs7607270 | 0.213 | -0.431 | -2.277 | -2.76 | -0.004 | -0.112 | -0.112 | 0 | 0.7 | 0.44 | 0.71 | 0.95 | 0.69 | 0.79 |
| 2 | 107807743 | rs7566414 | 2.378 | -0.489 | -2.232 | -2.732 | -0.013 | -0.017 | -0.017 | 0 | 0.07 | 0.11 | 0.02 | 0.02 | 0.02 | 0.06 |
| 2 | 107812973 | rs3922303 | -0.367 | -0.45 | -2.194 | -2.746 | 0.008 | -0.133 | -0.133 | 0.13 | 0.8 | 0.54 | 0.79 | 0.97 | 0.8 | 0.89 |
| 2 | 107813837 | rs1558398 | 1.005 | -0.695 | -2.283 | -2.949 | -0.064 | 0.209 | 0.209 | 0 | 0.43 | 0.15 | 0.59 | 0.86 | 0.5 | 0.61 |
| 2 | 107813952 | rs10496412 | 0.514 | -0.437 | -2.086 | -2.797 | -0.023 | -0.122 | -0.122 | 1.919 | 0.18 | 0.46 | 0.14 | 0.02 | 0.13 | 0.06 |
| 2 | 107816413 | rs2191588 | 1.506 | -0.488 | -2.181 | -2.822 | -0.009 | 0.011 | 0.011 | 5.214 | 0.14 | 0.12 | 0.09 | 0.07 | 0.18 | 0.12 |
| 2 | 107820073 | rs2191590 | 0.539 | -0.544 | -2.158 | -2.831 | 0.023 | 0.048 | 0.048 | 5.379 | 0.62 | 0.28 | 0.69 | 0.93 | 0.68 | 0.73 |
| 2 | 107822307 | rs17719622 | 1.023 | -0.543 | -2.114 | -2.782 | 0.018 | 0.056 | 0.056 | 0.068 | 0.17 | 0.04 | 0.09 | 0.04 | 0.13 | 0.15 |
| 2 | 107823219 | rs17033988 | 0.708 | -0.164 | -0.732 | -2.492 | 0.03 | -0.089 | -0.089 | 0 | 0.14 | 0.36 | 0.13 | 0.02 | 0.11 | 0.07 |
| 2 | 107830992 | rs4676327 | -0.749 | -0.076 | -0.698 | -2.338 | 0.024 | -0.064 | -0.064 | 0 | 0.87 | 0.51 | 0.82 | 0.96 | 0.87 | 0.9 |
| 2 | 107835048 | rs715379 | 0.821 | -0.046 | -1.191 | -2.418 | -0.037 | 0.098 | 0.098 | 0 | 0.36 | 0.13 | 0.29 | 0.16 | 0.42 | 0.28 |
| 2 | 107836419 | rs13420797 | 2.167 | -0.093 | -1.281 | -2.515 | -0.02 | -0.038 | -0.038 | 0 | 0.22 | 0.05 | 0.42 | 0.72 | 0.22 | 0.43 |
| 2 | 107836502 | rs7584779 | -0.004 | -0.101 | -1.117 | -2.548 | -0.011 | -0.05 | -0.05 | 0.372 | 0.69 | 0.38 | 0.79 | 0.88 | 0.76 | 0.74 |
| 2 | 107839788 | rs13035614 | 2.458 | -0.024 | -1.155 | -2.15 | -0.058 | -0.125 | -0.125 | 0 | 0.21 | 0.01 | 0.23 | 0.61 | 0.19 | 0.38 |
| 2 | 107848693 | rs10167469 | 0.409 | 0.041 | -0.854 | -2.362 | -0.017 | -0.078 | -0.078 | 0.006 | 0.2 | 0.56 | 0.11 | 0.08 | 0.15 | 0.15 |
| 2 | 107850627 | rs6755833 | 1.295 | 0.063 | -0.896 | -2.351 | -0.011 | 0.008 | 0.008 | 0.177 | 0.28 | 0.17 | 0.47 | 0.72 | 0.27 | 0.46 |
| 2 | 107857740 | rs12617850 | 1.404 | 0.066 | -0.912 | -2.351 | -0.048 | -0.01 | -0.01 | 0 | 0.28 | 0.06 | 0.46 | 0.72 | 0.27 | 0.46 |
| 2 | 107858218 | rs7560461 | -0.144 | 0.034 | -0.818 | -2.392 | 0.01 | -0.078 | -0.078 | 0 | 0.77 | 0.44 | 0.8 | 0.91 | 0.76 | 0.79 |
| 2 | 107862323 | rs6753056 | 1.291 | -0.013 | -0.848 | -1.738 | -0.01 | -0.008 | -0.008 | 1.26 | 0.29 | 0.17 | 0.47 | 0.69 | 0.27 | 0.46 |
| 2 | 107862691 | rs6725836 | 0.723 | -0.073 | -0.884 | -1.781 | 0.008 | 0.016 | 0.016 | 0 | 0.65 | 0.3 | 0.76 | 0.88 | 0.7 | 0.73 |
| 2 | 107862905 | rs13401181 | 1.999 | -0.044 | -0.481 | -1.809 | 0.002 | -0.011 | -0.011 | 0 | 0.11 | 0.16 | 0.06 | 0.06 | 0.12 | 0.1 |
| 2 | 107862981 | rs2191595 | -0.51 | -0.052 | -0.481 | -1.808 | -0.006 | 0.055 | 0.055 | 0 | 0.58 | 0.76 | 0.39 | 0.27 | 0.57 | 0.43 |
| 2 | 107864763 | rs7563746 | 1.405 | -0.127 | -0.516 | -1.795 | -0.008 | -0.034 | -0.034 | 0 | 0.1 | 0.4 | 0.05 | 0.02 | 0.02 | 0.05 |
| 2 | 107869470 | rs12478937 | 0.024 | 0.019 | -1.031 | -1.743 | -0.001 | -0.014 | -0.014 | 0 | 0.75 | 0.25 | 0.8 | 0.91 | 0.78 | 0.79 |
| 2 | 107870390 | rs10166265 | 0.3 | -0.025 | -1.018 | -1.729 | -0.015 | -0.051 | -0.051 | 0 | 0.22 | 0.7 | 0.12 | 0.08 | 0.14 | 0.15 |
| 2 | 107870934 | rs12612007 | 0.309 | 0.206 | -1.166 | -1.473 | -0.006 | 0.045 | 0.045 | 0 | 0.38 | 0.13 | 0.51 | 0.72 | 0.36 | 0.52 |
| 2 | 107883105 | rs7587131 | -0.352 | 0.279 | -0.379 | -1.334 | -0.003 | -0.032 | -0.032 | 0 | 0.85 | 0.81 | 0.87 | 0.96 | 0.86 | 0.88 |
| 2 | 107906065 | rs758822 | 0.837 | -0.152 | 0.297 | -1.6 | -0.018 | -0.095 | -0.095 | 0 | 0.69 | 0.45 | 0.79 | 0.9 | 0.73 | 0.74 |
| 2 | 107906659 | rs992634 | 1.592 | -0.094 | -0.292 | -1.545 | -0.002 | -0.023 | -0.023 | 0 | 0.06 | 0.09 | 0.04 | 0 | 0.06 | 0.01 |
| 2 | 107908592 | rs6747330 | -0.528 | -0.067 | -0.337 | -1.516 | -0.013 | -0.078 | -0.078 | 0.542 | 0.84 | 0.65 | 0.85 | 0.96 | 0.86 | 0.86 |
| 2 | 107929052 | rs7597900 | 0.332 | 0.07 | -1.178 | -1.369 | 0.031 | 0.099 | 0.099 | 0 | 0.64 | 0.2 | 0.76 | 0.89 | 0.72 | 0.73 |
| 2 | 107943371 | rs17683291 | 0.614 | 0.448 | -1.558 | -1.256 | -0.065 | 0.017 | 0.017 | 0.482 | 0.19 | 0.02 | 0.16 | 0.1 | 0.21 | 0.13 |
| 2 | 107950290 | rs4676120 | 1.221 | 0.419 | -2.348 | -1.106 | -0.072 | -0.119 | -0.119 | 0 | 0.23 | 0.01 | 0.33 | 0.67 | 0.23 | 0.34 |
| 2 | 107955058 | rs6753182 | 1.333 | -0.049 | -2.245 | -1.292 | -0.007 | -0.011 | -0.011 | 0 | 0.11 | 0 | 0.08 | 0.09 | 0.11 | 0.07 |
| 2 | 107962440 | rs6709735 | -0.422 | -0.332 | -1.981 | -1.638 | 0.025 | 0.04 | 0.04 | 0 | 0.26 | 0.2 | 0.23 | 0.13 | 0.24 | 0.17 |
| 2 | 107974838 | rs17827020 | 1.31 | -0.4 | -1.972 | -1.674 | -0.001 | 0.001 | 0.001 | 0 | 0.09 | 0.07 | 0.15 | 0 | 0.11 | 0.04 |
| 2 | 107975072 | rs17230424 | -0.897 | -0.428 | -1.934 | -1.687 | 0.012 | 0.015 | 0.015 | 0 | 0.28 | 0.28 | 0.24 | 0.14 | 0.24 | 0.18 |
| 2 | 107981795 | rs17230670 | 0.38 | -0.416 | -1.972 | -1.676 | 0.026 | 0.009 | 0.009 | 0 | 0.17 | 0.12 | 0.09 | 0.14 | 0.13 | 0.15 |
| 2 | 107987035 | rs6746088 | 1.442 | -0.386 | -1.394 | -1.907 | 0.064 | 0.131 | 0.131 | 0 | 0.53 | 0.15 | 0.58 | 0.76 | 0.59 | 0.5 |
| 2 | 107989333 | rs12999789 | 1.392 | -0.325 | -1.137 | -1.545 | -0.002 | -0.006 | -0.006 | 0 | 0.11 | 0.01 | 0.08 | 0.09 | 0.11 | 0.08 |
| 2 | 107990409 | rs2889765 | -0.698 | -0.322 | -0.847 | -1.54 | 0.007 | -0.042 | -0.042 | 0 | 0.82 | 0.52 | 0.83 | 0.9 | 0.83 | 0.68 |
| 2 | 107999545 | rs13006861 | -0.041 | -0.373 | -1.091 | -1.25 | 0.025 | 0.008 | 0.008 | 0 | 0.17 | 0.12 | 0.09 | 0.14 | 0.13 | 0.14 |
| 2 | 108000932 | rs10427352 | 0.552 | -0.374 | -1.091 | -1.249 | -0.018 | -0.002 | -0.002 | 0.012 | 0.06 | 0.11 | 0.01 | 0.05 | 0.01 | 0.07 |
| 2 | 108002088 | rs17663778 | 0.085 | -0.35 | -1.079 | -1.242 | 0.025 | 0.008 | 0.008 | 0 | 0.17 | 0.12 | 0.09 | 0.14 | 0.13 | 0.14 |
| 2 | 108018641 | rs11123663 | 0.196 | -0.121 | -0.13 | -1.187 | -0.009 | -0.076 | -0.076 | 0 | 0.71 | 0.43 | 0.77 | 0.91 | 0.71 | 0.56 |
| 2 | 108019522 | rs972881 | 1.442 | -0.019 | -0.402 | -0.951 | -0.087 | -0.166 | -0.166 | 1.744 | 0.27 | 0.01 | 0.41 | 0.57 | 0.29 | 0.25 |
| 2 | 108033742 | rs266243 | 1.066 | 0.176 | 0.995 | -0.96 | 0.008 | 0.027 | 0.027 | 0.017 | 0.66 | 0.27 | 0.75 | 0.86 | 0.69 | 0.44 |
| 2 | 108036641 | rs266247 | 0.615 | 0.127 | 1.329 | -0.986 | 0 | -0.003 | -0.003 | 0 | 0.71 | 0.63 | 0.78 | 0.86 | 0.69 | 0.52 |

| | | | | | | | | | | | | | | | | |
|---|---|---|---|---|---|---|---|---|---|---|---|---|---|---|---|---|
| 2 | 108038761 | rs2561445 | 1.336 | 0.111 | 1.269 | -0.994 | 0.018 | 0.059 | 0.059 | 0.097 | 0.65 | 0.22 | 0.75 | 0.86 | 0.68 | 0.44 |
| 2 | 108040409 | rs266253 | -0.037 | 0.217 | 1.171 | -0.987 | 0 | 0.014 | 0.014 | 0.017 | 0.78 | 0.78 | 0.81 | 0.92 | 0.74 | 0.68 |
| 2 | 108043628 | rs266168 | 1.192 | 0.161 | 1.211 | -0.959 | 0.019 | 0.054 | 0.054 | 0.103 | 0.65 | 0.24 | 0.75 | 0.89 | 0.7 | 0.46 |
| 2 | 108047544 | rs266170 | 1.328 | 0.216 | 1.263 | -0.922 | 0.019 | 0.054 | 0.054 | 0 | 0.66 | 0.24 | 0.75 | 0.85 | 0.69 | 0.46 |
| 2 | 108048120 | rs10496415 | 0.196 | 0.323 | 1.223 | -0.919 | -0.001 | 0 | 0 | 1.417 | 0.06 | 0.08 | 0.02 | 0.06 | 0.05 | 0.16 |
| 2 | 108051515 | rs266177 | 0.642 | 0.326 | 1.298 | -0.921 | 0 | 0.01 | 0.01 | 0 | 0.72 | 0.67 | 0.79 | 0.85 | 0.69 | 0.53 |
| 2 | 108071524 | rs266216 | 1.228 | -0.24 | 1.367 | -0.884 | 0.007 | 0.023 | 0.023 | 0 | 0.67 | 0.29 | 0.76 | 0.85 | 0.68 | 0.45 |
| 2 | 108074510 | rs266229 | 1.252 | -0.244 | 1.363 | -0.886 | 0.008 | 0.027 | 0.027 | 0 | 0.67 | 0.28 | 0.75 | 0.85 | 0.68 | 0.45 |
| 2 | 108076452 | rs2552312 | 0.716 | -0.247 | 1.396 | -0.888 | -0.004 | -0.037 | -0.037 | 0.143 | 0.7 | 0.56 | 0.79 | 0.85 | 0.68 | 0.45 |
| 2 | 108079648 | rs2715913 | 1.235 | -0.235 | 1.125 | -0.882 | 0.008 | 0.027 | 0.027 | 0 | 0.67 | 0.28 | 0.75 | 0.85 | 0.68 | 0.45 |
| 2 | 108081665 | rs959014 | 1.258 | -0.234 | 1.126 | -0.841 | 0.009 | 0.029 | 0.029 | 0 | 0.66 | 0.28 | 0.75 | 0.85 | 0.68 | 0.45 |
| 2 | 108100980 | rs266141 | 1.108 | -0.228 | 1.159 | -0.838 | -0.008 | -0.037 | -0.037 | 0 | 0.67 | 0.38 | 0.76 | 0.86 | 0.68 | 0.45 |
| 2 | 108107550 | rs10496413 | 0.144 | -0.152 | 0.424 | -0.772 | -0.001 | 0.001 | 0.001 | 0.126 | 0.06 | 0.08 | 0.02 | 0.06 | 0.06 | 0.16 |
| 2 | 108114085 | rs166464 | 0.864 | -0.111 | 0.299 | -0.782 | -0.009 | -0.045 | -0.045 | 0 | 0.68 | 0.42 | 0.77 | 0.86 | 0.68 | 0.46 |
| 2 | 108123370 | rs187862 | 0.908 | -0.103 | 0.081 | -0.778 | -0.006 | -0.03 | -0.03 | 0 | 0.68 | 0.39 | 0.77 | 0.86 | 0.68 | 0.46 |
| 2 | 108140880 | rs266273 | 0.1 | 0.262 | -0.836 | -0.42 | 0 | 0.023 | 0.023 | 0 | 0.79 | 0.8 | 0.82 | 0.92 | 0.75 | 0.69 |
| 2 | 108157513 | rs6744070 | -0.577 | 0.251 | -0.488 | 0.18 | -0.002 | 0.058 | 0.058 | 0 | 0.3 | 0.51 | 0.48 | 0.54 | 0.32 | 0.31 |
| 2 | 108160811 | rs7604787 | -0.425 | 0.168 | -0.149 | 0.143 | 0.003 | 0.042 | 0.042 | 0 | 0.28 | 0.4 | 0.46 | 0.54 | 0.32 | 0.26 |
| 2 | 108162684 | rs4299335 | -0.715 | -0.22 | -0.862 | -0.501 | -0.042 | 0.053 | 0.053 | 0 | 0.66 | 0.95 | 0.74 | 0.78 | 0.66 | 0.69 |
| 2 | 108167041 | rs4676023 | -0.18 | -0.53 | -0.259 | -0.395 | -0.059 | 0.228 | 0.228 | 0 | 0.11 | 0.32 | 0.26 | 0.54 | 0.06 | 0.2 |
| 2 | 108167800 | rs12468034 | 0.048 | -0.545 | -0.192 | -0.401 | -0.048 | 0.202 | 0.202 | 0 | 0.1 | 0.26 | 0.25 | 0.54 | 0.06 | 0.2 |
| 2 | 108169377 | rs1345961 | -1.026 | -0.539 | -0.171 | -0.384 | -0.002 | 0.061 | 0.061 | 0 | 0.3 | 0.56 | 0.48 | 0.56 | 0.32 | 0.31 |
| 2 | 108176718 | rs6721257 | 0.754 | -0.428 | -0.334 | -0.443 | -0.055 | 0.058 | 0.058 | 0 | 0.16 | 0.04 | 0.2 | 0 | 0.25 | 0.06 |
| 2 | 108190233 | rs1108853 | 0.876 | -0.348 | 0.015 | -0.319 | 0.001 | -0.022 | -0.022 | 1.103 | 0.13 | 0.12 | 0.19 | 0 | 0.22 | 0.06 |
| 2 | 108198802 | rs6542726 | 0.996 | -0.604 | 0.311 | -0.44 | -0.002 | -0.003 | -0.003 | 1.811 | 0.64 | 0.35 | 0.51 | 0.36 | 0.64 | 0.57 |
| 2 | 108199394 | rs6542727 | 0.09 | -1.106 | 0.935 | 0.216 | -0.044 | 0.175 | 0.175 | 0 | 0.9 | 0.76 | 0.76 | 0.46 | 0.94 | 0.8 |
| 2 | 108207995 | rs10184114 | 0.107 | -1.395 | 0.655 | -0.064 | -0.047 | 0.023 | 0.023 | 0 | 0.18 | 0.28 | 0.09 | 0.24 | 0.07 | 0.08 |
| 2 | 108212321 | rs908436 | -0.465 | -0.578 | 1.089 | 0.396 | -0.002 | -0.009 | -0.009 | 0 | 0.08 | 0.03 | 0.03 | 0.08 | 0.07 | 0.12 |
| 2 | 108241786 | rs1516217 | -1.777 | -0.663 | 1.567 | 0.392 | -0.045 | 0.045 | 0.045 | 0 | 0.21 | 0.43 | 0.28 | 0.65 | 0.14 | 0.39 |
| 2 | 108286602 | rs11123669 | -0.526 | -0.486 | 1.569 | 0.475 | -0.018 | 0.178 | 0.178 | 0 | 0.09 | 0.19 | 0.24 | 0.55 | 0.08 | 0.2 |
| 2 | 108290772 | rs10865014 | -1.605 | -0.482 | 1.512 | 0.359 | -0.035 | 0.102 | 0.102 | 0 | 0.2 | 0.46 | 0.31 | 0.63 | 0.15 | 0.38 |
| 2 | 108353629 | rs832013 | -0.686 | -0.064 | 0.623 | 0.763 | -0.016 | 0.295 | 0.295 | 0 | 0.09 | 0.26 | 0.27 | 0.54 | 0.06 | 0.21 |
| 2 | 108368694 | rs700855 | -1.108 | 0.091 | -0.119 | 0.189 | 0.024 | 0.025 | 0.025 | 0.734 | 0.39 | 0.79 | 0.44 | 0.66 | 0.29 | 0.42 |
| 2 | 108378250 | rs831984 | -0.736 | -0.07 | 0.685 | 0.162 | -0.019 | 0.068 | 0.068 | 0.729 | 0.3 | 0.46 | 0.37 | 0.54 | 0.25 | 0.33 |
| 2 | 108379520 | rs831983 | -0.98 | -0.087 | 0.796 | 0.153 | -0.004 | 0.006 | 0.006 | 0.551 | 0.33 | 0.64 | 0.38 | 0.54 | 0.25 | 0.33 |
| 2 | 108380249 | rs17021733 | -0.077 | 0.12 | 0.896 | 0.215 | 0 | -0.066 | -0.066 | 0.01 | 0.22 | 0.23 | 0.12 | 0 | 0.2 | 0.15 |
| 2 | 108390945 | rs6725400 | -1.368 | 0.333 | 1.081 | 0.368 | 0.072 | 0.074 | 0.074 | 1.874 | 0.05 | 0.52 | 0.04 | 0.01 | 0 | 0 |
| 2 | 108404052 | rs328035 | 0.083 | 1.169 | -0.125 | 0.059 | -0.103 | 0.047 | 0.047 | 0.185 | 0.75 | 0.9 | 0.69 | 0.83 | 0.61 | 0.67 |
| 2 | 108415702 | rs6542736 | 0.95 | 0.766 | -0.596 | 0.119 | 0.046 | 0.031 | 0.031 | 0.394 | 0.39 | 0.17 | 0.35 | 0.34 | 0.35 | 0.52 |
| 2 | 108422810 | rs4972265 | 0.049 | 0.775 | -0.673 | 0.32 | -0.017 | 0.032 | 0.032 | 0.679 | 0.34 | 0.43 | 0.32 | 0.49 | 0.26 | 0.19 |
| 2 | 108423990 | rs328032 | 0.271 | 2.037 | -0.734 | 0.302 | -0.056 | 0.025 | 0.025 | 0.012 | 0.79 | 0.87 | 0.73 | 0.83 | 0.71 | 0.74 |
| 2 | 108427959 | rs167869 | 0.336 | 1.897 | -0.876 | 0.369 | -0.106 | 0.025 | 0.025 | 0.068 | 0.78 | 0.93 | 0.71 | 0.83 | 0.66 | 0.72 |
| 2 | 108429937 | rs831973 | 0.675 | 1.784 | -0.871 | 1.281 | -0.057 | -0.083 | -0.083 | 0 | 0.85 | 0.94 | 0.66 | 0.44 | 0.77 | 0.81 |
| 2 | 108431941 | rs831971 | 1.008 | 2.19 | -0.988 | 1.33 | -0.059 | -0.07 | -0.07 | 0.738 | 0.84 | 0.94 | 0.64 | 0.37 | 0.77 | 0.85 |
| 2 | 108433169 | rs1659885 | 1.812 | 2.014 | 0.249 | 1.037 | 0.055 | 0.028 | 0.028 | 0 | 0.75 | 0.55 | 0.5 | 0.33 | 0.7 | 0.71 |
| 2 | 108437881 | rs831991 | 1.628 | 1.951 | 0.656 | 1.004 | -0.007 | -0.006 | -0.006 | 0.08 | 0.78 | 0.79 | 0.51 | 0.34 | 0.7 | 0.71 |
| 2 | 108438922 | rs831990 | 2.019 | 1.92 | 0.801 | 0.989 | 0.058 | 0.031 | 0.031 | 0 | 0.74 | 0.52 | 0.5 | 0.34 | 0.7 | 0.71 |
| 2 | 108440138 | rs831989 | 1.667 | 1.923 | 0.84 | 1.02 | 0.002 | 0 | 0 | 0 | 0.78 | 0.74 | 0.51 | 0.34 | 0.7 | 0.71 |
| 2 | 108545409 | rs6711763 | 0.818 | 1.685 | 1.494 | 0.945 | -0.018 | -0.021 | -0.021 | 0 | 0.83 | 0.87 | 0.64 | 0.37 | 0.76 | 0.85 |
| 2 | 108549097 | rs7604041 | 0.419 | 1.033 | 0.616 | -0.405 | 0.002 | 0.087 | 0.087 | 0 | 0.43 | 0.22 | 0.32 | 0.03 | 0.41 | 0.38 |
| 2 | 108556791 | rs1863688 | -0.997 | 1.302 | 1.071 | -0.311 | 0.031 | 0.017 | 0.017 | 0 | 0.08 | 0.23 | 0.15 | 0.06 | 0.08 | 0.17 |
| 2 | 108559419 | rs4383289 | 1.448 | 1.286 | 1.103 | -0.243 | 0.03 | 0.031 | 0.031 | 0 | 0.78 | 0.66 | 0.52 | 0.34 | 0.72 | 0.73 |
| 2 | 108575574 | rs10170368 | -1.186 | 1.202 | 1.03 | -0.208 | 0.016 | 0.02 | 0.02 | 0 | 0.09 | 0.22 | 0.15 | 0.07 | 0.08 | 0.17 |
| 2 | 108588343 | rs13416160 | 0.066 | 1.198 | 0.926 | -0.212 | 0.001 | -0.008 | -0.008 | 0 | 0.06 | 0.09 | 0.13 | 0.02 | 0.06 | 0.17 |
| 2 | 108593468 | rs6721036 | -0.054 | 1.375 | 0.798 | 0.307 | -0.042 | -0.085 | -0.085 | 0 | 0.88 | 0.99 | 0.83 | 0.54 | 0.88 | 0.92 |
| 2 | 108597499 | rs10184700 | 1.666 | 1.312 | 1.173 | 0.108 | 0.046 | 0.035 | 0.035 | 0 | 0.75 | 0.55 | 0.51 | 0.33 | 0.71 | 0.73 |
| 2 | 108599710 | rs13020302 | 1.023 | 0.705 | 0.315 | -0.743 | -0.122 | 0.173 | 0.173 | 0 | 0.26 | 0.01 | 0.25 | 0.03 | 0.33 | 0.28 |
| 2 | 108602145 | rs1021247 | 1.269 | 0.778 | 0.478 | -0.722 | 0.042 | 0.043 | 0.043 | 0 | 0.76 | 0.5 | 0.6 | 0.33 | 0.73 | 0.74 |
| 2 | 108602755 | rs3806536 | 0.912 | 0.708 | 0.38 | -0.711 | 0.062 | 0.097 | 0.097 | 0.013 | 0.48 | 0.27 | 0.33 | 0.29 | 0.4 | 0.47 |
| 2 | 108603431 | rs2433718 | -0.154 | 0.804 | 0.417 | -0.634 | -0.02 | -0.032 | -0.032 | 0 | 0.86 | 0.89 | 0.67 | 0.39 | 0.81 | 0.9 |
| 2 | 108605836 | rs6542746 | -0.344 | 0.235 | 0.092 | -0.976 | -0.006 | 0.02 | 0.02 | 0.201 | 0.56 | 0.59 | 0.49 | 0.32 | 0.51 | 0.77 |
| 2 | 108607100 | rs333235 | -0.497 | 0.24 | 0.053 | -0.973 | -0.026 | -0.036 | -0.036 | 0 | 0.85 | 0.89 | 0.67 | 0.39 | 0.8 | 0.9 |
| 2 | 108607394 | rs333234 | 1.273 | 0.207 | 0.125 | -0.972 | -0.001 | -0.012 | -0.012 | 0.56 | 0.3 | 0.3 | 0.17 | 0.07 | 0.28 | 0.13 |
| 2 | 108609099 | rs10195701 | 0.295 | 0.352 | -0.325 | -0.655 | 0.001 | -0.001 | -0.001 | 2.825 | 0.24 | 0.26 | 0.19 | 0.27 | 0.15 | 0.34 |
| 2 | 108610264 | rs333217 | -0.092 | 0.92 | -0.288 | -0.341 | -0.019 | -0.02 | -0.02 | 0 | 0.87 | 0.89 | 0.67 | 0.39 | 0.81 | 0.9 |
| 2 | 108610555 | rs333216 | -0.554 | 1.214 | -0.021 | -1.029 | 0.019 | -0.127 | -0.127 | 0.477 | 0.23 | 0.43 | 0.25 | 0.05 | 0.29 | 0.26 |
| 2 | 108618540 | rs3731683 | -0.984 | 1.274 | -0.572 | -0.744 | 0.001 | -0.002 | -0.002 | 1.032 | 0.33 | 0.6 | 0.45 | 0.26 | 0.38 | 0.24 |
| 2 | 108623666 | rs333224 | -0.701 | 1.27 | -1.369 | -1.01 | -0.051 | 0.091 | 0.091 | 1.944 | 0.85 | 0.99 | 0.94 | 0.99 | 0.87 | 0.88 |
| 2 | 108626468 | rs6753886 | 1.388 | 1.532 | -1.504 | -1.126 | 0.042 | 0.061 | 0.061 | 0 | 0.48 | 0.3 | 0.3 | 0.69 | 0.39 | 0.51 |

| | | | | | | | | | | | | | | | | |
|---|---|---|---|---|---|---|---|---|---|---|---|---|---|---|---|---|
| 2 | 108627917 | rs333226 | -0.895 | 1.361 | -1.159 | -0.729 | -0.052 | 0.094 | 0.094 | 1.257 | 0.85 | 0.99 | 0.94 | 0.99 | 0.87 | 0.87 |
| 2 | 108629104 | rs333227 | -0.822 | 1.362 | -1.073 | -0.728 | -0.03 | 0.074 | 0.074 | 1.447 | 0.84 | 0.96 | 0.93 | 0.99 | 0.87 | 0.87 |
| 2 | 108629260 | rs2046510 | 0.682 | 1.388 | -0.612 | -0.663 | 0.001 | -0.003 | -0.003 | 1.838 | 0.63 | 0.48 | 0.47 | 0.72 | 0.58 | 0.6 |
| 2 | 108631729 | rs10496417 | 0.508 | 1.388 | -0.675 | -0.651 | 0.005 | -0.033 | -0.033 | 0 | 0.66 | 0.52 | 0.65 | 0.85 | 0.62 | 0.65 |
| 2 | 108645033 | rs12712007 | -0.344 | 1.122 | -0.609 | -1.543 | 0.007 | 0.059 | 0.059 | 0.014 | 0.51 | 0.58 | 0.47 | 0.12 | 0.5 | 0.35 |
| 2 | 108651021 | rs6542747 | 0.674 | 1.017 | -0.161 | -1.89 | -0.002 | -0.052 | -0.052 | 0.294 | 0.32 | 0.29 | 0.38 | 0.08 | 0.37 | 0.23 |
| 2 | 108656351 | rs12617914 | 0.926 | 1.216 | -0.204 | -2.098 | -0.011 | 0.153 | 0.153 | 0.185 | 0.39 | 0.23 | 0.42 | 0.87 | 0.36 | 0.57 |
| 2 | 108657625 | rs7560566 | -0.731 | 1.242 | -0.014 | -2.063 | 0.015 | -0.029 | -0.029 | 0.812 | 0.91 | 0.88 | 0.88 | 0.99 | 0.85 | 0.92 |
| 2 | 108671330 | rs1542665 | 0.481 | 0.991 | 0.08 | -2.115 | -0.011 | -0.085 | -0.085 | 0.178 | 0.36 | 0.44 | 0.39 | 0.11 | 0.37 | 0.25 |
| 2 | 108674165 | rs1038533 | 0.928 | 1.033 | 0.51 | -2.088 | 0.001 | 0.016 | 0.016 | 0 | 0.38 | 0.34 | 0.42 | 0.87 | 0.35 | 0.54 |
| 2 | 108699104 | rs10865017 | 0.195 | 1.201 | -0.491 | -1.976 | -0.003 | -0.027 | -0.027 | 0.298 | 0.75 | 0.75 | 0.81 | 0.98 | 0.73 | 0.8 |
| 2 | 108703884 | rs832658 | 0.728 | 1.298 | -0.778 | -1.944 | -0.022 | 0.024 | 0.024 | 0 | 0.08 | 0.05 | 0.1 | 0.01 | 0.13 | 0.06 |
| 2 | 108711926 | rs2378089 | 0.687 | 1.077 | -0.577 | -2.008 | 0.005 | 0.057 | 0.057 | 0 | 0.4 | 0.32 | 0.43 | 0.87 | 0.36 | 0.54 |
| 2 | 108717242 | rs1385956 | 0.423 | 1.21 | -0.735 | -1.44 | -0.017 | 0.042 | 0.042 | 0 | 0.73 | 0.81 | 0.82 | 0.95 | 0.72 | 0.77 |
| 2 | 108722210 | rs1718024 | 0.422 | 0.717 | -1.162 | -1.598 | 0.019 | 0.182 | 0.182 | 0 | 0.55 | 0.18 | 0.6 | 0.17 | 0.59 | 0.45 |
| 2 | 108723751 | rs1027003 | -0.137 | 0.536 | -0.362 | -1.461 | 0.001 | -0.017 | -0.017 | 0 | 0.84 | 0.7 | 0.82 | 0.88 | 0.84 | 0.67 |
| 2 | 108724001 | rs1027002 | 0.114 | 0.363 | -0.191 | -1.7 | 0.003 | 0.013 | 0.013 | 0 | 0.3 | 0.22 | 0.34 | 0.73 | 0.28 | 0.27 |
| 2 | 108725337 | rs2378093 | 0.574 | 0.344 | 0.27 | -1.582 | 0.016 | 0.161 | 0.161 | 0 | 0.5 | 0.27 | 0.45 | 0.14 | 0.53 | 0.38 |
| 2 | 108731604 | rs832665 | 0.891 | 0.348 | 0.039 | -1.612 | -0.041 | 0.272 | 0.272 | 0 | 0.46 | 0.12 | 0.44 | 0.11 | 0.52 | 0.35 |
| 2 | 108735612 | rs1601548 | -0.327 | 0.324 | 0.158 | -1.639 | 0.003 | 0.051 | 0.051 | 0 | 0.41 | 0.29 | 0.41 | 0.74 | 0.38 | 0.37 |
| 2 | 108736675 | rs931326 | -1.046 | 0.335 | 0.415 | -2.305 | 0.003 | 0.031 | 0.031 | 0 | 0.51 | 0.63 | 0.54 | 0.89 | 0.48 | 0.65 |
| 2 | 108742857 | rs4676177 | -0.268 | 0.283 | 0.612 | -2.287 | 0.006 | 0.071 | 0.071 | 0 | 0.4 | 0.29 | 0.42 | 0.76 | 0.37 | 0.36 |
| 2 | 108746221 | rs750523 | -0.313 | 0.279 | 0.575 | -2.287 | 0.007 | 0.064 | 0.064 | 0 | 0.41 | 0.3 | 0.46 | 0.78 | 0.37 | 0.36 |
| 2 | 108760643 | rs1003019 | -0.664 | 0.292 | 0.608 | -2.281 | -0.002 | -0.027 | -0.027 | 0.732 | 0.46 | 0.58 | 0.49 | 0.78 | 0.4 | 0.38 |
| 2 | 108763528 | rs832668 | 0.59 | 0.591 | -0.106 | -2.722 | 0.032 | -0.071 | -0.071 | 0 | 0.22 | 0.35 | 0.24 | 0.1 | 0.24 | 0.17 |
| 2 | 108780731 | rs1522028 | -0.358 | 0.383 | -0.283 | -2.8 | 0.002 | -0.002 | -0.002 | 0 | 0.45 | 0.62 | 0.5 | 0.8 | 0.42 | 0.44 |
| 2 | 108791642 | rs1718031 | 0.592 | 0.334 | -0.384 | -2.627 | 0.006 | 0.082 | 0.082 | 0 | 0.36 | 0.25 | 0.29 | 0.1 | 0.37 | 0.42 |
| 2 | 108795460 | rs855017 | 0.768 | 0.323 | -0.563 | -2.611 | -0.013 | 0.151 | 0.151 | 0 | 0.34 | 0.14 | 0.28 | 0.1 | 0.37 | 0.42 |
| 2 | 108808424 | rs855034 | 0.615 | 0.209 | 0.03 | -2.647 | 0.008 | 0.056 | 0.056 | 0 | 0.36 | 0.26 | 0.29 | 0.11 | 0.37 | 0.43 |
| 2 | 108808752 | rs855035 | 0.437 | 0.206 | 0.032 | -2.648 | 0.006 | 0.009 | 0.009 | 0 | 0.38 | 0.34 | 0.3 | 0.11 | 0.37 | 0.43 |
| 2 | 108816273 | rs855044 | 1.447 | 0.18 | -0.064 | -2.712 | -0.055 | 0.205 | 0.205 | 0 | 0.29 | 0.02 | 0.16 | 0.01 | 0.29 | 0.2 |
| 2 | 108816885 | rs6761157 | 0.022 | 0.282 | 0.17 | -2.56 | -0.015 | -0.019 | -0.019 | 0.113 | 0.34 | 0.47 | 0.45 | 0.7 | 0.27 | 0.3 |
| 2 | 108818916 | rs9631035 | 0.339 | 0.286 | 0.146 | -2.558 | -0.003 | -0.005 | -0.005 | 0 | 0.32 | 0.36 | 0.44 | 0.7 | 0.27 | 0.3 |
| 2 | 108830233 | rs855047 | -0.909 | 0.316 | 0.045 | -2.611 | -0.055 | 0.086 | 0.086 | 0 | 0.75 | 0.93 | 0.79 | 0.9 | 0.72 | 0.78 |
| 2 | 108840613 | rs4676183 | -0.424 | 0.123 | 0.161 | -2.548 | 0.003 | -0.006 | -0.006 | 0.657 | 0.7 | 0.71 | 0.73 | 0.81 | 0.64 | 0.76 |
| 2 | 108852039 | rs2198472 | 0.872 | 0.215 | 0.02 | -2.517 | 0.005 | 0.047 | 0.047 | 0 | 0.37 | 0.27 | 0.29 | 0.15 | 0.37 | 0.46 |
| 2 | 108855168 | rs2123693 | 0.77 | 0.235 | -0.276 | -2.614 | -0.045 | 0.241 | 0.241 | 0 | 0.42 | 0.11 | 0.31 | 0.11 | 0.41 | 0.47 |
| 2 | 108857505 | rs2123694 | 0.998 | 0.221 | -0.095 | -2.559 | -0.002 | 0.002 | 0.002 | 0 | 0.23 | 0.27 | 0.38 | 0.68 | 0.19 | 0.21 |
| 2 | 108863650 | rs9646947 | 1.56 | 0.214 | -0.151 | -2.603 | -0.089 | 0.169 | 0.169 | 0 | 0.33 | 0.03 | 0.26 | 0.1 | 0.36 | 0.41 |
| 2 | 108863898 | rs10209928 | 0.509 | -0.14 | 0.451 | -2.086 | -0.008 | 0.005 | 0.005 | 0 | 0.24 | 0.32 | 0.39 | 0.7 | 0.2 | 0.24 |
| 2 | 108872012 | rs13392744 | 0.634 | -0.056 | 0.265 | -2.089 | 0.002 | -0.004 | -0.004 | 0 | 0.08 | 0.07 | 0.05 | 0 | 0.05 | 0.06 |
| 2 | 108875244 | rs6722745 | -0.927 | -0.062 | 0.367 | -2.075 | 0.014 | 0.015 | 0.015 | 0 | 0.38 | 0.67 | 0.47 | 0.71 | 0.28 | 0.35 |
| 2 | 108883866 | rs13388627 | 0.41 | 0.111 | 0.221 | -2.108 | -0.011 | 0.004 | 0.004 | 0 | 0.23 | 0.3 | 0.4 | 0.69 | 0.2 | 0.25 |
| 2 | 108884042 | rs4638749 | 0.424 | 0.112 | 0.226 | -2.094 | -0.01 | 0.013 | 0.013 | 0 | 0.23 | 0.3 | 0.4 | 0.66 | 0.19 | 0.21 |
| 2 | 108885526 | rs11891937 | 0.45 | 0.344 | 0.23 | -2.237 | 0.01 | -0.026 | -0.026 | 0 | 0.07 | 0.15 | 0.06 | 0.02 | 0.08 | 0.09 |
| 2 | 108885895 | rs6542759 | -0.231 | 0.079 | 0.298 | -2.488 | -0.048 | -0.027 | -0.027 | 0 | 0.31 | 0.45 | 0.43 | 0.71 | 0.22 | 0.23 |
| 2 | 108886435 | rs17820651 | 1.072 | 0.449 | -0.721 | -2.984 | -0.152 | 0.082 | 0.082 | 0 | 0.22 | 0.03 | 0.29 | 0.1 | 0.41 | 0.35 |
| 2 | 108893909 | rs6542761 | -1.222 | 0.245 | -0.519 | -3.003 | 0.028 | 0.079 | 0.079 | 0 | 0.61 | 0.78 | 0.79 | 0.83 | 0.71 | 0.68 |
| 2 | 108899288 | rs11684742 | 0.939 | 0.255 | -0.474 | -3.019 | -0.164 | 0.09 | 0.09 | 0 | 0.25 | 0.03 | 0.3 | 0.1 | 0.43 | 0.34 |
| 2 | 108900509 | rs2198465 | -1.135 | 0.121 | -0.116 | -3.029 | 0.019 | 0.044 | 0.044 | 0 | 0.61 | 0.73 | 0.79 | 0.82 | 0.71 | 0.68 |
| 2 | 108901868 | rs6713582 | -0.407 | -0.021 | -0.076 | -3.134 | 0.013 | 0.013 | 0.013 | 0.515 | 0.5 | 0.45 | 0.75 | 0.8 | 0.65 | 0.58 |
| 2 | 108902219 | rs6745104 | -1.204 | -0.002 | -0.217 | -3.124 | 0.004 | 0.003 | 0.003 | 0 | 0.6 | 0.65 | 0.79 | 0.81 | 0.71 | 0.68 |
| 2 | 108902718 | rs7568215 | 0.376 | 0.168 | 0.05 | -2.768 | 0.002 | 0.005 | 0.005 | 0 | 0.7 | 0.7 | 0.83 | 0.81 | 0.74 | 0.64 |
| 2 | 108905030 | rs2305484 | -0.641 | 0.173 | 0.162 | -2.703 | -0.001 | -0.039 | -0.039 | 0 | 0.39 | 0.57 | 0.54 | 0.83 | 0.41 | 0.43 |
| 2 | 108905048 | rs2305485 | 0.522 | 0.697 | 0.408 | -2.418 | 0.002 | 0.014 | 0.014 | 0 | 0.2 | 0.29 | 0.41 | 0.73 | 0.21 | 0.21 |
| 2 | 108906495 | rs17821152 | 0.782 | -0.081 | 0.246 | -2.628 | -0.063 | 0.314 | 0.314 | 0.124 | 0.45 | 0.08 | 0.38 | 0.05 | 0.53 | 0.37 |
| 2 | 108907361 | rs10173554 | 0.228 | -0.081 | 0.36 | -2.589 | -0.004 | -0.061 | -0.061 | 0 | 0.55 | 0.51 | 0.44 | 0.16 | 0.6 | 0.45 |
| 2 | 108911809 | rs17036061 | 0.139 | -0.094 | 0.367 | -2.583 | -0.004 | -0.06 | -0.06 | 0 | 0.55 | 0.51 | 0.44 | 0.16 | 0.59 | 0.46 |
| 2 | 108915136 | rs7590599 | -0.462 | -0.123 | 0.39 | -2.837 | -0.025 | 0.114 | 0.114 | 0.111 | 0.65 | 0.87 | 0.79 | 0.81 | 0.66 | 0.6 |
| 2 | 108915263 | rs4149422 | -0.599 | -0.123 | 0.39 | -2.838 | -0.03 | 0.183 | 0.183 | 0.072 | 0.66 | 0.87 | 0.79 | 0.92 | 0.66 | 0.62 |
| 2 | 108915428 | rs7576937 | -0.509 | -0.114 | 0.433 | -2.834 | -0.001 | 0.11 | 0.11 | 0.005 | 0.66 | 0.79 | 0.79 | 0.92 | 0.66 | 0.61 |
| 2 | 108915608 | rs9646948 | 1.119 | -0.15 | 0.304 | -2.888 | -0.057 | 0.116 | 0.116 | 0.19 | 0.36 | 0.19 | 0.35 | 0.04 | 0.44 | 0.29 |
| 2 | 108916044 | rs4149423 | 0.436 | -0.207 | 0.232 | -2.519 | -0.013 | -0.011 | -0.011 | 0 | 0.2 | 0.35 | 0.39 | 0.76 | 0.16 | 0.28 |
| 2 | 108917596 | rs4149425 | -0.136 | -0.147 | -0.072 | -2.021 | 0.011 | -0.016 | -0.016 | 1.549 | 0.07 | 0.03 | 0.03 | 0.12 | 0.06 | 0.04 |
| 2 | 108921391 | rs11885032 | -0.246 | 0.023 | -1.136 | -1.933 | 0.014 | -0.018 | -0.018 | 0 | 0.08 | 0.03 | 0.03 | 0.12 | 0.06 | 0.05 |
| 2 | 108921495 | rs11885078 | 0.148 | 0.022 | -1.136 | -1.933 | 0.01 | 0.021 | 0.021 | 0 | 0.07 | 0.03 | 0.03 | 0 | 0.05 | 0.03 |
| 2 | 108921795 | rs11890318 | 0.148 | 0.022 | -1.136 | -1.933 | 0.01 | 0.021 | 0.021 | 0 | 0.07 | 0.03 | 0.03 | 0.01 | 0.05 | 0.03 |
| 2 | 108922036 | rs17036104 | -0.096 | 0.032 | -1.131 | -1.93 | 0.014 | -0.017 | -0.017 | 0.139 | 0.08 | 0.03 | 0.03 | 0.12 | 0.06 | 0.05 |
| 2 | 108925220 | rs3731680 | -0.096 | 0.051 | -1.706 | -1.885 | 0.014 | -0.018 | -0.018 | 0 | 0.08 | 0.03 | 0.03 | 0.12 | 0.06 | 0.05 |

| | | | | | | | | | | | | | | | | |
|---|---|---|---|---|---|---|---|---|---|---|---|---|---|---|---|---|
| 2 | 108927768 | rs4676193 | 0.191 | 0.294 | -1.456 | -2.265 | 0.002 | 0.108 | 0.108 | 0.216 | 0.68 | 0.83 | 0.81 | 0.91 | 0.7 | 0.61 |
| 2 | 108932436 | rs1989096 | -0.505 | -0.399 | -0.758 | -2.907 | -0.022 | -0.093 | -0.093 | 0 | 0.28 | 0.38 | 0.15 | 0.04 | 0.24 | 0.29 |
| 2 | 108938552 | rs1879495 | -0.365 | 0.247 | -1.298 | -3.283 | -0.07 | 0.062 | 0.062 | 0.227 | 0.74 | 0.84 | 0.77 | 1 | 0.58 | 0.79 |
| 2 | 108938735 | rs12477830 | 0.904 | -0.347 | -1.745 | -3.679 | 0.013 | 0.05 | 0.05 | 2.01 | 0.19 | 0.05 | 0.39 | 0.86 | 0.16 | 0.1 |
| 2 | 108939940 | rs7595791 | 0.349 | -0.591 | -1.262 | -3.812 | 0.022 | 0.063 | 0.063 | 0 | 0.51 | 0.62 | 0.36 | 0.13 | 0.41 | 0.68 |
| 2 | 108940336 | rs12476238 | 0.808 | -0.562 | -1.487 | -3.607 | 0.018 | 0.052 | 0.052 | 1.658 | 0.19 | 0.06 | 0.39 | 0.86 | 0.16 | 0.1 |
| 2 | 108946069 | rs17036146 | 0.031 | -0.304 | -0.848 | -3.356 | -0.01 | -0.068 | -0.068 | 0.301 | 0.24 | 0.31 | 0.13 | 0.06 | 0.23 | 0.24 |
| 2 | 108946178 | rs6709159 | -0.385 | -0.218 | -0.773 | -3.516 | -0.007 | -0.028 | -0.028 | 0 | 0.69 | 0.68 | 0.74 | 0.96 | 0.55 | 0.65 |
| 2 | 108946860 | rs6709978 | 0.314 | -0.177 | -0.864 | -3.133 | 0.012 | -0.002 | -0.002 | 0 | 0.46 | 0.38 | 0.62 | 0.91 | 0.33 | 0.41 |
| 2 | 108950832 | rs730631 | -0.853 | -0.43 | -0.772 | -2.912 | 0.016 | 0.013 | 0.013 | 1.012 | 0.24 | 0.76 | 0.12 | 0.07 | 0.19 | 0.26 |
| 2 | 108968224 | rs7577165 | -0.426 | -0.496 | 1.002 | -2.952 | -0.018 | -0.098 | -0.098 | 0 | 0.17 | 0.37 | 0.09 | 0.06 | 0.17 | 0.24 |
| 2 | 108971351 | rs1402464 | 0.577 | -0.963 | 0.535 | -3.47 | -0.004 | -0.023 | -0.023 | 0 | 0.16 | 0.18 | 0.07 | 0.03 | 0.08 | 0.18 |
| 2 | 108986048 | rs1105109 | 0.609 | -0.965 | 0.534 | -3.482 | -0.004 | -0.026 | -0.026 | 0 | 0.15 | 0.17 | 0.07 | 0.03 | 0.08 | 0.18 |
| 2 | 108988625 | rs10167564 | -1.35 | -1.048 | 0.644 | -3.541 | 0.073 | 0.144 | 0.144 | 0.063 | 0.37 | 0.83 | 0.18 | 0.11 | 0.26 | 0.44 |
| 2 | 108990797 | rs6716875 | 0.108 | -1.046 | 0.767 | -3.54 | -0.018 | -0.094 | -0.094 | 0.257 | 0.16 | 0.35 | 0.09 | 0.06 | 0.16 | 0.23 |
| 2 | 108995325 | rs4149432 | -1.322 | -1.072 | 0.65 | -3.553 | 0.077 | 0.151 | 0.151 | 0.016 | 0.37 | 0.83 | 0.17 | 0.11 | 0.26 | 0.45 |
| 2 | 108999786 | rs4149436 | -1.623 | -1.082 | 0.653 | -3.558 | 0.067 | 0.26 | 0.26 | 0.268 | 0.39 | 0.95 | 0.19 | 0.11 | 0.26 | 0.45 |
| 2 | 109001623 | rs4149437 | -0.514 | -1.087 | 0.826 | -3.559 | -0.025 | -0.08 | -0.08 | 0 | 0.2 | 0.6 | 0.1 | 0.07 | 0.18 | 0.26 |
| 2 | 109001646 | rs6739402 | 0.144 | -1.086 | 0.826 | -3.559 | -0.017 | -0.091 | -0.091 | 0 | 0.16 | 0.33 | 0.09 | 0.06 | 0.16 | 0.23 |
| 2 | 109002048 | rs4149438 | -1.318 | -1.086 | 0.794 | -3.559 | 0.077 | 0.151 | 0.151 | 0.016 | 0.37 | 0.83 | 0.17 | 0.11 | 0.26 | 0.45 |
| 2 | 109006665 | rs13021399 | 1.477 | -0.652 | -0.202 | -4.264 | -0.073 | 0.127 | 0.127 | 0 | 0.26 | 0.02 | 0.53 | 0.89 | 0.26 | 0.34 |
| 2 | 109012055 | rs11673744 | 1.025 | -0.68 | 0.031 | -4.13 | -0.005 | -0.025 | -0.025 | 0 | 0.15 | 0.19 | 0.07 | 0.03 | 0.08 | 0.19 |
| 2 | 109025768 | rs10196756 | 0.846 | -0.742 | 0.239 | -4.156 | -0.02 | -0.048 | -0.048 | 0 | 0.16 | 0.25 | 0.08 | 0.03 | 0.08 | 0.19 |
| 2 | 109028656 | rs6755756 | -0.995 | -0.623 | 0.099 | -4.219 | 0.066 | 0.184 | 0.184 | 0.107 | 0.38 | 0.88 | 0.19 | 0.11 | 0.28 | 0.45 |
| 2 | 109034995 | rs4395219 | 0.677 | -0.585 | 0.371 | -4.245 | 0.004 | -0.078 | -0.078 | 0 | 0.16 | 0.3 | 0.09 | 0.07 | 0.17 | 0.22 |
| 2 | 109045009 | rs7596199 | -0.173 | -0.169 | 0.189 | -4.532 | -0.001 | -0.083 | -0.083 | 2.369 | 0.52 | 0.47 | 0.68 | 0.96 | 0.55 | 0.61 |
| 2 | 109055717 | rs10203795 | 0.839 | -0.129 | 0.136 | -4.515 | -0.029 | 0.119 | 0.119 | 0 | 0.34 | 0.14 | 0.58 | 0.89 | 0.37 | 0.39 |
| 2 | 109055908 | rs10169264 | 0.637 | -0.126 | 0.13 | -4.513 | -0.014 | 0.089 | 0.089 | 0 | 0.36 | 0.17 | 0.59 | 0.89 | 0.38 | 0.39 |
| 2 | 109065004 | rs2465956 | 0.296 | 0.045 | 0.147 | -4.581 | 0.005 | -0.077 | -0.077 | 2.724 | 0.19 | 0.35 | 0.12 | 0.08 | 0.21 | 0.24 |
| 2 | 109066424 | rs2378113 | 1.175 | 0.03 | 0.008 | -4.592 | -0.088 | 0.18 | 0.18 | 0 | 0.33 | 0.06 | 0.57 | 0.88 | 0.36 | 0.38 |
| 2 | 109099477 | rs2577613 | -0.458 | -0.004 | 0.114 | -4.573 | -0.002 | -0.095 | -0.095 | 0.29 | 0.55 | 0.52 | 0.7 | 0.97 | 0.59 | 0.63 |
| 2 | 109105194 | rs2577594 | -1.411 | -0.013 | -0.161 | -4.579 | -0.114 | 0.191 | 0.191 | 0 | 0.73 | 0.97 | 0.8 | 1 | 0.67 | 0.83 |
| 2 | 109150164 | rs11123706 | 1.735 | 0.138 | 0.016 | -4.426 | -0.049 | 0.104 | 0.104 | 0 | 0.25 | 0.02 | 0.5 | 0.88 | 0.23 | 0.33 |
| 2 | 109156275 | rs6754683 | 0.929 | -0.072 | 0.433 | -4.208 | -0.03 | 0.105 | 0.105 | 0 | 0.34 | 0.15 | 0.59 | 0.89 | 0.38 | 0.39 |
| 2 | 109158237 | rs2718740 | -0.417 | -0.106 | 0.639 | -4.239 | 0.005 | -0.023 | -0.023 | 0.008 | 0.55 | 0.58 | 0.71 | 0.97 | 0.59 | 0.63 |
| 2 | 109170054 | rs6747243 | 0.465 | -0.077 | 0.605 | -4.234 | -0.015 | 0.03 | 0.03 | 0 | 0.38 | 0.25 | 0.62 | 0.96 | 0.41 | 0.56 |
| 2 | 109170731 | rs1469964 | 0.61 | -0.075 | 0.602 | -4.232 | -0.022 | 0.063 | 0.063 | 0 | 0.37 | 0.19 | 0.61 | 0.95 | 0.4 | 0.55 |
| 2 | 109182594 | rs7568838 | -0.257 | -0.068 | 0.556 | -4.236 | 0.004 | -0.014 | -0.014 | 0 | 0.05 | 0.05 | 0.04 | 0.06 | 0.06 | 0.17 |
| 2 | 109197856 | rs2718721 | -0.582 | -0.102 | 0.702 | -4.265 | -0.001 | 0.144 | 0.144 | 0 | 0.57 | 0.75 | 0.71 | 0.98 | 0.59 | 0.63 |
| 2 | 109198530 | rs13413437 | 1.221 | -0.053 | 0.6 | -4.217 | -0.045 | 0.127 | 0.127 | 0 | 0.32 | 0.1 | 0.58 | 0.88 | 0.37 | 0.38 |
| 2 | 109232388 | rs826682 | 0.68 | -0.313 | 0.935 | -4.075 | -0.003 | -0.084 | -0.084 | 0.866 | 0.14 | 0.3 | 0.08 | 0.01 | 0.15 | 0.06 |
| 2 | 109233079 | rs6744135 | 0.623 | -0.312 | 0.642 | -4.118 | -0.039 | 0.11 | 0.11 | 0 | 0.38 | 0.15 | 0.62 | 0.95 | 0.43 | 0.55 |
| 2 | 109260259 | rs10170690 | 0.617 | -0.312 | 0.643 | -4.118 | -0.039 | 0.11 | 0.11 | 0 | 0.38 | 0.15 | 0.62 | 0.95 | 0.43 | 0.55 |
| 2 | 109301074 | rs826590 | -0.264 | -0.339 | 0.955 | -3.768 | 0.008 | 0.135 | 0.135 | 0 | 0.45 | 0.27 | 0.29 | 0.02 | 0.41 | 0.37 |
| 2 | 109401259 | rs826556 | -1.216 | -0.56 | 2.116 | -3.235 | 0.022 | 0.025 | 0.025 | 0.161 | 0.33 | 0.76 | 0.22 | 0.08 | 0.24 | 0.37 |
| 2 | 109408208 | rs6740879 | -0.933 | -0.574 | 2.301 | -3.218 | -0.019 | -0.107 | -0.107 | 2.715 | 0.28 | 0.36 | 0.19 | 0.09 | 0.23 | 0.37 |
| 2 | 109412959 | rs4387742 | 0.696 | -0.608 | 2.223 | -3.207 | -0.019 | -0.107 | -0.107 | 0 | 0.72 | 0.64 | 0.81 | 0.91 | 0.77 | 0.63 |
| 2 | 109419600 | rs10190045 | 0.357 | -0.447 | 1.633 | -3.143 | 0.009 | -0.061 | -0.061 | 0.975 | 0.84 | 0.75 | 0.88 | 0.94 | 0.82 | 0.79 |
| 2 | 109434315 | rs1997120 | 0.239 | -0.442 | 1.629 | -3.14 | -0.002 | -0.009 | -0.009 | 0.613 | 0.85 | 0.85 | 0.88 | 0.94 | 0.82 | 0.79 |
| 2 | 109442014 | rs10190977 | 0.239 | -0.442 | 1.629 | -3.14 | -0.002 | -0.009 | -0.009 | 0 | 0.85 | 0.85 | 0.88 | 0.94 | 0.82 | 0.78 |
| 2 | 109472671 | rs10865023 | 0.244 | -0.589 | 1.935 | -2.818 | 0.002 | -0.02 | -0.02 | 0 | 0.93 | 0.9 | 0.93 | 0.99 | 0.9 | 0.92 |
| 2 | 109491020 | rs10195050 | 0.418 | -0.691 | 2.118 | -2.892 | -0.012 | -0.032 | -0.032 | 0 | 0.13 | 0.19 | 0.06 | 0.03 | 0.06 | 0.16 |
| 2 | 109498997 | rs765089 | 1.424 | -0.744 | 2.363 | -2.916 | -0.039 | -0.096 | -0.096 | 0 | 0.75 | 0.35 | 0.84 | 0.95 | 0.84 | 0.73 |
| 2 | 109501066 | rs1478516 | 1.437 | -0.758 | 2.392 | -2.888 | -0.025 | -0.05 | -0.05 | 1.571 | 0.74 | 0.28 | 0.83 | 0.89 | 0.84 | 0.62 |
| 2 | 109501748 | rs17037027 | 0.625 | -0.738 | 2.304 | -2.88 | 0.007 | -0.005 | -0.005 | 0 | 0.12 | 0.11 | 0.06 | 0.03 | 0.06 | 0.16 |
| 2 | 109507073 | rs12712027 | 0.066 | -0.737 | 2.299 | -2.88 | 0 | -0.012 | -0.012 | 0 | 0.93 | 0.91 | 0.94 | 0.99 | 0.9 | 0.92 |
| 2 | 109512234 | rs13001699 | 1.246 | -0.738 | 2.296 | -2.881 | -0.04 | -0.113 | -0.113 | 0.022 | 0.76 | 0.35 | 0.85 | 0.96 | 0.84 | 0.76 |
| 2 | 109512428 | rs3749096 | -0.028 | -0.738 | 2.358 | -2.88 | -0.055 | -0.093 | -0.093 | 0.217 | 0.15 | 0.38 | 0.08 | 0.03 | 0.06 | 0.16 |
| 2 | 109512606 | rs3749097 | 0.497 | -0.737 | 2.203 | -2.879 | -0.016 | -0.037 | -0.037 | 0.069 | 0.13 | 0.19 | 0.06 | 0.03 | 0.06 | 0.16 |
| 2 | 109513654 | rs12623957 | 1.369 | -0.729 | 2.04 | -2.875 | -0.039 | -0.096 | -0.096 | 0 | 0.75 | 0.35 | 0.84 | 0.95 | 0.84 | 0.73 |
| 2 | 109525530 | rs260633 | 1.584 | -0.634 | 1.964 | -2.932 | -0.021 | -0.079 | -0.079 | 0 | 0.71 | 0.37 | 0.8 | 0.9 | 0.76 | 0.69 |
| 2 | 109527373 | rs260630 | 0.332 | -0.58 | 1.805 | -2.706 | 0.004 | -0.025 | -0.025 | 2.706 | 0.93 | 0.9 | 0.93 | 0.99 | 0.91 | 0.92 |
| 2 | 109531958 | rs17037099 | 0.579 | -0.568 | 1.676 | -2.673 | 0.02 | 0.016 | 0.016 | 0.078 | 0.12 | 0.06 | 0.06 | 0.03 | 0.05 | 0.1 |
| 2 | 109536784 | rs260642 | 1.179 | -0.497 | 1.682 | -2.568 | -0.024 | -0.087 | -0.087 | 0 | 0.75 | 0.55 | 0.85 | 0.9 | 0.82 | 0.65 |
| 2 | 109539430 | rs749810 | -0.6 | -0.764 | 1.116 | -2.188 | -0.084 | -0.102 | -0.102 | 0 | 0.17 | 0.57 | 0.09 | 0.05 | 0.05 | 0.13 |
| 2 | 109584599 | rs10779899 | 1.31 | -0.311 | 1.318 | -2.366 | -0.019 | -0.102 | -0.102 | 0 | 0.82 | 0.61 | 0.89 | 0.97 | 0.84 | 0.85 |
| 2 | 109586437 | rs11691107 | -1.419 | -0.324 | 1.815 | -2.165 | 0.036 | 0.069 | 0.069 | 0.323 | 0.06 | 0.33 | 0.04 | 0.12 | 0.03 | 0.06 |
| 2 | 109590380 | rs12992554 | 0.882 | -0.996 | 1.22 | -2.656 | 0.037 | 0.23 | 0.23 | 4.384 | 0.54 | 0.12 | 0.38 | 0.12 | 0.6 | 0.74 |
| 2 | 109599256 | rs260674 | 0.126 | -1.269 | 1.483 | -2.051 | -0.032 | -0.029 | -0.029 | 0 | 0.8 | 0.66 | 0.56 | 0.2 | 0.85 | 0.87 |

| | | | | | | | | | | | | | | | | |
|---|---|---|---|---|---|---|---|---|---|---|---|---|---|---|---|---|
| 2 | 109599670 | rs17269487 | 1.779 | -1.118 | 0.985 | -2.057 | 0.045 | 0.295 | 0.295 | 2.197 | 0.5 | 0.12 | 0.37 | 0.06 | 0.59 | 0.69 |
| 2 | 109603847 | rs393987 | 0.161 | -1.08 | 1.008 | -2.032 | -0.009 | 0.045 | 0.045 | 1.513 | 0.22 | 0.14 | 0.15 | 0 | 0.23 | 0.1 |
| 2 | 109607256 | rs6749207 | 0.095 | -1.097 | 1.123 | -1.952 | -0.021 | -0.024 | -0.024 | 0 | 0.19 | 0.34 | 0.44 | 0.81 | 0.15 | 0.15 |
| 2 | 109609765 | rs4676225 | -0.037 | -1.03 | 0.931 | -1.965 | 0 | 0 | 0 | 2.581 | 0.75 | 0.69 | 0.83 | 0.87 | 0.76 | 0.86 |
| 2 | 109610788 | rs6746808 | 1.723 | -1.008 | 0.844 | -1.958 | 0.036 | 0.336 | 0.336 | 1.86 | 0.5 | 0.09 | 0.37 | 0.03 | 0.58 | 0.64 |
| 2 | 109611097 | rs6750059 | 0.282 | -0.847 | 0.766 | -1.796 | -0.003 | -0.039 | -0.039 | 0 | 0.73 | 0.59 | 0.82 | 0.87 | 0.75 | 0.85 |
| 2 | 109622445 | rs6716645 | -0.026 | -0.849 | 0.51 | -1.041 | -0.017 | 0.086 | 0.086 | 0.795 | 0.18 | 0.44 | 0.36 | 0.71 | 0.14 | 0.17 |
| 2 | 109623949 | rs260617 | -0.107 | -0.862 | 0.255 | -1.555 | 0.008 | 0.047 | 0.047 | 1.977 | 0.26 | 0.21 | 0.16 | 0.1 | 0.23 | 0.1 |
| 2 | 109626157 | rs171856 | -0.933 | -0.627 | -0.927 | -1.428 | -0.001 | 0.376 | 0.376 | 1.684 | 0.52 | 0.87 | 0.67 | 0.96 | 0.43 | 0.4 |
| 2 | 109635257 | rs4676049 | -0.973 | -0.8 | -0.082 | -0.768 | -0.02 | 0.038 | 0.038 | 0.015 | 0.12 | 0.33 | 0.09 | 0.15 | 0.07 | 0.08 |
| 2 | 109635905 | rs17034806 | -1.221 | -0.863 | 0.051 | -0.649 | -0.025 | 0.055 | 0.055 | 0.068 | 0.13 | 0.36 | 0.12 | 0.2 | 0.07 | 0.1 |
| 2 | 109637508 | rs921876 | 0.534 | -0.575 | -0.496 | -1.039 | 0.012 | 0.087 | 0.087 | 6.273 | 0.52 | 0.43 | 0.67 | 0.78 | 0.56 | 0.69 |
| 2 | 109638248 | rs4477868 | 0.429 | -0.54 | -0.499 | -1.125 | 0.01 | 0.056 | 0.056 | 6.546 | 0.53 | 0.44 | 0.71 | 0.84 | 0.58 | 0.71 |
| 2 | 109644873 | rs13422589 | -0.064 | -0.762 | 0.07 | -1.376 | -0.004 | -0.066 | -0.066 | 0 | 0.2 | 0.21 | 0.12 | 0 | 0.16 | 0.09 |
| 2 | 109646359 | rs9653456 | 0.372 | -0.728 | 0.294 | -1.148 | -0.01 | -0.052 | -0.052 | 0 | 0.71 | 0.64 | 0.82 | 0.86 | 0.73 | 0.8 |
| 2 | 109648715 | rs4676051 | 0.639 | -0.652 | 0.022 | -1.24 | -0.019 | 0.046 | 0.046 | 0.304 | 0.73 | 0.84 | 0.82 | 0.87 | 0.69 | 0.73 |
| 2 | 109652644 | rs12998605 | 1.554 | -1.12 | 1.099 | -0.545 | -0.036 | 0.091 | 0.091 | 0 | 0.91 | 0.47 | 0.93 | 0.87 | 0.96 | 0.92 |
| 2 | 109662043 | rs4676060 | 1.492 | -1.155 | 1.14 | -0.485 | -0.025 | 0.144 | 0.144 | 0 | 0.91 | 0.54 | 0.93 | 0.86 | 0.96 | 0.91 |
| 2 | 109668527 | rs17269557 | -0.407 | -0.96 | 0.531 | -0.882 | -0.017 | 0.06 | 0.06 | 0 | 0.17 | 0.07 | 0.18 | 0 | 0.15 | 0.08 |
| 2 | 109669494 | rs6737482 | -1.034 | -1.076 | 0.877 | -0.45 | -0.042 | 0.008 | 0.008 | 0.399 | 0.19 | 0.26 | 0.35 | 0.76 | 0.08 | 0.15 |
| 2 | 109677619 | rs425238 | -0.986 | -0.646 | 0.168 | -0.199 | 0.01 | 0.004 | 0.004 | 0.479 | 0.23 | 0.2 | 0.11 | 0.17 | 0.16 | 0.16 |
| 2 | 109678210 | rs417999 | -0.261 | -0.53 | 0.055 | -0.154 | 0.001 | 0.003 | 0.003 | 0.379 | 0.17 | 0.07 | 0.09 | 0.12 | 0.14 | 0.07 |
| 2 | 109680352 | rs11689524 | -0.257 | -0.222 | -0.115 | -0.521 | -0.02 | 0.025 | 0.025 | 0 | 0.23 | 0.15 | 0.17 | 0.11 | 0.31 | 0.1 |
| 2 | 109686568 | rs260652 | 0.004 | -0.337 | 0.027 | -0.498 | 0 | 0 | 0 | 0 | 0.18 | 0.16 | 0.1 | 0.12 | 0.14 | 0.07 |
| 2 | 109688373 | rs260651 | -1.387 | -0.366 | 0.524 | -0.564 | 0.002 | -0.011 | -0.011 | 0 | 0.44 | 0.45 | 0.3 | 0.26 | 0.47 | 0.21 |
| 2 | 109697513 | rs11688442 | -0.531 | 0.012 | 0.327 | -0.297 | -0.053 | 0.017 | 0.017 | 0 | 0.23 | 0.08 | 0.16 | 0.14 | 0.29 | 0.13 |
| 2 | 109716551 | rs11674275 | -0.597 | -0.145 | 0.512 | -0.43 | -0.069 | 0.052 | 0.052 | 0 | 0.24 | 0.03 | 0.16 | 0.1 | 0.28 | 0.15 |
| 2 | 109742307 | rs6542797 | 1.367 | -0.145 | 0.985 | -0.351 | 0 | -0.015 | -0.015 | 0 | 0.61 | 0.39 | 0.4 | 0.25 | 0.62 | 0.65 |
| 2 | 109749303 | rs6740122 | -0.451 | 0.024 | 0.597 | -0.279 | -0.078 | 0.044 | 0.044 | 0 | 0.24 | 0.03 | 0.16 | 0.1 | 0.28 | 0.15 |
| 2 | 109765916 | rs10188630 | 0.639 | 0.035 | 0.801 | -0.26 | 0.015 | 0.003 | 0.003 | 0 | 0.79 | 0.75 | 0.46 | 0.21 | 0.76 | 0.71 |
| 2 | 109767173 | rs11676676 | -0.348 | -0.086 | 0.978 | -0.433 | 0.003 | 0.003 | 0.003 | 0 | 0.2 | 0.16 | 0.12 | 0.1 | 0.19 | 0.14 |
| 2 | 109769146 | rs737478 | -0.063 | -0.053 | 0.73 | -0.415 | 0.01 | 0.021 | 0.021 | 0 | 0.18 | 0.06 | 0.12 | 0.09 | 0.18 | 0.14 |
| 2 | 109769979 | rs6739562 | 0.755 | 0.122 | 0.821 | -0.236 | -0.006 | 0 | 0 | 0 | 0.79 | 0.8 | 0.46 | 0.18 | 0.76 | 0.71 |
| 2 | 109773689 | rs6752599 | -0.111 | -0.03 | 0.884 | -0.203 | -0.002 | -0.013 | -0.013 | 0 | 0.82 | 0.78 | 0.87 | 0.9 | 0.82 | 0.89 |
| 2 | 109776566 | rs891421 | -0.102 | -0.031 | 0.967 | -0.203 | -0.003 | -0.017 | -0.017 | 0.725 | 0.82 | 0.75 | 0.87 | 0.9 | 0.82 | 0.89 |
| 2 | 109798726 | rs4676258 | 1.305 | -0.036 | 1.585 | 0.756 | 0.001 | -0.013 | -0.013 | 0 | 0.6 | 0.28 | 0.45 | 0.4 | 0.63 | 0.63 |
| 2 | 109799833 | rs6749764 | -0.185 | -0.006 | 1.472 | 0.702 | -0.011 | -0.016 | -0.016 | 0 | 0.82 | 0.76 | 0.88 | 0.9 | 0.84 | 0.89 |
| 2 | 109809695 | rs6542803 | -0.157 | 0.003 | 1.476 | 0.75 | -0.012 | -0.007 | -0.007 | 0.235 | 0.81 | 0.76 | 0.87 | 0.86 | 0.84 | 0.87 |
| 2 | 109819331 | rs17035137 | -0.373 | -0.017 | 1.617 | 0.841 | 0.007 | 0.038 | 0.038 | 0 | 0.21 | 0.24 | 0.41 | 0.52 | 0.21 | 0.24 |
| 2 | 109822524 | rs17269661 | 0.99 | 0.047 | 0.867 | 0.12 | -0.022 | 0.159 | 0.159 | 0 | 0.38 | 0.13 | 0.21 | 0.09 | 0.41 | 0.37 |
| 2 | 109825516 | rs12328666 | -0.751 | 0.027 | 1.372 | 0.256 | 0.013 | 0.114 | 0.114 | 0 | 0.65 | 0.9 | 0.78 | 0.9 | 0.67 | 0.77 |
| 2 | 109827020 | rs12619389 | -0.092 | -0.004 | 1.306 | 0.256 | 0.008 | 0.054 | 0.054 | 0 | 0.23 | 0.34 | 0.41 | 0.49 | 0.22 | 0.28 |
| 2 | 109829904 | rs11691266 | 0.973 | 0.043 | 1.174 | 0.264 | -0.041 | 0.193 | 0.193 | 0 | 0.38 | 0.11 | 0.21 | 0.09 | 0.41 | 0.39 |
| 2 | 109856486 | rs6760953 | -1.295 | 0.247 | 1.221 | 0.417 | 0.104 | 0.526 | 0.526 | 0.154 | 0.06 | 0.49 | 0.16 | 0.33 | 0.04 | 0.12 |
| 2 | 109859519 | rs1864474 | -0.744 | 0.078 | 1.006 | 0.266 | 0.022 | 0.115 | 0.115 | 0 | 0.63 | 0.87 | 0.79 | 0.9 | 0.66 | 0.75 |
| 2 | 109861063 | rs7586264 | 0.159 | 0.038 | 0.802 | 0.238 | -0.004 | -0.014 | -0.014 | 1.842 | 0.23 | 0.2 | 0.4 | 0.48 | 0.21 | 0.26 |
| 2 | 109874379 | rs891423 | -0.846 | 0.029 | -0.501 | -0.085 | 0.017 | 0.123 | 0.123 | 0 | 0.65 | 0.9 | 0.8 | 0.9 | 0.68 | 0.76 |
| 2 | 109881236 | rs1465630 | 0.865 | 0.137 | -0.535 | 0.002 | -0.047 | 0.145 | 0.145 | 0.063 | 0.36 | 0.12 | 0.23 | 0.1 | 0.43 | 0.38 |
| 2 | 109890276 | rs6719705 | -0.146 | -0.165 | -0.195 | 0.231 | -0.006 | 0.013 | 0.013 | 0 | 0.33 | 0.38 | 0.41 | 0.48 | 0.25 | 0.28 |
| 2 | 109893210 | rs891425 | -0.116 | -0.158 | -0.323 | 0.249 | 0.002 | -0.006 | -0.006 | 0 | 0.33 | 0.31 | 0.41 | 0.48 | 0.25 | 0.31 |
| 2 | 109893879 | rs744898 | 1.13 | 0.188 | -0.345 | 0.15 | -0.004 | -0.004 | -0.004 | 0.704 | 0.27 | 0.25 | 0.34 | 0.38 | 0.39 | 0.43 |
| 2 | 109895831 | rs2163261 | -0.649 | 0.196 | -0.365 | 0.091 | -0.005 | -0.034 | -0.034 | 0 | 0.6 | 0.56 | 0.75 | 0.86 | 0.64 | 0.74 |
| 2 | 109897005 | rs4504023 | -0.803 | 0.222 | -0.632 | 0.051 | 0.005 | 0.049 | 0.049 | 0 | 0.62 | 0.74 | 0.77 | 0.9 | 0.65 | 0.76 |
| 2 | 109901013 | rs730491 | 0.18 | 0.787 | -0.741 | 0.42 | 0.022 | 0.039 | 0.039 | 0 | 0.31 | 0.54 | 0.38 | 0.42 | 0.4 | 0.46 |
| 2 | 109917069 | rs7597710 | 0.845 | 0.885 | -1.656 | 0.027 | -0.083 | 0.06 | 0.06 | 0 | 0.22 | 0.06 | 0.18 | 0.09 | 0.33 | 0.34 |
| 2 | 109930463 | rs12472919 | 0.54 | 0.806 | -1.361 | 0.513 | -0.024 | -0.034 | -0.034 | 0 | 0.09 | 0.03 | 0.2 | 0.24 | 0.12 | 0.1 |
| 2 | 109930525 | rs4632376 | 0.318 | 0.255 | -1.467 | 0.247 | -0.007 | -0.004 | -0.004 | 0 | 0.32 | 0.6 | 0.31 | 0.22 | 0.23 | 0.26 |
| 2 | 109931911 | rs1025378 | 0.016 | 0.201 | -1.588 | 0.096 | -0.014 | -0.03 | -0.03 | 0 | 0.24 | 0.15 | 0.39 | 0.64 | 0.26 | 0.22 |
| 2 | 109932096 | rs17035400 | 0.488 | 0.205 | -1.65 | 0.104 | -0.007 | -0.004 | -0.004 | 0 | 0.32 | 0.6 | 0.31 | 0.22 | 0.23 | 0.26 |
| 2 | 109935914 | rs891428 | -0.074 | 0.213 | -1.62 | 0.117 | -0.001 | 0.001 | 0.001 | 0 | 0.24 | 0.23 | 0.39 | 0.64 | 0.26 | 0.23 |
| 2 | 109937740 | rs4676273 | -0.961 | 0.247 | -2.18 | 0.074 | -0.044 | 0.201 | 0.201 | 0 | 0.56 | 0.82 | 0.7 | 0.85 | 0.49 | 0.48 |
| 2 | 109939843 | rs11689995 | 0.898 | 0.236 | -2.11 | 0.117 | -0.036 | 0.138 | 0.138 | 0.018 | 0.39 | 0.2 | 0.29 | 0.15 | 0.47 | 0.47 |
| 2 | 109941554 | rs11123741 | 1.06 | 0.26 | -2.211 | 0.22 | -0.019 | -0.028 | -0.028 | 0 | 0.09 | 0.04 | 0.2 | 0.22 | 0.12 | 0.1 |
| 2 | 109942206 | rs891415 | -0.938 | 0.227 | -2.299 | 0.191 | -0.036 | -0.125 | -0.125 | 1.323 | 0.8 | 0.98 | 0.81 | 0.58 | 0.82 | 0.82 |
| 2 | 109948465 | rs4676078 | -0.042 | 0.705 | -1.856 | -0.153 | 0.018 | 0.077 | 0.077 | 0 | 0.48 | 0.24 | 0.48 | 0.38 | 0.6 | 0.57 |
| 2 | 109948720 | rs4676274 | 0.524 | 0.78 | -2.126 | -0.519 | -0.143 | 0.184 | 0.184 | 0.083 | 0.37 | 0.04 | 0.28 | 0.11 | 0.47 | 0.44 |
| 2 | 109960595 | rs12470895 | 0.574 | 0.469 | -1.281 | -0.497 | -0.001 | -0.048 | -0.048 | 0.034 | 0.2 | 0.21 | 0.11 | 0 | 0.17 | 0.11 |
| 2 | 109962283 | rs6710178 | 0.384 | 0.328 | -0.642 | -0.473 | -0.005 | -0.102 | -0.102 | 0 | 0.22 | 0.37 | 0.13 | 0.04 | 0.18 | 0.14 |
| 2 | 109962344 | rs6754115 | -1.229 | 0.531 | -1.131 | -0.817 | -0.064 | 0.139 | 0.139 | 0 | 0.69 | 0.91 | 0.75 | 0.89 | 0.61 | 0.61 |

| | | | | | | | | | | | | | | | | |
|---|---|---|---|---|---|---|---|---|---|---|---|---|---|---|---|---|
| 2 | 109966153 | rs13407916 | 0.176 | -0.15 | -0.661 | -0.245 | -0.019 | 0.014 | 0.014 | 0.215 | 0.29 | 0.37 | 0.35 | 0.61 | 0.21 | 0.29 |
| 2 | 109972407 | rs7605935 | 0.786 | -0.278 | -0.532 | -0.46 | -0.001 | -0.032 | -0.032 | 0 | 0.19 | 0.19 | 0.08 | 0 | 0.12 | 0.11 |
| 2 | 109974444 | rs6716113 | -0.07 | -0.135 | 0.023 | -0.76 | -0.001 | -0.032 | -0.032 | 0 | 0.51 | 0.49 | 0.33 | 0.16 | 0.52 | 0.53 |
| 2 | 109992208 | rs10192569 | 0.598 | -0.044 | 0.241 | -0.404 | 0.018 | -0.074 | -0.074 | 0 | 0.34 | 0.5 | 0.27 | 0.15 | 0.41 | 0.41 |
| 2 | 109994532 | rs11687726 | 1.289 | -0.144 | -0.021 | -0.462 | -0.072 | 0.111 | 0.111 | 0 | 0.29 | 0.07 | 0.22 | 0.1 | 0.38 | 0.35 |
| 2 | 109995814 | rs6730477 | 0.002 | -0.484 | 0.121 | -0.108 | -0.006 | 0.066 | 0.066 | 0 | 0.54 | 0.69 | 0.44 | 0.27 | 0.54 | 0.55 |
| 2 | 110002052 | rs7605559 | 0.279 | -0.47 | -0.013 | -0.15 | -0.001 | -0.005 | -0.005 | 1.674 | 0.51 | 0.43 | 0.42 | 0.22 | 0.52 | 0.48 |
| 2 | 110004170 | rs3860481 | -0.254 | -0.524 | 0.123 | 0.19 | -0.011 | 0.007 | 0.007 | 0 | 0.57 | 0.63 | 0.3 | 0.4 | 0.53 | 0.63 |
| 2 | 110004196 | rs3860482 | -0.006 | -0.623 | 0.523 | 0.167 | -0.008 | 0.038 | 0.038 | 0 | 0.53 | 0.34 | 0.27 | 0.34 | 0.52 | 0.54 |
| 2 | 110021006 | rs11123744 | -0.128 | -1.132 | 0.644 | 0.464 | -0.028 | 0.036 | 0.036 | 0 | 0.23 | 0.24 | 0.08 | 0.5 | 0.11 | 0.21 |
| 2 | 110024153 | rs3922817 | 0.338 | -0.92 | 0.98 | 0.399 | 0.017 | 0.134 | 0.134 | 0 | 0.48 | 0.23 | 0.26 | 0.16 | 0.46 | 0.42 |
| 2 | 110026219 | rs4234110 | -0.517 | -0.778 | 1.293 | 0.251 | -0.017 | -0.019 | -0.019 | 0 | 0.64 | 0.7 | 0.34 | 0.49 | 0.55 | 0.64 |
| 2 | 110026466 | rs11684909 | -0.166 | -0.776 | 1.23 | 0.249 | 0.009 | 0.003 | 0.003 | 0 | 0.61 | 0.55 | 0.31 | 0.44 | 0.52 | 0.54 |
| 2 | 110030964 | rs10202141 | -0.358 | -0.646 | 0.866 | 0.262 | 0.006 | 0.008 | 0.008 | 0 | 0.63 | 0.57 | 0.33 | 0.5 | 0.55 | 0.64 |
| 2 | 110042294 | rs10207608 | -0.949 | -0.775 | 0.235 | -0.102 | -0.011 | -0.017 | -0.017 | 0.772 | 0.57 | 0.67 | 0.63 | 0.5 | 0.5 | 0.55 |
| 2 | 110048555 | rs7568694 | -0.322 | -0.174 | 0.36 | 0.065 | -0.003 | -0.025 | -0.025 | 0 | 0.12 | 0.12 | 0.09 | 0.01 | 0.1 | 0.09 |
| 2 | 110053209 | rs7370446 | -0.218 | -0.204 | 0.197 | -0.004 | -0.006 | -0.016 | -0.016 | 0.112 | 0.54 | 0.2 | 0.63 | 0.44 | 0.59 | 0.56 |
| 2 | 110053248 | rs7369523 | 0.598 | -0.098 | -0.023 | -0.083 | -0.082 | -0.106 | -0.106 | 0 | 0.18 | 0.04 | 0.44 | 0.39 | 0.31 | 0.23 |
| 2 | 110057303 | rs7568486 | -0.383 | -0.141 | 0.682 | 0.224 | -0.014 | -0.049 | -0.049 | 0 | 0.1 | 0.19 | 0.07 | 0.01 | 0.06 | 0.09 |
| 2 | 110067876 | rs7600602 | -0.848 | -0.015 | 0.287 | 0.339 | -0.024 | -0.057 | -0.057 | 0 | 0.66 | 0.79 | 0.7 | 0.5 | 0.63 | 0.65 |
| 2 | 110077153 | rs6542832 | -0.348 | 0.369 | -0.029 | 0.705 | 0 | 0.001 | 0.001 | 0.218 | 0.53 | 0.58 | 0.63 | 0.44 | 0.55 | 0.55 |
| 2 | 110086491 | rs7369007 | -0.11 | 0.72 | -1.393 | 0.863 | -0.037 | -0.015 | -0.015 | 0.175 | 0.48 | 0.16 | 0.6 | 0.44 | 0.54 | 0.49 |
| 2 | 110102339 | rs9646952 | -1.011 | 1.599 | -0.187 | 1.487 | 0.012 | 0.019 | 0.019 | 0 | 0.16 | 0.21 | 0.4 | 0.29 | 0.23 | 0.21 |
| 2 | 111705427 | rs1398948 | -0.68 | 0.06 | -0.41 | 0.732 | -0.002 | 0.029 | 0.029 | 0.192 | 0.72 | 0.35 | 0.49 | 0.59 | 0.69 | 0.58 |
| 2 | 111712703 | rs1877655 | -0.935 | 0.121 | -0.097 | 1.036 | 0.002 | -0.002 | -0.002 | 0.103 | 0.63 | 0.54 | 0.7 | 0.71 | 0.56 | 0.52 |
| 2 | 111713046 | rs4577288 | -0.351 | 0.167 | -0.156 | 0.859 | -0.002 | 0 | 0 | 0 | 0.56 | 0.41 | 0.53 | 0.54 | 0.54 | 0.48 |
| 2 | 111717100 | rs3789109 | -0.427 | 0.282 | -0.564 | 0.629 | 0.001 | -0.007 | -0.007 | 0 | 0.57 | 0.45 | 0.69 | 0.76 | 0.49 | 0.67 |
| 2 | 111720884 | rs3789106 | -0.446 | 0.846 | -0.951 | 0.462 | -0.101 | 0.175 | 0.175 | 0 | 0.38 | 0.11 | 0.42 | 0.15 | 0.44 | 0.32 |
| 2 | 111722118 | rs13386443 | -0.724 | 1.026 | -0.836 | 0.756 | -0.061 | 0.009 | 0.009 | 0 | 0.39 | 0.14 | 0.57 | 0.34 | 0.44 | 0.36 |
| 2 | 111723230 | rs2341912 | -0.172 | 0.715 | -0.323 | 0.654 | -0.046 | -0.034 | -0.034 | 0.195 | 0.13 | 0.34 | 0.07 | 0.05 | 0.05 | 0.22 |
| 2 | 111723551 | rs1905356 | -1.509 | 0.776 | -0.274 | 0.95 | -0.004 | 0.037 | 0.037 | 0 | 0.55 | 0.65 | 0.73 | 0.83 | 0.53 | 0.66 |
| 2 | 111728482 | rs4849303 | -1.23 | 0.77 | -0.497 | 0.607 | 0 | 0 | 0 | 0 | 0.51 | 0.51 | 0.63 | 0.39 | 0.48 | 0.57 |
| 2 | 111737275 | rs17041670 | -1.085 | 0.638 | -0.384 | 0.864 | -0.019 | 0.036 | 0.036 | 0 | 0.29 | 0.44 | 0.37 | 0.39 | 0.2 | 0.41 |
| 2 | 111740221 | rs11123271 | -0.611 | 0.909 | -0.635 | 0.647 | 0.012 | 0.023 | 0.023 | 0 | 0.42 | 0.62 | 0.51 | 0.25 | 0.38 | 0.41 |
| 2 | 111740671 | rs17481486 | 1.765 | 0.957 | -0.885 | 0.718 | -0.016 | 0.194 | 0.194 | 0 | 0.35 | 0.02 | 0.16 | 0.06 | 0.31 | 0.23 |
| 2 | 111746872 | rs7589645 | -0.443 | 0.977 | -0.159 | 0.854 | 0.013 | 0.025 | 0.025 | 0 | 0.37 | 0.6 | 0.33 | 0.06 | 0.32 | 0.36 |
| 2 | 111749200 | rs10496436 | 0.932 | 1.094 | -0.101 | 1.451 | -0.007 | 0.046 | 0.046 | 0 | 0.53 | 0.35 | 0.37 | 0.39 | 0.52 | 0.49 |
| 2 | 111749254 | rs3789096 | 0.929 | 1.088 | -0.104 | 1.48 | -0.008 | 0.003 | 0.003 | 0 | 0.53 | 0.34 | 0.37 | 0.47 | 0.52 | 0.49 |
| 2 | 111759540 | rs1483534 | -0.283 | 1.027 | 0.81 | 1.487 | -0.01 | -0.053 | -0.053 | 3.873 | 0.34 | 0.51 | 0.32 | 0.13 | 0.32 | 0.37 |
| 2 | 111766119 | rs1384792 | 0.818 | 0.963 | 0.8 | 1.794 | 0.014 | 0.092 | 0.092 | 0 | 0.88 | 0.84 | 0.69 | 0.6 | 0.84 | 0.85 |
| 2 | 111767152 | rs2341658 | 0.772 | 0.964 | 0.801 | 1.795 | 0.015 | 0.095 | 0.095 | 0 | 0.88 | 0.84 | 0.69 | 0.6 | 0.84 | 0.85 |
| 2 | 111775107 | rs6738187 | -1.039 | 0.797 | 0.989 | 1.668 | -0.042 | 0.106 | 0.106 | 0 | 0.13 | 0.29 | 0.12 | 0.31 | 0.07 | 0.28 |
| 2 | 111779171 | rs2306583 | 0.014 | 0.829 | 0.911 | 1.517 | -0.044 | 0.001 | 0.001 | 0 | 0.1 | 0.2 | 0.04 | 0.06 | 0.02 | 0.18 |
| 2 | 111789092 | rs3789090 | 0.887 | 0.929 | 0.96 | 1.72 | 0.001 | 0.001 | 0.001 | 0 | 0.49 | 0.5 | 0.26 | 0.36 | 0.38 | 0.48 |
| 2 | 111789791 | rs13012948 | 0.021 | 0.979 | 0.731 | 1.708 | -0.027 | -0.023 | -0.023 | 0 | 0.18 | 0.1 | 0.14 | 0.23 | 0.17 | 0.19 |
| 2 | 111794388 | rs3789085 | 0.734 | 1.135 | 0.836 | 1.717 | 0 | 0.001 | 0.001 | 0 | 0.55 | 0.43 | 0.25 | 0.43 | 0.45 | 0.4 |
| 2 | 111795509 | rs7602030 | 1.917 | 1.075 | 0.785 | 1.656 | 0.025 | 0.083 | 0.083 | 0.182 | 0.36 | 0.22 | 0.17 | 0.14 | 0.31 | 0.22 |
| 2 | 111796039 | rs11123312 | 0.278 | 1.073 | 0.807 | 1.706 | -0.002 | -0.034 | -0.034 | 0 | 0.17 | 0.07 | 0.07 | 0.24 | 0.14 | 0.18 |
| 2 | 111796195 | rs17549227 | 0.966 | 1.381 | 0.656 | 1.497 | -0.008 | 0.007 | 0.007 | 0 | 0.16 | 0.14 | 0.2 | 0.01 | 0.21 | 0.14 |
| 2 | 111796833 | rs6706577 | 0.962 | 1.458 | 0.792 | 1.62 | -0.004 | 0.004 | 0.004 | 0 | 0.56 | 0.62 | 0.44 | 0.4 | 0.49 | 0.54 |
| 2 | 111797458 | rs17483466 | 1.048 | 1.472 | 0.683 | 1.639 | -0.038 | 0.068 | 0.068 | 0 | 0.15 | 0.04 | 0.2 | 0.01 | 0.21 | 0.14 |
| 2 | 111797531 | rs3789080 | -0.986 | 1.403 | 0.759 | 1.747 | -0.03 | -0.049 | -0.049 | 0 | 0.89 | 0.97 | 0.71 | 0.69 | 0.85 | 0.86 |
| 2 | 111797565 | rs1992263 | -1.264 | 1.309 | 0.886 | 1.625 | 0.024 | 0.056 | 0.056 | 0 | 0.19 | 0.47 | 0.27 | 0.25 | 0.18 | 0.33 |
| 2 | 111797679 | rs3789078 | 0.299 | 1.305 | 0.675 | 1.505 | 0.003 | 0.004 | 0.004 | 0 | 0.1 | 0.2 | 0.12 | 0.06 | 0.06 | 0.2 |
| 2 | 111801402 | rs967895 | 0.002 | 1.424 | 0.225 | 1.619 | -0.028 | -0.038 | -0.038 | 0.166 | 0.73 | 0.81 | 0.49 | 0.63 | 0.64 | 0.71 |
| 2 | 111802105 | rs3789071 | 1.605 | 1.102 | 0.071 | 1.378 | 0.021 | 0.021 | 0.021 | 0 | 0.37 | 0.28 | 0.21 | 0.34 | 0.35 | 0.25 |
| 2 | 111807027 | rs3789069 | -1.022 | 1.496 | 0.477 | 1.671 | -0.002 | 0.027 | 0.027 | 0 | 0.31 | 0.4 | 0.28 | 0.46 | 0.26 | 0.42 |
| 2 | 111807303 | rs3827538 | 0.811 | 1.71 | 0.528 | 1.606 | 0 | 0.021 | 0.021 | 0 | 0.69 | 0.64 | 0.65 | 0.54 | 0.73 | 0.74 |
| 2 | 111808558 | rs12986962 | -0.98 | 1.791 | 0.369 | 1.38 | 0.007 | 0.014 | 0.014 | 0 | 0.3 | 0.25 | 0.34 | 0.49 | 0.38 | 0.45 |
| 2 | 111810281 | rs13000374 | 1.144 | 1.701 | 0.316 | 1.21 | 0.004 | 0.004 | 0.004 | 0.167 | 0.72 | 0.56 | 0.66 | 0.73 | 0.73 | 0.79 |
| 2 | 111812617 | rs10187034 | -0.773 | 1.802 | -0.107 | 1.333 | -0.029 | -0.064 | -0.064 | 0 | 0.88 | 0.99 | 0.76 | 0.8 | 0.88 | 0.88 |
| 2 | 111813085 | rs875063 | -1.012 | 1.795 | 0.21 | 1.356 | 0.002 | 0.029 | 0.029 | 0 | 0.34 | 0.36 | 0.33 | 0.47 | 0.31 | 0.45 |
| 2 | 111818527 | rs11692337 | 1.564 | 1.833 | -0.286 | 1.198 | 0.012 | 0.046 | 0.046 | 0 | 0.39 | 0.28 | 0.28 | 0.26 | 0.38 | 0.29 |
| 2 | 111825459 | rs17041842 | 0.243 | 1.86 | -0.326 | 1.386 | -0.072 | -0.101 | -0.101 | 1.119 | 0.18 | 0.03 | 0.25 | 0.42 | 0.24 | 0.18 |
| 2 | 111830273 | rs7576069 | 1.806 | 1.948 | -0.169 | 1.366 | 0.025 | 0.03 | 0.03 | 0.01 | 0.34 | 0.26 | 0.23 | 0.25 | 0.27 | 0.25 |
| 2 | 111836538 | rs7578982 | -0.535 | 2.517 | 0.514 | 1.541 | 0.02 | 0.036 | 0.036 | 0.987 | 0.17 | 0.18 | 0.29 | 0.43 | 0.29 | 0.22 |
| 2 | 111849659 | rs10207392 | -1.652 | 2.225 | 0.318 | 1.35 | 0.004 | 0.004 | 0.004 | 0.573 | 0.38 | 0.53 | 0.42 | 0.53 | 0.47 | 0.5 |
| 2 | 111851499 | rs7558938 | -1.51 | 2.025 | -0.093 | 1.094 | 0.024 | 0.026 | 0.026 | 0.507 | 0.36 | 0.44 | 0.38 | 0.54 | 0.44 | 0.49 |
| 2 | 111851616 | rs7559055 | -0.427 | 2.312 | -0.186 | 1.126 | 0.015 | -0.025 | -0.025 | 0 | 0.18 | 0.23 | 0.12 | 0.12 | 0.2 | 0.28 |

| | | | | | | | | | | | | | | | | |
|---|---|---|---|---|---|---|---|---|---|---|---|---|---|---|---|---|
| 2 | 111854524 | rs11691193 | -0.568 | 2.168 | -0.462 | 1.483 | -0.02 | -0.023 | -0.023 | 0 | 0.15 | 0.08 | 0.25 | 0.42 | 0.22 | 0.18 |
| 2 | 111856057 | rs4533473 | -0.828 | 1.496 | -0.055 | 0.632 | 0 | -0.067 | -0.067 | 0.697 | 0.19 | 0.39 | 0.15 | 0.09 | 0.19 | 0.13 |
| 2 | 111863438 | rs7567444 | -1.193 | 1.77 | -0.878 | 0.914 | -0.011 | -0.031 | -0.031 | 0.12 | 0.4 | 0.81 | 0.35 | 0.57 | 0.54 | 0.3 |
| 2 | 111868010 | rs11681263 | -0.305 | 3.044 | 0.575 | 1.584 | 0.028 | 0.026 | 0.026 | 0 | 0.13 | 0.2 | 0.11 | 0.17 | 0.24 | 0.11 |
| 2 | 111870220 | rs2289321 | -0.124 | 3.255 | 1.172 | 1.494 | 0.002 | 0 | 0 | 0 | 0.09 | 0.11 | 0.08 | 0.11 | 0.07 | 0.08 |
| 2 | 111871897 | rs1439287 | -1.63 | 3.176 | 1.163 | 1.441 | 0.04 | 0.035 | 0.035 | 0.002 | 0.27 | 0.38 | 0.24 | 0.46 | 0.47 | 0.19 |
| 2 | 111876148 | rs1054873 | -1.583 | 3.202 | 1.53 | 1.446 | 0.062 | 0.056 | 0.056 | 0.385 | 0.27 | 0.4 | 0.25 | 0.46 | 0.47 | 0.21 |
| 2 | 111896243 | rs17041869 | -0.619 | 3.342 | 1.515 | 1.446 | 0.009 | 0.031 | 0.031 | 0 | 0.07 | 0.09 | 0.08 | 0.29 | 0.1 | 0.06 |
| 2 | 111897506 | rs12613243 | -0.507 | 3.297 | 1.677 | 1.444 | 0.01 | 0.006 | 0.006 | 0 | 0.09 | 0.16 | 0.09 | 0.11 | 0.07 | 0.07 |
| 2 | 111903527 | rs3761705 | -0.829 | 3.361 | 1.579 | 1.457 | 0.012 | 0.003 | 0.003 | 0.403 | 0.17 | 0.2 | 0.14 | 0.17 | 0.3 | 0.15 |
| 2 | 111907691 | rs724710 | -2.664 | 2.663 | 1.44 | 1.128 | 0.088 | 0.154 | 0.154 | 0 | 0.55 | 0.8 | 0.5 | 0.85 | 0.67 | 0.54 |
| 2 | 111908262 | rs9308731 | -2.627 | 2.814 | 1.398 | 1.304 | -0.019 | -0.042 | -0.042 | 0 | 0.39 | 0.82 | 0.36 | 0.56 | 0.53 | 0.31 |
| 2 | 111909742 | rs17041883 | 0.51 | 3.217 | 1.408 | 1.318 | 0.027 | 0.065 | 0.065 | 0 | 0.05 | 0.09 | 0.07 | 0.29 | 0.11 | 0.05 |
| 2 | 111912729 | rs6727356 | -1.614 | 2.924 | 1.494 | 1.306 | 0.088 | 0.043 | 0.043 | 0 | 0.23 | 0.56 | 0.23 | 0.37 | 0.31 | 0.18 |
| 2 | 111915946 | rs1470053 | 0.575 | 2.628 | 0.805 | 1.359 | -0.05 | -0.065 | -0.065 | 2.896 | 0.11 | 0.01 | 0.1 | 0.18 | 0.18 | 0.13 |
| 2 | 111918472 | rs10189015 | -0.812 | 1.805 | 1.065 | 1.055 | 0.037 | 0.035 | 0.035 | 0.719 | 0.26 | 0.34 | 0.25 | 0.47 | 0.46 | 0.22 |
| 2 | 111919346 | rs686952 | -2.167 | 1.464 | 0.013 | 0.902 | 0.051 | 0.175 | 0.175 | 0.006 | 0.57 | 0.97 | 0.59 | 0.85 | 0.69 | 0.56 |
| 2 | 111920741 | rs3827536 | -1.55 | 1.527 | 0.198 | 1.056 | -0.022 | -0.035 | -0.035 | 0.687 | 0.39 | 0.82 | 0.36 | 0.57 | 0.54 | 0.31 |
| 2 | 111934107 | rs590097 | -1.795 | 1.564 | -0.068 | 0.688 | 0.066 | 0.251 | 0.251 | 0 | 0.53 | 0.93 | 0.5 | 0.86 | 0.62 | 0.5 |
| 2 | 111935939 | rs12615741 | 0.547 | 1.606 | 0.091 | 0.679 | 0.01 | 0.01 | 0.01 | 0 | 0.05 | 0.1 | 0.08 | 0.11 | 0.07 | 0.06 |
| 2 | 111943200 | rs604126 | -2.011 | 1.629 | -0.046 | 0.584 | 0.042 | 0.245 | 0.245 | 0.003 | 0.55 | 0.95 | 0.57 | 0.85 | 0.65 | 0.52 |
| 2 | 111948809 | rs938095 | -1.38 | 1.795 | 0.767 | 1.006 | -0.033 | 0.007 | 0.007 | 0 | 0.37 | 0.74 | 0.31 | 0.38 | 0.5 | 0.25 |
| 2 | 111949096 | rs6755484 | -1.168 | 1.804 | 0.832 | 1.008 | 0.025 | -0.002 | -0.002 | 0 | 0.35 | 0.54 | 0.29 | 0.38 | 0.5 | 0.25 |
| 2 | 111950594 | rs11123407 | -0.85 | 1.189 | 0.427 | 0.773 | -0.001 | 0 | 0 | 0.285 | 0.46 | 0.48 | 0.32 | 0.45 | 0.54 | 0.3 |
| 2 | 111952208 | rs880892 | -1.411 | 1.283 | 0.123 | 0.907 | 0.057 | -0.028 | -0.028 | 0 | 0.54 | 0.77 | 0.47 | 0.49 | 0.64 | 0.36 |
| 2 | 111953117 | rs12620756 | 0.575 | 1.529 | 0.069 | 1.037 | 0.017 | 0.067 | 0.067 | 0 | 0.07 | 0.12 | 0.15 | 0.16 | 0.1 | 0.09 |
| 2 | 111953254 | rs1107480 | 0.067 | 1.85 | -0.143 | 0.899 | 0.003 | 0.004 | 0.004 | 0 | 0.08 | 0.08 | 0.09 | 0.28 | 0.18 | 0.06 |
| 2 | 111955485 | rs622274 | 0.547 | 1.021 | -0.423 | 0.316 | 0.023 | 0.021 | 0.021 | 0.082 | 0.54 | 0.44 | 0.45 | 0.24 | 0.32 | 0.47 |
| 2 | 111957073 | rs11123410 | 0.918 | 0.913 | -0.446 | 0.283 | 0.023 | 0.009 | 0.009 | 0 | 0.45 | 0.37 | 0.5 | 0.42 | 0.35 | 0.64 |
| 2 | 111957313 | rs938101 | -1.387 | 0.755 | -0.327 | 0.25 | -0.032 | -0.022 | -0.022 | 0 | 0.88 | 0.81 | 0.92 | 0.98 | 0.92 | 0.94 |
| 2 | 111957594 | rs13431908 | 0.628 | 1.102 | -0.427 | 0.266 | 0.013 | 0.045 | 0.045 | 0 | 0.13 | 0.17 | 0.25 | 0.47 | 0.24 | 0.13 |
| 2 | 111957894 | rs6749625 | 1.212 | 0.977 | -0.498 | 0.205 | 0.037 | 0.007 | 0.007 | 0 | 0.44 | 0.22 | 0.49 | 0.42 | 0.35 | 0.64 |
| 2 | 111966222 | rs1398028 | -1.28 | 1.428 | -0.182 | 0.164 | -0.013 | 0.016 | 0.016 | 0 | 0.74 | 0.74 | 0.82 | 0.68 | 0.81 | 0.63 |
| 2 | 111966540 | rs3747653 | 1.434 | 1.488 | -0.357 | 0.209 | 0.035 | 0.025 | 0.025 | 1.009 | 0.3 | 0.16 | 0.39 | 0.19 | 0.24 | 0.33 |
| 2 | 111970330 | rs4374378 | 1.436 | 1.481 | -0.322 | 0.205 | 0.03 | 0.057 | 0.057 | 0.951 | 0.3 | 0.2 | 0.4 | 0.13 | 0.24 | 0.33 |
| 2 | 111972005 | rs1398030 | 0.532 | 1.623 | -0.276 | 0.242 | 0.022 | 0.033 | 0.033 | 0 | 0.13 | 0.2 | 0.26 | 0.4 | 0.24 | 0.13 |
| 2 | 111976515 | rs13411196 | 0.445 | 1.612 | -0.282 | 0.022 | 0 | 0.017 | 0.017 | 0 | 0.14 | 0.15 | 0.26 | 0.46 | 0.24 | 0.13 |
| 2 | 111979815 | rs6542368 | -0.096 | 1.587 | -0.319 | -0.01 | -0.002 | 0.004 | 0.004 | 0 | 0.58 | 0.43 | 0.55 | 0.22 | 0.57 | 0.5 |
| 2 | 111988016 | rs6749633 | 0.067 | 1.346 | -0.285 | 0.353 | 0.013 | -0.015 | -0.015 | 0 | 0.19 | 0.22 | 0.14 | 0.09 | 0.3 | 0.15 |
| 2 | 111989372 | rs6720394 | 1.156 | 1.421 | -0.301 | 0.335 | 0.019 | -0.032 | -0.032 | 0 | 0.07 | 0.14 | 0.07 | 0 | 0.13 | 0.05 |
| 2 | 111991133 | rs6716091 | -0.13 | 1.431 | -0.423 | 0.339 | 0.021 | -0.022 | -0.022 | 0.161 | 0.19 | 0.26 | 0.14 | 0.09 | 0.29 | 0.15 |
| 2 | 111992975 | rs1464023 | -1.073 | 1.548 | -0.89 | 0.642 | 0.001 | -0.005 | -0.005 | 0 | 0.69 | 0.8 | 0.75 | 0.68 | 0.7 | 0.61 |
| 2 | 111996785 | rs10197366 | 0.167 | 1.581 | -0.864 | 0.663 | 0.039 | 0.087 | 0.087 | 0 | 0.14 | 0.23 | 0.26 | 0.46 | 0.24 | 0.13 |
| 2 | 112000058 | rs13395957 | -0.23 | 1.588 | -0.959 | 0.651 | -0.005 | -0.002 | -0.002 | 0 | 0.57 | 0.73 | 0.68 | 0.59 | 0.54 | 0.52 |
| 2 | 112003867 | rs2271404 | 0.747 | 1.957 | -0.615 | 1.395 | 0.016 | 0.033 | 0.033 | 0 | 0.06 | 0.08 | 0.1 | 0.29 | 0.14 | 0.04 |
| 2 | 112004205 | rs3817185 | 0.38 | 1.067 | -0.649 | 0.613 | 0.011 | 0.049 | 0.049 | 0 | 0.08 | 0.14 | 0.16 | 0.2 | 0.1 | 0.09 |
| 2 | 112010486 | rs4308124 | -0.055 | 0.736 | -0.834 | -0.12 | -0.006 | -0.026 | -0.026 | 0 | 0.45 | 0.54 | 0.47 | 0.22 | 0.43 | 0.48 |
| 2 | 112143413 | rs2139376 | -0.362 | 0.712 | -1.133 | 0.179 | 0.011 | 0.032 | 0.032 | 0 | 0.52 | 0.55 | 0.4 | 0.21 | 0.42 | 0.39 |
| 2 | 112156243 | rs4849696 | -0.922 | 0.803 | -1.328 | 0.254 | -0.005 | 0.008 | 0.008 | 0 | 0.13 | 0.07 | 0.21 | 0.07 | 0.12 | 0.08 |
| 2 | 112220359 | rs1448187 | 0.076 | 0.811 | -1.068 | 0.31 | -0.008 | -0.005 | -0.005 | 0.972 | 0.67 | 0.47 | 0.49 | 0.27 | 0.73 | 0.57 |
| 2 | 112237234 | rs6721071 | 0.06 | 1.46 | -1.708 | 0.137 | -0.04 | -0.001 | -0.001 | 0 | 0.74 | 0.82 | 0.72 | 0.77 | 0.59 | 0.61 |
| 2 | 112253302 | rs1463118 | -0.195 | 1.276 | -2.493 | -0.065 | -0.055 | -0.003 | -0.003 | 0.077 | 0.3 | 0.07 | 0.21 | 0.22 | 0.35 | 0.27 |
| 2 | 112264992 | rs12611870 | -0.081 | 1.266 | -2.556 | -0.085 | -0.106 | -0.05 | -0.05 | 0 | 0.28 | 0.02 | 0.21 | 0.22 | 0.34 | 0.27 |
| 2 | 112270857 | rs6731210 | 0.009 | 0.648 | -1.894 | -0.416 | 0.017 | 0.053 | 0.053 | 0 | 0.52 | 0.6 | 0.52 | 0.26 | 0.62 | 0.39 |
| 2 | 112277340 | rs11123551 | -0.058 | 0.542 | -1.799 | -0.411 | 0.022 | 0.08 | 0.08 | 0 | 0.53 | 0.67 | 0.53 | 0.25 | 0.62 | 0.39 |
| 2 | 112277492 | rs17842119 | 0.198 | 0.284 | -2.252 | -0.749 | -0.034 | -0.035 | -0.035 | 0 | 0.3 | 0.06 | 0.38 | 0.74 | 0.24 | 0.48 |
| 2 | 112278999 | rs4849784 | 0.611 | 0.349 | -2.05 | -0.724 | 0.019 | 0.051 | 0.051 | 0 | 0.5 | 0.55 | 0.52 | 0.25 | 0.62 | 0.37 |
| 2 | 112295182 | rs935598 | 0.343 | 0.234 | -1.832 | 0.128 | -0.039 | -0.064 | -0.064 | 0 | 0.13 | 0.04 | 0.11 | 0.28 | 0.12 | 0.15 |
| 2 | 112295707 | rs1666714 | 0.183 | -0.15 | -1.184 | 0.107 | -0.029 | -0.039 | -0.039 | 0.186 | 0.91 | 0.71 | 0.96 | 0.99 | 0.96 | 0.92 |
| 2 | 112305653 | rs2218659 | 0.546 | -0.001 | -0.981 | 0.236 | -0.008 | 0.023 | 0.023 | 0 | 0.66 | 0.57 | 0.71 | 0.48 | 0.72 | 0.62 |
| 2 | 112323703 | rs3109453 | 0.582 | -0.314 | -0.168 | -0.199 | -0.02 | -0.024 | -0.024 | 0 | 0.91 | 0.59 | 0.95 | 0.99 | 0.98 | 0.9 |
| 2 | 112373169 | rs10171109 | -0.466 | -0.039 | -1.075 | -0.254 | 0 | -0.007 | -0.007 | 0 | 0.12 | 0.16 | 0.08 | 0.03 | 0.1 | 0.1 |
| 2 | 112373660 | rs13011755 | -0.062 | -0.11 | -1.209 | -0.289 | -0.002 | -0.02 | -0.02 | 0 | 0.74 | 0.38 | 0.77 | 0.81 | 0.77 | 0.83 |
| 2 | 112373702 | rs13006364 | 0.426 | 0.079 | -1.443 | -0.274 | 0.026 | 0.111 | 0.111 | 0 | 0.5 | 0.3 | 0.58 | 0.26 | 0.62 | 0.58 |
| 2 | 112375450 | rs11686181 | 0.387 | -0.2 | -1.722 | -0.654 | -0.038 | 0.191 | 0.191 | 0.007 | 0.38 | 0.1 | 0.43 | 0.12 | 0.61 | 0.46 |
| 2 | 112375963 | rs1448190 | -0.233 | -0.873 | -0.537 | -0.65 | -0.019 | -0.035 | -0.035 | 0 | 0.63 | 0.51 | 0.66 | 0.25 | 0.75 | 0.63 |
| 2 | 112376294 | rs4849903 | 0.829 | -1.037 | -0.702 | -0.77 | 0.01 | 0.212 | 0.212 | 0 | 0.46 | 0.16 | 0.46 | 0.15 | 0.61 | 0.47 |
| 2 | 112383896 | rs2121914 | 0.234 | -1.442 | -0.01 | -0.014 | -0.02 | 0.065 | 0.065 | 0 | 0.73 | 0.58 | 0.68 | 0.6 | 0.79 | 0.68 |
| 2 | 112388378 | rs12619327 | 0.907 | -1.535 | -0.302 | -0.023 | 0.003 | 0.2 | 0.2 | 0 | 0.07 | 0.19 | 0.12 | 0.32 | 0.05 | 0.11 |

| | | | | | | | | | | | | | | | | |
|---|---|---|---|---|---|---|---|---|---|---|---|---|---|---|---|---|
| 2 | 112395534 | rs7593992 | 0.128 | -1.774 | -0.116 | -0.077 | 0.019 | 0.037 | 0.037 | 0 | 0.64 | 0.75 | 0.71 | 0.73 | 0.74 | 0.63 |
| 2 | 112396165 | rs11693743 | -0.221 | -1.773 | -0.062 | -0.079 | -0.021 | -0.019 | -0.019 | 0 | 0.78 | 0.7 | 0.78 | 0.83 | 0.87 | 0.72 |
| 2 | 112397505 | rs9308627 | 0.063 | -1.78 | -0.517 | -0.083 | 0.013 | 0.059 | 0.059 | 0 | 0.28 | 0.15 | 0.16 | 0.19 | 0.2 | 0.26 |
| 2 | 112398498 | rs17776994 | 1.7 | -1.389 | -0.656 | -0.187 | 0 | -0.013 | -0.013 | 0.172 | 0.05 | 0.05 | 0.03 | 0.01 | 0.05 | 0.01 |
| 2 | 112400597 | rs7596295 | 0.871 | -1.423 | -0.799 | -0.148 | 0.021 | -0.013 | -0.013 | 2.535 | 0.47 | 0.32 | 0.54 | 0.46 | 0.6 | 0.55 |
| 2 | 112409842 | rs2311552 | -0.556 | -1.272 | -0.651 | -0.167 | 0.032 | 0.189 | 0.189 | 0.208 | 0.86 | 0.53 | 0.83 | 0.73 | 0.86 | 0.93 |
| 2 | 112411416 | rs12328538 | 0.544 | -0.993 | -1.214 | -0.786 | 0.009 | 0.008 | 0.008 | 0 | 0.21 | 0.16 | 0.2 | 0.17 | 0.13 | 0.28 |
| 2 | 112413845 | rs10208216 | 0.53 | -0.965 | -1.413 | -0.773 | 0.018 | 0.017 | 0.017 | 0 | 0.21 | 0.13 | 0.2 | 0.16 | 0.13 | 0.28 |
| 2 | 112414382 | rs13432469 | 0.598 | -0.932 | -1.395 | -0.757 | 0.016 | 0.015 | 0.015 | 0 | 0.2 | 0.13 | 0.2 | 0.16 | 0.13 | 0.28 |
| 2 | 112415726 | rs12612210 | 0.863 | -0.933 | -1.447 | -0.767 | -0.146 | 0.169 | 0.169 | 0.108 | 0.39 | 0.05 | 0.42 | 0.14 | 0.55 | 0.44 |
| 2 | 112417919 | rs4848137 | -0.399 | -0.476 | -1.226 | -0.584 | 0.007 | 0.04 | 0.04 | 0 | 0.6 | 0.31 | 0.6 | 0.3 | 0.63 | 0.71 |
| 2 | 112421566 | rs10210313 | 0.153 | -0.483 | -1.196 | -0.434 | 0.017 | 0.016 | 0.016 | 0 | 0.2 | 0.14 | 0.2 | 0.16 | 0.13 | 0.28 |
| 2 | 112422223 | rs13034716 | 0.016 | 0.16 | -0.931 | -0.257 | -0.001 | -0.011 | -0.011 | 0 | 0.55 | 0.61 | 0.47 | 0.21 | 0.54 | 0.69 |
| 2 | 112425540 | rs1399665 | -0.456 | 0.063 | -1.079 | 0.407 | -0.016 | -0.026 | -0.026 | 0 | 0.71 | 0.86 | 0.68 | 0.67 | 0.67 | 0.72 |
| 2 | 112428409 | rs6754001 | -0.336 | -0.345 | -0.543 | 0.558 | -0.021 | 0.048 | 0.048 | 0.125 | 0.25 | 0.32 | 0.33 | 0.42 | 0.18 | 0.3 |
| 2 | 112433668 | rs17777313 | 0.749 | -0.536 | -0.404 | 0.766 | -0.003 | 0.007 | 0.007 | 2.114 | 0.06 | 0.05 | 0.02 | 0.1 | 0.03 | 0.12 |
| 2 | 112435483 | rs17017403 | -0.798 | -0.636 | -0.387 | 0.912 | -0.052 | 0.047 | 0.047 | 0 | 0.19 | 0.34 | 0.16 | 0.25 | 0.09 | 0.21 |
| 2 | 112437140 | rs1464095 | 0.133 | -0.772 | -0.036 | 0.819 | 0.015 | 0.012 | 0.012 | 0 | 0.51 | 0.58 | 0.51 | 0.43 | 0.56 | 0.51 |
| 2 | 112444459 | rs10178648 | -0.814 | -0.808 | -0.097 | 0.817 | 0.004 | -0.026 | -0.026 | 0.041 | 0.64 | 0.77 | 0.65 | 0.58 | 0.62 | 0.6 |
| 2 | 112447888 | rs10174353 | 0.235 | -0.821 | -0.038 | 0.829 | -0.001 | -0.001 | -0.001 | 0 | 0.5 | 0.52 | 0.51 | 0.43 | 0.57 | 0.5 |
| 2 | 112458934 | rs3898028 | 0.385 | -0.632 | -0.038 | 0.624 | -0.008 | 0.011 | 0.011 | 0 | 0.09 | 0.12 | 0.13 | 0.15 | 0.05 | 0.09 |
| 2 | 112464137 | rs12619033 | 0.631 | -0.842 | -0.323 | 0.41 | -0.088 | 0.132 | 0.132 | 0 | 0.38 | 0.07 | 0.26 | 0.14 | 0.41 | 0.28 |
| 2 | 112464175 | rs17842577 | 0.653 | -0.737 | 0.337 | 0.399 | 0.022 | 0.024 | 0.024 | 0 | 0.09 | 0.14 | 0.23 | 0.18 | 0.17 | 0.11 |
| 2 | 112466265 | rs12373588 | -0.003 | -0.679 | 0.113 | 0.315 | -0.013 | 0.075 | 0.075 | 0.043 | 0.48 | 0.21 | 0.39 | 0.29 | 0.46 | 0.47 |
| 2 | 112473872 | rs7605443 | -0.278 | -0.69 | 0.086 | 0.346 | -0.009 | 0.008 | 0.008 | 0 | 0.53 | 0.3 | 0.41 | 0.39 | 0.5 | 0.58 |
| 2 | 112581082 | rs6742309 | 0.247 | -0.375 | 0.113 | 0.421 | 0.001 | 0.002 | 0.002 | 0 | 0.12 | 0.12 | 0.24 | 0.17 | 0.17 | 0.12 |
| 2 | 112591026 | rs7568001 | 0.648 | -0.806 | 0.145 | 0.25 | 0.013 | -0.005 | -0.005 | 0 | 0.11 | 0.04 | 0.13 | 0.12 | 0.07 | 0.2 |
| 2 | 112598119 | rs12617909 | 0.545 | -0.587 | 0.141 | 0.047 | -0.003 | 0.034 | 0.034 | 0 | 0.28 | 0.08 | 0.17 | 0.16 | 0.23 | 0.19 |
| 2 | 112625575 | rs6742515 | -0.349 | -0.534 | 0.948 | 0.195 | 0 | -0.002 | -0.002 | 0.06 | 0.64 | 0.36 | 0.66 | 0.56 | 0.66 | 0.67 |
| 2 | 112627223 | rs10197222 | -0.349 | -0.534 | 0.901 | 0.195 | -0.001 | 0 | 0 | 0 | 0.64 | 0.37 | 0.66 | 0.56 | 0.66 | 0.67 |
| 2 | 112634339 | rs1516637 | -0.402 | -0.511 | 0.721 | 0.19 | 0.001 | -0.008 | -0.008 | 0.25 | 0.66 | 0.74 | 0.7 | 0.63 | 0.66 | 0.68 |
| 2 | 112634737 | rs9653414 | -0.44 | -0.363 | -1.256 | 0.186 | 0.001 | -0.003 | -0.003 | 0 | 0.64 | 0.35 | 0.67 | 0.57 | 0.66 | 0.67 |
| 2 | 112645250 | rs4369862 | -0.128 | -0.695 | -1.56 | -0.231 | 0.074 | 0.057 | 0.057 | 0 | 0.3 | 0.15 | 0.11 | 0.16 | 0.16 | 0.26 |
| 2 | 112649472 | rs13016143 | -0.094 | 0.287 | -1.481 | -0.736 | -0.002 | 0.03 | 0.03 | 0 | 0.32 | 0.22 | 0.29 | 0.13 | 0.32 | 0.29 |
| 2 | 112649794 | rs12329119 | -0.1 | 0.293 | -1.381 | -0.973 | -0.001 | -0.022 | -0.022 | 0 | 0.78 | 0.79 | 0.84 | 0.98 | 0.8 | 0.94 |
| 2 | 112655242 | rs6736093 | 0.666 | 0.685 | -1.918 | -1.609 | -0.051 | 0.077 | 0.077 | 0.527 | 0.27 | 0.11 | 0.19 | 0.17 | 0.28 | 0.23 |
| 2 | 112665201 | rs17174870 | 0.307 | 0.816 | -0.415 | -1.914 | 0 | -0.004 | -0.004 | 0 | 0.2 | 0.18 | 0.35 | 0.12 | 0.26 | 0.36 |
| 2 | 112669950 | rs7568632 | 0.48 | 0.807 | -0.416 | -1.918 | -0.022 | 0.032 | 0.032 | 0.339 | 0.19 | 0.1 | 0.3 | 0.08 | 0.26 | 0.36 |
| 2 | 112670461 | rs1400323 | 0.344 | 0.54 | 0.002 | -2.016 | 0.005 | 0.009 | 0.009 | 0 | 0.5 | 0.46 | 0.45 | 0.71 | 0.46 | 0.41 |
| 2 | 112686988 | rs13027171 | 1.284 | 0.57 | -0.215 | -2.012 | -0.1 | 0.059 | 0.059 | 0.969 | 0.25 | 0.02 | 0.18 | 0.17 | 0.29 | 0.23 |
| 2 | 112690175 | rs6711987 | -0.528 | 0.478 | 0.275 | -2.064 | -0.001 | -0.013 | -0.013 | 1.709 | 0.5 | 0.54 | 0.54 | 0.27 | 0.54 | 0.59 |
| 2 | 112712870 | rs17175331 | 0.458 | 0.465 | 0.452 | -2.049 | -0.003 | 0.001 | 0.001 | 0.377 | 0.2 | 0.18 | 0.34 | 0.1 | 0.26 | 0.36 |
| 2 | 112723117 | rs4848908 | 0.551 | 0.268 | 0.54 | -2.453 | 0.013 | 0.025 | 0.025 | 0.404 | 0.15 | 0.18 | 0.26 | 0.69 | 0.19 | 0.16 |
| 2 | 112738562 | rs6725192 | -0.178 | -0.391 | -0.072 | -2.83 | 0.002 | 0.002 | 0.002 | 0 | 0.61 | 0.67 | 0.6 | 0.28 | 0.62 | 0.62 |
| 2 | 112745481 | rs870763 | 1.833 | -0.391 | -0.066 | -2.831 | -0.038 | 0.071 | 0.071 | 1.338 | 0.27 | 0.12 | 0.19 | 0.17 | 0.29 | 0.23 |
| 2 | 112751928 | rs7604639 | -0.176 | -0.389 | -0.093 | -2.827 | 0.002 | 0.002 | 0.002 | 0.001 | 0.61 | 0.67 | 0.6 | 0.28 | 0.62 | 0.62 |
| 2 | 112753950 | rs13003919 | 1.974 | -0.293 | -0.504 | -2.772 | -0.094 | 0.066 | 0.066 | 1.42 | 0.26 | 0.03 | 0.18 | 0.17 | 0.29 | 0.23 |
| 2 | 112754828 | rs3811632 | 0.528 | -0.292 | -0.482 | -2.774 | 0.02 | -0.052 | -0.052 | 0 | 0.24 | 0.36 | 0.38 | 0.11 | 0.3 | 0.38 |
| 2 | 112754943 | rs3811634 | 1.891 | -0.246 | -0.633 | -2.705 | -0.056 | 0.078 | 0.078 | 1.292 | 0.27 | 0.1 | 0.19 | 0.17 | 0.29 | 0.23 |
| 2 | 112762623 | rs7580261 | 0.999 | 0.01 | -0.507 | -1.912 | -0.007 | -0.035 | -0.035 | 0 | 0.12 | 0.02 | 0.09 | 0.23 | 0.14 | 0.07 |
| 2 | 112765562 | rs10195619 | 0.72 | 0.04 | -0.047 | -2.461 | -0.008 | 0.069 | 0.069 | 2.078 | 0.38 | 0.28 | 0.25 | 0.17 | 0.37 | 0.26 |
| 2 | 112767406 | rs11884641 | 1.009 | -0.005 | 0.062 | -2.533 | 0.002 | -0.01 | -0.01 | 0 | 0.2 | 0.22 | 0.35 | 0.1 | 0.26 | 0.36 |
| 2 | 112770770 | rs4374383 | -0.035 | -0.173 | -0.073 | -2.537 | -0.006 | -0.027 | -0.027 | 7.473 | 0.61 | 0.5 | 0.6 | 0.29 | 0.62 | 0.62 |
| 2 | 112773153 | rs4627569 | 0.17 | -0.071 | 0.037 | -2.53 | 0.003 | 0.003 | 0.003 | 0 | 0.57 | 0.42 | 0.59 | 0.28 | 0.62 | 0.6 |
| 2 | 112781917 | rs13419523 | 0.503 | 0.031 | -0.175 | -2.462 | -0.005 | -0.052 | -0.052 | 0 | 0.1 | 0.22 | 0.06 | 0 | 0.08 | 0.03 |
| 2 | 112783957 | rs6730521 | 0.98 | 0.095 | -0.11 | -2.44 | -0.014 | 0.018 | 0.018 | 0.857 | 0.19 | 0.16 | 0.33 | 0.1 | 0.26 | 0.35 |
| 2 | 112790562 | rs4848973 | 1.528 | 0.202 | -0.289 | -2.347 | -0.052 | 0.081 | 0.081 | 1.248 | 0.27 | 0.1 | 0.2 | 0.19 | 0.28 | 0.23 |
| 2 | 112795908 | rs11897014 | 1.241 | 0.195 | -0.065 | -2.331 | -0.012 | 0.043 | 0.043 | 0.468 | 0.3 | 0.23 | 0.23 | 0.19 | 0.32 | 0.24 |
| 2 | 112803874 | rs6748256 | 0.53 | -0.03 | 0.551 | -2.155 | -0.001 | -0.001 | -0.001 | 0.596 | 0.53 | 0.53 | 0.36 | 0.22 | 0.47 | 0.45 |
| 2 | 112816047 | rs4848978 | -0.381 | 0.089 | 0.002 | -2.115 | -0.009 | 0.063 | 0.063 | 0.006 | 0.72 | 0.8 | 0.65 | 0.91 | 0.68 | 0.64 |
| 2 | 112843244 | rs10165845 | 0.383 | -0.174 | 0.547 | -1.931 | 0.001 | 0.048 | 0.048 | 0.951 | 0.49 | 0.43 | 0.48 | 0.88 | 0.5 | 0.4 |
| 2 | 112844091 | rs4613287 | -0.839 | -0.202 | 0.297 | -2.037 | -0.082 | 0.151 | 0.151 | 2.971 | 0.69 | 0.96 | 0.59 | 0.91 | 0.62 | 0.6 |
| 2 | 112847164 | rs11123128 | 1.58 | -0.203 | 0.325 | -1.978 | -0.022 | 0.068 | 0.068 | 0.746 | 0.36 | 0.21 | 0.3 | 0.19 | 0.4 | 0.24 |
| 2 | 112850581 | rs2067928 | 0.091 | -0.189 | 0.492 | -1.956 | -0.024 | 0.022 | 0.022 | 0 | 0.14 | 0.28 | 0.19 | 0.69 | 0.1 | 0.17 |
| 2 | 112856165 | rs4849007 | 0.02 | -0.258 | 0.763 | -1.976 | -0.013 | 0.061 | 0.061 | 2.259 | 0.55 | 0.72 | 0.5 | 0.88 | 0.49 | 0.41 |
| 2 | 112857642 | rs9653422 | 0.405 | -0.261 | 0.787 | -1.98 | 0 | -0.008 | -0.008 | 0 | 0.49 | 0.5 | 0.48 | 0.88 | 0.49 | 0.41 |
| 2 | 112857865 | rs9653401 | 0.004 | -0.246 | 0.688 | -1.967 | -0.01 | 0.059 | 0.059 | 2.204 | 0.55 | 0.72 | 0.5 | 0.88 | 0.51 | 0.42 |
| 2 | 112858485 | rs7583755 | -0.032 | -0.217 | 0.589 | -1.808 | -0.029 | 0.023 | 0.023 | 0 | 0.14 | 0.28 | 0.18 | 0.69 | 0.09 | 0.17 |
| 2 | 112866558 | rs954521 | 0.175 | 0.081 | 0.286 | -1.835 | 0.01 | -0.036 | -0.036 | 5.419 | 0.08 | 0.15 | 0.05 | 0 | 0.06 | 0.01 |

| | | | | | | | | | | | | | | | | |
|---|---|---|---|---|---|---|---|---|---|---|---|---|---|---|---|---|
| 2 | 112868426 | rs10180018 | -0.197 | -0.127 | 0.553 | -1.731 | 0.017 | -0.045 | -0.045 | 0 | 0.09 | 0.25 | 0.05 | 0 | 0.05 | 0.02 |
| 2 | 112869041 | rs10864898 | 0.009 | -0.145 | 0.161 | -1.828 | -0.023 | 0.146 | 0.146 | 3.053 | 0.57 | 0.83 | 0.52 | 0.88 | 0.53 | 0.42 |
| 2 | 112901019 | rs7607362 | 0.156 | 0.371 | 0.028 | -1.331 | -0.001 | 0.011 | 0.011 | 0.174 | 0.52 | 0.51 | 0.37 | 0.25 | 0.5 | 0.44 |
| 2 | 112902721 | rs6755850 | 0.183 | 0.417 | -0.167 | -1.295 | 0 | -0.021 | -0.021 | 0 | 0.52 | 0.45 | 0.37 | 0.23 | 0.5 | 0.43 |
| 2 | 112920219 | rs4849047 | -0.653 | 0.2 | -1.249 | -1.184 | -0.014 | 0.041 | 0.041 | 0.03 | 0.64 | 0.74 | 0.45 | 0.26 | 0.55 | 0.48 |
| 2 | 112924973 | rs10197919 | 0.54 | 0.446 | -1.636 | -0.948 | -0.073 | 0.146 | 0.146 | 0.301 | 0.37 | 0.09 | 0.3 | 0.22 | 0.42 | 0.28 |
| 2 | 112928815 | rs10200201 | -0.238 | 0.567 | -0.915 | -0.697 | 0.001 | -0.012 | -0.012 | 0.531 | 0.6 | 0.56 | 0.42 | 0.26 | 0.54 | 0.46 |
| 2 | 112933521 | rs12711735 | -1.007 | 0.596 | -1.018 | 0.422 | -0.003 | -0.009 | -0.009 | 0 | 0.11 | 0.07 | 0.18 | 0.32 | 0.14 | 0.09 |
| 2 | 112939548 | rs4849050 | -1.466 | 0.446 | -0.442 | 0.371 | -0.002 | 0.074 | 0.074 | 0 | 0.13 | 0.19 | 0.18 | 0.33 | 0.14 | 0.1 |
| 2 | 112945220 | rs11123130 | -0.653 | 0.121 | -0.359 | -0.424 | -0.001 | -0.042 | -0.042 | 0.069 | 0.21 | 0.22 | 0.13 | 0.01 | 0.21 | 0.13 |
| 2 | 112949713 | rs12620553 | 1.547 | 0.173 | -1.294 | -0.579 | -0.085 | 0.001 | 0.001 | 1.255 | 0.29 | 0.02 | 0.21 | 0.25 | 0.32 | 0.26 |
| 2 | 112953542 | rs10183121 | 0.284 | -0.089 | -1.073 | -0.889 | -0.037 | 0.16 | 0.16 | 0 | 0.47 | 0.12 | 0.31 | 0.28 | 0.52 | 0.46 |
| 2 | 112966509 | rs4849056 | 1.816 | 0.2 | -1.292 | -1.014 | -0.096 | 0.003 | 0.003 | 1.147 | 0.31 | 0.02 | 0.22 | 0.26 | 0.34 | 0.26 |
| 2 | 113001329 | rs2312459 | -1.338 | -0.058 | -0.735 | -0.869 | 0.003 | 0.01 | 0.01 | 0 | 0.25 | 0.33 | 0.27 | 0.32 | 0.27 | 0.17 |
| 2 | 113007757 | rs17781007 | 0.944 | -0.05 | -0.926 | -0.875 | -0.002 | -0.03 | -0.03 | 0 | 0.1 | 0.14 | 0.07 | 0 | 0.07 | 0.01 |
| 2 | 113013979 | rs11691431 | 1.911 | 0.344 | -1.301 | -0.918 | -0.12 | 0.023 | 0.023 | 1.283 | 0.31 | 0.02 | 0.23 | 0.26 | 0.37 | 0.27 |
| 2 | 113014605 | rs10169925 | -1.032 | 0.129 | -1.161 | -1.025 | -0.016 | -0.096 | -0.096 | 0.006 | 0.7 | 0.92 | 0.59 | 0.59 | 0.68 | 0.54 |
| 2 | 113015512 | rs6542045 | 0.786 | 0.168 | -0.969 | -1.153 | -0.052 | 0.124 | 0.124 | 0.417 | 0.43 | 0.19 | 0.3 | 0.26 | 0.52 | 0.39 |
| 2 | 113053061 | rs4430968 | -1.169 | 0.185 | -1.169 | -1.228 | 0.005 | 0.146 | 0.146 | 0.477 | 0.45 | 0.8 | 0.57 | 0.74 | 0.39 | 0.5 |
| 2 | 113089267 | rs7601200 | -2.053 | 0.186 | -1.183 | -1.241 | -0.131 | 0.125 | 0.125 | 0.802 | 0.57 | 0.97 | 0.65 | 0.74 | 0.55 | 0.61 |
| 2 | 113090047 | rs4578863 | -2.03 | 0.187 | -1.183 | -1.24 | -0.13 | 0.128 | 0.128 | 0.833 | 0.57 | 0.97 | 0.65 | 0.74 | 0.55 | 0.62 |
| 2 | 113091340 | rs6758110 | -1.24 | 0.339 | -1.071 | -1.132 | 0.005 | 0.146 | 0.146 | 0.477 | 0.45 | 0.8 | 0.57 | 0.74 | 0.39 | 0.5 |
| 2 | 113091685 | rs6761599 | -1.24 | 0.339 | -1.071 | -1.132 | 0.005 | 0.146 | 0.146 | 0.477 | 0.45 | 0.8 | 0.57 | 0.74 | 0.39 | 0.5 |
| 2 | 113094793 | rs7591439 | -1.24 | 0.347 | -1.071 | -1.132 | 0.008 | 0.146 | 0.146 | 0.477 | 0.45 | 0.8 | 0.57 | 0.74 | 0.39 | 0.5 |
| 2 | 113192136 | rs6542056 | -1.253 | 0.45 | -0.261 | -0.671 | 0.001 | 0.012 | 0.012 | 0.325 | 0.14 | 0.07 | 0.08 | 0 | 0.16 | 0.14 |
| 2 | 113232433 | rs11890681 | -0.033 | 0.375 | -0.064 | -0.251 | -0.01 | 0.005 | 0.005 | 0 | 0.09 | 0.06 | 0.09 | 0 | 0.1 | 0.02 |
| 2 | 113238530 | rs6747985 | -0.826 | 0.347 | 0.154 | -0.193 | 0.01 | -0.038 | -0.038 | 0 | 0.14 | 0.26 | 0.13 | 0.1 | 0.1 | 0.05 |
| 2 | 113246011 | rs6726169 | -0.56 | 0.189 | 0.059 | -0.313 | -0.01 | 0.031 | 0.031 | 0 | 0.12 | 0.04 | 0.05 | 0 | 0.13 | 0.13 |
| 2 | 113251000 | rs7569407 | -1.781 | 0.152 | 0.348 | 0.195 | -0.002 | 0.009 | 0.009 | 0 | 0.38 | 0.52 | 0.37 | 0.55 | 0.39 | 0.48 |
| 2 | 113270599 | rs7576952 | -0.984 | 0.163 | 0.334 | 0.767 | 0 | 0.085 | 0.085 | 0.937 | 0.05 | 0.04 | 0.2 | 0.27 | 0.05 | 0.12 |
| 2 | 113276716 | rs17043158 | -1.4 | 0.161 | 0.593 | 0.563 | 0.01 | 0.025 | 0.025 | 0 | 0.32 | 0.47 | 0.36 | 0.55 | 0.38 | 0.47 |
| 2 | 113280880 | rs6718489 | -0.558 | 0.209 | 0.721 | 0.777 | -0.009 | -0.02 | -0.02 | 0 | 0.18 | 0.16 | 0.3 | 0.54 | 0.26 | 0.32 |
| 2 | 113289701 | rs7506 | 0.851 | 0.25 | 0.866 | 0.587 | 0.025 | 0.017 | 0.017 | 0.125 | 0.95 | 0.62 | 0.97 | 1 | 1 | 0.99 |
| 2 | 113294298 | rs10153801 | -1.487 | 0.304 | 0.701 | 0.725 | -0.001 | -0.001 | -0.001 | 0.028 | 0.33 | 0.39 | 0.38 | 0.57 | 0.39 | 0.49 |
| 2 | 113309473 | rs1545133 | 1.686 | 0.17 | 0.719 | 0.644 | 0 | 0.001 | 0.001 | 0 | 0.58 | 0.34 | 0.5 | 0.4 | 0.57 | 0.47 |
| 2 | 113313794 | rs3789724 | -1.129 | 0.197 | 0.687 | 0.823 | 0.016 | 0.176 | 0.176 | 0 | 0.09 | 0.26 | 0.23 | 0.3 | 0.08 | 0.16 |
| 2 | 113318239 | rs6760906 | -0.429 | 0.19 | 0.759 | 0.751 | 0.002 | 0.014 | 0.014 | 0.015 | 0.84 | 0.77 | 0.79 | 0.71 | 0.78 | 0.8 |
| 2 | 113319186 | rs7586668 | 1.956 | -0.072 | 0.711 | 0.387 | 0.016 | 0.059 | 0.059 | 0 | 0.51 | 0.39 | 0.61 | 0.7 | 0.55 | 0.6 |
| 2 | 113322346 | rs3816141 | 2.879 | -0.076 | 0.722 | 0.437 | -0.046 | -0.003 | -0.003 | 0 | 0.42 | 0.22 | 0.39 | 0.42 | 0.47 | 0.44 |
| 2 | 113329522 | rs13395779 | -0.427 | -0.053 | 0.635 | 0.46 | 0 | 0.124 | 0.124 | 0 | 0.08 | 0.14 | 0.2 | 0.29 | 0.07 | 0.16 |
| 2 | 113330638 | rs6734172 | -0.994 | -0.072 | 0.52 | 0.43 | -0.005 | -0.085 | -0.085 | 1.968 | 0.29 | 0.24 | 0.16 | 0 | 0.22 | 0.2 |
| 2 | 113333410 | rs10205187 | -0.992 | 0.088 | 0.322 | 0.784 | 0 | 0.154 | 0.154 | 0 | 0.1 | 0.27 | 0.21 | 0.29 | 0.07 | 0.16 |
| 2 | 113339719 | rs11691023 | -0.506 | 0.134 | -0.492 | 0.2 | 0.001 | 0.008 | 0.008 | 0 | 0.84 | 0.78 | 0.79 | 0.71 | 0.78 | 0.8 |
| 2 | 113342682 | rs719667 | 1.31 | 0.127 | -0.934 | -0.035 | -0.027 | 0.188 | 0.188 | 0 | 0.52 | 0.06 | 0.41 | 0.26 | 0.58 | 0.57 |
| 2 | 113351218 | rs7594007 | 1.141 | -0.319 | -0.19 | -0.03 | 0.012 | 0.092 | 0.092 | 0.021 | 0.54 | 0.26 | 0.53 | 0.71 | 0.55 | 0.59 |
| 2 | 113355181 | rs11694460 | 0.819 | -0.362 | 0.326 | -0.062 | -0.001 | -0.004 | -0.004 | 1.031 | 0.59 | 0.62 | 0.56 | 0.69 | 0.55 | 0.59 |
| 2 | 113355320 | rs10171686 | 1.158 | -0.262 | -0.732 | -0.013 | 0.012 | 0.086 | 0.086 | 0.085 | 0.54 | 0.21 | 0.53 | 0.69 | 0.54 | 0.59 |
| 2 | 113373950 | rs3849317 | -0.11 | -0.626 | -0.636 | -0.017 | 0.004 | -0.009 | -0.009 | 0 | 0.75 | 0.79 | 0.59 | 0.69 | 0.75 | 0.77 |
| 2 | 113377202 | rs17647292 | 0.295 | -0.222 | -1.001 | -0.938 | -0.048 | 0.055 | 0.055 | 0.035 | 0.1 | 0 | 0.05 | 0 | 0.15 | 0.08 |
| 2 | 113389232 | rs908545 | 1.322 | -0.509 | -1.247 | -0.9 | -0.09 | 0.096 | 0.096 | 0 | 0.45 | 0.03 | 0.5 | 0.25 | 0.55 | 0.58 |
| 2 | 113393165 | rs17042199 | -0.73 | -0.333 | 0.162 | -0.863 | -0.011 | -0.134 | -0.134 | 0.084 | 0.23 | 0.53 | 0.12 | 0 | 0.18 | 0.14 |
| 2 | 113394119 | rs4849086 | -0.132 | -0.374 | 0.115 | -0.893 | -0.006 | -0.002 | -0.002 | 0 | 0.75 | 0.64 | 0.59 | 0.25 | 0.75 | 0.75 |
| 2 | 113395241 | rs12996187 | -0.703 | -0.231 | 0.555 | -0.909 | -0.006 | -0.165 | -0.165 | 0 | 0.23 | 0.5 | 0.12 | 0 | 0.18 | 0.14 |
| 2 | 113402530 | rs1061254 | 0.278 | -0.128 | 0.341 | -0.858 | 0.028 | -0.144 | -0.144 | 2.946 | 0.81 | 0.58 | 0.9 | 1 | 0.83 | 0.88 |
| 2 | 113404708 | rs4849091 | 1.323 | -0.253 | 0.197 | -0.923 | 0.014 | 0.16 | 0.16 | 0 | 0.5 | 0.16 | 0.45 | 0.22 | 0.55 | 0.52 |
| 2 | 113410015 | rs10758 | -0.204 | -0.22 | 0.166 | -0.922 | 0.005 | 0.005 | 0.005 | 0 | 0.29 | 0.26 | 0.44 | 0.78 | 0.28 | 0.36 |
| 2 | 113421334 | rs6793 | 0.382 | -0.044 | 0.073 | -0.695 | 0.022 | -0.139 | -0.139 | 0.097 | 0.81 | 0.6 | 0.9 | 1 | 0.83 | 0.88 |
| 2 | 113431092 | rs6705487 | 0.528 | -0.242 | -1.048 | -0.687 | -0.076 | 0.109 | 0.109 | 0 | 0.43 | 0.08 | 0.34 | 0.21 | 0.48 | 0.5 |
| 2 | 113442808 | rs6728935 | -0.036 | -0.326 | -0.136 | -0.65 | -0.001 | 0 | 0 | 0 | 0.6 | 0.63 | 0.5 | 0.24 | 0.55 | 0.56 |
| 2 | 113444837 | rs4849101 | 0.005 | -0.372 | -0.156 | -0.74 | -0.001 | 0.006 | 0.006 | 0 | 0.59 | 0.64 | 0.5 | 0.18 | 0.56 | 0.54 |
| 2 | 113451498 | rs6746908 | 0.271 | -0.119 | 0.086 | -0.942 | 0 | -0.028 | -0.028 | 0 | 0.53 | 0.54 | 0.41 | 0.17 | 0.49 | 0.49 |
| 2 | 113457049 | rs10211073 | -0.108 | -0.393 | -0.085 | -1.027 | -0.021 | 0.136 | 0.136 | 0 | 0.56 | 0.77 | 0.41 | 0.17 | 0.5 | 0.52 |
| 2 | 113463525 | rs1018317 | 0.222 | -0.389 | 0.366 | -1.025 | -0.001 | -0.024 | -0.024 | 0 | 0.52 | 0.56 | 0.39 | 0.17 | 0.5 | 0.49 |
| 2 | 113467333 | rs2048876 | 0.071 | -0.419 | 0.107 | -0.801 | 0.015 | 0.037 | 0.037 | 0 | 0.51 | 0.59 | 0.48 | 0.26 | 0.46 | 0.56 |
| 2 | 113470159 | rs908547 | 0.066 | -0.418 | -0.179 | -0.738 | -0.001 | -0.001 | -0.001 | 1.13 | 0.5 | 0.52 | 0.6 | 0.25 | 0.56 | 0.57 |
| 2 | 113475835 | rs17042270 | -1.015 | -0.78 | -0.48 | 0.395 | -0.018 | 0.075 | 0.075 | 0.004 | 0.11 | 0.15 | 0.2 | 0.52 | 0.04 | 0.16 |
| 2 | 113476084 | rs17042273 | 0.125 | -1.529 | -0.562 | -0.124 | -0.006 | -0.026 | -0.026 | 0 | 0.17 | 0.15 | 0.05 | 0 | 0.06 | 0.12 |
| 2 | 113476126 | rs4849110 | -0.784 | -1.259 | -0.482 | -0.286 | -0.019 | 0.034 | 0.034 | 0.1 | 0.57 | 0.76 | 0.68 | 0.7 | 0.6 | 0.69 |
| 2 | 113476596 | rs4849113 | 0.283 | -0.991 | -0.274 | -0.525 | -0.016 | 0.015 | 0.015 | 0 | 0.33 | 0.28 | 0.45 | 0.18 | 0.51 | 0.4 |

| | | | | | | | | | | | | | | | | |
|---|---|---|---|---|---|---|---|---|---|---|---|---|---|---|---|---|
| 2 | 113478830 | rs17042278 | -0.281 | -1.245 | -0.059 | -0.645 | -0.031 | -0.012 | -0.012 | 0 | 0.06 | 0.11 | 0.12 | 0.04 | 0.01 | 0.05 |
| 2 | 113479614 | rs13386452 | 0.101 | -1.036 | -0.281 | -0.701 | 0.02 | 0.024 | 0.024 | 0.293 | 0.55 | 0.36 | 0.35 | 0.26 | 0.57 | 0.52 |
| 2 | 113486079 | rs908551 | 0.166 | -0.652 | -0.456 | -0.628 | -0.031 | -0.003 | -0.003 | 0.024 | 0.37 | 0.2 | 0.53 | 0.69 | 0.46 | 0.44 |
| 2 | 113489255 | rs12622683 | -0.007 | -0.557 | 0.27 | -0.743 | 0.005 | -0.004 | -0.004 | 0.556 | 0.42 | 0.4 | 0.57 | 0.69 | 0.51 | 0.46 |
| 2 | 113491565 | rs13019249 | 0.079 | -0.601 | -0.02 | -0.754 | -0.167 | -0.064 | -0.064 | 0 | 0.4 | 0.06 | 0.55 | 0.69 | 0.51 | 0.46 |
| 2 | 113494250 | rs4260249 | -1.194 | -0.652 | -0.049 | -0.921 | 0 | -0.001 | -0.001 | 0 | 0.95 | 0.93 | 0.96 | 1 | 0.95 | 0.99 |
| 2 | 113495785 | rs10209160 | -1.194 | -0.652 | -0.049 | -0.921 | 0 | -0.001 | -0.001 | 0 | 0.95 | 0.93 | 0.96 | 1 | 0.95 | 0.99 |
| 2 | 113498566 | rs3811040 | 0.215 | -0.662 | 0.07 | -0.925 | -0.166 | -0.063 | -0.063 | 0 | 0.4 | 0.06 | 0.55 | 0.69 | 0.51 | 0.46 |
| 2 | 113506001 | rs6721044 | -0.013 | -0.653 | 0.162 | -0.918 | 0.001 | 0.032 | 0.032 | 0 | 0.54 | 0.67 | 0.4 | 0.31 | 0.44 | 0.52 |
| 2 | 113513825 | rs6731822 | -0.987 | -0.48 | 0.368 | -0.869 | 0 | -0.001 | -0.001 | 0 | 0.95 | 0.93 | 0.96 | 1 | 0.95 | 0.98 |
| 2 | 113514180 | rs17042342 | 0.044 | -0.483 | 0.394 | -0.871 | 0.004 | -0.036 | -0.036 | 0 | 0.07 | 0.21 | 0.05 | 0 | 0.05 | 0.01 |
| 2 | 113523130 | rs7592947 | 0.272 | -0.872 | 0.598 | -1.265 | -0.056 | -0.101 | -0.101 | 0.011 | 0.19 | 0.38 | 0.17 | 0.06 | 0.1 | 0.19 |
| 2 | 113523967 | rs2048874 | 0.555 | -0.817 | 0.525 | -1.303 | 0.022 | 0.078 | 0.078 | 0 | 0.17 | 0.07 | 0.07 | 0 | 0.12 | 0.14 |
| 2 | 113527906 | rs4848300 | 0.264 | -0.353 | 0.092 | -1.352 | -0.021 | -0.087 | -0.087 | 0.069 | 0.37 | 0.4 | 0.28 | 0.07 | 0.29 | 0.33 |
| 2 | 113537223 | rs17561 | 0.422 | -0.357 | 0.123 | -1.354 | 0.043 | 0.118 | 0.118 | 0 | 0.36 | 0.17 | 0.27 | 0.07 | 0.29 | 0.33 |
| 2 | 113541925 | rs2856837 | 0.415 | -0.358 | 0.211 | -1.355 | 0.036 | 0.084 | 0.084 | 0 | 0.36 | 0.2 | 0.27 | 0.07 | 0.29 | 0.33 |
| 2 | 113542960 | rs1800587 | 0.269 | -0.347 | 0.22 | -1.349 | -0.018 | -0.079 | -0.079 | 0 | 0.37 | 0.4 | 0.28 | 0.07 | 0.29 | 0.31 |
| 2 | 113552737 | rs4848304 | 0.415 | -0.319 | 0.044 | -1.336 | 0.04 | 0.103 | 0.103 | 0 | 0.36 | 0.17 | 0.26 | 0.07 | 0.29 | 0.31 |
| 2 | 113553426 | rs6746923 | -0.224 | -0.302 | -0.008 | -1.276 | 0.001 | -0.001 | -0.001 | 0.663 | 0.64 | 0.59 | 0.75 | 0.78 | 0.61 | 0.61 |
| 2 | 113553668 | rs10496444 | 1.277 | -0.346 | -0.266 | -1.207 | -0.008 | -0.013 | -0.013 | 0 | 0.27 | 0.2 | 0.48 | 0.7 | 0.32 | 0.3 |
| 2 | 113555592 | rs11687624 | -0.432 | -0.206 | -0.255 | -0.998 | -0.001 | -0.004 | -0.004 | 0 | 0.53 | 0.5 | 0.43 | 0.29 | 0.55 | 0.55 |
| 2 | 113558914 | rs17042407 | 0.808 | 0.583 | -0.389 | -0.367 | 0.002 | 0.003 | 0.003 | 0 | 0.17 | 0.18 | 0.15 | 0.21 | 0.24 | 0.24 |
| 2 | 113569039 | rs4849123 | 0.53 | 0.446 | -0.36 | -0.387 | 0.006 | 0.003 | 0.003 | 0 | 0.19 | 0.25 | 0.18 | 0.22 | 0.3 | 0.24 |
| 2 | 113570581 | rs6542097 | -0.704 | 0.022 | -0.271 | -0.843 | 0.001 | -0.032 | -0.032 | 0.369 | 0.81 | 0.81 | 0.9 | 1 | 0.87 | 0.86 |
| 2 | 113570809 | rs11680809 | 0.126 | -0.148 | 0.072 | -0.717 | 0.016 | -0.024 | -0.024 | 0 | 0.61 | 0.45 | 0.71 | 0.78 | 0.57 | 0.62 |
| 2 | 113576902 | rs4849124 | 0.395 | -0.799 | -0.591 | -1.225 | 0.104 | 0.215 | 0.215 | 0 | 0.46 | 0.2 | 0.67 | 0.92 | 0.69 | 0.56 |
| 2 | 113577957 | rs7596461 | -0.836 | -1.005 | 0.199 | -1.343 | -0.057 | -0.115 | -0.115 | 0 | 0.83 | 0.68 | 0.94 | 1 | 0.92 | 0.87 |
| 2 | 113578203 | rs7596684 | 0.36 | -0.897 | 0.002 | -1.192 | 0.029 | 0.05 | 0.05 | 0 | 0.33 | 0.23 | 0.13 | 0.07 | 0.24 | 0.17 |
| 2 | 113582782 | rs3917368 | 0.989 | -0.5 | 0.042 | -0.176 | -0.04 | -0.05 | -0.05 | 0 | 0.22 | 0.14 | 0.28 | 0.5 | 0.34 | 0.25 |
| 2 | 113583880 | rs3917366 | 0.464 | -0.931 | 0.026 | -0.546 | 0.034 | 0.089 | 0.089 | 0.279 | 0.33 | 0.18 | 0.13 | 0.02 | 0.25 | 0.15 |
| 2 | 113587121 | rs2853550 | -0.755 | -0.936 | 0.4 | -0.362 | -0.065 | -0.047 | -0.047 | 0 | 0.82 | 0.62 | 0.83 | 0.92 | 0.92 | 0.76 |
| 2 | 113590390 | rs1143634 | 0.404 | -0.661 | -0.519 | -0.372 | 0.032 | 0.132 | 0.132 | 0.1 | 0.32 | 0.12 | 0.13 | 0.02 | 0.25 | 0.15 |
| 2 | 113590467 | rs1143633 | 0.834 | -0.509 | -0.432 | -0.097 | -0.025 | -0.029 | -0.029 | 0.007 | 0.23 | 0.17 | 0.29 | 0.54 | 0.33 | 0.25 |
| 2 | 113591275 | rs3136558 | 0.352 | -0.936 | -0.458 | 0.47 | 0.024 | -0.037 | -0.037 | 0.077 | 0.29 | 0.16 | 0.28 | 0.35 | 0.25 | 0.17 |
| 2 | 113594387 | rs1143627 | -0.511 | -0.458 | 0.253 | 0.107 | 0.011 | -0.003 | -0.003 | 0 | 0.56 | 0.37 | 0.45 | 0.52 | 0.65 | 0.4 |
| 2 | 113594867 | rs16944 | -0.608 | -0.454 | 0.301 | 0.109 | 0.002 | -0.001 | -0.001 | 0 | 0.57 | 0.43 | 0.45 | 0.53 | 0.65 | 0.4 |
| 2 | 113598255 | rs12621220 | 1.812 | -0.664 | 0.213 | 0.046 | -0.048 | -0.084 | -0.084 | 0 | 0.24 | 0.08 | 0.47 | 0.4 | 0.29 | 0.34 |
| 2 | 113602559 | rs4849127 | -0.989 | -0.742 | 0.784 | -0.01 | -0.054 | -0.09 | -0.09 | 0.075 | 0.83 | 0.72 | 0.94 | 1 | 0.94 | 0.85 |
| 2 | 113603728 | rs10169916 | -0.487 | -0.733 | 0.781 | 0.01 | -0.008 | 0.003 | 0.003 | 0 | 0.58 | 0.5 | 0.46 | 0.52 | 0.65 | 0.4 |
| 2 | 113605957 | rs4447608 | 0.819 | -1.065 | 0.795 | -0.154 | 0.001 | 0.001 | 0.001 | 0.109 | 0.28 | 0.27 | 0.34 | 0.45 | 0.44 | 0.27 |
| 2 | 113606645 | rs4369864 | -0.73 | -1.032 | 0.774 | -0.122 | -0.075 | -0.075 | -0.075 | 1.472 | 0.81 | 0.61 | 0.93 | 0.93 | 0.94 | 0.75 |
| 2 | 113610780 | rs12053091 | 1.868 | -0.838 | 0.511 | -0.057 | -0.01 | -0.023 | -0.023 | 0 | 0.24 | 0.15 | 0.47 | 0.42 | 0.28 | 0.35 |
| 2 | 113631525 | rs11674397 | 0.57 | -0.96 | 0.145 | -0.351 | -0.002 | 0.169 | 0.169 | 0.362 | 0.27 | 0.04 | 0.1 | 0.06 | 0.21 | 0.13 |
| 2 | 113633009 | rs17042509 | 1.474 | -0.932 | 0.392 | -0.296 | 0.013 | 0.001 | 0.001 | 0 | 0.12 | 0.05 | 0.03 | 0 | 0.06 | 0.14 |
| 2 | 113641631 | rs2708919 | -0.11 | -0.781 | 0.793 | 0.157 | 0.002 | 0.003 | 0.003 | 0.768 | 0.45 | 0.5 | 0.4 | 0.46 | 0.52 | 0.38 |
| 2 | 113645942 | rs2708924 | 0.606 | -0.642 | 0.862 | 0.068 | 0.13 | 0.025 | 0.025 | 0 | 0.06 | 0.36 | 0.06 | 0 | 0.09 | 0.02 |
| 2 | 113654849 | rs12472089 | 1.638 | -0.584 | 0.51 | 0.063 | -0.09 | -0.082 | -0.082 | 0 | 0.25 | 0.06 | 0.47 | 0.36 | 0.33 | 0.37 |
| 2 | 113658015 | rs11690539 | 1.586 | -0.634 | 0.513 | 0.063 | -0.08 | -0.074 | -0.074 | 0 | 0.25 | 0.09 | 0.47 | 0.35 | 0.33 | 0.38 |
| 2 | 113658075 | rs2723154 | 0.006 | -0.698 | 0.931 | -0.641 | -0.058 | -0.058 | -0.058 | 1.15 | 0.68 | 0.45 | 0.87 | 0.82 | 0.85 | 0.77 |
| 2 | 113671378 | rs3811046 | 0.624 | -0.661 | 0.994 | -0.595 | 0.047 | 0.137 | 0.137 | 2.233 | 0.6 | 0.18 | 0.8 | 0.81 | 0.7 | 0.71 |
| 2 | 113671410 | rs3811047 | 0.594 | -0.649 | 1.008 | -0.596 | 0.041 | 0.092 | 0.092 | 2.279 | 0.6 | 0.24 | 0.81 | 0.81 | 0.7 | 0.71 |
| 2 | 113678856 | rs4241122 | 0.635 | -0.647 | 0.457 | -0.587 | 0.047 | 0.142 | 0.142 | 2.125 | 0.6 | 0.16 | 0.8 | 0.81 | 0.7 | 0.71 |
| 2 | 113680436 | rs13023984 | 0.176 | -0.484 | 0.798 | -0.539 | 0.06 | 0.085 | 0.085 | 0 | 0.91 | 0.47 | 0.92 | 0.87 | 0.92 | 0.88 |
| 2 | 113683875 | rs10864907 | 0.297 | -0.555 | 0.462 | -0.418 | -0.094 | 0.056 | 0.056 | 0.839 | 0.52 | 0.09 | 0.57 | 0.33 | 0.49 | 0.52 |
| 2 | 113690077 | rs2723196 | 0.012 | -0.559 | 0.739 | -0.405 | 0.062 | 0.082 | 0.082 | 0 | 0.91 | 0.48 | 0.92 | 0.87 | 0.92 | 0.88 |
| 2 | 113691288 | rs4387792 | -0.876 | -0.495 | 0.367 | -0.491 | 0.002 | 0.008 | 0.008 | 0 | 0.67 | 0.44 | 0.46 | 0.58 | 0.65 | 0.53 |
| 2 | 113691499 | rs12987491 | 0.34 | -0.534 | 0.283 | -0.49 | -0.063 | -0.054 | -0.054 | 0 | 0.38 | 0.13 | 0.34 | 0.56 | 0.43 | 0.35 |
| 2 | 113694145 | rs11689621 | 0.06 | -0.529 | 0.332 | -0.651 | -0.005 | -0.056 | -0.056 | 0 | 0.27 | 0.33 | 0.12 | 0.01 | 0.22 | 0.19 |
| 2 | 113694339 | rs2723163 | 0.405 | -0.521 | -0.064 | -0.576 | -0.056 | -0.051 | -0.051 | 1.091 | 0.38 | 0.14 | 0.34 | 0.56 | 0.43 | 0.33 |
| 2 | 113698078 | rs11696039 | 1.381 | -0.509 | -0.16 | -0.65 | -0.011 | -0.025 | -0.025 | 7.697 | 0.28 | 0.14 | 0.47 | 0.31 | 0.27 | 0.36 |
| 2 | 113699617 | rs7602576 | 0.127 | -0.426 | -0.068 | -0.791 | 0.007 | 0.024 | 0.024 | 0.559 | 0.25 | 0.2 | 0.11 | 0.01 | 0.22 | 0.18 |
| 2 | 113715676 | rs10188765 | -0.043 | -0.475 | 0.252 | -0.527 | -0.005 | 0.006 | 0.006 | 0.793 | 0.54 | 0.61 | 0.59 | 0.42 | 0.48 | 0.63 |
| 2 | 113716222 | rs879710 | 0.019 | -0.488 | 0.416 | -0.537 | -0.001 | 0.001 | 0.001 | 0.657 | 0.54 | 0.56 | 0.59 | 0.41 | 0.48 | 0.63 |
| 2 | 113718077 | rs7578034 | -0.328 | -0.465 | 0.676 | -0.591 | -0.015 | -0.009 | -0.009 | 0.078 | 0.58 | 0.77 | 0.66 | 0.42 | 0.57 | 0.67 |
| 2 | 113725172 | rs11692384 | 1.179 | -0.601 | 1.325 | -0.748 | -0.001 | -0.003 | -0.003 | 0 | 0.32 | 0.26 | 0.49 | 0.41 | 0.34 | 0.48 |
| 2 | 113732560 | rs6542107 | 0.281 | -0.526 | 1.025 | -0.752 | 0.007 | 0.035 | 0.035 | 0 | 0.19 | 0.11 | 0.1 | 0.02 | 0.16 | 0.16 |
| 2 | 113738023 | rs13410389 | -0.52 | -0.615 | 1.409 | -0.731 | -0.045 | -0.091 | -0.091 | 0.034 | 0.25 | 0.5 | 0.12 | 0.1 | 0.16 | 0.26 |
| 2 | 113742734 | rs7584409 | -1.137 | -0.535 | 0.972 | -1.245 | -0.011 | -0.014 | -0.014 | 0 | 0.8 | 0.86 | 0.82 | 0.99 | 0.72 | 0.93 |
| 2 | 113745157 | rs10173618 | -0.291 | -0.614 | 1.092 | -1.233 | -0.043 | -0.093 | -0.093 | 0.303 | 0.25 | 0.5 | 0.12 | 0.09 | 0.16 | 0.26 |

| | | | | | | | | | | | | | | | | |
|---|---|---|---|---|---|---|---|---|---|---|---|---|---|---|---|---|
| 2 | 113745428 | rs13407838 | -0.226 | -0.601 | 1.101 | -1.226 | -0.043 | -0.095 | -0.095 | 0.059 | 0.25 | 0.49 | 0.12 | 0.09 | 0.16 | 0.26 |
| 2 | 113751613 | rs1562305 | 0.74 | -0.643 | 0.91 | -1.182 | 0.006 | -0.082 | -0.082 | 0 | 0.25 | 0.11 | 0.22 | 0.57 | 0.23 | 0.27 |
| 2 | 113755190 | rs6542108 | 0.784 | -0.604 | 0.99 | -1.232 | -0.002 | 0.066 | 0.066 | 0 | 0.55 | 0.36 | 0.7 | 0.9 | 0.57 | 0.7 |
| 2 | 113757973 | rs1446514 | -0.876 | -0.6 | 1.037 | -1.234 | 0.025 | -0.115 | -0.115 | 0 | 0.78 | 0.68 | 0.82 | 0.99 | 0.73 | 0.93 |
| 2 | 113759330 | rs6714534 | 0.164 | -0.642 | 0.883 | -1.201 | -0.009 | -0.026 | -0.026 | 0 | 0.23 | 0.26 | 0.11 | 0.09 | 0.16 | 0.22 |
| 2 | 113759950 | rs28938790 | 0.164 | -0.651 | 0.88 | -1.204 | -0.009 | -0.026 | -0.026 | 0 | 0.23 | 0.24 | 0.1 | 0.09 | 0.16 | 0.15 |
| 2 | 113761473 | rs1020404 | -1.045 | -0.407 | 0.697 | -1.073 | -0.011 | -0.014 | -0.014 | 0 | 0.8 | 0.86 | 0.82 | 0.99 | 0.73 | 0.93 |
| 2 | 113762543 | rs1446511 | -1.06 | -0.407 | 0.648 | -1.073 | -0.026 | 0.021 | 0.021 | 0.238 | 0.8 | 0.9 | 0.82 | 0.99 | 0.73 | 0.93 |
| 2 | 113763575 | rs895497 | -0.791 | -0.421 | 0.829 | -1.081 | 0.008 | -0.067 | -0.067 | 0 | 0.78 | 0.76 | 0.82 | 0.99 | 0.72 | 0.93 |
| 2 | 113768620 | rs1867828 | 0.237 | -0.414 | 0.833 | -0.978 | 0.001 | -0.002 | -0.002 | 0.046 | 0.27 | 0.32 | 0.23 | 0.57 | 0.23 | 0.28 |
| 2 | 113773561 | rs1867831 | -1.039 | -0.135 | 0.858 | -0.568 | -0.029 | 0.019 | 0.019 | 0.043 | 0.83 | 0.92 | 0.83 | 0.93 | 0.73 | 0.93 |
| 2 | 113779206 | rs13033574 | 1.351 | -0.319 | 0.671 | -0.629 | -0.037 | 0.064 | 0.064 | 0.496 | 0.56 | 0.14 | 0.57 | 0.72 | 0.57 | 0.7 |
| 2 | 113779533 | rs11894902 | -0.933 | -0.324 | 0.748 | -0.635 | -0.011 | 0.008 | 0.008 | 0 | 0.83 | 0.88 | 0.83 | 0.93 | 0.73 | 0.93 |
| 2 | 113779875 | rs4849142 | 1.276 | -0.273 | 0.386 | -0.675 | -0.037 | 0.063 | 0.063 | 0.471 | 0.56 | 0.14 | 0.57 | 0.72 | 0.57 | 0.7 |
| 2 | 113782059 | rs10496447 | 0.013 | -0.335 | 0.471 | -0.666 | 0.009 | 0.048 | 0.048 | 0 | 0.19 | 0.11 | 0.09 | 0.01 | 0.15 | 0.12 |
| 2 | 113784802 | rs34862832 | -0.235 | -0.327 | 0.358 | -0.659 | 0.001 | -0.02 | -0.02 | 0 | 0.2 | 0.2 | 0.1 | 0.01 | 0.15 | 0.12 |
| 2 | 113785202 | rs11673918 | 0.949 | -0.197 | -0.134 | -0.579 | -0.048 | 0.062 | 0.062 | 0.49 | 0.56 | 0.15 | 0.57 | 0.72 | 0.57 | 0.7 |
| 2 | 113789641 | rs6724667 | -0.752 | 0.185 | -0.233 | -0.531 | -0.084 | 0.033 | 0.033 | 0 | 0.83 | 0.97 | 0.82 | 0.95 | 0.7 | 0.92 |
| 2 | 113790127 | rs17042721 | 0.395 | 0.266 | 0.018 | -0.693 | 0.006 | -0.033 | -0.033 | 0 | 0.07 | 0.12 | 0.02 | 0 | 0.08 | 0.05 |
| 2 | 113790553 | rs2197578 | -0.458 | 0.177 | 0.691 | -0.728 | 0.01 | -0.015 | -0.015 | 0 | 0.8 | 0.79 | 0.81 | 0.95 | 0.7 | 0.92 |
| 2 | 113791193 | rs6758965 | 1.313 | -0.036 | 0.879 | -0.768 | -0.001 | -0.001 | -0.001 | 1.503 | 0.58 | 0.46 | 0.58 | 0.72 | 0.51 | 0.67 |
| 2 | 113796666 | rs1374280 | 1.891 | -0.136 | 0.258 | -0.79 | -0.064 | 0.215 | 0.215 | 0.768 | 0.44 | 0.07 | 0.42 | 0.17 | 0.43 | 0.54 |
| 2 | 113797677 | rs2862774 | -0.16 | 0.105 | 0.598 | -0.788 | 0.001 | -0.037 | -0.037 | 0.089 | 0.17 | 0.2 | 0.1 | 0.01 | 0.19 | 0.13 |
| 2 | 113798388 | rs7570159 | -0.626 | 0.231 | 0.606 | -0.555 | -0.005 | 0.003 | 0.003 | 0 | 0.82 | 0.85 | 0.82 | 0.95 | 0.7 | 0.93 |
| 2 | 113800049 | rs4849144 | -0.643 | 0.238 | 0.541 | -0.551 | -0.005 | 0.003 | 0.003 | 0 | 0.82 | 0.85 | 0.82 | 0.95 | 0.7 | 0.93 |
| 2 | 113811057 | rs11123158 | -0.063 | 0.319 | 0.808 | -0.556 | 0.037 | 0.034 | 0.034 | 0 | 0.78 | 0.68 | 0.69 | 0.73 | 0.7 | 0.83 |
| 2 | 113815997 | rs2515391 | 0.312 | 0.303 | 0.813 | -0.557 | 0.066 | 0.028 | 0.028 | 0 | 0.75 | 0.62 | 0.67 | 0.73 | 0.62 | 0.81 |
| 2 | 113816942 | rs2278717 | 1.819 | 0.045 | 0.714 | -0.603 | -0.008 | 0.196 | 0.196 | 0.184 | 0.46 | 0.16 | 0.34 | 0.19 | 0.42 | 0.58 |
| 2 | 113817566 | rs1530551 | -0.19 | 0.314 | 0.651 | -0.542 | -0.009 | 0.006 | 0.006 | 0 | 0.78 | 0.8 | 0.69 | 0.94 | 0.62 | 0.91 |
| 2 | 113819614 | rs2252007 | -0.165 | 0.306 | 0.615 | -0.546 | -0.017 | 0.013 | 0.013 | 0 | 0.78 | 0.81 | 0.69 | 0.94 | 0.62 | 0.91 |
| 2 | 113820476 | rs2515401 | 0.216 | 0.255 | 0.683 | -0.516 | 0.019 | 0.007 | 0.007 | 0 | 0.75 | 0.72 | 0.68 | 0.73 | 0.62 | 0.82 |
| 2 | 113820530 | rs1800930 | 0.216 | 0.255 | 0.684 | -0.516 | 0.018 | 0.006 | 0.006 | 0 | 0.75 | 0.72 | 0.68 | 0.73 | 0.62 | 0.82 |
| 2 | 113820580 | rs2515402 | 0.216 | 0.255 | 0.681 | -0.516 | 0.019 | 0.007 | 0.007 | 0 | 0.75 | 0.72 | 0.68 | 0.73 | 0.62 | 0.82 |
| 2 | 113820672 | rs3180235 | 0.216 | 0.255 | 0.681 | -0.516 | 0.019 | 0.007 | 0.007 | 0 | 0.75 | 0.72 | 0.68 | 0.73 | 0.62 | 0.81 |
| 2 | 113820732 | rs2515403 | 0.216 | 0.255 | 0.681 | -0.516 | 0.019 | 0.007 | 0.007 | 0 | 0.75 | 0.72 | 0.68 | 0.73 | 0.62 | 0.82 |
| 2 | 113821117 | rs2515404 | -0.661 | 0.315 | 0.66 | -0.522 | -0.108 | 0.032 | 0.032 | 0 | 0.81 | 0.98 | 0.71 | 0.94 | 0.7 | 0.93 |
| 2 | 113821336 | rs957200 | 0.104 | 0.075 | 0.798 | -0.389 | -0.021 | -0.042 | -0.042 | 0 | 0.16 | 0.32 | 0.18 | 0.05 | 0.11 | 0.1 |
| 2 | 113821653 | rs768627 | 0.087 | 0.085 | 0.793 | -0.385 | 0.019 | 0.007 | 0.007 | 0 | 0.75 | 0.72 | 0.68 | 0.73 | 0.62 | 0.82 |
| 2 | 113821887 | rs2515405 | 0.07 | 0.057 | 0.799 | -0.37 | -0.022 | -0.042 | -0.042 | 0 | 0.16 | 0.32 | 0.18 | 0.05 | 0.11 | 0.1 |
| 2 | 113822135 | rs996878 | 0.07 | 0.057 | 0.799 | -0.37 | -0.022 | -0.042 | -0.042 | 0 | 0.16 | 0.32 | 0.18 | 0.05 | 0.11 | 0.1 |
| 2 | 113822168 | rs2515406 | 0.052 | 0.049 | 0.795 | -0.374 | -0.022 | -0.042 | -0.042 | 0.533 | 0.16 | 0.32 | 0.18 | 0.05 | 0.11 | 0.11 |
| 2 | 113822241 | rs996879 | 0.07 | 0.058 | 0.799 | -0.37 | -0.022 | -0.042 | -0.042 | 1.023 | 0.16 | 0.32 | 0.18 | 0.05 | 0.11 | 0.1 |
| 2 | 113825966 | rs921065 | 1.001 | 0.086 | 0.48 | -0.374 | -0.007 | 0.003 | 0.003 | 0 | 0.59 | 0.41 | 0.5 | 0.67 | 0.51 | 0.71 |
| 2 | 113827845 | rs1156701 | 0.554 | 0.011 | 0.309 | -0.421 | 0.006 | 0.014 | 0.014 | 0 | 0.13 | 0.09 | 0.15 | 0.03 | 0.11 | 0.09 |
| 2 | 113828425 | rs9308682 | 0.174 | 0.04 | 0.457 | -0.161 | -0.019 | 0.114 | 0.114 | 0.139 | 0.14 | 0.23 | 0.15 | 0.51 | 0.07 | 0.13 |
| 2 | 113828450 | rs10165797 | 0.86 | 0.104 | 0.425 | -0.249 | 0.015 | -0.007 | -0.007 | 0.374 | 0.63 | 0.49 | 0.52 | 0.7 | 0.53 | 0.73 |
| 2 | 113829709 | rs12711749 | 0.353 | 0.246 | 0.152 | -0.219 | -0.012 | -0.027 | -0.027 | 0 | 0.67 | 0.75 | 0.56 | 0.43 | 0.63 | 0.8 |
| 2 | 113829869 | rs13019891 | 1.619 | 0.191 | -0.081 | -0.262 | -0.047 | 0.216 | 0.216 | 0.712 | 0.48 | 0.09 | 0.37 | 0.19 | 0.45 | 0.59 |
| 2 | 113830173 | rs3811050 | -0.206 | 0.654 | 0.728 | 0.014 | 0.005 | 0.065 | 0.065 | 0 | 0.14 | 0.27 | 0.3 | 0.24 | 0.15 | 0.19 |
| 2 | 113830688 | rs4145013 | 0.851 | 1.026 | 1.039 | 0.496 | 0.004 | 0.008 | 0.008 | 0 | 0.74 | 0.34 | 0.67 | 0.67 | 0.68 | 0.8 |
| 2 | 113831289 | rs3827763 | 0.662 | 1.027 | 0.789 | 0.347 | -0.006 | -0.025 | -0.025 | 0 | 0.74 | 0.77 | 0.73 | 0.49 | 0.7 | 0.81 |
| 2 | 113832312 | rs6761276 | 1.238 | 1.257 | 1.028 | 0.332 | 0.025 | -0.024 | -0.024 | 0 | 0.7 | 0.58 | 0.58 | 0.18 | 0.61 | 0.67 |
| 2 | 113832333 | rs6743376 | 0.363 | 1.315 | 1.078 | 0.366 | 0.03 | 0.032 | 0.032 | 0 | 0.81 | 0.73 | 0.71 | 0.68 | 0.69 | 0.8 |
| 2 | 113832981 | rs13005572 | 1.639 | 1.43 | 0.715 | 0.065 | -0.013 | 0.031 | 0.031 | 0.321 | 0.2 | 0.1 | 0.29 | 0.1 | 0.24 | 0.3 |
| 2 | 113833570 | rs13011842 | 1.632 | 1.423 | 0.676 | 0.075 | -0.014 | 0.032 | 0.032 | 0.327 | 0.2 | 0.1 | 0.29 | 0.1 | 0.24 | 0.3 |
| 2 | 113834739 | rs28928309 | 0.108 | 1.426 | 0.769 | 0.363 | -0.004 | 0.099 | 0.099 | 0.005 | 0.1 | 0.16 | 0.13 | 0.51 | 0.08 | 0.14 |
| 2 | 113836944 | rs10186133 | 0.996 | 1.535 | 0.741 | 0.167 | 0.026 | -0.028 | -0.028 | 0.538 | 0.71 | 0.57 | 0.59 | 0.17 | 0.62 | 0.67 |
| 2 | 113837416 | rs7574787 | -0.763 | 1.651 | 0.625 | 0.501 | -0.017 | 0.003 | 0.003 | 0 | 0.92 | 0.94 | 0.91 | 0.97 | 0.83 | 0.98 |
| 2 | 113840539 | rs11899198 | -1.02 | 1.552 | 1.175 | 0.649 | 0.118 | 0.348 | 0.348 | 0 | 0.06 | 0.36 | 0.16 | 0.28 | 0.08 | 0.12 |
| 2 | 113848574 | rs4849152 | 1.368 | 1.598 | 1.449 | 0.577 | 0.062 | 0.108 | 0.108 | 0.071 | 0.82 | 0.42 | 0.73 | 0.68 | 0.74 | 0.84 |
| 2 | 113849837 | rs1542176 | 0.221 | 1.609 | 0.918 | 0.277 | -0.052 | 0.004 | 0.004 | 0.226 | 0.59 | 0.82 | 0.49 | 0.35 | 0.49 | 0.47 |
| 2 | 113850114 | rs11123162 | 0.161 | 1.527 | 0.893 | 0.325 | -0.05 | -0.091 | -0.091 | 0.2 | 0.33 | 0.09 | 0.42 | 0.61 | 0.34 | 0.51 |
| 2 | 113865808 | rs11677140 | 0.476 | 1.507 | 0.562 | 0.386 | 0.008 | -0.073 | -0.073 | 0 | 0.28 | 0.39 | 0.33 | 0.11 | 0.28 | 0.36 |
| 2 | 113869347 | rs1688072 | 0.511 | 1.293 | 0.345 | 0.478 | -0.003 | 0.001 | 0.001 | 0 | 0.08 | 0.09 | 0.1 | 0.04 | 0.17 | 0.02 |
| 2 | 113869843 | rs315931 | -0.115 | 1.042 | 0.399 | 0.571 | -0.005 | 0.026 | 0.026 | 0.004 | 0.66 | 0.48 | 0.56 | 0.38 | 0.67 | 0.53 |
| 2 | 113870663 | rs17042917 | -0.588 | 1.318 | 0.196 | 0.931 | 0.025 | 0.117 | 0.117 | 0 | 0.08 | 0.2 | 0.17 | 0.28 | 0.11 | 0.12 |
| 2 | 113873018 | rs315920 | -0.081 | 1.587 | -0.165 | 0.544 | -0.017 | 0.016 | 0.016 | 0 | 0.1 | 0.07 | 0.1 | 0.02 | 0.18 | 0.03 |
| 2 | 113875509 | rs2234677 | 1.014 | 1.483 | -0.607 | 0.428 | -0.098 | 0.053 | 0.053 | 0.578 | 0.21 | 0.02 | 0.32 | 0.11 | 0.29 | 0.29 |
| 2 | 113875565 | rs2234678 | 1.014 | 1.524 | -0.607 | 0.428 | -0.095 | 0.053 | 0.053 | 0.578 | 0.21 | 0.02 | 0.32 | 0.11 | 0.29 | 0.29 |

| | | | | | | | | | | | | | | | | |
|---|---|---|---|---|---|---|---|---|---|---|---|---|---|---|---|---|
| 2 | 113875584 | rs2234679 | 1.014 | 1.474 | -0.607 | 0.428 | -0.095 | 0.053 | 0.053 | 0.578 | 0.21 | 0.02 | 0.32 | 0.11 | 0.29 | 0.29 |
| 2 | 113876213 | rs315919 | 0.344 | 0.997 | 0.277 | 0.705 | 0.003 | 0.009 | 0.009 | 0.144 | 0.62 | 0.5 | 0.52 | 0.41 | 0.57 | 0.52 |
| 2 | 113877413 | rs4251985 | 1.081 | 1.172 | 0.564 | 0.601 | -0.092 | 0.056 | 0.056 | 0.543 | 0.21 | 0.02 | 0.32 | 0.11 | 0.29 | 0.29 |
| 2 | 113879297 | rs3213448 | -0.495 | 1.145 | 0.86 | 0.785 | 0.003 | 0.192 | 0.192 | 0 | 0.13 | 0.27 | 0.14 | 0.56 | 0.14 | 0.14 |
| 2 | 113886049 | rs3181052 | -0.347 | 1.141 | 0.84 | 0.783 | -0.001 | 0.115 | 0.115 | 0 | 0.12 | 0.19 | 0.14 | 0.56 | 0.14 | 0.14 |
| 2 | 113887207 | rs419598 | 1.042 | 1.278 | 0.359 | 0.619 | -0.082 | 0.073 | 0.073 | 0.34 | 0.21 | 0.03 | 0.33 | 0.09 | 0.29 | 0.3 |
| 2 | 113887262 | rs423904 | 1.042 | 1.278 | 0.359 | 0.619 | -0.082 | 0.073 | 0.073 | 0.34 | 0.21 | 0.03 | 0.33 | 0.09 | 0.29 | 0.3 |
| 2 | 113889006 | rs431726 | 1.054 | 1.28 | 0.606 | 0.597 | -0.033 | 0.047 | 0.047 | 0.071 | 0.2 | 0.11 | 0.33 | 0.09 | 0.28 | 0.29 |
| 2 | 113889100 | rs3087266 | -0.758 | 1.128 | 1.1 | 0.644 | 0.025 | 0.27 | 0.27 | 0 | 0.14 | 0.51 | 0.15 | 0.57 | 0.16 | 0.13 |
| 2 | 113890304 | rs315952 | -1.016 | 1.222 | 1.06 | 0.782 | 0.04 | 0.187 | 0.187 | 0 | 0.21 | 0.47 | 0.21 | 0.56 | 0.28 | 0.15 |
| 2 | 113890586 | rs315951 | -0.982 | 1.2 | 1.152 | 0.702 | 0.041 | 0.168 | 0.168 | 0 | 0.21 | 0.46 | 0.21 | 0.59 | 0.28 | 0.15 |
| 2 | 113891412 | rs9005 | 0.678 | 1.382 | 0.713 | 0.683 | -0.022 | -0.018 | -0.018 | 0 | 0.24 | 0.18 | 0.51 | 0.33 | 0.31 | 0.38 |
| 2 | 113893256 | rs315948 | -0.075 | 1.28 | 0.657 | -0.12 | 0.005 | -0.025 | -0.025 | 6.508 | 0.1 | 0.19 | 0.09 | 0.02 | 0.12 | 0.03 |
| 2 | 113899025 | rs895496 | -0.838 | 1.111 | 0.296 | -0.253 | -0.001 | -0.023 | -0.023 | 0 | 0.47 | 0.51 | 0.74 | 0.9 | 0.58 | 0.52 |
| 2 | 113900177 | rs11885498 | 0.793 | 1.132 | 0.128 | -0.242 | 0.02 | 0.12 | 0.12 | 0 | 0.5 | 0.31 | 0.24 | 0.1 | 0.42 | 0.48 |
| 2 | 113901313 | rs315957 | -0.678 | 1.242 | 0.266 | 0.203 | 0.007 | -0.014 | -0.014 | 0 | 0.25 | 0.41 | 0.36 | 0.81 | 0.31 | 0.24 |
| 2 | 113914312 | rs6739883 | -0.264 | 0.992 | -0.573 | 0.171 | -0.002 | 0.009 | 0.009 | 0 | 0.19 | 0.15 | 0.09 | 0.04 | 0.18 | 0.26 |
| 2 | 113925061 | rs4849159 | -0.578 | 1.023 | -0.55 | 0.872 | -0.001 | -0.005 | -0.005 | 0.011 | 0.24 | 0.21 | 0.46 | 0.51 | 0.23 | 0.22 |
| 2 | 113930867 | rs12614131 | -0.08 | 0.881 | -0.625 | 1.06 | 0.002 | 0.051 | 0.051 | 0 | 0.07 | 0.03 | 0.24 | 0.5 | 0.06 | 0.08 |
| 2 | 113932431 | rs6760120 | -1.243 | 0.574 | -0.924 | 0.225 | -0.007 | 0.024 | 0.024 | 2.569 | 0.55 | 0.69 | 0.52 | 0.62 | 0.45 | 0.51 |
| 2 | 113933804 | rs17043051 | 0.571 | 0.488 | -0.896 | 0.31 | 0.001 | 0.056 | 0.056 | 0 | 0.07 | 0.03 | 0.24 | 0.5 | 0.06 | 0.08 |
| 2 | 113935028 | rs4849163 | -0.465 | 0.535 | -0.297 | 0.122 | -0.021 | -0.001 | -0.001 | 1.853 | 0.37 | 0.57 | 0.42 | 0.62 | 0.23 | 0.36 |
| 2 | 113935222 | rs10864910 | 0.026 | 0.421 | 0.199 | 0.03 | 0.024 | -0.033 | -0.033 | 0 | 0.29 | 0.21 | 0.37 | 0.52 | 0.18 | 0.24 |
| 2 | 113937563 | rs3762494 | -0.818 | 0.465 | 0.496 | 0.044 | 0.027 | 0.062 | 0.062 | 0 | 0.07 | 0.34 | 0.05 | 0.08 | 0.05 | 0.12 |
| 2 | 113940681 | rs3748914 | -1.212 | 0.594 | -0.076 | 0.051 | -0.005 | 0.02 | 0.02 | 1.688 | 0.55 | 0.74 | 0.52 | 0.6 | 0.44 | 0.5 |
| 2 | 113941769 | rs3791334 | -1.759 | 1.074 | -0.864 | -0.375 | -0.05 | 0.083 | 0.083 | 1.314 | 0.66 | 0.95 | 0.68 | 0.87 | 0.68 | 0.73 |
| 2 | 113945258 | rs17624793 | 0.476 | 1.157 | -0.273 | -0.082 | -0.012 | 0.004 | 0.004 | 0 | 0.17 | 0.1 | 0.1 | 0 | 0.21 | 0.16 |
| 2 | 113945334 | rs3791336 | -0.682 | 1.142 | 0.104 | 0.227 | 0.008 | 0.034 | 0.034 | 0.56 | 0.48 | 0.36 | 0.5 | 0.78 | 0.43 | 0.58 |
| 2 | 113948159 | rs6753019 | 1.576 | 1.421 | 0.971 | 0.802 | 0.093 | 0.138 | 0.138 | 0.343 | 0.89 | 0.56 | 0.87 | 0.75 | 0.82 | 0.8 |
| 2 | 113953508 | rs6755077 | -2.054 | 1.469 | 0.87 | 0.558 | 0.076 | 0.029 | 0.029 | 0 | 0.14 | 0.49 | 0.18 | 0.13 | 0.23 | 0.17 |
| 2 | 113954879 | rs902696 | 1.668 | 1.47 | 0.311 | 0.361 | -0.016 | 0.041 | 0.041 | 0 | 0.53 | 0.18 | 0.53 | 0.65 | 0.47 | 0.49 |
| 2 | 113967075 | rs11123169 | 0.702 | 1.514 | 0.393 | 0.276 | 0.002 | -0.007 | -0.007 | 0 | 0.67 | 0.62 | 0.68 | 0.78 | 0.67 | 0.65 |
| 2 | 113967990 | rs1015754 | 0.548 | 1.348 | 0.442 | 0.328 | 0.001 | -0.011 | -0.011 | 0 | 0.68 | 0.63 | 0.7 | 0.79 | 0.68 | 0.66 |
| 2 | 113973964 | rs895412 | 1.599 | 1.385 | 0.36 | 0.327 | -0.007 | 0.04 | 0.04 | 0 | 0.54 | 0.23 | 0.54 | 0.65 | 0.47 | 0.49 |
| 2 | 113975066 | rs1478 | -1.659 | 1.404 | 0.532 | 0.332 | 0.103 | 0.019 | 0.019 | 0 | 0.14 | 0.47 | 0.17 | 0.1 | 0.22 | 0.16 |
| 2 | 113985170 | rs4849179 | -1.449 | 1.401 | 0.541 | 0.297 | -0.001 | 0 | 0 | 0 | 0.38 | 0.36 | 0.37 | 0.33 | 0.39 | 0.47 |
| 2 | 113989236 | rs2863242 | 0.314 | 1.394 | 0.143 | 0.258 | -0.084 | 0.031 | 0.031 | 0 | 0.18 | 0.02 | 0.23 | 0.07 | 0.23 | 0.18 |
| 2 | 113993344 | rs7589901 | -1.625 | 1.327 | 0.273 | 0.29 | -0.007 | 0.001 | 0.001 | 0 | 0.37 | 0.26 | 0.36 | 0.33 | 0.38 | 0.46 |
| 2 | 113994417 | rs1110839 | 1.928 | 1.179 | 0.342 | 0.339 | -0.016 | 0.024 | 0.024 | 0 | 0.55 | 0.28 | 0.56 | 0.65 | 0.49 | 0.5 |
| 2 | 114000866 | rs2863244 | -2.009 | 0.703 | 0.338 | 0.374 | -0.003 | 0.015 | 0.015 | 0 | 0.44 | 0.59 | 0.44 | 0.33 | 0.46 | 0.5 |
| 2 | 114003237 | rs11886472 | -1.096 | 0.484 | 0.325 | 0.241 | -0.042 | -0.017 | -0.017 | 0.158 | 0.18 | 0.22 | 0.11 | 0.06 | 0.08 | 0.12 |
| 2 | 114010791 | rs4848320 | 0.05 | 0.65 | 0.04 | 0.191 | -0.065 | -0.004 | -0.004 | 1.653 | 0.22 | 0.05 | 0.25 | 0.2 | 0.25 | 0.35 |
| 2 | 114013468 | rs1374163 | -1.783 | 0.771 | 0.219 | 0.259 | -0.015 | -0.026 | -0.026 | 0 | 0.26 | 0.55 | 0.22 | 0.11 | 0.25 | 0.17 |
| 2 | 114017029 | rs6734610 | -1.918 | 0.845 | 0.136 | 0.293 | -0.011 | 0.037 | 0.037 | 0 | 0.47 | 0.59 | 0.46 | 0.31 | 0.51 | 0.52 |
| 2 | 114018826 | rs6542125 | -0.475 | 1.173 | 0.337 | 0.305 | 0.187 | 0.061 | 0.061 | 0.051 | 0.06 | 0.26 | 0.09 | 0.05 | 0.15 | 0.05 |
| 2 | 114022856 | rs11691437 | -0.527 | 1.048 | 0.286 | 0.216 | -0.003 | 0.048 | 0.048 | 0 | 0.26 | 0.16 | 0.23 | 0.07 | 0.25 | 0.23 |
| 2 | 114023217 | rs895415 | -1.058 | 1.038 | 0.1 | 0.237 | 0.04 | -0.074 | -0.074 | 0.069 | 0.32 | 0.43 | 0.33 | 0.12 | 0.41 | 0.27 |
| 2 | 114024505 | rs708633 | -1.058 | 1.038 | 0.1 | 0.237 | 0.04 | -0.074 | -0.074 | 0 | 0.32 | 0.43 | 0.33 | 0.12 | 0.41 | 0.27 |
| 2 | 114032767 | rs4848323 | 0.945 | 0.976 | 0.049 | 0.287 | 0.026 | -0.041 | -0.041 | 0 | 0.65 | 0.53 | 0.65 | 0.77 | 0.55 | 0.57 |
| 2 | 114050116 | rs10209795 | 1.391 | 0.348 | -0.579 | 0.404 | -0.002 | -0.008 | -0.008 | 0 | 0.56 | 0.51 | 0.59 | 0.73 | 0.59 | 0.51 |
| 2 | 114050323 | rs12476056 | 0.008 | 0.427 | -0.943 | 0.414 | -0.074 | 0.07 | 0.07 | 0 | 0.19 | 0.03 | 0.23 | 0.07 | 0.24 | 0.2 |
| 2 | 114057929 | rs4849195 | 0.398 | 0.547 | -0.774 | 0.507 | -0.02 | 0.026 | 0.026 | 0.96 | 0.83 | 0.69 | 0.86 | 0.79 | 0.87 | 0.76 |
| 2 | 114060028 | rs10864913 | -0.307 | 0.566 | -0.854 | 0.627 | -0.043 | 0.081 | 0.081 | 0 | 0.27 | 0.06 | 0.25 | 0.16 | 0.28 | 0.25 |
| 2 | 114062391 | rs13388160 | 0.615 | 0.642 | -0.648 | 0.815 | 0.001 | 0.007 | 0.007 | 0 | 0.57 | 0.57 | 0.61 | 0.63 | 0.58 | 0.51 |
| 2 | 114084836 | rs1191694 | -0.183 | 0.303 | -0.309 | 1.337 | 0 | -0.002 | -0.002 | 0.068 | 0.64 | 0.32 | 0.53 | 0.51 | 0.66 | 0.58 |
| 2 | 114085509 | rs1191691 | 0.249 | 0.284 | -0.867 | 1.104 | -0.034 | 0.025 | 0.025 | 0 | 0.39 | 0.21 | 0.38 | 0.34 | 0.45 | 0.31 |
| 2 | 114095279 | rs1191684 | -0.111 | 0.105 | -0.945 | 0.991 | 0.022 | 0.086 | 0.086 | 0.032 | 0.47 | 0.21 | 0.69 | 0.73 | 0.56 | 0.5 |
| 2 | 114096780 | rs1191682 | -0.091 | 0.13 | -0.944 | 0.88 | 0.016 | 0.112 | 0.112 | 0.201 | 0.47 | 0.2 | 0.73 | 0.78 | 0.56 | 0.5 |
| 2 | 114099346 | rs2121243 | -0.136 | 0.241 | -1.227 | 0.96 | 0.01 | -0.051 | -0.051 | 0 | 0.35 | 0.19 | 0.32 | 0.49 | 0.33 | 0.45 |
| 2 | 114107284 | rs13032628 | 0.742 | 0.548 | -0.515 | 0.941 | -0.004 | -0.028 | -0.028 | 0 | 0.24 | 0.27 | 0.25 | 0.1 | 0.22 | 0.24 |
| 2 | 114109977 | rs2004323 | -0.486 | 0.507 | -0.54 | 1.249 | -0.001 | 0.05 | 0.05 | 0 | 0.11 | 0.09 | 0.2 | 0.37 | 0.11 | 0.18 |
| 2 | 114119809 | rs4849199 | -0.341 | 0.83 | -0.985 | 0.359 | 0.035 | 0.074 | 0.074 | 1.432 | 0.52 | 0.79 | 0.54 | 0.34 | 0.62 | 0.47 |
| 2 | 114120785 | rs13424459 | -0.315 | 1.169 | -0.129 | 0.272 | 0.071 | -0.11 | -0.11 | 0.282 | 0.19 | 0.47 | 0.19 | 0.06 | 0.27 | 0.12 |
| 2 | 114138771 | rs4439970 | 0.349 | 0.986 | -0.843 | 0.74 | 0.016 | 0.015 | 0.015 | 0 | 0.61 | 0.49 | 0.38 | 0.42 | 0.57 | 0.59 |
| 2 | 114142218 | rs2121374 | 0.333 | 0.995 | -0.877 | 0.744 | 0.019 | 0.018 | 0.018 | 0 | 0.61 | 0.49 | 0.38 | 0.42 | 0.57 | 0.59 |
| 2 | 114162983 | rs1867868 | 0.277 | 1.041 | -1.294 | 0.764 | -0.003 | -0.003 | -0.003 | 0.623 | 0.62 | 0.63 | 0.4 | 0.41 | 0.57 | 0.59 |
| 2 | 114273105 | rs761647 | -0.533 | 0.924 | -2.688 | 0.568 | -0.109 | -0.026 | -0.026 | 0.063 | 0.41 | 0.03 | 0.55 | 0.43 | 0.5 | 0.38 |
| 2 | 114311513 | rs3982819 | -0.111 | 0.51 | -1.967 | 0.884 | 0.01 | -0.009 | -0.009 | 0 | 0.06 | 0.02 | 0.06 | 0.16 | 0.03 | 0.11 |
| 2 | 114329388 | rs10174451 | 0.36 | 0.885 | -2.063 | 0.536 | -0.009 | 0.036 | 0.036 | 0.225 | 0.09 | 0.01 | 0.03 | 0 | 0.1 | 0.09 |

| | | | | | | | | | | | | | | | | |
|---|---|---|---|---|---|---|---|---|---|---|---|---|---|---|---|---|
| 2 | 114331775 | rs2921879 | 0.049 | 0.974 | -2.919 | 0.671 | -0.051 | -0.102 | -0.102 | 0 | 0.61 | 0.86 | 0.46 | 0.45 | 0.62 | 0.48 |
| 2 | 114404126 | rs3982465 | -0.363 | 1.167 | -2.387 | 0.6 | -0.02 | 0.036 | 0.036 | 1.134 | 0.14 | 0.07 | 0.07 | 0.02 | 0.17 | 0.14 |
| 2 | 114430089 | rs12620183 | 0.866 | 1.527 | -2.519 | 1.306 | -0.029 | -0.015 | -0.015 | 0.587 | 0.06 | 0.01 | 0.05 | 0.18 | 0.13 | 0.12 |
| 2 | 114432293 | rs17046611 | -0.535 | 0.613 | -1.576 | 1.401 | -0.068 | 0.112 | 0.112 | 0.319 | 0.15 | 0.56 | 0.22 | 0.32 | 0.08 | 0.16 |
| 2 | 114432890 | rs7575011 | 0.482 | 0.775 | -2.129 | 1.2 | -0.098 | -0.172 | -0.172 | 0.099 | 0.78 | 0.97 | 0.59 | 0.56 | 0.68 | 0.64 |
| 2 | 114439648 | rs4544450 | 2.089 | 0.74 | -1.908 | 1.088 | 0.006 | 0.121 | 0.121 | 0 | 0.49 | 0.28 | 0.3 | 0.23 | 0.44 | 0.34 |
| 2 | 114447184 | rs11123205 | 1.101 | 0.89 | -1.884 | 1.144 | -0.072 | -0.071 | -0.071 | 0 | 0.64 | 0.84 | 0.52 | 0.55 | 0.52 | 0.51 |
| 2 | 114451884 | rs10187723 | -0.187 | 0.304 | -1.006 | 1.331 | -0.054 | 0.095 | 0.095 | 0.078 | 0.17 | 0.02 | 0.07 | 0.01 | 0.19 | 0.16 |
| 2 | 114453880 | rs6748854 | -0.271 | 0.613 | -0.283 | 1.576 | -0.043 | 0.058 | 0.058 | 0.209 | 0.2 | 0.43 | 0.23 | 0.32 | 0.17 | 0.23 |
| 2 | 114464777 | rs1044836 | 0.535 | 0.885 | -0.344 | 1.136 | -0.001 | 0 | 0 | 0 | 0.07 | 0.08 | 0.08 | 0.05 | 0.11 | 0.08 |
| 2 | 114465556 | rs17048299 | -1.315 | 0.831 | -0.204 | 1.12 | -0.003 | 0.004 | 0.004 | 0 | 0.11 | 0.11 | 0.05 | 0.02 | 0.11 | 0.09 |
| 2 | 114470778 | rs2290108 | 0.236 | 0.844 | -0.206 | 1.294 | -0.022 | 0.122 | 0.122 | 0.028 | 0.74 | 0.54 | 0.51 | 0.4 | 0.75 | 0.69 |
| 2 | 114485126 | rs17632109 | 0.692 | 0.868 | -0.285 | 1.254 | -0.017 | 0.031 | 0.031 | 0 | 0.06 | 0.01 | 0.1 | 0.01 | 0.07 | 0.16 |
| 2 | 114491636 | rs4848343 | 0.301 | 0.899 | 0.581 | 1.332 | -0.013 | 0.076 | 0.076 | 0.042 | 0.75 | 0.65 | 0.52 | 0.4 | 0.75 | 0.69 |
| 2 | 114493221 | rs10172842 | 0.248 | 0.907 | 0.695 | 1.336 | -0.009 | 0.053 | 0.053 | 0.016 | 0.75 | 0.66 | 0.52 | 0.4 | 0.75 | 0.69 |
| 2 | 114493315 | rs10199515 | -1.555 | 1.003 | 0.448 | 1.101 | 0.009 | -0.03 | -0.03 | 0 | 0.12 | 0.19 | 0.06 | 0.02 | 0.12 | 0.09 |
| 2 | 114500473 | rs3748905 | -0.16 | 1.134 | 0.279 | 1.145 | -0.018 | 0.018 | 0.018 | 0 | 0.09 | 0.05 | 0.08 | 0.04 | 0.14 | 0.1 |
| 2 | 114529665 | rs4848348 | -0.058 | 1.122 | 0.301 | 1.108 | 0.011 | -0.023 | -0.023 | 0 | 0.12 | 0.14 | 0.13 | 0.04 | 0.18 | 0.11 |
| 2 | 114541449 | rs10209330 | 1.545 | 0.228 | 0.869 | 1.621 | 0.001 | 0.007 | 0.007 | 0.916 | 0.64 | 0.37 | 0.47 | 0.41 | 0.62 | 0.59 |
| 2 | 114543194 | rs2687627 | 0.587 | 0.227 | 1.088 | 1.644 | -0.009 | 0.06 | 0.06 | 0.017 | 0.75 | 0.65 | 0.52 | 0.41 | 0.74 | 0.67 |
| 2 | 114556845 | rs3877611 | -1.335 | 0.382 | 0.347 | 1.759 | -0.039 | -0.053 | -0.053 | 0 | 0.14 | 0.05 | 0.15 | 0.21 | 0.17 | 0.31 |
| 2 | 114569119 | rs10208204 | -1.356 | 0.102 | 1.145 | 2.025 | -0.017 | 0.048 | 0.048 | 0 | 0.2 | 0.28 | 0.43 | 0.43 | 0.18 | 0.23 |
| 2 | 114570255 | rs6542174 | -0.49 | 0.154 | 1.684 | 2.022 | 0.021 | 0.086 | 0.086 | 0 | 0.89 | 0.57 | 0.93 | 0.84 | 0.91 | 0.89 |
| 2 | 114579876 | rs4295010 | -1.178 | 0.189 | 1.689 | 1.903 | -0.014 | 0.06 | 0.06 | 0 | 0.18 | 0.25 | 0.4 | 0.44 | 0.16 | 0.25 |
| 2 | 114597466 | rs10864924 | -1.391 | 0.175 | 1.734 | 1.863 | 0 | 0.004 | 0.004 | 0 | 0.2 | 0.2 | 0.41 | 0.4 | 0.19 | 0.25 |
| 2 | 114601242 | rs1028246 | 1.486 | 0.379 | 2.03 | 1.56 | 0.086 | 0.002 | 0.002 | 0 | 0.95 | 0.76 | 0.95 | 1 | 0.93 | 0.97 |
| 2 | 114609336 | rs4848357 | 1.502 | 0.325 | 2.185 | 1.58 | 0.123 | 0.066 | 0.066 | 0.366 | 0.94 | 0.6 | 0.92 | 0.97 | 0.94 | 0.96 |
| 2 | 114637231 | rs6720853 | -1.509 | 0.157 | 2.29 | 2.15 | 0.012 | 0.264 | 0.264 | 0 | 0.09 | 0.32 | 0.15 | 0.38 | 0.05 | 0.22 |
| 2 | 114644053 | rs17197611 | 2.332 | 0.034 | 1.626 | 1.726 | -0.101 | 0.181 | 0.181 | 0.026 | 0.4 | 0.03 | 0.25 | 0.12 | 0.44 | 0.22 |
| 2 | 114655601 | rs11681425 | 0.914 | 0.033 | 2.282 | 2.367 | -0.002 | -0.002 | -0.002 | 0.266 | 0.1 | 0.08 | 0.04 | 0.17 | 0.1 | 0.1 |
| 2 | 114672304 | rs12618683 | -1.631 | 0.02 | 2.29 | 2.472 | 0.014 | 0.295 | 0.295 | 0 | 0.1 | 0.45 | 0.16 | 0.38 | 0.05 | 0.23 |
| 2 | 114684848 | rs3748997 | 0.805 | 0.151 | 1.199 | 2.357 | -0.004 | -0.012 | -0.012 | 0 | 0.1 | 0.06 | 0.04 | 0.17 | 0.1 | 0.1 |
| 2 | 114712977 | rs974057 | 0.777 | 0.08 | 1.448 | 2.133 | 0.008 | -0.041 | -0.041 | 0.293 | 0.84 | 0.9 | 0.69 | 0.7 | 0.85 | 0.88 |
| 2 | 114739110 | rs7577695 | -1.881 | 0.21 | 1.234 | 1.499 | -0.002 | 0.046 | 0.046 | 0.151 | 0.28 | 0.8 | 0.35 | 0.1 | 0.27 | 0.36 |
| 2 | 114752390 | rs276821 | 1.028 | 0.192 | 0.925 | 2.061 | 0.003 | -0.01 | -0.01 | 0.897 | 0.83 | 0.85 | 0.67 | 0.49 | 0.83 | 0.76 |
| 2 | 114754877 | rs276318 | -1.215 | 0.196 | 0.8 | 2.063 | 0.005 | -0.037 | -0.037 | 0.14 | 0.17 | 0.12 | 0.32 | 0.51 | 0.17 | 0.23 |
| 2 | 114756395 | rs156712 | 0.371 | 0.32 | 0.683 | 1.508 | -0.044 | 0.052 | 0.052 | 2.857 | 0.86 | 0.96 | 0.82 | 0.95 | 0.82 | 0.74 |
| 2 | 114765717 | rs11893923 | -1.273 | 0.256 | 1.497 | 1.995 | -0.004 | 0.14 | 0.14 | 0.481 | 0.14 | 0.31 | 0.26 | 0.4 | 0.13 | 0.14 |
| 2 | 114766249 | rs17046870 | -0.654 | 0.134 | 1.615 | 1.869 | -0.008 | 0.065 | 0.065 | 1.273 | 0.16 | 0.22 | 0.27 | 0.38 | 0.14 | 0.18 |
| 2 | 114767730 | rs13016677 | 1.616 | 0.135 | 1.575 | 1.777 | -0.011 | -0.071 | -0.071 | 0 | 0.58 | 0.09 | 0.43 | 0.44 | 0.59 | 0.45 |
| 2 | 114784936 | rs465316 | 1.011 | 0.078 | 2.108 | 1.769 | 0.006 | -0.013 | -0.013 | 0 | 0.68 | 0.76 | 0.55 | 0.58 | 0.67 | 0.57 |
| 2 | 114800662 | rs11123222 | 1.384 | -0.315 | 1.098 | 1.294 | 0.012 | -0.04 | -0.04 | 0 | 0.62 | 0.06 | 0.47 | 0.61 | 0.66 | 0.58 |
| 2 | 114802570 | rs275414 | 0.709 | -0.318 | 1.488 | 1.28 | -0.006 | 0.013 | 0.013 | 0 | 0.73 | 0.62 | 0.55 | 0.64 | 0.73 | 0.67 |
| 2 | 114810707 | rs4848361 | 1.317 | -0.32 | 1.231 | 1.279 | 0.024 | -0.007 | -0.007 | 0 | 0.64 | 0.16 | 0.49 | 0.61 | 0.66 | 0.58 |
| 2 | 114820819 | rs275413 | 0.549 | -0.263 | 1.111 | 1.322 | -0.003 | 0.011 | 0.011 | 0 | 0.75 | 0.64 | 0.56 | 0.66 | 0.75 | 0.67 |
| 2 | 114830803 | rs923229 | 1.322 | -0.259 | 0.179 | 1.322 | 0.009 | -0.037 | -0.037 | 0 | 0.63 | 0.06 | 0.47 | 0.63 | 0.68 | 0.59 |
| 2 | 114844433 | rs7594713 | 0.816 | -0.012 | 0.447 | 1.382 | -0.005 | 0.007 | 0.007 | 0.012 | 0.73 | 0.38 | 0.53 | 0.66 | 0.74 | 0.67 |
| 2 | 114870510 | rs6733541 | -0.858 | -0.08 | -0.348 | 0.962 | 0.022 | 0.029 | 0.029 | 0.025 | 0.41 | 0.72 | 0.36 | 0.3 | 0.33 | 0.4 |
| 2 | 114874553 | rs11123232 | -0.773 | -0.199 | -0.144 | 1.268 | 0.007 | -0.059 | -0.059 | 0 | 0.38 | 0.19 | 0.36 | 0.44 | 0.33 | 0.46 |
| 2 | 114876417 | rs3132069 | 0.053 | 0.195 | -0.014 | 1.56 | 0.006 | 0.069 | 0.069 | 0 | 0.84 | 0.79 | 0.68 | 0.62 | 0.82 | 0.8 |
| 2 | 114888742 | rs4849308 | -0.937 | -0.084 | 0.15 | 1.2 | -0.001 | -0.001 | -0.001 | 1.825 | 0.2 | 0.19 | 0.22 | 0.17 | 0.19 | 0.3 |
| 2 | 114896263 | rs2062374 | -0.94 | -0.09 | 0.243 | 1.197 | 0.001 | 0.002 | 0.002 | 0 | 0.2 | 0.19 | 0.21 | 0.17 | 0.19 | 0.31 |
| 2 | 114907835 | rs7587746 | -1.07 | -0.584 | 0.778 | 1.21 | -0.032 | 0.017 | 0.017 | 0.096 | 0.19 | 0.39 | 0.15 | 0.19 | 0.1 | 0.15 |
| 2 | 114909198 | rs3860407 | -2.086 | -0.631 | 1.247 | 1.344 | -0.022 | 0.012 | 0.012 | 0.109 | 0.35 | 0.56 | 0.31 | 0.4 | 0.23 | 0.37 |
| 2 | 114912069 | rs3105488 | 0.788 | -0.58 | -0.046 | 0.742 | -0.06 | -0.004 | -0.004 | 0 | 0.29 | 0.03 | 0.18 | 0.26 | 0.29 | 0.32 |
| 2 | 114918233 | rs6754886 | -0.242 | -0.514 | 0.284 | 0.732 | -0.007 | 0.007 | 0.007 | 0 | 0.15 | 0.16 | 0.16 | 0.21 | 0.13 | 0.23 |
| 2 | 114924533 | rs17040427 | 0.996 | -0.377 | 0.209 | 0.573 | -0.003 | 0.008 | 0.008 | 0 | 0.05 | 0.04 | 0.06 | 0 | 0.07 | 0.1 |
| 2 | 114928223 | rs1118559 | -1.622 | -0.656 | 0.603 | 0.663 | 0.035 | -0.021 | -0.021 | 0 | 0.35 | 0.74 | 0.33 | 0.39 | 0.24 | 0.37 |
| 2 | 114934218 | rs3132078 | -1.366 | -0.69 | 0.812 | 0.593 | -0.007 | -0.012 | -0.012 | 0.001 | 0.32 | 0.58 | 0.3 | 0.18 | 0.22 | 0.29 |
| 2 | 114937429 | rs17759541 | -0.232 | -0.728 | 0.938 | 0.575 | 0.004 | -0.001 | -0.001 | 0 | 0.2 | 0.09 | 0.21 | 0.18 | 0.15 | 0.19 |
| 2 | 114939981 | rs17048216 | -1.036 | -0.704 | 0.986 | 0.616 | -0.015 | -0.03 | -0.03 | 0.51 | 0.28 | 0.5 | 0.29 | 0.18 | 0.22 | 0.29 |
| 2 | 114949094 | rs34616215 | -0.576 | -0.662 | 0.448 | 0.636 | 0.004 | 0.014 | 0.014 | 0.077 | 0.24 | 0.13 | 0.26 | 0.18 | 0.21 | 0.29 |
| 2 | 114961651 | rs3132025 | 0.71 | -0.469 | -0.039 | 0.468 | -0.072 | 0.003 | 0.003 | 1.475 | 0.3 | 0.03 | 0.18 | 0.25 | 0.3 | 0.33 |
| 2 | 114968227 | rs1549775 | -1.45 | -0.414 | 0.749 | 0.386 | 0.002 | -0.009 | -0.009 | 0 | 0.31 | 0.64 | 0.27 | 0.1 | 0.2 | 0.27 |
| 2 | 114976978 | rs3105495 | 1.522 | -0.364 | 1.122 | 1.266 | -0.046 | 0.103 | 0.103 | 0.127 | 0.8 | 0.48 | 0.85 | 0.61 | 0.9 | 0.83 |
| 2 | 114979501 | rs3105493 | -0.713 | -0.116 | 0.708 | 0.694 | -0.022 | -0.005 | -0.005 | 0.094 | 0.12 | 0.27 | 0.04 | 0.1 | 0.06 | 0.07 |
| 2 | 114980457 | rs2419782 | -0.588 | -0.104 | 0.473 | 0.7 | -0.021 | -0.007 | -0.007 | 0 | 0.12 | 0.22 | 0.04 | 0.11 | 0.06 | 0.07 |
| 2 | 114987326 | rs11123233 | 0.84 | 0.257 | -0.121 | 0.454 | -0.002 | 0.041 | 0.041 | 0.223 | 0.62 | 0.28 | 0.76 | 0.73 | 0.63 | 0.64 |
| 2 | 114992849 | rs4849321 | -0.553 | 0.326 | 0.628 | 0.282 | 0.001 | 0.001 | 0.001 | 0.024 | 0.2 | 0.22 | 0.1 | 0.23 | 0.19 | 0.15 |

| | | | | | | | | | | | | | | | | |
|---|---|---|---|---|---|---|---|---|---|---|---|---|---|---|---|---|
| 2 | 114996082 | rs3132038 | -1.482 | 0.333 | 1.381 | 0.397 | 0 | -0.003 | -0.003 | 0.016 | 0.32 | 0.4 | 0.19 | 0.23 | 0.34 | 0.32 |
| 2 | 114999941 | rs1827746 | -2.102 | 0.414 | 1.679 | 0.384 | -0.001 | 0.003 | 0.003 | 0 | 0.37 | 0.65 | 0.23 | 0.28 | 0.36 | 0.35 |
| 2 | 115010406 | rs1001131 | -0.214 | 0.406 | 1.567 | 0.4 | -0.034 | 0.011 | 0.011 | 0 | 0.08 | 0.01 | 0.07 | 0.04 | 0.11 | 0.14 |
| 2 | 115018255 | rs882575 | -1.622 | 0.321 | 2.158 | 0.464 | -0.015 | 0.003 | 0.003 | 0 | 0.29 | 0.22 | 0.17 | 0.28 | 0.32 | 0.33 |
| 2 | 115025412 | rs3132058 | 2.043 | 0.6 | 2.165 | 0.512 | -0.022 | 0.028 | 0.028 | 0.233 | 0.59 | 0.21 | 0.58 | 0.65 | 0.57 | 0.6 |
| 2 | 115043178 | rs6542199 | 2.086 | 0.589 | 2.542 | 0.513 | -0.04 | 0.028 | 0.028 | 0.435 | 0.59 | 0.15 | 0.58 | 0.64 | 0.58 | 0.6 |
| 2 | 115050713 | rs7559290 | -2.375 | 0.604 | 2.47 | 0.521 | 0.008 | -0.01 | -0.01 | 0 | 0.39 | 0.53 | 0.41 | 0.36 | 0.42 | 0.4 |
| 2 | 115063581 | rs7340228 | -2.215 | 0.753 | 2.358 | 0.523 | 0.007 | -0.006 | -0.006 | 0 | 0.38 | 0.43 | 0.4 | 0.36 | 0.43 | 0.4 |
| 2 | 115080639 | rs952596 | -1.092 | 0.637 | 2.11 | 0.471 | -0.012 | -0.01 | -0.01 | 0 | 0.27 | 0.16 | 0.3 | 0.32 | 0.3 | 0.25 |
| 2 | 115088587 | rs4491737 | -0.536 | 0.658 | 1.902 | 0.384 | 0.005 | -0.018 | -0.018 | 0 | 0.09 | 0.13 | 0.08 | 0.04 | 0.1 | 0.13 |
| 2 | 115103743 | rs6719696 | 1.86 | 0.733 | 1.555 | 1.159 | -0.009 | -0.015 | -0.015 | 0 | 0.59 | 0.35 | 0.59 | 0.46 | 0.57 | 0.6 |
| 2 | 115104954 | rs13014575 | -1.06 | 0.589 | 1.068 | 0.771 | -0.009 | 0.034 | 0.034 | 0 | 0.16 | 0.12 | 0.11 | 0.03 | 0.18 | 0.17 |
| 2 | 115109616 | rs4849329 | 1.509 | 0.769 | 1.614 | 0.786 | 0.041 | -0.075 | -0.075 | 0 | 0.85 | 0.6 | 0.88 | 0.97 | 0.84 | 0.83 |
| 2 | 115112543 | rs7593179 | 1.552 | 0.765 | 1.6 | 0.788 | 0.035 | -0.068 | -0.068 | 0 | 0.86 | 0.64 | 0.89 | 0.97 | 0.85 | 0.84 |
| 2 | 115118072 | rs12618741 | -1.061 | 0.576 | 1.351 | 0.968 | -0.002 | -0.022 | -0.022 | 1.099 | 0.27 | 0.2 | 0.32 | 0.48 | 0.28 | 0.32 |
| 2 | 115131208 | rs13023816 | -1.325 | 0.602 | 1.378 | 0.919 | 0 | -0.001 | -0.001 | 0.199 | 0.17 | 0.14 | 0.33 | 0.11 | 0.29 | 0.22 |
| 2 | 115141616 | rs10864931 | 2.115 | 0.522 | 1.53 | 1.036 | -0.034 | -0.05 | -0.05 | 0 | 0.58 | 0.18 | 0.55 | 0.48 | 0.58 | 0.53 |
| 2 | 115143068 | rs7568000 | -2.133 | 0.727 | 1.884 | 1.092 | 0.049 | 0.037 | 0.037 | 0.459 | 0.24 | 0.52 | 0.38 | 0.34 | 0.32 | 0.34 |
| 2 | 115181049 | rs11123242 | 1.511 | 0.822 | 1.26 | 1.003 | 0.03 | 0.046 | 0.046 | 0.004 | 0.84 | 0.61 | 0.87 | 0.75 | 0.83 | 0.77 |
| 2 | 115185121 | rs11898348 | -0.784 | 0.687 | 0.5 | 1.062 | 0.005 | 0.117 | 0.117 | 0.385 | 0.16 | 0.3 | 0.44 | 0.41 | 0.18 | 0.17 |
| 2 | 115196754 | rs1430094 | 0.247 | 0.446 | -0.122 | 0.913 | -0.001 | 0 | 0 | 0 | 0.81 | 0.81 | 0.48 | 0.68 | 0.69 | 0.78 |
| 2 | 115198886 | rs13392783 | 0.342 | 0.404 | -0.285 | 0.035 | 0.019 | 0.009 | 0.009 | 0 | 0.73 | 0.66 | 0.43 | 0.34 | 0.63 | 0.64 |
| 2 | 115202129 | rs1036022 | 0.279 | 0.388 | -0.358 | 0.006 | 0.009 | 0.001 | 0.001 | 0 | 0.7 | 0.66 | 0.42 | 0.33 | 0.62 | 0.63 |
| 2 | 115203917 | rs10208402 | 0.382 | 0.485 | -0.464 | 0.103 | 0.005 | -0.012 | -0.012 | 2.784 | 0.59 | 0.56 | 0.4 | 0.33 | 0.55 | 0.54 |
| 2 | 115208681 | rs10192393 | 0.189 | 0.603 | -1.213 | -0.628 | 0.003 | 0.032 | 0.032 | 0 | 0.42 | 0.31 | 0.23 | 0.05 | 0.38 | 0.35 |
| 2 | 115209729 | rs4849333 | -0.448 | 0.495 | -1.675 | -0.476 | 0.01 | 0.071 | 0.071 | 0 | 0.33 | 0.14 | 0.16 | 0.12 | 0.27 | 0.32 |
| 2 | 115210642 | rs1430090 | -0.606 | 0.471 | -1.775 | 0.428 | -0.027 | -0.035 | -0.035 | 0 | 0.31 | 0.11 | 0.2 | 0.38 | 0.31 | 0.32 |
| 2 | 115213675 | rs6737251 | -0.468 | 0.041 | -1.314 | 0.396 | 0 | 0 | 0 | 0 | 0.65 | 0.66 | 0.68 | 0.71 | 0.67 | 0.54 |
| 2 | 115213756 | rs17048372 | 0.736 | -0.235 | -1.414 | 0.261 | 0.016 | 0.034 | 0.034 | 0 | 0.21 | 0.12 | 0.09 | 0.13 | 0.19 | 0.21 |
| 2 | 115214509 | rs17763180 | 2.229 | -0.526 | -1.738 | 0.238 | -0.13 | 0.038 | 0.038 | 0.887 | 0.25 | 0.02 | 0.25 | 0.1 | 0.34 | 0.13 |
| 2 | 115218975 | rs4849337 | 0.533 | -0.893 | -0.92 | 0.097 | 0.101 | 0.09 | 0.09 | 1.491 | 0.48 | 0.22 | 0.61 | 0.69 | 0.62 | 0.44 |
| 2 | 115219702 | rs7609142 | 1.252 | -0.861 | -0.328 | 0.108 | -0.014 | -0.029 | -0.029 | 0.194 | 0.07 | 0.22 | 0.03 | 0 | 0.02 | 0.04 |
| 2 | 115228540 | rs13011555 | 2.213 | -0.844 | -0.439 | 0.181 | -0.131 | -0.122 | -0.122 | 0.019 | 0.27 | 0.03 | 0.43 | 0.33 | 0.34 | 0.19 |
| 2 | 115240680 | rs1430097 | 0.373 | -1.298 | -0.088 | -0.311 | 0.101 | 0.081 | 0.081 | 2.528 | 0.55 | 0.19 | 0.63 | 0.69 | 0.65 | 0.5 |
| 2 | 115241769 | rs17048411 | 1.155 | -1.201 | 0.261 | -0.233 | -0.02 | -0.033 | -0.033 | 0 | 0.06 | 0.1 | 0.02 | 0 | 0.02 | 0.04 |
| 2 | 115248223 | rs7558334 | 1.266 | -1.18 | 0.535 | -0.223 | -0.013 | -0.027 | -0.027 | 0 | 0.06 | 0.13 | 0.02 | 0 | 0.02 | 0.04 |
| 2 | 115249736 | rs6761461 | 0.39 | -1.184 | 0.645 | -0.225 | 0.028 | 0.006 | 0.006 | 1.373 | 0.08 | 0.45 | 0.04 | 0 | 0.02 | 0.04 |
| 2 | 115255344 | rs13432035 | 0.536 | -1.167 | 0.3 | -0.217 | 0.015 | -0.004 | -0.004 | 0 | 0.07 | 0.4 | 0.03 | 0 | 0.02 | 0.04 |
| 2 | 115258063 | rs10496465 | 0.706 | -1.131 | 0.063 | -0.25 | -0.013 | 0.046 | 0.046 | 0 | 0.17 | 0.06 | 0.09 | 0.06 | 0.15 | 0.21 |
| 2 | 115259063 | rs9308696 | 0.197 | -1.146 | 0.78 | 0.012 | 0.027 | 0.005 | 0.005 | 0.971 | 0.07 | 0.44 | 0.04 | 0 | 0.02 | 0.05 |
| 2 | 115270207 | rs1820924 | -0.401 | -1.152 | 0.28 | 0.092 | -0.053 | -0.012 | -0.012 | 0 | 0.3 | 0.44 | 0.17 | 0.28 | 0.2 | 0.37 |
| 2 | 115271789 | rs6750402 | -0.862 | -0.986 | 0.274 | 0.124 | -0.011 | 0.03 | 0.03 | 0 | 0.3 | 0.55 | 0.24 | 0.41 | 0.28 | 0.31 |
| 2 | 115271841 | rs10187644 | -0.896 | -1.131 | 0.055 | -0.227 | 0 | 0.005 | 0.005 | 0 | 0.47 | 0.42 | 0.33 | 0.46 | 0.47 | 0.46 |
| 2 | 115272561 | rs1346762 | 2.404 | -1.085 | -0.723 | -0.063 | -0.051 | -0.017 | -0.017 | 0.324 | 0.23 | 0.09 | 0.45 | 0.21 | 0.31 | 0.16 |
| 2 | 115272713 | rs1346763 | 1.13 | -1.041 | -0.701 | -0.015 | -0.05 | -0.028 | -0.028 | 0 | 0.36 | 0.19 | 0.56 | 0.38 | 0.42 | 0.34 |
| 2 | 115278738 | rs1430101 | -0.657 | -0.57 | -0.599 | 0.475 | -0.001 | 0.003 | 0.003 | 0 | 0.8 | 0.78 | 0.9 | 0.66 | 0.82 | 0.71 |
| 2 | 115280624 | rs2420165 | 0.16 | -0.174 | -0.768 | 0.277 | -0.002 | -0.011 | -0.011 | 0.51 | 0.13 | 0.07 | 0.11 | 0.17 | 0.12 | 0.19 |
| 2 | 115281036 | rs11674031 | 0.257 | 0.399 | -0.671 | 0.102 | -0.004 | 0.044 | 0.044 | 0 | 0.21 | 0.09 | 0.1 | 0.05 | 0.2 | 0.17 |
| 2 | 115285693 | rs6746254 | -0.103 | 0.404 | -0.171 | 0.852 | 0.008 | 0.038 | 0.038 | 0 | 0.12 | 0.23 | 0.27 | 0.24 | 0.15 | 0.13 |
| 2 | 115292826 | rs1430108 | -0.635 | 0.848 | -0.629 | 0.52 | -0.005 | 0.004 | 0.004 | 0 | 0.41 | 0.37 | 0.47 | 0.68 | 0.48 | 0.54 |
| 2 | 115294661 | rs6542214 | -0.675 | 0.365 | -0.731 | 0.775 | -0.004 | -0.007 | -0.007 | 0 | 0.43 | 0.52 | 0.46 | 0.69 | 0.41 | 0.49 |
| 2 | 115303050 | rs7590021 | -0.652 | 0.056 | -0.754 | 1.139 | 0.002 | -0.002 | -0.002 | 3.786 | 0.25 | 0.23 | 0.27 | 0.52 | 0.21 | 0.45 |
| 2 | 115305711 | rs1430112 | -1.185 | 0.763 | 0.215 | 1.013 | 0.059 | 0.063 | 0.063 | 0 | 0.18 | 0.5 | 0.13 | 0.26 | 0.18 | 0.16 |
| 2 | 115306432 | rs6738538 | -1.276 | 0.713 | 0.41 | 0.906 | 0.04 | 0.02 | 0.02 | 0.024 | 0.18 | 0.55 | 0.14 | 0.24 | 0.18 | 0.13 |
| 2 | 115341833 | rs6729737 | 1.08 | 1.019 | 0.23 | 0.46 | 0.006 | -0.009 | -0.009 | 0 | 0.67 | 0.64 | 0.65 | 0.71 | 0.65 | 0.63 |
| 2 | 115345516 | rs996135 | -1.031 | 1.124 | 0.297 | 0.371 | 0.008 | 0.001 | 0.001 | 0 | 0.22 | 0.22 | 0.21 | 0.25 | 0.25 | 0.27 |
| 2 | 115345817 | rs17048489 | -1.031 | 1.124 | 0.297 | 0.371 | 0.008 | 0.001 | 0.001 | 0 | 0.22 | 0.22 | 0.21 | 0.25 | 0.25 | 0.27 |
| 2 | 115349969 | rs12999482 | -0.543 | 0.909 | 0.466 | 0.578 | -0.01 | 0.034 | 0.034 | 0.031 | 0.08 | 0 | 0.12 | 0.03 | 0.09 | 0.1 |
| 2 | 115352728 | rs17048497 | -0.776 | 0.94 | 0.709 | 0.591 | 0.033 | 0.093 | 0.093 | 0 | 0.11 | 0.26 | 0.13 | 0.21 | 0.11 | 0.13 |
| 2 | 115360811 | rs17782840 | 0.327 | 0.888 | 0.548 | 0.282 | -0.002 | -0.012 | -0.012 | 0 | 0.09 | 0.05 | 0.06 | 0 | 0.13 | 0.08 |
| 2 | 115365883 | rs2195607 | -1.339 | 0.65 | 0.849 | 0.491 | 0 | 0.001 | 0.001 | 0 | 0.14 | 0.11 | 0.16 | 0.12 | 0.13 | 0.09 |
| 2 | 115370635 | rs785041 | -0.551 | 0.741 | 1.534 | 0.603 | 0.068 | 0.205 | 0.205 | 0 | 0.06 | 0.24 | 0.07 | 0.2 | 0.06 | 0.03 |
| 2 | 115378901 | rs6726651 | -2.111 | 0.711 | 1.689 | 0.551 | 0.004 | 0.046 | 0.046 | 0 | 0.22 | 0.54 | 0.21 | 0.29 | 0.19 | 0.14 |
| 2 | 115380251 | rs10186236 | -2.01 | 0.704 | 1.623 | 0.547 | 0.002 | 0.029 | 0.029 | 0 | 0.21 | 0.62 | 0.21 | 0.29 | 0.19 | 0.14 |
| 2 | 115386455 | rs17048536 | -1.135 | 0.225 | 1.483 | 0.58 | 0.102 | 0.205 | 0.205 | 0 | 0.07 | 0.41 | 0.08 | 0.17 | 0.06 | 0.03 |
| 2 | 115388286 | rs12711800 | -2.131 | 0.212 | 1.387 | 0.574 | 0.001 | 0.037 | 0.037 | 0 | 0.22 | 0.55 | 0.21 | 0.29 | 0.19 | 0.14 |
| 2 | 115390606 | rs7557857 | -1.052 | 0.023 | 0.481 | 0.391 | -0.015 | 0.016 | 0.016 | 0 | 0.13 | 0.02 | 0.11 | 0.09 | 0.12 | 0.1 |
| 2 | 115394527 | rs12996109 | -1.16 | 0.075 | 0.572 | 0.396 | -0.006 | -0.009 | -0.009 | 0 | 0.15 | 0.25 | 0.13 | 0.11 | 0.13 | 0.12 |
| 2 | 115405471 | rs1435874 | -1.675 | -0.378 | 0.444 | 0.506 | -0.023 | 0.019 | 0.019 | 0.009 | 0.21 | 0.68 | 0.35 | 0.27 | 0.17 | 0.13 |

| | | | | | | | | | | | | | | | | |
|---|---|---|---|---|---|---|---|---|---|---|---|---|---|---|---|---|
| 2 | 115406526 | rs10496470 | -0.153 | -0.323 | 0.262 | 0.569 | -0.015 | -0.042 | -0.042 | 0 | 0.08 | 0.02 | 0.23 | 0.16 | 0.1 | 0.07 |
| 2 | 115410964 | rs13022574 | -0.302 | -0.298 | 0.437 | 0.588 | -0.001 | -0.004 | -0.004 | 0.044 | 0.06 | 0.05 | 0.08 | 0.11 | 0.06 | 0.06 |
| 2 | 115413975 | rs7607402 | -1.548 | -0.581 | 0.41 | 0.325 | -0.032 | 0.007 | 0.007 | 0 | 0.25 | 0.61 | 0.37 | 0.27 | 0.18 | 0.21 |
| 2 | 115414557 | rs10167941 | -0.923 | -0.365 | -0.163 | 0.417 | -0.007 | 0.01 | 0.01 | 0 | 0.21 | 0.28 | 0.36 | 0.29 | 0.19 | 0.19 |
| 2 | 115418679 | rs1561070 | 0.006 | -0.986 | -1.244 | 0.709 | 0.069 | -0.091 | -0.091 | 4.472 | 0.18 | 0.03 | 0.17 | 0.33 | 0.12 | 0.19 |
| 2 | 115430912 | rs713370 | -0.637 | -0.192 | -0.762 | 0.359 | -0.002 | -0.006 | -0.006 | 0 | 0.48 | 0.51 | 0.48 | 0.37 | 0.41 | 0.56 |
| 2 | 115432648 | rs718062 | -0.121 | -0.443 | -0.548 | 0.051 | 0 | 0.001 | 0.001 | 0 | 0.34 | 0.27 | 0.37 | 0.35 | 0.31 | 0.47 |
| 2 | 115434678 | rs12478101 | -0.316 | -0.515 | -0.509 | -0.009 | -0.009 | -0.019 | -0.019 | 0 | 0.07 | 0.08 | 0.06 | 0.02 | 0.03 | 0.08 |
| 2 | 115445395 | rs17043357 | -0.407 | -0.515 | -0.528 | -0.027 | -0.006 | -0.019 | -0.019 | 0.922 | 0.07 | 0.08 | 0.06 | 0.02 | 0.04 | 0.09 |
| 2 | 115453185 | rs13032740 | -1.556 | -0.587 | -0.165 | -0.08 | -0.014 | -0.088 | -0.088 | 0.3 | 0.14 | 0.31 | 0.11 | 0.02 | 0.11 | 0.09 |
| 2 | 115472604 | rs7605779 | -1.338 | -0.493 | -0.024 | -0.225 | -0.015 | -0.101 | -0.101 | 0 | 0.15 | 0.35 | 0.11 | 0.02 | 0.12 | 0.11 |
| 2 | 115473521 | rs17043407 | 0.081 | -0.546 | -0.184 | -0.085 | -0.037 | -0.019 | -0.019 | 0 | 0.39 | 0.12 | 0.42 | 0.39 | 0.38 | 0.47 |
| 2 | 115490198 | rs7606549 | -0.92 | -0.392 | -0.176 | -0.036 | 0 | -0.003 | -0.003 | 1.217 | 0.53 | 0.46 | 0.52 | 0.41 | 0.49 | 0.56 |
| 2 | 115500323 | rs2117851 | -1.188 | 0.536 | -0.577 | 0.476 | -0.076 | -0.07 | -0.07 | 0 | 0.89 | 0.99 | 0.83 | 0.79 | 0.76 | 0.84 |
| 2 | 115508903 | rs6542220 | -0.751 | 0.295 | -0.258 | 0.011 | -0.011 | -0.029 | -0.029 | 0.042 | 0.08 | 0.11 | 0.05 | 0.02 | 0.04 | 0.1 |
| 2 | 115511716 | rs2082875 | 0.796 | 0.182 | -0.191 | -0.204 | -0.004 | 0.098 | 0.098 | 0.121 | 0.36 | 0.17 | 0.23 | 0.13 | 0.34 | 0.41 |
| 2 | 115524203 | rs11890570 | -0.764 | 0.25 | 0.061 | -0.135 | 0.061 | 0.03 | 0.03 | 0 | 0.06 | 0.37 | 0.04 | 0 | 0.03 | 0.01 |
| 2 | 115526908 | rs319853 | -0.886 | 0.171 | 0.331 | -0.045 | -0.019 | -0.001 | -0.001 | 2.776 | 0.59 | 0.68 | 0.56 | 0.44 | 0.53 | 0.6 |
| 2 | 115527730 | rs319855 | 0.181 | 0.178 | 0.288 | -0.058 | 0.001 | 0.013 | 0.013 | 0 | 0.45 | 0.3 | 0.46 | 0.42 | 0.46 | 0.49 |
| 2 | 115530841 | rs319858 | 0.166 | 0.225 | 0.293 | 0.142 | -0.002 | -0.01 | -0.01 | 0 | 0.46 | 0.31 | 0.46 | 0.49 | 0.47 | 0.49 |
| 2 | 115536791 | rs6754133 | -1.257 | 0.158 | 0.26 | -0.668 | -0.008 | -0.057 | -0.057 | 0.751 | 0.1 | 0.18 | 0.08 | 0.02 | 0.07 | 0.12 |
| 2 | 115543481 | rs12478225 | -0.754 | 0.196 | 0.281 | -0.647 | -0.018 | -0.059 | -0.059 | 0 | 0.11 | 0.2 | 0.07 | 0.02 | 0.06 | 0.11 |
| 2 | 115546549 | rs388581 | -0.688 | 0.228 | 0.035 | -0.703 | -0.007 | 0.031 | 0.031 | 0.249 | 0.55 | 0.61 | 0.35 | 0.22 | 0.47 | 0.51 |
| 2 | 115551783 | rs12711805 | -0.56 | 0.348 | 0.044 | -0.653 | -0.003 | -0.081 | -0.081 | 0 | 0.59 | 0.5 | 0.74 | 0.86 | 0.61 | 0.63 |
| 2 | 115557174 | rs12472049 | -0.61 | 0.317 | -0.007 | -0.658 | 0.012 | 0.03 | 0.03 | 0 | 0.11 | 0.04 | 0.07 | 0.01 | 0.07 | 0.1 |
| 2 | 115560289 | rs2139412 | 0.145 | 0.394 | 0.07 | -0.57 | 0 | 0.002 | 0.002 | 0.56 | 0.48 | 0.47 | 0.67 | 0.77 | 0.54 | 0.54 |
| 2 | 115568614 | rs11904812 | 0.212 | 0.79 | 0.021 | -0.56 | 0.004 | 0.046 | 0.046 | 0.705 | 0.48 | 0.39 | 0.66 | 0.78 | 0.49 | 0.52 |
| 2 | 115569596 | rs12711807 | -0.948 | 0.629 | 0.051 | -0.561 | 0.013 | -0.054 | -0.054 | 2.252 | 0.13 | 0.24 | 0.09 | 0.01 | 0.13 | 0.1 |
| 2 | 115573050 | rs2420449 | -0.651 | 0.958 | -0.523 | -0.643 | -0.002 | -0.021 | -0.021 | 0 | 0.6 | 0.59 | 0.73 | 0.82 | 0.61 | 0.61 |
| 2 | 115573499 | rs908931 | -0.029 | 0.909 | -0.521 | -0.514 | 0 | -0.003 | -0.003 | 0 | 0.49 | 0.53 | 0.63 | 0.79 | 0.47 | 0.52 |
| 2 | 115577204 | rs1519674 | -0.056 | 1.694 | -1.022 | 0.07 | -0.088 | -0.053 | -0.053 | 0.072 | 0.14 | 0.01 | 0.17 | 0.21 | 0.23 | 0.2 |
| 2 | 115578181 | rs7570960 | -1.077 | 0.963 | -0.547 | -0.211 | 0 | -0.015 | -0.015 | 2.051 | 0.12 | 0.15 | 0.08 | 0.02 | 0.11 | 0.09 |
| 2 | 115586939 | rs6732028 | -0.616 | 0.951 | -0.75 | -0.645 | -0.009 | 0.06 | 0.06 | 0.644 | 0.42 | 0.33 | 0.3 | 0.14 | 0.47 | 0.35 |
| 2 | 115588066 | rs7597658 | 0.074 | 0.569 | -0.222 | -1.006 | -0.001 | -0.02 | -0.02 | 0.808 | 0.58 | 0.57 | 0.71 | 0.87 | 0.59 | 0.73 |
| 2 | 115591358 | rs7558870 | 0.556 | 0.336 | -0.185 | -0.174 | 0.019 | -0.012 | -0.012 | 0.486 | 0.6 | 0.17 | 0.59 | 0.63 | 0.61 | 0.68 |
| 2 | 115599013 | rs10496476 | 0.435 | 0.424 | -0.014 | -0.172 | -0.004 | -0.037 | -0.037 | 0.208 | 0.23 | 0.12 | 0.27 | 0.35 | 0.26 | 0.22 |
| 2 | 115605062 | rs6755195 | -0.751 | 0.368 | 1.084 | -0.287 | 0.026 | -0.055 | -0.055 | 0 | 0.13 | 0.45 | 0.11 | 0.03 | 0.12 | 0.09 |
| 2 | 115605265 | rs6745105 | -0.932 | 0.364 | 0.838 | -0.289 | -0.015 | -0.059 | -0.059 | 0 | 0.16 | 0.69 | 0.12 | 0.03 | 0.12 | 0.09 |
| 2 | 115613314 | rs7604124 | 0.004 | 0.4 | -0.299 | -0.69 | -0.003 | 0.113 | 0.113 | 0 | 0.33 | 0.17 | 0.2 | 0.1 | 0.3 | 0.41 |
| 2 | 115613465 | rs4353658 | -0.129 | 0.684 | -0.658 | -1.009 | -0.014 | 0.194 | 0.194 | 0 | 0.47 | 0.26 | 0.29 | 0.15 | 0.45 | 0.51 |
| 2 | 115613577 | rs1879124 | 0.899 | 0.909 | -0.477 | -0.736 | 0.003 | -0.007 | -0.007 | 0 | 0.28 | 0.19 | 0.3 | 0.31 | 0.27 | 0.27 |
| 2 | 115618939 | rs11684641 | 1.143 | 0.742 | -0.746 | -0.823 | 0.013 | 0.088 | 0.088 | 0 | 0.26 | 0.08 | 0.23 | 0.12 | 0.22 | 0.22 |
| 2 | 115622199 | rs13007523 | 0.535 | 0.451 | -1.143 | -1.051 | -0.032 | 0.04 | 0.04 | 0 | 0.41 | 0.22 | 0.3 | 0.29 | 0.43 | 0.49 |
| 2 | 115626029 | rs6542227 | 1.006 | 0.487 | -0.49 | -1.079 | 0.022 | 0.107 | 0.107 | 0 | 0.22 | 0.44 | 0.4 | 0.59 | 0.28 | 0.23 |
| 2 | 115627410 | rs9308704 | 0.031 | 0.48 | -0.485 | -1.085 | 0.003 | 0.016 | 0.016 | 0 | 0.48 | 0.53 | 0.64 | 0.7 | 0.49 | 0.45 |
| 2 | 115627690 | rs12471830 | 0.37 | 0.286 | -1.067 | -1.209 | -0.044 | 0.043 | 0.043 | 0 | 0.38 | 0.13 | 0.26 | 0.26 | 0.36 | 0.48 |
| 2 | 115647583 | rs998429 | 0.715 | 0.26 | -0.906 | -1.132 | -0.025 | 0.148 | 0.148 | 0.496 | 0.34 | 0.04 | 0.27 | 0.12 | 0.28 | 0.27 |
| 2 | 115649874 | rs1879123 | 0.025 | 0.397 | -0.625 | -0.962 | 0.005 | 0.068 | 0.068 | 0 | 0.51 | 0.68 | 0.67 | 0.71 | 0.51 | 0.48 |
| 2 | 115665948 | rs13032069 | 1.282 | 0.773 | -0.443 | 0.391 | 0.023 | 0.035 | 0.035 | 0 | 0.16 | 0.28 | 0.23 | 0.28 | 0.24 | 0.17 |
| 2 | 115676090 | rs970882 | 0.003 | 0.853 | -0.995 | 0.369 | -0.073 | 0.035 | 0.035 | 0.041 | 0.42 | 0.15 | 0.46 | 0.6 | 0.46 | 0.55 |
| 2 | 115679311 | rs7559740 | -0.005 | 1.612 | -0.53 | 0.723 | -0.012 | 0.05 | 0.05 | 0.449 | 0.06 | 0.03 | 0.2 | 0.32 | 0.12 | 0.07 |
| 2 | 115683104 | rs11123259 | 1.054 | 1.483 | -0.192 | 0.604 | 0.01 | 0.037 | 0.037 | 0 | 0.18 | 0.19 | 0.2 | 0.36 | 0.25 | 0.21 |
| 2 | 115687956 | rs4849358 | 1.366 | 1.493 | -0.492 | 0.776 | -0.007 | -0.013 | -0.013 | 0.098 | 0.13 | 0.12 | 0.23 | 0.22 | 0.16 | 0.23 |
| 2 | 115688530 | rs1914973 | 0.422 | 1.685 | -0.451 | 0.528 | 0.003 | -0.023 | -0.023 | 0.844 | 0.09 | 0.11 | 0.07 | 0 | 0.14 | 0.04 |
| 2 | 115697773 | rs10496481 | 1.439 | 1.656 | -0.388 | 0.582 | 0.004 | 0.013 | 0.013 | 0 | 0.19 | 0.19 | 0.23 | 0.36 | 0.26 | 0.23 |
| 2 | 115710826 | rs980316 | 0.993 | 1.293 | -0.472 | 0.193 | 0.061 | -0.004 | -0.004 | 0 | 0.43 | 0.14 | 0.37 | 0.41 | 0.35 | 0.44 |
| 2 | 115718524 | rs10210557 | -1.452 | 1.215 | 0.117 | 0.302 | 0.058 | 0.073 | 0.073 | 0 | 0.78 | 0.58 | 0.75 | 0.65 | 0.69 | 0.77 |
| 2 | 115720456 | rs2136544 | 1.76 | 1.309 | 0.141 | 0.25 | -0.001 | 0.002 | 0.002 | 0 | 0.14 | 0.15 | 0.25 | 0.25 | 0.17 | 0.24 |
| 2 | 115720896 | rs2175176 | 0.514 | 1.292 | 0.302 | 0.238 | 0.005 | 0.002 | 0.002 | 0.506 | 0.33 | 0.35 | 0.49 | 0.59 | 0.47 | 0.46 |
| 2 | 115721233 | rs6740064 | 1.377 | 0.943 | 0.174 | 0.133 | 0.035 | 0.026 | 0.026 | 0.015 | 0.37 | 0.26 | 0.32 | 0.3 | 0.25 | 0.39 |
| 2 | 115722009 | rs2420589 | 0.081 | 0.853 | -0.23 | 0.131 | -0.006 | -0.027 | -0.027 | 0.102 | 0.31 | 0.22 | 0.37 | 0.35 | 0.32 | 0.35 |
| 2 | 115730919 | rs6745631 | 0.504 | 0.897 | -0.239 | 0.157 | -0.004 | -0.003 | -0.003 | 0 | 0.25 | 0.25 | 0.31 | 0.25 | 0.26 | 0.29 |
| 2 | 115735755 | rs2036779 | 0.857 | 0.943 | -0.511 | 0.065 | 0.003 | -0.023 | -0.023 | 0 | 0.1 | 0.11 | 0.09 | 0 | 0.16 | 0.05 |
| 2 | 115741770 | rs1396933 | -0.117 | 0.911 | -0.4 | 0.094 | -0.004 | -0.006 | -0.006 | 0.011 | 0.44 | 0.47 | 0.56 | 0.6 | 0.57 | 0.52 |
| 2 | 115744112 | rs17355236 | 1.154 | 0.607 | -0.811 | 0.128 | 0.087 | 0.033 | 0.033 | 0.008 | 0.34 | 0.11 | 0.27 | 0.29 | 0.22 | 0.38 |
| 2 | 115745224 | rs17043876 | 1.141 | 1.398 | -0.392 | 0.151 | 0.002 | 0.004 | 0.004 | 0 | 0.19 | 0.18 | 0.25 | 0.35 | 0.31 | 0.23 |
| 2 | 115745317 | rs17452458 | -0.127 | 1.095 | -0.502 | -0.255 | -0.024 | 0.017 | 0.017 | 0 | 0.2 | 0.12 | 0.15 | 0 | 0.25 | 0.12 |
| 2 | 115756457 | rs1980007 | -0.216 | -0.244 | -1.036 | 0.328 | 0.031 | -0.002 | -0.002 | 0 | 0.62 | 0.38 | 0.61 | 0.65 | 0.8 | 0.55 |
| 2 | 115757276 | rs1876860 | -0.118 | 0.298 | -1.847 | 0.233 | -0.049 | 0.021 | 0.021 | 0.124 | 0.26 | 0.08 | 0.17 | 0.16 | 0.33 | 0.23 |

| | | | | | | | | | | | | | | | | |
|---|---|---|---|---|---|---|---|---|---|---|---|---|---|---|---|---|
| 2 | 115767705 | rs1396921 | 2.081 | 0.348 | -1.825 | 0.478 | -0.07 | -0.063 | -0.063 | 0 | 0.22 | 0.08 | 0.31 | 0.32 | 0.32 | 0.22 |
| 2 | 115817986 | rs1024114 | -0.752 | -0.092 | -1.033 | 0.123 | 0 | -0.001 | -0.001 | 0 | 0.48 | 0.54 | 0.57 | 0.53 | 0.45 | 0.6 |
| 2 | 115822755 | rs17452616 | -1.375 | -0.423 | -0.58 | -0.212 | -0.007 | 0.023 | 0.023 | 0 | 0.54 | 0.34 | 0.55 | 0.35 | 0.5 | 0.6 |
| 2 | 115826561 | rs2901277 | 0.4 | -0.405 | -0.465 | -0.093 | -0.001 | -0.001 | -0.001 | 0 | 0.8 | 0.73 | 0.91 | 0.83 | 0.82 | 0.85 |
| 2 | 115827788 | rs1396919 | 0.657 | -0.299 | -0.537 | -0.012 | 0.004 | 0.005 | 0.005 | 0 | 0.13 | 0.14 | 0.08 | 0.17 | 0.17 | 0.14 |
| 2 | 115832574 | rs10864934 | 0.112 | -0.357 | -0.656 | -0.024 | -0.012 | 0.045 | 0.045 | 0.781 | 0.22 | 0.27 | 0.34 | 0.52 | 0.16 | 0.21 |
| 2 | 115834653 | rs17043985 | 0.322 | -0.386 | -0.638 | -0.038 | -0.021 | 0.025 | 0.025 | 1.49 | 0.21 | 0.25 | 0.28 | 0.35 | 0.11 | 0.18 |
| 2 | 115845875 | rs12987050 | 2.506 | -0.326 | -1.196 | -0.153 | -0.035 | -0.018 | -0.018 | 2.117 | 0.28 | 0.14 | 0.32 | 0.3 | 0.35 | 0.24 |
| 2 | 115848781 | rs11695180 | 1.117 | -0.282 | -0.911 | -0.173 | -0.002 | 0.008 | 0.008 | 0 | 0.43 | 0.32 | 0.46 | 0.63 | 0.57 | 0.42 |
| 2 | 115861277 | rs11123279 | 0.376 | -0.297 | -0.933 | -0.177 | -0.057 | 0.097 | 0.097 | 1.237 | 0.21 | 0.36 | 0.29 | 0.35 | 0.11 | 0.18 |
| 2 | 115866745 | rs10201854 | -0.726 | -0.294 | -0.929 | -0.221 | -0.001 | -0.01 | -0.01 | 0.177 | 0.67 | 0.74 | 0.76 | 0.98 | 0.68 | 0.6 |
| 2 | 115867786 | rs17044073 | 0.687 | -0.175 | -0.985 | -0.177 | -0.022 | -0.016 | -0.016 | 5.079 | 0.14 | 0.07 | 0.09 | 0.16 | 0.2 | 0.16 |
| 2 | 115872796 | rs4277531 | 1.16 | -0.211 | -0.978 | -0.239 | -0.003 | -0.006 | -0.006 | 0 | 0.47 | 0.48 | 0.59 | 0.64 | 0.43 | 0.4 |
| 2 | 115874542 | rs4459743 | 1.726 | -0.151 | -1.223 | -0.263 | -0.071 | 0.014 | 0.014 | 0 | 0.35 | 0.14 | 0.36 | 0.3 | 0.45 | 0.29 |
| 2 | 115881852 | rs7579207 | -0.463 | -0.5 | -0.979 | -0.519 | 0.003 | -0.009 | -0.009 | 0.637 | 0.68 | 0.74 | 0.77 | 0.98 | 0.69 | 0.61 |
| 2 | 115889896 | rs4560123 | -1.335 | -2.015 | -0.409 | 0.114 | -0.003 | 0.006 | 0.006 | 0 | 0.16 | 0.2 | 0.3 | 0.29 | 0.09 | 0.17 |
| 2 | 115890104 | rs4491738 | -0.149 | -2.07 | -0.396 | 0.115 | -0.008 | 0.025 | 0.025 | 0 | 0.11 | 0.15 | 0.27 | 0.29 | 0.05 | 0.14 |
| 2 | 115891114 | rs10199796 | -0.028 | -2.092 | -0.416 | 0.139 | -0.005 | -0.009 | -0.009 | 0 | 0.12 | 0.09 | 0.07 | 0.15 | 0.14 | 0.11 |
| 2 | 115893956 | rs983829 | -0.688 | -2.227 | -0.444 | 0.162 | -0.001 | 0.001 | 0.001 | 0 | 0.16 | 0.18 | 0.28 | 0.28 | 0.07 | 0.15 |
| 2 | 115897734 | rs958457 | -0.195 | -2.274 | -0.589 | -0.507 | 0.039 | 0.047 | 0.047 | 0 | 0.64 | 0.78 | 0.8 | 0.96 | 0.76 | 0.61 |
| 2 | 115900885 | rs13416481 | -0.056 | -1.837 | 0.004 | -0.275 | 0.052 | -0.073 | -0.073 | 1.501 | 0.12 | 0.38 | 0.07 | 0 | 0.12 | 0.05 |
| 2 | 115917470 | rs4131302 | 0.384 | -2.179 | -0.331 | -0.232 | 0.044 | 0.072 | 0.072 | 0 | 0.55 | 0.71 | 0.54 | 0.76 | 0.68 | 0.53 |
| 2 | 115922821 | rs12711818 | 1.385 | -2.157 | -0.254 | -0.604 | 0.006 | 0.005 | 0.005 | 0 | 0.46 | 0.44 | 0.46 | 0.36 | 0.64 | 0.43 |
| 2 | 115932634 | rs10195710 | 2.26 | -2.024 | 0.086 | -0.54 | -0.13 | -0.071 | -0.071 | 4.767 | 0.33 | 0.07 | 0.38 | 0.36 | 0.52 | 0.39 |
| 2 | 115943835 | rs10192079 | 0.581 | -2.45 | 0.849 | -0.336 | -0.01 | -0.029 | -0.029 | 0 | 0.65 | 0.67 | 0.82 | 0.88 | 0.8 | 0.78 |
| 2 | 115946323 | rs7558702 | 0.58 | -2.457 | 1.154 | -0.348 | -0.016 | 0.001 | 0.001 | 0 | 0.72 | 0.6 | 0.83 | 0.76 | 0.77 | 0.7 |
| 2 | 115948786 | rs7566796 | -0.932 | -2.661 | 1.179 | -0.418 | 0.003 | -0.005 | -0.005 | 0 | 0.25 | 0.27 | 0.38 | 0.4 | 0.2 | 0.26 |
| 2 | 115949774 | rs9653426 | 2.031 | -2.639 | 1.089 | -0.419 | 0.028 | 0.01 | 0.01 | 0 | 0.57 | 0.29 | 0.49 | 0.37 | 0.66 | 0.51 |
| 2 | 115955900 | rs6722872 | 0.544 | -2.131 | 1.379 | -0.181 | 0.008 | 0.01 | 0.01 | 0 | 0.08 | 0.04 | 0.04 | 0 | 0.05 | 0.03 |
| 2 | 115958079 | rs7581057 | 0.847 | -2.157 | 1.428 | -0.198 | -0.022 | -0.051 | -0.051 | 0 | 0.11 | 0.18 | 0.06 | 0.03 | 0.07 | 0.05 |
| 2 | 115958505 | rs17355616 | 0.137 | -2.145 | 1.358 | -0.191 | 0.008 | 0.002 | 0.002 | 0 | 0.09 | 0.07 | 0.04 | 0 | 0.05 | 0.03 |
| 2 | 115960540 | rs956534 | 2.054 | -2.157 | 1.344 | -0.194 | 0.006 | 0.075 | 0.075 | 0 | 0.48 | 0.17 | 0.42 | 0.32 | 0.56 | 0.44 |
| 2 | 115961802 | rs9308710 | -0.254 | -2.673 | 1.512 | -0.265 | 0.016 | 0.254 | 0.254 | 0 | 0.08 | 0.3 | 0.3 | 0.38 | 0.07 | 0.16 |
| 2 | 115964006 | rs11123288 | -0.564 | -2.672 | 1.553 | -0.376 | 0.012 | 0.144 | 0.144 | 0 | 0.22 | 0.43 | 0.38 | 0.54 | 0.25 | 0.35 |
| 2 | 115964611 | rs12472611 | 2.693 | -2.669 | 1.306 | -0.397 | -0.18 | -0.077 | -0.077 | 0.636 | 0.37 | 0.05 | 0.39 | 0.33 | 0.49 | 0.39 |
| 2 | 115964965 | rs17355679 | 1.377 | -2.693 | 1.387 | -0.353 | 0.005 | 0.006 | 0.006 | 0 | 0.58 | 0.38 | 0.5 | 0.37 | 0.62 | 0.48 |
| 2 | 115969880 | rs4849387 | -0.225 | -2.675 | 1.278 | -0.377 | 0.003 | 0.045 | 0.045 | 0.283 | 0.8 | 0.98 | 0.89 | 0.91 | 0.88 | 0.85 |
| 2 | 115973833 | rs10204084 | 1.799 | -2.615 | 1.345 | -0.363 | 0.028 | 0.037 | 0.037 | 0 | 0.49 | 0.33 | 0.46 | 0.37 | 0.58 | 0.47 |
| 2 | 116003178 | rs4849389 | 0.192 | -2.507 | 1.423 | -0.395 | -0.003 | -0.023 | -0.023 | 0 | 0.17 | 0.14 | 0.1 | 0.03 | 0.1 | 0.07 |
| 2 | 116009839 | rs10496482 | 2.375 | -2.525 | 1.456 | -0.373 | -0.013 | 0.01 | 0.01 | 0 | 0.41 | 0.24 | 0.42 | 0.35 | 0.57 | 0.41 |
| 2 | 116023110 | rs13386690 | -1.278 | -2.559 | 1.533 | -0.36 | 0.009 | -0.035 | -0.035 | 0 | 0.29 | 0.24 | 0.4 | 0.54 | 0.26 | 0.43 |
| 2 | 116026002 | rs11123292 | 2.579 | -2.476 | 1.504 | -0.324 | -0.123 | -0.077 | -0.077 | 0 | 0.38 | 0.05 | 0.41 | 0.34 | 0.57 | 0.42 |
| 2 | 116028220 | rs12466330 | 0.108 | -2.465 | 1.566 | -0.312 | -0.003 | 0.006 | 0.006 | 0 | 0.17 | 0.2 | 0.33 | 0.36 | 0.14 | 0.26 |
| 2 | 116029766 | rs4849390 | 1.237 | -2.295 | 1.764 | -0.229 | -0.024 | -0.001 | -0.001 | 0 | 0.66 | 0.56 | 0.8 | 0.71 | 0.77 | 0.71 |
| 2 | 116045282 | rs7565854 | 1.41 | -2.276 | 1.578 | -0.283 | -0.024 | -0.054 | -0.054 | 0 | 0.61 | 0.49 | 0.78 | 0.88 | 0.75 | 0.79 |
| 2 | 116086464 | rs925849 | 1.297 | -1.906 | 1.758 | -0.446 | -0.032 | -0.007 | -0.007 | 0 | 0.63 | 0.55 | 0.77 | 0.73 | 0.76 | 0.73 |
| 2 | 116094446 | rs7569681 | -0.255 | -1.9 | 1.881 | -0.452 | -0.016 | -0.05 | -0.05 | 0 | 0.09 | 0.19 | 0.06 | 0 | 0.07 | 0.04 |
| 2 | 116097036 | rs4849392 | 0.056 | -1.917 | 1.811 | -0.458 | 0.004 | -0.016 | -0.016 | 0 | 0.83 | 0.87 | 0.88 | 0.76 | 0.88 | 0.83 |
| 2 | 116098665 | rs11123298 | 1.138 | -1.844 | 1.698 | -0.431 | -0.015 | 0.006 | 0.006 | 0 | 0.67 | 0.56 | 0.79 | 0.7 | 0.75 | 0.66 |
| 2 | 116130709 | rs2421100 | 1.422 | -1.801 | 1.639 | -0.422 | -0.021 | -0.011 | -0.011 | 0 | 0.59 | 0.54 | 0.76 | 0.71 | 0.71 | 0.65 |
| 2 | 116152030 | rs7578789 | 1.791 | -1.511 | 1.765 | -0.271 | -0.001 | 0.01 | 0.01 | 0 | 0.42 | 0.27 | 0.4 | 0.37 | 0.57 | 0.42 |
| 2 | 116156643 | rs949888 | 0.23 | -1.312 | 1.515 | -0.264 | 0.014 | -0.006 | -0.006 | 3.024 | 0.24 | 0.2 | 0.38 | 0.34 | 0.17 | 0.25 |
| 2 | 116158650 | rs17453235 | -0.071 | -1.083 | 1.187 | -0.377 | -0.034 | 0.035 | 0.035 | 0 | 0.07 | 0.01 | 0.05 | 0.03 | 0.11 | 0.13 |
| 2 | 116162614 | rs11683441 | 1.099 | -1.298 | 1.541 | -0.279 | 0.028 | 0.021 | 0.021 | 0 | 0.53 | 0.27 | 0.43 | 0.4 | 0.61 | 0.5 |
| 2 | 116173796 | rs843393 | -0.03 | -1.213 | 1.611 | -0.251 | -0.007 | -0.05 | -0.05 | 0.76 | 0.1 | 0.23 | 0.07 | 0.02 | 0.07 | 0.11 |
| 2 | 116174484 | rs34796827 | 0.145 | -1.213 | 1.633 | -0.251 | 0.005 | 0.005 | 0.005 | 0.342 | 0.09 | 0.06 | 0.06 | 0.02 | 0.07 | 0.11 |
| 2 | 116177579 | rs843385 | -0.02 | -1.218 | 1.796 | -0.254 | 0.004 | -0.049 | -0.049 | 0.152 | 0.1 | 0.33 | 0.07 | 0.02 | 0.07 | 0.11 |
| 2 | 116185335 | rs843399 | 0.465 | -1.238 | 1.709 | -0.251 | -0.014 | 0.008 | 0.008 | 0 | 0.62 | 0.49 | 0.49 | 0.41 | 0.68 | 0.61 |
| 2 | 116212878 | rs1070872 | 0.338 | -1.247 | 1.78 | -0.128 | -0.025 | 0.177 | 0.177 | 0.657 | 0.89 | 0.42 | 0.93 | 0.76 | 0.96 | 0.89 |
| 2 | 116238748 | rs4530380 | 0.3 | -1.242 | 1.79 | -0.136 | -0.023 | 0.186 | 0.186 | 0.743 | 0.89 | 0.42 | 0.93 | 0.76 | 0.96 | 0.9 |
| 2 | 116241186 | rs1375144 | 1.081 | -1.263 | 1.753 | -0.169 | 0.037 | 0.014 | 0.014 | 0 | 0.55 | 0.34 | 0.46 | 0.38 | 0.67 | 0.54 |
| 2 | 116243747 | rs1530689 | 1.25 | -1.289 | 1.611 | -0.182 | 0.088 | 0.028 | 0.028 | 0 | 0.53 | 0.15 | 0.44 | 0.38 | 0.67 | 0.53 |
| 2 | 116247721 | rs4849399 | -0.547 | -1.267 | 1.455 | -0.169 | 0.024 | -0.006 | -0.006 | 2.377 | 0.34 | 0.12 | 0.48 | 0.38 | 0.29 | 0.36 |
| 2 | 116249606 | rs7606086 | 0.017 | -1.261 | 1.656 | -0.163 | -0.017 | 0.138 | 0.138 | 0 | 0.09 | 0.28 | 0.05 | 0.24 | 0.03 | 0.1 |
| 2 | 116254100 | rs17362720 | 0.291 | -1.259 | 1.435 | -0.176 | 0.003 | 0.047 | 0.047 | 0.451 | 0.09 | 0.01 | 0.05 | 0.02 | 0.07 | 0.11 |
| 2 | 116260989 | rs4241135 | 0.356 | -1.251 | 1.446 | -0.166 | 0.008 | -0.047 | -0.047 | 0 | 0.24 | 0.1 | 0.42 | 0.37 | 0.22 | 0.25 |
| 2 | 116274213 | rs7600647 | -0.434 | -1.539 | 1.498 | -0.146 | -0.022 | 0.201 | 0.201 | 0.23 | 0.11 | 0.38 | 0.06 | 0.24 | 0.03 | 0.1 |
| 2 | 116279799 | rs13429901 | -0.831 | -1.485 | 1.425 | -0.123 | -0.046 | 0.071 | 0.071 | 0.414 | 0.2 | 0.45 | 0.12 | 0.26 | 0.1 | 0.21 |
| 2 | 116296595 | rs1971190 | 0.198 | -1.479 | 1.349 | -0.143 | -0.039 | -0.079 | -0.079 | 0 | 0.26 | 0.03 | 0.44 | 0.37 | 0.25 | 0.27 |

| | | | | | | | | | | | | | | | | |
|---|---|---|---|---|---|---|---|---|---|---|---|---|---|---|---|---|
| 2 | 116298589 | rs10210316 | -0.925 | -2.102 | 1.587 | -0.021 | -0.046 | 0.071 | 0.071 | 0.414 | 0.2 | 0.45 | 0.12 | 0.26 | 0.1 | 0.21 |
| 2 | 116302600 | rs17044656 | 0.406 | -2.029 | 1.55 | 0.031 | -0.01 | 0.091 | 0.091 | 0 | 0.08 | 0.14 | 0.04 | 0.24 | 0.03 | 0.1 |
| 2 | 116316515 | rs17044697 | 0.422 | -1.715 | 1.247 | 0.155 | -0.005 | -0.051 | -0.051 | 0 | 0.22 | 0.11 | 0.42 | 0.38 | 0.23 | 0.23 |
| 2 | 116320675 | rs1545396 | -0.177 | -1.84 | 1.271 | 0.064 | -0.063 | -0.108 | -0.108 | 0 | 0.84 | 0.53 | 0.91 | 0.98 | 0.92 | 0.91 |
| 2 | 116323717 | rs1550985 | -0.795 | -1.829 | 1.251 | 0.071 | -0.03 | -0.009 | -0.009 | 0 | 0.37 | 0.46 | 0.5 | 0.4 | 0.3 | 0.33 |
| 2 | 116332653 | rs17044733 | 0.538 | -1.754 | 1.141 | 0.057 | -0.003 | -0.015 | -0.015 | 0 | 0.13 | 0.09 | 0.05 | 0.03 | 0.08 | 0.11 |
| 2 | 116336847 | rs11890841 | 0.241 | -1.696 | 0.656 | 0.086 | -0.053 | -0.089 | -0.089 | 0 | 0.15 | 0.36 | 0.08 | 0.03 | 0.08 | 0.11 |
| 2 | 116337401 | rs11123313 | 1.478 | -1.685 | 0.193 | 0.082 | -0.081 | -0.05 | -0.05 | 0 | 0.42 | 0.03 | 0.38 | 0.34 | 0.56 | 0.45 |
| 2 | 116362694 | rs10205952 | 0.477 | -1.58 | -0.013 | 0.138 | -0.011 | -0.054 | -0.054 | 0 | 0.22 | 0.09 | 0.42 | 0.37 | 0.24 | 0.23 |
| 2 | 116369034 | rs7570085 | -1.493 | -1.554 | -0.147 | 0.178 | 0.01 | 0.004 | 0.004 | 0 | 0.56 | 0.87 | 0.61 | 0.64 | 0.43 | 0.5 |
| 2 | 116372286 | rs7576503 | -0.764 | -1.73 | -0.102 | 0.184 | -0.03 | -0.015 | -0.015 | 0.881 | 0.37 | 0.51 | 0.5 | 0.41 | 0.31 | 0.34 |
| 2 | 116373937 | rs10496499 | -0.465 | -1.494 | 0.726 | 0.132 | -0.036 | 0.065 | 0.065 | 0 | 0.19 | 0.37 | 0.11 | 0.25 | 0.11 | 0.16 |
| 2 | 116384329 | rs6740354 | -0.699 | -1.063 | -0.223 | 0.34 | 0.015 | 0.002 | 0.002 | 0.211 | 0.34 | 0.66 | 0.18 | 0.28 | 0.19 | 0.27 |
| 2 | 116415803 | rs11681542 | 0.577 | -0.193 | -0.285 | -0.053 | -0.021 | -0.044 | -0.044 | 0 | 0.73 | 0.35 | 0.83 | 0.92 | 0.83 | 0.7 |
| 2 | 116417861 | rs4849416 | 0.721 | -0.461 | 1.445 | -0.187 | 0.022 | 0.023 | 0.023 | 0.895 | 0.93 | 0.54 | 0.97 | 0.99 | 1 | 0.9 |
| 2 | 116419064 | rs7589670 | -0.697 | 0.31 | 1.638 | -0.262 | 0.026 | 0.007 | 0.007 | 1.091 | 0.06 | 0.1 | 0.11 | 0.07 | 0.16 | 0.15 |
| 2 | 116422188 | rs6734985 | 1.775 | 0.219 | 1.748 | -0.312 | -0.007 | -0.046 | -0.046 | 0 | 0.72 | 0.38 | 0.8 | 0.92 | 0.75 | 0.68 |
| 2 | 116426293 | rs4599118 | -1.383 | 0.2 | 2.28 | -0.318 | 0.065 | 0.065 | 0.065 | 1.19 | 0.08 | 0.38 | 0.11 | 0.06 | 0.09 | 0.14 |
| 2 | 116438276 | rs13032384 | 0.691 | 0.252 | 1.105 | -0.599 | -0.001 | 0.001 | 0.001 | 0 | 0.3 | 0.24 | 0.38 | 0.27 | 0.28 | 0.27 |
| 2 | 116453722 | rs6542289 | -0.061 | 0.25 | 1.173 | -0.564 | 0.018 | 0.02 | 0.02 | 0.031 | 0.94 | 0.69 | 0.98 | 0.99 | 1 | 0.89 |
| 2 | 116463021 | rs12991458 | 1.267 | 0.268 | 1.151 | -0.575 | -0.003 | -0.027 | -0.027 | 0 | 0.71 | 0.32 | 0.79 | 0.91 | 0.75 | 0.68 |
| 2 | 116467283 | rs10173996 | 0.431 | 0.232 | 1.086 | -0.594 | 0.029 | 0.023 | 0.023 | 2.462 | 0.92 | 0.78 | 0.91 | 0.93 | 0.91 | 0.86 |
| 2 | 116470425 | rs2421277 | 0.31 | 0.221 | 1.098 | -0.599 | -0.038 | -0.089 | -0.089 | 0 | 0.82 | 0.71 | 0.93 | 0.99 | 0.91 | 0.83 |
| 2 | 116481940 | rs13019180 | 0.764 | -0.043 | 1.037 | -0.892 | -0.042 | 0.003 | 0.003 | 0 | 0.31 | 0.06 | 0.38 | 0.27 | 0.32 | 0.27 |
| 2 | 116484040 | rs12621324 | 1.085 | 0.093 | 1.247 | -0.929 | -0.042 | -0.015 | -0.015 | 0.324 | 0.39 | 0.1 | 0.4 | 0.65 | 0.44 | 0.42 |
| 2 | 116484698 | rs967418 | -0.803 | 0.07 | 1.457 | -0.937 | -0.02 | -0.078 | -0.078 | 1.022 | 0.87 | 0.79 | 0.95 | 1 | 0.91 | 0.93 |
| 2 | 116486762 | rs12711833 | -0.365 | 0.062 | 1.491 | -0.95 | 0.002 | 0.027 | 0.027 | 0 | 0.45 | 0.71 | 0.51 | 0.34 | 0.4 | 0.5 |
| 2 | 116487245 | rs7608947 | -0.39 | 0.097 | 1.475 | -0.956 | 0.005 | 0.032 | 0.032 | 0 | 0.46 | 0.72 | 0.51 | 0.34 | 0.4 | 0.5 |
| 2 | 116488520 | rs13024934 | -0.721 | 0.116 | 1.697 | -0.947 | 0.056 | 0.009 | 0.009 | 2.02 | 0.09 | 0.27 | 0.13 | 0.07 | 0.15 | 0.14 |
| 2 | 116491246 | rs4308128 | 0.993 | 0.028 | 1.635 | -0.942 | -0.125 | -0.03 | -0.03 | 0.022 | 0.4 | 0.08 | 0.43 | 0.65 | 0.51 | 0.43 |
| 2 | 116499896 | rs1446492 | 0.469 | 0.077 | 1.84 | -1.001 | 0.03 | 0.09 | 0.09 | 0 | 0.48 | 0.35 | 0.75 | 0.73 | 0.63 | 0.58 |
| 2 | 116507473 | rs1807396 | 0.575 | 0.077 | 1.763 | -0.985 | 0.01 | 0.134 | 0.134 | 0 | 0.47 | 0.15 | 0.74 | 0.74 | 0.63 | 0.58 |
| 2 | 116521409 | rs10496504 | 0.209 | 0.059 | 1.868 | -0.932 | 0.003 | 0.01 | 0.01 | 0 | 0.36 | 0.29 | 0.18 | 0.28 | 0.29 | 0.36 |
| 2 | 116522596 | rs7566591 | 0.179 | 0.059 | 1.918 | -0.898 | 0.101 | 0.101 | 0.101 | 1.659 | 0.92 | 0.55 | 0.89 | 0.93 | 0.91 | 0.88 |
| 2 | 116527831 | rs12476345 | 0.808 | -0.039 | 1.769 | -0.963 | -0.053 | -0.004 | -0.004 | 0 | 0.3 | 0.05 | 0.17 | 0.27 | 0.29 | 0.24 |
| 2 | 116529255 | rs7603307 | 0.866 | 0.032 | 1.97 | -0.928 | -0.057 | 0.025 | 0.025 | 0 | 0.43 | 0.19 | 0.66 | 0.65 | 0.54 | 0.46 |
| 2 | 116548326 | rs1374268 | 1.036 | 0.127 | 1.943 | -0.935 | -0.106 | -0.003 | -0.003 | 0 | 0.42 | 0.1 | 0.65 | 0.66 | 0.53 | 0.45 |
| 2 | 116548526 | rs6735355 | 0.1 | 0.08 | 2.153 | -0.926 | -0.002 | -0.019 | -0.019 | 0 | 0.82 | 0.78 | 0.86 | 0.93 | 0.82 | 0.81 |
| 2 | 116551040 | rs13005041 | 0.57 | 0.088 | 2.144 | -0.905 | -0.005 | -0.005 | -0.005 | 0 | 0.05 | 0.05 | 0.08 | 0.07 | 0.09 | 0.12 |
| 2 | 116551962 | rs4849427 | 0.674 | 0.075 | 2.238 | -0.919 | -0.015 | -0.062 | -0.062 | 1.116 | 0.78 | 0.41 | 0.84 | 0.93 | 0.82 | 0.81 |
| 2 | 116564254 | rs7605007 | 0.103 | 0.087 | 2.244 | -0.903 | 0.025 | 0.026 | 0.026 | 0.719 | 0.06 | 0.19 | 0.09 | 0.07 | 0.09 | 0.12 |
| 2 | 116564960 | rs13416281 | -0.653 | 0.67 | 2.231 | -0.934 | 0.002 | 0.022 | 0.022 | 0 | 0.52 | 0.57 | 0.26 | 0.27 | 0.43 | 0.43 |
| 2 | 116576057 | rs1037131 | -0.45 | 0.555 | 2.104 | -1.037 | 0.003 | 0.033 | 0.033 | 0 | 0.47 | 0.36 | 0.23 | 0.27 | 0.4 | 0.41 |
| 2 | 116581067 | rs17718031 | 0.854 | 0.526 | 2.056 | -1.031 | 0.007 | 0.061 | 0.061 | 0.365 | 0.12 | 0.02 | 0.05 | 0 | 0.09 | 0.07 |
| 2 | 116625784 | rs272043 | 0.114 | 0.526 | 2.048 | -0.889 | 0.004 | -0.009 | -0.009 | 6.609 | 0.57 | 0.48 | 0.71 | 0.66 | 0.57 | 0.53 |
| 2 | 116628321 | rs272042 | -0.273 | 0.541 | 1.711 | -0.89 | 0 | 0.003 | 0.003 | 0 | 0.62 | 0.42 | 0.79 | 0.73 | 0.68 | 0.64 |
| 2 | 116632989 | rs272014 | 1.213 | 0.428 | 1.588 | 0.216 | -0.094 | 0.073 | 0.073 | 0 | 0.39 | 0.08 | 0.54 | 0.19 | 0.47 | 0.42 |
| 2 | 116634049 | rs12992776 | 0.563 | 0.425 | 1.596 | 0.479 | 0.002 | 0.004 | 0.004 | 0.414 | 0.06 | 0.08 | 0.08 | 0.1 | 0.1 | 0.11 |
| 2 | 116638503 | rs272034 | 1.205 | 0.411 | 1.585 | 0.471 | -0.071 | 0.091 | 0.091 | 0 | 0.41 | 0.1 | 0.55 | 0.19 | 0.47 | 0.44 |
| 2 | 116638647 | rs4848401 | 0.765 | 0.386 | 1.58 | 0.474 | 0.007 | -0.075 | -0.075 | 3.383 | 0.13 | 0.02 | 0.14 | 0.44 | 0.1 | 0.09 |
| 2 | 116640033 | rs272032 | 0.189 | 0.393 | 1.613 | 0.471 | -0.006 | 0.02 | 0.02 | 0.12 | 0.55 | 0.31 | 0.7 | 0.63 | 0.57 | 0.53 |
| 2 | 116641667 | rs183886 | 0.474 | 0.389 | 1.586 | 0.466 | 0.007 | 0.042 | 0.042 | 0 | 0.48 | 0.38 | 0.65 | 0.3 | 0.57 | 0.55 |
| 2 | 116644188 | rs272021 | 1.463 | 0.18 | 0.698 | -0.106 | -0.043 | 0.058 | 0.058 | 0 | 0.25 | 0.04 | 0.42 | 0.13 | 0.27 | 0.24 |
| 2 | 116654040 | rs272002 | 0.679 | 0.067 | 0.82 | 0.007 | -0.077 | 0.097 | 0.097 | 0 | 0.41 | 0.1 | 0.55 | 0.19 | 0.47 | 0.44 |
| 2 | 116654179 | rs272001 | -0.662 | 0.115 | 1.235 | 0.029 | -0.002 | -0.009 | -0.009 | 0 | 0.62 | 0.45 | 0.79 | 0.73 | 0.68 | 0.64 |
| 2 | 116658046 | rs775336 | -0.566 | 0.13 | 1.026 | 0.037 | 0.002 | 0.009 | 0.009 | 0 | 0.61 | 0.38 | 0.78 | 0.73 | 0.68 | 0.64 |
| 2 | 116668062 | rs11890985 | -0.461 | -0.163 | 0.94 | -0.229 | -0.001 | -0.028 | -0.028 | 0.05 | 0.56 | 0.5 | 0.44 | 0.81 | 0.53 | 0.57 |
| 2 | 116669314 | rs7569451 | 0.211 | -0.181 | 0.877 | 0.28 | 0.003 | -0.004 | -0.004 | 0 | 0.42 | 0.47 | 0.3 | 0.38 | 0.41 | 0.48 |
| 2 | 116673688 | rs7567890 | -0.554 | -0.228 | 0.904 | 0.315 | -0.005 | 0.038 | 0.038 | 0 | 0.58 | 0.7 | 0.46 | 0.85 | 0.53 | 0.59 |
| 2 | 116677020 | rs7579538 | -0.62 | -0.237 | 1.466 | 0.263 | -0.016 | 0.067 | 0.067 | 0 | 0.59 | 0.73 | 0.44 | 0.83 | 0.51 | 0.61 |
| 2 | 116679009 | rs12711838 | -0.063 | -0.426 | 1.487 | 0.479 | 0.002 | 0.005 | 0.005 | 0 | 0.51 | 0.55 | 0.33 | 0.44 | 0.47 | 0.58 |
| 2 | 116687240 | rs10198403 | 0.023 | -0.435 | 1.446 | 0.47 | -0.002 | 0.005 | 0.005 | 0 | 0.48 | 0.37 | 0.32 | 0.44 | 0.46 | 0.58 |
| 2 | 116696434 | rs1447241 | -0.091 | -0.354 | 0.967 | 0.717 | 0.023 | 0.049 | 0.049 | 0 | 0.53 | 0.75 | 0.37 | 0.7 | 0.39 | 0.5 |
| 2 | 116697363 | rs6542297 | -0.737 | -0.491 | -0.065 | 0.528 | -0.092 | 0.122 | 0.122 | 0 | 0.66 | 0.92 | 0.55 | 0.91 | 0.53 | 0.72 |
| 2 | 116709850 | rs10178229 | -0.762 | -0.917 | 0.719 | 0.647 | -0.026 | 0.096 | 0.096 | 0.687 | 0.1 | 0.42 | 0.04 | 0.14 | 0.01 | 0.17 |
| 2 | 116716621 | rs13387046 | -0.801 | -0.952 | 0.598 | 0.673 | -0.04 | 0.048 | 0.048 | 0.192 | 0.11 | 0.33 | 0.03 | 0.14 | 0.01 | 0.17 |
| 2 | 116733337 | rs1374597 | -1.4 | -0.99 | 0.51 | 0.773 | -0.046 | 0.066 | 0.066 | 0.802 | 0.19 | 0.71 | 0.17 | 0.48 | 0.06 | 0.19 |
| 2 | 116743703 | rs10173690 | -1.479 | -0.939 | 0.515 | 0.718 | -0.052 | 0.018 | 0.018 | 2.411 | 0.13 | 0.64 | 0.05 | 0.14 | 0.02 | 0.17 |
| 2 | 116761542 | rs2084307 | 0.523 | -0.843 | -0.534 | 0.728 | -0.065 | -0.022 | -0.022 | 3.323 | 0.39 | 0.04 | 0.28 | 0.33 | 0.36 | 0.41 |

| | | | | | | | | | | | | | | | | |
|---|---|---|---|---|---|---|---|---|---|---|---|---|---|---|---|---|
| 2 | 116770267 | rs17045365 | 0.569 | -0.625 | -0.023 | 0.958 | 0.014 | 0.002 | 0.002 | 0 | 0.06 | 0.03 | 0.12 | 0.35 | 0.04 | 0.03 |
| 2 | 116776554 | rs11686430 | -1.302 | -0.209 | 0.104 | 0.466 | -0.001 | 0.004 | 0.004 | 0 | 0.26 | 0.29 | 0.26 | 0.61 | 0.24 | 0.38 |
| 2 | 116781809 | rs17045399 | -0.507 | -0.071 | 0.053 | 0.447 | -0.004 | -0.065 | -0.065 | 0 | 0.19 | 0.1 | 0.18 | 0.55 | 0.18 | 0.2 |
| 2 | 116800903 | rs6715118 | -1.851 | -0.305 | 1.6 | 0.284 | -0.013 | 0.031 | 0.031 | 0 | 0.3 | 0.6 | 0.29 | 0.59 | 0.24 | 0.42 |
| 2 | 116801679 | rs10176215 | -0.468 | -0.326 | 1.509 | 0.274 | 0 | 0.005 | 0.005 | 0 | 0.19 | 0.16 | 0.19 | 0.54 | 0.17 | 0.19 |
| 2 | 116804730 | rs11692828 | -2.41 | -0.667 | 1.524 | 0.053 | 0.033 | 0.054 | 0.054 | 0 | 0.41 | 0.82 | 0.38 | 0.67 | 0.32 | 0.58 |
| 2 | 116813932 | rs10170412 | -1.589 | -0.669 | 1.684 | 0.042 | -0.003 | 0 | 0 | 0 | 0.3 | 0.34 | 0.28 | 0.62 | 0.25 | 0.36 |
| 2 | 116815289 | rs6542299 | -2.279 | -0.633 | 1.522 | 0.059 | 0.015 | 0.021 | 0.021 | 0 | 0.4 | 0.71 | 0.37 | 0.67 | 0.32 | 0.58 |
| 2 | 116819705 | rs4849436 | -0.173 | -0.583 | 1.122 | 0.062 | 0.002 | -0.002 | -0.002 | 0.003 | 0.1 | 0.12 | 0.09 | 0.08 | 0.08 | 0.16 |
| 2 | 116841792 | rs10181163 | -2.141 | -0.613 | 1.307 | 0.036 | -0.032 | -0.026 | -0.026 | 0.75 | 0.21 | 0.59 | 0.19 | 0.13 | 0.14 | 0.36 |
| 2 | 116864243 | rs7557744 | -2.279 | -0.476 | 1.074 | 0.143 | 0.013 | 0.015 | 0.015 | 0 | 0.39 | 0.69 | 0.37 | 0.63 | 0.32 | 0.57 |
| 2 | 116867596 | rs6542304 | -1.155 | -0.461 | 1.197 | 0.193 | -0.001 | 0.026 | 0.026 | 0 | 0.26 | 0.34 | 0.26 | 0.5 | 0.24 | 0.4 |
| 2 | 116872975 | rs6707419 | -1.717 | 0.008 | -0.094 | -0.542 | -0.104 | 0.181 | 0.181 | 0 | 0.72 | 0.97 | 0.76 | 1 | 0.66 | 0.92 |
| 2 | 116883963 | rs10196779 | -1.572 | -0.201 | 0.371 | -0.212 | 0.025 | 0.037 | 0.037 | 0 | 0.46 | 0.64 | 0.4 | 0.63 | 0.36 | 0.61 |
| 2 | 116891472 | rs1480488 | -0.898 | -0.246 | -0.094 | -0.178 | 0.033 | 0.019 | 0.019 | 0 | 0.4 | 0.3 | 0.35 | 0.59 | 0.31 | 0.55 |
| 2 | 116894086 | rs7579948 | 0.183 | -0.298 | -0.341 | -0.032 | 0.024 | -0.099 | -0.099 | 0 | 0.26 | 0.13 | 0.19 | 0.49 | 0.19 | 0.24 |
| 2 | 116897625 | rs17045530 | -0.148 | -0.189 | -0.186 | -0.189 | 0.016 | -0.01 | -0.01 | 0.919 | 0.09 | 0.03 | 0.08 | 0.08 | 0.06 | 0.14 |
| 2 | 116900138 | rs11675755 | 0.659 | -0.131 | -0.137 | -0.219 | 0.013 | -0.056 | -0.056 | 0 | 0.26 | 0.16 | 0.21 | 0.55 | 0.23 | 0.23 |
| 2 | 116905270 | rs11903740 | 0.334 | -0.084 | 0.709 | -0.127 | -0.015 | 0.032 | 0.032 | 0 | 0.29 | 0.36 | 0.13 | 0.54 | 0.23 | 0.22 |
| 2 | 116908221 | rs17045597 | 0.042 | 0.032 | 0.82 | -0.289 | 0.005 | 0.005 | 0.005 | 0 | 0.08 | 0.1 | 0.11 | 0.08 | 0.1 | 0.16 |
| 2 | 116909251 | rs11123333 | -1.444 | -0.085 | 1.141 | -0.415 | 0.039 | 0.064 | 0.064 | 0 | 0.48 | 0.72 | 0.33 | 0.63 | 0.41 | 0.59 |
| 2 | 116923465 | rs2127701 | 0.253 | -0.078 | 0.968 | -0.398 | 0.006 | -0.01 | -0.01 | 0 | 0.31 | 0.32 | 0.14 | 0.55 | 0.25 | 0.22 |
| 2 | 116950492 | rs10179985 | -1.248 | -0.067 | 1.023 | -0.453 | 0 | -0.001 | -0.001 | 0 | 0.46 | 0.47 | 0.32 | 0.63 | 0.41 | 0.59 |
| 2 | 116955767 | rs4849449 | -1.287 | 0.015 | 0.968 | -0.527 | 0.002 | 0.004 | 0.004 | 0 | 0.47 | 0.46 | 0.34 | 0.65 | 0.42 | 0.61 |
| 2 | 116963387 | rs12105693 | 0.85 | -0.001 | 0.814 | -0.518 | 0.002 | -0.122 | -0.122 | 0.6 | 0.27 | 0.06 | 0.12 | 0.57 | 0.23 | 0.21 |
| 2 | 116976066 | rs2010823 | -0.516 | -0.035 | 0.973 | -0.49 | 0.01 | 0.003 | 0.003 | 0 | 0.1 | 0.2 | 0.11 | 0.08 | 0.11 | 0.16 |
| 2 | 116980641 | rs2901453 | -1.296 | 0.091 | 0.959 | -0.52 | -0.002 | -0.006 | -0.006 | 0 | 0.49 | 0.49 | 0.35 | 0.65 | 0.44 | 0.65 |
| 2 | 117007403 | rs4849451 | -0.149 | 0.177 | 0.86 | -0.444 | 0.011 | 0.012 | 0.012 | 0 | 0.08 | 0.12 | 0.1 | 0.07 | 0.1 | 0.15 |
| 2 | 117019964 | rs10199125 | -1.309 | 0.05 | 1.208 | -0.558 | 0.024 | 0.076 | 0.076 | 0 | 0.51 | 0.71 | 0.37 | 0.64 | 0.45 | 0.62 |
| 2 | 117034216 | rs7599159 | 0.849 | -0.001 | 0.907 | -0.545 | 0.004 | -0.095 | -0.095 | 0.534 | 0.28 | 0.08 | 0.13 | 0.58 | 0.25 | 0.22 |
| 2 | 117054882 | rs6735305 | -2.371 | 0.512 | 0.392 | -1.007 | -0.051 | 0.137 | 0.137 | 0.325 | 0.77 | 0.98 | 0.84 | 1 | 0.79 | 0.96 |
| 2 | 117064215 | rs6750553 | -2.099 | 0.335 | 0.526 | -0.826 | -0.062 | 0.029 | 0.029 | 0.14 | 0.74 | 0.93 | 0.77 | 0.8 | 0.72 | 0.9 |
| 2 | 117072368 | rs17045782 | 0.354 | 0.366 | 0.693 | -0.906 | 0.044 | 0.045 | 0.045 | 0 | 0.08 | 0.2 | 0.1 | 0.08 | 0.09 | 0.15 |
| 2 | 117072522 | rs1472780 | 1.81 | 0.401 | 0.427 | -0.971 | -0.024 | 0.038 | 0.038 | 0 | 0.22 | 0.08 | 0.38 | 0.15 | 0.27 | 0.27 |
| 2 | 117078901 | rs1505758 | -0.512 | 0.084 | 0.737 | -0.677 | -0.01 | -0.018 | -0.018 | 0 | 0.42 | 0.5 | 0.26 | 0.64 | 0.37 | 0.39 |
| 2 | 117080187 | rs7586235 | -1.128 | 0.15 | 0.486 | -0.659 | 0.005 | -0.045 | -0.045 | 0.467 | 0.19 | 0.38 | 0.22 | 0.08 | 0.2 | 0.36 |
| 2 | 117080769 | rs17045802 | -0.334 | 0.258 | 0.047 | -0.551 | 0.015 | -0.001 | -0.001 | 0 | 0.11 | 0.24 | 0.12 | 0.07 | 0.12 | 0.16 |
| 2 | 117082368 | rs13024154 | 0.654 | 0.029 | -0.429 | -0.399 | 0.01 | -0.071 | -0.071 | 0 | 0.28 | 0.08 | 0.13 | 0.6 | 0.24 | 0.22 |
| 2 | 117104789 | rs1566994 | -0.608 | -0.074 | -0.039 | -0.551 | -0.002 | -0.004 | -0.004 | 0 | 0.48 | 0.49 | 0.36 | 0.67 | 0.42 | 0.48 |
| 2 | 117115097 | rs11681662 | 0.384 | -0.204 | -0.323 | -0.482 | 0.045 | -0.023 | -0.023 | 0 | 0.35 | 0.2 | 0.17 | 0.61 | 0.28 | 0.26 |
| 2 | 117126557 | rs11123348 | 0.744 | -0.267 | -0.59 | -0.484 | 0.04 | -0.086 | -0.086 | 0 | 0.32 | 0.07 | 0.15 | 0.61 | 0.26 | 0.22 |
| 2 | 117130037 | rs7558789 | 1.266 | 0.074 | -0.614 | -0.757 | -0.008 | 0.021 | 0.021 | 0 | 0.2 | 0.17 | 0.27 | 0.09 | 0.28 | 0.26 |
| 2 | 117130417 | rs1394994 | 1.616 | -0.152 | -0.29 | -0.482 | -0.003 | -0.002 | -0.002 | 0 | 0.06 | 0.04 | 0.08 | 0 | 0.07 | 0.16 |
| 2 | 117132173 | rs4849464 | 1.568 | -0.157 | -0.292 | -0.485 | -0.003 | -0.002 | -0.002 | 0 | 0.06 | 0.04 | 0.08 | 0 | 0.07 | 0.16 |
| 2 | 117132704 | rs13023630 | -0.833 | -0.24 | 0.136 | -0.568 | 0.016 | 0.023 | 0.023 | 0 | 0.49 | 0.57 | 0.38 | 0.65 | 0.42 | 0.46 |
| 2 | 117133635 | rs1505753 | -0.776 | -0.26 | 0.139 | -0.621 | 0.015 | 0.021 | 0.021 | 0 | 0.49 | 0.56 | 0.37 | 0.67 | 0.41 | 0.46 |
| 2 | 117134847 | rs13015577 | 0.654 | -0.276 | 0.1 | -0.552 | 0.047 | -0.062 | -0.062 | 0 | 0.33 | 0.08 | 0.15 | 0.63 | 0.26 | 0.22 |
| 2 | 117134904 | rs17045852 | -0.042 | -0.281 | 0.264 | -0.703 | -0.019 | -0.017 | -0.017 | 0 | 0.4 | 0.39 | 0.29 | 0.68 | 0.32 | 0.29 |
| 2 | 117146367 | rs13432983 | -0.092 | -0.047 | 0.296 | -0.639 | 0.007 | 0.013 | 0.013 | 0 | 0.44 | 0.39 | 0.28 | 0.66 | 0.39 | 0.4 |
| 2 | 117154934 | rs10864945 | 0.384 | -0.079 | 0.284 | -0.673 | -0.027 | 0.012 | 0.012 | 0 | 0.4 | 0.07 | 0.26 | 0.66 | 0.38 | 0.39 |
| 2 | 117155498 | rs6737541 | -0.085 | -0.03 | 0.426 | -0.652 | 0.004 | 0.008 | 0.008 | 0 | 0.46 | 0.42 | 0.29 | 0.66 | 0.4 | 0.4 |
| 2 | 117160547 | rs7605652 | -0.096 | -0.023 | 0.468 | -0.648 | 0.009 | 0.022 | 0.022 | 0 | 0.45 | 0.33 | 0.28 | 0.66 | 0.4 | 0.41 |
| 2 | 117181551 | rs2029580 | -0.1 | -0.099 | 0.842 | -0.596 | 0.008 | 0.037 | 0.037 | 0.03 | 0.93 | 0.8 | 0.89 | 0.93 | 0.95 | 0.88 |
| 2 | 117182880 | rs11681836 | 0.233 | -0.135 | 0.75 | -0.711 | -0.007 | 0.036 | 0.036 | 0 | 0.42 | 0.11 | 0.26 | 0.66 | 0.39 | 0.4 |
| 2 | 117193056 | rs6706985 | 0.321 | -0.194 | 0.812 | -0.749 | 0.012 | 0.087 | 0.087 | 0 | 0.49 | 0.33 | 0.61 | 0.27 | 0.56 | 0.48 |
| 2 | 117200788 | rs7604982 | 0.473 | -0.331 | 0.662 | -0.673 | 0 | 0.005 | 0.005 | 0 | 0.16 | 0.15 | 0.36 | 0.12 | 0.22 | 0.32 |
| 2 | 117207537 | rs1388402 | 0.127 | -0.337 | 0.797 | -0.54 | -0.003 | 0.026 | 0.026 | 0.076 | 0.92 | 0.8 | 0.9 | 0.93 | 0.95 | 0.88 |
| 2 | 117213613 | rs6719461 | 0.597 | -0.381 | 0.324 | -0.94 | 0.001 | 0.132 | 0.132 | 0.058 | 0.44 | 0.22 | 0.47 | 0.2 | 0.5 | 0.5 |
| 2 | 117217244 | rs4848420 | -0.801 | -0.391 | 0.235 | -0.849 | -0.001 | -0.008 | -0.008 | 0.882 | 0.84 | 0.87 | 0.75 | 0.93 | 0.83 | 0.79 |
| 2 | 117221756 | rs12619029 | 1.097 | -0.458 | 0.027 | -0.841 | 0.011 | -0.022 | -0.022 | 0 | 0.28 | 0.11 | 0.15 | 0.66 | 0.25 | 0.18 |
| 2 | 117224730 | rs6742479 | -0.541 | -0.478 | 0.396 | -0.863 | -0.019 | -0.062 | -0.062 | 0 | 0.83 | 0.7 | 0.74 | 0.93 | 0.83 | 0.79 |
| 2 | 117226672 | rs6740625 | 0.173 | -0.384 | 0.062 | -0.807 | 0.038 | 0.078 | 0.078 | 0 | 0.49 | 0.36 | 0.57 | 0.26 | 0.55 | 0.59 |
| 2 | 117230574 | rs7604612 | 0.045 | -0.262 | -0.079 | -0.729 | 0.016 | 0.032 | 0.032 | 0 | 0.53 | 0.4 | 0.58 | 0.26 | 0.56 | 0.6 |
| 2 | 117235005 | rs7609533 | -0.712 | -0.234 | -0.313 | -0.333 | 0.026 | -0.001 | -0.001 | 0.057 | 0.67 | 0.8 | 0.82 | 0.33 | 0.72 | 0.78 |
| 2 | 117249923 | rs2130980 | 0.061 | -0.03 | -0.159 | -0.512 | 0.016 | 0.101 | 0.101 | 0 | 0.49 | 0.25 | 0.45 | 0.25 | 0.5 | 0.54 |
| 2 | 117251750 | rs1002233 | 0.98 | -0.102 | 0.096 | -0.425 | -0.012 | -0.032 | -0.032 | 0 | 0.35 | 0.11 | 0.32 | 0.6 | 0.39 | 0.26 |
| 2 | 117252338 | rs7583510 | 0.125 | -0.09 | 0.135 | -0.422 | 0.016 | 0.08 | 0.08 | 0 | 0.49 | 0.27 | 0.45 | 0.27 | 0.5 | 0.54 |
| 2 | 117254847 | rs11683338 | 0.986 | -0.097 | 0.043 | -0.202 | -0.012 | -0.058 | -0.058 | 0.368 | 0.35 | 0.11 | 0.31 | 0.53 | 0.39 | 0.26 |
| 2 | 117257001 | rs11691675 | 0.411 | -0.117 | 0.123 | -0.213 | -0.029 | 0.002 | 0.002 | 0.296 | 0.84 | 0.38 | 0.77 | 0.88 | 0.89 | 0.81 |

| | | | | | | | | | | | | | | | | |
|---|---|---|---|---|---|---|---|---|---|---|---|---|---|---|---|---|
| 2 | 117258775 | rs10200752 | 0.134 | -0.091 | 0.124 | -0.216 | 0.019 | 0.07 | 0.07 | 0 | 0.49 | 0.25 | 0.46 | 0.35 | 0.52 | 0.54 |
| 2 | 117261493 | rs957549 | 0.395 | -0.107 | 0.133 | -0.364 | -0.028 | 0.007 | 0.007 | 0.258 | 0.84 | 0.38 | 0.77 | 0.88 | 0.89 | 0.8 |
| 2 | 117266851 | rs2172185 | 0.983 | -0.089 | 0.217 | -0.363 | 0.006 | 0.029 | 0.029 | 0 | 0.07 | 0.16 | 0.18 | 0.12 | 0.09 | 0.17 |
| 2 | 117277447 | rs12471941 | -0.104 | -0.025 | -0.295 | -0.312 | 0.017 | 0.085 | 0.085 | 0 | 0.49 | 0.25 | 0.46 | 0.3 | 0.5 | 0.53 |
| 2 | 117284895 | rs7565856 | 0.871 | 0.071 | -0.688 | -0.521 | -0.091 | 0.139 | 0.139 | 0 | 0.33 | 0.08 | 0.36 | 0.12 | 0.37 | 0.3 |
| 2 | 117293709 | rs1388410 | 0.065 | 0.432 | -0.809 | -0.465 | -0.001 | 0.132 | 0.132 | 0 | 0.49 | 0.23 | 0.42 | 0.75 | 0.55 | 0.5 |
| 2 | 117304779 | rs1986494 | -0.535 | 0.312 | -0.566 | -0.489 | -0.004 | 0.024 | 0.024 | 0 | 0.84 | 0.47 | 0.79 | 0.87 | 0.88 | 0.8 |
| 2 | 117304901 | rs13009572 | 0.502 | 0.294 | -0.65 | -0.536 | -0.114 | 0.105 | 0.105 | 0 | 0.46 | 0.05 | 0.41 | 0.75 | 0.52 | 0.48 |
| 2 | 117309557 | rs4423607 | 0.759 | 0.289 | -0.621 | -0.488 | 0.002 | -0.025 | -0.025 | 0 | 0.36 | 0.31 | 0.38 | 0.19 | 0.35 | 0.31 |
| 2 | 117316473 | rs7605495 | 0.84 | 0.299 | -0.697 | -0.484 | -0.002 | -0.011 | -0.011 | 0 | 0.35 | 0.26 | 0.38 | 0.12 | 0.35 | 0.31 |
| 2 | 117319154 | rs1986685 | 1.222 | 0.354 | -0.858 | -0.292 | -0.062 | -0.103 | -0.103 | 2.179 | 0.34 | 0.06 | 0.31 | 0.56 | 0.39 | 0.27 |
| 2 | 117320047 | rs6742579 | -0.266 | 0.353 | -0.887 | -0.295 | 0 | -0.003 | -0.003 | 0 | 0.51 | 0.42 | 0.48 | 0.31 | 0.5 | 0.53 |
| 2 | 117321526 | rs12467804 | 1.582 | 0.353 | -0.889 | -0.301 | -0.021 | -0.021 | -0.021 | 0 | 0.12 | 0.02 | 0.09 | 0.12 | 0.13 | 0.21 |
| 2 | 117322280 | rs10210242 | -0.226 | 0.361 | -0.862 | -0.283 | 0 | -0.007 | -0.007 | 0 | 0.51 | 0.42 | 0.47 | 0.24 | 0.5 | 0.53 |
| 2 | 117324347 | rs2901477 | 1.159 | 0.406 | -1.009 | -0.246 | -0.068 | -0.111 | -0.111 | 0.029 | 0.34 | 0.05 | 0.32 | 0.56 | 0.39 | 0.27 |
| 2 | 117336732 | rs17553584 | 1.068 | 0.433 | -0.959 | -0.436 | -0.039 | 0.003 | 0.003 | 2.097 | 0.17 | 0.02 | 0.1 | 0.12 | 0.18 | 0.24 |
| 2 | 117337817 | rs10496521 | -0.531 | 0.453 | -0.606 | -0.014 | 0.008 | 0.031 | 0.031 | 0 | 0.81 | 0.56 | 0.61 | 0.75 | 0.81 | 0.71 |
| 2 | 117341109 | rs2584995 | 1.594 | 0.429 | -0.729 | -0.062 | 0.007 | -0.091 | -0.091 | 1.019 | 0.26 | 0.1 | 0.15 | 0.44 | 0.28 | 0.16 |
| 2 | 117351488 | rs2584964 | -0.31 | 0.482 | -0.626 | -0.023 | 0.01 | 0.074 | 0.074 | 0 | 0.81 | 0.5 | 0.61 | 0.67 | 0.81 | 0.71 |
| 2 | 117356414 | rs2099878 | 1.253 | 0.486 | -0.972 | 0.008 | 0.01 | -0.047 | -0.047 | 0 | 0.27 | 0.16 | 0.16 | 0.44 | 0.28 | 0.17 |
| 2 | 117364078 | rs2584969 | -0.175 | 0.569 | -0.9 | 0.03 | 0.009 | 0.036 | 0.036 | 0 | 0.8 | 0.51 | 0.6 | 0.75 | 0.81 | 0.7 |
| 2 | 117367385 | rs2709267 | 0.505 | 0.613 | -1.489 | -0.235 | -0.001 | 0.009 | 0.009 | 0.007 | 0.45 | 0.19 | 0.27 | 0.55 | 0.47 | 0.4 |
| 2 | 117373537 | rs6735366 | 1.858 | 0.531 | -1.964 | -0.466 | -0.079 | 0.013 | 0.013 | 0 | 0.19 | 0.03 | 0.23 | 0.13 | 0.26 | 0.07 |
| 2 | 117375212 | rs7598642 | 0.589 | 0.624 | -1.613 | -0.23 | 0.008 | -0.052 | -0.052 | 8.566 | 0.35 | 0.38 | 0.34 | 0.2 | 0.34 | 0.3 |
| 2 | 117375251 | rs2584980 | 0.128 | 0.549 | -1.956 | 0.114 | -0.026 | -0.035 | -0.035 | 0 | 0.42 | 0.12 | 0.26 | 0.54 | 0.46 | 0.39 |
| 2 | 117375672 | rs2584981 | 0.788 | 0.362 | -1.554 | 0.355 | 0.002 | -0.106 | -0.106 | 0.119 | 0.26 | 0.09 | 0.16 | 0.43 | 0.29 | 0.17 |
| 2 | 117407288 | rs2952738 | 0.465 | -0.115 | -0.877 | -0.453 | 0 | -0.003 | -0.003 | 0 | 0.37 | 0.36 | 0.37 | 0.63 | 0.37 | 0.4 |
| 2 | 117410934 | rs12999723 | -0.401 | -0.255 | -0.614 | -0.525 | 0.019 | 0.005 | 0.005 | 0 | 0.57 | 0.24 | 0.59 | 0.37 | 0.61 | 0.58 |
| 2 | 117412520 | rs1837986 | 0.29 | 0.127 | -0.569 | -0.463 | -0.059 | -0.017 | -0.017 | 0 | 0.17 | 0.01 | 0.11 | 0.12 | 0.22 | 0.22 |
| 2 | 117426163 | rs4525709 | -1.292 | 0.001 | -0.239 | -0.51 | -0.001 | 0 | 0 | 0.434 | 0.45 | 0.54 | 0.45 | 0.36 | 0.53 | 0.37 |
| 2 | 117444585 | rs1822314 | 0.39 | 0.124 | -0.182 | -0.528 | -0.06 | -0.008 | -0.008 | 0 | 0.17 | 0.02 | 0.13 | 0.14 | 0.28 | 0.22 |
| 2 | 117445623 | rs17046267 | -0.916 | 0.062 | 0.016 | -0.575 | -0.016 | -0.044 | -0.044 | 0.016 | 0.08 | 0.19 | 0.03 | 0 | 0.04 | 0.04 |
| 2 | 117451223 | rs10165591 | -1.178 | -0.036 | 0.082 | -0.532 | -0.004 | -0.001 | -0.001 | 0 | 0.39 | 0.34 | 0.43 | 0.37 | 0.52 | 0.35 |
| 2 | 117455795 | rs17046287 | 0.055 | -0.198 | -0.02 | -0.5 | -0.005 | 0.009 | 0.009 | 0 | 0.22 | 0.25 | 0.16 | 0.22 | 0.2 | 0.14 |
| 2 | 117463256 | rs10167828 | -1.403 | -0.252 | 0.047 | -0.454 | 0.006 | 0.03 | 0.03 | 0.02 | 0.48 | 0.67 | 0.33 | 0.37 | 0.5 | 0.39 |
| 2 | 117468479 | rs7568311 | -0.736 | 0.043 | 0.05 | -0.333 | 0.001 | 0 | 0 | 0 | 0.38 | 0.38 | 0.3 | 0.36 | 0.48 | 0.37 |
| 2 | 117476586 | rs7594984 | -1.154 | 0.115 | -0.154 | -0.29 | -0.001 | 0.008 | 0.008 | 1.542 | 0.45 | 0.55 | 0.32 | 0.36 | 0.5 | 0.39 |
| 2 | 117482174 | rs13386927 | -0.842 | -0.203 | -0.083 | -0.01 | -0.016 | -0.044 | -0.044 | 0.484 | 0.08 | 0.16 | 0.03 | 0 | 0.05 | 0.04 |
| 2 | 117516618 | rs17732795 | 0.71 | 0.272 | -0.809 | 0.116 | -0.072 | -0.017 | -0.017 | 0.426 | 0.12 | 0.01 | 0.21 | 0.11 | 0.21 | 0.25 |
| 2 | 117523594 | rs908254 | -0.594 | -0.082 | -0.127 | 0.193 | 0.021 | 0.023 | 0.023 | 0.449 | 0.31 | 0.42 | 0.37 | 0.35 | 0.39 | 0.37 |
| 2 | 117526462 | rs9308732 | -1.331 | -0.581 | -0.828 | -0.323 | -0.072 | -0.117 | -0.117 | 0.045 | 0.62 | 0.95 | 0.6 | 0.4 | 0.59 | 0.6 |
| 2 | 117537339 | rs7579812 | -1.159 | -0.463 | -0.405 | -0.294 | -0.012 | 0.001 | 0.001 | 5.273 | 0.41 | 0.7 | 0.41 | 0.35 | 0.44 | 0.41 |
| 2 | 117542629 | rs1515126 | -1.491 | -0.724 | -0.622 | -0.145 | -0.117 | -0.145 | -0.145 | 0 | 0.69 | 0.99 | 0.62 | 0.51 | 0.62 | 0.75 |
| 2 | 117554931 | rs1983308 | -1.273 | -0.522 | -0.126 | -0.094 | -0.013 | -0.002 | -0.002 | 2.588 | 0.41 | 0.71 | 0.41 | 0.35 | 0.44 | 0.41 |
| 2 | 117565888 | rs1515128 | -0.818 | -0.526 | -0.047 | -0.217 | 0.004 | 0.001 | 0.001 | 3.189 | 0.35 | 0.57 | 0.39 | 0.33 | 0.4 | 0.37 |
| 2 | 117575925 | rs6723540 | -1.11 | -0.405 | -0.172 | -0.152 | -0.011 | 0.005 | 0.005 | 4.515 | 0.4 | 0.68 | 0.4 | 0.33 | 0.44 | 0.4 |
| 2 | 117577139 | rs17046388 | 0.846 | -0.507 | -0.181 | -0.183 | -0.038 | -0.053 | -0.053 | 0 | 0.11 | 0.02 | 0.12 | 0.19 | 0.16 | 0.11 |
| 2 | 117580737 | rs10496528 | -0.827 | -0.865 | 0.089 | -0.192 | -0.011 | 0.007 | 0.007 | 4.509 | 0.4 | 0.68 | 0.4 | 0.33 | 0.44 | 0.4 |
| 2 | 117590751 | rs1515138 | -0.945 | -0.853 | 0.179 | -0.201 | -0.006 | 0.002 | 0.002 | 4.38 | 0.4 | 0.62 | 0.4 | 0.33 | 0.44 | 0.4 |
| 2 | 117606351 | rs7558397 | -0.825 | -0.853 | 0.175 | -0.194 | -0.012 | 0.007 | 0.007 | 4.726 | 0.4 | 0.68 | 0.41 | 0.33 | 0.44 | 0.4 |
| 2 | 117623172 | rs12620252 | -0.038 | -0.868 | 0.324 | -0.211 | 0.028 | 0.022 | 0.022 | 1.38 | 0.31 | 0.42 | 0.37 | 0.33 | 0.39 | 0.38 |
| 2 | 117623762 | rs1356054 | -0.929 | -0.865 | 0.19 | -0.206 | -0.004 | 0.001 | 0.001 | 3.495 | 0.4 | 0.6 | 0.4 | 0.33 | 0.44 | 0.4 |
| 2 | 117637447 | rs12104589 | -0.822 | -0.865 | 0.083 | -0.207 | -0.01 | 0.007 | 0.007 | 4.018 | 0.4 | 0.67 | 0.4 | 0.33 | 0.44 | 0.4 |
| 2 | 117644685 | rs2421760 | -0.017 | -0.807 | 0.231 | -0.19 | 0.016 | 0.013 | 0.013 | 1.476 | 0.3 | 0.36 | 0.37 | 0.33 | 0.39 | 0.38 |
| 2 | 117649859 | rs13412286 | -0.774 | -0.797 | 0.173 | -0.133 | -0.009 | 0.01 | 0.01 | 5.121 | 0.41 | 0.69 | 0.4 | 0.33 | 0.42 | 0.41 |
| 2 | 117664241 | rs6752687 | -0.709 | -0.787 | 0.431 | -0.129 | -0.004 | 0.009 | 0.009 | 4.166 | 0.41 | 0.62 | 0.4 | 0.3 | 0.42 | 0.39 |
| 2 | 117675916 | rs17046478 | -0.774 | -0.805 | 0.536 | -0.12 | -0.004 | 0.001 | 0.001 | 5.544 | 0.39 | 0.61 | 0.4 | 0.33 | 0.42 | 0.39 |
| 2 | 117686719 | rs2421821 | -0.78 | -0.807 | 0.478 | -0.121 | -0.004 | 0.001 | 0.001 | 5.597 | 0.39 | 0.61 | 0.39 | 0.33 | 0.42 | 0.4 |
| 2 | 117693287 | rs10496530 | 0 | -0.841 | 0.641 | -0.142 | 0.024 | 0.016 | 0.016 | 3.767 | 0.3 | 0.39 | 0.37 | 0.33 | 0.39 | 0.37 |
| 2 | 117695712 | rs9653433 | -1.175 | -0.957 | -0.361 | -0.206 | -0.13 | -0.173 | -0.173 | 0.06 | 0.7 | 0.98 | 0.61 | 0.47 | 0.6 | 0.72 |
| 2 | 117716628 | rs2118392 | -1.193 | -0.461 | 0.641 | 0.048 | -0.003 | -0.003 | -0.003 | 0 | 0.4 | 0.61 | 0.4 | 0.36 | 0.42 | 0.43 |
| 2 | 117721104 | rs17040558 | -0.043 | -0.484 | -0.197 | 0.053 | -0.073 | -0.069 | -0.069 | 0 | 0.27 | 0.12 | 0.35 | 0.33 | 0.36 | 0.37 |
| 2 | 117721773 | rs4305291 | -1.421 | -0.496 | -0.183 | 0.045 | -0.013 | -0.008 | -0.008 | 0 | 0.43 | 0.77 | 0.42 | 0.36 | 0.42 | 0.43 |
| 2 | 117724192 | rs7608004 | -1.361 | -0.175 | -0.269 | 0.192 | -0.003 | -0.003 | -0.003 | 0 | 0.4 | 0.61 | 0.39 | 0.36 | 0.42 | 0.43 |
| 2 | 117725092 | rs4453704 | -0.468 | 0.695 | -1.055 | 0.603 | -0.073 | -0.07 | -0.07 | 0 | 0.27 | 0.12 | 0.34 | 0.33 | 0.36 | 0.37 |
| 2 | 117737432 | rs6542345 | -0.535 | 0.155 | -0.693 | 0.386 | 0.018 | -0.015 | -0.015 | 0.134 | 0.38 | 0.8 | 0.39 | 0.6 | 0.36 | 0.34 |
| 2 | 117772652 | rs4848430 | 0.34 | 0.1 | 0.419 | 0.168 | 0.014 | 0.012 | 0.012 | 0 | 0.58 | 0.19 | 0.58 | 0.38 | 0.59 | 0.64 |
| 2 | 117784322 | rs11901513 | -0.635 | -0.097 | 0.897 | 0.2 | 0.034 | 0.012 | 0.012 | 0.067 | 0.05 | 0.34 | 0.04 | 0 | 0.03 | 0.01 |
| 2 | 117785135 | rs1583594 | 0.951 | -0.205 | 1.215 | 0.146 | -0.024 | -0.044 | -0.044 | 0 | 0.92 | 0.28 | 0.93 | 1 | 0.97 | 0.99 |

| | | | | | | | | | | | | | | | | |
|---|---|---|---|---|---|---|---|---|---|---|---|---|---|---|---|---|
| 2 | 117819728 | rs12463859 | 0.642 | -0.165 | 0.482 | 0.179 | 0.024 | 0.004 | 0.004 | 1.576 | 0.93 | 0.43 | 0.95 | 1 | 0.97 | 0.99 |
| 2 | 117820115 | rs4999658 | 0.152 | -0.052 | -0.068 | 0.277 | 0.013 | -0.003 | -0.003 | 0 | 0.63 | 0.33 | 0.62 | 0.35 | 0.66 | 0.63 |
| 2 | 117820944 | rs13023828 | 0.209 | -0.024 | -0.942 | -0.057 | 0.012 | 0.008 | 0.008 | 1.263 | 0.25 | 0.08 | 0.37 | 0.2 | 0.22 | 0.26 |
| 2 | 117821875 | rs1437433 | 0.42 | 0.171 | -0.703 | -0.384 | 0.007 | -0.014 | -0.014 | 0 | 0.35 | 0.33 | 0.31 | 0.14 | 0.4 | 0.28 |
| 2 | 117823299 | rs17046609 | -1.183 | -0.28 | -0.491 | 0.302 | -0.011 | 0.1 | 0.1 | 0 | 0.14 | 0.21 | 0.22 | 0.48 | 0.09 | 0.21 |
| 2 | 117823797 | rs4407257 | 0.227 | -0.395 | -1.514 | -0.526 | 0.021 | 0.08 | 0.08 | 4.834 | 0.63 | 0.21 | 0.8 | 0.79 | 0.68 | 0.7 |
| 2 | 117836737 | rs13406669 | 0.113 | -0.51 | -0.83 | -0.699 | 0 | 0 | 0 | 7.104 | 0.23 | 0.2 | 0.13 | 0.19 | 0.23 | 0.09 |
| 2 | 117838220 | rs6727107 | -0.118 | -0.505 | -0.869 | -0.635 | -0.004 | 0.006 | 0.006 | 0 | 0.12 | 0.07 | 0.22 | 0.49 | 0.12 | 0.22 |
| 2 | 117838706 | rs17798906 | 0.397 | -0.542 | -0.964 | -0.69 | -0.036 | -0.009 | -0.009 | 0 | 0.17 | 0.01 | 0.09 | 0.19 | 0.16 | 0.07 |
| 2 | 117844244 | rs1437434 | 0.333 | -0.108 | -1.199 | -0.988 | -0.006 | 0.017 | 0.017 | 0.442 | 0.71 | 0.78 | 0.81 | 0.81 | 0.67 | 0.86 |
| 2 | 117844549 | rs332895 | 0.913 | -0.319 | -0.83 | -1.03 | -0.006 | -0.003 | -0.003 | 0 | 0.64 | 0.55 | 0.65 | 0.32 | 0.66 | 0.69 |
| 2 | 117847914 | rs729670 | -1.187 | -0.373 | -0.807 | -1.024 | -0.028 | 0.158 | 0.158 | 0 | 0.22 | 0.45 | 0.26 | 0.48 | 0.15 | 0.34 |
| 2 | 117848641 | rs744832 | 0.48 | -0.244 | -1.652 | -0.942 | -0.024 | 0.016 | 0.016 | 0 | 0.07 | 0 | 0.21 | 0.47 | 0.08 | 0.16 |
| 2 | 117850546 | rs2615828 | 1.469 | -0.22 | -1.519 | -0.977 | -0.013 | 0.049 | 0.049 | 0.689 | 0.51 | 0.07 | 0.59 | 0.32 | 0.6 | 0.5 |
| 2 | 117859507 | rs332906 | -0.087 | -0.373 | -0.82 | -0.569 | -0.043 | -0.004 | -0.004 | 0.157 | 0.22 | 0.01 | 0.19 | 0.15 | 0.21 | 0.22 |
| 2 | 117899586 | rs332925 | -0.534 | -0.079 | 0.493 | -0.264 | 0.003 | -0.011 | -0.011 | 0 | 0.26 | 0.29 | 0.17 | 0.22 | 0.3 | 0.16 |
| 2 | 117900052 | rs10205198 | -1.14 | -0.206 | 0.98 | -0.415 | 0.005 | -0.093 | -0.093 | 0 | 0.14 | 0.43 | 0.07 | 0.02 | 0.1 | 0.21 |
| 2 | 117908078 | rs1545123 | 1.723 | -0.251 | 1.011 | -0.431 | -0.011 | 0.172 | 0.172 | 0 | 0.58 | 0.12 | 0.75 | 0.84 | 0.6 | 0.63 |
| 2 | 117911357 | rs2035171 | 1.621 | -0.206 | 1.048 | -0.408 | -0.01 | 0.151 | 0.151 | 0 | 0.59 | 0.14 | 0.76 | 0.84 | 0.6 | 0.63 |
| 2 | 117912256 | rs10192798 | 0.236 | -0.211 | 1.065 | -0.41 | 0.012 | -0.041 | -0.041 | 0.531 | 0.74 | 0.6 | 0.83 | 0.86 | 0.7 | 0.84 |
| 2 | 117931948 | rs2007087 | -1.418 | -0.331 | 1.256 | -0.411 | -0.005 | -0.105 | -0.105 | 0.593 | 0.15 | 0.45 | 0.07 | 0.02 | 0.1 | 0.21 |
| 2 | 117942921 | rs17800371 | 1.512 | -0.289 | 0.919 | -0.369 | -0.009 | 0.137 | 0.137 | 0 | 0.59 | 0.13 | 0.76 | 0.78 | 0.61 | 0.65 |
| 2 | 117945566 | rs7585507 | -1.299 | -0.096 | 0.885 | 0.013 | -0.006 | -0.056 | -0.056 | 0 | 0.12 | 0.24 | 0.05 | 0.02 | 0.08 | 0.18 |
| 2 | 117973154 | rs333852 | 1.543 | -0.608 | 1.541 | -0.075 | -0.04 | -0.09 | -0.09 | 0 | 0.81 | 0.31 | 0.89 | 0.96 | 0.92 | 0.78 |
| 2 | 117986078 | rs432349 | 1.641 | -0.625 | 1.178 | -0.083 | 0 | -0.03 | -0.03 | 0 | 0.79 | 0.24 | 0.89 | 0.96 | 0.92 | 0.78 |
| 2 | 117994723 | rs1074282 | 0.408 | 0.094 | 0.819 | 1.064 | -0.014 | 0.034 | 0.034 | 0 | 0.08 | 0.06 | 0.28 | 0.38 | 0.15 | 0.31 |
| 2 | 118005365 | rs333885 | 1.83 | -0.121 | 1.224 | 1.028 | -0.005 | 0.01 | 0.01 | 3.288 | 0.73 | 0.3 | 0.62 | 0.62 | 0.76 | 0.46 |
| 2 | 118006384 | rs71470 | 1.338 | -0.142 | 1.535 | 0.952 | -0.067 | -0.138 | -0.138 | 0 | 0.81 | 0.44 | 0.84 | 0.96 | 0.92 | 0.8 |
| 2 | 118029195 | rs1453234 | -0.6 | -0.159 | 1.606 | 0.945 | -0.008 | -0.03 | -0.03 | 0.017 | 0.09 | 0.15 | 0.03 | 0.01 | 0.05 | 0.13 |
| 2 | 118033253 | rs7609039 | 0.503 | -0.163 | 1.724 | 0.943 | -0.017 | -0.047 | -0.047 | 0 | 0.9 | 0.8 | 0.96 | 0.99 | 0.95 | 0.86 |
| 2 | 118043647 | rs6726585 | 0.515 | -0.138 | 1.735 | 0.97 | -0.013 | -0.027 | -0.027 | 0.895 | 0.94 | 0.89 | 0.88 | 0.97 | 0.97 | 0.93 |
| 2 | 118056771 | rs1377407 | 1.266 | -0.11 | 1.823 | 1.069 | 0.003 | 0.022 | 0.022 | 0.077 | 0.86 | 0.8 | 0.66 | 0.58 | 0.81 | 0.61 |
| 2 | 118075351 | rs13032220 | -1.156 | -0.136 | 2.368 | 1.115 | 0.017 | 0.081 | 0.081 | 0 | 0.16 | 0.32 | 0.26 | 0.43 | 0.2 | 0.44 |
| 2 | 118081209 | rs10496537 | -1.013 | -0.126 | 2.623 | 1.126 | 0.058 | 0.205 | 0.205 | 1.456 | 0.12 | 0.4 | 0.23 | 0.42 | 0.15 | 0.31 |
| 2 | 118099393 | rs334855 | 0.561 | -0.135 | 1.487 | 0.758 | -0.073 | 0.019 | 0.019 | 0 | 0.22 | 0.02 | 0.14 | 0.15 | 0.27 | 0.13 |
| 2 | 118121536 | rs334835 | 0.294 | -0.25 | 1.291 | 0.767 | -0.001 | 0.003 | 0.003 | 0 | 0.59 | 0.55 | 0.58 | 0.36 | 0.63 | 0.32 |
| 2 | 118139446 | rs1996291 | -0.918 | 0.017 | 1.416 | 0.954 | 0.004 | 0.097 | 0.097 | 1.012 | 0.22 | 0.45 | 0.36 | 0.48 | 0.23 | 0.47 |
| 2 | 118156840 | rs2421932 | -0.844 | 0.035 | 1.216 | 0.991 | 0.012 | 0.112 | 0.112 | 0.847 | 0.22 | 0.48 | 0.37 | 0.5 | 0.22 | 0.47 |
| 2 | 118157713 | rs6757570 | 0.31 | -0.035 | 0.483 | 0.675 | 0.012 | 0.045 | 0.045 | 3.021 | 0.39 | 0.16 | 0.21 | 0.3 | 0.35 | 0.31 |
| 2 | 118168045 | rs1996295 | 1.492 | -0.292 | 1.061 | 1.22 | -0.002 | 0 | 0 | 2.815 | 0.68 | 0.24 | 0.58 | 0.59 | 0.69 | 0.48 |
| 2 | 118178568 | rs17046973 | 0.337 | -0.046 | 1.233 | 1.461 | 0.028 | 0.066 | 0.066 | 0 | 0.08 | 0.13 | 0.22 | 0.36 | 0.17 | 0.27 |
| 2 | 118178623 | rs6727686 | -1.264 | -0.083 | 1.223 | 1.118 | 0.007 | -0.001 | -0.001 | 0 | 0.93 | 0.92 | 0.93 | 0.94 | 0.9 | 0.92 |
| 2 | 118180359 | rs10864955 | 0.641 | -0.354 | 0.92 | 0.999 | 0.009 | 0.018 | 0.018 | 2.383 | 0.21 | 0.1 | 0.1 | 0.12 | 0.17 | 0.1 |
| 2 | 118187490 | rs6760213 | 1.264 | -0.021 | 1.717 | 1.742 | 0.018 | 0.03 | 0.03 | 0 | 0.81 | 0.75 | 0.7 | 0.56 | 0.71 | 0.7 |
| 2 | 118190338 | rs2090340 | 1.941 | -0.106 | 1.297 | 1.604 | -0.002 | -0.011 | -0.011 | 0 | 0.66 | 0.26 | 0.54 | 0.5 | 0.64 | 0.51 |
| 2 | 118194106 | rs5025876 | 0.027 | 0.071 | 0.682 | 0.737 | -0.012 | 0.01 | 0.01 | 0 | 0.9 | 0.93 | 0.93 | 1 | 0.87 | 0.94 |
| 2 | 118224170 | rs10182096 | 0.073 | 0.076 | 0.649 | 0.799 | -0.005 | -0.005 | -0.005 | 0.257 | 0.9 | 0.88 | 0.89 | 0.86 | 0.87 | 0.94 |
| 2 | 118247119 | rs4849560 | 0.545 | 0.198 | 0.514 | 0.774 | -0.038 | -0.022 | -0.022 | 0.376 | 0.06 | 0.01 | 0.16 | 0.11 | 0.15 | 0.07 |
| 2 | 118250637 | rs10164651 | -1.415 | 0.142 | 0.951 | 0.68 | 0.034 | -0.002 | -0.002 | 0 | 0.07 | 0.39 | 0.05 | 0 | 0.03 | 0.01 |
| 2 | 118258797 | rs290876 | -1.551 | 0.14 | 0.821 | 0.679 | 0.018 | -0.017 | -0.017 | 0 | 0.08 | 0.41 | 0.05 | 0 | 0.03 | 0.01 |
| 2 | 118274002 | rs4849561 | 0.09 | 0.216 | -0.131 | 0.703 | 0.011 | -0.05 | -0.05 | 0 | 0.89 | 0.79 | 0.91 | 1 | 0.87 | 0.94 |
| 2 | 118303189 | rs6724956 | 1.022 | 0.092 | -0.706 | 0.821 | -0.002 | 0.02 | 0.02 | 0.894 | 0.64 | 0.11 | 0.53 | 0.57 | 0.65 | 0.53 |
| 2 | 118316309 | rs17047088 | 0.394 | 0.314 | -1.121 | 1.088 | -0.041 | -0.096 | -0.096 | 0.468 | 0.13 | 0 | 0.24 | 0.33 | 0.16 | 0.23 |
| 2 | 118318731 | rs13390020 | -1.128 | 0.074 | -0.302 | 0.546 | -0.004 | 0.001 | 0.001 | 0.242 | 0.26 | 0.64 | 0.38 | 0.43 | 0.19 | 0.42 |
| 2 | 118335509 | rs6749073 | -1.016 | 0.136 | 0.21 | 0.577 | -0.018 | 0.044 | 0.044 | 0 | 0.24 | 0.48 | 0.37 | 0.43 | 0.19 | 0.42 |
| 2 | 118354342 | rs4849584 | -1.023 | 0.283 | -0.148 | 0.5 | -0.007 | -0.001 | -0.001 | 0 | 0.17 | 0.18 | 0.27 | 0.24 | 0.21 | 0.17 |
| 2 | 118355754 | rs17818667 | 0.156 | 0.309 | -0.308 | 0.555 | -0.007 | -0.007 | -0.007 | 0 | 0.07 | 0.07 | 0.2 | 0.24 | 0.12 | 0.12 |
| 2 | 118362380 | rs11123431 | 0.66 | 0.229 | -0.274 | 0.523 | 0.042 | 0.003 | 0.003 | 1.822 | 0.83 | 0.61 | 0.74 | 0.75 | 0.79 | 0.83 |
| 2 | 118362441 | rs12472414 | 1.099 | 0.226 | -1.292 | 0.595 | -0.013 | -0.003 | -0.003 | 0.06 | 0.63 | 0.08 | 0.52 | 0.56 | 0.68 | 0.51 |
| 2 | 118364695 | rs13034275 | 1.117 | 0.447 | -1.595 | 0.215 | -0.067 | -0.043 | -0.043 | 0.466 | 0.43 | 0.05 | 0.33 | 0.54 | 0.43 | 0.47 |
| 2 | 118364722 | rs10187234 | 0.207 | 0.193 | -0.455 | -0.024 | -0.018 | -0.055 | -0.055 | 0 | 0.78 | 0.68 | 0.86 | 0.9 | 0.81 | 0.89 |
| 2 | 118365400 | rs17047157 | -0.639 | 0.366 | -0.71 | 0.332 | 0 | -0.001 | -0.001 | 0 | 0.09 | 0.11 | 0.18 | 0.19 | 0.1 | 0.16 |
| 2 | 118369298 | rs9973888 | -1.096 | 0.187 | -0.713 | 0.119 | 0.007 | 0.003 | 0.003 | 0 | 0.08 | 0.08 | 0.03 | 0 | 0.05 | 0.01 |
| 2 | 118369880 | rs7600929 | -1.096 | 0.187 | -0.713 | 0.119 | 0.007 | 0.003 | 0.003 | 0 | 0.08 | 0.08 | 0.03 | 0 | 0.05 | 0.01 |
| 2 | 118376620 | rs10439399 | -1.169 | 0.06 | 0.845 | -0.003 | -0.004 | -0.015 | -0.015 | 0.05 | 0.08 | 0.1 | 0.04 | 0 | 0.05 | 0.01 |
| 2 | 118389164 | rs1516006 | 0.706 | 0.559 | 1.109 | -0.248 | 0.099 | -0.071 | -0.071 | 0 | 0.89 | 0.56 | 0.88 | 1 | 0.82 | 0.94 |
| 2 | 118396819 | rs962315 | -0.45 | 0.358 | 0.732 | -0.248 | 0 | -0.001 | -0.001 | 0 | 0.24 | 0.21 | 0.3 | 0.32 | 0.2 | 0.27 |
| 2 | 118403937 | rs13402352 | 1.806 | 0.34 | 0.972 | -0.304 | 0 | -0.107 | -0.107 | 0 | 0.82 | 0.44 | 0.9 | 0.97 | 0.84 | 0.89 |
| 2 | 118413757 | rs13032870 | 2.228 | -0.023 | 0.151 | -0.237 | -0.053 | 0.014 | 0.014 | 1.846 | 0.51 | 0.04 | 0.54 | 0.52 | 0.52 | 0.53 |

| | | | | | | | | | | | | | | | | |
|---|---|---|---|---|---|---|---|---|---|---|---|---|---|---|---|---|
| 2 | 118414432 | rs6705449 | 1.634 | 0.074 | 1.254 | -0.226 | -0.003 | -0.091 | -0.091 | 0 | 0.83 | 0.38 | 0.89 | 0.97 | 0.84 | 0.89 |
| 2 | 118422277 | rs1035602 | 2.405 | 0.276 | 0.983 | -0.454 | -0.048 | 0.095 | 0.095 | 0.698 | 0.51 | 0.08 | 0.42 | 0.6 | 0.48 | 0.58 |
| 2 | 118428425 | rs10186120 | -1.424 | 0.365 | 1.547 | -0.314 | -0.025 | -0.035 | -0.035 | 0 | 0.1 | 0.3 | 0.05 | 0.03 | 0.06 | 0.07 |
| 2 | 118434604 | rs12474388 | 1.05 | 0.371 | 1.687 | -0.32 | 0.065 | -0.047 | -0.047 | 0 | 0.88 | 0.55 | 0.92 | 0.97 | 0.85 | 0.9 |
| 2 | 118434667 | rs12466589 | -0.709 | 0.392 | 1.623 | -0.321 | -0.001 | 0.002 | 0.002 | 0 | 0.26 | 0.24 | 0.34 | 0.36 | 0.28 | 0.28 |
| 2 | 118442596 | rs7577373 | -0.645 | 0.389 | 1.529 | -0.325 | 0.009 | 0.003 | 0.003 | 1.873 | 0.06 | 0.24 | 0.03 | 0.03 | 0.05 | 0.06 |
| 2 | 118448025 | rs12475536 | 0.279 | 0.585 | 1.478 | -0.382 | -0.017 | 0.005 | 0.005 | 0 | 0.09 | 0.03 | 0.25 | 0.09 | 0.17 | 0.11 |
| 2 | 118448486 | rs1477450 | 2.56 | 0.543 | 1.507 | -0.385 | -0.027 | 0.071 | 0.071 | 2.797 | 0.55 | 0.09 | 0.53 | 0.63 | 0.53 | 0.6 |
| 2 | 118455405 | rs918860 | -1.146 | 0.527 | 1.786 | -0.371 | -0.007 | -0.007 | -0.007 | 0 | 0.25 | 0.18 | 0.36 | 0.34 | 0.31 | 0.28 |
| 2 | 118466558 | rs2010944 | 0.407 | 0.572 | 1.807 | -0.349 | 0.062 | -0.021 | -0.021 | 0 | 0.91 | 0.65 | 0.94 | 0.97 | 0.89 | 0.91 |
| 2 | 118468843 | rs999497 | 1.857 | 0.494 | 1.332 | -0.393 | -0.02 | 0.044 | 0.044 | 2.829 | 0.59 | 0.17 | 0.55 | 0.63 | 0.56 | 0.61 |
| 2 | 118474016 | rs4849613 | -0.709 | 1.111 | 1.046 | -0.074 | -0.002 | -0.002 | -0.002 | 0 | 0.06 | 0.05 | 0.1 | 0.13 | 0.1 | 0.14 |
| 2 | 118477333 | rs7568324 | -0.974 | 1.039 | 1.025 | -0.175 | -0.004 | -0.02 | -0.02 | 0 | 0.06 | 0.08 | 0.05 | 0 | 0.06 | 0.03 |
| 2 | 118484594 | rs17498503 | 1.998 | 1.159 | 0.983 | 0.142 | -0.023 | -0.021 | -0.021 | 0.14 | 0.56 | 0.17 | 0.49 | 0.39 | 0.48 | 0.46 |
| 2 | 118489676 | rs4849620 | -0.771 | 1.243 | 0.563 | 0.177 | 0 | -0.006 | -0.006 | 0.102 | 0.06 | 0.07 | 0.1 | 0.03 | 0.1 | 0.14 |
| 2 | 118493077 | rs10210418 | 1.424 | 1.075 | 0.502 | 0.314 | -0.016 | -0.008 | -0.008 | 0 | 0.6 | 0.3 | 0.52 | 0.51 | 0.54 | 0.55 |
| 2 | 118504764 | rs417651 | -0.276 | 1.149 | -0.054 | 0.422 | -0.028 | -0.039 | -0.039 | 0 | 0.92 | 0.99 | 0.82 | 0.84 | 0.91 | 0.91 |
| 2 | 118506345 | rs367201 | -0.985 | 1.266 | -0.802 | -0.299 | -0.033 | 0.151 | 0.151 | 0.217 | 0.52 | 0.78 | 0.46 | 0.23 | 0.53 | 0.53 |
| 2 | 118514931 | rs17499330 | -0.716 | 1.418 | -0.3 | -0.584 | -0.004 | -0.006 | -0.006 | 1.857 | 0.14 | 0.11 | 0.1 | 0 | 0.15 | 0.07 |
| 2 | 118517860 | rs370828 | 1.373 | 1.221 | -0.406 | -0.504 | 0.004 | -0.003 | -0.003 | 0 | 0.69 | 0.62 | 0.56 | 0.71 | 0.66 | 0.76 |
| 2 | 118520639 | rs6728975 | 1.399 | 1.203 | -0.372 | -0.511 | 0.006 | -0.004 | -0.004 | 0 | 0.68 | 0.6 | 0.55 | 0.71 | 0.66 | 0.76 |
| 2 | 118529856 | rs6542393 | 1.47 | 1.133 | -0.473 | -0.606 | -0.008 | 0.007 | 0.007 | 0 | 0.42 | 0.34 | 0.51 | 0.59 | 0.47 | 0.42 |
| 2 | 118534260 | rs17047392 | 2.319 | 1.128 | -0.493 | -0.61 | -0.05 | -0.088 | -0.088 | 0 | 0.35 | 0.11 | 0.33 | 0.53 | 0.38 | 0.34 |
| 2 | 118542042 | rs13416893 | 0.369 | 1.065 | -0.356 | -0.697 | -0.015 | -0.065 | -0.065 | 0.359 | 0.3 | 0.46 | 0.19 | 0.17 | 0.28 | 0.39 |
| 2 | 118550719 | rs204855 | 1.025 | 1.108 | -0.427 | -0.728 | 0.011 | 0.034 | 0.034 | 0 | 0.5 | 0.37 | 0.54 | 0.59 | 0.54 | 0.54 |
| 2 | 118551825 | rs2060332 | 0.08 | 1.142 | -0.426 | -0.7 | -0.019 | -0.061 | -0.061 | 0 | 0.33 | 0.49 | 0.23 | 0.17 | 0.28 | 0.42 |
| 2 | 118555803 | rs6757293 | 2.375 | 1.123 | -0.666 | -0.716 | -0.109 | -0.154 | -0.154 | 0.218 | 0.33 | 0.03 | 0.32 | 0.53 | 0.38 | 0.34 |
| 2 | 118569196 | rs17507637 | 0.577 | 1.166 | -0.402 | -0.957 | 0.011 | 0.006 | 0.006 | 0.109 | 0.19 | 0.12 | 0.1 | 0.01 | 0.14 | 0.18 |
| 2 | 118573396 | rs11689073 | 1.637 | 1.342 | -0.331 | -0.727 | -0.041 | -0.033 | -0.033 | 0 | 0.42 | 0.17 | 0.36 | 0.53 | 0.45 | 0.39 |
| 2 | 118578114 | rs2432721 | 0.577 | 1.347 | -0.139 | -0.747 | 0.01 | 0.01 | 0.01 | 0.584 | 0.53 | 0.4 | 0.57 | 0.59 | 0.61 | 0.46 |
| 2 | 118579085 | rs10490632 | 0.392 | 1.355 | -0.165 | -0.748 | -0.003 | -0.036 | -0.036 | 0.146 | 0.09 | 0.13 | 0.04 | 0 | 0.08 | 0.05 |
| 2 | 118580670 | rs6715763 | -0.313 | 1.371 | -0.181 | -0.696 | 0.001 | -0.026 | -0.026 | 0.141 | 0.08 | 0.12 | 0.07 | 0.01 | 0.08 | 0.02 |
| 2 | 118583940 | rs2303348 | 2.382 | 1.373 | -0.209 | -0.715 | -0.108 | -0.149 | -0.149 | 2.295 | 0.33 | 0.04 | 0.32 | 0.53 | 0.38 | 0.34 |
| 2 | 118589029 | rs13030497 | -0.083 | 1.322 | -0.238 | -0.695 | 0 | -0.017 | -0.017 | 0 | 0.07 | 0.09 | 0.07 | 0 | 0.08 | 0.02 |
| 2 | 118602560 | rs4849629 | 1.454 | 1.355 | -0.184 | -0.65 | -0.019 | 0.015 | 0.015 | 1.495 | 0.43 | 0.23 | 0.53 | 0.59 | 0.54 | 0.41 |
| 2 | 118606236 | rs2570207 | 0.499 | 1.286 | -0.15 | -0.514 | 0.005 | 0.053 | 0.053 | 0.017 | 0.82 | 0.68 | 0.65 | 0.71 | 0.82 | 0.83 |
| 2 | 118608769 | rs4849630 | 0.285 | 1.249 | -0.542 | -0.502 | -0.038 | -0.051 | -0.051 | 0 | 0.8 | 0.97 | 0.76 | 0.76 | 0.83 | 0.83 |
| 2 | 118611789 | rs6712766 | -1.287 | 1.367 | -0.082 | -0.315 | 0.045 | 0.033 | 0.033 | 2.108 | 0.07 | 0.28 | 0.13 | 0.05 | 0.1 | 0.06 |
| 2 | 118617542 | rs7593686 | -0.149 | 1.585 | -0.384 | -0.459 | -0.006 | -0.07 | -0.07 | 0 | 0.34 | 0.48 | 0.25 | 0.16 | 0.29 | 0.41 |
| 2 | 118618718 | rs1433526 | 1.916 | 1.63 | -0.503 | -0.47 | -0.102 | -0.128 | -0.128 | 0.056 | 0.39 | 0.04 | 0.35 | 0.54 | 0.43 | 0.4 |
| 2 | 118618928 | rs6749268 | -0.108 | 1.623 | -0.449 | -0.442 | -0.006 | -0.07 | -0.07 | 0.053 | 0.34 | 0.48 | 0.25 | 0.16 | 0.29 | 0.41 |
| 2 | 118622990 | rs2570211 | -1.127 | 1.629 | -0.782 | -0.329 | -0.015 | -0.068 | -0.068 | 0.067 | 0.88 | 0.98 | 0.8 | 0.75 | 0.91 | 0.88 |
| 2 | 118627100 | rs9677816 | 0.26 | 1.676 | -0.572 | -0.292 | 0.004 | -0.02 | -0.02 | 0 | 0.05 | 0.1 | 0.06 | 0 | 0.07 | 0.01 |
| 2 | 118632862 | rs6542406 | 2.008 | 1.613 | -0.752 | -0.31 | -0.066 | -0.146 | -0.146 | 0 | 0.34 | 0.04 | 0.32 | 0.55 | 0.36 | 0.35 |
| 2 | 118639481 | rs13035453 | 0.021 | 1.654 | -0.357 | -0.271 | -0.001 | -0.027 | -0.027 | 0.053 | 0.06 | 0.11 | 0.06 | 0 | 0.06 | 0.01 |
| 2 | 118643492 | rs967252 | 0.432 | 1.732 | -0.198 | -0.336 | -0.001 | -0.003 | -0.003 | 0 | 0.56 | 0.44 | 0.6 | 0.61 | 0.62 | 0.44 |
| 2 | 118646201 | rs1433531 | 1.284 | 1.639 | -0.565 | -0.348 | -0.006 | -0.024 | -0.024 | 0 | 0.44 | 0.15 | 0.46 | 0.57 | 0.46 | 0.34 |
| 2 | 118672017 | rs6757088 | -1.392 | 1.794 | -0.314 | -0.01 | 0.032 | 0.034 | 0.034 | 1.048 | 0.52 | 0.66 | 0.49 | 0.34 | 0.6 | 0.58 |
| 2 | 118672529 | rs7591195 | -1.564 | 1.757 | -0.194 | -0.028 | 0.043 | 0.047 | 0.047 | 1.481 | 0.54 | 0.77 | 0.5 | 0.34 | 0.6 | 0.58 |
| 2 | 118673663 | rs1046330 | -1.642 | 1.718 | -0.214 | -0.047 | 0.04 | 0.043 | 0.043 | 1.129 | 0.54 | 0.78 | 0.5 | 0.34 | 0.6 | 0.61 |
| 2 | 118673738 | rs1046328 | -0.464 | 1.862 | -0.114 | -0.129 | 0.009 | -0.055 | -0.055 | 0 | 0.35 | 0.56 | 0.25 | 0.14 | 0.35 | 0.31 |
| 2 | 118676429 | rs11556267 | 0.245 | 1.927 | -0.16 | -0.124 | -0.03 | 0.015 | 0.015 | 0 | 0.09 | 0.02 | 0.14 | 0.04 | 0.17 | 0.11 |
| 2 | 118676836 | rs1863040 | -1.852 | 1.903 | -0.158 | -0.12 | 0.043 | 0.047 | 0.047 | 1.481 | 0.54 | 0.77 | 0.5 | 0.34 | 0.6 | 0.58 |
| 2 | 118679906 | rs3771933 | 0.737 | 2.081 | 0.307 | -0.191 | -0.002 | 0.009 | 0.009 | 0 | 0.25 | 0.2 | 0.13 | 0.13 | 0.23 | 0.28 |
| 2 | 118701764 | rs3732320 | -0.419 | 2.04 | 0.147 | -0.115 | -0.01 | -0.013 | -0.013 | 0 | 0.1 | 0.08 | 0.21 | 0.2 | 0.17 | 0.23 |
| 2 | 118710482 | rs3755498 | 0.796 | 2.027 | 0.21 | -0.201 | -0.011 | 0.038 | 0.038 | 0 | 0.57 | 0.31 | 0.71 | 0.86 | 0.57 | 0.64 |
| 2 | 118755038 | rs6750836 | -0.812 | 1.978 | -0.085 | -0.176 | 0.011 | -0.055 | -0.055 | 0 | 0.4 | 0.5 | 0.27 | 0.14 | 0.43 | 0.36 |
| 2 | 118755669 | rs17512698 | 1.03 | 1.987 | -0.087 | -0.173 | 0 | 0.041 | 0.041 | 0 | 0.55 | 0.33 | 0.61 | 0.82 | 0.48 | 0.58 |
| 2 | 118763068 | rs11684454 | 2.29 | 1.896 | -0.192 | 0.035 | 0.026 | -0.018 | -0.018 | 0 | 0.34 | 0.17 | 0.3 | 0.39 | 0.31 | 0.23 |
| 2 | 118786833 | rs9646905 | 0.244 | 2.035 | -0.113 | -0.107 | 0.009 | -0.055 | -0.055 | 2.482 | 0.28 | 0.44 | 0.17 | 0.14 | 0.26 | 0.29 |
| 2 | 118791400 | rs11690896 | 0.433 | 1.899 | -0.151 | -0.029 | -0.001 | -0.039 | -0.039 | 0 | 0.92 | 0.83 | 0.92 | 1 | 0.91 | 0.99 |
| 2 | 118803518 | rs6721751 | 0.355 | 1.9 | -0.155 | -0.028 | -0.001 | -0.037 | -0.037 | 0 | 0.92 | 0.83 | 0.92 | 1 | 0.91 | 0.99 |
| 2 | 118809451 | rs1385925 | 1.557 | 2.035 | -0.135 | 0.011 | 0.016 | 0.004 | 0.004 | 6.108 | 0.82 | 0.69 | 0.71 | 0.8 | 0.75 | 0.76 |
| 2 | 118814868 | rs2276695 | -0.572 | 2.008 | -0.161 | 0.101 | -0.001 | -0.002 | -0.002 | 1.81 | 0.42 | 0.59 | 0.4 | 0.43 | 0.36 | 0.42 |
| 2 | 118819855 | rs1541714 | -0.696 | 2.067 | -0.205 | 0.098 | -0.005 | -0.003 | -0.003 | 0 | 0.16 | 0.14 | 0.24 | 0.22 | 0.24 | 0.27 |
| 2 | 118822403 | rs11687724 | 0.679 | 1.719 | -0.342 | 0.194 | -0.011 | -0.02 | -0.02 | 1.536 | 0.07 | 0.03 | 0.09 | 0.18 | 0.07 | 0.16 |
| 2 | 118827810 | rs17047697 | 2.011 | 1.466 | -0.316 | 0.221 | 0.041 | -0.031 | -0.031 | 0.047 | 0.34 | 0.1 | 0.29 | 0.35 | 0.31 | 0.29 |
| 2 | 118827859 | rs4849674 | 0.656 | 1.565 | -0.308 | 0.282 | -0.005 | 0.029 | 0.029 | 0 | 0.13 | 0.15 | 0.22 | 0.28 | 0.09 | 0.15 |
| 2 | 118836025 | rs7566605 | -2.1 | 1.509 | -0.538 | 0.101 | 0.037 | -0.008 | -0.008 | 0 | 0.64 | 0.8 | 0.7 | 0.65 | 0.69 | 0.71 |

| | | | | | | | | | | | | | | | | |
|---|---|---|---|---|---|---|---|---|---|---|---|---|---|---|---|---|
| 2 | 118838281 | rs2161830 | -1.517 | 1.5 | -0.457 | 0.202 | 0.036 | 0.018 | 0.018 | 2.015 | 0.57 | 0.72 | 0.62 | 0.65 | 0.6 | 0.69 |
| 2 | 118841325 | rs17047733 | 0.177 | 1.341 | 0.109 | 0.624 | -0.014 | 0.127 | 0.127 | 0 | 0.14 | 0.23 | 0.21 | 0.33 | 0.09 | 0.15 |
| 2 | 118855927 | rs1559509 | 0.251 | 2.003 | 0.243 | 0.259 | 0.003 | -0.022 | -0.022 | 0.818 | 0.25 | 0.3 | 0.15 | 0.12 | 0.24 | 0.25 |
| 2 | 118856914 | rs2161829 | 0.799 | 1.907 | 0.139 | 0.311 | -0.004 | -0.004 | -0.004 | 0.283 | 0.59 | 0.4 | 0.52 | 0.52 | 0.57 | 0.55 |
| 2 | 118860471 | rs889904 | -1.271 | 1.922 | -0.189 | -0.306 | -0.002 | 0.014 | 0.014 | 0 | 0.47 | 0.65 | 0.4 | 0.13 | 0.49 | 0.37 |
| 2 | 118861435 | rs17047757 | 1.619 | 1.78 | -0.445 | -0.092 | -0.022 | -0.071 | -0.071 | 2.358 | 0.11 | 0.01 | 0.19 | 0.26 | 0.09 | 0.1 |
| 2 | 118862492 | rs10490624 | -0.861 | 1.825 | -0.341 | -0.303 | 0.005 | -0.016 | -0.016 | 0 | 0.07 | 0.1 | 0.15 | 0.02 | 0.11 | 0.06 |
| 2 | 118863874 | rs9308762 | -0.423 | 1.86 | -0.457 | -0.258 | -0.001 | 0.015 | 0.015 | 0.331 | 0.81 | 0.84 | 0.71 | 0.5 | 0.83 | 0.69 |
| 2 | 118872427 | rs3849327 | -1.07 | 1.905 | -0.386 | -0.685 | 0.037 | -0.116 | -0.116 | 0 | 0.16 | 0.53 | 0.21 | 0.02 | 0.17 | 0.11 |
| 2 | 118873587 | rs12151787 | -1.31 | 1.847 | -0.478 | -0.764 | 0.013 | -0.165 | -0.165 | 0.117 | 0.3 | 0.56 | 0.3 | 0.02 | 0.34 | 0.16 |
| 2 | 118873636 | rs11681903 | 0.796 | 1.723 | -0.401 | -0.49 | -0.016 | 0.036 | 0.036 | 0 | 0.18 | 0.08 | 0.1 | 0.12 | 0.17 | 0.2 |
| 2 | 118876072 | rs11673894 | -1.209 | 1.729 | -0.115 | -0.388 | 0.023 | -0.07 | -0.07 | 4.01 | 0.21 | 0.51 | 0.15 | 0.12 | 0.19 | 0.18 |
| 2 | 118876116 | rs11673900 | -0.503 | 1.769 | -0.278 | -0.341 | 0.002 | -0.002 | -0.002 | 1.586 | 0.12 | 0.15 | 0.1 | 0.12 | 0.12 | 0.14 |
| 2 | 118876384 | rs11675138 | -0.666 | 1.949 | -0.299 | -0.351 | 0 | 0 | 0 | 0 | 0.13 | 0.16 | 0.11 | 0.12 | 0.16 | 0.16 |
| 2 | 118881001 | rs13400438 | 1.794 | 1.588 | -0.452 | -0.271 | 0.006 | -0.005 | -0.005 | 0.629 | 0.38 | 0.24 | 0.4 | 0.37 | 0.42 | 0.35 |
| 2 | 118881068 | rs17047787 | -0.054 | 1.456 | -0.05 | -0.407 | 0.025 | -0.041 | -0.041 | 0.012 | 0.09 | 0.37 | 0.06 | 0 | 0.07 | 0.04 |
| 2 | 118889525 | rs4848493 | -0.507 | 1.517 | -0.415 | -0.38 | -0.012 | 0.03 | 0.03 | 0 | 0.22 | 0.13 | 0.14 | 0.11 | 0.22 | 0.19 |
| 2 | 118891996 | rs17047805 | 0.131 | 1.212 | 0.127 | -0.174 | 0.03 | -0.031 | -0.031 | 0 | 0.08 | 0.33 | 0.05 | 0 | 0.07 | 0.03 |
| 2 | 118892712 | rs6740055 | 0.224 | 1.308 | -0.054 | 0.128 | -0.007 | 0.055 | 0.055 | 0 | 0.69 | 0.46 | 0.56 | 0.56 | 0.72 | 0.67 |
| 2 | 118894127 | rs7593540 | -1.055 | 1.388 | 0.086 | -0.25 | 0.051 | -0.074 | -0.074 | 0 | 0.2 | 0.43 | 0.23 | 0.05 | 0.23 | 0.17 |
| 2 | 118903295 | rs6720263 | 1.225 | 1.129 | -0.466 | -0.237 | -0.01 | -0.015 | -0.015 | 0.451 | 0.4 | 0.24 | 0.36 | 0.43 | 0.47 | 0.3 |
| 2 | 118904460 | rs12616545 | 0.286 | 1.009 | -0.643 | -0.135 | 0.003 | -0.004 | -0.004 | 0 | 0.56 | 0.6 | 0.44 | 0.43 | 0.57 | 0.46 |
| 2 | 118908358 | rs3849331 | -0.112 | 0.895 | -1.085 | -0.075 | 0.01 | -0.046 | -0.046 | 0 | 0.37 | 0.22 | 0.5 | 0.49 | 0.36 | 0.4 |
| 2 | 118926955 | rs13025020 | 1.383 | 0.761 | -0.989 | 0.018 | 0.001 | 0.023 | 0.023 | 1.088 | 0.49 | 0.19 | 0.5 | 0.43 | 0.51 | 0.34 |
| 2 | 118927760 | rs11123477 | -0.683 | 0.812 | 0.169 | -0.452 | 0.001 | -0.02 | -0.02 | 0.546 | 0.14 | 0.15 | 0.17 | 0.04 | 0.15 | 0.21 |
| 2 | 118928678 | rs4848495 | 0.831 | 0.703 | 0.703 | -0.479 | 0.036 | -0.032 | -0.032 | 0 | 0.89 | 0.49 | 0.91 | 0.92 | 0.93 | 0.82 |
| 2 | 118930311 | rs4088767 | 0.001 | 0.705 | 0.675 | -0.478 | 0 | -0.007 | -0.007 | 0 | 0.92 | 0.88 | 0.95 | 0.92 | 0.93 | 0.83 |
| 2 | 118934355 | rs3849332 | 0.595 | 0.701 | 1.001 | -0.48 | 0.01 | -0.03 | -0.03 | 0 | 0.91 | 0.75 | 0.94 | 0.92 | 0.93 | 0.83 |
| 2 | 118946643 | rs4849683 | 1.763 | 0.598 | 0.446 | -0.487 | -0.008 | 0.07 | 0.07 | 2.178 | 0.66 | 0.21 | 0.7 | 0.88 | 0.7 | 0.51 |
| 2 | 118946854 | rs4848497 | 1.587 | 0.612 | 0.453 | -0.48 | 0 | -0.03 | -0.03 | 0 | 0.68 | 0.3 | 0.71 | 0.88 | 0.7 | 0.51 |
| 2 | 118959792 | rs12711897 | 1.186 | 0.167 | 0.899 | 0.068 | -0.011 | -0.048 | -0.048 | 2.826 | 0.81 | 0.2 | 0.91 | 1 | 0.92 | 0.91 |
| 2 | 118964860 | rs17047865 | -1.58 | 0.138 | 1.442 | 0.066 | -0.024 | -0.043 | -0.043 | 0.023 | 0.1 | 0.61 | 0.07 | 0 | 0.03 | 0.04 |
| 2 | 118971945 | rs12711898 | 1.067 | 0.137 | 1.658 | 0.055 | -0.044 | -0.114 | -0.114 | 1.162 | 0.83 | 0.37 | 0.92 | 1 | 0.92 | 0.91 |
| 2 | 118978890 | rs3914668 | 1.386 | 0.132 | 1.646 | 0.052 | -0.022 | -0.065 | -0.065 | 0.885 | 0.81 | 0.23 | 0.91 | 1 | 0.92 | 0.91 |
| 2 | 118984208 | rs7566010 | 1.284 | 0.145 | 1.962 | 0.074 | -0.006 | -0.028 | -0.028 | 0 | 0.91 | 0.46 | 0.94 | 1 | 0.97 | 0.96 |
| 2 | 118989619 | rs4849688 | 1.477 | 0.145 | 1.833 | 0.074 | -0.038 | -0.056 | -0.056 | 0 | 0.9 | 0.33 | 0.94 | 1 | 0.97 | 0.96 |
| 2 | 118997824 | rs10185114 | 1.133 | 0.147 | 2.063 | 0.059 | -0.045 | -0.118 | -0.118 | 0 | 0.83 | 0.39 | 0.93 | 1 | 0.92 | 0.91 |
| 2 | 118998844 | rs6733786 | 0.57 | 0.173 | 1.745 | 0.084 | -0.006 | -0.022 | -0.022 | 0 | 0.93 | 0.74 | 0.97 | 1 | 0.97 | 0.96 |
| 2 | 119000584 | rs17047898 | 0.716 | 0.22 | 1.604 | 0.103 | -0.002 | -0.026 | -0.026 | 0 | 0.08 | 0.09 | 0.02 | 0 | 0.05 | 0.06 |
| 2 | 119000652 | rs10864971 | -0.014 | 0.297 | 0.786 | 0.325 | -0.022 | -0.024 | -0.024 | 0 | 0.31 | 0.09 | 0.27 | 0.4 | 0.34 | 0.39 |
| 2 | 119004158 | rs7573503 | 0.501 | 0.106 | 1.066 | 0.287 | 0 | -0.063 | -0.063 | 0.699 | 0.11 | 0.4 | 0.05 | 0 | 0.05 | 0.05 |
| 2 | 119006422 | rs10186364 | 0.867 | -0.002 | 0.591 | 0.302 | -0.045 | -0.051 | -0.051 | 0.072 | 0.74 | 0.4 | 0.86 | 0.82 | 0.85 | 0.79 |
| 2 | 119007927 | rs12711901 | 0.626 | 0.002 | 0.42 | 0.309 | -0.047 | -0.009 | -0.009 | 0.042 | 0.81 | 0.4 | 0.88 | 0.81 | 0.89 | 0.84 |
| 2 | 119009388 | rs7604243 | 0.621 | 0.162 | -0.33 | 0.408 | -0.008 | 0.002 | 0.002 | 0 | 0.23 | 0.09 | 0.2 | 0.22 | 0.24 | 0.28 |
| 2 | 119010201 | rs13005524 | 1.035 | -0.199 | -0.185 | 0.399 | 0.01 | 0.011 | 0.011 | 0.397 | 0.72 | 0.22 | 0.84 | 0.82 | 0.84 | 0.79 |
| 2 | 119010598 | rs17770539 | 1.655 | 0.146 | -0.843 | 0.848 | -0.08 | -0.098 | -0.098 | 1.603 | 0.4 | 0.05 | 0.44 | 0.43 | 0.42 | 0.42 |
| 2 | 119012578 | rs11690269 | 0.176 | -0.157 | -0.551 | 0.847 | 0.024 | -0.026 | -0.026 | 0 | 0.6 | 0.17 | 0.65 | 0.53 | 0.69 | 0.67 |
| 2 | 119013032 | rs11123481 | 1.657 | -0.312 | 0.35 | 0.812 | 0.025 | 0.016 | 0.016 | 5.494 | 0.69 | 0.17 | 0.8 | 0.72 | 0.8 | 0.73 |
| 2 | 119023101 | rs1517531 | 1.463 | -0.065 | 1.032 | 0.925 | -0.015 | 0.032 | 0.032 | 0.806 | 0.8 | 0.39 | 0.84 | 0.74 | 0.84 | 0.78 |
| 2 | 119026322 | rs1122021 | 1.432 | -0.061 | 1.075 | 0.91 | -0.012 | 0.038 | 0.038 | 0.893 | 0.8 | 0.41 | 0.84 | 0.74 | 0.84 | 0.78 |
| 2 | 119029737 | rs13399825 | 0.243 | 0.231 | 0.655 | 0.953 | -0.025 | 0.038 | 0.038 | 0.058 | 0.18 | 0.08 | 0.27 | 0.09 | 0.3 | 0.25 |
| 2 | 119033518 | rs6732183 | 1.07 | 0.074 | 1.05 | 0.968 | 0 | -0.008 | -0.008 | 3.35 | 0.67 | 0.34 | 0.62 | 0.72 | 0.62 | 0.66 |
| 2 | 119034265 | rs4849690 | 1.742 | -0.04 | 0.866 | 1.097 | 0.009 | -0.001 | -0.001 | 0 | 0.52 | 0.28 | 0.54 | 0.55 | 0.51 | 0.48 |
| 2 | 119038598 | rs1878526 | -1.154 | -0.308 | 1.139 | 1.155 | -0.008 | 0.019 | 0.019 | 0 | 0.25 | 0.35 | 0.16 | 0.35 | 0.18 | 0.28 |
| 2 | 119039841 | rs7595438 | -1.734 | -0.277 | 1.558 | 0.775 | 0.009 | 0.014 | 0.014 | 0 | 0.39 | 0.71 | 0.37 | 0.26 | 0.37 | 0.4 |
| 2 | 119040942 | rs17771321 | 0.308 | -0.272 | 1.469 | 0.699 | -0.023 | 0.059 | 0.059 | 0.38 | 0.18 | 0.08 | 0.26 | 0.08 | 0.26 | 0.22 |
| 2 | 119042079 | rs6542441 | -1.719 | -0.217 | 1.67 | 0.747 | 0 | -0.002 | -0.002 | 0 | 0.38 | 0.66 | 0.37 | 0.23 | 0.37 | 0.41 |
| 2 | 119042700 | rs13420052 | -0.693 | -0.582 | 2.062 | 1.191 | -0.019 | 0.118 | 0.118 | 0 | 0.1 | 0.34 | 0.06 | 0.17 | 0.04 | 0.13 |
| 2 | 119043209 | rs9308765 | -1.218 | -0.585 | 2.055 | 1.185 | -0.019 | 0.017 | 0.017 | 0 | 0.18 | 0.35 | 0.09 | 0.17 | 0.11 | 0.19 |
| 2 | 119043311 | rs10432590 | -0.712 | -0.556 | 2.094 | 1.076 | -0.019 | 0.089 | 0.089 | 0 | 0.1 | 0.34 | 0.06 | 0.14 | 0.04 | 0.13 |
| 2 | 119052905 | rs4564774 | -0.585 | -0.516 | 2.406 | 1.004 | 0.002 | -0.081 | -0.081 | 0 | 0.12 | 0.43 | 0.06 | 0 | 0.06 | 0.05 |
| 2 | 119056234 | rs10170466 | 2.175 | -0.643 | 2.303 | 1.134 | 0.02 | -0.004 | -0.004 | 0.892 | 0.61 | 0.26 | 0.65 | 0.58 | 0.73 | 0.65 |
| 2 | 119072376 | rs4309580 | 1.054 | -0.509 | 2.409 | 0.852 | 0.003 | -0.002 | -0.002 | 0 | 0.8 | 0.83 | 0.87 | 0.8 | 0.84 | 0.78 |
| 2 | 119083739 | rs7564201 | -1.911 | -0.2 | 2.989 | 0.825 | 0.047 | 0.06 | 0.06 | 1.12 | 0.08 | 0.33 | 0.07 | 0.06 | 0.06 | 0.07 |
| 2 | 119085908 | rs7562732 | -2.048 | -0.19 | 2.881 | 0.829 | 0.045 | 0.058 | 0.058 | 2.354 | 0.08 | 0.43 | 0.08 | 0.06 | 0.06 | 0.07 |
| 2 | 119090434 | rs6542449 | -2.069 | -0.233 | 2.404 | 0.785 | 0.007 | 0.023 | 0.023 | 5.217 | 0.25 | 0.82 | 0.26 | 0.19 | 0.17 | 0.2 |
| 2 | 119092344 | rs13426749 | -0.998 | -0.136 | 2.603 | 0.753 | -0.016 | -0.097 | -0.097 | 0.095 | 0.14 | 0.44 | 0.06 | 0 | 0.07 | 0.06 |
| 2 | 119099822 | rs4849694 | 0.618 | -0.204 | 2.403 | 0.942 | -0.007 | 0.007 | 0.007 | 0 | 0.83 | 0.81 | 0.87 | 0.79 | 0.86 | 0.8 |
| 2 | 119104699 | rs13400406 | -0.956 | -0.157 | 2.591 | 0.921 | 0.019 | -0.011 | -0.011 | 0.61 | 0.16 | 0.34 | 0.18 | 0.15 | 0.12 | 0.12 |

| | | | | | | | | | | | | | | | | |
|---|---|---|---|---|---|---|---|---|---|---|---|---|---|---|---|---|
| 2 | 119104977 | rs4276037 | 2.016 | -0.215 | 2.677 | 0.925 | 0.013 | -0.005 | -0.005 | 0.475 | 0.64 | 0.35 | 0.68 | 0.64 | 0.75 | 0.68 |
| 2 | 119109623 | rs12616543 | -1.322 | -0.175 | 2.701 | 0.909 | 0.023 | -0.019 | -0.019 | 0.047 | 0.17 | 0.39 | 0.18 | 0.15 | 0.12 | 0.12 |
| 2 | 119124673 | rs13383360 | -0.472 | -0.097 | 2.465 | 0.941 | 0.047 | 0.066 | 0.066 | 0.02 | 0.05 | 0.11 | 0.13 | 0.15 | 0.06 | 0.06 |
| 2 | 119129071 | rs12467546 | 1.92 | -0.168 | 2.226 | 0.904 | -0.001 | -0.001 | -0.001 | 0 | 0.65 | 0.41 | 0.68 | 0.65 | 0.75 | 0.68 |
| 2 | 119129814 | rs7576666 | 1.528 | -0.129 | 2.13 | 1.013 | -0.028 | 0.086 | 0.086 | 0 | 0.75 | 0.55 | 0.72 | 0.65 | 0.81 | 0.73 |
| 2 | 119134468 | rs4849699 | -1.09 | 0.02 | 2.156 | 0.802 | 0.07 | 0.076 | 0.076 | 2.013 | 0.07 | 0.28 | 0.15 | 0.12 | 0.05 | 0.06 |
| 2 | 119138025 | rs6542456 | 0.615 | -0.164 | 2.009 | 1.076 | 0.018 | -0.028 | -0.028 | 0 | 0.83 | 0.89 | 0.89 | 0.76 | 0.86 | 0.8 |
| 2 | 119141118 | rs4313994 | 1.75 | -0.209 | 2.284 | 0.938 | 0.03 | -0.033 | -0.033 | 1.648 | 0.65 | 0.23 | 0.69 | 0.64 | 0.74 | 0.68 |
| 2 | 119141494 | rs6753124 | 1.059 | -0.212 | 2.472 | 0.944 | -0.055 | 0.104 | 0.104 | 0 | 0.8 | 0.6 | 0.87 | 0.76 | 0.86 | 0.8 |
| 2 | 119147509 | rs6708595 | -0.897 | -0.104 | 1.918 | 0.834 | 0.015 | -0.031 | -0.031 | 0 | 0.16 | 0.35 | 0.18 | 0.12 | 0.12 | 0.13 |
| 2 | 119161638 | rs4073566 | 1.529 | -0.141 | 1.364 | 0.911 | 0.043 | -0.031 | -0.031 | 0 | 0.66 | 0.13 | 0.72 | 0.69 | 0.79 | 0.71 |
| 2 | 119174985 | rs6741265 | -1.654 | -0.259 | 2.056 | 0.866 | 0.006 | -0.002 | -0.002 | 0.566 | 0.31 | 0.68 | 0.17 | 0.22 | 0.21 | 0.26 |
| 2 | 119178877 | rs7605250 | -1.138 | -0.284 | 2.288 | 0.852 | -0.044 | 0.012 | 0.012 | 0 | 0.25 | 0.4 | 0.11 | 0.18 | 0.16 | 0.19 |
| 2 | 119182917 | rs6542461 | -1.706 | -0.254 | 1.722 | 0.828 | 0.027 | -0.006 | -0.006 | 0.065 | 0.32 | 0.73 | 0.18 | 0.22 | 0.21 | 0.26 |
| 2 | 119188451 | rs6718717 | -0.837 | 0.17 | 1.699 | 0.143 | -0.004 | -0.064 | -0.064 | 1.917 | 0.22 | 0.26 | 0.12 | 0 | 0.16 | 0.18 |
| 2 | 119190301 | rs6739488 | 1.373 | 0.674 | 1.034 | -0.339 | 0.01 | 0.056 | 0.056 | 0 | 0.52 | 0.57 | 0.48 | 0.78 | 0.45 | 0.57 |
| 2 | 119194992 | rs1370512 | 0.343 | 0.729 | 1.116 | -0.188 | -0.008 | 0.009 | 0.009 | 0 | 0.72 | 0.72 | 0.6 | 0.77 | 0.64 | 0.72 |
| 2 | 119198753 | rs17048101 | -1.013 | 0.552 | 1.386 | -0.275 | 0.02 | 0.044 | 0.044 | 0 | 0.06 | 0.24 | 0.07 | 0.06 | 0.06 | 0.07 |
| 2 | 119199259 | rs17048104 | -1.182 | 0.547 | 1.375 | -0.277 | 0.031 | 0.058 | 0.058 | 0 | 0.07 | 0.29 | 0.07 | 0.06 | 0.06 | 0.07 |
| 2 | 119204784 | rs10165720 | 0.162 | 0.587 | 0.593 | -0.688 | -0.009 | 0.034 | 0.034 | 0 | 0.7 | 0.76 | 0.59 | 0.77 | 0.65 | 0.72 |
| 2 | 119205841 | rs10192071 | -1.039 | 1.032 | 0.899 | -0.654 | -0.013 | -0.081 | -0.081 | 0 | 0.31 | 0.46 | 0.2 | 0.06 | 0.25 | 0.28 |
| 2 | 119218881 | rs1437690 | -1.726 | 1.866 | 0.486 | -0.919 | 0.03 | -0.064 | -0.064 | 0 | 0.34 | 0.54 | 0.34 | 0.18 | 0.39 | 0.38 |
| 2 | 119220906 | rs1965130 | 0.016 | 1.677 | 0.089 | -0.865 | -0.057 | 0.141 | 0.141 | 0 | 0.21 | 0.01 | 0.12 | 0 | 0.19 | 0.16 |
| 2 | 119224568 | rs17016813 | -0.85 | 1.494 | 0.415 | -0.644 | 0.011 | -0.001 | -0.001 | 0 | 0.12 | 0.22 | 0.09 | 0.1 | 0.13 | 0.11 |
| 2 | 119227798 | rs6542463 | -2 | 1.273 | 0.244 | -0.377 | 0.029 | 0.001 | 0.001 | 0 | 0.61 | 0.81 | 0.64 | 0.4 | 0.69 | 0.56 |
| 2 | 119231292 | rs6542464 | -1.283 | 1.432 | 0.987 | -0.221 | 0.035 | -0.086 | -0.086 | 0.984 | 0.16 | 0.49 | 0.11 | 0.01 | 0.14 | 0.08 |
| 2 | 119235336 | rs17838804 | 0.221 | 1.336 | 0.569 | -0.181 | -0.005 | -0.002 | -0.002 | 0 | 0.12 | 0.04 | 0.15 | 0.12 | 0.09 | 0.17 |
| 2 | 119236996 | rs17824540 | 0.483 | 1.362 | 0.568 | -0.191 | -0.004 | 0.036 | 0.036 | 0 | 0.08 | 0.02 | 0.05 | 0 | 0.08 | 0.03 |
| 2 | 119239524 | rs6542466 | -2.395 | 1.742 | 0.495 | -0.156 | 0.048 | 0.037 | 0.037 | 0 | 0.51 | 0.77 | 0.55 | 0.41 | 0.58 | 0.48 |
| 2 | 119240369 | rs17016802 | -0.466 | 1.746 | 0.534 | -0.156 | -0.001 | -0.008 | -0.008 | 0.077 | 0.1 | 0.15 | 0.06 | 0.06 | 0.09 | 0.12 |
| 2 | 119249559 | rs12988135 | 0.006 | 1.84 | 0.587 | -0.145 | -0.015 | -0.01 | -0.01 | 0 | 0.16 | 0.12 | 0.25 | 0.23 | 0.27 | 0.13 |
| 2 | 119250430 | rs1344896 | -2.036 | 1.735 | 0.783 | -0.219 | -0.002 | 0.001 | 0.001 | 0 | 0.38 | 0.69 | 0.4 | 0.29 | 0.46 | 0.28 |
| 2 | 119250862 | rs7558656 | -0.259 | 1.835 | 0.861 | -0.032 | -0.002 | -0.053 | -0.053 | 2.507 | 0.1 | 0.26 | 0.07 | 0 | 0.08 | 0.03 |
| 2 | 119253438 | rs934753 | -0.191 | 1.84 | 0.856 | -0.029 | -0.001 | -0.052 | -0.052 | 0 | 0.1 | 0.25 | 0.07 | 0 | 0.08 | 0.03 |
| 2 | 119255493 | rs7557274 | -1.086 | 1.799 | 0.442 | 0.038 | 0.053 | -0.026 | -0.026 | 0.022 | 0.28 | 0.53 | 0.34 | 0.23 | 0.37 | 0.16 |
| 2 | 119263092 | rs10496548 | -0.571 | 1.889 | -0.081 | 0.256 | 0.002 | 0.009 | 0.009 | 0 | 0.17 | 0.17 | 0.3 | 0.3 | 0.2 | 0.3 |
| 2 | 119270317 | rs934756 | -1.314 | 1.822 | 0.283 | 0.049 | 0.017 | 0.019 | 0.019 | 0 | 0.26 | 0.41 | 0.37 | 0.3 | 0.29 | 0.35 |
| 2 | 119271253 | rs17825502 | 0.433 | 1.889 | 0.092 | -0.07 | -0.012 | 0.031 | 0.031 | 0 | 0.08 | 0 | 0.06 | 0.01 | 0.08 | 0.03 |
| 2 | 119271830 | rs6704674 | -0.053 | 1.801 | 0.737 | -0.14 | -0.005 | -0.042 | -0.042 | 0 | 0.1 | 0.16 | 0.07 | 0 | 0.07 | 0.03 |
| 2 | 119274053 | rs11690936 | 0.121 | 1.914 | 0.584 | -0.256 | -0.05 | 0.018 | 0.018 | 0 | 0.14 | 0.05 | 0.12 | 0.12 | 0.25 | 0.1 |
| 2 | 119279230 | rs1000950 | -2.193 | 1.564 | 0.938 | -0.251 | 0.089 | 0.021 | 0.021 | 0.05 | 0.52 | 0.82 | 0.57 | 0.41 | 0.63 | 0.48 |
| 2 | 119279471 | rs12478798 | 0.546 | 1.782 | 0.698 | -0.025 | -0.015 | -0.016 | -0.016 | 0 | 0.07 | 0.04 | 0.13 | 0.16 | 0.12 | 0.14 |
| 2 | 119280794 | rs2008776 | -1.499 | 1.585 | 0.712 | -0.135 | -0.001 | -0.001 | -0.001 | 0 | 0.41 | 0.64 | 0.5 | 0.41 | 0.53 | 0.45 |
| 2 | 119288813 | rs17009143 | -0.362 | 1.364 | 0.053 | -0.106 | 0.011 | -0.038 | -0.038 | 0 | 0.07 | 0.19 | 0.07 | 0 | 0.08 | 0.05 |
| 2 | 119292114 | rs4849706 | -1.058 | 1.382 | -0.267 | -0.229 | -0.008 | 0.01 | 0.01 | 0 | 0.18 | 0.14 | 0.25 | 0.02 | 0.24 | 0.16 |
| 2 | 119296554 | rs7571800 | 0.004 | 0.695 | 0.439 | 0.027 | -0.002 | -0.012 | -0.012 | 0 | 0.32 | 0.25 | 0.45 | 0.37 | 0.34 | 0.29 |
| 2 | 119298887 | rs1437683 | -0.973 | 0.64 | 0.299 | -0.038 | 0.002 | -0.001 | -0.001 | 0 | 0.45 | 0.64 | 0.53 | 0.38 | 0.44 | 0.32 |
| 2 | 119312232 | rs6719023 | -0.954 | 0.612 | -0.026 | -0.359 | -0.001 | -0.003 | -0.003 | 1.047 | 0.66 | 0.68 | 0.73 | 0.77 | 0.74 | 0.53 |
| 2 | 119314299 | rs747725 | 0.036 | 0.673 | -0.059 | -0.445 | -0.026 | 0.023 | 0.023 | 0 | 0.13 | 0.1 | 0.19 | 0.02 | 0.21 | 0.11 |
| 2 | 119318751 | rs12994932 | -0.276 | 0.499 | 0.435 | -0.486 | 0.003 | 0.006 | 0.006 | 0.04 | 0.57 | 0.4 | 0.65 | 0.76 | 0.65 | 0.53 |
| 2 | 119320083 | rs2118379 | -1.335 | 0.527 | 0.309 | -0.452 | 0.013 | 0.041 | 0.041 | 0.179 | 0.69 | 0.83 | 0.73 | 0.77 | 0.74 | 0.58 |
| 2 | 119325355 | rs7581452 | 0.336 | 0.613 | 0.421 | -0.413 | 0.03 | -0.036 | -0.036 | 0 | 0.08 | 0.25 | 0.07 | 0 | 0.08 | 0.01 |
| 2 | 119341661 | rs11888583 | -1.738 | 0.461 | 0.156 | -0.454 | 0 | 0.037 | 0.037 | 2.316 | 0.74 | 0.83 | 0.74 | 0.83 | 0.77 | 0.62 |
| 2 | 119344750 | rs13017945 | 1.151 | 0.566 | 0.43 | -0.034 | 0.002 | 0.01 | 0.01 | 0 | 0.22 | 0.25 | 0.16 | 0.25 | 0.23 | 0.17 |
| 2 | 119349987 | rs10496550 | 1.307 | 0.634 | 0.234 | -0.175 | 0 | -0.009 | -0.009 | 2.01 | 0.06 | 0.1 | 0.05 | 0 | 0.08 | 0.01 |
| 2 | 119371196 | rs1451206 | -0.182 | 0.501 | 0.208 | -0.231 | 0.001 | 0.008 | 0.008 | 0 | 0.5 | 0.41 | 0.47 | 0.75 | 0.52 | 0.42 |
| 2 | 119377442 | rs563449 | -0.787 | 0.508 | -0.018 | -0.257 | -0.043 | -0.024 | -0.024 | 0 | 0.9 | 0.99 | 0.87 | 0.93 | 0.85 | 0.85 |
| 2 | 119379115 | rs17776428 | 0.867 | 0.562 | -0.171 | -0.469 | -0.004 | 0.017 | 0.017 | 0 | 0.09 | 0.04 | 0.07 | 0 | 0.11 | 0.05 |
| 2 | 119395691 | rs651477 | 1.439 | 0.538 | -0.085 | -0.367 | 0.003 | 0.042 | 0.042 | 0.843 | 0.32 | 0.23 | 0.33 | 0.18 | 0.25 | 0.42 |
| 2 | 119401093 | rs6736836 | 0.269 | 0.501 | -0.031 | 0.122 | 0.001 | 0 | 0 | 0 | 0.46 | 0.36 | 0.5 | 0.52 | 0.48 | 0.36 |
| 2 | 119414453 | rs563697 | 1.462 | 0.544 | -0.139 | 0.101 | 0.003 | 0.05 | 0.05 | 1.076 | 0.31 | 0.2 | 0.27 | 0.17 | 0.25 | 0.42 |
| 2 | 119424298 | rs11891602 | 1.323 | 0.539 | -0.179 | 0.162 | 0.001 | -0.01 | -0.01 | 0.554 | 0.06 | 0.09 | 0.05 | 0 | 0.08 | 0.01 |
| 2 | 119435628 | rs10204148 | 0.625 | 0.634 | -0.089 | 0.404 | 0.001 | 0.005 | 0.005 | 0 | 0.26 | 0.27 | 0.27 | 0.29 | 0.27 | 0.18 |
| 2 | 119444229 | rs332105 | -0.079 | 0.529 | -0.189 | 0.129 | 0 | 0.017 | 0.017 | 0 | 0.53 | 0.56 | 0.58 | 0.71 | 0.56 | 0.38 |
| 2 | 119446030 | rs332104 | 0.23 | 0.69 | -0.424 | 0.25 | -0.045 | -0.06 | -0.06 | 0 | 0.86 | 0.99 | 0.85 | 0.81 | 0.82 | 0.74 |
| 2 | 119449172 | rs332103 | 0.526 | 0.79 | -0.345 | 0.205 | -0.004 | -0.003 | -0.003 | 0 | 0.83 | 0.84 | 0.81 | 0.8 | 0.71 | 0.72 |
| 2 | 119449992 | rs45140 | 0.432 | 0.841 | -0.299 | 0.23 | 0.031 | 0.009 | 0.009 | 2.198 | 0.86 | 0.82 | 0.82 | 0.8 | 0.72 | 0.75 |
| 2 | 119450163 | rs512681 | 1.276 | 0.291 | -0.424 | -0.009 | 0.017 | 0.014 | 0.014 | 1.133 | 0.3 | 0.26 | 0.23 | 0.09 | 0.15 | 0.34 |
| 2 | 119457212 | rs332657 | -0.27 | 0.412 | -0.29 | -0.12 | 0.045 | 0.065 | 0.065 | 0 | 0.53 | 0.63 | 0.69 | 0.84 | 0.7 | 0.57 |

| | | | | | | | | | | | | | | | | |
|---|---|---|---|---|---|---|---|---|---|---|---|---|---|---|---|---|
| 2 | 119465312 | rs458340 | -0.787 | 0.339 | -0.326 | -0.154 | 0.024 | 0.024 | 0.024 | 0 | 0.59 | 0.64 | 0.74 | 0.87 | 0.78 | 0.58 |
| 2 | 119472709 | rs11676636 | 1.141 | 0.781 | -0.262 | 0.209 | 0.029 | 0.013 | 0.013 | 0 | 0.07 | 0.09 | 0.15 | 0.19 | 0.2 | 0.12 |
| 2 | 119474785 | rs332088 | -0.944 | 0.578 | -0.206 | 0.136 | 0.073 | 0.09 | 0.09 | 0 | 0.61 | 0.78 | 0.75 | 0.87 | 0.78 | 0.58 |
| 2 | 119475432 | rs332089 | 0.159 | 0.574 | -0.092 | 0.16 | 0 | -0.004 | -0.004 | 1.159 | 0.45 | 0.46 | 0.53 | 0.66 | 0.5 | 0.46 |
| 2 | 119476578 | rs332092 | 0.158 | 0.577 | -0.115 | 0.162 | -0.001 | 0.007 | 0.007 | 0 | 0.46 | 0.55 | 0.54 | 0.66 | 0.5 | 0.46 |
| 2 | 119481109 | rs332100 | 0.693 | 0.373 | 0.065 | 0.219 | -0.007 | -0.018 | -0.018 | 0 | 0.4 | 0.25 | 0.49 | 0.52 | 0.48 | 0.36 |
| 2 | 119482773 | rs332705 | -0.805 | 0.324 | 0.041 | 0.136 | 0.019 | 0.02 | 0.02 | 0 | 0.61 | 0.66 | 0.75 | 0.87 | 0.79 | 0.6 |
| 2 | 119486759 | rs332702 | -1.051 | 0.332 | 0.059 | 0.132 | 0.058 | 0.075 | 0.075 | 0 | 0.63 | 0.78 | 0.76 | 0.87 | 0.79 | 0.6 |
| 2 | 119497419 | rs332668 | -2.302 | -0.02 | -0.437 | -0.153 | 0.051 | 0.083 | 0.083 | 0.169 | 0.76 | 0.95 | 0.81 | 0.91 | 0.87 | 0.69 |
| 2 | 119498667 | rs332665 | 0.046 | 0.237 | -0.177 | 0.009 | 0.004 | 0.023 | 0.023 | 0 | 0.49 | 0.59 | 0.57 | 0.84 | 0.51 | 0.48 |
| 2 | 119505210 | rs332080 | 0.81 | 0.317 | -0.521 | -0.053 | -0.008 | -0.001 | -0.001 | 0 | 0.37 | 0.45 | 0.47 | 0.63 | 0.37 | 0.4 |
| 2 | 119507984 | rs332073 | 0.11 | 0.42 | -0.574 | -0.054 | -0.002 | 0.013 | 0.013 | 0 | 0.48 | 0.51 | 0.54 | 0.66 | 0.49 | 0.43 |
| 2 | 119511582 | rs332683 | 0.272 | 0.645 | -0.979 | 0.017 | -0.008 | 0.003 | 0.003 | 0 | 0.84 | 0.83 | 0.71 | 0.85 | 0.67 | 0.78 |
| 2 | 119514587 | rs332675 | 2.18 | 0.267 | -1.214 | 0.347 | 0.001 | -0.013 | -0.013 | 0 | 0.15 | 0.1 | 0.26 | 0.22 | 0.15 | 0.2 |
| 2 | 119520620 | rs17006295 | 0.314 | 0.1 | -0.872 | 0.438 | 0.076 | 0.068 | 0.068 | 0 | 0.36 | 0.2 | 0.15 | 0.16 | 0.18 | 0.36 |
| 2 | 119522156 | rs4848521 | 0.737 | 0.881 | -0.471 | 0.641 | 0.007 | 0.013 | 0.013 | 0 | 0.49 | 0.29 | 0.51 | 0.56 | 0.51 | 0.42 |
| 2 | 119522687 | rs10177490 | -0.761 | 0.928 | -0.471 | 0.764 | -0.007 | 0.002 | 0.002 | 0 | 0.46 | 0.65 | 0.39 | 0.42 | 0.43 | 0.52 |
| 2 | 119526970 | rs11679318 | 0.215 | 1.772 | -0.414 | 0.729 | 0.008 | -0.004 | -0.004 | 0 | 0.07 | 0.1 | 0.16 | 0.16 | 0.2 | 0.2 |
| 2 | 119533754 | rs10196575 | -0.223 | 0.894 | -0.172 | 0.171 | 0.047 | -0.03 | -0.03 | 0.122 | 0.12 | 0.37 | 0.11 | 0.03 | 0.12 | 0.03 |
| 2 | 119536884 | rs1551133 | 1.131 | -0.161 | -1.096 | 0.172 | 0.072 | 0.014 | 0.014 | 0 | 0.2 | 0.04 | 0.26 | 0.09 | 0.07 | 0.19 |
| 2 | 119539832 | rs1527417 | 0.015 | 0.495 | -0.875 | -0.048 | 0.002 | 0.022 | 0.022 | 0 | 0.48 | 0.54 | 0.45 | 0.22 | 0.52 | 0.41 |
| 2 | 119540832 | rs1527418 | -0.615 | 0.41 | -1.035 | 0.307 | -0.011 | 0.004 | 0.004 | 0 | 0.66 | 0.74 | 0.57 | 0.71 | 0.66 | 0.53 |
| 2 | 119544022 | rs4849721 | 1.682 | 0.612 | -1.289 | 0.089 | -0.056 | 0.026 | 0.026 | 0.086 | 0.11 | 0.01 | 0.14 | 0.02 | 0.19 | 0.04 |
| 2 | 119547211 | rs10221632 | -0.008 | 0.679 | -1.203 | 0.037 | 0.009 | 0.103 | 0.103 | 0 | 0.5 | 0.69 | 0.45 | 0.22 | 0.52 | 0.4 |
| 2 | 119549440 | rs17006621 | 1.258 | 0.695 | -1.604 | 0.097 | 0.011 | -0.124 | -0.124 | 0 | 0.16 | 0.02 | 0.12 | 0.49 | 0.14 | 0.12 |
| 2 | 119551291 | rs1527420 | 1.535 | 0.751 | -1.442 | 0.073 | -0.056 | 0.026 | 0.026 | 0.086 | 0.11 | 0.01 | 0.14 | 0.02 | 0.19 | 0.04 |
| 2 | 119560844 | rs6749677 | -0.7 | 0.746 | -1.412 | 0.083 | -0.015 | 0.003 | 0.003 | 0 | 0.67 | 0.78 | 0.58 | 0.72 | 0.66 | 0.53 |
| 2 | 119561237 | rs12988660 | 0.035 | 0.707 | -1.374 | 0.186 | 0.015 | 0.006 | 0.006 | 0 | 0.49 | 0.51 | 0.4 | 0.59 | 0.43 | 0.45 |
| 2 | 119566431 | rs12993402 | 0.995 | 0.579 | -1.702 | -0.418 | 0 | -0.047 | -0.047 | 0.011 | 0.3 | 0.62 | 0.2 | 0.04 | 0.25 | 0.25 |
| 2 | 119566559 | rs10191670 | -0.573 | 0.56 | -2.039 | -0.186 | -0.014 | -0.019 | -0.019 | 0 | 0.61 | 0.67 | 0.46 | 0.51 | 0.58 | 0.44 |
| 2 | 119567815 | rs11694004 | -0.425 | 0.743 | -2.333 | -0.155 | 0.068 | 0.104 | 0.104 | 0 | 0.55 | 0.73 | 0.53 | 0.18 | 0.67 | 0.55 |
| 2 | 119575277 | rs972143 | 1.016 | 0.7 | -1.867 | -0.171 | 0 | 0.013 | 0.013 | 0 | 0.31 | 0.27 | 0.2 | 0.11 | 0.29 | 0.28 |
| 2 | 119577465 | rs6718150 | -1.417 | 0.654 | -1.805 | -0.142 | 0.014 | -0.005 | -0.005 | 0 | 0.73 | 0.75 | 0.65 | 0.71 | 0.82 | 0.7 |
| 2 | 119579558 | rs11687120 | -0.303 | 0.908 | -1.604 | -0.181 | 0.091 | -0.023 | -0.023 | 4.659 | 0.26 | 0.47 | 0.34 | 0.15 | 0.42 | 0.3 |
| 2 | 119580473 | rs11684772 | 0.65 | 0.812 | -1.302 | -0.049 | 0.164 | 0.101 | 0.101 | 0.804 | 0.11 | 0.48 | 0.18 | 0.12 | 0.2 | 0.24 |
| 2 | 119580634 | rs17177826 | 0.977 | 0.034 | -1.694 | -0.271 | 0.009 | 0.089 | 0.089 | 0 | 0.26 | 0.12 | 0.22 | 0.04 | 0.2 | 0.23 |
| 2 | 119581251 | rs10186271 | -0.262 | 0.013 | -1.355 | 0.486 | 0.019 | -0.018 | -0.018 | 0 | 0.71 | 0.76 | 0.69 | 0.64 | 0.8 | 0.67 |
| 2 | 119582484 | rs4849726 | -0.206 | 0.668 | -1.054 | 0.38 | 0.001 | -0.002 | -0.002 | 0 | 0.17 | 0.19 | 0.17 | 0.12 | 0.26 | 0.08 |
| 2 | 119582505 | rs17006964 | 0.215 | -0.123 | -1.782 | 0.706 | 0.052 | -0.138 | -0.138 | 0 | 0.27 | 0.01 | 0.11 | 0.44 | 0.11 | 0.2 |
| 2 | 119587710 | rs4848526 | 0.361 | 0.511 | -0.809 | 0.584 | 0 | 0 | 0 | 0.699 | 0.6 | 0.49 | 0.66 | 0.38 | 0.56 | 0.72 |
| 2 | 119592977 | rs7567463 | -0.12 | 0.496 | -0.835 | 0.643 | -0.016 | 0.019 | 0.019 | 0 | 0.72 | 0.46 | 0.83 | 0.3 | 0.82 | 0.75 |
| 2 | 119598854 | rs13023152 | 1.122 | 0.514 | -1.09 | 0.346 | 0.088 | 0.055 | 0.055 | 0.537 | 0.41 | 0.21 | 0.33 | 0.2 | 0.24 | 0.35 |
| 2 | 119608582 | rs3754855 | -0.451 | 0.635 | -0.78 | 0.488 | 0.003 | 0.017 | 0.017 | 0 | 0.38 | 0.44 | 0.28 | 0.61 | 0.4 | 0.25 |
| 2 | 119610043 | rs6719822 | 1.09 | 0.581 | -0.187 | 0.561 | 0.001 | 0.002 | 0.002 | 0 | 0.22 | 0.16 | 0.08 | 0.49 | 0.13 | 0.21 |
| 2 | 119615001 | rs17007326 | 1.212 | 0.641 | -0.373 | 0.438 | -0.017 | -0.159 | -0.159 | 0 | 0.21 | 0.01 | 0.07 | 0.52 | 0.13 | 0.22 |
| 2 | 119617076 | rs6726762 | 1.105 | 0.576 | -0.408 | -0.039 | 0.067 | 0.065 | 0.065 | 0 | 0.39 | 0.22 | 0.3 | 0.17 | 0.21 | 0.34 |
| 2 | 119618696 | rs4849728 | -0.074 | 1.718 | -0.043 | -0.061 | 0.043 | 0.004 | 0.004 | 0.564 | 0.78 | 0.64 | 0.6 | 0.78 | 0.63 | 0.61 |
| 2 | 119619074 | rs7558638 | 0.308 | 0.819 | 0.179 | 0.147 | 0 | 0.02 | 0.02 | 0 | 0.63 | 0.63 | 0.4 | 0.78 | 0.35 | 0.57 |
| 2 | 119636505 | rs7602200 | 0.574 | 1.09 | -0.194 | 0.738 | 0.002 | 0 | 0 | 0 | 0.7 | 0.37 | 0.58 | 0.73 | 0.6 | 0.59 |
| 2 | 119639040 | rs6759783 | -0.173 | 0.182 | -0.298 | -0.041 | 0.02 | 0.062 | 0.062 | 0.412 | 0.45 | 0.38 | 0.35 | 0.2 | 0.39 | 0.42 |
| 2 | 119639054 | rs7557460 | 0.184 | -0.058 | 0.135 | 0.224 | -0.006 | 0.076 | 0.076 | 0 | 0.17 | 0.28 | 0.17 | 0.59 | 0.11 | 0.32 |
| 2 | 119640441 | rs1438846 | -1.484 | -0.117 | 0.101 | 0.026 | -0.064 | 0.11 | 0.11 | 3.182 | 0.69 | 0.88 | 0.67 | 0.91 | 0.55 | 0.88 |
| 2 | 119642078 | rs7565844 | 0.261 | -0.142 | 0.273 | 0.419 | 0.021 | 0.163 | 0.163 | 0 | 0.16 | 0.44 | 0.18 | 0.57 | 0.1 | 0.26 |
| 2 | 119642949 | rs10169004 | 0.262 | -0.138 | 0.261 | 0.421 | 0.024 | 0.172 | 0.172 | 0 | 0.16 | 0.45 | 0.18 | 0.57 | 0.1 | 0.26 |
| 2 | 119645055 | rs10200064 | 0.407 | -0.122 | 0.377 | 0.426 | 0.019 | 0.153 | 0.153 | 0 | 0.15 | 0.39 | 0.17 | 0.57 | 0.1 | 0.26 |
| 2 | 119650598 | rs6742106 | 0.354 | -0.275 | 0.074 | 0.24 | 0.009 | 0.029 | 0.029 | 0.004 | 0.41 | 0.38 | 0.33 | 0.2 | 0.35 | 0.4 |
| 2 | 119661546 | rs4849731 | -0.661 | -0.144 | 0.597 | 0.853 | -0.004 | 0.045 | 0.045 | 0 | 0.24 | 0.3 | 0.32 | 0.4 | 0.2 | 0.27 |
| 2 | 119663173 | rs4849732 | -0.517 | -0.169 | 0.797 | 0.964 | -0.008 | 0.041 | 0.041 | 0.111 | 0.1 | 0.17 | 0.14 | 0.14 | 0.08 | 0.17 |
| 2 | 119664339 | rs4849733 | -1.298 | -0.222 | 0.715 | 0.941 | -0.014 | -0.021 | -0.021 | 0 | 0.27 | 0.54 | 0.41 | 0.72 | 0.21 | 0.44 |
| 2 | 119664399 | rs17008480 | 0.545 | -0.15 | 0.542 | 0.87 | -0.008 | -0.039 | -0.039 | 1.076 | 0.14 | 0.03 | 0.18 | 0.4 | 0.12 | 0.16 |
| 2 | 119665911 | rs1898706 | -0.337 | -0.193 | 0.698 | 0.477 | -0.001 | -0.02 | -0.02 | 5.419 | 0.1 | 0.09 | 0.12 | 0 | 0.08 | 0.08 |
| 2 | 119668520 | rs920331 | 0.389 | -0.182 | 0.551 | 0.484 | -0.001 | 0.02 | 0.02 | 0 | 0.06 | 0.04 | 0.08 | 0.26 | 0.04 | 0.21 |
| 2 | 119671090 | rs12615639 | 0.27 | -0.319 | 1.09 | 0.433 | 0.048 | 0.298 | 0.298 | 0 | 0.14 | 0.39 | 0.18 | 0.47 | 0.14 | 0.13 |
| 2 | 119675587 | rs11691838 | 1.299 | -0.426 | 0.672 | 0.107 | -0.067 | 0.062 | 0.062 | 3.335 | 0.34 | 0.1 | 0.27 | 0.18 | 0.44 | 0.23 |
| 2 | 119676029 | rs17180039 | 1.32 | -0.425 | 0.671 | 0.107 | -0.07 | 0.056 | 0.056 | 3.405 | 0.34 | 0.08 | 0.29 | 0.18 | 0.44 | 0.23 |
| 2 | 119694552 | rs2119110 | -0.226 | -0.526 | 0.514 | 0.179 | 0.003 | 0.007 | 0.007 | 0 | 0.59 | 0.6 | 0.6 | 0.65 | 0.65 | 0.44 |
| 2 | 119698574 | rs3806496 | 0.558 | -0.511 | 0.568 | 0.232 | 0.003 | -0.004 | -0.004 | 0 | 0.47 | 0.54 | 0.48 | 0.52 | 0.57 | 0.22 |
| 2 | 119710400 | rs1985119 | 0.365 | -0.471 | 0.357 | 0.344 | -0.005 | -0.013 | -0.013 | 0.01 | 0.12 | 0.07 | 0.19 | 0.24 | 0.13 | 0.28 |
| 2 | 119718028 | rs4491733 | -1.624 | -0.44 | 0.745 | 0.138 | -0.002 | 0.01 | 0.01 | 0.231 | 0.3 | 0.69 | 0.28 | 0.47 | 0.24 | 0.63 |

| | | | | | | | | | | | | | | | | |
|---|---|---|---|---|---|---|---|---|---|---|---|---|---|---|---|---|
| 2 | 119724969 | rs7559955 | -1.289 | -0.423 | 0.948 | 0.001 | -0.013 | 0.032 | 0.032 | 0 | 0.17 | 0.36 | 0.09 | 0.22 | 0.11 | 0.28 |
| 2 | 119725151 | rs17009716 | -1.289 | -0.423 | 0.948 | 0.001 | -0.013 | 0.032 | 0.032 | 0 | 0.17 | 0.36 | 0.09 | 0.23 | 0.11 | 0.35 |
| 2 | 119731188 | rs4849743 | 0.532 | -0.842 | 0.748 | 0.069 | -0.001 | -0.002 | -0.002 | 0 | 0.13 | 0.08 | 0.19 | 0.21 | 0.13 | 0.3 |
| 2 | 119732042 | rs11693199 | 1.873 | -0.845 | 0.441 | -0.15 | -0.114 | 0.159 | 0.159 | 3.569 | 0.29 | 0.04 | 0.28 | 0.07 | 0.44 | 0.08 |
| 2 | 119732948 | rs1371562 | 1.421 | -0.884 | 0.573 | -0.135 | -0.064 | 0.107 | 0.107 | 1.39 | 0.33 | 0.1 | 0.29 | 0.17 | 0.46 | 0.21 |
| 2 | 119735251 | rs2278590 | -0.673 | -0.991 | 0.923 | -0.209 | -0.013 | 0.092 | 0.092 | 0 | 0.1 | 0.35 | 0.05 | 0.12 | 0.04 | 0.19 |
| 2 | 119740290 | rs7599352 | 0.891 | -0.967 | -0.178 | -0.308 | 0.03 | 0.034 | 0.034 | 0.671 | 0.46 | 0.12 | 0.49 | 0.66 | 0.62 | 0.35 |
| 2 | 119742584 | rs2011839 | 1.293 | -0.625 | -0.13 | -0.036 | -0.1 | 0.195 | 0.195 | 4.269 | 0.33 | 0.05 | 0.29 | 0.05 | 0.45 | 0.11 |
| 2 | 119742865 | rs3731612 | -1.153 | -0.193 | 0.535 | 0.418 | -0.01 | 0.065 | 0.065 | 0.826 | 0.22 | 0.62 | 0.22 | 0.34 | 0.14 | 0.45 |
| 2 | 119747502 | rs979387 | -0.574 | -0.652 | 0.423 | 0.46 | -0.026 | -0.025 | -0.025 | 0 | 0.22 | 0.54 | 0.32 | 0.12 | 0.16 | 0.28 |
| 2 | 119749611 | rs4848530 | 0.992 | -0.578 | -0.441 | 0.58 | -0.113 | 0.042 | 0.042 | 0 | 0.37 | 0.1 | 0.31 | 0.28 | 0.5 | 0.33 |
| 2 | 119749858 | rs3765035 | -0.137 | -0.811 | -0.136 | 0.663 | -0.007 | 0 | 0 | 0 | 0.56 | 0.65 | 0.62 | 0.4 | 0.68 | 0.57 |
| 2 | 119751867 | rs11678719 | -0.13 | -0.646 | 0.022 | 0.65 | -0.012 | 0.003 | 0.003 | 1.84 | 0.18 | 0.46 | 0.29 | 0.11 | 0.17 | 0.23 |
| 2 | 119765899 | rs6749056 | -0.328 | -0.697 | -0.339 | 0.719 | 0.017 | -0.019 | -0.019 | 0 | 0.61 | 0.78 | 0.64 | 0.45 | 0.7 | 0.68 |
| 2 | 119768536 | rs4848532 | 0.311 | -0.37 | -0.559 | 0.589 | -0.088 | 0.04 | 0.04 | 0 | 0.38 | 0.09 | 0.33 | 0.31 | 0.51 | 0.33 |
| 2 | 119773399 | rs6712180 | -0.437 | -0.129 | -0.649 | 1.258 | 0.002 | -0.003 | -0.003 | 0 | 0.23 | 0.22 | 0.28 | 0.46 | 0.16 | 0.26 |
| 2 | 119783031 | rs13391263 | -0.625 | -0.205 | -0.296 | 0.705 | -0.019 | -0.026 | -0.026 | 0 | 0.21 | 0.52 | 0.22 | 0.14 | 0.18 | 0.24 |
| 2 | 119783186 | rs13403020 | -0.874 | -0.769 | -1.424 | 0.635 | -0.04 | 0.008 | 0.008 | 0 | 0.63 | 0.87 | 0.62 | 0.67 | 0.48 | 0.64 |
| 2 | 119784193 | rs729735 | -0.712 | -1.019 | -1.482 | 0.19 | 0.065 | 0.166 | 0.166 | 0 | 0.5 | 0.79 | 0.63 | 0.24 | 0.41 | 0.4 |
| 2 | 119793795 | rs2422460 | -0.345 | -1.028 | -0.203 | 0.186 | -0.001 | -0.013 | -0.013 | 0 | 0.41 | 0.38 | 0.59 | 0.24 | 0.38 | 0.37 |
| 2 | 119810661 | rs2083384 | -0.983 | -1.076 | -0.338 | 0.677 | 0.025 | 0.044 | 0.044 | 0 | 0.52 | 0.75 | 0.66 | 0.63 | 0.42 | 0.48 |
| 2 | 119811352 | rs12465239 | 0.212 | -0.968 | -0.508 | -0.184 | 0 | -0.01 | -0.01 | 0 | 0.29 | 0.28 | 0.44 | 0.12 | 0.27 | 0.19 |
| 2 | 119813161 | rs13411411 | 0.442 | -0.576 | -0.561 | 0.07 | -0.001 | 0.018 | 0.018 | 0 | 0.13 | 0.16 | 0.15 | 0.22 | 0.12 | 0.19 |
| 2 | 119814585 | rs12711919 | -0.501 | -0.7 | -0.827 | -0.607 | 0.002 | -0.014 | -0.014 | 4.973 | 0.37 | 0.64 | 0.48 | 0.13 | 0.29 | 0.22 |
| 2 | 119819515 | rs12472071 | 0.149 | -0.224 | -1.456 | -0.855 | -0.037 | 0.234 | 0.234 | 0 | 0.42 | 0.11 | 0.38 | 0.05 | 0.48 | 0.43 |
| 2 | 119823534 | rs10177450 | -0.24 | -0.763 | -0.17 | -1.009 | -0.038 | -0.052 | -0.052 | 1.644 | 0.1 | 0.4 | 0.03 | 0 | 0.02 | 0.04 |
| 2 | 119826093 | rs13023588 | 0.683 | -0.828 | -0.109 | -0.477 | -0.029 | 0.007 | 0.007 | 0 | 0.74 | 0.34 | 0.8 | 0.77 | 0.87 | 0.78 |
| 2 | 119827462 | rs1439950 | -0.441 | -0.801 | 0.002 | -0.462 | -0.004 | 0.024 | 0.024 | 0 | 0.85 | 0.81 | 0.84 | 0.77 | 0.89 | 0.82 |
| 2 | 119827616 | rs1439951 | 0.383 | -0.644 | -0.158 | -0.053 | -0.071 | -0.076 | -0.076 | 0 | 0.33 | 0.08 | 0.35 | 0.49 | 0.36 | 0.34 |
| 2 | 119833296 | rs330770 | -1.03 | -0.666 | 0.258 | -0.088 | 0 | -0.012 | -0.012 | 0.142 | 0.54 | 0.59 | 0.59 | 0.94 | 0.49 | 0.59 |
| 2 | 119836043 | rs330773 | 0.575 | -0.634 | 0.172 | -0.057 | 0.001 | 0.044 | 0.044 | 0 | 0.13 | 0.22 | 0.16 | 0.23 | 0.11 | 0.18 |
| 2 | 119840084 | rs893769 | 0.598 | -0.671 | 0.194 | -0.068 | 0.003 | 0.098 | 0.098 | 0 | 0.48 | 0.21 | 0.45 | 0.31 | 0.52 | 0.48 |
| 2 | 119844557 | rs6759463 | 0.605 | -1.025 | 0.003 | -0.353 | 0.001 | 0.05 | 0.05 | 0.918 | 0.46 | 0.31 | 0.33 | 0.31 | 0.49 | 0.39 |
| 2 | 119845034 | rs11674178 | 1.209 | -0.667 | -0.359 | -0.197 | -0.084 | -0.101 | -0.101 | 0.274 | 0.24 | 0.06 | 0.32 | 0.46 | 0.33 | 0.29 |
| 2 | 119861103 | rs330758 | -0.259 | -0.887 | 0.462 | -0.038 | -0.013 | 0.013 | 0.013 | 0 | 0.23 | 0.37 | 0.33 | 0.22 | 0.18 | 0.3 |
| 2 | 119864163 | rs330763 | 1.296 | -0.69 | 0.697 | -0.152 | 0.002 | 0.141 | 0.141 | 0 | 0.92 | 0.73 | 0.92 | 0.75 | 0.97 | 0.93 |
| 2 | 119873176 | rs7603104 | -0.703 | -0.621 | 0.33 | -0.24 | -0.016 | -0.044 | -0.044 | 0.854 | 0.1 | 0.14 | 0.03 | 0 | 0.03 | 0.05 |
| 2 | 119874129 | rs4849753 | 0.23 | -0.635 | 0.23 | -0.5 | 0 | -0.003 | -0.003 | 0 | 0.42 | 0.37 | 0.29 | 0.31 | 0.42 | 0.33 |
| 2 | 119874248 | rs4849754 | -0.037 | -0.521 | 0.1 | -0.399 | -0.004 | -0.015 | -0.015 | 0.512 | 0.47 | 0.45 | 0.29 | 0.31 | 0.43 | 0.37 |
| 2 | 119879588 | rs10496555 | 0.229 | -0.792 | -0.166 | -0.685 | 0.005 | -0.012 | -0.012 | 0 | 0.33 | 0.27 | 0.26 | 0.05 | 0.26 | 0.34 |
| 2 | 119879640 | rs11674898 | 0.764 | -0.137 | -0.638 | -0.694 | -0.082 | -0.025 | -0.025 | 0.239 | 0.3 | 0.1 | 0.31 | 0.27 | 0.41 | 0.31 |
| 2 | 119880803 | rs12469532 | 0.11 | -0.37 | -0.794 | -0.486 | -0.014 | -0.01 | -0.01 | 0 | 0.57 | 0.28 | 0.59 | 0.48 | 0.57 | 0.61 |
| 2 | 119883331 | rs7570552 | 0.212 | -0.244 | -0.205 | -0.396 | -0.001 | 0.01 | 0.01 | 0 | 0.32 | 0.43 | 0.42 | 0.7 | 0.33 | 0.33 |
| 2 | 119886918 | rs9636365 | 0.037 | -0.243 | -0.339 | -0.395 | -0.004 | 0.007 | 0.007 | 0 | 0.33 | 0.58 | 0.42 | 0.7 | 0.33 | 0.32 |
| 2 | 119887435 | rs10496557 | 0.756 | -0.015 | -0.284 | -0.474 | -0.018 | -0.034 | -0.034 | 0 | 0.27 | 0.14 | 0.35 | 0.46 | 0.33 | 0.27 |
| 2 | 119893632 | rs17012263 | 0.406 | -0.424 | -0.225 | -0.913 | 0 | -0.035 | -0.035 | 0.985 | 0.32 | 0.27 | 0.25 | 0.03 | 0.24 | 0.33 |
| 2 | 119894965 | rs7557802 | -0.163 | 0.069 | -0.398 | -0.79 | 0.002 | -0.011 | -0.011 | 0 | 0.19 | 0.18 | 0.21 | 0.1 | 0.2 | 0.25 |
| 2 | 119896763 | rs6542505 | -0.753 | 0.189 | -0.606 | -1.212 | 0.002 | 0.009 | 0.009 | 0 | 0.76 | 0.8 | 0.81 | 0.94 | 0.74 | 0.82 |
| 2 | 119896848 | rs6728751 | 0.15 | 0.659 | -0.533 | -1.356 | 0.009 | -0.04 | -0.04 | 0 | 0.28 | 0.26 | 0.36 | 0.06 | 0.31 | 0.29 |
| 2 | 119898769 | rs12711922 | 1.166 | 0.061 | -0.308 | -0.866 | -0.006 | 0.024 | 0.024 | 0 | 0.12 | 0.15 | 0.14 | 0.22 | 0.09 | 0.15 |
| 2 | 119899119 | rs12711923 | 0.972 | 0.039 | -0.254 | -0.877 | -0.011 | 0.039 | 0.039 | 0 | 0.13 | 0.2 | 0.14 | 0.22 | 0.09 | 0.15 |
| 2 | 119899176 | rs13017443 | 1.399 | -0.131 | -0.369 | -1.002 | -0.044 | -0.067 | -0.067 | 0.143 | 0.31 | 0.09 | 0.28 | 0.66 | 0.34 | 0.36 |
| 2 | 119899933 | rs10205049 | -0.286 | 0.004 | -0.212 | -1.366 | 0.004 | 0.021 | 0.021 | 0.152 | 0.49 | 0.33 | 0.53 | 0.12 | 0.56 | 0.47 |
| 2 | 119900093 | rs5018771 | 0.178 | 0.06 | -0.28 | -1.317 | 0.004 | 0.013 | 0.013 | 0 | 0.47 | 0.49 | 0.45 | 0.88 | 0.43 | 0.53 |
| 2 | 119905653 | rs10207133 | 0.876 | 0.158 | -0.415 | -1.353 | -0.034 | 0.061 | 0.061 | 0 | 0.24 | 0.13 | 0.25 | 0.07 | 0.33 | 0.18 |
| 2 | 119908054 | rs4849760 | -0.924 | -0.645 | -0.241 | -1.371 | -0.022 | -0.036 | -0.036 | 0 | 0.91 | 0.87 | 0.95 | 1 | 0.96 | 0.98 |
| 2 | 119910314 | rs745888 | 1.76 | -0.864 | -0.835 | -1.124 | -0.065 | -0.129 | -0.129 | 0 | 0.25 | 0.04 | 0.2 | 0.61 | 0.31 | 0.32 |
| 2 | 119911161 | rs970013 | -0.414 | -0.829 | -0.438 | -1.095 | -0.015 | -0.008 | -0.008 | 1.414 | 0.58 | 0.48 | 0.71 | 0.35 | 0.64 | 0.57 |
| 2 | 119912814 | rs6542507 | -0.296 | -0.792 | -0.382 | -1.089 | 0.001 | 0 | 0 | 0.309 | 0.56 | 0.4 | 0.71 | 0.35 | 0.62 | 0.55 |
| 2 | 119914101 | rs6758051 | -0.444 | -0.813 | -0.473 | -1.017 | 0.001 | 0 | 0 | 1.187 | 0.6 | 0.41 | 0.73 | 0.34 | 0.67 | 0.57 |
| 2 | 119920455 | rs1867750 | 0.238 | -0.616 | -0.635 | -1.072 | -0.003 | -0.006 | -0.006 | 0 | 0.31 | 0.23 | 0.37 | 0.3 | 0.26 | 0.35 |
| 2 | 119926403 | rs7564432 | 1.262 | -0.689 | -0.691 | -1.233 | -0.004 | -0.012 | -0.012 | 0.227 | 0.16 | 0.1 | 0.23 | 0 | 0.15 | 0.15 |
| 2 | 119928438 | rs11680207 | 0.203 | -0.497 | -0.855 | -1.151 | 0 | 0.008 | 0.008 | 3.475 | 0.45 | 0.41 | 0.52 | 0.34 | 0.56 | 0.43 |
| 2 | 119935802 | rs935361 | 1.155 | -0.315 | -1.149 | -0.343 | -0.039 | -0.135 | -0.135 | 0.201 | 0.23 | 0.02 | 0.18 | 0.61 | 0.26 | 0.31 |
| 2 | 119936683 | rs12623976 | 0.128 | -0.327 | -0.333 | -0.148 | -0.022 | 0.087 | 0.087 | 0.18 | 0.18 | 0.36 | 0.15 | 0.28 | 0.12 | 0.21 |
| 2 | 119939585 | rs6715243 | 0.611 | -0.212 | -0.75 | -0.25 | 0.003 | -0.019 | -0.019 | 0 | 0.26 | 0.27 | 0.17 | 0.59 | 0.25 | 0.38 |
| 2 | 119942569 | rs7584975 | -0.033 | 0.052 | -0.755 | -0.847 | -0.001 | -0.025 | -0.025 | 0.02 | 0.19 | 0.13 | 0.28 | 0 | 0.18 | 0.23 |
| 2 | 119953966 | rs4849768 | 0.222 | 0.057 | -0.749 | -1.744 | -0.005 | -0.002 | -0.002 | 0 | 0.16 | 0.11 | 0.26 | 0 | 0.16 | 0.15 |
| 2 | 119957026 | rs4849770 | -0.632 | -0.281 | -0.367 | -1.816 | -0.007 | -0.007 | -0.007 | 0 | 0.34 | 0.39 | 0.37 | 0.25 | 0.31 | 0.28 |

| | | | | | | | | | | | | | | | | |
|---|---|---|---|---|---|---|---|---|---|---|---|---|---|---|---|---|
| 2 | 119965951 | rs4848543 | 0.776 | -0.759 | -1.258 | -2.033 | -0.096 | -0.089 | -0.089 | 0 | 0.33 | 0.02 | 0.5 | 0.68 | 0.33 | 0.42 |
| 2 | 119968031 | rs6754799 | 0.266 | -0.918 | -0.371 | -2.21 | -0.001 | -0.011 | -0.011 | 0 | 0.42 | 0.45 | 0.55 | 0.68 | 0.37 | 0.54 |
| 2 | 119971173 | rs1562256 | -0.192 | -1.009 | -0.453 | -2.405 | 0.005 | 0.052 | 0.052 | 1.047 | 0.49 | 0.37 | 0.39 | 0.3 | 0.55 | 0.42 |
| 2 | 119971475 | rs1446125 | 0.813 | -1.023 | -0.178 | -2.416 | -0.004 | -0.011 | -0.011 | 0.008 | 0.37 | 0.42 | 0.52 | 0.7 | 0.33 | 0.42 |
| 2 | 119972145 | rs11677262 | 0.934 | -0.891 | -0.544 | -2.356 | -0.003 | -0.021 | -0.021 | 0 | 0.24 | 0.24 | 0.23 | 0.07 | 0.34 | 0.2 |
| 2 | 119977331 | rs838065 | 0.146 | -0.827 | -0.566 | -2.119 | 0.033 | -0.028 | -0.028 | 0.049 | 0.25 | 0.09 | 0.16 | 0.23 | 0.21 | 0.21 |
| 2 | 119984674 | rs838103 | -0.321 | -1.093 | 0.015 | -2.116 | 0.009 | 0.026 | 0.026 | 0.466 | 0.52 | 0.64 | 0.39 | 0.35 | 0.57 | 0.53 |
| 2 | 119984733 | rs838102 | 0.758 | -0.885 | -0.348 | -1.996 | -0.016 | -0.013 | -0.013 | 0 | 0.26 | 0.19 | 0.28 | 0.34 | 0.42 | 0.23 |
| 2 | 119987237 | rs708670 | 0.726 | -0.924 | -0.676 | -1.982 | -0.006 | -0.006 | -0.006 | 0 | 0.26 | 0.24 | 0.27 | 0.34 | 0.41 | 0.24 |
| 2 | 119988565 | rs838100 | 0.816 | -0.857 | -0.817 | -1.948 | -0.038 | -0.032 | -0.032 | 0 | 0.26 | 0.16 | 0.26 | 0.34 | 0.42 | 0.24 |
| 2 | 119989636 | rs838095 | 0.655 | -0.932 | -0.819 | -1.978 | -0.01 | -0.008 | -0.008 | 0 | 0.27 | 0.22 | 0.27 | 0.34 | 0.42 | 0.24 |
| 2 | 119997159 | rs838090 | 0.241 | -1.073 | -1.275 | -2.117 | -0.072 | -0.015 | -0.015 | 0 | 0.34 | 0.12 | 0.27 | 0.3 | 0.45 | 0.31 |
| 2 | 119999893 | rs17804991 | 0.704 | -1.457 | -1.118 | -2.171 | -0.007 | -0.026 | -0.026 | 0 | 0.09 | 0.15 | 0.05 | 0 | 0.05 | 0.05 |
| 2 | 120000611 | rs6728660 | 0.988 | -1.565 | -1.471 | -2.412 | -0.051 | -0.042 | -0.042 | 0 | 0.33 | 0.09 | 0.53 | 0.68 | 0.34 | 0.48 |
| 2 | 120001502 | rs838086 | 0.787 | -1.568 | -1.614 | -2.79 | 0.071 | 0.196 | 0.196 | 0 | 0.56 | 0.19 | 0.79 | 0.97 | 0.72 | 0.64 |
| 2 | 120003780 | rs10188946 | 0.91 | -1.292 | -1.158 | -2.251 | -0.019 | 0.018 | 0.018 | 0.571 | 0.38 | 0.15 | 0.57 | 0.71 | 0.38 | 0.48 |
| 2 | 120005312 | rs17013371 | 0.741 | -1.422 | -0.834 | -2.241 | -0.034 | -0.049 | -0.049 | 0 | 0.09 | 0.23 | 0.03 | 0 | 0.01 | 0.07 |
| 2 | 120008556 | rs734846 | 0.199 | -1.449 | -0.931 | -2.339 | -0.024 | -0.074 | -0.074 | 0 | 0.2 | 0.29 | 0.08 | 0 | 0.1 | 0.12 |
| 2 | 120009220 | rs895406 | 0.022 | -1.356 | -1.932 | -2.323 | -0.052 | 0.053 | 0.053 | 0.674 | 0.43 | 0.18 | 0.33 | 0.31 | 0.53 | 0.33 |
| 2 | 120012171 | rs12711924 | 1.263 | -1.234 | -1.981 | -2.283 | -0.014 | -0.007 | -0.007 | 0.008 | 0.33 | 0.12 | 0.53 | 0.69 | 0.32 | 0.45 |
| 2 | 120012888 | rs838075 | 0.875 | -1.118 | -1.821 | -2.18 | -0.004 | -0.001 | -0.001 | 0 | 0.07 | 0.06 | 0.04 | 0 | 0.08 | 0.03 |
| 2 | 120014609 | rs838073 | -0.482 | -1.16 | -2.009 | -2.226 | -0.047 | 0.026 | 0.026 | 0.064 | 0.61 | 0.81 | 0.68 | 0.7 | 0.5 | 0.69 |
| 2 | 120021785 | rs7581339 | -1.018 | -0.957 | -2.467 | -1.745 | -0.005 | 0 | 0 | 0 | 0.94 | 0.95 | 0.95 | 1 | 0.92 | 0.97 |
| 2 | 120021917 | rs3731603 | 1.265 | -1.011 | -2.594 | -1.784 | -0.006 | -0.037 | -0.037 | 0.126 | 0.28 | 0.13 | 0.52 | 0.67 | 0.29 | 0.46 |
| 2 | 120023188 | rs4849771 | -0.424 | -0.316 | -2.039 | -1.213 | 0.002 | -0.021 | -0.021 | 2.489 | 0.93 | 0.91 | 0.95 | 1 | 0.92 | 0.97 |
| 2 | 120023260 | rs11694139 | 0.937 | -0.2 | -2.379 | -1.186 | -0.11 | 0.09 | 0.09 | 0 | 0.23 | 0.06 | 0.22 | 0.08 | 0.38 | 0.17 |
| 2 | 120024442 | rs1530561 | 0.582 | -0.031 | -2.029 | -0.777 | 0.012 | 0.018 | 0.018 | 1.914 | 0.4 | 0.18 | 0.57 | 0.67 | 0.39 | 0.6 |
| 2 | 120025472 | rs6707849 | 1.188 | 0.001 | -1.449 | -0.522 | 0.021 | 0.026 | 0.026 | 0 | 0.79 | 0.83 | 0.91 | 0.99 | 0.88 | 0.81 |
| 2 | 120026899 | rs1545360 | 0.683 | 0.559 | -1.053 | 0.073 | 0.001 | -0.001 | -0.001 | 0.058 | 0.91 | 0.9 | 0.98 | 0.99 | 0.96 | 0.83 |
| 2 | 120035899 | rs2166504 | -0.538 | 1.42 | -1.047 | 0.503 | -0.007 | -0.004 | -0.004 | 0 | 0.25 | 0.22 | 0.4 | 0.29 | 0.36 | 0.34 |
| 2 | 120043214 | rs3795891 | -1.105 | 0.938 | -0.179 | 0.692 | 0.022 | 0.028 | 0.028 | 0 | 0.36 | 0.46 | 0.45 | 0.42 | 0.41 | 0.48 |
| 2 | 120046905 | rs6711245 | -0.453 | 0.011 | -0.63 | -0.271 | -0.028 | -0.004 | -0.004 | 0 | 0.67 | 0.81 | 0.58 | 0.31 | 0.62 | 0.6 |
| 2 | 120051015 | rs4848546 | -0.279 | -0.055 | -0.422 | -0.292 | 0 | -0.001 | -0.001 | 0 | 0.65 | 0.65 | 0.57 | 0.31 | 0.62 | 0.55 |
| 2 | 120057962 | rs7424214 | 0.618 | -0.412 | 0.345 | -0.513 | 0.04 | 0.031 | 0.031 | 0 | 0.38 | 0.23 | 0.15 | 0.01 | 0.28 | 0.21 |
| 2 | 120060082 | rs1052500 | -1.031 | -0.412 | 0.335 | -0.496 | -0.065 | -0.11 | -0.11 | 0 | 0.71 | 0.92 | 0.6 | 0.43 | 0.65 | 0.62 |
| 2 | 120061461 | rs13015640 | 1.205 | -0.492 | 0.292 | -0.542 | 0.049 | 0.058 | 0.058 | 0.005 | 0.31 | 0.15 | 0.12 | 0.01 | 0.23 | 0.13 |
| 2 | 120068172 | rs11123515 | 1.002 | -0.465 | 0.339 | -0.522 | -0.032 | -0.019 | -0.019 | 1.306 | 0.23 | 0.14 | 0.39 | 0.3 | 0.34 | 0.34 |
| 2 | 120073602 | rs4369868 | 0.527 | -0.58 | 0.455 | -0.601 | -0.033 | -0.115 | -0.115 | 0.987 | 0.37 | 0.59 | 0.15 | 0.01 | 0.24 | 0.13 |
| 2 | 120074190 | rs7560740 | -1.053 | -0.528 | 0.294 | -0.567 | -0.025 | -0.05 | -0.05 | 0 | 0.7 | 0.84 | 0.59 | 0.43 | 0.65 | 0.62 |
| 2 | 120076903 | rs6721130 | 1.193 | -0.576 | 0.581 | -0.63 | 0.068 | 0.147 | 0.147 | 0.193 | 0.3 | 0.07 | 0.11 | 0.01 | 0.23 | 0.13 |
| 2 | 120087676 | rs12711926 | -0.153 | 0.068 | 0.364 | -0.858 | -0.005 | 0.006 | 0.006 | 0 | 0.58 | 0.63 | 0.39 | 0.29 | 0.61 | 0.47 |
| 2 | 120096359 | rs7578714 | -1.469 | 0.39 | 0.228 | -0.87 | -0.028 | -0.082 | -0.082 | 2.45 | 0.79 | 0.9 | 0.63 | 0.43 | 0.77 | 0.69 |
| 2 | 120096939 | rs6711445 | 0.528 | 0.708 | 0.544 | -0.923 | 0.011 | 0.011 | 0.011 | 0.002 | 0.08 | 0.14 | 0.04 | 0.08 | 0.07 | 0.12 |
| 2 | 120104458 | rs6542524 | 0.743 | 0.583 | 0.591 | -1.03 | 0.004 | 0.026 | 0.026 | 0.205 | 0.45 | 0.44 | 0.33 | 0.21 | 0.51 | 0.31 |
| 2 | 120108230 | rs935394 | -0.201 | 1.016 | 0.712 | -0.984 | 0.013 | 0.015 | 0.015 | 0 | 0.12 | 0.16 | 0.16 | 0.21 | 0.23 | 0.18 |
| 2 | 120122340 | rs12613135 | -1.129 | 1.063 | 0.824 | -0.93 | 0.023 | 0.033 | 0.033 | 0 | 0.22 | 0.33 | 0.21 | 0.3 | 0.3 | 0.28 |
| 2 | 120123277 | rs11693629 | -1.129 | 1.063 | 0.824 | -0.93 | 0.023 | 0.033 | 0.033 | 0 | 0.22 | 0.33 | 0.21 | 0.3 | 0.3 | 0.28 |
| 2 | 120123704 | rs6756192 | 0.372 | 1.076 | 0.815 | -0.923 | 0.014 | 0.026 | 0.026 | 0 | 0.09 | 0.12 | 0.15 | 0.22 | 0.2 | 0.12 |
| 2 | 120125936 | rs2244135 | -1.596 | 1.248 | 0.759 | -0.956 | -0.021 | -0.016 | -0.016 | 0 | 0.86 | 0.96 | 0.8 | 0.87 | 0.85 | 0.81 |
| 2 | 120129405 | rs3754854 | -0.315 | 0.731 | 1.022 | -1.116 | -0.002 | 0.009 | 0.009 | 0 | 0.54 | 0.62 | 0.37 | 0.31 | 0.55 | 0.41 |
| 2 | 120130796 | rs6727485 | -0.311 | 0.919 | 0.901 | -0.888 | -0.007 | 0.009 | 0.009 | 0 | 0.14 | 0.2 | 0.06 | 0.13 | 0.1 | 0.16 |
| 2 | 120135491 | rs11898177 | -1.381 | 0.994 | 0.814 | -0.901 | 0.034 | 0.093 | 0.093 | 0 | 0.24 | 0.46 | 0.22 | 0.35 | 0.29 | 0.27 |
| 2 | 120145637 | rs6759748 | 0.266 | 1.041 | 0.555 | -0.662 | 0.043 | 0.093 | 0.093 | 0 | 0.09 | 0.2 | 0.16 | 0.27 | 0.19 | 0.12 |
| 2 | 120145655 | rs1374313 | -0.028 | 0.798 | 0.212 | -0.791 | 0.013 | 0.052 | 0.052 | 0 | 0.52 | 0.81 | 0.38 | 0.36 | 0.52 | 0.37 |
| 2 | 120159868 | rs12994683 | 1.106 | 0.156 | 0.171 | -1.014 | 0.017 | -0.001 | -0.001 | 0 | 0.29 | 0.23 | 0.17 | 0.01 | 0.25 | 0.13 |
| 2 | 120160459 | rs12469579 | 0.621 | 0.136 | 0.422 | -1.072 | 0 | -0.057 | -0.057 | 0 | 0.34 | 0.35 | 0.19 | 0.01 | 0.27 | 0.17 |
| 2 | 120161698 | rs4849779 | 0.858 | 0.452 | -0.005 | -0.955 | 0 | -0.104 | -0.104 | 0 | 0.27 | 0.09 | 0.34 | 0.54 | 0.21 | 0.36 |
| 2 | 120165021 | rs10427304 | 0.21 | 0.15 | 0.645 | -0.953 | -0.036 | -0.057 | -0.057 | 0 | 0.38 | 0.54 | 0.22 | 0.14 | 0.31 | 0.2 |
| 2 | 120176240 | rs6705957 | 0.662 | 0.094 | 0.711 | -1.114 | -0.038 | -0.15 | -0.15 | 0 | 0.32 | 0.52 | 0.19 | 0.01 | 0.25 | 0.13 |
| 2 | 120187344 | rs11123523 | 0.466 | -0.104 | -0.537 | 0.035 | 0.033 | 0.004 | 0.004 | 0 | 0.61 | 0.29 | 0.72 | 0.56 | 0.69 | 0.57 |
| 2 | 120197670 | rs1042656 | 0.336 | -0.222 | 0.695 | -0.103 | 0.022 | 0.011 | 0.011 | 0 | 0.6 | 0.37 | 0.71 | 0.56 | 0.69 | 0.54 |
| 2 | 120198927 | rs2587663 | 1.879 | -0.259 | 0.132 | -0.757 | -0.012 | 0.09 | 0.09 | 0 | 0.25 | 0.14 | 0.18 | 0 | 0.29 | 0.11 |
| 2 | 120204052 | rs11123526 | -1.888 | -0.711 | 0.6 | -0.203 | -0.016 | -0.004 | -0.004 | 0 | 0.9 | 0.97 | 0.84 | 0.91 | 0.87 | 0.9 |
| 2 | 120207853 | rs12479137 | 0.519 | -0.73 | 0.767 | -0.171 | 0.005 | 0.002 | 0.002 | 0 | 0.6 | 0.41 | 0.72 | 0.58 | 0.69 | 0.54 |
| 2 | 120217053 | rs10165398 | 0.669 | -0.802 | 0.605 | -0.2 | 0.014 | -0.013 | -0.013 | 0 | 0.21 | 0.16 | 0.09 | 0.33 | 0.17 | 0.31 |
| 2 | 120219797 | rs2254122 | 0.793 | -0.833 | 0.518 | -0.223 | 0.045 | 0.025 | 0.025 | 0 | 0.59 | 0.3 | 0.7 | 0.58 | 0.68 | 0.54 |
| 2 | 120219940 | rs2579644 | 1.657 | -0.756 | 0.204 | -0.195 | 0.003 | -0.002 | -0.002 | 0 | 0.47 | 0.19 | 0.62 | 0.53 | 0.57 | 0.46 |
| 2 | 120222696 | rs6750751 | 0.135 | -0.832 | 0.826 | -0.082 | -0.004 | 0.001 | 0.001 | 0 | 0.22 | 0.23 | 0.1 | 0.33 | 0.17 | 0.31 |
| 2 | 120226922 | rs2918939 | 0.29 | -0.725 | 0.98 | 0.075 | -0.002 | 0.055 | 0.055 | 0.029 | 0.16 | 0.28 | 0.09 | 0.33 | 0.13 | 0.28 |

| | | | | | | | | | | | | | | | | |
|---|---|---|---|---|---|---|---|---|---|---|---|---|---|---|---|---|
| 2 | 120230901 | rs2579640 | 0.555 | -0.693 | 0.456 | -0.429 | 0.011 | 0.015 | 0.015 | 1.406 | 0.78 | 0.28 | 0.76 | 0.86 | 0.83 | 0.83 |
| 2 | 120231303 | rs2587702 | -1.211 | -0.465 | 0.254 | -0.306 | 0 | 0 | 0 | 0.018 | 0.89 | 0.92 | 0.82 | 0.86 | 0.86 | 0.86 |
| 2 | 120244580 | rs6710114 | 0.855 | -0.321 | -0.372 | -0.357 | -0.043 | -0.023 | -0.023 | 0 | 0.16 | 0.01 | 0.28 | 0.12 | 0.15 | 0.3 |
| 2 | 120244703 | rs6757604 | 0.927 | -0.509 | 0.445 | -0.414 | 0.039 | 0.034 | 0.034 | 0 | 0.59 | 0.27 | 0.69 | 0.6 | 0.68 | 0.55 |
| 2 | 120247486 | rs6751578 | -0.388 | -0.348 | 0.684 | -0.328 | -0.038 | -0.017 | -0.017 | 0 | 0.26 | 0.36 | 0.09 | 0.27 | 0.17 | 0.33 |
| 2 | 120248482 | rs2579654 | 1.003 | 0.184 | 0.942 | -0.311 | -0.013 | -0.053 | -0.053 | 0 | 0.79 | 0.61 | 0.77 | 0.86 | 0.83 | 0.83 |
| 2 | 120248802 | rs2587677 | 0.083 | 0.749 | -0.082 | 0.027 | -0.015 | 0.001 | 0.001 | 0 | 0.87 | 0.96 | 0.81 | 0.87 | 0.85 | 0.87 |
| 2 | 120261127 | rs11687793 | 0.297 | 1.016 | -0.456 | 0.197 | -0.003 | 0.001 | 0.001 | 0 | 0.6 | 0.34 | 0.45 | 0.41 | 0.57 | 0.61 |
| 2 | 120261891 | rs2579600 | 0.568 | 1.003 | -0.325 | -0.474 | 0.007 | 0.015 | 0.015 | 0 | 0.55 | 0.44 | 0.37 | 0.85 | 0.42 | 0.56 |
| 2 | 120264850 | rs2579604 | 0.135 | 1.569 | -0.538 | -0.373 | -0.005 | -0.002 | -0.002 | 0 | 0.74 | 0.8 | 0.69 | 0.92 | 0.68 | 0.8 |
| 2 | 120278171 | rs4383344 | -0.304 | 1.715 | -0.873 | 0.148 | -0.074 | 0.016 | 0.016 | 0 | 0.16 | 0.07 | 0.25 | 0.09 | 0.27 | 0.28 |
| 2 | 120285695 | rs2587686 | 0.277 | 1.407 | -0.39 | 0.106 | -0.002 | 0.021 | 0.021 | 3.175 | 0.56 | 0.6 | 0.58 | 0.72 | 0.53 | 0.61 |
| 2 | 120293582 | rs2165061 | 0.532 | 1.408 | 0.216 | 0.186 | 0.022 | 0.038 | 0.038 | 3.859 | 0.52 | 0.32 | 0.51 | 0.65 | 0.48 | 0.58 |
| 2 | 120294700 | rs2579621 | 0.158 | 1.387 | 0.134 | 0.128 | -0.027 | 0.086 | 0.086 | 3.363 | 0.57 | 0.81 | 0.59 | 0.73 | 0.54 | 0.6 |
| 2 | 120321817 | rs2587695 | 0.166 | 1.358 | 0.198 | 0.124 | -0.007 | 0.046 | 0.046 | 3.278 | 0.57 | 0.71 | 0.59 | 0.73 | 0.54 | 0.61 |
| 2 | 120337094 | rs17049444 | -0.347 | 1.883 | 0.375 | 0.047 | 0.038 | 0.022 | 0.022 | 0 | 0.13 | 0.27 | 0.16 | 0.13 | 0.18 | 0.14 |
| 2 | 120343832 | rs10187631 | -0.702 | 1.807 | 0.901 | 0.104 | 0.061 | 0.087 | 0.087 | 0 | 0.16 | 0.49 | 0.19 | 0.19 | 0.19 | 0.15 |
| 2 | 120357089 | rs12373810 | 0.467 | 1.147 | 1.026 | 0.029 | 0.011 | 0.032 | 0.032 | 0 | 0.23 | 0.11 | 0.3 | 0.12 | 0.23 | 0.33 |
| 2 | 120364042 | rs2860875 | 0.263 | 1.235 | 1.269 | 0.066 | 0.031 | 0.052 | 0.052 | 0 | 0.83 | 0.48 | 0.8 | 0.82 | 0.81 | 0.85 |
| 2 | 120368359 | rs7570863 | 0.571 | 1.233 | 0.675 | 0.535 | -0.103 | -0.018 | -0.018 | 1.65 | 0.28 | 0.05 | 0.28 | 0.27 | 0.33 | 0.31 |
| 2 | 120369105 | rs17660087 | 0.177 | 0.973 | 1.552 | 0.603 | 0.001 | 0.002 | 0.002 | 0 | 0.13 | 0.14 | 0.12 | 0.12 | 0.15 | 0.13 |
| 2 | 120408400 | rs939775 | 0.005 | 0.994 | 1.364 | 0.583 | 0.031 | 0.018 | 0.018 | 0 | 0.85 | 0.69 | 0.83 | 0.87 | 0.82 | 0.84 |
| 2 | 120409616 | rs11686014 | 0.093 | 0.977 | 1.162 | 0.598 | -0.009 | 0.016 | 0.016 | 0 | 0.13 | 0.22 | 0.12 | 0.13 | 0.13 | 0.13 |
| 2 | 120414083 | rs6761357 | 0.509 | 0.98 | 0.893 | 0.714 | 0 | 0.006 | 0.006 | 0 | 0.64 | 0.48 | 0.59 | 0.61 | 0.64 | 0.68 |
| 2 | 120416787 | rs11678053 | -0.046 | 0.916 | 1.533 | 0.647 | 0.016 | 0.046 | 0.046 | 0 | 0.15 | 0.29 | 0.16 | 0.17 | 0.16 | 0.15 |
| 2 | 120418222 | rs7596716 | 0.188 | 0.931 | 1.794 | 0.636 | 0.021 | 0.047 | 0.047 | 0 | 0.83 | 0.52 | 0.8 | 0.83 | 0.82 | 0.82 |
| 2 | 120432637 | rs11895880 | -0.006 | 0.936 | 1.824 | 0.626 | 0.029 | 0.051 | 0.051 | 0 | 0.84 | 0.62 | 0.81 | 0.83 | 0.82 | 0.84 |
| 2 | 120435393 | rs1568539 | 0.156 | 0.935 | 1.795 | 0.626 | 0.02 | 0.042 | 0.042 | 0 | 0.83 | 0.5 | 0.8 | 0.83 | 0.82 | 0.84 |
| 2 | 120436773 | rs2272053 | 0.666 | 0.934 | 1.579 | 0.627 | -0.029 | -0.03 | -0.03 | 0 | 0.12 | 0.02 | 0.12 | 0.12 | 0.15 | 0.13 |
| 2 | 120439459 | rs4849797 | -0.156 | 0.928 | 1.734 | 0.616 | 0.021 | 0.062 | 0.062 | 0 | 0.84 | 0.63 | 0.83 | 0.81 | 0.84 | 0.86 |
| 2 | 120443014 | rs2177095 | 0.105 | 0.911 | 1.318 | 0.618 | 0.009 | 0.048 | 0.048 | 0 | 0.83 | 0.52 | 0.83 | 0.81 | 0.84 | 0.86 |
| 2 | 120469993 | rs6542545 | 0.399 | 0.907 | 0.526 | 0.333 | -0.012 | -0.023 | -0.023 | 0 | 0.27 | 0.38 | 0.24 | 0.21 | 0.24 | 0.34 |
| 2 | 120475953 | rs11123535 | -0.543 | 0.984 | 0.344 | 0.366 | 0 | -0.009 | -0.009 | 0 | 0.41 | 0.44 | 0.38 | 0.32 | 0.42 | 0.5 |
| 2 | 120485053 | rs11693193 | 0.681 | 1.047 | -0.114 | 0.468 | -0.007 | -0.011 | -0.011 | 0 | 0.13 | 0.05 | 0.12 | 0.11 | 0.14 | 0.12 |
| 2 | 120489760 | rs12620790 | 1.074 | 1.179 | -0.927 | 0.487 | -0.059 | -0.141 | -0.141 | 0 | 0.24 | 0.02 | 0.28 | 0.4 | 0.26 | 0.26 |
| 2 | 120498760 | rs971651 | 0.714 | 1.148 | -0.95 | 0.089 | -0.063 | 0.023 | 0.023 | 0.351 | 0.27 | 0.06 | 0.26 | 0.23 | 0.29 | 0.23 |
| 2 | 120507425 | rs10174596 | 1.064 | 0.909 | -0.139 | 0.126 | -0.005 | -0.008 | -0.008 | 0 | 0.29 | 0.32 | 0.26 | 0.2 | 0.27 | 0.36 |
| 2 | 120540224 | rs12990769 | -0.322 | 1.224 | -0.204 | 0.073 | -0.02 | 0.059 | 0.059 | 0 | 0.6 | 0.22 | 0.56 | 0.77 | 0.53 | 0.53 |
| 2 | 120573539 | rs7606911 | -1.221 | 0.913 | 0.431 | 0.109 | 0.002 | 0.003 | 0.003 | 0 | 0.7 | 0.71 | 0.73 | 0.79 | 0.72 | 0.65 |
| 2 | 120578206 | rs3106246 | 0.518 | 0.915 | 0.517 | 0.1 | 0 | 0 | 0 | 0.849 | 0.34 | 0.65 | 0.32 | 0.2 | 0.31 | 0.37 |
| 2 | 120592580 | rs6542550 | 1.107 | 0.934 | 0.484 | 0.094 | -0.01 | -0.02 | -0.02 | 0.614 | 0.3 | 0.39 | 0.27 | 0.21 | 0.28 | 0.35 |
| 2 | 120617488 | rs949530 | 0.838 | 1.016 | -0.295 | -0.13 | -0.123 | -0.146 | -0.146 | 0 | 0.27 | 0.02 | 0.3 | 0.44 | 0.36 | 0.32 |
| 2 | 120628463 | rs1995916 | 0.211 | 1.02 | -0.244 | -0.116 | 0.037 | 0.041 | 0.041 | 0 | 0.49 | 0.75 | 0.44 | 0.34 | 0.42 | 0.49 |
| 2 | 120661001 | rs3106242 | 0.897 | 1.038 | 0.101 | -0.103 | 0.013 | 0.016 | 0.016 | 0 | 0.33 | 0.72 | 0.32 | 0.23 | 0.28 | 0.34 |
| 2 | 120682525 | rs6730447 | -0.961 | 1.016 | 0.066 | -0.107 | 0.001 | 0.008 | 0.008 | 0.806 | 0.77 | 0.89 | 0.85 | 0.78 | 0.82 | 0.82 |
| 2 | 120686847 | rs4404280 | -0.956 | 0.865 | 0.319 | -0.042 | -0.012 | -0.034 | -0.034 | 0.402 | 0.68 | 0.51 | 0.7 | 0.79 | 0.69 | 0.63 |
| 2 | 120737825 | rs1046433 | 0.764 | 0.87 | 0.322 | -0.099 | -0.012 | -0.043 | -0.043 | 0.545 | 0.32 | 0.48 | 0.3 | 0.19 | 0.31 | 0.37 |
| 2 | 120752342 | rs7578989 | -0.854 | 0.898 | 0.061 | -0.041 | -0.02 | -0.03 | -0.03 | 0 | 0.66 | 0.48 | 0.71 | 0.79 | 0.71 | 0.65 |
| 2 | 120764058 | rs12622262 | -0.506 | 0.845 | -0.479 | -0.072 | 0.023 | 0.051 | 0.051 | 0 | 0.62 | 0.23 | 0.65 | 0.79 | 0.65 | 0.62 |
| 2 | 120766790 | rs4849811 | 0.962 | 0.988 | -0.937 | -0.052 | -0.019 | 0.003 | 0.003 | 0.165 | 0.19 | 0.06 | 0.22 | 0.21 | 0.18 | 0.17 |
| 2 | 120769787 | rs1627943 | 1.041 | 0.659 | -0.171 | -0.065 | -0.006 | -0.008 | -0.008 | 0.029 | 0.28 | 0.32 | 0.24 | 0.2 | 0.23 | 0.34 |
| 2 | 120776988 | rs17669093 | 1.415 | 0.635 | -0.675 | -0.308 | -0.029 | 0.058 | 0.058 | 0 | 0.19 | 0.01 | 0.09 | 0.04 | 0.17 | 0.12 |
| 2 | 120798425 | rs12476760 | 0.948 | 0.429 | -0.516 | 0.183 | -0.051 | -0.139 | -0.139 | 0 | 0.18 | 0.01 | 0.23 | 0.44 | 0.22 | 0.16 |
| 2 | 120800573 | rs12466598 | -0.988 | 0.323 | 0.266 | 0.11 | -0.025 | 0 | 0 | 0 | 0.7 | 0.4 | 0.71 | 0.68 | 0.8 | 0.63 |
| 2 | 120855940 | rs4849821 | -0.555 | 0.332 | 0.283 | 0.106 | 0.018 | 0.001 | 0.001 | 0 | 0.65 | 0.28 | 0.67 | 0.65 | 0.74 | 0.63 |
| 2 | 120864763 | rs13029879 | -0.862 | 0.408 | 0.335 | 0.152 | -0.014 | 0 | 0 | 0.048 | 0.69 | 0.4 | 0.67 | 0.68 | 0.73 | 0.6 |
| 2 | 120885310 | rs1034489 | -0.771 | 0.396 | 0.554 | 0.15 | -0.011 | -0.002 | -0.002 | 0 | 0.68 | 0.38 | 0.66 | 0.67 | 0.73 | 0.6 |
| 2 | 120942349 | rs7340249 | -1.503 | 0.408 | -0.576 | 0.238 | 0.019 | -0.028 | -0.028 | 0 | 0.75 | 0.87 | 0.75 | 0.66 | 0.8 | 0.63 |
| 2 | 120948573 | rs12618213 | -0.406 | 0.289 | -0.109 | 0.106 | 0.002 | 0 | 0 | 2.112 | 0.63 | 0.35 | 0.63 | 0.65 | 0.67 | 0.61 |
| 2 | 120949109 | rs34520190 | 1.056 | 0.389 | -0.47 | 0.11 | -0.051 | -0.133 | -0.133 | 0 | 0.18 | 0.01 | 0.24 | 0.43 | 0.22 | 0.16 |
| 2 | 120949126 | rs3961960 | -0.208 | 0.408 | 0.073 | 0.226 | -0.003 | -0.001 | -0.001 | 0 | 0.64 | 0.38 | 0.63 | 0.65 | 0.67 | 0.61 |
| 2 | 120975831 | rs1437421 | 0.081 | 0.218 | 0.195 | 0.298 | 0.004 | 0.002 | 0.002 | 0.374 | 0.64 | 0.35 | 0.65 | 0.67 | 0.68 | 0.57 |
| 2 | 120986708 | rs6542572 | -0.849 | 0.219 | 0.05 | 0.483 | -0.032 | -0.021 | -0.021 | 0.009 | 0.73 | 0.91 | 0.71 | 0.71 | 0.73 | 0.6 |
| 2 | 120986872 | rs6542573 | -0.714 | 0.14 | -0.103 | 0.521 | -0.136 | -0.018 | -0.018 | 0 | 0.73 | 0.99 | 0.68 | 0.79 | 0.63 | 0.6 |
| 2 | 120993920 | rs6733204 | -0.535 | 0.129 | 0.767 | 0.531 | 0 | 0 | 0 | 0 | 0.7 | 0.71 | 0.69 | 0.67 | 0.72 | 0.59 |
| 2 | 121018537 | rs11893187 | -0.434 | 0.126 | 0.459 | -0.039 | -0.019 | -0.014 | -0.014 | 0 | 0.68 | 0.77 | 0.52 | 0.4 | 0.58 | 0.62 |
| 2 | 121019800 | rs11690576 | 0.178 | -0.463 | 0.95 | 0.104 | -0.001 | 0 | 0 | 0 | 0.62 | 0.39 | 0.49 | 0.4 | 0.59 | 0.54 |
| 2 | 121037018 | rs3896630 | 0.593 | -0.445 | 0.542 | 0.116 | 0.007 | 0.056 | 0.056 | 0.534 | 0.57 | 0.19 | 0.45 | 0.33 | 0.58 | 0.48 |
| 2 | 121041347 | rs3931840 | -0.724 | -0.408 | 0.591 | 0.158 | -0.004 | -0.012 | -0.012 | 0.273 | 0.72 | 0.78 | 0.58 | 0.4 | 0.71 | 0.63 |

| | | | | | | | | | | | | | | | | |
|---|---|---|---|---|---|---|---|---|---|---|---|---|---|---|---|---|
| 2 | 121052324 | rs10496561 | 0.696 | -0.372 | -0.528 | 0.078 | 0.007 | -0.008 | -0.008 | 0 | 0.13 | 0.11 | 0.16 | 0.17 | 0.07 | 0.16 |
| 2 | 121058412 | rs2028147 | 0.726 | -0.124 | -0.012 | 0.433 | 0.005 | 0.052 | 0.052 | 0.461 | 0.57 | 0.21 | 0.46 | 0.33 | 0.58 | 0.48 |
| 2 | 121066311 | rs2043722 | -0.327 | 0.243 | -0.212 | 0.497 | 0.003 | -0.05 | -0.05 | 0 | 0.1 | 0.26 | 0.08 | 0.01 | 0.12 | 0.09 |
| 2 | 121070811 | rs11673874 | 0.018 | 0.376 | -0.828 | 1.007 | -0.002 | -0.006 | -0.006 | 0.16 | 0.73 | 0.74 | 0.58 | 0.42 | 0.72 | 0.61 |
| 2 | 121077184 | rs13018516 | 0.446 | -0.064 | -1.015 | 0.812 | 0.028 | -0.012 | -0.012 | 0.933 | 0.52 | 0.27 | 0.65 | 0.54 | 0.62 | 0.55 |
| 2 | 121081469 | rs934721 | 0.457 | -0.682 | -0.594 | 1.002 | -0.038 | 0.065 | 0.065 | 0 | 0.79 | 0.74 | 0.85 | 0.5 | 0.91 | 0.73 |
| 2 | 121086679 | rs7593535 | -0.937 | -0.408 | -0.716 | 0.926 | -0.004 | 0.007 | 0.007 | 0 | 0.28 | 0.39 | 0.35 | 0.41 | 0.23 | 0.3 |
| 2 | 121089731 | rs17625845 | -0.266 | -0.157 | -1.294 | 0.686 | -0.049 | 0.013 | 0.013 | 0 | 0.13 | 0.05 | 0.21 | 0.07 | 0.22 | 0.1 |
| 2 | 121094955 | rs10864990 | -0.812 | -0.285 | -1.582 | 0.276 | -0.002 | 0.127 | 0.127 | 0.145 | 0.54 | 0.83 | 0.49 | 0.29 | 0.42 | 0.57 |
| 2 | 121095920 | rs7589138 | -0.763 | -0.283 | -1.574 | 0.312 | -0.01 | 0.13 | 0.13 | 0.223 | 0.54 | 0.84 | 0.49 | 0.3 | 0.43 | 0.58 |
| 2 | 121100836 | rs11897635 | -0.877 | -0.429 | -0.585 | 0.984 | -0.018 | 0.043 | 0.043 | 0 | 0.09 | 0.24 | 0.05 | 0.16 | 0.04 | 0.24 |
| 2 | 121102572 | rs13419301 | -1.257 | -0.503 | -0.512 | 0.896 | -0.025 | 0.022 | 0.022 | 0 | 0.13 | 0.29 | 0.06 | 0.16 | 0.06 | 0.24 |
| 2 | 121106003 | rs11902591 | -1.119 | -0.267 | -0.308 | 1.754 | -0.004 | 0.188 | 0.188 | 0 | 0.08 | 0.16 | 0.19 | 0.48 | 0.08 | 0.06 |
| 2 | 121114263 | rs7576183 | -0.836 | -0.266 | -0.878 | 0.751 | 0.025 | 0.071 | 0.071 | 0 | 0.5 | 0.64 | 0.4 | 0.74 | 0.41 | 0.74 |
| 2 | 121118124 | rs7579169 | -0.166 | -0.455 | -1.276 | 0.803 | -0.056 | 0.014 | 0.014 | 0 | 0.65 | 0.85 | 0.67 | 0.75 | 0.63 | 0.59 |
| 2 | 121120552 | rs7589683 | -1.148 | -0.613 | -0.495 | 1.024 | -0.014 | 0.238 | 0.238 | 1.807 | 0.17 | 0.41 | 0.25 | 0.6 | 0.15 | 0.39 |
| 2 | 121122730 | rs11123550 | 1.703 | -1.028 | -1.011 | 0.409 | -0.019 | 0.035 | 0.035 | 0 | 0.34 | 0.24 | 0.37 | 0.03 | 0.4 | 0.17 |
| 2 | 121123383 | rs12711941 | -0.136 | -0.728 | -0.676 | 0.768 | -0.054 | 0.052 | 0.052 | 0 | 0.64 | 0.86 | 0.67 | 0.81 | 0.63 | 0.59 |
| 2 | 121124076 | rs11887035 | -1.081 | -0.834 | -0.48 | 0.944 | 0.004 | -0.007 | -0.007 | 0.187 | 0.1 | 0.09 | 0.05 | 0.18 | 0.07 | 0.32 |
| 2 | 121125072 | rs13410055 | 0.665 | -0.807 | -0.09 | 0.367 | -0.001 | -0.012 | -0.012 | 0 | 0.55 | 0.45 | 0.65 | 0.82 | 0.61 | 0.54 |
| 2 | 121126493 | rs12616090 | -0.832 | -0.415 | -0.654 | 0.738 | -0.001 | 0.057 | 0.057 | 1.209 | 0.09 | 0.08 | 0.19 | 0.61 | 0.12 | 0.35 |
| 2 | 121133245 | rs4076867 | -0.494 | -0.686 | -1.033 | 0.519 | -0.059 | 0.069 | 0.069 | 0 | 0.32 | 0.15 | 0.27 | 0.14 | 0.35 | 0.38 |
| 2 | 121135655 | rs4366914 | -1.97 | -1.063 | -1.026 | 0.363 | -0.082 | 0.061 | 0.061 | 0.02 | 0.59 | 0.87 | 0.54 | 0.75 | 0.47 | 0.79 |
| 2 | 121141060 | rs12479311 | 0.77 | -0.893 | -1.055 | 0.446 | -0.062 | 0.034 | 0.034 | 0 | 0.16 | 0.06 | 0.23 | 0.07 | 0.21 | 0.16 |
| 2 | 121145352 | rs10177102 | -2.059 | -1.16 | -1.03 | 0.357 | -0.08 | 0.064 | 0.064 | 0.012 | 0.58 | 0.91 | 0.51 | 0.74 | 0.46 | 0.78 |
| 2 | 121152814 | rs10168508 | 0.564 | -1.555 | 0.008 | 0.034 | -0.019 | -0.075 | -0.075 | 1.533 | 0.2 | 0.28 | 0.08 | 0 | 0.12 | 0.1 |
| 2 | 121154330 | rs13393184 | -0.033 | -1.461 | 0.637 | 0.508 | -0.024 | -0.006 | -0.006 | 0 | 0.3 | 0.45 | 0.2 | 0.25 | 0.21 | 0.28 |
| 2 | 121155187 | rs6542592 | 0.366 | -1.486 | 0.707 | 0.507 | 0.009 | -0.033 | -0.033 | 0 | 0.16 | 0.43 | 0.2 | 0.09 | 0.15 | 0.13 |
| 2 | 121155352 | rs6735848 | 0.001 | -1.402 | 0.56 | 0.564 | -0.031 | -0.008 | -0.008 | 0 | 0.3 | 0.47 | 0.2 | 0.25 | 0.2 | 0.28 |
| 2 | 121160145 | rs10206874 | -2.401 | -1.317 | 0.372 | 0.482 | -0.049 | 0.027 | 0.027 | 0 | 0.65 | 0.84 | 0.54 | 0.77 | 0.53 | 0.78 |
| 2 | 121161402 | rs12623934 | -1.505 | -1.269 | 0.286 | 0.556 | -0.013 | 0.015 | 0.015 | 0 | 0.26 | 0.38 | 0.25 | 0.69 | 0.2 | 0.55 |
| 2 | 121164027 | rs7608411 | -0.84 | -1.21 | 0.128 | 0.662 | -0.008 | 0.021 | 0.021 | 2.317 | 0.14 | 0.24 | 0.14 | 0.25 | 0.1 | 0.35 |
| 2 | 121171678 | rs9308780 | -2.091 | -1.231 | 0.622 | 0.969 | -0.003 | 0.034 | 0.034 | 0 | 0.55 | 0.72 | 0.48 | 0.73 | 0.47 | 0.6 |
| 2 | 121171926 | rs10432608 | -1.729 | -1.226 | 0.645 | 0.973 | -0.001 | -0.001 | -0.001 | 0 | 0.49 | 0.52 | 0.38 | 0.51 | 0.43 | 0.42 |
| 2 | 121175888 | rs4456710 | -0.301 | -0.98 | -0.079 | 0.915 | -0.004 | 0.058 | 0.058 | 0 | 0.11 | 0.16 | 0.09 | 0.42 | 0.1 | 0.19 |
| 2 | 121178252 | rs7562511 | -0.756 | -0.864 | -0.438 | 0.019 | -0.006 | -0.004 | -0.004 | 0 | 0.33 | 0.66 | 0.39 | 0.72 | 0.29 | 0.5 |
| 2 | 121184220 | rs6731937 | -0.665 | -0.711 | -0.059 | 0.503 | -0.002 | 0.03 | 0.03 | 0 | 0.54 | 0.7 | 0.48 | 0.73 | 0.46 | 0.6 |
| 2 | 121187063 | rs7592778 | -1.686 | -0.613 | 1.329 | 1.279 | -0.008 | 0.274 | 0.274 | 0.365 | 0.13 | 0.43 | 0.12 | 0.62 | 0.07 | 0.3 |
| 2 | 121189187 | rs13408457 | -1.187 | -0.854 | 1.62 | -0.023 | 0.024 | -0.07 | -0.07 | 1.665 | 0.21 | 0.37 | 0.19 | 0.05 | 0.22 | 0.12 |
| 2 | 121192652 | rs11890441 | -2.124 | -1.531 | 1.98 | 0.067 | -0.018 | -0.022 | -0.022 | 0.416 | 0.29 | 0.62 | 0.32 | 0.18 | 0.24 | 0.26 |
| 2 | 121197733 | rs7578773 | -1.278 | -1.485 | 2.311 | 0.079 | -0.054 | 0.028 | 0.028 | 0 | 0.12 | 0.33 | 0.16 | 0.12 | 0.08 | 0.15 |
| 2 | 121202152 | rs4280462 | -2.438 | -1.736 | 2.205 | 0.037 | -0.043 | -0.016 | -0.016 | 0.016 | 0.38 | 0.56 | 0.28 | 0.31 | 0.2 | 0.51 |
| 2 | 121202941 | rs10198258 | -2.853 | -1.794 | 1.49 | -0.013 | 0.095 | 0.074 | 0.074 | 0 | 0.46 | 0.92 | 0.41 | 0.36 | 0.31 | 0.6 |
| 2 | 121203511 | rs10186456 | -1.629 | -0.81 | 2.079 | 0.647 | 0.001 | -0.001 | -0.001 | 0 | 0.14 | 0.11 | 0.07 | 0.25 | 0.06 | 0.33 |
| 2 | 121214402 | rs4504000 | -2.51 | -0.504 | 2.504 | 0.577 | 0.023 | 0.009 | 0.009 | 1.614 | 0.19 | 0.58 | 0.31 | 0.17 | 0.22 | 0.29 |
| 2 | 121214469 | rs11681505 | -1.056 | -0.431 | 2.157 | 0.657 | 0 | -0.001 | -0.001 | 0 | 0.09 | 0.08 | 0.04 | 0.19 | 0.04 | 0.31 |
| 2 | 121216828 | rs6542598 | -1.61 | -0.34 | 2.346 | 0.656 | -0.001 | 0.022 | 0.022 | 0 | 0.12 | 0.22 | 0.2 | 0.13 | 0.13 | 0.24 |
| 2 | 121217568 | rs4599120 | -1.736 | -0.266 | 2.191 | 0.608 | 0.057 | 0.01 | 0.01 | 0 | 0.07 | 0.16 | 0.1 | 0.05 | 0.09 | 0.05 |
| 2 | 121223274 | rs17050160 | -1.705 | -0.672 | 1.764 | 0.677 | -0.02 | 0.021 | 0.021 | 0.114 | 0.1 | 0.23 | 0.07 | 0.16 | 0.05 | 0.3 |
| 2 | 121225558 | rs2140778 | 1.308 | -0.827 | 1.594 | 0.64 | -0.04 | 0.019 | 0.019 | 0 | 0.82 | 0.72 | 0.9 | 0.76 | 0.91 | 0.67 |
| 2 | 121228177 | rs6542599 | 1.566 | -0.813 | 1.412 | 0.467 | -0.009 | 0 | 0 | 0 | 0.63 | 0.45 | 0.64 | 0.59 | 0.7 | 0.42 |
| 2 | 121229487 | rs3941990 | -2.026 | -0.807 | 1.777 | 0.272 | -0.011 | -0.005 | -0.005 | 0 | 0.25 | 0.38 | 0.11 | 0.26 | 0.18 | 0.4 |
| 2 | 121229807 | rs4849877 | 1.974 | -0.902 | 2.08 | 0.155 | -0.012 | -0.109 | -0.109 | 0 | 0.85 | 0.53 | 0.91 | 1 | 0.91 | 0.97 |
| 2 | 121231515 | rs4848598 | 2.171 | -0.847 | 1.789 | 0.233 | -0.017 | 0.003 | 0.003 | 0 | 0.75 | 0.62 | 0.84 | 0.68 | 0.8 | 0.54 |
| 2 | 121234860 | rs10204717 | 1.389 | -0.733 | 1.581 | 0.344 | 0.01 | 0.015 | 0.015 | 0.16 | 0.91 | 0.8 | 0.94 | 0.89 | 0.91 | 0.81 |
| 2 | 121235399 | rs940677 | 2.325 | -0.743 | 1.362 | 0.359 | -0.024 | 0.022 | 0.022 | 0 | 0.8 | 0.68 | 0.88 | 0.68 | 0.87 | 0.58 |
| 2 | 121235551 | rs2310899 | 2.221 | -0.717 | 1.317 | 0.195 | -0.022 | 0.005 | 0.005 | 0 | 0.81 | 0.7 | 0.88 | 0.72 | 0.87 | 0.58 |
| 2 | 121237547 | rs940678 | 1.454 | -1.057 | 1.221 | 0.165 | -0.039 | 0.028 | 0.028 | 0 | 0.89 | 0.72 | 0.93 | 0.8 | 0.96 | 0.77 |
| 2 | 121244947 | rs4849884 | 1.764 | -0.898 | 1.056 | 0.093 | -0.014 | -0.003 | -0.003 | 0.028 | 0.85 | 0.7 | 0.87 | 0.79 | 0.9 | 0.75 |
| 2 | 121257128 | rs17050219 | -1.839 | -1.363 | 1.212 | 0.133 | -0.055 | -0.007 | -0.007 | 0.113 | 0.19 | 0.35 | 0.08 | 0.21 | 0.07 | 0.32 |
| 2 | 121257342 | rs940671 | 1.383 | -1.114 | 1.28 | 0.18 | -0.027 | -0.007 | -0.007 | 0 | 0.91 | 0.65 | 0.93 | 0.93 | 0.98 | 0.92 |
| 2 | 121264471 | rs874816 | 1.637 | -0.919 | 0.234 | 0.189 | 0.015 | 0.01 | 0.01 | 0.28 | 0.76 | 0.22 | 0.82 | 0.76 | 0.92 | 0.73 |
| 2 | 121267560 | rs4848605 | 1.475 | -0.873 | 0.243 | 0.271 | 0.005 | 0.004 | 0.004 | 0.014 | 0.75 | 0.28 | 0.84 | 0.76 | 0.88 | 0.75 |
| 2 | 121271649 | rs4849900 | -1.356 | -0.604 | 0.366 | -0.199 | 0.003 | -0.052 | -0.052 | 0 | 0.17 | 0.17 | 0.11 | 0.02 | 0.16 | 0.13 |
| 2 | 121271677 | rs10188887 | -2.063 | -1.253 | 0.86 | -0.424 | -0.039 | -0.084 | -0.084 | 0 | 0.27 | 0.43 | 0.16 | 0.07 | 0.16 | 0.22 |
| 2 | 121280603 | rs7585627 | -2.343 | -1.287 | 1.021 | -0.112 | -0.02 | -0.039 | -0.039 | 0 | 0.29 | 0.51 | 0.18 | 0.19 | 0.21 | 0.41 |
| 2 | 121283724 | rs4848607 | 2.016 | -1.774 | 1.044 | 0.023 | 0.027 | 0.025 | 0.025 | 0.028 | 0.64 | 0.28 | 0.8 | 0.7 | 0.78 | 0.53 |
| 2 | 121289107 | rs2311302 | -1.661 | -2.04 | 1.116 | -0.09 | -0.031 | -0.038 | -0.038 | 2.486 | 0.28 | 0.51 | 0.12 | 0.26 | 0.19 | 0.28 |
| 2 | 121289392 | rs17050253 | -0.763 | -2.046 | 1.273 | -0.096 | -0.001 | -0.042 | -0.042 | 0 | 0.19 | 0.2 | 0.07 | 0 | 0.15 | 0.07 |

| | | | | | | | | | | | | | | | | |
|---|---|---|---|---|---|---|---|---|---|---|---|---|---|---|---|---|
| 2 | 121290666 | rs11903416 | -1.911 | -1.189 | 0.906 | -0.348 | -0.003 | -0.046 | -0.046 | 0 | 0.37 | 0.51 | 0.18 | 0.26 | 0.31 | 0.34 |
| 2 | 121290925 | rs13427257 | -0.983 | -1.077 | 0.92 | -0.391 | 0.009 | -0.085 | -0.085 | 0 | 0.27 | 0.37 | 0.13 | 0.03 | 0.25 | 0.2 |
| 2 | 121296485 | rs6758414 | -1.429 | -0.957 | 0.688 | 0.066 | -0.013 | 0.038 | 0.038 | 0 | 0.1 | 0.3 | 0.06 | 0.21 | 0.06 | 0.09 |
| 2 | 121301760 | rs6721532 | -1.624 | -1.039 | 0.62 | 0.091 | -0.047 | 0.04 | 0.04 | 0 | 0.15 | 0.53 | 0.09 | 0.24 | 0.08 | 0.23 |
| 2 | 121302719 | rs2140779 | -1.044 | -0.663 | -0.615 | 1.219 | -0.03 | -0.068 | -0.068 | 1.097 | 0.11 | 0.01 | 0.23 | 0.43 | 0.14 | 0.22 |
| 2 | 121303240 | rs17389324 | 0.905 | -0.546 | -1.704 | 0.46 | -0.066 | -0.046 | -0.046 | 0.188 | 0.42 | 0.04 | 0.33 | 0.4 | 0.45 | 0.34 |
| 2 | 121303783 | rs17050267 | -0.874 | -0.841 | -1.196 | 1.213 | 0.005 | -0.003 | -0.003 | 0 | 0.08 | 0.07 | 0.07 | 0.13 | 0.05 | 0.13 |
| 2 | 121306440 | rs17050272 | 0.38 | -0.359 | -2.151 | 1.179 | -0.113 | -0.146 | -0.146 | 0 | 0.33 | 0.05 | 0.39 | 0.46 | 0.45 | 0.31 |
| 2 | 121309488 | rs2030746 | -0.349 | -0.404 | -0.873 | 1.172 | -0.002 | 0 | 0 | 0 | 0.45 | 0.53 | 0.43 | 0.5 | 0.4 | 0.37 |
| 2 | 121309759 | rs12617659 | 0.814 | 0.022 | -0.316 | 1.523 | 0.044 | 0.03 | 0.03 | 0 | 0.09 | 0.17 | 0.09 | 0.2 | 0.16 | 0.1 |
| 2 | 121322038 | rs7568633 | -0.712 | -0.119 | 0.024 | 1.376 | -0.001 | -0.004 | -0.004 | 0.63 | 0.43 | 0.62 | 0.43 | 0.53 | 0.4 | 0.53 |
| 2 | 121327514 | rs940670 | -0.537 | 0.093 | 0.065 | 1.362 | -0.004 | 0.013 | 0.013 | 0 | 0.4 | 0.53 | 0.47 | 0.52 | 0.38 | 0.55 |
| 2 | 121332030 | rs1467326 | -0.611 | 0.232 | 0.261 | 0.913 | -0.004 | -0.058 | -0.058 | 0.071 | 0.8 | 0.79 | 0.94 | 1 | 0.89 | 0.91 |
| 2 | 121337196 | rs9308614 | 0.423 | 0.752 | 0.536 | 1.636 | 0.003 | -0.001 | -0.001 | 0 | 0.09 | 0.1 | 0.08 | 0.1 | 0.15 | 0.09 |
| 2 | 121341281 | rs1869026 | -0.205 | 0.034 | 1.198 | 1.817 | 0.002 | 0 | 0 | 0 | 0.58 | 0.56 | 0.52 | 0.44 | 0.49 | 0.63 |
| 2 | 121343284 | rs6726523 | 0.206 | -0.047 | 0.62 | 1.555 | 0 | 0 | 0 | 0 | 0.56 | 0.46 | 0.43 | 0.45 | 0.51 | 0.65 |
| 2 | 121348224 | rs3860374 | -0.112 | -0.146 | 0.104 | 1.549 | -0.001 | -0.012 | -0.012 | 0 | 0.59 | 0.52 | 0.48 | 0.67 | 0.55 | 0.68 |
| 2 | 121350385 | rs896499 | 0.175 | 0.241 | -0.96 | 0.946 | 0.003 | 0.009 | 0.009 | 0 | 0.55 | 0.22 | 0.42 | 0.6 | 0.52 | 0.67 |
| 2 | 121357095 | rs1568836 | -1.431 | -0.025 | 0.176 | 1.171 | -0.009 | 0.145 | 0.145 | 1.64 | 0.12 | 0.54 | 0.08 | 0.23 | 0.07 | 0.03 |
| 2 | 121358255 | rs4849909 | -0.522 | -0.081 | -0.316 | 0.298 | 0.001 | -0.003 | -0.003 | 0 | 0.44 | 0.54 | 0.62 | 0.82 | 0.38 | 0.64 |
| 2 | 121358941 | rs4849911 | 1.144 | -0.854 | -0.596 | 0.677 | 0.037 | 0.01 | 0.01 | 1.378 | 0.73 | 0.19 | 0.84 | 0.8 | 0.84 | 0.93 |
| 2 | 121379349 | rs1349561 | 1.22 | -1.512 | -0.756 | 1.009 | 0.1 | -0.017 | -0.017 | 0 | 0.55 | 0.22 | 0.64 | 0.54 | 0.72 | 0.71 |
| 2 | 121380247 | rs7573498 | -1.201 | -1.749 | -0.125 | 1.199 | -0.011 | 0.055 | 0.055 | 1.238 | 0.14 | 0.19 | 0.08 | 0.24 | 0.09 | 0.07 |
| 2 | 121380361 | rs6541729 | -2.044 | -1.896 | 0.414 | 1.142 | -0.008 | 0 | 0 | 1.015 | 0.25 | 0.62 | 0.18 | 0.24 | 0.17 | 0.08 |
| 2 | 121393032 | rs4848608 | 0.087 | -1.913 | 0.398 | 1.159 | 0.011 | -0.012 | -0.012 | 0 | 0.79 | 0.74 | 0.81 | 0.66 | 0.88 | 0.79 |
| 2 | 121395161 | rs10201680 | -0.446 | -1.901 | 0.954 | 1.139 | -0.002 | 0.13 | 0.13 | 0 | 0.1 | 0.42 | 0.11 | 0.13 | 0.08 | 0.02 |
| 2 | 121400276 | rs4848094 | 1.955 | -1.905 | 0.479 | 1.035 | 0.123 | -0.021 | -0.021 | 0 | 0.55 | 0.1 | 0.62 | 0.42 | 0.72 | 0.71 |
| 2 | 121411430 | rs13412688 | -1.189 | -1.576 | 1.398 | 0.732 | 0 | -0.09 | -0.09 | 0 | 0.13 | 0.41 | 0.12 | 0 | 0.1 | 0.01 |
| 2 | 121416218 | rs7598132 | 0.343 | -1.444 | 0.74 | 1.04 | 0.01 | -0.014 | -0.014 | 0 | 0.8 | 0.79 | 0.82 | 0.63 | 0.89 | 0.8 |
| 2 | 121433891 | rs11674661 | 1.511 | -0.307 | 0.569 | 0.557 | -0.002 | 0.015 | 0.015 | 0.276 | 0.69 | 0.19 | 0.55 | 0.78 | 0.72 | 0.9 |
| 2 | 121439840 | rs11903904 | -0.749 | -0.177 | 1.178 | 0.517 | 0.025 | 0.013 | 0.013 | 0 | 0.14 | 0.17 | 0.3 | 0.14 | 0.24 | 0.07 |
| 2 | 121445220 | rs1979165 | 0.332 | -0.693 | 1.427 | 0.159 | -0.052 | -0.034 | -0.034 | 0 | 0.84 | 0.58 | 0.92 | 0.92 | 0.94 | 0.97 |
| 2 | 121456859 | rs965959 | 1.84 | -0.604 | 0.7 | 0.109 | 0.057 | 0.12 | 0.12 | 0.237 | 0.66 | 0.1 | 0.54 | 0.87 | 0.75 | 0.91 |
| 2 | 121460980 | rs1109002 | 0.406 | -1.036 | 0.035 | -0.277 | 0.052 | 0.079 | 0.079 | 4.718 | 0.24 | 0.08 | 0.07 | 0.08 | 0.1 | 0.06 |
| 2 | 121464750 | rs4848101 | -0.495 | -0.904 | 0.64 | -0.223 | -0.013 | -0.055 | -0.055 | 0 | 0.14 | 0.34 | 0.23 | 0.05 | 0.16 | 0.05 |
| 2 | 121465230 | rs2310938 | -0.163 | -0.893 | 0.481 | -0.218 | -0.011 | -0.044 | -0.044 | 0 | 0.13 | 0.25 | 0.23 | 0.05 | 0.16 | 0.05 |
| 2 | 121469965 | rs10496562 | -0.101 | -0.883 | 0.376 | -0.213 | -0.011 | -0.038 | -0.038 | 0.866 | 0.13 | 0.22 | 0.22 | 0.05 | 0.15 | 0.05 |
| 2 | 121470971 | rs12463908 | -0.299 | -0.868 | 0.359 | -0.222 | -0.001 | -0.042 | -0.042 | 0 | 0.15 | 0.22 | 0.24 | 0.05 | 0.2 | 0.07 |
| 2 | 121471046 | rs11685627 | 1.697 | -0.361 | -0.828 | -0.255 | -0.113 | -0.096 | -0.096 | 0.296 | 0.33 | 0.08 | 0.3 | 0.4 | 0.5 | 0.57 |
| 2 | 121472844 | rs13410771 | 0.265 | -0.646 | -0.217 | -0.495 | 0.032 | 0.037 | 0.037 | 5.475 | 0.26 | 0.18 | 0.07 | 0.08 | 0.1 | 0.07 |
| 2 | 121475836 | rs4848111 | -0.69 | -0.089 | -0.495 | -0.131 | 0.021 | 0.02 | 0.02 | 3.236 | 0.45 | 0.61 | 0.47 | 0.13 | 0.3 | 0.15 |
| 2 | 121485425 | rs2084802 | -0.018 | 1.06 | -0.653 | 0.543 | 0.015 | 0.02 | 0.02 | 0 | 0.18 | 0.22 | 0.23 | 0.34 | 0.29 | 0.25 |
| 2 | 121491552 | rs707482 | -0.288 | 0.46 | -1.769 | -0.117 | 0.073 | 0.186 | 0.186 | 0.582 | 0.53 | 0.9 | 0.55 | 0.24 | 0.36 | 0.3 |
| 2 | 121503661 | rs707485 | -0.577 | -0.035 | -0.856 | -0.175 | 0.001 | 0.009 | 0.009 | 0.052 | 0.56 | 0.59 | 0.49 | 0.33 | 0.51 | 0.47 |
| 2 | 121507873 | rs17389904 | 1.188 | 0.068 | -1.068 | -0.154 | -0.072 | -0.048 | -0.048 | 1.029 | 0.16 | 0.04 | 0.17 | 0.19 | 0.26 | 0.24 |
| 2 | 121509378 | rs776501 | -0.521 | -0.007 | -1.121 | -0.144 | 0.062 | 0.097 | 0.097 | 0.533 | 0.53 | 0.87 | 0.58 | 0.38 | 0.37 | 0.29 |
| 2 | 121513197 | rs776504 | -1.536 | -0.149 | -0.993 | -0.029 | -0.088 | -0.114 | -0.114 | 0 | 0.67 | 0.96 | 0.73 | 0.59 | 0.6 | 0.53 |
| 2 | 121513993 | rs11901641 | 0.791 | -0.087 | -0.896 | 0.012 | -0.019 | -0.01 | -0.01 | 0 | 0.19 | 0.11 | 0.18 | 0.19 | 0.26 | 0.26 |
| 2 | 121515390 | rs13034141 | 1.425 | -0.956 | -0.897 | -0.202 | -0.004 | 0.028 | 0.028 | 4.072 | 0.19 | 0.04 | 0.09 | 0.13 | 0.15 | 0.14 |
| 2 | 121515982 | rs707489 | -0.218 | -0.988 | -0.544 | -0.214 | -0.058 | 0.076 | 0.076 | 0.006 | 0.16 | 0.5 | 0.28 | 0.26 | 0.12 | 0.05 |
| 2 | 121524842 | rs17050378 | 0.577 | -1.143 | -0.292 | -0.169 | -0.039 | 0.082 | 0.082 | 0.073 | 0.11 | 0.21 | 0.17 | 0.25 | 0.09 | 0.04 |
| 2 | 121526871 | rs4848115 | 1.236 | -1.174 | -0.285 | -0.254 | -0.002 | -0.012 | -0.012 | 0 | 0.2 | 0.19 | 0.1 | 0.13 | 0.15 | 0.16 |
| 2 | 121530262 | rs4848627 | -0.33 | -1.094 | -0.635 | -0.214 | 0.011 | 0.002 | 0.002 | 1.433 | 0.35 | 0.61 | 0.3 | 0.38 | 0.24 | 0.2 |
| 2 | 121530351 | rs11677381 | 0.703 | -1.014 | -0.62 | -0.267 | 0.004 | 0.037 | 0.037 | 0 | 0.1 | 0.04 | 0.16 | 0 | 0.07 | 0.06 |
| 2 | 121532932 | rs2121992 | -0.264 | -0.916 | -0.534 | -0.176 | 0.007 | 0.005 | 0.005 | 2.478 | 0.34 | 0.58 | 0.28 | 0.3 | 0.24 | 0.2 |
| 2 | 121536569 | rs4848628 | 0.087 | -0.609 | -0.232 | -0.339 | -0.014 | -0.024 | -0.024 | 0 | 0.27 | 0.57 | 0.15 | 0.19 | 0.17 | 0.18 |
| 2 | 121544316 | rs2053989 | -1.483 | -0.362 | -0.889 | -0.296 | -0.137 | -0.043 | -0.043 | 0 | 0.58 | 0.95 | 0.5 | 0.38 | 0.47 | 0.49 |
| 2 | 121547526 | rs10170242 | -1.148 | -0.216 | 0.225 | 0.06 | -0.002 | -0.015 | -0.015 | 0 | 0.32 | 0.45 | 0.35 | 0.19 | 0.3 | 0.31 |
| 2 | 121551745 | rs7565812 | -1.453 | -0.217 | -0.293 | 0.087 | 0.011 | -0.011 | -0.011 | 0.014 | 0.36 | 0.71 | 0.41 | 0.38 | 0.31 | 0.34 |
| 2 | 121551822 | rs7605011 | -1.346 | -0.468 | 0.229 | -0.209 | -0.007 | -0.065 | -0.065 | 0 | 0.13 | 0.46 | 0.2 | 0.05 | 0.08 | 0.07 |
| 2 | 121553965 | rs4848630 | -1.095 | -0.162 | -0.568 | -0.052 | -0.015 | -0.071 | -0.071 | 0 | 0.11 | 0.66 | 0.2 | 0.02 | 0.06 | 0.02 |
| 2 | 121554317 | rs7569100 | -1.233 | -0.167 | -0.736 | -0.035 | -0.019 | -0.038 | -0.038 | 0 | 0.13 | 0.77 | 0.22 | 0.04 | 0.06 | 0.02 |
| 2 | 121555948 | rs735557 | 0.192 | -0.213 | -1.528 | 0.113 | 0.019 | 0.019 | 0.019 | 1.12 | 0.65 | 0.22 | 0.59 | 0.71 | 0.69 | 0.81 |
| 2 | 121562218 | rs746345 | 0.333 | -0.024 | -2.07 | 0.19 | 0.06 | 0.084 | 0.084 | 0.636 | 0.55 | 0.08 | 0.55 | 0.69 | 0.65 | 0.74 |
| 2 | 121562595 | rs4848632 | 0.176 | 0.061 | -1.897 | 0.217 | 0.051 | 0.072 | 0.072 | 0.038 | 0.58 | 0.19 | 0.55 | 0.7 | 0.65 | 0.74 |
| 2 | 121570665 | rs895485 | -0.247 | 0.592 | -1.914 | 0.478 | -0.055 | -0.024 | -0.024 | 0.824 | 0.37 | 0.23 | 0.5 | 0.48 | 0.49 | 0.56 |
| 2 | 121575289 | rs17005238 | -0.207 | 0.259 | -1.539 | 0.489 | -0.01 | 0.01 | 0.01 | 0 | 0.11 | 0.14 | 0.07 | 0.13 | 0.09 | 0.08 |
| 2 | 121578827 | rs895491 | 0.228 | 0.024 | -0.767 | 0.352 | -0.041 | -0.028 | -0.028 | 2.782 | 0.34 | 0.21 | 0.48 | 0.48 | 0.48 | 0.55 |
| 2 | 121579245 | rs4848637 | 0.856 | -0.332 | -0.767 | -0.227 | 0.023 | -0.021 | -0.021 | 1.689 | 0.36 | 0.16 | 0.32 | 0.39 | 0.3 | 0.3 |

| | | | | | | | | | | | | | | | | |
|---|---|---|---|---|---|---|---|---|---|---|---|---|---|---|---|---|
| 2 | 121580431 | rs7561607 | 0.183 | 0.333 | -0.352 | -0.121 | -0.003 | 0.027 | 0.027 | 0 | 0.52 | 0.36 | 0.41 | 0.29 | 0.48 | 0.37 |
| 2 | 121582095 | rs3943553 | -1.048 | 0.237 | -0.123 | -0.098 | -0.012 | 0.016 | 0.016 | 0.014 | 0.13 | 0.28 | 0.1 | 0.13 | 0.09 | 0.08 |
| 2 | 121585366 | rs4848640 | 1.348 | 0.181 | 0.176 | -0.166 | 0.012 | -0.008 | -0.008 | 0.045 | 0.95 | 0.72 | 0.95 | 1 | 0.98 | 1 |
| 2 | 121588250 | rs12185756 | -0.549 | 0.124 | -0.399 | -0.216 | -0.003 | -0.003 | -0.003 | 0 | 0.2 | 0.18 | 0.22 | 0.19 | 0.18 | 0.1 |
| 2 | 121588532 | rs1992900 | 0.442 | 0.348 | -0.979 | -0.487 | -0.11 | -0.135 | -0.135 | 0.123 | 0.32 | 0.03 | 0.41 | 0.62 | 0.42 | 0.51 |
| 2 | 121589440 | rs895479 | -0.827 | 0.448 | -0.444 | -0.413 | -0.001 | -0.001 | -0.001 | 0 | 0.22 | 0.24 | 0.22 | 0.19 | 0.18 | 0.1 |
| 2 | 121592523 | rs17005273 | -0.535 | 0.276 | -0.081 | -0.904 | -0.009 | -0.073 | -0.073 | 0 | 0.21 | 0.27 | 0.17 | 0 | 0.18 | 0.06 |
| 2 | 121595616 | rs7559777 | -1.405 | 0.374 | 0.038 | -0.993 | -0.003 | -0.013 | -0.013 | 0 | 0.55 | 0.51 | 0.65 | 0.75 | 0.6 | 0.47 |
| 2 | 121598768 | rs7557372 | -1.183 | 0.015 | 0.661 | -0.861 | -0.034 | -0.095 | -0.095 | 1.514 | 0.27 | 0.49 | 0.21 | 0.11 | 0.22 | 0.09 |
| 2 | 121600223 | rs17005295 | -0.481 | -0.015 | 0.684 | -0.872 | -0.033 | -0.139 | -0.139 | 0.079 | 0.22 | 0.4 | 0.16 | 0 | 0.18 | 0.05 |
| 2 | 121603584 | rs4848641 | -1.155 | 0.075 | 0.235 | -0.83 | -0.013 | -0.043 | -0.043 | 0.047 | 0.29 | 0.64 | 0.23 | 0.1 | 0.23 | 0.09 |
| 2 | 121612659 | rs2166898 | -0.375 | 0.714 | -0.319 | -0.62 | -0.003 | 0 | 0 | 0 | 0.12 | 0.15 | 0.15 | 0.13 | 0.15 | 0.1 |
| 2 | 121620828 | rs4143116 | 0.049 | 0.981 | -0.432 | -0.354 | 0.006 | 0.029 | 0.029 | 0 | 0.15 | 0.18 | 0.32 | 0.64 | 0.23 | 0.21 |
| 2 | 121627175 | rs4848126 | -0.576 | 0.508 | -0.633 | -0.831 | -0.001 | 0.057 | 0.057 | 0.164 | 0.46 | 0.32 | 0.59 | 0.77 | 0.53 | 0.41 |
| 2 | 121647802 | rs6541743 | -0.132 | 0.216 | -0.652 | -1.028 | 0.033 | 0.076 | 0.076 | 0 | 0.5 | 0.65 | 0.41 | 0.26 | 0.39 | 0.58 |
| 2 | 121648806 | rs12478509 | -0.955 | -0.267 | -0.029 | -1.256 | -0.037 | -0.138 | -0.138 | 0.653 | 0.19 | 0.5 | 0.17 | 0.01 | 0.13 | 0.09 |
| 2 | 121649279 | rs10199310 | -0.913 | -0.275 | 0.097 | -1.259 | -0.035 | -0.137 | -0.137 | 0 | 0.18 | 0.51 | 0.17 | 0.01 | 0.13 | 0.09 |
| 2 | 121651457 | rs277534 | 0.222 | 0.101 | -0.524 | -1.21 | 0.002 | -0.032 | -0.032 | 0 | 0.77 | 0.65 | 0.77 | 0.92 | 0.79 | 0.89 |
| 2 | 121652747 | rs277536 | -0.127 | -0.701 | -0.744 | -0.858 | 0.101 | 0.068 | 0.068 | 0 | 0.48 | 0.71 | 0.43 | 0.34 | 0.34 | 0.53 |
| 2 | 121656429 | rs277541 | -0.699 | -0.55 | -0.513 | -0.78 | -0.018 | -0.032 | -0.032 | 0.122 | 0.07 | 0.12 | 0.04 | 0 | 0.04 | 0.03 |
| 2 | 121668836 | rs6724298 | 0.359 | -0.861 | -0.682 | -0.91 | 0.015 | 0.017 | 0.017 | 0 | 0.37 | 0.24 | 0.41 | 0.24 | 0.27 | 0.52 |
| 2 | 121669113 | rs1187936 | -0.254 | -0.621 | -1.184 | -0.826 | 0.102 | 0.147 | 0.147 | 0.048 | 0.51 | 0.86 | 0.5 | 0.26 | 0.35 | 0.58 |
| 2 | 121670602 | rs1992902 | -1.236 | -0.512 | -0.057 | -0.865 | -0.028 | -0.143 | -0.143 | 1.159 | 0.18 | 0.53 | 0.15 | 0 | 0.13 | 0.08 |
| 2 | 121670669 | rs1992901 | 0.416 | -0.352 | -0.247 | -0.395 | 0.004 | -0.01 | -0.01 | 0.357 | 0.57 | 0.64 | 0.57 | 0.49 | 0.61 | 0.64 |
| 2 | 121670757 | rs277524 | 0.301 | -0.248 | 0.194 | 0.147 | 0.005 | 0.04 | 0.04 | 0 | 0.74 | 0.69 | 0.68 | 0.47 | 0.7 | 0.66 |
| 2 | 121676655 | rs895547 | 1.462 | -0.834 | 0.527 | -0.225 | -0.011 | -0.021 | -0.021 | 0 | 0.9 | 0.89 | 0.95 | 1 | 0.96 | 0.95 |
| 2 | 121678973 | rs7592746 | 2.334 | -0.985 | 1.3 | -0.299 | -0.061 | -0.084 | -0.084 | 0 | 0.87 | 0.6 | 0.92 | 1 | 0.95 | 0.94 |
| 2 | 121686439 | rs4848648 | 0.832 | -0.785 | 1.099 | -0.213 | -0.009 | -0.02 | -0.02 | 0 | 0.94 | 0.92 | 0.97 | 1 | 0.98 | 0.98 |
| 2 | 121696149 | rs2871874 | 2.178 | -0.994 | 1.866 | -0.311 | -0.028 | -0.036 | -0.036 | 0 | 0.91 | 0.41 | 0.94 | 1 | 0.99 | 0.99 |
| 2 | 121721467 | rs895545 | 2.697 | -1.128 | 1.883 | -0.415 | -0.069 | -0.101 | -0.101 | 0.279 | 0.82 | 0.3 | 0.9 | 1 | 0.95 | 0.95 |
| 2 | 121727565 | rs2592594 | 2.474 | -0.91 | 1.993 | -0.224 | -0.051 | -0.059 | -0.059 | 0 | 0.9 | 0.41 | 0.94 | 1 | 0.98 | 0.98 |
| 2 | 121727842 | rs1867906 | 1.578 | -0.772 | 1.624 | -0.157 | 0.01 | 0.002 | 0.002 | 0 | 0.94 | 0.73 | 0.96 | 1 | 0.98 | 0.99 |
| 2 | 121729360 | rs2592592 | 1.91 | -0.784 | 1.615 | -0.163 | 0.015 | 0.006 | 0.006 | 0.063 | 0.93 | 0.59 | 0.95 | 1 | 0.98 | 0.98 |
| 2 | 121733476 | rs280199 | 0.112 | -1.056 | 0.592 | 0.09 | 0.052 | -0.033 | -0.033 | 0.015 | 0.59 | 0.23 | 0.58 | 0.47 | 0.74 | 0.5 |
| 2 | 121735335 | rs7588213 | -0.05 | -1.266 | 1.129 | 0.021 | 0.009 | -0.016 | -0.016 | 0 | 0.35 | 0.25 | 0.37 | 0.53 | 0.25 | 0.48 |
| 2 | 121738820 | rs280194 | 0.117 | -1.222 | 1.387 | 0.038 | -0.025 | 0.023 | 0.023 | 0 | 0.61 | 0.52 | 0.61 | 0.49 | 0.74 | 0.51 |
| 2 | 121741811 | rs3820720 | 0.13 | -1.309 | 1.124 | 0.05 | 0.014 | -0.012 | -0.012 | 0 | 0.6 | 0.35 | 0.6 | 0.48 | 0.72 | 0.47 |
| 2 | 121746956 | rs3738880 | 0.217 | -1.368 | 1.123 | 0.114 | 0.013 | -0.01 | -0.01 | 0 | 0.6 | 0.36 | 0.58 | 0.47 | 0.72 | 0.48 |
| 2 | 121762093 | rs735370 | 0.887 | -0.66 | -0.141 | -0.308 | -0.13 | -0.148 | -0.148 | 0 | 0.3 | 0.03 | 0.38 | 0.44 | 0.41 | 0.27 |
| 2 | 121786101 | rs280210 | -0.053 | -1.26 | 0.955 | 0.048 | 0.009 | -0.016 | -0.016 | 0 | 0.36 | 0.27 | 0.38 | 0.53 | 0.3 | 0.52 |
| 2 | 121788164 | rs775806 | -0.459 | -1.302 | 1.505 | 0.028 | 0.018 | -0.015 | -0.015 | 0 | 0.39 | 0.65 | 0.41 | 0.53 | 0.3 | 0.53 |
| 2 | 121788630 | rs280213 | -0.459 | -1.302 | 1.54 | 0.049 | 0.015 | -0.013 | -0.013 | 2.997 | 0.39 | 0.63 | 0.41 | 0.54 | 0.3 | 0.52 |
| 2 | 121793815 | rs1878516 | -0.024 | -1.435 | 1.717 | 0.204 | 0.025 | -0.046 | -0.046 | 0 | 0.35 | 0.18 | 0.38 | 0.49 | 0.3 | 0.52 |
| 2 | 121797319 | rs11693648 | 0.15 | -1.439 | 1.714 | 0.219 | 0.031 | -0.016 | -0.016 | 0.054 | 0.6 | 0.32 | 0.58 | 0.51 | 0.69 | 0.47 |
| 2 | 121806835 | rs12613541 | 0.406 | -1.114 | 1.181 | 0.456 | 0.047 | -0.02 | -0.02 | 0 | 0.57 | 0.2 | 0.55 | 0.52 | 0.66 | 0.45 |
| 2 | 121807052 | rs4848142 | 0.349 | -1.12 | 1.194 | 0.585 | 0.042 | -0.044 | -0.044 | 0 | 0.57 | 0.22 | 0.56 | 0.5 | 0.66 | 0.45 |
| 2 | 121807776 | rs2249979 | -0.437 | -1.146 | 1.626 | 0.572 | -0.004 | 0.005 | 0.005 | 0 | 0.39 | 0.53 | 0.41 | 0.5 | 0.34 | 0.53 |
| 2 | 121816852 | rs280224 | -0.703 | -1.304 | 1.01 | 0.691 | -0.027 | -0.005 | -0.005 | 1.765 | 0.4 | 0.44 | 0.39 | 0.61 | 0.3 | 0.57 |
| 2 | 121824898 | rs2677510 | 0.862 | -1.323 | 0.288 | 0.772 | 0.044 | -0.059 | -0.059 | 0 | 0.66 | 0.2 | 0.64 | 0.49 | 0.76 | 0.52 |
| 2 | 121829977 | rs13384487 | -0.386 | -1.51 | 0.466 | 0.904 | -0.005 | 0.015 | 0.015 | 0 | 0.22 | 0.2 | 0.23 | 0.48 | 0.12 | 0.35 |
| 2 | 121835283 | rs4467269 | -0.659 | -1.89 | 0.33 | 0.552 | 0.011 | -0.011 | -0.011 | 0 | 0.35 | 0.66 | 0.39 | 0.46 | 0.19 | 0.39 |
| 2 | 121837519 | rs12711539 | 1.22 | -1.64 | -0.594 | 0.021 | -0.068 | -0.11 | -0.11 | 0 | 0.36 | 0.07 | 0.42 | 0.54 | 0.59 | 0.39 |
| 2 | 121840183 | rs4076328 | -0.14 | -1.328 | 0.834 | 0.619 | 0.013 | -0.024 | -0.024 | 1.678 | 0.27 | 0.19 | 0.34 | 0.45 | 0.19 | 0.37 |
| 2 | 121853648 | rs7602673 | -1.139 | -1.261 | 1.357 | 0.43 | 0.046 | -0.019 | -0.019 | 0.151 | 0.39 | 0.76 | 0.4 | 0.5 | 0.24 | 0.4 |
| 2 | 121853817 | rs7602881 | -1.069 | -1.263 | 1.333 | 0.429 | 0.071 | -0.027 | -0.027 | 0 | 0.4 | 0.8 | 0.4 | 0.5 | 0.24 | 0.42 |
| 2 | 121856858 | rs7561272 | -0.824 | -1.311 | 1.71 | 0.499 | 0.016 | -0.003 | -0.003 | 3.836 | 0.05 | 0.24 | 0.03 | 0 | 0.01 | 0.01 |
| 2 | 121859148 | rs17005869 | -0.692 | -1.3 | 1.636 | 0.52 | -0.028 | 0.023 | 0.023 | 0 | 0.32 | 0.42 | 0.35 | 0.49 | 0.19 | 0.36 |
| 2 | 121860596 | rs11122844 | -0.321 | -1.107 | 1.588 | 0.638 | -0.026 | 0.031 | 0.031 | 0 | 0.28 | 0.31 | 0.33 | 0.5 | 0.17 | 0.32 |
| 2 | 121868436 | rs7571982 | -0.351 | -1.1 | 1.579 | 0.69 | -0.038 | 0.059 | 0.059 | 0 | 0.28 | 0.34 | 0.28 | 0.47 | 0.17 | 0.33 |
| 2 | 121883746 | rs10169502 | -0.446 | -0.763 | 0.863 | 0.523 | -0.03 | 0.046 | 0.046 | 0 | 0.3 | 0.39 | 0.3 | 0.45 | 0.19 | 0.35 |
| 2 | 121886826 | rs6709558 | -0.266 | -0.616 | 0.982 | 0.419 | -0.02 | 0.039 | 0.039 | 0 | 0.27 | 0.35 | 0.28 | 0.49 | 0.18 | 0.34 |
| 2 | 121899272 | rs875175 | -0.617 | -0.568 | 0.531 | 0.436 | -0.029 | 0.054 | 0.054 | 0 | 0.31 | 0.49 | 0.35 | 0.51 | 0.21 | 0.37 |
| 2 | 121901126 | rs11885558 | -0.438 | -0.387 | -0.38 | 0.519 | 0.012 | -0.03 | -0.03 | 0 | 0.29 | 0.21 | 0.27 | 0.49 | 0.22 | 0.35 |
| 2 | 121909486 | rs4271771 | -0.528 | 0.188 | -0.252 | 0.058 | -0.005 | -0.002 | -0.002 | 0 | 0.54 | 0.54 | 0.54 | 0.52 | 0.4 | 0.37 |
| 2 | 121911207 | rs12616919 | 0.379 | 0.413 | -0.263 | 0.024 | 0 | 0 | 0 | 0 | 0.44 | 0.4 | 0.45 | 0.48 | 0.55 | 0.62 |
| 2 | 121911251 | rs2677533 | 0.175 | -0.457 | -0.03 | -0.319 | -0.02 | -0.119 | -0.119 | 1.092 | 0.77 | 0.56 | 0.88 | 0.97 | 0.85 | 0.91 |
| 2 | 121913022 | rs2677532 | 0.248 | 0.19 | -0.208 | 0.006 | 0.003 | 0.003 | 0.003 | 0 | 0.46 | 0.37 | 0.34 | 0.52 | 0.48 | 0.53 |
| 2 | 121914455 | rs13018265 | 0.128 | -0.014 | -0.418 | -0.126 | 0.019 | 0.024 | 0.024 | 0.239 | 0.2 | 0.14 | 0.1 | 0.03 | 0.09 | 0.11 |
| 2 | 121916540 | rs12617448 | -0.038 | 1.024 | -1.01 | 0.725 | -0.084 | -0.133 | -0.133 | 0.118 | 0.29 | 0.06 | 0.49 | 0.43 | 0.36 | 0.3 |

| | | | | | | | | | | | | | | | | |
|---|---|---|---|---|---|---|---|---|---|---|---|---|---|---|---|---|
| 2 | 121918815 | rs4848685 | -0.296 | 0.903 | -0.839 | 0.687 | -0.04 | -0.039 | -0.039 | 0 | 0.37 | 0.19 | 0.54 | 0.5 | 0.46 | 0.46 |
| 2 | 121925398 | rs4488664 | -0.328 | 0.661 | -1.636 | 0.62 | -0.003 | 0.037 | 0.037 | 0 | 0.49 | 0.18 | 0.49 | 0.36 | 0.51 | 0.51 |
| 2 | 121937079 | rs10171378 | -0.428 | -0.037 | -1.145 | 0.464 | -0.003 | -0.001 | -0.001 | 0.073 | 0.36 | 0.35 | 0.25 | 0.29 | 0.27 | 0.53 |
| 2 | 121940001 | rs4848692 | 0.72 | -0.152 | -1.558 | 0.23 | -0.052 | -0.063 | -0.063 | 0 | 0.4 | 0.14 | 0.6 | 0.52 | 0.53 | 0.37 |
| 2 | 121942213 | rs4848694 | -0.339 | -0.45 | -0.773 | 0.163 | 0 | -0.009 | -0.009 | 0 | 0.82 | 0.79 | 0.89 | 0.86 | 0.82 | 0.91 |
| 2 | 121943196 | rs10864869 | 0.557 | -0.763 | -0.404 | -0.154 | 0.011 | -0.003 | -0.003 | 0 | 0.47 | 0.33 | 0.65 | 0.52 | 0.58 | 0.37 |
| 2 | 121944266 | rs11890342 | 0.088 | -0.841 | -0.335 | -0.167 | -0.021 | 0.007 | 0.007 | 0 | 0.34 | 0.42 | 0.24 | 0.34 | 0.24 | 0.55 |
| 2 | 121947629 | rs10189866 | -0.726 | -1.212 | -0.376 | -0.149 | 0.003 | 0 | 0 | 0 | 0.49 | 0.53 | 0.35 | 0.5 | 0.41 | 0.55 |
| 2 | 121947761 | rs12463998 | -0.417 | -1.319 | -0.567 | -0.102 | -0.019 | -0.058 | -0.058 | 2.999 | 0.39 | 0.16 | 0.3 | 0.52 | 0.42 | 0.53 |
| 2 | 121948307 | rs4622731 | -0.179 | -0.79 | -0.418 | 0.025 | 0.001 | 0.032 | 0.032 | 0 | 0.48 | 0.35 | 0.39 | 0.38 | 0.53 | 0.64 |
| 2 | 121948846 | rs17006117 | 0.326 | -0.533 | -0.967 | 0.199 | -0.001 | -0.018 | -0.018 | 0.883 | 0.41 | 0.27 | 0.5 | 0.48 | 0.4 | 0.33 |
| 2 | 121955160 | rs4848147 | 0.42 | -0.337 | -1.792 | 0.283 | -0.044 | -0.093 | -0.093 | 0 | 0.37 | 0.06 | 0.37 | 0.45 | 0.34 | 0.25 |
| 2 | 121957702 | rs17006161 | -0.195 | -0.638 | -1.229 | 0.466 | 0.018 | -0.007 | -0.007 | 0.459 | 0.35 | 0.18 | 0.27 | 0.38 | 0.32 | 0.59 |
| 2 | 121962947 | rs6753743 | 1.06 | -0.526 | -0.02 | 0.639 | -0.031 | 0.032 | 0.032 | 0.122 | 0.84 | 0.4 | 0.82 | 0.79 | 0.89 | 0.89 |
| 2 | 121965134 | rs2002067 | -1.657 | -0.545 | 0.627 | 0.631 | 0.039 | 0.187 | 0.187 | 0.955 | 0.09 | 0.49 | 0.17 | 0.18 | 0.06 | 0.06 |
| 2 | 121968032 | rs4076519 | 1.526 | -0.732 | 0.45 | 0.541 | -0.054 | 0.009 | 0.009 | 0.075 | 0.78 | 0.56 | 0.77 | 0.78 | 0.85 | 0.82 |
| 2 | 121968414 | rs4073806 | 1.241 | -0.342 | 0.333 | 0.678 | 0 | 0.051 | 0.051 | 0 | 0.9 | 0.74 | 0.84 | 0.79 | 0.91 | 0.89 |
| 2 | 121969468 | rs9653393 | -1.388 | -0.387 | 0.227 | 0.505 | -0.03 | -0.015 | -0.015 | 0 | 0.16 | 0.43 | 0.11 | 0.13 | 0.11 | 0.16 |
| 2 | 121976642 | rs17794420 | 0.348 | 0.083 | -1.139 | 0.616 | -0.095 | -0.05 | -0.05 | 0.007 | 0.34 | 0.11 | 0.38 | 0.34 | 0.46 | 0.37 |
| 2 | 121977438 | rs17006183 | 0.084 | -0.565 | -0.511 | 0.568 | -0.008 | -0.013 | -0.013 | 2.139 | 0.34 | 0.33 | 0.29 | 0.26 | 0.3 | 0.46 |
| 2 | 121981210 | rs10200387 | -0.06 | -0.506 | -0.287 | 0.436 | -0.017 | -0.028 | -0.028 | 0 | 0.36 | 0.43 | 0.3 | 0.26 | 0.31 | 0.48 |
| 2 | 121981608 | rs3795853 | 0.076 | -0.456 | -0.275 | 0.435 | -0.012 | -0.023 | -0.023 | 0 | 0.36 | 0.42 | 0.3 | 0.26 | 0.31 | 0.46 |
| 2 | 121981950 | rs11890430 | -0.067 | -0.451 | -0.3 | 0.45 | -0.016 | -0.029 | -0.029 | 0 | 0.36 | 0.44 | 0.3 | 0.26 | 0.31 | 0.46 |
| 2 | 121989489 | rs2304667 | 0.924 | -0.322 | -0.66 | 0.398 | -0.15 | -0.041 | -0.041 | 0 | 0.4 | 0.07 | 0.41 | 0.35 | 0.5 | 0.37 |
| 2 | 121991967 | rs11677172 | 0.773 | -0.477 | -0.458 | 0.302 | 0.006 | -0.023 | -0.023 | 0 | 0.08 | 0.07 | 0.1 | 0 | 0.15 | 0.06 |
| 2 | 121992083 | rs2713195 | -0.351 | -0.449 | -0.214 | 0.331 | -0.003 | -0.004 | -0.004 | 0 | 0.62 | 0.49 | 0.69 | 0.74 | 0.69 | 0.54 |
| 2 | 121992208 | rs3768904 | 1 | -0.479 | -0.31 | 0.394 | -0.09 | 0.001 | 0.001 | 0 | 0.42 | 0.15 | 0.42 | 0.35 | 0.5 | 0.37 |
| 2 | 121992921 | rs2250092 | 0.325 | -0.704 | -0.044 | 0.605 | -0.037 | 0.093 | 0.093 | 0.286 | 0.12 | 0.25 | 0.17 | 0.39 | 0.04 | 0.09 |
| 2 | 121996388 | rs2713203 | -0.148 | -0.754 | 0.21 | 0.24 | 0.032 | -0.034 | -0.034 | 0 | 0.78 | 0.66 | 0.78 | 0.9 | 0.71 | 0.77 |
| 2 | 121997698 | rs750416 | 1.254 | -1.139 | 0.2 | 0.417 | 0.011 | 0.013 | 0.013 | 0 | 0.51 | 0.26 | 0.54 | 0.54 | 0.54 | 0.43 |
| 2 | 122000796 | rs10496564 | -0.296 | -1.623 | 0.099 | 0.424 | -0.022 | -0.011 | -0.011 | 0 | 0.06 | 0.12 | 0.03 | 0.02 | 0 | 0.03 |
| 2 | 122003073 | rs882759 | -2.004 | -1.545 | 0.113 | 0.691 | 0.004 | 0.036 | 0.036 | 0 | 0.54 | 0.9 | 0.46 | 0.65 | 0.4 | 0.6 |
| 2 | 122003356 | rs3754633 | -0.529 | -1.638 | 0.424 | 0.757 | 0.003 | -0.003 | -0.003 | 0 | 0.22 | 0.24 | 0.19 | 0.24 | 0.18 | 0.37 |
| 2 | 122003442 | rs2713209 | 1.424 | -1.634 | 0.32 | 0.753 | 0.012 | 0.002 | 0.002 | 1.216 | 0.08 | 0.07 | 0.07 | 0.29 | 0.03 | 0.04 |
| 2 | 122005928 | rs1530124 | -0.535 | -1.582 | 0.563 | 0.704 | -0.002 | 0.001 | 0.001 | 0 | 0.22 | 0.27 | 0.19 | 0.24 | 0.18 | 0.37 |
| 2 | 122007858 | rs2713207 | -0.491 | -1.149 | 0.407 | 0.78 | -0.008 | 0.011 | 0.011 | 0.657 | 0.23 | 0.3 | 0.18 | 0.33 | 0.17 | 0.09 |
| 2 | 122016946 | rs2953080 | -1.248 | -1.121 | 0.048 | 0.551 | 0.064 | 0.089 | 0.089 | 0 | 0.49 | 0.76 | 0.44 | 0.71 | 0.37 | 0.51 |
| 2 | 122036871 | rs2953083 | 0.927 | -1.29 | -0.331 | 0.493 | -0.015 | 0.094 | 0.094 | 0.36 | 0.48 | 0.18 | 0.55 | 0.4 | 0.52 | 0.43 |
| 2 | 122037415 | rs11688688 | 0.235 | -2.19 | 0.547 | 0.36 | -0.006 | -0.046 | -0.046 | 0 | 0.14 | 0.15 | 0.1 | 0 | 0.11 | 0.05 |
| 2 | 122037433 | rs6714806 | -0.784 | -2.354 | 0.937 | 0.735 | -0.042 | 0.01 | 0.01 | 0 | 0.27 | 0.48 | 0.22 | 0.27 | 0.17 | 0.16 |
| 2 | 122039095 | rs7596689 | -0.587 | -1.885 | 1.079 | 0.459 | -0.03 | -0.037 | -0.037 | 0.24 | 0.13 | 0.27 | 0.05 | 0.04 | 0.06 | 0.11 |
| 2 | 122043586 | rs6717937 | 0.241 | -1.887 | 0.732 | 0.449 | -0.021 | -0.052 | -0.052 | 0 | 0.1 | 0.14 | 0.03 | 0 | 0.05 | 0.07 |
| 2 | 122046743 | rs2713245 | -1.093 | -1.982 | -0.338 | 0.399 | -0.161 | -0.038 | -0.038 | 0 | 0.71 | 0.96 | 0.55 | 0.78 | 0.51 | 0.76 |
| 2 | 122047352 | rs2580344 | 0.432 | -2.004 | 0.211 | 0.575 | -0.002 | -0.001 | -0.001 | 0 | 0.46 | 0.48 | 0.45 | 0.64 | 0.38 | 0.41 |
| 2 | 122051230 | rs753727 | -2.083 | -1.921 | 1.378 | 0.778 | -0.007 | -0.001 | -0.001 | 0.211 | 0.24 | 0.69 | 0.14 | 0.23 | 0.11 | 0.37 |
| 2 | 122055684 | rs6760235 | -0.991 | -1.712 | 1.328 | 0.613 | -0.035 | 0.12 | 0.12 | 0.014 | 0.13 | 0.44 | 0.09 | 0.16 | 0.06 | 0.13 |
| 2 | 122063844 | rs2580359 | -1.401 | -1.75 | 0.831 | 0.376 | 0.016 | 0.006 | 0.006 | 0.314 | 0.23 | 0.78 | 0.17 | 0.12 | 0.08 | 0.14 |
| 2 | 122064537 | rs2580358 | -0.929 | -1.56 | 0.503 | -0.076 | -0.084 | -0.133 | -0.133 | 3.478 | 0.18 | 0.42 | 0.08 | 0 | 0.07 | 0.13 |
| 2 | 122078698 | rs2580373 | -0.824 | -1.512 | -0.724 | 0.07 | 0.129 | 0.084 | 0.084 | 0.046 | 0.43 | 0.88 | 0.34 | 0.22 | 0.27 | 0.4 |
| 2 | 122083042 | rs2580352 | -0.722 | -1.538 | -0.248 | 0.069 | 0.068 | 0.046 | 0.046 | 0 | 0.41 | 0.7 | 0.33 | 0.22 | 0.27 | 0.41 |
| 2 | 122084428 | rs2580347 | -0.795 | -1.507 | 0.025 | 0.075 | -0.025 | -0.016 | -0.016 | 4.795 | 0.4 | 0.46 | 0.31 | 0.22 | 0.27 | 0.42 |
| 2 | 122095640 | rs2919051 | -1.035 | -1.686 | -0.215 | -0.141 | 0.016 | 0.015 | 0.015 | 0.194 | 0.26 | 0.7 | 0.19 | 0.13 | 0.1 | 0.23 |
| 2 | 122115494 | rs12615868 | -0.691 | -1.856 | 0.889 | -0.242 | -0.032 | 0 | 0 | 0 | 0.15 | 0.27 | 0.11 | 0.13 | 0.07 | 0.2 |
| 2 | 122117738 | rs12479320 | -1.318 | -2.551 | -0.386 | -0.32 | -0.107 | -0.007 | -0.007 | 0 | 0.66 | 0.98 | 0.65 | 0.78 | 0.43 | 0.67 |
| 2 | 122123637 | rs6735895 | 0.032 | -2.522 | 0.075 | -0.299 | -0.036 | 0 | 0 | 0 | 0.15 | 0.26 | 0.11 | 0.13 | 0.06 | 0.2 |
| 2 | 122145157 | rs2015439 | 0.321 | -2.098 | 0.316 | -0.346 | -0.013 | -0.015 | -0.015 | 0 | 0.12 | 0.14 | 0.09 | 0.05 | 0.05 | 0.08 |
| 2 | 122148174 | rs10496565 | 1.329 | -2.617 | -0.07 | -0.026 | -0.011 | -0.101 | -0.101 | 0.023 | 0.13 | 0.01 | 0.27 | 0.5 | 0.14 | 0.24 |
| 2 | 122161850 | rs9807995 | -0.557 | -2.59 | 0.819 | -0.032 | -0.064 | -0.008 | -0.008 | 0 | 0.18 | 0.52 | 0.14 | 0.13 | 0.07 | 0.2 |
| 2 | 122164469 | rs10208849 | -0.942 | -2.588 | 0.954 | -0.075 | -0.04 | 0.054 | 0.054 | 0.161 | 0.2 | 0.56 | 0.2 | 0.28 | 0.1 | 0.25 |
| 2 | 122167101 | rs6725314 | -0.04 | -2.593 | 0.922 | -0.01 | -0.039 | 0.017 | 0.017 | 0 | 0.15 | 0.3 | 0.11 | 0.15 | 0.07 | 0.2 |
| 2 | 122176722 | rs17006463 | 0.022 | -2.594 | 0.809 | -0.047 | -0.032 | 0.001 | 0.001 | 0 | 0.15 | 0.27 | 0.11 | 0.13 | 0.07 | 0.2 |
| 2 | 122193423 | rs11685153 | 1.437 | -2.775 | 0.715 | -0.157 | 0.011 | 0.108 | 0.108 | 0 | 0.22 | 0.05 | 0.11 | 0 | 0.16 | 0.14 |
| 2 | 122207005 | rs1975305 | -1.217 | -2.809 | 0.836 | -0.153 | 0.137 | 0.126 | 0.126 | 0.203 | 0.5 | 0.84 | 0.36 | 0.27 | 0.29 | 0.43 |
| 2 | 122212519 | rs17006496 | -0.889 | -2.325 | 1.257 | 0.154 | -0.043 | -0.031 | -0.031 | 0.167 | 0.19 | 0.64 | 0.14 | 0.1 | 0.06 | 0.2 |
| 2 | 122216419 | rs2304560 | -0.771 | -2.32 | 1.286 | 0.14 | -0.053 | -0.013 | -0.013 | 0 | 0.18 | 0.6 | 0.14 | 0.13 | 0.07 | 0.2 |
| 2 | 122224357 | rs7587659 | 0.544 | -2.35 | 1.232 | 0.155 | 0.049 | -0.043 | -0.043 | 0 | 0.38 | 0.23 | 0.4 | 0.51 | 0.3 | 0.37 |
| 2 | 122252266 | rs10185275 | -1.751 | -2.227 | 1.39 | 0.064 | 0.045 | -0.008 | -0.008 | 0 | 0.28 | 0.75 | 0.24 | 0.27 | 0.13 | 0.3 |
| 2 | 122275058 | rs12614801 | -1.043 | -2.077 | 1.423 | 0.14 | -0.031 | 0.031 | 0.031 | 0.305 | 0.2 | 0.64 | 0.2 | 0.27 | 0.1 | 0.25 |
| 2 | 122288029 | rs2289192 | -0.783 | -2.079 | 1.439 | 0.133 | -0.053 | -0.011 | -0.011 | 0.878 | 0.18 | 0.59 | 0.14 | 0.13 | 0.07 | 0.2 |

| | | | | | | | | | | | | | | | | |
|---|---|---|---|---|---|---|---|---|---|---|---|---|---|---|---|---|
| 2 | 122319019 | rs7557276 | -1.735 | -1.208 | 0.987 | -0.292 | -0.157 | 0.15 | 0.15 | 0 | 0.76 | 0.98 | 0.79 | 0.99 | 0.57 | 0.89 |
| 2 | 122324025 | rs4482489 | -1.009 | -1.32 | 1.498 | 0.181 | -0.031 | 0.03 | 0.03 | 0.337 | 0.2 | 0.64 | 0.2 | 0.28 | 0.1 | 0.25 |
| 2 | 122326718 | rs6541785 | -1.039 | -1.325 | 1.495 | 0.178 | -0.029 | 0.027 | 0.027 | 0.632 | 0.2 | 0.64 | 0.2 | 0.28 | 0.1 | 0.25 |
| 2 | 122343704 | rs6541786 | -0.944 | -1.313 | 1.506 | 0.203 | -0.038 | 0.076 | 0.076 | 0.013 | 0.19 | 0.56 | 0.19 | 0.28 | 0.1 | 0.25 |
| 2 | 122345071 | rs6746090 | -0.597 | -1.314 | 1.509 | 0.205 | -0.055 | -0.003 | -0.003 | 3.11 | 0.17 | 0.54 | 0.13 | 0.13 | 0.07 | 0.2 |
| 2 | 122346821 | rs10205258 | -0.136 | -1.32 | 1.609 | 0.201 | -0.011 | 0 | 0 | 0.629 | 0.14 | 0.18 | 0.1 | 0.13 | 0.07 | 0.2 |
| 2 | 122349120 | rs10496574 | -1.134 | -1.32 | 1.598 | 0.202 | -0.038 | 0.053 | 0.053 | 0.317 | 0.2 | 0.59 | 0.2 | 0.28 | 0.1 | 0.25 |
| 2 | 122354655 | rs6707415 | -1.101 | -1.319 | 1.602 | 0.202 | -0.038 | 0.056 | 0.056 | 0.209 | 0.19 | 0.58 | 0.2 | 0.28 | 0.1 | 0.25 |
| 2 | 122375371 | rs7571169 | -0.982 | -1.319 | 1.602 | 0.202 | -0.038 | 0.076 | 0.076 | 0.013 | 0.19 | 0.56 | 0.19 | 0.28 | 0.1 | 0.25 |
| 2 | 122376857 | rs287790 | -1.669 | -1.378 | 1.627 | 0.17 | 0.048 | -0.016 | -0.016 | 0 | 0.27 | 0.78 | 0.24 | 0.28 | 0.13 | 0.29 |
| 2 | 122377347 | rs10173198 | -1.112 | -0.987 | 1.55 | 0.417 | -0.038 | 0.076 | 0.076 | 0.013 | 0.19 | 0.56 | 0.19 | 0.28 | 0.1 | 0.25 |
| 2 | 122386658 | rs12613842 | -0.295 | -0.987 | 1.556 | 0.496 | -0.023 | 0.007 | 0.007 | 0 | 0.14 | 0.22 | 0.11 | 0.14 | 0.07 | 0.2 |
| 2 | 122400728 | rs2034138 | -1.167 | -0.986 | 1.227 | 0.497 | -0.037 | 0.059 | 0.059 | 0.337 | 0.2 | 0.59 | 0.2 | 0.28 | 0.1 | 0.25 |
| 2 | 122401005 | rs2034136 | -0.536 | -0.966 | 1.229 | 0.505 | -0.054 | 0.026 | 0.026 | 0 | 0.17 | 0.46 | 0.13 | 0.14 | 0.07 | 0.2 |
| 2 | 122423386 | rs6717212 | -0.885 | -0.963 | 0.916 | 0.474 | -0.038 | 0.086 | 0.086 | 0.013 | 0.19 | 0.56 | 0.19 | 0.28 | 0.1 | 0.25 |
| 2 | 122453962 | rs10210011 | -0.762 | -0.803 | 0.905 | 0.51 | -0.023 | 0.108 | 0.108 | 0.129 | 0.17 | 0.56 | 0.19 | 0.28 | 0.1 | 0.23 |
| 2 | 122473940 | rs12999360 | 0.317 | 0.412 | 0.048 | 0.487 | -0.155 | -0.064 | -0.064 | 0 | 0.24 | 0.01 | 0.31 | 0.21 | 0.42 | 0.27 |
| 2 | 122482142 | rs7581005 | -0.869 | 0.279 | 0.602 | 0.553 | 0.062 | 0.21 | 0.21 | 1.06 | 0.06 | 0.41 | 0.07 | 0.14 | 0.02 | 0.13 |
| 2 | 122484622 | rs17006765 | -0.829 | 0.279 | 0.602 | 0.553 | 0.064 | 0.213 | 0.213 | 1.042 | 0.06 | 0.41 | 0.07 | 0.14 | 0.02 | 0.13 |
| 2 | 122489374 | rs12616209 | -1.316 | 0.295 | 0.62 | 0.751 | 0.06 | 0.281 | 0.281 | 1.097 | 0.09 | 0.45 | 0.14 | 0.27 | 0.06 | 0.17 |
| 2 | 122489932 | rs6726677 | -1.335 | 0.383 | 0.62 | 0.736 | 0.065 | 0.275 | 0.275 | 1.228 | 0.09 | 0.46 | 0.15 | 0.27 | 0.06 | 0.17 |
| 2 | 122492869 | rs11676686 | 0.063 | 0.44 | 0.139 | 0.627 | -0.172 | -0.066 | -0.066 | 0 | 0.27 | 0.01 | 0.32 | 0.23 | 0.44 | 0.28 |
| 2 | 122509570 | rs11689101 | 0.936 | 0.641 | 0.133 | 0.539 | -0.034 | 0.029 | 0.029 | 0 | 0.06 | 0.01 | 0.07 | 0 | 0.14 | 0.11 |
| 2 | 122515214 | rs11122867 | -0.029 | 0.343 | 0.245 | 0.561 | -0.164 | -0.064 | -0.064 | 0.871 | 0.27 | 0.01 | 0.32 | 0.22 | 0.41 | 0.28 |
| 2 | 122547210 | rs7573614 | 0.294 | -0.032 | 0.651 | 0.446 | 0.001 | 0.008 | 0.008 | 0 | 0.83 | 0.86 | 0.9 | 0.9 | 0.84 | 0.77 |
| 2 | 122562188 | rs6734147 | 1.166 | -0.102 | 1.108 | 0.526 | -0.012 | 0.025 | 0.025 | 0.843 | 0.74 | 0.41 | 0.75 | 0.65 | 0.79 | 0.69 |
| 2 | 122562779 | rs1037366 | 0.575 | -0.142 | 1.399 | 0.513 | 0.017 | 0.049 | 0.049 | 0.811 | 0.83 | 0.63 | 0.8 | 0.75 | 0.82 | 0.88 |
| 2 | 122566566 | rs1026224 | 1.533 | -0.246 | 1.315 | 0.567 | -0.018 | 0.035 | 0.035 | 2.544 | 0.73 | 0.44 | 0.75 | 0.65 | 0.78 | 0.69 |
| 2 | 122567150 | rs11904524 | -0.506 | -0.239 | 1.097 | 0.382 | 0.014 | -0.006 | -0.006 | 0.173 | 0.09 | 0.05 | 0.03 | 0.1 | 0.03 | 0.19 |
| 2 | 122574017 | rs10864871 | 0.71 | -0.434 | 0.374 | 0.143 | 0.017 | -0.003 | -0.003 | 0 | 0.45 | 0.44 | 0.4 | 0.61 | 0.3 | 0.31 |
| 2 | 122574851 | rs4522608 | 0.396 | 0.39 | 0.183 | -0.069 | 0.005 | -0.007 | -0.007 | 0 | 0.27 | 0.24 | 0.32 | 0.11 | 0.41 | 0.33 |
| 2 | 122576458 | rs6541804 | 0.638 | 0.314 | 0.003 | 0.036 | -0.004 | 0.004 | 0.004 | 0 | 0.7 | 0.64 | 0.72 | 0.71 | 0.72 | 0.62 |
| 2 | 122577163 | rs13018495 | 0.272 | 0.287 | -0.474 | -0.54 | 0.009 | 0.083 | 0.083 | 0.004 | 0.18 | 0.03 | 0.09 | 0 | 0.14 | 0.14 |
| 2 | 122578292 | rs7567579 | -0.031 | 0.025 | -0.693 | -1.284 | 0.001 | -0.018 | -0.018 | 0 | 0.55 | 0.63 | 0.32 | 0.07 | 0.42 | 0.44 |
| 2 | 122578634 | rs4848725 | 0.791 | -0.011 | -0.861 | -1.281 | -0.012 | 0.103 | 0.103 | 0.407 | 0.2 | 0.02 | 0.11 | 0 | 0.18 | 0.18 |
| 2 | 122579512 | rs7571542 | -0.946 | -0.06 | -0.716 | -1.011 | -0.002 | 0.003 | 0.003 | 0 | 0.79 | 0.74 | 0.59 | 0.19 | 0.79 | 0.73 |
| 2 | 122586232 | rs6541806 | 1.247 | -0.061 | -0.75 | -1.056 | -0.025 | -0.099 | -0.099 | 0 | 0.29 | 0.4 | 0.16 | 0 | 0.15 | 0.11 |
| 2 | 122588385 | rs4490191 | -0.127 | 0.237 | -1.136 | -0.774 | -0.011 | -0.035 | -0.035 | 0 | 0.42 | 0.17 | 0.49 | 0.59 | 0.39 | 0.41 |
| 2 | 122590318 | rs4390770 | 1.191 | 0.301 | -0.739 | -1.118 | 0.053 | 0.019 | 0.019 | 0 | 0.38 | 0.26 | 0.43 | 0.71 | 0.23 | 0.35 |
| 2 | 122591071 | rs9808536 | 0.235 | 0.667 | -0.393 | -1.33 | -0.002 | -0.023 | -0.023 | 0.722 | 0.58 | 0.58 | 0.59 | 0.79 | 0.43 | 0.58 |
| 2 | 122598583 | rs12614797 | 0.25 | 0.786 | -0.623 | -1.167 | -0.032 | 0.032 | 0.032 | 0 | 0.14 | 0.04 | 0.11 | 0.08 | 0.19 | 0.23 |
| 2 | 122599806 | rs10460462 | 1.531 | 0.699 | -0.378 | -1.055 | 0 | 0.013 | 0.013 | 0 | 0.05 | 0.06 | 0.11 | 0.12 | 0.09 | 0.06 |
| 2 | 122601302 | rs10460453 | 1.149 | 0.68 | -0.219 | -1.064 | -0.003 | 0.044 | 0.044 | 0 | 0.06 | 0.12 | 0.11 | 0.12 | 0.09 | 0.06 |
| 2 | 122607245 | rs6731977 | 0.195 | 1.348 | -0.164 | -1.256 | 0.009 | -0.05 | -0.05 | 0 | 0.73 | 0.45 | 0.75 | 0.91 | 0.69 | 0.69 |
| 2 | 122608702 | rs6710409 | 0.757 | 1.386 | -0.253 | -1.33 | -0.004 | -0.015 | -0.015 | 0 | 0.57 | 0.53 | 0.58 | 0.72 | 0.4 | 0.43 |
| 2 | 122612746 | rs11122875 | 0.183 | 1.669 | -0.666 | -1.263 | -0.13 | 0.092 | 0.092 | 0 | 0.26 | 0.04 | 0.2 | 0.1 | 0.37 | 0.3 |
| 2 | 122615926 | rs4267513 | -0.583 | 1.185 | -0.184 | -1.163 | 0.002 | 0.032 | 0.032 | 0 | 0.41 | 0.25 | 0.4 | 0.27 | 0.6 | 0.56 |
| 2 | 122616143 | rs4316942 | -0.633 | 1.145 | 0.065 | -1.181 | 0 | 0.004 | 0.004 | 0 | 0.41 | 0.41 | 0.41 | 0.27 | 0.6 | 0.56 |
| 2 | 122616990 | rs4848730 | 0.75 | 1.154 | -0.376 | -1.227 | 0.002 | -0.042 | -0.042 | 0 | 0.15 | 0.2 | 0.11 | 0 | 0.17 | 0.09 |
| 2 | 122620193 | rs4430963 | -1.423 | 0.875 | -0.322 | -1.103 | 0.04 | 0.025 | 0.025 | 0.019 | 0.59 | 0.66 | 0.52 | 0.28 | 0.77 | 0.66 |
| 2 | 122628361 | rs11889169 | -0.796 | 1.878 | -0.306 | -0.985 | -0.001 | -0.014 | -0.014 | 0 | 0.35 | 0.41 | 0.29 | 0.15 | 0.47 | 0.4 |
| 2 | 122630611 | rs9679491 | -0.415 | 2.101 | -0.142 | -0.774 | 0.043 | -0.032 | -0.032 | 0 | 0.16 | 0.33 | 0.16 | 0.09 | 0.25 | 0.21 |
| 2 | 122656947 | rs4337480 | 0.291 | 2.033 | -0.162 | -0.806 | 0.03 | -0.054 | -0.054 | 0.657 | 0.11 | 0.26 | 0.11 | 0 | 0.16 | 0.09 |
| 2 | 122694063 | rs7604413 | -1.143 | 2.484 | 0.006 | -0.18 | 0.083 | 0.06 | 0.06 | 1.711 | 0.17 | 0.41 | 0.33 | 0.26 | 0.27 | 0.24 |
| 2 | 122696902 | rs1002411 | -0.645 | 2.449 | -0.218 | -0.81 | -0.022 | 0.08 | 0.08 | 1.073 | 0.8 | 0.94 | 0.88 | 0.93 | 0.79 | 0.79 |
| 2 | 122708759 | rs17778385 | 0.741 | 2.344 | -0.121 | -0.817 | -0.012 | 0.002 | 0.002 | 0.104 | 0.09 | 0.06 | 0.1 | 0 | 0.16 | 0.09 |
| 2 | 122747975 | rs1219913 | -0.803 | 2.311 | -0.105 | -0.772 | 0.062 | 0.01 | 0.01 | 0.303 | 0.16 | 0.34 | 0.17 | 0.17 | 0.27 | 0.19 |
| 2 | 122756270 | rs9646911 | 0.95 | 2.39 | 0.322 | -0.825 | 0.021 | -0.023 | -0.023 | 0.061 | 0.63 | 0.44 | 0.71 | 0.74 | 0.55 | 0.61 |
| 2 | 122761849 | rs1712865 | 0.14 | 2.327 | 0.295 | -0.872 | 0.031 | -0.061 | -0.061 | 0 | 0.12 | 0.26 | 0.12 | 0 | 0.16 | 0.09 |
| 2 | 122777669 | rs1878592 | 0.502 | 2.25 | 0.21 | -0.834 | 0.019 | -0.119 | -0.119 | 0.934 | 0.68 | 0.49 | 0.77 | 0.9 | 0.65 | 0.7 |
| 2 | 122797678 | rs6741948 | 0.811 | 2.258 | 0.127 | -0.788 | 0.043 | -0.032 | -0.032 | 0 | 0.65 | 0.5 | 0.72 | 0.73 | 0.55 | 0.63 |
| 2 | 122833317 | rs908677 | 1.265 | 0.996 | -0.445 | -1.372 | 0.023 | -0.012 | -0.012 | 0 | 0.3 | 0.23 | 0.4 | 0.64 | 0.13 | 0.35 |
| 2 | 122837193 | rs1733820 | -1.293 | 1.37 | -0.83 | -1.532 | -0.007 | 0.048 | 0.048 | 0.061 | 0.81 | 0.93 | 0.89 | 0.9 | 0.82 | 0.79 |
| 2 | 122870285 | rs1560603 | -1.258 | 1.596 | -0.696 | -1.543 | -0.015 | 0.006 | 0.006 | 0 | 0.91 | 0.95 | 0.9 | 1 | 0.84 | 0.91 |
| 2 | 122873914 | rs2902343 | 1.471 | 1.841 | -0.933 | -1.794 | -0.024 | 0.052 | 0.052 | 0.322 | 0.21 | 0.13 | 0.15 | 0 | 0.25 | 0.11 |
| 2 | 122884720 | rs13029963 | 1.605 | 1.706 | -0.906 | -1.835 | -0.002 | 0.014 | 0.014 | 0 | 0.1 | 0.08 | 0.14 | 0.04 | 0.14 | 0.16 |
| 2 | 122892378 | rs4848747 | 1.262 | 1.954 | -0.898 | -1.852 | -0.04 | 0.061 | 0.061 | 0.064 | 0.23 | 0.13 | 0.16 | 0 | 0.28 | 0.12 |
| 2 | 122898265 | rs13005471 | 0.632 | 1.63 | -0.701 | -1.851 | -0.025 | -0.014 | -0.014 | 0 | 0.43 | 0.49 | 0.56 | 0.71 | 0.28 | 0.53 |
| 2 | 122918215 | rs4848752 | 1.916 | 2.039 | -0.861 | -1.83 | -0.022 | 0.115 | 0.115 | 0 | 0.18 | 0.01 | 0.1 | 0 | 0.15 | 0.06 |

| | | | | | | | | | | | | | | | | |
|---|---|---|---|---|---|---|---|---|---|---|---|---|---|---|---|---|
| 2 | 122926268 | rs12612050 | 0.242 | 1.809 | -0.797 | -1.94 | -0.002 | -0.017 | -0.017 | 0 | 0.58 | 0.56 | 0.67 | 0.83 | 0.46 | 0.72 |
| 2 | 122932022 | rs1545016 | -0.212 | 2.036 | -0.735 | -2.045 | 0.016 | -0.104 | -0.104 | 0 | 0.66 | 0.59 | 0.73 | 1 | 0.57 | 0.79 |
| 2 | 122936070 | rs10192552 | 1.24 | 1.803 | -0.95 | -1.91 | 0.001 | 0.012 | 0.012 | 0.123 | 0.18 | 0.08 | 0.11 | 0.14 | 0.18 | 0.2 |
| 2 | 122936260 | rs10206024 | 0.155 | 1.977 | -0.883 | -1.896 | -0.04 | 0.133 | 0.133 | 0.002 | 0.41 | 0.22 | 0.27 | 0.14 | 0.46 | 0.33 |
| 2 | 122936737 | rs1432312 | -0.388 | 1.828 | -0.741 | -1.859 | 0.032 | 0.069 | 0.069 | 0.358 | 0.5 | 0.34 | 0.33 | 0.29 | 0.56 | 0.39 |
| 2 | 122943468 | rs4848174 | 0.292 | 1.433 | -0.982 | -1.767 | 0.012 | 0.105 | 0.105 | 0.013 | 0.37 | 0.2 | 0.21 | 0.14 | 0.33 | 0.26 |
| 2 | 122945745 | rs9678809 | -0.298 | 1.569 | -0.473 | -1.644 | -0.001 | -0.013 | -0.013 | 0.055 | 0.45 | 0.48 | 0.32 | 0.14 | 0.52 | 0.32 |
| 2 | 122947107 | rs9679739 | 0.223 | 1.183 | -0.294 | -1.308 | 0.019 | 0.044 | 0.044 | 0 | 0.09 | 0.36 | 0.11 | 0.08 | 0.06 | 0.1 |
| 2 | 122952356 | rs7597014 | 1.227 | 1.316 | -0.749 | -1.466 | -0.049 | -0.01 | -0.01 | 0 | 0.18 | 0.03 | 0.12 | 0.13 | 0.2 | 0.24 |
| 2 | 122953150 | rs1432302 | 0.97 | 1.617 | -0.602 | -1.502 | -0.104 | 0.164 | 0.164 | 0 | 0.26 | 0.03 | 0.18 | 0.01 | 0.34 | 0.2 |
| 2 | 122953533 | rs10169452 | -0.055 | 0.967 | 0.112 | -0.96 | 0.027 | 0.048 | 0.048 | 0.064 | 0.09 | 0.4 | 0.11 | 0.07 | 0.06 | 0.09 |
| 2 | 122961410 | rs10207028 | 0.55 | 1.014 | 0.158 | -0.938 | 0.014 | 0.031 | 0.031 | 0 | 0.07 | 0.25 | 0.09 | 0.07 | 0.06 | 0.09 |
| 2 | 122965688 | rs13018000 | -1.032 | 1.09 | 0.156 | -0.985 | 0.035 | 0.073 | 0.073 | 0 | 0.16 | 0.47 | 0.22 | 0.24 | 0.19 | 0.15 |
| 2 | 122966157 | rs6751865 | -1.038 | 1.212 | 0.139 | -0.974 | 0.06 | 0.077 | 0.077 | 0 | 0.16 | 0.48 | 0.23 | 0.24 | 0.21 | 0.16 |
| 2 | 122966970 | rs1432303 | 0.056 | 1.225 | 0.205 | -0.961 | 0.036 | 0.044 | 0.044 | 0 | 0.08 | 0.18 | 0.11 | 0.16 | 0.13 | 0.05 |
| 2 | 122970146 | rs10193471 | -0.916 | 1.072 | 0.214 | -0.993 | 0.006 | 0.082 | 0.082 | 0 | 0.16 | 0.5 | 0.17 | 0.24 | 0.14 | 0.14 |
| 2 | 122971252 | rs1432307 | -0.754 | 1.021 | -0.593 | -1.633 | -0.094 | 0.171 | 0.171 | 0 | 0.65 | 0.89 | 0.77 | 0.98 | 0.54 | 0.75 |
| 2 | 122973151 | rs765886 | 0.033 | 0.018 | -0.133 | -1.618 | -0.015 | -0.018 | -0.018 | 0.133 | 0.41 | 0.54 | 0.56 | 0.8 | 0.29 | 0.47 |
| 2 | 122976173 | rs934388 | -0.078 | -0.057 | -0.155 | -1.631 | 0.028 | 0.03 | 0.03 | 0 | 0.42 | 0.64 | 0.58 | 0.8 | 0.29 | 0.45 |
| 2 | 122976844 | rs6745986 | 1.875 | 0.054 | -0.274 | -1.706 | -0.05 | -0.045 | -0.045 | 1.194 | 0.16 | 0.02 | 0.14 | 0.18 | 0.18 | 0.24 |
| 2 | 122979880 | rs4848758 | 1.259 | -0.14 | -0.031 | -1.457 | 0.025 | -0.018 | -0.018 | 0.307 | 0.28 | 0.19 | 0.42 | 0.62 | 0.14 | 0.36 |
| 2 | 122982362 | rs10200535 | -0.514 | 0.063 | 0.019 | -1.414 | 0.03 | 0.084 | 0.084 | 0.031 | 0.13 | 0.46 | 0.13 | 0.18 | 0.13 | 0.09 |
| 2 | 122986481 | rs1432309 | 1.122 | 0.126 | -0.069 | -1.368 | 0.014 | 0.047 | 0.047 | 0 | 0.07 | 0.15 | 0.06 | 0.15 | 0.09 | 0.04 |
| 2 | 122992346 | rs11690406 | 1.555 | 0.419 | -0.461 | -1.632 | -0.147 | 0.189 | 0.189 | 0 | 0.28 | 0.04 | 0.22 | 0.01 | 0.43 | 0.21 |
| 2 | 122994633 | rs7603857 | -0.825 | 0.319 | -0.197 | -1.463 | -0.044 | 0.231 | 0.231 | 0 | 0.59 | 0.89 | 0.7 | 0.99 | 0.44 | 0.68 |
| 2 | 123028173 | rs1560601 | -1.39 | 0.145 | 0.298 | -1.002 | 0.041 | 0.075 | 0.075 | 0 | 0.93 | 0.85 | 0.87 | 0.84 | 0.9 | 0.95 |
| 2 | 123028642 | rs934392 | 1.054 | -0.17 | 0.065 | -1.085 | 0.014 | -0.016 | -0.016 | 0 | 0.28 | 0.2 | 0.38 | 0.62 | 0.13 | 0.35 |
| 2 | 123030939 | rs1320871 | -0.4 | 0.177 | 0.429 | -0.928 | 0.097 | 0.122 | 0.122 | 0 | 0.91 | 0.63 | 0.85 | 0.84 | 0.88 | 0.94 |
| 2 | 123039255 | rs1344759 | -0.731 | 0.214 | 0.43 | -0.924 | 0.122 | 0.132 | 0.132 | 0 | 0.92 | 0.64 | 0.85 | 0.87 | 0.88 | 0.94 |
| 2 | 123047991 | rs13421929 | -0.58 | 0.524 | -0.096 | -0.984 | 0.034 | -0.018 | -0.018 | 0.42 | 0.09 | 0.26 | 0.08 | 0.03 | 0.13 | 0.05 |
| 2 | 123051775 | rs17007488 | 0.28 | 0.12 | -0.414 | -1.09 | 0.002 | -0.001 | -0.001 | 0.283 | 0.35 | 0.32 | 0.43 | 0.65 | 0.22 | 0.39 |
| 2 | 123053585 | rs12612483 | 1.157 | 0.368 | -0.739 | -1.062 | -0.051 | -0.041 | -0.041 | 0 | 0.21 | 0.07 | 0.17 | 0.26 | 0.24 | 0.28 |
| 2 | 123058365 | rs12466028 | 1.389 | 0.065 | -0.646 | -1.068 | 0.006 | -0.015 | -0.015 | 0.296 | 0.26 | 0.18 | 0.37 | 0.59 | 0.11 | 0.33 |
| 2 | 123065343 | rs12467940 | 0.861 | 0.023 | -0.485 | -1.017 | -0.069 | 0.015 | 0.015 | 0.297 | 0.3 | 0.35 | 0.39 | 0.64 | 0.12 | 0.39 |
| 2 | 123065521 | rs1406747 | 1.469 | 0.335 | -0.928 | -1.098 | -0.07 | -0.036 | -0.036 | 0 | 0.19 | 0.04 | 0.16 | 0.21 | 0.23 | 0.22 |
| 2 | 123065622 | rs934393 | 1.693 | 0.443 | -0.894 | -1.274 | -0.03 | 0.143 | 0.143 | 0 | 0.28 | 0.06 | 0.22 | 0.01 | 0.31 | 0.24 |
| 2 | 123092005 | rs1881887 | -0.094 | 0.349 | -0.365 | -1.009 | 0.007 | -0.028 | -0.028 | 0 | 0.12 | 0.25 | 0.08 | 0.06 | 0.11 | 0.12 |
| 2 | 123099547 | rs10496581 | 0.038 | 0.039 | -0.274 | -1.071 | -0.05 | -0.005 | -0.005 | 0 | 0.39 | 0.46 | 0.46 | 0.64 | 0.22 | 0.46 |
| 2 | 123102720 | rs12615635 | 0.887 | -0.178 | -0.253 | -0.901 | -0.015 | -0.002 | -0.002 | 0 | 0.29 | 0.22 | 0.38 | 0.64 | 0.12 | 0.39 |
| 2 | 123118913 | rs10496583 | 1.123 | -0.148 | -0.254 | -0.457 | -0.011 | -0.003 | -0.003 | 0 | 0.27 | 0.2 | 0.36 | 0.55 | 0.11 | 0.34 |
| 2 | 123120079 | rs6541823 | -0.448 | 0.614 | -0.151 | -0.538 | 0.011 | 0.001 | 0.001 | 0 | 0.88 | 0.83 | 0.83 | 0.86 | 0.77 | 0.92 |
| 2 | 123122558 | rs11689992 | 0.333 | 0.52 | -0.335 | -0.809 | -0.006 | 0.051 | 0.051 | 0 | 0.38 | 0.26 | 0.28 | 0.1 | 0.43 | 0.35 |
| 2 | 123126062 | rs7580362 | 0.35 | 0.62 | 0.122 | -0.635 | 0.045 | -0.039 | -0.039 | 0.047 | 0.83 | 0.6 | 0.83 | 0.92 | 0.78 | 0.88 |
| 2 | 123134412 | rs1320869 | -1.22 | 0.604 | 0.232 | -0.592 | 0.005 | 0.001 | 0.001 | 0.074 | 0.93 | 0.92 | 0.91 | 0.92 | 0.9 | 0.91 |
| 2 | 123135010 | rs33920762 | 1.092 | 0.01 | -0.338 | -0.021 | -0.007 | 0.029 | 0.029 | 0 | 0.2 | 0.16 | 0.31 | 0.46 | 0.13 | 0.22 |
| 2 | 123139842 | rs17007644 | 0.582 | -0.331 | -0.573 | -0.406 | 0.007 | 0.003 | 0.003 | 0 | 0.36 | 0.25 | 0.28 | 0.37 | 0.32 | 0.38 |
| 2 | 123139879 | rs1453675 | 2.257 | -0.473 | -0.69 | -0.419 | -0.051 | 0 | 0 | 0 | 0.24 | 0.04 | 0.23 | 0.2 | 0.26 | 0.26 |
| 2 | 123140603 | rs1453678 | -0.607 | -0.49 | -0.641 | -0.429 | 0.001 | -0.023 | -0.023 | 0 | 0.57 | 0.49 | 0.6 | 0.82 | 0.44 | 0.6 |
| 2 | 123141545 | rs4848768 | 1.078 | -0.49 | -0.638 | -0.422 | -0.032 | 0.085 | 0.085 | 0 | 0.22 | 0.29 | 0.32 | 0.46 | 0.13 | 0.22 |
| 2 | 123143224 | rs6756499 | -0.602 | -0.41 | -0.755 | -0.395 | -0.002 | 0.006 | 0.006 | 0 | 0.57 | 0.54 | 0.6 | 0.82 | 0.44 | 0.6 |
| 2 | 123144514 | rs6735231 | 0.51 | -0.208 | -0.813 | -0.495 | -0.01 | 0.005 | 0.005 | 0 | 0.11 | 0.15 | 0.05 | 0.17 | 0.06 | 0.12 |
| 2 | 123151264 | rs6705997 | 0.287 | -0.013 | -0.795 | -0.446 | 0.008 | 0.005 | 0.005 | 0 | 0.37 | 0.3 | 0.29 | 0.37 | 0.31 | 0.38 |
| 2 | 123160373 | rs6721174 | 0.548 | 0.124 | -0.861 | -1.283 | 0.015 | -0.109 | -0.109 | 0 | 0.36 | 0.49 | 0.31 | 0.02 | 0.46 | 0.26 |
| 2 | 123161474 | rs13400671 | 1.688 | 0.168 | -1.123 | -1.309 | -0.062 | 0.115 | 0.115 | 0 | 0.24 | 0.06 | 0.22 | 0.02 | 0.34 | 0.23 |
| 2 | 123169027 | rs12711583 | -0.924 | -0.19 | -0.972 | -0.902 | 0.005 | -0.018 | -0.018 | 0 | 0.75 | 0.88 | 0.64 | 0.66 | 0.77 | 0.7 |
| 2 | 123170459 | rs7589494 | 0.355 | 0.007 | -0.773 | -1.189 | 0.018 | -0.106 | -0.106 | 0.174 | 0.37 | 0.5 | 0.32 | 0.02 | 0.51 | 0.27 |
| 2 | 123171461 | rs6715671 | -0.247 | -0.278 | -0.538 | -0.651 | -0.033 | 0.033 | 0.033 | 0 | 0.13 | 0.28 | 0.07 | 0.17 | 0.07 | 0.08 |
| 2 | 123171791 | rs6716061 | 0.001 | 0.003 | -1.089 | -0.767 | 0.006 | -0.05 | -0.05 | 0 | 0.87 | 0.73 | 0.81 | 0.94 | 0.87 | 0.89 |
| 2 | 123174260 | rs12471936 | 1.154 | 0.073 | -2.52 | -0.839 | -0.055 | 0.019 | 0.019 | 0 | 0.33 | 0.02 | 0.31 | 0.27 | 0.3 | 0.37 |
| 2 | 123176322 | rs1881881 | 0.231 | 0.129 | -2.153 | -0.8 | -0.001 | 0.009 | 0.009 | 0 | 0.5 | 0.5 | 0.43 | 0.44 | 0.48 | 0.46 |
| 2 | 123182738 | rs4848773 | -0.001 | 0.653 | -2.389 | -0.872 | 0.066 | -0.117 | -0.117 | 0 | 0.73 | 0.56 | 0.55 | 0.93 | 0.6 | 0.69 |
| 2 | 123186585 | rs10496586 | 1.015 | 0.491 | -2.378 | -0.723 | 0.013 | -0.057 | -0.057 | 0 | 0.29 | 0.17 | 0.25 | 0.47 | 0.26 | 0.29 |
| 2 | 123192216 | rs17007730 | 0.874 | 0.619 | -2.12 | -0.606 | 0.087 | -0.085 | -0.085 | 0 | 0.12 | 0.4 | 0.1 | 0 | 0.16 | 0.04 |
| 2 | 123194367 | rs12053379 | 1.984 | 0.466 | -3.091 | -0.709 | -0.031 | 0.03 | 0.03 | 0 | 0.24 | 0.03 | 0.19 | 0.17 | 0.24 | 0.22 |
| 2 | 123196967 | rs17007740 | 0.685 | 0.199 | -1.792 | -0.35 | -0.008 | 0.014 | 0.014 | 0 | 0.22 | 0.2 | 0.19 | 0.45 | 0.11 | 0.25 |
| 2 | 123200413 | rs1580069 | -1.104 | 0.277 | -1.806 | -0.352 | -0.029 | 0.014 | 0.014 | 0 | 0.93 | 1 | 0.85 | 0.97 | 0.9 | 0.9 |
| 2 | 123201835 | rs11122903 | 1.3 | 0.268 | -1.784 | -0.366 | -0.017 | 0.014 | 0.014 | 0 | 0.51 | 0.3 | 0.36 | 0.49 | 0.47 | 0.48 |
| 2 | 123202227 | rs1598022 | -0.238 | 0.316 | -1.808 | -0.359 | -0.016 | 0.006 | 0.006 | 0 | 0.3 | 0.39 | 0.27 | 0.62 | 0.21 | 0.32 |
| 2 | 123204730 | rs1453681 | 0.274 | 0.328 | -1.813 | -0.356 | 0.007 | -0.023 | -0.023 | 1.132 | 0.63 | 0.76 | 0.47 | 0.49 | 0.62 | 0.52 |

| | | | | | | | | | | | | | | | | |
|---|---|---|---|---|---|---|---|---|---|---|---|---|---|---|---|---|
| 2 | 123210344 | rs10169108 | -0.311 | 0.553 | -1.807 | -0.238 | -0.068 | 0.043 | 0.043 | 0 | 0.84 | 0.97 | 0.66 | 0.94 | 0.73 | 0.77 |
| 2 | 123214056 | rs11903734 | -0.388 | 0.75 | -1.432 | -0.07 | 0.003 | 0.017 | 0.017 | 0 | 0.09 | 0.19 | 0.08 | 0.17 | 0.1 | 0.07 |
| 2 | 123214283 | rs12471629 | 1.092 | 0.74 | -1.433 | -0.131 | -0.071 | 0.014 | 0.014 | 0 | 0.39 | 0.06 | 0.26 | 0.33 | 0.35 | 0.4 |
| 2 | 123216847 | rs6713772 | -0.528 | 0.086 | -0.369 | -0.003 | 0.07 | -0.082 | -0.082 | 0 | 0.12 | 0.44 | 0.11 | 0 | 0.15 | 0.04 |
| 2 | 123218071 | rs6718275 | 0.338 | -0.381 | -0.5 | 0.228 | -0.008 | 0.001 | 0.001 | 0 | 0.62 | 0.5 | 0.59 | 0.44 | 0.69 | 0.59 |
| 2 | 123222183 | rs6541829 | -1.189 | -0.056 | 0.051 | -0.062 | 0.01 | -0.009 | -0.009 | 0 | 0.2 | 0.6 | 0.19 | 0.16 | 0.25 | 0.1 |
| 2 | 123224400 | rs4848780 | -1.365 | -1.368 | 0.724 | -0.199 | -0.037 | -0.031 | -0.031 | 0 | 0.27 | 0.44 | 0.3 | 0.24 | 0.21 | 0.24 |
| 2 | 123225623 | rs17007761 | -0.965 | -1.404 | 0.492 | -0.162 | 0.002 | -0.002 | -0.002 | 0 | 0.06 | 0.02 | 0.16 | 0.11 | 0.05 | 0.18 |
| 2 | 123233702 | rs1982463 | -0.575 | -1.527 | 0.779 | -0.463 | -0.01 | -0.041 | -0.041 | 0 | 0.08 | 0.17 | 0.11 | 0 | 0.04 | 0 |
| 2 | 123234578 | rs896902 | 2.041 | -1.848 | 0.177 | -0.655 | 0.104 | 0.146 | 0.146 | 0.848 | 0.55 | 0.12 | 0.49 | 0.73 | 0.72 | 0.59 |
| 2 | 123234677 | rs753620 | 1.476 | -1.711 | 0.878 | -0.617 | -0.028 | -0.072 | -0.072 | 0.459 | 0.68 | 0.47 | 0.69 | 0.88 | 0.8 | 0.81 |
| 2 | 123235520 | rs1219701 | 1.598 | -1.581 | 0.912 | -0.528 | -0.004 | -0.015 | -0.015 | 0 | 0.66 | 0.42 | 0.55 | 0.78 | 0.77 | 0.68 |
| 2 | 123238843 | rs4459739 | -1.613 | -0.995 | 0.672 | -0.014 | -0.033 | 0.025 | 0.025 | 0.162 | 0.15 | 0.3 | 0.33 | 0.16 | 0.08 | 0.19 |
| 2 | 123244252 | rs1474470 | -0.783 | -1.195 | -0.445 | -0.143 | -0.02 | 0.049 | 0.049 | 0 | 0.19 | 0.01 | 0.12 | 0.12 | 0.15 | 0.22 |
| 2 | 123249909 | rs10205651 | 1.635 | -1.406 | -0.041 | -0.169 | 0.058 | 0.008 | 0.008 | 1.602 | 0.76 | 0.15 | 0.65 | 0.82 | 0.87 | 0.78 |
| 2 | 123274766 | rs893742 | -1.364 | -1.28 | 0.848 | -0.196 | -0.032 | -0.072 | -0.072 | 0 | 0.13 | 0.23 | 0.13 | 0 | 0.07 | 0.03 |
| 2 | 123276791 | rs11122908 | 1.607 | -1.291 | 1.452 | -0.202 | -0.062 | -0.11 | -0.11 | 0 | 0.82 | 0.37 | 0.83 | 1 | 0.93 | 0.97 |
| 2 | 123281539 | rs11679684 | 2.095 | -1.445 | 1.003 | -0.19 | 0.062 | 0.015 | 0.015 | 1.434 | 0.75 | 0.15 | 0.65 | 0.83 | 0.86 | 0.79 |
| 2 | 123291022 | rs1527243 | 1.974 | -1.346 | 0.998 | -0.07 | -0.001 | 0 | 0 | 0 | 0.76 | 0.28 | 0.72 | 0.82 | 0.84 | 0.77 |
| 2 | 123297807 | rs17039776 | -1.057 | -0.929 | 0.817 | 0.149 | -0.002 | -0.006 | -0.006 | 0 | 0.11 | 0.09 | 0.2 | 0.17 | 0.12 | 0.2 |
| 2 | 123302906 | rs10188484 | 1.198 | -1.042 | 0.825 | 0.128 | -0.013 | 0.059 | 0.059 | 0.018 | 0.84 | 0.49 | 0.77 | 0.82 | 0.87 | 0.79 |
| 2 | 123304053 | rs10168196 | -0.762 | -1.512 | 0.118 | -0.517 | 0.009 | 0.01 | 0.01 | 0 | 0.31 | 0.3 | 0.22 | 0.15 | 0.2 | 0.32 |
| 2 | 123305740 | rs992725 | 0.998 | -1.681 | -1.146 | -0.555 | -0.042 | -0.032 | -0.032 | 0.018 | 0.4 | 0.1 | 0.4 | 0.6 | 0.6 | 0.35 |
| 2 | 123320715 | rs766757 | 0.642 | -2.313 | -1.02 | -0.863 | 0.058 | 0.077 | 0.077 | 0 | 0.55 | 0.24 | 0.53 | 0.67 | 0.68 | 0.42 |
| 2 | 123327494 | rs17008069 | -0.457 | -2.548 | -0.244 | -1.026 | 0.023 | 0.019 | 0.019 | 4.179 | 0.07 | 0.57 | 0.05 | 0 | 0.01 | 0 |
| 2 | 123332354 | rs999492 | 0.779 | -2.654 | 0.538 | -0.811 | 0.043 | 0.053 | 0.053 | 0 | 0.53 | 0.76 | 0.55 | 0.65 | 0.67 | 0.44 |
| 2 | 123339341 | rs1157873 | 0.942 | -2.785 | 0.455 | -0.972 | 0.006 | 0.008 | 0.008 | 0.433 | 0.55 | 0.64 | 0.47 | 0.64 | 0.65 | 0.38 |
| 2 | 123343577 | rs1439842 | -0.781 | -2.858 | 0.346 | -1.075 | -0.015 | -0.018 | -0.018 | 0 | 0.84 | 0.82 | 0.91 | 0.86 | 0.9 | 0.8 |
| 2 | 123354282 | rs1405604 | 1.375 | -2.765 | 0.711 | -1.018 | 0.001 | 0 | 0 | 0 | 0.51 | 0.58 | 0.46 | 0.64 | 0.66 | 0.41 |
| 2 | 123356151 | rs6727132 | -1.008 | -2.757 | 0.662 | -1.017 | -0.039 | -0.086 | -0.086 | 0 | 0.13 | 0.55 | 0.06 | 0 | 0.04 | 0.05 |
| 2 | 123356656 | rs6541836 | -1.008 | -2.757 | 0.662 | -1.017 | -0.039 | -0.086 | -0.086 | 0 | 0.13 | 0.55 | 0.06 | 0 | 0.04 | 0.05 |
| 2 | 123360311 | rs6746132 | 2.158 | -2.748 | 0.505 | -1.012 | -0.019 | 0.031 | 0.031 | 0 | 0.41 | 0.13 | 0.41 | 0.64 | 0.62 | 0.37 |
| 2 | 123366899 | rs2043940 | 1.177 | -2.699 | 0.478 | -0.992 | 0.021 | 0.025 | 0.025 | 0 | 0.53 | 0.66 | 0.54 | 0.65 | 0.67 | 0.46 |
| 2 | 123368169 | rs4848184 | -0.073 | -2.341 | 0.037 | -0.859 | -0.05 | -0.043 | -0.043 | 0 | 0.22 | 0.01 | 0.26 | 0.21 | 0.21 | 0.31 |
| 2 | 123377942 | rs7566539 | 0.615 | -1.825 | 0.547 | -0.771 | -0.006 | -0.028 | -0.028 | 1.636 | 0.06 | 0.1 | 0.02 | 0 | 0.04 | 0.05 |
| 2 | 123386902 | rs1371492 | 1.027 | -1.833 | 0.616 | -0.902 | -0.016 | -0.1 | -0.1 | 0 | 0.79 | 0.34 | 0.67 | 0.99 | 0.83 | 0.81 |
| 2 | 123388562 | rs10192183 | 1.448 | -2.212 | 0.067 | -0.886 | 0.013 | 0.045 | 0.045 | 0.093 | 0.44 | 0.14 | 0.51 | 0.62 | 0.61 | 0.43 |
| 2 | 123389473 | rs13428951 | 0.988 | -2.257 | 0.717 | -0.91 | 0.02 | 0.015 | 0.015 | 0 | 0.52 | 0.46 | 0.73 | 0.63 | 0.74 | 0.51 |
| 2 | 123390288 | rs12474979 | -0.181 | -2.015 | 0.44 | -0.783 | 0.018 | -0.053 | -0.053 | 0 | 0.09 | 0.27 | 0.21 | 0 | 0.12 | 0.09 |
| 2 | 123392303 | rs987707 | -0.409 | -1.951 | 0.147 | -0.653 | 0.03 | 0.023 | 0.023 | 0 | 0.4 | 0.14 | 0.31 | 0.37 | 0.32 | 0.51 |
| 2 | 123395522 | rs6725236 | 0.776 | -1.131 | 0.171 | -0.772 | 0.031 | -0.004 | -0.004 | 0.017 | 0.36 | 0.41 | 0.46 | 0.62 | 0.51 | 0.43 |
| 2 | 123395877 | rs13400652 | 0.834 | -0.936 | -0.098 | -0.025 | -0.072 | -0.048 | -0.048 | 1.299 | 0.34 | 0.16 | 0.44 | 0.39 | 0.52 | 0.4 |
| 2 | 123395921 | rs13400673 | 0.273 | -0.127 | 0.556 | -0.043 | -0.002 | 0.003 | 0.003 | 0 | 0.42 | 0.62 | 0.43 | 0.36 | 0.46 | 0.45 |
| 2 | 123396005 | rs7575942 | -0.739 | 0.176 | 1.204 | 0.106 | 0 | -0.007 | -0.007 | 0 | 0.46 | 0.49 | 0.54 | 0.4 | 0.5 | 0.53 |
| 2 | 123397360 | rs7582698 | -2.148 | 0.223 | 2.443 | 0.484 | -0.014 | -0.094 | -0.094 | 0 | 0.18 | 0.45 | 0.13 | 0.05 | 0.12 | 0.17 |
| 2 | 123399137 | rs17008278 | -0.639 | 0.225 | 2.033 | 0.483 | -0.007 | -0.033 | -0.033 | 0 | 0.07 | 0.14 | 0.04 | 0 | 0.05 | 0.03 |
| 2 | 123401043 | rs7594595 | -2.354 | 0.234 | 1.673 | 0.485 | -0.02 | -0.086 | -0.086 | 0 | 0.23 | 0.59 | 0.15 | 0.05 | 0.16 | 0.21 |
| 2 | 123407317 | rs10179725 | -2.307 | 0.254 | 1.671 | 0.494 | -0.019 | -0.086 | -0.086 | 6.231 | 0.23 | 0.59 | 0.15 | 0.05 | 0.17 | 0.21 |
| 2 | 123407739 | rs10203851 | -1.273 | 0.633 | 0.791 | 1.045 | 0 | -0.011 | -0.011 | 0 | 0.09 | 0.1 | 0.07 | 0.05 | 0.08 | 0.14 |
| 2 | 123414629 | rs11893130 | -1.788 | 0.423 | 1.166 | 1.415 | 0.007 | 0.104 | 0.104 | 0 | 0.06 | 0.23 | 0.1 | 0.17 | 0.04 | 0.09 |
| 2 | 123415817 | rs11894795 | -2.509 | 0.256 | 1.66 | 1.183 | -0.015 | 0.034 | 0.034 | 0 | 0.13 | 0.34 | 0.13 | 0.16 | 0.08 | 0.16 |
| 2 | 123416609 | rs17008320 | -2.87 | 0.089 | 1.901 | 1.696 | 0.018 | -0.03 | -0.03 | 0.368 | 0.31 | 0.78 | 0.37 | 0.48 | 0.22 | 0.33 |
| 2 | 123420627 | rs1371493 | -2.734 | 0.09 | 2.187 | 1.838 | -0.021 | 0.034 | 0.034 | 2.528 | 0.21 | 0.7 | 0.19 | 0.37 | 0.13 | 0.17 |
| 2 | 123423297 | rs715054 | -1.745 | 0.041 | 2.01 | 1.316 | 0.003 | -0.008 | -0.008 | 0 | 0.11 | 0.07 | 0.17 | 0.12 | 0.09 | 0.15 |
| 2 | 123428804 | rs11122923 | -2.82 | -0.004 | 2.667 | 1.339 | -0.01 | 0.097 | 0.097 | 0 | 0.21 | 0.56 | 0.2 | 0.34 | 0.17 | 0.2 |
| 2 | 123431555 | rs10201125 | 1.923 | -0.032 | 2.993 | 1.152 | -0.016 | -0.077 | -0.077 | 3.033 | 0.88 | 0.51 | 0.94 | 0.99 | 0.96 | 0.92 |
| 2 | 123431631 | rs10177647 | 3.211 | -0.035 | 3.019 | 1.166 | 0.017 | -0.012 | -0.012 | 0.038 | 0.71 | 0.23 | 0.79 | 0.66 | 0.83 | 0.72 |
| 2 | 123433978 | rs11691961 | 3.379 | -0.035 | 2.972 | 1.165 | 0.029 | -0.021 | -0.021 | 0 | 0.68 | 0.13 | 0.76 | 0.65 | 0.79 | 0.72 |
| 2 | 123437061 | rs13013395 | -2.063 | -0.046 | 2.985 | 1.16 | -0.033 | -0.065 | -0.065 | 0.027 | 0.12 | 0.03 | 0.12 | 0.33 | 0.12 | 0.19 |
| 2 | 123438127 | rs4848791 | 3.412 | -0.045 | 2.98 | 1.165 | 0.029 | -0.028 | -0.028 | 0 | 0.68 | 0.11 | 0.76 | 0.65 | 0.79 | 0.72 |
| 2 | 123440075 | rs4848792 | 3.352 | -0.045 | 3.022 | 1.18 | 0.028 | -0.001 | -0.001 | 0 | 0.69 | 0.14 | 0.76 | 0.68 | 0.8 | 0.73 |
| 2 | 123442764 | rs9784127 | -1.608 | 0.03 | 3.131 | 1.131 | -0.004 | -0.044 | -0.044 | 4.376 | 0.1 | 0.13 | 0.03 | 0 | 0.05 | 0.08 |
| 2 | 123444468 | rs12052515 | -2.395 | -0.035 | 3.238 | 1.154 | 0 | 0.004 | 0.004 | 0 | 0.14 | 0.15 | 0.13 | 0.31 | 0.11 | 0.2 |
| 2 | 123446216 | rs12052977 | -3.205 | 0.024 | 3.432 | 1.147 | -0.022 | 0.054 | 0.054 | 0 | 0.22 | 0.49 | 0.19 | 0.32 | 0.16 | 0.2 |
| 2 | 123447262 | rs17008455 | -2.951 | 0.023 | 3.417 | 1.151 | -0.023 | 0.066 | 0.066 | 0.009 | 0.19 | 0.41 | 0.15 | 0.31 | 0.11 | 0.2 |
| 2 | 123463049 | rs11690346 | -2.119 | 0.068 | 2.899 | 1.237 | -0.004 | -0.035 | -0.035 | 0 | 0.13 | 0.06 | 0.11 | 0.37 | 0.12 | 0.21 |
| 2 | 123467108 | rs4241113 | 3.165 | 0.052 | 2.934 | 1.214 | 0.029 | -0.022 | -0.022 | 0.033 | 0.67 | 0.12 | 0.76 | 0.65 | 0.79 | 0.71 |
| 2 | 123470077 | rs6718776 | -2.644 | 0.114 | 2.988 | 1.251 | -0.001 | 0.035 | 0.035 | 0 | 0.17 | 0.24 | 0.16 | 0.35 | 0.15 | 0.21 |
| 2 | 123488984 | rs17008539 | -2.456 | 0.139 | 1.877 | 1.001 | -0.001 | 0.053 | 0.053 | 0.51 | 0.14 | 0.23 | 0.13 | 0.29 | 0.12 | 0.21 |

| | | | | | | | | | | | | | | | | |
|---|---|---|---|---|---|---|---|---|---|---|---|---|---|---|---|---|
| 2 | 123489337 | rs1400967 | 2.888 | 0.129 | 1.648 | 0.987 | 0.027 | -0.008 | -0.008 | 0 | 0.67 | 0.13 | 0.77 | 0.7 | 0.78 | 0.71 |
| 2 | 123490744 | rs1518057 | -2.053 | 0.243 | 1.464 | 1.286 | 0 | -0.002 | -0.002 | 0 | 0.16 | 0.18 | 0.16 | 0.29 | 0.14 | 0.22 |
| 2 | 123492420 | rs6541851 | 1.551 | 0.363 | 1.532 | 1.069 | -0.029 | -0.013 | -0.013 | 0 | 0.79 | 0.41 | 0.77 | 0.8 | 0.83 | 0.83 |
| 2 | 123502497 | rs10496595 | -2.328 | 0.655 | 1.36 | 1.259 | 0.002 | 0.012 | 0.012 | 0 | 0.16 | 0.2 | 0.19 | 0.24 | 0.15 | 0.17 |
| 2 | 123503566 | rs6749889 | -3.155 | 0.514 | 2.079 | 1.098 | -0.006 | -0.032 | -0.032 | 0 | 0.2 | 0.39 | 0.19 | 0.17 | 0.16 | 0.25 |
| 2 | 123512526 | rs12327981 | -1.606 | 0.556 | 1.929 | 1.025 | 0.001 | -0.006 | -0.006 | 0 | 0.07 | 0.06 | 0.12 | 0.03 | 0.1 | 0.09 |
| 2 | 123520650 | rs6725154 | -2.594 | 0.81 | 2.221 | 1.215 | 0.052 | 0.144 | 0.144 | 0.23 | 0.2 | 0.42 | 0.29 | 0.38 | 0.29 | 0.24 |
| 2 | 123524253 | rs6541859 | -2.572 | 0.831 | 2.229 | 1.19 | 0.032 | 0.028 | 0.028 | 0.221 | 0.14 | 0.3 | 0.17 | 0.2 | 0.17 | 0.11 |
| 2 | 123524663 | rs12711597 | -2.429 | 0.83 | 2.189 | 1.19 | 0.004 | 0.003 | 0.003 | 0.095 | 0.13 | 0.14 | 0.16 | 0.2 | 0.17 | 0.11 |
| 2 | 123525820 | rs7600772 | -3.574 | 0.797 | 2.181 | 1.178 | 0.006 | -0.006 | -0.006 | 0 | 0.22 | 0.43 | 0.22 | 0.24 | 0.22 | 0.25 |
| 2 | 123538727 | rs17008648 | -1.369 | 0.881 | 1.723 | 1.188 | 0.006 | 0.047 | 0.047 | 0.117 | 0.06 | 0.09 | 0.09 | 0.14 | 0.09 | 0.07 |
| 2 | 123539121 | rs2136316 | -1.77 | 0.786 | 1.605 | 0.984 | -0.006 | -0.002 | -0.002 | 0.061 | 0.1 | 0.07 | 0.15 | 0.17 | 0.16 | 0.1 |
| 2 | 123542070 | rs10181686 | -1.997 | 0.796 | 1.548 | 0.983 | 0.001 | 0.001 | 0.001 | 0.116 | 0.06 | 0.04 | 0.09 | 0.04 | 0.05 | 0.13 |
| 2 | 123544134 | rs6754547 | -1.821 | 0.877 | 1.517 | 0.973 | -0.018 | -0.006 | -0.006 | 0 | 0.1 | 0.04 | 0.15 | 0.17 | 0.17 | 0.1 |
| 2 | 123548703 | rs10496599 | -1.896 | 0.818 | 1.522 | 0.973 | 0.002 | 0 | 0 | 0.538 | 0.06 | 0.03 | 0.09 | 0.04 | 0.05 | 0.13 |
| 2 | 123556174 | rs6724312 | -2.529 | 0.508 | 2.179 | 0.815 | -0.015 | -0.031 | -0.031 | 0 | 0.22 | 0.4 | 0.21 | 0.19 | 0.19 | 0.23 |
| 2 | 123564927 | rs167164 | 1.158 | 0.584 | 2.236 | 0.869 | 0 | -0.001 | -0.001 | 0 | 0.88 | 0.88 | 0.82 | 0.82 | 0.82 | 0.9 |
| 2 | 123566021 | rs294671 | 2.824 | 1.271 | 2.804 | 0.966 | 0.004 | 0.002 | 0.002 | 0 | 0.68 | 0.39 | 0.67 | 0.67 | 0.61 | 0.72 |
| 2 | 123574925 | rs296039 | 2.152 | 1.241 | 3.079 | 0.964 | 0.009 | -0.001 | -0.001 | 0 | 0.81 | 0.61 | 0.77 | 0.79 | 0.8 | 0.82 |
| 2 | 123582223 | rs159395 | 1.037 | 1.238 | 3.062 | 0.961 | 0.008 | 0.002 | 0.002 | 0 | 0.87 | 0.85 | 0.8 | 0.82 | 0.81 | 0.89 |
| 2 | 123582300 | rs10169855 | -1.266 | 1.218 | 3.202 | 0.949 | -0.003 | -0.014 | -0.014 | 0 | 0.05 | 0.22 | 0.01 | 0 | 0.01 | 0.03 |
| 2 | 123588921 | rs296019 | 3.017 | 1.395 | 3.289 | 0.934 | -0.024 | -0.016 | -0.016 | 0 | 0.65 | 0.31 | 0.57 | 0.62 | 0.54 | 0.64 |
| 2 | 123595281 | rs10178940 | -1.849 | 1.417 | 3.252 | 0.947 | 0.016 | 0.002 | 0.002 | 0.074 | 0.07 | 0.33 | 0.03 | 0 | 0.01 | 0.03 |
| 2 | 123598445 | rs296027 | 3.212 | 1.213 | 2.634 | 0.48 | -0.082 | -0.029 | -0.029 | 0.093 | 0.5 | 0.03 | 0.5 | 0.61 | 0.46 | 0.54 |
| 2 | 123600418 | rs296031 | 2.79 | 1.472 | 2.853 | 0.664 | -0.004 | -0.002 | -0.002 | 0 | 0.67 | 0.35 | 0.66 | 0.67 | 0.59 | 0.73 |
| 2 | 123604856 | rs190917 | 2.152 | 1.542 | 3.035 | 0.692 | 0.077 | 0.064 | 0.064 | 0 | 0.74 | 0.61 | 0.61 | 0.63 | 0.56 | 0.74 |
| 2 | 123610375 | rs7609162 | -0.862 | 1.668 | 2.376 | 0.714 | 0.012 | 0.012 | 0.012 | 0 | 0.08 | 0.09 | 0.1 | 0.13 | 0.21 | 0.07 |
| 2 | 123627808 | rs1962159 | -1.54 | 1.495 | 2.721 | 0.571 | -0.035 | -0.129 | -0.129 | 0.084 | 0.17 | 0.5 | 0.08 | 0.01 | 0.09 | 0.11 |
| 2 | 123634829 | rs1406007 | -2.978 | 1.56 | 2.644 | 0.639 | -0.02 | 0.011 | 0.011 | 1.658 | 0.35 | 0.82 | 0.35 | 0.26 | 0.38 | 0.36 |
| 2 | 123636499 | rs4142668 | -1.591 | 1.775 | 2.587 | 0.746 | 0.036 | 0.022 | 0.022 | 0 | 0.12 | 0.18 | 0.21 | 0.17 | 0.25 | 0.14 |
| 2 | 123646963 | rs296068 | 2.413 | 1.691 | 2.756 | 0.779 | 0.005 | 0 | 0 | 0 | 0.64 | 0.44 | 0.61 | 0.63 | 0.55 | 0.61 |
| 2 | 123647424 | rs296069 | 1.921 | 1.697 | 2.689 | 0.774 | -0.019 | -0.029 | -0.029 | 2.594 | 0.76 | 0.63 | 0.81 | 0.8 | 0.79 | 0.75 |
| 2 | 123653197 | rs17367160 | -0.455 | 1.716 | 2.519 | 0.79 | -0.047 | -0.04 | -0.04 | 2.012 | 0.07 | 0.01 | 0.1 | 0.12 | 0.21 | 0.06 |
| 2 | 123662993 | rs1527973 | 2.494 | 1.76 | 2.753 | 0.781 | 0.068 | 0.031 | 0.031 | 0.067 | 0.7 | 0.58 | 0.65 | 0.64 | 0.53 | 0.7 |
| 2 | 123669787 | rs17008910 | 2.767 | 1.74 | 2.765 | 0.786 | -0.002 | -0.001 | -0.001 | 0.006 | 0.61 | 0.44 | 0.62 | 0.63 | 0.5 | 0.62 |
| 2 | 123670708 | rs12471361 | 1.388 | 1.766 | 2.819 | 0.788 | 0.011 | -0.003 | -0.003 | 0 | 0.84 | 0.85 | 0.84 | 0.83 | 0.78 | 0.89 |
| 2 | 123671811 | rs6541865 | 2.143 | 1.746 | 2.837 | 0.789 | 0.014 | 0.003 | 0.003 | 0 | 0.76 | 0.79 | 0.72 | 0.69 | 0.58 | 0.81 |
| 2 | 123674046 | rs7574508 | -0.234 | 1.718 | 2.825 | 0.781 | -0.023 | -0.037 | -0.037 | 0 | 0.09 | 0.14 | 0.03 | 0.01 | 0.04 | 0.09 |
| 2 | 123678650 | rs17008932 | -1.45 | 1.74 | 2.756 | 0.803 | 0.007 | 0.038 | 0.038 | 0 | 0.06 | 0.18 | 0.07 | 0.05 | 0.04 | 0.11 |
| 2 | 123680257 | rs951528 | 1.309 | 1.778 | 2.764 | 0.798 | 0.014 | -0.002 | -0.002 | 0 | 0.86 | 0.85 | 0.85 | 0.83 | 0.81 | 0.91 |
| 2 | 123681660 | rs2090539 | -1.448 | 1.765 | 2.728 | 0.802 | 0.007 | 0.038 | 0.038 | 0 | 0.06 | 0.18 | 0.07 | 0.05 | 0.04 | 0.11 |
| 2 | 123688145 | rs2139011 | 0.089 | 1.78 | 2.585 | 0.802 | 0.005 | 0.006 | 0.006 | 0 | 0.91 | 0.92 | 0.88 | 0.83 | 0.85 | 0.94 |
| 2 | 123694920 | rs297470 | 3.089 | 1.768 | 2.604 | 0.801 | -0.018 | -0.014 | -0.014 | 1.509 | 0.57 | 0.25 | 0.52 | 0.58 | 0.46 | 0.56 |
| 2 | 123705845 | rs297481 | 2.272 | 1.97 | 2.002 | 0.898 | 0.076 | 0.049 | 0.049 | 0 | 0.76 | 0.7 | 0.68 | 0.64 | 0.56 | 0.81 |
| 2 | 123720746 | rs17008996 | -1.833 | 2.057 | 1.967 | 0.943 | 0.064 | 0.064 | 0.064 | 0 | 0.11 | 0.24 | 0.18 | 0.2 | 0.25 | 0.1 |
| 2 | 123726800 | rs7349363 | -0.405 | 2.058 | 1.952 | 0.928 | 0.01 | 0 | 0 | 0 | 0.07 | 0.06 | 0.11 | 0.12 | 0.21 | 0.07 |
| 2 | 123730872 | rs297478 | 2.295 | 2.06 | 1.998 | 0.929 | 0.093 | 0.057 | 0.057 | 0.049 | 0.76 | 0.61 | 0.67 | 0.64 | 0.55 | 0.82 |
| 2 | 123742223 | rs1358423 | 2.703 | 1.944 | 1.898 | 0.912 | 0.016 | 0.001 | 0.001 | 0 | 0.62 | 0.52 | 0.64 | 0.63 | 0.49 | 0.62 |
| 2 | 123742974 | rs11674052 | -0.474 | 2.321 | 1.164 | 1.027 | 0.005 | 0.001 | 0.001 | 0 | 0.1 | 0.05 | 0.04 | 0.01 | 0.06 | 0.11 |
| 2 | 123749705 | rs2247120 | 2.407 | 2.658 | 1.033 | 1.34 | 0.019 | 0.015 | 0.015 | 0.296 | 0.77 | 0.74 | 0.64 | 0.6 | 0.55 | 0.81 |
| 2 | 123752100 | rs1395930 | 2.447 | 2.66 | 1.081 | 1.336 | 0.102 | 0.074 | 0.074 | 0.032 | 0.73 | 0.55 | 0.59 | 0.6 | 0.52 | 0.75 |
| 2 | 123760784 | rs999672 | 0.582 | 2.659 | 1.061 | 1.332 | 0.01 | 0.005 | 0.005 | 0 | 0.9 | 0.91 | 0.82 | 0.84 | 0.81 | 0.92 |
| 2 | 123763312 | rs1507672 | -1.961 | 2.568 | 1.004 | 1.343 | 0.021 | 0.014 | 0.014 | 0 | 0.16 | 0.22 | 0.21 | 0.23 | 0.24 | 0.12 |
| 2 | 123766356 | rs2696233 | -1.748 | 2.135 | 0.659 | 1.018 | 0.049 | 0.016 | 0.016 | 0.065 | 0.25 | 0.3 | 0.4 | 0.33 | 0.48 | 0.24 |
| 2 | 123768369 | rs10203485 | -1.994 | 1.196 | 0.033 | 0.698 | 0.017 | -0.018 | -0.018 | 0 | 0.07 | 0.13 | 0.03 | 0.03 | 0.05 | 0.03 |
| 2 | 123769845 | rs10207388 | -1.19 | 1.315 | -0.436 | 0.684 | 0.002 | -0.012 | -0.012 | 0 | 0.05 | 0.04 | 0.02 | 0 | 0.05 | 0.03 |
| 2 | 123774660 | rs12613332 | -0.194 | 1.106 | -1.123 | 0.801 | -0.028 | -0.028 | -0.028 | 0 | 0.2 | 0.08 | 0.19 | 0.29 | 0.23 | 0.15 |
| 2 | 123777605 | rs3113834 | -0.656 | 2.007 | -0.869 | 0.99 | 0.034 | 0.013 | 0.013 | 0 | 0.07 | 0.14 | 0.19 | 0.09 | 0.15 | 0.05 |
| 2 | 123780262 | rs17009204 | 0.146 | 0.899 | -0.273 | 1.272 | -0.042 | 0.057 | 0.057 | 0 | 0.1 | 0.21 | 0.03 | 0.16 | 0.02 | 0.09 |
| 2 | 123795919 | rs13013898 | 0.844 | -0.001 | -1.041 | 0.796 | 0.028 | 0.035 | 0.035 | 0 | 0.19 | 0.08 | 0.31 | 0.08 | 0.16 | 0.1 |
| 2 | 123796040 | rs7571273 | -1.588 | -0.863 | -1.061 | 0.674 | -0.071 | -0.001 | -0.001 | 3.356 | 0.67 | 0.85 | 0.77 | 0.7 | 0.57 | 0.52 |
| 2 | 123797042 | rs7574923 | 1.607 | -0.843 | -1.03 | 0.668 | -0.024 | -0.026 | -0.026 | 0 | 0.2 | 0.09 | 0.2 | 0.29 | 0.23 | 0.15 |
| 2 | 123799416 | rs17465431 | 1.137 | -1.083 | -1.197 | 0.507 | 0.028 | 0.021 | 0.021 | 0.299 | 0.14 | 0.04 | 0.29 | 0.07 | 0.12 | 0.1 |
| 2 | 123801083 | rs9678754 | 1.471 | -1.054 | -1.057 | 0.654 | -0.034 | 0.023 | 0.023 | 0 | 0.09 | 0.16 | 0.03 | 0.12 | 0.01 | 0.08 |
| 2 | 123804013 | rs299572 | 1.644 | -1.031 | -1.341 | 0.554 | -0.073 | -0.01 | -0.01 | 0 | 0.32 | 0.13 | 0.22 | 0.3 | 0.41 | 0.47 |
| 2 | 123804915 | rs953374 | 0.912 | -1.128 | -0.405 | 0.5 | -0.039 | 0.12 | 0.12 | 0.024 | 0.11 | 0.39 | 0.04 | 0.15 | 0.02 | 0.09 |
| 2 | 123805126 | rs299573 | 1.239 | -1.078 | -0.341 | 0.553 | 0.06 | 0.013 | 0.013 | 0 | 0.54 | 0.3 | 0.44 | 0.63 | 0.66 | 0.63 |
| 2 | 123808961 | rs299575 | 0.319 | -0.533 | -0.504 | 0.44 | -0.001 | 0.001 | 0.001 | 0 | 0.74 | 0.76 | 0.5 | 0.83 | 0.7 | 0.85 |
| 2 | 123809708 | rs299576 | -0.511 | -0.027 | -0.393 | 0.578 | 0.025 | 0.007 | 0.007 | 0 | 0.93 | 0.9 | 0.82 | 0.91 | 0.85 | 0.95 |

| | | | | | | | | | | | | | | | | |
|---|---|---|---|---|---|---|---|---|---|---|---|---|---|---|---|---|
| 2 | 123818296 | rs299544 | 0.332 | 0.094 | -0.812 | 0.33 | -0.014 | -0.024 | -0.024 | 0.138 | 0.43 | 0.22 | 0.32 | 0.46 | 0.48 | 0.42 |
| 2 | 123818496 | rs6728117 | 0.622 | -0.09 | -0.251 | 0.176 | -0.008 | -0.011 | -0.011 | 1.003 | 0.15 | 0.32 | 0.09 | 0.13 | 0.12 | 0.2 |
| 2 | 123818682 | rs17368508 | 1.654 | -0.204 | -0.451 | 0.033 | -0.018 | -0.058 | -0.058 | 0.153 | 0.2 | 0.03 | 0.22 | 0.32 | 0.15 | 0.13 |
| 2 | 123818973 | rs1395931 | -0.778 | -0.222 | -0.223 | 0.175 | 0 | 0.006 | 0.006 | 0 | 0.51 | 0.52 | 0.56 | 0.48 | 0.5 | 0.53 |
| 2 | 123823730 | rs6541877 | 0.2 | -0.199 | -0.057 | 0.152 | -0.062 | 0.012 | 0.012 | 0 | 0.35 | 0.11 | 0.47 | 0.35 | 0.37 | 0.33 |
| 2 | 123825356 | rs17039820 | 0.953 | -0.206 | -0.003 | 0.156 | -0.029 | -0.005 | -0.005 | 2.243 | 0.14 | 0.23 | 0.05 | 0.13 | 0.07 | 0.13 |
| 2 | 123826562 | rs299541 | 0.654 | -0.193 | 0.062 | 0.161 | 0.071 | 0.044 | 0.044 | 0 | 0.94 | 0.71 | 0.85 | 0.94 | 0.93 | 0.88 |
| 2 | 123828854 | rs13011089 | 0.019 | -0.219 | 0.039 | 0.149 | -0.022 | -0.024 | -0.024 | 0 | 0.81 | 0.36 | 0.88 | 0.87 | 0.88 | 0.8 |
| 2 | 123834466 | rs2673185 | 1.19 | -0.18 | 0.037 | 0.192 | -0.039 | -0.028 | -0.028 | 0.292 | 0.42 | 0.11 | 0.3 | 0.38 | 0.48 | 0.42 |
| 2 | 123840759 | rs2661038 | 0.853 | -0.163 | 0.087 | 0.153 | 0.127 | 0.054 | 0.054 | 2.039 | 0.95 | 0.58 | 0.93 | 1 | 0.95 | 0.93 |
| 2 | 123842381 | rs7608426 | -0.676 | -0.157 | 0.059 | 0.165 | 0.001 | 0.029 | 0.029 | 0 | 0.5 | 0.33 | 0.53 | 0.48 | 0.46 | 0.47 |
| 2 | 123846339 | rs6761846 | -0.315 | -0.169 | 0.082 | 0.16 | 0.001 | -0.015 | -0.015 | 0 | 0.28 | 0.3 | 0.3 | 0.16 | 0.3 | 0.33 |
| 2 | 123860377 | rs299562 | 0.823 | -0.208 | -0.249 | 0.212 | -0.02 | -0.028 | -0.028 | 0 | 0.44 | 0.23 | 0.4 | 0.43 | 0.49 | 0.46 |
| 2 | 123878592 | rs10171294 | 1.57 | -0.241 | -0.344 | 0.268 | -0.021 | -0.031 | -0.031 | 0 | 0.2 | 0.05 | 0.24 | 0.32 | 0.19 | 0.13 |
| 2 | 123900548 | rs7602084 | 1.483 | -0.21 | -0.311 | 0.277 | -0.022 | -0.025 | -0.025 | 1.33 | 0.21 | 0.06 | 0.25 | 0.32 | 0.19 | 0.14 |
| 2 | 123908725 | rs6749937 | -0.027 | -0.284 | -0.215 | 0.195 | -0.032 | 0.009 | 0.009 | 0 | 0.2 | 0.61 | 0.09 | 0.21 | 0.1 | 0.22 |
| 2 | 123910170 | rs10171011 | -0.244 | -0.295 | -0.207 | 0.2 | 0.027 | -0.012 | -0.012 | 0 | 0.41 | 0.69 | 0.34 | 0.54 | 0.29 | 0.36 |
| 2 | 123912534 | rs1040134 | 1.298 | -0.32 | -0.282 | 0.163 | -0.076 | -0.079 | -0.079 | 0.138 | 0.41 | 0.06 | 0.28 | 0.37 | 0.47 | 0.39 |
| 2 | 123916216 | rs7605383 | -0.207 | -0.321 | -0.231 | 0.187 | 0.027 | -0.012 | -0.012 | 0 | 0.41 | 0.69 | 0.34 | 0.54 | 0.29 | 0.36 |
| 2 | 123918648 | rs17009469 | 0.653 | -0.194 | 0.167 | 0.287 | -0.05 | -0.005 | -0.005 | 0.097 | 0.15 | 0.31 | 0.05 | 0.13 | 0.06 | 0.13 |
| 2 | 123925634 | rs13010499 | 0.253 | 0.094 | 0.045 | 0.253 | -0.057 | 0.044 | 0.044 | 0 | 0.17 | 0.37 | 0.07 | 0.21 | 0.08 | 0.15 |
| 2 | 123927679 | rs13401808 | 0.366 | 0.322 | -0.31 | 0.122 | -0.006 | 0.003 | 0.003 | 0 | 0.15 | 0.13 | 0.26 | 0.04 | 0.24 | 0.2 |
| 2 | 123928492 | rs1515158 | -0.258 | 0.116 | -0.093 | 0.19 | -0.08 | 0.018 | 0.018 | 0 | 0.81 | 0.56 | 0.79 | 0.81 | 0.93 | 0.77 |
| 2 | 123929100 | rs1515160 | -0.235 | 0.115 | -0.093 | 0.189 | -0.081 | 0.017 | 0.017 | 0 | 0.81 | 0.56 | 0.79 | 0.81 | 0.93 | 0.77 |
| 2 | 123929932 | rs6706214 | 0.062 | 0.091 | -0.025 | 0.25 | -0.08 | 0.043 | 0.043 | 0 | 0.8 | 0.46 | 0.79 | 0.76 | 0.93 | 0.76 |
| 2 | 123930041 | rs6735307 | 1.217 | 0.129 | -0.591 | -0.262 | -0.02 | 0.094 | 0.094 | 0 | 0.52 | 0.12 | 0.48 | 0.72 | 0.58 | 0.44 |
| 2 | 123935107 | rs4411698 | 1.406 | 0.19 | -0.876 | -0.254 | -0.047 | -0.102 | -0.102 | 2.916 | 0.28 | 0.06 | 0.23 | 0.37 | 0.35 | 0.3 |
| 2 | 123936217 | rs2048768 | 0.458 | 0.273 | -0.738 | -0.406 | -0.011 | 0.021 | 0.021 | 0 | 0.18 | 0.13 | 0.24 | 0.04 | 0.23 | 0.22 |
| 2 | 123937555 | rs2419554 | 1.195 | 0.318 | -0.67 | -0.316 | 0.011 | -0.002 | -0.002 | 0 | 0.08 | 0.11 | 0.05 | 0.05 | 0.1 | 0.02 |
| 2 | 123943220 | rs10198199 | -0.141 | 0.168 | -0.601 | -0.357 | -0.017 | -0.045 | -0.045 | 0 | 0.75 | 0.42 | 0.86 | 0.86 | 0.82 | 0.78 |
| 2 | 123951832 | rs1519293 | 0.622 | 0.255 | -0.965 | -0.314 | -0.078 | 0.027 | 0.027 | 0 | 0.15 | 0.01 | 0.22 | 0.05 | 0.22 | 0.23 |
| 2 | 123952420 | rs6541885 | 1.089 | -0.181 | -1.085 | -0.335 | 0.011 | 0.009 | 0.009 | 0 | 0.1 | 0.04 | 0.09 | 0.06 | 0.07 | 0.1 |
| 2 | 123955792 | rs1913660 | 1.831 | -0.113 | -1.156 | -0.005 | -0.017 | -0.046 | -0.046 | 1.732 | 0.25 | 0.06 | 0.26 | 0.37 | 0.22 | 0.13 |
| 2 | 123958889 | rs12623767 | 1.822 | -0.109 | -1.135 | 0.012 | -0.014 | -0.04 | -0.04 | 0 | 0.25 | 0.06 | 0.26 | 0.37 | 0.22 | 0.18 |
| 2 | 123963717 | rs6744585 | 0.861 | -0.089 | -1.014 | 0.026 | 0.025 | -0.013 | -0.013 | 0.019 | 0.36 | 0.14 | 0.35 | 0.44 | 0.29 | 0.24 |
| 2 | 123963834 | rs11892911 | 1.458 | 0.046 | -0.811 | 0.014 | 0.01 | -0.035 | -0.035 | 0 | 0.08 | 0.15 | 0.05 | 0 | 0.09 | 0.01 |
| 2 | 123967985 | rs1033294 | -1.19 | -0.183 | -0.663 | 0.004 | -0.059 | -0.007 | -0.007 | 1.184 | 0.84 | 0.65 | 0.93 | 0.86 | 0.93 | 0.88 |
| 2 | 123970983 | rs7567104 | 1.934 | -0.091 | -0.89 | -0.005 | -0.092 | -0.101 | -0.101 | 0 | 0.2 | 0.01 | 0.15 | 0.26 | 0.3 | 0.28 |
| 2 | 123972573 | rs1018332 | -1.115 | -0.312 | -0.564 | -0.086 | -0.066 | 0 | 0 | 0 | 0.84 | 0.57 | 0.93 | 0.86 | 0.93 | 0.88 |
| 2 | 123981311 | rs1850810 | -0.328 | -0.241 | -0.63 | -0.056 | -0.031 | -0.046 | -0.046 | 0 | 0.76 | 0.46 | 0.88 | 0.86 | 0.84 | 0.87 |
| 2 | 124006345 | rs12477571 | 0.485 | -0.237 | -0.806 | -0.053 | -0.098 | -0.111 | -0.111 | 0 | 0.37 | 0.13 | 0.51 | 0.41 | 0.53 | 0.57 |
| 2 | 124007505 | rs7607381 | -0.141 | -0.275 | -0.533 | -0.019 | -0.023 | -0.033 | -0.033 | 0 | 0.75 | 0.38 | 0.87 | 0.86 | 0.84 | 0.87 |
| 2 | 124018896 | rs2139300 | -0.442 | -0.141 | -0.831 | 0.058 | -0.003 | -0.001 | -0.001 | 0 | 0.54 | 0.58 | 0.64 | 0.49 | 0.63 | 0.77 |
| 2 | 124024096 | rs1996588 | 0.704 | 0.201 | -0.633 | -0.045 | -0.007 | -0.002 | -0.002 | 3.287 | 0.07 | 0.02 | 0.12 | 0.05 | 0.07 | 0.12 |
| 2 | 124033771 | rs4848836 | -0.165 | 0.47 | -0.189 | -0.092 | 0.09 | 0.016 | 0.016 | 0.136 | 0.18 | 0.48 | 0.32 | 0.14 | 0.26 | 0.33 |
| 2 | 124035542 | rs11884534 | 0.329 | 0.152 | -0.196 | -0.156 | 0.017 | -0.012 | -0.012 | 0 | 0.44 | 0.29 | 0.31 | 0.47 | 0.42 | 0.51 |
| 2 | 124049457 | rs13002629 | 0.878 | 0.206 | -0.181 | -0.148 | 0.011 | -0.02 | -0.02 | 0 | 0.37 | 0.22 | 0.19 | 0.42 | 0.35 | 0.4 |
| 2 | 124057939 | rs955303 | 0.344 | 0.123 | -0.039 | -0.236 | 0.014 | -0.004 | -0.004 | 0 | 0.46 | 0.33 | 0.33 | 0.48 | 0.42 | 0.52 |
| 2 | 124061531 | rs1519283 | -0.794 | 0.055 | -0.082 | -0.221 | 0.012 | 0.018 | 0.018 | 0.08 | 0.63 | 0.76 | 0.64 | 0.61 | 0.68 | 0.85 |
| 2 | 124062001 | rs12711636 | 0.896 | 0.057 | -0.177 | -0.22 | -0.003 | 0.011 | 0.011 | 0.361 | 0.35 | 0.09 | 0.35 | 0.39 | 0.32 | 0.15 |
| 2 | 124069523 | rs12612635 | -0.554 | 0.142 | -0.256 | -0.167 | -0.01 | -0.007 | -0.007 | 0 | 0.6 | 0.46 | 0.61 | 0.58 | 0.68 | 0.78 |
| 2 | 124082249 | rs12615507 | 1.023 | 0.1 | -0.51 | -0.237 | -0.008 | -0.061 | -0.061 | 0 | 0.31 | 0.07 | 0.4 | 0.44 | 0.28 | 0.27 |
| 2 | 124088863 | rs1356669 | 0.267 | 0.564 | -0.103 | 0.21 | 0.004 | 0 | 0 | 0 | 0.15 | 0.18 | 0.29 | 0.1 | 0.26 | 0.26 |
| 2 | 124099844 | rs6541893 | -1.012 | 0.512 | -0.051 | 0.181 | 0.012 | 0.019 | 0.019 | 0.041 | 0.62 | 0.76 | 0.64 | 0.61 | 0.68 | 0.84 |
| 2 | 124107356 | rs970066 | -0.815 | 0.559 | -0.13 | 0.218 | -0.008 | -0.006 | -0.006 | 0 | 0.6 | 0.56 | 0.6 | 0.58 | 0.68 | 0.78 |
| 2 | 124109213 | rs993934 | 0.132 | 0.57 | -0.131 | 0.216 | 0.01 | -0.005 | -0.005 | 0 | 0.45 | 0.34 | 0.31 | 0.48 | 0.42 | 0.51 |
| 2 | 124116694 | rs12711638 | -0.821 | 0.555 | -0.291 | 0.09 | -0.001 | -0.001 | -0.001 | 0.18 | 0.61 | 0.62 | 0.61 | 0.61 | 0.68 | 0.84 |
| 2 | 124130763 | rs17246066 | 0.905 | 0.684 | -0.908 | 0.012 | -0.063 | 0.037 | 0.037 | 0 | 0.12 | 0.03 | 0.14 | 0.04 | 0.25 | 0.21 |
| 2 | 124135616 | rs9636352 | -0.78 | 0.599 | -0.805 | 0.027 | -0.01 | -0.004 | -0.004 | 0 | 0.59 | 0.54 | 0.48 | 0.55 | 0.67 | 0.78 |
| 2 | 124144379 | rs10201655 | 0.401 | 0.754 | -1.35 | 0.072 | -0.012 | 0.01 | 0.01 | 0 | 0.13 | 0.13 | 0.15 | 0.04 | 0.25 | 0.21 |
| 2 | 124144870 | rs10178547 | -0.487 | 0.542 | -1.147 | 0.238 | 0.008 | 0.003 | 0.003 | 0 | 0.58 | 0.39 | 0.47 | 0.55 | 0.68 | 0.72 |
| 2 | 124148965 | rs6709341 | 0.066 | 0.244 | -1.161 | 0.547 | 0.001 | -0.02 | -0.02 | 0 | 0.33 | 0.24 | 0.18 | 0.44 | 0.35 | 0.39 |
| 2 | 124155237 | rs4848220 | -0.385 | 0.397 | -1.571 | 0.708 | -0.048 | -0.076 | -0.076 | 0 | 0.3 | 0.15 | 0.46 | 0.42 | 0.52 | 0.41 |
| 2 | 124169595 | rs1405093 | 0.444 | -0.922 | -1.106 | -0.141 | 0.015 | 0 | 0 | 6.618 | 0.49 | 0.42 | 0.42 | 0.58 | 0.35 | 0.55 |
| 2 | 124171272 | rs12052848 | 0.218 | -0.984 | -0.69 | -0.173 | -0.013 | -0.022 | -0.022 | 0.057 | 0.31 | 0.29 | 0.47 | 0.42 | 0.53 | 0.43 |
| 2 | 124179505 | rs12619549 | 0.764 | -1.808 | -0.945 | -0.266 | -0.013 | -0.014 | -0.014 | 0 | 0.31 | 0.33 | 0.26 | 0.25 | 0.24 | 0.36 |
| 2 | 124222386 | rs11683811 | -0.177 | -1.731 | -0.481 | -0.263 | 0.014 | 0 | 0 | 0.219 | 0.49 | 0.51 | 0.43 | 0.59 | 0.35 | 0.54 |
| 2 | 124235774 | rs2419751 | -0.199 | -1.688 | -0.518 | -0.24 | -0.004 | 0 | 0 | 0 | 0.49 | 0.48 | 0.41 | 0.59 | 0.36 | 0.54 |
| 2 | 124250217 | rs11122985 | -0.038 | -1.725 | -0.418 | -0.259 | 0.021 | -0.001 | -0.001 | 0.002 | 0.48 | 0.39 | 0.4 | 0.59 | 0.36 | 0.53 |

| | | | | | | | | | | | | | | | |
|---|---|---|---|---|---|---|---|---|---|---|---|---|---|---|---|
| 2 | 124251487 | rs4429478 | -0.106 | -1.687 | -0.494 | -0.24 | 0.029 | 0.002 | 0.002 | 0 | 0.49 | 0.45 | 0.59 | 0.41 | 0.64 | 0.47 |
| 2 | 124260724 | rs12471979 | 0.75 | -1.85 | -0.838 | -0.426 | -0.07 | -0.091 | -0.091 | 0 | 0.3 | 0.12 | 0.48 | 0.44 | 0.52 | 0.44 |
| 2 | 124263199 | rs6737383 | -1.315 | -2.226 | -1.086 | -0.603 | -0.002 | -0.006 | -0.006 | 0 | 0.82 | 0.79 | 0.83 | 0.86 | 0.85 | 0.79 |
| 2 | 124269785 | rs7598708 | 0.064 | -2.129 | -0.914 | -0.492 | -0.045 | 0.021 | 0.021 | 0 | 0.33 | 0.46 | 0.26 | 0.42 | 0.21 | 0.32 |
| 2 | 124271902 | rs7579595 | 1.615 | -1.822 | -0.972 | -0.603 | 0.014 | 0.004 | 0.004 | 1.338 | 0.17 | 0.1 | 0.08 | 0 | 0.12 | 0.02 |
| 2 | 124276444 | rs10496614 | 0.779 | -1.851 | -0.641 | -0.569 | -0.004 | -0.033 | -0.033 | 2.2 | 0.14 | 0.28 | 0.09 | 0.06 | 0.1 | 0.14 |
| 2 | 124278828 | rs6707592 | -1.139 | -1.299 | -1.233 | -0.941 | 0.001 | 0.009 | 0.009 | 0 | 0.77 | 0.79 | 0.8 | 0.92 | 0.77 | 0.73 |
| 2 | 124278997 | rs13385327 | 0.326 | -0.797 | -1.383 | -0.691 | -0.005 | -0.016 | -0.016 | 0 | 0.24 | 0.51 | 0.2 | 0.18 | 0.22 | 0.27 |
| 2 | 124279468 | rs10496615 | 1.802 | -0.864 | -1.548 | -0.49 | -0.049 | -0.053 | -0.053 | 1.131 | 0.16 | 0.08 | 0.28 | 0.38 | 0.31 | 0.27 |
| 2 | 124279775 | rs12616423 | -0.59 | -1.062 | -1.471 | -0.436 | 0.036 | 0.018 | 0.018 | 0 | 0.59 | 0.69 | 0.63 | 0.56 | 0.66 | 0.55 |
| 2 | 124280149 | rs2419976 | 1.309 | -0.783 | -1.56 | -0.109 | 0.039 | -0.048 | -0.048 | 0 | 0.18 | 0.09 | 0.17 | 0.35 | 0.11 | 0.18 |
| 2 | 124280718 | rs17195650 | 0.383 | -0.389 | -1.381 | -0.054 | -0.035 | -0.027 | -0.027 | 0.031 | 0.34 | 0.22 | 0.46 | 0.38 | 0.44 | 0.31 |
| 2 | 124286924 | rs1344228 | 1.072 | -0.817 | -1.331 | 0.219 | 0.016 | -0.028 | -0.028 | 0 | 0.16 | 0.11 | 0.17 | 0.34 | 0.12 | 0.16 |
| 2 | 124287002 | rs6719156 | 0.963 | 0.051 | -1.434 | -0.324 | -0.03 | 0.017 | 0.017 | 0 | 0.22 | 0.16 | 0.25 | 0.02 | 0.39 | 0.15 |
| 2 | 124287245 | rs2419967 | -0.825 | -0.421 | -1.69 | -0.467 | 0.006 | 0.02 | 0.02 | 0.649 | 0.61 | 0.37 | 0.73 | 0.79 | 0.67 | 0.66 |
| 2 | 124288380 | rs6726405 | -0.067 | -0.535 | -1.802 | -0.572 | 0.061 | 0.094 | 0.094 | 0.122 | 0.51 | 0.3 | 0.56 | 0.78 | 0.38 | 0.52 |
| 2 | 124301185 | rs7556991 | 2.63 | -0.756 | -1.311 | -0.566 | -0.005 | -0.032 | -0.032 | 2.252 | 0.13 | 0.13 | 0.08 | 0 | 0.09 | 0.02 |
| 2 | 124306147 | rs11684317 | 0.779 | -0.712 | -1.28 | -0.572 | 0 | -0.005 | -0.005 | 0 | 0.36 | 0.37 | 0.24 | 0.2 | 0.33 | 0.31 |
| 2 | 124319428 | rs6737930 | 1.118 | -0.706 | -1.465 | -0.523 | -0.002 | -0.031 | -0.031 | 0 | 0.38 | 0.09 | 0.32 | 0.43 | 0.33 | 0.35 |
| 2 | 124330320 | rs11904509 | 1.157 | -0.848 | -1.459 | -0.558 | 0.013 | 0.016 | 0.016 | 0 | 0.21 | 0.18 | 0.2 | 0.08 | 0.16 | 0.1 |
| 2 | 124331148 | rs2901018 | 0.597 | -0.601 | -1.289 | -0.347 | -0.021 | -0.023 | -0.023 | 0 | 0.6 | 0.27 | 0.52 | 0.51 | 0.49 | 0.44 |
| 2 | 124336412 | rs2419986 | 1.16 | -0.647 | -1.168 | -0.217 | 0.002 | -0.014 | -0.014 | 0 | 0.08 | 0.02 | 0.14 | 0.08 | 0.08 | 0.08 |
| 2 | 124356455 | rs13032568 | 0.883 | -0.656 | -1.063 | -0.207 | 0.001 | -0.008 | -0.008 | 0 | 0.1 | 0.09 | 0.05 | 0.12 | 0.09 | 0.11 |
| 2 | 124363708 | rs2419979 | -0.615 | -0.184 | -0.626 | -0.397 | 0.189 | -0.05 | -0.05 | 0 | 0.89 | 0.72 | 0.81 | 0.98 | 0.71 | 0.87 |
| 2 | 124368356 | rs11677528 | 1.26 | -0.637 | -0.792 | -0.466 | 0 | 0.005 | 0.005 | 0 | 0.39 | 0.31 | 0.35 | 0.41 | 0.38 | 0.38 |
| 2 | 124379041 | rs2670616 | -2.012 | -0.873 | -0.868 | -0.536 | 0.007 | -0.01 | -0.01 | 0 | 0.87 | 0.93 | 0.82 | 0.63 | 0.91 | 0.74 |
| 2 | 124382362 | rs2037033 | -0.482 | -0.911 | -0.858 | -0.597 | 0.031 | -0.059 | -0.059 | 0 | 0.74 | 0.87 | 0.66 | 0.56 | 0.83 | 0.66 |
| 2 | 124382907 | rs13403945 | 0.227 | -0.953 | -0.785 | -0.608 | -0.056 | -0.057 | -0.057 | 0 | 0.27 | 0.52 | 0.15 | 0.13 | 0.16 | 0.13 |
| 2 | 124399883 | rs7565459 | 0.343 | -0.862 | -0.842 | -0.564 | -0.057 | -0.063 | -0.063 | 3.812 | 0.66 | 0.84 | 0.5 | 0.54 | 0.54 | 0.51 |
| 2 | 124401476 | rs6735319 | 0.507 | -0.818 | -0.699 | -0.507 | -0.014 | 0.02 | 0.02 | 0.012 | 0.14 | 0.33 | 0.08 | 0.13 | 0.09 | 0.12 |
| 2 | 124408674 | rs2901024 | 0.318 | -0.795 | -0.775 | -0.496 | -0.01 | 0.026 | 0.026 | 0.297 | 0.14 | 0.39 | 0.09 | 0.13 | 0.1 | 0.14 |
| 2 | 124408746 | rs2419997 | 1.231 | -0.525 | -0.983 | -0.503 | -0.045 | 0.029 | 0.029 | 0 | 0.08 | 0.01 | 0.16 | 0.02 | 0.29 | 0.15 |
| 2 | 124409054 | rs6729527 | 1.479 | -0.789 | -0.967 | -0.519 | -0.025 | -0.012 | -0.012 | 0 | 0.37 | 0.14 | 0.33 | 0.41 | 0.38 | 0.35 |
| 2 | 124409144 | rs6742867 | 1.427 | -0.793 | -0.938 | -0.522 | -0.01 | 0 | 0 | 2.356 | 0.38 | 0.18 | 0.34 | 0.41 | 0.38 | 0.35 |
| 2 | 124428936 | rs17010366 | 2.497 | -0.831 | -0.766 | -0.538 | -0.029 | -0.066 | -0.066 | 0 | 0.13 | 0.27 | 0.09 | 0 | 0.07 | 0.01 |
| 2 | 124431915 | rs1156077 | -0.068 | -0.746 | -0.89 | -0.49 | -0.001 | -0.001 | -0.001 | 0.002 | 0.65 | 0.68 | 0.59 | 0.61 | 0.57 | 0.59 |
| 2 | 124432181 | rs2203293 | 1.581 | -0.849 | -1.12 | -0.524 | -0.016 | -0.008 | -0.008 | 0.58 | 0.38 | 0.16 | 0.34 | 0.41 | 0.39 | 0.36 |
| 2 | 124434749 | rs1467136 | -1.804 | -1.006 | -1.141 | -0.407 | 0.017 | -0.038 | -0.038 | 0.164 | 0.86 | 0.94 | 0.82 | 0.63 | 0.92 | 0.74 |
| 2 | 124437197 | rs9653411 | -0.594 | -0.792 | -0.742 | -0.457 | -0.033 | -0.013 | -0.013 | 0 | 0.26 | 0.42 | 0.24 | 0.21 | 0.18 | 0.22 |
| 2 | 124454519 | rs12990473 | 0.338 | -0.864 | -0.999 | -0.314 | 0.009 | -0.013 | -0.013 | 0 | 0.25 | 0.23 | 0.25 | 0.5 | 0.15 | 0.29 |
| 2 | 124454811 | rs2707551 | -1.239 | -0.603 | -1.361 | -0.454 | 0.002 | 0.071 | 0.071 | 0.006 | 0.48 | 0.78 | 0.58 | 0.59 | 0.51 | 0.49 |
| 2 | 124457999 | rs12467497 | -0.159 | -1.281 | -1.228 | -0.353 | -0.007 | -0.001 | -0.001 | 0.943 | 0.31 | 0.66 | 0.29 | 0.5 | 0.15 | 0.29 |
| 2 | 124469758 | rs17010412 | 0.593 | -0.324 | -1.246 | -0.389 | 0.019 | 0.009 | 0.009 | 0 | 0.1 | 0.05 | 0.14 | 0.08 | 0.09 | 0.07 |
| 2 | 124476871 | rs1159139 | 1.034 | 0.215 | -1.045 | -0.521 | -0.018 | 0.043 | 0.043 | 0 | 0.1 | 0.02 | 0.1 | 0.01 | 0.1 | 0.11 |
| 2 | 124477143 | rs17010427 | 0.016 | 0.171 | -0.696 | 0.018 | -0.014 | 0.051 | 0.051 | 0 | 0.14 | 0.18 | 0.08 | 0.32 | 0.07 | 0.18 |
| 2 | 124479731 | rs17198253 | 0.664 | 0.467 | -1.317 | -0.164 | -0.017 | 0.037 | 0.037 | 0.001 | 0.4 | 0.16 | 0.32 | 0.36 | 0.39 | 0.34 |
| 2 | 124481347 | rs7593488 | -2.023 | 0.376 | -1.006 | -0.139 | -0.003 | -0.016 | -0.016 | 0 | 0.89 | 0.86 | 0.92 | 0.95 | 0.92 | 0.87 |
| 2 | 124481367 | rs12620260 | 0.214 | 0.531 | -1.198 | -0.214 | 0.107 | 0.094 | 0.094 | 0 | 0.48 | 0.21 | 0.48 | 0.36 | 0.68 | 0.48 |
| 2 | 124483400 | rs13009072 | -0.085 | 1.279 | -0.624 | -0.405 | -0.068 | 0.036 | 0.036 | 0.041 | 0.1 | 0.02 | 0.15 | 0.02 | 0.29 | 0.14 |
| 2 | 124484294 | rs12476179 | 0.057 | 1.42 | -0.217 | -0.378 | -0.002 | 0.11 | 0.11 | 1.511 | 0.23 | 0.44 | 0.17 | 0.37 | 0.18 | 0.32 |
| 2 | 124488139 | rs1040049 | 1.39 | 1.322 | -0.472 | -0.597 | 0.068 | -0.06 | -0.06 | 0 | 0.39 | 0.13 | 0.47 | 0.6 | 0.31 | 0.32 |
| 2 | 124488562 | rs1513789 | 0.643 | 1.376 | -0.393 | -0.566 | -0.015 | -0.01 | -0.01 | 0 | 0.65 | 0.35 | 0.66 | 0.62 | 0.52 | 0.54 |
| 2 | 124508051 | rs10496618 | 0.793 | 1.331 | -0.449 | -0.597 | 0.008 | 0.055 | 0.055 | 0 | 0.26 | 0.17 | 0.19 | 0.02 | 0.21 | 0.22 |
| 2 | 124509138 | rs2037034 | -0.44 | 1.727 | 0.236 | 0.014 | -0.005 | 0.09 | 0.09 | 2.489 | 0.25 | 0.43 | 0.19 | 0.36 | 0.22 | 0.34 |
| 2 | 124523311 | rs1355834 | 0.888 | 1.362 | -0.291 | 0.074 | -0.019 | -0.013 | -0.013 | 0 | 0.64 | 0.31 | 0.65 | 0.62 | 0.52 | 0.54 |
| 2 | 124531331 | rs12987158 | -0.747 | 1.884 | -0.215 | -0.156 | 0.019 | -0.01 | -0.01 | 0 | 0.11 | 0.12 | 0.16 | 0.02 | 0.28 | 0.15 |
| 2 | 124531480 | rs3981253 | -0.207 | 0.843 | 0.438 | 0.142 | -0.017 | 0.12 | 0.12 | 0.193 | 0.11 | 0.24 | 0.07 | 0.31 | 0.07 | 0.16 |
| 2 | 124538116 | rs1554001 | 0.012 | 1.146 | 0.369 | 0.092 | -0.008 | 0.011 | 0.011 | 0 | 0.07 | 0.03 | 0.09 | 0.02 | 0.11 | 0.13 |
| 2 | 124538974 | rs782380 | -0.624 | 0.819 | 0.54 | -0.343 | 0.021 | 0.016 | 0.016 | 1.007 | 0.51 | 0.6 | 0.35 | 0.38 | 0.6 | 0.56 |
| 2 | 124540542 | rs782377 | -0.732 | 0.815 | 0.621 | -0.345 | 0.031 | 0.024 | 0.024 | 0.865 | 0.52 | 0.65 | 0.36 | 0.38 | 0.6 | 0.57 |
| 2 | 124565374 | rs13028007 | 0.522 | 0.712 | 0.524 | -0.264 | -0.029 | 0.015 | 0.015 | 0 | 0.07 | 0.01 | 0.09 | 0.02 | 0.11 | 0.13 |
| 2 | 124591112 | rs6717076 | 1.438 | 0.747 | 0.573 | -0.318 | 0.039 | -0.025 | -0.025 | 0 | 0.4 | 0.15 | 0.53 | 0.59 | 0.31 | 0.31 |
| 2 | 124600797 | rs1588939 | -0.415 | 0.772 | 0.732 | -0.309 | 0.023 | -0.027 | -0.027 | 0.011 | 0.08 | 0.22 | 0.11 | 0.02 | 0.11 | 0.13 |
| 2 | 124608981 | rs2138431 | -1.089 | 0.847 | 0.364 | -0.442 | -0.08 | 0.123 | 0.123 | 0 | 0.75 | 0.92 | 0.82 | 0.98 | 0.62 | 0.75 |
| 2 | 124621037 | rs6541929 | 1.162 | 0.847 | 0.411 | -0.347 | 0.049 | -0.014 | -0.014 | 0 | 0.42 | 0.21 | 0.54 | 0.59 | 0.3 | 0.3 |
| 2 | 124631703 | rs12624214 | 1.42 | 0.886 | 0.483 | -0.381 | -0.005 | 0.004 | 0.004 | 0.393 | 0.12 | 0.1 | 0.09 | 0.05 | 0.14 | 0.15 |
| 2 | 124663350 | rs17267326 | 1.441 | 0.782 | 0.332 | -0.336 | 0.017 | -0.075 | -0.075 | 0 | 0.39 | 0.09 | 0.53 | 0.58 | 0.3 | 0.31 |
| 2 | 124665077 | rs925796 | -0.764 | 0.827 | 0.544 | -0.357 | -0.003 | 0.024 | 0.024 | 0 | 0.47 | 0.67 | 0.38 | 0.38 | 0.57 | 0.54 |
| 2 | 124670783 | rs1516080 | 1.336 | 0.792 | 0.436 | -0.443 | -0.004 | -0.037 | -0.037 | 0 | 0.39 | 0.08 | 0.55 | 0.61 | 0.34 | 0.35 |

| | | | | | | | | | | | | | | | | |
|---|---|---|---|---|---|---|---|---|---|---|---|---|---|---|---|---|
| 2 | 124675291 | rs10496621 | 0.439 | 0.638 | 0.843 | -0.368 | -0.002 | 0.023 | 0.023 | 0 | 0.1 | 0.1 | 0.05 | 0.3 | 0.06 | 0.15 |
| 2 | 124676369 | rs871475 | -0.406 | 0.671 | 0.974 | -0.405 | 0.001 | -0.002 | -0.002 | 0 | 0.38 | 0.41 | 0.25 | 0.32 | 0.43 | 0.37 |
| 2 | 124678253 | rs12469604 | 0.991 | 0.662 | 0.688 | -0.413 | -0.014 | 0.032 | 0.032 | 0.159 | 0.14 | 0.07 | 0.08 | 0.05 | 0.14 | 0.15 |
| 2 | 124683913 | rs1214131 | -0.633 | 0.72 | 0.657 | -0.36 | -0.002 | 0.003 | 0.003 | 0 | 0.4 | 0.6 | 0.27 | 0.32 | 0.44 | 0.37 |
| 2 | 124685686 | rs1214132 | 0.648 | 0.802 | 0.137 | -0.338 | -0.022 | 0.029 | 0.029 | 0 | 0.27 | 0.18 | 0.19 | 0.02 | 0.37 | 0.21 |
| 2 | 124698518 | rs1214123 | 0.055 | 0.966 | -0.069 | -0.085 | 0.004 | -0.006 | -0.006 | 0 | 0.38 | 0.44 | 0.25 | 0.32 | 0.44 | 0.37 |
| 2 | 124701856 | rs6760710 | 0.754 | 0.944 | -0.073 | -0.116 | 0.011 | 0.022 | 0.022 | 0.25 | 0.44 | 0.25 | 0.64 | 0.63 | 0.42 | 0.45 |
| 2 | 124710795 | rs1214081 | -0.491 | 1.044 | -0.271 | 0.387 | 0.003 | -0.006 | -0.006 | 0 | 0.38 | 0.44 | 0.25 | 0.32 | 0.44 | 0.37 |
| 2 | 124739775 | rs13415885 | -0.006 | 0.986 | -0.207 | 0.903 | 0 | 0.042 | 0.042 | 0 | 0.16 | 0.22 | 0.18 | 0.26 | 0.15 | 0.25 |
| 2 | 124740888 | rs1921827 | 0.455 | 0.957 | -0.188 | 0.602 | -0.001 | -0.005 | -0.005 | 0.97 | 0.23 | 0.24 | 0.35 | 0.24 | 0.26 | 0.21 |
| 2 | 124741766 | rs6722224 | 0.355 | 0.892 | -0.046 | 0.57 | -0.002 | -0.015 | -0.015 | 0.733 | 0.24 | 0.33 | 0.36 | 0.24 | 0.26 | 0.2 |
| 2 | 124748246 | rs13022471 | 0.745 | 0.836 | -0.093 | 0.57 | -0.001 | -0.008 | -0.008 | 0 | 0.15 | 0.1 | 0.18 | 0.26 | 0.15 | 0.25 |
| 2 | 124749564 | rs1921819 | -0.666 | 1.387 | -0.142 | 0.281 | 0 | 0 | 0 | 0 | 0.44 | 0.56 | 0.59 | 0.51 | 0.51 | 0.5 |
| 2 | 124749962 | rs1921817 | 0.662 | 1.49 | -1.136 | -0.042 | -0.035 | 0.044 | 0.044 | 0.293 | 0.21 | 0.09 | 0.14 | 0.09 | 0.25 | 0.22 |
| 2 | 124753418 | rs4848904 | -0.054 | 1.317 | -1.257 | -0.07 | -0.016 | -0.035 | -0.035 | 0.115 | 0.45 | 0.18 | 0.34 | 0.49 | 0.44 | 0.42 |
| 2 | 124754742 | rs13431665 | 1.173 | 0.824 | -1.353 | 0.26 | 0.025 | -0.058 | -0.058 | 0 | 0.22 | 0.07 | 0.12 | 0.32 | 0.14 | 0.15 |
| 2 | 124759480 | rs17010715 | 1.231 | 0.828 | -1.368 | 0.261 | 0.022 | -0.068 | -0.068 | 0 | 0.21 | 0.06 | 0.12 | 0.31 | 0.14 | 0.11 |
| 2 | 124771097 | rs6746763 | 1.925 | 1.148 | -1.38 | 0.436 | -0.04 | -0.039 | -0.039 | 0 | 0.07 | 0 | 0.05 | 0.12 | 0.1 | 0.09 |
| 2 | 124775660 | rs6745116 | -0.448 | 0.946 | -1.076 | 0.137 | -0.001 | 0.015 | 0.015 | 0.173 | 0.61 | 0.3 | 0.52 | 0.76 | 0.59 | 0.62 |
| 2 | 124779084 | rs4848910 | -0.157 | 1.006 | -0.973 | 0.122 | -0.002 | -0.021 | -0.021 | 0 | 0.45 | 0.24 | 0.34 | 0.5 | 0.44 | 0.42 |
| 2 | 124785753 | rs1439131 | -0.565 | 1.002 | -0.918 | -0.048 | -0.011 | 0.018 | 0.018 | 0 | 0.86 | 0.91 | 0.91 | 1 | 0.85 | 0.87 |
| 2 | 124797837 | rs17010743 | 1.031 | 0.596 | -1.103 | 0.248 | 0.032 | -0.041 | -0.041 | 0 | 0.2 | 0.06 | 0.12 | 0.27 | 0.14 | 0.11 |
| 2 | 124801531 | rs1213952 | 0.417 | 0.956 | -1.165 | -0.111 | -0.041 | 0.058 | 0.058 | 0 | 0.16 | 0.07 | 0.14 | 0.01 | 0.24 | 0.15 |
| 2 | 124815809 | rs7566987 | 1.126 | 1.039 | -0.759 | -0.06 | -0.005 | 0.006 | 0.006 | 0 | 0.21 | 0.21 | 0.39 | 0.19 | 0.3 | 0.22 |
| 2 | 124819487 | rs1213940 | -1.796 | 0.861 | -0.472 | 0.266 | -0.009 | 0.016 | 0.016 | 0 | 0.84 | 0.8 | 0.84 | 0.8 | 0.9 | 0.76 |
| 2 | 124819530 | rs17010807 | 2.599 | 0.885 | -0.694 | 0.3 | -0.014 | -0.018 | -0.018 | 0 | 0.07 | 0 | 0.04 | 0.08 | 0.06 | 0.09 |
| 2 | 124819540 | rs17010809 | 1.842 | 0.673 | -0.809 | 0.308 | 0.033 | -0.087 | -0.087 | 0.371 | 0.18 | 0.04 | 0.12 | 0.33 | 0.13 | 0.11 |
| 2 | 124821380 | rs17271568 | 1.288 | 0.643 | -0.48 | 0.219 | 0.004 | -0.015 | -0.015 | 0.722 | 0.23 | 0.3 | 0.4 | 0.19 | 0.31 | 0.22 |
| 2 | 124821430 | rs17010817 | 1.865 | 0.625 | -0.506 | 0.267 | 0.037 | -0.07 | -0.07 | 0 | 0.18 | 0.04 | 0.12 | 0.29 | 0.13 | 0.11 |
| 2 | 124822388 | rs1213946 | 0.475 | 0.884 | -0.074 | 0.082 | -0.007 | 0.008 | 0.008 | 0 | 0.28 | 0.18 | 0.23 | 0.2 | 0.3 | 0.33 |
| 2 | 124822591 | rs980374 | 1.434 | 0.912 | -0.116 | 0.087 | -0.001 | -0.002 | -0.002 | 0 | 0.22 | 0.24 | 0.4 | 0.19 | 0.3 | 0.22 |
| 2 | 124826460 | rs4473382 | 0.493 | 0.857 | 0.122 | 0.049 | 0.015 | -0.032 | -0.032 | 1.731 | 0.29 | 0.39 | 0.44 | 0.19 | 0.41 | 0.24 |
| 2 | 124832503 | rs3902861 | 1.404 | 0.87 | -0.019 | 0.064 | 0 | -0.003 | -0.003 | 0 | 0.22 | 0.25 | 0.39 | 0.19 | 0.31 | 0.22 |
| 2 | 124833536 | rs1816698 | 0.586 | 0.828 | -0.009 | 0.068 | -0.027 | 0.023 | 0.023 | 0 | 0.27 | 0.14 | 0.23 | 0.18 | 0.3 | 0.3 |
| 2 | 124836896 | rs17212552 | 1.798 | 1.005 | -0.162 | -0.042 | -0.016 | 0.015 | 0.015 | 0 | 0.06 | 0.02 | 0.09 | 0.01 | 0.13 | 0.07 |
| 2 | 124837809 | rs2420441 | 0.539 | 0.816 | -0.029 | 0 | -0.008 | 0.007 | 0.007 | 0.091 | 0.27 | 0.26 | 0.43 | 0.19 | 0.41 | 0.23 |
| 2 | 124841261 | rs7571072 | -0.254 | 0.922 | -0.186 | -0.054 | 0.001 | 0.017 | 0.017 | 0 | 0.38 | 0.34 | 0.29 | 0.28 | 0.34 | 0.43 |
| 2 | 124847337 | rs1347876 | 0.223 | 0.898 | -0.267 | -0.066 | -0.002 | 0.004 | 0.004 | 0 | 0.29 | 0.3 | 0.23 | 0.2 | 0.3 | 0.32 |
| 2 | 124849293 | rs13032868 | 1.34 | 0.891 | -0.321 | -0.041 | -0.011 | 0.016 | 0.016 | 0 | 0.21 | 0.15 | 0.39 | 0.17 | 0.29 | 0.22 |
| 2 | 124856339 | rs2043885 | 0.498 | 0.732 | -0.491 | 0.015 | -0.015 | -0.016 | -0.016 | 0.872 | 0.4 | 0.19 | 0.53 | 0.47 | 0.45 | 0.3 |
| 2 | 124858946 | rs12105596 | -0.263 | 0.723 | -0.401 | -0.045 | -0.001 | 0.001 | 0.001 | 0.12 | 0.4 | 0.44 | 0.3 | 0.28 | 0.43 | 0.45 |
| 2 | 124864686 | rs13422063 | 1.906 | 0.713 | -0.488 | 0.049 | 0.003 | -0.006 | -0.006 | 0 | 0.17 | 0.14 | 0.16 | 0.21 | 0.11 | 0.25 |
| 2 | 124869709 | rs7577916 | -0.285 | 0.71 | -0.678 | 0.037 | -0.003 | 0.004 | 0.004 | 0 | 0.6 | 0.45 | 0.66 | 0.52 | 0.64 | 0.67 |
| 2 | 124875366 | rs4449139 | 0.269 | 0.77 | -0.93 | 0.012 | 0.008 | 0.009 | 0.009 | 0 | 0.52 | 0.3 | 0.61 | 0.46 | 0.52 | 0.63 |
| 2 | 124876342 | rs10496625 | 2.186 | 0.78 | -0.875 | 0.031 | 0.031 | -0.073 | -0.073 | 1.469 | 0.14 | 0.04 | 0.12 | 0.34 | 0.11 | 0.06 |
| 2 | 124879107 | rs7592198 | 0.364 | 0.906 | -0.849 | -0.01 | -0.013 | 0.017 | 0.017 | 0 | 0.43 | 0.29 | 0.57 | 0.38 | 0.45 | 0.47 |
| 2 | 124905952 | rs17391285 | 0.638 | 1.005 | -1.338 | 0.024 | -0.014 | 0.019 | 0.019 | 0.197 | 0.21 | 0.14 | 0.2 | 0.13 | 0.24 | 0.27 |
| 2 | 124909452 | rs1919839 | 0.159 | 0.254 | -1.589 | -0.162 | -0.02 | 0.082 | 0.082 | 0 | 0.31 | 0.13 | 0.43 | 0.18 | 0.39 | 0.38 |
| 2 | 124909788 | rs17010934 | 1.817 | -0.02 | -1.706 | 0.167 | 0.025 | -0.06 | -0.06 | 0 | 0.15 | 0.06 | 0.13 | 0.35 | 0.1 | 0.06 |
| 2 | 124910001 | rs2420559 | -0.392 | 0.001 | -1.333 | 0.016 | 0 | -0.019 | -0.019 | 0.396 | 0.68 | 0.73 | 0.57 | 0.82 | 0.61 | 0.62 |
| 2 | 124913517 | rs960537 | 2.33 | 0.036 | -1.478 | 0.142 | -0.031 | -0.052 | -0.052 | 1.138 | 0.08 | 0 | 0.05 | 0.24 | 0.08 | 0.15 |
| 2 | 124915643 | rs1529289 | 0.347 | -0.275 | -1.449 | 0.153 | -0.02 | 0.085 | 0.085 | 0 | 0.31 | 0.13 | 0.43 | 0.18 | 0.39 | 0.37 |
| 2 | 124917657 | rs1881793 | -1.902 | -0.592 | -1.275 | 0.648 | -0.002 | 0.105 | 0.105 | 0 | 0.91 | 0.85 | 0.95 | 0.76 | 0.94 | 0.85 |
| 2 | 124919174 | rs2579824 | -0.399 | -0.61 | -1.291 | 0.525 | -0.041 | 0.048 | 0.048 | 0.143 | 0.73 | 0.62 | 0.8 | 0.42 | 0.84 | 0.76 |
| 2 | 124925587 | rs4848917 | 1.385 | -0.282 | -1.888 | 0.582 | -0.017 | -0.062 | -0.062 | 0 | 0.16 | 0.05 | 0.14 | 0.18 | 0.11 | 0.18 |
| 2 | 124926513 | rs17319464 | 0.592 | -0.064 | -1.489 | 0.618 | -0.02 | 0.026 | 0.026 | 0 | 0.08 | 0.02 | 0.09 | 0.02 | 0.15 | 0.18 |
| 2 | 124928508 | rs1529294 | 0.996 | -0.535 | -1.237 | 1.013 | 0.024 | -0.047 | -0.047 | 0 | 0.16 | 0.06 | 0.1 | 0.32 | 0.1 | 0.05 |
| 2 | 124929266 | rs2699381 | 0.206 | -0.41 | -1.293 | 1.014 | 0.041 | -0.064 | -0.064 | 0.119 | 0.24 | 0.05 | 0.12 | 0.32 | 0.16 | 0.1 |
| 2 | 124929288 | rs11123025 | 0.039 | -0.251 | -1.035 | 0.522 | 0.003 | -0.002 | -0.002 | 0 | 0.65 | 0.73 | 0.78 | 0.42 | 0.78 | 0.72 |
| 2 | 124940432 | rs2699376 | -0.573 | -0.207 | -0.786 | 0.37 | -0.005 | -0.002 | -0.002 | 1.567 | 0.75 | 0.71 | 0.84 | 0.83 | 0.85 | 0.8 |
| 2 | 124946945 | rs7569367 | 0.398 | 0.149 | -0.724 | 0.315 | -0.005 | 0.001 | 0.001 | 0 | 0.41 | 0.29 | 0.33 | 0.45 | 0.39 | 0.49 |
| 2 | 124947363 | rs17011058 | 0.941 | 0.511 | -0.512 | -0.061 | 0.015 | -0.074 | -0.074 | 0 | 0.15 | 0.27 | 0.16 | 0.01 | 0.21 | 0.05 |
| 2 | 124948463 | rs1406662 | -0.91 | 0.712 | -0.685 | -0.042 | 0.001 | 0.006 | 0.006 | 0 | 0.7 | 0.71 | 0.81 | 0.83 | 0.74 | 0.79 |
| 2 | 124953763 | rs12711674 | 0.554 | 0.677 | -0.72 | -0.095 | 0.042 | 0.011 | 0.011 | 0 | 0.45 | 0.32 | 0.31 | 0.4 | 0.3 | 0.55 |
| 2 | 124954598 | rs4848919 | -1.526 | 0.763 | -0.216 | -0.19 | 0.009 | -0.068 | -0.068 | 0 | 0.85 | 0.77 | 0.84 | 1 | 0.79 | 0.94 |
| 2 | 124970109 | rs12711675 | 1.891 | 0.66 | -0.36 | -0.286 | 0.002 | -0.015 | -0.015 | 0.021 | 0.1 | 0.12 | 0.11 | 0.03 | 0.1 | 0.21 |
| 2 | 124971023 | rs12711676 | 0.83 | 0.672 | -0.046 | -0.004 | 0.045 | 0.003 | 0.003 | 0 | 0.44 | 0.24 | 0.29 | 0.4 | 0.3 | 0.55 |
| 2 | 124976131 | rs17011129 | 1.385 | 0.576 | -0.113 | -0.006 | 0.015 | -0.03 | -0.03 | 0 | 0.1 | 0.05 | 0.05 | 0.2 | 0.05 | 0.14 |
| 2 | 124976163 | rs11123030 | 0.182 | 0.804 | 0.254 | 0.142 | -0.007 | -0.002 | -0.002 | 0 | 0.64 | 0.35 | 0.48 | 0.64 | 0.54 | 0.69 |

| | | | | | | | | | | | | | | | | |
|---|---|---|---|---|---|---|---|---|---|---|---|---|---|---|---|---|
| 2 | 124976279 | rs1529292 | 1.073 | 0.824 | 0.109 | 0.195 | -0.021 | -0.003 | -0.003 | 0 | 0.2 | 0.08 | 0.16 | 0.22 | 0.24 | 0.12 |
| 2 | 124981162 | rs6541945 | 0.26 | 0.801 | 0.459 | 0.19 | 0.025 | 0.008 | 0.008 | 1.137 | 0.65 | 0.49 | 0.49 | 0.64 | 0.53 | 0.69 |
| 2 | 124982477 | rs1170611 | -0.995 | 0.69 | 0.442 | 0.225 | 0.013 | -0.092 | -0.092 | 0 | 0.84 | 0.68 | 0.83 | 1 | 0.79 | 0.94 |
| 2 | 124986440 | rs1170617 | 1.447 | 0.791 | 0.135 | 0.4 | 0 | 0.008 | 0.008 | 0.481 | 0.14 | 0.18 | 0.31 | 0.36 | 0.15 | 0.25 |
| 2 | 124990440 | rs1170624 | 1.447 | 0.791 | 0.122 | 0.4 | 0 | 0.012 | 0.012 | 0.481 | 0.14 | 0.19 | 0.31 | 0.36 | 0.15 | 0.25 |
| 2 | 124994170 | rs1170628 | 1.728 | 0.79 | 0.108 | 0.4 | -0.009 | -0.051 | -0.051 | 0 | 0.13 | 0.06 | 0.29 | 0.36 | 0.15 | 0.25 |
| 2 | 124996309 | rs779994 | 1.297 | 0.789 | 0.108 | 0.399 | 0 | 0.013 | 0.013 | 0.584 | 0.15 | 0.2 | 0.31 | 0.36 | 0.15 | 0.25 |
| 2 | 124998081 | rs779992 | -0.077 | 0.808 | 0.092 | 0.326 | -0.001 | -0.003 | -0.003 | 0 | 0.36 | 0.35 | 0.51 | 0.6 | 0.39 | 0.38 |
| 2 | 125004689 | rs779985 | 0.4 | 0.824 | 0.101 | 0.368 | -0.002 | -0.005 | -0.005 | 0 | 0.27 | 0.23 | 0.43 | 0.37 | 0.37 | 0.3 |
| 2 | 125005099 | rs1170585 | -1.104 | 0.644 | 0.305 | 0.393 | 0.005 | -0.004 | -0.004 | 0 | 0.8 | 0.81 | 0.84 | 0.81 | 0.76 | 0.9 |
| 2 | 125009858 | rs1919840 | 0.547 | 0.701 | 0.158 | 0.35 | 0.01 | 0.002 | 0.002 | 0.519 | 0.48 | 0.59 | 0.38 | 0.44 | 0.29 | 0.58 |
| 2 | 125022152 | rs812952 | -0.9 | 0.707 | 0.026 | 0.404 | -0.003 | 0 | 0 | 0 | 0.8 | 0.83 | 0.84 | 0.81 | 0.76 | 0.9 |
| 2 | 125024950 | rs1186066 | -0.299 | 0.608 | 0.452 | 0.198 | 0.022 | 0.01 | 0.01 | 0.337 | 0.35 | 0.48 | 0.5 | 0.39 | 0.45 | 0.34 |
| 2 | 125027173 | rs780029 | 0.513 | 0.695 | 0.572 | 0.248 | -0.01 | -0.018 | -0.018 | 0 | 0.27 | 0.12 | 0.4 | 0.35 | 0.34 | 0.3 |
| 2 | 125032306 | rs961304 | 0.985 | 0.651 | 0.676 | 0.54 | 0.032 | 0.001 | 0.001 | 0 | 0.44 | 0.26 | 0.32 | 0.44 | 0.31 | 0.58 |
| 2 | 125033303 | rs780049 | -1.113 | 0.657 | 0.597 | 0.555 | 0.001 | 0 | 0 | 0.063 | 0.8 | 0.83 | 0.84 | 0.82 | 0.77 | 0.92 |
| 2 | 125036924 | rs6755697 | 1.26 | 0.288 | 0.071 | 0.53 | 0.017 | -0.017 | -0.017 | 0 | 0.09 | 0.06 | 0.05 | 0.17 | 0.04 | 0.11 |
| 2 | 125042374 | rs780012 | -0.424 | 0.462 | 0.324 | 0.454 | 0.036 | -0.007 | -0.007 | 0 | 0.74 | 0.55 | 0.74 | 0.79 | 0.65 | 0.88 |
| 2 | 125044519 | rs796643 | 0.453 | 0.432 | 0.344 | 0.42 | -0.002 | -0.004 | -0.004 | 0.791 | 0.29 | 0.23 | 0.41 | 0.35 | 0.34 | 0.3 |
| 2 | 125044669 | rs17011309 | 1.797 | 0.484 | 0.029 | 0.337 | -0.028 | -0.03 | -0.03 | 0 | 0.12 | 0 | 0.08 | 0.1 | 0.14 | 0.02 |
| 2 | 125050005 | rs11123033 | 0.832 | 0.366 | 0.188 | 0.342 | 0.024 | 0.002 | 0.002 | 0 | 0.45 | 0.32 | 0.33 | 0.44 | 0.32 | 0.58 |
| 2 | 125054254 | rs17011381 | 0.499 | 0.635 | 0.492 | 0.181 | 0.002 | -0.067 | -0.067 | 0.164 | 0.13 | 0.19 | 0.13 | 0 | 0.15 | 0.04 |
| 2 | 125057421 | rs2901264 | 0.36 | 0.467 | 0.365 | 0.172 | 0.005 | -0.022 | -0.022 | 0 | 0.71 | 0.67 | 0.75 | 0.88 | 0.66 | 0.82 |
| 2 | 125058227 | rs2115890 | -0.304 | 0.485 | 0.385 | 0.154 | -0.002 | -0.003 | -0.003 | 0 | 0.77 | 0.8 | 0.81 | 0.91 | 0.71 | 0.86 |
| 2 | 125061838 | rs733112 | -1.106 | 0.013 | 0.216 | 0.182 | 0.002 | 0.002 | 0.002 | 0.228 | 0.51 | 0.51 | 0.4 | 0.43 | 0.58 | 0.3 |
| 2 | 125064801 | rs1430263 | 0.471 | 0.139 | -0.268 | 0.322 | 0 | -0.003 | -0.003 | 0 | 0.22 | 0.17 | 0.14 | 0.32 | 0.24 | 0.13 |
| 2 | 125065631 | rs17011394 | 0.498 | 0.143 | -0.266 | 0.324 | 0 | -0.003 | -0.003 | 0 | 0.22 | 0.17 | 0.14 | 0.32 | 0.24 | 0.13 |
| 2 | 125066751 | rs780010 | -1.46 | 0.079 | -0.514 | 0.476 | -0.014 | 0.028 | 0.028 | 0 | 0.6 | 0.79 | 0.62 | 0.68 | 0.64 | 0.48 |
| 2 | 125066976 | rs17011399 | -0.543 | 0.11 | -0.002 | 0.229 | 0.005 | -0.011 | -0.011 | 0 | 0.31 | 0.42 | 0.39 | 0.27 | 0.35 | 0.3 |
| 2 | 125076612 | rs17724018 | -0.317 | -0.044 | -0.118 | 0.101 | -0.001 | 0 | 0 | 0 | 0.08 | 0.09 | 0.03 | 0.12 | 0.06 | 0.12 |
| 2 | 125078104 | rs779979 | -1.149 | -0.27 | -0.16 | -0.514 | -0.004 | 0.104 | 0.104 | 0.243 | 0.52 | 0.75 | 0.31 | 0.18 | 0.47 | 0.31 |
| 2 | 125080181 | rs2553625 | 0.602 | 0.034 | 0.159 | -0.721 | 0 | -0.052 | -0.052 | 0.701 | 0.26 | 0.27 | 0.17 | 0 | 0.27 | 0.1 |
| 2 | 125080866 | rs17011420 | -0.421 | 0.01 | 0.238 | -0.738 | 0 | -0.045 | -0.045 | 0 | 0.36 | 0.42 | 0.23 | 0.05 | 0.36 | 0.18 |
| 2 | 125083298 | rs2553628 | -0.743 | 0.004 | 0.219 | -0.739 | -0.001 | -0.034 | -0.034 | 1.043 | 0.38 | 0.42 | 0.23 | 0.05 | 0.41 | 0.19 |
| 2 | 125085600 | rs17011429 | 0.62 | -0.024 | 0.155 | -0.732 | -0.002 | -0.028 | -0.028 | 0 | 0.25 | 0.2 | 0.16 | 0 | 0.27 | 0.1 |
| 2 | 125091035 | rs7574198 | 0.572 | 0.058 | 0.253 | -0.587 | 0.002 | -0.053 | -0.053 | 0 | 0.25 | 0.28 | 0.2 | 0.03 | 0.25 | 0.12 |
| 2 | 125099924 | rs12475383 | -0.063 | 0.028 | 0.342 | -0.52 | 0.012 | -0.056 | -0.056 | 0 | 0.35 | 0.41 | 0.37 | 0.19 | 0.4 | 0.31 |
| 2 | 125107948 | rs7567224 | -0.456 | -0.137 | 0.08 | -0.441 | 0.004 | 0.03 | 0.03 | 0 | 0.41 | 0.34 | 0.31 | 0.23 | 0.39 | 0.21 |
| 2 | 125108025 | rs779999 | -2.193 | -0.122 | 0.134 | -0.307 | -0.022 | -0.082 | -0.082 | 0 | 0.67 | 0.92 | 0.56 | 0.51 | 0.69 | 0.48 |
| 2 | 125108191 | rs796641 | -2.077 | -0.19 | 0.226 | -0.314 | 0.004 | -0.057 | -0.057 | 0 | 0.66 | 0.84 | 0.55 | 0.51 | 0.71 | 0.48 |
| 2 | 125108364 | rs7582767 | 0.545 | -0.185 | 0.196 | -0.369 | -0.001 | -0.003 | -0.003 | 0 | 0.32 | 0.33 | 0.27 | 0.23 | 0.3 | 0.19 |
| 2 | 125109738 | rs7560392 | 1.167 | 0.018 | 0.039 | -0.238 | 0.002 | 0.004 | 0.004 | 0 | 0.11 | 0.1 | 0.09 | 0.16 | 0.16 | 0.19 |
| 2 | 125126822 | rs779982 | 0.136 | 0.13 | -0.202 | -0.251 | 0.009 | 0.006 | 0.006 | 0 | 0.11 | 0.25 | 0.1 | 0.1 | 0.15 | 0.07 |
| 2 | 125127498 | rs720477 | -1.823 | 0.077 | -0.306 | -0.332 | -0.014 | -0.1 | -0.1 | 0.165 | 0.66 | 0.93 | 0.56 | 0.51 | 0.71 | 0.47 |
| 2 | 125131007 | rs11123037 | -0.564 | 0.104 | -0.29 | -0.375 | 0.001 | 0.028 | 0.028 | 0 | 0.5 | 0.64 | 0.44 | 0.42 | 0.51 | 0.4 |
| 2 | 125132013 | rs780058 | -0.708 | 0.109 | -0.279 | -0.284 | 0.004 | 0 | 0 | 0 | 0.89 | 0.83 | 0.9 | 0.91 | 0.85 | 0.93 |
| 2 | 125133276 | rs4848240 | -1.215 | -0.186 | -0.202 | -0.182 | -0.006 | 0.01 | 0.01 | 0 | 0.54 | 0.66 | 0.46 | 0.42 | 0.56 | 0.4 |
| 2 | 125135589 | rs13384799 | 0.412 | 0.157 | -0.223 | -0.408 | 0.005 | -0.08 | -0.08 | 0 | 0.26 | 0.28 | 0.16 | 0 | 0.3 | 0.11 |
| 2 | 125143225 | rs2565762 | 0.45 | 0.331 | -1.227 | 0.223 | -0.008 | -0.034 | -0.034 | 0 | 0.43 | 0.21 | 0.52 | 0.53 | 0.37 | 0.55 |
| 2 | 125147162 | rs780005 | 0.476 | -0.5 | -0.603 | 0.015 | -0.011 | -0.069 | -0.069 | 0.697 | 0.75 | 0.44 | 0.72 | 0.9 | 0.75 | 0.79 |
| 2 | 125156841 | rs2043072 | 1.158 | -0.703 | -1.144 | 0.192 | 0.036 | 0.004 | 0.004 | 0 | 0.55 | 0.25 | 0.59 | 0.62 | 0.61 | 0.6 |
| 2 | 125158776 | rs1820931 | 1.34 | -0.654 | -1.191 | 0.266 | 0.044 | -0.005 | -0.005 | 0.366 | 0.52 | 0.18 | 0.54 | 0.53 | 0.56 | 0.58 |
| 2 | 125159292 | rs2584353 | 0.124 | -0.267 | -0.764 | 0.558 | 0.004 | -0.04 | -0.04 | 0 | 0.11 | 0.06 | 0.06 | 0.24 | 0.09 | 0.1 |
| 2 | 125160773 | rs1834442 | 0.341 | -0.25 | -0.189 | -0.22 | -0.015 | -0.06 | -0.06 | 0 | 0.81 | 0.49 | 0.83 | 0.9 | 0.82 | 0.8 |
| 2 | 125163827 | rs1835353 | -0.216 | 0.121 | -0.618 | -0.282 | -0.006 | 0.005 | 0.005 | 0 | 0.12 | 0.11 | 0.11 | 0.04 | 0.18 | 0.14 |
| 2 | 125177190 | rs11123043 | -0.125 | 0.126 | -0.628 | -0.279 | -0.013 | 0.021 | 0.021 | 0 | 0.11 | 0.07 | 0.1 | 0.04 | 0.18 | 0.14 |
| 2 | 125183450 | rs17011501 | 0.306 | -0.153 | -0.054 | 0.004 | -0.025 | 0.06 | 0.06 | 0 | 0.25 | 0.36 | 0.31 | 0.48 | 0.21 | 0.22 |
| 2 | 125189434 | rs1543901 | 1.07 | -0.045 | -1.194 | -0.363 | -0.133 | 0.038 | 0.038 | 0.124 | 0.44 | 0.03 | 0.33 | 0.34 | 0.48 | 0.42 |
| 2 | 125197319 | rs12053037 | 0.242 | 1.211 | -0.218 | 0.327 | -0.026 | -0.026 | -0.026 | 0 | 0.15 | 0.09 | 0.22 | 0.33 | 0.22 | 0.31 |
| 2 | 125202316 | rs2003039 | -0.789 | 1.242 | -0.091 | 0.184 | -0.034 | 0.043 | 0.043 | 0 | 0.15 | 0.08 | 0.12 | 0 | 0.25 | 0.1 |
| 2 | 125204264 | rs6715476 | 0.121 | 1.181 | 0.218 | 0.272 | -0.027 | 0.009 | 0.009 | 0 | 0.1 | 0.24 | 0.14 | 0.13 | 0.04 | 0.18 |
| 2 | 125213719 | rs2420858 | -0.07 | 1.219 | 0.214 | 0.267 | 0.017 | -0.027 | -0.027 | 1.426 | 0.95 | 0.87 | 0.98 | 1 | 0.93 | 0.98 |
| 2 | 125232609 | rs4411700 | 1.126 | 1.203 | 0.293 | 0.272 | -0.02 | 0.002 | 0.002 | 0.282 | 0.83 | 0.54 | 0.81 | 0.83 | 0.89 | 0.72 |
| 2 | 125246092 | rs2421083 | -1.528 | 1.294 | 0.117 | 0.383 | 0.018 | 0.031 | 0.031 | 0 | 0.3 | 0.43 | 0.46 | 0.52 | 0.33 | 0.62 |
| 2 | 125250587 | rs6745541 | 0.223 | 1.284 | 0.065 | 0.401 | -0.033 | -0.034 | -0.034 | 0 | 0.15 | 0.08 | 0.28 | 0.35 | 0.23 | 0.34 |
| 2 | 125256925 | rs728049 | 0.145 | 1.152 | 0.021 | 0.369 | -0.022 | 0.011 | 0.011 | 0 | 0.09 | 0.23 | 0.14 | 0.13 | 0.03 | 0.19 |
| 2 | 125260743 | rs1504001 | 0.111 | 1.118 | 0.02 | 0.312 | -0.025 | 0.003 | 0.003 | 0 | 0.09 | 0.23 | 0.14 | 0.11 | 0.03 | 0.19 |
| 2 | 125265936 | rs869760 | -0.425 | 1.242 | 0.051 | 0.371 | 0.006 | -0.033 | -0.033 | 0 | 0.89 | 0.89 | 0.9 | 1 | 0.82 | 0.93 |
| 2 | 125272371 | rs884376 | 1.221 | 1.179 | -0.254 | 0.368 | 0.003 | 0.001 | 0.001 | 0 | 0.58 | 0.46 | 0.44 | 0.5 | 0.47 | 0.31 |

| | | | | | | | | | | | | | | | | |
|---|---|---|---|---|---|---|---|---|---|---|---|---|---|---|---|---|
| 2 | 125278326 | rs6722179 | -0.459 | 1.194 | -1.02 | 0.485 | -0.04 | -0.059 | -0.059 | 0 | 0.25 | 0.16 | 0.35 | 0.42 | 0.27 | 0.55 |
| 2 | 125278565 | rs6722312 | -0.962 | 1.243 | -0.924 | 0.455 | -0.023 | -0.015 | -0.015 | 1.832 | 0.3 | 0.24 | 0.37 | 0.4 | 0.34 | 0.57 |
| 2 | 125290839 | rs12612022 | -0.482 | 1.328 | -0.711 | 0.784 | -0.021 | -0.03 | -0.03 | 0 | 0.26 | 0.19 | 0.32 | 0.47 | 0.32 | 0.48 |
| 2 | 125291034 | rs13015323 | -1.1 | 0.642 | -0.445 | 0.456 | -0.004 | 0.004 | 0.004 | 0 | 0.44 | 0.51 | 0.47 | 0.5 | 0.57 | 0.58 |
| 2 | 125299661 | rs12470517 | 0.56 | -0.281 | -0.301 | -0.077 | 0 | 0 | 0 | 0 | 0.56 | 0.59 | 0.54 | 0.61 | 0.3 | 0.35 |
| 2 | 125303307 | rs7574777 | -0.277 | -0.33 | -0.222 | -0.22 | -0.013 | -0.014 | -0.014 | 0 | 0.13 | 0.14 | 0.07 | 0.08 | 0.12 | 0.17 |
| 2 | 125303786 | rs987656 | -0.826 | 0.649 | -0.163 | -0.289 | 0 | 0.006 | 0.006 | 0 | 0.88 | 0.9 | 0.86 | 0.97 | 0.69 | 0.85 |
| 2 | 125304035 | rs12711680 | -0.977 | 0.555 | -0.001 | -0.349 | 0.009 | 0.01 | 0.01 | 0.012 | 0.32 | 0.37 | 0.33 | 0.36 | 0.39 | 0.5 |
| 2 | 125304779 | rs11886998 | 1.335 | -0.85 | -0.32 | -1.075 | -0.01 | -0.075 | -0.075 | 0 | 0.33 | 0.37 | 0.16 | 0 | 0.13 | 0.11 |
| 2 | 125305213 | rs10172675 | 0.927 | -0.613 | -0.357 | -0.732 | -0.003 | -0.001 | -0.001 | 0 | 0.54 | 0.51 | 0.52 | 0.51 | 0.29 | 0.33 |
| 2 | 125309057 | rs2047876 | 0.841 | -0.705 | -0.436 | -1.115 | -0.009 | -0.003 | -0.003 | 0.573 | 0.54 | 0.51 | 0.55 | 0.59 | 0.34 | 0.35 |
| 2 | 125312883 | rs13021400 | 0.525 | -0.466 | -0.979 | -0.871 | -0.09 | -0.089 | -0.089 | 1.148 | 0.16 | 0.02 | 0.26 | 0.29 | 0.27 | 0.33 |
| 2 | 125313021 | rs7601703 | 0.031 | -0.679 | -0.646 | -0.754 | -0.019 | -0.007 | -0.007 | 1.447 | 0.1 | 0.12 | 0.04 | 0.09 | 0.07 | 0.09 |
| 2 | 125315748 | rs6707944 | 1.052 | -0.765 | -0.952 | -0.712 | 0.036 | -0.095 | -0.095 | 0 | 0.21 | 0.06 | 0.37 | 0.59 | 0.17 | 0.26 |
| 2 | 125315822 | rs4848246 | -0.463 | -0.834 | -0.342 | -1.077 | 0.009 | 0.035 | 0.035 | 0.709 | 0.29 | 0.46 | 0.33 | 0.36 | 0.32 | 0.38 |
| 2 | 125317108 | rs2047877 | 1.14 | -0.712 | -0.479 | -0.921 | 0.018 | -0.093 | -0.093 | 0.555 | 0.21 | 0.06 | 0.38 | 0.6 | 0.2 | 0.23 |
| 2 | 125318315 | rs6748545 | -0.599 | -0.393 | -0.212 | -1.172 | 0.005 | 0.001 | 0.001 | 0 | 0.39 | 0.3 | 0.6 | 0.95 | 0.37 | 0.53 |
| 2 | 125322481 | rs17320237 | 1.359 | -0.392 | -0.628 | -1.442 | -0.092 | 0.077 | 0.077 | 0 | 0.13 | 0.01 | 0.12 | 0 | 0.29 | 0.12 |
| 2 | 125322835 | rs6541961 | 0.377 | -0.418 | -0.059 | -1.28 | -0.014 | -0.143 | -0.143 | 0 | 0.59 | 0.55 | 0.39 | 0.02 | 0.63 | 0.47 |
| 2 | 125327962 | rs17320299 | 0.449 | -0.112 | -0.958 | -0.97 | -0.041 | -0.059 | -0.059 | 0.272 | 0.11 | 0.02 | 0.15 | 0.3 | 0.15 | 0.26 |
| 2 | 125328441 | rs7420022 | 0.043 | -0.161 | -0.986 | -1.059 | -0.03 | 0.134 | 0.134 | 0.401 | 0.32 | 0.07 | 0.53 | 0.89 | 0.35 | 0.48 |
| 2 | 125330439 | rs17011723 | 1.292 | -0.198 | -1.043 | -0.82 | -0.005 | -0.137 | -0.137 | 0 | 0.18 | 0.02 | 0.37 | 0.58 | 0.17 | 0.22 |
| 2 | 125331126 | rs10445861 | -0.059 | -0.594 | -0.515 | -0.861 | -0.002 | 0.006 | 0.006 | 0 | 0.47 | 0.5 | 0.62 | 0.67 | 0.5 | 0.46 |
| 2 | 125335411 | rs13008208 | 0.32 | -0.477 | -0.999 | -0.608 | 0 | -0.021 | -0.021 | 0.055 | 0.17 | 0.23 | 0.14 | 0.02 | 0.18 | 0.24 |
| 2 | 125337929 | rs4402762 | -0.501 | -0.464 | -0.813 | -0.399 | 0 | -0.006 | -0.006 | 0 | 0.34 | 0.41 | 0.34 | 0.41 | 0.38 | 0.49 |
| 2 | 125341075 | rs6753520 | -0.242 | -0.35 | -0.757 | -0.371 | -0.027 | 0.021 | 0.021 | 0 | 0.42 | 0.28 | 0.54 | 0.66 | 0.47 | 0.43 |
| 2 | 125341886 | rs12619786 | -0.06 | -0.286 | -0.811 | -0.965 | -0.031 | 0.032 | 0.032 | 0 | 0.22 | 0.18 | 0.16 | 0.02 | 0.3 | 0.18 |
| 2 | 125342637 | rs4848941 | 0.537 | -0.755 | -0.179 | -0.676 | -0.017 | 0.016 | 0.016 | 1.016 | 0.22 | 0.24 | 0.41 | 0.69 | 0.16 | 0.2 |
| 2 | 125344627 | rs4364025 | -0.973 | -0.857 | 0.538 | -0.794 | -0.001 | 0.022 | 0.022 | 0 | 0.46 | 0.58 | 0.59 | 0.72 | 0.46 | 0.38 |
| 2 | 125346934 | rs1504017 | -0.038 | -0.799 | 0.458 | -0.829 | 0.038 | -0.093 | -0.093 | 0.021 | 0.24 | 0.37 | 0.18 | 0.03 | 0.31 | 0.18 |
| 2 | 125347357 | rs4848944 | 0.914 | -1.003 | 0.246 | -1.02 | -0.032 | 0.089 | 0.089 | 0 | 0.47 | 0.22 | 0.35 | 0.25 | 0.46 | 0.62 |
| 2 | 125348819 | rs2421092 | 1.257 | -1 | 1.07 | -1.008 | 0.001 | -0.046 | -0.046 | 0 | 0.9 | 0.53 | 0.93 | 1 | 0.95 | 0.99 |
| 2 | 125358226 | rs12990799 | 0.999 | -1.255 | 0.209 | -1.2 | -0.026 | 0.119 | 0.119 | 0 | 0.45 | 0.2 | 0.38 | 0.19 | 0.47 | 0.45 |
| 2 | 125362837 | rs11123053 | 0.1 | -1.428 | 0.494 | -1.334 | 0.001 | -0.011 | -0.011 | 0 | 0.59 | 0.62 | 0.44 | 0.19 | 0.55 | 0.46 |
| 2 | 125373757 | rs7578835 | -0.68 | -1.579 | 0.3 | -1.272 | -0.001 | -0.001 | -0.001 | 0.298 | 0.39 | 0.39 | 0.51 | 0.75 | 0.4 | 0.48 |
| 2 | 125379346 | rs7573997 | 0.254 | -1.201 | -0.004 | -1.352 | -0.009 | -0.035 | -0.035 | 0.107 | 0.07 | 0.16 | 0.03 | 0 | 0.05 | 0.01 |
| 2 | 125394795 | rs11900792 | 0.749 | -0.957 | -0.406 | -1.181 | -0.002 | -0.11 | -0.11 | 0.703 | 0.2 | 0.02 | 0.34 | 0.73 | 0.16 | 0.36 |
| 2 | 125415526 | rs6541966 | -0.027 | -0.76 | 0 | -1.337 | -0.006 | 0.014 | 0.014 | 0.051 | 0.61 | 0.43 | 0.47 | 0.21 | 0.57 | 0.45 |
| 2 | 125415995 | rs10202736 | -0.112 | -0.761 | 0.102 | -1.338 | 0.002 | -0.011 | -0.011 | 0 | 0.62 | 0.49 | 0.48 | 0.21 | 0.57 | 0.46 |
| 2 | 125416607 | rs6739399 | -0.789 | 0.185 | 0.153 | -1.439 | -0.005 | -0.007 | -0.007 | 0 | 0.22 | 0.33 | 0.14 | 0.14 | 0.2 | 0.18 |
| 2 | 125422631 | rs12619819 | -0.041 | 0.305 | -0.177 | -1.39 | -0.012 | -0.018 | -0.018 | 0 | 0.15 | 0.08 | 0.11 | 0.17 | 0.18 | 0.19 |
| 2 | 125425762 | rs768318 | 1.217 | -0.008 | -0.526 | -1.759 | -0.058 | 0.094 | 0.094 | 0 | 0.25 | 0.08 | 0.26 | 0 | 0.32 | 0.2 |
| 2 | 125427543 | rs11684957 | -0.685 | 0.025 | -0.33 | -1.733 | -0.006 | 0.068 | 0.068 | 0 | 0.48 | 0.64 | 0.41 | 0.13 | 0.5 | 0.38 |
| 2 | 125431247 | rs12467127 | 0.805 | 0.094 | -0.702 | -1.76 | -0.048 | 0.084 | 0.084 | 0 | 0.29 | 0.12 | 0.29 | 0.01 | 0.37 | 0.21 |
| 2 | 125435930 | rs7567761 | -0.469 | -0.457 | -0.35 | -1.44 | -0.019 | -0.019 | -0.019 | 0 | 0.21 | 0.37 | 0.15 | 0.12 | 0.16 | 0.17 |
| 2 | 125437743 | rs13413315 | -0.916 | -0.453 | -0.353 | -1.442 | -0.017 | -0.021 | -0.021 | 0.084 | 0.23 | 0.37 | 0.18 | 0.12 | 0.19 | 0.18 |
| 2 | 125443155 | rs10496638 | -0.174 | -0.22 | -0.932 | -1.282 | 0.009 | 0.009 | 0.009 | 0.983 | 0.18 | 0.1 | 0.12 | 0.12 | 0.16 | 0.19 |
| 2 | 125450693 | rs7603818 | -0.666 | 0.061 | -0.86 | -1.446 | 0.01 | 0.01 | 0.01 | 0 | 0.6 | 0.65 | 0.47 | 0.16 | 0.69 | 0.54 |
| 2 | 125457240 | rs7580383 | 0.115 | 0.998 | -0.731 | -1.704 | 0.01 | -0.144 | -0.144 | 0.051 | 0.42 | 0.5 | 0.29 | 0.01 | 0.45 | 0.24 |
| 2 | 125460161 | rs17321135 | -1.119 | 1.876 | -0.259 | -0.891 | 0 | 0 | 0 | 0 | 0.33 | 0.36 | 0.27 | 0.28 | 0.43 | 0.31 |
| 2 | 125465194 | rs17393209 | -0.605 | 2.21 | -0.206 | -0.918 | 0.037 | 0.029 | 0.029 | 0 | 0.14 | 0.22 | 0.16 | 0.25 | 0.21 | 0.25 |
| 2 | 125468195 | rs13021629 | -1.343 | 2.16 | -0.284 | -0.918 | 0.003 | -0.002 | -0.002 | 0 | 0.33 | 0.38 | 0.27 | 0.29 | 0.43 | 0.31 |
| 2 | 125491192 | rs17393410 | -0.265 | 2.243 | -0.2 | -1.14 | -0.041 | -0.001 | -0.001 | 0.706 | 0.11 | 0.06 | 0.18 | 0.13 | 0.21 | 0.25 |
| 2 | 125499930 | rs10194544 | -1.438 | 2.467 | -0.074 | -1.378 | 0.008 | -0.033 | -0.033 | 0 | 0.39 | 0.47 | 0.4 | 0.19 | 0.51 | 0.35 |
| 2 | 125503362 | rs12468133 | -0.295 | 1.974 | -0.413 | -1.79 | -0.016 | 0.036 | 0.036 | 0.001 | 0.24 | 0.16 | 0.21 | 0.04 | 0.31 | 0.1 |
| 2 | 125511904 | rs7588987 | 0.094 | 1.421 | 0.232 | -1.981 | 0.004 | -0.07 | -0.07 | 0.276 | 0.63 | 0.62 | 0.72 | 0.96 | 0.6 | 0.89 |
| 2 | 125524699 | rs10166480 | -0.109 | 1.389 | 0.464 | -1.995 | -0.015 | 0.024 | 0.024 | 0.912 | 0.65 | 0.71 | 0.74 | 0.96 | 0.6 | 0.89 |
| 2 | 125526506 | rs2168024 | 0.388 | 1.361 | 0.431 | -1.98 | 0.007 | -0.052 | -0.052 | 0 | 0.58 | 0.5 | 0.6 | 0.83 | 0.51 | 0.78 |
| 2 | 125533425 | rs17011975 | -0.991 | 0.989 | 0.851 | -1.65 | 0.068 | -0.008 | -0.008 | 0 | 0.2 | 0.47 | 0.26 | 0.17 | 0.22 | 0.15 |
| 2 | 125533656 | rs17393704 | 1.131 | 0.405 | 0.481 | -2.191 | 0.033 | 0.083 | 0.083 | 0.102 | 0.26 | 0.08 | 0.11 | 0.01 | 0.19 | 0.21 |
| 2 | 125543111 | rs10202684 | 0.305 | 0.43 | 1.122 | -1.915 | -0.065 | -0.077 | -0.077 | 0 | 0.16 | 0.42 | 0.07 | 0.03 | 0.04 | 0.15 |
| 2 | 125544987 | rs12474831 | -1.409 | 0.45 | 1.03 | -1.932 | -0.002 | -0.015 | -0.015 | 0.302 | 0.34 | 0.61 | 0.31 | 0.21 | 0.26 | 0.3 |
| 2 | 125548699 | rs10198988 | 0.571 | 0.469 | 1.051 | -1.898 | -0.04 | -0.048 | -0.048 | 0 | 0.14 | 0.26 | 0.06 | 0.03 | 0.04 | 0.15 |
| 2 | 125553842 | rs922862 | -0.174 | 0.489 | 0.813 | -1.949 | 0.002 | -0.014 | -0.014 | 0 | 0.22 | 0.19 | 0.15 | 0.02 | 0.15 | 0.09 |
| 2 | 125562545 | rs2602642 | 0.581 | 0.706 | 1 | -2.081 | 0.075 | -0.035 | -0.035 | 0 | 0.8 | 0.63 | 0.75 | 0.85 | 0.7 | 0.85 |
| 2 | 125564823 | rs314718 | 0.17 | 0.728 | 0.895 | -2.074 | 0.028 | -0.01 | -0.01 | 0 | 0.82 | 0.79 | 0.76 | 0.85 | 0.7 | 0.85 |
| 2 | 125567543 | rs314712 | 0.527 | 0.549 | 1.064 | -2.062 | 0.012 | -0.048 | -0.048 | 0 | 0.77 | 0.48 | 0.81 | 0.85 | 0.79 | 0.76 |
| 2 | 125575440 | rs7556875 | -1.049 | 0.56 | 0.93 | -2.055 | -0.02 | -0.072 | -0.072 | 2.25 | 0.3 | 0.61 | 0.19 | 0.06 | 0.16 | 0.21 |
| 2 | 125576665 | rs314710 | 1.13 | 0.589 | 0.913 | -2.066 | -0.001 | -0.004 | -0.004 | 0 | 0.66 | 0.34 | 0.69 | 0.81 | 0.67 | 0.7 |

| | | | | | | | | | | | | | | | | |
|---|---|---|---|---|---|---|---|---|---|---|---|---|---|---|---|---|
| 2 | 125579839 | rs35374656 | 0.449 | 0.758 | 1.476 | -1.616 | -0.019 | 0.034 | 0.034 | 0 | 0.12 | 0.06 | 0.11 | 0 | 0.12 | 0.06 |
| 2 | 125592972 | rs314700 | 0.465 | 0.559 | 1.572 | -1.81 | 0.002 | 0.178 | 0.178 | 1.427 | 0.49 | 0.24 | 0.59 | 0.83 | 0.5 | 0.56 |
| 2 | 125605170 | rs7561882 | -0.477 | 0.535 | 2.293 | -1.063 | 0.012 | -0.076 | -0.076 | 0 | 0.16 | 0.39 | 0.15 | 0.03 | 0.12 | 0.06 |
| 2 | 125610827 | rs17012066 | -0.368 | 0.553 | 2.151 | -1.055 | 0.009 | -0.055 | -0.055 | 0.817 | 0.14 | 0.31 | 0.14 | 0.03 | 0.12 | 0.06 |
| 2 | 125641511 | rs2602652 | -0.608 | 0.483 | -0.329 | -0.661 | -0.01 | -0.033 | -0.033 | 2.228 | 0.86 | 0.94 | 0.62 | 0.24 | 0.85 | 0.68 |
| 2 | 125643441 | rs2249026 | -0.53 | 0.224 | -0.156 | -1.346 | 0.005 | 0.066 | 0.066 | 0.665 | 0.49 | 0.66 | 0.38 | 0.17 | 0.47 | 0.47 |
| 2 | 125664021 | rs2602647 | -0.205 | 0.132 | -0.043 | -1.458 | 0 | -0.046 | -0.046 | 0.086 | 0.45 | 0.54 | 0.34 | 0.17 | 0.47 | 0.44 |
| 2 | 125669565 | rs730668 | 0.048 | 0.291 | -0.269 | -1.542 | -0.03 | -0.036 | -0.036 | 0 | 0.07 | 0.01 | 0.09 | 0.13 | 0.11 | 0.17 |
| 2 | 125676313 | rs2247279 | -0.513 | 0.055 | 0.004 | -1.458 | 0.007 | 0.124 | 0.124 | 0.108 | 0.48 | 0.75 | 0.37 | 0.16 | 0.47 | 0.44 |
| 2 | 125683957 | rs2602644 | -0.431 | 0.055 | 0.028 | -1.458 | 0.004 | 0.086 | 0.086 | 0.019 | 0.48 | 0.7 | 0.36 | 0.16 | 0.47 | 0.44 |
| 2 | 125722442 | rs11123069 | -0.104 | 0.064 | -0.152 | -1.426 | 0 | -0.033 | -0.033 | 0 | 0.54 | 0.46 | 0.61 | 0.83 | 0.53 | 0.57 |
| 2 | 125735879 | rs17637939 | 0.057 | 0.296 | -1.233 | -1.312 | 0.002 | 0.181 | 0.181 | 0 | 0.49 | 0.19 | 0.59 | 0.83 | 0.53 | 0.56 |
| 2 | 125741700 | rs10190995 | -0.462 | -0.051 | -1.694 | -1.455 | 0.003 | 0.033 | 0.033 | 0 | 0.24 | 0.04 | 0.25 | 0.11 | 0.22 | 0.21 |
| 2 | 125745721 | rs4459740 | 0.318 | -0.183 | -1.235 | -2.482 | -0.001 | -0.024 | -0.024 | 0.036 | 0.45 | 0.46 | 0.56 | 0.89 | 0.53 | 0.56 |
| 2 | 125752937 | rs2421353 | 0.786 | -0.374 | -1.159 | -2.571 | 0 | -0.007 | -0.007 | 0 | 0.07 | 0.04 | 0.04 | 0 | 0.08 | 0.02 |
| 2 | 125760078 | rs2421356 | 0.773 | -0.469 | -0.942 | -2.633 | -0.011 | 0.032 | 0.032 | 0.464 | 0.41 | 0.31 | 0.51 | 0.89 | 0.47 | 0.56 |
| 2 | 125769695 | rs7578686 | 0.046 | -0.642 | -0.874 | -2.654 | 0.014 | 0.021 | 0.021 | 0 | 0.49 | 0.41 | 0.57 | 0.89 | 0.57 | 0.58 |
| 2 | 125792078 | rs6733261 | 0.771 | -0.762 | -0.653 | -2.715 | -0.011 | 0.07 | 0.07 | 0.538 | 0.41 | 0.26 | 0.51 | 0.89 | 0.49 | 0.56 |
| 2 | 125792747 | rs7560137 | 0.493 | -0.754 | -0.451 | -2.94 | -0.005 | -0.004 | -0.004 | 0 | 0.09 | 0.08 | 0.08 | 0 | 0.16 | 0.09 |
| 2 | 125798306 | rs13034713 | -0.122 | -0.751 | -0.44 | -2.923 | -0.008 | -0.056 | -0.056 | 0 | 0.1 | 0.25 | 0.06 | 0 | 0.08 | 0.02 |
| 2 | 125803157 | rs2421342 | -0.506 | -0.812 | -0.017 | -3.073 | 0.06 | 0.158 | 0.158 | 1.182 | 0.62 | 0.9 | 0.69 | 0.9 | 0.7 | 0.65 |
| 2 | 125806843 | rs4622728 | 0.22 | -0.727 | 0.426 | -2.913 | 0.006 | 0 | 0 | 0 | 0.5 | 0.54 | 0.58 | 0.86 | 0.55 | 0.57 |
| 2 | 125843421 | rs7572346 | 0.904 | -0.854 | 0.386 | -3.078 | -0.002 | -0.011 | -0.011 | 0.021 | 0.41 | 0.37 | 0.52 | 0.89 | 0.45 | 0.55 |
| 2 | 125859777 | rs4497887 | 1.851 | -1.245 | -0.117 | -3.541 | 0.035 | -0.001 | -0.001 | 0 | 0.16 | 0.03 | 0.33 | 0.87 | 0.08 | 0.37 |
| 2 | 125860398 | rs12465385 | -0.385 | -1.345 | -0.078 | -3.537 | 0.01 | 0.005 | 0.005 | 0.008 | 0.5 | 0.55 | 0.58 | 0.89 | 0.54 | 0.58 |
| 2 | 125864242 | rs4468815 | 0.324 | -1.309 | -0.167 | -3.51 | 0.001 | -0.026 | -0.026 | 0 | 0.4 | 0.36 | 0.52 | 0.89 | 0.45 | 0.56 |
| 2 | 125895045 | rs4338967 | 0.388 | -1.31 | -0.195 | -3.511 | -0.021 | 0.069 | 0.069 | 0.23 | 0.39 | 0.25 | 0.52 | 0.89 | 0.45 | 0.55 |
| 2 | 125896861 | rs13019609 | 1.459 | -1.32 | -0.179 | -3.529 | 0 | -0.006 | -0.006 | 0 | 0.08 | 0.05 | 0.04 | 0 | 0.1 | 0.02 |
| 2 | 125919832 | rs13007822 | 0.621 | -1.334 | 0.138 | -3.535 | -0.007 | -0.058 | -0.058 | 0.078 | 0.1 | 0.19 | 0.06 | 0 | 0.1 | 0.02 |
| 2 | 125926130 | rs4625907 | 0.981 | -1.326 | 0.088 | -3.543 | -0.006 | -0.001 | -0.001 | 0 | 0.08 | 0.07 | 0.08 | 0 | 0.16 | 0.09 |
| 2 | 125944407 | rs4396725 | 0.209 | -1.152 | -0.165 | -3.436 | -0.005 | 0.003 | 0.003 | 0.184 | 0.4 | 0.35 | 0.52 | 0.89 | 0.45 | 0.56 |
| 2 | 125966466 | rs2573111 | -1.4 | -1.157 | -0.173 | -3.451 | 0.002 | -0.033 | -0.033 | 0 | 0.77 | 0.81 | 0.81 | 0.98 | 0.62 | 0.82 |
| 2 | 125975601 | rs314915 | -0.215 | -1.077 | -0.047 | -3.284 | 0.023 | 0.069 | 0.069 | 0 | 0.52 | 0.62 | 0.43 | 0.12 | 0.46 | 0.42 |
| 2 | 126005563 | rs314964 | -0.792 | -0.996 | -0.079 | -3.266 | -0.025 | 0.144 | 0.144 | 0.003 | 0.58 | 0.82 | 0.46 | 0.1 | 0.52 | 0.41 |
| 2 | 126007559 | rs17628820 | 1.557 | -0.97 | -0.145 | -3.187 | -0.002 | 0.021 | 0.021 | 0 | 0.08 | 0.03 | 0.05 | 0 | 0.09 | 0.04 |
| 2 | 126013778 | rs314948 | 0.136 | -0.977 | 0.091 | -3.231 | -0.014 | -0.062 | -0.062 | 0 | 0.42 | 0.49 | 0.35 | 0.1 | 0.36 | 0.32 |
| 2 | 126015125 | rs314946 | -0.394 | -0.965 | 0.058 | -3.224 | 0.025 | 0.055 | 0.055 | 0.143 | 0.51 | 0.64 | 0.41 | 0.1 | 0.45 | 0.36 |
| 2 | 126021552 | rs11692066 | 0.02 | -0.857 | -0.156 | -3.115 | -0.001 | -0.026 | -0.026 | 0 | 0.42 | 0.46 | 0.35 | 0.1 | 0.36 | 0.32 |
| 2 | 126024238 | rs6739515 | 0.144 | -0.517 | -0.029 | -2.961 | -0.008 | -0.047 | -0.047 | 0.011 | 0.32 | 0.38 | 0.25 | 0.1 | 0.27 | 0.23 |
| 2 | 126034009 | rs1459166 | -0.884 | -0.054 | -0.423 | -2.735 | -0.115 | 0.111 | 0.111 | 0.741 | 0.74 | 0.89 | 0.82 | 0.97 | 0.59 | 0.83 |
| 2 | 126038422 | rs6735587 | -0.255 | -0.285 | -0.002 | -2.826 | -0.013 | 0.004 | 0.004 | 2.271 | 0.67 | 0.65 | 0.75 | 0.97 | 0.51 | 0.78 |
| 2 | 126042867 | rs11123082 | 0.444 | -0.622 | 0.463 | -2.838 | 0.025 | 0.042 | 0.042 | 3.517 | 0.54 | 0.33 | 0.66 | 0.9 | 0.39 | 0.72 |
| 2 | 126054305 | rs11903126 | 0.071 | -0.937 | 0.479 | -2.656 | -0.013 | -0.021 | -0.021 | 0.018 | 0.14 | 0.38 | 0.1 | 0.07 | 0.11 | 0.07 |
| 2 | 126070876 | rs1898216 | 1.432 | -1.369 | 0.102 | -2.659 | 0.088 | 0.008 | 0.008 | 0.103 | 0.24 | 0.08 | 0.4 | 0.8 | 0.12 | 0.45 |
| 2 | 126077250 | rs313295 | -0.68 | -0.159 | 0.307 | -2.63 | -0.02 | 0.057 | 0.057 | 4.232 | 0.42 | 0.72 | 0.29 | 0.1 | 0.5 | 0.25 |
| 2 | 126095439 | rs314980 | -0.119 | -0.146 | -0.219 | -2.675 | 0.004 | -0.028 | -0.028 | 0 | 0.33 | 0.66 | 0.23 | 0.1 | 0.41 | 0.19 |
| 2 | 126109030 | rs10496641 | 1.47 | -0.141 | -0.331 | -2.667 | -0.005 | 0.046 | 0.046 | 0 | 0.09 | 0.01 | 0.05 | 0 | 0.09 | 0.06 |
| 2 | 126115362 | rs13426616 | -0.011 | -0.141 | -0.269 | -2.669 | 0.023 | -0.093 | -0.093 | 0 | 0.32 | 0.59 | 0.22 | 0.06 | 0.4 | 0.19 |
| 2 | 126125141 | rs6541985 | -0.732 | -0.149 | -0.754 | -2.644 | -0.039 | 0.137 | 0.137 | 4.619 | 0.43 | 0.8 | 0.29 | 0.1 | 0.5 | 0.25 |
| 2 | 126128433 | rs13028050 | 0.165 | 0.08 | -0.933 | -2.561 | -0.026 | 0.058 | 0.058 | 0.973 | 0.31 | 0.21 | 0.19 | 0.06 | 0.4 | 0.2 |
| 2 | 126132036 | rs10451603 | 1.082 | 0.257 | -1.014 | -2.419 | -0.049 | 0.075 | 0.075 | 1.541 | 0.21 | 0.11 | 0.13 | 0.02 | 0.29 | 0.13 |
| 2 | 126137939 | rs1435578 | 0.193 | 0.181 | -0.89 | -2.415 | -0.002 | 0.096 | 0.096 | 0 | 0.55 | 0.3 | 0.53 | 0.86 | 0.51 | 0.65 |
| 2 | 126152664 | rs4493268 | -0.471 | 0.272 | -1.275 | -2.307 | -0.063 | 0.101 | 0.101 | 0.889 | 0.67 | 0.87 | 0.64 | 0.9 | 0.62 | 0.71 |
| 2 | 126158145 | rs1019442 | -0.158 | 0.193 | -1.335 | -2.359 | -0.056 | 0.182 | 0.182 | 0.72 | 0.58 | 0.88 | 0.59 | 0.9 | 0.51 | 0.64 |
| 2 | 126159556 | rs1836858 | 0.987 | -0.492 | -0.473 | -2.446 | -0.097 | -0.037 | -0.037 | 0 | 0.28 | 0.36 | 0.37 | 0.84 | 0.12 | 0.46 |
| 2 | 126166731 | rs1436304 | 1.027 | -0.487 | -0.393 | -2.453 | -0.08 | -0.035 | -0.035 | 0 | 0.27 | 0.35 | 0.38 | 0.85 | 0.12 | 0.46 |
| 2 | 126171392 | rs975150 | -0.09 | 0.287 | -0.666 | -2.74 | 0.017 | 0.149 | 0.149 | 0.266 | 0.55 | 0.83 | 0.56 | 0.86 | 0.42 | 0.61 |
| 2 | 126172619 | rs1436298 | 1.258 | 0.242 | -0.432 | -2.757 | -0.098 | -0.036 | -0.036 | 0 | 0.28 | 0.38 | 0.38 | 0.84 | 0.12 | 0.46 |
| 2 | 126173324 | rs1436301 | -0.56 | 0.356 | -0.659 | -2.857 | -0.118 | 0.134 | 0.134 | 0.483 | 0.66 | 0.91 | 0.62 | 0.9 | 0.54 | 0.68 |
| 2 | 126182771 | rs13036195 | 1.72 | 0.779 | -0.767 | -2.888 | -0.025 | 0.054 | 0.054 | 0 | 0.09 | 0 | 0.05 | 0 | 0.11 | 0.07 |
| 2 | 126186444 | rs2421538 | 0.133 | 0.737 | -0.412 | -2.922 | 0.019 | 0.106 | 0.106 | 0.122 | 0.55 | 0.77 | 0.53 | 0.86 | 0.42 | 0.61 |
| 2 | 126187262 | rs6724173 | -1.246 | 0.635 | -0.733 | -3.161 | -0.084 | 0.003 | 0.003 | 0 | 0.86 | 0.99 | 0.8 | 0.93 | 0.78 | 0.89 |
| 2 | 126188449 | rs7605018 | 0.759 | 0.361 | -0.526 | -3.119 | 0.014 | -0.111 | -0.111 | 6.82 | 0.28 | 0.45 | 0.17 | 0.02 | 0.3 | 0.15 |
| 2 | 126191031 | rs2193915 | 1.786 | 0.434 | -0.541 | -3.218 | -0.033 | 0.083 | 0.083 | 0.204 | 0.19 | 0.08 | 0.09 | 0.02 | 0.2 | 0.11 |
| 2 | 126192895 | rs6732625 | 1.284 | 0.426 | -0.469 | -3.196 | 0.059 | -0.037 | -0.037 | 4.088 | 0.09 | 0.37 | 0.08 | 0 | 0.1 | 0.04 |
| 2 | 126193197 | rs10490483 | 0.91 | 0.374 | -0.625 | -3.213 | 0.022 | 0.021 | 0.021 | 0.016 | 0.36 | 0.69 | 0.44 | 0.84 | 0.22 | 0.5 |
| 2 | 126193494 | rs9308661 | 0.91 | 0.374 | -0.625 | -3.213 | 0.022 | 0.021 | 0.021 | 0.016 | 0.36 | 0.69 | 0.44 | 0.84 | 0.22 | 0.5 |
| 2 | 126195411 | rs17531922 | 0.645 | 0.411 | -0.638 | -3.213 | -0.026 | 0.096 | 0.096 | 0.063 | 0.3 | 0.15 | 0.18 | 0.06 | 0.32 | 0.18 |
| 2 | 126207171 | rs12469322 | -0.043 | 0.447 | -0.654 | -3.177 | -0.046 | 0.064 | 0.064 | 0.365 | 0.65 | 0.79 | 0.62 | 0.9 | 0.54 | 0.68 |

| | | | | | | | | | | | | | | | | |
|---|---|---|---|---|---|---|---|---|---|---|---|---|---|---|---|---|
| 2 | 126224987 | rs12328873 | 2.2 | 0.163 | -0.565 | -3.129 | 0.088 | 0.005 | 0.005 | 0 | 0.24 | 0.08 | 0.33 | 0.83 | 0.11 | 0.45 |
| 2 | 126229399 | rs7581261 | 1.605 | 0.439 | -0.029 | -3.093 | 0.025 | -0.037 | -0.037 | 0 | 0.07 | 0.17 | 0.07 | 0 | 0.1 | 0.05 |
| 2 | 126239051 | rs7595772 | 1.33 | 0.431 | 0.108 | -3.096 | 0.043 | -0.04 | -0.04 | 0 | 0.08 | 0.27 | 0.07 | 0 | 0.1 | 0.05 |
| 2 | 126247087 | rs1377122 | 1.094 | 0.482 | 0.077 | -3.208 | 0.03 | 0.041 | 0.041 | 0 | 0.47 | 0.37 | 0.46 | 0.86 | 0.32 | 0.58 |
| 2 | 126252232 | rs10490485 | -0.062 | 0.455 | 0.127 | -3.127 | -0.067 | 0.087 | 0.087 | 0.161 | 0.67 | 0.86 | 0.63 | 0.9 | 0.54 | 0.7 |
| 2 | 126268453 | rs10187811 | 1.761 | 0.32 | 0.209 | -3.132 | -0.085 | -0.027 | -0.027 | 0.072 | 0.28 | 0.38 | 0.37 | 0.84 | 0.12 | 0.46 |
| 2 | 126271749 | rs13430283 | 0.113 | 0.349 | 0.219 | -3.131 | -0.003 | -0.025 | -0.025 | 0.111 | 0.37 | 0.4 | 0.26 | 0.05 | 0.42 | 0.23 |
| 2 | 126272972 | rs2016244 | -0.057 | 0.388 | 0.065 | -3.113 | -0.067 | 0.086 | 0.086 | 0.349 | 0.66 | 0.86 | 0.63 | 0.9 | 0.54 | 0.7 |
| 2 | 126273115 | rs13001190 | 1.863 | 0.333 | -0.153 | -3.159 | -0.055 | 0.057 | 0.057 | 0 | 0.13 | 0 | 0.13 | 0.03 | 0.16 | 0.18 |
| 2 | 126275143 | rs1116286 | 1.81 | 0.229 | 0.012 | -3.101 | -0.067 | -0.018 | -0.018 | 0 | 0.28 | 0.32 | 0.37 | 0.84 | 0.12 | 0.46 |
| 2 | 126280007 | rs2084869 | 1.955 | 0.218 | 0.337 | -3.108 | 0.027 | 0.014 | 0.014 | 0 | 0.27 | 0.18 | 0.35 | 0.84 | 0.12 | 0.46 |
| 2 | 126290376 | rs1820556 | 1.907 | 0.108 | 0.17 | -2.774 | -0.051 | -0.009 | -0.009 | 0.138 | 0.26 | 0.24 | 0.36 | 0.84 | 0.11 | 0.46 |
| 2 | 126293367 | rs1017627 | -0.034 | 0.501 | 0.037 | -2.872 | -0.097 | 0.105 | 0.105 | 0.746 | 0.67 | 0.88 | 0.65 | 0.9 | 0.55 | 0.7 |
| 2 | 126299170 | rs4146084 | 0.188 | 0.415 | 0.113 | -2.88 | -0.054 | 0.071 | 0.071 | 0.205 | 0.64 | 0.8 | 0.6 | 0.86 | 0.53 | 0.69 |
| 2 | 126303416 | rs2198318 | 0.069 | 0.415 | 0.256 | -2.882 | -0.06 | 0.085 | 0.085 | 0.22 | 0.65 | 0.82 | 0.63 | 0.89 | 0.54 | 0.69 |
| 2 | 126310056 | rs12464056 | 2 | 0.309 | 0.159 | -2.867 | -0.05 | -0.008 | -0.008 | 0 | 0.26 | 0.25 | 0.36 | 0.84 | 0.11 | 0.46 |
| 2 | 126310110 | rs10427350 | -1.639 | 0.576 | 0.137 | -2.767 | -0.064 | 0.007 | 0.007 | 0.025 | 0.87 | 0.98 | 0.82 | 0.93 | 0.77 | 0.9 |
| 2 | 126319594 | rs11675893 | 0.705 | 0.41 | 0.258 | -2.75 | -0.004 | -0.039 | -0.039 | 0 | 0.56 | 0.46 | 0.55 | 0.89 | 0.44 | 0.64 |
| 2 | 126327505 | rs7571928 | 0.86 | 0.403 | 0.338 | -2.566 | 0.026 | -0.048 | -0.048 | 0 | 0.09 | 0.25 | 0.07 | 0 | 0.1 | 0.05 |
| 2 | 126331343 | rs7586797 | 1.323 | 0.102 | 0.296 | -2.073 | -0.017 | 0 | 0 | 0 | 0.3 | 0.26 | 0.29 | 0.75 | 0.19 | 0.48 |
| 2 | 126332264 | rs1452515 | 0.152 | 0.582 | 0.122 | -2.031 | -0.029 | 0.015 | 0.015 | 0.022 | 0.67 | 0.76 | 0.57 | 0.8 | 0.6 | 0.69 |
| 2 | 126334309 | rs13402931 | 0.836 | 0.279 | 0.381 | -1.945 | 0.034 | 0.033 | 0.033 | 0 | 0.35 | 0.23 | 0.46 | 0.86 | 0.24 | 0.56 |
| 2 | 126335543 | rs17630886 | 0.559 | 0.586 | 0.425 | -2.071 | -0.031 | 0.13 | 0.13 | 0 | 0.36 | 0.21 | 0.27 | 0.06 | 0.36 | 0.26 |
| 2 | 126338255 | rs1452512 | -1.025 | 0.641 | 0.678 | -2.038 | 0.047 | -0.011 | -0.011 | 5.626 | 0.85 | 0.76 | 0.83 | 0.91 | 0.78 | 0.89 |
| 2 | 126346332 | rs1365788 | 0.638 | 0.583 | 0.54 | -2.067 | -0.033 | 0.125 | 0.125 | 0 | 0.35 | 0.17 | 0.26 | 0.06 | 0.36 | 0.26 |
| 2 | 126349761 | rs12465056 | -0.465 | 0.812 | 0.801 | -1.862 | 0.127 | -0.018 | -0.018 | 0.061 | 0.84 | 0.54 | 0.81 | 0.91 | 0.75 | 0.89 |
| 2 | 126363407 | rs11123096 | -0.359 | 0.637 | 0.56 | -1.84 | -0.01 | 0.01 | 0.01 | 0 | 0.75 | 0.8 | 0.78 | 0.91 | 0.66 | 0.83 |
| 2 | 126367137 | rs1452510 | -0.027 | 0.653 | 0.605 | -1.843 | 0.053 | -0.072 | -0.072 | 0.858 | 0.73 | 0.54 | 0.75 | 0.91 | 0.64 | 0.83 |
| 2 | 126368399 | rs6715359 | 0.318 | 0.333 | 0.508 | -1.898 | -0.009 | 0.041 | 0.041 | 0 | 0.4 | 0.35 | 0.46 | 0.18 | 0.4 | 0.29 |
| 2 | 126370871 | rs1452509 | 0.87 | 0.293 | 0.515 | -1.733 | -0.003 | -0.001 | -0.001 | 0 | 0.34 | 0.26 | 0.3 | 0.69 | 0.24 | 0.55 |
| 2 | 126395158 | rs12618696 | 0.99 | 0.255 | 0.375 | -1.692 | 0.024 | -0.009 | -0.009 | 0.287 | 0.33 | 0.19 | 0.29 | 0.69 | 0.24 | 0.55 |
| 2 | 126398345 | rs1863680 | 0.271 | 0.327 | 0.249 | -1.687 | -0.015 | 0.051 | 0.051 | 0 | 0.4 | 0.3 | 0.46 | 0.18 | 0.4 | 0.28 |
| 2 | 126401516 | rs3980656 | 0.9 | 0.253 | 0.099 | -1.594 | -0.001 | -0.001 | -0.001 | 0 | 0.32 | 0.25 | 0.31 | 0.69 | 0.27 | 0.56 |
| 2 | 126408611 | rs2162486 | 1.092 | 0.105 | -0.103 | -1.475 | 0.034 | -0.032 | -0.032 | 0.306 | 0.28 | 0.16 | 0.28 | 0.7 | 0.22 | 0.51 |
| 2 | 126411247 | rs12053356 | -0.263 | 0.228 | 0.127 | -1.595 | 0.016 | -0.016 | -0.016 | 0.193 | 0.73 | 0.61 | 0.77 | 0.86 | 0.66 | 0.83 |
| 2 | 126415098 | rs6746555 | -0.287 | 0.284 | 0.113 | -1.656 | -0.004 | -0.032 | -0.032 | 0 | 0.57 | 0.48 | 0.56 | 0.16 | 0.58 | 0.4 |
| 2 | 126423834 | rs12624172 | 1.477 | 0.173 | -0.037 | -1.643 | 0.07 | -0.056 | -0.056 | 0 | 0.28 | 0.1 | 0.27 | 0.7 | 0.18 | 0.49 |
| 2 | 126425287 | rs12998726 | 0.36 | 0.109 | 0.031 | -1.763 | -0.002 | 0.002 | 0.002 | 0 | 0.42 | 0.34 | 0.48 | 0.16 | 0.44 | 0.32 |
| 2 | 126429993 | rs1427308 | -0.504 | 0.114 | 0.126 | -1.769 | 0.028 | 0 | 0 | 0 | 0.83 | 0.68 | 0.83 | 0.86 | 0.77 | 0.89 |
| 2 | 126440151 | rs2052815 | 1.288 | 0.111 | 0.143 | -1.681 | 0.025 | -0.017 | -0.017 | 0 | 0.3 | 0.2 | 0.28 | 0.69 | 0.2 | 0.51 |
| 2 | 126440697 | rs2114601 | 0.052 | 0.165 | 0.083 | -1.654 | 0.011 | -0.013 | -0.013 | 0 | 0.72 | 0.62 | 0.77 | 0.86 | 0.64 | 0.84 |
| 2 | 126441311 | rs3943891 | -1.321 | 0.305 | 0.069 | -1.548 | 0.029 | 0.012 | 0.012 | 0 | 0.87 | 0.8 | 0.85 | 0.86 | 0.81 | 0.91 |
| 2 | 126442332 | rs933964 | 0.165 | 0.125 | -0.139 | -1.598 | -0.002 | 0.002 | 0.002 | 0 | 0.42 | 0.34 | 0.48 | 0.17 | 0.44 | 0.32 |
| 2 | 126455657 | rs7563352 | 0.632 | 0.116 | -0.017 | -1.794 | 0.012 | -0.028 | -0.028 | 0 | 0.62 | 0.41 | 0.62 | 0.75 | 0.55 | 0.77 |
| 2 | 126456344 | rs6542014 | 0.659 | -0.198 | 0.113 | -1.822 | -0.046 | -0.005 | -0.005 | 0 | 0.87 | 0.62 | 0.83 | 0.91 | 0.93 | 0.95 |
| 2 | 126461407 | rs4444536 | 0.27 | -0.043 | 0.088 | -1.842 | -0.02 | -0.037 | -0.037 | 0.348 | 0.77 | 0.6 | 0.77 | 0.91 | 0.8 | 0.88 |
| 2 | 126466408 | rs10165470 | 0.888 | -0.073 | -0.002 | -1.801 | 0.014 | 0.002 | 0.002 | 0 | 0.15 | 0.19 | 0.13 | 0.16 | 0.22 | 0.11 |
| 2 | 126468675 | rs17012921 | 0.335 | -0.243 | 0.005 | -1.815 | -0.003 | 0 | 0 | 0 | 0.08 | 0.09 | 0.14 | 0.09 | 0.07 | 0.05 |
| 2 | 126468706 | rs10496646 | 0.335 | -0.243 | 0.005 | -1.815 | -0.003 | 0 | 0 | 0.07 | 0.08 | 0.09 | 0.14 | 0.09 | 0.07 | 0.05 |
| 2 | 126476559 | rs3962035 | -0.81 | -0.274 | 0.004 | -1.837 | 0.008 | -0.024 | -0.024 | 0 | 0.35 | 0.5 | 0.33 | 0.24 | 0.42 | 0.28 |
| 2 | 126477788 | rs13429984 | 0.165 | -0.205 | -0.317 | -1.831 | -0.006 | 0.006 | 0.006 | 0 | 0.25 | 0.22 | 0.22 | 0.15 | 0.35 | 0.23 |
| 2 | 126488824 | rs933967 | 0.322 | -0.253 | -0.76 | -2.145 | -0.028 | 0.019 | 0.019 | 0 | 0.32 | 0.18 | 0.25 | 0.02 | 0.37 | 0.26 |
| 2 | 126491789 | rs1872374 | -0.163 | -0.362 | -0.824 | -1.709 | 0.022 | -0.008 | -0.008 | 0 | 0.09 | 0.06 | 0.1 | 0.09 | 0.07 | 0.05 |
| 2 | 126503735 | rs2421680 | -1.322 | -0.39 | -0.698 | -1.541 | -0.007 | -0.002 | -0.002 | 0 | 0.69 | 0.68 | 0.56 | 0.26 | 0.77 | 0.55 |
| 2 | 126508647 | rs2901496 | 0.94 | -0.098 | -0.631 | -1.802 | 0.015 | -0.017 | -0.017 | 0 | 0.17 | 0.21 | 0.16 | 0.09 | 0.23 | 0.16 |
| 2 | 126513548 | rs7589808 | -0.724 | -0.205 | -0.918 | -1.837 | -0.008 | -0.01 | -0.01 | 0 | 0.6 | 0.57 | 0.44 | 0.18 | 0.69 | 0.49 |
| 2 | 126520538 | rs17633694 | 1.892 | -0.125 | -1.412 | -1.917 | -0.013 | 0.081 | 0.081 | 0 | 0.14 | 0.01 | 0.07 | 0 | 0.15 | 0.07 |
| 2 | 126523149 | rs7565438 | 0.374 | -0.231 | -1.183 | -1.736 | -0.002 | -0.001 | -0.001 | 0 | 0.09 | 0.1 | 0.12 | 0.08 | 0.07 | 0.06 |
| 2 | 126523177 | rs2421684 | 0.633 | -0.048 | -1.088 | -1.674 | -0.03 | 0.033 | 0.033 | 0 | 0.27 | 0.15 | 0.19 | 0.09 | 0.32 | 0.27 |
| 2 | 126535687 | rs17685310 | 1.032 | 0.295 | -1.33 | -1.567 | -0.051 | -0.021 | -0.021 | 0 | 0.09 | 0.01 | 0.08 | 0.07 | 0.13 | 0.1 |
| 2 | 126549789 | rs4536659 | -1.339 | -0.613 | -0.981 | -1.429 | 0.02 | 0 | 0 | 0.195 | 0.7 | 0.73 | 0.55 | 0.29 | 0.76 | 0.53 |
| 2 | 126574928 | rs1319922 | 0.7 | -0.341 | -1.426 | -1.54 | -0.009 | 0.019 | 0.019 | 0 | 0.23 | 0.12 | 0.12 | 0.09 | 0.2 | 0.2 |
| 2 | 126575349 | rs13421098 | 0.023 | -1.34 | -1.282 | -1.238 | 0 | -0.001 | -0.001 | 0 | 0.12 | 0.13 | 0.12 | 0.08 | 0.09 | 0.06 |
| 2 | 126575750 | rs6720122 | -0.016 | -1.338 | -1.278 | -1.227 | -0.001 | -0.001 | -0.001 | 0 | 0.12 | 0.14 | 0.14 | 0.09 | 0.09 | 0.06 |
| 2 | 126580416 | rs6726563 | 0.787 | -1.508 | -1.773 | -1.562 | -0.01 | 0.162 | 0.162 | 0 | 0.41 | 0.07 | 0.36 | 0.15 | 0.54 | 0.45 |
| 2 | 126590327 | rs17635031 | -0.093 | -1.314 | -1.514 | -1.43 | 0.008 | 0.086 | 0.086 | 0 | 0.25 | 0.11 | 0.15 | 0.03 | 0.27 | 0.11 |
| 2 | 126602160 | rs10201523 | 0.301 | -1.255 | -1.409 | -1.385 | 0.004 | 0.045 | 0.045 | 0 | 0.21 | 0.11 | 0.13 | 0.01 | 0.22 | 0.07 |
| 2 | 126602658 | rs10186786 | 0.398 | -1.841 | -0.977 | -1.797 | -0.035 | -0.11 | -0.11 | 0.125 | 0.18 | 0.37 | 0.11 | 0 | 0.12 | 0.06 |
| 2 | 126625561 | rs4263115 | 1.441 | -1.895 | -1.212 | -1.813 | -0.041 | 0.089 | 0.089 | 0 | 0.14 | 0.01 | 0.09 | 0 | 0.19 | 0.06 |

| | | | | | | | | | | | | | | | | |
|---|---|---|---|---|---|---|---|---|---|---|---|---|---|---|---|---|
| 2 | 126627665 | rs10496650 | 1.443 | -1.866 | -1.156 | -1.791 | -0.01 | -0.027 | -0.027 | 0 | 0.07 | 0.12 | 0.04 | 0.01 | 0.03 | 0.01 |
| 2 | 126631551 | rs1317407 | -0.985 | -1.905 | -0.667 | -1.79 | 0.024 | -0.003 | -0.003 | 0 | 0.42 | 0.68 | 0.26 | 0.01 | 0.35 | 0.13 |
| 2 | 126633803 | rs13396428 | 0.266 | -1.36 | -0.607 | -1.25 | -0.006 | -0.065 | -0.065 | 0.203 | 0.23 | 0.26 | 0.15 | 0.01 | 0.22 | 0.07 |
| 2 | 126668449 | rs17013142 | 0.251 | -1.044 | -1.13 | -0.296 | 0.004 | 0 | 0 | 0 | 0.22 | 0.16 | 0.42 | 0.79 | 0.13 | 0.38 |
| 2 | 126677957 | rs1531078 | -0.414 | -1.081 | -1.117 | -0.003 | -0.007 | 0.003 | 0.003 | 0 | 0.3 | 0.31 | 0.41 | 0.58 | 0.2 | 0.35 |
| 2 | 126679224 | rs7607959 | -0.307 | -0.719 | -1.821 | -0.167 | -0.006 | -0.122 | -0.122 | 0 | 0.73 | 0.98 | 0.68 | 0.34 | 0.77 | 0.49 |
| 2 | 126693321 | rs11694158 | 0.917 | -0.486 | -2.059 | -0.781 | -0.017 | 0.056 | 0.056 | 0 | 0.14 | 0.02 | 0.08 | 0 | 0.16 | 0.03 |
| 2 | 126711616 | rs12994681 | 0.198 | -0.061 | -1.973 | -1.406 | -0.014 | -0.006 | -0.006 | 0.878 | 0.43 | 0.34 | 0.28 | 0 | 0.46 | 0.17 |
| 2 | 126712081 | rs12711718 | 0.15 | -0.084 | -1.638 | -1.417 | 0 | -0.097 | -0.097 | 0 | 0.43 | 0.46 | 0.29 | 0 | 0.46 | 0.17 |
| 2 | 126715716 | rs13009198 | 0.953 | -0.05 | -1.628 | -1.373 | 0.004 | -0.081 | -0.081 | 7.588 | 0.29 | 0.33 | 0.2 | 0 | 0.29 | 0.13 |
| 2 | 126719125 | rs1455627 | -0.686 | -0.464 | -1.546 | -1.545 | -0.019 | -0.04 | -0.04 | 0.878 | 0.64 | 0.61 | 0.47 | 0.13 | 0.74 | 0.46 |
| 2 | 126722908 | rs874770 | 0.391 | -0.428 | -1.541 | -1.58 | 0.006 | -0.146 | -0.146 | 0 | 0.44 | 0.51 | 0.29 | 0 | 0.46 | 0.17 |
| 2 | 126728027 | rs17013251 | 2.299 | -0.71 | -1.392 | -1.373 | -0.011 | 0.056 | 0.056 | 0.04 | 0.06 | 0.06 | 0.18 | 0.22 | 0.03 | 0.03 |
| 2 | 126728274 | rs1455608 | 0.857 | -0.539 | -1.489 | -1.328 | -0.036 | 0.007 | 0.007 | 0.419 | 0.19 | 0.06 | 0.17 | 0.14 | 0.25 | 0.28 |
| 2 | 126728541 | rs17689012 | 0.658 | -0.506 | -1.425 | -1.348 | -0.001 | 0.001 | 0.001 | 0 | 0.2 | 0.19 | 0.27 | 0.22 | 0.19 | 0.06 |
| 2 | 126729166 | rs1563338 | 0.203 | -0.235 | -1.33 | -1.146 | 0.009 | 0.031 | 0.031 | 0.317 | 0.5 | 0.43 | 0.48 | 0.22 | 0.51 | 0.2 |
| 2 | 126736245 | rs1902241 | 0.632 | -0.073 | -1.604 | -1.25 | -0.006 | 0.102 | 0.102 | 1.98 | 0.4 | 0.77 | 0.35 | 0.13 | 0.34 | 0.17 |
| 2 | 126736747 | rs10496652 | -0.323 | 0.245 | -1.609 | -1.107 | -0.048 | -0.001 | -0.001 | 0 | 0.8 | 0.92 | 0.83 | 0.81 | 0.7 | 0.68 |
| 2 | 126742136 | rs1455611 | -0.946 | 0.061 | -1.427 | -1.31 | -0.015 | -0.066 | -0.066 | 0 | 0.72 | 0.96 | 0.6 | 0.26 | 0.75 | 0.47 |
| 2 | 126752687 | rs17013262 | -0.073 | 0.365 | -0.647 | -1.429 | -0.013 | -0.066 | -0.066 | 0.034 | 0.11 | 0.33 | 0.06 | 0 | 0.04 | 0.01 |
| 2 | 126764347 | rs17013287 | -1.265 | 0.654 | -0.605 | -1.345 | 0.015 | -0.059 | -0.059 | 2.289 | 0.31 | 0.52 | 0.29 | 0.13 | 0.39 | 0.36 |
| 2 | 126766945 | rs12475137 | -0.232 | 0.752 | -0.687 | -1.315 | -0.025 | 0.007 | 0.007 | 0.043 | 0.15 | 0.1 | 0.16 | 0.13 | 0.23 | 0.3 |
| 2 | 126768087 | rs12990876 | -1.039 | 0.667 | -0.616 | -1.38 | 0.03 | -0.084 | -0.084 | 0.712 | 0.28 | 0.39 | 0.25 | 0.1 | 0.37 | 0.24 |
| 2 | 126768155 | rs17013295 | 0.064 | 0.703 | -0.525 | -1.307 | 0.094 | 0.022 | 0.022 | 0 | 0.07 | 0.27 | 0.06 | 0.06 | 0.11 | 0.07 |
| 2 | 126795430 | rs17695513 | 0.951 | 0.691 | -0.765 | -1.322 | -0.012 | 0.016 | 0.016 | 0 | 0.11 | 0.06 | 0.09 | 0 | 0.16 | 0.07 |
| 2 | 126797090 | rs11687421 | -1.646 | 0.6 | -0.736 | -1.468 | -0.005 | 0.012 | 0.012 | 0 | 0.4 | 0.6 | 0.32 | 0.13 | 0.45 | 0.37 |
| 2 | 126799433 | rs17695598 | -0.372 | 0.681 | -1.045 | -1.406 | -0.011 | 0.066 | 0.066 | 0 | 0.18 | 0.03 | 0.12 | 0.04 | 0.16 | 0.12 |
| 2 | 126800645 | rs1455618 | -1.157 | 0.685 | -0.797 | -1.393 | 0.017 | -0.065 | -0.065 | 0 | 0.25 | 0.36 | 0.22 | 0.14 | 0.27 | 0.19 |
| 2 | 126802800 | rs10496654 | 0.448 | 0.702 | -0.74 | -1.361 | 0.111 | 0.043 | 0.043 | 0 | 0.07 | 0.24 | 0.09 | 0.09 | 0.11 | 0.08 |
| 2 | 126806674 | rs13030845 | 0.23 | 0.707 | -0.88 | -1.359 | 0.118 | 0.046 | 0.046 | 0 | 0.07 | 0.29 | 0.09 | 0.09 | 0.11 | 0.08 |
| 2 | 126807931 | rs13014869 | 0.23 | 0.707 | -0.88 | -1.359 | 0.118 | 0.046 | 0.046 | 0 | 0.07 | 0.29 | 0.09 | 0.09 | 0.11 | 0.08 |
| 2 | 126808935 | rs6753603 | -1.678 | 0.59 | -1.146 | -1.392 | 0.001 | -0.025 | -0.025 | 0 | 0.4 | 0.53 | 0.34 | 0.17 | 0.45 | 0.37 |
| 2 | 126826913 | rs11885991 | -1.103 | 0.535 | -1.178 | -0.975 | 0.017 | -0.065 | -0.065 | 1.018 | 0.25 | 0.36 | 0.22 | 0.14 | 0.27 | 0.19 |
| 2 | 126833044 | rs4296429 | -0.41 | 0.506 | -1.621 | -0.76 | 0.013 | -0.034 | -0.034 | 0 | 0.66 | 0.98 | 0.71 | 0.31 | 0.73 | 0.54 |
| 2 | 126858204 | rs28715475 | -1.232 | 1.124 | -0.283 | -0.496 | 0.028 | -0.06 | -0.06 | 0.171 | 0.23 | 0.36 | 0.22 | 0.14 | 0.26 | 0.16 |
| 2 | 126886084 | rs35448183 | -0.464 | 1.344 | -0.175 | -0.191 | 0.055 | 0.058 | 0.058 | 0 | 0.06 | 0.14 | 0.09 | 0.16 | 0.11 | 0.06 |
| 2 | 126904601 | rs4271774 | -0.353 | 1.253 | -0.196 | -0.262 | -0.005 | 0.008 | 0.008 | 0 | 0.07 | 0.06 | 0.1 | 0.02 | 0.1 | 0.08 |
| 2 | 126906956 | rs4311069 | -0.935 | 1.546 | -0.195 | -0.352 | 0.048 | 0.029 | 0.029 | 0 | 0.08 | 0.16 | 0.1 | 0.16 | 0.14 | 0.08 |
| 2 | 126943392 | rs28703141 | -1.057 | 1.515 | 0.275 | -0.389 | 0.07 | 0.042 | 0.042 | 0 | 0.08 | 0.2 | 0.1 | 0.16 | 0.16 | 0.08 |
| 2 | 126947592 | rs10496655 | -1.023 | 1.52 | 0.277 | -0.386 | 0.071 | 0.043 | 0.043 | 0 | 0.08 | 0.2 | 0.1 | 0.16 | 0.16 | 0.08 |
| 2 | 126962441 | rs1484448 | -2.306 | 1.262 | 0.129 | -0.923 | 0.006 | -0.04 | -0.04 | 0.019 | 0.4 | 0.5 | 0.35 | 0.21 | 0.46 | 0.35 |
| 2 | 126965458 | rs17013415 | -1.969 | 1.223 | 0.104 | -0.958 | 0.008 | -0.011 | -0.011 | 0 | 0.25 | 0.31 | 0.24 | 0.19 | 0.31 | 0.17 |
| 2 | 126966164 | rs4500895 | -2.179 | 1.196 | 0.136 | -0.948 | 0.008 | -0.05 | -0.05 | 0 | 0.4 | 0.54 | 0.35 | 0.21 | 0.46 | 0.35 |
| 2 | 126984196 | rs17013438 | -1.953 | 0.898 | 0.514 | -0.799 | 0.004 | -0.048 | -0.048 | 0 | 0.32 | 0.52 | 0.24 | 0.25 | 0.34 | 0.15 |
| 2 | 126986871 | rs2404205 | 0.525 | 0.907 | 0.368 | -0.805 | -0.012 | 0.057 | 0.057 | 0 | 0.11 | 0.03 | 0.07 | 0 | 0.14 | 0.06 |
| 2 | 126987912 | rs17013442 | -1.624 | 0.991 | 0.503 | -0.748 | 0.024 | -0.01 | -0.01 | 0.336 | 0.2 | 0.4 | 0.16 | 0.25 | 0.2 | 0.09 |
| 2 | 126992626 | rs523147 | -1.341 | 1.027 | 0.551 | -0.623 | 0.077 | 0.067 | 0.067 | 0 | 0.12 | 0.35 | 0.14 | 0.28 | 0.16 | 0.09 |
| 2 | 127008196 | rs771078 | -1.714 | 0.905 | 0.506 | -0.701 | 0.034 | -0.028 | -0.028 | 0 | 0.23 | 0.43 | 0.2 | 0.28 | 0.3 | 0.11 |
| 2 | 127009760 | rs729446 | 0.109 | 0.856 | 0.312 | -0.734 | -0.001 | -0.028 | -0.028 | 0 | 0.11 | 0.11 | 0.07 | 0 | 0.14 | 0.02 |
| 2 | 127035725 | rs10929011 | -0.988 | 0.882 | 0.435 | -0.442 | 0.068 | 0.092 | 0.092 | 0 | 0.1 | 0.3 | 0.16 | 0.31 | 0.16 | 0.1 |
| 2 | 127041555 | rs360234 | -2.368 | 0.982 | -0.513 | -0.753 | 0.036 | -0.115 | -0.115 | 0 | 0.62 | 0.99 | 0.65 | 0.46 | 0.74 | 0.51 |
| 2 | 127046316 | rs10928762 | -1.585 | 1.495 | 0.369 | -0.584 | 0.011 | -0.008 | -0.008 | 0.008 | 0.28 | 0.36 | 0.25 | 0.33 | 0.35 | 0.13 |
| 2 | 127048167 | rs360390 | -2.539 | 1.602 | 0.582 | -0.643 | -0.001 | 0.002 | 0.002 | 0 | 0.38 | 0.62 | 0.33 | 0.35 | 0.44 | 0.19 |
| 2 | 127056534 | rs360259 | -1.262 | 1.57 | 0.579 | -0.524 | 0.032 | 0.043 | 0.043 | 0.86 | 0.2 | 0.47 | 0.21 | 0.33 | 0.2 | 0.1 |
| 2 | 127082310 | rs360308 | -0.031 | 1.68 | 0.183 | -0.528 | 0.017 | -0.046 | -0.046 | 0 | 0.1 | 0.14 | 0.08 | 0 | 0.18 | 0.04 |
| 2 | 127083389 | rs360314 | 0.33 | 1.681 | 0.187 | -0.528 | -0.008 | -0.001 | -0.001 | 0.031 | 0.09 | 0.07 | 0.06 | 0 | 0.16 | 0.04 |
| 2 | 127083960 | rs360317 | -0.038 | 1.68 | 0.183 | -0.528 | 0.017 | -0.046 | -0.046 | 0 | 0.1 | 0.14 | 0.08 | 0 | 0.18 | 0.04 |
| 2 | 127085797 | rs360323 | -1.54 | 1.672 | 0.32 | -0.536 | 0.013 | -0.01 | -0.01 | 0 | 0.29 | 0.36 | 0.23 | 0.34 | 0.36 | 0.14 |
| 2 | 127102573 | rs359683 | -1.759 | 1.654 | 0.311 | -0.545 | 0.023 | -0.024 | -0.024 | 0 | 0.31 | 0.46 | 0.26 | 0.34 | 0.37 | 0.15 |
| 2 | 127115202 | rs359687 | -1.197 | 1.904 | 0.377 | -0.447 | 0.077 | 0.049 | 0.049 | 0 | 0.16 | 0.34 | 0.2 | 0.34 | 0.29 | 0.11 |
| 2 | 127120893 | rs360385 | 0.672 | 1.849 | 0.406 | -0.426 | 0.104 | 0.212 | 0.212 | 0 | 0.93 | 0.72 | 0.87 | 0.67 | 0.89 | 0.93 |
| 2 | 127123647 | rs168604 | 1.796 | 2.1 | 0.281 | -0.531 | 0.054 | 0.085 | 0.085 | 0 | 0.86 | 0.67 | 0.83 | 0.67 | 0.82 | 0.88 |
| 2 | 127124505 | rs359697 | 0.956 | 1.877 | 0.445 | -0.44 | 0.082 | 0.056 | 0.056 | 0 | 0.84 | 0.67 | 0.81 | 0.67 | 0.72 | 0.9 |
| 2 | 127126224 | rs243417 | -0.584 | 1.869 | 0.42 | -0.449 | -0.019 | 0.015 | 0.015 | 0 | 0.92 | 0.95 | 0.94 | 1 | 0.84 | 0.96 |
| 2 | 127131081 | rs359706 | -0.484 | 1.868 | 0.42 | -0.45 | -0.004 | -0.01 | -0.01 | 0 | 0.91 | 0.93 | 0.94 | 1 | 0.84 | 0.96 |
| 2 | 127133165 | rs7600474 | -0.798 | 1.887 | 0.452 | -0.417 | 0.099 | 0.207 | 0.207 | 0 | 0.06 | 0.24 | 0.13 | 0.33 | 0.12 | 0.07 |
| 2 | 127137005 | rs359688 | -0.56 | 1.885 | 0.444 | -0.421 | -0.014 | 0.008 | 0.008 | 0 | 0.92 | 0.95 | 0.94 | 1 | 0.84 | 0.96 |
| 2 | 127139060 | rs359690 | 1.156 | 1.879 | 0.573 | -0.423 | 0.113 | 0.081 | 0.081 | 0 | 0.84 | 0.58 | 0.79 | 0.66 | 0.72 | 0.89 |
| 2 | 127145800 | rs7567897 | -0.799 | 1.888 | 0.513 | -0.517 | 0.099 | 0.16 | 0.16 | 0 | 0.06 | 0.24 | 0.08 | 0.28 | 0.11 | 0.06 |

| | | | | | | | | | | | | | | | | |
|---|---|---|---|---|---|---|---|---|---|---|---|---|---|---|---|---|
| 2 | 127148459 | rs437866 | 1.927 | 1.896 | 0.495 | -0.681 | -0.001 | -0.003 | -0.003 | 0 | 0.6 | 0.39 | 0.7 | 0.7 | 0.53 | 0.77 |
| 2 | 127149467 | rs7602532 | -1.108 | 1.969 | 0.797 | -0.076 | 0.094 | 0.037 | 0.037 | 0 | 0.15 | 0.31 | 0.15 | 0.28 | 0.28 | 0.1 |
| 2 | 127167247 | rs12998429 | -1.232 | 1.861 | 0.687 | 0.013 | 0.055 | 0.033 | 0.033 | 0 | 0.14 | 0.31 | 0.13 | 0.28 | 0.24 | 0.08 |
| 2 | 127218915 | rs10928857 | -1.184 | 1.998 | 1.021 | -0.166 | 0.058 | -0.001 | -0.001 | 0 | 0.15 | 0.25 | 0.14 | 0.21 | 0.29 | 0.09 |
| 2 | 127223373 | rs13019936 | -0.667 | 2.077 | 1.052 | -0.187 | 0.044 | -0.064 | -0.064 | 0 | 0.11 | 0.23 | 0.08 | 0 | 0.18 | 0.05 |
| 2 | 127228805 | rs6754762 | -2.389 | 1.981 | 1.291 | -0.513 | -0.003 | -0.004 | -0.004 | 0 | 0.3 | 0.64 | 0.27 | 0.13 | 0.42 | 0.24 |
| 2 | 127229413 | rs13024895 | -0.272 | 1.491 | 1.25 | -0.528 | 0.025 | -0.052 | -0.052 | 0.982 | 0.11 | 0.17 | 0.07 | 0 | 0.18 | 0.05 |
| 2 | 127230124 | rs10201795 | -0.997 | 1.53 | 1.235 | -0.527 | 0.006 | -0.043 | -0.043 | 1.346 | 0.13 | 0.21 | 0.12 | 0.02 | 0.13 | 0.13 |
| 2 | 127238737 | rs6753696 | -2.022 | 1.552 | 0.679 | -0.534 | 0.003 | -0.045 | -0.045 | 3.03 | 0.26 | 0.64 | 0.21 | 0.02 | 0.31 | 0.18 |
| 2 | 127240800 | rs7606197 | -1.882 | 1.45 | 0.063 | -0.356 | 0.104 | -0.039 | -0.039 | 0.622 | 0.54 | 0.93 | 0.58 | 0.45 | 0.72 | 0.59 |
| 2 | 127265612 | rs13425595 | -1.585 | 1.665 | 0.681 | -0.45 | -0.007 | -0.049 | -0.049 | 0 | 0.17 | 0.56 | 0.13 | 0.11 | 0.13 | 0.13 |
| 2 | 127267188 | rs4577234 | -2.073 | 1.666 | 0.659 | -0.472 | 0.016 | -0.061 | -0.061 | 0 | 0.26 | 0.57 | 0.19 | 0.12 | 0.31 | 0.18 |
| 2 | 127279139 | rs13019920 | -0.366 | 1.678 | 0.519 | -0.496 | -0.026 | 0.02 | 0.02 | 0 | 0.1 | 0.05 | 0.06 | 0 | 0.18 | 0.04 |
| 2 | 127296122 | rs11883942 | -0.305 | 1.671 | 0.496 | -0.499 | -0.017 | 0.006 | 0.006 | 0 | 0.1 | 0.05 | 0.07 | 0 | 0.18 | 0.04 |
| 2 | 127310729 | rs7565286 | -1.946 | 1.439 | 0.828 | -0.353 | 0.027 | -0.073 | -0.073 | 0 | 0.17 | 0.6 | 0.14 | 0.02 | 0.19 | 0.1 |
| 2 | 127322284 | rs17013910 | -1.01 | 1.418 | 0.885 | -0.328 | 0.032 | -0.043 | -0.043 | 0 | 0.11 | 0.32 | 0.08 | 0.02 | 0.1 | 0.07 |
| 2 | 127325786 | rs2165222 | -1.365 | 0.932 | -0.392 | -0.078 | 0.011 | -0.122 | -0.122 | 0.007 | 0.69 | 0.96 | 0.62 | 0.51 | 0.79 | 0.63 |
| 2 | 127329197 | rs1025788 | -1.741 | 1.027 | 0.253 | -0.528 | 0.02 | -0.084 | -0.084 | 0 | 0.21 | 0.59 | 0.16 | 0.04 | 0.22 | 0.12 |
| 2 | 127335267 | rs10175150 | -1.718 | 0.955 | 0.204 | -0.719 | 0.021 | -0.083 | -0.083 | 0.479 | 0.17 | 0.6 | 0.14 | 0.03 | 0.19 | 0.07 |
| 2 | 127336615 | rs6728481 | -1.1 | 0.87 | 0.336 | -0.724 | 0.031 | -0.054 | -0.054 | 0.02 | 0.12 | 0.4 | 0.11 | 0.03 | 0.12 | 0.05 |
| 2 | 127340075 | rs7596310 | 0.949 | 0.481 | 0.156 | -0.777 | 0.033 | -0.022 | -0.022 | 0 | 0.28 | 0.1 | 0.39 | 0.63 | 0.2 | 0.36 |
| 2 | 127342981 | rs6707983 | -2.249 | 1.116 | 0.996 | -0.832 | 0.032 | -0.113 | -0.113 | 0 | 0.18 | 0.48 | 0.14 | 0.02 | 0.19 | 0.1 |
| 2 | 127351413 | rs6713953 | -1.163 | 1.108 | 1.079 | -0.834 | 0.019 | -0.044 | -0.044 | 0 | 0.11 | 0.34 | 0.09 | 0.02 | 0.11 | 0.04 |
| 2 | 127353125 | rs6710365 | -2.236 | 1.133 | 0.865 | -0.826 | 0.009 | -0.093 | -0.093 | 0 | 0.18 | 0.58 | 0.14 | 0.02 | 0.19 | 0.1 |
| 2 | 127359823 | rs10183848 | -1.412 | 0.842 | 0.474 | -0.564 | 0.015 | -0.049 | -0.049 | 0.046 | 0.12 | 0.42 | 0.09 | 0.03 | 0.11 | 0.04 |
| 2 | 127372728 | rs6747825 | -0.71 | 1.038 | -0.344 | -0.691 | -0.001 | -0.003 | -0.003 | 0 | 0.33 | 0.32 | 0.2 | 0.14 | 0.33 | 0.17 |
| 2 | 127374341 | rs6712439 | 0.49 | 1.156 | -0.275 | -0.968 | 0 | -0.04 | -0.04 | 0 | 0.58 | 0.51 | 0.71 | 0.88 | 0.56 | 0.73 |
| 2 | 127376895 | rs6724089 | -0.335 | 1.297 | -0.437 | -1.174 | -0.035 | 0.041 | 0.041 | 0 | 0.27 | 0.22 | 0.2 | 0.14 | 0.36 | 0.2 |
| 2 | 127378841 | rs6714256 | -0.619 | 1.104 | 0.016 | -1.263 | 0.004 | -0.021 | -0.021 | 0 | 0.3 | 0.37 | 0.22 | 0.14 | 0.36 | 0.2 |
| 2 | 127378997 | rs9287557 | -1.044 | 0.957 | 0.185 | -1.513 | 0.005 | -0.054 | -0.054 | 0 | 0.4 | 0.57 | 0.3 | 0.14 | 0.52 | 0.25 |
| 2 | 127383914 | rs6714079 | -1.025 | 0.431 | 0.251 | -1.667 | -0.073 | -0.096 | -0.096 | 0.628 | 0.23 | 0.51 | 0.13 | 0.04 | 0.15 | 0.12 |
| 2 | 127384735 | rs1899038 | 0.055 | 0.481 | 0.198 | -1.688 | -0.045 | 0.044 | 0.044 | 0.11 | 0.28 | 0.15 | 0.23 | 0.13 | 0.44 | 0.22 |
| 2 | 127387469 | rs12692111 | -1.305 | 0.439 | -0.149 | -1.575 | 0.014 | 0.083 | 0.083 | 0 | 0.51 | 0.68 | 0.37 | 0.18 | 0.58 | 0.34 |
| 2 | 127396231 | rs6741528 | -0.312 | 0.312 | -0.384 | -1.515 | -0.046 | -0.048 | -0.048 | 0 | 0.16 | 0.26 | 0.08 | 0.03 | 0.07 | 0.1 |
| 2 | 127398038 | rs7563897 | -0.909 | 0.769 | -0.933 | -1.392 | 0.022 | 0.144 | 0.144 | 0 | 0.51 | 0.74 | 0.36 | 0.16 | 0.59 | 0.34 |
| 2 | 127402833 | rs6431169 | 0.04 | 0.875 | -1.033 | -0.315 | -0.022 | -0.027 | -0.027 | 0 | 0.78 | 0.83 | 0.63 | 0.66 | 0.69 | 0.7 |
| 2 | 127421701 | rs12468957 | -0.408 | -0.029 | -0.975 | -0.449 | 0.011 | 0.004 | 0.004 | 0 | 0.42 | 0.62 | 0.62 | 0.57 | 0.37 | 0.53 |
| 2 | 127422573 | rs4662685 | -0.818 | -0.097 | -0.992 | -0.658 | 0.027 | 0.041 | 0.041 | 0 | 0.51 | 0.72 | 0.68 | 0.57 | 0.47 | 0.56 |
| 2 | 127422593 | rs7589096 | 0.634 | -0.561 | -0.984 | -0.747 | 0.042 | 0.046 | 0.046 | 0 | 0.18 | 0.11 | 0.06 | 0.01 | 0.09 | 0.15 |
| 2 | 127423066 | rs4663038 | 0.291 | -0.569 | -0.397 | -0.746 | 0.003 | 0.012 | 0.012 | 0 | 0.21 | 0.19 | 0.53 | 0.56 | 0.28 | 0.38 |
| 2 | 127424849 | rs11694376 | -0.315 | -0.596 | -0.914 | -1.059 | 0.051 | 0.037 | 0.037 | 0 | 0.51 | 0.34 | 0.72 | 0.64 | 0.69 | 0.62 |
| 2 | 127426034 | rs6761722 | -1.609 | -0.527 | -1.237 | -1.032 | 0.004 | -0.013 | -0.013 | 0 | 0.73 | 0.78 | 0.82 | 0.65 | 0.78 | 0.77 |
| 2 | 127426697 | rs17741574 | 0.459 | -0.317 | -1.336 | -1.007 | -0.074 | 0.127 | 0.127 | 0 | 0.3 | 0.1 | 0.19 | 0.08 | 0.41 | 0.23 |
| 2 | 127429676 | rs12990356 | 0.585 | -0.521 | -0.843 | -0.858 | -0.009 | -0.03 | -0.03 | 0 | 0.19 | 0.2 | 0.07 | 0.01 | 0.09 | 0.15 |
| 2 | 127430211 | rs6736203 | -0.347 | 0.24 | -1.167 | -0.137 | -0.035 | -0.069 | -0.069 | 0.173 | 0.15 | 0.06 | 0.33 | 0.42 | 0.25 | 0.29 |
| 2 | 127430549 | rs17014125 | 0.65 | 0.279 | -1.105 | -0.073 | -0.012 | 0.023 | 0.023 | 0 | 0.09 | 0.05 | 0.28 | 0.42 | 0.14 | 0.26 |
| 2 | 127431312 | rs17741655 | 1.128 | 0.265 | -1.263 | -0.322 | -0.092 | 0.097 | 0.097 | 0 | 0.25 | 0.05 | 0.16 | 0.08 | 0.33 | 0.21 |
| 2 | 127432998 | rs908641 | 0.975 | 0.39 | -1.182 | -0.179 | -0.006 | 0.051 | 0.051 | 0 | 0.08 | 0.06 | 0.28 | 0.43 | 0.14 | 0.24 |
| 2 | 127433923 | rs6711745 | -0.293 | 0.238 | -0.76 | -0.322 | 0 | -0.015 | -0.015 | 0 | 0.42 | 0.29 | 0.51 | 0.51 | 0.58 | 0.47 |
| 2 | 127436724 | rs4143021 | 1.054 | 0.409 | -0.883 | -0.202 | -0.016 | 0.031 | 0.031 | 0 | 0.08 | 0.05 | 0.29 | 0.43 | 0.14 | 0.26 |
| 2 | 127437540 | rs6724592 | 0.923 | -0.342 | -0.576 | -0.409 | -0.023 | -0.06 | -0.06 | 0 | 0.14 | 0.24 | 0.06 | 0 | 0.08 | 0.08 |
| 2 | 127437880 | rs28387166 | 0.572 | -0.331 | -0.625 | -0.402 | -0.017 | 0.037 | 0.037 | 0 | 0.07 | 0 | 0.05 | 0 | 0.11 | 0.03 |
| 2 | 127440524 | rs12990441 | 0.44 | -0.284 | -0.496 | -0.31 | -0.004 | -0.045 | -0.045 | 0 | 0.19 | 0.23 | 0.1 | 0 | 0.17 | 0.11 |
| 2 | 127441158 | rs1866657 | -0.881 | -0.248 | -0.662 | -0.087 | 0.002 | 0 | 0 | 0 | 0.65 | 0.68 | 0.77 | 0.66 | 0.76 | 0.68 |
| 2 | 127441605 | rs13425954 | -0.161 | 0.488 | -1.05 | 0.262 | -0.026 | -0.028 | -0.028 | 2.239 | 0.39 | 0.3 | 0.46 | 0.51 | 0.48 | 0.44 |
| 2 | 127443592 | rs4662686 | 0.08 | 0.359 | -0.945 | 0.541 | -0.003 | -0.002 | -0.002 | 0 | 0.55 | 0.56 | 0.45 | 0.42 | 0.41 | 0.54 |
| 2 | 127443627 | rs1866658 | 0.655 | 1.25 | -1.353 | 0.878 | -0.036 | -0.039 | -0.039 | 0.355 | 0.66 | 0.17 | 0.45 | 0.57 | 0.64 | 0.6 |
| 2 | 127443771 | rs7599354 | 0.389 | 1.608 | -0.462 | 1.343 | 0.059 | 0.058 | 0.058 | 0 | 0.79 | 0.71 | 0.68 | 0.67 | 0.66 | 0.84 |
| 2 | 127445050 | rs755516 | -0.949 | 1.049 | -0.661 | 0.689 | 0.017 | 0.081 | 0.081 | 0 | 0.54 | 0.75 | 0.6 | 0.68 | 0.55 | 0.47 |
| 2 | 127453664 | rs1050967 | -0.955 | 1.005 | -0.65 | 0.666 | 0.014 | 0.077 | 0.077 | 0 | 0.54 | 0.75 | 0.57 | 0.68 | 0.55 | 0.47 |
| 2 | 127454230 | rs6568 | -0.21 | 1.447 | -0.165 | 1.145 | 0.129 | 0.337 | 0.337 | 0.391 | 0.15 | 0.44 | 0.37 | 0.43 | 0.22 | 0.13 |
| 2 | 127456021 | rs6732528 | -0.8 | 1.298 | -0.068 | 1.046 | 0.069 | 0.153 | 0.153 | 0.334 | 0.21 | 0.56 | 0.41 | 0.35 | 0.29 | 0.17 |
| 2 | 127456098 | rs6431172 | -1.285 | 1.189 | -0.203 | 0.899 | 0.01 | 0.041 | 0.041 | 0 | 0.55 | 0.68 | 0.56 | 0.67 | 0.56 | 0.47 |
| 2 | 127457468 | rs17014263 | -0.467 | 1.091 | -0.041 | 0.651 | 0.032 | -0.054 | -0.054 | 1.443 | 0.1 | 0.37 | 0.07 | 0 | 0.11 | 0.05 |
| 2 | 127459321 | rs1000858 | -0.044 | 0.958 | -0.698 | 0.857 | 0.009 | 0.012 | 0.012 | 0.181 | 0.33 | 0.13 | 0.15 | 0.32 | 0.26 | 0.28 |
| 2 | 127463952 | rs10187088 | -0.531 | 0.851 | -0.678 | 0.462 | -0.009 | 0 | 0 | 0 | 0.41 | 0.7 | 0.38 | 0.32 | 0.45 | 0.5 |
| 2 | 127464740 | rs2404962 | -1.515 | 0.726 | -0.733 | 0.377 | 0.005 | 0.025 | 0.025 | 0 | 0.69 | 0.81 | 0.81 | 0.78 | 0.68 | 0.76 |
| 2 | 127465648 | rs881944 | -0.22 | 0.701 | -0.578 | 0.43 | 0.015 | -0.085 | -0.085 | 0 | 0.38 | 0.46 | 0.32 | 0.11 | 0.43 | 0.47 |
| 2 | 127469104 | rs4662687 | -0.021 | 0.899 | -0.559 | 1.26 | -0.006 | -0.066 | -0.066 | 1.568 | 0.28 | 0.11 | 0.43 | 0.46 | 0.23 | 0.26 |

| | | | | | | | | | | | | | | | | |
|---|---|---|---|---|---|---|---|---|---|---|---|---|---|---|---|---|
| 2 | 127473880 | rs10164664 | -0.266 | 0.934 | -0.582 | 0.921 | 0.004 | 0.001 | 0.001 | 0 | 0.3 | 0.27 | 0.47 | 0.67 | 0.23 | 0.29 |
| 2 | 127497136 | rs10210965 | -0.641 | 0.802 | -0.848 | 0.563 | -0.006 | -0.004 | -0.004 | 0 | 0.43 | 0.64 | 0.55 | 0.53 | 0.42 | 0.44 |
| 2 | 127503457 | rs11674717 | -0.141 | 0.203 | -1.169 | 0.17 | 0.031 | 0.075 | 0.075 | 0.861 | 0.24 | 0.1 | 0.14 | 0.05 | 0.19 | 0.17 |
| 2 | 127503533 | rs4663044 | 1.03 | 0.323 | -0.849 | 0.219 | 0 | -0.001 | -0.001 | 0 | 0.36 | 0.37 | 0.23 | 0.29 | 0.36 | 0.42 |
| 2 | 127511804 | rs4663045 | 0.955 | 0.253 | -0.729 | 0.194 | 0.01 | 0.016 | 0.016 | 0 | 0.37 | 0.32 | 0.23 | 0.29 | 0.36 | 0.42 |
| 2 | 127512967 | rs4663046 | -0.198 | 0.649 | -0.617 | 0.544 | 0.026 | 0.018 | 0.018 | 0.56 | 0.66 | 0.52 | 0.45 | 0.51 | 0.62 | 0.63 |
| 2 | 127514060 | rs12613274 | -0.901 | 0.79 | -1.053 | 0.511 | -0.091 | -0.092 | -0.092 | 0 | 0.74 | 0.95 | 0.5 | 0.52 | 0.67 | 0.66 |
| 2 | 127515846 | rs1914458 | -0.385 | 0.702 | -0.034 | 0.586 | -0.011 | 0.002 | 0.002 | 0 | 0.32 | 0.41 | 0.23 | 0.21 | 0.26 | 0.21 |
| 2 | 127516506 | rs6732888 | -0.878 | 0.662 | -0.378 | 0.569 | -0.007 | -0.002 | -0.002 | 0 | 0.35 | 0.54 | 0.26 | 0.22 | 0.31 | 0.24 |
| 2 | 127524084 | rs7581770 | 0.129 | 0.444 | -1.018 | 0.268 | 0.033 | 0.06 | 0.06 | 0.797 | 0.25 | 0.12 | 0.16 | 0.08 | 0.19 | 0.19 |
| 2 | 127524572 | rs12692118 | -0.438 | 1.003 | -1.339 | 0.578 | -0.12 | -0.109 | -0.109 | 0.306 | 0.77 | 0.98 | 0.52 | 0.61 | 0.71 | 0.69 |
| 2 | 127524945 | rs11690189 | 0.171 | 0.469 | -1.25 | 0.36 | -0.029 | -0.024 | -0.024 | 0 | 0.63 | 0.8 | 0.41 | 0.52 | 0.52 | 0.43 |
| 2 | 127525025 | rs2176427 | 0.31 | 0.272 | -1.225 | 0.203 | -0.004 | 0.002 | 0.002 | 0.323 | 0.6 | 0.81 | 0.4 | 0.52 | 0.5 | 0.43 |
| 2 | 127527100 | rs10210850 | -0.296 | 1.082 | 0.014 | -0.111 | 0.125 | -0.071 | -0.071 | 0 | 0.21 | 0.5 | 0.43 | 0.11 | 0.29 | 0.37 |
| 2 | 127528413 | rs13013047 | 0.621 | 1.316 | 0.279 | -0.121 | -0.014 | 0.012 | 0.012 | 0 | 0.15 | 0.12 | 0.36 | 0.09 | 0.2 | 0.33 |
| 2 | 127529784 | rs4344883 | -0.33 | 1.318 | 0.304 | -0.127 | 0.153 | -0.073 | -0.073 | 0.717 | 0.21 | 0.45 | 0.42 | 0.11 | 0.3 | 0.36 |
| 2 | 127533079 | rs6431176 | 0.243 | 0.533 | 0.057 | -0.151 | 0 | -0.001 | -0.001 | 0 | 0.58 | 0.55 | 0.63 | 0.51 | 0.6 | 0.63 |
| 2 | 127540119 | rs12373775 | 1.328 | 0.437 | -0.34 | -0.275 | 0.061 | -0.012 | -0.012 | 0.166 | 0.41 | 0.12 | 0.2 | 0.37 | 0.38 | 0.3 |
| 2 | 127548764 | rs4366852 | -0.152 | 0.457 | 0.24 | -0.292 | 0.01 | 0.028 | 0.028 | 0.801 | 0.63 | 0.53 | 0.33 | 0.48 | 0.53 | 0.43 |
| 2 | 127550278 | rs7601320 | -0.216 | 0.475 | 0.196 | -0.293 | -0.002 | -0.005 | -0.005 | 0 | 0.64 | 0.63 | 0.35 | 0.48 | 0.53 | 0.43 |
| 2 | 127552872 | rs4075997 | 0.052 | 0.681 | 0.444 | -0.155 | 0 | -0.003 | -0.003 | 0 | 0.54 | 0.67 | 0.32 | 0.47 | 0.51 | 0.44 |
| 2 | 127555838 | rs12998207 | 0.37 | 0.712 | 0.458 | -0.149 | 0.03 | 0.004 | 0.004 | 0 | 0.51 | 0.63 | 0.29 | 0.48 | 0.44 | 0.41 |
| 2 | 127556299 | rs4402698 | -0.917 | 0.752 | 0.593 | -0.196 | 0.016 | -0.03 | -0.03 | 2.814 | 0.83 | 0.81 | 0.63 | 0.93 | 0.77 | 0.67 |
| 2 | 127556418 | rs4413104 | 0.566 | 0.285 | 0.914 | 0.25 | 0.053 | 0.007 | 0.007 | 0.42 | 0.41 | 0.13 | 0.21 | 0.35 | 0.38 | 0.31 |
| 2 | 127556548 | rs4322791 | -0.249 | 0.325 | 0.881 | 0.253 | 0.016 | 0.003 | 0.003 | 0 | 0.51 | 0.61 | 0.29 | 0.47 | 0.43 | 0.41 |
| 2 | 127565232 | rs4405698 | -0.414 | 0.35 | 0.814 | 0.241 | -0.001 | -0.001 | -0.001 | 0 | 0.52 | 0.57 | 0.3 | 0.47 | 0.5 | 0.44 |
| 2 | 127576875 | rs10181296 | 0.274 | 0.659 | 0.771 | 0.438 | 0.008 | -0.001 | -0.001 | 0 | 0.34 | 0.24 | 0.19 | 0.33 | 0.32 | 0.31 |
| 2 | 127577393 | rs7557097 | -0.414 | 0.67 | 0.615 | 0.433 | 0.007 | -0.002 | -0.002 | 0 | 0.43 | 0.59 | 0.24 | 0.45 | 0.37 | 0.4 |
| 2 | 127577888 | rs11891357 | -0.494 | 0.688 | 0.473 | 0.437 | 0.024 | -0.004 | -0.004 | 0 | 0.44 | 0.67 | 0.25 | 0.45 | 0.37 | 0.4 |
| 2 | 127578744 | rs10167103 | -0.593 | 0.468 | 0.476 | 0.487 | 0.003 | 0.004 | 0.004 | 0 | 0.46 | 0.65 | 0.28 | 0.45 | 0.45 | 0.44 |
| 2 | 127582332 | rs10177866 | 0.175 | 0.308 | 0.556 | 0.508 | -0.009 | -0.003 | -0.003 | 0 | 0.34 | 0.36 | 0.19 | 0.32 | 0.33 | 0.28 |
| 2 | 127583622 | rs4072704 | 0.073 | -0.129 | 1.014 | 0.388 | -0.003 | 0.002 | 0.002 | 0 | 0.4 | 0.34 | 0.23 | 0.32 | 0.44 | 0.32 |
| 2 | 127587337 | rs10197495 | -0.666 | -0.143 | 1.198 | 0.423 | 0.003 | -0.002 | -0.002 | 0 | 0.48 | 0.7 | 0.3 | 0.46 | 0.5 | 0.41 |
| 2 | 127593168 | rs7592735 | -0.048 | -0.165 | 1.131 | 0.064 | 0 | -0.002 | -0.002 | 0 | 0.41 | 0.42 | 0.25 | 0.32 | 0.44 | 0.32 |
| 2 | 127594176 | rs13405490 | -0.12 | -0.215 | 1.059 | 0.054 | 0.003 | -0.007 | -0.007 | 0 | 0.42 | 0.45 | 0.25 | 0.32 | 0.44 | 0.32 |
| 2 | 127597911 | rs13419545 | 0.553 | -0.16 | 0.823 | -0.171 | -0.092 | -0.009 | -0.009 | 0.055 | 0.26 | 0.08 | 0.19 | 0.22 | 0.39 | 0.19 |
| 2 | 127604576 | rs7564109 | -2.275 | -1.166 | 1.921 | -0.145 | -0.001 | -0.033 | -0.033 | 0 | 0.13 | 0.44 | 0.08 | 0.03 | 0.06 | 0.11 |
| 2 | 127609718 | rs11677887 | -1.23 | -1.299 | 1.483 | -0.313 | -0.016 | -0.094 | -0.094 | 0 | 0.17 | 0.29 | 0.1 | 0.01 | 0.14 | 0.14 |
| 2 | 127610281 | rs11678173 | -1.064 | -1.299 | 1.106 | -0.277 | -0.006 | -0.051 | -0.051 | 0 | 0.15 | 0.21 | 0.09 | 0.01 | 0.14 | 0.13 |
| 2 | 127619117 | rs4637050 | 1.125 | -1.438 | 1.359 | -0.296 | -0.044 | -0.104 | -0.104 | 0 | 0.82 | 0.62 | 0.91 | 1 | 0.88 | 0.9 |
| 2 | 127620861 | rs4616428 | 2.04 | -1.818 | 1.732 | -0.43 | -0.027 | -0.05 | -0.05 | 0 | 0.72 | 0.33 | 0.85 | 0.93 | 0.84 | 0.77 |
| 2 | 127621089 | rs12692119 | 1.276 | -1.814 | 1.845 | -0.428 | -0.057 | -0.127 | -0.127 | 0 | 0.81 | 0.46 | 0.89 | 1 | 0.88 | 0.9 |
| 2 | 127622469 | rs13398472 | -0.467 | -1.704 | 1.636 | -0.348 | 0.005 | 0.004 | 0.004 | 0 | 0.08 | 0.09 | 0.03 | 0.07 | 0.04 | 0.13 |
| 2 | 127631859 | rs7423827 | 1.55 | -1.723 | 1.662 | -0.363 | -0.048 | -0.056 | -0.056 | 0 | 0.86 | 0.54 | 0.93 | 0.93 | 0.95 | 0.85 |
| 2 | 127635553 | rs7425789 | 0.845 | -1.761 | 1.823 | -0.391 | -0.049 | -0.115 | -0.115 | 0 | 0.84 | 0.54 | 0.9 | 1 | 0.89 | 0.92 |
| 2 | 127639995 | rs4663054 | 1.693 | -1.874 | 2.081 | 0.058 | -0.061 | 0.047 | 0.047 | 1.225 | 0.86 | 0.36 | 0.89 | 0.83 | 0.95 | 0.86 |
| 2 | 127644785 | rs10928992 | 1.138 | -2.233 | 1.135 | -0.067 | -0.104 | 0.001 | 0.001 | 0 | 0.34 | 0.09 | 0.37 | 0.31 | 0.52 | 0.37 |
| 2 | 127645411 | rs12329197 | -0.259 | -2.217 | 2.109 | -0.14 | -0.012 | -0.005 | -0.005 | 0.048 | 0.08 | 0.14 | 0.04 | 0.07 | 0.04 | 0.13 |
| 2 | 127646013 | rs4663057 | 2.387 | -2.282 | 2.113 | -0.187 | 0.02 | 0.012 | 0.012 | 0.404 | 0.72 | 0.26 | 0.81 | 0.83 | 0.85 | 0.77 |
| 2 | 127653076 | rs13393833 | -0.258 | -2.246 | 2.012 | -0.101 | -0.012 | 0.002 | 0.002 | 0 | 0.08 | 0.13 | 0.04 | 0.09 | 0.04 | 0.13 |
| 2 | 127653376 | rs12469732 | 1.495 | -2.238 | 2.004 | -0.124 | -0.048 | -0.011 | -0.011 | 0.537 | 0.82 | 0.57 | 0.86 | 0.9 | 0.89 | 0.9 |
| 2 | 127659845 | rs17743968 | -0.737 | -2.262 | 1.26 | -0.076 | 0.016 | 0.056 | 0.056 | 0 | 0.13 | 0.03 | 0.08 | 0.01 | 0.1 | 0.09 |
| 2 | 127662386 | rs6733255 | 1.988 | -2.303 | 1.289 | -0.055 | 0.018 | -0.012 | -0.012 | 0 | 0.84 | 0.17 | 0.88 | 0.82 | 0.95 | 0.85 |
| 2 | 127665049 | rs951816 | -0.721 | -2.275 | 1.479 | -0.041 | -0.026 | 0.084 | 0.084 | 0.015 | 0.09 | 0.22 | 0.06 | 0.18 | 0.04 | 0.14 |
| 2 | 127665641 | rs6737070 | -2.367 | -2.303 | 1.515 | -0.069 | 0.033 | 0.011 | 0.011 | 0.431 | 0.27 | 0.71 | 0.19 | 0.19 | 0.15 | 0.26 |
| 2 | 127666335 | rs13411741 | -1.099 | -1.979 | 1.44 | 0.142 | -0.026 | 0.076 | 0.076 | 0 | 0.1 | 0.23 | 0.06 | 0.18 | 0.04 | 0.16 |
| 2 | 127673527 | rs11693919 | -0.994 | -2.089 | 1.027 | -0.059 | 0.005 | 0.004 | 0.004 | 0 | 0.22 | 0.21 | 0.12 | 0.14 | 0.16 | 0.24 |
| 2 | 127679248 | rs4663061 | 1.323 | -2.069 | 0.187 | -0.076 | 0.025 | 0.024 | 0.024 | 1.571 | 0.7 | 0.28 | 0.83 | 0.87 | 0.84 | 0.75 |
| 2 | 127688836 | rs13385899 | -1.379 | -1.949 | 0.082 | -0.001 | -0.006 | -0.002 | -0.002 | 0 | 0.28 | 0.64 | 0.18 | 0.18 | 0.17 | 0.26 |
| 2 | 127690797 | rs12616704 | -0.85 | -2.267 | -0.035 | -0.142 | 0.013 | -0.003 | -0.003 | 1.675 | 0.22 | 0.15 | 0.19 | 0.16 | 0.15 | 0.2 |
| 2 | 127696077 | rs4662692 | 0.872 | -2.345 | 0.773 | -0.325 | -0.077 | -0.074 | -0.074 | 2.16 | 0.75 | 0.45 | 0.86 | 0.9 | 0.86 | 0.71 |
| 2 | 127710392 | rs11887389 | -0.724 | -2.317 | 0.772 | -0.315 | 0.009 | -0.01 | -0.01 | 0.91 | 0.2 | 0.13 | 0.1 | 0 | 0.14 | 0.07 |
| 2 | 127724610 | rs4662695 | 0.705 | -2.204 | 0.94 | -0.222 | -0.066 | -0.039 | -0.039 | 0 | 0.81 | 0.67 | 0.89 | 0.9 | 0.91 | 0.73 |
| 2 | 127724850 | rs4663077 | 0.705 | -2.204 | 0.94 | -0.222 | -0.066 | -0.039 | -0.039 | 0 | 0.81 | 0.67 | 0.89 | 0.9 | 0.91 | 0.73 |
| 2 | 127732462 | rs12999411 | -0.305 | -1.712 | 0.155 | -0.361 | 0.015 | 0.035 | 0.035 | 0 | 0.18 | 0.07 | 0.11 | 0 | 0.17 | 0.09 |
| 2 | 127738513 | rs574015 | -0.09 | -1.191 | 0.109 | 0.136 | -0.011 | -0.004 | -0.004 | 0 | 0.44 | 0.54 | 0.52 | 0.64 | 0.66 | 0.37 |
| 2 | 127740315 | rs544587 | 0.133 | -0.929 | -0.879 | 0.253 | 0.017 | -0.009 | -0.009 | 0 | 0.39 | 0.15 | 0.48 | 0.65 | 0.63 | 0.36 |
| 2 | 127741821 | rs556902 | 0.204 | -0.942 | -0.989 | 0.286 | 0.018 | 0.003 | 0.003 | 0 | 0.4 | 0.24 | 0.49 | 0.65 | 0.63 | 0.37 |
| 2 | 127756813 | rs9287564 | -0.957 | -0.531 | -0.78 | 0.552 | -0.006 | 0.001 | 0.001 | 0 | 0.15 | 0.15 | 0.18 | 0.2 | 0.08 | 0.25 |

| | | | | | | | | | | | | | | | | |
|---|---|---|---|---|---|---|---|---|---|---|---|---|---|---|---|---|
| 2 | 127756954 | rs10178199 | -0.824 | -0.537 | -0.765 | 0.548 | 0.004 | -0.006 | -0.006 | 0 | 0.14 | 0.11 | 0.18 | 0.2 | 0.08 | 0.25 |
| 2 | 127757411 | rs754737 | -1.778 | 0.638 | -0.739 | -0.098 | -0.012 | 0.016 | 0.016 | 0 | 0.37 | 0.7 | 0.34 | 0.21 | 0.39 | 0.41 |
| 2 | 127758448 | rs12993006 | -0.531 | 0.962 | -0.678 | -0.113 | 0.2 | -0.089 | -0.089 | 0.016 | 0.13 | 0.55 | 0.11 | 0.01 | 0.22 | 0.05 |
| 2 | 127762988 | rs10194509 | -1.313 | 0.706 | -1.293 | 0.188 | -0.003 | 0 | 0 | 0.188 | 0.34 | 0.7 | 0.32 | 0.31 | 0.38 | 0.41 |
| 2 | 127771006 | rs9287568 | -1.144 | 0.921 | -1.239 | 0.227 | -0.026 | 0.031 | 0.031 | 0 | 0.36 | 0.8 | 0.41 | 0.23 | 0.39 | 0.4 |
| 2 | 127773880 | rs11899372 | -1.015 | 0.925 | -0.958 | 0.462 | 0 | -0.001 | -0.001 | 0 | 0.06 | 0.06 | 0.04 | 0.08 | 0.05 | 0.15 |
| 2 | 127777100 | rs934827 | -1.215 | 1.145 | -0.957 | 0.889 | -0.002 | -0.002 | -0.002 | 0 | 0.23 | 0.24 | 0.32 | 0.35 | 0.23 | 0.35 |
| 2 | 127778235 | rs7573824 | -0.323 | 1.042 | -1.079 | 0.89 | -0.005 | -0.003 | -0.003 | 2.399 | 0.38 | 0.54 | 0.43 | 0.38 | 0.37 | 0.47 |
| 2 | 127780904 | rs6713701 | -1.144 | 0.861 | 0.058 | 0.825 | -0.001 | -0.001 | -0.001 | 0 | 0.53 | 0.49 | 0.4 | 0.41 | 0.52 | 0.53 |
| 2 | 127783891 | rs2404175 | 0.721 | 0.791 | 0.567 | 0.539 | -0.002 | 0.005 | 0.005 | 0 | 0.2 | 0.23 | 0.22 | 0.23 | 0.2 | 0.14 |
| 2 | 127785403 | rs2404176 | -1.639 | 0.793 | 0.504 | 0.538 | 0 | 0.009 | 0.009 | 0.406 | 0.53 | 0.6 | 0.41 | 0.41 | 0.52 | 0.53 |
| 2 | 127789641 | rs17694125 | 0.672 | 0.859 | 0.459 | 0.541 | -0.006 | -0.02 | -0.02 | 0 | 0.19 | 0.1 | 0.21 | 0.23 | 0.2 | 0.14 |
| 2 | 127796020 | rs12476995 | -1.981 | 0.79 | 0.522 | 0.534 | -0.008 | -0.018 | -0.018 | 0 | 0.58 | 0.75 | 0.56 | 0.56 | 0.56 | 0.71 |
| 2 | 127797114 | rs7605172 | -0.202 | 0.717 | 0.797 | 0.474 | 0.007 | -0.004 | -0.004 | 0 | 0.19 | 0.29 | 0.13 | 0.16 | 0.18 | 0.2 |
| 2 | 127797656 | rs11902275 | -0.954 | 0.692 | 0.933 | 0.878 | -0.004 | 0.005 | 0.005 | 0.998 | 0.37 | 0.4 | 0.43 | 0.44 | 0.38 | 0.51 |
| 2 | 127800981 | rs4662703 | -0.905 | 0.668 | 0.911 | 0.906 | 0.003 | -0.013 | -0.013 | 0 | 0.37 | 0.31 | 0.43 | 0.46 | 0.38 | 0.51 |
| 2 | 127804699 | rs12476339 | -0.984 | 0.686 | 0.892 | 1.108 | 0 | -0.001 | -0.001 | 0 | 0.38 | 0.37 | 0.43 | 0.41 | 0.38 | 0.5 |
| 2 | 127807019 | rs3768865 | -2.092 | 0.722 | 0.863 | 1.124 | -0.009 | -0.011 | -0.011 | 0.133 | 0.58 | 0.77 | 0.55 | 0.56 | 0.56 | 0.71 |
| 2 | 127810697 | rs17014818 | -0.949 | 0.732 | 0.743 | 1.118 | 0.004 | -0.003 | -0.003 | 0 | 0.37 | 0.34 | 0.43 | 0.41 | 0.37 | 0.49 |
| 2 | 127814781 | rs3754618 | -0.794 | 0.405 | 0.516 | 0.958 | 0.001 | 0.008 | 0.008 | 0.496 | 0.25 | 0.72 | 0.15 | 0.15 | 0.18 | 0.2 |
| 2 | 127814835 | rs3754617 | 0.884 | 0.426 | 0.459 | 1.004 | -0.024 | -0.054 | -0.054 | 0 | 0.18 | 0.05 | 0.21 | 0.24 | 0.19 | 0.15 |
| 2 | 127816424 | rs2276582 | -1.554 | 0.14 | 0.298 | 0.845 | 0.003 | 0.01 | 0.01 | 0.275 | 0.49 | 0.81 | 0.49 | 0.55 | 0.41 | 0.5 |
| 2 | 127821085 | rs2071268 | -1.011 | 0.1 | 0.205 | 0.861 | -0.001 | -0.017 | -0.017 | 0.269 | 0.38 | 0.15 | 0.4 | 0.39 | 0.37 | 0.5 |
| 2 | 127826533 | rs1060743 | -0.682 | 0.156 | 0.419 | 0.957 | -0.002 | -0.027 | -0.027 | 0 | 0.31 | 0.24 | 0.37 | 0.45 | 0.29 | 0.46 |
| 2 | 127829282 | rs3768857 | 0.78 | 0.134 | 0.21 | 1.039 | -0.032 | -0.058 | -0.058 | 0 | 0.17 | 0.05 | 0.19 | 0.3 | 0.19 | 0.13 |
| 2 | 127830299 | rs753431 | -1.21 | 0.011 | 0.723 | 0.875 | 0 | 0.003 | 0.003 | 0.457 | 0.5 | 0.49 | 0.48 | 0.46 | 0.51 | 0.64 |
| 2 | 127834136 | rs2276575 | -0.885 | 0.405 | 0.781 | 0.617 | -0.005 | -0.019 | -0.019 | 0 | 0.17 | 0.22 | 0.12 | 0.08 | 0.16 | 0.24 |
| 2 | 127834459 | rs7562227 | -1.631 | -0.064 | 0.339 | 0.612 | -0.025 | -0.015 | -0.015 | 1.526 | 0.57 | 0.82 | 0.48 | 0.36 | 0.47 | 0.7 |
| 2 | 127835005 | rs6750960 | -0.301 | 0.058 | 0.379 | 0.537 | -0.001 | -0.003 | -0.003 | 0 | 0.16 | 0.16 | 0.11 | 0.08 | 0.16 | 0.24 |
| 2 | 127837041 | rs13430599 | -0.51 | 0.063 | 0.313 | 0.539 | -0.005 | -0.02 | -0.02 | 2.803 | 0.17 | 0.23 | 0.12 | 0.08 | 0.16 | 0.24 |
| 2 | 127839781 | rs10194375 | -0.717 | 0.22 | 0.369 | 0.622 | -0.009 | 0.011 | 0.011 | 0 | 0.3 | 0.24 | 0.31 | 0.21 | 0.36 | 0.27 |
| 2 | 127841337 | rs13426725 | -0.102 | 0.187 | 0.366 | 0.644 | -0.003 | -0.017 | -0.017 | 0 | 0.17 | 0.21 | 0.12 | 0.09 | 0.16 | 0.21 |
| 2 | 127841769 | rs10200967 | -2.337 | 0.348 | 0.223 | 0.587 | 0.002 | -0.024 | -0.024 | 0 | 0.72 | 0.79 | 0.71 | 0.5 | 0.73 | 0.73 |
| 2 | 127841930 | rs17014923 | 0.814 | 0.331 | 0.136 | 0.538 | -0.004 | -0.015 | -0.015 | 0 | 0.24 | 0.29 | 0.17 | 0.09 | 0.17 | 0.28 |
| 2 | 127842383 | rs11691237 | 0.394 | 0.506 | -0.114 | 0.558 | -0.031 | 0.025 | 0.025 | 0 | 0.19 | 0.06 | 0.22 | 0.12 | 0.25 | 0.1 |
| 2 | 127844735 | rs6709337 | -1.02 | 0.503 | 0.117 | 0.755 | 0.002 | -0.005 | -0.005 | 0.051 | 0.57 | 0.66 | 0.6 | 0.49 | 0.57 | 0.59 |
| 2 | 127851697 | rs11678252 | -0.414 | 0.546 | 0.216 | 0.673 | -0.004 | 0.028 | 0.028 | 0 | 0.44 | 0.39 | 0.37 | 0.23 | 0.43 | 0.37 |
| 2 | 127855409 | rs13019394 | -1.134 | 0.604 | 0.118 | 0.698 | 0.001 | -0.028 | -0.028 | 0 | 0.58 | 0.75 | 0.6 | 0.49 | 0.57 | 0.57 |
| 2 | 127859441 | rs749008 | 0.058 | 0.436 | -0.019 | 0.571 | -0.005 | -0.002 | -0.002 | 0 | 0.36 | 0.45 | 0.4 | 0.32 | 0.33 | 0.38 |
| 2 | 127865694 | rs6751498 | 0.225 | 0.607 | -0.009 | 0.626 | -0.014 | -0.017 | -0.017 | 0 | 0.09 | 0.17 | 0.05 | 0.06 | 0.05 | 0.12 |
| 2 | 127868435 | rs6743470 | 0.761 | 0.653 | -0.093 | 0.614 | 0.005 | 0.009 | 0.009 | 0 | 0.29 | 0.28 | 0.17 | 0.13 | 0.2 | 0.27 |
| 2 | 127873035 | rs4663098 | -1.81 | 1.172 | -0.458 | 0.932 | 0.01 | -0.022 | -0.022 | 0 | 0.74 | 0.84 | 0.7 | 0.61 | 0.75 | 0.8 |
| 2 | 127878839 | rs10929009 | -1.713 | 1.168 | -0.405 | 0.847 | 0.002 | -0.003 | -0.003 | 0 | 0.73 | 0.76 | 0.7 | 0.61 | 0.75 | 0.8 |
| 2 | 127887482 | rs4663101 | 0.278 | 1.091 | -0.19 | 0.595 | -0.001 | -0.005 | -0.005 | 0 | 0.49 | 0.42 | 0.6 | 0.7 | 0.52 | 0.57 |
| 2 | 127889637 | rs7561528 | -0.914 | 1.457 | 0.1 | 1.016 | -0.006 | 0.005 | 0.005 | 0 | 0.23 | 0.19 | 0.26 | 0.12 | 0.32 | 0.13 |
| 2 | 127889932 | rs6710467 | 0.08 | 1.227 | -0.043 | 1.158 | 0.018 | 0.085 | 0.085 | 0 | 0.11 | 0.2 | 0.19 | 0.22 | 0.13 | 0.12 |
| 2 | 127894615 | rs744373 | -0.477 | 1.227 | -0.32 | 0.928 | 0.024 | 0.038 | 0.038 | 0.047 | 0.25 | 0.52 | 0.39 | 0.34 | 0.27 | 0.22 |
| 2 | 127895573 | rs1866235 | 0.577 | 1.129 | -1.211 | 0.718 | -0.017 | -0.051 | -0.051 | 0 | 0.3 | 0.15 | 0.47 | 0.58 | 0.34 | 0.32 |
| 2 | 127895655 | rs1866236 | -0.114 | 0.732 | -0.756 | 0.018 | -0.009 | -0.032 | -0.032 | 0 | 0.68 | 0.59 | 0.8 | 0.89 | 0.72 | 0.7 |
| 2 | 127896232 | rs1469979 | -0.544 | 0.797 | -1.223 | -0.082 | -0.001 | 0.024 | 0.024 | 0 | 0.25 | 0.23 | 0.15 | 0.08 | 0.25 | 0.21 |
| 2 | 127897320 | rs3943703 | 0.397 | 0.429 | -1.825 | 0.31 | 0.02 | -0.055 | -0.055 | 0 | 0.3 | 0.15 | 0.28 | 0.46 | 0.26 | 0.35 |
| 2 | 127898467 | rs17701375 | 0.736 | 0.789 | -2.342 | 0.233 | -0.069 | -0.02 | -0.02 | 0 | 0.28 | 0.09 | 0.23 | 0.25 | 0.42 | 0.21 |
| 2 | 127901657 | rs2028211 | 0.856 | 0.872 | -2.358 | 0.234 | -0.051 | -0.022 | -0.022 | 0.014 | 0.27 | 0.11 | 0.22 | 0.25 | 0.39 | 0.22 |
| 2 | 127903692 | rs2118507 | -0.198 | 0.714 | -1.793 | -0.065 | -0.001 | 0 | 0 | 0 | 0.7 | 0.36 | 0.57 | 0.71 | 0.75 | 0.71 |
| 2 | 127905440 | rs6761675 | 0.57 | 0.824 | -1.355 | 0.211 | 0.02 | 0.075 | 0.075 | 0 | 0.29 | 0.13 | 0.13 | 0.07 | 0.22 | 0.33 |
| 2 | 127907061 | rs6758116 | -1.423 | 0.866 | -1.304 | 0.252 | -0.014 | -0.013 | -0.013 | 0 | 0.92 | 0.95 | 0.71 | 0.83 | 0.89 | 0.91 |
| 2 | 127909259 | rs2118508 | 1.596 | 0.776 | -1.291 | 0.656 | 0.002 | -0.025 | -0.025 | 0 | 0.27 | 0.07 | 0.24 | 0.31 | 0.25 | 0.2 |
| 2 | 127910264 | rs2896885 | -0.171 | 0.85 | -1.172 | 0.678 | -0.014 | 0.049 | 0.049 | 0.45 | 0.63 | 0.82 | 0.59 | 0.76 | 0.67 | 0.58 |
| 2 | 127913044 | rs10496660 | -1.066 | 0.914 | -1.421 | 0.659 | -0.018 | -0.049 | -0.049 | 0.173 | 0.75 | 0.93 | 0.66 | 0.7 | 0.77 | 0.81 |
| 2 | 127913835 | rs729666 | 0.521 | 0.982 | -1.257 | 0.595 | -0.012 | -0.047 | -0.047 | 1.059 | 0.36 | 0.6 | 0.25 | 0.13 | 0.33 | 0.35 |
| 2 | 127915119 | rs11680864 | 2.016 | 1.052 | -1.283 | 0.673 | -0.017 | -0.058 | -0.058 | 0 | 0.16 | 0.04 | 0.19 | 0.31 | 0.15 | 0.17 |
| 2 | 127915335 | rs7422269 | -1.258 | 1.033 | -1.324 | 0.649 | -0.027 | -0.064 | -0.064 | 0.55 | 0.75 | 0.95 | 0.66 | 0.7 | 0.77 | 0.8 |
| 2 | 127919620 | rs905656 | 1.093 | 0.942 | -1.162 | 0.603 | 0.021 | 0.008 | 0.008 | 0 | 0.23 | 0.27 | 0.23 | 0.26 | 0.3 | 0.28 |
| 2 | 127919641 | rs11682330 | 0.324 | 0.471 | -0.886 | 0.335 | -0.019 | -0.063 | -0.063 | 0 | 0.36 | 0.53 | 0.24 | 0.13 | 0.32 | 0.35 |
| 2 | 127924714 | rs6730476 | 0.991 | 1.459 | -0.537 | 0.633 | 0.04 | 0.03 | 0.03 | 0.581 | 0.17 | 0.24 | 0.23 | 0.31 | 0.3 | 0.18 |
| 2 | 127924981 | rs6751149 | -0.375 | 1.19 | -0.61 | 0.507 | 0.009 | -0.002 | -0.002 | 0.147 | 0.4 | 0.49 | 0.39 | 0.42 | 0.49 | 0.28 |
| 2 | 127926992 | rs6738800 | 0.511 | 0.612 | -0.752 | 0.194 | -0.002 | -0.011 | -0.011 | 0 | 0.22 | 0.19 | 0.16 | 0.11 | 0.19 | 0.1 |
| 2 | 127927990 | rs6431231 | -0.097 | 0.456 | -0.821 | 0.394 | 0 | -0.005 | -0.005 | 0 | 0.42 | 0.35 | 0.47 | 0.48 | 0.41 | 0.6 |
| 2 | 127932213 | rs17015010 | 1.148 | 0.32 | -1.349 | 0.063 | -0.114 | -0.17 | -0.17 | 0 | 0.25 | 0.02 | 0.5 | 0.54 | 0.34 | 0.28 |

| | | | | | | | | | | | | | | | | |
|---|---|---|---|---|---|---|---|---|---|---|---|---|---|---|---|---|
| 2 | 127936284 | rs4510165 | 1.532 | 0.112 | -1.065 | -0.02 | 0.014 | 0.055 | 0.055 | 0 | 0.16 | 0.05 | 0.04 | 0 | 0.12 | 0.07 |
| 2 | 127936624 | rs2404529 | -0.764 | -0.211 | -0.874 | -0.337 | -0.01 | -0.014 | -0.014 | 0 | 0.75 | 0.84 | 0.72 | 0.66 | 0.71 | 0.63 |
| 2 | 127937551 | rs13035268 | 0.541 | -0.359 | -0.512 | -0.407 | -0.007 | -0.008 | -0.008 | 0 | 0.52 | 0.43 | 0.64 | 0.6 | 0.54 | 0.5 |
| 2 | 127940615 | rs7355659 | -0.534 | 0.217 | -0.605 | 0.013 | -0.086 | -0.109 | -0.109 | 2.789 | 0.76 | 0.93 | 0.73 | 0.66 | 0.7 | 0.64 |
| 2 | 127949198 | rs7606194 | -0.511 | 0.212 | -0.397 | 0.012 | -0.041 | -0.056 | -0.056 | 0.169 | 0.75 | 0.86 | 0.72 | 0.66 | 0.7 | 0.61 |
| 2 | 127951797 | rs12477559 | 0.877 | -0.18 | -0.314 | -0.287 | -0.014 | -0.024 | -0.024 | 0 | 0.34 | 0.34 | 0.57 | 0.56 | 0.43 | 0.3 |
| 2 | 127953054 | rs7568070 | 0.192 | -0.536 | 0.034 | -0.613 | -0.013 | -0.032 | -0.032 | 0.011 | 0.4 | 0.46 | 0.15 | 0.11 | 0.27 | 0.3 |
| 2 | 127956543 | rs11683427 | 0.449 | -0.464 | -0.148 | -0.743 | 0.043 | 0.088 | 0.088 | 0 | 0.36 | 0.15 | 0.12 | 0.09 | 0.26 | 0.29 |
| 2 | 127962493 | rs7585314 | -0.652 | -0.583 | 0.12 | -0.917 | 0.008 | 0 | 0 | 0 | 0.6 | 0.67 | 0.83 | 0.91 | 0.71 | 0.65 |
| 2 | 127962635 | rs7585198 | 0.904 | -0.341 | 0.238 | -0.716 | -0.03 | -0.028 | -0.028 | 0 | 0.35 | 0.25 | 0.62 | 0.61 | 0.47 | 0.31 |
| 2 | 127977118 | rs1504136 | -0.662 | -0.084 | 0.292 | -0.627 | 0 | -0.021 | -0.021 | 5.333 | 0.59 | 0.64 | 0.83 | 0.91 | 0.7 | 0.65 |
| 2 | 127987049 | rs4359587 | 0.806 | -0.053 | -0.169 | -0.594 | -0.053 | -0.065 | -0.065 | 0 | 0.16 | 0.05 | 0.18 | 0.3 | 0.23 | 0.33 |
| 2 | 127987391 | rs1566822 | 0.724 | -0.136 | -0.117 | -0.602 | -0.027 | -0.026 | -0.026 | 0 | 0.37 | 0.24 | 0.61 | 0.61 | 0.47 | 0.32 |
| 2 | 128002572 | rs4536600 | -0.469 | -0.128 | 0.104 | -0.798 | 0.046 | 0.103 | 0.103 | 0 | 0.55 | 0.35 | 0.77 | 0.89 | 0.67 | 0.65 |
| 2 | 128006078 | rs2174270 | 0.636 | -0.1 | 0.205 | -0.679 | -0.014 | -0.017 | -0.017 | 0 | 0.38 | 0.34 | 0.61 | 0.61 | 0.46 | 0.29 |
| 2 | 128006991 | rs6430935 | -0.157 | -0.252 | 0.552 | -0.834 | 0.041 | 0.103 | 0.103 | 1.26 | 0.25 | 0.44 | 0.24 | 0.39 | 0.29 | 0.48 |
| 2 | 128007686 | rs6430936 | -1.258 | -0.287 | 0.456 | -0.863 | 0.032 | 0.035 | 0.035 | 0 | 0.65 | 0.81 | 0.86 | 1 | 0.76 | 0.76 |
| 2 | 128015367 | rs4662718 | -1.088 | -0.225 | 0.28 | -0.856 | 0.003 | -0.066 | -0.066 | 1.029 | 0.63 | 0.73 | 0.86 | 1 | 0.75 | 0.77 |
| 2 | 128018192 | rs1566823 | 1.148 | -0.019 | 0.162 | -0.906 | -0.009 | -0.029 | -0.029 | 0 | 0.33 | 0.23 | 0.6 | 0.6 | 0.41 | 0.27 |
| 2 | 128023884 | rs1143407 | 0.823 | -0.018 | 0.184 | -0.901 | -0.031 | -0.042 | -0.042 | 0.164 | 0.18 | 0.1 | 0.19 | 0.29 | 0.23 | 0.38 |
| 2 | 128029503 | rs4150496 | 1.225 | -0.003 | 0.177 | -0.872 | -0.006 | -0.041 | -0.041 | 0 | 0.33 | 0.2 | 0.56 | 0.6 | 0.38 | 0.26 |
| 2 | 128058627 | rs13431911 | 0.894 | 0.085 | 0.328 | -0.742 | 0 | 0.048 | 0.048 | 0 | 0.06 | 0.19 | 0.04 | 0.11 | 0.05 | 0.11 |
| 2 | 128062048 | rs3088374 | 0.648 | 0.111 | 0.304 | -0.785 | 0 | 0 | 0 | 0 | 0.38 | 0.39 | 0.6 | 0.7 | 0.43 | 0.34 |
| 2 | 128063945 | rs10850 | -1.343 | 0.1 | 0.151 | -0.809 | 0.015 | -0.024 | -0.024 | 0 | 0.64 | 0.77 | 0.86 | 1 | 0.75 | 0.76 |
| 2 | 128074433 | rs13429925 | 0.64 | 0.117 | 0.148 | -0.804 | -0.029 | -0.041 | -0.041 | 0 | 0.19 | 0.14 | 0.19 | 0.29 | 0.23 | 0.38 |
| 2 | 128079806 | rs3732209 | 0.897 | 0.123 | 0.165 | -0.831 | 0.006 | -0.045 | -0.045 | 0 | 0.37 | 0.27 | 0.17 | 0.02 | 0.28 | 0.27 |
| 2 | 128090144 | rs10496661 | 1.031 | 0.114 | 0.153 | -0.833 | 0.015 | -0.024 | -0.024 | 0.698 | 0.36 | 0.23 | 0.14 | 0 | 0.25 | 0.24 |
| 2 | 128118553 | rs12479003 | 1.157 | 0.029 | 0.044 | -0.727 | -0.005 | -0.041 | -0.041 | 1.817 | 0.32 | 0.2 | 0.56 | 0.6 | 0.38 | 0.26 |
| 2 | 128151003 | rs11679414 | 0.785 | -0.059 | 0.144 | -0.951 | 0.005 | -0.054 | -0.054 | 2.538 | 0.37 | 0.27 | 0.17 | 0 | 0.28 | 0.24 |
| 2 | 128162623 | rs6430938 | 0.713 | -0.03 | -0.049 | -0.938 | -0.008 | -0.111 | -0.111 | 0 | 0.38 | 0.37 | 0.18 | 0 | 0.3 | 0.25 |
| 2 | 128169899 | rs6753288 | -0.549 | -0.065 | -0.157 | -0.868 | 0.026 | 0.154 | 0.154 | 0 | 0.53 | 0.74 | 0.28 | 0.13 | 0.42 | 0.38 |
| 2 | 128173417 | rs878461 | 0.105 | 0.003 | 0.072 | -0.685 | -0.013 | 0.056 | 0.056 | 0.066 | 0.09 | 0.26 | 0.04 | 0.12 | 0.05 | 0.08 |
| 2 | 128175862 | rs1799808 | 1.294 | -0.108 | -0.413 | -0.629 | -0.01 | -0.072 | -0.072 | 0 | 0.29 | 0.17 | 0.54 | 0.55 | 0.37 | 0.22 |
| 2 | 128175875 | rs1799809 | 0.073 | -0.122 | -0.44 | -0.702 | 0.024 | 0.162 | 0.162 | 0 | 0.47 | 0.26 | 0.71 | 0.82 | 0.59 | 0.62 |
| 2 | 128177377 | rs1158867 | 0.068 | -0.119 | -0.432 | -0.7 | 0.025 | 0.16 | 0.16 | 0 | 0.47 | 0.25 | 0.71 | 0.82 | 0.58 | 0.62 |
| 2 | 128178191 | rs2069912 | -0.282 | -0.091 | -0.403 | -0.687 | -0.004 | -0.013 | -0.013 | 0 | 0.27 | 0.29 | 0.2 | 0.45 | 0.26 | 0.51 |
| 2 | 128184770 | rs5937 | 0.94 | -0.234 | -0.346 | -0.857 | 0.001 | -0.071 | -0.071 | 0 | 0.38 | 0.35 | 0.21 | 0 | 0.31 | 0.24 |
| 2 | 128185688 | rs2069933 | 0.432 | 0.097 | -0.502 | -0.567 | -0.029 | -0.05 | -0.05 | 0.11 | 0.7 | 0.8 | 0.78 | 0.58 | 0.67 | 0.48 |
| 2 | 128189308 | rs1401296 | 0.462 | 0.097 | -0.491 | -0.567 | -0.025 | -0.044 | -0.044 | 0 | 0.7 | 0.78 | 0.77 | 0.58 | 0.67 | 0.48 |
| 2 | 128190055 | rs1568277 | 0.941 | 0.479 | -0.645 | -0.444 | -0.019 | -0.045 | -0.045 | 0 | 0.15 | 0.07 | 0.1 | 0.25 | 0.15 | 0.39 |
| 2 | 128193928 | rs10928766 | 0.978 | 0.498 | -0.625 | -0.371 | -0.013 | -0.027 | -0.027 | 0 | 0.14 | 0.1 | 0.1 | 0.28 | 0.15 | 0.38 |
| 2 | 128195387 | rs6754999 | 1.518 | 0.349 | -0.583 | -0.5 | -0.007 | -0.032 | -0.032 | 0 | 0.26 | 0.19 | 0.55 | 0.57 | 0.36 | 0.23 |
| 2 | 128195442 | rs6755028 | -0.304 | 0.386 | -0.544 | -0.55 | 0.036 | 0.064 | 0.064 | 0 | 0.48 | 0.42 | 0.74 | 0.85 | 0.65 | 0.63 |
| 2 | 128196424 | rs6749002 | 0.581 | 0.451 | -0.602 | -0.566 | -0.034 | -0.035 | -0.035 | 1.288 | 0.34 | 0.27 | 0.64 | 0.57 | 0.5 | 0.25 |
| 2 | 128196930 | rs13408910 | 0.21 | 0.313 | -0.459 | -0.551 | -0.001 | -0.004 | -0.004 | 0 | 0.41 | 0.47 | 0.65 | 0.79 | 0.52 | 0.58 |
| 2 | 128211219 | rs11679431 | 0.874 | 0.446 | -0.471 | -0.283 | -0.022 | -0.042 | -0.042 | 0 | 0.14 | 0.05 | 0.1 | 0.23 | 0.16 | 0.33 |
| 2 | 128212588 | rs777567 | 0.49 | 0.154 | -0.428 | -0.321 | -0.005 | 0.005 | 0.005 | 0 | 0.08 | 0.09 | 0.03 | 0.15 | 0.04 | 0.14 |
| 2 | 128225228 | rs7590705 | -1.027 | 0.042 | -0.75 | -0.792 | -0.009 | -0.023 | -0.023 | 0.522 | 0.55 | 0.78 | 0.41 | 0.43 | 0.5 | 0.59 |
| 2 | 128237754 | rs10928769 | 1.179 | -0.303 | -0.689 | -0.638 | -0.037 | -0.062 | -0.062 | 0.017 | 0.14 | 0.03 | 0.09 | 0.19 | 0.15 | 0.34 |
| 2 | 128238166 | rs11694581 | 0.587 | -0.31 | -0.751 | -0.725 | 0.034 | -0.016 | -0.016 | 0 | 0.53 | 0.22 | 0.67 | 0.58 | 0.62 | 0.43 |
| 2 | 128243139 | rs12476558 | 0.587 | -0.31 | -0.751 | -0.725 | 0.034 | -0.016 | -0.016 | 0 | 0.53 | 0.22 | 0.67 | 0.58 | 0.62 | 0.43 |
| 2 | 128259371 | rs1549767 | -0.202 | -0.347 | -0.547 | -0.739 | 0.002 | 0.002 | 0.002 | 0 | 0.67 | 0.34 | 0.78 | 0.77 | 0.77 | 0.77 |
| 2 | 128264246 | rs2078028 | 0.399 | -0.454 | -0.157 | -0.785 | -0.027 | 0.13 | 0.13 | 0.98 | 0.1 | 0.32 | 0.11 | 0.23 | 0.06 | 0.16 |
| 2 | 128264431 | rs334162 | -0.887 | -0.512 | -0.085 | -0.814 | -0.064 | 0.112 | 0.112 | 2.501 | 0.14 | 0.55 | 0.13 | 0.23 | 0.06 | 0.16 |
| 2 | 128265139 | rs12620037 | 1.304 | -0.424 | -0.28 | -0.776 | -0.037 | -0.062 | -0.062 | 1.203 | 0.14 | 0.03 | 0.1 | 0.19 | 0.15 | 0.34 |
| 2 | 128266533 | rs17015347 | 0.483 | -0.624 | -0.206 | -0.751 | -0.026 | 0.089 | 0.089 | 0.77 | 0.1 | 0.22 | 0.1 | 0.23 | 0.06 | 0.16 |
| 2 | 128279047 | rs777554 | -1.45 | -1.172 | -0.446 | -0.924 | 0 | 0.016 | 0.016 | 0 | 0.81 | 0.88 | 0.9 | 1 | 0.83 | 0.93 |
| 2 | 128288850 | rs6758364 | -0.43 | -1.149 | -0.459 | -0.906 | -0.009 | 0.018 | 0.018 | 0 | 0.69 | 0.84 | 0.82 | 0.81 | 0.7 | 0.58 |
| 2 | 128289963 | rs7570609 | -0.161 | -1.283 | -0.282 | -0.907 | -0.036 | 0.11 | 0.11 | 1.124 | 0.14 | 0.37 | 0.12 | 0.23 | 0.07 | 0.18 |
| 2 | 128292280 | rs2082182 | 1.279 | -1.306 | -0.554 | -0.91 | -0.033 | -0.067 | -0.067 | 0 | 0.13 | 0.01 | 0.08 | 0.19 | 0.13 | 0.35 |
| 2 | 128303003 | rs10460269 | 0.953 | -1.396 | -0.513 | -0.988 | 0.057 | -0.008 | -0.008 | 0 | 0.51 | 0.27 | 0.68 | 0.56 | 0.64 | 0.39 |
| 2 | 128303364 | rs6707186 | 0.108 | -1.379 | -0.515 | -0.962 | -0.037 | 0.078 | 0.078 | 1.067 | 0.13 | 0.28 | 0.1 | 0.24 | 0.07 | 0.17 |
| 2 | 128305196 | rs1019842 | -1.666 | -1.366 | -0.717 | -0.965 | -0.007 | 0.061 | 0.061 | 0 | 0.82 | 0.92 | 0.91 | 1 | 0.84 | 0.93 |
| 2 | 128308167 | rs6727506 | 0.385 | -1.288 | -0.775 | -0.97 | 0.013 | 0.017 | 0.017 | 0 | 0.61 | 0.35 | 0.74 | 0.76 | 0.71 | 0.75 |
| 2 | 128310381 | rs2116026 | 1.083 | -1.158 | -1.087 | -0.758 | -0.031 | -0.067 | -0.067 | 0 | 0.13 | 0.01 | 0.08 | 0.19 | 0.14 | 0.34 |
| 2 | 128319195 | rs13403811 | -0.429 | -0.966 | -0.675 | -1.301 | -0.044 | 0.119 | 0.119 | 1.154 | 0.14 | 0.35 | 0.11 | 0.24 | 0.07 | 0.17 |
| 2 | 128328375 | rs10803588 | 1.45 | -0.75 | -1.615 | -1.708 | -0.09 | -0.035 | -0.035 | 0.163 | 0.27 | 0.08 | 0.56 | 0.71 | 0.45 | 0.22 |
| 2 | 128328836 | rs10177052 | -1.111 | -0.819 | -1.804 | -1.946 | -0.062 | 0.146 | 0.146 | 0.011 | 0.78 | 0.98 | 0.89 | 1 | 0.78 | 0.91 |
| 2 | 128330458 | rs10803589 | -0.823 | -0.788 | -1.653 | -1.909 | -0.004 | 0.114 | 0.114 | 0 | 0.73 | 0.89 | 0.88 | 1 | 0.78 | 0.9 |

| | | | | | | | | | | | | | | | | |
|---|---|---|---|---|---|---|---|---|---|---|---|---|---|---|---|---|
| 2 | 128334413 | rs12999486 | 1.323 | -0.894 | -1.565 | -1.758 | -0.046 | -0.005 | -0.005 | 0 | 0.14 | 0.05 | 0.11 | 0.11 | 0.17 | 0.31 |
| 2 | 128334470 | rs968756 | 0.922 | -1.069 | -1.068 | -1.878 | -0.047 | -0.071 | -0.071 | 0 | 0.19 | 0.32 | 0.15 | 0.04 | 0.11 | 0.16 |
| 2 | 128336189 | rs6709113 | -1.487 | -0.807 | -1.337 | -1.779 | -0.032 | 0.143 | 0.143 | 0 | 0.79 | 0.99 | 0.92 | 1 | 0.83 | 0.92 |
| 2 | 128337512 | rs4611601 | -0.545 | -0.735 | -1.122 | -1.712 | 0.003 | 0.006 | 0.006 | 1.294 | 0.49 | 0.54 | 0.35 | 0.31 | 0.4 | 0.71 |
| 2 | 128340855 | rs11693296 | 0.747 | -0.543 | -1.301 | -1.497 | -0.038 | -0.12 | -0.12 | 0 | 0.24 | 0.35 | 0.15 | 0.01 | 0.16 | 0.22 |
| 2 | 128341603 | rs10928772 | 0.912 | 0.145 | -2.029 | -1.031 | -0.1 | -0.068 | -0.068 | 0.168 | 0.26 | 0.1 | 0.53 | 0.68 | 0.43 | 0.13 |
| 2 | 128350808 | rs4662743 | -0.979 | 0.009 | -2.171 | -0.933 | -0.129 | -0.053 | -0.053 | 0.043 | 0.71 | 0.93 | 0.45 | 0.31 | 0.53 | 0.81 |
| 2 | 128364684 | rs11691981 | 0.892 | 0.089 | -2.152 | -1.059 | -0.058 | -0.009 | -0.009 | 0 | 0.15 | 0.05 | 0.12 | 0.13 | 0.2 | 0.34 |
| 2 | 128366272 | rs17261971 | 0.921 | -0.306 | -2.475 | -1.425 | -0.007 | 0.108 | 0.108 | 0.059 | 0.21 | 0.06 | 0.08 | 0 | 0.16 | 0.11 |
| 2 | 128368235 | rs777433 | -0.485 | -0.317 | -2.09 | -1.495 | -0.012 | 0.065 | 0.065 | 0 | 0.4 | 0.37 | 0.25 | 0.13 | 0.38 | 0.44 |
| 2 | 128378563 | rs2245711 | -0.774 | -0.456 | -2.002 | -1.472 | -0.005 | 0.068 | 0.068 | 0 | 0.45 | 0.28 | 0.26 | 0.27 | 0.4 | 0.61 |
| 2 | 128387937 | rs13417920 | -0.047 | -0.457 | -2.243 | -1.512 | 0.022 | 0.062 | 0.062 | 0.032 | 0.3 | 0.11 | 0.14 | 0.14 | 0.22 | 0.23 |
| 2 | 128388790 | rs13025791 | 1.004 | -0.121 | -2.003 | -1.253 | -0.053 | -0.016 | -0.016 | 0.012 | 0.16 | 0.05 | 0.12 | 0.13 | 0.2 | 0.36 |
| 2 | 128388862 | rs13025959 | 1.02 | -0.122 | -2.003 | -1.253 | -0.053 | -0.016 | -0.016 | 0 | 0.16 | 0.05 | 0.12 | 0.13 | 0.2 | 0.36 |
| 2 | 128392547 | rs4662750 | -1.795 | -0.06 | -2.107 | -1.289 | -0.119 | -0.124 | -0.124 | 0 | 0.74 | 0.97 | 0.52 | 0.35 | 0.64 | 0.82 |
| 2 | 128393289 | rs2255177 | -1.157 | 0.281 | -1.429 | -0.893 | -0.001 | 0 | 0 | 0 | 0.33 | 0.32 | 0.27 | 0.28 | 0.34 | 0.55 |
| 2 | 128393800 | rs2255161 | 0.382 | 0.359 | -1.193 | -0.683 | 0.001 | -0.001 | -0.001 | 0 | 0.18 | 0.19 | 0.14 | 0.15 | 0.2 | 0.42 |
| 2 | 128394955 | rs11686946 | 0.455 | 0.378 | -1.402 | -0.624 | -0.017 | -0.004 | -0.004 | 1.203 | 0.18 | 0.11 | 0.12 | 0.19 | 0.2 | 0.42 |
| 2 | 128395498 | rs2288657 | 0.027 | 0.233 | -1.191 | -0.37 | -0.001 | 0.054 | 0.054 | 0 | 0.06 | 0.09 | 0.04 | 0.14 | 0.05 | 0.12 |
| 2 | 128398840 | rs2244871 | -0.605 | 0.19 | -1.368 | -0.648 | -0.001 | -0.002 | -0.002 | 0 | 0.5 | 0.5 | 0.31 | 0.33 | 0.46 | 0.62 |
| 2 | 128399414 | rs13419955 | -0.188 | 0.062 | -1.321 | -0.906 | -0.013 | -0.042 | -0.042 | 1.535 | 0.29 | 0.36 | 0.14 | 0.14 | 0.22 | 0.14 |
| 2 | 128400992 | rs2244513 | 0.527 | 0.191 | -1.772 | -0.756 | 0 | 0.009 | 0.009 | 0 | 0.21 | 0.12 | 0.12 | 0.16 | 0.17 | 0.16 |
| 2 | 128401091 | rs2244507 | -0.956 | 0.31 | -2.254 | -0.726 | -0.052 | -0.055 | -0.055 | 0.89 | 0.68 | 0.93 | 0.48 | 0.34 | 0.61 | 0.8 |
| 2 | 128403997 | rs11687066 | 0.217 | 0.64 | -1.678 | -0.766 | 0.005 | -0.005 | -0.005 | 0.009 | 0.21 | 0.28 | 0.18 | 0.15 | 0.25 | 0.4 |
| 2 | 128404998 | rs2244042 | -0.661 | 0.247 | -2.269 | -0.767 | -0.037 | 0.001 | 0.001 | 0 | 0.61 | 0.94 | 0.47 | 0.3 | 0.57 | 0.76 |
| 2 | 128405921 | rs2243934 | -0.381 | 0.209 | -1.437 | -0.54 | 0.005 | 0.017 | 0.017 | 0 | 0.06 | 0.08 | 0.05 | 0.12 | 0.07 | 0.14 |
| 2 | 128409419 | rs13021001 | -0.641 | 0.217 | -1.772 | -0.65 | -0.018 | -0.03 | -0.03 | 0 | 0.14 | 0.03 | 0.12 | 0.16 | 0.16 | 0.27 |
| 2 | 128422745 | rs10185637 | -0.79 | -0.483 | -1.389 | -1.349 | 0.015 | 0.024 | 0.024 | 0 | 0.44 | 0.41 | 0.24 | 0.17 | 0.36 | 0.34 |
| 2 | 128433897 | rs3806586 | -0.145 | -0.353 | -0.656 | -1.236 | -0.001 | -0.001 | -0.001 | 0 | 0.14 | 0.15 | 0.16 | 0.15 | 0.16 | 0.25 |
| 2 | 128451894 | rs6430945 | -0.874 | -1.166 | -0.091 | -1.272 | 0.01 | 0.006 | 0.006 | 0 | 0.45 | 0.61 | 0.29 | 0.32 | 0.33 | 0.55 |
| 2 | 128457941 | rs7580688 | -0.94 | -1.239 | -0.084 | -1.3 | 0.009 | 0.005 | 0.005 | 0 | 0.45 | 0.61 | 0.31 | 0.32 | 0.33 | 0.56 |
| 2 | 128465547 | rs11681053 | 0.359 | -1.323 | 0.004 | -1.345 | 0 | 0 | 0 | 0 | 0.15 | 0.11 | 0.14 | 0.15 | 0.15 | 0.28 |
| 2 | 128468363 | rs2270160 | -1.221 | -1.051 | 0.22 | -1.552 | 0.02 | 0.025 | 0.025 | 0 | 0.5 | 0.62 | 0.35 | 0.36 | 0.4 | 0.62 |
| 2 | 128476484 | rs7595381 | -0.968 | -1.246 | 0.13 | -1.631 | -0.002 | -0.01 | -0.01 | 0 | 0.27 | 0.3 | 0.21 | 0.2 | 0.24 | 0.35 |
| 2 | 128480973 | rs17599575 | 1.9 | -1.176 | 0.1 | -1.589 | 0.008 | -0.013 | -0.013 | 0 | 0.05 | 0.06 | 0.05 | 0.04 | 0.07 | 0.06 |
| 2 | 128486506 | rs6430947 | 0.947 | -1.2 | 0.152 | -1.584 | -0.01 | -0.003 | -0.003 | 0 | 0.22 | 0.27 | 0.14 | 0.16 | 0.14 | 0.26 |
| 2 | 128495645 | rs6754215 | -1.618 | -1.331 | 0.104 | -1.698 | 0.005 | 0.014 | 0.014 | 0 | 0.56 | 0.66 | 0.37 | 0.36 | 0.44 | 0.63 |
| 2 | 128496524 | rs7603285 | 1.074 | -1.33 | 0.111 | -1.695 | -0.01 | -0.005 | -0.005 | 0 | 0.22 | 0.25 | 0.12 | 0.13 | 0.13 | 0.23 |
| 2 | 128498986 | rs800860 | -2.278 | -1.731 | -0.646 | -2.141 | -0.176 | -0.128 | -0.128 | 0 | 0.72 | 0.97 | 0.46 | 0.38 | 0.57 | 0.81 |
| 2 | 128516543 | rs7576459 | 0.814 | -1.682 | -0.392 | -2.043 | -0.002 | -0.002 | -0.002 | 0.152 | 0.16 | 0.22 | 0.15 | 0.15 | 0.15 | 0.28 |
| 2 | 128522207 | rs17534123 | 2.399 | -1.647 | -0.407 | -2.028 | 0.008 | -0.013 | -0.013 | 0 | 0.05 | 0.06 | 0.05 | 0.04 | 0.07 | 0.06 |
| 2 | 128522383 | rs11557685 | 2.399 | -1.647 | -0.407 | -2.028 | 0.008 | -0.013 | -0.013 | 0 | 0.05 | 0.06 | 0.05 | 0.04 | 0.07 | 0.06 |
| 2 | 128549177 | rs6750523 | -0.142 | -1.855 | -0.49 | -2.125 | -0.024 | -0.014 | -0.014 | 0.027 | 0.38 | 0.54 | 0.27 | 0.27 | 0.28 | 0.51 |
| 2 | 128552970 | rs6722353 | 0.669 | -1.731 | -0.483 | -2.084 | -0.006 | -0.011 | -0.011 | 0.005 | 0.27 | 0.3 | 0.17 | 0.17 | 0.2 | 0.29 |
| 2 | 128567755 | rs335136 | 1.161 | -1.76 | -0.331 | -2.137 | -0.012 | -0.056 | -0.056 | 0 | 0.16 | 0.22 | 0.09 | 0.02 | 0.13 | 0.19 |
| 2 | 128569248 | rs7581360 | 0.738 | -1.608 | -0.529 | -2.025 | -0.019 | -0.032 | -0.032 | 0 | 0.27 | 0.39 | 0.18 | 0.17 | 0.2 | 0.28 |
| 2 | 128572853 | rs335128 | -0.865 | -1.479 | -0.553 | -1.98 | -0.019 | -0.055 | -0.055 | 0.636 | 0.37 | 0.43 | 0.26 | 0.15 | 0.33 | 0.46 |
| 2 | 128575580 | rs10515925 | 0.785 | -1.408 | -0.902 | -1.922 | 0 | 0.001 | 0.001 | 0 | 0.17 | 0.06 | 0.15 | 0.13 | 0.16 | 0.32 |
| 2 | 128576820 | rs10200640 | -0.822 | -1.487 | -0.569 | -1.902 | -0.008 | 0 | 0 | 0 | 0.31 | 0.38 | 0.27 | 0.31 | 0.28 | 0.54 |
| 2 | 128577892 | rs335130 | -2.091 | -0.338 | -1.032 | -1.921 | -0.059 | 0.01 | 0.01 | 0 | 0.91 | 0.99 | 0.8 | 0.97 | 0.76 | 0.92 |
| 2 | 128581226 | rs6743767 | -0.776 | -0.825 | -0.859 | -1.938 | -0.002 | -0.003 | -0.003 | 1.545 | 0.52 | 0.5 | 0.34 | 0.35 | 0.42 | 0.61 |
| 2 | 128587317 | rs7562358 | -1.056 | -0.525 | -0.155 | -1.538 | 0.005 | -0.004 | -0.004 | 0 | 0.57 | 0.51 | 0.39 | 0.36 | 0.49 | 0.76 |
| 2 | 128591365 | rs11890668 | 1.485 | -0.473 | -0.383 | -1.427 | 0.017 | -0.046 | -0.046 | 2.858 | 0.17 | 0.01 | 0.09 | 0.13 | 0.1 | 0.16 |
| 2 | 128600609 | rs17600636 | -0.531 | -0.558 | -0.52 | -1.528 | 0 | -0.006 | -0.006 | 7.093 | 0.41 | 0.4 | 0.34 | 0.22 | 0.41 | 0.59 |
| 2 | 128611806 | rs3816127 | 1.085 | -0.574 | -0.811 | -1.475 | 0.045 | -0.015 | -0.015 | 0.238 | 0.2 | 0.05 | 0.1 | 0.14 | 0.1 | 0.16 |
| 2 | 128613156 | rs3771290 | -1.578 | -0.446 | -0.617 | -1.48 | -0.017 | -0.006 | -0.006 | 0.007 | 0.63 | 0.66 | 0.47 | 0.36 | 0.51 | 0.75 |
| 2 | 128618446 | rs3755305 | -0.471 | -0.189 | -0.737 | -1.223 | -0.003 | -0.073 | -0.073 | 0 | 0.36 | 0.07 | 0.34 | 0.32 | 0.33 | 0.66 |
| 2 | 128620372 | rs2261682 | 1.607 | -0.007 | -0.889 | -1.196 | -0.182 | -0.126 | -0.126 | 0 | 0.37 | 0.03 | 0.5 | 0.62 | 0.49 | 0.26 |
| 2 | 128621094 | rs1045832 | -1.219 | 0.474 | 0.037 | -0.671 | 0.005 | 0 | 0 | 0 | 0.42 | 0.71 | 0.42 | 0.37 | 0.4 | 0.7 |
| 2 | 128624998 | rs13397060 | -0.044 | 0.226 | 0.514 | -0.589 | 0.153 | 0.031 | 0.031 | 1.506 | 0.05 | 0.26 | 0.07 | 0.05 | 0.09 | 0.04 |
| 2 | 128628225 | rs13018398 | -0.877 | 0.063 | 0.16 | -0.513 | -0.018 | 0.004 | 0.004 | 0 | 0.37 | 0.45 | 0.36 | 0.32 | 0.3 | 0.66 |
| 2 | 128641031 | rs13398811 | -0.225 | -0.187 | 0.464 | -0.31 | 0.001 | 0.194 | 0.194 | 0.776 | 0.06 | 0.32 | 0.13 | 0.14 | 0 | 0.13 |
| 2 | 128645618 | rs2304518 | 0.101 | 0.122 | -0.511 | -0.57 | 0.002 | 0.054 | 0.054 | 0 | 0.24 | 0.04 | 0.26 | 0.13 | 0.21 | 0.44 |
| 2 | 128649042 | rs6749005 | -0.653 | -0.172 | -0.062 | -0.618 | 0.055 | -0.043 | -0.043 | 0 | 0.14 | 0.54 | 0.1 | 0.06 | 0.12 | 0.08 |
| 2 | 128653761 | rs9677367 | -0.896 | -0.104 | -0.254 | -0.385 | 0.003 | 0.003 | 0.003 | 0 | 0.39 | 0.62 | 0.4 | 0.32 | 0.31 | 0.64 |
| 2 | 128660326 | rs6715338 | -0.958 | 0.012 | -0.689 | -0.292 | 0.037 | 0.015 | 0.015 | 0 | 0.43 | 0.75 | 0.41 | 0.35 | 0.31 | 0.66 |
| 2 | 128699833 | rs1370229 | 1.166 | 0.149 | -0.933 | 0.228 | -0.146 | -0.167 | -0.167 | 0.442 | 0.36 | 0.03 | 0.48 | 0.5 | 0.53 | 0.26 |
| 2 | 128706637 | rs17015719 | 0.815 | -0.172 | -0.203 | 0.134 | -0.029 | 0.025 | 0.025 | 2.085 | 0.07 | 0.25 | 0.06 | 0.09 | 0.02 | 0.01 |
| 2 | 128706690 | rs2438039 | 0.16 | -0.268 | -0.081 | -0.011 | 0.052 | 0.056 | 0.056 | 0 | 0.89 | 0.67 | 0.86 | 0.86 | 0.89 | 0.76 |

| | | | | | | | | | | | | | | | | |
|---|---|---|---|---|---|---|---|---|---|---|---|---|---|---|---|---|
| 2 | 128716817 | rs11682273 | 0.754 | -0.59 | -0.068 | -0.203 | 0.023 | 0.086 | 0.086 | 0 | 0.29 | 0.06 | 0.19 | 0.12 | 0.21 | 0.41 |
| 2 | 128726382 | rs2917657 | 0.124 | -0.197 | -0.103 | -0.103 | 0.052 | 0.056 | 0.056 | 0 | 0.89 | 0.67 | 0.86 | 0.86 | 0.89 | 0.76 |
| 2 | 128726477 | rs2461443 | 1.603 | -0.04 | -0.297 | -0.107 | -0.102 | -0.041 | -0.041 | 0 | 0.37 | 0.07 | 0.5 | 0.64 | 0.53 | 0.28 |
| 2 | 128742474 | rs2438028 | 1.565 | 0.386 | 0.587 | 0.234 | -0.01 | -0.044 | -0.044 | 0 | 0.74 | 0.5 | 0.77 | 0.85 | 0.77 | 0.7 |
| 2 | 128748238 | rs11680146 | 1.632 | 0.546 | 0.566 | 0.23 | -0.009 | 0.004 | 0.004 | 0 | 0.73 | 0.45 | 0.74 | 0.71 | 0.77 | 0.69 |
| 2 | 128780173 | rs10166905 | -0.915 | 0.542 | 0.125 | -0.391 | -0.013 | -0.057 | -0.057 | 0 | 0.15 | 0.23 | 0.09 | 0.01 | 0.14 | 0.05 |
| 2 | 128801224 | rs4533428 | -0.451 | 0.844 | -0.53 | -0.332 | -0.048 | 0.046 | 0.046 | 0 | 0.11 | 0.04 | 0.09 | 0 | 0.22 | 0.06 |
| 2 | 128818557 | rs6430975 | 1.053 | 0.613 | -0.625 | -0.517 | -0.001 | 0.065 | 0.065 | 0 | 0.49 | 0.24 | 0.58 | 0.69 | 0.51 | 0.34 |
| 2 | 128822702 | rs13022015 | 0.566 | 0.631 | -1.201 | -0.719 | 0.009 | 0.067 | 0.067 | 0 | 0.25 | 0.03 | 0.13 | 0.12 | 0.18 | 0.38 |
| 2 | 128822741 | rs6728375 | -1.22 | 0.789 | -0.779 | -0.634 | 0.007 | 0 | 0 | 0.035 | 0.14 | 0.18 | 0.21 | 0.12 | 0.28 | 0.24 |
| 2 | 128831582 | rs7589262 | -0.003 | 0.43 | -0.478 | -0.616 | 0.026 | 0.009 | 0.009 | 0 | 0.68 | 0.78 | 0.79 | 0.73 | 0.81 | 0.47 |
| 2 | 128832809 | rs11683368 | -0.302 | 0.667 | -0.709 | -0.618 | -0.045 | 0.042 | 0.042 | 0 | 0.08 | 0.03 | 0.1 | 0 | 0.2 | 0.05 |
| 2 | 128838634 | rs12617108 | 1.318 | 0.959 | -0.48 | -0.578 | 0.019 | 0.03 | 0.03 | 0 | 0.5 | 0.55 | 0.59 | 0.67 | 0.45 | 0.36 |
| 2 | 128849114 | rs4476299 | -0.141 | 0.737 | -0.472 | -0.573 | -0.008 | 0.144 | 0.144 | 0 | 0.93 | 0.7 | 0.91 | 0.85 | 0.98 | 0.8 |
| 2 | 128851760 | rs2609 | -0.181 | 0.858 | -0.684 | -0.764 | -0.073 | 0.127 | 0.127 | 0 | 0.35 | 0.08 | 0.28 | 0.17 | 0.39 | 0.41 |
| 2 | 128870808 | rs4662775 | 0.162 | 0.849 | -0.56 | -0.764 | -0.008 | 0.156 | 0.156 | 0 | 0.92 | 0.65 | 0.9 | 0.85 | 0.98 | 0.8 |
| 2 | 128891067 | rs11903649 | 1.665 | 0.776 | -0.334 | -0.814 | 0.011 | 0.017 | 0.017 | 0 | 0.47 | 0.42 | 0.52 | 0.67 | 0.43 | 0.32 |
| 2 | 128897455 | rs10194565 | 0.048 | 0.836 | -0.367 | -0.785 | 0.049 | -0.034 | -0.034 | 0 | 0.07 | 0.09 | 0.09 | 0.01 | 0.15 | 0.08 |
| 2 | 128920741 | rs13005793 | 0.695 | 0.84 | -0.509 | -0.748 | 0.017 | 0.055 | 0.055 | 0 | 0.27 | 0.05 | 0.12 | 0.17 | 0.18 | 0.34 |
| 2 | 128932760 | rs10196158 | -1.056 | 0.757 | -0.459 | -0.749 | 0.001 | 0.009 | 0.009 | 0 | 0.43 | 0.43 | 0.38 | 0.32 | 0.41 | 0.6 |
| 2 | 128939695 | rs17793615 | 0.735 | 0.801 | -0.48 | -0.751 | -0.001 | 0.054 | 0.054 | 0 | 0.27 | 0.04 | 0.13 | 0.17 | 0.2 | 0.34 |
| 2 | 128939817 | rs1699 | -0.945 | 0.782 | -0.43 | -0.737 | 0.001 | 0.012 | 0.012 | 0 | 0.43 | 0.42 | 0.38 | 0.32 | 0.41 | 0.6 |
| 2 | 128947505 | rs11542865 | -0.987 | 0.783 | -0.45 | -0.736 | 0.001 | 0.006 | 0.006 | 0 | 0.43 | 0.44 | 0.38 | 0.32 | 0.41 | 0.6 |
| 2 | 128959560 | rs2405983 | -0.151 | 0.865 | -0.894 | -0.697 | -0.105 | 0.036 | 0.036 | 0 | 0.33 | 0.05 | 0.27 | 0.22 | 0.39 | 0.43 |
| 2 | 128971040 | rs13384737 | -0.896 | 0.114 | -1.017 | -0.991 | 0.023 | -0.062 | -0.062 | 1.866 | 0.33 | 0.52 | 0.3 | 0.11 | 0.41 | 0.18 |
| 2 | 128972833 | rs6725404 | -0.079 | 0.146 | -0.484 | -0.956 | 0.009 | 0.152 | 0.152 | 0 | 0.08 | 0.4 | 0.1 | 0.14 | 0.02 | 0.18 |
| 2 | 128974065 | rs6716830 | 0.025 | 0.146 | -0.676 | -0.967 | -0.038 | 0.006 | 0.006 | 0 | 0.33 | 0.52 | 0.24 | 0.33 | 0.23 | 0.52 |
| 2 | 128996070 | rs10167836 | 0.14 | 0.212 | -0.739 | -0.948 | -0.031 | 0.014 | 0.014 | 0 | 0.32 | 0.45 | 0.24 | 0.35 | 0.24 | 0.5 |
| 2 | 129004419 | rs13430943 | 0.243 | 0.346 | -0.832 | -1.093 | -0.004 | -0.021 | -0.021 | 0.521 | 0.2 | 0.15 | 0.12 | 0.01 | 0.21 | 0.13 |
| 2 | 129005979 | rs10191718 | 0.243 | 0.346 | -0.832 | -1.093 | -0.004 | -0.021 | -0.021 | 0 | 0.2 | 0.15 | 0.12 | 0.01 | 0.21 | 0.13 |
| 2 | 129007373 | rs6742117 | -0.517 | 0.287 | -1.054 | -1.195 | 0.019 | -0.003 | -0.003 | 0.277 | 0.18 | 0.38 | 0.24 | 0.14 | 0.21 | 0.25 |
| 2 | 129007461 | rs6732469 | -0.993 | 0.457 | -1.446 | -1.021 | -0.004 | 0.005 | 0.005 | 0 | 0.38 | 0.61 | 0.36 | 0.15 | 0.42 | 0.37 |
| 2 | 129007898 | rs11689623 | 0.823 | 0.064 | -1.47 | -0.887 | -0.003 | -0.006 | -0.006 | 0 | 0.25 | 0.05 | 0.14 | 0.21 | 0.22 | 0.34 |
| 2 | 129012073 | rs13414154 | -1.184 | 0.083 | -0.855 | -1.165 | -0.007 | -0.024 | -0.024 | 0 | 0.33 | 0.58 | 0.29 | 0.15 | 0.3 | 0.36 |
| 2 | 129013157 | rs6731871 | -0.355 | 0.037 | -1.172 | -1.13 | -0.046 | -0.023 | -0.023 | 0 | 0.22 | 0.39 | 0.16 | 0.14 | 0.16 | 0.31 |
| 2 | 129013461 | rs6431004 | -1.243 | 0.307 | -2.146 | -1.126 | 0.049 | 0.171 | 0.171 | 0 | 0.47 | 0.88 | 0.4 | 0.21 | 0.39 | 0.49 |
| 2 | 129015968 | rs1868394 | -1.226 | 0.665 | -1.816 | -1.362 | -0.074 | 0.147 | 0.147 | 2.177 | 0.61 | 0.94 | 0.72 | 0.84 | 0.57 | 0.5 |
| 2 | 129016127 | rs4429419 | -1.253 | 1.089 | -1.415 | -1.038 | -0.071 | 0.223 | 0.223 | 0.258 | 0.54 | 0.94 | 0.7 | 0.84 | 0.52 | 0.5 |
| 2 | 129020968 | rs6727849 | -1.411 | 1.094 | -0.041 | -0.427 | -0.032 | 0.001 | 0.001 | 3.065 | 0.1 | 0.68 | 0.07 | 0.05 | 0.03 | 0.05 |
| 2 | 129027235 | rs734193 | -1.403 | 1.715 | 0.041 | -0.643 | 0 | -0.018 | -0.018 | 0 | 0.47 | 0.52 | 0.7 | 0.77 | 0.48 | 0.32 |
| 2 | 129028201 | rs6756336 | 0.018 | 1.995 | 0.227 | -0.644 | 0 | 0.007 | 0.007 | 0 | 0.19 | 0.2 | 0.43 | 0.67 | 0.18 | 0.13 |
| 2 | 129028302 | rs13428208 | -0.223 | 2.121 | 0.262 | -0.599 | -0.013 | 0.008 | 0.008 | 0 | 0.07 | 0.04 | 0.07 | 0.02 | 0.15 | 0.14 |
| 2 | 129030337 | rs4662784 | -1.261 | 2.117 | 0.313 | -0.623 | 0.007 | -0.018 | -0.018 | 0 | 0.38 | 0.44 | 0.65 | 0.77 | 0.36 | 0.28 |
| 2 | 129030554 | rs4662785 | -0.842 | 1.758 | 0.454 | -0.75 | 0.013 | -0.031 | -0.031 | 0 | 0.08 | 0.1 | 0.08 | 0.02 | 0.14 | 0.14 |
| 2 | 129031073 | rs6717511 | -0.634 | 1.776 | 0.353 | -0.741 | 0.001 | -0.014 | -0.014 | 0 | 0.08 | 0.06 | 0.08 | 0.02 | 0.14 | 0.14 |
| 2 | 129035083 | rs17016070 | -1.396 | 1.795 | 0.359 | -0.748 | -0.001 | -0.003 | -0.003 | 0 | 0.09 | 0.24 | 0.12 | 0.09 | 0.07 | 0.09 |
| 2 | 129035892 | rs4662786 | -1.561 | 1.807 | 0.321 | -0.776 | 0.008 | -0.041 | -0.041 | 0.196 | 0.12 | 0.15 | 0.1 | 0.02 | 0.16 | 0.16 |
| 2 | 129039724 | rs17794012 | -0.591 | 1.83 | 0.349 | -0.752 | 0.02 | -0.033 | -0.033 | 0.32 | 0.08 | 0.1 | 0.08 | 0.02 | 0.15 | 0.14 |
| 2 | 129043010 | rs6711447 | -2.021 | 1.818 | 0.725 | -0.776 | 0.052 | -0.062 | -0.062 | 0 | 0.18 | 0.48 | 0.21 | 0.11 | 0.22 | 0.28 |
| 2 | 129044265 | rs6431007 | -1.36 | 1.853 | -0.159 | -1.107 | -0.046 | 0.128 | 0.128 | 0.016 | 0.44 | 0.79 | 0.7 | 0.79 | 0.44 | 0.32 |
| 2 | 129045765 | rs4143782 | -0.1 | 1.703 | 0.064 | -1.016 | -0.008 | 0.002 | 0.002 | 0 | 0.2 | 0.13 | 0.47 | 0.78 | 0.18 | 0.21 |
| 2 | 129050909 | rs1375347 | -0.548 | 1.834 | 0.418 | -0.872 | -0.001 | -0.049 | -0.049 | 1.481 | 0.3 | 0.48 | 0.56 | 0.8 | 0.29 | 0.28 |
| 2 | 129052655 | rs4662791 | -1.498 | 2.075 | -0.025 | -0.921 | 0.01 | 0.127 | 0.127 | 0.195 | 0.47 | 0.73 | 0.74 | 0.82 | 0.54 | 0.47 |
| 2 | 129052860 | rs7566627 | -0.5 | 2.103 | -0.011 | -0.832 | -0.004 | -0.031 | -0.031 | 0 | 0.31 | 0.57 | 0.56 | 0.8 | 0.3 | 0.28 |
| 2 | 129061452 | rs714129 | -0.74 | 2.198 | -0.553 | -0.359 | -0.078 | 0.081 | 0.081 | 0.085 | 0.14 | 0.01 | 0.17 | 0.01 | 0.24 | 0.19 |
| 2 | 129082949 | rs6733432 | -1.304 | 1.437 | 0.562 | -0.09 | 0.115 | 0.041 | 0.041 | 0.486 | 0.05 | 0.36 | 0.06 | 0 | 0.06 | 0.02 |
| 2 | 129086164 | rs2579663 | 0.58 | 1.424 | 0.285 | 0.149 | 0 | 0 | 0 | 0 | 0.7 | 0.35 | 0.44 | 0.29 | 0.75 | 0.72 |
| 2 | 129090109 | rs11890277 | -1.189 | 0.929 | 0.013 | 0.634 | -0.003 | 0.005 | 0.005 | 0 | 0.11 | 0.11 | 0.28 | 0.17 | 0.12 | 0.2 |
| 2 | 129104653 | rs17016151 | 0.485 | 1.291 | 0.023 | 0.499 | -0.008 | -0.026 | -0.026 | 0 | 0.59 | 0.29 | 0.55 | 0.43 | 0.62 | 0.77 |
| 2 | 129106110 | rs4662799 | 1.251 | 1.642 | -0.224 | 0.278 | -0.113 | 0.095 | 0.095 | 0 | 0.49 | 0.04 | 0.31 | 0.26 | 0.51 | 0.58 |
| 2 | 129110365 | rs11888763 | -0.36 | 1.509 | -0.034 | 0.288 | -0.012 | -0.024 | -0.024 | 0.092 | 0.06 | 0.07 | 0.04 | 0 | 0.06 | 0.02 |
| 2 | 129116922 | rs750785 | 0.29 | 1.621 | 0.285 | 0.302 | 0.005 | 0.034 | 0.034 | 0 | 0.66 | 0.52 | 0.5 | 0.37 | 0.72 | 0.78 |
| 2 | 129118215 | rs10204956 | 0.326 | 1.65 | -0.034 | 0.359 | 0 | -0.007 | -0.007 | 0 | 0.64 | 0.32 | 0.48 | 0.38 | 0.72 | 0.78 |
| 2 | 129123452 | rs2897293 | 1.101 | 1.81 | -0.7 | 0.035 | 0.062 | 0.042 | 0.042 | 0 | 0.47 | 0.12 | 0.33 | 0.35 | 0.4 | 0.63 |
| 2 | 129124530 | rs936127 | -1.522 | 1.891 | -0.288 | 0.149 | -0.001 | -0.002 | -0.002 | 0.089 | 0.18 | 0.19 | 0.14 | 0.13 | 0.18 | 0.27 |
| 2 | 129125972 | rs882275 | -0.001 | 1.366 | -0.711 | -0.406 | 0 | -0.001 | -0.001 | 0 | 0.55 | 0.58 | 0.45 | 0.4 | 0.5 | 0.63 |
| 2 | 129128066 | rs4662800 | -0.21 | 1.203 | -0.732 | -0.434 | 0.004 | 0.002 | 0.002 | 0.005 | 0.59 | 0.58 | 0.49 | 0.41 | 0.57 | 0.66 |
| 2 | 129128239 | rs4662801 | 1.353 | 1.162 | -1.177 | -0.466 | -0.084 | 0.016 | 0.016 | 0 | 0.37 | 0.02 | 0.31 | 0.23 | 0.38 | 0.36 |
| 2 | 129134764 | rs17016212 | -0.564 | 0.598 | -0.423 | -0.332 | -0.011 | -0.022 | -0.022 | 0.013 | 0.1 | 0.2 | 0.12 | 0.03 | 0.06 | 0.12 |

| | | | | | | | | | | | | | | | | |
|---|---|---|---|---|---|---|---|---|---|---|---|---|---|---|---|---|
| 2 | 129135431 | rs934915 | 1.298 | 0.459 | -0.939 | -0.344 | -0.083 | -0.031 | -0.031 | 0 | 0.39 | 0.05 | 0.27 | 0.3 | 0.41 | 0.41 |
| 2 | 129136550 | rs7560558 | 1.274 | 0.439 | -0.889 | -0.348 | -0.063 | -0.011 | -0.011 | 0.092 | 0.39 | 0.07 | 0.27 | 0.3 | 0.42 | 0.41 |
| 2 | 129137161 | rs12473414 | 0.247 | 0.55 | -1.047 | -0.03 | -0.103 | -0.16 | -0.16 | 0 | 0.26 | 0.02 | 0.34 | 0.5 | 0.33 | 0.11 |
| 2 | 129140084 | rs9287544 | -1.141 | 0.556 | -0.604 | -0.023 | -0.016 | -0.004 | -0.004 | 0 | 0.15 | 0.33 | 0.15 | 0.07 | 0.12 | 0.24 |
| 2 | 129143954 | rs7565310 | -0.689 | 0.39 | -0.642 | -0.116 | -0.004 | -0.015 | -0.015 | 0.133 | 0.37 | 0.32 | 0.43 | 0.53 | 0.41 | 0.15 |
| 2 | 129147159 | rs6726600 | -1.359 | -0.297 | -0.221 | -0.327 | 0 | -0.01 | -0.01 | 0 | 0.47 | 0.5 | 0.4 | 0.75 | 0.51 | 0.3 |
| 2 | 129153267 | rs13002365 | -0.311 | -0.335 | 0.312 | -0.538 | -0.012 | -0.038 | -0.038 | 0.126 | 0.22 | 0.19 | 0.13 | 0.02 | 0.14 | 0.2 |
| 2 | 129153304 | rs880721 | 0.127 | -0.342 | 0.253 | -0.455 | -0.049 | -0.068 | -0.068 | 0.183 | 0.31 | 0.16 | 0.32 | 0.55 | 0.42 | 0.13 |
| 2 | 129161203 | rs17216354 | 0.438 | 0.041 | 0.179 | -0.203 | -0.005 | -0.022 | -0.022 | 0.43 | 0.14 | 0.14 | 0.12 | 0.02 | 0.11 | 0.16 |
| 2 | 129163372 | rs1866461 | -0.963 | 0.272 | 0.03 | -0.018 | -0.002 | -0.002 | -0.002 | 0.923 | 0.54 | 0.62 | 0.48 | 0.58 | 0.56 | 0.33 |
| 2 | 129171305 | rs9783995 | -0.923 | 0.092 | -0.179 | -0.51 | 0.005 | 0.027 | 0.027 | 0 | 0.56 | 0.62 | 0.52 | 0.74 | 0.61 | 0.35 |
| 2 | 129179239 | rs4662808 | -1.548 | -0.292 | 0.594 | -0.656 | -0.07 | -0.028 | -0.028 | 0 | 0.23 | 0.43 | 0.13 | 0.2 | 0.12 | 0.23 |
| 2 | 129180470 | rs745071 | -1.324 | -0.286 | 0.606 | -0.65 | -0.063 | -0.118 | -0.118 | 0.948 | 0.22 | 0.43 | 0.13 | 0.02 | 0.12 | 0.23 |
| 2 | 129184166 | rs2124432 | -2.394 | -0.131 | 0.659 | -0.848 | 0.009 | 0.02 | 0.02 | 0 | 0.37 | 0.66 | 0.33 | 0.22 | 0.31 | 0.31 |
| 2 | 129184866 | rs1540493 | -0.142 | -0.198 | 0.907 | -0.866 | -0.012 | -0.045 | -0.045 | 0 | 0.12 | 0.13 | 0.1 | 0 | 0.08 | 0.1 |
| 2 | 129185316 | rs10191637 | -0.92 | -0.096 | 0.839 | -0.801 | -0.049 | -0.019 | -0.019 | 2.237 | 0.22 | 0.38 | 0.13 | 0.2 | 0.13 | 0.24 |
| 2 | 129193361 | rs6748470 | -2.113 | -0.05 | 0.595 | -0.799 | 0.015 | 0.026 | 0.026 | 0 | 0.37 | 0.67 | 0.33 | 0.22 | 0.3 | 0.3 |
| 2 | 129196612 | rs4662818 | -0.578 | 0.035 | 0.459 | -0.666 | 0.012 | 0.01 | 0.01 | 0 | 0.15 | 0.21 | 0.19 | 0.2 | 0.18 | 0.07 |
| 2 | 129202441 | rs11690893 | 1.219 | 0.001 | -0.002 | -0.724 | 0.019 | 0.043 | 0.043 | 0 | 0.57 | 0.34 | 0.64 | 0.78 | 0.63 | 0.66 |
| 2 | 129203736 | rs7567587 | -0.986 | 0.127 | 0.251 | -0.516 | -0.026 | -0.073 | -0.073 | 0 | 0.19 | 0.32 | 0.18 | 0.02 | 0.12 | 0.09 |
| 2 | 129211017 | rs4517924 | -0.093 | 0.287 | -0.168 | -0.565 | -0.003 | -0.03 | -0.03 | 0.045 | 0.17 | 0.15 | 0.14 | 0.03 | 0.13 | 0.12 |
| 2 | 129212869 | rs10186968 | -0.241 | 0.441 | 0.124 | -0.598 | 0.005 | 0 | 0 | 0 | 0.08 | 0.08 | 0.07 | 0.07 | 0.1 | 0.14 |
| 2 | 129213567 | rs10175328 | -0.391 | 0.532 | -0.055 | -0.854 | 0.005 | 0.009 | 0.009 | 0 | 0.67 | 0.48 | 0.63 | 0.4 | 0.65 | 0.74 |
| 2 | 129215649 | rs10181968 | 1.103 | 0.498 | 0.135 | -1.308 | -0.011 | 0.157 | 0.157 | 0 | 0.61 | 0.19 | 0.69 | 0.9 | 0.65 | 0.71 |
| 2 | 129222649 | rs17283348 | 0.347 | 0.503 | 0.388 | -1.067 | -0.012 | -0.002 | -0.002 | 0 | 0.07 | 0.04 | 0.08 | 0.07 | 0.11 | 0.13 |
| 2 | 129222802 | rs1378716 | 1.073 | 0.522 | 0.416 | -1.075 | 0.007 | -0.05 | -0.05 | 1.14 | 0.79 | 0.34 | 0.84 | 0.93 | 0.79 | 0.84 |
| 2 | 129226743 | rs4662827 | -0.289 | 0.587 | 0.034 | -0.987 | 0.016 | -0.003 | -0.003 | 0.048 | 0.08 | 0.15 | 0.08 | 0.07 | 0.11 | 0.14 |
| 2 | 129235440 | rs935846 | -0.212 | 0.61 | 0.701 | -0.979 | 0.024 | -0.061 | -0.061 | 0 | 0.9 | 0.73 | 0.94 | 1 | 0.9 | 0.98 |
| 2 | 129245547 | rs705055 | -0.805 | 0.352 | -0.25 | -1.177 | 0.032 | 0.164 | 0.164 | 0 | 0.42 | 0.75 | 0.3 | 0.1 | 0.34 | 0.32 |
| 2 | 129259855 | rs11686500 | 0.09 | 0.552 | 0.242 | -0.82 | 0 | 0 | 0 | 0 | 0.12 | 0.12 | 0.17 | 0.09 | 0.09 | 0.18 |
| 2 | 129260444 | rs11896212 | -1.131 | 0.39 | 0.922 | -1.047 | 0.003 | -0.063 | -0.063 | 0 | 0.23 | 0.58 | 0.25 | 0.09 | 0.2 | 0.21 |
| 2 | 129274119 | rs11681913 | 1.023 | 0.564 | 0.676 | -0.998 | 0.003 | -0.001 | -0.001 | 0 | 0.09 | 0.08 | 0.17 | 0.07 | 0.11 | 0.11 |
| 2 | 129274601 | rs11683275 | 0.749 | 0.553 | 1.209 | -1.004 | 0.044 | -0.007 | -0.007 | 0 | 0.09 | 0.18 | 0.18 | 0.07 | 0.11 | 0.11 |
| 2 | 129276328 | rs11681298 | 0.719 | 0.549 | 1.17 | -1.006 | 0.002 | -0.002 | -0.002 | 0 | 0.09 | 0.09 | 0.17 | 0.07 | 0.11 | 0.1 |
| 2 | 129277705 | rs13390327 | -0.552 | 0.411 | 1.27 | -1.057 | -0.022 | -0.028 | -0.028 | 0 | 0.06 | 0.19 | 0.03 | 0 | 0.01 | 0 |
| 2 | 129278276 | rs841095 | -0.292 | 0.399 | 1.652 | -1.082 | 0.017 | -0.044 | -0.044 | 0 | 0.1 | 0.35 | 0.07 | 0.03 | 0.1 | 0.1 |
| 2 | 129280640 | rs841078 | -1.703 | 0.37 | 1.011 | -1.19 | -0.006 | 0.107 | 0.107 | 1.511 | 0.28 | 0.84 | 0.29 | 0.1 | 0.23 | 0.21 |
| 2 | 129284155 | rs841080 | -0.839 | 0.439 | 0.348 | -0.744 | -0.089 | -0.19 | -0.19 | 0 | 0.73 | 0.98 | 0.72 | 0.5 | 0.7 | 0.62 |
| 2 | 129284521 | rs841081 | -1.148 | 0.409 | 1.435 | -0.592 | 0.002 | -0.022 | -0.022 | 0 | 0.23 | 0.65 | 0.26 | 0.09 | 0.23 | 0.21 |
| 2 | 129291445 | rs2445231 | -1.646 | 0.442 | 0.785 | -0.618 | 0 | 0.102 | 0.102 | 0.789 | 0.29 | 0.81 | 0.29 | 0.09 | 0.23 | 0.21 |
| 2 | 129294941 | rs11687419 | 0.007 | 0.591 | 0.994 | -0.202 | 0.032 | -0.03 | -0.03 | 0 | 0.12 | 0.18 | 0.18 | 0.06 | 0.12 | 0.11 |
| 2 | 129300580 | rs7564226 | -0.262 | 0.451 | -0.104 | -0.007 | 0.005 | -0.001 | -0.001 | 0.359 | 0.43 | 0.37 | 0.48 | 0.43 | 0.42 | 0.42 |
| 2 | 129314229 | rs2438162 | 0.703 | 0.652 | 0.471 | 0.394 | 0.031 | 0.053 | 0.053 | 0 | 0.81 | 0.72 | 0.75 | 0.34 | 0.75 | 0.56 |
| 2 | 129316382 | rs705071 | -1.261 | -0.062 | 1.592 | -0.372 | -0.059 | -0.046 | -0.046 | 0 | 0.14 | 0.4 | 0.18 | 0.05 | 0.05 | 0.02 |
| 2 | 129317105 | rs841302 | -1.991 | -0.288 | 1.469 | -0.556 | -0.055 | -0.063 | -0.063 | 0 | 0.22 | 0.66 | 0.23 | 0.06 | 0.11 | 0.13 |
| 2 | 129319031 | rs697363 | -0.388 | -0.284 | 1.976 | -0.544 | -0.03 | -0.02 | -0.02 | 0 | 0.11 | 0.26 | 0.16 | 0.05 | 0.05 | 0.02 |
| 2 | 129326003 | rs841292 | -1.485 | -0.27 | 2.08 | -0.533 | -0.057 | -0.039 | -0.039 | 0 | 0.14 | 0.57 | 0.2 | 0.05 | 0.05 | 0.02 |
| 2 | 129355260 | rs841463 | 2.174 | -0.242 | 1.957 | -0.546 | -0.068 | -0.085 | -0.085 | 0 | 0.78 | 0.41 | 0.86 | 0.94 | 0.89 | 0.81 |
| 2 | 129360694 | rs841452 | -1.917 | -0.192 | 2.031 | -0.553 | 0 | -0.052 | -0.052 | 2.29 | 0.1 | 0.41 | 0.07 | 0.01 | 0.07 | 0.17 |
| 2 | 129370246 | rs841470 | -1.815 | -0.207 | 2.063 | -0.424 | -0.069 | -0.073 | -0.073 | 0 | 0.17 | 0.55 | 0.12 | 0.06 | 0.06 | 0.11 |
| 2 | 129394742 | rs13386062 | -0.138 | -0.042 | 1.737 | -0.492 | -0.021 | -0.023 | -0.023 | 0 | 0.79 | 0.93 | 0.68 | 0.25 | 0.77 | 0.77 |
| 2 | 129395115 | rs10204926 | -2.74 | -0.122 | 2.443 | -0.374 | -0.009 | -0.099 | -0.099 | 0 | 0.14 | 0.46 | 0.11 | 0.01 | 0.12 | 0.12 |
| 2 | 129404992 | rs17048656 | -2.335 | -0.206 | 2.044 | -0.355 | -0.013 | -0.06 | -0.06 | 0.746 | 0.12 | 0.26 | 0.1 | 0.01 | 0.13 | 0.12 |
| 2 | 129408831 | rs7573450 | -1.683 | 0.022 | 2.045 | -0.312 | -0.001 | -0.016 | -0.016 | 0.067 | 0.12 | 0.11 | 0.08 | 0.01 | 0.13 | 0.12 |
| 2 | 129413217 | rs6729786 | -1.8 | 0.015 | 2.007 | -0.308 | -0.003 | -0.084 | -0.084 | 0 | 0.15 | 0.32 | 0.1 | 0.01 | 0.12 | 0.12 |
| 2 | 129416319 | rs6738894 | -2.01 | -0.01 | 2.213 | -0.32 | -0.015 | -0.09 | -0.09 | 0.246 | 0.17 | 0.65 | 0.13 | 0.01 | 0.12 | 0.12 |
| 2 | 129421376 | rs6723103 | -1.851 | 0.075 | 1.442 | -0.278 | -0.003 | -0.089 | -0.089 | 0.115 | 0.15 | 0.34 | 0.1 | 0.01 | 0.12 | 0.13 |
| 2 | 129427325 | rs13010732 | 0.396 | 0.097 | -0.353 | -0.029 | -0.019 | 0.018 | 0.018 | 0 | 0.42 | 0.2 | 0.53 | 0.37 | 0.53 | 0.64 |
| 2 | 129435535 | rs10176331 | -0.577 | -0.268 | -0.544 | -0.496 | 0 | -0.04 | -0.04 | 0 | 0.76 | 0.9 | 0.74 | 0.38 | 0.77 | 0.84 |
| 2 | 129441341 | rs3843310 | -0.176 | -0.14 | 0.117 | -0.749 | -0.02 | -0.125 | -0.125 | 0 | 0.31 | 0.44 | 0.18 | 0.01 | 0.24 | 0.2 |
| 2 | 129448422 | rs3845649 | 0.593 | -0.007 | -0.206 | -0.118 | 0.01 | 0.006 | 0.006 | 0.23 | 0.59 | 0.21 | 0.66 | 0.37 | 0.63 | 0.72 |
| 2 | 129451446 | rs11893223 | -0.965 | -0.075 | 0.13 | -0.145 | 0.004 | -0.027 | -0.027 | 0.655 | 0.12 | 0.23 | 0.07 | 0.09 | 0.13 | 0.13 |
| 2 | 129451757 | rs3845650 | -0.454 | -0.139 | -0.174 | -0.175 | -0.023 | -0.113 | -0.113 | 0 | 0.75 | 0.93 | 0.78 | 0.46 | 0.75 | 0.86 |
| 2 | 129453152 | rs17287302 | 0.681 | -0.101 | -0.188 | -0.224 | 0.018 | 0.012 | 0.012 | 1.053 | 0.58 | 0.15 | 0.66 | 0.37 | 0.63 | 0.72 |
| 2 | 129457798 | rs10928827 | 1.242 | -0.185 | -0.602 | -0.321 | -0.106 | 0.186 | 0.186 | 0.408 | 0.48 | 0.03 | 0.55 | 0.21 | 0.53 | 0.55 |
| 2 | 129461606 | rs2405985 | 0.757 | -0.072 | -0.147 | -0.129 | 0.01 | 0.007 | 0.007 | 0.947 | 0.58 | 0.13 | 0.66 | 0.38 | 0.63 | 0.72 |
| 2 | 129467773 | rs7570769 | 0.48 | 0.21 | 0.044 | 0.131 | 0.003 | -0.004 | -0.004 | 0 | 0.66 | 0.28 | 0.71 | 0.37 | 0.68 | 0.75 |
| 2 | 129471297 | rs3845654 | 0.299 | 0.316 | -0.376 | 0.183 | -0.001 | 0.007 | 0.007 | 0 | 0.67 | 0.55 | 0.74 | 0.37 | 0.68 | 0.74 |
| 2 | 129478842 | rs4662834 | -0.903 | 0.843 | -1.018 | 0.322 | 0.058 | 0.042 | 0.042 | 0 | 0.49 | 0.78 | 0.58 | 0.46 | 0.6 | 0.41 |

| | | | | | | | | | | | | | | | | |
|---|---|---|---|---|---|---|---|---|---|---|---|---|---|---|---|---|
| 2 | 129483795 | rs11684311 | -0.85 | 0.927 | -0.847 | 0.322 | 0.04 | 0.034 | 0.034 | 0 | 0.48 | 0.74 | 0.57 | 0.46 | 0.58 | 0.41 |
| 2 | 129488506 | rs7594747 | -0.108 | 1.788 | -0.487 | 0.273 | 0.088 | 0.136 | 0.136 | 0.25 | 0.15 | 0.51 | 0.23 | 0.28 | 0.2 | 0.14 |
| 2 | 129489556 | rs6744496 | -0.387 | 1.882 | -0.751 | 0.225 | -0.045 | 0.083 | 0.083 | 0 | 0.34 | 0.18 | 0.36 | 0.18 | 0.41 | 0.28 |
| 2 | 129490984 | rs12991269 | -1.272 | 1.873 | -1.154 | 0.32 | 0.063 | 0.041 | 0.041 | 0 | 0.49 | 0.74 | 0.59 | 0.46 | 0.62 | 0.42 |
| 2 | 129493277 | rs17222130 | -0.123 | 2.164 | -1.306 | 0.573 | -0.027 | 0.007 | 0.007 | 0.075 | 0.14 | 0.09 | 0.13 | 0.13 | 0.16 | 0.08 |
| 2 | 129497314 | rs2406261 | -1.017 | 1.568 | -1.368 | 0.759 | -0.025 | 0.003 | 0.003 | 0 | 0.43 | 0.78 | 0.56 | 0.46 | 0.46 | 0.31 |
| 2 | 129505570 | rs3907872 | -0.746 | 0.726 | -0.421 | 1.354 | -0.051 | 0.009 | 0.009 | 0 | 0.26 | 0.44 | 0.23 | 0.26 | 0.17 | 0.18 |
| 2 | 129507314 | rs6722639 | -0.759 | 0.712 | -0.428 | 1.421 | -0.051 | 0.014 | 0.014 | 0 | 0.26 | 0.44 | 0.23 | 0.27 | 0.17 | 0.21 |
| 2 | 129509766 | rs2897394 | -1.53 | 0.839 | -0.211 | 1.38 | 0.017 | -0.07 | -0.07 | 0.967 | 0.37 | 0.86 | 0.47 | 0.52 | 0.33 | 0.32 |
| 2 | 129514127 | rs13390906 | -1.197 | 0.715 | 0.673 | 1.465 | -0.015 | 0.059 | 0.059 | 0.068 | 0.29 | 0.66 | 0.4 | 0.51 | 0.26 | 0.28 |
| 2 | 129536443 | rs7355382 | -0.798 | 0.301 | 0.812 | 1.258 | -0.056 | 0.015 | 0.015 | 0.008 | 0.26 | 0.64 | 0.24 | 0.29 | 0.18 | 0.23 |
| 2 | 129537700 | rs7578952 | -0.798 | 0.301 | 0.812 | 1.258 | -0.056 | 0.015 | 0.015 | 0.008 | 0.26 | 0.64 | 0.24 | 0.29 | 0.18 | 0.23 |
| 2 | 129538149 | rs9287545 | -1.107 | 0.22 | 0.899 | 1.296 | 0.054 | 0.289 | 0.289 | 2.455 | 0.06 | 0.54 | 0.11 | 0.16 | 0.02 | 0.04 |
| 2 | 129539213 | rs10209450 | -0.316 | 0.363 | 0.096 | 1.113 | 0.028 | 0.039 | 0.039 | 0 | 0.2 | 0.09 | 0.13 | 0.12 | 0.15 | 0.19 |
| 2 | 129541542 | rs12473285 | -1.074 | -0.013 | 0.653 | 1.691 | -0.055 | 0.2 | 0.2 | 0.018 | 0.13 | 0.39 | 0.2 | 0.38 | 0.1 | 0.17 |
| 2 | 129542173 | rs7565335 | -0.684 | 0.094 | -0.654 | 0.824 | -0.006 | 0.089 | 0.089 | 0 | 0.63 | 0.83 | 0.71 | 0.82 | 0.68 | 0.58 |
| 2 | 129542735 | rs11690223 | 0.838 | 0.618 | -1.181 | 0.719 | -0.085 | 0.029 | 0.029 | 0 | 0.23 | 0.08 | 0.2 | 0.13 | 0.36 | 0.31 |
| 2 | 129542746 | rs11696078 | -0.346 | -0.074 | -0.699 | 0.49 | -0.004 | 0.017 | 0.017 | 0 | 0.57 | 0.59 | 0.71 | 0.84 | 0.59 | 0.7 |
| 2 | 129542829 | rs17048785 | -0.375 | -0.086 | -0.171 | 0.952 | -0.002 | 0.035 | 0.035 | 0 | 0.17 | 0.18 | 0.33 | 0.54 | 0.14 | 0.11 |
| 2 | 129545520 | rs3845668 | -1.458 | -0.013 | -0.335 | 0.809 | -0.019 | 0.044 | 0.044 | 0 | 0.91 | 0.99 | 0.97 | 0.99 | 0.92 | 0.9 |
| 2 | 129545568 | rs3845669 | 0.79 | -0.275 | 0.171 | 0.71 | -0.046 | 0 | 0 | 0 | 0.22 | 0.36 | 0.22 | 0.2 | 0.14 | 0.34 |
| 2 | 129549283 | rs12692083 | -0.228 | -0.202 | 0.226 | 0.754 | -0.028 | -0.022 | -0.022 | 0 | 0.69 | 0.62 | 0.75 | 0.79 | 0.78 | 0.57 |
| 2 | 129555886 | rs1317473 | 0.867 | -0.004 | -0.198 | 0.788 | 0.024 | 0.104 | 0.104 | 0 | 0.5 | 0.28 | 0.34 | 0.28 | 0.54 | 0.41 |
| 2 | 129557610 | rs3922666 | 0.682 | -0.027 | -0.13 | 1.201 | 0.011 | -0.008 | -0.008 | 0 | 0.54 | 0.33 | 0.38 | 0.47 | 0.57 | 0.44 |
| 2 | 129559317 | rs1344242 | -1.038 | 0.042 | -0.03 | 1.209 | 0.001 | -0.001 | -0.001 | 0 | 0.24 | 0.7 | 0.42 | 0.52 | 0.22 | 0.16 |
| 2 | 129563143 | rs1898779 | -1.271 | 0.083 | 0.434 | 0.972 | 0.005 | -0.05 | -0.05 | 0 | 0.89 | 0.79 | 0.96 | 0.99 | 0.92 | 0.92 |
| 2 | 129567011 | rs12478682 | 0.931 | 0.185 | 0.337 | 1.013 | 0.007 | 0.088 | 0.088 | 0.215 | 0.53 | 0.31 | 0.37 | 0.29 | 0.52 | 0.45 |
| 2 | 129577382 | rs12615148 | -0.141 | 0.344 | 1.56 | 1.67 | 0.046 | 0.141 | 0.141 | 0.015 | 0.15 | 0.46 | 0.31 | 0.31 | 0.15 | 0.06 |
| 2 | 129589480 | rs4662848 | 0.391 | 0.322 | 1.517 | 1.686 | 0 | 0.003 | 0.003 | 0 | 0.21 | 0.25 | 0.29 | 0.38 | 0.23 | 0.4 |
| 2 | 129590826 | rs6734640 | -0.907 | 0.329 | 1.751 | 1.661 | 0 | -0.001 | -0.001 | 0.76 | 0.34 | 0.38 | 0.57 | 0.7 | 0.38 | 0.46 |
| 2 | 129593844 | rs1866472 | -0.438 | 0.323 | 1.647 | 1.73 | -0.011 | -0.035 | -0.035 | 0.053 | 0.3 | 0.17 | 0.52 | 0.5 | 0.34 | 0.38 |
| 2 | 129599075 | rs6748903 | 0.503 | 0.299 | 1.846 | 1.712 | -0.007 | 0.013 | 0.013 | 0 | 0.19 | 0.36 | 0.27 | 0.19 | 0.2 | 0.33 |
| 2 | 129627236 | rs17048850 | 1.114 | 1.065 | 1.569 | 1.757 | -0.007 | -0.022 | -0.022 | 0 | 0.17 | 0.06 | 0.25 | 0.19 | 0.2 | 0.32 |
| 2 | 129627445 | rs17048852 | 0.29 | 1.073 | 1.738 | 1.748 | 0.008 | 0.029 | 0.029 | 0 | 0.13 | 0.14 | 0.27 | 0.32 | 0.14 | 0.06 |
| 2 | 129628406 | rs1562319 | 0.568 | 1.091 | 1.659 | 1.686 | -0.004 | 0.048 | 0.048 | 0 | 0.2 | 0.22 | 0.31 | 0.38 | 0.23 | 0.36 |
| 2 | 129633203 | rs2252663 | 0.712 | 1.067 | 1.854 | 1.738 | 0.026 | 0.174 | 0.174 | 0.095 | 0.82 | 0.5 | 0.67 | 0.57 | 0.83 | 0.9 |
| 2 | 129635464 | rs1562320 | 0.61 | 1.023 | 1.955 | 1.644 | -0.001 | 0.004 | 0.004 | 0 | 0.64 | 0.42 | 0.42 | 0.37 | 0.63 | 0.58 |
| 2 | 129635987 | rs3910997 | -0.383 | 1.027 | 1.821 | 1.65 | -0.004 | -0.016 | -0.016 | 0 | 0.31 | 0.28 | 0.52 | 0.5 | 0.34 | 0.38 |
| 2 | 129647350 | rs10803597 | -0.227 | 1.031 | 1.732 | 1.648 | -0.014 | -0.045 | -0.045 | 0 | 0.3 | 0.19 | 0.51 | 0.5 | 0.34 | 0.38 |
| 2 | 129651161 | rs2251907 | 0.569 | 1.057 | 1.672 | 1.567 | 0.001 | 0.011 | 0.011 | 0 | 0.65 | 0.5 | 0.44 | 0.37 | 0.63 | 0.58 |
| 2 | 129655682 | rs2100079 | -0.716 | 1.168 | 1.706 | 1.538 | 0.006 | -0.008 | -0.008 | 0 | 0.34 | 0.5 | 0.52 | 0.41 | 0.35 | 0.42 |
| 2 | 129669656 | rs4662863 | 0.476 | 1.077 | 1.764 | 1.595 | -0.005 | -0.015 | -0.015 | 0 | 0.22 | 0.34 | 0.27 | 0.21 | 0.22 | 0.36 |
| 2 | 129691750 | rs2263458 | 0.575 | 1.092 | 1.82 | 1.673 | 0.002 | -0.026 | -0.026 | 4.102 | 0.64 | 0.28 | 0.57 | 0.47 | 0.67 | 0.55 |
| 2 | 129696104 | rs4331451 | 0.476 | 1.087 | 1.801 | 1.561 | 0.02 | 0.009 | 0.009 | 0 | 0.11 | 0.19 | 0.25 | 0.2 | 0.13 | 0.06 |
| 2 | 129700617 | rs2246289 | 0.633 | 1.018 | 1.919 | 1.664 | 0.002 | -0.002 | -0.002 | 0 | 0.65 | 0.3 | 0.58 | 0.56 | 0.69 | 0.6 |
| 2 | 129708945 | rs10168295 | 0.336 | 1.016 | 1.966 | 1.663 | -0.025 | 0.004 | 0.004 | 0 | 0.23 | 0.42 | 0.16 | 0.24 | 0.18 | 0.34 |
| 2 | 129725930 | rs7601699 | 0.502 | 1.029 | 1.851 | 1.64 | 0.019 | 0.008 | 0.008 | 0 | 0.11 | 0.18 | 0.25 | 0.2 | 0.13 | 0.06 |
| 2 | 129728083 | rs2251444 | -0.502 | 1.019 | 1.857 | 1.647 | -0.024 | -0.002 | -0.002 | 0 | 0.76 | 0.56 | 0.84 | 0.76 | 0.82 | 0.66 |
| 2 | 129728567 | rs2251354 | 0.774 | 1.014 | 1.848 | 1.698 | 0.002 | -0.057 | -0.057 | 0 | 0.63 | 0.17 | 0.56 | 0.48 | 0.67 | 0.55 |
| 2 | 129738917 | rs1979046 | 0.565 | 1.045 | 1.676 | 1.692 | 0.005 | -0.001 | -0.001 | 0 | 0.22 | 0.22 | 0.15 | 0.24 | 0.18 | 0.34 |
| 2 | 129740917 | rs17048931 | 0.381 | 1.068 | 1.59 | 1.694 | 0.041 | 0.022 | 0.022 | 0 | 0.12 | 0.25 | 0.26 | 0.2 | 0.13 | 0.06 |
| 2 | 129741337 | rs10928856 | 0.379 | 1.054 | 1.024 | 1.701 | 0.002 | -0.001 | -0.001 | 0 | 0.65 | 0.31 | 0.57 | 0.56 | 0.7 | 0.6 |
| 2 | 129753114 | rs876647 | 0.761 | 1.022 | 0.265 | 1.777 | 0.005 | -0.031 | -0.031 | 0.067 | 0.21 | 0.1 | 0.14 | 0.27 | 0.18 | 0.36 |
| 2 | 129755276 | rs1107024 | 1.501 | 1.06 | -0.048 | 1.62 | -0.005 | -0.02 | -0.02 | 0 | 0.11 | 0.1 | 0.18 | 0.27 | 0.15 | 0.19 |
| 2 | 129762245 | rs10190944 | 0.723 | 1.156 | 0.189 | 1.396 | 0.026 | 0.026 | 0.026 | 0.059 | 0.11 | 0.24 | 0.24 | 0.2 | 0.1 | 0.05 |
| 2 | 129762727 | rs4662627 | 0.104 | 0.908 | -0.895 | 0.985 | -0.012 | 0.007 | 0.007 | 0 | 0.4 | 0.16 | 0.4 | 0.58 | 0.42 | 0.64 |
| 2 | 129766470 | rs11693887 | 1.484 | 0.107 | -0.569 | 1.346 | -0.006 | -0.022 | -0.022 | 3.81 | 0.1 | 0.07 | 0.18 | 0.26 | 0.14 | 0.2 |
| 2 | 129769189 | rs4662878 | 0.358 | 0.102 | -0.521 | 1.08 | -0.027 | -0.008 | -0.008 | 0.325 | 0.39 | 0.3 | 0.38 | 0.54 | 0.51 | 0.59 |
| 2 | 129772622 | rs4662881 | 1.859 | 0.125 | -0.618 | 1.147 | -0.026 | -0.059 | -0.059 | 0 | 0.1 | 0.01 | 0.17 | 0.27 | 0.14 | 0.21 |
| 2 | 129773278 | rs4662883 | 1.528 | 0.127 | -0.675 | 1.112 | -0.11 | -0.075 | -0.075 | 0 | 0.26 | 0.01 | 0.18 | 0.26 | 0.36 | 0.38 |
| 2 | 129773352 | rs4662630 | -0.567 | 0.049 | -0.351 | 0.527 | 0.045 | 0.124 | 0.124 | 0.034 | 0.55 | 0.79 | 0.66 | 0.84 | 0.66 | 0.68 |
| 2 | 129774095 | rs10928858 | -0.606 | 0.352 | 0.432 | 0.73 | 0.05 | 0.073 | 0.073 | 0.737 | 0.17 | 0.49 | 0.27 | 0.3 | 0.16 | 0.09 |
| 2 | 129774475 | rs10928860 | -0.426 | 0.362 | 0.459 | 0.735 | 0.055 | 0.078 | 0.078 | 0.377 | 0.17 | 0.42 | 0.27 | 0.3 | 0.16 | 0.09 |
| 2 | 129777555 | rs7584643 | 0.038 | -0.158 | 0.144 | 0.296 | -0.018 | -0.046 | -0.046 | 0.314 | 0.22 | 0.34 | 0.11 | 0.1 | 0.15 | 0.14 |
| 2 | 129780529 | rs6712840 | 0.099 | 0.057 | -0.125 | 0.33 | -0.036 | -0.013 | -0.013 | 0.02 | 0.13 | 0.05 | 0.27 | 0.19 | 0.17 | 0.07 |
| 2 | 129811792 | rs4662634 | 1.985 | -0.26 | -0.659 | 0.097 | -0.095 | 0.048 | 0.048 | 0 | 0.33 | 0.02 | 0.22 | 0.19 | 0.42 | 0.47 |
| 2 | 129812129 | rs4146904 | 0.509 | -0.408 | -0.309 | 0.103 | -0.008 | -0.026 | -0.026 | 2.602 | 0.19 | 0.28 | 0.1 | 0.11 | 0.15 | 0.14 |
| 2 | 129816397 | rs11679295 | 0.669 | -0.407 | -0.401 | 0.2 | 0.01 | -0.008 | -0.008 | 0 | 0.19 | 0.18 | 0.23 | 0.23 | 0.12 | 0.12 |
| 2 | 129817704 | rs7602203 | 0.488 | -0.42 | -0.156 | 0.163 | 0.004 | 0 | 0 | 0 | 0.58 | 0.51 | 0.5 | 0.42 | 0.58 | 0.59 |

| | | | | | | | | | | | | | | | | |
|---|---|---|---|---|---|---|---|---|---|---|---|---|---|---|---|---|
| 2 | 129819469 | rs11682667 | 0.561 | -0.78 | -0.058 | 0.522 | 0.014 | -0.026 | -0.026 | 0 | 0.18 | 0.09 | 0.22 | 0.23 | 0.12 | 0.12 |
| 2 | 129823158 | rs4662637 | 1.769 | -0.829 | -0.162 | 0.439 | -0.058 | 0.11 | 0.11 | 0 | 0.39 | 0.1 | 0.25 | 0.18 | 0.48 | 0.5 |
| 2 | 129826412 | rs11692464 | 1.134 | -0.875 | 0.683 | 0.914 | 0 | -0.002 | -0.002 | 0 | 0.6 | 0.38 | 0.5 | 0.42 | 0.63 | 0.6 |
| 2 | 129826843 | rs13031494 | 1.167 | -0.856 | 0.65 | 0.924 | 0.002 | -0.003 | -0.003 | 0 | 0.6 | 0.36 | 0.5 | 0.42 | 0.63 | 0.6 |
| 2 | 129830093 | rs10928871 | -0.104 | -0.586 | 0.559 | 0.949 | 0.008 | 0.014 | 0.014 | 0 | 0.13 | 0.17 | 0.16 | 0.2 | 0.13 | 0.05 |
| 2 | 129830473 | rs11684678 | 0.528 | -0.7 | 0.454 | 1.051 | 0.005 | -0.007 | -0.007 | 0 | 0.1 | 0.08 | 0.21 | 0.27 | 0.05 | 0.06 |
| 2 | 129835103 | rs908848 | -2.35 | -0.746 | 0.277 | 0.781 | -0.14 | 0.17 | 0.17 | 1.216 | 0.57 | 0.95 | 0.7 | 0.87 | 0.49 | 0.53 |
| 2 | 129838543 | rs17049081 | -0.491 | -0.536 | 0.913 | 0.983 | 0.018 | 0.03 | 0.03 | 0 | 0.13 | 0.27 | 0.17 | 0.2 | 0.13 | 0.05 |
| 2 | 129846840 | rs1519448 | -1.15 | -0.582 | 0.884 | 0.692 | -0.006 | -0.03 | -0.03 | 0 | 0.38 | 0.44 | 0.56 | 0.76 | 0.33 | 0.36 |
| 2 | 129860589 | rs6715935 | -0.615 | -0.589 | 1.1 | 1.406 | 0.017 | 0.164 | 0.164 | 0 | 0.13 | 0.36 | 0.2 | 0.4 | 0.09 | 0.08 |
| 2 | 129861592 | rs10189325 | -1.549 | -0.61 | 0.785 | 1.395 | -0.011 | 0.076 | 0.076 | 0.053 | 0.19 | 0.61 | 0.26 | 0.42 | 0.14 | 0.1 |
| 2 | 129861630 | rs17049106 | -0.366 | -0.353 | -0.377 | 1.463 | 0.007 | 0.104 | 0.104 | 0 | 0.12 | 0.23 | 0.2 | 0.4 | 0.09 | 0.08 |
| 2 | 129864132 | rs13428187 | -0.996 | -0.28 | -1.385 | 1.275 | 0.025 | 0.065 | 0.065 | 0.066 | 0.49 | 0.91 | 0.44 | 0.67 | 0.37 | 0.4 |
| 2 | 129866605 | rs2406579 | 0.913 | -0.145 | 0.683 | 0.629 | -0.016 | -0.07 | -0.07 | 0 | 0.88 | 0.65 | 0.93 | 1 | 0.93 | 0.96 |
| 2 | 129875814 | rs11693521 | -0.77 | 0.431 | 0.538 | 1.006 | -0.021 | -0.028 | -0.028 | 0 | 0.22 | 0.16 | 0.21 | 0.4 | 0.26 | 0.33 |
| 2 | 129881317 | rs2139347 | -0.77 | 0.404 | 0.527 | 0.996 | -0.02 | -0.028 | -0.028 | 5.625 | 0.22 | 0.16 | 0.21 | 0.4 | 0.26 | 0.33 |
| 2 | 129881863 | rs1607576 | -0.935 | 0.323 | 0.83 | 0.993 | -0.005 | -0.006 | -0.006 | 0.008 | 0.23 | 0.22 | 0.22 | 0.38 | 0.26 | 0.33 |
| 2 | 129884769 | rs9287550 | -1.051 | 0.205 | 2.3 | 0.909 | 0.012 | 0.022 | 0.022 | 0 | 0.25 | 0.34 | 0.23 | 0.4 | 0.26 | 0.33 |
| 2 | 129886603 | rs2090647 | 1.118 | 0.165 | 2.284 | 0.881 | -0.003 | -0.008 | -0.008 | 0 | 0.72 | 0.28 | 0.73 | 0.6 | 0.74 | 0.67 |
| 2 | 129891257 | rs10928882 | -0.873 | 0.224 | 1.87 | 0.911 | 0.006 | 0.01 | 0.01 | 0 | 0.24 | 0.29 | 0.22 | 0.4 | 0.26 | 0.33 |
| 2 | 129899376 | rs13024819 | -0.294 | -0.378 | 0.39 | 0.557 | 0.033 | 0.052 | 0.052 | 1.588 | 0.2 | 0.11 | 0.09 | 0.05 | 0.16 | 0.07 |
| 2 | 129903894 | rs2056281 | 1.12 | -0.252 | 0.471 | 0.646 | 0.027 | 0.023 | 0.023 | 0.604 | 0.52 | 0.27 | 0.65 | 0.54 | 0.58 | 0.6 |
| 2 | 129911352 | rs6715410 | 0.774 | -0.305 | 1.001 | 0.84 | -0.02 | 0.013 | 0.013 | 0 | 0.84 | 0.52 | 0.84 | 0.89 | 0.89 | 0.81 |
| 2 | 129912236 | rs4641887 | -0.472 | -0.038 | 0.396 | 1.092 | -0.023 | -0.035 | -0.035 | 0.183 | 0.12 | 0.07 | 0.12 | 0.26 | 0.16 | 0.23 |
| 2 | 129912509 | rs1113307 | -0.836 | -0.085 | 0.515 | 1.069 | -0.001 | 0 | 0 | 0 | 0.14 | 0.17 | 0.13 | 0.26 | 0.16 | 0.24 |
| 2 | 129914648 | rs10496675 | -0.791 | 0.07 | 0.067 | 0.615 | -0.015 | 0.057 | 0.057 | 0 | 0.13 | 0.03 | 0.07 | 0 | 0.14 | 0.05 |
| 2 | 129914893 | rs6713126 | 1.28 | 0.053 | 0.777 | 0.785 | -0.004 | 0.02 | 0.02 | 0 | 0.71 | 0.38 | 0.7 | 0.65 | 0.73 | 0.58 |
| 2 | 129915151 | rs10496676 | -1.717 | -0.121 | 1.51 | 1.046 | 0.012 | 0.052 | 0.052 | 0.015 | 0.26 | 0.35 | 0.25 | 0.36 | 0.28 | 0.34 |
| 2 | 129915467 | rs17049177 | -1.484 | 0.124 | 1.586 | 0.996 | 0.004 | 0.004 | 0.004 | 0 | 0.13 | 0.28 | 0.29 | 0.07 | 0.12 | 0.12 |
| 2 | 129915740 | rs964343 | 1.659 | 0.061 | 1.843 | 0.97 | 0.007 | 0.036 | 0.036 | 0 | 0.85 | 0.48 | 0.76 | 0.89 | 0.88 | 0.88 |
| 2 | 129918146 | rs10496677 | -1.017 | -0.137 | 1.779 | 0.844 | 0.003 | -0.012 | -0.012 | 1.717 | 0.14 | 0.12 | 0.04 | 0 | 0.04 | 0.1 |
| 2 | 129921407 | rs908849 | 2.083 | -0.191 | 1.787 | 0.83 | 0.052 | 0.049 | 0.049 | 1.286 | 0.7 | 0.19 | 0.65 | 0.88 | 0.84 | 0.77 |
| 2 | 129930092 | rs2707808 | 2.096 | -0.247 | 1.757 | 0.845 | 0.065 | 0.054 | 0.054 | 2.033 | 0.7 | 0.15 | 0.65 | 0.88 | 0.84 | 0.78 |
| 2 | 129939764 | rs2204904 | -3.161 | -0.342 | 1.454 | 1.121 | -0.038 | -0.105 | -0.105 | 0.346 | 0.45 | 0.96 | 0.49 | 0.37 | 0.35 | 0.46 |
| 2 | 129943346 | rs10171432 | -1.895 | -0.284 | 1.633 | 1.186 | 0.058 | 0.103 | 0.103 | 0 | 0.24 | 0.48 | 0.41 | 0.33 | 0.3 | 0.27 |
| 2 | 129947153 | rs17049228 | -1.258 | -0.325 | 1.617 | 0.89 | 0.011 | 0.012 | 0.012 | 0 | 0.2 | 0.16 | 0.08 | 0.01 | 0.13 | 0.14 |
| 2 | 129954793 | rs6754324 | -0.506 | -0.364 | 2.02 | 0.971 | 0.032 | 0.188 | 0.188 | 0.322 | 0.07 | 0.35 | 0.25 | 0.12 | 0.02 | 0.08 |
| 2 | 129961812 | rs2222901 | -2.307 | -0.398 | 1.437 | 1.364 | 0.004 | 0.001 | 0.001 | 0 | 0.46 | 0.61 | 0.46 | 0.51 | 0.42 | 0.54 |
| 2 | 129962060 | rs2204903 | -2.178 | -0.399 | 1.465 | 1.363 | -0.001 | -0.001 | -0.001 | 0 | 0.45 | 0.49 | 0.45 | 0.51 | 0.42 | 0.54 |
| 2 | 129964946 | rs2952619 | -2.841 | -0.409 | 1.349 | 1.353 | -0.091 | 0.094 | 0.094 | 0.143 | 0.53 | 0.96 | 0.71 | 0.63 | 0.44 | 0.62 |
| 2 | 129975956 | rs997005 | -1.877 | -0.288 | 1.536 | 1.323 | -0.009 | -0.026 | -0.026 | 0 | 0.39 | 0.3 | 0.43 | 0.47 | 0.4 | 0.41 |
| 2 | 129978391 | rs1441139 | -0.376 | -0.196 | 0.963 | 1.359 | -0.046 | -0.09 | -0.09 | 2.481 | 0.19 | 0.09 | 0.34 | 0.5 | 0.26 | 0.27 |
| 2 | 129980879 | rs6741167 | -1.717 | -0.18 | 1.024 | 1.432 | 0.004 | 0.01 | 0.01 | 0 | 0.46 | 0.61 | 0.46 | 0.65 | 0.41 | 0.53 |
| 2 | 129983541 | rs1251210 | 1.447 | 0.299 | 0.778 | 0.951 | 0.002 | 0.035 | 0.035 | 0 | 0.46 | 0.42 | 0.32 | 0.16 | 0.51 | 0.34 |
| 2 | 129984659 | rs1251213 | 0.067 | 0.216 | 0.86 | 0.863 | -0.026 | -0.019 | -0.019 | 0 | 0.92 | 0.84 | 0.98 | 0.96 | 0.97 | 0.84 |
| 2 | 129990869 | rs13033217 | -0.944 | 0.286 | 1.012 | 1.085 | -0.098 | 0.091 | 0.091 | 0 | 0.43 | 0.11 | 0.51 | 0.71 | 0.43 | 0.4 |
| 2 | 129991704 | rs751070 | 0.044 | 0.274 | 1.038 | 1.098 | -0.026 | -0.019 | -0.019 | 0 | 0.92 | 0.84 | 0.98 | 0.96 | 0.97 | 0.84 |
| 2 | 129998443 | rs2952621 | 1.392 | 0.28 | 1.244 | 1.087 | -0.001 | 0.083 | 0.083 | 0 | 0.48 | 0.67 | 0.47 | 0.25 | 0.54 | 0.44 |
| 2 | 130002588 | rs1372117 | -0.723 | 0.463 | 0.764 | 1.308 | -0.079 | -0.011 | -0.011 | 0.085 | 0.4 | 0.08 | 0.48 | 0.62 | 0.38 | 0.38 |
| 2 | 130007750 | rs2197022 | 1.577 | 0.467 | 1.151 | 1.135 | 0.004 | 0.134 | 0.134 | 0 | 0.46 | 0.31 | 0.32 | 0.16 | 0.51 | 0.35 |
| 2 | 130024033 | rs837899 | 0.75 | 0.206 | 1.956 | 2.063 | -0.019 | 0.014 | 0.014 | 0.116 | 0.86 | 0.75 | 0.92 | 0.82 | 0.91 | 0.83 |
| 2 | 130024589 | rs837898 | -0.816 | 0.49 | 1.922 | 1.985 | 0.016 | -0.015 | -0.015 | 0.51 | 0.37 | 0.51 | 0.58 | 0.66 | 0.4 | 0.45 |
| 2 | 130031647 | rs4662908 | -0.292 | 0.459 | 1.975 | 2.075 | -0.079 | -0.088 | -0.088 | 0 | 0.29 | 0.11 | 0.34 | 0.41 | 0.35 | 0.25 |
| 2 | 130034012 | rs837841 | -0.952 | 0.361 | 1.813 | 2.046 | 0.02 | -0.015 | -0.015 | 5.367 | 0.36 | 0.49 | 0.58 | 0.66 | 0.38 | 0.45 |
| 2 | 130034326 | rs7567687 | 1.651 | 0.347 | 1.821 | 2.005 | 0.002 | 0.216 | 0.216 | 0.187 | 0.5 | 0.24 | 0.33 | 0.16 | 0.53 | 0.37 |
| 2 | 130048509 | rs837855 | -0.524 | 0.513 | 1.915 | 2.027 | 0.037 | 0.041 | 0.041 | 0 | 0.31 | 0.39 | 0.48 | 0.49 | 0.37 | 0.36 |
| 2 | 130051445 | rs1251193 | 0.768 | 0.507 | 1.552 | 1.623 | -0.029 | 0.026 | 0.026 | 0.174 | 0.85 | 0.73 | 0.92 | 0.79 | 0.91 | 0.81 |
| 2 | 130057509 | rs4662909 | -0.62 | 0.642 | 1.447 | 1.769 | 0.027 | 0.019 | 0.019 | 0 | 0.31 | 0.36 | 0.47 | 0.31 | 0.36 | 0.34 |
| 2 | 130060157 | rs17049328 | -0.176 | 1.047 | 0.653 | 1.75 | -0.006 | -0.034 | -0.034 | 0 | 0.26 | 0.07 | 0.28 | 0.22 | 0.22 | 0.22 |
| 2 | 130060598 | rs4662910 | -0.737 | 0.598 | 0.545 | 1.544 | 0.028 | 0.016 | 0.016 | 0 | 0.33 | 0.51 | 0.48 | 0.31 | 0.37 | 0.35 |
| 2 | 130065486 | rs7349444 | 0.977 | 0.409 | 0.246 | 1.47 | -0.01 | -0.009 | -0.009 | 0 | 0.56 | 0.41 | 0.5 | 0.62 | 0.59 | 0.44 |
| 2 | 130066172 | rs10928902 | 1.48 | -0.498 | 1.097 | 1.799 | -0.109 | 0.18 | 0.18 | 3.205 | 0.82 | 0.49 | 0.82 | 0.54 | 0.95 | 0.72 |
| 2 | 130066311 | rs12463402 | 1.455 | -0.701 | 1.184 | 1.683 | -0.057 | 0.078 | 0.078 | 0.1 | 0.82 | 0.71 | 0.81 | 0.52 | 0.94 | 0.71 |
| 2 | 130071251 | rs4446032 | -1.017 | -1.032 | 0.152 | 0.328 | 0.036 | 0.083 | 0.083 | 0 | 0.5 | 0.62 | 0.61 | 0.84 | 0.38 | 0.53 |
| 2 | 130076139 | rs7599712 | 0.216 | -1.366 | -0.097 | 0.092 | -0.056 | -0.063 | -0.063 | 2.727 | 0.86 | 0.68 | 0.93 | 0.96 | 0.95 | 0.9 |
| 2 | 130076319 | rs3940590 | 0.541 | -1.636 | -0.837 | 0.595 | 0.033 | -0.028 | -0.028 | 0 | 0.29 | 0.08 | 0.3 | 0.31 | 0.21 | 0.16 |
| 2 | 130077270 | rs755569 | 1.57 | -1.069 | -0.646 | 0.107 | -0.121 | 0.132 | 0.132 | 0.926 | 0.36 | 0.04 | 0.31 | 0.12 | 0.57 | 0.37 |
| 2 | 130082114 | rs837868 | 0.377 | -1.248 | 0.115 | 0.301 | 0.007 | 0.006 | 0.006 | 0 | 0.55 | 0.58 | 0.56 | 0.32 | 0.71 | 0.65 |
| 2 | 130083785 | rs13433028 | -0.526 | -0.565 | 0.7 | 0.673 | 0.035 | 0.031 | 0.031 | 0.048 | 0.13 | 0.25 | 0.22 | 0.16 | 0.15 | 0.09 |

| | | | | | | | | | | | | | | | | |
|---|---|---|---|---|---|---|---|---|---|---|---|---|---|---|---|---|
| 2 | 130087664 | rs10169498 | -0.849 | -0.574 | 0.771 | 0.666 | 0.045 | 0.036 | 0.036 | 0.301 | 0.14 | 0.33 | 0.23 | 0.16 | 0.15 | 0.09 |
| 2 | 130088091 | rs10169910 | -0.83 | -0.537 | 0.766 | 0.669 | 0.049 | 0.037 | 0.037 | 0.403 | 0.14 | 0.33 | 0.23 | 0.16 | 0.15 | 0.09 |
| 2 | 130088717 | rs10210526 | -0.456 | -0.54 | 0.453 | 0.676 | 0.018 | 0.005 | 0.005 | 0.006 | 0.12 | 0.2 | 0.2 | 0.12 | 0.14 | 0.09 |
| 2 | 130091066 | rs7590490 | 0.997 | -0.426 | -0.442 | 0.181 | -0.011 | 0.116 | 0.116 | 0 | 0.36 | 0.08 | 0.21 | 0.15 | 0.29 | 0.35 |
| 2 | 130094203 | rs837870 | 0.888 | -1.242 | 1.085 | 0.565 | -0.026 | -0.03 | -0.03 | 0 | 0.88 | 0.52 | 0.9 | 0.96 | 0.95 | 0.92 |
| 2 | 130110882 | rs13397518 | 0.519 | -1.242 | 1.375 | 1.375 | 0.002 | -0.007 | -0.007 | 0 | 0.67 | 0.73 | 0.63 | 0.55 | 0.7 | 0.78 |
| 2 | 130111078 | rs4662917 | 0.742 | -1.172 | 1.471 | 1.293 | -0.019 | 0.001 | 0.001 | 0 | 0.64 | 0.55 | 0.55 | 0.32 | 0.69 | 0.65 |
| 2 | 130111979 | rs10196948 | 0.417 | -1.145 | 1.263 | 1.213 | -0.005 | -0.033 | -0.033 | 0.261 | 0.9 | 0.83 | 0.92 | 0.96 | 0.92 | 0.92 |
| 2 | 130124417 | rs6711309 | 1.081 | -1.207 | 1.327 | 1.241 | -0.002 | -0.001 | -0.001 | 0 | 0.63 | 0.39 | 0.54 | 0.32 | 0.69 | 0.65 |
| 2 | 130126993 | rs944906 | 0.497 | -1.312 | 0.672 | 1.221 | -0.038 | -0.148 | -0.148 | 0.279 | 0.19 | 0.02 | 0.29 | 0.42 | 0.22 | 0.13 |
| 2 | 130130648 | rs1404074 | 0.842 | -1.461 | 1.242 | 0.987 | 0.004 | -0.037 | -0.037 | 0 | 0.89 | 0.76 | 0.91 | 0.96 | 0.91 | 0.91 |
| 2 | 130132291 | rs11681622 | 1.109 | -1.457 | 1.152 | 0.997 | 0.012 | -0.043 | -0.043 | 0 | 0.88 | 0.65 | 0.9 | 0.96 | 0.91 | 0.91 |
| 2 | 130133365 | rs2030664 | 1.062 | -1.602 | 0.273 | 1.145 | 0.048 | -0.075 | -0.075 | 0.398 | 0.62 | 0.08 | 0.48 | 0.36 | 0.69 | 0.63 |
| 2 | 130140383 | rs1097704 | -0.378 | -1.649 | 0.547 | 1.348 | 0.011 | -0.037 | -0.037 | 0 | 0.28 | 0.17 | 0.36 | 0.39 | 0.28 | 0.21 |
| 2 | 130140883 | rs2123522 | 1.082 | -1.617 | 0.29 | 1.182 | 0.047 | -0.072 | -0.072 | 0.21 | 0.62 | 0.08 | 0.48 | 0.36 | 0.69 | 0.64 |
| 2 | 130154906 | rs4662923 | 1.081 | -1.612 | 0.334 | 1.169 | 0.054 | -0.062 | -0.062 | 0.011 | 0.62 | 0.11 | 0.49 | 0.36 | 0.69 | 0.64 |
| 2 | 130159699 | rs6431075 | 1.09 | -1.469 | 1.061 | 1.24 | -0.02 | 0.03 | 0.03 | 1.557 | 0.92 | 0.45 | 0.92 | 0.93 | 0.96 | 1 |
| 2 | 130167351 | rs985162 | -1.474 | -1.634 | 1.307 | 1.26 | 0.042 | 0.006 | 0.006 | 0 | 0.37 | 0.77 | 0.52 | 0.69 | 0.31 | 0.41 |
| 2 | 130167765 | rs13387772 | -1.671 | -1.654 | 1.579 | 1.352 | -0.028 | 0.163 | 0.163 | 1.365 | 0.09 | 0.58 | 0.16 | 0.24 | 0.03 | 0.15 |
| 2 | 130175746 | rs7588843 | -0.458 | -1.647 | 1.532 | 1.354 | 0.014 | -0.065 | -0.065 | 0 | 0.28 | 0.19 | 0.37 | 0.45 | 0.28 | 0.27 |
| 2 | 130186163 | rs13427948 | -0.709 | -1.468 | 1.431 | 1.024 | 0.044 | 0.117 | 0.117 | 0.031 | 0.06 | 0.36 | 0.07 | 0.07 | 0.03 | 0 |
| 2 | 130191184 | rs10205941 | 1.047 | -1.477 | 0.968 | 0.878 | -0.002 | -0.01 | -0.01 | 0 | 0.56 | 0.51 | 0.35 | 0.25 | 0.58 | 0.39 |
| 2 | 130191927 | rs17518713 | 0.256 | -1.518 | 0.833 | 0.952 | -0.037 | -0.03 | -0.03 | 0 | 0.15 | 0.03 | 0.24 | 0.14 | 0.16 | 0.23 |
| 2 | 130197500 | rs1540552 | -0.441 | -1.546 | 0.996 | 0.924 | -0.001 | -0.019 | -0.019 | 0 | 0.07 | 0.16 | 0.04 | 0.02 | 0.04 | 0.11 |
| 2 | 130200925 | rs10174187 | 0.707 | -1.555 | 1.239 | 0.901 | 0 | 0 | 0 | 0 | 0.6 | 0.63 | 0.39 | 0.25 | 0.62 | 0.43 |
| 2 | 130201273 | rs7570620 | -1.319 | -1.605 | 1.278 | 0.874 | 0.007 | 0.007 | 0.007 | 0.162 | 0.12 | 0.49 | 0.1 | 0.07 | 0.08 | 0.04 |
| 2 | 130201986 | rs17049483 | -0.55 | -1.614 | 1.259 | 0.889 | -0.001 | -0.001 | -0.001 | 0 | 0.19 | 0.19 | 0.26 | 0.16 | 0.16 | 0.32 |
| 2 | 130205544 | rs921622 | -1.833 | -1.602 | 1.03 | 0.894 | 0.003 | 0.002 | 0.002 | 0 | 0.31 | 0.68 | 0.36 | 0.23 | 0.24 | 0.36 |
| 2 | 130208064 | rs2406820 | -0.939 | -1.5 | 0.964 | 0.964 | 0.001 | -0.001 | -0.001 | 1.124 | 0.79 | 0.82 | 0.65 | 0.42 | 0.78 | 0.75 |
| 2 | 130216043 | rs4662945 | 0.522 | -1.445 | 0.983 | 0.899 | 0.003 | 0.023 | 0.023 | 0.104 | 0.63 | 0.75 | 0.4 | 0.27 | 0.6 | 0.5 |
| 2 | 130217485 | rs10210419 | -0.571 | -1.41 | 1.472 | 0.901 | -0.016 | 0.046 | 0.046 | 0.051 | 0.77 | 0.65 | 0.61 | 0.35 | 0.76 | 0.75 |
| 2 | 130218178 | rs2406822 | 2.222 | -1.762 | 1.109 | 0.675 | -0.072 | 0.164 | 0.164 | 0 | 0.44 | 0.04 | 0.28 | 0.18 | 0.51 | 0.36 |
| 2 | 130219822 | rs6705917 | 0.31 | -1.767 | 1.552 | 0.681 | -0.023 | 0.067 | 0.067 | 0.117 | 0.24 | 0.4 | 0.39 | 0.65 | 0.24 | 0.24 |
| 2 | 130221148 | rs4427966 | 1.99 | -1.782 | 1.652 | 0.758 | 0.012 | 0.093 | 0.093 | 0 | 0.47 | 0.23 | 0.35 | 0.27 | 0.52 | 0.44 |
| 2 | 130222522 | rs12373608 | 0.937 | -1.735 | 1.653 | 0.965 | 0.004 | -0.022 | -0.022 | 0.192 | 0.2 | 0.14 | 0.31 | 0.4 | 0.24 | 0.14 |
| 2 | 130222801 | rs13394229 | 0.252 | -1.734 | 1.707 | 0.95 | -0.028 | 0.138 | 0.138 | 0.157 | 0.24 | 0.45 | 0.37 | 0.56 | 0.24 | 0.24 |
| 2 | 130231345 | rs10185238 | 1.81 | -1.61 | 1.731 | 0.713 | 0.027 | -0.001 | -0.001 | 0 | 0.65 | 0.17 | 0.63 | 0.68 | 0.74 | 0.52 |
| 2 | 130231480 | rs7607327 | 1.374 | -1.51 | 1.739 | 0.791 | 0.083 | 0.31 | 0.31 | 1.849 | 0.95 | 0.56 | 0.9 | 0.84 | 0.99 | 0.9 |
| 2 | 130240631 | rs12478202 | -0.051 | -1.532 | 1.627 | 0.726 | -0.01 | -0.011 | -0.011 | 0 | 0.17 | 0.06 | 0.24 | 0.16 | 0.16 | 0.25 |
| 2 | 130245943 | rs11680033 | -0.535 | -1.414 | 0.823 | 0.532 | 0 | 0.023 | 0.023 | 0 | 0.1 | 0.06 | 0.03 | 0 | 0.09 | 0.06 |
| 2 | 130250978 | rs2123518 | -0.421 | -1.066 | 1.029 | 0.718 | 0.003 | 0.067 | 0.067 | 0 | 0.2 | 0.35 | 0.24 | 0.28 | 0.16 | 0.37 |
| 2 | 130258186 | rs10200446 | 0.44 | -1.221 | 1.028 | 0.822 | -0.013 | 0.063 | 0.063 | 0 | 0.21 | 0.37 | 0.32 | 0.34 | 0.22 | 0.18 |
| 2 | 130262209 | rs10496678 | 2.182 | -1.518 | 0.658 | 0.579 | -0.114 | 0.008 | 0.008 | 0 | 0.43 | 0.04 | 0.36 | 0.37 | 0.51 | 0.39 |
| 2 | 130269829 | rs7601707 | -1.146 | -1.514 | 0.713 | 0.341 | 0 | -0.006 | -0.006 | 0 | 0.3 | 0.35 | 0.28 | 0.23 | 0.25 | 0.42 |
| 2 | 130271510 | rs1453147 | 2.239 | -1.516 | 0.75 | 0.34 | -0.124 | -0.057 | -0.057 | 0 | 0.43 | 0.03 | 0.36 | 0.43 | 0.51 | 0.38 |
| 2 | 130297109 | rs4519460 | 0.158 | -1.582 | 0.839 | 0.449 | 0.002 | 0.036 | 0.036 | 0 | 0.8 | 0.66 | 0.76 | 0.77 | 0.85 | 0.63 |
| 2 | 130306785 | rs10203768 | 0.521 | -1.515 | 0.657 | 0.462 | -0.012 | 0.05 | 0.05 | 0 | 0.2 | 0.33 | 0.32 | 0.35 | 0.22 | 0.18 |
| 2 | 130313935 | rs12617480 | 0.962 | -1.401 | 0.706 | 0.418 | -0.01 | -0.007 | -0.007 | 0 | 0.7 | 0.42 | 0.72 | 0.77 | 0.79 | 0.54 |
| 2 | 130315974 | rs2406983 | 0.575 | -1.746 | 0.485 | 0.26 | 0.016 | -0.027 | -0.027 | 0 | 0.23 | 0.12 | 0.3 | 0.34 | 0.23 | 0.16 |
| 2 | 130316023 | rs2003288 | 1.346 | -1.762 | 0.517 | 0.242 | 0.028 | 0.023 | 0.023 | 0 | 0.49 | 0.3 | 0.42 | 0.43 | 0.58 | 0.38 |
| 2 | 130316931 | rs4662959 | 0.795 | -1.537 | 0.541 | 0.324 | -0.012 | -0.012 | -0.012 | 0 | 0.69 | 0.5 | 0.72 | 0.77 | 0.77 | 0.54 |
| 2 | 130317048 | rs4613217 | 0.382 | -1.5 | 0.7 | 0.417 | 0.004 | 0.013 | 0.013 | 0 | 0.76 | 0.62 | 0.74 | 0.77 | 0.82 | 0.55 |
| 2 | 130318908 | rs10153725 | -0.52 | -1.336 | 0.355 | 0.46 | 0.013 | 0.002 | 0.002 | 0 | 0.33 | 0.27 | 0.35 | 0.34 | 0.29 | 0.22 |
| 2 | 130319985 | rs11685256 | -0.52 | -1.336 | 0.354 | 0.46 | 0.012 | 0.002 | 0.002 | 0 | 0.33 | 0.27 | 0.35 | 0.34 | 0.29 | 0.21 |
| 2 | 130323107 | rs2214271 | -1.068 | -0.607 | 0.759 | 0.761 | 0 | 0.003 | 0.003 | 0 | 0.31 | 0.3 | 0.43 | 0.49 | 0.36 | 0.33 |
| 2 | 130334543 | rs2521951 | -1.645 | -0.707 | 1.21 | 0.653 | 0.004 | 0.008 | 0.008 | 0 | 0.48 | 0.55 | 0.48 | 0.46 | 0.45 | 0.44 |
| 2 | 130336029 | rs2214270 | -1.134 | -0.68 | 1.283 | 0.669 | -0.002 | -0.002 | -0.002 | 0 | 0.42 | 0.36 | 0.46 | 0.46 | 0.43 | 0.43 |
| 2 | 130344802 | rs886263 | -1.015 | -0.683 | 1.241 | 0.689 | -0.026 | 0.035 | 0.035 | 0 | 0.2 | 0.14 | 0.21 | 0.07 | 0.24 | 0.27 |
| 2 | 130346959 | rs11900978 | 0.48 | -0.959 | 1.391 | 0.762 | -0.008 | 0.018 | 0.018 | 0 | 0.21 | 0.21 | 0.25 | 0.39 | 0.19 | 0.16 |
| 2 | 130354239 | rs2188907 | -0.769 | -0.818 | 1.388 | 0.604 | 0.001 | -0.004 | -0.004 | 4.325 | 0.13 | 0.15 | 0.17 | 0.07 | 0.16 | 0.21 |
| 2 | 130355440 | rs2704533 | -1.446 | -0.824 | 1.382 | 0.598 | -0.011 | 0.012 | 0.012 | 0 | 0.2 | 0.19 | 0.22 | 0.07 | 0.24 | 0.27 |
| 2 | 130355506 | rs3109303 | 0.335 | -0.882 | 1.429 | 0.609 | -0.025 | -0.035 | -0.035 | 0 | 0.06 | 0.14 | 0.01 | 0 | 0.02 | 0.01 |
| 2 | 130362926 | rs3109306 | -1.052 | -0.916 | 1.522 | 0.646 | -0.05 | 0.1 | 0.1 | 0 | 0.11 | 0.38 | 0.07 | 0.15 | 0.02 | 0.2 |
| 2 | 130363260 | rs1034646 | -2.334 | -0.937 | 1.413 | 0.585 | 0.007 | 0.016 | 0.016 | 0 | 0.34 | 0.71 | 0.31 | 0.22 | 0.26 | 0.48 |
| 2 | 130367080 | rs3094419 | -0.861 | -1.112 | 1.501 | 0.547 | -0.029 | 0.023 | 0.023 | 0 | 0.33 | 0.54 | 0.36 | 0.55 | 0.28 | 0.42 |
| 2 | 130368088 | rs3094417 | -0.255 | -1.048 | 1.608 | 0.56 | -0.006 | -0.05 | -0.05 | 0.029 | 0.91 | 0.8 | 0.94 | 1 | 0.92 | 0.94 |
| 2 | 130376011 | rs1012734 | 2.077 | -1.024 | 1.397 | 0.56 | -0.008 | -0.013 | -0.013 | 0.007 | 0.46 | 0.14 | 0.44 | 0.41 | 0.55 | 0.36 |
| 2 | 130384917 | rs3109015 | 0.007 | -1.153 | 1.66 | 0.489 | -0.013 | 0.068 | 0.068 | 0 | 0.91 | 0.75 | 0.92 | 0.86 | 0.97 | 0.75 |
| 2 | 130385050 | rs2704555 | -1.772 | -1.189 | 1.83 | 0.436 | -0.002 | -0.093 | -0.093 | 0 | 0.25 | 0.38 | 0.25 | 0.06 | 0.22 | 0.22 |

| | | | | | | | | | | | | | | | | |
|---|---|---|---|---|---|---|---|---|---|---|---|---|---|---|---|---|
| 2 | 130387174 | rs2521930 | -1.362 | -1.169 | 1.721 | 0.43 | 0.009 | 0.001 | 0.001 | 0 | 0.18 | 0.31 | 0.24 | 0.17 | 0.16 | 0.4 |
| 2 | 130392796 | rs2704553 | -1.147 | -1.104 | 1.531 | 0.465 | 0.006 | -0.068 | -0.068 | 0 | 0.16 | 0.26 | 0.19 | 0.04 | 0.14 | 0.17 |
| 2 | 130392975 | rs2704552 | -0.19 | -1.042 | 1.186 | 0.498 | 0.015 | -0.025 | -0.025 | 1.118 | 0.09 | 0.14 | 0.17 | 0.04 | 0.13 | 0.17 |
| 2 | 130393136 | rs2521920 | 1.607 | -1.06 | 1.2 | 0.49 | 0.005 | 0.011 | 0.011 | 0 | 0.66 | 0.35 | 0.68 | 0.82 | 0.75 | 0.52 |
| 2 | 130396140 | rs3109320 | 0.795 | -0.675 | 0.587 | 0.431 | 0.079 | -0.002 | -0.002 | 0 | 0.55 | 0.29 | 0.64 | 0.46 | 0.68 | 0.52 |
| 2 | 130400730 | rs2023803 | -0.706 | -0.384 | 0.64 | 0.462 | 0.051 | -0.05 | -0.05 | 0 | 0.12 | 0.29 | 0.18 | 0.04 | 0.12 | 0.17 |
| 2 | 130403009 | rs2521933 | 1.32 | -0.548 | 0.158 | 0.482 | 0.008 | 0.01 | 0.01 | 0 | 0.47 | 0.19 | 0.48 | 0.43 | 0.55 | 0.35 |
| 2 | 130405842 | rs2406998 | -1.418 | -0.541 | 0.186 | 0.396 | 0.013 | 0.075 | 0.075 | 0 | 0.53 | 0.8 | 0.54 | 0.62 | 0.45 | 0.65 |
| 2 | 130410719 | rs4146382 | -0.399 | -0.656 | 0.484 | 0.378 | 0.005 | 0 | 0 | 0 | 0.36 | 0.38 | 0.34 | 0.59 | 0.31 | 0.48 |
| 2 | 130412891 | rs3109009 | 1.206 | -0.56 | 0.173 | 0.42 | 0.002 | 0.079 | 0.079 | 0 | 0.47 | 0.17 | 0.45 | 0.38 | 0.55 | 0.35 |
| 2 | 130419906 | rs2407013 | 1.262 | -0.075 | -0.019 | 0.245 | -0.07 | 0.042 | 0.042 | 0 | 0.41 | 0.17 | 0.39 | 0.36 | 0.42 | 0.32 |
| 2 | 130427167 | rs12479404 | -0.131 | 0.044 | 0.831 | 0.193 | 0.053 | -0.012 | -0.012 | 0 | 0.07 | 0.2 | 0.16 | 0.04 | 0.09 | 0.12 |
| 2 | 130428389 | rs3109276 | -0.758 | -0.231 | 0.999 | 0.249 | -0.005 | -0.013 | -0.013 | 0 | 0.26 | 0.32 | 0.13 | 0.15 | 0.18 | 0.38 |
| 2 | 130429646 | rs7572865 | 0.197 | -0.199 | 1.165 | 0.265 | 0.013 | -0.015 | -0.015 | 0 | 0.07 | 0.09 | 0.15 | 0.04 | 0.09 | 0.12 |
| 2 | 130430091 | rs3108980 | -0.894 | -0.269 | 1.132 | 0.331 | 0.008 | 0.02 | 0.02 | 0 | 0.46 | 0.41 | 0.43 | 0.6 | 0.47 | 0.57 |
| 2 | 130432067 | rs2407015 | -0.138 | -0.282 | 1.1 | 0.318 | -0.001 | 0.011 | 0.011 | 0 | 0.2 | 0.2 | 0.11 | 0.15 | 0.17 | 0.37 |
| 2 | 130433313 | rs6725384 | 0.641 | -0.288 | 1.071 | 0.321 | 0.007 | -0.033 | -0.033 | 0.698 | 0.25 | 0.18 | 0.31 | 0.45 | 0.3 | 0.19 |
| 2 | 130437195 | rs10928920 | 0.26 | -0.504 | 0.898 | 0.35 | -0.003 | -0.017 | -0.017 | 0 | 0.79 | 0.68 | 0.88 | 0.85 | 0.83 | 0.63 |
| 2 | 130456202 | rs12994016 | 1.453 | 0.199 | 0.248 | 0.24 | -0.017 | 0.02 | 0.02 | 0 | 0.4 | 0.21 | 0.39 | 0.37 | 0.39 | 0.3 |
| 2 | 130457465 | rs12473869 | 0.135 | 0.343 | 0.471 | 0.781 | -0.009 | -0.009 | -0.009 | 0 | 0.31 | 0.24 | 0.46 | 0.34 | 0.43 | 0.31 |
| 2 | 130457993 | rs17049692 | -0.954 | -0.194 | 0.454 | 0.819 | 0.002 | -0.005 | -0.005 | 0 | 0.22 | 0.22 | 0.18 | 0.33 | 0.16 | 0.42 |
| 2 | 130458709 | rs6745475 | -1.065 | -0.193 | 0.374 | 0.818 | -0.016 | 0.025 | 0.025 | 0 | 0.23 | 0.36 | 0.19 | 0.32 | 0.16 | 0.42 |
| 2 | 130460096 | rs4662980 | -0.41 | -0.222 | -0.106 | 0.393 | -0.013 | -0.011 | -0.011 | 0 | 0.45 | 0.28 | 0.54 | 0.44 | 0.5 | 0.41 |
| 2 | 130460115 | rs4662981 | -0.823 | -0.456 | 0.528 | 0.186 | 0.004 | 0.015 | 0.015 | 1.223 | 0.46 | 0.44 | 0.47 | 0.36 | 0.48 | 0.45 |
| 2 | 130461143 | rs10928926 | 0.076 | -0.645 | 0.738 | 0.454 | 0 | -0.006 | -0.006 | 0 | 0.66 | 0.52 | 0.56 | 0.67 | 0.58 | 0.82 |
| 2 | 130469908 | rs4662986 | 1.829 | -1.284 | 1.229 | 0.455 | -0.01 | -0.002 | -0.002 | 0 | 0.75 | 0.35 | 0.72 | 0.7 | 0.87 | 0.65 |
| 2 | 130477704 | rs7601871 | -1.404 | -1.005 | 1.97 | 0.178 | 0.011 | -0.056 | -0.056 | 2 | 0.1 | 0.34 | 0.06 | 0 | 0.07 | 0.05 |
| 2 | 130483910 | rs4233595 | 1.385 | -1.017 | 1.479 | 0.173 | -0.027 | -0.069 | -0.069 | 0 | 0.86 | 0.34 | 0.9 | 1 | 0.93 | 0.94 |
| 2 | 130491622 | rs6710206 | 1.383 | -1.012 | 1 | 0.172 | -0.024 | -0.065 | -0.065 | 0 | 0.86 | 0.34 | 0.9 | 1 | 0.93 | 0.94 |
| 2 | 130507364 | rs7589671 | 0.748 | -0.966 | 0.17 | 0.68 | -0.08 | 0.034 | 0.034 | 0 | 0.77 | 0.51 | 0.73 | 0.7 | 0.91 | 0.62 |
| 2 | 130527538 | rs4622658 | 0.138 | -0.371 | -0.292 | 0.538 | -0.011 | -0.003 | -0.003 | 0.152 | 0.6 | 0.5 | 0.43 | 0.31 | 0.63 | 0.39 |
| 2 | 130534281 | rs7578253 | -0.37 | -0.307 | -0.059 | 0.054 | 0.004 | -0.028 | -0.028 | 0 | 0.35 | 0.57 | 0.4 | 0.29 | 0.34 | 0.27 |
| 2 | 130543746 | rs2952605 | -0.16 | -0.35 | 0.684 | -0.236 | 0.001 | -0.014 | -0.014 | 0 | 0.79 | 0.77 | 0.89 | 0.89 | 0.79 | 0.89 |
| 2 | 130544098 | rs2707645 | 0.449 | -0.836 | 1.385 | -0.54 | -0.01 | -0.045 | -0.045 | 0 | 0.7 | 0.45 | 0.85 | 0.89 | 0.77 | 0.88 |
| 2 | 130550135 | rs2464786 | 0.895 | -1.175 | 1.341 | -0.522 | -0.026 | -0.004 | -0.004 | 0.475 | 0.37 | 0.15 | 0.46 | 0.6 | 0.44 | 0.58 |
| 2 | 130551925 | rs13013015 | 0.57 | -1.185 | 1.181 | -0.522 | -0.073 | 0.007 | 0.007 | 0 | 0.32 | 0.09 | 0.36 | 0.3 | 0.34 | 0.29 |
| 2 | 130552008 | rs13011607 | 0.438 | -1.178 | 1.6 | -0.52 | -0.008 | 0.016 | 0.016 | 0 | 0.34 | 0.28 | 0.38 | 0.3 | 0.34 | 0.29 |
| 2 | 130553849 | rs2464792 | -0.369 | -1.241 | 1.598 | -0.371 | -0.011 | 0.03 | 0.03 | 0 | 0.81 | 0.9 | 0.9 | 0.9 | 0.81 | 0.87 |
| 2 | 130568652 | rs4662665 | -0.144 | -1.699 | 1.639 | -0.593 | 0.008 | 0.017 | 0.017 | 0 | 0.23 | 0.15 | 0.09 | 0.06 | 0.16 | 0.14 |
| 2 | 130571830 | rs2464776 | 0.832 | -1.754 | 1.132 | -0.622 | -0.079 | 0.003 | 0.003 | 2.123 | 0.38 | 0.15 | 0.51 | 0.63 | 0.57 | 0.5 |
| 2 | 130576290 | rs7584759 | 0.126 | -2.012 | 1.44 | -0.492 | 0.023 | 0.022 | 0.022 | 0 | 0.36 | 0.27 | 0.35 | 0.33 | 0.27 | 0.36 |
| 2 | 130579222 | rs2464777 | 0.764 | -1.949 | 1.281 | -0.602 | -0.001 | 0.048 | 0.048 | 1.893 | 0.41 | 0.19 | 0.55 | 0.65 | 0.63 | 0.49 |
| 2 | 130583503 | rs2707652 | 0.754 | -1.905 | 0.922 | -0.58 | 0.003 | 0.044 | 0.044 | 1.823 | 0.41 | 0.23 | 0.56 | 0.64 | 0.63 | 0.48 |
| 2 | 130590169 | rs4662666 | -0.531 | -1.848 | 0.96 | -0.182 | 0.031 | 0.056 | 0.056 | 0.207 | 0.18 | 0.07 | 0.05 | 0.06 | 0.1 | 0.13 |
| 2 | 130603995 | rs7421552 | -0.634 | -1.846 | 1.037 | -0.181 | 0.008 | 0.006 | 0.006 | 0 | 0.19 | 0.15 | 0.06 | 0.06 | 0.1 | 0.13 |
| 2 | 130618373 | rs7566086 | -0.769 | -1.597 | 0.282 | 0.76 | -0.021 | 0.033 | 0.033 | 0 | 0.14 | 0.17 | 0.25 | 0.35 | 0.08 | 0.2 |
| 2 | 130621840 | rs4470286 | -0.802 | -2.487 | 0.732 | 0.896 | -0.097 | -0.019 | -0.019 | 0 | 0.19 | 0.49 | 0.08 | 0.19 | 0.06 | 0.23 |
| 2 | 130622888 | rs1955394 | -1.675 | -2.449 | 0.55 | 0.913 | -0.013 | -0.002 | -0.002 | 0.3 | 0.33 | 0.66 | 0.33 | 0.54 | 0.13 | 0.43 |
| 2 | 130630344 | rs6720670 | -1.635 | -2.538 | 0.396 | 0.938 | -0.047 | 0.009 | 0.009 | 0 | 0.35 | 0.57 | 0.32 | 0.51 | 0.14 | 0.33 |
| 2 | 130643612 | rs4662667 | 1.049 | -1.949 | -0.47 | 1.183 | -0.022 | 0.016 | 0.016 | 0 | 0.66 | 0.57 | 0.66 | 0.46 | 0.76 | 0.67 |
| 2 | 130645159 | rs6726811 | 0.483 | -1.91 | -0.544 | 1.294 | 0.013 | -0.022 | -0.022 | 0.145 | 0.75 | 0.85 | 0.69 | 0.6 | 0.8 | 0.78 |
| 2 | 130645784 | rs11898081 | 1.236 | -1.972 | -0.558 | 1.12 | -0.004 | 0 | 0 | 0 | 0.6 | 0.55 | 0.64 | 0.34 | 0.73 | 0.54 |
| 2 | 130646173 | rs4233600 | 1.223 | -1.974 | -0.559 | 1.119 | -0.009 | 0.001 | 0.001 | 0 | 0.6 | 0.54 | 0.64 | 0.34 | 0.73 | 0.54 |
| 2 | 130647012 | rs1316468 | 0.633 | -1.29 | -1.107 | 1.231 | 0.009 | -0.019 | -0.019 | 0 | 0.75 | 0.81 | 0.68 | 0.51 | 0.79 | 0.79 |
| 2 | 130650828 | rs4663004 | 0.007 | -0.357 | -1.945 | 0.825 | -0.014 | 0.04 | 0.04 | 0 | 0.61 | 0.86 | 0.51 | 0.29 | 0.61 | 0.55 |
| 2 | 130658715 | rs2407072 | -0.013 | -0.532 | -2.145 | 1.255 | 0.021 | -0.056 | -0.056 | 0.036 | 0.8 | 0.88 | 0.72 | 0.48 | 0.84 | 0.65 |
| 2 | 130658908 | rs2407073 | 0.231 | -0.183 | -2.252 | 0.667 | -0.051 | -0.02 | -0.02 | 0 | 0.37 | 0.21 | 0.28 | 0.33 | 0.47 | 0.31 |
| 2 | 130659897 | rs4663006 | 0.462 | -0.067 | -2.521 | 0.126 | -0.069 | 0.019 | 0.019 | 0 | 0.22 | 0.03 | 0.26 | 0.1 | 0.26 | 0.15 |
| 2 | 130662519 | rs2407074 | 0.824 | 0.39 | -2.08 | 0.09 | 0.033 | 0.024 | 0.024 | 0 | 0.5 | 0.58 | 0.44 | 0.62 | 0.38 | 0.49 |
| 2 | 130681447 | rs11897873 | 0.29 | 0.041 | -1.443 | 0.488 | 0.004 | -0.001 | -0.001 | 0 | 0.7 | 0.77 | 0.69 | 0.67 | 0.73 | 0.67 |
| 2 | 130683344 | rs10928942 | 0.364 | 0.038 | -1.351 | 0.487 | 0.007 | -0.005 | -0.005 | 0 | 0.71 | 0.84 | 0.69 | 0.67 | 0.73 | 0.66 |
| 2 | 130701189 | rs1922633 | 0.532 | -0.138 | -0.339 | 0.611 | -0.027 | 0.002 | 0.002 | 0.78 | 0.66 | 0.47 | 0.69 | 0.6 | 0.7 | 0.68 |
| 2 | 130718827 | rs6752853 | -1.437 | -0.185 | 0.482 | 0.472 | 0.024 | 0.009 | 0.009 | 2.151 | 0.07 | 0.48 | 0.05 | 0.01 | 0.02 | 0 |
| 2 | 130731130 | rs11886689 | -1.539 | -0.227 | 0.351 | 0.548 | 0.01 | 0.129 | 0.129 | 3.372 | 0.08 | 0.58 | 0.06 | 0.14 | 0.01 | 0.02 |
| 2 | 131685875 | rs4247444 | 1.631 | -0.513 | 2.007 | 1.182 | 0.118 | -0.036 | -0.036 | 0.052 | 0.9 | 0.51 | 0.86 | 1 | 0.88 | 0.93 |
| 2 | 131755154 | rs10185789 | -1.78 | 0.029 | 1.425 | 1.335 | -0.016 | 0.002 | 0.002 | 0.031 | 0.27 | 0.56 | 0.2 | 0.29 | 0.23 | 0.43 |
| 2 | 131756501 | rs13399132 | -2.07 | 0.126 | 1.323 | 1.493 | 0 | -0.001 | -0.001 | 0 | 0.33 | 0.69 | 0.33 | 0.36 | 0.29 | 0.46 |
| 2 | 131762450 | rs10196530 | -1.336 | -0.14 | 1.2 | 1.573 | -0.046 | 0.068 | 0.068 | 0.62 | 0.22 | 0.59 | 0.18 | 0.36 | 0.15 | 0.4 |
| 2 | 131762905 | rs7579742 | -1.449 | -0.019 | 0.655 | 0.985 | 0.013 | 0.008 | 0.008 | 0 | 0.11 | 0.2 | 0.17 | 0.11 | 0.13 | 0.09 |

| | | | | | | | | | | | | | | | | |
|---|---|---|---|---|---|---|---|---|---|---|---|---|---|---|---|---|
| 2 | 131763953 | rs13032375 | -0.783 | -0.301 | 0.253 | 0.842 | -0.005 | -0.001 | -0.001 | 0.851 | 0.15 | 0.16 | 0.16 | 0.13 | 0.14 | 0.22 |
| 2 | 131768531 | rs6430514 | 1.118 | -0.59 | 0.455 | 0.578 | -0.06 | -0.029 | -0.029 | 0 | 0.83 | 0.35 | 0.92 | 0.89 | 0.97 | 0.77 |
| 2 | 131771762 | rs2403238 | 1.366 | -0.157 | 1.082 | 0.549 | -0.025 | -0.056 | -0.056 | 0.261 | 0.83 | 0.43 | 0.89 | 0.91 | 0.92 | 0.78 |
| 2 | 131774186 | rs12466662 | -0.143 | 0.107 | 0.169 | 0.902 | -0.048 | -0.067 | -0.067 | 0 | 0.14 | 0.02 | 0.17 | 0.22 | 0.25 | 0.17 |
| 2 | 131781750 | rs6706055 | 0.603 | -0.094 | 1.322 | 0.471 | -0.055 | -0.022 | -0.022 | 0 | 0.87 | 0.69 | 0.95 | 0.92 | 0.98 | 0.78 |
| 2 | 131797367 | rs2278115 | -0.971 | -1.112 | 1.637 | 0.027 | -0.06 | -0.036 | -0.036 | 0 | 0.23 | 0.42 | 0.13 | 0.13 | 0.08 | 0.23 |
| 2 | 131799563 | rs2241461 | -1 | -1.17 | 1.838 | -0.011 | -0.076 | -0.048 | -0.048 | 0 | 0.23 | 0.47 | 0.13 | 0.13 | 0.08 | 0.23 |
| 2 | 131802332 | rs10928516 | -0.962 | -0.953 | 1.946 | -0.048 | -0.009 | -0.008 | -0.008 | 0.312 | 0.23 | 0.28 | 0.12 | 0.13 | 0.11 | 0.24 |
| 2 | 131819651 | rs6712737 | -0.059 | -1.47 | 1.333 | -0.494 | -0.011 | 0.069 | 0.069 | 0 | 0.78 | 0.96 | 0.86 | 0.91 | 0.75 | 0.8 |
| 2 | 131833758 | rs11900545 | -1.042 | -1.437 | 2.224 | -0.592 | -0.008 | -0.008 | -0.008 | 0 | 0.26 | 0.72 | 0.16 | 0.13 | 0.12 | 0.24 |
| 2 | 131852643 | rs7597771 | -0.754 | -1.458 | 2.322 | -0.6 | -0.06 | -0.055 | -0.055 | 0 | 0.23 | 0.5 | 0.14 | 0.13 | 0.11 | 0.23 |
| 2 | 131854373 | rs11685313 | -0.641 | -1.459 | 2.322 | -0.601 | -0.022 | -0.02 | -0.02 | 0 | 0.23 | 0.32 | 0.12 | 0.13 | 0.11 | 0.23 |
| 2 | 131855231 | rs11892761 | -0.816 | -1.459 | 2.31 | -0.601 | -0.065 | -0.059 | -0.059 | 0 | 0.24 | 0.51 | 0.15 | 0.13 | 0.11 | 0.23 |
| 2 | 131865064 | rs11901794 | -0.458 | -1.489 | 2.296 | -0.638 | 0.003 | 0.003 | 0.003 | 0 | 0.22 | 0.23 | 0.12 | 0.13 | 0.11 | 0.25 |
| 2 | 131869747 | rs918958 | -0.706 | -1.336 | 2.11 | -0.727 | -0.018 | 0.031 | 0.031 | 0 | 0.84 | 0.96 | 0.89 | 0.91 | 0.78 | 0.8 |
| 2 | 131889179 | rs10928539 | -1.016 | -1.402 | 2.445 | -0.833 | -0.027 | -0.029 | -0.029 | 0 | 0.24 | 0.34 | 0.13 | 0.13 | 0.11 | 0.25 |
| 2 | 131895230 | rs11895872 | -1.095 | -1.401 | 2.459 | -0.832 | -0.068 | -0.072 | -0.072 | 8.167 | 0.25 | 0.51 | 0.15 | 0.13 | 0.11 | 0.25 |
| 2 | 131896476 | rs11897491 | -1.393 | -1.409 | 2.665 | -0.838 | -0.008 | -0.01 | -0.01 | 0 | 0.27 | 0.68 | 0.17 | 0.13 | 0.12 | 0.26 |
| 2 | 131898811 | rs11899277 | -1.027 | -1.431 | 2.554 | -0.829 | -0.027 | -0.029 | -0.029 | 0 | 0.24 | 0.34 | 0.13 | 0.13 | 0.11 | 0.25 |
| 2 | 131945159 | rs6761005 | -0.955 | -1.364 | 2.806 | -0.788 | -0.024 | -0.017 | -0.017 | 0.023 | 0.23 | 0.34 | 0.13 | 0.15 | 0.11 | 0.25 |
| 2 | 131951961 | rs1834274 | -1.194 | -1.361 | 2.541 | -0.786 | -0.036 | -0.027 | -0.027 | 0 | 0.25 | 0.59 | 0.15 | 0.15 | 0.11 | 0.25 |
| 2 | 132077379 | rs6414023 | 0.987 | -1.799 | 1.876 | -1.124 | 0.024 | 0.016 | 0.016 | 0 | 0.73 | 0.26 | 0.86 | 0.84 | 0.87 | 0.74 |
| 2 | 132096387 | rs2313725 | 0.531 | -1.77 | 1.924 | -1.021 | -0.041 | -0.008 | -0.008 | 0.002 | 0.77 | 0.58 | 0.88 | 0.83 | 0.89 | 0.75 |
| 2 | 132098142 | rs6430737 | 0.491 | -1.761 | 1.923 | -1.064 | -0.037 | -0.016 | -0.016 | 0 | 0.77 | 0.6 | 0.89 | 0.84 | 0.89 | 0.75 |
| 2 | 132140614 | rs1997219 | 0.295 | -1.233 | 1.692 | -1.214 | -0.007 | -0.015 | -0.015 | 0 | 0.74 | 0.68 | 0.88 | 0.84 | 0.83 | 0.73 |
| 2 | 132141767 | rs13425822 | 0.565 | -1.26 | 1.587 | -1.227 | -0.024 | -0.043 | -0.043 | 0 | 0.72 | 0.52 | 0.86 | 0.84 | 0.83 | 0.73 |
| 2 | 132151248 | rs4850191 | 0.559 | -1.23 | 1.111 | -1.215 | -0.027 | -0.043 | -0.043 | 0.033 | 0.72 | 0.53 | 0.86 | 0.84 | 0.83 | 0.71 |
| 2 | 132159846 | rs2313693 | 0.765 | -1.128 | 0.933 | -1.242 | -0.014 | -0.017 | -0.017 | 0 | 0.69 | 0.4 | 0.84 | 0.83 | 0.82 | 0.72 |
| 2 | 132161304 | rs7573771 | 0.443 | -0.996 | 1.287 | -1.282 | -0.019 | -0.098 | -0.098 | 0 | 0.83 | 0.61 | 0.86 | 0.98 | 0.89 | 0.88 |
| 2 | 132164854 | rs2272355 | -0.11 | -0.416 | 0.694 | -1.318 | -0.005 | 0.006 | 0.006 | 0 | 0.33 | 0.32 | 0.58 | 0.74 | 0.35 | 0.42 |
| 2 | 132166387 | rs7573518 | 0.153 | -0.36 | 0.144 | -1.29 | -0.039 | -0.014 | -0.014 | 0.09 | 0.3 | 0.09 | 0.52 | 0.73 | 0.31 | 0.4 |
| 2 | 132170888 | rs7589111 | -0.975 | -0.581 | 0.079 | -1.531 | 0.03 | 0.128 | 0.128 | 0 | 0.48 | 0.85 | 0.66 | 0.74 | 0.38 | 0.43 |
| 2 | 132171917 | rs6739473 | -0.206 | -0.458 | 1.109 | -1.117 | -0.004 | -0.008 | -0.008 | 0 | 0.3 | 0.47 | 0.55 | 0.74 | 0.28 | 0.38 |
| 2 | 132177438 | rs11695582 | 0.208 | -0.522 | 0.89 | -0.684 | -0.02 | -0.011 | -0.011 | 0 | 0.71 | 0.53 | 0.52 | 0.26 | 0.75 | 0.66 |
| 2 | 132182159 | rs3730148 | 0.097 | -0.342 | 0.412 | -0.584 | -0.007 | -0.023 | -0.023 | 0.702 | 0.23 | 0.13 | 0.51 | 0.75 | 0.23 | 0.31 |
| 2 | 132187123 | rs870943 | -0.39 | -0.424 | 0.778 | -0.678 | -0.001 | 0 | 0 | 0 | 0.28 | 0.33 | 0.54 | 0.76 | 0.23 | 0.31 |
| 2 | 132196434 | rs13411754 | -0.245 | -0.422 | 0.594 | -0.677 | 0.012 | 0.001 | 0.001 | 0 | 0.27 | 0.2 | 0.53 | 0.76 | 0.23 | 0.31 |
| 2 | 132234986 | rs6731616 | -0.755 | -0.31 | 1.212 | -0.786 | 0.051 | -0.071 | -0.071 | 0.364 | 0.12 | 0.46 | 0.1 | 0 | 0.09 | 0.06 |
| 2 | 132248876 | rs6709245 | -1.525 | -0.582 | 0.91 | -0.896 | 0.025 | 0.046 | 0.046 | 0 | 0.39 | 0.73 | 0.63 | 0.76 | 0.31 | 0.41 |
| 2 | 132255853 | rs12469056 | -0.117 | -0.559 | 0.77 | -0.732 | 0.03 | -0.041 | -0.041 | 0 | 0.26 | 0.13 | 0.43 | 0.68 | 0.22 | 0.3 |
| 2 | 132262229 | rs10177008 | -0.117 | -0.559 | 0.77 | -0.732 | 0.03 | -0.041 | -0.041 | 0 | 0.26 | 0.13 | 0.43 | 0.68 | 0.22 | 0.3 |
| 2 | 132266537 | rs7572535 | 0.151 | -0.441 | 1.043 | -0.742 | 0.027 | -0.047 | -0.047 | 0 | 0.09 | 0.2 | 0.08 | 0 | 0.1 | 0.06 |
| 2 | 132276359 | rs7608954 | -0.151 | -0.48 | 0.672 | -0.652 | 0.031 | -0.041 | -0.041 | 0 | 0.26 | 0.13 | 0.43 | 0.68 | 0.22 | 0.3 |
| 2 | 132298468 | rs12472079 | -1.039 | -0.043 | 0.816 | -0.811 | 0.001 | -0.007 | -0.007 | 0 | 0.42 | 0.57 | 0.6 | 0.71 | 0.48 | 0.46 |
| 2 | 132312325 | rs11900136 | 0.774 | -0.124 | 0.577 | -0.862 | 0 | 0.023 | 0.023 | 0 | 0.55 | 0.34 | 0.4 | 0.29 | 0.52 | 0.55 |
| 2 | 132317186 | rs4850213 | 0.181 | -0.394 | -0.453 | -0.762 | 0.012 | -0.043 | -0.043 | 0 | 0.09 | 0.02 | 0.14 | 0.18 | 0.06 | 0.2 |
| 2 | 132323011 | rs1396837 | -0.231 | -0.199 | -0.268 | -0.766 | 0.002 | -0.01 | -0.01 | 0 | 0.3 | 0.22 | 0.47 | 0.71 | 0.37 | 0.32 |
| 2 | 132323205 | rs1828293 | -0.204 | -0.186 | -0.382 | -0.759 | 0 | -0.015 | -0.015 | 0 | 0.3 | 0.18 | 0.47 | 0.71 | 0.37 | 0.32 |
| 2 | 132339510 | rs11674634 | 0.063 | 0.023 | -0.05 | -0.504 | 0.013 | 0.019 | 0.019 | 0 | 0.47 | 0.38 | 0.38 | 0.36 | 0.42 | 0.47 |
| 2 | 132349648 | rs13029900 | 0.825 | -0.068 | 0.214 | -0.963 | -0.031 | 0.071 | 0.071 | 0 | 0.34 | 0.13 | 0.21 | 0.15 | 0.35 | 0.27 |
| 2 | 133224391 | rs6753618 | -0.195 | -1.485 | 0.407 | -0.734 | 0.007 | 0.133 | 0.133 | 0 | 0.42 | 0.13 | 0.52 | 0.78 | 0.29 | 0.5 |
| 2 | 133225803 | rs11689323 | 1.253 | -1.556 | 1.013 | -0.946 | 0.007 | -0.055 | -0.055 | 0 | 0.36 | 0.33 | 0.35 | 0.07 | 0.56 | 0.31 |
| 2 | 133226267 | rs10177527 | 1.256 | -1.564 | 1.006 | -0.95 | 0.006 | -0.053 | -0.053 | 2.092 | 0.36 | 0.33 | 0.35 | 0.07 | 0.56 | 0.31 |
| 2 | 133227542 | rs10803531 | -0.114 | -1.596 | 1.189 | -0.836 | -0.025 | -0.095 | -0.095 | 0 | 0.81 | 0.62 | 0.88 | 0.94 | 0.86 | 0.83 |
| 2 | 133234905 | rs13034310 | 0.557 | -1.647 | 0.103 | -0.998 | -0.062 | 0.188 | 0.188 | 0 | 0.39 | 0.04 | 0.35 | 0.07 | 0.62 | 0.36 |
| 2 | 133249282 | rs13001535 | 0.111 | -2.066 | 1.035 | -0.938 | 0.004 | 0.002 | 0.002 | 0 | 0.51 | 0.59 | 0.46 | 0.2 | 0.67 | 0.42 |
| 2 | 133253928 | rs10803532 | 0.836 | -2.016 | 1.379 | -1.063 | 0.016 | -0.001 | -0.001 | 0 | 0.85 | 0.5 | 0.88 | 0.88 | 0.9 | 0.86 |
| 2 | 133254834 | rs13389584 | 0.002 | -2.117 | 1.088 | -1.189 | 0.074 | 0.166 | 0.166 | 0.019 | 0.49 | 0.17 | 0.51 | 0.91 | 0.34 | 0.6 |
| 2 | 133265427 | rs2315593 | -0.987 | -2.143 | 1.176 | -1.218 | 0.007 | -0.086 | -0.086 | 0 | 0.62 | 0.53 | 0.6 | 0.96 | 0.44 | 0.7 |
| 2 | 133266673 | rs12691821 | -0.062 | -2.202 | 1.374 | -1.183 | 0.019 | 0.032 | 0.032 | 0 | 0.47 | 0.47 | 0.46 | 0.21 | 0.65 | 0.39 |
| 2 | 133267181 | rs10210934 | 1.01 | -2.201 | 1.349 | -1.189 | -0.041 | 0.068 | 0.068 | 0.063 | 0.35 | 0.26 | 0.37 | 0.04 | 0.56 | 0.27 |
| 2 | 133268418 | rs12691822 | 1.051 | -2.537 | 0.459 | -1.243 | -0.012 | -0.015 | -0.015 | 0.374 | 0.37 | 0.37 | 0.38 | 0.04 | 0.56 | 0.27 |
| 2 | 133269261 | rs10496687 | -0.055 | -2.503 | 0.384 | -1.197 | -0.003 | -0.014 | -0.014 | 0 | 0.11 | 0.07 | 0.07 | 0.17 | 0.09 | 0.12 |
| 2 | 133271657 | rs6750271 | 0.914 | -2.412 | 0.333 | -1.438 | -0.014 | -0.009 | -0.009 | 0 | 0.36 | 0.38 | 0.38 | 0.04 | 0.56 | 0.27 |
| 2 | 133272901 | rs2315595 | -0.158 | -2.993 | -0.39 | -1.249 | -0.02 | -0.039 | -0.039 | 0 | 0.55 | 0.58 | 0.49 | 0.16 | 0.72 | 0.45 |
| 2 | 133273008 | rs2315596 | 0.546 | -3.085 | -1.16 | -1.343 | 0.009 | 0.26 | 0.26 | 0.361 | 0.43 | 0.08 | 0.53 | 0.07 | 0.65 | 0.38 |
| 2 | 133274642 | rs4954337 | 0.187 | -2.361 | -0.49 | -1.474 | 0.041 | 0.05 | 0.05 | 0 | 0.5 | 0.54 | 0.77 | 0.75 | 0.68 | 0.51 |
| 2 | 133275867 | rs4606980 | 0.416 | -2.123 | -0.458 | -1.581 | -0.012 | 0.024 | 0.024 | 0.323 | 0.37 | 0.33 | 0.5 | 0.18 | 0.54 | 0.44 |
| 2 | 133276480 | rs11889041 | 0.177 | -2.147 | -0.033 | -1.61 | 0.002 | -0.023 | -0.023 | 0 | 0.39 | 0.46 | 0.52 | 0.18 | 0.56 | 0.44 |

| | | | | | | | | | | | | | | | | |
|---|---|---|---|---|---|---|---|---|---|---|---|---|---|---|---|---|
| 2 | 133277257 | rs6430365 | -0.763 | -2.227 | 0.211 | -1.577 | -0.005 | -0.026 | -0.026 | 0 | 0.57 | 0.53 | 0.74 | 0.8 | 0.68 | 0.68 |
| 2 | 133277683 | rs4395295 | 0.218 | -2.246 | -0.262 | -1.499 | -0.048 | 0.076 | 0.076 | 0.294 | 0.22 | 0.45 | 0.26 | 0.62 | 0.13 | 0.24 |
| 2 | 133278691 | rs4954338 | 0.429 | -2.198 | -0.318 | -1.532 | -0.012 | 0.039 | 0.039 | 0.392 | 0.38 | 0.3 | 0.51 | 0.18 | 0.56 | 0.44 |
| 2 | 133282008 | rs10178456 | -0.647 | -2.155 | -0.603 | -1.739 | 0.043 | 0.091 | 0.091 | 0 | 0.58 | 0.74 | 0.77 | 0.8 | 0.7 | 0.67 |
| 2 | 133283943 | rs6710643 | -0.202 | -2.069 | -0.563 | -1.704 | 0.032 | 0.059 | 0.059 | 0 | 0.4 | 0.29 | 0.39 | 0.72 | 0.32 | 0.38 |
| 2 | 133285727 | rs3109140 | -1.696 | -1.855 | -0.744 | -1.826 | -0.086 | 0.076 | 0.076 | 0.407 | 0.75 | 0.9 | 0.56 | 0.9 | 0.52 | 0.67 |
| 2 | 133299230 | rs12478654 | 0.855 | -2.072 | -0.718 | -1.718 | 0.047 | 0.021 | 0.021 | 0.148 | 0.23 | 0.04 | 0.09 | 0.18 | 0.14 | 0.23 |
| 2 | 133300624 | rs3115031 | -0.083 | -1.922 | -0.374 | -1.7 | -0.004 | -0.009 | -0.009 | 0.948 | 0.41 | 0.43 | 0.39 | 0.72 | 0.32 | 0.38 |
| 2 | 133302257 | rs3115027 | 0.697 | -2.05 | -0.422 | -1.715 | -0.02 | -0.033 | -0.033 | 0 | 0.26 | 0.38 | 0.18 | 0.13 | 0.19 | 0.17 |
| 2 | 133302558 | rs12476183 | 0.711 | -2.156 | -0.062 | -1.68 | 0.001 | 0.002 | 0.002 | 0 | 0.27 | 0.27 | 0.12 | 0.23 | 0.14 | 0.24 |
| 2 | 133303203 | rs714546 | 1.085 | -2.081 | -0.244 | -1.641 | -0.026 | -0.102 | -0.102 | 0 | 0.14 | 0.01 | 0.22 | 0.54 | 0.13 | 0.2 |
| 2 | 133304228 | rs3109134 | -0.48 | -2.094 | -0.218 | -1.653 | 0.041 | 0.03 | 0.03 | 0 | 0.54 | 0.68 | 0.3 | 0.36 | 0.33 | 0.42 |
| 2 | 133306684 | rs3109137 | -0.441 | -2.143 | -0.167 | -1.648 | 0.015 | 0.011 | 0.011 | 0 | 0.53 | 0.61 | 0.3 | 0.36 | 0.32 | 0.42 |
| 2 | 133307332 | rs6741093 | -0.757 | -2.156 | -0.123 | -1.662 | 0.003 | -0.076 | -0.076 | 0 | 0.61 | 0.53 | 0.49 | 0.87 | 0.43 | 0.53 |
| 2 | 133307991 | rs10198735 | 0.163 | -2.163 | -0.132 | -1.658 | 0.018 | 0.018 | 0.018 | 0 | 0.47 | 0.45 | 0.27 | 0.33 | 0.3 | 0.33 |
| 2 | 133308005 | rs10166090 | 0.163 | -2.163 | -0.132 | -1.658 | 0.018 | 0.018 | 0.018 | 0 | 0.47 | 0.45 | 0.27 | 0.33 | 0.3 | 0.33 |
| 2 | 133310237 | rs10173080 | -1.068 | -1.749 | -0.166 | -1.7 | 0.021 | -0.085 | -0.085 | 0.575 | 0.67 | 0.63 | 0.51 | 0.9 | 0.46 | 0.62 |
| 2 | 133313845 | rs9287421 | 1.471 | -1.77 | -0.131 | -1.63 | 0.006 | 0.021 | 0.021 | 0 | 0.19 | 0.1 | 0.14 | 0.1 | 0.16 | 0.09 |
| 2 | 133314973 | rs10928428 | -1.09 | -1.767 | -0.149 | -1.686 | 0 | -0.039 | -0.039 | 0 | 0.65 | 0.67 | 0.53 | 0.92 | 0.44 | 0.56 |
| 2 | 133320414 | rs3109144 | 0.975 | -1.599 | -0.432 | -1.445 | 0.002 | -0.012 | -0.012 | 0.076 | 0.21 | 0.2 | 0.2 | 0.13 | 0.22 | 0.15 |
| 2 | 133320593 | rs11684083 | 0.832 | -1.818 | -0.782 | -1.307 | 0.044 | -0.043 | -0.043 | 0.251 | 0.23 | 0.05 | 0.08 | 0.28 | 0.13 | 0.25 |
| 2 | 133322919 | rs12624113 | -0.049 | -1.508 | -0.439 | -1.827 | 0.025 | 0.034 | 0.034 | 0 | 0.38 | 0.28 | 0.32 | 0.79 | 0.26 | 0.45 |
| 2 | 133331994 | rs17396006 | 1.271 | -1.443 | -1.298 | -1.707 | 0.026 | 0.012 | 0.012 | 0 | 0.2 | 0.03 | 0.09 | 0.18 | 0.13 | 0.15 |
| 2 | 133333289 | rs3109152 | -1.092 | -1.302 | -1.404 | -1.843 | -0.118 | 0.186 | 0.186 | 0.523 | 0.64 | 0.92 | 0.61 | 0.92 | 0.53 | 0.64 |
| 2 | 133334386 | rs3115011 | -0.779 | -1.156 | -1.447 | -1.769 | -0.01 | 0.001 | 0.001 | 0.565 | 0.6 | 0.33 | 0.54 | 0.92 | 0.52 | 0.64 |
| 2 | 133342529 | rs3109154 | -0.511 | -0.298 | -1.212 | -1.138 | -0.003 | -0.02 | -0.02 | 0 | 0.41 | 0.45 | 0.47 | 0.74 | 0.4 | 0.47 |
| 2 | 133342658 | rs4378857 | 0.49 | 0.188 | -1.714 | -0.704 | 0.003 | -0.103 | -0.103 | 0.086 | 0.16 | 0.02 | 0.26 | 0.6 | 0.17 | 0.3 |
| 2 | 133344452 | rs3109155 | -0.148 | 1.181 | -1.208 | -1.037 | 0.019 | -0.052 | -0.052 | 0 | 0.26 | 0.26 | 0.2 | 0.03 | 0.42 | 0.34 |
| 2 | 133344676 | rs9287422 | 0.415 | 1.05 | -2.174 | -0.826 | -0.119 | 0.115 | 0.115 | 1.064 | 0.21 | 0.03 | 0.16 | 0.01 | 0.36 | 0.29 |
| 2 | 133346235 | rs3109156 | -0.163 | -0.223 | -1.849 | -0.537 | 0.061 | 0.058 | 0.058 | 0 | 0.64 | 0.81 | 0.83 | 0.75 | 0.77 | 0.76 |
| 2 | 133346735 | rs6712250 | -0.625 | 0.36 | -1.366 | -0.932 | 0 | -0.076 | -0.076 | 0.103 | 0.35 | 0.59 | 0.25 | 0.13 | 0.32 | 0.18 |
| 2 | 133348083 | rs3109157 | -0.377 | -0.029 | -1.623 | -0.83 | 0.023 | 0.072 | 0.072 | 0 | 0.4 | 0.72 | 0.45 | 0.19 | 0.29 | 0.4 |
| 2 | 133350137 | rs4954345 | -0.643 | -0.098 | -2.311 | -1.134 | -0.056 | 0.215 | 0.215 | 0 | 0.47 | 0.13 | 0.47 | 0.81 | 0.46 | 0.63 |
| 2 | 133351415 | rs16837640 | 0.354 | -0.144 | -2.346 | -0.781 | -0.004 | -0.045 | -0.045 | 0 | 0.21 | 0.07 | 0.3 | 0.69 | 0.21 | 0.36 |
| 2 | 133351703 | rs12616464 | -0.595 | -0.172 | -2.009 | -0.094 | -0.009 | 0.056 | 0.056 | 0 | 0.48 | 0.29 | 0.47 | 0.64 | 0.44 | 0.62 |
| 2 | 133353129 | rs17324208 | 0.382 | 0.295 | -1.626 | 0.143 | -0.007 | -0.003 | -0.003 | 0.9 | 0.16 | 0.11 | 0.15 | 0.18 | 0.27 | 0.15 |
| 2 | 133353827 | rs16837705 | 0.494 | 0.163 | -0.948 | -0.299 | -0.003 | -0.029 | -0.029 | 0.212 | 0.24 | 0.31 | 0.25 | 0.12 | 0.17 | 0.17 |
| 2 | 133362943 | rs11680742 | 0.364 | -0.015 | -1.068 | 0.033 | 0.001 | 0.01 | 0.01 | 0 | 0.13 | 0.15 | 0.13 | 0.29 | 0.08 | 0.24 |
| 2 | 133363129 | rs12622406 | 0.619 | 0.149 | -0.99 | 0.305 | 0.002 | 0.009 | 0.009 | 0.196 | 0.23 | 0.21 | 0.24 | 0.11 | 0.16 | 0.17 |
| 2 | 133367209 | rs4449192 | -0.874 | 0.211 | -1.133 | 0.193 | -0.007 | 0.053 | 0.053 | 0 | 0.51 | 0.35 | 0.57 | 0.67 | 0.49 | 0.47 |
| 2 | 133368556 | rs2315523 | -0.273 | 0.26 | -1.347 | 0.624 | -0.041 | -0.134 | -0.134 | 0 | 0.26 | 0.06 | 0.32 | 0.54 | 0.32 | 0.3 |
| 2 | 133377826 | rs6760538 | -0.73 | 0.271 | -1.262 | 0.014 | -0.07 | -0.047 | -0.047 | 0 | 0.61 | 0.81 | 0.56 | 0.48 | 0.51 | 0.39 |
| 2 | 133381436 | rs2315536 | 0.19 | 0.512 | -1.331 | 0.055 | 0.002 | 0.009 | 0.009 | 0.046 | 0.26 | 0.24 | 0.28 | 0.34 | 0.31 | 0.22 |
| 2 | 133383855 | rs6719342 | -0.031 | 0.265 | -1.609 | -0.383 | 0.03 | 0.067 | 0.067 | 0.046 | 0.31 | 0.19 | 0.18 | 0.1 | 0.2 | 0.17 |
| 2 | 133392633 | rs10208154 | -0.013 | 0.81 | -1.869 | -0.135 | -0.08 | 0.145 | 0.145 | 0.365 | 0.42 | 0.13 | 0.46 | 0.3 | 0.47 | 0.41 |
| 2 | 133396179 | rs17324515 | 0.073 | 1.037 | -2.007 | 0.002 | -0.068 | 0.082 | 0.082 | 0 | 0.41 | 0.08 | 0.5 | 0.32 | 0.39 | 0.38 |
| 2 | 133398177 | rs2315521 | -0.105 | 1.029 | -2.169 | -0.049 | -0.112 | 0.115 | 0.115 | 0.333 | 0.44 | 0.1 | 0.47 | 0.33 | 0.48 | 0.52 |
| 2 | 133400992 | rs7558730 | -0.381 | 0.618 | -2.138 | -0.349 | -0.051 | 0.176 | 0.176 | 0 | 0.49 | 0.88 | 0.53 | 0.34 | 0.47 | 0.51 |
| 2 | 133402607 | rs1131671 | -0.074 | 0.598 | -0.558 | -0.357 | 0 | -0.012 | -0.012 | 0 | 0.46 | 0.55 | 0.5 | 0.34 | 0.46 | 0.5 |
| 2 | 133410440 | rs9973717 | 0.048 | 0.607 | 0.285 | -0.353 | -0.002 | 0.027 | 0.027 | 0 | 0.47 | 0.38 | 0.51 | 0.33 | 0.5 | 0.57 |
| 2 | 133413724 | rs713535 | -0.27 | 0.457 | 0.595 | -0.266 | 0.008 | 0.043 | 0.043 | 0 | 0.54 | 0.34 | 0.49 | 0.66 | 0.58 | 0.47 |
| 2 | 133415170 | rs964842 | 0.931 | 0.259 | -0.176 | -0.151 | 0.01 | 0 | 0 | 3.164 | 0.13 | 0.08 | 0.07 | 0.13 | 0.09 | 0.07 |
| 2 | 133422262 | rs2566535 | -1.067 | 0.247 | -0.2 | -0.158 | -0.063 | 0.078 | 0.078 | 0 | 0.59 | 0.91 | 0.55 | 0.66 | 0.59 | 0.47 |
| 2 | 133437172 | rs2566530 | -1.066 | 0.249 | 0.817 | -0.202 | -0.042 | 0.091 | 0.091 | 0 | 0.59 | 0.9 | 0.52 | 0.67 | 0.58 | 0.47 |
| 2 | 133440227 | rs2709549 | -0.453 | 0.194 | 0.659 | -0.215 | -0.015 | 0.159 | 0.159 | 0 | 0.52 | 0.12 | 0.45 | 0.66 | 0.58 | 0.47 |
| 2 | 133452455 | rs9287423 | 0.429 | 0.437 | 0.325 | 0.256 | -0.103 | -0.061 | -0.061 | 0 | 0.26 | 0.02 | 0.17 | 0.23 | 0.31 | 0.3 |
| 2 | 133468426 | rs2250481 | -1.219 | 0.332 | 0.37 | 0.03 | 0.001 | 0 | 0 | 0 | 0.59 | 0.62 | 0.62 | 0.84 | 0.61 | 0.58 |
| 2 | 133470543 | rs2566520 | -1.247 | 0.763 | 0.193 | -0.036 | -0.001 | -0.024 | -0.024 | 0 | 0.6 | 0.62 | 0.62 | 0.85 | 0.64 | 0.59 |
| 2 | 133471789 | rs10166009 | 0.526 | 0.995 | -0.434 | 0.423 | -0.083 | -0.076 | -0.076 | 0 | 0.24 | 0.01 | 0.15 | 0.22 | 0.26 | 0.25 |
| 2 | 133472002 | rs6708476 | -0.714 | 0.909 | 0.2 | 0.165 | 0 | -0.013 | -0.013 | 0.592 | 0.46 | 0.46 | 0.56 | 0.74 | 0.52 | 0.53 |
| 2 | 133472054 | rs11686423 | -0.508 | 0.869 | 0.244 | 0.499 | -0.002 | -0.009 | -0.009 | 0.992 | 0.38 | 0.58 | 0.5 | 0.63 | 0.36 | 0.4 |
| 2 | 133474412 | rs17325076 | 0.674 | 1.005 | 0.182 | 0.774 | 0.001 | 0.001 | 0.001 | 0.283 | 0.11 | 0.06 | 0.05 | 0.12 | 0.08 | 0.05 |
| 2 | 133478977 | rs6736839 | -1.026 | 1.301 | 0.253 | 1.203 | -0.012 | -0.024 | -0.024 | 2.051 | 0.58 | 0.77 | 0.36 | 0.42 | 0.58 | 0.41 |
| 2 | 133482699 | rs4953986 | -1.665 | 1.354 | -0.024 | 1.171 | -0.052 | 0.132 | 0.132 | 0.151 | 0.67 | 0.99 | 0.68 | 0.86 | 0.68 | 0.62 |
| 2 | 133484851 | rs13025233 | 1.104 | 1.435 | 0.071 | 1.455 | -0.015 | -0.016 | -0.016 | 0 | 0.1 | 0.07 | 0.09 | 0.15 | 0.16 | 0.07 |
| 2 | 133487103 | rs7596376 | 0.06 | 1.404 | 0.116 | 1.56 | 0.001 | 0.08 | 0.08 | 0 | 0.06 | 0.03 | 0.3 | 0.45 | 0.1 | 0.22 |
| 2 | 133491553 | rs17397163 | 0.913 | 1.339 | 0.052 | 1.496 | -0.028 | 0.034 | 0.034 | 0 | 0.27 | 0.05 | 0.15 | 0.1 | 0.26 | 0.25 |
| 2 | 133492206 | rs12619898 | 0.864 | 1.335 | 0.35 | 1.494 | 0.004 | 0.043 | 0.043 | 0 | 0.28 | 0.11 | 0.15 | 0.1 | 0.26 | 0.25 |
| 2 | 133494689 | rs2010511 | -0.183 | 1.309 | 0.356 | 1.5 | -0.013 | 0.025 | 0.025 | 0.272 | 0.39 | 0.17 | 0.21 | 0.25 | 0.41 | 0.31 |

| | | | | | | | | | | | | | | | | |
|---|---|---|---|---|---|---|---|---|---|---|---|---|---|---|---|---|
| 2 | 133499279 | rs16840590 | -0.619 | 1.381 | 0.727 | 1.548 | 0.054 | 0.15 | 0.15 | 0 | 0.19 | 0.3 | 0.38 | 0.59 | 0.24 | 0.27 |
| 2 | 133503784 | rs11890169 | 0.705 | 1.364 | 0.098 | 1.36 | 0.004 | 0.005 | 0.005 | 0.105 | 0.12 | 0.13 | 0.07 | 0.15 | 0.15 | 0.06 |
| 2 | 133511761 | rs7595365 | -1.091 | 1.304 | -0.252 | 1.135 | -0.088 | 0.256 | 0.256 | 0.203 | 0.56 | 0.93 | 0.62 | 0.86 | 0.5 | 0.59 |
| 2 | 133513086 | rs6430373 | -1.041 | 1.299 | -0.222 | 1.226 | -0.073 | 0.229 | 0.229 | 0.439 | 0.54 | 0.9 | 0.58 | 0.84 | 0.5 | 0.58 |
| 2 | 133514933 | rs1019841 | -2.189 | 1.238 | -0.692 | 1.066 | -0.077 | 0.116 | 0.116 | 0 | 0.71 | 0.98 | 0.7 | 0.87 | 0.67 | 0.74 |
| 2 | 133515838 | rs11687264 | 0.969 | 1.654 | -0.469 | 1.369 | -0.014 | 0.047 | 0.047 | 0 | 0.27 | 0.06 | 0.15 | 0.11 | 0.26 | 0.23 |
| 2 | 133515862 | rs1430626 | 0.127 | 1.728 | -0.387 | 1.811 | -0.008 | -0.005 | -0.005 | 0 | 0.09 | 0.06 | 0.08 | 0.27 | 0.12 | 0.13 |
| 2 | 133516599 | rs17397380 | 0.741 | 1.343 | -0.179 | 1.94 | -0.014 | 0.05 | 0.05 | 0 | 0.27 | 0.06 | 0.15 | 0.11 | 0.26 | 0.22 |
| 2 | 133517783 | rs13006029 | 0.325 | 1.037 | 0.364 | 1.808 | 0.004 | 0.009 | 0.009 | 0 | 0.11 | 0.11 | 0.09 | 0.17 | 0.15 | 0.14 |
| 2 | 133519504 | rs11884795 | -0.87 | 1.007 | 0.634 | 1.853 | 0.011 | 0.041 | 0.041 | 0.99 | 0.22 | 0.28 | 0.17 | 0.32 | 0.25 | 0.21 |
| 2 | 133520889 | rs17325495 | 0.681 | 1.04 | 0.304 | 1.875 | -0.033 | -0.04 | -0.04 | 0 | 0.1 | 0.02 | 0.08 | 0.17 | 0.15 | 0.13 |
| 2 | 133521231 | rs10496692 | -0.703 | 0.75 | 0.527 | 1.798 | -0.003 | 0.02 | 0.02 | 4.317 | 0.13 | 0.22 | 0.09 | 0.15 | 0.11 | 0.09 |
| 2 | 133530556 | rs16841075 | 0.609 | 1.05 | -0.152 | 1.399 | -0.091 | -0.053 | -0.053 | 0 | 0.28 | 0.04 | 0.25 | 0.24 | 0.32 | 0.26 |
| 2 | 133532709 | rs10496693 | 0.439 | 0.549 | 0.667 | 1.456 | 0 | 0.022 | 0.022 | 0 | 0.28 | 0.34 | 0.35 | 0.37 | 0.24 | 0.33 |
| 2 | 133533325 | rs6735149 | 0.739 | 0.627 | 0.526 | 1.588 | -0.012 | -0.009 | -0.009 | 0 | 0.07 | 0.04 | 0.14 | 0.2 | 0.09 | 0.1 |
| 2 | 133539566 | rs6730282 | -1.136 | 0.404 | 0.849 | 1.347 | 0 | -0.018 | -0.018 | 0.134 | 0.51 | 0.5 | 0.48 | 0.75 | 0.47 | 0.48 |
| 2 | 133540605 | rs13016342 | 0.219 | 0.371 | 1.156 | 1.315 | -0.001 | -0.011 | -0.011 | 0 | 0.16 | 0.17 | 0.12 | 0.07 | 0.14 | 0.09 |
| 2 | 133541454 | rs12691830 | -1.525 | 0.337 | 1.057 | 1.271 | -0.039 | 0.195 | 0.195 | 0.068 | 0.57 | 0.83 | 0.67 | 0.87 | 0.49 | 0.61 |
| 2 | 133541575 | rs12611515 | -1.264 | 0.413 | 1.045 | 1.347 | -0.008 | 0.083 | 0.083 | 0.016 | 0.29 | 0.46 | 0.31 | 0.5 | 0.3 | 0.25 |
| 2 | 133544048 | rs16841335 | -0.499 | 0.306 | 1.109 | 1.421 | 0.023 | 0.155 | 0.155 | 0 | 0.22 | 0.44 | 0.3 | 0.49 | 0.25 | 0.24 |
| 2 | 133554288 | rs1868076 | -1.784 | 0.243 | 1.079 | 1.07 | -0.01 | 0.052 | 0.052 | 0 | 0.61 | 0.71 | 0.67 | 0.89 | 0.54 | 0.62 |
| 2 | 133569227 | rs6430375 | -1.021 | 0.094 | 1.16 | 1.296 | -0.002 | 0.098 | 0.098 | 0.06 | 0.27 | 0.52 | 0.31 | 0.51 | 0.27 | 0.23 |
| 2 | 133577404 | rs10199393 | -1.125 | 0.134 | 1.135 | 1.348 | -0.032 | -0.001 | -0.001 | 0 | 0.3 | 0.43 | 0.18 | 0.31 | 0.22 | 0.18 |
| 2 | 133577528 | rs10199481 | -1.607 | 0.149 | 1.018 | 1.352 | -0.005 | 0.023 | 0.023 | 0 | 0.34 | 0.55 | 0.32 | 0.51 | 0.33 | 0.25 |
| 2 | 133581889 | rs2320155 | -1.015 | 0.084 | 0.996 | 1.336 | -0.022 | -0.02 | -0.02 | 0 | 0.29 | 0.32 | 0.17 | 0.19 | 0.21 | 0.17 |
| 2 | 133584471 | rs1374740 | 1.113 | 0.18 | 0.771 | 1.37 | -0.011 | -0.064 | -0.064 | 0 | 0.25 | 0.06 | 0.17 | 0.26 | 0.21 | 0.23 |
| 2 | 133607540 | rs7571473 | -0.62 | 0.06 | 1.24 | 1.37 | -0.021 | 0.094 | 0.094 | 0 | 0.24 | 0.44 | 0.27 | 0.39 | 0.2 | 0.2 |
| 2 | 133609463 | rs6430377 | -1.754 | 0.01 | 0.96 | 1.212 | -0.044 | 0.131 | 0.131 | 0 | 0.62 | 0.77 | 0.67 | 0.92 | 0.54 | 0.59 |
| 2 | 133610357 | rs11680119 | 1.167 | -0.011 | 0.725 | 1.296 | -0.014 | -0.068 | -0.068 | 3.276 | 0.25 | 0.07 | 0.17 | 0.28 | 0.21 | 0.21 |
| 2 | 133615057 | rs1447553 | 0.9 | 0.008 | 0.917 | 1.157 | -0.002 | 0.01 | 0.01 | 0 | 0.27 | 0.3 | 0.34 | 0.41 | 0.22 | 0.33 |
| 2 | 133616690 | rs6736014 | 0.615 | 0.047 | 0.856 | 1.221 | 0 | 0.002 | 0.002 | 0 | 0.29 | 0.34 | 0.47 | 0.61 | 0.26 | 0.38 |
| 2 | 133617800 | rs1447550 | -1.089 | -0.002 | 0.892 | 1.14 | -0.011 | 0.007 | 0.007 | 0 | 0.33 | 0.3 | 0.29 | 0.38 | 0.28 | 0.25 |
| 2 | 133618220 | rs1447549 | 0.671 | 0.185 | 0.655 | 1.163 | -0.005 | 0.024 | 0.024 | 0 | 0.28 | 0.37 | 0.35 | 0.41 | 0.22 | 0.33 |
| 2 | 133627136 | rs12464177 | -1.392 | 0.232 | 0.628 | 1.118 | 0.004 | -0.008 | -0.008 | 3.103 | 0.56 | 0.54 | 0.62 | 0.67 | 0.49 | 0.53 |
| 2 | 133629289 | rs6709364 | -1.081 | 0.195 | 0.555 | 1.099 | 0 | 0 | 0 | 0 | 0.48 | 0.48 | 0.61 | 0.79 | 0.46 | 0.49 |
| 2 | 133629401 | rs2320159 | -0.351 | 0.187 | 0.139 | 1.211 | -0.003 | 0.009 | 0.009 | 0 | 0.22 | 0.22 | 0.4 | 0.39 | 0.21 | 0.27 |
| 2 | 133629564 | rs1037175 | -0.351 | 0.187 | 0.139 | 1.211 | -0.003 | 0.009 | 0.009 | 0 | 0.22 | 0.23 | 0.39 | 0.39 | 0.21 | 0.28 |
| 2 | 133633125 | rs10177329 | 0.371 | 0.256 | -0.058 | 1.136 | -0.008 | 0.061 | 0.061 | 0 | 0.3 | 0.5 | 0.38 | 0.53 | 0.25 | 0.33 |
| 2 | 133636696 | rs12987460 | -0.176 | 0.25 | -0.591 | 0.753 | 0.008 | -0.001 | -0.001 | 0.007 | 0.38 | 0.3 | 0.28 | 0.32 | 0.33 | 0.28 |
| 2 | 133637592 | rs6720847 | -0.553 | 0.297 | -0.46 | 0.879 | 0.016 | 0.029 | 0.029 | 0 | 0.47 | 0.3 | 0.33 | 0.33 | 0.41 | 0.32 |
| 2 | 133638721 | rs10186393 | -1.198 | 0.455 | -1.245 | 0.943 | -0.011 | 0.03 | 0.03 | 0 | 0.6 | 0.71 | 0.66 | 0.79 | 0.54 | 0.56 |
| 2 | 133639374 | rs10928436 | -0.081 | 0.493 | -0.999 | 1.495 | -0.018 | -0.074 | -0.074 | 0 | 0.24 | 0.08 | 0.18 | 0.27 | 0.21 | 0.2 |
| 2 | 133643197 | rs6752610 | -0.376 | 0.859 | -0.209 | 1.634 | 0.012 | 0.005 | 0.005 | 0.233 | 0.21 | 0.28 | 0.38 | 0.3 | 0.32 | 0.22 |
| 2 | 133648157 | rs1348960 | 0.575 | 0.082 | -0.166 | 1.419 | 0.044 | 0.045 | 0.045 | 0 | 0.44 | 0.29 | 0.38 | 0.27 | 0.36 | 0.46 |
| 2 | 133650465 | rs1374744 | 0.365 | 0.245 | -0.305 | 1.263 | 0.008 | 0.028 | 0.028 | 0.009 | 0.53 | 0.58 | 0.44 | 0.66 | 0.46 | 0.57 |
| 2 | 133657593 | rs11883458 | 0.064 | -0.417 | -0.377 | 1.355 | -0.004 | 0.001 | 0.001 | 0 | 0.23 | 0.28 | 0.49 | 0.31 | 0.2 | 0.19 |
| 2 | 133664437 | rs4953862 | -0.741 | -0.319 | -0.178 | 1.491 | 0.027 | 0.238 | 0.238 | 0 | 0.05 | 0.28 | 0.37 | 0.26 | 0.04 | 0.02 |
| 2 | 133683658 | rs2577634 | -0.626 | -0.64 | -1.311 | 1.428 | -0.032 | -0.03 | -0.03 | 0.016 | 0.59 | 0.98 | 0.83 | 0.69 | 0.68 | 0.7 |
| 2 | 133688972 | rs1370602 | -0.471 | -0.309 | -0.435 | 1.684 | 0.031 | -0.042 | -0.042 | 0 | 0.3 | 0.8 | 0.57 | 0.51 | 0.2 | 0.21 |
| 2 | 133693415 | rs1020430 | -0.585 | -0.042 | -0.431 | 1.768 | 0.001 | 0.011 | 0.011 | 0 | 0.07 | 0.07 | 0.37 | 0.25 | 0.11 | 0.03 |
| 2 | 133696554 | rs10207480 | 0.091 | -0.324 | 0.048 | 1.824 | -0.026 | 0.001 | 0.001 | 0 | 0.28 | 0.45 | 0.53 | 0.32 | 0.21 | 0.19 |
| 2 | 133697627 | rs6747820 | -0.587 | -0.429 | -0.619 | 1.58 | -0.174 | 0.158 | 0.158 | 1.379 | 0.68 | 0.97 | 0.74 | 0.93 | 0.54 | 0.66 |
| 2 | 133700309 | rs281575 | 1.051 | -0.583 | 0.578 | 2.274 | 0.057 | -0.044 | -0.044 | 0 | 0.6 | 0.17 | 0.38 | 0.51 | 0.72 | 0.76 |
| 2 | 133707706 | rs281583 | 0.631 | -0.506 | 0.177 | 1.547 | 0.025 | 0.044 | 0.044 | 0 | 0.68 | 0.83 | 0.48 | 0.73 | 0.74 | 0.79 |
| 2 | 133708257 | rs6752737 | -0.762 | -0.291 | 0.481 | 2.084 | -0.044 | -0.01 | -0.01 | 0 | 0.28 | 0.55 | 0.54 | 0.31 | 0.2 | 0.19 |
| 2 | 133726458 | rs12619661 | -1.414 | 0.364 | 0.506 | 2.878 | 0.013 | 0.299 | 0.299 | 0 | 0.11 | 0.33 | 0.43 | 0.49 | 0.11 | 0.06 |
| 2 | 133739616 | rs6727803 | -0.71 | -0.154 | -0.102 | 2.347 | -0.009 | 0.008 | 0.008 | 0 | 0.28 | 0.68 | 0.19 | 0.36 | 0.21 | 0.34 |
| 2 | 133746273 | rs4954014 | 0.142 | 0.978 | -0.409 | 2.476 | -0.004 | 0.009 | 0.009 | 0 | 0.89 | 0.9 | 0.62 | 0.61 | 0.76 | 0.83 |
| 2 | 133748769 | rs4954015 | 0.678 | 0.616 | -0.424 | 2.753 | -0.019 | -0.027 | -0.027 | 0.132 | 0.85 | 0.9 | 0.62 | 0.59 | 0.75 | 0.81 |
| 2 | 133754272 | rs6735163 | -0.167 | 0.632 | -0.085 | 2.197 | 0.005 | 0.043 | 0.043 | 0.009 | 0.45 | 0.36 | 0.37 | 0.19 | 0.38 | 0.34 |
| 2 | 133754360 | rs12475645 | 0.732 | 0.506 | -0.3 | 1.937 | 0.021 | 0.028 | 0.028 | 2.395 | 0.33 | 0.19 | 0.19 | 0.22 | 0.27 | 0.44 |
| 2 | 133759852 | rs6708646 | 0.417 | 0.818 | 0.508 | 2.169 | 0.04 | 0.056 | 0.056 | 0 | 0.78 | 0.65 | 0.55 | 0.38 | 0.64 | 0.76 |
| 2 | 133761794 | rs1437897 | -0.151 | 0.892 | -0.043 | 1.765 | -0.026 | -0.023 | -0.023 | 0.082 | 0.83 | 0.89 | 0.7 | 0.69 | 0.69 | 0.84 |
| 2 | 133768995 | rs11683464 | 0.081 | 0.14 | 0.204 | 1.41 | 0 | 0.015 | 0.015 | 0 | 0.5 | 0.56 | 0.55 | 0.63 | 0.45 | 0.5 |
| 2 | 133769673 | rs4953868 | 0.448 | 0.051 | 0.312 | 1.628 | 0 | 0.002 | 0.002 | 0 | 0.46 | 0.42 | 0.38 | 0.26 | 0.4 | 0.39 |
| 2 | 133771658 | rs1437903 | -0.383 | 0.094 | -0.353 | 0.912 | -0.061 | -0.091 | -0.091 | 0 | 0.83 | 0.96 | 0.76 | 0.64 | 0.75 | 0.82 |
| 2 | 133772581 | rs12988086 | 0.023 | -0.151 | 0.127 | 0.374 | 0.007 | 0.008 | 0.008 | 2.107 | 0.35 | 0.28 | 0.2 | 0.05 | 0.3 | 0.35 |
| 2 | 133776904 | rs7572356 | 0.252 | -0.586 | -0.146 | 0.403 | -0.008 | -0.019 | -0.019 | 0 | 0.16 | 0.14 | 0.09 | 0.04 | 0.09 | 0.08 |
| 2 | 133778855 | rs35615951 | -0.186 | -0.195 | -0.697 | 0.016 | -0.006 | -0.006 | -0.006 | 0.057 | 0.41 | 0.5 | 0.36 | 0.36 | 0.38 | 0.34 |

| | | | | | | | | | | | | | | | | |
|---|---|---|---|---|---|---|---|---|---|---|---|---|---|---|---|---|
| 2 | 133781914 | rs6705690 | -0.668 | -0.016 | -1.537 | -0.511 | -0.008 | 0.058 | 0.058 | 0 | 0.47 | 0.16 | 0.49 | 0.38 | 0.53 | 0.4 |
| 2 | 133783103 | rs6730963 | -0.911 | -0.168 | -0.735 | -0.582 | 0.006 | 0.023 | 0.023 | 0 | 0.52 | 0.62 | 0.54 | 0.38 | 0.55 | 0.41 |
| 2 | 133784184 | rs16845737 | 1.517 | 0.062 | -0.8 | -0.7 | -0.007 | 0.026 | 0.026 | 0 | 0.08 | 0.06 | 0.2 | 0.35 | 0.12 | 0.04 |
| 2 | 133785054 | rs2196672 | -1.063 | 0.078 | -1.047 | -0.895 | -0.002 | 0 | 0 | 3.329 | 0.35 | 0.61 | 0.37 | 0.37 | 0.37 | 0.3 |
| 2 | 133787580 | rs1006444 | 0.315 | -0.121 | -1.75 | -1.116 | -0.092 | 0.212 | 0.212 | 0 | 0.31 | 0.03 | 0.22 | 0.02 | 0.36 | 0.29 |
| 2 | 133787995 | rs17733147 | 0.793 | 0.106 | -1.355 | -0.949 | -0.017 | 0.019 | 0.019 | 0 | 0.24 | 0.15 | 0.2 | 0.02 | 0.31 | 0.26 |
| 2 | 133788260 | rs6430386 | 1.155 | 0.142 | -1.545 | -0.626 | -0.002 | -0.014 | -0.014 | 0 | 0.08 | 0.06 | 0.06 | 0.01 | 0.05 | 0.07 |
| 2 | 133792119 | rs1898562 | 1.183 | 0.399 | -1.611 | 0.047 | -0.013 | 0.026 | 0.026 | 0 | 0.08 | 0.05 | 0.2 | 0.38 | 0.11 | 0.03 |
| 2 | 133797169 | rs1437909 | -0.152 | 0.373 | -1.46 | -0.754 | -0.002 | 0.111 | 0.111 | 0 | 0.45 | 0.73 | 0.34 | 0.03 | 0.43 | 0.42 |
| 2 | 133806124 | rs13395693 | 0.057 | 0.574 | -1.564 | 0.125 | -0.03 | 0.003 | 0.003 | 0 | 0.34 | 0.15 | 0.31 | 0.32 | 0.38 | 0.5 |
| 2 | 133806432 | rs4954023 | 1.358 | 0.538 | -1.118 | 0.395 | 0.005 | 0.034 | 0.034 | 1.358 | 0.08 | 0.06 | 0.21 | 0.4 | 0.05 | 0.02 |
| 2 | 133806508 | rs13431190 | 0.808 | 0.882 | -1.101 | -0.184 | -0.009 | -0.005 | -0.005 | 0.147 | 0.06 | 0.04 | 0.06 | 0.08 | 0.09 | 0.13 |
| 2 | 133811842 | rs1550243 | -0.92 | 0.843 | -0.954 | -0.596 | -0.043 | 0.218 | 0.218 | 0.37 | 0.55 | 0.83 | 0.63 | 0.96 | 0.52 | 0.69 |
| 2 | 133816484 | rs1595416 | -1.641 | 1.066 | -1.166 | -0.795 | 0.001 | 0.086 | 0.086 | 0 | 0.7 | 0.85 | 0.77 | 0.97 | 0.73 | 0.81 |
| 2 | 133826358 | rs7588373 | 0.208 | 1.321 | -0.713 | -0.538 | -0.013 | 0.011 | 0.011 | 0 | 0.21 | 0.13 | 0.18 | 0 | 0.26 | 0.17 |
| 2 | 133830790 | rs1437904 | -0.822 | 1.183 | 0.008 | -0.391 | 0.013 | 0.038 | 0.038 | 0.018 | 0.27 | 0.53 | 0.35 | 0.62 | 0.31 | 0.33 |
| 2 | 133838236 | rs6430390 | -0.221 | 1.378 | 0.299 | -0.473 | 0.016 | -0.075 | -0.075 | 0 | 0.25 | 0.33 | 0.18 | 0.01 | 0.3 | 0.19 |
| 2 | 133838788 | rs6712510 | -0.21 | 1.374 | 0.319 | -0.475 | 0.015 | -0.072 | -0.072 | 0.026 | 0.25 | 0.31 | 0.18 | 0.01 | 0.3 | 0.2 |
| 2 | 133841008 | rs16847299 | 0.51 | 1.069 | 0.597 | -0.237 | -0.013 | -0.018 | -0.018 | 0 | 0.15 | 0.12 | 0.28 | 0.6 | 0.23 | 0.26 |
| 2 | 133841199 | rs16847317 | 0.292 | 1.137 | 0.598 | -0.297 | 0.004 | 0.016 | 0.016 | 0.022 | 0.07 | 0.08 | 0.1 | 0.23 | 0.18 | 0.23 |
| 2 | 133844409 | rs765609 | 1.445 | 0.94 | 0.894 | -0.636 | -0.005 | -0.036 | -0.036 | 0.081 | 0.07 | 0.18 | 0.05 | 0 | 0.05 | 0.04 |
| 2 | 133851032 | rs2217747 | -0.244 | 0.8 | 0.755 | -0.786 | -0.009 | 0.152 | 0.152 | 0 | 0.39 | 0.17 | 0.54 | 0.96 | 0.43 | 0.63 |
| 2 | 133862100 | rs16848001 | 0.731 | 0.824 | 1.423 | -0.077 | 0.002 | -0.036 | -0.036 | 0 | 0.08 | 0.26 | 0.05 | 0 | 0.05 | 0.04 |
| 2 | 133863624 | rs12052290 | 0.197 | 0.968 | 1.491 | 0.142 | 0 | 0.034 | 0.034 | 0 | 0.14 | 0.16 | 0.29 | 0.54 | 0.16 | 0.17 |
| 2 | 133864452 | rs4437017 | -1.531 | 0.99 | 1.517 | -0.075 | -0.007 | 0.002 | 0.002 | 0 | 0.44 | 0.73 | 0.51 | 0.55 | 0.45 | 0.38 |
| 2 | 133869449 | rs6712988 | -0.967 | 0.898 | 1.504 | 0.36 | 0.009 | 0.169 | 0.169 | 0.197 | 0.18 | 0.43 | 0.32 | 0.55 | 0.18 | 0.2 |
| 2 | 133875148 | rs3795847 | -0.848 | 0.876 | 1.418 | 0.248 | 0.005 | 0.12 | 0.12 | 0 | 0.2 | 0.4 | 0.34 | 0.57 | 0.18 | 0.22 |
| 2 | 133881219 | rs2278456 | -0.03 | 0.933 | 1.198 | 0.419 | -0.01 | 0.042 | 0.042 | 0.025 | 0.15 | 0.21 | 0.22 | 0.37 | 0.11 | 0.1 |
| 2 | 133883098 | rs2217748 | -0.113 | 0.93 | 1.242 | 0.417 | -0.026 | 0.1 | 0.1 | 0.141 | 0.16 | 0.3 | 0.23 | 0.37 | 0.12 | 0.1 |
| 2 | 133884188 | rs7592507 | 0.171 | 0.929 | 1.078 | 0.396 | 0.013 | -0.03 | -0.03 | 0 | 0.14 | 0.1 | 0.2 | 0.38 | 0.11 | 0.1 |
| 2 | 133891025 | rs4954029 | -1.573 | 0.962 | 0.973 | 0.067 | 0.006 | 0.004 | 0.004 | 0 | 0.43 | 0.69 | 0.51 | 0.57 | 0.4 | 0.42 |
| 2 | 133892268 | rs10194253 | 0.69 | 0.957 | 0.538 | 0.063 | -0.005 | 0.006 | 0.006 | 0 | 0.06 | 0.04 | 0.1 | 0.19 | 0.07 | 0.16 |
| 2 | 133892552 | rs2117307 | 0.887 | 0.925 | 0.451 | 0.053 | -0.003 | 0.025 | 0.025 | 0.142 | 0.12 | 0.11 | 0.21 | 0.37 | 0.06 | 0.04 |
| 2 | 133893981 | rs10198384 | -1.643 | 0.936 | 0.356 | 0.043 | 0.005 | 0 | 0 | 0 | 0.44 | 0.82 | 0.51 | 0.58 | 0.41 | 0.42 |
| 2 | 133899078 | rs1434228 | -0.602 | 1.002 | 0.39 | 0.078 | -0.026 | 0.058 | 0.058 | 0.02 | 0.23 | 0.37 | 0.29 | 0.38 | 0.18 | 0.16 |
| 2 | 133902333 | rs13000201 | 0.479 | 1.02 | 0.347 | 0.024 | 0.001 | 0.013 | 0.013 | 0 | 0.18 | 0.12 | 0.13 | 0 | 0.2 | 0.16 |
| 2 | 133903318 | rs13432172 | 0.713 | 1.056 | -0.07 | 0.072 | -0.012 | -0.005 | -0.005 | 1.083 | 0.06 | 0.02 | 0.1 | 0.19 | 0.07 | 0.16 |
| 2 | 133906887 | rs17793104 | 0.998 | 1.407 | -0.174 | -0.047 | -0.002 | 0.003 | 0.003 | 0.164 | 0.05 | 0.03 | 0.03 | 0.01 | 0.12 | 0.07 |
| 2 | 133919427 | rs6716343 | -1.337 | 1.154 | -0.034 | 0.002 | -0.011 | 0.03 | 0.03 | 0 | 0.3 | 0.57 | 0.39 | 0.57 | 0.25 | 0.32 |
| 2 | 133921317 | rs6430397 | 0.305 | 0.852 | -0.382 | 0.114 | 0.001 | 0.007 | 0.007 | 0 | 0.15 | 0.14 | 0.21 | 0.38 | 0.13 | 0.1 |
| 2 | 133925939 | rs1434231 | -0.071 | 1.119 | -0.268 | -0.768 | 0.006 | -0.014 | -0.014 | 0 | 0.25 | 0.32 | 0.25 | 0.19 | 0.27 | 0.32 |
| 2 | 133933462 | rs10174389 | -0.056 | 0.413 | -0.949 | -0.211 | 0.034 | 0.02 | 0.02 | 0 | 0.31 | 0.2 | 0.44 | 0.73 | 0.28 | 0.36 |
| 2 | 133934181 | rs6733829 | -0.236 | 0.868 | -1.186 | -0.064 | 0.025 | -0.02 | -0.02 | 2.32 | 0.12 | 0.17 | 0.06 | 0.06 | 0.18 | 0.18 |
| 2 | 133934367 | rs1344809 | -0.905 | 0.965 | -1.809 | 0.013 | -0.076 | 0.178 | 0.178 | 0 | 0.55 | 0.96 | 0.61 | 0.79 | 0.51 | 0.61 |
| 2 | 133951544 | rs10176380 | -0.56 | 1.433 | -0.636 | 1.179 | 0 | 0.009 | 0.009 | 0 | 0.24 | 0.28 | 0.43 | 0.52 | 0.3 | 0.33 |
| 2 | 133952040 | rs12691834 | -0.958 | 1.257 | -0.305 | 0.805 | 0 | -0.002 | -0.002 | 2.716 | 0.25 | 0.31 | 0.3 | 0.76 | 0.29 | 0.31 |
| 2 | 133957689 | rs13029552 | -0.565 | 1.662 | 1.014 | 1.121 | -0.036 | -0.034 | -0.034 | 0 | 0.06 | 0.01 | 0.1 | 0.14 | 0.09 | 0.17 |
| 2 | 133962422 | rs1560891 | -1.353 | 2.064 | 0.766 | 1.384 | 0.02 | 0.019 | 0.019 | 0.679 | 0.33 | 0.53 | 0.57 | 0.45 | 0.47 | 0.35 |
| 2 | 133964894 | rs1821421 | -1.382 | 2.051 | 0.76 | 1.57 | 0.019 | 0.022 | 0.022 | 0.652 | 0.33 | 0.53 | 0.57 | 0.5 | 0.47 | 0.36 |
| 2 | 133968768 | rs4953880 | -1.301 | 2.081 | 0.231 | 1.591 | 0.022 | 0.025 | 0.025 | 0.522 | 0.33 | 0.37 | 0.56 | 0.5 | 0.47 | 0.35 |
| 2 | 133969611 | rs17738103 | -1.155 | 2.077 | -0.248 | 1.478 | -0.071 | -0.07 | -0.07 | 0 | 0.31 | 0.13 | 0.54 | 0.45 | 0.47 | 0.35 |
| 2 | 133978555 | rs12477764 | -0.682 | 1.879 | -0.075 | 1.229 | -0.053 | -0.014 | -0.014 | 0 | 0.25 | 0.12 | 0.44 | 0.31 | 0.38 | 0.19 |
| 2 | 133981057 | rs10469554 | -1.431 | 1.376 | -0.585 | 1.084 | -0.066 | 0.027 | 0.027 | 0 | 0.43 | 0.88 | 0.55 | 0.36 | 0.47 | 0.31 |
| 2 | 133985708 | rs1430171 | -0.803 | 2.189 | -0.856 | 1.459 | -0.071 | -0.037 | -0.037 | 0 | 0.25 | 0.06 | 0.41 | 0.32 | 0.34 | 0.18 |
| 2 | 134000064 | rs6749561 | -1.735 | 2.31 | 0.21 | 1.778 | 0.097 | 0.117 | 0.117 | 0 | 0.12 | 0.38 | 0.24 | 0.18 | 0.18 | 0.26 |
| 2 | 134005569 | rs10496702 | -1.485 | 2.295 | 0.444 | 1.836 | 0.065 | 0.089 | 0.089 | 0 | 0.09 | 0.33 | 0.2 | 0.13 | 0.11 | 0.14 |
| 2 | 134009296 | rs13413333 | -0.819 | 2.303 | 0.415 | 1.835 | -0.003 | 0 | 0 | 0 | 0.06 | 0.04 | 0.16 | 0.14 | 0.11 | 0.14 |
| 2 | 134014961 | rs7600871 | -1.824 | 2.285 | 0.474 | 1.839 | 0.043 | 0.11 | 0.11 | 0 | 0.09 | 0.25 | 0.18 | 0.19 | 0.11 | 0.22 |
| 2 | 134033399 | rs1430170 | -1.531 | 2.104 | 0.129 | 1.955 | 0.006 | 0.038 | 0.038 | 0 | 0.23 | 0.36 | 0.3 | 0.3 | 0.28 | 0.26 |
| 2 | 134034022 | rs16823003 | -0.227 | 2.41 | 0.04 | 2.486 | -0.01 | -0.008 | -0.008 | 0 | 0.07 | 0.03 | 0.08 | 0.28 | 0.12 | 0.11 |
| 2 | 134040519 | rs7577925 | -1.752 | 2.308 | 0.229 | 2.015 | 0.028 | 0.019 | 0.019 | 0 | 0.26 | 0.45 | 0.34 | 0.26 | 0.36 | 0.3 |
| 2 | 134041922 | rs4589811 | -1.014 | 1.954 | 0.246 | 2.067 | 0.018 | 0.011 | 0.011 | 0 | 0.08 | 0.14 | 0.18 | 0.12 | 0.14 | 0.13 |
| 2 | 134044131 | rs10496704 | -1.09 | 1.958 | 0.237 | 2.099 | 0.024 | 0.049 | 0.049 | 0 | 0.09 | 0.18 | 0.19 | 0.17 | 0.14 | 0.14 |
| 2 | 134046163 | rs1344726 | -1.46 | 1.783 | -0.039 | 1.705 | 0.003 | 0.008 | 0.008 | 1.948 | 0.26 | 0.46 | 0.32 | 0.27 | 0.3 | 0.19 |
| 2 | 134051863 | rs1529570 | -0.514 | 1.811 | -0.583 | 1.422 | -0.04 | 0.005 | 0.005 | 0 | 0.14 | 0.01 | 0.09 | 0.06 | 0.19 | 0.06 |
| 2 | 134060719 | rs7604084 | -0.807 | 1.512 | 0.138 | 1.421 | -0.019 | 0.005 | 0.005 | 0 | 0.16 | 0.32 | 0.1 | 0.12 | 0.1 | 0.14 |
| 2 | 134070417 | rs934265 | -0.785 | 1.701 | 0.196 | 1.367 | 0.001 | -0.003 | -0.003 | 0 | 0.12 | 0.15 | 0.07 | 0.07 | 0.12 | 0.15 |
| 2 | 134081421 | rs13409478 | -1.53 | 1.679 | 0.689 | 1.695 | 0.007 | 0.031 | 0.031 | 0.08 | 0.19 | 0.32 | 0.15 | 0.27 | 0.2 | 0.17 |
| 2 | 134086271 | rs10928450 | -1.413 | 2.086 | 0.633 | 1.4 | -0.011 | 0.017 | 0.017 | 0 | 0.2 | 0.15 | 0.2 | 0.14 | 0.25 | 0.18 |

| | | | | | | | | | | | | | | | | |
|---|---|---|---|---|---|---|---|---|---|---|---|---|---|---|---|---|
| 2 | 134088150 | rs10165555 | -0.041 | 2.894 | 1.089 | 1.8 | 0.1 | 0.148 | 0.148 | 0.21 | 0.09 | 0.25 | 0.27 | 0.33 | 0.14 | 0.23 |
| 2 | 134089004 | rs1430166 | -1.423 | 2.577 | 1.419 | 1.644 | 0.011 | -0.062 | -0.062 | 0 | 0.2 | 0.42 | 0.19 | 0.13 | 0.19 | 0.15 |
| 2 | 134103374 | rs13432602 | -1.069 | 2.567 | 1.446 | 1.625 | -0.003 | 0.008 | 0.008 | 0 | 0.18 | 0.17 | 0.17 | 0.13 | 0.19 | 0.16 |
| 2 | 134110924 | rs13407503 | -1.663 | 2.56 | 1.485 | 1.623 | 0.01 | -0.073 | -0.073 | 0 | 0.22 | 0.46 | 0.23 | 0.13 | 0.23 | 0.21 |
| 2 | 134116904 | rs10175744 | -0.985 | 2.557 | 1.446 | 1.623 | -0.005 | 0.01 | 0.01 | 0 | 0.17 | 0.14 | 0.16 | 0.12 | 0.18 | 0.18 |
| 2 | 134117177 | rs1435556 | -1.058 | 2.557 | 1.465 | 1.623 | -0.003 | 0.005 | 0.005 | 0 | 0.18 | 0.17 | 0.22 | 0.13 | 0.19 | 0.18 |
| 2 | 134128159 | rs2289076 | 0.131 | 2.279 | 0.824 | 1.403 | 0.018 | 0.015 | 0.015 | 0 | 0.18 | 0.26 | 0.26 | 0.31 | 0.21 | 0.22 |
| 2 | 134151457 | rs13417230 | -0.229 | 1.98 | 0.361 | 0.996 | -0.017 | 0.027 | 0.027 | 0 | 0.12 | 0.07 | 0.08 | 0 | 0.14 | 0.04 |
| 2 | 134152610 | rs7572861 | -0.678 | 1.716 | -0.036 | 0.8 | -0.064 | -0.076 | -0.076 | 1.012 | 0.83 | 0.98 | 0.75 | 0.81 | 0.75 | 0.8 |
| 2 | 134154923 | rs16824299 | -1.525 | 1.254 | 0.745 | 0.437 | 0 | -0.007 | -0.007 | 0.555 | 0.07 | 0.04 | 0.11 | 0.13 | 0.09 | 0.18 |
| 2 | 134157295 | rs17802295 | -1.349 | 1.222 | 0.798 | 0.468 | 0.007 | -0.046 | -0.046 | 0 | 0.2 | 0.33 | 0.18 | 0.12 | 0.19 | 0.18 |
| 2 | 134162470 | rs9287426 | -0.829 | 1.226 | 0.747 | 0.47 | 0.001 | -0.007 | -0.007 | 0.204 | 0.17 | 0.18 | 0.16 | 0.12 | 0.18 | 0.18 |
| 2 | 134168015 | rs12327930 | -1.243 | 1.24 | 0.62 | 0.429 | -0.019 | 0.04 | 0.04 | 0 | 0.21 | 0.1 | 0.19 | 0.12 | 0.24 | 0.23 |
| 2 | 134172551 | rs10184630 | -0.997 | 1.258 | 0.649 | 0.441 | 0.003 | -0.018 | -0.018 | 0 | 0.18 | 0.23 | 0.16 | 0.12 | 0.18 | 0.18 |
| 2 | 134175132 | rs16824573 | -1.421 | 1.266 | 0.459 | 0.44 | 0.017 | -0.066 | -0.066 | 0 | 0.22 | 0.43 | 0.22 | 0.12 | 0.24 | 0.21 |
| 2 | 134175855 | rs1369510 | -1.03 | 1.336 | -0.611 | 0.647 | -0.015 | 0.03 | 0.03 | 0 | 0.19 | 0.1 | 0.16 | 0.12 | 0.19 | 0.19 |
| 2 | 134184420 | rs4954050 | -0.67 | 0.917 | -0.779 | 0.8 | 0.005 | -0.003 | -0.003 | 0.153 | 0.41 | 0.32 | 0.35 | 0.51 | 0.36 | 0.31 |
| 2 | 134187067 | rs6430417 | -0.371 | 0.056 | -0.927 | 0.425 | 0.006 | 0.004 | 0.004 | 0.892 | 0.42 | 0.62 | 0.33 | 0.36 | 0.28 | 0.3 |
| 2 | 134188405 | rs11883588 | -0.316 | 0.001 | -0.71 | 0.483 | -0.006 | -0.003 | -0.003 | 0.356 | 0.41 | 0.56 | 0.33 | 0.38 | 0.28 | 0.3 |
| 2 | 134193885 | rs2117737 | -1.14 | 0.633 | -1.513 | -0.051 | -0.093 | 0.079 | 0.079 | 1.136 | 0.76 | 0.96 | 0.66 | 0.91 | 0.66 | 0.63 |
| 2 | 134194530 | rs11689516 | -0.558 | 0.579 | -1.482 | -0.51 | 0.043 | 0.034 | 0.034 | 0 | 0.24 | 0.52 | 0.27 | 0.32 | 0.25 | 0.27 |
| 2 | 134195776 | rs6414013 | 0.304 | 0.529 | -0.924 | -0.532 | -0.019 | 0.073 | 0.073 | 0 | 0.1 | 0.24 | 0.08 | 0.27 | 0.03 | 0.12 |
| 2 | 134195981 | rs10199285 | -0.043 | 0.662 | -0.863 | -0.572 | 0.091 | -0.024 | -0.024 | 0.032 | 0.09 | 0.24 | 0.09 | 0.04 | 0.18 | 0.13 |
| 2 | 134196632 | rs13422893 | -0.476 | 0.336 | -1.019 | -0.799 | 0.006 | -0.069 | -0.069 | 0 | 0.33 | 0.39 | 0.27 | 0.05 | 0.35 | 0.2 |
| 2 | 134197301 | rs12623281 | 1.628 | 0.222 | -1.17 | -0.724 | 0.01 | 0.048 | 0.048 | 0 | 0.19 | 0.05 | 0.08 | 0.08 | 0.18 | 0.13 |
| 2 | 134197843 | rs11896860 | -0.733 | 0.426 | -0.512 | -0.229 | 0.06 | -0.043 | -0.043 | 0.13 | 0.14 | 0.27 | 0.18 | 0.05 | 0.22 | 0.14 |
| 2 | 134204245 | rs7596517 | -0.957 | 0.391 | -0.579 | -0.233 | -0.001 | -0.003 | -0.003 | 0 | 0.41 | 0.63 | 0.32 | 0.32 | 0.32 | 0.29 |
| 2 | 134206763 | rs7604549 | -1.074 | 0.331 | -0.549 | -0.231 | -0.001 | 0.002 | 0.002 | 0 | 0.42 | 0.63 | 0.35 | 0.29 | 0.39 | 0.32 |
| 2 | 134207106 | rs6712629 | -0.16 | 0.76 | -0.252 | 0.448 | 0.059 | -0.009 | -0.009 | 0 | 0.07 | 0.24 | 0.06 | 0.04 | 0.11 | 0.1 |
| 2 | 134213436 | rs16825174 | -0.329 | 0.576 | -0.657 | 0.648 | 0.002 | 0.084 | 0.084 | 0 | 0.05 | 0.14 | 0.07 | 0.24 | 0.03 | 0.1 |
| 2 | 134221398 | rs6430423 | -0.659 | 0.652 | -0.208 | 0.921 | 0.024 | 0.185 | 0.185 | 0.256 | 0.14 | 0.3 | 0.21 | 0.41 | 0.16 | 0.23 |
| 2 | 134222450 | rs4953882 | -0.544 | 0.657 | -0.19 | 0.925 | 0.024 | 0.173 | 0.173 | 0.215 | 0.13 | 0.29 | 0.21 | 0.38 | 0.16 | 0.23 |
| 2 | 134228810 | rs1369506 | -0.953 | 0.316 | -1.119 | 0.146 | -0.043 | 0.159 | 0.159 | 0.914 | 0.54 | 0.85 | 0.6 | 0.82 | 0.48 | 0.57 |
| 2 | 134229008 | rs13419826 | -0.653 | 0.228 | -0.676 | 0.258 | 0.015 | 0.055 | 0.055 | 0.48 | 0.44 | 0.72 | 0.48 | 0.7 | 0.37 | 0.43 |
| 2 | 134230217 | rs1821625 | -0.677 | 0.607 | 0.565 | 1.494 | 0.187 | 0.333 | 0.333 | 0 | 0.07 | 0.44 | 0.11 | 0.3 | 0.1 | 0.04 |
| 2 | 134230426 | rs1821624 | -0.296 | 0.826 | 0.27 | 1.231 | -0.003 | -0.013 | -0.013 | 0 | 0.32 | 0.19 | 0.35 | 0.39 | 0.34 | 0.33 |
| 2 | 134231234 | rs13408209 | -0.722 | 0.873 | 1.124 | 1.222 | 0.064 | 0.142 | 0.142 | 1.344 | 0.08 | 0.49 | 0.08 | 0.12 | 0.06 | 0.17 |
| 2 | 134232010 | rs10206425 | -0.722 | 0.873 | 1.157 | 1.222 | 0.062 | 0.138 | 0.138 | 1.344 | 0.08 | 0.49 | 0.08 | 0.12 | 0.06 | 0.17 |
| 2 | 134234914 | rs16825505 | 0.013 | 0.888 | 1.309 | 1.218 | 0.022 | 0.039 | 0.039 | 0.036 | 0.06 | 0.2 | 0.06 | 0.06 | 0.06 | 0.17 |
| 2 | 134242471 | rs1836669 | -0.172 | 0.742 | 1.125 | 1.499 | 0.007 | 0.003 | 0.003 | 0.47 | 0.22 | 0.19 | 0.27 | 0.3 | 0.29 | 0.33 |
| 2 | 134245403 | rs2874399 | -0.839 | 0.48 | 1.163 | 1.352 | 0.041 | 0.009 | 0.009 | 0.099 | 0.3 | 0.43 | 0.33 | 0.32 | 0.37 | 0.38 |
| 2 | 134250093 | rs1017368 | -1.115 | 0.295 | 1.2 | 1.127 | 0.005 | 0.003 | 0.003 | 0 | 0.36 | 0.57 | 0.44 | 0.4 | 0.42 | 0.39 |
| 2 | 134253001 | rs4953883 | -1.642 | 0.256 | 1.03 | 1.11 | -0.043 | -0.026 | -0.026 | 0 | 0.43 | 0.81 | 0.5 | 0.45 | 0.47 | 0.57 |
| 2 | 134253973 | rs7559239 | -1.305 | 0.302 | 1.041 | 1.105 | -0.027 | -0.016 | -0.016 | 0 | 0.37 | 0.75 | 0.46 | 0.4 | 0.42 | 0.39 |
| 2 | 134259612 | rs2163654 | -0.835 | 0.509 | 0.988 | 1.158 | 0.011 | 0.002 | 0.002 | 0.035 | 0.29 | 0.26 | 0.34 | 0.28 | 0.4 | 0.38 |
| 2 | 134261816 | rs6736284 | 0.061 | 0.417 | 1.014 | 1.165 | 0.056 | 0.181 | 0.181 | 0.193 | 0.13 | 0.32 | 0.19 | 0.44 | 0.15 | 0.09 |
| 2 | 134262118 | rs10928453 | -0.455 | 0.428 | 0.998 | 1.156 | 0.041 | 0.124 | 0.124 | 0.437 | 0.15 | 0.33 | 0.21 | 0.41 | 0.19 | 0.1 |
| 2 | 134274625 | rs1966628 | -1.149 | 0.553 | 0.874 | 1.065 | 0.001 | -0.001 | -0.001 | 0 | 0.3 | 0.22 | 0.38 | 0.61 | 0.37 | 0.46 |
| 2 | 134287762 | rs12612615 | -1.169 | 0.918 | 0.903 | 1.399 | 0.028 | 0.301 | 0.301 | 1.472 | 0.16 | 0.48 | 0.22 | 0.45 | 0.13 | 0.29 |
| 2 | 134289974 | rs1347044 | -1.109 | 0.935 | 0.889 | 1.347 | 0.03 | 0.306 | 0.306 | 1.316 | 0.15 | 0.46 | 0.22 | 0.48 | 0.12 | 0.29 |
| 2 | 134311186 | rs16826387 | 0.447 | 1.045 | 0.33 | 1.459 | 0.002 | 0.08 | 0.08 | 0 | 0.06 | 0.08 | 0.09 | 0.3 | 0.05 | 0.14 |
| 2 | 134311713 | rs1835818 | -0.669 | 0.626 | -0.096 | 0.958 | -0.005 | -0.012 | -0.012 | 0.191 | 0.3 | 0.16 | 0.29 | 0.35 | 0.3 | 0.31 |
| 2 | 134312960 | rs1368091 | -0.733 | 1.007 | -0.286 | 0.703 | 0.001 | -0.034 | -0.034 | 0 | 0.83 | 0.88 | 0.84 | 1 | 0.75 | 0.86 |
| 2 | 134316522 | rs1897433 | 0.682 | 0.397 | -0.484 | 0.425 | 0.014 | -0.023 | -0.023 | 0 | 0.38 | 0.2 | 0.22 | 0.39 | 0.34 | 0.43 |
| 2 | 134319270 | rs1816382 | 0.263 | 0.59 | -0.156 | 0.413 | 0.001 | 0.004 | 0.004 | 0 | 0.47 | 0.33 | 0.45 | 0.51 | 0.53 | 0.63 |
| 2 | 134325713 | rs4143759 | 0.959 | 0.48 | -0.045 | 0.434 | -0.007 | 0.003 | 0.003 | 0 | 0.05 | 0.07 | 0.01 | 0.09 | 0.02 | 0.07 |
| 2 | 134326523 | rs13421559 | 0.469 | 0.356 | 0.251 | 0.223 | -0.001 | -0.001 | -0.001 | 0 | 0.29 | 0.24 | 0.29 | 0.37 | 0.18 | 0.19 |
| 2 | 134332096 | rs891821 | 0.494 | 0.765 | 0.099 | 0.301 | -0.013 | -0.009 | -0.009 | 0 | 0.43 | 0.32 | 0.44 | 0.49 | 0.52 | 0.6 |
| 2 | 134332761 | rs7588354 | -1.112 | 0.591 | -0.302 | 0.336 | -0.013 | 0.061 | 0.061 | 2.47 | 0.76 | 0.9 | 0.86 | 0.97 | 0.76 | 0.89 |
| 2 | 134337990 | rs13027438 | -0.967 | 0.981 | -0.206 | 0.647 | -0.001 | 0.049 | 0.049 | 0.462 | 0.71 | 0.91 | 0.85 | 0.82 | 0.73 | 0.82 |
| 2 | 134340162 | rs12465334 | -0.258 | 1.743 | 0.885 | 1.656 | 0.009 | -0.013 | -0.013 | 0 | 0.23 | 0.17 | 0.28 | 0.32 | 0.16 | 0.16 |
| 2 | 134350570 | rs6715224 | -0.363 | 1.941 | 0.263 | 1.637 | 0.057 | 0.015 | 0.015 | 0 | 0.08 | 0.11 | 0.18 | 0.09 | 0.19 | 0.19 |
| 2 | 134356109 | rs2436193 | 0.328 | 0.056 | -0.464 | 1.969 | 0.007 | -0.009 | -0.009 | 0 | 0.76 | 0.82 | 0.71 | 0.68 | 0.83 | 0.84 |
| 2 | 134363032 | rs7588567 | -0.632 | 0.652 | -0.295 | 1.4 | -0.001 | -0.003 | -0.003 | 0 | 0.42 | 0.41 | 0.47 | 0.65 | 0.38 | 0.44 |
| 2 | 134363401 | rs11676278 | -1.287 | 1.194 | 0.498 | 1.907 | 0.048 | 0.054 | 0.054 | 0.708 | 0.2 | 0.59 | 0.29 | 0.26 | 0.26 | 0.31 |
| 2 | 134367842 | rs1669470 | 0.955 | 1.173 | 0.994 | 2.359 | 0.015 | 0.127 | 0.127 | 0 | 0.79 | 0.54 | 0.57 | 0.46 | 0.74 | 0.74 |
| 2 | 134369949 | rs1653794 | 1.732 | 1.033 | 1.364 | 3.16 | 0.011 | 0.459 | 0.459 | 0 | 0.95 | 0.7 | 0.78 | 0.54 | 0.97 | 0.89 |
| 2 | 134370162 | rs7564515 | -0.908 | 1.037 | 1.413 | 3.093 | -0.002 | -0.002 | -0.002 | 0 | 0.18 | 0.16 | 0.24 | 0.32 | 0.24 | 0.23 |
| 2 | 134379150 | rs12328892 | -1.22 | 1.634 | 1.843 | 2.848 | 0.08 | 0.015 | 0.015 | 0.013 | 0.05 | 0.11 | 0.06 | 0.03 | 0.1 | 0.06 |

| | | | | | | | | | | | | | | | | |
|---|---|---|---|---|---|---|---|---|---|---|---|---|---|---|---|---|
| 2 | 134382097 | rs13393385 | -1.523 | 1.607 | 2.167 | 2.821 | 0.166 | 0.062 | 0.062 | 0.199 | 0.06 | 0.28 | 0.07 | 0.02 | 0.1 | 0.06 |
| 2 | 134392926 | rs13382906 | -2.22 | 1.562 | 2.03 | 2.876 | 0.004 | 0.44 | 0.44 | 0 | 0.05 | 0.3 | 0.22 | 0.46 | 0.03 | 0.11 |
| 2 | 134397457 | rs6748388 | -1.55 | 1.588 | 1.866 | 2.923 | 0.046 | 0.127 | 0.127 | 0 | 0.07 | 0.1 | 0.09 | 0.26 | 0.1 | 0.15 |
| 2 | 134426649 | rs7607128 | -0.487 | 0.921 | 0.069 | 1.655 | 0.042 | 0.215 | 0.215 | 0 | 0.5 | 0.8 | 0.54 | 0.8 | 0.45 | 0.46 |
| 2 | 134427773 | rs721270 | -0.754 | 1.329 | -0.044 | 1.458 | -0.057 | 0.19 | 0.19 | 0 | 0.56 | 0.8 | 0.64 | 0.86 | 0.54 | 0.56 |
| 2 | 134427974 | rs721269 | -0.11 | 1.179 | 0.105 | 1.5 | -0.006 | 0.001 | 0.001 | 0 | 0.38 | 0.42 | 0.27 | 0.39 | 0.34 | 0.37 |
| 2 | 134431078 | rs10928458 | -0.306 | 1.508 | -1.326 | 1.201 | -0.001 | -0.018 | -0.018 | 0 | 0.46 | 0.14 | 0.44 | 0.58 | 0.55 | 0.57 |
| 2 | 134431443 | rs1901435 | -0.43 | 2.084 | -1.626 | 0.678 | -0.001 | 0.096 | 0.096 | 0 | 0.49 | 0.18 | 0.41 | 0.67 | 0.54 | 0.55 |
| 2 | 134434824 | rs7567288 | -0.627 | 2.374 | -1.085 | 0.399 | 0.017 | -0.023 | -0.023 | 0 | 0.16 | 0.32 | 0.17 | 0.1 | 0.19 | 0.18 |
| 2 | 134437102 | rs10439201 | -1.476 | 2.363 | -0.848 | 0.152 | -0.009 | -0.019 | -0.019 | 1.394 | 0.64 | 0.47 | 0.65 | 0.8 | 0.7 | 0.67 |
| 2 | 134439898 | rs7586145 | 0.087 | 2.804 | -0.295 | 0.392 | 0.045 | 0.029 | 0.029 | 0 | 0.13 | 0.24 | 0.17 | 0.17 | 0.17 | 0.17 |
| 2 | 134440433 | rs17767492 | 0.297 | 2.809 | -0.519 | 0.395 | -0.054 | -0.042 | -0.042 | 0 | 0.12 | 0.04 | 0.15 | 0.17 | 0.17 | 0.17 |
| 2 | 134440486 | rs1901441 | -1.431 | 2.779 | -0.396 | 0.353 | 0.044 | -0.01 | -0.01 | 0 | 0.3 | 0.59 | 0.34 | 0.26 | 0.36 | 0.35 |
| 2 | 134441766 | rs16827937 | 1.16 | 2.581 | -0.455 | 0.334 | 0.003 | -0.02 | -0.02 | 0 | 0.3 | 0.18 | 0.34 | 0.55 | 0.32 | 0.28 |
| 2 | 134442532 | rs6430443 | -1.858 | 2.19 | -0.478 | 0.264 | 0.065 | 0.046 | 0.046 | 0 | 0.69 | 0.85 | 0.76 | 0.8 | 0.81 | 0.68 |
| 2 | 134458912 | rs11896655 | 0.083 | 2.399 | 0.008 | 0.353 | 0.063 | -0.047 | -0.047 | 0 | 0.12 | 0.4 | 0.12 | 0.04 | 0.14 | 0.11 |
| 2 | 134461021 | rs4305321 | -0.778 | 2.57 | -0.206 | 0.52 | 0.026 | -0.066 | -0.066 | 0 | 0.66 | 0.87 | 0.7 | 0.48 | 0.69 | 0.69 |
| 2 | 134467013 | rs12474261 | -1.79 | 2.291 | -0.191 | 0.336 | 0.081 | 0.063 | 0.063 | 0 | 0.67 | 0.87 | 0.76 | 0.79 | 0.8 | 0.67 |
| 2 | 134467871 | rs6738353 | -0.649 | 2.374 | -0.157 | 0.393 | 0.044 | 0.07 | 0.07 | 0 | 0.54 | 0.29 | 0.63 | 0.76 | 0.66 | 0.56 |
| 2 | 134473174 | rs6712538 | -0.218 | 2.64 | 0.023 | 0.659 | 0.086 | 0.011 | 0.011 | 0.693 | 0.05 | 0.11 | 0.15 | 0.05 | 0.16 | 0.06 |
| 2 | 134473360 | rs6430446 | -1.131 | 2.754 | 0.057 | 0.512 | 0.053 | 0.087 | 0.087 | 0 | 0.57 | 0.71 | 0.59 | 0.8 | 0.67 | 0.62 |
| 2 | 134487163 | rs4522658 | -1.221 | 2.96 | 0.244 | 0.685 | 0.017 | 0.002 | 0.002 | 0.374 | 0.28 | 0.3 | 0.41 | 0.27 | 0.46 | 0.32 |
| 2 | 134491266 | rs1992784 | 0.207 | 3.016 | 0.131 | 0.711 | -0.022 | -0.022 | -0.022 | 0 | 0.13 | 0.08 | 0.16 | 0.19 | 0.17 | 0.16 |
| 2 | 134495945 | rs1374405 | -0.046 | 2.829 | 0.195 | 0.645 | 0.002 | -0.019 | -0.019 | 0 | 0.12 | 0.21 | 0.12 | 0.04 | 0.14 | 0.12 |
| 2 | 134501131 | rs12614438 | 1.187 | 2.958 | 0.206 | 0.656 | 0.038 | -0.077 | -0.077 | 0.003 | 0.29 | 0.12 | 0.17 | 0.52 | 0.22 | 0.31 |
| 2 | 134501207 | rs12614459 | 1.221 | 2.955 | 0.182 | 0.707 | 0.021 | -0.167 | -0.167 | 0 | 0.28 | 0.03 | 0.16 | 0.52 | 0.22 | 0.31 |
| 2 | 134507967 | rs895539 | 0.192 | 2.965 | 0.219 | 0.676 | 0.024 | -0.027 | -0.027 | 0 | 0.4 | 0.33 | 0.27 | 0.55 | 0.34 | 0.41 |
| 2 | 134509873 | rs7568496 | 0.194 | 2.951 | 0.228 | 0.635 | -0.001 | -0.006 | -0.006 | 0 | 0.11 | 0.19 | 0.11 | 0.04 | 0.12 | 0.11 |
| 2 | 134512727 | rs7559696 | -0.66 | 2.994 | 0.398 | 0.952 | 0.007 | 0.007 | 0.007 | 2.632 | 0.16 | 0.34 | 0.16 | 0.19 | 0.15 | 0.14 |
| 2 | 134513100 | rs6414015 | -1.307 | 3.123 | 0.367 | 0.912 | 0.046 | 0.105 | 0.105 | 0 | 0.59 | 0.83 | 0.61 | 0.79 | 0.66 | 0.62 |
| 2 | 134514960 | rs6731434 | -0.245 | 2.932 | 0.365 | 1.115 | 0.003 | 0.011 | 0.011 | 0 | 0.45 | 0.6 | 0.36 | 0.71 | 0.39 | 0.47 |
| 2 | 134519547 | rs1867897 | 0.633 | 2.864 | 0.34 | 1.135 | -0.006 | 0.006 | 0.006 | 0 | 0.09 | 0.09 | 0.09 | 0.04 | 0.12 | 0.11 |
| 2 | 134528954 | rs746838 | -1.249 | 2.902 | 0.389 | 1.207 | 0.064 | 0.064 | 0.064 | 0 | 0.27 | 0.52 | 0.32 | 0.38 | 0.32 | 0.3 |
| 2 | 134529098 | rs1446740 | -0.958 | 2.895 | 0.301 | 1.185 | -0.004 | 0.05 | 0.05 | 0.048 | 0.54 | 0.71 | 0.46 | 0.75 | 0.51 | 0.57 |
| 2 | 134529356 | rs731972 | 1.568 | 2.314 | 0.349 | 1.182 | 0.047 | -0.084 | -0.084 | 0 | 0.25 | 0.04 | 0.12 | 0.36 | 0.19 | 0.28 |
| 2 | 134531412 | rs9917166 | 0.418 | 2.34 | 0.494 | 1.163 | 0.049 | -0.035 | -0.035 | 0.106 | 0.1 | 0.33 | 0.11 | 0.04 | 0.12 | 0.11 |
| 2 | 134532317 | rs1446743 | -0.101 | 2.357 | 0.356 | 1.204 | 0.009 | -0.007 | -0.007 | 0 | 0.41 | 0.67 | 0.3 | 0.56 | 0.34 | 0.41 |
| 2 | 134537665 | rs16829308 | 1.452 | 2.262 | 0.042 | 1.224 | 0.049 | -0.082 | -0.082 | 0.175 | 0.25 | 0.04 | 0.12 | 0.37 | 0.19 | 0.27 |
| 2 | 134539036 | rs4423652 | -0.577 | 2.971 | 0.304 | 1.211 | 0.017 | 0.091 | 0.091 | 0 | 0.49 | 0.61 | 0.52 | 0.76 | 0.58 | 0.52 |
| 2 | 134541742 | rs16829379 | -0.97 | 3.052 | 0.514 | 1.274 | 0.114 | 0.125 | 0.125 | 0 | 0.09 | 0.31 | 0.21 | 0.19 | 0.18 | 0.07 |
| 2 | 134544190 | rs10496712 | 1.381 | 2.929 | 0.424 | 1.222 | 0.05 | -0.04 | -0.04 | 0 | 0.27 | 0.13 | 0.12 | 0.37 | 0.2 | 0.31 |
| 2 | 134546481 | rs10496713 | 0.59 | 3.002 | 0.415 | 1.162 | -0.004 | 0.002 | 0.002 | 0 | 0.1 | 0.11 | 0.1 | 0.04 | 0.12 | 0.09 |
| 2 | 134546741 | rs2321287 | -0.028 | 3.039 | 0.416 | 1.17 | -0.047 | -0.046 | -0.046 | 4.895 | 0.14 | 0.04 | 0.18 | 0.2 | 0.21 | 0.14 |
| 2 | 134548819 | rs6430452 | -1.38 | 3.043 | 0.46 | 1.162 | 0.033 | 0.08 | 0.08 | 0 | 0.6 | 0.79 | 0.64 | 0.79 | 0.71 | 0.61 |
| 2 | 134549860 | rs4589813 | 0.079 | 3.024 | 0.5 | 1.158 | 0.003 | 0 | 0 | 0.106 | 0.42 | 0.39 | 0.33 | 0.57 | 0.41 | 0.45 |
| 2 | 134552026 | rs744129 | 0.303 | 2.979 | 0.516 | 1.165 | -0.011 | -0.001 | -0.001 | 0 | 0.38 | 0.47 | 0.25 | 0.41 | 0.32 | 0.41 |
| 2 | 134559550 | rs7603190 | 0.497 | 2.978 | 0.532 | 1.175 | -0.001 | 0.001 | 0.001 | 0 | 0.36 | 0.56 | 0.33 | 0.56 | 0.37 | 0.38 |
| 2 | 134559761 | rs6430455 | 1.289 | 2.903 | 0.539 | 1.19 | -0.023 | 0.009 | 0.009 | 0 | 0.28 | 0.34 | 0.14 | 0.37 | 0.2 | 0.31 |
| 2 | 134560999 | rs981668 | -0.952 | 2.893 | 0.487 | 1.145 | -0.024 | -0.005 | -0.005 | 0 | 0.24 | 0.23 | 0.3 | 0.23 | 0.34 | 0.22 |
| 2 | 134561145 | rs6711922 | -1.505 | 2.717 | 0.22 | 1.072 | 0.022 | 0.048 | 0.048 | 0 | 0.64 | 0.79 | 0.67 | 0.79 | 0.74 | 0.68 |
| 2 | 134561352 | rs13429336 | 0 | 2.854 | 0.225 | 1.156 | -0.043 | -0.042 | -0.042 | 5.873 | 0.13 | 0.04 | 0.18 | 0.2 | 0.21 | 0.13 |
| 2 | 134563241 | rs7570764 | -1.264 | 2.543 | 0.083 | 1.21 | 0.021 | 0.091 | 0.091 | 0 | 0.6 | 0.81 | 0.63 | 0.79 | 0.66 | 0.66 |
| 2 | 134563476 | rs16829491 | 0.598 | 2.502 | -0.058 | 1.22 | -0.031 | -0.039 | -0.039 | 0.91 | 0.1 | 0.03 | 0.14 | 0.21 | 0.13 | 0.13 |
| 2 | 134565505 | rs895535 | -0.219 | 2.337 | 0.095 | 0.911 | 0.001 | 0.003 | 0.003 | 0 | 0.47 | 0.55 | 0.39 | 0.62 | 0.45 | 0.53 |
| 2 | 134566099 | rs1374399 | -0.74 | 2.332 | -0.034 | 0.944 | 0.02 | -0.01 | -0.01 | 0 | 0.08 | 0.06 | 0.18 | 0.03 | 0.19 | 0.12 |
| 2 | 134568892 | rs17279673 | 0.352 | 2.073 | 0.193 | 1.124 | 0 | 0.001 | 0.001 | 0 | 0.44 | 0.53 | 0.6 | 0.39 | 0.57 | 0.4 |
| 2 | 134569297 | rs7584077 | -0.568 | 2.066 | 0.275 | 0.877 | 0 | -0.046 | -0.046 | 0 | 0.88 | 0.78 | 0.87 | 0.97 | 0.89 | 0.89 |
| 2 | 134574681 | rs4954079 | 0.29 | 2.399 | 0.356 | 1.032 | 0 | 0.001 | 0.001 | 0 | 0.52 | 0.51 | 0.4 | 0.41 | 0.53 | 0.46 |
| 2 | 134575885 | rs6735017 | 1.302 | 2.125 | 0.349 | 1.116 | 0.002 | -0.054 | -0.054 | 0.88 | 0.37 | 0.53 | 0.23 | 0.15 | 0.38 | 0.24 |
| 2 | 134576249 | rs20349 | 1.444 | 2.129 | 0.274 | 1.118 | 0.005 | -0.042 | -0.042 | 0 | 0.35 | 0.4 | 0.21 | 0.15 | 0.38 | 0.24 |
| 2 | 134579078 | rs6750345 | 0.193 | 1.918 | -0.017 | 1.14 | -0.019 | -0.068 | -0.068 | 2.947 | 0.16 | 0.02 | 0.16 | 0.25 | 0.14 | 0.24 |
| 2 | 134579389 | rs17821164 | 1.508 | 1.885 | 0.124 | 1.053 | -0.024 | 0.061 | 0.061 | 0 | 0.34 | 0.18 | 0.21 | 0.15 | 0.39 | 0.24 |
| 2 | 134580522 | rs4954080 | -1.543 | 1.63 | 0.01 | 1.115 | 0.041 | -0.159 | -0.159 | 0.098 | 0.74 | 0.98 | 0.9 | 0.54 | 0.83 | 0.67 |
| 2 | 134583314 | rs6430460 | 0.093 | 1.008 | 0.312 | 1.105 | -0.003 | 0 | 0 | 0 | 0.55 | 0.5 | 0.72 | 0.56 | 0.69 | 0.47 |
| 2 | 134584046 | rs4954083 | -1.569 | 1.06 | -0.388 | 1.168 | 0.04 | -0.025 | -0.025 | 0 | 0.75 | 0.94 | 0.87 | 0.75 | 0.83 | 0.71 |
| 2 | 134584179 | rs17770239 | 0.212 | 1.053 | -0.264 | 1.12 | -0.007 | -0.011 | -0.011 | 0.012 | 0.17 | 0.1 | 0.13 | 0.17 | 0.16 | 0.25 |
| 2 | 134589401 | rs7602946 | 1.456 | 1.257 | -0.078 | 1.026 | -0.034 | -0.002 | -0.002 | 0.045 | 0.35 | 0.24 | 0.28 | 0.34 | 0.46 | 0.25 |
| 2 | 134589967 | rs1348837 | 0.137 | 1.268 | -0.103 | 1.023 | 0.028 | 0.006 | 0.006 | 0 | 0.53 | 0.67 | 0.66 | 0.48 | 0.62 | 0.43 |
| 2 | 134591340 | rs6739279 | -1.039 | 0.968 | 0.149 | 1.057 | 0.004 | -0.014 | -0.014 | 0 | 0.23 | 0.6 | 0.47 | 0.24 | 0.23 | 0.21 |

| | | | | | | | | | | | | | | | | |
|---|---|---|---|---|---|---|---|---|---|---|---|---|---|---|---|---|
| 2 | 134591987 | rs12618948 | -0.42 | 0.966 | 0.085 | 1.097 | 0.006 | -0.001 | -0.001 | 0 | 0.17 | 0.36 | 0.22 | 0.2 | 0.16 | 0.14 |
| 2 | 134594110 | rs11691129 | -0.525 | 0.937 | 0.001 | 1.051 | 0.016 | 0.002 | 0.002 | 0.231 | 0.18 | 0.36 | 0.42 | 0.24 | 0.2 | 0.17 |
| 2 | 134595521 | rs13020966 | 0.108 | 0.789 | 0.094 | 0.897 | -0.009 | -0.021 | -0.021 | 0 | 0.14 | 0.32 | 0.15 | 0.12 | 0.11 | 0.12 |
| 2 | 134600686 | rs7569670 | -0.775 | 0.924 | 0.009 | 1.034 | -0.004 | 0.007 | 0.007 | 0 | 0.35 | 0.56 | 0.54 | 0.41 | 0.32 | 0.42 |
| 2 | 134603157 | rs1562345 | 0.554 | 1.479 | -0.002 | 1.177 | -0.109 | -0.004 | -0.004 | 0 | 0.36 | 0.12 | 0.29 | 0.29 | 0.49 | 0.25 |
| 2 | 134606584 | rs10928462 | -0.682 | 0.238 | 0.364 | 1.371 | 0 | 0.029 | 0.029 | 0.802 | 0.14 | 0.46 | 0.23 | 0.16 | 0.13 | 0.15 |
| 2 | 134607680 | rs1900346 | -0.733 | 0.248 | -0.304 | 1.43 | 0.022 | 0.062 | 0.062 | 0 | 0.51 | 0.85 | 0.52 | 0.6 | 0.47 | 0.39 |
| 2 | 134608321 | rs7579501 | -0.576 | 0.293 | -0.406 | 0.926 | -0.005 | -0.001 | -0.001 | 1.112 | 0.45 | 0.5 | 0.42 | 0.42 | 0.43 | 0.34 |
| 2 | 134617840 | rs16829656 | -0.977 | 0.546 | 0.102 | 0.983 | 0.02 | 0.028 | 0.028 | 0 | 0.13 | 0.35 | 0.23 | 0.16 | 0.15 | 0.16 |
| 2 | 134624276 | rs4603824 | -0.095 | 0.222 | -1.176 | 0.891 | 0.03 | -0.014 | -0.014 | 0 | 0.57 | 0.25 | 0.51 | 0.61 | 0.63 | 0.7 |
| 2 | 134626686 | rs10208557 | -0.537 | -0.359 | -0.692 | 0.478 | 0.001 | -0.031 | -0.031 | 0.036 | 0.17 | 0.28 | 0.21 | 0.07 | 0.18 | 0.14 |
| 2 | 134632112 | rs2084229 | 0.69 | -1.156 | -0.306 | 0.162 | -0.001 | -0.007 | -0.007 | 0 | 0.32 | 0.25 | 0.19 | 0.12 | 0.24 | 0.23 |
| 2 | 134632157 | rs13431659 | 0.048 | -1.184 | -0.062 | 0.157 | -0.003 | 0.009 | 0.009 | 0 | 0.16 | 0.11 | 0.19 | 0.03 | 0.19 | 0.13 |
| 2 | 134633660 | rs2321299 | 0.73 | -0.924 | -0.396 | 0.26 | -0.028 | 0 | 0 | 0 | 0.17 | 0.12 | 0.16 | 0.19 | 0.35 | 0.21 |
| 2 | 134635194 | rs10185444 | -0.865 | -0.971 | -0.16 | 0.448 | 0.01 | -0.01 | -0.01 | 0.093 | 0.36 | 0.52 | 0.54 | 0.35 | 0.55 | 0.38 |
| 2 | 134637112 | rs10928465 | -0.093 | -0.95 | -0.182 | 0.444 | 0.019 | 0.033 | 0.033 | 0.11 | 0.49 | 0.51 | 0.39 | 0.15 | 0.43 | 0.36 |
| 2 | 134638499 | rs1530572 | -1.08 | -0.92 | -0.405 | 0.566 | 0.035 | -0.041 | -0.041 | 0.702 | 0.68 | 0.76 | 0.71 | 0.47 | 0.78 | 0.6 |
| 2 | 134639305 | rs12691850 | -0.167 | -0.963 | -0.298 | 0.384 | 0.007 | -0.052 | -0.052 | 0.339 | 0.14 | 0.25 | 0.19 | 0.03 | 0.15 | 0.13 |
| 2 | 134641137 | rs12618404 | 0.163 | -1.236 | -0.874 | 0.584 | 0.082 | 0.003 | 0.003 | 0 | 0.41 | 0.08 | 0.33 | 0.37 | 0.33 | 0.49 |
| 2 | 134642665 | rs12476265 | -0.504 | -0.911 | -0.508 | 0.617 | -0.002 | -0.008 | -0.008 | 0 | 0.56 | 0.49 | 0.51 | 0.38 | 0.48 | 0.62 |
| 2 | 134644096 | rs6711228 | 0.594 | -0.782 | -1.095 | 0.401 | 0.023 | -0.037 | -0.037 | 0 | 0.21 | 0.06 | 0.18 | 0.17 | 0.12 | 0.15 |
| 2 | 134657740 | rs1446732 | 1.054 | -0.571 | -0.469 | -0.02 | -0.004 | -0.004 | -0.004 | 0.257 | 0.39 | 0.37 | 0.39 | 0.4 | 0.44 | 0.35 |
| 2 | 134659485 | rs6714405 | 0.079 | -0.874 | -0.258 | -0.289 | -0.014 | -0.048 | -0.048 | 0.252 | 0.28 | 0.28 | 0.11 | 0.06 | 0.15 | 0.21 |
| 2 | 134666625 | rs1446738 | 0.191 | -0.077 | 0.489 | 0.591 | -0.014 | -0.028 | -0.028 | 0 | 0.84 | 0.86 | 0.66 | 0.55 | 0.77 | 0.68 |
| 2 | 134676982 | rs17317784 | 1.14 | -0.071 | 0.413 | 0.572 | -0.022 | -0.01 | -0.01 | 0.023 | 0.06 | 0 | 0.05 | 0.05 | 0.08 | 0.04 |
| 2 | 134678907 | rs10803543 | 1.197 | 0.105 | 0.55 | 0.601 | -0.014 | -0.027 | -0.027 | 0 | 0.66 | 0.72 | 0.5 | 0.45 | 0.58 | 0.52 |
| 2 | 134685937 | rs13399984 | 0.893 | 0.033 | 0.403 | 0.591 | 0.009 | 0.05 | 0.05 | 1.071 | 0.78 | 0.73 | 0.62 | 0.52 | 0.73 | 0.6 |
| 2 | 134686681 | rs17318092 | -0.05 | 0.07 | 0.247 | 0.594 | -0.005 | 0.04 | 0.04 | 0 | 0.18 | 0.07 | 0.15 | 0.09 | 0.19 | 0.16 |
| 2 | 134687715 | rs13382243 | 0.022 | 0.071 | 0.247 | 0.595 | -0.006 | 0.038 | 0.038 | 0 | 0.17 | 0.06 | 0.15 | 0.09 | 0.19 | 0.16 |
| 2 | 134698086 | rs11884370 | 1.411 | 0.089 | 0.328 | 0.634 | -0.006 | -0.006 | -0.006 | 0 | 0.6 | 0.67 | 0.49 | 0.4 | 0.53 | 0.46 |
| 2 | 134703352 | rs12691854 | 0.064 | 0.152 | -0.016 | 0.723 | -0.068 | -0.112 | -0.112 | 0 | 0.85 | 0.99 | 0.7 | 0.58 | 0.79 | 0.75 |
| 2 | 134713030 | rs11897842 | 1.016 | 0.128 | -0.013 | 0.719 | -0.073 | -0.116 | -0.116 | 0.794 | 0.67 | 0.93 | 0.56 | 0.47 | 0.58 | 0.53 |
| 2 | 134713735 | rs2087006 | 0.94 | 0.026 | 0.192 | 0.712 | 0.027 | 0.125 | 0.125 | 0.686 | 0.78 | 0.53 | 0.62 | 0.51 | 0.74 | 0.71 |
| 2 | 134716160 | rs6748729 | 1.527 | 0.047 | 0.24 | 0.676 | 0.006 | 0.006 | 0.006 | 0.121 | 0.6 | 0.49 | 0.46 | 0.4 | 0.52 | 0.5 |
| 2 | 134716952 | rs6736758 | -0.098 | 0.085 | 0.176 | 0.638 | -0.065 | -0.106 | -0.106 | 1.422 | 0.86 | 0.99 | 0.68 | 0.59 | 0.79 | 0.76 |
| 2 | 134717120 | rs6752844 | 1.439 | 0.078 | 0.385 | 0.546 | -0.003 | -0.005 | -0.005 | 0 | 0.58 | 0.35 | 0.45 | 0.41 | 0.52 | 0.5 |
| 2 | 134717465 | rs10207061 | -0.177 | 0.287 | -0.047 | 0.624 | -0.006 | 0.03 | 0.03 | 0 | 0.19 | 0.06 | 0.14 | 0.12 | 0.22 | 0.23 |
| 2 | 134719933 | rs6430476 | 0.142 | 0.306 | -0.145 | 0.613 | 0.023 | 0.119 | 0.119 | 0.153 | 0.8 | 0.57 | 0.6 | 0.5 | 0.75 | 0.72 |
| 2 | 134721888 | rs4429522 | -0.2 | 0.414 | -0.976 | 0.639 | -0.011 | 0.039 | 0.039 | 0 | 0.19 | 0.06 | 0.14 | 0.09 | 0.22 | 0.22 |
| 2 | 134724568 | rs2101331 | 0.656 | 0.368 | -0.341 | 1.133 | 0.015 | 0.092 | 0.092 | 0.162 | 0.79 | 0.62 | 0.62 | 0.5 | 0.74 | 0.72 |
| 2 | 134727858 | rs1542656 | -0.009 | -0.561 | -0.776 | 0.342 | 0.019 | 0.019 | 0.019 | 0.35 | 0.44 | 0.53 | 0.39 | 0.24 | 0.27 | 0.29 |
| 2 | 134731059 | rs12477481 | 0.144 | -0.689 | -1.282 | -0.142 | 0.023 | 0.177 | 0.177 | 0.032 | 0.42 | 0.11 | 0.35 | 0.18 | 0.39 | 0.28 |
| 2 | 134734490 | rs2321537 | -0.056 | -1.083 | -0.766 | 0.976 | 0.038 | 0.031 | 0.031 | 0 | 0.51 | 0.34 | 0.48 | 0.57 | 0.43 | 0.41 |
| 2 | 134736340 | rs16829877 | -0.001 | -1.126 | -0.667 | 1.208 | -0.014 | 0.154 | 0.154 | 0 | 0.07 | 0.14 | 0.14 | 0.33 | 0.04 | 0.11 |
| 2 | 134737301 | rs12691856 | 0.252 | -1.149 | -0.226 | 1.591 | 0.039 | 0.081 | 0.081 | 0 | 0.46 | 0.23 | 0.42 | 0.3 | 0.42 | 0.34 |
| 2 | 134738411 | rs4411763 | 0.392 | -1.128 | -0.265 | 1.571 | 0.048 | 0.132 | 0.132 | 0.111 | 0.45 | 0.2 | 0.35 | 0.23 | 0.39 | 0.3 |
| 2 | 134741635 | rs7606427 | 0.687 | -0.595 | -0.273 | 1.551 | -0.013 | 0.002 | 0.002 | 0 | 0.1 | 0.07 | 0.14 | 0.09 | 0.14 | 0.21 |
| 2 | 134745702 | rs2321550 | -0.998 | -0.839 | 0.515 | 1.459 | -0.032 | 0.214 | 0.214 | 0.139 | 0.13 | 0.41 | 0.17 | 0.32 | 0.07 | 0.14 |
| 2 | 134758642 | rs12612011 | -1.041 | -0.847 | 0.524 | 1.448 | -0.03 | 0.207 | 0.207 | 0.126 | 0.13 | 0.41 | 0.17 | 0.33 | 0.07 | 0.15 |
| 2 | 134765869 | rs11683312 | 0.769 | -0.345 | -0.084 | 0.754 | 0 | 0.001 | 0.001 | 0.234 | 0.48 | 0.47 | 0.54 | 0.5 | 0.56 | 0.45 |
| 2 | 134766833 | rs6717488 | -0.046 | -0.512 | -0.009 | 0.742 | -0.008 | -0.001 | -0.001 | 0 | 0.7 | 0.62 | 0.62 | 0.36 | 0.73 | 0.62 |
| 2 | 134768717 | rs11891791 | -1.014 | -0.271 | -0.07 | 0.664 | -0.034 | 0.006 | 0.006 | 0.097 | 0.31 | 0.58 | 0.4 | 0.64 | 0.21 | 0.36 |
| 2 | 134772837 | rs11685957 | 0.833 | -0.278 | 0.027 | 1.153 | -0.001 | -0.004 | -0.004 | 0 | 0.12 | 0.12 | 0.19 | 0.08 | 0.11 | 0.24 |
| 2 | 134775529 | rs1160118 | -1.34 | -0.301 | -0.059 | 1.317 | -0.015 | 0.025 | 0.025 | 0 | 0.81 | 0.93 | 0.87 | 0.89 | 0.81 | 0.84 |
| 2 | 134775918 | rs7594310 | 0.27 | -0.303 | -0.013 | 1.295 | -0.016 | -0.011 | -0.011 | 0 | 0.2 | 0.28 | 0.14 | 0.19 | 0.14 | 0.15 |
| 2 | 134778035 | rs6740690 | 1.862 | -0.298 | -0.145 | 1.249 | -0.167 | 0.163 | 0.163 | 0.649 | 0.36 | 0.05 | 0.23 | 0.13 | 0.49 | 0.23 |
| 2 | 134788709 | rs6719360 | 0.639 | -0.352 | 0.024 | 1.282 | -0.001 | -0.013 | -0.013 | 0 | 0.12 | 0.16 | 0.19 | 0.08 | 0.11 | 0.23 |
| 2 | 134790276 | rs1038511 | -0.633 | -0.408 | 0.144 | 1.424 | -0.025 | 0.348 | 0.348 | 0 | 0.11 | 0.39 | 0.27 | 0.44 | 0.07 | 0.21 |
| 2 | 134791174 | rs6736502 | 1.572 | -0.388 | -0.04 | 1.076 | -0.037 | 0.062 | 0.062 | 0.349 | 0.38 | 0.25 | 0.25 | 0.05 | 0.49 | 0.23 |
| 2 | 134792531 | rs6430487 | -0.866 | -0.431 | -0.112 | 1.143 | 0.002 | -0.002 | -0.002 | 0 | 0.4 | 0.58 | 0.6 | 0.76 | 0.32 | 0.6 |
| 2 | 134793274 | rs1466569 | 0.213 | -0.263 | 0.006 | 1.297 | -0.002 | -0.022 | -0.022 | 0 | 0.76 | 0.57 | 0.78 | 0.76 | 0.81 | 0.82 |
| 2 | 134806823 | rs4953899 | 1.339 | -0.379 | -0.169 | 1.487 | 0.014 | -0.034 | -0.034 | 0 | 0.16 | 0.04 | 0.1 | 0.21 | 0.14 | 0.14 |
| 2 | 134813786 | rs1519305 | 1.082 | -0.428 | 0.09 | 1.736 | 0.016 | 0.037 | 0.037 | 0 | 0.56 | 0.38 | 0.51 | 0.63 | 0.64 | 0.58 |
| 2 | 134817016 | rs1715008 | 1.025 | 0.038 | 0.255 | 1.942 | 0.011 | 0.052 | 0.052 | 0 | 0.71 | 0.62 | 0.58 | 0.56 | 0.69 | 0.56 |
| 2 | 134830363 | rs11684241 | -0.069 | -0.356 | -0.248 | 1.594 | 0.008 | 0.034 | 0.034 | 0 | 0.19 | 0.08 | 0.22 | 0.08 | 0.18 | 0.22 |
| 2 | 134832158 | rs10175819 | -0.908 | -0.381 | -0.056 | 1.489 | -0.033 | 0.068 | 0.068 | 0 | 0.3 | 0.53 | 0.4 | 0.46 | 0.23 | 0.45 |
| 2 | 134837980 | rs13410504 | -1.549 | -0.201 | -0.002 | 1.172 | -0.003 | -0.033 | -0.033 | 0 | 0.1 | 0.2 | 0.09 | 0.02 | 0.09 | 0.06 |
| 2 | 134840155 | rs4954106 | -1.389 | -0.197 | -0.03 | 1.174 | 0 | 0 | 0 | 0 | 0.09 | 0.13 | 0.07 | 0.02 | 0.09 | 0.06 |
| 2 | 134840967 | rs6430491 | -0.739 | -0.546 | 0.263 | 1.571 | -0.016 | 0.045 | 0.045 | 0 | 0.29 | 0.51 | 0.32 | 0.39 | 0.2 | 0.18 |

| | | | | | | | | | | | | | | | | |
|---|---|---|---|---|---|---|---|---|---|---|---|---|---|---|---|---|
| 2 | 134845085 | rs764686 | -1.156 | -0.679 | 0.392 | 1.834 | -0.006 | -0.005 | -0.005 | 0.001 | 0.37 | 0.56 | 0.41 | 0.63 | 0.28 | 0.32 |
| 2 | 134848764 | rs13382580 | -1.897 | -0.669 | 0.277 | 1.798 | 0.028 | 0.075 | 0.075 | 0.958 | 0.45 | 0.85 | 0.49 | 0.65 | 0.35 | 0.35 |
| 2 | 134853516 | rs9287434 | -1.214 | -0.714 | 0.912 | 1.921 | -0.007 | 0.059 | 0.059 | 0 | 0.15 | 0.32 | 0.14 | 0.26 | 0.14 | 0.17 |
| 2 | 134855665 | rs7598485 | -0.859 | -0.731 | 0.89 | 2.242 | -0.011 | -0.047 | -0.047 | 0.914 | 0.32 | 0.54 | 0.17 | 0.17 | 0.24 | 0.19 |
| 2 | 134856414 | rs16830053 | 0.384 | -0.728 | 0.733 | 2.264 | 0 | 0.022 | 0.022 | 0 | 0.08 | 0.07 | 0.08 | 0.24 | 0.08 | 0.13 |
| 2 | 134856737 | rs6754184 | -1.804 | -0.738 | 0.729 | 2.255 | 0.024 | -0.03 | -0.03 | 0 | 0.41 | 0.85 | 0.27 | 0.41 | 0.32 | 0.31 |
| 2 | 134858756 | rs6718163 | -1.265 | -0.773 | 0.897 | 2.288 | -0.011 | -0.025 | -0.025 | 0 | 0.3 | 0.66 | 0.16 | 0.17 | 0.2 | 0.15 |
| 2 | 134859017 | rs16830058 | -0.009 | -0.756 | 0.887 | 2.256 | -0.011 | -0.013 | -0.013 | 0 | 0.22 | 0.33 | 0.09 | 0.15 | 0.14 | 0.11 |
| 2 | 134861320 | rs12466056 | -1.812 | -0.723 | 0.703 | 1.994 | 0.028 | 0.032 | 0.032 | 2.395 | 0.38 | 0.85 | 0.49 | 0.66 | 0.29 | 0.3 |
| 2 | 134862678 | rs16830064 | -2.476 | -0.853 | 0.827 | 1.773 | -0.008 | -0.042 | -0.042 | 0 | 0.1 | 0.36 | 0.07 | 0.03 | 0.07 | 0.05 |
| 2 | 134864344 | rs10179986 | -1.911 | -0.952 | 0.496 | 1.859 | 0.06 | 0.075 | 0.075 | 4.107 | 0.44 | 0.85 | 0.41 | 0.69 | 0.3 | 0.31 |
| 2 | 134865196 | rs1996589 | -1.922 | -0.928 | 0.436 | 1.542 | 0.022 | -0.006 | -0.006 | 0.373 | 0.45 | 0.68 | 0.25 | 0.44 | 0.31 | 0.3 |
| 2 | 134872037 | rs1879019 | -0.573 | -0.889 | 0.332 | 1.477 | -0.008 | -0.042 | -0.042 | 0.262 | 0.14 | 0.25 | 0.08 | 0.03 | 0.12 | 0.06 |
| 2 | 134876049 | rs4953902 | -0.728 | -0.602 | 0.198 | 1.785 | -0.014 | 0.03 | 0.03 | 1.574 | 0.29 | 0.35 | 0.15 | 0.39 | 0.19 | 0.24 |
| 2 | 134880830 | rs1111961 | -1.147 | -0.591 | 0.199 | 1.804 | -0.008 | 0.011 | 0.011 | 1.546 | 0.33 | 0.37 | 0.17 | 0.4 | 0.23 | 0.28 |
| 2 | 134889971 | rs4600717 | -1.505 | -0.834 | -0.591 | 1.471 | 0.067 | 0.017 | 0.017 | 1.282 | 0.43 | 0.85 | 0.4 | 0.59 | 0.29 | 0.35 |
| 2 | 134891588 | rs10200776 | -0.416 | -0.887 | 0.078 | 1.776 | -0.014 | 0.073 | 0.073 | 0 | 0.17 | 0.34 | 0.13 | 0.22 | 0.14 | 0.23 |
| 2 | 134893447 | rs7601756 | -1.078 | -0.843 | 0.012 | 1.817 | -0.013 | 0.03 | 0.03 | 0 | 0.31 | 0.51 | 0.36 | 0.56 | 0.26 | 0.27 |
| 2 | 134895113 | rs12464954 | 0.688 | -0.832 | -0.164 | 1.625 | -0.006 | -0.039 | -0.039 | 0 | 0.12 | 0.05 | 0.12 | 0.19 | 0.14 | 0.18 |
| 2 | 134899485 | rs11890697 | -0.087 | -0.795 | 0.094 | 1.395 | -0.01 | -0.052 | -0.052 | 0 | 0.19 | 0.49 | 0.13 | 0.1 | 0.12 | 0.08 |
| 2 | 134901659 | rs11886886 | -1.473 | -0.535 | -0.528 | 1.492 | 0.017 | -0.048 | -0.048 | 0.157 | 0.33 | 0.85 | 0.4 | 0.53 | 0.31 | 0.34 |
| 2 | 134912114 | rs2139311 | 0.359 | -0.415 | -0.674 | 0.918 | 0.001 | 0.018 | 0.018 | 1.846 | 0.39 | 0.7 | 0.18 | 0.18 | 0.36 | 0.23 |
| 2 | 134913756 | rs11900787 | -0.487 | -0.455 | -0.804 | 1.093 | -0.027 | 0.035 | 0.035 | 0.449 | 0.57 | 0.88 | 0.46 | 0.65 | 0.58 | 0.48 |
| 2 | 134913954 | rs11897287 | -1.595 | -0.53 | -1.009 | 0.905 | -0.03 | 0.034 | 0.034 | 0 | 0.74 | 0.95 | 0.67 | 0.82 | 0.71 | 0.65 |
| 2 | 134917005 | rs11692586 | 0.896 | -0.347 | -0.266 | 0.869 | -0.009 | 0.107 | 0.107 | 0.882 | 0.34 | 0.17 | 0.11 | 0.13 | 0.32 | 0.25 |
| 2 | 134920164 | rs10928484 | 0.207 | 0.065 | 0.787 | 1.76 | -0.006 | -0.025 | -0.025 | 0.177 | 0.16 | 0.1 | 0.15 | 0.2 | 0.22 | 0.25 |
| 2 | 134922529 | rs10167538 | -2.302 | 0.032 | 0.784 | 1.742 | -0.01 | -0.014 | -0.014 | 0 | 0.1 | 0.68 | 0.08 | 0.07 | 0.04 | 0.07 |
| 2 | 134923031 | rs12473666 | 0.405 | 0.051 | 0.633 | 1.761 | -0.013 | -0.031 | -0.031 | 0 | 0.15 | 0.08 | 0.14 | 0.19 | 0.22 | 0.25 |
| 2 | 134929915 | rs16830149 | -0.688 | 0.054 | 0.985 | 1.696 | -0.014 | 0.007 | 0.007 | 0 | 0.22 | 0.4 | 0.18 | 0.2 | 0.22 | 0.3 |
| 2 | 134932432 | rs10496720 | -0.338 | 0.183 | 0.662 | 1.703 | 0.001 | 0.001 | 0.001 | 0.234 | 0.2 | 0.26 | 0.17 | 0.2 | 0.25 | 0.31 |
| 2 | 134932707 | rs2139307 | 0.333 | 0.205 | 0.187 | 1.738 | -0.021 | -0.054 | -0.054 | 0 | 0.16 | 0.05 | 0.14 | 0.19 | 0.21 | 0.24 |
| 2 | 134934400 | rs4954112 | -2.375 | -0.036 | 1 | 1.869 | -0.007 | 0.071 | 0.071 | 2.243 | 0.12 | 0.71 | 0.32 | 0.24 | 0.07 | 0.11 |
| 2 | 134935746 | rs10928485 | -2.061 | -0.018 | 0.76 | 1.773 | -0.009 | 0.091 | 0.091 | 5.195 | 0.17 | 0.66 | 0.36 | 0.37 | 0.14 | 0.11 |
| 2 | 134935979 | rs9808181 | -1.391 | 0.092 | 0.274 | 1.421 | 0.005 | 0.011 | 0.011 | 0.151 | 0.11 | 0.27 | 0.26 | 0.14 | 0.11 | 0.09 |
| 2 | 134945535 | rs1257174 | 0.362 | -0.087 | -0.402 | 1.268 | 0.018 | 0.015 | 0.015 | 1.155 | 0.5 | 0.43 | 0.36 | 0.43 | 0.37 | 0.46 |
| 2 | 134945868 | rs4954117 | -0.971 | 0.037 | -0.262 | 1.678 | 0.01 | 0.004 | 0.004 | 0.725 | 0.11 | 0.21 | 0.27 | 0.13 | 0.11 | 0.12 |
| 2 | 134955210 | rs4953906 | -1.042 | 0.016 | -0.276 | 1.981 | 0.007 | 0.076 | 0.076 | 0.861 | 0.11 | 0.21 | 0.34 | 0.37 | 0.11 | 0.15 |
| 2 | 134961089 | rs13017917 | 0.173 | -0.284 | -1.095 | 1.69 | 0.042 | -0.04 | -0.04 | 0 | 0.21 | 0.05 | 0.15 | 0.25 | 0.11 | 0.21 |
| 2 | 134962379 | rs9646694 | 0.158 | -0.25 | -1.088 | 1.691 | 0.039 | -0.056 | -0.056 | 0.171 | 0.21 | 0.05 | 0.15 | 0.3 | 0.11 | 0.22 |
| 2 | 134965464 | rs3748896 | -0.675 | -0.354 | -1.088 | 0.81 | 0.07 | 0.093 | 0.093 | 0.158 | 0.44 | 0.84 | 0.44 | 0.75 | 0.29 | 0.38 |
| 2 | 134973418 | rs999717 | -0.482 | 0.08 | -0.704 | 0.724 | -0.015 | 0.032 | 0.032 | 0.796 | 0.63 | 0.72 | 0.62 | 0.81 | 0.57 | 0.74 |
| 2 | 134984070 | rs1257208 | -0.476 | 0.598 | 0.042 | 0.793 | 0 | -0.001 | -0.001 | 0.535 | 0.85 | 0.8 | 0.73 | 0.85 | 0.83 | 0.81 |
| 2 | 134985914 | rs1257210 | 0.873 | 0.432 | 0.726 | 1.215 | 0.012 | 0.074 | 0.074 | 0.542 | 0.72 | 0.47 | 0.6 | 0.51 | 0.71 | 0.65 |
| 2 | 134991024 | rs1257192 | -0.051 | 0.776 | 0.528 | 1.117 | 0 | -0.003 | -0.003 | 0 | 0.82 | 0.74 | 0.7 | 0.8 | 0.81 | 0.8 |
| 2 | 134992922 | rs1401568 | 0.868 | 0.39 | 0.375 | 0.79 | 0.012 | -0.062 | -0.062 | 0.042 | 0.3 | 0.34 | 0.16 | 0.08 | 0.3 | 0.14 |
| 2 | 134993328 | rs6752163 | -0.183 | 0.444 | 0.366 | 0.786 | -0.004 | 0.002 | 0.002 | 2.857 | 0.38 | 0.6 | 0.25 | 0.3 | 0.38 | 0.25 |
| 2 | 134993718 | rs1257193 | 0.021 | 0.061 | -0.296 | 0.512 | 0.002 | -0.003 | -0.003 | 0 | 0.68 | 0.62 | 0.61 | 0.69 | 0.66 | 0.71 |
| 2 | 134994825 | rs1313363 | -0.111 | -0.486 | -0.119 | 0.758 | 0.004 | 0.007 | 0.007 | 0.078 | 0.46 | 0.5 | 0.49 | 0.69 | 0.4 | 0.62 |
| 2 | 134997741 | rs1257184 | 1.011 | -0.739 | 0.3 | 1.565 | 0.033 | -0.026 | -0.026 | 0.104 | 0.65 | 0.17 | 0.57 | 0.37 | 0.73 | 0.63 |
| 2 | 135005832 | rs1257196 | 0.918 | -0.447 | 0.717 | 1.708 | -0.001 | 0.001 | 0.001 | 0.081 | 0.69 | 0.27 | 0.62 | 0.5 | 0.73 | 0.66 |
| 2 | 135007736 | rs1257197 | -0.04 | 0.078 | 0.077 | 1.132 | 0.002 | -0.007 | -0.007 | 0 | 0.82 | 0.89 | 0.7 | 0.78 | 0.76 | 0.76 |
| 2 | 135014116 | rs2460382 | 0.096 | 0.105 | -0.312 | 1.164 | -0.001 | 0 | 0 | 0 | 0.8 | 0.79 | 0.67 | 0.78 | 0.75 | 0.76 |
| 2 | 135017074 | rs16830277 | -0.929 | 0.122 | -0.121 | 1.36 | 0.006 | 0.003 | 0.003 | 0 | 0.06 | 0.07 | 0.04 | 0.05 | 0.11 | 0.03 |
| 2 | 135022414 | rs3791271 | -0.697 | -0.593 | 0.74 | 2.124 | -0.007 | 0.031 | 0.031 | 0.185 | 0.08 | 0.13 | 0.09 | 0.14 | 0.05 | 0.1 |
| 2 | 135022906 | rs6735741 | -1.203 | -0.593 | 0.731 | 2.113 | -0.002 | 0.242 | 0.242 | 0.523 | 0.09 | 0.32 | 0.23 | 0.34 | 0.06 | 0.2 |
| 2 | 135024766 | rs935471 | -1.358 | -1.164 | 0.642 | 1.656 | -0.056 | 0.056 | 0.056 | 0 | 0.25 | 0.43 | 0.27 | 0.34 | 0.12 | 0.29 |
| 2 | 135028858 | rs7607690 | -0.499 | -0.984 | 0.521 | 1.4 | -0.045 | 0.019 | 0.019 | 0.924 | 0.17 | 0.33 | 0.18 | 0.21 | 0.06 | 0.19 |
| 2 | 135038848 | rs11686546 | -0.501 | -1.325 | 1.854 | 1.929 | 0.024 | 0.054 | 0.054 | 0 | 0.21 | 0.08 | 0.08 | 0.07 | 0.13 | 0.27 |
| 2 | 135040234 | rs11903405 | -1.178 | -1.294 | 2.01 | 1.974 | 0.043 | 0.573 | 0.573 | 0.674 | 0.06 | 0.41 | 0.17 | 0.36 | 0.02 | 0.1 |
| 2 | 135052071 | rs4954126 | 0.682 | -1.381 | 1.836 | 1.737 | -0.015 | 0.056 | 0.056 | 0.438 | 0.95 | 0.83 | 0.88 | 0.87 | 0.99 | 0.97 |
| 2 | 135054058 | rs2166736 | -0.354 | -2.394 | 1.304 | 0.931 | 0.001 | 0.001 | 0.001 | 2.446 | 0.53 | 0.48 | 0.33 | 0.26 | 0.32 | 0.41 |
| 2 | 135056095 | rs6430504 | 0.145 | -2.44 | 1.43 | 0.851 | -0.023 | -0.06 | -0.06 | 1.346 | 0.23 | 0.3 | 0.1 | 0.06 | 0.13 | 0.26 |
| 2 | 135057424 | rs11887041 | -0.085 | -2.344 | 1.328 | 0.856 | 0.026 | 0.009 | 0.009 | 1.769 | 0.95 | 0.69 | 0.96 | 1 | 0.99 | 1 |
| 2 | 135058732 | rs4954130 | 1.874 | -0.909 | 0.87 | 1.231 | -0.022 | -0.038 | -0.038 | 0 | 0.07 | 0.01 | 0.15 | 0.18 | 0.26 | 0.09 |
| 2 | 135061348 | rs6430506 | 0.538 | -2.38 | 1.177 | 0.916 | 0.035 | -0.081 | -0.081 | 0 | 0.39 | 0.15 | 0.59 | 0.55 | 0.66 | 0.54 |
| 2 | 135062598 | rs1439112 | 0.394 | -2.388 | 1.498 | 0.94 | 0.026 | 0.021 | 0.021 | 4.133 | 0.42 | 0.33 | 0.69 | 0.69 | 0.67 | 0.58 |
| 2 | 135065943 | rs7573104 | 0.5 | -2.38 | 1.434 | 0.941 | 0.026 | -0.037 | -0.037 | 0 | 0.4 | 0.24 | 0.59 | 0.55 | 0.65 | 0.54 |
| 2 | 135067289 | rs749048 | 0.558 | -2.376 | 1.475 | 0.953 | 0.021 | -0.02 | -0.02 | 0.65 | 0.41 | 0.31 | 0.6 | 0.56 | 0.65 | 0.54 |
| 2 | 135068007 | rs755586 | 0.185 | -2.41 | 1.333 | 0.568 | -0.011 | 0.001 | 0.001 | 0 | 0.24 | 0.23 | 0.13 | 0.25 | 0.14 | 0.31 |

| | | | | | | | | | | | | | | | | |
|---|---|---|---|---|---|---|---|---|---|---|---|---|---|---|---|---|
| 2 | 135068656 | rs17715420 | 1.586 | -2.512 | 1.04 | 0.461 | 0.034 | 0.068 | 0.068 | 4.259 | 0.25 | 0.09 | 0.13 | 0.05 | 0.19 | 0.11 |
| 2 | 135070460 | rs17715461 | 0.975 | -2.648 | 1.072 | 0.463 | 0.062 | 0.085 | 0.085 | 8.497 | 0.29 | 0.14 | 0.14 | 0.05 | 0.18 | 0.1 |
| 2 | 135075621 | rs3791281 | 0.334 | -2.731 | 1.457 | 0.565 | -0.051 | 0.024 | 0.024 | 0 | 0.22 | 0.31 | 0.06 | 0.26 | 0.12 | 0.27 |
| 2 | 135080139 | rs935473 | -1.167 | -2.849 | 1.703 | 0.512 | -0.034 | -0.05 | -0.05 | 0.376 | 0.7 | 0.59 | 0.84 | 0.92 | 0.81 | 0.88 |
| 2 | 135080569 | rs17715961 | 1.645 | -2.837 | 1.727 | 0.504 | 0.04 | 0.059 | 0.059 | 3.537 | 0.25 | 0.11 | 0.12 | 0.05 | 0.18 | 0.1 |
| 2 | 135082444 | rs11674550 | 0.159 | -2.695 | 1.716 | 0.591 | 0.075 | 0.034 | 0.034 | 3.223 | 0.48 | 0.3 | 0.78 | 0.66 | 0.69 | 0.61 |
| 2 | 135096302 | rs935480 | 0.142 | -2.552 | 1.615 | 0.576 | 0.044 | 0.023 | 0.023 | 2.08 | 0.48 | 0.36 | 0.78 | 0.66 | 0.67 | 0.6 |
| 2 | 135099144 | rs2290047 | 0.821 | -2.473 | 1.171 | 0.964 | 0.006 | -0.003 | -0.003 | 0 | 0.07 | 0.03 | 0.22 | 0.22 | 0 | 0.06 |
| 2 | 135105378 | rs6707591 | 0.182 | -2.822 | 0.882 | 0.68 | 0.071 | 0.018 | 0.018 | 1.542 | 0.43 | 0.18 | 0.74 | 0.66 | 0.63 | 0.57 |
| 2 | 135106510 | rs2028296 | 0.215 | -2.773 | 0.843 | 0.669 | 0.059 | 0.008 | 0.008 | 1.312 | 0.43 | 0.17 | 0.73 | 0.66 | 0.62 | 0.57 |
| 2 | 135111093 | rs1439107 | 0.858 | -2.621 | 1.543 | 0.868 | -0.012 | -0.002 | -0.002 | 0 | 0.3 | 0.23 | 0.15 | 0.25 | 0.23 | 0.17 |
| 2 | 135111439 | rs3791289 | -0.41 | -2.617 | 1.449 | 0.847 | -0.019 | -0.008 | -0.008 | 0 | 0.65 | 0.49 | 0.81 | 0.76 | 0.75 | 0.83 |
| 2 | 135114889 | rs2290049 | 0.403 | -2.57 | 0.816 | 0.916 | 0.136 | -0.062 | -0.062 | 0 | 0.56 | 0.16 | 0.55 | 0.51 | 0.74 | 0.74 |
| 2 | 135128486 | rs4953915 | 0.535 | -2.214 | 1.247 | 0.48 | -0.033 | -0.007 | -0.007 | 0.585 | 0.31 | 0.47 | 0.17 | 0.25 | 0.23 | 0.16 |
| 2 | 135130826 | rs16830502 | 0.771 | -2.266 | 0.825 | 0.141 | -0.034 | -0.05 | -0.05 | 0.366 | 0.29 | 0.34 | 0.15 | 0.08 | 0.2 | 0.09 |
| 2 | 135142222 | rs6714520 | -0.257 | -2.267 | 0.128 | -0.141 | 0.027 | 0.021 | 0.021 | 0 | 0.61 | 0.71 | 0.86 | 0.91 | 0.78 | 0.74 |
| 2 | 135144454 | rs755503 | 0.115 | -1.894 | -0.351 | 0.404 | 0 | -0.015 | -0.015 | 0 | 0.33 | 0.33 | 0.63 | 0.58 | 0.58 | 0.59 |
| 2 | 135157953 | rs999925 | 0.993 | -1.761 | -0.32 | 0.36 | -0.032 | 0.017 | 0.017 | 4.844 | 0.27 | 0.28 | 0.21 | 0.31 | 0.17 | 0.15 |
| 2 | 135158128 | rs13012494 | 0.186 | -2.004 | -0.5 | -0.093 | 0.043 | 0.078 | 0.078 | 0.26 | 0.36 | 0.17 | 0.16 | 0.08 | 0.27 | 0.28 |
| 2 | 135158382 | rs2165064 | 0.518 | -1.984 | -0.414 | 0.141 | -0.008 | -0.001 | -0.001 | 1.105 | 0.41 | 0.33 | 0.27 | 0.32 | 0.28 | 0.29 |
| 2 | 135161934 | rs4954142 | 0.244 | -0.939 | -0.497 | 0.546 | 0.002 | 0.008 | 0.008 | 8.445 | 0.61 | 0.58 | 0.58 | 0.51 | 0.6 | 0.35 |
| 2 | 135162755 | rs3791295 | -0.233 | -1.107 | -0.495 | 0.701 | -0.003 | -0.005 | -0.005 | 0.094 | 0.71 | 0.75 | 0.63 | 0.59 | 0.65 | 0.53 |
| 2 | 135163759 | rs3791298 | 1.52 | -0.161 | -0.587 | 1.036 | 0.085 | 0.033 | 0.033 | 0 | 0.06 | 0.08 | 0.18 | 0.17 | 0.27 | 0.09 |
| 2 | 135167600 | rs2118844 | 0.35 | -0.663 | -0.524 | 0.821 | -0.039 | -0.073 | -0.073 | 0 | 0.27 | 0.09 | 0.32 | 0.38 | 0.32 | 0.47 |
| 2 | 135167653 | rs2118843 | 0.719 | -1.651 | 0.398 | 0.335 | -0.043 | -0.019 | -0.019 | 0 | 0.85 | 0.5 | 0.91 | 0.89 | 0.94 | 0.82 |
| 2 | 135171675 | rs3791306 | 1.54 | -2.723 | -0.222 | 0.325 | -0.01 | -0.001 | -0.001 | 0 | 0.26 | 0.22 | 0.13 | 0.29 | 0.08 | 0.2 |
| 2 | 135178756 | rs7588521 | -0.088 | -3.263 | 0.095 | 0.076 | 0.011 | 0.01 | 0.01 | 0.096 | 0.48 | 0.62 | 0.33 | 0.45 | 0.22 | 0.37 |
| 2 | 135180526 | rs2289466 | 1.251 | -3.2 | 0.193 | 0.06 | -0.005 | 0.002 | 0.002 | 0 | 0.06 | 0.07 | 0.04 | 0.13 | 0.04 | 0.07 |
| 2 | 135180550 | rs2289467 | 0.967 | -3.314 | 0.133 | 0.013 | 0 | -0.001 | -0.001 | 0 | 0.08 | 0.06 | 0.04 | 0.09 | 0.05 | 0.14 |
| 2 | 135181515 | rs16830596 | 0.116 | -3.356 | 0.199 | -0.008 | -0.02 | 0.016 | 0.016 | 0.42 | 0.1 | 0.24 | 0.12 | 0.15 | 0.06 | 0.17 |
| 2 | 135183045 | rs3791312 | -0.495 | -3.363 | 0.172 | -0.063 | -0.027 | -0.029 | -0.029 | 0 | 0.21 | 0.3 | 0.19 | 0.14 | 0.14 | 0.18 |
| 2 | 135185391 | rs16830611 | 0.282 | -3.451 | -0.055 | -0.043 | -0.076 | -0.028 | -0.028 | 0 | 0.4 | 0.55 | 0.29 | 0.33 | 0.18 | 0.24 |
| 2 | 135189585 | rs3791322 | -0.83 | -3.314 | -0.035 | 0.015 | -0.053 | 0.046 | 0.046 | 0.021 | 0.17 | 0.5 | 0.18 | 0.29 | 0.1 | 0.23 |
| 2 | 135195182 | rs1866867 | 0.521 | -3.171 | -0.601 | -0.063 | 0.031 | 0.063 | 0.063 | 0 | 0.14 | 0.02 | 0.13 | 0 | 0.1 | 0.11 |
| 2 | 135199564 | rs2289468 | 3.617 | -3.291 | -0.607 | 0.073 | 0.095 | -0.05 | -0.05 | 0 | 0.16 | 0 | 0.01 | 0.17 | 0.02 | 0.05 |
| 2 | 135202455 | rs535900 | 1.014 | -3.122 | -0.858 | 0.146 | -0.103 | 0.017 | 0.017 | 1.761 | 0.34 | 0.12 | 0.55 | 0.3 | 0.52 | 0.46 |
| 2 | 135203150 | rs3791324 | -0.299 | -2.784 | 0.121 | 0.639 | -0.043 | 0.061 | 0.061 | 0 | 0.14 | 0.3 | 0.11 | 0.25 | 0.07 | 0.21 |
| 2 | 135203162 | rs661899 | -0.392 | -2.741 | 0.359 | 0.568 | 0.133 | 0.102 | 0.102 | 0 | 0.82 | 0.61 | 0.81 | 0.78 | 0.71 | 0.9 |
| 2 | 135203537 | rs566964 | 1.132 | -2.84 | 0.271 | 0.497 | -0.063 | 0.007 | 0.007 | 4.403 | 0.34 | 0.61 | 0.17 | 0.46 | 0.1 | 0.28 |
| 2 | 135205735 | rs629794 | -0.112 | -2.775 | 0.116 | 0.406 | 0.083 | 0.076 | 0.076 | 0 | 0.79 | 0.66 | 0.81 | 0.74 | 0.69 | 0.85 |
| 2 | 135209154 | rs636975 | -0.126 | -2.492 | -0.592 | 0.232 | -0.02 | 0.079 | 0.079 | 3.704 | 0.45 | 0.23 | 0.64 | 0.31 | 0.56 | 0.57 |
| 2 | 135210390 | rs2290483 | 0.591 | -4.7 | -0.507 | 0.228 | 0.088 | 0.031 | 0.031 | 11.659 | 0.4 | 0.69 | 0.25 | 0.6 | 0.11 | 0.2 |
| 2 | 135212329 | rs539588 | -0.367 | -3.858 | -0.62 | -0.244 | -0.012 | 0.001 | 0.001 | 0.277 | 0.76 | 0.79 | 0.66 | 0.97 | 0.44 | 0.82 |
| 2 | 135212421 | rs10185078 | 0.134 | -4.061 | -0.363 | 0.09 | -0.067 | 0.09 | 0.09 | 0 | 0.15 | 0.3 | 0.09 | 0.25 | 0.06 | 0.09 |
| 2 | 135212784 | rs543944 | 0.033 | -4.175 | -0.137 | 0.05 | 0.004 | -0.091 | -0.091 | 0.063 | 0.69 | 0.56 | 0.53 | 0.88 | 0.4 | 0.66 |
| 2 | 135213256 | rs609164 | 0.998 | -4.024 | -0.036 | 0.165 | 0.082 | 0.017 | 0.017 | 3.588 | 0.49 | 0.26 | 0.4 | 0.58 | 0.33 | 0.54 |
| 2 | 135213762 | rs610624 | 0.846 | -3.859 | -0.557 | 0.137 | 0.002 | 0.001 | 0.001 | 1.458 | 0.53 | 0.55 | 0.46 | 0.67 | 0.34 | 0.59 |
| 2 | 135214259 | rs2289470 | -0.487 | -2.979 | -0.497 | -0.165 | -0.013 | -0.079 | -0.079 | 0 | 0.18 | 0.31 | 0.19 | 0.07 | 0.15 | 0.21 |
| 2 | 135215941 | rs547177 | -0.623 | -3.445 | -1.718 | 0.04 | 0.059 | 0.002 | 0.002 | 8.84 | 0.4 | 0.08 | 0.46 | 0.39 | 0.32 | 0.64 |
| 2 | 135218605 | rs6430516 | 1.146 | -4.437 | -1.286 | -0.117 | -0.063 | 0.011 | 0.011 | 10.228 | 0.34 | 0.59 | 0.18 | 0.56 | 0.11 | 0.26 |
| 2 | 135219050 | rs11904278 | -1.009 | -4.788 | -1.161 | -0.088 | 0.038 | -0.024 | -0.024 | 0 | 0.32 | 0.75 | 0.26 | 0.4 | 0.22 | 0.31 |
| 2 | 135219539 | rs583446 | -2.065 | -4.698 | -1.213 | -0.084 | -0.074 | 0.062 | 0.062 | 11.209 | 0.67 | 0.89 | 0.7 | 0.81 | 0.52 | 0.83 |
| 2 | 135221136 | rs16830673 | 0.937 | -4.882 | -1.202 | 0.154 | 0.048 | -0.017 | -0.017 | 0 | 0.16 | 0.07 | 0.17 | 0.17 | 0.07 | 0.19 |
| 2 | 135221749 | rs7565308 | -0.803 | -4.127 | -0.718 | 0.586 | 0.006 | 0.385 | 0.385 | 1.523 | 0.09 | 0.49 | 0.13 | 0.33 | 0.05 | 0.09 |
| 2 | 135223497 | rs2289471 | 2.698 | -4.122 | -0.898 | 0.376 | -0.003 | -0.001 | -0.001 | 0 | 0.21 | 0.21 | 0.08 | 0.22 | 0.05 | 0.1 |
| 2 | 135224353 | rs587201 | 1.438 | -3.866 | -1.364 | 0.15 | -0.06 | -0.063 | -0.063 | 0 | 0.15 | 0.01 | 0.25 | 0.24 | 0.22 | 0.33 |
| 2 | 135237274 | rs480368 | 0.261 | -3.677 | -1.254 | -0.089 | 0.096 | 0.126 | 0.126 | 0.324 | 0.49 | 0.72 | 0.44 | 0.82 | 0.33 | 0.59 |
| 2 | 135240324 | rs494905 | 1.035 | -3.107 | -1.708 | -0.158 | -0.008 | -0.01 | -0.01 | 0.09 | 0.17 | 0.08 | 0.16 | 0.16 | 0.2 | 0.3 |
| 2 | 135242989 | rs673260 | 1.182 | -3.527 | -1.912 | -0.107 | 0.074 | -0.038 | -0.038 | 0 | 0.34 | 0.11 | 0.17 | 0.33 | 0.21 | 0.32 |
| 2 | 135243291 | rs685113 | 0.297 | -3.733 | -1.892 | -0.334 | 0.059 | 0.024 | 0.024 | 0.647 | 0.48 | 0.29 | 0.27 | 0.4 | 0.37 | 0.52 |
| 2 | 135243753 | rs7580799 | 0.622 | -3.444 | -1.884 | -0.308 | -0.023 | 0.045 | 0.045 | 0.452 | 0.13 | 0.03 | 0.09 | 0.07 | 0.16 | 0.21 |
| 2 | 135244569 | rs12473509 | 0.575 | -3.441 | -1.882 | -0.306 | -0.023 | 0.045 | 0.045 | 0.475 | 0.14 | 0.03 | 0.09 | 0.07 | 0.16 | 0.21 |
| 2 | 135248544 | rs598668 | -0.111 | -3.45 | -1.871 | -0.296 | -0.021 | 0.031 | 0.031 | 5.513 | 0.31 | 0.15 | 0.25 | 0.23 | 0.36 | 0.5 |
| 2 | 135258363 | rs522227 | -0.69 | -3.484 | -1.837 | -0.42 | 0.123 | 0.147 | 0.147 | 4.659 | 0.52 | 0.83 | 0.51 | 0.74 | 0.34 | 0.62 |
| 2 | 135260903 | rs662438 | 0.927 | -3.557 | -1.637 | -0.342 | -0.005 | -0.001 | -0.001 | 0 | 0.24 | 0.29 | 0.23 | 0.25 | 0.23 | 0.4 |
| 2 | 135261737 | rs540053 | 1.123 | -3.549 | -1.71 | -0.338 | 0.007 | -0.012 | -0.012 | 0 | 0.23 | 0.1 | 0.22 | 0.25 | 0.23 | 0.4 |
| 2 | 135266560 | rs579076 | 0.753 | -2.711 | -1.229 | -0.35 | 0.015 | -0.005 | -0.005 | 0.317 | 0.55 | 0.4 | 0.34 | 0.48 | 0.4 | 0.65 |
| 2 | 135268006 | rs523905 | 0.1 | -2.624 | -1.676 | -0.241 | 0.006 | 0.065 | 0.065 | 0 | 0.68 | 0.79 | 0.52 | 0.9 | 0.36 | 0.64 |
| 2 | 135268255 | rs749460 | -0.551 | -1.925 | -1.05 | 0.24 | -0.046 | 0.044 | 0.044 | 1.061 | 0.26 | 0.64 | 0.29 | 0.49 | 0.12 | 0.21 |

| | | | | | | | | | | | | | | | | |
|---|---|---|---|---|---|---|---|---|---|---|---|---|---|---|---|---|
| 2 | 135278207 | rs624817 | -0.139 | -0.493 | -1.219 | -0.169 | -0.044 | 0.026 | 0.026 | 0 | 0.79 | 0.87 | 0.63 | 0.97 | 0.52 | 0.84 |
| 2 | 135280476 | rs12614292 | -0.594 | -0.957 | -0.703 | 0.614 | -0.005 | 0.035 | 0.035 | 0 | 0.13 | 0.21 | 0.16 | 0.22 | 0.1 | 0.12 |
| 2 | 135285678 | rs515983 | -0.883 | -0.798 | -0.708 | 0.368 | -0.012 | 0.015 | 0.015 | 0 | 0.35 | 0.51 | 0.43 | 0.56 | 0.34 | 0.38 |
| 2 | 135285736 | rs6730241 | 2.803 | -2.413 | -1.291 | 0.007 | 0.031 | -0.007 | -0.007 | 0 | 0.19 | 0.09 | 0.01 | 0.18 | 0.01 | 0.12 |
| 2 | 135286448 | rs4953921 | 0.206 | -1.703 | -1.45 | -0.496 | 0.016 | 0.019 | 0.019 | 0.348 | 0.31 | 0.23 | 0.36 | 0.25 | 0.3 | 0.38 |
| 2 | 135288913 | rs4954152 | -0.642 | -1.934 | -1.506 | -0.679 | 0.057 | 0.087 | 0.087 | 0.135 | 0.49 | 0.33 | 0.43 | 0.23 | 0.39 | 0.44 |
| 2 | 135292926 | rs542851 | 0.313 | -1.869 | -1.465 | -0.968 | 0 | -0.037 | -0.037 | 0 | 0.17 | 0.15 | 0.19 | 0.04 | 0.21 | 0.24 |
| 2 | 135298713 | rs613828 | -1.81 | -2.315 | -1.138 | -0.678 | 0.077 | -0.037 | -0.037 | 0 | 0.73 | 0.6 | 0.79 | 0.83 | 0.64 | 0.8 |
| 2 | 135300482 | rs2924943 | -0.466 | -2.223 | -1.889 | -0.659 | -0.026 | 0.101 | 0.101 | 0.253 | 0.27 | 0.01 | 0.24 | 0.11 | 0.28 | 0.21 |
| 2 | 135338640 | rs6730778 | 0.119 | -2.397 | -1.074 | -0.128 | -0.01 | 0.012 | 0.012 | 0 | 0.06 | 0.03 | 0.13 | 0 | 0.11 | 0.04 |
| 2 | 135358298 | rs16830847 | -0.426 | -2.486 | -0.592 | 0.13 | -0.002 | -0.008 | -0.008 | 0 | 0.12 | 0.1 | 0.09 | 0.06 | 0.1 | 0.17 |
| 2 | 135389197 | rs6759065 | -1.82 | -2.727 | 0.038 | 0.238 | 0.007 | -0.035 | -0.035 | 4.17 | 0.29 | 0.53 | 0.53 | 0.27 | 0.57 | 0.47 |
| 2 | 135397569 | rs6705916 | -0.627 | -2.64 | 0.095 | 0.264 | 0.113 | 0.012 | 0.012 | 4.077 | 0.17 | 0.37 | 0.44 | 0.21 | 0.48 | 0.3 |
| 2 | 135400321 | rs6745632 | -1.346 | -3.335 | -0.233 | 0.397 | 0.105 | -0.064 | -0.064 | 0.345 | 0.65 | 0.99 | 0.73 | 0.35 | 0.8 | 0.63 |
| 2 | 135402521 | rs4953922 | 1.054 | -3.141 | 0.039 | 0.155 | 0.083 | 0.155 | 0.155 | 0.144 | 0.32 | 0.09 | 0.16 | 0.03 | 0.23 | 0.13 |
| 2 | 135414882 | rs7422711 | -0.551 | -2.997 | 0.273 | 0.349 | -0.038 | -0.05 | -0.05 | 2.23 | 0.14 | 0.36 | 0.11 | 0.07 | 0.09 | 0.17 |
| 2 | 135419163 | rs7560738 | -0.392 | -2.875 | -0.177 | 0.915 | -0.021 | -0.058 | -0.058 | 0.004 | 0.91 | 0.99 | 0.83 | 0.74 | 0.9 | 0.76 |
| 2 | 135437405 | rs4440018 | -0.111 | -2.446 | 0.297 | 0.067 | 0.016 | -0.007 | -0.007 | 0 | 0.45 | 0.63 | 0.44 | 0.05 | 0.59 | 0.24 |
| 2 | 135466520 | rs16830921 | 0.225 | -2.515 | 0.491 | 0.242 | 0.002 | 0 | 0 | 0.688 | 0.11 | 0.09 | 0.08 | 0.1 | 0.1 | 0.18 |
| 2 | 135468385 | rs10164986 | -0.044 | -2.177 | 0.574 | 0.441 | 0.016 | 0.015 | 0.015 | 0.134 | 0.42 | 0.63 | 0.27 | 0.24 | 0.23 | 0.19 |
| 2 | 135469769 | rs6747870 | 0.71 | -2.227 | 0.792 | 0.418 | 0.006 | -0.023 | -0.023 | 0 | 0.34 | 0.27 | 0.15 | 0.03 | 0.2 | 0.13 |
| 2 | 135472847 | rs4953926 | 0.014 | -2.206 | 0.776 | 0.43 | 0.016 | 0.015 | 0.015 | 0.134 | 0.42 | 0.63 | 0.27 | 0.24 | 0.23 | 0.19 |
| 2 | 135483381 | rs3739034 | 0.555 | -3.031 | 0.881 | 0.49 | 0.029 | 0.007 | 0.007 | 0 | 0.36 | 0.26 | 0.21 | 0.29 | 0.21 | 0.18 |
| 2 | 135483413 | rs948583 | 0.245 | -3.028 | 0.878 | 0.492 | -0.035 | -0.011 | -0.011 | 0.094 | 0.41 | 0.5 | 0.25 | 0.29 | 0.23 | 0.21 |
| 2 | 135483960 | rs3886740 | 0.235 | -3.028 | 0.861 | 0.492 | -0.051 | -0.017 | -0.017 | 0 | 0.41 | 0.53 | 0.26 | 0.29 | 0.23 | 0.21 |
| 2 | 135493304 | rs10469555 | -1.469 | -2.293 | 1.027 | 0.676 | 0.038 | 0.005 | 0.005 | 1.806 | 0.06 | 0.36 | 0.06 | 0 | 0.02 | 0.02 |
| 2 | 135539967 | rs6430538 | -0.589 | -2.134 | 0.177 | 0.572 | -0.088 | 0.065 | 0.065 | 2.136 | 0.88 | 0.98 | 0.7 | 0.98 | 0.5 | 0.8 |
| 2 | 135545747 | rs6734966 | -0.182 | -2.552 | 0.833 | 0.621 | -0.027 | -0.046 | -0.046 | 0 | 0.2 | 0.3 | 0.15 | 0.1 | 0.18 | 0.24 |
| 2 | 135549730 | rs6430539 | -0.251 | -1.845 | -0.397 | 1.16 | 0.072 | -0.004 | -0.004 | 0 | 0.59 | 0.77 | 0.36 | 0.37 | 0.38 | 0.43 |
| 2 | 135574583 | rs1942050 | -0.425 | -1.824 | -0.755 | 1.25 | 0.026 | -0.095 | -0.095 | 0 | 0.62 | 0.87 | 0.47 | 0.49 | 0.41 | 0.48 |
| 2 | 135594699 | rs1954874 | -0.463 | -1.621 | -1.256 | 1.094 | 0.021 | -0.123 | -0.123 | 0 | 0.63 | 0.93 | 0.47 | 0.48 | 0.41 | 0.47 |
| 2 | 135599876 | rs10198552 | -1.087 | -1.006 | -0.071 | 1.194 | -0.005 | -0.035 | -0.035 | 0.899 | 0.07 | 0.14 | 0.04 | 0 | 0.04 | 0.04 |
| 2 | 135673387 | rs12151827 | -1.105 | -1.375 | 1.053 | 1.98 | -0.006 | 0.023 | 0.023 | 1.683 | 0.22 | 0.25 | 0.3 | 0.39 | 0.22 | 0.4 |
| 2 | 135677161 | rs3769025 | -0.444 | -1.253 | 0.912 | 2.205 | 0.006 | 0.006 | 0.006 | 1.223 | 0.14 | 0.04 | 0.21 | 0.11 | 0.17 | 0.17 |
| 2 | 135694848 | rs4953938 | -0.412 | -1.052 | 1.211 | 2.181 | 0.098 | -0.003 | -0.003 | 0.734 | 0.9 | 0.8 | 0.65 | 0.88 | 0.55 | 0.79 |
| 2 | 135711516 | rs3814355 | -0.357 | -1.059 | 1.209 | 2.177 | 0.108 | -0.004 | -0.004 | 0 | 0.9 | 0.77 | 0.64 | 0.88 | 0.55 | 0.79 |
| 2 | 135711657 | rs3814354 | 0.167 | -1.092 | 1.421 | 2.161 | 0.15 | -0.013 | -0.013 | 0 | 0.87 | 0.66 | 0.62 | 0.88 | 0.55 | 0.79 |
| 2 | 135713703 | rs3769023 | 0.167 | -1.092 | 1.363 | 2.184 | 0.131 | -0.011 | -0.011 | 0 | 0.87 | 0.68 | 0.62 | 0.88 | 0.55 | 0.79 |
| 2 | 135719056 | rs12463963 | -1.767 | -1.567 | 0.943 | 2.577 | -0.013 | 0.071 | 0.071 | 1.588 | 0.23 | 0.32 | 0.32 | 0.39 | 0.26 | 0.43 |
| 2 | 135721175 | rs16831203 | -0.98 | -1.433 | 0.292 | 2.556 | -0.003 | 0.133 | 0.133 | 0 | 0.06 | 0.1 | 0.07 | 0.29 | 0.05 | 0.23 |
| 2 | 135728958 | rs12621721 | -1.139 | -1.466 | -0.084 | 2.101 | -0.025 | 0.336 | 0.336 | 0.263 | 0.06 | 0.22 | 0.05 | 0.39 | 0.01 | 0.04 |
| 2 | 135745129 | rs16831235 | -0.974 | -1.352 | -0.71 | 2.03 | 0.008 | 0.025 | 0.025 | 0.016 | 0.18 | 0.04 | 0.23 | 0.12 | 0.2 | 0.13 |
| 2 | 135759095 | rs6706490 | -1.15 | -1.41 | -0.354 | 1.434 | -0.015 | 0 | 0 | 2.72 | 0.18 | 0.22 | 0.25 | 0.26 | 0.11 | 0.37 |
| 2 | 135762344 | rs16831243 | 0.091 | -3.251 | -0.477 | 1.329 | -0.102 | 0.027 | 0.027 | 0.217 | 0.33 | 0.49 | 0.23 | 0.38 | 0.18 | 0.34 |
| 2 | 135763318 | rs7582173 | -0.155 | -3.33 | -0.503 | 1.141 | -0.029 | 0.002 | 0.002 | 0.587 | 0.37 | 0.56 | 0.27 | 0.39 | 0.25 | 0.38 |
| 2 | 135818907 | rs2874739 | -1.366 | -3.48 | 0.661 | 0.902 | 0.005 | 0.002 | 0.002 | 3.75 | 0.27 | 0.72 | 0.33 | 0.31 | 0.2 | 0.42 |
| 2 | 135829309 | rs6736412 | -1.739 | -3.484 | 0.676 | 0.894 | 0.031 | 0.033 | 0.033 | 0 | 0.08 | 0.4 | 0.08 | 0.05 | 0.09 | 0.06 |
| 2 | 135839805 | rs6728452 | -0.213 | -3.601 | 0.651 | 0.852 | 0.021 | -0.018 | -0.018 | 1.142 | 0.14 | 0.05 | 0.21 | 0.24 | 0.1 | 0.26 |
| 2 | 135876308 | rs2034840 | -0.823 | -3.61 | 0.763 | 0.847 | 0.022 | -0.006 | -0.006 | 0 | 0.17 | 0.1 | 0.23 | 0.26 | 0.11 | 0.36 |
| 2 | 135887893 | rs2305595 | -0.9 | -3.66 | 0.989 | 0.819 | 0.022 | -0.005 | -0.005 | 0 | 0.17 | 0.11 | 0.23 | 0.26 | 0.11 | 0.36 |
| 2 | 135893372 | rs10445686 | 0.376 | -3.882 | 0.864 | 0.747 | 0.139 | -0.136 | -0.136 | 0 | 0.3 | 0.02 | 0.2 | 0.39 | 0.19 | 0.33 |
| 2 | 135914653 | rs4954223 | -1.131 | -3.704 | 0.936 | 0.339 | -0.044 | 0.006 | 0.006 | 3.027 | 0.18 | 0.3 | 0.24 | 0.26 | 0.11 | 0.36 |
| 2 | 135925002 | rs6705610 | -2.141 | -3.771 | 0.89 | 0.31 | 0.001 | 0.001 | 0.001 | 4.052 | 0.27 | 0.7 | 0.33 | 0.3 | 0.2 | 0.43 |
| 2 | 135965376 | rs7578111 | -1.49 | -3.865 | 0.963 | 0.204 | -0.043 | -0.008 | -0.008 | 0 | 0.2 | 0.35 | 0.27 | 0.22 | 0.14 | 0.37 |
| 2 | 135972093 | rs10928532 | -1.519 | -3.866 | 0.974 | 0.204 | -0.058 | -0.014 | -0.014 | 0 | 0.21 | 0.56 | 0.28 | 0.22 | 0.13 | 0.37 |
| 2 | 135988416 | rs935615 | -1.438 | -3.863 | 0.92 | 0.205 | -0.039 | -0.006 | -0.006 | 0 | 0.19 | 0.33 | 0.27 | 0.22 | 0.13 | 0.37 |
| 2 | 136002500 | rs1900741 | -1.265 | -4.472 | 0.628 | -0.04 | 0.147 | 0.147 | 0.147 | 3.11 | 0.48 | 0.72 | 0.51 | 0.84 | 0.3 | 0.57 |
| 2 | 136081193 | rs6758362 | -1.995 | -4.289 | 0.981 | 0.257 | -0.043 | -0.012 | -0.012 | 0 | 0.2 | 0.34 | 0.26 | 0.21 | 0.13 | 0.37 |
| 2 | 136123848 | rs4954246 | -2.189 | -4.28 | 0.953 | 0.261 | -0.04 | -0.015 | -0.015 | 1.22 | 0.22 | 0.66 | 0.28 | 0.21 | 0.13 | 0.37 |
| 2 | 136151774 | rs16831601 | -1.393 | -4.286 | 0.943 | 0.257 | 0.031 | 0.019 | 0.019 | 3.722 | 0.17 | 0.08 | 0.22 | 0.17 | 0.11 | 0.36 |
| 2 | 136169799 | rs6742013 | -2.687 | -4.32 | 0.926 | 0.238 | 0.008 | 0.008 | 0.008 | 4.726 | 0.27 | 0.72 | 0.32 | 0.22 | 0.2 | 0.42 |
| 2 | 136226626 | rs10167001 | -2.202 | -4.32 | 0.883 | 0.238 | -0.042 | -0.016 | -0.016 | 0 | 0.22 | 0.66 | 0.28 | 0.21 | 0.13 | 0.37 |
| 2 | 136264631 | rs16831751 | -0.735 | -4.315 | 0.545 | 0.24 | 0.023 | 0.01 | 0.01 | 1.346 | 0.14 | 0.05 | 0.21 | 0.15 | 0.1 | 0.26 |
| 2 | 136288273 | rs3806502 | -2.523 | -4.291 | 0.548 | 0.254 | 0.006 | 0.006 | 0.006 | 4.726 | 0.27 | 0.72 | 0.32 | 0.22 | 0.2 | 0.42 |
| 2 | 136324225 | rs4954265 | -2.335 | -4.081 | 0.594 | 0.361 | -0.052 | -0.011 | -0.011 | 0 | 0.2 | 0.66 | 0.26 | 0.21 | 0.11 | 0.37 |
| 2 | 136360455 | rs13397889 | -2.245 | -4.079 | 0.385 | 0.362 | -0.052 | -0.013 | -0.013 | 0 | 0.19 | 0.66 | 0.26 | 0.19 | 0.11 | 0.33 |
| 2 | 136385683 | rs6761268 | -1.237 | -4.057 | -0.083 | 0.452 | 0.027 | 0.008 | 0.008 | 3.918 | 0.14 | 0.08 | 0.21 | 0.17 | 0.08 | 0.36 |
| 2 | 136388473 | rs10187054 | -1.467 | -4.874 | -0.52 | 0.533 | 0.2 | 0.087 | 0.087 | 5.336 | 0.46 | 0.81 | 0.38 | 0.37 | 0.24 | 0.57 |
| 2 | 136393157 | rs10928541 | 1.294 | -5.614 | -0.024 | 0.18 | -0.039 | -0.03 | -0.03 | 0 | 0.38 | 0.33 | 0.24 | 0.16 | 0.13 | 0.36 |

| | | | | | | | | | | | | | | | | |
|---|---|---|---|---|---|---|---|---|---|---|---|---|---|---|---|---|
| 2 | 136393658 | rs961360 | -0.305 | -5.906 | 0.005 | 0.48 | 0.028 | -0.022 | -0.022 | 0 | 0.34 | 0.25 | 0.27 | 0.47 | 0.23 | 0.33 |
| 2 | 136396300 | rs2278731 | -0.287 | -5.908 | 0.003 | 0.499 | 0.027 | -0.019 | -0.019 | 0 | 0.34 | 0.25 | 0.27 | 0.46 | 0.23 | 0.33 |
| 2 | 136402117 | rs4954276 | -0.591 | -5.912 | 0.018 | 0.663 | -0.01 | 0.07 | 0.07 | 3.067 | 0.77 | 0.9 | 0.58 | 0.87 | 0.4 | 0.75 |
| 2 | 136416533 | rs1900306 | 0.109 | -5.869 | 0.081 | 0.686 | 0.053 | -0.042 | -0.042 | 0 | 0.3 | 0.19 | 0.23 | 0.46 | 0.2 | 0.31 |
| 2 | 136423904 | rs2289959 | -0.454 | -5.857 | 0.253 | 0.735 | -0.024 | -0.003 | -0.003 | 0 | 0.35 | 0.36 | 0.31 | 0.65 | 0.23 | 0.38 |
| 2 | 136436317 | rs6753232 | -0.333 | -5.85 | 0.217 | 0.696 | 0.027 | -0.019 | -0.019 | 0.099 | 0.34 | 0.25 | 0.27 | 0.46 | 0.23 | 0.33 |
| 2 | 136462661 | rs2117511 | -1.229 | -5.805 | 0.088 | 0.695 | 0.063 | -0.007 | -0.007 | 0.051 | 0.56 | 0.36 | 0.68 | 0.58 | 0.84 | 0.57 |
| 2 | 136488978 | rs6739713 | 0.337 | -5.674 | 0.322 | 0.764 | -0.048 | 0.024 | 0.024 | 5.711 | 0.7 | 0.5 | 0.5 | 0.68 | 0.36 | 0.68 |
| 2 | 136494186 | rs10202489 | 0.151 | -5.626 | 0.033 | 1.076 | 0.003 | -0.077 | -0.077 | 0 | 0.25 | 0.12 | 0.45 | 0.38 | 0.61 | 0.34 |
| 2 | 136499166 | rs1438307 | 0.162 | -5.604 | 0.038 | 1.054 | -0.001 | -0.075 | -0.075 | 0 | 0.25 | 0.12 | 0.45 | 0.39 | 0.61 | 0.34 |
| 2 | 136511817 | rs1050115 | 0.181 | -5.751 | 0.149 | 0.961 | 0.046 | 0.033 | 0.033 | 0.144 | 0.3 | 0.17 | 0.21 | 0.19 | 0.21 | 0.23 |
| 2 | 136522710 | rs9287442 | -0.362 | -5.775 | 0.28 | 0.996 | 0.023 | -0.024 | -0.024 | 0 | 0.3 | 0.29 | 0.48 | 0.44 | 0.65 | 0.4 |
| 2 | 136539513 | rs10188066 | -0.426 | -5.756 | 0.276 | 0.956 | 0.023 | -0.004 | -0.004 | 0 | 0.3 | 0.29 | 0.51 | 0.61 | 0.65 | 0.41 |
| 2 | 136543147 | rs16832006 | -0.704 | -5.773 | 0.495 | 1.022 | -0.034 | 0.003 | 0.003 | 0.012 | 0.11 | 0.17 | 0.16 | 0.17 | 0.06 | 0.25 |
| 2 | 136552188 | rs3213890 | -0.01 | -5.386 | 1.048 | 1.173 | 0.01 | 0.011 | 0.011 | 0.137 | 0.31 | 0.24 | 0.21 | 0.19 | 0.21 | 0.23 |
| 2 | 136554890 | rs746857 | 0.031 | -5.384 | 1.117 | 1.199 | -0.034 | -0.017 | -0.017 | 0 | 0.68 | 0.65 | 0.78 | 0.79 | 0.79 | 0.72 |
| 2 | 136555659 | rs2322659 | -0.645 | -5.496 | 1.079 | 1.247 | 0.017 | -0.022 | -0.022 | 0 | 0.29 | 0.3 | 0.47 | 0.46 | 0.64 | 0.4 |
| 2 | 136556182 | rs3769013 | -0.068 | -5.471 | 1.168 | 1.193 | -0.015 | -0.006 | -0.006 | 0 | 0.68 | 0.68 | 0.78 | 0.82 | 0.78 | 0.75 |
| 2 | 136557319 | rs2322660 | 0.038 | -5.426 | 1.102 | 1.223 | 0.008 | -0.048 | -0.048 | 0.893 | 0.24 | 0.18 | 0.44 | 0.4 | 0.6 | 0.33 |
| 2 | 136561557 | rs2304371 | 0.153 | -5.459 | 1.469 | 1.265 | -0.013 | -0.002 | -0.002 | 0 | 0.64 | 0.4 | 0.75 | 0.78 | 0.75 | 0.73 |
| 2 | 136562472 | rs3739022 | 1.594 | -5.455 | 1.466 | 1.29 | 0.05 | 0.051 | 0.051 | 13.293 | 0.38 | 0.22 | 0.28 | 0.22 | 0.15 | 0.37 |
| 2 | 136569848 | rs12988076 | 0.194 | -5.398 | 1.415 | 1.3 | -0.007 | -0.08 | -0.08 | 0.669 | 0.23 | 0.12 | 0.43 | 0.4 | 0.6 | 0.33 |
| 2 | 136583192 | rs745500 | 0.194 | -5.398 | 1.416 | 1.292 | -0.007 | -0.078 | -0.078 | 0 | 0.23 | 0.12 | 0.43 | 0.4 | 0.6 | 0.34 |
| 2 | 136586183 | rs3769008 | -0.183 | -5.397 | 1.528 | 1.234 | -0.048 | -0.034 | -0.034 | 0 | 0.33 | 0.34 | 0.21 | 0.17 | 0.22 | 0.26 |
| 2 | 136589278 | rs9636213 | 0.087 | -5.403 | 1.514 | 1.247 | -0.001 | -0.005 | -0.005 | 0.669 | 0.65 | 0.37 | 0.73 | 0.63 | 0.78 | 0.71 |
| 2 | 136590558 | rs3754690 | 1.599 | -5.415 | 1.478 | 1.264 | 0.057 | 0.056 | 0.056 | 12.369 | 0.38 | 0.22 | 0.31 | 0.19 | 0.15 | 0.34 |
| 2 | 136590746 | rs3754689 | -0.428 | -5.273 | 1.552 | 1.326 | -0.048 | 0.013 | 0.013 | 0.421 | 0.34 | 0.54 | 0.26 | 0.37 | 0.22 | 0.29 |
| 2 | 136615222 | rs3213871 | 1.476 | -5.165 | 1.687 | 1.259 | 0.072 | 0.069 | 0.069 | 9.361 | 0.4 | 0.22 | 0.33 | 0.19 | 0.16 | 0.38 |
| 2 | 136643900 | rs309126 | 1.118 | -5.371 | 1.253 | 1.216 | 0.037 | 0.005 | 0.005 | 7.739 | 0.44 | 0.34 | 0.37 | 0.36 | 0.18 | 0.42 |
| 2 | 136648077 | rs12472293 | -0.4 | -5.277 | 0.874 | 1.25 | 0.071 | 0.055 | 0.055 | 0.626 | 0.33 | 0.18 | 0.21 | 0.21 | 0.23 | 0.31 |
| 2 | 136657252 | rs309152 | -0.514 | -5.324 | 1.033 | 1.296 | -0.072 | 0.019 | 0.019 | 0.887 | 0.34 | 0.44 | 0.23 | 0.38 | 0.21 | 0.3 |
| 2 | 136674028 | rs309172 | -0.345 | -5.345 | 1.138 | 1.286 | 0.047 | -0.026 | -0.026 | 0 | 0.33 | 0.23 | 0.22 | 0.38 | 0.22 | 0.29 |
| 2 | 136685228 | rs309160 | -0.477 | -5.449 | 1.089 | 1.296 | -0.02 | -0.086 | -0.086 | 0 | 0.79 | 0.92 | 0.63 | 0.62 | 0.41 | 0.71 |
| 2 | 136688749 | rs309161 | -0.412 | -5.45 | 1.15 | 1.294 | 0 | -0.007 | -0.007 | 1.819 | 0.34 | 0.34 | 0.23 | 0.38 | 0.21 | 0.31 |
| 2 | 136707480 | rs6750549 | -0.212 | -5.45 | 1.159 | 1.319 | 0.094 | 0.066 | 0.066 | 0.593 | 0.32 | 0.15 | 0.22 | 0.21 | 0.22 | 0.28 |
| 2 | 136721603 | rs12615624 | 0.016 | -5.224 | 1.452 | 1.33 | -0.005 | 0.024 | 0.024 | 0.718 | 0.77 | 0.7 | 0.59 | 0.57 | 0.38 | 0.69 |
| 2 | 136743490 | rs2278683 | 1.241 | -5.022 | 0.202 | 1.09 | 0.079 | 0.075 | 0.075 | 8.756 | 0.42 | 0.26 | 0.36 | 0.19 | 0.17 | 0.38 |
| 2 | 136765951 | rs309137 | 0.4 | -2.504 | -0.627 | 1.963 | -0.033 | -0.068 | -0.068 | 0.116 | 0.79 | 0.92 | 0.63 | 0.65 | 0.42 | 0.69 |
| 2 | 136785688 | rs7585965 | -1.398 | -2.141 | -0.036 | 1.904 | -0.028 | 0.133 | 0.133 | 0.019 | 0.06 | 0.14 | 0.11 | 0.27 | 0.02 | 0.16 |
| 2 | 136788978 | rs7585506 | -0.753 | -2.273 | -0.086 | 1.287 | -0.024 | -0.011 | -0.011 | 1.975 | 0.12 | 0.14 | 0.04 | 0.02 | 0.05 | 0.07 |
| 2 | 136790283 | rs7578528 | -1.301 | -2.073 | 0.126 | 1.969 | 0.006 | 0.014 | 0.014 | 2.518 | 0.09 | 0.06 | 0.18 | 0.28 | 0.05 | 0.2 |
| 2 | 136793114 | rs4954555 | -1.826 | -2.404 | 0.758 | 1.875 | -0.071 | 0.203 | 0.203 | 1.486 | 0.18 | 0.41 | 0.25 | 0.42 | 0.08 | 0.33 |
| 2 | 136817088 | rs749873 | -0.174 | -1.514 | 0 | 1.087 | -0.074 | 0.014 | 0.014 | 0.233 | 0.11 | 0.03 | 0.27 | 0.02 | 0.53 | 0.16 |
| 2 | 136822223 | rs1879087 | -1.478 | -2.2 | 0.942 | 1.523 | -0.073 | 0.125 | 0.125 | 0.201 | 0.18 | 0.4 | 0.23 | 0.69 | 0.08 | 0.32 |
| 2 | 136836365 | rs6430600 | -1.663 | -1.292 | 0.901 | 0.768 | -0.008 | -0.03 | -0.03 | 0 | 0.3 | 0.62 | 0.4 | 0.18 | 0.27 | 0.35 |
| 2 | 136840424 | rs11890513 | -1.203 | -1.146 | 0.92 | 0.73 | -0.002 | -0.008 | -0.008 | 0.111 | 0.09 | 0.11 | 0.07 | 0.03 | 0.08 | 0.1 |
| 2 | 136849515 | rs10189863 | -1.689 | -1.146 | 1.37 | 0.716 | 0.017 | -0.035 | -0.035 | 0 | 0.23 | 0.47 | 0.37 | 0.15 | 0.25 | 0.32 |
| 2 | 136864703 | rs16832731 | -1.476 | -1.338 | 1.434 | 0.878 | 0.023 | 0.021 | 0.021 | 3.677 | 0.18 | 0.53 | 0.34 | 0.13 | 0.19 | 0.28 |
| 2 | 136873084 | rs2228014 | 0.328 | -1.408 | 1.072 | 1.145 | 0.015 | -0.017 | -0.017 | 3.064 | 0.07 | 0.01 | 0.03 | 0.15 | 0.04 | 0.08 |
| 2 | 136873549 | rs2680880 | -0.421 | 0.47 | 1.156 | 1.076 | -0.073 | 0.063 | 0.063 | 1.978 | 0.12 | 0.03 | 0.25 | 0 | 0.42 | 0.23 |
| 2 | 136880474 | rs12691874 | -0.419 | -1.905 | 0.49 | 0.751 | -0.058 | -0.002 | -0.002 | 0.391 | 0.32 | 0.15 | 0.31 | 0.29 | 0.52 | 0.35 |
| 2 | 136883823 | rs4954391 | 0.35 | -1.952 | 0.97 | 0.714 | 0.061 | -0.072 | -0.072 | 0 | 0.84 | 0.47 | 0.66 | 0.97 | 0.8 | 0.77 |
| 2 | 136884679 | rs10191360 | -0.288 | -2.021 | 1.063 | 0.729 | -0.056 | -0.015 | -0.015 | 0 | 0.31 | 0.12 | 0.31 | 0.3 | 0.52 | 0.34 |
| 2 | 136886830 | rs4278980 | 0.509 | -1.848 | 1.008 | 0.849 | 0.038 | -0.078 | -0.078 | 0 | 0.84 | 0.41 | 0.65 | 0.97 | 0.8 | 0.77 |
| 2 | 136895508 | rs4954564 | 0.57 | -2.099 | 1.256 | 0.853 | 0.004 | -0.063 | -0.063 | 0.091 | 0.75 | 0.57 | 0.6 | 0.86 | 0.43 | 0.58 |
| 2 | 136900469 | rs6726457 | 0.25 | -2.322 | 1.257 | 0.707 | -0.004 | 0.001 | 0.001 | 0 | 0.21 | 0.21 | 0.11 | 0.12 | 0.12 | 0.12 |
| 2 | 136901587 | rs4954567 | 0.658 | -2.243 | 1.229 | 0.738 | 0.004 | -0.043 | -0.043 | 0 | 0.75 | 0.57 | 0.6 | 0.81 | 0.43 | 0.58 |
| 2 | 136917165 | rs4954393 | -0.282 | -2.69 | 0.324 | 1.086 | 0.001 | -0.002 | -0.002 | 0 | 0.15 | 0.12 | 0.14 | 0.22 | 0.12 | 0.17 |
| 2 | 136917330 | rs4954394 | 1.268 | -1.878 | 0.65 | 1.535 | 0.097 | 0.038 | 0.038 | 7.56 | 0.54 | 0.28 | 0.34 | 0.66 | 0.31 | 0.45 |
| 2 | 136922672 | rs6711997 | 1.45 | -1.721 | 0.635 | 0.988 | 0.074 | 0.026 | 0.026 | 11.639 | 0.5 | 0.35 | 0.28 | 0.45 | 0.27 | 0.35 |
| 2 | 136926146 | rs11687597 | -0.399 | -1.286 | 0.592 | 0.836 | -0.002 | -0.022 | -0.022 | 0 | 0.13 | 0.14 | 0.16 | 0 | 0.21 | 0.23 |
| 2 | 136934449 | rs1519523 | -0.283 | -1.221 | 0.628 | 1.124 | 0.191 | 0.023 | 0.023 | 2.216 | 0.88 | 0.74 | 0.76 | 0.8 | 0.62 | 0.8 |
| 2 | 136950383 | rs6734803 | 0.039 | -1.793 | 0.154 | 0.982 | 0.04 | -0.043 | -0.043 | 0 | 0.23 | 0.12 | 0.16 | 0.34 | 0.14 | 0.21 |
| 2 | 136956733 | rs7578292 | -0.027 | -1.795 | 0.152 | 0.981 | 0.042 | -0.044 | -0.044 | 0 | 0.23 | 0.12 | 0.16 | 0.34 | 0.14 | 0.22 |
| 2 | 136963494 | rs6716987 | 1.051 | -0.71 | -0.043 | 0.692 | 0 | -0.026 | -0.026 | 1.311 | 0.74 | 0.66 | 0.61 | 0.79 | 0.42 | 0.57 |
| 2 | 136976255 | rs882300 | 0.297 | 0.031 | 0.007 | 1.2 | 0.202 | 0.032 | 0.032 | 2.259 | 0.87 | 0.73 | 0.77 | 0.79 | 0.63 | 0.8 |
| 2 | 136976914 | rs17467077 | -0.622 | -1.194 | -0.951 | 1.395 | -0.009 | -0.071 | -0.071 | 0.466 | 0.34 | 0.14 | 0.43 | 0.43 | 0.38 | 0.49 |
| 2 | 136976936 | rs7371043 | -1.017 | -1.816 | -0.492 | 1.712 | -0.013 | 0.07 | 0.07 | 0 | 0.11 | 0.16 | 0.22 | 0.26 | 0.09 | 0.09 |
| 2 | 136986303 | rs4954580 | -0.966 | -2.856 | -0.421 | 1.062 | -0.026 | -0.001 | -0.001 | 0 | 0.61 | 0.64 | 0.75 | 0.55 | 0.86 | 0.71 |

| | | | | | | | | | | | | | | | | |
|---|---|---|---|---|---|---|---|---|---|---|---|---|---|---|---|---|
| 2 | 136989136 | rs6430608 | -0.948 | -2.957 | 0.173 | 1.011 | -0.048 | -0.003 | -0.003 | 0.158 | 0.59 | 0.47 | 0.74 | 0.55 | 0.86 | 0.68 |
| 2 | 136990327 | rs4440020 | -0.867 | -3.019 | 0.466 | 0.98 | 0.011 | 0.003 | 0.003 | 14.19 | 0.58 | 0.41 | 0.74 | 0.55 | 0.86 | 0.69 |
| 2 | 136998168 | rs10202310 | -0.986 | -3.032 | 0.401 | 0.973 | -0.06 | -0.003 | -0.003 | 0 | 0.6 | 0.58 | 0.75 | 0.56 | 0.86 | 0.71 |
| 2 | 136998394 | rs4954585 | -0.135 | -3.238 | 0.038 | 0.81 | 0.01 | 0.022 | 0.022 | 3.257 | 0.37 | 0.35 | 0.51 | 0.16 | 0.6 | 0.39 |
| 2 | 137003145 | rs4075810 | 1.512 | -3.341 | 0.21 | 0.805 | 0.155 | -0.028 | -0.028 | 0 | 0.37 | 0.1 | 0.22 | 0.45 | 0.14 | 0.29 |
| 2 | 137006198 | rs6430612 | 0.24 | -3.087 | 0.552 | 0.784 | 0.166 | -0.028 | -0.028 | 0.558 | 0.11 | 0.28 | 0.26 | 0.05 | 0.47 | 0.22 |
| 2 | 137007076 | rs6756490 | -0.505 | -3.117 | 0.459 | 0.78 | -0.006 | 0.029 | 0.029 | 0.988 | 0.23 | 0.27 | 0.24 | 0.39 | 0.26 | 0.32 |
| 2 | 137007943 | rs7608128 | -0.807 | -3.24 | 0.504 | 0.727 | -0.022 | 0.061 | 0.061 | 2.97 | 0.26 | 0.41 | 0.39 | 0.48 | 0.26 | 0.4 |
| 2 | 137009006 | rs4477975 | -0.474 | -3.178 | 0.438 | 0.724 | 0 | 0.005 | 0.005 | 1.065 | 0.23 | 0.24 | 0.24 | 0.4 | 0.26 | 0.32 |
| 2 | 137013525 | rs4954592 | 0.381 | -3.353 | -0.235 | 0.559 | 0.096 | -0.019 | -0.019 | 0 | 0.3 | 0.07 | 0.25 | 0.32 | 0.18 | 0.26 |
| 2 | 137013606 | rs10221893 | 0.58 | -2.607 | -0.06 | 0.394 | -0.053 | 0.03 | 0.03 | 1.96 | 0.09 | 0.03 | 0.23 | 0.03 | 0.47 | 0.22 |
| 2 | 137032560 | rs4389374 | 0.011 | -2.913 | -0.985 | 0.861 | -0.011 | -0.035 | -0.035 | 0.073 | 0.09 | 0 | 0.07 | 0.09 | 0.11 | 0.06 |
| 2 | 137035789 | rs13426918 | 1.644 | -3.184 | -0.879 | 0.306 | 0.018 | 0.028 | 0.028 | 20.632 | 0.26 | 0.2 | 0.07 | 0.13 | 0.06 | 0.1 |
| 2 | 137035909 | rs13387850 | 1.635 | -3.183 | -0.879 | 0.307 | 0.018 | 0.028 | 0.028 | 20.682 | 0.26 | 0.2 | 0.08 | 0.13 | 0.06 | 0.1 |
| 2 | 137037119 | rs13420371 | -0.149 | -3.31 | -0.951 | 0.558 | -0.014 | -0.14 | -0.14 | 0 | 0.26 | 0.07 | 0.23 | 0.45 | 0.25 | 0.31 |
| 2 | 137037386 | rs4954599 | 0.016 | -3.306 | -0.931 | 0.552 | 0.012 | 0.011 | 0.011 | 9.939 | 0.46 | 0.57 | 0.53 | 0.37 | 0.68 | 0.56 |
| 2 | 137039230 | rs10206655 | 1.14 | -2.88 | -0.253 | 0.513 | -0.019 | 0.002 | 0.002 | 21.013 | 0.26 | 0.23 | 0.09 | 0.13 | 0.06 | 0.1 |
| 2 | 137039910 | rs12623165 | -0.601 | -2.218 | -0.866 | 0.531 | -0.064 | -0.028 | -0.028 | 0 | 0.67 | 0.47 | 0.72 | 0.74 | 0.84 | 0.72 |
| 2 | 137055404 | rs4516492 | 0.756 | -1.438 | -1.599 | 0.42 | 0.075 | 0.18 | 0.18 | 2.712 | 0.53 | 0.11 | 0.44 | 0.17 | 0.62 | 0.51 |
| 2 | 137057148 | rs13008975 | 0.474 | -1.719 | -1.335 | 0.81 | 0.109 | 0.037 | 0.037 | 1.008 | 0.59 | 0.2 | 0.5 | 0.32 | 0.77 | 0.64 |
| 2 | 137059906 | rs10167166 | -0.759 | -2.021 | -0.22 | 1.609 | -0.06 | 0.071 | 0.071 | 2.648 | 0.23 | 0.42 | 0.31 | 0.3 | 0.14 | 0.16 |
| 2 | 137061077 | rs4954603 | 1.536 | -1.767 | 1.102 | 2.069 | 0.163 | 0.206 | 0.206 | 1.138 | 0.92 | 0.55 | 0.93 | 0.77 | 0.89 | 0.83 |
| 2 | 137062927 | rs12616701 | -0.663 | -1.617 | 0.544 | 2.168 | 0.003 | -0.011 | -0.011 | 2.134 | 0.14 | 0.08 | 0.32 | 0.42 | 0.15 | 0.23 |
| 2 | 137069775 | rs6758115 | -0.784 | -1.527 | 0.491 | 2.165 | -0.003 | -0.02 | -0.02 | 0 | 0.15 | 0.08 | 0.34 | 0.41 | 0.2 | 0.21 |
| 2 | 137072784 | rs6745602 | 1.175 | -2.025 | 1.092 | 2.005 | 0.006 | -0.014 | -0.014 | 0 | 0.8 | 0.45 | 0.85 | 0.79 | 0.79 | 0.73 |
| 2 | 137074132 | rs6708702 | -1.762 | -2.391 | 1.104 | 1.928 | -0.002 | -0.005 | -0.005 | 0 | 0.34 | 0.35 | 0.56 | 0.73 | 0.38 | 0.38 |
| 2 | 137074850 | rs12465599 | 2.384 | -2.396 | 1.406 | 1.935 | 0.009 | 0.019 | 0.019 | 0.891 | 0.52 | 0.54 | 0.36 | 0.28 | 0.53 | 0.45 |
| 2 | 137075288 | rs13028760 | 2.729 | -2.482 | 1.124 | 1.796 | 0.026 | 0.277 | 0.277 | 2.688 | 0.48 | 0.1 | 0.31 | 0.12 | 0.52 | 0.41 |
| 2 | 137076425 | rs4954411 | 2.541 | -2.479 | 1.197 | 1.797 | 0.059 | 0.24 | 0.24 | 2.125 | 0.49 | 0.16 | 0.33 | 0.12 | 0.52 | 0.44 |
| 2 | 137084682 | rs13006092 | 2.846 | -2.516 | 1.023 | 1.778 | -0.019 | 0.282 | 0.282 | 3.781 | 0.46 | 0.06 | 0.3 | 0.12 | 0.5 | 0.4 |
| 2 | 137084999 | rs6719065 | 0.471 | -2.441 | 1.124 | 2.305 | -0.001 | -0.001 | -0.001 | 0.99 | 0.12 | 0.08 | 0.28 | 0.23 | 0.16 | 0.15 |
| 2 | 137085598 | rs7570134 | -1.192 | -2.468 | 1.22 | 2.256 | 0.005 | -0.024 | -0.024 | 0.663 | 0.22 | 0.15 | 0.29 | 0.32 | 0.23 | 0.23 |
| 2 | 137087603 | rs7589775 | -0.135 | -2.513 | 1.147 | 2.279 | 0.031 | 0.013 | 0.013 | 0 | 0.15 | 0.09 | 0.17 | 0.09 | 0.06 | 0.07 |
| 2 | 137090865 | rs6739956 | -0.603 | -2.472 | 1.162 | 2.268 | 0.018 | -0.018 | -0.018 | 0 | 0.18 | 0.12 | 0.25 | 0.25 | 0.12 | 0.14 |
| 2 | 137093578 | rs6430628 | 0.166 | -1.944 | 1.419 | 2.07 | -0.001 | -0.004 | -0.004 | 0 | 0.12 | 0.09 | 0.28 | 0.21 | 0.16 | 0.15 |
| 2 | 137102171 | rs10197003 | 1.631 | -2.064 | 1.396 | 2.045 | 0.014 | -0.008 | -0.008 | 0 | 0.68 | 0.28 | 0.59 | 0.64 | 0.74 | 0.65 |
| 2 | 137111399 | rs6740091 | -1.744 | -2.087 | 1.908 | 2.115 | 0.004 | 0.038 | 0.038 | 1.166 | 0.26 | 0.55 | 0.47 | 0.37 | 0.31 | 0.3 |
| 2 | 137112562 | rs1515069 | -0.57 | -2.079 | 2.236 | 2.123 | 0.012 | 0.017 | 0.017 | 0 | 0.16 | 0.19 | 0.32 | 0.27 | 0.27 | 0.23 |
| 2 | 137114910 | rs16833821 | -1.102 | -2.079 | 2.151 | 2.062 | 0 | 0.074 | 0.074 | 0.312 | 0.08 | 0.33 | 0.15 | 0.09 | 0.03 | 0.06 |
| 2 | 137116544 | rs16833840 | -0.052 | -2.171 | 1.798 | 1.979 | 0.025 | -0.015 | -0.015 | 0 | 0.14 | 0.03 | 0.09 | 0.15 | 0.09 | 0.14 |
| 2 | 137120582 | rs12615012 | 2.381 | -2.117 | 1.843 | 2.044 | 0.039 | 0.006 | 0.006 | 0.929 | 0.56 | 0.14 | 0.42 | 0.31 | 0.6 | 0.49 |
| 2 | 137131321 | rs1849873 | 0.498 | -2.084 | 1.87 | 1.876 | -0.007 | -0.006 | -0.006 | 0 | 0.82 | 0.8 | 0.9 | 0.9 | 0.91 | 0.8 |
| 2 | 137137913 | rs16833925 | -1.642 | -1.883 | 1.856 | 2.006 | 0.003 | 0.039 | 0.039 | 0.852 | 0.25 | 0.27 | 0.34 | 0.53 | 0.31 | 0.3 |
| 2 | 137139548 | rs6738835 | -1.577 | -1.908 | 1.932 | 1.993 | 0.001 | 0.029 | 0.029 | 1.344 | 0.25 | 0.27 | 0.34 | 0.53 | 0.31 | 0.3 |
| 2 | 137141711 | rs16833946 | -0.767 | -1.838 | 1.485 | 1.907 | -0.005 | -0.007 | -0.007 | 0 | 0.18 | 0.14 | 0.32 | 0.35 | 0.28 | 0.24 |
| 2 | 137146324 | rs16833964 | -1.416 | -1.823 | 1.544 | 1.816 | -0.001 | -0.003 | -0.003 | 0 | 0.25 | 0.22 | 0.34 | 0.51 | 0.31 | 0.3 |
| 2 | 137149716 | rs4588246 | 1.803 | -1.451 | 2.078 | 1.152 | 0.013 | -0.03 | -0.03 | 0.323 | 0.92 | 0.61 | 0.95 | 1 | 0.96 | 0.96 |
| 2 | 137152765 | rs6430636 | 0.603 | -1.414 | 1.788 | 1.172 | -0.005 | -0.03 | -0.03 | 0.173 | 0.94 | 0.85 | 0.97 | 1 | 0.97 | 0.96 |
| 2 | 137155352 | rs16834018 | -0.33 | -1.537 | 1.784 | 1.163 | -0.02 | -0.011 | -0.011 | 0 | 0.14 | 0.24 | 0.07 | 0.1 | 0.06 | 0.13 |
| 2 | 137155850 | rs2048756 | 1.649 | 0.275 | 1.359 | 1.146 | 0.048 | -0.008 | -0.008 | 0 | 0.71 | 0.37 | 0.73 | 0.71 | 0.62 | 0.71 |
| 2 | 137159383 | rs10496744 | -1.417 | 0.847 | 0.669 | 1.262 | 0.014 | -0.014 | -0.014 | 0 | 0.1 | 0.1 | 0.08 | 0.05 | 0.15 | 0.11 |
| 2 | 137163071 | rs6738782 | -0.941 | -2.569 | -0.397 | 0.83 | 0.036 | -0.002 | -0.002 | 0 | 0.55 | 0.63 | 0.64 | 0.48 | 0.79 | 0.67 |
| 2 | 137163904 | rs6729725 | 0.149 | -2.583 | -0.743 | 0.474 | 0.028 | 0.027 | 0.027 | 0.029 | 0.15 | 0.05 | 0.08 | 0.11 | 0.09 | 0.19 |
| 2 | 137164476 | rs9646660 | 0.512 | -2.737 | -0.605 | 0.244 | 0.024 | 0.018 | 0.018 | 0 | 0.16 | 0.15 | 0.06 | 0.01 | 0.08 | 0.07 |
| 2 | 137166151 | rs527620 | -0.665 | -2.92 | -0.359 | 0.238 | 0.028 | 0.035 | 0.035 | 0 | 0.23 | 0.17 | 0.12 | 0.06 | 0.13 | 0.21 |
| 2 | 137169495 | rs4954609 | 1.378 | -3.079 | 0.336 | 0.264 | -0.006 | 0.001 | 0.001 | 3.642 | 0.43 | 0.38 | 0.37 | 0.63 | 0.22 | 0.35 |
| 2 | 137170151 | rs13394842 | 0.633 | -3.339 | 0.401 | 0.101 | 0.015 | 0.003 | 0.003 | 0.142 | 0.06 | 0.26 | 0.04 | 0 | 0.02 | 0.02 |
| 2 | 137173367 | rs12691887 | 0.567 | -3.387 | 0.257 | 0.391 | -0.001 | -0.083 | -0.083 | 0 | 0.3 | 0.13 | 0.52 | 0.41 | 0.64 | 0.44 |
| 2 | 137173571 | rs17382822 | -0.026 | -3.363 | 0.287 | 0.331 | 0.047 | 0.075 | 0.075 | 0 | 0.22 | 0.11 | 0.12 | 0.04 | 0.12 | 0.2 |
| 2 | 137177456 | rs13401318 | 1.385 | -3.366 | 0.332 | 0.386 | -0.014 | 0 | 0 | 0 | 0.43 | 0.39 | 0.37 | 0.63 | 0.21 | 0.35 |
| 2 | 137178771 | rs11687599 | 0.652 | -3.366 | 0.251 | 0.344 | -0.014 | -0.016 | -0.016 | 0 | 0.29 | 0.08 | 0.47 | 0.23 | 0.64 | 0.41 |
| 2 | 137179433 | rs16834111 | 1.125 | -3.177 | 0.764 | 0.416 | -0.049 | 0.004 | 0.004 | 3.56 | 0.43 | 0.43 | 0.33 | 0.53 | 0.21 | 0.34 |
| 2 | 137182038 | rs10180839 | -0.955 | -3.137 | 0.333 | 0.436 | 0.133 | -0.006 | -0.006 | 0 | 0.53 | 0.29 | 0.65 | 0.46 | 0.77 | 0.65 |
| 2 | 137199326 | rs603666 | -0.669 | -3.38 | 0.182 | 0.151 | -0.007 | -0.01 | -0.01 | 0 | 0.9 | 0.89 | 0.96 | 0.97 | 0.95 | 0.92 |
| 2 | 137200214 | rs4954614 | 2.213 | -4.082 | 0.093 | 0.373 | 0.046 | -0.035 | -0.035 | 0 | 0.27 | 0.14 | 0.33 | 0.37 | 0.14 | 0.2 |
| 2 | 137207082 | rs10496746 | 1.647 | -4.425 | -0.261 | -0.003 | -0.034 | 0.092 | 0.092 | 0.158 | 0.22 | 0.08 | 0.38 | 0.02 | 0.6 | 0.26 |
| 2 | 137209221 | rs7603072 | 1.958 | -4.494 | 0.589 | 0.058 | -0.076 | 0.037 | 0.037 | 3.371 | 0.31 | 0.43 | 0.36 | 0.41 | 0.14 | 0.25 |
| 2 | 137210991 | rs578935 | 0.064 | -4.596 | 0.536 | 0.085 | 0.088 | 0.068 | 0.068 | 0 | 0.29 | 0.1 | 0.12 | 0.22 | 0.16 | 0.25 |
| 2 | 137214747 | rs12612658 | 2.52 | -4.568 | 0.734 | 0.17 | 0.049 | -0.037 | -0.037 | 1.516 | 0.27 | 0.14 | 0.33 | 0.37 | 0.14 | 0.21 |

| | | | | | | | | | | | | | | | | |
|---|---|---|---|---|---|---|---|---|---|---|---|---|---|---|---|---|
| 2 | 137219133 | rs563709 | -1.315 | -4.564 | 0.766 | 0.101 | -0.016 | -0.021 | -0.021 | 0 | 0.9 | 0.87 | 0.96 | 0.97 | 0.95 | 0.92 |
| 2 | 137220743 | rs47150 | -0.5 | -4.528 | 0.893 | 0.191 | 0.01 | -0.005 | -0.005 | 0.236 | 0.64 | 0.67 | 0.44 | 0.64 | 0.32 | 0.51 |
| 2 | 137233178 | rs2885003 | 1.57 | -4.402 | 1.139 | 0.253 | -0.086 | 0.033 | 0.033 | 3.56 | 0.34 | 0.53 | 0.38 | 0.5 | 0.16 | 0.27 |
| 2 | 137241454 | rs643821 | -0.381 | -4.174 | 0.872 | 0.295 | 0.009 | 0.004 | 0.004 | 2.647 | 0.64 | 0.67 | 0.49 | 0.72 | 0.32 | 0.52 |
| 2 | 137254205 | rs581054 | -1.312 | -3.958 | 0.755 | 0.331 | 0.001 | 0.006 | 0.006 | 0.301 | 0.77 | 0.75 | 0.49 | 0.67 | 0.38 | 0.6 |
| 2 | 137259124 | rs10200731 | 1.137 | -4.349 | 0.762 | 0.406 | -0.02 | -0.04 | -0.04 | 0 | 0.25 | 0.09 | 0.47 | 0.25 | 0.63 | 0.39 |
| 2 | 137263270 | rs16834591 | 2.068 | -4.577 | 1.019 | 0.349 | -0.023 | 0.002 | 0.002 | 5.879 | 0.3 | 0.32 | 0.29 | 0.41 | 0.14 | 0.25 |
| 2 | 137265745 | rs16834638 | 1.47 | -4.615 | 1.135 | 0.329 | -0.077 | 0.029 | 0.029 | 3.789 | 0.35 | 0.53 | 0.37 | 0.53 | 0.15 | 0.29 |
| 2 | 137270625 | rs10208178 | 1.094 | -5.732 | 0.632 | 0.398 | 0 | -0.007 | -0.007 | 0 | 0.26 | 0.18 | 0.47 | 0.25 | 0.63 | 0.39 |
| 2 | 137282346 | rs479300 | 1.887 | -5.787 | 0.619 | 0.387 | -0.04 | 0.023 | 0.023 | 4.349 | 0.31 | 0.32 | 0.34 | 0.49 | 0.15 | 0.25 |
| 2 | 137283019 | rs562370 | -0.983 | -5.844 | 0.647 | 0.381 | -0.006 | 0.003 | 0.003 | 0.471 | 0.45 | 0.57 | 0.19 | 0.32 | 0.23 | 0.37 |
| 2 | 137283342 | rs10496747 | 2.178 | -5.744 | -0.089 | 0.387 | 0.06 | -0.007 | -0.007 | 0 | 0.29 | 0.14 | 0.28 | 0.31 | 0.14 | 0.22 |
| 2 | 137288459 | rs881748 | 2.272 | -5.713 | -0.079 | 0.401 | 0.069 | -0.012 | -0.012 | 0 | 0.28 | 0.12 | 0.27 | 0.31 | 0.13 | 0.22 |
| 2 | 137295709 | rs667607 | 0.424 | -5.852 | -0.301 | 0.057 | 0.087 | 0.065 | 0.065 | 0 | 0.28 | 0.07 | 0.12 | 0.2 | 0.16 | 0.24 |
| 2 | 137300091 | rs10205768 | 1.517 | -5.523 | -0.432 | 0.139 | -0.04 | 0.09 | 0.09 | 0.376 | 0.22 | 0.08 | 0.39 | 0.02 | 0.61 | 0.25 |
| 2 | 137301605 | rs10496748 | 1.811 | -5 | -0.229 | 0.831 | 0.056 | 0.043 | 0.043 | 6.58 | 0.28 | 0.14 | 0.16 | 0.17 | 0.13 | 0.21 |
| 2 | 137304516 | rs1427683 | 1.811 | -4.895 | -0.757 | 0.491 | 0.056 | -0.038 | -0.038 | 0 | 0.28 | 0.14 | 0.33 | 0.39 | 0.14 | 0.23 |
| 2 | 137312484 | rs1374109 | 0.614 | -4.432 | -0.754 | 0.462 | 0.05 | 0.008 | 0.008 | 0 | 0.18 | 0.08 | 0.1 | 0.17 | 0.1 | 0.18 |
| 2 | 137312731 | rs766674 | -1.191 | -5.339 | -0.538 | 0.18 | 0.016 | 0.007 | 0.007 | 1.291 | 0.49 | 0.57 | 0.48 | 0.59 | 0.34 | 0.56 |
| 2 | 137314139 | rs580879 | -0.478 | -5.365 | -0.153 | -0.047 | -0.052 | -0.096 | -0.096 | 4.112 | 0.35 | 0.48 | 0.22 | 0.11 | 0.25 | 0.35 |
| 2 | 137314240 | rs12612204 | 0.962 | -5.629 | -0.557 | -0.005 | -0.006 | -0.051 | -0.051 | 0.597 | 0.27 | 0.15 | 0.64 | 0.53 | 0.62 | 0.4 |
| 2 | 137315739 | rs12691893 | 0.782 | -5.698 | -0.416 | 0.003 | 0.126 | 0.083 | 0.083 | 3.697 | 0.5 | 0.28 | 0.54 | 0.35 | 0.68 | 0.46 |
| 2 | 137321430 | rs13005339 | 0.617 | -4.867 | -0.556 | 0.45 | -0.004 | -0.051 | -0.051 | 0 | 0.28 | 0.14 | 0.64 | 0.54 | 0.63 | 0.4 |
| 2 | 137325995 | rs11686763 | 1.156 | -4.848 | -0.771 | 0.396 | -0.057 | -0.069 | -0.069 | 0.104 | 0.23 | 0.07 | 0.56 | 0.3 | 0.6 | 0.26 |
| 2 | 137332558 | rs13412518 | -0.052 | -5.015 | -0.403 | 0.341 | -0.005 | 0.032 | 0.032 | 0.722 | 0.16 | 0.15 | 0.3 | 0.36 | 0.15 | 0.24 |
| 2 | 137338721 | rs934298 | 0.415 | -4.939 | -0.322 | 0.276 | -0.008 | -0.011 | -0.011 | 0 | 0.13 | 0.15 | 0.12 | 0.1 | 0.13 | 0.2 |
| 2 | 137340092 | rs4270400 | -0.922 | -5.267 | -0.339 | 0.099 | 0.054 | 0.04 | 0.04 | 5.762 | 0.56 | 0.38 | 0.84 | 0.64 | 0.85 | 0.69 |
| 2 | 137340497 | rs841435 | -0.376 | -4.336 | 0.257 | 0.57 | -0.029 | 0.013 | 0.013 | 0.053 | 0.94 | 0.88 | 0.98 | 0.91 | 0.99 | 0.93 |
| 2 | 137341712 | rs10193021 | 0.558 | -4.206 | -0.11 | 0.604 | -0.023 | -0.029 | -0.029 | 0 | 0.27 | 0.09 | 0.47 | 0.27 | 0.62 | 0.38 |
| 2 | 137342808 | rs10496750 | -0.354 | -4.324 | -0.007 | 0.094 | 0.061 | 0.03 | 0.03 | 3.851 | 0.4 | 0.25 | 0.76 | 0.64 | 0.75 | 0.6 |
| 2 | 137348398 | rs1446099 | -1 | -3.207 | 0.808 | 0.145 | -0.024 | -0.026 | -0.026 | 0 | 0.14 | 0.21 | 0.11 | 0.1 | 0.11 | 0.21 |
| 2 | 137351216 | rs841443 | -0.52 | -2.843 | 1.112 | 0.306 | -0.017 | -0.026 | -0.026 | 0 | 0.08 | 0.35 | 0.04 | 0 | 0.02 | 0.06 |
| 2 | 137364202 | rs574516 | -0.671 | -2.832 | 1.295 | 0.405 | 0.013 | 0.032 | 0.032 | 2.712 | 0.08 | 0.51 | 0.06 | 0.02 | 0.03 | 0.06 |
| 2 | 137372841 | rs4954630 | -0.346 | -3.594 | 0.168 | 0.291 | 0.088 | -0.009 | -0.009 | 0 | 0.37 | 0.21 | 0.34 | 0.65 | 0.22 | 0.44 |
| 2 | 137374734 | rs13388535 | 0.419 | -3.424 | 0.511 | 0.249 | -0.04 | -0.048 | -0.048 | 1.068 | 0.09 | 0.3 | 0.04 | 0 | 0.02 | 0.06 |
| 2 | 137382887 | rs7608688 | -0.617 | -3.343 | 0.306 | 0.183 | -0.038 | -0.047 | -0.047 | 0 | 0.14 | 0.29 | 0.07 | 0.04 | 0.08 | 0.12 |
| 2 | 137385578 | rs1374112 | -0.607 | -3.286 | -0.629 | 0.248 | 0.089 | 0.046 | 0.046 | 2.643 | 0.43 | 0.26 | 0.37 | 0.75 | 0.28 | 0.54 |
| 2 | 137389497 | rs10210687 | 0.54 | -3.079 | -0.763 | 0.016 | 0.183 | 0.122 | 0.122 | 0 | 0.53 | 0.19 | 0.75 | 0.69 | 0.76 | 0.52 |
| 2 | 137391962 | rs7576758 | 0.209 | -3.127 | -1.092 | -0.084 | 0.119 | 0.104 | 0.104 | 0 | 0.58 | 0.22 | 0.77 | 0.78 | 0.8 | 0.64 |
| 2 | 137398249 | rs7588497 | 0.167 | -1.97 | -0.72 | 0.396 | -0.008 | 0.004 | 0.004 | 0 | 0.05 | 0.08 | 0.02 | 0.08 | 0.04 | 0.12 |
| 2 | 137404246 | rs7607648 | 0.104 | -2.764 | -1.193 | -0.024 | 0.084 | 0.05 | 0.05 | 2.451 | 0.24 | 0.11 | 0.14 | 0.17 | 0.12 | 0.17 |
| 2 | 137405571 | rs1823084 | -0.027 | -3.086 | -1.332 | 0.129 | -0.002 | 0.003 | 0.003 | 0 | 0.19 | 0.17 | 0.26 | 0.59 | 0.15 | 0.38 |
| 2 | 137406682 | rs1446100 | 0.485 | -3.229 | -0.453 | -0.236 | -0.037 | -0.051 | -0.051 | 0 | 0.73 | 0.56 | 0.65 | 0.22 | 0.79 | 0.53 |
| 2 | 137409714 | rs17654195 | 1.714 | -0.555 | -0.811 | -0.468 | -0.096 | 0.027 | 0.027 | 1.228 | 0.13 | 0.02 | 0.18 | 0.09 | 0.25 | 0.11 |
| 2 | 137413062 | rs9789419 | -0.705 | -0.417 | -0.402 | -0.372 | 0 | -0.001 | -0.001 | 0 | 0.28 | 0.24 | 0.34 | 0.76 | 0.23 | 0.48 |
| 2 | 137414562 | rs10192163 | 1.057 | -0.535 | -0.608 | -0.515 | 0.022 | -0.054 | -0.054 | 0.026 | 0.64 | 0.54 | 0.55 | 0.2 | 0.55 | 0.41 |
| 2 | 137416035 | rs1530518 | 0.422 | -0.777 | -0.345 | -0.468 | -0.034 | -0.033 | -0.033 | 0 | 0.72 | 0.63 | 0.66 | 0.24 | 0.8 | 0.52 |
| 2 | 137431134 | rs13392503 | -1.178 | -1.345 | 0.315 | -0.881 | 0.045 | 0.06 | 0.06 | 0 | 0.56 | 0.62 | 0.5 | 0.88 | 0.33 | 0.71 |
| 2 | 137433902 | rs1374111 | 1.581 | -1.355 | 0.209 | -0.948 | 0.012 | 0.092 | 0.092 | 0 | 0.34 | 0.2 | 0.46 | 0.03 | 0.6 | 0.21 |
| 2 | 137434487 | rs12466458 | -0.173 | -1.477 | 0.378 | -0.854 | 0.002 | -0.009 | -0.009 | 0.438 | 0.11 | 0.06 | 0.09 | 0.18 | 0.11 | 0.16 |
| 2 | 137435455 | rs4954638 | -1.019 | -1.533 | 0.693 | -0.888 | -0.011 | -0.019 | -0.019 | 1.927 | 0.56 | 0.57 | 0.44 | 0.8 | 0.3 | 0.64 |
| 2 | 137435625 | rs4954639 | 1.672 | -1.656 | 0.778 | -0.883 | 0.065 | 0.066 | 0.066 | 0.034 | 0.21 | 0.09 | 0.08 | 0.1 | 0.09 | 0.18 |
| 2 | 137438726 | rs4954640 | -0.593 | -1.645 | 0.833 | -0.878 | 0.021 | 0.019 | 0.019 | 0 | 0.51 | 0.48 | 0.42 | 0.79 | 0.26 | 0.63 |
| 2 | 137447448 | rs4419275 | 0.116 | -1.128 | 0.835 | -0.753 | -0.004 | 0.011 | 0.011 | 0 | 0.6 | 0.35 | 0.76 | 0.78 | 0.64 | 0.62 |
| 2 | 137452607 | rs4434047 | 0.966 | -0.937 | 0.828 | -0.653 | 0.179 | 0.007 | 0.007 | 1.067 | 0.84 | 0.67 | 0.81 | 0.84 | 0.7 | 0.8 |
| 2 | 137465622 | rs16836162 | 0.779 | -1.22 | 0.705 | 0.012 | 0.017 | -0.059 | -0.059 | 0 | 0.12 | 0.03 | 0.21 | 0.39 | 0.11 | 0.26 |
| 2 | 137474142 | rs1427611 | 0.768 | -1.197 | 0.777 | -0.07 | 0.081 | -0.015 | -0.015 | 0 | 0.74 | 0.56 | 0.78 | 0.73 | 0.65 | 0.72 |
| 2 | 137480048 | rs2361173 | -0.609 | -1.112 | 0.752 | -0.031 | -0.005 | 0.013 | 0.013 | 0.524 | 0.13 | 0.21 | 0.1 | 0.18 | 0.11 | 0.16 |
| 2 | 137488316 | rs9287454 | 0.954 | -1.275 | 0.838 | 0.008 | 0.106 | 0.087 | 0.087 | 0 | 0.75 | 0.59 | 0.61 | 0.46 | 0.63 | 0.65 |
| 2 | 137491121 | rs1582861 | 0.878 | -1.377 | 0.744 | -0.069 | 0.001 | 0.008 | 0.008 | 0 | 0.52 | 0.51 | 0.52 | 0.33 | 0.51 | 0.37 |
| 2 | 137492022 | rs7340271 | 1.466 | -1.415 | 0.792 | 0.019 | 0.002 | 0.045 | 0.045 | 0 | 0.42 | 0.3 | 0.48 | 0.24 | 0.47 | 0.3 |
| 2 | 137492648 | rs1427587 | -0.334 | -1.366 | 0.854 | 0.052 | -0.003 | 0.017 | 0.017 | 0 | 0.19 | 0.16 | 0.28 | 0.54 | 0.17 | 0.31 |
| 2 | 137493598 | rs12476718 | -0.654 | -1.37 | 0.869 | 0.048 | -0.016 | 0.039 | 0.039 | 0 | 0.21 | 0.23 | 0.3 | 0.55 | 0.18 | 0.32 |
| 2 | 137493893 | rs12475945 | 0.704 | -1.317 | 0.703 | 0.086 | -0.011 | -0.035 | -0.035 | 0.017 | 0.07 | 0.01 | 0.07 | 0.16 | 0.08 | 0.07 |
| 2 | 137498009 | rs4473441 | -1.013 | -1.321 | 0.751 | -0.007 | -0.002 | -0.006 | -0.006 | 0.011 | 0.48 | 0.45 | 0.47 | 0.65 | 0.49 | 0.63 |
| 2 | 137506263 | rs1559972 | -1.653 | -1.213 | 0.762 | -0.038 | 0.01 | 0.005 | 0.005 | 0 | 0.9 | 0.88 | 0.96 | 0.91 | 0.95 | 0.92 |
| 2 | 137506665 | rs10182108 | 1.194 | -1.185 | 0.764 | -0.072 | 0.003 | 0.074 | 0.074 | 0 | 0.47 | 0.3 | 0.5 | 0.24 | 0.51 | 0.34 |
| 2 | 137508544 | rs1896477 | 0.026 | -1.35 | 0.861 | -0.122 | -0.007 | 0.044 | 0.044 | 0 | 0.17 | 0.19 | 0.29 | 0.55 | 0.15 | 0.28 |
| 2 | 137509045 | rs10496754 | -0.44 | -1.356 | 0.869 | -0.148 | -0.061 | -0.048 | -0.048 | 0 | 0.67 | 0.64 | 0.88 | 0.82 | 0.86 | 0.73 |

| | | | | | | | | | | | | | | | | |
|---|---|---|---|---|---|---|---|---|---|---|---|---|---|---|---|---|
| 2 | 137514418 | rs1427591 | 1.158 | -1.292 | 0.787 | -0.139 | 0.079 | 0.068 | 0.068 | 0 | 0.47 | 0.31 | 0.57 | 0.25 | 0.68 | 0.38 |
| 2 | 137515382 | rs1427592 | -0.38 | -1.122 | 0.843 | 0.084 | 0.05 | 0.014 | 0.014 | 0 | 0.65 | 0.78 | 0.82 | 0.63 | 0.81 | 0.69 |
| 2 | 137518086 | rs2253551 | -1.503 | -0.568 | 0.653 | 0.096 | 0.026 | 0.008 | 0.008 | 0 | 0.9 | 0.98 | 0.97 | 0.9 | 0.96 | 0.93 |
| 2 | 137527377 | rs16855705 | 1.281 | -0.58 | 0.516 | 0.224 | 0.024 | -0.054 | -0.054 | 0 | 0.1 | 0 | 0.03 | 0.24 | 0.04 | 0.07 |
| 2 | 137527965 | rs2710184 | 0.645 | -0.74 | 0.716 | 0.154 | -0.04 | -0.01 | -0.01 | 0 | 0.27 | 0.33 | 0.16 | 0.27 | 0.15 | 0.23 |
| 2 | 137528204 | rs6752506 | 0.634 | -0.406 | 0.144 | 0.001 | -0.019 | -0.071 | -0.071 | 2.321 | 0.17 | 0.05 | 0.26 | 0.38 | 0.22 | 0.33 |
| 2 | 137529247 | rs1427602 | -1.228 | -0.144 | 0.237 | -0.118 | 0.017 | 0.049 | 0.049 | 0 | 0.57 | 0.7 | 0.55 | 0.9 | 0.61 | 0.71 |
| 2 | 137531892 | rs2558095 | -0.455 | -0.201 | 0.302 | -0.051 | 0 | 0 | 0 | 0 | 0.44 | 0.61 | 0.44 | 0.64 | 0.38 | 0.6 |
| 2 | 137535866 | rs13016946 | 1.47 | -0.323 | -0.054 | -0.634 | -0.003 | 0.081 | 0.081 | 0 | 0.33 | 0.16 | 0.34 | 0.03 | 0.33 | 0.16 |
| 2 | 137536904 | rs2465095 | -1.484 | -0.211 | 0.308 | -0.588 | -0.002 | -0.014 | -0.014 | 0 | 0.64 | 0.68 | 0.57 | 0.93 | 0.47 | 0.8 |
| 2 | 137543318 | rs2114750 | 0.182 | -0.387 | 0.803 | -0.316 | -0.015 | 0.112 | 0.112 | 0.004 | 0.13 | 0.3 | 0.11 | 0.22 | 0.08 | 0.15 |
| 2 | 137547542 | rs1427599 | 1.442 | -0.687 | 0.587 | -0.466 | 0.082 | 0.083 | 0.083 | 0 | 0.23 | 0.09 | 0.1 | 0.11 | 0.12 | 0.2 |
| 2 | 137548444 | rs1427598 | -1.936 | -0.761 | 0.571 | -0.416 | 0.02 | -0.017 | -0.017 | 0.937 | 0.9 | 0.93 | 0.97 | 0.81 | 0.96 | 0.93 |
| 2 | 137551335 | rs1593760 | -1.271 | -0.75 | 0.613 | -0.443 | 0.021 | 0.062 | 0.062 | 0 | 0.56 | 0.7 | 0.62 | 0.77 | 0.62 | 0.76 |
| 2 | 137552254 | rs1593759 | 0.324 | -0.733 | 0.628 | -0.427 | 0.01 | 0.01 | 0.01 | 0 | 0.2 | 0.26 | 0.4 | 0.45 | 0.42 | 0.4 |
| 2 | 137553966 | rs2710188 | 1.237 | -0.763 | 0.488 | -0.573 | -0.003 | -0.003 | -0.003 | 0 | 0.24 | 0.21 | 0.11 | 0.11 | 0.12 | 0.21 |
| 2 | 137554697 | rs2710190 | -1.824 | -0.645 | 0.218 | -0.513 | 0 | 0.047 | 0.047 | 0 | 0.67 | 0.79 | 0.65 | 0.97 | 0.67 | 0.84 |
| 2 | 137559630 | rs4954438 | -0.489 | -0.616 | 0.411 | -0.134 | 0.013 | -0.005 | -0.005 | 0 | 0.39 | 0.43 | 0.28 | 0.37 | 0.39 | 0.4 |
| 2 | 137573885 | rs778199 | 0.825 | -1.042 | -0.138 | -0.542 | -0.006 | 0.004 | 0.004 | 3.405 | 0.32 | 0.22 | 0.25 | 0.02 | 0.36 | 0.24 |
| 2 | 137580234 | rs10496758 | 1.133 | -1.355 | -0.526 | -0.651 | 0.136 | 0.135 | 0.135 | 12.836 | 0.33 | 0.09 | 0.16 | 0.13 | 0.17 | 0.21 |
| 2 | 137583444 | rs12617926 | 0.44 | -0.826 | -0.397 | -0.18 | -0.035 | -0.031 | -0.031 | 0 | 0.11 | 0.04 | 0.36 | 0.4 | 0.17 | 0.26 |
| 2 | 137591618 | rs778192 | -0.415 | -0.986 | 0.145 | -0.161 | 0.002 | 0.002 | 0.002 | 2.375 | 0.55 | 0.45 | 0.74 | 0.58 | 0.63 | 0.71 |
| 2 | 137592114 | rs809012 | 1.403 | -1.065 | 0.101 | -0.298 | -0.014 | 0.054 | 0.054 | 1.649 | 0.31 | 0.16 | 0.26 | 0.03 | 0.36 | 0.25 |
| 2 | 137592304 | rs778189 | 1.308 | -1.025 | -0.077 | -0.271 | -0.008 | 0.006 | 0.006 | 1.647 | 0.32 | 0.21 | 0.26 | 0.03 | 0.36 | 0.25 |
| 2 | 137595545 | rs778187 | -1.408 | -0.739 | 0.269 | -0.017 | 0.063 | 0.061 | 0.061 | 9.213 | 0.69 | 0.83 | 0.86 | 0.85 | 0.86 | 0.81 |
| 2 | 137597032 | rs778195 | -1.12 | -0.702 | 0.082 | 0.063 | -0.117 | -0.04 | -0.04 | 2.158 | 0.67 | 0.54 | 0.83 | 0.75 | 0.86 | 0.81 |
| 2 | 137599649 | rs778193 | 0.835 | -0.631 | -0.082 | 0.061 | -0.004 | 0.047 | 0.047 | 1.377 | 0.41 | 0.26 | 0.34 | 0.2 | 0.44 | 0.39 |
| 2 | 137601781 | rs12613655 | -1.156 | -0.475 | -0.57 | 0.149 | -0.052 | -0.039 | -0.039 | 0.326 | 0.68 | 0.65 | 0.85 | 0.86 | 0.86 | 0.81 |
| 2 | 137602926 | rs1961273 | -1.71 | 1.215 | -0.657 | 1.215 | -0.029 | -0.05 | -0.05 | 0 | 0.17 | 0.12 | 0.47 | 0.55 | 0.27 | 0.39 |
| 2 | 137610035 | rs13015892 | -0.428 | 1.431 | -0.492 | 1.399 | 0.001 | 0.055 | 0.055 | 0 | 0.08 | 0.1 | 0.36 | 0.37 | 0.16 | 0.23 |
| 2 | 137612229 | rs778202 | 0.643 | 0.931 | -0.341 | 1.352 | 0.022 | 0.017 | 0.017 | 0 | 0.84 | 0.81 | 0.86 | 0.69 | 0.75 | 0.76 |
| 2 | 137616423 | rs1649540 | -0.068 | 0.983 | -0.355 | 1.364 | 0.171 | 0.053 | 0.053 | 0 | 0.94 | 0.83 | 0.89 | 0.89 | 0.79 | 0.89 |
| 2 | 137621738 | rs1346733 | 0.581 | 0.976 | -0.364 | 1.337 | 0.023 | 0.017 | 0.017 | 0 | 0.84 | 0.82 | 0.86 | 0.71 | 0.75 | 0.76 |
| 2 | 137621910 | rs1652435 | 1.069 | 0.96 | -0.384 | 1.347 | 0.005 | 0.011 | 0.011 | 0.009 | 0.81 | 0.82 | 0.8 | 0.55 | 0.69 | 0.7 |
| 2 | 137623006 | rs16837192 | -0.229 | 0.674 | -0.208 | 1.468 | -0.002 | 0.047 | 0.047 | 0 | 0.08 | 0.1 | 0.29 | 0.37 | 0.12 | 0.22 |
| 2 | 137623359 | rs10496760 | 0.26 | 0.4 | -0.262 | 1.013 | 0.019 | -0.013 | -0.013 | 4.927 | 0.11 | 0.03 | 0.05 | 0.18 | 0.07 | 0.11 |
| 2 | 137629019 | rs813570 | 1.231 | 0.48 | -0.277 | 0.985 | 0.023 | 0.005 | 0.005 | 0.8 | 0.76 | 0.69 | 0.59 | 0.32 | 0.63 | 0.57 |
| 2 | 137629273 | rs16837222 | 0.033 | -1.08 | -0.792 | 0.577 | 0.021 | -0.025 | -0.025 | 2.476 | 0.11 | 0.02 | 0.05 | 0.17 | 0.07 | 0.11 |
| 2 | 137632620 | rs1649546 | 0.493 | -1.708 | -0.554 | 0.085 | 0.018 | 0.017 | 0.017 | 0 | 0.59 | 0.35 | 0.54 | 0.2 | 0.7 | 0.48 |
| 2 | 137634079 | rs1652443 | 0.401 | -1.755 | 0.228 | 0.061 | -0.021 | -0.032 | -0.032 | 0 | 0.6 | 0.5 | 0.56 | 0.2 | 0.71 | 0.49 |
| 2 | 137638965 | rs1540422 | -1.003 | -2.184 | 0.495 | 0.494 | -0.031 | -0.002 | -0.002 | 0 | 0.79 | 0.75 | 0.92 | 0.82 | 0.89 | 0.89 |
| 2 | 137641815 | rs2059729 | -0.523 | -2.165 | 0.533 | 0.511 | -0.026 | 0.063 | 0.063 | 0 | 0.21 | 0.39 | 0.37 | 0.62 | 0.19 | 0.41 |
| 2 | 137643190 | rs10194354 | -0.458 | -2.168 | 0.63 | 0.509 | -0.02 | 0.048 | 0.048 | 0 | 0.2 | 0.34 | 0.36 | 0.62 | 0.18 | 0.4 |
| 2 | 137643850 | rs12691900 | -1.579 | -2.279 | 0.666 | 0.488 | -0.009 | -0.004 | -0.004 | 0 | 0.84 | 0.84 | 0.93 | 0.91 | 0.91 | 0.93 |
| 2 | 137644809 | rs16837261 | 0.392 | -2.605 | 0.312 | 0.163 | 0.017 | -0.023 | -0.023 | 4.212 | 0.13 | 0.02 | 0.07 | 0.19 | 0.09 | 0.14 |
| 2 | 137647008 | rs12474381 | 1.045 | -2.615 | 0.33 | 0.185 | -0.007 | 0.018 | 0.018 | 0 | 0.06 | 0 | 0.03 | 0.03 | 0.04 | 0.08 |
| 2 | 137648914 | rs1346735 | 2.04 | -2.884 | 0.201 | -0.013 | -0.018 | -0.14 | -0.14 | 0.073 | 0.43 | 0.45 | 0.49 | 0.03 | 0.6 | 0.34 |
| 2 | 137649203 | rs7588481 | -0.292 | -2.945 | 0.111 | 0.603 | 0.023 | -0.065 | -0.065 | 0 | 0.18 | 0.13 | 0.31 | 0.4 | 0.16 | 0.25 |
| 2 | 137651500 | rs550612 | 0.338 | -2.718 | 0.536 | 0.468 | -0.014 | -0.013 | -0.013 | 0 | 0.63 | 0.49 | 0.69 | 0.73 | 0.72 | 0.74 |
| 2 | 137651678 | rs548794 | 0.338 | -2.718 | 0.536 | 0.451 | -0.014 | -0.017 | -0.017 | 0.102 | 0.63 | 0.49 | 0.69 | 0.75 | 0.72 | 0.74 |
| 2 | 137651892 | rs554259 | 0.361 | -2.766 | 0.573 | 0.438 | -0.011 | -0.031 | -0.031 | 1.296 | 0.63 | 0.49 | 0.68 | 0.81 | 0.71 | 0.72 |
| 2 | 137652051 | rs4954446 | 0.595 | -2.741 | 0.519 | 0.45 | -0.003 | -0.014 | -0.014 | 0.766 | 0.15 | 0.02 | 0.08 | 0.19 | 0.13 | 0.18 |
| 2 | 137652747 | rs524958 | 2.306 | -2.806 | 0.568 | 0.434 | -0.008 | 0.057 | 0.057 | 0 | 0.38 | 0.25 | 0.34 | 0.16 | 0.51 | 0.3 |
| 2 | 137652942 | rs17722726 | 2.572 | -2.803 | 0.547 | 0.411 | -0.043 | 0.142 | 0.142 | 0 | 0.35 | 0.12 | 0.32 | 0.03 | 0.5 | 0.29 |
| 2 | 137658112 | rs487311 | 0.679 | -2.835 | 0.873 | 0.415 | -0.031 | -0.018 | -0.018 | 0 | 0.61 | 0.52 | 0.58 | 0.29 | 0.74 | 0.56 |
| 2 | 137659635 | rs543309 | 2.026 | -2.694 | 0.801 | 0.364 | -0.005 | 0.112 | 0.112 | 0 | 0.4 | 0.21 | 0.34 | 0.12 | 0.52 | 0.33 |
| 2 | 137668190 | rs1835313 | 2.09 | -2.512 | 1.011 | 0.741 | -0.013 | 0.159 | 0.159 | 0 | 0.39 | 0.16 | 0.32 | 0.06 | 0.52 | 0.32 |
| 2 | 137678829 | rs567375 | 1.626 | -2.539 | 1.403 | 0.77 | 0.019 | 0.039 | 0.039 | 0 | 0.46 | 0.34 | 0.35 | 0.21 | 0.56 | 0.37 |
| 2 | 137679501 | rs765336 | -0.125 | -2.632 | 1.409 | 0.99 | 0.024 | -0.041 | -0.041 | 0.008 | 0.18 | 0.13 | 0.32 | 0.42 | 0.14 | 0.28 |
| 2 | 137685287 | rs1344714 | 1.094 | -2.664 | 1.503 | 0.965 | -0.02 | -0.005 | -0.005 | 2.123 | 0.2 | 0.35 | 0.24 | 0.2 | 0.17 | 0.2 |
| 2 | 137695294 | rs576902 | 2.278 | -2.641 | 1.345 | 0.898 | -0.026 | 0.184 | 0.184 | 0 | 0.39 | 0.13 | 0.32 | 0.07 | 0.52 | 0.32 |
| 2 | 137707025 | rs515093 | 0.812 | -2.627 | 1.403 | 0.927 | -0.008 | -0.002 | -0.002 | 0 | 0.59 | 0.39 | 0.43 | 0.37 | 0.69 | 0.52 |
| 2 | 137708631 | rs17724217 | 0.712 | -2.569 | 1.35 | 0.907 | -0.029 | -0.027 | -0.027 | 0 | 0.13 | 0.01 | 0.07 | 0.17 | 0.13 | 0.15 |
| 2 | 137718197 | rs13431367 | -0.437 | -2.581 | 1.368 | 0.93 | -0.004 | 0.007 | 0.007 | 0 | 0.2 | 0.18 | 0.1 | 0.32 | 0.17 | 0.2 |
| 2 | 137719211 | rs11884290 | 0.448 | -2.541 | 1.402 | 0.908 | -0.005 | 0.088 | 0.088 | 0 | 0.07 | 0.17 | 0.03 | 0.14 | 0.04 | 0.05 |
| 2 | 137722985 | rs10803553 | -1.197 | -2.559 | 1.439 | 0.905 | -0.019 | -0.005 | -0.005 | 0 | 0.4 | 0.6 | 0.57 | 0.63 | 0.31 | 0.48 |
| 2 | 137723026 | rs543086 | -0.594 | -2.525 | 1.398 | 0.909 | 0 | 0.001 | 0.001 | 0.01 | 0.86 | 0.9 | 0.92 | 0.82 | 0.87 | 0.85 |
| 2 | 137740120 | rs574135 | 1.666 | -2.534 | 1.432 | 0.889 | 0.029 | 0.064 | 0.064 | 0 | 0.46 | 0.3 | 0.35 | 0.19 | 0.57 | 0.38 |
| 2 | 137743384 | rs7564933 | -2.254 | -2.542 | 1.401 | 0.846 | 0.003 | 0.104 | 0.104 | 0 | 0.59 | 0.79 | 0.67 | 0.94 | 0.47 | 0.68 |

| | | | | | | | | | | | | | | | | |
|---|---|---|---|---|---|---|---|---|---|---|---|---|---|---|---|---|
| 2 | 137745116 | rs1367128 | -0.583 | -2.601 | 1.305 | 0.988 | 0.024 | -0.059 | -0.059 | 1.071 | 0.25 | 0.14 | 0.37 | 0.57 | 0.22 | 0.34 |
| 2 | 137747337 | rs766687 | 1.669 | -2.143 | 1.108 | 1.078 | 0.012 | -0.007 | -0.007 | 0 | 0.15 | 0.11 | 0.09 | 0.2 | 0.07 | 0.15 |
| 2 | 137756717 | rs1897419 | -1.011 | -1.311 | 1.318 | 1.396 | 0.019 | -0.037 | -0.037 | 1.878 | 0.26 | 0.2 | 0.37 | 0.57 | 0.22 | 0.34 |
| 2 | 137757933 | rs4954368 | -1.272 | -1.093 | 1.066 | 0.877 | 0.014 | -0.049 | -0.049 | 0.174 | 0.26 | 0.4 | 0.27 | 0.2 | 0.28 | 0.28 |
| 2 | 137759233 | rs4426576 | -0.262 | -0.763 | 0.576 | 1.064 | 0.019 | -0.038 | -0.038 | 3.267 | 0.11 | 0.21 | 0.18 | 0.03 | 0.14 | 0.13 |
| 2 | 137764620 | rs13391444 | -0.735 | -0.83 | 0.366 | 1.725 | -0.063 | 0.213 | 0.213 | 0.221 | 0.15 | 0.37 | 0.28 | 0.43 | 0.09 | 0.27 |
| 2 | 137767548 | rs6750400 | 0.758 | 0.41 | -0.442 | 1.265 | -0.022 | -0.001 | -0.001 | 0.097 | 0.75 | 0.86 | 0.74 | 0.74 | 0.66 | 0.79 |
| 2 | 137768012 | rs10928585 | -0.448 | 0.068 | -0.197 | 1.505 | 0 | 0.01 | 0.01 | 0 | 0.05 | 0.06 | 0.12 | 0.14 | 0.07 | 0.06 |
| 2 | 137768702 | rs979976 | 1.232 | 0.248 | 0.186 | 1.38 | -0.002 | 0.001 | 0.001 | 0.333 | 0.67 | 0.35 | 0.63 | 0.64 | 0.7 | 0.75 |
| 2 | 137769420 | rs2116538 | -0.148 | 0.28 | 0.361 | 1.303 | 0.022 | 0.044 | 0.044 | 0.567 | 0.8 | 0.46 | 0.67 | 0.66 | 0.8 | 0.86 |
| 2 | 137771279 | rs4954460 | -0.556 | -0.064 | 0.247 | 1.904 | -0.028 | 0.097 | 0.097 | 0 | 0.08 | 0.21 | 0.04 | 0.24 | 0.02 | 0.14 |
| 2 | 137772007 | rs1347033 | 2.322 | -0.154 | 0.155 | 1.85 | -0.049 | -0.104 | -0.104 | 0 | 0.59 | 0.04 | 0.55 | 0.4 | 0.67 | 0.61 |
| 2 | 137785783 | rs1368056 | -0.062 | -0.139 | 0.731 | 1.811 | -0.004 | 0.051 | 0.051 | 0 | 0.06 | 0.12 | 0.12 | 0.14 | 0.06 | 0.06 |
| 2 | 137787676 | rs7581350 | -0.775 | -0.079 | 0.837 | 1.705 | -0.029 | 0.082 | 0.082 | 0 | 0.08 | 0.2 | 0.04 | 0.23 | 0.03 | 0.13 |
| 2 | 137795717 | rs1432234 | -1.081 | -0.039 | 0.981 | 1.709 | 0.107 | 0.05 | 0.05 | 4.025 | 0.86 | 0.62 | 0.8 | 0.8 | 0.87 | 0.92 |
| 2 | 137799664 | rs1432232 | 1.638 | -0.01 | 0.363 | 1.696 | -0.002 | -0.049 | -0.049 | 0.462 | 0.64 | 0.11 | 0.67 | 0.55 | 0.73 | 0.67 |
| 2 | 137805397 | rs13026531 | -1.303 | 0.591 | 1.41 | 2.041 | 0.025 | 0.052 | 0.052 | 0 | 0.15 | 0.3 | 0.23 | 0.24 | 0.16 | 0.15 |
| 2 | 137808068 | rs16837950 | -1.756 | 0.597 | 1.571 | 2.044 | 0.12 | 0.098 | 0.098 | 0 | 0.06 | 0.32 | 0.15 | 0.07 | 0.06 | 0.03 |
| 2 | 137808507 | rs1432237 | -1.843 | 0.742 | 2.046 | 2.661 | 0.018 | 0.17 | 0.17 | 0 | 0.18 | 0.39 | 0.25 | 0.48 | 0.21 | 0.25 |
| 2 | 137811629 | rs11888486 | -0.786 | 0.738 | 2.045 | 2.564 | 0.008 | -0.049 | -0.049 | 0.009 | 0.12 | 0.04 | 0.11 | 0.22 | 0.11 | 0.13 |
| 2 | 137811955 | rs9784168 | -1.592 | 0.987 | 2.345 | 2.347 | 0.135 | 0.025 | 0.025 | 0 | 0.06 | 0.32 | 0.13 | 0.02 | 0.06 | 0.03 |
| 2 | 137846562 | rs6430676 | -1.753 | 0.949 | 1.973 | 2.423 | 0.167 | 0.482 | 0.482 | 0.055 | 0.07 | 0.47 | 0.15 | 0.29 | 0.1 | 0.14 |
| 2 | 137852698 | rs7561482 | -2.808 | 0.656 | 1.969 | 2.475 | 0.039 | 0.216 | 0.216 | 2.116 | 0.14 | 0.58 | 0.31 | 0.4 | 0.12 | 0.15 |
| 2 | 137854851 | rs1432229 | -1.775 | 0.649 | 1.542 | 1.924 | 0.014 | 0.032 | 0.032 | 0 | 0.07 | 0.22 | 0.1 | 0.1 | 0.06 | 0.07 |
| 2 | 137865422 | rs10496767 | 1.932 | 1.23 | 1.34 | 1.835 | 0.043 | 0.011 | 0.011 | 2.403 | 0.8 | 0.39 | 0.7 | 0.78 | 0.78 | 0.8 |
| 2 | 137866726 | rs4954466 | 2.295 | 0.846 | 1.316 | 2.112 | -0.004 | 0.005 | 0.005 | 5.517 | 0.84 | 0.24 | 0.73 | 0.83 | 0.89 | 0.88 |
| 2 | 137868740 | rs1432220 | 0.159 | 0.102 | 0.678 | 1.743 | -0.014 | -0.008 | -0.008 | 0 | 0.63 | 0.46 | 0.64 | 0.63 | 0.74 | 0.53 |
| 2 | 137868769 | rs1368061 | 0.708 | 0.165 | 0.533 | 1.686 | 0.03 | 0.034 | 0.034 | 1.081 | 0.55 | 0.29 | 0.56 | 0.56 | 0.66 | 0.47 |
| 2 | 137868836 | rs1432219 | 1.31 | -0.161 | 0.3 | 1.78 | 0.002 | -0.035 | -0.035 | 0 | 0.46 | 0.11 | 0.45 | 0.41 | 0.6 | 0.34 |
| 2 | 137870582 | rs974528 | 0.676 | -0.09 | 0.265 | 1.521 | 0.028 | 0.039 | 0.039 | 1.105 | 0.55 | 0.3 | 0.56 | 0.59 | 0.67 | 0.47 |
| 2 | 137872961 | rs745907 | -1.301 | -0.159 | 0.417 | 1.262 | 0.008 | -0.042 | -0.042 | 0.064 | 0.19 | 0.31 | 0.19 | 0.17 | 0.22 | 0.11 |
| 2 | 137872992 | rs745908 | 1.054 | -0.015 | -0.402 | 1.395 | -0.002 | 0.014 | 0.014 | 1.016 | 0.46 | 0.22 | 0.46 | 0.52 | 0.58 | 0.4 |
| 2 | 137874930 | rs10490742 | -0.351 | -0.021 | -0.497 | 1.361 | -0.018 | 0.01 | 0.01 | 0 | 0.13 | 0.08 | 0.16 | 0.15 | 0.21 | 0.1 |
| 2 | 137875875 | rs1469622 | 1.028 | -0.013 | -0.379 | 1.395 | 0.003 | 0.016 | 0.016 | 0.96 | 0.46 | 0.24 | 0.44 | 0.53 | 0.58 | 0.4 |
| 2 | 137875955 | rs1469621 | 1.472 | -0.281 | -0.878 | 0.979 | -0.13 | 0.097 | 0.097 | 2.916 | 0.34 | 0.04 | 0.29 | 0.13 | 0.43 | 0.25 |
| 2 | 137879819 | rs1347030 | -0.654 | -0.446 | -0.559 | 0.7 | -0.013 | -0.011 | -0.011 | 0 | 0.57 | 0.45 | 0.65 | 0.73 | 0.78 | 0.51 |
| 2 | 137886196 | rs10803554 | 0.249 | -0.45 | -0.953 | 0.727 | -0.002 | 0.022 | 0.022 | 0.639 | 0.43 | 0.29 | 0.49 | 0.56 | 0.57 | 0.42 |
| 2 | 137896264 | rs1432205 | 0.539 | -0.345 | -1.202 | 0.769 | -0.026 | -0.027 | -0.027 | 0 | 0.39 | 0.25 | 0.42 | 0.5 | 0.55 | 0.37 |
| 2 | 137898753 | rs984295 | -0.132 | -0.667 | -0.935 | 0.531 | 0.024 | 0.023 | 0.023 | 0.367 | 0.56 | 0.4 | 0.64 | 0.71 | 0.77 | 0.51 |
| 2 | 137901019 | rs16838223 | -0.176 | -0.659 | -1.159 | 0.428 | -0.01 | 0 | 0 | 0.526 | 0.46 | 0.52 | 0.41 | 0.52 | 0.28 | 0.32 |
| 2 | 137901254 | rs892874 | -0.233 | -0.411 | -1.204 | 0.029 | -0.025 | 0.05 | 0.05 | 0.138 | 0.63 | 0.77 | 0.55 | 0.72 | 0.43 | 0.62 |
| 2 | 137903383 | rs10490738 | 0.698 | -0.303 | -1.373 | 0.206 | -0.026 | -0.138 | -0.138 | 0.45 | 0.22 | 0.03 | 0.21 | 0.4 | 0.2 | 0.47 |
| 2 | 137911174 | rs1432227 | -0.461 | -0.418 | -0.08 | 0 | -0.001 | 0.035 | 0.035 | 3.06 | 0.75 | 0.68 | 0.67 | 0.58 | 0.73 | 0.49 |
| 2 | 137911275 | rs1432226 | 0.49 | -0.455 | -0.399 | -0.491 | -0.085 | 0.117 | 0.117 | 0 | 0.35 | 0.15 | 0.34 | 0.17 | 0.47 | 0.34 |
| 2 | 137915407 | rs10190090 | -0.914 | -0.322 | -0.195 | -0.343 | 0 | 0.002 | 0.002 | 2.427 | 0.76 | 0.75 | 0.7 | 0.6 | 0.77 | 0.52 |
| 2 | 137916325 | rs7573893 | -0.437 | -0.142 | -0.046 | -0.944 | -0.002 | 0 | 0 | 0 | 0.22 | 0.21 | 0.24 | 0.18 | 0.34 | 0.14 |
| 2 | 137916344 | rs17795220 | 0.68 | -0.334 | 0.219 | -0.891 | 0.011 | 0.027 | 0.027 | 4.616 | 0.5 | 0.32 | 0.43 | 0.38 | 0.39 | 0.36 |
| 2 | 137917103 | rs13413874 | 0.144 | -0.38 | 0.171 | -1.11 | -0.054 | 0.154 | 0.154 | 0 | 0.39 | 0.16 | 0.37 | 0.14 | 0.51 | 0.36 |
| 2 | 137921051 | rs6737431 | -0.839 | -0.517 | 0.09 | -1.117 | 0.069 | 0.113 | 0.113 | 0.376 | 0.57 | 0.79 | 0.63 | 0.26 | 0.73 | 0.47 |
| 2 | 137937793 | rs6430686 | 0.526 | -0.175 | 0.476 | -0.887 | 0 | -0.014 | -0.014 | 0.027 | 0.12 | 0.23 | 0.14 | 0.13 | 0.15 | 0.05 |
| 2 | 137939891 | rs4954477 | 0.361 | -0.119 | -0.025 | -0.938 | -0.003 | 0.281 | 0.281 | 1.312 | 0.51 | 0.09 | 0.42 | 0.78 | 0.43 | 0.6 |
| 2 | 137956006 | rs7424822 | 0.817 | 0.383 | 0.762 | -0.584 | 0.078 | 0.042 | 0.042 | 1.273 | 0.92 | 0.61 | 0.81 | 0.91 | 0.92 | 0.97 |
| 2 | 137964683 | rs6735885 | -0.097 | 0.313 | 0.477 | -0.545 | 0.006 | -0.001 | -0.001 | 0 | 0.54 | 0.7 | 0.51 | 0.46 | 0.53 | 0.74 |
| 2 | 137980318 | rs4610107 | 1.089 | 0.159 | 0.308 | -0.686 | -0.041 | 0.079 | 0.079 | 0.057 | 0.25 | 0.15 | 0.24 | 0.07 | 0.31 | 0.28 |
| 2 | 137981047 | rs6722499 | 0.081 | 0.176 | 0.214 | -0.619 | -0.011 | 0.006 | 0.006 | 2.677 | 0.43 | 0.18 | 0.41 | 0.46 | 0.44 | 0.23 |
| 2 | 137981675 | rs2196346 | -0.053 | 0.159 | 0.519 | -0.63 | 0.001 | 0.004 | 0.004 | 1.974 | 0.45 | 0.41 | 0.43 | 0.45 | 0.47 | 0.25 |
| 2 | 137983759 | rs6430699 | -0.096 | 0.129 | 0.582 | -0.656 | 0.003 | 0.008 | 0.008 | 2.801 | 0.46 | 0.57 | 0.44 | 0.46 | 0.45 | 0.24 |
| 2 | 137987412 | rs12466293 | 0.278 | 0.144 | 0.369 | -0.652 | 0.013 | -0.02 | -0.02 | 0 | 0.4 | 0.15 | 0.3 | 0.45 | 0.39 | 0.19 |
| 2 | 137993414 | rs10195563 | -0.085 | 0.237 | -0.989 | -0.45 | 0.017 | 0.007 | 0.007 | 2.948 | 0.44 | 0.62 | 0.41 | 0.45 | 0.39 | 0.23 |
| 2 | 137999060 | rs4954489 | 0.63 | 0.166 | -2.243 | -0.087 | -0.08 | -0.177 | -0.177 | 0.367 | 0.27 | 0.03 | 0.26 | 0.39 | 0.27 | 0.46 |
| 2 | 138001680 | rs4954491 | -0.126 | -0.236 | -1.726 | 0.195 | 0.042 | -0.042 | -0.042 | 0.194 | 0.36 | 0.14 | 0.26 | 0.45 | 0.31 | 0.18 |
| 2 | 138003520 | rs10211290 | 0.313 | -0.144 | -1.575 | 0.161 | 0.065 | -0.034 | -0.034 | 0.605 | 0.58 | 0.23 | 0.61 | 0.45 | 0.65 | 0.78 |
| 2 | 138011809 | rs17346940 | 0.586 | 0.113 | -1.517 | 0.175 | -0.009 | 0.004 | 0.004 | 0 | 0.06 | 0.03 | 0.08 | 0.09 | 0.12 | 0.08 |
| 2 | 138012183 | rs10490734 | -0.021 | -0.434 | -0.834 | 0.128 | -0.004 | -0.001 | -0.001 | 0 | 0.17 | 0.28 | 0.17 | 0.2 | 0.17 | 0.13 |
| 2 | 138015427 | rs1865590 | 1.255 | -0.34 | -1.452 | -0.478 | 0.021 | -0.042 | -0.042 | 0.052 | 0.37 | 0.15 | 0.34 | 0.34 | 0.31 | 0.47 |
| 2 | 138018893 | rs16838484 | 0.777 | -0.517 | -1.749 | -0.627 | 0.041 | -0.119 | -0.119 | 0.479 | 0.19 | 0.02 | 0.2 | 0.42 | 0.11 | 0.12 |
| 2 | 138022356 | rs11680942 | 1.431 | -0.384 | -1.714 | -0.618 | -0.009 | 0.007 | 0.007 | 0 | 0.06 | 0.02 | 0.08 | 0.07 | 0.13 | 0.08 |
| 2 | 138030234 | rs12622896 | 3.24 | -0.483 | -1.657 | -0.613 | -0.009 | -0.001 | -0.001 | 0 | 0.07 | 0.08 | 0.02 | 0.06 | 0.02 | 0.05 |
| 2 | 138038158 | rs11895622 | 0.211 | -0.195 | -1.517 | -0.503 | 0.017 | -0.027 | -0.027 | 0.821 | 0.23 | 0.18 | 0.23 | 0.48 | 0.14 | 0.12 |

| | | | | | | | | | | | | | | | | |
|---|---|---|---|---|---|---|---|---|---|---|---|---|---|---|---|---|
| 2 | 138045048 | rs2602347 | -0.327 | -0.34 | -0.824 | -0.866 | 0.069 | -0.122 | -0.122 | 0 | 0.73 | 0.57 | 0.67 | 0.95 | 0.57 | 0.7 |
| 2 | 138048340 | rs11688662 | 1.372 | -0.294 | -1.159 | -0.827 | -0.009 | 0.005 | 0.005 | 0 | 0.06 | 0.02 | 0.08 | 0.09 | 0.12 | 0.08 |
| 2 | 138049087 | rs6728076 | 1.046 | -0.364 | -0.709 | -0.746 | 0.003 | 0 | 0 | 0 | 0.51 | 0.5 | 0.46 | 0.48 | 0.45 | 0.58 |
| 2 | 138050346 | rs1202281 | 0.217 | 0.483 | -0.362 | -1.255 | 0.019 | -0.059 | -0.059 | 0 | 0.73 | 0.67 | 0.67 | 0.95 | 0.56 | 0.7 |
| 2 | 138073500 | rs6749316 | -0.421 | 0.445 | -0.176 | -1.206 | 0.029 | -0.076 | -0.076 | 0 | 0.27 | 0.37 | 0.31 | 0.06 | 0.4 | 0.26 |
| 2 | 138077603 | rs13415724 | -0.456 | 0.273 | -0.364 | -0.887 | -0.021 | 0.044 | 0.044 | 1.615 | 0.24 | 0.12 | 0.15 | 0.15 | 0.26 | 0.34 |
| 2 | 138078236 | rs10198627 | -0.537 | 0.194 | -0.006 | -0.91 | -0.002 | 0.015 | 0.015 | 1.689 | 0.24 | 0.22 | 0.16 | 0.15 | 0.26 | 0.33 |
| 2 | 138080695 | rs1453301 | 1.232 | -0.384 | 0.245 | -1.1 | 0.016 | -0.008 | -0.008 | 0.127 | 0.22 | 0.09 | 0.17 | 0.19 | 0.16 | 0.33 |
| 2 | 138084741 | rs6715610 | 0.888 | -0.282 | 1.168 | -1.344 | 0.024 | -0.053 | -0.053 | 0 | 0.11 | 0.17 | 0.09 | 0 | 0.17 | 0.03 |
| 2 | 138090607 | rs2123613 | 1.11 | -1.117 | 1.965 | -0.844 | 0.013 | 0.04 | 0.04 | 0.537 | 0.64 | 0.24 | 0.66 | 0.79 | 0.68 | 0.58 |
| 2 | 138091057 | rs6740175 | -0.987 | -1.1 | 2.005 | -0.839 | -0.002 | -0.035 | -0.035 | 0 | 0.3 | 0.42 | 0.2 | 0.15 | 0.31 | 0.33 |
| 2 | 138095149 | rs1597937 | 0 | -1.039 | 1.856 | -0.852 | 0.042 | 0.033 | 0.033 | 2.003 | 0.21 | 0.09 | 0.1 | 0.13 | 0.15 | 0.28 |
| 2 | 138107683 | rs13395172 | -0.647 | -1.005 | 1.876 | -0.805 | -0.042 | 0.026 | 0.026 | 1.542 | 0.26 | 0.1 | 0.22 | 0.15 | 0.31 | 0.34 |
| 2 | 138116294 | rs1453294 | 1.332 | -0.954 | 2.159 | -0.743 | 0.072 | 0.092 | 0.092 | 2.65 | 0.94 | 0.53 | 0.86 | 0.94 | 0.98 | 0.91 |
| 2 | 138116333 | rs1453293 | 1.456 | -1.011 | 2.056 | -0.804 | 0.011 | 0.065 | 0.065 | 0.99 | 0.66 | 0.15 | 0.62 | 0.79 | 0.68 | 0.58 |
| 2 | 138135771 | rs10180601 | -0.347 | -0.946 | 1.51 | -0.68 | 0.033 | 0.01 | 0.01 | 2.546 | 0.23 | 0.09 | 0.17 | 0.15 | 0.18 | 0.29 |
| 2 | 138166061 | rs13428514 | -2.147 | -0.973 | 1.99 | -0.768 | -0.014 | 0.032 | 0.032 | 1.548 | 0.11 | 0.49 | 0.17 | 0.09 | 0.06 | 0.11 |
| 2 | 138166938 | rs6741735 | -1.518 | -0.957 | 1.997 | -0.825 | 0.008 | 0.03 | 0.03 | 0.831 | 0.33 | 0.76 | 0.38 | 0.22 | 0.32 | 0.41 |
| 2 | 138167717 | rs4954505 | 1.404 | -0.978 | 1.993 | -0.835 | 0.013 | 0.051 | 0.051 | 1.695 | 0.66 | 0.19 | 0.62 | 0.78 | 0.68 | 0.59 |
| 2 | 138170561 | rs9287473 | -0.608 | -0.933 | 2.02 | -0.817 | 0.006 | -0.064 | -0.064 | 0 | 0.25 | 0.46 | 0.23 | 0.13 | 0.26 | 0.3 |
| 2 | 138196578 | rs9989745 | -1.115 | -0.95 | 2.076 | -0.815 | 0.001 | -0.052 | -0.052 | 0 | 0.29 | 0.53 | 0.26 | 0.15 | 0.3 | 0.32 |
| 2 | 138197269 | rs10496771 | 0.024 | -0.928 | 1.993 | -0.804 | 0 | -0.002 | -0.002 | 0 | 0.21 | 0.21 | 0.2 | 0.13 | 0.26 | 0.34 |
| 2 | 138201585 | rs10190462 | 1.843 | -0.965 | 1.909 | -0.938 | -0.002 | 0.045 | 0.045 | 1.182 | 0.9 | 0.55 | 0.83 | 0.91 | 0.94 | 0.88 |
| 2 | 138207269 | rs719790 | 0.065 | -0.915 | 1.691 | -0.935 | -0.005 | 0.012 | 0.012 | 1.653 | 0.21 | 0.16 | 0.19 | 0.11 | 0.26 | 0.29 |
| 2 | 138208631 | rs1453302 | 0.722 | -1.057 | 1.819 | -1.019 | -0.001 | -0.017 | -0.017 | 0 | 0.68 | 0.64 | 0.76 | 0.86 | 0.68 | 0.68 |
| 2 | 138211966 | rs10182199 | 1.093 | -0.995 | 1.454 | -0.987 | 0.065 | 0.11 | 0.11 | 4.476 | 0.94 | 0.57 | 0.85 | 0.92 | 0.98 | 0.87 |
| 2 | 138219404 | rs1453290 | 1.219 | -1.026 | 0.767 | -0.982 | 0.028 | 0.104 | 0.104 | 1.162 | 0.61 | 0.15 | 0.61 | 0.79 | 0.66 | 0.59 |
| 2 | 138225792 | rs3943484 | 1.53 | -0.656 | 1.447 | -0.706 | -0.02 | 0.02 | 0.02 | 11.695 | 0.92 | 0.38 | 0.84 | 0.93 | 0.98 | 0.9 |
| 2 | 138240046 | rs4954516 | -0.178 | -0.898 | 1.099 | -0.853 | -0.009 | -0.007 | -0.007 | 0 | 0.7 | 0.36 | 0.66 | 0.76 | 0.82 | 0.58 |
| 2 | 138245997 | rs4954378 | 1.327 | -0.977 | 0.805 | -0.903 | -0.031 | 0.056 | 0.056 | 0.205 | 0.44 | 0.17 | 0.44 | 0.64 | 0.5 | 0.28 |
| 2 | 138262443 | rs3748877 | 0.423 | -0.964 | 0.831 | -0.902 | -0.009 | 0.01 | 0.01 | 0.139 | 0.16 | 0.14 | 0.18 | 0.1 | 0.23 | 0.3 |
| 2 | 138266571 | rs10928615 | 0.966 | -1.014 | 1.046 | -0.889 | 0 | -0.003 | -0.003 | 0.075 | 0.5 | 0.45 | 0.58 | 0.7 | 0.54 | 0.39 |
| 2 | 138283751 | rs10196011 | -1.576 | -0.984 | 0.403 | -0.869 | -0.035 | 0.044 | 0.044 | 0.009 | 0.56 | 0.83 | 0.56 | 0.37 | 0.5 | 0.72 |
| 2 | 138289249 | rs10496773 | 0.846 | -1.029 | 0.163 | -0.828 | 0.02 | 0.023 | 0.023 | 0 | 0.23 | 0.13 | 0.18 | 0.15 | 0.16 | 0.28 |
| 2 | 138289558 | rs10496774 | 0.612 | -1.005 | 0.053 | -0.817 | 0.004 | 0.005 | 0.005 | 0 | 0.09 | 0.09 | 0.05 | 0.03 | 0.07 | 0.03 |
| 2 | 138290299 | rs9653194 | 0.101 | -0.828 | -0.027 | -0.718 | -0.012 | 0.016 | 0.016 | 0 | 0.17 | 0.13 | 0.16 | 0.11 | 0.22 | 0.3 |
| 2 | 138310150 | rs1448917 | -0.624 | -0.771 | -0.007 | -0.663 | 0.005 | 0.061 | 0.061 | 1.526 | 0.4 | 0.23 | 0.34 | 0.26 | 0.38 | 0.58 |
| 2 | 138319054 | rs1562661 | -0.687 | -0.423 | 0.577 | -0.572 | 0.019 | -0.068 | -0.068 | 0 | 0.22 | 0.51 | 0.19 | 0.12 | 0.24 | 0.31 |
| 2 | 138323267 | rs11900859 | -0.278 | -0.942 | 0.575 | -0.884 | 0.032 | 0.024 | 0.024 | 0 | 0.34 | 0.22 | 0.34 | 0.25 | 0.24 | 0.37 |
| 2 | 138328342 | rs3919758 | -0.08 | -0.942 | 0.614 | -0.884 | 0.038 | 0.027 | 0.027 | 0 | 0.33 | 0.17 | 0.33 | 0.25 | 0.24 | 0.37 |
| 2 | 138332365 | rs4954379 | -1.47 | -0.913 | 0.586 | -0.903 | -0.004 | 0.057 | 0.057 | 0 | 0.54 | 0.65 | 0.47 | 0.29 | 0.48 | 0.68 |
| 2 | 138337790 | rs13430689 | -0.124 | -1.051 | 0.802 | -1.064 | 0 | -0.041 | -0.041 | 0.008 | 0.08 | 0.24 | 0.07 | 0.01 | 0.07 | 0.18 |
| 2 | 138353736 | rs352188 | -1.635 | -1.191 | 0.204 | -1.138 | -0.03 | 0.193 | 0.193 | 0 | 0.53 | 0.94 | 0.44 | 0.23 | 0.41 | 0.63 |
| 2 | 138355734 | rs1375357 | -0.69 | -1.089 | 0.479 | -1.14 | -0.005 | -0.024 | -0.024 | 0 | 0.36 | 0.38 | 0.25 | 0.12 | 0.23 | 0.41 |
| 2 | 138370085 | rs352192 | -0.437 | -0.925 | 0.081 | -1.064 | -0.013 | 0.025 | 0.025 | 0.322 | 0.19 | 0.11 | 0.12 | 0.05 | 0.24 | 0.31 |
| 2 | 138374461 | rs10496782 | 0.849 | -1.073 | -0.187 | -1.132 | 0.014 | 0.053 | 0.053 | 0 | 0.19 | 0.06 | 0.13 | 0.07 | 0.13 | 0.21 |
| 2 | 138376210 | rs16839162 | 0.741 | -0.981 | -0.182 | -1.002 | 0.026 | 0.002 | 0.002 | 0.124 | 0.07 | 0 | 0.03 | 0.03 | 0.02 | 0.02 |
| 2 | 138379559 | rs6730607 | -1.258 | -0.981 | 0.146 | -0.73 | 0.006 | 0.014 | 0.014 | 0 | 0.32 | 0.7 | 0.29 | 0.18 | 0.29 | 0.43 |
| 2 | 138386595 | rs1448919 | -1.258 | -0.981 | 0.146 | -0.932 | 0.006 | 0.014 | 0.014 | 0 | 0.32 | 0.7 | 0.29 | 0.12 | 0.29 | 0.43 |
| 2 | 138398906 | rs881639 | -1.087 | -0.371 | -0.893 | -0.921 | 0.004 | 0.067 | 0.067 | 0.195 | 0.54 | 0.3 | 0.39 | 0.23 | 0.51 | 0.54 |
| 2 | 138399937 | rs3828170 | 0.588 | -0.896 | -1.119 | -1.149 | 0.004 | 0.003 | 0.003 | 1.415 | 0.18 | 0.12 | 0.16 | 0.07 | 0.1 | 0.15 |
| 2 | 138400685 | rs10178306 | -1.266 | -0.611 | -1.245 | -1.167 | 0.017 | 0.086 | 0.086 | 0.275 | 0.53 | 0.28 | 0.34 | 0.22 | 0.39 | 0.51 |
| 2 | 138401390 | rs5010976 | 0.476 | -0.563 | -1.478 | -0.736 | 0 | 0.045 | 0.045 | 2.177 | 0.17 | 0.01 | 0.15 | 0.07 | 0.1 | 0.15 |
| 2 | 138412985 | rs10496784 | 0.42 | -0.545 | -1.359 | -0.727 | 0.015 | 0.057 | 0.057 | 0.639 | 0.15 | 0.03 | 0.06 | 0.05 | 0.11 | 0.11 |
| 2 | 138413585 | rs7591709 | 0.279 | -0.61 | -1.036 | -0.727 | 0.005 | 0 | 0 | 0 | 0.16 | 0.14 | 0.07 | 0.05 | 0.12 | 0.14 |
| 2 | 138418155 | rs16839281 | -0.784 | -0.713 | -1.086 | -0.106 | 0 | -0.001 | -0.001 | 0 | 0.31 | 0.3 | 0.17 | 0.3 | 0.28 | 0.24 |
| 2 | 138418875 | rs12994131 | 0.457 | -0.627 | -1.896 | -0.233 | -0.06 | 0.025 | 0.025 | 0 | 0.27 | 0.03 | 0.27 | 0.21 | 0.27 | 0.31 |
| 2 | 138420996 | rs10206850 | -1.265 | -1.314 | -1.19 | -0.097 | -0.001 | -0.001 | -0.001 | 0.001 | 0.66 | 0.66 | 0.54 | 0.55 | 0.56 | 0.59 |
| 2 | 138425728 | rs998288 | -0.238 | -0.753 | -1.458 | -0.257 | -0.092 | 0.093 | 0.093 | 0.006 | 0.36 | 0.15 | 0.35 | 0.23 | 0.42 | 0.39 |
| 2 | 138425911 | rs1446222 | -0.365 | -0.757 | -1.457 | -0.296 | -0.088 | 0.121 | 0.121 | 0 | 0.38 | 0.15 | 0.36 | 0.2 | 0.42 | 0.39 |
| 2 | 138427264 | rs2044433 | 0.583 | -0.949 | -0.526 | 0.169 | 0.028 | -0.006 | -0.006 | 0 | 0.57 | 0.37 | 0.67 | 0.62 | 0.69 | 0.79 |
| 2 | 138435014 | rs1048821 | 1.002 | -0.725 | 0.305 | -0.244 | 0.002 | -0.025 | -0.025 | 0 | 0.66 | 0.39 | 0.67 | 0.89 | 0.65 | 0.68 |
| 2 | 138436339 | rs11686818 | 1.32 | -0.772 | 0.294 | -0.284 | 0.023 | -0.009 | -0.009 | 0 | 0.86 | 0.41 | 0.87 | 0.91 | 0.9 | 0.81 |
| 2 | 138440691 | rs2218539 | 1.451 | -0.771 | 0.945 | -0.304 | 0.006 | -0.07 | -0.07 | 0 | 0.65 | 0.47 | 0.67 | 0.89 | 0.66 | 0.68 |
| 2 | 138440922 | rs2197551 | 0.947 | -0.122 | 1.796 | 0.295 | 0.061 | 0.034 | 0.034 | 0.305 | 0.89 | 0.61 | 0.89 | 0.91 | 0.91 | 0.81 |
| 2 | 138441476 | rs12993553 | 1.29 | -0.126 | 1.615 | 0.3 | 0.066 | 0.022 | 0.022 | 1.265 | 0.87 | 0.5 | 0.87 | 0.91 | 0.9 | 0.81 |
| 2 | 138446791 | rs13002302 | -1.539 | -1.096 | 1.502 | 0.398 | 0.054 | 0.051 | 0.051 | 0.189 | 0.05 | 0.28 | 0.05 | 0.02 | 0.02 | 0.02 |
| 2 | 138447427 | rs1348739 | 0.672 | -1.16 | 1.665 | 0.42 | -0.045 | 0.022 | 0.022 | 0.27 | 0.91 | 0.78 | 0.92 | 0.91 | 0.99 | 0.89 |
| 2 | 138447831 | rs1446232 | 1.2 | -1.355 | 1.286 | 0.437 | 0.051 | -0.011 | -0.011 | 0 | 0.64 | 0.27 | 0.58 | 0.45 | 0.79 | 0.69 |

| | | | | | | | | | | | | | | | | |
|---|---|---|---|---|---|---|---|---|---|---|---|---|---|---|---|---|
| 2 | 138448951 | rs12052327 | 0.974 | -1.121 | 0.946 | 0.559 | -0.058 | 0.037 | 0.037 | 0 | 0.69 | 0.55 | 0.63 | 0.47 | 0.8 | 0.71 |
| 2 | 138451457 | rs13388919 | 1.098 | -0.876 | 0.405 | 0.555 | 0.064 | -0.037 | -0.037 | 0 | 0.64 | 0.16 | 0.58 | 0.42 | 0.76 | 0.74 |
| 2 | 138451475 | rs13415092 | 0.735 | -0.929 | 1.37 | 0.836 | -0.034 | 0.351 | 0.351 | 0 | 0.89 | 0.62 | 0.73 | 0.53 | 0.97 | 0.88 |
| 2 | 138465850 | rs2121234 | 0.09 | -1.003 | 1.728 | 0.721 | -0.046 | 0.012 | 0.012 | 0 | 0.13 | 0.3 | 0.05 | 0.1 | 0.05 | 0.05 |
| 2 | 138471717 | rs11680805 | 1.391 | -0.988 | 1.615 | 0.789 | 0.06 | -0.041 | -0.041 | 0 | 0.65 | 0.16 | 0.58 | 0.42 | 0.76 | 0.74 |
| 2 | 138483702 | rs7600086 | -0.975 | -0.632 | 1.706 | 0.899 | -0.017 | -0.058 | -0.058 | 0 | 0.23 | 0.42 | 0.15 | 0.12 | 0.21 | 0.13 |
| 2 | 138492290 | rs7592123 | -0.816 | -0.631 | 1.676 | 0.888 | -0.011 | -0.047 | -0.047 | 0.808 | 0.22 | 0.35 | 0.15 | 0.11 | 0.21 | 0.13 |
| 2 | 138495892 | rs10207996 | 0.847 | -0.619 | 1.424 | 0.894 | -0.021 | -0.066 | -0.066 | 0 | 0.76 | 0.51 | 0.84 | 0.88 | 0.79 | 0.87 |
| 2 | 138522393 | rs1374154 | 0.026 | 0.191 | 0.268 | 0.496 | -0.041 | 0.008 | 0.008 | 0.009 | 0.07 | 0.01 | 0.09 | 0.09 | 0.16 | 0.04 |
| 2 | 138537633 | rs1374153 | 0.808 | -0.54 | 0.924 | 0.623 | -0.074 | 0.104 | 0.104 | 0 | 0.76 | 0.62 | 0.7 | 0.45 | 0.91 | 0.76 |
| 2 | 138563356 | rs10181206 | -0.169 | 0.439 | 0.834 | 0.88 | -0.089 | 0.034 | 0.034 | 0.833 | 0.13 | 0.01 | 0.16 | 0.08 | 0.32 | 0.14 |
| 2 | 138566028 | rs974033 | 1.057 | -0.741 | 1.687 | 0.996 | 0.024 | -0.031 | -0.031 | 0.707 | 0.69 | 0.27 | 0.6 | 0.45 | 0.78 | 0.69 |
| 2 | 138591741 | rs13028477 | -0.576 | -0.589 | 1.841 | 1.097 | 0 | 0.044 | 0.044 | 0 | 0.17 | 0.2 | 0.34 | 0.42 | 0.19 | 0.25 |
| 2 | 138600801 | rs6730857 | -0.321 | -0.747 | 1.402 | 0.967 | -0.062 | -0.009 | -0.009 | 0 | 0.85 | 0.71 | 0.93 | 0.88 | 0.95 | 0.95 |
| 2 | 138605815 | rs6759022 | -0.685 | -0.692 | 1.383 | 0.998 | 0.013 | 0.161 | 0.161 | 0 | 0.18 | 0.42 | 0.35 | 0.43 | 0.19 | 0.25 |
| 2 | 138612416 | rs1973076 | -0.077 | -0.843 | 1.625 | 0.491 | -0.054 | -0.007 | -0.007 | 0 | 0.86 | 0.78 | 0.93 | 0.88 | 0.95 | 0.95 |
| 2 | 138619715 | rs6430748 | 0.68 | -0.839 | 1.054 | 0.422 | -0.098 | 0.061 | 0.061 | 0 | 0.8 | 0.48 | 0.72 | 0.65 | 0.95 | 0.82 |
| 2 | 138624319 | rs1456005 | 0.342 | 0.015 | 1.865 | 0.243 | 0.005 | 0.047 | 0.047 | 0 | 0.87 | 0.84 | 0.7 | 0.69 | 0.84 | 0.83 |
| 2 | 138626889 | rs10928633 | 0.923 | -0.019 | 1.21 | -0.338 | 0.003 | 0.118 | 0.118 | 0 | 0.5 | 0.17 | 0.35 | 0.62 | 0.43 | 0.6 |
| 2 | 138631472 | rs7591286 | 1.203 | -0.161 | 1.08 | -0.493 | 0.018 | 0.044 | 0.044 | 2.274 | 0.1 | 0.02 | 0.05 | 0.01 | 0.08 | 0.01 |
| 2 | 138636040 | rs10496793 | 0.663 | -0.117 | 0.87 | -0.478 | -0.028 | 0.03 | 0.03 | 0 | 0.23 | 0.15 | 0.27 | 0.06 | 0.33 | 0.23 |
| 2 | 138637196 | rs17642263 | 0.268 | -0.106 | 0.866 | -0.478 | 0.029 | -0.054 | -0.054 | 0.026 | 0.26 | 0.35 | 0.32 | 0.07 | 0.41 | 0.23 |
| 2 | 138638997 | rs11691636 | 0.4 | -0.124 | 0.699 | -0.495 | 0.051 | -0.105 | -0.105 | 0.206 | 0.25 | 0.44 | 0.28 | 0.06 | 0.33 | 0.23 |
| 2 | 138644649 | rs6430755 | 0.789 | -0.241 | 0.663 | -0.466 | -0.026 | 0.011 | 0.011 | 0 | 0.76 | 0.57 | 0.63 | 0.67 | 0.76 | 0.83 |
| 2 | 138645684 | rs6738271 | 1.235 | -0.186 | 0.515 | -0.502 | -0.017 | 0.12 | 0.12 | 0 | 0.5 | 0.09 | 0.33 | 0.62 | 0.42 | 0.6 |
| 2 | 138650882 | rs7590072 | 0.862 | -0.149 | 0.622 | -0.398 | -0.027 | 0.01 | 0.01 | 0 | 0.76 | 0.51 | 0.63 | 0.67 | 0.76 | 0.83 |
| 2 | 138650963 | rs1979243 | 0.763 | -0.145 | 0.664 | -0.396 | -0.024 | 0.014 | 0.014 | 0 | 0.76 | 0.59 | 0.63 | 0.67 | 0.76 | 0.83 |
| 2 | 138656057 | rs1840646 | 0.786 | -0.125 | 0.648 | -0.386 | -0.027 | 0.014 | 0.014 | 0 | 0.76 | 0.54 | 0.63 | 0.67 | 0.76 | 0.83 |
| 2 | 138678381 | rs1840656 | 0.862 | -0.135 | 0.546 | -0.391 | -0.031 | 0.009 | 0.009 | 0 | 0.75 | 0.55 | 0.62 | 0.67 | 0.76 | 0.82 |
| 2 | 138683670 | rs4954547 | 0.508 | 0.294 | 0.377 | -0.545 | 0.109 | 0.069 | 0.069 | 3.393 | 0.95 | 0.48 | 0.92 | 0.99 | 0.92 | 0.99 |
| 2 | 138685102 | rs1840641 | 0.085 | 0.251 | -0.714 | -0.516 | -0.011 | 0.024 | 0.024 | 0.421 | 0.62 | 0.34 | 0.63 | 0.93 | 0.58 | 0.76 |
| 2 | 138699716 | rs6430763 | -0.091 | -0.356 | -0.905 | 0.04 | 0.036 | 0.019 | 0.019 | 0 | 0.51 | 0.42 | 0.38 | 0.41 | 0.4 | 0.57 |
| 2 | 138700958 | rs3100714 | -0.153 | -0.363 | -0.909 | 0.038 | 0.036 | 0.02 | 0.02 | 0 | 0.49 | 0.58 | 0.62 | 0.59 | 0.6 | 0.43 |
| 2 | 138701957 | rs10928638 | 0.18 | -1.076 | -1.188 | -0.087 | 0.007 | -0.007 | -0.007 | 0 | 0.13 | 0.1 | 0.09 | 0.18 | 0.06 | 0.25 |
| 2 | 138702469 | rs3100713 | -0.577 | -1.582 | -1.575 | -0.416 | -0.009 | -0.005 | -0.005 | 0.28 | 0.81 | 0.7 | 0.78 | 0.81 | 0.79 | 0.85 |
| 2 | 138703598 | rs16839937 | -0.527 | -1.592 | -1.58 | -0.42 | 0.067 | 0.032 | 0.032 | 0 | 0.32 | 0.12 | 0.16 | 0.22 | 0.19 | 0.42 |
| 2 | 138703723 | rs13023679 | 0.634 | -1.647 | -1.578 | -0.443 | -0.01 | -0.02 | -0.02 | 0 | 0.17 | 0.03 | 0.19 | 0.19 | 0.21 | 0.15 |
| 2 | 138704004 | rs3113209 | 0.771 | -1.73 | -1.617 | -0.489 | -0.015 | 0.003 | 0.003 | 0 | 0.66 | 0.6 | 0.69 | 0.63 | 0.73 | 0.6 |
| 2 | 138706592 | rs3113227 | 0.41 | -1.646 | -0.585 | -0.665 | -0.006 | -0.024 | -0.024 | 0 | 0.18 | 0.17 | 0.07 | 0.04 | 0.13 | 0.17 |
| 2 | 138708730 | rs3113237 | -0.71 | -1.629 | -0.581 | -0.661 | 0.023 | 0.041 | 0.041 | 0 | 0.35 | 0.2 | 0.26 | 0.23 | 0.34 | 0.32 |
| 2 | 138717595 | rs13432464 | 0.659 | -1.902 | -0.105 | -0.515 | 0.041 | 0.024 | 0.024 | 0 | 0.46 | 0.22 | 0.58 | 0.56 | 0.54 | 0.56 |
| 2 | 138718534 | rs12475378 | 0.895 | -1.799 | -0.434 | -0.49 | -0.013 | -0.026 | -0.026 | 0 | 0.16 | 0.03 | 0.19 | 0.19 | 0.21 | 0.15 |
| 2 | 138720423 | rs6430764 | 0.489 | -1.985 | -0.343 | -0.642 | 0.033 | 0.051 | 0.051 | 0 | 0.48 | 0.36 | 0.65 | 0.64 | 0.55 | 0.56 |
| 2 | 138721597 | rs2071048 | 0.447 | -1.994 | 0.157 | -0.647 | 0.005 | 0.006 | 0.006 | 0 | 0.5 | 0.54 | 0.68 | 0.65 | 0.55 | 0.56 |
| 2 | 138727607 | rs3100702 | -0.613 | -2.022 | 0.332 | -0.668 | 0.013 | 0.02 | 0.02 | 0 | 0.49 | 0.37 | 0.32 | 0.34 | 0.45 | 0.43 |
| 2 | 138729113 | rs1982494 | 0.522 | -2.042 | 0.36 | -0.681 | 0.04 | 0.071 | 0.071 | 0 | 0.48 | 0.33 | 0.66 | 0.68 | 0.55 | 0.57 |
| 2 | 138730349 | rs3100701 | -0.628 | -1.967 | 0.03 | -0.648 | 0.016 | 0.027 | 0.027 | 0 | 0.49 | 0.35 | 0.32 | 0.32 | 0.45 | 0.43 |
| 2 | 138731765 | rs3791244 | 0.749 | -1.693 | -0.296 | -0.676 | -0.017 | 0.004 | 0.004 | 0 | 0.15 | 0.03 | 0.12 | 0.12 | 0.2 | 0.16 |
| 2 | 138743166 | rs3100725 | -0.117 | -1.938 | 0.049 | -0.899 | -0.005 | -0.003 | -0.003 | 0.539 | 0.31 | 0.3 | 0.32 | 0.24 | 0.22 | 0.28 |
| 2 | 138745144 | rs1349992 | 1.267 | -0.995 | -0.295 | -1.352 | -0.043 | -0.011 | -0.011 | 0.657 | 0.31 | 0.15 | 0.37 | 0.6 | 0.38 | 0.37 |
| 2 | 138746039 | rs17583889 | 1.15 | -0.991 | -0.499 | -1.422 | -0.022 | 0.022 | 0.022 | 0 | 0.18 | 0.03 | 0.11 | 0.13 | 0.2 | 0.21 |
| 2 | 138749057 | rs1580111 | -0.293 | -0.917 | -0.178 | -1.385 | 0.014 | 0.018 | 0.018 | 0 | 0.5 | 0.45 | 0.43 | 0.36 | 0.42 | 0.49 |
| 2 | 138763546 | rs2737385 | 0.385 | -0.961 | 0.659 | -1.356 | 0.011 | 0.006 | 0.006 | 0 | 0.29 | 0.24 | 0.31 | 0.26 | 0.21 | 0.33 |
| 2 | 138767527 | rs993891 | -0.091 | -0.976 | 0.694 | -1.362 | 0.017 | 0.012 | 0.012 | 0 | 0.67 | 0.3 | 0.66 | 0.74 | 0.78 | 0.67 |
| 2 | 138769951 | rs4954940 | 0.471 | -0.951 | 0.393 | -1.35 | 0.039 | 0.013 | 0.013 | 0 | 0.28 | 0.16 | 0.31 | 0.26 | 0.21 | 0.33 |
| 2 | 138772694 | rs4245861 | 0.471 | -0.951 | 0.393 | -1.35 | 0.039 | 0.013 | 0.013 | 0 | 0.28 | 0.16 | 0.31 | 0.26 | 0.21 | 0.33 |
| 2 | 138773229 | rs4646333 | 0.471 | -0.951 | 0.393 | -1.35 | 0.039 | 0.013 | 0.013 | 0 | 0.28 | 0.16 | 0.31 | 0.26 | 0.21 | 0.33 |
| 2 | 138773638 | rs1455158 | -0.023 | -0.953 | 0.394 | -1.351 | -0.039 | -0.022 | -0.022 | 0 | 0.32 | 0.58 | 0.33 | 0.26 | 0.22 | 0.33 |
| 2 | 138773752 | rs1455157 | -0.023 | -0.953 | 0.394 | -1.351 | -0.039 | -0.022 | -0.022 | 0.266 | 0.32 | 0.58 | 0.33 | 0.26 | 0.22 | 0.33 |
| 2 | 138776597 | rs16840158 | 0.325 | -0.95 | 0.498 | -1.353 | 0.015 | 0.008 | 0.008 | 0 | 0.29 | 0.24 | 0.3 | 0.26 | 0.22 | 0.33 |
| 2 | 138780932 | rs17645700 | 0.216 | -0.889 | 0.001 | -1.398 | -0.038 | 0.086 | 0.086 | 0 | 0.18 | 0.04 | 0.17 | 0.04 | 0.2 | 0.12 |
| 2 | 138787927 | rs4954961 | 0.409 | -0.77 | -0.724 | -1.232 | 0.001 | 0.004 | 0.004 | 0 | 0.25 | 0.09 | 0.3 | 0.27 | 0.26 | 0.29 |
| 2 | 138789029 | rs11687448 | 0.044 | -0.764 | -0.955 | -1.391 | 0.003 | 0.089 | 0.089 | 0 | 0.54 | 0.19 | 0.45 | 0.36 | 0.49 | 0.42 |
| 2 | 138804673 | rs10496795 | -0.235 | -0.256 | -0.235 | -1.063 | 0.01 | -0.028 | -0.028 | 0 | 0.13 | 0.25 | 0.15 | 0.06 | 0.13 | 0.14 |
| 2 | 138810921 | rs12463785 | -0.55 | 0.456 | -0.65 | -0.529 | -0.016 | -0.014 | -0.014 | 0.729 | 0.21 | 0.12 | 0.2 | 0.28 | 0.25 | 0.36 |
| 2 | 138815778 | rs17585698 | 1.071 | 0.021 | -1.197 | -0.947 | 0.031 | 0.111 | 0.111 | 0.256 | 0.23 | 0.06 | 0.12 | 0.07 | 0.17 | 0.12 |
| 2 | 138846608 | rs4273272 | -0.454 | 0.038 | -0.342 | -0.826 | 0.012 | -0.015 | -0.015 | 0 | 0.95 | 0.79 | 0.97 | 1 | 0.98 | 0.97 |
| 2 | 138848169 | rs1980293 | -1.141 | -0.039 | -0.383 | -0.907 | -0.005 | 0.008 | 0.008 | 0 | 0.61 | 0.63 | 0.6 | 0.39 | 0.6 | 0.53 |
| 2 | 138851111 | rs6747709 | 1.482 | -0.034 | -0.481 | -0.852 | -0.044 | -0.039 | -0.039 | 0.021 | 0.33 | 0.12 | 0.36 | 0.56 | 0.38 | 0.38 |

| | | | | | | | | | | | | | | | | |
|---|---|---|---|---|---|---|---|---|---|---|---|---|---|---|---|---|
| 2 | 138860501 | rs13031368 | 0.354 | 0.181 | -0.123 | -0.833 | -0.008 | 0.017 | 0.017 | 0 | 0.16 | 0.22 | 0.15 | 0.25 | 0.13 | 0.21 |
| 2 | 138865871 | rs1568648 | 0.288 | 0.581 | -0.945 | -0.817 | 0.005 | 0.036 | 0.036 | 0 | 0.3 | 0.2 | 0.19 | 0.07 | 0.3 | 0.2 |
| 2 | 138868450 | rs16840318 | -0.062 | 0.89 | -0.245 | 0.261 | -0.025 | -0.011 | -0.011 | 0 | 0.09 | 0.02 | 0.18 | 0.07 | 0.11 | 0.08 |
| 2 | 138881079 | rs6756311 | -0.998 | 0.692 | 0.093 | 0.443 | 0.004 | -0.008 | -0.008 | 0.169 | 0.36 | 0.44 | 0.43 | 0.28 | 0.39 | 0.28 |
| 2 | 138888963 | rs7597777 | -0.936 | 0.549 | 0.965 | 1.397 | -0.004 | 0.026 | 0.026 | 0.269 | 0.11 | 0.11 | 0.25 | 0.5 | 0.13 | 0.12 |
| 2 | 138898229 | rs2204181 | -0.586 | 0.365 | 0.705 | 1.314 | 0.001 | -0.04 | -0.04 | 0.59 | 0.15 | 0.03 | 0.12 | 0.21 | 0.12 | 0.19 |
| 2 | 138900983 | rs6713969 | 1.918 | 0.297 | 1.304 | 1.263 | -0.004 | -0.003 | -0.003 | 0 | 0.71 | 0.36 | 0.6 | 0.29 | 0.74 | 0.69 |
| 2 | 138909352 | rs10185017 | 1.33 | 0.106 | 1.35 | 1.175 | -0.01 | 0.041 | 0.041 | 0 | 0.81 | 0.62 | 0.85 | 0.78 | 0.86 | 0.77 |
| 2 | 138936400 | rs6758531 | 0.092 | 0.034 | 1.167 | 1.197 | 0.01 | -0.02 | -0.02 | 0 | 0.09 | 0.03 | 0.06 | 0.19 | 0.05 | 0.2 |
| 2 | 138954903 | rs1357407 | -0.406 | 0.032 | 1.21 | 1.183 | 0.004 | -0.006 | -0.006 | 0 | 0.11 | 0.08 | 0.08 | 0.19 | 0.07 | 0.21 |
| 2 | 138990082 | rs13000810 | -1.819 | 0.155 | 1.533 | 1.2 | -0.009 | 0.019 | 0.019 | 0.004 | 0.23 | 0.44 | 0.39 | 0.72 | 0.19 | 0.31 |
| 2 | 139008811 | rs28569024 | -1.338 | -0.511 | 0.824 | 0.491 | 0.033 | 0.053 | 0.053 | 3.061 | 0.68 | 0.8 | 0.83 | 0.93 | 0.74 | 0.81 |
| 2 | 139010773 | rs6734558 | -0.252 | -0.441 | 0.929 | 0.543 | 0.008 | 0.209 | 0.209 | 0 | 0.13 | 0.3 | 0.32 | 0.53 | 0.12 | 0.1 |
| 2 | 139020862 | rs4643591 | 0.837 | -0.536 | 1.017 | 0.517 | -0.009 | 0.115 | 0.115 | 0.019 | 0.42 | 0.21 | 0.4 | 0.17 | 0.52 | 0.49 |
| 2 | 139022658 | rs11885046 | 0.533 | -0.521 | 0.862 | 0.527 | -0.006 | 0.051 | 0.051 | 0.46 | 0.1 | 0.14 | 0.08 | 0.2 | 0.07 | 0.2 |
| 2 | 139029250 | rs6710702 | -1.074 | -0.516 | 0.477 | 0.473 | 0.023 | 0.032 | 0.032 | 0.958 | 0.65 | 0.73 | 0.8 | 0.9 | 0.71 | 0.79 |
| 2 | 139045852 | rs35211371 | 0.96 | -0.417 | 0.283 | 0.61 | -0.018 | -0.051 | -0.051 | 0 | 0.08 | 0 | 0.09 | 0.17 | 0.07 | 0.13 |
| 2 | 139045987 | rs13033503 | -0.043 | -0.395 | 0.398 | 0.77 | 0.006 | 0.154 | 0.154 | 0 | 0.12 | 0.25 | 0.29 | 0.49 | 0.12 | 0.1 |
| 2 | 139046385 | rs13382964 | -1.176 | -0.38 | -0.099 | 0.744 | 0.037 | 0.055 | 0.055 | 1.739 | 0.63 | 0.76 | 0.81 | 0.93 | 0.71 | 0.79 |
| 2 | 139049380 | rs4399769 | 0.913 | 0.174 | -0.171 | 0.609 | -0.125 | 0.131 | 0.131 | 0.002 | 0.35 | 0.07 | 0.37 | 0.13 | 0.45 | 0.38 |
| 2 | 139052164 | rs4311107 | -0.645 | -0.294 | -0.49 | 0.828 | 0.04 | -0.025 | -0.025 | 0 | 0.6 | 0.75 | 0.62 | 0.45 | 0.68 | 0.74 |
| 2 | 139054836 | rs10211175 | -0.907 | -0.344 | -0.455 | 0.396 | 0.037 | 0.056 | 0.056 | 1.708 | 0.63 | 0.75 | 0.81 | 0.93 | 0.71 | 0.79 |
| 2 | 139057105 | rs4328686 | -0.354 | -0.126 | -0.417 | 0.679 | 0.024 | 0.021 | 0.021 | 1.319 | 0.53 | 0.64 | 0.7 | 0.69 | 0.63 | 0.58 |
| 2 | 139057484 | rs13403593 | 0.479 | 0.309 | -1.053 | 0.15 | -0.05 | 0.066 | 0.066 | 0.291 | 0.36 | 0.19 | 0.32 | 0.22 | 0.5 | 0.45 |
| 2 | 139060160 | rs11682582 | 0.825 | -0.469 | -1.369 | 0.118 | 0.05 | 0.108 | 0.108 | 0.926 | 0.29 | 0.05 | 0.15 | 0.14 | 0.22 | 0.22 |
| 2 | 139063638 | rs13432566 | 0.258 | -0.447 | -0.996 | 0.134 | 0.022 | 0.082 | 0.082 | 0.148 | 0.44 | 0.25 | 0.45 | 0.23 | 0.58 | 0.51 |
| 2 | 139065123 | rs7564491 | -0.836 | -0.509 | -0.402 | 0.025 | -0.003 | -0.044 | -0.044 | 0.283 | 0.89 | 0.86 | 0.94 | 1 | 0.89 | 0.95 |
| 2 | 139065411 | rs4954919 | -0.676 | -0.55 | -0.423 | 0.015 | -0.004 | -0.039 | -0.039 | 0.036 | 0.88 | 0.86 | 0.94 | 1 | 0.89 | 0.95 |
| 2 | 139080891 | rs11885193 | 0.77 | -0.795 | 0.007 | 0.522 | -0.016 | 0.057 | 0.057 | 0 | 0.76 | 0.68 | 0.67 | 0.59 | 0.78 | 0.84 |
| 2 | 139085690 | rs11900575 | 1.041 | -1.155 | 0.283 | 0.385 | -0.007 | 0.021 | 0.021 | 0 | 0.7 | 0.68 | 0.62 | 0.54 | 0.7 | 0.77 |
| 2 | 139102704 | rs13393072 | 1.234 | -1.204 | 0.632 | 0.937 | -0.01 | 0.02 | 0.02 | 0.329 | 0.68 | 0.39 | 0.6 | 0.54 | 0.7 | 0.77 |
| 2 | 139104124 | rs4550720 | 1.293 | -1.023 | 0.5 | 0.953 | -0.027 | -0.031 | -0.031 | 0.039 | 0.08 | 0.01 | 0.06 | 0.11 | 0.12 | 0.05 |
| 2 | 139108517 | rs13423279 | -0.425 | -0.615 | 0.312 | 0.235 | -0.006 | 0.002 | 0.002 | 0.02 | 0.42 | 0.64 | 0.37 | 0.09 | 0.49 | 0.47 |
| 2 | 139112953 | rs10188086 | -0.825 | -1.031 | -0.119 | 0.145 | 0.013 | 0.022 | 0.022 | 0 | 0.51 | 0.74 | 0.6 | 0.39 | 0.58 | 0.54 |
| 2 | 139130069 | rs7598286 | -0.37 | -0.671 | 0.095 | 0.222 | 0.02 | 0.011 | 0.011 | 0 | 0.06 | 0.21 | 0.07 | 0.05 | 0.08 | 0.07 |
| 2 | 139144543 | rs7577825 | -0.339 | -0.735 | 0.336 | 0.129 | -0.003 | -0.045 | -0.045 | 1.389 | 0.64 | 0.67 | 0.69 | 0.97 | 0.53 | 0.59 |
| 2 | 139149053 | rs7576870 | 0.37 | -0.735 | 0.193 | 0.13 | -0.007 | 0.034 | 0.034 | 0.133 | 0.57 | 0.34 | 0.6 | 0.92 | 0.45 | 0.52 |
| 2 | 139177661 | rs11896100 | -0.235 | -0.657 | -0.015 | 0.09 | -0.005 | -0.043 | -0.043 | 0.509 | 0.65 | 0.72 | 0.69 | 0.99 | 0.53 | 0.66 |
| 2 | 139185789 | rs9287504 | 0.879 | -0.028 | -0.942 | 0.495 | -0.035 | 0.015 | 0.015 | 0 | 0.41 | 0.15 | 0.33 | 0.4 | 0.5 | 0.33 |
| 2 | 139190391 | rs13024237 | 0.534 | -0.821 | -0.185 | 0.757 | 0.009 | 0.024 | 0.024 | 0.615 | 0.52 | 0.2 | 0.4 | 0.62 | 0.43 | 0.47 |
| 2 | 139205853 | rs4560178 | -0.275 | -1.003 | -0.192 | 0.594 | 0.031 | 0.013 | 0.013 | 0.987 | 0.36 | 0.51 | 0.47 | 0.4 | 0.43 | 0.42 |
| 2 | 139211959 | rs10171464 | -0.465 | -0.981 | -0.199 | 0.631 | -0.001 | -0.003 | -0.003 | 0 | 0.43 | 0.55 | 0.55 | 0.43 | 0.53 | 0.45 |
| 2 | 139225571 | rs11679023 | 1.087 | -1.322 | -0.128 | 0.38 | 0.003 | -0.006 | -0.006 | 0 | 0.35 | 0.32 | 0.32 | 0.55 | 0.31 | 0.4 |
| 2 | 139230947 | rs10184636 | -0.534 | -1.362 | 0.044 | 0.336 | 0.074 | 0.02 | 0.02 | 0 | 0.49 | 0.27 | 0.57 | 0.45 | 0.61 | 0.52 |
| 2 | 139232021 | rs12612989 | 0.887 | -1.139 | -0.012 | 0.679 | 0 | -0.012 | -0.012 | 0.566 | 0.07 | 0 | 0.17 | 0.38 | 0.05 | 0.05 |
| 2 | 139238623 | rs12472370 | 1.025 | -1.187 | 0 | 0.609 | 0.002 | -0.012 | -0.012 | 1.944 | 0.07 | 0 | 0.28 | 0.36 | 0.05 | 0.06 |
| 2 | 139247616 | rs4256008 | 0.877 | -1.251 | 0.048 | 0.469 | -0.03 | -0.122 | -0.122 | 0 | 0.21 | 0.36 | 0.11 | 0 | 0.14 | 0.1 |
| 2 | 139251957 | rs11691283 | 1.195 | -1.364 | -0.07 | 0.621 | 0.055 | -0.062 | -0.062 | 0 | 0.5 | 0.04 | 0.41 | 0.37 | 0.65 | 0.54 |
| 2 | 139258603 | rs13405108 | 1.19 | -1.35 | -0.063 | 0.627 | 0.054 | -0.063 | -0.063 | 0.029 | 0.5 | 0.04 | 0.41 | 0.37 | 0.66 | 0.55 |
| 2 | 139278922 | rs10170310 | -0.166 | -1.35 | 0.072 | 0.626 | -0.01 | 0.008 | 0.008 | 0.937 | 0.8 | 0.62 | 0.82 | 0.75 | 0.84 | 0.76 |
| 2 | 139289801 | rs4954934 | -1.308 | -1.359 | 0.081 | 0.603 | -0.033 | -0.073 | -0.073 | 0 | 0.86 | 0.76 | 0.91 | 1 | 0.92 | 0.92 |
| 2 | 139291103 | rs11892957 | 0.759 | -1.366 | 0.084 | 0.601 | -0.042 | -0.148 | -0.148 | 0 | 0.23 | 0.4 | 0.11 | 0.02 | 0.14 | 0.16 |
| 2 | 139303548 | rs13004492 | 0.221 | -1.3 | 0.119 | 0.642 | 0.048 | 0.108 | 0.108 | 0.093 | 0.07 | 0.2 | 0.1 | 0.25 | 0.08 | 0.16 |
| 2 | 139327055 | rs7598316 | 1.002 | -2.426 | 0.126 | 0.553 | -0.036 | -0.08 | -0.08 | 0 | 0.14 | 0.28 | 0.09 | 0 | 0.08 | 0.08 |
| 2 | 139328264 | rs12470170 | 1.288 | -2.327 | -0.05 | 0.615 | 0.004 | 0.002 | 0.002 | 0 | 0.06 | 0.01 | 0.28 | 0.36 | 0.05 | 0.05 |
| 2 | 139340051 | rs7574224 | 1.01 | -2.239 | 0.076 | 0.247 | -0.017 | -0.086 | -0.086 | 0 | 0.2 | 0.28 | 0.11 | 0 | 0.14 | 0.1 |
| 2 | 139368389 | rs2139990 | -0.041 | -1.662 | -0.622 | 0.767 | -0.023 | -0.026 | -0.026 | 0 | 0.41 | 0.48 | 0.48 | 0.2 | 0.27 | 0.24 |
| 2 | 139379303 | rs13429541 | 0.483 | -1.59 | -0.738 | 0.542 | -0.024 | -0.117 | -0.117 | 0 | 0.33 | 0.39 | 0.2 | 0 | 0.22 | 0.18 |
| 2 | 139380666 | rs6759330 | -0.249 | -0.963 | -1.252 | 1.034 | 0.146 | -0.001 | -0.001 | 0.051 | 0.48 | 0.86 | 0.58 | 0.46 | 0.3 | 0.39 |
| 2 | 139393369 | rs6705862 | -0.35 | -0.987 | -1.553 | 1.06 | 0.123 | -0.045 | -0.045 | 0 | 0.49 | 0.94 | 0.59 | 0.48 | 0.29 | 0.45 |
| 2 | 139394256 | rs16840903 | 1.557 | -0.817 | -1.312 | 1.097 | 0.001 | -0.037 | -0.037 | 0 | 0.07 | 0 | 0.02 | 0.22 | 0.04 | 0.06 |
| 2 | 139404163 | rs4954942 | 0.708 | -0.164 | -1.092 | 0.536 | -0.157 | 0.023 | 0.023 | 2.233 | 0.18 | 0.02 | 0.2 | 0.11 | 0.4 | 0.28 |
| 2 | 139406389 | rs12612384 | 0.24 | -0.833 | -0.104 | 0.857 | -0.029 | -0.025 | -0.025 | 0.863 | 0.58 | 0.5 | 0.54 | 0.76 | 0.73 | 0.74 |
| 2 | 139409318 | rs4954943 | 0.036 | -0.747 | -1.137 | 0.955 | 0.026 | 0.109 | 0.109 | 0 | 0.05 | 0.13 | 0.11 | 0.23 | 0.06 | 0.15 |
| 2 | 139409648 | rs1521756 | 0.51 | -0.595 | -1.848 | 0.633 | 0.115 | 0.006 | 0.006 | 0.982 | 0.51 | 0.15 | 0.41 | 0.51 | 0.67 | 0.58 |
| 2 | 139414418 | rs7608950 | 0.342 | -0.287 | -1.264 | 0.264 | 0.01 | -0.01 | -0.01 | 0 | 0.83 | 0.86 | 0.9 | 1 | 0.89 | 0.9 |
| 2 | 139415944 | rs1521754 | 0.106 | -0.303 | -1.953 | 1.093 | 0.034 | -0.057 | -0.057 | 0 | 0.65 | 0.88 | 0.64 | 0.5 | 0.68 | 0.41 |
| 2 | 139416228 | rs6711947 | 0.435 | 0.225 | -2.176 | 0.64 | -0.012 | -0.001 | -0.001 | 0 | 0.18 | 0.09 | 0.17 | 0.22 | 0.19 | 0.14 |
| 2 | 139418739 | rs11682262 | -0.637 | 0.424 | -2.631 | 0.507 | 0.019 | -0.032 | -0.032 | 0 | 0.63 | 0.85 | 0.65 | 0.44 | 0.66 | 0.41 |

| | | | | | | | | | | | | | | | | |
|---|---|---|---|---|---|---|---|---|---|---|---|---|---|---|---|---|
| 2 | 139424201 | rs1521753 | -0.076 | 0.331 | -2.411 | 0.359 | 0.026 | 0.038 | 0.038 | 0.12 | 0.56 | 0.8 | 0.56 | 0.39 | 0.57 | 0.35 |
| 2 | 139427669 | rs10496799 | 1.355 | -0.122 | -1.519 | -0.215 | 0.043 | -0.012 | -0.012 | 3.481 | 0.21 | 0.34 | 0.32 | 0.17 | 0.3 | 0.19 |
| 2 | 139429068 | rs10803570 | -1.295 | -0.158 | -1.587 | -0.253 | 0.039 | -0.048 | -0.048 | 1.796 | 0.66 | 0.89 | 0.69 | 0.45 | 0.72 | 0.48 |
| 2 | 139433750 | rs10197504 | 0.815 | -0.284 | -1.127 | -0.332 | 0.013 | -0.011 | -0.011 | 0.976 | 0.22 | 0.31 | 0.21 | 0.22 | 0.25 | 0.15 |
| 2 | 139434161 | rs10197712 | -1.437 | -0.348 | -0.544 | -0.359 | 0.016 | -0.009 | -0.009 | 0 | 0.64 | 0.73 | 0.68 | 0.45 | 0.71 | 0.42 |
| 2 | 139434880 | rs7579695 | 0.167 | -0.337 | -0.601 | -0.373 | 0 | -0.049 | -0.049 | 1.318 | 0.35 | 0.52 | 0.36 | 0.17 | 0.33 | 0.22 |
| 2 | 139434892 | rs7566008 | 1.502 | -0.34 | -0.411 | -0.376 | -0.032 | 0.008 | 0.008 | 0 | 0.2 | 0.13 | 0.31 | 0.17 | 0.3 | 0.19 |
| 2 | 139442287 | rs10928660 | 0.589 | -0.491 | -0.385 | -0.304 | -0.028 | -0.011 | -0.011 | 0 | 0.24 | 0.08 | 0.24 | 0.28 | 0.29 | 0.17 |
| 2 | 139443559 | rs17598060 | 1.55 | -0.476 | -0.374 | -0.291 | -0.013 | -0.01 | -0.01 | 1.125 | 0.17 | 0.07 | 0.15 | 0.23 | 0.17 | 0.13 |
| 2 | 139454756 | rs4954948 | 0.577 | -0.507 | -0.371 | -0.305 | -0.005 | 0.089 | 0.089 | 0 | 0.56 | 0.2 | 0.61 | 0.71 | 0.59 | 0.79 |
| 2 | 139472180 | rs10206559 | 0.863 | -0.788 | -0.114 | -0.28 | -0.063 | 0.026 | 0.026 | 0 | 0.25 | 0.15 | 0.35 | 0.17 | 0.42 | 0.2 |
| 2 | 139472210 | rs6724976 | 1.005 | -0.928 | -0.074 | -0.076 | 0.003 | 0.007 | 0.007 | 0 | 0.19 | 0.24 | 0.17 | 0.24 | 0.17 | 0.15 |
| 2 | 139473896 | rs4531866 | -1.725 | -0.923 | -0.442 | -0.06 | -0.028 | -0.043 | -0.043 | 0.296 | 0.88 | 0.86 | 0.96 | 1 | 0.97 | 0.99 |
| 2 | 139489417 | rs10928663 | -1.249 | -1.004 | -0.418 | -0.1 | -0.055 | -0.073 | -0.073 | 0 | 0.86 | 0.78 | 0.96 | 1 | 0.97 | 0.98 |
| 2 | 139489497 | rs16841084 | 0.518 | -0.656 | -1.123 | 0.073 | 0.006 | -0.025 | -0.025 | 0 | 0.07 | 0.13 | 0.11 | 0.01 | 0.12 | 0.04 |
| 2 | 139492011 | rs4130252 | 0.713 | -0.732 | -1.875 | 0.039 | 0.026 | -0.032 | -0.032 | 0 | 0.27 | 0.21 | 0.27 | 0.53 | 0.17 | 0.46 |
| 2 | 139497309 | rs11680550 | -0.108 | 0.298 | -1.957 | 0.157 | -0.149 | 0.021 | 0.021 | 0 | 0.3 | 0.05 | 0.38 | 0.21 | 0.48 | 0.34 |
| 2 | 139503337 | rs16841109 | -0.867 | 0.243 | -1.625 | 0.859 | 0 | -0.039 | -0.039 | 0.015 | 0.17 | 0.1 | 0.18 | 0.35 | 0.14 | 0.22 |
| 2 | 139505708 | rs4322786 | -0.021 | 0.226 | -1.602 | 1.278 | 0 | 0.042 | 0.042 | 0 | 0.12 | 0.11 | 0.18 | 0.46 | 0.11 | 0.28 |
| 2 | 139505868 | rs17599153 | 0.251 | 0.983 | -2.219 | 0.886 | -0.064 | -0.032 | -0.032 | 0 | 0.16 | 0.04 | 0.19 | 0.2 | 0.19 | 0.15 |
| 2 | 139507550 | rs17657900 | 0.251 | 0.983 | -2.239 | 0.886 | -0.068 | -0.036 | -0.036 | 0 | 0.16 | 0.03 | 0.19 | 0.2 | 0.19 | 0.15 |
| 2 | 139509236 | rs16841114 | 0.385 | 1.311 | -2.3 | 0.482 | -0.033 | -0.053 | -0.053 | 3.648 | 0.27 | 0.01 | 0.35 | 0.25 | 0.23 | 0.35 |
| 2 | 139510272 | rs11681160 | 0.337 | 1.373 | -1.552 | -0.285 | 0.013 | -0.048 | -0.048 | 0.038 | 0.18 | 0.22 | 0.2 | 0 | 0.32 | 0.12 |
| 2 | 139512134 | rs10193564 | 0.038 | 1.027 | -1.848 | -0.087 | 0.006 | 0.03 | 0.03 | 0.112 | 0.38 | 0.18 | 0.5 | 0.69 | 0.35 | 0.63 |
| 2 | 139513373 | rs10928667 | 0.054 | 1.564 | -1.472 | -0.28 | -0.013 | -0.001 | -0.001 | 0 | 0.18 | 0.16 | 0.19 | 0 | 0.31 | 0.12 |
| 2 | 139515243 | rs13000377 | -0.377 | 1.194 | -1.646 | -0.346 | -0.012 | 0.039 | 0.039 | 0.022 | 0.09 | 0.02 | 0.06 | 0 | 0.09 | 0.05 |
| 2 | 139518044 | rs4524069 | 0.636 | 1.186 | -2.453 | -0.562 | 0 | -0.021 | -0.021 | 0 | 0.3 | 0.15 | 0.33 | 0.32 | 0.28 | 0.23 |
| 2 | 139530239 | rs10200922 | -0.054 | 0.917 | -1.958 | -0.627 | 0.016 | 0.028 | 0.028 | 2.485 | 0.44 | 0.28 | 0.41 | 0.32 | 0.37 | 0.3 |
| 2 | 139531083 | rs16841174 | 0.808 | 0.963 | -2.068 | -0.487 | 0.011 | 0.008 | 0.008 | 0 | 0.21 | 0.11 | 0.15 | 0.18 | 0.15 | 0.16 |
| 2 | 139535928 | rs10188629 | 0.257 | 0.907 | -1.082 | -0.52 | 0.002 | 0.027 | 0.027 | 0.549 | 0.51 | 0.59 | 0.49 | 0.32 | 0.44 | 0.4 |
| 2 | 139537261 | rs6742749 | -0.461 | 0.914 | -0.97 | -0.413 | 0.005 | 0.016 | 0.016 | 0 | 0.71 | 0.7 | 0.63 | 0.5 | 0.59 | 0.56 |
| 2 | 139552197 | rs4561602 | 0.248 | 0.952 | -1.399 | -0.433 | -0.046 | -0.021 | -0.021 | 0 | 0.27 | 0.05 | 0.18 | 0.22 | 0.23 | 0.29 |
| 2 | 139555034 | rs4257340 | -1.003 | 0.742 | -1.113 | -0.529 | 0.038 | 0.033 | 0.033 | 0.199 | 0.63 | 0.24 | 0.71 | 0.69 | 0.74 | 0.75 |
| 2 | 139573272 | rs12474367 | 0.405 | 0.738 | -0.477 | -0.56 | -0.042 | -0.027 | -0.027 | 0 | 0.27 | 0.04 | 0.18 | 0.22 | 0.23 | 0.29 |
| 2 | 139573631 | rs9636230 | -0.006 | 0.68 | -0.417 | -0.604 | -0.007 | -0.008 | -0.008 | 0 | 0.6 | 0.35 | 0.44 | 0.52 | 0.5 | 0.54 |
| 2 | 139575344 | rs4347759 | 0.906 | 0.644 | 0.025 | -0.561 | 0.01 | 0.004 | 0.004 | 0 | 0.35 | 0.29 | 0.35 | 0.34 | 0.33 | 0.26 |
| 2 | 139577089 | rs10174953 | 0.074 | 0.599 | 0.293 | -0.66 | -0.007 | 0.035 | 0.035 | 0 | 0.61 | 0.71 | 0.73 | 0.78 | 0.67 | 0.66 |
| 2 | 139585106 | rs12470031 | 0.694 | 0.603 | 0.303 | -0.649 | -0.021 | -0.065 | -0.065 | 0 | 0.26 | 0.13 | 0.35 | 0.44 | 0.34 | 0.4 |
| 2 | 139593079 | rs4611595 | 1.118 | 0.549 | 0.392 | -0.67 | 0.018 | 0.002 | 0.002 | 0 | 0.34 | 0.23 | 0.35 | 0.34 | 0.33 | 0.26 |
| 2 | 139599513 | rs12470293 | 0.347 | 0.675 | 0.404 | -0.685 | 0.003 | -0.012 | -0.012 | 0 | 0.29 | 0.39 | 0.22 | 0.22 | 0.25 | 0.28 |
| 2 | 139600035 | rs10496801 | 0.429 | 0.678 | 0.317 | -0.684 | -0.016 | 0.019 | 0.019 | 0 | 0.29 | 0.16 | 0.2 | 0.22 | 0.25 | 0.28 |
| 2 | 139603269 | rs6430811 | -1.179 | 0.638 | 0.404 | -0.653 | 0.01 | -0.055 | -0.055 | 0.421 | 0.9 | 0.84 | 0.93 | 1 | 0.92 | 0.93 |
| 2 | 139607947 | rs4245869 | -1.208 | 0.665 | 0.348 | -0.652 | 0.008 | -0.05 | -0.05 | 0 | 0.9 | 0.85 | 0.93 | 1 | 0.91 | 0.93 |
| 2 | 139615924 | rs1905738 | 0.828 | 0.665 | 0.272 | -0.652 | 0.061 | -0.057 | -0.057 | 0 | 0.1 | 0.37 | 0.09 | 0 | 0.09 | 0.07 |
| 2 | 139619228 | rs13020961 | 0.948 | 0.625 | -0.544 | -0.744 | -0.04 | 0.088 | 0.088 | 0 | 0.14 | 0.02 | 0.09 | 0 | 0.14 | 0.11 |
| 2 | 139620096 | rs1351919 | -0.827 | 0.524 | -0.441 | -0.545 | 0.018 | 0.038 | 0.038 | 0.02 | 0.55 | 0.34 | 0.55 | 0.66 | 0.59 | 0.67 |
| 2 | 139624610 | rs11885104 | 0.211 | 0.656 | -0.216 | -0.134 | -0.005 | -0.04 | -0.04 | 0 | 0.24 | 0.16 | 0.32 | 0.44 | 0.27 | 0.38 |
| 2 | 139641542 | rs1485066 | 0.174 | 0.411 | -0.238 | -0.301 | 0 | -0.025 | -0.025 | 0 | 0.71 | 0.74 | 0.86 | 0.93 | 0.72 | 0.71 |
| 2 | 139642496 | rs13425763 | 0.135 | 0.215 | -0.037 | -0.007 | -0.023 | 0.171 | 0.171 | 0 | 0.07 | 0.17 | 0.22 | 0.38 | 0.02 | 0.17 |
| 2 | 139644839 | rs12612135 | 1.635 | 0.023 | -0.688 | -0.266 | -0.019 | -0.044 | -0.044 | 0.139 | 0.21 | 0.09 | 0.34 | 0.28 | 0.27 | 0.17 |
| 2 | 139645564 | rs10432381 | 1.635 | 0.023 | -0.688 | -0.266 | -0.019 | -0.044 | -0.044 | 0 | 0.21 | 0.09 | 0.34 | 0.28 | 0.27 | 0.17 |
| 2 | 139646468 | rs719259 | -0.382 | 0.03 | -0.544 | -0.307 | 0 | -0.021 | -0.021 | 0 | 0.71 | 0.75 | 0.85 | 0.93 | 0.71 | 0.7 |
| 2 | 139659274 | rs12470151 | 0.875 | -0.046 | -0.23 | -0.405 | -0.022 | 0.034 | 0.034 | 0 | 0.36 | 0.23 | 0.28 | 0.24 | 0.38 | 0.29 |
| 2 | 139659849 | rs6713376 | -0.105 | -0.045 | -0.144 | -0.293 | -0.035 | 0.258 | 0.258 | 0 | 0.14 | 0.43 | 0.24 | 0.41 | 0.06 | 0.25 |
| 2 | 139659956 | rs17660836 | 1.197 | 0.043 | -0.284 | -0.512 | -0.011 | 0.021 | 0.021 | 0 | 0.17 | 0.11 | 0.09 | 0.07 | 0.19 | 0.22 |
| 2 | 139666757 | rs2889267 | 0.182 | -0.006 | -0.169 | -0.477 | -0.016 | -0.005 | -0.005 | 0 | 0.35 | 0.49 | 0.57 | 0.69 | 0.33 | 0.41 |
| 2 | 139667920 | rs1948019 | 0.988 | 0.292 | -0.306 | -0.535 | -0.011 | 0.02 | 0.02 | 0 | 0.17 | 0.11 | 0.09 | 0.07 | 0.19 | 0.21 |
| 2 | 139677323 | rs6757538 | -1.401 | -0.477 | -0.107 | -0.507 | -0.008 | -0.062 | -0.062 | 3.213 | 0.88 | 0.84 | 0.94 | 1 | 0.9 | 0.92 |
| 2 | 139687078 | rs11889428 | 0.035 | -0.784 | 0.228 | -0.678 | -0.064 | -0.053 | -0.053 | 0 | 0.21 | 0.35 | 0.07 | 0.09 | 0.1 | 0.24 |
| 2 | 139687146 | rs11889468 | 1.14 | -0.802 | 0.306 | -0.5 | -0.026 | -0.012 | -0.012 | 0 | 0.13 | 0.16 | 0.05 | 0.06 | 0.05 | 0.16 |
| 2 | 139687946 | rs1485053 | 0.543 | -0.945 | 0.119 | -0.422 | -0.01 | 0.004 | 0.004 | 0 | 0.39 | 0.29 | 0.47 | 0.6 | 0.4 | 0.42 |
| 2 | 139692323 | rs1564835 | 1.438 | -0.864 | 0.032 | -0.408 | -0.096 | -0.032 | -0.032 | 0 | 0.27 | 0.1 | 0.39 | 0.31 | 0.4 | 0.26 |
| 2 | 139703024 | rs1485091 | 1.377 | -0.871 | 0.037 | -0.43 | -0.051 | -0.004 | -0.004 | 0 | 0.27 | 0.21 | 0.39 | 0.3 | 0.4 | 0.26 |
| 2 | 139704410 | rs1485090 | -0.647 | -0.893 | 0.071 | -0.447 | -0.01 | 0.011 | 0.011 | 0 | 0.61 | 0.64 | 0.54 | 0.7 | 0.5 | 0.67 |
| 2 | 139709252 | rs4954642 | -1.679 | -0.855 | 0.055 | -0.454 | -0.005 | -0.042 | -0.042 | 0 | 0.88 | 0.88 | 0.94 | 1 | 0.9 | 0.92 |
| 2 | 139715737 | rs6751856 | 1.557 | -0.831 | 0.098 | -0.406 | -0.002 | -0.008 | -0.008 | 0 | 0.06 | 0.11 | 0.02 | 0.03 | 0.05 | 0.07 |
| 2 | 139717085 | rs13009652 | 1.5 | -0.798 | -0.33 | -0.639 | -0.084 | 0.189 | 0.189 | 1.329 | 0.27 | 0.02 | 0.16 | 0 | 0.27 | 0.15 |
| 2 | 139721734 | rs1485088 | -0.432 | -0.619 | 0.173 | -0.259 | 0 | 0.002 | 0.002 | 0.301 | 0.5 | 0.55 | 0.49 | 0.39 | 0.54 | 0.49 |
| 2 | 139725320 | rs16841431 | -0.08 | -0.709 | 0.357 | -0.29 | -0.059 | -0.066 | -0.066 | 0 | 0.23 | 0.41 | 0.1 | 0.09 | 0.14 | 0.24 |

| | | | | | | | | | | | | | | | | |
|---|---|---|---|---|---|---|---|---|---|---|---|---|---|---|---|---|
| 2 | 139730483 | rs1351921 | -0.161 | -0.707 | 0.296 | -0.277 | -0.052 | -0.057 | -0.057 | 0.187 | 0.23 | 0.42 | 0.1 | 0.09 | 0.14 | 0.24 |
| 2 | 139737015 | rs16841454 | 0.761 | -0.714 | 0.289 | -0.276 | -0.072 | -0.048 | -0.048 | 0 | 0.15 | 0.35 | 0.07 | 0.06 | 0.06 | 0.16 |
| 2 | 139748069 | rs2101294 | 0.761 | -0.559 | 0.303 | -0.209 | 0.004 | -0.023 | -0.023 | 0.186 | 0.09 | 0.17 | 0.05 | 0.03 | 0.09 | 0.08 |
| 2 | 139757823 | rs1872386 | 0.509 | -0.677 | 0.186 | -0.239 | 0.006 | 0.018 | 0.018 | 0 | 0.43 | 0.31 | 0.47 | 0.62 | 0.4 | 0.45 |
| 2 | 139758906 | rs3922799 | 0.481 | -0.387 | 0.226 | -0.109 | -0.024 | 0.003 | 0.003 | 0 | 0.34 | 0.29 | 0.42 | 0.35 | 0.41 | 0.4 |
| 2 | 139778282 | rs1505846 | 1.348 | -0.365 | 0.132 | -0.133 | -0.098 | -0.025 | -0.025 | 0.048 | 0.26 | 0.1 | 0.38 | 0.29 | 0.4 | 0.27 |
| 2 | 139780817 | rs6714843 | 0.821 | -0.292 | 0.186 | -0.112 | 0.004 | -0.02 | -0.02 | 0 | 0.09 | 0.17 | 0.05 | 0.03 | 0.09 | 0.08 |
| 2 | 139782825 | rs1505842 | 1.06 | -0.356 | 0.006 | 0.312 | -0.01 | 0.018 | 0.018 | 0 | 0.32 | 0.26 | 0.26 | 0.2 | 0.33 | 0.25 |
| 2 | 139783751 | rs4954732 | 1.077 | -0.349 | -0.106 | 0.315 | -0.018 | 0.028 | 0.028 | 0 | 0.32 | 0.21 | 0.26 | 0.2 | 0.33 | 0.25 |
| 2 | 139797216 | rs1395010 | 0.592 | 0.375 | 0.274 | 0.578 | -0.004 | 0.009 | 0.009 | 0.65 | 0.55 | 0.38 | 0.47 | 0.6 | 0.5 | 0.53 |
| 2 | 139800877 | rs10803575 | -0.774 | 0.554 | -0.416 | 0.685 | 0.005 | -0.007 | -0.007 | 0 | 0.43 | 0.48 | 0.51 | 0.4 | 0.5 | 0.47 |
| 2 | 139801479 | rs16841529 | 0.503 | 0.456 | -1.028 | 0.816 | -0.032 | -0.005 | -0.005 | 0 | 0.08 | 0.16 | 0.09 | 0.07 | 0.02 | 0.13 |
| 2 | 139804141 | rs1605136 | -0.514 | 0.65 | -1.086 | 0.981 | -0.009 | -0.007 | -0.007 | 0 | 0.34 | 0.32 | 0.48 | 0.43 | 0.45 | 0.39 |
| 2 | 139808906 | rs11688290 | 0.459 | 0.598 | -1.22 | 1.054 | -0.121 | -0.091 | -0.091 | 0 | 0.24 | 0.07 | 0.36 | 0.35 | 0.4 | 0.24 |
| 2 | 139810224 | rs13395791 | -0.079 | 0.767 | -1.139 | 0.974 | -0.009 | -0.011 | -0.011 | 0 | 0.31 | 0.31 | 0.45 | 0.43 | 0.41 | 0.37 |
| 2 | 139813797 | rs4144773 | 0.136 | 0.203 | -1.366 | 1.255 | 0.018 | -0.042 | -0.042 | 0 | 0.75 | 0.8 | 0.68 | 0.58 | 0.86 | 0.52 |
| 2 | 139820383 | rs1121728 | 0.239 | -0.112 | -1.729 | 1.018 | 0.05 | 0.008 | 0.008 | 0 | 0.54 | 0.7 | 0.52 | 0.37 | 0.7 | 0.42 |
| 2 | 139823059 | rs16841575 | 0.049 | 0.394 | -1.978 | 0.774 | -0.034 | -0.063 | -0.063 | 0 | 0.19 | 0.09 | 0.26 | 0.6 | 0.29 | 0.22 |
| 2 | 139823945 | rs12617255 | 1.162 | 0.404 | -2.058 | 1.147 | -0.001 | -0.056 | -0.056 | 8.835 | 0.09 | 0 | 0.04 | 0.18 | 0.1 | 0.04 |
| 2 | 139825194 | rs16841580 | 2.08 | 0.453 | -1.301 | 0.306 | 0 | -0.062 | -0.062 | 0.909 | 0.12 | 0.2 | 0.06 | 0 | 0.1 | 0.04 |
| 2 | 139826398 | rs1437314 | -1.137 | -0.205 | -1.359 | -0.028 | -0.02 | -0.088 | -0.088 | 1.128 | 0.76 | 0.61 | 0.81 | 0.94 | 0.85 | 0.76 |
| 2 | 139826907 | rs1437316 | 0.943 | 0.025 | -1.541 | -0.183 | 0.002 | 0.007 | 0.007 | 0 | 0.36 | 0.23 | 0.41 | 0.34 | 0.33 | 0.43 |
| 2 | 139836023 | rs963629 | 1.081 | 0.028 | -0.859 | -0.185 | -0.003 | -0.01 | -0.01 | 0 | 0.34 | 0.38 | 0.4 | 0.27 | 0.31 | 0.43 |
| 2 | 139836688 | rs6430823 | -1.336 | 0.034 | -0.958 | -0.311 | 0.007 | 0.014 | 0.014 | 0.308 | 0.77 | 0.77 | 0.82 | 0.94 | 0.86 | 0.76 |
| 2 | 139843788 | rs1437321 | -1.412 | 0.018 | -0.989 | -0.307 | 0.007 | 0.02 | 0.02 | 0.249 | 0.77 | 0.79 | 0.82 | 0.94 | 0.85 | 0.76 |
| 2 | 139848110 | rs12470704 | 1.24 | 0.042 | -1.047 | -0.278 | -0.001 | -0.003 | -0.003 | 0 | 0.32 | 0.21 | 0.39 | 0.33 | 0.31 | 0.43 |
| 2 | 139859202 | rs4954647 | -0.437 | 0.162 | -1.018 | 0.039 | -0.022 | -0.027 | -0.027 | 0 | 0.29 | 0.2 | 0.34 | 0.6 | 0.42 | 0.29 |
| 2 | 139866554 | rs2099575 | -0.318 | 0.175 | -1.2 | 0.049 | -0.042 | -0.053 | -0.053 | 0 | 0.28 | 0.15 | 0.34 | 0.6 | 0.42 | 0.29 |
| 2 | 139876065 | rs12613183 | 0.038 | 0.273 | -1.294 | 0.049 | -0.054 | -0.074 | -0.074 | 0 | 0.23 | 0.1 | 0.32 | 0.6 | 0.38 | 0.24 |
| 2 | 139878422 | rs10178021 | 0.609 | 0.181 | -1.284 | -0.057 | 0.013 | 0.076 | 0.076 | 0 | 0.49 | 0.66 | 0.48 | 0.33 | 0.43 | 0.47 |
| 2 | 139879838 | rs11680642 | -0.362 | -0.505 | -0.959 | -0.362 | 0.014 | 0.038 | 0.038 | 0 | 0.72 | 0.76 | 0.81 | 0.94 | 0.81 | 0.71 |
| 2 | 139886095 | rs1437329 | 0.499 | -0.441 | -0.467 | 0.069 | -0.004 | 0.027 | 0.027 | 0 | 0.06 | 0.1 | 0.03 | 0.08 | 0.03 | 0.12 |
| 2 | 139895553 | rs1437330 | 0.383 | -0.564 | -0.459 | -0.001 | 0.014 | 0.023 | 0.023 | 0 | 0.55 | 0.36 | 0.52 | 0.71 | 0.63 | 0.46 |
| 2 | 139897941 | rs6716603 | 1.199 | -0.415 | -0.367 | 0.037 | 0.002 | 0.043 | 0.043 | 0 | 0.06 | 0.11 | 0.09 | 0.12 | 0.06 | 0.08 |
| 2 | 139908499 | rs12691976 | -0.625 | -0.415 | -0.162 | -0.341 | -0.001 | 0.046 | 0.046 | 0 | 0.7 | 0.82 | 0.66 | 0.91 | 0.73 | 0.68 |
| 2 | 139911372 | rs934706 | -0.37 | -0.369 | -0.199 | -0.574 | -0.025 | 0.101 | 0.101 | 0.408 | 0.56 | 0.35 | 0.54 | 0.88 | 0.5 | 0.74 |
| 2 | 139914782 | rs13029794 | -0.344 | -0.502 | -0.15 | -0.526 | -0.02 | 0.108 | 0.108 | 0.049 | 0.54 | 0.34 | 0.53 | 0.85 | 0.5 | 0.71 |
| 2 | 139930003 | rs1370315 | 0.708 | -0.501 | 0.164 | -0.538 | 0.014 | -0.065 | -0.065 | 0 | 0.38 | 0.4 | 0.3 | 0.08 | 0.46 | 0.22 |
| 2 | 139933602 | rs13013702 | 0.717 | -0.702 | -0.312 | -0.31 | 0.002 | -0.028 | -0.028 | 0 | 0.32 | 0.27 | 0.28 | 0.51 | 0.28 | 0.46 |
| 2 | 139943589 | rs4954752 | -0.07 | -0.699 | -0.287 | -0.322 | -0.001 | 0 | 0 | 0.11 | 0.57 | 0.54 | 0.54 | 0.41 | 0.65 | 0.43 |
| 2 | 139952914 | rs10205677 | 0.817 | -0.715 | -0.058 | -0.357 | -0.012 | 0.015 | 0.015 | 0 | 0.39 | 0.31 | 0.31 | 0.08 | 0.47 | 0.22 |
| 2 | 139954090 | rs2118269 | 0.787 | -0.724 | -0.041 | -0.34 | -0.013 | 0.044 | 0.044 | 2.283 | 0.19 | 0.3 | 0.24 | 0.33 | 0.18 | 0.2 |
| 2 | 139957431 | rs9646664 | 0.748 | -0.619 | -0.772 | -0.578 | -0.002 | -0.036 | -0.036 | 0 | 0.31 | 0.2 | 0.27 | 0.52 | 0.28 | 0.46 |
| 2 | 139961825 | rs1437305 | -0.664 | -0.457 | -1.418 | -0.71 | -0.019 | 0.025 | 0.025 | 0.026 | 0.93 | 0.85 | 0.84 | 0.92 | 0.96 | 0.93 |
| 2 | 139964259 | rs10496809 | 0.818 | -0.445 | -1.416 | -0.727 | -0.07 | 0.125 | 0.125 | 0.034 | 0.36 | 0.13 | 0.29 | 0 | 0.47 | 0.15 |
| 2 | 139988814 | rs17706564 | 1.283 | -0.143 | -1.956 | -0.673 | -0.141 | 0.163 | 0.163 | 0 | 0.24 | 0.01 | 0.22 | 0 | 0.37 | 0.12 |
| 2 | 139994754 | rs7566877 | -0.663 | -0.164 | -1.769 | 0.053 | -0.015 | 0.004 | 0.004 | 0 | 0.94 | 0.86 | 0.84 | 0.94 | 0.96 | 0.94 |
| 2 | 140000301 | rs4954758 | 0.027 | 0.013 | -2.075 | 0.354 | -0.025 | -0.104 | -0.104 | 0 | 0.29 | 0.09 | 0.26 | 0.53 | 0.27 | 0.46 |
| 2 | 140002534 | rs7567555 | 0.132 | 0.151 | -2.099 | -0.489 | -0.076 | 0.159 | 0.159 | 0 | 0.36 | 0.16 | 0.31 | 0.04 | 0.52 | 0.23 |
| 2 | 140002855 | rs7577179 | -0.129 | -0.524 | -1.568 | 0.017 | 0.023 | 0.003 | 0.003 | 0 | 0.41 | 0.36 | 0.46 | 0.64 | 0.31 | 0.45 |
| 2 | 140004659 | rs6721816 | 0.09 | -0.401 | -0.906 | 0.195 | -0.004 | 0.029 | 0.029 | 0 | 0.06 | 0.16 | 0.05 | 0.09 | 0.05 | 0.1 |
| 2 | 140007493 | rs12373816 | 0.385 | -0.417 | -0.977 | -0.119 | -0.001 | -0.014 | -0.014 | 0 | 0.19 | 0.2 | 0.17 | 0.05 | 0.16 | 0.17 |
| 2 | 140019620 | rs7558498 | -0.865 | -0.145 | -0.463 | -0.304 | 0.006 | -0.04 | -0.04 | 0.121 | 0.3 | 0.38 | 0.29 | 0.14 | 0.29 | 0.32 |
| 2 | 140020932 | rs10928703 | 0.515 | -0.158 | -0.366 | -0.276 | -0.005 | -0.026 | -0.026 | 0 | 0.19 | 0.2 | 0.17 | 0.05 | 0.16 | 0.17 |
| 2 | 140026707 | rs17758500 | 0.81 | -0.239 | -0.385 | -0.399 | -0.006 | 0.062 | 0.062 | 0 | 0.17 | 0.06 | 0.15 | 0.04 | 0.16 | 0.12 |
| 2 | 140027282 | rs16855914 | 0.932 | -0.513 | -0.883 | 0.279 | 0.01 | -0.136 | -0.136 | 0 | 0.2 | 0.02 | 0.1 | 0.42 | 0.16 | 0.27 |
| 2 | 140030968 | rs13416582 | 1.111 | -0.408 | -0.827 | -0.173 | -0.003 | 0.008 | 0.008 | 0 | 0.06 | 0.09 | 0.06 | 0.09 | 0.05 | 0.1 |
| 2 | 140031202 | rs736666 | -0.093 | -0.403 | -0.491 | -0.153 | 0.03 | -0.008 | -0.008 | 0 | 0.32 | 0.26 | 0.23 | 0.72 | 0.26 | 0.5 |
| 2 | 140036852 | rs6742944 | -0.456 | -0.446 | -0.386 | -0.239 | 0.004 | -0.042 | -0.042 | 0 | 0.31 | 0.39 | 0.29 | 0.14 | 0.29 | 0.32 |
| 2 | 140037016 | rs9287512 | 0.432 | -0.362 | -0.583 | -0.135 | 0.01 | -0.001 | -0.001 | 0.319 | 0.11 | 0.16 | 0.11 | 0.09 | 0.13 | 0.15 |
| 2 | 140049207 | rs1347861 | 1.545 | -0.459 | -0.943 | 0.111 | -0.005 | -0.024 | -0.024 | 0 | 0.15 | 0 | 0.07 | 0.11 | 0.13 | 0.16 |
| 2 | 140050693 | rs12992680 | 0.843 | -0.247 | -0.715 | -0.075 | 0.003 | 0.011 | 0.011 | 0 | 0.21 | 0.15 | 0.2 | 0.05 | 0.19 | 0.2 |
| 2 | 140056052 | rs1371110 | 1.085 | 0.301 | -0.64 | 0.142 | -0.025 | 0.043 | 0.043 | 0.23 | 0.34 | 0.25 | 0.44 | 0.11 | 0.42 | 0.15 |
| 2 | 140056305 | rs13401409 | -0.325 | 0.61 | -0.391 | 0.599 | -0.001 | 0 | 0 | 0 | 0.11 | 0.12 | 0.12 | 0.09 | 0.13 | 0.15 |
| 2 | 140062182 | rs10496816 | -0.268 | 0.541 | -0.264 | 0.629 | -0.003 | 0.003 | 0.003 | 0 | 0.37 | 0.41 | 0.28 | 0.6 | 0.37 | 0.52 |
| 2 | 140064644 | rs1439080 | -0.927 | 0.604 | -0.179 | 0.529 | -0.002 | 0.001 | 0.001 | 0 | 0.93 | 0.93 | 0.94 | 0.97 | 0.92 | 0.88 |
| 2 | 140069737 | rs4954763 | 1.773 | 0.553 | -0.056 | 0.532 | -0.029 | 0.027 | 0.027 | 0 | 0.08 | 0.02 | 0.04 | 0.04 | 0.14 | 0.03 |
| 2 | 140071800 | rs1371112 | -0.528 | 0.501 | 0.2 | 0.534 | 0.036 | -0.013 | -0.013 | 0 | 0.92 | 0.79 | 0.93 | 0.97 | 0.92 | 0.88 |
| 2 | 140074659 | rs344914 | 1.071 | 1.142 | -0.08 | 0.318 | 0.004 | 0.005 | 0.005 | 0 | 0.36 | 0.33 | 0.45 | 0.23 | 0.3 | 0.21 |

| | | | | | | | | | | | | | | | | |
|---|---|---|---|---|---|---|---|---|---|---|---|---|---|---|---|---|
| 2 | 140074943 | rs16842051 | 0.689 | 1.104 | 0.243 | 0.322 | 0.014 | -0.028 | -0.028 | 0.256 | 0.28 | 0.27 | 0.23 | 0.56 | 0.23 | 0.49 |
| 2 | 140077038 | rs344907 | -1.201 | 1.104 | 0.23 | 0.245 | 0.014 | -0.009 | -0.009 | 0.212 | 0.93 | 0.87 | 0.93 | 0.97 | 0.92 | 0.88 |
| 2 | 140077355 | rs344905 | -0.778 | 1.106 | 0.163 | 0.246 | 0.05 | -0.008 | -0.008 | 0 | 0.91 | 0.74 | 0.92 | 0.97 | 0.92 | 0.88 |
| 2 | 140082554 | rs344891 | 0.573 | 0.911 | -0.149 | 0.306 | 0 | -0.003 | -0.003 | 0.007 | 0.08 | 0.07 | 0.09 | 0.08 | 0.07 | 0.07 |
| 2 | 140087319 | rs13416465 | -0.667 | 1.069 | -0.12 | 0.368 | 0 | 0.003 | 0.003 | 0 | 0.57 | 0.57 | 0.48 | 0.74 | 0.61 | 0.67 |
| 2 | 140095800 | rs344868 | 1.279 | 1.158 | -0.67 | 0.335 | -0.046 | -0.004 | -0.004 | 0 | 0.19 | 0.05 | 0.2 | 0.17 | 0.26 | 0.15 |
| 2 | 140100106 | rs344689 | 1.323 | 0.932 | -0.744 | 0.322 | -0.049 | -0.027 | -0.027 | 0 | 0.19 | 0.06 | 0.2 | 0.24 | 0.24 | 0.15 |
| 2 | 140102096 | rs10186150 | 1.218 | 0.426 | -0.573 | 0.35 | -0.001 | 0.006 | 0.006 | 0 | 0.09 | 0.07 | 0.04 | 0.05 | 0.12 | 0.05 |
| 2 | 140102691 | rs1877937 | 0.581 | 0.303 | -0.446 | 0.544 | 0.001 | 0 | 0 | 0 | 0.26 | 0.24 | 0.17 | 0.25 | 0.21 | 0.21 |
| 2 | 140126129 | rs4517921 | 1.164 | 0.23 | -0.904 | 0.176 | 0.017 | 0.113 | 0.113 | 0.309 | 0.2 | 0.02 | 0.11 | 0.03 | 0.16 | 0.14 |
| 2 | 140127528 | rs1367905 | 0.696 | 0.476 | -0.267 | 0.718 | -0.001 | -0.001 | -0.001 | 0.276 | 0.1 | 0.11 | 0.06 | 0.08 | 0.12 | 0.14 |
| 2 | 140133117 | rs17710801 | 0.941 | 0.525 | -0.354 | 0.671 | -0.006 | 0.027 | 0.027 | 0 | 0.09 | 0.04 | 0.04 | 0.01 | 0.1 | 0.1 |
| 2 | 140138681 | rs11884896 | -0.578 | 0.493 | -0.376 | 0.865 | -0.011 | -0.068 | -0.068 | 0 | 0.18 | 0.07 | 0.14 | 0.34 | 0.19 | 0.41 |
| 2 | 140159379 | rs4954772 | 1.001 | 0.607 | -0.251 | 0.71 | -0.001 | 0 | 0 | 0 | 0.09 | 0.08 | 0.05 | 0.08 | 0.13 | 0.09 |
| 2 | 140166857 | rs6741585 | -0.619 | 0.668 | -0.254 | 0.607 | -0.012 | 0.004 | 0.004 | 0.305 | 0.54 | 0.69 | 0.6 | 0.53 | 0.53 | 0.59 |
| 2 | 140178191 | rs1473242 | -0.366 | 0.674 | -0.255 | 0.572 | 0.005 | 0.006 | 0.006 | 0.141 | 0.48 | 0.6 | 0.55 | 0.53 | 0.46 | 0.56 |
| 2 | 140179200 | rs12614356 | -0.366 | 0.674 | -0.255 | 0.564 | 0.005 | 0.007 | 0.007 | 0.141 | 0.48 | 0.6 | 0.55 | 0.53 | 0.46 | 0.56 |
| 2 | 140180201 | rs6718997 | 0.868 | 0.685 | -0.349 | 0.415 | 0 | 0.011 | 0.011 | 0.224 | 0.28 | 0.25 | 0.38 | 0.15 | 0.27 | 0.11 |
| 2 | 140197851 | rs16842293 | 1.111 | 0.661 | -0.366 | 0.465 | -0.002 | 0.003 | 0.003 | 0 | 0.06 | 0.05 | 0.04 | 0 | 0.06 | 0.03 |
| 2 | 140200435 | rs11688725 | 1.078 | 0.745 | -0.422 | 0.322 | -0.002 | 0.016 | 0.016 | 0.688 | 0.26 | 0.2 | 0.36 | 0.15 | 0.25 | 0.1 |
| 2 | 140204444 | rs16855933 | -0.137 | 0.59 | 0.239 | 0.512 | -0.004 | 0.009 | 0.009 | 0.246 | 0.38 | 0.58 | 0.5 | 0.45 | 0.32 | 0.46 |
| 2 | 140208503 | rs13384348 | -0.558 | 0.648 | -0.058 | 0.436 | 0.013 | -0.009 | -0.009 | 0 | 0.46 | 0.19 | 0.34 | 0.58 | 0.44 | 0.66 |
| 2 | 140211403 | rs4538141 | -0.565 | 0.647 | -0.059 | 0.435 | 0.013 | -0.009 | -0.009 | 1.31 | 0.46 | 0.19 | 0.34 | 0.58 | 0.44 | 0.66 |
| 2 | 140211435 | rs10205939 | 1.952 | 0.539 | -0.226 | 0.251 | -0.013 | 0.031 | 0.031 | 0 | 0.1 | 0.01 | 0.07 | 0.05 | 0.13 | 0.05 |
| 2 | 140218663 | rs1431873 | 1.076 | 0.526 | -0.085 | 0.28 | -0.004 | -0.032 | -0.032 | 2.239 | 0.31 | 0.56 | 0.42 | 0.15 | 0.28 | 0.1 |
| 2 | 140220527 | rs11890914 | 0.85 | 0.498 | -0.201 | 0.305 | 0.004 | 0.012 | 0.012 | 1.488 | 0.33 | 0.69 | 0.44 | 0.18 | 0.28 | 0.18 |
| 2 | 140224777 | rs10496820 | 1.087 | 0.461 | -0.094 | 0.442 | -0.015 | -0.037 | -0.037 | 0 | 0.23 | 0.04 | 0.15 | 0.24 | 0.2 | 0.22 |
| 2 | 140235838 | rs1515177 | 0.638 | 0.527 | 0.127 | 0.688 | 0.003 | -0.008 | -0.008 | 1.431 | 0.07 | 0.06 | 0.07 | 0.3 | 0.05 | 0.35 |
| 2 | 140238458 | rs11900853 | -1.753 | 0.638 | 0.105 | 0.6 | -0.051 | -0.015 | -0.015 | 7.734 | 0.79 | 0.91 | 0.78 | 0.77 | 0.72 | 0.85 |
| 2 | 140244750 | rs2860047 | -1.74 | 0.639 | 0.114 | 0.6 | -0.048 | -0.013 | -0.013 | 8.451 | 0.79 | 0.92 | 0.78 | 0.77 | 0.73 | 0.85 |
| 2 | 140250506 | rs13408369 | 0.157 | 0.561 | 0.213 | 0.562 | 0.009 | 0.035 | 0.035 | 0 | 0.34 | 0.13 | 0.23 | 0.26 | 0.34 | 0.29 |
| 2 | 140263382 | rs4600589 | -0.478 | 0.536 | 0.204 | 0.552 | 0.022 | 0.039 | 0.039 | 0 | 0.41 | 0.65 | 0.3 | 0.28 | 0.37 | 0.37 |
| 2 | 140265440 | rs1121136 | 0.095 | 0.623 | -0.047 | 0.548 | 0.002 | 0.022 | 0.022 | 0 | 0.34 | 0.1 | 0.22 | 0.29 | 0.34 | 0.29 |
| 2 | 140279603 | rs4549018 | 0.369 | 0.685 | 0.028 | 0.778 | 0 | 0.041 | 0.041 | 0 | 0.08 | 0.09 | 0.09 | 0.35 | 0.05 | 0.42 |
| 2 | 140301222 | rs2376446 | 0.946 | 0.812 | 0.122 | 0.708 | 0.026 | -0.008 | -0.008 | 0.285 | 0.24 | 0.38 | 0.25 | 0.24 | 0.28 | 0.15 |
| 2 | 140313269 | rs2376447 | 1.113 | 0.982 | -0.424 | 0.712 | 0.056 | 0.144 | 0.144 | 0.469 | 0.05 | 0.2 | 0.27 | 0.16 | 0.08 | 0.03 |
| 2 | 140314266 | rs10167444 | -0.035 | 0.576 | -1.109 | 0.572 | -0.014 | -0.004 | -0.004 | 0 | 0.34 | 0.05 | 0.22 | 0.28 | 0.34 | 0.3 |
| 2 | 140318199 | rs6430841 | 0.932 | 0.517 | -1.149 | 0.678 | -0.034 | -0.027 | -0.027 | 0 | 0.2 | 0.01 | 0.22 | 0.19 | 0.18 | 0.21 |
| 2 | 140320186 | rs838042 | -1.551 | 0.691 | -1.392 | 0.498 | -0.106 | -0.055 | -0.055 | 0 | 0.78 | 0.98 | 0.77 | 0.76 | 0.69 | 0.83 |
| 2 | 140323265 | rs1347729 | -0.065 | 0.422 | -1.419 | 0.446 | 0.015 | -0.015 | -0.015 | 0 | 0.38 | 0.08 | 0.52 | 0.66 | 0.33 | 0.63 |
| 2 | 140340859 | rs6742261 | 0.378 | 0.282 | -0.395 | -0.542 | -0.001 | -0.052 | -0.052 | 0 | 0.36 | 0.41 | 0.19 | 0.1 | 0.33 | 0.2 |
| 2 | 140348800 | rs16848205 | 1.227 | 0.394 | -0.271 | -0.538 | -0.005 | 0.02 | 0.02 | 0 | 0.27 | 0.2 | 0.14 | 0.06 | 0.29 | 0.12 |
| 2 | 140356067 | rs1370651 | 1.259 | 0.411 | -0.28 | -0.568 | -0.034 | 0.088 | 0.088 | 0 | 0.27 | 0.12 | 0.14 | 0.05 | 0.32 | 0.13 |
| 2 | 140360216 | rs16842634 | 0.295 | 0.381 | 0.248 | -0.57 | 0.041 | 0.028 | 0.028 | 0 | 0.07 | 0.31 | 0.05 | 0.04 | 0.03 | 0.08 |
| 2 | 140364648 | rs1595435 | -0.088 | 0.332 | 0.418 | -0.505 | 0.031 | 0.025 | 0.025 | 0.005 | 0.39 | 0.22 | 0.53 | 0.69 | 0.31 | 0.62 |
| 2 | 140372174 | rs7564459 | -1.204 | 0.362 | 0.292 | -0.496 | 0.002 | -0.004 | -0.004 | 0 | 0.73 | 0.67 | 0.72 | 0.79 | 0.67 | 0.83 |
| 2 | 140380671 | rs7567706 | -0.037 | 0.332 | 0.122 | -0.457 | 0.033 | 0.021 | 0.021 | 0.009 | 0.38 | 0.09 | 0.52 | 0.7 | 0.31 | 0.61 |
| 2 | 140395703 | rs16842707 | 0.719 | 0.37 | 0.274 | -0.409 | 0.015 | 0.009 | 0.009 | 0 | 0.06 | 0.21 | 0.05 | 0.04 | 0.04 | 0.08 |
| 2 | 140423871 | rs1837161 | -1.156 | 0.281 | 0.163 | -0.683 | 0.009 | -0.012 | -0.012 | 0 | 0.73 | 0.64 | 0.72 | 0.79 | 0.67 | 0.83 |
| 2 | 140426628 | rs1837162 | -0.018 | 0.253 | 0.147 | -0.714 | 0.037 | 0.028 | 0.028 | 0 | 0.38 | 0.15 | 0.52 | 0.7 | 0.31 | 0.62 |
| 2 | 140432189 | rs12692010 | -0.366 | 0.257 | 0.197 | -0.722 | -0.001 | 0 | 0 | 0.823 | 0.66 | 0.32 | 0.67 | 0.75 | 0.63 | 0.74 |
| 2 | 140448156 | rs7574941 | 0.165 | 0.376 | 0.193 | -0.469 | 0.01 | -0.081 | -0.081 | 0.199 | 0.34 | 0.49 | 0.21 | 0.12 | 0.36 | 0.21 |
| 2 | 140464009 | rs6430852 | 0.053 | -0.244 | 0.259 | -0.034 | 0.029 | 0.035 | 0.035 | 0 | 0.06 | 0.25 | 0.06 | 0.06 | 0.03 | 0.09 |
| 2 | 140473877 | rs4954797 | -0.12 | -0.336 | -0.124 | -0.057 | -0.006 | 0.002 | 0.002 | 0.106 | 0.69 | 0.65 | 0.53 | 0.34 | 0.74 | 0.42 |
| 2 | 140484883 | rs1595432 | 0.23 | -0.301 | -0.962 | -0.781 | -0.04 | 0.149 | 0.149 | 0 | 0.34 | 0.12 | 0.21 | 0.06 | 0.42 | 0.19 |
| 2 | 140494663 | rs10193630 | 0.344 | -0.643 | -1.074 | -0.86 | 0.089 | 0.152 | 0.152 | 0 | 0.52 | 0.2 | 0.4 | 0.22 | 0.68 | 0.31 |
| 2 | 140500673 | rs12464074 | 0.997 | -0.912 | -1.215 | -0.657 | -0.018 | -0.008 | -0.008 | 0 | 0.14 | 0.03 | 0.13 | 0.14 | 0.17 | 0.08 |
| 2 | 140509516 | rs7596826 | 0.975 | -0.825 | -1.327 | -0.757 | -0.01 | 0.088 | 0.088 | 0 | 0.32 | 0.15 | 0.18 | 0.07 | 0.35 | 0.11 |
| 2 | 140518301 | rs6723612 | -0.875 | -1.226 | -0.715 | -0.862 | -0.016 | 0.039 | 0.039 | 0 | 0.82 | 0.73 | 0.6 | 0.4 | 0.89 | 0.49 |
| 2 | 140518725 | rs2376761 | 0.094 | -1.185 | -0.111 | -0.878 | -0.009 | -0.009 | -0.009 | 0.116 | 0.27 | 0.28 | 0.17 | 0.18 | 0.19 | 0.17 |
| 2 | 140532933 | rs13384228 | 1.806 | -1.18 | -0.015 | -0.967 | -0.033 | 0.025 | 0.025 | 2.839 | 0.21 | 0.14 | 0.22 | 0.15 | 0.32 | 0.2 |
| 2 | 140533235 | rs938827 | 0.136 | -1.183 | 0.039 | -0.904 | -0.001 | -0.002 | -0.002 | 0.02 | 0.27 | 0.28 | 0.17 | 0.17 | 0.19 | 0.17 |
| 2 | 140544481 | rs16842866 | 1.82 | -1.195 | 0.086 | -0.84 | -0.002 | 0 | 0 | 0 | 0.13 | 0.1 | 0.03 | 0.04 | 0.02 | 0.09 |
| 2 | 140549533 | rs13023556 | 0.862 | -0.887 | 0.267 | -0.756 | 0 | -0.01 | -0.01 | 0.928 | 0.33 | 0.27 | 0.19 | 0.05 | 0.36 | 0.12 |
| 2 | 140551751 | rs9808190 | -1.068 | -0.911 | 0.22 | -0.677 | 0.006 | -0.004 | -0.004 | 0 | 0.84 | 0.87 | 0.64 | 0.4 | 0.89 | 0.51 |
| 2 | 140553141 | rs10928725 | 0.837 | -0.821 | -0.039 | -0.771 | -0.001 | 0.031 | 0.031 | 0 | 0.32 | 0.23 | 0.19 | 0.05 | 0.37 | 0.12 |
| 2 | 140562342 | rs10165615 | 0.033 | -0.696 | 0.097 | -0.643 | 0.005 | 0.033 | 0.033 | 0 | 0.51 | 0.42 | 0.43 | 0.32 | 0.51 | 0.4 |
| 2 | 140562947 | rs2376762 | 0.463 | -0.626 | 0.088 | -0.604 | -0.004 | -0.008 | -0.008 | 0 | 0.28 | 0.31 | 0.2 | 0.18 | 0.2 | 0.17 |
| 2 | 140562994 | rs2376763 | -0.083 | -0.668 | 0.033 | -0.736 | -0.004 | 0.046 | 0.046 | 0 | 0.74 | 0.57 | 0.54 | 0.4 | 0.73 | 0.49 |

| | | | | | | | | | | | | | | | | |
|---|---|---|---|---|---|---|---|---|---|---|---|---|---|---|---|---|
| 2 | 140565448 | rs4395203 | 0.739 | -0.581 | 0.023 | -0.5 | -0.01 | 0.032 | 0.032 | 0.598 | 0.44 | 0.2 | 0.33 | 0.36 | 0.52 | 0.32 |
| 2 | 140565694 | rs1401935 | 1.797 | -0.571 | -0.083 | -0.512 | -0.066 | -0.029 | -0.029 | 0 | 0.22 | 0.06 | 0.23 | 0.3 | 0.32 | 0.24 |
| 2 | 140566297 | rs10179028 | 0.015 | -0.694 | 0.223 | -0.266 | 0.005 | 0 | 0 | 0.001 | 0.6 | 0.39 | 0.51 | 0.43 | 0.69 | 0.45 |
| 2 | 140567666 | rs1356799 | 1.112 | -0.328 | -0.474 | -0.31 | 0.003 | 0.007 | 0.007 | 0 | 0.23 | 0.16 | 0.11 | 0.06 | 0.22 | 0.09 |
| 2 | 140567880 | rs17787272 | 0.641 | 0.13 | -0.594 | 0.243 | 0.007 | -0.006 | -0.006 | 0 | 0.09 | 0.16 | 0.11 | 0.07 | 0.13 | 0.06 |
| 2 | 140577118 | rs10197954 | -0.912 | -0.062 | -0.859 | 0.607 | 0.011 | -0.04 | -0.04 | 0 | 0.88 | 0.94 | 0.66 | 0.52 | 0.91 | 0.53 |
| 2 | 140579673 | rs6724734 | -0.461 | -0.066 | -0.514 | 0.201 | 0.023 | -0.049 | -0.049 | 0.754 | 0.21 | 0.32 | 0.22 | 0.09 | 0.31 | 0.13 |
| 2 | 140579813 | rs6724964 | 0.887 | -0.047 | -0.562 | 0.174 | -0.052 | 0.054 | 0.054 | 0.074 | 0.1 | 0.01 | 0.09 | 0.01 | 0.18 | 0.06 |
| 2 | 140580235 | rs7581253 | -0.819 | -0.051 | -0.61 | 0.179 | 0.004 | -0.073 | -0.073 | 0 | 0.86 | 0.95 | 0.66 | 0.52 | 0.89 | 0.53 |
| 2 | 140580413 | rs7590643 | 0.843 | -0.417 | -0.531 | -0.247 | 0.009 | 0.004 | 0.004 | 1.423 | 0.4 | 0.37 | 0.19 | 0.12 | 0.27 | 0.17 |
| 2 | 140580795 | rs16842932 | 0.987 | -0.293 | -0.665 | -0.155 | 0.024 | -0.005 | -0.005 | 0 | 0.11 | 0.29 | 0.13 | 0.08 | 0.13 | 0.07 |
| 2 | 140586311 | rs9989822 | 1.67 | -0.319 | -0.778 | -0.104 | -0.119 | -0.07 | -0.07 | 1.026 | 0.23 | 0.03 | 0.22 | 0.31 | 0.31 | 0.23 |
| 2 | 140586741 | rs2889460 | 0.613 | -0.413 | -0.74 | 0.353 | 0.003 | 0.006 | 0.006 | 1.923 | 0.67 | 0.63 | 0.43 | 0.57 | 0.59 | 0.47 |
| 2 | 140588462 | rs10206439 | 1.267 | -0.101 | -0.847 | 0.53 | -0.073 | -0.1 | -0.1 | 0.432 | 0.26 | 0.1 | 0.23 | 0.45 | 0.32 | 0.3 |
| 2 | 140596960 | rs13034655 | -0.283 | -0.421 | -0.584 | 0.243 | -0.002 | -0.026 | -0.026 | 0 | 0.37 | 0.38 | 0.29 | 0.14 | 0.35 | 0.19 |
| 2 | 140599522 | rs2376764 | 0.866 | -0.508 | -0.382 | 0.416 | 0.007 | 0.071 | 0.071 | 0.37 | 0.49 | 0.31 | 0.35 | 0.32 | 0.53 | 0.33 |
| 2 | 140603682 | rs16842968 | 1.54 | -0.764 | -0.134 | 0.399 | 0.016 | 0.018 | 0.018 | 2.504 | 0.19 | 0.15 | 0.1 | 0.06 | 0.08 | 0.09 |
| 2 | 140604808 | rs17731461 | 2.234 | -0.658 | -0.158 | 0.405 | 0.03 | 0.016 | 0.016 | 7.285 | 0.06 | 0.15 | 0.09 | 0.08 | 0.1 | 0.04 |
| 2 | 140605306 | rs6758413 | 1.546 | -0.514 | -0.542 | 0.167 | -0.01 | -0.039 | -0.039 | 0.176 | 0.23 | 0.28 | 0.11 | 0.05 | 0.21 | 0.08 |
| 2 | 140608231 | rs10176558 | 1.433 | -0.325 | -0.883 | 0.217 | -0.136 | 0.007 | 0.007 | 1.947 | 0.28 | 0.03 | 0.27 | 0.26 | 0.39 | 0.29 |
| 2 | 140616483 | rs4954665 | -1.685 | -0.154 | -0.754 | 0.33 | 0.004 | 0.006 | 0.006 | 0 | 0.9 | 0.94 | 0.67 | 0.64 | 0.91 | 0.59 |
| 2 | 140629092 | rs10928728 | 1.132 | -0.121 | -0.77 | 0.167 | -0.029 | 0.045 | 0.045 | 0 | 0.08 | 0 | 0.05 | 0.01 | 0.11 | 0.02 |
| 2 | 140629263 | rs10928729 | -1.336 | -0.033 | -0.633 | 0.403 | -0.005 | -0.023 | -0.023 | 0.057 | 0.85 | 0.92 | 0.6 | 0.67 | 0.81 | 0.57 |
| 2 | 140629964 | rs2176805 | 0.605 | -0.213 | -0.457 | 0.171 | -0.018 | -0.047 | -0.047 | 0 | 0.28 | 0.36 | 0.17 | 0.13 | 0.18 | 0.11 |
| 2 | 140651329 | rs6722928 | 0.834 | 0.03 | -0.582 | 0.27 | -0.011 | -0.01 | -0.01 | 0 | 0.32 | 0.27 | 0.31 | 0.41 | 0.4 | 0.36 |
| 2 | 140651778 | rs4588123 | 0.841 | -0.255 | -0.575 | -0.185 | -0.003 | -0.02 | -0.02 | 0 | 0.23 | 0.26 | 0.11 | 0.05 | 0.21 | 0.08 |
| 2 | 140652285 | rs13035654 | -0.051 | -0.323 | -0.403 | -0.18 | -0.001 | -0.015 | -0.015 | 0 | 0.28 | 0.32 | 0.17 | 0.08 | 0.18 | 0.11 |
| 2 | 140652826 | rs16843017 | 0.706 | -0.298 | -0.311 | -0.118 | -0.004 | -0.01 | -0.01 | 0 | 0.19 | 0.25 | 0.11 | 0.07 | 0.08 | 0.1 |
| 2 | 140654531 | rs7575680 | 0.82 | -0.365 | -0.135 | -0.2 | -0.003 | -0.016 | -0.016 | 0 | 0.19 | 0.18 | 0.1 | 0.04 | 0.08 | 0.06 |
| 2 | 140657370 | rs11901615 | -0.666 | 0.495 | -0.794 | 0.071 | -0.036 | -0.07 | -0.07 | 0 | 0.77 | 0.89 | 0.48 | 0.5 | 0.66 | 0.46 |
| 2 | 140657911 | rs12692022 | 0.382 | 0.228 | -0.644 | -0.032 | -0.009 | 0.039 | 0.039 | 0.501 | 0.44 | 0.33 | 0.31 | 0.28 | 0.53 | 0.31 |
| 2 | 140661931 | rs3953980 | 0.289 | 0.044 | -0.616 | 0.195 | -0.007 | 0.006 | 0.006 | 0 | 0.08 | 0.1 | 0.03 | 0.1 | 0.07 | 0.07 |
| 2 | 140665986 | rs13009101 | 0.693 | 0.313 | -0.907 | 0.005 | -0.042 | 0.042 | 0.042 | 0 | 0.08 | 0.01 | 0.06 | 0.01 | 0.11 | 0.02 |
| 2 | 140666027 | rs7606774 | -1.664 | -0.088 | -1.619 | -0.343 | -0.077 | -0.031 | -0.031 | 0.534 | 0.62 | 0.92 | 0.42 | 0.32 | 0.53 | 0.43 |
| 2 | 140666340 | rs11675023 | 0.83 | -0.015 | -1.694 | -0.549 | -0.051 | -0.003 | -0.003 | 0 | 0.19 | 0.03 | 0.14 | 0.13 | 0.21 | 0.21 |
| 2 | 140680731 | rs16843069 | 0.695 | -0.137 | -1.025 | -0.535 | -0.062 | -0.074 | -0.074 | 0 | 0.24 | 0.52 | 0.15 | 0.08 | 0.1 | 0.12 |
| 2 | 140683305 | rs7568693 | 0.997 | -0.11 | -1.084 | -0.569 | -0.001 | -0.005 | -0.005 | 0 | 0.46 | 0.45 | 0.64 | 0.78 | 0.53 | 0.64 |
| 2 | 140699470 | rs12479038 | -0.206 | 0.222 | -1.558 | -0.343 | -0.112 | 0.04 | 0.04 | 0.021 | 0.27 | 0.03 | 0.2 | 0.14 | 0.32 | 0.23 |
| 2 | 140709142 | rs13035203 | -0.097 | 0.213 | -1.165 | -0.299 | -0.016 | 0.025 | 0.025 | 0.036 | 0.12 | 0.05 | 0.08 | 0.01 | 0.15 | 0.05 |
| 2 | 140713355 | rs10174598 | 1.796 | 0.129 | -1.704 | -0.095 | -0.095 | -0.145 | -0.145 | 0 | 0.32 | 0.04 | 0.49 | 0.64 | 0.39 | 0.58 |
| 2 | 140714614 | rs7607195 | -1.326 | -0.182 | -1.023 | -0.512 | -0.007 | -0.041 | -0.041 | 0.125 | 0.8 | 0.79 | 0.87 | 0.99 | 0.9 | 0.9 |
| 2 | 140723694 | rs3856354 | 0.723 | 0.043 | -1.152 | -0.485 | -0.007 | 0.061 | 0.061 | 0 | 0.46 | 0.69 | 0.56 | 0.81 | 0.48 | 0.68 |
| 2 | 140724573 | rs1920526 | -0.685 | 0.192 | -1.405 | -0.38 | -0.079 | 0.044 | 0.044 | 1.775 | 0.34 | 0.09 | 0.31 | 0.19 | 0.42 | 0.22 |
| 2 | 140733326 | rs1920519 | 0.454 | 0.575 | -1.036 | -0.17 | -0.083 | -0.036 | -0.036 | 0.086 | 0.21 | 0.02 | 0.21 | 0.17 | 0.27 | 0.18 |
| 2 | 140737439 | rs2380747 | -1.146 | 0.487 | -0.776 | -0.304 | -0.002 | -0.09 | -0.09 | 0 | 0.83 | 0.69 | 0.85 | 0.98 | 0.85 | 0.88 |
| 2 | 140739894 | rs12986848 | -0.447 | 0.457 | -0.61 | 0.053 | 0.015 | 0.182 | 0.182 | 0 | 0.17 | 0.38 | 0.22 | 0.35 | 0.18 | 0.31 |
| 2 | 140743217 | rs2141652 | 0.682 | -0.252 | -0.537 | 0.066 | -0.001 | 0.008 | 0.008 | 0 | 0.53 | 0.54 | 0.62 | 0.79 | 0.54 | 0.69 |
| 2 | 140752290 | rs4577229 | 0.94 | -0.227 | -0.712 | 0.153 | 0.067 | -0.048 | -0.048 | 0 | 0.58 | 0.12 | 0.63 | 0.61 | 0.7 | 0.56 |
| 2 | 140753902 | rs7580389 | 0.492 | -0.191 | -0.346 | 0.381 | -0.005 | 0.035 | 0.035 | 0 | 0.1 | 0.13 | 0.17 | 0.27 | 0.09 | 0.27 |
| 2 | 140756534 | rs7579130 | -1.069 | -0.151 | -0.056 | 0.181 | -0.046 | 0.102 | 0.102 | 0 | 0.24 | 0.45 | 0.22 | 0.36 | 0.16 | 0.26 |
| 2 | 140768891 | rs12162312 | -0.362 | -0.148 | -0.054 | 0.104 | -0.029 | 0.112 | 0.112 | 0 | 0.15 | 0.27 | 0.19 | 0.34 | 0.12 | 0.26 |
| 2 | 140773470 | rs1569097 | -1.309 | -0.143 | -0.002 | -0.039 | -0.016 | -0.093 | -0.093 | 0 | 0.84 | 0.67 | 0.87 | 0.98 | 0.9 | 0.87 |
| 2 | 140774793 | rs10175740 | 0.948 | -0.101 | -0.178 | 0.003 | 0.033 | -0.004 | -0.004 | 0.262 | 0.6 | 0.33 | 0.66 | 0.63 | 0.73 | 0.6 |
| 2 | 140786028 | rs1527847 | 0.436 | 0.082 | -0.471 | 0.028 | -0.084 | -0.048 | -0.048 | 3.59 | 0.21 | 0.02 | 0.21 | 0.2 | 0.28 | 0.18 |
| 2 | 140790828 | rs12052525 | 0.536 | 0.016 | -0.215 | 0.123 | -0.005 | 0.046 | 0.046 | 0 | 0.11 | 0.13 | 0.18 | 0.3 | 0.12 | 0.24 |
| 2 | 140791757 | rs1015739 | -1.276 | 0.111 | -0.176 | -0.166 | -0.011 | 0.012 | 0.012 | 0.303 | 0.37 | 0.47 | 0.37 | 0.52 | 0.37 | 0.45 |
| 2 | 140794290 | rs10928736 | -0.712 | 0.105 | -0.186 | -0.162 | -0.003 | 0.004 | 0.004 | 0.087 | 0.33 | 0.32 | 0.36 | 0.51 | 0.32 | 0.42 |
| 2 | 140794737 | rs10221883 | 1.938 | 0.101 | -0.288 | -0.219 | -0.101 | -0.079 | -0.079 | 0 | 0.36 | 0.07 | 0.41 | 0.43 | 0.4 | 0.4 |
| 2 | 140797961 | rs4359577 | -1.279 | -0.005 | -0.137 | -0.198 | 0.003 | -0.094 | -0.094 | 2.071 | 0.81 | 0.69 | 0.82 | 0.98 | 0.81 | 0.85 |
| 2 | 140809596 | rs16843261 | 0.9 | 0.155 | -0.349 | -0.123 | -0.003 | 0.007 | 0.007 | 0.538 | 0.48 | 0.45 | 0.46 | 0.47 | 0.47 | 0.4 |
| 2 | 140813276 | rs12105000 | 1.524 | 0.249 | -0.848 | 0.01 | -0.012 | -0.017 | -0.017 | 0.457 | 0.37 | 0.18 | 0.41 | 0.43 | 0.38 | 0.37 |
| 2 | 140819890 | rs12053320 | 0.465 | 0.388 | -0.982 | 0.18 | -0.06 | -0.039 | -0.039 | 0 | 0.18 | 0.02 | 0.17 | 0.15 | 0.2 | 0.17 |
| 2 | 140821180 | rs1527849 | -0.862 | 0.362 | -0.678 | -0.039 | 0.001 | -0.004 | -0.004 | 0 | 0.33 | 0.3 | 0.36 | 0.51 | 0.33 | 0.46 |
| 2 | 140829061 | rs6739604 | -1.205 | 0.724 | -1.281 | 0.098 | -0.061 | 0.023 | 0.023 | 0.05 | 0.58 | 0.9 | 0.56 | 0.63 | 0.55 | 0.56 |
| 2 | 140829708 | rs1405933 | -1.334 | 0.728 | -1.369 | 0.091 | -0.071 | -0.026 | -0.026 | 0 | 0.6 | 0.95 | 0.58 | 0.63 | 0.61 | 0.58 |
| 2 | 140841941 | rs2380750 | -1.284 | 0.537 | -1.192 | 0.166 | -0.03 | 0.011 | 0.011 | 0 | 0.58 | 0.87 | 0.56 | 0.63 | 0.6 | 0.57 |
| 2 | 140843027 | rs1569103 | -1.189 | 0.502 | -1.207 | 0.26 | -0.091 | 0.018 | 0.018 | 0 | 0.56 | 0.93 | 0.55 | 0.63 | 0.55 | 0.53 |
| 2 | 140847560 | rs2380752 | 0.3 | 0.743 | -0.732 | -0.205 | -0.009 | -0.009 | -0.009 | 0 | 0.11 | 0.09 | 0.16 | 0.22 | 0.15 | 0.15 |
| 2 | 140858451 | rs16843311 | 0.607 | 0.55 | -0.452 | -0.257 | -0.03 | -0.062 | -0.062 | 0.25 | 0.22 | 0.48 | 0.17 | 0.13 | 0.18 | 0.09 |

| | | | | | | | | | | | | | | | | |
|---|---|---|---|---|---|---|---|---|---|---|---|---|---|---|---|---|
| 2 | 140868186 | rs1405930 | -0.077 | 0.322 | -0.824 | -0.374 | 0.03 | 0.036 | 0.036 | 0 | 0.77 | 0.88 | 0.91 | 0.93 | 0.84 | 0.86 |
| 2 | 140870484 | rs6718792 | 1.377 | 0.507 | -0.87 | -0.323 | -0.054 | -0.06 | -0.06 | 0 | 0.4 | 0.14 | 0.46 | 0.57 | 0.43 | 0.63 |
| 2 | 140875531 | rs11889562 | 0.681 | -0.192 | -0.817 | -0.236 | -0.007 | -0.013 | -0.013 | 0 | 0.11 | 0.12 | 0.06 | 0.07 | 0.1 | 0.12 |
| 2 | 140880629 | rs6430871 | -0.108 | -0.283 | -0.808 | -0.255 | -0.005 | -0.018 | -0.018 | 0 | 0.68 | 0.73 | 0.81 | 0.88 | 0.75 | 0.88 |
| 2 | 140884146 | rs4954669 | 1.503 | 0.174 | -1.939 | -0.187 | -0.058 | -0.101 | -0.101 | 0.021 | 0.36 | 0.1 | 0.38 | 0.58 | 0.36 | 0.54 |
| 2 | 140896080 | rs1969503 | 1.401 | 1.632 | -0.137 | 0.993 | 0.002 | 0 | 0 | 4.926 | 0.71 | 0.7 | 0.59 | 0.7 | 0.61 | 0.66 |
| 2 | 140900604 | rs2966395 | 0.689 | 1.478 | -0.35 | 1.082 | -0.02 | -0.015 | -0.015 | 0 | 0.83 | 0.88 | 0.76 | 0.81 | 0.75 | 0.79 |
| 2 | 140910687 | rs7568724 | -0.274 | 0.631 | -0.089 | 0.849 | 0.002 | 0.025 | 0.025 | 0 | 0.48 | 0.52 | 0.44 | 0.36 | 0.49 | 0.38 |
| 2 | 140915674 | rs1405923 | 0.514 | 0.627 | 0.273 | 1.293 | 0.055 | 0.028 | 0.028 | 0 | 0.41 | 0.28 | 0.38 | 0.35 | 0.36 | 0.38 |
| 2 | 140922911 | rs825714 | 0.674 | 0.596 | 0.868 | 1.002 | 0.026 | 0.013 | 0.013 | 2.392 | 0.9 | 0.62 | 0.96 | 0.92 | 0.95 | 0.92 |
| 2 | 140923936 | rs10174088 | -0.081 | 0.876 | 0.652 | 1.286 | -0.008 | 0.021 | 0.021 | 0 | 0.37 | 0.32 | 0.44 | 0.2 | 0.51 | 0.4 |
| 2 | 140926018 | rs884460 | 0.105 | 0.871 | 0.246 | 1.284 | -0.045 | 0.045 | 0.045 | 0 | 0.35 | 0.21 | 0.42 | 0.21 | 0.47 | 0.36 |
| 2 | 140930736 | rs1527860 | -0.322 | 0.71 | -0.22 | 1.169 | 0 | 0 | 0 | 0 | 0.43 | 0.55 | 0.44 | 0.47 | 0.5 | 0.44 |
| 2 | 140943640 | rs9287533 | -0.123 | -0.093 | -0.791 | 0.799 | 0.05 | 0.03 | 0.03 | 0.434 | 0.48 | 0.68 | 0.48 | 0.45 | 0.33 | 0.44 |
| 2 | 140945164 | rs6715982 | -0.022 | -0.159 | -0.428 | 0.857 | 0.013 | 0.01 | 0.01 | 0.056 | 0.45 | 0.56 | 0.42 | 0.39 | 0.31 | 0.42 |
| 2 | 140948967 | rs10197964 | -0.636 | 0.191 | -0.848 | 0.743 | -0.04 | 0.015 | 0.015 | 0 | 0.74 | 0.82 | 0.69 | 0.82 | 0.58 | 0.65 |
| 2 | 140949078 | rs1492390 | 0.294 | 0.064 | -0.099 | 0.705 | 0.009 | -0.016 | -0.016 | 2.039 | 0.29 | 0.26 | 0.27 | 0.42 | 0.25 | 0.22 |
| 2 | 140956836 | rs923739 | 0.117 | 0.127 | -0.092 | 0.703 | -0.012 | 0.018 | 0.018 | 0 | 0.3 | 0.39 | 0.29 | 0.42 | 0.26 | 0.22 |
| 2 | 140961449 | rs6725195 | 0.414 | 0.124 | -0.057 | 0.717 | 0.039 | 0.007 | 0.007 | 0.154 | 0.47 | 0.6 | 0.42 | 0.5 | 0.32 | 0.43 |
| 2 | 140962137 | rs10200179 | 0.704 | 0.12 | -0.083 | 0.662 | -0.009 | -0.006 | -0.006 | 0 | 0.43 | 0.4 | 0.4 | 0.4 | 0.31 | 0.42 |
| 2 | 140966549 | rs4954670 | 0.512 | -0.118 | -0.406 | 0.451 | 0 | -0.017 | -0.017 | 0 | 0.09 | 0.09 | 0.06 | 0 | 0.07 | 0.07 |
| 2 | 140968308 | rs1008695 | 0.929 | -0.036 | -0.435 | 0.475 | 0.011 | -0.065 | -0.065 | 0.007 | 0.18 | 0.1 | 0.21 | 0.32 | 0.19 | 0.16 |
| 2 | 140973503 | rs745021 | -1.093 | 0.639 | -0.478 | 0.272 | -0.029 | 0.016 | 0.016 | 0 | 0.79 | 0.86 | 0.76 | 0.91 | 0.63 | 0.73 |
| 2 | 140978431 | rs4954671 | 0.458 | 0.387 | -0.39 | 0.285 | 0.002 | -0.008 | -0.008 | 0 | 0.09 | 0.06 | 0.07 | 0 | 0.07 | 0.07 |
| 2 | 140982918 | rs2029629 | 0.384 | -0.079 | -0.451 | 0.248 | 0.007 | 0.004 | 0.004 | 0 | 0.47 | 0.52 | 0.44 | 0.46 | 0.34 | 0.46 |
| 2 | 140988796 | rs12463968 | 1.509 | -0.717 | -0.34 | -0.125 | 0.01 | 0.003 | 0.003 | 0 | 0.26 | 0.21 | 0.31 | 0.28 | 0.21 | 0.14 |
| 2 | 140989843 | rs1047874 | 0 | -0.739 | -0.22 | -0.152 | 0.058 | 0.034 | 0.034 | 0 | 0.51 | 0.4 | 0.54 | 0.54 | 0.64 | 0.53 |
| 2 | 140992337 | rs13387241 | 0.234 | -0.731 | -0.205 | -0.186 | 0.02 | 0.023 | 0.023 | 0 | 0.46 | 0.32 | 0.53 | 0.51 | 0.59 | 0.45 |
| 2 | 140998670 | rs1388809 | 0.72 | -0.749 | -0.197 | -0.199 | -0.027 | -0.024 | -0.024 | 0.006 | 0.37 | 0.39 | 0.37 | 0.31 | 0.27 | 0.37 |
| 2 | 140999818 | rs12464830 | 1.583 | -0.751 | -0.242 | -0.191 | 0.01 | 0.003 | 0.003 | 0 | 0.26 | 0.21 | 0.3 | 0.28 | 0.21 | 0.15 |
| 2 | 141000302 | rs16843644 | 0.741 | -0.491 | -0.27 | 0.044 | 0.003 | -0.034 | -0.034 | 0.361 | 0.23 | 0.2 | 0.23 | 0.34 | 0.24 | 0.23 |
| 2 | 141005223 | rs17385521 | 0.938 | -0.454 | -0.384 | 0.042 | -0.151 | -0.026 | -0.026 | 0.06 | 0.26 | 0.03 | 0.31 | 0.2 | 0.4 | 0.27 |
| 2 | 141013919 | rs13021003 | -0.597 | -0.536 | -0.293 | 0.03 | 0.023 | 0.038 | 0.038 | 0 | 0.57 | 0.4 | 0.61 | 0.65 | 0.68 | 0.53 |
| 2 | 141023643 | rs10469593 | 0.238 | -0.526 | -0.299 | 0.051 | 0.037 | 0.013 | 0.013 | 0.028 | 0.45 | 0.71 | 0.42 | 0.49 | 0.34 | 0.5 |
| 2 | 141023886 | rs13004107 | 0.139 | -0.424 | -0.35 | 0.028 | -0.105 | 0.061 | 0.061 | 0 | 0.34 | 0.1 | 0.37 | 0.19 | 0.45 | 0.31 |
| 2 | 141024545 | rs16843732 | 1.636 | -0.422 | -0.341 | 0.036 | -0.014 | -0.117 | -0.117 | 0 | 0.17 | 0.01 | 0.2 | 0.31 | 0.19 | 0.18 |
| 2 | 141032476 | rs1492393 | 0.17 | -0.354 | -0.271 | 0.072 | 0.021 | 0.037 | 0.037 | 0 | 0.44 | 0.64 | 0.42 | 0.35 | 0.33 | 0.48 |
| 2 | 141040236 | rs2171322 | 0.725 | -0.369 | -0.228 | 0.071 | -0.039 | -0.032 | -0.032 | 0 | 0.37 | 0.47 | 0.37 | 0.32 | 0.26 | 0.38 |
| 2 | 141043913 | rs1486969 | -0.048 | -0.3 | -0.207 | 0.091 | 0.041 | 0.106 | 0.106 | 0 | 0.93 | 0.73 | 0.97 | 0.84 | 0.95 | 0.88 |
| 2 | 141049869 | rs10496839 | 1.552 | -0.597 | -0.296 | -0.304 | 0.019 | 0.078 | 0.078 | 0 | 0.09 | 0.15 | 0.28 | 0.29 | 0.12 | 0.1 |
| 2 | 141050608 | rs10496840 | 0.564 | -0.174 | -0.33 | 0.064 | -0.001 | -0.001 | -0.001 | 1.398 | 0.4 | 0.59 | 0.44 | 0.32 | 0.28 | 0.39 |
| 2 | 141060408 | rs12692042 | 0.601 | -0.046 | -0.73 | -0.007 | -0.053 | 0.005 | 0.005 | 0 | 0.24 | 0.15 | 0.31 | 0.2 | 0.39 | 0.27 |
| 2 | 141083159 | rs654051 | -1.299 | -0.164 | -0.601 | -0.027 | -0.039 | -0.033 | -0.033 | 0 | 0.63 | 0.58 | 0.58 | 0.69 | 0.73 | 0.63 |
| 2 | 141084395 | rs11894060 | -0.055 | -0.173 | -0.637 | 0.027 | 0.024 | -0.054 | -0.054 | 0 | 0.33 | 0.17 | 0.29 | 0.37 | 0.31 | 0.25 |
| 2 | 141090153 | rs16843791 | 0.789 | -0.202 | -0.632 | -0.122 | 0.003 | -0.004 | -0.004 | 0 | 0.09 | 0.08 | 0.08 | 0 | 0.08 | 0.04 |
| 2 | 141090216 | rs10195473 | 1.155 | -0.265 | -0.388 | -0.131 | -0.008 | -0.007 | -0.007 | 0.864 | 0.36 | 0.35 | 0.41 | 0.31 | 0.27 | 0.37 |
| 2 | 141098446 | rs600802 | -0.694 | 0.023 | -0.555 | -0.056 | 0.007 | -0.018 | -0.018 | 0 | 0.38 | 0.46 | 0.42 | 0.21 | 0.5 | 0.32 |
| 2 | 141101586 | rs13019002 | 0.229 | 0.048 | -0.665 | -0.054 | 0.003 | 0.002 | 0.002 | 0 | 0.11 | 0.11 | 0.08 | 0.02 | 0.11 | 0.05 |
| 2 | 141101864 | rs13002228 | -0.535 | 0.046 | -0.662 | -0.056 | -0.024 | 0.032 | 0.032 | 0 | 0.36 | 0.31 | 0.4 | 0.21 | 0.5 | 0.32 |
| 2 | 141102038 | rs35699793 | 0.746 | 0.064 | -0.676 | -0.043 | -0.061 | -0.002 | -0.002 | 0 | 0.23 | 0.1 | 0.31 | 0.2 | 0.37 | 0.26 |
| 2 | 141110677 | rs13034302 | 0.559 | 0.074 | -0.511 | 0.035 | 0.004 | 0.004 | 0.004 | 0 | 0.09 | 0.05 | 0.08 | 0 | 0.08 | 0.04 |
| 2 | 141123496 | rs4954830 | 0.488 | 0.078 | -0.508 | 0.033 | -0.001 | -0.019 | -0.019 | 0 | 0.09 | 0.11 | 0.07 | 0 | 0.07 | 0.04 |
| 2 | 141126089 | rs12997262 | -0.418 | 0.082 | -0.498 | 0.03 | -0.001 | -0.021 | -0.021 | 0 | 0.13 | 0.2 | 0.1 | 0.02 | 0.12 | 0.06 |
| 2 | 141130097 | rs2931465 | -0.186 | 0.081 | -0.422 | 0.087 | 0.045 | 0.134 | 0.134 | 0 | 0.06 | 0.25 | 0.03 | 0.17 | 0.05 | 0.1 |
| 2 | 141130695 | rs16843864 | -0.4 | 0.083 | -0.427 | 0.078 | 0 | -0.019 | -0.019 | 0 | 0.13 | 0.2 | 0.1 | 0.02 | 0.12 | 0.06 |
| 2 | 141133405 | rs670617 | -0.242 | 0.101 | -0.459 | 0.105 | 0.048 | 0.132 | 0.132 | 0 | 0.06 | 0.25 | 0.03 | 0.17 | 0.05 | 0.1 |
| 2 | 141141060 | rs13028094 | 0.367 | 0.102 | -0.595 | -0.013 | 0.002 | -0.012 | -0.012 | 0 | 0.1 | 0.12 | 0.08 | 0 | 0.09 | 0.04 |
| 2 | 141151319 | rs16843900 | -0.336 | 0.108 | -0.586 | -0.017 | 0 | -0.012 | -0.012 | 0 | 0.13 | 0.17 | 0.1 | 0.02 | 0.12 | 0.06 |
| 2 | 141153229 | rs10460268 | -0.731 | 0.108 | -0.727 | -0.022 | 0.003 | -0.002 | -0.002 | 0.554 | 0.36 | 0.36 | 0.26 | 0.36 | 0.35 | 0.31 |
| 2 | 141166435 | rs1568256 | -0.378 | 0.14 | -0.597 | -0.327 | 0 | -0.012 | -0.012 | 0.229 | 0.13 | 0.17 | 0.1 | 0.02 | 0.13 | 0.06 |
| 2 | 141173769 | rs4954834 | 0.118 | 0.13 | -0.579 | -0.325 | -0.01 | -0.064 | -0.064 | 0.163 | 0.11 | 0.23 | 0.09 | 0 | 0.09 | 0.04 |
| 2 | 141180816 | rs4954835 | -0.672 | 0.886 | -0.835 | -0.38 | -0.006 | -0.004 | -0.004 | 0 | 0.46 | 0.45 | 0.36 | 0.49 | 0.5 | 0.4 |
| 2 | 141190436 | rs11897270 | -0.134 | 1.334 | -0.592 | -0.349 | 0.022 | -0.084 | -0.084 | 1.256 | 0.23 | 0.31 | 0.19 | 0.02 | 0.32 | 0.26 |
| 2 | 141194627 | rs17387292 | 1.316 | 1.407 | -0.752 | -0.317 | -0.056 | 0.06 | 0.06 | 0 | 0.13 | 0.04 | 0.11 | 0.02 | 0.2 | 0.18 |
| 2 | 141197873 | rs1518444 | 0.468 | 1.403 | -0.731 | -0.326 | -0.02 | 0.006 | 0.006 | 0 | 0.17 | 0.18 | 0.13 | 0.02 | 0.26 | 0.21 |
| 2 | 141202644 | rs7420069 | 0.563 | 1.575 | -0.578 | -0.054 | -0.002 | 0.007 | 0.007 | 0 | 0.1 | 0.06 | 0.18 | 0.23 | 0.11 | 0.16 |
| 2 | 141204257 | rs7422845 | 1.128 | 1.431 | -0.632 | -0.637 | -0.032 | 0.029 | 0.029 | 0 | 0.13 | 0.09 | 0.11 | 0.02 | 0.2 | 0.18 |
| 2 | 141239598 | rs3923350 | 1.228 | 0.767 | -0.351 | -0.825 | 0.039 | 0.041 | 0.041 | 0 | 0.32 | 0.19 | 0.42 | 0.24 | 0.27 | 0.15 |
| 2 | 141245486 | rs7563696 | 0.847 | 1.389 | 0.023 | -0.246 | -0.002 | -0.029 | -0.029 | 0.193 | 0.57 | 0.46 | 0.54 | 0.28 | 0.46 | 0.46 |

| | | | | | | | | | | | | | | | | |
|---|---|---|---|---|---|---|---|---|---|---|---|---|---|---|---|---|
| 2 | 141253408 | rs11683334 | -1.078 | 1.38 | 0.068 | -0.256 | -0.001 | 0.006 | 0.006 | 0 | 0.42 | 0.46 | 0.45 | 0.72 | 0.54 | 0.53 |
| 2 | 141258419 | rs13422213 | -0.196 | 1.901 | -0.076 | -0.04 | 0.037 | -0.086 | -0.086 | 0 | 0.18 | 0.3 | 0.14 | 0.02 | 0.27 | 0.23 |
| 2 | 141269323 | rs6430904 | 0.753 | 1.916 | -0.137 | 1.245 | 0.019 | 0.038 | 0.038 | 0 | 0.81 | 0.73 | 0.73 | 0.44 | 0.71 | 0.6 |
| 2 | 141269874 | rs1463615 | 0.647 | 1.901 | -0.081 | 1.238 | -0.012 | -0.013 | -0.013 | 0 | 0.82 | 0.83 | 0.74 | 0.44 | 0.71 | 0.6 |
| 2 | 141270001 | rs1463614 | 0.594 | 1.913 | -0.09 | 1.647 | -0.02 | -0.026 | -0.026 | 0 | 0.82 | 0.84 | 0.74 | 0.53 | 0.71 | 0.61 |
| 2 | 141274576 | rs4954672 | 0.843 | 2.044 | 0.218 | 1.651 | 0.039 | 0.079 | 0.079 | 0.475 | 0.83 | 0.69 | 0.74 | 0.42 | 0.75 | 0.63 |
| 2 | 141281486 | rs6715219 | 0.352 | 2.408 | 0.496 | 0.951 | 0.001 | -0.022 | -0.022 | 0 | 0.13 | 0.14 | 0.13 | 0.01 | 0.21 | 0.18 |
| 2 | 141285404 | rs1352046 | -0.249 | 2.332 | 0.442 | 1.007 | 0.008 | -0.001 | -0.001 | 0 | 0.86 | 0.82 | 0.87 | 0.9 | 0.79 | 0.8 |
| 2 | 141285694 | rs1385769 | -0.016 | 2.322 | 0.556 | 0.999 | 0.053 | -0.009 | -0.009 | 1.621 | 0.85 | 0.67 | 0.85 | 0.9 | 0.79 | 0.8 |
| 2 | 141287873 | rs1485848 | -1.302 | 1.972 | 0.404 | 0.873 | 0.008 | -0.001 | -0.001 | 0.322 | 0.37 | 0.44 | 0.45 | 0.61 | 0.51 | 0.62 |
| 2 | 141288260 | rs10496845 | -0.339 | 1.836 | 0.437 | 0.82 | -0.017 | -0.023 | -0.023 | 0 | 0.08 | 0.18 | 0.03 | 0.03 | 0.03 | 0.01 |
| 2 | 141295248 | rs2380885 | 0.591 | 2.302 | 0.072 | 0.87 | -0.032 | -0.025 | -0.025 | 0 | 0.05 | 0.01 | 0.23 | 0.17 | 0.14 | 0.05 |
| 2 | 141297317 | rs16844206 | 0.729 | 2.145 | 0.183 | 0.815 | -0.001 | -0.011 | -0.011 | 0 | 0.06 | 0.04 | 0.03 | 0 | 0.03 | 0 |
| 2 | 141298467 | rs6729619 | -0.692 | 2.183 | 0.265 | 0.848 | 0.005 | 0.002 | 0.002 | 0 | 0.31 | 0.35 | 0.42 | 0.58 | 0.48 | 0.62 |
| 2 | 141300273 | rs7598849 | -0.341 | 2.634 | 0.285 | 1.804 | -0.068 | -0.053 | -0.053 | 0 | 0.26 | 0.12 | 0.38 | 0.35 | 0.4 | 0.48 |
| 2 | 141300595 | rs6430913 | -0.814 | 2.619 | 0.281 | 1.734 | -0.043 | -0.024 | -0.024 | 0 | 0.29 | 0.25 | 0.42 | 0.58 | 0.48 | 0.61 |
| 2 | 141307263 | rs6430918 | -1.473 | 2.639 | 0.267 | 1.736 | -0.006 | -0.002 | -0.002 | 0.28 | 0.36 | 0.37 | 0.44 | 0.58 | 0.49 | 0.62 |
| 2 | 141312831 | rs13032617 | 1.558 | 2.856 | 0.152 | 2.143 | -0.059 | -0.048 | -0.048 | 0 | 0.09 | 0.01 | 0.11 | 0.12 | 0.17 | 0.16 |
| 2 | 141321183 | rs2380887 | 1.171 | 1.799 | -0.238 | 1.33 | -0.005 | -0.008 | -0.008 | 0.598 | 0.55 | 0.47 | 0.51 | 0.36 | 0.43 | 0.28 |
| 2 | 141331966 | rs11686165 | 0.387 | 1.687 | -0.394 | 1.359 | -0.001 | 0.009 | 0.009 | 0 | 0.15 | 0.2 | 0.09 | 0.22 | 0.14 | 0.19 |
| 2 | 141334108 | rs10496849 | -0.276 | 1.827 | -0.521 | 1.361 | -0.062 | -0.046 | -0.046 | 1.067 | 0.29 | 0.14 | 0.38 | 0.38 | 0.42 | 0.53 |
| 2 | 141334737 | rs6732761 | 1.162 | 1.592 | -0.316 | 1.247 | -0.001 | -0.002 | -0.002 | 0 | 0.54 | 0.43 | 0.52 | 0.35 | 0.44 | 0.28 |
| 2 | 141336597 | rs10170926 | -0.196 | 1.083 | -0.394 | 1.254 | 0.004 | 0.004 | 0.004 | 0.503 | 0.18 | 0.22 | 0.29 | 0.25 | 0.24 | 0.39 |
| 2 | 141338918 | rs7600406 | 0.119 | 0.124 | -0.393 | 0.779 | -0.005 | 0.001 | 0.001 | 1.374 | 0.56 | 0.58 | 0.71 | 0.57 | 0.63 | 0.6 |
| 2 | 141340626 | rs1564933 | 0.547 | -0.175 | -0.624 | 0.658 | 0 | 0.004 | 0.004 | 1.763 | 0.45 | 0.39 | 0.62 | 0.47 | 0.55 | 0.55 |
| 2 | 141343100 | rs16844281 | 0.495 | -0.254 | -0.282 | 0.669 | -0.007 | 0.002 | 0.002 | 0 | 0.12 | 0.3 | 0.1 | 0.09 | 0.09 | 0.05 |
| 2 | 141346007 | rs1485845 | -0.542 | -0.829 | -0.877 | 0.588 | 0.006 | 0 | 0 | 0 | 0.37 | 0.39 | 0.21 | 0.33 | 0.25 | 0.27 |
| 2 | 141348478 | rs6752743 | 1.852 | -0.238 | -0.515 | 0.881 | 0.048 | 0.084 | 0.084 | 0 | 0.12 | 0.16 | 0.19 | 0.3 | 0.2 | 0.2 |
| 2 | 141348712 | rs693474 | -0.536 | -0.901 | -0.692 | 0.322 | -0.014 | -0.053 | -0.053 | 0 | 0.36 | 0.59 | 0.25 | 0.09 | 0.31 | 0.17 |
| 2 | 141349306 | rs483128 | 0.016 | -0.871 | -1.01 | 0.395 | -0.006 | 0.004 | 0.004 | 0 | 0.6 | 0.68 | 0.55 | 0.33 | 0.56 | 0.33 |
| 2 | 141350100 | rs511625 | 1.465 | -1.128 | -0.871 | 0.202 | -0.001 | 0 | 0 | 0 | 0.35 | 0.37 | 0.36 | 0.24 | 0.35 | 0.26 |
| 2 | 141354396 | rs1372256 | 0.525 | -1.145 | -1.031 | 0.272 | -0.002 | -0.058 | -0.058 | 0 | 0.25 | 0.11 | 0.25 | 0.37 | 0.21 | 0.45 |
| 2 | 141355107 | rs492190 | 2.234 | -1.226 | -0.905 | 0.202 | -0.012 | -0.035 | -0.035 | 0.15 | 0.07 | 0.21 | 0.03 | 0 | 0.05 | 0.08 |
| 2 | 141355954 | rs1025832 | 3.194 | -1.134 | -1.052 | 0.254 | -0.034 | -0.022 | -0.022 | 0.477 | 0.21 | 0.09 | 0.29 | 0.24 | 0.25 | 0.15 |
| 2 | 141361688 | rs1441467 | 1.864 | -1.212 | -1.081 | 0.322 | 0.041 | -0.036 | -0.036 | 0 | 0.14 | 0.06 | 0.04 | 0.21 | 0.06 | 0.16 |
| 2 | 141361747 | rs1441468 | -1.136 | -1.206 | -0.959 | 0.318 | 0.025 | 0.016 | 0.016 | 0 | 0.5 | 0.41 | 0.44 | 0.46 | 0.42 | 0.52 |
| 2 | 141361994 | rs16844322 | 1.943 | -0.867 | -0.758 | 0.548 | -0.05 | -0.038 | -0.038 | 0.037 | 0.08 | 0.01 | 0.21 | 0.16 | 0.15 | 0.28 |
| 2 | 141363377 | rs12470296 | 1.335 | -0.882 | -0.53 | 0.375 | -0.016 | -0.011 | -0.011 | 0 | 0.14 | 0.26 | 0.13 | 0.09 | 0.12 | 0.04 |
| 2 | 141363453 | rs502443 | 0.317 | -0.896 | -0.556 | 0.344 | 0.003 | -0.002 | -0.002 | 0 | 0.58 | 0.58 | 0.61 | 0.54 | 0.64 | 0.48 |
| 2 | 141365171 | rs10153880 | 2.64 | -0.58 | -0.525 | 0.582 | -0.026 | -0.003 | -0.003 | 0 | 0.26 | 0.11 | 0.31 | 0.24 | 0.28 | 0.18 |
| 2 | 141367562 | rs11674356 | 1 | -0.47 | 0.246 | 1.106 | -0.002 | -0.002 | -0.002 | 0 | 0.13 | 0.18 | 0.12 | 0.09 | 0.12 | 0.04 |
| 2 | 141367579 | rs554360 | 0.934 | -0.538 | 0.582 | 1.158 | 0.052 | 0.083 | 0.083 | 0 | 0.17 | 0.34 | 0.22 | 0.3 | 0.25 | 0.26 |
| 2 | 141369826 | rs544510 | -1.371 | -0.684 | 0.509 | 1.078 | 0.005 | 0 | 0 | 0 | 0.54 | 0.66 | 0.45 | 0.46 | 0.44 | 0.6 |
| 2 | 141370160 | rs531021 | -0.851 | -0.452 | 0.612 | 1.32 | -0.005 | 0 | 0 | 0 | 0.45 | 0.57 | 0.41 | 0.46 | 0.38 | 0.55 |
| 2 | 141391918 | rs506281 | -1.095 | -0.684 | 1.353 | 1.652 | -0.018 | 0.03 | 0.03 | 0 | 0.13 | 0.27 | 0.28 | 0.19 | 0.1 | 0.15 |
| 2 | 141398489 | rs10166217 | 1.798 | -0.783 | 1.263 | 1.83 | 0.034 | -0.048 | -0.048 | 0 | 0.59 | 0.16 | 0.46 | 0.44 | 0.68 | 0.44 |
| 2 | 141399837 | rs1564934 | 1.865 | -0.873 | 1.438 | 1.489 | 0.008 | -0.007 | -0.007 | 0 | 0.6 | 0.33 | 0.48 | 0.49 | 0.69 | 0.45 |
| 2 | 141407101 | rs4954847 | -0.919 | -0.622 | 1.351 | 1.477 | -0.017 | 0.02 | 0.02 | 0 | 0.12 | 0.22 | 0.28 | 0.19 | 0.1 | 0.14 |
| 2 | 141407182 | rs6709373 | 0.214 | -0.422 | 1.106 | 1.317 | 0.013 | -0.013 | -0.013 | 0 | 0.16 | 0.24 | 0.22 | 0.1 | 0.19 | 0.23 |
| 2 | 141409166 | rs514478 | -0.341 | -0.422 | 1.056 | 1.318 | 0.026 | -0.021 | -0.021 | 0.091 | 0.84 | 0.7 | 0.78 | 0.9 | 0.81 | 0.78 |
| 2 | 141419779 | rs552747 | -1.392 | -0.772 | 0.812 | 1.748 | -0.041 | 0.03 | 0.03 | 0 | 0.17 | 0.32 | 0.16 | 0.25 | 0.08 | 0.24 |
| 2 | 141423316 | rs1441470 | 1.698 | -0.721 | 0.948 | 1.966 | -0.062 | 0.093 | 0.093 | 0 | 0.78 | 0.58 | 0.71 | 0.59 | 0.88 | 0.69 |
| 2 | 141426726 | rs16844450 | -1.27 | -0.663 | 0.832 | 2.156 | -0.004 | 0.014 | 0.014 | 0 | 0.14 | 0.15 | 0.05 | 0.33 | 0.06 | 0.23 |
| 2 | 141428751 | rs6737361 | 1.431 | -0.719 | 0.859 | 2.063 | -0.044 | 0.03 | 0.03 | 0 | 0.8 | 0.67 | 0.73 | 0.67 | 0.89 | 0.69 |
| 2 | 141434988 | rs6711580 | 0.31 | -0.705 | 0.767 | 2.008 | -0.013 | 0.045 | 0.045 | 0 | 0.91 | 0.86 | 0.97 | 0.71 | 0.98 | 0.89 |
| 2 | 141436934 | rs6748249 | 1.429 | -0.718 | 0.681 | 1.924 | -0.051 | 0.093 | 0.093 | 0 | 0.78 | 0.63 | 0.66 | 0.43 | 0.88 | 0.69 |
| 2 | 141448312 | rs2165506 | 1.07 | -0.436 | 0.662 | 1.41 | 0.011 | -0.019 | -0.019 | 0.239 | 0.63 | 0.3 | 0.63 | 0.48 | 0.7 | 0.52 |
| 2 | 141467526 | rs13383867 | 0.319 | -0.299 | 0.512 | 1.332 | -0.04 | -0.001 | -0.001 | 0.822 | 0.06 | 0.01 | 0.15 | 0.06 | 0.13 | 0.19 |
| 2 | 141469911 | rs12691557 | -1.171 | -0.378 | 1.179 | 1.642 | 0.063 | -0.013 | -0.013 | 0.247 | 0.1 | 0.29 | 0.17 | 0.09 | 0.14 | 0.25 |
| 2 | 141471725 | rs6746848 | 1.231 | -0.639 | 1.146 | 1.283 | 0.003 | -0.003 | -0.003 | 0.41 | 0.61 | 0.66 | 0.6 | 0.47 | 0.72 | 0.64 |
| 2 | 141478468 | rs436323 | -0.407 | -0.593 | 1.26 | 1.203 | 0.067 | -0.012 | -0.012 | 0 | 0.09 | 0.25 | 0.16 | 0.06 | 0.13 | 0.19 |
| 2 | 141487897 | rs288097 | -0.532 | -0.682 | 1.143 | 1.236 | -0.018 | -0.002 | -0.002 | 0.016 | 0.25 | 0.27 | 0.13 | 0.29 | 0.16 | 0.22 |
| 2 | 141495311 | rs10439180 | 2.126 | -0.493 | 1.178 | 1.21 | 0.012 | 0.017 | 0.017 | 0 | 0.48 | 0.22 | 0.4 | 0.38 | 0.56 | 0.39 |
| 2 | 141495719 | rs288122 | -0.49 | -0.534 | 1.245 | 1.21 | -0.012 | -0.018 | -0.018 | 2.407 | 0.93 | 0.85 | 0.82 | 0.96 | 0.95 | 0.92 |
| 2 | 141496596 | rs288125 | -0.683 | -0.32 | 1.033 | 1.308 | -0.006 | -0.002 | -0.002 | 0 | 0.28 | 0.28 | 0.17 | 0.29 | 0.22 | 0.3 |
| 2 | 141500692 | rs16844617 | -1.291 | -0.244 | 1.128 | 1.412 | 0.042 | 0.166 | 0.166 | 0.623 | 0.05 | 0.16 | 0.16 | 0.28 | 0.07 | 0.12 |
| 2 | 141504719 | rs387927 | -1.31 | -0.407 | 0.818 | 1.28 | 0.001 | 0.013 | 0.013 | 0 | 0.38 | 0.3 | 0.3 | 0.35 | 0.36 | 0.52 |
| 2 | 141517652 | rs185959 | -1.368 | -0.36 | 0.344 | 1.104 | 0.004 | 0.001 | 0.001 | 0 | 0.41 | 0.62 | 0.51 | 0.59 | 0.37 | 0.51 |
| 2 | 141521573 | rs1486126 | 1.64 | -0.203 | -0.768 | 0.684 | -0.084 | -0.068 | -0.068 | 0.096 | 0.4 | 0.06 | 0.35 | 0.38 | 0.46 | 0.36 |

| | | | | | | | | | | | | | | | | |
|---|---|---|---|---|---|---|---|---|---|---|---|---|---|---|---|---|
| 2 | 141525130 | rs6738178 | 1.212 | -0.307 | 0.288 | 0.32 | -0.004 | 0.016 | 0.016 | 0 | 0.58 | 0.6 | 0.64 | 0.76 | 0.57 | 0.49 |
| 2 | 141526325 | rs6727622 | 0.137 | -0.454 | 0.298 | 0.178 | -0.027 | -0.07 | -0.07 | 0 | 0.84 | 0.75 | 0.91 | 1 | 0.9 | 0.93 |
| 2 | 141532786 | rs9287300 | -0.845 | -0.468 | 0.956 | 0.317 | 0.07 | 0.112 | 0.112 | 0.813 | 0.21 | 0.38 | 0.34 | 0.41 | 0.3 | 0.33 |
| 2 | 141533662 | rs996361 | 0.915 | -0.544 | 0.873 | 0.262 | -0.007 | -0.053 | -0.053 | 0 | 0.76 | 0.75 | 0.9 | 1 | 0.87 | 0.81 |
| 2 | 141536876 | rs13030928 | -0.596 | -0.405 | 0.778 | 0.388 | -0.033 | -0.019 | -0.019 | 0 | 0.19 | 0.14 | 0.26 | 0.24 | 0.29 | 0.32 |
| 2 | 141537758 | rs12691568 | 0.135 | -0.392 | 0.801 | 0.403 | -0.015 | -0.01 | -0.01 | 0 | 0.14 | 0.1 | 0.21 | 0.2 | 0.22 | 0.29 |
| 2 | 141546140 | rs12621209 | 2.386 | -0.618 | 0.82 | 0.267 | -0.012 | 0.018 | 0.018 | 0 | 0.5 | 0.13 | 0.4 | 0.58 | 0.54 | 0.47 |
| 2 | 141551350 | rs6735489 | -1.169 | -0.514 | 1.201 | 0.403 | 0.054 | 0.122 | 0.122 | 0.461 | 0.22 | 0.45 | 0.34 | 0.42 | 0.28 | 0.34 |
| 2 | 141560538 | rs7606602 | 2.165 | -0.579 | 1.015 | 0.384 | 0 | 0.059 | 0.059 | 0 | 0.53 | 0.19 | 0.42 | 0.66 | 0.56 | 0.47 |
| 2 | 141561255 | rs2890537 | 0.337 | -0.57 | 1.066 | 0.367 | -0.034 | -0.082 | -0.082 | 0.016 | 0.83 | 0.72 | 0.77 | 0.99 | 0.87 | 0.92 |
| 2 | 141566421 | rs873727 | -0.245 | -0.533 | 1.006 | 0.426 | -0.001 | -0.001 | -0.001 | 0.201 | 0.05 | 0.05 | 0.05 | 0.04 | 0.05 | 0.04 |
| 2 | 141570937 | rs3828265 | 2.264 | -0.611 | 0.891 | 0.386 | -0.003 | 0.025 | 0.025 | 0 | 0.51 | 0.22 | 0.41 | 0.57 | 0.54 | 0.45 |
| 2 | 141577113 | rs1429331 | 0.774 | -0.444 | 1.61 | 0.998 | -0.027 | -0.079 | -0.079 | 3.987 | 0.84 | 0.7 | 0.77 | 0.99 | 0.88 | 0.92 |
| 2 | 141578414 | rs1835164 | 1.804 | -0.532 | 1.693 | 1.067 | -0.019 | -0.01 | -0.01 | 0 | 0.76 | 0.38 | 0.68 | 0.81 | 0.86 | 0.86 |
| 2 | 141582377 | rs1429328 | 1.767 | -0.498 | 1.914 | 0.905 | -0.05 | -0.064 | -0.064 | 0 | 0.77 | 0.53 | 0.7 | 0.9 | 0.86 | 0.87 |
| 2 | 141587797 | rs4954859 | 1.584 | -0.472 | 1.475 | 0.542 | -0.03 | -0.093 | -0.093 | 0 | 0.78 | 0.65 | 0.77 | 0.99 | 0.86 | 0.87 |
| 2 | 141592252 | rs6709738 | -0.877 | -0.229 | 1.208 | 0.657 | 0.002 | 0.005 | 0.005 | 0 | 0.07 | 0.11 | 0.08 | 0.09 | 0.05 | 0.07 |
| 2 | 141594886 | rs6718790 | -0.776 | -0.204 | 1.093 | 0.54 | 0.003 | 0.005 | 0.005 | 0.02 | 0.08 | 0.04 | 0.05 | 0.05 | 0.06 | 0.04 |
| 2 | 141600541 | rs4461204 | 0.082 | -0.216 | 1.217 | 0.484 | -0.002 | -0.06 | -0.06 | 0 | 0.89 | 0.69 | 0.93 | 1 | 0.92 | 0.95 |
| 2 | 141602546 | rs17576906 | -1.131 | -0.058 | 0.803 | 0.64 | -0.004 | -0.01 | -0.01 | 0 | 0.09 | 0.02 | 0.07 | 0.09 | 0.09 | 0.03 |
| 2 | 141604286 | rs10166257 | -1.433 | -0.106 | 1.147 | 0.326 | -0.018 | -0.081 | -0.081 | 0 | 0.18 | 0.27 | 0.17 | 0 | 0.11 | 0.14 |
| 2 | 141604697 | rs2220257 | 0.649 | 0.112 | 1.385 | 0.854 | 0.019 | 0.131 | 0.131 | 0.018 | 0.85 | 0.67 | 0.74 | 0.65 | 0.84 | 0.79 |
| 2 | 141605983 | rs11901880 | 2.19 | -0.023 | 1.191 | 0.733 | 0.007 | 0.009 | 0.009 | 0 | 0.59 | 0.25 | 0.52 | 0.63 | 0.62 | 0.59 |
| 2 | 141623459 | rs6429850 | 0.189 | -0.005 | 1.238 | 0.853 | -0.019 | 0.099 | 0.099 | 0 | 0.87 | 0.73 | 0.8 | 0.7 | 0.88 | 0.81 |
| 2 | 141623557 | rs7565456 | 1.912 | -0.051 | 1.402 | 0.796 | -0.015 | 0.001 | 0.001 | 0.001 | 0.67 | 0.38 | 0.7 | 0.68 | 0.77 | 0.67 |
| 2 | 141625410 | rs12987572 | -0.047 | 0.079 | 1.326 | 0.786 | -0.022 | 0.005 | 0.005 | 0.077 | 0.08 | 0.05 | 0.2 | 0.07 | 0.17 | 0.08 |
| 2 | 141627898 | rs13416050 | 2.346 | 0.088 | 1.354 | 0.786 | 0.001 | 0.005 | 0.005 | 0 | 0.59 | 0.24 | 0.51 | 0.61 | 0.61 | 0.59 |
| 2 | 141632103 | rs12999130 | -0.182 | 0.19 | 1.197 | 0.546 | -0.023 | 0.006 | 0.006 | 0 | 0.08 | 0.06 | 0.2 | 0.07 | 0.17 | 0.08 |
| 2 | 141639645 | rs10496858 | -2.468 | 0.191 | 1.461 | 0.659 | 0.019 | -0.044 | -0.044 | 0 | 0.33 | 0.52 | 0.36 | 0.21 | 0.36 | 0.25 |
| 2 | 141640478 | rs12691579 | -2.323 | 0.212 | 1.299 | 0.669 | 0.015 | -0.034 | -0.034 | 0 | 0.32 | 0.39 | 0.35 | 0.19 | 0.36 | 0.24 |
| 2 | 141641717 | rs16844912 | -0.522 | 0.115 | 1.017 | 0.716 | 0 | 0.005 | 0.005 | 0 | 0.06 | 0.04 | 0.06 | 0.12 | 0.09 | 0.03 |
| 2 | 141655736 | rs12987986 | -2.033 | 0.124 | 1.07 | 0.543 | 0.002 | -0.034 | -0.034 | 0 | 0.28 | 0.29 | 0.31 | 0.08 | 0.31 | 0.27 |
| 2 | 141659527 | rs1549709 | -1.593 | 0.114 | 1.011 | 0.527 | -0.002 | 0.016 | 0.016 | 0 | 0.22 | 0.18 | 0.27 | 0.08 | 0.25 | 0.15 |
| 2 | 141660215 | rs1366802 | -2.824 | 0.061 | 1.44 | 0.738 | 0.001 | 0.001 | 0.001 | 0 | 0.44 | 0.46 | 0.47 | 0.48 | 0.47 | 0.44 |
| 2 | 141662215 | rs10170474 | 2.556 | 0.087 | 1.343 | 0.747 | -0.003 | -0.01 | -0.01 | 0 | 0.55 | 0.27 | 0.52 | 0.53 | 0.53 | 0.56 |
| 2 | 141668678 | rs10496859 | 2.59 | 0.17 | 1.308 | 0.751 | -0.011 | -0.009 | -0.009 | 0 | 0.55 | 0.24 | 0.52 | 0.54 | 0.53 | 0.53 |
| 2 | 141669902 | rs16844936 | 1.273 | 0.061 | 0.977 | 1.013 | -0.01 | 0.01 | 0.01 | 2.294 | 0.1 | 0 | 0.07 | 0.06 | 0.09 | 0.01 |
| 2 | 141671250 | rs10496860 | -1.223 | 0.141 | 1.963 | 1.333 | -0.007 | 0.104 | 0.104 | 0 | 0.15 | 0.29 | 0.16 | 0.37 | 0.16 | 0.17 |
| 2 | 141674190 | rs1429343 | -1.85 | 0.138 | 1.975 | 1.322 | -0.009 | 0.087 | 0.087 | 0 | 0.19 | 0.35 | 0.19 | 0.39 | 0.2 | 0.23 |
| 2 | 141677131 | rs7576000 | -1.223 | 0.186 | 1.983 | 1.355 | 0.005 | 0.181 | 0.181 | 0 | 0.15 | 0.39 | 0.17 | 0.37 | 0.16 | 0.17 |
| 2 | 141677631 | rs7562670 | 0.381 | 0.32 | 1.613 | 0.97 | 0.036 | -0.01 | -0.01 | 0 | 0.91 | 0.83 | 0.79 | 0.93 | 0.84 | 0.92 |
| 2 | 141688383 | rs17519490 | -1.068 | -0.637 | 1.604 | 1.091 | -0.009 | 0 | 0 | 0.042 | 0.1 | 0.03 | 0.06 | 0.09 | 0.12 | 0.05 |
| 2 | 141689095 | rs10166782 | -2.759 | -0.808 | 1.953 | 1.193 | -0.027 | 0.002 | 0.002 | 0 | 0.32 | 0.46 | 0.3 | 0.35 | 0.26 | 0.24 |
| 2 | 141690863 | rs10928079 | -0.682 | -0.845 | 1.985 | 1.32 | -0.026 | 0.102 | 0.102 | 0.05 | 0.11 | 0.29 | 0.18 | 0.25 | 0.06 | 0.14 |
| 2 | 141695517 | rs6721069 | -0.578 | -0.844 | 1.944 | 1.251 | -0.025 | 0.074 | 0.074 | 0.019 | 0.1 | 0.27 | 0.17 | 0.23 | 0.06 | 0.14 |
| 2 | 141698709 | rs1546785 | 1.327 | -0.844 | 1.781 | 1.228 | -0.016 | -0.027 | -0.027 | 0.703 | 0.85 | 0.8 | 0.93 | 0.91 | 0.9 | 0.92 |
| 2 | 141707610 | rs6719711 | -2.114 | -0.665 | 1.373 | 1.451 | -0.034 | 0.052 | 0.052 | 0 | 0.24 | 0.49 | 0.26 | 0.34 | 0.2 | 0.24 |
| 2 | 141709406 | rs1991540 | -0.898 | -0.624 | 0.816 | 1.556 | -0.02 | 0.051 | 0.051 | 0 | 0.1 | 0.18 | 0.16 | 0.23 | 0.06 | 0.14 |
| 2 | 141713655 | rs16845014 | -1.004 | -0.506 | 0.821 | 1.564 | -0.011 | 0.027 | 0.027 | 0.022 | 0.1 | 0.14 | 0.12 | 0.21 | 0.07 | 0.14 |
| 2 | 141713836 | rs16845015 | -1.178 | -0.548 | 1.061 | 1.623 | -0.019 | 0.06 | 0.06 | 0.032 | 0.1 | 0.22 | 0.12 | 0.22 | 0.06 | 0.15 |
| 2 | 141717646 | rs13016717 | -0.602 | -1.034 | 1.784 | 1.978 | -0.001 | 0 | 0 | 0 | 0.19 | 0.16 | 0.11 | 0.24 | 0.18 | 0.21 |
| 2 | 141721159 | rs10928082 | -1.704 | -1.109 | 1.489 | 1.913 | -0.006 | 0.003 | 0.003 | 0 | 0.19 | 0.22 | 0.19 | 0.25 | 0.17 | 0.27 |
| 2 | 141722532 | rs16845035 | -1.206 | -1.054 | 1.33 | 1.729 | 0 | -0.011 | -0.011 | 0.192 | 0.27 | 0.13 | 0.15 | 0.24 | 0.28 | 0.26 |
| 2 | 141729597 | rs7586607 | -1.435 | -1.016 | 1.243 | 1.492 | -0.006 | 0.016 | 0.016 | 0 | 0.13 | 0.1 | 0.1 | 0.04 | 0.13 | 0.08 |
| 2 | 141731683 | rs980288 | 1.912 | -1.058 | 1.389 | 1.491 | -0.016 | -0.01 | -0.01 | 0 | 0.73 | 0.6 | 0.77 | 0.73 | 0.77 | 0.72 |
| 2 | 141737664 | rs1429362 | -1.457 | -1.071 | 1.538 | 1.995 | -0.034 | 0.041 | 0.041 | 0 | 0.33 | 0.43 | 0.29 | 0.54 | 0.28 | 0.46 |
| 2 | 141743077 | rs16845089 | -0.02 | -0.996 | 1.509 | 1.818 | 0.003 | -0.02 | -0.02 | 0.345 | 0.07 | 0.11 | 0.05 | 0.02 | 0.07 | 0.01 |
| 2 | 141751592 | rs13007735 | -2.282 | -1.105 | 1.725 | 1.848 | 0.019 | 0.01 | 0.01 | 0 | 0.46 | 0.61 | 0.38 | 0.61 | 0.37 | 0.5 |
| 2 | 141752588 | rs2380919 | 2.1 | -1.061 | 1.782 | 1.844 | 0.029 | 0.024 | 0.024 | 0 | 0.54 | 0.33 | 0.6 | 0.38 | 0.59 | 0.46 |
| 2 | 141756334 | rs1949871 | 1.522 | -1.052 | 1.678 | 1.867 | -0.012 | 0.002 | 0.002 | 0 | 0.62 | 0.48 | 0.66 | 0.4 | 0.66 | 0.47 |
| 2 | 141757246 | rs1474407 | -1.822 | -1.121 | 1.63 | 1.874 | -0.025 | 0.005 | 0.005 | 0 | 0.37 | 0.44 | 0.33 | 0.6 | 0.31 | 0.49 |
| 2 | 141762830 | rs13413663 | 1.893 | -1.061 | 1.791 | 1.926 | 0.008 | 0.007 | 0.007 | 0 | 0.55 | 0.44 | 0.6 | 0.38 | 0.55 | 0.44 |
| 2 | 141764693 | rs13012143 | -0.191 | -1.104 | 1.813 | 2.018 | 0.007 | -0.016 | -0.016 | 2.746 | 0.2 | 0.11 | 0.16 | 0.21 | 0.22 | 0.23 |
| 2 | 141771962 | rs6758426 | -1.385 | -1.059 | 1.955 | 2.441 | 0 | -0.002 | -0.002 | 0 | 0.32 | 0.3 | 0.38 | 0.85 | 0.33 | 0.49 |
| 2 | 141778702 | rs1525599 | -1.436 | -0.918 | 1.755 | 2.06 | -0.005 | 0.011 | 0.011 | 0 | 0.14 | 0.19 | 0.14 | 0.24 | 0.12 | 0.23 |
| 2 | 141779955 | rs6429856 | 1.47 | -0.904 | 1.237 | 1.846 | 0.002 | -0.004 | -0.004 | 0 | 0.57 | 0.43 | 0.64 | 0.55 | 0.55 | 0.51 |
| 2 | 141782073 | rs1880620 | 1.594 | -0.871 | 1.209 | 1.567 | 0.008 | 0.015 | 0.015 | 0 | 0.55 | 0.35 | 0.59 | 0.38 | 0.54 | 0.45 |
| 2 | 141784262 | rs11682851 | 2.012 | -0.918 | 1.011 | 1.547 | 0.016 | 0.05 | 0.05 | 0 | 0.5 | 0.2 | 0.57 | 0.38 | 0.53 | 0.43 |
| 2 | 141790694 | rs12468894 | -1.542 | -0.574 | 0.952 | 1.733 | 0.001 | -0.002 | -0.002 | 0 | 0.41 | 0.45 | 0.38 | 0.61 | 0.44 | 0.53 |

| | | | | | | | | | | | | | | | | |
|---|---|---|---|---|---|---|---|---|---|---|---|---|---|---|---|---|
| 2 | 141795620 | rs10210358 | 1.347 | -1.084 | 0.74 | 1.67 | -0.018 | 0.029 | 0.029 | 0 | 0.77 | 0.42 | 0.72 | 0.65 | 0.82 | 0.66 |
| 2 | 141796109 | rs17588066 | -0.296 | -0.916 | 0.445 | 1.143 | 0.001 | 0.003 | 0.003 | 1.31 | 0.09 | 0.07 | 0.02 | 0.06 | 0.05 | 0.1 |
| 2 | 141796256 | rs10202342 | 1.356 | -1.22 | 0.18 | 1.146 | 0.007 | 0.011 | 0.011 | 0 | 0.65 | 0.34 | 0.77 | 0.72 | 0.74 | 0.72 |
| 2 | 141799879 | rs2091302 | 1.937 | -1.29 | 0.106 | 1.303 | 0.074 | 0.066 | 0.066 | 0 | 0.49 | 0.2 | 0.57 | 0.31 | 0.62 | 0.46 |
| 2 | 141801011 | rs7603711 | 1.571 | -0.924 | 0.453 | 0.681 | -0.006 | -0.009 | -0.009 | 0.249 | 0.66 | 0.4 | 0.77 | 0.74 | 0.77 | 0.74 |
| 2 | 141803496 | rs7606917 | 0.688 | -0.826 | 0.541 | 0.052 | -0.018 | -0.085 | -0.085 | 0 | 0.8 | 0.63 | 0.9 | 0.94 | 0.88 | 0.88 |
| 2 | 141803602 | rs7594687 | -1.092 | -0.356 | -0.04 | 0.612 | -0.005 | -0.034 | -0.034 | 0 | 0.11 | 0.16 | 0.06 | 0 | 0.09 | 0.09 |
| 2 | 141810244 | rs2380922 | 0.824 | -0.465 | 0.108 | 0.79 | -0.025 | -0.011 | -0.011 | 0 | 0.83 | 0.75 | 0.85 | 0.88 | 0.91 | 0.78 |
| 2 | 141813216 | rs16845233 | -0.1 | -0.658 | -0.025 | 0.703 | -0.002 | -0.028 | -0.028 | 0 | 0.07 | 0 | 0.1 | 0.1 | 0.04 | 0.15 |
| 2 | 141819422 | rs1525583 | 1.911 | -0.57 | 0.175 | 0.528 | -0.022 | -0.025 | -0.025 | 1.533 | 0.78 | 0.69 | 0.83 | 0.89 | 0.87 | 0.77 |
| 2 | 141823300 | rs4954875 | 0.296 | -0.15 | -0.1 | 0.875 | -0.011 | -0.016 | -0.016 | 1.167 | 0.2 | 0.18 | 0.34 | 0.44 | 0.25 | 0.21 |
| 2 | 141824403 | rs1473662 | -0.101 | -0.151 | -0.09 | 0.863 | -0.016 | -0.037 | -0.037 | 0 | 0.89 | 0.75 | 0.95 | 0.99 | 0.94 | 0.93 |
| 2 | 141826948 | rs4954876 | 1.525 | -0.154 | 0.306 | 0.826 | 0.002 | 0.008 | 0.008 | 0 | 0.57 | 0.45 | 0.49 | 0.45 | 0.62 | 0.56 |
| 2 | 141833787 | rs1525589 | 1.566 | -0.601 | 0.452 | 1.203 | -0.014 | -0.01 | -0.01 | 0 | 0.73 | 0.68 | 0.82 | 0.84 | 0.84 | 0.69 |
| 2 | 141840541 | rs1474408 | 1.81 | -0.593 | 0.631 | 1.165 | 0 | -0.002 | -0.002 | 0.06 | 0.52 | 0.5 | 0.47 | 0.39 | 0.59 | 0.48 |
| 2 | 141841860 | rs2140872 | 0.964 | -0.615 | 0.621 | 1.129 | -0.01 | -0.012 | -0.012 | 0 | 0.8 | 0.75 | 0.84 | 0.89 | 0.87 | 0.77 |
| 2 | 141844434 | rs7601662 | 1.031 | -0.353 | 0.6 | 1.133 | -0.01 | -0.017 | -0.017 | 0 | 0.79 | 0.72 | 0.82 | 0.89 | 0.86 | 0.77 |
| 2 | 141845251 | rs3856356 | 0.521 | -0.247 | 0.563 | 1.295 | -0.029 | -0.05 | -0.05 | 0 | 0.18 | 0.09 | 0.33 | 0.51 | 0.22 | 0.2 |
| 2 | 141845679 | rs3845640 | 1.952 | -0.421 | 0.635 | 1.025 | -0.005 | 0.057 | 0.057 | 0 | 0.48 | 0.31 | 0.4 | 0.23 | 0.55 | 0.4 |
| 2 | 141846346 | rs4954880 | 0.464 | -0.442 | 0.705 | 1.012 | -0.018 | -0.021 | -0.021 | 0.013 | 0.82 | 0.63 | 0.84 | 0.89 | 0.88 | 0.8 |
| 2 | 141847207 | rs7573798 | -1.03 | -0.383 | 0.728 | 1.096 | -0.006 | -0.016 | -0.016 | 0.002 | 0.33 | 0.28 | 0.44 | 0.66 | 0.34 | 0.4 |
| 2 | 141852424 | rs1882628 | -0.322 | -0.337 | 0.706 | 1.185 | -0.001 | 0.004 | 0.004 | 0 | 0.25 | 0.26 | 0.35 | 0.64 | 0.26 | 0.31 |
| 2 | 141857090 | rs12996731 | 2.087 | -0.08 | 0.66 | 1.116 | 0.011 | 0.061 | 0.061 | 0 | 0.49 | 0.33 | 0.39 | 0.22 | 0.51 | 0.4 |
| 2 | 141861910 | rs1882642 | 0.546 | -0.1 | 0.691 | 1.105 | 0.004 | -0.019 | -0.019 | 2.801 | 0.82 | 0.62 | 0.82 | 0.88 | 0.85 | 0.81 |
| 2 | 141863233 | rs12623287 | 0.815 | -0.04 | 0.587 | 1.17 | -0.029 | -0.051 | -0.051 | 0 | 0.18 | 0.09 | 0.31 | 0.5 | 0.22 | 0.21 |
| 2 | 141867448 | rs1922704 | -0.976 | -0.052 | 0.586 | 1.135 | -0.009 | -0.021 | -0.021 | 0 | 0.32 | 0.26 | 0.43 | 0.66 | 0.34 | 0.41 |
| 2 | 141871495 | rs16845474 | 0.152 | 0.054 | 0.788 | 1.335 | 0.007 | 0.036 | 0.036 | 0 | 0.21 | 0.25 | 0.33 | 0.65 | 0.22 | 0.28 |
| 2 | 141872505 | rs1922697 | 0.623 | 0.016 | 0.701 | 1.306 | 0.001 | 0.003 | 0.003 | 0 | 0.69 | 0.37 | 0.6 | 0.34 | 0.7 | 0.65 |
| 2 | 141877201 | rs1882638 | 0.325 | -0.073 | 0.465 | 1.387 | -0.005 | -0.001 | -0.001 | 0 | 0.87 | 0.79 | 0.87 | 0.9 | 0.91 | 0.87 |
| 2 | 141877890 | rs1882637 | 0.434 | 0.3 | 0.355 | 1.58 | -0.039 | -0.058 | -0.058 | 0.037 | 0.18 | 0.08 | 0.31 | 0.51 | 0.25 | 0.23 |
| 2 | 141882963 | rs13013197 | 1.241 | -0.361 | 0.093 | 0.954 | -0.002 | 0.007 | 0.007 | 0 | 0.41 | 0.34 | 0.45 | 0.31 | 0.47 | 0.24 |
| 2 | 141888035 | rs11677400 | -0.64 | -0.4 | -0.053 | 1.342 | -0.011 | -0.012 | -0.012 | 0.281 | 0.82 | 0.88 | 0.69 | 0.57 | 0.76 | 0.75 |
| 2 | 141890746 | rs2141921 | 1.81 | -0.587 | -0.166 | 1.502 | 0.012 | -0.007 | -0.007 | 0 | 0.06 | 0.02 | 0.03 | 0.17 | 0.02 | 0.1 |
| 2 | 141891149 | rs13002102 | 1.399 | -1.027 | 0.413 | 1.159 | 0.003 | 0.047 | 0.047 | 0 | 0.51 | 0.34 | 0.53 | 0.31 | 0.52 | 0.41 |
| 2 | 141893894 | rs12476507 | 2.298 | -1.148 | 0.243 | 1.017 | -0.04 | 0.025 | 0.025 | 0 | 0.35 | 0.14 | 0.42 | 0.24 | 0.45 | 0.22 |
| 2 | 141895211 | rs11694803 | 0.69 | -1.221 | 0.184 | 0.748 | -0.033 | 0.01 | 0.01 | 0.163 | 0.14 | 0.01 | 0.06 | 0.05 | 0.13 | 0.15 |
| 2 | 141897042 | rs7573569 | -0.609 | -1.204 | 0.962 | 0.743 | 0.067 | 0.023 | 0.023 | 0.216 | 0.08 | 0.41 | 0.06 | 0.03 | 0.07 | 0.03 |
| 2 | 141916520 | rs6718638 | 1.734 | -1.357 | 0.331 | 0.721 | -0.029 | 0.205 | 0.205 | 0 | 0.44 | 0.15 | 0.45 | 0.06 | 0.51 | 0.28 |
| 2 | 141919280 | rs13025185 | 1.862 | -1.128 | 0.655 | 0.899 | -0.013 | 0.23 | 0.23 | 0 | 0.49 | 0.14 | 0.42 | 0.1 | 0.58 | 0.31 |
| 2 | 141919943 | rs2222233 | -0.109 | -0.69 | 1.271 | 1.25 | 0.037 | 0.074 | 0.074 | 0 | 0.77 | 0.73 | 0.61 | 0.33 | 0.72 | 0.65 |
| 2 | 141920218 | rs16845527 | 1.102 | -0.403 | 0.909 | 1.359 | -0.027 | -0.027 | -0.027 | 0 | 0.09 | 0.03 | 0.14 | 0.24 | 0.16 | 0.15 |
| 2 | 141929225 | rs1249458 | -0.367 | -0.695 | 1.112 | 0.518 | 0.032 | 0.062 | 0.062 | 0 | 0.24 | 0.28 | 0.39 | 0.66 | 0.29 | 0.38 |
| 2 | 141942235 | rs7424161 | 1.528 | -0.788 | 1.298 | 0.389 | -0.007 | 0.057 | 0.057 | 0 | 0.56 | 0.26 | 0.37 | 0.24 | 0.6 | 0.41 |
| 2 | 141945852 | rs1249433 | -1.071 | -0.475 | 1.157 | 0.549 | 0.005 | 0.002 | 0.002 | 0 | 0.33 | 0.29 | 0.57 | 0.72 | 0.35 | 0.49 |
| 2 | 141955000 | rs17592904 | -0.607 | -0.383 | 1.492 | 0.568 | 0.017 | 0.006 | 0.006 | 0.007 | 0.3 | 0.33 | 0.48 | 0.71 | 0.39 | 0.43 |
| 2 | 141956759 | rs16845593 | 0.739 | -0.528 | 0.94 | 0.032 | 0.001 | -0.009 | -0.009 | 0 | 0.09 | 0.06 | 0.09 | 0.15 | 0.09 | 0.1 |
| 2 | 141962644 | rs971251 | 0.403 | -0.491 | 1.039 | 0.006 | 0.004 | 0.015 | 0.015 | 0 | 0.18 | 0.18 | 0.42 | 0.67 | 0.27 | 0.28 |
| 2 | 141967308 | rs300353 | 0.373 | -0.637 | 1.157 | -0.077 | 0.013 | 0.101 | 0.101 | 0 | 0.86 | 0.57 | 0.87 | 0.79 | 0.87 | 0.8 |
| 2 | 141971994 | rs406689 | -1.409 | 0.121 | 0.934 | -0.322 | 0.003 | 0.016 | 0.016 | 0 | 0.47 | 0.42 | 0.64 | 0.75 | 0.51 | 0.62 |
| 2 | 141972943 | rs11902930 | -0.24 | -0.095 | 0.9 | -0.233 | 0.066 | 0.062 | 0.062 | 0.304 | 0.06 | 0.35 | 0.04 | 0.06 | 0.04 | 0.11 |
| 2 | 141975850 | rs300374 | -0.48 | -0.074 | 0.401 | -0.26 | 0.01 | 0.053 | 0.053 | 0 | 0.18 | 0.09 | 0.17 | 0.04 | 0.13 | 0.21 |
| 2 | 141983765 | rs300388 | -0.009 | -0.003 | -0.048 | -0.192 | -0.001 | 0.047 | 0.047 | 0 | 0.15 | 0.07 | 0.16 | 0.04 | 0.11 | 0.19 |
| 2 | 141988708 | rs10496872 | 0.947 | 0.086 | 0.139 | -0.064 | 0.004 | 0.002 | 0.002 | 0 | 0.11 | 0.09 | 0.05 | 0.09 | 0.11 | 0.13 |
| 2 | 141997870 | rs1401308 | 0.509 | -0.027 | 0.416 | 0.002 | -0.007 | 0.032 | 0.032 | 0 | 0.11 | 0.16 | 0.11 | 0.16 | 0.1 | 0.1 |
| 2 | 141998656 | rs2049153 | 1.73 | -0.061 | 0.836 | -0.137 | 0.003 | 0.001 | 0.001 | 0 | 0.57 | 0.35 | 0.64 | 0.62 | 0.59 | 0.49 |
| 2 | 142003921 | rs2679452 | 0.485 | -0.07 | 0.733 | -0.126 | 0 | -0.001 | -0.001 | 0 | 0.14 | 0.11 | 0.08 | 0.15 | 0.16 | 0.24 |
| 2 | 142005960 | rs10496874 | 1.221 | -0.07 | 0.731 | -0.128 | 0.006 | 0.003 | 0.003 | 0 | 0.12 | 0.08 | 0.05 | 0.09 | 0.11 | 0.13 |
| 2 | 142006135 | rs1521108 | 1.13 | -0.099 | 0.924 | -0.147 | -0.02 | -0.01 | -0.01 | 0 | 0.71 | 0.48 | 0.72 | 0.77 | 0.75 | 0.7 |
| 2 | 142006662 | rs9287315 | 2.109 | 0.128 | 0.609 | -0.354 | 0.019 | 0.071 | 0.071 | 0.244 | 0.39 | 0.2 | 0.18 | 0.06 | 0.33 | 0.22 |
| 2 | 142009056 | rs380870 | -0.599 | 0.288 | 0.032 | -0.21 | 0.003 | 0 | 0 | 0 | 0.24 | 0.22 | 0.21 | 0.19 | 0.22 | 0.25 |
| 2 | 142010268 | rs418443 | -0.599 | 0.288 | 0.032 | -0.21 | 0.003 | 0 | 0 | 0 | 0.24 | 0.22 | 0.21 | 0.19 | 0.22 | 0.25 |
| 2 | 142013695 | rs2714211 | 0.536 | 0.27 | 0.042 | -0.222 | 0.001 | 0 | 0 | 0 | 0.16 | 0.15 | 0.09 | 0.15 | 0.16 | 0.24 |
| 2 | 142014406 | rs373908 | -1.833 | 0.274 | 0.012 | -0.073 | 0.002 | -0.002 | -0.002 | 0 | 0.4 | 0.39 | 0.4 | 0.39 | 0.37 | 0.49 |
| 2 | 142018926 | rs969575 | 1.5 | 0.389 | -0.197 | -0.033 | -0.052 | -0.068 | -0.068 | 0.252 | 0.15 | 0.07 | 0.46 | 0.57 | 0.26 | 0.24 |
| 2 | 142025228 | rs300399 | -0.27 | 0.362 | -0.133 | -0.03 | -0.019 | 0.011 | 0.011 | 0 | 0.17 | 0.08 | 0.27 | 0.14 | 0.18 | 0.28 |
| 2 | 142032671 | rs2090806 | 1.665 | 0.359 | -0.177 | -0.042 | 0.003 | -0.041 | -0.041 | 0 | 0.07 | 0 | 0.08 | 0.15 | 0.07 | 0.07 |
| 2 | 142047759 | rs955681 | -1.163 | 0.332 | 0.018 | -0.109 | -0.014 | 0.013 | 0.013 | 0.29 | 0.29 | 0.37 | 0.21 | 0.31 | 0.27 | 0.36 |
| 2 | 142054467 | rs13400044 | 1.435 | 0.536 | -0.433 | 0.029 | -0.051 | -0.065 | -0.065 | 0 | 0.15 | 0.06 | 0.34 | 0.51 | 0.26 | 0.24 |
| 2 | 142063085 | rs12465020 | 0.351 | 0.322 | -0.564 | 0.065 | -0.041 | 0.002 | 0.002 | 0 | 0.13 | 0.01 | 0.22 | 0.08 | 0.13 | 0.17 |

| | | | | | | | | | | | | | | | | |
|---|---|---|---|---|---|---|---|---|---|---|---|---|---|---|---|---|
| 2 | 142073871 | rs17543266 | 0.903 | 0.357 | -0.322 | 0.225 | 0.008 | 0.004 | 0.004 | 0 | 0.12 | 0.07 | 0.05 | 0.09 | 0.11 | 0.14 |
| 2 | 142076913 | rs16845771 | 0.191 | 0.39 | -0.093 | 0.17 | -0.019 | 0.058 | 0.058 | 0 | 0.11 | 0.2 | 0.1 | 0.19 | 0.09 | 0.09 |
| 2 | 142077028 | rs983810 | -0.249 | 0.677 | -0.553 | -0.351 | -0.009 | -0.004 | -0.004 | 0 | 0.89 | 0.93 | 0.75 | 0.86 | 0.85 | 0.78 |
| 2 | 142077308 | rs17602834 | -0.601 | 0.308 | -0.33 | -0.308 | -0.022 | -0.005 | -0.005 | 0 | 0.34 | 0.32 | 0.44 | 0.65 | 0.42 | 0.43 |
| 2 | 142077946 | rs2381140 | -0.147 | 0.391 | 0.301 | -0.529 | 0.031 | 0.008 | 0.008 | 0.005 | 0.27 | 0.53 | 0.41 | 0.69 | 0.34 | 0.33 |
| 2 | 142078003 | rs2683815 | -1.666 | -0.14 | 0.508 | -0.716 | 0.029 | -0.008 | -0.008 | 0 | 0.08 | 0.26 | 0.09 | 0.05 | 0.1 | 0.13 |
| 2 | 142078226 | rs16855038 | -1.103 | -0.782 | 0.437 | -1.036 | -0.045 | -0.086 | -0.086 | 0 | 0.23 | 0.47 | 0.24 | 0.05 | 0.13 | 0.21 |
| 2 | 142082244 | rs975607 | -0.789 | -0.424 | 0.179 | -0.957 | 0.008 | -0.002 | -0.002 | 0.05 | 0.75 | 0.87 | 0.59 | 0.3 | 0.77 | 0.6 |
| 2 | 142088764 | rs2714185 | 0.558 | -0.739 | -0.042 | -1.42 | 0.045 | 0.104 | 0.104 | 0.092 | 0.52 | 0.75 | 0.34 | 0.21 | 0.6 | 0.4 |
| 2 | 142092565 | rs2683854 | 0.606 | -0.744 | 0.026 | -1.423 | 0.04 | 0.086 | 0.086 | 0.057 | 0.51 | 0.71 | 0.34 | 0.21 | 0.6 | 0.4 |
| 2 | 142094193 | rs4954898 | 0.694 | -0.74 | -0.004 | -1.451 | 0.035 | 0.086 | 0.086 | 0.023 | 0.49 | 0.71 | 0.32 | 0.21 | 0.59 | 0.38 |
| 2 | 142099985 | rs1033311 | -0.074 | -0.886 | -0.066 | -1.528 | 0.014 | 0.037 | 0.037 | 1.022 | 0.59 | 0.74 | 0.37 | 0.23 | 0.63 | 0.45 |
| 2 | 142102998 | rs2714217 | 0.288 | -0.97 | 0.414 | -1.547 | -0.003 | -0.015 | -0.015 | 0 | 0.55 | 0.61 | 0.34 | 0.23 | 0.61 | 0.44 |
| 2 | 142107016 | rs2714214 | 0.662 | -1.006 | 0.543 | -1.58 | 0 | -0.011 | -0.011 | 1.741 | 0.5 | 0.59 | 0.34 | 0.23 | 0.61 | 0.43 |
| 2 | 142108725 | rs11893168 | 0.786 | -0.969 | 0.422 | -1.558 | 0.007 | 0.004 | 0.004 | 2.029 | 0.08 | 0.05 | 0.03 | 0.01 | 0.02 | 0.06 |
| 2 | 142108997 | rs2683824 | 1.106 | -0.946 | 0.62 | -1.574 | 0.018 | 0.131 | 0.131 | 0 | 0.44 | 0.18 | 0.27 | 0.21 | 0.58 | 0.38 |
| 2 | 142116085 | rs2683834 | 0.619 | -1.03 | 0.772 | -1.624 | 0.003 | -0.004 | -0.004 | 0 | 0.52 | 0.45 | 0.33 | 0.21 | 0.65 | 0.42 |
| 2 | 142118351 | rs16845926 | 0.209 | -0.95 | 0.571 | -1.566 | 0 | -0.02 | -0.02 | 0 | 0.05 | 0.11 | 0.06 | 0.02 | 0.04 | 0.07 |
| 2 | 142119402 | rs1402470 | 1.056 | -1.192 | -0.115 | -2.006 | -0.072 | 0.132 | 0.132 | 0 | 0.35 | 0.12 | 0.23 | 0.13 | 0.54 | 0.25 |
| 2 | 142124008 | rs2714177 | 1 | -1.34 | 0.254 | -1.58 | 0.016 | 0.129 | 0.129 | 0 | 0.43 | 0.18 | 0.27 | 0.22 | 0.58 | 0.38 |
| 2 | 142134407 | rs10208287 | -0.89 | -1.254 | 0.667 | -1.104 | -0.028 | -0.064 | -0.064 | 0 | 0.12 | 0.24 | 0.06 | 0.01 | 0.05 | 0.07 |
| 2 | 142148511 | rs7560169 | 0.202 | -1.352 | 0.303 | -1.057 | 0.026 | 0.008 | 0.008 | 0 | 0.3 | 0.19 | 0.42 | 0.69 | 0.23 | 0.4 |
| 2 | 142149328 | rs4245855 | 0.771 | -1.571 | 0.231 | -1.169 | 0.039 | 0.042 | 0.042 | 0 | 0.55 | 0.29 | 0.48 | 0.29 | 0.71 | 0.51 |
| 2 | 142156180 | rs12052880 | -0.239 | -1.29 | -0.662 | -1.007 | 0.043 | 0.023 | 0.023 | 0.183 | 0.33 | 0.09 | 0.43 | 0.71 | 0.26 | 0.45 |
| 2 | 142157532 | rs4258750 | 0.253 | -1.243 | -0.199 | -0.875 | 0.008 | -0.037 | -0.037 | 0.009 | 0.91 | 0.53 | 0.93 | 1 | 0.96 | 0.94 |
| 2 | 142158186 | rs11687134 | 0.034 | -1.254 | -0.424 | -0.681 | 0.042 | -0.016 | -0.016 | 0 | 0.66 | 0.87 | 0.62 | 0.4 | 0.73 | 0.68 |
| 2 | 142158967 | rs4485511 | 0.477 | -0.828 | -0.325 | -0.452 | -0.021 | -0.006 | -0.006 | 0 | 0.62 | 0.42 | 0.56 | 0.38 | 0.7 | 0.6 |
| 2 | 142166561 | rs6429872 | -0.397 | -0.341 | -1.444 | 0.183 | 0.024 | -0.047 | -0.047 | 0.189 | 0.77 | 0.91 | 0.69 | 0.4 | 0.81 | 0.69 |
| 2 | 142172366 | rs4954702 | -0.029 | 0.175 | -1.783 | 0.128 | 0.014 | 0.069 | 0.069 | 0.427 | 0.53 | 0.15 | 0.62 | 0.68 | 0.57 | 0.53 |
| 2 | 142176624 | rs4954906 | 0.766 | 0.528 | -1.64 | 0.582 | 0.015 | 0.015 | 0.015 | 0.001 | 0.19 | 0.25 | 0.33 | 0.21 | 0.3 | 0.16 |
| 2 | 142179191 | rs12468848 | 0.766 | 0.533 | -1.64 | 0.582 | 0.016 | 0.015 | 0.015 | 0.001 | 0.19 | 0.26 | 0.33 | 0.21 | 0.3 | 0.17 |
| 2 | 142180729 | rs11892609 | 0.467 | 0.212 | -1.743 | 0.484 | 0.034 | -0.077 | -0.077 | 0 | 0.3 | 0.07 | 0.27 | 0.47 | 0.2 | 0.3 |
| 2 | 142180831 | rs6717169 | -0.39 | 0.428 | -1.055 | 0.111 | 0.011 | -0.027 | -0.027 | 1.782 | 0.83 | 0.68 | 0.82 | 0.93 | 0.81 | 0.82 |
| 2 | 142184803 | rs12691606 | 0.84 | 0.236 | -1.089 | 0.121 | 0.031 | -0.077 | -0.077 | 0 | 0.3 | 0.07 | 0.27 | 0.47 | 0.21 | 0.31 |
| 2 | 142188957 | rs4507038 | 0.142 | -0.026 | -1.337 | 0.012 | 0.014 | 0.007 | 0.007 | 0.389 | 0.59 | 0.42 | 0.47 | 0.63 | 0.48 | 0.61 |
| 2 | 142191011 | rs7585927 | 0.879 | -0.044 | -1.237 | -0.003 | -0.003 | -0.059 | -0.059 | 1.083 | 0.3 | 0.37 | 0.2 | 0.16 | 0.27 | 0.3 |
| 2 | 142194933 | rs6726240 | 0.368 | -0.189 | -1.354 | -0.071 | -0.008 | -0.019 | -0.019 | 0 | 0.36 | 0.36 | 0.22 | 0.26 | 0.3 | 0.35 |
| 2 | 142196011 | rs10185384 | -0.099 | -0.505 | -1.425 | -0.238 | 0.022 | -0.007 | -0.007 | 0.286 | 0.62 | 0.43 | 0.49 | 0.72 | 0.51 | 0.65 |
| 2 | 142199346 | rs4309525 | 1.002 | -0.438 | -1.633 | -0.188 | -0.006 | -0.142 | -0.142 | 0 | 0.29 | 0.02 | 0.27 | 0.47 | 0.21 | 0.33 |
| 2 | 142208793 | rs13029426 | 1.609 | -0.581 | -1.223 | -0.41 | -0.011 | 0.028 | 0.028 | 0.403 | 0.2 | 0.05 | 0.13 | 0.14 | 0.21 | 0.26 |
| 2 | 142210528 | rs6753877 | -0.509 | -0.291 | -0.687 | -0.367 | 0.034 | -0.033 | -0.033 | 0 | 0.94 | 0.87 | 0.93 | 1 | 0.91 | 0.98 |
| 2 | 142210946 | rs12993942 | 0.608 | -0.856 | -0.827 | -0.324 | 0.009 | -0.049 | -0.049 | 1.101 | 0.33 | 0.11 | 0.31 | 0.47 | 0.27 | 0.38 |
| 2 | 142213886 | rs11902970 | -0.014 | -0.602 | 0.055 | -0.264 | -0.022 | 0.015 | 0.015 | 0.043 | 0.08 | 0.16 | 0.02 | 0.11 | 0.03 | 0.04 |
| 2 | 142216515 | rs7578144 | 1.236 | -0.433 | 0.605 | -0.254 | 0.035 | -0.032 | -0.032 | 0 | 0.91 | 0.49 | 0.92 | 1 | 0.96 | 0.95 |
| 2 | 142216767 | rs7605083 | 1.035 | -0.951 | 0.355 | -0.285 | -0.016 | -0.005 | -0.005 | 0 | 0.71 | 0.33 | 0.79 | 0.78 | 0.87 | 0.65 |
| 2 | 142218508 | rs10928103 | 1.057 | -0.673 | 0.494 | -0.343 | -0.019 | -0.071 | -0.071 | 0.013 | 0.71 | 0.48 | 0.79 | 0.88 | 0.81 | 0.65 |
| 2 | 142219033 | rs4622654 | 0.725 | -0.808 | -0.047 | -0.326 | 0.005 | 0.004 | 0.004 | 0 | 0.33 | 0.34 | 0.45 | 0.34 | 0.52 | 0.38 |
| 2 | 142219810 | rs6429882 | -1.612 | -0.854 | 0.168 | -0.405 | -0.016 | -0.034 | -0.034 | 0 | 0.87 | 0.67 | 0.93 | 0.97 | 0.89 | 0.85 |
| 2 | 142222972 | rs4456644 | 2.147 | -0.792 | 0.104 | -0.369 | 0.007 | 0.002 | 0.002 | 0 | 0.19 | 0.16 | 0.36 | 0.21 | 0.39 | 0.2 |
| 2 | 142226832 | rs6719026 | 1.002 | -0.937 | 0.046 | -0.37 | 0.07 | 0.036 | 0.036 | 0.125 | 0.27 | 0.34 | 0.44 | 0.31 | 0.48 | 0.25 |
| 2 | 142227602 | rs4401160 | 0.434 | -1.092 | 0.035 | -0.435 | 0.022 | 0.014 | 0.014 | 0 | 0.19 | 0.15 | 0.13 | 0.12 | 0.07 | 0.31 |
| 2 | 142228509 | rs12472911 | 0.471 | -1.124 | -0.024 | -0.459 | -0.027 | -0.034 | -0.034 | 0 | 0.64 | 0.4 | 0.77 | 0.85 | 0.78 | 0.52 |
| 2 | 142230158 | rs13432585 | -0.317 | -1.062 | -0.083 | -0.431 | -0.024 | -0.015 | -0.015 | 0.123 | 0.6 | 0.3 | 0.49 | 0.66 | 0.41 | 0.61 |
| 2 | 142231898 | rs13030335 | 1.235 | -1.034 | -0.161 | -0.392 | -0.022 | -0.118 | -0.118 | 0 | 0.37 | 0.07 | 0.33 | 0.54 | 0.3 | 0.28 |
| 2 | 142234165 | rs6729073 | 0.43 | -0.899 | -0.034 | -0.453 | 0.026 | 0.038 | 0.038 | 0.024 | 0.52 | 0.28 | 0.42 | 0.66 | 0.43 | 0.46 |
| 2 | 142238800 | rs10173806 | -0.014 | -0.919 | -0.017 | -0.471 | 0.004 | -0.005 | -0.005 | 1.815 | 0.42 | 0.43 | 0.53 | 0.34 | 0.52 | 0.52 |
| 2 | 142244994 | rs9287319 | 0.281 | -1.007 | -0.054 | -0.54 | -0.029 | -0.066 | -0.066 | 0 | 0.82 | 0.52 | 0.87 | 0.94 | 0.92 | 0.78 |
| 2 | 142245717 | rs4547471 | 0.003 | -0.996 | -0.085 | -0.534 | -0.019 | -0.049 | -0.049 | 0 | 0.84 | 0.68 | 0.88 | 0.94 | 0.92 | 0.78 |
| 2 | 142245947 | rs10496882 | -0.314 | -1.044 | -0.16 | -0.562 | -0.075 | 0.022 | 0.022 | 3.771 | 0.61 | 0.2 | 0.48 | 0.72 | 0.49 | 0.64 |
| 2 | 142248347 | rs16846251 | 0.729 | -1.017 | 0.066 | -0.473 | 0.005 | 0.005 | 0.005 | 0 | 0.11 | 0.11 | 0.08 | 0.06 | 0.05 | 0.2 |
| 2 | 142253278 | rs7557886 | 0.749 | -0.988 | -0.122 | -0.43 | 0.014 | 0.013 | 0.013 | 0 | 0.11 | 0.07 | 0.07 | 0.06 | 0.04 | 0.19 |
| 2 | 142261821 | rs4591293 | 0.839 | -0.792 | 0.024 | -0.296 | -0.001 | -0.001 | -0.001 | 0 | 0.3 | 0.26 | 0.43 | 0.42 | 0.43 | 0.35 |
| 2 | 142263979 | rs4260183 | 2.412 | -0.995 | -0.199 | -0.387 | -0.02 | -0.016 | -0.016 | 0.573 | 0.11 | 0.01 | 0.1 | 0.11 | 0.11 | 0.1 |
| 2 | 142266867 | rs4954914 | -0.069 | -1.15 | -0.186 | -0.484 | -0.021 | -0.045 | -0.045 | 0 | 0.74 | 0.65 | 0.84 | 0.91 | 0.82 | 0.64 |
| 2 | 142268315 | rs13027965 | 2.448 | -1.207 | -0.406 | -0.481 | -0.053 | -0.032 | -0.032 | 0 | 0.18 | 0.1 | 0.35 | 0.2 | 0.38 | 0.18 |
| 2 | 142268997 | rs16846362 | 0.867 | -1.086 | -0.201 | -0.399 | 0 | -0.002 | -0.002 | 0 | 0.14 | 0.16 | 0.09 | 0.03 | 0.14 | 0.16 |
| 2 | 142277783 | rs1905291 | -0.426 | -1.106 | -0.146 | -0.407 | -0.018 | -0.038 | -0.038 | 0.614 | 0.27 | 0.39 | 0.17 | 0.09 | 0.18 | 0.37 |
| 2 | 142285767 | rs10496885 | -0.245 | -1.328 | 0.006 | -0.306 | 0.011 | 0.009 | 0.009 | 0 | 0.25 | 0.22 | 0.18 | 0.15 | 0.17 | 0.44 |
| 2 | 142286840 | rs12617263 | 1.793 | -1.537 | -0.275 | -0.343 | -0.018 | -0.105 | -0.105 | 0 | 0.23 | 0.01 | 0.22 | 0.37 | 0.2 | 0.17 |

| | | | | | | | | | | | | | | | | |
|---|---|---|---|---|---|---|---|---|---|---|---|---|---|---|---|---|
| 2 | 142287827 | rs1483141 | -0.237 | -1.167 | 0.381 | -0.352 | 0.016 | 0.017 | 0.017 | 0 | 0.51 | 0.34 | 0.54 | 0.36 | 0.59 | 0.66 |
| 2 | 142290687 | rs16846416 | 1.274 | -1.138 | 0.379 | -0.34 | -0.005 | -0.001 | -0.001 | 3.97 | 0.07 | 0.11 | 0.09 | 0.06 | 0.03 | 0.17 |
| 2 | 142292571 | rs1483137 | 1.533 | -1.122 | 0.287 | -0.349 | -0.066 | -0.009 | -0.009 | 0 | 0.27 | 0.13 | 0.36 | 0.2 | 0.41 | 0.22 |
| 2 | 142302196 | rs6719851 | -0.982 | -1.137 | 0.377 | -0.323 | -0.001 | 0 | 0 | 0 | 0.31 | 0.29 | 0.26 | 0.32 | 0.27 | 0.49 |
| 2 | 142302920 | rs2890615 | 1.054 | -1.04 | 0.078 | -0.31 | 0.028 | -0.024 | -0.024 | 0 | 0.37 | 0.23 | 0.34 | 0.48 | 0.3 | 0.27 |
| 2 | 142307179 | rs16855058 | 1.172 | -0.782 | 0.08 | -0.189 | -0.023 | 0.017 | 0.017 | 0.443 | 0.09 | 0.03 | 0.09 | 0.07 | 0.1 | 0.13 |
| 2 | 142321708 | rs7602707 | 0.243 | -0.852 | 0.357 | -0.195 | 0.001 | -0.04 | -0.04 | 0 | 0.77 | 0.63 | 0.83 | 0.89 | 0.78 | 0.73 |
| 2 | 142324536 | rs12621499 | 0.896 | -1.049 | 0.064 | -0.262 | 0.016 | -0.051 | -0.051 | 0.016 | 0.37 | 0.16 | 0.36 | 0.63 | 0.32 | 0.3 |
| 2 | 142329146 | rs7592987 | 1.69 | -0.962 | 0.044 | -0.235 | -0.002 | 0.003 | 0.003 | 0 | 0.1 | 0.02 | 0.06 | 0.04 | 0.11 | 0.16 |
| 2 | 142331556 | rs10196328 | 2.565 | -0.957 | 0.047 | -0.23 | 0.015 | -0.013 | -0.013 | 0 | 0.07 | 0 | 0.02 | 0.07 | 0.03 | 0.09 |
| 2 | 142336104 | rs4363973 | 0.182 | -0.97 | 0.164 | -0.371 | 0.008 | 0.005 | 0.005 | 0.022 | 0.45 | 0.35 | 0.45 | 0.7 | 0.36 | 0.49 |
| 2 | 142345100 | rs10188788 | 0.758 | -1.008 | -0.489 | -0.398 | 0.031 | 0.003 | 0.003 | 0 | 0.17 | 0.03 | 0.09 | 0.09 | 0.14 | 0.26 |
| 2 | 142346445 | rs12612624 | 1.454 | -1.002 | -0.558 | -0.334 | -0.062 | -0.149 | -0.149 | 0 | 0.32 | 0.03 | 0.33 | 0.51 | 0.3 | 0.26 |
| 2 | 142347455 | rs6429912 | 1.243 | -0.923 | -0.464 | -0.232 | -0.066 | 0.012 | 0.012 | 0 | 0.31 | 0.13 | 0.36 | 0.2 | 0.42 | 0.23 |
| 2 | 142349495 | rs10178963 | -0.192 | -0.939 | -0.066 | -0.067 | -0.001 | 0.005 | 0.005 | 0 | 0.55 | 0.53 | 0.53 | 0.76 | 0.43 | 0.6 |
| 2 | 142358209 | rs7575223 | -1.233 | -1.163 | 0.1 | 0.302 | 0.026 | -0.006 | -0.006 | 0.83 | 0.33 | 0.74 | 0.23 | 0.34 | 0.2 | 0.5 |
| 2 | 142359850 | rs13001145 | 1.231 | -0.452 | -0.644 | 0.467 | -0.063 | -0.009 | -0.009 | 0 | 0.25 | 0.09 | 0.32 | 0.2 | 0.37 | 0.22 |
| 2 | 142363473 | rs16855061 | -0.222 | -0.704 | 0.162 | 0.56 | -0.013 | 0.014 | 0.014 | 0 | 0.09 | 0.16 | 0.1 | 0.1 | 0.04 | 0.2 |
| 2 | 142368156 | rs7606738 | 0.809 | -0.566 | -0.214 | 0.465 | -0.004 | -0.001 | -0.001 | 0 | 0.08 | 0.13 | 0.03 | 0.06 | 0.03 | 0.1 |
| 2 | 142374378 | rs7568970 | -1.19 | -0.716 | -0.004 | 0.526 | -0.064 | 0.035 | 0.035 | 0 | 0.24 | 0.44 | 0.17 | 0.29 | 0.1 | 0.35 |
| 2 | 142375798 | rs446401 | 0.882 | -0.623 | 0.01 | 0.171 | -0.003 | -0.003 | -0.003 | 0 | 0.7 | 0.68 | 0.77 | 0.76 | 0.79 | 0.53 |
| 2 | 142376540 | rs11681602 | -0.116 | -0.712 | -0.113 | 0.145 | 0.005 | 0.001 | 0.001 | 0.065 | 0.17 | 0.09 | 0.18 | 0.15 | 0.13 | 0.27 |
| 2 | 142376873 | rs6755592 | 0.829 | -0.635 | -0.149 | -0.028 | -0.002 | -0.005 | -0.005 | 0 | 0.36 | 0.36 | 0.46 | 0.53 | 0.37 | 0.27 |
| 2 | 142378380 | rs355576 | 0.489 | -0.715 | 0.183 | 0.04 | -0.029 | -0.002 | -0.002 | 0.076 | 0.1 | 0.13 | 0.02 | 0.09 | 0.02 | 0.19 |
| 2 | 142381045 | rs355582 | -0.687 | -0.819 | 0.439 | -0.078 | -0.043 | -0.03 | -0.03 | 0 | 0.25 | 0.35 | 0.15 | 0.17 | 0.13 | 0.37 |
| 2 | 142381172 | rs12992601 | 0.732 | -0.77 | 0.525 | -0.095 | -0.025 | -0.044 | -0.044 | 0.143 | 0.72 | 0.6 | 0.83 | 0.82 | 0.8 | 0.62 |
| 2 | 142383174 | rs16846697 | 0.786 | -0.739 | 0.832 | -0.109 | -0.021 | -0.042 | -0.042 | 0.012 | 0.72 | 0.46 | 0.81 | 0.82 | 0.8 | 0.62 |
| 2 | 142384387 | rs10928113 | 0.707 | -0.753 | 1.06 | -0.116 | -0.026 | -0.056 | -0.056 | 0 | 0.72 | 0.53 | 0.81 | 0.82 | 0.8 | 0.61 |
| 2 | 142384798 | rs10190730 | 0.379 | -0.844 | 0.728 | -0.066 | -0.011 | -0.017 | -0.017 | 0.015 | 0.14 | 0.2 | 0.11 | 0.09 | 0.1 | 0.18 |
| 2 | 142386991 | rs1370324 | 1.082 | -0.707 | 0.199 | -0.252 | -0.003 | -0.022 | -0.022 | 0 | 0.26 | 0.23 | 0.37 | 0.54 | 0.26 | 0.25 |
| 2 | 142391367 | rs355597 | -1.051 | -0.829 | 0.28 | -0.465 | -0.005 | 0.006 | 0.006 | 0 | 0.16 | 0.29 | 0.14 | 0.16 | 0.11 | 0.28 |
| 2 | 142392337 | rs355600 | -0.948 | -0.925 | 0.141 | -0.58 | -0.013 | -0.068 | -0.068 | 0 | 0.85 | 0.73 | 0.93 | 0.99 | 0.89 | 0.82 |
| 2 | 142397669 | rs6747694 | 0.182 | -0.974 | -0.277 | -0.719 | -0.004 | 0.042 | 0.042 | 0 | 0.53 | 0.66 | 0.48 | 0.31 | 0.55 | 0.51 |
| 2 | 142398540 | rs6751890 | 1.256 | -1.003 | -0.061 | -0.781 | -0.008 | 0.019 | 0.019 | 0 | 0.4 | 0.29 | 0.33 | 0.21 | 0.47 | 0.29 |
| 2 | 142403763 | rs7560183 | -0.432 | -0.991 | -0.052 | -0.709 | -0.017 | 0.01 | 0.01 | 0.06 | 0.12 | 0.33 | 0.13 | 0.11 | 0.06 | 0.21 |
| 2 | 142408609 | rs1437352 | -0.209 | -0.949 | -0.347 | -0.704 | -0.004 | 0 | 0 | 0 | 0.58 | 0.7 | 0.56 | 0.36 | 0.58 | 0.6 |
| 2 | 142412195 | rs17737803 | 2.171 | -1.169 | -1.077 | -1.083 | -0.08 | 0.103 | 0.103 | 0 | 0.21 | 0.05 | 0.13 | 0.06 | 0.33 | 0.17 |
| 2 | 142414531 | rs2196598 | 1.849 | -1.565 | -1.164 | -0.8 | -0.04 | -0.138 | -0.138 | 0 | 0.17 | 0.01 | 0.24 | 0.44 | 0.17 | 0.16 |
| 2 | 142423793 | rs11898305 | 0.574 | -1.432 | -0.76 | -0.817 | 0.033 | 0.007 | 0.007 | 0 | 0.07 | 0.16 | 0.12 | 0.06 | 0.1 | 0.11 |
| 2 | 142428166 | rs12691618 | 0.39 | -1.574 | -0.958 | -0.892 | 0.006 | -0.049 | -0.049 | 0 | 0.4 | 0.16 | 0.41 | 0.64 | 0.35 | 0.4 |
| 2 | 142428940 | rs6752277 | -0.72 | -1.593 | -0.673 | -0.754 | -0.011 | 0.046 | 0.046 | 0.278 | 0.16 | 0.36 | 0.21 | 0.19 | 0.1 | 0.3 |
| 2 | 142431539 | rs16846822 | 2.391 | -1.735 | -0.922 | -0.772 | 0.016 | -0.046 | -0.046 | 0.173 | 0.11 | 0.03 | 0.07 | 0.12 | 0.06 | 0.07 |
| 2 | 142434892 | rs12995584 | 1.88 | -1.677 | -0.847 | -0.709 | -0.011 | 0.003 | 0.003 | 0.008 | 0.12 | 0.01 | 0.09 | 0.09 | 0.1 | 0.15 |
| 2 | 142436459 | rs164979 | -1.087 | -1.576 | -0.653 | -0.713 | 0.015 | -0.03 | -0.03 | 0 | 0.16 | 0.51 | 0.18 | 0.09 | 0.14 | 0.24 |
| 2 | 142440110 | rs1437351 | -0.411 | -1.604 | -0.728 | -0.716 | 0.004 | 0 | 0 | 0 | 0.44 | 0.61 | 0.49 | 0.61 | 0.39 | 0.49 |
| 2 | 142443923 | rs4954922 | -0.158 | -2.059 | -0.616 | -0.559 | -0.001 | -0.001 | -0.001 | 0 | 0.41 | 0.56 | 0.37 | 0.59 | 0.29 | 0.42 |
| 2 | 142446465 | rs9653179 | 1.113 | -2.023 | -0.628 | -0.528 | 0.006 | -0.045 | -0.045 | 0 | 0.28 | 0.21 | 0.32 | 0.51 | 0.25 | 0.25 |
| 2 | 142456242 | rs16846884 | 0.509 | -2.039 | -0.452 | -0.54 | -0.049 | -0.001 | -0.001 | 0 | 0.11 | 0.2 | 0.03 | 0.09 | 0.03 | 0.15 |
| 2 | 142456528 | rs12613378 | -0.067 | -2.008 | -0.526 | -0.531 | -0.027 | 0.01 | 0.01 | 0 | 0.39 | 0.44 | 0.35 | 0.59 | 0.27 | 0.4 |
| 2 | 142465872 | rs16846918 | -0.227 | -1.955 | -0.999 | -0.505 | -0.055 | -0.014 | -0.014 | 0 | 0.14 | 0.25 | 0.05 | 0.09 | 0.05 | 0.18 |
| 2 | 142467595 | rs925193 | -1.087 | -2.048 | -1.091 | -0.582 | -0.024 | -0.004 | -0.004 | 0 | 0.34 | 0.43 | 0.34 | 0.31 | 0.25 | 0.45 |
| 2 | 142468442 | rs1370328 | 0.276 | -1.922 | -1.088 | -0.562 | -0.007 | -0.004 | -0.004 | 0 | 0.21 | 0.18 | 0.29 | 0.22 | 0.22 | 0.28 |
| 2 | 142468472 | rs17801458 | 1.342 | -1.875 | -1.106 | -0.546 | -0.017 | -0.03 | -0.03 | 0 | 0.05 | 0.01 | 0.09 | 0.1 | 0.04 | 0.12 |
| 2 | 142468592 | rs1370327 | -1.135 | -1.835 | -1.01 | -0.982 | 0.012 | -0.034 | -0.034 | 0 | 0.59 | 0.52 | 0.62 | 0.81 | 0.49 | 0.69 |
| 2 | 142471029 | rs9677566 | 1.051 | -1.655 | -1.189 | -0.828 | -0.004 | 0.007 | 0.007 | 0 | 0.15 | 0.15 | 0.18 | 0.12 | 0.17 | 0.17 |
| 2 | 142475665 | rs16846962 | 0.142 | -1.457 | -0.752 | -0.69 | -0.047 | -0.012 | -0.012 | 0 | 0.13 | 0.2 | 0.04 | 0.09 | 0.04 | 0.16 |
| 2 | 142477274 | rs1437349 | -0.357 | -1.004 | -0.924 | -0.606 | 0 | 0 | 0 | 0 | 0.49 | 0.49 | 0.45 | 0.69 | 0.33 | 0.55 |
| 2 | 142481583 | rs13016120 | 0.77 | -1.196 | -1.078 | -0.669 | 0.009 | -0.05 | -0.05 | 0 | 0.26 | 0.08 | 0.27 | 0.25 | 0.22 | 0.24 |
| 2 | 142482165 | rs12477186 | -1.205 | -0.929 | -0.809 | -0.701 | 0.103 | -0.117 | -0.117 | 0 | 0.74 | 0.55 | 0.69 | 0.93 | 0.53 | 0.77 |
| 2 | 142486253 | rs13024316 | 0.2 | -0.807 | -1.172 | 0.105 | 0.002 | -0.07 | -0.07 | 0 | 0.25 | 0.07 | 0.32 | 0.28 | 0.22 | 0.22 |
| 2 | 142486771 | rs2118272 | 0.62 | -0.727 | -0.729 | 0.09 | 0.006 | 0.077 | 0.077 | 0 | 0.17 | 0.27 | 0.26 | 0.53 | 0.17 | 0.21 |
| 2 | 142486875 | rs906651 | 0.607 | -1.374 | -0.357 | 0.294 | 0.064 | -0.03 | -0.03 | 0 | 0.62 | 0.75 | 0.66 | 0.59 | 0.76 | 0.52 |
| 2 | 142487449 | rs906649 | 0.542 | -1.356 | -0.209 | 0.224 | -0.008 | -0.002 | -0.002 | 0 | 0.83 | 0.83 | 0.89 | 0.89 | 0.93 | 0.67 |
| 2 | 142488529 | rs906648 | 1.52 | -1.173 | 0.122 | 0.152 | 0.032 | 0.024 | 0.024 | 0 | 0.49 | 0.32 | 0.53 | 0.6 | 0.63 | 0.45 |
| 2 | 142488678 | rs754943 | -0.287 | -1.139 | 0.115 | 0.158 | -0.006 | 0.005 | 0.005 | 0 | 0.31 | 0.19 | 0.32 | 0.29 | 0.3 | 0.22 |
| 2 | 142489456 | rs6713237 | 1.393 | -0.725 | 1.017 | 0.061 | -0.024 | -0.018 | -0.018 | 0 | 0.79 | 0.74 | 0.85 | 0.88 | 0.88 | 0.64 |
| 2 | 142489811 | rs1813194 | 1.829 | -0.677 | 0.706 | 0.055 | -0.019 | -0.046 | -0.046 | 0 | 0.7 | 0.38 | 0.76 | 0.88 | 0.78 | 0.63 |
| 2 | 142490499 | rs10172574 | 1.829 | -0.677 | 0.706 | 0.062 | -0.019 | -0.044 | -0.044 | 0 | 0.7 | 0.39 | 0.76 | 0.88 | 0.78 | 0.63 |
| 2 | 142495019 | rs1898422 | -1.727 | -0.439 | 0.868 | 0.278 | -0.021 | -0.042 | -0.042 | 0 | 0.23 | 0.53 | 0.19 | 0.12 | 0.19 | 0.3 |

| | | | | | | | | | | | | | | | | |
|---|---|---|---|---|---|---|---|---|---|---|---|---|---|---|---|---|
| 2 | 142496207 | rs6713253 | -1.422 | -0.36 | 0.735 | 0.348 | 0.037 | -0.006 | -0.006 | 0.558 | 0.07 | 0.3 | 0.05 | 0 | 0.06 | 0.02 |
| 2 | 142498632 | rs6708353 | -1.153 | -0.777 | 0.009 | 0.927 | -0.06 | 0.069 | 0.069 | 0 | 0.22 | 0.57 | 0.13 | 0.37 | 0.11 | 0.29 |
| 2 | 142500134 | rs6758126 | 0.524 | 0.157 | -0.363 | 0.774 | -0.003 | -0.009 | -0.009 | 0 | 0.74 | 0.78 | 0.67 | 0.69 | 0.72 | 0.68 |
| 2 | 142506366 | rs10496894 | -0.427 | -0.191 | -0.469 | 0.645 | 0.01 | 0.001 | 0.001 | 0.105 | 0.19 | 0.12 | 0.32 | 0.16 | 0.12 | 0.18 |
| 2 | 142509570 | rs1437347 | 0.513 | 0.222 | -0.687 | 0.461 | 0.005 | -0.018 | -0.018 | 1.939 | 0.41 | 0.33 | 0.45 | 0.47 | 0.4 | 0.38 |
| 2 | 142512880 | rs11679435 | 0.879 | 0.48 | -0.066 | 0.226 | -0.003 | -0.001 | -0.001 | 0.005 | 0.71 | 0.5 | 0.62 | 0.74 | 0.73 | 0.61 |
| 2 | 142513131 | rs10928120 | 0.873 | 0.483 | -0.101 | 0.2 | -0.003 | -0.004 | -0.004 | 0 | 0.71 | 0.51 | 0.62 | 0.74 | 0.73 | 0.61 |
| 2 | 142517668 | rs10193328 | 0.345 | 0.504 | -0.416 | -0.149 | 0.014 | 0.032 | 0.032 | 0.278 | 0.64 | 0.34 | 0.56 | 0.83 | 0.75 | 0.76 |
| 2 | 142518602 | rs16847064 | -1.627 | 0.908 | 0.556 | 0.191 | 0.064 | 0.02 | 0.02 | 1.858 | 0.08 | 0.38 | 0.09 | 0.06 | 0.08 | 0.04 |
| 2 | 142524373 | rs1866024 | -1.192 | 1.03 | 0.548 | -0.098 | -0.001 | -0.018 | -0.018 | 0 | 0.12 | 0.1 | 0.09 | 0 | 0.15 | 0.08 |
| 2 | 142530120 | rs10928122 | -1.23 | 0.767 | 0.848 | -0.446 | -0.005 | -0.06 | -0.06 | 0 | 0.26 | 0.46 | 0.23 | 0.13 | 0.27 | 0.25 |
| 2 | 142530945 | rs16847120 | 0.469 | 1.57 | 0.363 | -0.077 | -0.034 | -0.034 | -0.034 | 0 | 0.16 | 0.07 | 0.34 | 0.45 | 0.35 | 0.28 |
| 2 | 142531733 | rs7604400 | -0.812 | 1.37 | 0.388 | -0.054 | -0.005 | -0.06 | -0.06 | 0 | 0.26 | 0.46 | 0.23 | 0.13 | 0.27 | 0.25 |
| 2 | 142533158 | rs16847133 | -0.053 | 1.368 | 0.394 | -0.053 | -0.02 | -0.025 | -0.025 | 0 | 0.2 | 0.36 | 0.18 | 0.13 | 0.17 | 0.23 |
| 2 | 142533719 | rs7558916 | -0.706 | 1.307 | 0.178 | -0.312 | -0.054 | -0.064 | -0.064 | 0 | 0.31 | 0.15 | 0.57 | 0.51 | 0.51 | 0.45 |
| 2 | 142533965 | rs7598314 | -1.301 | 1.333 | 0.198 | -0.319 | 0 | -0.016 | -0.016 | 0 | 0.37 | 0.27 | 0.61 | 0.51 | 0.59 | 0.47 |
| 2 | 142535388 | rs16847151 | 0.566 | 1.242 | 0.62 | -0.374 | 0.001 | 0.047 | 0.047 | 0.053 | 0.07 | 0.23 | 0.04 | 0.1 | 0.06 | 0.11 |
| 2 | 142536819 | rs10496896 | -0.191 | 1.27 | 0.665 | -0.376 | -0.054 | -0.048 | -0.048 | 0 | 0.25 | 0.15 | 0.52 | 0.51 | 0.43 | 0.39 |
| 2 | 142561219 | rs2164698 | -0.998 | 1.131 | 0.694 | -0.413 | -0.002 | -0.029 | -0.029 | 0 | 0.2 | 0.33 | 0.15 | 0.11 | 0.23 | 0.19 |
| 2 | 142562218 | rs1009398 | -0.044 | 1.158 | 0.683 | -0.421 | -0.031 | -0.027 | -0.027 | 0 | 0.75 | 0.82 | 0.48 | 0.5 | 0.56 | 0.61 |
| 2 | 142565292 | rs972485 | 2.139 | 0.876 | 0.675 | -0.453 | 0.119 | -0.048 | -0.048 | 2.136 | 0.39 | 0.07 | 0.17 | 0.35 | 0.22 | 0.29 |
| 2 | 142567010 | rs974974 | 1.019 | 0.872 | 0.76 | -0.464 | 0.017 | -0.009 | -0.009 | 0 | 0.56 | 0.66 | 0.3 | 0.46 | 0.38 | 0.43 |
| 2 | 142567218 | rs974973 | 1.615 | 0.881 | 0.808 | -0.464 | 0.01 | 0.005 | 0.005 | 0.202 | 0.47 | 0.45 | 0.25 | 0.35 | 0.32 | 0.31 |
| 2 | 142567436 | rs16847202 | 0.26 | 0.835 | 0.691 | -0.452 | -0.012 | 0.021 | 0.021 | 0.213 | 0.09 | 0.22 | 0.06 | 0.11 | 0.07 | 0.12 |
| 2 | 142568206 | rs17747853 | 0.652 | 0.844 | 0.64 | -0.459 | -0.002 | -0.01 | -0.01 | 0 | 0.12 | 0.12 | 0.13 | 0.03 | 0.1 | 0.12 |
| 2 | 142568465 | rs1866026 | 1.546 | 0.731 | 0.764 | -0.377 | 0.062 | 0.023 | 0.023 | 0.45 | 0.46 | 0.31 | 0.23 | 0.35 | 0.31 | 0.31 |
| 2 | 142572590 | rs7584919 | -0.734 | 1.159 | 1.138 | -0.045 | -0.011 | 0.032 | 0.032 | 1.461 | 0.11 | 0.4 | 0.07 | 0.11 | 0.07 | 0.12 |
| 2 | 142574431 | rs10206213 | -0.777 | 1.274 | 0.912 | -0.065 | 0.049 | -0.024 | -0.024 | 0 | 0.06 | 0.17 | 0.05 | 0 | 0.09 | 0.02 |
| 2 | 142576915 | rs1465759 | -1.041 | 1.929 | 1.159 | 0.105 | 0.098 | 0.099 | 0.099 | 0.205 | 0.26 | 0.42 | 0.37 | 0.54 | 0.44 | 0.39 |
| 2 | 142580764 | rs7602631 | -0.354 | 1.746 | 1.043 | -0.068 | 0 | -0.019 | -0.019 | 0 | 0.13 | 0.16 | 0.15 | 0.03 | 0.15 | 0.17 |
| 2 | 142585975 | rs7561165 | -0.373 | 1.447 | 0.908 | -0.326 | 0.002 | 0.005 | 0.005 | 0.557 | 0.3 | 0.34 | 0.35 | 0.21 | 0.25 | 0.34 |
| 2 | 142586200 | rs352980 | -0.711 | 1.827 | 0.744 | -0.179 | 0.006 | 0.017 | 0.017 | 0 | 0.85 | 0.88 | 0.74 | 0.56 | 0.67 | 0.77 |
| 2 | 142591851 | rs6721469 | -0.978 | 1.767 | 0.886 | -0.295 | 0 | -0.005 | -0.005 | 0.254 | 0.41 | 0.48 | 0.51 | 0.24 | 0.4 | 0.48 |
| 2 | 142591936 | rs6721397 | -1.148 | 1.705 | 0.835 | -0.325 | 0.002 | 0.005 | 0.005 | 1.669 | 0.43 | 0.64 | 0.53 | 0.24 | 0.4 | 0.48 |
| 2 | 142592088 | rs173291 | 1.18 | 1.671 | 0.813 | -0.346 | -0.001 | 0 | 0 | 0.193 | 0.61 | 0.42 | 0.51 | 0.43 | 0.45 | 0.42 |
| 2 | 142592560 | rs11884813 | 0.638 | 1.571 | 0.786 | -0.277 | 0 | 0.012 | 0.012 | 0 | 0.2 | 0.22 | 0.3 | 0.1 | 0.18 | 0.14 |
| 2 | 142593683 | rs352985 | -0.886 | 1.525 | 0.442 | -0.166 | 0.003 | 0.036 | 0.036 | 0 | 0.86 | 0.89 | 0.81 | 0.56 | 0.77 | 0.79 |
| 2 | 142601992 | rs16847336 | 1.192 | 1.773 | 0.47 | 0.309 | -0.009 | -0.004 | -0.004 | 0.048 | 0.11 | 0.05 | 0.16 | 0.28 | 0.21 | 0.12 |
| 2 | 142602144 | rs11902392 | -0.591 | 1.317 | 0.239 | -0.267 | -0.002 | -0.04 | -0.04 | 0.306 | 0.34 | 0.45 | 0.4 | 0.21 | 0.34 | 0.35 |
| 2 | 142602912 | rs1541976 | 0.407 | 1.372 | 0.057 | -0.274 | 0.019 | 0.028 | 0.028 | 0 | 0.72 | 0.67 | 0.61 | 0.53 | 0.59 | 0.61 |
| 2 | 142608783 | rs10496903 | 0.652 | 1.337 | -0.079 | -0.448 | 0.011 | -0.051 | -0.051 | 0 | 0.22 | 0.33 | 0.34 | 0.1 | 0.25 | 0.15 |
| 2 | 142610271 | rs13031421 | 1.839 | 1.311 | -0.912 | -0.256 | 0.043 | -0.023 | -0.023 | 0 | 0.37 | 0.04 | 0.2 | 0.3 | 0.25 | 0.26 |
| 2 | 142611767 | rs6429925 | -0.866 | 1.354 | -0.857 | -0.224 | 0.027 | 0.093 | 0.093 | 0 | 0.85 | 0.82 | 0.8 | 0.54 | 0.76 | 0.79 |
| 2 | 142617164 | rs13385804 | 0.619 | 0.292 | -1.438 | -0.19 | 0.097 | 0.018 | 0.018 | 2.427 | 0.47 | 0.17 | 0.25 | 0.41 | 0.33 | 0.44 |
| 2 | 142618914 | rs16847412 | 0.005 | 0.223 | -1.216 | -0.149 | -0.012 | -0.001 | -0.001 | 0 | 0.1 | 0.14 | 0.05 | 0.11 | 0.07 | 0.21 |
| 2 | 142624753 | rs13007371 | 0.358 | 0.266 | -1.7 | -0.131 | 0.002 | 0.046 | 0.046 | 3.529 | 0.47 | 0.07 | 0.35 | 0.35 | 0.41 | 0.37 |
| 2 | 142625386 | rs4662269 | 0.371 | 0.271 | -1.735 | -0.117 | -0.006 | 0.034 | 0.034 | 3.456 | 0.47 | 0.06 | 0.35 | 0.35 | 0.41 | 0.37 |
| 2 | 142634018 | rs2053175 | -0.93 | 0.625 | -0.691 | -0.157 | -0.007 | 0.013 | 0.013 | 0.002 | 0.14 | 0.13 | 0.19 | 0.04 | 0.18 | 0.21 |
| 2 | 142634758 | rs10178875 | -0.28 | 0.796 | -0.879 | -0.073 | -0.033 | 0.044 | 0.044 | 0 | 0.11 | 0.03 | 0.18 | 0.04 | 0.18 | 0.14 |
| 2 | 142636460 | rs10496905 | -0.871 | -0.422 | -0.262 | -0.25 | -0.09 | -0.061 | -0.061 | 0 | 0.22 | 0.39 | 0.11 | 0.12 | 0.08 | 0.33 |
| 2 | 142636523 | rs12466165 | -0.944 | -0.56 | 0.021 | -0.296 | -0.042 | -0.015 | -0.015 | 0 | 0.16 | 0.25 | 0.16 | 0.13 | 0.09 | 0.23 |
| 2 | 142636907 | rs12613827 | 0.868 | -0.753 | -1.111 | -0.2 | 0.006 | -0.07 | -0.07 | 3.48 | 0.36 | 0.13 | 0.35 | 0.42 | 0.3 | 0.24 |
| 2 | 142637466 | rs1369529 | -0.613 | -0.288 | -1.073 | -0.351 | -0.005 | 0.012 | 0.012 | 0 | 0.9 | 0.89 | 0.9 | 1 | 0.86 | 0.88 |
| 2 | 142639313 | rs11693445 | -0.657 | 0.049 | -0.567 | -0.817 | 0.003 | 0.012 | 0.012 | 0.934 | 0.37 | 0.42 | 0.35 | 0.56 | 0.37 | 0.56 |
| 2 | 142639394 | rs7565791 | -0.76 | 0.133 | -0.537 | -0.798 | 0.001 | -0.037 | -0.037 | 0.995 | 0.89 | 0.81 | 0.89 | 1 | 0.86 | 0.88 |
| 2 | 142639632 | rs16847499 | 0.48 | 0.379 | -0.491 | -0.726 | 0.001 | -0.014 | -0.014 | 0 | 0.07 | 0.14 | 0.08 | 0.06 | 0.06 | 0.07 |
| 2 | 142646363 | rs1991997 | 1.073 | 0.34 | -0.471 | -0.62 | -0.004 | -0.002 | -0.002 | 0.311 | 0.54 | 0.41 | 0.55 | 0.46 | 0.49 | 0.33 |
| 2 | 142646622 | rs13408365 | -0.712 | 0.352 | -0.388 | -0.625 | 0.004 | 0.02 | 0.02 | 1.793 | 0.35 | 0.38 | 0.34 | 0.54 | 0.36 | 0.56 |
| 2 | 142662928 | rs4662559 | -0.812 | 0.42 | -0.337 | -0.577 | 0.007 | -0.047 | -0.047 | 0 | 0.91 | 0.79 | 0.92 | 1 | 0.88 | 0.89 |
| 2 | 142665064 | rs10496907 | 0.993 | 0.207 | -0.872 | -0.757 | -0.053 | -0.077 | -0.077 | 0 | 0.2 | 0.05 | 0.22 | 0.37 | 0.27 | 0.29 |
| 2 | 142668576 | rs1529869 | -0.877 | 0.009 | -0.832 | -0.925 | 0 | -0.019 | -0.019 | 0.51 | 0.44 | 0.3 | 0.38 | 0.54 | 0.41 | 0.56 |
| 2 | 142671167 | rs1898018 | 1.428 | -0.028 | -0.854 | -0.99 | -0.039 | -0.044 | -0.044 | 0 | 0.45 | 0.14 | 0.51 | 0.46 | 0.47 | 0.32 |
| 2 | 142681720 | rs10928131 | -1.433 | 0.014 | -0.687 | -0.989 | 0.006 | -0.042 | -0.042 | 0 | 0.91 | 0.8 | 0.91 | 1 | 0.88 | 0.88 |
| 2 | 142685324 | rs13013316 | 2.07 | -0.247 | -1.137 | -1.294 | -0.005 | -0.066 | -0.066 | 0 | 0.27 | 0.08 | 0.31 | 0.41 | 0.26 | 0.24 |
| 2 | 142699457 | rs1435585 | -0.853 | -0.039 | -0.729 | -1.045 | 0.04 | -0.068 | -0.068 | 0.052 | 0.89 | 0.45 | 0.89 | 1 | 0.88 | 0.88 |
| 2 | 142699984 | rs10179103 | -1.582 | -0.014 | -0.782 | -1.032 | 0.011 | -0.043 | -0.043 | 0 | 0.92 | 0.8 | 0.91 | 1 | 0.88 | 0.88 |
| 2 | 142702456 | rs16847711 | 1.568 | -0.133 | -1.011 | -1.051 | 0.01 | 0.015 | 0.015 | 0 | 0.07 | 0.01 | 0.05 | 0.04 | 0.06 | 0.05 |
| 2 | 142702491 | rs10205836 | 2.194 | -0.394 | -1.405 | -0.778 | 0.037 | -0.02 | -0.02 | 0 | 0.21 | 0.07 | 0.23 | 0.24 | 0.1 | 0.11 |
| 2 | 142707328 | rs12471235 | 0.833 | 0.427 | -0.824 | -0.434 | 0 | 0 | 0 | 0 | 0.55 | 0.47 | 0.55 | 0.45 | 0.5 | 0.33 |

| | | | | | | | | | | | | | | | | |
|---|---|---|---|---|---|---|---|---|---|---|---|---|---|---|---|---|
| 2 | 142716216 | rs16847729 | 1.015 | 0.71 | -1.713 | -0.359 | -0.049 | -0.053 | -0.053 | 0 | 0.15 | 0.06 | 0.2 | 0.29 | 0.25 | 0.24 |
| 2 | 142717631 | rs7421755 | 0.988 | 0.543 | -1.669 | -0.259 | 0.01 | -0.004 | -0.004 | 0 | 0.49 | 0.12 | 0.51 | 0.53 | 0.46 | 0.34 |
| 2 | 142739121 | rs7564502 | 0.184 | 0.978 | -1.401 | -0.046 | -0.001 | -0.001 | -0.001 | 0.063 | 0.69 | 0.68 | 0.68 | 0.65 | 0.57 | 0.58 |
| 2 | 142744474 | rs10496909 | 0.071 | 0.904 | -1.598 | -0.017 | -0.013 | 0.013 | 0.013 | 0 | 0.17 | 0.09 | 0.16 | 0.12 | 0.22 | 0.26 |
| 2 | 142746083 | rs7568221 | 0.612 | 1.084 | -1.742 | 0.095 | -0.057 | -0.067 | -0.067 | 0.853 | 0.17 | 0.04 | 0.19 | 0.29 | 0.26 | 0.15 |
| 2 | 142750477 | rs1435609 | 1.356 | 1.392 | -1.752 | -0.112 | 0.045 | -0.003 | -0.003 | 0.042 | 0.52 | 0.65 | 0.53 | 0.51 | 0.37 | 0.37 |
| 2 | 142754084 | rs720755 | 0.555 | 1.681 | -1.631 | 0.072 | -0.058 | -0.034 | -0.034 | 0.053 | 0.72 | 0.83 | 0.69 | 0.64 | 0.6 | 0.66 |
| 2 | 142756717 | rs10928132 | 0.156 | 1.169 | -1.247 | -0.102 | 0.003 | -0.013 | -0.013 | 0 | 0.87 | 0.84 | 0.88 | 0.94 | 0.86 | 0.81 |
| 2 | 142759003 | rs1191970 | -1.146 | 0.466 | -1.256 | -0.454 | -0.002 | -0.012 | -0.012 | 0 | 0.44 | 0.49 | 0.3 | 0.19 | 0.36 | 0.45 |
| 2 | 142763365 | rs1191972 | -0.613 | 0.545 | -1.339 | 0.011 | -0.001 | -0.001 | -0.001 | 0 | 0.44 | 0.49 | 0.58 | 0.58 | 0.51 | 0.62 |
| 2 | 142765788 | rs1191973 | -1.133 | 0.5 | -1.457 | -0.053 | 0.011 | 0.006 | 0.006 | 5.455 | 0.5 | 0.5 | 0.69 | 0.61 | 0.6 | 0.66 |
| 2 | 142768518 | rs1191974 | -0.6 | -0.316 | -0.862 | -0.062 | 0.03 | -0.007 | -0.007 | 0 | 0.93 | 0.75 | 0.88 | 0.98 | 0.91 | 0.96 |
| 2 | 142776441 | rs7597320 | 0.359 | -0.798 | -1.039 | -0.066 | 0.049 | 0.026 | 0.026 | 10.161 | 0.47 | 0.17 | 0.26 | 0.4 | 0.35 | 0.31 |
| 2 | 142785189 | rs13394203 | -0.316 | -0.948 | -0.705 | 0.315 | -0.006 | 0.004 | 0.004 | 0 | 0.11 | 0.15 | 0.04 | 0.15 | 0.06 | 0.16 |
| 2 | 142787997 | rs13430581 | 1.024 | -1.347 | -1.156 | 0.204 | 0.038 | -0.008 | -0.008 | 0.006 | 0.31 | 0.08 | 0.16 | 0.31 | 0.19 | 0.18 |
| 2 | 142788546 | rs12623786 | -0.764 | -0.925 | -0.154 | 0.148 | 0.036 | -0.038 | -0.038 | 0 | 0.09 | 0.03 | 0.04 | 0.18 | 0.02 | 0.16 |
| 2 | 142791459 | rs12617185 | -0.892 | -0.923 | 0.057 | -0.434 | -0.05 | 0.004 | 0.004 | 0 | 0.11 | 0.27 | 0.03 | 0.09 | 0.02 | 0.16 |
| 2 | 142794718 | rs11677303 | 0.265 | -0.79 | 0.158 | -0.978 | -0.013 | -0.113 | -0.113 | 1.025 | 0.75 | 0.54 | 0.88 | 0.98 | 0.79 | 0.85 |
| 2 | 142801609 | rs17772688 | 0.38 | -1.161 | -0.441 | -1.205 | 0.013 | 0.053 | 0.053 | 0 | 0.15 | 0 | 0.05 | 0.07 | 0.09 | 0.19 |
| 2 | 142819607 | rs11688838 | 1.528 | -0.961 | -0.034 | -1.138 | 0.011 | -0.007 | -0.007 | 0 | 0.37 | 0.35 | 0.61 | 0.68 | 0.48 | 0.39 |
| 2 | 142822193 | rs7586417 | -0.434 | -1.119 | -0.203 | -1.19 | 0.071 | 0.042 | 0.042 | 0 | 0.36 | 0.18 | 0.24 | 0.28 | 0.25 | 0.47 |
| 2 | 142824137 | rs4662219 | 1 | -1.185 | 0.257 | -1.321 | 0.005 | 0.006 | 0.006 | 0 | 0.49 | 0.45 | 0.65 | 0.72 | 0.59 | 0.46 |
| 2 | 142831758 | rs13396415 | 0.47 | -1.158 | 0.505 | -1.306 | 0.011 | -0.067 | -0.067 | 0.425 | 0.13 | 0.19 | 0.09 | 0 | 0.17 | 0.07 |
| 2 | 142835779 | rs1449480 | 0.224 | -1.197 | 0.37 | -1.306 | 0.025 | 0.018 | 0.018 | 0 | 0.62 | 0.64 | 0.74 | 0.72 | 0.75 | 0.53 |
| 2 | 142839970 | rs2044731 | -0.506 | -1.234 | 0.468 | -1.366 | 0 | -0.032 | -0.032 | 0 | 0.26 | 0.28 | 0.13 | 0.02 | 0.27 | 0.14 |
| 2 | 142855291 | rs7583748 | -1.641 | -1.239 | 0.511 | -1.331 | 0 | -0.007 | -0.007 | 0.628 | 0.91 | 0.82 | 0.95 | 0.94 | 0.91 | 0.93 |
| 2 | 142856523 | rs11680286 | 1.909 | -1.204 | 0.507 | -1.318 | 0.017 | -0.009 | -0.009 | 0.056 | 0.36 | 0.37 | 0.61 | 0.7 | 0.49 | 0.4 |
| 2 | 142862830 | rs10181071 | -0.449 | -1.22 | 0.378 | -1.298 | -0.002 | -0.055 | -0.055 | 0 | 0.18 | 0.29 | 0.11 | 0.02 | 0.19 | 0.1 |
| 2 | 142868147 | rs1001628 | -0.335 | -1.211 | 0.359 | -1.294 | -0.001 | -0.049 | -0.049 | 0 | 0.18 | 0.27 | 0.1 | 0.02 | 0.19 | 0.1 |
| 2 | 142876051 | rs2290140 | 0.861 | -1.386 | 0.266 | -1.231 | -0.001 | -0.003 | -0.003 | 0.584 | 0.5 | 0.46 | 0.68 | 0.73 | 0.64 | 0.47 |
| 2 | 142878411 | rs10193058 | -1.077 | -1.289 | 0.106 | -1.122 | 0.048 | 0.066 | 0.066 | 0 | 0.51 | 0.36 | 0.3 | 0.24 | 0.36 | 0.52 |
| 2 | 142882593 | rs10496915 | -0.423 | -1.322 | 0.279 | -1.04 | -0.049 | -0.046 | -0.046 | 0 | 0.28 | 0.51 | 0.33 | 0.16 | 0.17 | 0.18 |
| 2 | 142883177 | rs1449487 | -1.856 | -1.353 | 0.129 | -1.281 | 0.019 | 0.125 | 0.125 | 0.283 | 0.55 | 0.79 | 0.49 | 0.22 | 0.43 | 0.42 |
| 2 | 142885880 | rs11904731 | 0.092 | -1.36 | 0.515 | -1.246 | -0.034 | -0.031 | -0.031 | 0 | 0.25 | 0.37 | 0.31 | 0.16 | 0.17 | 0.18 |
| 2 | 142889137 | rs10164457 | -0.346 | -1.687 | 0.317 | -1.257 | -0.01 | -0.033 | -0.033 | 0 | 0.19 | 0.25 | 0.12 | 0.05 | 0.14 | 0.17 |
| 2 | 142889154 | rs10211565 | -2.434 | -1.745 | 0.042 | -1.273 | -0.031 | 0.001 | 0.001 | 0 | 0.63 | 0.8 | 0.46 | 0.33 | 0.48 | 0.65 |
| 2 | 142896437 | rs7586590 | -0.975 | -1.701 | 0.446 | -1.238 | -0.024 | -0.071 | -0.071 | 0.504 | 0.23 | 0.58 | 0.16 | 0.05 | 0.18 | 0.19 |
| 2 | 142901507 | rs17832157 | 0.33 | -1.647 | 0.214 | -1.232 | 0.005 | 0.035 | 0.035 | 0 | 0.14 | 0.09 | 0.05 | 0.01 | 0.12 | 0.06 |
| 2 | 142909146 | rs4662234 | 0.395 | -1.732 | 0.181 | -1.249 | 0.035 | 0.026 | 0.026 | 0 | 0.22 | 0.11 | 0.22 | 0.15 | 0.14 | 0.17 |
| 2 | 142910683 | rs10183140 | -0.485 | -1.587 | 0.417 | -1.214 | -0.01 | -0.055 | -0.055 | 0.139 | 0.2 | 0.29 | 0.15 | 0.05 | 0.19 | 0.19 |
| 2 | 142925716 | rs11695507 | -0.191 | -1.55 | 0.404 | -1.071 | 0.014 | 0.008 | 0.008 | 0 | 0.25 | 0.2 | 0.27 | 0.2 | 0.18 | 0.24 |
| 2 | 142927676 | rs7590590 | -0.761 | -1.636 | 0.783 | -1.115 | -0.048 | -0.034 | -0.034 | 0 | 0.29 | 0.53 | 0.3 | 0.2 | 0.2 | 0.25 |
| 2 | 142928238 | rs4662239 | -0.274 | -1.633 | 0.654 | -1.114 | 0.02 | 0.014 | 0.014 | 0 | 0.26 | 0.2 | 0.29 | 0.2 | 0.2 | 0.25 |
| 2 | 142931146 | rs17776169 | 1.478 | -1.565 | 0.359 | -1.184 | -0.004 | 0.001 | 0.001 | 0 | 0.09 | 0.08 | 0.03 | 0 | 0.09 | 0.05 |
| 2 | 142934735 | rs7557962 | -1.343 | -1.621 | 0.277 | -1.162 | 0.047 | 0.061 | 0.061 | 0.004 | 0.43 | 0.67 | 0.35 | 0.23 | 0.31 | 0.32 |
| 2 | 142938021 | rs4662423 | -1.284 | -1.627 | 0.341 | -1.218 | 0.034 | 0.048 | 0.048 | 0 | 0.42 | 0.65 | 0.35 | 0.19 | 0.31 | 0.32 |
| 2 | 142940335 | rs2177416 | -1.67 | -1.727 | 0.48 | -1.258 | -0.001 | -0.001 | -0.001 | 0 | 0.53 | 0.57 | 0.42 | 0.28 | 0.39 | 0.45 |
| 2 | 142942795 | rs17832932 | 1.022 | -1.671 | 0.398 | -1.223 | 0.009 | -0.046 | -0.046 | 2.765 | 0.11 | 0.22 | 0.07 | 0.03 | 0.11 | 0.19 |
| 2 | 142942925 | rs10172452 | -0.622 | -1.773 | 0.154 | -1.246 | 0.034 | 0.02 | 0.02 | 0 | 0.26 | 0.1 | 0.14 | 0.16 | 0.2 | 0.25 |
| 2 | 142947513 | rs13408409 | 0.19 | -1.79 | 0.127 | -1.292 | 0.011 | 0.016 | 0.016 | 0 | 0.13 | 0.08 | 0.07 | 0.02 | 0.1 | 0.07 |
| 2 | 142952096 | rs13406850 | -1.565 | -1.845 | 0.281 | -1.326 | 0.07 | 0.067 | 0.067 | 0 | 0.48 | 0.29 | 0.38 | 0.28 | 0.36 | 0.45 |
| 2 | 142954841 | rs7579516 | -2.37 | -1.813 | 0.349 | -1.33 | 0.004 | -0.005 | -0.005 | 0 | 0.63 | 0.58 | 0.46 | 0.31 | 0.49 | 0.64 |
| 2 | 142963846 | rs1523706 | -0.502 | -1.86 | 0.371 | -1.41 | -0.039 | -0.045 | -0.045 | 0 | 0.26 | 0.37 | 0.28 | 0.12 | 0.16 | 0.34 |
| 2 | 142963860 | rs1523705 | 0.418 | -1.88 | 0.28 | -1.419 | 0.032 | 0.036 | 0.036 | 0 | 0.2 | 0.1 | 0.2 | 0.1 | 0.12 | 0.19 |
| 2 | 142963976 | rs1523704 | 0.944 | -2.047 | 0.173 | -1.477 | 0.006 | 0.018 | 0.018 | 0 | 0.14 | 0.01 | 0.17 | 0.08 | 0.11 | 0.17 |
| 2 | 142967224 | rs10496917 | 0.437 | -1.887 | 0.241 | -1.478 | 0.027 | 0.045 | 0.045 | 0 | 0.2 | 0.1 | 0.21 | 0.08 | 0.13 | 0.19 |
| 2 | 142969568 | rs10203049 | 0.25 | -1.878 | 0.119 | -1.524 | 0.017 | 0.038 | 0.038 | 0 | 0.18 | 0.09 | 0.08 | 0.02 | 0.13 | 0.08 |
| 2 | 142969782 | rs11890556 | -0.44 | -1.91 | 0.457 | -1.52 | -0.047 | -0.084 | -0.084 | 0 | 0.27 | 0.44 | 0.29 | 0.11 | 0.18 | 0.34 |
| 2 | 142976147 | rs1628975 | 2.432 | -1.853 | 0.105 | -1.474 | -0.034 | 0.019 | 0.019 | 0 | 0.36 | 0.21 | 0.52 | 0.72 | 0.5 | 0.36 |
| 2 | 142976841 | rs1665164 | 0.754 | -1.742 | 0.193 | -1.502 | 0.026 | 0.043 | 0.043 | 0.224 | 0.61 | 0.3 | 0.66 | 0.89 | 0.69 | 0.6 |
| 2 | 142980375 | rs10174738 | -0.088 | -1.436 | 0.128 | -1.213 | 0.012 | 0.018 | 0.018 | 0 | 0.18 | 0.12 | 0.07 | 0.02 | 0.13 | 0.08 |
| 2 | 142989440 | rs4473327 | -0.275 | -1.067 | 0.285 | -1.101 | -0.002 | 0.004 | 0.004 | 0 | 0.19 | 0.28 | 0.13 | 0.18 | 0.17 | 0.19 |
| 2 | 142993458 | rs6752143 | 0.453 | -0.915 | 0.126 | -1.326 | 0.011 | 0.035 | 0.035 | 0 | 0.2 | 0.1 | 0.22 | 0.09 | 0.16 | 0.19 |
| 2 | 142994328 | rs1446355 | -0.277 | -0.764 | 0.469 | -1.304 | -0.005 | -0.046 | -0.046 | 0 | 0.22 | 0.2 | 0.12 | 0.02 | 0.2 | 0.11 |
| 2 | 142998565 | rs13026179 | -0.794 | -0.674 | 0.314 | -1.227 | -0.025 | -0.031 | -0.031 | 0.155 | 0.32 | 0.48 | 0.22 | 0.21 | 0.28 | 0.25 |
| 2 | 142999394 | rs7596606 | -0.658 | -0.645 | 0.034 | -1.213 | -0.009 | -0.011 | -0.011 | 0.594 | 0.31 | 0.36 | 0.21 | 0.21 | 0.28 | 0.25 |
| 2 | 143008971 | rs1584201 | 1.746 | -0.34 | -0.362 | -1.185 | -0.05 | 0.014 | 0.014 | 0 | 0.36 | 0.18 | 0.47 | 0.68 | 0.44 | 0.36 |
| 2 | 143011761 | rs16857018 | 0.6 | -0.337 | -0.265 | -1.064 | 0.009 | 0.017 | 0.017 | 0 | 0.15 | 0.05 | 0.16 | 0.1 | 0.11 | 0.17 |
| 2 | 143018256 | rs6429957 | -0.948 | -0.386 | 0.198 | -1.189 | -0.002 | -0.022 | -0.022 | 0 | 0.3 | 0.3 | 0.24 | 0.1 | 0.25 | 0.23 |

| | | | | | | | | | | | | | | | | |
|---|---|---|---|---|---|---|---|---|---|---|---|---|---|---|---|---|
| 2 | 143043285 | rs2121279 | 1.502 | -0.249 | 0.16 | -1.091 | -0.029 | 0.04 | 0.04 | 0.004 | 0.08 | 0.01 | 0.05 | 0 | 0.13 | 0.05 |
| 2 | 143047409 | rs1732605 | -0.186 | -0.53 | 0.693 | -1.155 | -0.017 | -0.085 | -0.085 | 0 | 0.14 | 0.28 | 0.06 | 0 | 0.09 | 0.07 |
| 2 | 143059566 | rs13431378 | -0.094 | 0.146 | 0.228 | -0.989 | -0.054 | -0.045 | -0.045 | 0 | 0.32 | 0.19 | 0.27 | 0.38 | 0.43 | 0.24 |
| 2 | 143076629 | rs4662472 | 1.05 | 0.254 | 0.195 | -0.974 | -0.052 | 0.029 | 0.029 | 0 | 0.18 | 0.06 | 0.11 | 0.11 | 0.27 | 0.09 |
| 2 | 143077372 | rs13034522 | -0.758 | -0.215 | 0.816 | -1.125 | 0.005 | 0.002 | 0.002 | 0 | 0.48 | 0.35 | 0.42 | 0.47 | 0.52 | 0.41 |
| 2 | 143077695 | rs6720779 | 1.83 | -0.451 | 0.883 | -1.154 | -0.003 | 0.009 | 0.009 | 0 | 0.08 | 0.05 | 0.03 | 0 | 0.08 | 0.05 |
| 2 | 143079431 | rs1446343 | -1.95 | -0.456 | 0.934 | -1.197 | 0.008 | -0.036 | -0.036 | 0.034 | 0.65 | 0.79 | 0.53 | 0.47 | 0.66 | 0.47 |
| 2 | 143093585 | rs17175240 | 0.921 | -0.451 | 1.14 | -1.186 | -0.001 | -0.028 | -0.028 | 0.052 | 0.12 | 0.12 | 0.07 | 0 | 0.12 | 0.06 |
| 2 | 143107213 | rs7578744 | -1.874 | -0.484 | 1.196 | -1.211 | 0.008 | -0.014 | -0.014 | 0 | 0.63 | 0.71 | 0.52 | 0.47 | 0.64 | 0.45 |
| 2 | 143110338 | rs4662491 | -0.741 | -0.387 | 0.831 | -1.408 | 0.041 | -0.002 | -0.002 | 1.413 | 0.54 | 0.36 | 0.59 | 0.47 | 0.68 | 0.57 |
| 2 | 143118816 | rs960549 | -2.281 | -0.475 | 0.819 | -1.101 | -0.005 | -0.075 | -0.075 | 0.249 | 0.67 | 0.86 | 0.54 | 0.47 | 0.64 | 0.46 |
| 2 | 143121279 | rs11682012 | -1.114 | -0.316 | 0.906 | -1.107 | -0.03 | -0.081 | -0.081 | 0.33 | 0.33 | 0.45 | 0.24 | 0.09 | 0.21 | 0.22 |
| 2 | 143126665 | rs4662498 | 0.726 | -0.167 | 0.815 | -1.06 | 0.001 | 0.001 | 0.001 | 0 | 0.16 | 0.13 | 0.15 | 0.09 | 0.09 | 0.15 |
| 2 | 143133296 | rs7590398 | -2.117 | -0.238 | 0.728 | -1.027 | 0.005 | -0.018 | -0.018 | 0.144 | 0.64 | 0.7 | 0.52 | 0.47 | 0.64 | 0.46 |
| 2 | 143136026 | rs1160251 | -0.168 | -0.33 | 0.785 | -0.826 | -0.018 | -0.015 | -0.015 | 0 | 0.32 | 0.25 | 0.28 | 0.38 | 0.42 | 0.24 |
| 2 | 143149884 | rs10206016 | -1.641 | -0.893 | 1.047 | -0.505 | -0.004 | 0 | 0 | 0 | 0.56 | 0.64 | 0.47 | 0.49 | 0.53 | 0.44 |
| 2 | 143154021 | rs961109 | 0.665 | -0.972 | 0.711 | -0.532 | -0.001 | -0.012 | -0.012 | 0 | 0.1 | 0.12 | 0.05 | 0.01 | 0.09 | 0.05 |
| 2 | 143157016 | rs12691651 | -0.657 | -0.988 | 0.682 | -0.523 | 0.002 | 0.019 | 0.019 | 0 | 0.35 | 0.43 | 0.3 | 0.43 | 0.37 | 0.26 |
| 2 | 143160364 | rs4662289 | 0.759 | -0.902 | -0.138 | -0.489 | 0.012 | 0.017 | 0.017 | 0 | 0.12 | 0.05 | 0.13 | 0.05 | 0.07 | 0.14 |
| 2 | 143165531 | rs4273167 | 1.447 | -0.779 | -0.17 | -0.389 | -0.033 | -0.021 | -0.021 | 0 | 0.43 | 0.2 | 0.51 | 0.52 | 0.48 | 0.56 |
| 2 | 143174049 | rs10928143 | -0.308 | -0.536 | -0.246 | -0.255 | -0.011 | -0.021 | -0.021 | 0 | 0.29 | 0.28 | 0.29 | 0.42 | 0.37 | 0.24 |
| 2 | 143177484 | rs4327191 | 1.169 | -0.446 | -0.661 | -0.16 | -0.064 | -0.039 | -0.039 | 0.141 | 0.43 | 0.14 | 0.5 | 0.5 | 0.47 | 0.56 |
| 2 | 143220907 | rs4311004 | 0.485 | -0.57 | -0.347 | -0.225 | 0.018 | 0.013 | 0.013 | 0 | 0.65 | 0.53 | 0.67 | 0.58 | 0.59 | 0.74 |
| 2 | 143222005 | rs13416800 | 1.418 | -0.863 | -0.307 | -0.325 | -0.004 | -0.007 | -0.007 | 0.086 | 0.41 | 0.32 | 0.5 | 0.5 | 0.47 | 0.54 |
| 2 | 143223382 | rs13407187 | 0.698 | -0.822 | -0.312 | -0.386 | -0.049 | -0.061 | -0.061 | 0 | 0.21 | 0.11 | 0.34 | 0.42 | 0.32 | 0.2 |
| 2 | 143227120 | rs6751887 | 0.045 | -0.802 | -0.487 | -0.132 | -0.004 | -0.003 | -0.003 | 0 | 0.24 | 0.21 | 0.1 | 0.08 | 0.12 | 0.19 |
| 2 | 143230267 | rs4550612 | 0.445 | 0 | -1.779 | 0.234 | -0.03 | -0.04 | -0.04 | 0 | 0.39 | 0.2 | 0.48 | 0.5 | 0.47 | 0.54 |
| 2 | 143237865 | rs7604603 | -0.263 | 0.492 | -1.668 | 0.625 | -0.035 | -0.069 | -0.069 | 0.027 | 0.66 | 0.81 | 0.5 | 0.53 | 0.63 | 0.67 |
| 2 | 143239105 | rs7572373 | -1.001 | 0.164 | -0.619 | 0.679 | 0.002 | 0.006 | 0.006 | 0 | 0.13 | 0.26 | 0.1 | 0.13 | 0.13 | 0.12 |
| 2 | 143239532 | rs7558786 | -0.686 | 0.175 | -0.996 | 0.612 | -0.004 | 0.005 | 0.005 | 0.303 | 0.08 | 0.04 | 0.05 | 0 | 0.12 | 0.07 |
| 2 | 143244498 | rs10199004 | 0.752 | -0.031 | -0.758 | 0.889 | -0.002 | -0.005 | -0.005 | 0 | 0.91 | 0.85 | 0.95 | 0.92 | 0.94 | 0.88 |
| 2 | 143261959 | rs6747453 | -1.257 | -0.656 | -1.495 | 1.288 | 0.09 | -0.156 | -0.156 | 3.897 | 0.33 | 0.94 | 0.21 | 0.49 | 0.18 | 0.43 |
| 2 | 143268255 | rs13003495 | -1.279 | -0.764 | -1.021 | 1.663 | 0.098 | -0.155 | -0.155 | 0 | 0.34 | 0.94 | 0.21 | 0.49 | 0.18 | 0.44 |
| 2 | 143269307 | rs4344876 | -1.061 | -0.887 | 0.352 | 1.75 | 0.009 | 0.238 | 0.238 | 2.005 | 0.07 | 0.38 | 0.06 | 0.23 | 0.04 | 0.08 |
| 2 | 143269355 | rs16857354 | -2.135 | -0.935 | 0.677 | 1.727 | -0.04 | 0.284 | 0.284 | 10.315 | 0.13 | 0.57 | 0.11 | 0.36 | 0.06 | 0.1 |
| 2 | 143272903 | rs1020426 | -0.849 | -0.974 | 0.653 | 1.621 | -0.046 | -0.078 | -0.078 | 1.246 | 0.22 | 0.37 | 0.1 | 0.13 | 0.13 | 0.35 |
| 2 | 143276280 | rs13400749 | -0.754 | -0.959 | 0.661 | 1.625 | -0.052 | -0.074 | -0.074 | 0 | 0.21 | 0.37 | 0.1 | 0.13 | 0.12 | 0.34 |
| 2 | 143278394 | rs16857378 | -1.285 | -0.891 | 0.807 | 1.663 | 0.005 | 0.138 | 0.138 | 4.24 | 0.05 | 0.22 | 0.05 | 0.13 | 0.02 | 0.02 |
| 2 | 143291667 | rs6712159 | -1.504 | -0.467 | 0.704 | 1.536 | -0.014 | 0 | 0 | 0 | 0.28 | 0.66 | 0.16 | 0.35 | 0.15 | 0.42 |
| 2 | 143300106 | rs13032123 | -1.524 | -0.433 | 1.023 | 1.535 | -0.01 | 0 | 0 | 0 | 0.28 | 0.66 | 0.17 | 0.35 | 0.17 | 0.42 |
| 2 | 143301347 | rs7594541 | -1.532 | -0.44 | 1.019 | 1.532 | -0.01 | 0 | 0 | 0 | 0.28 | 0.66 | 0.17 | 0.35 | 0.17 | 0.42 |
| 2 | 143312024 | rs4662556 | -0.049 | 0.309 | 0.36 | 2.009 | -0.111 | -0.079 | -0.079 | 0 | 0.16 | 0.01 | 0.45 | 0.21 | 0.32 | 0.3 |
| 2 | 143320073 | rs6429977 | -1.882 | 0.084 | 1.464 | 1.954 | -0.005 | 0.153 | 0.153 | 8.731 | 0.11 | 0.52 | 0.1 | 0.21 | 0.08 | 0.08 |
| 2 | 143320815 | rs11897421 | -1.302 | 0.112 | 1.336 | 1.969 | 0.023 | 0.175 | 0.175 | 2.163 | 0.09 | 0.36 | 0.09 | 0.21 | 0.07 | 0.08 |
| 2 | 143356378 | rs752661 | -2.018 | -0.256 | 1.18 | 1.9 | 0.05 | -0.06 | -0.06 | 0 | 0.3 | 0.88 | 0.2 | 0.34 | 0.19 | 0.24 |
| 2 | 143356687 | rs4146077 | -2.131 | -0.283 | 1.474 | 1.976 | -0.002 | 0.148 | 0.148 | 0.888 | 0.11 | 0.24 | 0.1 | 0.32 | 0.11 | 0.1 |
| 2 | 143387612 | rs4662294 | -0.516 | -0.186 | 1.227 | 1.99 | -0.055 | -0.041 | -0.041 | 2.995 | 0.22 | 0.06 | 0.41 | 0.3 | 0.31 | 0.5 |
| 2 | 143389735 | rs6715407 | -1.433 | -0.363 | 1.481 | 1.994 | 0.006 | 0.166 | 0.166 | 0 | 0.07 | 0.15 | 0.08 | 0.31 | 0.09 | 0.1 |
| 2 | 143408555 | rs7578423 | -0.434 | -0.431 | 1.687 | 1.627 | -0.017 | -0.03 | -0.03 | 0 | 0.06 | 0.18 | 0.04 | 0 | 0.03 | 0.05 |
| 2 | 143420408 | rs7603249 | -2.577 | -0.703 | 1.715 | 1.856 | 0.041 | -0.033 | -0.033 | 11.225 | 0.3 | 0.76 | 0.19 | 0.42 | 0.18 | 0.39 |
| 2 | 143436591 | rs13431986 | -2.316 | -0.608 | 1.637 | 1.935 | 0.007 | -0.01 | -0.01 | 0 | 0.21 | 0.76 | 0.16 | 0.39 | 0.13 | 0.3 |
| 2 | 143453889 | rs7568557 | -1.49 | -0.807 | 2.194 | 1.433 | -0.003 | -0.003 | -0.003 | 0.109 | 0.4 | 0.36 | 0.5 | 0.54 | 0.44 | 0.54 |
| 2 | 143458993 | rs6738045 | -0.989 | -0.794 | 2.204 | 1.351 | -0.03 | -0.056 | -0.056 | 0 | 0.12 | 0.32 | 0.06 | 0.03 | 0.07 | 0.1 |
| 2 | 143475728 | rs7598800 | -2.522 | -0.865 | 2.139 | 1.503 | -0.033 | 0.028 | 0.028 | 0 | 0.58 | 0.82 | 0.65 | 0.67 | 0.53 | 0.81 |
| 2 | 143508243 | rs12477181 | -2.162 | -0.906 | 2.494 | 1.499 | 0 | 0.003 | 0.003 | 0 | 0.52 | 0.52 | 0.61 | 0.61 | 0.52 | 0.68 |
| 2 | 143519238 | rs4662565 | 0.124 | -0.862 | 1.678 | 1.893 | -0.1 | -0.097 | -0.097 | 0 | 0.17 | 0.01 | 0.38 | 0.29 | 0.27 | 0.31 |
| 2 | 143533594 | rs12987477 | -2.737 | -0.832 | 1.815 | 1.877 | -0.049 | 0.027 | 0.027 | 0 | 0.58 | 0.85 | 0.65 | 0.66 | 0.53 | 0.81 |
| 2 | 143550868 | rs6712291 | -1.203 | -0.561 | 2.235 | 1.954 | -0.004 | 0 | 0 | 0 | 0.15 | 0.21 | 0.1 | 0.18 | 0.11 | 0.16 |
| 2 | 143553642 | rs1490949 | -1.894 | -0.708 | 2.266 | 1.94 | -0.029 | -0.03 | -0.03 | 0.023 | 0.23 | 0.59 | 0.17 | 0.15 | 0.21 | 0.29 |
| 2 | 143554042 | rs13032886 | -1.606 | -0.727 | 2.311 | 1.949 | -0.028 | -0.026 | -0.026 | 0.504 | 0.21 | 0.49 | 0.16 | 0.15 | 0.2 | 0.29 |
| 2 | 143554417 | rs4662298 | -2.278 | -0.611 | 1.909 | 2.258 | -0.008 | 0 | 0 | 0 | 0.46 | 0.66 | 0.45 | 0.56 | 0.49 | 0.63 |
| 2 | 143558760 | rs4662299 | -2.296 | -0.616 | 1.893 | 2.304 | -0.009 | 0.018 | 0.018 | 0 | 0.46 | 0.67 | 0.59 | 0.61 | 0.49 | 0.64 |
| 2 | 143560569 | rs13402740 | -1.946 | -0.754 | 1.779 | 2.457 | -0.004 | 0 | 0 | 0 | 0.24 | 0.71 | 0.33 | 0.22 | 0.2 | 0.29 |
| 2 | 143572404 | rs12691656 | -1.946 | -0.754 | 1.779 | 2.457 | -0.004 | 0 | 0 | 0 | 0.24 | 0.71 | 0.33 | 0.22 | 0.2 | 0.29 |
| 2 | 143574859 | rs4662300 | -1.946 | -0.754 | 1.779 | 2.457 | -0.004 | 0 | 0 | 0 | 0.24 | 0.71 | 0.33 | 0.22 | 0.2 | 0.29 |
| 2 | 143580262 | rs12992599 | -0.502 | -0.72 | 1.452 | 2.333 | -0.088 | -0.039 | -0.039 | 0.281 | 0.23 | 0.01 | 0.25 | 0.24 | 0.25 | 0.36 |
| 2 | 143582624 | rs2381381 | -1.926 | -0.742 | 1.738 | 2.431 | -0.01 | 0.017 | 0.017 | 0 | 0.24 | 0.71 | 0.33 | 0.38 | 0.2 | 0.3 |
| 2 | 143583508 | rs11902906 | -2.414 | -0.754 | 1.91 | 2.489 | -0.01 | 0.021 | 0.021 | 0 | 0.26 | 0.72 | 0.34 | 0.43 | 0.19 | 0.37 |
| 2 | 143592109 | rs966624 | -2.145 | -0.837 | 2.123 | 2.474 | 0.001 | 0.003 | 0.003 | 0.006 | 0.42 | 0.28 | 0.39 | 0.6 | 0.42 | 0.65 |

| | | | | | | | | | | | | | | | | |
|---|---|---|---|---|---|---|---|---|---|---|---|---|---|---|---|---|
| 2 | 143594676 | rs12470820 | -0.628 | -0.763 | 1.977 | 2.196 | -0.044 | -0.016 | -0.016 | 0.887 | 0.21 | 0.05 | 0.25 | 0.26 | 0.24 | 0.35 |
| 2 | 143602503 | rs13027801 | -1.531 | -0.855 | 2.203 | 2.123 | 0.003 | -0.014 | -0.014 | 0.017 | 0.3 | 0.58 | 0.46 | 0.31 | 0.28 | 0.46 |
| 2 | 143603794 | rs10198899 | -2.525 | -0.861 | 2.197 | 2.121 | -0.002 | 0.075 | 0.075 | 4.73 | 0.08 | 0.57 | 0.21 | 0.08 | 0.02 | 0.09 |
| 2 | 143607317 | rs1017994 | -0.679 | -0.749 | 1.593 | 2.084 | -0.098 | -0.026 | -0.026 | 0.12 | 0.22 | 0.01 | 0.24 | 0.23 | 0.26 | 0.37 |
| 2 | 143610406 | rs13007757 | -1.743 | -0.886 | 2.267 | 2.279 | -0.064 | 0.067 | 0.067 | 0 | 0.14 | 0.36 | 0.12 | 0.22 | 0.07 | 0.18 |
| 2 | 143612415 | rs6748749 | -2.185 | -0.889 | 2.6 | 2.302 | 0.005 | -0.016 | -0.016 | 0.195 | 0.38 | 0.5 | 0.51 | 0.34 | 0.39 | 0.56 |
| 2 | 143615226 | rs1473288 | -2.113 | -0.891 | 2.686 | 2.301 | 0.004 | -0.01 | -0.01 | 0.002 | 0.37 | 0.44 | 0.51 | 0.34 | 0.39 | 0.56 |
| 2 | 143620691 | rs7574678 | -1.752 | -0.897 | 2.73 | 2.447 | -0.02 | 0.009 | 0.009 | 1.63 | 0.35 | 0.21 | 0.39 | 0.34 | 0.39 | 0.57 |
| 2 | 143627604 | rs7581038 | -0.657 | -0.927 | 2.596 | 2.462 | -0.032 | 0.051 | 0.051 | 0 | 0.1 | 0.21 | 0.08 | 0.21 | 0.05 | 0.12 |
| 2 | 143638569 | rs1439875 | -1.101 | -0.921 | 2.773 | 2.498 | -0.034 | 0.132 | 0.132 | 0 | 0.12 | 0.35 | 0.27 | 0.25 | 0.07 | 0.13 |
| 2 | 143648031 | rs17805075 | -0.731 | -0.908 | 2.609 | 2.471 | -0.094 | -0.029 | -0.029 | 0 | 0.24 | 0.02 | 0.3 | 0.25 | 0.27 | 0.4 |
| 2 | 143652818 | rs7562371 | -2.262 | -0.94 | 2.757 | 2.459 | 0.003 | -0.006 | -0.006 | 0 | 0.37 | 0.47 | 0.56 | 0.38 | 0.38 | 0.55 |
| 2 | 143667100 | rs164732 | -2.97 | -0.938 | 2.569 | 2.406 | 0.018 | 0.02 | 0.02 | 0.164 | 0.49 | 0.66 | 0.66 | 0.59 | 0.47 | 0.69 |
| 2 | 143674826 | rs3768853 | -2.267 | -0.815 | 2.501 | 2.335 | 0.005 | 0.013 | 0.013 | 0 | 0.11 | 0.27 | 0.08 | 0.1 | 0.11 | 0.16 |
| 2 | 143683066 | rs6755074 | -2.04 | -0.636 | 2.443 | 2.35 | -0.004 | 0.002 | 0.002 | 1.57 | 0.35 | 0.31 | 0.39 | 0.34 | 0.39 | 0.56 |
| 2 | 143694424 | rs13030111 | -1.102 | -0.511 | 2.272 | 2.19 | -0.133 | -0.004 | -0.004 | 0 | 0.23 | 0.03 | 0.34 | 0.17 | 0.35 | 0.44 |
| 2 | 143699250 | rs9917201 | -1.284 | -0.589 | 2.565 | 2.361 | -0.056 | 0.083 | 0.083 | 0 | 0.18 | 0.33 | 0.14 | 0.32 | 0.11 | 0.18 |
| 2 | 143701255 | rs902924 | -1.319 | -0.59 | 2.564 | 2.361 | -0.063 | 0.097 | 0.097 | 0 | 0.18 | 0.35 | 0.14 | 0.32 | 0.11 | 0.18 |
| 2 | 143705417 | rs164735 | -1.994 | -0.595 | 2.52 | 2.357 | 0.021 | 0.109 | 0.109 | 2.236 | 0.06 | 0.32 | 0.18 | 0.1 | 0.01 | 0.08 |
| 2 | 143709091 | rs12997823 | -1.907 | -0.614 | 1.989 | 2.352 | -0.034 | -0.075 | -0.075 | 0 | 0.35 | 0.13 | 0.38 | 0.45 | 0.34 | 0.53 |
| 2 | 143711005 | rs13024113 | -1.913 | -0.654 | 1.958 | 2.342 | -0.048 | -0.097 | -0.097 | 0 | 0.34 | 0.07 | 0.39 | 0.45 | 0.36 | 0.53 |
| 2 | 143711265 | rs351676 | -2.451 | -0.896 | 2.369 | 2.475 | -0.008 | -0.002 | -0.002 | 0 | 0.25 | 0.71 | 0.33 | 0.42 | 0.11 | 0.25 |
| 2 | 143719772 | rs187090 | -2.715 | -0.859 | 2.359 | 2.434 | -0.019 | 0.063 | 0.063 | 8.034 | 0.08 | 0.57 | 0.22 | 0.1 | 0.01 | 0.08 |
| 2 | 143720342 | rs351685 | -2.178 | -0.855 | 2.188 | 2.422 | -0.054 | 0.045 | 0.045 | 0 | 0.24 | 0.63 | 0.32 | 0.42 | 0.12 | 0.26 |
| 2 | 143722839 | rs7561260 | -2.111 | -0.707 | 1.964 | 2.322 | -0.032 | 0.056 | 0.056 | 0 | 0.25 | 0.37 | 0.18 | 0.41 | 0.22 | 0.36 |
| 2 | 143729878 | rs10928161 | -1.008 | -0.551 | 1.558 | 2.231 | 0.058 | -0.048 | -0.048 | 0.156 | 0.16 | 0.02 | 0.09 | 0.19 | 0.08 | 0.18 |
| 2 | 143730019 | rs16855223 | -1.008 | -0.551 | 1.558 | 2.231 | 0.058 | -0.048 | -0.048 | 0 | 0.16 | 0.02 | 0.09 | 0.19 | 0.08 | 0.18 |
| 2 | 143738648 | rs11686073 | -0.697 | -0.024 | 0.732 | 2.142 | -0.105 | -0.027 | -0.027 | 0 | 0.24 | 0.06 | 0.28 | 0.28 | 0.32 | 0.46 |
| 2 | 143743012 | rs6734849 | -2.209 | -0.074 | 1.498 | 2.161 | 0.001 | 0.003 | 0.003 | 0.233 | 0.47 | 0.43 | 0.42 | 0.69 | 0.42 | 0.73 |
| 2 | 143744183 | rs960822 | -2.382 | -0.106 | 1.528 | 2.165 | 0.012 | 0.028 | 0.028 | 0.408 | 0.48 | 0.58 | 0.44 | 0.69 | 0.41 | 0.73 |
| 2 | 143745800 | rs7607734 | -0.869 | 0.044 | 1.495 | 1.796 | 0.004 | 0.059 | 0.059 | 0 | 0.09 | 0.12 | 0.07 | 0.27 | 0.12 | 0.12 |
| 2 | 143746849 | rs1050950 | -2.096 | -0.017 | 1.548 | 1.838 | -0.04 | 0.04 | 0.04 | 0 | 0.29 | 0.55 | 0.2 | 0.47 | 0.2 | 0.3 |
| 2 | 143749073 | rs1371514 | -1.871 | -0.002 | 1.638 | 1.838 | -0.036 | 0.038 | 0.038 | 0 | 0.27 | 0.37 | 0.18 | 0.46 | 0.19 | 0.31 |
| 2 | 143751071 | rs12466715 | -1.939 | 0.073 | 1.596 | 1.816 | -0.051 | 0.054 | 0.054 | 0 | 0.27 | 0.41 | 0.18 | 0.46 | 0.19 | 0.3 |
| 2 | 143754303 | rs10184722 | -1.904 | 0.079 | 1.558 | 1.636 | -0.047 | -0.017 | -0.017 | 0 | 0.25 | 0.58 | 0.18 | 0.22 | 0.15 | 0.29 |
| 2 | 143756755 | rs2452477 | -1.188 | 0.09 | 1.591 | 1.68 | -0.034 | 0.022 | 0.022 | 0 | 0.2 | 0.35 | 0.19 | 0.25 | 0.15 | 0.24 |
| 2 | 143759457 | rs352890 | -0.767 | 0.087 | 1.345 | 1.682 | 0.006 | -0.007 | -0.007 | 0 | 0.17 | 0.17 | 0.15 | 0.23 | 0.08 | 0.19 |
| 2 | 143759691 | rs352889 | -2.348 | 0.035 | 1.247 | 1.622 | 0.012 | 0.003 | 0.003 | 0 | 0.48 | 0.59 | 0.45 | 0.51 | 0.4 | 0.71 |
| 2 | 143760056 | rs352888 | -0.631 | 0.064 | 1.156 | 1.617 | 0.006 | -0.007 | -0.007 | 0 | 0.17 | 0.18 | 0.15 | 0.23 | 0.08 | 0.2 |
| 2 | 143760384 | rs352887 | -2.808 | 0.054 | 1.044 | 1.636 | -0.034 | 0.052 | 0.052 | 1.299 | 0.59 | 0.78 | 0.52 | 0.76 | 0.52 | 0.81 |
| 2 | 143781731 | rs13008636 | -0.181 | -0.147 | 0.867 | 1.738 | -0.065 | -0.046 | -0.046 | 0 | 0.21 | 0.01 | 0.2 | 0.25 | 0.19 | 0.42 |
| 2 | 143783182 | rs407867 | -2.078 | -0.322 | 1.215 | 1.749 | -0.021 | 0.051 | 0.051 | 0 | 0.3 | 0.55 | 0.27 | 0.51 | 0.28 | 0.36 |
| 2 | 143791014 | rs2304700 | -0.771 | 0.065 | 1.02 | 1.791 | 0.001 | 0.041 | 0.041 | 0 | 0.09 | 0.12 | 0.06 | 0.26 | 0.13 | 0.12 |
| 2 | 143817718 | rs354690 | 1.669 | 0.055 | 1.116 | 1.225 | 0.012 | 0.049 | 0.049 | 0.477 | 0.52 | 0.37 | 0.53 | 0.23 | 0.56 | 0.24 |
| 2 | 143825002 | rs1095654 | 0.593 | -0.277 | 0.877 | 1.321 | -0.022 | 0.055 | 0.055 | 0 | 0.77 | 0.72 | 0.77 | 0.52 | 0.79 | 0.7 |
| 2 | 143838780 | rs10202197 | -0.351 | -0.081 | 0.004 | 1.36 | -0.019 | -0.024 | -0.024 | 0.029 | 0.21 | 0.07 | 0.19 | 0.27 | 0.16 | 0.41 |
| 2 | 143844480 | rs2099716 | -0.351 | -0.081 | 0.004 | 1.36 | -0.019 | -0.024 | -0.024 | 0.948 | 0.21 | 0.07 | 0.19 | 0.27 | 0.16 | 0.41 |
| 2 | 143868223 | rs6750969 | -0.281 | -0.081 | -0.05 | 1.361 | -0.026 | -0.033 | -0.033 | 0 | 0.21 | 0.04 | 0.19 | 0.28 | 0.16 | 0.42 |
| 2 | 143868915 | rs1561541 | -0.98 | -0.318 | 0.123 | 1.194 | -0.058 | 0.059 | 0.059 | 0 | 0.28 | 0.45 | 0.21 | 0.44 | 0.22 | 0.33 |
| 2 | 143886216 | rs354706 | 1.073 | 0.109 | -0.468 | 0.723 | 0.043 | 0.077 | 0.077 | 0 | 0.49 | 0.28 | 0.53 | 0.25 | 0.61 | 0.25 |
| 2 | 143893576 | rs6430004 | -1.216 | 0.328 | -0.275 | 0.384 | -0.009 | -0.041 | -0.041 | 0 | 0.25 | 0.66 | 0.23 | 0.09 | 0.24 | 0.3 |
| 2 | 143898781 | rs10173270 | 0.053 | 0.318 | -1.1 | 0.684 | -0.047 | -0.04 | -0.04 | 0.493 | 0.25 | 0.08 | 0.21 | 0.32 | 0.26 | 0.22 |
| 2 | 143899318 | rs3845643 | -0.242 | 0.236 | -0.706 | 0.836 | -0.007 | -0.029 | -0.029 | 0 | 0.29 | 0.2 | 0.23 | 0.47 | 0.27 | 0.25 |
| 2 | 143902663 | rs3856363 | -0.941 | -0.082 | -0.005 | 1.176 | -0.008 | 0.005 | 0.005 | 0 | 0.36 | 0.47 | 0.28 | 0.65 | 0.28 | 0.57 |
| 2 | 143909719 | rs827228 | 0.048 | 0.345 | -0.461 | 0.295 | 0.001 | -0.043 | -0.043 | 0 | 0.7 | 0.65 | 0.54 | 0.93 | 0.65 | 0.7 |
| 2 | 143920386 | rs187133 | 1.309 | 0.474 | -0.545 | 0.395 | -0.002 | 0.005 | 0.005 | 1.509 | 0.41 | 0.42 | 0.31 | 0.45 | 0.37 | 0.46 |
| 2 | 143922613 | rs827226 | 0.612 | 0.628 | -0.459 | 0.124 | -0.003 | 0.001 | 0.001 | 0 | 0.72 | 0.78 | 0.55 | 0.93 | 0.65 | 0.71 |
| 2 | 143925517 | rs354694 | 0.483 | 1.006 | -0.431 | 0.234 | -0.015 | 0.012 | 0.012 | 1.576 | 0.78 | 0.82 | 0.56 | 0.92 | 0.63 | 0.74 |
| 2 | 143927574 | rs6713802 | -0.698 | 0.395 | -0.418 | 0.393 | 0.012 | 0.003 | 0.003 | 0 | 0.38 | 0.31 | 0.34 | 0.55 | 0.34 | 0.5 |
| 2 | 143929917 | rs10176686 | -0.232 | 0.247 | -0.514 | 0.483 | 0.006 | -0.013 | -0.013 | 0.123 | 0.2 | 0.17 | 0.17 | 0.34 | 0.18 | 0.29 |
| 2 | 143931505 | rs354696 | 0.823 | 0.204 | 0.274 | 0.567 | -0.015 | 0.019 | 0.019 | 0 | 0.61 | 0.53 | 0.65 | 0.54 | 0.66 | 0.51 |
| 2 | 143944775 | rs354728 | 0.153 | 0.474 | 0.323 | 0.044 | 0.009 | -0.019 | -0.019 | 0 | 0.21 | 0.16 | 0.39 | 0.06 | 0.32 | 0.17 |
| 2 | 143949956 | rs6716653 | 0.757 | 0.365 | 0.558 | 0.032 | -0.004 | -0.014 | -0.014 | 0 | 0.18 | 0.25 | 0.18 | 0.13 | 0.17 | 0.2 |
| 2 | 143951268 | rs7607443 | 0.791 | 0.364 | 0.558 | 0.031 | -0.004 | -0.014 | -0.014 | 0 | 0.18 | 0.25 | 0.18 | 0.13 | 0.17 | 0.19 |
| 2 | 143951325 | rs7595079 | 2.067 | 0.066 | 0.703 | 0.052 | 0.005 | -0.019 | -0.019 | 0 | 0.36 | 0.28 | 0.26 | 0.43 | 0.34 | 0.23 |
| 2 | 143951865 | rs16858783 | 0.598 | -0.076 | 0.689 | 0.49 | 0.018 | -0.014 | -0.014 | 0.812 | 0.12 | 0.05 | 0.04 | 0.18 | 0.05 | 0.18 |
| 2 | 143960933 | rs6711340 | 0.789 | 0.029 | 0.684 | 0.464 | -0.004 | -0.014 | -0.014 | 3.755 | 0.18 | 0.25 | 0.18 | 0.13 | 0.17 | 0.19 |
| 2 | 143981258 | rs7580937 | 1.045 | 0.028 | 0.684 | 0.464 | -0.004 | 0.015 | 0.015 | 0 | 0.16 | 0.08 | 0.16 | 0.13 | 0.17 | 0.19 |
| 2 | 143988118 | rs10195682 | -0.566 | 0.001 | 0.729 | 0.499 | 0.017 | -0.034 | -0.034 | 0 | 0.22 | 0.14 | 0.16 | 0.33 | 0.18 | 0.29 |

| | | | | | | | | | | | | | | | | |
|---|---|---|---|---|---|---|---|---|---|---|---|---|---|---|---|---|
| 2 | 143988492 | rs7607384 | 0.689 | 0.04 | 0.864 | 0.468 | -0.002 | -0.01 | -0.01 | 0.077 | 0.18 | 0.22 | 0.18 | 0.12 | 0.17 | 0.19 |
| 2 | 143998944 | rs10496945 | 0.902 | 0.021 | 0.821 | 0.46 | -0.001 | -0.007 | -0.007 | 0.305 | 0.17 | 0.2 | 0.17 | 0.12 | 0.17 | 0.19 |
| 2 | 144000004 | rs10178148 | -0.891 | 0.279 | 0.569 | 0.393 | -0.088 | 0.025 | 0.025 | 0 | 0.86 | 0.99 | 0.65 | 0.94 | 0.75 | 0.85 |
| 2 | 144007986 | rs6430017 | 0.752 | 0.201 | 0.73 | 0.458 | -0.003 | -0.013 | -0.013 | 0 | 0.18 | 0.24 | 0.17 | 0.12 | 0.17 | 0.19 |
| 2 | 144012112 | rs6714864 | -0.522 | 0.156 | 0.111 | 0.19 | 0.01 | -0.033 | -0.033 | 0 | 0.3 | 0.15 | 0.23 | 0.33 | 0.26 | 0.35 |
| 2 | 144013418 | rs6719577 | -0.541 | -0.043 | 0.774 | 0.648 | -0.009 | -0.001 | -0.001 | 0 | 0.3 | 0.33 | 0.22 | 0.32 | 0.21 | 0.38 |
| 2 | 144015093 | rs13022541 | 0.594 | -0.034 | 0.453 | 0.652 | 0.005 | 0.026 | 0.026 | 0 | 0.18 | 0.09 | 0.16 | 0.11 | 0.16 | 0.19 |
| 2 | 144016529 | rs4662318 | 0.157 | 0.064 | 0.424 | 0.694 | 0.002 | 0.012 | 0.012 | 0 | 0.1 | 0.13 | 0.12 | 0.17 | 0.13 | 0.11 |
| 2 | 144026991 | rs6751190 | -1.493 | 0.077 | -0.062 | 0.665 | 0.044 | -0.023 | -0.023 | 0 | 0.43 | 0.8 | 0.38 | 0.54 | 0.35 | 0.58 |
| 2 | 144032962 | rs12993643 | -0.606 | 0.575 | -0.592 | 0.447 | -0.005 | -0.054 | -0.054 | 0.166 | 0.33 | 0.11 | 0.53 | 0.37 | 0.32 | 0.42 |
| 2 | 144033069 | rs7602120 | -1.716 | 0.601 | -0.43 | 0.383 | 0.001 | 0.004 | 0.004 | 0 | 0.5 | 0.51 | 0.64 | 0.43 | 0.49 | 0.68 |
| 2 | 144034099 | rs17228139 | 0.12 | 0.905 | -0.881 | 0.208 | -0.032 | -0.028 | -0.028 | 0.072 | 0.05 | 0.01 | 0.25 | 0.11 | 0.15 | 0.12 |
| 2 | 144035442 | rs2381404 | 0.236 | 0.792 | -0.838 | 0.193 | -0.025 | 0.023 | 0.023 | 0 | 0.22 | 0.1 | 0.33 | 0.15 | 0.25 | 0.21 |
| 2 | 144039139 | rs16858984 | -1.289 | 0.776 | -0.779 | 0.183 | -0.004 | -0.005 | -0.005 | 0.876 | 0.45 | 0.52 | 0.55 | 0.37 | 0.4 | 0.55 |
| 2 | 144040731 | rs7579046 | 0.566 | 0.8 | -0.82 | 0.201 | 0.001 | 0.01 | 0.01 | 0 | 0.19 | 0.1 | 0.29 | 0.15 | 0.17 | 0.19 |
| 2 | 144042333 | rs2381407 | -2.053 | 0.842 | -0.788 | 0.261 | 0 | -0.004 | -0.004 | 0.134 | 0.56 | 0.61 | 0.68 | 0.5 | 0.6 | 0.65 |
| 2 | 144043421 | rs7596229 | 0.189 | 0.362 | -0.604 | 0.905 | 0.001 | 0.002 | 0.002 | 0 | 0.08 | 0.08 | 0.08 | 0.13 | 0.11 | 0.08 |
| 2 | 144047727 | rs4662198 | -0.815 | 0.32 | -0.892 | 0.891 | 0.031 | -0.056 | -0.056 | 0.145 | 0.31 | 0.13 | 0.45 | 0.36 | 0.24 | 0.39 |
| 2 | 144053578 | rs10928169 | -0.741 | 0.892 | -0.423 | 0.167 | 0.001 | -0.005 | -0.005 | 0.533 | 0.34 | 0.55 | 0.48 | 0.33 | 0.38 | 0.31 |
| 2 | 144056248 | rs12328672 | -0.697 | 1.118 | -0.321 | -0.155 | 0.02 | -0.052 | -0.052 | 0 | 0.12 | 0.26 | 0.09 | 0.02 | 0.13 | 0.07 |
| 2 | 144057444 | rs11681284 | 0.771 | 1.135 | -1.278 | -0.408 | -0.017 | -0.064 | -0.064 | 0.283 | 0.37 | 0.12 | 0.47 | 0.46 | 0.33 | 0.31 |
| 2 | 144063836 | rs4662320 | 0.351 | 1.43 | -1.083 | -0.604 | -0.014 | 0.024 | 0.024 | 0 | 0.09 | 0.04 | 0.11 | 0 | 0.16 | 0.09 |
| 2 | 144067374 | rs2381461 | -0.468 | 1.27 | -1.231 | -1.003 | -0.006 | -0.087 | -0.087 | 0 | 0.77 | 0.89 | 0.8 | 0.49 | 0.76 | 0.57 |
| 2 | 144070983 | rs6430022 | -0.155 | 1.121 | -1.67 | -1.211 | 0.027 | -0.02 | -0.02 | 0.206 | 0.53 | 0.89 | 0.69 | 0.58 | 0.6 | 0.5 |
| 2 | 144076740 | rs2381408 | 0 | 1.235 | -1.656 | -1.224 | -0.01 | -0.007 | -0.007 | 0 | 0.52 | 0.87 | 0.68 | 0.58 | 0.55 | 0.5 |
| 2 | 144095579 | rs10803491 | 1.82 | 1.306 | -1.644 | -1.203 | -0.035 | -0.092 | -0.092 | 0.004 | 0.27 | 0.09 | 0.25 | 0.46 | 0.28 | 0.26 |
| 2 | 144096234 | rs17229439 | 0.712 | 1.625 | -1.705 | -1.041 | -0.064 | 0.183 | 0.183 | 0 | 0.26 | 0.01 | 0.12 | 0.01 | 0.25 | 0.19 |
| 2 | 144101819 | rs13406291 | -0.036 | 1.556 | -1.631 | -1.047 | -0.005 | -0.01 | -0.01 | 0 | 0.13 | 0.14 | 0.12 | 0 | 0.22 | 0.11 |
| 2 | 144108930 | rs2381464 | 0.826 | 1.527 | -1.492 | -1.077 | -0.002 | -0.049 | -0.049 | 0 | 0.1 | 0.25 | 0.08 | 0.01 | 0.08 | 0.05 |
| 2 | 144111366 | rs7587606 | -0.162 | 1.716 | -1.893 | -0.886 | -0.011 | 0.181 | 0.181 | 0 | 0.67 | 0.92 | 0.84 | 0.99 | 0.71 | 0.78 |
| 2 | 144112684 | rs10205708 | -0.913 | 1.56 | -2.124 | -0.795 | 0.029 | 0.112 | 0.112 | 0 | 0.75 | 0.93 | 0.89 | 0.99 | 0.82 | 0.81 |
| 2 | 144118862 | rs11890035 | -0.24 | 1.667 | -2.092 | -0.786 | -0.004 | 0.01 | 0.01 | 0 | 0.15 | 0.13 | 0.14 | 0 | 0.23 | 0.11 |
| 2 | 144125853 | rs10208099 | 0.473 | 1.552 | -2.125 | -0.787 | -0.016 | -0.011 | -0.011 | 0 | 0.55 | 0.83 | 0.68 | 0.57 | 0.56 | 0.49 |
| 2 | 144127612 | rs2890714 | -0.8 | 1.527 | -2.151 | -0.823 | -0.007 | 0.123 | 0.123 | 0 | 0.74 | 0.93 | 0.88 | 0.99 | 0.77 | 0.82 |
| 2 | 144128251 | rs16822400 | 0.818 | 1.422 | -1.881 | -0.546 | 0.002 | 0.028 | 0.028 | 0 | 0.1 | 0.08 | 0.15 | 0.41 | 0.1 | 0.28 |
| 2 | 144129055 | rs6729300 | 0.433 | 1.616 | -1.436 | -0.196 | 0.001 | -0.003 | -0.003 | 0 | 0.44 | 0.67 | 0.57 | 0.58 | 0.4 | 0.4 |
| 2 | 144131703 | rs7597018 | -0.107 | 1.515 | -1.588 | -0.064 | -0.006 | -0.004 | -0.004 | 0 | 0.55 | 0.77 | 0.68 | 0.57 | 0.56 | 0.44 |
| 2 | 144135819 | rs2381467 | -0.169 | 1.029 | -1.536 | -0.022 | 0.028 | -0.001 | -0.001 | 0.044 | 0.56 | 0.77 | 0.69 | 0.58 | 0.61 | 0.44 |
| 2 | 144135930 | rs2890715 | -0.169 | 1.029 | -1.536 | -0.022 | 0.028 | -0.001 | -0.001 | 0 | 0.56 | 0.76 | 0.69 | 0.58 | 0.61 | 0.44 |
| 2 | 144139538 | rs13409533 | -0.664 | 1.359 | -1.159 | 0.18 | -0.03 | -0.006 | -0.006 | 0 | 0.11 | 0.05 | 0.32 | 0.11 | 0.2 | 0.17 |
| 2 | 144142507 | rs7591447 | -1.527 | 0.806 | -0.22 | -0.053 | -0.003 | -0.05 | -0.05 | 0 | 0.09 | 0.35 | 0.07 | 0 | 0.04 | 0.01 |
| 2 | 144142749 | rs10754918 | 0.079 | 0.68 | -0.713 | -0.257 | -0.003 | 0.018 | 0.018 | 0 | 0.81 | 0.91 | 0.9 | 0.89 | 0.83 | 0.85 |
| 2 | 144146850 | rs7580625 | -0.68 | -0.209 | -0.485 | 0.273 | 0.007 | -0.004 | -0.004 | 1.242 | 0.3 | 0.3 | 0.4 | 0.3 | 0.19 | 0.37 |
| 2 | 144147475 | rs16822430 | 1.49 | 0.118 | -1.266 | 0.256 | -0.041 | -0.099 | -0.099 | 2.236 | 0.23 | 0.06 | 0.13 | 0.47 | 0.24 | 0.17 |
| 2 | 144155919 | rs7608439 | -0.601 | -0.101 | -0.792 | 0.09 | -0.001 | -0.032 | -0.032 | 0.512 | 0.81 | 0.76 | 0.9 | 0.94 | 0.81 | 0.83 |
| 2 | 144157738 | rs16822448 | 1.341 | -0.203 | -0.809 | -0.126 | -0.001 | 0.014 | 0.014 | 0 | 0.06 | 0.06 | 0.11 | 0.16 | 0.06 | 0.15 |
| 2 | 144159751 | rs4662327 | 1.43 | -0.167 | -0.671 | -0.486 | -0.048 | 0.085 | 0.085 | 0.275 | 0.26 | 0.08 | 0.24 | 0 | 0.36 | 0.14 |
| 2 | 144166523 | rs6430030 | -0.721 | -0.183 | -0.491 | -0.378 | -0.01 | -0.016 | -0.016 | 6.234 | 0.57 | 0.7 | 0.77 | 0.46 | 0.58 | 0.62 |
| 2 | 144168037 | rs10048784 | -1.173 | -0.1 | -0.234 | -0.334 | 0.002 | 0.014 | 0.014 | 0 | 0.7 | 0.67 | 0.8 | 0.47 | 0.7 | 0.71 |
| 2 | 144169816 | rs6723640 | 0.392 | -0.287 | -0.153 | -0.372 | -0.009 | 0.018 | 0.018 | 0.138 | 0.21 | 0.29 | 0.39 | 0.3 | 0.16 | 0.3 |
| 2 | 144178400 | rs12612084 | 0.298 | -0.216 | 0.042 | -0.414 | 0.001 | 0.004 | 0.004 | 0 | 0.45 | 0.47 | 0.54 | 0.79 | 0.38 | 0.48 |
| 2 | 144180499 | rs7608859 | -0.709 | -0.174 | 0.096 | -0.401 | 0.062 | 0.045 | 0.045 | 0.024 | 0.95 | 0.76 | 0.94 | 0.94 | 0.93 | 0.92 |
| 2 | 144184279 | rs12105739 | 1.595 | -0.218 | -0.19 | -0.441 | -0.013 | 0.099 | 0.099 | 0 | 0.16 | 0.03 | 0.05 | 0 | 0.14 | 0.13 |
| 2 | 144186113 | rs10928176 | 0.336 | -0.286 | -0.162 | -0.414 | -0.028 | 0.051 | 0.051 | 3.518 | 0.21 | 0.4 | 0.4 | 0.3 | 0.16 | 0.3 |
| 2 | 144194565 | rs17230746 | 1.2 | 0.029 | -0.309 | -0.62 | -0.035 | 0.119 | 0.119 | 0 | 0.19 | 0.03 | 0.09 | 0 | 0.21 | 0.15 |
| 2 | 144200043 | rs13010737 | 0.322 | -0.011 | -0.189 | -0.495 | -0.001 | -0.004 | -0.004 | 0.537 | 0.41 | 0.44 | 0.49 | 0.78 | 0.31 | 0.48 |
| 2 | 144205842 | rs10803494 | 0.748 | 0.006 | -0.206 | -0.146 | -0.011 | -0.002 | -0.002 | 0 | 0.19 | 0.29 | 0.31 | 0.22 | 0.16 | 0.29 |
| 2 | 144226581 | rs12478952 | 1.111 | 0 | -0.035 | -0.195 | -0.003 | 0.008 | 0.008 | 0.018 | 0.27 | 0.15 | 0.28 | 0.71 | 0.21 | 0.29 |
| 2 | 144236866 | rs4260182 | 1.217 | 0.029 | -0.321 | -0.032 | -0.001 | 0.065 | 0.065 | 0 | 0.07 | 0.08 | 0.18 | 0.23 | 0.06 | 0.14 |
| 2 | 144240994 | rs11690735 | 0.782 | 0.058 | -0.357 | -0.028 | -0.01 | 0.017 | 0.017 | 0.007 | 0.3 | 0.15 | 0.32 | 0.71 | 0.28 | 0.32 |
| 2 | 144255201 | rs12465778 | 0.033 | 0.033 | -0.509 | -0.142 | 0.004 | 0.06 | 0.06 | 0 | 0.51 | 0.29 | 0.51 | 0.24 | 0.55 | 0.44 |
| 2 | 144263033 | rs10180461 | 1.109 | 0.15 | -0.476 | -0.345 | -0.022 | 0.02 | 0.02 | 0.121 | 0.28 | 0.16 | 0.26 | 0 | 0.37 | 0.14 |
| 2 | 144263416 | rs13020925 | 0.635 | 0.189 | -0.613 | 0.029 | -0.009 | 0.028 | 0.028 | 0 | 0.3 | 0.14 | 0.32 | 0.72 | 0.27 | 0.37 |
| 2 | 144266489 | rs12621600 | 0.607 | 0.229 | -0.821 | 0.025 | -0.039 | -0.065 | -0.065 | 0.636 | 0.27 | 0.09 | 0.29 | 0.64 | 0.26 | 0.31 |
| 2 | 144266504 | rs7561228 | 0.028 | 0.081 | -0.111 | -0.027 | -0.023 | 0.047 | 0.047 | 2.537 | 0.25 | 0.45 | 0.37 | 0.37 | 0.22 | 0.43 |
| 2 | 144266623 | rs7561163 | 0.099 | 0.056 | -0.121 | 0.327 | -0.01 | 0.022 | 0.022 | 3.023 | 0.25 | 0.36 | 0.34 | 0.42 | 0.2 | 0.42 |
| 2 | 144270623 | rs10928182 | -1.049 | 0.288 | -0.328 | 0.387 | -0.012 | -0.016 | -0.016 | 0 | 0.81 | 0.86 | 0.86 | 0.58 | 0.76 | 0.75 |
| 2 | 144272138 | rs6727288 | -0.406 | -0.162 | -0.255 | 0.086 | 0.004 | 0.016 | 0.016 | 0 | 0.51 | 0.49 | 0.49 | 0.77 | 0.44 | 0.69 |
| 2 | 144272229 | rs4233566 | 0.636 | -0.196 | -0.273 | 0.008 | 0.013 | -0.004 | -0.004 | 0.831 | 0.36 | 0.33 | 0.46 | 0.22 | 0.45 | 0.24 |

| | | | | | | | | | | | | | | | | |
|---|---|---|---|---|---|---|---|---|---|---|---|---|---|---|---|---|
| 2 | 144272299 | rs4563166 | 0.139 | -0.291 | -0.139 | 0.112 | -0.013 | 0.003 | 0.003 | 0.032 | 0.31 | 0.41 | 0.35 | 0.32 | 0.19 | 0.44 |
| 2 | 144275787 | rs12691682 | -0.012 | 0.497 | -0.373 | -0.06 | 0.008 | -0.007 | -0.007 | 0 | 0.61 | 0.44 | 0.54 | 0.42 | 0.6 | 0.53 |
| 2 | 144278534 | rs6736741 | -0.823 | 0.955 | -0.25 | -0.092 | -0.004 | -0.003 | -0.003 | 0.042 | 0.87 | 0.89 | 0.75 | 0.8 | 0.82 | 0.86 |
| 2 | 144282078 | rs11898020 | 1.466 | 1.124 | -0.471 | -0.404 | -0.003 | -0.061 | -0.061 | 0 | 0.24 | 0.2 | 0.19 | 0.01 | 0.25 | 0.1 |
| 2 | 144282793 | rs6732131 | 0.421 | 1.197 | -0.991 | -0.069 | 0.006 | -0.012 | -0.012 | 0 | 0.36 | 0.23 | 0.34 | 0.45 | 0.33 | 0.44 |
| 2 | 144290246 | rs12614491 | 0.301 | 1.136 | -0.97 | -0.1 | 0.001 | 0.016 | 0.016 | 0.428 | 0.51 | 0.6 | 0.39 | 0.35 | 0.49 | 0.42 |
| 2 | 144303318 | rs10084340 | 0.363 | 1.111 | -1.14 | -0.052 | 0 | -0.033 | -0.033 | 0.885 | 0.36 | 0.16 | 0.34 | 0.45 | 0.34 | 0.45 |
| 2 | 144318832 | rs4662339 | -0.902 | 1.257 | -0.957 | 0.017 | 0.014 | 0.021 | 0.021 | 0 | 0.87 | 0.85 | 0.74 | 0.81 | 0.82 | 0.88 |
| 2 | 144347994 | rs4594385 | -0.85 | 1.218 | -0.996 | 0.016 | 0.023 | 0.038 | 0.038 | 0 | 0.87 | 0.81 | 0.74 | 0.8 | 0.82 | 0.85 |
| 2 | 144349620 | rs13386631 | 0.785 | 1.067 | -0.938 | -0.258 | -0.003 | 0.044 | 0.044 | 0 | 0.28 | 0.52 | 0.21 | 0.35 | 0.23 | 0.31 |
| 2 | 144352350 | rs10803495 | -0.698 | 1.276 | -1.047 | -0.284 | 0.032 | 0.032 | 0.032 | 0 | 0.86 | 0.77 | 0.73 | 0.8 | 0.78 | 0.77 |
| 2 | 144352423 | rs13415550 | 1.59 | 1.142 | -1.811 | -0.469 | -0.028 | -0.089 | -0.089 | 0 | 0.23 | 0.04 | 0.28 | 0.44 | 0.21 | 0.3 |
| 2 | 144362349 | rs16822700 | 0.941 | 1.246 | -1.153 | -0.145 | -0.001 | 0.012 | 0.012 | 0 | 0.27 | 0.31 | 0.2 | 0.35 | 0.23 | 0.33 |
| 2 | 144363902 | rs12052670 | -1.336 | 0.838 | -2.034 | -0.557 | 0.006 | 0.035 | 0.035 | 1.146 | 0.77 | 0.87 | 0.82 | 0.99 | 0.76 | 0.9 |
| 2 | 144364876 | rs13389286 | 0.006 | 0.814 | -2.015 | -0.008 | 0.009 | -0.027 | -0.027 | 0.63 | 0.36 | 0.13 | 0.35 | 0.44 | 0.34 | 0.44 |
| 2 | 144370273 | rs1533526 | -0.775 | 1.151 | -1.665 | -0.575 | -0.009 | 0.022 | 0.022 | 0.016 | 0.7 | 0.79 | 0.77 | 0.99 | 0.67 | 0.78 |
| 2 | 144371034 | rs2969584 | -0.017 | 1.29 | -1.721 | -0.371 | -0.015 | 0.119 | 0.119 | 0 | 0.54 | 0.23 | 0.59 | 0.8 | 0.49 | 0.61 |
| 2 | 144371604 | rs938815 | 0.897 | 1.04 | -2.263 | -0.295 | -0.008 | 0.002 | 0.002 | 0 | 0.23 | 0.09 | 0.3 | 0.25 | 0.24 | 0.18 |
| 2 | 144374203 | rs2678290 | 0.24 | 1.01 | -2.172 | -0.16 | 0.026 | -0.056 | -0.056 | 0 | 0.32 | 0.15 | 0.28 | 0.56 | 0.26 | 0.44 |
| 2 | 144376033 | rs10928190 | 0.682 | 1.189 | -1.639 | -0.354 | 0.001 | 0.071 | 0.071 | 0.083 | 0.5 | 0.32 | 0.53 | 0.25 | 0.52 | 0.3 |
| 2 | 144385104 | rs2731561 | 0.433 | 0.655 | -1.857 | 0.104 | -0.014 | 0.004 | 0.004 | 0 | 0.23 | 0.15 | 0.23 | 0.23 | 0.26 | 0.34 |
| 2 | 144386869 | rs950013 | -0.206 | 0.879 | -1.964 | 0.029 | -0.08 | 0.055 | 0.055 | 0 | 0.37 | 0.12 | 0.48 | 0.71 | 0.44 | 0.56 |
| 2 | 144391132 | rs2048955 | -0.348 | 0.725 | -1.964 | -0.03 | 0.014 | 0.014 | 0.014 | 0.424 | 0.88 | 0.48 | 0.86 | 0.91 | 0.92 | 0.95 |
| 2 | 144394101 | rs7559052 | 0.778 | 0.876 | -2.104 | 0.161 | -0.019 | -0.068 | -0.068 | 0.063 | 0.1 | 0 | 0.07 | 0.33 | 0.09 | 0.2 |
| 2 | 144394398 | rs11679600 | 1.079 | 0.857 | -2.14 | 0.069 | -0.1 | -0.106 | -0.106 | 0.212 | 0.24 | 0.02 | 0.36 | 0.33 | 0.31 | 0.27 |
| 2 | 144403649 | rs12691685 | -0.311 | 0.664 | -1.751 | -0.308 | 0.006 | 0.007 | 0.007 | 0.41 | 0.88 | 0.45 | 0.86 | 0.91 | 0.92 | 0.95 |
| 2 | 144403796 | rs13409480 | 0.568 | 0.726 | -1.839 | -0.341 | 0.008 | 0.067 | 0.067 | 0.083 | 0.51 | 0.38 | 0.34 | 0.2 | 0.48 | 0.39 |
| 2 | 144409949 | rs13430960 | 1.775 | 0.775 | -1.769 | -0.36 | 0.003 | -0.034 | -0.034 | 0 | 0.29 | 0.25 | 0.22 | 0.01 | 0.28 | 0.14 |
| 2 | 144415159 | rs10211220 | -0.412 | 0.526 | -1.446 | 0.085 | -0.013 | -0.046 | -0.046 | 0.041 | 0.83 | 0.5 | 0.82 | 0.9 | 0.85 | 0.91 |
| 2 | 144415905 | rs6750487 | 0.463 | 0.623 | -1.785 | 0.249 | -0.004 | -0.004 | -0.004 | 0 | 0.15 | 0.11 | 0.08 | 0.2 | 0.13 | 0.21 |
| 2 | 144428480 | rs11685813 | 0.681 | 0.447 | -1.744 | 0.149 | 0.005 | 0.011 | 0.011 | 0 | 0.62 | 0.34 | 0.66 | 0.68 | 0.68 | 0.65 |
| 2 | 144429770 | rs1517912 | -0.141 | 0.38 | -1.58 | 0.067 | -0.037 | -0.055 | -0.055 | 0.065 | 0.82 | 0.46 | 0.82 | 0.91 | 0.88 | 0.91 |
| 2 | 144442150 | rs6708276 | 1.052 | 0.326 | -2.04 | -0.443 | 0.004 | 0.001 | 0.001 | 0 | 0.33 | 0.31 | 0.31 | 0.1 | 0.33 | 0.21 |
| 2 | 144450181 | rs1996447 | 0.999 | 0.263 | -2.413 | -0.046 | -0.121 | -0.114 | -0.114 | 0 | 0.3 | 0.02 | 0.39 | 0.36 | 0.36 | 0.31 |
| 2 | 144450292 | rs16822931 | 0.281 | 0.283 | -2.267 | 0.19 | -0.013 | -0.006 | -0.006 | 0 | 0.2 | 0.08 | 0.15 | 0.23 | 0.2 | 0.32 |
| 2 | 144463315 | rs10469562 | 0.615 | 0.232 | -2.163 | 0.182 | 0 | -0.007 | -0.007 | 0 | 0.64 | 0.32 | 0.69 | 0.45 | 0.69 | 0.56 |
| 2 | 144477751 | rs7606067 | 0.16 | 0.269 | -2.362 | 0.293 | -0.016 | 0.064 | 0.064 | 0 | 0.47 | 0.26 | 0.64 | 0.62 | 0.53 | 0.49 |
| 2 | 144484493 | rs10189240 | 1.015 | 0.298 | -2.493 | 0.36 | 0.012 | 0.14 | 0.14 | 0 | 0.36 | 0.12 | 0.32 | 0.15 | 0.35 | 0.21 |
| 2 | 144486169 | rs2292834 | 0.859 | 0.331 | -2.354 | 0.7 | 0.003 | 0.026 | 0.026 | 0 | 0.06 | 0.05 | 0.09 | 0.26 | 0.05 | 0.09 |
| 2 | 144498693 | rs12989517 | 0.682 | 0.322 | -2.462 | 0.574 | -0.118 | -0.008 | -0.008 | 0 | 0.3 | 0.02 | 0.25 | 0.21 | 0.37 | 0.34 |
| 2 | 144503257 | rs4662214 | 0.545 | 0.521 | -1.785 | 0.86 | -0.001 | -0.025 | -0.025 | 1.916 | 0.72 | 0.25 | 0.65 | 0.6 | 0.78 | 0.64 |
| 2 | 144503694 | rs16822986 | 0.523 | 0.65 | -1.569 | 0.528 | -0.021 | -0.024 | -0.024 | 0.128 | 0.12 | 0 | 0.14 | 0.13 | 0.13 | 0.09 |
| 2 | 144505198 | rs1712957 | 0.941 | 0.857 | -0.65 | 0.697 | -0.008 | 0.106 | 0.106 | 0 | 0.53 | 0.29 | 0.5 | 0.2 | 0.56 | 0.37 |
| 2 | 144505690 | rs938817 | -0.23 | 0.816 | -0.358 | 0.972 | 0.001 | 0.025 | 0.025 | 0 | 0.19 | 0.16 | 0.35 | 0.49 | 0.18 | 0.35 |
| 2 | 144509425 | rs6745506 | 0.269 | 0.941 | -0.353 | 1.053 | -0.006 | -0.004 | -0.004 | 0 | 0.16 | 0.11 | 0.34 | 0.46 | 0.18 | 0.25 |
| 2 | 144511229 | rs11898334 | -0.91 | 1.073 | -0.488 | 0.309 | -0.001 | -0.01 | -0.01 | 0 | 0.45 | 0.52 | 0.57 | 0.83 | 0.48 | 0.61 |
| 2 | 144517069 | rs1517915 | 0.745 | 1.255 | 0.038 | 1.311 | 0.036 | 0.089 | 0.089 | 0.017 | 0.76 | 0.56 | 0.78 | 0.64 | 0.74 | 0.71 |
| 2 | 144523686 | rs7568511 | -0.24 | 1.147 | 0.085 | 1.157 | 0.002 | 0.01 | 0.01 | 0 | 0.17 | 0.15 | 0.22 | 0.25 | 0.17 | 0.23 |
| 2 | 144523872 | rs7581136 | -1.099 | 0.861 | 0.739 | 0.955 | -0.004 | 0.034 | 0.034 | 0 | 0.27 | 0.48 | 0.27 | 0.31 | 0.21 | 0.28 |
| 2 | 144529508 | rs12995474 | -0.368 | 1.343 | 0.767 | 0.514 | 0.037 | -0.052 | -0.052 | 0.041 | 0.14 | 0.29 | 0.15 | 0.04 | 0.17 | 0.08 |
| 2 | 144535175 | rs6729239 | -0.979 | 1.536 | 0.772 | 0.365 | 0.014 | -0.024 | -0.024 | 2.255 | 0.21 | 0.29 | 0.23 | 0.09 | 0.32 | 0.14 |
| 2 | 144554309 | rs2381558 | 0.841 | 1.197 | 1.159 | 0.843 | 0.006 | 0.145 | 0.145 | 0.474 | 0.94 | 0.73 | 0.92 | 0.81 | 0.97 | 0.92 |
| 2 | 144555529 | rs11680552 | -0.44 | 1.655 | 1.161 | 1.378 | 0.027 | 0.015 | 0.015 | 0 | 0.18 | 0.26 | 0.36 | 0.24 | 0.34 | 0.12 |
| 2 | 144562877 | rs13029549 | 0.829 | 1.6 | 1.105 | 1.391 | -0.006 | -0.021 | -0.021 | 0 | 0.16 | 0.09 | 0.19 | 0.25 | 0.15 | 0.24 |
| 2 | 144578152 | rs10928200 | -0.598 | 1.495 | 1.109 | 1.241 | 0.018 | 0.008 | 0.008 | 0 | 0.2 | 0.27 | 0.38 | 0.24 | 0.37 | 0.13 |
| 2 | 144579820 | rs7573324 | -1.458 | 1.426 | 1.107 | 1.148 | -0.02 | -0.008 | -0.008 | 0 | 0.41 | 0.67 | 0.59 | 0.51 | 0.52 | 0.38 |
| 2 | 144582568 | rs2381511 | 0.929 | 1.249 | 1.191 | 1.073 | 0.06 | 0.119 | 0.119 | 0 | 0.76 | 0.51 | 0.55 | 0.55 | 0.6 | 0.79 |
| 2 | 144587571 | rs10185589 | 0.953 | 1.26 | 1.119 | 1.057 | 0.056 | 0.095 | 0.095 | 0 | 0.76 | 0.47 | 0.55 | 0.57 | 0.6 | 0.8 |
| 2 | 144624649 | rs1447560 | 0.411 | 1.673 | 1.042 | 1.052 | 0.007 | 0.001 | 0.001 | 0 | 0.21 | 0.26 | 0.17 | 0.21 | 0.23 | 0.26 |
| 2 | 144624936 | rs1257346 | -1.848 | 1.724 | 0.963 | 0.95 | 0.083 | 0.078 | 0.078 | 1.36 | 0.52 | 0.79 | 0.65 | 0.66 | 0.68 | 0.51 |
| 2 | 144632311 | rs10928203 | 0.253 | 1.902 | 1.054 | 1.02 | 0.025 | 0.033 | 0.033 | 0 | 0.14 | 0.16 | 0.31 | 0.25 | 0.24 | 0.1 |
| 2 | 144634670 | rs1868078 | -1.468 | 0.681 | 0.354 | 0.403 | 0.004 | 0 | 0 | 0 | 0.84 | 0.88 | 0.91 | 0.81 | 0.93 | 0.91 |
| 2 | 144638809 | rs17834780 | -0.149 | 0.654 | 0.141 | 0.234 | -0.005 | 0.006 | 0.006 | 0 | 0.14 | 0.1 | 0.1 | 0.01 | 0.2 | 0.08 |
| 2 | 144639420 | rs12477840 | 0.971 | 0.91 | 0.257 | 0.309 | 0.083 | 0.102 | 0.102 | 0 | 0.11 | 0.29 | 0.28 | 0.21 | 0.19 | 0.1 |
| 2 | 144639630 | rs12469850 | 1.08 | 0.916 | 0.235 | 0.31 | 0.04 | 0.066 | 0.066 | 0 | 0.11 | 0.19 | 0.28 | 0.23 | 0.19 | 0.1 |
| 2 | 144645286 | rs787159 | 1.649 | 0.657 | -0.072 | 0.134 | -0.02 | -0.059 | -0.059 | 0 | 0.32 | 0.09 | 0.3 | 0.35 | 0.24 | 0.38 |
| 2 | 144653631 | rs787163 | 0.672 | 0.534 | -0.314 | 0.2 | 0.001 | -0.019 | -0.019 | 0 | 0.47 | 0.5 | 0.65 | 0.77 | 0.46 | 0.63 |
| 2 | 144655236 | rs2593713 | 1.734 | 0.208 | -0.874 | -0.127 | -0.01 | -0.067 | -0.067 | 0 | 0.24 | 0.09 | 0.27 | 0.35 | 0.19 | 0.34 |
| 2 | 144655691 | rs787152 | 2.178 | 0.094 | -1.163 | -0.492 | -0.043 | 0.037 | 0.037 | 0 | 0.19 | 0.02 | 0.12 | 0.12 | 0.16 | 0.22 |

| | | | | | | | | | | | | | | | | |
|---|---|---|---|---|---|---|---|---|---|---|---|---|---|---|---|---|
| 2 | 144655836 | rs787151 | -0.364 | 0.294 | -0.904 | -0.763 | 0.008 | 0 | 0 | 0 | 0.72 | 0.76 | 0.86 | 0.96 | 0.74 | 0.92 |
| 2 | 144656743 | rs12998885 | 0.481 | 1.096 | -0.537 | -0.292 | 0.001 | -0.001 | -0.001 | 0 | 0.25 | 0.24 | 0.21 | 0.22 | 0.28 | 0.29 |
| 2 | 144660336 | rs10803501 | -0.518 | 1.101 | -0.546 | -0.268 | 0.006 | -0.004 | -0.004 | 0 | 0.32 | 0.32 | 0.4 | 0.25 | 0.45 | 0.23 |
| 2 | 144662334 | rs13402786 | 0.848 | 1.232 | -0.384 | -0.111 | 0.028 | 0.017 | 0.017 | 0 | 0.19 | 0.29 | 0.31 | 0.21 | 0.24 | 0.15 |
| 2 | 144664940 | rs1019389 | -1.025 | 1.203 | -0.474 | -0.102 | 0.051 | -0.015 | -0.015 | 4.738 | 0.54 | 0.68 | 0.6 | 0.47 | 0.7 | 0.5 |
| 2 | 144669586 | rs10496954 | -0.832 | 1.442 | -0.488 | -0.347 | 0.007 | -0.025 | -0.025 | 0 | 0.15 | 0.21 | 0.12 | 0.04 | 0.23 | 0.08 |
| 2 | 144670651 | rs11888667 | -0.272 | 1.247 | -0.338 | -0.021 | 0.024 | 0 | 0 | 0 | 0.22 | 0.34 | 0.2 | 0.22 | 0.27 | 0.27 |
| 2 | 144675354 | rs787143 | 0.305 | 1.229 | -0.77 | 0.161 | 0 | 0 | 0 | 0 | 0.69 | 0.67 | 0.78 | 0.62 | 0.69 | 0.61 |
| 2 | 144684875 | rs768145 | -0.255 | 1.579 | -0.732 | -0.441 | -0.001 | -0.005 | -0.005 | 0 | 0.34 | 0.25 | 0.32 | 0.16 | 0.53 | 0.32 |
| 2 | 144689662 | rs787166 | -0.096 | 1.314 | -0.949 | -0.517 | -0.036 | 0.062 | 0.062 | 0 | 0.35 | 0.17 | 0.32 | 0.19 | 0.47 | 0.42 |
| 2 | 144689725 | rs4662216 | -0.224 | 0.656 | -0.602 | -0.466 | -0.003 | -0.015 | -0.015 | 0.377 | 0.14 | 0.17 | 0.13 | 0.03 | 0.1 | 0.07 |
| 2 | 144690117 | rs1011397 | 0.334 | 0.591 | -0.6 | -0.344 | -0.003 | -0.013 | -0.013 | 0 | 0.79 | 0.55 | 0.79 | 0.8 | 0.85 | 0.8 |
| 2 | 144691346 | rs973543 | 0.146 | 1.014 | -1.397 | 0.145 | -0.043 | -0.031 | -0.031 | 0 | 0.4 | 0.19 | 0.46 | 0.55 | 0.43 | 0.4 |
| 2 | 144697113 | rs787172 | -0.594 | 0.918 | -1.357 | 0.245 | -0.02 | -0.018 | -0.018 | 0 | 0.82 | 0.95 | 0.88 | 0.83 | 0.79 | 0.79 |
| 2 | 144712417 | rs787175 | -0.505 | 0.892 | -0.971 | 0.23 | -0.014 | 0.096 | 0.096 | 0 | 0.58 | 0.82 | 0.72 | 0.78 | 0.56 | 0.63 |
| 2 | 144718006 | rs16823373 | -0.523 | 0.842 | -0.337 | 0.366 | 0.006 | 0.05 | 0.05 | 0 | 0.07 | 0.19 | 0.1 | 0.16 | 0.04 | 0.14 |
| 2 | 144720437 | rs12472227 | 1.102 | 0.622 | -0.575 | 0.231 | -0.018 | 0.03 | 0.03 | 0.09 | 0.06 | 0 | 0.03 | 0 | 0.07 | 0.12 |
| 2 | 144728879 | rs16823382 | -0.557 | 0.539 | -0.399 | 0.32 | -0.014 | 0.042 | 0.042 | 0.116 | 0.06 | 0.18 | 0.09 | 0.16 | 0.02 | 0.13 |
| 2 | 144732454 | rs787182 | -0.149 | 0.632 | 0.221 | 0.382 | -0.002 | -0.005 | -0.005 | 0 | 0.47 | 0.26 | 0.54 | 0.53 | 0.47 | 0.44 |
| 2 | 144778389 | rs10928205 | 1.249 | 0.361 | 0.139 | 1.139 | 0.016 | -0.027 | -0.027 | 0 | 0.13 | 0.04 | 0.16 | 0.23 | 0.05 | 0.06 |
| 2 | 144788389 | rs6719835 | -1.034 | 0.341 | 0.904 | 1.143 | 0.046 | 0.209 | 0.209 | 0.323 | 0.08 | 0.42 | 0.16 | 0.26 | 0.05 | 0.15 |
| 2 | 144795011 | rs959766 | -0.591 | 0.336 | 1.033 | 1.152 | 0.013 | 0.11 | 0.11 | 0 | 0.06 | 0.2 | 0.14 | 0.27 | 0.05 | 0.15 |
| 2 | 144803955 | rs1560222 | -0.508 | 0.183 | 0.96 | 0.892 | 0.002 | 0.05 | 0.05 | 0.178 | 0.5 | 0.21 | 0.54 | 0.59 | 0.49 | 0.42 |
| 2 | 144816988 | rs17666845 | 0.802 | 0.386 | 0.804 | 0.938 | -0.09 | -0.115 | -0.115 | 0 | 0.24 | 0.02 | 0.29 | 0.38 | 0.33 | 0.29 |
| 2 | 144826531 | rs10496956 | -1.143 | 0.265 | 1.213 | 0.724 | -0.017 | 0.106 | 0.106 | 0 | 0.57 | 0.75 | 0.7 | 0.84 | 0.53 | 0.57 |
| 2 | 144859463 | rs16823489 | 1.338 | 0.134 | 1.211 | 0.896 | 0.016 | -0.003 | -0.003 | 0 | 0.13 | 0.05 | 0.15 | 0.13 | 0.04 | 0.06 |
| 2 | 144869044 | rs1560224 | 0.146 | 0.569 | 1.496 | 0.708 | -0.015 | 0.027 | 0.027 | 0.193 | 0.48 | 0.2 | 0.53 | 0.57 | 0.5 | 0.42 |
| 2 | 144899559 | rs3731958 | -0.514 | 0.504 | 0.881 | 0.17 | -0.006 | 0.108 | 0.108 | 0 | 0.57 | 0.74 | 0.71 | 0.84 | 0.54 | 0.56 |
| 2 | 144918226 | rs16823541 | -1.95 | 0.536 | 1.291 | 0.88 | 0.073 | 0.142 | 0.142 | 1.145 | 0.07 | 0.48 | 0.1 | 0.09 | 0.03 | 0.07 |
| 2 | 144931112 | rs6707579 | -1.6 | 0.617 | 0.24 | 0.893 | 0.028 | 0.076 | 0.076 | 0.365 | 0.06 | 0.3 | 0.09 | 0.09 | 0.03 | 0.06 |
| 2 | 144950248 | rs16823566 | -1.403 | 0.506 | -0.01 | 0.641 | -0.006 | -0.008 | -0.008 | 0 | 0.1 | 0.29 | 0.1 | 0.07 | 0.07 | 0.14 |
| 2 | 144967929 | rs13392253 | -1.086 | 0.428 | 0.564 | 0.601 | 0.01 | 0.007 | 0.007 | 0 | 0.08 | 0.18 | 0.09 | 0.07 | 0.07 | 0.14 |
| 2 | 144969618 | rs16823583 | -1.938 | 0.428 | 0.567 | 0.601 | 0.007 | 0.003 | 0.003 | 0 | 0.1 | 0.35 | 0.1 | 0.07 | 0.07 | 0.15 |
| 2 | 144993201 | rs1012272 | -0.979 | 0.433 | 0.564 | 0.604 | 0.011 | 0.009 | 0.009 | 0 | 0.08 | 0.17 | 0.09 | 0.07 | 0.07 | 0.14 |
| 2 | 145027454 | rs7589628 | 0.288 | 0.37 | -0.207 | 0.452 | -0.022 | -0.006 | -0.006 | 0.486 | 0.43 | 0.16 | 0.64 | 0.69 | 0.53 | 0.41 |
| 2 | 145057036 | rs12615626 | -0.345 | 0.898 | -0.482 | 0.458 | -0.026 | -0.019 | -0.019 | 0 | 0.06 | 0.02 | 0.14 | 0.19 | 0.1 | 0.13 |
| 2 | 145058077 | rs997840 | -0.593 | 0.319 | -0.33 | 0.341 | 0.021 | -0.017 | -0.017 | 0 | 0.08 | 0.34 | 0.09 | 0.02 | 0.05 | 0.09 |
| 2 | 145079065 | rs10496963 | 0.937 | 0.05 | -1.565 | 0.061 | -0.015 | 0.049 | 0.049 | 2.181 | 0.21 | 0.02 | 0.09 | 0.07 | 0.18 | 0.19 |
| 2 | 145126711 | rs12185711 | -0.45 | 0.387 | -1.012 | 0.462 | -0.004 | -0.017 | -0.017 | 0.044 | 0.06 | 0.01 | 0.1 | 0.14 | 0.06 | 0.11 |
| 2 | 145134588 | rs3928425 | 0.988 | 0.316 | -1.009 | 0.629 | 0 | -0.011 | -0.011 | 0 | 0.81 | 0.85 | 0.67 | 0.71 | 0.82 | 0.71 |
| 2 | 145141541 | rs12991836 | 1.126 | 0.32 | -1.319 | 0.291 | -0.066 | -0.093 | -0.093 | 0.087 | 0.29 | 0.12 | 0.36 | 0.52 | 0.37 | 0.22 |
| 2 | 145147758 | rs16855324 | -0.528 | -0.69 | -0.331 | -0.382 | 0.008 | -0.039 | -0.039 | 0 | 0.2 | 0.36 | 0.14 | 0.12 | 0.2 | 0.22 |
| 2 | 145155731 | rs10185359 | 0.545 | -1.523 | -0.201 | -0.737 | 0.022 | 0.02 | 0.02 | 0.4 | 0.6 | 0.41 | 0.82 | 0.83 | 0.78 | 0.68 |
| 2 | 145173029 | rs16823675 | -0.687 | -1.364 | -0.143 | -0.617 | 0.04 | 0.017 | 0.017 | 0 | 0.17 | 0.42 | 0.22 | 0.23 | 0.22 | 0.35 |
| 2 | 145177708 | rs6748886 | -0.15 | -1.493 | -0.285 | -0.691 | -0.022 | -0.04 | -0.04 | 5.129 | 0.21 | 0.36 | 0.14 | 0.12 | 0.17 | 0.23 |
| 2 | 145178110 | rs7600752 | 1.756 | -1.554 | -0.623 | -0.685 | -0.087 | -0.028 | -0.028 | 0 | 0.41 | 0.08 | 0.59 | 0.57 | 0.54 | 0.33 |
| 2 | 145184316 | rs10192562 | 1.25 | -1.523 | -0.251 | -0.776 | 0.012 | 0.014 | 0.014 | 2.959 | 0.2 | 0.11 | 0.03 | 0.08 | 0.05 | 0.08 |
| 2 | 145184666 | rs12996983 | 1.452 | -1.434 | -0.339 | -0.773 | -0.003 | 0.002 | 0.002 | 0 | 0.07 | 0.03 | 0.06 | 0.06 | 0.1 | 0.16 |
| 2 | 145186749 | rs12614546 | 2.145 | -1.477 | -0.501 | -0.807 | -0.191 | -0.073 | -0.073 | 0 | 0.36 | 0.02 | 0.53 | 0.57 | 0.46 | 0.32 |
| 2 | 145187783 | rs2288353 | -0.77 | -1.54 | 0.106 | -0.618 | -0.012 | -0.015 | -0.015 | 0 | 0.36 | 0.52 | 0.26 | 0.31 | 0.27 | 0.44 |
| 2 | 145202375 | rs13013418 | 0.126 | -1.415 | 0.045 | -0.666 | 0.037 | -0.035 | -0.035 | 0.948 | 0.13 | 0.28 | 0.12 | 0.07 | 0.18 | 0.23 |
| 2 | 145204758 | rs16823732 | 0.889 | -1.575 | -0.14 | -0.715 | 0.046 | 0.039 | 0.039 | 3.718 | 0.2 | 0.09 | 0.03 | 0.08 | 0.05 | 0.09 |
| 2 | 145204976 | rs7568133 | -0.458 | -1.785 | -0.142 | -0.878 | -0.004 | -0.008 | -0.008 | 0 | 0.24 | 0.35 | 0.25 | 0.27 | 0.22 | 0.35 |
| 2 | 145208193 | rs12105918 | 0.971 | -1.763 | -0.624 | -0.752 | 0.062 | 0.049 | 0.049 | 3.023 | 0.19 | 0.07 | 0.03 | 0.08 | 0.05 | 0.09 |
| 2 | 145209076 | rs7597006 | 1.251 | -1.67 | -0.673 | -0.703 | -0.045 | 0.011 | 0.011 | 0.952 | 0.42 | 0.1 | 0.59 | 0.57 | 0.57 | 0.33 |
| 2 | 145214607 | rs7599224 | -1.516 | -1.306 | -0.007 | 0.013 | 0.013 | 0.03 | 0.03 | 0.106 | 0.51 | 0.56 | 0.37 | 0.43 | 0.47 | 0.66 |
| 2 | 145216048 | rs13389578 | 0.074 | -1.486 | 0.381 | -0.049 | -0.085 | -0.071 | -0.071 | 0 | 0.23 | 0.35 | 0.06 | 0.09 | 0.08 | 0.11 |
| 2 | 145218605 | rs12327962 | -1.621 | -0.67 | 0.85 | 0.583 | -0.022 | -0.014 | -0.014 | 1.102 | 0.16 | 0.41 | 0.15 | 0.1 | 0.14 | 0.08 |
| 2 | 145220163 | rs13002663 | -0.767 | -0.04 | 0.047 | 0.745 | 0.013 | -0.005 | -0.005 | 0 | 0.3 | 0.36 | 0.24 | 0.37 | 0.39 | 0.61 |
| 2 | 145220327 | rs10172508 | -0.75 | -0.074 | -0.178 | 1.039 | -0.001 | -0.007 | -0.007 | 0.127 | 0.06 | 0.06 | 0.07 | 0 | 0.09 | 0.03 |
| 2 | 145222038 | rs13403907 | -1.588 | -0.047 | -0.244 | 1.769 | -0.014 | 0.027 | 0.027 | 0 | 0.05 | 0.03 | 0.25 | 0.31 | 0.12 | 0.13 |
| 2 | 145223620 | rs13382811 | -2.372 | -0.127 | 0.602 | 1.711 | 0.08 | 0.178 | 0.178 | 0 | 0.08 | 0.31 | 0.32 | 0.25 | 0.14 | 0.12 |
| 2 | 145223732 | rs17677601 | -0.441 | -0.94 | -0.156 | 1.503 | 0.081 | 0.045 | 0.045 | 0.931 | 0.31 | 0.03 | 0.14 | 0.2 | 0.17 | 0.38 |
| 2 | 145227258 | rs13410341 | -1.428 | -0.786 | 0.087 | 1.625 | 0.044 | 0.177 | 0.177 | 0 | 0.06 | 0.18 | 0.28 | 0.28 | 0.11 | 0.13 |
| 2 | 145235108 | rs4662223 | 0.416 | -1.001 | 0.673 | 1.622 | -0.006 | 0 | 0 | 0 | 0.67 | 0.66 | 0.83 | 0.76 | 0.84 | 0.62 |
| 2 | 145236142 | rs7600781 | 0.859 | -0.946 | 0.322 | 1.349 | -0.021 | 0.015 | 0.015 | 0.019 | 0.6 | 0.48 | 0.51 | 0.47 | 0.7 | 0.48 |
| 2 | 145242122 | rs6755392 | 0.755 | -0.954 | 0.345 | 1.324 | -0.015 | 0.008 | 0.008 | 0.004 | 0.6 | 0.44 | 0.51 | 0.46 | 0.7 | 0.48 |
| 2 | 145255778 | rs1035822 | 1.742 | -0.363 | 0.318 | 0.949 | -0.002 | -0.026 | -0.026 | 0.082 | 0.86 | 0.34 | 0.87 | 0.94 | 0.89 | 0.81 |
| 2 | 145270592 | rs6740731 | -0.714 | -0.916 | 0.295 | 1.182 | -0.037 | 0.035 | 0.035 | 1.535 | 0.26 | 0.39 | 0.38 | 0.42 | 0.16 | 0.26 |

| | | | | | | | | | | | | | | | | |
|---|---|---|---|---|---|---|---|---|---|---|---|---|---|---|---|---|
| 2 | 145282685 | rs12691694 | -1.059 | -0.709 | -0.233 | 0.885 | 0.009 | -0.081 | -0.081 | 0.24 | 0.38 | 0.11 | 0.44 | 0.56 | 0.34 | 0.38 |
| 2 | 145287079 | rs1469346 | 1.213 | -0.705 | 0.197 | 0.673 | -0.041 | -0.073 | -0.073 | 1.125 | 0.84 | 0.56 | 0.93 | 0.96 | 0.89 | 0.85 |
| 2 | 145288341 | rs16823835 | -0.094 | -0.496 | -0.644 | 0.817 | -0.002 | -0.11 | -0.11 | 0 | 0.26 | 0.07 | 0.43 | 0.58 | 0.2 | 0.26 |
| 2 | 145292497 | rs10173242 | -1.223 | -0.006 | -0.343 | 0.796 | 0.003 | 0.001 | 0.001 | 0.561 | 0.44 | 0.47 | 0.54 | 0.61 | 0.41 | 0.44 |
| 2 | 145296263 | rs2052807 | -1.201 | 0 | -0.323 | 0.801 | 0.002 | 0.001 | 0.001 | 0.694 | 0.44 | 0.47 | 0.5 | 0.62 | 0.4 | 0.43 |
| 2 | 145298323 | rs17689833 | -0.791 | 0.68 | -0.519 | 0.525 | -0.021 | 0.034 | 0.034 | 0 | 0.11 | 0.05 | 0.08 | 0.01 | 0.19 | 0.09 |
| 2 | 145305277 | rs1365778 | -0.054 | 1.195 | -0.982 | 0.976 | -0.052 | -0.037 | -0.037 | 0 | 0.25 | 0.1 | 0.33 | 0.35 | 0.33 | 0.2 |
| 2 | 145315492 | rs17741144 | -0.082 | 1.213 | -0.533 | 0.976 | -0.052 | -0.037 | -0.037 | 0 | 0.26 | 0.1 | 0.34 | 0.34 | 0.33 | 0.2 |
| 2 | 145316407 | rs4662380 | -0.107 | 0.795 | -0.55 | 0.8 | 0.001 | 0.001 | 0.001 | 0 | 0.2 | 0.17 | 0.11 | 0.23 | 0.19 | 0.38 |
| 2 | 145317284 | rs6746122 | 1.001 | 0.46 | -0.298 | 0.436 | 0.035 | -0.012 | -0.012 | 0 | 0.4 | 0.21 | 0.42 | 0.38 | 0.33 | 0.31 |
| 2 | 145319073 | rs6722387 | 0.104 | 0.394 | 0.008 | 0.342 | 0 | 0 | 0 | 0 | 0.21 | 0.25 | 0.12 | 0.22 | 0.19 | 0.38 |
| 2 | 145324977 | rs16823866 | 0.359 | 0.341 | -0.098 | 0.314 | -0.041 | -0.035 | -0.035 | 0 | 0.19 | 0.01 | 0.1 | 0.23 | 0.19 | 0.34 |
| 2 | 145328876 | rs13413446 | -0.132 | 0.332 | 0.146 | 0.31 | -0.011 | -0.014 | -0.014 | 0 | 0.22 | 0.34 | 0.12 | 0.23 | 0.19 | 0.38 |
| 2 | 145334304 | rs4563167 | -0.473 | 0.39 | 0.222 | 0.3 | 0 | -0.001 | -0.001 | 0 | 0.9 | 0.84 | 0.9 | 0.95 | 0.85 | 0.89 |
| 2 | 145341259 | rs16823886 | 0.028 | 0.129 | -0.076 | 0.199 | -0.029 | -0.025 | -0.025 | 0.145 | 0.14 | 0.02 | 0.06 | 0.17 | 0.14 | 0.27 |
| 2 | 145345343 | rs12467963 | 0.218 | 0.352 | -0.171 | 0.3 | -0.102 | -0.037 | -0.037 | 0.443 | 0.32 | 0.08 | 0.36 | 0.36 | 0.4 | 0.31 |
| 2 | 145362155 | rs17741930 | -0.037 | 0.749 | 0.317 | -0.227 | -0.06 | -0.016 | -0.016 | 0.408 | 0.16 | 0.02 | 0.09 | 0.17 | 0.19 | 0.27 |
| 2 | 145368852 | rs2162571 | 0.922 | 0.587 | 1.558 | -0.306 | -0.007 | -0.037 | -0.037 | 0 | 0.78 | 0.38 | 0.84 | 0.83 | 0.81 | 0.71 |
| 2 | 145386594 | rs4143010 | 1.405 | 0.613 | 1.546 | -0.266 | 0.039 | 0.035 | 0.035 | 5.509 | 0.94 | 0.43 | 0.93 | 1 | 1 | 1 |
| 2 | 145400317 | rs1427499 | 0.894 | 0.791 | 1.274 | -0.605 | -0.004 | -0.045 | -0.045 | 0 | 0.69 | 0.4 | 0.76 | 0.83 | 0.69 | 0.68 |
| 2 | 145416400 | rs1427497 | 1.147 | 0.738 | 1.545 | -0.491 | 0.08 | 0.07 | 0.07 | 4.613 | 0.95 | 0.48 | 0.94 | 1 | 0.99 | 0.99 |
| 2 | 145442190 | rs980772 | 0.919 | 0.772 | 1.205 | -0.883 | -0.001 | -0.02 | -0.02 | 0 | 0.64 | 0.4 | 0.72 | 0.78 | 0.65 | 0.68 |
| 2 | 145446205 | rs2195115 | -0.173 | 0.839 | 1.138 | -0.935 | -0.062 | 0.009 | 0.009 | 0 | 0.24 | 0.03 | 0.2 | 0.22 | 0.27 | 0.28 |
| 2 | 145463764 | rs2195114 | 1.042 | 0.844 | 1.229 | -0.951 | -0.001 | -0.057 | -0.057 | 0.404 | 0.64 | 0.47 | 0.73 | 0.78 | 0.65 | 0.69 |
| 2 | 145504693 | rs7603516 | 0.267 | 0.778 | 1.367 | -0.917 | 0.004 | -0.046 | -0.046 | 0 | 0.08 | 0.2 | 0.03 | 0 | 0.08 | 0.03 |
| 2 | 145506108 | rs13032840 | 0.712 | 0.778 | 1.341 | -0.925 | 0 | -0.03 | -0.03 | 0.234 | 0.66 | 0.57 | 0.73 | 0.77 | 0.64 | 0.7 |
| 2 | 145519623 | rs11883829 | 0.662 | 0.779 | 1.324 | -0.925 | -0.001 | -0.02 | -0.02 | 0 | 0.65 | 0.4 | 0.72 | 0.77 | 0.64 | 0.69 |
| 2 | 145527778 | rs11689575 | 0.536 | 0.786 | 1.173 | -0.941 | 0.001 | -0.033 | -0.033 | 0.226 | 0.08 | 0.13 | 0.03 | 0 | 0.08 | 0.03 |
| 2 | 145531555 | rs13413613 | 0.023 | 0.763 | 1.092 | -0.875 | -0.027 | 0.028 | 0.028 | 0 | 0.25 | 0.09 | 0.2 | 0.22 | 0.27 | 0.29 |
| 2 | 145584570 | rs786490 | 0.467 | 0.783 | 1.314 | -0.959 | -0.006 | -0.045 | -0.045 | 0 | 0.68 | 0.46 | 0.74 | 0.78 | 0.7 | 0.69 |
| 2 | 145590819 | rs13028678 | 0.068 | 0.655 | 1.464 | -1.027 | -0.024 | 0.03 | 0.03 | 0 | 0.26 | 0.11 | 0.21 | 0.22 | 0.3 | 0.29 |
| 2 | 145592070 | rs13018654 | 0.652 | 0.644 | 1.548 | -1.033 | 0.005 | 0.023 | 0.023 | 0 | 0.66 | 0.29 | 0.72 | 0.78 | 0.7 | 0.68 |
| 2 | 145595571 | rs786500 | 0.253 | 0.676 | 1.555 | -1.016 | 0 | -0.037 | -0.037 | 0 | 0.7 | 0.57 | 0.75 | 0.78 | 0.7 | 0.71 |
| 2 | 145598913 | rs786480 | -0.227 | 0.231 | 1.157 | -1.298 | -0.029 | -0.046 | -0.046 | 0 | 0.91 | 0.65 | 0.95 | 1 | 0.98 | 0.97 |
| 2 | 145599194 | rs10928230 | 0.879 | 0.283 | 1.029 | -1.33 | -0.002 | -0.022 | -0.022 | 0 | 0.58 | 0.41 | 0.71 | 0.78 | 0.62 | 0.65 |
| 2 | 145602015 | rs17479071 | 1.587 | 0.28 | 0.198 | -1.437 | -0.063 | 0.077 | 0.077 | 2.174 | 0.3 | 0.05 | 0.21 | 0.12 | 0.35 | 0.24 |
| 2 | 145603306 | rs11690035 | -0.215 | 0.272 | 1.163 | -1.199 | -0.001 | -0.049 | -0.049 | 0 | 0.09 | 0.21 | 0.04 | 0.01 | 0.06 | 0.03 |
| 2 | 145603911 | rs1406120 | -1.199 | 0.355 | 1.29 | -1.256 | -0.023 | -0.111 | -0.111 | 0 | 0.16 | 0.42 | 0.1 | 0 | 0.11 | 0.06 |
| 2 | 145616899 | rs7560871 | 1.147 | 0.391 | 1.084 | -1.24 | -0.008 | 0.027 | 0.027 | 0 | 0.06 | 0.01 | 0.02 | 0 | 0.06 | 0.02 |
| 2 | 145619967 | rs10188324 | -0.685 | 0.386 | 1.095 | -1.228 | -0.023 | -0.066 | -0.066 | 0.378 | 0.11 | 0.36 | 0.07 | 0 | 0.05 | 0.05 |
| 2 | 145627927 | rs6430068 | -1.19 | 0.415 | 0.646 | -1.218 | -0.04 | -0.131 | -0.131 | 0 | 0.19 | 0.58 | 0.11 | 0 | 0.11 | 0.06 |
| 2 | 145630762 | rs17479267 | 0.987 | 0.301 | -0.686 | -1.428 | 0.005 | 0.12 | 0.12 | 0 | 0.31 | 0.11 | 0.18 | 0.09 | 0.31 | 0.33 |
| 2 | 145632077 | rs7591518 | 0.257 | 0.695 | -0.752 | -1.713 | -0.025 | 0.087 | 0.087 | 0 | 0.31 | 0.14 | 0.19 | 0.08 | 0.31 | 0.27 |
| 2 | 145634153 | rs6738560 | -0.451 | 0.793 | 0.012 | -1.436 | 0.029 | -0.009 | -0.009 | 0.236 | 0.77 | 0.55 | 0.68 | 0.83 | 0.72 | 0.83 |
| 2 | 145634861 | rs1438900 | 0.769 | 0.742 | 0.279 | -1.456 | 0.026 | 0.056 | 0.056 | 0 | 0.45 | 0.3 | 0.48 | 0.76 | 0.41 | 0.56 |
| 2 | 145639443 | rs11677207 | -0.33 | 0.745 | 0.312 | -1.483 | -0.008 | 0.022 | 0.022 | 0 | 0.41 | 0.31 | 0.46 | 0.81 | 0.42 | 0.55 |
| 2 | 145639577 | rs11677257 | 1.268 | 0.737 | 0.309 | -1.473 | -0.032 | -0.045 | -0.045 | 4.234 | 0.08 | 0.01 | 0.25 | 0.74 | 0.11 | 0.28 |
| 2 | 145640663 | rs2118775 | -0.923 | 0.635 | 0.289 | -1.562 | 0.021 | 0.04 | 0.04 | 0.391 | 0.56 | 0.33 | 0.66 | 0.9 | 0.63 | 0.69 |
| 2 | 145646072 | rs1438896 | -0.201 | 0.691 | 0.213 | -1.525 | 0.038 | -0.01 | -0.01 | 0.462 | 0.77 | 0.51 | 0.67 | 0.83 | 0.72 | 0.83 |
| 2 | 145661823 | rs786280 | -1.616 | 0.674 | 0.262 | -1.54 | 0.025 | 0.056 | 0.056 | 0.018 | 0.65 | 0.79 | 0.8 | 0.98 | 0.7 | 0.73 |
| 2 | 145672413 | rs786274 | -1.103 | 0.502 | 0.784 | -1.48 | 0 | -0.062 | -0.062 | 0.059 | 0.57 | 0.44 | 0.71 | 0.98 | 0.57 | 0.7 |
| 2 | 145674075 | rs2579022 | -1.763 | 0.532 | 0.491 | -1.464 | 0.004 | 0.04 | 0.04 | 0.277 | 0.64 | 0.75 | 0.76 | 0.98 | 0.63 | 0.73 |
| 2 | 145677006 | rs813684 | -1.734 | 0.541 | 0.491 | -1.445 | 0.002 | 0.054 | 0.054 | 0.277 | 0.63 | 0.75 | 0.73 | 0.96 | 0.62 | 0.72 |
| 2 | 145684278 | rs11684042 | 1.041 | 0.687 | 0.334 | -1.284 | -0.03 | -0.017 | -0.017 | 0 | 0.17 | 0.07 | 0.27 | 0.17 | 0.22 | 0.15 |
| 2 | 145704055 | rs10201023 | 0.675 | 0.542 | 0.318 | -1.387 | -0.004 | 0.054 | 0.054 | 0 | 0.28 | 0.14 | 0.17 | 0 | 0.25 | 0.2 |
| 2 | 145714354 | rs1438898 | 1.342 | 0.514 | 0.699 | -1.355 | -0.073 | -0.058 | -0.058 | 1.212 | 0.16 | 0.04 | 0.27 | 0.19 | 0.22 | 0.15 |
| 2 | 145721472 | rs786260 | -1.842 | 0.499 | 0.855 | -1.425 | 0.012 | 0.046 | 0.046 | 0.232 | 0.63 | 0.78 | 0.77 | 0.99 | 0.63 | 0.72 |
| 2 | 145721524 | rs13010313 | 0.732 | 0.534 | 0.451 | -1.404 | -0.01 | -0.052 | -0.052 | 0 | 0.11 | 0.02 | 0.26 | 0.8 | 0.1 | 0.35 |
| 2 | 145725530 | rs11674165 | 0.633 | 0.55 | 0.502 | -1.393 | -0.038 | -0.011 | -0.011 | 2.175 | 0.2 | 0.12 | 0.3 | 0.18 | 0.28 | 0.17 |
| 2 | 145732884 | rs1881266 | 0.615 | 0.563 | 0.585 | -1.45 | 0 | -0.049 | -0.049 | 0.174 | 0.29 | 0.3 | 0.18 | 0 | 0.25 | 0.2 |
| 2 | 145749357 | rs1528188 | -0.748 | 0.482 | 0.267 | -1.296 | -0.026 | 0.064 | 0.064 | 2.617 | 0.32 | 0.23 | 0.56 | 0.98 | 0.38 | 0.51 |
| 2 | 145759899 | rs13386034 | 1.013 | 0.583 | 0.161 | -1.093 | -0.046 | 0.023 | 0.023 | 0.225 | 0.17 | 0.04 | 0.2 | 0.11 | 0.23 | 0.15 |
| 2 | 145764034 | rs7583863 | 0.903 | 0.473 | 0.124 | -1.252 | -0.028 | 0.046 | 0.046 | 0 | 0.22 | 0.1 | 0.22 | 0.08 | 0.25 | 0.19 |
| 2 | 145764746 | rs12691702 | -0.586 | 0.379 | 0.108 | -1.198 | -0.025 | 0.016 | 0.016 | 0 | 0.78 | 0.92 | 0.79 | 0.82 | 0.76 | 0.77 |
| 2 | 145770621 | rs2250546 | -1.171 | 0.399 | 0.639 | -1.111 | 0 | -0.039 | -0.039 | 0 | 0.43 | 0.56 | 0.53 | 0.87 | 0.38 | 0.54 |
| 2 | 145775399 | rs1006923 | 2.052 | 0.461 | 0.504 | -1.122 | -0.062 | 0.185 | 0.185 | 0.224 | 0.32 | 0.03 | 0.2 | 0.03 | 0.35 | 0.29 |
| 2 | 145784885 | rs2246142 | -0.815 | 0.771 | 1.119 | -0.763 | 0.004 | -0.016 | -0.016 | 0 | 0.36 | 0.47 | 0.51 | 0.79 | 0.35 | 0.46 |
| 2 | 145793076 | rs2243647 | -1.011 | 0.769 | 1.164 | -0.734 | 0.012 | -0.036 | -0.036 | 0 | 0.37 | 0.55 | 0.52 | 0.78 | 0.38 | 0.46 |
| 2 | 145794405 | rs12616953 | 0.85 | 0.794 | 1.075 | -0.689 | -0.004 | -0.019 | -0.019 | 0 | 0.09 | 0.05 | 0.25 | 0.7 | 0.06 | 0.26 |

| | | | | | | | | | | | | | | | | |
|---|---|---|---|---|---|---|---|---|---|---|---|---|---|---|---|---|
| 2 | 145797829 | rs1406225 | 0.171 | 0.766 | 1.133 | -0.716 | 0.01 | -0.043 | -0.043 | 0 | 0.26 | 0.41 | 0.25 | 0.08 | 0.29 | 0.19 |
| 2 | 145798956 | rs4662413 | 0.787 | 0.439 | 0.847 | -0.626 | -0.001 | -0.016 | -0.016 | 0.408 | 0.09 | 0.05 | 0.25 | 0.71 | 0.06 | 0.26 |
| 2 | 145801113 | rs2252654 | 1.214 | 0.37 | 1.13 | -0.707 | 0.011 | 0.014 | 0.014 | 0.361 | 0.43 | 0.59 | 0.25 | 0.1 | 0.34 | 0.38 |
| 2 | 145801461 | rs2252641 | 0.121 | 0.411 | 0.74 | -0.634 | -0.042 | 0.106 | 0.106 | 0.32 | 0.58 | 0.86 | 0.55 | 0.81 | 0.46 | 0.68 |
| 2 | 145820196 | rs12618061 | 0.743 | 0.599 | 0.769 | -0.512 | 0 | -0.012 | -0.012 | 0 | 0.09 | 0.03 | 0.24 | 0.7 | 0.06 | 0.24 |
| 2 | 145822778 | rs12691704 | 1.364 | 0.531 | 1.059 | -0.619 | 0.009 | 0.033 | 0.033 | 0.011 | 0.39 | 0.24 | 0.22 | 0.03 | 0.35 | 0.36 |
| 2 | 145822995 | rs7561039 | 0.866 | 0.53 | 1.149 | -0.609 | -0.003 | 0.076 | 0.076 | 0 | 0.08 | 0.09 | 0.25 | 0.7 | 0.05 | 0.23 |
| 2 | 145825555 | rs1830321 | 0.793 | 0.536 | 1.102 | -0.616 | 0.009 | 0.024 | 0.024 | 0 | 0.47 | 0.33 | 0.47 | 0.74 | 0.4 | 0.59 |
| 2 | 145835757 | rs6740528 | 0.527 | 0.82 | 0.745 | -0.212 | -0.004 | 0.024 | 0.024 | 0 | 0.09 | 0.07 | 0.22 | 0.53 | 0.08 | 0.24 |
| 2 | 145837441 | rs1125128 | -0.862 | 0.841 | 0.888 | -0.523 | 0.025 | 0.181 | 0.181 | 0.25 | 0.56 | 0.25 | 0.72 | 0.97 | 0.62 | 0.64 |
| 2 | 145854691 | rs17408733 | -0.509 | 1.593 | 1.77 | 0.423 | -0.015 | 0.041 | 0.041 | 0 | 0.22 | 0.12 | 0.2 | 0.06 | 0.26 | 0.14 |
| 2 | 145868177 | rs17408803 | 0.007 | 1.354 | 1.756 | -0.445 | -0.055 | 0.111 | 0.111 | 0 | 0.18 | 0.03 | 0.13 | 0 | 0.21 | 0.12 |
| 2 | 145888369 | rs787435 | 0.804 | 1.369 | 2.22 | -0.453 | -0.001 | -0.081 | -0.081 | 1.112 | 0.73 | 0.26 | 0.78 | 0.98 | 0.73 | 0.81 |
| 2 | 145892427 | rs2697329 | 0.836 | 1.37 | 2.318 | -0.455 | 0 | -0.104 | -0.104 | 0 | 0.73 | 0.28 | 0.78 | 0.98 | 0.72 | 0.79 |
| 2 | 145908283 | rs1085641 | 1 | 1.373 | 2.059 | -0.454 | -0.001 | -0.078 | -0.078 | 0 | 0.73 | 0.26 | 0.78 | 0.98 | 0.72 | 0.8 |
| 2 | 145951277 | rs6718620 | -0.377 | 1.213 | 1.739 | -0.466 | 0 | -0.043 | -0.043 | 0 | 0.2 | 0.2 | 0.17 | 0.02 | 0.21 | 0.17 |
| 2 | 145963547 | rs959809 | 1.409 | 0.822 | 1.317 | 0.2 | 0.04 | -0.033 | -0.033 | 0.223 | 0.91 | 0.44 | 0.93 | 0.99 | 0.93 | 0.97 |
| 2 | 145983262 | rs13398905 | -1.379 | 0.978 | 1.139 | 0.488 | 0.048 | 0.056 | 0.056 | 0 | 0.11 | 0.38 | 0.12 | 0.14 | 0.12 | 0.08 |
| 2 | 145984088 | rs16824634 | -1.558 | 0.953 | 1.616 | 0.475 | 0.067 | 0.077 | 0.077 | 0 | 0.11 | 0.44 | 0.12 | 0.14 | 0.12 | 0.08 |
| 2 | 146003188 | rs17409319 | -0.614 | 0.936 | 1.748 | 0.514 | -0.013 | -0.011 | -0.011 | 0 | 0.07 | 0.05 | 0.08 | 0.1 | 0.11 | 0.07 |
| 2 | 146006855 | rs17409360 | 0.027 | 1.515 | 1.071 | -0.227 | -0.077 | 0.1 | 0.1 | 0 | 0.21 | 0.06 | 0.19 | 0 | 0.32 | 0.11 |
| 2 | 146011003 | rs10203763 | -1.223 | 1.211 | 1.543 | -0.216 | 0.001 | -0.012 | -0.012 | 0 | 0.34 | 0.63 | 0.33 | 0.15 | 0.43 | 0.19 |
| 2 | 146015334 | rs6756883 | -0.945 | 1.167 | 1.325 | -0.233 | 0.026 | -0.054 | -0.054 | 0 | 0.32 | 0.42 | 0.31 | 0.13 | 0.43 | 0.22 |
| 2 | 146028558 | rs13003259 | 0.842 | 0.545 | 0.092 | -0.204 | -0.047 | -0.025 | -0.025 | 0.001 | 0.29 | 0.02 | 0.12 | 0.32 | 0.24 | 0.3 |
| 2 | 146035130 | rs2381710 | 0.881 | 0.602 | 0.547 | -0.21 | -0.002 | -0.01 | -0.01 | 1.78 | 0.34 | 0.31 | 0.32 | 0.1 | 0.31 | 0.32 |
| 2 | 146035401 | rs4662238 | -0.424 | 0.606 | 0.261 | 0.025 | -0.003 | -0.006 | -0.006 | 0 | 0.63 | 0.61 | 0.6 | 0.24 | 0.73 | 0.53 |
| 2 | 146044052 | rs1822881 | 0.729 | 0.217 | -0.098 | -0.004 | 0.003 | 0.004 | 0.004 | 0 | 0.31 | 0.24 | 0.28 | 0.1 | 0.3 | 0.33 |
| 2 | 146045060 | rs17409597 | 0.122 | 0.299 | -0.29 | 0.332 | 0 | 0.075 | 0.075 | 0 | 0.48 | 0.18 | 0.42 | 0.35 | 0.5 | 0.34 |
| 2 | 146046344 | rs1445304 | 0.633 | 0.341 | 0.179 | 0.807 | -0.008 | 0.249 | 0.249 | 0.853 | 0.87 | 0.54 | 0.81 | 0.68 | 0.94 | 0.75 |
| 2 | 146068096 | rs7609091 | 0.502 | 0.718 | -0.618 | -0.078 | -0.043 | 0.107 | 0.107 | 0 | 0.25 | 0.08 | 0.18 | 0.01 | 0.3 | 0.16 |
| 2 | 146068496 | rs17409653 | 0.469 | 0.652 | 0.101 | -0.111 | -0.04 | 0.105 | 0.105 | 0 | 0.25 | 0.11 | 0.18 | 0.01 | 0.3 | 0.16 |
| 2 | 146071853 | rs10180522 | 1.496 | 0.423 | 0.145 | -0.249 | 0.019 | 0.084 | 0.084 | 0 | 0.31 | 0.12 | 0.28 | 0.1 | 0.29 | 0.33 |
| 2 | 146078530 | rs12615682 | 0.3 | 0.774 | 0.623 | 0.421 | 0.017 | 0.08 | 0.08 | 0 | 0.11 | 0.15 | 0.17 | 0.33 | 0.13 | 0.29 |
| 2 | 146082440 | rs10188215 | 1.187 | 0.608 | 0.686 | 0.399 | -0.009 | 0.053 | 0.053 | 0 | 0.17 | 0.2 | 0.23 | 0.38 | 0.11 | 0.1 |
| 2 | 146083333 | rs10496977 | 0.164 | 0.598 | 0.892 | 0.494 | 0.001 | -0.023 | -0.023 | 0 | 0.05 | 0.08 | 0.05 | 0.01 | 0.08 | 0.06 |
| 2 | 146087653 | rs7590923 | 1.279 | 0.488 | 0.738 | 0.387 | 0.007 | 0.033 | 0.033 | 0 | 0.69 | 0.38 | 0.67 | 0.48 | 0.68 | 0.56 |
| 2 | 146089267 | rs1822882 | -0.633 | 0.514 | 0.927 | 0.413 | 0.035 | 0.135 | 0.135 | 0 | 0.89 | 0.77 | 0.82 | 0.67 | 0.87 | 0.71 |
| 2 | 146090174 | rs10496978 | 0.177 | 0.372 | 0.618 | 0.322 | -0.008 | -0.001 | -0.001 | 0 | 0.38 | 0.21 | 0.4 | 0.38 | 0.38 | 0.23 |
| 2 | 146095070 | rs1516144 | -1.525 | 0.251 | 0.726 | 0.379 | -0.001 | -0.022 | -0.022 | 0 | 0.19 | 0.37 | 0.15 | 0.19 | 0.19 | 0.16 |
| 2 | 146095468 | rs12373773 | -0.022 | 0.313 | 0.571 | 0.578 | 0.023 | 0.034 | 0.034 | 0 | 0.08 | 0.2 | 0.07 | 0.17 | 0.09 | 0.06 |
| 2 | 146105254 | rs10445672 | -1.042 | 0.168 | 0.622 | 0.364 | -0.013 | -0.053 | -0.053 | 0.059 | 0.19 | 0.39 | 0.15 | 0.12 | 0.16 | 0.18 |
| 2 | 146108380 | rs1996289 | 0.315 | -0.408 | -0.358 | 0.69 | 0.021 | -0.028 | -0.028 | 6.857 | 0.18 | 0.11 | 0.2 | 0.36 | 0.08 | 0.09 |
| 2 | 146111971 | rs10928245 | -0.319 | 0.06 | -0.137 | 0.703 | -0.002 | 0.01 | 0.01 | 0 | 0.2 | 0.19 | 0.3 | 0.32 | 0.19 | 0.15 |
| 2 | 146114898 | rs4422110 | -0.37 | -0.085 | -0.129 | 0.244 | -0.011 | -0.006 | -0.006 | 0 | 0.59 | 0.62 | 0.63 | 0.56 | 0.5 | 0.55 |
| 2 | 146120511 | rs1040173 | -0.1 | -0.393 | -0.331 | -0.076 | 0.002 | 0.002 | 0.002 | 0 | 0.48 | 0.42 | 0.46 | 0.47 | 0.43 | 0.32 |
| 2 | 146134774 | rs17410058 | 1.425 | 0.039 | -0.822 | -0.842 | -0.047 | 0.137 | 0.137 | 0.049 | 0.27 | 0.05 | 0.18 | 0 | 0.32 | 0.14 |
| 2 | 146138077 | rs10496979 | 1.484 | -0.101 | -0.377 | -0.555 | -0.012 | -0.003 | -0.003 | 0 | 0.11 | 0.16 | 0.17 | 0.09 | 0.07 | 0.23 |
| 2 | 146141390 | rs10928247 | 0.255 | 0.031 | -0.517 | -0.504 | 0.011 | -0.008 | -0.008 | 0.058 | 0.12 | 0.14 | 0.17 | 0.06 | 0.19 | 0.14 |
| 2 | 146153407 | rs16824898 | 1.601 | -0.177 | -0.756 | -0.559 | -0.014 | 0.037 | 0.037 | 0 | 0.26 | 0.12 | 0.15 | 0.16 | 0.22 | 0.17 |
| 2 | 146153841 | rs10928249 | 1.439 | -0.042 | -0.652 | -0.533 | 0 | -0.014 | -0.014 | 0 | 0.07 | 0.06 | 0.09 | 0 | 0.08 | 0.07 |
| 2 | 146154475 | rs10204857 | -0.573 | -0.031 | -0.362 | -0.542 | 0.004 | -0.006 | -0.006 | 0 | 0.58 | 0.61 | 0.52 | 0.3 | 0.49 | 0.54 |
| 2 | 146156197 | rs6721116 | 0.392 | -0.04 | -0.334 | -0.544 | -0.006 | 0.006 | 0.006 | 0 | 0.42 | 0.31 | 0.47 | 0.7 | 0.51 | 0.44 |
| 2 | 146158880 | rs13389677 | 1.489 | -0.124 | -0.81 | -0.364 | -0.015 | 0.031 | 0.031 | 0 | 0.25 | 0.12 | 0.15 | 0.16 | 0.22 | 0.17 |
| 2 | 146168208 | rs16824949 | 0.521 | -0.124 | -0.654 | -0.24 | -0.068 | -0.005 | -0.005 | 0.525 | 0.39 | 0.12 | 0.46 | 0.66 | 0.56 | 0.43 |
| 2 | 146171033 | rs1533427 | -0.415 | -0.292 | 0.023 | -0.372 | 0.003 | 0.014 | 0.014 | 0 | 0.54 | 0.45 | 0.65 | 0.76 | 0.63 | 0.7 |
| 2 | 146178188 | rs6738331 | 0.704 | -0.245 | 0.18 | -0.199 | -0.047 | -0.001 | -0.001 | 0.421 | 0.14 | 0.35 | 0.19 | 0.11 | 0.07 | 0.28 |
| 2 | 146217925 | rs13008797 | 1.251 | -0.378 | -0.31 | -0.364 | -0.097 | -0.013 | -0.013 | 0 | 0.27 | 0.02 | 0.19 | 0.21 | 0.28 | 0.19 |
| 2 | 146238529 | rs983115 | 0.016 | -0.211 | 0.456 | -0.179 | -0.015 | -0.005 | -0.005 | 0 | 0.74 | 0.35 | 0.76 | 0.86 | 0.89 | 0.69 |
| 2 | 146247003 | rs6430083 | 0.182 | -0.217 | 0.418 | -0.193 | -0.005 | 0 | 0 | 1.72 | 0.73 | 0.32 | 0.8 | 0.86 | 0.89 | 0.67 |
| 2 | 146249965 | rs1385367 | -2.156 | -0.086 | 0.505 | -0.13 | -0.021 | -0.02 | -0.02 | 0 | 0.9 | 0.77 | 0.95 | 0.97 | 0.96 | 0.95 |
| 2 | 146253714 | rs12614884 | 2.381 | -0.043 | 0.197 | -0.109 | 0.002 | 0.002 | 0.002 | 0.137 | 0.09 | 0.06 | 0.04 | 0.03 | 0.04 | 0.03 |
| 2 | 146260496 | rs1351907 | 0.939 | -0.004 | 0.31 | -0.051 | -0.04 | -0.004 | -0.004 | 1.724 | 0.13 | 0.22 | 0.13 | 0.11 | 0.07 | 0.27 |
| 2 | 146276286 | rs4662243 | 1.076 | 0.54 | -0.228 | 0.552 | -0.028 | 0.02 | 0.02 | 0 | 0.06 | 0.01 | 0.21 | 0.36 | 0.18 | 0.16 |
| 2 | 146294931 | rs10180063 | 0.808 | -0.045 | 0.282 | 0.329 | -0.052 | 0.005 | 0.005 | 0.053 | 0.14 | 0.27 | 0.13 | 0.1 | 0.07 | 0.27 |
| 2 | 146300691 | rs10199699 | 0.512 | -0.151 | 0.237 | 0.202 | 0.01 | 0.009 | 0.009 | 1.45 | 0.71 | 0.32 | 0.74 | 0.85 | 0.86 | 0.64 |
| 2 | 146308114 | rs10496982 | 0.864 | -0.212 | 0.151 | 0.18 | -0.054 | 0.008 | 0.008 | 0.018 | 0.14 | 0.31 | 0.13 | 0.1 | 0.07 | 0.27 |
| 2 | 146309946 | rs17410535 | 1.246 | -0.307 | -1.007 | -0.138 | -0.083 | 0.054 | 0.054 | 0 | 0.3 | 0.02 | 0.22 | 0.17 | 0.28 | 0.18 |
| 2 | 146316319 | rs10193706 | 0.477 | -0.165 | -0.728 | 0.251 | -0.086 | -0.071 | -0.071 | 0.896 | 0.38 | 0.12 | 0.5 | 0.62 | 0.55 | 0.41 |
| 2 | 146323219 | rs900980 | -0.08 | -0.657 | 0.11 | 0.398 | -0.08 | 0.032 | 0.032 | 0.053 | 0.79 | 0.59 | 0.85 | 0.84 | 0.89 | 0.72 |

| | | | | | | | | | | | | | | | | |
|---|---|---|---|---|---|---|---|---|---|---|---|---|---|---|---|---|
| 2 | 146337062 | rs16825205 | 1.212 | -0.518 | -0.917 | 0.506 | 0.011 | -0.04 | -0.04 | 2.73 | 0.07 | 0 | 0.1 | 0.21 | 0.05 | 0.1 |
| 2 | 146344233 | rs10186882 | 0.393 | -0.712 | -0.714 | 0.302 | -0.003 | 0 | 0 | 0.932 | 0.74 | 0.33 | 0.75 | 0.83 | 0.85 | 0.66 |
| 2 | 146346331 | rs16825228 | -0.087 | -0.021 | -0.916 | 0.299 | -0.085 | -0.082 | -0.082 | 0 | 0.35 | 0.14 | 0.46 | 0.59 | 0.47 | 0.38 |
| 2 | 146348863 | rs7573795 | 0.101 | -0.197 | -0.46 | 0.205 | -0.025 | -0.001 | -0.001 | 0.018 | 0.13 | 0.16 | 0.13 | 0.17 | 0.06 | 0.09 |
| 2 | 146349854 | rs1484917 | -1.561 | -0.348 | -0.513 | 0.173 | 0.012 | -0.03 | -0.03 | 0 | 0.57 | 0.82 | 0.67 | 0.61 | 0.61 | 0.51 |
| 2 | 146354458 | rs12476415 | 0.57 | -0.293 | -0.573 | 0.204 | -0.085 | -0.036 | -0.036 | 0.041 | 0.28 | 0.14 | 0.39 | 0.38 | 0.46 | 0.32 |
| 2 | 146355456 | rs4662248 | -1.277 | -0.163 | -0.499 | 0.363 | 0.004 | -0.014 | -0.014 | 0 | 0.53 | 0.82 | 0.65 | 0.62 | 0.58 | 0.49 |
| 2 | 146371961 | rs13396091 | -0.763 | -0.191 | -0.379 | 0.379 | 0.006 | 0.012 | 0.012 | 0 | 0.46 | 0.67 | 0.49 | 0.43 | 0.56 | 0.43 |
| 2 | 146381260 | rs10496984 | 0.521 | -0.703 | 0.206 | 0.74 | -0.036 | 0.04 | 0.04 | 0 | 0.08 | 0.14 | 0.16 | 0.18 | 0.01 | 0.07 |
| 2 | 146382236 | rs2381750 | 1.377 | -0.462 | -0.054 | 0.251 | 0.003 | 0.01 | 0.01 | 0.117 | 0.11 | 0.11 | 0.11 | 0.05 | 0.08 | 0.08 |
| 2 | 146392350 | rs12619445 | -1.221 | -0.332 | -0.056 | 0.2 | 0.003 | -0.006 | -0.006 | 0.193 | 0.26 | 0.69 | 0.36 | 0.52 | 0.18 | 0.31 |
| 2 | 146393483 | rs2381755 | -1.34 | -0.093 | -0.182 | 0.569 | 0.001 | -0.002 | -0.002 | 0 | 0.89 | 0.89 | 0.94 | 0.88 | 0.88 | 0.92 |
| 2 | 146404349 | rs4632277 | -0.424 | -0.016 | -0.642 | 0.704 | -0.002 | 0.055 | 0.055 | 0 | 0.14 | 0.2 | 0.28 | 0.5 | 0.11 | 0.25 |
| 2 | 146407820 | rs16825330 | -0.376 | 0.05 | -0.812 | 0.726 | 0 | 0.024 | 0.024 | 0 | 0.14 | 0.16 | 0.28 | 0.51 | 0.11 | 0.22 |
| 2 | 146408408 | rs10183073 | 0.477 | 0.122 | -0.731 | -0.088 | 0.001 | -0.004 | -0.004 | 0.101 | 0.11 | 0.09 | 0.13 | 0.03 | 0.11 | 0.11 |
| 2 | 146416386 | rs11896112 | 0.336 | 0.135 | -0.526 | 0.073 | -0.011 | -0.003 | -0.003 | 0 | 0.13 | 0.17 | 0.06 | 0.12 | 0.11 | 0.08 |
| 2 | 146422643 | rs2381757 | 0.628 | 0.228 | -1.203 | -0.397 | -0.007 | 0.022 | 0.022 | 0 | 0.3 | 0.19 | 0.29 | 0.16 | 0.32 | 0.23 |
| 2 | 146423803 | rs11689535 | 0.959 | 0.443 | -1.112 | -0.4 | -0.001 | -0.009 | -0.009 | 0.009 | 0.07 | 0.07 | 0.09 | 0.01 | 0.11 | 0.07 |
| 2 | 146425531 | rs16825349 | 0.365 | 0.535 | -1.165 | -0.712 | -0.001 | 0.03 | 0.03 | 0 | 0.19 | 0.16 | 0.1 | 0.09 | 0.19 | 0.1 |
| 2 | 146426796 | rs11683083 | 0.383 | 0.417 | -1.162 | -0.708 | 0.005 | 0.028 | 0.028 | 0 | 0.19 | 0.16 | 0.1 | 0.1 | 0.18 | 0.1 |
| 2 | 146429247 | rs10170496 | 0.5 | 0.55 | -0.188 | -0.372 | 0.036 | -0.037 | -0.037 | 0.298 | 0.08 | 0.22 | 0.1 | 0.01 | 0.1 | 0.11 |
| 2 | 146442306 | rs6709699 | -0.311 | 0.239 | -0.78 | -0.317 | -0.026 | 0.036 | 0.036 | 0 | 0.63 | 0.78 | 0.5 | 0.28 | 0.62 | 0.5 |
| 2 | 146446673 | rs12691712 | 0.082 | -0.169 | -0.074 | 0.35 | -0.015 | 0.067 | 0.067 | 0.059 | 0.82 | 0.7 | 0.63 | 0.4 | 0.88 | 0.79 |
| 2 | 146450099 | rs2381762 | 0.315 | -0.217 | 0.429 | 0.335 | -0.026 | 0.136 | 0.136 | 0.063 | 0.8 | 0.57 | 0.61 | 0.4 | 0.88 | 0.79 |
| 2 | 146455861 | rs2381766 | 0.442 | -0.448 | -0.404 | -0.396 | -0.027 | 0.239 | 0.239 | 0 | 0.48 | 0.11 | 0.35 | 0.13 | 0.55 | 0.48 |
| 2 | 146461554 | rs7592820 | -0.044 | -0.536 | 0.138 | -0.4 | 0.002 | -0.045 | -0.045 | 0.101 | 0.57 | 0.48 | 0.46 | 0.16 | 0.57 | 0.48 |
| 2 | 146479241 | rs6737874 | -0.038 | -0.546 | 0.119 | -0.415 | 0 | -0.01 | -0.01 | 0 | 0.57 | 0.52 | 0.46 | 0.16 | 0.57 | 0.48 |
| 2 | 146483034 | rs10197322 | 0.486 | -0.654 | 0.342 | -0.477 | 0.011 | 0.096 | 0.096 | 0 | 0.5 | 0.28 | 0.42 | 0.14 | 0.55 | 0.47 |
| 2 | 146490561 | rs2246147 | -0.708 | -0.664 | 0.222 | -0.302 | 0.001 | 0.004 | 0.004 | 0 | 0.86 | 0.81 | 0.76 | 0.38 | 0.9 | 0.78 |
| 2 | 146499959 | rs2704783 | 0.616 | -0.663 | 0.26 | -0.391 | -0.024 | 0.031 | 0.031 | 0.009 | 0.28 | 0.16 | 0.28 | 0.22 | 0.32 | 0.3 |
| 2 | 146501581 | rs11891563 | 1.251 | -0.567 | 0.487 | -0.387 | -0.018 | -0.016 | -0.016 | 3.842 | 0.06 | 0.1 | 0.04 | 0.02 | 0.02 | 0.01 |
| 2 | 146512376 | rs2577215 | -0.966 | -0.924 | 0.174 | -0.597 | -0.003 | 0.016 | 0.016 | 0 | 0.55 | 0.46 | 0.59 | 0.87 | 0.48 | 0.55 |
| 2 | 146512861 | rs16825450 | 0.84 | -0.585 | -0.068 | 0.456 | -0.034 | -0.072 | -0.072 | 0 | 0.12 | 0.02 | 0.06 | 0.24 | 0.1 | 0.16 |
| 2 | 146513307 | rs2704789 | -0.328 | -0.753 | 0.083 | 0.71 | 0.007 | -0.005 | -0.005 | 0 | 0.39 | 0.31 | 0.43 | 0.58 | 0.37 | 0.39 |
| 2 | 146516811 | rs1437711 | 0.729 | -0.539 | -0.311 | 0.658 | -0.019 | 0.013 | 0.013 | 0.75 | 0.28 | 0.12 | 0.3 | 0.3 | 0.3 | 0.27 |
| 2 | 146522726 | rs1370523 | -0.768 | -0.376 | -0.14 | 0.518 | -0.006 | 0.015 | 0.015 | 0 | 0.55 | 0.67 | 0.52 | 0.82 | 0.49 | 0.56 |
| 2 | 146528869 | rs16825477 | 0.991 | -0.057 | -0.364 | 0.97 | -0.014 | -0.067 | -0.067 | 0 | 0.11 | 0.02 | 0.05 | 0.23 | 0.08 | 0.16 |
| 2 | 146548018 | rs10427396 | 0.526 | -0.015 | 0.072 | 0.937 | -0.006 | 0.057 | 0.057 | 0.058 | 0.14 | 0.23 | 0.07 | 0.26 | 0.09 | 0.18 |
| 2 | 146548704 | rs10496990 | -0.894 | -0.296 | 0.324 | 0.984 | -0.027 | 0.046 | 0.046 | 3.412 | 0.28 | 0.58 | 0.28 | 0.57 | 0.19 | 0.32 |
| 2 | 146551147 | rs1437714 | -0.802 | -0.277 | 0.106 | 0.993 | -0.024 | 0.047 | 0.047 | 3.525 | 0.27 | 0.46 | 0.27 | 0.57 | 0.19 | 0.32 |
| 2 | 146553275 | rs16825539 | 0.725 | -0.148 | -0.143 | 1.005 | -0.002 | 0.025 | 0.025 | 0.431 | 0.13 | 0.18 | 0.07 | 0.26 | 0.09 | 0.18 |
| 2 | 146556096 | rs4591296 | 1.336 | 0.161 | -0.965 | 0.259 | -0.071 | -0.018 | -0.018 | 0.425 | 0.21 | 0.01 | 0.2 | 0.11 | 0.25 | 0.3 |
| 2 | 146578234 | rs1228061 | 0.594 | -0.167 | -0.444 | 0.587 | -0.037 | 0.017 | 0.017 | 0.11 | 0.23 | 0.1 | 0.26 | 0.23 | 0.28 | 0.18 |
| 2 | 146584312 | rs6735536 | -0.564 | -0.393 | 0.384 | 1.009 | -0.015 | 0.01 | 0.01 | 3.623 | 0.3 | 0.4 | 0.27 | 0.62 | 0.2 | 0.35 |
| 2 | 146595034 | rs1437713 | -0.594 | -0.405 | 0.266 | 0.973 | -0.039 | 0.014 | 0.014 | 2.545 | 0.31 | 0.49 | 0.33 | 0.65 | 0.19 | 0.35 |
| 2 | 146602191 | rs7608036 | 0.141 | -0.065 | 0.288 | 1.297 | 0.006 | 0.258 | 0.258 | 0 | 0.05 | 0.21 | 0.14 | 0.3 | 0.01 | 0.06 |
| 2 | 146618875 | rs1228091 | 0.599 | -0.326 | -0.187 | 1.203 | 0.02 | -0.01 | -0.01 | 3.725 | 0.71 | 0.25 | 0.72 | 0.34 | 0.82 | 0.65 |
| 2 | 146639739 | rs1922035 | 0.305 | 0.046 | -0.627 | 1.146 | -0.028 | -0.073 | -0.073 | 0 | 0.18 | 0.09 | 0.12 | 0.28 | 0.18 | 0.18 |
| 2 | 146644346 | rs11902459 | -0.261 | -0.13 | -0.591 | 0.788 | -0.011 | 0.035 | 0.035 | 0 | 0.33 | 0.51 | 0.44 | 0.55 | 0.3 | 0.24 |
| 2 | 146658324 | rs2890798 | 0.303 | -0.069 | -1.431 | 0.315 | -0.009 | 0.077 | 0.077 | 0.131 | 0.31 | 0.09 | 0.17 | 0.1 | 0.31 | 0.28 |
| 2 | 146660741 | rs1317530 | 0.054 | 0.277 | -0.892 | 0.763 | -0.003 | -0.003 | -0.003 | 2.126 | 0.21 | 0.19 | 0.38 | 0.62 | 0.24 | 0.29 |
| 2 | 146666257 | rs10496992 | 0.796 | -0.172 | -1.46 | 0.338 | -0.104 | -0.088 | -0.088 | 0.106 | 0.33 | 0.05 | 0.26 | 0.32 | 0.38 | 0.29 |
| 2 | 146672778 | rs997033 | -0.813 | -0.325 | -0.901 | 0.207 | -0.006 | 0.085 | 0.085 | 0 | 0.65 | 0.79 | 0.69 | 0.91 | 0.64 | 0.65 |
| 2 | 146679403 | rs1515238 | -0.091 | -0.373 | -0.278 | 0.869 | -0.001 | -0.004 | -0.004 | 0 | 0.21 | 0.19 | 0.38 | 0.59 | 0.23 | 0.3 |
| 2 | 146689981 | rs12612543 | -0.322 | -0.312 | 0.204 | 0.376 | 0.002 | 0.023 | 0.023 | 0 | 0.23 | 0.32 | 0.31 | 0.32 | 0.23 | 0.29 |
| 2 | 146691431 | rs10200377 | 0.88 | -0.562 | -0.038 | 0.235 | 0.025 | 0.008 | 0.008 | 0 | 0.66 | 0.24 | 0.63 | 0.68 | 0.74 | 0.64 |
| 2 | 146708258 | rs2381864 | 0.56 | -0.585 | 0.75 | 0.087 | 0.009 | -0.017 | -0.017 | 0.192 | 0.93 | 0.73 | 0.96 | 1 | 0.97 | 0.96 |
| 2 | 146715742 | rs12617212 | -0.154 | -0.724 | 0.763 | 0.167 | 0.001 | 0.004 | 0.004 | 0 | 0.2 | 0.12 | 0.12 | 0.19 | 0.2 | 0.15 |
| 2 | 146719809 | rs2381866 | 1.188 | -1.019 | -0.2 | -0.318 | -0.14 | -0.084 | -0.084 | 0 | 0.35 | 0.06 | 0.38 | 0.59 | 0.42 | 0.36 |
| 2 | 146736345 | rs10173102 | -0.222 | -1.067 | 0.43 | -0.33 | -0.004 | -0.003 | -0.003 | 0 | 0.26 | 0.27 | 0.23 | 0.24 | 0.22 | 0.25 |
| 2 | 146737198 | rs16825854 | 0.257 | -1.116 | 0.338 | -0.535 | -0.001 | -0.001 | -0.001 | 0.13 | 0.19 | 0.16 | 0.08 | 0.17 | 0.12 | 0.14 |
| 2 | 146740793 | rs1515241 | 0.653 | -1.216 | 0.435 | -0.582 | 0.044 | 0.045 | 0.045 | 0.518 | 0.72 | 0.26 | 0.86 | 0.82 | 0.85 | 0.82 |
| 2 | 146742030 | rs16825861 | 0.768 | -1.343 | 0.325 | -0.686 | -0.033 | 0.073 | 0.073 | 0 | 0.32 | 0.16 | 0.38 | 0.17 | 0.38 | 0.36 |
| 2 | 146742779 | rs13010057 | 0.605 | -1.261 | 0.46 | -0.613 | 0.002 | -0.002 | -0.002 | 0 | 0.05 | 0.04 | 0.11 | 0.06 | 0.06 | 0.1 |
| 2 | 146752161 | rs2130858 | 1.098 | -1.279 | 0.422 | -0.605 | 0.076 | 0.067 | 0.067 | 0.7 | 0.66 | 0.22 | 0.75 | 0.77 | 0.8 | 0.72 |
| 2 | 146774636 | rs6714304 | 0.02 | -1.212 | 0.9 | -0.603 | 0.003 | -0.025 | -0.025 | 0.014 | 0.92 | 0.7 | 0.96 | 1 | 0.97 | 0.95 |
| 2 | 146775353 | rs13404625 | 0.577 | -1.119 | 0.836 | -0.627 | -0.002 | 0.001 | 0.001 | 0 | 0.05 | 0.04 | 0.11 | 0.04 | 0.07 | 0.1 |
| 2 | 146776699 | rs16825929 | 0.52 | -1.262 | 1.071 | -0.692 | -0.03 | -0.014 | -0.014 | 0.938 | 0.18 | 0.24 | 0.08 | 0.15 | 0.1 | 0.13 |
| 2 | 146779243 | rs1515252 | 0.651 | -1.589 | 0.81 | -0.816 | -0.026 | 0.075 | 0.075 | 0 | 0.33 | 0.18 | 0.38 | 0.17 | 0.38 | 0.36 |

| | | | | | | | | | | | | | | | | |
|---|---|---|---|---|---|---|---|---|---|---|---|---|---|---|---|---|
| 2 | 146779528 | rs2035786 | 0.697 | -1.534 | 0.939 | -0.751 | 0.047 | 0.043 | 0.043 | 0.525 | 0.71 | 0.29 | 0.87 | 0.84 | 0.85 | 0.82 |
| 2 | 146789314 | rs4662448 | -0.045 | -1.36 | 1.041 | -0.741 | 0.012 | -0.022 | -0.022 | 0 | 0.92 | 0.67 | 0.96 | 1 | 0.97 | 0.95 |
| 2 | 146815737 | rs7594453 | -0.761 | -1.603 | 0.905 | -0.868 | 0.01 | -0.054 | -0.054 | 0 | 0.68 | 0.51 | 0.63 | 0.83 | 0.64 | 0.65 |
| 2 | 146819721 | rs7583107 | -0.867 | -1.599 | 0.914 | -0.866 | 0.009 | -0.04 | -0.04 | 0 | 0.69 | 0.57 | 0.63 | 0.83 | 0.64 | 0.65 |
| 2 | 146822909 | rs12612023 | 1.53 | -1.592 | 0.677 | -0.804 | -0.185 | -0.07 | -0.07 | 0 | 0.35 | 0.03 | 0.38 | 0.64 | 0.43 | 0.36 |
| 2 | 146824868 | rs16825985 | 0.363 | -1.582 | 0.766 | -0.794 | -0.035 | -0.02 | -0.02 | 5.102 | 0.19 | 0.27 | 0.09 | 0.15 | 0.12 | 0.14 |
| 2 | 146835908 | rs1515246 | 0.906 | -1.565 | 0.702 | -0.788 | -0.061 | 0.054 | 0.054 | 0.003 | 0.41 | 0.14 | 0.51 | 0.67 | 0.49 | 0.46 |
| 2 | 146835932 | rs10496995 | 0.631 | -1.489 | 0.798 | -0.762 | -0.03 | -0.015 | -0.015 | 0 | 0.17 | 0.22 | 0.09 | 0.15 | 0.11 | 0.13 |
| 2 | 146853654 | rs11902246 | 0.196 | -1.35 | 1.037 | -0.734 | 0.017 | 0.026 | 0.026 | 0 | 0.06 | 0.17 | 0.12 | 0.05 | 0.06 | 0.1 |
| 2 | 146857571 | rs13025579 | 0.196 | -1.35 | 1.037 | -0.787 | 0.017 | 0.008 | 0.008 | 0 | 0.06 | 0.17 | 0.12 | 0.04 | 0.06 | 0.1 |
| 2 | 146858890 | rs1602445 | -0.224 | -1.45 | 0.994 | -0.865 | -0.002 | -0.035 | -0.035 | 0.886 | 0.62 | 0.53 | 0.61 | 0.84 | 0.61 | 0.59 |
| 2 | 146889404 | rs11680349 | 0.725 | -1.697 | 0.57 | -0.849 | 0.007 | 0.009 | 0.009 | 0 | 0.71 | 0.29 | 0.76 | 0.81 | 0.86 | 0.73 |
| 2 | 146891701 | rs222810 | -0.245 | -1.673 | 0.48 | -0.816 | -0.09 | -0.065 | -0.065 | 0 | 0.23 | 0.51 | 0.12 | 0.16 | 0.1 | 0.17 |
| 2 | 146897275 | rs4407186 | -0.135 | -1.693 | 0.316 | -0.809 | -0.078 | -0.053 | -0.053 | 0.067 | 0.22 | 0.56 | 0.13 | 0.16 | 0.11 | 0.19 |
| 2 | 146906634 | rs12473657 | 0.755 | -1.817 | -0.056 | -0.976 | -0.069 | 0.086 | 0.086 | 0 | 0.28 | 0.07 | 0.33 | 0.14 | 0.38 | 0.36 |
| 2 | 146921487 | rs10185692 | -0.139 | -1.857 | 0.21 | -0.994 | 0.064 | 0.103 | 0.103 | 1.768 | 0.05 | 0.45 | 0.05 | 0.05 | 0 | 0.06 |
| 2 | 146926010 | rs2381906 | 0.88 | -1.842 | -0.028 | -0.994 | -0.058 | 0.074 | 0.074 | 0 | 0.28 | 0.09 | 0.33 | 0.14 | 0.38 | 0.36 |
| 2 | 146931550 | rs10490592 | 0.595 | -1.919 | 0.195 | -0.89 | 0.019 | 0.018 | 0.018 | 0.532 | 0.16 | 0.1 | 0.08 | 0.11 | 0.11 | 0.13 |
| 2 | 146932243 | rs2194675 | 1.323 | -1.719 | 0.369 | -0.813 | -0.012 | -0.015 | -0.015 | 0.046 | 0.8 | 0.24 | 0.79 | 0.9 | 0.91 | 0.82 |
| 2 | 146961859 | rs16826097 | 0.147 | -1.458 | -0.004 | -0.676 | 0 | 0 | 0 | 0 | 0.06 | 0.05 | 0.02 | 0 | 0.06 | 0.03 |
| 2 | 146963650 | rs2059419 | 0.614 | -1.62 | -0.332 | -0.902 | -0.06 | 0.073 | 0.073 | 0 | 0.28 | 0.08 | 0.32 | 0.14 | 0.37 | 0.39 |
| 2 | 146964115 | rs7597570 | 0.939 | -1.732 | -0.403 | -1.507 | -0.004 | 0.198 | 0.198 | 0.449 | 0.5 | 0.14 | 0.48 | 0.75 | 0.54 | 0.44 |
| 2 | 146971415 | rs12997609 | 1.423 | -1.792 | -0.651 | -1.663 | -0.01 | 0.033 | 0.033 | 0.101 | 0.07 | 0 | 0.02 | 0.02 | 0.05 | 0.14 |
| 2 | 146972549 | rs17735680 | 1.703 | -1.896 | -0.881 | -1.704 | -0.189 | -0.049 | -0.049 | 0 | 0.33 | 0.02 | 0.39 | 0.66 | 0.43 | 0.32 |
| 2 | 146991064 | rs2016340 | 0.919 | -1.642 | -0.726 | -1.738 | -0.001 | 0.189 | 0.189 | 0.527 | 0.51 | 0.15 | 0.48 | 0.75 | 0.54 | 0.44 |
| 2 | 146992350 | rs10928284 | -0.021 | -1.418 | -0.69 | -1.673 | -0.001 | -0.004 | -0.004 | 2.861 | 0.65 | 0.39 | 0.63 | 0.8 | 0.66 | 0.56 |
| 2 | 146997640 | rs11692798 | 1.359 | -1.063 | -0.869 | -1.419 | -0.127 | -0.102 | -0.102 | 3.252 | 0.31 | 0.03 | 0.38 | 0.59 | 0.39 | 0.28 |
| 2 | 146998758 | rs6745791 | 0.453 | -1.132 | -0.424 | -1.418 | -0.013 | 0.024 | 0.024 | 5.186 | 0.92 | 0.35 | 0.92 | 0.93 | 0.98 | 0.96 |
| 2 | 147000282 | rs2194684 | 0.833 | -1.213 | -0.724 | -1.642 | -0.005 | 0.067 | 0.067 | 0.04 | 0.22 | 0.09 | 0.18 | 0.05 | 0.21 | 0.28 |
| 2 | 147009206 | rs1559608 | 0.199 | -1.317 | -0.406 | -1.47 | 0.022 | 0.068 | 0.068 | 4.254 | 0.92 | 0.44 | 0.93 | 0.93 | 0.98 | 0.96 |
| 2 | 147013450 | rs12614416 | 0.199 | -1.317 | -0.406 | -1.47 | 0.022 | 0.068 | 0.068 | 4.254 | 0.92 | 0.44 | 0.93 | 0.93 | 0.98 | 0.96 |
| 2 | 147024137 | rs2381911 | -0.05 | -1.277 | -0.633 | -1.465 | -0.006 | -0.028 | -0.028 | 0 | 0.69 | 0.35 | 0.74 | 0.88 | 0.76 | 0.66 |
| 2 | 147026018 | rs13013481 | -0.505 | -1.143 | -0.502 | -1.394 | 0.006 | 0.017 | 0.017 | 0.016 | 0.39 | 0.32 | 0.37 | 0.3 | 0.37 | 0.39 |
| 2 | 147045525 | rs2552538 | 0.309 | -0.831 | -0.015 | -1.011 | 0.019 | 0.256 | 0.256 | 0.029 | 0.92 | 0.54 | 0.82 | 0.79 | 0.99 | 0.86 |
| 2 | 147054952 | rs2552546 | -0.024 | -0.909 | -0.338 | -1.22 | -0.002 | 0.065 | 0.065 | 0 | 0.52 | 0.34 | 0.47 | 0.19 | 0.55 | 0.56 |
| 2 | 147076910 | rs6733867 | 0.356 | -0.604 | -0.755 | -1.074 | -0.083 | 0.008 | 0.008 | 0 | 0.41 | 0.18 | 0.41 | 0.6 | 0.45 | 0.3 |
| 2 | 147083675 | rs4662459 | 0.541 | -0.581 | -0.749 | -1.047 | -0.086 | -0.017 | -0.017 | 0 | 0.38 | 0.16 | 0.4 | 0.59 | 0.43 | 0.29 |
| 2 | 147087949 | rs2381914 | 0.842 | -0.838 | -0.938 | -1.255 | -0.127 | -0.061 | -0.061 | 1.042 | 0.29 | 0.07 | 0.38 | 0.65 | 0.44 | 0.32 |
| 2 | 147094385 | rs2890845 | 0.98 | -0.184 | 0.23 | -0.506 | 0.002 | 0.041 | 0.041 | 1.945 | 0.49 | 0.61 | 0.39 | 0.11 | 0.47 | 0.45 |
| 2 | 147100152 | rs6752130 | 0.695 | -0.172 | 0.256 | -0.514 | -0.031 | 0.052 | 0.052 | 0.164 | 0.25 | 0.1 | 0.26 | 0.15 | 0.35 | 0.26 |
| 2 | 147131153 | rs7584296 | 1.197 | -0.252 | 0.328 | -0.53 | 0.002 | -0.003 | -0.003 | 0 | 0.51 | 0.54 | 0.39 | 0.11 | 0.45 | 0.44 |
| 2 | 147137675 | rs4488597 | -0.692 | 0.05 | 0.453 | -0.282 | -0.018 | 0.07 | 0.07 | 0 | 0.94 | 0.84 | 0.9 | 0.86 | 0.99 | 0.89 |
| 2 | 147186982 | rs7559270 | -0.175 | 0.143 | -0.202 | -0.246 | -0.006 | -0.058 | -0.058 | 2.541 | 0.15 | 0.07 | 0.24 | 0.59 | 0.14 | 0.14 |
| 2 | 147190164 | rs12611768 | -1.501 | 0.844 | 0.168 | 0.139 | -0.006 | 0.019 | 0.019 | 4.154 | 0.45 | 0.38 | 0.6 | 0.89 | 0.52 | 0.55 |
| 2 | 147193915 | rs4522547 | -1.512 | 0.838 | 0.267 | 0.136 | -0.001 | -0.036 | -0.036 | 1.851 | 0.46 | 0.53 | 0.61 | 0.89 | 0.52 | 0.55 |
| 2 | 147219964 | rs11897013 | -0.917 | 1.001 | -0.576 | 0.225 | -0.015 | 0.025 | 0.025 | 0 | 0.11 | 0.24 | 0.11 | 0.14 | 0.04 | 0.16 |
| 2 | 147232091 | rs10928295 | -1.085 | 1.182 | -0.565 | 0.388 | -0.005 | 0.034 | 0.034 | 3.91 | 0.45 | 0.38 | 0.54 | 0.79 | 0.53 | 0.52 |
| 2 | 147242774 | rs6715004 | -0.348 | 0.911 | -0.389 | 0.673 | -0.002 | 0.003 | 0.003 | 2.238 | 0.31 | 0.25 | 0.36 | 0.29 | 0.35 | 0.38 |
| 2 | 147292648 | rs13429725 | -1.337 | 1.22 | 0.985 | 1.845 | 0.058 | 0.278 | 0.278 | 0.623 | 0.16 | 0.48 | 0.26 | 0.39 | 0.19 | 0.22 |
| 2 | 147318243 | rs12988074 | -1.521 | 1.088 | 1.436 | 2.103 | 0.029 | 0.211 | 0.211 | 1.21 | 0.14 | 0.39 | 0.23 | 0.37 | 0.16 | 0.24 |
| 2 | 147318659 | rs9653189 | -1.606 | 1.079 | 1.515 | 2.099 | 0.046 | 0.286 | 0.286 | 1.957 | 0.15 | 0.5 | 0.24 | 0.37 | 0.16 | 0.24 |
| 2 | 147356327 | rs4433946 | -0.47 | 1.021 | 0.848 | 1.296 | 0.001 | 0.004 | 0.004 | 1.739 | 0.33 | 0.26 | 0.35 | 0.22 | 0.31 | 0.43 |
| 2 | 147365709 | rs6720072 | -1.153 | 0.69 | 0.885 | 1.155 | -0.02 | 0.006 | 0.006 | 0 | 0.11 | 0.23 | 0.1 | 0.14 | 0.05 | 0.22 |
| 2 | 147380204 | rs10928303 | -1.138 | 0.674 | 0.876 | 1.462 | -0.019 | 0.037 | 0.037 | 0 | 0.11 | 0.24 | 0.12 | 0.2 | 0.04 | 0.23 |
| 2 | 147406770 | rs13421774 | 0.255 | 0.937 | 1.215 | 2.06 | -0.002 | -0.003 | -0.003 | 0 | 0.26 | 0.15 | 0.29 | 0.23 | 0.3 | 0.3 |
| 2 | 147422813 | rs4662263 | 1.15 | 0.814 | 1.048 | 1.941 | -0.032 | 0.056 | 0.056 | 0.002 | 0.87 | 0.66 | 0.88 | 0.8 | 0.95 | 0.76 |
| 2 | 147451283 | rs4275971 | -0.4 | 0.971 | 0.797 | 1.657 | -0.004 | 0.005 | 0.005 | 0.082 | 0.4 | 0.54 | 0.57 | 0.69 | 0.53 | 0.52 |
| 2 | 147492614 | rs4233577 | -0.35 | 1.162 | 0.785 | 1.948 | -0.001 | 0 | 0 | 1.683 | 0.41 | 0.56 | 0.45 | 0.52 | 0.56 | 0.49 |
| 2 | 147497999 | rs4467208 | 0.708 | 1.209 | 0.456 | 1.804 | 0.044 | 0.099 | 0.099 | 0.872 | 0.48 | 0.13 | 0.5 | 0.4 | 0.39 | 0.41 |
| 2 | 147504133 | rs4133034 | 0.48 | 1.291 | 0.365 | 1.682 | 0.038 | 0.08 | 0.08 | 0.254 | 0.5 | 0.31 | 0.5 | 0.33 | 0.39 | 0.34 |
| 2 | 147527155 | rs4280392 | 0.766 | 1.634 | 0.462 | 2.232 | 0.069 | 0.158 | 0.158 | 0 | 0.87 | 0.73 | 0.73 | 0.5 | 0.76 | 0.75 |
| 2 | 147529677 | rs12473513 | -0.312 | 1.219 | 0.008 | 1.867 | -0.001 | -0.017 | -0.017 | 0 | 0.08 | 0.05 | 0.01 | 0.08 | 0.04 | 0.09 |
| 2 | 147537807 | rs4459683 | 0.918 | 1.226 | -0.521 | 1.663 | 0.005 | 0.12 | 0.12 | 0.184 | 0.42 | 0.12 | 0.39 | 0.25 | 0.4 | 0.24 |
| 2 | 147539571 | rs6430137 | -0.524 | 1.434 | -0.21 | 1.919 | 0.025 | 0.03 | 0.03 | 0 | 0.48 | 0.67 | 0.39 | 0.59 | 0.38 | 0.58 |
| 2 | 147541992 | rs6430138 | 0.928 | 1.393 | -0.027 | 1.851 | 0.037 | 0.035 | 0.035 | 0 | 0.85 | 0.69 | 0.76 | 0.76 | 0.77 | 0.71 |
| 2 | 147558579 | rs9287403 | -0.633 | 1.008 | 0.313 | 1.869 | -0.001 | -0.001 | -0.001 | 0 | 0.45 | 0.51 | 0.33 | 0.37 | 0.39 | 0.54 |
| 2 | 147559341 | rs10193880 | -1.333 | 0.481 | 0.601 | 1.571 | -0.027 | 0.001 | 0.001 | 0 | 0.08 | 0.18 | 0.05 | 0.08 | 0.02 | 0.12 |
| 2 | 147586507 | rs6729729 | -1.073 | 0.381 | 1.025 | 1.584 | -0.021 | 0.008 | 0.008 | 0 | 0.11 | 0.28 | 0.06 | 0.09 | 0.06 | 0.1 |
| 2 | 147591712 | rs4491670 | -1.601 | 0.185 | 0.374 | 1.273 | -0.015 | 0.009 | 0.009 | 0 | 0.57 | 0.7 | 0.62 | 0.69 | 0.49 | 0.58 |

| | | | | | | | | | | | | | | | | |
|---|---|---|---|---|---|---|---|---|---|---|---|---|---|---|---|---|
| 2 | 147609864 | rs6746781 | 0.746 | -0.295 | -0.444 | 0.445 | -0.047 | 0.016 | 0.016 | 0 | 0.2 | 0.02 | 0.31 | 0.2 | 0.21 | 0.16 |
| 2 | 147612107 | rs7349350 | 0.472 | -0.306 | -0.32 | 0.436 | -0.003 | 0.025 | 0.025 | 0 | 0.21 | 0.09 | 0.32 | 0.22 | 0.21 | 0.16 |
| 2 | 147612734 | rs4128364 | -0.824 | -0.125 | -0.168 | 0.695 | -0.004 | -0.01 | -0.01 | 2.149 | 0.38 | 0.41 | 0.48 | 0.35 | 0.36 | 0.33 |
| 2 | 147674203 | rs13389478 | -1.641 | -0.797 | -0.125 | 0.979 | 0.023 | 0.018 | 0.018 | 0 | 0.48 | 0.65 | 0.52 | 0.47 | 0.41 | 0.45 |
| 2 | 147691096 | rs4524072 | -0.615 | -0.765 | 0.095 | 1.136 | 0.038 | 0.081 | 0.081 | 0 | 0.08 | 0.26 | 0.14 | 0.14 | 0.12 | 0.15 |
| 2 | 147722211 | rs2176056 | -0.381 | -0.918 | -0.074 | 1.026 | -0.014 | 0.012 | 0.012 | 0.069 | 0.1 | 0.28 | 0.07 | 0.11 | 0.06 | 0.11 |
| 2 | 147731153 | rs12468849 | -0.298 | -0.88 | -0.405 | 1.071 | 0.015 | -0.035 | -0.035 | 1.589 | 0.28 | 0.05 | 0.29 | 0.32 | 0.2 | 0.19 |
| 2 | 147745593 | rs6733155 | -1.305 | -0.847 | -0.093 | 0.978 | 0.004 | 0.003 | 0.003 | 0.358 | 0.44 | 0.42 | 0.38 | 0.4 | 0.31 | 0.27 |
| 2 | 147766421 | rs1546602 | -1.967 | -1.033 | 0.017 | 0.823 | 0.002 | 0.04 | 0.04 | 0.014 | 0.53 | 0.69 | 0.56 | 0.73 | 0.45 | 0.53 |
| 2 | 147777020 | rs1112719 | -1.151 | -1.377 | 0.873 | 0.896 | -0.037 | 0 | 0 | 0 | 0.13 | 0.32 | 0.07 | 0.11 | 0.06 | 0.05 |
| 2 | 147777746 | rs1606809 | -1.24 | -1.376 | 0.866 | 0.897 | -0.041 | 0.002 | 0.002 | 0.185 | 0.13 | 0.37 | 0.07 | 0.11 | 0.06 | 0.05 |
| 2 | 147787283 | rs6759721 | -0.525 | -1.259 | 0.757 | 0.972 | 0.026 | 0.007 | 0.007 | 0 | 0.06 | 0.19 | 0.12 | 0.05 | 0.09 | 0.11 |
| 2 | 147795457 | rs1515265 | -0.87 | -1.318 | 0.545 | 0.95 | -0.024 | -0.005 | -0.005 | 3.685 | 0.14 | 0.2 | 0.06 | 0.11 | 0.06 | 0.04 |
| 2 | 147804074 | rs2034526 | -1.706 | -1.482 | 0.667 | 0.926 | -0.045 | -0.024 | -0.024 | 0 | 0.23 | 0.56 | 0.2 | 0.16 | 0.15 | 0.16 |
| 2 | 147807932 | rs1376382 | 0.631 | -1.784 | 0.848 | 1.079 | -0.013 | -0.001 | -0.001 | 0 | 0.81 | 0.76 | 0.74 | 0.75 | 0.86 | 0.8 |
| 2 | 147812704 | rs2706035 | 2.007 | -2.183 | 1.076 | 1.331 | 0.037 | -0.01 | -0.01 | 0 | 0.57 | 0.3 | 0.55 | 0.46 | 0.69 | 0.56 |
| 2 | 147819724 | rs1020522 | -1.423 | -2.211 | 1.145 | 1.307 | -0.035 | 0.035 | 0.035 | 0 | 0.24 | 0.43 | 0.19 | 0.3 | 0.18 | 0.24 |
| 2 | 147824480 | rs10928337 | 0.001 | -2.21 | 1.144 | 1.299 | 0.008 | -0.018 | -0.018 | 0 | 0.16 | 0.06 | 0.25 | 0.25 | 0.14 | 0.18 |
| 2 | 147825689 | rs16826827 | -0.759 | -2.243 | 1.284 | 1.376 | -0.002 | 0.003 | 0.003 | 0 | 0.14 | 0.16 | 0.12 | 0.18 | 0.12 | 0.19 |
| 2 | 147826517 | rs6726059 | -0.346 | -2.201 | 1.163 | 1.337 | 0.004 | -0.001 | -0.001 | 0.008 | 0.18 | 0.16 | 0.25 | 0.26 | 0.14 | 0.18 |
| 2 | 147835933 | rs4662486 | -2.031 | -2.127 | 1.132 | 1.333 | -0.017 | 0.004 | 0.004 | 0 | 0.42 | 0.59 | 0.45 | 0.55 | 0.31 | 0.42 |
| 2 | 147843645 | rs373970 | 2.064 | -1.768 | 1.039 | 1.328 | 0.03 | 0.017 | 0.017 | 0.101 | 0.51 | 0.3 | 0.54 | 0.42 | 0.61 | 0.51 |
| 2 | 147855299 | rs7559672 | 0.214 | -1.804 | 0.929 | 1.327 | 0.003 | -0.018 | -0.018 | 0 | 0.16 | 0.07 | 0.25 | 0.27 | 0.15 | 0.19 |
| 2 | 147863014 | rs2650020 | 1.341 | -1.819 | 1.134 | 1.311 | -0.021 | 0.007 | 0.007 | 0.041 | 0.68 | 0.44 | 0.79 | 0.7 | 0.76 | 0.7 |
| 2 | 147868345 | rs16826971 | -0.249 | -1.823 | 0.958 | 1.345 | -0.001 | 0.005 | 0.005 | 0 | 0.1 | 0.12 | 0.07 | 0.18 | 0.08 | 0.18 |
| 2 | 147869810 | rs10221820 | -0.797 | -1.778 | 1.021 | 1.188 | 0.003 | -0.009 | -0.009 | 0 | 0.28 | 0.25 | 0.18 | 0.31 | 0.24 | 0.3 |
| 2 | 147880646 | rs2258224 | 1.968 | -1.864 | 0.999 | 1.446 | 0.047 | 0.128 | 0.128 | 0.01 | 0.49 | 0.23 | 0.52 | 0.23 | 0.61 | 0.36 |
| 2 | 147887551 | rs422792 | 0.413 | -1.84 | 1.1 | 1.474 | -0.013 | 0.004 | 0.004 | 0 | 0.81 | 0.75 | 0.74 | 0.73 | 0.85 | 0.68 |
| 2 | 147890980 | rs4662278 | -1.109 | -1.734 | 1.054 | 1.56 | -0.016 | 0.045 | 0.045 | 0 | 0.31 | 0.47 | 0.24 | 0.5 | 0.24 | 0.33 |
| 2 | 147905651 | rs6711477 | -0.144 | -1.657 | 0.311 | 1.477 | 0.008 | 0 | 0 | 0 | 0.21 | 0.11 | 0.26 | 0.27 | 0.19 | 0.21 |
| 2 | 147916442 | rs778087 | 0.548 | -1.755 | 0.533 | 1.474 | -0.016 | 0.04 | 0.04 | 0 | 0.73 | 0.51 | 0.8 | 0.69 | 0.77 | 0.62 |
| 2 | 147918261 | rs1083455 | 2.051 | -1.847 | 0.55 | 1.491 | 0.009 | 0.119 | 0.119 | 0.057 | 0.46 | 0.19 | 0.51 | 0.23 | 0.57 | 0.34 |
| 2 | 147922777 | rs706629 | 1.422 | -1.82 | 1.068 | 1.626 | 0.009 | -0.012 | -0.012 | 2.502 | 0.66 | 0.3 | 0.75 | 0.5 | 0.76 | 0.55 |
| 2 | 147927239 | rs365549 | 0.664 | -1.752 | 1.049 | 1.663 | -0.01 | 0.037 | 0.037 | 0 | 0.74 | 0.56 | 0.8 | 0.69 | 0.77 | 0.6 |
| 2 | 147933935 | rs16827083 | -0.31 | -1.699 | 0.835 | 1.691 | -0.005 | 0.151 | 0.151 | 1.142 | 0.06 | 0.21 | 0.05 | 0.19 | 0.01 | 0.05 |
| 2 | 147936676 | rs431727 | 0.826 | -1.43 | 0.502 | 1.144 | -0.014 | 0.037 | 0.037 | 0 | 0.73 | 0.54 | 0.8 | 0.69 | 0.77 | 0.62 |
| 2 | 147962263 | rs10497010 | -1.096 | -2.116 | 0.532 | 0.996 | -0.066 | 0.017 | 0.017 | 0.504 | 0.29 | 0.49 | 0.35 | 0.34 | 0.17 | 0.4 |
| 2 | 147992491 | rs16827115 | 0.046 | -1.934 | 0.479 | 1.1 | -0.014 | 0.025 | 0.025 | 0 | 0.06 | 0.06 | 0.16 | 0.14 | 0.02 | 0.11 |
| 2 | 147993969 | rs12476865 | 1.519 | -2.142 | -0.012 | 0.961 | -0.09 | 0.025 | 0.025 | 0 | 0.35 | 0.19 | 0.42 | 0.28 | 0.5 | 0.36 |
| 2 | 148014073 | rs16827139 | -0.357 | -1.762 | 0.727 | 1.065 | -0.019 | 0.04 | 0.04 | 0 | 0.06 | 0.11 | 0.19 | 0.14 | 0.02 | 0.12 |
| 2 | 148014862 | rs16827142 | -0.187 | -1.761 | 0.688 | 1.066 | -0.018 | 0.06 | 0.06 | 0 | 0.06 | 0.11 | 0.19 | 0.14 | 0.02 | 0.12 |
| 2 | 148029505 | rs4662497 | -0.03 | -1.862 | 0.502 | 1.041 | -0.003 | -0.024 | -0.024 | 0 | 0.17 | 0.04 | 0.13 | 0.19 | 0.16 | 0.22 |
| 2 | 148048320 | rs16827173 | -0.511 | -1.828 | 0.746 | 1.078 | -0.01 | 0.106 | 0.106 | 0.036 | 0.07 | 0.18 | 0.18 | 0.14 | 0.02 | 0.12 |
| 2 | 148048910 | rs16827175 | -1.312 | -1.849 | 0.583 | 1.008 | -0.042 | 0.019 | 0.019 | 0.019 | 0.28 | 0.5 | 0.34 | 0.34 | 0.19 | 0.4 |
| 2 | 148091377 | rs17720710 | -0.052 | -1.664 | 0.085 | 0.91 | -0.01 | -0.002 | -0.002 | 0 | 0.16 | 0.04 | 0.14 | 0.15 | 0.17 | 0.22 |
| 2 | 148103267 | rs939984 | -0.1 | -1.674 | 0.539 | 0.798 | 0.031 | -0.016 | -0.016 | 0 | 0.93 | 0.86 | 0.93 | 0.98 | 0.91 | 0.92 |
| 2 | 148104717 | rs1403263 | -0.899 | -1.795 | 0.917 | 0.903 | -0.01 | 0.006 | 0.006 | 0 | 0.23 | 0.23 | 0.31 | 0.27 | 0.19 | 0.32 |
| 2 | 148105737 | rs16827224 | -0.272 | -1.789 | 0.966 | 0.936 | -0.019 | 0.063 | 0.063 | 0 | 0.06 | 0.14 | 0.17 | 0.13 | 0.02 | 0.09 |
| 2 | 148114271 | rs4662500 | 1.357 | -2.397 | 0.789 | 0.879 | 0.011 | 0 | 0 | 0 | 0.56 | 0.42 | 0.55 | 0.59 | 0.7 | 0.47 |
| 2 | 148121393 | rs6430168 | 0.328 | -2.396 | 0.509 | 0.61 | 0.006 | 0.012 | 0.012 | 0 | 0.7 | 0.8 | 0.76 | 0.85 | 0.75 | 0.74 |
| 2 | 148122467 | rs12475208 | 0.227 | -2.393 | 0.372 | 0.71 | 0.018 | -0.006 | -0.006 | 0 | 0.22 | 0.04 | 0.15 | 0.21 | 0.16 | 0.18 |
| 2 | 148123250 | rs12476278 | -0.373 | -2.472 | 0.953 | 0.887 | 0.007 | 0 | 0 | 0 | 0.31 | 0.28 | 0.3 | 0.29 | 0.18 | 0.26 |
| 2 | 148131947 | rs16827260 | 0.274 | -2.456 | 0.725 | 0.888 | 0.042 | -0.006 | -0.006 | 0.577 | 0.24 | 0.04 | 0.15 | 0.21 | 0.14 | 0.21 |
| 2 | 148137464 | rs7567739 | 0.147 | -2.215 | 1.114 | 0.907 | 0.03 | -0.023 | -0.023 | 0 | 0.92 | 0.83 | 0.91 | 0.98 | 0.91 | 0.93 |
| 2 | 148148960 | rs16827299 | 0.273 | -2.427 | 1.344 | 0.923 | -0.022 | 0.041 | 0.041 | 0 | 0.08 | 0.23 | 0.06 | 0.11 | 0.01 | 0.04 |
| 2 | 148153368 | rs1439678 | 1.514 | -2.463 | 1.314 | 0.926 | -0.014 | 0.003 | 0.003 | 0.012 | 0.79 | 0.37 | 0.7 | 0.79 | 0.86 | 0.82 |
| 2 | 148154682 | rs7591837 | 1.365 | -2.291 | 0.484 | 1.208 | -0.121 | 0.027 | 0.027 | 0 | 0.4 | 0.11 | 0.43 | 0.29 | 0.51 | 0.48 |
| 2 | 148155161 | rs6746117 | 0.067 | -2.07 | 1.325 | 1.454 | -0.022 | 0.015 | 0.015 | 0 | 0.07 | 0.16 | 0.05 | 0.1 | 0.01 | 0.04 |
| 2 | 148165153 | rs1371040 | 1.074 | -2.019 | 1.203 | 1.439 | -0.02 | 0.033 | 0.033 | 0 | 0.81 | 0.62 | 0.7 | 0.79 | 0.86 | 0.82 |
| 2 | 148183865 | rs1550312 | 0.592 | -2.162 | 0.558 | 1.46 | 0.151 | 0.063 | 0.063 | 0.212 | 0.53 | 0.29 | 0.6 | 0.36 | 0.79 | 0.59 |
| 2 | 148192027 | rs934965 | 0.834 | -2.067 | 0.946 | 1.685 | -0.028 | 0.079 | 0.079 | 0.005 | 0.79 | 0.4 | 0.75 | 0.54 | 0.87 | 0.77 |
| 2 | 148198672 | rs2382041 | 0.203 | -1.975 | 0.837 | 1.736 | -0.023 | 0.193 | 0.193 | 0 | 0.84 | 0.7 | 0.82 | 0.56 | 0.9 | 0.79 |
| 2 | 148222783 | rs10210722 | -0.727 | -1.991 | 0.487 | 1.775 | 0.019 | 0.137 | 0.137 | 0.09 | 0.06 | 0.26 | 0.11 | 0.11 | 0.03 | 0.03 |
| 2 | 148226355 | rs1550322 | 0.459 | -1.875 | -0.559 | 1.507 | -0.017 | 0.038 | 0.038 | 0 | 0.8 | 0.78 | 0.85 | 0.63 | 0.84 | 0.76 |
| 2 | 148227502 | rs16827454 | 0.605 | -1.628 | -1.141 | 1.045 | 0.088 | 0.029 | 0.029 | 1.811 | 0.26 | 0.07 | 0.18 | 0.17 | 0.08 | 0.17 |
| 2 | 148241782 | rs6430189 | 1.049 | -0.47 | -0.635 | 1.848 | 0.02 | -0.026 | -0.026 | 0 | 0.71 | 0.27 | 0.7 | 0.56 | 0.82 | 0.62 |
| 2 | 148243073 | rs7580064 | 1.394 | 0.029 | -0.765 | 1.791 | 0.004 | -0.024 | -0.024 | 0 | 0.62 | 0.25 | 0.63 | 0.49 | 0.62 | 0.51 |
| 2 | 148245219 | rs877616 | -1.367 | -0.542 | -0.36 | 1.924 | -0.005 | 0.003 | 0.003 | 1.356 | 0.29 | 0.66 | 0.29 | 0.44 | 0.19 | 0.39 |
| 2 | 148259029 | rs13031932 | -1.028 | 0.108 | -1.004 | 1.786 | 0.012 | -0.055 | -0.055 | 0 | 0.11 | 0.01 | 0.1 | 0.15 | 0.11 | 0.18 |

| | | | | | | | | | | | | | | | | |
|---|---|---|---|---|---|---|---|---|---|---|---|---|---|---|---|---|
| 2 | 148261932 | rs17666990 | -1.144 | 1.211 | -1.328 | 1.815 | -0.082 | -0.048 | -0.048 | 0 | 0.21 | 0.05 | 0.29 | 0.23 | 0.32 | 0.39 |
| 2 | 148267476 | rs4662515 | 1.641 | 1.493 | -0.691 | 1.869 | -0.008 | -0.005 | -0.005 | 0.037 | 0.71 | 0.7 | 0.63 | 0.64 | 0.6 | 0.55 |
| 2 | 148276923 | rs723529 | 1.023 | 1.227 | -0.428 | 1.804 | 0.036 | 0.04 | 0.04 | 0 | 0.93 | 0.76 | 0.91 | 0.87 | 0.92 | 0.94 |
| 2 | 148282050 | rs10197459 | 1.622 | 1.088 | 0.129 | 2.355 | -0.016 | 0.114 | 0.114 | 2.217 | 0.84 | 0.34 | 0.8 | 0.64 | 0.87 | 0.86 |
| 2 | 148286117 | rs934881 | 0.826 | 1.507 | -0.07 | 2.348 | -0.013 | -0.017 | -0.017 | 0 | 0.82 | 0.84 | 0.61 | 0.65 | 0.73 | 0.66 |
| 2 | 148287033 | rs1438873 | 0.986 | 0.992 | -0.272 | 1.609 | -0.009 | 0.016 | 0.016 | 0 | 0.61 | 0.69 | 0.5 | 0.64 | 0.55 | 0.48 |
| 2 | 148287410 | rs12992493 | -0.901 | 0.67 | 0.183 | 1.872 | -0.011 | 0.029 | 0.029 | 0 | 0.12 | 0.16 | 0.11 | 0.25 | 0.07 | 0.15 |
| 2 | 148300228 | rs934880 | 1.624 | 0.549 | 0.956 | 1.828 | -0.003 | -0.023 | -0.023 | 0 | 0.66 | 0.3 | 0.46 | 0.44 | 0.65 | 0.53 |
| 2 | 148304568 | rs13390757 | 1.899 | 0.526 | 0.679 | 1.79 | -0.024 | -0.005 | -0.005 | 0 | 0.46 | 0.23 | 0.36 | 0.38 | 0.47 | 0.33 |
| 2 | 148305357 | rs16827511 | -0.254 | 0.52 | 0.381 | 1.796 | 0.023 | -0.022 | -0.022 | 0 | 0.09 | 0.04 | 0.06 | 0.15 | 0.02 | 0.12 |
| 2 | 148310480 | rs7576755 | 0.078 | 0.599 | 0.669 | 1.737 | 0.013 | 0.01 | 0.01 | 0 | 0.82 | 0.78 | 0.6 | 0.68 | 0.72 | 0.68 |
| 2 | 148313839 | rs10184962 | 1.753 | 0.37 | 0.424 | 1.953 | -0.071 | 0.063 | 0.063 | 0.189 | 0.45 | 0.1 | 0.31 | 0.3 | 0.49 | 0.31 |
| 2 | 148318521 | rs12464146 | -0.363 | 0.256 | 0.41 | 2.027 | 0.032 | -0.019 | -0.019 | 0.032 | 0.11 | 0.04 | 0.07 | 0.14 | 0.02 | 0.11 |
| 2 | 148321813 | rs7578749 | -0.014 | 0.262 | 0.424 | 2.103 | -0.043 | -0.043 | -0.043 | 0.03 | 0.14 | 0.06 | 0.38 | 0.33 | 0.25 | 0.32 |
| 2 | 148327589 | rs12327971 | 2.001 | 0.2 | 0.529 | 2.056 | -0.04 | 0.073 | 0.073 | 0.373 | 0.46 | 0.15 | 0.32 | 0.29 | 0.49 | 0.31 |
| 2 | 148329807 | rs12691759 | 1.174 | 0.206 | 0.516 | 2.052 | -0.03 | -0.021 | -0.021 | 0 | 0.56 | 0.19 | 0.39 | 0.43 | 0.51 | 0.43 |
| 2 | 148336830 | rs1550316 | 1.925 | 0.3 | 0.388 | 1.923 | -0.056 | 0.083 | 0.083 | 0.302 | 0.45 | 0.13 | 0.31 | 0.3 | 0.48 | 0.3 |
| 2 | 148337163 | rs10181805 | -0.401 | 0.341 | 1.22 | 1.671 | -0.029 | -0.025 | -0.025 | 0.189 | 0.26 | 0.38 | 0.15 | 0.18 | 0.21 | 0.23 |
| 2 | 148347811 | rs2118769 | -0.881 | 1.094 | 0.912 | 2.067 | -0.015 | -0.01 | -0.01 | 0 | 0.16 | 0.11 | 0.43 | 0.48 | 0.31 | 0.39 |
| 2 | 148353164 | rs11883765 | -1.836 | 0.691 | 1.14 | 1.618 | -0.006 | 0.016 | 0.016 | 1.373 | 0.46 | 0.71 | 0.66 | 0.58 | 0.53 | 0.63 |
| 2 | 148356683 | rs6719751 | 1.547 | 0.932 | 1.439 | 1.673 | 0.003 | 0.024 | 0.024 | 0.442 | 0.55 | 0.45 | 0.37 | 0.29 | 0.46 | 0.32 |
| 2 | 148356872 | rs4586588 | -1.647 | 0.946 | 1.562 | 1.69 | -0.001 | 0.003 | 0.003 | 0.554 | 0.43 | 0.35 | 0.58 | 0.71 | 0.54 | 0.68 |
| 2 | 148365343 | rs4591297 | -0.366 | 0.986 | 1.548 | 1.649 | -0.002 | -0.012 | -0.012 | 0.555 | 0.25 | 0.2 | 0.14 | 0.08 | 0.21 | 0.21 |
| 2 | 148365565 | rs7557760 | -0.439 | 1.084 | 1.495 | 1.758 | -0.048 | -0.058 | -0.058 | 0.057 | 0.16 | 0.07 | 0.42 | 0.48 | 0.3 | 0.39 |
| 2 | 148368803 | rs13405066 | -0.67 | 0.877 | 1.727 | 1.613 | -0.005 | -0.012 | -0.012 | 0 | 0.27 | 0.28 | 0.18 | 0.23 | 0.23 | 0.29 |
| 2 | 148373440 | rs7594106 | 1.236 | 0.681 | 1.076 | 1.185 | 0.031 | 0.102 | 0.102 | 0 | 0.5 | 0.3 | 0.28 | 0.27 | 0.41 | 0.27 |
| 2 | 148374771 | rs4662532 | 1.429 | 1.035 | 1.165 | 0.846 | -0.017 | -0.017 | -0.017 | 1.314 | 0.84 | 0.6 | 0.83 | 0.9 | 0.91 | 0.94 |
| 2 | 148379602 | rs7590392 | 1.573 | 0.587 | 0.898 | 0.738 | 0.008 | 0.056 | 0.056 | 0 | 0.44 | 0.27 | 0.23 | 0.2 | 0.42 | 0.28 |
| 2 | 148387591 | rs12611929 | 2.04 | 0.565 | 0.661 | 0.661 | -0.08 | 0.101 | 0.101 | 0 | 0.38 | 0.03 | 0.18 | 0.15 | 0.37 | 0.26 |
| 2 | 148394342 | rs6709045 | 2.007 | 0.354 | 0.751 | 0.663 | -0.043 | 0.121 | 0.121 | 0 | 0.38 | 0.1 | 0.18 | 0.15 | 0.39 | 0.25 |
| 2 | 148398301 | rs1919443 | -1.121 | 0.404 | 1.084 | 0.718 | -0.003 | 0.059 | 0.059 | 0 | 0.12 | 0.26 | 0.16 | 0.15 | 0.11 | 0.08 |
| 2 | 148402109 | rs10199422 | 1.449 | 0.405 | 0.772 | 0.681 | 0.007 | 0.081 | 0.081 | 0 | 0.46 | 0.36 | 0.24 | 0.15 | 0.42 | 0.27 |
| 2 | 148406827 | rs1406448 | -0.676 | 0.623 | 0.719 | 0.938 | -0.054 | -0.06 | -0.06 | 0.055 | 0.87 | 0.96 | 0.63 | 0.41 | 0.72 | 0.59 |
| 2 | 148409679 | rs17676417 | 1.508 | 0.422 | 1.078 | 0.586 | 0.003 | 0.028 | 0.028 | 0 | 0.45 | 0.37 | 0.25 | 0.2 | 0.44 | 0.27 |
| 2 | 148409737 | rs17676429 | 1.659 | 0.413 | 1.105 | 0.583 | -0.017 | 0.115 | 0.115 | 0 | 0.43 | 0.16 | 0.22 | 0.18 | 0.44 | 0.27 |
| 2 | 148412107 | rs6710158 | 1.026 | 0.428 | 1.095 | 0.602 | 0 | 0.016 | 0.016 | 0 | 0.52 | 0.45 | 0.28 | 0.2 | 0.47 | 0.29 |
| 2 | 148418820 | rs10188716 | -1.021 | 0.446 | 0.945 | 0.617 | 0.031 | 0.089 | 0.089 | 0.363 | 0.08 | 0.31 | 0.14 | 0.12 | 0.05 | 0.06 |
| 2 | 148423372 | rs7583258 | -0.509 | 0.466 | 0.83 | 0.627 | 0.009 | 0.049 | 0.049 | 0.014 | 0.07 | 0.15 | 0.11 | 0.12 | 0.05 | 0.06 |
| 2 | 148432679 | rs4662539 | 0.428 | 0.493 | 0.324 | 0.659 | 0.008 | -0.01 | -0.01 | 0 | 0.58 | 0.5 | 0.39 | 0.3 | 0.53 | 0.34 |
| 2 | 148439796 | rs13394781 | 0.279 | 0.517 | -0.148 | 0.672 | -0.007 | 0.004 | 0.004 | 0 | 0.6 | 0.62 | 0.4 | 0.3 | 0.53 | 0.34 |
| 2 | 148453079 | rs16827689 | 1.105 | 0.792 | -1.197 | 0.621 | -0.017 | 0.087 | 0.087 | 0 | 0.39 | 0.11 | 0.2 | 0.18 | 0.34 | 0.26 |
| 2 | 148455301 | rs1547596 | 0.205 | 1.367 | -0.686 | 1.135 | -0.04 | -0.048 | -0.048 | 0 | 0.79 | 0.9 | 0.49 | 0.43 | 0.62 | 0.47 |
| 2 | 148460036 | rs6730140 | -0.579 | 1.006 | -0.681 | 1.183 | 0.006 | -0.008 | -0.008 | 0 | 0.12 | 0.13 | 0.19 | 0.17 | 0.11 | 0.07 |
| 2 | 148460947 | rs6733995 | -0.443 | 0.409 | 0.66 | 0.774 | -0.006 | -0.014 | -0.014 | 0 | 0.73 | 0.72 | 0.86 | 0.89 | 0.75 | 0.84 |
| 2 | 148461491 | rs1528838 | 0.312 | 0.503 | 0.772 | 0.804 | -0.002 | -0.002 | -0.002 | 1.294 | 0.18 | 0.13 | 0.4 | 0.36 | 0.31 | 0.5 |
| 2 | 148467554 | rs6711640 | 0.312 | 0.503 | 0.772 | 0.804 | -0.002 | -0.002 | -0.002 | 0 | 0.18 | 0.13 | 0.4 | 0.36 | 0.31 | 0.5 |
| 2 | 148473247 | rs1406444 | -0.29 | 0.41 | 0.938 | 0.75 | -0.033 | -0.063 | -0.063 | 0 | 0.71 | 0.57 | 0.84 | 0.89 | 0.75 | 0.84 |
| 2 | 148478767 | rs6706259 | 1.889 | 0.463 | 0.901 | 0.768 | 0.008 | 0.002 | 0.002 | 0 | 0.41 | 0.32 | 0.25 | 0.36 | 0.33 | 0.27 |
| 2 | 148480979 | rs1113958 | 2.164 | 0.48 | 0.632 | 0.782 | -0.017 | -0.047 | -0.047 | 0 | 0.37 | 0.11 | 0.22 | 0.36 | 0.31 | 0.27 |
| 2 | 148495285 | rs1406449 | -1.138 | 0.522 | 0.677 | 0.787 | -0.015 | -0.019 | -0.019 | 0 | 0.3 | 0.25 | 0.58 | 0.54 | 0.42 | 0.57 |
| 2 | 148496821 | rs13004499 | 2.089 | 0.426 | 0.801 | 0.765 | 0.002 | -0.006 | -0.006 | 0 | 0.38 | 0.17 | 0.25 | 0.35 | 0.32 | 0.27 |
| 2 | 148500974 | rs10928364 | 0.341 | 0.483 | 0.779 | 0.799 | 0.007 | 0.007 | 0.007 | 0 | 0.18 | 0.2 | 0.42 | 0.37 | 0.3 | 0.49 |
| 2 | 148505544 | rs12691767 | 1.39 | 0.163 | 1.021 | 0.676 | 0.015 | 0.017 | 0.017 | 0 | 0.47 | 0.38 | 0.29 | 0.35 | 0.37 | 0.29 |
| 2 | 148505928 | rs7607970 | -0.503 | 0.142 | 0.777 | 0.686 | 0.008 | 0.011 | 0.011 | 0 | 0.73 | 0.77 | 0.87 | 0.89 | 0.76 | 0.84 |
| 2 | 148515565 | rs4662293 | 1.365 | -0.003 | 0.983 | 0.706 | 0.001 | 0.001 | 0.001 | 0 | 0.43 | 0.38 | 0.27 | 0.35 | 0.33 | 0.27 |
| 2 | 148517019 | rs13026223 | 1.721 | 0.025 | 0.967 | 0.719 | 0.008 | 0.004 | 0.004 | 0 | 0.4 | 0.23 | 0.26 | 0.35 | 0.32 | 0.27 |
| 2 | 148538149 | rs1364660 | 1.873 | -0.086 | 0.632 | 0.748 | -0.032 | -0.007 | -0.007 | 0 | 0.37 | 0.08 | 0.25 | 0.32 | 0.32 | 0.25 |
| 2 | 148542963 | rs1424949 | 1.35 | -0.204 | 0.876 | 0.773 | 0.015 | -0.007 | -0.007 | 0 | 0.44 | 0.34 | 0.39 | 0.46 | 0.33 | 0.3 |
| 2 | 148543509 | rs4972316 | 1.767 | -0.191 | 0.715 | 0.771 | -0.019 | -0.025 | -0.025 | 1.49 | 0.39 | 0.09 | 0.33 | 0.38 | 0.32 | 0.28 |
| 2 | 148563890 | rs13016400 | 0.95 | -0.15 | 0.651 | 0.774 | -0.017 | -0.006 | -0.006 | 0.229 | 0.65 | 0.32 | 0.48 | 0.57 | 0.54 | 0.45 |
| 2 | 148582309 | rs4972356 | 2.291 | -0.199 | 0.545 | 0.51 | -0.075 | -0.117 | -0.117 | 0.088 | 0.37 | 0.02 | 0.36 | 0.43 | 0.32 | 0.26 |
| 2 | 148596760 | rs7566230 | 0.125 | -0.108 | 0.886 | 0.512 | -0.04 | -0.033 | -0.033 | 0.704 | 0.23 | 0.15 | 0.34 | 0.4 | 0.4 | 0.54 |
| 2 | 148611528 | rs6747792 | 0.91 | -0.129 | 0.871 | 0.502 | -0.019 | 0.008 | 0.008 | 0 | 0.22 | 0.04 | 0.1 | 0.11 | 0.21 | 0.15 |
| 2 | 148619746 | rs1895694 | -0.056 | -0.124 | 1.061 | 0.503 | 0.011 | 0.007 | 0.007 | 0 | 0.25 | 0.27 | 0.36 | 0.41 | 0.4 | 0.55 |
| 2 | 148651458 | rs4972314 | 0.156 | -0.134 | 1.068 | 0.51 | 0.075 | 0.065 | 0.065 | 4.14 | 0.95 | 0.63 | 0.95 | 1 | 0.99 | 1 |
| 2 | 148657117 | rs1128919 | 1.883 | -0.116 | 0.681 | 0.561 | 0.018 | -0.008 | -0.008 | 0 | 0.42 | 0.29 | 0.46 | 0.47 | 0.33 | 0.28 |
| 2 | 148657589 | rs2288190 | -0.99 | 0.028 | 0.157 | 0.642 | 0.015 | -0.024 | -0.024 | 0 | 0.45 | 0.19 | 0.44 | 0.52 | 0.61 | 0.71 |
| 2 | 148687731 | rs17692648 | 0.858 | 0.056 | -0.028 | 0.584 | -0.011 | 0 | 0 | 0.966 | 0.22 | 0.04 | 0.09 | 0.11 | 0.2 | 0.15 |
| 2 | 148692776 | rs3768686 | 0.601 | 0.354 | 0.018 | 0.653 | -0.1 | -0.097 | -0.097 | 0 | 0.17 | 0.03 | 0.29 | 0.41 | 0.35 | 0.51 |

| | | | | | | | | | | | | | | | |
|---|---|---|---|---|---|---|---|---|---|---|---|---|---|---|---|
| 2 | 148711175 | rs6731371 | -0.198 | 0.131 | 0.479 | 0.634 | -0.001 | -0.002 | -0.002 | 0 | 0.76 | 0.79 | 0.65 | 0.59 | 0.59 | 0.46 |
| 2 | 148716428 | rs2307394 | 1.862 | -0.099 | 0.857 | 0.523 | 0.018 | -0.013 | -0.013 | 0.006 | 0.42 | 0.29 | 0.46 | 0.48 | 0.33 | 0.28 |
| 2 | 148746693 | rs2100469 | -0.565 | -0.119 | 0.876 | 0.516 | -0.018 | -0.015 | -0.015 | 0 | 0.78 | 0.84 | 0.65 | 0.59 | 0.59 | 0.46 |
| 2 | 148758479 | rs7598361 | 2.138 | -0.153 | 0.797 | 0.55 | -0.024 | -0.063 | -0.063 | 0 | 0.37 | 0.08 | 0.37 | 0.43 | 0.33 | 0.26 |
| 2 | 148778272 | rs897172 | 1.486 | -0.133 | 0.802 | 0.55 | 0.017 | -0.018 | -0.018 | 0 | 0.42 | 0.3 | 0.46 | 0.49 | 0.33 | 0.28 |
| 2 | 148817543 | rs1234428 | 0.609 | -0.243 | 1.047 | 0.586 | 0.007 | 0.004 | 0.004 | 0 | 0.19 | 0.15 | 0.1 | 0.1 | 0.12 | 0.14 |
| 2 | 148818437 | rs10497026 | 1.729 | -0.23 | 0.695 | 0.59 | -0.036 | -0.027 | -0.027 | 0 | 0.42 | 0.09 | 0.38 | 0.43 | 0.41 | 0.27 |
| 2 | 148892754 | rs11685710 | -0.22 | 0.074 | 1.043 | 0.675 | -0.029 | -0.021 | -0.021 | 0 | 0.25 | 0.2 | 0.36 | 0.4 | 0.41 | 0.55 |
| 2 | 148953669 | rs6430301 | -0.323 | 0.093 | 0.975 | 0.681 | -0.034 | -0.016 | -0.016 | 0 | 0.78 | 0.66 | 0.87 | 0.89 | 0.87 | 0.86 |
| 2 | 148954077 | rs7578002 | -0.104 | 0.119 | 0.408 | 0.694 | -0.066 | -0.036 | -0.036 | 1.152 | 0.76 | 0.53 | 0.85 | 0.89 | 0.87 | 0.86 |
| 2 | 148955901 | rs13395141 | -0.574 | 0.585 | -0.019 | 0.665 | -0.017 | -0.013 | -0.013 | 0 | 0.25 | 0.25 | 0.36 | 0.4 | 0.41 | 0.53 |
| 2 | 148965489 | rs1982387 | -1.273 | 0.304 | 0.21 | 0.171 | 0.014 | -0.011 | -0.011 | 0.134 | 0.1 | 0.19 | 0.19 | 0.05 | 0.13 | 0.07 |
| 2 | 148970147 | rs11694397 | 1.038 | -0.379 | 0.129 | -0.075 | 0.017 | 0.021 | 0.021 | 0.114 | 0.66 | 0.26 | 0.56 | 0.8 | 0.71 | 0.75 |
| 2 | 148970615 | rs1840140 | 0.447 | -0.414 | 0.009 | -0.088 | 0.016 | -0.014 | -0.014 | 0 | 0.17 | 0.08 | 0.18 | 0.15 | 0.12 | 0.15 |
| 2 | 148970738 | rs1840139 | -0.764 | -0.337 | 0.075 | -0.148 | -0.001 | 0 | 0 | 0 | 0.09 | 0.09 | 0.19 | 0.05 | 0.13 | 0.07 |
| 2 | 148977189 | rs2382215 | -0.402 | -0.319 | 0.474 | -0.029 | 0.001 | 0.002 | 0.002 | 0 | 0.2 | 0.25 | 0.37 | 0.18 | 0.24 | 0.22 |
| 2 | 148977583 | rs2382214 | 0.917 | -0.314 | 0.438 | -0.024 | 0.005 | -0.027 | -0.027 | 0 | 0.13 | 0.05 | 0.18 | 0.13 | 0.11 | 0.15 |
| 2 | 148996824 | rs13397840 | -0.613 | -0.241 | 0.532 | -0.064 | 0.027 | 0.007 | 0.007 | 0 | 0.06 | 0.12 | 0.19 | 0.05 | 0.13 | 0.07 |
| 2 | 149016812 | rs13032996 | 0.298 | -0.427 | -0.09 | -0.376 | -0.032 | 0.025 | 0.025 | 0 | 0.3 | 0.15 | 0.37 | 0.24 | 0.31 | 0.41 |
| 2 | 149022674 | rs991554 | 0.476 | -0.71 | -0.248 | -0.631 | 0.008 | 0.08 | 0.08 | 0 | 0.51 | 0.38 | 0.5 | 0.74 | 0.51 | 0.55 |
| 2 | 149037012 | rs7605678 | 0.316 | -0.62 | 0.019 | -0.612 | 0.007 | -0.033 | -0.033 | 0.601 | 0.64 | 0.48 | 0.51 | 0.82 | 0.57 | 0.74 |
| 2 | 149039379 | rs4972277 | 0.757 | -0.7 | -0.718 | -0.652 | -0.047 | 0.021 | 0.021 | 0 | 0.28 | 0.1 | 0.36 | 0.24 | 0.29 | 0.33 |
| 2 | 149055810 | rs13393889 | -0.335 | -0.634 | -0.749 | -0.635 | -0.012 | 0.019 | 0.019 | 0.017 | 0.33 | 0.28 | 0.46 | 0.18 | 0.44 | 0.26 |
| 2 | 149057121 | rs6713264 | 1.107 | -0.875 | -1.164 | -0.854 | 0.005 | 0.015 | 0.015 | 0 | 0.15 | 0.11 | 0.18 | 0.06 | 0.12 | 0.12 |
| 2 | 149079674 | rs11677584 | 1.176 | -0.697 | -0.963 | -0.734 | 0.001 | -0.001 | -0.001 | 0.221 | 0.18 | 0.22 | 0.22 | 0.15 | 0.22 | 0.15 |
| 2 | 149087536 | rs17276501 | 0.92 | -0.714 | -1.035 | -0.746 | -0.072 | 0.014 | 0.014 | 0 | 0.28 | 0.07 | 0.36 | 0.24 | 0.29 | 0.33 |
| 2 | 149091754 | rs4972279 | 1.269 | -0.709 | -1.091 | -0.752 | 0.029 | -0.087 | -0.087 | 0 | 0.31 | 0.12 | 0.13 | 0.55 | 0.25 | 0.4 |
| 2 | 149097747 | rs16828564 | 1.587 | -0.256 | -0.744 | -0.457 | 0 | 0 | 0 | 0 | 0.15 | 0.14 | 0.21 | 0.12 | 0.21 | 0.15 |
| 2 | 149109237 | rs16828574 | 0.541 | -0.26 | -0.705 | -0.459 | -0.007 | 0.007 | 0.007 | 0 | 0.08 | 0.18 | 0.17 | 0.08 | 0.07 | 0.08 |
| 2 | 149110873 | rs6744194 | 0.347 | -0.257 | -0.708 | -0.458 | 0.002 | 0.002 | 0.002 | 0 | 0.23 | 0.32 | 0.38 | 0.21 | 0.27 | 0.23 |
| 2 | 149119241 | rs7567204 | -0.239 | -0.169 | -0.787 | -0.478 | -0.006 | 0.013 | 0.013 | 0 | 0.57 | 0.45 | 0.5 | 0.76 | 0.53 | 0.63 |
| 2 | 149136540 | rs10205611 | 0.405 | -0.153 | -0.795 | -0.47 | 0.018 | 0.069 | 0.069 | 0 | 0.48 | 0.26 | 0.34 | 0.67 | 0.46 | 0.56 |
| 2 | 149154823 | rs13026304 | 0.356 | -0.148 | -0.766 | -0.456 | 0.015 | 0.072 | 0.072 | 0 | 0.48 | 0.26 | 0.35 | 0.67 | 0.47 | 0.56 |
| 2 | 149159429 | rs10199719 | -0.347 | -0.152 | -0.7 | -0.471 | 0.005 | -0.026 | -0.026 | 0 | 0.59 | 0.5 | 0.51 | 0.76 | 0.54 | 0.69 |
| 2 | 149164048 | rs12617242 | 1.127 | -0.118 | -0.578 | 0.042 | 0.01 | -0.019 | -0.019 | 0 | 0.15 | 0.11 | 0.04 | 0.29 | 0.1 | 0.18 |
| 2 | 149167661 | rs13430177 | 0.575 | 0.105 | -0.607 | -0.111 | 0 | 0.002 | 0.002 | 0 | 0.06 | 0.08 | 0.04 | 0.03 | 0.06 | 0.04 |
| 2 | 149167935 | rs10196996 | 0.775 | 0.132 | -0.621 | -0.152 | 0 | -0.024 | -0.024 | 0 | 0.31 | 0.22 | 0.26 | 0.42 | 0.31 | 0.33 |
| 2 | 149170115 | rs2103127 | -0.816 | 0.296 | -0.597 | -0.084 | -0.028 | -0.018 | -0.018 | 0 | 0.59 | 0.84 | 0.56 | 0.51 | 0.58 | 0.5 |
| 2 | 149178974 | rs11885393 | -0.954 | 0.243 | 0.304 | -0.225 | 0.003 | -0.084 | -0.084 | 0 | 0.22 | 0.48 | 0.25 | 0.06 | 0.21 | 0.13 |
| 2 | 149192933 | rs6706461 | -0.941 | 0.249 | -0.121 | -0.222 | 0 | -0.07 | -0.07 | 0 | 0.22 | 0.42 | 0.25 | 0.06 | 0.21 | 0.13 |
| 2 | 149206943 | rs4246608 | 0.771 | -0.142 | -0.421 | -0.326 | -0.015 | -0.061 | -0.061 | 0.564 | 0.82 | 0.77 | 0.94 | 1 | 0.91 | 0.97 |
| 2 | 149208646 | rs11690040 | 0.061 | 1 | -1.306 | -0.653 | -0.093 | 0.013 | 0.013 | 0 | 0.16 | 0.03 | 0.23 | 0.08 | 0.31 | 0.16 |
| 2 | 149210898 | rs1979038 | 0.869 | -0.285 | -0.902 | -0.927 | 0.032 | 0.028 | 0.028 | 0.045 | 0.75 | 0.2 | 0.87 | 0.98 | 0.9 | 0.92 |
| 2 | 149224187 | rs6755778 | 0.563 | -0.266 | 0.363 | -0.755 | -0.012 | -0.06 | -0.06 | 0 | 0.87 | 0.51 | 0.93 | 0.97 | 0.94 | 0.99 |
| 2 | 149229721 | rs13406161 | 0.916 | -0.321 | -0.486 | -0.201 | 0.02 | -0.05 | -0.05 | 0 | 0.62 | 0.21 | 0.56 | 0.42 | 0.7 | 0.57 |
| 2 | 149231816 | rs12475202 | 0.187 | -0.494 | 0.209 | -0.217 | -0.008 | -0.058 | -0.058 | 0 | 0.87 | 0.54 | 0.93 | 0.97 | 0.94 | 0.99 |
| 2 | 149234991 | rs1543255 | 1.02 | -0.774 | 0.072 | -0.395 | -0.04 | -0.076 | -0.076 | 0.242 | 0.81 | 0.39 | 0.89 | 0.97 | 0.91 | 0.96 |
| 2 | 149236792 | rs1867844 | 0.466 | -0.677 | 0.14 | -0.325 | -0.034 | -0.067 | -0.067 | 0 | 0.85 | 0.39 | 0.92 | 0.97 | 0.94 | 0.99 |
| 2 | 149243260 | rs2121344 | -0.246 | -0.671 | -0.409 | -0.288 | 0.049 | 0.081 | 0.081 | 0.003 | 0.54 | 0.29 | 0.77 | 0.72 | 0.64 | 0.74 |
| 2 | 149245727 | rs2166485 | 0.09 | -0.955 | 0.551 | 0.614 | -0.034 | 0.043 | 0.043 | 0.474 | 0.9 | 0.82 | 0.89 | 0.85 | 0.98 | 0.98 |
| 2 | 149250722 | rs4265953 | 1.331 | -1.345 | 0.301 | -0.258 | 0.055 | 0.137 | 0.137 | 0 | 0.56 | 0.26 | 0.52 | 0.09 | 0.72 | 0.44 |
| 2 | 149251702 | rs10928398 | -0.687 | -1.024 | 1.242 | 0.418 | -0.032 | 0.042 | 0.042 | 0.161 | 0.18 | 0.51 | 0.09 | 0.19 | 0.1 | 0.15 |
| 2 | 149263673 | rs12619515 | -1.04 | -1.037 | 1.342 | 0.463 | -0.033 | 0.017 | 0.017 | 1.708 | 0.19 | 0.6 | 0.1 | 0.22 | 0.1 | 0.15 |
| 2 | 149268931 | rs1404183 | 0.221 | -0.751 | 0.267 | 0.603 | -0.006 | 0.01 | 0.01 | 0 | 0.12 | 0.12 | 0.05 | 0.2 | 0.08 | 0.09 |
| 2 | 149271144 | rs16828718 | 0.242 | -0.686 | 0.205 | 0.492 | 0.001 | -0.007 | -0.007 | 0 | 0.1 | 0.07 | 0.05 | 0.16 | 0.08 | 0.07 |
| 2 | 149271642 | rs16828726 | -0.745 | -0.715 | 1.08 | 0.403 | 0.015 | 0.021 | 0.021 | 0 | 0.14 | 0.36 | 0.1 | 0.16 | 0.12 | 0.09 |
| 2 | 149277896 | rs10928399 | -0.797 | -0.458 | 1.173 | 0.655 | 0.012 | 0.044 | 0.044 | 0 | 0.15 | 0.19 | 0.39 | 0.65 | 0.17 | 0.39 |
| 2 | 149283070 | rs881184 | -0.666 | -0.935 | 1.037 | 0.602 | -0.015 | -0.016 | -0.016 | 0.184 | 0.13 | 0.15 | 0.06 | 0.05 | 0.05 | 0.08 |
| 2 | 149284068 | rs1108567 | 1.038 | -0.971 | 1.199 | 0.626 | 0.001 | -0.003 | -0.003 | 0 | 0.76 | 0.78 | 0.59 | 0.29 | 0.82 | 0.53 |
| 2 | 149284724 | rs1446554 | 1.435 | -1.275 | 1.449 | 0.69 | -0.01 | -0.001 | -0.001 | 0 | 0.71 | 0.36 | 0.85 | 0.7 | 0.87 | 0.82 |
| 2 | 149293831 | rs1550858 | -0.094 | -1.232 | 1.666 | 0.852 | -0.015 | 0.064 | 0.064 | 0 | 0.13 | 0.21 | 0.06 | 0.26 | 0.08 | 0.11 |
| 2 | 149306071 | rs3936439 | 1.771 | -1.238 | 1.606 | 0.733 | 0.015 | 0.039 | 0.039 | 0 | 0.59 | 0.35 | 0.49 | 0.09 | 0.7 | 0.45 |
| 2 | 149306450 | rs748519 | 1.395 | -1.239 | 1.71 | 0.804 | -0.037 | 0.013 | 0.013 | 0 | 0.71 | 0.47 | 0.84 | 0.68 | 0.82 | 0.8 |
| 2 | 149310384 | rs6712173 | 0.272 | -1.219 | 1.579 | 0.624 | 0.01 | 0.01 | 0.01 | 0.127 | 0.88 | 0.9 | 0.94 | 0.94 | 0.93 | 0.93 |
| 2 | 149312470 | rs7578059 | 0.32 | -1.221 | 1.558 | 0.662 | -0.035 | -0.069 | -0.069 | 0.124 | 0.11 | 0.03 | 0.34 | 0.62 | 0.12 | 0.34 |
| 2 | 149312728 | rs16828836 | 0.32 | -1.221 | 1.558 | 0.666 | -0.035 | -0.069 | -0.069 | 0 | 0.11 | 0.03 | 0.34 | 0.63 | 0.12 | 0.34 |
| 2 | 149314037 | rs10164839 | 0.42 | -1.258 | 1.624 | 0.643 | -0.005 | -0.006 | -0.006 | 0 | 0.87 | 0.84 | 0.93 | 0.94 | 0.93 | 0.93 |
| 2 | 149325251 | rs7606475 | -0.259 | -1.268 | 1.586 | 0.682 | -0.002 | -0.005 | -0.005 | 0 | 0.13 | 0.11 | 0.35 | 0.62 | 0.12 | 0.34 |
| 2 | 149325798 | rs16828910 | -2.237 | -1.332 | 1.742 | 0.603 | -0.01 | -0.06 | -0.06 | 0 | 0.4 | 0.53 | 0.51 | 0.91 | 0.3 | 0.54 |

| | | | | | | | | | | | | | | | | |
|---|---|---|---|---|---|---|---|---|---|---|---|---|---|---|---|---|
| 2 | 149326328 | rs13008779 | -1.431 | -1.303 | 1.544 | 0.648 | -0.001 | -0.001 | -0.001 | 0 | 0.26 | 0.28 | 0.42 | 0.68 | 0.19 | 0.41 |
| 2 | 149326704 | rs13010027 | 0.194 | -1.217 | 1.22 | 0.705 | -0.029 | -0.065 | -0.065 | 0 | 0.12 | 0.04 | 0.35 | 0.62 | 0.12 | 0.34 |
| 2 | 149330251 | rs921243 | 1.741 | -1.305 | 1.457 | 0.537 | 0.03 | 0.088 | 0.088 | 0 | 0.58 | 0.31 | 0.48 | 0.1 | 0.7 | 0.46 |
| 2 | 149337111 | rs7592501 | -0.686 | -1.321 | 1.541 | 0.551 | 0.014 | 0.015 | 0.015 | 0.022 | 0.13 | 0.1 | 0.06 | 0.06 | 0.07 | 0.07 |
| 2 | 149344429 | rs4972382 | 0.033 | -1.327 | 1.88 | 0.604 | -0.019 | 0.062 | 0.062 | 0 | 0.86 | 0.69 | 0.93 | 0.78 | 0.92 | 0.88 |
| 2 | 149350128 | rs720844 | -0.188 | -1.186 | 1.656 | 0.733 | 0.002 | 0.035 | 0.035 | 0 | 0.13 | 0.18 | 0.38 | 0.62 | 0.15 | 0.34 |
| 2 | 149361998 | rs6430331 | -0.928 | -1.238 | 1.549 | 0.745 | -0.015 | -0.017 | -0.017 | 0 | 0.13 | 0.17 | 0.07 | 0.05 | 0.07 | 0.07 |
| 2 | 149365347 | rs16829052 | -0.646 | -1.219 | 1.378 | 0.762 | -0.01 | -0.009 | -0.009 | 0 | 0.12 | 0.15 | 0.05 | 0.05 | 0.06 | 0.07 |
| 2 | 149372769 | rs16829057 | -1.178 | -1.207 | 1.238 | 0.739 | 0.007 | -0.038 | -0.038 | 0 | 0.23 | 0.14 | 0.4 | 0.67 | 0.18 | 0.39 |
| 2 | 149383708 | rs16829091 | -1.074 | -0.945 | 1.785 | 0.613 | 0.043 | 0.016 | 0.016 | 0.781 | 0.05 | 0.21 | 0.03 | 0.03 | 0.03 | 0.02 |
| 2 | 149394197 | rs7600461 | 1.369 | -1.658 | 1.242 | 0.473 | 0.043 | 0.007 | 0.007 | 5.006 | 0.61 | 0.31 | 0.54 | 0.41 | 0.79 | 0.59 |
| 2 | 149414439 | rs1530720 | 0.681 | -1.784 | 1.675 | 0.566 | -0.046 | 0.075 | 0.075 | 2.672 | 0.72 | 0.53 | 0.59 | 0.5 | 0.84 | 0.65 |
| 2 | 149417458 | rs16829136 | -0.596 | -1.778 | 1.293 | 0.568 | 0.03 | -0.068 | -0.068 | 0 | 0.25 | 0.1 | 0.38 | 0.5 | 0.16 | 0.35 |
| 2 | 149423647 | rs10497039 | 2.005 | -2.031 | 0.379 | -0.271 | -0.019 | 0.118 | 0.118 | 0 | 0.21 | 0.01 | 0.14 | 0.02 | 0.23 | 0.17 |
| 2 | 149449185 | rs6724787 | 1.675 | -1.887 | 0.4 | -0.118 | 0.106 | 0.231 | 0.231 | 1.461 | 0.51 | 0.23 | 0.48 | 0.11 | 0.71 | 0.44 |
| 2 | 149469713 | rs7558075 | 1.216 | -2.016 | 0.209 | -0.121 | 0.056 | 0.014 | 0.014 | 7.25 | 0.6 | 0.31 | 0.54 | 0.4 | 0.78 | 0.55 |
| 2 | 149487344 | rs4972294 | 0.502 | -1.822 | -0.131 | 0.004 | 0.037 | -0.104 | -0.104 | 0 | 0.21 | 0.05 | 0.36 | 0.47 | 0.13 | 0.33 |
| 2 | 149510282 | rs12614977 | 1.656 | -1.812 | -0.189 | 0.005 | -0.015 | -0.064 | -0.064 | 0 | 0.1 | 0.01 | 0.06 | 0.29 | 0.06 | 0.11 |
| 2 | 149529212 | rs16829255 | 0.17 | -1.57 | 0.189 | 0.192 | 0.041 | -0.131 | -0.131 | 1.758 | 0.22 | 0.04 | 0.36 | 0.47 | 0.13 | 0.33 |
| 2 | 149551882 | rs2121432 | -0.304 | -1.56 | 0.225 | 0.197 | 0.017 | -0.034 | -0.034 | 1.793 | 0.26 | 0.15 | 0.38 | 0.5 | 0.17 | 0.39 |
| 2 | 149557825 | rs2121434 | 0.619 | -0.963 | 0.135 | 0.312 | -0.01 | 0.013 | 0.013 | 0.356 | 0.68 | 0.36 | 0.55 | 0.49 | 0.77 | 0.57 |
| 2 | 149558625 | rs1374441 | -0.297 | -0.836 | -0.58 | 0.364 | -0.001 | -0.006 | -0.006 | 4.427 | 0.77 | 0.82 | 0.59 | 0.53 | 0.79 | 0.59 |
| 2 | 149566472 | rs12467520 | 1.003 | -0.508 | -1.487 | -0.175 | -0.171 | 0.196 | 0.196 | 2.446 | 0.35 | 0.05 | 0.35 | 0.08 | 0.46 | 0.25 |
| 2 | 149568396 | rs4499362 | -0.573 | -0.474 | -0.447 | 1.041 | -0.01 | 0.022 | 0.022 | 0 | 0.27 | 0.35 | 0.36 | 0.36 | 0.22 | 0.42 |
| 2 | 149568729 | rs10171238 | 0.566 | -0.576 | -1.555 | 0.221 | 0.039 | 0.054 | 0.054 | 0.651 | 0.63 | 0.29 | 0.61 | 0.16 | 0.77 | 0.46 |
| 2 | 149584946 | rs6430345 | -0.256 | -0.407 | -1.961 | 1.398 | 0.029 | -0.071 | -0.071 | 0.028 | 0.11 | 0.01 | 0.1 | 0.34 | 0.05 | 0.25 |
| 2 | 149587457 | rs1001644 | 0.174 | 0.857 | -2.58 | 0.779 | -0.088 | 0.071 | 0.071 | 0 | 0.24 | 0.07 | 0.51 | 0.12 | 0.38 | 0.26 |
| 2 | 149590270 | rs16829355 | -0.111 | 0.723 | -2.262 | 1.411 | -0.027 | -0.092 | -0.092 | 4.238 | 0.15 | 0.01 | 0.06 | 0.3 | 0.12 | 0.05 |
| 2 | 149604300 | rs12691814 | -0.373 | 1.59 | -2.455 | 1.314 | -0.117 | -0.125 | -0.125 | 0 | 0.17 | 0.02 | 0.21 | 0.38 | 0.32 | 0.28 |
| 2 | 149606989 | rs2166561 | -0.107 | 0.358 | -1.25 | 0.315 | -0.037 | -0.106 | -0.106 | 0.617 | 0.84 | 0.63 | 0.88 | 0.99 | 0.9 | 0.94 |
| 2 | 149624472 | rs3923181 | 0.276 | 0.13 | -0.577 | 0.203 | -0.052 | -0.122 | -0.122 | 1.902 | 0.81 | 0.44 | 0.87 | 0.99 | 0.9 | 0.94 |
| 2 | 149645404 | rs11899232 | -0.745 | -0.573 | -0.168 | -0.163 | -0.075 | -0.138 | -0.138 | 1.132 | 0.26 | 0.53 | 0.12 | 0.02 | 0.11 | 0.08 |
| 2 | 149647205 | rs12616265 | 1.105 | -0.504 | -1 | -0.108 | -0.058 | -0.095 | -0.095 | 0.191 | 0.17 | 0.06 | 0.18 | 0.38 | 0.23 | 0.12 |
| 2 | 149673362 | rs11885388 | -0.572 | -0.996 | -0.018 | -0.473 | -0.088 | -0.14 | -0.14 | 0 | 0.25 | 0.43 | 0.11 | 0.03 | 0.11 | 0.08 |
| 2 | 149797754 | rs35668735 | 0.314 | -0.921 | 0.343 | -0.451 | -0.022 | -0.079 | -0.079 | 0.005 | 0.14 | 0.22 | 0.09 | 0.01 | 0.09 | 0.05 |
| 2 | 149816918 | rs6710001 | -0.514 | -0.905 | 0.409 | -0.398 | -0.073 | -0.103 | -0.103 | 0 | 0.25 | 0.35 | 0.1 | 0.05 | 0.11 | 0.08 |
| 2 | 149830944 | rs6755213 | -0.496 | -0.944 | 0.788 | -0.417 | -0.084 | -0.118 | -0.118 | 0 | 0.26 | 0.42 | 0.11 | 0.05 | 0.11 | 0.08 |
| 2 | 149853960 | rs6738277 | -0.048 | -0.449 | 0.286 | -0.317 | -0.042 | -0.013 | -0.013 | 0 | 0.71 | 0.82 | 0.65 | 0.31 | 0.66 | 0.66 |
| 2 | 149858906 | rs6721330 | 1.288 | -0.599 | 0.391 | -0.446 | -0.049 | 0.124 | 0.124 | 1.236 | 0.45 | 0.15 | 0.53 | 0.25 | 0.56 | 0.57 |
| 2 | 149863406 | rs4452104 | 0.89 | -0.482 | 0.662 | -0.333 | 0.009 | 0.079 | 0.079 | 2.255 | 0.56 | 0.18 | 0.58 | 0.29 | 0.64 | 0.62 |
| 2 | 149865389 | rs7603107 | 0.791 | -0.54 | 0.882 | -0.33 | -0.058 | -0.048 | -0.048 | 1.002 | 0.11 | 0.17 | 0.02 | 0.02 | 0.02 | 0.04 |
| 2 | 149875961 | rs7577450 | 0.21 | -0.191 | 0.656 | -0.205 | -0.015 | -0.001 | -0.001 | 7.529 | 0.69 | 0.78 | 0.64 | 0.31 | 0.63 | 0.63 |
| 2 | 149886591 | rs1568853 | 0.447 | -0.007 | 0.465 | 0.244 | -0.074 | -0.08 | -0.08 | 0.143 | 0.22 | 0.05 | 0.24 | 0.37 | 0.27 | 0.22 |
| 2 | 149888184 | rs17347194 | 1.551 | -0.127 | 0.084 | -0.231 | -0.036 | -0.015 | -0.015 | 0 | 0.11 | 0.01 | 0.24 | 0.11 | 0.14 | 0.16 |
| 2 | 149892352 | rs10165688 | 0.392 | 0.114 | 0.607 | 0.062 | -0.02 | 0.015 | 0.015 | 0.092 | 0.63 | 0.29 | 0.59 | 0.31 | 0.63 | 0.63 |
| 2 | 149906958 | rs922860 | -0.683 | 0.306 | 0.229 | -0.069 | 0.016 | -0.072 | -0.072 | 0 | 0.87 | 0.67 | 0.89 | 0.97 | 0.87 | 0.94 |
| 2 | 149910003 | rs9636278 | 1.252 | -0.188 | -0.07 | 0.342 | -0.009 | -0.035 | -0.035 | 0 | 0.08 | 0.02 | 0.1 | 0.32 | 0.07 | 0.08 |
| 2 | 149912550 | rs1525171 | 0.36 | -0.187 | -0.399 | 0.188 | -0.034 | -0.083 | -0.083 | 0 | 0.23 | 0.08 | 0.22 | 0.38 | 0.22 | 0.23 |
| 2 | 149914069 | rs12622062 | -0.649 | -0.327 | -0.365 | -0.177 | 0.003 | -0.042 | -0.042 | 0 | 0.58 | 0.53 | 0.63 | 0.83 | 0.51 | 0.6 |
| 2 | 149915786 | rs1471649 | 0.386 | 0.152 | 0.11 | 0.19 | 0.036 | -0.007 | -0.007 | 1.206 | 0.53 | 0.3 | 0.5 | 0.51 | 0.62 | 0.55 |
| 2 | 149915977 | rs6723935 | -0.991 | 0.169 | -0.022 | 0.191 | -0.001 | -0.004 | -0.004 | 0 | 0.74 | 0.72 | 0.69 | 0.87 | 0.78 | 0.68 |
| 2 | 149916356 | rs6724307 | -0.651 | 0.473 | 0.216 | 1.064 | -0.016 | 0.047 | 0.047 | 0 | 0.19 | 0.29 | 0.18 | 0.38 | 0.16 | 0.13 |
| 2 | 149919348 | rs2222597 | -0.015 | 0.56 | 0.125 | 0.713 | -0.028 | -0.056 | -0.056 | 0 | 0.86 | 0.72 | 0.88 | 0.95 | 0.92 | 0.95 |
| 2 | 149919888 | rs11682788 | -0.599 | 1.224 | -0.62 | 1.249 | -0.027 | -0.032 | -0.032 | 0 | 0.14 | 0.08 | 0.2 | 0.35 | 0.19 | 0.21 |
| 2 | 149927066 | rs4667324 | -0.842 | 0.06 | -0.459 | 1.028 | -0.014 | 0.031 | 0.031 | 0.04 | 0.2 | 0.25 | 0.21 | 0.32 | 0.14 | 0.22 |
| 2 | 149927914 | rs17347516 | 0.256 | -0.477 | -1.409 | 0.19 | 0.041 | 0.118 | 0.118 | 1.16 | 0.31 | 0.07 | 0.13 | 0.08 | 0.2 | 0.12 |
| 2 | 149928090 | rs4667360 | -0.121 | -0.299 | -1.066 | 0.371 | 0.008 | -0.021 | -0.021 | 0.483 | 0.18 | 0.14 | 0.19 | 0.32 | 0.13 | 0.22 |
| 2 | 149931224 | rs10803787 | 0.236 | 0.307 | -0.699 | -0.104 | -0.003 | 0.006 | 0.006 | 0 | 0.38 | 0.37 | 0.35 | 0.17 | 0.4 | 0.46 |
| 2 | 149934100 | rs4622681 | 0.251 | 0.34 | -0.539 | -0.212 | -0.009 | 0.03 | 0.03 | 0 | 0.38 | 0.34 | 0.35 | 0.17 | 0.4 | 0.46 |
| 2 | 149936767 | rs2377510 | 0.025 | 0.448 | -0.498 | 0.409 | -0.081 | -0.1 | -0.1 | 0.749 | 0.53 | 0.07 | 0.47 | 0.5 | 0.44 | 0.38 |
| 2 | 149938538 | rs1525172 | 0.143 | 0.381 | 0.017 | 0.357 | 0.003 | -0.036 | -0.036 | 2.977 | 0.4 | 0.49 | 0.39 | 0.19 | 0.42 | 0.49 |
| 2 | 149940220 | rs6432716 | 0.066 | 0.229 | 0.06 | 0.105 | -0.001 | 0 | 0 | 0 | 0.53 | 0.52 | 0.45 | 0.48 | 0.55 | 0.59 |
| 2 | 149942519 | rs11900448 | -0.204 | 0.606 | 0.441 | -0.293 | 0.014 | 0.005 | 0.005 | 0 | 0.5 | 0.4 | 0.7 | 0.59 | 0.68 | 0.47 |
| 2 | 149946198 | rs16826516 | 1.247 | 0.413 | -0.029 | -0.295 | 0.004 | -0.107 | -0.107 | 2.334 | 0.18 | 0.04 | 0.21 | 0.39 | 0.14 | 0.19 |
| 2 | 149946582 | rs13005197 | 0.761 | 0.895 | 0.39 | -0.429 | -0.03 | -0.16 | -0.16 | 0 | 0.34 | 0.05 | 0.38 | 0.51 | 0.27 | 0.29 |
| 2 | 149948063 | rs2170606 | -0.323 | 1.02 | 0.557 | -0.569 | 0.137 | -0.035 | -0.035 | 0 | 0.14 | 0.32 | 0.27 | 0.08 | 0.34 | 0.16 |
| 2 | 149949244 | rs10930135 | -0.537 | 0.863 | 0.679 | -0.595 | 0.026 | 0.021 | 0.021 | 0 | 0.53 | 0.65 | 0.69 | 0.62 | 0.62 | 0.48 |
| 2 | 149958933 | rs11688597 | 0.343 | 0.65 | 0.873 | -0.612 | 0.028 | -0.025 | -0.025 | 0 | 0.38 | 0.24 | 0.43 | 0.55 | 0.28 | 0.37 |
| 2 | 149959855 | rs7592055 | 0.475 | 0.638 | 0.947 | -0.715 | 0.014 | 0.032 | 0.032 | 0.747 | 0.37 | 0.29 | 0.43 | 0.86 | 0.27 | 0.42 |

| | | | | | | | | | | | | | | | | |
|---|---|---|---|---|---|---|---|---|---|---|---|---|---|---|---|---|
| 2 | 149960273 | rs9784080 | -0.19 | 1.018 | 0.963 | -0.932 | 0.052 | -0.076 | -0.076 | 0 | 0.25 | 0.38 | 0.31 | 0.08 | 0.4 | 0.18 |
| 2 | 149972869 | rs7425814 | -0.588 | 0.801 | 0.981 | -0.973 | 0.001 | 0.042 | 0.042 | 0 | 0.61 | 0.68 | 0.73 | 0.93 | 0.63 | 0.6 |
| 2 | 149979291 | rs6720513 | -0.069 | 0.889 | 1.439 | -0.797 | 0.061 | 0.088 | 0.088 | 1.161 | 0.24 | 0.46 | 0.32 | 0.36 | 0.36 | 0.23 |
| 2 | 149979727 | rs7577550 | 0.133 | 0.726 | 1.153 | -1.084 | 0.007 | 0.14 | 0.14 | 0.048 | 0.82 | 0.48 | 0.85 | 0.63 | 0.87 | 0.75 |
| 2 | 149981513 | rs10803829 | 0.474 | 0.843 | 0.972 | -1.292 | 0.018 | -0.073 | -0.073 | 0 | 0.62 | 0.48 | 0.59 | 0.88 | 0.51 | 0.64 |
| 2 | 149981717 | rs12621827 | 1.361 | 0.785 | 0.632 | -1.296 | -0.003 | -0.124 | -0.124 | 0 | 0.33 | 0.05 | 0.39 | 0.54 | 0.25 | 0.36 |
| 2 | 149983918 | rs1026889 | 0.51 | 0.489 | 0.892 | -1.298 | 0.008 | -0.035 | -0.035 | 0 | 0.22 | 0.22 | 0.28 | 0.05 | 0.35 | 0.17 |
| 2 | 149985179 | rs17432490 | 1.05 | 0.58 | 0.959 | -1.341 | -0.002 | 0.001 | 0.001 | 0 | 0.24 | 0.21 | 0.17 | 0 | 0.26 | 0.22 |
| 2 | 149985897 | rs7600157 | -0.479 | 0.573 | 0.978 | -1.33 | -0.012 | -0.028 | -0.028 | 0 | 0.85 | 0.75 | 0.89 | 0.93 | 0.89 | 0.92 |
| 2 | 149986664 | rs6432964 | 0.122 | 0.593 | 1.105 | -1.341 | -0.002 | -0.056 | -0.056 | 3.591 | 0.61 | 0.58 | 0.7 | 0.93 | 0.63 | 0.71 |
| 2 | 149988911 | rs1404683 | 0.934 | 0.736 | 1.029 | -1.281 | -0.008 | -0.011 | -0.011 | 0.494 | 0.41 | 0.32 | 0.54 | 0.55 | 0.45 | 0.53 |
| 2 | 149999742 | rs678915 | 0.445 | 0.747 | 0.835 | -1.275 | 0.009 | 0.037 | 0.037 | 2.247 | 0.51 | 0.32 | 0.51 | 0.62 | 0.4 | 0.55 |
| 2 | 150002543 | rs7570723 | 0.657 | 0.928 | 1.176 | -1.163 | -0.003 | -0.007 | -0.007 | 0 | 0.61 | 0.38 | 0.59 | 0.56 | 0.55 | 0.69 |
| 2 | 150005013 | rs421468 | 0.4 | 0.855 | 1.105 | -0.956 | -0.007 | -0.033 | -0.033 | 0.81 | 0.12 | 0.29 | 0.12 | 0.07 | 0.09 | 0.03 |
| 2 | 150011247 | rs170672 | -0.923 | 1.06 | 1.101 | -0.868 | 0 | 0.029 | 0.029 | 0.058 | 0.81 | 0.79 | 0.85 | 0.63 | 0.83 | 0.82 |
| 2 | 150016500 | rs229333 | -0.845 | 1.059 | 1.143 | -0.869 | 0 | 0.061 | 0.061 | 0.028 | 0.8 | 0.74 | 0.84 | 0.63 | 0.83 | 0.82 |
| 2 | 150018835 | rs11695116 | 1.161 | 0.999 | 0.684 | -1.04 | -0.071 | -0.11 | -0.11 | 0 | 0.43 | 0.02 | 0.41 | 0.56 | 0.33 | 0.52 |
| 2 | 150020329 | rs229335 | -0.555 | 0.877 | 1.059 | -1.083 | -0.01 | 0.034 | 0.034 | 0 | 0.37 | 0.75 | 0.44 | 0.08 | 0.5 | 0.3 |
| 2 | 150023463 | rs10803869 | -0.874 | 0.517 | 1.226 | -0.642 | 0.019 | -0.004 | -0.004 | 0 | 0.89 | 0.64 | 0.87 | 0.93 | 0.91 | 0.97 |
| 2 | 150033334 | rs10183565 | 0.613 | 0.53 | 0.825 | -0.597 | -0.001 | -0.013 | -0.013 | 0.367 | 0.19 | 0.13 | 0.15 | 0.36 | 0.17 | 0.18 |
| 2 | 150036071 | rs4667348 | -0.929 | 0.549 | 0.908 | -0.621 | 0.019 | 0.001 | 0.001 | 0 | 0.9 | 0.65 | 0.87 | 0.93 | 0.91 | 0.97 |
| 2 | 150049955 | rs7576669 | -0.507 | 0.53 | 0.99 | -0.631 | 0.019 | -0.002 | -0.002 | 0 | 0.89 | 0.54 | 0.87 | 0.93 | 0.91 | 0.97 |
| 2 | 150052713 | rs10490386 | 0.472 | 0.512 | 0.908 | -0.607 | -0.001 | 0.015 | 0.015 | 0.522 | 0.2 | 0.21 | 0.16 | 0.36 | 0.17 | 0.19 |
| 2 | 150058128 | rs11686575 | -0.944 | 0.575 | 0.73 | -0.626 | 0.027 | 0.011 | 0.011 | 0 | 0.9 | 0.62 | 0.88 | 0.93 | 0.92 | 0.96 |
| 2 | 150065692 | rs1023632 | 0.002 | 0.559 | 0.239 | -0.602 | 0.003 | 0.04 | 0.04 | 0.879 | 0.24 | 0.37 | 0.29 | 0.37 | 0.28 | 0.24 |
| 2 | 150067228 | rs4667345 | 0.92 | 0.51 | -0.323 | -0.919 | -0.001 | -0.029 | -0.029 | 0 | 0.19 | 0.21 | 0.24 | 0 | 0.29 | 0.25 |
| 2 | 150071949 | rs6715376 | -0.594 | 0.486 | -0.33 | -0.906 | 0.003 | 0.004 | 0.004 | 4.6 | 0.43 | 0.43 | 0.41 | 0.37 | 0.46 | 0.43 |
| 2 | 150074246 | rs10207105 | 0.677 | 0.41 | -0.731 | -1.024 | -0.002 | 0.027 | 0.027 | 1.156 | 0.21 | 0.24 | 0.17 | 0.37 | 0.17 | 0.18 |
| 2 | 150077123 | rs10514634 | 0.702 | 0.212 | -1.483 | -0.916 | 0.015 | 0.05 | 0.05 | 1.309 | 0.2 | 0.09 | 0.12 | 0 | 0.18 | 0.19 |
| 2 | 150077190 | rs1035588 | 0.241 | 0.482 | -1.219 | -0.441 | -0.134 | -0.119 | -0.119 | 0.473 | 0.42 | 0.03 | 0.42 | 0.55 | 0.38 | 0.53 |
| 2 | 150092739 | rs7605137 | -0.78 | 0.315 | 0.504 | -0.701 | 0.024 | -0.003 | -0.003 | 1.789 | 0.08 | 0.53 | 0.05 | 0 | 0.02 | 0.01 |
| 2 | 150097468 | rs7584841 | -0.713 | 0.039 | -0.027 | -1.242 | 0.013 | 0.013 | 0.013 | 0 | 0.37 | 0.67 | 0.31 | 0.09 | 0.33 | 0.28 |
| 2 | 150133728 | rs16826803 | 0.247 | 0.117 | 0.463 | -1.3 | 0 | -0.011 | -0.011 | 2.448 | 0.26 | 0.25 | 0.24 | 0.09 | 0.21 | 0.2 |
| 2 | 150140202 | rs6706233 | -1.228 | 0.247 | -0.298 | -1.316 | 0.024 | 0.068 | 0.068 | 0 | 0.48 | 0.62 | 0.54 | 0.24 | 0.43 | 0.36 |
| 2 | 150146800 | rs751869 | 0.458 | 0.493 | -1.226 | -1.027 | 0.024 | 0.091 | 0.091 | 0 | 0.25 | 0.09 | 0.22 | 0.09 | 0.21 | 0.2 |
| 2 | 150147278 | rs10432496 | -0.243 | 0.962 | -0.691 | -0.396 | -0.024 | -0.032 | -0.032 | 0 | 0.12 | 0.02 | 0.16 | 0.15 | 0.11 | 0.12 |
| 2 | 150167831 | rs11689839 | -0.737 | 1.341 | -0.357 | -0.281 | 0.023 | 0.026 | 0.026 | 0 | 0.08 | 0.13 | 0.16 | 0.15 | 0.14 | 0.09 |
| 2 | 150176769 | rs12464726 | -0.461 | 1.083 | -0.768 | -0.584 | 0 | -0.001 | -0.001 | 0 | 0.63 | 0.72 | 0.66 | 0.4 | 0.71 | 0.51 |
| 2 | 150178701 | rs6717230 | -0.456 | 1.08 | -0.769 | -0.585 | 0 | -0.001 | -0.001 | 0 | 0.63 | 0.72 | 0.64 | 0.4 | 0.71 | 0.51 |
| 2 | 150184523 | rs10208058 | 0.035 | 1.107 | -0.698 | -0.704 | 0 | 0.047 | 0.047 | 0 | 0.56 | 0.66 | 0.51 | 0.26 | 0.65 | 0.45 |
| 2 | 150186142 | rs4667440 | 0.55 | 0.885 | -0.96 | -0.959 | 0.001 | 0.105 | 0.105 | 0 | 0.33 | 0.13 | 0.33 | 0.11 | 0.36 | 0.34 |
| 2 | 150188932 | rs1420356 | 0.307 | 0.925 | -0.693 | -0.745 | 0.001 | 0.099 | 0.099 | 0 | 0.47 | 0.26 | 0.41 | 0.11 | 0.47 | 0.43 |
| 2 | 150193025 | rs6735761 | -0.427 | 0.698 | -0.612 | -0.173 | -0.002 | 0.013 | 0.013 | 0 | 0.76 | 0.72 | 0.71 | 0.41 | 0.81 | 0.59 |
| 2 | 150194002 | rs10206766 | 0.091 | 0.639 | -0.245 | -0.162 | 0.004 | 0.028 | 0.028 | 0.283 | 0.25 | 0.41 | 0.28 | 0.35 | 0.3 | 0.15 |
| 2 | 150196099 | rs1990649 | -0.529 | 0.61 | -0.142 | -0.527 | -0.003 | 0.027 | 0.027 | 0 | 0.23 | 0.36 | 0.32 | 0.61 | 0.19 | 0.48 |
| 2 | 150196498 | rs17348064 | 0.32 | 0.399 | -0.917 | -1.053 | 0.002 | 0.011 | 0.011 | 0 | 0.34 | 0.22 | 0.29 | 0.28 | 0.37 | 0.15 |
| 2 | 150204378 | rs16826885 | 0.503 | 0.384 | -1.441 | -0.798 | 0.009 | -0.055 | -0.055 | 0 | 0.17 | 0.11 | 0.28 | 0.56 | 0.12 | 0.39 |
| 2 | 150206753 | rs1196696 | -1.083 | 0.32 | -1.426 | -0.98 | -0.059 | 0.146 | 0.146 | 0.094 | 0.6 | 0.84 | 0.63 | 0.89 | 0.54 | 0.66 |
| 2 | 150209892 | rs11897524 | -0.097 | 0.42 | -1.304 | -0.502 | 0.007 | 0.011 | 0.011 | 0 | 0.33 | 0.19 | 0.36 | 0.28 | 0.36 | 0.16 |
| 2 | 150212988 | rs1196699 | -0.381 | 0.047 | -1.947 | -1.632 | 0.057 | 0.285 | 0.285 | 0.068 | 0.5 | 0.86 | 0.68 | 0.9 | 0.38 | 0.59 |
| 2 | 150225950 | rs1196714 | -0.497 | -0.06 | -1.185 | -1.728 | 0.037 | 0.203 | 0.203 | 0 | 0.52 | 0.79 | 0.69 | 0.9 | 0.39 | 0.59 |
| 2 | 150227436 | rs888103 | 0.032 | -0.499 | -0.786 | -1.807 | -0.001 | -0.009 | -0.009 | 0 | 0.23 | 0.22 | 0.1 | 0.15 | 0.2 | 0.13 |
| 2 | 150230301 | rs4667437 | 0.675 | -0.618 | -0.401 | -2.005 | 0.01 | -0.001 | -0.001 | 0 | 0.23 | 0.18 | 0.52 | 0.75 | 0.18 | 0.44 |
| 2 | 150233741 | rs13027838 | 1.315 | -0.389 | -1.098 | -2.307 | -0.042 | 0.101 | 0.101 | 0.107 | 0.16 | 0.01 | 0.06 | 0 | 0.17 | 0.23 |
| 2 | 150237153 | rs2377376 | 1.652 | -0.504 | -0.999 | -2.187 | 0.016 | -0.037 | -0.037 | 0.306 | 0.18 | 0.1 | 0.49 | 0.71 | 0.12 | 0.42 |
| 2 | 150242490 | rs1196686 | 0.135 | -0.291 | -0.556 | -2.198 | -0.001 | -0.006 | -0.006 | 0 | 0.23 | 0.24 | 0.19 | 0.21 | 0.21 | 0.13 |
| 2 | 150247476 | rs1196687 | 0.962 | -0.329 | -1.046 | -2.31 | 0.02 | 0.196 | 0.196 | 0 | 0.53 | 0.15 | 0.67 | 0.79 | 0.61 | 0.6 |
| 2 | 150248022 | rs1196688 | -0.593 | -0.046 | -0.533 | -2.152 | -0.005 | 0.063 | 0.063 | 0.016 | 0.81 | 0.92 | 0.91 | 1 | 0.81 | 0.77 |
| 2 | 150270689 | rs7556835 | -0.359 | 0.021 | 0.068 | -2.03 | 0.009 | -0.034 | -0.034 | 0 | 0.23 | 0.5 | 0.23 | 0.14 | 0.2 | 0.21 |
| 2 | 150271975 | rs9288269 | 1.578 | -0.001 | -0.286 | -2.435 | -0.014 | 0.072 | 0.072 | 0.016 | 0.34 | 0.08 | 0.43 | 0.82 | 0.39 | 0.44 |
| 2 | 150274458 | rs6721513 | -0.618 | 0.014 | 0.456 | -1.881 | -0.003 | -0.029 | -0.029 | 0 | 0.24 | 0.6 | 0.24 | 0.14 | 0.2 | 0.21 |
| 2 | 150278005 | rs6434827 | -0.322 | 0.02 | 0.576 | -1.878 | 0.015 | -0.034 | -0.034 | 0 | 0.22 | 0.47 | 0.22 | 0.14 | 0.2 | 0.21 |
| 2 | 150305703 | rs11681264 | -0.043 | 0.046 | 0.678 | -1.86 | 0.004 | -0.005 | -0.005 | 0 | 0.2 | 0.22 | 0.2 | 0.14 | 0.2 | 0.19 |
| 2 | 150311503 | rs17433067 | -0.399 | 0.002 | 0.699 | -1.887 | 0 | -0.002 | -0.002 | 0 | 0.23 | 0.22 | 0.21 | 0.14 | 0.21 | 0.2 |
| 2 | 150313107 | rs1196642 | 1.308 | 0.049 | 0.635 | -1.917 | 0.006 | 0.039 | 0.039 | 0 | 0.59 | 0.31 | 0.7 | 0.85 | 0.64 | 0.65 |
| 2 | 150318916 | rs1196652 | -0.153 | 0.141 | 1.013 | -1.476 | -0.002 | -0.001 | -0.001 | 2.877 | 0.83 | 0.87 | 0.94 | 0.99 | 0.83 | 0.86 |
| 2 | 150321233 | rs1196656 | 1.214 | 0.058 | 0.932 | -1.444 | -0.01 | -0.034 | -0.034 | 0 | 0.7 | 0.36 | 0.76 | 0.87 | 0.76 | 0.74 |
| 2 | 150329973 | rs871 | -0.502 | 0.343 | 0.804 | -1.036 | -0.001 | -0.017 | -0.017 | 0.433 | 0.87 | 0.88 | 0.96 | 0.99 | 0.88 | 0.86 |
| 2 | 150334751 | rs7600746 | -1.058 | 0.371 | 1.061 | -1.018 | 0.035 | 0.105 | 0.105 | 0.901 | 0.1 | 0.47 | 0.25 | 0.13 | 0.1 | 0.13 |

| | | | | | | | | | | | | | | | | |
|---|---|---|---|---|---|---|---|---|---|---|---|---|---|---|---|---|
| 2 | 150341051 | rs1196677 | -0.521 | 0.374 | 1.014 | -1.047 | -0.001 | -0.017 | -0.017 | 0 | 0.87 | 0.88 | 0.96 | 0.99 | 0.88 | 0.86 |
| 2 | 150342476 | rs1196678 | 1.286 | 0.326 | 1.046 | -1.078 | -0.009 | -0.025 | -0.025 | 0.277 | 0.74 | 0.36 | 0.69 | 0.86 | 0.76 | 0.74 |
| 2 | 150344508 | rs10497046 | 0.137 | 0.519 | 1.049 | -0.94 | 0.026 | 0.087 | 0.087 | 0 | 0.07 | 0.22 | 0.14 | 0.13 | 0.08 | 0.11 |
| 2 | 150350535 | rs6716959 | 0.855 | 0.574 | 0.823 | -0.947 | 0.009 | -0.04 | -0.04 | 0 | 0.78 | 0.51 | 0.72 | 0.87 | 0.78 | 0.75 |
| 2 | 150373254 | rs7369134 | 0.671 | 0.592 | 0.593 | -0.939 | 0.006 | -0.027 | -0.027 | 0.012 | 0.79 | 0.63 | 0.8 | 0.86 | 0.78 | 0.76 |
| 2 | 150382926 | rs13422595 | -0.006 | 0.599 | 0.501 | -0.943 | 0.002 | 0.003 | 0.003 | 0 | 0.14 | 0.13 | 0.06 | 0.01 | 0.13 | 0.13 |
| 2 | 150384023 | rs6729581 | 0.07 | 0.639 | 0.41 | -0.852 | 0.061 | 0.133 | 0.133 | 0 | 0.07 | 0.31 | 0.14 | 0.13 | 0.08 | 0.1 |
| 2 | 150393466 | rs4667426 | 0.634 | 0.623 | -0.386 | -0.88 | -0.006 | -0.03 | -0.03 | 3.793 | 0.77 | 0.39 | 0.71 | 0.86 | 0.78 | 0.76 |
| 2 | 150425265 | rs10932440 | 0.701 | 1.061 | -1.761 | -1.287 | -0.038 | 0.151 | 0.151 | 0 | 0.59 | 0.2 | 0.52 | 0.86 | 0.6 | 0.68 |
| 2 | 150425584 | rs13020214 | 1.128 | 1.076 | -1.794 | -1.347 | -0.048 | 0.085 | 0.085 | 0.516 | 0.17 | 0.03 | 0.15 | 0 | 0.17 | 0.07 |
| 2 | 150425917 | rs10173317 | 0.679 | 1.095 | -1.705 | -1.35 | -0.018 | 0.105 | 0.105 | 0 | 0.6 | 0.26 | 0.53 | 0.86 | 0.62 | 0.68 |
| 2 | 150445221 | rs4530332 | 0.613 | 1.026 | -1.299 | -0.923 | 0 | 0.001 | 0.001 | 0.148 | 0.76 | 0.29 | 0.68 | 0.87 | 0.77 | 0.76 |
| 2 | 150450828 | rs4667337 | 0.613 | 1.026 | -1.299 | -0.923 | 0 | 0.001 | 0.001 | 0.148 | 0.76 | 0.3 | 0.68 | 0.86 | 0.77 | 0.76 |
| 2 | 150459870 | rs10490073 | 0.29 | 0.987 | -1.721 | -0.656 | -0.042 | 0.075 | 0.075 | 0.398 | 0.37 | 0.08 | 0.4 | 0.81 | 0.42 | 0.5 |
| 2 | 150468495 | rs759438 | 0.73 | 1.099 | -0.391 | 0.57 | 0.009 | 0.007 | 0.007 | 4.133 | 0.86 | 0.31 | 0.77 | 0.87 | 0.86 | 0.88 |
| 2 | 150491670 | rs17737841 | -0.675 | 1.274 | -0.385 | 1.463 | 0.005 | -0.005 | -0.005 | 0 | 0.2 | 0.17 | 0.28 | 0.17 | 0.26 | 0.2 |
| 2 | 150552390 | rs848223 | -0.092 | 0.735 | 0.003 | 1.271 | 0.004 | 0.001 | 0.001 | 0.278 | 0.06 | 0.05 | 0.03 | 0.05 | 0.05 | 0.05 |
| 2 | 150553984 | rs6727151 | -0.284 | 1.105 | 0.031 | 1.252 | 0.014 | 0.177 | 0.177 | 0.088 | 0.06 | 0.1 | 0.16 | 0.47 | 0.1 | 0.29 |
| 2 | 150572567 | rs7560430 | 1.13 | 1.305 | -0.364 | 0.375 | 0.008 | 0.002 | 0.002 | 0.077 | 0.66 | 0.57 | 0.67 | 0.36 | 0.56 | 0.47 |
| 2 | 150578924 | rs1526278 | -0.614 | 0.974 | -0.225 | 0.577 | -0.022 | -0.003 | -0.003 | 0 | 0.08 | 0.17 | 0.04 | 0.06 | 0.05 | 0.07 |
| 2 | 150583820 | rs848260 | -0.71 | 0.969 | -0.187 | 0.575 | -0.024 | -0.002 | -0.002 | 3.551 | 0.08 | 0.2 | 0.04 | 0.06 | 0.05 | 0.07 |
| 2 | 150588829 | rs1100185 | -0.584 | 0.97 | -0.197 | 0.575 | -0.021 | -0.003 | -0.003 | 0 | 0.08 | 0.16 | 0.04 | 0.06 | 0.05 | 0.07 |
| 2 | 150592070 | rs4667404 | 0.668 | 0.726 | -0.57 | -0.073 | -0.002 | 0.01 | 0.01 | 0 | 0.56 | 0.38 | 0.58 | 0.33 | 0.51 | 0.43 |
| 2 | 150597927 | rs4667402 | -0.596 | 0.771 | -0.852 | 0.228 | -0.005 | -0.01 | -0.01 | 0.013 | 0.16 | 0.13 | 0.22 | 0.48 | 0.21 | 0.26 |
| 2 | 150603067 | rs968871 | -0.717 | 0.818 | -0.885 | 0.062 | -0.007 | -0.011 | -0.011 | 0.146 | 0.35 | 0.3 | 0.34 | 0.54 | 0.4 | 0.45 |
| 2 | 150604671 | rs4667400 | -0.204 | 0.788 | -0.934 | 0.067 | -0.021 | -0.045 | -0.045 | 0 | 0.28 | 0.19 | 0.29 | 0.49 | 0.35 | 0.42 |
| 2 | 150604789 | rs4667399 | 0.375 | 0.821 | -0.958 | 0.102 | -0.033 | -0.032 | -0.032 | 0 | 0.1 | 0.03 | 0.18 | 0.42 | 0.15 | 0.23 |
| 2 | 150605349 | rs1641389 | -0.653 | 0.758 | -0.808 | -0.033 | -0.009 | -0.014 | -0.014 | 0 | 0.34 | 0.27 | 0.34 | 0.54 | 0.4 | 0.43 |
| 2 | 150606904 | rs1641394 | -1.531 | 0.678 | -0.758 | -0.41 | 0 | 0 | 0 | 0 | 0.57 | 0.6 | 0.53 | 0.63 | 0.73 | 0.69 |
| 2 | 150607967 | rs11687398 | -0.094 | 0.678 | -0.582 | -0.442 | 0.038 | -0.059 | -0.059 | 0 | 0.23 | 0.35 | 0.21 | 0.11 | 0.33 | 0.27 |
| 2 | 150609339 | rs1405224 | -0.045 | 0.658 | -0.582 | -0.451 | 0.042 | -0.059 | -0.059 | 0 | 0.22 | 0.35 | 0.21 | 0.11 | 0.33 | 0.27 |
| 2 | 150609764 | rs1724885 | -1.698 | 0.615 | -0.577 | -0.447 | 0.016 | 0.007 | 0.007 | 0 | 0.58 | 0.67 | 0.54 | 0.63 | 0.72 | 0.69 |
| 2 | 150619995 | rs1641417 | -0.178 | 0.634 | -0.667 | -0.563 | 0.05 | -0.075 | -0.075 | 0.001 | 0.23 | 0.41 | 0.22 | 0.1 | 0.32 | 0.25 |
| 2 | 150620074 | rs2160339 | 1.47 | 0.642 | -0.818 | -0.504 | -0.06 | 0.04 | 0.04 | 0.025 | 0.17 | 0.01 | 0.1 | 0.04 | 0.19 | 0.16 |
| 2 | 150621743 | rs1724836 | -0.374 | 0.544 | -0.784 | -0.493 | -0.012 | -0.007 | -0.007 | 0 | 0.35 | 0.29 | 0.33 | 0.54 | 0.42 | 0.44 |
| 2 | 150623050 | rs1724875 | -0.374 | 0.544 | -0.784 | -0.493 | -0.012 | -0.007 | -0.007 | 0.909 | 0.35 | 0.29 | 0.33 | 0.54 | 0.42 | 0.44 |
| 2 | 150624073 | rs10172203 | -0.374 | 0.544 | -0.784 | -0.493 | -0.012 | -0.007 | -0.007 | 0.025 | 0.35 | 0.29 | 0.33 | 0.54 | 0.42 | 0.44 |
| 2 | 150625294 | rs730098 | -0.374 | 0.544 | -0.784 | -0.493 | -0.012 | -0.007 | -0.007 | 0 | 0.35 | 0.29 | 0.33 | 0.54 | 0.42 | 0.44 |
| 2 | 150634825 | rs848242 | -0.598 | 0.098 | -0.891 | -0.715 | 0.001 | 0.006 | 0.006 | 0 | 0.43 | 0.37 | 0.35 | 0.54 | 0.43 | 0.5 |
| 2 | 150642505 | rs6437067 | -1.794 | 0.142 | -0.908 | -0.742 | -0.024 | -0.011 | -0.011 | 0 | 0.64 | 0.61 | 0.55 | 0.65 | 0.73 | 0.75 |
| 2 | 150656433 | rs2058686 | 0.926 | 0.304 | -0.975 | -0.78 | -0.057 | 0.073 | 0.073 | 0 | 0.15 | 0.03 | 0.13 | 0.03 | 0.21 | 0.12 |
| 2 | 150663824 | rs4667395 | -1.747 | 0.298 | -0.929 | -0.774 | -0.03 | -0.015 | -0.015 | 0 | 0.63 | 0.54 | 0.55 | 0.65 | 0.73 | 0.74 |
| 2 | 150668256 | rs17348700 | 0.894 | 0.323 | -1.045 | -0.769 | -0.057 | 0.07 | 0.07 | 0 | 0.15 | 0.03 | 0.13 | 0.03 | 0.21 | 0.11 |
| 2 | 150670406 | rs13429279 | -0.229 | 0.599 | -0.862 | -0.48 | -0.03 | -0.174 | -0.174 | 0.09 | 0.23 | 0.03 | 0.23 | 0.5 | 0.24 | 0.33 |
| 2 | 150670765 | rs1405214 | -0.944 | 0.635 | -0.823 | -0.483 | -0.03 | 0.061 | 0.061 | 0 | 0.45 | 0.23 | 0.4 | 0.62 | 0.52 | 0.62 |
| 2 | 150673889 | rs13018598 | 0.808 | 0.652 | -0.842 | -0.476 | -0.03 | 0.021 | 0.021 | 0 | 0.22 | 0.15 | 0.17 | 0.12 | 0.28 | 0.29 |
| 2 | 150675936 | rs7420486 | -0.842 | 0.668 | -0.699 | -0.399 | 0.001 | 0.017 | 0.017 | 0 | 0.3 | 0.39 | 0.32 | 0.58 | 0.33 | 0.45 |
| 2 | 150683374 | rs9288699 | -1.688 | 0.305 | -1.312 | -0.67 | 0 | 0.018 | 0.018 | 0 | 0.77 | 0.72 | 0.65 | 0.64 | 0.77 | 0.74 |
| 2 | 150685360 | rs4130046 | -1.305 | 0.118 | -1.739 | -0.719 | 0.02 | 0.044 | 0.044 | 0 | 0.72 | 0.58 | 0.57 | 0.56 | 0.69 | 0.61 |
| 2 | 150686397 | rs9710887 | 0.87 | 0.094 | -2.483 | -0.717 | -0.02 | 0.132 | 0.132 | 0 | 0.25 | 0.07 | 0.17 | 0.02 | 0.24 | 0.14 |
| 2 | 150688553 | rs12616417 | -0.113 | 0.104 | -2.613 | -0.652 | 0.008 | -0.015 | -0.015 | 0.482 | 0.41 | 0.41 | 0.25 | 0.08 | 0.29 | 0.24 |
| 2 | 150689164 | rs4132254 | 0.343 | 1.046 | -2.332 | -0.271 | -0.003 | -0.064 | -0.064 | 0.938 | 0.28 | 0.31 | 0.2 | 0.02 | 0.25 | 0.14 |
| 2 | 150698290 | rs9751098 | 0.368 | 1.618 | -2.392 | 0.865 | -0.069 | 0.033 | 0.033 | 0 | 0.16 | 0.01 | 0.1 | 0.03 | 0.19 | 0.15 |
| 2 | 150701094 | rs9710823 | -0.826 | 1.213 | -1.593 | 1.477 | -0.024 | -0.033 | -0.033 | 0 | 0.11 | 0.05 | 0.17 | 0.45 | 0.18 | 0.21 |
| 2 | 150714840 | rs4667393 | -0.846 | 1.279 | -1.729 | 1.359 | -0.056 | -0.101 | -0.101 | 1.03 | 0.14 | 0.01 | 0.22 | 0.36 | 0.18 | 0.23 |
| 2 | 150727326 | rs13016092 | -0.809 | 1.381 | -1.734 | 1.354 | -0.059 | -0.099 | -0.099 | 0.122 | 0.13 | 0.01 | 0.22 | 0.36 | 0.18 | 0.23 |
| 2 | 150727944 | rs10176816 | -0.093 | 1.515 | -1.484 | 0.926 | 0.019 | -0.017 | -0.017 | 0 | 0.32 | 0.48 | 0.29 | 0.34 | 0.35 | 0.35 |
| 2 | 150729153 | rs1437966 | -1.287 | 1.112 | -1.592 | 0.497 | 0 | -0.023 | -0.023 | 0 | 0.81 | 0.76 | 0.89 | 0.94 | 0.82 | 0.84 |
| 2 | 150729453 | rs10171287 | 0.809 | 1.6 | -1.326 | 0.478 | -0.001 | -0.003 | -0.003 | 0 | 0.21 | 0.12 | 0.18 | 0.26 | 0.21 | 0.18 |
| 2 | 150739256 | rs6710538 | -0.485 | 0.874 | -1.726 | 0.189 | 0 | 0.001 | 0.001 | 0 | 0.54 | 0.49 | 0.56 | 0.58 | 0.5 | 0.5 |
| 2 | 150740751 | rs7573044 | 0.477 | 0.814 | -1.704 | 0.148 | 0.023 | -0.008 | -0.008 | 0.928 | 0.3 | 0.44 | 0.28 | 0.34 | 0.34 | 0.31 |
| 2 | 150742467 | rs7588167 | 1.099 | 0.367 | -1.944 | 0.25 | 0.035 | -0.06 | -0.06 | 0 | 0.21 | 0.01 | 0.27 | 0.24 | 0.14 | 0.16 |
| 2 | 150748536 | rs11690195 | -0.148 | 0.796 | -1.288 | 0.238 | 0.005 | -0.001 | -0.001 | 0 | 0.49 | 0.58 | 0.59 | 0.58 | 0.57 | 0.46 |
| 2 | 150751907 | rs17363367 | -0.754 | 1.055 | -0.85 | 0.699 | 0.004 | 0.029 | 0.029 | 0.143 | 0.19 | 0.21 | 0.18 | 0.34 | 0.22 | 0.26 |
| 2 | 150761377 | rs2218027 | 0.678 | 1.039 | -0.49 | 0.421 | 0.011 | -0.018 | -0.018 | 1.424 | 0.68 | 0.4 | 0.66 | 0.81 | 0.64 | 0.63 |
| 2 | 150766179 | rs6735540 | -0.988 | 1.14 | 0.048 | 0.734 | 0.008 | 0.002 | 0.002 | 0 | 0.32 | 0.43 | 0.39 | 0.31 | 0.34 | 0.45 |
| 2 | 150777288 | rs4667329 | -1.092 | 0.985 | -0.269 | 0.892 | -0.005 | -0.002 | -0.002 | 0.099 | 0.65 | 0.61 | 0.72 | 0.71 | 0.7 | 0.67 |
| 2 | 150782216 | rs10804400 | 0.885 | 1.149 | -0.303 | 0.872 | -0.011 | -0.009 | -0.009 | 0.172 | 0.33 | 0.18 | 0.32 | 0.39 | 0.36 | 0.21 |
| 2 | 150787550 | rs1561313 | -0.303 | 1.079 | 0.118 | 0.765 | -0.004 | 0 | 0 | 0 | 0.56 | 0.42 | 0.6 | 0.57 | 0.63 | 0.52 |

| | | | | | | | | | | | | | | | | |
|---|---|---|---|---|---|---|---|---|---|---|---|---|---|---|---|---|
| 2 | 150796369 | rs17781497 | 0.108 | 1.08 | 0.246 | 0.77 | -0.004 | 0.002 | 0.002 | 0.004 | 0.24 | 0.23 | 0.27 | 0.17 | 0.27 | 0.3 |
| 2 | 150812317 | rs6709313 | 0.605 | 0.981 | 0.278 | 0.766 | -0.013 | 0.005 | 0.005 | 0 | 0.38 | 0.45 | 0.36 | 0.44 | 0.29 | 0.41 |
| 2 | 150818921 | rs6728165 | 0.993 | 1.174 | 0.073 | 0.717 | -0.017 | -0.01 | -0.01 | 0.068 | 0.15 | 0.01 | 0.19 | 0.1 | 0.13 | 0.09 |
| 2 | 150828833 | rs1020088 | 0.251 | 1.388 | 0.303 | 0.705 | 0.02 | -0.023 | -0.023 | 0 | 0.73 | 0.61 | 0.71 | 0.82 | 0.71 | 0.67 |
| 2 | 150843084 | rs10171121 | 1.126 | 1.44 | 0.254 | 0.741 | 0.009 | -0.003 | -0.003 | 0.038 | 0.32 | 0.24 | 0.26 | 0.3 | 0.27 | 0.32 |
| 2 | 150852395 | rs331118 | 0.886 | 1.433 | 0.328 | 1.032 | -0.005 | 0 | 0 | 0 | 0.32 | 0.23 | 0.33 | 0.35 | 0.35 | 0.21 |
| 2 | 150853996 | rs6735775 | 1.105 | 1.15 | 0.366 | 1.017 | 0.048 | 0.157 | 0.157 | 0 | 0.35 | 0.11 | 0.18 | 0.08 | 0.25 | 0.24 |
| 2 | 150855837 | rs331112 | -0.908 | 1.095 | 0.461 | 0.954 | -0.011 | -0.006 | -0.006 | 0 | 0.63 | 0.53 | 0.7 | 0.71 | 0.68 | 0.66 |
| 2 | 150858540 | rs745720 | 0.093 | 0.832 | 0.442 | 1.028 | -0.005 | -0.002 | -0.002 | 0 | 0.18 | 0.12 | 0.24 | 0.17 | 0.2 | 0.28 |
| 2 | 150877992 | rs732227 | -0.005 | 0.549 | 0.373 | 1.097 | 0.006 | -0.003 | -0.003 | 0.062 | 0.42 | 0.42 | 0.39 | 0.44 | 0.36 | 0.47 |
| 2 | 150886421 | rs16827723 | 0.554 | 0.655 | 0.102 | 1.055 | 0.007 | 0.005 | 0.005 | 0 | 0.33 | 0.24 | 0.27 | 0.27 | 0.3 | 0.32 |
| 2 | 150890233 | rs330650 | -0.235 | 0.747 | 0.348 | 0.802 | -0.002 | 0.005 | 0.005 | 1.211 | 0.54 | 0.39 | 0.45 | 0.44 | 0.5 | 0.5 |
| 2 | 150904671 | rs330648 | -0.365 | 0.842 | 0.979 | 0.814 | 0 | -0.001 | -0.001 | 0 | 0.56 | 0.53 | 0.59 | 0.61 | 0.56 | 0.62 |
| 2 | 150911159 | rs330625 | -0.407 | 0.797 | 0.849 | 0.774 | 0 | -0.001 | -0.001 | 0.087 | 0.56 | 0.57 | 0.59 | 0.59 | 0.56 | 0.59 |
| 2 | 150913894 | rs330623 | -0.898 | 0.805 | 0.488 | 0.779 | -0.034 | -0.041 | -0.041 | 0.621 | 0.6 | 0.88 | 0.63 | 0.6 | 0.59 | 0.63 |
| 2 | 150917809 | rs16827761 | 1.084 | 0.732 | 0.263 | 0.749 | 0.007 | 0.007 | 0.007 | 2.755 | 0.16 | 0.12 | 0.06 | 0.05 | 0.11 | 0.1 |
| 2 | 150922856 | rs330613 | 0.397 | 0.741 | 0.051 | 0.759 | -0.002 | 0 | 0 | 0.199 | 0.34 | 0.35 | 0.38 | 0.37 | 0.39 | 0.33 |
| 2 | 150940963 | rs330594 | -0.55 | 1.002 | -0.099 | 0.759 | -0.009 | 0.025 | 0.025 | 0.388 | 0.58 | 0.69 | 0.7 | 0.68 | 0.54 | 0.57 |
| 2 | 150943313 | rs570148 | -1.001 | 1.104 | -0.653 | 0.66 | -0.055 | 0.139 | 0.139 | 0.047 | 0.63 | 0.9 | 0.73 | 0.85 | 0.62 | 0.74 |
| 2 | 150945777 | rs330633 | 0.031 | 0.91 | -0.567 | 0.996 | 0.025 | 0.005 | 0.005 | 0 | 0.45 | 0.29 | 0.51 | 0.44 | 0.41 | 0.39 |
| 2 | 150948703 | rs330630 | -0.548 | 1.15 | -0.479 | 1.365 | 0.021 | 0.14 | 0.14 | 0 | 0.12 | 0.21 | 0.37 | 0.52 | 0.17 | 0.3 |
| 2 | 150958446 | rs1446243 | 1.115 | 1.289 | -1.444 | 0.641 | -0.069 | 0.088 | 0.088 | 0.079 | 0.31 | 0.03 | 0.22 | 0.14 | 0.32 | 0.24 |
| 2 | 150958611 | rs330634 | 1.2 | 1.714 | -1.339 | 1.022 | -0.013 | 0.032 | 0.032 | 0 | 0.05 | 0.03 | 0.16 | 0.33 | 0.11 | 0.18 |
| 2 | 150964274 | rs17349216 | 1.324 | 0.757 | -0.619 | -0.091 | 0.008 | 0.011 | 0.011 | 0 | 0.16 | 0.08 | 0.17 | 0.03 | 0.14 | 0.1 |
| 2 | 150970181 | rs179357 | -0.429 | 0.817 | -0.499 | -0.219 | 0 | 0 | 0 | 0 | 0.83 | 0.82 | 0.88 | 0.89 | 0.82 | 0.86 |
| 2 | 150975854 | rs502957 | 0.957 | 0.806 | -0.161 | -0.228 | -0.049 | 0.025 | 0.025 | 1.749 | 0.43 | 0.19 | 0.58 | 0.67 | 0.5 | 0.53 |
| 2 | 150977222 | rs973103 | -0.454 | 0.772 | -0.171 | -0.289 | 0.013 | 0.053 | 0.053 | 2.248 | 0.69 | 0.81 | 0.83 | 0.89 | 0.73 | 0.83 |
| 2 | 150977649 | rs554545 | 0.845 | 0.776 | -0.212 | -0.146 | -0.049 | 0.025 | 0.025 | 1.749 | 0.43 | 0.19 | 0.58 | 0.67 | 0.5 | 0.53 |
| 2 | 150985765 | rs330609 | 1.318 | 0.855 | -0.136 | -0.286 | -0.053 | 0.024 | 0.024 | 1.886 | 0.4 | 0.15 | 0.57 | 0.69 | 0.48 | 0.5 |
| 2 | 150987418 | rs16827839 | -0.348 | 0.695 | 0.664 | -0.292 | 0.041 | -0.025 | -0.025 | 0 | 0.08 | 0.33 | 0.04 | 0.01 | 0.05 | 0.11 |
| 2 | 150993329 | rs6705368 | -0.535 | 0.725 | 0.228 | -0.414 | -0.001 | 0.008 | 0.008 | 0 | 0.34 | 0.68 | 0.29 | 0.2 | 0.32 | 0.36 |
| 2 | 150994953 | rs330582 | -0.689 | 0.405 | 0.604 | -0.071 | 0.014 | 0.032 | 0.032 | 4.755 | 0.75 | 0.83 | 0.87 | 0.91 | 0.81 | 0.87 |
| 2 | 151000108 | rs330585 | 0.889 | 0.445 | 0.483 | 0.012 | 0.052 | 0.071 | 0.071 | 3.945 | 0.56 | 0.13 | 0.76 | 0.72 | 0.73 | 0.66 |
| 2 | 151001420 | rs330586 | -0.194 | 0.306 | 0.676 | -0.284 | -0.023 | -0.042 | -0.042 | 0 | 0.67 | 0.4 | 0.84 | 0.9 | 0.83 | 0.85 |
| 2 | 151007043 | rs476837 | 1.132 | 0.834 | 0.95 | -0.225 | 0.052 | -0.111 | -0.111 | 0.125 | 0.84 | 0.56 | 0.81 | 0.99 | 0.85 | 0.81 |
| 2 | 151010323 | rs10174406 | 0.738 | 0.238 | 0.552 | -0.099 | 0.036 | 0.052 | 0.052 | 2.016 | 0.22 | 0.11 | 0.09 | 0.06 | 0.12 | 0.09 |
| 2 | 151016041 | rs6727767 | -0.194 | 0.394 | 0.906 | 0.17 | -0.002 | -0.001 | -0.001 | 0 | 0.31 | 0.31 | 0.16 | 0.26 | 0.2 | 0.22 |
| 2 | 151016074 | rs6727863 | -0.955 | 0.364 | 0.88 | 0.133 | -0.001 | -0.001 | -0.001 | 0 | 0.37 | 0.59 | 0.3 | 0.26 | 0.25 | 0.28 |
| 2 | 151019241 | rs4667379 | -0.71 | 0.54 | 0.875 | 0.228 | 0 | 0 | 0 | 0.148 | 0.36 | 0.63 | 0.36 | 0.27 | 0.25 | 0.29 |
| 2 | 151025381 | rs1374158 | 0.418 | 0.584 | 0.552 | 0.197 | 0.018 | 0.018 | 0.018 | 3.168 | 0.24 | 0.22 | 0.1 | 0.06 | 0.11 | 0.08 |
| 2 | 151033471 | rs7602256 | 0.54 | 1.211 | 0.916 | 0.593 | 0.039 | -0.015 | -0.015 | 0 | 0.05 | 0.17 | 0.13 | 0 | 0.06 | 0.04 |
| 2 | 151043023 | rs12692632 | -0.346 | 0.88 | 0.482 | 0.775 | 0.004 | -0.004 | -0.004 | 0.136 | 0.55 | 0.56 | 0.65 | 0.48 | 0.59 | 0.45 |
| 2 | 151044800 | rs7580941 | -0.621 | 0.821 | 0.583 | 0.744 | 0.007 | -0.025 | -0.025 | 0.268 | 0.58 | 0.74 | 0.67 | 0.48 | 0.58 | 0.47 |
| 2 | 151063726 | rs13013440 | 1.385 | 0.899 | 0.44 | 1.036 | -0.041 | -0.042 | -0.042 | 2.112 | 0.1 | 0 | 0.06 | 0.16 | 0.1 | 0.29 |
| 2 | 151064165 | rs971810 | 1.58 | 1.161 | 0.376 | 0.643 | -0.022 | 0.024 | 0.024 | 0 | 0.06 | 0.02 | 0.07 | 0 | 0.08 | 0.08 |
| 2 | 151076984 | rs4260269 | 0.131 | 0.321 | 0.755 | 0.82 | 0.032 | -0.012 | -0.012 | 0 | 0.53 | 0.3 | 0.62 | 0.47 | 0.58 | 0.44 |
| 2 | 151077809 | rs13024803 | 0.348 | 0.37 | 0.652 | 0.841 | 0.019 | 0.13 | 0.13 | 0.996 | 0.5 | 0.21 | 0.44 | 0.25 | 0.52 | 0.32 |
| 2 | 151090995 | rs10177101 | 1.331 | 0.209 | -0.152 | 0.534 | -0.044 | 0.064 | 0.064 | 1.116 | 0.28 | 0.01 | 0.17 | 0.12 | 0.29 | 0.14 |
| 2 | 151112873 | rs4318407 | -0.384 | 0.239 | 0.254 | 0.521 | 0.002 | -0.01 | -0.01 | 0 | 0.57 | 0.67 | 0.57 | 0.51 | 0.56 | 0.44 |
| 2 | 151131188 | rs10190378 | 0.767 | -0.14 | -0.07 | 0.355 | 0.022 | -0.011 | -0.011 | 0 | 0.23 | 0.02 | 0.15 | 0.18 | 0.16 | 0.13 |
| 2 | 151135525 | rs4233802 | 0.859 | 0.137 | 0.75 | 0.29 | 0.039 | -0.016 | -0.016 | 0 | 0.92 | 0.35 | 0.83 | 0.97 | 0.95 | 0.92 |
| 2 | 151175668 | rs13030734 | 0.27 | 0.206 | -0.383 | 0.368 | -0.063 | -0.083 | -0.083 | 0 | 0.36 | 0.08 | 0.3 | 0.44 | 0.39 | 0.43 |
| 2 | 151201049 | rs10177059 | 1.085 | 0.478 | -0.397 | 0.334 | -0.014 | 0.026 | 0.026 | 0 | 0.24 | 0.16 | 0.35 | 0.14 | 0.31 | 0.17 |
| 2 | 151201834 | rs10180142 | 1.32 | -0.092 | -0.94 | 0.44 | 0.041 | -0.05 | -0.05 | 1.263 | 0.22 | 0.03 | 0.1 | 0.21 | 0.13 | 0.19 |
| 2 | 151205806 | rs983231 | 0.967 | -0.166 | -1.102 | 0.691 | -0.003 | -0.053 | -0.053 | 0.135 | 0.09 | 0 | 0.12 | 0.31 | 0.08 | 0.09 |
| 2 | 151210898 | rs12994436 | 1.236 | -0.245 | 0.155 | 0.613 | -0.001 | 0.003 | 0.003 | 0.289 | 0.11 | 0.08 | 0.06 | 0.1 | 0.1 | 0.26 |
| 2 | 151218033 | rs6760903 | 0.915 | -0.28 | 0.557 | 0.615 | 0.002 | -0.01 | -0.01 | 0 | 0.12 | 0.21 | 0.07 | 0.1 | 0.1 | 0.26 |
| 2 | 151218776 | rs7600018 | 0.931 | -0.344 | 0.659 | 0.596 | -0.003 | 0.061 | 0.061 | 0 | 0.05 | 0.09 | 0.1 | 0.21 | 0.03 | 0.07 |
| 2 | 151219089 | rs10930090 | 0.459 | -0.306 | 0.479 | 0.502 | -0.006 | 0.034 | 0.034 | 0.026 | 0.51 | 0.21 | 0.43 | 0.54 | 0.52 | 0.46 |
| 2 | 151227364 | rs1866979 | -0.883 | -0.321 | 0.123 | 0.48 | 0.014 | -0.058 | -0.058 | 0 | 0.68 | 0.5 | 0.59 | 0.85 | 0.63 | 0.8 |
| 2 | 151237876 | rs7590701 | 0.701 | -0.194 | -0.389 | 0.571 | -0.002 | -0.013 | -0.013 | 0.018 | 0.15 | 0.11 | 0.15 | 0.31 | 0.15 | 0.32 |
| 2 | 151247366 | rs4287765 | -1.837 | -0.186 | -0.759 | 0.547 | -0.045 | 0.028 | 0.028 | 0 | 0.76 | 0.89 | 0.64 | 0.86 | 0.67 | 0.83 |
| 2 | 151249628 | rs10187222 | 1.298 | -0.188 | -0.859 | 0.68 | 0.039 | -0.059 | -0.059 | 0 | 0.22 | 0.03 | 0.09 | 0.23 | 0.12 | 0.19 |
| 2 | 151250458 | rs11684751 | 0.976 | 0.11 | -0.739 | 0.223 | -0.02 | -0.006 | -0.006 | 0 | 0.28 | 0.18 | 0.32 | 0.31 | 0.36 | 0.29 |
| 2 | 151251552 | rs976396 | -0.449 | 0.239 | -0.43 | 0.582 | -0.011 | 0.005 | 0.005 | 0 | 0.4 | 0.52 | 0.27 | 0.55 | 0.27 | 0.52 |
| 2 | 151259604 | rs6432753 | 0.42 | 0.755 | -0.881 | 0.297 | -0.001 | -0.016 | -0.016 | 0 | 0.52 | 0.46 | 0.49 | 0.75 | 0.47 | 0.51 |
| 2 | 151259655 | rs6432754 | 0.427 | 0.895 | -0.221 | 0.868 | 0.04 | -0.039 | -0.039 | 0 | 0.84 | 0.61 | 0.88 | 0.88 | 0.82 | 0.71 |
| 2 | 151263268 | rs12692704 | 0.026 | 0.937 | -1.155 | 1.311 | -0.023 | -0.069 | -0.069 | 0.459 | 0.12 | 0.02 | 0.13 | 0.26 | 0.12 | 0.1 |
| 2 | 151263611 | rs12474944 | -0.771 | 0.764 | -0.559 | 1.054 | 0 | 0.004 | 0.004 | 0.837 | 0.26 | 0.25 | 0.2 | 0.37 | 0.25 | 0.35 |

| | | | | | | | | | | | | | | | | |
|---|---|---|---|---|---|---|---|---|---|---|---|---|---|---|---|---|
| 2 | 151273111 | rs1166856 | -0.327 | 0.583 | -0.305 | 1.093 | -0.015 | 0.142 | 0.142 | 0.029 | 0.16 | 0.26 | 0.08 | 0.44 | 0.11 | 0.18 |
| 2 | 151279247 | rs3108514 | 0.387 | 0.628 | -0.118 | -0.148 | -0.01 | -0.065 | -0.065 | 0.613 | 0.73 | 0.61 | 0.84 | 0.96 | 0.76 | 0.79 |
| 2 | 151283141 | rs2053495 | 0.123 | 0.456 | -0.298 | -0.248 | 0.013 | 0.073 | 0.073 | 0 | 0.15 | 0.03 | 0.08 | 0.04 | 0.12 | 0.15 |
| 2 | 151296205 | rs17350459 | 0.075 | 0.521 | -0.27 | -0.232 | 0.01 | 0.064 | 0.064 | 0 | 0.14 | 0.03 | 0.09 | 0.04 | 0.12 | 0.15 |
| 2 | 151296330 | rs11679729 | 0.216 | 1.307 | -0.354 | -0.465 | 0.027 | -0.007 | -0.007 | 1.99 | 0.18 | 0.16 | 0.17 | 0.13 | 0.32 | 0.3 |
| 2 | 151297117 | rs816895 | 0.903 | 0.609 | -0.334 | -0.236 | -0.026 | -0.116 | -0.116 | 1.442 | 0.73 | 0.52 | 0.84 | 0.96 | 0.77 | 0.79 |
| 2 | 151297852 | rs17435056 | -0.798 | 0.721 | -0.776 | -0.045 | 0.014 | 0.025 | 0.025 | 0 | 0.22 | 0.21 | 0.14 | 0.04 | 0.2 | 0.19 |
| 2 | 151311055 | rs16828194 | -0.655 | 0.723 | -0.633 | 0.135 | -0.015 | -0.05 | -0.05 | 0 | 0.19 | 0.32 | 0.11 | 0.04 | 0.14 | 0.19 |
| 2 | 151316465 | rs10930121 | -0.049 | 1.307 | -1.249 | 0.506 | -0.004 | 0.073 | 0.073 | 0 | 0.49 | 0.24 | 0.55 | 0.6 | 0.51 | 0.58 |
| 2 | 151317118 | rs2697338 | 0.917 | 1.093 | -0.297 | 1.47 | 0.012 | 0.341 | 0.341 | 0 | 0.93 | 0.8 | 0.9 | 0.67 | 0.95 | 0.89 |
| 2 | 151317140 | rs2697339 | -0.522 | 1.357 | -0.177 | 0.66 | 0.006 | -0.049 | -0.049 | 1.348 | 0.14 | 0.14 | 0.17 | 0.01 | 0.19 | 0.09 |
| 2 | 151324400 | rs6432774 | -0.744 | 1.188 | 0.741 | 0.227 | 0.015 | -0.041 | -0.041 | 0 | 0.15 | 0.25 | 0.11 | 0.04 | 0.12 | 0.07 |
| 2 | 151324688 | rs1372438 | -0.549 | 1.189 | 0.805 | 0.227 | 0.019 | -0.031 | -0.031 | 0 | 0.13 | 0.21 | 0.11 | 0.04 | 0.12 | 0.07 |
| 2 | 151325295 | rs816889 | -0.536 | 0.301 | 0.451 | 0.363 | 0.03 | -0.044 | -0.044 | 0 | 0.74 | 0.8 | 0.8 | 0.51 | 0.83 | 0.85 |
| 2 | 151325303 | rs10185950 | -0.255 | 0.553 | 0.462 | 0.143 | 0.019 | -0.033 | -0.033 | 0 | 0.14 | 0.21 | 0.11 | 0.04 | 0.12 | 0.07 |
| 2 | 151335648 | rs7579153 | 0.136 | 0.522 | 0.357 | 0.12 | 0.007 | -0.016 | -0.016 | 0.382 | 0.11 | 0.2 | 0.1 | 0.05 | 0.11 | 0.07 |
| 2 | 151336842 | rs1528429 | 0.58 | 0.752 | -0.455 | -0.26 | -0.041 | 0.027 | 0.027 | 0 | 0.24 | 0.07 | 0.44 | 0.18 | 0.33 | 0.21 |
| 2 | 151344984 | rs3814395 | -1.163 | 0.315 | -0.745 | -0.24 | -0.007 | -0.034 | -0.034 | 0 | 0.69 | 0.77 | 0.59 | 0.5 | 0.64 | 0.46 |
| 2 | 151345113 | rs2222760 | 0.103 | -0.098 | -1.362 | -0.814 | -0.034 | 0.08 | 0.08 | 0 | 0.27 | 0.14 | 0.2 | 0.15 | 0.26 | 0.12 |
| 2 | 151346377 | rs1528431 | -1.058 | 0.342 | -0.717 | -0.307 | -0.002 | 0 | 0 | 0.625 | 0.65 | 0.73 | 0.57 | 0.39 | 0.62 | 0.42 |
| 2 | 151347639 | rs16828256 | 0.61 | 0.289 | -1.473 | -0.958 | 0.007 | -0.032 | -0.032 | 1.521 | 0.08 | 0.13 | 0.06 | 0 | 0.11 | 0.05 |
| 2 | 151352154 | rs12692720 | -0.312 | 0.231 | -1.954 | -1.008 | 0.004 | 0.11 | 0.11 | 3.422 | 0.45 | 0.29 | 0.29 | 0.17 | 0.39 | 0.24 |
| 2 | 151375161 | rs10497061 | 0.184 | 0.474 | -1.796 | -0.788 | -0.004 | -0.004 | -0.004 | 0 | 0.07 | 0.08 | 0.06 | 0 | 0.11 | 0.05 |
| 2 | 151375551 | rs11681459 | -0.052 | 0.465 | -1.782 | -0.646 | -0.015 | 0.122 | 0.122 | 4.38 | 0.29 | 0.1 | 0.17 | 0.01 | 0.25 | 0.14 |
| 2 | 151392059 | rs6743187 | -0.05 | 0.55 | -1.441 | -0.128 | 0.013 | 0.081 | 0.081 | 3.39 | 0.36 | 0.14 | 0.17 | 0.13 | 0.27 | 0.17 |
| 2 | 151401771 | rs6759010 | -0.067 | 0.582 | -1.403 | -0.112 | 0.017 | 0.073 | 0.073 | 3.569 | 0.36 | 0.17 | 0.17 | 0.14 | 0.28 | 0.17 |
| 2 | 151409434 | rs7592790 | 0.074 | 0.775 | -1.9 | 0.016 | 0.003 | -0.015 | -0.015 | 0.01 | 0.12 | 0.26 | 0.09 | 0.06 | 0.14 | 0.25 |
| 2 | 151413170 | rs1406242 | 0.04 | 0.94 | -2.334 | 0.326 | -0.014 | 0.146 | 0.146 | 0.683 | 0.5 | 0.07 | 0.71 | 0.81 | 0.59 | 0.58 |
| 2 | 151429183 | rs4635532 | 0.53 | 1.467 | -1.292 | 0.906 | 0.01 | 0.009 | 0.009 | 0 | 0.82 | 0.8 | 0.58 | 0.75 | 0.77 | 0.82 |
| 2 | 151430525 | rs2340857 | 1.045 | 1.547 | -1.905 | 0.927 | -0.019 | -0.013 | -0.013 | 0 | 0.64 | 0.71 | 0.5 | 0.55 | 0.59 | 0.68 |
| 2 | 151436367 | rs10469691 | 1.324 | 1.897 | -0.633 | 0.83 | -0.002 | -0.001 | -0.001 | 0.98 | 0.6 | 0.34 | 0.44 | 0.53 | 0.53 | 0.62 |
| 2 | 151457605 | rs2340861 | 0.468 | 2.024 | -0.247 | 0.852 | 0.003 | 0.001 | 0.001 | 0 | 0.77 | 0.78 | 0.54 | 0.74 | 0.72 | 0.77 |
| 2 | 151462867 | rs16828372 | -0.438 | 1.957 | -0.057 | 0.816 | -0.001 | -0.001 | -0.001 | 0 | 0.14 | 0.17 | 0.07 | 0.17 | 0.14 | 0.1 |
| 2 | 151465458 | rs10930176 | 1.068 | 1.188 | -0.585 | 0.602 | -0.01 | 0.045 | 0.045 | 0 | 0.12 | 0.19 | 0.08 | 0.22 | 0.12 | 0.28 |
| 2 | 151469069 | rs2340864 | 1.194 | 1.451 | -0.179 | 1.101 | -0.014 | 0.008 | 0.008 | 0 | 0.57 | 0.25 | 0.4 | 0.41 | 0.51 | 0.6 |
| 2 | 151470729 | rs10202222 | -0.501 | 1.592 | -0.21 | 0.825 | 0.014 | 0.008 | 0.008 | 0 | 0.81 | 0.79 | 0.56 | 0.76 | 0.78 | 0.81 |
| 2 | 151485519 | rs2341068 | -0.067 | 1.429 | -0.184 | 0.904 | 0.026 | 0.019 | 0.019 | 0.333 | 0.9 | 0.8 | 0.94 | 0.86 | 0.88 | 0.96 |
| 2 | 151487613 | rs4355108 | 1.088 | 1.311 | -0.151 | 0.802 | 0.031 | 0.011 | 0.011 | 0 | 0.67 | 0.49 | 0.47 | 0.58 | 0.58 | 0.68 |
| 2 | 151494381 | rs4271766 | -0.546 | 1.405 | -0.07 | 0.865 | -0.001 | -0.001 | -0.001 | 0 | 0.24 | 0.3 | 0.48 | 0.28 | 0.31 | 0.28 |
| 2 | 151494809 | rs10497066 | -0.392 | 1.402 | 0.03 | 0.896 | -0.02 | -0.008 | -0.008 | 0 | 0.23 | 0.22 | 0.48 | 0.28 | 0.31 | 0.28 |
| 2 | 151496161 | rs7562436 | 0.046 | 1.391 | 0.035 | 0.863 | 0.049 | 0.042 | 0.042 | 0 | 0.91 | 0.71 | 0.94 | 0.86 | 0.89 | 0.96 |
| 2 | 151496524 | rs7603668 | 0.887 | 1.394 | 0.016 | 0.865 | 0 | -0.001 | -0.001 | 0 | 0.69 | 0.62 | 0.48 | 0.6 | 0.58 | 0.69 |
| 2 | 151498770 | rs4408725 | -0.797 | 1.436 | -0.021 | 0.774 | -0.003 | -0.003 | -0.003 | 0.244 | 0.93 | 0.93 | 0.96 | 0.86 | 0.89 | 0.96 |
| 2 | 151503700 | rs2879932 | -0.886 | 1.372 | -0.018 | 0.797 | -0.003 | -0.002 | -0.002 | 0 | 0.93 | 0.93 | 0.96 | 0.89 | 0.91 | 0.96 |
| 2 | 151504694 | rs10803803 | -0.81 | 1.352 | -0.021 | 0.906 | -0.003 | -0.001 | -0.001 | 1.132 | 0.93 | 0.93 | 0.97 | 0.85 | 0.91 | 0.96 |
| 2 | 151506626 | rs2341092 | 0.733 | 1.296 | 0.03 | 0.865 | 0.037 | 0.061 | 0.061 | 0 | 0.88 | 0.56 | 0.93 | 0.85 | 0.88 | 0.93 |
| 2 | 151509486 | rs12463905 | 0.104 | 1.341 | -0.532 | 0.886 | -0.126 | -0.027 | -0.027 | 0 | 0.2 | 0.01 | 0.45 | 0.23 | 0.29 | 0.25 |
| 2 | 151535966 | rs4664722 | -0.755 | 1.446 | -0.454 | 0.943 | -0.08 | -0.052 | -0.052 | 0 | 0.25 | 0.12 | 0.48 | 0.41 | 0.4 | 0.32 |
| 2 | 151538472 | rs16828577 | 0.847 | 1.097 | -0.224 | 0.77 | -0.007 | -0.005 | -0.005 | 0 | 0.05 | 0.03 | 0.03 | 0.13 | 0.08 | 0.04 |
| 2 | 151548341 | rs1533664 | 1.225 | 0.937 | 0.208 | 0.41 | 0.023 | -0.015 | -0.015 | 0 | 0.89 | 0.63 | 0.93 | 0.95 | 0.9 | 0.92 |
| 2 | 151556877 | rs13412348 | -0.37 | 0.954 | 0.057 | 0.406 | 0.006 | -0.021 | -0.021 | 0 | 0.07 | 0.18 | 0.04 | 0.03 | 0.07 | 0.05 |
| 2 | 151584332 | rs2139425 | 1.111 | 0.846 | 0.03 | 0.328 | 0.044 | 0.095 | 0.095 | 0 | 0.48 | 0.29 | 0.38 | 0.71 | 0.42 | 0.57 |
| 2 | 151588181 | rs13400288 | 0.349 | 0.884 | 0.423 | 0.657 | 0.017 | -0.01 | -0.01 | 0.338 | 0.05 | 0.16 | 0.03 | 0.03 | 0.07 | 0.03 |
| 2 | 151607040 | rs12477164 | 1.146 | 0.672 | 0.655 | 0.504 | -0.005 | -0.05 | -0.05 | 1.024 | 0.85 | 0.6 | 0.9 | 0.94 | 0.89 | 0.9 |
| 2 | 151608765 | rs10497071 | -0.28 | 1.196 | 0.339 | 0.916 | 0 | 0 | 0 | 0 | 0.22 | 0.25 | 0.47 | 0.32 | 0.28 | 0.26 |
| 2 | 151611392 | rs1519790 | 0.455 | 0.693 | 0.581 | 0.822 | -0.006 | -0.021 | -0.021 | 0 | 0.89 | 0.81 | 0.94 | 0.95 | 0.92 | 0.94 |
| 2 | 151619345 | rs970307 | 0.87 | 0.707 | 0.821 | 1.002 | -0.004 | 0.028 | 0.028 | 0 | 0.87 | 0.79 | 0.93 | 0.79 | 0.89 | 0.88 |
| 2 | 151619693 | rs1519754 | 1.334 | 0.724 | 0.688 | 0.959 | 0.001 | 0.003 | 0.003 | 0 | 0.58 | 0.45 | 0.68 | 0.57 | 0.56 | 0.56 |
| 2 | 151622783 | rs1519756 | 0.555 | 0.638 | 0.987 | 1.165 | -0.002 | -0.019 | -0.019 | 0 | 0.7 | 0.59 | 0.75 | 0.75 | 0.64 | 0.65 |
| 2 | 151622877 | rs1519757 | 0.085 | 0.653 | 0.95 | 1.178 | 0.001 | -0.002 | -0.002 | 0.459 | 0.1 | 0.05 | 0.05 | 0.18 | 0.08 | 0.08 |
| 2 | 151624882 | rs6734792 | 1.363 | 0.65 | 0.929 | 1.171 | 0.004 | 0.008 | 0.008 | 0 | 0.58 | 0.41 | 0.68 | 0.56 | 0.56 | 0.57 |
| 2 | 151627106 | rs9789673 | -0.525 | 0.645 | 0.746 | 1.159 | -0.003 | 0.023 | 0.023 | 0 | 0.28 | 0.21 | 0.22 | 0.23 | 0.33 | 0.31 |
| 2 | 151633204 | rs1519761 | 1.062 | 0.69 | 0.713 | 1.2 | -0.006 | -0.012 | -0.012 | 0.286 | 0.59 | 0.53 | 0.7 | 0.58 | 0.56 | 0.57 |
| 2 | 151635832 | rs6757804 | 1.054 | 0.735 | 0.712 | 1.202 | -0.004 | -0.008 | -0.008 | 3.011 | 0.59 | 0.52 | 0.7 | 0.57 | 0.56 | 0.57 |
| 2 | 151645865 | rs10515926 | -1.289 | 0.547 | 0.615 | 1.21 | -0.005 | 0.006 | 0.006 | 0 | 0.41 | 0.44 | 0.27 | 0.6 | 0.43 | 0.46 |
| 2 | 151648276 | rs1519768 | 1.105 | 0.562 | 0.761 | 1.201 | -0.002 | 0.001 | 0.001 | 0 | 0.58 | 0.49 | 0.72 | 0.4 | 0.57 | 0.54 |
| 2 | 151650314 | rs2190375 | 0.536 | 0.439 | 0.83 | 1.204 | -0.025 | 0.08 | 0.08 | 0 | 0.82 | 0.68 | 0.91 | 0.66 | 0.84 | 0.84 |
| 2 | 151653870 | rs6728560 | 0.554 | 0.737 | 0.745 | 1.311 | 0.005 | 0.021 | 0.021 | 0 | 0.73 | 0.69 | 0.77 | 0.59 | 0.71 | 0.65 |
| 2 | 151655513 | rs7580162 | -0.763 | 0.597 | 0.563 | 1.166 | -0.004 | 0.01 | 0.01 | 0 | 0.28 | 0.23 | 0.22 | 0.26 | 0.29 | 0.31 |

| | | | | | | | | | | | | | | | | |
|---|---|---|---|---|---|---|---|---|---|---|---|---|---|---|---|---|
| 2 | 151663105 | rs4141070 | 0.338 | 0.152 | 0.815 | 1.194 | 0.019 | 0.208 | 0.208 | 0 | 0.86 | 0.53 | 0.92 | 0.66 | 0.85 | 0.82 |
| 2 | 151669375 | rs1519775 | 0.392 | 0.145 | 0.841 | 1.19 | 0.014 | 0.19 | 0.19 | 0 | 0.86 | 0.53 | 0.91 | 0.65 | 0.85 | 0.83 |
| 2 | 151669661 | rs6753674 | 0.375 | 0.147 | 0.892 | 1.189 | 0.006 | 0.139 | 0.139 | 0 | 0.86 | 0.62 | 0.92 | 0.65 | 0.85 | 0.83 |
| 2 | 151673960 | rs1401800 | -0.082 | 0.146 | 0.838 | 1.202 | 0.013 | 0.022 | 0.022 | 0 | 0.88 | 0.71 | 0.93 | 0.81 | 0.85 | 0.88 |
| 2 | 151676174 | rs12998083 | -0.524 | 0.163 | 0.843 | 1.202 | 0.005 | 0.01 | 0.01 | 0 | 0.89 | 0.8 | 0.94 | 0.81 | 0.86 | 0.88 |
| 2 | 151677478 | rs11683080 | 0.539 | 0.132 | 0.566 | 1.206 | -0.005 | 0.123 | 0.123 | 0 | 0.85 | 0.45 | 0.9 | 0.65 | 0.85 | 0.83 |
| 2 | 151681039 | rs7564415 | 0.396 | 0.098 | 0.556 | 1.189 | 0.017 | 0.188 | 0.188 | 0 | 0.86 | 0.61 | 0.92 | 0.65 | 0.85 | 0.82 |
| 2 | 151685251 | rs6729783 | 0.476 | 0.095 | 0.413 | 1.188 | 0 | 0.148 | 0.148 | 0 | 0.85 | 0.46 | 0.91 | 0.66 | 0.85 | 0.83 |
| 2 | 151693284 | rs1519781 | -0.429 | 0.186 | 0.391 | 1.107 | 0 | -0.001 | -0.001 | 0.002 | 0.88 | 0.85 | 0.94 | 0.81 | 0.85 | 0.89 |
| 2 | 151698157 | rs1850879 | -0.132 | 0.124 | 0.234 | 1.059 | 0.007 | 0.013 | 0.013 | 0 | 0.87 | 0.68 | 0.93 | 0.81 | 0.85 | 0.88 |
| 2 | 151704756 | rs929388 | 0.923 | 0.169 | -0.507 | 0.792 | 0.005 | 0.006 | 0.006 | 0 | 0.3 | 0.25 | 0.37 | 0.13 | 0.19 | 0.22 |
| 2 | 151711700 | rs7579851 | 0.175 | -0.151 | -0.721 | 0.762 | -0.058 | -0.038 | -0.038 | 0 | 0.73 | 0.58 | 0.88 | 0.84 | 0.81 | 0.74 |
| 2 | 151714618 | rs767118 | 0.026 | 0.032 | -0.633 | 0.903 | -0.048 | -0.025 | -0.025 | 0 | 0.76 | 0.56 | 0.89 | 0.84 | 0.82 | 0.76 |
| 2 | 151714880 | rs1114978 | 0.341 | 0.059 | -0.938 | 1.007 | -0.02 | 0.009 | 0.009 | 0.006 | 0.66 | 0.42 | 0.8 | 0.58 | 0.74 | 0.72 |
| 2 | 151715691 | rs4664779 | 0.25 | 0.848 | -1.465 | 1.136 | -0.061 | -0.043 | -0.043 | 0 | 0.22 | 0.01 | 0.18 | 0.25 | 0.23 | 0.2 |
| 2 | 151718133 | rs6433010 | 0.826 | 0.698 | -0.13 | 1.628 | -0.024 | 0.259 | 0.259 | 0 | 0.85 | 0.54 | 0.9 | 0.59 | 0.9 | 0.91 |
| 2 | 151726326 | rs13004017 | -0.197 | 1.425 | -0.888 | 1.259 | -0.044 | 0.044 | 0.044 | 0 | 0.16 | 0.09 | 0.12 | 0.05 | 0.27 | 0.3 |
| 2 | 151726959 | rs12467192 | 1.319 | 0.65 | -0.465 | 1.505 | 0.023 | -0.034 | -0.034 | 0 | 0.77 | 0.24 | 0.85 | 0.58 | 0.86 | 0.78 |
| 2 | 151728377 | rs1005274 | 0.98 | 0.918 | -0.057 | 1.683 | -0.015 | 0.077 | 0.077 | 0.576 | 0.85 | 0.31 | 0.87 | 0.58 | 0.9 | 0.91 |
| 2 | 151729571 | rs6433018 | -0.673 | 0.819 | -0.021 | 1.432 | 0 | 0.222 | 0.222 | 0 | 0.11 | 0.3 | 0.09 | 0.3 | 0.07 | 0.03 |
| 2 | 151731930 | rs10189619 | -0.735 | 1.062 | -0.759 | 0.917 | -0.011 | 0.014 | 0.014 | 1.077 | 0.44 | 0.4 | 0.41 | 0.39 | 0.5 | 0.58 |
| 2 | 151731950 | rs1990149 | 0.761 | 1.125 | -0.611 | 0.969 | 0.023 | 0.148 | 0.148 | 0 | 0.92 | 0.79 | 0.84 | 0.68 | 0.89 | 0.95 |
| 2 | 151733201 | rs960615 | 0.582 | 0.04 | -1.808 | 0.183 | 0.079 | 0.095 | 0.095 | 3.362 | 0.31 | 0.06 | 0.2 | 0.13 | 0.21 | 0.2 |
| 2 | 151735155 | rs6723323 | 0.073 | 0.349 | -1.993 | 0.351 | 0.002 | -0.01 | -0.01 | 1.095 | 0.76 | 0.76 | 0.7 | 0.58 | 0.75 | 0.83 |
| 2 | 151738700 | rs757921 | -1.316 | 0.342 | -2.004 | 0.363 | 0.015 | -0.019 | -0.019 | 0.418 | 0.92 | 0.88 | 0.93 | 0.97 | 0.88 | 0.93 |
| 2 | 151739579 | rs10210872 | -0.341 | 0.742 | -2.204 | 0.208 | 0.006 | -0.021 | -0.021 | 0.002 | 0.24 | 0.22 | 0.3 | 0.51 | 0.21 | 0.22 |
| 2 | 151741365 | rs7585613 | 0.966 | 0.795 | -2.341 | 0.25 | -0.007 | -0.022 | -0.022 | 0 | 0.07 | 0.03 | 0.12 | 0.23 | 0.07 | 0.05 |
| 2 | 151741406 | rs7558214 | 0.294 | 0.753 | -2.093 | 0.229 | -0.002 | 0.078 | 0.078 | 0 | 0.09 | 0.21 | 0.14 | 0.23 | 0.07 | 0.05 |
| 2 | 151742522 | rs1859724 | -1.336 | 0.459 | -2.049 | 0.036 | -0.003 | -0.01 | -0.01 | 0 | 0.61 | 0.46 | 0.71 | 0.81 | 0.68 | 0.73 |
| 2 | 151744426 | rs10209774 | 1.073 | 0.386 | -2.665 | -0.002 | 0.007 | -0.014 | -0.014 | 0 | 0.11 | 0.06 | 0.22 | 0.4 | 0.1 | 0.09 |
| 2 | 151746876 | rs2214892 | 0.923 | 1.34 | -2.775 | -0.101 | -0.019 | 0.073 | 0.073 | 0 | 0.26 | 0.11 | 0.25 | 0.08 | 0.28 | 0.44 |
| 2 | 151750506 | rs6730185 | 1.02 | 0.74 | -2.717 | -0.042 | 0.009 | -0.024 | -0.024 | 0 | 0.11 | 0.07 | 0.22 | 0.4 | 0.1 | 0.09 |
| 2 | 151750639 | rs10930331 | 0.521 | 1.069 | -2.672 | -0.12 | -0.083 | 0.092 | 0.092 | 0 | 0.29 | 0.13 | 0.3 | 0.08 | 0.4 | 0.47 |
| 2 | 151753903 | rs10188669 | -0.306 | 0.986 | -2.688 | -0.127 | -0.037 | 0.128 | 0.128 | 0.156 | 0.41 | 0.16 | 0.36 | 0.2 | 0.51 | 0.59 |
| 2 | 151754978 | rs12996110 | 0.247 | 0.317 | -2.778 | -0.631 | -0.07 | 0.223 | 0.223 | 0 | 0.43 | 0.15 | 0.35 | 0.08 | 0.47 | 0.48 |
| 2 | 151756935 | rs10198677 | -0.244 | 1.13 | -1.984 | 0.043 | -0.034 | 0.009 | 0.009 | 0.559 | 0.37 | 0.25 | 0.43 | 0.31 | 0.47 | 0.52 |
| 2 | 151760808 | rs740274 | -0.801 | 1.515 | -1.666 | 0.586 | -0.027 | -0.031 | -0.031 | 0.002 | 0.32 | 0.27 | 0.52 | 0.45 | 0.38 | 0.35 |
| 2 | 151762741 | rs10200704 | -0.429 | 1.336 | -1.692 | -0.003 | -0.026 | 0.06 | 0.06 | 0.505 | 0.25 | 0.12 | 0.31 | 0.12 | 0.33 | 0.18 |
| 2 | 151764512 | rs917237 | 0.031 | 1.128 | -1.208 | -0.039 | -0.001 | -0.023 | -0.023 | 0.109 | 0.81 | 0.79 | 0.76 | 0.95 | 0.79 | 0.89 |
| 2 | 151765163 | rs13010870 | -0.065 | 0.687 | -0.931 | 0.17 | -0.016 | -0.008 | -0.008 | 0 | 0.3 | 0.38 | 0.36 | 0.68 | 0.25 | 0.48 |
| 2 | 151773732 | rs1859720 | 0.781 | 0.903 | -1.735 | 0.358 | -0.084 | -0.02 | -0.02 | 0 | 0.34 | 0.05 | 0.25 | 0.26 | 0.34 | 0.26 |
| 2 | 151773871 | rs13003656 | 0.578 | 0.566 | -1.036 | 0.785 | -0.006 | 0.016 | 0.016 | 0 | 0.21 | 0.23 | 0.31 | 0.35 | 0.17 | 0.35 |
| 2 | 151774093 | rs2190373 | 0.949 | 0.441 | -1.203 | 0.48 | -0.024 | 0.043 | 0.043 | 0.194 | 0.54 | 0.14 | 0.45 | 0.34 | 0.56 | 0.44 |
| 2 | 151774124 | rs11673872 | 0.185 | 0.152 | -0.999 | 0.296 | -0.002 | 0.006 | 0.006 | 0 | 0.28 | 0.3 | 0.35 | 0.33 | 0.26 | 0.38 |
| 2 | 151774643 | rs12328269 | -0.222 | 0.299 | -0.55 | 0.273 | -0.014 | 0.016 | 0.016 | 0 | 0.37 | 0.3 | 0.37 | 0.31 | 0.42 | 0.32 |
| 2 | 151776216 | rs1548632 | -0.091 | 0.115 | -0.329 | 0.265 | -0.003 | -0.018 | -0.018 | 0 | 0.58 | 0.54 | 0.67 | 0.81 | 0.63 | 0.5 |
| 2 | 151781053 | rs12692888 | -0.369 | 0.037 | -0.214 | 0.318 | -0.06 | -0.03 | -0.03 | 0 | 0.79 | 0.54 | 0.89 | 0.86 | 0.89 | 0.68 |
| 2 | 151785370 | rs1465234 | -0.874 | 0.026 | 0.113 | 0.287 | -0.077 | -0.077 | -0.077 | 0 | 0.83 | 0.58 | 0.91 | 0.95 | 0.92 | 0.72 |
| 2 | 151793627 | rs10930342 | -0.362 | 0.009 | 0.162 | 0.308 | 0.015 | 0.089 | 0.089 | 0 | 0.16 | 0.39 | 0.15 | 0.28 | 0.15 | 0.17 |
| 2 | 151798095 | rs10201074 | -1.032 | -0.041 | 0.254 | 0.299 | -0.057 | -0.057 | -0.057 | 0.953 | 0.83 | 0.69 | 0.92 | 0.95 | 0.92 | 0.72 |
| 2 | 151802790 | rs1385430 | 0.259 | 0.203 | 0.112 | 0.562 | 0.007 | 0.031 | 0.031 | 0 | 0.14 | 0.18 | 0.3 | 0.35 | 0.19 | 0.13 |
| 2 | 151802844 | rs1019035 | 0.357 | 0.2 | -0.11 | 0.563 | -0.001 | -0.003 | -0.003 | 0 | 0.47 | 0.4 | 0.39 | 0.49 | 0.45 | 0.39 |
| 2 | 151816753 | rs13003635 | 1.211 | 0.116 | -0.177 | 0.275 | 0.005 | 0.002 | 0.002 | 0 | 0.35 | 0.26 | 0.27 | 0.32 | 0.33 | 0.26 |
| 2 | 151816764 | rs12998436 | 1.211 | 0.116 | -0.177 | 0.275 | 0.005 | 0.002 | 0.002 | 0 | 0.35 | 0.26 | 0.27 | 0.32 | 0.33 | 0.26 |
| 2 | 151821242 | rs6716456 | -0.845 | 0.183 | -0.224 | 0.292 | 0.009 | 0.015 | 0.015 | 0 | 0.27 | 0.31 | 0.43 | 0.58 | 0.33 | 0.27 |
| 2 | 151825296 | rs12992982 | 1.438 | 0.181 | -0.536 | 0.306 | -0.014 | 0.008 | 0.008 | 0 | 0.33 | 0.11 | 0.25 | 0.26 | 0.31 | 0.24 |
| 2 | 151828436 | rs12995627 | -0.537 | 0.134 | -0.221 | 0.164 | -0.008 | -0.057 | -0.057 | 0 | 0.62 | 0.57 | 0.7 | 0.9 | 0.65 | 0.54 |
| 2 | 151832511 | rs16829095 | 1.324 | 0.144 | -0.071 | 0.256 | 0.005 | 0.003 | 0.003 | 0 | 0.36 | 0.28 | 0.27 | 0.32 | 0.33 | 0.27 |
| 2 | 151834334 | rs6708093 | -0.597 | 0.15 | -0.019 | 0.187 | -0.007 | -0.052 | -0.052 | 0 | 0.62 | 0.58 | 0.7 | 0.9 | 0.65 | 0.54 |
| 2 | 151836383 | rs6713372 | 1.158 | 0.212 | -0.063 | 0.55 | -0.002 | 0.001 | 0.001 | 0 | 0.09 | 0.08 | 0.11 | 0.22 | 0.11 | 0.07 |
| 2 | 151836787 | rs6433099 | -0.568 | 0.185 | -0.019 | 0.457 | -0.014 | -0.081 | -0.081 | 0 | 0.62 | 0.5 | 0.69 | 0.9 | 0.65 | 0.54 |
| 2 | 151839884 | rs1118980 | -0.461 | 0.225 | -0.08 | 0.493 | -0.01 | -0.044 | -0.044 | 0 | 0.61 | 0.54 | 0.69 | 0.84 | 0.64 | 0.58 |
| 2 | 151862198 | rs10211364 | 1.271 | 0.092 | -0.16 | 0.369 | -0.011 | 0.045 | 0.045 | 0.295 | 0.23 | 0.14 | 0.23 | 0.05 | 0.28 | 0.19 |
| 2 | 151865022 | rs10206535 | 0.139 | 0.073 | 0.052 | 0.523 | -0.05 | 0.008 | 0.008 | 0 | 0.75 | 0.51 | 0.82 | 0.74 | 0.82 | 0.72 |
| 2 | 151870187 | rs7598311 | 0.295 | 0.211 | 0.013 | 0.668 | 0.022 | 0.041 | 0.041 | 0 | 0.14 | 0.22 | 0.32 | 0.24 | 0.19 | 0.13 |
| 2 | 151871888 | rs4664875 | 0.482 | 0.229 | -0.086 | 0.694 | -0.026 | 0.03 | 0.03 | 0 | 0.71 | 0.48 | 0.79 | 0.63 | 0.78 | 0.69 |
| 2 | 151873098 | rs12464767 | 1.05 | 0.214 | -0.019 | 0.675 | 0.001 | 0.019 | 0.019 | 0 | 0.09 | 0.1 | 0.09 | 0.22 | 0.11 | 0.08 |
| 2 | 151873828 | rs10186209 | 0.753 | 0.195 | -0.163 | 0.709 | -0.002 | 0.018 | 0.018 | 0.021 | 0.63 | 0.45 | 0.74 | 0.54 | 0.64 | 0.66 |
| 2 | 151874500 | rs6735085 | 0.097 | 0.01 | -0.048 | 0.604 | -0.042 | 0.014 | 0.014 | 0 | 0.75 | 0.48 | 0.82 | 0.72 | 0.81 | 0.72 |

| | | | | | | | | | | | | | | | | |
|---|---|---|---|---|---|---|---|---|---|---|---|---|---|---|---|---|
| 2 | 151885556 | rs7603077 | 0.654 | 0.137 | 0.013 | 0.469 | -0.001 | -0.02 | -0.02 | 0 | 0.27 | 0.33 | 0.47 | 0.15 | 0.29 | 0.28 |
| 2 | 151890227 | rs16829311 | 0.733 | 0.117 | 0.024 | 0.575 | -0.019 | 0 | 0 | 2.664 | 0.06 | 0.13 | 0.04 | 0.06 | 0.02 | 0.06 |
| 2 | 151890819 | rs6433123 | -0.757 | 0.06 | 0.096 | 0.511 | -0.002 | -0.002 | -0.002 | 1.055 | 0.51 | 0.54 | 0.68 | 0.64 | 0.63 | 0.45 |
| 2 | 151892897 | rs1485127 | -0.595 | 0.962 | 0.354 | 0.572 | 0.01 | 0.01 | 0.01 | 0 | 0.2 | 0.22 | 0.18 | 0.43 | 0.35 | 0.23 |
| 2 | 151892910 | rs1485126 | -0.537 | 0.305 | 0.313 | 0.161 | -0.022 | -0.057 | -0.057 | 0 | 0.59 | 0.54 | 0.48 | 0.88 | 0.66 | 0.58 |
| 2 | 151895088 | rs1463609 | 0.881 | 0.305 | 0.378 | 0.071 | 0.007 | 0.002 | 0.002 | 0 | 0.32 | 0.24 | 0.46 | 0.06 | 0.3 | 0.27 |
| 2 | 151895659 | rs16829328 | 0.149 | 0.794 | 0.452 | 0.384 | 0.057 | 0.087 | 0.087 | 0 | 0.08 | 0.22 | 0.11 | 0.24 | 0.11 | 0.1 |
| 2 | 151896995 | rs10497076 | -0.228 | 0.608 | 0.584 | 0.233 | -0.008 | 0.003 | 0.003 | 0 | 0.09 | 0.38 | 0.07 | 0.05 | 0.04 | 0.1 |
| 2 | 151897448 | rs6728040 | 0.036 | 0.811 | 0.313 | 0.328 | -0.031 | -0.029 | -0.029 | 0 | 0.13 | 0.05 | 0.1 | 0.23 | 0.24 | 0.13 |
| 2 | 151910669 | rs4664900 | 1.536 | 0.647 | -0.02 | 0.112 | -0.011 | -0.123 | -0.123 | 0 | 0.31 | 0.03 | 0.22 | 0.36 | 0.26 | 0.24 |
| 2 | 151918816 | rs12692945 | -0.071 | 0.811 | 0.191 | 0.057 | 0.023 | 0.079 | 0.079 | 0 | 0.49 | 0.22 | 0.34 | 0.63 | 0.51 | 0.58 |
| 2 | 151922077 | rs11686669 | -0.094 | 0.513 | 0.166 | 0.089 | 0.069 | 0.002 | 0.002 | 0.274 | 0.51 | 0.25 | 0.64 | 0.52 | 0.65 | 0.46 |
| 2 | 151924098 | rs2341849 | 1.12 | 0.996 | 0.43 | 0.418 | 0.011 | 0.024 | 0.024 | 0 | 0.06 | 0.08 | 0.1 | 0.24 | 0.11 | 0.04 |
| 2 | 151930856 | rs17197919 | 0.22 | 1.061 | 0.266 | 0.291 | -0.056 | -0.06 | -0.06 | 0 | 0.16 | 0.02 | 0.09 | 0.22 | 0.21 | 0.12 |
| 2 | 151931005 | rs11675356 | -1.29 | 1.024 | 0.523 | 0.245 | 0.003 | 0 | 0 | 0 | 0.28 | 0.4 | 0.17 | 0.27 | 0.28 | 0.3 |
| 2 | 151931439 | rs13026867 | 1.973 | 0.913 | 0.23 | 0.134 | -0.041 | -0.131 | -0.131 | 0 | 0.23 | 0.03 | 0.2 | 0.36 | 0.23 | 0.23 |
| 2 | 151932232 | rs7585884 | 0.897 | 0.818 | 0.328 | 0.108 | -0.019 | 0.061 | 0.061 | 0.574 | 0.43 | 0.22 | 0.57 | 0.32 | 0.47 | 0.38 |
| 2 | 151936086 | rs2046421 | 0.505 | 0.234 | 0.641 | 0.228 | -0.028 | -0.023 | -0.023 | 0 | 0.87 | 0.49 | 0.89 | 0.95 | 0.93 | 0.88 |
| 2 | 151938118 | rs2046422 | -0.834 | 0.407 | 0.619 | 0.29 | -0.018 | -0.042 | -0.042 | 0.571 | 0.22 | 0.18 | 0.15 | 0.52 | 0.28 | 0.29 |
| 2 | 151942632 | rs17197926 | 1.121 | 0.414 | 0.642 | 0.203 | 0.009 | 0.012 | 0.012 | 0.746 | 0.37 | 0.26 | 0.48 | 0.06 | 0.34 | 0.33 |
| 2 | 151946149 | rs720066 | 1.222 | 0.413 | 0.63 | 0.203 | 0.016 | 0.059 | 0.059 | 0.349 | 0.36 | 0.21 | 0.48 | 0.06 | 0.34 | 0.33 |
| 2 | 151954166 | rs749691 | -1.096 | 0.411 | 0.675 | 0.209 | -0.01 | -0.047 | -0.047 | 0 | 0.61 | 0.58 | 0.5 | 0.89 | 0.66 | 0.59 |
| 2 | 151959513 | rs16829500 | -0.154 | 0.452 | 0.476 | 0.272 | 0.009 | 0.1 | 0.1 | 0 | 0.06 | 0.17 | 0.07 | 0.25 | 0.06 | 0.03 |
| 2 | 151960452 | rs1485099 | -0.604 | 0.438 | 0.477 | 0.241 | 0.033 | 0.081 | 0.081 | 0.829 | 0.53 | 0.31 | 0.41 | 0.88 | 0.59 | 0.55 |
| 2 | 151962991 | rs4664951 | -0.221 | 0.442 | 0.42 | 0.249 | 0.025 | 0.081 | 0.081 | 1.158 | 0.48 | 0.18 | 0.35 | 0.63 | 0.53 | 0.51 |
| 2 | 151970688 | rs4297880 | -0.478 | 0.564 | 0.3 | 0.381 | -0.023 | -0.033 | -0.033 | 0.058 | 0.21 | 0.08 | 0.12 | 0.28 | 0.23 | 0.29 |
| 2 | 151971783 | rs7584360 | 0.511 | 0.652 | 0.412 | 0.405 | -0.009 | -0.015 | -0.015 | 0 | 0.42 | 0.43 | 0.56 | 0.36 | 0.4 | 0.43 |
| 2 | 151972837 | rs10167013 | 0.053 | 0.614 | 0.185 | 0.52 | 0.046 | 0.245 | 0.245 | 0 | 0.06 | 0.21 | 0.07 | 0.3 | 0.06 | 0.11 |
| 2 | 151983855 | rs816709 | -0.327 | 0.607 | -0.526 | 0.244 | -0.012 | 0.08 | 0.08 | 0.031 | 0.48 | 0.85 | 0.42 | 0.72 | 0.45 | 0.55 |
| 2 | 151987931 | rs6711018 | -0.064 | 0.398 | -0.271 | 0.497 | 0.001 | 0.007 | 0.007 | 0 | 0.2 | 0.26 | 0.14 | 0.2 | 0.23 | 0.14 |
| 2 | 151996692 | rs13001465 | 0.133 | 0.456 | -0.347 | 0.603 | -0.018 | -0.025 | -0.025 | 0 | 0.36 | 0.24 | 0.29 | 0.45 | 0.36 | 0.42 |
| 2 | 151997571 | rs289946 | 0.146 | 0.556 | -0.564 | 0.61 | -0.01 | -0.009 | -0.009 | 0 | 0.34 | 0.32 | 0.46 | 0.36 | 0.35 | 0.29 |
| 2 | 152000346 | rs289955 | -1.545 | -0.137 | 0.273 | 0.527 | 0.035 | 0.064 | 0.064 | 9.615 | 0.06 | 0.39 | 0.04 | 0.05 | 0 | 0.13 |
| 2 | 152006025 | rs289932 | -0.826 | -0.281 | -0.118 | 0.437 | -0.013 | 0.032 | 0.032 | 2.576 | 0.33 | 0.5 | 0.33 | 0.58 | 0.36 | 0.51 |
| 2 | 152007227 | rs403973 | 0.503 | -0.522 | -1.034 | 0.189 | 0.018 | 0.012 | 0.012 | 0 | 0.5 | 0.4 | 0.56 | 0.61 | 0.54 | 0.58 |
| 2 | 152008089 | rs172342 | -0.169 | -0.798 | -0.643 | 0.321 | -0.022 | 0.036 | 0.036 | 0 | 0.15 | 0.37 | 0.08 | 0.24 | 0.13 | 0.28 |
| 2 | 152015683 | rs1949463 | 0.488 | -0.774 | -0.593 | 0.663 | -0.01 | 0.005 | 0.005 | 2.54 | 0.18 | 0.11 | 0.21 | 0.16 | 0.24 | 0.19 |
| 2 | 152021778 | rs10193255 | -0.242 | -1.01 | 0.158 | 0.393 | -0.012 | -0.007 | -0.007 | 0 | 0.36 | 0.53 | 0.2 | 0.34 | 0.28 | 0.47 |
| 2 | 152024889 | rs289848 | -1.542 | -0.95 | 0.047 | 0.33 | -0.02 | -0.004 | -0.004 | 0 | 0.58 | 0.77 | 0.44 | 0.47 | 0.52 | 0.68 |
| 2 | 152030598 | rs10930471 | -0.325 | -1 | 0.321 | 0.258 | -0.004 | 0.009 | 0.009 | 0 | 0.17 | 0.24 | 0.09 | 0.27 | 0.16 | 0.33 |
| 2 | 152049608 | rs289880 | 0.538 | -1.065 | 0.506 | 0.283 | -0.03 | 0.086 | 0.086 | 1.247 | 0.87 | 0.63 | 0.9 | 0.85 | 0.94 | 0.9 |
| 2 | 152050243 | rs289854 | 1.114 | -1.08 | 0.533 | 0.101 | 0.007 | -0.004 | -0.004 | 0.093 | 0.2 | 0.17 | 0.11 | 0.04 | 0.13 | 0.16 |
| 2 | 152055452 | rs289862 | -1.029 | -1.052 | 0.515 | 0.145 | 0.004 | 0.005 | 0.005 | 5.1 | 0.49 | 0.48 | 0.38 | 0.45 | 0.45 | 0.56 |
| 2 | 152070336 | rs289827 | 0.739 | -1.041 | 0.152 | 0.116 | -0.01 | -0.039 | -0.039 | 0.076 | 0.22 | 0.23 | 0.13 | 0.04 | 0.14 | 0.17 |
| 2 | 152085643 | rs289904 | -1.304 | -1.046 | -0.323 | 0.039 | 0.006 | -0.004 | -0.004 | 1.642 | 0.58 | 0.49 | 0.44 | 0.47 | 0.54 | 0.6 |
| 2 | 152101800 | rs289890 | -0.431 | -1.065 | -0.31 | 0.088 | -0.013 | 0.04 | 0.04 | 0 | 0.95 | 0.83 | 0.98 | 0.94 | 0.97 | 0.97 |
| 2 | 152103077 | rs4665114 | 2.03 | -1.126 | -0.408 | 0.123 | -0.168 | -0.117 | -0.117 | 0 | 0.33 | 0.05 | 0.49 | 0.39 | 0.43 | 0.34 |
| 2 | 152106653 | rs289883 | 0.987 | -1.26 | -0.322 | 0.016 | -0.042 | -0.098 | -0.098 | 0.176 | 0.21 | 0.34 | 0.12 | 0.03 | 0.12 | 0.17 |
| 2 | 152108405 | rs4665120 | -0.308 | -1.382 | -0.422 | -0.039 | -0.066 | -0.077 | -0.077 | 0 | 0.75 | 0.49 | 0.86 | 0.9 | 0.86 | 0.8 |
| 2 | 152111965 | rs11683285 | 1.306 | -1.308 | -0.554 | 0.002 | -0.021 | 0.034 | 0.034 | 0 | 0.07 | 0 | 0.04 | 0.02 | 0.07 | 0.1 |
| 2 | 152117780 | rs2271811 | 1.947 | -1.191 | -0.425 | 0.076 | -0.091 | -0.049 | -0.049 | 0.134 | 0.36 | 0.14 | 0.5 | 0.38 | 0.44 | 0.34 |
| 2 | 152117957 | rs2271813 | 0.531 | -1.249 | -0.484 | 0.175 | -0.052 | -0.052 | -0.052 | 0.3 | 0.14 | 0 | 0.08 | 0.22 | 0.18 | 0.26 |
| 2 | 152126603 | rs453226 | 0.24 | -1.312 | -0.234 | 0.067 | -0.056 | -0.103 | -0.103 | 0 | 0.22 | 0.39 | 0.12 | 0.05 | 0.12 | 0.17 |
| 2 | 152127011 | rs11730 | 2.08 | -1.275 | -0.416 | 0.03 | -0.167 | -0.106 | -0.106 | 0 | 0.35 | 0.05 | 0.49 | 0.39 | 0.44 | 0.33 |
| 2 | 152129879 | rs446791 | -0.358 | -1.21 | -0.202 | 0.059 | 0.012 | 0.012 | 0.012 | 0.061 | 0.26 | 0.69 | 0.17 | 0.11 | 0.14 | 0.23 |
| 2 | 152132130 | rs289834 | 0.449 | -1.185 | -0.237 | 0.028 | -0.042 | -0.081 | -0.081 | 0.05 | 0.21 | 0.32 | 0.12 | 0.05 | 0.12 | 0.17 |
| 2 | 152135415 | rs11551174 | 2.049 | -1.217 | -0.536 | -0.014 | 0.002 | 0 | 0 | 0 | 0.06 | 0.05 | 0.12 | 0 | 0.05 | 0.09 |
| 2 | 152138522 | rs3771886 | 0.629 | -1.195 | -0.568 | 0.038 | -0.001 | -0.004 | -0.004 | 0.141 | 0.6 | 0.65 | 0.63 | 0.44 | 0.56 | 0.5 |
| 2 | 152139416 | rs1048135 | 2.084 | -1.196 | -0.803 | 0.034 | -0.166 | -0.104 | -0.104 | 0 | 0.36 | 0.05 | 0.49 | 0.38 | 0.44 | 0.34 |
| 2 | 152143524 | rs13383563 | -0.031 | -1.115 | -0.645 | -0.079 | -0.018 | -0.04 | -0.04 | 0 | 0.23 | 0.68 | 0.15 | 0.05 | 0.12 | 0.17 |
| 2 | 152146672 | rs2278089 | 0.324 | -1.098 | -0.706 | -0.01 | -0.042 | -0.017 | -0.017 | 0.188 | 0.64 | 0.9 | 0.69 | 0.59 | 0.59 | 0.58 |
| 2 | 152159880 | rs6726634 | 1.464 | -1 | -0.529 | 0.174 | -0.022 | 0.013 | 0.013 | 0.604 | 0.43 | 0.24 | 0.55 | 0.55 | 0.49 | 0.41 |
| 2 | 152186147 | rs10490522 | 0.289 | -1.032 | -0.39 | 0.155 | -0.019 | -0.018 | -0.018 | 0 | 0.69 | 0.81 | 0.78 | 0.6 | 0.62 | 0.55 |
| 2 | 152193359 | rs4664349 | 1.605 | -1.041 | -0.343 | 0.23 | -0.008 | 0.005 | 0.005 | 0.11 | 0.43 | 0.4 | 0.51 | 0.4 | 0.46 | 0.34 |
| 2 | 152195436 | rs6725961 | 0.148 | -1.001 | -0.33 | 0.253 | -0.044 | -0.024 | -0.024 | 0 | 0.23 | 0.31 | 0.23 | 0.11 | 0.14 | 0.17 |
| 2 | 152195934 | rs4664356 | 0.896 | -0.913 | -0.636 | 0.189 | 0.012 | -0.007 | -0.007 | 0 | 0.54 | 0.42 | 0.6 | 0.49 | 0.57 | 0.48 |
| 2 | 152201213 | rs12471886 | 1.587 | -0.653 | -1.187 | 0.322 | -0.138 | -0.061 | -0.061 | 0 | 0.4 | 0.07 | 0.5 | 0.39 | 0.47 | 0.36 |
| 2 | 152201469 | rs6715902 | -0.222 | -0.702 | -1.023 | 0.295 | -0.006 | -0.035 | -0.035 | 0.026 | 0.72 | 0.82 | 0.79 | 0.59 | 0.67 | 0.55 |
| 2 | 152205741 | rs6433371 | -0.904 | -0.592 | -0.892 | 0.219 | 0.07 | -0.037 | -0.037 | 0 | 0.9 | 0.6 | 0.83 | 0.99 | 0.9 | 0.93 |

| | | | | | | | | | | | | | | | | |
|---|---|---|---|---|---|---|---|---|---|---|---|---|---|---|---|---|
| 2 | 152206979 | rs12466578 | 1.801 | -0.162 | -1.556 | 0.308 | -0.074 | -0.058 | -0.058 | 0 | 0.38 | 0.13 | 0.47 | 0.42 | 0.38 | 0.35 |
| 2 | 152210713 | rs2342909 | 0.232 | 0.172 | -1.267 | 0.818 | -0.026 | -0.059 | -0.059 | 0 | 0.23 | 0.12 | 0.19 | 0.4 | 0.3 | 0.37 |
| 2 | 152218231 | rs3771892 | -0.327 | 0.628 | -1.057 | 1.046 | 0.041 | 0.075 | 0.075 | 0 | 0.12 | 0.31 | 0.11 | 0.13 | 0.14 | 0.14 |
| 2 | 152224339 | rs3771895 | -1.105 | 0.604 | -1.077 | 1.05 | 0.007 | 0.019 | 0.019 | 4.299 | 0.5 | 0.42 | 0.35 | 0.57 | 0.51 | 0.59 |
| 2 | 152226570 | rs1046668 | -1.075 | 0.925 | 0.275 | 1.077 | 0.043 | 0.078 | 0.078 | 0 | 0.11 | 0.3 | 0.11 | 0.13 | 0.14 | 0.14 |
| 2 | 152231294 | rs3845844 | -0.956 | 0.965 | -0.113 | 0.777 | 0.025 | 0.018 | 0.018 | 0 | 0.11 | 0.21 | 0.11 | 0.08 | 0.14 | 0.1 |
| 2 | 152236398 | rs1046675 | -0.956 | 0.965 | -0.113 | 0.777 | 0.025 | 0.018 | 0.018 | 0 | 0.11 | 0.21 | 0.11 | 0.08 | 0.14 | 0.1 |
| 2 | 152253359 | rs12618981 | 0.424 | 1.106 | -1.049 | 0.641 | -0.113 | -0.056 | -0.056 | 0 | 0.26 | 0.02 | 0.28 | 0.25 | 0.31 | 0.19 |
| 2 | 152253918 | rs10197940 | -0.407 | 0.94 | -0.285 | 0.833 | -0.002 | -0.003 | -0.003 | 0 | 0.38 | 0.37 | 0.44 | 0.49 | 0.4 | 0.44 |
| 2 | 152266311 | rs4664423 | -0.405 | 0.932 | -0.183 | 0.803 | 0.002 | 0.002 | 0.002 | 0 | 0.38 | 0.42 | 0.45 | 0.49 | 0.4 | 0.44 |
| 2 | 152266473 | rs2290368 | -0.405 | 0.932 | -0.196 | 0.803 | 0.002 | 0.003 | 0.003 | 0 | 0.38 | 0.42 | 0.45 | 0.49 | 0.4 | 0.44 |
| 2 | 152267359 | rs16830019 | 0.686 | 0.984 | -0.875 | 0.78 | -0.046 | -0.111 | -0.111 | 0 | 0.3 | 0.02 | 0.36 | 0.31 | 0.3 | 0.18 |
| 2 | 152271907 | rs4315520 | 0.129 | 0.751 | -0.887 | 0.631 | 0.012 | -0.017 | -0.017 | 0 | 0.3 | 0.42 | 0.29 | 0.25 | 0.32 | 0.48 |
| 2 | 152287897 | rs2432957 | 0.852 | 0.835 | -0.658 | 0.524 | 0.009 | -0.027 | -0.027 | 0 | 0.35 | 0.45 | 0.35 | 0.25 | 0.37 | 0.2 |
| 2 | 152311570 | rs2444263 | -0.245 | 0.825 | -0.659 | 0.52 | 0.013 | -0.025 | -0.025 | 0 | 0.7 | 0.58 | 0.71 | 0.76 | 0.68 | 0.52 |
| 2 | 152313539 | rs7422009 | 0.945 | 0.808 | -0.656 | 0.548 | -0.015 | -0.085 | -0.085 | 0 | 0.3 | 0.06 | 0.36 | 0.32 | 0.3 | 0.22 |
| 2 | 152320118 | rs2123465 | -0.244 | 0.816 | -0.644 | 0.534 | 0.012 | -0.025 | -0.025 | 0 | 0.7 | 0.58 | 0.71 | 0.76 | 0.68 | 0.52 |
| 2 | 152331745 | rs16830067 | 1.019 | 0.822 | -0.404 | 0.607 | -0.03 | -0.093 | -0.093 | 0.768 | 0.29 | 0.04 | 0.35 | 0.31 | 0.29 | 0.18 |
| 2 | 152336428 | rs3815851 | -0.58 | 0.838 | -0.385 | 0.587 | 0.01 | -0.024 | -0.024 | 0 | 0.73 | 0.58 | 0.74 | 0.76 | 0.73 | 0.53 |
| 2 | 152346979 | rs1061305 | 0.736 | 0.805 | -0.261 | 0.682 | 0.009 | -0.025 | -0.025 | 0 | 0.35 | 0.46 | 0.35 | 0.25 | 0.37 | 0.2 |
| 2 | 152351126 | rs4994327 | -0.45 | 0.649 | -0.051 | 0.775 | 0.009 | -0.022 | -0.022 | 0 | 0.69 | 0.59 | 0.71 | 0.77 | 0.66 | 0.52 |
| 2 | 152382199 | rs7585717 | -0.527 | 0.707 | 0.161 | 0.819 | 0.001 | -0.005 | -0.005 | 0 | 0.69 | 0.64 | 0.7 | 0.75 | 0.66 | 0.49 |
| 2 | 152387553 | rs4664475 | 0.098 | 0.742 | 0.268 | 0.865 | -0.001 | 0.002 | 0.002 | 0.611 | 0.61 | 0.35 | 0.66 | 0.56 | 0.62 | 0.36 |
| 2 | 152401311 | rs9287983 | -0.078 | 0.743 | 0.197 | 0.861 | 0.002 | -0.004 | -0.004 | 0 | 0.61 | 0.46 | 0.66 | 0.56 | 0.63 | 0.36 |
| 2 | 152402171 | rs4664055 | -0.518 | 0.74 | 0.213 | 0.862 | 0.002 | -0.01 | -0.01 | 0 | 0.64 | 0.61 | 0.69 | 0.76 | 0.63 | 0.47 |
| 2 | 152412203 | rs4664058 | 0.955 | 0.654 | 0.074 | 0.92 | -0.037 | -0.111 | -0.111 | 0 | 0.24 | 0.03 | 0.35 | 0.31 | 0.28 | 0.16 |
| 2 | 152417999 | rs7424331 | -0.631 | 0.737 | 0.155 | 0.854 | 0 | -0.016 | -0.016 | 0 | 0.65 | 0.61 | 0.7 | 0.77 | 0.68 | 0.53 |
| 2 | 152419855 | rs16830228 | 0.859 | 0.954 | -0.236 | 0.852 | -0.065 | -0.105 | -0.105 | 0.52 | 0.25 | 0.04 | 0.38 | 0.3 | 0.33 | 0.18 |
| 2 | 152421937 | rs7602715 | 0.992 | 0.933 | -0.167 | 0.85 | 0.006 | -0.012 | -0.012 | 1.24 | 0.31 | 0.22 | 0.25 | 0.4 | 0.27 | 0.26 |
| 2 | 152422087 | rs2288211 | 0.683 | 1.035 | -0.417 | 0.903 | -0.065 | -0.105 | -0.105 | 0 | 0.26 | 0.04 | 0.38 | 0.3 | 0.33 | 0.18 |
| 2 | 152467001 | rs11902616 | -1.227 | 0.641 | 0.109 | 1.031 | 0.036 | 0.2 | 0.2 | 0.066 | 0.08 | 0.38 | 0.04 | 0.18 | 0.03 | 0.16 |
| 2 | 152476130 | rs6717213 | -0.676 | 0.717 | -0.184 | 1.069 | 0.008 | 0.131 | 0.131 | 0.289 | 0.06 | 0.25 | 0.04 | 0.18 | 0.03 | 0.13 |
| 2 | 152485754 | rs6750342 | 0.78 | 0.843 | -1.06 | 0.948 | -0.048 | 0.019 | 0.019 | 1.036 | 0.32 | 0.14 | 0.28 | 0.25 | 0.36 | 0.21 |
| 2 | 152486782 | rs4664492 | 0.704 | 0.955 | -1.052 | 0.874 | -0.041 | -0.099 | -0.099 | 0 | 0.22 | 0.03 | 0.34 | 0.29 | 0.26 | 0.16 |
| 2 | 152487574 | rs4233653 | 0.466 | 0.873 | -0.972 | 0.929 | -0.057 | -0.084 | -0.084 | 0 | 0.25 | 0.07 | 0.38 | 0.3 | 0.32 | 0.16 |
| 2 | 152492324 | rs4664493 | 0.121 | 0.484 | -0.087 | 0.713 | -0.004 | -0.049 | -0.049 | 0 | 0.61 | 0.49 | 0.68 | 0.77 | 0.65 | 0.5 |
| 2 | 152496526 | rs6713162 | -2.041 | 0.078 | 0.751 | 1.715 | 0.013 | -0.023 | -0.023 | 1.162 | 0.26 | 0.72 | 0.16 | 0.63 | 0.17 | 0.36 |
| 2 | 152517979 | rs10192459 | -1.338 | 0.116 | 1.513 | 1.727 | 0.007 | -0.032 | -0.032 | 0.84 | 0.09 | 0.24 | 0.05 | 0.05 | 0.06 | 0.06 |
| 2 | 152521096 | rs10170273 | -2.766 | 0.08 | 1.799 | 1.779 | 0.008 | -0.015 | -0.015 | 0 | 0.25 | 0.68 | 0.16 | 0.63 | 0.18 | 0.36 |
| 2 | 152526082 | rs6712533 | -1.772 | 0.097 | 1.94 | 1.802 | -0.019 | 0.227 | 0.227 | 0.293 | 0.16 | 0.42 | 0.11 | 0.58 | 0.12 | 0.31 |
| 2 | 152531077 | rs6711382 | 1.513 | 0.087 | 2.314 | 2.052 | -0.004 | 0.238 | 0.238 | 0.156 | 0.85 | 0.59 | 0.92 | 0.58 | 0.88 | 0.7 |
| 2 | 152551210 | rs10193188 | -2.733 | 0.239 | 2.236 | 1.967 | -0.003 | 0.011 | 0.011 | 0.246 | 0.25 | 0.63 | 0.15 | 0.62 | 0.19 | 0.37 |
| 2 | 152567908 | rs6433575 | 2.451 | 0.2 | 2.197 | 1.981 | 0.006 | -0.02 | -0.02 | 0 | 0.75 | 0.27 | 0.84 | 0.38 | 0.81 | 0.63 |
| 2 | 152573981 | rs4611637 | 2.425 | 0.197 | 2.252 | 2.01 | 0.002 | -0.006 | -0.006 | 2.315 | 0.75 | 0.27 | 0.85 | 0.47 | 0.81 | 0.63 |
| 2 | 152574284 | rs4413159 | 2.425 | 0.197 | 2.252 | 2.01 | 0.002 | -0.006 | -0.006 | 0 | 0.75 | 0.27 | 0.85 | 0.47 | 0.81 | 0.63 |
| 2 | 152584153 | rs4664499 | 0.879 | 0.207 | 2.398 | 2.046 | 0.017 | 0.331 | 0.331 | 0 | 0.9 | 0.67 | 0.94 | 0.58 | 0.91 | 0.73 |
| 2 | 152585020 | rs13025954 | -1.481 | 0.214 | 2.453 | 1.953 | 0.015 | 0.007 | 0.007 | 0 | 0.1 | 0.2 | 0.05 | 0.13 | 0.08 | 0.07 |
| 2 | 152587344 | rs10165921 | 1.868 | 0.22 | 2.278 | 1.837 | -0.012 | 0.294 | 0.294 | 1.204 | 0.85 | 0.46 | 0.9 | 0.53 | 0.89 | 0.7 |
| 2 | 152587814 | rs6760384 | 2.455 | 0.175 | 1.851 | 1.828 | -0.001 | 0.006 | 0.006 | 0.335 | 0.75 | 0.27 | 0.85 | 0.48 | 0.81 | 0.63 |
| 2 | 152593795 | rs4510224 | 0.895 | 0.237 | 1.869 | 1.844 | 0.013 | 0.308 | 0.308 | 0.003 | 0.9 | 0.65 | 0.94 | 0.58 | 0.91 | 0.73 |
| 2 | 152609432 | rs4274593 | 2.519 | 0.245 | 1.796 | 1.85 | -0.003 | 0.019 | 0.019 | 0.312 | 0.75 | 0.28 | 0.85 | 0.47 | 0.81 | 0.63 |
| 2 | 152632406 | rs4664066 | 0.815 | 0.254 | 1.48 | 2.097 | 0.009 | 0.281 | 0.281 | 0 | 0.91 | 0.7 | 0.94 | 0.58 | 0.91 | 0.73 |
| 2 | 152651592 | rs11682909 | 0.643 | 0.26 | 1.399 | 2.093 | 0.009 | 0.274 | 0.274 | 0 | 0.91 | 0.72 | 0.95 | 0.58 | 0.91 | 0.73 |
| 2 | 152652644 | rs7594616 | -2.194 | 0.36 | 1.505 | 1.921 | 0.013 | -0.01 | -0.01 | 0.157 | 0.14 | 0.38 | 0.09 | 0.11 | 0.09 | 0.1 |
| 2 | 152655652 | rs2293222 | -2.243 | 0.356 | 1.474 | 1.919 | 0.01 | -0.013 | -0.013 | 0 | 0.14 | 0.37 | 0.09 | 0.11 | 0.09 | 0.1 |
| 2 | 152672843 | rs10182533 | -2.204 | 0.365 | 1.479 | 1.923 | 0.011 | -0.012 | -0.012 | 0 | 0.14 | 0.37 | 0.09 | 0.1 | 0.09 | 0.1 |
| 2 | 152694448 | rs13410490 | -1.525 | 0.043 | -0.194 | 0.888 | -0.008 | -0.04 | -0.04 | 0 | 0.1 | 0.21 | 0.06 | 0.03 | 0.08 | 0.09 |
| 2 | 152716628 | rs3768653 | 0.66 | -0.456 | -1.234 | 0.326 | -0.042 | 0.103 | 0.103 | 0.555 | 0.23 | 0.02 | 0.13 | 0.04 | 0.19 | 0.16 |
| 2 | 152725937 | rs3768652 | -1.177 | -0.43 | -0.816 | 0.554 | -0.003 | 0.001 | 0.001 | 0 | 0.1 | 0.16 | 0.07 | 0.13 | 0.09 | 0.1 |
| 2 | 152736391 | rs6742375 | -0.275 | -0.862 | -0.731 | 0.147 | 0.027 | 0.068 | 0.068 | 0.532 | 0.34 | 0.2 | 0.16 | 0.07 | 0.23 | 0.12 |
| 2 | 152741725 | rs1401780 | 0.88 | -1.637 | -0.391 | 0.405 | 0.037 | 0.129 | 0.129 | 1.038 | 0.48 | 0.04 | 0.49 | 0.2 | 0.66 | 0.45 |
| 2 | 152760078 | rs11695363 | 0.784 | -2.048 | -0.983 | -0.093 | 0.01 | 0.13 | 0.13 | 0 | 0.21 | 0.01 | 0.07 | 0.02 | 0.13 | 0.17 |
| 2 | 152763157 | rs17270290 | 0.145 | -1.643 | 0.07 | 0.322 | 0.014 | 0.025 | 0.025 | 0.453 | 0.34 | 0.23 | 0.16 | 0.07 | 0.24 | 0.12 |
| 2 | 152775308 | rs12471646 | 0.958 | -1.645 | -0.2 | 0.369 | -0.041 | 0.094 | 0.094 | 0.341 | 0.46 | 0.05 | 0.48 | 0.2 | 0.63 | 0.42 |
| 2 | 152775388 | rs956745 | -0.895 | -1.68 | 0.24 | 0.365 | 0.071 | 0.107 | 0.107 | 0 | 0.49 | 0.76 | 0.48 | 0.8 | 0.33 | 0.55 |
| 2 | 152788303 | rs11895837 | 0.137 | -1.778 | 0.266 | 0.115 | 0.015 | 0.027 | 0.027 | 2.375 | 0.35 | 0.23 | 0.17 | 0.07 | 0.24 | 0.11 |
| 2 | 152790117 | rs12623883 | 0.211 | -1.908 | 0.286 | 0.409 | 0.014 | -0.011 | -0.011 | 0 | 0.09 | 0.04 | 0.05 | 0.57 | 0.07 | 0.42 |
| 2 | 152794904 | rs1356732 | 0.878 | -1.941 | 0.839 | 0.4 | 0.082 | 0.311 | 0.311 | 3.581 | 0.94 | 0.5 | 0.72 | 0.82 | 0.97 | 0.94 |
| 2 | 152794981 | rs1356733 | -0.518 | -1.897 | 0.505 | 0.395 | 0.001 | -0.039 | -0.039 | 0.079 | 0.13 | 0.08 | 0.06 | 0.57 | 0.11 | 0.44 |

| | | | | | | | | | | | | | | | | |
|---|---|---|---|---|---|---|---|---|---|---|---|---|---|---|---|---|
| 2 | 152806958 | rs12618445 | 0.241 | -1.913 | 0.518 | 0.246 | 0.013 | -0.014 | -0.014 | 0.165 | 0.09 | 0.04 | 0.06 | 0.61 | 0.07 | 0.41 |
| 2 | 152807966 | rs7571637 | 1.039 | -1.954 | 0.445 | 0.155 | 0.044 | 0.182 | 0.182 | 1.298 | 0.47 | 0.1 | 0.49 | 0.18 | 0.65 | 0.42 |
| 2 | 152808468 | rs7597915 | 1.263 | -2.026 | 0.692 | 0.208 | 0.012 | 0.123 | 0.123 | 5.058 | 0.93 | 0.43 | 0.73 | 0.88 | 0.97 | 0.94 |
| 2 | 152809065 | rs6725269 | 0.017 | -1.925 | 0.571 | 0.257 | 0.014 | -0.009 | -0.009 | 0.433 | 0.1 | 0.08 | 0.07 | 0.61 | 0.07 | 0.41 |
| 2 | 152811852 | rs4664510 | -0.278 | -1.884 | 0.718 | 0.053 | 0.02 | 0.04 | 0.04 | 1.135 | 0.39 | 0.3 | 0.19 | 0.08 | 0.28 | 0.14 |
| 2 | 152826896 | rs6725537 | -0.268 | -1.88 | 0.648 | 0.05 | 0.027 | 0.071 | 0.071 | 1.071 | 0.39 | 0.26 | 0.19 | 0.1 | 0.28 | 0.14 |
| 2 | 152844074 | rs17270394 | 1.349 | -1.991 | 0.572 | -0.098 | -0.009 | 0.1 | 0.1 | 1.346 | 0.22 | 0.05 | 0.12 | 0.06 | 0.17 | 0.06 |
| 2 | 152849322 | rs4664517 | -0.518 | -1.786 | 0.949 | 0.393 | 0.017 | 0.132 | 0.132 | 0 | 0.05 | 0.12 | 0.26 | 0.26 | 0.07 | 0.12 |
| 2 | 152849625 | rs7564177 | 0.956 | -2.087 | 1.511 | 0.338 | 0.053 | 0.093 | 0.093 | 0.294 | 0.49 | 0.33 | 0.5 | 0.18 | 0.65 | 0.42 |
| 2 | 152855708 | rs1879138 | -0.437 | -2.115 | 1.649 | 0.307 | 0.012 | 0.01 | 0.01 | 1.149 | 0.41 | 0.35 | 0.43 | 0.36 | 0.32 | 0.23 |
| 2 | 152856062 | rs1879140 | 0.626 | -2.091 | 1.812 | 0.309 | -0.036 | 0.319 | 0.319 | 0.059 | 0.86 | 0.58 | 0.91 | 0.54 | 0.94 | 0.62 |
| 2 | 152859992 | rs12693199 | -0.597 | -2.055 | 1.724 | 0.375 | 0.001 | -0.001 | -0.001 | 0.694 | 0.42 | 0.42 | 0.43 | 0.43 | 0.33 | 0.24 |
| 2 | 152860003 | rs12693200 | 0.109 | -2.158 | 1.507 | 0.157 | 0.038 | 0.057 | 0.057 | 1.639 | 0.34 | 0.22 | 0.15 | 0.13 | 0.21 | 0.09 |
| 2 | 152873157 | rs6727569 | 1.388 | -2.06 | 1.289 | 0.098 | -0.022 | 0.118 | 0.118 | 0 | 0.42 | 0.16 | 0.44 | 0.18 | 0.55 | 0.37 |
| 2 | 152877234 | rs7565396 | 0.149 | -2.062 | 1.382 | 0.044 | 0.016 | 0.023 | 0.023 | 1.891 | 0.34 | 0.27 | 0.18 | 0.1 | 0.23 | 0.09 |
| 2 | 152879315 | rs4664073 | -0.038 | -2.175 | 1.3 | -0.039 | 0.009 | 0.007 | 0.007 | 2.943 | 0.34 | 0.29 | 0.18 | 0.06 | 0.25 | 0.1 |
| 2 | 152880509 | rs16830608 | -0.108 | -2.106 | 1.259 | 0.012 | 0.002 | -0.008 | -0.008 | 1.457 | 0.07 | 0.11 | 0.04 | 0.03 | 0.06 | 0.05 |
| 2 | 152892240 | rs9784082 | 0.172 | -2.138 | 1.443 | -0.012 | -0.005 | -0.023 | -0.023 | 0.578 | 0.61 | 0.62 | 0.8 | 0.86 | 0.7 | 0.87 |
| 2 | 152899057 | rs7582231 | 1.522 | -2.061 | 1.589 | 0.01 | -0.037 | 0.206 | 0.206 | 0 | 0.42 | 0.14 | 0.45 | 0.11 | 0.55 | 0.35 |
| 2 | 152911430 | rs10930892 | -0.213 | -2.048 | 1.479 | 0.204 | 0.002 | 0.061 | 0.061 | 0.084 | 0.07 | 0.09 | 0.26 | 0.29 | 0.1 | 0.15 |
| 2 | 152912368 | rs4664523 | -1.613 | -2.073 | 1.402 | 0.091 | -0.025 | 0.125 | 0.125 | 0 | 0.57 | 0.86 | 0.55 | 0.81 | 0.44 | 0.64 |
| 2 | 152919055 | rs12327907 | 0.089 | -2.044 | 1.391 | 0.042 | 0.015 | 0.008 | 0.008 | 2.525 | 0.34 | 0.25 | 0.18 | 0.04 | 0.25 | 0.09 |
| 2 | 152923904 | rs7588958 | 0.192 | -1.914 | 1.379 | 0.266 | -0.023 | 0.21 | 0.21 | 0 | 0.87 | 0.67 | 0.89 | 0.52 | 0.93 | 0.59 |
| 2 | 152926830 | rs6741737 | -1.789 | -2.027 | 0.698 | -0.605 | -0.062 | 0.126 | 0.126 | 0 | 0.63 | 0.85 | 0.58 | 0.86 | 0.48 | 0.67 |
| 2 | 152930618 | rs10930910 | -1.812 | -1.756 | 0.545 | -0.634 | -0.137 | 0.135 | 0.135 | 0 | 0.65 | 0.89 | 0.59 | 0.86 | 0.51 | 0.68 |
| 2 | 152931734 | rs10803938 | -0.967 | -1.839 | 0.894 | -0.693 | 0.019 | 0.091 | 0.091 | 0 | 0.52 | 0.64 | 0.54 | 0.86 | 0.44 | 0.64 |
| 2 | 152934475 | rs10190546 | -1.359 | -1.718 | 0.694 | -0.56 | 0.011 | 0.041 | 0.041 | 0 | 0.29 | 0.14 | 0.38 | 0.82 | 0.28 | 0.61 |
| 2 | 152947703 | rs13429172 | -0.18 | -1.735 | 0.846 | -0.565 | 0.007 | 0.05 | 0.05 | 0.008 | 0.4 | 0.26 | 0.43 | 0.3 | 0.39 | 0.27 |
| 2 | 152948293 | rs13416727 | -0.566 | -1.731 | 1.285 | -0.468 | 0.01 | -0.006 | -0.006 | 0.127 | 0.06 | 0.24 | 0.04 | 0 | 0.02 | 0.02 |
| 2 | 152955805 | rs13390828 | -0.246 | -1.724 | 1.236 | -0.464 | 0.006 | -0.009 | -0.009 | 0 | 0.05 | 0.18 | 0.03 | 0 | 0.02 | 0.02 |
| 2 | 152957411 | rs3820706 | -0.596 | -1.782 | 1.257 | -0.447 | -0.02 | 0.076 | 0.076 | 0 | 0.16 | 0.23 | 0.1 | 0.57 | 0.11 | 0.35 |
| 2 | 152963360 | rs6725180 | -0.773 | -1.769 | 0.987 | -0.501 | 0.079 | 0.042 | 0.042 | 2.248 | 0.47 | 0.67 | 0.29 | 0.59 | 0.3 | 0.4 |
| 2 | 152981335 | rs16830728 | -0.518 | -2.002 | 1.227 | -0.352 | -0.025 | 0.073 | 0.073 | 0 | 0.14 | 0.2 | 0.07 | 0.58 | 0.08 | 0.34 |
| 2 | 152996974 | rs17398329 | 1.654 | -1.951 | 0.988 | -0.604 | -0.122 | 0.164 | 0.164 | 0.116 | 0.37 | 0.07 | 0.43 | 0.13 | 0.5 | 0.36 |
| 2 | 152998467 | rs16830744 | -0.457 | -1.884 | 1.118 | -0.545 | -0.03 | 0.089 | 0.089 | 0 | 0.14 | 0.22 | 0.07 | 0.58 | 0.08 | 0.34 |
| 2 | 153005844 | rs10184862 | -1.706 | -1.981 | 1.568 | -0.709 | -0.009 | -0.064 | -0.064 | 0 | 0.32 | 0.43 | 0.37 | 0.84 | 0.31 | 0.57 |
| 2 | 153009409 | rs10930958 | -1.786 | -1.959 | 1.24 | -0.701 | -0.072 | 0.153 | 0.153 | 0.085 | 0.61 | 0.86 | 0.56 | 0.87 | 0.5 | 0.64 |
| 2 | 153029067 | rs12328733 | -0.396 | -1.795 | 1.303 | -0.697 | 0.002 | 0.002 | 0.002 | 0 | 0.16 | 0.14 | 0.29 | 0.26 | 0.22 | 0.23 |
| 2 | 153045036 | rs4664089 | -0.62 | -1.723 | 1.228 | -1.134 | -0.008 | 0.041 | 0.041 | 0.148 | 0.23 | 0.1 | 0.31 | 0.85 | 0.25 | 0.52 |
| 2 | 153092375 | rs4664544 | 0.457 | -1.704 | 1.405 | -1.123 | -0.001 | -0.008 | -0.008 | 0 | 0.06 | 0.09 | 0.04 | 0.04 | 0.06 | 0.05 |
| 2 | 153105583 | rs4233657 | -1.056 | -1.789 | 1.492 | -1.209 | 0.021 | 0.029 | 0.029 | 1.227 | 0.74 | 0.82 | 0.84 | 0.97 | 0.82 | 0.94 |
| 2 | 153115431 | rs6722815 | -0.744 | -1.806 | 1.724 | -1.21 | 0.048 | 0.039 | 0.039 | 0 | 0.06 | 0.4 | 0.04 | 0.01 | 0.01 | 0.01 |
| 2 | 153119272 | rs2345816 | -0.245 | -1.815 | 1.543 | -1.227 | -0.003 | -0.054 | -0.054 | 0.173 | 0.67 | 0.32 | 0.79 | 0.97 | 0.8 | 0.92 |
| 2 | 153129366 | rs12988834 | 0.816 | -1.803 | 1.509 | -1.218 | 0 | -0.002 | -0.002 | 0 | 0.06 | 0.06 | 0.04 | 0.04 | 0.06 | 0.05 |
| 2 | 153141491 | rs4664558 | -0.311 | -1.799 | 1.514 | -1.221 | -0.024 | -0.103 | -0.103 | 0 | 0.67 | 0.4 | 0.8 | 0.97 | 0.8 | 0.92 |
| 2 | 153153159 | rs4308089 | -1.105 | -2.055 | 1.375 | -1.269 | 0.001 | -0.006 | -0.006 | 0 | 0.3 | 0.27 | 0.37 | 0.94 | 0.3 | 0.56 |
| 2 | 153153555 | rs17326775 | 1.161 | -2.054 | 1.6 | -1.319 | 0.026 | 0.027 | 0.027 | 2.717 | 0.24 | 0.14 | 0.15 | 0.03 | 0.18 | 0.06 |
| 2 | 153154612 | rs7568759 | -1.665 | -2.047 | 1.479 | -1.297 | -0.013 | -0.108 | -0.108 | 0 | 0.35 | 0.48 | 0.39 | 0.92 | 0.31 | 0.57 |
| 2 | 153164068 | rs17326796 | 1.64 | -2.041 | 1.197 | -1.294 | -0.098 | 0.233 | 0.233 | 0.398 | 0.38 | 0.07 | 0.44 | 0.05 | 0.51 | 0.37 |
| 2 | 153164514 | rs2345876 | -1.117 | -2.003 | 1.204 | -1.268 | 0.001 | 0.007 | 0.007 | 0 | 0.3 | 0.27 | 0.37 | 0.92 | 0.3 | 0.56 |
| 2 | 153183646 | rs6761021 | -0.371 | -2.048 | 1.245 | -1.15 | -0.022 | -0.087 | -0.087 | 0 | 0.69 | 0.41 | 0.81 | 0.97 | 0.81 | 0.92 |
| 2 | 153190223 | rs4664100 | -0.737 | -1.994 | 1.389 | -1.12 | 0.022 | 0.041 | 0.041 | 0.063 | 0.47 | 0.35 | 0.47 | 0.9 | 0.39 | 0.51 |
| 2 | 153193698 | rs6722740 | 0.916 | -1.863 | 0.207 | -1.044 | 0.04 | 0.168 | 0.168 | 0.013 | 0.47 | 0.25 | 0.49 | 0.09 | 0.59 | 0.47 |
| 2 | 153197805 | rs28394649 | -0.811 | -1.598 | 0.205 | -0.77 | 0.001 | 0.028 | 0.028 | 0 | 0.22 | 0.16 | 0.32 | 0.88 | 0.22 | 0.45 |
| 2 | 153208493 | rs16830964 | -0.257 | -1.744 | 0.39 | -0.741 | -0.01 | -0.011 | -0.011 | 0 | 0.07 | 0.15 | 0.05 | 0.04 | 0.05 | 0.11 |
| 2 | 153208633 | rs12468749 | -0.218 | -1.848 | 0.254 | -0.428 | 0.008 | -0.053 | -0.053 | 0.338 | 0.1 | 0.01 | 0.25 | 0.67 | 0.07 | 0.27 |
| 2 | 153211284 | rs6704965 | 0.778 | -2.029 | 0.766 | -0.701 | -0.011 | -0.022 | -0.022 | 0 | 0.78 | 0.53 | 0.9 | 0.81 | 0.83 | 0.84 |
| 2 | 153219836 | rs4254462 | 1.306 | -2.314 | 0.368 | -0.998 | 0.043 | 0.089 | 0.089 | 6.649 | 0.22 | 0.09 | 0.13 | 0.03 | 0.15 | 0.05 |
| 2 | 153221287 | rs4664574 | 0.139 | -2.247 | 0.468 | -0.941 | -0.01 | -0.011 | -0.011 | 0 | 0.08 | 0.13 | 0.05 | 0.04 | 0.05 | 0.11 |
| 2 | 153224567 | rs13432002 | 0.008 | -2.28 | 0.788 | -0.937 | 0.01 | 0.022 | 0.022 | 0 | 0.13 | 0.2 | 0.07 | 0.21 | 0.15 | 0.15 |
| 2 | 153225649 | rs11885624 | 1.828 | -2.392 | 0.641 | -1.011 | -0.015 | 0.062 | 0.062 | 1.444 | 0.38 | 0.22 | 0.47 | 0.06 | 0.55 | 0.38 |
| 2 | 153228568 | rs4544376 | -0.107 | -2.491 | 0.53 | -0.843 | 0.07 | 0.005 | 0.005 | 3.666 | 0.34 | 0.14 | 0.39 | 0.69 | 0.23 | 0.32 |
| 2 | 153232345 | rs16831009 | 0.183 | -2.401 | 0.64 | -0.908 | -0.005 | -0.015 | -0.015 | 0 | 0.07 | 0.09 | 0.04 | 0.02 | 0.06 | 0.08 |
| 2 | 153233059 | rs16831013 | 0.096 | -2.404 | 0.559 | -0.895 | 0.057 | 0.009 | 0.009 | 2.297 | 0.32 | 0.18 | 0.38 | 0.7 | 0.22 | 0.32 |
| 2 | 153236814 | rs10931075 | 0.784 | -2.244 | 1.045 | -0.436 | -0.012 | -0.006 | -0.006 | 0 | 0.57 | 0.44 | 0.55 | 0.61 | 0.64 | 0.68 |
| 2 | 153236960 | rs12474369 | 0.784 | -2.244 | 1.045 | -0.426 | -0.012 | -0.006 | -0.006 | 0 | 0.57 | 0.44 | 0.55 | 0.6 | 0.64 | 0.68 |
| 2 | 153238858 | rs16831021 | 1.312 | -2.243 | 0.968 | -0.379 | 0.047 | 0.074 | 0.074 | 8.727 | 0.2 | 0.1 | 0.12 | 0.05 | 0.13 | 0.05 |
| 2 | 153242179 | rs4664106 | -1.161 | -2.179 | 1.184 | -0.331 | -0.002 | 0.009 | 0.009 | 0 | 0.19 | 0.31 | 0.11 | 0.24 | 0.18 | 0.25 |
| 2 | 153248404 | rs2345873 | 0.06 | -2.17 | 1.217 | -0.325 | 0.013 | 0.039 | 0.039 | 0 | 0.11 | 0.23 | 0.07 | 0.2 | 0.12 | 0.13 |

| | | | | | | | | | | | | | | | | |
|---|---|---|---|---|---|---|---|---|---|---|---|---|---|---|---|---|
| 2 | 153248943 | rs6761178 | -0.339 | -2.341 | 1.476 | -0.45 | 0.001 | -0.018 | -0.018 | 0.064 | 0.71 | 0.72 | 0.87 | 0.95 | 0.84 | 0.93 |
| 2 | 153249105 | rs6707033 | -0.565 | -2.318 | 1.483 | -0.436 | -0.023 | -0.044 | -0.044 | 0 | 0.92 | 0.82 | 0.98 | 1 | 0.97 | 0.97 |
| 2 | 153251742 | rs6751576 | 0.41 | -2.356 | 1.433 | -0.456 | -0.013 | -0.044 | -0.044 | 0.87 | 0.62 | 0.58 | 0.82 | 0.91 | 0.77 | 0.82 |
| 2 | 153257617 | rs12614465 | 1.517 | -2.317 | 1.194 | -0.417 | 0.043 | 0.029 | 0.029 | 4.292 | 0.47 | 0.23 | 0.52 | 0.59 | 0.6 | 0.65 |
| 2 | 153268611 | rs6751778 | -0.874 | -1.998 | 1.373 | -0.372 | 0 | -0.001 | -0.001 | 0 | 0.38 | 0.48 | 0.39 | 0.22 | 0.26 | 0.2 |
| 2 | 153268746 | rs6741728 | -1.161 | -2.026 | 1.397 | -0.402 | -0.048 | -0.054 | -0.054 | 0.265 | 0.14 | 0.33 | 0.04 | 0.04 | 0.05 | 0.12 |
| 2 | 153271524 | rs3811577 | 0.206 | -1.965 | 1.381 | -0.291 | 0.018 | 0.055 | 0.055 | 0 | 0.11 | 0.2 | 0.07 | 0.2 | 0.12 | 0.14 |
| 2 | 153273219 | rs2164402 | -0.551 | -2.002 | 1.352 | -0.326 | -0.014 | -0.017 | -0.017 | 0 | 0.36 | 0.41 | 0.22 | 0.28 | 0.31 | 0.29 |
| 2 | 153275316 | rs9917302 | -1.41 | -1.875 | 1.411 | -0.207 | 0.018 | 0.095 | 0.095 | 1.123 | 0.07 | 0.24 | 0.26 | 0.19 | 0.07 | 0.14 |
| 2 | 153275850 | rs6761815 | -0.073 | -1.915 | 1.682 | -0.17 | 0.061 | 0.159 | 0.159 | 0 | 0.11 | 0.37 | 0.08 | 0.21 | 0.12 | 0.14 |
| 2 | 153276774 | rs7587450 | 0.039 | -2.05 | 1.316 | -0.325 | 0.055 | 0.09 | 0.09 | 6.993 | 0.3 | 0.19 | 0.14 | 0.05 | 0.19 | 0.08 |
| 2 | 153281714 | rs4664583 | 0.911 | -2.133 | 1.404 | -0.308 | 0.054 | -0.008 | -0.008 | 0 | 0.56 | 0.22 | 0.54 | 0.55 | 0.63 | 0.66 |
| 2 | 153282375 | rs7600624 | -0.569 | -2.03 | 1.63 | -0.243 | 0.014 | 0.109 | 0.109 | 0.007 | 0.15 | 0.25 | 0.3 | 0.39 | 0.18 | 0.27 |
| 2 | 153282883 | rs3811575 | -1.444 | -2.023 | 1.196 | -0.242 | 0.09 | 0.019 | 0.019 | 2.001 | 0.5 | 0.83 | 0.47 | 0.45 | 0.38 | 0.36 |
| 2 | 153286715 | rs13388095 | -0.065 | -1.867 | 1.074 | -0.392 | 0.024 | 0.022 | 0.022 | 5.869 | 0.31 | 0.29 | 0.15 | 0.05 | 0.19 | 0.09 |
| 2 | 153292174 | rs6727037 | -0.599 | -1.857 | 1.007 | -0.341 | -0.045 | -0.048 | -0.048 | 3.138 | 0.31 | 0.51 | 0.37 | 0.21 | 0.22 | 0.11 |
| 2 | 153296210 | rs7582905 | -1.611 | -1.462 | 1.371 | -0.003 | 0.068 | 0.22 | 0.22 | 1.537 | 0.07 | 0.4 | 0.26 | 0.18 | 0.05 | 0.07 |
| 2 | 153299488 | rs11684450 | 1.025 | -2.082 | 0.704 | -0.818 | 0.002 | 0.17 | 0.17 | 3.989 | 0.41 | 0.13 | 0.46 | 0.03 | 0.55 | 0.36 |
| 2 | 153299943 | rs10179418 | -0.42 | -2.228 | 0.771 | -0.501 | 0.063 | 0.048 | 0.048 | 7.56 | 0.45 | 0.72 | 0.46 | 0.68 | 0.3 | 0.39 |
| 2 | 153315366 | rs10432493 | -0.388 | -2.313 | 0.741 | -0.489 | -0.001 | 0.001 | 0.001 | 4.054 | 0.43 | 0.64 | 0.44 | 0.67 | 0.28 | 0.38 |
| 2 | 153323026 | rs10166703 | 0.303 | -2.357 | 0.772 | -0.433 | 0.044 | -0.016 | -0.016 | 0 | 0.36 | 0.3 | 0.2 | 0.54 | 0.25 | 0.31 |
| 2 | 153323135 | rs13424038 | -0.119 | -2.406 | 0.939 | -0.554 | 0.001 | 0.056 | 0.056 | 0 | 0.14 | 0.22 | 0.08 | 0.31 | 0.14 | 0.26 |
| 2 | 153323158 | rs1878632 | -1.331 | -2.433 | 0.977 | -0.582 | 0.023 | 0.196 | 0.196 | 2.046 | 0.58 | 0.89 | 0.53 | 0.98 | 0.44 | 0.63 |
| 2 | 153324632 | rs7559217 | -0.275 | -2.412 | 1.13 | -0.556 | 0.003 | 0.098 | 0.098 | 0 | 0.14 | 0.29 | 0.09 | 0.34 | 0.14 | 0.26 |
| 2 | 153331400 | rs4664587 | -0.216 | -2.527 | 1.117 | -0.633 | 0.015 | 0.01 | 0.01 | 1.913 | 0.43 | 0.45 | 0.46 | 0.63 | 0.32 | 0.39 |
| 2 | 153334614 | rs6744391 | -1.334 | -2.579 | 0.52 | -0.784 | 0.032 | 0.173 | 0.173 | 0.305 | 0.6 | 0.9 | 0.52 | 0.98 | 0.4 | 0.64 |
| 2 | 153346845 | rs3811574 | -1.069 | -2.279 | 0.591 | -0.593 | -0.003 | 0.125 | 0.125 | 0.445 | 0.64 | 0.89 | 0.56 | 0.97 | 0.45 | 0.65 |
| 2 | 153354994 | rs10173398 | 0.499 | -2.317 | 0.634 | -0.443 | 0.065 | 0.002 | 0.002 | 1.741 | 0.43 | 0.35 | 0.23 | 0.54 | 0.28 | 0.33 |
| 2 | 153358005 | rs2346205 | -0.256 | -2.352 | 0.712 | -0.45 | -0.007 | 0.138 | 0.138 | 0 | 0.88 | 0.69 | 0.93 | 0.69 | 0.91 | 0.75 |
| 2 | 153358329 | rs11892232 | 0.579 | -2.316 | 0.655 | -0.414 | 0.087 | -0.006 | -0.006 | 0 | 0.42 | 0.28 | 0.22 | 0.53 | 0.29 | 0.32 |
| 2 | 153361084 | rs12992900 | -1.122 | -1.3 | 1.33 | 0.278 | 0.02 | 0.081 | 0.081 | 0.37 | 0.06 | 0.25 | 0.25 | 0.13 | 0.05 | 0.06 |
| 2 | 153366129 | rs16831246 | -1.228 | -1.306 | 1.223 | 0.29 | 0.025 | 0.096 | 0.096 | 0.771 | 0.07 | 0.3 | 0.25 | 0.13 | 0.05 | 0.06 |
| 2 | 153368897 | rs6434078 | 1.168 | -1.341 | 0.673 | 0.309 | -0.025 | 0.112 | 0.112 | 0 | 0.82 | 0.43 | 0.68 | 0.58 | 0.86 | 0.7 |
| 2 | 153371042 | rs4664113 | 0.778 | -0.666 | 0.631 | 0.379 | 0.008 | 0.037 | 0.037 | 0.012 | 0.79 | 0.63 | 0.66 | 0.7 | 0.79 | 0.69 |
| 2 | 153373267 | rs6434081 | -0.809 | 0.147 | 0.113 | -0.305 | 0.01 | 0.107 | 0.107 | 0 | 0.43 | 0.26 | 0.46 | 0.1 | 0.53 | 0.41 |
| 2 | 153373875 | rs7591606 | 0.153 | 0.173 | 0.599 | 0.026 | 0.051 | 0.275 | 0.275 | 0 | 0.92 | 0.71 | 0.72 | 0.65 | 0.89 | 0.77 |
| 2 | 153374122 | rs17399537 | 0.674 | 0.841 | 0.676 | -0.148 | 0.004 | -0.008 | -0.008 | 0 | 0.16 | 0.13 | 0.27 | 0.06 | 0.27 | 0.23 |
| 2 | 153377986 | rs13427485 | 0.704 | 0.728 | 1.116 | -0.174 | 0.001 | 0.007 | 0.007 | 0 | 0.64 | 0.47 | 0.53 | 0.61 | 0.63 | 0.59 |
| 2 | 153378459 | rs4664114 | 0.473 | 0.735 | 1.163 | -0.081 | 0 | -0.008 | -0.008 | 0 | 0.68 | 0.64 | 0.77 | 0.75 | 0.67 | 0.66 |
| 2 | 153378652 | rs4664593 | 0.761 | 0.713 | 1.628 | -0.093 | -0.001 | -0.002 | -0.002 | 0 | 0.64 | 0.38 | 0.52 | 0.61 | 0.63 | 0.59 |
| 2 | 153379368 | rs6760321 | -0.164 | 0.711 | 1.503 | -0.099 | -0.004 | 0.054 | 0.054 | 0 | 0.27 | 0.17 | 0.19 | 0.03 | 0.26 | 0.18 |
| 2 | 153384255 | rs10209069 | 1.623 | 0.667 | 1.07 | -0.106 | 0.096 | 0.015 | 0.015 | 0 | 0.48 | 0.25 | 0.26 | 0.55 | 0.36 | 0.37 |
| 2 | 153386833 | rs9288108 | 1.401 | 0.911 | 1.014 | -0.172 | 0.045 | 0.023 | 0.023 | 0 | 0.53 | 0.29 | 0.27 | 0.59 | 0.41 | 0.39 |
| 2 | 153388487 | rs17327752 | -0.179 | 0.954 | 0.733 | -0.327 | -0.011 | 0.092 | 0.092 | 0 | 0.27 | 0.14 | 0.19 | 0.03 | 0.26 | 0.18 |
| 2 | 153391282 | rs13002771 | 0.137 | 0.71 | 0.76 | -0.362 | 0.001 | -0.024 | -0.024 | 0.233 | 0.19 | 0.21 | 0.16 | 0.07 | 0.22 | 0.12 |
| 2 | 153391582 | rs12474927 | 1.594 | 0.789 | 0.406 | -0.337 | -0.059 | 0.045 | 0.045 | 0 | 0.1 | 0.02 | 0.24 | 0.03 | 0.21 | 0.2 |
| 2 | 153393747 | rs2577182 | 1.402 | 0.768 | 0.596 | -0.334 | 0.046 | 0.014 | 0.014 | 0 | 0.49 | 0.39 | 0.27 | 0.55 | 0.36 | 0.41 |
| 2 | 153399594 | rs2678301 | -1.424 | 1.168 | 0.099 | -0.378 | 0.028 | -0.019 | -0.019 | 0.3 | 0.89 | 0.88 | 0.76 | 0.96 | 0.79 | 0.8 |
| 2 | 153402571 | rs17327856 | 0.596 | 0.885 | -0.152 | -0.382 | -0.04 | 0.073 | 0.073 | 0 | 0.12 | 0.01 | 0.05 | 0 | 0.12 | 0.08 |
| 2 | 153405594 | rs1155779 | -0.773 | 0.128 | -0.191 | -0.496 | -0.002 | -0.013 | -0.013 | 0.037 | 0.52 | 0.49 | 0.67 | 0.8 | 0.59 | 0.78 |
| 2 | 153407808 | rs2083098 | 0.414 | 0.227 | -0.318 | -0.349 | 0.012 | -0.011 | -0.011 | 0.061 | 0.62 | 0.32 | 0.5 | 0.58 | 0.64 | 0.6 |
| 2 | 153408076 | rs10497104 | 1.599 | 0.603 | -0.454 | -0.49 | -0.008 | 0.003 | 0.003 | 0 | 0.12 | 0.1 | 0.24 | 0.02 | 0.2 | 0.21 |
| 2 | 153409344 | rs10497105 | 0.384 | 0.557 | -0.271 | -0.447 | 0.004 | -0.006 | -0.006 | 0 | 0.61 | 0.36 | 0.5 | 0.58 | 0.6 | 0.6 |
| 2 | 153409828 | rs10497106 | 0.653 | 0.705 | 0.084 | -0.395 | 0.052 | 0.101 | 0.101 | 0.189 | 0.16 | 0.46 | 0.15 | 0.24 | 0.17 | 0.14 |
| 2 | 153415664 | rs7599894 | 1.239 | 0.561 | 0.147 | -0.69 | 0.005 | -0.028 | -0.028 | 0.04 | 0.13 | 0.18 | 0.12 | 0.04 | 0.13 | 0.07 |
| 2 | 153417325 | rs6713836 | -0.467 | 0.488 | -0.482 | -0.721 | -0.001 | -0.001 | -0.001 | 0.005 | 0.78 | 0.76 | 0.65 | 0.82 | 0.77 | 0.74 |
| 2 | 153421718 | rs6724909 | 2.353 | 0.75 | -0.659 | -0.639 | -0.002 | 0.007 | 0.007 | 0 | 0.08 | 0.06 | 0.24 | 0.03 | 0.19 | 0.15 |
| 2 | 153435288 | rs12693407 | 0.892 | 0.039 | -0.269 | -0.563 | 0.014 | 0.046 | 0.046 | 0.906 | 0.6 | 0.37 | 0.72 | 0.9 | 0.63 | 0.69 |
| 2 | 153435434 | rs16831399 | 0.668 | 0.026 | -0.161 | -0.576 | 0.01 | -0.043 | -0.043 | 0 | 0.14 | 0.29 | 0.13 | 0.04 | 0.13 | 0.08 |
| 2 | 153436318 | rs7608585 | 0.009 | 0.059 | -0.268 | -0.47 | -0.013 | -0.063 | -0.063 | 0 | 0.75 | 0.66 | 0.84 | 0.93 | 0.76 | 0.76 |
| 2 | 153444836 | rs1437679 | -0.599 | 0.132 | -1.435 | -0.448 | -0.007 | 0.021 | 0.021 | 0 | 0.36 | 0.26 | 0.42 | 0.27 | 0.36 | 0.29 |
| 2 | 153453241 | rs4664597 | 0.891 | 0.638 | -1.209 | 0.173 | 0 | -0.048 | -0.048 | 0 | 0.6 | 0.21 | 0.5 | 0.6 | 0.64 | 0.53 |
| 2 | 153454707 | rs16831418 | -0.461 | 0.938 | -0.26 | 0.049 | 0.067 | 0.019 | 0.019 | 0.056 | 0.11 | 0.35 | 0.11 | 0.06 | 0.11 | 0.06 |
| 2 | 153464448 | rs1370497 | 0.104 | 0.83 | -1.688 | -0.242 | -0.017 | 0.154 | 0.154 | 0 | 0.53 | 0.24 | 0.44 | 0.83 | 0.46 | 0.53 |
| 2 | 153464499 | rs727327 | -0.484 | 0.829 | -1.68 | -0.734 | -0.011 | 0.068 | 0.068 | 0 | 0.25 | 0.13 | 0.12 | 0.05 | 0.25 | 0.15 |
| 2 | 153464580 | rs727326 | 0.577 | 0.714 | -1.084 | -0.49 | 0 | 0.003 | 0.003 | 0 | 0.5 | 0.49 | 0.36 | 0.64 | 0.49 | 0.46 |
| 2 | 153466602 | rs6741131 | 0.887 | 0.641 | -1.237 | -0.391 | 0.002 | -0.007 | -0.007 | 0.335 | 0.14 | 0.07 | 0.25 | 0.14 | 0.14 | 0.16 |
| 2 | 153466691 | rs6734194 | 0.197 | 0.293 | -0.766 | -0.961 | -0.001 | 0 | 0 | 5.327 | 0.6 | 0.4 | 0.57 | 0.76 | 0.55 | 0.58 |
| 2 | 153468187 | rs2304558 | 0.191 | -0.015 | -0.599 | -0.781 | 0.001 | 0 | 0 | 0 | 0.13 | 0.12 | 0.06 | 0.09 | 0.09 | 0.1 |

| | | | | | | | | | | | | | | | | |
|---|---|---|---|---|---|---|---|---|---|---|---|---|---|---|---|---|
| 2 | 153470115 | rs16831440 | 0.632 | 0.171 | -0.393 | -0.953 | -0.003 | -0.052 | -0.052 | 0.505 | 0.18 | 0.26 | 0.26 | 0.03 | 0.16 | 0.21 |
| 2 | 153473521 | rs12693415 | -0.664 | 0.068 | -0.142 | -1.076 | 0.024 | 0.039 | 0.039 | 0 | 0.42 | 0.68 | 0.41 | 0.27 | 0.33 | 0.42 |
| 2 | 153473794 | rs2304556 | -0.546 | 0.016 | 0.164 | -1.012 | -0.012 | -0.014 | -0.014 | 0 | 0.4 | 0.46 | 0.37 | 0.28 | 0.31 | 0.42 |
| 2 | 153478066 | rs16831456 | 1.414 | 0.006 | 0.168 | -1.089 | -0.002 | -0.037 | -0.037 | 0 | 0.17 | 0.18 | 0.25 | 0.02 | 0.16 | 0.2 |
| 2 | 153478642 | rs1370499 | 1.587 | 0.007 | 0.061 | -1.088 | -0.009 | 0.076 | 0.076 | 0.115 | 0.16 | 0.03 | 0.24 | 0.02 | 0.16 | 0.2 |
| 2 | 153484685 | rs3811570 | 0.619 | 0.068 | -0.127 | -1.675 | -0.001 | -0.023 | -0.023 | 0 | 0.5 | 0.51 | 0.37 | 0.82 | 0.46 | 0.47 |
| 2 | 153485865 | rs4146903 | 0.088 | 0.108 | -0.205 | -1.697 | 0.003 | -0.016 | -0.016 | 0 | 0.7 | 0.68 | 0.63 | 0.85 | 0.63 | 0.68 |
| 2 | 153488963 | rs10185455 | -0.151 | 0.007 | -0.278 | -1.695 | -0.019 | -0.054 | -0.054 | 2.226 | 0.67 | 0.59 | 0.67 | 0.84 | 0.73 | 0.66 |
| 2 | 153492910 | rs1561267 | 0.18 | -0.004 | -0.243 | -1.579 | -0.027 | -0.012 | -0.012 | 0.466 | 0.84 | 0.67 | 0.91 | 0.85 | 0.89 | 0.87 |
| 2 | 153504552 | rs12693429 | 0.251 | 0.069 | -0.131 | -1.711 | 0.022 | -0.147 | -0.147 | 0.423 | 0.36 | 0.53 | 0.56 | 0.02 | 0.43 | 0.39 |
| 2 | 153505003 | rs1065266 | -0.23 | -0.113 | -0.16 | -1.497 | 0.017 | -0.033 | -0.033 | 0 | 0.77 | 0.91 | 0.89 | 0.73 | 0.82 | 0.77 |
| 2 | 153507483 | rs9288121 | -0.004 | -0.122 | -0.147 | -1.501 | 0.012 | -0.009 | -0.009 | 0 | 0.75 | 0.81 | 0.89 | 0.74 | 0.82 | 0.8 |
| 2 | 153523111 | rs9288125 | 0.062 | -0.124 | -0.07 | -1.502 | -0.02 | 0.01 | 0.01 | 0 | 0.74 | 0.69 | 0.88 | 0.73 | 0.82 | 0.78 |
| 2 | 153528136 | rs1437677 | 1.845 | 0.014 | -0.288 | -1.603 | -0.071 | 0.07 | 0.07 | 0.368 | 0.14 | 0.02 | 0.28 | 0.01 | 0.27 | 0.19 |
| 2 | 153534527 | rs4664601 | 1.074 | -0.008 | -0.301 | -1.627 | -0.013 | 0.017 | 0.017 | 0 | 0.38 | 0.15 | 0.31 | 0.71 | 0.37 | 0.37 |
| 2 | 153543165 | rs2346537 | 1.806 | -0.33 | -0.976 | -1.554 | -0.033 | -0.115 | -0.115 | 1.689 | 0.16 | 0.01 | 0.18 | 0.56 | 0.15 | 0.26 |
| 2 | 153545639 | rs6744923 | 0.492 | -0.281 | -0.883 | -1.292 | -0.013 | -0.019 | -0.019 | 0 | 0.38 | 0.15 | 0.31 | 0.67 | 0.37 | 0.37 |
| 2 | 153557883 | rs6728189 | 2.316 | -0.79 | -0.713 | -1.196 | -0.002 | 0.045 | 0.045 | 0.076 | 0.17 | 0.09 | 0.24 | 0.02 | 0.16 | 0.2 |
| 2 | 153564230 | rs1962313 | 0.071 | -0.789 | -0.597 | -1.186 | -0.012 | -0.015 | -0.015 | 0 | 0.57 | 0.53 | 0.62 | 0.72 | 0.66 | 0.57 |
| 2 | 153572156 | rs3811567 | 1.623 | -0.766 | -0.91 | -1.157 | -0.036 | -0.1 | -0.1 | 0 | 0.14 | 0.01 | 0.18 | 0.57 | 0.15 | 0.25 |
| 2 | 153573528 | rs4664605 | 0.318 | -0.747 | -0.186 | -1.055 | 0.02 | 0.033 | 0.033 | 0 | 0.58 | 0.72 | 0.64 | 0.72 | 0.65 | 0.57 |
| 2 | 153577786 | rs10497113 | 1.947 | -0.898 | -0.179 | -1.048 | -0.012 | 0.076 | 0.076 | 0.03 | 0.16 | 0.04 | 0.24 | 0.02 | 0.16 | 0.2 |
| 2 | 153578900 | rs2346549 | 0.238 | -0.832 | -0.065 | -0.984 | 0.019 | 0.025 | 0.025 | 0 | 0.58 | 0.69 | 0.65 | 0.72 | 0.66 | 0.58 |
| 2 | 153586899 | rs1986743 | 0.254 | -0.753 | -0.645 | -0.9 | 0.002 | 0.002 | 0.002 | 0 | 0.58 | 0.63 | 0.65 | 0.73 | 0.66 | 0.57 |
| 2 | 153587296 | rs2346550 | 0.478 | -0.614 | -0.59 | -0.482 | 0 | -0.001 | -0.001 | 0.312 | 0.76 | 0.76 | 0.89 | 0.74 | 0.82 | 0.78 |
| 2 | 153593671 | rs16831728 | 0.233 | -0.718 | -0.912 | -0.516 | -0.01 | -0.034 | -0.034 | 0.337 | 0.1 | 0 | 0.05 | 0.14 | 0.07 | 0.09 |
| 2 | 153607110 | rs10497116 | 0.285 | -0.693 | -0.811 | -0.473 | 0.004 | -0.034 | -0.034 | 0 | 0.15 | 0.02 | 0.04 | 0.12 | 0.1 | 0.13 |
| 2 | 153617420 | rs1128804 | 0.467 | -0.681 | -0.588 | -0.461 | 0.001 | -0.002 | -0.002 | 0.027 | 0.75 | 0.77 | 0.89 | 0.74 | 0.82 | 0.78 |
| 2 | 153622532 | rs6710313 | -0.418 | -0.357 | -0.402 | -0.756 | -0.002 | 0.002 | 0.002 | 0 | 0.24 | 0.25 | 0.43 | 0.81 | 0.27 | 0.4 |
| 2 | 153627907 | rs3791626 | -0.614 | -0.94 | -0.112 | -1.158 | -0.025 | -0.039 | -0.039 | 0 | 0.66 | 0.63 | 0.88 | 0.9 | 0.83 | 0.81 |
| 2 | 153640839 | rs890590 | 0.454 | -0.994 | -0.126 | -1.034 | 0.041 | 0.046 | 0.046 | 0 | 0.34 | 0.22 | 0.11 | 0.1 | 0.18 | 0.2 |
| 2 | 153647254 | rs10931290 | 0.254 | -0.724 | -0.387 | -0.939 | 0 | 0.002 | 0.002 | 0 | 0.19 | 0.23 | 0.39 | 0.7 | 0.23 | 0.32 |
| 2 | 153648587 | rs2114653 | 1.497 | -0.326 | -0.572 | -1.098 | 0.014 | -0.008 | -0.008 | 0 | 0.14 | 0.13 | 0.33 | 0.06 | 0.33 | 0.19 |
| 2 | 153649490 | rs11692707 | 1.993 | -0.538 | -0.593 | -1.01 | -0.007 | 0.065 | 0.065 | 0 | 0.2 | 0.09 | 0.11 | 0.03 | 0.2 | 0.21 |
| 2 | 153654720 | rs12478462 | 0.345 | -0.471 | -0.507 | -0.839 | 0.003 | 0.008 | 0.008 | 0 | 0.2 | 0.26 | 0.41 | 0.7 | 0.25 | 0.35 |
| 2 | 153657990 | rs6434203 | 1.074 | -0.656 | -0.903 | -0.967 | -0.023 | 0.073 | 0.073 | 0.207 | 0.28 | 0.08 | 0.14 | 0.14 | 0.24 | 0.26 |
| 2 | 153670186 | rs7593238 | 0.079 | -0.396 | -0.245 | -0.628 | 0.005 | 0.01 | 0.01 | 0 | 0.21 | 0.27 | 0.42 | 0.71 | 0.25 | 0.35 |
| 2 | 153675592 | rs1427405 | -0.336 | -0.515 | 0.138 | -0.679 | -0.033 | -0.029 | -0.029 | 0 | 0.15 | 0.43 | 0.07 | 0.1 | 0.06 | 0.14 |
| 2 | 153683524 | rs12621161 | -0.321 | -0.491 | 0.044 | -0.135 | 0.024 | 0.035 | 0.035 | 0 | 0.31 | 0.61 | 0.54 | 0.5 | 0.4 | 0.31 |
| 2 | 153683596 | rs17401154 | -0.279 | -0.481 | 0.004 | -0.147 | 0.012 | -0.023 | -0.023 | 0 | 0.32 | 0.63 | 0.43 | 0.16 | 0.5 | 0.32 |
| 2 | 153685066 | rs17329272 | 1.318 | -0.281 | -0.387 | 0.005 | -0.051 | 0.044 | 0.044 | 0 | 0.17 | 0.08 | 0.29 | 0.06 | 0.33 | 0.17 |
| 2 | 153685992 | rs11883777 | 1.312 | -0.276 | -0.385 | 0.007 | -0.051 | 0.045 | 0.045 | 0 | 0.17 | 0.08 | 0.29 | 0.06 | 0.33 | 0.17 |
| 2 | 153719002 | rs6434230 | -0.298 | -0.397 | -0.158 | 0.161 | 0.002 | 0.123 | 0.123 | 0 | 0.13 | 0.43 | 0.15 | 0.24 | 0.1 | 0.25 |
| 2 | 153719421 | rs6760678 | 0.918 | -0.414 | -0.511 | 0.151 | 0.013 | -0.015 | -0.015 | 0 | 0.1 | 0.05 | 0.2 | 0.42 | 0.06 | 0.11 |
| 2 | 153721223 | rs17401286 | 1.256 | -0.245 | -0.476 | -0.067 | -0.03 | 0.022 | 0.022 | 0 | 0.17 | 0.13 | 0.3 | 0.06 | 0.33 | 0.17 |
| 2 | 153729824 | rs10209901 | -0.489 | -0.315 | -0.244 | -0.019 | 0.013 | 0 | 0 | 0 | 0.75 | 0.71 | 0.6 | 0.71 | 0.59 | 0.62 |
| 2 | 153731644 | rs10931352 | 0.016 | -0.504 | -0.162 | 0.059 | -0.016 | 0.076 | 0.076 | 0 | 0.14 | 0.2 | 0.24 | 0.44 | 0.07 | 0.14 |
| 2 | 153734205 | rs755306 | 0.615 | -0.457 | -0.138 | 0.014 | 0 | 0.075 | 0.075 | 0 | 0.1 | 0.2 | 0.11 | 0.24 | 0.08 | 0.21 |
| 2 | 153735193 | rs1427396 | 0.836 | -0.633 | -0.211 | -0.191 | 0.036 | 0.02 | 0.02 | 0 | 0.13 | 0.26 | 0.11 | 0.13 | 0.18 | 0.13 |
| 2 | 153740617 | rs13035127 | 2.613 | -1.001 | -0.712 | -0.492 | -0.035 | 0.077 | 0.077 | 0 | 0.19 | 0.01 | 0.12 | 0.04 | 0.18 | 0.15 |
| 2 | 153742470 | rs13410800 | 1.395 | -1.036 | -0.705 | -0.473 | 0.046 | 0.145 | 0.145 | 0 | 0.28 | 0.01 | 0.09 | 0.07 | 0.16 | 0.18 |
| 2 | 153746585 | rs7578801 | -0.421 | -0.948 | -0.528 | -0.343 | -0.022 | 0.099 | 0.099 | 0 | 0.86 | 0.76 | 0.78 | 0.55 | 0.93 | 0.86 |
| 2 | 153750286 | rs1427398 | 0.38 | -0.873 | -0.386 | -0.391 | 0.07 | 0.013 | 0.013 | 0 | 0.26 | 0.41 | 0.42 | 0.31 | 0.41 | 0.4 |
| 2 | 153752923 | rs11695096 | 0.226 | -0.919 | -0.372 | -0.33 | 0.003 | 0.086 | 0.086 | 0 | 0.76 | 0.55 | 0.66 | 0.43 | 0.77 | 0.74 |
| 2 | 153760605 | rs12693506 | 1.294 | -1.057 | -0.848 | -0.303 | 0.049 | 0.148 | 0.148 | 0 | 0.27 | 0.01 | 0.1 | 0.07 | 0.16 | 0.18 |
| 2 | 153761031 | rs1582842 | -1.064 | -0.85 | -0.501 | -0.026 | -0.011 | 0.102 | 0.102 | 0.036 | 0.89 | 0.81 | 0.78 | 0.56 | 0.94 | 0.88 |
| 2 | 153767696 | rs7581172 | 0.755 | -0.71 | -0.488 | -0.056 | 0.006 | 0.094 | 0.094 | 0 | 0.1 | 0.22 | 0.11 | 0.23 | 0.08 | 0.21 |
| 2 | 153776863 | rs1365841 | 1.035 | -0.686 | -0.837 | -0.008 | -0.004 | -0.023 | -0.023 | 0 | 0.09 | 0.04 | 0.1 | 0.16 | 0.08 | 0.18 |
| 2 | 153781648 | rs1025246 | -1.325 | -1.126 | -1.019 | -0.3 | 0.055 | 0.086 | 0.086 | 0 | 0.55 | 0.67 | 0.79 | 0.88 | 0.69 | 0.69 |
| 2 | 153782249 | rs6731128 | -0.11 | -0.439 | -0.915 | -0.208 | 0.031 | -0.02 | -0.02 | 0.604 | 0.35 | 0.49 | 0.45 | 0.25 | 0.5 | 0.35 |
| 2 | 153787485 | rs10931387 | -0.756 | -0.026 | -0.771 | -0.1 | 0.002 | -0.015 | -0.015 | 0 | 0.68 | 0.56 | 0.74 | 0.76 | 0.69 | 0.64 |
| 2 | 153789391 | rs6730152 | 0.423 | -0.039 | -1.065 | 0.481 | -0.123 | -0.023 | -0.023 | 0 | 0.23 | 0.06 | 0.41 | 0.25 | 0.41 | 0.39 |
| 2 | 153811011 | rs4664633 | 0.12 | -0.283 | -0.998 | 0.456 | -0.091 | -0.061 | -0.061 | 0 | 0.26 | 0.09 | 0.58 | 0.61 | 0.43 | 0.43 |
| 2 | 153814233 | rs7601781 | -0.164 | -0.344 | -0.835 | 0.429 | -0.041 | -0.024 | -0.024 | 0 | 0.28 | 0.2 | 0.6 | 0.62 | 0.44 | 0.45 |
| 2 | 153817179 | rs7599452 | 0.727 | -0.658 | -0.456 | 0.267 | -0.034 | 0.02 | 0.02 | 0.078 | 0.15 | 0.27 | 0.08 | 0.17 | 0.09 | 0.12 |
| 2 | 153829171 | rs12476976 | 1.352 | -0.824 | -0.672 | -0.013 | -0.004 | 0.028 | 0.028 | 0 | 0.3 | 0.24 | 0.34 | 0.05 | 0.3 | 0.24 |
| 2 | 153833730 | rs17810828 | 0.101 | -0.668 | -1.219 | 0.158 | -0.045 | -0.034 | -0.034 | 0.022 | 0.28 | 0.19 | 0.47 | 0.6 | 0.44 | 0.44 |
| 2 | 153843128 | rs1435022 | -0.071 | -0.848 | -1.217 | 0.171 | 0.078 | 0.032 | 0.032 | 0 | 0.58 | 0.32 | 0.8 | 0.66 | 0.74 | 0.71 |
| 2 | 153845053 | rs13008832 | 0.68 | -0.594 | -1.297 | -0.147 | 0.037 | 0.152 | 0.152 | 0 | 0.46 | 0.29 | 0.65 | 0.15 | 0.59 | 0.43 |

| | | | | | | | | | | | | | | | | |
|---|---|---|---|---|---|---|---|---|---|---|---|---|---|---|---|---|
| 2 | 153848663 | rs17604102 | 1.981 | -0.427 | -1.52 | -0.283 | -0.079 | 0.054 | 0.054 | 0 | 0.13 | 0.01 | 0.26 | 0.05 | 0.27 | 0.17 |
| 2 | 153850952 | rs13013816 | 0.982 | -0.911 | -1.077 | 0.054 | 0.028 | 0.194 | 0.194 | 0 | 0.46 | 0.23 | 0.65 | 0.11 | 0.58 | 0.41 |
| 2 | 153859292 | rs10196299 | 0.541 | -0.682 | -1.034 | 0.338 | -0.016 | -0.035 | -0.035 | 0.206 | 0.26 | 0.28 | 0.11 | 0.13 | 0.16 | 0.16 |
| 2 | 153871336 | rs1435035 | 0.239 | 0.289 | -1.119 | 1.446 | -0.034 | -0.041 | -0.041 | 0 | 0.08 | 0.01 | 0.12 | 0.16 | 0.13 | 0.18 |
| 2 | 153872617 | rs6753428 | -0.54 | 0.285 | -1.112 | 1.141 | -0.01 | -0.015 | -0.015 | 0 | 0.13 | 0.09 | 0.22 | 0.61 | 0.16 | 0.3 |
| 2 | 153880653 | rs1435031 | -0.488 | 0.31 | -1.117 | 1.204 | -0.012 | -0.015 | -0.015 | 0 | 0.13 | 0.09 | 0.22 | 0.59 | 0.16 | 0.3 |
| 2 | 153881453 | rs10931434 | 0.735 | -0.527 | -1.3 | 1.15 | 0.035 | -0.019 | -0.019 | 0 | 0.72 | 0.25 | 0.76 | 0.39 | 0.83 | 0.73 |
| 2 | 153882052 | rs6717712 | 0.049 | -0.652 | -0.687 | 0.705 | -0.016 | -0.007 | -0.007 | 0 | 0.4 | 0.31 | 0.46 | 0.62 | 0.45 | 0.46 |
| 2 | 153889619 | rs7595323 | -0.206 | -0.719 | 0.173 | 0.689 | -0.037 | -0.077 | -0.077 | 0 | 0.87 | 0.73 | 0.94 | 1 | 0.95 | 0.94 |
| 2 | 153891830 | rs7583048 | -0.189 | -0.76 | -0.15 | 0.498 | 0.024 | 0.046 | 0.046 | 0 | 0.28 | 0.6 | 0.43 | 0.51 | 0.35 | 0.33 |
| 2 | 153892382 | rs7570033 | 0.339 | -0.759 | -0.082 | 0.447 | 0.051 | -0.045 | -0.045 | 4.285 | 0.23 | 0.55 | 0.33 | 0.11 | 0.31 | 0.19 |
| 2 | 153895168 | rs1897920 | 0.28 | -0.892 | -0.202 | 0.377 | -0.032 | -0.019 | -0.019 | 0 | 0.78 | 0.68 | 0.89 | 0.85 | 0.88 | 0.83 |
| 2 | 153897536 | rs9636312 | -0.619 | -0.854 | -0.103 | 0.409 | -0.03 | 0.009 | 0.009 | 0.806 | 0.85 | 0.71 | 0.92 | 0.85 | 0.91 | 0.87 |
| 2 | 153899603 | rs4664132 | -0.143 | -0.754 | -0.167 | 0.481 | 0.053 | 0.095 | 0.095 | 0 | 0.27 | 0.56 | 0.43 | 0.48 | 0.35 | 0.32 |
| 2 | 153905669 | rs6714254 | 1.412 | -0.751 | -0.384 | 0.01 | -0.035 | 0.105 | 0.105 | 0 | 0.51 | 0.18 | 0.47 | 0.36 | 0.51 | 0.5 |
| 2 | 153906947 | rs6728620 | 0.549 | -0.585 | -0.161 | -0.569 | 0.013 | -0.011 | -0.011 | 0 | 0.19 | 0.2 | 0.29 | 0.09 | 0.31 | 0.18 |
| 2 | 153912273 | rs16832068 | 0.819 | -0.513 | -0.121 | -0.504 | 0.06 | -0.027 | -0.027 | 0 | 0.17 | 0.28 | 0.29 | 0.09 | 0.31 | 0.18 |
| 2 | 153918127 | rs4664643 | 0.454 | -0.902 | -0.235 | -0.393 | -0.039 | -0.085 | -0.085 | 1.223 | 0.81 | 0.72 | 0.93 | 0.99 | 0.93 | 0.91 |
| 2 | 153921800 | rs6746077 | 0.031 | -0.986 | -0.255 | -0.619 | -0.056 | -0.101 | -0.101 | 0 | 0.78 | 0.62 | 0.91 | 0.97 | 0.91 | 0.86 |
| 2 | 153922805 | rs1946677 | 1.071 | -0.65 | -1.558 | -0.236 | -0.074 | -0.055 | -0.055 | 0 | 0.46 | 0.03 | 0.48 | 0.59 | 0.52 | 0.51 |
| 2 | 153925229 | rs2347397 | 0.633 | -0.641 | -1.257 | -0.207 | -0.028 | 0.012 | 0.012 | 0 | 0.57 | 0.15 | 0.54 | 0.62 | 0.61 | 0.61 |
| 2 | 153927273 | rs1435023 | 0.367 | -0.598 | -0.736 | -0.478 | -0.044 | -0.083 | -0.083 | 0 | 0.74 | 0.42 | 0.88 | 0.93 | 0.9 | 0.86 |
| 2 | 153929960 | rs7572009 | 1.663 | -0.234 | -0.922 | -0.312 | -0.018 | 0.009 | 0.009 | 0 | 0.12 | 0.07 | 0.25 | 0.1 | 0.25 | 0.14 |
| 2 | 153935493 | rs13001527 | 1.355 | -0.33 | -1.003 | -0.386 | 0 | 0.003 | 0.003 | 0 | 0.1 | 0.08 | 0.05 | 0.03 | 0.07 | 0.09 |
| 2 | 153938665 | rs7559920 | 1.215 | -0.244 | -0.841 | -0.265 | 0.035 | -0.005 | -0.005 | 2.79 | 0.14 | 0.22 | 0.26 | 0.13 | 0.26 | 0.14 |
| 2 | 153943546 | rs2347399 | -0.883 | -0.578 | -0.338 | -0.32 | -0.058 | -0.076 | -0.076 | 0 | 0.88 | 0.69 | 0.95 | 1 | 0.98 | 0.94 |
| 2 | 153945439 | rs4643488 | -0.124 | -0.377 | -1.333 | -0.275 | -0.061 | 0.071 | 0.071 | 0 | 0.3 | 0.14 | 0.33 | 0.17 | 0.4 | 0.24 |
| 2 | 153950651 | rs10167782 | -0.336 | -0.337 | -1.358 | -0.254 | -0.015 | 0.02 | 0.02 | 0 | 0.32 | 0.31 | 0.34 | 0.19 | 0.4 | 0.24 |
| 2 | 153957064 | rs4664645 | 0.368 | -0.444 | -1.857 | 0.21 | 0.022 | -0.104 | -0.104 | 0 | 0.25 | 0.05 | 0.2 | 0.57 | 0.25 | 0.36 |
| 2 | 153960362 | rs13427732 | 0.383 | -0.128 | -1.374 | -0.049 | 0.009 | 0.014 | 0.014 | 0 | 0.16 | 0.11 | 0.06 | 0.03 | 0.1 | 0.15 |
| 2 | 153968637 | rs16832124 | -0.188 | -0.148 | -1.572 | 0.124 | -0.011 | -0.034 | -0.034 | 1.651 | 0.13 | 0.05 | 0.12 | 0.26 | 0.11 | 0.16 |
| 2 | 153971405 | rs10804036 | 1.398 | 0.166 | -1.697 | 0.079 | -0.027 | -0.002 | -0.002 | 0.269 | 0.49 | 0.16 | 0.64 | 0.45 | 0.57 | 0.59 |
| 2 | 153972564 | rs1865465 | 1.352 | 0.143 | -1.549 | 0.019 | -0.018 | 0.009 | 0.009 | 0.166 | 0.52 | 0.25 | 0.66 | 0.44 | 0.59 | 0.63 |
| 2 | 153982792 | rs10931475 | -0.175 | 0.568 | -1.022 | 0.107 | 0.022 | 0.002 | 0.002 | 0 | 0.14 | 0.2 | 0.1 | 0.16 | 0.18 | 0.11 |
| 2 | 153986336 | rs7582602 | -1.157 | 0.5 | -0.869 | 0.108 | 0.008 | 0.022 | 0.022 | 0 | 0.26 | 0.32 | 0.2 | 0.36 | 0.24 | 0.23 |
| 2 | 153987318 | rs1347393 | 0.942 | 0.411 | -0.941 | 0.039 | 0.004 | 0.047 | 0.047 | 1.161 | 0.54 | 0.29 | 0.73 | 0.65 | 0.61 | 0.72 |
| 2 | 153992778 | rs10931477 | 0.782 | 0.441 | -0.953 | -0.01 | 0.001 | 0.046 | 0.046 | 0 | 0.22 | 0.11 | 0.11 | 0.03 | 0.19 | 0.16 |
| 2 | 153993149 | rs1435019 | 0.748 | 0.509 | -0.877 | 0.214 | 0.018 | -0.003 | -0.003 | 0 | 0.53 | 0.22 | 0.69 | 0.52 | 0.63 | 0.68 |
| 2 | 153993808 | rs1369256 | -1.173 | 0.325 | -0.848 | 0.262 | 0.005 | 0.105 | 0.105 | 0.016 | 0.91 | 0.65 | 0.94 | 0.84 | 0.96 | 0.97 |
| 2 | 153997058 | rs10497126 | 0.559 | 0.29 | -1.237 | 0.174 | -0.002 | 0.002 | 0.002 | 0 | 0.09 | 0.05 | 0.05 | 0.07 | 0.07 | 0.07 |
| 2 | 153999220 | rs1369260 | 0.45 | 0.349 | -1.171 | 0.202 | 0.018 | 0.008 | 0.008 | 0 | 0.5 | 0.42 | 0.7 | 0.54 | 0.67 | 0.64 |
| 2 | 154003081 | rs7606384 | -0.208 | 0.444 | -1.213 | -0.047 | -0.007 | -0.051 | -0.051 | 0 | 0.77 | 0.74 | 0.76 | 0.96 | 0.79 | 0.77 |
| 2 | 154003158 | rs7562599 | -0.367 | 0.515 | -1.529 | 0.552 | 0.001 | -0.003 | -0.003 | 0.359 | 0.22 | 0.19 | 0.15 | 0.32 | 0.14 | 0.21 |
| 2 | 154003392 | rs11689662 | -0.764 | 0.916 | -1.741 | 0.488 | -0.017 | -0.026 | -0.026 | 0 | 0.11 | 0.04 | 0.1 | 0.16 | 0.16 | 0.08 |
| 2 | 154003962 | rs12996693 | -0.427 | 0.961 | -2.399 | -0.277 | -0.073 | 0.116 | 0.116 | 0 | 0.35 | 0.11 | 0.39 | 0.17 | 0.41 | 0.2 |
| 2 | 154006439 | rs6434405 | -0.641 | 0.622 | -2.362 | -0.72 | -0.015 | 0.046 | 0.046 | 0 | 0.66 | 0.78 | 0.69 | 0.85 | 0.63 | 0.55 |
| 2 | 154013361 | rs16832173 | -0.26 | -0.036 | -2.629 | -0.497 | 0.011 | -0.068 | -0.068 | 0.011 | 0.19 | 0.06 | 0.32 | 0.61 | 0.15 | 0.15 |
| 2 | 154018552 | rs12616616 | -0.426 | 0.051 | -2.699 | -0.962 | 0.02 | 0.129 | 0.129 | 0.437 | 0.51 | 0.79 | 0.63 | 0.75 | 0.55 | 0.45 |
| 2 | 154018799 | rs1435041 | -0.846 | -0.172 | -2.178 | -1.18 | -0.003 | 0.01 | 0.01 | 0 | 0.6 | 0.6 | 0.67 | 0.84 | 0.6 | 0.5 |
| 2 | 154019476 | rs1435042 | 0.798 | 0.1 | -1.724 | -1.174 | -0.001 | -0.096 | -0.096 | 0 | 0.22 | 0.34 | 0.16 | 0.05 | 0.22 | 0.21 |
| 2 | 154025701 | rs6434419 | -1.571 | 0.192 | -1.984 | -1.057 | -0.047 | 0.052 | 0.052 | 0 | 0.72 | 0.93 | 0.74 | 0.83 | 0.7 | 0.57 |
| 2 | 154029978 | rs13414678 | -0.729 | 0.339 | -1.264 | -0.558 | 0.008 | 0.021 | 0.021 | 0 | 0.48 | 0.38 | 0.51 | 0.67 | 0.45 | 0.39 |
| 2 | 154031682 | rs2881691 | 0.653 | 0.961 | -0.912 | -0.737 | -0.001 | -0.001 | -0.001 | 6.498 | 0.13 | 0.11 | 0.11 | 0.11 | 0.14 | 0.11 |
| 2 | 154032484 | rs892747 | -0.422 | 0.809 | -0.997 | -0.785 | -0.006 | 0.012 | 0.012 | 0.021 | 0.34 | 0.32 | 0.48 | 0.74 | 0.36 | 0.36 |
| 2 | 154052591 | rs1074750 | -0.178 | 1.165 | -0.676 | -0.677 | 0.018 | -0.038 | -0.038 | 0 | 0.1 | 0.17 | 0.11 | 0.01 | 0.17 | 0.07 |
| 2 | 154054061 | rs2013235 | 0.217 | 1.281 | -1.135 | -0.771 | -0.052 | 0.051 | 0.051 | 0 | 0.1 | 0.02 | 0.11 | 0 | 0.17 | 0.06 |
| 2 | 154058372 | rs4664649 | 0.614 | 1.948 | -0.806 | -0.801 | 0.039 | -0.005 | -0.005 | 0.454 | 0.11 | 0.17 | 0.18 | 0.1 | 0.18 | 0.14 |
| 2 | 154059622 | rs1550376 | -0.487 | 1.404 | -1.687 | -1.515 | -0.05 | 0.174 | 0.174 | 0 | 0.33 | 0.07 | 0.23 | 0.08 | 0.39 | 0.26 |
| 2 | 154061615 | rs13405095 | -0.048 | 1.271 | -1.125 | -1.633 | 0 | -0.018 | -0.018 | 0.122 | 0.23 | 0.23 | 0.12 | 0.07 | 0.18 | 0.18 |
| 2 | 154067245 | rs4535003 | -1.737 | 0.733 | -1.696 | -1.746 | 0.017 | -0.079 | -0.079 | 0.051 | 0.8 | 0.94 | 0.67 | 0.39 | 0.85 | 0.6 |
| 2 | 154074197 | rs1371394 | -0.407 | 0.748 | -1.116 | -1.858 | -0.001 | -0.043 | -0.043 | 0 | 0.35 | 0.39 | 0.27 | 0.09 | 0.38 | 0.25 |
| 2 | 154084126 | rs1439638 | 1.766 | 0.894 | -0.839 | -1.711 | 0.034 | 0.001 | 0.001 | 0 | 0.11 | 0.17 | 0.19 | 0.12 | 0.17 | 0.14 |
| 2 | 154084200 | rs1015802 | 0.929 | 0.798 | -0.988 | -1.859 | -0.002 | -0.023 | -0.023 | 0 | 0.41 | 0.39 | 0.27 | 0.16 | 0.38 | 0.23 |
| 2 | 154085028 | rs2881710 | -0.776 | 0.838 | -1.047 | -1.804 | -0.015 | 0.022 | 0.022 | 0.359 | 0.7 | 0.78 | 0.48 | 0.26 | 0.68 | 0.47 |
| 2 | 154130843 | rs11680628 | 0.864 | 0.719 | -1.02 | -1.895 | -0.006 | -0.046 | -0.046 | 0 | 0.43 | 0.4 | 0.26 | 0.15 | 0.37 | 0.23 |
| 2 | 154131206 | rs17230094 | 1.614 | 0.739 | -1.071 | -1.925 | -0.05 | 0.064 | 0.064 | 0 | 0.12 | 0.02 | 0.13 | 0 | 0.2 | 0.08 |
| 2 | 154131374 | rs1439630 | 0.155 | 0.695 | -0.991 | -1.913 | -0.001 | -0.009 | -0.009 | 0 | 0.55 | 0.51 | 0.39 | 0.2 | 0.57 | 0.31 |
| 2 | 154135071 | rs10195227 | 1.217 | 0.623 | -1.065 | -1.858 | 0.004 | 0.028 | 0.028 | 0 | 0.12 | 0.04 | 0.06 | 0.08 | 0.1 | 0.16 |
| 2 | 154141087 | rs9288196 | 0.079 | 0.572 | -0.812 | -1.917 | -0.005 | 0.039 | 0.039 | 0.04 | 0.56 | 0.63 | 0.34 | 0.23 | 0.47 | 0.38 |

| | | | | | | | | | | | | | | | | |
|---|---|---|---|---|---|---|---|---|---|---|---|---|---|---|---|---|
| 2 | 154141994 | rs7556698 | 0.598 | 0.595 | -1.17 | -1.897 | 0.029 | 0.144 | 0.144 | 0.095 | 0.45 | 0.73 | 0.3 | 0.2 | 0.37 | 0.23 |
| 2 | 154143853 | rs4410241 | -0.433 | 0.706 | -1.268 | -1.89 | -0.148 | 0.098 | 0.098 | 0 | 0.34 | 0.09 | 0.36 | 0.17 | 0.47 | 0.36 |
| 2 | 154144551 | rs13387164 | 0.532 | 0.659 | -1.057 | -1.847 | -0.021 | -0.04 | -0.04 | 0.01 | 0.15 | 0.34 | 0.1 | 0.07 | 0.1 | 0.15 |
| 2 | 154147124 | rs10203000 | 0.061 | 0.755 | -1.185 | -1.847 | -0.006 | 0.007 | 0.007 | 0 | 0.24 | 0.26 | 0.26 | 0.17 | 0.26 | 0.3 |
| 2 | 154151887 | rs7574077 | -1.555 | 0.543 | -1.341 | -1.709 | 0.021 | -0.052 | -0.052 | 0 | 0.8 | 0.92 | 0.67 | 0.34 | 0.84 | 0.61 |
| 2 | 154154347 | rs1020146 | 0.165 | 0.961 | -1.099 | -1.591 | -0.059 | 0.095 | 0.095 | 0 | 0.23 | 0.06 | 0.19 | 0.08 | 0.31 | 0.22 |
| 2 | 154156259 | rs1439629 | 1.425 | 0.936 | -1.073 | -1.558 | -0.061 | -0.022 | -0.022 | 0 | 0.1 | 0.01 | 0.18 | 0.11 | 0.16 | 0.16 |
| 2 | 154156666 | rs16832680 | 0.739 | 0.906 | -0.668 | -1.57 | 0.034 | -0.008 | -0.008 | 0 | 0.13 | 0.26 | 0.21 | 0.11 | 0.16 | 0.16 |
| 2 | 154158793 | rs1921768 | -0.533 | 0.329 | -0.483 | -1.762 | 0.001 | -0.036 | -0.036 | 0 | 0.4 | 0.5 | 0.45 | 0.19 | 0.54 | 0.38 |
| 2 | 154163944 | rs1347927 | -0.266 | 0.085 | -0.662 | -1.661 | -0.006 | -0.014 | -0.014 | 0 | 0.52 | 0.57 | 0.52 | 0.3 | 0.61 | 0.46 |
| 2 | 154171024 | rs6757880 | 0.223 | 0.166 | -0.878 | -1.705 | -0.039 | 0.107 | 0.107 | 0.112 | 0.41 | 0.21 | 0.38 | 0.23 | 0.43 | 0.29 |
| 2 | 154171376 | rs10183071 | -1.495 | -0.248 | -0.98 | -1.961 | 0.013 | -0.037 | -0.037 | 0 | 0.83 | 0.92 | 0.69 | 0.37 | 0.86 | 0.66 |
| 2 | 154171712 | rs1579684 | 0.273 | -0.216 | -0.12 | -2.057 | -0.001 | 0.023 | 0.023 | 0 | 0.47 | 0.44 | 0.46 | 0.24 | 0.52 | 0.31 |
| 2 | 154173881 | rs13021542 | 0.999 | -0.226 | -0.07 | -2.217 | -0.004 | -0.022 | -0.022 | 0.357 | 0.32 | 0.3 | 0.32 | 0.04 | 0.36 | 0.18 |
| 2 | 154178478 | rs1439622 | 0.117 | -0.352 | 0.631 | -2.023 | -0.024 | -0.032 | -0.032 | 0 | 0.66 | 0.49 | 0.54 | 0.25 | 0.71 | 0.52 |
| 2 | 154178949 | rs6434548 | -1.357 | -0.357 | 0.504 | -1.952 | 0.007 | -0.012 | -0.012 | 0.042 | 0.83 | 0.85 | 0.69 | 0.38 | 0.87 | 0.67 |
| 2 | 154185954 | rs7579834 | 0.19 | -0.358 | 0.495 | -2.148 | 0 | -0.009 | -0.009 | 4.286 | 0.62 | 0.59 | 0.5 | 0.16 | 0.66 | 0.44 |
| 2 | 154189126 | rs1074166 | -0.01 | -0.387 | 0.524 | -2.118 | -0.004 | -0.007 | -0.007 | 3.083 | 0.66 | 0.6 | 0.55 | 0.24 | 0.74 | 0.46 |
| 2 | 154190197 | rs10204385 | 0.783 | -0.375 | 0.435 | -2.242 | 0.009 | 0.011 | 0.011 | 0 | 0.17 | 0.15 | 0.08 | 0.06 | 0.14 | 0.17 |
| 2 | 154195839 | rs2165156 | -1.751 | -0.306 | 0.31 | -2.146 | 0.009 | -0.037 | -0.037 | 0 | 0.84 | 0.92 | 0.69 | 0.37 | 0.86 | 0.66 |
| 2 | 154199490 | rs10192439 | 0.766 | -0.258 | 0.696 | -2.341 | -0.002 | -0.043 | -0.043 | 0.285 | 0.4 | 0.35 | 0.39 | 0.04 | 0.44 | 0.2 |
| 2 | 154204271 | rs2178593 | 0.469 | -0.358 | 0.536 | -2.276 | 0.021 | 0.15 | 0.15 | 1.635 | 0.52 | 0.22 | 0.48 | 0.24 | 0.54 | 0.41 |
| 2 | 154205978 | rs4372831 | -0.277 | -0.284 | 0.976 | -2.203 | -0.003 | -0.032 | -0.032 | 0.034 | 0.69 | 0.45 | 0.56 | 0.23 | 0.68 | 0.57 |
| 2 | 154207962 | rs7578875 | -1.475 | -0.285 | 1.037 | -2.209 | -0.011 | 0.03 | 0.03 | 0 | 0.82 | 0.75 | 0.66 | 0.37 | 0.85 | 0.66 |
| 2 | 154211122 | rs1921772 | 0.23 | -0.227 | 0.928 | -2.305 | 0.002 | -0.007 | -0.007 | 0 | 0.21 | 0.43 | 0.2 | 0.2 | 0.19 | 0.22 |
| 2 | 154214176 | rs6434567 | 0.353 | -0.197 | 0.761 | -2.251 | 0 | -0.044 | -0.044 | 0 | 0.56 | 0.45 | 0.44 | 0.1 | 0.57 | 0.37 |
| 2 | 154223319 | rs4664671 | -1.873 | -0.194 | 0.332 | -2.14 | 0.009 | -0.045 | -0.045 | 0 | 0.84 | 0.93 | 0.69 | 0.37 | 0.86 | 0.66 |
| 2 | 154224516 | rs13004477 | 0.218 | -0.143 | 0.43 | -2.133 | 0.002 | -0.001 | -0.001 | 0 | 0.19 | 0.29 | 0.19 | 0.21 | 0.18 | 0.23 |
| 2 | 154226249 | rs10167225 | 0.878 | -0.154 | -0.151 | -2.239 | 0.009 | 0.018 | 0.018 | 0 | 0.16 | 0.13 | 0.08 | 0.06 | 0.14 | 0.18 |
| 2 | 154231628 | rs6745205 | 0.825 | -0.043 | -0.324 | -2.337 | -0.017 | 0.059 | 0.059 | 0.686 | 0.39 | 0.24 | 0.35 | 0.04 | 0.42 | 0.19 |
| 2 | 154232697 | rs13415443 | 0.64 | -0.059 | -0.458 | -1.968 | 0.012 | 0.052 | 0.052 | 0 | 0.16 | 0.06 | 0.07 | 0.05 | 0.15 | 0.19 |
| 2 | 154244198 | rs11686661 | -0.069 | -0.168 | -0.36 | -1.527 | 0.001 | 0.001 | 0.001 | 0 | 0.68 | 0.72 | 0.56 | 0.27 | 0.71 | 0.48 |
| 2 | 154262090 | rs13022448 | 0.861 | -0.249 | -0.569 | -1.513 | -0.013 | -0.008 | -0.008 | 0 | 0.14 | 0.05 | 0.1 | 0.14 | 0.12 | 0.23 |
| 2 | 154264858 | rs10179200 | 0.719 | -0.359 | -0.981 | -1.976 | -0.029 | 0.042 | 0.042 | 0.631 | 0.28 | 0.16 | 0.29 | 0.04 | 0.35 | 0.31 |
| 2 | 154267276 | rs7597597 | -0.539 | -0.536 | -0.62 | -1.795 | 0.006 | 0.037 | 0.037 | 0 | 0.4 | 0.74 | 0.28 | 0.24 | 0.41 | 0.27 |
| 2 | 154273475 | rs11673716 | -1.043 | -0.37 | -0.237 | -1.218 | 0.007 | -0.019 | -0.019 | 0 | 0.81 | 0.89 | 0.62 | 0.36 | 0.85 | 0.6 |
| 2 | 154278240 | rs3863911 | 0.843 | -0.357 | 0.152 | -1.305 | -0.001 | -0.002 | -0.002 | 0 | 0.18 | 0.2 | 0.14 | 0.1 | 0.28 | 0.09 |
| 2 | 154287246 | rs11886199 | 0.623 | -0.308 | 0.462 | -1.286 | -0.007 | -0.04 | -0.04 | 0 | 0.1 | 0.23 | 0.07 | 0.03 | 0.07 | 0.16 |
| 2 | 154310368 | rs11676810 | 0.84 | -0.227 | -0.198 | -1.262 | 0.006 | -0.011 | -0.011 | 0.083 | 0.18 | 0.18 | 0.13 | 0.1 | 0.28 | 0.08 |
| 2 | 154311343 | rs1658551 | -1.477 | -0.375 | 0.256 | -0.611 | -0.028 | -0.015 | -0.015 | 0 | 0.38 | 0.5 | 0.47 | 0.67 | 0.28 | 0.58 |
| 2 | 154313125 | rs4286235 | 0.973 | -0.21 | -0.251 | -0.516 | -0.018 | -0.007 | -0.007 | 0 | 0.63 | 0.6 | 0.53 | 0.34 | 0.73 | 0.41 |
| 2 | 154318002 | rs7583243 | -0.35 | 0.264 | -0.429 | 0.027 | 0.006 | -0.038 | -0.038 | 0 | 0.82 | 0.92 | 0.65 | 0.41 | 0.85 | 0.61 |
| 2 | 154318295 | rs11692573 | 0.869 | 0.295 | -0.574 | 0.001 | -0.021 | 0.004 | 0.004 | 1.045 | 0.7 | 0.64 | 0.55 | 0.33 | 0.77 | 0.44 |
| 2 | 154331653 | rs10190938 | -0.107 | 0.051 | -1.56 | 0.223 | 0.009 | -0.028 | -0.028 | 0.13 | 0.09 | 0.03 | 0.08 | 0.14 | 0.07 | 0.16 |
| 2 | 154338317 | rs1344655 | 0.008 | 0.093 | -1.432 | 0.366 | 0.008 | -0.06 | -0.06 | 0.393 | 0.83 | 0.93 | 0.66 | 0.4 | 0.85 | 0.65 |
| 2 | 154348863 | rs12466830 | -0.27 | 0.473 | -1.595 | 0.194 | -0.019 | 0.012 | 0.012 | 0 | 0.2 | 0.09 | 0.33 | 0.16 | 0.2 | 0.31 |
| 2 | 154352501 | rs1221839 | -0.672 | 0.101 | -1.802 | 0.134 | 0.004 | 0.113 | 0.113 | 1.042 | 0.53 | 0.78 | 0.54 | 0.74 | 0.4 | 0.63 |
| 2 | 154365426 | rs4035233 | 0.413 | 0.03 | -1.589 | 0.558 | -0.004 | -0.009 | -0.009 | 0.036 | 0.09 | 0.07 | 0.06 | 0.15 | 0.08 | 0.17 |
| 2 | 154369594 | rs13430459 | 0.417 | 0.004 | -1.598 | 0.594 | -0.002 | -0.01 | -0.01 | 0 | 0.09 | 0.07 | 0.06 | 0.21 | 0.09 | 0.17 |
| 2 | 154370972 | rs7603221 | 0.904 | -0.548 | -0.66 | 0.172 | -0.018 | -0.087 | -0.087 | 0 | 0.17 | 0.3 | 0.09 | 0.02 | 0.1 | 0.07 |
| 2 | 154372766 | rs1221754 | 0.054 | -0.593 | -0.226 | 0.178 | 0.006 | -0.021 | -0.021 | 0.865 | 0.3 | 0.26 | 0.48 | 0.55 | 0.24 | 0.39 |
| 2 | 154374874 | rs9784007 | 1.283 | -0.546 | -0.084 | 0.122 | -0.008 | -0.057 | -0.057 | 0 | 0.15 | 0.21 | 0.09 | 0.02 | 0.1 | 0.07 |
| 2 | 154384644 | rs2155471 | -0.953 | -0.531 | -0.429 | 0.197 | -0.003 | 0.096 | 0.096 | 0.733 | 0.52 | 0.33 | 0.6 | 0.76 | 0.42 | 0.63 |
| 2 | 154390780 | rs1595455 | -1.162 | -0.544 | -0.488 | 0.178 | -0.035 | 0.12 | 0.12 | 1.079 | 0.55 | 0.83 | 0.65 | 0.76 | 0.42 | 0.62 |
| 2 | 154397112 | rs1595457 | -0.197 | -0.569 | -0.894 | 0.335 | -0.002 | -0.005 | -0.005 | 0 | 0.3 | 0.31 | 0.38 | 0.37 | 0.28 | 0.48 |
| 2 | 154401748 | rs1347740 | -1.08 | -0.238 | -1.097 | 0.239 | -0.091 | 0.117 | 0.117 | 0.33 | 0.58 | 0.88 | 0.66 | 0.78 | 0.43 | 0.64 |
| 2 | 154405645 | rs4641895 | -1.176 | 0.261 | -0.768 | 0.565 | -0.006 | -0.015 | -0.015 | 0 | 0.89 | 0.94 | 0.9 | 0.82 | 0.88 | 0.84 |
| 2 | 154411685 | rs16833593 | 0.326 | 0.282 | -0.41 | 0.525 | 0.016 | 0.014 | 0.014 | 0 | 0.11 | 0.21 | 0.19 | 0.14 | 0.15 | 0.24 |
| 2 | 154413874 | rs4664144 | -0.462 | 0.103 | -0.526 | 0.341 | 0.008 | 0.014 | 0.014 | 0 | 0.5 | 0.62 | 0.6 | 0.58 | 0.41 | 0.49 |
| 2 | 154414033 | rs16833611 | -0.14 | 0.068 | -0.021 | 0.546 | 0.011 | 0.047 | 0.047 | 0.153 | 0.1 | 0.27 | 0.07 | 0.12 | 0.08 | 0.15 |
| 2 | 154414404 | rs7570293 | -0.559 | -0.097 | -0.494 | 0.093 | -0.004 | 0.006 | 0.006 | 0.755 | 0.57 | 0.62 | 0.6 | 0.65 | 0.49 | 0.64 |
| 2 | 154414456 | rs10931682 | -0.175 | -0.216 | -0.323 | -0.018 | -0.004 | -0.027 | -0.027 | 0 | 0.37 | 0.46 | 0.33 | 0.17 | 0.36 | 0.4 |
| 2 | 154416171 | rs1940266 | 0.06 | -0.298 | -0.13 | 0.508 | 0.02 | -0.002 | -0.002 | 0 | 0.41 | 0.66 | 0.54 | 0.51 | 0.35 | 0.31 |
| 2 | 154417270 | rs7564934 | -0.247 | -0.466 | 0.335 | 0.821 | -0.03 | 0.013 | 0.013 | 0.331 | 0.15 | 0.23 | 0.13 | 0.17 | 0.09 | 0.18 |
| 2 | 154418706 | rs2053272 | -0.69 | -0.109 | 0.389 | 0.441 | -0.001 | -0.002 | -0.002 | 0 | 0.48 | 0.61 | 0.33 | 0.51 | 0.46 | 0.49 |
| 2 | 154419821 | rs16833642 | 0.853 | -0.074 | 0.456 | 0.516 | -0.002 | -0.001 | -0.001 | 0 | 0.11 | 0.09 | 0.08 | 0.1 | 0.09 | 0.18 |
| 2 | 154420012 | rs7589528 | 0.087 | -0.144 | 0.452 | 0.451 | -0.006 | 0.109 | 0.109 | 0.118 | 0.12 | 0.32 | 0.08 | 0.36 | 0.11 | 0.24 |
| 2 | 154420175 | rs1438093 | -0.065 | -0.166 | 0.309 | 0.363 | -0.005 | 0.04 | 0.04 | 0 | 0.37 | 0.25 | 0.25 | 0.15 | 0.36 | 0.26 |
| 2 | 154421703 | rs4533431 | -1.121 | -0.227 | 0.037 | 0.538 | -0.001 | 0.006 | 0.006 | 0 | 0.74 | 0.8 | 0.71 | 0.86 | 0.72 | 0.63 |

| | | | | | | | | | | | | | | | | |
|---|---|---|---|---|---|---|---|---|---|---|---|---|---|---|---|---|
| 2 | 154423181 | rs12693760 | -1.032 | -0.169 | -0.443 | 0.569 | 0.007 | 0 | 0 | 0.151 | 0.89 | 0.94 | 0.92 | 0.9 | 0.91 | 0.79 |
| 2 | 154423961 | rs766767 | 0.72 | 0.348 | -0.458 | 0.76 | -0.011 | 0.049 | 0.049 | 0 | 0.26 | 0.11 | 0.27 | 0.15 | 0.28 | 0.34 |
| 2 | 154424279 | rs1370676 | 0.622 | 0.272 | -0.018 | 1.433 | -0.001 | -0.018 | -0.018 | 0 | 0.24 | 0.18 | 0.38 | 0.35 | 0.25 | 0.12 |
| 2 | 154429323 | rs12472086 | 0.648 | 0.345 | -0.26 | 1.331 | 0.001 | -0.033 | -0.033 | 0 | 0.25 | 0.11 | 0.38 | 0.33 | 0.26 | 0.14 |
| 2 | 154431399 | rs4664146 | 0.879 | 0.44 | 0.435 | 1.478 | 0.016 | 0.194 | 0.194 | 0 | 0.94 | 0.74 | 0.93 | 0.8 | 0.93 | 0.82 |
| 2 | 154435087 | rs7565204 | -0.724 | 0.317 | 0.541 | 0.9 | 0.002 | -0.034 | -0.034 | 0.034 | 0.07 | 0.21 | 0.05 | 0 | 0.05 | 0.01 |
| 2 | 154436100 | rs4449085 | -1.124 | 0.332 | -0.225 | 0.826 | 0.042 | -0.011 | -0.011 | 0.062 | 0.48 | 0.84 | 0.38 | 0.63 | 0.41 | 0.64 |
| 2 | 154436676 | rs1940269 | -0.748 | 0.113 | 0.424 | 0.887 | -0.02 | 0.019 | 0.019 | 0 | 0.3 | 0.55 | 0.18 | 0.47 | 0.26 | 0.48 |
| 2 | 154442130 | rs1940255 | 0.307 | 0.181 | 0.433 | 0.806 | -0.001 | -0.001 | -0.001 | 0 | 0.17 | 0.22 | 0.11 | 0.15 | 0.14 | 0.16 |
| 2 | 154442520 | rs1940254 | -0.09 | 0.094 | 0.754 | 0.877 | -0.013 | 0.036 | 0.036 | 0 | 0.23 | 0.38 | 0.18 | 0.44 | 0.21 | 0.46 |
| 2 | 154450248 | rs746759 | -1.766 | 0.072 | 0.552 | 0.866 | 0.028 | -0.066 | -0.066 | 1.827 | 0.48 | 0.9 | 0.35 | 0.59 | 0.4 | 0.63 |
| 2 | 154452223 | rs10469747 | -1.186 | 0.119 | 0.729 | 0.893 | 0.001 | -0.001 | -0.001 | 0 | 0.41 | 0.59 | 0.29 | 0.58 | 0.35 | 0.63 |
| 2 | 154457104 | rs10497133 | -0.037 | 0.089 | 0.744 | 0.865 | -0.01 | -0.005 | -0.005 | 0 | 0.19 | 0.31 | 0.12 | 0.15 | 0.15 | 0.17 |
| 2 | 154461799 | rs4594396 | -1.645 | 0.09 | 0.511 | 0.866 | 0.031 | -0.025 | -0.025 | 0 | 0.46 | 0.74 | 0.37 | 0.6 | 0.4 | 0.64 |
| 2 | 154467433 | rs10199044 | -0.584 | 0.606 | -0.099 | 0.648 | 0 | -0.003 | -0.003 | 0 | 0.91 | 0.92 | 0.87 | 0.97 | 0.86 | 0.91 |
| 2 | 154469734 | rs1401741 | -0.541 | 0.379 | -0.2 | 0.814 | 0.001 | 0.001 | 0.001 | 0 | 0.23 | 0.27 | 0.14 | 0.17 | 0.19 | 0.18 |
| 2 | 154472066 | rs10197706 | -0.488 | 0.39 | -0.195 | 0.819 | 0 | 0.001 | 0.001 | 0 | 0.23 | 0.28 | 0.14 | 0.17 | 0.19 | 0.18 |
| 2 | 154479664 | rs6719847 | -0.002 | 0.466 | -0.259 | 0.973 | 0.001 | 0 | 0 | 0.384 | 0.18 | 0.21 | 0.11 | 0.16 | 0.15 | 0.17 |
| 2 | 154491513 | rs11693958 | -0.667 | 0.733 | -0.238 | 0.968 | 0.004 | 0.015 | 0.015 | 1.144 | 0.23 | 0.33 | 0.16 | 0.41 | 0.26 | 0.47 |
| 2 | 154494454 | rs10497135 | -1.174 | 0.553 | -0.749 | 1.076 | 0.013 | -0.024 | -0.024 | 0 | 0.34 | 0.32 | 0.52 | 0.55 | 0.32 | 0.4 |
| 2 | 154496150 | rs2009834 | 0.041 | 0.731 | -0.1 | 1.278 | 0 | 0.144 | 0.144 | 0.32 | 0.12 | 0.23 | 0.08 | 0.47 | 0.11 | 0.3 |
| 2 | 154507409 | rs7571900 | -0.193 | 0.934 | -0.17 | 1.038 | 0 | 0 | 0 | 0.452 | 0.93 | 0.92 | 0.95 | 0.92 | 0.93 | 0.82 |
| 2 | 154509239 | rs1546676 | 1.887 | 0.344 | -0.933 | 0.245 | -0.099 | 0.227 | 0.227 | 0 | 0.42 | 0.08 | 0.33 | 0.12 | 0.52 | 0.29 |
| 2 | 154510972 | rs13397382 | -0.314 | 0.111 | -0.48 | 0.21 | -0.014 | 0.076 | 0.076 | 0 | 0.19 | 0.32 | 0.4 | 0.29 | 0.16 | 0.15 |
| 2 | 154512950 | rs16834011 | -0.742 | 0.11 | -0.414 | 0.209 | -0.019 | 0.036 | 0.036 | 0 | 0.26 | 0.37 | 0.43 | 0.29 | 0.22 | 0.18 |
| 2 | 154513849 | rs13035707 | 2.953 | 0.559 | -0.697 | 0.002 | -0.077 | 0.152 | 0.152 | 0 | 0.34 | 0.03 | 0.17 | 0.12 | 0.33 | 0.2 |
| 2 | 154515483 | rs4664695 | 1.282 | 0.607 | -0.659 | 0.018 | -0.021 | 0.02 | 0.02 | 0.239 | 0.08 | 0.06 | 0.16 | 0 | 0.18 | 0.1 |
| 2 | 154516655 | rs6723606 | 1.203 | 0.543 | -0.593 | 0.016 | 0.006 | 0.007 | 0.007 | 0.002 | 0.61 | 0.45 | 0.61 | 0.41 | 0.55 | 0.37 |
| 2 | 154519277 | rs908919 | 1.302 | 0.649 | -0.948 | 0.078 | -0.021 | 0.019 | 0.019 | 0.225 | 0.08 | 0.06 | 0.15 | 0 | 0.19 | 0.1 |
| 2 | 154519448 | rs1519643 | 2.686 | 0.551 | -0.963 | 0.106 | -0.076 | 0.149 | 0.149 | 0 | 0.34 | 0.03 | 0.17 | 0.12 | 0.33 | 0.2 |
| 2 | 154521363 | rs4233663 | 0.438 | 0.481 | -0.568 | 0.397 | 0.002 | 0.015 | 0.015 | 0 | 0.08 | 0.06 | 0.31 | 0.23 | 0.05 | 0.07 |
| 2 | 154536942 | rs6750051 | 0.688 | 0.492 | -0.525 | -0.25 | -0.001 | -0.008 | -0.008 | 0 | 0.86 | 0.81 | 0.92 | 0.93 | 0.88 | 0.76 |
| 2 | 154540201 | rs1519640 | -1.547 | 0.364 | -0.607 | -0.396 | 0 | -0.004 | -0.004 | 0 | 0.4 | 0.54 | 0.53 | 0.22 | 0.44 | 0.32 |
| 2 | 154541088 | rs2203890 | 0.222 | 0.267 | -0.621 | -0.603 | -0.007 | 0.05 | 0.05 | 0 | 0.78 | 0.65 | 0.73 | 0.34 | 0.8 | 0.51 |
| 2 | 154545453 | rs12995533 | 2.359 | 0.29 | -1.056 | -0.632 | -0.027 | 0.167 | 0.167 | 0 | 0.39 | 0.14 | 0.2 | 0.11 | 0.38 | 0.2 |
| 2 | 154546923 | rs1401743 | 1.545 | 0.153 | -0.523 | -0.295 | 0.003 | -0.001 | -0.001 | 0 | 0.62 | 0.39 | 0.54 | 0.33 | 0.58 | 0.36 |
| 2 | 154549105 | rs4664152 | -0.583 | 0.406 | -0.705 | -1.34 | -0.013 | 0.008 | 0.008 | 0 | 0.14 | 0.13 | 0.17 | 0 | 0.22 | 0.15 |
| 2 | 154553429 | rs13013240 | 0.735 | -0.057 | -0.433 | -1.184 | -0.035 | -0.012 | -0.012 | 0 | 0.7 | 0.48 | 0.61 | 0.32 | 0.77 | 0.7 |
| 2 | 154561625 | rs16834075 | -0.751 | 0.058 | -0.236 | -1.142 | 0.049 | 0.137 | 0.137 | 0.455 | 0.09 | 0.41 | 0.06 | 0.21 | 0.05 | 0.25 |
| 2 | 154574231 | rs12479422 | -0.089 | 0.107 | -0.426 | -1.093 | -0.003 | 0.008 | 0.008 | 0 | 0.08 | 0.1 | 0.05 | 0.02 | 0.08 | 0.05 |
| 2 | 154582806 | rs1022386 | 0.82 | -0.126 | -0.479 | -1.277 | -0.015 | -0.02 | -0.02 | 0.845 | 0.65 | 0.53 | 0.59 | 0.32 | 0.69 | 0.66 |
| 2 | 154592770 | rs10164711 | -0.293 | -0.193 | -0.59 | -1.173 | -0.002 | -0.021 | -0.021 | 0 | 0.87 | 0.89 | 0.8 | 0.36 | 0.86 | 0.76 |
| 2 | 154605456 | rs13035809 | 2.201 | -0.251 | -1.392 | -1.769 | -0.116 | 0.161 | 0.161 | 0 | 0.35 | 0.02 | 0.19 | 0.08 | 0.34 | 0.19 |
| 2 | 154605760 | rs11889857 | 0.2 | -0.452 | -0.261 | -1.679 | -0.03 | -0.026 | -0.026 | 0.438 | 0.13 | 0.3 | 0.15 | 0.15 | 0.09 | 0.14 |
| 2 | 154609331 | rs1596111 | -1.05 | -0.374 | -0.172 | -1.843 | -0.025 | -0.083 | -0.083 | 1.274 | 0.28 | 0.44 | 0.46 | 0.18 | 0.26 | 0.25 |
| 2 | 154614114 | rs6713577 | -0.702 | -0.459 | 0.123 | -1.917 | -0.002 | -0.086 | -0.086 | 1.03 | 0.31 | 0.37 | 0.34 | 0.04 | 0.33 | 0.19 |
| 2 | 154619808 | rs10183136 | 1.668 | -0.481 | -0.252 | -1.974 | 0.01 | 0.284 | 0.284 | 0 | 0.49 | 0.15 | 0.5 | 0.1 | 0.52 | 0.31 |
| 2 | 154623397 | rs7579033 | -1.398 | -0.425 | 0.063 | -1.916 | 0.002 | -0.003 | -0.003 | 0.11 | 0.36 | 0.65 | 0.36 | 0.13 | 0.36 | 0.36 |
| 2 | 154650420 | rs16834236 | -0.541 | -0.356 | 0.172 | -1.785 | -0.042 | -0.067 | -0.067 | 0.009 | 0.19 | 0.4 | 0.16 | 0.09 | 0.12 | 0.22 |
| 2 | 154661248 | rs16834313 | -0.077 | -0.239 | 0.143 | -1.784 | 0.004 | 0.036 | 0.036 | 2.595 | 0.19 | 0.1 | 0.41 | 0.04 | 0.2 | 0.08 |
| 2 | 154664602 | rs1020223 | -1.117 | -0.24 | 0.472 | -1.864 | -0.003 | -0.008 | -0.008 | 0.736 | 0.35 | 0.32 | 0.6 | 0.08 | 0.44 | 0.21 |
| 2 | 154664805 | rs10187119 | -0.709 | -0.241 | 0.455 | -1.728 | -0.005 | 0.031 | 0.031 | 1.868 | 0.87 | 0.86 | 0.9 | 0.41 | 0.9 | 0.76 |
| 2 | 154667317 | rs4664703 | -0.168 | -0.271 | 0.555 | -1.741 | -0.018 | 0.066 | 0.066 | 3.643 | 0.85 | 0.77 | 0.89 | 0.42 | 0.9 | 0.75 |
| 2 | 154669222 | rs10931792 | 0.351 | -0.189 | 0.593 | -1.772 | 0.005 | -0.031 | -0.031 | 1.082 | 0.17 | 0.22 | 0.2 | 0.05 | 0.24 | 0.13 |
| 2 | 154671218 | rs6434925 | -1.561 | -0.213 | 0.571 | -1.797 | -0.002 | -0.094 | -0.094 | 0 | 0.44 | 0.53 | 0.66 | 0.09 | 0.5 | 0.3 |
| 2 | 154674451 | rs11695673 | 2.341 | -0.284 | 0.329 | -1.989 | -0.029 | 0.066 | 0.066 | 0.104 | 0.38 | 0.17 | 0.21 | 0.19 | 0.35 | 0.38 |
| 2 | 154688728 | rs1406084 | 0.959 | -0.187 | 0.454 | -1.946 | -0.02 | -0.003 | -0.003 | 0.307 | 0.74 | 0.48 | 0.81 | 0.27 | 0.78 | 0.58 |
| 2 | 154690220 | rs11899213 | 0.218 | -0.154 | 0.329 | -1.828 | -0.009 | 0.032 | 0.032 | 0 | 0.81 | 0.72 | 0.86 | 0.41 | 0.84 | 0.68 |
| 2 | 154691177 | rs10497137 | 0.084 | -0.177 | 0.318 | -1.862 | -0.007 | -0.053 | -0.053 | 0 | 0.11 | 0.21 | 0.06 | 0.01 | 0.08 | 0.13 |
| 2 | 154703431 | rs1372391 | 0.588 | -0.184 | 0.145 | -1.852 | -0.011 | 0.031 | 0.031 | 0 | 0.81 | 0.71 | 0.86 | 0.41 | 0.84 | 0.68 |
| 2 | 154704737 | rs1441872 | 1.261 | 0.079 | -0.06 | -1.461 | 0.002 | 0.01 | 0.01 | 0 | 0.53 | 0.43 | 0.31 | 0.35 | 0.49 | 0.6 |
| 2 | 154725568 | rs1373582 | -0.254 | 0.351 | -0.594 | -1.242 | -0.003 | 0.004 | 0.004 | 0 | 0.17 | 0.17 | 0.19 | 0.08 | 0.24 | 0.13 |
| 2 | 154731055 | rs10221630 | 1.197 | 0 | -1.14 | -1.684 | -0.011 | 0.15 | 0.15 | 0 | 0.39 | 0.17 | 0.22 | 0.07 | 0.36 | 0.32 |
| 2 | 154737638 | rs1444684 | -0.19 | -0.005 | -0.913 | -1.943 | -0.003 | -0.007 | -0.007 | 0.37 | 0.76 | 0.83 | 0.57 | 0.96 | 0.74 | 0.9 |
| 2 | 154739376 | rs1955152 | 0.707 | -0.086 | -0.649 | -2.123 | -0.006 | -0.024 | -0.024 | 0 | 0.61 | 0.62 | 0.38 | 0.94 | 0.48 | 0.72 |
| 2 | 154742427 | rs13412310 | 0.549 | -0.041 | -0.639 | -2.093 | 0.004 | -0.022 | -0.022 | 0.205 | 0.16 | 0.21 | 0.19 | 0.02 | 0.26 | 0.17 |
| 2 | 154743167 | rs17585462 | 0.199 | -0.328 | -0.136 | -1.888 | -0.034 | 0.013 | 0.013 | 0 | 0.16 | 0.25 | 0.08 | 0.27 | 0.1 | 0.25 |
| 2 | 154746555 | rs17585476 | 0.199 | -0.316 | -0.136 | -1.884 | -0.031 | 0.018 | 0.018 | 0 | 0.16 | 0.25 | 0.09 | 0.28 | 0.1 | 0.25 |
| 2 | 154748983 | rs1373581 | 0.868 | -0.466 | -0.421 | -2.118 | 0.003 | -0.013 | -0.013 | 0 | 0.58 | 0.62 | 0.48 | 0.81 | 0.63 | 0.67 |

| | | | | | | | | | | | | | | | | |
|---|---|---|---|---|---|---|---|---|---|---|---|---|---|---|---|---|
| 2 | 154750594 | rs9636320 | -0.306 | -0.46 | -0.419 | -2.058 | -0.006 | -0.015 | -0.015 | 4.372 | 0.18 | 0.23 | 0.1 | 0.15 | 0.12 | 0.23 |
| 2 | 154751502 | rs12612456 | -0.306 | -0.46 | -0.419 | -2.058 | -0.006 | -0.015 | -0.015 | 0.351 | 0.18 | 0.23 | 0.1 | 0.15 | 0.12 | 0.23 |
| 2 | 154767281 | rs10931826 | 0.293 | -0.444 | -0.487 | -2.102 | -0.003 | -0.063 | -0.063 | 1.408 | 0.68 | 0.65 | 0.54 | 0.91 | 0.71 | 0.77 |
| 2 | 154769276 | rs10210440 | 2.679 | -0.428 | -0.775 | -2.147 | -0.086 | 0.133 | 0.133 | 0.29 | 0.31 | 0.01 | 0.17 | 0.06 | 0.3 | 0.17 |
| 2 | 154783869 | rs17511012 | 0.438 | -0.398 | -0.597 | -2.149 | 0.009 | -0.033 | -0.033 | 3.153 | 0.17 | 0.24 | 0.2 | 0.02 | 0.27 | 0.17 |
| 2 | 154808697 | rs6714832 | 2.052 | -0.427 | -0.76 | -2.203 | -0.097 | 0.23 | 0.23 | 1.402 | 0.41 | 0.03 | 0.34 | 0.07 | 0.55 | 0.3 |
| 2 | 154815432 | rs2680680 | 1.336 | -0.396 | -0.655 | -2.212 | -0.041 | 0.025 | 0.025 | 0.081 | 0.07 | 0.01 | 0.3 | 0.03 | 0.12 | 0.06 |
| 2 | 154826491 | rs10196189 | -0.667 | -0.548 | -0.351 | -2.139 | -0.017 | -0.057 | -0.057 | 1.288 | 0.23 | 0.38 | 0.12 | 0.12 | 0.14 | 0.28 |
| 2 | 154830754 | rs10931862 | 1.611 | -0.576 | -0.219 | -2.113 | -0.023 | 0.127 | 0.127 | 0.143 | 0.07 | 0.18 | 0.08 | 0.62 | 0.04 | 0.18 |
| 2 | 154833426 | rs6748364 | -1.694 | -0.529 | -0.399 | -2.229 | 0.001 | -0.034 | -0.034 | 0.063 | 0.38 | 0.57 | 0.36 | 0.83 | 0.42 | 0.67 |
| 2 | 154836803 | rs10931864 | 1.023 | -0.547 | -0.43 | -2.254 | 0.036 | 0.105 | 0.105 | 0 | 0.58 | 0.29 | 0.43 | 0.21 | 0.65 | 0.46 |
| 2 | 154844785 | rs7575630 | 0.24 | -0.444 | -0.401 | -2.233 | -0.026 | 0.011 | 0.011 | 0 | 0.66 | 0.78 | 0.41 | 0.96 | 0.51 | 0.77 |
| 2 | 154849098 | rs7590906 | 0.153 | -0.431 | -0.4 | -2.218 | 0.009 | -0.029 | -0.029 | 0.225 | 0.16 | 0.21 | 0.19 | 0.02 | 0.27 | 0.18 |
| 2 | 154861176 | rs1013693 | -0.698 | -0.45 | -0.412 | -2.219 | 0 | -0.038 | -0.038 | 0 | 0.76 | 0.8 | 0.57 | 0.96 | 0.75 | 0.9 |
| 2 | 154865481 | rs12624082 | -0.015 | -0.403 | -0.453 | -2.194 | -0.024 | 0.015 | 0.015 | 0 | 0.17 | 0.13 | 0.19 | 0.01 | 0.28 | 0.24 |
| 2 | 154875613 | rs13400898 | -0.275 | -0.501 | -0.384 | -2.159 | -0.018 | -0.07 | -0.07 | 0.345 | 0.13 | 0.33 | 0.06 | 0.02 | 0.07 | 0.17 |
| 2 | 154886245 | rs13427314 | 2.666 | -0.529 | -0.617 | -2.173 | -0.082 | 0.104 | 0.104 | 0.039 | 0.32 | 0.01 | 0.18 | 0.11 | 0.31 | 0.18 |
| 2 | 154900325 | rs1863086 | 0.766 | -0.514 | -0.428 | -2.18 | -0.003 | -0.031 | -0.031 | 0 | 0.59 | 0.64 | 0.37 | 0.95 | 0.47 | 0.65 |
| 2 | 154903010 | rs1519213 | 1.129 | -0.623 | -0.451 | -2.208 | -0.015 | 0.119 | 0.119 | 0 | 0.55 | 0.31 | 0.29 | 0.24 | 0.45 | 0.45 |
| 2 | 154908367 | rs16835732 | 1.689 | -0.661 | -0.289 | -2.083 | -0.021 | 0.129 | 0.129 | 0.161 | 0.06 | 0.13 | 0.07 | 0.6 | 0.04 | 0.18 |
| 2 | 154910880 | rs6435029 | -0.897 | -0.676 | -0.284 | -2.186 | 0.003 | -0.063 | -0.063 | 0 | 0.77 | 0.76 | 0.56 | 0.96 | 0.75 | 0.89 |
| 2 | 154912289 | rs10490537 | 1.718 | -0.658 | -0.291 | -2.125 | -0.02 | 0.129 | 0.129 | 0.15 | 0.06 | 0.13 | 0.07 | 0.6 | 0.04 | 0.18 |
| 2 | 154937930 | rs10490534 | -0.455 | -0.754 | -0.342 | -2.233 | -0.074 | -0.039 | -0.039 | 0 | 0.23 | 0.44 | 0.17 | 0.83 | 0.14 | 0.43 |
| 2 | 154938172 | rs16836014 | 1.41 | -0.688 | -0.797 | -2.207 | 0.016 | 0.027 | 0.027 | 0.984 | 0.11 | 0.05 | 0.06 | 0.11 | 0.08 | 0.11 |
| 2 | 154944691 | rs10175190 | -1.153 | -0.673 | -0.904 | -2.243 | -0.012 | 0.025 | 0.025 | 0 | 0.34 | 0.33 | 0.28 | 0.14 | 0.38 | 0.4 |
| 2 | 154946026 | rs7587580 | 2.568 | -1.029 | -0.941 | -2.02 | -0.09 | 0.133 | 0.133 | 0.388 | 0.31 | 0.01 | 0.17 | 0.06 | 0.31 | 0.17 |
| 2 | 154984566 | rs16836088 | -0.408 | -1.22 | -0.007 | -1.294 | 0.008 | 0.007 | 0.007 | 0.854 | 0.18 | 0.16 | 0.09 | 0.12 | 0.12 | 0.23 |
| 2 | 154994668 | rs7592932 | 0.987 | -1.224 | 0.011 | -1.254 | -0.018 | 0.093 | 0.093 | 0.056 | 0.06 | 0.11 | 0.07 | 0.6 | 0.04 | 0.18 |
| 2 | 155012888 | rs6744657 | -2.475 | -1.23 | 0.191 | -1.332 | 0.029 | 0.032 | 0.032 | 0.901 | 0.36 | 0.71 | 0.23 | 0.85 | 0.2 | 0.6 |
| 2 | 155021606 | rs16836152 | 0.685 | -0.84 | -0.398 | -1.048 | 0.001 | 0.025 | 0.025 | 1.153 | 0.11 | 0.01 | 0.05 | 0.11 | 0.08 | 0.12 |
| 2 | 155024641 | rs1160471 | -0.264 | -0.827 | -0.273 | -1.016 | 0.002 | -0.007 | -0.007 | 0 | 0.18 | 0.18 | 0.1 | 0.11 | 0.12 | 0.24 |
| 2 | 155026455 | rs7558032 | 0.802 | -0.617 | -0.291 | -0.757 | -0.018 | 0.091 | 0.091 | 0.15 | 0.06 | 0.11 | 0.08 | 0.61 | 0.04 | 0.18 |
| 2 | 155027447 | rs12620249 | -0.759 | -0.533 | -0.324 | -1.089 | -0.017 | -0.047 | -0.047 | 0 | 0.24 | 0.33 | 0.12 | 0.13 | 0.14 | 0.29 |
| 2 | 155028927 | rs7580788 | 0.608 | -0.436 | -0.358 | -0.416 | 0 | 0 | 0 | 0 | 0.61 | 0.43 | 0.71 | 0.65 | 0.66 | 0.52 |
| 2 | 155038788 | rs11677129 | -0.281 | -0.533 | -0.199 | -0.507 | -0.006 | -0.033 | -0.033 | 0 | 0.27 | 0.31 | 0.16 | 0.11 | 0.21 | 0.24 |
| 2 | 155053140 | rs7592907 | 0.782 | -0.246 | -0.151 | 0.299 | 0.016 | 0.106 | 0.106 | 0.867 | 0.09 | 0.19 | 0.06 | 0.34 | 0.12 | 0.22 |
| 2 | 155058065 | rs13432170 | 1.09 | -0.306 | -0.329 | 0.252 | -0.006 | -0.033 | -0.033 | 0.121 | 0.4 | 0.44 | 0.24 | 0.13 | 0.31 | 0.16 |
| 2 | 155072003 | rs888581 | -1.026 | -0.207 | -0.619 | 0.213 | -0.029 | -0.051 | -0.051 | 0 | 0.85 | 0.97 | 0.8 | 0.74 | 0.82 | 0.74 |
| 2 | 155072139 | rs10211548 | 1.073 | -0.113 | -0.471 | 0.227 | -0.001 | -0.008 | -0.008 | 0 | 0.17 | 0.17 | 0.1 | 0.09 | 0.15 | 0.19 |
| 2 | 155084230 | rs10189484 | 1.928 | 0.028 | -0.492 | 0.255 | -0.006 | 0.081 | 0.081 | 0 | 0.3 | 0.08 | 0.2 | 0.12 | 0.26 | 0.12 |
| 2 | 155099537 | rs12477393 | 1.627 | 0.064 | -0.58 | 0.309 | -0.042 | 0.023 | 0.023 | 0.198 | 0.16 | 0.03 | 0.36 | 0.08 | 0.21 | 0.19 |
| 2 | 155100980 | rs11686394 | 1.032 | -0.176 | -0.388 | 0.17 | -0.002 | -0.016 | -0.016 | 1.018 | 0.18 | 0.18 | 0.1 | 0.06 | 0.15 | 0.19 |
| 2 | 155131619 | rs4664160 | -0.525 | -0.075 | -0.195 | 0.277 | -0.005 | 0.016 | 0.016 | 0.1 | 0.42 | 0.36 | 0.61 | 0.77 | 0.53 | 0.64 |
| 2 | 155151484 | rs4233668 | -1.745 | -0.008 | -0.315 | 0.268 | 0.004 | 0.01 | 0.01 | 0 | 0.69 | 0.72 | 0.78 | 0.88 | 0.74 | 0.87 |
| 2 | 155164091 | rs16836440 | 0.039 | -0.04 | -0.269 | 0.48 | -0.006 | -0.02 | -0.02 | 0 | 0.31 | 0.27 | 0.41 | 0.48 | 0.36 | 0.4 |
| 2 | 155166873 | rs7571016 | -0.861 | 0.181 | -0.589 | 0.564 | -0.002 | 0.003 | 0.003 | 0 | 0.58 | 0.61 | 0.58 | 0.65 | 0.56 | 0.64 |
| 2 | 155170857 | rs7566164 | 0.102 | 0.465 | -1.023 | 0.269 | -0.077 | 0.035 | 0.035 | 0 | 0.27 | 0.13 | 0.54 | 0.22 | 0.43 | 0.31 |
| 2 | 155171139 | rs12693973 | 1.518 | 0.153 | -1.226 | 0.271 | 0.003 | 0.04 | 0.04 | 0.174 | 0.32 | 0.07 | 0.23 | 0.17 | 0.26 | 0.17 |
| 2 | 155171476 | rs16836481 | 0.647 | 0.37 | -1.059 | 0.567 | 0.003 | 0.106 | 0.106 | 1.644 | 0.11 | 0.21 | 0.05 | 0.34 | 0.1 | 0.23 |
| 2 | 155177873 | rs10193792 | 0.795 | 0.108 | -0.766 | -0.114 | -0.005 | -0.003 | -0.003 | 0 | 0.36 | 0.34 | 0.29 | 0.29 | 0.25 | 0.22 |
| 2 | 155178214 | rs10183389 | 1.282 | 0.174 | -1.297 | -0.4 | 0.002 | 0.004 | 0.004 | 0 | 0.16 | 0.12 | 0.09 | 0.09 | 0.14 | 0.17 |
| 2 | 155179331 | rs13393190 | 1.912 | 0.232 | -1.976 | -0.396 | 0.03 | 0.017 | 0.017 | 0 | 0.24 | 0.07 | 0.2 | 0.17 | 0.19 | 0.18 |
| 2 | 155181150 | rs13032599 | 1.562 | 0.55 | -2.06 | -0.446 | -0.044 | 0.114 | 0.114 | 0.972 | 0.21 | 0.06 | 0.32 | 0.02 | 0.27 | 0.23 |
| 2 | 155182433 | rs7576163 | 1.554 | 0.585 | -2.02 | -0.369 | 0.019 | 0.252 | 0.252 | 0.019 | 0.07 | 0.2 | 0.05 | 0.5 | 0.07 | 0.18 |
| 2 | 155188523 | rs707025 | -0.951 | 0.625 | -2.953 | -0.562 | -0.019 | 0.001 | 0.001 | 0 | 0.65 | 0.84 | 0.58 | 0.71 | 0.62 | 0.66 |
| 2 | 155188665 | rs16836531 | 0.597 | 0.926 | -2.486 | -0.077 | -0.006 | 0.105 | 0.105 | 2.303 | 0.2 | 0.62 | 0.14 | 0.57 | 0.13 | 0.22 |
| 2 | 155197361 | rs808948 | 0.406 | 1.37 | -2.578 | -0.101 | -0.049 | 0.117 | 0.117 | 1.149 | 0.21 | 0.05 | 0.32 | 0.02 | 0.27 | 0.23 |
| 2 | 155204133 | rs17811332 | -0.328 | 1.151 | -1.557 | 0.996 | -0.058 | -0.047 | -0.047 | 0.129 | 0.09 | 0.01 | 0.21 | 0.16 | 0.2 | 0.18 |
| 2 | 155206097 | rs13431051 | -0.87 | 0.067 | -1.032 | 0.679 | -0.002 | -0.006 | -0.006 | 0 | 0.16 | 0.14 | 0.09 | 0.1 | 0.15 | 0.17 |
| 2 | 155211698 | rs16836643 | -1.164 | -0.108 | -1.312 | 1.209 | 0.034 | -0.049 | -0.049 | 1.395 | 0.12 | 0.04 | 0.04 | 0.24 | 0.07 | 0.2 |
| 2 | 155213618 | rs1527865 | 0.497 | -0.797 | -0.881 | 0.387 | -0.044 | -0.046 | -0.046 | 1.401 | 0.65 | 0.51 | 0.7 | 0.81 | 0.71 | 0.6 |
| 2 | 155214199 | rs13001607 | 0.03 | -1.184 | -1.09 | 0.273 | 0.003 | 0.001 | 0.001 | 0.028 | 0.1 | 0.08 | 0.08 | 0.11 | 0.07 | 0.06 |
| 2 | 155219508 | rs1524915 | 0.377 | -0.886 | -0.764 | 0.522 | -0.026 | -0.037 | -0.037 | 0 | 0.65 | 0.54 | 0.7 | 0.81 | 0.68 | 0.61 |
| 2 | 155222220 | rs17838576 | 0.262 | -0.964 | -0.134 | 0.482 | 0.003 | 0.003 | 0.003 | 0 | 0.67 | 0.68 | 0.72 | 0.81 | 0.71 | 0.61 |
| 2 | 155225035 | rs1527867 | 0.502 | 0.158 | -0.202 | 0.982 | 0 | 0.001 | 0.001 | 0 | 0.58 | 0.58 | 0.65 | 0.66 | 0.54 | 0.48 |
| 2 | 155226644 | rs707039 | -0.374 | 0.16 | -0.015 | 1.022 | -0.015 | -0.013 | -0.013 | 0.015 | 0.35 | 0.42 | 0.3 | 0.24 | 0.32 | 0.39 |
| 2 | 155230428 | rs1527869 | 1.341 | 0.204 | -0.012 | 1.039 | 0.002 | 0.003 | 0.003 | 0 | 0.46 | 0.33 | 0.56 | 0.58 | 0.49 | 0.4 |
| 2 | 155232409 | rs2037660 | 0.639 | 0.127 | 0.011 | 1.018 | 0.007 | 0.005 | 0.005 | 0 | 0.54 | 0.44 | 0.62 | 0.65 | 0.65 | 0.54 |
| 2 | 155232423 | rs2037659 | 1.403 | 0.122 | -0.016 | 1.013 | -0.004 | -0.001 | -0.001 | 0 | 0.46 | 0.33 | 0.55 | 0.58 | 0.48 | 0.4 |

| | | | | | | | | | | | | | | | | |
|---|---|---|---|---|---|---|---|---|---|---|---|---|---|---|---|---|
| 2 | 155234318 | rs12621256 | 0.4 | 0.195 | -0.038 | 0.999 | -0.01 | -0.009 | -0.009 | 0 | 0.08 | 0.08 | 0.07 | 0.1 | 0.17 | 0.13 |
| 2 | 155236334 | rs1154759 | 0.732 | 0.11 | 0.094 | 0.967 | -0.003 | -0.002 | -0.002 | 0 | 0.54 | 0.44 | 0.61 | 0.65 | 0.62 | 0.53 |
| 2 | 155241841 | rs799793 | -0.943 | 0.015 | 0.302 | 0.884 | -0.011 | -0.011 | -0.011 | 0 | 0.44 | 0.5 | 0.38 | 0.32 | 0.37 | 0.45 |
| 2 | 155243685 | rs707043 | -1.007 | -0.051 | 0.556 | 0.81 | -0.007 | -0.012 | -0.012 | 0 | 0.45 | 0.54 | 0.35 | 0.25 | 0.38 | 0.44 |
| 2 | 155248839 | rs707051 | -0.82 | -0.369 | 0.485 | 0.561 | -0.044 | -0.035 | -0.035 | 0 | 0.34 | 0.48 | 0.27 | 0.25 | 0.24 | 0.4 |
| 2 | 155254734 | rs16837121 | 1.642 | -0.451 | 0.597 | 0.586 | -0.022 | 0.034 | 0.034 | 0 | 0.47 | 0.21 | 0.57 | 0.64 | 0.45 | 0.42 |
| 2 | 155255152 | rs993412 | 1.717 | -0.431 | 0.605 | 0.59 | -0.02 | 0.029 | 0.029 | 0 | 0.46 | 0.23 | 0.57 | 0.65 | 0.46 | 0.42 |
| 2 | 155268668 | rs17814066 | 2.027 | -0.121 | -0.668 | -0.056 | -0.087 | 0.097 | 0.097 | 0.778 | 0.23 | 0.01 | 0.12 | 0.09 | 0.23 | 0.16 |
| 2 | 155272829 | rs1012721 | 1.175 | -0.335 | -0.377 | 0.689 | -0.032 | 0.006 | 0.006 | 0 | 0.44 | 0.2 | 0.48 | 0.44 | 0.45 | 0.39 |
| 2 | 155275025 | rs799761 | 0.799 | -0.292 | -0.326 | 0.36 | 0.007 | 0.019 | 0.019 | 0 | 0.65 | 0.47 | 0.76 | 0.58 | 0.57 | 0.63 |
| 2 | 155279037 | rs707057 | -1.006 | -0.653 | -0.077 | 0.001 | -0.004 | -0.003 | -0.003 | 0 | 0.35 | 0.4 | 0.28 | 0.22 | 0.26 | 0.35 |
| 2 | 155281705 | rs707059 | -0.921 | -0.172 | -0.343 | -0.3 | 0.011 | -0.013 | -0.013 | 0 | 0.32 | 0.31 | 0.25 | 0.01 | 0.27 | 0.28 |
| 2 | 155287778 | rs10185435 | 1.244 | 0.122 | 0.059 | -0.076 | 0.008 | -0.049 | -0.049 | 0.555 | 0.1 | 0.17 | 0.08 | 0.01 | 0.11 | 0.07 |
| 2 | 155289620 | rs10932026 | -0.175 | 0.354 | -0.07 | 0.341 | 0.008 | 0.036 | 0.036 | 0.107 | 0.16 | 0.14 | 0.15 | 0.49 | 0.22 | 0.2 |
| 2 | 155291579 | rs799770 | -0.152 | 0.16 | -0.314 | 0.346 | -0.012 | -0.075 | -0.075 | 0 | 0.16 | 0.01 | 0.13 | 0.24 | 0.12 | 0.19 |
| 2 | 155294795 | rs1527862 | 0.169 | 0.187 | -0.133 | 0.361 | 0.009 | -0.017 | -0.017 | 0.699 | 0.15 | 0.11 | 0.1 | 0.22 | 0.14 | 0.11 |
| 2 | 155298880 | rs799822 | 0.403 | 0.143 | -0.006 | 0.005 | 0.005 | -0.002 | -0.002 | 0 | 0.12 | 0.12 | 0.12 | 0.01 | 0.08 | 0.19 |
| 2 | 155300526 | rs799825 | -0.029 | 0.226 | 0.136 | 0.055 | 0.013 | 0.073 | 0.073 | 0.728 | 0.8 | 0.47 | 0.85 | 0.67 | 0.78 | 0.75 |
| 2 | 155303560 | rs13419570 | 1.433 | 0.192 | 0.269 | -0.008 | 0.01 | -0.05 | -0.05 | 0.148 | 0.1 | 0.17 | 0.08 | 0.01 | 0.11 | 0.07 |
| 2 | 155303655 | rs707073 | 2.022 | 0.239 | 0.091 | -0.125 | -0.036 | 0.246 | 0.246 | 0.151 | 0.54 | 0.18 | 0.58 | 0.16 | 0.45 | 0.52 |
| 2 | 155303969 | rs13032771 | -0.384 | 0.313 | 0.005 | 0.027 | -0.001 | 0 | 0 | 0 | 0.25 | 0.23 | 0.26 | 0.51 | 0.32 | 0.22 |
| 2 | 155304170 | rs707075 | -0.733 | 0.165 | 0.357 | 0.206 | 0.038 | 0.223 | 0.223 | 0.147 | 0.9 | 0.65 | 0.94 | 0.69 | 0.89 | 0.81 |
| 2 | 155307617 | rs707081 | -0.871 | 0.331 | -0.025 | 0.108 | -0.007 | -0.019 | -0.019 | 0.195 | 0.33 | 0.24 | 0.32 | 0.5 | 0.35 | 0.3 |
| 2 | 155307832 | rs707082 | -0.4 | 0.375 | -0.059 | 0.324 | 0.037 | 0.245 | 0.245 | 0.166 | 0.89 | 0.62 | 0.9 | 0.67 | 0.89 | 0.82 |
| 2 | 155308061 | rs707084 | -1.549 | 0.385 | -0.495 | 0.24 | 0 | 0.018 | 0.018 | 0 | 0.48 | 0.65 | 0.43 | 0.51 | 0.47 | 0.52 |
| 2 | 155308672 | rs7582614 | 1.845 | 0.389 | -0.53 | 0.22 | -0.021 | 0.221 | 0.221 | 0 | 0.43 | 0.13 | 0.48 | 0.16 | 0.41 | 0.33 |
| 2 | 155308898 | rs707085 | -1.64 | 0.392 | -0.345 | 0.363 | -0.001 | 0.011 | 0.011 | 0 | 0.48 | 0.62 | 0.43 | 0.51 | 0.47 | 0.52 |
| 2 | 155313749 | rs707093 | -0.51 | 0.566 | -0.137 | 0.97 | 0.001 | 0.016 | 0.016 | 0 | 0.16 | 0.17 | 0.16 | 0.28 | 0.16 | 0.24 |
| 2 | 155314294 | rs17815112 | -0.299 | 0.447 | -0.2 | 0.678 | -0.003 | -0.031 | -0.031 | 0 | 0.4 | 0.28 | 0.32 | 0.53 | 0.45 | 0.35 |
| 2 | 155314404 | rs6435200 | 1.782 | 0.394 | 0.302 | 0.593 | -0.001 | -0.043 | -0.043 | 0.041 | 0.4 | 0.41 | 0.5 | 0.08 | 0.38 | 0.35 |
| 2 | 155315021 | rs13000699 | 0.854 | 0.684 | 0.595 | 0.927 | 0.07 | 0.036 | 0.036 | 0 | 0.08 | 0.21 | 0.08 | 0.07 | 0.14 | 0.09 |
| 2 | 155315938 | rs741602 | -0.76 | 0.444 | 0.336 | 0.991 | 0.005 | 0.033 | 0.033 | 0.235 | 0.43 | 0.33 | 0.33 | 0.73 | 0.45 | 0.42 |
| 2 | 155316574 | rs10932050 | -0.494 | 0.446 | 0.32 | 1.128 | -0.001 | -0.021 | -0.021 | 0 | 0.41 | 0.3 | 0.32 | 0.5 | 0.45 | 0.35 |
| 2 | 155320829 | rs10490525 | -0.916 | 0.16 | 0.442 | 0.986 | 0.001 | 0.003 | 0.003 | 0.337 | 0.39 | 0.37 | 0.28 | 0.74 | 0.39 | 0.36 |
| 2 | 155321482 | rs1157233 | -0.853 | 0.195 | 0.736 | 1.162 | 0.011 | -0.007 | -0.007 | 0 | 0.4 | 0.48 | 0.32 | 0.64 | 0.48 | 0.42 |
| 2 | 155322083 | rs707104 | 1.347 | 0.163 | 0.936 | 1.106 | -0.001 | -0.011 | -0.011 | 0.197 | 0.53 | 0.5 | 0.64 | 0.18 | 0.48 | 0.53 |
| 2 | 155322559 | rs10932055 | 0.838 | 0.242 | 1.164 | 1.089 | -0.001 | 0.029 | 0.029 | 0.34 | 0.19 | 0.22 | 0.13 | 0.4 | 0.17 | 0.18 |
| 2 | 155322696 | rs707107 | -0.289 | 0.217 | 1.186 | 1.073 | 0 | -0.002 | -0.002 | 0.993 | 0.74 | 0.72 | 0.82 | 0.6 | 0.76 | 0.81 |
| 2 | 155323278 | rs697651 | 0.504 | 0.267 | 1.147 | 1.074 | 0.014 | 0.044 | 0.044 | 0 | 0.67 | 0.53 | 0.73 | 0.52 | 0.63 | 0.71 |
| 2 | 155323352 | rs6750412 | -1.317 | 0.168 | 1.111 | 0.751 | 0.022 | 0.032 | 0.032 | 0 | 0.2 | 0.29 | 0.14 | 0.26 | 0.24 | 0.24 |
| 2 | 155330735 | rs1066376 | -0.202 | 0.344 | 0.782 | 0.837 | 0.045 | 0.019 | 0.019 | 0.148 | 0.91 | 0.74 | 0.91 | 0.92 | 0.87 | 0.9 |
| 2 | 155332023 | rs1609150 | -0.686 | 0.054 | 0.638 | 0.812 | -0.02 | 0.014 | 0.014 | 0 | 0.15 | 0.18 | 0.07 | 0.2 | 0.12 | 0.1 |
| 2 | 155340061 | rs992587 | 1.099 | 0.21 | 0.217 | 0.374 | -0.006 | -0.03 | -0.03 | 0 | 0.16 | 0.08 | 0.23 | 0.4 | 0.17 | 0.15 |
| 2 | 155341520 | rs11675853 | 0.328 | 0.203 | 0.476 | 0.274 | 0.046 | 0.013 | 0.013 | 0 | 0.11 | 0.16 | 0.14 | 0.12 | 0.2 | 0.18 |
| 2 | 155342678 | rs1429261 | 0.061 | -0.052 | 0.311 | 0.183 | 0.012 | -0.016 | -0.016 | 0 | 0.14 | 0.08 | 0.03 | 0.25 | 0.08 | 0.17 |
| 2 | 155345947 | rs12617045 | -1.312 | 0.016 | 0.612 | 0.086 | 0.003 | -0.038 | -0.038 | 0.761 | 0.43 | 0.43 | 0.4 | 0.77 | 0.45 | 0.5 |
| 2 | 155346569 | rs249694 | -0.944 | -0.082 | 0.745 | -0.007 | -0.014 | -0.034 | -0.034 | 0.131 | 0.92 | 0.81 | 0.91 | 0.99 | 0.91 | 0.97 |
| 2 | 155346958 | rs1003743 | 0.067 | -0.073 | 0.58 | 0.01 | 0.011 | 0.002 | 0.002 | 0.097 | 0.31 | 0.17 | 0.26 | 0.64 | 0.25 | 0.33 |
| 2 | 155347161 | rs17553946 | 2.29 | -0.18 | 0.041 | -0.436 | -0.007 | 0.1 | 0.1 | 0 | 0.24 | 0.03 | 0.23 | 0.15 | 0.21 | 0.2 |
| 2 | 155353711 | rs249703 | 1.51 | -0.095 | 0.744 | 0.258 | -0.003 | 0.107 | 0.107 | 0 | 0.49 | 0.36 | 0.52 | 0.21 | 0.47 | 0.47 |
| 2 | 155359252 | rs10211143 | -0.685 | 0.071 | 0.825 | 0.359 | 0.028 | -0.011 | -0.011 | 0 | 0.17 | 0.29 | 0.21 | 0.13 | 0.29 | 0.2 |
| 2 | 155362422 | rs1366748 | -1.256 | -0.026 | 0.661 | 0.355 | 0.004 | -0.01 | -0.01 | 0 | 0.43 | 0.44 | 0.41 | 0.64 | 0.45 | 0.43 |
| 2 | 155364263 | rs13033023 | 0.146 | 0.271 | 0.288 | 0.078 | 0.06 | 0.021 | 0.021 | 0 | 0.11 | 0.19 | 0.14 | 0.13 | 0.2 | 0.16 |
| 2 | 155366693 | rs774516 | -1.801 | -0.098 | -0.035 | 0.099 | -0.011 | 0.074 | 0.074 | 0 | 0.57 | 0.68 | 0.53 | 0.79 | 0.56 | 0.6 |
| 2 | 155366857 | rs16837770 | 0.632 | -0.681 | -0.384 | 0.556 | -0.047 | -0.131 | -0.131 | 0.027 | 0.18 | 0.02 | 0.27 | 0.45 | 0.22 | 0.17 |
| 2 | 155371557 | rs16837788 | -1.666 | -0.619 | 0.263 | 0.615 | -0.043 | 0.009 | 0.009 | 0 | 0.22 | 0.28 | 0.08 | 0.21 | 0.11 | 0.25 |
| 2 | 155372611 | rs16837790 | 0.195 | -0.688 | 0.077 | 0.655 | 0.016 | 0.022 | 0.022 | 0 | 0.05 | 0.12 | 0.03 | 0.06 | 0.05 | 0.05 |
| 2 | 155373390 | rs16837794 | 0.251 | -0.614 | -0.328 | 0.674 | 0.002 | -0.003 | -0.003 | 0 | 0.08 | 0.06 | 0.02 | 0.14 | 0.03 | 0.14 |
| 2 | 155381331 | rs1896831 | -0.994 | -0.115 | -0.085 | 0.392 | -0.046 | -0.046 | -0.046 | 0 | 0.15 | 0.36 | 0.07 | 0.07 | 0.07 | 0.09 |
| 2 | 155390620 | rs1429258 | -0.407 | 0.327 | -0.912 | 0.611 | 0.023 | -0.01 | -0.01 | 0.818 | 0.42 | 0.28 | 0.42 | 0.54 | 0.4 | 0.33 |
| 2 | 155390631 | rs1429259 | -0.247 | 1.226 | -0.855 | 0.29 | -0.008 | 0.005 | 0.005 | 0 | 0.81 | 0.81 | 0.77 | 0.94 | 0.72 | 0.84 |
| 2 | 155391005 | rs2695412 | 0.687 | 0.804 | -0.782 | 0.905 | -0.002 | -0.005 | -0.005 | 0 | 0.86 | 0.9 | 0.82 | 0.74 | 0.85 | 0.82 |
| 2 | 155391797 | rs2695415 | -1.049 | 0.669 | -0.22 | 0.592 | 0.001 | -0.026 | -0.026 | 0 | 0.14 | 0.22 | 0.11 | 0.07 | 0.12 | 0.11 |
| 2 | 155392530 | rs13432931 | -0.683 | 0.698 | -0.006 | 0.953 | -0.004 | 0.022 | 0.022 | 0.711 | 0.3 | 0.43 | 0.28 | 0.57 | 0.25 | 0.37 |
| 2 | 155392642 | rs1527452 | -0.277 | 0.495 | 0.126 | 0.631 | -0.004 | -0.034 | -0.034 | 2.514 | 0.14 | 0.18 | 0.08 | 0.04 | 0.13 | 0.07 |
| 2 | 155393553 | rs774500 | -1.279 | 0.295 | 0.401 | 1.23 | 0.002 | 0.008 | 0.008 | 0 | 0.29 | 0.61 | 0.28 | 0.3 | 0.3 | 0.29 |
| 2 | 155399506 | rs10202391 | 0.318 | 0.209 | 0.056 | 1.572 | -0.002 | -0.049 | -0.049 | 0.629 | 0.09 | 0 | 0.07 | 0.12 | 0.04 | 0.09 |
| 2 | 155399735 | rs774503 | -1.401 | 0.208 | -0.686 | 1.07 | 0.017 | 0.052 | 0.052 | 0 | 0.72 | 0.92 | 0.65 | 0.77 | 0.78 | 0.71 |
| 2 | 155402812 | rs1366749 | 1.071 | 0.623 | 0.054 | 1.278 | -0.009 | 0.005 | 0.005 | 0 | 0.29 | 0.28 | 0.42 | 0.16 | 0.45 | 0.28 |

| | | | | | | | | | | | | | | | | |
|---|---|---|---|---|---|---|---|---|---|---|---|---|---|---|---|---|
| 2 | 155406625 | rs16837822 | 0.473 | 0.693 | 0.59 | 1.164 | 0.002 | -0.047 | -0.047 | 0 | 0.11 | 0.23 | 0.1 | 0.01 | 0.1 | 0.16 |
| 2 | 155408097 | rs1918367 | -0.044 | 0.503 | 0.878 | 1.457 | -0.004 | 0.002 | 0.002 | 0 | 0.43 | 0.59 | 0.62 | 0.4 | 0.63 | 0.49 |
| 2 | 155418877 | rs11687614 | 0.968 | 0.45 | 0.367 | 1.411 | -0.001 | 0.012 | 0.012 | 0.143 | 0.05 | 0.03 | 0.01 | 0.2 | 0.03 | 0.07 |
| 2 | 155423435 | rs4461221 | 0.012 | 0.529 | 0.562 | 1.18 | 0.005 | -0.022 | -0.022 | 0.042 | 0.4 | 0.54 | 0.53 | 0.17 | 0.56 | 0.44 |
| 2 | 155439118 | rs16837852 | -0.689 | 0.34 | 0.711 | 1.346 | 0.048 | 0.127 | 0.127 | 0.387 | 0.13 | 0.44 | 0.18 | 0.24 | 0.16 | 0.21 |
| 2 | 155448977 | rs774509 | 0.025 | -0.053 | 0.469 | 0.938 | -0.002 | -0.016 | -0.016 | 0 | 0.56 | 0.46 | 0.53 | 0.75 | 0.4 | 0.64 |
| 2 | 155449286 | rs13414552 | 0.784 | 0.213 | 0.271 | 0.827 | -0.076 | 0.124 | 0.124 | 0 | 0.31 | 0.13 | 0.42 | 0.04 | 0.54 | 0.28 |
| 2 | 155450180 | rs808935 | 0.223 | 0.176 | 0.439 | 0.938 | 0.033 | 0.071 | 0.071 | 0.32 | 0.55 | 0.31 | 0.49 | 0.72 | 0.38 | 0.63 |
| 2 | 155452433 | rs1835153 | 1.44 | -0.257 | -0.11 | 1.151 | -0.014 | -0.004 | -0.004 | 0 | 0.34 | 0.04 | 0.45 | 0.35 | 0.29 | 0.37 |
| 2 | 155463709 | rs10185725 | 0.879 | 0.067 | 0.183 | 1.182 | -0.101 | 0.118 | 0.118 | 0 | 0.3 | 0.13 | 0.3 | 0.04 | 0.49 | 0.25 |
| 2 | 155471901 | rs774505 | -0.055 | -0.091 | 0.317 | 1.33 | -0.015 | 0.002 | 0.002 | 0 | 0.11 | 0.19 | 0.08 | 0.12 | 0.06 | 0.12 |
| 2 | 155474355 | rs13422673 | 0.85 | -0.099 | 0.19 | 1.435 | -0.081 | 0.041 | 0.041 | 0 | 0.31 | 0.16 | 0.3 | 0.16 | 0.5 | 0.26 |
| 2 | 155482937 | rs6728625 | -1.658 | -0.29 | -0.12 | 1.483 | -0.009 | 0 | 0 | 0 | 0.62 | 0.97 | 0.53 | 0.64 | 0.68 | 0.62 |
| 2 | 155494606 | rs1429254 | -1.34 | -0.305 | -0.084 | 1.508 | -0.022 | 0.028 | 0.028 | 0 | 0.58 | 0.95 | 0.51 | 0.63 | 0.64 | 0.58 |
| 2 | 155496698 | rs1835154 | -0.238 | -0.06 | 0.568 | 1.354 | -0.013 | 0.04 | 0.04 | 0 | 0.11 | 0.23 | 0.04 | 0.2 | 0.06 | 0.08 |
| 2 | 155499463 | rs12329336 | -0.312 | -0.064 | 0.472 | 1.415 | 0.006 | 0.002 | 0.002 | 0 | 0.45 | 0.62 | 0.38 | 0.4 | 0.58 | 0.37 |
| 2 | 155527139 | rs6760772 | 0.774 | 0.304 | -0.029 | 0.94 | -0.08 | 0.037 | 0.037 | 0 | 0.29 | 0.1 | 0.27 | 0.17 | 0.47 | 0.24 |
| 2 | 155543535 | rs10210023 | 0.713 | 0.484 | 0.171 | 0.287 | -0.08 | 0.099 | 0.099 | 0 | 0.27 | 0.12 | 0.28 | 0.03 | 0.44 | 0.27 |
| 2 | 155545670 | rs4571015 | -0.329 | 0.387 | 0.421 | 0.468 | 0.001 | 0.042 | 0.042 | 0 | 0.11 | 0.2 | 0.1 | 0.18 | 0.1 | 0.12 |
| 2 | 155547853 | rs3106666 | 0.819 | 0.391 | 0.42 | 0.342 | -0.075 | 0.074 | 0.074 | 0 | 0.27 | 0.11 | 0.29 | 0.05 | 0.46 | 0.28 |
| 2 | 155548559 | rs3111037 | -0.734 | 0.224 | 0.407 | 0.379 | 0.007 | 0.005 | 0.005 | 0 | 0.44 | 0.63 | 0.47 | 0.35 | 0.61 | 0.55 |
| 2 | 155556576 | rs11690166 | -0.518 | 0.113 | -0.248 | 0.25 | 0 | 0.02 | 0.02 | 0 | 0.43 | 0.29 | 0.43 | 0.32 | 0.58 | 0.38 |
| 2 | 155567389 | rs3111020 | -0.891 | 0.463 | -0.108 | 0.368 | -0.017 | -0.02 | -0.02 | 0 | 0.42 | 0.73 | 0.42 | 0.46 | 0.49 | 0.45 |
| 2 | 155571123 | rs11895478 | 0.656 | 0.493 | -0.563 | 0.328 | -0.077 | -0.012 | -0.012 | 0 | 0.22 | 0.07 | 0.26 | 0.15 | 0.32 | 0.26 |
| 2 | 155575560 | rs3106653 | 0.178 | 0.682 | -0.389 | 0.332 | -0.023 | -0.036 | -0.036 | 0 | 0.26 | 0.21 | 0.27 | 0.27 | 0.29 | 0.28 |
| 2 | 155581638 | rs3111017 | -0.423 | 0.677 | -0.403 | 0.298 | -0.028 | -0.024 | -0.024 | 0 | 0.3 | 0.22 | 0.3 | 0.27 | 0.35 | 0.31 |
| 2 | 155590902 | rs16838062 | 0.096 | 0.622 | -0.039 | 0.407 | 0.006 | -0.002 | -0.002 | 0.619 | 0.08 | 0.11 | 0.05 | 0.09 | 0.08 | 0.05 |
| 2 | 155594099 | rs3111006 | -0.476 | 0.238 | -0.241 | 0.32 | -0.011 | -0.007 | -0.007 | 1.493 | 0.37 | 0.32 | 0.34 | 0.36 | 0.43 | 0.33 |
| 2 | 155596137 | rs12471193 | -0.01 | 0.118 | -0.231 | 0.144 | -0.003 | 0.005 | 0.005 | 0.279 | 0.32 | 0.31 | 0.27 | 0.21 | 0.38 | 0.26 |
| 2 | 155596438 | rs6711727 | -0.788 | -0.248 | -0.605 | -0.196 | 0 | 0.113 | 0.113 | 0.179 | 0.51 | 0.83 | 0.43 | 0.3 | 0.5 | 0.38 |
| 2 | 155605132 | rs1823002 | 0.506 | -0.619 | -0.377 | -0.05 | -0.027 | 0.045 | 0.045 | 0 | 0.76 | 0.7 | 0.8 | 0.6 | 0.86 | 0.84 |
| 2 | 155605737 | rs2652443 | 1.203 | -0.489 | -0.725 | -0.174 | -0.034 | -0.067 | -0.067 | 2.042 | 0.4 | 0.06 | 0.47 | 0.47 | 0.36 | 0.52 |
| 2 | 155606965 | rs2591161 | 1.658 | -0.485 | -0.9 | -0.216 | -0.04 | -0.107 | -0.107 | 0.072 | 0.35 | 0.03 | 0.45 | 0.46 | 0.31 | 0.48 |
| 2 | 155607148 | rs7574878 | 0.151 | -0.48 | -0.835 | -0.283 | -0.056 | -0.041 | -0.041 | 0 | 0.34 | 0.17 | 0.36 | 0.43 | 0.49 | 0.36 |
| 2 | 155607966 | rs2591163 | 0.988 | -0.204 | -0.205 | -0.215 | 0 | -0.001 | -0.001 | 0.053 | 0.45 | 0.43 | 0.54 | 0.67 | 0.34 | 0.63 |
| 2 | 155610205 | rs2591164 | 0.279 | -0.07 | -0.226 | -0.092 | -0.022 | -0.038 | -0.038 | 0.781 | 0.79 | 0.6 | 0.88 | 0.89 | 0.85 | 0.9 |
| 2 | 155617896 | rs2591167 | 0.304 | -0.069 | -0.413 | -0.053 | -0.033 | -0.04 | -0.04 | 0.049 | 0.75 | 0.5 | 0.83 | 0.81 | 0.83 | 0.88 |
| 2 | 155618326 | rs2961957 | 0.063 | 0.228 | -0.361 | 0.046 | 0.035 | 0.023 | 0.023 | 0.178 | 0.33 | 0.47 | 0.34 | 0.36 | 0.48 | 0.36 |
| 2 | 155619314 | rs1867648 | 0.014 | 0.18 | -0.358 | -0.312 | 0.024 | -0.044 | -0.044 | 1.11 | 0.33 | 0.44 | 0.32 | 0.14 | 0.49 | 0.3 |
| 2 | 155619395 | rs17623227 | 0.762 | 0.37 | -1.243 | -0.567 | -0.077 | 0.094 | 0.094 | 1.875 | 0.26 | 0.11 | 0.26 | 0.03 | 0.44 | 0.27 |
| 2 | 155634565 | rs10804161 | 0.01 | -0.078 | -1.089 | -0.301 | -0.001 | -0.009 | -0.009 | 0 | 0.48 | 0.74 | 0.46 | 0.51 | 0.58 | 0.48 |
| 2 | 155634982 | rs1445657 | 0.405 | -0.346 | -0.405 | -0.739 | -0.011 | -0.054 | -0.054 | 0 | 0.83 | 0.8 | 0.9 | 1 | 0.9 | 0.92 |
| 2 | 155637238 | rs2053671 | 0.313 | -0.268 | -0.274 | -0.623 | -0.043 | -0.109 | -0.109 | 0.363 | 0.82 | 0.69 | 0.93 | 1 | 0.93 | 0.93 |
| 2 | 155640626 | rs6719455 | -0.465 | 0.461 | -0.546 | -0.692 | 0.005 | -0.039 | -0.039 | 0 | 0.33 | 0.36 | 0.28 | 0.08 | 0.41 | 0.31 |
| 2 | 155647666 | rs2591158 | 0.166 | 0.654 | -1.005 | -0.996 | -0.056 | 0.077 | 0.077 | 0 | 0.74 | 0.88 | 0.78 | 0.95 | 0.65 | 0.79 |
| 2 | 155647819 | rs17640676 | -0.567 | 0.669 | -0.804 | -1.261 | -0.031 | 0.041 | 0.041 | 0 | 0.29 | 0.23 | 0.25 | 0.05 | 0.41 | 0.21 |
| 2 | 155648521 | rs2652442 | 0.522 | 0.209 | -0.961 | -0.427 | -0.001 | 0.008 | 0.008 | 0 | 0.16 | 0.14 | 0.21 | 0.32 | 0.14 | 0.24 |
| 2 | 155649979 | rs2591156 | 0.486 | 0.168 | -1.02 | -0.389 | -0.002 | 0.005 | 0.005 | 0 | 0.17 | 0.15 | 0.2 | 0.31 | 0.12 | 0.21 |
| 2 | 155651371 | rs1838675 | -1.026 | -0.206 | -0.936 | -0.782 | -0.006 | -0.009 | -0.009 | 0 | 0.75 | 0.77 | 0.61 | 0.46 | 0.66 | 0.67 |
| 2 | 155651541 | rs1838674 | 0.613 | -0.421 | -0.962 | -0.355 | -0.002 | 0.01 | 0.01 | 0 | 0.18 | 0.16 | 0.21 | 0.37 | 0.12 | 0.23 |
| 2 | 155652357 | rs1037091 | -0.006 | 0.225 | -0.656 | -0.67 | 0.014 | 0.045 | 0.045 | 0 | 0.32 | 0.43 | 0.44 | 0.52 | 0.34 | 0.37 |
| 2 | 155654692 | rs6716140 | 0.847 | 0.201 | -0.585 | -0.745 | -0.004 | -0.036 | -0.036 | 0.043 | 0.07 | 0.16 | 0.03 | 0.01 | 0.06 | 0.08 |
| 2 | 155655308 | rs2591153 | -0.481 | -0.182 | -0.754 | -1.172 | -0.028 | -0.089 | -0.089 | 0 | 0.72 | 0.56 | 0.79 | 0.9 | 0.81 | 0.79 |
| 2 | 155656548 | rs1037094 | 1.634 | 0.026 | -0.94 | -0.763 | -0.027 | -0.04 | -0.04 | 0 | 0.12 | 0.07 | 0.26 | 0.46 | 0.21 | 0.23 |
| 2 | 155660800 | rs991801 | -0.534 | -0.066 | -0.808 | -0.928 | -0.029 | -0.123 | -0.123 | 0.019 | 0.71 | 0.54 | 0.78 | 0.94 | 0.81 | 0.79 |
| 2 | 155661432 | rs1445653 | 0.169 | 0.036 | -0.924 | -0.998 | -0.001 | -0.001 | -0.001 | 0 | 0.56 | 0.44 | 0.52 | 0.44 | 0.59 | 0.55 |
| 2 | 155663723 | rs2652450 | 0.449 | 0.005 | -0.941 | -0.627 | 0.008 | 0.002 | 0.002 | 0 | 0.52 | 0.4 | 0.51 | 0.42 | 0.57 | 0.47 |
| 2 | 155665752 | rs17641121 | 1.446 | 0.258 | -1.346 | -0.698 | 0.015 | -0.025 | -0.025 | 0 | 0.2 | 0.23 | 0.22 | 0.04 | 0.35 | 0.12 |
| 2 | 155666016 | rs2652451 | 1.932 | 0.109 | -1.467 | -0.698 | -0.011 | -0.129 | -0.129 | 0 | 0.25 | 0.02 | 0.26 | 0.39 | 0.17 | 0.26 |
| 2 | 155668615 | rs1445651 | -0.543 | 0.033 | -1.065 | -0.973 | -0.013 | -0.071 | -0.071 | 0 | 0.65 | 0.43 | 0.75 | 0.93 | 0.75 | 0.68 |
| 2 | 155682556 | rs1445649 | 0.218 | 0.282 | -1.195 | -0.599 | -0.005 | 0.02 | 0.02 | 0 | 0.48 | 0.24 | 0.48 | 0.55 | 0.52 | 0.39 |
| 2 | 155685078 | rs2591140 | 0.424 | 0.279 | -1.241 | -0.695 | -0.031 | -0.004 | -0.004 | 0 | 0.45 | 0.18 | 0.47 | 0.54 | 0.51 | 0.38 |
| 2 | 155688576 | rs2971898 | 1.251 | 0.684 | -1.055 | -0.572 | -0.012 | -0.02 | -0.02 | 0 | 0.14 | 0.12 | 0.27 | 0.35 | 0.22 | 0.23 |
| 2 | 155703068 | rs1900132 | 0.374 | 0.605 | -0.537 | -0.754 | 0.02 | 0.04 | 0.04 | 0 | 0.24 | 0.36 | 0.34 | 0.39 | 0.31 | 0.25 |
| 2 | 155703266 | rs2971906 | 1.756 | 0.643 | -0.381 | -0.874 | 0.017 | -0.049 | -0.049 | 0 | 0.09 | 0.26 | 0.07 | 0 | 0.08 | 0.03 |
| 2 | 155711357 | rs17642086 | 1.244 | 0.944 | -0.768 | -0.949 | -0.023 | 0.022 | 0.022 | 0 | 0.2 | 0.13 | 0.21 | 0.04 | 0.35 | 0.12 |
| 2 | 155716084 | rs2591139 | -0.575 | 0.396 | -0.59 | -1.134 | -0.022 | -0.083 | -0.083 | 0.733 | 0.66 | 0.41 | 0.76 | 0.93 | 0.8 | 0.73 |
| 2 | 155739562 | rs7565079 | 2.026 | 0.532 | -0.701 | -0.971 | -0.002 | -0.02 | -0.02 | 0.184 | 0.06 | 0.06 | 0.02 | 0 | 0.06 | 0.01 |
| 2 | 155743470 | rs13000959 | 1.276 | 0.742 | -0.621 | -0.879 | -0.02 | 0.014 | 0.014 | 0 | 0.2 | 0.16 | 0.22 | 0.07 | 0.36 | 0.13 |

| | | | | | | | | | | | | | | | | |
|---|---|---|---|---|---|---|---|---|---|---|---|---|---|---|---|---|
| 2 | 155750882 | rs765439 | -0.154 | 0.669 | -0.483 | -0.991 | 0 | -0.016 | -0.016 | 0.307 | 0.54 | 0.43 | 0.6 | 0.86 | 0.5 | 0.62 |
| 2 | 155757249 | rs977869 | -0.154 | 0.669 | -0.515 | -1.007 | -0.001 | -0.009 | -0.009 | 0 | 0.54 | 0.43 | 0.6 | 0.86 | 0.5 | 0.62 |
| 2 | 155765902 | rs9973321 | 1.105 | 0.41 | -0.591 | -0.827 | -0.002 | 0.001 | 0.001 | 0.673 | 0.32 | 0.26 | 0.28 | 0.47 | 0.21 | 0.3 |
| 2 | 155774933 | rs10177522 | -1.876 | 0.039 | -0.413 | -1.039 | -0.006 | -0.052 | -0.052 | 0 | 0.82 | 0.79 | 0.9 | 1 | 0.87 | 0.92 |
| 2 | 155778187 | rs4479389 | -0.397 | 0.72 | -0.651 | -0.832 | -0.001 | -0.025 | -0.025 | 0 | 0.58 | 0.44 | 0.65 | 0.86 | 0.62 | 0.64 |
| 2 | 155795999 | rs4664750 | -0.067 | 0.644 | -1.037 | -0.987 | 0.008 | 0.033 | 0.033 | 0.239 | 0.55 | 0.36 | 0.65 | 0.91 | 0.62 | 0.57 |
| 2 | 155799304 | rs7425016 | -0.385 | 0.655 | -1.029 | -0.981 | 0.003 | 0.006 | 0.006 | 0 | 0.58 | 0.38 | 0.66 | 0.92 | 0.63 | 0.63 |
| 2 | 155807051 | rs4426483 | 0.28 | 0.64 | -1.086 | -1.006 | 0.024 | 0.139 | 0.139 | 0 | 0.5 | 0.3 | 0.64 | 0.9 | 0.58 | 0.56 |
| 2 | 155807489 | rs10182711 | 0.28 | 0.64 | -1.086 | -1.006 | 0.024 | 0.139 | 0.139 | 0 | 0.5 | 0.3 | 0.64 | 0.9 | 0.58 | 0.56 |
| 2 | 155815731 | rs10179618 | 0.375 | 0.612 | -1.053 | -0.975 | -0.015 | 0.094 | 0.094 | 0.408 | 0.45 | 0.29 | 0.6 | 0.88 | 0.51 | 0.55 |
| 2 | 155818294 | rs4274557 | -1.17 | 0.57 | -1.001 | -1.049 | -0.037 | -0.165 | -0.165 | 0 | 0.69 | 0.5 | 0.78 | 0.98 | 0.8 | 0.74 |
| 2 | 155879752 | rs16838515 | 1.603 | 0.628 | -0.965 | -0.949 | -0.008 | -0.022 | -0.022 | 0 | 0.07 | 0.08 | 0.02 | 0.02 | 0.05 | 0.07 |
| 2 | 155882144 | rs2350121 | -0.634 | 0.784 | -1.064 | -0.829 | -0.011 | -0.011 | -0.011 | 0 | 0.39 | 0.36 | 0.46 | 0.52 | 0.51 | 0.46 |
| 2 | 155883358 | rs10432523 | 1.76 | 0.754 | -1.069 | -0.841 | -0.012 | -0.009 | -0.009 | 0 | 0.14 | 0.13 | 0.28 | 0.42 | 0.23 | 0.25 |
| 2 | 155889400 | rs16838536 | -0.461 | 0.742 | -1.131 | -0.875 | 0.001 | 0.005 | 0.005 | 0 | 0.55 | 0.37 | 0.48 | 0.57 | 0.57 | 0.5 |
| 2 | 155919140 | rs10203676 | 1.274 | 0.669 | -1.14 | -0.792 | 0 | -0.049 | -0.049 | 0 | 0.3 | 0.15 | 0.3 | 0.45 | 0.28 | 0.3 |
| 2 | 155937322 | rs13409316 | -0.518 | 0.701 | -1.178 | -0.801 | -0.018 | -0.013 | -0.013 | 2.234 | 0.38 | 0.34 | 0.46 | 0.54 | 0.51 | 0.45 |
| 2 | 155944539 | rs10195519 | 0.148 | 0.74 | -1.253 | -0.787 | -0.037 | -0.041 | -0.041 | 0 | 0.31 | 0.26 | 0.44 | 0.52 | 0.46 | 0.38 |
| 2 | 155953495 | rs1482320 | 1.231 | 0.719 | -1.31 | -0.705 | -0.018 | -0.097 | -0.097 | 7.481 | 0.29 | 0.06 | 0.29 | 0.45 | 0.29 | 0.3 |
| 2 | 155982775 | rs13419149 | 1.721 | 0.881 | -1.236 | -0.643 | -0.012 | -0.003 | -0.003 | 0 | 0.14 | 0.12 | 0.28 | 0.44 | 0.23 | 0.25 |
| 2 | 155986195 | rs6744254 | -0.034 | 0.755 | -1.173 | -0.86 | -0.002 | 0.012 | 0.012 | 0 | 0.49 | 0.27 | 0.46 | 0.54 | 0.52 | 0.49 |
| 2 | 155990263 | rs13393958 | 1.375 | 0.803 | -1.185 | -0.898 | -0.006 | -0.071 | -0.071 | 0 | 0.3 | 0.11 | 0.29 | 0.45 | 0.29 | 0.29 |
| 2 | 155994114 | rs1882459 | -1.217 | 0.772 | -1.11 | -0.934 | -0.035 | -0.16 | -0.16 | 0 | 0.68 | 0.49 | 0.76 | 0.98 | 0.79 | 0.75 |
| 2 | 155999124 | rs13000491 | 1.343 | 0.821 | -1.231 | -0.905 | -0.014 | 0.016 | 0.016 | 0.569 | 0.16 | 0.11 | 0.16 | 0.07 | 0.23 | 0.14 |
| 2 | 156003177 | rs4664757 | 1.892 | 0.816 | -1.196 | -0.873 | -0.02 | -0.022 | -0.022 | 0.031 | 0.14 | 0.1 | 0.28 | 0.44 | 0.23 | 0.29 |
| 2 | 156011900 | rs11893828 | 2.045 | 0.798 | -1.175 | -0.845 | -0.005 | -0.028 | -0.028 | 0 | 0.06 | 0.1 | 0.02 | 0 | 0.05 | 0.01 |
| 2 | 156023165 | rs1994247 | -0.117 | 0.887 | -1.407 | -0.739 | -0.008 | 0.004 | 0.004 | 0 | 0.48 | 0.26 | 0.46 | 0.54 | 0.51 | 0.48 |
| 2 | 156038786 | rs958209 | 1.178 | 0.878 | -1.44 | -0.744 | -0.006 | -0.072 | -0.072 | 0 | 0.3 | 0.13 | 0.29 | 0.46 | 0.29 | 0.34 |
| 2 | 156050016 | rs900990 | 1.178 | 0.878 | -1.44 | -0.744 | -0.006 | -0.072 | -0.072 | 0 | 0.3 | 0.13 | 0.29 | 0.46 | 0.29 | 0.34 |
| 2 | 156070516 | rs13010269 | 0.919 | 0.892 | -1.427 | -0.741 | -0.014 | 0.021 | 0.021 | 0.142 | 0.18 | 0.11 | 0.16 | 0.08 | 0.23 | 0.14 |
| 2 | 156091492 | rs1484974 | -0.14 | 0.901 | -1.429 | -0.729 | -0.004 | 0.006 | 0.006 | 0 | 0.48 | 0.28 | 0.46 | 0.54 | 0.52 | 0.48 |
| 2 | 156103249 | rs10210535 | 1.415 | 1.101 | -1.479 | -0.338 | -0.019 | -0.024 | -0.024 | 0.691 | 0.14 | 0.1 | 0.13 | 0.38 | 0.22 | 0.25 |
| 2 | 156114451 | rs1484972 | 0.942 | 1.072 | -1.501 | -0.356 | -0.028 | -0.104 | -0.104 | 0 | 0.29 | 0.07 | 0.29 | 0.45 | 0.29 | 0.28 |
| 2 | 156116188 | rs12052956 | 1.133 | 1.051 | -1.208 | -0.109 | -0.019 | -0.023 | -0.023 | 0.043 | 0.14 | 0.1 | 0.13 | 0.39 | 0.22 | 0.25 |
| 2 | 156130909 | rs9646758 | 1.44 | 0.65 | -1.534 | -0.263 | -0.048 | -0.124 | -0.124 | 0.842 | 0.18 | 0.01 | 0.21 | 0.32 | 0.14 | 0.2 |
| 2 | 156160747 | rs7589312 | -2.04 | 0.231 | -0.859 | -0.534 | 0.008 | -0.041 | -0.041 | 0 | 0.92 | 0.77 | 0.91 | 1 | 0.93 | 0.97 |
| 2 | 156161150 | rs10191533 | -0.569 | 0.131 | -0.856 | -0.563 | -0.01 | 0.012 | 0.012 | 0 | 0.85 | 0.68 | 0.86 | 0.83 | 0.86 | 0.82 |
| 2 | 156177947 | rs4664171 | -0.767 | 0.124 | -0.86 | -0.55 | -0.005 | 0.01 | 0.01 | 0 | 0.86 | 0.68 | 0.86 | 0.86 | 0.86 | 0.89 |
| 2 | 156183489 | rs1385902 | 0.51 | 0.171 | -0.397 | -0.786 | -0.003 | 0.018 | 0.018 | 0 | 0.69 | 0.37 | 0.67 | 0.43 | 0.67 | 0.59 |
| 2 | 156213347 | rs4664759 | 1.853 | 0.191 | -0.291 | -0.758 | 0.024 | 0.105 | 0.105 | 0 | 0.11 | 0.16 | 0.17 | 0.4 | 0.18 | 0.22 |
| 2 | 156220537 | rs12476912 | 2.004 | 0.414 | -0.446 | -0.842 | -0.071 | -0.113 | -0.113 | 0 | 0.18 | 0.02 | 0.22 | 0.32 | 0.2 | 0.23 |
| 2 | 156237750 | rs7605630 | 1.516 | 0.3 | -0.466 | -1.177 | 0.014 | 0.022 | 0.022 | 0.308 | 0.36 | 0.28 | 0.36 | 0.09 | 0.29 | 0.29 |
| 2 | 156246016 | rs1486062 | 2.094 | 0.363 | -0.46 | -1.133 | 0.011 | 0.067 | 0.067 | 0 | 0.11 | 0.12 | 0.17 | 0.4 | 0.18 | 0.22 |
| 2 | 156253110 | rs17287295 | 2.403 | 0.384 | -0.522 | -1.185 | -0.041 | 0.058 | 0.058 | 0 | 0.1 | 0.01 | 0.06 | 0.02 | 0.13 | 0.06 |
| 2 | 156261310 | rs1601527 | -0.327 | 0.355 | -0.454 | -1.204 | -0.033 | -0.01 | -0.01 | 0 | 0.81 | 0.58 | 0.84 | 0.83 | 0.85 | 0.83 |
| 2 | 156266988 | rs13031664 | 0.85 | 0.349 | -0.485 | -1.215 | 0.019 | 0.107 | 0.107 | 0 | 0.47 | 0.29 | 0.45 | 0.11 | 0.42 | 0.35 |
| 2 | 156267449 | rs11678977 | 1.68 | 0.367 | -0.112 | -0.849 | 0.022 | 0.101 | 0.101 | 0 | 0.11 | 0.17 | 0.17 | 0.4 | 0.18 | 0.23 |
| 2 | 156277532 | rs11691512 | 1.708 | 0.369 | -0.167 | -0.848 | 0.011 | 0.066 | 0.066 | 0 | 0.11 | 0.14 | 0.17 | 0.4 | 0.18 | 0.23 |
| 2 | 156283312 | rs2350797 | 2.125 | 0.342 | -0.237 | -0.865 | -0.041 | 0.058 | 0.058 | 0 | 0.1 | 0.01 | 0.06 | 0.02 | 0.13 | 0.06 |
| 2 | 156312118 | rs1505727 | -1.942 | 0.199 | 0.145 | -0.529 | 0.023 | -0.018 | -0.018 | 0 | 0.91 | 0.69 | 0.9 | 0.98 | 0.92 | 0.97 |
| 2 | 156314691 | rs2135269 | -0.418 | 0.158 | 0.055 | -0.596 | -0.005 | 0.007 | 0.007 | 0 | 0.83 | 0.64 | 0.84 | 0.83 | 0.84 | 0.78 |
| 2 | 156315179 | rs16839095 | 1.525 | -0.095 | 0.13 | -0.762 | -0.013 | -0.006 | -0.006 | 0 | 0.08 | 0.16 | 0.19 | 0.07 | 0.06 | 0.07 |
| 2 | 156322314 | rs4490128 | -2.024 | 0.27 | 0.29 | -0.547 | 0.006 | -0.044 | -0.044 | 0 | 0.92 | 0.78 | 0.91 | 1 | 0.92 | 0.97 |
| 2 | 156338251 | rs1876110 | 1.091 | 0.32 | -0.171 | -0.511 | -0.001 | -0.01 | -0.01 | 0 | 0.62 | 0.32 | 0.63 | 0.43 | 0.6 | 0.54 |
| 2 | 156344073 | rs4664764 | 1.727 | 0.356 | -0.307 | -0.479 | 0.009 | 0.062 | 0.062 | 0 | 0.1 | 0.1 | 0.17 | 0.4 | 0.18 | 0.23 |
| 2 | 156348737 | rs4664767 | 1.121 | 0.33 | -0.382 | -0.482 | 0 | -0.017 | -0.017 | 0 | 0.61 | 0.29 | 0.63 | 0.43 | 0.6 | 0.54 |
| 2 | 156351432 | rs1524900 | -0.568 | 0.478 | -0.395 | -0.294 | 0 | 0.002 | 0.002 | 0 | 0.84 | 0.64 | 0.84 | 0.86 | 0.84 | 0.85 |
| 2 | 156356200 | rs1404495 | 0.474 | 0.619 | -0.627 | -0.104 | 0.005 | 0.045 | 0.045 | 0 | 0.14 | 0.17 | 0.18 | 0.39 | 0.19 | 0.23 |
| 2 | 156359533 | rs1394976 | 1.369 | 0.601 | -0.629 | -0.113 | 0.025 | 0.1 | 0.1 | 0 | 0.1 | 0.16 | 0.17 | 0.39 | 0.18 | 0.23 |
| 2 | 156360692 | rs12611450 | 0.383 | 0.596 | -0.632 | -0.115 | 0.005 | 0.045 | 0.045 | 0 | 0.14 | 0.17 | 0.18 | 0.39 | 0.19 | 0.23 |
| 2 | 156365796 | rs11886299 | 0.499 | 0.617 | -0.736 | -0.124 | -0.006 | -0.005 | -0.005 | 0 | 0.14 | 0.11 | 0.17 | 0.39 | 0.19 | 0.23 |
| 2 | 156367484 | rs13030468 | 0.739 | 0.462 | -0.524 | -0.282 | 0.004 | 0.046 | 0.046 | 0 | 0.68 | 0.46 | 0.66 | 0.43 | 0.65 | 0.55 |
| 2 | 156374550 | rs11891913 | 0.527 | 0.515 | -0.695 | -0.22 | -0.004 | -0.002 | -0.002 | 0 | 0.14 | 0.12 | 0.17 | 0.39 | 0.18 | 0.23 |
| 2 | 156381959 | rs1524909 | 0.935 | 0.482 | -1.435 | -0.723 | -0.11 | -0.11 | -0.11 | 2.13 | 0.28 | 0.03 | 0.28 | 0.34 | 0.32 | 0.29 |
| 2 | 156402518 | rs1021929 | 1.822 | 0.462 | -1.293 | -0.639 | -0.018 | -0.004 | -0.004 | 0 | 0.1 | 0.06 | 0.15 | 0.39 | 0.18 | 0.23 |
| 2 | 156404460 | rs1524912 | 0.475 | 0.193 | -1.068 | -0.899 | 0.004 | 0.002 | 0.002 | 0 | 0.48 | 0.42 | 0.53 | 0.48 | 0.46 | 0.48 |
| 2 | 156404781 | rs16839226 | 1.853 | 0.186 | -0.987 | -0.916 | -0.001 | -0.031 | -0.031 | 2.342 | 0.06 | 0.09 | 0.02 | 0 | 0.06 | 0.01 |
| 2 | 156408483 | rs1524907 | 0.766 | 0.192 | -1.049 | -0.882 | 0 | -0.001 | -0.001 | 0.781 | 0.44 | 0.41 | 0.52 | 0.48 | 0.46 | 0.47 |
| 2 | 156409123 | rs6435845 | 0.454 | 0.331 | -0.98 | -0.815 | 0.004 | 0.002 | 0.002 | 0 | 0.48 | 0.42 | 0.53 | 0.48 | 0.46 | 0.48 |

| | | | | | | | | | | | | | | | | |
|---|---|---|---|---|---|---|---|---|---|---|---|---|---|---|---|---|
| 2 | 156412732 | rs1524908 | 0.513 | 0.337 | -0.977 | -0.812 | 0.003 | 0.002 | 0.002 | 0 | 0.48 | 0.42 | 0.53 | 0.48 | 0.46 | 0.47 |
| 2 | 156420492 | rs278865 | 0.609 | 0.513 | -0.87 | -1.415 | -0.001 | -0.045 | -0.045 | 0 | 0.66 | 0.45 | 0.75 | 0.8 | 0.66 | 0.72 |
| 2 | 156422048 | rs13394504 | 0.862 | 0.801 | -0.789 | -1.619 | -0.001 | -0.003 | -0.003 | 0 | 0.07 | 0.05 | 0.04 | 0.1 | 0.09 | 0.11 |
| 2 | 156423437 | rs843198 | 0.157 | 0.826 | -0.806 | -1.732 | 0.037 | -0.001 | -0.001 | 0 | 0.71 | 0.59 | 0.64 | 0.7 | 0.57 | 0.58 |
| 2 | 156431249 | rs2691782 | 0.141 | 0.84 | -0.759 | -1.725 | 0.038 | -0.001 | -0.001 | 0 | 0.71 | 0.59 | 0.64 | 0.7 | 0.57 | 0.58 |
| 2 | 156434090 | rs2691775 | 0.574 | 0.772 | -0.826 | -1.718 | 0.045 | -0.018 | -0.018 | 4.083 | 0.66 | 0.5 | 0.61 | 0.7 | 0.52 | 0.56 |
| 2 | 156436247 | rs10180851 | 1.089 | 0.83 | -0.89 | -1.564 | -0.003 | -0.004 | -0.004 | 0 | 0.07 | 0.05 | 0.04 | 0.1 | 0.09 | 0.11 |
| 2 | 156440448 | rs6706687 | 0.948 | 0.762 | -0.99 | -1.67 | -0.008 | -0.015 | -0.015 | 0 | 0.2 | 0.16 | 0.24 | 0.58 | 0.24 | 0.24 |
| 2 | 156446586 | rs278871 | 0.698 | 0.648 | -1.017 | -1.87 | 0.045 | 0.057 | 0.057 | 0 | 0.49 | 0.34 | 0.4 | 0.13 | 0.34 | 0.34 |
| 2 | 156447246 | rs278872 | 1.087 | 0.632 | -1.036 | -1.886 | 0.036 | 0.027 | 0.027 | 0 | 0.45 | 0.34 | 0.37 | 0.12 | 0.28 | 0.31 |
| 2 | 156461568 | rs2677475 | 0.705 | 0.663 | -1.006 | -1.861 | 0.045 | 0.057 | 0.057 | 0 | 0.49 | 0.34 | 0.4 | 0.13 | 0.34 | 0.34 |
| 2 | 156461906 | rs10432531 | 1.923 | 0.736 | -0.984 | -1.809 | -0.018 | -0.053 | -0.053 | 0 | 0.16 | 0.07 | 0.2 | 0.58 | 0.18 | 0.23 |
| 2 | 156463461 | rs10432447 | 1.905 | 0.732 | -0.986 | -1.811 | -0.018 | -0.053 | -0.053 | 0.037 | 0.16 | 0.07 | 0.2 | 0.58 | 0.18 | 0.23 |
| 2 | 156465014 | rs11679712 | 1.467 | 0.371 | -0.917 | -2.089 | -0.016 | -0.017 | -0.017 | 0 | 0.09 | 0.15 | 0.19 | 0.06 | 0.07 | 0.05 |
| 2 | 156465700 | rs12465886 | -0.809 | 0.677 | -0.795 | -1.851 | 0.062 | -0.043 | -0.043 | 0.392 | 0.9 | 0.69 | 0.86 | 0.98 | 0.84 | 0.94 |
| 2 | 156466403 | rs2691787 | 0.335 | 0.808 | -0.739 | -1.793 | 0.001 | -0.037 | -0.037 | 0 | 0.54 | 0.43 | 0.43 | 0.13 | 0.39 | 0.35 |
| 2 | 156467786 | rs843212 | -0.015 | 1.381 | -0.059 | -1.121 | 0.05 | -0.004 | -0.004 | 0 | 0.78 | 0.64 | 0.67 | 0.81 | 0.67 | 0.7 |
| 2 | 156468897 | rs707121 | 0.837 | 0.978 | -0.196 | -1.096 | 0.02 | -0.102 | -0.102 | 0 | 0.74 | 0.55 | 0.81 | 0.94 | 0.71 | 0.83 |
| 2 | 156470796 | rs843205 | 0.614 | 1.176 | -0.09 | -0.933 | 0.075 | -0.123 | -0.123 | 1.035 | 0.79 | 0.52 | 0.8 | 0.97 | 0.68 | 0.91 |
| 2 | 156473057 | rs12694373 | 0.638 | 0.886 | -0.125 | -0.925 | 0.041 | -0.045 | -0.045 | 0 | 0.65 | 0.46 | 0.62 | 0.78 | 0.5 | 0.65 |
| 2 | 156473598 | rs843209 | 0.245 | 0.862 | 0.177 | -0.827 | 0.092 | -0.072 | -0.072 | 0.121 | 0.86 | 0.55 | 0.86 | 0.97 | 0.78 | 0.95 |
| 2 | 156476895 | rs10199474 | 0.14 | 0.735 | -0.145 | -0.643 | 0 | 0.001 | 0.001 | 0 | 0.05 | 0.05 | 0.03 | 0.13 | 0.05 | 0.1 |
| 2 | 156483829 | rs843236 | 0.204 | 0.789 | 0.264 | -0.69 | 0.056 | -0.038 | -0.038 | 0 | 0.86 | 0.63 | 0.88 | 0.95 | 0.79 | 0.94 |
| 2 | 156485345 | rs1917440 | 0.489 | 0.759 | -0.099 | -0.756 | 0.009 | -0.009 | -0.009 | 0 | 0.27 | 0.27 | 0.45 | 0.21 | 0.34 | 0.35 |
| 2 | 156489245 | rs843249 | 0.842 | 0.413 | -0.45 | -0.693 | 0.048 | 0.025 | 0.025 | 0.053 | 0.46 | 0.16 | 0.36 | 0.57 | 0.35 | 0.4 |
| 2 | 156490877 | rs843247 | -0.11 | 0.686 | 0.064 | -0.763 | -0.003 | 0.002 | 0.002 | 0 | 0.37 | 0.34 | 0.52 | 0.25 | 0.46 | 0.37 |
| 2 | 156493455 | rs1400702 | 0.106 | 0.7 | 0.147 | -0.771 | 0.006 | -0.013 | -0.013 | 0 | 0.1 | 0.2 | 0.08 | 0.06 | 0.12 | 0.05 |
| 2 | 156496507 | rs707120 | -1.111 | 0.728 | -0.116 | -0.769 | -0.03 | 0.02 | 0.02 | 0 | 0.74 | 0.85 | 0.56 | 0.8 | 0.66 | 0.68 |
| 2 | 156498720 | rs8179654 | 0.939 | 0.619 | -0.352 | -0.815 | 0.029 | -0.007 | -0.007 | 0 | 0.44 | 0.12 | 0.34 | 0.56 | 0.34 | 0.43 |
| 2 | 156501487 | rs13431999 | 0.382 | 0.64 | -0.066 | -0.76 | 0.008 | -0.007 | -0.007 | 0 | 0.16 | 0.25 | 0.11 | 0.15 | 0.2 | 0.2 |
| 2 | 156506662 | rs10174401 | -0.643 | 0.63 | -0.278 | -0.77 | 0.028 | -0.024 | -0.024 | 0.02 | 0.25 | 0.44 | 0.17 | 0.22 | 0.32 | 0.25 |
| 2 | 156508589 | rs2048932 | 1.305 | 0.633 | -0.553 | -0.793 | -0.114 | -0.018 | -0.018 | 0.468 | 0.24 | 0.02 | 0.43 | 0.2 | 0.33 | 0.3 |
| 2 | 156513815 | rs1533563 | 0.74 | 0.756 | -0.437 | -0.727 | 0.004 | -0.033 | -0.033 | 0 | 0.44 | 0.06 | 0.34 | 0.56 | 0.36 | 0.41 |
| 2 | 156516166 | rs1018320 | 0.74 | 0.756 | -0.437 | -0.727 | 0.004 | -0.033 | -0.033 | 0 | 0.44 | 0.06 | 0.34 | 0.56 | 0.36 | 0.41 |
| 2 | 156521982 | rs1033254 | 0.627 | 0.749 | -0.228 | -0.747 | 0.023 | -0.027 | -0.027 | 0 | 0.74 | 0.44 | 0.82 | 0.78 | 0.67 | 0.74 |
| 2 | 156523141 | rs16839401 | 0.713 | 0.73 | -0.364 | -0.704 | 0.004 | -0.033 | -0.033 | 0 | 0.44 | 0.06 | 0.34 | 0.56 | 0.36 | 0.41 |
| 2 | 156524611 | rs1283773 | -0.052 | 0.777 | -0.086 | -0.658 | 0.024 | -0.012 | -0.012 | 0 | 0.9 | 0.71 | 0.91 | 0.93 | 0.88 | 0.94 |
| 2 | 156533969 | rs12466169 | 0.591 | 0.714 | -0.065 | -0.611 | 0.007 | 0.006 | 0.006 | 0 | 0.06 | 0.16 | 0.05 | 0.07 | 0.09 | 0.05 |
| 2 | 156540276 | rs2677469 | 0.165 | 0.742 | -0.145 | -0.635 | 0.017 | -0.014 | -0.014 | 0 | 0.18 | 0.27 | 0.11 | 0.15 | 0.23 | 0.2 |
| 2 | 156541170 | rs2677468 | -0.394 | 0.743 | -0.105 | -0.647 | 0.006 | -0.014 | -0.014 | 0.362 | 0.63 | 0.46 | 0.46 | 0.71 | 0.6 | 0.61 |
| 2 | 156546655 | rs1283766 | -0.268 | 0.771 | -0.327 | -0.634 | -0.003 | 0.008 | 0.008 | 0.265 | 0.62 | 0.34 | 0.45 | 0.71 | 0.6 | 0.61 |
| 2 | 156550602 | rs1283763 | -0.438 | 0.757 | -0.749 | -0.709 | 0.023 | -0.035 | -0.035 | 0 | 0.27 | 0.47 | 0.17 | 0.22 | 0.34 | 0.25 |
| 2 | 156553448 | rs1283760 | -0.882 | 0.72 | -1.357 | -0.603 | 0.004 | -0.047 | -0.047 | 0 | 0.59 | 0.93 | 0.66 | 0.44 | 0.66 | 0.59 |
| 2 | 156564805 | rs1914648 | -0.833 | 0.751 | -1.361 | -0.629 | 0.015 | -0.026 | -0.026 | 0 | 0.59 | 0.92 | 0.65 | 0.43 | 0.66 | 0.57 |
| 2 | 156568446 | rs1283754 | -0.187 | 0.859 | -0.906 | -0.522 | 0.027 | -0.06 | -0.06 | 0 | 0.24 | 0.47 | 0.14 | 0.15 | 0.26 | 0.21 |
| 2 | 156571307 | rs4664773 | 0.187 | 0.939 | -1.047 | -0.215 | 0.001 | -0.006 | -0.006 | 0 | 0.29 | 0.31 | 0.46 | 0.21 | 0.31 | 0.31 |
| 2 | 156582825 | rs10200621 | 0.155 | 0.949 | -1.578 | 0.485 | -0.034 | -0.098 | -0.098 | 0 | 0.15 | 0.03 | 0.17 | 0.46 | 0.18 | 0.24 |
| 2 | 156586568 | rs1980211 | -0.336 | 0.867 | -1.241 | -0.536 | 0 | -0.031 | -0.031 | 0 | 0.63 | 0.36 | 0.7 | 0.88 | 0.64 | 0.78 |
| 2 | 156587414 | rs6758770 | 0.11 | 0.89 | -1.279 | -0.369 | 0 | 0 | 0 | 0 | 0.22 | 0.17 | 0.11 | 0.16 | 0.18 | 0.21 |
| 2 | 156589930 | rs17306659 | 0.397 | 1.178 | -1.371 | -0.536 | 0.002 | -0.011 | -0.011 | 0 | 0.1 | 0.11 | 0.09 | 0.05 | 0.15 | 0.07 |
| 2 | 156597010 | rs12477257 | 0.631 | 1.228 | -1.741 | 0.06 | -0.033 | -0.082 | -0.082 | 0 | 0.12 | 0.01 | 0.16 | 0.4 | 0.18 | 0.24 |
| 2 | 156604837 | rs13035923 | -0.439 | 1.114 | -0.945 | -0.251 | 0.038 | -0.08 | -0.08 | 0.866 | 0.17 | 0.32 | 0.15 | 0.08 | 0.23 | 0.11 |
| 2 | 156625848 | rs2882513 | -0.392 | 1.006 | -1.147 | -0.548 | 0.009 | -0.083 | -0.083 | 0 | 0.34 | 0.39 | 0.25 | 0.09 | 0.38 | 0.18 |
| 2 | 156647746 | rs10192440 | 1.549 | 0.921 | -1.273 | -0.532 | -0.01 | -0.034 | -0.034 | 0 | 0.11 | 0.04 | 0.1 | 0.22 | 0.12 | 0.13 |
| 2 | 156652388 | rs13005153 | 0.095 | 1.018 | -1.223 | -0.669 | 0.002 | -0.062 | -0.062 | 0 | 0.29 | 0.33 | 0.2 | 0.08 | 0.3 | 0.16 |
| 2 | 156663824 | rs7608383 | -0.409 | 0.908 | -0.959 | -0.441 | 0.033 | -0.042 | -0.042 | 0.341 | 0.11 | 0.37 | 0.1 | 0.04 | 0.11 | 0.06 |
| 2 | 156667228 | rs1520342 | -0.794 | 0.536 | -0.727 | -0.625 | -0.002 | -0.015 | -0.015 | 0.259 | 0.58 | 0.44 | 0.47 | 0.68 | 0.62 | 0.56 |
| 2 | 156682657 | rs1518284 | -1.275 | 0.757 | -0.926 | -0.48 | -0.001 | 0.006 | 0.006 | 0 | 0.66 | 0.68 | 0.54 | 0.72 | 0.68 | 0.61 |
| 2 | 156696136 | rs16839640 | 0.189 | 0.295 | -0.755 | 0.119 | -0.005 | 0.013 | 0.013 | 0 | 0.17 | 0.18 | 0.17 | 0.32 | 0.14 | 0.29 |
| 2 | 156697282 | rs9288534 | -0.603 | 0.282 | -0.737 | -0.047 | -0.017 | 0.004 | 0.004 | 0 | 0.36 | 0.55 | 0.33 | 0.58 | 0.28 | 0.45 |
| 2 | 156704216 | rs7573309 | -0.934 | 0.439 | -0.807 | -0.114 | -0.003 | -0.001 | -0.001 | 0 | 0.38 | 0.57 | 0.35 | 0.59 | 0.32 | 0.48 |
| 2 | 156706397 | rs884799 | -1.331 | 0.912 | -0.803 | -0.026 | 0.015 | 0.011 | 0.011 | 0.206 | 0.88 | 0.85 | 0.91 | 0.86 | 0.84 | 0.94 |
| 2 | 156721846 | rs12471133 | 0.232 | 0.696 | -0.725 | 0.024 | -0.009 | 0.024 | 0.024 | 0 | 0.18 | 0.21 | 0.18 | 0.32 | 0.15 | 0.29 |
| 2 | 156730260 | rs13382244 | 1.08 | 0.593 | -0.833 | 0.023 | 0.019 | -0.058 | -0.058 | 0 | 0.12 | 0.03 | 0.14 | 0.31 | 0.11 | 0.23 |
| 2 | 156751921 | rs1520339 | -0.565 | 0.984 | -1.11 | -0.216 | -0.043 | 0.03 | 0.03 | 0 | 0.73 | 0.86 | 0.82 | 0.82 | 0.68 | 0.88 |
| 2 | 156752459 | rs17236969 | 1.757 | 0.468 | -1.06 | -0.12 | -0.05 | 0.073 | 0.073 | 0 | 0.14 | 0.01 | 0.09 | 0.04 | 0.14 | 0.07 |
| 2 | 156752588 | rs7594921 | -1.366 | -0.02 | -0.531 | 0.247 | 0.018 | 0.007 | 0.007 | 0 | 0.43 | 0.66 | 0.39 | 0.59 | 0.41 | 0.48 |
| 2 | 156753946 | rs10202061 | 0.042 | 0.422 | -0.674 | 0.037 | -0.017 | 0.019 | 0.019 | 0 | 0.21 | 0.38 | 0.18 | 0.26 | 0.18 | 0.18 |
| 2 | 156755591 | rs13420750 | 0.567 | 0.38 | -0.577 | 0.027 | -0.014 | 0.023 | 0.023 | 0 | 0.17 | 0.24 | 0.13 | 0.26 | 0.14 | 0.16 |

| | | | | | | | | | | | | | | | | |
|---|---|---|---|---|---|---|---|---|---|---|---|---|---|---|---|---|
| 2 | 156759291 | rs1402071 | -0.077 | 0.36 | -1.002 | 0.114 | 0.006 | -0.001 | -0.001 | 0 | 0.25 | 0.43 | 0.23 | 0.26 | 0.27 | 0.19 |
| 2 | 156765002 | rs13383584 | 0.372 | 0.684 | -2.065 | 0.062 | 0.043 | 0.126 | 0.126 | 0.356 | 0.49 | 0.22 | 0.56 | 0.33 | 0.54 | 0.48 |
| 2 | 156765630 | rs16839725 | 0.126 | 0.682 | -2.622 | 0.091 | -0.024 | -0.085 | -0.085 | 0 | 0.32 | 0.1 | 0.45 | 0.39 | 0.34 | 0.35 |
| 2 | 156765776 | rs16839730 | 0.619 | 0.294 | -2.312 | 0.276 | 0.005 | -0.065 | -0.065 | 0 | 0.16 | 0.1 | 0.36 | 0.34 | 0.16 | 0.29 |
| 2 | 156765867 | rs7573784 | 0.451 | 0.355 | -1.96 | 0.645 | 0 | -0.024 | -0.024 | 0 | 0.09 | 0.06 | 0.09 | 0.21 | 0.08 | 0.12 |
| 2 | 156766219 | rs17309678 | 0.628 | 0.731 | -2.267 | 0.126 | -0.08 | 0.033 | 0.033 | 0 | 0.44 | 0.09 | 0.5 | 0.33 | 0.45 | 0.35 |
| 2 | 156768868 | rs13432856 | -0.063 | 0.851 | -1.998 | 0.042 | 0.001 | -0.01 | -0.01 | 0 | 0.09 | 0.11 | 0.06 | 0.01 | 0.12 | 0.13 |
| 2 | 156785138 | rs4664187 | 0.427 | 0.694 | -1.116 | -0.074 | -0.002 | -0.002 | -0.002 | 0 | 0.59 | 0.43 | 0.63 | 0.36 | 0.64 | 0.44 |
| 2 | 156786929 | rs10189214 | 1.288 | 0.635 | -0.604 | 0.104 | -0.009 | 0.085 | 0.085 | 0 | 0.14 | 0.23 | 0.18 | 0.34 | 0.13 | 0.32 |
| 2 | 156802816 | rs7607309 | 0.584 | 0.611 | -0.627 | 0.088 | 0.002 | -0.013 | -0.013 | 0 | 0.08 | 0.11 | 0.05 | 0.01 | 0.1 | 0.13 |
| 2 | 156808365 | rs10198113 | 0.162 | 0.698 | -0.513 | 0.161 | -0.005 | 0.003 | 0.003 | 0 | 0.26 | 0.29 | 0.17 | 0.29 | 0.22 | 0.25 |
| 2 | 156813500 | rs769353 | 1.538 | 0.808 | -0.192 | 0.093 | 0.027 | -0.02 | -0.02 | 0.237 | 0.08 | 0.17 | 0.09 | 0.04 | 0.11 | 0.03 |
| 2 | 156814699 | rs13421080 | 0.021 | 0.403 | -0.466 | 0.245 | -0.003 | 0.002 | 0.002 | 0 | 0.26 | 0.29 | 0.18 | 0.29 | 0.23 | 0.25 |
| 2 | 156829816 | rs13031378 | 0.135 | -0.107 | -0.56 | 0.111 | -0.004 | -0.002 | -0.002 | 0.195 | 0.6 | 0.43 | 0.63 | 0.36 | 0.65 | 0.45 |
| 2 | 156833208 | rs6436095 | -0.516 | -0.084 | -0.788 | 0.122 | -0.014 | -0.007 | -0.007 | 0.176 | 0.4 | 0.48 | 0.36 | 0.63 | 0.34 | 0.55 |
| 2 | 156842199 | rs1527754 | 0.52 | 0.115 | -1.39 | 0.176 | -0.097 | -0.137 | -0.137 | 0 | 0.33 | 0.04 | 0.43 | 0.4 | 0.34 | 0.36 |
| 2 | 156866619 | rs1949991 | -0.154 | -0.172 | -0.79 | 0.035 | -0.009 | 0 | 0 | 0 | 0.46 | 0.35 | 0.55 | 0.44 | 0.49 | 0.41 |
| 2 | 156869592 | rs1405869 | 1.211 | -0.297 | -0.485 | 0.168 | -0.002 | -0.001 | -0.001 | 0 | 0.27 | 0.24 | 0.27 | 0.31 | 0.32 | 0.37 |
| 2 | 156877774 | rs6748811 | -0.664 | -0.75 | 0.253 | -0.091 | -0.037 | 0.031 | 0.031 | 0 | 0.84 | 0.69 | 0.89 | 0.76 | 0.92 | 0.92 |
| 2 | 156877890 | rs11695569 | -0.154 | -0.359 | -0.058 | -0.148 | -0.033 | -0.042 | -0.042 | 1.397 | 0.11 | 0.27 | 0.07 | 0.06 | 0.06 | 0.03 |
| 2 | 156885750 | rs7570959 | 0.509 | -0.513 | -0.274 | -0.227 | -0.011 | -0.011 | -0.011 | 0 | 0.67 | 0.36 | 0.77 | 0.75 | 0.75 | 0.79 |
| 2 | 156886978 | rs7572191 | 0.445 | -0.699 | 0.107 | -0.208 | -0.021 | -0.023 | -0.023 | 0.304 | 0.08 | 0.21 | 0.05 | 0.06 | 0.05 | 0.03 |
| 2 | 156893605 | rs1918173 | 0.656 | -0.84 | 0.115 | -0.297 | -0.012 | -0.002 | -0.002 | 0.021 | 0.68 | 0.35 | 0.81 | 0.75 | 0.85 | 0.84 |
| 2 | 156893877 | rs2882618 | -0.515 | -0.623 | 0.019 | -0.169 | -0.053 | -0.01 | -0.01 | 0 | 0.8 | 0.66 | 0.87 | 0.8 | 0.91 | 0.85 |
| 2 | 156903540 | rs2177951 | 0.477 | 0.213 | -0.43 | -0.274 | 0.001 | -0.008 | -0.008 | 0 | 0.46 | 0.45 | 0.65 | 0.63 | 0.48 | 0.64 |
| 2 | 156911208 | rs1358227 | 0.493 | 0.501 | -0.83 | 0.292 | -0.001 | -0.01 | -0.01 | 0.122 | 0.13 | 0.06 | 0.18 | 0.51 | 0.1 | 0.19 |
| 2 | 156913082 | rs6705303 | 0.487 | 0.537 | -0.792 | 0.3 | 0.001 | 0.041 | 0.041 | 0.052 | 0.1 | 0.05 | 0.16 | 0.51 | 0.07 | 0.17 |
| 2 | 156916479 | rs16839935 | 0.368 | 0.906 | -0.78 | 0.32 | -0.006 | 0.028 | 0.028 | 0.013 | 0.11 | 0.06 | 0.2 | 0.52 | 0.15 | 0.19 |
| 2 | 156917835 | rs7578557 | -1.277 | 0.645 | -0.825 | -0.157 | -0.012 | -0.004 | -0.004 | 0 | 0.39 | 0.36 | 0.4 | 0.58 | 0.48 | 0.29 |
| 2 | 156918327 | rs16839941 | -0.248 | 0.782 | -0.976 | -0.179 | -0.034 | -0.075 | -0.075 | 0 | 0.23 | 0.11 | 0.29 | 0.52 | 0.31 | 0.21 |
| 2 | 156919338 | rs2204584 | -1.207 | 0.798 | -0.992 | -0.203 | -0.034 | -0.017 | -0.017 | 0 | 0.34 | 0.27 | 0.37 | 0.57 | 0.48 | 0.29 |
| 2 | 156924698 | rs16839959 | -0.74 | 0.626 | -0.681 | 0.102 | -0.004 | -0.003 | -0.003 | 0 | 0.19 | 0.19 | 0.24 | 0.55 | 0.25 | 0.24 |
| 2 | 156926963 | rs13032427 | -0.799 | 0.637 | -0.449 | 0.187 | 0 | 0.023 | 0.023 | 0.092 | 0.15 | 0.16 | 0.31 | 0.55 | 0.21 | 0.23 |
| 2 | 156928039 | rs17185498 | 0.288 | 0.568 | -0.172 | -0.388 | -0.004 | 0.064 | 0.064 | 0 | 0.51 | 0.34 | 0.31 | 0.31 | 0.5 | 0.43 |
| 2 | 156928056 | rs16839972 | -0.354 | 0.496 | 0.092 | -0.725 | 0.011 | -0.05 | -0.05 | 0 | 0.13 | 0.24 | 0.2 | 0.03 | 0.17 | 0.07 |
| 2 | 156928123 | rs6730147 | -0.928 | 0.508 | 0.141 | -0.634 | -0.009 | -0.045 | -0.045 | 0 | 0.7 | 0.62 | 0.64 | 0.87 | 0.74 | 0.7 |
| 2 | 156928496 | rs6748197 | -0.328 | 0.478 | 0.021 | -0.601 | 0.007 | -0.057 | -0.057 | 0 | 0.62 | 0.5 | 0.47 | 0.83 | 0.57 | 0.65 |
| 2 | 156933553 | rs10177924 | 0.697 | 0.396 | -0.167 | -0.536 | 0 | 0.001 | 0.001 | 0 | 0.28 | 0.25 | 0.16 | 0.13 | 0.28 | 0.34 |
| 2 | 156935758 | rs10932860 | -0.797 | 0.265 | -0.089 | -0.115 | 0.017 | 0.015 | 0.015 | 0 | 0.38 | 0.49 | 0.63 | 0.54 | 0.49 | 0.29 |
| 2 | 156948047 | rs1113060 | -0.383 | 0.1 | -0.315 | -0.332 | -0.019 | -0.003 | -0.003 | 0 | 0.68 | 0.61 | 0.66 | 0.69 | 0.8 | 0.77 |
| 2 | 156949425 | rs11885399 | -0.881 | 0.193 | -0.162 | -0.238 | -0.043 | 0.02 | 0.02 | 0 | 0.75 | 0.62 | 0.69 | 0.69 | 0.89 | 0.8 |
| 2 | 156950640 | rs11692441 | 0.053 | 0.549 | -0.472 | 0.096 | 0.028 | 0.01 | 0.01 | 0 | 0.14 | 0.25 | 0.31 | 0.19 | 0.22 | 0.11 |
| 2 | 156950824 | rs1519797 | -0.786 | 0.392 | -0.723 | 0.166 | 0.019 | -0.014 | -0.014 | 0.022 | 0.75 | 0.82 | 0.7 | 0.69 | 0.88 | 0.8 |
| 2 | 156954563 | rs10497173 | -0.307 | 0.413 | -0.679 | -0.275 | 0.03 | 0.03 | 0.03 | 0 | 0.59 | 0.72 | 0.59 | 0.3 | 0.73 | 0.61 |
| 2 | 156970611 | rs10196668 | 0.326 | 0.409 | 0.007 | -0.186 | 0.071 | -0.015 | -0.015 | 0 | 0.07 | 0.29 | 0.05 | 0 | 0.08 | 0.03 |
| 2 | 156975459 | rs6736241 | 0.37 | 0.284 | -0.191 | -0.211 | -0.011 | 0.057 | 0.057 | 0 | 0.08 | 0.17 | 0.12 | 0.19 | 0.09 | 0.15 |
| 2 | 156981694 | rs1519805 | -0.318 | 0.307 | -0.553 | -0.201 | 0.012 | 0.053 | 0.053 | 0 | 0.15 | 0.45 | 0.17 | 0.19 | 0.17 | 0.19 |
| 2 | 156984088 | rs13011491 | 0.53 | 0.433 | -1.228 | -0.345 | -0.04 | 0.095 | 0.095 | 0 | 0.4 | 0.25 | 0.37 | 0.1 | 0.48 | 0.39 |
| 2 | 156990306 | rs12694551 | -0.039 | 0.02 | -1.455 | -0.201 | 0.041 | 0.081 | 0.081 | 0 | 0.5 | 0.72 | 0.51 | 0.3 | 0.63 | 0.57 |
| 2 | 156990320 | rs16840042 | 0.565 | -0.041 | -1.836 | 0.165 | -0.05 | -0.089 | -0.089 | 0 | 0.14 | 0.02 | 0.1 | 0.31 | 0.15 | 0.18 |
| 2 | 156990715 | rs17186802 | -0.277 | -0.453 | -2.142 | -0.297 | 0.026 | 0.153 | 0.153 | 0 | 0.43 | 0.11 | 0.53 | 0.84 | 0.32 | 0.45 |
| 2 | 156991900 | rs11896224 | 0.43 | -0.424 | -1.776 | 0.055 | 0.001 | -0.055 | -0.055 | 0.031 | 0.18 | 0.07 | 0.09 | 0.32 | 0.13 | 0.21 |
| 2 | 157009370 | rs1519794 | 0.195 | -0.24 | -0.36 | -0.182 | 0.018 | 0.001 | 0.001 | 1.759 | 0.94 | 0.74 | 0.95 | 1 | 0.98 | 1 |
| 2 | 157011778 | rs4664800 | 1.376 | -0.515 | -0.667 | -0.237 | 0.028 | -0.063 | -0.063 | 0 | 0.18 | 0.1 | 0.37 | 0.53 | 0.1 | 0.28 |
| 2 | 157012321 | rs10189160 | 0.817 | -0.694 | -0.682 | -0.527 | 0.035 | 0.082 | 0.082 | 0 | 0.47 | 0.34 | 0.34 | 0.11 | 0.64 | 0.36 |
| 2 | 157023248 | rs1818991 | 0.098 | -0.643 | -0.942 | -0.424 | 0.011 | -0.048 | -0.048 | 0 | 0.27 | 0.13 | 0.22 | 0.36 | 0.24 | 0.36 |
| 2 | 157030835 | rs2049582 | 1.283 | -0.459 | -1.349 | -0.541 | -0.084 | 0.153 | 0.153 | 0 | 0.37 | 0.15 | 0.26 | 0.09 | 0.49 | 0.3 |
| 2 | 157031032 | rs12997268 | 1.361 | -0.502 | -1.369 | -0.562 | -0.132 | 0.171 | 0.171 | 0.023 | 0.36 | 0.06 | 0.24 | 0.09 | 0.49 | 0.3 |
| 2 | 157032115 | rs2037341 | 0.843 | -0.285 | -1.03 | -0.44 | 0.003 | 0.004 | 0.004 | 0 | 0.56 | 0.24 | 0.63 | 0.63 | 0.59 | 0.59 |
| 2 | 157034020 | rs10932909 | 0.824 | -0.266 | -1.02 | -0.43 | 0.001 | 0.002 | 0.002 | 0 | 0.56 | 0.24 | 0.63 | 0.63 | 0.59 | 0.59 |
| 2 | 157053380 | rs7567451 | 0.267 | -0.367 | -0.902 | -0.365 | 0.031 | -0.019 | -0.019 | 0.615 | 0.64 | 0.27 | 0.69 | 0.64 | 0.73 | 0.64 |
| 2 | 157071176 | rs2351534 | -0.581 | -0.324 | -0.486 | -0.38 | 0 | -0.011 | -0.011 | 0 | 0.34 | 0.58 | 0.29 | 0.28 | 0.33 | 0.37 |
| 2 | 157094848 | rs2351535 | 0.19 | -0.294 | -0.18 | -0.373 | 0.12 | 0 | 0 | 0.019 | 0.07 | 0.32 | 0.05 | 0 | 0.09 | 0.02 |
| 2 | 157105605 | rs1234876 | 1.517 | -0.577 | -0.645 | -0.592 | -0.133 | 0.169 | 0.169 | 0 | 0.36 | 0.06 | 0.25 | 0.09 | 0.49 | 0.3 |
| 2 | 157108393 | rs4664804 | 1.589 | -0.71 | -0.505 | -0.544 | 0.014 | -0.022 | -0.022 | 0 | 0.19 | 0.15 | 0.38 | 0.62 | 0.1 | 0.29 |
| 2 | 157122035 | rs2697368 | -0.198 | -0.585 | -0.368 | -0.542 | -0.001 | -0.001 | -0.001 | 0 | 0.72 | 0.74 | 0.78 | 0.73 | 0.79 | 0.66 |
| 2 | 157126321 | rs16840135 | -0.385 | -0.309 | -0.15 | -0.433 | -0.001 | -0.001 | -0.001 | 0.202 | 0.39 | 0.59 | 0.56 | 0.64 | 0.33 | 0.37 |
| 2 | 157141280 | rs1030599 | 1.19 | -0.317 | -0.321 | -0.413 | -0.004 | -0.13 | -0.13 | 0 | 0.25 | 0.01 | 0.43 | 0.63 | 0.18 | 0.3 |
| 2 | 157143309 | rs7589499 | 1.039 | -0.27 | -0.269 | -0.389 | 0.034 | -0.071 | -0.071 | 0.183 | 0.26 | 0.08 | 0.43 | 0.63 | 0.18 | 0.3 |

| | | | | | | | | | | | | | | | | |
|---|---|---|---|---|---|---|---|---|---|---|---|---|---|---|---|---|
| 2 | 157145030 | rs13417852 | 0.033 | -0.195 | -0.147 | -0.358 | 0.116 | -0.002 | -0.002 | 0 | 0.07 | 0.32 | 0.06 | 0 | 0.09 | 0.03 |
| 2 | 157148002 | rs6436290 | -0.2 | -0.207 | -0.659 | -0.295 | -0.003 | -0.002 | -0.002 | 0 | 0.39 | 0.57 | 0.56 | 0.64 | 0.33 | 0.37 |
| 2 | 157178503 | rs1150144 | -0.114 | -0.094 | -0.561 | -0.172 | 0.02 | -0.02 | -0.02 | 0 | 0.6 | 0.28 | 0.4 | 0.47 | 0.72 | 0.57 |
| 2 | 157180255 | rs1150146 | -0.164 | -0.042 | -0.527 | -0.162 | 0.019 | -0.021 | -0.021 | 0 | 0.61 | 0.29 | 0.4 | 0.47 | 0.72 | 0.58 |
| 2 | 157181925 | rs12803 | 0.851 | -0.154 | -0.956 | -0.36 | -0.064 | 0.1 | 0.1 | 0 | 0.41 | 0.12 | 0.23 | 0.19 | 0.53 | 0.4 |
| 2 | 157188145 | rs834835 | -0.455 | -0.122 | -0.914 | -0.336 | 0.017 | -0.018 | -0.018 | 0.267 | 0.61 | 0.3 | 0.41 | 0.47 | 0.72 | 0.56 |
| 2 | 157192613 | rs13428968 | 0.256 | 0.031 | -1.03 | -0.325 | -0.02 | -0.073 | -0.073 | 0.598 | 0.16 | 0.03 | 0.15 | 0.28 | 0.18 | 0.17 |
| 2 | 157197931 | rs1107005 | -0.141 | 0.083 | -0.442 | -0.359 | 0.049 | -0.015 | -0.015 | 0.787 | 0.08 | 0.37 | 0.13 | 0 | 0.06 | 0.07 |
| 2 | 157214370 | rs707132 | 0.502 | 0.333 | -1.079 | -0.247 | -0.081 | 0.074 | 0.074 | 0.102 | 0.41 | 0.1 | 0.23 | 0.21 | 0.53 | 0.4 |
| 2 | 157216692 | rs10166471 | -0.05 | 0.435 | -0.864 | 0.006 | -0.027 | -0.079 | -0.079 | 0.308 | 0.17 | 0.02 | 0.15 | 0.28 | 0.18 | 0.16 |
| 2 | 157244794 | rs298329 | -0.211 | 0.895 | -0.744 | -0.045 | -0.018 | -0.034 | -0.034 | 0 | 0.18 | 0.08 | 0.16 | 0.27 | 0.26 | 0.17 |
| 2 | 157252095 | rs6436373 | 0.464 | 0.797 | -0.239 | -0.098 | 0.003 | 0.02 | 0.02 | 0 | 0.78 | 0.62 | 0.79 | 0.73 | 0.74 | 0.83 |
| 2 | 157280645 | rs298291 | -0.229 | 0.829 | -0.444 | -0.021 | -0.021 | -0.04 | -0.04 | 0.222 | 0.19 | 0.06 | 0.16 | 0.27 | 0.26 | 0.17 |
| 2 | 157285151 | rs6721605 | 0.063 | 1.075 | -0.345 | -0.414 | 0.001 | -0.006 | -0.006 | 0 | 0.6 | 0.58 | 0.7 | 0.74 | 0.53 | 0.77 |
| 2 | 157315748 | rs4664206 | 0.742 | 1.153 | -0.779 | -0.486 | -0.092 | -0.055 | -0.055 | 0 | 0.24 | 0.01 | 0.14 | 0.2 | 0.3 | 0.33 |
| 2 | 157324825 | rs298259 | -0.752 | 0.808 | -0.497 | -0.476 | 0.079 | -0.073 | -0.073 | 0 | 0.6 | 0.14 | 0.41 | 0.44 | 0.76 | 0.56 |
| 2 | 157388702 | rs692187 | -0.111 | 0.774 | -0.06 | -0.634 | 0 | -0.007 | -0.007 | 0 | 0.73 | 0.58 | 0.72 | 0.78 | 0.69 | 0.7 |
| 2 | 157391734 | rs1368249 | 0.017 | 0.748 | 0.073 | -0.646 | -0.004 | -0.013 | -0.013 | 0 | 0.72 | 0.52 | 0.71 | 0.79 | 0.69 | 0.7 |
| 2 | 157392032 | rs1821227 | -0.42 | 0.745 | 0.117 | -0.652 | 0 | -0.003 | -0.003 | 0 | 0.75 | 0.66 | 0.73 | 0.79 | 0.7 | 0.7 |
| 2 | 157400888 | rs4629116 | 1.068 | 0.731 | -0.079 | -0.653 | -0.047 | -0.026 | -0.026 | 0.067 | 0.2 | 0.03 | 0.15 | 0.16 | 0.27 | 0.27 |
| 2 | 157401269 | rs1878769 | -0.29 | 0.786 | 0.041 | -0.62 | 0 | -0.005 | -0.005 | 0.066 | 0.73 | 0.65 | 0.72 | 0.78 | 0.69 | 0.7 |
| 2 | 157403130 | rs3754770 | 1.449 | 0.808 | -0.194 | -0.632 | 0.003 | -0.006 | -0.006 | 0 | 0.32 | 0.28 | 0.43 | 0.55 | 0.22 | 0.42 |
| 2 | 157404713 | rs16840461 | 0.262 | 1.096 | -0.684 | -0.74 | 0 | -0.008 | -0.008 | 0 | 0.13 | 0.16 | 0.1 | 0.04 | 0.14 | 0.12 |
| 2 | 157404968 | rs1560628 | 0.5 | 1.123 | -0.787 | -0.602 | -0.097 | 0.018 | 0.018 | 0 | 0.27 | 0.05 | 0.17 | 0.2 | 0.33 | 0.15 |
| 2 | 157437583 | rs17200880 | 0.428 | 1.127 | -0.785 | -0.602 | -0.096 | 0.012 | 0.012 | 0 | 0.27 | 0.04 | 0.17 | 0.21 | 0.33 | 0.15 |
| 2 | 157448166 | rs10497165 | 0.639 | 1.195 | -0.783 | -0.677 | -0.095 | 0.01 | 0.01 | 0 | 0.25 | 0.04 | 0.17 | 0.18 | 0.31 | 0.14 |
| 2 | 157449118 | rs1368246 | -0.499 | 1.168 | -0.477 | -0.796 | -0.005 | 0.004 | 0.004 | 0 | 0.2 | 0.2 | 0.14 | 0.05 | 0.2 | 0.15 |
| 2 | 157456236 | rs7561693 | 0.863 | 0.932 | -0.363 | -0.744 | -0.001 | -0.001 | -0.001 | 2.863 | 0.23 | 0.23 | 0.21 | 0.2 | 0.3 | 0.3 |
| 2 | 157456540 | rs6713675 | 1.028 | 0.735 | -0.116 | -0.623 | 0.019 | -0.021 | -0.021 | 0 | 0.34 | 0.27 | 0.46 | 0.57 | 0.22 | 0.45 |
| 2 | 157463796 | rs12694677 | -1.268 | 0.846 | 0.03 | -0.738 | -0.001 | -0.001 | -0.001 | 0.196 | 0.65 | 0.66 | 0.52 | 0.46 | 0.77 | 0.56 |
| 2 | 157470262 | rs7597171 | 1.001 | 0.888 | -0.039 | -0.752 | 0.019 | -0.019 | -0.019 | 0.447 | 0.35 | 0.27 | 0.47 | 0.55 | 0.23 | 0.45 |
| 2 | 157478890 | rs10195414 | 0.053 | 1.163 | -0.479 | -0.806 | 0 | -0.006 | -0.006 | 0 | 0.82 | 0.83 | 0.88 | 0.95 | 0.82 | 0.86 |
| 2 | 157479462 | rs10497164 | 0.233 | 1.125 | -0.783 | -0.717 | -0.09 | 0.012 | 0.012 | 0.314 | 0.41 | 0.11 | 0.36 | 0.35 | 0.55 | 0.39 |
| 2 | 157488277 | rs6436555 | -0.486 | 1.076 | -0.555 | -0.638 | 0.004 | -0.019 | -0.019 | 0 | 0.43 | 0.48 | 0.36 | 0.24 | 0.51 | 0.45 |
| 2 | 157515560 | rs6706146 | 0.715 | 0.817 | -0.406 | -0.619 | 0.02 | -0.003 | -0.003 | 0 | 0.43 | 0.36 | 0.55 | 0.55 | 0.34 | 0.47 |
| 2 | 157516519 | rs11689150 | 0.004 | 1.027 | -0.438 | -0.676 | -0.003 | -0.023 | -0.023 | 0 | 0.33 | 0.46 | 0.25 | 0.23 | 0.3 | 0.38 |
| 2 | 157519736 | rs7575920 | -0.438 | 0.982 | -0.45 | -0.687 | 0.004 | -0.025 | -0.025 | 0.019 | 0.38 | 0.47 | 0.28 | 0.24 | 0.41 | 0.41 |
| 2 | 157531815 | rs1432572 | 0.703 | 0.838 | -0.147 | -0.673 | 0.014 | 0.025 | 0.025 | 0 | 0.5 | 0.25 | 0.6 | 0.55 | 0.42 | 0.52 |
| 2 | 157533704 | rs11678196 | -0.066 | 0.822 | -0.158 | -0.672 | -0.009 | -0.017 | -0.017 | 0.014 | 0.32 | 0.47 | 0.25 | 0.24 | 0.35 | 0.39 |
| 2 | 157534037 | rs17268385 | 1.075 | 0.829 | -0.26 | -0.658 | -0.071 | -0.033 | -0.033 | 0 | 0.15 | 0.02 | 0.12 | 0.21 | 0.23 | 0.08 |
| 2 | 157539961 | rs10497163 | 0.419 | 0.936 | 0.286 | -0.453 | -0.004 | -0.001 | -0.001 | 0.206 | 0.26 | 0.3 | 0.2 | 0.23 | 0.26 | 0.37 |
| 2 | 157548367 | rs6740709 | -0.127 | 1.135 | -0.484 | -0.325 | -0.009 | -0.014 | -0.014 | 0.043 | 0.32 | 0.51 | 0.25 | 0.24 | 0.35 | 0.39 |
| 2 | 157552860 | rs7601713 | -0.61 | 0.755 | -1.09 | -0.294 | -0.01 | 0.036 | 0.036 | 0 | 0.73 | 0.88 | 0.85 | 0.79 | 0.74 | 0.84 |
| 2 | 157558583 | rs7580515 | -0.262 | 0.895 | -0.745 | -0.095 | -0.005 | 0.002 | 0.002 | 0 | 0.45 | 0.48 | 0.62 | 0.55 | 0.48 | 0.46 |
| 2 | 157565192 | rs1543932 | -0.578 | 0.559 | -1.374 | -0.58 | -0.018 | 0.037 | 0.037 | 0 | 0.55 | 0.85 | 0.55 | 0.34 | 0.53 | 0.56 |
| 2 | 157566204 | rs1870898 | -0.171 | 0.52 | -0.601 | -0.779 | -0.008 | 0.032 | 0.032 | 0 | 0.45 | 0.34 | 0.44 | 0.31 | 0.44 | 0.55 |
| 2 | 157583475 | rs10202702 | -0.417 | 0.439 | -0.012 | -0.743 | -0.004 | 0.021 | 0.021 | 0 | 0.45 | 0.6 | 0.31 | 0.31 | 0.43 | 0.51 |
| 2 | 157588429 | rs4664212 | -0.005 | 0.509 | -0.04 | -0.918 | 0.02 | -0.022 | -0.022 | 0 | 0.7 | 0.41 | 0.77 | 0.8 | 0.64 | 0.71 |
| 2 | 157591413 | rs1031808 | -0.55 | 0.578 | -0.394 | -0.829 | -0.007 | 0.002 | 0.002 | 0 | 0.77 | 0.8 | 0.84 | 0.8 | 0.71 | 0.72 |
| 2 | 157592784 | rs1381693 | -0.338 | 0.449 | -0.663 | -0.961 | 0.025 | 0.057 | 0.057 | 0 | 0.22 | 0.08 | 0.12 | 0.08 | 0.15 | 0.23 |
| 2 | 157593363 | rs6716032 | 0.417 | 1.024 | -0.736 | -0.927 | -0.046 | 0.046 | 0.046 | 0 | 0.12 | 0.04 | 0.13 | 0 | 0.23 | 0.04 |
| 2 | 157596697 | rs10167258 | -0.618 | 0.049 | -0.34 | -0.803 | -0.003 | 0.061 | 0.061 | 0 | 0.57 | 0.77 | 0.37 | 0.27 | 0.54 | 0.5 |
| 2 | 157598678 | rs8179646 | 0.82 | -0.01 | -0.451 | -0.865 | -0.011 | 0.016 | 0.016 | 0 | 0.36 | 0.14 | 0.57 | 0.68 | 0.38 | 0.46 |
| 2 | 157599502 | rs6743360 | -1.256 | -0.052 | -0.331 | -0.837 | 0 | -0.016 | -0.016 | 0 | 0.95 | 0.93 | 0.95 | 1 | 0.92 | 0.98 |
| 2 | 157617191 | rs6712741 | -0.42 | -0.174 | 0.068 | -0.95 | -0.005 | 0.039 | 0.039 | 0 | 0.56 | 0.68 | 0.34 | 0.26 | 0.49 | 0.51 |
| 2 | 157617340 | rs6716024 | -0.395 | -0.375 | 0.971 | -1.017 | 0 | -0.004 | -0.004 | 0.523 | 0.54 | 0.46 | 0.34 | 0.19 | 0.52 | 0.52 |
| 2 | 157622147 | rs12620493 | 0.512 | -0.449 | 0.673 | -1.213 | -0.013 | 0.033 | 0.033 | 0 | 0.36 | 0.27 | 0.23 | 0.11 | 0.39 | 0.32 |
| 2 | 157634151 | rs6436638 | -0.9 | -0.434 | 0.504 | -1.158 | -0.023 | 0.071 | 0.071 | 0 | 0.58 | 0.8 | 0.36 | 0.26 | 0.52 | 0.52 |
| 2 | 157642017 | rs7565809 | 1.053 | -0.485 | 0.514 | -1.109 | -0.033 | 0.035 | 0.035 | 0 | 0.36 | 0.11 | 0.58 | 0.74 | 0.4 | 0.46 |
| 2 | 157649957 | rs6436652 | -0.196 | -0.758 | 0.618 | -0.639 | -0.012 | 0.048 | 0.048 | 0 | 0.4 | 0.26 | 0.24 | 0.22 | 0.39 | 0.3 |
| 2 | 157650780 | rs9288624 | -1.076 | -0.615 | 0.596 | -0.534 | -0.029 | 0.013 | 0.013 | 0.09 | 0.54 | 0.83 | 0.33 | 0.37 | 0.45 | 0.47 |
| 2 | 157654171 | rs2131801 | 0.653 | -0.668 | 0.723 | -0.443 | -0.003 | 0.005 | 0.005 | 0.172 | 0.3 | 0.25 | 0.17 | 0.21 | 0.28 | 0.23 |
| 2 | 157677493 | rs6753202 | -1.325 | -1.1 | 1.3 | -0.119 | -0.04 | -0.04 | -0.04 | 0 | 0.25 | 0.59 | 0.25 | 0.13 | 0.13 | 0.21 |
| 2 | 157680572 | rs1874064 | -1.428 | -1.078 | 1.06 | -0.045 | 0.033 | 0.033 | 0.033 | 0.768 | 0.26 | 0.77 | 0.27 | 0.13 | 0.13 | 0.2 |
| 2 | 157708966 | rs6757549 | -1.44 | -1.33 | 1.221 | -0.06 | -0.083 | -0.028 | -0.028 | 0 | 0.25 | 0.57 | 0.24 | 0.19 | 0.1 | 0.22 |
| 2 | 157724633 | rs7609000 | -1.588 | -1.053 | 1.201 | -0.082 | -0.075 | -0.06 | -0.06 | 0.109 | 0.2 | 0.52 | 0.22 | 0.12 | 0.1 | 0.16 |
| 2 | 157731677 | rs295830 | 1.85 | -1.319 | 1.405 | -0.215 | 0.049 | 0.045 | 0.045 | 5.24 | 0.75 | 0.17 | 0.72 | 0.85 | 0.85 | 0.81 |
| 2 | 157731753 | rs7565121 | -0.099 | -1.335 | 2.268 | -0.23 | 0.016 | -0.027 | -0.027 | 0.081 | 0.07 | 0.23 | 0.05 | 0 | 0.06 | 0 |
| 2 | 157751825 | rs295793 | 2.347 | -1.35 | 2.625 | -0.244 | 0.017 | 0.017 | 0.017 | 4.815 | 0.75 | 0.22 | 0.73 | 0.85 | 0.85 | 0.81 |

| | | | | | | | | | | | | | | | | |
|---|---|---|---|---|---|---|---|---|---|---|---|---|---|---|---|---|
| 2 | 157758173 | rs10204885 | -0.56 | -1.272 | 2.628 | -0.229 | 0.028 | -0.025 | -0.025 | 0 | 0.08 | 0.34 | 0.07 | 0 | 0.07 | 0 |
| 2 | 157774494 | rs13394736 | -1.893 | -1.351 | 2.799 | -0.169 | -0.076 | -0.049 | -0.049 | 0 | 0.2 | 0.52 | 0.22 | 0.13 | 0.11 | 0.16 |
| 2 | 157778161 | rs11900160 | -0.698 | -1.346 | 2.98 | -0.166 | -0.048 | 0 | 0 | 0 | 0.12 | 0.24 | 0.15 | 0.13 | 0.04 | 0.16 |
| 2 | 157785456 | rs10497159 | -0.071 | -1.316 | 2.921 | -0.162 | 0.026 | -0.022 | -0.022 | 0 | 0.07 | 0.24 | 0.05 | 0 | 0.07 | 0.01 |
| 2 | 157791556 | rs9973369 | -1.936 | -1.329 | 2.907 | -0.25 | 0.023 | -0.065 | -0.065 | 0.048 | 0.11 | 0.41 | 0.1 | 0 | 0.1 | 0.02 |
| 2 | 157797614 | rs1871780 | -2.714 | -1.402 | 2.66 | -0.228 | 0.035 | 0.032 | 0.032 | 3.718 | 0.26 | 0.8 | 0.31 | 0.17 | 0.19 | 0.21 |
| 2 | 157805435 | rs995283 | -2.008 | -1.392 | 2.761 | -0.217 | -0.064 | -0.04 | -0.04 | 0.008 | 0.21 | 0.61 | 0.26 | 0.13 | 0.11 | 0.18 |
| 2 | 157818436 | rs16841124 | -0.215 | -1.393 | 1.521 | -0.614 | -0.006 | 0.014 | 0.014 | 0 | 0.69 | 0.84 | 0.81 | 0.25 | 0.69 | 0.57 |
| 2 | 157856323 | rs6732936 | 1.166 | -1.477 | 1.902 | -0.655 | 0.002 | -0.012 | -0.012 | 0 | 0.54 | 0.4 | 0.6 | 0.14 | 0.63 | 0.39 |
| 2 | 157856605 | rs11888555 | -0.614 | -1.359 | 2.14 | -0.376 | -0.048 | -0.001 | -0.001 | 0 | 0.12 | 0.22 | 0.17 | 0.11 | 0.03 | 0.16 |
| 2 | 157863647 | rs6740075 | -1.195 | -1.376 | 2.36 | -0.384 | -0.077 | -0.006 | -0.006 | 1.619 | 0.14 | 0.38 | 0.2 | 0.11 | 0.04 | 0.18 |
| 2 | 157873775 | rs6747313 | -2.313 | -1.414 | 2.061 | -0.405 | -0.06 | -0.016 | -0.016 | 0 | 0.19 | 0.66 | 0.25 | 0.14 | 0.08 | 0.18 |
| 2 | 157875846 | rs10497155 | -1.095 | -1.42 | 1.849 | -0.385 | -0.079 | 0.001 | 0.001 | 0.232 | 0.14 | 0.39 | 0.2 | 0.11 | 0.03 | 0.17 |
| 2 | 157878707 | rs16841206 | -2.296 | -1.358 | 1.309 | -0.361 | -0.017 | -0.018 | -0.018 | 0.697 | 0.23 | 0.65 | 0.17 | 0.13 | 0.18 | 0.21 |
| 2 | 157885769 | rs6717435 | 0.4 | -2.027 | 0.763 | 0.503 | -0.017 | 0.014 | 0.014 | 0 | 0.67 | 0.61 | 0.84 | 0.56 | 0.76 | 0.68 |
| 2 | 157891197 | rs1481388 | 0.486 | -2.056 | 0.763 | 0.73 | -0.018 | 0.037 | 0.037 | 0 | 0.66 | 0.58 | 0.83 | 0.46 | 0.74 | 0.66 |
| 2 | 157906208 | rs1481385 | -0.76 | -1.84 | 0.278 | 0.651 | -0.023 | -0.109 | -0.109 | 0 | 0.79 | 0.99 | 0.91 | 0.62 | 0.84 | 0.8 |
| 2 | 157915086 | rs17254856 | 0.746 | -1.719 | 0.696 | 0.614 | 0.009 | -0.034 | -0.034 | 0 | 0.06 | 0.11 | 0.05 | 0 | 0.07 | 0.01 |
| 2 | 157921236 | rs1481234 | -0.772 | -1.9 | 0.712 | 0.596 | 0.01 | -0.005 | -0.005 | 0.021 | 0.34 | 0.27 | 0.18 | 0.41 | 0.3 | 0.29 |
| 2 | 157923137 | rs16841278 | 0.253 | -1.674 | 0.918 | 0.816 | -0.009 | 0.027 | 0.027 | 0 | 0.1 | 0.19 | 0.18 | 0.19 | 0.07 | 0.18 |
| 2 | 157924492 | rs7355634 | 2.064 | -1.756 | 0.343 | 0.763 | -0.046 | -0.002 | -0.002 | 0 | 0.48 | 0.05 | 0.56 | 0.3 | 0.56 | 0.49 |
| 2 | 157947400 | rs13401572 | -1.032 | -1.493 | 0.782 | 0.997 | 0.019 | 0.038 | 0.038 | 5.159 | 0.1 | 0.4 | 0.07 | 0.11 | 0.11 | 0.02 |
| 2 | 157990064 | rs12694882 | 0.159 | -1.436 | 0.343 | 0.982 | 0.008 | -0.028 | -0.028 | 0.415 | 0.71 | 0.82 | 0.86 | 0.49 | 0.76 | 0.73 |
| 2 | 157993888 | rs1481235 | 0.033 | -1.428 | 0.195 | 0.901 | 0.019 | -0.064 | -0.064 | 0.196 | 0.72 | 0.83 | 0.76 | 0.45 | 0.79 | 0.73 |
| 2 | 158004920 | rs12477460 | 1.794 | -1.414 | 0.261 | 0.88 | 0.024 | -0.001 | -0.001 | 0 | 0.77 | 0.19 | 0.82 | 0.74 | 0.82 | 0.81 |
| 2 | 158005419 | rs16841331 | -1.43 | -1.4 | 0.287 | 0.867 | -0.018 | -0.017 | -0.017 | 1.086 | 0.2 | 0.67 | 0.16 | 0.16 | 0.15 | 0.18 |
| 2 | 158009296 | rs962286 | -0.226 | -1.416 | 0.126 | 0.923 | 0.01 | -0.074 | -0.074 | 1.507 | 0.74 | 0.93 | 0.77 | 0.55 | 0.82 | 0.74 |
| 2 | 158015425 | rs10497177 | 0.344 | -1.409 | 0.121 | 0.683 | -0.038 | -0.149 | -0.149 | 0 | 0.23 | 0.02 | 0.22 | 0.47 | 0.18 | 0.25 |
| 2 | 158015603 | rs263326 | -0.065 | -1.426 | 0.191 | 0.672 | 0.012 | -0.062 | -0.062 | 0.202 | 0.26 | 0.13 | 0.24 | 0.58 | 0.21 | 0.27 |
| 2 | 158024652 | rs263317 | -1.536 | -1.252 | 0.315 | 0.626 | -0.001 | -0.002 | -0.002 | 0 | 0.46 | 0.53 | 0.38 | 0.69 | 0.41 | 0.46 |
| 2 | 158041480 | rs17279165 | 0.101 | -1.272 | -0.211 | 0.615 | 0.005 | -0.035 | -0.035 | 0 | 0.07 | 0.13 | 0.05 | 0 | 0.09 | 0.01 |
| 2 | 158044578 | rs2623041 | -2.106 | -1.315 | -0.259 | 0.631 | 0.009 | 0.014 | 0.014 | 7.286 | 0.23 | 0.8 | 0.19 | 0.2 | 0.21 | 0.19 |
| 2 | 158057860 | rs10177010 | -0.095 | -1.011 | 0.629 | 1.203 | 0.033 | -0.066 | -0.066 | 0 | 0.24 | 0.11 | 0.22 | 0.46 | 0.13 | 0.22 |
| 2 | 158066572 | rs834128 | -2.45 | -0.891 | 0.606 | 1.422 | 0.007 | -0.037 | -0.037 | 0 | 0.24 | 0.81 | 0.2 | 0.43 | 0.22 | 0.21 |
| 2 | 158068785 | rs263310 | -1.723 | -0.872 | 0.679 | 1.352 | 0.034 | 0.041 | 0.041 | 1.403 | 0.42 | 0.7 | 0.36 | 0.32 | 0.29 | 0.4 |
| 2 | 158075906 | rs834140 | -1.8 | -1.324 | 1.35 | 1.261 | 0.019 | -0.02 | -0.02 | 0 | 0.36 | 0.73 | 0.35 | 0.47 | 0.27 | 0.25 |
| 2 | 158080876 | rs263305 | -1.566 | -1.173 | 1.443 | 1.171 | 0.02 | -0.014 | -0.014 | 0 | 0.08 | 0.34 | 0.06 | 0 | 0.03 | 0.04 |
| 2 | 158091611 | rs834132 | -3.042 | -0.983 | 1.177 | 1.395 | 0.003 | 0.003 | 0.003 | 12.107 | 0.17 | 0.79 | 0.15 | 0.11 | 0.13 | 0.03 |
| 2 | 158095057 | rs707137 | -2.159 | -0.842 | 1.209 | 1.466 | 0.021 | -0.061 | -0.061 | 0 | 0.38 | 0.82 | 0.37 | 0.57 | 0.31 | 0.3 |
| 2 | 158098935 | rs6706447 | -1.371 | -0.594 | 1.335 | 1.515 | 0.003 | 0.013 | 0.013 | 0.015 | 0.29 | 0.4 | 0.31 | 0.45 | 0.28 | 0.19 |
| 2 | 158105322 | rs11891616 | -2.516 | -0.422 | 0.684 | 1.391 | 0.008 | 0.015 | 0.015 | 11.394 | 0.18 | 0.85 | 0.16 | 0.12 | 0.17 | 0.06 |
| 2 | 158136933 | rs6756958 | -1.58 | -0.813 | 1.184 | 1.73 | 0.031 | 0.017 | 0.017 | 8.194 | 0.07 | 0.51 | 0.07 | 0 | 0.02 | 0.02 |
| 2 | 158140717 | rs2289942 | -0.552 | -0.849 | 0.89 | 1.807 | -0.002 | 0.007 | 0.007 | 0 | 0.24 | 0.32 | 0.24 | 0.48 | 0.17 | 0.16 |
| 2 | 158141753 | rs2121347 | -1.205 | -0.767 | 0.186 | 1.722 | 0.04 | -0.081 | -0.081 | 0.187 | 0.33 | 0.86 | 0.33 | 0.58 | 0.23 | 0.23 |
| 2 | 158144195 | rs6437044 | 0.979 | -0.1 | 0.616 | 1.468 | 0.05 | 0.074 | 0.074 | 0.03 | 0.54 | 0.15 | 0.47 | 0.32 | 0.66 | 0.62 |
| 2 | 158144782 | rs11903808 | -2.017 | 0.255 | 2.604 | 1.546 | 0.063 | 0.219 | 0.219 | 0.014 | 0.07 | 0.4 | 0.06 | 0.14 | 0.04 | 0.04 |
| 2 | 158150015 | rs11890241 | -2.206 | 0.253 | 2.566 | 1.501 | 0.049 | 0.224 | 0.224 | 0.475 | 0.07 | 0.39 | 0.06 | 0.16 | 0.04 | 0.04 |
| 2 | 158157361 | rs3214040 | 2.249 | 0.239 | 2.594 | 1.49 | 0.006 | 0.108 | 0.108 | 0 | 0.88 | 0.57 | 0.8 | 0.84 | 0.93 | 0.93 |
| 2 | 158157794 | rs11895393 | -1.714 | 0.239 | 2.711 | 1.489 | 0.021 | 0.008 | 0.008 | 0.234 | 0.06 | 0.32 | 0.05 | 0 | 0.01 | 0.01 |
| 2 | 158157899 | rs4664867 | 0.51 | 0.242 | 2.167 | 1.491 | -0.002 | 0.003 | 0.003 | 0 | 0.95 | 0.96 | 0.86 | 0.84 | 0.94 | 0.94 |
| 2 | 158162033 | rs10183902 | 2.405 | 0.24 | 1.999 | 1.493 | 0.001 | 0 | 0 | 0 | 0.82 | 0.22 | 0.77 | 0.84 | 0.92 | 0.91 |
| 2 | 158165004 | rs6728956 | 1.624 | 0.682 | 1.642 | 1.608 | -0.005 | 0.199 | 0.199 | 0.044 | 0.88 | 0.56 | 0.81 | 0.74 | 0.93 | 0.93 |
| 2 | 158165401 | rs2305965 | 1.336 | 0.745 | 1.377 | 1.639 | 0.007 | 0.208 | 0.208 | 0 | 0.89 | 0.62 | 0.82 | 0.74 | 0.93 | 0.93 |
| 2 | 158177489 | rs16841611 | -2.247 | 0.679 | 1.794 | 1.039 | -0.029 | 0.005 | 0.005 | 1.337 | 0.13 | 0.68 | 0.1 | 0.09 | 0.04 | 0.12 |
| 2 | 158186653 | rs1867846 | -1.046 | 1.626 | 1.472 | 1.037 | -0.04 | -0.007 | -0.007 | 0 | 0.07 | 0.01 | 0.08 | 0.08 | 0.19 | 0.16 |
| 2 | 158219122 | rs155619 | 2.159 | 0.679 | 2.398 | 1.056 | -0.029 | 0.025 | 0.025 | 3.251 | 0.87 | 0.36 | 0.79 | 0.89 | 0.96 | 0.88 |
| 2 | 158270142 | rs155594 | 1.981 | 0.711 | 2.439 | 1.064 | -0.024 | -0.006 | -0.006 | 0 | 0.89 | 0.43 | 0.92 | 0.97 | 0.96 | 0.93 |
| 2 | 158270960 | rs198688 | 2.255 | 0.71 | 2.407 | 1.119 | -0.024 | 0.049 | 0.049 | 2.68 | 0.88 | 0.37 | 0.8 | 0.87 | 0.96 | 0.88 |
| 2 | 158271512 | rs267992 | 2.08 | 0.736 | 2.258 | 1.131 | -0.026 | -0.009 | -0.009 | 0.025 | 0.88 | 0.43 | 0.92 | 0.96 | 0.96 | 0.92 |
| 2 | 158278740 | rs267991 | 1.511 | 0.833 | 1.267 | 0.729 | -0.007 | -0.035 | -0.035 | 0 | 0.92 | 0.81 | 0.95 | 1 | 0.96 | 0.94 |
| 2 | 158283299 | rs155615 | 2.052 | 0.88 | 1.46 | 0.957 | 0.01 | 0.055 | 0.055 | 0.481 | 0.89 | 0.64 | 0.83 | 0.83 | 0.91 | 0.89 |
| 2 | 158284056 | rs155613 | 1.604 | 0.81 | 1.549 | 0.993 | -0.026 | 0.04 | 0.04 | 0.021 | 0.93 | 0.8 | 0.96 | 0.9 | 0.98 | 0.92 |
| 2 | 158291539 | rs10497185 | -2.471 | 0.83 | 1.542 | 0.803 | 0 | -0.026 | -0.026 | 0 | 0.08 | 0.24 | 0.14 | 0.01 | 0.06 | 0.05 |
| 2 | 158310166 | rs155597 | 1.17 | 0.875 | 1.308 | 0.884 | 0.005 | 0.009 | 0.009 | 0 | 0.81 | 0.77 | 0.79 | 0.78 | 0.8 | 0.77 |
| 2 | 158311691 | rs267989 | 0.703 | 0.845 | 1.189 | 0.895 | -0.013 | -0.019 | -0.019 | 0.05 | 0.84 | 0.94 | 0.91 | 0.9 | 0.85 | 0.8 |
| 2 | 158312472 | rs2123992 | 1.424 | 1.274 | 2.167 | 0.693 | 0.008 | 0.008 | 0.008 | 0 | 0.79 | 0.65 | 0.84 | 0.79 | 0.77 | 0.76 |
| 2 | 158314418 | rs267993 | 0.579 | 0.993 | 1.04 | -0.439 | -0.031 | 0.101 | 0.101 | 0 | 0.64 | 0.96 | 0.8 | 0.9 | 0.66 | 0.66 |
| 2 | 158324190 | rs16841822 | -3.255 | 0.803 | 2.137 | -0.273 | 0.122 | 0.035 | 0.035 | 0.315 | 0.07 | 0.49 | 0.15 | 0 | 0.07 | 0.02 |
| 2 | 158329471 | rs2198581 | -2.819 | 0.846 | 2.195 | -0.282 | 0.132 | -0.027 | -0.027 | 0 | 0.09 | 0.45 | 0.18 | 0 | 0.13 | 0.04 |

| | | | | | | | | | | | | | | | | |
|---|---|---|---|---|---|---|---|---|---|---|---|---|---|---|---|---|
| 2 | 158341335 | rs7579654 | -2.995 | 0.806 | 2.705 | -0.298 | 0.13 | 0.05 | 0.05 | 0.033 | 0.06 | 0.46 | 0.14 | 0 | 0.07 | 0.03 |
| 2 | 158342497 | rs7597165 | -2.285 | 0.816 | 2.766 | -0.295 | 0.136 | -0.01 | -0.01 | 0 | 0.08 | 0.41 | 0.16 | 0 | 0.11 | 0.04 |
| 2 | 158352326 | rs6713245 | -2.977 | 0.707 | 2.709 | -0.044 | 0.09 | 0.035 | 0.035 | 0.464 | 0.11 | 0.55 | 0.2 | 0.11 | 0.13 | 0.03 |
| 2 | 158364734 | rs16841871 | 1.668 | 0.406 | 2.682 | -0.435 | -0.024 | -0.061 | -0.061 | 0 | 0.71 | 0.43 | 0.73 | 0.91 | 0.78 | 0.72 |
| 2 | 158371626 | rs7579893 | 1.113 | 0.572 | 2.501 | -0.405 | 0.004 | 0.007 | 0.007 | 0 | 0.76 | 0.77 | 0.87 | 0.91 | 0.83 | 0.73 |
| 2 | 158388779 | rs12993800 | 1.611 | 0.392 | 2.79 | -0.341 | -0.031 | -0.047 | -0.047 | 0.623 | 0.77 | 0.39 | 0.78 | 0.91 | 0.84 | 0.75 |
| 2 | 158391031 | rs4664888 | 0.998 | 0.493 | 2.467 | -0.308 | -0.006 | -0.004 | -0.004 | 0 | 0.82 | 0.77 | 0.91 | 0.92 | 0.88 | 0.74 |
| 2 | 158405375 | rs16826016 | 1.503 | 0.307 | 2.354 | -0.45 | -0.047 | -0.075 | -0.075 | 0.334 | 0.76 | 0.49 | 0.78 | 0.93 | 0.84 | 0.74 |
| 2 | 158409131 | rs4299296 | 1.829 | 0.283 | 2.184 | -0.531 | -0.005 | -0.022 | -0.022 | 0 | 0.7 | 0.36 | 0.72 | 0.93 | 0.77 | 0.75 |
| 2 | 158410196 | rs6437106 | -2.656 | 0.479 | 2.4 | -0.486 | 0.017 | -0.057 | -0.057 | 0.096 | 0.12 | 0.57 | 0.09 | 0.01 | 0.11 | 0.05 |
| 2 | 158417363 | rs11691149 | -0.842 | 0.747 | 1.731 | -1.465 | 0.004 | -0.068 | -0.068 | 0 | 0.36 | 0.34 | 0.27 | 0.01 | 0.41 | 0.21 |
| 2 | 158432280 | rs3934681 | -0.843 | 0.743 | 1.854 | -1.317 | -0.012 | 0.003 | 0.003 | 0 | 0.36 | 0.26 | 0.27 | 0.06 | 0.41 | 0.21 |
| 2 | 158442005 | rs10184376 | -1.212 | 0.621 | 2.062 | -0.553 | -0.016 | 0.096 | 0.096 | 0 | 0.09 | 0.21 | 0.18 | 0.26 | 0.06 | 0.09 |
| 2 | 158443889 | rs4556933 | -1.529 | 1.076 | 2.346 | -0.67 | 0.011 | 0.044 | 0.044 | 0.439 | 0.44 | 0.7 | 0.43 | 0.32 | 0.39 | 0.27 |
| 2 | 158444844 | rs6743822 | -0.488 | 1.125 | 2.26 | -0.669 | -0.001 | -0.013 | -0.013 | 0 | 0.33 | 0.26 | 0.23 | 0.06 | 0.33 | 0.18 |
| 2 | 158451825 | rs11899550 | -1.213 | 0.626 | 2.043 | -0.584 | -0.013 | 0.106 | 0.106 | 0 | 0.08 | 0.21 | 0.18 | 0.26 | 0.06 | 0.09 |
| 2 | 158455980 | rs13383985 | -1.151 | 0.638 | 2.036 | -0.572 | -0.01 | 0.113 | 0.113 | 0 | 0.08 | 0.21 | 0.18 | 0.26 | 0.06 | 0.09 |
| 2 | 158460413 | rs2603276 | 1.222 | 0.735 | 1.716 | -0.588 | -0.003 | 0.097 | 0.097 | 0.164 | 0.51 | 0.24 | 0.52 | 0.68 | 0.51 | 0.66 |
| 2 | 158497658 | rs1220123 | 1.022 | 0.192 | 2.146 | -0.422 | 0 | -0.005 | -0.005 | 0 | 0.9 | 0.93 | 0.9 | 0.99 | 0.87 | 0.94 |
| 2 | 158499868 | rs1220127 | 2.593 | -0.565 | 2.054 | -0.584 | 0.019 | 0.023 | 0.023 | 1.294 | 0.68 | 0.27 | 0.78 | 0.87 | 0.79 | 0.86 |
| 2 | 158503739 | rs2444770 | 2.116 | -0.614 | 2.342 | -0.618 | -0.045 | -0.069 | -0.069 | 0.175 | 0.76 | 0.41 | 0.84 | 0.93 | 0.85 | 0.89 |
| 2 | 158508271 | rs11900786 | 1.832 | -0.557 | 2.008 | -0.547 | -0.049 | -0.064 | -0.064 | 1.166 | 0.79 | 0.63 | 0.86 | 0.93 | 0.86 | 0.92 |
| 2 | 158513202 | rs16841926 | 0.124 | -0.588 | 1.73 | -0.533 | -0.007 | 0 | 0 | 1.676 | 0.08 | 0.08 | 0.01 | 0.06 | 0.01 | 0.01 |
| 2 | 158514510 | rs2509903 | 1.67 | -0.562 | 1.726 | -0.545 | -0.032 | -0.051 | -0.051 | 0 | 0.79 | 0.67 | 0.87 | 0.93 | 0.86 | 0.92 |
| 2 | 158520905 | rs10195713 | 1.257 | -0.31 | 1.037 | -0.532 | -0.002 | 0 | 0 | 0 | 0.88 | 0.9 | 0.89 | 0.99 | 0.86 | 0.93 |
| 2 | 158525226 | rs12694933 | 1.959 | -0.669 | 1.582 | -0.497 | -0.041 | -0.057 | -0.057 | 0.167 | 0.78 | 0.65 | 0.87 | 0.9 | 0.86 | 0.89 |
| 2 | 158526978 | rs2848647 | 2.022 | -0.443 | 1.668 | -0.527 | -0.015 | -0.084 | -0.084 | 0.3 | 0.79 | 0.66 | 0.82 | 0.97 | 0.77 | 0.9 |
| 2 | 158530857 | rs897085 | 1.963 | -0.425 | 1.647 | -0.473 | -0.01 | -0.079 | -0.079 | 0 | 0.8 | 0.69 | 0.83 | 0.97 | 0.79 | 0.9 |
| 2 | 158531739 | rs10497187 | 1.96 | -0.832 | 1.715 | -0.466 | -0.05 | -0.066 | -0.066 | 0 | 0.76 | 0.4 | 0.84 | 0.89 | 0.85 | 0.86 |
| 2 | 158531809 | rs10933439 | 2.061 | -1.029 | 0.832 | -0.492 | 0.124 | 0.163 | 0.163 | 0.497 | 0.62 | 0.13 | 0.77 | 0.87 | 0.8 | 0.76 |
| 2 | 158536555 | rs1220121 | -2.152 | -0.563 | 1.824 | 0.17 | -0.023 | -0.016 | -0.016 | 0 | 0.29 | 0.57 | 0.3 | 0.23 | 0.21 | 0.19 |
| 2 | 158549371 | rs16841991 | -1.091 | -0.562 | 2.839 | 0.241 | 0.002 | 0.056 | 0.056 | 0 | 0.06 | 0.27 | 0.03 | 0.1 | 0.02 | 0.02 |
| 2 | 158549742 | rs747282 | -2.375 | -0.604 | 3.094 | 0.347 | -0.015 | 0.006 | 0.006 | 0 | 0.28 | 0.59 | 0.2 | 0.32 | 0.22 | 0.16 |
| 2 | 158555607 | rs1220141 | -2.994 | -0.582 | 3.134 | 0.474 | 0.031 | 0.002 | 0.002 | 0.106 | 0.36 | 0.73 | 0.23 | 0.37 | 0.25 | 0.23 |
| 2 | 158574077 | rs6756623 | -2.949 | -0.218 | 3.043 | 0.404 | 0.015 | 0.014 | 0.014 | 0.007 | 0.35 | 0.71 | 0.25 | 0.28 | 0.28 | 0.19 |
| 2 | 158593357 | rs12997 | -3.17 | -0.004 | 3.12 | 0.514 | -0.012 | -0.016 | -0.016 | 0.096 | 0.35 | 0.94 | 0.25 | 0.28 | 0.27 | 0.19 |
| 2 | 158596197 | rs17182166 | -1.648 | 0.108 | 3.471 | 0.579 | 0.005 | -0.023 | -0.023 | 0.865 | 0.12 | 0.19 | 0.08 | 0.05 | 0.15 | 0.09 |
| 2 | 158603316 | rs1220133 | -2.716 | 0.1 | 2.905 | 0.584 | 0.015 | 0.01 | 0.01 | 0.266 | 0.31 | 0.76 | 0.23 | 0.26 | 0.24 | 0.16 |
| 2 | 158626980 | rs1146031 | 1.198 | 0.325 | 2.342 | 0.541 | -0.007 | -0.005 | -0.005 | 0 | 0.94 | 0.74 | 0.96 | 0.97 | 0.99 | 0.98 |
| 2 | 158634289 | rs1146037 | 1.668 | 0.769 | 1.665 | 0.621 | 0.006 | -0.006 | -0.006 | 0 | 0.84 | 0.81 | 0.84 | 0.92 | 0.79 | 0.83 |
| 2 | 158646738 | rs6719924 | 1.828 | 0.714 | 1.861 | 0.609 | 0.023 | -0.022 | -0.022 | 0.019 | 0.84 | 0.73 | 0.84 | 0.92 | 0.79 | 0.85 |
| 2 | 158657433 | rs12694937 | -1.789 | 0.372 | 1.903 | 0.535 | -0.024 | -0.029 | -0.029 | 0.254 | 0.06 | 0.21 | 0.04 | 0.03 | 0.02 | 0.02 |
| 2 | 158667217 | rs10497191 | 2.953 | 0.416 | 1.515 | 0.448 | 0.021 | 0.069 | 0.069 | 5.086 | 0.78 | 0.08 | 0.81 | 0.9 | 0.88 | 0.9 |
| 2 | 158668445 | rs4380178 | 1.601 | 0.208 | 1.675 | 1.138 | 0 | 0.005 | 0.005 | 0.263 | 0.86 | 0.79 | 0.88 | 0.8 | 0.84 | 0.86 |
| 2 | 158671700 | rs10497192 | 3.202 | 0.343 | 1.348 | 1.645 | -0.069 | -0.158 | -0.158 | 0 | 0.67 | 0.05 | 0.71 | 0.41 | 0.72 | 0.71 |
| 2 | 158675263 | rs4664898 | 1.718 | 0.359 | 1.463 | 1.373 | 0.007 | 0.014 | 0.014 | 0.085 | 0.86 | 0.76 | 0.86 | 0.81 | 0.82 | 0.83 |
| 2 | 158691926 | rs4233672 | 2.188 | 0.34 | 1.179 | 1.338 | 0.009 | 0 | 0 | 7.108 | 0.75 | 0.21 | 0.79 | 0.79 | 0.81 | 0.81 |
| 2 | 158694121 | rs10933443 | 2.03 | 0.234 | 1.349 | 1.235 | -0.001 | -0.003 | -0.003 | 0 | 0.72 | 0.28 | 0.75 | 0.44 | 0.73 | 0.76 |
| 2 | 158694380 | rs13398650 | -2.116 | -0.097 | 1.559 | 0.596 | -0.04 | -0.07 | -0.07 | 0 | 0.18 | 0.64 | 0.14 | 0.06 | 0.1 | 0.12 |
| 2 | 158705117 | rs7563276 | -1.3 | 0.005 | 1.195 | 0.509 | -0.001 | -0.025 | -0.025 | 0.792 | 0.11 | 0.1 | 0.07 | 0 | 0.07 | 0.09 |
| 2 | 158709678 | rs10497193 | -2.296 | 0.116 | 1.852 | 0.358 | 0.026 | -0.127 | -0.127 | 3.177 | 0.22 | 0.48 | 0.17 | 0.05 | 0.18 | 0.16 |
| 2 | 158712538 | rs4664901 | 2.325 | -0.239 | 1.443 | 0.975 | 0.043 | -0.075 | -0.075 | 0 | 0.62 | 0.16 | 0.73 | 0.47 | 0.75 | 0.59 |
| 2 | 158717966 | rs10201181 | 2.203 | -0.682 | 1.787 | 1.063 | 0.071 | -0.078 | -0.078 | 0.177 | 0.69 | 0.22 | 0.82 | 0.49 | 0.83 | 0.68 |
| 2 | 158724552 | rs16842215 | -1.013 | -0.628 | 2.024 | 1.087 | 0.014 | -0.008 | -0.008 | 0 | 0.06 | 0.13 | 0.05 | 0.02 | 0.07 | 0.05 |
| 2 | 158735265 | rs9288700 | 2.136 | -0.647 | 2.674 | 1.088 | -0.014 | -0.053 | -0.053 | 0.018 | 0.86 | 0.7 | 0.93 | 0.95 | 0.9 | 0.92 |
| 2 | 158737086 | rs16842230 | -1.21 | -0.6 | 2.636 | 1.117 | 0.018 | -0.007 | -0.007 | 0 | 0.06 | 0.13 | 0.05 | 0.02 | 0.06 | 0.04 |
| 2 | 158750454 | rs4438436 | 1.529 | -0.79 | 2.479 | 1.039 | -0.034 | 0.018 | 0.018 | 0.228 | 0.71 | 0.39 | 0.87 | 0.5 | 0.88 | 0.7 |
| 2 | 158765030 | rs12472133 | 1.672 | -0.35 | 0.495 | 0.96 | 0.083 | -0.069 | -0.069 | 2.118 | 0.65 | 0.12 | 0.79 | 0.6 | 0.78 | 0.71 |
| 2 | 158772058 | rs7578180 | 1.643 | 0.417 | 0.477 | 0.872 | 0.003 | -0.079 | -0.079 | 1.179 | 0.65 | 0.18 | 0.77 | 0.49 | 0.67 | 0.66 |
| 2 | 158772233 | rs16842263 | -1.497 | 0.294 | 1.455 | 0.559 | -0.013 | -0.04 | -0.04 | 0 | 0.09 | 0.33 | 0.04 | 0.02 | 0.03 | 0.04 |
| 2 | 158780204 | rs10183640 | -1.391 | 1.064 | 1.137 | 0.279 | 0.006 | 0.006 | 0.006 | 0 | 0.27 | 0.29 | 0.18 | 0.38 | 0.3 | 0.26 |
| 2 | 158781445 | rs12997150 | -0.666 | 1.133 | 0.89 | 0.136 | -0.023 | 0.047 | 0.047 | 0 | 0.11 | 0.02 | 0.04 | 0.02 | 0.1 | 0.07 |
| 2 | 158784567 | rs4441418 | 0.496 | 1.228 | 1.049 | -0.067 | -0.002 | -0.018 | -0.018 | 0 | 0.87 | 0.87 | 0.91 | 0.99 | 0.8 | 0.9 |
| 2 | 158786869 | rs6737672 | 2.8 | 0.855 | 0.848 | -0.064 | -0.074 | -0.01 | -0.01 | 1.205 | 0.52 | 0.05 | 0.67 | 0.6 | 0.49 | 0.63 |
| 2 | 158789851 | rs10190823 | -0.526 | 1.294 | 1.147 | -0.185 | 0 | 0.005 | 0.005 | 0 | 0.17 | 0.15 | 0.15 | 0.35 | 0.25 | 0.15 |
| 2 | 158791560 | rs6720264 | 1.716 | 1.205 | 1.893 | -0.252 | 0.017 | -0.135 | -0.135 | 0 | 0.72 | 0.44 | 0.82 | 0.95 | 0.69 | 0.79 |
| 2 | 158797105 | rs7595677 | 0.292 | 1.183 | 1.496 | -0.2 | -0.002 | -0.018 | -0.018 | 0 | 0.87 | 0.88 | 0.91 | 0.99 | 0.8 | 0.91 |
| 2 | 158800569 | rs6741915 | 2.653 | 0.482 | 0.773 | -0.561 | -0.048 | -0.053 | -0.053 | 0 | 0.43 | 0.05 | 0.63 | 0.6 | 0.41 | 0.61 |
| 2 | 158802021 | rs13419857 | -1.717 | 0.428 | 1.573 | -0.669 | 0.002 | -0.088 | -0.088 | 0 | 0.19 | 0.56 | 0.12 | 0.05 | 0.15 | 0.08 |

| | | | | | | | | | | | | | | | | |
|---|---|---|---|---|---|---|---|---|---|---|---|---|---|---|---|---|
| 2 | 158806924 | rs12995606 | -0.925 | 0.422 | 1.412 | -0.453 | 0.005 | -0.033 | -0.033 | 0 | 0.24 | 0.14 | 0.16 | 0.42 | 0.22 | 0.22 |
| 2 | 158813048 | rs10182041 | -1.595 | 0.479 | 1.943 | -0.468 | 0.003 | 0.039 | 0.039 | 0 | 0.29 | 0.44 | 0.22 | 0.44 | 0.3 | 0.25 |
| 2 | 158813398 | rs7560416 | -2.342 | 0.517 | 1.758 | -0.483 | -0.007 | -0.009 | -0.009 | 0 | 0.41 | 0.71 | 0.28 | 0.47 | 0.4 | 0.3 |
| 2 | 158817388 | rs13391200 | -1.608 | 0.284 | 1.849 | -0.614 | 0.014 | -0.082 | -0.082 | 0 | 0.18 | 0.47 | 0.11 | 0.05 | 0.15 | 0.08 |
| 2 | 158821440 | rs12694947 | -1.67 | 0.254 | 1.836 | -0.628 | 0.013 | -0.086 | -0.086 | 0 | 0.18 | 0.48 | 0.11 | 0.05 | 0.15 | 0.08 |
| 2 | 158839672 | rs4664914 | 0.474 | 0.6 | 1.543 | -0.677 | -0.022 | 0.019 | 0.019 | 0 | 0.75 | 0.83 | 0.81 | 0.99 | 0.61 | 0.88 |
| 2 | 158846569 | rs11892485 | -1.574 | -0.028 | 2.001 | -0.334 | -0.04 | 0.131 | 0.131 | 0 | 0.19 | 0.48 | 0.13 | 0.41 | 0.12 | 0.2 |
| 2 | 158848642 | rs2108811 | 0.946 | -0.204 | 1.911 | -0.573 | 0.002 | -0.097 | -0.097 | 0 | 0.64 | 0.64 | 0.79 | 0.97 | 0.63 | 0.84 |
| 2 | 158848804 | rs720159 | 2.103 | -0.252 | 2.033 | -0.47 | 0.007 | 0.075 | 0.075 | 0.196 | 0.51 | 0.37 | 0.71 | 0.77 | 0.55 | 0.73 |
| 2 | 158854828 | rs6742818 | 0.937 | -0.187 | 2.027 | -0.461 | 0.001 | -0.074 | -0.074 | 0.007 | 0.65 | 0.64 | 0.8 | 0.97 | 0.62 | 0.84 |
| 2 | 158857396 | rs2355163 | 2.17 | -0.19 | 1.897 | -0.361 | 0.014 | 0.035 | 0.035 | 0.553 | 0.51 | 0.32 | 0.69 | 0.56 | 0.53 | 0.68 |
| 2 | 158858365 | rs10804388 | 0.912 | -0.161 | 1.633 | -0.392 | -0.017 | 0.068 | 0.068 | 1.099 | 0.66 | 0.8 | 0.82 | 0.97 | 0.63 | 0.84 |
| 2 | 158860293 | rs758279 | 2.096 | -0.335 | 1.496 | -0.108 | 0.034 | 0.013 | 0.013 | 1.845 | 0.56 | 0.18 | 0.74 | 0.56 | 0.64 | 0.7 |
| 2 | 158865831 | rs6437126 | -1.982 | -0.362 | 1.512 | -0.199 | 0.022 | -0.024 | -0.024 | 0.167 | 0.27 | 0.8 | 0.17 | 0.4 | 0.18 | 0.21 |
| 2 | 158870618 | rs1860163 | 2.129 | -0.308 | 1.257 | -0.149 | 0.033 | 0.043 | 0.043 | 1.273 | 0.53 | 0.25 | 0.73 | 0.57 | 0.61 | 0.67 |
| 2 | 158874478 | rs4664919 | 2.307 | -0.368 | 1.197 | 0.085 | 0.011 | -0.002 | -0.002 | 0 | 0.51 | 0.12 | 0.71 | 0.5 | 0.58 | 0.65 |
| 2 | 158876018 | rs11895999 | -0.683 | -0.384 | 1.401 | -0.155 | -0.028 | 0.036 | 0.036 | 0 | 0.15 | 0.31 | 0.1 | 0.21 | 0.09 | 0.13 |
| 2 | 158879580 | rs4664920 | 1.851 | -0.205 | 0.473 | -0.2 | 0.027 | 0.04 | 0.04 | 1.216 | 0.51 | 0.26 | 0.73 | 0.56 | 0.6 | 0.66 |
| 2 | 158885519 | rs1160638 | -0.882 | -0.287 | 0.155 | -0.08 | -0.058 | 0.08 | 0.08 | 0.745 | 0.19 | 0.56 | 0.11 | 0.37 | 0.1 | 0.15 |
| 2 | 158886433 | rs10755053 | 0.531 | 0.072 | -0.606 | 0.004 | -0.011 | -0.005 | -0.005 | 0.446 | 0.63 | 0.42 | 0.78 | 0.63 | 0.71 | 0.74 |
| 2 | 158888256 | rs13024527 | -0.792 | 0.295 | -1.067 | -0.046 | -0.026 | 0.047 | 0.047 | 0 | 0.1 | 0.01 | 0.03 | 0.02 | 0.09 | 0.11 |
| 2 | 158890762 | rs6731028 | -1.187 | -0.133 | -1.122 | 0.04 | 0.013 | 0.017 | 0.017 | 0 | 0.23 | 0.16 | 0.11 | 0.14 | 0.15 | 0.19 |
| 2 | 158893323 | rs7605276 | -0.174 | -0.12 | -1.002 | 0.21 | -0.01 | 0.017 | 0.017 | 0.009 | 0.07 | 0.11 | 0.01 | 0.12 | 0.03 | 0.07 |
| 2 | 158895094 | rs2079656 | -0.641 | -0.063 | -1.583 | -0.702 | -0.048 | 0.085 | 0.085 | 0 | 0.65 | 0.77 | 0.53 | 0.95 | 0.54 | 0.7 |
| 2 | 158895546 | rs4664238 | -0.603 | -0.529 | -0.871 | -0.218 | 0.005 | 0.017 | 0.017 | 0.029 | 0.28 | 0.46 | 0.23 | 0.34 | 0.28 | 0.17 |
| 2 | 158895655 | rs13007683 | 1.448 | -0.995 | -1.242 | -0.065 | -0.048 | -0.167 | -0.167 | 0.703 | 0.21 | 0.01 | 0.26 | 0.47 | 0.21 | 0.41 |
| 2 | 158898944 | rs13023991 | -0.58 | -0.974 | -0.977 | -0.093 | -0.005 | 0.005 | 0.005 | 0.251 | 0.31 | 0.52 | 0.24 | 0.34 | 0.28 | 0.17 |
| 2 | 158900762 | rs6707195 | -1.571 | -0.643 | -0.987 | -0.182 | -0.088 | 0.171 | 0.171 | 0 | 0.67 | 0.91 | 0.57 | 0.97 | 0.58 | 0.7 |
| 2 | 158902989 | rs1860165 | 0.392 | -1.246 | 0.018 | -0.026 | 0.044 | 0.034 | 0.034 | 0 | 0.52 | 0.4 | 0.63 | 0.35 | 0.69 | 0.45 |
| 2 | 158903284 | rs2190959 | -0.237 | -1.373 | 0.632 | 0 | -0.013 | 0.01 | 0.01 | 0 | 0.61 | 0.56 | 0.66 | 0.52 | 0.72 | 0.65 |
| 2 | 158906795 | rs981727 | -0.296 | -1.395 | 0.713 | 0.096 | 0.03 | -0.026 | -0.026 | 0 | 0.62 | 0.75 | 0.67 | 0.54 | 0.72 | 0.58 |
| 2 | 158910156 | rs16842499 | -0.106 | -1.375 | 0.767 | 0.101 | -0.026 | 0.015 | 0.015 | 0.002 | 0.1 | 0.12 | 0.02 | 0.17 | 0.03 | 0.13 |
| 2 | 158915735 | rs13398821 | 0.543 | -1.383 | 0.709 | 0.142 | 0.032 | 0.015 | 0.015 | 0 | 0.52 | 0.42 | 0.73 | 0.39 | 0.7 | 0.52 |
| 2 | 158918573 | rs768364 | -0.889 | -1.341 | 0.357 | 0.247 | -0.039 | -0.166 | -0.166 | 1.07 | 0.69 | 0.98 | 0.79 | 0.54 | 0.73 | 0.59 |
| 2 | 158922575 | rs4664922 | 0.177 | -1.363 | 0.498 | 0.177 | 0.053 | 0.018 | 0.018 | 0.027 | 0.55 | 0.73 | 0.75 | 0.39 | 0.7 | 0.53 |
| 2 | 158935854 | rs6712771 | -0.862 | -1.193 | 0.483 | 0.039 | 0.004 | -0.092 | -0.092 | 0.387 | 0.69 | 0.92 | 0.7 | 0.48 | 0.72 | 0.55 |
| 2 | 158950622 | rs16842547 | -0.604 | -0.926 | 1.081 | 0.045 | 0.018 | 0.058 | 0.058 | 0 | 0.15 | 0.31 | 0.14 | 0.27 | 0.17 | 0.13 |
| 2 | 158958378 | rs6739665 | -1.107 | -1.057 | 1.607 | -0.05 | -0.069 | 0.031 | 0.031 | 0.203 | 0.15 | 0.3 | 0.04 | 0.16 | 0.04 | 0.12 |
| 2 | 158958605 | rs6710480 | -1.289 | -1.081 | 1.623 | -0.059 | -0.087 | 0.021 | 0.021 | 0.279 | 0.17 | 0.37 | 0.06 | 0.16 | 0.04 | 0.12 |
| 2 | 158959335 | rs7606259 | -1.267 | -1.069 | 1.63 | -0.078 | -0.086 | 0.018 | 0.018 | 0.58 | 0.17 | 0.37 | 0.06 | 0.16 | 0.04 | 0.12 |
| 2 | 158966186 | rs6707705 | 0.399 | -1.068 | 1.619 | -0.188 | 0.005 | 0.057 | 0.057 | 0 | 0.52 | 0.62 | 0.57 | 0.21 | 0.56 | 0.4 |
| 2 | 158968794 | rs10174157 | 0.173 | -1.071 | 1.623 | -0.19 | -0.006 | 0.075 | 0.075 | 0 | 0.55 | 0.69 | 0.58 | 0.21 | 0.56 | 0.44 |
| 2 | 158971237 | rs13418976 | 1.325 | -1.059 | 1.863 | -0.211 | -0.047 | 0.101 | 0.101 | 0 | 0.39 | 0.24 | 0.51 | 0.05 | 0.51 | 0.31 |
| 2 | 158972966 | rs17603277 | -0.597 | -1.012 | 1.961 | -0.087 | 0.006 | -0.014 | -0.014 | 0.319 | 0.1 | 0.05 | 0.09 | 0.26 | 0.09 | 0.1 |
| 2 | 158977794 | rs2074955 | -1.099 | -0.898 | 2.098 | -0.015 | 0.028 | 0.095 | 0.095 | 0 | 0.13 | 0.3 | 0.13 | 0.27 | 0.14 | 0.11 |
| 2 | 158980689 | rs6437137 | 0.554 | -0.861 | 1.512 | -0.068 | 0.008 | 0.1 | 0.1 | 0 | 0.52 | 0.31 | 0.55 | 0.21 | 0.56 | 0.4 |
| 2 | 158987439 | rs13419905 | 1.22 | -0.857 | 1.4 | -0.105 | -0.03 | 0.125 | 0.125 | 0 | 0.42 | 0.24 | 0.52 | 0.05 | 0.55 | 0.33 |
| 2 | 158991525 | rs2074954 | -0.227 | -0.742 | 1.7 | 0.195 | 0 | 0.026 | 0.026 | 0 | 0.08 | 0.09 | 0.08 | 0.26 | 0.06 | 0.08 |
| 2 | 158992725 | rs6437140 | -0.547 | -0.712 | 2.14 | 0.144 | 0.174 | 0.065 | 0.065 | 0.017 | 0.05 | 0.33 | 0.07 | 0.01 | 0.07 | 0.03 |
| 2 | 158994950 | rs17603340 | -0.039 | -0.711 | 1.55 | 0.167 | -0.006 | 0.044 | 0.044 | 0 | 0.07 | 0.11 | 0.08 | 0.25 | 0.05 | 0.08 |
| 2 | 158994990 | rs16842635 | -2.17 | -0.824 | 1.557 | 0.185 | 0.013 | -0.017 | -0.017 | 0 | 0.29 | 0.73 | 0.19 | 0.43 | 0.17 | 0.22 |
| 2 | 158995999 | rs736671 | -0.675 | -0.697 | 1.297 | 0.205 | -0.015 | 0.113 | 0.113 | 0 | 0.09 | 0.25 | 0.09 | 0.26 | 0.05 | 0.08 |
| 2 | 159008513 | rs1357184 | 0.911 | -0.481 | 0.924 | -0.014 | 0 | 0.034 | 0.034 | 0 | 0.47 | 0.44 | 0.59 | 0.28 | 0.56 | 0.42 |
| 2 | 159009069 | rs13027877 | 0.124 | -0.24 | -0.21 | 0.134 | 0.033 | -0.032 | -0.032 | 3.998 | 0.1 | 0.01 | 0.02 | 0.15 | 0.04 | 0.05 |
| 2 | 159026540 | rs1402866 | 0.546 | -0.305 | -0.273 | 0.161 | -0.051 | -0.149 | -0.149 | 0.1 | 0.29 | 0.02 | 0.27 | 0.57 | 0.29 | 0.45 |
| 2 | 159027173 | rs10804390 | -0.163 | -0.051 | -0.012 | -0.155 | 0.028 | 0.035 | 0.035 | 0.219 | 0.4 | 0.08 | 0.3 | 0.71 | 0.32 | 0.5 |
| 2 | 159029330 | rs16842681 | -1.168 | 0.44 | 1.063 | 0.096 | 0.028 | -0.023 | -0.023 | 0 | 0.07 | 0.14 | 0.08 | 0.01 | 0.11 | 0.09 |
| 2 | 159032245 | rs1357182 | 0.833 | 0.674 | -0.139 | 0.23 | 0.016 | -0.018 | -0.018 | 0 | 0.93 | 0.86 | 0.92 | 0.99 | 0.89 | 0.91 |
| 2 | 159036321 | rs956986 | -0.261 | 0.654 | -0.532 | 0.39 | 0.004 | -0.011 | -0.011 | 0.169 | 0.38 | 0.29 | 0.42 | 0.58 | 0.34 | 0.6 |
| 2 | 159037050 | rs16842710 | -0.757 | 0.165 | -0.155 | 0.218 | -0.013 | 0.001 | 0.001 | 0 | 0.11 | 0.2 | 0.05 | 0.11 | 0.07 | 0.06 |
| 2 | 159037227 | rs1010279 | -1.752 | 0.398 | -0.046 | 0.233 | -0.003 | 0.01 | 0.01 | 0 | 0.13 | 0.2 | 0.05 | 0.19 | 0.08 | 0.09 |
| 2 | 159044007 | rs926013 | 1.067 | 0.139 | 0.515 | -0.51 | -0.027 | -0.124 | -0.124 | 0.062 | 0.8 | 0.41 | 0.89 | 1 | 0.87 | 0.95 |
| 2 | 159047406 | rs7592105 | -1.519 | 0.289 | 0.999 | -0.887 | -0.003 | -0.022 | -0.022 | 0 | 0.36 | 0.64 | 0.26 | 0.21 | 0.33 | 0.14 |
| 2 | 159051128 | rs1357186 | -0.357 | 0.591 | 0.055 | -1.264 | 0.016 | -0.052 | -0.052 | 0 | 0.69 | 0.51 | 0.75 | 0.84 | 0.65 | 0.74 |
| 2 | 159051791 | rs12612640 | 1.29 | 0.644 | -0.32 | -1.155 | 0.014 | 0.008 | 0.008 | 0 | 0.38 | 0.29 | 0.51 | 0.64 | 0.31 | 0.64 |
| 2 | 159053112 | rs12466131 | 1.004 | 0.478 | -0.501 | -0.906 | -0.039 | 0.013 | 0.013 | 0 | 0.14 | 0.02 | 0.14 | 0.13 | 0.19 | 0.03 |
| 2 | 159058679 | rs4233674 | -0.745 | 0.342 | -0.155 | -1.063 | 0.004 | -0.011 | -0.011 | 0.108 | 0.45 | 0.45 | 0.37 | 0.33 | 0.55 | 0.29 |
| 2 | 159059679 | rs7567129 | 0.401 | 0.2 | -0.468 | -1.07 | -0.003 | 0.01 | 0.01 | 0.729 | 0.2 | 0.14 | 0.13 | 0.1 | 0.24 | 0.2 |
| 2 | 159061258 | rs7604580 | -0.489 | 0.197 | -0.362 | -0.998 | -0.011 | 0.021 | 0.021 | 0.071 | 0.28 | 0.18 | 0.2 | 0.17 | 0.34 | 0.23 |

| | | | | | | | | | | | | | | | | |
|---|---|---|---|---|---|---|---|---|---|---|---|---|---|---|---|---|
| 2 | 159061535 | rs7579360 | -0.504 | 0.197 | -0.318 | -0.998 | -0.003 | 0.007 | 0.007 | 0.065 | 0.28 | 0.24 | 0.21 | 0.17 | 0.34 | 0.23 |
| 2 | 159063290 | rs13432858 | 1.397 | -0.053 | -0.644 | -1.089 | 0 | 0.058 | 0.058 | 0 | 0.53 | 0.08 | 0.69 | 0.71 | 0.6 | 0.68 |
| 2 | 159068798 | rs6718403 | -0.905 | -0.015 | -0.396 | -0.957 | 0.005 | 0.015 | 0.015 | 0 | 0.05 | 0.16 | 0.04 | 0.08 | 0.04 | 0.03 |
| 2 | 159072395 | rs4664935 | -1.404 | -0.116 | 0.097 | -0.99 | 0.003 | 0.024 | 0.024 | 0.55 | 0.95 | 0.78 | 0.98 | 0.99 | 1 | 0.99 |
| 2 | 159085899 | rs4315473 | 0.463 | -0.092 | -0.113 | -1.096 | -0.009 | -0.024 | -0.024 | 0.084 | 0.68 | 0.43 | 0.77 | 0.88 | 0.69 | 0.73 |
| 2 | 159088394 | rs2030487 | 0.73 | -0.048 | -0.319 | -1.139 | 0.003 | 0.019 | 0.019 | 1.53 | 0.55 | 0.45 | 0.65 | 0.74 | 0.5 | 0.71 |
| 2 | 159088963 | rs11693922 | 1.664 | 0.04 | -1.21 | -1.088 | -0.136 | -0.036 | -0.036 | 0 | 0.4 | 0.07 | 0.56 | 0.63 | 0.43 | 0.67 |
| 2 | 159089280 | rs10192799 | 1.064 | 0.247 | -0.901 | -0.879 | -0.041 | 0.014 | 0.014 | 0 | 0.14 | 0.02 | 0.13 | 0.14 | 0.19 | 0.03 |
| 2 | 159092141 | rs12994903 | 0.137 | 0.225 | -1.398 | -1.204 | -0.078 | 0.042 | 0.042 | 0.517 | 0.25 | 0.02 | 0.38 | 0.09 | 0.34 | 0.24 |
| 2 | 159093577 | rs1608037 | 1.777 | 0.032 | -1.62 | -1.31 | -0.091 | -0.098 | -0.098 | 0 | 0.35 | 0.04 | 0.22 | 0.58 | 0.36 | 0.61 |
| 2 | 159095185 | rs13413279 | 0.287 | -0.028 | -0.803 | -1.162 | -0.004 | 0.02 | 0.02 | 0 | 0.21 | 0.15 | 0.27 | 0.21 | 0.23 | 0.05 |
| 2 | 159101213 | rs7559893 | 0.289 | -0.222 | 0.436 | -1.219 | -0.024 | 0.064 | 0.064 | 1.772 | 0.9 | 0.55 | 0.94 | 0.88 | 0.96 | 0.95 |
| 2 | 159103164 | rs4664944 | 1.682 | -0.311 | -0.416 | -1.31 | -0.04 | 0.045 | 0.045 | 0.12 | 0.4 | 0.14 | 0.47 | 0.68 | 0.41 | 0.66 |
| 2 | 159105196 | rs1402867 | 1.419 | -0.35 | 0.039 | -1.324 | 0.002 | 0.018 | 0.018 | 0 | 0.43 | 0.4 | 0.5 | 0.68 | 0.41 | 0.66 |
| 2 | 159105867 | rs1521865 | 1.804 | -0.309 | 0.016 | -1.304 | -0.098 | -0.082 | -0.082 | 0 | 0.38 | 0.06 | 0.46 | 0.57 | 0.39 | 0.65 |
| 2 | 159106259 | rs10209732 | 1.035 | -0.333 | -0.037 | -1.344 | -0.036 | 0.031 | 0.031 | 0.555 | 0.19 | 0.01 | 0.22 | 0.13 | 0.21 | 0.03 |
| 2 | 159106959 | rs1521867 | 0.647 | -0.265 | -0.056 | -1.414 | -0.002 | 0.007 | 0.007 | 0 | 0.55 | 0.48 | 0.59 | 0.69 | 0.53 | 0.74 |
| 2 | 159107659 | rs10933464 | 0.647 | -0.265 | -0.136 | -1.414 | -0.002 | 0.014 | 0.014 | 0 | 0.55 | 0.44 | 0.59 | 0.69 | 0.53 | 0.74 |
| 2 | 159109369 | rs7573649 | 0.647 | -0.265 | -0.221 | -1.414 | -0.005 | 0.033 | 0.033 | 0 | 0.55 | 0.38 | 0.58 | 0.68 | 0.53 | 0.74 |
| 2 | 159109925 | rs12328335 | -0.392 | -0.136 | -0.053 | -1.294 | -0.018 | -0.031 | -0.031 | 1.627 | 0.17 | 0.35 | 0.13 | 0.11 | 0.14 | 0.08 |
| 2 | 159111281 | rs965432 | 0.55 | -0.004 | -0.34 | -1.143 | -0.011 | 0.008 | 0.008 | 0 | 0.58 | 0.64 | 0.62 | 0.62 | 0.53 | 0.75 |
| 2 | 159114394 | rs10497198 | 0.828 | -0.026 | -0.987 | -1.142 | -0.035 | 0.019 | 0.019 | 0.43 | 0.18 | 0.03 | 0.22 | 0.14 | 0.21 | 0.03 |
| 2 | 159117099 | rs1960641 | 1.595 | -0.033 | -0.814 | -1.106 | -0.007 | -0.001 | -0.001 | 0.878 | 0.41 | 0.18 | 0.58 | 0.58 | 0.4 | 0.68 |
| 2 | 159120721 | rs3936747 | -0.709 | 0.175 | -1.246 | -1.166 | -0.054 | 0.102 | 0.102 | 0 | 0.39 | 0.06 | 0.29 | 0.2 | 0.41 | 0.25 |
| 2 | 159127158 | rs4664946 | 1.903 | -0.2 | -0.493 | -1.215 | -0.009 | 0 | 0 | 0.505 | 0.39 | 0.31 | 0.57 | 0.62 | 0.41 | 0.67 |
| 2 | 159141224 | rs2602189 | -0.016 | -0.299 | -0.515 | -1.225 | 0.001 | -0.001 | -0.001 | 0 | 0.16 | 0.12 | 0.11 | 0.17 | 0.14 | 0.14 |
| 2 | 159159916 | rs4514833 | -1.239 | -0.323 | -0.485 | -1.261 | -0.002 | -0.015 | -0.015 | 0.044 | 0.93 | 0.93 | 0.94 | 1 | 0.94 | 0.97 |
| 2 | 159160404 | rs11897570 | 0.018 | -0.345 | -0.487 | -1.239 | 0.001 | -0.001 | -0.001 | 0.147 | 0.16 | 0.12 | 0.11 | 0.17 | 0.14 | 0.14 |
| 2 | 159170978 | rs6746288 | 1.119 | -0.387 | 0.044 | -1.204 | -0.015 | -0.022 | -0.022 | 0 | 0.06 | 0.19 | 0.02 | 0 | 0.01 | 0 |
| 2 | 159173779 | rs4581847 | 0.94 | -0.425 | 0.079 | -1.261 | -0.051 | -0.082 | -0.082 | 4.802 | 0.82 | 0.64 | 0.88 | 0.97 | 0.9 | 0.92 |
| 2 | 159180748 | rs12617658 | -1.052 | -0.487 | 0.135 | -1.205 | -0.059 | 0.064 | 0.064 | 0.54 | 0.16 | 0.43 | 0.1 | 0.21 | 0.08 | 0.07 |
| 2 | 159181502 | rs10188238 | -1.272 | -0.442 | -0.149 | -1.407 | 0.002 | 0.006 | 0.006 | 0 | 0.46 | 0.52 | 0.29 | 0.38 | 0.42 | 0.23 |
| 2 | 159190540 | rs2356086 | 0.493 | -0.36 | -0.183 | -1.475 | -0.002 | -0.006 | -0.006 | 0 | 0.85 | 0.84 | 0.85 | 0.97 | 0.83 | 0.9 |
| 2 | 159195225 | rs11686762 | 1.321 | -0.302 | -0.214 | -1.495 | -0.067 | 0.019 | 0.019 | 1.157 | 0.2 | 0.02 | 0.12 | 0.14 | 0.25 | 0.06 |
| 2 | 159195282 | rs16842890 | -0.835 | -0.336 | -0.207 | -1.431 | -0.059 | 0.075 | 0.075 | 0.413 | 0.17 | 0.44 | 0.09 | 0.22 | 0.08 | 0.07 |
| 2 | 159196771 | rs12620556 | -0.835 | -0.336 | -0.207 | -1.431 | -0.059 | 0.075 | 0.075 | 0.413 | 0.17 | 0.44 | 0.09 | 0.22 | 0.08 | 0.07 |
| 2 | 159204593 | rs9917145 | 0.699 | -0.317 | -0.275 | -1.424 | -0.015 | 0.051 | 0.051 | 0.018 | 0.76 | 0.84 | 0.77 | 0.95 | 0.72 | 0.8 |
| 2 | 159213880 | rs13399899 | -0.604 | -0.3 | -0.159 | -1.384 | -0.011 | -0.006 | -0.006 | 0 | 0.36 | 0.49 | 0.21 | 0.36 | 0.33 | 0.13 |
| 2 | 159222561 | rs13401538 | 0.128 | -0.277 | -0.083 | -1.308 | -0.013 | 0.022 | 0.022 | 0.248 | 0.1 | 0.35 | 0.06 | 0.07 | 0.04 | 0.04 |
| 2 | 159237284 | rs10167316 | 2.126 | -0.192 | -0.578 | -1.316 | -0.01 | -0.003 | -0.003 | 0 | 0.39 | 0.29 | 0.55 | 0.59 | 0.4 | 0.66 |
| 2 | 159239052 | rs2883854 | 0.866 | -0.13 | -0.633 | -1.307 | -0.004 | 0.005 | 0.005 | 0.441 | 0.23 | 0.19 | 0.16 | 0.2 | 0.27 | 0.09 |
| 2 | 159249741 | rs10190089 | 1.626 | -0.16 | -0.986 | -1.457 | -0.009 | -0.041 | -0.041 | 0 | 0.62 | 0.54 | 0.68 | 0.8 | 0.66 | 0.74 |
| 2 | 159250362 | rs2602205 | -0.145 | -0.239 | -0.821 | -1.309 | -0.002 | -0.013 | -0.013 | 1.665 | 0.71 | 0.67 | 0.83 | 0.74 | 0.71 | 0.91 |
| 2 | 159259303 | rs2602191 | -0.404 | -0.25 | -0.775 | -1.255 | 0.008 | 0.121 | 0.121 | 0 | 0.91 | 0.68 | 0.95 | 0.86 | 0.96 | 0.97 |
| 2 | 159261308 | rs9711224 | 0.964 | -0.264 | -0.877 | -1.288 | -0.031 | 0.019 | 0.019 | 0 | 0.1 | 0.01 | 0.07 | 0.02 | 0.11 | 0.08 |
| 2 | 159263645 | rs10933480 | 1.586 | -0.261 | -0.87 | -1.32 | -0.013 | -0.036 | -0.036 | 0 | 0.72 | 0.54 | 0.83 | 0.82 | 0.75 | 0.84 |
| 2 | 159266933 | rs2193700 | 1.026 | -0.259 | -0.829 | -1.296 | -0.021 | -0.004 | -0.004 | 0 | 0.81 | 0.55 | 0.9 | 0.85 | 0.86 | 0.92 |
| 2 | 159274227 | rs2193701 | 1.023 | 0.062 | -1.61 | -1.232 | -0.058 | 0.054 | 0.054 | 1.078 | 0.2 | 0.02 | 0.12 | 0.11 | 0.25 | 0.06 |
| 2 | 159282923 | rs7579690 | -0.848 | 0.055 | -1.624 | -1.261 | -0.01 | 0.048 | 0.048 | 0 | 0.2 | 0.14 | 0.13 | 0.03 | 0.21 | 0.14 |
| 2 | 159289746 | rs11675870 | -0.907 | -0.088 | -1.577 | -1.197 | -0.068 | 0.177 | 0.177 | 0.443 | 0.38 | 0.08 | 0.22 | 0.14 | 0.41 | 0.16 |
| 2 | 159300522 | rs16842933 | 0.485 | -0.125 | -0.93 | -0.751 | 0 | -0.005 | -0.005 | 0 | 0.07 | 0.05 | 0.02 | 0.01 | 0.03 | 0.03 |
| 2 | 159308789 | rs2193705 | -0.249 | 0.079 | -1.078 | -0.517 | -0.017 | -0.044 | -0.044 | 0 | 0.9 | 0.97 | 0.93 | 0.82 | 0.88 | 0.9 |
| 2 | 159316583 | rs1125661 | -0.192 | -0.02 | -0.781 | -1.053 | -0.016 | 0.064 | 0.064 | 0.078 | 0.69 | 0.8 | 0.79 | 0.89 | 0.62 | 0.87 |
| 2 | 159320951 | rs2193707 | 0.214 | -0.004 | -0.935 | -1.165 | -0.023 | 0.097 | 0.097 | 0 | 0.63 | 0.77 | 0.67 | 0.87 | 0.56 | 0.76 |
| 2 | 159325925 | rs4664962 | 1.654 | -0.527 | -0.446 | -1.33 | -0.016 | 0.015 | 0.015 | 0.009 | 0.38 | 0.26 | 0.53 | 0.68 | 0.39 | 0.61 |
| 2 | 159329196 | rs17492981 | 0.053 | -0.516 | 0.052 | -1.325 | -0.014 | -0.03 | -0.03 | 0 | 0.21 | 0.22 | 0.12 | 0.05 | 0.15 | 0.13 |
| 2 | 159330659 | rs16842963 | -0.592 | -0.49 | 0.065 | -1.311 | -0.002 | -0.011 | -0.011 | 0 | 0.26 | 0.24 | 0.23 | 0.07 | 0.22 | 0.23 |
| 2 | 159333765 | rs6717437 | -1.265 | -0.432 | 0.016 | -1.394 | 0.013 | -0.012 | -0.012 | 1.086 | 0.94 | 0.85 | 0.93 | 0.98 | 0.91 | 0.95 |
| 2 | 159339424 | rs2111880 | 1.21 | -0.455 | 0.154 | -1.409 | -0.015 | -0.06 | -0.06 | 0 | 0.64 | 0.43 | 0.68 | 0.9 | 0.69 | 0.69 |
| 2 | 159341805 | rs4664964 | -0.325 | -0.449 | 0.102 | -1.408 | -0.004 | 0.006 | 0.006 | 0.017 | 0.68 | 0.71 | 0.78 | 0.75 | 0.61 | 0.86 |
| 2 | 159350135 | rs4664966 | 2.093 | -0.43 | 0.006 | -1.58 | -0.021 | 0.015 | 0.015 | 0.008 | 0.38 | 0.22 | 0.53 | 0.68 | 0.39 | 0.61 |
| 2 | 159364472 | rs2356186 | 0.132 | -0.523 | 0.03 | -1.565 | -0.011 | 0.019 | 0.019 | 0.017 | 0.68 | 0.74 | 0.79 | 0.75 | 0.61 | 0.86 |
| 2 | 159368196 | rs1861984 | 2.304 | -0.611 | 0.041 | -1.628 | -0.034 | 0.008 | 0.008 | 0.225 | 0.35 | 0.22 | 0.53 | 0.68 | 0.41 | 0.61 |
| 2 | 159385972 | rs7589768 | -0.957 | -0.482 | -0.055 | -1.54 | 0.052 | 0.087 | 0.087 | 0 | 0.93 | 0.77 | 0.92 | 0.84 | 0.89 | 0.94 |
| 2 | 159391780 | rs6728605 | -0.383 | -0.568 | -0.157 | -1.551 | -0.01 | -0.024 | -0.024 | 0 | 0.25 | 0.24 | 0.22 | 0.07 | 0.2 | 0.23 |
| 2 | 159393449 | rs3771595 | 0.355 | -0.424 | -0.467 | -1.468 | -0.011 | 0.022 | 0.022 | 0.635 | 0.26 | 0.13 | 0.14 | 0.22 | 0.3 | 0.09 |
| 2 | 159411420 | rs3771603 | 1.405 | -0.116 | -0.984 | -1.669 | -0.036 | 0.045 | 0.045 | 0 | 0.33 | 0.22 | 0.55 | 0.81 | 0.4 | 0.51 |
| 2 | 159412448 | rs4664250 | 0.282 | -0.975 | -0.226 | -1.36 | -0.028 | -0.077 | -0.077 | 0 | 0.73 | 0.57 | 0.82 | 0.93 | 0.8 | 0.82 |
| 2 | 159414979 | rs3771604 | 1.013 | -1.09 | -0.21 | -0.694 | 0.013 | -0.011 | -0.011 | 0 | 0.44 | 0.4 | 0.71 | 0.52 | 0.6 | 0.66 |

| | | | | | | | | | | | | | | | | |
|---|---|---|---|---|---|---|---|---|---|---|---|---|---|---|---|---|
| 2 | 159415520 | rs2069100 | 0.86 | -0.962 | -0.33 | -1.184 | -0.045 | 0.102 | 0.102 | 0.522 | 0.27 | 0.02 | 0.13 | 0.13 | 0.28 | 0.08 |
| 2 | 159426661 | rs3821290 | 0.816 | -0.958 | -0.26 | -1.179 | 0 | -0.023 | -0.023 | 0 | 0.46 | 0.54 | 0.68 | 0.79 | 0.51 | 0.74 |
| 2 | 159429406 | rs2214903 | 2.03 | -0.964 | -0.237 | -1.129 | -0.079 | -0.043 | -0.043 | 0.045 | 0.33 | 0.11 | 0.53 | 0.63 | 0.41 | 0.6 |
| 2 | 159433065 | rs1990155 | 0.381 | -0.821 | -0.002 | -0.971 | -0.02 | -0.005 | -0.005 | 0 | 0.73 | 0.49 | 0.81 | 0.8 | 0.78 | 0.82 |
| 2 | 159442447 | rs13417157 | 1.15 | -0.881 | -0.138 | -1.012 | -0.018 | 0.008 | 0.008 | 0 | 0.12 | 0.01 | 0.06 | 0.05 | 0.11 | 0.12 |
| 2 | 159444459 | rs7566874 | -1.042 | -0.846 | -0.02 | -1.002 | 0.01 | -0.01 | -0.01 | 0 | 0.93 | 0.84 | 0.93 | 0.97 | 0.91 | 0.95 |
| 2 | 159446818 | rs2108215 | 1.138 | -0.877 | -0.083 | -1.014 | -0.005 | 0.013 | 0.013 | 0 | 0.45 | 0.42 | 0.67 | 0.66 | 0.49 | 0.73 |
| 2 | 159450163 | rs3771635 | 0.643 | -0.85 | -0.278 | -0.98 | -0.043 | 0.1 | 0.1 | 0.424 | 0.28 | 0.03 | 0.13 | 0.14 | 0.28 | 0.09 |
| 2 | 159467006 | rs3771644 | 0.793 | -0.856 | -0.297 | -1.015 | -0.041 | 0.09 | 0.09 | 0.811 | 0.27 | 0.02 | 0.13 | 0.13 | 0.27 | 0.08 |
| 2 | 159469920 | rs7607589 | 1.346 | -0.821 | -0.119 | -0.874 | -0.033 | 0.003 | 0.003 | 0.872 | 0.42 | 0.27 | 0.66 | 0.61 | 0.49 | 0.7 |
| 2 | 159485213 | rs7597080 | 1.513 | -0.803 | 0.197 | -0.822 | -0.007 | -0.014 | -0.014 | 0 | 0.07 | 0.22 | 0.04 | 0 | 0.01 | 0.01 |
| 2 | 159489481 | rs886770 | 0.753 | -0.834 | 0.045 | -0.947 | -0.028 | -0.073 | -0.073 | 0 | 0.82 | 0.41 | 0.86 | 0.97 | 0.9 | 0.92 |
| 2 | 159491583 | rs976426 | 0.073 | -0.818 | 0.104 | -0.938 | 0.003 | -0.06 | -0.06 | 0.038 | 0.86 | 0.56 | 0.88 | 0.98 | 0.9 | 0.94 |
| 2 | 159494988 | rs2356140 | 1.343 | -0.802 | -0.163 | -0.962 | -0.021 | 0.058 | 0.058 | 1.2 | 0.42 | 0.29 | 0.66 | 0.79 | 0.5 | 0.72 |
| 2 | 159512538 | rs2528584 | 0.861 | -0.8 | -0.111 | -0.945 | 0.023 | 0.051 | 0.051 | 0.616 | 0.68 | 0.25 | 0.79 | 0.92 | 0.77 | 0.8 |
| 2 | 159512997 | rs1010332 | 0.639 | -0.769 | -0.043 | -0.928 | -0.01 | -0.037 | -0.037 | 0.169 | 0.72 | 0.38 | 0.8 | 0.92 | 0.78 | 0.82 |
| 2 | 159515697 | rs1010334 | -0.759 | -0.692 | -0.207 | -0.882 | 0.009 | -0.01 | -0.01 | 0 | 0.93 | 0.85 | 0.93 | 0.97 | 0.91 | 0.95 |
| 2 | 159518352 | rs2286250 | 0.704 | -0.781 | -0.178 | -0.889 | 0.002 | 0.008 | 0.008 | 0 | 0.14 | 0.13 | 0.07 | 0.05 | 0.12 | 0.12 |
| 2 | 159527936 | rs2528591 | 1.376 | -0.867 | -0.096 | -0.893 | -0.041 | 0.001 | 0.001 | 0.779 | 0.41 | 0.23 | 0.65 | 0.61 | 0.49 | 0.71 |
| 2 | 159528642 | rs7573340 | 1.333 | -0.77 | -0.005 | -1.094 | -0.009 | -0.017 | -0.017 | 1.066 | 0.07 | 0.24 | 0.05 | 0 | 0.01 | 0.01 |
| 2 | 159534889 | rs2711040 | -0.116 | -0.769 | -0.771 | -1.169 | -0.021 | 0.105 | 0.105 | 0 | 0.64 | 0.82 | 0.78 | 0.84 | 0.6 | 0.85 |
| 2 | 159538716 | rs2711042 | 0.492 | -0.471 | -0.411 | -1.154 | 0.014 | -0.005 | -0.005 | 0 | 0.74 | 0.57 | 0.84 | 0.78 | 0.71 | 0.8 |
| 2 | 159539434 | rs1055579 | 0.239 | -0.672 | -0.483 | -1.183 | -0.014 | -0.033 | -0.033 | 0 | 0.3 | 0.28 | 0.17 | 0.14 | 0.22 | 0.1 |
| 2 | 159539677 | rs2711043 | 0.359 | -0.589 | -0.441 | -1.148 | -0.014 | -0.019 | -0.019 | 0 | 0.6 | 0.48 | 0.76 | 0.7 | 0.66 | 0.82 |
| 2 | 159548035 | rs2711083 | 0.254 | -0.643 | -1.174 | -0.958 | 0.019 | 0.019 | 0.019 | 0 | 0.63 | 0.7 | 0.7 | 0.68 | 0.63 | 0.71 |
| 2 | 159548100 | rs6727441 | 0.963 | -0.562 | -1.231 | -0.993 | -0.041 | 0.033 | 0.033 | 0 | 0.13 | 0.03 | 0.08 | 0.05 | 0.17 | 0.11 |
| 2 | 159552465 | rs2528603 | 0.352 | -0.58 | -1.228 | -0.991 | 0.015 | 0.027 | 0.027 | 0 | 0.62 | 0.69 | 0.7 | 0.68 | 0.61 | 0.71 |
| 2 | 159553686 | rs2528604 | 1.107 | -0.657 | -1.103 | -1.024 | 0 | -0.001 | -0.001 | 0 | 0.47 | 0.47 | 0.71 | 0.65 | 0.54 | 0.72 |
| 2 | 159558658 | rs2528609 | 0.502 | -0.638 | -1.109 | -0.99 | 0.014 | 0.013 | 0.013 | 0 | 0.86 | 0.74 | 0.83 | 0.81 | 0.8 | 0.81 |
| 2 | 159559624 | rs2254443 | 0.026 | -0.653 | -1.066 | -1.006 | 0.016 | 0.036 | 0.036 | 0.06 | 0.91 | 0.76 | 0.93 | 0.84 | 0.88 | 0.93 |
| 2 | 159560048 | rs2528611 | 1.463 | -0.695 | -1.115 | -1.005 | 0.008 | -0.007 | -0.007 | 0 | 0.42 | 0.45 | 0.6 | 0.62 | 0.45 | 0.6 |
| 2 | 159563930 | rs2078507 | 0.81 | -0.657 | -1.355 | -0.992 | -0.018 | -0.014 | -0.014 | 0 | 0.09 | 0.03 | 0.14 | 0.07 | 0.14 | 0.13 |
| 2 | 159564276 | rs2528614 | -0.479 | -0.667 | -1.073 | -1.221 | 0.029 | 0.05 | 0.05 | 0 | 0.92 | 0.76 | 0.94 | 0.85 | 0.89 | 0.94 |
| 2 | 159564450 | rs10489982 | -0.066 | -0.882 | -1.363 | -1.396 | -0.002 | 0.105 | 0.105 | 0 | 0.25 | 0.07 | 0.11 | 0.06 | 0.21 | 0.13 |
| 2 | 159567817 | rs2711047 | 0.482 | -0.868 | -0.933 | -0.886 | -0.013 | -0.02 | -0.02 | 0 | 0.67 | 0.6 | 0.83 | 0.81 | 0.75 | 0.87 |
| 2 | 159570350 | rs2158832 | 1.646 | -0.8 | -1.112 | -0.865 | 0.017 | -0.006 | -0.006 | 0 | 0.11 | 0.01 | 0.06 | 0.13 | 0.09 | 0.06 |
| 2 | 159572270 | rs725806 | -0.034 | -0.462 | -0.834 | -0.722 | -0.028 | -0.009 | -0.009 | 0 | 0.65 | 0.48 | 0.83 | 0.69 | 0.74 | 0.84 |
| 2 | 159574883 | rs2711053 | 0.324 | 0.654 | -0.393 | -0.258 | 0.018 | 0.009 | 0.009 | 0 | 0.32 | 0.36 | 0.54 | 0.62 | 0.4 | 0.58 |
| 2 | 159575554 | rs1990153 | 0.498 | 0.325 | -0.356 | -0.456 | 0.015 | 0.005 | 0.005 | 0.812 | 0.93 | 0.76 | 0.94 | 0.95 | 0.91 | 0.95 |
| 2 | 159577307 | rs16843269 | -0.692 | 0.105 | -0.493 | -0.777 | -0.001 | -0.03 | -0.03 | 0 | 0.12 | 0.16 | 0.11 | 0.02 | 0.09 | 0.09 |
| 2 | 159577391 | rs757928 | 0.759 | -1.085 | -0.299 | -1.469 | -0.03 | -0.072 | -0.072 | 0.019 | 0.61 | 0.47 | 0.75 | 0.9 | 0.79 | 0.74 |
| 2 | 159579618 | rs6744767 | -0.182 | -1.132 | -0.436 | -1.242 | 0.034 | 0.021 | 0.021 | 0 | 0.2 | 0.09 | 0.1 | 0.12 | 0.12 | 0.18 |
| 2 | 159582975 | rs11674976 | -0.574 | -0.52 | -0.18 | -1.874 | 0 | -0.027 | -0.027 | 0 | 0.43 | 0.44 | 0.29 | 0.15 | 0.43 | 0.29 |
| 2 | 159583610 | rs2528626 | 1.083 | -0.636 | -0.16 | -1.988 | -0.035 | 0.081 | 0.081 | 1.988 | 0.4 | 0.26 | 0.57 | 0.83 | 0.47 | 0.59 |
| 2 | 159585082 | rs2528627 | 0.361 | -0.636 | -0.102 | -2.117 | -0.004 | -0.016 | -0.016 | 0 | 0.58 | 0.37 | 0.67 | 0.95 | 0.58 | 0.78 |
| 2 | 159586005 | rs4665001 | 0.771 | -0.6 | -0.167 | -1.989 | 0.011 | 0.022 | 0.022 | 0 | 0.16 | 0.11 | 0.1 | 0.05 | 0.13 | 0.08 |
| 2 | 159586721 | rs17828333 | 0.843 | -0.301 | -0.602 | -2.281 | -0.128 | 0.119 | 0.119 | 0.429 | 0.22 | 0.01 | 0.15 | 0.01 | 0.3 | 0.06 |
| 2 | 159587170 | rs2711048 | 0.032 | -0.381 | -0.393 | -2.323 | -0.075 | 0.186 | 0.186 | 0 | 0.63 | 0.86 | 0.76 | 0.94 | 0.63 | 0.85 |
| 2 | 159587256 | rs13415523 | 0.83 | -0.373 | -0.207 | -2.16 | 0.025 | 0.033 | 0.033 | 0 | 0.16 | 0.09 | 0.08 | 0.05 | 0.07 | 0.11 |
| 2 | 159588589 | rs13430918 | -0.241 | -0.715 | 0.253 | -2.346 | -0.019 | -0.042 | -0.042 | 1.112 | 0.25 | 0.3 | 0.13 | 0.05 | 0.11 | 0.11 |
| 2 | 159588969 | rs2711086 | 1.496 | -0.912 | 0.409 | -2.515 | -0.018 | 0.033 | 0.033 | 0.407 | 0.34 | 0.27 | 0.51 | 0.8 | 0.4 | 0.52 |
| 2 | 159589457 | rs13407077 | 0.022 | -0.977 | 0.626 | -2.546 | -0.036 | -0.066 | -0.066 | 1.85 | 0.26 | 0.35 | 0.14 | 0.05 | 0.11 | 0.11 |
| 2 | 159589475 | rs2528632 | 0.169 | -0.938 | 0.646 | -2.543 | 0.007 | -0.046 | -0.046 | 0.455 | 0.58 | 0.56 | 0.65 | 0.85 | 0.5 | 0.63 |
| 2 | 159589828 | rs2528633 | 0.436 | -0.433 | 0.822 | -2.648 | -0.003 | 0.02 | 0.02 | 0.236 | 0.53 | 0.38 | 0.67 | 0.9 | 0.58 | 0.82 |
| 2 | 159591062 | rs13007462 | -1.51 | -0.343 | 0.59 | -2.088 | -0.003 | 0.002 | 0.002 | 0 | 0.69 | 0.66 | 0.49 | 0.29 | 0.61 | 0.57 |
| 2 | 159593394 | rs4664262 | -1.504 | -0.34 | 0.599 | -2.087 | 0.001 | -0.001 | -0.001 | 0 | 0.69 | 0.68 | 0.49 | 0.29 | 0.61 | 0.57 |
| 2 | 159597415 | rs908405 | -1.055 | -0.435 | 0.678 | -1.948 | 0.004 | -0.014 | -0.014 | 3.787 | 0.63 | 0.52 | 0.46 | 0.32 | 0.62 | 0.56 |
| 2 | 159598188 | rs6750583 | -0.013 | -0.399 | 0.496 | -2.13 | 0.002 | -0.008 | -0.008 | 0 | 0.25 | 0.27 | 0.21 | 0.21 | 0.3 | 0.36 |
| 2 | 159599779 | rs1996274 | 0.366 | -0.406 | 0.052 | -2.082 | -0.01 | 0.133 | 0.133 | 0 | 0.36 | 0.11 | 0.23 | 0.12 | 0.32 | 0.19 |
| 2 | 159602186 | rs6705792 | -1.339 | -0.273 | 0.472 | -1.935 | 0.01 | 0.001 | 0.001 | 0 | 0.7 | 0.61 | 0.46 | 0.32 | 0.65 | 0.57 |
| 2 | 159606437 | rs1040166 | 0.439 | -0.591 | 0.38 | -2.321 | -0.017 | -0.027 | -0.027 | 0 | 0.64 | 0.56 | 0.71 | 0.75 | 0.69 | 0.59 |
| 2 | 159613837 | rs6729674 | 0.064 | -0.75 | 0.526 | -2.488 | 0.012 | 0.003 | 0.003 | 0 | 0.78 | 0.81 | 0.84 | 0.77 | 0.85 | 0.72 |
| 2 | 159616934 | rs10497202 | 0.465 | -0.722 | 0.859 | -2.494 | 0.007 | -0.055 | -0.055 | 0 | 0.11 | 0.2 | 0.12 | 0 | 0.16 | 0.05 |
| 2 | 159622208 | rs1515926 | 1.606 | -0.663 | 0.544 | -2.639 | -0.017 | 0.018 | 0.018 | 0.071 | 0.3 | 0.21 | 0.51 | 0.75 | 0.36 | 0.54 |
| 2 | 159628474 | rs4665004 | 0.776 | -0.786 | 0.406 | -2.585 | 0.009 | -0.006 | -0.006 | 0.109 | 0.07 | 0.06 | 0.06 | 0.12 | 0.03 | 0.05 |
| 2 | 159630705 | rs11689479 | 0.221 | -0.777 | 0.704 | -2.753 | -0.003 | 0.014 | 0.014 | 0.898 | 0.47 | 0.45 | 0.67 | 0.85 | 0.54 | 0.63 |
| 2 | 159632662 | rs4665006 | -0.263 | -0.727 | 0.631 | -2.692 | -0.012 | -0.017 | -0.017 | 0 | 0.83 | 0.73 | 0.87 | 0.88 | 0.88 | 0.78 |
| 2 | 159639345 | rs2356507 | 1.568 | -0.713 | 0.232 | -2.735 | -0.023 | 0.033 | 0.033 | 0 | 0.32 | 0.21 | 0.51 | 0.78 | 0.36 | 0.5 |
| 2 | 159644006 | rs2356508 | -0.538 | -0.777 | 0.307 | -2.598 | -0.001 | 0 | 0 | 0 | 0.9 | 0.92 | 0.91 | 0.86 | 0.92 | 0.78 |

| | | | | | | | | | | | | | | | | |
|---|---|---|---|---|---|---|---|---|---|---|---|---|---|---|---|---|
| 2 | 159645941 | rs4254463 | 0.722 | -0.847 | 0.247 | -2.727 | -0.005 | 0.106 | 0.106 | 0 | 0.46 | 0.31 | 0.64 | 0.79 | 0.55 | 0.54 |
| 2 | 159649340 | rs1515924 | 1.02 | -0.949 | -0.302 | -2.87 | -0.11 | 0.13 | 0.13 | 0.202 | 0.22 | 0.02 | 0.14 | 0.03 | 0.3 | 0.12 |
| 2 | 159650386 | rs4561615 | 0.99 | -0.945 | -0.152 | -2.864 | -0.005 | 0.005 | 0.005 | 0 | 0.11 | 0.04 | 0.11 | 0 | 0.16 | 0.04 |
| 2 | 159651734 | rs925781 | -0.112 | -0.891 | 0.24 | -2.896 | 0.011 | -0.05 | -0.05 | 0 | 0.64 | 0.59 | 0.75 | 0.83 | 0.63 | 0.66 |
| 2 | 159660870 | rs17810398 | 0.988 | -0.802 | 0.116 | -2.758 | -0.03 | 0.061 | 0.061 | 0 | 0.1 | 0.02 | 0.1 | 0 | 0.16 | 0.04 |
| 2 | 159661485 | rs3914570 | 1.248 | -0.879 | 0.329 | -2.813 | -0.001 | -0.015 | -0.015 | 0.109 | 0.43 | 0.47 | 0.61 | 0.83 | 0.43 | 0.59 |
| 2 | 159663599 | rs9869 | 0.954 | -0.933 | 0.279 | -2.713 | -0.003 | -0.038 | -0.038 | 0.169 | 0.45 | 0.54 | 0.62 | 0.8 | 0.43 | 0.58 |
| 2 | 159664703 | rs17810584 | 0.853 | -0.605 | 0.082 | -2.549 | -0.003 | -0.009 | -0.009 | 0 | 0.1 | 0.06 | 0.1 | 0 | 0.16 | 0.04 |
| 2 | 159667328 | rs6716178 | 1.442 | -0.885 | -0.479 | -2.809 | -0.036 | 0.011 | 0.011 | 0 | 0.31 | 0.11 | 0.48 | 0.73 | 0.3 | 0.43 |
| 2 | 159667731 | rs908402 | -1.019 | -0.733 | -0.017 | -2.712 | -0.004 | 0.014 | 0.014 | 0 | 0.9 | 0.88 | 0.91 | 0.84 | 0.92 | 0.76 |
| 2 | 159668758 | rs4665011 | 0.643 | -0.857 | 0.168 | -2.86 | -0.017 | -0.029 | -0.029 | 0 | 0.63 | 0.5 | 0.68 | 0.77 | 0.64 | 0.6 |
| 2 | 159669573 | rs6437203 | 0.139 | -0.617 | -0.13 | -2.71 | -0.001 | -0.009 | -0.009 | 0 | 0.79 | 0.77 | 0.8 | 0.84 | 0.75 | 0.72 |
| 2 | 159670449 | rs16843372 | -0.995 | -0.172 | -0.422 | -2.895 | 0 | -0.006 | -0.006 | 0 | 0.54 | 0.56 | 0.44 | 0.12 | 0.6 | 0.33 |
| 2 | 159671992 | rs17810816 | 1.008 | -0.115 | -0.756 | -2.753 | 0 | 0.08 | 0.08 | 0 | 0.18 | 0.04 | 0.12 | 0 | 0.19 | 0.07 |
| 2 | 159674805 | rs7603880 | -0.929 | 0.276 | -0.856 | -2.746 | -0.004 | -0.04 | -0.04 | 1.051 | 0.57 | 0.53 | 0.46 | 0.13 | 0.62 | 0.39 |
| 2 | 159680230 | rs10196949 | -0.753 | 0.254 | -0.966 | -2.819 | 0.035 | 0.163 | 0.163 | 0 | 0.53 | 0.26 | 0.42 | 0.11 | 0.62 | 0.39 |
| 2 | 159689830 | rs2356510 | -0.745 | 0.251 | -0.984 | -2.82 | 0.036 | 0.164 | 0.164 | 0 | 0.53 | 0.26 | 0.42 | 0.11 | 0.62 | 0.39 |
| 2 | 159700075 | rs12694971 | -0.388 | 0.008 | -1.357 | -2.633 | 0.021 | 0.084 | 0.084 | 0.026 | 0.38 | 0.24 | 0.29 | 0.08 | 0.34 | 0.24 |
| 2 | 159723506 | rs6757197 | -0.482 | 0.1 | -1.528 | -2.62 | 0.004 | -0.027 | -0.027 | 0 | 0.47 | 0.48 | 0.36 | 0.08 | 0.6 | 0.32 |
| 2 | 159726920 | rs11687201 | -0.585 | 0.784 | -1.256 | -1.999 | 0.016 | 0.138 | 0.138 | 0 | 0.47 | 0.23 | 0.37 | 0.1 | 0.57 | 0.5 |
| 2 | 159735081 | rs13033775 | -0.249 | -0.086 | -1.146 | -1.779 | 0.01 | -0.002 | -0.002 | 0 | 0.34 | 0.29 | 0.21 | 0.06 | 0.27 | 0.2 |
| 2 | 159736419 | rs2884003 | -0.388 | -0.472 | -1.111 | -1.783 | 0.005 | -0.02 | -0.02 | 0 | 0.4 | 0.37 | 0.23 | 0.06 | 0.28 | 0.28 |
| 2 | 159755842 | rs7587737 | -0.055 | -0.071 | -1.095 | -1.946 | -0.001 | 0.105 | 0.105 | 1.206 | 0.45 | 0.71 | 0.33 | 0.08 | 0.38 | 0.29 |
| 2 | 159760442 | rs11685166 | 0.885 | -0.275 | -0.858 | -2.17 | -0.002 | -0.026 | -0.026 | 0.117 | 0.24 | 0.26 | 0.2 | 0.02 | 0.31 | 0.23 |
| 2 | 159760687 | rs6719412 | -0.77 | -0.803 | -0.797 | -2.624 | 0.008 | 0.017 | 0.017 | 0.07 | 0.5 | 0.59 | 0.37 | 0.07 | 0.52 | 0.35 |
| 2 | 159761682 | rs17716892 | 1.694 | -0.721 | -0.963 | -2.585 | 0.015 | -0.039 | -0.039 | 0 | 0.21 | 0.26 | 0.2 | 0.02 | 0.31 | 0.23 |
| 2 | 159762132 | rs17727768 | 0.69 | -0.817 | -1.129 | -2.609 | 0.011 | 0.074 | 0.074 | 0 | 0.23 | 0.06 | 0.13 | 0.06 | 0.2 | 0.11 |
| 2 | 159764389 | rs1522699 | 0.287 | -0.661 | -0.804 | -2.415 | -0.05 | 0.182 | 0.182 | 0 | 0.43 | 0.17 | 0.32 | 0.07 | 0.48 | 0.29 |
| 2 | 159770970 | rs939998 | 0.243 | -0.979 | -0.939 | -1.851 | 0 | 0.007 | 0.007 | 0 | 0.31 | 0.39 | 0.41 | 0.68 | 0.34 | 0.33 |
| 2 | 159772066 | rs4665018 | -1.383 | -0.947 | -0.911 | -1.512 | 0.031 | -0.005 | -0.005 | 0.501 | 0.79 | 0.93 | 0.77 | 0.79 | 0.86 | 0.69 |
| 2 | 159778783 | rs11686680 | 0.582 | -0.834 | -0.618 | -1.384 | -0.002 | -0.003 | -0.003 | 0 | 0.33 | 0.31 | 0.42 | 0.63 | 0.38 | 0.32 |
| 2 | 159785388 | rs2037506 | 0.11 | -1.182 | -0.007 | -1.139 | 0.073 | 0.139 | 0.139 | 0.094 | 0.06 | 0.38 | 0.05 | 0.09 | 0.03 | 0.06 |
| 2 | 159787793 | rs998896 | 0.059 | -1.24 | -0.197 | -1.288 | -0.035 | -0.004 | -0.004 | 0 | 0.68 | 0.46 | 0.71 | 0.7 | 0.81 | 0.58 |
| 2 | 159792033 | rs11894687 | 0.441 | -1.018 | 0.051 | -1.356 | -0.008 | -0.033 | -0.033 | 0.101 | 0.07 | 0.32 | 0.05 | 0 | 0.02 | 0.02 |
| 2 | 159809860 | rs6437215 | 0.223 | -1.037 | -0.062 | -1.417 | -0.005 | -0.022 | -0.022 | 0 | 0.11 | 0.31 | 0.09 | 0.05 | 0.11 | 0.1 |
| 2 | 159810691 | rs7565830 | 0.407 | -1.043 | -0.274 | -1.493 | -0.008 | 0.02 | 0.02 | 0 | 0.27 | 0.33 | 0.51 | 0.62 | 0.29 | 0.27 |
| 2 | 159836227 | rs842067 | 0.051 | -1.083 | -0.356 | -1.39 | 0.006 | -0.016 | -0.016 | 0 | 0.32 | 0.25 | 0.28 | 0.49 | 0.27 | 0.49 |
| 2 | 159840743 | rs842064 | 0.826 | -1.171 | -0.285 | -1.414 | -0.005 | -0.027 | -0.027 | 0 | 0.24 | 0.15 | 0.14 | 0.33 | 0.23 | 0.49 |
| 2 | 159851549 | rs16843534 | 1.167 | -1.178 | -0.114 | -1.301 | -0.006 | 0.005 | 0.005 | 0 | 0.07 | 0.09 | 0.14 | 0.17 | 0.04 | 0.02 |
| 2 | 159857642 | rs16843536 | 0.881 | -1.18 | -0.11 | -1.302 | -0.019 | 0.025 | 0.025 | 0 | 0.07 | 0.13 | 0.14 | 0.17 | 0.04 | 0.02 |
| 2 | 159859820 | rs6716681 | -0.224 | -1.267 | 0.003 | -1.371 | -0.019 | 0.032 | 0.032 | 0 | 0.33 | 0.53 | 0.31 | 0.5 | 0.27 | 0.51 |
| 2 | 159865053 | rs1365907 | -0.883 | -1.374 | -0.225 | -1.281 | 0.003 | -0.003 | -0.003 | 0 | 0.43 | 0.6 | 0.37 | 0.56 | 0.37 | 0.6 |
| 2 | 159870038 | rs6735689 | 0.701 | -1.371 | -0.232 | -1.233 | -0.023 | 0.026 | 0.026 | 0 | 0.08 | 0.14 | 0.15 | 0.17 | 0.04 | 0.02 |
| 2 | 159886836 | rs264604 | 1.361 | -1.185 | -0.415 | -1.182 | 0.013 | -0.029 | -0.029 | 0 | 0.19 | 0.11 | 0.1 | 0.33 | 0.14 | 0.45 |
| 2 | 159900278 | rs447106 | -0.813 | -1.354 | -0.412 | -1.304 | -0.005 | 0.063 | 0.063 | 0 | 0.73 | 0.65 | 0.62 | 0.56 | 0.72 | 0.72 |
| 2 | 159908922 | rs17812009 | 2.817 | -1.029 | -0.863 | -1.491 | -0.098 | 0.117 | 0.117 | 0.046 | 0.18 | 0.02 | 0.13 | 0 | 0.25 | 0.04 |
| 2 | 159913886 | rs16843639 | -0.731 | -2.011 | -0.131 | -0.953 | -0.04 | 0.024 | 0.024 | 0.402 | 0.17 | 0.45 | 0.23 | 0.23 | 0.15 | 0.11 |
| 2 | 159926849 | rs264653 | -0.273 | -1.846 | -0.148 | -0.942 | -0.006 | 0.025 | 0.025 | 0 | 0.71 | 0.67 | 0.61 | 0.56 | 0.72 | 0.72 |
| 2 | 159933898 | rs16843649 | 0.855 | -1.995 | -0.356 | -0.882 | -0.014 | -0.154 | -0.154 | 0 | 0.2 | 0.01 | 0.13 | 0.39 | 0.21 | 0.22 |
| 2 | 159937019 | rs2032757 | 0.797 | -1.89 | -0.186 | -1.03 | -0.007 | -0.036 | -0.036 | 0.023 | 0.23 | 0.13 | 0.14 | 0.33 | 0.22 | 0.4 |
| 2 | 159948101 | rs264626 | -0.765 | -1.71 | 0.207 | -1.19 | -0.009 | 0.055 | 0.055 | 0.862 | 0.09 | 0.37 | 0.16 | 0.16 | 0.04 | 0.13 |
| 2 | 159950898 | rs10497204 | 1.422 | -1.668 | -0.31 | -1.276 | -0.078 | 0.141 | 0.141 | 0.047 | 0.37 | 0.14 | 0.31 | 0.06 | 0.45 | 0.16 |
| 2 | 159950952 | rs169073 | -0.251 | -1.104 | 0.346 | -0.622 | -0.009 | 0.003 | 0.003 | 0 | 0.07 | 0.1 | 0.14 | 0.16 | 0.05 | 0.11 |
| 2 | 159953466 | rs10167864 | 0.81 | -1.099 | 0.252 | -0.609 | -0.017 | -0.02 | -0.02 | 0 | 0.77 | 0.46 | 0.54 | 0.75 | 0.76 | 0.76 |
| 2 | 159955575 | rs10198413 | 0.097 | -0.753 | -0.164 | -0.747 | -0.001 | -0.001 | -0.001 | 0.21 | 0.24 | 0.21 | 0.18 | 0.3 | 0.25 | 0.42 |
| 2 | 159957945 | rs4665022 | -0.002 | -0.719 | -0.151 | -0.737 | 0 | -0.01 | -0.01 | 0 | 0.29 | 0.22 | 0.31 | 0.35 | 0.29 | 0.42 |
| 2 | 159958620 | rs264639 | -0.063 | -0.881 | 0.267 | -0.736 | -0.005 | 0.199 | 0.199 | 0 | 0.09 | 0.17 | 0.06 | 0.59 | 0.1 | 0.22 |
| 2 | 159964552 | rs448513 | -0.586 | -1.097 | -0.224 | -1.085 | -0.006 | -0.001 | -0.001 | 0.092 | 0.36 | 0.59 | 0.43 | 0.64 | 0.35 | 0.45 |
| 2 | 159964866 | rs17207648 | 0.888 | -1.135 | -0.151 | -1.14 | 0.019 | 0.04 | 0.04 | 0 | 0.22 | 0.15 | 0.12 | 0.04 | 0.2 | 0.2 |
| 2 | 159965140 | rs6714340 | 0.832 | -1.128 | -0.211 | -1.137 | 0.008 | -0.003 | -0.003 | 0 | 0.22 | 0.18 | 0.13 | 0.04 | 0.2 | 0.2 |
| 2 | 159970939 | rs13021373 | 1.545 | -0.973 | -1.046 | -1.004 | -0.158 | 0.217 | 0.217 | 0.014 | 0.3 | 0.02 | 0.23 | 0.01 | 0.34 | 0.13 |
| 2 | 159971363 | rs264581 | -0.48 | -1.133 | -0.233 | -0.268 | -0.036 | 0.018 | 0.018 | 0 | 0.32 | 0.45 | 0.19 | 0.62 | 0.3 | 0.43 |
| 2 | 159985511 | rs6724143 | 0.339 | -1.015 | -0.521 | -0.641 | 0.012 | -0.001 | -0.001 | 0 | 0.25 | 0.21 | 0.13 | 0.03 | 0.22 | 0.25 |
| 2 | 159986806 | rs3732285 | 0.041 | -0.959 | -0.287 | -0.45 | 0.016 | 0.029 | 0.029 | 0 | 0.35 | 0.29 | 0.19 | 0.12 | 0.3 | 0.27 |
| 2 | 159990664 | rs6754934 | 0.236 | -0.97 | -0.491 | -0.158 | 0.007 | -0.027 | -0.027 | 0 | 0.23 | 0.17 | 0.27 | 0.42 | 0.22 | 0.41 |
| 2 | 159994048 | rs1421385 | 0.956 | -1.056 | -0.216 | -0.242 | -0.035 | -0.042 | -0.042 | 0.041 | 0.33 | 0.24 | 0.5 | 0.46 | 0.46 | 0.26 |
| 2 | 160002919 | rs4413115 | 0.687 | -0.947 | -0.343 | -0.305 | 0.007 | 0.271 | 0.271 | 0 | 0.07 | 0.18 | 0.26 | 0.46 | 0.08 | 0.17 |
| 2 | 160005259 | rs12463769 | 1.179 | -1.006 | -0.544 | -0.479 | -0.007 | 0.034 | 0.034 | 0.067 | 0.12 | 0.17 | 0.2 | 0.3 | 0.07 | 0.23 |
| 2 | 160006143 | rs7586743 | 1.196 | -0.746 | -0.905 | -0.782 | -0.101 | 0.124 | 0.124 | 0 | 0.29 | 0.07 | 0.25 | 0.06 | 0.37 | 0.1 |

| | | | | | | | | | | | | | | | | |
|---|---|---|---|---|---|---|---|---|---|---|---|---|---|---|---|---|
| 2 | 160016914 | rs10200189 | 0.992 | -0.753 | -0.862 | -0.77 | -0.098 | 0.117 | 0.117 | 0 | 0.31 | 0.1 | 0.27 | 0.07 | 0.4 | 0.13 |
| 2 | 160032007 | rs6721087 | -0.992 | -0.775 | -0.668 | -0.781 | -0.045 | 0.055 | 0.055 | 0 | 0.68 | 0.83 | 0.72 | 0.93 | 0.59 | 0.84 |
| 2 | 160035195 | rs12988603 | 1.207 | -0.715 | -0.858 | -0.754 | -0.101 | 0.123 | 0.123 | 0 | 0.28 | 0.07 | 0.25 | 0.06 | 0.37 | 0.1 |
| 2 | 160041993 | rs17494000 | -0.258 | -1.031 | -0.212 | -0.095 | -0.001 | 0 | 0 | 0 | 0.07 | 0.06 | 0.04 | 0.07 | 0.06 | 0.06 |
| 2 | 160042430 | rs13389867 | 0.169 | -1.147 | -0.207 | -0.153 | 0.034 | 0.071 | 0.071 | 0.35 | 0.33 | 0.22 | 0.15 | 0.05 | 0.26 | 0.25 |
| 2 | 160048036 | rs10191093 | -0.036 | -0.52 | -0.425 | 0.28 | -0.009 | 0.017 | 0.017 | 0 | 0.05 | 0.12 | 0.03 | 0.06 | 0.03 | 0.05 |
| 2 | 160054932 | rs6432515 | -0.436 | -0.472 | -0.609 | 0.273 | 0.026 | 0.03 | 0.03 | 0 | 0.51 | 0.61 | 0.39 | 0.33 | 0.37 | 0.52 |
| 2 | 160058776 | rs10175176 | -0.225 | -0.248 | -0.543 | 0.727 | -0.001 | 0.005 | 0.005 | 0.725 | 0.28 | 0.29 | 0.19 | 0.46 | 0.31 | 0.31 |
| 2 | 160064299 | rs6709296 | -0.782 | -0.774 | -0.788 | -0.016 | 0.028 | 0.133 | 0.133 | 0.205 | 0.54 | 0.68 | 0.66 | 0.87 | 0.45 | 0.77 |
| 2 | 160065027 | rs12692542 | -0.326 | -1.329 | -0.296 | 0.572 | -0.05 | 0.017 | 0.017 | 0.133 | 0.35 | 0.36 | 0.17 | 0.53 | 0.26 | 0.52 |
| 2 | 160065816 | rs4665040 | -0.478 | -1.53 | -0.496 | 0.329 | -0.033 | 0 | 0 | 0.635 | 0.9 | 0.71 | 0.94 | 0.91 | 0.96 | 0.95 |
| 2 | 160069021 | rs2159874 | -0.123 | -1.803 | 0.05 | 0.702 | -0.008 | 0.006 | 0.006 | 0 | 0.37 | 0.51 | 0.6 | 0.56 | 0.33 | 0.47 |
| 2 | 160071704 | rs12465443 | -0.061 | -1.996 | -0.203 | 0.825 | -0.014 | -0.021 | -0.021 | 0 | 0.36 | 0.11 | 0.57 | 0.37 | 0.35 | 0.46 |
| 2 | 160072181 | rs6432518 | 0.934 | -2.088 | 0.102 | 0.815 | 0 | -0.014 | -0.014 | 0.292 | 0.47 | 0.49 | 0.35 | 0.14 | 0.55 | 0.23 |
| 2 | 160072555 | rs9287784 | -0.334 | -2.087 | 0.099 | 0.839 | -0.074 | 0.276 | 0.276 | 0.231 | 0.17 | 0.4 | 0.08 | 0.49 | 0.09 | 0.31 |
| 2 | 160073035 | rs4665041 | 0.277 | -2.036 | -0.02 | 0.816 | -0.05 | -0.046 | -0.046 | 0 | 0.33 | 0.04 | 0.55 | 0.31 | 0.34 | 0.39 |
| 2 | 160078959 | rs12692543 | -0.043 | -2.057 | 0.125 | 0.866 | -0.014 | -0.021 | -0.021 | 0 | 0.36 | 0.11 | 0.56 | 0.37 | 0.35 | 0.46 |
| 2 | 160082724 | rs11685555 | 1.658 | -1.309 | -0.372 | 0.382 | -0.061 | 0.004 | 0.004 | 0.423 | 0.11 | 0 | 0.07 | 0.04 | 0.15 | 0.08 |
| 2 | 160084263 | rs2042495 | 0.053 | -1.494 | -0.154 | 0.321 | -0.065 | 0.046 | 0.046 | 0.013 | 0.78 | 0.65 | 0.9 | 0.66 | 0.89 | 0.72 |
| 2 | 160084473 | rs2288110 | -0.06 | -1.483 | -0.164 | 0.326 | 0.006 | -0.002 | -0.002 | 0 | 0.4 | 0.35 | 0.6 | 0.62 | 0.37 | 0.53 |
| 2 | 160086692 | rs3821296 | -0.06 | -1.483 | -0.164 | 0.326 | 0.006 | -0.002 | -0.002 | 0 | 0.4 | 0.35 | 0.6 | 0.62 | 0.37 | 0.53 |
| 2 | 160087535 | rs3821301 | 0.059 | -1.488 | -0.164 | 0.334 | -0.063 | 0.044 | 0.044 | 0.013 | 0.78 | 0.65 | 0.9 | 0.66 | 0.89 | 0.72 |
| 2 | 160088793 | rs1065594 | 0.818 | -1.472 | -0.218 | 0.339 | -0.062 | -0.076 | -0.076 | 0 | 0.33 | 0.02 | 0.55 | 0.33 | 0.34 | 0.39 |
| 2 | 160096387 | rs11690351 | 1.208 | -1.454 | -0.116 | 0.267 | -0.023 | 0.103 | 0.103 | 0 | 0.37 | 0.18 | 0.3 | 0.05 | 0.5 | 0.18 |
| 2 | 160107072 | rs7603170 | 1.108 | -1.443 | -0.181 | 0.272 | -0.006 | 0.02 | 0.02 | 0.19 | 0.38 | 0.3 | 0.3 | 0.05 | 0.51 | 0.18 |
| 2 | 160109798 | rs6723106 | 0.841 | -1.447 | -0.163 | 0.301 | 0 | 0.002 | 0.002 | 0.262 | 0.42 | 0.54 | 0.34 | 0.27 | 0.52 | 0.26 |
| 2 | 160112881 | rs7591849 | 0.287 | -1.617 | -0.083 | 0.212 | 0.053 | 0.115 | 0.115 | 0.156 | 0.51 | 0.74 | 0.37 | 0.27 | 0.54 | 0.32 |
| 2 | 160120618 | rs6743012 | 0.81 | -1.529 | -0.22 | 0.238 | -0.025 | -0.038 | -0.038 | 0 | 0.07 | 0.13 | 0.02 | 0 | 0.02 | 0.06 |
| 2 | 160123080 | rs6761921 | 0.883 | -1.505 | -0.492 | 0.271 | -0.063 | -0.077 | -0.077 | 0 | 0.32 | 0.02 | 0.54 | 0.33 | 0.34 | 0.38 |
| 2 | 160125205 | rs17494490 | 2.307 | -1.445 | -0.496 | 0.194 | -0.138 | 0.177 | 0.177 | 0 | 0.25 | 0.02 | 0.21 | 0.01 | 0.35 | 0.1 |
| 2 | 160132089 | rs16843852 | -0.68 | -1.946 | -0.014 | 0.597 | -0.051 | 0.13 | 0.13 | 0.064 | 0.13 | 0.44 | 0.07 | 0.23 | 0.03 | 0.14 |
| 2 | 160137738 | rs52848 | -0.161 | -1.86 | -0.213 | 0.301 | 0.001 | -0.053 | -0.053 | 0.03 | 0.39 | 0.15 | 0.6 | 0.43 | 0.39 | 0.44 |
| 2 | 160146177 | rs13026048 | 1.299 | -1.964 | -0.065 | 0.367 | -0.023 | 0.005 | 0.005 | 0.572 | 0.7 | 0.3 | 0.85 | 0.38 | 0.86 | 0.57 |
| 2 | 160147145 | rs174261 | 0.438 | -1.49 | 0.199 | -0.061 | 0.03 | 0.051 | 0.051 | 0 | 0.51 | 0.4 | 0.38 | 0.13 | 0.58 | 0.29 |
| 2 | 160148610 | rs10169221 | 0.471 | -1.528 | -0.27 | 0.122 | -0.011 | 0.011 | 0.011 | 0 | 0.22 | 0.01 | 0.11 | 0.13 | 0.17 | 0.22 |
| 2 | 160158746 | rs174253 | -0.704 | -1.824 | 0.385 | 0.301 | 0.003 | 0.001 | 0.001 | 0 | 0.42 | 0.45 | 0.56 | 0.66 | 0.33 | 0.62 |
| 2 | 160163375 | rs11676419 | 1.5 | -1.864 | 0.326 | 0.236 | -0.039 | 0.075 | 0.075 | 0 | 0.37 | 0.11 | 0.29 | 0.2 | 0.52 | 0.22 |
| 2 | 160164187 | rs174244 | 0.029 | -1.468 | 1.5 | 0.917 | -0.036 | 0.047 | 0.047 | 0 | 0.78 | 0.56 | 0.85 | 0.71 | 0.84 | 0.76 |
| 2 | 160164716 | rs174240 | -0.239 | -1.464 | 1.463 | 0.892 | -0.031 | -0.026 | -0.026 | 0.25 | 0.79 | 0.63 | 0.87 | 0.86 | 0.85 | 0.84 |
| 2 | 160182302 | rs415793 | -0.87 | -1.459 | 1.502 | 0.898 | 0.005 | 0.004 | 0.004 | 0 | 0.3 | 0.33 | 0.18 | 0.23 | 0.21 | 0.26 |
| 2 | 160185226 | rs11688519 | 0.879 | -1.415 | 1.462 | 0.907 | 0.005 | -0.002 | -0.002 | 0 | 0.67 | 0.34 | 0.78 | 0.61 | 0.77 | 0.66 |
| 2 | 160189755 | rs16843897 | -1.351 | -1.431 | 1.594 | 0.943 | -0.006 | 0.105 | 0.105 | 0 | 0.08 | 0.3 | 0.05 | 0.15 | 0.02 | 0.13 |
| 2 | 160200749 | rs9678906 | -0.828 | -1.389 | 0.555 | 0.818 | 0.002 | 0.002 | 0.002 | 0 | 0.3 | 0.33 | 0.19 | 0.24 | 0.21 | 0.26 |
| 2 | 160201955 | rs11884022 | -0.921 | -1.271 | 0.598 | 0.938 | -0.029 | 0.026 | 0.026 | 0 | 0.11 | 0.15 | 0.06 | 0.17 | 0.06 | 0.14 |
| 2 | 160203810 | rs10172339 | -0.651 | -1.402 | -0.463 | 1.126 | 0.038 | -0.053 | -0.053 | 0.013 | 0.3 | 0.15 | 0.18 | 0.46 | 0.22 | 0.46 |
| 2 | 160220164 | rs3771686 | 0.46 | -1.542 | 0.053 | 1.07 | -0.04 | 0.066 | 0.066 | 0 | 0.73 | 0.43 | 0.51 | 0.56 | 0.8 | 0.7 |
| 2 | 160220745 | rs10181248 | 0.992 | -1.956 | -0.171 | 0.971 | 0.024 | -0.017 | -0.017 | 0 | 0.61 | 0.3 | 0.47 | 0.48 | 0.76 | 0.6 |
| 2 | 160224542 | rs11686677 | 0.806 | -1.982 | -0.023 | 1.197 | -0.025 | -0.012 | -0.012 | 0 | 0.64 | 0.36 | 0.48 | 0.67 | 0.79 | 0.63 |
| 2 | 160227307 | rs34459426 | -0.022 | -1.782 | -0.793 | 0.901 | -0.001 | -0.003 | -0.003 | 0 | 0.32 | 0.27 | 0.21 | 0.39 | 0.32 | 0.48 |
| 2 | 160232059 | rs3771698 | -0.032 | -2.001 | -0.712 | 0.737 | 0.05 | 0.049 | 0.049 | 0 | 0.21 | 0.11 | 0.06 | 0.1 | 0.08 | 0.18 |
| 2 | 160233383 | rs768072 | 0.568 | -1.982 | -0.724 | 0.766 | -0.025 | -0.012 | -0.012 | 0 | 0.64 | 0.36 | 0.48 | 0.68 | 0.79 | 0.64 |
| 2 | 160238292 | rs13005337 | 0.675 | -1.949 | -0.837 | 0.724 | 0.027 | 0.025 | 0.025 | 0 | 0.59 | 0.31 | 0.45 | 0.68 | 0.74 | 0.59 |
| 2 | 160239378 | rs2302927 | 1.865 | -1.74 | -1.001 | 0.824 | -0.182 | -0.05 | -0.05 | 0 | 0.32 | 0.03 | 0.27 | 0.29 | 0.48 | 0.16 |
| 2 | 160242424 | rs3771706 | -1.215 | -1.682 | -0.388 | 0.857 | -0.022 | 0.027 | 0.027 | 0 | 0.87 | 0.7 | 0.57 | 0.78 | 0.88 | 0.82 |
| 2 | 160243726 | rs3771709 | -0.791 | -1.717 | -0.536 | 0.857 | -0.04 | 0.064 | 0.064 | 0 | 0.84 | 0.47 | 0.54 | 0.71 | 0.88 | 0.74 |
| 2 | 160244080 | rs3771711 | -0.756 | -1.726 | -0.543 | 0.852 | -0.041 | 0.061 | 0.061 | 0 | 0.84 | 0.43 | 0.54 | 0.71 | 0.88 | 0.74 |
| 2 | 160245001 | rs12692547 | -0.369 | -1.886 | -0.811 | 0.722 | -0.002 | -0.027 | -0.027 | 3.779 | 0.33 | 0.22 | 0.2 | 0.41 | 0.32 | 0.49 |
| 2 | 160247075 | rs7560385 | -0.363 | -1.888 | -0.838 | 0.721 | -0.002 | -0.027 | -0.027 | 0.697 | 0.33 | 0.22 | 0.2 | 0.41 | 0.32 | 0.49 |
| 2 | 160259258 | rs13028985 | 1.557 | -1.951 | -1.093 | 0.567 | -0.104 | -0.002 | -0.002 | 0 | 0.39 | 0.12 | 0.29 | 0.29 | 0.5 | 0.18 |
| 2 | 160262693 | rs2032777 | 0.058 | -1.979 | -0.802 | 0.771 | -0.037 | -0.014 | -0.014 | 0.912 | 0.73 | 0.46 | 0.49 | 0.72 | 0.8 | 0.75 |
| 2 | 160269937 | rs6730764 | -0.313 | -1.956 | -0.816 | 0.738 | -0.017 | 0.016 | 0.016 | 0 | 0.14 | 0.22 | 0.05 | 0.23 | 0.07 | 0.13 |
| 2 | 160288062 | rs10180868 | 0.568 | -1.91 | -0.996 | 0.764 | 0.01 | -0.002 | -0.002 | 0 | 0.57 | 0.34 | 0.36 | 0.41 | 0.58 | 0.38 |
| 2 | 160290519 | rs10188117 | 0.411 | -1.967 | -1.07 | 0.353 | 0.054 | 0.053 | 0.053 | 0 | 0.24 | 0.17 | 0.07 | 0.11 | 0.11 | 0.23 |
| 2 | 160291593 | rs10195878 | 0.641 | -1.625 | -1.078 | 0.348 | 0.001 | 0.002 | 0.002 | 0.179 | 0.55 | 0.25 | 0.35 | 0.41 | 0.57 | 0.38 |
| 2 | 160294842 | rs3213790 | 0.389 | -1.72 | -1.053 | 0.341 | 0.053 | 0.054 | 0.054 | 0 | 0.24 | 0.17 | 0.07 | 0.11 | 0.11 | 0.23 |
| 2 | 160298123 | rs2193923 | 1.859 | -1.559 | -1.063 | 0.272 | -0.108 | -0.069 | -0.069 | 0.348 | 0.32 | 0.12 | 0.28 | 0.39 | 0.47 | 0.22 |
| 2 | 160310246 | rs10202670 | 0.57 | -1.516 | -0.95 | 0.132 | 0.045 | 0.145 | 0.145 | 0 | 0.22 | 0.03 | 0.06 | 0.01 | 0.12 | 0.15 |
| 2 | 160315563 | rs6749118 | 0.335 | -1.6 | -0.839 | 0.088 | -0.01 | -0.03 | -0.03 | 0 | 0.1 | 0.13 | 0.05 | 0 | 0.05 | 0.06 |
| 2 | 160318403 | rs12471541 | 0.734 | -1.675 | -0.871 | 0.139 | 0.012 | 0 | 0 | 0 | 0.57 | 0.36 | 0.35 | 0.41 | 0.58 | 0.36 |

| | | | | | | | | | | | | | | | | |
|---|---|---|---|---|---|---|---|---|---|---|---|---|---|---|---|---|
| 2 | 160319338 | rs6736104 | 0.692 | -1.587 | -1.164 | 0.159 | -0.016 | -0.039 | -0.039 | 0 | 0.1 | 0.15 | 0.05 | 0 | 0.05 | 0.07 |
| 2 | 160321788 | rs7576462 | -0.508 | -1.316 | -0.88 | 0.333 | 0.001 | 0.009 | 0.009 | 0 | 0.34 | 0.35 | 0.58 | 0.59 | 0.37 | 0.56 |
| 2 | 160321840 | rs7564053 | 0.223 | -1.213 | -1.079 | 0.466 | 0.008 | 0.047 | 0.047 | 0 | 0.19 | 0.19 | 0.15 | 0.42 | 0.23 | 0.42 |
| 2 | 160322656 | rs16844045 | -0.466 | -1.304 | -1.488 | -0.097 | 0.06 | 0.064 | 0.064 | 0 | 0.32 | 0.2 | 0.11 | 0.01 | 0.17 | 0.22 |
| 2 | 160323691 | rs4665072 | 0.179 | 0.065 | -1.823 | 0.475 | -0.01 | 0.014 | 0.014 | 1.892 | 0.46 | 0.22 | 0.6 | 0.42 | 0.45 | 0.37 |
| 2 | 160323884 | rs964176 | 0.672 | 1.037 | -2.288 | 0.854 | -0.192 | -0.125 | -0.125 | 0 | 0.27 | 0.03 | 0.3 | 0.27 | 0.48 | 0.14 |
| 2 | 160325182 | rs10170119 | 0.035 | 0.202 | -1.169 | 1.028 | 0.019 | 0.096 | 0.096 | 0 | 0.09 | 0.23 | 0.09 | 0.2 | 0.09 | 0.1 |
| 2 | 160334363 | rs6725557 | -0.536 | 0.351 | -0.864 | 0.893 | 0.003 | 0.003 | 0.003 | 0 | 0.6 | 0.63 | 0.32 | 0.49 | 0.47 | 0.64 |
| 2 | 160363669 | rs10205689 | 0.5 | -0.206 | -0.673 | 0.51 | 0.013 | 0.018 | 0.018 | 0.342 | 0.37 | 0.3 | 0.16 | 0.28 | 0.28 | 0.37 |
| 2 | 160364747 | rs1425045 | 0.6 | -0.19 | -0.832 | 0.518 | 0.021 | 0.033 | 0.033 | 0.528 | 0.36 | 0.24 | 0.15 | 0.28 | 0.28 | 0.37 |
| 2 | 160368787 | rs1863218 | -0.168 | 0.092 | -0.985 | 0.457 | 0.005 | -0.001 | -0.001 | 0 | 0.48 | 0.47 | 0.41 | 0.47 | 0.56 | 0.42 |
| 2 | 160386260 | rs1030328 | 0.155 | 0.157 | -1.144 | 0.492 | -0.038 | -0.046 | -0.046 | 0 | 0.42 | 0.22 | 0.37 | 0.48 | 0.53 | 0.32 |
| 2 | 160386903 | rs11690478 | 0.327 | 0.049 | -1.015 | 0.652 | 0.005 | -0.037 | -0.037 | 0 | 0.17 | 0.06 | 0.11 | 0.2 | 0.16 | 0.18 |
| 2 | 160403400 | rs4665079 | -0.544 | 0.078 | -0.452 | 0.564 | 0 | 0.028 | 0.028 | 0 | 0.86 | 0.71 | 0.87 | 0.79 | 0.87 | 0.82 |
| 2 | 160405498 | rs10489997 | 0.982 | -0.009 | -1.147 | 0.371 | -0.159 | -0.025 | -0.025 | 0 | 0.32 | 0.03 | 0.51 | 0.25 | 0.44 | 0.17 |
| 2 | 160406890 | rs7585513 | 0.592 | -0.389 | -0.764 | 0.266 | -0.005 | 0 | 0 | 0 | 0.32 | 0.47 | 0.23 | 0.32 | 0.28 | 0.45 |
| 2 | 160413623 | rs6432535 | 0.946 | -0.327 | -0.753 | 0.292 | -0.005 | -0.03 | -0.03 | 0.441 | 0.05 | 0.13 | 0.03 | 0 | 0.03 | 0.1 |
| 2 | 160417561 | rs6755927 | -0.299 | -0.633 | -0.954 | 0.082 | 0.015 | -0.007 | -0.007 | 3.552 | 0.21 | 0.16 | 0.09 | 0 | 0.11 | 0.19 |
| 2 | 160428587 | rs11693108 | 0.112 | -0.327 | -0.994 | 0.259 | 0.006 | 0.112 | 0.112 | 0 | 0.5 | 0.72 | 0.42 | 0.75 | 0.49 | 0.65 |
| 2 | 160430127 | rs4665083 | 0.903 | -0.436 | -0.67 | 0.436 | -0.001 | 0.001 | 0.001 | 0 | 0.38 | 0.36 | 0.26 | 0.39 | 0.37 | 0.49 |
| 2 | 160432819 | rs17605928 | 0.786 | -0.097 | -0.949 | 0.482 | -0.109 | -0.137 | -0.137 | 0 | 0.34 | 0.06 | 0.49 | 0.44 | 0.39 | 0.22 |
| 2 | 160457075 | rs7563368 | 0.75 | -0.127 | -0.654 | 0.558 | -0.002 | 0 | 0 | 0 | 0.39 | 0.54 | 0.23 | 0.38 | 0.37 | 0.49 |
| 2 | 160491675 | rs357027 | -0.33 | -0.085 | -0.707 | 0.698 | -0.002 | 0.091 | 0.091 | 0 | 0.11 | 0.18 | 0.15 | 0.36 | 0.12 | 0.17 |
| 2 | 160499483 | rs4461223 | -1.546 | -0.071 | -0.925 | 0.721 | -0.008 | -0.005 | -0.005 | 4.008 | 0.94 | 0.97 | 0.95 | 0.78 | 0.93 | 0.9 |
| 2 | 160514739 | rs7594159 | -1.43 | -0.078 | -0.949 | 0.735 | -0.008 | -0.009 | -0.009 | 0.132 | 0.94 | 0.97 | 0.95 | 0.78 | 0.93 | 0.9 |
| 2 | 160540261 | rs7556888 | 0.205 | -0.144 | -1.151 | 0.561 | -0.119 | 0.106 | 0.106 | 0 | 0.44 | 0.07 | 0.59 | 0.26 | 0.49 | 0.25 |
| 2 | 160567537 | rs1549576 | -1.506 | -0.111 | -1.056 | 0.652 | -0.006 | -0.005 | -0.005 | 0 | 0.94 | 0.96 | 0.95 | 0.78 | 0.93 | 0.9 |
| 2 | 160569276 | rs10173538 | 0.83 | -0.195 | -0.74 | 0.482 | -0.002 | -0.001 | -0.001 | 0 | 0.39 | 0.54 | 0.23 | 0.38 | 0.37 | 0.48 |
| 2 | 160575226 | rs17813538 | 1.419 | -0.274 | -0.939 | 0.391 | -0.104 | 0.009 | 0.009 | 0 | 0.28 | 0.03 | 0.49 | 0.27 | 0.33 | 0.12 |
| 2 | 160580867 | rs7578537 | 0.665 | -0.292 | -0.642 | 0.376 | 0.042 | 0.001 | 0.001 | 2.319 | 0.07 | 0.39 | 0.05 | 0 | 0.03 | 0.1 |
| 2 | 160582184 | rs6761145 | 1.074 | -0.323 | -0.693 | 0.336 | -0.002 | 0 | 0 | 0 | 0.37 | 0.37 | 0.19 | 0.34 | 0.35 | 0.46 |
| 2 | 160584761 | rs7607143 | 0.487 | -0.333 | -0.716 | 0.33 | 0.016 | 0.024 | 0.024 | 0 | 0.44 | 0.76 | 0.25 | 0.34 | 0.38 | 0.56 |
| 2 | 160595127 | rs4477859 | -0.26 | -0.327 | -0.753 | 0.294 | -0.02 | 0.018 | 0.018 | 0 | 0.57 | 0.8 | 0.29 | 0.34 | 0.45 | 0.64 |
| 2 | 160601870 | rs5002908 | 0.006 | -0.317 | -0.766 | 0.311 | 0.001 | 0.06 | 0.06 | 0 | 0.88 | 0.83 | 0.83 | 0.63 | 0.86 | 0.81 |
| 2 | 160604452 | rs11555185 | 1.524 | -0.3 | -0.759 | 0.316 | -0.102 | 0.008 | 0.008 | 0 | 0.28 | 0.03 | 0.49 | 0.27 | 0.33 | 0.12 |
| 2 | 160604812 | rs1427329 | 0.989 | -0.426 | -0.622 | 0.356 | -0.002 | 0 | 0 | 0.003 | 0.37 | 0.37 | 0.2 | 0.34 | 0.35 | 0.46 |
| 2 | 160604936 | rs13024801 | 1.553 | -0.469 | -0.645 | 0.334 | 0.015 | 0.043 | 0.043 | 0.011 | 0.1 | 0.03 | 0.05 | 0 | 0.07 | 0.08 |
| 2 | 160605268 | rs1365803 | -1.566 | -0.448 | -0.644 | 0.381 | -0.012 | -0.003 | -0.003 | 1.422 | 0.94 | 0.97 | 0.93 | 0.73 | 0.92 | 0.9 |
| 2 | 160621924 | rs16844315 | 0.499 | -0.628 | -0.432 | 0.218 | -0.022 | -0.074 | -0.074 | 0.297 | 0.16 | 0.24 | 0.08 | 0 | 0.1 | 0.18 |
| 2 | 160623366 | rs4664301 | 0.765 | -0.464 | -0.403 | 0.385 | -0.002 | 0 | 0 | 0 | 0.37 | 0.37 | 0.19 | 0.34 | 0.35 | 0.46 |
| 2 | 160639970 | rs2114626 | 0.783 | -0.432 | -0.675 | 0.401 | -0.002 | 0 | 0 | 0 | 0.37 | 0.37 | 0.19 | 0.34 | 0.35 | 0.46 |
| 2 | 160647090 | rs3764968 | 0.838 | -0.407 | -0.78 | 0.414 | -0.019 | -0.029 | -0.029 | 0 | 0.35 | 0.11 | 0.16 | 0.34 | 0.35 | 0.46 |
| 2 | 160653720 | rs7588230 | -0.49 | -0.396 | -0.74 | 0.438 | -0.001 | -0.003 | -0.003 | 0 | 0.57 | 0.6 | 0.74 | 0.39 | 0.58 | 0.43 |
| 2 | 160658739 | rs1896270 | -1.333 | -0.36 | -0.739 | 0.625 | -0.006 | -0.016 | -0.016 | 0.275 | 0.93 | 0.96 | 0.93 | 0.73 | 0.93 | 0.89 |
| 2 | 160662058 | rs2042772 | 0.478 | -0.552 | -0.494 | 0.609 | -0.018 | 0.003 | 0.003 | 0 | 0.67 | 0.59 | 0.76 | 0.39 | 0.77 | 0.57 |
| 2 | 160664907 | rs2556088 | -0.71 | -0.541 | -0.357 | 0.512 | -0.007 | 0.18 | 0.18 | 2.936 | 0.12 | 0.46 | 0.08 | 0.24 | 0.06 | 0.2 |
| 2 | 160665936 | rs2729722 | 0.673 | -0.471 | -0.601 | 0.553 | -0.011 | 0.149 | 0.149 | 1.318 | 0.07 | 0.18 | 0.03 | 0.24 | 0.03 | 0.16 |
| 2 | 160668108 | rs2729726 | 0.347 | -0.508 | -0.942 | 0.405 | 0.002 | 0.002 | 0.002 | 0 | 0.41 | 0.54 | 0.22 | 0.33 | 0.37 | 0.54 |
| 2 | 160670424 | rs6432565 | 0.47 | -0.152 | -0.986 | 0.67 | -0.031 | 0.039 | 0.039 | 0 | 0.34 | 0.25 | 0.57 | 0.25 | 0.42 | 0.19 |
| 2 | 160673123 | rs2729715 | 0.304 | -0.288 | -1.289 | 0.786 | -0.015 | 0.128 | 0.128 | 0.942 | 0.08 | 0.16 | 0.03 | 0.25 | 0.03 | 0.17 |
| 2 | 160673526 | rs2059696 | 0.885 | 0.009 | -1.974 | 0.319 | -0.098 | 0.051 | 0.051 | 0 | 0.28 | 0.03 | 0.48 | 0.22 | 0.33 | 0.11 |
| 2 | 160676427 | rs12692566 | -1.348 | -0.514 | -2.059 | 0.063 | 0.017 | 0.061 | 0.061 | 0 | 0.75 | 0.9 | 0.88 | 0.91 | 0.8 | 0.73 |
| 2 | 160685431 | rs1344632 | 0.584 | -0.666 | -2.368 | 0.114 | -0.084 | -0.09 | -0.09 | 0 | 0.34 | 0.13 | 0.31 | 0.51 | 0.51 | 0.35 |
| 2 | 160685529 | rs2556094 | -1.701 | -0.742 | -1.703 | 0.074 | -0.011 | -0.029 | -0.029 | 0 | 0.74 | 0.7 | 0.87 | 0.92 | 0.81 | 0.73 |
| 2 | 160685769 | rs13023761 | 0.634 | -1.123 | -1.706 | -0.038 | -0.064 | -0.075 | -0.075 | 0 | 0.23 | 0.44 | 0.15 | 0.11 | 0.13 | 0.18 |
| 2 | 160686816 | rs2556096 | 0.718 | -0.916 | -1.659 | 0.08 | -0.084 | -0.09 | -0.09 | 0.181 | 0.34 | 0.13 | 0.31 | 0.51 | 0.51 | 0.35 |
| 2 | 160688557 | rs16844380 | 0.785 | -0.695 | -1.373 | 0.178 | -0.002 | -0.035 | -0.035 | 0.046 | 0.16 | 0.1 | 0.37 | 0.3 | 0.16 | 0.2 |
| 2 | 160691494 | rs7609477 | 0.381 | -0.528 | -1.515 | 0.333 | -0.049 | -0.05 | -0.05 | 0 | 0.08 | 0.01 | 0.14 | 0.27 | 0.17 | 0.15 |
| 2 | 160692461 | rs6750782 | -0.553 | -0.947 | -1.723 | 0.133 | 0.035 | 0.005 | 0.005 | 0 | 0.49 | 0.57 | 0.71 | 0.57 | 0.69 | 0.41 |
| 2 | 160695800 | rs7574420 | -1.748 | -1.102 | -1.826 | 0.284 | -0.035 | 0.008 | 0.008 | 0 | 0.72 | 0.64 | 0.89 | 0.71 | 0.85 | 0.69 |
| 2 | 160701198 | rs7573748 | 0.081 | -1.095 | -1.746 | -0.025 | -0.04 | -0.063 | -0.063 | 0 | 0.25 | 0.46 | 0.18 | 0.13 | 0.17 | 0.28 |
| 2 | 160701224 | rs1469350 | -1.141 | -1.102 | -1.886 | 0.068 | 0.066 | 0.008 | 0.008 | 0.005 | 0.6 | 0.75 | 0.55 | 0.4 | 0.72 | 0.49 |
| 2 | 160710922 | rs2729705 | -0.615 | -0.423 | -1.426 | 0.556 | 0.016 | -0.019 | -0.019 | 0 | 0.41 | 0.74 | 0.44 | 0.46 | 0.57 | 0.3 |
| 2 | 160711556 | rs975490 | -0.582 | -0.285 | -1.188 | 0.366 | 0.047 | -0.007 | -0.007 | 0 | 0.92 | 0.58 | 0.94 | 1 | 0.96 | 0.98 |
| 2 | 160725948 | rs10497205 | 1.849 | -0.128 | -1.538 | 0.34 | -0.13 | 0.116 | 0.116 | 1.199 | 0.19 | 0.03 | 0.17 | 0 | 0.36 | 0.06 |
| 2 | 160726544 | rs17231342 | 1.43 | -0.33 | -1.526 | 0.365 | -0.168 | 0.1 | 0.1 | 0.718 | 0.22 | 0.03 | 0.21 | 0.03 | 0.45 | 0.12 |
| 2 | 160729186 | rs10168271 | 0.216 | -0.573 | -1.416 | 0.472 | -0.012 | 0.054 | 0.054 | 1.694 | 0.62 | 0.16 | 0.55 | 0.71 | 0.43 | 0.79 |
| 2 | 160734722 | rs2292387 | 0.991 | -0.927 | -1.57 | 0.583 | -0.004 | -0.036 | -0.036 | 0 | 0.17 | 0.1 | 0.37 | 0.35 | 0.16 | 0.19 |
| 2 | 160738677 | rs16822581 | 0.958 | -0.927 | -1.545 | 0.675 | -0.002 | -0.021 | -0.021 | 0.162 | 0.17 | 0.11 | 0.37 | 0.32 | 0.16 | 0.19 |

| | | | | | | | | | | | | | | | | |
|---|---|---|---|---|---|---|---|---|---|---|---|---|---|---|---|---|
| 2 | 160742771 | rs2666986 | 0.597 | -1.011 | -1.399 | 0.615 | 0.098 | 0.022 | 0.022 | 0 | 0.49 | 0.3 | 0.25 | 0.4 | 0.26 | 0.6 |
| 2 | 160743793 | rs12692572 | 0.111 | -0.784 | -1.302 | 0.749 | -0.002 | -0.012 | -0.012 | 0 | 0.63 | 0.33 | 0.58 | 0.51 | 0.43 | 0.7 |
| 2 | 160754050 | rs7601374 | 0.718 | -0.593 | -1.463 | 0.818 | -0.111 | -0.127 | -0.127 | 0.37 | 0.28 | 0.09 | 0.31 | 0.49 | 0.54 | 0.28 |
| 2 | 160759168 | rs2292391 | 0.401 | -0.825 | -1.454 | 0.78 | 0 | -0.004 | -0.004 | 0.022 | 0.24 | 0.23 | 0.44 | 0.32 | 0.23 | 0.28 |
| 2 | 160761808 | rs3806603 | 1.048 | -0.944 | -1.548 | 0.815 | -0.122 | -0.036 | -0.036 | 1.684 | 0.24 | 0.04 | 0.31 | 0.29 | 0.54 | 0.19 |
| 2 | 160769588 | rs6735923 | 0.825 | -0.958 | -1.515 | 0.836 | -0.062 | -0.075 | -0.075 | 0 | 0.26 | 0.12 | 0.31 | 0.49 | 0.54 | 0.32 |
| 2 | 160770031 | rs12692575 | -0.85 | -1.1 | -1.266 | 0.77 | -0.001 | 0 | 0 | 0.196 | 0.51 | 0.52 | 0.78 | 0.8 | 0.78 | 0.6 |
| 2 | 160780926 | rs7576140 | -0.983 | -1.157 | -1.392 | 0.631 | 0.065 | 0.062 | 0.062 | 0 | 0.54 | 0.35 | 0.76 | 0.8 | 0.78 | 0.6 |
| 2 | 160782196 | rs12692577 | 0.789 | -1.061 | -1.549 | 0.736 | -0.012 | -0.052 | -0.052 | 0 | 0.23 | 0.08 | 0.43 | 0.32 | 0.23 | 0.24 |
| 2 | 160782335 | rs12472767 | 1.209 | -1.129 | -1.529 | 0.526 | -0.011 | -0.009 | -0.009 | 0 | 0.39 | 0.39 | 0.22 | 0.19 | 0.22 | 0.42 |
| 2 | 160802644 | rs1511223 | 1.687 | -0.483 | -1.818 | 0.691 | -0.121 | -0.021 | -0.021 | 3.719 | 0.21 | 0.04 | 0.24 | 0.27 | 0.42 | 0.11 |
| 2 | 160805582 | rs10929959 | -0.475 | -1.029 | -1.711 | 0.538 | -0.029 | -0.041 | -0.041 | 0 | 0.92 | 0.38 | 0.93 | 1 | 0.99 | 0.98 |
| 2 | 160808075 | rs3828323 | 1.096 | -0.938 | -2.047 | 0.541 | -0.134 | -0.024 | -0.024 | 0 | 0.24 | 0.04 | 0.28 | 0.29 | 0.52 | 0.19 |
| 2 | 160811184 | rs949753 | 1.067 | -1.096 | -1.92 | 0.4 | 0.118 | 0.099 | 0.099 | 0 | 0.39 | 0.07 | 0.18 | 0.19 | 0.21 | 0.4 |
| 2 | 160833188 | rs2715918 | -1.47 | -1.232 | -2.05 | 0.297 | 0.125 | 0.17 | 0.17 | 0.597 | 0.63 | 0.91 | 0.86 | 0.86 | 0.83 | 0.62 |
| 2 | 160840151 | rs979129 | 0.994 | -1.081 | -1.6 | 0.796 | -0.131 | -0.087 | -0.087 | 0 | 0.22 | 0.04 | 0.26 | 0.35 | 0.5 | 0.23 |
| 2 | 160841307 | rs1546512 | 0.067 | -1.289 | -1.471 | 0.872 | -0.034 | 0.12 | 0.12 | 3.196 | 0.1 | 0.61 | 0.1 | 0.18 | 0.03 | 0.11 |
| 2 | 160841828 | rs2715921 | -2.099 | -1.331 | -1.814 | 0.726 | 0.083 | 0.134 | 0.134 | 0 | 0.67 | 0.95 | 0.87 | 0.86 | 0.83 | 0.64 |
| 2 | 160857482 | rs12692581 | -0.855 | -0.694 | -1.682 | 1.049 | 0.177 | 0.019 | 0.019 | 1.085 | 0.48 | 0.14 | 0.74 | 0.62 | 0.76 | 0.51 |
| 2 | 160861094 | rs1397711 | 0.422 | -0.49 | -1.555 | 1.034 | -0.012 | -0.005 | -0.005 | 0 | 0.26 | 0.1 | 0.48 | 0.28 | 0.27 | 0.28 |
| 2 | 160861579 | rs17829933 | 0.323 | 0.99 | -1.503 | 1.258 | -0.079 | -0.048 | -0.048 | 0 | 0.11 | 0.02 | 0.16 | 0.13 | 0.3 | 0.12 |
| 2 | 160863209 | rs1355418 | 0.43 | -0.736 | -1.411 | 1.055 | 0.003 | -0.019 | -0.019 | 3.743 | 0.22 | 0.09 | 0.42 | 0.27 | 0.2 | 0.24 |
| 2 | 160867349 | rs2248403 | 0.65 | -0.892 | -0.903 | 1.055 | -0.024 | 0.163 | 0.163 | 0.587 | 0.09 | 0.29 | 0.06 | 0.18 | 0.02 | 0.08 |
| 2 | 160870601 | rs1039931 | -1.537 | -0.928 | -1.185 | 0.661 | -0.002 | -0.018 | -0.018 | 0.243 | 0.58 | 0.62 | 0.78 | 0.86 | 0.78 | 0.59 |
| 2 | 160870944 | rs2221808 | 0.742 | -0.478 | -1.419 | 1.048 | -0.143 | -0.097 | -0.097 | 0.045 | 0.21 | 0.04 | 0.27 | 0.35 | 0.5 | 0.22 |
| 2 | 160873270 | rs2667040 | -1.521 | -1.065 | -1.193 | 0.637 | 0.009 | -0.003 | -0.003 | 0.207 | 0.56 | 0.6 | 0.76 | 0.86 | 0.75 | 0.6 |
| 2 | 160874354 | rs7594498 | 0.069 | -0.892 | -0.945 | 1.014 | -0.023 | 0.133 | 0.133 | 0 | 0.11 | 0.33 | 0.08 | 0.21 | 0.06 | 0.12 |
| 2 | 160878364 | rs17830558 | 0.45 | -0.693 | -1.279 | 0.766 | -0.124 | -0.085 | -0.085 | 0 | 0.24 | 0.04 | 0.29 | 0.34 | 0.53 | 0.25 |
| 2 | 160879651 | rs883423 | -1.207 | -1.195 | -1.065 | 0.226 | -0.014 | -0.043 | -0.043 | 0 | 0.56 | 0.54 | 0.76 | 0.86 | 0.73 | 0.66 |
| 2 | 160885442 | rs3749117 | -0.079 | -0.414 | -0.73 | 0.905 | -0.142 | -0.058 | -0.058 | 0.388 | 0.25 | 0.06 | 0.27 | 0.35 | 0.47 | 0.2 |
| 2 | 160885872 | rs3792189 | -1.018 | -0.602 | -0.558 | 1.063 | 0.022 | -0.011 | -0.011 | 0.037 | 0.42 | 0.28 | 0.41 | 0.57 | 0.61 | 0.38 |
| 2 | 160891009 | rs17241282 | 0.616 | -0.294 | -0.661 | 1.084 | -0.127 | -0.076 | -0.076 | 0.563 | 0.19 | 0.03 | 0.24 | 0.35 | 0.42 | 0.16 |
| 2 | 160891055 | rs10196882 | 0.705 | -0.882 | -0.534 | 1.136 | -0.006 | 0.013 | 0.013 | 0.014 | 0.16 | 0.15 | 0.13 | 0.22 | 0.14 | 0.17 |
| 2 | 160893260 | rs6722275 | 0.284 | -0.651 | -0.752 | 1.163 | -0.143 | -0.073 | -0.073 | 0 | 0.22 | 0.04 | 0.28 | 0.35 | 0.47 | 0.21 |
| 2 | 160893796 | rs6740901 | -1.432 | -1.479 | -0.36 | 0.874 | -0.043 | 0 | 0 | 0 | 0.7 | 0.61 | 0.87 | 0.74 | 0.84 | 0.65 |
| 2 | 160897655 | rs16844706 | 0.02 | -1.134 | -0.11 | 0.917 | -0.047 | 0.13 | 0.13 | 0.55 | 0.17 | 0.43 | 0.15 | 0.35 | 0.11 | 0.18 |
| 2 | 160902420 | rs17341301 | 0.902 | -0.613 | -0.45 | 0.615 | -0.039 | -0.025 | -0.025 | 0 | 0.16 | 0.08 | 0.22 | 0.34 | 0.41 | 0.2 |
| 2 | 160903759 | rs877199 | 1.094 | -0.878 | -0.861 | 0.375 | 0.005 | 0.028 | 0.028 | 0 | 0.11 | 0.04 | 0.36 | 0.02 | 0.09 | 0.12 |
| 2 | 160907869 | rs4665145 | -1.6 | -1.162 | -0.9 | 0.18 | 0.062 | 0.108 | 0.108 | 0 | 0.61 | 0.81 | 0.86 | 0.95 | 0.76 | 0.61 |
| 2 | 160912747 | rs6759924 | 0.698 | 0.031 | -0.628 | 1.192 | -0.106 | -0.045 | -0.045 | 0 | 0.16 | 0.03 | 0.2 | 0.27 | 0.4 | 0.2 |
| 2 | 160913599 | rs2715928 | -1.503 | -0.232 | -0.543 | 1.036 | 0.046 | 0.047 | 0.047 | 0.786 | 0.46 | 0.7 | 0.7 | 0.4 | 0.61 | 0.46 |
| 2 | 160915106 | rs16844715 | 0.496 | 0.142 | -0.843 | 1.317 | -0.114 | -0.091 | -0.091 | 0 | 0.17 | 0.03 | 0.23 | 0.42 | 0.42 | 0.2 |
| 2 | 160918386 | rs16844726 | 0.005 | -0.378 | -0.172 | 1.023 | -0.031 | 0.111 | 0.111 | 0.295 | 0.17 | 0.34 | 0.1 | 0.28 | 0.09 | 0.25 |
| 2 | 160921555 | rs2667026 | -1.247 | -0.559 | -0.53 | 0.783 | 0.043 | -0.003 | -0.003 | 0 | 0.34 | 0.71 | 0.19 | 0.3 | 0.16 | 0.33 |
| 2 | 160935582 | rs7598162 | -1.729 | -0.613 | -0.846 | 0.699 | 0.098 | -0.096 | -0.096 | 0.03 | 0.47 | 0.94 | 0.6 | 0.46 | 0.3 | 0.47 |
| 2 | 160949387 | rs7573119 | 0.751 | -0.491 | -0.644 | 0.659 | 0.007 | 0.01 | 0.01 | 0 | 0.09 | 0.04 | 0.05 | 0 | 0.07 | 0.07 |
| 2 | 160955896 | rs4665154 | 0.564 | -0.557 | -0.472 | 0.806 | -0.005 | 0.012 | 0.012 | 0.155 | 0.13 | 0.22 | 0.4 | 0.18 | 0.13 | 0.14 |
| 2 | 160963372 | rs13403826 | 0.265 | -0.683 | -0.145 | 0.823 | -0.005 | 0.14 | 0.14 | 0 | 0.12 | 0.26 | 0.07 | 0.28 | 0.07 | 0.17 |
| 2 | 160963603 | rs10497213 | -0.065 | -0.373 | -0.612 | 0.884 | -0.103 | -0.021 | -0.021 | 0 | 0.25 | 0.07 | 0.28 | 0.32 | 0.48 | 0.24 |
| 2 | 160965146 | rs6432593 | -2.218 | -0.401 | -0.503 | 0.828 | 0.019 | 0.04 | 0.04 | 0.925 | 0.61 | 0.98 | 0.83 | 0.78 | 0.71 | 0.64 |
| 2 | 160968628 | rs16844790 | 0.534 | -0.206 | -0.256 | 0.798 | -0.004 | 0.002 | 0.002 | 1.78 | 0.16 | 0.22 | 0.42 | 0.18 | 0.16 | 0.15 |
| 2 | 160986542 | rs4580348 | -0.724 | -0.189 | -0.37 | 0.824 | 0.012 | -0.013 | -0.013 | 0 | 0.4 | 0.3 | 0.71 | 0.5 | 0.64 | 0.39 |
| 2 | 160989296 | rs16844824 | -1.411 | -0.216 | -0.308 | 0.817 | 0.03 | 0.007 | 0.007 | 2.448 | 0.48 | 0.63 | 0.75 | 0.5 | 0.66 | 0.47 |
| 2 | 161015066 | rs983916 | 1.642 | 1.094 | -0.5 | 0.182 | -0.033 | 0.032 | 0.032 | 0 | 0.09 | 0.03 | 0.08 | 0 | 0.24 | 0.02 |
| 2 | 161019869 | rs6730023 | -0.08 | 0.743 | -0.295 | 0.283 | -0.055 | 0.203 | 0.203 | 2.802 | 0.84 | 0.35 | 0.89 | 0.57 | 0.95 | 0.83 |
| 2 | 161026336 | rs2290706 | 0.47 | 0.753 | -0.173 | 0.285 | -0.033 | 0.252 | 0.252 | 0.267 | 0.13 | 0.33 | 0.08 | 0.42 | 0.05 | 0.17 |
| 2 | 161029218 | rs2305818 | 0.343 | 1.066 | -0.338 | 0.321 | -0.102 | -0.072 | -0.072 | 0 | 0.16 | 0.03 | 0.26 | 0.33 | 0.42 | 0.16 |
| 2 | 161051363 | rs3111397 | -0.805 | 0.766 | -0.541 | 0.264 | -0.002 | 0.001 | 0.001 | 0.489 | 0.82 | 0.77 | 0.57 | 0.79 | 0.82 | 0.85 |
| 2 | 161055315 | rs16844863 | 0.465 | 0.799 | -0.642 | 0.186 | -0.044 | -0.029 | -0.029 | 0.284 | 0.07 | 0.01 | 0.17 | 0.33 | 0.18 | 0.14 |
| 2 | 161060315 | rs2925754 | -0.798 | 0.715 | -0.538 | 0.21 | -0.001 | -0.001 | -0.001 | 0 | 0.82 | 0.78 | 0.58 | 0.81 | 0.82 | 0.86 |
| 2 | 161064585 | rs7588696 | 0.326 | 0.882 | -0.49 | 0.226 | 0.055 | 0.119 | 0.119 | 0 | 0.22 | 0.38 | 0.19 | 0.44 | 0.3 | 0.21 |
| 2 | 161065568 | rs13014884 | 1.168 | 0.8 | -1.475 | 0.183 | 0.003 | -0.005 | -0.005 | 0 | 0.07 | 0.04 | 0.04 | 0 | 0.04 | 0.07 |
| 2 | 161070078 | rs4665164 | -1.254 | 0.764 | -1.515 | 0.182 | -0.005 | 0.109 | 0.109 | 0 | 0.44 | 0.77 | 0.44 | 0.77 | 0.54 | 0.49 |
| 2 | 161072585 | rs2925755 | 0.415 | 1.045 | -0.766 | 0.524 | -0.004 | -0.008 | -0.008 | 0 | 0.64 | 0.7 | 0.36 | 0.49 | 0.57 | 0.57 |
| 2 | 161079435 | rs3101043 | 0.367 | 0.871 | -0.604 | -0.273 | 0.01 | -0.086 | -0.086 | 0.159 | 0.59 | 0.48 | 0.65 | 0.08 | 0.51 | 0.58 |
| 2 | 161079450 | rs3111398 | 0.274 | 0.419 | -0.449 | -0.116 | 0.011 | 0.002 | 0.002 | 0 | 0.59 | 0.49 | 0.69 | 0.4 | 0.53 | 0.68 |
| 2 | 161079873 | rs13031645 | 0.402 | 0.576 | -0.06 | -0.303 | 0.007 | -0.031 | -0.031 | 0 | 0.73 | 0.45 | 0.68 | 0.8 | 0.69 | 0.71 |
| 2 | 161085348 | rs7560257 | -0.96 | 0.653 | -0.245 | -0.117 | -0.001 | 0.005 | 0.005 | 0 | 0.09 | 0.07 | 0.08 | 0.21 | 0.15 | 0.16 |
| 2 | 161090873 | rs6711375 | 0.147 | -0.218 | -0.213 | -0.53 | 0.006 | 0.006 | 0.006 | 0 | 0.59 | 0.59 | 0.76 | 0.82 | 0.68 | 0.52 |

| | | | | | | | | | | | | | | | | |
|---|---|---|---|---|---|---|---|---|---|---|---|---|---|---|---|---|
| 2 | 161096189 | rs1563574 | -0.297 | -0.542 | -0.136 | -0.727 | -0.023 | 0.328 | 0.328 | 0.116 | 0.11 | 0.43 | 0.17 | 0.6 | 0.06 | 0.18 |
| 2 | 161105876 | rs10803762 | 0.021 | -0.456 | -0.091 | -0.782 | -0.011 | -0.022 | -0.022 | 0.269 | 0.59 | 0.57 | 0.76 | 0.82 | 0.68 | 0.52 |
| 2 | 161118687 | rs10170600 | -0.362 | -0.791 | -0.25 | -0.963 | -0.014 | -0.029 | -0.029 | 0.105 | 0.69 | 0.58 | 0.79 | 0.82 | 0.73 | 0.63 |
| 2 | 161119392 | rs733659 | 2.304 | -0.695 | -0.43 | -1.267 | 0.018 | 0.1 | 0.1 | 0 | 0.28 | 0.1 | 0.38 | 0.01 | 0.19 | 0.16 |
| 2 | 161124182 | rs12692588 | 0.181 | -0.542 | -0.23 | -1.185 | 0.001 | 0 | 0 | 0 | 0.6 | 0.48 | 0.69 | 0.6 | 0.57 | 0.46 |
| 2 | 161131762 | rs3772069 | 0.502 | -0.37 | -0.03 | -0.903 | -0.003 | 0.004 | 0.004 | 0 | 0.2 | 0.23 | 0.25 | 0.53 | 0.33 | 0.2 |
| 2 | 161133471 | rs3772070 | -0.401 | -0.299 | -0.1 | -0.889 | -0.022 | -0.035 | -0.035 | 0 | 0.69 | 0.58 | 0.77 | 0.83 | 0.74 | 0.65 |
| 2 | 161163559 | rs9917155 | 0.337 | -0.077 | 0.044 | -0.977 | -0.001 | 0.001 | 0.001 | 0 | 0.56 | 0.48 | 0.69 | 0.54 | 0.55 | 0.4 |
| 2 | 161166234 | rs4077463 | -1.018 | -0.072 | -0.04 | -1.022 | 0.014 | 0.022 | 0.022 | 0.058 | 0.82 | 0.57 | 0.81 | 0.85 | 0.79 | 0.79 |
| 2 | 161177594 | rs10201614 | 1.979 | -0.115 | -0.128 | -1.108 | 0.029 | 0.143 | 0.143 | 0 | 0.31 | 0.1 | 0.42 | 0.01 | 0.21 | 0.16 |
| 2 | 161180007 | rs6708182 | 1.572 | 0.011 | 0 | -1.028 | -0.001 | 0.021 | 0.021 | 0 | 0.14 | 0.12 | 0.21 | 0.53 | 0.3 | 0.17 |
| 2 | 161192070 | rs4410242 | -0.97 | -0.099 | 0.174 | -0.991 | 0.013 | 0.02 | 0.02 | 0.037 | 0.82 | 0.6 | 0.81 | 0.85 | 0.79 | 0.79 |
| 2 | 161202588 | rs10168504 | -0.937 | -0.101 | 0.225 | -0.992 | 0.011 | 0.018 | 0.018 | 0.055 | 0.81 | 0.57 | 0.8 | 0.85 | 0.79 | 0.79 |
| 2 | 161203395 | rs7425274 | -0.866 | -0.146 | 0.014 | -1.078 | 0.001 | 0.001 | 0.001 | 0 | 0.45 | 0.4 | 0.32 | 0.57 | 0.45 | 0.44 |
| 2 | 161214175 | rs6718526 | -1.443 | -0.135 | 0.188 | -1.046 | -0.001 | -0.003 | -0.003 | 0 | 0.84 | 0.87 | 0.84 | 0.87 | 0.81 | 0.8 |
| 2 | 161214857 | rs10929979 | 1.579 | -0.134 | -0.3 | -1.117 | 0.002 | 0.025 | 0.025 | 0 | 0.14 | 0.1 | 0.21 | 0.53 | 0.3 | 0.17 |
| 2 | 161219032 | rs6751676 | 2.194 | -0.134 | -0.502 | -1.089 | -0.015 | 0.225 | 0.225 | 0 | 0.31 | 0.03 | 0.42 | 0.01 | 0.21 | 0.17 |
| 2 | 161220126 | rs6755940 | -0.036 | 0.029 | -0.05 | -0.779 | 0.045 | 0.096 | 0.096 | 0 | 0.76 | 0.45 | 0.73 | 0.58 | 0.66 | 0.61 |
| 2 | 161242295 | rs12998587 | -0.113 | 0.299 | -0.082 | -1.194 | -0.014 | -0.052 | -0.052 | 0 | 0.62 | 0.56 | 0.75 | 0.89 | 0.71 | 0.59 |
| 2 | 161246317 | rs4544378 | 0.359 | 0.53 | -0.149 | -1.08 | -0.003 | -0.009 | -0.009 | 0 | 0.56 | 0.48 | 0.69 | 0.62 | 0.58 | 0.41 |
| 2 | 161265099 | rs13009374 | -0.225 | 0.428 | -0.125 | -1 | 0.011 | 0.016 | 0.016 | 0 | 0.63 | 0.69 | 0.75 | 0.83 | 0.71 | 0.59 |
| 2 | 161265910 | rs12692596 | 0.994 | 0.715 | -0.598 | -1.103 | -0.014 | 0.033 | 0.033 | 0 | 0.37 | 0.21 | 0.48 | 0.3 | 0.36 | 0.34 |
| 2 | 161277644 | rs12463865 | 0.907 | 0.831 | -0.352 | -0.946 | 0.063 | 0.101 | 0.101 | 0 | 0.16 | 0.21 | 0.22 | 0.59 | 0.31 | 0.17 |
| 2 | 161281691 | rs4664325 | 0.522 | 0.607 | -0.483 | -1.133 | 0.008 | 0.026 | 0.026 | 0 | 0.48 | 0.31 | 0.66 | 0.62 | 0.55 | 0.34 |
| 2 | 161292830 | rs10929986 | -1.234 | 0.472 | -0.05 | -1.065 | -0.014 | -0.03 | -0.03 | 0.514 | 0.94 | 0.84 | 0.98 | 1 | 0.98 | 0.97 |
| 2 | 161317460 | rs6742799 | 0.739 | 0.236 | -0.157 | -1.285 | -0.003 | -0.044 | -0.044 | 0 | 0.17 | 0.32 | 0.19 | 0.08 | 0.19 | 0.21 |
| 2 | 161324155 | rs13017450 | 1.869 | 0.249 | -0.674 | -1.451 | -0.002 | 0.15 | 0.15 | 0 | 0.32 | 0.1 | 0.44 | 0.03 | 0.26 | 0.16 |
| 2 | 161324786 | rs7425305 | 0.168 | 0.254 | -0.617 | -1.392 | 0.013 | 0.009 | 0.009 | 0 | 0.66 | 0.64 | 0.88 | 0.7 | 0.77 | 0.55 |
| 2 | 161329644 | rs7423892 | 0.709 | 0.167 | -0.443 | -1.451 | 0.007 | 0.016 | 0.016 | 0 | 0.51 | 0.54 | 0.65 | 0.11 | 0.45 | 0.38 |
| 2 | 161332702 | rs7585160 | -1.267 | -0.046 | -0.782 | -1.34 | -0.016 | -0.031 | -0.031 | 0 | 0.94 | 0.84 | 0.98 | 1 | 0.98 | 0.97 |
| 2 | 161333557 | rs13020444 | 1.455 | 0.03 | -0.9 | -1.309 | 0.002 | 0.15 | 0.15 | 0 | 0.32 | 0.1 | 0.44 | 0.03 | 0.26 | 0.18 |
| 2 | 161351779 | rs10929989 | -0.635 | 0.422 | -0.438 | -0.865 | -0.009 | 0.006 | 0.006 | 0.096 | 0.38 | 0.54 | 0.46 | 0.67 | 0.53 | 0.37 |
| 2 | 161368905 | rs4664334 | -0.647 | 0.565 | -0.336 | -0.906 | 0 | 0 | 0 | 0.013 | 0.34 | 0.63 | 0.45 | 0.73 | 0.46 | 0.36 |
| 2 | 161375040 | rs6432626 | -0.266 | 0.323 | -0.239 | -1.341 | 0.017 | -0.098 | -0.098 | 0 | 0.64 | 0.55 | 0.7 | 0.92 | 0.56 | 0.7 |
| 2 | 161380888 | rs10192369 | 0.477 | 0.373 | -0.106 | -1.344 | -0.028 | 0.055 | 0.055 | 0.585 | 0.57 | 0.21 | 0.64 | 0.69 | 0.52 | 0.58 |
| 2 | 161386729 | rs13382644 | 0.775 | 0.014 | 0.102 | -1.278 | -0.008 | -0.08 | -0.08 | 0 | 0.36 | 0.38 | 0.41 | 0.02 | 0.23 | 0.22 |
| 2 | 161395188 | rs6432628 | 1.197 | 0.028 | 0.101 | -1.27 | 0.017 | 0.033 | 0.033 | 0.711 | 0.32 | 0.18 | 0.38 | 0.02 | 0.22 | 0.2 |
| 2 | 161398208 | rs7423194 | 0.552 | 0.059 | -0.006 | -1.196 | -0.01 | -0.015 | -0.015 | 0 | 0.38 | 0.41 | 0.42 | 0.25 | 0.26 | 0.32 |
| 2 | 161406793 | rs7598938 | 0.586 | 0.05 | -0.444 | -1.314 | -0.007 | -0.006 | -0.006 | 0 | 0.35 | 0.61 | 0.42 | 0.24 | 0.23 | 0.31 |
| 2 | 161410542 | rs13410476 | 0.265 | 0.087 | -0.28 | -1.254 | 0.016 | 0.023 | 0.023 | 0.316 | 0.4 | 0.64 | 0.45 | 0.24 | 0.26 | 0.33 |
| 2 | 161414622 | rs6729770 | 0.873 | 0 | -0.358 | -1.359 | 0.028 | 0.086 | 0.086 | 0.12 | 0.34 | 0.17 | 0.38 | 0.02 | 0.23 | 0.23 |
| 2 | 161421950 | rs7589022 | 0.778 | 0.01 | -0.251 | -1.333 | 0.014 | 0.008 | 0.008 | 0.871 | 0.35 | 0.26 | 0.4 | 0.02 | 0.23 | 0.21 |
| 2 | 161423570 | rs11888274 | 0.286 | -0.035 | -0.253 | -1.204 | 0.015 | 0.021 | 0.021 | 0.513 | 0.41 | 0.63 | 0.46 | 0.26 | 0.27 | 0.32 |
| 2 | 161424684 | rs11681327 | 0.71 | 0.08 | -1.027 | -1.136 | 0.021 | 0.051 | 0.051 | 0.949 | 0.35 | 0.22 | 0.4 | 0.02 | 0.23 | 0.21 |
| 2 | 161426931 | rs13407070 | 1.035 | 0.546 | -1.542 | -1.158 | -0.046 | -0.092 | -0.092 | 0.059 | 0.15 | 0.03 | 0.22 | 0.65 | 0.22 | 0.18 |
| 2 | 161431604 | rs7603576 | 0.74 | 0.366 | -1.142 | -1.322 | 0.008 | 0.125 | 0.125 | 0.41 | 0.32 | 0.12 | 0.39 | 0.02 | 0.23 | 0.22 |
| 2 | 161440007 | rs4341947 | -0.778 | 0.399 | -0.471 | -0.117 | -0.008 | 0.115 | 0.115 | 0 | 0.93 | 0.88 | 0.94 | 0.66 | 0.93 | 0.9 |
| 2 | 161440199 | rs7557611 | -0.007 | 0.205 | -0.42 | -0.365 | 0.002 | -0.012 | -0.012 | 0 | 0.31 | 0.26 | 0.3 | 0.45 | 0.34 | 0.45 |
| 2 | 161464707 | rs6737230 | -0.961 | 0.069 | 0.2 | -0.272 | 0.007 | 0.25 | 0.25 | 0 | 0.92 | 0.73 | 0.92 | 0.66 | 0.94 | 0.88 |
| 2 | 161477970 | rs4268941 | 1.108 | 0.1 | 0.11 | -0.453 | 0.007 | -0.016 | -0.016 | 0 | 0.18 | 0.18 | 0.21 | 0.44 | 0.19 | 0.22 |
| 2 | 161490982 | rs12991101 | 1.009 | 0.572 | 0.309 | -0.511 | 0 | -0.049 | -0.049 | 0 | 0.29 | 0.22 | 0.15 | 0.02 | 0.29 | 0.25 |
| 2 | 161503223 | rs6432641 | 0.374 | 0.44 | 0.053 | -0.428 | 0.023 | -0.037 | -0.037 | 0 | 0.35 | 0.25 | 0.3 | 0.46 | 0.34 | 0.43 |
| 2 | 161515983 | rs4664346 | -0.341 | 0.436 | 0.221 | -0.474 | -0.022 | -0.043 | -0.043 | 3.617 | 0.59 | 0.45 | 0.64 | 0.23 | 0.63 | 0.6 |
| 2 | 161523902 | rs13011975 | 1.196 | 0.541 | 0.165 | -0.611 | -0.007 | -0.01 | -0.01 | 0 | 0.29 | 0.19 | 0.15 | 0.02 | 0.28 | 0.24 |
| 2 | 161525601 | rs13390172 | 0.787 | 0.494 | 0.045 | -0.456 | 0.021 | -0.039 | -0.039 | 0 | 0.32 | 0.22 | 0.27 | 0.43 | 0.3 | 0.27 |
| 2 | 161542328 | rs12053303 | -0.484 | 0.705 | 0.57 | 0.004 | 0.023 | -0.041 | -0.041 | 0 | 0.31 | 0.4 | 0.51 | 0.22 | 0.37 | 0.39 |
| 2 | 161547697 | rs6752228 | 0.946 | 0.695 | 0.55 | 0.101 | -0.006 | -0.016 | -0.016 | 0 | 0.59 | 0.33 | 0.41 | 0.47 | 0.57 | 0.52 |
| 2 | 161550899 | rs12618371 | 1.603 | 0.693 | 0.545 | 0.103 | -0.007 | 0.005 | 0.005 | 0 | 0.06 | 0 | 0.05 | 0.31 | 0.06 | 0.08 |
| 2 | 161553284 | rs4664022 | 0.411 | 0.961 | 0.588 | 0.086 | -0.012 | -0.003 | -0.003 | 0.336 | 0.14 | 0.11 | 0.38 | 0.18 | 0.21 | 0.1 |
| 2 | 161555610 | rs6718138 | 0.78 | 0.844 | 0.61 | 0.095 | -0.014 | 0.023 | 0.023 | 0.022 | 0.73 | 0.44 | 0.79 | 0.64 | 0.78 | 0.64 |
| 2 | 161560869 | rs1451826 | -0.417 | 0.893 | 0.34 | 0.034 | -0.024 | 0.071 | 0.071 | 0.08 | 0.83 | 0.96 | 0.89 | 0.95 | 0.84 | 0.74 |
| 2 | 161572781 | rs6732189 | 1.188 | 0.968 | 0.336 | 0.109 | -0.002 | -0.004 | -0.004 | 0 | 0.59 | 0.41 | 0.33 | 0.46 | 0.56 | 0.52 |
| 2 | 161598210 | rs1480577 | -0.248 | 0.973 | 0.205 | 0.12 | 0.038 | 0.012 | 0.012 | 0.013 | 0.16 | 0.4 | 0.41 | 0.18 | 0.21 | 0.13 |
| 2 | 161600175 | rs17219937 | 0.574 | 1.202 | 0.043 | 0.099 | -0.051 | 0.049 | 0.049 | 0.209 | 0.15 | 0.04 | 0.1 | 0.07 | 0.17 | 0.12 |
| 2 | 161600547 | rs4664366 | -0.157 | 1.2 | 0.123 | 0.1 | 0.02 | -0.001 | -0.001 | 0 | 0.17 | 0.28 | 0.42 | 0.16 | 0.26 | 0.26 |
| 2 | 161603829 | rs584526 | -0.749 | 1.016 | 0.189 | 0.118 | 0.019 | -0.016 | -0.016 | 0 | 0.32 | 0.42 | 0.51 | 0.23 | 0.42 | 0.38 |
| 2 | 161604844 | rs616635 | -1.308 | 0.984 | 0.168 | 0.215 | -0.007 | -0.006 | -0.006 | 0.365 | 0.4 | 0.4 | 0.65 | 0.56 | 0.49 | 0.47 |
| 2 | 161605747 | rs270919 | -1.288 | 1.046 | 0.144 | 0.234 | -0.001 | -0.001 | -0.001 | 0 | 0.41 | 0.62 | 0.67 | 0.56 | 0.47 | 0.47 |
| 2 | 161606567 | rs175238 | 0.912 | 0.962 | -0.319 | -0.034 | -0.015 | 0.055 | 0.055 | 0 | 0.56 | 0.66 | 0.43 | 0.81 | 0.53 | 0.53 |

| | | | | | | | | | | | | | | | | |
|---|---|---|---|---|---|---|---|---|---|---|---|---|---|---|---|---|
| 2 | 161622729 | rs648120 | 0.544 | 1.2 | 0.037 | 0.652 | 0.024 | 0.075 | 0.075 | 0.108 | 0.78 | 0.63 | 0.46 | 0.63 | 0.72 | 0.69 |
| 2 | 161622892 | rs532146 | -0.77 | 1.22 | 0.02 | 0.691 | -0.017 | -0.011 | -0.011 | 6.259 | 0.88 | 0.9 | 0.63 | 0.84 | 0.79 | 0.78 |
| 2 | 161631419 | rs7574698 | 0.536 | 1.234 | 0.079 | 0.538 | 0.004 | -0.007 | -0.007 | 1.902 | 0.17 | 0.14 | 0.13 | 0.1 | 0.2 | 0.14 |
| 2 | 161637237 | rs270924 | 0.482 | 1.234 | 0.114 | 0.566 | 0.013 | 0.046 | 0.046 | 0.108 | 0.78 | 0.66 | 0.46 | 0.63 | 0.72 | 0.69 |
| 2 | 161646558 | rs10190870 | 1.312 | 0.887 | 0.05 | 0.479 | 0 | -0.005 | -0.005 | 0 | 0.55 | 0.41 | 0.3 | 0.53 | 0.48 | 0.54 |
| 2 | 161661244 | rs565736 | 0.852 | 0.902 | 0.035 | 0.49 | 0.019 | 0.022 | 0.022 | 0.185 | 0.61 | 0.51 | 0.33 | 0.53 | 0.52 | 0.55 |
| 2 | 161670760 | rs270996 | 1.093 | 0.815 | 0.013 | 0.438 | -0.01 | -0.028 | -0.028 | 0 | 0.06 | 0.1 | 0.02 | 0 | 0.05 | 0.01 |
| 2 | 161685696 | rs270982 | -0.856 | 1.033 | 0.032 | 0.475 | -0.004 | -0.003 | -0.003 | 0 | 0.88 | 0.87 | 0.63 | 0.84 | 0.79 | 0.8 |
| 2 | 161688392 | rs270965 | -0.913 | 0.969 | 0.084 | 0.549 | -0.001 | 0.011 | 0.011 | 0 | 0.32 | 0.48 | 0.33 | 0.32 | 0.31 | 0.26 |
| 2 | 161690170 | rs7585907 | 0.28 | 0.97 | 0.081 | 0.55 | 0.021 | -0.065 | -0.065 | 0 | 0.2 | 0.39 | 0.16 | 0.1 | 0.2 | 0.16 |
| 2 | 161693170 | rs270975 | -0.886 | 1.011 | 0.143 | 0.57 | -0.019 | -0.01 | -0.01 | 0 | 0.88 | 0.9 | 0.63 | 0.84 | 0.79 | 0.8 |
| 2 | 161694980 | rs270979 | 1.175 | 0.952 | 0.107 | 0.558 | -0.001 | 0.002 | 0.002 | 0 | 0.55 | 0.5 | 0.31 | 0.52 | 0.48 | 0.54 |
| 2 | 161715638 | rs270958 | -0.829 | 0.866 | 0.059 | 0.544 | -0.002 | 0.007 | 0.007 | 0 | 0.33 | 0.43 | 0.32 | 0.32 | 0.31 | 0.26 |
| 2 | 161725523 | rs270955 | -0.598 | 0.974 | 0.032 | 1.035 | -0.007 | -0.003 | -0.003 | 0 | 0.87 | 0.87 | 0.62 | 0.68 | 0.78 | 0.8 |
| 2 | 161739740 | rs6729946 | 1.304 | 0.943 | 0.066 | 0.63 | -0.02 | -0.012 | -0.012 | 0 | 0.06 | 0 | 0.15 | 0.22 | 0.07 | 0.09 |
| 2 | 161739776 | rs6730050 | -0.178 | 0.811 | 0.255 | 0.516 | 0.005 | 0.023 | 0.023 | 0.026 | 0.27 | 0.36 | 0.29 | 0.32 | 0.27 | 0.22 |
| 2 | 161742051 | rs17287105 | 1.554 | 0.432 | -0.923 | 0.016 | -0.082 | 0.144 | 0.144 | 0 | 0.25 | 0.05 | 0.13 | 0.04 | 0.25 | 0.2 |
| 2 | 161749220 | rs10497221 | 1.239 | 1.126 | -0.088 | 0.777 | -0.006 | -0.007 | -0.007 | 0 | 0.57 | 0.38 | 0.31 | 0.52 | 0.48 | 0.56 |
| 2 | 161750217 | rs1722650 | 0.22 | 1.228 | -0.422 | 0.813 | 0.008 | 0.002 | 0.002 | 0.161 | 0.13 | 0.14 | 0.37 | 0.16 | 0.21 | 0.2 |
| 2 | 161751552 | rs13014133 | -0.455 | 1.186 | -0.374 | 0.82 | -0.024 | -0.045 | -0.045 | 0 | 0.27 | 0.12 | 0.28 | 0.32 | 0.31 | 0.25 |
| 2 | 161752484 | rs16845296 | 1.023 | 1.184 | -0.409 | 0.785 | -0.007 | -0.022 | -0.022 | 0.393 | 0.06 | 0.07 | 0.02 | 0 | 0.04 | 0.04 |
| 2 | 161756280 | rs1615929 | -1.02 | 1.247 | -0.616 | 0.856 | -0.015 | -0.013 | -0.013 | 0 | 0.35 | 0.31 | 0.64 | 0.48 | 0.48 | 0.41 |
| 2 | 161759570 | rs1710649 | -0.083 | 1.455 | -0.764 | 1.049 | 0.02 | 0.01 | 0.01 | 0.161 | 0.13 | 0.17 | 0.38 | 0.19 | 0.21 | 0.2 |
| 2 | 161761464 | rs1710648 | -0.012 | 1.46 | -0.931 | 1.197 | -0.002 | 0 | 0 | 1.14 | 0.13 | 0.13 | 0.37 | 0.24 | 0.21 | 0.19 |
| 2 | 161766458 | rs12620236 | 0.058 | 1.489 | -1.2 | 0.923 | -0.026 | -0.005 | -0.005 | 0.005 | 0.13 | 0.09 | 0.37 | 0.14 | 0.21 | 0.15 |
| 2 | 161768312 | rs1710647 | 0.264 | 1.05 | -1.067 | 0.901 | -0.002 | -0.003 | -0.003 | 0.046 | 0.55 | 0.41 | 0.28 | 0.57 | 0.47 | 0.58 |
| 2 | 161775489 | rs12470772 | -0.288 | 1.473 | -1.598 | 1.128 | -0.065 | -0.07 | -0.07 | 0.002 | 0.27 | 0.11 | 0.31 | 0.54 | 0.42 | 0.24 |
| 2 | 161776405 | rs13015721 | 0.271 | 1.779 | -1.35 | 1.447 | 0.006 | -0.029 | -0.029 | 0 | 0.19 | 0.12 | 0.13 | 0.34 | 0.2 | 0.17 |
| 2 | 161776746 | rs1710630 | -1.369 | 1.911 | -1.171 | 1.013 | -0.003 | -0.012 | -0.012 | 0.334 | 0.49 | 0.49 | 0.51 | 0.71 | 0.65 | 0.55 |
| 2 | 161781773 | rs1722636 | -0.089 | 2.076 | -0.892 | 1.037 | 0.023 | -0.003 | -0.003 | 0 | 0.22 | 0.44 | 0.22 | 0.17 | 0.23 | 0.31 |
| 2 | 161782375 | rs1516100 | 0.384 | 2.147 | -1.003 | 1.154 | -0.053 | -0.051 | -0.051 | 2.315 | 0.08 | 0.01 | 0.16 | 0.21 | 0.22 | 0.07 |
| 2 | 161783717 | rs1710627 | -1.312 | 1.623 | -1.252 | 0.758 | 0.01 | 0.001 | 0.001 | 0.029 | 0.39 | 0.42 | 0.41 | 0.37 | 0.46 | 0.44 |
| 2 | 161784195 | rs1516099 | -0.453 | 1.513 | -1.182 | 0.528 | 0 | -0.049 | -0.049 | 0.702 | 0.84 | 0.83 | 0.9 | 0.99 | 0.86 | 0.87 |
| 2 | 161784214 | rs16845318 | -0.877 | 1.189 | -1.257 | 0.757 | 0.01 | -0.017 | -0.017 | 0 | 0.06 | 0.02 | 0.09 | 0.17 | 0.06 | 0.23 |
| 2 | 161784702 | rs1722634 | 0.663 | 1.345 | -1.161 | 0.856 | 0.005 | 0.081 | 0.081 | 0.823 | 0.89 | 0.82 | 0.94 | 0.65 | 0.91 | 0.88 |
| 2 | 161785169 | rs1829975 | -0.209 | 1.042 | -0.973 | 0.732 | -0.002 | 0.024 | 0.024 | 0.012 | 0.16 | 0.2 | 0.22 | 0.32 | 0.17 | 0.26 |
| 2 | 161802019 | rs2357973 | 0.03 | 1.08 | -0.993 | 0.469 | -0.007 | -0.076 | -0.076 | 0 | 0.23 | 0.29 | 0.17 | 0.01 | 0.18 | 0.17 |
| 2 | 161808966 | rs1722642 | 0.833 | 1.034 | -0.943 | 0.735 | 0.005 | 0.188 | 0.188 | 0 | 0.94 | 0.85 | 0.97 | 0.66 | 0.96 | 0.89 |
| 2 | 161816754 | rs9287801 | 0.87 | 1.363 | -1.105 | 0.637 | 0.011 | 0.011 | 0.011 | 0 | 0.63 | 0.55 | 0.7 | 0.51 | 0.57 | 0.61 |
| 2 | 161822765 | rs960595 | 0.873 | 1.345 | -1.091 | 0.626 | 0.011 | 0.011 | 0.011 | 0 | 0.63 | 0.55 | 0.7 | 0.5 | 0.57 | 0.61 |
| 2 | 161825001 | rs1722659 | 0.3 | 1.716 | -1.303 | 0.942 | -0.004 | 0.031 | 0.031 | 0 | 0.9 | 0.92 | 0.89 | 0.51 | 0.76 | 0.78 |
| 2 | 161840835 | rs1850178 | 0.844 | 1.661 | -1.27 | 0.767 | 0.012 | 0.009 | 0.009 | 0 | 0.87 | 0.83 | 0.88 | 0.78 | 0.77 | 0.85 |
| 2 | 161842590 | rs2048827 | 1.56 | 1.073 | -1.917 | 0.039 | -0.031 | 0.162 | 0.162 | 0 | 0.24 | 0.01 | 0.39 | 0.01 | 0.23 | 0.25 |
| 2 | 161849209 | rs17289782 | 0.455 | 1.828 | -1.312 | 0.74 | -0.037 | -0.041 | -0.041 | 0 | 0.07 | 0.01 | 0.1 | 0.17 | 0.21 | 0.08 |
| 2 | 161863052 | rs10208511 | -0.181 | 1.216 | -0.529 | 0.769 | -0.004 | -0.001 | -0.001 | 0.717 | 0.24 | 0.34 | 0.17 | 0.27 | 0.21 | 0.23 |
| 2 | 161866727 | rs2102759 | 0.472 | 1.439 | -0.373 | 0.719 | 0.013 | -0.008 | -0.008 | 0 | 0.69 | 0.51 | 0.75 | 0.73 | 0.64 | 0.69 |
| 2 | 161877135 | rs4664030 | -0.625 | 1.457 | -0.463 | 0.74 | -0.018 | -0.005 | -0.005 | 0 | 0.28 | 0.23 | 0.22 | 0.26 | 0.36 | 0.28 |
| 2 | 161878831 | rs13004784 | 1.359 | 0.842 | -0.931 | 0.5 | -0.016 | -0.067 | -0.067 | 0 | 0.2 | 0.02 | 0.09 | 0.3 | 0.17 | 0.12 |
| 2 | 161889193 | rs2551691 | -0.776 | 0.843 | -0.678 | 0.364 | -0.017 | -0.08 | -0.08 | 0.248 | 0.81 | 0.7 | 0.87 | 1 | 0.85 | 0.85 |
| 2 | 161891382 | rs6757645 | 1.227 | 1.064 | -0.189 | 0.467 | -0.003 | -0.009 | -0.009 | 0 | 0.37 | 0.44 | 0.56 | 0.31 | 0.34 | 0.36 |
| 2 | 161894663 | rs197273 | 0.258 | 1.301 | -0.42 | 0.567 | -0.023 | -0.01 | -0.01 | 0 | 0.54 | 0.72 | 0.7 | 0.45 | 0.49 | 0.61 |
| 2 | 161899538 | rs10490561 | -0.183 | 1.821 | -0.244 | 0.854 | 0.002 | 0.018 | 0.018 | 0 | 0.07 | 0.08 | 0.09 | 0.25 | 0.19 | 0.12 |
| 2 | 161901650 | rs4664396 | 0.288 | 1.416 | 0.031 | 0.532 | 0.002 | 0.004 | 0.004 | 0.057 | 0.48 | 0.53 | 0.68 | 0.57 | 0.56 | 0.5 |
| 2 | 161902484 | rs197265 | -0.349 | 1.471 | 0.288 | 0.088 | -0.007 | -0.07 | -0.07 | 0 | 0.8 | 0.74 | 0.86 | 1 | 0.81 | 0.86 |
| 2 | 161902546 | rs4664398 | -0.75 | 1.664 | 0.587 | 0.417 | 0.139 | 0.174 | 0.174 | 0 | 0.07 | 0.28 | 0.13 | 0.22 | 0.17 | 0.14 |
| 2 | 161913992 | rs9287806 | 0.04 | 1.389 | 0.851 | 0.257 | -0.044 | -0.152 | -0.152 | 0 | 0.78 | 0.46 | 0.83 | 1 | 0.82 | 0.85 |
| 2 | 161915361 | rs9287807 | 0.643 | 1.254 | 0.585 | 0.279 | 0.019 | -0.015 | -0.015 | 0.136 | 0.3 | 0.15 | 0.21 | 0.37 | 0.27 | 0.24 |
| 2 | 161916409 | rs747003 | 1.242 | 1.302 | 0.465 | 0.251 | -0.013 | -0.014 | -0.014 | 0 | 0.39 | 0.2 | 0.51 | 0.41 | 0.36 | 0.48 |
| 2 | 161916743 | rs1114337 | -0.032 | 1.307 | 0.374 | 0.227 | 0.014 | 0.028 | 0.028 | 0 | 0.05 | 0.06 | 0.11 | 0.22 | 0.18 | 0.12 |
| 2 | 161917192 | rs889922 | -0.193 | 1.211 | 0.448 | 0.147 | -0.032 | -0.119 | -0.119 | 0.436 | 0.78 | 0.58 | 0.85 | 1 | 0.82 | 0.85 |
| 2 | 161931112 | rs13013777 | 1.66 | 1.139 | -0.09 | 0.3 | -0.018 | -0.053 | -0.053 | 0 | 0.21 | 0.02 | 0.09 | 0.25 | 0.16 | 0.13 |
| 2 | 161938839 | rs6432670 | -0.091 | 1.107 | 0.347 | 0.133 | -0.02 | -0.083 | -0.083 | 0.132 | 0.78 | 0.66 | 0.86 | 1 | 0.82 | 0.84 |
| 2 | 161942911 | rs1991110 | -1.015 | 1.066 | 0.329 | 0.091 | 0.014 | 0.055 | 0.055 | 0 | 0.86 | 0.96 | 0.91 | 1 | 0.86 | 0.9 |
| 2 | 161949299 | rs6750514 | 0.789 | 1.29 | 0.656 | 0.177 | 0.001 | -0.012 | -0.012 | 0 | 0.69 | 0.36 | 0.72 | 0.78 | 0.64 | 0.73 |
| 2 | 161952938 | rs4664405 | -0.428 | 1.32 | 0.679 | 0.164 | -0.006 | -0.049 | -0.049 | 0 | 0.8 | 0.74 | 0.87 | 1 | 0.82 | 0.85 |
| 2 | 161965564 | rs16845639 | -0.367 | 1.441 | 0.548 | 0.299 | 0.113 | 0.152 | 0.152 | 0 | 0.05 | 0.16 | 0.12 | 0.22 | 0.17 | 0.11 |
| 2 | 161977647 | rs2884364 | 1.064 | 1.566 | 0.441 | 0.301 | -0.01 | -0.013 | -0.013 | 0 | 0.39 | 0.21 | 0.52 | 0.41 | 0.36 | 0.49 |
| 2 | 161979394 | rs13391342 | -1.1 | 1.551 | 0.347 | 0.317 | 0.12 | 0.164 | 0.164 | 0.027 | 0.08 | 0.3 | 0.14 | 0.22 | 0.18 | 0.11 |
| 2 | 161991197 | rs17705608 | 1.616 | 1.035 | -0.121 | -0.096 | -0.127 | 0.185 | 0.185 | 0 | 0.3 | 0.06 | 0.42 | 0.01 | 0.38 | 0.22 |

| | | | | | | | | | | | | | | | | |
|---|---|---|---|---|---|---|---|---|---|---|---|---|---|---|---|---|
| 2 | 162012135 | rs1267068 | 0.161 | 0.928 | 0.29 | 0.316 | 0.016 | -0.02 | -0.02 | 0 | 0.33 | 0.14 | 0.24 | 0.39 | 0.3 | 0.24 |
| 2 | 162018643 | rs1267059 | 0.256 | 0.935 | 0.511 | 0.327 | -0.028 | -0.015 | -0.015 | 0 | 0.74 | 0.4 | 0.79 | 0.81 | 0.77 | 0.74 |
| 2 | 162059817 | rs1267075 | -0.375 | -0.207 | 0.605 | 0.113 | -0.005 | -0.056 | -0.056 | 0 | 0.81 | 0.73 | 0.87 | 1 | 0.81 | 0.85 |
| 2 | 162069234 | rs1267032 | -0.439 | -0.154 | 0.248 | 0.14 | -0.002 | -0.042 | -0.042 | 0 | 0.79 | 0.75 | 0.87 | 1 | 0.81 | 0.85 |
| 2 | 162070325 | rs10930013 | 0.665 | 0.299 | -0.158 | 0.532 | -0.013 | -0.008 | -0.008 | 1.787 | 0.41 | 0.25 | 0.55 | 0.42 | 0.47 | 0.5 |
| 2 | 162072738 | rs1267034 | 0.012 | 0.226 | -0.44 | 0.564 | -0.009 | 0.004 | 0.004 | 0 | 0.1 | 0.05 | 0.11 | 0.08 | 0.13 | 0.09 |
| 2 | 162092640 | rs7309 | 0.355 | 0.373 | -1.011 | 0.645 | -0.028 | -0.014 | -0.014 | 0 | 0.41 | 0.22 | 0.56 | 0.42 | 0.46 | 0.5 |
| 2 | 162107295 | rs3731769 | 0.701 | -0.099 | -0.567 | 0.706 | -0.006 | 0.007 | 0.007 | 0 | 0.67 | 0.4 | 0.72 | 0.51 | 0.68 | 0.59 |
| 2 | 162114060 | rs1882364 | 0.59 | -0.161 | 0.104 | 0.673 | -0.012 | 0.029 | 0.029 | 0 | 0.7 | 0.63 | 0.76 | 0.52 | 0.69 | 0.59 |
| 2 | 162127611 | rs1267044 | -0.337 | -0.318 | 0.117 | 0.84 | -0.051 | 0.094 | 0.094 | 0.154 | 0.15 | 0.31 | 0.12 | 0.3 | 0.1 | 0.21 |
| 2 | 162169631 | rs4664044 | 0.015 | -0.316 | 0.081 | 0.816 | -0.024 | -0.014 | -0.014 | 2.306 | 0.85 | 0.95 | 0.87 | 0.81 | 0.79 | 0.8 |
| 2 | 162173778 | rs10490567 | -0.395 | -0.336 | 0.186 | 0.808 | -0.054 | 0.106 | 0.106 | 0.22 | 0.16 | 0.33 | 0.12 | 0.3 | 0.1 | 0.21 |
| 2 | 162176044 | rs2013879 | 0.618 | -0.353 | 0.405 | 0.762 | 0.008 | -0.008 | -0.008 | 0.04 | 0.78 | 0.79 | 0.85 | 0.7 | 0.81 | 0.72 |
| 2 | 162183507 | rs10469684 | 1.301 | -0.344 | 0.393 | 0.783 | -0.013 | 0.025 | 0.025 | 0 | 0.69 | 0.63 | 0.75 | 0.51 | 0.69 | 0.58 |
| 2 | 162186610 | rs2113480 | 0.918 | -0.373 | 0.399 | 0.76 | 0.008 | -0.005 | -0.005 | 0.077 | 0.74 | 0.75 | 0.82 | 0.7 | 0.77 | 0.68 |
| 2 | 162198993 | rs10197817 | 1.324 | -0.349 | 0.321 | 0.758 | -0.017 | 0.035 | 0.035 | 0 | 0.69 | 0.62 | 0.75 | 0.51 | 0.69 | 0.58 |
| 2 | 162207510 | rs3769963 | -0.326 | -0.351 | 0.342 | 0.758 | -0.033 | 0.056 | 0.056 | 0.11 | 0.14 | 0.24 | 0.12 | 0.3 | 0.1 | 0.2 |
| 2 | 162208985 | rs1111417 | -0.548 | -0.349 | 0.326 | 0.755 | -0.007 | 0.036 | 0.036 | 0 | 0.89 | 0.97 | 0.94 | 1 | 0.87 | 0.88 |
| 2 | 162211730 | rs6709356 | 1.551 | -0.362 | 0.401 | 0.751 | -0.01 | 0.014 | 0.014 | 0.005 | 0.68 | 0.4 | 0.72 | 0.51 | 0.68 | 0.58 |
| 2 | 162224495 | rs2303318 | -0.432 | -0.355 | -0.072 | 0.691 | -0.047 | 0.083 | 0.083 | 0.16 | 0.16 | 0.29 | 0.12 | 0.3 | 0.11 | 0.21 |
| 2 | 162226838 | rs1116173 | 0.828 | -0.359 | -0.13 | 0.693 | -0.003 | 0 | 0 | 0 | 0.75 | 0.74 | 0.82 | 0.7 | 0.77 | 0.68 |
| 2 | 162251437 | rs11886577 | 0.787 | -0.439 | -1.093 | 0.017 | 0.022 | -0.063 | -0.063 | 0.548 | 0.38 | 0.17 | 0.29 | 0.52 | 0.31 | 0.38 |
| 2 | 162255778 | rs7574882 | 1.659 | -0.678 | -1.946 | 0.213 | -0.044 | -0.006 | -0.006 | 1.7 | 0.26 | 0.02 | 0.14 | 0.16 | 0.25 | 0.11 |
| 2 | 162264215 | rs2018759 | 1.191 | -0.314 | -1.69 | -0.123 | -0.172 | 0.199 | 0.199 | 0 | 0.28 | 0.03 | 0.41 | 0.01 | 0.37 | 0.2 |
| 2 | 162265270 | rs16845844 | -0.493 | -0.214 | -0.69 | 0.794 | -0.053 | 0.11 | 0.11 | 0.167 | 0.16 | 0.33 | 0.12 | 0.3 | 0.1 | 0.21 |
| 2 | 162276981 | rs10175058 | 0.617 | 1.003 | -0.375 | 0.811 | 0.059 | 0.041 | 0.041 | 0 | 0.69 | 0.57 | 0.67 | 0.56 | 0.48 | 0.6 |
| 2 | 162277301 | rs3769956 | -0.247 | 0.501 | -0.011 | 0.782 | -0.013 | 0.028 | 0.028 | 0.089 | 0.08 | 0.14 | 0.07 | 0.11 | 0.08 | 0.08 |
| 2 | 162280748 | rs890076 | 0.748 | 0.708 | 0.765 | 0.695 | 0.02 | 0.042 | 0.042 | 0.247 | 0.85 | 0.55 | 0.83 | 0.84 | 0.81 | 0.85 |
| 2 | 162295445 | rs2358139 | 0.664 | 0.693 | 0.815 | 0.619 | -0.004 | -0.054 | -0.054 | 0 | 0.88 | 0.61 | 0.93 | 0.99 | 0.9 | 0.91 |
| 2 | 162297519 | rs768835 | 0.46 | 0.694 | 0.901 | 0.62 | 0.007 | -0.039 | -0.039 | 0 | 0.9 | 0.69 | 0.94 | 0.99 | 0.9 | 0.91 |
| 2 | 162325200 | rs6432686 | 0.668 | -0.162 | 0.221 | 0.084 | 0.009 | -0.026 | -0.026 | 0 | 0.35 | 0.27 | 0.33 | 0.49 | 0.27 | 0.3 |
| 2 | 162326199 | rs2194734 | -0.245 | 0.045 | -0.143 | 0.141 | -0.005 | -0.029 | -0.029 | 0.016 | 0.91 | 0.85 | 0.95 | 0.99 | 0.9 | 0.91 |
| 2 | 162350772 | rs3849350 | -0.106 | 0.045 | -0.124 | 0.141 | 0.006 | -0.035 | -0.035 | 0.016 | 0.91 | 0.75 | 0.94 | 0.99 | 0.9 | 0.91 |
| 2 | 162365202 | rs7588198 | 0.997 | -0.134 | -0.187 | 0.146 | 0.003 | 0.019 | 0.019 | 0 | 0.84 | 0.55 | 0.86 | 0.84 | 0.82 | 0.81 |
| 2 | 162379760 | rs6432689 | -0.973 | -0.133 | -0.494 | 0.141 | 0.002 | 0.025 | 0.025 | 0 | 0.93 | 0.99 | 0.97 | 0.99 | 0.93 | 0.93 |
| 2 | 162404452 | rs10182681 | 0.37 | -0.14 | -0.496 | 0.139 | -0.013 | -0.031 | -0.031 | 0 | 0.89 | 0.8 | 0.95 | 0.99 | 0.92 | 0.9 |
| 2 | 162407930 | rs3911925 | 1.315 | -0.542 | -1.007 | -0.3 | -0.194 | 0.237 | 0.237 | 0 | 0.32 | 0.03 | 0.42 | 0.01 | 0.43 | 0.25 |
| 2 | 162412380 | rs10191709 | -0.802 | -0.484 | -0.944 | -0.244 | -0.003 | -0.013 | -0.013 | 0 | 0.92 | 0.9 | 0.96 | 0.99 | 0.92 | 0.91 |
| 2 | 162458401 | rs2389550 | 1.47 | -0.496 | -0.995 | -0.362 | -0.18 | 0.213 | 0.213 | 0 | 0.3 | 0.03 | 0.42 | 0.01 | 0.41 | 0.25 |
| 2 | 162468888 | rs979375 | 1.799 | -0.301 | -0.455 | 0.174 | -0.03 | 0.087 | 0.087 | 0.021 | 0.73 | 0.53 | 0.8 | 0.52 | 0.74 | 0.59 |
| 2 | 162473284 | rs1397087 | -0.387 | 0.015 | -0.46 | -0.212 | -0.003 | -0.026 | -0.026 | 0.977 | 0.92 | 0.9 | 0.95 | 1 | 0.92 | 0.91 |
| 2 | 162502191 | rs6759390 | 0.887 | -0.013 | -0.454 | -0.223 | -0.034 | -0.084 | -0.084 | 0 | 0.87 | 0.69 | 0.94 | 1 | 0.91 | 0.9 |
| 2 | 162508594 | rs1876816 | 0.908 | -0.003 | -0.454 | -0.218 | -0.034 | -0.085 | -0.085 | 2.588 | 0.87 | 0.68 | 0.94 | 1 | 0.91 | 0.9 |
| 2 | 162512296 | rs4408743 | 0.908 | -0.003 | -0.454 | -0.218 | -0.034 | -0.085 | -0.085 | 0 | 0.87 | 0.68 | 0.94 | 1 | 0.91 | 0.9 |
| 2 | 162532520 | rs1158463 | 0.371 | 0.089 | -0.778 | -0.172 | -0.005 | -0.029 | -0.029 | 0 | 0.89 | 0.85 | 0.95 | 1 | 0.92 | 0.9 |
| 2 | 162549321 | rs1040135 | 1.388 | -0.166 | -1.089 | -0.077 | -0.062 | -0.008 | -0.008 | 1.721 | 0.2 | 0.02 | 0.12 | 0.12 | 0.21 | 0.05 |
| 2 | 162551718 | rs1399650 | 1.185 | -0.412 | -0.374 | 0.22 | -0.003 | -0.003 | -0.003 | 0 | 0.84 | 0.69 | 0.87 | 0.85 | 0.82 | 0.81 |
| 2 | 162558490 | rs4664436 | 0.233 | -0.284 | -0.592 | 0.093 | 0.001 | -0.013 | -0.013 | 0 | 0.91 | 0.88 | 0.96 | 1 | 0.93 | 0.94 |
| 2 | 162580384 | rs3903713 | 1.164 | -0.245 | -0.458 | 0.193 | -0.001 | 0 | 0 | 0 | 0.86 | 0.77 | 0.88 | 0.85 | 0.83 | 0.83 |
| 2 | 162615115 | rs7581747 | -1.774 | -0.808 | 0.07 | -0.105 | -0.044 | -0.027 | -0.027 | 0 | 0.21 | 0.55 | 0.15 | 0.15 | 0.17 | 0.2 |
| 2 | 162621531 | rs7597134 | 1.233 | -0.376 | -0.733 | -0.38 | -0.171 | -0.019 | -0.019 | 0.305 | 0.2 | 0.02 | 0.19 | 0.12 | 0.4 | 0.06 |
| 2 | 162626768 | rs13006199 | -1.484 | -0.526 | -0.295 | -0.26 | -0.04 | -0.023 | -0.023 | 0 | 0.21 | 0.43 | 0.14 | 0.16 | 0.17 | 0.2 |
| 2 | 162632256 | rs16846107 | -1.074 | -0.746 | -0.178 | -0.36 | -0.012 | -0.005 | -0.005 | 0 | 0.17 | 0.23 | 0.11 | 0.15 | 0.16 | 0.18 |
| 2 | 162635439 | rs1449636 | -0.176 | -0.751 | -0.256 | -0.365 | -0.015 | -0.033 | -0.033 | 0 | 0.11 | 0.12 | 0.04 | 0.01 | 0.06 | 0.07 |
| 2 | 162636614 | rs12692642 | 0.937 | -1.463 | -1.12 | -0.953 | -0.139 | 0.103 | 0.103 | 0 | 0.36 | 0.04 | 0.27 | 0.12 | 0.46 | 0.19 |
| 2 | 162639726 | rs2389533 | -1.517 | -1.217 | -0.555 | -1.272 | -0.039 | -0.024 | -0.024 | 0.344 | 0.21 | 0.6 | 0.14 | 0.15 | 0.16 | 0.18 |
| 2 | 162640725 | rs2892792 | -1.543 | -1.558 | -1.09 | -1.52 | -0.057 | 0.201 | 0.201 | 0 | 0.72 | 0.97 | 0.62 | 0.99 | 0.72 | 0.74 |
| 2 | 162685991 | rs1667597 | -1.6 | -1.617 | -0.318 | -1.176 | -0.033 | -0.019 | -0.019 | 0 | 0.22 | 0.52 | 0.14 | 0.17 | 0.19 | 0.21 |
| 2 | 162695741 | rs1227919 | -0.14 | -1.272 | -0.696 | -0.949 | 0.043 | 0.084 | 0.084 | 0 | 0.56 | 0.73 | 0.57 | 0.76 | 0.66 | 0.48 |
| 2 | 162697503 | rs961324 | -0.351 | -1.124 | -0.466 | -1.406 | 0.015 | -0.013 | -0.013 | 0.36 | 0.6 | 0.66 | 0.57 | 0.98 | 0.68 | 0.65 |
| 2 | 162720101 | rs1227929 | -0.122 | -1.111 | -0.357 | -1.379 | 0.002 | 0 | 0 | 0 | 0.57 | 0.59 | 0.51 | 0.85 | 0.6 | 0.56 |
| 2 | 162737714 | rs6707956 | -0.407 | -1.172 | -0.348 | -1.415 | 0.002 | -0.007 | -0.007 | 0 | 0.47 | 0.47 | 0.51 | 0.15 | 0.42 | 0.44 |
| 2 | 162758123 | rs6726796 | -0.187 | -1.203 | -0.487 | -1.398 | -0.012 | -0.021 | -0.021 | 0 | 0.07 | 0.07 | 0.02 | 0 | 0.03 | 0.07 |
| 2 | 162760195 | rs6734760 | 0.508 | -1.127 | -0.089 | -1.368 | 0.016 | 0.08 | 0.08 | 0 | 0.47 | 0.33 | 0.46 | 0.84 | 0.55 | 0.54 |
| 2 | 162760974 | rs7594396 | -0.04 | -0.921 | -0.612 | -1.464 | 0.009 | -0.035 | -0.035 | 0 | 0.39 | 0.35 | 0.44 | 0.02 | 0.32 | 0.36 |
| 2 | 162772897 | rs6715439 | -0.019 | -0.943 | -0.605 | -1.475 | 0.008 | -0.037 | -0.037 | 0 | 0.39 | 0.35 | 0.44 | 0.02 | 0.32 | 0.36 |
| 2 | 162785323 | rs10930036 | -0.023 | -0.904 | -0.897 | -1.455 | 0.021 | 0.025 | 0.025 | 0 | 0.39 | 0.29 | 0.44 | 0.02 | 0.32 | 0.36 |
| 2 | 162801454 | rs2176135 | -0.608 | 0.005 | -1.126 | -1.286 | -0.066 | 0.107 | 0.107 | 0.05 | 0.81 | 0.97 | 0.86 | 0.99 | 0.77 | 0.9 |
| 2 | 162802993 | rs12474587 | 0.431 | 0.012 | -1.369 | -1.567 | -0.077 | 0.132 | 0.132 | 0 | 0.34 | 0.1 | 0.46 | 0.11 | 0.4 | 0.41 |

| | | | | | | | | | | | | | | | | |
|---|---|---|---|---|---|---|---|---|---|---|---|---|---|---|---|---|
| 2 | 162816167 | rs13011013 | 0.544 | 0.051 | -1.392 | -1.644 | -0.077 | 0.172 | 0.172 | 0 | 0.33 | 0.09 | 0.44 | 0.01 | 0.39 | 0.38 |
| 2 | 162820220 | rs16846221 | 1.066 | -0.191 | -1.292 | -1.256 | -0.016 | 0.025 | 0.025 | 0 | 0.09 | 0.15 | 0.01 | 0.25 | 0.03 | 0.24 |
| 2 | 162831523 | rs10497223 | 0.748 | -0.223 | -0.77 | -1.272 | -0.035 | 0.113 | 0.113 | 0 | 0.1 | 0.29 | 0.02 | 0.24 | 0.03 | 0.24 |
| 2 | 162834595 | rs4316948 | -1.136 | -0.17 | -0.673 | -1.475 | -0.003 | 0 | 0 | 0 | 0.8 | 0.87 | 0.85 | 0.96 | 0.79 | 0.9 |
| 2 | 162843668 | rs2052400 | -1.443 | -0.082 | -0.83 | -1.471 | -0.124 | 0.191 | 0.191 | 0 | 0.72 | 0.98 | 0.57 | 0.99 | 0.64 | 0.7 |
| 2 | 162854908 | rs2287509 | -0.79 | -0.15 | -0.76 | -1.389 | -0.078 | 0.061 | 0.061 | 0 | 0.66 | 0.95 | 0.57 | 0.78 | 0.62 | 0.48 |
| 2 | 162859000 | rs7565794 | 1.304 | -0.296 | -0.138 | -1.424 | 0.001 | -0.001 | -0.001 | 0 | 0.32 | 0.36 | 0.35 | 0.65 | 0.35 | 0.25 |
| 2 | 162864475 | rs2909447 | 1.07 | -0.328 | -0.253 | -1.446 | -0.018 | -0.022 | -0.022 | 0 | 0.07 | 0.07 | 0.02 | 0.04 | 0.01 | 0.06 |
| 2 | 162866255 | rs2909448 | -0.256 | -0.092 | -0.178 | -1.342 | -0.008 | 0.073 | 0.073 | 0 | 0.57 | 0.77 | 0.54 | 0.74 | 0.59 | 0.39 |
| 2 | 162867791 | rs2909450 | 1.425 | -0.047 | -0.257 | -1.323 | 0 | -0.04 | -0.04 | 0 | 0.2 | 0.19 | 0.15 | 0.04 | 0.22 | 0.11 |
| 2 | 162872492 | rs2268890 | -0.803 | -0.04 | -0.478 | -1.265 | -0.061 | 0.026 | 0.026 | 0 | 0.64 | 0.93 | 0.56 | 0.74 | 0.61 | 0.47 |
| 2 | 162878769 | rs3788980 | 1.008 | -0.364 | 0.011 | -0.798 | 0.011 | 0.001 | 0.001 | 0 | 0.08 | 0.04 | 0.01 | 0.26 | 0.03 | 0.22 |
| 2 | 162883962 | rs2300755 | -0.28 | 0.08 | 0.472 | -0.599 | 0.005 | 0.017 | 0.017 | 0.083 | 0.41 | 0.69 | 0.39 | 0.68 | 0.39 | 0.36 |
| 2 | 162885110 | rs2268889 | 0.213 | 0.415 | 0.196 | -0.227 | 0 | -0.004 | -0.004 | 5.273 | 0.33 | 0.37 | 0.35 | 0.68 | 0.35 | 0.27 |
| 2 | 162901327 | rs16822665 | 0.425 | 0.5 | -0.383 | -0.157 | -0.003 | -0.007 | -0.007 | 0.027 | 0.3 | 0.26 | 0.33 | 0.64 | 0.33 | 0.24 |
| 2 | 162921135 | rs3788976 | -0.381 | 0.348 | -0.288 | 0.087 | 0.003 | -0.001 | -0.001 | 0 | 0.36 | 0.61 | 0.23 | 0.36 | 0.25 | 0.42 |
| 2 | 162929300 | rs1861975 | 0.233 | 1.106 | -0.287 | -0.409 | -0.005 | 0.003 | 0.003 | 1.084 | 0.3 | 0.21 | 0.17 | 0.05 | 0.35 | 0.28 |
| 2 | 162939467 | rs10930042 | 0.261 | 0.899 | -0.355 | 0.05 | 0.024 | 0.111 | 0.111 | 0.166 | 0.48 | 0.64 | 0.31 | 0.07 | 0.4 | 0.29 |
| 2 | 162952349 | rs4420712 | -0.197 | 0.911 | -0.606 | 0.598 | -0.034 | -0.017 | -0.017 | 0 | 0.58 | 0.7 | 0.43 | 0.56 | 0.46 | 0.6 |
| 2 | 162959748 | rs2270447 | -0.22 | 0.643 | -0.268 | 0.934 | -0.002 | 0.108 | 0.108 | 0 | 0.11 | 0.19 | 0.14 | 0.36 | 0.06 | 0.3 |
| 2 | 162960873 | rs7564465 | -0.107 | 0.774 | -0.743 | 0.727 | -0.009 | -0.034 | -0.034 | 0 | 0.61 | 0.71 | 0.74 | 0.52 | 0.55 | 0.63 |
| 2 | 162961929 | rs2080712 | -0.645 | 0.631 | -0.529 | 0.452 | 0.003 | 0.071 | 0.071 | 0 | 0.55 | 0.66 | 0.39 | 0.8 | 0.54 | 0.64 |
| 2 | 162969662 | rs11897585 | -0.267 | 0.334 | 0.024 | 0.672 | 0.006 | 0.037 | 0.037 | 0 | 0.47 | 0.32 | 0.33 | 0.65 | 0.4 | 0.54 |
| 2 | 162978811 | rs12692647 | -0.096 | 0.297 | 0.105 | 0.665 | -0.011 | 0.047 | 0.047 | 0 | 0.45 | 0.19 | 0.31 | 0.64 | 0.4 | 0.54 |
| 2 | 162983749 | rs2052352 | -0.682 | 0.406 | 0.029 | 0.714 | 0 | 0.072 | 0.072 | 0 | 0.53 | 0.69 | 0.38 | 0.67 | 0.43 | 0.58 |
| 2 | 162984829 | rs13431489 | -0.473 | 0.585 | -0.484 | 0.801 | 0.007 | 0.034 | 0.034 | 0 | 0.47 | 0.34 | 0.33 | 0.67 | 0.4 | 0.55 |
| 2 | 163011235 | rs3761656 | -1.114 | -0.036 | 0.118 | 1.362 | -0.003 | 0 | 0 | 0 | 0.13 | 0.19 | 0.18 | 0.14 | 0.06 | 0.08 |
| 2 | 163030336 | rs3764917 | -0.997 | 0.116 | 0.773 | 1.754 | -0.063 | -0.089 | -0.089 | 0 | 0.13 | 0.01 | 0.1 | 0.35 | 0.19 | 0.44 |
| 2 | 163076146 | rs2075302 | 1.356 | -0.181 | 0.243 | 0.645 | -0.069 | 0.213 | 0.213 | 0.17 | 0.42 | 0.12 | 0.34 | 0.15 | 0.45 | 0.27 |
| 2 | 163082395 | rs11679244 | 1.362 | -0.184 | 0.216 | 0.781 | -0.07 | 0.212 | 0.212 | 0.186 | 0.42 | 0.12 | 0.34 | 0.15 | 0.45 | 0.26 |
| 2 | 163110536 | rs2111485 | 1.379 | -0.637 | 0.854 | 0.909 | 0.039 | 0.264 | 0.264 | 0 | 0.51 | 0.18 | 0.39 | 0.17 | 0.6 | 0.43 |
| 2 | 163124051 | rs1990760 | 1.289 | -0.612 | 1.312 | 0.969 | 0.016 | 0.25 | 0.25 | 0 | 0.53 | 0.13 | 0.39 | 0.19 | 0.61 | 0.56 |
| 2 | 163128824 | rs3747517 | 1.179 | -0.766 | 1.295 | 0.951 | -0.019 | -0.005 | -0.005 | 0 | 0.69 | 0.58 | 0.77 | 0.31 | 0.75 | 0.6 |
| 2 | 163207480 | rs3788964 | -0.705 | -0.274 | 0.489 | 1.205 | -0.024 | -0.105 | -0.105 | 0 | 0.15 | 0.01 | 0.11 | 0.36 | 0.15 | 0.3 |
| 2 | 163208893 | rs17783344 | 0.317 | -0.016 | 0.536 | 0.406 | -0.044 | 0.134 | 0.134 | 9.266 | 0.21 | 0.01 | 0.07 | 0 | 0.16 | 0.04 |
| 2 | 163224521 | rs984971 | -1.286 | 0.212 | 1.964 | 1.319 | 0.004 | 0.115 | 0.115 | 0.811 | 0.41 | 0.75 | 0.52 | 0.83 | 0.36 | 0.43 |
| 2 | 163311202 | rs6734436 | -1.862 | -0.2 | 1.862 | 1.845 | 0.057 | 0.145 | 0.145 | 0.148 | 0.07 | 0.58 | 0.28 | 0.16 | 0.06 | 0.03 |
| 2 | 163337649 | rs888274 | -1.131 | -0.305 | 1.191 | 2.004 | 0.005 | -0.064 | -0.064 | 0 | 0.31 | 0.83 | 0.53 | 0.56 | 0.27 | 0.27 |
| 2 | 163343148 | rs7572827 | -1.932 | -0.226 | 1.954 | 2.429 | 0.087 | 0.262 | 0.262 | 0.017 | 0.05 | 0.43 | 0.15 | 0.11 | 0.01 | 0.12 |
| 2 | 163358537 | rs3736200 | -0.713 | -0.436 | 0.427 | 2.6 | 0.019 | -0.147 | -0.147 | 0.123 | 0.18 | 0 | 0.08 | 0.51 | 0.12 | 0.3 |
| 2 | 163362430 | rs918159 | 1.732 | 0.049 | 1.802 | 2.585 | 0.042 | 0.262 | 0.262 | 0.035 | 0.9 | 0.46 | 0.61 | 0.76 | 0.92 | 0.82 |
| 2 | 163396349 | rs13404874 | -1.75 | 0.065 | 2.096 | 2.302 | 0.051 | 0.246 | 0.246 | 0.339 | 0.09 | 0.54 | 0.15 | 0.19 | 0.07 | 0.06 |
| 2 | 163415640 | rs7557730 | -1.514 | -0.227 | 1.213 | 2.238 | 0.006 | 0.328 | 0.328 | 1.548 | 0.08 | 0.56 | 0.14 | 0.21 | 0.01 | 0.04 |
| 2 | 163416873 | rs867824 | -0.363 | -0.127 | 0.4 | 2.02 | -0.006 | -0.006 | -0.006 | 0.576 | 0.42 | 0.53 | 0.57 | 0.37 | 0.36 | 0.55 |
| 2 | 163429955 | rs985309 | -0.475 | 0.156 | 0.622 | 2.249 | -0.011 | -0.022 | -0.022 | 0 | 0.56 | 0.72 | 0.72 | 0.48 | 0.49 | 0.59 |
| 2 | 163465698 | rs12692651 | -0.602 | 0.115 | 0.611 | 2.513 | -0.065 | 0.056 | 0.056 | 0.798 | 0.58 | 0.86 | 0.73 | 0.67 | 0.5 | 0.65 |
| 2 | 163466991 | rs2200475 | -0.605 | 0.12 | 0.616 | 2.515 | -0.087 | 0.049 | 0.049 | 0.987 | 0.58 | 0.88 | 0.73 | 0.67 | 0.5 | 0.64 |
| 2 | 163479034 | rs2389739 | -0.198 | -0.124 | 0.451 | 2.507 | 0.008 | 0.009 | 0.009 | 0.015 | 0.51 | 0.53 | 0.5 | 0.38 | 0.42 | 0.52 |
| 2 | 163509615 | rs11884055 | -1.437 | -0.452 | 1.037 | 2.835 | -0.041 | 0.085 | 0.085 | 0.115 | 0.21 | 0.6 | 0.11 | 0.49 | 0.13 | 0.21 |
| 2 | 163513014 | rs12613181 | -1.151 | -0.52 | 1.809 | 3.026 | -0.014 | 0.05 | 0.05 | 0 | 0.15 | 0.18 | 0.06 | 0.34 | 0.12 | 0.13 |
| 2 | 163515417 | rs4667811 | -0.72 | -0.178 | 1.796 | 3.529 | 0 | 0.031 | 0.031 | 0 | 0.05 | 0.02 | 0.05 | 0.25 | 0.1 | 0.14 |
| 2 | 163517449 | rs13390563 | -1.725 | -0.38 | 2.005 | 3.435 | -0.043 | 0.051 | 0.051 | 0.114 | 0.2 | 0.57 | 0.1 | 0.34 | 0.13 | 0.15 |
| 2 | 163517482 | rs13390414 | -1.679 | -0.412 | 1.999 | 3.221 | 0.05 | 0.046 | 0.046 | 0.651 | 0.05 | 0.4 | 0.04 | 0 | 0 | 0.02 |
| 2 | 163519902 | rs13022851 | -1.667 | -0.526 | 2.003 | 3.715 | -0.016 | 0.052 | 0.052 | 0 | 0.19 | 0.21 | 0.1 | 0.5 | 0.14 | 0.25 |
| 2 | 163520048 | rs13400734 | -2.272 | -0.666 | 2.252 | 3.972 | 0.003 | 0.128 | 0.128 | 3.249 | 0.06 | 0.68 | 0.07 | 0.09 | 0 | 0.03 |
| 2 | 163537420 | rs1435009 | -2.398 | -0.64 | 2.32 | 3.951 | 0.047 | -0.06 | -0.06 | 0.475 | 0.25 | 0.82 | 0.16 | 0.59 | 0.15 | 0.28 |
| 2 | 163571544 | rs10199249 | -1.469 | -0.723 | 3.093 | 4.014 | -0.013 | 0.024 | 0.024 | 0 | 0.18 | 0.14 | 0.06 | 0.35 | 0.13 | 0.17 |
| 2 | 163573790 | rs6736615 | -2.242 | -0.73 | 2.898 | 3.894 | 0.04 | -0.047 | -0.047 | 0.003 | 0.25 | 0.77 | 0.17 | 0.59 | 0.15 | 0.26 |
| 2 | 163576941 | rs9677443 | -2.115 | -0.728 | 3.027 | 3.901 | -0.032 | 0.048 | 0.048 | 0 | 0.23 | 0.61 | 0.15 | 0.57 | 0.14 | 0.25 |
| 2 | 163577551 | rs13390307 | -1.56 | -0.721 | 3.06 | 3.911 | -0.043 | 0.168 | 0.168 | 0 | 0.19 | 0.43 | 0.1 | 0.42 | 0.13 | 0.17 |
| 2 | 163595967 | rs11682423 | -2.033 | -0.661 | 2.052 | 3.638 | 0.055 | -0.066 | -0.066 | 0.009 | 0.25 | 0.83 | 0.17 | 0.59 | 0.14 | 0.26 |
| 2 | 163611203 | rs305697 | 1.198 | -0.745 | 1.78 | 3.133 | -0.06 | 0.168 | 0.168 | 0 | 0.84 | 0.72 | 0.87 | 0.48 | 0.92 | 0.91 |
| 2 | 163627077 | rs1434999 | -1.099 | -0.024 | 1.515 | 2.422 | 0.01 | 0.035 | 0.035 | 0 | 0.22 | 0.5 | 0.29 | 0.29 | 0.21 | 0.14 |
| 2 | 163636185 | rs1273201 | -1.149 | 0.111 | 2.269 | 2.267 | 0.012 | 0.005 | 0.005 | 0 | 0.1 | 0.35 | 0.15 | 0.09 | 0.11 | 0.14 |
| 2 | 163638249 | rs16847319 | 1.453 | 0.07 | 2.712 | 2.328 | 0.006 | 0.027 | 0.027 | 0 | 0.87 | 0.48 | 0.82 | 0.88 | 0.89 | 0.83 |
| 2 | 163639107 | rs1032412 | 2.045 | 0.049 | 2.45 | 2.991 | 0.001 | -0.086 | -0.086 | 0 | 0.65 | 0.13 | 0.55 | 0.43 | 0.7 | 0.71 |
| 2 | 163639601 | rs6729147 | -0.901 | 0.038 | 2.576 | 3.082 | 0.001 | 0.008 | 0.008 | 0 | 0.21 | 0.3 | 0.26 | 0.28 | 0.19 | 0.1 |
| 2 | 163643407 | rs1017406 | 2.035 | 0.038 | 2.483 | 3.051 | 0.005 | -0.074 | -0.074 | 0.122 | 0.65 | 0.16 | 0.56 | 0.43 | 0.7 | 0.71 |
| 2 | 163661011 | rs12622595 | -1.322 | 0.477 | 2.099 | 2.515 | 0.002 | -0.003 | -0.003 | 0.472 | 0.28 | 0.32 | 0.32 | 0.28 | 0.31 | 0.11 |

| | | | | | | | | | | | | | | | |
|---|---|---|---|---|---|---|---|---|---|---|---|---|---|---|---|
| 2 | 163671035 | rs16847359 | -0.318 | 0.416 | 2.512 | 2.489 | 0.001 | -0.002 | -0.002 | 0 | 0.07 | 0.07 | 0.11 | 0.01 | 0.06 | 0.11 |
| 2 | 163692347 | rs10169170 | 1.56 | 0.45 | 2.568 | 2.673 | -0.007 | -0.018 | -0.018 | 0 | 0.67 | 0.33 | 0.62 | 0.53 | 0.65 | 0.86 |
| 2 | 163697314 | rs13031375 | -0.3 | 0.522 | 2.801 | 2.642 | -0.006 | -0.007 | -0.007 | 0 | 0.1 | 0.07 | 0.2 | 0.24 | 0.16 | 0.04 |
| 2 | 163708978 | rs11903092 | 2.107 | 0.512 | 2.68 | 2.611 | -0.013 | -0.006 | -0.006 | 0 | 0.61 | 0.26 | 0.53 | 0.6 | 0.59 | 0.75 |
| 2 | 163710056 | rs12617678 | 1.624 | 0.52 | 2.684 | 2.616 | -0.004 | -0.004 | -0.004 | 1.346 | 0.68 | 0.37 | 0.63 | 0.6 | 0.65 | 0.86 |
| 2 | 163723667 | rs10174959 | 1.76 | 0.515 | 2.597 | 2.566 | 0.001 | 0 | 0 | 0 | 0.64 | 0.44 | 0.55 | 0.6 | 0.6 | 0.75 |
| 2 | 163738632 | rs2014527 | 1.604 | 0.461 | 2.624 | 2.665 | -0.005 | 0.023 | 0.023 | 0.268 | 0.75 | 0.39 | 0.7 | 0.61 | 0.77 | 0.89 |
| 2 | 163738883 | rs1834216 | 2.727 | 0.403 | 2.545 | 2.641 | -0.02 | -0.058 | -0.058 | 0 | 0.58 | 0.12 | 0.5 | 0.43 | 0.59 | 0.69 |
| 2 | 163739071 | rs10451557 | -1.64 | 0.368 | 3.04 | 2.757 | -0.002 | 0.033 | 0.033 | 0.145 | 0.2 | 0.33 | 0.22 | 0.3 | 0.19 | 0.08 |
| 2 | 163739189 | rs993283 | 2.009 | 0.368 | 2.905 | 2.625 | -0.003 | 0.013 | 0.013 | 0.791 | 0.73 | 0.35 | 0.69 | 0.45 | 0.76 | 0.86 |
| 2 | 163767608 | rs307864 | -2.152 | 0.361 | 3.013 | 2.691 | -0.013 | 0.095 | 0.095 | 0.272 | 0.26 | 0.5 | 0.28 | 0.56 | 0.23 | 0.15 |
| 2 | 163775203 | rs6432738 | 2.447 | -0.235 | 1.599 | 2.007 | -0.016 | -0.006 | -0.006 | 0 | 0.47 | 0.17 | 0.51 | 0.45 | 0.52 | 0.64 |
| 2 | 163781234 | rs16847639 | -0.669 | -0.106 | 1.926 | 2.172 | -0.013 | 0.015 | 0.015 | 0.166 | 0.06 | 0.1 | 0.03 | 0.16 | 0.01 | 0.07 |
| 2 | 163781563 | rs307862 | -1.428 | -0.15 | 2.058 | 2.083 | 0.004 | 0.057 | 0.057 | 0.126 | 0.25 | 0.43 | 0.27 | 0.37 | 0.24 | 0.07 |
| 2 | 163781787 | rs1424760 | 2.094 | -0.158 | 1.835 | 2.042 | -0.047 | -0.059 | -0.059 | 0 | 0.65 | 0.16 | 0.61 | 0.44 | 0.62 | 0.85 |
| 2 | 163789057 | rs307857 | 1.461 | -0.262 | 1.801 | 1.842 | 0.031 | -0.065 | -0.065 | 0.444 | 0.83 | 0.64 | 0.87 | 0.96 | 0.81 | 0.88 |
| 2 | 163793048 | rs9653278 | -0.798 | -0.242 | 1.934 | 1.795 | 0.075 | -0.037 | -0.037 | 1.123 | 0.93 | 0.85 | 0.91 | 1 | 0.86 | 0.95 |
| 2 | 163797893 | rs7568706 | -0.69 | -0.11 | 2.125 | 1.888 | 0.043 | 0.087 | 0.087 | 0 | 0.07 | 0.23 | 0.09 | 0.14 | 0.08 | 0.05 |
| 2 | 163798127 | rs2081364 | -1.48 | -0.101 | 1.509 | 1.79 | 0.004 | 0.067 | 0.067 | 0.282 | 0.26 | 0.53 | 0.28 | 0.37 | 0.23 | 0.07 |
| 2 | 163811155 | rs10183854 | 1.058 | -0.051 | 1.623 | 1.625 | 0.023 | -0.093 | -0.093 | 0 | 0.84 | 0.68 | 0.89 | 1 | 0.84 | 0.91 |
| 2 | 163815994 | rs12692662 | 1.153 | -0.043 | 1.616 | 1.659 | 0.016 | -0.093 | -0.093 | 0 | 0.83 | 0.68 | 0.89 | 1 | 0.84 | 0.91 |
| 2 | 163831487 | rs7599823 | -1.082 | -0.256 | 0.002 | 1.592 | -0.003 | 0.009 | 0.009 | 0.773 | 0.29 | 0.7 | 0.27 | 0.47 | 0.26 | 0.25 |
| 2 | 163841985 | rs1834220 | -0.967 | -0.475 | 0.091 | 1.447 | -0.013 | 0.031 | 0.031 | 0.382 | 0.31 | 0.64 | 0.27 | 0.47 | 0.27 | 0.24 |
| 2 | 163848125 | rs10194641 | -2.056 | -0.446 | -0.64 | 1.083 | -0.014 | 0.01 | 0.01 | 0 | 0.62 | 0.8 | 0.52 | 0.65 | 0.58 | 0.41 |
| 2 | 163851763 | rs12990598 | 0.291 | -0.518 | -0.816 | 0.951 | -0.113 | 0.075 | 0.075 | 0 | 0.21 | 0.02 | 0.16 | 0.09 | 0.27 | 0.08 |
| 2 | 163851855 | rs10930087 | -0.635 | -0.621 | -0.778 | 1.013 | 0.008 | -0.018 | -0.018 | 0 | 0.32 | 0.74 | 0.27 | 0.47 | 0.27 | 0.22 |
| 2 | 163858149 | rs13033137 | 0.216 | -0.532 | -0.718 | 0.874 | -0.046 | 0.059 | 0.059 | 0 | 0.21 | 0.08 | 0.17 | 0.09 | 0.27 | 0.08 |
| 2 | 163867002 | rs6740318 | -0.191 | -0.681 | -0.166 | 1.18 | -0.007 | -0.019 | -0.019 | 0 | 0.25 | 0.44 | 0.18 | 0.18 | 0.24 | 0.14 |
| 2 | 163885183 | rs10200294 | -2.1 | -0.591 | -0.253 | 1.201 | -0.003 | 0.009 | 0.009 | 0.213 | 0.58 | 0.72 | 0.45 | 0.39 | 0.55 | 0.32 |
| 2 | 163894302 | rs918932 | -0.282 | -0.58 | -0.434 | 1.084 | -0.034 | 0.068 | 0.068 | 0 | 0.24 | 0.14 | 0.17 | 0.08 | 0.27 | 0.07 |
| 2 | 163909113 | rs13413840 | 0.26 | -0.572 | -1.026 | 0.99 | -0.068 | 0.078 | 0.078 | 0 | 0.21 | 0.07 | 0.17 | 0.08 | 0.27 | 0.08 |
| 2 | 163921145 | rs16848037 | 0.137 | -0.547 | -0.453 | 0.937 | -0.002 | -0.003 | -0.003 | 0 | 0.22 | 0.33 | 0.14 | 0.21 | 0.23 | 0.14 |
| 2 | 163933929 | rs9287826 | -0.713 | -0.39 | -0.46 | 1.026 | 0.004 | 0.005 | 0.005 | 0 | 0.28 | 0.34 | 0.25 | 0.37 | 0.28 | 0.14 |
| 2 | 163943304 | rs2389844 | 0.655 | -0.546 | -0.638 | 1.062 | 0.038 | -0.028 | -0.028 | 0.037 | 0.09 | 0.02 | 0.09 | 0.1 | 0.04 | 0.11 |
| 2 | 163944063 | rs11896390 | -0.145 | -0.55 | -0.585 | 1.079 | -0.026 | 0.062 | 0.062 | 0.093 | 0.11 | 0.15 | 0.15 | 0.36 | 0.04 | 0.17 |
| 2 | 163965160 | rs10167931 | -1.169 | -0.49 | -0.89 | 0.948 | 0 | 0.025 | 0.025 | 0 | 0.35 | 0.28 | 0.28 | 0.18 | 0.31 | 0.18 |
| 2 | 163968169 | rs2451280 | -2.305 | -0.489 | -1.526 | 0.617 | -0.04 | -0.001 | -0.001 | 0.18 | 0.66 | 0.87 | 0.55 | 0.68 | 0.62 | 0.41 |
| 2 | 163977186 | rs1586627 | 0.422 | -0.429 | -1.463 | 0.947 | -0.11 | -0.017 | -0.017 | 0.238 | 0.21 | 0.02 | 0.19 | 0.15 | 0.28 | 0.08 |
| 2 | 163987112 | rs2451281 | -1.943 | -0.537 | -1.435 | 0.828 | -0.083 | 0.105 | 0.105 | 0 | 0.61 | 0.9 | 0.54 | 0.74 | 0.56 | 0.37 |
| 2 | 163988314 | rs1906891 | -2.763 | -0.556 | -1.636 | 0.896 | -0.118 | 0.053 | 0.053 | 0.358 | 0.72 | 0.98 | 0.61 | 0.83 | 0.65 | 0.42 |
| 2 | 163999139 | rs6432750 | -1.243 | -1.048 | 0.077 | 1.928 | -0.001 | 0 | 0 | 0 | 0.36 | 0.37 | 0.35 | 0.51 | 0.31 | 0.26 |
| 2 | 164003374 | rs6755666 | -1.222 | -1.038 | -0.011 | 1.904 | 0.001 | -0.004 | -0.004 | 0.116 | 0.35 | 0.34 | 0.39 | 0.55 | 0.31 | 0.25 |
| 2 | 164005790 | rs6738745 | -0.917 | -1.77 | 0.019 | 1.685 | 0.02 | 0.027 | 0.027 | 0 | 0.33 | 0.23 | 0.23 | 0.2 | 0.29 | 0.23 |
| 2 | 164009870 | rs12619849 | -1.101 | -1.786 | 0.008 | 1.773 | 0.03 | -0.038 | -0.038 | 0.007 | 0.36 | 0.24 | 0.32 | 0.48 | 0.31 | 0.27 |
| 2 | 164015688 | rs1387588 | -2.096 | -1.659 | -0.016 | 1.776 | -0.012 | 0.023 | 0.023 | 0 | 0.57 | 0.34 | 0.51 | 0.63 | 0.53 | 0.34 |
| 2 | 164017775 | rs12995161 | -0.069 | -1.767 | 0.443 | 1.713 | -0.025 | 0.004 | 0.004 | 0 | 0.21 | 0.08 | 0.19 | 0.15 | 0.22 | 0.07 |
| 2 | 164026233 | rs10497232 | 0.059 | -2.422 | 0.45 | 1.741 | 0.026 | -0.011 | -0.011 | 0 | 0.09 | 0 | 0.08 | 0.07 | 0.05 | 0.11 |
| 2 | 164038636 | rs16848118 | -1.109 | -2.425 | 0.602 | 1.738 | -0.038 | -0.009 | -0.009 | 0.166 | 0.13 | 0.24 | 0.11 | 0.1 | 0.06 | 0.13 |
| 2 | 164041560 | rs16848135 | -1.229 | -2.422 | 0.648 | 1.742 | -0.043 | -0.013 | -0.013 | 0 | 0.14 | 0.25 | 0.11 | 0.1 | 0.06 | 0.13 |
| 2 | 164048659 | rs13389923 | -0.604 | -2.406 | 1.047 | 1.885 | -0.055 | 0.059 | 0.059 | 1.508 | 0.22 | 0.51 | 0.14 | 0.29 | 0.12 | 0.07 |
| 2 | 164056323 | rs12476131 | -1.094 | -2.113 | 0.667 | 1.901 | -0.031 | -0.008 | -0.008 | 0 | 0.13 | 0.22 | 0.11 | 0.09 | 0.06 | 0.12 |
| 2 | 164091503 | rs1018151 | 1.61 | -1.782 | 0.416 | 2.448 | 0.094 | 0.052 | 0.052 | 0.049 | 0.58 | 0.13 | 0.68 | 0.35 | 0.75 | 0.75 |
| 2 | 164096043 | rs1587803 | 1.201 | -1.385 | 0.752 | 2.609 | -0.022 | 0.102 | 0.102 | 0 | 0.94 | 0.8 | 0.96 | 0.81 | 0.98 | 0.93 |
| 2 | 164114775 | rs4144827 | -1.054 | -1.585 | 0.836 | 1.915 | -0.089 | 0.121 | 0.121 | 0 | 0.21 | 0.47 | 0.15 | 0.42 | 0.07 | 0.12 |
| 2 | 164133258 | rs6740640 | -2.038 | -1.723 | 0.856 | 1.682 | 0.066 | 0.03 | 0.03 | 0 | 0.42 | 0.79 | 0.32 | 0.63 | 0.28 | 0.24 |
| 2 | 164145180 | rs7596375 | -0.011 | -1.734 | 0.939 | 1.628 | 0.046 | 0.051 | 0.051 | 0.096 | 0.14 | 0.03 | 0.03 | 0 | 0.05 | 0.02 |
| 2 | 164149690 | rs6752189 | -1.651 | -1.732 | 1.541 | 1.956 | 0.012 | -0.008 | -0.008 | 0.927 | 0.32 | 0.28 | 0.15 | 0.41 | 0.15 | 0.14 |
| 2 | 164180977 | rs1015797 | -0.026 | -2.145 | 1.207 | 1.974 | -0.013 | 0.024 | 0.024 | 0 | 0.13 | 0.04 | 0.12 | 0 | 0.19 | 0.08 |
| 2 | 164184329 | rs1032843 | -1.978 | -2.496 | 1.535 | 2.567 | -0.096 | 0.043 | 0.043 | 0 | 0.26 | 0.5 | 0.16 | 0.62 | 0.08 | 0.16 |
| 2 | 164187575 | rs988990 | -2.033 | -1.813 | 1.486 | 2.782 | -0.035 | 0.186 | 0.186 | 0.023 | 0.08 | 0.34 | 0.1 | 0.21 | 0.01 | 0.07 |
| 2 | 164192642 | rs1504262 | -0.594 | -1.687 | 0.357 | 2.751 | -0.015 | -0.044 | -0.044 | 0 | 0.09 | 0.02 | 0.09 | 0.31 | 0.1 | 0.09 |
| 2 | 164193154 | rs1221518 | -2.358 | -1.906 | 0.575 | 2.747 | -0.018 | 0.012 | 0.012 | 0.097 | 0.27 | 0.34 | 0.15 | 0.59 | 0.16 | 0.23 |
| 2 | 164196683 | rs1221522 | -0.926 | -1.889 | 0.321 | 2.825 | -0.029 | 0.004 | 0.004 | 0 | 0.12 | 0.2 | 0.05 | 0.1 | 0.05 | 0.07 |
| 2 | 164197701 | rs4144828 | -1.951 | -1.387 | -0.338 | 2.806 | 0.014 | -0.051 | -0.051 | 0 | 0.15 | 0.07 | 0.1 | 0.49 | 0.11 | 0.17 |
| 2 | 164209580 | rs1922092 | -0.667 | -1.191 | -0.44 | 2.363 | -0.007 | -0.043 | -0.043 | 0 | 0.09 | 0.02 | 0.09 | 0.29 | 0.09 | 0.09 |
| 2 | 164212362 | rs16848323 | -0.603 | -1.189 | -0.355 | 2.262 | 0 | 0 | 0 | 0.797 | 0.08 | 0.12 | 0.05 | 0.09 | 0.09 | 0.04 |
| 2 | 164213746 | rs16848326 | -0.603 | -1.189 | -0.335 | 2.262 | -0.003 | 0.004 | 0.004 | 0.02 | 0.08 | 0.14 | 0.06 | 0.09 | 0.09 | 0.04 |
| 2 | 164250311 | rs1676038 | -1.605 | -1.93 | -0.716 | 2.064 | 0.082 | -0.058 | -0.058 | 0 | 0.27 | 0.79 | 0.22 | 0.58 | 0.08 | 0.14 |
| 2 | 164263699 | rs2952381 | -1.954 | -1.868 | -0.572 | 1.599 | 0.049 | -0.009 | -0.009 | 1.136 | 0.22 | 0.86 | 0.16 | 0.2 | 0.08 | 0.16 |

| | | | | | | | | | | | | | | | | |
|---|---|---|---|---|---|---|---|---|---|---|---|---|---|---|---|---|
| 2 | 164275935 | rs13402758 | -1.538 | -1.896 | 0.078 | 1.364 | -0.046 | -0.056 | -0.056 | 0.653 | 0.12 | 0.18 | 0.04 | 0.01 | 0.02 | 0.06 |
| 2 | 164276423 | rs13428507 | -1.684 | -1.899 | 0.075 | 1.494 | -0.043 | -0.008 | -0.008 | 0 | 0.13 | 0.18 | 0.04 | 0.08 | 0.02 | 0.1 |
| 2 | 164279151 | rs13413641 | -2.55 | -2.007 | 0.636 | 1.514 | -0.019 | 0 | 0 | 1.636 | 0.18 | 0.76 | 0.13 | 0.15 | 0.03 | 0.14 |
| 2 | 164282402 | rs7587898 | -1.716 | -1.87 | 0.449 | 1.379 | -0.043 | -0.008 | -0.008 | 0 | 0.13 | 0.19 | 0.04 | 0.08 | 0.02 | 0.09 |
| 2 | 164291227 | rs13401220 | -1.449 | -1.802 | 0.345 | 1.323 | -0.002 | -0.008 | -0.008 | 0 | 0.05 | 0.08 | 0.03 | 0.03 | 0.03 | 0.02 |
| 2 | 164295200 | rs7589756 | -0.544 | -1.959 | -0.556 | 0.902 | 0.002 | 0.065 | 0.065 | 0.543 | 0.35 | 0.15 | 0.43 | 0.22 | 0.32 | 0.12 |
| 2 | 164296921 | rs1462167 | 0.102 | -1.91 | -0.571 | 0.886 | -0.039 | 0.031 | 0.031 | 1.461 | 0.29 | 0.05 | 0.39 | 0.19 | 0.29 | 0.11 |
| 2 | 164297858 | rs7574166 | -0.666 | -1.963 | -0.135 | 0.99 | 0.011 | 0.028 | 0.028 | 0.468 | 0.36 | 0.25 | 0.44 | 0.29 | 0.33 | 0.16 |
| 2 | 164299829 | rs17190104 | 0.092 | -1.938 | -0.148 | 0.987 | -0.01 | 0.043 | 0.043 | 1.461 | 0.29 | 0.09 | 0.4 | 0.19 | 0.29 | 0.11 |
| 2 | 164305897 | rs12991587 | 0.14 | -1.959 | -0.156 | 0.986 | -0.011 | 0.033 | 0.033 | 1.78 | 0.28 | 0.1 | 0.4 | 0.2 | 0.29 | 0.11 |
| 2 | 164317722 | rs4668348 | -1.367 | -1.807 | 0.599 | 0.961 | -0.003 | -0.005 | -0.005 | 0 | 0.47 | 0.52 | 0.51 | 0.38 | 0.37 | 0.21 |
| 2 | 164317876 | rs12476047 | 0.015 | -1.516 | 0.076 | 0.928 | -0.034 | 0.042 | 0.042 | 1.534 | 0.3 | 0.1 | 0.42 | 0.2 | 0.32 | 0.13 |
| 2 | 164328182 | rs1870871 | -0.192 | -1.521 | 0.243 | 1.021 | -0.014 | -0.002 | -0.002 | 0 | 0.33 | 0.18 | 0.44 | 0.31 | 0.33 | 0.16 |
| 2 | 164330382 | rs6732555 | -0.059 | -1.522 | 0.21 | 1.045 | -0.06 | -0.003 | -0.003 | 0 | 0.31 | 0.07 | 0.4 | 0.25 | 0.33 | 0.13 |
| 2 | 164333146 | rs1462163 | -1.254 | -1.847 | 0.455 | 0.634 | 0 | 0.003 | 0.003 | 0 | 0.53 | 0.56 | 0.56 | 0.43 | 0.45 | 0.28 |
| 2 | 164340544 | rs1551825 | -0.8 | -1.946 | 0.628 | 0.554 | 0.005 | 0.004 | 0.004 | 0.408 | 0.46 | 0.54 | 0.51 | 0.39 | 0.39 | 0.23 |
| 2 | 164348579 | rs6713957 | -0.856 | -2.036 | 1.29 | 0.652 | 0.004 | 0.004 | 0.004 | 0 | 0.48 | 0.46 | 0.49 | 0.39 | 0.4 | 0.22 |
| 2 | 164351562 | rs2615330 | -1.403 | -2.067 | 1.376 | 0.646 | -0.004 | -0.005 | -0.005 | 0 | 0.54 | 0.75 | 0.6 | 0.5 | 0.47 | 0.28 |
| 2 | 164370435 | rs10193051 | -1.481 | -2.039 | 1.398 | 0.663 | 0.062 | 0.083 | 0.083 | 0.107 | 0.07 | 0.34 | 0.1 | 0.12 | 0.06 | 0.06 |
| 2 | 164372976 | rs16848517 | -1.068 | -2.112 | 1.458 | 0.641 | -0.029 | 0.008 | 0.008 | 0 | 0.18 | 0.42 | 0.14 | 0.18 | 0.12 | 0.14 |
| 2 | 164375814 | rs13000422 | -1.175 | -2.157 | 1.354 | 0.625 | 0.003 | 0.005 | 0.005 | 0.017 | 0.52 | 0.46 | 0.56 | 0.47 | 0.45 | 0.29 |
| 2 | 164377868 | rs1899112 | -1.328 | -1.99 | 1.354 | 0.623 | 0.056 | 0.032 | 0.032 | 0.008 | 0.07 | 0.35 | 0.07 | 0.03 | 0.05 | 0.03 |
| 2 | 164394511 | rs893945 | -0.585 | -1.91 | 1.145 | 0.72 | -0.058 | -0.012 | -0.012 | 0 | 0.14 | 0.55 | 0.09 | 0.09 | 0.06 | 0.08 |
| 2 | 164402710 | rs2704358 | -0.564 | -1.888 | 1.123 | 0.722 | -0.053 | -0.011 | -0.011 | 0 | 0.14 | 0.55 | 0.09 | 0.09 | 0.06 | 0.08 |
| 2 | 164403334 | rs7421396 | 0.146 | -1.602 | 0.431 | 0.664 | -0.043 | -0.024 | -0.024 | 0 | 0.28 | 0.08 | 0.4 | 0.28 | 0.3 | 0.12 |
| 2 | 164408314 | rs1882347 | 0.432 | -1.247 | 0.225 | 0.83 | -0.002 | 0.012 | 0.012 | 0 | 0.08 | 0.08 | 0.04 | 0.23 | 0.07 | 0.09 |
| 2 | 164409218 | rs7424428 | -1.079 | -1.498 | 0.409 | 0.466 | -0.015 | 0.011 | 0.011 | 0 | 0.3 | 0.39 | 0.2 | 0.33 | 0.23 | 0.27 |
| 2 | 164415389 | rs995711 | -1.433 | -1.372 | 1.007 | 0.069 | 0.034 | -0.025 | -0.025 | 0 | 0.12 | 0.53 | 0.34 | 0.06 | 0.11 | 0.08 |
| 2 | 164417751 | rs7608146 | -1.072 | -1.723 | 1.337 | -0.195 | -0.02 | -0.119 | -0.119 | 0.026 | 0.19 | 0.39 | 0.38 | 0.01 | 0.14 | 0.14 |
| 2 | 164421747 | rs4668420 | -0.158 | -1.663 | 1.181 | -0.162 | 0.008 | -0.029 | -0.029 | 0 | 0.06 | 0.14 | 0.29 | 0 | 0.07 | 0.04 |
| 2 | 164430333 | rs1036697 | -0.158 | -1.663 | 1.197 | -0.162 | 0.007 | -0.028 | -0.028 | 0 | 0.06 | 0.13 | 0.29 | 0 | 0.07 | 0.04 |
| 2 | 164430356 | rs16848641 | 0.038 | -2.104 | 0.752 | -0.076 | 0.013 | -0.043 | -0.043 | 0 | 0.13 | 0.04 | 0.04 | 0.22 | 0.08 | 0.15 |
| 2 | 164432175 | rs10199735 | -2.383 | -2.398 | 0.996 | -0.241 | 0.02 | 0.019 | 0.019 | 0 | 0.4 | 0.68 | 0.5 | 0.27 | 0.28 | 0.34 |
| 2 | 164435125 | rs16848651 | 0.084 | -2.306 | 0.779 | -0.168 | 0.013 | -0.037 | -0.037 | 0.017 | 0.13 | 0.07 | 0.04 | 0.22 | 0.07 | 0.15 |
| 2 | 164436645 | rs7597888 | -0.743 | -2.31 | 0.186 | -0.383 | 0.01 | -0.028 | -0.028 | 0 | 0.73 | 0.81 | 0.8 | 0.45 | 0.75 | 0.8 |
| 2 | 164436807 | rs10169055 | -0.242 | -2.026 | 0.361 | -0.37 | 0.024 | -0.032 | -0.032 | 0.299 | 0.07 | 0.22 | 0.3 | 0.01 | 0.07 | 0.05 |
| 2 | 164439331 | rs7568213 | 1.132 | -2.169 | -0.207 | -0.427 | -0.017 | 0.026 | 0.026 | 1.352 | 0.34 | 0.25 | 0.35 | 0.26 | 0.55 | 0.53 |
| 2 | 164440431 | rs4668423 | -0.445 | -2.337 | -0.095 | -0.606 | -0.002 | -0.002 | -0.002 | 0 | 0.32 | 0.33 | 0.37 | 0.24 | 0.22 | 0.21 |
| 2 | 164441947 | rs12692697 | -1.176 | -2.312 | -0.371 | -0.581 | 0.047 | 0.05 | 0.05 | 0 | 0.48 | 0.62 | 0.45 | 0.25 | 0.29 | 0.29 |
| 2 | 164444305 | rs12692700 | 0.83 | -2.185 | -0.576 | -0.458 | -0.016 | -0.03 | -0.03 | 0 | 0.6 | 0.49 | 0.58 | 0.79 | 0.74 | 0.6 |
| 2 | 164445182 | rs16848671 | 0.496 | -2.205 | -0.678 | -0.27 | -0.001 | 0.051 | 0.051 | 0 | 0.16 | 0.22 | 0.16 | 0.53 | 0.16 | 0.09 |
| 2 | 164447403 | rs10185395 | -0.647 | -2.445 | -0.793 | -0.411 | 0.018 | -0.034 | -0.034 | 1.09 | 0.28 | 0.23 | 0.21 | 0.53 | 0.21 | 0.16 |
| 2 | 164449900 | rs10930107 | 0.334 | -2.449 | -0.487 | -0.31 | -0.059 | -0.105 | -0.105 | 0 | 0.65 | 0.53 | 0.6 | 0.87 | 0.8 | 0.81 |
| 2 | 164453389 | rs11904552 | -1.595 | -1.75 | 0.369 | 0.365 | -0.036 | 0.067 | 0.067 | 0 | 0.27 | 0.51 | 0.21 | 0.6 | 0.21 | 0.32 |
| 2 | 164454604 | rs4668424 | 0.252 | -1.644 | 0.184 | 0.469 | 0 | 0.054 | 0.054 | 0 | 0.13 | 0.17 | 0.13 | 0.52 | 0.12 | 0.07 |
| 2 | 164454947 | rs2704371 | 0.67 | -1.659 | 0.373 | 0.524 | -0.006 | 0.017 | 0.017 | 0 | 0.9 | 0.74 | 0.93 | 0.9 | 0.93 | 0.78 |
| 2 | 164457461 | rs1561606 | -0.386 | -1.614 | -0.253 | 0.551 | 0.01 | -0.022 | -0.022 | 0.33 | 0.23 | 0.22 | 0.17 | 0.53 | 0.17 | 0.12 |
| 2 | 164464777 | rs10209354 | -0.987 | -1.5 | -0.449 | 0.423 | -0.026 | 0.006 | 0.006 | 0 | 0.35 | 0.55 | 0.25 | 0.62 | 0.23 | 0.37 |
| 2 | 164468360 | rs2231899 | -1.131 | -1.181 | -0.228 | 0.738 | -0.036 | 0.041 | 0.041 | 0.059 | 0.26 | 0.48 | 0.4 | 0.44 | 0.19 | 0.19 |
| 2 | 164469186 | rs12692701 | 1.481 | -1.518 | -1.091 | 0.402 | -0.058 | -0.149 | -0.149 | 0.214 | 0.41 | 0.03 | 0.35 | 0.45 | 0.61 | 0.52 |
| 2 | 164470044 | rs16848716 | -0.13 | -1.045 | -0.112 | 0.878 | -0.017 | 0.134 | 0.134 | 0 | 0.07 | 0.18 | 0.07 | 0.32 | 0.03 | 0.06 |
| 2 | 164472105 | rs2028406 | 0.428 | -1.17 | 0.008 | 0.494 | -0.043 | -0.055 | -0.055 | 0 | 0.69 | 0.52 | 0.77 | 0.82 | 0.79 | 0.7 |
| 2 | 164477772 | rs7589668 | -1.681 | -0.708 | 0.636 | 0.921 | -0.024 | 0.079 | 0.079 | 0.158 | 0.18 | 0.34 | 0.15 | 0.45 | 0.12 | 0.29 |
| 2 | 164485597 | rs1440472 | -1.654 | -0.061 | 1.129 | 0.206 | 0.084 | -0.063 | -0.063 | 1.536 | 0.17 | 0.52 | 0.37 | 0.09 | 0.23 | 0.18 |
| 2 | 164489014 | rs13003002 | -1.358 | 0.105 | 1.002 | 0.135 | 0.049 | -0.07 | -0.07 | 0.723 | 0.13 | 0.29 | 0.34 | 0.01 | 0.23 | 0.13 |
| 2 | 164499515 | rs1561609 | 1.466 | -0.624 | 0.945 | 0.524 | -0.012 | 0.011 | 0.011 | 0 | 0.86 | 0.83 | 0.87 | 0.79 | 0.89 | 0.82 |
| 2 | 164505303 | rs1371826 | 0.865 | -0.429 | 0.743 | 0.482 | -0.001 | 0.001 | 0.001 | 0 | 0.89 | 0.9 | 0.92 | 0.85 | 0.9 | 0.88 |
| 2 | 164506159 | rs10177481 | 1.816 | -0.448 | 0.095 | 0.168 | -0.032 | 0.086 | 0.086 | 1.105 | 0.59 | 0.06 | 0.68 | 0.73 | 0.67 | 0.62 |
| 2 | 164534434 | rs1239137 | 1.823 | -0.3 | 1.748 | 0.588 | 0 | 0 | 0 | 0.404 | 0.68 | 0.35 | 0.71 | 0.54 | 0.68 | 0.67 |
| 2 | 164543036 | rs11896989 | -2.076 | -0.258 | 2.083 | 0.548 | 0.053 | 0.029 | 0.029 | 0.342 | 0.05 | 0.31 | 0.06 | 0.01 | 0.04 | 0.01 |
| 2 | 164576841 | rs13026329 | 2.101 | -0.139 | 1.507 | 0.503 | -0.016 | -0.055 | -0.055 | 2.641 | 0.66 | 0.14 | 0.69 | 0.58 | 0.68 | 0.68 |
| 2 | 164577569 | rs728700 | -0.507 | -0.059 | 1.394 | 0.565 | -0.031 | -0.007 | -0.007 | 0 | 0.05 | 0.01 | 0.1 | 0.06 | 0.11 | 0.11 |
| 2 | 164579999 | rs16848859 | 0.384 | -0.661 | 1.545 | 0.449 | -0.019 | -0.026 | -0.026 | 0 | 0.06 | 0.1 | 0.02 | 0 | 0.03 | 0.01 |
| 2 | 164581241 | rs10192119 | -1.852 | -0.154 | 1.594 | 0.731 | -0.018 | 0.036 | 0.036 | 0.192 | 0.24 | 0.59 | 0.18 | 0.36 | 0.17 | 0.15 |
| 2 | 164581991 | rs1531360 | -1.339 | 0.05 | 0.513 | 0.833 | -0.005 | 0.027 | 0.027 | 0 | 0.21 | 0.27 | 0.13 | 0.36 | 0.17 | 0.15 |
| 2 | 164609151 | rs10197470 | -1.33 | 0.437 | 1.016 | 0.354 | 0.009 | -0.062 | -0.062 | 0 | 0.27 | 0.49 | 0.51 | 0.12 | 0.28 | 0.22 |
| 2 | 164611028 | rs1380893 | 1.625 | 0.43 | 1.124 | 0.508 | -0.008 | -0.013 | -0.013 | 0.052 | 0.61 | 0.24 | 0.39 | 0.58 | 0.58 | 0.6 |
| 2 | 164611921 | rs13383835 | -0.994 | 0.414 | 1.732 | 0.367 | -0.004 | 0.002 | 0.002 | 0.151 | 0.07 | 0.26 | 0.09 | 0.01 | 0.04 | 0.02 |
| 2 | 164614747 | rs17809516 | -1.34 | 0.329 | 2.021 | 0.35 | -0.026 | 0.027 | 0.027 | 0.147 | 0.24 | 0.2 | 0.45 | 0.14 | 0.32 | 0.32 |

| | | | | | | | | | | | | | | | | |
|---|---|---|---|---|---|---|---|---|---|---|---|---|---|---|---|---|
| 2 | 164614897 | rs1031736 | -2.314 | 0.287 | 2.425 | 0.427 | -0.002 | -0.004 | -0.004 | 0.404 | 0.34 | 0.68 | 0.57 | 0.33 | 0.36 | 0.34 |
| 2 | 164615800 | rs17203738 | -0.374 | 0.36 | 2.369 | 0.377 | -0.05 | 0.015 | 0.015 | 0 | 0.15 | 0.07 | 0.39 | 0.09 | 0.24 | 0.2 |
| 2 | 164616028 | rs16848906 | -0.838 | 0.413 | 2.595 | 0.438 | -0.016 | -0.01 | -0.01 | 0 | 0.07 | 0.18 | 0.08 | 0.01 | 0.04 | 0.02 |
| 2 | 164616458 | rs6750731 | -0.313 | 0.394 | 1.737 | 0.218 | 0.013 | -0.029 | -0.029 | 0 | 0.1 | 0.23 | 0.06 | 0.04 | 0.09 | 0.13 |
| 2 | 164619926 | rs1460669 | -2.287 | 0.227 | 0.977 | 0.3 | -0.019 | 0 | 0 | 0 | 0.44 | 0.81 | 0.63 | 0.38 | 0.44 | 0.5 |
| 2 | 164630062 | rs4667702 | -1.332 | 0.235 | 0.798 | 0.338 | -0.034 | 0.027 | 0.027 | 0.437 | 0.19 | 0.61 | 0.18 | 0.2 | 0.14 | 0.14 |
| 2 | 164638458 | rs2169219 | -1.889 | 0.428 | 0.383 | 0.346 | -0.011 | -0.01 | -0.01 | 0 | 0.43 | 0.73 | 0.61 | 0.4 | 0.44 | 0.5 |
| 2 | 164645126 | rs4146580 | -1.78 | 0.511 | -0.244 | 0.231 | -0.067 | -0.003 | -0.003 | 0 | 0.42 | 0.86 | 0.61 | 0.34 | 0.46 | 0.43 |
| 2 | 164648662 | rs17810175 | -1.034 | 0.637 | 0.086 | 0.17 | 0 | -0.024 | -0.024 | 0 | 0.16 | 0.2 | 0.15 | 0.01 | 0.2 | 0.16 |
| 2 | 164653712 | rs16848928 | -1.004 | 0.326 | -0.017 | 0.556 | 0 | 0.002 | 0.002 | 0.368 | 0.14 | 0.16 | 0.1 | 0.25 | 0.13 | 0.17 |
| 2 | 164663695 | rs16848934 | -0.875 | 0.327 | 0.125 | 0.144 | 0.001 | 0.009 | 0.009 | 0.06 | 0.13 | 0.14 | 0.09 | 0.05 | 0.13 | 0.16 |
| 2 | 164665096 | rs6432773 | -1.44 | -0.1 | -0.157 | -0.544 | -0.028 | 0.183 | 0.183 | 0 | 0.54 | 0.84 | 0.64 | 0.15 | 0.54 | 0.55 |
| 2 | 164668038 | rs6734152 | -0.637 | -0.094 | 0.117 | -0.235 | 0.001 | 0.049 | 0.049 | 0 | 0.4 | 0.68 | 0.55 | 0.12 | 0.41 | 0.39 |
| 2 | 164670064 | rs17810529 | -0.311 | -0.13 | 0.578 | -0.338 | -0.011 | 0.026 | 0.026 | 0 | 0.32 | 0.25 | 0.49 | 0.08 | 0.39 | 0.37 |
| 2 | 164683893 | rs4131081 | -0.624 | -0.062 | 0.446 | -0.304 | 0.003 | -0.054 | -0.054 | 0 | 0.37 | 0.58 | 0.53 | 0.09 | 0.4 | 0.38 |
| 2 | 164700130 | rs13027834 | -0.42 | 0.579 | 0.322 | 0.123 | 0.047 | 0.006 | 0.006 | 0 | 0.28 | 0.43 | 0.26 | 0.24 | 0.36 | 0.29 |
| 2 | 164703935 | rs2921842 | -0.212 | -0.007 | -0.421 | -0.605 | -0.01 | -0.046 | -0.046 | 0 | 0.63 | 0.62 | 0.59 | 0.91 | 0.7 | 0.62 |
| 2 | 164707825 | rs438326 | 1.166 | -0.012 | -0.532 | -0.195 | -0.012 | -0.08 | -0.08 | 0 | 0.36 | 0.15 | 0.34 | 0.6 | 0.33 | 0.33 |
| 2 | 164714987 | rs429340 | -0.41 | -0.142 | -0.188 | -0.394 | 0.011 | 0.022 | 0.022 | 0 | 0.68 | 0.72 | 0.63 | 0.91 | 0.71 | 0.64 |
| 2 | 164715908 | rs4145309 | 0.223 | -0.187 | -0.297 | -0.381 | -0.015 | -0.051 | -0.051 | 0.251 | 0.59 | 0.41 | 0.51 | 0.89 | 0.69 | 0.61 |
| 2 | 164718561 | rs16848967 | 1.411 | -0.33 | 0.222 | 0.195 | -0.014 | -0.027 | -0.027 | 0 | 0.32 | 0.15 | 0.26 | 0.48 | 0.32 | 0.31 |
| 2 | 164720199 | rs12994524 | 0.727 | -0.046 | 0.34 | 0.187 | -0.038 | -0.076 | -0.076 | 0 | 0.24 | 0.08 | 0.19 | 0.34 | 0.3 | 0.28 |
| 2 | 164720735 | rs1446851 | 0.194 | -0.122 | 0.459 | 0.011 | -0.014 | -0.025 | -0.025 | 0 | 0.58 | 0.41 | 0.49 | 0.83 | 0.68 | 0.59 |
| 2 | 164729019 | rs1122116 | 0.171 | -0.227 | 0.607 | -0.046 | -0.006 | -0.031 | -0.031 | 0 | 0.59 | 0.44 | 0.51 | 0.88 | 0.69 | 0.61 |
| 2 | 164730245 | rs1123172 | 0.268 | -0.239 | 0.72 | -0.032 | 0.021 | 0.026 | 0.026 | 0 | 0.58 | 0.37 | 0.46 | 0.82 | 0.68 | 0.6 |
| 2 | 164731394 | rs717366 | 1.14 | -0.181 | 0.865 | 0.201 | 0.02 | 0.035 | 0.035 | 0 | 0.34 | 0.41 | 0.3 | 0.59 | 0.37 | 0.31 |
| 2 | 164747772 | rs1821849 | 0.178 | -0.634 | 0.894 | -0.19 | -0.013 | -0.028 | -0.028 | 0 | 0.62 | 0.48 | 0.5 | 0.83 | 0.72 | 0.58 |
| 2 | 164752462 | rs10497248 | 0.129 | -0.627 | 0.919 | -0.184 | -0.024 | -0.084 | -0.084 | 0 | 0.62 | 0.54 | 0.58 | 0.9 | 0.73 | 0.6 |
| 2 | 164774492 | rs13390293 | 1.765 | -0.461 | 0.573 | -0.116 | -0.092 | -0.086 | -0.086 | 0 | 0.29 | 0.04 | 0.25 | 0.65 | 0.35 | 0.29 |
| 2 | 164787621 | rs7574118 | 0.853 | -0.515 | 0.972 | -0.152 | 0.003 | -0.003 | -0.003 | 0 | 0.25 | 0.24 | 0.21 | 0.19 | 0.34 | 0.26 |
| 2 | 164800264 | rs357295 | 1.886 | -0.566 | 0.817 | 0.063 | -0.081 | -0.142 | -0.142 | 0 | 0.28 | 0.04 | 0.25 | 0.55 | 0.35 | 0.28 |
| 2 | 164800581 | rs995614 | 0.708 | -0.466 | 1.062 | 0.283 | -0.113 | 0.008 | 0.008 | 0 | 0.25 | 0.07 | 0.19 | 0.19 | 0.33 | 0.26 |
| 2 | 164808567 | rs3860432 | -0.585 | -0.584 | 1.257 | 0.027 | -0.002 | -0.014 | -0.014 | 0 | 0.37 | 0.53 | 0.27 | 0.3 | 0.34 | 0.29 |
| 2 | 164814041 | rs6710706 | -0.524 | -0.581 | 0.959 | 0.039 | -0.003 | -0.008 | -0.008 | 0.167 | 0.34 | 0.71 | 0.36 | 0.38 | 0.33 | 0.3 |
| 2 | 164833680 | rs6748298 | 0.44 | -0.28 | 0.546 | 0.141 | -0.044 | -0.035 | -0.035 | 0.367 | 0.24 | 0.17 | 0.2 | 0.31 | 0.31 | 0.26 |
| 2 | 164837081 | rs165226 | -0.292 | -0.425 | 0.521 | -0.021 | 0.017 | 0.063 | 0.063 | 0 | 0.68 | 0.86 | 0.65 | 0.92 | 0.72 | 0.61 |
| 2 | 164837656 | rs165223 | 1.166 | -0.417 | 0.373 | 0.04 | -0.012 | -0.078 | -0.078 | 2.123 | 0.39 | 0.15 | 0.31 | 0.53 | 0.4 | 0.32 |
| 2 | 164840529 | rs165221 | 0.771 | -0.407 | 0.501 | -0.141 | 0.003 | 0.003 | 0.003 | 0 | 0.44 | 0.46 | 0.43 | 0.61 | 0.4 | 0.35 |
| 2 | 164843556 | rs3901479 | 0.861 | -0.21 | 0.092 | -0.052 | -0.107 | -0.099 | -0.099 | 0 | 0.22 | 0.06 | 0.25 | 0.33 | 0.32 | 0.25 |
| 2 | 164846937 | rs3910511 | 0.377 | -0.148 | 0.215 | -0.037 | 0.003 | 0.002 | 0.002 | 0 | 0.24 | 0.29 | 0.27 | 0.33 | 0.32 | 0.24 |
| 2 | 164858188 | rs653310 | 1.172 | -0.566 | -0.483 | -0.246 | 0 | -0.029 | -0.029 | 0 | 0.39 | 0.24 | 0.32 | 0.59 | 0.4 | 0.35 |
| 2 | 164866669 | rs357309 | 1.238 | -0.538 | -0.624 | -0.196 | -0.019 | -0.089 | -0.089 | 0.655 | 0.39 | 0.11 | 0.31 | 0.55 | 0.4 | 0.36 |
| 2 | 164879136 | rs1516296 | 0.224 | -0.93 | 0.128 | -0.272 | 0.004 | -0.064 | -0.064 | 2.572 | 0.19 | 0.26 | 0.33 | 0.05 | 0.21 | 0.26 |
| 2 | 164885707 | rs2176241 | 1.038 | -0.951 | -0.048 | -0.331 | -0.002 | 0.03 | 0.03 | 0 | 0.11 | 0.04 | 0.25 | 0.02 | 0.11 | 0.11 |
| 2 | 164889562 | rs13017068 | -1.048 | -0.842 | 0.092 | -0.219 | 0.002 | 0.005 | 0.005 | 0.109 | 0.5 | 0.51 | 0.59 | 0.49 | 0.51 | 0.63 |
| 2 | 164894561 | rs12692709 | -1.498 | -0.748 | -0.699 | -0.194 | -0.075 | -0.06 | -0.06 | 0.313 | 0.57 | 0.89 | 0.65 | 0.49 | 0.52 | 0.63 |
| 2 | 164899430 | rs17351368 | 0.155 | -0.259 | -0.433 | 0.153 | -0.006 | -0.018 | -0.018 | 0 | 0.11 | 0.12 | 0.05 | 0.03 | 0.08 | 0.08 |
| 2 | 164915279 | rs13002621 | 0.641 | -0.105 | -0.636 | 0.327 | -0.018 | -0.049 | -0.049 | 0 | 0.2 | 0.12 | 0.2 | 0.37 | 0.22 | 0.24 |
| 2 | 164936940 | rs1546631 | -0.436 | -0.812 | -0.824 | -0.361 | 0.008 | 0.05 | 0.05 | 0.024 | 0.22 | 0.15 | 0.37 | 0.05 | 0.2 | 0.26 |
| 2 | 164940807 | rs268268 | 1.419 | -0.875 | -0.763 | -0.176 | -0.021 | -0.025 | -0.025 | 0 | 0.43 | 0.14 | 0.35 | 0.52 | 0.55 | 0.43 |
| 2 | 164941552 | rs268271 | -0.591 | -0.926 | -0.197 | -0.211 | -0.007 | -0.034 | -0.034 | 0 | 0.9 | 0.85 | 0.95 | 0.99 | 0.92 | 0.95 |
| 2 | 164952015 | rs171985 | -0.069 | -0.954 | 0.438 | -0.225 | -0.004 | -0.07 | -0.07 | 0 | 0.88 | 0.62 | 0.92 | 0.99 | 0.92 | 0.95 |
| 2 | 164952970 | rs268264 | 1.212 | -0.967 | 0.206 | -0.22 | 0.011 | -0.018 | -0.018 | 0 | 0.45 | 0.13 | 0.37 | 0.52 | 0.59 | 0.46 |
| 2 | 164961382 | rs7566116 | -0.027 | -0.674 | 0.625 | -0.291 | -0.002 | 0.004 | 0.004 | 1.494 | 0.15 | 0.16 | 0.34 | 0.06 | 0.16 | 0.22 |
| 2 | 164965571 | rs1545358 | -1.092 | -0.621 | 0.665 | -0.184 | 0.01 | -0.017 | -0.017 | 0.077 | 0.39 | 0.66 | 0.57 | 0.44 | 0.33 | 0.5 |
| 2 | 164973051 | rs2121315 | -0.966 | -0.571 | 0.735 | -0.133 | -0.005 | 0.011 | 0.011 | 0.222 | 0.37 | 0.48 | 0.56 | 0.44 | 0.33 | 0.49 |
| 2 | 164973229 | rs13014943 | 0.42 | -0.673 | 0.361 | -0.266 | -0.015 | 0.055 | 0.055 | 0 | 0.12 | 0.02 | 0.05 | 0.01 | 0.14 | 0.07 |
| 2 | 164976416 | rs13009997 | 1.266 | -0.618 | 0.198 | -0.183 | -0.034 | -0.134 | -0.134 | 1.912 | 0.17 | 0.01 | 0.17 | 0.37 | 0.17 | 0.23 |
| 2 | 164994653 | rs7603599 | 0.179 | -0.403 | 0.353 | -0.066 | -0.002 | 0.003 | 0.003 | 1.682 | 0.15 | 0.16 | 0.37 | 0.11 | 0.17 | 0.26 |
| 2 | 164999883 | rs1374264 | -1.548 | -0.436 | 0.411 | -0.055 | 0.01 | 0.012 | 0.012 | 0.418 | 0.48 | 0.64 | 0.63 | 0.49 | 0.48 | 0.57 |
| 2 | 165009070 | rs16849320 | 0.647 | -0.313 | 0.347 | -0.121 | 0.007 | -0.04 | -0.04 | 0 | 0.19 | 0.14 | 0.18 | 0.35 | 0.17 | 0.24 |
| 2 | 165022831 | rs4667455 | -0.136 | -0.249 | 0.621 | -0.572 | -0.004 | -0.067 | -0.067 | 0.441 | 0.2 | 0.37 | 0.36 | 0.03 | 0.17 | 0.26 |
| 2 | 165026230 | rs2034256 | 0.112 | -0.221 | 0.366 | -0.559 | -0.011 | 0.054 | 0.054 | 0.194 | 0.18 | 0.07 | 0.34 | 0.03 | 0.17 | 0.25 |
| 2 | 165026936 | rs6738278 | -0.095 | -0.217 | 0.392 | -0.557 | -0.001 | -0.015 | -0.015 | 0.663 | 0.19 | 0.17 | 0.35 | 0.03 | 0.17 | 0.25 |
| 2 | 165043460 | rs6712094 | -0.017 | -0.371 | 0.212 | -0.515 | 0.01 | -0.039 | -0.039 | 0 | 0.3 | 0.12 | 0.22 | 0.31 | 0.3 | 0.27 |
| 2 | 165047345 | rs4667731 | 0.12 | -0.408 | 0.455 | -0.63 | -0.001 | -0.031 | -0.031 | 0.074 | 0.19 | 0.2 | 0.36 | 0.03 | 0.17 | 0.25 |
| 2 | 165048706 | rs10172929 | 1.885 | 0.169 | 0.14 | -0.818 | -0.031 | 0.025 | 0.025 | 0.207 | 0.05 | 0 | 0.07 | 0 | 0.11 | 0.01 |
| 2 | 165049684 | rs6711015 | -1.497 | -0.427 | 0.667 | -0.62 | 0.008 | 0.012 | 0.012 | 0 | 0.53 | 0.7 | 0.62 | 0.38 | 0.47 | 0.53 |
| 2 | 165051004 | rs400022 | -0.035 | -0.335 | 0.452 | -0.964 | -0.005 | -0.099 | -0.099 | 0 | 0.79 | 0.56 | 0.62 | 0.96 | 0.83 | 0.75 |

| | | | | | | | | | | | | | | | | |
|---|---|---|---|---|---|---|---|---|---|---|---|---|---|---|---|---|
| 2 | 165056704 | rs1595642 | 0.881 | -0.445 | 0.03 | -0.842 | 0.006 | -0.083 | -0.083 | 0 | 0.19 | 0.07 | 0.18 | 0.27 | 0.16 | 0.26 |
| 2 | 165072927 | rs13035163 | -0.628 | -0.475 | 0.112 | -0.927 | 0.035 | 0.014 | 0.014 | 0 | 0.36 | 0.27 | 0.24 | 0.28 | 0.3 | 0.31 |
| 2 | 165074029 | rs1439210 | 0.679 | 0.051 | 0.025 | -0.965 | -0.017 | -0.06 | -0.06 | 0.014 | 0.23 | 0.1 | 0.2 | 0.27 | 0.23 | 0.26 |
| 2 | 165074774 | rs4667732 | 0.358 | -0.424 | 0.189 | -0.866 | -0.013 | 0.057 | 0.057 | 0.022 | 0.18 | 0.07 | 0.34 | 0.04 | 0.17 | 0.26 |
| 2 | 165078704 | rs13027527 | -0.336 | -0.396 | 0.215 | -0.797 | 0.023 | 0.01 | 0.01 | 0 | 0.36 | 0.36 | 0.25 | 0.28 | 0.3 | 0.31 |
| 2 | 165090463 | rs13398947 | 0.245 | -0.226 | 0.26 | -0.775 | 0.006 | -0.051 | -0.051 | 0.121 | 0.17 | 0.25 | 0.36 | 0.04 | 0.17 | 0.25 |
| 2 | 165096178 | rs12463826 | -0.977 | -0.335 | 0.369 | -0.772 | 0.003 | 0.008 | 0.008 | 0 | 0.51 | 0.58 | 0.61 | 0.31 | 0.46 | 0.51 |
| 2 | 165100379 | rs4667736 | -1.229 | -0.477 | 0.13 | -0.776 | 0.002 | 0.031 | 0.031 | 0 | 0.56 | 0.74 | 0.64 | 0.31 | 0.47 | 0.5 |
| 2 | 165107298 | rs9287843 | -1.209 | -0.466 | 0.106 | -0.771 | 0.003 | 0.024 | 0.024 | 0 | 0.56 | 0.7 | 0.64 | 0.31 | 0.47 | 0.5 |
| 2 | 165109326 | rs6707461 | 0.091 | -0.336 | -0.295 | -0.787 | -0.023 | 0.062 | 0.062 | 0.017 | 0.21 | 0.08 | 0.36 | 0.03 | 0.23 | 0.31 |
| 2 | 165130868 | rs16849404 | -0.348 | -0.779 | -0.028 | -0.619 | -0.019 | -0.004 | -0.004 | 1.726 | 0.35 | 0.41 | 0.25 | 0.28 | 0.24 | 0.19 |
| 2 | 165133462 | rs4260259 | -0.852 | -1.013 | 0.024 | -1.05 | -0.017 | -0.072 | -0.072 | 0 | 0.28 | 0.39 | 0.34 | 0.07 | 0.21 | 0.29 |
| 2 | 165134402 | rs4267526 | -0.598 | -1.684 | -0.183 | -1.347 | -0.073 | -0.066 | -0.066 | 0.346 | 0.23 | 0.31 | 0.09 | 0.03 | 0.08 | 0.08 |
| 2 | 165142629 | rs16849420 | -0.779 | -1.345 | 0.238 | -1.179 | -0.017 | 0.032 | 0.032 | 0.314 | 0.09 | 0.39 | 0.05 | 0.03 | 0.02 | 0.04 |
| 2 | 165161878 | rs4667744 | -0.037 | -1.346 | 0.085 | -1.188 | 0.008 | -0.022 | -0.022 | 0.038 | 0.17 | 0.22 | 0.29 | 0.05 | 0.19 | 0.2 |
| 2 | 165166534 | rs1371185 | -0.994 | -1.339 | -0.212 | -1.164 | 0.001 | -0.003 | -0.003 | 1.182 | 0.26 | 0.74 | 0.39 | 0.06 | 0.21 | 0.24 |
| 2 | 165178853 | rs4667458 | -0.912 | -1.327 | -0.329 | -1.175 | -0.004 | -0.027 | -0.027 | 0 | 0.26 | 0.7 | 0.36 | 0.06 | 0.21 | 0.23 |
| 2 | 165180232 | rs6432792 | -0.958 | -1.318 | -0.358 | -1.171 | 0.001 | -0.002 | -0.002 | 0 | 0.26 | 0.74 | 0.4 | 0.06 | 0.21 | 0.24 |
| 2 | 165180407 | rs13033944 | 0.748 | -1.226 | -0.458 | -1.163 | 0.001 | -0.004 | -0.004 | 0 | 0.11 | 0.1 | 0.24 | 0.05 | 0.11 | 0.14 |
| 2 | 165183257 | rs10167913 | -1.508 | -1.319 | -1.216 | -1.002 | -0.045 | -0.059 | -0.059 | 0 | 0.58 | 0.97 | 0.7 | 0.34 | 0.42 | 0.48 |
| 2 | 165195314 | rs1530897 | -0.727 | -0.312 | -0.474 | -0.672 | 0.007 | -0.036 | -0.036 | 0 | 0.25 | 0.62 | 0.4 | 0.09 | 0.24 | 0.24 |
| 2 | 165205260 | rs2196802 | -0.502 | -0.4 | -1.258 | -0.577 | -0.045 | -0.048 | -0.048 | 0.235 | 0.72 | 0.87 | 0.76 | 0.33 | 0.67 | 0.56 |
| 2 | 165213471 | rs13405823 | 0.901 | -1.376 | -1.743 | -1.45 | -0.07 | 0.125 | 0.125 | 0 | 0.28 | 0.07 | 0.24 | 0.02 | 0.35 | 0.18 |
| 2 | 165218517 | rs13423685 | 0.183 | -2.113 | -0.904 | -1.836 | -0.112 | -0.121 | -0.121 | 0.122 | 0.19 | 0.46 | 0.1 | 0.01 | 0.03 | 0.05 |
| 2 | 165219140 | rs10171285 | -0.786 | -2.284 | -0.499 | -1.97 | -0.019 | 0.036 | 0.036 | 0 | 0.62 | 0.69 | 0.52 | 0.07 | 0.51 | 0.43 |
| 2 | 165230397 | rs1839883 | 1.202 | -2.105 | -0.476 | -1.837 | 0.001 | 0.013 | 0.013 | 0 | 0.14 | 0.09 | 0.19 | 0.04 | 0.13 | 0.19 |
| 2 | 165231635 | rs6731651 | 0.337 | -2.066 | -0.419 | -1.904 | -0.026 | 0.097 | 0.097 | 0 | 0.43 | 0.22 | 0.43 | 0.06 | 0.48 | 0.38 |
| 2 | 165246738 | rs1025750 | 1.208 | -2.243 | -0.684 | -1.599 | 0 | -0.073 | -0.073 | 0 | 0.15 | 0.01 | 0.18 | 0.23 | 0.15 | 0.18 |
| 2 | 165252345 | rs6724372 | -0.273 | -2.493 | -0.125 | -1.634 | -0.058 | -0.064 | -0.064 | 0.014 | 0.29 | 0.51 | 0.27 | 0.08 | 0.15 | 0.23 |
| 2 | 165263217 | rs2100323 | 1.766 | -2.363 | -0.287 | -1.605 | -0.095 | -0.121 | -0.121 | 0.037 | 0.21 | 0.05 | 0.19 | 0.6 | 0.32 | 0.35 |
| 2 | 165275535 | rs10192863 | 0.149 | -2.384 | -0.004 | -1.722 | 0.003 | -0.037 | -0.037 | 0 | 0.42 | 0.48 | 0.48 | 0.9 | 0.48 | 0.56 |
| 2 | 165275993 | rs13002029 | 1.211 | -2.359 | -0.156 | -1.677 | 0.003 | 0.004 | 0.004 | 0 | 0.16 | 0.12 | 0.2 | 0.07 | 0.13 | 0.22 |
| 2 | 165285147 | rs17352892 | 0.918 | -2.371 | -0.078 | -1.683 | -0.006 | -0.008 | -0.008 | 0 | 0.18 | 0.19 | 0.2 | 0.07 | 0.13 | 0.22 |
| 2 | 165286833 | rs510806 | -1.385 | -2.108 | -0.433 | -1.598 | -0.034 | 0.057 | 0.057 | 0 | 0.71 | 0.83 | 0.72 | 0.98 | 0.63 | 0.82 |
| 2 | 165290258 | rs1947326 | 0.005 | -1.921 | -0.426 | -1.475 | -0.015 | 0.056 | 0.056 | 0 | 0.41 | 0.33 | 0.47 | 0.9 | 0.48 | 0.54 |
| 2 | 165312183 | rs13426015 | 0.672 | -1.513 | -0.097 | -1.085 | -0.009 | 0.045 | 0.045 | 0 | 0.16 | 0.28 | 0.28 | 0.29 | 0.18 | 0.19 |
| 2 | 165312786 | rs473532 | -0.971 | -1.685 | -0.017 | -1.232 | 0.043 | 0.008 | 0.008 | 0 | 0.45 | 0.68 | 0.51 | 0.36 | 0.34 | 0.47 |
| 2 | 165320390 | rs12466514 | 0.239 | -1.226 | -0.005 | -1.439 | -0.015 | 0.024 | 0.024 | 0 | 0.4 | 0.38 | 0.5 | 0.28 | 0.51 | 0.33 |
| 2 | 165323774 | rs6737277 | 0.285 | -1.159 | -0.131 | -1.724 | -0.006 | -0.022 | -0.022 | 0 | 0.43 | 0.4 | 0.49 | 0.03 | 0.5 | 0.29 |
| 2 | 165326140 | rs7580761 | 1.259 | -1.096 | -0.114 | -1.704 | -0.031 | 0.033 | 0.033 | 0 | 0.27 | 0.2 | 0.26 | 0.01 | 0.37 | 0.15 |
| 2 | 165332055 | rs10930127 | 0.562 | -1.235 | -0.075 | -1.267 | -0.011 | 0.03 | 0.03 | 0 | 0.16 | 0.19 | 0.27 | 0.36 | 0.16 | 0.2 |
| 2 | 165344029 | rs1402700 | -0.737 | -1.248 | 0.064 | -1.118 | -0.031 | -0.033 | -0.033 | 0 | 0.89 | 0.82 | 0.98 | 0.98 | 0.97 | 0.96 |
| 2 | 165345783 | rs10930129 | -0.761 | -1.242 | 0.072 | -1.113 | -0.032 | -0.033 | -0.033 | 2.253 | 0.89 | 0.82 | 0.98 | 0.98 | 0.97 | 0.96 |
| 2 | 165346592 | rs12474679 | 0.082 | -1.284 | -0.003 | -1.24 | 0.008 | 0.015 | 0.015 | 0.103 | 0.56 | 0.39 | 0.7 | 0.37 | 0.65 | 0.48 |
| 2 | 165349902 | rs3828191 | 0.028 | -1.298 | -0.029 | -1.221 | -0.001 | -0.002 | -0.002 | 0 | 0.57 | 0.43 | 0.7 | 0.37 | 0.66 | 0.48 |
| 2 | 165352165 | rs670757 | 0.377 | -1.068 | -0.126 | -1.122 | 0.002 | 0.008 | 0.008 | 0 | 0.32 | 0.44 | 0.28 | 0.61 | 0.31 | 0.48 |
| 2 | 165353323 | rs17353146 | 0.066 | -1.933 | -0.726 | -1.32 | -0.001 | -0.019 | -0.019 | 0 | 0.17 | 0.18 | 0.24 | 0.01 | 0.17 | 0.15 |
| 2 | 165356343 | rs570657 | -0.472 | -1.539 | -1.24 | -1.05 | 0.022 | -0.052 | -0.052 | 0 | 0.71 | 0.84 | 0.82 | 0.4 | 0.78 | 0.56 |
| 2 | 165364887 | rs8192674 | 0.488 | -1.696 | -1.347 | -1.319 | 0.016 | 0.072 | 0.072 | 0.653 | 0.19 | 0.07 | 0.19 | 0.01 | 0.13 | 0.11 |
| 2 | 165394580 | rs12464825 | 0.766 | -2.022 | -1.593 | -1.464 | -0.047 | 0.199 | 0.199 | 0 | 0.4 | 0.14 | 0.38 | 0.02 | 0.55 | 0.32 |
| 2 | 165395437 | rs1106257 | -0.696 | -1.628 | -1.72 | -1.116 | 0.009 | -0.045 | -0.045 | 0 | 0.73 | 0.86 | 0.82 | 0.39 | 0.77 | 0.56 |
| 2 | 165424575 | rs1402698 | -0.033 | -0.849 | -1.438 | -0.701 | 0.012 | 0.022 | 0.022 | 0.634 | 0.45 | 0.36 | 0.57 | 0.38 | 0.61 | 0.4 |
| 2 | 165424994 | rs16849597 | -0.114 | -0.291 | -1.152 | 0.401 | -0.005 | 0.033 | 0.033 | 0 | 0.11 | 0.14 | 0.19 | 0.34 | 0.09 | 0.13 |
| 2 | 165437722 | rs13027837 | -0.082 | -0.125 | -1.467 | 0.365 | -0.017 | -0.07 | -0.07 | 0.014 | 0.8 | 0.91 | 0.83 | 0.4 | 0.8 | 0.63 |
| 2 | 165438230 | rs4423615 | 0.409 | -0.496 | -1.57 | -0.411 | 0 | 0.087 | 0.087 | 0 | 0.47 | 0.24 | 0.51 | 0.35 | 0.56 | 0.44 |
| 2 | 165449620 | rs7568923 | -0.763 | -1.055 | -1.163 | -0.554 | 0.029 | 0.029 | 0.029 | 0.487 | 0.27 | 0.84 | 0.33 | 0.14 | 0.17 | 0.15 |
| 2 | 165451353 | rs11885631 | -0.482 | -1.08 | -0.002 | -0.684 | -0.029 | -0.109 | -0.109 | 0 | 0.23 | 0.37 | 0.28 | 0.01 | 0.16 | 0.12 |
| 2 | 165452742 | rs4319945 | -0.508 | -1.472 | -0.867 | -0.636 | 0.036 | -0.041 | -0.041 | 0 | 0.63 | 0.86 | 0.86 | 0.47 | 0.73 | 0.55 |
| 2 | 165458333 | rs4130269 | -0.393 | -1.449 | -0.591 | -0.856 | 0.091 | 0.05 | 0.05 | 0 | 0.6 | 0.79 | 0.87 | 0.36 | 0.78 | 0.46 |
| 2 | 165466449 | rs11895467 | 0.526 | -1.347 | -0.088 | -0.836 | -0.003 | -0.009 | -0.009 | 0.003 | 0.45 | 0.47 | 0.62 | 0.34 | 0.63 | 0.36 |
| 2 | 165468435 | rs4667765 | -0.153 | -1.403 | -0.779 | -0.463 | 0.105 | 0.02 | 0.02 | 0 | 0.6 | 0.84 | 0.87 | 0.38 | 0.79 | 0.47 |
| 2 | 165474413 | rs6754749 | -0.226 | -1.323 | -0.911 | -0.366 | 0.096 | 0.014 | 0.014 | 0 | 0.6 | 0.82 | 0.86 | 0.39 | 0.78 | 0.46 |
| 2 | 165487381 | rs6713688 | 0.105 | -0.914 | -0.781 | -0.373 | 0.001 | -0.006 | -0.006 | 0.92 | 0.11 | 0.12 | 0.08 | 0.03 | 0.11 | 0.07 |
| 2 | 165491225 | rs4425091 | -0.082 | -1.073 | -1.023 | -0.312 | 0.091 | 0.039 | 0.039 | 0 | 0.54 | 0.74 | 0.8 | 0.37 | 0.72 | 0.45 |
| 2 | 165509579 | rs7607886 | -0.877 | -0.635 | -0.466 | -0.658 | -0.005 | -0.03 | -0.03 | 0 | 0.32 | 0.62 | 0.18 | 0.04 | 0.25 | 0.15 |
| 2 | 165525147 | rs6723985 | -0.119 | -0.798 | 0.066 | -0.774 | -0.011 | -0.008 | -0.008 | 0.064 | 0.12 | 0.19 | 0.06 | 0.04 | 0.08 | 0.06 |
| 2 | 165526338 | rs7592412 | -2.038 | 0.344 | -0.214 | -0.849 | -0.006 | 0.087 | 0.087 | 0 | 0.4 | 0.76 | 0.25 | 0.1 | 0.4 | 0.21 |
| 2 | 165540800 | rs12328675 | 0.149 | 0.469 | 0.475 | -0.869 | -0.004 | -0.062 | -0.062 | 1.281 | 0.18 | 0.19 | 0.08 | 0 | 0.16 | 0.08 |
| 2 | 165548569 | rs10490694 | 0.149 | 0.496 | 0.472 | -0.88 | -0.006 | -0.066 | -0.066 | 0 | 0.18 | 0.2 | 0.08 | 0 | 0.16 | 0.08 |

| | | | | | | | | | | | | | | | | |
|---|---|---|---|---|---|---|---|---|---|---|---|---|---|---|---|---|
| 2 | 165551201 | rs7607980 | 0.149 | 0.496 | 0.45 | -0.88 | -0.005 | -0.059 | -0.059 | 5.321 | 0.18 | 0.19 | 0.08 | 0 | 0.15 | 0.08 |
| 2 | 165555207 | rs7609045 | -1.443 | 0.437 | 0.088 | -0.843 | -0.011 | 0.044 | 0.044 | 0.01 | 0.37 | 0.72 | 0.24 | 0.07 | 0.41 | 0.16 |
| 2 | 165558252 | rs12692738 | -1.304 | 0.407 | 0.181 | -0.848 | 0.006 | 0.006 | 0.006 | 0 | 0.34 | 0.68 | 0.21 | 0.06 | 0.28 | 0.16 |
| 2 | 165562893 | rs13017482 | -1.607 | 0.834 | -0.053 | -0.738 | -0.041 | 0.108 | 0.108 | 0.275 | 0.39 | 0.81 | 0.32 | 0.13 | 0.45 | 0.21 |
| 2 | 165565548 | rs3769872 | -1.872 | 0.75 | 0.514 | -0.756 | 0.029 | 0.075 | 0.075 | 0 | 0.05 | 0.25 | 0.05 | 0.09 | 0.06 | 0.04 |
| 2 | 165567695 | rs6705646 | -1.79 | 0.456 | -0.584 | -0.491 | -0.038 | -0.071 | -0.071 | 0 | 0.6 | 0.89 | 0.51 | 0.4 | 0.6 | 0.47 |
| 2 | 165568585 | rs3769874 | 0.813 | 0.406 | -0.477 | -0.427 | 0.012 | -0.02 | -0.02 | 0 | 0.21 | 0.12 | 0.19 | 0.29 | 0.15 | 0.25 |
| 2 | 165569779 | rs3769877 | -1.041 | 0.562 | -0.848 | 0.177 | -0.029 | -0.066 | -0.066 | 0.522 | 0.89 | 0.97 | 0.78 | 0.58 | 0.89 | 0.75 |
| 2 | 165573702 | rs7586858 | -0.396 | 0.315 | -0.253 | -0.093 | -0.004 | -0.002 | -0.002 | 0 | 0.17 | 0.22 | 0.1 | 0.11 | 0.16 | 0.11 |
| 2 | 165579065 | rs10167778 | -0.221 | 0.325 | 0.082 | -0.032 | 0.005 | 0.015 | 0.015 | 0 | 0.15 | 0.34 | 0.11 | 0.11 | 0.16 | 0.09 |
| 2 | 165580775 | rs13410987 | 1.139 | 0.378 | 0.231 | -0.205 | -0.002 | -0.052 | -0.052 | 1.156 | 0.17 | 0.16 | 0.08 | 0 | 0.14 | 0.09 |
| 2 | 165581707 | rs12619501 | -1.876 | 0.298 | -0.006 | -0.198 | -0.038 | -0.077 | -0.077 | 0.003 | 0.6 | 0.89 | 0.51 | 0.4 | 0.6 | 0.47 |
| 2 | 165584049 | rs3769884 | 0.095 | 0.303 | 0.105 | -0.174 | 0.011 | 0.016 | 0.016 | 0 | 0.14 | 0.33 | 0.11 | 0.08 | 0.14 | 0.06 |
| 2 | 165586466 | rs13392281 | -0.06 | 0.306 | 0.107 | -0.173 | 0.005 | 0.014 | 0.014 | 0 | 0.15 | 0.33 | 0.11 | 0.11 | 0.16 | 0.09 |
| 2 | 165592390 | rs3769885 | -1.123 | 0.299 | 0.101 | -0.198 | -0.007 | 0.049 | 0.049 | 0.448 | 0.48 | 0.68 | 0.43 | 0.35 | 0.5 | 0.4 |
| 2 | 165594756 | rs1074545 | 1.191 | 0.097 | -0.023 | -0.388 | 0.03 | 0.045 | 0.045 | 0.434 | 0.15 | 0.06 | 0.16 | 0.01 | 0.08 | 0.15 |
| 2 | 165600957 | rs3769889 | 0.508 | 0.17 | 0.024 | -0.143 | 0.017 | -0.009 | -0.009 | 0 | 0.2 | 0.12 | 0.17 | 0.25 | 0.14 | 0.18 |
| 2 | 165602181 | rs3769891 | -0.604 | 0.28 | 0.196 | -0.271 | -0.004 | -0.022 | -0.022 | 0 | 0.32 | 0.59 | 0.21 | 0.13 | 0.3 | 0.18 |
| 2 | 165612039 | rs355825 | 0.228 | 0.267 | 0.267 | -0.3 | -0.002 | -0.037 | -0.037 | 0 | 0.26 | 0.25 | 0.15 | 0.04 | 0.22 | 0.14 |
| 2 | 165616471 | rs4667772 | -0.642 | 0.096 | 0.091 | -0.022 | -0.021 | 0.016 | 0.016 | 0.725 | 0.31 | 0.38 | 0.25 | 0.39 | 0.24 | 0.24 |
| 2 | 165619163 | rs355838 | -2.271 | 0.257 | -0.136 | 0.047 | -0.08 | -0.168 | -0.168 | 0.697 | 0.62 | 0.97 | 0.61 | 0.51 | 0.64 | 0.42 |
| 2 | 165625830 | rs355849 | 1.373 | 0.278 | 0.186 | -0.148 | -0.001 | -0.031 | -0.031 | 0 | 0.16 | 0.13 | 0.08 | 0 | 0.14 | 0.1 |
| 2 | 165631433 | rs355870 | 0.19 | 0.308 | 0.143 | -0.095 | -0.006 | -0.092 | -0.092 | 0.859 | 0.26 | 0.41 | 0.17 | 0.04 | 0.23 | 0.14 |
| 2 | 165635869 | rs355895 | -1.799 | 0.215 | -0.324 | -0.079 | -0.049 | -0.059 | -0.059 | 0 | 0.59 | 0.85 | 0.42 | 0.44 | 0.46 | 0.41 |
| 2 | 165637285 | rs3820989 | 0.535 | 0.029 | -0.372 | -0.142 | 0.017 | -0.035 | -0.035 | 0 | 0.2 | 0.11 | 0.17 | 0.38 | 0.15 | 0.23 |
| 2 | 165638070 | rs355900 | -2.278 | 0.114 | -0.362 | -0.18 | -0.095 | -0.177 | -0.177 | 0.368 | 0.65 | 0.98 | 0.61 | 0.53 | 0.64 | 0.46 |
| 2 | 165647332 | rs355910 | -1.048 | 0.137 | 0.045 | -0.148 | -0.018 | 0.008 | 0.008 | 0 | 0.44 | 0.69 | 0.41 | 0.44 | 0.47 | 0.31 |
| 2 | 165647341 | rs355911 | -0.064 | 0.31 | 0.138 | -0.079 | -0.007 | 0.005 | 0.005 | 0 | 0.32 | 0.43 | 0.19 | 0.34 | 0.27 | 0.26 |
| 2 | 165665490 | rs10173063 | -0.221 | 0.229 | 0.015 | -0.133 | -0.014 | -0.018 | -0.018 | 0 | 0.24 | 0.25 | 0.2 | 0.1 | 0.18 | 0.15 |
| 2 | 165677808 | rs1509103 | -0.167 | 0.266 | 0.187 | -0.238 | 0.055 | -0.035 | -0.035 | 0 | 0.17 | 0.33 | 0.27 | 0.1 | 0.26 | 0.1 |
| 2 | 165682946 | rs355808 | -1.712 | 0.183 | 0.178 | -0.165 | -0.018 | 0.089 | 0.089 | 0 | 0.43 | 0.77 | 0.49 | 0.24 | 0.45 | 0.26 |
| 2 | 165685646 | rs355810 | -2.288 | 0.136 | -0.254 | -0.086 | -0.1 | -0.156 | -0.156 | 0 | 0.65 | 0.98 | 0.6 | 0.57 | 0.64 | 0.47 |
| 2 | 165701577 | rs6724963 | -0.438 | 0.119 | 0.421 | -0.592 | -0.011 | -0.009 | -0.009 | 0 | 0.24 | 0.25 | 0.2 | 0.13 | 0.19 | 0.15 |
| 2 | 165705829 | rs355819 | -0.414 | 0.094 | 0.668 | -0.618 | -0.022 | -0.084 | -0.084 | 0 | 0.16 | 0.43 | 0.1 | 0.03 | 0.11 | 0.08 |
| 2 | 165710110 | rs7569506 | -0.545 | 0.032 | 0.646 | -0.638 | -0.006 | -0.004 | -0.004 | 0 | 0.25 | 0.26 | 0.22 | 0.14 | 0.18 | 0.15 |
| 2 | 165712301 | rs2969609 | -0.108 | -0.139 | 0.604 | -0.658 | -0.011 | -0.088 | -0.088 | 0 | 0.14 | 0.43 | 0.09 | 0 | 0.09 | 0.06 |
| 2 | 165719734 | rs355801 | 1.68 | -0.005 | 0.447 | -0.647 | -0.012 | 0.104 | 0.104 | 0 | 0.56 | 0.2 | 0.35 | 0.76 | 0.56 | 0.74 |
| 2 | 165726586 | rs11694908 | -0.298 | 0.226 | 0.543 | -0.521 | 0 | 0 | 0 | 0.049 | 0.19 | 0.2 | 0.4 | 0.22 | 0.29 | 0.14 |
| 2 | 165727448 | rs2390124 | 0.482 | 0.249 | 0.396 | -0.517 | 0.047 | 0.001 | 0.001 | 0 | 0.77 | 0.58 | 0.59 | 0.76 | 0.7 | 0.83 |
| 2 | 165730246 | rs16849871 | 0.139 | 0.195 | 0.009 | -0.523 | -0.008 | -0.062 | -0.062 | 6.351 | 0.13 | 0.28 | 0.09 | 0 | 0.09 | 0.06 |
| 2 | 165732418 | rs1396598 | 0.529 | 0.457 | -0.012 | -0.453 | 0.065 | -0.005 | -0.005 | 0 | 0.78 | 0.48 | 0.58 | 0.78 | 0.7 | 0.86 |
| 2 | 165737931 | rs1509097 | -0.828 | 0.512 | -1.01 | -0.747 | -0.079 | 0.023 | 0.023 | 0 | 0.26 | 0.17 | 0.52 | 0.23 | 0.48 | 0.14 |
| 2 | 165739014 | rs10497251 | 2.533 | 0.213 | -1.735 | -1.521 | -0.023 | 0.149 | 0.149 | 0 | 0.22 | 0.01 | 0.14 | 0 | 0.2 | 0.11 |
| 2 | 165741394 | rs7600482 | 0.029 | 0.119 | -1.404 | -0.951 | 0.079 | -0.014 | -0.014 | 0 | 0.66 | 0.26 | 0.81 | 0.65 | 0.83 | 0.72 |
| 2 | 165747260 | rs1509104 | -0.304 | 0.087 | -1.081 | -0.957 | -0.065 | 0.012 | 0.012 | 0 | 0.69 | 0.51 | 0.87 | 0.67 | 0.84 | 0.74 |
| 2 | 165749160 | rs11679541 | -0.391 | 0.07 | -0.913 | -0.966 | -0.039 | 0.008 | 0.008 | 0 | 0.7 | 0.66 | 0.88 | 0.67 | 0.84 | 0.74 |
| 2 | 165749532 | rs12990122 | 1.258 | 0.114 | -0.963 | -0.942 | 0.001 | 0.022 | 0.022 | 0 | 0.07 | 0.03 | 0.07 | 0.01 | 0.06 | 0.07 |
| 2 | 165752007 | rs16849912 | 1.853 | 0.159 | -0.993 | -0.972 | -0.007 | -0.087 | -0.087 | 0 | 0.34 | 0.09 | 0.22 | 0.46 | 0.3 | 0.52 |
| 2 | 165753118 | rs3817135 | 1.403 | 0.191 | -0.933 | -0.932 | 0.003 | -0.06 | -0.06 | 0.586 | 0.41 | 0.14 | 0.3 | 0.46 | 0.36 | 0.58 |
| 2 | 165769051 | rs9287852 | 0.297 | 0.532 | -0.982 | -0.964 | 0.002 | 0.006 | 0.006 | 0 | 0.58 | 0.65 | 0.68 | 0.68 | 0.6 | 0.65 |
| 2 | 165773955 | rs1454406 | 0.365 | 0.65 | -0.923 | -0.924 | -0.006 | -0.005 | -0.005 | 0.012 | 0.12 | 0.13 | 0.31 | 0.19 | 0.23 | 0.06 |
| 2 | 165776177 | rs10202959 | -1.225 | 0.646 | -0.853 | -1.122 | 0.099 | -0.06 | -0.06 | 0.57 | 0.89 | 0.75 | 0.79 | 1 | 0.8 | 0.92 |
| 2 | 165785586 | rs1840328 | 0.732 | 0.743 | -1.198 | -0.986 | 0 | 0.006 | 0.006 | 0 | 0.49 | 0.53 | 0.59 | 0.7 | 0.51 | 0.56 |
| 2 | 165791732 | rs3067 | 0.721 | 0.846 | -1.304 | -0.884 | 0.001 | 0.028 | 0.028 | 0 | 0.09 | 0.05 | 0.08 | 0.01 | 0.07 | 0.1 |
| 2 | 165792339 | rs1902054 | 0.633 | 0.961 | -1.34 | -0.75 | -0.013 | -0.012 | -0.012 | 0 | 0.11 | 0.07 | 0.31 | 0.32 | 0.19 | 0.06 |
| 2 | 165794477 | rs1840329 | 0.73 | 0.957 | -1.322 | -0.765 | 0.001 | 0.028 | 0.028 | 0 | 0.09 | 0.05 | 0.08 | 0.01 | 0.07 | 0.1 |
| 2 | 165794594 | rs1840330 | -0.369 | 0.897 | -1.359 | -0.914 | 0.037 | -0.116 | -0.116 | 0.969 | 0.79 | 0.65 | 0.74 | 1 | 0.71 | 0.85 |
| 2 | 165797110 | rs4468831 | 0.607 | 0.914 | -1.375 | -0.89 | 0.014 | -0.107 | -0.107 | 0.149 | 0.7 | 0.6 | 0.66 | 0.99 | 0.64 | 0.76 |
| 2 | 165804660 | rs1869543 | 1.004 | -0.241 | -1.348 | -0.786 | 0 | 0.032 | 0.032 | 0 | 0.58 | 0.35 | 0.5 | 0.66 | 0.43 | 0.7 |
| 2 | 165805055 | rs10205696 | 1.013 | -0.24 | -1.38 | -0.773 | 0 | 0.03 | 0.03 | 0 | 0.57 | 0.35 | 0.5 | 0.62 | 0.43 | 0.7 |
| 2 | 165808057 | rs4667775 | 0.464 | 0.185 | -1.593 | -0.491 | 0.048 | -0.071 | -0.071 | 0 | 0.26 | 0.08 | 0.26 | 0.29 | 0.15 | 0.23 |
| 2 | 165812428 | rs1463342 | -0.209 | 1.296 | -0.781 | -0.824 | 0.01 | -0.086 | -0.086 | 0 | 0.85 | 0.62 | 0.78 | 0.95 | 0.85 | 0.87 |
| 2 | 165812719 | rs1454408 | -0.313 | 1.557 | -1.576 | -0.186 | -0.019 | -0.022 | -0.022 | 0 | 0.12 | 0.07 | 0.13 | 0.28 | 0.3 | 0.07 |
| 2 | 165826943 | rs10184126 | -0.818 | 0.999 | -1.587 | -0.252 | -0.057 | 0.029 | 0.029 | 0 | 0.36 | 0.19 | 0.26 | 0.31 | 0.47 | 0.24 |
| 2 | 165829017 | rs10803794 | -0.61 | 0.122 | -0.989 | -0.751 | 0.002 | -0.037 | -0.037 | 0 | 0.35 | 0.32 | 0.18 | 0.08 | 0.28 | 0.16 |
| 2 | 165834236 | rs10190215 | 0.444 | -0.25 | -1.055 | -0.52 | 0.111 | 0.106 | 0.106 | 0 | 0.47 | 0.17 | 0.52 | 0.7 | 0.33 | 0.62 |
| 2 | 165838776 | rs10497255 | 0.259 | -0.015 | -1.051 | -0.568 | 0.104 | 0.128 | 0.128 | 0 | 0.49 | 0.18 | 0.53 | 0.71 | 0.37 | 0.64 |
| 2 | 165840077 | rs10186138 | 0.013 | 0.106 | -0.933 | -0.711 | 0.003 | -0.001 | -0.001 | 0 | 0.69 | 0.29 | 0.6 | 0.71 | 0.52 | 0.77 |
| 2 | 165841197 | rs6721075 | 0.13 | -0.465 | 0.12 | -0.374 | 0.054 | 0.169 | 0.169 | 0.338 | 0.95 | 0.77 | 0.98 | 0.84 | 0.96 | 0.93 |

| | | | | | | | | | | | | | | | | |
|---|---|---|---|---|---|---|---|---|---|---|---|---|---|---|---|---|
| 2 | 165843619 | rs1993308 | 1.015 | -0.555 | 0.254 | -0.417 | 0.023 | 0.169 | 0.169 | 1.376 | 0.9 | 0.59 | 0.94 | 0.76 | 0.95 | 0.92 |
| 2 | 165858997 | rs10497256 | 0.543 | -1.043 | -0.042 | -0.684 | 0.003 | 0.11 | 0.11 | 0.313 | 0.17 | 0.01 | 0.05 | 0 | 0.13 | 0.12 |
| 2 | 165862603 | rs16850017 | 0.43 | -1.999 | -0.353 | -0.094 | 0.015 | -0.009 | -0.009 | 0 | 0.14 | 0.06 | 0.25 | 0.47 | 0.07 | 0.31 |
| 2 | 165864012 | rs11895069 | -0.391 | -2.146 | 0.074 | -0.158 | -0.067 | 0.089 | 0.089 | 0 | 0.21 | 0.37 | 0.31 | 0.53 | 0.08 | 0.35 |
| 2 | 165864123 | rs11900639 | -0.391 | -2.146 | 0.074 | -0.158 | -0.067 | 0.089 | 0.089 | 0 | 0.21 | 0.37 | 0.31 | 0.53 | 0.08 | 0.35 |
| 2 | 165868477 | rs6759510 | 0.695 | -2.225 | -0.25 | -0.192 | 0.111 | 0.035 | 0.035 | 0.062 | 0.58 | 0.22 | 0.6 | 0.34 | 0.78 | 0.44 |
| 2 | 165881866 | rs6730413 | -0.149 | -1.967 | -0.38 | 0.007 | 0.03 | -0.041 | -0.041 | 0 | 0.18 | 0.1 | 0.28 | 0.51 | 0.08 | 0.32 |
| 2 | 165902214 | rs1077617 | 0.629 | -2.091 | -0.592 | -0.214 | 0.016 | -0.006 | -0.006 | 0 | 0.12 | 0.08 | 0.12 | 0.17 | 0.08 | 0.07 |
| 2 | 165903749 | rs10930147 | -0.56 | -2.385 | -0.927 | -0.595 | 0.14 | 0.153 | 0.153 | 0 | 0.56 | 0.86 | 0.55 | 0.84 | 0.3 | 0.63 |
| 2 | 165918124 | rs11676673 | -0.065 | -0.753 | -1.25 | -0.136 | -0.013 | -0.031 | -0.031 | 0 | 0.25 | 0.05 | 0.18 | 0.27 | 0.23 | 0.31 |
| 2 | 165925668 | rs1117794 | -0.605 | -0.86 | -0.917 | -0.37 | 0.003 | 0.007 | 0.007 | 0 | 0.41 | 0.37 | 0.3 | 0.29 | 0.39 | 0.43 |
| 2 | 165926663 | rs6432808 | -1.065 | 0.365 | -1.649 | -0.29 | -0.044 | -0.054 | -0.054 | 0.348 | 0.74 | 0.89 | 0.47 | 0.4 | 0.66 | 0.6 |
| 2 | 165956508 | rs3816195 | 0.583 | 0.008 | -1.709 | -0.238 | -0.032 | -0.047 | -0.047 | 0 | 0.23 | 0.03 | 0.17 | 0.26 | 0.22 | 0.29 |
| 2 | 165972870 | rs10930150 | 0.79 | 0.202 | -1.636 | -0.36 | -0.005 | -0.004 | -0.004 | 0.366 | 0.08 | 0.11 | 0.07 | 0.1 | 0.17 | 0.03 |
| 2 | 165972903 | rs10930151 | 0.584 | 0.046 | -1.424 | -0.439 | 0.01 | -0.014 | -0.014 | 0 | 0.25 | 0.16 | 0.21 | 0.27 | 0.22 | 0.31 |
| 2 | 165980313 | rs1158135 | -1.703 | 0.615 | -1.495 | -0.437 | -0.018 | 0.033 | 0.033 | 0 | 0.77 | 0.88 | 0.72 | 0.89 | 0.74 | 0.73 |
| 2 | 165991766 | rs1439806 | 1.363 | 0.491 | -1.155 | -0.586 | -0.008 | 0.118 | 0.118 | 0 | 0.27 | 0.45 | 0.36 | 0.52 | 0.22 | 0.29 |
| 2 | 165995621 | rs13015737 | 1.048 | 0.763 | -1.082 | -0.2 | -0.002 | 0 | 0 | 0 | 0.07 | 0.11 | 0.06 | 0.07 | 0.16 | 0.03 |
| 2 | 165999868 | rs1439807 | 0.097 | 0.477 | -1.158 | -0.212 | 0.008 | -0.004 | -0.004 | 0.074 | 0.28 | 0.27 | 0.21 | 0.29 | 0.2 | 0.29 |
| 2 | 166003479 | rs16850131 | 0.118 | 0.483 | -1.153 | -0.214 | 0.006 | -0.005 | -0.005 | 0 | 0.28 | 0.27 | 0.2 | 0.31 | 0.2 | 0.29 |
| 2 | 166008216 | rs7596422 | -0.103 | 0.615 | -1.238 | -0.23 | 0.013 | 0.03 | 0.03 | 0 | 0.5 | 0.74 | 0.53 | 0.61 | 0.54 | 0.43 |
| 2 | 166010069 | rs4667786 | -0.254 | 0.89 | -0.736 | -0.031 | 0 | 0.006 | 0.006 | 0 | 0.69 | 0.7 | 0.68 | 0.85 | 0.71 | 0.74 |
| 2 | 166018354 | rs4667792 | -0.186 | 0.74 | -0.632 | -0.093 | -0.039 | 0.051 | 0.051 | 0 | 0.67 | 0.79 | 0.63 | 0.81 | 0.54 | 0.7 |
| 2 | 166020893 | rs7579000 | -1.009 | 0.224 | 0.003 | 0.453 | -0.013 | 0.184 | 0.184 | 0.742 | 0.15 | 0.39 | 0.29 | 0.55 | 0.12 | 0.2 |
| 2 | 166041895 | rs11898238 | -1.031 | 0.323 | -0.473 | 0.422 | 0.001 | 0.019 | 0.019 | 0.365 | 0.15 | 0.2 | 0.29 | 0.51 | 0.14 | 0.2 |
| 2 | 166059919 | rs2390165 | -0.649 | 0.447 | -0.723 | -0.093 | -0.044 | 0.043 | 0.043 | 0.328 | 0.61 | 0.83 | 0.64 | 0.73 | 0.55 | 0.53 |
| 2 | 166068283 | rs6727857 | -1.369 | 0.913 | 0.29 | -0.059 | 0.006 | 0 | 0 | 0 | 0.2 | 0.73 | 0.18 | 0.16 | 0.28 | 0.13 |
| 2 | 166076177 | rs7561455 | -0.617 | 0.428 | 0.495 | -0.107 | -0.002 | 0.059 | 0.059 | 3.336 | 0.06 | 0.3 | 0.07 | 0.03 | 0.01 | 0 |
| 2 | 166079757 | rs7576138 | -1.026 | 0.507 | 0.425 | -0.118 | 0.02 | -0.013 | -0.013 | 0 | 0.12 | 0.4 | 0.09 | 0.03 | 0.11 | 0.09 |
| 2 | 166088989 | rs1866603 | -0.396 | 0.458 | -0.495 | -0.118 | -0.011 | 0.038 | 0.038 | 0 | 0.56 | 0.71 | 0.57 | 0.66 | 0.5 | 0.59 |
| 2 | 166095728 | rs6718960 | 0.906 | 1.204 | -0.652 | -0.169 | -0.016 | -0.004 | -0.004 | 0 | 0.1 | 0.07 | 0.13 | 0.12 | 0.27 | 0.15 |
| 2 | 166102076 | rs1439804 | 1.517 | 0.497 | -0.712 | -0.461 | 0.02 | 0.058 | 0.058 | 0 | 0.2 | 0.1 | 0.14 | 0.1 | 0.15 | 0.21 |
| 2 | 166106258 | rs12997787 | -2.514 | 0.187 | -1.014 | -0.728 | 0.009 | -0.03 | -0.03 | 0 | 0.85 | 0.95 | 0.8 | 0.79 | 0.89 | 0.78 |
| 2 | 166108771 | rs6736704 | 1.727 | 0.168 | -1.013 | -0.72 | 0.021 | 0.06 | 0.06 | 0 | 0.2 | 0.09 | 0.14 | 0.1 | 0.15 | 0.22 |
| 2 | 166109634 | rs10182570 | 0.613 | -0.104 | -0.644 | -0.667 | -0.001 | 0.05 | 0.05 | 0 | 0.32 | 0.47 | 0.39 | 0.43 | 0.33 | 0.34 |
| 2 | 166116345 | rs1898970 | -0.482 | -0.035 | -0.655 | -0.707 | -0.008 | 0.018 | 0.018 | 0 | 0.58 | 0.66 | 0.57 | 0.66 | 0.5 | 0.58 |
| 2 | 166117860 | rs7600082 | -1.7 | -0.06 | -0.658 | -0.647 | 0.005 | -0.003 | -0.003 | 0 | 0.8 | 0.84 | 0.77 | 0.78 | 0.84 | 0.75 |
| 2 | 166125219 | rs10174400 | 0.413 | -0.215 | -0.553 | -0.626 | -0.004 | 0.042 | 0.042 | 0 | 0.33 | 0.49 | 0.41 | 0.48 | 0.33 | 0.33 |
| 2 | 166126863 | rs7573433 | 1.467 | -0.039 | -0.45 | -0.483 | -0.037 | -0.037 | -0.037 | 5.363 | 0.18 | 0.28 | 0.09 | 0.09 | 0.09 | 0.08 |
| 2 | 166127283 | rs2119067 | -1.925 | 0.326 | -0.974 | -0.553 | -0.019 | -0.01 | -0.01 | 0 | 0.79 | 0.91 | 0.78 | 0.76 | 0.77 | 0.73 |
| 2 | 166128186 | rs10207911 | 0.341 | 0.22 | -0.658 | -0.661 | 0.001 | -0.013 | -0.013 | 0.471 | 0.36 | 0.37 | 0.29 | 0.14 | 0.35 | 0.31 |
| 2 | 166129323 | rs353121 | -1.298 | 0.266 | -0.797 | -0.357 | -0.004 | 0.006 | 0.006 | 0 | 0.73 | 0.77 | 0.73 | 0.75 | 0.71 | 0.69 |
| 2 | 166133632 | rs353116 | 0.347 | 0.598 | -0.69 | -0.062 | 0.017 | 0.018 | 0.018 | 0 | 0.42 | 0.29 | 0.5 | 0.6 | 0.38 | 0.51 |
| 2 | 166134322 | rs353115 | 1.276 | 0.662 | -0.821 | -0.245 | -0.003 | 0.031 | 0.031 | 0 | 0.3 | 0.19 | 0.26 | 0.23 | 0.31 | 0.34 |
| 2 | 166138549 | rs9287856 | 1.042 | 1.049 | -0.534 | -0.514 | -0.008 | -0.002 | -0.002 | 0 | 0.13 | 0.1 | 0.12 | 0.03 | 0.2 | 0.07 |
| 2 | 166138608 | rs353128 | 0.61 | 1.086 | -0.311 | -0.324 | 0.009 | 0.016 | 0.016 | 0 | 0.42 | 0.35 | 0.52 | 0.64 | 0.38 | 0.51 |
| 2 | 166140020 | rs6740895 | -0.152 | 1.368 | -0.006 | 0.201 | 0.019 | -0.011 | -0.011 | 0.408 | 0.24 | 0.32 | 0.2 | 0.21 | 0.29 | 0.22 |
| 2 | 166146520 | rs4387807 | -0.238 | 2.031 | 0.263 | 0.229 | 0.038 | -0.012 | -0.012 | 0.001 | 0.17 | 0.26 | 0.18 | 0.14 | 0.3 | 0.17 |
| 2 | 166148647 | rs1371466 | -0.713 | 1.633 | 0.191 | -0.008 | 0.008 | 0.017 | 0.017 | 0 | 0.54 | 0.61 | 0.47 | 0.29 | 0.58 | 0.47 |
| 2 | 166151876 | rs2075703 | 0.954 | 1.72 | 0.148 | 0.682 | 0.001 | 0.026 | 0.026 | 0 | 0.12 | 0.12 | 0.31 | 0.36 | 0.25 | 0.22 |
| 2 | 166152389 | rs17183814 | 1.31 | 1.528 | 0.082 | 0.639 | -0.007 | -0.019 | -0.019 | 0 | 0.07 | 0.02 | 0.08 | 0.14 | 0.06 | 0.14 |
| 2 | 166164887 | rs1965757 | -1.373 | 1.346 | 0.237 | 0.717 | 0.019 | -0.003 | -0.003 | 1.028 | 0.67 | 0.76 | 0.78 | 0.65 | 0.83 | 0.69 |
| 2 | 166179357 | rs1816918 | -1.487 | 1.374 | 0.287 | 0.72 | 0.024 | 0.018 | 0.018 | 0 | 0.68 | 0.79 | 0.8 | 0.74 | 0.83 | 0.7 |
| 2 | 166179650 | rs1867864 | -0.51 | 1.527 | 0.335 | 0.75 | 0.008 | 0.016 | 0.016 | 0 | 0.55 | 0.61 | 0.46 | 0.34 | 0.56 | 0.49 |
| 2 | 166189640 | rs7589614 | -1.237 | 1.688 | 0.662 | 1.03 | 0.009 | 0.008 | 0.008 | 0 | 0.61 | 0.72 | 0.73 | 0.69 | 0.71 | 0.64 |
| 2 | 166192733 | rs4303727 | 0.944 | 1.752 | 0.681 | 1.006 | 0 | -0.002 | -0.002 | 0 | 0.1 | 0.12 | 0.12 | 0.14 | 0.18 | 0.14 |
| 2 | 166193965 | rs13387970 | -0.658 | 1.722 | 0.834 | 1.017 | 0.001 | 0.011 | 0.011 | 0 | 0.5 | 0.61 | 0.44 | 0.39 | 0.46 | 0.48 |
| 2 | 166208049 | rs6744911 | 0.153 | 1.715 | 0.818 | 0.839 | 0.04 | 0.13 | 0.13 | 0.248 | 0.15 | 0.32 | 0.34 | 0.37 | 0.25 | 0.19 |
| 2 | 166213297 | rs764660 | 0.202 | 1.458 | 0.773 | 0.703 | -0.002 | -0.003 | -0.003 | 0 | 0.37 | 0.34 | 0.29 | 0.22 | 0.28 | 0.34 |
| 2 | 166224047 | rs2304010 | -0.454 | 1.546 | 0.598 | 0.74 | 0.06 | 0.065 | 0.065 | 1.74 | 0.25 | 0.44 | 0.46 | 0.51 | 0.43 | 0.33 |
| 2 | 166248047 | rs1007722 | -0.132 | 1.633 | 0.324 | 0.784 | 0.017 | 0.019 | 0.019 | 0.255 | 0.22 | 0.26 | 0.44 | 0.52 | 0.43 | 0.33 |
| 2 | 166250129 | rs2390258 | 0.023 | 1.169 | 0.072 | 0.672 | 0.018 | 0.025 | 0.025 | 0 | 0.37 | 0.28 | 0.28 | 0.21 | 0.28 | 0.31 |
| 2 | 166257941 | rs6705474 | 0.274 | 1.575 | 0.147 | 0.699 | 0.014 | -0.005 | -0.005 | 0 | 0.15 | 0.21 | 0.14 | 0.16 | 0.2 | 0.18 |
| 2 | 166260790 | rs1432546 | -0.567 | 1.219 | 0.621 | 0.501 | 0.029 | 0.013 | 0.013 | 0 | 0.35 | 0.45 | 0.25 | 0.41 | 0.38 | 0.43 |
| 2 | 166263130 | rs6432823 | -1.481 | 0.847 | 0.721 | 0.146 | -0.001 | -0.029 | -0.029 | 0.489 | 0.83 | 0.83 | 0.87 | 1 | 0.9 | 0.95 |
| 2 | 166264307 | rs12052859 | 1.469 | 1.014 | 0.545 | 0.27 | 0.044 | 0.063 | 0.063 | 0 | 0.35 | 0.14 | 0.31 | 0.25 | 0.27 | 0.31 |
| 2 | 166265016 | rs16850592 | 1.232 | 1.104 | 0.509 | 0.497 | -0.023 | -0.025 | -0.025 | 0 | 0.12 | 0.07 | 0.27 | 0.27 | 0.25 | 0.18 |
| 2 | 166266805 | rs10207716 | 0.406 | 1.207 | 0.934 | 0.456 | -0.006 | 0.002 | 0.002 | 0.531 | 0.57 | 0.45 | 0.52 | 0.63 | 0.5 | 0.67 |
| 2 | 166268797 | rs6728727 | 0.86 | 1.174 | 1.16 | 0.352 | 0.052 | -0.079 | -0.079 | 0 | 0.15 | 0.39 | 0.09 | 0.06 | 0.15 | 0.1 |

| | | | | | | | | | | | | | | | | |
|---|---|---|---|---|---|---|---|---|---|---|---|---|---|---|---|---|
| 2 | 166274594 | rs1355995 | -1.091 | 1.127 | 1.112 | 0.418 | 0.022 | 0.029 | 0.029 | 0 | 0.47 | 0.56 | 0.55 | 0.72 | 0.63 | 0.63 |
| 2 | 166274638 | rs667884 | 1.306 | 1.217 | 1.199 | 0.574 | 0.035 | 0.056 | 0.056 | 0 | 0.36 | 0.27 | 0.32 | 0.23 | 0.27 | 0.32 |
| 2 | 166277393 | rs6711980 | -0.269 | 1.324 | 1.172 | 0.584 | -0.063 | -0.006 | -0.006 | 0.27 | 0.32 | 0.16 | 0.46 | 0.64 | 0.48 | 0.53 |
| 2 | 166278012 | rs1849811 | 0.737 | 1.278 | 0.882 | 0.68 | -0.007 | -0.014 | -0.014 | 0 | 0.21 | 0.18 | 0.2 | 0.36 | 0.23 | 0.35 |
| 2 | 166278856 | rs601400 | -0.835 | 1.273 | 0.926 | 0.677 | 0.01 | 0.029 | 0.029 | 0 | 0.79 | 0.72 | 0.79 | 0.64 | 0.77 | 0.65 |
| 2 | 166281132 | rs6706169 | 0.736 | 1.384 | 0.555 | 0.477 | 0.11 | 0.193 | 0.193 | 0.113 | 0.13 | 0.38 | 0.3 | 0.29 | 0.25 | 0.22 |
| 2 | 166288740 | rs11683781 | 0.369 | 0.923 | -0.043 | -0.196 | 0.009 | 0.032 | 0.032 | 4.518 | 0.3 | 0.23 | 0.18 | 0.05 | 0.24 | 0.12 |
| 2 | 166290534 | rs615122 | 0.692 | 0.774 | -0.9 | 0.259 | 0.03 | 0.068 | 0.068 | 0 | 0.34 | 0.11 | 0.29 | 0.24 | 0.28 | 0.31 |
| 2 | 166295400 | rs1541901 | 0.831 | 0.691 | -0.877 | 0.201 | 0.018 | 0.065 | 0.065 | 0 | 0.32 | 0.11 | 0.21 | 0.21 | 0.28 | 0.31 |
| 2 | 166295822 | rs581972 | 0.823 | 0.686 | -0.542 | 0.205 | 0.02 | 0.047 | 0.047 | 0 | 0.32 | 0.17 | 0.21 | 0.21 | 0.28 | 0.31 |
| 2 | 166295835 | rs7567296 | 0.521 | 0.725 | 0.064 | -0.344 | -0.001 | -0.052 | -0.052 | 0.767 | 0.28 | 0.34 | 0.17 | 0.05 | 0.24 | 0.13 |
| 2 | 166299635 | rs1514755 | 0.953 | 0.898 | -0.035 | 0.243 | -0.034 | -0.038 | -0.038 | 0 | 0.13 | 0.08 | 0.28 | 0.29 | 0.25 | 0.18 |
| 2 | 166303284 | rs7600830 | 0.044 | 1.215 | 0.246 | -0.332 | 0.006 | -0.067 | -0.067 | 1.65 | 0.3 | 0.36 | 0.19 | 0.05 | 0.37 | 0.2 |
| 2 | 166303598 | rs687415 | -1.311 | 0.998 | 0.723 | -0.094 | -0.034 | -0.034 | -0.034 | 0 | 0.83 | 0.72 | 0.89 | 0.94 | 0.92 | 0.95 |
| 2 | 166306826 | rs6726130 | 1.704 | 1.018 | 0.768 | -0.084 | -0.003 | -0.003 | -0.003 | 0 | 0.15 | 0.12 | 0.08 | 0.06 | 0.08 | 0.05 |
| 2 | 166310177 | rs670663 | -1.542 | 1.134 | 0.759 | -0.094 | -0.024 | -0.023 | -0.023 | 0 | 0.84 | 0.8 | 0.91 | 0.94 | 0.92 | 0.95 |
| 2 | 166314930 | rs2943460 | 1.606 | 1.152 | 0.762 | -0.086 | -0.018 | -0.017 | -0.017 | 0 | 0.15 | 0.17 | 0.09 | 0.06 | 0.08 | 0.05 |
| 2 | 166317355 | rs16850708 | 1.191 | 1.312 | 0.857 | -0.154 | 0.058 | -0.067 | -0.067 | 0.12 | 0.15 | 0.34 | 0.13 | 0.05 | 0.23 | 0.08 |
| 2 | 166318788 | rs4667825 | 0.21 | 1.315 | 0.874 | 0.19 | 0.001 | 0.007 | 0.007 | 1.379 | 0.29 | 0.27 | 0.36 | 0.4 | 0.33 | 0.24 |
| 2 | 166319297 | rs11687370 | -1.591 | 1.161 | 0.822 | 0.085 | 0.023 | -0.055 | -0.055 | 0 | 0.62 | 0.89 | 0.59 | 0.45 | 0.7 | 0.47 |
| 2 | 166326888 | rs10497261 | 0.246 | 1.185 | 0.952 | 0.34 | -0.002 | -0.003 | -0.003 | 0 | 0.28 | 0.24 | 0.35 | 0.35 | 0.33 | 0.24 |
| 2 | 166332166 | rs1514758 | 0.582 | 1.448 | 1.011 | 0.562 | 0.019 | 0.061 | 0.061 | 0 | 0.15 | 0.19 | 0.28 | 0.39 | 0.25 | 0.19 |
| 2 | 166336696 | rs4438496 | 0.561 | 1.443 | 1.008 | 0.559 | 0.018 | 0.06 | 0.06 | 0 | 0.15 | 0.19 | 0.28 | 0.39 | 0.25 | 0.19 |
| 2 | 166355411 | rs10930169 | 0.633 | 1.44 | 1.165 | 0.555 | 0.028 | 0.086 | 0.086 | 0.011 | 0.15 | 0.27 | 0.29 | 0.39 | 0.25 | 0.19 |
| 2 | 166368788 | rs6756041 | -0.464 | 1.521 | 1.438 | 0.417 | 0.011 | 0.006 | 0.006 | 0 | 0.31 | 0.67 | 0.45 | 0.43 | 0.47 | 0.3 |
| 2 | 166369996 | rs16850819 | 0.868 | 1.573 | 1.556 | 0.354 | 0.038 | -0.062 | -0.062 | 0 | 0.15 | 0.27 | 0.13 | 0.04 | 0.22 | 0.07 |
| 2 | 166374737 | rs1514759 | 0.876 | 1.54 | 1.681 | 0.469 | 0.042 | 0.082 | 0.082 | 0 | 0.14 | 0.3 | 0.29 | 0.33 | 0.25 | 0.19 |
| 2 | 166377674 | rs17196365 | 0.915 | 1.539 | 1.568 | 0.412 | 0.013 | -0.034 | -0.034 | 0.013 | 0.14 | 0.19 | 0.13 | 0.04 | 0.22 | 0.07 |
| 2 | 166386181 | rs2175990 | 0.331 | 1.289 | 1.249 | 0.449 | 0.001 | 0.025 | 0.025 | 0 | 0.11 | 0.09 | 0.22 | 0.36 | 0.08 | 0.23 |
| 2 | 166395803 | rs2292609 | 0.98 | 1.424 | 1.285 | 0.4 | 0.06 | -0.028 | -0.028 | 0 | 0.16 | 0.32 | 0.16 | 0.09 | 0.21 | 0.07 |
| 2 | 166395881 | rs930792 | -1.35 | 0.924 | 0.697 | 0.192 | 0.04 | 0.097 | 0.097 | 0 | 0.58 | 0.85 | 0.75 | 0.74 | 0.68 | 0.61 |
| 2 | 166396406 | rs4667489 | -0.796 | 0.783 | 0.768 | 0.179 | -0.001 | -0.002 | -0.002 | 0 | 0.44 | 0.52 | 0.46 | 0.47 | 0.44 | 0.42 |
| 2 | 166396485 | rs10198543 | -0.842 | 0.79 | 0.687 | 0.188 | 0 | -0.005 | -0.005 | 0 | 0.46 | 0.58 | 0.47 | 0.47 | 0.43 | 0.43 |
| 2 | 166398634 | rs1020626 | -0.047 | 0.12 | 0.447 | 0.057 | 0.013 | 0.032 | 0.032 | 0.001 | 0.51 | 0.39 | 0.55 | 0.74 | 0.6 | 0.5 |
| 2 | 166399158 | rs10427253 | 0.177 | -0.043 | 0.696 | -0.075 | -0.007 | -0.076 | -0.076 | 0 | 0.12 | 0.34 | 0.06 | 0.01 | 0.08 | 0.12 |
| 2 | 166399467 | rs10930170 | -0.309 | -0.108 | 0.493 | -0.113 | -0.008 | -0.021 | -0.021 | 1.043 | 0.63 | 0.54 | 0.64 | 0.77 | 0.71 | 0.57 |
| 2 | 166401496 | rs10174164 | 0.452 | 0.056 | 0.236 | 0.366 | -0.008 | -0.01 | -0.01 | 0 | 0.22 | 0.15 | 0.19 | 0.48 | 0.27 | 0.28 |
| 2 | 166415099 | rs6432836 | -0.737 | -0.309 | 0.331 | 0.292 | 0.011 | 0.008 | 0.008 | 0 | 0.74 | 0.82 | 0.7 | 0.77 | 0.8 | 0.69 |
| 2 | 166421414 | rs16850912 | 0.086 | -0.354 | 0.578 | 0.273 | -0.014 | -0.008 | -0.008 | 0.099 | 0.3 | 0.31 | 0.29 | 0.25 | 0.19 | 0.29 |
| 2 | 166422010 | rs16850914 | 0.335 | -0.336 | 0.162 | 0.288 | 0.028 | 0.012 | 0.012 | 0 | 0.28 | 0.11 | 0.27 | 0.25 | 0.19 | 0.29 |
| 2 | 166425732 | rs1514760 | 0.226 | -0.869 | 0.556 | 0.166 | -0.022 | -0.018 | -0.018 | 9.884 | 0.66 | 0.52 | 0.73 | 0.72 | 0.72 | 0.78 |
| 2 | 166426989 | rs7557669 | -0.986 | -0.823 | 0.549 | 0.205 | 0.006 | -0.008 | -0.008 | 0.074 | 0.78 | 0.85 | 0.81 | 0.74 | 0.82 | 0.8 |
| 2 | 166428188 | rs10181227 | 1.42 | -0.774 | 0.14 | 0.18 | -0.043 | -0.04 | -0.04 | 0 | 0.27 | 0.08 | 0.46 | 0.23 | 0.35 | 0.26 |
| 2 | 166432227 | rs10200396 | 1.138 | -0.789 | 0.506 | 0.171 | -0.002 | -0.002 | -0.002 | 0 | 0.29 | 0.29 | 0.48 | 0.23 | 0.35 | 0.3 |
| 2 | 166439804 | rs12621964 | 1.376 | -0.749 | 0.405 | 0.179 | -0.029 | -0.027 | -0.027 | 0 | 0.27 | 0.1 | 0.46 | 0.23 | 0.35 | 0.29 |
| 2 | 166450268 | rs4667490 | -0.472 | -0.829 | 0.693 | 0.14 | -0.001 | 0.006 | 0.006 | 0 | 0.49 | 0.53 | 0.34 | 0.5 | 0.47 | 0.5 |
| 2 | 166455078 | rs13389345 | 1.525 | -0.68 | 0.788 | 0.247 | -0.009 | -0.008 | -0.008 | 0 | 0.25 | 0.21 | 0.44 | 0.23 | 0.33 | 0.25 |
| 2 | 166458111 | rs7595846 | -0.52 | -0.51 | 1.094 | 0.37 | -0.001 | -0.001 | -0.001 | 1.434 | 0.52 | 0.5 | 0.36 | 0.49 | 0.5 | 0.55 |
| 2 | 166459298 | rs1007732 | -0.52 | -0.51 | 1.078 | 0.37 | -0.001 | 0.007 | 0.007 | 0 | 0.52 | 0.54 | 0.37 | 0.49 | 0.5 | 0.55 |
| 2 | 166460479 | rs1551336 | 0.179 | -0.599 | 1.137 | 0.335 | -0.006 | 0.002 | 0.002 | 0 | 0.42 | 0.28 | 0.3 | 0.49 | 0.41 | 0.53 |
| 2 | 166467351 | rs2034710 | 0.325 | -0.57 | 1.075 | 0.311 | 0.002 | 0.004 | 0.004 | 0 | 0.44 | 0.39 | 0.63 | 0.62 | 0.6 | 0.34 |
| 2 | 166470623 | rs12987780 | 0.893 | -0.566 | 1.094 | 0.301 | -0.007 | 0.011 | 0.011 | 0 | 0.36 | 0.28 | 0.49 | 0.23 | 0.43 | 0.28 |
| 2 | 166474235 | rs11899256 | 0.488 | -0.586 | 1.024 | 0.237 | 0.017 | 0.014 | 0.014 | 0.506 | 0.34 | 0.26 | 0.17 | 0.11 | 0.23 | 0.46 |
| 2 | 166486320 | rs729867 | -1.499 | -0.321 | 0.939 | 0.189 | -0.002 | 0.009 | 0.009 | 0 | 0.81 | 0.8 | 0.82 | 0.73 | 0.85 | 0.82 |
| 2 | 166487713 | rs10180533 | 0.554 | -0.309 | 0.749 | 0.29 | -0.041 | -0.043 | -0.043 | 0 | 0.34 | 0.15 | 0.38 | 0.61 | 0.38 | 0.5 |
| 2 | 166490343 | rs10211296 | 0.854 | -0.339 | 0.884 | 0.24 | -0.007 | 0.024 | 0.024 | 0 | 0.34 | 0.26 | 0.37 | 0.11 | 0.43 | 0.26 |
| 2 | 166491207 | rs1869473 | 1.6 | -0.104 | 0.988 | 0.469 | 0.019 | 0.045 | 0.045 | 0 | 0.09 | 0.1 | 0.14 | 0.36 | 0.17 | 0.06 |
| 2 | 166492582 | rs10207753 | 0.599 | -0.171 | 0.62 | 0.296 | -0.049 | 0.035 | 0.035 | 2.291 | 0.53 | 0.13 | 0.55 | 0.35 | 0.55 | 0.71 |
| 2 | 166496811 | rs12467436 | 0.069 | -0.161 | 0.624 | 0.287 | -0.003 | 0.001 | 0.001 | 0.758 | 0.6 | 0.33 | 0.61 | 0.35 | 0.64 | 0.71 |
| 2 | 166503040 | rs6720974 | 0.455 | 0.021 | 0.85 | 0.062 | 0.015 | -0.048 | -0.048 | 0 | 0.08 | 0.23 | 0.07 | 0 | 0.09 | 0.02 |
| 2 | 166509827 | rs10167479 | 0.763 | 0.05 | 0.36 | 0.071 | -0.03 | 0.075 | 0.075 | 0 | 0.28 | 0.07 | 0.31 | 0.11 | 0.34 | 0.28 |
| 2 | 166511783 | rs6432838 | -0.599 | 0.084 | 0.447 | 0.207 | -0.013 | 0.041 | 0.041 | 0 | 0.78 | 0.66 | 0.8 | 0.7 | 0.83 | 0.81 |
| 2 | 166512004 | rs6432839 | 0.298 | 0.178 | -0.181 | 0.225 | -0.003 | -0.021 | -0.021 | 0 | 0.33 | 0.39 | 0.34 | 0.14 | 0.34 | 0.29 |
| 2 | 166518516 | rs2390289 | 0.581 | 0.404 | -0.437 | 0.22 | 0.003 | 0.002 | 0.002 | 0 | 0.47 | 0.46 | 0.45 | 0.59 | 0.4 | 0.55 |
| 2 | 166521260 | rs10497266 | 0.685 | 0.572 | -1.242 | 0.486 | -0.04 | 0.055 | 0.055 | 0 | 0.25 | 0.07 | 0.31 | 0.12 | 0.34 | 0.24 |
| 2 | 166526250 | rs12692774 | 0.739 | 0.62 | -1.356 | 0.505 | -0.061 | 0.055 | 0.055 | 0 | 0.24 | 0.04 | 0.31 | 0.12 | 0.34 | 0.24 |
| 2 | 166526941 | rs10189153 | 0.068 | 0.249 | -0.6 | 0.973 | 0 | -0.01 | -0.01 | 2.712 | 0.81 | 0.85 | 0.83 | 0.73 | 0.84 | 0.81 |
| 2 | 166534206 | rs1820325 | -0.378 | 0.254 | -0.142 | 1.13 | 0.004 | 0.094 | 0.094 | 0.025 | 0.13 | 0.27 | 0.14 | 0.29 | 0.11 | 0.15 |
| 2 | 166535918 | rs777346 | 0.267 | 0.056 | -0.132 | 0.959 | -0.001 | 0 | 0 | 0 | 0.37 | 0.34 | 0.31 | 0.37 | 0.5 | 0.31 |

| | | | | | | | | | | | | | | | | |
|---|---|---|---|---|---|---|---|---|---|---|---|---|---|---|---|---|
| 2 | 166551808 | rs777354 | -0.701 | -0.961 | -0.494 | 0.445 | 0.031 | -0.029 | -0.029 | 0 | 0.61 | 0.81 | 0.58 | 0.59 | 0.73 | 0.48 |
| 2 | 166555672 | rs777358 | 0.338 | -0.816 | -0.056 | 0.491 | 0.036 | -0.007 | -0.007 | 0.422 | 0.35 | 0.46 | 0.34 | 0.38 | 0.5 | 0.32 |
| 2 | 166576956 | rs12185748 | 0.835 | -1.075 | 0.204 | 0.411 | -0.049 | -0.016 | -0.016 | 0 | 0.28 | 0.18 | 0.32 | 0.36 | 0.49 | 0.31 |
| 2 | 166577489 | rs7586085 | 0.863 | -1.096 | 0.263 | 0.4 | -0.073 | -0.026 | -0.026 | 0 | 0.28 | 0.15 | 0.31 | 0.36 | 0.49 | 0.31 |
| 2 | 166583244 | rs6710518 | 0.863 | -1.096 | 0.263 | 0.4 | -0.073 | -0.026 | -0.026 | 0 | 0.28 | 0.15 | 0.31 | 0.36 | 0.49 | 0.31 |
| 2 | 166598478 | rs1974924 | -0.203 | -1.149 | 0.635 | 0.418 | 0.007 | 0.014 | 0.014 | 0 | 0.95 | 0.72 | 0.88 | 0.98 | 0.99 | 0.99 |
| 2 | 166613243 | rs3791848 | -0.383 | -1.034 | 0.245 | 0.303 | 0.006 | -0.002 | -0.002 | 0.327 | 0.63 | 0.41 | 0.57 | 0.62 | 0.49 | 0.63 |
| 2 | 166621236 | rs6721582 | 0.489 | -1.43 | -0.092 | 0.121 | 0.018 | -0.02 | -0.02 | 6.006 | 0.34 | 0.3 | 0.32 | 0.41 | 0.23 | 0.34 |
| 2 | 166626606 | rs4667836 | 0.159 | -1.01 | -0.11 | 0.544 | 0.047 | 0.086 | 0.086 | 0.211 | 0.48 | 0.27 | 0.38 | 0.31 | 0.38 | 0.4 |
| 2 | 166630707 | rs16851009 | 0.86 | -1.045 | 0.097 | 0.768 | -0.004 | 0.034 | 0.034 | 0 | 0.12 | 0.13 | 0.24 | 0.3 | 0.09 | 0.17 |
| 2 | 166647689 | rs13032536 | 0.288 | -1.158 | -0.051 | 0.774 | 0.045 | 0.074 | 0.074 | 0.651 | 0.49 | 0.32 | 0.28 | 0.34 | 0.38 | 0.45 |
| 2 | 166648792 | rs4621175 | 0.284 | -0.98 | 0.641 | 0.831 | -0.043 | 0.11 | 0.11 | 0 | 0.82 | 0.65 | 0.64 | 0.68 | 0.9 | 0.82 |
| 2 | 166656155 | rs12151689 | 0.491 | -0.537 | 0.362 | 0.105 | 0.01 | 0.026 | 0.026 | 0.036 | 0.6 | 0.37 | 0.4 | 0.19 | 0.65 | 0.53 |
| 2 | 166658586 | rs6745114 | 1.824 | -0.491 | 0.424 | 0.338 | 0.01 | -0.018 | -0.018 | 0 | 0.14 | 0.05 | 0.05 | 0.28 | 0.11 | 0.19 |
| 2 | 166659242 | rs16851057 | 0.954 | -0.482 | 0.644 | 0.466 | 0.006 | -0.023 | -0.023 | 1.299 | 0.14 | 0.06 | 0.29 | 0.33 | 0.12 | 0.21 |
| 2 | 166659327 | rs10930179 | 1.568 | -0.453 | 0.825 | 0.436 | 0 | -0.002 | -0.002 | 0 | 0.15 | 0.15 | 0.06 | 0.28 | 0.11 | 0.19 |
| 2 | 166662119 | rs4667494 | -1.552 | -0.438 | 1.013 | 0.492 | 0 | 0.001 | 0.001 | 0 | 0.91 | 0.9 | 0.92 | 0.86 | 0.88 | 0.92 |
| 2 | 166670105 | rs10497270 | 1.74 | -0.465 | 1.122 | 0.446 | 0.005 | -0.01 | -0.01 | 0 | 0.14 | 0.08 | 0.18 | 0.34 | 0.11 | 0.19 |
| 2 | 166672949 | rs13413443 | 1.726 | -0.536 | 0.393 | 0.071 | -0.128 | 0.068 | 0.068 | 0.615 | 0.19 | 0.03 | 0.17 | 0.1 | 0.38 | 0.31 |
| 2 | 166679148 | rs4667840 | -1.605 | -0.521 | 0.499 | 0.086 | 0.013 | 0.013 | 0.013 | 0 | 0.91 | 0.82 | 0.91 | 0.87 | 0.88 | 0.92 |
| 2 | 166686414 | rs717027 | 1.668 | -0.543 | 0.484 | 0.06 | -0.075 | 0.059 | 0.059 | 0.568 | 0.2 | 0.1 | 0.17 | 0.1 | 0.38 | 0.31 |
| 2 | 166688188 | rs10930182 | -0.615 | -0.61 | 0.715 | 0.068 | -0.011 | 0.048 | 0.048 | 0 | 0.75 | 0.58 | 0.71 | 0.53 | 0.77 | 0.73 |
| 2 | 166688858 | rs13400882 | 1.052 | -0.648 | 0.373 | 0.061 | 0.052 | 0.088 | 0.088 | 0 | 0.4 | 0.26 | 0.23 | 0.09 | 0.27 | 0.21 |
| 2 | 166689342 | rs2060167 | -0.464 | -0.583 | 0.31 | 0.248 | -0.014 | 0.044 | 0.044 | 0 | 0.74 | 0.48 | 0.71 | 0.53 | 0.77 | 0.73 |
| 2 | 166694257 | rs1432274 | 1.824 | -0.996 | -0.938 | -0.171 | 0.014 | 0.02 | 0.02 | 0 | 0.22 | 0.02 | 0.15 | 0.09 | 0.15 | 0.17 |
| 2 | 166699798 | rs1560594 | -0.722 | -0.435 | -0.065 | 0.304 | -0.008 | -0.041 | -0.041 | 0 | 0.78 | 0.92 | 0.88 | 0.59 | 0.77 | 0.71 |
| 2 | 166700939 | rs16851139 | 1.823 | -0.439 | -0.074 | 0.311 | 0.008 | -0.033 | -0.033 | 0.005 | 0.14 | 0.03 | 0.05 | 0.28 | 0.11 | 0.19 |
| 2 | 166702446 | rs10930184 | -1.466 | -0.375 | -0.05 | 0.326 | -0.009 | -0.008 | -0.008 | 1.795 | 0.91 | 0.95 | 0.93 | 0.87 | 0.88 | 0.89 |
| 2 | 166714081 | rs2304003 | -0.803 | -0.377 | -0.319 | 0.324 | -0.001 | -0.01 | -0.01 | 0 | 0.78 | 0.84 | 0.87 | 0.59 | 0.78 | 0.73 |
| 2 | 166715857 | rs9677856 | 1.373 | 0.957 | -0.348 | 0.251 | -0.04 | 0.039 | 0.039 | 0.095 | 0.19 | 0.14 | 0.16 | 0.1 | 0.33 | 0.27 |
| 2 | 166720619 | rs16851176 | 0.843 | 0.689 | -0.306 | 0.62 | 0.006 | -0.032 | -0.032 | 0.465 | 0.14 | 0.06 | 0.25 | 0.3 | 0.11 | 0.17 |
| 2 | 166721251 | rs962501 | -0.44 | 1.222 | -0.227 | 0.48 | -0.015 | -0.012 | -0.012 | 0 | 0.77 | 0.83 | 0.82 | 0.59 | 0.66 | 0.66 |
| 2 | 166728820 | rs10204478 | -1.463 | 1.05 | -0.205 | 0.531 | -0.009 | -0.009 | -0.009 | 0 | 0.92 | 0.96 | 0.94 | 0.87 | 0.88 | 0.89 |
| 2 | 166731930 | rs10185938 | -1.175 | 1.008 | -0.098 | 0.536 | 0.001 | 0 | 0 | 0.037 | 0.91 | 0.92 | 0.91 | 0.87 | 0.85 | 0.9 |
| 2 | 166741358 | rs9332423 | 0.139 | 0.522 | 0.358 | 0.37 | -0.009 | 0.004 | 0.004 | 0 | 0.55 | 0.31 | 0.55 | 0.44 | 0.5 | 0.43 |
| 2 | 166743340 | rs891819 | 0.447 | 0.507 | 0.356 | 0.403 | 0.011 | 0.082 | 0.082 | 0 | 0.15 | 0.22 | 0.29 | 0.31 | 0.17 | 0.19 |
| 2 | 166748913 | rs6432851 | 0.232 | 0.432 | 1.074 | 0.37 | -0.016 | 0.004 | 0.004 | 0 | 0.41 | 0.33 | 0.28 | 0.5 | 0.49 | 0.57 |
| 2 | 166754152 | rs12692787 | 0.106 | 0.413 | 1.166 | 0.376 | -0.018 | 0.005 | 0.005 | 0 | 0.55 | 0.24 | 0.54 | 0.44 | 0.5 | 0.43 |
| 2 | 166760357 | rs1036147 | 0.468 | 0.425 | 1.2 | 0.397 | 0.004 | 0.045 | 0.045 | 0 | 0.15 | 0.16 | 0.29 | 0.31 | 0.17 | 0.18 |
| 2 | 166770120 | rs10176588 | 0.837 | 0.399 | 1.417 | 0.356 | -0.013 | 0.011 | 0.011 | 0.524 | 0.27 | 0.26 | 0.23 | 0.2 | 0.36 | 0.33 |
| 2 | 166771663 | rs7590559 | 0.635 | 0.395 | 1.615 | 0.36 | 0.021 | -0.011 | -0.011 | 0 | 0.28 | 0.46 | 0.38 | 0.26 | 0.37 | 0.33 |
| 2 | 166773971 | rs6750044 | 0.81 | 0.397 | 1.637 | 0.358 | -0.003 | 0.002 | 0.002 | 0.59 | 0.27 | 0.31 | 0.23 | 0.2 | 0.37 | 0.33 |
| 2 | 166774101 | rs3749030 | 1.395 | 0.384 | 1.64 | 0.355 | -0.007 | 0.013 | 0.013 | 0 | 0.16 | 0.27 | 0.07 | 0.3 | 0.13 | 0.24 |
| 2 | 166777119 | rs10206075 | 0.839 | 0.398 | 1.528 | 0.356 | -0.013 | 0.011 | 0.011 | 0.524 | 0.27 | 0.26 | 0.22 | 0.2 | 0.37 | 0.33 |
| 2 | 166786351 | rs1560596 | 1.529 | 0.422 | 1.656 | 0.352 | -0.003 | 0.008 | 0.008 | 0 | 0.16 | 0.23 | 0.07 | 0.3 | 0.13 | 0.24 |
| 2 | 166796552 | rs919683 | -0.866 | 0.348 | 1.396 | 0.334 | 0.026 | -0.024 | -0.024 | 0 | 0.73 | 0.55 | 0.75 | 0.8 | 0.63 | 0.67 |
| 2 | 166810570 | rs6432854 | 0.299 | 0.277 | 1.458 | 0.501 | -0.008 | 0.01 | 0.01 | 0 | 0.2 | 0.7 | 0.28 | 0.36 | 0.16 | 0.25 |
| 2 | 166821406 | rs2390319 | 0.243 | 0.361 | 1.567 | 0.461 | 0.069 | 0.31 | 0.31 | 0.199 | 0.11 | 0.37 | 0.29 | 0.3 | 0.15 | 0.14 |
| 2 | 166846730 | rs10497275 | 1.87 | 0.479 | 0.884 | 0.562 | 0.001 | 0.084 | 0.084 | 0 | 0.06 | 0.05 | 0.27 | 0.3 | 0.15 | 0.15 |
| 2 | 166873723 | rs10182473 | 0.515 | 0.415 | 0.976 | 0.474 | 0.024 | 0.053 | 0.053 | 0 | 0.22 | 0.33 | 0.35 | 0.61 | 0.3 | 0.39 |
| 2 | 166882828 | rs478389 | -1.114 | 0.399 | 0.524 | 0.253 | -0.041 | 0.015 | 0.015 | 0.351 | 0.74 | 0.88 | 0.78 | 0.79 | 0.62 | 0.67 |
| 2 | 166898249 | rs6731591 | 1.302 | 0.274 | 0.685 | 0.25 | 0.004 | 0 | 0 | 0 | 0.18 | 0.2 | 0.09 | 0.3 | 0.21 | 0.29 |
| 2 | 166905375 | rs1542484 | -0.207 | 0.268 | 0.765 | 0.183 | -0.01 | -0.013 | -0.013 | 2.94 | 0.42 | 0.34 | 0.48 | 0.6 | 0.54 | 0.58 |
| 2 | 166913475 | rs16851382 | 0.252 | -0.347 | 0.183 | -0.076 | -0.01 | 0.064 | 0.064 | 0 | 0.18 | 0.26 | 0.31 | 0.3 | 0.17 | 0.21 |
| 2 | 166913962 | rs545331 | 1.711 | -0.3 | -0.343 | -0.152 | -0.034 | 0.102 | 0.102 | 0.02 | 0.31 | 0.07 | 0.2 | 0.09 | 0.28 | 0.19 |
| 2 | 166915897 | rs10188577 | 0.918 | -0.056 | -0.23 | 0.038 | -0.031 | 0.007 | 0.007 | 0.194 | 0.24 | 0.1 | 0.19 | 0.21 | 0.32 | 0.28 |
| 2 | 166919528 | rs16851400 | 1.934 | 0.09 | 0 | 0.158 | 0.023 | 0.142 | 0.142 | 0 | 0.06 | 0.08 | 0.27 | 0.31 | 0.15 | 0.15 |
| 2 | 166919976 | rs2217198 | 0.024 | -0.195 | 0.396 | -0.055 | 0.026 | 0.008 | 0.008 | 0.004 | 0.31 | 0.43 | 0.54 | 0.7 | 0.37 | 0.45 |
| 2 | 166920430 | rs1841548 | 1.54 | -0.214 | 0.26 | -0.062 | -0.009 | -0.017 | -0.017 | 0 | 0.21 | 0.15 | 0.25 | 0.39 | 0.22 | 0.3 |
| 2 | 166920909 | rs557222 | -0.655 | -0.355 | 0.285 | -0.224 | 0 | 0.02 | 0.02 | 0.93 | 0.71 | 0.75 | 0.77 | 0.79 | 0.67 | 0.7 |
| 2 | 166921698 | rs492299 | 1.734 | -0.386 | 0.057 | -0.266 | -0.029 | 0.09 | 0.09 | 0.059 | 0.31 | 0.08 | 0.2 | 0.09 | 0.28 | 0.2 |
| 2 | 166922028 | rs13397210 | 1.524 | -0.101 | 0.253 | -0.042 | -0.009 | -0.015 | -0.015 | 0 | 0.21 | 0.15 | 0.25 | 0.39 | 0.22 | 0.3 |
| 2 | 166924971 | rs1824549 | 0.138 | -0.011 | 0.384 | -0.066 | 0.034 | 0.013 | 0.013 | 0 | 0.3 | 0.43 | 0.54 | 0.7 | 0.37 | 0.45 |
| 2 | 166926428 | rs11692675 | 0.966 | -0.138 | 0.411 | -0.153 | -0.01 | 0.012 | 0.012 | 0.324 | 0.24 | 0.19 | 0.21 | 0.19 | 0.31 | 0.28 |
| 2 | 166927659 | rs1020852 | 1.325 | 0.052 | 0.737 | 0.132 | -0.001 | 0 | 0 | 0 | 0.22 | 0.21 | 0.25 | 0.39 | 0.22 | 0.3 |
| 2 | 166931326 | rs6735544 | 0.476 | -0.012 | 0.714 | 0.11 | -0.032 | -0.043 | -0.043 | 0 | 0.08 | 0.28 | 0.04 | 0.01 | 0.01 | 0.01 |
| 2 | 166931526 | rs6722462 | -0.728 | 0.252 | 0.155 | 0.2 | -0.007 | 0.015 | 0.015 | 0 | 0.39 | 0.71 | 0.57 | 0.72 | 0.41 | 0.47 |
| 2 | 166933023 | rs12613942 | 0.31 | 0.457 | 0.24 | 0.797 | 0.066 | 0.262 | 0.262 | 0.001 | 0.08 | 0.22 | 0.28 | 0.31 | 0.15 | 0.15 |
| 2 | 166937368 | rs13406905 | 0.594 | 0.372 | -0.414 | 0.451 | -0.017 | -0.007 | -0.007 | 0 | 0.18 | 0.15 | 0.17 | 0.19 | 0.26 | 0.24 |

| | | | | | | | | | | | | | | | | |
|---|---|---|---|---|---|---|---|---|---|---|---|---|---|---|---|---|
| 2 | 166940925 | rs487374 | 0.507 | 0.278 | -0.445 | 0.576 | -0.001 | 0.023 | 0.023 | 0 | 0.16 | 0.14 | 0.49 | 0.42 | 0.25 | 0.21 |
| 2 | 166949372 | rs1030445 | 0.759 | 0.335 | -0.458 | 0.404 | 0 | -0.007 | -0.007 | 0 | 0.08 | 0.07 | 0.08 | 0.21 | 0.2 | 0.3 |
| 2 | 166950510 | rs580041 | 0.512 | -0.847 | -0.024 | 0.424 | 0.004 | -0.002 | -0.002 | 0 | 0.19 | 0.24 | 0.18 | 0.23 | 0.26 | 0.16 |
| 2 | 166951160 | rs498918 | -0.833 | -1.311 | 0.163 | 0.226 | -0.028 | -0.008 | -0.008 | 0.181 | 0.33 | 0.51 | 0.43 | 0.37 | 0.29 | 0.27 |
| 2 | 166955439 | rs10930204 | 0.886 | -1.146 | -0.006 | 0.345 | -0.044 | -0.086 | -0.086 | 0 | 0.16 | 0.03 | 0.28 | 0.32 | 0.24 | 0.26 |
| 2 | 166978750 | rs7587026 | 0.507 | -1.314 | 0.265 | -0.001 | 0.005 | -0.008 | -0.008 | 0 | 0.19 | 0.26 | 0.18 | 0.18 | 0.25 | 0.16 |
| 2 | 166983181 | rs533202 | -0.805 | -1.831 | -0.186 | -0.342 | 0.026 | 0.02 | 0.02 | 0.352 | 0.74 | 0.77 | 0.9 | 0.97 | 0.87 | 0.85 |
| 2 | 166987347 | rs519110 | 0.453 | -1.666 | -0.043 | -0.228 | 0.035 | 0.047 | 0.047 | 0 | 0.49 | 0.37 | 0.51 | 0.63 | 0.61 | 0.68 |
| 2 | 166992750 | rs11681250 | -0.204 | -1.353 | 0.277 | 0.051 | -0.074 | 0.012 | 0.012 | 0.067 | 0.15 | 0.36 | 0.38 | 0.19 | 0.04 | 0.09 |
| 2 | 166993034 | rs11896706 | 0.67 | -0.667 | 0.065 | 0.291 | 0.006 | -0.004 | -0.004 | 0 | 0.16 | 0.21 | 0.17 | 0.18 | 0.24 | 0.18 |
| 2 | 166996841 | rs4233806 | -0.817 | -0.408 | -0.295 | 0.66 | -0.06 | -0.1 | -0.1 | 0.908 | 0.84 | 0.98 | 0.84 | 0.68 | 0.75 | 0.76 |
| 2 | 167004685 | rs6749736 | -0.784 | -0.408 | -0.313 | 0.544 | -0.06 | -0.1 | -0.1 | 0 | 0.84 | 0.96 | 0.52 | 0.52 | 0.74 | 0.75 |
| 2 | 167005694 | rs16851666 | 0.331 | -0.64 | -0.069 | 0.334 | -0.002 | -0.014 | -0.014 | 0 | 0.48 | 0.48 | 0.27 | 0.3 | 0.39 | 0.5 |
| 2 | 167011230 | rs720087 | -0.207 | -0.149 | -0.155 | 0.46 | -0.017 | -0.014 | -0.014 | 0.011 | 0.34 | 0.22 | 0.22 | 0.34 | 0.4 | 0.28 |
| 2 | 167011681 | rs4386334 | -0.207 | -0.153 | -0.143 | 0.46 | -0.014 | -0.013 | -0.013 | 0 | 0.34 | 0.23 | 0.25 | 0.34 | 0.4 | 0.28 |
| 2 | 167021197 | rs6715117 | 0.581 | -0.471 | -0.07 | -0.042 | -0.01 | -0.035 | -0.035 | 0 | 0.56 | 0.51 | 0.49 | 0.76 | 0.67 | 0.66 |
| 2 | 167027454 | rs11897214 | 0.775 | -0.874 | -0.249 | -0.134 | 0.03 | 0.03 | 0.03 | 0.856 | 0.23 | 0.11 | 0.06 | 0.02 | 0.11 | 0.11 |
| 2 | 167028525 | rs10210369 | 0.617 | -0.762 | -0.073 | -0.01 | -0.022 | -0.029 | -0.029 | 0.01 | 0.68 | 0.81 | 0.4 | 0.46 | 0.6 | 0.57 |
| 2 | 167035346 | rs6722166 | 0.326 | -0.762 | -0.067 | -0.008 | -0.014 | -0.046 | -0.046 | 0 | 0.72 | 0.84 | 0.7 | 0.63 | 0.67 | 0.59 |
| 2 | 167041305 | rs7556913 | -0.585 | -0.76 | -0.119 | -0.014 | 0.005 | 0.035 | 0.035 | 0 | 0.85 | 0.96 | 0.88 | 0.95 | 0.87 | 0.8 |
| 2 | 167042887 | rs7563811 | 0.456 | -0.821 | -0.131 | -0.04 | -0.005 | 0.005 | 0.005 | 0 | 0.54 | 0.8 | 0.38 | 0.49 | 0.57 | 0.53 |
| 2 | 167044714 | rs4667511 | 0.348 | -0.802 | -0.142 | -0.022 | -0.014 | -0.049 | -0.049 | 0 | 0.3 | 0.15 | 0.24 | 0.32 | 0.3 | 0.27 |
| 2 | 167048572 | rs12473643 | 1.267 | -0.822 | 0.132 | -0.037 | -0.012 | 0.003 | 0.003 | 0 | 0.43 | 0.37 | 0.31 | 0.48 | 0.49 | 0.46 |
| 2 | 167053386 | rs16851751 | -0.437 | -0.721 | 0.099 | 0.016 | -0.01 | -0.024 | -0.024 | 0 | 0.1 | 0.16 | 0.05 | 0.03 | 0.08 | 0.07 |
| 2 | 167056428 | rs4303728 | -0.48 | -0.681 | 0.127 | 0.051 | -0.01 | 0.004 | 0.004 | 0 | 0.1 | 0.16 | 0.31 | 0.16 | 0.08 | 0.07 |
| 2 | 167061309 | rs16851778 | 1.241 | -0.668 | -0.081 | 0.049 | -0.032 | -0.041 | -0.041 | 2.123 | 0.12 | 0.03 | 0.18 | 0.3 | 0.19 | 0.21 |
| 2 | 167072851 | rs7589835 | -1.317 | -1.091 | -0.584 | -0.344 | -0.026 | 0.003 | 0.003 | 0 | 0.91 | 0.99 | 0.91 | 0.96 | 0.9 | 0.84 |
| 2 | 167077032 | rs7605012 | 1.382 | -1.008 | -0.505 | -0.299 | 0.018 | 0.043 | 0.043 | 4.648 | 0.19 | 0.09 | 0.06 | 0.02 | 0.11 | 0.06 |
| 2 | 167086414 | rs16851799 | 1.233 | -0.792 | -0.333 | -0.007 | -0.021 | 0.017 | 0.017 | 0.048 | 0.34 | 0.26 | 0.26 | 0.27 | 0.41 | 0.41 |
| 2 | 167095109 | rs6758728 | -0.472 | -0.422 | -0.75 | 0.147 | -0.005 | -0.019 | -0.019 | 1.676 | 0.85 | 0.82 | 0.87 | 0.96 | 0.87 | 0.81 |
| 2 | 167097547 | rs2155877 | -0.247 | -0.259 | -0.726 | 0.228 | 0.009 | 0.009 | 0.009 | 0 | 0.8 | 0.83 | 0.87 | 0.95 | 0.87 | 0.76 |
| 2 | 167113336 | rs7607967 | 0.435 | -0.213 | -1.345 | 0.235 | 0.012 | 0.008 | 0.008 | 0 | 0.49 | 0.58 | 0.37 | 0.5 | 0.59 | 0.49 |
| 2 | 167120956 | rs4426541 | 0.691 | 0.018 | -1.015 | 0.124 | 0.003 | -0.002 | -0.002 | 0.018 | 0.76 | 0.74 | 0.83 | 0.94 | 0.79 | 0.76 |
| 2 | 167136589 | rs3924001 | -0.434 | 0.871 | -0.801 | 0.771 | 0.01 | -0.003 | -0.003 | 1.653 | 0.28 | 0.35 | 0.32 | 0.3 | 0.4 | 0.37 |
| 2 | 167137261 | rs13007020 | 0.463 | 0.344 | -0.71 | 0.148 | -0.013 | -0.015 | -0.015 | 0 | 0.13 | 0.18 | 0.19 | 0.08 | 0.08 | 0.05 |
| 2 | 167140838 | rs7607896 | -1.079 | -0.122 | -0.371 | -0.085 | -0.012 | 0.055 | 0.055 | 0.061 | 0.46 | 0.8 | 0.54 | 0.38 | 0.48 | 0.43 |
| 2 | 167143210 | rs9917401 | 0.643 | -0.093 | -0.647 | -0.071 | -0.01 | -0.047 | -0.047 | 0 | 0.13 | 0.17 | 0.05 | 0.02 | 0.08 | 0.04 |
| 2 | 167143255 | rs9917250 | 0.519 | -0.084 | -0.642 | -0.064 | -0.014 | -0.015 | -0.015 | 0 | 0.13 | 0.19 | 0.19 | 0.08 | 0.08 | 0.05 |
| 2 | 167144748 | rs7596100 | -1.135 | -0.39 | -1.633 | -0.19 | -0.103 | 0.069 | 0.069 | 0 | 0.71 | 0.97 | 0.69 | 0.85 | 0.69 | 0.79 |
| 2 | 167144995 | rs13402180 | 0.723 | -0.336 | -1.566 | -0.156 | 0.011 | -0.003 | -0.003 | 0 | 0.28 | 0.36 | 0.32 | 0.31 | 0.4 | 0.37 |
| 2 | 167149203 | rs4128577 | -0.828 | 0.031 | -1.498 | 0.086 | -0.106 | 0.042 | 0.042 | 0.221 | 0.72 | 0.97 | 0.69 | 0.84 | 0.69 | 0.79 |
| 2 | 167153789 | rs11688164 | -0.189 | -0.205 | -0.951 | 0.88 | -0.01 | 0.179 | 0.179 | 0 | 0.18 | 0.42 | 0.28 | 0.4 | 0.15 | 0.26 |
| 2 | 167159946 | rs13017637 | -0.145 | 0.785 | -1.701 | 0.06 | -0.049 | 0.083 | 0.083 | 0 | 0.33 | 0.19 | 0.28 | 0.17 | 0.43 | 0.41 |
| 2 | 167160029 | rs12994338 | -0.759 | 0.38 | -1.894 | 0.273 | -0.02 | 0 | 0 | 6.334 | 0.74 | 0.9 | 0.73 | 0.74 | 0.73 | 0.81 |
| 2 | 167161420 | rs6432896 | 0.178 | 0.786 | -1.124 | 0.466 | 0.028 | -0.011 | -0.011 | 0 | 0.63 | 0.49 | 0.45 | 0.7 | 0.52 | 0.6 |
| 2 | 167161598 | rs4605385 | 0.006 | 0.904 | -1.351 | 0.159 | -0.008 | -0.003 | -0.003 | 0 | 0.87 | 0.9 | 0.77 | 0.88 | 0.77 | 0.81 |
| 2 | 167162649 | rs10171225 | -0.326 | 0.745 | -0.768 | -0.132 | -0.02 | -0.124 | -0.124 | 0.017 | 0.27 | 0.47 | 0.15 | 0.1 | 0.21 | 0.23 |
| 2 | 167163043 | rs9646771 | -1.216 | 0.465 | -1.307 | 0.414 | 0.004 | -0.015 | -0.015 | 0 | 0.58 | 0.86 | 0.61 | 0.46 | 0.63 | 0.57 |
| 2 | 167168093 | rs6432901 | -0.893 | 0.476 | -1.137 | 0.214 | 0.024 | 0.027 | 0.027 | 0 | 0.54 | 0.76 | 0.46 | 0.41 | 0.61 | 0.54 |
| 2 | 167178155 | rs4455169 | -1.153 | 0.355 | -1.29 | 0.174 | 0.013 | -0.013 | -0.013 | 0 | 0.57 | 0.93 | 0.52 | 0.42 | 0.67 | 0.55 |
| 2 | 167179540 | rs7558866 | 0.144 | 0.83 | -0.967 | 0.217 | 0.016 | 0.031 | 0.031 | 0 | 0.09 | 0.15 | 0.17 | 0.14 | 0.18 | 0.16 |
| 2 | 167191814 | rs1528487 | 0.778 | 0.755 | -1.279 | -0.05 | -0.121 | 0.018 | 0.018 | 0.269 | 0.22 | 0.02 | 0.19 | 0.16 | 0.33 | 0.24 |
| 2 | 167196794 | rs2893013 | -0.196 | 0.636 | 0.045 | 0.203 | 0.059 | 0.207 | 0.207 | 1.13 | 0.1 | 0.39 | 0.18 | 0.22 | 0.13 | 0.16 |
| 2 | 167200916 | rs16851943 | 0.599 | 0.142 | 0.184 | -0.157 | -0.033 | -0.077 | -0.077 | 0 | 0.22 | 0.39 | 0.11 | 0.1 | 0.15 | 0.12 |
| 2 | 167204366 | rs1919177 | 0.399 | -0.594 | -0.595 | -0.148 | -0.034 | 0.084 | 0.084 | 0 | 0.45 | 0.15 | 0.42 | 0.3 | 0.46 | 0.42 |
| 2 | 167209218 | rs6750593 | -1.409 | -0.691 | -0.584 | -0.197 | -0.04 | -0.061 | -0.061 | 0 | 0.69 | 0.85 | 0.57 | 0.5 | 0.62 | 0.56 |
| 2 | 167210004 | rs952462 | -1.161 | -0.637 | -0.396 | -0.197 | -0.011 | -0.012 | -0.012 | 0.054 | 0.66 | 0.68 | 0.52 | 0.49 | 0.55 | 0.54 |
| 2 | 167210495 | rs13000233 | 1.271 | -0.676 | -0.278 | -0.241 | -0.011 | -0.031 | -0.031 | 0.136 | 0.19 | 0.22 | 0.09 | 0.11 | 0.15 | 0.12 |
| 2 | 167211444 | rs17766243 | 0.315 | -0.593 | -0.606 | -0.191 | -0.019 | 0.079 | 0.079 | 0 | 0.45 | 0.22 | 0.43 | 0.31 | 0.46 | 0.42 |
| 2 | 167223238 | rs16852031 | 1.688 | -0.763 | -0.718 | -0.281 | 0.074 | -0.034 | -0.034 | 0 | 0.23 | 0.03 | 0.12 | 0.22 | 0.11 | 0.18 |
| 2 | 167223512 | rs6721003 | -1.541 | -0.263 | -0.23 | -0.558 | -0.026 | 0.156 | 0.156 | 0 | 0.48 | 0.83 | 0.46 | 0.28 | 0.51 | 0.38 |
| 2 | 167224823 | rs6714902 | -0.024 | -0.214 | -0.145 | -0.593 | -0.011 | -0.013 | -0.013 | 0 | 0.27 | 0.65 | 0.17 | 0.18 | 0.19 | 0.18 |
| 2 | 167227872 | rs12692796 | 0.01 | -0.31 | -0.098 | -0.498 | -0.015 | -0.021 | -0.021 | 0 | 0.27 | 0.63 | 0.15 | 0.17 | 0.19 | 0.19 |
| 2 | 167228744 | rs10930218 | 1.073 | -0.151 | -0.596 | -0.445 | -0.11 | 0.047 | 0.047 | 0.136 | 0.18 | 0.01 | 0.29 | 0.11 | 0.29 | 0.22 |
| 2 | 167232463 | rs1881440 | 0.326 | -0.107 | -0.333 | -0.415 | -0.019 | -0.095 | -0.095 | 0 | 0.24 | 0.4 | 0.13 | 0.09 | 0.19 | 0.17 |
| 2 | 167235607 | rs2091545 | -0.594 | -0.105 | -0.474 | -0.323 | 0.006 | -0.036 | -0.036 | 0 | 0.42 | 0.47 | 0.4 | 0.29 | 0.45 | 0.37 |
| 2 | 167242799 | rs6738209 | 0.252 | -0.302 | -0.319 | -0.07 | -0.006 | -0.062 | -0.062 | 1.192 | 0.14 | 0.29 | 0.08 | 0.05 | 0.11 | 0.04 |
| 2 | 167253015 | rs2031002 | 0.775 | -0.801 | -1.148 | -0.065 | 0.066 | 0.014 | 0.014 | 0 | 0.19 | 0.03 | 0.03 | 0.11 | 0.08 | 0.12 |
| 2 | 167261996 | rs3579 | 0.895 | -0.826 | -1.161 | -0.088 | 0.062 | 0.085 | 0.085 | 0 | 0.18 | 0.03 | 0.03 | 0.02 | 0.08 | 0.12 |

| | | | | | | | | | | | | | | | | |
|---|---|---|---|---|---|---|---|---|---|---|---|---|---|---|---|---|
| 2 | 167262088 | rs16852112 | 0.611 | -0.871 | -0.732 | -0.093 | 0.014 | 0.009 | 0.009 | 0 | 0.19 | 0.14 | 0.04 | 0.11 | 0.08 | 0.13 |
| 2 | 167264978 | rs1406275 | -1.201 | -0.651 | -0.468 | 0.09 | 0.014 | 0.018 | 0.018 | 0 | 0.73 | 0.71 | 0.62 | 0.63 | 0.6 | 0.63 |
| 2 | 167265040 | rs1406274 | -0.157 | -0.867 | -0.062 | 0.204 | 0.004 | 0.022 | 0.022 | 0 | 0.49 | 0.45 | 0.48 | 0.39 | 0.51 | 0.41 |
| 2 | 167267987 | rs7562525 | -0.465 | -1.003 | -0.04 | 0.007 | -0.004 | -0.048 | -0.048 | 0 | 0.09 | 0.22 | 0.06 | 0 | 0.06 | 0.04 |
| 2 | 167273603 | rs7561953 | -0.406 | -0.934 | -0.343 | -0.279 | -0.006 | -0.01 | -0.01 | 0 | 0.33 | 0.4 | 0.22 | 0.28 | 0.29 | 0.19 |
| 2 | 167279922 | rs6738031 | -2.116 | -0.661 | -0.77 | -0.295 | -0.092 | -0.121 | -0.121 | 0 | 0.78 | 0.96 | 0.68 | 0.65 | 0.66 | 0.68 |
| 2 | 167281710 | rs5024297 | 1.991 | -0.596 | -0.836 | -0.322 | -0.089 | 0.014 | 0.014 | 0 | 0.17 | 0.01 | 0.25 | 0.11 | 0.21 | 0.23 |
| 2 | 167284923 | rs13404431 | 1 | -0.622 | -0.535 | -0.282 | 0.002 | 0.074 | 0.074 | 0.369 | 0.06 | 0.17 | 0.12 | 0.14 | 0.03 | 0.11 |
| 2 | 167286044 | rs3791253 | 0.305 | -0.823 | -0.432 | -0.355 | -0.024 | 0.001 | 0.001 | 0 | 0.21 | 0.34 | 0.15 | 0.23 | 0.19 | 0.15 |
| 2 | 167288172 | rs7597971 | -1.932 | -0.756 | -0.216 | -0.293 | -0.003 | -0.004 | -0.004 | 0 | 0.76 | 0.77 | 0.65 | 0.65 | 0.66 | 0.68 |
| 2 | 167307460 | rs16852148 | 0.861 | -0.728 | -0.218 | -0.277 | -0.006 | 0 | 0 | 0 | 0.18 | 0.17 | 0.11 | 0.23 | 0.13 | 0.13 |
| 2 | 167307529 | rs9967700 | -0.458 | -0.741 | -0.222 | -0.323 | 0 | 0.012 | 0.012 | 0 | 0.48 | 0.4 | 0.45 | 0.5 | 0.45 | 0.41 |
| 2 | 167313451 | rs11888208 | 1.623 | -0.773 | -0.186 | -0.345 | 0 | -0.017 | -0.017 | 0.013 | 0.2 | 0.18 | 0.07 | 0.02 | 0.14 | 0.16 |
| 2 | 167313904 | rs13014266 | -1.057 | -0.76 | -0.178 | -0.336 | 0.022 | 0.034 | 0.034 | 0 | 0.69 | 0.57 | 0.52 | 0.51 | 0.58 | 0.56 |
| 2 | 167321931 | rs13417885 | 0.293 | -0.743 | -0.115 | -0.328 | 0.032 | 0.07 | 0.07 | 0.003 | 0.07 | 0.18 | 0.14 | 0.14 | 0.08 | 0.12 |
| 2 | 167322522 | rs7593452 | 0.748 | -0.773 | -0.15 | -0.349 | -0.059 | 0.083 | 0.083 | 0 | 0.29 | 0.14 | 0.33 | 0.16 | 0.32 | 0.27 |
| 2 | 167326652 | rs13017982 | 1.631 | -0.765 | -0.149 | -0.333 | 0 | -0.018 | -0.018 | 0 | 0.2 | 0.17 | 0.07 | 0.02 | 0.14 | 0.16 |
| 2 | 167337367 | rs2893022 | -1.016 | -0.747 | -0.138 | -0.267 | 0.02 | 0.035 | 0.035 | 0 | 0.68 | 0.57 | 0.52 | 0.52 | 0.58 | 0.56 |
| 2 | 167337735 | rs11893208 | 0.975 | -0.752 | -0.13 | -0.261 | -0.007 | 0.001 | 0.001 | 0 | 0.17 | 0.18 | 0.11 | 0.22 | 0.13 | 0.13 |
| 2 | 167340424 | rs11687273 | 0.859 | -0.747 | -0.13 | -0.298 | -0.025 | 0.031 | 0.031 | 0 | 0.18 | 0.22 | 0.11 | 0.33 | 0.13 | 0.14 |
| 2 | 167342641 | rs1172384 | 1.736 | -0.699 | -0.19 | -0.277 | -0.002 | -0.002 | -0.002 | 0.249 | 0.25 | 0.25 | 0.35 | 0.34 | 0.34 | 0.32 |
| 2 | 167345058 | rs13411795 | 0.199 | -0.663 | -0.52 | -0.302 | 0.027 | 0.069 | 0.069 | 0.175 | 0.07 | 0.2 | 0.13 | 0.14 | 0.08 | 0.12 |
| 2 | 167348363 | rs17769186 | 1.735 | -0.772 | -0.647 | -0.343 | -0.093 | 0.025 | 0.025 | 0 | 0.19 | 0.01 | 0.26 | 0.11 | 0.22 | 0.24 |
| 2 | 167348635 | rs13022308 | 1.215 | -1.184 | 0.242 | 0.04 | 0.028 | 0.048 | 0.048 | 0 | 0.2 | 0.12 | 0.06 | 0.02 | 0.13 | 0.16 |
| 2 | 167352856 | rs2049711 | -0.547 | -0.855 | 0.383 | 0.389 | 0.012 | 0.017 | 0.017 | 0 | 0.5 | 0.37 | 0.46 | 0.53 | 0.51 | 0.43 |
| 2 | 167356561 | rs10188223 | -0.162 | -0.727 | 1.066 | 1.264 | 0.002 | 0.08 | 0.08 | 0.577 | 0.05 | 0.15 | 0.12 | 0.11 | 0.02 | 0.1 |
| 2 | 167370260 | rs1374592 | -0.56 | -0.615 | 0.494 | 1.024 | -0.02 | 0.028 | 0.028 | 0 | 0.18 | 0.19 | 0.11 | 0.36 | 0.13 | 0.14 |
| 2 | 167372934 | rs10497287 | -0.505 | -0.584 | 0.243 | 0.663 | 0 | -0.002 | -0.002 | 0 | 0.17 | 0.15 | 0.11 | 0.21 | 0.13 | 0.13 |
| 2 | 167376605 | rs10497288 | 0.559 | -0.56 | 0.618 | 0.605 | -0.008 | 0.034 | 0.034 | 0.056 | 0.1 | 0.12 | 0.09 | 0.21 | 0.08 | 0.08 |
| 2 | 167377995 | rs7422259 | 1.034 | -0.598 | 0.522 | 0.06 | 0.007 | 0.173 | 0.173 | 0 | 0.46 | 0.24 | 0.45 | 0.13 | 0.49 | 0.48 |
| 2 | 167379259 | rs13426300 | -0.181 | -0.774 | 0.505 | -0.01 | 0.013 | -0.001 | -0.001 | 0.567 | 0.15 | 0.09 | 0.07 | 0 | 0.06 | 0.14 |
| 2 | 167393816 | rs4667514 | 0.499 | -0.853 | 0.671 | 0.2 | -0.048 | 0.073 | 0.073 | 0 | 0.88 | 0.7 | 0.91 | 0.78 | 0.96 | 0.86 |
| 2 | 167401072 | rs6432918 | -0.203 | -0.851 | 0.819 | 0.204 | 0.01 | 0.043 | 0.043 | 0 | 0.68 | 0.58 | 0.63 | 0.47 | 0.59 | 0.69 |
| 2 | 167401890 | rs16852287 | 1.154 | -0.891 | 0.786 | 0.191 | -0.014 | 0.054 | 0.054 | 0.107 | 0.1 | 0.16 | 0.1 | 0.2 | 0.08 | 0.08 |
| 2 | 167403939 | rs10930228 | 1.94 | -0.801 | 0.733 | 0.035 | -0.097 | 0.219 | 0.219 | 0 | 0.38 | 0.07 | 0.38 | 0.12 | 0.39 | 0.31 |
| 2 | 167414424 | rs16852300 | 0.053 | -0.884 | 0.801 | 0.102 | -0.021 | 0.069 | 0.069 | 0 | 0.14 | 0.21 | 0.15 | 0.33 | 0.09 | 0.15 |
| 2 | 167422236 | rs10196943 | -0.031 | -0.782 | 0.749 | 0.125 | -0.03 | -0.049 | -0.049 | 0 | 0.08 | 0.24 | 0.05 | 0.02 | 0.03 | 0.06 |
| 2 | 167430697 | rs10208695 | 1.169 | -0.78 | 0.776 | 0.116 | -0.01 | -0.015 | -0.015 | 0 | 0.22 | 0.18 | 0.31 | 0.43 | 0.37 | 0.21 |
| 2 | 167435558 | rs6717308 | 0.229 | -0.668 | 0.583 | 0.088 | -0.023 | -0.047 | -0.047 | 0.02 | 0.9 | 0.6 | 0.93 | 0.99 | 0.97 | 0.94 |
| 2 | 167448639 | rs7592457 | 0.184 | -0.667 | 0.494 | 0.062 | 0 | -0.005 | -0.005 | 0 | 0.58 | 0.43 | 0.53 | 0.26 | 0.51 | 0.61 |
| 2 | 167450282 | rs12470129 | 0.184 | -0.667 | 0.494 | 0.062 | 0 | -0.005 | -0.005 | 0 | 0.58 | 0.43 | 0.53 | 0.26 | 0.51 | 0.61 |
| 2 | 167453461 | rs1596996 | -0.306 | -0.674 | 0.567 | 0.106 | 0 | -0.002 | -0.002 | 0 | 0.41 | 0.41 | 0.45 | 0.64 | 0.48 | 0.38 |
| 2 | 167456786 | rs7591369 | 1.785 | -0.712 | 0.52 | 0.017 | -0.097 | 0.215 | 0.215 | 0 | 0.37 | 0.07 | 0.38 | 0.13 | 0.39 | 0.31 |
| 2 | 167459550 | rs1584286 | 0.816 | -0.718 | 0.551 | 0.019 | 0.022 | 0.172 | 0.172 | 0.257 | 0.49 | 0.2 | 0.48 | 0.25 | 0.49 | 0.55 |
| 2 | 167460233 | rs1597004 | -0.311 | -0.609 | 0.208 | 0.082 | 0 | -0.007 | -0.007 | 0 | 0.41 | 0.41 | 0.45 | 0.74 | 0.49 | 0.39 |
| 2 | 167463166 | rs6745751 | -0.822 | -0.937 | 0.203 | 0.113 | -0.032 | -0.026 | -0.026 | 1.002 | 0.2 | 0.35 | 0.15 | 0.14 | 0.11 | 0.3 |
| 2 | 167464366 | rs11903255 | 0.269 | -0.736 | -0.488 | 0.193 | -0.012 | -0.015 | -0.015 | 0 | 0.31 | 0.25 | 0.36 | 0.43 | 0.44 | 0.31 |
| 2 | 167472743 | rs2218603 | 0.591 | -0.662 | -0.597 | -0.497 | 0.015 | -0.005 | -0.005 | 0 | 0.32 | 0.37 | 0.4 | 0.64 | 0.5 | 0.33 |
| 2 | 167472916 | rs10183984 | -0.707 | -1.204 | 0.419 | 0.362 | -0.003 | -0.028 | -0.028 | 0.107 | 0.11 | 0.13 | 0.07 | 0 | 0.08 | 0.19 |
| 2 | 167475237 | rs4667911 | 0.572 | -1.213 | 0.383 | 0.353 | -0.076 | 0.013 | 0.013 | 0 | 0.8 | 0.6 | 0.85 | 0.77 | 0.94 | 0.82 |
| 2 | 167475482 | rs4233808 | 0.703 | -0.954 | -0.085 | 0.497 | -0.054 | -0.029 | -0.029 | 0 | 0.69 | 0.44 | 0.78 | 0.77 | 0.85 | 0.63 |
| 2 | 167490384 | rs2390381 | 0.991 | -0.561 | 0.183 | 0.874 | -0.011 | 0.046 | 0.046 | 0.173 | 0.86 | 0.69 | 0.87 | 0.79 | 0.9 | 0.83 |
| 2 | 167493988 | rs6432937 | 1.024 | -1.287 | -0.45 | 0.938 | 0.055 | -0.001 | -0.001 | 0 | 0.61 | 0.18 | 0.61 | 0.34 | 0.72 | 0.41 |
| 2 | 167494012 | rs6432938 | 0.467 | -1.684 | -1.111 | 0.499 | -0.001 | 0.008 | 0.008 | 0.601 | 0.14 | 0.14 | 0.12 | 0.21 | 0.12 | 0.14 |
| 2 | 167496600 | rs1447237 | 1.403 | -1.776 | -0.456 | 0.998 | -0.001 | 0.306 | 0.306 | 0 | 0.47 | 0.04 | 0.49 | 0.13 | 0.6 | 0.27 |
| 2 | 167506317 | rs9287871 | 0.368 | -1.87 | -0.474 | 0.659 | -0.054 | -0.036 | -0.036 | 3.018 | 0.74 | 0.44 | 0.88 | 0.85 | 0.88 | 0.67 |
| 2 | 167515013 | rs10497293 | -0.177 | -2.01 | 0.065 | 0.604 | 0.01 | 0.009 | 0.009 | 0 | 0.2 | 0.16 | 0.08 | 0.12 | 0.12 | 0.34 |
| 2 | 167525345 | rs16852468 | 0.855 | -2.013 | 0.022 | 0.587 | 0.009 | -0.027 | -0.027 | 0.381 | 0.13 | 0.08 | 0.11 | 0.21 | 0.12 | 0.14 |
| 2 | 167534187 | rs4667919 | 2.121 | -2.096 | -0.102 | 0.493 | -0.012 | 0.294 | 0.294 | 0 | 0.46 | 0.04 | 0.49 | 0.13 | 0.59 | 0.27 |
| 2 | 167540475 | rs10167856 | 0.285 | -2.006 | 0.322 | 0.796 | -0.053 | 0.113 | 0.113 | 0.818 | 0.81 | 0.56 | 0.85 | 0.75 | 0.88 | 0.86 |
| 2 | 167549363 | rs12619000 | 1.212 | -2.044 | 0.165 | 0.787 | 0.062 | -0.033 | -0.033 | 0.129 | 0.6 | 0.26 | 0.76 | 0.59 | 0.75 | 0.52 |
| 2 | 167575861 | rs925285 | 0.282 | -2.051 | 0.302 | 0.806 | -0.056 | 0.102 | 0.102 | 0.893 | 0.81 | 0.51 | 0.85 | 0.76 | 0.88 | 0.85 |
| 2 | 167584926 | rs10205288 | -0.923 | -2.079 | 0.389 | 0.806 | -0.011 | 0.001 | 0.001 | 0 | 0.28 | 0.41 | 0.38 | 0.7 | 0.28 | 0.39 |
| 2 | 167586125 | rs10497301 | 0.171 | -2.08 | 0.389 | 0.749 | 0.011 | 0.131 | 0.131 | 0 | 0.14 | 0.24 | 0.27 | 0.47 | 0.17 | 0.25 |
| 2 | 167601454 | rs1588350 | -2.04 | -2.131 | 0.391 | 0.732 | 0.018 | 0.033 | 0.033 | 0 | 0.49 | 0.58 | 0.46 | 0.82 | 0.41 | 0.73 |
| 2 | 167601485 | rs1588351 | -0.888 | -2.12 | 0.357 | 0.958 | -0.011 | 0.03 | 0.03 | 0 | 0.28 | 0.41 | 0.38 | 0.64 | 0.28 | 0.39 |
| 2 | 167614459 | rs16852630 | -0.197 | -2.104 | 0.243 | 0.93 | 0.015 | -0.004 | -0.004 | 0 | 0.21 | 0.13 | 0.07 | 0.18 | 0.13 | 0.34 |
| 2 | 167618745 | rs9646729 | 0.917 | -2.012 | 0.122 | 0.929 | -0.032 | 0.017 | 0.017 | 0 | 0.09 | 0.01 | 0.06 | 0.06 | 0.11 | 0.1 |
| 2 | 167620935 | rs13399002 | -0.521 | -2.021 | 0.177 | 0.963 | 0 | -0.001 | -0.001 | 0.602 | 0.23 | 0.25 | 0.18 | 0.23 | 0.23 | 0.25 |

| | | | | | | | | | | | | | | | | |
|---|---|---|---|---|---|---|---|---|---|---|---|---|---|---|---|---|
| 2 | 167629579 | rs2664016 | -2.035 | -2.313 | -0.243 | 0.546 | -0.009 | 0.163 | 0.163 | 0 | 0.59 | 0.84 | 0.5 | 0.84 | 0.44 | 0.75 |
| 2 | 167647580 | rs11887203 | -1.139 | -2.748 | -0.1 | 0.455 | 0.045 | -0.007 | -0.007 | 1.42 | 0.42 | 0.72 | 0.22 | 0.36 | 0.24 | 0.47 |
| 2 | 167649537 | rs12993796 | 0.601 | -2.856 | 0.298 | 0.46 | 0.014 | -0.005 | -0.005 | 0 | 0.19 | 0.19 | 0.13 | 0.15 | 0.14 | 0.19 |
| 2 | 167650494 | rs12616334 | 0.183 | -2.89 | 0.464 | 0.451 | -0.004 | 0.001 | 0.001 | 0 | 0.16 | 0.16 | 0.05 | 0.16 | 0.11 | 0.28 |
| 2 | 167652463 | rs2390485 | -1.277 | -2.888 | 0.424 | 0.452 | 0.039 | -0.008 | -0.008 | 0 | 0.41 | 0.72 | 0.22 | 0.36 | 0.24 | 0.47 |
| 2 | 167652676 | rs13392463 | 0.712 | -2.824 | 0.229 | 0.463 | 0.024 | -0.012 | -0.012 | 0 | 0.19 | 0.13 | 0.12 | 0.15 | 0.14 | 0.19 |
| 2 | 167656920 | rs17775012 | 0.308 | -3.415 | -0.319 | 0.062 | 0 | 0.099 | 0.099 | 1.127 | 0.21 | 0.01 | 0.12 | 0.05 | 0.18 | 0.18 |
| 2 | 167659346 | rs1316477 | -1.371 | -3.489 | -0.239 | 0.053 | 0 | 0 | 0 | 0 | 0.54 | 0.43 | 0.28 | 0.37 | 0.4 | 0.64 |
| 2 | 167669165 | rs13032769 | -1.815 | -3.559 | -0.656 | -0.328 | -0.015 | 0.167 | 0.167 | 0 | 0.6 | 0.9 | 0.51 | 0.83 | 0.4 | 0.75 |
| 2 | 167671324 | rs13009938 | 0.15 | -3.438 | -0.626 | -0.329 | 0.001 | 0.1 | 0.1 | 1.365 | 0.21 | 0.01 | 0.12 | 0.05 | 0.18 | 0.18 |
| 2 | 167671649 | rs9287876 | -1.094 | -3.164 | -0.324 | -0.168 | 0.014 | 0.013 | 0.013 | 0 | 0.41 | 0.61 | 0.2 | 0.26 | 0.26 | 0.45 |
| 2 | 167719966 | rs6723198 | -1.068 | -2.514 | -1.117 | -0.051 | 0.084 | 0.144 | 0.144 | 2.198 | 0.43 | 0.81 | 0.4 | 0.74 | 0.3 | 0.55 |
| 2 | 167742601 | rs7566386 | -0.28 | -1.67 | -1.012 | -0.18 | -0.009 | 0.034 | 0.034 | 0.792 | 0.24 | 0.18 | 0.33 | 0.11 | 0.24 | 0.22 |
| 2 | 167746159 | rs7566782 | -0.142 | -1.666 | -1.078 | -0.178 | -0.02 | 0.064 | 0.064 | 0.827 | 0.24 | 0.15 | 0.33 | 0.11 | 0.24 | 0.22 |
| 2 | 167753994 | rs6755120 | -0.056 | -1.768 | -0.954 | -0.235 | -0.008 | 0.048 | 0.048 | 0.303 | 0.25 | 0.19 | 0.33 | 0.1 | 0.24 | 0.23 |
| 2 | 167757572 | rs13406651 | 0.942 | -2.227 | -1.33 | -0.1 | 0.03 | -0.025 | -0.025 | 0.035 | 0.18 | 0.01 | 0.09 | 0.12 | 0.12 | 0.19 |
| 2 | 167758908 | rs7595556 | 0.394 | -1.959 | -1.255 | -0.038 | -0.025 | 0.116 | 0.116 | 0.97 | 0.22 | 0.03 | 0.3 | 0.05 | 0.2 | 0.21 |
| 2 | 167759948 | rs7599152 | -1.906 | -2.273 | -1.057 | 0.177 | 0.005 | 0.044 | 0.044 | 0 | 0.58 | 0.77 | 0.65 | 0.7 | 0.41 | 0.54 |
| 2 | 167772293 | rs1567534 | -0.587 | -1.838 | -1.035 | 0.436 | -0.02 | -0.085 | -0.085 | 0.012 | 0.28 | 0.17 | 0.49 | 0.54 | 0.29 | 0.34 |
| 2 | 167806610 | rs16852806 | -1.454 | -1.353 | -0.806 | 0.908 | -0.001 | 0.007 | 0.007 | 0 | 0.09 | 0.11 | 0.06 | 0.25 | 0.14 | 0.1 |
| 2 | 167808487 | rs11896680 | -1.12 | -0.962 | -0.819 | 1.094 | 0.001 | -0.002 | -0.002 | 0 | 0.09 | 0.11 | 0.08 | 0.12 | 0.17 | 0.11 |
| 2 | 167816291 | rs13383718 | -1.399 | -1.918 | -0.421 | 1.092 | 0.017 | 0.006 | 0.006 | 0 | 0.53 | 0.63 | 0.44 | 0.43 | 0.32 | 0.51 |
| 2 | 167818552 | rs1473041 | -1.977 | -1.965 | -0.37 | 1.23 | -0.01 | 0.076 | 0.076 | 0 | 0.6 | 0.77 | 0.63 | 0.8 | 0.37 | 0.63 |
| 2 | 167839415 | rs12623180 | -1.329 | -1.973 | -0.131 | 1.297 | 0.009 | 0.048 | 0.048 | 0 | 0.09 | 0.14 | 0.08 | 0.28 | 0.09 | 0.1 |
| 2 | 167844513 | rs4493277 | -0.394 | -2.007 | -0.028 | 1.269 | -0.026 | -0.049 | -0.049 | 0.134 | 0.44 | 0.47 | 0.37 | 0.14 | 0.23 | 0.4 |
| 2 | 167855303 | rs6715263 | -0.719 | -2.105 | -0.072 | 1.752 | -0.002 | 0.19 | 0.19 | 0 | 0.06 | 0.14 | 0.18 | 0.37 | 0.05 | 0.12 |
| 2 | 167857670 | rs16852842 | -1.619 | -2.098 | -0.069 | 1.747 | -0.005 | 0.116 | 0.116 | 0 | 0.11 | 0.21 | 0.19 | 0.46 | 0.08 | 0.2 |
| 2 | 167862974 | rs869000 | -1.667 | -2.095 | -0.081 | 1.773 | -0.007 | 0.149 | 0.149 | 0 | 0.12 | 0.25 | 0.2 | 0.46 | 0.08 | 0.21 |
| 2 | 167863420 | rs1877192 | -2.38 | -2.096 | -0.081 | 1.594 | 0.011 | 0.048 | 0.048 | 0 | 0.14 | 0.26 | 0.25 | 0.65 | 0.14 | 0.23 |
| 2 | 167904042 | rs1511199 | -0.437 | -2.539 | 0.17 | 1.076 | 0.058 | 0.044 | 0.044 | 0 | 0.37 | 0.21 | 0.12 | 0.19 | 0.15 | 0.29 |
| 2 | 167907699 | rs13017014 | -0.876 | -2.448 | 0.263 | 1.184 | -0.037 | -0.015 | -0.015 | 0 | 0.42 | 0.47 | 0.19 | 0.32 | 0.21 | 0.34 |
| 2 | 167911826 | rs1511202 | -1.469 | -2.386 | 0.116 | 1.197 | 0.067 | 0.009 | 0.009 | 0 | 0.5 | 0.66 | 0.23 | 0.45 | 0.24 | 0.49 |
| 2 | 167912604 | rs35472201 | -1.659 | -2.38 | 0.124 | 1.112 | 0.035 | 0.015 | 0.015 | 0 | 0.5 | 0.59 | 0.23 | 0.37 | 0.24 | 0.47 |
| 2 | 167923974 | rs7565761 | 0.028 | -1.864 | 0.421 | 1.18 | -0.014 | -0.012 | -0.012 | 0 | 0.05 | 0.08 | 0.03 | 0.02 | 0.01 | 0.08 |
| 2 | 167925854 | rs10497306 | 0.182 | -2.149 | 0.017 | 0.948 | 0.022 | 0.052 | 0.052 | 0.007 | 0.17 | 0.03 | 0.07 | 0.06 | 0.09 | 0.15 |
| 2 | 167925954 | rs7599374 | -2.548 | -2.395 | 0.35 | 1.255 | -0.039 | 0.051 | 0.051 | 1.018 | 0.2 | 0.62 | 0.15 | 0.31 | 0.08 | 0.22 |
| 2 | 167926359 | rs16852935 | -2.18 | -2.399 | 0.501 | 1.238 | -0.048 | 0.07 | 0.07 | 0.155 | 0.17 | 0.57 | 0.11 | 0.26 | 0.06 | 0.2 |
| 2 | 167933831 | rs6733384 | -2.218 | -2.391 | 0.513 | 1.225 | -0.005 | 0 | 0 | 0 | 0.35 | 0.66 | 0.17 | 0.32 | 0.15 | 0.35 |
| 2 | 167961457 | rs7607028 | -2.461 | -2.603 | 0.569 | 1.276 | 0.008 | 0.003 | 0.003 | 0 | 0.39 | 0.63 | 0.2 | 0.35 | 0.15 | 0.39 |
| 2 | 167976035 | rs2303877 | -0.34 | -2.717 | 1.001 | 1.345 | 0.018 | 0.136 | 0.136 | 0 | 0.07 | 0.26 | 0.05 | 0.13 | 0.03 | 0.12 |
| 2 | 167991933 | rs16852998 | -2.13 | -2.978 | 0.694 | 1.257 | 0.013 | 0.019 | 0.019 | 1.122 | 0.06 | 0.15 | 0.07 | 0.05 | 0.08 | 0.02 |
| 2 | 167997309 | rs17551389 | 0.204 | -3.053 | 0.617 | 1.23 | 0.03 | 0.057 | 0.057 | 0 | 0.17 | 0.03 | 0.07 | 0.06 | 0.08 | 0.15 |
| 2 | 167997978 | rs1004767 | -0.151 | -3.065 | 0.785 | 1.254 | 0.02 | 0.02 | 0.02 | 0 | 0.18 | 0.15 | 0.08 | 0.06 | 0.09 | 0.15 |
| 2 | 168005852 | rs6755389 | -0.133 | -3.102 | 1.459 | 1.557 | 0.022 | 0.023 | 0.023 | 0 | 0.18 | 0.13 | 0.07 | 0.06 | 0.08 | 0.15 |
| 2 | 168036615 | rs16853128 | -1.839 | -3.113 | 1.374 | 1.554 | 0.148 | 0.28 | 0.28 | 0.016 | 0.06 | 0.36 | 0.07 | 0.14 | 0.09 | 0.12 |
| 2 | 168039548 | rs16853166 | -2.166 | -3.114 | 1.451 | 1.519 | 0.124 | 0.34 | 0.34 | 0 | 0.07 | 0.35 | 0.1 | 0.23 | 0.09 | 0.13 |
| 2 | 168046215 | rs10451542 | -1.416 | -3.115 | 1.149 | 1.492 | -0.029 | 0.028 | 0.028 | 1.657 | 0.27 | 0.56 | 0.2 | 0.34 | 0.19 | 0.29 |
| 2 | 168055739 | rs2390548 | 0.333 | -3.076 | 0.804 | 1.393 | 0.044 | 0.055 | 0.055 | 0 | 0.17 | 0.06 | 0.07 | 0.07 | 0.08 | 0.15 |
| 2 | 168062616 | rs6432984 | -2.578 | -3.025 | 0.91 | 1.458 | -0.017 | 0.014 | 0.014 | 2.821 | 0.22 | 0.65 | 0.19 | 0.29 | 0.12 | 0.25 |
| 2 | 168066066 | rs7585061 | -2.211 | -2.947 | 1.152 | 1.493 | 0.038 | -0.002 | -0.002 | 0.094 | 0.39 | 0.66 | 0.28 | 0.38 | 0.23 | 0.41 |
| 2 | 168066491 | rs16853221 | -0.428 | -2.904 | 1.506 | 1.461 | -0.026 | 0.05 | 0.05 | 0.245 | 0.22 | 0.39 | 0.18 | 0.3 | 0.14 | 0.29 |
| 2 | 168073830 | rs6731283 | -2.588 | -3.152 | 1.11 | 1.719 | 0.111 | -0.005 | -0.005 | 0 | 0.43 | 0.84 | 0.31 | 0.39 | 0.25 | 0.49 |
| 2 | 168099796 | rs16853306 | -2.279 | -3.111 | 1.596 | 1.805 | -0.052 | -0.001 | -0.001 | 0 | 0.21 | 0.39 | 0.17 | 0.22 | 0.13 | 0.24 |
| 2 | 168103925 | rs7607246 | -1.938 | -3.146 | 1.852 | 1.985 | 0.014 | 0.028 | 0.028 | 0 | 0.11 | 0.29 | 0.12 | 0.12 | 0.11 | 0.17 |
| 2 | 168104199 | rs7581190 | -1.938 | -3.146 | 1.852 | 1.985 | 0.014 | 0.028 | 0.028 | 0 | 0.11 | 0.29 | 0.12 | 0.12 | 0.11 | 0.17 |
| 2 | 168106085 | rs16853328 | -1.938 | -3.146 | 1.852 | 1.985 | 0.014 | 0.028 | 0.028 | 0 | 0.11 | 0.29 | 0.12 | 0.12 | 0.11 | 0.17 |
| 2 | 168106246 | rs16853330 | -2.723 | -3.14 | 1.835 | 1.99 | -0.05 | 0 | 0 | 0 | 0.21 | 0.36 | 0.17 | 0.22 | 0.13 | 0.24 |
| 2 | 168107155 | rs16853331 | -2.448 | -3.14 | 1.831 | 1.992 | -0.042 | -0.035 | -0.035 | 0 | 0.2 | 0.33 | 0.13 | 0.12 | 0.12 | 0.23 |
| 2 | 168107491 | rs3749004 | -2.448 | -3.14 | 1.831 | 1.988 | -0.042 | -0.035 | -0.035 | 0 | 0.2 | 0.33 | 0.13 | 0.12 | 0.12 | 0.23 |
| 2 | 168108357 | rs1842233 | -3.239 | -3.138 | 1.746 | 1.976 | -0.029 | 0.044 | 0.044 | 0.171 | 0.24 | 0.55 | 0.25 | 0.33 | 0.17 | 0.34 |
| 2 | 168115710 | rs3749006 | -2.053 | -3.084 | 1.579 | 1.996 | 0.015 | 0.08 | 0.08 | 0 | 0.11 | 0.32 | 0.13 | 0.2 | 0.11 | 0.18 |
| 2 | 168117790 | rs1479588 | -2.464 | -3.332 | 1.452 | 1.871 | 0.115 | 0.022 | 0.022 | 0.176 | 0.44 | 0.79 | 0.32 | 0.37 | 0.25 | 0.49 |
| 2 | 168117941 | rs1479587 | -2.037 | -3.005 | 1.573 | 1.827 | 0.017 | 0.001 | 0.001 | 0 | 0.11 | 0.32 | 0.12 | 0.11 | 0.11 | 0.17 |
| 2 | 168127837 | rs7578175 | -2.644 | -3.058 | 1.558 | 1.8 | -0.003 | 0.038 | 0.038 | 0 | 0.13 | 0.33 | 0.16 | 0.2 | 0.12 | 0.18 |
| 2 | 168161260 | rs1403894 | 1.993 | -3.034 | 1.635 | 2.109 | -0.022 | 0.087 | 0.087 | 0 | 0.85 | 0.58 | 0.86 | 0.78 | 0.89 | 0.8 |
| 2 | 168178179 | rs2177440 | 3.148 | -3.033 | 1.619 | 2.232 | -0.01 | 0.094 | 0.094 | 0.233 | 0.81 | 0.43 | 0.79 | 0.65 | 0.84 | 0.72 |
| 2 | 168187606 | rs1608518 | 2.159 | -3.007 | 1.635 | 2.241 | -0.023 | 0.047 | 0.047 | 0 | 0.84 | 0.5 | 0.86 | 0.83 | 0.89 | 0.79 |
| 2 | 168187747 | rs1600148 | 1.551 | -3.29 | 1.456 | 2.444 | 0.115 | -0.033 | -0.033 | 0 | 0.63 | 0.18 | 0.73 | 0.63 | 0.81 | 0.63 |
| 2 | 168194827 | rs1523879 | 1.32 | -3.376 | 0.736 | 2.431 | 0.005 | -0.07 | -0.07 | 0 | 0.17 | 0.05 | 0.11 | 0.33 | 0.14 | 0.19 |

| | | | | | | | | | | | | | | | | |
|---|---|---|---|---|---|---|---|---|---|---|---|---|---|---|---|---|
| 2 | 168196591 | rs1357602 | 2.301 | -3.104 | 0.973 | 1.636 | -0.061 | 0.06 | 0.06 | 1.294 | 0.84 | 0.42 | 0.84 | 0.76 | 0.95 | 0.79 |
| 2 | 168203116 | rs2140501 | 1.321 | -3.059 | 0.82 | 1.147 | 0.008 | 0.054 | 0.054 | 0 | 0.9 | 0.77 | 0.89 | 0.84 | 0.9 | 0.83 |
| 2 | 168207901 | rs2617367 | 1.601 | -3.191 | 0.392 | 0.881 | -0.093 | 0.159 | 0.159 | 0 | 0.38 | 0.04 | 0.48 | 0.15 | 0.6 | 0.27 |
| 2 | 168216090 | rs899185 | 1.919 | -3.27 | 1.005 | 0.921 | -0.025 | -0.017 | -0.017 | 0.133 | 0.85 | 0.64 | 0.91 | 0.94 | 0.9 | 0.83 |
| 2 | 168217204 | rs2617391 | 1.919 | -3.254 | 1.005 | 0.929 | -0.021 | -0.005 | -0.005 | 0 | 0.85 | 0.64 | 0.91 | 0.93 | 0.9 | 0.83 |
| 2 | 168222414 | rs2250619 | 1.869 | -3.267 | 0.977 | 0.84 | -0.026 | -0.014 | -0.014 | 0 | 0.85 | 0.63 | 0.91 | 0.93 | 0.9 | 0.81 |
| 2 | 168240431 | rs1357604 | 0.628 | -3.244 | 0.882 | 0.829 | -0.025 | 0.02 | 0.02 | 0 | 0.93 | 0.85 | 0.94 | 0.94 | 0.98 | 0.92 |
| 2 | 168255670 | rs1159502 | 0.803 | -3.26 | 0.782 | 0.904 | 0.015 | -0.103 | -0.103 | 0 | 0.15 | 0 | 0.1 | 0.35 | 0.09 | 0.15 |
| 2 | 168257411 | rs1880171 | 1.145 | -2.422 | 1.495 | 0.189 | -0.033 | 0.028 | 0.028 | 2.717 | 0.93 | 0.83 | 0.95 | 0.94 | 0.99 | 0.92 |
| 2 | 168287130 | rs1403895 | 2.417 | -2.425 | 1.272 | 0.185 | -0.05 | -0.026 | -0.026 | 0 | 0.83 | 0.47 | 0.9 | 0.91 | 0.9 | 0.81 |
| 2 | 168330745 | rs4387746 | 1.809 | -2.216 | 0.653 | 0.032 | -0.047 | -0.085 | -0.085 | 0.038 | 0.81 | 0.51 | 0.92 | 0.92 | 0.88 | 0.86 |
| 2 | 168338520 | rs7567362 | 0.198 | -2.269 | 0.328 | -0.152 | 0.013 | 0.015 | 0.015 | 0 | 0.48 | 0.46 | 0.46 | 0.81 | 0.35 | 0.62 |
| 2 | 168340426 | rs1919854 | 0.191 | -2.255 | 0.326 | -0.083 | 0.01 | 0.013 | 0.013 | 0 | 0.48 | 0.47 | 0.4 | 0.83 | 0.35 | 0.61 |
| 2 | 168356194 | rs1730725 | 0.025 | -2.394 | -0.111 | -0.464 | 0.051 | 0.127 | 0.127 | 0 | 0.52 | 0.69 | 0.49 | 0.86 | 0.35 | 0.62 |
| 2 | 168403680 | rs1227136 | -0.487 | -1.978 | 0.213 | 0.321 | -0.022 | 0.055 | 0.055 | 0.346 | 0.06 | 0.12 | 0.09 | 0.18 | 0.02 | 0.05 |
| 2 | 168405906 | rs11890435 | -0.382 | -1.71 | -0.024 | 0.193 | -0.012 | -0.014 | -0.014 | 0 | 0.06 | 0.22 | 0.02 | 0 | 0 | 0.07 |
| 2 | 168413249 | rs1236837 | 0.326 | -1.959 | -1.023 | -0.842 | 0.011 | 0.035 | 0.035 | 0.013 | 0.42 | 0.31 | 0.45 | 0.84 | 0.33 | 0.6 |
| 2 | 168426623 | rs1227139 | -0.609 | -1.038 | -1.499 | -0.386 | -0.174 | 0.173 | 0.173 | 0 | 0.74 | 0.98 | 0.64 | 0.99 | 0.56 | 0.86 |
| 2 | 168434795 | rs988843 | -1.095 | -2.072 | -0.117 | 0.698 | -0.072 | -0.019 | -0.019 | 0 | 0.14 | 0.34 | 0.05 | 0.05 | 0.03 | 0.12 |
| 2 | 168434845 | rs988842 | -0.931 | -1.54 | -0.614 | 1.067 | 0.017 | -0.002 | -0.002 | 0.716 | 0.12 | 0.08 | 0.01 | 0.1 | 0.03 | 0.19 |
| 2 | 168438360 | rs986697 | -1.103 | -1.663 | -0.435 | 1.023 | 0.004 | 0.006 | 0.006 | 0.381 | 0.3 | 0.67 | 0.19 | 0.17 | 0.12 | 0.32 |
| 2 | 168448561 | rs6743050 | 0.927 | -0.968 | -0.013 | 1.359 | 0.001 | 0.001 | 0.001 | 0 | 0.79 | 0.84 | 0.89 | 0.84 | 0.83 | 0.82 |
| 2 | 168461161 | rs734284 | 0.733 | -0.954 | 0.175 | 1.368 | -0.007 | -0.002 | -0.002 | 0.793 | 0.82 | 0.76 | 0.91 | 0.81 | 0.85 | 0.83 |
| 2 | 168468707 | rs1367919 | 1.534 | -1.106 | 1.399 | 1.197 | -0.016 | -0.028 | -0.028 | 0 | 0.75 | 0.41 | 0.86 | 0.8 | 0.8 | 0.79 |
| 2 | 168470528 | rs4667959 | 1.42 | -1.125 | 1.526 | 1.187 | -0.016 | -0.029 | -0.029 | 0 | 0.75 | 0.38 | 0.86 | 0.8 | 0.8 | 0.79 |
| 2 | 168477472 | rs1431900 | 0.896 | -1.093 | 1.328 | 1.21 | -0.016 | -0.008 | -0.008 | 0 | 0.75 | 0.62 | 0.84 | 0.78 | 0.79 | 0.72 |
| 2 | 168479374 | rs1406665 | 1.008 | -1.079 | 1.365 | 1.198 | 0.011 | 0.008 | 0.008 | 0 | 0.74 | 0.82 | 0.84 | 0.77 | 0.79 | 0.72 |
| 2 | 168485936 | rs2052981 | 1.626 | -1.13 | 1.354 | 1.592 | -0.016 | 0.007 | 0.007 | 0.099 | 0.67 | 0.41 | 0.71 | 0.65 | 0.76 | 0.68 |
| 2 | 168497830 | rs1147148 | -1.063 | -0.957 | 0.765 | 1.678 | -0.009 | -0.005 | -0.005 | 0 | 0.25 | 0.26 | 0.16 | 0.23 | 0.19 | 0.26 |
| 2 | 168500389 | rs1226906 | -1.166 | -0.882 | 0.609 | 1.54 | 0.007 | -0.013 | -0.013 | 0 | 0.28 | 0.25 | 0.19 | 0.34 | 0.22 | 0.29 |
| 2 | 168513346 | rs935640 | 1.077 | -0.792 | 1.387 | 1.927 | -0.013 | 0.266 | 0.266 | 4.166 | 0.9 | 0.49 | 0.93 | 0.71 | 0.96 | 0.95 |
| 2 | 168513854 | rs935641 | -0.245 | -0.708 | 0.395 | 1.01 | -0.028 | 0.084 | 0.084 | 1.542 | 0.19 | 0.05 | 0.19 | 0.01 | 0.2 | 0.1 |
| 2 | 168515362 | rs2390601 | 0.982 | -1.335 | 0.057 | 1.265 | -0.01 | 0.084 | 0.084 | 1.599 | 0.43 | 0.17 | 0.48 | 0.25 | 0.59 | 0.51 |
| 2 | 168518957 | rs1839191 | 0.637 | -1.561 | 0.882 | 1.666 | -0.054 | 0.115 | 0.115 | 0 | 0.87 | 0.43 | 0.81 | 0.75 | 0.96 | 0.88 |
| 2 | 168538351 | rs2029084 | 1.026 | -0.543 | -0.637 | 0.529 | -0.044 | 0.191 | 0.191 | 2.994 | 0.3 | 0.06 | 0.13 | 0.04 | 0.3 | 0.23 |
| 2 | 168538458 | rs7591667 | -1.143 | -0.642 | 0.238 | 0.938 | -0.016 | 0.244 | 0.244 | 0.28 | 0.12 | 0.34 | 0.15 | 0.39 | 0.07 | 0.11 |
| 2 | 168539669 | rs1463152 | 0.532 | 0.312 | -0.274 | 0.988 | -0.04 | -0.035 | -0.035 | 0 | 0.12 | 0.05 | 0.37 | 0.2 | 0.3 | 0.29 |
| 2 | 168542174 | rs2029083 | 0.811 | -0.001 | -0.96 | 0.363 | -0.052 | 0.112 | 0.112 | 0 | 0.19 | 0.04 | 0.23 | 0 | 0.21 | 0.14 |
| 2 | 168544279 | rs2029079 | 0.581 | 0.278 | -1.085 | 0.501 | -0.056 | 0.011 | 0.011 | 0 | 0.16 | 0.05 | 0.3 | 0.11 | 0.28 | 0.28 |
| 2 | 168544776 | rs7605048 | -0.053 | -0.305 | -1.122 | 0.225 | 0.008 | 0.032 | 0.032 | 0.006 | 0.29 | 0.18 | 0.39 | 0.19 | 0.26 | 0.25 |
| 2 | 168551879 | rs1448843 | -2.1 | -0.417 | -0.619 | -0.226 | -0.008 | -0.054 | -0.054 | 0.008 | 0.84 | 0.72 | 0.91 | 0.94 | 0.89 | 0.88 |
| 2 | 168553125 | rs11690497 | 0.781 | -0.223 | -0.485 | 0.024 | -0.067 | 0.003 | 0.003 | 0 | 0.18 | 0.06 | 0.32 | 0.14 | 0.3 | 0.27 |
| 2 | 168558146 | rs6718232 | 0.401 | -0.309 | -0.768 | 0.127 | -0.029 | 0.055 | 0.055 | 0 | 0.09 | 0.22 | 0.16 | 0.16 | 0.03 | 0.11 |
| 2 | 168559995 | rs6726109 | 1.106 | -0.213 | -1.015 | 0.076 | -0.078 | -0.001 | -0.001 | 0 | 0.16 | 0.06 | 0.3 | 0.12 | 0.29 | 0.28 |
| 2 | 168560206 | rs1947192 | -0.494 | -0.754 | -1.558 | 0.331 | 0.007 | -0.011 | -0.011 | 0 | 0.28 | 0.26 | 0.27 | 0.58 | 0.2 | 0.34 |
| 2 | 168563180 | rs12151513 | 0.609 | -0.49 | -1.878 | 0.608 | -0.022 | -0.051 | -0.051 | 0 | 0.13 | 0.05 | 0.09 | 0.36 | 0.15 | 0.16 |
| 2 | 168564354 | rs17581506 | 1.84 | -0.11 | -2.263 | 0.067 | -0.009 | 0.088 | 0.088 | 1.399 | 0.18 | 0.05 | 0.05 | 0 | 0.16 | 0.1 |
| 2 | 168565693 | rs13408808 | 0.347 | 0.083 | -2.117 | 0.367 | -0.016 | -0.061 | -0.061 | 0 | 0.15 | 0.05 | 0.1 | 0.38 | 0.16 | 0.17 |
| 2 | 168567310 | rs17581611 | 1.052 | -0.054 | -2.16 | 0.034 | -0.038 | 0.079 | 0.079 | 0 | 0.33 | 0.14 | 0.32 | 0.16 | 0.36 | 0.25 |
| 2 | 168574523 | rs2044682 | -0.658 | -0.051 | -1.143 | 0.324 | 0.014 | -0.006 | -0.006 | 0 | 0.37 | 0.51 | 0.56 | 0.68 | 0.44 | 0.46 |
| 2 | 168577470 | rs1868382 | -0.462 | -0.157 | -0.874 | 0.588 | -0.027 | -0.01 | -0.01 | 0 | 0.69 | 0.48 | 0.58 | 0.72 | 0.78 | 0.82 |
| 2 | 168577771 | rs16854089 | 0.29 | 0.496 | -0.98 | 0.738 | -0.007 | -0.013 | -0.013 | 1.753 | 0.21 | 0.19 | 0.16 | 0.5 | 0.23 | 0.26 |
| 2 | 168581320 | rs7582020 | -1.057 | 0.341 | -0.788 | 0.717 | -0.024 | 0.003 | 0.003 | 0 | 0.72 | 0.46 | 0.6 | 0.72 | 0.79 | 0.82 |
| 2 | 168583981 | rs1900833 | -1.587 | 0.346 | -0.742 | 0.699 | -0.026 | 0.027 | 0.027 | 0 | 0.76 | 0.52 | 0.63 | 0.74 | 0.83 | 0.84 |
| 2 | 168585251 | rs11676513 | 1.237 | 0.549 | -1.461 | 0.547 | -0.019 | 0.037 | 0.037 | 0 | 0.23 | 0.1 | 0.2 | 0.11 | 0.22 | 0.18 |
| 2 | 168585553 | rs11687831 | -0.117 | 0.676 | -1.264 | 0.734 | -0.021 | 0.002 | 0.002 | 0 | 0.56 | 0.15 | 0.44 | 0.32 | 0.62 | 0.65 |
| 2 | 168586216 | rs6752240 | 0.532 | 0.943 | -0.755 | 0.989 | 0.031 | 0.078 | 0.078 | 0 | 0.15 | 0.28 | 0.14 | 0.32 | 0.16 | 0.16 |
| 2 | 168587946 | rs10180263 | 0.275 | 1 | -0.715 | 0.999 | 0.04 | 0.088 | 0.088 | 0 | 0.16 | 0.31 | 0.16 | 0.41 | 0.19 | 0.17 |
| 2 | 168589319 | rs10173035 | 0.742 | 1.027 | -1.063 | 0.91 | 0.003 | 0.009 | 0.009 | 0 | 0.13 | 0.15 | 0.13 | 0.3 | 0.17 | 0.15 |
| 2 | 168591057 | rs9646774 | 0.688 | 0.686 | -1.379 | 0.475 | -0.009 | 0.048 | 0.048 | 0 | 0.25 | 0.1 | 0.22 | 0.11 | 0.24 | 0.21 |
| 2 | 168593889 | rs10930282 | 1.091 | 0.633 | -1.423 | 0.334 | 0.011 | 0.107 | 0.107 | 0 | 0.22 | 0.07 | 0.19 | 0.03 | 0.18 | 0.14 |
| 2 | 168596313 | rs4667970 | -0.399 | 0.609 | -0.918 | 0.589 | 0.001 | 0 | 0 | 0 | 0.54 | 0.47 | 0.6 | 0.43 | 0.57 | 0.59 |
| 2 | 168597986 | rs10168551 | 1.711 | 0.919 | -0.91 | 0.587 | 0.013 | -0.007 | -0.007 | 0 | 0.14 | 0.18 | 0.11 | 0.16 | 0.19 | 0.23 |
| 2 | 168598551 | rs13424531 | 1.16 | 0.779 | -0.917 | 0.801 | -0.01 | -0.027 | -0.027 | 0 | 0.14 | 0.06 | 0.09 | 0.32 | 0.15 | 0.15 |
| 2 | 168601735 | rs12479257 | -1.001 | 0.524 | -0.613 | 0.527 | -0.007 | 0.015 | 0.015 | 0 | 0.65 | 0.78 | 0.84 | 0.69 | 0.66 | 0.76 |
| 2 | 168602991 | rs7585307 | 0.708 | 0.543 | -0.363 | 0.554 | 0.024 | 0.029 | 0.029 | 0 | 0.31 | 0.36 | 0.43 | 0.42 | 0.42 | 0.46 |
| 2 | 168614346 | rs4667972 | 0.485 | 0.31 | -0.021 | 0.849 | 0.001 | 0 | 0 | 0 | 0.33 | 0.39 | 0.31 | 0.28 | 0.26 | 0.29 |
| 2 | 168616962 | rs6433011 | -0.636 | 0.276 | -0.082 | 0.981 | 0.029 | 0.176 | 0.176 | 0 | 0.51 | 0.76 | 0.47 | 0.85 | 0.54 | 0.56 |
| 2 | 168619462 | rs16854131 | 1.345 | 0.335 | -0.047 | 1.039 | -0.002 | 0.006 | 0.006 | 0 | 0.26 | 0.35 | 0.27 | 0.27 | 0.21 | 0.24 |

| | | | | | | | | | | | | | | | |
|---|---|---|---|---|---|---|---|---|---|---|---|---|---|---|---|
| 2 | 168621005 | rs1375311 | 1.972 | 0.411 | 0 | 1.251 | 0.002 | -0.001 | -0.001 | 0 | 0.12 | 0.12 | 0.11 | 0.09 | 0.2 | 0.22 |
| 2 | 168625881 | rs6741880 | 0.746 | 0.37 | -0.122 | 1.387 | -0.016 | -0.031 | -0.031 | 0 | 0.14 | 0.08 | 0.11 | 0.36 | 0.19 | 0.12 |
| 2 | 168628177 | rs12622054 | 0.851 | 0.311 | 0.061 | 1.318 | 0.008 | -0.01 | -0.01 | 0 | 0.33 | 0.35 | 0.16 | 0.22 | 0.34 | 0.3 |
| 2 | 168633681 | rs11692980 | 0.216 | 0.285 | -0.094 | 1.071 | -0.007 | 0.002 | 0.002 | 0 | 0.42 | 0.7 | 0.41 | 0.49 | 0.43 | 0.47 |
| 2 | 168635552 | rs17644183 | 0.173 | 0.284 | -0.088 | 1.052 | -0.006 | 0.022 | 0.022 | 0 | 0.42 | 0.69 | 0.41 | 0.59 | 0.43 | 0.47 |
| 2 | 168636999 | rs7600099 | 1.281 | 0.268 | 0.089 | 1.112 | -0.022 | 0.045 | 0.045 | 0.471 | 0.26 | 0.55 | 0.29 | 0.31 | 0.17 | 0.14 |
| 2 | 168639838 | rs11691258 | 0.886 | 0.579 | -0.184 | 1.198 | -0.016 | -0.02 | -0.02 | 0 | 0.13 | 0.07 | 0.1 | 0.27 | 0.17 | 0.11 |
| 2 | 168640233 | rs1004366 | 1.771 | 0.634 | -0.222 | 1.22 | -0.018 | -0.013 | -0.013 | 0.295 | 0.13 | 0.04 | 0.1 | 0.18 | 0.22 | 0.23 |
| 2 | 168641930 | rs1004369 | -0.839 | 0.097 | 0.483 | 1.296 | 0.026 | 0.081 | 0.081 | 0 | 0.22 | 0.51 | 0.18 | 0.36 | 0.23 | 0.27 |
| 2 | 168641970 | rs1004368 | -0.862 | 0.096 | 0.482 | 1.299 | 0.024 | 0.082 | 0.082 | 0 | 0.22 | 0.51 | 0.18 | 0.36 | 0.23 | 0.27 |
| 2 | 168645406 | rs836692 | -0.509 | -0.226 | 0.121 | 1.095 | 0.002 | 0.006 | 0.006 | 0.119 | 0.42 | 0.59 | 0.35 | 0.65 | 0.32 | 0.34 |
| 2 | 168649212 | rs12992085 | -0.967 | -0.203 | 0.27 | 1.022 | 0.008 | 0.013 | 0.013 | 0.127 | 0.08 | 0.21 | 0.05 | 0.07 | 0.06 | 0.17 |
| 2 | 168649399 | rs12992718 | -0.967 | -0.203 | 0.27 | 1.022 | 0.008 | 0.013 | 0.013 | 0.127 | 0.08 | 0.22 | 0.05 | 0.07 | 0.06 | 0.17 |
| 2 | 168649930 | rs12999150 | 0.689 | -0.251 | 0.089 | 0.858 | 0.003 | 0.016 | 0.016 | 0 | 0.27 | 0.21 | 0.1 | 0.07 | 0.2 | 0.26 |
| 2 | 168650226 | rs836684 | 0.157 | -0.208 | 0.517 | 1.313 | 0.008 | 0.057 | 0.057 | 0 | 0.14 | 0.19 | 0.09 | 0.37 | 0.14 | 0.12 |
| 2 | 168653425 | rs1446502 | -0.064 | -0.248 | 0.442 | 1.426 | -0.02 | 0.268 | 0.268 | 0 | 0.06 | 0.28 | 0.2 | 0.29 | 0.01 | 0.05 |
| 2 | 168655533 | rs836674 | -1.749 | -0.044 | -0.226 | 1.138 | -0.073 | -0.021 | -0.021 | 0 | 0.86 | 0.98 | 0.69 | 0.87 | 0.74 | 0.82 |
| 2 | 168656646 | rs836680 | -0.876 | -1.176 | 0.5 | 1.327 | 0.017 | 0.086 | 0.086 | 0 | 0.21 | 0.42 | 0.12 | 0.38 | 0.2 | 0.28 |
| 2 | 168658918 | rs836681 | -1.253 | -1.288 | 0.223 | 1.191 | 0.028 | 0.059 | 0.059 | 0 | 0.47 | 0.78 | 0.4 | 0.68 | 0.35 | 0.41 |
| 2 | 168659362 | rs4667977 | 0.131 | -1.241 | 0.484 | 1.271 | -0.017 | 0.035 | 0.035 | 0.367 | 0.29 | 0.43 | 0.3 | 0.37 | 0.2 | 0.24 |
| 2 | 168671309 | rs12614700 | 0.976 | -1.451 | -0.123 | 1.383 | -0.005 | -0.157 | -0.157 | 0.709 | 0.21 | 0 | 0.25 | 0.43 | 0.13 | 0.22 |
| 2 | 168671408 | rs836715 | -1.32 | -2.142 | 0.172 | 1.043 | 0.021 | 0.015 | 0.015 | 1.353 | 0.47 | 0.55 | 0.42 | 0.7 | 0.29 | 0.39 |
| 2 | 168673957 | rs836717 | -0.584 | -2.051 | 0.26 | 1.245 | 0.036 | -0.047 | -0.047 | 0 | 0.35 | 0.19 | 0.33 | 0.57 | 0.24 | 0.25 |
| 2 | 168675188 | rs836722 | -0.638 | -2.044 | 0.228 | 1.25 | 0.036 | -0.051 | -0.051 | 0.004 | 0.35 | 0.17 | 0.33 | 0.57 | 0.24 | 0.25 |
| 2 | 168679800 | rs6718868 | 0.461 | -2.213 | 0.14 | 1.216 | 0.036 | -0.1 | -0.1 | 0.42 | 0.26 | 0.04 | 0.26 | 0.4 | 0.16 | 0.28 |
| 2 | 168682277 | rs700546 | -0.516 | -2.093 | 0.359 | 1.105 | -0.008 | -0.003 | -0.003 | 0 | 0.17 | 0.18 | 0.09 | 0.21 | 0.13 | 0.14 |
| 2 | 168687153 | rs836696 | -1.218 | -2.084 | 0.633 | 0.894 | -0.021 | 0.007 | 0.007 | 0.263 | 0.43 | 0.41 | 0.35 | 0.55 | 0.29 | 0.36 |
| 2 | 168701171 | rs2034258 | -1.866 | -2.195 | 0.595 | 0.813 | 0.035 | 0.022 | 0.022 | 0.165 | 0.53 | 0.69 | 0.44 | 0.67 | 0.36 | 0.49 |
| 2 | 168710172 | rs10200961 | -0.799 | -1.675 | 1.212 | 1.147 | 0.012 | 0.126 | 0.126 | 0 | 0.15 | 0.25 | 0.19 | 0.38 | 0.13 | 0.17 |
| 2 | 168712041 | rs6734498 | -1.348 | -1.57 | 0.673 | 1.15 | -0.018 | -0.017 | -0.017 | 0 | 0.1 | 0.31 | 0.08 | 0.07 | 0.06 | 0.04 |
| 2 | 168712452 | rs10930293 | -0.591 | -1.389 | 0.175 | 1.138 | 0 | -0.003 | -0.003 | 0 | 0.16 | 0.11 | 0.18 | 0.4 | 0.14 | 0.17 |
| 2 | 168721539 | rs7598325 | -1.017 | -1.244 | 0.043 | 0.557 | 0.001 | -0.014 | -0.014 | 0 | 0.09 | 0.1 | 0.03 | 0 | 0.08 | 0.11 |
| 2 | 168723890 | rs12471704 | 1.105 | -1.557 | 0.391 | 0.724 | -0.041 | 0.03 | 0.03 | 0 | 0.71 | 0.53 | 0.68 | 0.59 | 0.78 | 0.67 |
| 2 | 168729982 | rs836708 | -1.545 | -1.777 | 0.023 | 0.675 | 0.1 | -0.018 | -0.018 | 0.135 | 0.49 | 0.83 | 0.53 | 0.53 | 0.32 | 0.42 |
| 2 | 168730580 | rs836709 | -1.393 | -1.74 | 0.142 | 0.405 | 0.066 | 0.099 | 0.099 | 0.635 | 0.47 | 0.78 | 0.46 | 0.33 | 0.34 | 0.34 |
| 2 | 168734817 | rs7577431 | -0.357 | -1.806 | 0.416 | 0.258 | -0.029 | -0.132 | -0.132 | 0 | 0.27 | 0.44 | 0.14 | 0 | 0.18 | 0.14 |
| 2 | 168737020 | rs1551824 | -0.218 | -1.603 | -0.476 | 0.442 | 0.007 | -0.032 | -0.032 | 0 | 0.2 | 0.06 | 0.28 | 0.31 | 0.17 | 0.2 |
| 2 | 168737748 | rs1374276 | -0.616 | -1.476 | -0.298 | 0.566 | 0.02 | 0.04 | 0.04 | 3.416 | 0.37 | 0.1 | 0.33 | 0.32 | 0.3 | 0.29 |
| 2 | 168749667 | rs10930295 | -1.433 | -1.595 | 0.088 | 0.711 | 0.079 | -0.002 | -0.002 | 0 | 0.49 | 0.7 | 0.52 | 0.54 | 0.31 | 0.49 |
| 2 | 168752887 | rs16854364 | -0.017 | -1.681 | 0.516 | 0.827 | 0.001 | 0.166 | 0.166 | 0.182 | 0.07 | 0.24 | 0.05 | 0.17 | 0.02 | 0.1 |
| 2 | 168759755 | rs836647 | -1.637 | -1.708 | -0.181 | 0.632 | 0.145 | -0.036 | -0.036 | 0 | 0.5 | 0.89 | 0.54 | 0.54 | 0.3 | 0.49 |
| 2 | 168761499 | rs836650 | -0.554 | -1.611 | 0.07 | 0.636 | 0.017 | 0.026 | 0.026 | 0 | 0.32 | 0.2 | 0.37 | 0.32 | 0.28 | 0.29 |
| 2 | 168764274 | rs1838836 | -1.092 | -1.576 | 0.392 | 0.733 | 0.012 | -0.011 | -0.011 | 0 | 0.39 | 0.4 | 0.5 | 0.53 | 0.28 | 0.46 |
| 2 | 168772522 | rs836664 | -0.858 | -1.232 | 0.249 | 0.388 | -0.038 | 0.078 | 0.078 | 0 | 0.17 | 0.43 | 0.23 | 0.3 | 0.12 | 0.16 |
| 2 | 168772574 | rs1947111 | 0.855 | -0.752 | -0.826 | 0.016 | -0.073 | 0.055 | 0.055 | 0.109 | 0.25 | 0.04 | 0.26 | 0.08 | 0.29 | 0.13 |
| 2 | 168775061 | rs7576460 | -0.449 | -0.738 | 0.117 | 0.539 | 0.008 | -0.062 | -0.062 | 0.001 | 0.14 | 0.03 | 0.19 | 0.3 | 0.12 | 0.16 |
| 2 | 168802437 | rs6729671 | -0.701 | -0.674 | -0.392 | 0.59 | -0.023 | 0.01 | 0.01 | 0 | 0.27 | 0.32 | 0.22 | 0.33 | 0.17 | 0.22 |
| 2 | 168814299 | rs13000880 | 0.113 | 0.192 | -0.796 | -0.004 | -0.026 | 0.019 | 0.019 | 0 | 0.29 | 0.23 | 0.34 | 0.03 | 0.4 | 0.22 |
| 2 | 168814408 | rs1446505 | -0.327 | 0.314 | -0.901 | 0.481 | -0.024 | 0.004 | 0.004 | 0 | 0.35 | 0.23 | 0.36 | 0.26 | 0.42 | 0.34 |
| 2 | 168817002 | rs10930299 | 0.324 | 0.053 | -1.495 | 0.923 | -0.029 | -0.091 | -0.091 | 0 | 0.13 | 0.04 | 0.22 | 0.39 | 0.16 | 0.15 |
| 2 | 168824134 | rs935381 | -1.127 | -0.719 | -1.468 | 0.437 | -0.002 | -0.001 | -0.001 | 0 | 0.37 | 0.62 | 0.4 | 0.66 | 0.33 | 0.41 |
| 2 | 168826850 | rs7579964 | -0.276 | -0.95 | -0.711 | 0.237 | 0.002 | 0.019 | 0.019 | 0 | 0.52 | 0.57 | 0.48 | 0.3 | 0.56 | 0.48 |
| 2 | 168833081 | rs6718279 | 0.489 | -1.016 | -0.753 | 0.218 | 0.031 | -0.021 | -0.021 | 0 | 0.07 | 0.01 | 0.02 | 0.22 | 0.03 | 0.12 |
| 2 | 168837207 | rs6734516 | 0.053 | -1.019 | -0.445 | 0.185 | -0.005 | 0.137 | 0.137 | 0 | 0.47 | 0.71 | 0.47 | 0.08 | 0.52 | 0.36 |
| 2 | 168840315 | rs10497332 | 1.243 | -1.026 | -0.116 | 0.083 | 0.011 | -0.057 | -0.057 | 0.396 | 0.3 | 0.29 | 0.33 | 0.03 | 0.37 | 0.22 |
| 2 | 168842047 | rs12986593 | 2.215 | -0.973 | -0.483 | -0.03 | -0.025 | 0.022 | 0.022 | 0.073 | 0.21 | 0.02 | 0.11 | 0.17 | 0.19 | 0.23 |
| 2 | 168848588 | rs4667994 | 0.123 | -1.022 | 0.04 | 0.1 | -0.005 | 0.079 | 0.079 | 0 | 0.46 | 0.65 | 0.46 | 0.03 | 0.52 | 0.29 |
| 2 | 168849447 | rs6712239 | 0.55 | -1.022 | 0.092 | 0.128 | 0.011 | -0.012 | -0.012 | 0.272 | 0.37 | 0.47 | 0.36 | 0.24 | 0.4 | 0.33 |
| 2 | 168849603 | rs13410387 | 0.861 | -0.947 | -0.013 | -0.048 | 0.003 | 0.013 | 0.013 | 0 | 0.14 | 0.08 | 0.11 | 0 | 0.15 | 0.07 |
| 2 | 168852219 | rs4667995 | 0.858 | -0.793 | 0.052 | 0.074 | 0.026 | -0.121 | -0.121 | 0 | 0.32 | 0.49 | 0.35 | 0.03 | 0.37 | 0.22 |
| 2 | 168855387 | rs7565986 | -1.037 | -0.853 | -0.062 | 0.459 | -0.015 | 0.014 | 0.014 | 0.061 | 0.69 | 0.85 | 0.73 | 0.69 | 0.72 | 0.63 |
| 2 | 168875155 | rs7574351 | -0.092 | -1.244 | 0.382 | 0.487 | -0.109 | 0.119 | 0.119 | 0 | 0.16 | 0.4 | 0.05 | 0.22 | 0.04 | 0.16 |
| 2 | 168888485 | rs2029076 | -0.993 | -1.271 | 0.383 | 0.478 | 0.012 | 0.026 | 0.026 | 5.02 | 0.64 | 0.8 | 0.74 | 0.67 | 0.73 | 0.58 |
| 2 | 168889664 | rs11891885 | 0.292 | -1.3 | 0.469 | 0.534 | -0.041 | 0.132 | 0.132 | 0 | 0.08 | 0.17 | 0.03 | 0.22 | 0.03 | 0.11 |
| 2 | 168892418 | rs9789702 | 0.602 | -1.358 | 0.391 | 0.36 | -0.017 | -0.011 | -0.011 | 0.089 | 0.4 | 0.3 | 0.56 | 0.45 | 0.54 | 0.4 |
| 2 | 168894649 | rs4668007 | -1.426 | -1.351 | 0.381 | 0.368 | -0.001 | 0.003 | 0.003 | 0.745 | 0.69 | 0.73 | 0.82 | 0.68 | 0.74 | 0.63 |
| 2 | 168896548 | rs10169506 | -0.711 | -1.276 | 0.319 | 0.456 | -0.045 | 0.018 | 0.018 | 0.022 | 0.24 | 0.43 | 0.17 | 0.21 | 0.19 | 0.18 |
| 2 | 168901077 | rs4668009 | 0.582 | -1.195 | -0.086 | 0.561 | -0.011 | 0 | 0 | 0 | 0.39 | 0.34 | 0.57 | 0.39 | 0.54 | 0.32 |
| 2 | 168906641 | rs715878 | 0.439 | -1.206 | -0.1 | 0.597 | -0.001 | -0.002 | -0.002 | 0 | 0.4 | 0.4 | 0.58 | 0.5 | 0.55 | 0.4 |

| | | | | | | | | | | | | | | | |
|---|---|---|---|---|---|---|---|---|---|---|---|---|---|---|---|
| 2 | 168912084 | rs12479070 | -0.202 | -1.267 | -0.507 | 0.651 | -0.044 | 0.068 | 0.068 | 0 | 0.1 | 0.14 | 0.13 | 0.2 | 0.03 | 0.12 |
| 2 | 168921669 | rs2290106 | 0.801 | -1.276 | -1.023 | 0.665 | 0.02 | -0.025 | -0.025 | 0.117 | 0.13 | 0.04 | 0.12 | 0.2 | 0.03 | 0.16 |
| 2 | 168924785 | rs17176633 | 1.007 | -0.864 | -1.117 | 1.005 | -0.032 | 0.013 | 0.013 | 0 | 0.31 | 0.19 | 0.38 | 0.3 | 0.43 | 0.25 |
| 2 | 168931610 | rs3739115 | 0.267 | -1.032 | -1.374 | 1.111 | 0.035 | -0.033 | -0.033 | 0 | 0.15 | 0.03 | 0.12 | 0.2 | 0.03 | 0.16 |
| 2 | 168932843 | rs4233815 | -1.025 | -0.417 | -1.319 | 0.76 | -0.029 | 0.009 | 0.009 | 1.013 | 0.71 | 0.89 | 0.85 | 0.7 | 0.74 | 0.63 |
| 2 | 168941044 | rs6750247 | 0.429 | -0.426 | -1.335 | 0.896 | 0.012 | -0.019 | -0.019 | 0 | 0.49 | 0.14 | 0.45 | 0.52 | 0.53 | 0.56 |
| 2 | 168943765 | rs4668014 | 0.595 | -0.191 | -1.503 | 1.123 | -0.029 | -0.026 | -0.026 | 0.012 | 0.06 | 0.02 | 0.14 | 0.24 | 0.11 | 0.17 |
| 2 | 168944228 | rs1992928 | 1.033 | -0.748 | -1.251 | 0.789 | -0.033 | 0.063 | 0.063 | 0 | 0.24 | 0.1 | 0.2 | 0.08 | 0.3 | 0.11 |
| 2 | 168950700 | rs11689980 | 0.014 | -0.906 | -0.656 | 0.773 | -0.001 | 0.009 | 0.009 | 0 | 0.33 | 0.29 | 0.32 | 0.16 | 0.33 | 0.22 |
| 2 | 168954815 | rs4667556 | 0.633 | -0.812 | -0.215 | 0.5 | -0.004 | 0.011 | 0.011 | 0 | 0.77 | 0.63 | 0.81 | 0.69 | 0.78 | 0.76 |
| 2 | 168962486 | rs6720684 | 0.787 | -0.826 | -0.088 | 0.522 | -0.007 | 0.018 | 0.018 | 0 | 0.76 | 0.6 | 0.64 | 0.69 | 0.77 | 0.76 |
| 2 | 168963731 | rs16854729 | 0.125 | -0.959 | 0.091 | 0.86 | -0.005 | -0.017 | -0.017 | 0 | 0.35 | 0.45 | 0.41 | 0.27 | 0.33 | 0.27 |
| 2 | 168969842 | rs1448828 | 0.248 | -0.962 | 0.169 | 0.879 | 0 | 0.005 | 0.005 | 0 | 0.33 | 0.3 | 0.4 | 0.25 | 0.33 | 0.22 |
| 2 | 168971139 | rs1813141 | 0.174 | -1.02 | 0.193 | 0.849 | -0.004 | -0.007 | -0.007 | 0.54 | 0.6 | 0.45 | 0.64 | 0.75 | 0.65 | 0.77 |
| 2 | 168975305 | rs6751710 | 1.422 | -0.938 | -0.037 | 0.76 | -0.031 | -0.017 | -0.017 | 0.012 | 0.4 | 0.16 | 0.29 | 0.41 | 0.43 | 0.48 |
| 2 | 168975320 | rs6433033 | 1.552 | -0.948 | -0.037 | 0.81 | -0.03 | 0.039 | 0.039 | 0.547 | 0.67 | 0.41 | 0.61 | 0.51 | 0.75 | 0.59 |
| 2 | 168975787 | rs1816977 | 0.094 | -0.877 | -0.148 | 0.853 | -0.011 | -0.011 | -0.011 | 0.037 | 0.81 | 0.86 | 0.78 | 0.77 | 0.81 | 0.78 |
| 2 | 168979569 | rs7562413 | -0.373 | -0.83 | -0.014 | 0.875 | -0.012 | -0.008 | -0.008 | 0 | 0.38 | 0.51 | 0.44 | 0.32 | 0.35 | 0.24 |
| 2 | 168984168 | rs1517334 | -0.004 | -0.692 | -0.249 | 0.974 | -0.007 | -0.007 | -0.007 | 0.071 | 0.82 | 0.82 | 0.78 | 0.77 | 0.79 | 0.78 |
| 2 | 168986767 | rs16854770 | -0.564 | -0.717 | 0.12 | 1.059 | -0.055 | 0.084 | 0.084 | 0.144 | 0.11 | 0.25 | 0.14 | 0.18 | 0.04 | 0.17 |
| 2 | 168990756 | rs2122080 | 0.541 | -0.491 | -0.08 | 0.885 | -0.001 | -0.001 | -0.001 | 0.065 | 0.48 | 0.47 | 0.45 | 0.5 | 0.46 | 0.49 |
| 2 | 168993929 | rs3769408 | -0.113 | -0.55 | -0.149 | 0.95 | -0.004 | -0.026 | -0.026 | 0 | 0.15 | 0.09 | 0.24 | 0.25 | 0.17 | 0.23 |
| 2 | 168998401 | rs3769405 | -1.417 | -0.73 | -0.074 | 0.751 | -0.005 | -0.011 | -0.011 | 0.253 | 0.38 | 0.57 | 0.44 | 0.71 | 0.29 | 0.45 |
| 2 | 169000293 | rs2166963 | -0.121 | -0.715 | -0.021 | 0.804 | -0.039 | 0.14 | 0.14 | 0.719 | 0.22 | 0.46 | 0.19 | 0.43 | 0.11 | 0.19 |
| 2 | 169002104 | rs1517328 | -1.353 | -0.711 | -0.017 | 0.781 | -0.001 | -0.001 | -0.001 | 0 | 0.37 | 0.58 | 0.44 | 0.69 | 0.29 | 0.42 |
| 2 | 169002591 | rs1517331 | 0.302 | -0.68 | 0.329 | 0.856 | -0.012 | 0.006 | 0.006 | 0.297 | 0.18 | 0.37 | 0.17 | 0.18 | 0.1 | 0.13 |
| 2 | 169002934 | rs3769396 | -0.365 | -0.685 | 0.202 | 0.834 | -0.028 | 0.08 | 0.08 | 1.193 | 0.22 | 0.55 | 0.2 | 0.43 | 0.11 | 0.19 |
| 2 | 169004125 | rs3769395 | -0.275 | -0.677 | 0.185 | 0.836 | -0.03 | 0.098 | 0.098 | 0.976 | 0.22 | 0.53 | 0.19 | 0.44 | 0.11 | 0.19 |
| 2 | 169012502 | rs1850440 | -0.02 | -0.404 | -0.311 | 0.487 | 0.006 | 0.013 | 0.013 | 0 | 0.65 | 0.74 | 0.8 | 0.67 | 0.65 | 0.76 |
| 2 | 169015673 | rs1517342 | -0.642 | -0.227 | -0.066 | 0.611 | -0.001 | -0.002 | -0.002 | 0 | 0.52 | 0.46 | 0.46 | 0.66 | 0.51 | 0.45 |
| 2 | 169016304 | rs3754774 | -0.642 | -0.227 | -0.066 | 0.611 | -0.001 | -0.002 | -0.002 | 0.046 | 0.52 | 0.46 | 0.46 | 0.66 | 0.51 | 0.45 |
| 2 | 169021742 | rs3769382 | -1.134 | -0.313 | 0.018 | 0.768 | -0.024 | 0.209 | 0.209 | 0.042 | 0.08 | 0.24 | 0.05 | 0.32 | 0.03 | 0.11 |
| 2 | 169022315 | rs3769381 | -1.673 | -0.335 | 0.022 | 0.759 | -0.026 | 0.247 | 0.247 | 0.309 | 0.09 | 0.32 | 0.06 | 0.33 | 0.04 | 0.11 |
| 2 | 169025379 | rs4668039 | -0.744 | -0.329 | -0.061 | 0.681 | -0.033 | 0.091 | 0.091 | 1.26 | 0.24 | 0.43 | 0.31 | 0.58 | 0.21 | 0.32 |
| 2 | 169027555 | rs6740826 | 0.758 | -0.334 | -0.027 | 0.606 | -0.01 | 0.022 | 0.022 | 0 | 0.17 | 0.34 | 0.16 | 0.2 | 0.1 | 0.17 |
| 2 | 169029951 | rs1400646 | 0.737 | -0.178 | -0.398 | 0.421 | -0.053 | 0.191 | 0.191 | 2.281 | 0.31 | 0.04 | 0.15 | 0.08 | 0.31 | 0.16 |
| 2 | 169040198 | rs1878486 | -0.44 | -0.177 | -0.105 | 0.588 | 0.003 | 0.241 | 0.241 | 1.004 | 0.05 | 0.25 | 0.03 | 0.25 | 0.01 | 0.05 |
| 2 | 169044118 | rs883406 | -0.502 | -0.212 | -0.201 | 0.479 | -0.011 | 0.021 | 0.021 | 0 | 0.35 | 0.4 | 0.42 | 0.46 | 0.27 | 0.39 |
| 2 | 169048087 | rs13426907 | -0.995 | -0.201 | -0.396 | 0.407 | 0.038 | 0.073 | 0.073 | 1.603 | 0.41 | 0.74 | 0.48 | 0.78 | 0.31 | 0.49 |
| 2 | 169062731 | rs6737602 | 1.624 | -0.061 | -0.366 | 0.451 | -0.024 | 0.022 | 0.022 | 1.378 | 0.29 | 0.21 | 0.37 | 0.14 | 0.39 | 0.35 |
| 2 | 169065468 | rs932167 | -0.647 | -0.293 | -0.424 | 0.571 | -0.011 | 0.024 | 0.024 | 0 | 0.35 | 0.4 | 0.42 | 0.46 | 0.27 | 0.38 |
| 2 | 169078534 | rs2271741 | 0.675 | -0.056 | -0.956 | 0.334 | -0.044 | 0.144 | 0.144 | 4.399 | 0.28 | 0.04 | 0.15 | 0.09 | 0.28 | 0.16 |
| 2 | 169082034 | rs2030162 | 1.159 | -0.119 | -0.721 | 0.319 | -0.044 | 0.073 | 0.073 | 2.108 | 0.35 | 0.23 | 0.27 | 0.12 | 0.47 | 0.34 |
| 2 | 169082242 | rs1517320 | 0.193 | 0.217 | 0.695 | 1.476 | -0.004 | 0.012 | 0.012 | 0.022 | 0.81 | 0.8 | 0.72 | 0.74 | 0.82 | 0.77 |
| 2 | 169089920 | rs16855160 | -0.224 | 0.321 | 0.434 | 1.548 | 0.024 | 0.064 | 0.064 | 0 | 0.1 | 0.21 | 0.1 | 0.2 | 0.1 | 0.15 |
| 2 | 169090562 | rs10194006 | 1.677 | 0.521 | -0.202 | 1.08 | -0.003 | 0.023 | 0.023 | 0.173 | 0.4 | 0.28 | 0.27 | 0.13 | 0.44 | 0.29 |
| 2 | 169090774 | rs13430810 | 1.669 | 0.518 | -0.204 | 1.079 | -0.003 | 0.024 | 0.024 | 0.195 | 0.4 | 0.28 | 0.27 | 0.13 | 0.44 | 0.29 |
| 2 | 169091942 | rs2390669 | 0.319 | 0.469 | -0.289 | 1.268 | 0.009 | -0.044 | -0.044 | 0.078 | 0.13 | 0.09 | 0.23 | 0.33 | 0.13 | 0.18 |
| 2 | 169095090 | rs10930316 | 0.417 | 0.462 | -0.385 | 1.093 | -0.07 | 0.154 | 0.154 | 0.707 | 0.26 | 0.04 | 0.13 | 0.04 | 0.27 | 0.21 |
| 2 | 169095854 | rs950535 | 1.988 | 0.534 | -0.298 | 1.133 | -0.01 | 0.093 | 0.093 | 0.39 | 0.39 | 0.2 | 0.27 | 0.12 | 0.44 | 0.29 |
| 2 | 169096765 | rs1517323 | 0.723 | 1.219 | 0.268 | 1.703 | -0.069 | 0.083 | 0.083 | 0 | 0.17 | 0.03 | 0.12 | 0.04 | 0.24 | 0.13 |
| 2 | 169100650 | rs1517325 | 1.052 | 1.023 | 0.602 | 1.998 | 0.016 | 0.046 | 0.046 | 1.47 | 0.81 | 0.47 | 0.84 | 0.73 | 0.83 | 0.78 |
| 2 | 169121841 | rs7569089 | -1.191 | 1.064 | 0.553 | 1.385 | 0.002 | 0.12 | 0.12 | 0.964 | 0.21 | 0.55 | 0.35 | 0.49 | 0.16 | 0.31 |
| 2 | 169125080 | rs11904323 | -1.141 | 1.07 | 0.541 | 1.363 | 0.012 | 0.19 | 0.19 | 0.67 | 0.21 | 0.46 | 0.35 | 0.49 | 0.16 | 0.31 |
| 2 | 169143035 | rs9917256 | 0.08 | 1.28 | -0.456 | 1.308 | -0.004 | -0.023 | -0.023 | 0.622 | 0.1 | 0.09 | 0.25 | 0.33 | 0.14 | 0.18 |
| 2 | 169143079 | rs6755949 | -1.01 | 1.56 | -0.6 | 1.157 | -0.02 | -0.008 | -0.008 | 0.117 | 0.33 | 0.31 | 0.38 | 0.38 | 0.42 | 0.34 |
| 2 | 169156907 | rs12616285 | -0.507 | 1.488 | -0.65 | 1.173 | 0.009 | 0.094 | 0.094 | 0.236 | 0.15 | 0.31 | 0.28 | 0.33 | 0.18 | 0.24 |
| 2 | 169157245 | rs7569877 | -0.082 | 1.658 | -0.891 | 0.915 | -0.072 | 0.083 | 0.083 | 0 | 0.21 | 0.08 | 0.13 | 0.06 | 0.29 | 0.16 |
| 2 | 169165476 | rs9646776 | 0.592 | 1.226 | -0.209 | 1.613 | 0.012 | 0.227 | 0.227 | 0.236 | 0.85 | 0.52 | 0.68 | 0.63 | 0.9 | 0.79 |
| 2 | 169166282 | rs4668059 | -0.468 | 1.369 | -0.901 | 1.632 | -0.01 | -0.046 | -0.046 | 0 | 0.1 | 0.03 | 0.25 | 0.22 | 0.12 | 0.21 |
| 2 | 169171760 | rs4668061 | -0.261 | 1.347 | -0.772 | 1.481 | 0.009 | -0.007 | -0.007 | 0 | 0.08 | 0.1 | 0.21 | 0.12 | 0.06 | 0.17 |
| 2 | 169173631 | rs4629128 | -0.079 | 2.24 | -0.737 | 1.266 | -0.11 | 0.035 | 0.035 | 0 | 0.18 | 0.02 | 0.14 | 0.06 | 0.31 | 0.16 |
| 2 | 169174367 | rs12618858 | -1.287 | 1.35 | 0.908 | 2.253 | 0.047 | 0.363 | 0.363 | 3.399 | 0.08 | 0.39 | 0.29 | 0.28 | 0.04 | 0.2 |
| 2 | 169187306 | rs4433965 | 0.281 | 0.882 | -1.023 | 1.052 | 0.089 | -0.022 | -0.022 | 0.262 | 0.37 | 0.08 | 0.23 | 0.46 | 0.2 | 0.35 |
| 2 | 169193891 | rs11676990 | 0.174 | 1.596 | -0.687 | 0.486 | 0.004 | -0.036 | -0.036 | 0 | 0.18 | 0.12 | 0.14 | 0.01 | 0.3 | 0.22 |
| 2 | 169193973 | rs6740553 | -0.81 | 1.63 | -0.24 | 0.976 | 0.03 | 0.007 | 0.007 | 0.272 | 0.29 | 0.42 | 0.36 | 0.28 | 0.43 | 0.37 |
| 2 | 169194601 | rs995761 | 1.309 | 1.685 | 0.196 | 1.05 | 0.033 | 0.007 | 0.007 | 0.868 | 0.36 | 0.17 | 0.25 | 0.44 | 0.24 | 0.34 |
| 2 | 169195767 | rs2601087 | -1.114 | 1.782 | 0.109 | 0.813 | 0 | -0.045 | -0.045 | 0 | 0.84 | 0.95 | 0.73 | 0.77 | 0.85 | 0.77 |
| 2 | 169196137 | rs2724178 | 1.592 | 1.766 | 0.068 | 0.801 | 0.025 | 0.011 | 0.011 | 0.766 | 0.38 | 0.13 | 0.25 | 0.45 | 0.28 | 0.36 |

| | | | | | | | | | | | | | | | | |
|---|---|---|---|---|---|---|---|---|---|---|---|---|---|---|---|---|
| 2 | 169196340 | rs7557093 | -0.313 | 1.859 | 0.223 | 0.769 | 0.045 | 0.024 | 0.024 | 0.032 | 0.27 | 0.46 | 0.36 | 0.28 | 0.4 | 0.36 |
| 2 | 169197854 | rs2724177 | -0.371 | 1.673 | 0.681 | 0.803 | -0.013 | 0.013 | 0.013 | 0 | 0.81 | 0.73 | 0.71 | 0.79 | 0.85 | 0.78 |
| 2 | 169202636 | rs7600566 | -1.533 | 1.746 | 0.919 | 0.876 | 0.004 | 0.003 | 0.003 | 0 | 0.46 | 0.62 | 0.46 | 0.32 | 0.57 | 0.4 |
| 2 | 169204872 | rs6719797 | -0.683 | 1.938 | 0.714 | 1.121 | -0.07 | 0.032 | 0.032 | 0 | 0.32 | 0.14 | 0.25 | 0.2 | 0.43 | 0.32 |
| 2 | 169204903 | rs2123194 | -1.108 | 1.935 | 0.915 | 1.099 | 0.011 | -0.044 | -0.044 | 0.896 | 0.08 | 0.17 | 0.05 | 0 | 0.09 | 0.04 |
| 2 | 169204918 | rs6748292 | 0.088 | 2.038 | 0.808 | 1.161 | -0.026 | 0.019 | 0.019 | 0 | 0.2 | 0.12 | 0.17 | 0.13 | 0.28 | 0.26 |
| 2 | 169205282 | rs7597374 | -1.562 | 1.904 | 0.769 | 1.213 | -0.001 | -0.001 | -0.001 | 0 | 0.43 | 0.51 | 0.44 | 0.32 | 0.55 | 0.4 |
| 2 | 169205599 | rs16855301 | -1.332 | 1.746 | 0.447 | 1.38 | 0.003 | 0.04 | 0.04 | 0 | 0.07 | 0.1 | 0.14 | 0.12 | 0.09 | 0.06 |
| 2 | 169214903 | rs7561814 | -1.327 | 1.918 | 0.479 | 1.081 | 0.011 | 0.008 | 0.008 | 0.01 | 0.24 | 0.32 | 0.25 | 0.19 | 0.28 | 0.15 |
| 2 | 169220101 | rs2044958 | -0.967 | 1.7 | 0.46 | 1.158 | 0.008 | 0.027 | 0.027 | 0.016 | 0.21 | 0.34 | 0.23 | 0.19 | 0.21 | 0.12 |
| 2 | 169222490 | rs6717927 | 0.887 | 1.558 | 0.235 | 1.004 | 0.01 | 0.009 | 0.009 | 0 | 0.52 | 0.46 | 0.47 | 0.47 | 0.39 | 0.41 |
| 2 | 169223035 | rs6736425 | -1.296 | 1.657 | 0.778 | 0.839 | 0.02 | -0.029 | -0.029 | 0 | 0.15 | 0.29 | 0.2 | 0.07 | 0.18 | 0.08 |
| 2 | 169223698 | rs7575325 | -1.628 | 1.759 | 0.636 | 0.886 | 0.006 | -0.009 | -0.009 | 0 | 0.22 | 0.27 | 0.25 | 0.15 | 0.25 | 0.1 |
| 2 | 169224527 | rs1466180 | -0.455 | 2.191 | 0.44 | 1.219 | -0.013 | -0.013 | -0.013 | 0 | 0.9 | 0.89 | 0.83 | 0.77 | 0.84 | 0.74 |
| 2 | 169234576 | rs11888499 | -0.438 | 2.23 | 0.472 | 1.173 | -0.011 | 0.015 | 0.015 | 0 | 0.09 | 0.03 | 0.07 | 0.02 | 0.13 | 0.07 |
| 2 | 169237675 | rs2601053 | 0.722 | 2.089 | 0.849 | 1.094 | 0.014 | 0.075 | 0.075 | 0 | 0.84 | 0.64 | 0.81 | 0.77 | 0.82 | 0.7 |
| 2 | 169243941 | rs2724144 | 1.676 | 2.091 | 1.136 | 1.01 | 0.015 | 0.018 | 0.018 | 0 | 0.67 | 0.54 | 0.68 | 0.68 | 0.61 | 0.61 |
| 2 | 169244775 | rs4503959 | 1.45 | 2.012 | 1.655 | 0.997 | 0 | 0.028 | 0.028 | 0 | 0.72 | 0.34 | 0.73 | 0.69 | 0.73 | 0.67 |
| 2 | 169250883 | rs7559622 | -2.073 | 1.95 | 1.94 | 0.893 | 0.051 | 0.068 | 0.068 | 0.045 | 0.08 | 0.46 | 0.17 | 0.07 | 0.07 | 0.06 |
| 2 | 169252417 | rs6433058 | -1.543 | 1.96 | 1.715 | 0.903 | 0.03 | 0.048 | 0.048 | 0 | 0.06 | 0.26 | 0.14 | 0.07 | 0.06 | 0.06 |
| 2 | 169261867 | rs6716086 | 1.553 | 1.97 | 1.653 | 1.055 | 0.028 | 0.041 | 0.041 | 0 | 0.67 | 0.46 | 0.54 | 0.6 | 0.58 | 0.58 |
| 2 | 169276227 | rs2724160 | -0.634 | 1.825 | 1.322 | 1.243 | 0.001 | 0.003 | 0.003 | 0 | 0.92 | 0.92 | 0.91 | 0.83 | 0.92 | 0.78 |
| 2 | 169277487 | rs2601084 | 0.856 | 1.722 | 2.337 | 1.175 | -0.035 | 0.088 | 0.088 | 0 | 0.85 | 0.51 | 0.86 | 0.83 | 0.89 | 0.71 |
| 2 | 169284018 | rs977171 | 0.148 | 1.826 | 2.257 | 1.221 | 0.009 | 0.065 | 0.065 | 0 | 0.89 | 0.78 | 0.77 | 0.77 | 0.88 | 0.95 |
| 2 | 169285488 | rs10497340 | -0.447 | 1.988 | 2.203 | 1.107 | -0.018 | 0.01 | 0.01 | 0 | 0.09 | 0.05 | 0.11 | 0 | 0.19 | 0.07 |
| 2 | 169287514 | rs1377502 | -2.169 | 2.273 | 1.211 | 0.757 | -0.037 | 0.035 | 0.035 | 0 | 0.42 | 0.79 | 0.37 | 0.29 | 0.52 | 0.55 |
| 2 | 169299532 | rs17806793 | -0.15 | 2.222 | 1.714 | 0.759 | 0.012 | -0.018 | -0.018 | 0 | 0.18 | 0.23 | 0.12 | 0.11 | 0.22 | 0.19 |
| 2 | 169300048 | rs1124893 | -1.475 | 2.213 | 1.482 | 0.808 | -0.002 | 0.001 | 0.001 | 0 | 0.32 | 0.66 | 0.26 | 0.29 | 0.32 | 0.48 |
| 2 | 169308231 | rs1453405 | 2.241 | 2.259 | 1.357 | 0.756 | 0.042 | 0.004 | 0.004 | 0 | 0.51 | 0.24 | 0.53 | 0.55 | 0.41 | 0.47 |
| 2 | 169323540 | rs1453406 | 0.214 | 2.06 | 1.007 | 0.169 | -0.013 | -0.011 | -0.011 | 0.007 | 0.83 | 0.88 | 0.85 | 1 | 0.72 | 0.89 |
| 2 | 169338829 | rs7557806 | 1.462 | 1.914 | 1.431 | 0.153 | 0.051 | -0.053 | -0.053 | 0 | 0.7 | 0.51 | 0.71 | 0.83 | 0.61 | 0.8 |
| 2 | 169340129 | rs16855379 | 0.903 | 1.894 | 1.107 | 0.519 | 0.005 | 0.046 | 0.046 | 0 | 0.07 | 0.09 | 0.11 | 0.2 | 0.08 | 0.02 |
| 2 | 169341723 | rs6733926 | 1.09 | 1.892 | 1.073 | 0.379 | 0.008 | 0.041 | 0.041 | 0 | 0.06 | 0.08 | 0.1 | 0.16 | 0.08 | 0.02 |
| 2 | 169351953 | rs13007801 | 1.409 | 1.775 | 0.707 | 0.298 | -0.149 | -0.072 | -0.072 | 0 | 0.67 | 0.06 | 0.58 | 0.65 | 0.58 | 0.71 |
| 2 | 169357139 | rs964987 | 0.336 | 0.928 | 0.244 | -0.433 | -0.013 | -0.013 | -0.013 | 0 | 0.55 | 0.71 | 0.64 | 0.58 | 0.51 | 0.48 |
| 2 | 169361542 | rs12479292 | 2.095 | 1.248 | 0.142 | -0.31 | -0.001 | 0.031 | 0.031 | 0 | 0.53 | 0.08 | 0.44 | 0.65 | 0.43 | 0.64 |
| 2 | 169366003 | rs17808106 | -0.754 | 1.192 | 0.048 | -0.35 | -0.107 | 0.004 | 0.004 | 0.296 | 0.28 | 0.04 | 0.24 | 0.13 | 0.34 | 0.34 |
| 2 | 169368019 | rs6710428 | -0.052 | 1.257 | 0.754 | -0.262 | 0.012 | 0.168 | 0.168 | 2.89 | 0.05 | 0.24 | 0.09 | 0.17 | 0.02 | 0.02 |
| 2 | 169371014 | rs1993224 | 0.2 | 1.276 | 0.893 | -0.419 | 0.024 | -0.011 | -0.011 | 0 | 0.19 | 0.28 | 0.29 | 0.15 | 0.29 | 0.19 |
| 2 | 169373229 | rs13419014 | -0.128 | 1.373 | -0.141 | -0.44 | -0.08 | 0.081 | 0.081 | 0 | 0.16 | 0.02 | 0.15 | 0.01 | 0.24 | 0.08 |
| 2 | 169377947 | rs10930327 | -1.118 | 1.51 | 0.304 | -0.545 | 0.053 | -0.125 | -0.125 | 0 | 0.23 | 0.44 | 0.21 | 0.01 | 0.3 | 0.17 |
| 2 | 169382342 | rs1020603 | 1.083 | 1.416 | -0.13 | -0.395 | -0.011 | 0 | 0 | 0 | 0.68 | 0.71 | 0.69 | 0.71 | 0.54 | 0.84 |
| 2 | 169386870 | rs13398999 | -0.283 | 1.3 | -0.771 | -0.406 | -0.028 | 0.036 | 0.036 | 0.031 | 0.17 | 0.08 | 0.15 | 0.01 | 0.24 | 0.07 |
| 2 | 169389091 | rs13393173 | -0.04 | 1.326 | -1.413 | -0.397 | -0.085 | 0.075 | 0.075 | 0.071 | 0.16 | 0.01 | 0.15 | 0.02 | 0.24 | 0.07 |
| 2 | 169401278 | rs11695350 | 0.602 | 1.049 | -1.695 | -0.453 | -0.085 | 0.033 | 0.033 | 0 | 0.2 | 0.02 | 0.17 | 0.11 | 0.28 | 0.08 |
| 2 | 169402127 | rs716335 | 2.181 | 0.764 | -1.868 | -0.37 | -0.057 | -0.15 | -0.15 | 0 | 0.37 | 0.03 | 0.37 | 0.54 | 0.36 | 0.34 |
| 2 | 169403721 | rs17202792 | 0.479 | 0.671 | -1.641 | -0.431 | -0.006 | 0.018 | 0.018 | 0 | 0.11 | 0.05 | 0.08 | 0.02 | 0.12 | 0.06 |
| 2 | 169414380 | rs12463569 | 1.903 | 0.288 | -1.602 | -0.138 | 0.028 | -0.165 | -0.165 | 0.168 | 0.35 | 0.03 | 0.33 | 0.54 | 0.27 | 0.32 |
| 2 | 169417219 | rs4667582 | -0.815 | 0.503 | -1.524 | -0.284 | -0.066 | 0.094 | 0.094 | 0.166 | 0.47 | 0.03 | 0.37 | 0.23 | 0.55 | 0.49 |
| 2 | 169427769 | rs1813595 | 1.049 | 0.802 | 0.403 | 0.143 | 0.187 | 0.093 | 0.093 | 0 | 0.95 | 0.67 | 0.84 | 0.95 | 0.9 | 0.97 |
| 2 | 169429220 | rs1869368 | 1.123 | 0.798 | 0.401 | 0.141 | 0.18 | 0.088 | 0.088 | 0 | 0.95 | 0.67 | 0.84 | 0.95 | 0.9 | 0.97 |
| 2 | 169448031 | rs719970 | 1.986 | 0.684 | 0.497 | 0.202 | 0.066 | 0.1 | 0.1 | 0.893 | 0.86 | 0.45 | 0.77 | 0.78 | 0.84 | 0.89 |
| 2 | 169465503 | rs1901834 | 1.826 | 0.145 | 0.8 | 0.211 | -0.03 | 0.044 | 0.044 | 4.866 | 0.88 | 0.28 | 0.87 | 0.8 | 0.94 | 0.89 |
| 2 | 169477814 | rs1002667 | -1.248 | -0.013 | 0.541 | -0.048 | -0.028 | -0.047 | -0.047 | 0 | 0.31 | 0.5 | 0.21 | 0.15 | 0.23 | 0.52 |
| 2 | 169485639 | rs4668079 | -1.207 | -0.023 | 0.534 | -0.019 | -0.023 | 0.009 | 0.009 | 0 | 0.3 | 0.51 | 0.23 | 0.29 | 0.23 | 0.54 |
| 2 | 169498189 | rs4668086 | -0.911 | 0.744 | 0.793 | 0.414 | 0.016 | 0.065 | 0.065 | 0 | 0.18 | 0.38 | 0.28 | 0.27 | 0.19 | 0.49 |
| 2 | 169498986 | rs6433082 | -1.207 | 0.992 | 1.63 | 0.43 | -0.01 | 0.024 | 0.024 | 0 | 0.23 | 0.61 | 0.36 | 0.27 | 0.19 | 0.5 |
| 2 | 169501848 | rs4668089 | -1.226 | 1.015 | 1.691 | 0.431 | 0.01 | 0.037 | 0.037 | 0 | 0.23 | 0.52 | 0.36 | 0.27 | 0.22 | 0.5 |
| 2 | 169503729 | rs2893097 | 0.136 | 0.975 | 1.316 | 0.428 | 0 | 0 | 0 | 0 | 0.13 | 0.12 | 0.26 | 0.13 | 0.15 | 0.39 |
| 2 | 169506879 | rs1806328 | 0.652 | 0.704 | 1.172 | 0.319 | -0.027 | -0.01 | -0.01 | 0 | 0.58 | 0.22 | 0.32 | 0.56 | 0.52 | 0.38 |
| 2 | 169511104 | rs16855658 | -0.677 | 0.417 | 0.619 | 0.286 | 0.022 | 0.033 | 0.033 | 0 | 0.17 | 0.11 | 0.1 | 0.05 | 0.12 | 0.1 |
| 2 | 169516830 | rs7565629 | -1.498 | 0.529 | 0.732 | 0.311 | 0.046 | -0.035 | -0.035 | 0 | 0.6 | 0.73 | 0.63 | 0.44 | 0.74 | 0.75 |
| 2 | 169516881 | rs13391857 | -0.688 | 0.764 | 0.629 | 0.118 | -0.015 | 0.035 | 0.035 | 0 | 0.17 | 0.13 | 0.13 | 0.03 | 0.2 | 0.09 |
| 2 | 169519631 | rs4668094 | -0.694 | 0.504 | 1.203 | 0.507 | 0.014 | -0.016 | -0.016 | 0.003 | 0.4 | 0.41 | 0.44 | 0.26 | 0.52 | 0.58 |
| 2 | 169528778 | rs10514622 | -0.442 | 0.383 | 0.715 | 0.605 | -0.037 | 0.013 | 0.013 | 0.736 | 0.37 | 0.22 | 0.43 | 0.27 | 0.5 | 0.51 |
| 2 | 169530154 | rs2042549 | 1.347 | -0.037 | 1.647 | 0.588 | -0.015 | 0.022 | 0.022 | 0 | 0.84 | 0.44 | 0.83 | 0.85 | 0.89 | 0.92 |
| 2 | 169530546 | rs10930335 | 1.161 | -0.551 | 1.069 | 0.282 | 0.046 | 0.051 | 0.051 | 0 | 0.82 | 0.93 | 0.96 | 0.98 | 0.89 | 0.88 |
| 2 | 169534683 | rs6715474 | -0.806 | -0.388 | 1.596 | 0.475 | 0.014 | -0.027 | -0.027 | 0 | 0.07 | 0.27 | 0.06 | 0.01 | 0.05 | 0.02 |
| 2 | 169536911 | rs918957 | 1.441 | -0.491 | 1.925 | 0.873 | -0.027 | 0.012 | 0.012 | 0 | 0.83 | 0.76 | 0.9 | 0.74 | 0.88 | 0.87 |

| | | | | | | | | | | | | | | | | |
|---|---|---|---|---|---|---|---|---|---|---|---|---|---|---|---|---|
| 2 | 169537411 | rs7601307 | -0.403 | -0.33 | 1.894 | 0.67 | -0.004 | -0.005 | -0.005 | 0.007 | 0.4 | 0.61 | 0.51 | 0.58 | 0.54 | 0.52 |
| 2 | 169546569 | rs2161977 | 1.434 | -0.385 | 1.577 | 0.557 | -0.05 | 0.049 | 0.049 | 0.105 | 0.78 | 0.38 | 0.84 | 0.61 | 0.85 | 0.81 |
| 2 | 169546710 | rs4668098 | 1.435 | -0.351 | 0.867 | 0.568 | 0 | -0.001 | -0.001 | 0 | 0.77 | 0.24 | 0.79 | 0.61 | 0.81 | 0.8 |
| 2 | 169554005 | rs11900907 | 1.015 | -0.035 | 0.267 | 0.657 | 0.013 | 0.021 | 0.021 | 1.381 | 0.45 | 0.23 | 0.45 | 0.4 | 0.38 | 0.36 |
| 2 | 169566184 | rs1424937 | 0.232 | 0.489 | -0.098 | 0.498 | -0.066 | -0.007 | -0.007 | 0 | 0.29 | 0.13 | 0.29 | 0.21 | 0.44 | 0.48 |
| 2 | 169580183 | rs7558531 | -1.225 | 0.133 | -0.29 | 0.54 | 0.037 | 0 | 0 | 2.539 | 0.5 | 0.6 | 0.45 | 0.45 | 0.63 | 0.61 |
| 2 | 169582484 | rs16855727 | -0.145 | 0.466 | -1.209 | -0.092 | -0.02 | -0.07 | -0.07 | 0 | 0.15 | 0.25 | 0.08 | 0 | 0.09 | 0.06 |
| 2 | 169583149 | rs6754773 | -0.301 | 1.178 | -1.646 | 0.579 | -0.186 | -0.052 | -0.052 | 0.244 | 0.31 | 0.02 | 0.3 | 0.2 | 0.47 | 0.49 |
| 2 | 169589698 | rs2390731 | 0.813 | 0.807 | -0.239 | 0.445 | 0.063 | 0.001 | 0.001 | 0.653 | 0.94 | 0.7 | 0.92 | 0.99 | 0.94 | 0.97 |
| 2 | 169597668 | rs4668102 | -0.082 | 0.969 | -1.336 | 0.546 | -0.092 | -0.098 | -0.098 | 0 | 0.34 | 0.08 | 0.6 | 0.37 | 0.54 | 0.46 |
| 2 | 169599429 | rs13390393 | 0.134 | 0.362 | -1.287 | 0.506 | 0.001 | -0.001 | -0.001 | 0 | 0.1 | 0.08 | 0.06 | 0.09 | 0.09 | 0.07 |
| 2 | 169605967 | rs2390732 | -0.053 | -0.459 | -1.822 | 0.048 | 0.022 | -0.035 | -0.035 | 0.086 | 0.54 | 0.82 | 0.71 | 0.43 | 0.58 | 0.52 |
| 2 | 169627271 | rs16855819 | 0.142 | -0.515 | -1.907 | -0.002 | -0.024 | -0.003 | -0.003 | 0 | 0.07 | 0.01 | 0.07 | 0.06 | 0.1 | 0.05 |
| 2 | 169630483 | rs1063729 | -0.502 | -0.707 | -1.943 | -0.047 | 0.026 | -0.027 | -0.027 | 0 | 0.61 | 0.79 | 0.79 | 0.5 | 0.68 | 0.58 |
| 2 | 169632725 | rs4233821 | -0.615 | -1.044 | -1.877 | -0.535 | 0.022 | 0.09 | 0.09 | 0.513 | 0.68 | 0.86 | 0.86 | 0.98 | 0.75 | 0.75 |
| 2 | 169636186 | rs12104858 | 0.326 | -1.028 | -0.935 | -0.634 | -0.012 | -0.027 | -0.027 | 0.656 | 0.14 | 0.24 | 0.15 | 0.07 | 0.11 | 0.04 |
| 2 | 169642621 | rs16855872 | -0.333 | -1.202 | -0.578 | -1.057 | 0.001 | -0.048 | -0.048 | 0 | 0.65 | 0.67 | 0.81 | 0.96 | 0.72 | 0.74 |
| 2 | 169643222 | rs7584086 | -1.145 | -1.477 | -0.355 | -0.911 | -0.033 | 0.059 | 0.059 | 0 | 0.27 | 0.43 | 0.22 | 0.48 | 0.2 | 0.17 |
| 2 | 169645524 | rs3856426 | -0.173 | -1.376 | -0.259 | -0.985 | 0.009 | 0.082 | 0.082 | 0.609 | 0.6 | 0.75 | 0.76 | 0.96 | 0.61 | 0.8 |
| 2 | 169647435 | rs16855886 | -0.631 | -1.285 | -0.089 | -0.535 | 0 | 0.078 | 0.078 | 0 | 0.07 | 0.06 | 0.06 | 0.42 | 0.04 | 0.17 |
| 2 | 169652568 | rs6749871 | 0.104 | -1.362 | 0.123 | -0.684 | -0.029 | -0.075 | -0.075 | 0 | 0.13 | 0.3 | 0.1 | 0 | 0.08 | 0.03 |
| 2 | 169652647 | rs16855893 | -0.86 | -1.255 | -0.001 | -0.526 | 0.002 | 0.068 | 0.068 | 0 | 0.08 | 0.07 | 0.07 | 0.43 | 0.05 | 0.19 |
| 2 | 169654055 | rs829950 | 1.027 | -0.614 | 0.194 | -1.025 | 0.006 | -0.011 | -0.011 | 0 | 0.36 | 0.49 | 0.32 | 0.67 | 0.39 | 0.51 |
| 2 | 169658231 | rs700547 | -2.249 | -0.959 | 0.118 | -1.101 | -0.01 | -0.026 | -0.026 | 0 | 0.92 | 0.95 | 0.98 | 1 | 0.96 | 0.97 |
| 2 | 169658783 | rs829958 | -0.807 | -0.901 | 0.235 | -1.328 | 0.009 | -0.071 | -0.071 | 0 | 0.14 | 0.25 | 0.11 | 0 | 0.16 | 0.09 |
| 2 | 169662024 | rs829955 | -1.131 | -0.588 | 0.058 | -1.324 | 0 | -0.016 | -0.016 | 0 | 0.86 | 0.88 | 0.84 | 0.99 | 0.86 | 0.94 |
| 2 | 169669961 | rs17200300 | -0.069 | -0.852 | -0.248 | -1.274 | 0.001 | -0.012 | -0.012 | 0.157 | 0.28 | 0.1 | 0.21 | 0.29 | 0.28 | 0.23 |
| 2 | 169673576 | rs2433680 | 1.632 | -0.75 | -0.247 | -1.147 | -0.027 | -0.023 | -0.023 | 0.012 | 0.07 | 0.03 | 0.14 | 0.12 | 0.1 | 0.04 |
| 2 | 169676045 | rs482435 | 0.625 | -0.916 | 0.002 | -1.313 | -0.005 | -0.073 | -0.073 | 0.172 | 0.68 | 0.59 | 0.7 | 0.89 | 0.67 | 0.84 |
| 2 | 169680122 | rs9287903 | 0.364 | -0.59 | 0.001 | -1.3 | -0.006 | -0.073 | -0.073 | 0.35 | 0.71 | 0.6 | 0.74 | 0.89 | 0.71 | 0.86 |
| 2 | 169683061 | rs10198693 | 0.052 | -0.746 | -0.422 | -1.393 | -0.015 | 0.141 | 0.141 | 0 | 0.29 | 0.09 | 0.21 | 0.01 | 0.29 | 0.12 |
| 2 | 169684457 | rs700548 | -2.322 | -0.703 | -0.275 | -1.077 | -0.006 | -0.021 | -0.021 | 0.664 | 0.91 | 0.93 | 0.98 | 1 | 0.96 | 0.98 |
| 2 | 169685756 | rs830020 | 1.931 | -0.266 | -0.073 | -1.002 | -0.003 | 0.001 | 0.001 | 0.23 | 0.39 | 0.36 | 0.47 | 0.62 | 0.36 | 0.62 |
| 2 | 169686559 | rs487756 | -0.199 | -0.447 | -0.587 | -1.019 | -0.057 | -0.008 | -0.008 | 0 | 0.33 | 0.09 | 0.34 | 0.3 | 0.41 | 0.28 |
| 2 | 169690089 | rs549582 | -1.807 | -0.394 | -0.665 | -0.986 | -0.034 | 0.037 | 0.037 | 0 | 0.88 | 0.98 | 0.84 | 0.97 | 0.84 | 0.96 |
| 2 | 169690729 | rs522893 | 0.749 | -0.45 | -0.354 | -0.944 | 0.006 | 0.018 | 0.018 | 0.839 | 0.5 | 0.6 | 0.6 | 0.7 | 0.54 | 0.69 |
| 2 | 169701054 | rs508423 | 0.25 | 0.041 | -1.067 | -0.817 | -0.022 | 0.016 | 0.016 | 0 | 0.65 | 0.84 | 0.63 | 0.71 | 0.58 | 0.73 |
| 2 | 169703896 | rs529002 | 0.168 | 0.075 | -0.996 | -0.738 | -0.023 | 0.009 | 0.009 | 0 | 0.67 | 0.87 | 0.64 | 0.7 | 0.57 | 0.75 |
| 2 | 169705037 | rs497808 | 1.163 | 0 | -1.239 | -0.769 | -0.037 | -0.027 | -0.027 | 0.337 | 0.36 | 0.12 | 0.43 | 0.61 | 0.35 | 0.58 |
| 2 | 169707428 | rs540652 | 0.539 | -0.086 | -1.178 | -0.703 | -0.036 | 0.031 | 0.031 | 2.026 | 0.45 | 0.18 | 0.5 | 0.61 | 0.48 | 0.64 |
| 2 | 169714893 | rs569997 | 0.559 | -0.111 | -1.119 | -0.692 | -0.012 | 0.043 | 0.043 | 2.044 | 0.46 | 0.22 | 0.5 | 0.61 | 0.49 | 0.64 |
| 2 | 169717624 | rs2685800 | -0.961 | -0.063 | -1.225 | -0.69 | -0.029 | 0.031 | 0.031 | 0 | 0.86 | 0.96 | 0.81 | 0.94 | 0.83 | 0.94 |
| 2 | 169718301 | rs10497348 | 1.214 | -0.626 | -0.649 | -0.965 | -0.096 | -0.084 | -0.084 | 0 | 0.21 | 0.54 | 0.1 | 0.06 | 0.06 | 0.08 |
| 2 | 169718981 | rs581363 | -0.775 | -0.558 | 0.127 | -0.929 | -0.002 | -0.027 | -0.027 | 0 | 0.84 | 0.75 | 0.8 | 0.93 | 0.83 | 0.9 |
| 2 | 169721381 | rs3931 | 1.088 | -0.416 | 0.083 | -0.934 | -0.02 | -0.015 | -0.015 | 0 | 0.19 | 0.09 | 0.2 | 0.27 | 0.29 | 0.23 |
| 2 | 169729849 | rs3821116 | -0.448 | -0.302 | 0.047 | -1.102 | 0.006 | -0.005 | -0.005 | 0 | 0.31 | 0.31 | 0.28 | 0.27 | 0.43 | 0.3 |
| 2 | 169734975 | rs509988 | -0.714 | -0.096 | -0.217 | -0.934 | -0.002 | -0.034 | -0.034 | 1.17 | 0.84 | 0.69 | 0.79 | 0.94 | 0.83 | 0.91 |
| 2 | 169742272 | rs830193 | 1.35 | -0.331 | -0.555 | -0.915 | -0.06 | -0.044 | -0.044 | 0 | 0.32 | 0.14 | 0.43 | 0.59 | 0.34 | 0.54 |
| 2 | 169748422 | rs574981 | 0.076 | -0.174 | 0.17 | -0.396 | -0.005 | 0.004 | 0.004 | 0 | 0.76 | 0.78 | 0.74 | 0.93 | 0.7 | 0.89 |
| 2 | 169749841 | rs853770 | 0.297 | -0.119 | 0.104 | -0.249 | -0.002 | 0.003 | 0.003 | 0 | 0.73 | 0.72 | 0.7 | 0.82 | 0.71 | 0.88 |
| 2 | 169750483 | rs477224 | 0.118 | 0.278 | 0.181 | 0.107 | -0.008 | -0.031 | -0.031 | 0 | 0.58 | 0.45 | 0.64 | 0.81 | 0.66 | 0.85 |
| 2 | 169756930 | rs540524 | 0.497 | 0.62 | -0.014 | 0.543 | 0.002 | -0.026 | -0.026 | 0 | 0.36 | 0.2 | 0.42 | 0.42 | 0.37 | 0.62 |
| 2 | 169757354 | rs1402837 | 0.63 | 0.665 | 0.5 | 0.302 | -0.003 | 0.004 | 0.004 | 3.475 | 0.32 | 0.33 | 0.19 | 0.41 | 0.23 | 0.21 |
| 2 | 169757541 | rs573225 | -1.142 | 0.745 | 0.331 | 0.19 | -0.108 | 0.085 | 0.085 | 0 | 0.8 | 0.97 | 0.8 | 0.94 | 0.68 | 0.84 |
| 2 | 169757562 | rs2232316 | 0.404 | 0.738 | 0.621 | 0.086 | -0.003 | -0.002 | -0.002 | 0 | 0.1 | 0.14 | 0.17 | 0.12 | 0.09 | 0.04 |
| 2 | 169760082 | rs16856161 | 0.227 | 0.745 | 0.589 | 0.067 | 0.006 | -0.052 | -0.052 | 0.511 | 0.11 | 0.31 | 0.17 | 0.02 | 0.09 | 0.02 |
| 2 | 169763148 | rs560887 | -1.574 | 0.799 | 0.421 | 0.126 | -0.107 | 0.103 | 0.103 | 0 | 0.83 | 0.99 | 0.81 | 0.98 | 0.7 | 0.9 |
| 2 | 169765737 | rs567243 | 0.13 | 0.683 | 0.727 | 0.231 | 0.009 | 0.002 | 0.002 | 0.029 | 0.46 | 0.58 | 0.59 | 0.43 | 0.44 | 0.66 |
| 2 | 169771420 | rs13419326 | -0.352 | -0.033 | 1.095 | 0.186 | 0.031 | 0.374 | 0.374 | 4.9 | 0.07 | 0.43 | 0.09 | 0.3 | 0.02 | 0.12 |
| 2 | 169771518 | rs4144858 | -0.352 | -0.033 | 1.095 | 0.186 | 0.031 | 0.374 | 0.374 | 4.9 | 0.07 | 0.42 | 0.09 | 0.3 | 0.02 | 0.12 |
| 2 | 169774071 | rs563694 | -1.057 | 0.332 | 0.687 | -0.019 | -0.086 | 0.089 | 0.089 | 0 | 0.8 | 0.93 | 0.78 | 0.97 | 0.64 | 0.83 |
| 2 | 169774646 | rs537183 | -1.057 | 0.332 | 0.687 | -0.019 | -0.086 | 0.089 | 0.089 | 0 | 0.8 | 0.93 | 0.78 | 0.97 | 0.64 | 0.83 |
| 2 | 169776746 | rs475612 | -1.049 | 0.324 | 0.79 | -0.007 | -0.042 | 0.037 | 0.037 | 0 | 0.79 | 0.85 | 0.74 | 0.96 | 0.64 | 0.82 |
| 2 | 169777595 | rs557462 | -1.085 | 0.332 | 0.711 | -0.006 | -0.133 | 0.113 | 0.113 | 0 | 0.8 | 0.99 | 0.79 | 0.97 | 0.64 | 0.83 |
| 2 | 169779156 | rs478333 | 0.594 | 0.062 | 0.944 | 0.065 | -0.034 | 0.03 | 0.03 | 0.613 | 0.61 | 0.22 | 0.5 | 0.66 | 0.53 | 0.67 |
| 2 | 169779205 | rs4148804 | 0.123 | 0.05 | 1.204 | 0.067 | 0.038 | 0.335 | 0.335 | 3.688 | 0.06 | 0.37 | 0.11 | 0.29 | 0.02 | 0.13 |
| 2 | 169779712 | rs496550 | 0.524 | 0.048 | 1.216 | -0.035 | -0.022 | 0.01 | 0.01 | 0.315 | 0.62 | 0.26 | 0.51 | 0.64 | 0.53 | 0.67 |
| 2 | 169779764 | rs495714 | 0.562 | 0.048 | 1.216 | -0.035 | -0.022 | 0.011 | 0.011 | 0.494 | 0.61 | 0.26 | 0.51 | 0.64 | 0.53 | 0.67 |
| 2 | 169779896 | rs473351 | -0.137 | 0.047 | 1.211 | -0.038 | -0.019 | 0.005 | 0.005 | 0 | 0.69 | 0.71 | 0.62 | 0.94 | 0.55 | 0.8 |

| | | | | | | | | | | | | | | | | |
|---|---|---|---|---|---|---|---|---|---|---|---|---|---|---|---|---|
| 2 | 169780366 | rs579275 | 0.548 | 0.054 | 1.226 | -0.032 | -0.033 | 0.014 | 0.014 | 0.353 | 0.61 | 0.22 | 0.5 | 0.64 | 0.53 | 0.68 |
| 2 | 169780818 | rs575671 | -0.199 | 0.05 | 1.259 | -0.035 | -0.018 | 0.001 | 0.001 | 0 | 0.7 | 0.73 | 0.62 | 0.96 | 0.55 | 0.77 |
| 2 | 169780885 | rs519887 | 0.548 | 0.052 | 1.212 | -0.033 | -0.033 | 0.014 | 0.014 | 0.353 | 0.61 | 0.22 | 0.5 | 0.64 | 0.53 | 0.68 |
| 2 | 169782149 | rs486981 | -1.194 | 0.058 | 1.209 | -0.036 | -0.067 | 0.066 | 0.066 | 0 | 0.79 | 0.86 | 0.78 | 0.98 | 0.64 | 0.83 |
| 2 | 169782574 | rs483234 | 1.015 | -0.044 | 1.028 | -0.067 | -0.046 | -0.166 | -0.166 | 0.371 | 0.34 | 0.01 | 0.39 | 0.52 | 0.32 | 0.58 |
| 2 | 169783039 | rs579060 | -1.286 | -0.042 | 1.043 | -0.096 | -0.067 | 0.066 | 0.066 | 0 | 0.79 | 0.86 | 0.78 | 0.98 | 0.64 | 0.83 |
| 2 | 169784493 | rs17540154 | 1.102 | -0.04 | 1.128 | -0.067 | 0.001 | 0.014 | 0.014 | 4.781 | 0.28 | 0.21 | 0.12 | 0.15 | 0.22 | 0.09 |
| 2 | 169784940 | rs7607790 | 0.832 | -0.152 | 1.356 | 0.073 | -0.002 | -0.007 | -0.007 | 0 | 0.09 | 0.09 | 0.16 | 0.02 | 0.08 | 0.02 |
| 2 | 169784955 | rs508506 | -1.262 | -0.15 | 1.339 | 0.057 | -0.112 | 0.113 | 0.113 | 0 | 0.8 | 0.95 | 0.81 | 0.99 | 0.64 | 0.83 |
| 2 | 169785449 | rs503931 | 0.585 | -0.152 | 1.343 | 0.062 | -0.033 | 0.014 | 0.014 | 0.372 | 0.61 | 0.22 | 0.51 | 0.64 | 0.53 | 0.67 |
| 2 | 169787686 | rs551754 | 0.594 | -0.152 | 1.343 | 0.062 | -0.034 | 0.015 | 0.015 | 0.44 | 0.61 | 0.22 | 0.51 | 0.64 | 0.53 | 0.67 |
| 2 | 169788474 | rs531772 | -0.074 | -0.152 | 1.346 | 0.06 | -0.031 | 0.031 | 0.031 | 0 | 0.69 | 0.72 | 0.62 | 0.94 | 0.55 | 0.8 |
| 2 | 169789016 | rs497692 | 0.51 | -0.085 | 1.382 | 0.041 | -0.029 | 0.017 | 0.017 | 0.416 | 0.61 | 0.23 | 0.51 | 0.64 | 0.53 | 0.67 |
| 2 | 169789306 | rs494874 | -1.078 | -0.016 | 1.434 | 0.055 | -0.051 | 0.046 | 0.046 | 0 | 0.79 | 0.87 | 0.8 | 0.99 | 0.64 | 0.83 |
| 2 | 169791438 | rs552976 | -0.482 | -0.03 | 0.909 | 0.023 | 0.055 | -0.131 | -0.131 | 1.524 | 0.76 | 0.58 | 0.77 | 0.99 | 0.64 | 0.82 |
| 2 | 169792171 | rs3755157 | 0.494 | -0.061 | 0.901 | 0.047 | -0.024 | 0.107 | 0.107 | 0 | 0.14 | 0.31 | 0.22 | 0.32 | 0.1 | 0.15 |
| 2 | 169792421 | rs472614 | -0.139 | 0.016 | 0.95 | 0.09 | 0.004 | -0.001 | -0.001 | 0.64 | 0.65 | 0.6 | 0.55 | 0.64 | 0.53 | 0.67 |
| 2 | 169794431 | rs567074 | -0.067 | 0.033 | 0.881 | 0.081 | 0.001 | 0 | 0 | 0 | 0.33 | 0.34 | 0.45 | 0.35 | 0.45 | 0.3 |
| 2 | 169796288 | rs2544367 | 0.304 | 0.027 | 0.856 | 0.093 | -0.015 | 0.014 | 0.014 | 0.087 | 0.61 | 0.28 | 0.52 | 0.65 | 0.54 | 0.68 |
| 2 | 169797652 | rs1581397 | 0.296 | -0.032 | 0.861 | 0.097 | -0.016 | 0.014 | 0.014 | 0.321 | 0.61 | 0.28 | 0.52 | 0.65 | 0.54 | 0.68 |
| 2 | 169798171 | rs2685812 | 0.289 | -0.034 | 0.859 | 0.096 | -0.016 | 0.014 | 0.014 | 0.327 | 0.61 | 0.28 | 0.52 | 0.65 | 0.53 | 0.68 |
| 2 | 169801628 | rs860510 | 0.337 | -0.04 | 0.948 | 0.093 | -0.03 | 0.025 | 0.025 | 0.646 | 0.6 | 0.22 | 0.55 | 0.65 | 0.54 | 0.67 |
| 2 | 169801905 | rs853788 | 0.22 | -0.061 | 0.822 | 0.114 | 0.005 | -0.004 | -0.004 | 0 | 0.61 | 0.43 | 0.58 | 0.65 | 0.54 | 0.67 |
| 2 | 169802252 | rs853787 | -1.362 | -0.065 | 0.836 | 0.104 | -0.041 | 0.029 | 0.029 | 0 | 0.8 | 0.88 | 0.79 | 0.98 | 0.65 | 0.85 |
| 2 | 169802310 | rs853786 | -0.413 | 0.086 | 0.96 | 0.542 | -0.017 | 0.014 | 0.014 | 0.351 | 0.61 | 0.3 | 0.56 | 0.65 | 0.54 | 0.67 |
| 2 | 169802329 | rs862662 | -0.812 | 0.087 | 0.926 | 0.542 | 0.017 | -0.003 | -0.003 | 0 | 0.66 | 0.6 | 0.6 | 0.65 | 0.55 | 0.7 |
| 2 | 169802594 | rs853785 | -0.466 | 0.089 | 0.913 | 0.543 | -0.004 | 0.002 | 0.002 | 0.233 | 0.61 | 0.36 | 0.57 | 0.65 | 0.54 | 0.67 |
| 2 | 169803674 | rs853784 | -0.378 | 0.062 | 0.913 | 0.555 | -0.026 | 0.022 | 0.022 | 0.564 | 0.6 | 0.24 | 0.52 | 0.65 | 0.52 | 0.67 |
| 2 | 169805511 | rs853783 | -0.489 | 0.067 | 0.728 | 0.558 | 0.007 | -0.007 | -0.007 | 0.051 | 0.61 | 0.41 | 0.54 | 0.65 | 0.53 | 0.67 |
| 2 | 169806321 | rs853781 | -0.869 | 0.08 | 0.718 | 0.561 | 0.009 | -0.001 | -0.001 | 0 | 0.66 | 0.64 | 0.55 | 0.65 | 0.53 | 0.7 |
| 2 | 169809672 | rs853779 | -0.639 | 0.3 | 0.271 | 0.74 | -0.005 | 0.005 | 0.005 | 0.228 | 0.6 | 0.37 | 0.52 | 0.66 | 0.52 | 0.67 |
| 2 | 169811224 | rs853778 | -0.84 | 0.335 | -0.475 | 0.738 | 0.023 | -0.016 | -0.016 | 0.185 | 0.62 | 0.46 | 0.54 | 0.65 | 0.53 | 0.67 |
| 2 | 169813396 | rs853774 | 0.435 | 0.19 | -1.748 | 0.737 | -0.07 | -0.192 | -0.192 | 0 | 0.29 | 0.01 | 0.39 | 0.49 | 0.3 | 0.47 |
| 2 | 169814655 | rs853772 | -0.409 | 0.586 | -1.099 | 0.791 | -0.03 | 0.033 | 0.033 | 0.535 | 0.6 | 0.23 | 0.51 | 0.66 | 0.51 | 0.67 |
| 2 | 169817713 | rs16823014 | -0.683 | 0.759 | -0.738 | 1.152 | 0.009 | -0.017 | -0.017 | 0 | 0.09 | 0.04 | 0.03 | 0.18 | 0.06 | 0.13 |
| 2 | 169818035 | rs4148797 | -0.248 | 0.847 | -0.993 | 1.272 | -0.055 | -0.058 | -0.058 | 0.281 | 0.23 | 0.07 | 0.38 | 0.24 | 0.27 | 0.45 |
| 2 | 169818302 | rs16856300 | -0.959 | 0.879 | -0.721 | 1.152 | -0.002 | -0.005 | -0.005 | 0 | 0.34 | 0.31 | 0.45 | 0.41 | 0.37 | 0.6 |
| 2 | 169820618 | rs4148796 | -0.622 | 0.477 | -1.155 | 1.23 | 0.004 | -0.075 | -0.075 | 0 | 0.25 | 0.09 | 0.37 | 0.36 | 0.21 | 0.51 |
| 2 | 169820619 | rs4148795 | -0.622 | 0.477 | -1.155 | 1.23 | 0.004 | -0.075 | -0.075 | 0.012 | 0.25 | 0.09 | 0.37 | 0.36 | 0.21 | 0.51 |
| 2 | 169823286 | rs4148790 | 0.158 | -0.122 | 0.001 | 2.139 | 0.018 | 0.038 | 0.038 | 0 | 0.64 | 0.81 | 0.47 | 0.78 | 0.63 | 0.62 |
| 2 | 169823408 | rs17267869 | -0.144 | 0.51 | -0.656 | 1.56 | -0.055 | 0.058 | 0.058 | 0 | 0.14 | 0.01 | 0.07 | 0.02 | 0.17 | 0.1 |
| 2 | 169823867 | rs3770585 | 0.925 | 0.103 | 0.169 | 1.396 | -0.004 | 0.013 | 0.013 | 0 | 0.43 | 0.35 | 0.33 | 0.29 | 0.44 | 0.5 |
| 2 | 169826877 | rs2193831 | -0.13 | -0.109 | 0.887 | 1.894 | -0.004 | -0.006 | -0.006 | 0 | 0.6 | 0.55 | 0.45 | 0.74 | 0.6 | 0.6 |
| 2 | 169827074 | rs4140387 | -0.085 | -0.222 | 0.828 | 1.837 | -0.004 | -0.005 | -0.005 | 0 | 0.59 | 0.55 | 0.45 | 0.74 | 0.6 | 0.6 |
| 2 | 169828325 | rs2241340 | -0.073 | -0.19 | 0.838 | 1.845 | -0.005 | -0.006 | -0.006 | 0 | 0.6 | 0.55 | 0.45 | 0.74 | 0.6 | 0.6 |
| 2 | 169828995 | rs3770586 | 0.698 | -0.069 | 0.852 | 1.779 | 0.003 | 0.02 | 0.02 | 0.014 | 0.48 | 0.39 | 0.35 | 0.44 | 0.49 | 0.5 |
| 2 | 169830155 | rs2287623 | -0.08 | -0.065 | 0.756 | 1.759 | -0.005 | -0.007 | -0.007 | 0.037 | 0.6 | 0.55 | 0.45 | 0.74 | 0.6 | 0.6 |
| 2 | 169830328 | rs2287622 | -0.061 | -0.065 | 0.756 | 1.756 | -0.005 | -0.006 | -0.006 | 0 | 0.6 | 0.55 | 0.45 | 0.7 | 0.6 | 0.6 |
| 2 | 169830515 | rs2287621 | -0.524 | -0.048 | 0.512 | 1.766 | 0.011 | 0.021 | 0.021 | 0 | 0.66 | 0.83 | 0.48 | 0.74 | 0.64 | 0.63 |
| 2 | 169833949 | rs2058996 | 0.076 | -0.136 | 0.487 | 1.648 | 0.011 | -0.019 | -0.019 | 0.306 | 0.59 | 0.74 | 0.41 | 0.57 | 0.58 | 0.56 |
| 2 | 169835090 | rs13384680 | 0.184 | -0.092 | 0.77 | 1.211 | -0.004 | -0.017 | -0.017 | 0 | 0.55 | 0.51 | 0.38 | 0.27 | 0.6 | 0.62 |
| 2 | 169836698 | rs2287620 | -0.021 | -0.135 | 0.978 | 1.209 | 0.02 | 0.07 | 0.07 | 0 | 0.58 | 0.71 | 0.39 | 0.27 | 0.6 | 0.64 |
| 2 | 169836730 | rs2287619 | 0.986 | -0.273 | 0.87 | 1.343 | -0.01 | -0.025 | -0.025 | 0 | 0.08 | 0.04 | 0.07 | 0.21 | 0.08 | 0.13 |
| 2 | 169837328 | rs3770589 | 0.56 | -0.336 | 1.191 | 1.385 | 0 | -0.008 | -0.008 | 0.268 | 0.52 | 0.53 | 0.36 | 0.27 | 0.59 | 0.64 |
| 2 | 169837520 | rs3770590 | 0.64 | -0.368 | 1.245 | 1.376 | 0.018 | 0.038 | 0.038 | 0 | 0.52 | 0.35 | 0.35 | 0.27 | 0.59 | 0.61 |
| 2 | 169837800 | rs3770591 | 0.654 | -0.321 | 1.184 | 1.364 | 0.001 | -0.004 | -0.004 | 0.049 | 0.52 | 0.53 | 0.35 | 0.27 | 0.58 | 0.63 |
| 2 | 169838997 | rs4148782 | 0.316 | -0.335 | 1.244 | 1.364 | 0.023 | 0.074 | 0.074 | 0 | 0.55 | 0.66 | 0.37 | 0.27 | 0.59 | 0.64 |
| 2 | 169842746 | rs7563233 | -1.728 | -0.321 | 1.379 | 1.45 | 0.041 | 0.021 | 0.021 | 2.421 | 0.06 | 0.4 | 0.05 | 0 | 0.02 | 0.03 |
| 2 | 169842809 | rs2287618 | -0.782 | -0.423 | 1.306 | 1.429 | 0.029 | 0.016 | 0.016 | 0 | 0.7 | 0.87 | 0.54 | 0.81 | 0.73 | 0.82 |
| 2 | 169844609 | rs6759156 | -0.709 | -0.428 | 1.367 | 1.43 | 0.018 | 0.02 | 0.02 | 0 | 0.69 | 0.8 | 0.51 | 0.81 | 0.71 | 0.81 |
| 2 | 169847758 | rs2287613 | -1.455 | -0.41 | 1.337 | 1.479 | 0 | -0.005 | -0.005 | 0 | 0.07 | 0.12 | 0.04 | 0.06 | 0.05 | 0.03 |
| 2 | 169851396 | rs10208831 | -0.77 | -0.445 | 1.44 | 1.451 | 0.013 | 0.016 | 0.016 | 0 | 0.7 | 0.88 | 0.52 | 0.81 | 0.71 | 0.81 |
| 2 | 169852183 | rs13407866 | -1.537 | -0.4 | 1.445 | 1.417 | 0.027 | 0.009 | 0.009 | 2.526 | 0.06 | 0.33 | 0.05 | 0 | 0.02 | 0.03 |
| 2 | 169856454 | rs7605199 | 1.527 | -1.17 | 0.99 | 0.938 | -0.004 | 0.015 | 0.015 | 0 | 0.43 | 0.33 | 0.3 | 0.26 | 0.51 | 0.56 |
| 2 | 169857566 | rs6733156 | -1.807 | -1.023 | 1.052 | 1.081 | 0.012 | 0 | 0 | 1.507 | 0.09 | 0.29 | 0.06 | 0.06 | 0.05 | 0.03 |
| 2 | 169863259 | rs10196426 | -1.646 | -1.011 | 1.326 | 1.074 | 0.019 | 0.002 | 0.002 | 1.982 | 0.09 | 0.32 | 0.06 | 0.06 | 0.04 | 0.03 |
| 2 | 169869331 | rs6759871 | -1.62 | -1.007 | 1.312 | 1.075 | 0.025 | 0.003 | 0.003 | 1.872 | 0.09 | 0.31 | 0.06 | 0.06 | 0.05 | 0.03 |
| 2 | 169870295 | rs2241339 | -1.9 | -1.013 | 1.34 | 1.241 | 0.021 | 0.33 | 0.33 | 1.924 | 0.09 | 0.38 | 0.1 | 0.34 | 0.05 | 0.07 |
| 2 | 169870882 | rs4148776 | -1.625 | -1.009 | 1.326 | 1.226 | 0.029 | 0.012 | 0.012 | 1.813 | 0.09 | 0.36 | 0.06 | 0.06 | 0.05 | 0.06 |

| | | | | | | | | | | | | | | | | |
|---|---|---|---|---|---|---|---|---|---|---|---|---|---|---|---|---|
| 2 | 169871543 | rs2389603 | -1.015 | -1.005 | 1.34 | 1.209 | 0.043 | 0.04 | 0.04 | 0.641 | 0.06 | 0.27 | 0.04 | 0.06 | 0.04 | 0.03 |
| 2 | 169872124 | rs10208608 | -1.015 | -1.005 | 1.334 | 1.209 | 0.04 | 0.037 | 0.037 | 0.641 | 0.06 | 0.28 | 0.04 | 0.06 | 0.04 | 0.03 |
| 2 | 169873205 | rs10199694 | -1.07 | -1.009 | 1.314 | 1.207 | 0.031 | 0.028 | 0.028 | 0.625 | 0.06 | 0.26 | 0.04 | 0.06 | 0.04 | 0.03 |
| 2 | 169873821 | rs4148773 | 0.76 | -1.206 | 1.184 | 1.153 | -0.001 | -0.029 | -0.029 | 0.039 | 0.11 | 0.04 | 0.08 | 0.21 | 0.08 | 0.13 |
| 2 | 169874238 | rs10202892 | -0.942 | -1.156 | 1.209 | 1.142 | 0.038 | 0.03 | 0.03 | 0.641 | 0.06 | 0.26 | 0.04 | 0.06 | 0.04 | 0.03 |
| 2 | 169875242 | rs3755161 | -0.942 | -1.167 | 1.214 | 1.142 | 0.033 | 0.03 | 0.03 | 0.641 | 0.06 | 0.26 | 0.04 | 0.06 | 0.04 | 0.03 |
| 2 | 169875736 | rs11892966 | -1.719 | -1.171 | 1.223 | 1.171 | 0.021 | 0.319 | 0.319 | 1.942 | 0.09 | 0.35 | 0.1 | 0.34 | 0.05 | 0.07 |
| 2 | 169880178 | rs7602171 | -0.406 | -1.298 | 0.98 | 0.846 | 0.001 | 0.007 | 0.007 | 0 | 0.74 | 0.88 | 0.53 | 0.82 | 0.72 | 0.82 |
| 2 | 169881691 | rs3770602 | -0.27 | -1.291 | 0.976 | 0.845 | 0.005 | 0.009 | 0.009 | 0 | 0.73 | 0.83 | 0.5 | 0.82 | 0.72 | 0.81 |
| 2 | 169883218 | rs3770603 | 0.558 | -1.325 | 0.966 | 1.405 | 0.003 | -0.005 | -0.005 | 1.184 | 0.12 | 0.15 | 0.08 | 0.2 | 0.08 | 0.15 |
| 2 | 169884648 | rs4668115 | -0.452 | -1.312 | 0.852 | 1.436 | 0.01 | 0.007 | 0.007 | 0 | 0.74 | 0.83 | 0.52 | 0.82 | 0.73 | 0.82 |
| 2 | 169887154 | rs4148768 | 0.809 | -1.263 | 0.79 | 1.481 | -0.001 | -0.025 | -0.025 | 0 | 0.11 | 0.05 | 0.07 | 0.2 | 0.08 | 0.13 |
| 2 | 169888842 | rs3829888 | -0.438 | -1.231 | 0.885 | 1.2 | 0.022 | 0.016 | 0.016 | 0.067 | 0.05 | 0.16 | 0.03 | 0.06 | 0.04 | 0.03 |
| 2 | 169889072 | rs3814381 | 0.029 | -1.303 | 0.602 | 1.151 | 0 | 0 | 0 | 0 | 0.27 | 0.3 | 0.5 | 0.19 | 0.27 | 0.18 |
| 2 | 169889579 | rs4148765 | 0.775 | -1.06 | 0.212 | 1.215 | -0.016 | -0.044 | -0.044 | 0 | 0.1 | 0 | 0.06 | 0.2 | 0.08 | 0.13 |
| 2 | 169891210 | rs7577650 | -1.115 | -1.7 | 0.29 | 0.992 | 0.082 | 0.039 | 0.039 | 0 | 0.49 | 0.72 | 0.38 | 0.38 | 0.35 | 0.33 |
| 2 | 169898016 | rs13403114 | -0.554 | -1.7 | 0.309 | 1.161 | 0.043 | 0.002 | 0.002 | 0 | 0.38 | 0.74 | 0.34 | 0.34 | 0.29 | 0.24 |
| 2 | 169902115 | rs9941622 | -0.556 | -1.671 | 0.321 | 1.087 | -0.011 | 0.002 | 0.002 | 0 | 0.16 | 0.27 | 0.14 | 0.16 | 0.13 | 0.13 |
| 2 | 169911189 | rs6433103 | -1.125 | -1.627 | -0.227 | 1.109 | 0.076 | 0.032 | 0.032 | 0 | 0.51 | 0.85 | 0.48 | 0.38 | 0.41 | 0.36 |
| 2 | 169925861 | rs2287611 | -0.139 | -1.183 | 0.383 | 1.182 | 0.001 | -0.049 | -0.049 | 0 | 0.13 | 0.24 | 0.09 | 0.03 | 0.13 | 0.11 |
| 2 | 169932975 | rs2892805 | -1.169 | -1.455 | -0.345 | 1.25 | 0.091 | 0.001 | 0.001 | 0 | 0.42 | 0.84 | 0.34 | 0.37 | 0.29 | 0.3 |
| 2 | 169933741 | rs1862069 | -1.442 | -0.93 | -0.55 | 1.588 | -0.156 | -0.101 | -0.101 | 2.654 | 0.63 | 0.98 | 0.5 | 0.63 | 0.52 | 0.49 |
| 2 | 169943259 | rs6753502 | 0.137 | -1.025 | -0.047 | 1.288 | -0.03 | 0.04 | 0.04 | 0 | 0.22 | 0.35 | 0.2 | 0.31 | 0.18 | 0.14 |
| 2 | 169952233 | rs1059261 | -0.101 | -0.967 | -0.393 | 1.299 | -0.045 | 0.036 | 0.036 | 0 | 0.25 | 0.42 | 0.25 | 0.32 | 0.18 | 0.14 |
| 2 | 169954185 | rs10515932 | -0.05 | -0.961 | -0.787 | 1.28 | -0.041 | 0.027 | 0.027 | 0 | 0.25 | 0.47 | 0.26 | 0.32 | 0.18 | 0.14 |
| 2 | 169962422 | rs12105342 | -0.628 | -0.102 | -1.6 | 1.032 | -0.026 | -0.016 | -0.016 | 0 | 0.61 | 0.87 | 0.54 | 0.63 | 0.48 | 0.53 |
| 2 | 169963330 | rs3815574 | -0.602 | -0.087 | -1.591 | 1.04 | -0.022 | -0.012 | -0.012 | 0 | 0.61 | 0.85 | 0.54 | 0.63 | 0.48 | 0.53 |
| 2 | 169974671 | rs6433105 | 0.769 | 0.027 | -0.714 | 1.232 | 0.001 | -0.008 | -0.008 | 0.078 | 0.73 | 0.77 | 0.78 | 0.52 | 0.74 | 0.77 |
| 2 | 169997051 | rs990626 | 1.112 | 0.177 | -0.746 | 1.06 | 0.003 | -0.011 | -0.011 | 0 | 0.72 | 0.27 | 0.69 | 0.48 | 0.73 | 0.78 |
| 2 | 169997621 | rs2268379 | 0.947 | 0.131 | -0.032 | 1.078 | -0.014 | 0.089 | 0.089 | 0 | 0.76 | 0.42 | 0.72 | 0.48 | 0.75 | 0.78 |
| 2 | 170003432 | rs4667591 | 1.411 | 0.156 | -0.552 | 0.749 | 0.022 | -0.052 | -0.052 | 0 | 0.74 | 0.15 | 0.73 | 0.49 | 0.79 | 0.81 |
| 2 | 170010985 | rs2075252 | 1.181 | 0.203 | -0.271 | 0.827 | -0.027 | -0.056 | -0.056 | 2.882 | 0.83 | 0.92 | 0.79 | 0.52 | 0.76 | 0.87 |
| 2 | 170016750 | rs6759013 | 0.721 | 0.069 | -0.2 | -0.078 | -0.002 | 0.043 | 0.043 | 0 | 0.48 | 0.64 | 0.64 | 0.13 | 0.48 | 0.47 |
| 2 | 170018638 | rs3821124 | 0.822 | 0.068 | 0.212 | -0.08 | 0 | -0.032 | -0.032 | 0 | 0.47 | 0.54 | 0.63 | 0.11 | 0.48 | 0.47 |
| 2 | 170020744 | rs2268375 | 0.966 | 0.041 | 0.4 | -0.094 | -0.005 | 0.023 | 0.023 | 0 | 0.46 | 0.45 | 0.62 | 0.11 | 0.48 | 0.46 |
| 2 | 170025340 | rs2239602 | -0.356 | 0.14 | -0.01 | 0.948 | -0.019 | -0.035 | -0.035 | 0 | 0.91 | 0.96 | 0.92 | 0.77 | 0.9 | 0.93 |
| 2 | 170036266 | rs2892803 | 0.403 | 0.132 | 0.701 | 0.442 | -0.002 | 0.032 | 0.032 | 0 | 0.54 | 0.59 | 0.68 | 0.14 | 0.52 | 0.53 |
| 2 | 170037070 | rs9789747 | 1.19 | 0.118 | 0.175 | 0.657 | 0.009 | -0.054 | -0.054 | 0 | 0.25 | 0.07 | 0.12 | 0.4 | 0.21 | 0.3 |
| 2 | 170039594 | rs10169879 | 0.712 | 0.067 | 0.495 | -0.225 | 0.006 | 0.035 | 0.035 | 0.056 | 0.76 | 0.75 | 0.76 | 0.49 | 0.69 | 0.81 |
| 2 | 170046850 | rs10210408 | 1.308 | -0.295 | 0.865 | -0.397 | 0.003 | -0.01 | -0.01 | 0 | 0.66 | 0.37 | 0.7 | 0.47 | 0.67 | 0.72 |
| 2 | 170047935 | rs7592152 | -1.464 | -0.34 | -0.172 | -1.016 | -0.109 | 0.133 | 0.133 | 0 | 0.7 | 0.97 | 0.48 | 0.91 | 0.65 | 0.63 |
| 2 | 170048929 | rs2389589 | -0.937 | -0.631 | 0.128 | -0.905 | 0.005 | -0.045 | -0.045 | 0.14 | 0.26 | 0.8 | 0.28 | 0.53 | 0.27 | 0.17 |
| 2 | 170049604 | rs3821125 | -0.49 | -0.6 | 0.649 | -0.901 | 0.016 | 0.122 | 0.122 | 0.736 | 0.22 | 0.5 | 0.23 | 0.51 | 0.26 | 0.16 |
| 2 | 170051167 | rs10490131 | 2.069 | -0.596 | 0.226 | -1.064 | -0.022 | -0.12 | -0.12 | 0.001 | 0.2 | 0.01 | 0.08 | 0.37 | 0.17 | 0.24 |
| 2 | 170053580 | rs2302696 | -0.793 | -0.429 | 0.868 | -0.966 | 0.009 | 0.135 | 0.135 | 0.705 | 0.24 | 0.45 | 0.23 | 0.53 | 0.26 | 0.17 |
| 2 | 170058497 | rs16856600 | -0.497 | -0.524 | 0.814 | -1.083 | 0 | 0.007 | 0.007 | 0 | 0.24 | 0.19 | 0.22 | 0.52 | 0.31 | 0.27 |
| 2 | 170058959 | rs2284675 | -0.291 | -0.53 | 0.726 | -1.1 | 0.024 | 0.145 | 0.145 | 0 | 0.22 | 0.45 | 0.25 | 0.54 | 0.27 | 0.17 |
| 2 | 170061466 | rs6725137 | -1.002 | -0.432 | 0.88 | -0.955 | 0.043 | -0.016 | -0.016 | 0 | 0.07 | 0.36 | 0.06 | 0 | 0.05 | 0.12 |
| 2 | 170070348 | rs11886219 | -1.359 | -0.46 | 0.907 | -1.006 | 0.033 | -0.022 | -0.022 | 2.473 | 0.09 | 0.49 | 0.1 | 0 | 0.05 | 0.03 |
| 2 | 170080596 | rs13389381 | -1.067 | -0.669 | 0.83 | -1.155 | -0.036 | -0.113 | -0.113 | 0 | 0.16 | 0.43 | 0.11 | 0 | 0.09 | 0.06 |
| 2 | 170083777 | rs6433109 | -1.13 | -0.212 | -0.087 | -1.497 | -0.018 | -0.013 | -0.013 | 0 | 0.64 | 0.78 | 0.42 | 0.48 | 0.5 | 0.78 |
| 2 | 170088046 | rs4668128 | -0.859 | -0.189 | -0.094 | -1.458 | -0.011 | -0.015 | -0.015 | 0 | 0.61 | 0.77 | 0.47 | 0.54 | 0.46 | 0.67 |
| 2 | 170088351 | rs2302694 | -0.859 | -0.04 | 0.084 | -1.389 | -0.023 | -0.084 | -0.084 | 0 | 0.16 | 0.42 | 0.19 | 0.05 | 0.11 | 0.07 |
| 2 | 170089070 | rs3821126 | 0.555 | -0.24 | 0.02 | -1.458 | 0.016 | -0.011 | -0.011 | 0.183 | 0.41 | 0.31 | 0.28 | 0.48 | 0.34 | 0.6 |
| 2 | 170089824 | rs2239596 | 0.184 | -0.19 | 0 | -1.484 | 0.029 | -0.003 | -0.003 | 0.892 | 0.44 | 0.41 | 0.51 | 0.47 | 0.33 | 0.62 |
| 2 | 170089851 | rs2239595 | 2.307 | -0.353 | -0.681 | -1.504 | 0.016 | -0.129 | -0.129 | 0 | 0.21 | 0.01 | 0.32 | 0.37 | 0.14 | 0.27 |
| 2 | 170099111 | rs10490130 | 0.342 | -0.182 | -0.455 | -1.473 | -0.002 | -0.045 | -0.045 | 0 | 0.1 | 0.2 | 0.08 | 0 | 0.09 | 0.04 |
| 2 | 170099446 | rs831041 | -0.961 | -0.063 | -0.454 | -1.476 | -0.004 | -0.005 | -0.005 | 0 | 0.59 | 0.72 | 0.64 | 0.48 | 0.46 | 0.77 |
| 2 | 170099473 | rs831042 | -0.676 | -0.08 | -0.436 | -1.484 | -0.001 | 0 | 0 | 0 | 0.55 | 0.61 | 0.62 | 0.48 | 0.46 | 0.75 |
| 2 | 170103336 | rs831043 | -0.945 | -0.025 | -0.528 | -1.443 | -0.005 | -0.006 | -0.006 | 0 | 0.59 | 0.72 | 0.64 | 0.48 | 0.46 | 0.77 |
| 2 | 170111945 | rs830956 | -0.912 | -0.025 | -0.496 | -1.467 | -0.001 | -0.001 | -0.001 | 0 | 0.59 | 0.68 | 0.63 | 0.48 | 0.46 | 0.77 |
| 2 | 170115588 | rs2241190 | -0.965 | -0.024 | -0.518 | -1.434 | -0.008 | -0.007 | -0.007 | 0.021 | 0.59 | 0.71 | 0.64 | 0.48 | 0.48 | 0.78 |
| 2 | 170116631 | rs830983 | -0.113 | 0.298 | -0.508 | -1.359 | 0.016 | -0.003 | -0.003 | 0.133 | 0.55 | 0.63 | 0.47 | 0.52 | 0.61 | 0.28 |
| 2 | 170117811 | rs12988804 | -0.951 | 0.516 | -0.072 | -1.887 | -0.012 | 0.072 | 0.072 | 0 | 0.28 | 0.18 | 0.22 | 0.07 | 0.27 | 0.3 |
| 2 | 170128337 | rs830992 | 1.762 | 1.188 | 0.424 | -1.288 | 0.004 | -0.051 | -0.051 | 2.222 | 0.9 | 0.64 | 0.72 | 0.97 | 0.91 | 0.81 |
| 2 | 170132243 | rs830997 | 0.669 | 1.332 | 0.282 | -1.323 | 0.001 | -0.044 | -0.044 | 0 | 0.75 | 0.63 | 0.76 | 0.92 | 0.75 | 0.58 |
| 2 | 170133086 | rs830998 | 0.21 | 1.408 | -0.299 | -1.267 | 0 | 0.018 | 0.018 | 0 | 0.79 | 0.79 | 0.8 | 0.92 | 0.81 | 0.72 |
| 2 | 170135215 | rs7568568 | -0.839 | 1.243 | -0.024 | -1.087 | 0.024 | -0.027 | -0.027 | 2.619 | 0.06 | 0.22 | 0.05 | 0 | 0.08 | 0.04 |
| 2 | 170135742 | rs831004 | 1.667 | 1.27 | -0.459 | -1.135 | 0.001 | -0.058 | -0.058 | 1.097 | 0.6 | 0.32 | 0.67 | 0.88 | 0.59 | 0.51 |

| | | | | | | | | | | | | | | | | |
|---|---|---|---|---|---|---|---|---|---|---|---|---|---|---|---|---|
| 2 | 170138506 | rs831007 | 0.896 | 0.602 | -1.445 | -0.618 | 0.005 | 0.009 | 0.009 | 0.007 | 0.75 | 0.71 | 0.79 | 0.88 | 0.78 | 0.61 |
| 2 | 170139346 | rs831010 | 0.029 | 0.626 | -1.479 | -0.541 | 0.008 | 0.021 | 0.021 | 0 | 0.83 | 0.83 | 0.83 | 0.9 | 0.85 | 0.75 |
| 2 | 170139862 | rs831012 | 0.493 | 0.683 | -1.39 | -0.409 | -0.001 | 0.044 | 0.044 | 0.31 | 0.71 | 0.72 | 0.75 | 0.88 | 0.69 | 0.56 |
| 2 | 170143494 | rs3770611 | 0.224 | 0.66 | -1.179 | 0.564 | 0.006 | 0.024 | 0.024 | 0.03 | 0.71 | 0.38 | 0.53 | 0.56 | 0.68 | 0.7 |
| 2 | 170145273 | rs2161039 | 0.767 | 0.334 | -1.786 | 0.125 | -0.022 | -0.001 | -0.001 | 0 | 0.47 | 0.22 | 0.37 | 0.48 | 0.46 | 0.52 |
| 2 | 170146592 | rs831019 | -0.725 | 0.032 | -1.464 | 0.207 | 0 | -0.002 | -0.002 | 0 | 0.42 | 0.56 | 0.52 | 0.45 | 0.41 | 0.36 |
| 2 | 170149613 | rs831022 | -0.016 | 0.155 | -1.3 | 0.41 | 0.018 | 0.068 | 0.068 | 0 | 0.27 | 0.54 | 0.46 | 0.45 | 0.3 | 0.28 |
| 2 | 170149854 | rs11887007 | 0.346 | -0.093 | -1.705 | 0.464 | 0.057 | 0.049 | 0.049 | 0 | 0.5 | 0.25 | 0.44 | 0.33 | 0.6 | 0.59 |
| 2 | 170153526 | rs2673164 | 0.388 | 0.466 | -1.082 | 0.983 | 0.004 | 0.058 | 0.058 | 0.003 | 0.73 | 0.37 | 0.46 | 0.56 | 0.72 | 0.76 |
| 2 | 170157559 | rs830965 | -0.579 | 0.233 | -0.628 | 0.805 | -0.002 | -0.009 | -0.009 | 0 | 0.36 | 0.43 | 0.25 | 0.17 | 0.27 | 0.33 |
| 2 | 170157956 | rs2892802 | 1.213 | 0.59 | -1.31 | 0.68 | -0.016 | -0.001 | -0.001 | 0 | 0.1 | 0.05 | 0.09 | 0.07 | 0.14 | 0.09 |
| 2 | 170158456 | rs700550 | -1.138 | 0.663 | -1.182 | 0.879 | 0.008 | 0.089 | 0.089 | 0 | 0.56 | 0.83 | 0.75 | 0.81 | 0.56 | 0.68 |
| 2 | 170159313 | rs830968 | -1.11 | 0.672 | -1.177 | 0.883 | 0.01 | 0.091 | 0.091 | 0 | 0.56 | 0.83 | 0.75 | 0.81 | 0.56 | 0.68 |
| 2 | 170160361 | rs853988 | -1.067 | 0.493 | -1.075 | 0.926 | -0.009 | 0.044 | 0.044 | 0.216 | 0.48 | 0.84 | 0.7 | 0.55 | 0.46 | 0.44 |
| 2 | 170163768 | rs2247506 | -0.174 | 0.277 | -0.271 | 0.687 | -0.011 | -0.048 | -0.048 | 0 | 0.23 | 0.33 | 0.16 | 0.07 | 0.2 | 0.21 |
| 2 | 170175109 | rs861239 | -0.617 | 0.765 | 0.233 | 0.733 | 0.02 | -0.02 | -0.02 | 0 | 0.25 | 0.48 | 0.21 | 0.28 | 0.28 | 0.2 |
| 2 | 170181305 | rs2673177 | -1.422 | 0.699 | 0.219 | 0.669 | 0.001 | 0.009 | 0.009 | 0 | 0.37 | 0.62 | 0.29 | 0.36 | 0.36 | 0.26 |
| 2 | 170196990 | rs10930352 | 0.065 | 0.843 | 0.28 | 0.681 | 0.003 | 0.013 | 0.013 | 0 | 0.18 | 0.3 | 0.14 | 0.27 | 0.18 | 0.16 |
| 2 | 170201843 | rs831037 | -1.282 | 0.869 | 0.03 | 0.935 | 0.001 | -0.004 | -0.004 | 0.01 | 0.37 | 0.62 | 0.31 | 0.46 | 0.36 | 0.27 |
| 2 | 170207896 | rs6713072 | 1.347 | 1.153 | -0.123 | 0.727 | 0.01 | 0.07 | 0.07 | 0 | 0.44 | 0.27 | 0.46 | 0.23 | 0.42 | 0.55 |
| 2 | 170208668 | rs12614394 | 0.171 | 1.033 | 0.073 | 1.12 | -0.07 | -0.017 | -0.017 | 0 | 0.21 | 0.08 | 0.2 | 0.17 | 0.3 | 0.2 |
| 2 | 170208975 | rs10199676 | 0.01 | 0.8 | 0.832 | 1.378 | -0.03 | 0.022 | 0.022 | 0 | 0.18 | 0.24 | 0.16 | 0.27 | 0.12 | 0.17 |
| 2 | 170211377 | rs2389557 | 1.22 | 0.859 | 0.936 | 1.324 | 0.003 | 0.002 | 0.002 | 0 | 0.55 | 0.46 | 0.57 | 0.55 | 0.52 | 0.59 |
| 2 | 170216267 | rs4668136 | 1.661 | 0.92 | 0.593 | 1.368 | -0.001 | 0.086 | 0.086 | 0 | 0.5 | 0.3 | 0.51 | 0.26 | 0.51 | 0.63 |
| 2 | 170216868 | rs3845732 | 0.796 | 0.879 | 0.598 | 1.456 | -0.01 | 0.005 | 0.005 | 0 | 0.7 | 0.36 | 0.71 | 0.4 | 0.81 | 0.77 |
| 2 | 170219881 | rs3755166 | 1.041 | 1.143 | 0.909 | 1.788 | -0.01 | -0.013 | -0.013 | 0 | 0.66 | 0.72 | 0.66 | 0.58 | 0.61 | 0.69 |
| 2 | 170223699 | rs830955 | 1.203 | 1.526 | 0.818 | 1.667 | -0.016 | 0.002 | 0.002 | 0 | 0.65 | 0.47 | 0.62 | 0.32 | 0.66 | 0.55 |
| 2 | 170227070 | rs2544371 | 1.655 | 1.602 | 0.814 | 1.616 | -0.001 | -0.02 | -0.02 | 0 | 0.55 | 0.42 | 0.51 | 0.18 | 0.46 | 0.38 |
| 2 | 170228677 | rs13002827 | 0.512 | 1.657 | 0.742 | 1.657 | 0 | -0.003 | -0.003 | 0 | 0.1 | 0.06 | 0.12 | 0.14 | 0.2 | 0.16 |
| 2 | 170230772 | rs12613663 | -1.029 | 1.19 | 0.545 | 0.93 | 0.032 | -0.002 | -0.002 | 0 | 0.3 | 0.41 | 0.34 | 0.75 | 0.4 | 0.37 |
| 2 | 170236380 | rs4667601 | -0.053 | 0.33 | 0.373 | 1.186 | -0.009 | 0.006 | 0.006 | 0 | 0.13 | 0 | 0.14 | 0.07 | 0.13 | 0.25 |
| 2 | 170238935 | rs4668139 | -0.007 | 0.42 | 0.418 | 1.158 | -0.032 | 0.016 | 0.016 | 0 | 0.15 | 0.03 | 0.16 | 0.1 | 0.2 | 0.25 |
| 2 | 170239158 | rs1606797 | 1.306 | 0.253 | 0.73 | 1.106 | 0.005 | -0.021 | -0.021 | 0 | 0.65 | 0.49 | 0.64 | 0.75 | 0.64 | 0.55 |
| 2 | 170249802 | rs12619539 | -0.757 | -0.236 | 1.031 | 1.719 | -0.045 | 0.164 | 0.164 | 0 | 0.16 | 0.33 | 0.16 | 0.5 | 0.07 | 0.11 |
| 2 | 170264277 | rs4668142 | 0.85 | -0.013 | -0.599 | 1.153 | -0.02 | 0.015 | 0.015 | 0 | 0.7 | 0.36 | 0.72 | 0.37 | 0.81 | 0.73 |
| 2 | 170271013 | rs4616447 | 0.715 | -0.143 | -0.608 | 1.122 | -0.04 | 0.058 | 0.058 | 0.279 | 0.72 | 0.41 | 0.78 | 0.42 | 0.83 | 0.77 |
| 2 | 170271626 | rs7591026 | 0.922 | -0.348 | -1.132 | 0.073 | -0.003 | -0.051 | -0.051 | 1.886 | 0.76 | 0.32 | 0.84 | 1 | 0.89 | 0.9 |
| 2 | 170272102 | rs10168193 | -0.009 | -0.054 | -2.273 | 0.218 | 0.052 | 0.112 | 0.112 | 0.595 | 0.49 | 0.12 | 0.49 | 0.7 | 0.58 | 0.69 |
| 2 | 170273127 | rs10930354 | 0.06 | 0.349 | -2.647 | 0.493 | -0.156 | -0.089 | -0.089 | 1.438 | 0.32 | 0.08 | 0.41 | 0.41 | 0.46 | 0.31 |
| 2 | 170278948 | rs6731329 | -0.487 | -0.537 | -1.766 | 0.488 | -0.004 | 0.003 | 0.003 | 0.224 | 0.3 | 0.67 | 0.28 | 0.3 | 0.29 | 0.31 |
| 2 | 170285339 | rs12465476 | -0.319 | -0.545 | -1.012 | 0.888 | 0.022 | 0.18 | 0.18 | 0 | 0.09 | 0.2 | 0.17 | 0.34 | 0.09 | 0.25 |
| 2 | 170290349 | rs13034809 | 0.754 | -0.693 | -0.241 | 0.729 | 0 | -0.002 | -0.002 | 0 | 0.65 | 0.36 | 0.64 | 0.53 | 0.67 | 0.47 |
| 2 | 170297339 | rs4491700 | 0.628 | -0.89 | 0.156 | 0.63 | 0.007 | -0.032 | -0.032 | 0 | 0.64 | 0.35 | 0.65 | 0.53 | 0.67 | 0.49 |
| 2 | 170298350 | rs12692903 | 0.991 | -1.515 | -0.439 | 0.262 | 0.027 | -0.004 | -0.004 | 1.117 | 0.45 | 0.32 | 0.46 | 0.47 | 0.58 | 0.39 |
| 2 | 170299584 | rs10185935 | 0.474 | -1.47 | -0.282 | 0.194 | -0.015 | -0.015 | -0.015 | 0.653 | 0.21 | 0.2 | 0.21 | 0.06 | 0.09 | 0.1 |
| 2 | 170303507 | rs6433120 | 0.371 | -1.357 | -0.263 | -0.048 | 0.008 | -0.004 | -0.004 | 0 | 0.56 | 0.57 | 0.66 | 0.9 | 0.7 | 0.68 |
| 2 | 170304533 | rs2883107 | -0.891 | -1.382 | -0.311 | -0.069 | 0.002 | 0.044 | 0.044 | 0 | 0.78 | 0.92 | 0.88 | 0.96 | 0.79 | 0.78 |
| 2 | 170305273 | rs2353182 | 0.41 | -1.357 | -0.182 | 0.114 | -0.018 | -0.027 | -0.027 | 1.537 | 0.55 | 0.46 | 0.64 | 0.79 | 0.68 | 0.66 |
| 2 | 170306991 | rs11692518 | 1.484 | -1.22 | -0.446 | 0.301 | -0.055 | -0.056 | -0.056 | 0 | 0.39 | 0.11 | 0.41 | 0.46 | 0.53 | 0.36 |
| 2 | 170325515 | rs9636291 | 0.149 | -1.155 | 0.174 | 0.255 | -0.023 | -0.044 | -0.044 | 1.185 | 0.71 | 0.43 | 0.83 | 0.85 | 0.77 | 0.76 |
| 2 | 170336659 | rs11901329 | -1.003 | -0.868 | 0.063 | 0.386 | -0.006 | -0.006 | -0.006 | 0 | 0.07 | 0.04 | 0.08 | 0.12 | 0.1 | 0.06 |
| 2 | 170336797 | rs16856972 | 1.273 | -0.885 | 0.062 | 0.601 | -0.01 | -0.012 | -0.012 | 0.144 | 0.06 | 0.01 | 0.1 | 0.2 | 0.06 | 0.25 |
| 2 | 170336811 | rs6736056 | 0.926 | -1.328 | 0.424 | 0.447 | -0.007 | 0.002 | 0.002 | 1.562 | 0.65 | 0.42 | 0.7 | 0.64 | 0.76 | 0.58 |
| 2 | 170340252 | rs12612150 | 0.704 | -1.289 | 0.384 | 0.467 | 0 | 0.03 | 0.03 | 0 | 0.07 | 0.05 | 0.12 | 0.32 | 0.1 | 0.28 |
| 2 | 170343083 | rs2353188 | 2.013 | -1.021 | 0.183 | 0.233 | -0.048 | -0.039 | -0.039 | 0 | 0.38 | 0.22 | 0.5 | 0.45 | 0.47 | 0.35 |
| 2 | 170344060 | rs6433124 | 0.874 | -1.029 | 0.259 | 0.247 | 0.022 | -0.028 | -0.028 | 0 | 0.64 | 0.31 | 0.68 | 0.52 | 0.72 | 0.55 |
| 2 | 170354790 | rs12619026 | -0.741 | -0.908 | 0.622 | 0.286 | 0.018 | 0.056 | 0.056 | 0 | 0.13 | 0.19 | 0.18 | 0.32 | 0.16 | 0.31 |
| 2 | 170359113 | rs11902340 | 1.837 | -0.851 | 0.445 | 0.274 | -0.017 | 0.015 | 0.015 | 0 | 0.06 | 0.03 | 0.06 | 0.02 | 0.12 | 0.1 |
| 2 | 170367099 | rs28763868 | 1.837 | -0.848 | 0.395 | 0.282 | -0.017 | 0.015 | 0.015 | 0 | 0.06 | 0.02 | 0.06 | 0.01 | 0.12 | 0.09 |
| 2 | 170373591 | rs13417370 | 0.874 | -0.553 | 0.657 | 0.26 | -0.005 | 0.048 | 0.048 | 0 | 0.48 | 0.68 | 0.64 | 0.7 | 0.54 | 0.43 |
| 2 | 170375686 | rs13387104 | 0.848 | -0.497 | 0.826 | 0.259 | 0.006 | -0.011 | -0.011 | 0 | 0.6 | 0.56 | 0.68 | 0.64 | 0.58 | 0.46 |
| 2 | 170384819 | rs16857010 | 2.005 | -0.679 | 0.711 | 0.26 | -0.069 | -0.109 | -0.109 | 0 | 0.37 | 0.09 | 0.48 | 0.46 | 0.43 | 0.35 |
| 2 | 170386678 | rs2166577 | 1.936 | -0.661 | 0.716 | 0.243 | -0.032 | -0.066 | -0.066 | 0 | 0.38 | 0.15 | 0.48 | 0.46 | 0.42 | 0.35 |
| 2 | 170390189 | rs16857020 | 1.416 | -0.583 | 0.907 | 0.333 | 0.014 | 0.058 | 0.058 | 0 | 0.06 | 0.09 | 0.11 | 0.2 | 0.07 | 0.24 |
| 2 | 170402752 | rs2290010 | 1.464 | -0.586 | 0.73 | 0.376 | -0.01 | -0.012 | -0.012 | 0 | 0.06 | 0.01 | 0.1 | 0.2 | 0.07 | 0.24 |
| 2 | 170403030 | rs2253680 | 0.66 | -0.671 | 0.91 | 0.28 | 0.003 | 0.021 | 0.021 | 0 | 0.63 | 0.49 | 0.67 | 0.51 | 0.62 | 0.48 |
| 2 | 170410637 | rs2441676 | 1.018 | -0.673 | 0.694 | 0.287 | -0.007 | -0.009 | -0.009 | 0 | 0.58 | 0.39 | 0.62 | 0.52 | 0.57 | 0.44 |
| 2 | 170411485 | rs2592823 | 1.842 | -0.608 | 0.551 | 0.36 | -0.019 | -0.043 | -0.043 | 4.951 | 0.39 | 0.22 | 0.49 | 0.46 | 0.44 | 0.34 |
| 2 | 170415441 | rs12052238 | 1.395 | -0.419 | 0.561 | 0.383 | -0.012 | -0.015 | -0.015 | 0.552 | 0.06 | 0.01 | 0.1 | 0.21 | 0.07 | 0.24 |

| | | | | | | | | | | | | | | | | |
|---|---|---|---|---|---|---|---|---|---|---|---|---|---|---|---|---|
| 2 | 170440746 | rs2592797 | 1.511 | -0.697 | 0.572 | 0.241 | -0.014 | 0.005 | 0.005 | 0.01 | 0.4 | 0.24 | 0.53 | 0.57 | 0.46 | 0.36 |
| 2 | 170447698 | rs2592804 | 1.653 | -0.691 | 0.584 | 0.244 | -0.015 | 0.005 | 0.005 | 0.014 | 0.4 | 0.24 | 0.53 | 0.57 | 0.46 | 0.36 |
| 2 | 170457994 | rs13422340 | 0.28 | -0.703 | 1.022 | 0.204 | 0.042 | 0.083 | 0.083 | 0 | 0.11 | 0.32 | 0.18 | 0.22 | 0.12 | 0.28 |
| 2 | 170460091 | rs1111534 | 1.701 | -0.711 | 1.003 | 0.21 | -0.013 | 0.007 | 0.007 | 0.02 | 0.4 | 0.24 | 0.54 | 0.57 | 0.46 | 0.36 |
| 2 | 170463551 | rs2592813 | 1.841 | -0.749 | 1.145 | 0.641 | -0.026 | -0.016 | -0.016 | 0 | 0.38 | 0.18 | 0.48 | 0.4 | 0.43 | 0.34 |
| 2 | 170469839 | rs16857073 | 0.913 | -0.733 | 1.135 | 0.261 | 0.008 | 0.011 | 0.011 | 0 | 0.19 | 0.2 | 0.13 | 0.05 | 0.13 | 0.11 |
| 2 | 170479662 | rs16857081 | 0.912 | -0.766 | 1.198 | 0.206 | -0.02 | -0.038 | -0.038 | 0.048 | 0.19 | 0.2 | 0.06 | 0.02 | 0.09 | 0.12 |
| 2 | 170481769 | rs13021966 | -0.221 | -0.731 | 1.128 | 0.204 | 0 | -0.03 | -0.03 | 0 | 0.29 | 0.27 | 0.15 | 0.05 | 0.25 | 0.23 |
| 2 | 170493863 | rs8207 | -0.251 | -0.811 | 1.093 | 0.163 | 0 | -0.024 | -0.024 | 1.222 | 0.3 | 0.27 | 0.15 | 0.05 | 0.25 | 0.23 |
| 2 | 170498773 | rs16857109 | 0.739 | -0.735 | 1.139 | 0.216 | -0.01 | -0.019 | -0.019 | 0 | 0.09 | 0.08 | 0.16 | 0.22 | 0.12 | 0.28 |
| 2 | 170505684 | rs3213946 | -1.311 | -0.794 | 1.165 | 0.211 | 0.017 | 0.088 | 0.088 | 0 | 0.51 | 0.62 | 0.3 | 0.21 | 0.42 | 0.36 |
| 2 | 170517503 | rs9287917 | 0.622 | -0.764 | 0.995 | 0.227 | -0.003 | -0.005 | -0.005 | 0 | 0.1 | 0.14 | 0.16 | 0.22 | 0.12 | 0.28 |
| 2 | 170517594 | rs7575494 | -0.397 | -0.851 | 0.921 | 0.183 | 0 | -0.002 | -0.002 | 0 | 0.3 | 0.27 | 0.15 | 0.16 | 0.28 | 0.26 |
| 2 | 170527791 | rs6433133 | -1.297 | -0.733 | 0.945 | 0.254 | 0.001 | -0.006 | -0.006 | 0 | 0.49 | 0.43 | 0.28 | 0.21 | 0.42 | 0.36 |
| 2 | 170531572 | rs11682248 | -1.297 | -0.733 | 0.945 | 0.254 | 0.001 | -0.006 | -0.006 | 0 | 0.49 | 0.43 | 0.28 | 0.21 | 0.42 | 0.36 |
| 2 | 170550727 | rs11690884 | 1.312 | -0.76 | 0.967 | 0.318 | -0.034 | 0.028 | 0.028 | 0 | 0.09 | 0.03 | 0.08 | 0.02 | 0.17 | 0.11 |
| 2 | 170557350 | rs3829806 | -1.216 | -0.854 | 0.989 | 0.324 | 0.001 | -0.005 | -0.005 | 0.038 | 0.49 | 0.43 | 0.29 | 0.21 | 0.42 | 0.37 |
| 2 | 170578701 | rs10164482 | -1.857 | -0.417 | 0.329 | 0.51 | -0.003 | -0.041 | -0.041 | 0 | 0.63 | 0.76 | 0.61 | 0.5 | 0.6 | 0.66 |
| 2 | 170583879 | rs7562456 | 0.915 | -0.309 | 0.282 | 0.588 | 0.005 | -0.002 | -0.002 | 0 | 0.11 | 0.13 | 0.1 | 0.09 | 0.18 | 0.1 |
| 2 | 170592709 | rs2288331 | 0.231 | -0.585 | 0.321 | 0.661 | 0.006 | 0.001 | 0.001 | 0 | 0.22 | 0.18 | 0.14 | 0.18 | 0.16 | 0.11 |
| 2 | 170594091 | rs12470910 | 0.033 | -0.649 | 0.272 | 0.575 | -0.003 | 0.016 | 0.016 | 0.697 | 0.28 | 0.08 | 0.34 | 0.26 | 0.25 | 0.47 |
| 2 | 170597835 | rs2303556 | 0.504 | -0.794 | 0.397 | 0.271 | 0.013 | 0.017 | 0.017 | 0 | 0.19 | 0.15 | 0.12 | 0.07 | 0.11 | 0.09 |
| 2 | 170599373 | rs6750738 | 1.328 | -1.109 | 0.609 | 0.228 | 0.044 | 0.043 | 0.043 | 0.087 | 0.51 | 0.32 | 0.74 | 0.67 | 0.67 | 0.69 |
| 2 | 170599690 | rs6725295 | 0.545 | -1.2 | 0.536 | 0.256 | -0.038 | -0.014 | -0.014 | 0 | 0.67 | 0.58 | 0.85 | 0.77 | 0.86 | 0.79 |
| 2 | 170602280 | rs7590503 | -1.32 | -0.569 | 0.452 | 0.267 | 0.007 | 0.034 | 0.034 | 0 | 0.44 | 0.68 | 0.28 | 0.27 | 0.38 | 0.27 |
| 2 | 170607750 | rs1344639 | -1.22 | -0.531 | 0.069 | 0.253 | -0.002 | -0.012 | -0.012 | 0 | 0.43 | 0.53 | 0.26 | 0.25 | 0.37 | 0.27 |
| 2 | 170613342 | rs7559061 | -0.217 | -0.183 | 0.403 | 0.445 | 0.031 | -0.021 | -0.021 | 0.873 | 0.25 | 0.51 | 0.17 | 0.18 | 0.27 | 0.18 |
| 2 | 170621358 | rs7583128 | 0.544 | -0.37 | 0.226 | 0.553 | -0.004 | -0.006 | -0.006 | 1.209 | 0.64 | 0.46 | 0.74 | 0.71 | 0.67 | 0.74 |
| 2 | 170623100 | rs7590347 | -0.717 | 0.009 | -0.062 | 1.164 | 0.003 | 0.002 | 0.002 | 0 | 0.3 | 0.33 | 0.55 | 0.57 | 0.4 | 0.41 |
| 2 | 170624221 | rs2114646 | 0.743 | 0.143 | -0.355 | 1.274 | -0.069 | -0.091 | -0.091 | 0 | 0.12 | 0.02 | 0.43 | 0.43 | 0.21 | 0.31 |
| 2 | 170626325 | rs6738458 | 1.448 | -0.301 | -0.041 | 0.89 | 0.003 | -0.022 | -0.022 | 0 | 0.4 | 0.44 | 0.31 | 0.28 | 0.43 | 0.35 |
| 2 | 170629301 | rs12692924 | 0.842 | -0.141 | -0.212 | 0.97 | -0.006 | -0.007 | -0.007 | 0 | 0.15 | 0.12 | 0.11 | 0.2 | 0.19 | 0.1 |
| 2 | 170630809 | rs2059705 | -0.511 | -0.648 | 0.145 | 1.045 | 0.001 | -0.002 | -0.002 | 0 | 0.3 | 0.29 | 0.37 | 0.4 | 0.28 | 0.45 |
| 2 | 170631928 | rs12692925 | 0.651 | -0.322 | 0.487 | 0.76 | 0.026 | 0.084 | 0.084 | 0 | 0.71 | 0.53 | 0.45 | 0.46 | 0.66 | 0.54 |
| 2 | 170632625 | rs10174880 | 0.179 | -0.439 | 0.644 | -0.102 | 0 | -0.001 | -0.001 | 0 | 0.8 | 0.75 | 0.87 | 0.9 | 0.83 | 0.85 |
| 2 | 170632807 | rs10930375 | -0.128 | -0.517 | 0.791 | 0.027 | 0.01 | 0.066 | 0.066 | 0 | 0.22 | 0.24 | 0.41 | 0.47 | 0.27 | 0.38 |
| 2 | 170639070 | rs4668163 | 1.125 | -0.559 | 0.316 | -0.011 | -0.006 | -0.018 | -0.018 | 0.108 | 0.58 | 0.27 | 0.46 | 0.43 | 0.59 | 0.48 |
| 2 | 170647382 | rs1559935 | -0.885 | -0.314 | 0.221 | 0.092 | 0.012 | 0.042 | 0.042 | 0 | 0.29 | 0.39 | 0.45 | 0.5 | 0.34 | 0.46 |
| 2 | 170657136 | rs7572691 | 0.919 | -0.133 | 0.719 | 0.038 | -0.015 | 0.005 | 0.005 | 0 | 0.78 | 0.56 | 0.89 | 0.84 | 0.84 | 0.87 |
| 2 | 170665108 | rs2556352 | 0.363 | -0.415 | 0.632 | -0.307 | -0.015 | 0.003 | 0.003 | 0 | 0.58 | 0.48 | 0.72 | 0.47 | 0.71 | 0.52 |
| 2 | 170672843 | rs10190853 | 0.754 | -0.417 | 0.547 | -0.231 | 0.028 | -0.018 | -0.018 | 0.383 | 0.25 | 0.21 | 0.15 | 0.38 | 0.17 | 0.37 |
| 2 | 170675765 | rs3769769 | 0.813 | -0.43 | 0.722 | -0.255 | 0.04 | 0.006 | 0.006 | 0 | 0.53 | 0.4 | 0.71 | 0.45 | 0.68 | 0.5 |
| 2 | 170676227 | rs1541776 | -0.769 | -0.309 | 0.861 | -0.193 | -0.008 | 0.002 | 0.002 | 0 | 0.74 | 0.69 | 0.79 | 0.6 | 0.83 | 0.62 |
| 2 | 170680879 | rs3769767 | -0.776 | 0.203 | 0.724 | -0.306 | 0.01 | 0.001 | 0.001 | 0 | 0.2 | 0.18 | 0.29 | 0.15 | 0.24 | 0.17 |
| 2 | 170733740 | rs6747239 | 0.794 | 0.101 | 0.728 | -0.325 | 0.025 | 0.049 | 0.049 | 1.976 | 0.84 | 0.53 | 0.72 | 0.87 | 0.86 | 0.89 |
| 2 | 170746427 | rs10803844 | 1.15 | 0.057 | 0.269 | -0.389 | -0.046 | -0.049 | -0.049 | 0.119 | 0.46 | 0.12 | 0.46 | 0.45 | 0.53 | 0.39 |
| 2 | 170754854 | rs17554448 | 1.648 | 0.17 | -0.229 | -0.826 | -0.009 | 0.037 | 0.037 | 0.003 | 0.14 | 0.07 | 0.08 | 0 | 0.16 | 0.1 |
| 2 | 170758703 | rs12991583 | 1.031 | -0.072 | -0.078 | -0.693 | -0.075 | -0.058 | -0.058 | 0 | 0.42 | 0.13 | 0.45 | 0.45 | 0.51 | 0.35 |
| 2 | 170771022 | rs1031773 | 1.984 | 0.284 | -0.119 | -0.603 | -0.024 | -0.142 | -0.142 | 0.029 | 0.2 | 0.03 | 0.27 | 0.42 | 0.16 | 0.18 |
| 2 | 170800441 | rs1461961 | 1.058 | 0.147 | 0.294 | -0.549 | -0.012 | 0.005 | 0.005 | 0.031 | 0.24 | 0.28 | 0.2 | 0.38 | 0.16 | 0.37 |
| 2 | 170817760 | rs1017771 | 0.368 | 0.291 | 0.526 | -0.619 | 0.04 | -0.024 | -0.024 | 9.694 | 0.06 | 0.18 | 0.07 | 0.01 | 0.08 | 0.06 |
| 2 | 170819774 | rs10200017 | 0.373 | 0.275 | 0.56 | -0.658 | -0.001 | 0 | 0 | 0.59 | 0.32 | 0.31 | 0.24 | 0.42 | 0.24 | 0.45 |
| 2 | 170835422 | rs7599582 | 0.811 | 0.274 | 0.595 | -0.708 | -0.012 | -0.002 | -0.002 | 0 | 0.49 | 0.2 | 0.46 | 0.45 | 0.53 | 0.39 |
| 2 | 170840224 | rs10184753 | 1.158 | 0.249 | 0.613 | -0.694 | -0.016 | 0.009 | 0.009 | 0.001 | 0.24 | 0.28 | 0.2 | 0.38 | 0.16 | 0.37 |
| 2 | 170858819 | rs13011286 | -0.043 | 0.21 | 0.769 | -0.708 | 0.014 | 0 | 0 | 0 | 0.6 | 0.35 | 0.68 | 0.56 | 0.67 | 0.49 |
| 2 | 170860765 | rs12987772 | 0.902 | 0.221 | 0.763 | -0.715 | -0.021 | -0.013 | -0.013 | 0 | 0.48 | 0.16 | 0.46 | 0.45 | 0.53 | 0.39 |
| 2 | 170860951 | rs4668190 | -0.073 | 0.227 | 0.771 | -0.696 | 0.012 | 0.013 | 0.013 | 0 | 0.12 | 0.21 | 0.23 | 0.11 | 0.14 | 0.11 |
| 2 | 170865808 | rs13389112 | 1.456 | 0.196 | 0.844 | -0.709 | 0.018 | -0.018 | -0.018 | 0.033 | 0.23 | 0.17 | 0.19 | 0.38 | 0.16 | 0.37 |
| 2 | 170866411 | rs6720533 | 0.019 | 0.221 | 0.895 | -0.691 | 0.02 | 0.023 | 0.023 | 0 | 0.87 | 0.74 | 0.78 | 0.89 | 0.86 | 0.89 |
| 2 | 170868888 | rs16857439 | 0.442 | 0.267 | 0.894 | -0.727 | 0.056 | -0.015 | -0.015 | 0.298 | 0.06 | 0.24 | 0.08 | 0.01 | 0.08 | 0.06 |
| 2 | 170882081 | rs10930391 | -0.026 | 0.235 | 0.695 | -0.708 | -0.001 | -0.001 | -0.001 | 0 | 0.37 | 0.35 | 0.43 | 0.39 | 0.36 | 0.54 |
| 2 | 170887864 | rs13430186 | 0.813 | 0.274 | 0.764 | -0.684 | 0.002 | -0.005 | -0.005 | 0 | 0.27 | 0.23 | 0.38 | 0.38 | 0.27 | 0.46 |
| 2 | 170894684 | rs1466400 | 0.564 | 0.345 | 0.617 | -0.674 | 0.004 | 0.032 | 0.032 | 0 | 0.85 | 0.6 | 0.9 | 0.88 | 0.89 | 0.93 |
| 2 | 170895909 | rs10189407 | 1.128 | 0.347 | 0.618 | -0.678 | 0.046 | -0.014 | -0.014 | 0 | 0.81 | 0.56 | 0.71 | 0.88 | 0.78 | 0.84 |
| 2 | 170901298 | rs13408281 | 0.17 | 0.351 | 0.386 | -0.753 | -0.006 | 0.022 | 0.022 | 0 | 0.56 | 0.38 | 0.65 | 0.83 | 0.52 | 0.71 |
| 2 | 170908633 | rs10930393 | 1.281 | 0.371 | 1.055 | -0.045 | -0.03 | -0.048 | -0.048 | 0 | 0.48 | 0.14 | 0.5 | 0.5 | 0.53 | 0.39 |
| 2 | 170910733 | rs7572721 | 0.028 | 0.39 | 1.122 | -0.043 | -0.002 | 0 | 0 | 0.029 | 0.32 | 0.31 | 0.19 | 0.37 | 0.25 | 0.45 |
| 2 | 170914784 | rs12465205 | 1.249 | 0.404 | 1.117 | -0.118 | -0.013 | -0.03 | -0.03 | 0 | 0.48 | 0.2 | 0.5 | 0.47 | 0.53 | 0.38 |
| 2 | 170917648 | rs10203212 | -0.012 | 0.413 | 1.142 | -0.035 | -0.005 | 0.001 | 0.001 | 0 | 0.32 | 0.32 | 0.19 | 0.41 | 0.25 | 0.45 |

| | | | | | | | | | | | | | | | | |
|---|---|---|---|---|---|---|---|---|---|---|---|---|---|---|---|---|
| 2 | 170932522 | rs10204475 | 0.83 | 0.474 | 1.093 | -0.039 | 0.055 | 0.058 | 0.058 | 0.384 | 0.88 | 0.63 | 0.75 | 0.88 | 0.85 | 0.88 |
| 2 | 170945200 | rs12619305 | -0.258 | 0.419 | 0.986 | 0.054 | -0.001 | 0 | 0 | 0 | 0.16 | 0.12 | 0.05 | 0.16 | 0.12 | 0.09 |
| 2 | 170947923 | rs10084272 | 1.022 | 0.442 | 1.011 | 0.052 | 0.003 | -0.004 | -0.004 | 0 | 0.09 | 0.09 | 0.04 | 0.04 | 0.08 | 0.08 |
| 2 | 170950113 | rs9287931 | 1.057 | 0.443 | 0.936 | 0.052 | -0.001 | 0 | 0 | 0 | 0.08 | 0.07 | 0.04 | 0.04 | 0.08 | 0.08 |
| 2 | 170952337 | rs6725973 | 0.217 | 0.467 | 1.017 | 0.078 | 0.004 | 0.091 | 0.091 | 0.395 | 0.9 | 0.68 | 0.94 | 0.88 | 0.94 | 0.95 |
| 2 | 170955247 | rs6738152 | 0.656 | 0.466 | 0.549 | 0.003 | 0.05 | -0.015 | -0.015 | 0 | 0.54 | 0.24 | 0.67 | 0.45 | 0.64 | 0.48 |
| 2 | 170960119 | rs6758094 | -0.758 | 0.617 | 0.884 | 0.074 | -0.007 | 0.002 | 0.002 | 0 | 0.94 | 0.85 | 0.98 | 0.96 | 0.97 | 0.99 |
| 2 | 170964822 | rs13384346 | -0.657 | 0.747 | 0.566 | -0.026 | 0.01 | -0.034 | -0.034 | 0 | 0.18 | 0.24 | 0.25 | 0.05 | 0.2 | 0.18 |
| 2 | 170966684 | rs11903124 | 0.782 | 0.905 | 0.293 | 0.078 | 0 | -0.004 | -0.004 | 0 | 0.71 | 0.66 | 0.71 | 0.82 | 0.72 | 0.79 |
| 2 | 170968227 | rs13395562 | -0.747 | 1.139 | -0.191 | -0.106 | -0.005 | 0.003 | 0.003 | 0 | 0.4 | 0.54 | 0.31 | 0.54 | 0.35 | 0.48 |
| 2 | 170968414 | rs4667617 | 0.506 | 0.767 | -0.289 | -0.013 | -0.004 | -0.002 | -0.002 | 0.011 | 0.56 | 0.34 | 0.49 | 0.46 | 0.54 | 0.45 |
| 2 | 170970994 | rs6731008 | -0.266 | 0.797 | -0.435 | -0.256 | -0.01 | -0.024 | -0.024 | 0 | 0.18 | 0.26 | 0.08 | 0.06 | 0.13 | 0.1 |
| 2 | 170971939 | rs13002765 | -0.626 | 1.313 | -0.895 | 0.119 | -0.027 | 0.036 | 0.036 | 0 | 0.14 | 0.04 | 0.1 | 0.03 | 0.19 | 0.1 |
| 2 | 170978340 | rs10180192 | -0.312 | 0.45 | -0.933 | 0.693 | 0.01 | -0.022 | -0.022 | 0 | 0.34 | 0.3 | 0.52 | 0.56 | 0.31 | 0.53 |
| 2 | 170988776 | rs6711421 | -0.648 | 0.54 | -0.807 | -0.139 | 0.008 | 0.011 | 0.011 | 0 | 0.61 | 0.55 | 0.7 | 0.59 | 0.56 | 0.7 |
| 2 | 170995274 | rs6756989 | -0.013 | 0.707 | -1 | -0.359 | -0.001 | 0.001 | 0.001 | 0 | 0.55 | 0.43 | 0.51 | 0.85 | 0.54 | 0.59 |
| 2 | 170996360 | rs10930404 | -1.034 | 0.85 | -0.9 | -0.38 | -0.014 | 0.015 | 0.015 | 0 | 0.78 | 0.82 | 0.68 | 0.97 | 0.75 | 0.81 |
| 2 | 170997061 | rs4667619 | 0.267 | 0.359 | -1.355 | -0.567 | -0.047 | -0.09 | -0.09 | 0 | 0.24 | 0.06 | 0.23 | 0.31 | 0.28 | 0.23 |
| 2 | 170997821 | rs13414831 | 0.385 | 0.549 | -1.071 | -0.558 | 0.017 | -0.022 | -0.022 | 0 | 0.36 | 0.3 | 0.36 | 0.57 | 0.32 | 0.42 |
| 2 | 170998121 | rs4668205 | 0.403 | 0.524 | -0.96 | -0.57 | 0.019 | -0.032 | -0.032 | 0.274 | 0.34 | 0.26 | 0.35 | 0.56 | 0.32 | 0.42 |
| 2 | 170998431 | rs12619735 | 0.973 | 0.469 | -1.053 | -0.613 | -0.016 | -0.049 | -0.049 | 0 | 0.18 | 0.1 | 0.16 | 0.29 | 0.24 | 0.16 |
| 2 | 170999176 | rs12474899 | -0.261 | 0.426 | -0.961 | -0.702 | -0.002 | 0.091 | 0.091 | 0 | 0.52 | 0.35 | 0.51 | 0.86 | 0.56 | 0.58 |
| 2 | 171000476 | rs13015810 | 1.299 | 0.413 | -1.069 | -0.859 | -0.023 | 0.042 | 0.042 | 0 | 0.21 | 0.11 | 0.11 | 0.02 | 0.22 | 0.18 |
| 2 | 171000799 | rs4667620 | -0.403 | 0.322 | -0.626 | -0.654 | -0.007 | -0.04 | -0.04 | 0 | 0.93 | 0.89 | 0.95 | 1 | 0.93 | 0.98 |
| 2 | 171012115 | rs16857804 | 0.257 | 0.38 | -0.65 | -0.547 | 0.013 | -0.026 | -0.026 | 0.862 | 0.35 | 0.23 | 0.36 | 0.59 | 0.33 | 0.49 |
| 2 | 171016042 | rs2883449 | 0.045 | 0.455 | -0.948 | -0.256 | -0.021 | -0.066 | -0.066 | 0.115 | 0.6 | 0.78 | 0.7 | 0.52 | 0.52 | 0.65 |
| 2 | 171017094 | rs11684893 | 1.047 | 0.617 | -0.499 | -0.219 | 0.005 | 0.018 | 0.018 | 0 | 0.17 | 0.07 | 0.29 | 0 | 0.12 | 0.13 |
| 2 | 171022073 | rs13007861 | 0.136 | 0.502 | 0.467 | 0.23 | -0.016 | 0.152 | 0.152 | 0 | 0.83 | 0.69 | 0.8 | 0.57 | 0.83 | 0.72 |
| 2 | 171024805 | rs13403741 | 0.585 | 0.475 | 0.983 | 0.456 | -0.007 | 0.106 | 0.106 | 0 | 0.41 | 0.24 | 0.23 | 0.11 | 0.39 | 0.31 |
| 2 | 171026184 | rs10930407 | 0.31 | 0.433 | 1.055 | 0.54 | 0.003 | 0.003 | 0.003 | 0 | 0.42 | 0.45 | 0.57 | 0.54 | 0.44 | 0.49 |
| 2 | 171029215 | rs11904398 | 0.71 | 0.511 | 0.8 | 0.419 | 0.001 | 0.035 | 0.035 | 0 | 0.4 | 0.34 | 0.24 | 0.11 | 0.39 | 0.34 |
| 2 | 171030988 | rs7607334 | -0.629 | 0.74 | 0.278 | 0.45 | 0.011 | -0.016 | -0.016 | 0 | 0.95 | 0.9 | 0.96 | 1 | 0.91 | 0.99 |
| 2 | 171032090 | rs4667623 | 0.219 | 0.728 | 0.337 | 0.489 | 0.02 | 0.022 | 0.022 | 0 | 0.93 | 0.83 | 0.95 | 0.88 | 0.91 | 0.89 |
| 2 | 171033501 | rs1035629 | 0.529 | 0.69 | 0.222 | 0.431 | -0.022 | -0.057 | -0.057 | 0 | 0.37 | 0.15 | 0.5 | 0.57 | 0.37 | 0.45 |
| 2 | 171053409 | rs1820293 | 0.345 | 0.62 | 0.446 | 0.735 | 0.005 | -0.016 | -0.016 | 0 | 0.41 | 0.64 | 0.37 | 0.46 | 0.37 | 0.42 |
| 2 | 171056994 | rs6759658 | 0.324 | 0.624 | 0.621 | 0.629 | -0.001 | -0.022 | -0.022 | 0 | 0.39 | 0.52 | 0.24 | 0.15 | 0.37 | 0.35 |
| 2 | 171060729 | rs6738892 | -0.327 | 0.69 | 0.335 | 0.337 | -0.006 | -0.003 | -0.003 | 0 | 0.55 | 0.57 | 0.64 | 0.54 | 0.54 | 0.56 |
| 2 | 171063461 | rs1424650 | 0.517 | 0.726 | 0.249 | 0.333 | 0 | -0.012 | -0.012 | 0 | 0.4 | 0.36 | 0.32 | 0.46 | 0.38 | 0.43 |
| 2 | 171064562 | rs2355343 | 0.695 | 0.952 | 0.104 | 0.333 | -0.003 | -0.031 | -0.031 | 0 | 0.37 | 0.17 | 0.53 | 0.42 | 0.35 | 0.34 |
| 2 | 171072781 | rs6724040 | 0.337 | 0.945 | -0.098 | 0.468 | 0.002 | -0.004 | -0.004 | 0.102 | 0.42 | 0.4 | 0.32 | 0.45 | 0.38 | 0.43 |
| 2 | 171073887 | rs2161916 | -0.196 | 1 | -0.158 | 0.544 | -0.001 | -0.001 | -0.001 | 0 | 0.57 | 0.63 | 0.42 | 0.57 | 0.56 | 0.63 |
| 2 | 171078637 | rs6730459 | -0.077 | 0.453 | -0.454 | -0.465 | 0.002 | -0.016 | -0.016 | 0 | 0.24 | 0.29 | 0.17 | 0 | 0.25 | 0.11 |
| 2 | 171082383 | rs4668224 | -0.089 | 0.007 | -0.647 | 0.152 | 0.01 | -0.043 | -0.043 | 0 | 0.63 | 0.82 | 0.7 | 0.53 | 0.65 | 0.46 |
| 2 | 171083808 | rs13021523 | 0.66 | -0.088 | 0.019 | 0.292 | 0.004 | 0.008 | 0.008 | 0 | 0.53 | 0.47 | 0.65 | 0.36 | 0.6 | 0.43 |
| 2 | 171087015 | rs7607331 | -0.683 | -0.375 | -0.265 | 0.37 | 0.001 | 0.001 | 0.001 | 0 | 0.84 | 0.86 | 0.89 | 0.93 | 0.94 | 0.7 |
| 2 | 171088262 | rs6748642 | 0.453 | -0.196 | -0.378 | 0.394 | 0.002 | 0.041 | 0.041 | 0 | 0.06 | 0.08 | 0.03 | 0.19 | 0.05 | 0.03 |
| 2 | 171091432 | rs7588868 | 0.756 | -0.385 | -0.556 | 0.052 | 0.016 | 0.003 | 0.003 | 0 | 0.52 | 0.42 | 0.64 | 0.55 | 0.6 | 0.43 |
| 2 | 171092865 | rs17578787 | -0.091 | -0.066 | -0.693 | 0.099 | -0.008 | -0.076 | -0.076 | 0.287 | 0.18 | 0.03 | 0.18 | 0.25 | 0.18 | 0.26 |
| 2 | 171096605 | rs7568852 | -0.204 | -0.043 | -0.038 | -0.463 | 0.002 | -0.004 | -0.004 | 0 | 0.64 | 0.64 | 0.73 | 0.72 | 0.59 | 0.66 |
| 2 | 171096766 | rs1424658 | -0.086 | 0.236 | -0.279 | -0.266 | 0.002 | -0.04 | -0.04 | 3.502 | 0.07 | 0.17 | 0.04 | 0 | 0.04 | 0.01 |
| 2 | 171101509 | rs2355351 | 0.024 | 0.287 | -0.919 | -0.163 | 0.004 | 0.077 | 0.077 | 0.374 | 0.5 | 0.31 | 0.66 | 0.72 | 0.51 | 0.61 |
| 2 | 171105318 | rs6745880 | -0.591 | 0.016 | -0.501 | -0.574 | -0.002 | -0.04 | -0.04 | 0 | 0.3 | 0.65 | 0.14 | 0.03 | 0.24 | 0.25 |
| 2 | 171106131 | rs10168221 | 0.291 | -0.016 | -0.48 | -0.184 | -0.048 | 0.124 | 0.124 | 0 | 0.48 | 0.11 | 0.26 | 0.3 | 0.45 | 0.62 |
| 2 | 171111774 | rs1364529 | 0.148 | 0.004 | 0.093 | -0.043 | -0.007 | 0.002 | 0.002 | 0 | 0.35 | 0.3 | 0.62 | 0.66 | 0.42 | 0.22 |
| 2 | 171125118 | rs3914752 | 1.151 | -0.023 | 1.267 | 0.059 | 0.041 | 0.042 | 0.042 | 2.407 | 0.92 | 0.64 | 0.96 | 0.99 | 0.98 | 0.91 |
| 2 | 171129618 | rs908784 | 1.051 | -0.033 | 1.122 | 0.034 | -0.001 | -0.008 | -0.008 | 0.031 | 0.91 | 0.74 | 0.91 | 0.97 | 0.91 | 0.95 |
| 2 | 171130071 | rs925921 | 1.142 | -0.027 | 1.042 | 0.075 | -0.001 | 0.004 | 0.004 | 0.832 | 0.89 | 0.68 | 0.92 | 0.96 | 0.91 | 0.9 |
| 2 | 171131199 | rs10930412 | 1.459 | -0.032 | 0.637 | 0.023 | -0.019 | -0.05 | -0.05 | 0.834 | 0.86 | 0.5 | 0.89 | 0.94 | 0.89 | 0.86 |
| 2 | 171134259 | rs10930413 | -0.426 | 0.066 | -0.418 | 0.661 | 0.007 | -0.034 | -0.034 | 0 | 0.33 | 0.76 | 0.57 | 0.53 | 0.34 | 0.25 |
| 2 | 171135046 | rs1401283 | 0.357 | 0.273 | -0.552 | 0.828 | -0.001 | -0.05 | -0.05 | 0 | 0.64 | 0.18 | 0.64 | 0.42 | 0.67 | 0.64 |
| 2 | 171135562 | rs4668231 | 0.147 | 0.267 | -0.217 | 0.217 | 0.001 | 0.01 | 0.01 | 0 | 0.45 | 0.38 | 0.54 | 0.27 | 0.43 | 0.62 |
| 2 | 171139256 | rs11884023 | -0.249 | -0.073 | 0.046 | -0.213 | -0.001 | -0.002 | -0.002 | 0 | 0.44 | 0.56 | 0.31 | 0.28 | 0.34 | 0.45 |
| 2 | 171139288 | rs10930416 | 0.805 | -0.131 | 0.066 | -0.296 | 0.037 | -0.026 | -0.026 | 0 | 0.91 | 0.75 | 0.72 | 0.97 | 0.88 | 0.84 |
| 2 | 171139494 | rs12611789 | 0.478 | -0.314 | 0.696 | -0.409 | -0.001 | 0.009 | 0.009 | 0 | 0.48 | 0.46 | 0.45 | 0.68 | 0.54 | 0.38 |
| 2 | 171140426 | rs1850685 | 0.582 | -0.342 | 0.652 | -0.527 | 0.016 | 0.01 | 0.01 | 0.046 | 0.94 | 0.79 | 0.97 | 0.97 | 0.96 | 0.9 |
| 2 | 171142750 | rs6722835 | 0.166 | -0.307 | 0.177 | -0.491 | 0.006 | 0.023 | 0.023 | 0 | 0.4 | 0.25 | 0.26 | 0.28 | 0.35 | 0.45 |
| 2 | 171153256 | rs3845739 | -0.249 | -0.29 | 0.243 | -0.468 | 0.022 | 0.031 | 0.031 | 0 | 0.57 | 0.68 | 0.73 | 0.72 | 0.66 | 0.56 |
| 2 | 171155823 | rs1518739 | -1.567 | -0.601 | 0.61 | -0.659 | -0.042 | -0.118 | -0.118 | 0.048 | 0.22 | 0.47 | 0.35 | 0.05 | 0.15 | 0.21 |
| 2 | 171169811 | rs6717173 | -0.982 | -0.187 | 1.459 | -0.539 | 0.13 | -0.004 | -0.004 | 5.29 | 0.07 | 0.34 | 0.3 | 0.01 | 0.11 | 0.14 |

| | | | | | | | | | | | | | | | | |
|---|---|---|---|---|---|---|---|---|---|---|---|---|---|---|---|---|
| 2 | 171169854 | rs16858142 | 0.255 | -0.289 | 0.984 | -0.706 | -0.009 | 0.004 | 0.004 | 0 | 0.32 | 0.2 | 0.24 | 0.27 | 0.34 | 0.43 |
| 2 | 171173261 | rs1568269 | 0.207 | -0.453 | 0.614 | -0.715 | -0.024 | 0.025 | 0.025 | 0.06 | 0.59 | 0.31 | 0.44 | 0.69 | 0.55 | 0.42 |
| 2 | 171173948 | rs11696033 | 0.483 | -0.433 | 0.2 | -0.714 | -0.046 | 0.053 | 0.053 | 0.111 | 0.57 | 0.2 | 0.43 | 0.69 | 0.55 | 0.41 |
| 2 | 171183010 | rs2176592 | 0.067 | -0.529 | 0.365 | -0.551 | -0.016 | 0.006 | 0.006 | 0 | 0.32 | 0.16 | 0.24 | 0.28 | 0.34 | 0.41 |
| 2 | 171192281 | rs6757443 | 0.439 | -0.673 | 0.555 | -0.643 | 0 | 0.001 | 0.001 | 0 | 0.57 | 0.6 | 0.72 | 0.68 | 0.61 | 0.53 |
| 2 | 171204750 | rs7580594 | 0.861 | -0.726 | 0.67 | -0.682 | 0.024 | 0.072 | 0.072 | 0 | 0.52 | 0.3 | 0.68 | 0.67 | 0.58 | 0.47 |
| 2 | 171205088 | rs16858186 | -1.245 | -0.923 | 0.726 | -0.495 | -0.052 | -0.078 | -0.078 | 0 | 0.14 | 0.25 | 0.05 | 0.01 | 0.06 | 0.11 |
| 2 | 171206891 | rs6723795 | -1.513 | -0.565 | 0.708 | -0.286 | 0.061 | 0.086 | 0.086 | 0 | 0.06 | 0.34 | 0.04 | 0.05 | 0.03 | 0.09 |
| 2 | 171208205 | rs12616969 | 0.602 | -0.262 | -0.772 | -0.205 | -0.023 | -0.078 | -0.078 | 2.079 | 0.32 | 0.12 | 0.46 | 0.56 | 0.34 | 0.32 |
| 2 | 171208441 | rs7578592 | -1.328 | -0.622 | 0.1 | 0.053 | -0.044 | -0.01 | -0.01 | 0 | 0.18 | 0.65 | 0.18 | 0.15 | 0.12 | 0.2 |
| 2 | 171209875 | rs6736110 | 0.503 | -0.239 | -0.522 | -0.085 | -0.013 | 0.021 | 0.021 | 0 | 0.2 | 0.06 | 0.13 | 0.14 | 0.19 | 0.21 |
| 2 | 171210531 | rs7559456 | 0.479 | -0.09 | -0.508 | -0.009 | -0.039 | 0.014 | 0.014 | 0 | 0.2 | 0.07 | 0.15 | 0.14 | 0.22 | 0.2 |
| 2 | 171210640 | rs17497248 | 0.84 | -0.052 | -0.502 | 0.062 | -0.014 | 0.003 | 0.003 | 0 | 0.09 | 0.04 | 0.07 | 0.05 | 0.12 | 0.1 |
| 2 | 171210691 | rs4296400 | 1.049 | -0.412 | -0.028 | 0.252 | -0.037 | 0.077 | 0.077 | 2.398 | 0.87 | 0.72 | 0.94 | 0.77 | 0.92 | 0.88 |
| 2 | 171212153 | rs17579090 | -0.247 | -0.368 | 0.023 | 0.263 | -0.012 | 0.072 | 0.072 | 0 | 0.06 | 0.15 | 0.04 | 0.15 | 0.05 | 0.08 |
| 2 | 171218574 | rs7596319 | -0.867 | -0.415 | 0.072 | 0.163 | 0 | -0.011 | -0.011 | 9.864 | 0.41 | 0.47 | 0.28 | 0.26 | 0.34 | 0.39 |
| 2 | 171219147 | rs1356589 | -0.321 | -0.43 | 0.115 | 0.147 | -0.015 | -0.053 | -0.053 | 0.052 | 0.31 | 0.58 | 0.21 | 0.16 | 0.2 | 0.24 |
| 2 | 171219176 | rs17083 | 1.306 | -0.381 | -0.22 | 0.166 | -0.039 | -0.019 | -0.019 | 0 | 0.09 | 0.01 | 0.07 | 0.07 | 0.12 | 0.08 |
| 2 | 171221115 | rs876751 | 0.782 | -0.333 | -0.058 | 0.265 | 0.003 | -0.008 | -0.008 | 0.908 | 0.65 | 0.33 | 0.77 | 0.61 | 0.72 | 0.65 |
| 2 | 171222398 | rs13004652 | 1.538 | -0.178 | -0.615 | 0.29 | -0.1 | -0.119 | -0.119 | 2.75 | 0.46 | 0.05 | 0.63 | 0.53 | 0.55 | 0.39 |
| 2 | 171223111 | rs6761096 | 0.985 | 0.276 | 0.381 | 0.64 | 0.003 | 0.132 | 0.132 | 0.032 | 0.92 | 0.72 | 0.94 | 0.81 | 0.94 | 0.9 |
| 2 | 171223154 | rs17425473 | -0.417 | 0.456 | -0.444 | 0.449 | 0.005 | -0.007 | -0.007 | 0 | 0.23 | 0.25 | 0.17 | 0.17 | 0.22 | 0.21 |
| 2 | 171224155 | rs1401282 | -0.486 | 0.352 | -1.31 | 0.406 | -0.01 | -0.013 | -0.013 | 0 | 0.25 | 0.14 | 0.14 | 0.26 | 0.23 | 0.26 |
| 2 | 171226982 | rs12471833 | -0.19 | 0.08 | -1.29 | -0.084 | 0.018 | 0.138 | 0.138 | 0 | 0.28 | 0.11 | 0.2 | 0.04 | 0.24 | 0.28 |
| 2 | 171229490 | rs16858239 | 0.617 | 0.12 | -0.398 | 0.507 | 0.013 | 0.156 | 0.156 | 1.082 | 0.88 | 0.42 | 0.91 | 0.74 | 0.92 | 0.84 |
| 2 | 171230013 | rs11690118 | 0.112 | 0.327 | -0.058 | 0.652 | 0.004 | 0.003 | 0.003 | 0 | 0.32 | 0.37 | 0.39 | 0.4 | 0.36 | 0.34 |
| 2 | 171230520 | rs11674021 | 0.333 | 0.288 | -0.086 | 0.662 | -0.005 | -0.029 | -0.029 | 0 | 0.3 | 0.22 | 0.37 | 0.42 | 0.33 | 0.29 |
| 2 | 171230560 | rs10176156 | -0.53 | -0.12 | 0.604 | 0.564 | -0.019 | -0.099 | -0.099 | 0.17 | 0.35 | 0.49 | 0.19 | 0.08 | 0.29 | 0.37 |
| 2 | 171231958 | rs17425581 | 0.26 | -0.224 | -0.084 | 0.613 | 0.042 | 0.086 | 0.086 | 0 | 0.27 | 0.14 | 0.12 | 0.05 | 0.2 | 0.28 |
| 2 | 171233990 | rs10207452 | -0.471 | -0.056 | -0.258 | 0.726 | -0.003 | -0.001 | -0.001 | 7.036 | 0.37 | 0.55 | 0.23 | 0.28 | 0.24 | 0.34 |
| 2 | 171236530 | rs10497365 | -0.541 | 0.728 | -0.553 | 1.403 | -0.015 | 0.092 | 0.092 | 0 | 0.05 | 0.04 | 0.17 | 0.46 | 0.08 | 0.14 |
| 2 | 171236596 | rs10497366 | 0.519 | 0.374 | -0.142 | 1.221 | 0.004 | 0.015 | 0.015 | 0.798 | 0.7 | 0.43 | 0.7 | 0.61 | 0.72 | 0.61 |
| 2 | 171236962 | rs6723790 | 0.55 | 0.373 | -0.141 | 1.127 | 0.005 | 0.013 | 0.013 | 0.748 | 0.69 | 0.43 | 0.6 | 0.4 | 0.72 | 0.56 |
| 2 | 171242127 | rs10169408 | -0.354 | 0.311 | 0.119 | 1.066 | -0.002 | -0.104 | -0.104 | 1.087 | 0.82 | 0.62 | 0.84 | 0.95 | 0.88 | 0.86 |
| 2 | 171242761 | rs17497636 | -0.63 | 0.842 | -0.569 | 1.06 | -0.023 | 0.034 | 0.034 | 0 | 0.13 | 0.05 | 0.1 | 0.08 | 0.16 | 0.2 |
| 2 | 171242878 | rs9646782 | 1.282 | 0.889 | -1.015 | 0.729 | -0.041 | -0.042 | -0.042 | 0 | 0.27 | 0.03 | 0.16 | 0.29 | 0.31 | 0.25 |
| 2 | 171243250 | rs17607321 | 0.439 | 0.842 | -1.089 | 0.644 | -0.007 | 0.045 | 0.045 | 0.01 | 0.06 | 0.04 | 0.19 | 0.45 | 0.08 | 0.18 |
| 2 | 171243363 | rs13004172 | 1.302 | 0.82 | -1.054 | 0.61 | -0.018 | -0.019 | -0.019 | 0 | 0.28 | 0.07 | 0.17 | 0.29 | 0.31 | 0.25 |
| 2 | 171244004 | rs1438627 | 0.19 | 0.755 | -0.962 | 0.577 | 0 | 0.014 | 0.014 | 0 | 0.39 | 0.33 | 0.26 | 0.34 | 0.41 | 0.33 |
| 2 | 171246544 | rs12613167 | -0.793 | 0.792 | -0.925 | 0.506 | 0.018 | 0.09 | 0.09 | 0 | 0.54 | 0.38 | 0.58 | 0.85 | 0.62 | 0.7 |
| 2 | 171247392 | rs6723362 | -0.571 | 0.887 | -0.941 | 0.577 | -0.017 | 0.018 | 0.018 | 0 | 0.14 | 0.14 | 0.1 | 0.09 | 0.16 | 0.15 |
| 2 | 171249093 | rs2013102 | 1.112 | 0.817 | -0.911 | 0.626 | -0.004 | -0.003 | -0.003 | 0 | 0.28 | 0.18 | 0.19 | 0.28 | 0.31 | 0.24 |
| 2 | 171252745 | rs908779 | 0.388 | 0.827 | -0.955 | 0.666 | -0.059 | -0.052 | -0.052 | 0.481 | 0.36 | 0.17 | 0.33 | 0.56 | 0.45 | 0.46 |
| 2 | 171256597 | rs7561268 | -0.141 | 0.766 | -0.84 | 0.674 | -0.008 | 0.023 | 0.023 | 0 | 0.45 | 0.34 | 0.34 | 0.62 | 0.54 | 0.54 |
| 2 | 171258964 | rs12623412 | -0.988 | 0.695 | -0.767 | 0.941 | -0.001 | 0.019 | 0.019 | 0 | 0.11 | 0.1 | 0.22 | 0.47 | 0.13 | 0.3 |
| 2 | 171260787 | rs6736609 | 0.809 | 0.685 | -0.217 | 0.897 | 0.013 | 0.151 | 0.151 | 0 | 0.9 | 0.68 | 0.89 | 0.73 | 0.9 | 0.76 |
| 2 | 171261484 | rs1344940 | 0.033 | 0.743 | -0.414 | 0.699 | 0 | 0.003 | 0.003 | 0 | 0.53 | 0.51 | 0.51 | 0.41 | 0.46 | 0.42 |
| 2 | 171262499 | rs10497368 | 0.522 | 0.703 | -0.575 | 0.674 | 0.004 | 0.009 | 0.009 | 0 | 0.46 | 0.54 | 0.4 | 0.16 | 0.4 | 0.27 |
| 2 | 171264727 | rs1370935 | 1.332 | 0.841 | -0.711 | 0.685 | -0.048 | -0.033 | -0.033 | 0 | 0.28 | 0.03 | 0.32 | 0.27 | 0.33 | 0.29 |
| 2 | 171266136 | rs17579806 | -1.384 | 0.848 | -0.55 | 0.817 | -0.012 | -0.013 | -0.013 | 0 | 0.14 | 0.13 | 0.18 | 0.55 | 0.17 | 0.35 |
| 2 | 171268878 | rs10497370 | -1.324 | 0.856 | -0.521 | 0.766 | -0.008 | -0.009 | -0.009 | 0 | 0.14 | 0.17 | 0.14 | 0.3 | 0.17 | 0.29 |
| 2 | 171269301 | rs10497371 | -1.248 | 0.863 | -0.522 | 0.77 | -0.016 | -0.026 | -0.026 | 0 | 0.13 | 0.11 | 0.13 | 0.33 | 0.17 | 0.29 |
| 2 | 171271317 | rs4667638 | -0.215 | 0.833 | -0.458 | 0.736 | 0.003 | -0.023 | -0.023 | 0.14 | 0.9 | 0.71 | 0.92 | 0.94 | 0.91 | 0.93 |
| 2 | 171274757 | rs11884349 | -0.718 | 0.863 | -0.479 | 0.761 | 0 | 0.024 | 0.024 | 0 | 0.08 | 0.1 | 0.08 | 0.27 | 0.11 | 0.19 |
| 2 | 171277794 | rs10930428 | 0.157 | 0.784 | -0.448 | 0.696 | 0.002 | -0.028 | -0.028 | 5.294 | 0.89 | 0.6 | 0.89 | 0.94 | 0.91 | 0.91 |
| 2 | 171278398 | rs10930429 | 0.263 | 0.795 | -0.712 | 0.651 | 0.006 | 0.032 | 0.032 | 0 | 0.49 | 0.57 | 0.63 | 0.36 | 0.44 | 0.35 |
| 2 | 171286303 | rs4668258 | 0.223 | 0.962 | -1.115 | 0.747 | -0.007 | -0.025 | -0.025 | 1.373 | 0.39 | 0.26 | 0.25 | 0.55 | 0.41 | 0.51 |
| 2 | 171287677 | rs12477031 | 1.212 | 0.9 | -1.237 | 0.749 | -0.024 | -0.016 | -0.016 | 0 | 0.3 | 0.03 | 0.25 | 0.27 | 0.31 | 0.31 |
| 2 | 171287710 | rs17426142 | -0.003 | 0.897 | -1.25 | 0.719 | -0.058 | 0.163 | 0.163 | 0 | 0.43 | 0.12 | 0.44 | 0.79 | 0.48 | 0.63 |
| 2 | 171293397 | rs13393912 | -0.709 | 0.955 | -1.193 | 0.705 | 0.026 | -0.004 | -0.004 | 0.005 | 0.08 | 0.11 | 0.11 | 0.09 | 0.13 | 0.2 |
| 2 | 171293484 | rs12620713 | 1.241 | 0.909 | -1.233 | 0.726 | -0.039 | -0.021 | -0.021 | 0.238 | 0.3 | 0.02 | 0.25 | 0.26 | 0.31 | 0.3 |
| 2 | 171295013 | rs10205129 | 0.31 | 0.638 | -1.066 | 0.745 | -0.003 | -0.015 | -0.015 | 0.022 | 0.07 | 0.07 | 0.05 | 0.03 | 0.07 | 0.17 |
| 2 | 171301387 | rs13403603 | 0.128 | 0.627 | -1.06 | 0.768 | 0.001 | 0.058 | 0.058 | 0 | 0.08 | 0.1 | 0.11 | 0.25 | 0.08 | 0.25 |
| 2 | 171302916 | rs17498117 | 1.047 | 0.515 | -1.013 | 0.503 | 0.003 | 0.032 | 0.032 | 0 | 0.31 | 0.22 | 0.39 | 0.1 | 0.31 | 0.16 |
| 2 | 171303727 | rs10170044 | 0.537 | 0.644 | -1.19 | 0.572 | -0.003 | -0.004 | -0.004 | 0 | 0.41 | 0.41 | 0.51 | 0.35 | 0.39 | 0.44 |
| 2 | 171307693 | rs1816670 | 0.123 | 0.551 | -0.547 | 0.709 | -0.014 | 0.003 | 0.003 | 0 | 0.07 | 0.01 | 0.06 | 0.06 | 0.1 | 0.07 |
| 2 | 171308909 | rs1946834 | 0.151 | 0.479 | -0.267 | 0.821 | 0.016 | 0.124 | 0.124 | 0.126 | 0.84 | 0.53 | 0.88 | 0.73 | 0.88 | 0.85 |
| 2 | 171319997 | rs12477518 | 0.097 | 0.216 | -1.502 | -0.027 | 0.057 | 0.177 | 0.177 | 0.48 | 0.53 | 0.21 | 0.43 | 0.16 | 0.61 | 0.43 |
| 2 | 171320948 | rs7559169 | 0.224 | 0 | -2.508 | 0.062 | -0.012 | 0.024 | 0.024 | 0 | 0.57 | 0.2 | 0.56 | 0.37 | 0.62 | 0.51 |

| | | | | | | | | | | | | | | | | |
|---|---|---|---|---|---|---|---|---|---|---|---|---|---|---|---|---|
| 2 | 171323416 | rs2164963 | 0.788 | -0.063 | -2.647 | 0.154 | -0.08 | 0 | 0 | 2.515 | 0.28 | 0.03 | 0.35 | 0.21 | 0.33 | 0.24 |
| 2 | 171325374 | rs12616150 | 0.839 | -0.44 | -1.528 | 0.563 | -0.01 | -0.023 | -0.023 | 0.302 | 0.32 | 0.37 | 0.21 | 0.23 | 0.29 | 0.3 |
| 2 | 171325695 | rs13005411 | 0.869 | -0.582 | -2.04 | 0.246 | -0.094 | 0.126 | 0.126 | 0.38 | 0.4 | 0.04 | 0.48 | 0.18 | 0.47 | 0.27 |
| 2 | 171326321 | rs2883655 | 0.481 | -1.197 | -2.377 | 0.09 | -0.096 | 0.105 | 0.105 | 0 | 0.36 | 0.06 | 0.45 | 0.19 | 0.41 | 0.3 |
| 2 | 171330363 | rs9287939 | -0.658 | -1.233 | -1.69 | 0.439 | 0 | 0.001 | 0.001 | 0 | 0.63 | 0.62 | 0.46 | 0.45 | 0.62 | 0.68 |
| 2 | 171340688 | rs934861 | 0.19 | -1.335 | -1.315 | 0.449 | -0.006 | -0.01 | -0.01 | 0 | 0.4 | 0.6 | 0.38 | 0.53 | 0.43 | 0.43 |
| 2 | 171343360 | rs7583952 | 2.129 | -1.518 | -1.291 | 0.28 | 0.009 | 0.024 | 0.024 | 1.699 | 0.14 | 0.13 | 0.11 | 0.02 | 0.14 | 0.06 |
| 2 | 171344289 | rs4668261 | 1.805 | -1.627 | -1.369 | 0.645 | 0.016 | -0.02 | -0.02 | 0 | 0.1 | 0.02 | 0.25 | 0.4 | 0.09 | 0.1 |
| 2 | 171345022 | rs12692953 | -0.013 | -2.001 | -1.164 | 0.302 | -0.017 | 0.034 | 0.034 | 2.258 | 0.48 | 0.21 | 0.55 | 0.62 | 0.51 | 0.49 |
| 2 | 171345197 | rs7578981 | -1.841 | -1.898 | -1.341 | 0.092 | -0.03 | 0.02 | 0.02 | 0.519 | 0.83 | 0.93 | 0.78 | 0.85 | 0.76 | 0.81 |
| 2 | 171345277 | rs10189587 | 0.756 | -1.927 | -1.007 | 0.151 | -0.051 | -0.123 | -0.123 | 0.066 | 0.22 | 0.51 | 0.15 | 0.02 | 0.15 | 0.1 |
| 2 | 171345915 | rs17498333 | 1.961 | -1.616 | -1.198 | 0.166 | -0.005 | 0.05 | 0.05 | 2.679 | 0.17 | 0.01 | 0.09 | 0.09 | 0.14 | 0.24 |
| 2 | 171348751 | rs11895735 | -0.53 | -1.108 | -0.738 | 0.345 | -0.006 | 0.015 | 0.015 | 0 | 0.64 | 0.73 | 0.7 | 0.72 | 0.63 | 0.62 |
| 2 | 171350680 | rs6716011 | 1.181 | -0.939 | 0.43 | 0.513 | 0.013 | 0.027 | 0.027 | 0 | 0.18 | 0.2 | 0.27 | 0.44 | 0.28 | 0.15 |
| 2 | 171351706 | rs4668263 | 1.676 | -1.026 | 0.394 | 0.415 | 0.006 | 0.12 | 0.12 | 0.177 | 0.26 | 0.02 | 0.14 | 0.09 | 0.23 | 0.25 |
| 2 | 171358785 | rs7596971 | -0.41 | -0.819 | 0.661 | 0.739 | 0.007 | 0.02 | 0.02 | 0 | 0.49 | 0.4 | 0.51 | 0.38 | 0.46 | 0.58 |
| 2 | 171360915 | rs4668268 | -1.23 | -0.382 | 0.45 | 1.038 | 0.06 | 0.09 | 0.09 | 0 | 0.84 | 0.73 | 0.7 | 0.56 | 0.72 | 0.91 |
| 2 | 171365134 | rs934858 | -0.26 | -0.46 | -0.032 | 0.274 | -0.031 | 0.046 | 0.046 | 0 | 0.27 | 0.53 | 0.16 | 0.31 | 0.2 | 0.32 |
| 2 | 171373446 | rs9967753 | 0.184 | -0.691 | -0.319 | 0.447 | -0.014 | -0.006 | -0.006 | 0 | 0.12 | 0.17 | 0.22 | 0.06 | 0.09 | 0.13 |
| 2 | 171373483 | rs13390244 | 0.928 | -0.711 | -0.522 | 0.364 | -0.017 | -0.048 | -0.048 | 0 | 0.2 | 0.13 | 0.27 | 0.4 | 0.21 | 0.19 |
| 2 | 171374662 | rs17271301 | 0.398 | -0.877 | -0.307 | -0.126 | -0.005 | -0.003 | -0.003 | 1.178 | 0.12 | 0.13 | 0.22 | 0.06 | 0.09 | 0.13 |
| 2 | 171375497 | rs10189774 | -1.572 | -0.743 | -0.903 | -0.372 | -0.028 | 0.129 | 0.129 | 0 | 0.72 | 0.97 | 0.84 | 0.89 | 0.7 | 0.75 |
| 2 | 171387595 | rs11686211 | -0.121 | -0.423 | -0.783 | -0.201 | 0.042 | 0.166 | 0.166 | 0 | 0.46 | 0.26 | 0.45 | 0.13 | 0.59 | 0.42 |
| 2 | 171395521 | rs6752204 | 0.457 | -0.468 | 0.093 | 0.808 | -0.066 | 0.038 | 0.038 | 0 | 0.82 | 0.49 | 0.74 | 0.86 | 0.91 | 0.86 |
| 2 | 171395578 | rs12477054 | 0.208 | -0.376 | -0.307 | 0.724 | -0.014 | 0.024 | 0.024 | 0 | 0.23 | 0.09 | 0.21 | 0.12 | 0.21 | 0.27 |
| 2 | 171402733 | rs724806 | 0.756 | 0.041 | -0.119 | 1.078 | 0.041 | 0.071 | 0.071 | 0 | 0.56 | 0.28 | 0.45 | 0.29 | 0.7 | 0.54 |
| 2 | 171405956 | rs13419943 | -0.385 | 0.097 | -0.068 | 1.163 | 0.071 | 0.276 | 0.276 | 3.992 | 0.05 | 0.42 | 0.08 | 0.1 | 0 | 0.08 |
| 2 | 171409269 | rs6730101 | -0.384 | 0.088 | -0.667 | 1.181 | 0.02 | -0.087 | -0.087 | 1.021 | 0.7 | 0.9 | 0.73 | 0.47 | 0.75 | 0.7 |
| 2 | 171417536 | rs6712261 | 0.076 | 0.163 | -0.571 | 1.095 | 0.037 | 0.208 | 0.208 | 0.972 | 0.07 | 0.39 | 0.09 | 0.11 | 0.04 | 0.1 |
| 2 | 171418907 | rs2192754 | -0.568 | 0.145 | -0.661 | 1.08 | 0.02 | -0.121 | -0.121 | 0.121 | 0.72 | 0.94 | 0.74 | 0.5 | 0.8 | 0.71 |
| 2 | 171421606 | rs6735844 | 0.81 | 0.703 | -0.563 | 1.136 | -0.045 | 0.031 | 0.031 | 0 | 0.13 | 0.05 | 0.17 | 0.08 | 0.25 | 0.06 |
| 2 | 171422060 | rs1012078 | -1.369 | -0.266 | 0.061 | 1.687 | 0.005 | 0.174 | 0.174 | 0.005 | 0.12 | 0.39 | 0.26 | 0.17 | 0.09 | 0.13 |
| 2 | 171422330 | rs1012077 | -1.285 | -0.274 | -0.582 | 1.52 | -0.019 | 0.238 | 0.238 | 0.114 | 0.14 | 0.53 | 0.29 | 0.26 | 0.09 | 0.19 |
| 2 | 171424479 | rs2355826 | 0.349 | -0.069 | -1.585 | 0.744 | 0.05 | 0.056 | 0.056 | 0 | 0.61 | 0.17 | 0.59 | 0.29 | 0.73 | 0.53 |
| 2 | 171429351 | rs16858767 | -1.157 | 0.104 | -1.245 | 1.369 | 0 | 0.002 | 0.002 | 0 | 0.07 | 0.04 | 0.07 | 0.26 | 0.04 | 0.18 |
| 2 | 171433825 | rs2883782 | -0.464 | 0.381 | -1.32 | 1.424 | -0.009 | -0.004 | -0.004 | 0 | 0.36 | 0.31 | 0.52 | 0.64 | 0.48 | 0.42 |
| 2 | 171435002 | rs6741612 | -0.066 | 0.355 | -1.207 | 1.295 | -0.009 | -0.006 | -0.006 | 6.826 | 0.31 | 0.26 | 0.49 | 0.44 | 0.48 | 0.27 |
| 2 | 171435131 | rs17279667 | 0.504 | 0.144 | -1.536 | 0.331 | -0.001 | -0.006 | -0.006 | 0 | 0.13 | 0.1 | 0.14 | 0.05 | 0.15 | 0.07 |
| 2 | 171438522 | rs17279771 | 0.18 | -0.338 | -1.377 | 0.39 | -0.021 | 0.161 | 0.161 | 2.288 | 0.4 | 0.13 | 0.45 | 0.01 | 0.59 | 0.31 |
| 2 | 171438669 | rs10490079 | -0.02 | -1.095 | -0.87 | 0.75 | 0.011 | 0.01 | 0.01 | 0 | 0.19 | 0.13 | 0.11 | 0.11 | 0.12 | 0.13 |
| 2 | 171440524 | rs972547 | -1.027 | -1.732 | -0.275 | 0.623 | -0.027 | -0.015 | -0.015 | 0 | 0.4 | 0.53 | 0.27 | 0.74 | 0.2 | 0.38 |
| 2 | 171440957 | rs741283 | -1.132 | -1.597 | 0.413 | 1.278 | 0 | 0.019 | 0.019 | 0 | 0.19 | 0.25 | 0.12 | 0.35 | 0.16 | 0.19 |
| 2 | 171441202 | rs735436 | -2.29 | -1.774 | 1.487 | 1.822 | -0.025 | 0.301 | 0.301 | 0.928 | 0.11 | 0.37 | 0.17 | 0.49 | 0.04 | 0.14 |
| 2 | 171442782 | rs4668285 | -1.667 | -2.104 | 1.301 | 1.214 | 0.008 | -0.004 | -0.004 | 0 | 0.44 | 0.59 | 0.26 | 0.5 | 0.22 | 0.4 |
| 2 | 171450161 | rs13425835 | -0.906 | -2.058 | 1.455 | 1.181 | -0.054 | -0.039 | -0.039 | 0 | 0.32 | 0.58 | 0.22 | 0.18 | 0.13 | 0.35 |
| 2 | 171450353 | rs4500910 | -1.344 | -2.068 | 1.42 | 1.187 | -0.036 | -0.03 | -0.03 | 0 | 0.36 | 0.6 | 0.24 | 0.19 | 0.15 | 0.36 |
| 2 | 171452334 | rs6734646 | -1.045 | -1.869 | 1.064 | 1.033 | -0.02 | 0.007 | 0.007 | 0 | 0.41 | 0.51 | 0.24 | 0.51 | 0.22 | 0.4 |
| 2 | 171458370 | rs6743969 | -0.678 | -1.126 | 1.449 | 0.556 | -0.051 | -0.146 | -0.146 | 0 | 0.2 | 0.45 | 0.11 | 0.01 | 0.09 | 0.13 |
| 2 | 171461132 | rs6749331 | -1.123 | -1.06 | 0.583 | 0.625 | -0.018 | -0.039 | -0.039 | 0 | 0.26 | 0.63 | 0.13 | 0.05 | 0.12 | 0.18 |
| 2 | 171461637 | rs6433223 | -1.295 | -0.948 | 0.428 | 0.629 | -0.006 | -0.029 | -0.029 | 0 | 0.28 | 0.64 | 0.16 | 0.05 | 0.15 | 0.19 |
| 2 | 171465659 | rs6708842 | -0.567 | 0.023 | 0.41 | 1.424 | -0.012 | 0.117 | 0.117 | 0 | 0.18 | 0.39 | 0.15 | 0.42 | 0.11 | 0.25 |
| 2 | 171469234 | rs6724616 | -1.21 | 0.555 | -0.373 | 1.268 | -0.012 | -0.038 | -0.038 | 0.668 | 0.29 | 0.78 | 0.38 | 0.52 | 0.29 | 0.35 |
| 2 | 171476133 | rs10191600 | 0.513 | 0.036 | -0.656 | 0.926 | 0.046 | 0.031 | 0.031 | 0 | 0.54 | 0.28 | 0.54 | 0.3 | 0.66 | 0.56 |
| 2 | 171485774 | rs724529 | 0.182 | 0.417 | -1.076 | 0.674 | -0.027 | 0.025 | 0.025 | 0 | 0.69 | 0.8 | 0.8 | 0.7 | 0.64 | 0.76 |
| 2 | 171492175 | rs12473121 | -0.293 | 0.103 | -0.739 | 0.683 | 0.016 | -0.037 | -0.037 | 0 | 0.19 | 0.08 | 0.19 | 0.3 | 0.14 | 0.2 |
| 2 | 171499887 | rs1861288 | -1.348 | -0.306 | -0.755 | 0.33 | 0.042 | 0.016 | 0.016 | 2.605 | 0.39 | 0.71 | 0.34 | 0.36 | 0.25 | 0.24 |
| 2 | 171500167 | rs10803851 | 0.236 | -0.466 | -0.649 | 0.34 | 0.005 | -0.026 | -0.026 | 0 | 0.17 | 0.08 | 0.15 | 0.22 | 0.16 | 0.14 |
| 2 | 171500624 | rs1990606 | -1.596 | 0.206 | -0.739 | 0.447 | 0.005 | 0.01 | 0.01 | 1.992 | 0.39 | 0.76 | 0.49 | 0.38 | 0.38 | 0.29 |
| 2 | 171508262 | rs6724627 | -1.269 | 0.289 | -0.49 | 0.374 | 0 | 0.003 | 0.003 | 2.191 | 0.33 | 0.69 | 0.41 | 0.29 | 0.32 | 0.25 |
| 2 | 171511791 | rs7569594 | 0.241 | 0.111 | -0.751 | 0.412 | 0.008 | -0.004 | -0.004 | 0 | 0.16 | 0.07 | 0.12 | 0.15 | 0.15 | 0.16 |
| 2 | 171522950 | rs7425766 | -0.378 | -0.369 | -0.986 | 0.102 | 0.006 | 0.006 | 0.006 | 0 | 0.25 | 0.18 | 0.17 | 0.19 | 0.16 | 0.28 |
| 2 | 171524901 | rs1157200 | -0.113 | -0.284 | -1.45 | -0.077 | 0.011 | 0.029 | 0.029 | 0.998 | 0.61 | 0.29 | 0.7 | 0.78 | 0.67 | 0.63 |
| 2 | 171526128 | rs2883785 | 0.3 | -0.189 | -1.582 | 0.041 | 0.01 | 0.021 | 0.021 | 0 | 0.18 | 0.07 | 0.11 | 0.13 | 0.14 | 0.2 |
| 2 | 171529969 | rs12619910 | 0.331 | 0.32 | -1.814 | 0.269 | -0.075 | -0.02 | -0.02 | 0.142 | 0.38 | 0.1 | 0.4 | 0.58 | 0.49 | 0.41 |
| 2 | 171534221 | rs10192049 | 0.161 | -0.126 | -1.38 | 0.518 | -0.008 | 0.002 | 0.002 | 0 | 0.68 | 0.66 | 0.59 | 0.73 | 0.72 | 0.58 |
| 2 | 171539261 | rs1035150 | -0.364 | -0.46 | -0.656 | 0.32 | -0.009 | 0.026 | 0.026 | 0.018 | 0.3 | 0.42 | 0.5 | 0.37 | 0.32 | 0.34 |
| 2 | 171540823 | rs2080401 | -0.765 | -0.382 | -0.19 | 0.056 | -0.005 | 0.001 | 0.001 | 0 | 0.36 | 0.38 | 0.57 | 0.28 | 0.34 | 0.4 |
| 2 | 171548493 | rs7574074 | -0.387 | -0.3 | -0.136 | 0.163 | -0.004 | 0 | 0 | 0 | 0.31 | 0.29 | 0.51 | 0.22 | 0.37 | 0.35 |
| 2 | 171554956 | rs6730170 | -1.493 | -0.44 | 0.535 | 0.07 | 0.038 | 0.058 | 0.058 | 5.396 | 0.11 | 0.44 | 0.21 | 0.14 | 0.06 | 0.07 |

| | | | | | | | | | | | | | | | | |
|---|---|---|---|---|---|---|---|---|---|---|---|---|---|---|---|---|
| 2 | 171557049 | rs2270336 | -0.732 | 0.101 | -0.298 | -0.319 | 0 | -0.007 | -0.007 | 0.793 | 0.37 | 0.58 | 0.46 | 0.28 | 0.36 | 0.32 |
| 2 | 171564929 | rs11694061 | 0.356 | 0.605 | -0.998 | -0.072 | -0.036 | 0.017 | 0.017 | 0 | 0.33 | 0.2 | 0.3 | 0.26 | 0.44 | 0.26 |
| 2 | 171565049 | rs1861465 | -0.143 | 0.011 | -0.757 | 0.065 | -0.04 | -0.019 | -0.019 | 0 | 0.67 | 0.52 | 0.52 | 0.8 | 0.73 | 0.56 |
| 2 | 171566925 | rs11680066 | 1.097 | 0.452 | -1.278 | 0.152 | -0.044 | -0.002 | -0.002 | 0 | 0.29 | 0.08 | 0.24 | 0.26 | 0.37 | 0.25 |
| 2 | 171566968 | rs10201752 | -0.506 | -0.046 | -0.585 | 0.181 | 0.005 | -0.005 | -0.005 | 0.042 | 0.77 | 0.8 | 0.7 | 0.8 | 0.77 | 0.65 |
| 2 | 171571401 | rs6723611 | -0.229 | -0.229 | -0.637 | 0.222 | 0.017 | -0.022 | -0.022 | 0.052 | 0.34 | 0.71 | 0.47 | 0.6 | 0.3 | 0.31 |
| 2 | 171576422 | rs7580460 | 0.222 | -0.282 | -0.15 | 0.2 | -0.016 | 0.051 | 0.051 | 0 | 0.31 | 0.44 | 0.32 | 0.6 | 0.29 | 0.28 |
| 2 | 171581341 | rs10930437 | -1.56 | 0.007 | -0.683 | 0.083 | 0.003 | 0.071 | 0.071 | 0 | 0.6 | 0.76 | 0.67 | 0.78 | 0.66 | 0.64 |
| 2 | 171606871 | rs10930438 | 0.191 | -0.495 | -0.507 | 0.127 | -0.002 | 0.001 | 0.001 | 0 | 0.34 | 0.58 | 0.33 | 0.6 | 0.3 | 0.29 |
| 2 | 171630631 | rs1420378 | 0.396 | -0.397 | -0.656 | 0.11 | -0.052 | -0.011 | -0.011 | 0 | 0.25 | 0.1 | 0.21 | 0.18 | 0.32 | 0.34 |
| 2 | 171635765 | rs1362490 | 0.112 | -0.413 | -0.601 | 0.158 | 0.012 | -0.015 | -0.015 | 0 | 0.35 | 0.66 | 0.46 | 0.59 | 0.3 | 0.3 |
| 2 | 171643177 | rs953680 | 0.55 | -0.402 | -0.612 | 0.166 | -0.017 | 0.04 | 0.04 | 0 | 0.32 | 0.41 | 0.31 | 0.59 | 0.29 | 0.28 |
| 2 | 171649402 | rs3764916 | 0.702 | -0.433 | -1.111 | 0.163 | -0.015 | -0.122 | -0.122 | 3.764 | 0.3 | 0.07 | 0.28 | 0.59 | 0.29 | 0.28 |
| 2 | 171652365 | rs7568115 | 0.076 | -0.494 | -1.363 | 0.073 | -0.001 | -0.002 | -0.002 | 0.112 | 0.7 | 0.64 | 0.64 | 0.81 | 0.67 | 0.65 |
| 2 | 171655723 | rs10181282 | 0.425 | -0.448 | -1.11 | 0.234 | -0.01 | 0.02 | 0.02 | 0 | 0.32 | 0.53 | 0.32 | 0.59 | 0.29 | 0.28 |
| 2 | 171662339 | rs6736129 | 0.541 | -0.479 | -1.274 | 0.228 | -0.027 | 0.023 | 0.023 | 0.641 | 0.3 | 0.59 | 0.27 | 0.6 | 0.22 | 0.27 |
| 2 | 171665709 | rs10803852 | -0.096 | -0.384 | -1.069 | 0.188 | -0.004 | -0.064 | -0.064 | 0 | 0.89 | 0.66 | 0.79 | 1 | 0.95 | 0.95 |
| 2 | 171666997 | rs6755814 | 0.697 | -0.319 | -1.201 | 0.153 | 0 | -0.003 | -0.003 | 0 | 0.66 | 0.63 | 0.59 | 0.81 | 0.63 | 0.62 |
| 2 | 171672191 | rs3791878 | -0.236 | -0.422 | -1.887 | 0.455 | -0.096 | -0.044 | -0.044 | 0.038 | 0.26 | 0.02 | 0.2 | 0.22 | 0.33 | 0.32 |
| 2 | 171673475 | rs3749034 | 0.873 | -0.763 | -1.648 | 1.149 | -0.002 | -0.053 | -0.053 | 0 | 0.23 | 0.01 | 0.2 | 0.27 | 0.21 | 0.24 |
| 2 | 171678379 | rs2241165 | -0.051 | -0.789 | -1.007 | 0.966 | 0.022 | 0.015 | 0.015 | 1.145 | 0.67 | 0.27 | 0.71 | 0.73 | 0.77 | 0.76 |
| 2 | 171682740 | rs3828275 | 1.304 | -0.394 | -1.161 | 0.867 | -0.044 | 0.1 | 0.1 | 0.165 | 0.41 | 0.17 | 0.39 | 0.21 | 0.45 | 0.34 |
| 2 | 171686559 | rs2241164 | 1.059 | -0.514 | -0.731 | 0.951 | -0.002 | 0.002 | 0.002 | 0.043 | 0.67 | 0.63 | 0.65 | 0.46 | 0.68 | 0.47 |
| 2 | 171702480 | rs701492 | -0.717 | -0.265 | -0.597 | 0.786 | 0 | 0 | 0 | 0.125 | 0.24 | 0.25 | 0.29 | 0.27 | 0.29 | 0.39 |
| 2 | 171702960 | rs16858988 | 0.026 | -0.137 | -1.251 | 0.832 | -0.04 | -0.048 | -0.048 | 0.976 | 0.17 | 0.05 | 0.11 | 0.24 | 0.23 | 0.35 |
| 2 | 171708100 | rs3791850 | 0.878 | -0.109 | -0.969 | 0.64 | 0.007 | 0.031 | 0.031 | 0.335 | 0.22 | 0.12 | 0.22 | 0.06 | 0.22 | 0.11 |
| 2 | 171716803 | rs769395 | 0.453 | -0.075 | -1.101 | 1.08 | -0.011 | -0.051 | -0.051 | 0 | 0.75 | 0.86 | 0.82 | 0.7 | 0.72 | 0.64 |
| 2 | 171719539 | rs4297845 | 0.297 | -0.061 | -0.241 | 0.573 | 0.008 | 0.014 | 0.014 | 0 | 0.53 | 0.35 | 0.5 | 0.38 | 0.53 | 0.49 |
| 2 | 171732267 | rs7602737 | 0.907 | 0.157 | 1.16 | 1.335 | 0.02 | 0.246 | 0.246 | 1.047 | 0.95 | 0.72 | 0.95 | 0.73 | 0.98 | 0.97 |
| 2 | 171745442 | rs4233833 | 0.475 | 0.252 | 0.649 | 0.044 | 0.001 | -0.048 | -0.048 | 0.006 | 0.58 | 0.56 | 0.71 | 0.96 | 0.52 | 0.8 |
| 2 | 171745475 | rs2356238 | 0.914 | 0.191 | 0.687 | 0.073 | -0.013 | 0.057 | 0.057 | 3.629 | 0.52 | 0.19 | 0.62 | 0.66 | 0.51 | 0.63 |
| 2 | 171774195 | rs4668343 | 0.881 | 0.205 | 0.947 | 0.312 | -0.006 | -0.015 | -0.015 | 0 | 0.62 | 0.18 | 0.62 | 0.65 | 0.66 | 0.69 |
| 2 | 171784896 | rs16859050 | 1.027 | 0.131 | 0.984 | 0.268 | 0.003 | -0.007 | -0.007 | 0 | 0.61 | 0.19 | 0.63 | 0.65 | 0.66 | 0.68 |
| 2 | 171831197 | rs2059723 | 2.04 | 0.123 | 1.016 | 0.253 | -0.013 | 0.028 | 0.028 | 7.231 | 0.84 | 0.22 | 0.76 | 0.7 | 0.92 | 0.8 |
| 2 | 171833910 | rs2723253 | 1.415 | 0.138 | 1.199 | 0.164 | 0.047 | -0.045 | -0.045 | 0 | 0.9 | 0.53 | 0.9 | 1 | 0.94 | 0.98 |
| 2 | 171849612 | rs1045327 | 0.143 | 0.217 | 1.027 | 0.163 | -0.052 | 0.082 | 0.082 | 0.452 | 0.23 | 0.06 | 0.16 | 0.05 | 0.28 | 0.12 |
| 2 | 171876212 | rs2676151 | -0.615 | 0.212 | 1.151 | 0.159 | 0.025 | -0.096 | -0.096 | 2.585 | 0.29 | 0.54 | 0.25 | 0.05 | 0.35 | 0.13 |
| 2 | 171893117 | rs754058 | 0.874 | 0.259 | 1.083 | 0.176 | -0.019 | -0.021 | -0.021 | 1.307 | 0.65 | 0.16 | 0.61 | 0.67 | 0.65 | 0.71 |
| 2 | 171897442 | rs10930452 | 1.164 | 0.197 | 1.15 | 0.21 | -0.02 | -0.028 | -0.028 | 0 | 0.93 | 0.7 | 0.96 | 1 | 1 | 0.99 |
| 2 | 171899813 | rs7584937 | 1.164 | 0.201 | 1.048 | 0.211 | -0.021 | -0.029 | -0.029 | 0.045 | 0.93 | 0.7 | 0.96 | 1 | 1 | 0.99 |
| 2 | 171912166 | rs13003218 | 0.343 | 0.188 | 0.979 | 0.195 | -0.065 | 0.088 | 0.088 | 0.975 | 0.23 | 0.02 | 0.15 | 0.04 | 0.28 | 0.12 |
| 2 | 171920785 | rs1035828 | 0.697 | 0.165 | 1.041 | 0.19 | -0.006 | 0.027 | 0.027 | 1.364 | 0.64 | 0.27 | 0.71 | 0.95 | 0.63 | 0.86 |
| 2 | 171934234 | rs7606822 | -0.316 | 0.086 | 0.888 | 0.203 | 0.03 | -0.081 | -0.081 | 0 | 0.28 | 0.43 | 0.25 | 0.05 | 0.36 | 0.13 |
| 2 | 171940975 | rs930035 | -1.367 | 0.086 | 0.819 | 0.243 | -0.01 | -0.01 | -0.01 | 0 | 0.38 | 0.8 | 0.39 | 0.34 | 0.37 | 0.31 |
| 2 | 171941018 | rs930036 | -1.33 | 0.079 | 0.773 | 0.214 | -0.005 | -0.005 | -0.005 | 0 | 0.38 | 0.77 | 0.38 | 0.34 | 0.37 | 0.31 |
| 2 | 171943820 | rs13016705 | -0.112 | 0.186 | 0.626 | 0.054 | -0.091 | 0.122 | 0.122 | 0.37 | 0.28 | 0.02 | 0.17 | 0.04 | 0.34 | 0.15 |
| 2 | 172034292 | rs1989613 | -1.473 | 0.191 | 0.478 | 0.667 | 0.021 | 0.303 | 0.303 | 4.424 | 0.1 | 0.5 | 0.18 | 0.29 | 0.06 | 0.22 |
| 2 | 172034918 | rs7582822 | -1.173 | 0.237 | 0.386 | 0.587 | -0.008 | -0.008 | -0.008 | 0 | 0.37 | 0.78 | 0.39 | 0.34 | 0.37 | 0.34 |
| 2 | 172043803 | rs16859206 | -1.715 | 0.671 | 0.547 | 1.038 | 0.049 | 0.311 | 0.311 | 4.332 | 0.1 | 0.5 | 0.2 | 0.29 | 0.08 | 0.22 |
| 2 | 172056351 | rs218307 | 0.687 | 0.616 | -0.801 | 0.775 | -0.026 | 0.06 | 0.06 | 0.757 | 0.56 | 0.2 | 0.57 | 0.72 | 0.54 | 0.7 |
| 2 | 172057864 | rs218311 | 0.188 | 0.563 | -0.543 | 1.023 | -0.018 | -0.012 | -0.012 | 0 | 0.67 | 0.22 | 0.52 | 0.68 | 0.66 | 0.69 |
| 2 | 172066068 | rs7568175 | -0.709 | 0.136 | 0.085 | 1.391 | -0.002 | -0.014 | -0.014 | 0 | 0.22 | 0.2 | 0.23 | 0.07 | 0.18 | 0.33 |
| 2 | 172069140 | rs1035250 | -0.956 | 0.177 | 0.292 | 2.076 | -0.025 | 0.071 | 0.071 | 0.254 | 0.27 | 0.47 | 0.36 | 0.53 | 0.21 | 0.45 |
| 2 | 172077806 | rs11674962 | 0.695 | 0.233 | 0.205 | 1.469 | -0.023 | 0.018 | 0.018 | 0.05 | 0.72 | 0.52 | 0.65 | 0.68 | 0.79 | 0.57 |
| 2 | 172092298 | rs12474993 | -0.097 | -0.401 | -0.343 | 1.366 | 0.026 | 0.037 | 0.037 | 0 | 0.31 | 0.18 | 0.25 | 0.14 | 0.26 | 0.29 |
| 2 | 172098069 | rs17283817 | 0.424 | -0.232 | -1.187 | 1.728 | -0.028 | -0.078 | -0.078 | 0 | 0.17 | 0.02 | 0.35 | 0.25 | 0.17 | 0.17 |
| 2 | 172104703 | rs10211242 | -0.002 | 0.543 | -0.637 | 1.614 | 0.017 | -0.004 | -0.004 | 0.861 | 0.12 | 0.2 | 0.11 | 0.1 | 0.15 | 0.23 |
| 2 | 172105380 | rs4667681 | -0.316 | 0.663 | -0.082 | 1.678 | 0.012 | 0.009 | 0.009 | 0 | 0.49 | 0.67 | 0.47 | 0.6 | 0.46 | 0.66 |
| 2 | 172107822 | rs7566096 | -0.188 | 0.948 | -0.275 | 1.098 | -0.01 | 0.016 | 0.016 | 0 | 0.66 | 0.8 | 0.57 | 0.75 | 0.65 | 0.76 |
| 2 | 172127920 | rs4667682 | 0.985 | 0.792 | -0.453 | 1.059 | 0.029 | -0.102 | -0.102 | 0.477 | 0.21 | 0.07 | 0.34 | 0.4 | 0.16 | 0.17 |
| 2 | 172129015 | rs13007471 | 1.28 | 0.883 | -0.272 | 0.604 | 0.027 | 0.036 | 0.036 | 0 | 0.32 | 0.24 | 0.25 | 0.11 | 0.25 | 0.26 |
| 2 | 172131682 | rs10930464 | 0.516 | 0.861 | -0.065 | 0.541 | 0.022 | 0.074 | 0.074 | 0.231 | 0.48 | 0.36 | 0.48 | 0.11 | 0.47 | 0.38 |
| 2 | 172142488 | rs10930467 | 0.339 | 1.08 | 0.205 | 0.733 | 0.003 | 0.015 | 0.015 | 0.038 | 0.52 | 0.47 | 0.54 | 0.2 | 0.58 | 0.49 |
| 2 | 172142609 | rs17218608 | -1.108 | 1.191 | 0.474 | 0.919 | 0.08 | 0.046 | 0.046 | 0.986 | 0.12 | 0.3 | 0.16 | 0.23 | 0.17 | 0.3 |
| 2 | 172143499 | rs888272 | 0.73 | 1.266 | 0.298 | 0.755 | 0.014 | 0.08 | 0.08 | 0.056 | 0.47 | 0.32 | 0.49 | 0.13 | 0.49 | 0.37 |
| 2 | 172143764 | rs17284660 | 1.306 | 1.105 | 0.2 | 0.93 | 0.015 | 0.007 | 0.007 | 0 | 0.18 | 0.02 | 0.19 | 0.11 | 0.14 | 0.12 |
| 2 | 172143822 | rs16859334 | -0.219 | 1.211 | 0.349 | 0.915 | 0.017 | 0.034 | 0.034 | 0 | 0.05 | 0.12 | 0.1 | 0.16 | 0.05 | 0.19 |
| 2 | 172160433 | rs7564482 | 0.127 | 0.57 | -0.021 | 0.056 | -0.002 | -0.03 | -0.03 | 0.153 | 0.6 | 0.51 | 0.61 | 0.8 | 0.56 | 0.68 |
| 2 | 172163971 | rs1019362 | 0.712 | 0.462 | 0.547 | 0.115 | 0.013 | -0.072 | -0.072 | 0 | 0.72 | 0.55 | 0.74 | 0.84 | 0.68 | 0.62 |

| | | | | | | | | | | | | | | | | |
|---|---|---|---|---|---|---|---|---|---|---|---|---|---|---|---|---|
| 2 | 172172864 | rs6722670 | -0.356 | 0.491 | 0.377 | 0.104 | 0.002 | -0.059 | -0.059 | 0 | 0.8 | 0.76 | 0.85 | 1 | 0.73 | 0.83 |
| 2 | 172174263 | rs2111459 | -0.636 | 0.479 | 0.368 | 0.089 | 0.002 | -0.061 | -0.061 | 0 | 0.83 | 0.78 | 0.87 | 1 | 0.76 | 0.84 |
| 2 | 172181222 | rs13000846 | 0 | 0.474 | 0.357 | 0.084 | -0.001 | -0.038 | -0.038 | 0 | 0.21 | 0.19 | 0.15 | 0 | 0.27 | 0.17 |
| 2 | 172182292 | rs13031794 | 0.406 | 0.415 | 0.337 | 0.025 | 0 | -0.038 | -0.038 | 0.724 | 0.16 | 0.18 | 0.13 | 0 | 0.24 | 0.17 |
| 2 | 172182957 | rs11691073 | 0.44 | 0.424 | 0.356 | 0.03 | 0.001 | -0.041 | -0.041 | 0 | 0.16 | 0.18 | 0.13 | 0 | 0.24 | 0.17 |
| 2 | 172188368 | rs13023973 | 0.556 | 0.427 | 0.349 | 0.032 | -0.003 | -0.019 | -0.019 | 0 | 0.16 | 0.14 | 0.13 | 0 | 0.24 | 0.17 |
| 2 | 172193808 | rs3731987 | -1 | 0.435 | 0.506 | -0.013 | 0.004 | -0.055 | -0.055 | 0 | 0.27 | 0.59 | 0.26 | 0.09 | 0.32 | 0.26 |
| 2 | 172212652 | rs12465868 | 1.557 | 0.651 | 0.217 | 0.018 | -0.061 | -0.081 | -0.081 | 1.408 | 0.31 | 0.07 | 0.38 | 0.65 | 0.34 | 0.35 |
| 2 | 172216969 | rs3731986 | -0.806 | 0.677 | 0.393 | -0.004 | 0.019 | -0.055 | -0.055 | 0.807 | 0.27 | 0.48 | 0.26 | 0.15 | 0.33 | 0.28 |
| 2 | 172218624 | rs888521 | -0.525 | 0.683 | 0.378 | 0.003 | 0.021 | -0.039 | -0.039 | 0.218 | 0.24 | 0.44 | 0.24 | 0.15 | 0.31 | 0.27 |
| 2 | 172239759 | rs17285688 | 0.266 | 0.675 | 0.146 | -0.048 | 0.01 | 0.053 | 0.053 | 0 | 0.4 | 0.23 | 0.35 | 0.23 | 0.33 | 0.37 |
| 2 | 172240163 | rs16859370 | 1.256 | 0.846 | 0.105 | -0.118 | -0.014 | 0.04 | 0.04 | 0 | 0.18 | 0.09 | 0.1 | 0.04 | 0.18 | 0.1 |
| 2 | 172259687 | rs7562674 | 1.613 | 0.809 | 0.07 | -0.019 | 0.013 | 0.006 | 0.006 | 0.046 | 0.2 | 0.12 | 0.19 | 0.13 | 0.14 | 0.11 |
| 2 | 172268061 | rs2884005 | 0.212 | 1.015 | 0.212 | -0.125 | -0.002 | -0.023 | -0.023 | 0.016 | 0.61 | 0.58 | 0.69 | 0.91 | 0.55 | 0.58 |
| 2 | 172272690 | rs10195398 | -0.088 | 0.815 | 0.471 | 0.024 | 0.038 | 0.076 | 0.076 | 0 | 0.06 | 0.2 | 0.09 | 0.13 | 0.07 | 0.11 |
| 2 | 172278694 | rs727231 | 2.179 | 0.821 | 0.174 | 0.051 | -0.006 | 0.059 | 0.059 | 0.181 | 0.26 | 0.01 | 0.32 | 0.1 | 0.22 | 0.21 |
| 2 | 172288175 | rs2160849 | 1.804 | 0.885 | 0.238 | -0.113 | 0.005 | -0.028 | -0.028 | 1.561 | 0.19 | 0.01 | 0.2 | 0.13 | 0.16 | 0.11 |
| 2 | 172291616 | rs6751956 | 0.298 | 0.91 | 0.475 | -0.135 | -0.01 | -0.005 | -0.005 | 0 | 0.29 | 0.38 | 0.34 | 0.27 | 0.23 | 0.31 |
| 2 | 172294792 | rs2356709 | 1.591 | 1.362 | 0.209 | -0.211 | -0.023 | -0.042 | -0.042 | 1.743 | 0.32 | 0.14 | 0.39 | 0.63 | 0.35 | 0.37 |
| 2 | 172295541 | rs6433291 | 0.364 | 1.026 | 0.39 | 0.007 | 0.001 | 0 | 0 | 1.102 | 0.27 | 0.24 | 0.31 | 0.27 | 0.23 | 0.31 |
| 2 | 172297434 | rs4522571 | 0.661 | 0.985 | 0.336 | -0.005 | 0.013 | 0.012 | 0.012 | 0 | 0.25 | 0.15 | 0.28 | 0.19 | 0.19 | 0.27 |
| 2 | 172303555 | rs10445762 | 1.656 | 0.974 | 0.227 | 0.014 | 0.005 | -0.025 | -0.025 | 0 | 0.19 | 0.01 | 0.2 | 0.14 | 0.16 | 0.11 |
| 2 | 172338194 | rs3795998 | 0.867 | 1.09 | 0.324 | 0.019 | 0.013 | 0.012 | 0.012 | 0 | 0.25 | 0.15 | 0.28 | 0.19 | 0.19 | 0.27 |
| 2 | 172338300 | rs3821084 | 0.867 | 1.09 | 0.324 | 0.019 | 0.013 | 0.012 | 0.012 | 0 | 0.25 | 0.15 | 0.28 | 0.19 | 0.19 | 0.27 |
| 2 | 172340419 | rs17221346 | 0.579 | 1.092 | 0.411 | 0.028 | -0.001 | -0.001 | -0.001 | 0 | 0.27 | 0.26 | 0.32 | 0.27 | 0.23 | 0.3 |
| 2 | 172341327 | rs9789572 | 0.864 | 0.997 | 0.298 | 0.026 | 0.013 | 0.012 | 0.012 | 0 | 0.25 | 0.15 | 0.28 | 0.19 | 0.19 | 0.27 |
| 2 | 172351233 | rs17287497 | 1.298 | 1.384 | 0.17 | -0.04 | -0.004 | 0.003 | 0.003 | 0 | 0.06 | 0.04 | 0.07 | 0.06 | 0.14 | 0.05 |
| 2 | 172353544 | rs2058926 | -0.366 | 1.296 | 0.412 | -0.13 | 0.003 | -0.058 | -0.058 | 0.1 | 0.88 | 0.78 | 0.9 | 1 | 0.87 | 0.89 |
| 2 | 172353787 | rs13384194 | -0.087 | 1.212 | 0.497 | -0.005 | -0.004 | -0.02 | -0.02 | 0 | 0.34 | 0.43 | 0.38 | 0.27 | 0.34 | 0.31 |
| 2 | 172366523 | rs7564847 | 0.885 | 1.771 | 0.463 | 0.067 | -0.003 | 0.003 | 0.003 | 0 | 0.79 | 0.79 | 0.79 | 0.85 | 0.7 | 0.76 |
| 2 | 172368120 | rs888427 | 0.826 | 1.669 | 0.805 | -0.05 | -0.001 | -0.021 | -0.021 | 0 | 0.64 | 0.42 | 0.69 | 0.81 | 0.62 | 0.67 |
| 2 | 172369108 | rs12052537 | 0.85 | 2.533 | 0.548 | 0.411 | 0.041 | 0.123 | 0.123 | 0.189 | 0.06 | 0.03 | 0.15 | 0.5 | 0.2 | 0.13 |
| 2 | 172370486 | rs7580094 | 0.407 | 2.343 | 0.661 | 0.011 | -0.001 | -0.001 | -0.001 | 0 | 0.83 | 0.84 | 0.82 | 0.79 | 0.76 | 0.78 |
| 2 | 172372593 | rs17221848 | 0.979 | 2.601 | 0.628 | 0.152 | 0.066 | 0 | 0 | 0.081 | 0.05 | 0.04 | 0.08 | 0.05 | 0.2 | 0.06 |
| 2 | 172373507 | rs1476110 | 0.865 | 1.956 | 0.894 | 0.076 | -0.004 | -0.012 | -0.012 | 2.133 | 0.65 | 0.4 | 0.71 | 0.74 | 0.62 | 0.63 |
| 2 | 172374395 | rs13406777 | 0.633 | 2.013 | 1.047 | 0.217 | 0.006 | 0.004 | 0.004 | 0 | 0.82 | 0.78 | 0.81 | 0.78 | 0.76 | 0.75 |
| 2 | 172375457 | rs11680268 | 0.759 | 1.782 | 0.594 | -0.185 | -0.002 | -0.049 | -0.049 | 1.62 | 0.15 | 0.23 | 0.23 | 0.02 | 0.12 | 0.2 |
| 2 | 172376785 | rs3806562 | -0.65 | 1.745 | 0.825 | 0.034 | 0.043 | 0.038 | 0.038 | 0.926 | 0.13 | 0.39 | 0.14 | 0.15 | 0.17 | 0.18 |
| 2 | 172379936 | rs960748 | -1.512 | 1.739 | 0.121 | -0.133 | -0.009 | 0.129 | 0.129 | 0.038 | 0.51 | 0.8 | 0.63 | 0.84 | 0.52 | 0.69 |
| 2 | 172395221 | rs12692966 | -0.735 | 1.675 | 0.29 | -0.487 | 0.035 | -0.034 | -0.034 | 0 | 0.33 | 0.44 | 0.41 | 0.23 | 0.41 | 0.51 |
| 2 | 172401763 | rs13392902 | 1.68 | 1.4 | -0.319 | -0.474 | -0.014 | 0.013 | 0.013 | 0.008 | 0.12 | 0.02 | 0.07 | 0.09 | 0.13 | 0.05 |
| 2 | 172402537 | rs13009270 | 0.206 | 1.473 | -0.088 | -0.543 | 0.019 | -0.01 | -0.01 | 0 | 0.11 | 0.17 | 0.12 | 0.06 | 0.18 | 0.19 |
| 2 | 172402667 | rs7596578 | 1.675 | 1.343 | -0.276 | -0.491 | -0.014 | 0.013 | 0.013 | 0 | 0.11 | 0.02 | 0.07 | 0.1 | 0.13 | 0.05 |
| 2 | 172405989 | rs11684782 | -0.64 | 1.427 | 0.236 | -0.721 | 0.011 | -0.021 | -0.021 | 0.118 | 0.34 | 0.37 | 0.43 | 0.24 | 0.46 | 0.51 |
| 2 | 172409721 | rs2542941 | -1.6 | 0.877 | -0.127 | -0.718 | 0.144 | 0.109 | 0.109 | 1.257 | 0.51 | 0.88 | 0.59 | 0.4 | 0.67 | 0.72 |
| 2 | 172411273 | rs10455 | -1.653 | 0.854 | -0.153 | -0.729 | 0.133 | 0.096 | 0.096 | 1.473 | 0.51 | 0.93 | 0.6 | 0.4 | 0.67 | 0.72 |
| 2 | 172411644 | rs1060248 | 0.628 | 1.044 | -0.061 | -0.755 | -0.014 | -0.01 | -0.01 | 0.261 | 0.09 | 0.04 | 0.07 | 0.15 | 0.15 | 0.15 |
| 2 | 172411737 | rs950163 | -1.704 | 1.047 | 0.008 | -0.763 | 0.133 | 0.102 | 0.102 | 1.293 | 0.49 | 0.81 | 0.59 | 0.4 | 0.67 | 0.72 |
| 2 | 172412510 | rs2542940 | -0.864 | 1.02 | 0.147 | -0.785 | -0.015 | 0.063 | 0.063 | 1.038 | 0.39 | 0.72 | 0.5 | 0.24 | 0.51 | 0.57 |
| 2 | 172412616 | rs7585194 | -0.254 | 1.662 | 0.794 | -0.885 | 0 | -0.056 | -0.056 | 5.662 | 0.29 | 0.39 | 0.37 | 0.19 | 0.27 | 0.49 |
| 2 | 172412907 | rs3821083 | 1.647 | 1.67 | 0.386 | -0.874 | -0.017 | 0.005 | 0.005 | 0 | 0.07 | 0.02 | 0.1 | 0.05 | 0.24 | 0.08 |
| 2 | 172413018 | rs2542939 | -1.664 | 1.384 | 0.405 | -0.854 | 0.119 | 0.106 | 0.106 | 1.128 | 0.48 | 0.8 | 0.58 | 0.38 | 0.67 | 0.72 |
| 2 | 172419265 | rs10490602 | -0.3 | 1.544 | 0.132 | -0.763 | -0.041 | -0.002 | -0.002 | 0 | 0.17 | 0.11 | 0.18 | 0.19 | 0.4 | 0.24 |
| 2 | 172422877 | rs2111720 | -1.81 | 1.103 | 0.316 | -0.845 | 0.007 | 0.005 | 0.005 | 0.064 | 0.58 | 0.41 | 0.61 | 0.4 | 0.77 | 0.76 |
| 2 | 172423811 | rs11674142 | 0.137 | 1.174 | 0.168 | -0.893 | -0.007 | 0.011 | 0.011 | 0 | 0.24 | 0.18 | 0.32 | 0.14 | 0.23 | 0.34 |
| 2 | 172424963 | rs2542943 | -1.622 | 0.983 | -0.137 | -0.883 | -0.05 | 0.096 | 0.096 | 0 | 0.52 | 0.89 | 0.56 | 0.38 | 0.53 | 0.57 |
| 2 | 172425464 | rs2249572 | 0.344 | 0.847 | 0.678 | -0.859 | -0.052 | -0.088 | -0.088 | 0 | 0.18 | 0.41 | 0.11 | 0.02 | 0.11 | 0.07 |
| 2 | 172428405 | rs6433299 | -0.42 | 1.015 | 0.609 | -0.944 | 0.074 | 0.08 | 0.08 | 0.043 | 0.1 | 0.28 | 0.13 | 0.19 | 0.19 | 0.18 |
| 2 | 172428729 | rs2356791 | 0.452 | 0.856 | 0.364 | -1.165 | 0.008 | 0.031 | 0.031 | 0 | 0.6 | 0.32 | 0.58 | 0.81 | 0.66 | 0.62 |
| 2 | 172429907 | rs6729179 | -1.007 | 0.786 | -0.063 | -1.148 | 0.008 | -0.017 | -0.017 | 2.482 | 0.33 | 0.38 | 0.35 | 0.33 | 0.32 | 0.42 |
| 2 | 172432306 | rs1399959 | -0.276 | 0.087 | -0.654 | -1.502 | 0.035 | 0.048 | 0.048 | 0 | 0.46 | 0.26 | 0.45 | 0.76 | 0.37 | 0.66 |
| 2 | 172450729 | rs12476299 | 0.49 | -0.229 | -0.231 | -1.29 | -0.014 | -0.002 | -0.002 | 0 | 0.42 | 0.46 | 0.42 | 0.57 | 0.3 | 0.44 |
| 2 | 172451719 | rs1399955 | -1.438 | 0.152 | -0.308 | -1.552 | -0.015 | 0.049 | 0.049 | 0 | 0.66 | 0.78 | 0.63 | 0.85 | 0.66 | 0.76 |
| 2 | 172454319 | rs4668394 | -0.279 | 0.097 | -0.261 | -1.506 | 0.006 | 0.007 | 0.007 | 0 | 0.52 | 0.46 | 0.63 | 0.58 | 0.43 | 0.54 |
| 2 | 172454933 | rs6745725 | -0.855 | 0.447 | -0.306 | -1.51 | -0.006 | 0.003 | 0.003 | 0 | 0.6 | 0.67 | 0.7 | 0.6 | 0.51 | 0.57 |
| 2 | 172455639 | rs2884028 | 0.473 | 0.5 | -0.321 | -1.201 | 0.021 | -0.036 | -0.036 | 0 | 0.23 | 0.13 | 0.35 | 0.56 | 0.21 | 0.37 |
| 2 | 172456061 | rs2884029 | -0.053 | 0.851 | -0.26 | -1.125 | 0.024 | 0.002 | 0.002 | 0 | 0.18 | 0.2 | 0.18 | 0.26 | 0.34 | 0.27 |
| 2 | 172462366 | rs10432422 | -0.23 | 0.503 | -0.614 | -1.029 | -0.003 | 0.002 | 0.002 | 0 | 0.31 | 0.35 | 0.43 | 0.58 | 0.28 | 0.4 |
| 2 | 172464008 | rs6433300 | 0.757 | 0.652 | -0.687 | -0.933 | -0.021 | 0.024 | 0.024 | 1.161 | 0.19 | 0.12 | 0.24 | 0.01 | 0.23 | 0.18 |

| | | | | | | | | | | | | | | | |
|---|---|---|---|---|---|---|---|---|---|---|---|---|---|---|---|
| 2 | 172468776 | rs3951105 | -1.993 | 0.344 | -0.575 | -0.941 | -0.054 | -0.057 | -0.057 | 0 | 0.65 | 0.56 | 0.81 | 0.86 | 0.82 | 0.85 |
| 2 | 172472584 | rs10930486 | -0.912 | 0.455 | -0.605 | -0.912 | 0.007 | 0.019 | 0.019 | 0 | 0.5 | 0.45 | 0.67 | 0.59 | 0.53 | 0.63 |
| 2 | 172482159 | rs6433305 | 0.683 | 0.343 | -0.694 | -0.632 | 0.011 | -0.039 | -0.039 | 0.005 | 0.22 | 0.13 | 0.35 | 0.55 | 0.22 | 0.37 |
| 2 | 172487577 | rs7602864 | -0.218 | 0.347 | -0.61 | -0.684 | 0.004 | -0.018 | -0.018 | 0 | 0.09 | 0.28 | 0.07 | 0.03 | 0.08 | 0.07 |
| 2 | 172490226 | rs10930488 | -0.416 | 0.239 | -0.634 | -0.775 | -0.004 | 0.004 | 0.004 | 0 | 0.33 | 0.41 | 0.46 | 0.58 | 0.33 | 0.44 |
| 2 | 172495507 | rs12995009 | -0.457 | 0.412 | -0.395 | -1.115 | 0.001 | -0.003 | -0.003 | 3.755 | 0.46 | 0.42 | 0.57 | 0.85 | 0.59 | 0.66 |
| 2 | 172510158 | rs1033217 | -1.562 | 0.509 | -0.52 | -1.152 | 0.012 | 0.012 | 0.012 | 0.264 | 0.67 | 0.77 | 0.84 | 0.87 | 0.83 | 0.84 |
| 2 | 172510689 | rs1399956 | -0.225 | 1.009 | -0.374 | -0.509 | -0.018 | 0.042 | 0.042 | 0 | 0.41 | 0.3 | 0.52 | 0.83 | 0.52 | 0.59 |
| 2 | 172511925 | rs951917 | -0.947 | 1.076 | -0.749 | -0.476 | 0.013 | 0.055 | 0.055 | 0 | 0.49 | 0.61 | 0.6 | 0.85 | 0.59 | 0.66 |
| 2 | 172512177 | rs7557017 | -0.598 | 1.164 | -0.234 | 0.421 | 0.008 | -0.021 | -0.021 | 0 | 0.07 | 0.2 | 0.06 | 0.02 | 0.07 | 0.07 |
| 2 | 172525884 | rs2102738 | 1.111 | 0.897 | -1.143 | 0.766 | 0.031 | -0.125 | -0.125 | 0 | 0.3 | 0.01 | 0.15 | 0.67 | 0.2 | 0.25 |
| 2 | 172528095 | rs979956 | 0.637 | 1.71 | -0.082 | 1.159 | 0.003 | 0.02 | 0.02 | 0 | 0.52 | 0.46 | 0.58 | 0.16 | 0.43 | 0.49 |
| 2 | 172535622 | rs11678892 | -0.238 | 1.85 | -0.13 | 1.114 | -0.007 | -0.015 | -0.015 | 0 | 0.15 | 0.44 | 0.29 | 0.11 | 0.11 | 0.15 |
| 2 | 172538530 | rs17580984 | 0.847 | 1.825 | -0.011 | 1.102 | 0.004 | -0.001 | -0.001 | 0.005 | 0.1 | 0.07 | 0.24 | 0.08 | 0.08 | 0.14 |
| 2 | 172541378 | rs17614941 | 1.212 | 2.304 | -0.061 | 0.958 | -0.065 | -0.016 | -0.016 | 0 | 0.64 | 0.11 | 0.39 | 0.69 | 0.59 | 0.68 |
| 2 | 172599615 | rs6738445 | 0.762 | 2.333 | 0.278 | 1.009 | -0.016 | 0.004 | 0.004 | 0.936 | 0.77 | 0.19 | 0.71 | 0.8 | 0.71 | 0.85 |
| 2 | 172601389 | rs6709804 | -0.733 | 1.981 | 0.352 | 0.937 | -0.005 | -0.005 | -0.005 | 0 | 0.05 | 0.06 | 0.01 | 0 | 0.01 | 0 |
| 2 | 172608956 | rs10184866 | -0.838 | 2.048 | 0.388 | 0.97 | 0.054 | 0.021 | 0.021 | 0 | 0.21 | 0.46 | 0.24 | 0.2 | 0.28 | 0.17 |
| 2 | 172626552 | rs1607254 | 1.345 | 2.063 | 0.396 | 0.977 | -0.053 | -0.008 | -0.008 | 3.195 | 0.64 | 0.14 | 0.39 | 0.7 | 0.6 | 0.68 |
| 2 | 172640734 | rs17499544 | 1.118 | 2.152 | 0.358 | 0.923 | -0.025 | 0.099 | 0.099 | 0.115 | 0.19 | 0.03 | 0.11 | 0.03 | 0.17 | 0.2 |
| 2 | 172646765 | rs7608403 | -0.898 | 1.992 | 0.488 | 1.059 | 0.054 | 0.027 | 0.027 | 0 | 0.21 | 0.47 | 0.25 | 0.21 | 0.28 | 0.17 |
| 2 | 172648592 | rs3770459 | 1.786 | 2.002 | 0.432 | 1.164 | -0.048 | 0.143 | 0.143 | 0 | 0.39 | 0.04 | 0.23 | 0.12 | 0.34 | 0.36 |
| 2 | 172649826 | rs10167419 | -0.835 | 1.999 | 0.45 | 1.163 | 0.071 | 0.035 | 0.035 | 0 | 0.2 | 0.45 | 0.24 | 0.2 | 0.28 | 0.17 |
| 2 | 172659376 | rs17581284 | 1.514 | 1.999 | 0.439 | 1.178 | -0.057 | -0.031 | -0.031 | 0.288 | 0.65 | 0.12 | 0.39 | 0.67 | 0.6 | 0.67 |
| 2 | 172673453 | rs6716901 | 0.822 | 2.171 | 0.407 | 1.111 | -0.038 | -0.017 | -0.017 | 0 | 0.06 | 0 | 0.07 | 0.08 | 0.12 | 0.11 |
| 2 | 172680678 | rs1400816 | -0.569 | 2.134 | 0.423 | 1.113 | -0.014 | 0.013 | 0.013 | 0.006 | 0.88 | 0.58 | 0.73 | 0.92 | 0.92 | 0.86 |
| 2 | 172683481 | rs17615262 | 0.533 | 2.137 | 0.419 | 1.116 | -0.006 | -0.105 | -0.105 | 0 | 0.26 | 0.07 | 0.16 | 0.57 | 0.26 | 0.31 |
| 2 | 172684403 | rs12692976 | 0.445 | 2.142 | 0.418 | 1.115 | -0.007 | -0.005 | -0.005 | 0.506 | 0.59 | 0.49 | 0.48 | 0.33 | 0.62 | 0.53 |
| 2 | 172693780 | rs3765166 | 0.668 | 2.14 | 0.405 | 1.11 | -0.014 | -0.1 | -0.1 | 0.583 | 0.25 | 0.07 | 0.16 | 0.57 | 0.26 | 0.31 |
| 2 | 172701157 | rs2271758 | 1.707 | 2.084 | 0.437 | 1.101 | -0.039 | -0.015 | -0.015 | 0 | 0.65 | 0.17 | 0.4 | 0.7 | 0.6 | 0.67 |
| 2 | 172704291 | rs4668414 | -0.714 | 2.175 | 0.499 | 1.133 | 0.064 | 0.043 | 0.043 | 0.016 | 0.17 | 0.3 | 0.23 | 0.21 | 0.27 | 0.17 |
| 2 | 172715401 | rs1105265 | 0.475 | 2.227 | 0.464 | 1.121 | -0.043 | -0.015 | -0.015 | 0.165 | 0.07 | 0 | 0.13 | 0.08 | 0.13 | 0.15 |
| 2 | 172725301 | rs1878583 | -0.5 | 1.643 | 0.525 | 1.148 | -0.014 | 0.013 | 0.013 | 0.006 | 0.88 | 0.58 | 0.73 | 0.92 | 0.92 | 0.86 |
| 2 | 172737981 | rs11675404 | 1.968 | 1.647 | 0.456 | 1.123 | -0.019 | 0.122 | 0.122 | 0 | 0.38 | 0.04 | 0.2 | 0.13 | 0.3 | 0.35 |
| 2 | 172754554 | rs6753859 | -0.659 | 1.553 | 0.743 | 1.169 | -0.016 | 0.005 | 0.005 | 0 | 0.89 | 0.71 | 0.73 | 0.92 | 0.92 | 0.86 |
| 2 | 172815510 | rs6757680 | 0.281 | 1.601 | 0.555 | 1.183 | -0.007 | 0.007 | 0.007 | 0 | 0.83 | 0.57 | 0.64 | 0.87 | 0.87 | 0.84 |
| 2 | 172820315 | rs10165126 | -1.169 | 1.638 | 0.578 | 1.197 | 0.046 | 0.064 | 0.064 | 0.04 | 0.2 | 0.36 | 0.26 | 0.73 | 0.3 | 0.26 |
| 2 | 172851552 | rs11695630 | 1.872 | 1.541 | 0.382 | 1.123 | 0.01 | 0.178 | 0.178 | 0 | 0.39 | 0.07 | 0.19 | 0.12 | 0.31 | 0.35 |
| 2 | 172858214 | rs10174055 | -0.861 | 1.575 | 0.162 | 1.217 | 0.039 | 0.018 | 0.018 | 0 | 0.18 | 0.26 | 0.25 | 0.21 | 0.31 | 0.18 |
| 2 | 172860402 | rs6433321 | -0.984 | 1.514 | 0.156 | 1.267 | 0.001 | -0.02 | -0.02 | 0.096 | 0.43 | 0.47 | 0.44 | 0.76 | 0.56 | 0.5 |
| 2 | 172873060 | rs6433323 | 1.223 | 1.491 | 0.104 | 1.159 | -0.005 | -0.003 | -0.003 | 0.364 | 0.68 | 0.31 | 0.48 | 0.7 | 0.62 | 0.69 |
| 2 | 172890588 | rs10930502 | -1.064 | 1.329 | -0.299 | 1.19 | 0.031 | 0.047 | 0.047 | 0.031 | 0.19 | 0.3 | 0.33 | 0.7 | 0.3 | 0.26 |
| 2 | 172895449 | rs7606918 | 0.427 | 1.28 | -0.223 | 1.241 | 0.043 | 0.077 | 0.077 | 0.036 | 0.08 | 0.23 | 0.14 | 0.13 | 0.15 | 0.03 |
| 2 | 172917076 | rs13013988 | 1.525 | 1.291 | -0.749 | 1.17 | -0.028 | 0.196 | 0.196 | 0 | 0.41 | 0.1 | 0.23 | 0.13 | 0.36 | 0.25 |
| 2 | 172921454 | rs17581715 | 0.591 | 1.246 | -0.639 | 1.481 | -0.006 | 0.017 | 0.017 | 0 | 0.1 | 0.08 | 0.23 | 0.17 | 0.08 | 0.28 |
| 2 | 172921832 | rs788171 | 0.399 | 1.313 | -0.937 | 1.283 | 0.003 | 0.014 | 0.014 | 0.307 | 0.69 | 0.41 | 0.51 | 0.6 | 0.63 | 0.56 |
| 2 | 172929901 | rs10497378 | -0.416 | 1.512 | 0.275 | 0.772 | 0.043 | 0.078 | 0.078 | 0 | 0.08 | 0.25 | 0.14 | 0.13 | 0.14 | 0.06 |
| 2 | 172931559 | rs788163 | -0.842 | 1.235 | -0.389 | 0.323 | 0 | -0.031 | -0.031 | 0 | 0.29 | 0.37 | 0.22 | 0.11 | 0.28 | 0.25 |
| 2 | 172942897 | rs788158 | -0.895 | 1.302 | -0.203 | 0.813 | 0.008 | 0.017 | 0.017 | 1.311 | 0.27 | 0.46 | 0.21 | 0.33 | 0.25 | 0.47 |
| 2 | 172943351 | rs16859964 | -0.749 | 1.895 | 0.004 | 1.047 | 0.019 | 0.026 | 0.026 | 0 | 0.15 | 0.26 | 0.26 | 0.18 | 0.19 | 0.16 |
| 2 | 172944650 | rs788159 | -1.471 | 2.326 | 0.382 | 0.669 | 0.009 | -0.011 | -0.011 | 0 | 0.4 | 0.47 | 0.4 | 0.32 | 0.42 | 0.45 |
| 2 | 172945529 | rs705869 | 0.129 | 2.195 | 0.762 | 0.633 | -0.001 | 0 | 0 | 0.271 | 0.85 | 0.85 | 0.85 | 0.89 | 0.82 | 0.86 |
| 2 | 172946527 | rs1047889 | -0.96 | 2.5 | 0.659 | 0.55 | -0.025 | -0.002 | -0.002 | 0 | 0.38 | 0.29 | 0.44 | 0.63 | 0.43 | 0.55 |
| 2 | 172953438 | rs788172 | -1.085 | 1.788 | 0.142 | -0.2 | 0.003 | 0.012 | 0.012 | 1.5 | 0.3 | 0.71 | 0.31 | 0.19 | 0.34 | 0.34 |
| 2 | 172953460 | rs788173 | -1.085 | 1.788 | 0.142 | -0.2 | 0.003 | 0.012 | 0.012 | 1.5 | 0.3 | 0.71 | 0.31 | 0.19 | 0.34 | 0.34 |
| 2 | 172955011 | rs10186317 | -0.628 | 1.807 | 0.16 | -0.323 | 0.021 | -0.031 | -0.031 | 0 | 0.2 | 0.41 | 0.22 | 0.13 | 0.29 | 0.25 |
| 2 | 172957643 | rs13390848 | 2.25 | 1.209 | -1.034 | -0.666 | 0.011 | 0.164 | 0.164 | 0 | 0.3 | 0.04 | 0.25 | 0.08 | 0.2 | 0.26 |
| 2 | 172963025 | rs1154721 | 0.506 | 1.482 | -0.824 | -0.426 | 0.016 | 0.027 | 0.027 | 0.922 | 0.08 | 0.02 | 0.05 | 0.01 | 0.03 | 0.02 |
| 2 | 172966250 | rs2228184 | -0.301 | 1.83 | -1.063 | -0.582 | -0.115 | -0.031 | -0.031 | 3.233 | 0.29 | 0.1 | 0.31 | 0.66 | 0.39 | 0.4 |
| 2 | 172972792 | rs13034296 | 0.334 | 1.373 | -0.724 | -0.475 | 0.012 | 0.027 | 0.027 | 0.673 | 0.08 | 0 | 0.06 | 0.01 | 0.04 | 0.02 |
| 2 | 172972971 | rs2016394 | -1.084 | 1.834 | -0.321 | -0.387 | -0.033 | 0.021 | 0.021 | 0 | 0.28 | 0.23 | 0.43 | 0.19 | 0.46 | 0.34 |
| 2 | 172984009 | rs10930506 | -0.882 | 0.938 | 0.056 | -0.351 | -0.007 | -0.011 | -0.011 | 0 | 0.13 | 0.21 | 0.23 | 0.11 | 0.13 | 0.12 |
| 2 | 172992016 | rs12692981 | 1.605 | -0.348 | -0.722 | -1.01 | 0.076 | 0.097 | 0.097 | 0 | 0.37 | 0.2 | 0.2 | 0.16 | 0.17 | 0.21 |
| 2 | 172994091 | rs4972623 | 0.756 | -0.657 | -0.26 | -1.047 | 0.067 | 0.072 | 0.072 | 0 | 0.49 | 0.38 | 0.48 | 0.26 | 0.3 | 0.37 |
| 2 | 173001058 | rs4972630 | -1.26 | -0.596 | -0.216 | -0.999 | -0.014 | 0 | 0 | 0 | 0.13 | 0.28 | 0.29 | 0.1 | 0.14 | 0.17 |
| 2 | 173008793 | rs6433331 | 0.534 | -0.101 | -0.054 | -1.242 | 0.001 | 0.002 | 0.002 | 0 | 0.56 | 0.49 | 0.57 | 0.38 | 0.43 | 0.59 |
| 2 | 173008906 | rs17218404 | -0.179 | -0.128 | -0.06 | -1.254 | 0.006 | -0.006 | -0.006 | 0 | 0.07 | 0.11 | 0.09 | 0.12 | 0.13 | 0.22 |
| 2 | 173009083 | rs1843756 | 0.952 | -0.284 | -0.195 | -1.286 | 0.072 | 0.078 | 0.078 | 0 | 0.49 | 0.38 | 0.48 | 0.26 | 0.31 | 0.37 |
| 2 | 173011537 | rs1386354 | -1.278 | -0.224 | -0.152 | -1.217 | -0.015 | -0.005 | -0.005 | 0 | 0.14 | 0.31 | 0.29 | 0.08 | 0.13 | 0.17 |

| | | | | | | | | | | | | | | | | |
|---|---|---|---|---|---|---|---|---|---|---|---|---|---|---|---|---|
| 2 | 173013373 | rs1486957 | 1.208 | 0.303 | -0.078 | -1.38 | 0.019 | 0.052 | 0.052 | 0 | 0.48 | 0.38 | 0.32 | 0.19 | 0.36 | 0.23 |
| 2 | 173015110 | rs1843755 | 0.528 | 0.102 | -0.346 | -1.339 | -0.021 | 0.048 | 0.048 | 0 | 0.59 | 0.68 | 0.47 | 0.82 | 0.47 | 0.59 |
| 2 | 173023346 | rs2054831 | -0.882 | -0.621 | 0.01 | -1.475 | 0.076 | 0.114 | 0.114 | 0.112 | 0.5 | 0.65 | 0.57 | 0.85 | 0.72 | 0.69 |
| 2 | 173027943 | rs3901602 | 0.252 | -1.287 | 0.436 | -1.443 | -0.048 | -0.085 | -0.085 | 0.286 | 0.41 | 0.51 | 0.19 | 0.16 | 0.21 | 0.43 |
| 2 | 173029863 | rs2171320 | -0.097 | -1.186 | 0.386 | -1.394 | -0.032 | -0.064 | -0.064 | 0.347 | 0.43 | 0.5 | 0.21 | 0.17 | 0.22 | 0.44 |
| 2 | 173034038 | rs13006876 | 1.23 | -1.003 | -0.193 | -1.515 | 0.041 | 0.168 | 0.168 | 1.379 | 0.28 | 0.04 | 0.12 | 0.02 | 0.16 | 0.16 |
| 2 | 173034649 | rs6433333 | -1.493 | -0.88 | 0.469 | -1.374 | -0.007 | -0.063 | -0.063 | 0 | 0.16 | 0.58 | 0.16 | 0.08 | 0.15 | 0.23 |
| 2 | 173046584 | rs1386351 | -0.35 | -0.937 | 0.568 | -1.196 | -0.007 | -0.013 | -0.013 | 0 | 0.45 | 0.56 | 0.33 | 0.26 | 0.29 | 0.38 |
| 2 | 173047982 | rs6730323 | -0.275 | -1.031 | 0.614 | -1.154 | 0.029 | 0.034 | 0.034 | 2.959 | 0.43 | 0.68 | 0.28 | 0.18 | 0.22 | 0.24 |
| 2 | 173060526 | rs1038565 | -0.278 | -1.088 | 0.567 | -1.148 | 0.052 | 0.063 | 0.063 | 3.385 | 0.43 | 0.72 | 0.28 | 0.19 | 0.2 | 0.25 |
| 2 | 173069236 | rs6759270 | -0.98 | -1.069 | 0.884 | -1.227 | -0.027 | -0.089 | -0.089 | 0 | 0.69 | 0.59 | 0.84 | 0.95 | 0.86 | 0.84 |
| 2 | 173070052 | rs10930529 | -0.439 | -1.01 | 1.08 | -1.237 | -0.038 | -0.057 | -0.057 | 0 | 0.63 | 0.5 | 0.75 | 0.8 | 0.81 | 0.8 |
| 2 | 173070217 | rs4972710 | -0.719 | -1.011 | 1.056 | -1.255 | -0.019 | -0.022 | -0.022 | 0 | 0.67 | 0.61 | 0.79 | 0.8 | 0.86 | 0.81 |
| 2 | 173070436 | rs4972713 | 0.427 | -0.964 | 0.794 | -1.236 | 0.087 | 0.133 | 0.133 | 3.629 | 0.54 | 0.2 | 0.68 | 0.77 | 0.76 | 0.77 |
| 2 | 173072719 | rs11893372 | -0.636 | -0.814 | 0.577 | -1.091 | 0.009 | -0.034 | -0.034 | 0 | 0.08 | 0.24 | 0.06 | 0.01 | 0.05 | 0.02 |
| 2 | 173076350 | rs4972717 | -0.204 | -0.8 | 0.171 | -1.076 | -0.024 | -0.039 | -0.039 | 0 | 0.59 | 0.44 | 0.74 | 0.79 | 0.82 | 0.79 |
| 2 | 173093268 | rs13396614 | -1.126 | -0.492 | 0.167 | -0.874 | 0.034 | -0.033 | -0.033 | 0 | 0.11 | 0.27 | 0.13 | 0.07 | 0.14 | 0.08 |
| 2 | 173093533 | rs13422459 | 0.438 | -1.008 | -0.412 | -0.951 | -0.006 | 0.03 | 0.03 | 0.399 | 0.22 | 0.12 | 0.22 | 0.07 | 0.23 | 0.26 |
| 2 | 173093926 | rs10204599 | -0.222 | -0.998 | -0.602 | -1.679 | -0.004 | 0.036 | 0.036 | 0 | 0.54 | 0.37 | 0.41 | 0.13 | 0.47 | 0.43 |
| 2 | 173096709 | rs4610020 | -0.667 | -0.935 | 0.419 | -1.176 | -0.014 | -0.013 | -0.013 | 0 | 0.32 | 0.5 | 0.33 | 0.31 | 0.3 | 0.35 |
| 2 | 173097848 | rs10194179 | -0.595 | -0.93 | 0.632 | -1.146 | -0.016 | 0.081 | 0.081 | 0.052 | 0.08 | 0.22 | 0.12 | 0.18 | 0.05 | 0.08 |
| 2 | 173104712 | rs4076083 | -0.362 | -0.924 | 0.592 | -1.178 | -0.002 | 0 | 0 | 0 | 0.28 | 0.36 | 0.3 | 0.31 | 0.28 | 0.34 |
| 2 | 173105479 | rs10930538 | 0.774 | -0.932 | 0.34 | -1.217 | 0.02 | 0.025 | 0.025 | 0.785 | 0.61 | 0.24 | 0.61 | 0.64 | 0.64 | 0.52 |
| 2 | 173108443 | rs4972740 | -0.313 | -0.712 | -0.152 | -1.349 | 0.006 | 0.028 | 0.028 | 1.257 | 0.32 | 0.25 | 0.43 | 0.93 | 0.29 | 0.5 |
| 2 | 173113143 | rs6757886 | -0.353 | -0.819 | -0.043 | -0.81 | -0.024 | 0.09 | 0.09 | 0 | 0.14 | 0.2 | 0.12 | 0.53 | 0.09 | 0.22 |
| 2 | 173114962 | rs6708374 | 0.68 | -0.65 | -0.208 | -0.889 | -0.004 | 0.008 | 0.008 | 0.237 | 0.22 | 0.25 | 0.17 | 0.33 | 0.2 | 0.27 |
| 2 | 173115214 | rs4435421 | 0.35 | -0.607 | -0.475 | -0.971 | 0.026 | -0.011 | -0.011 | 4.888 | 0.62 | 0.28 | 0.71 | 0.53 | 0.7 | 0.52 |
| 2 | 173116283 | rs4972462 | 0.643 | -0.605 | -0.47 | -0.83 | 0.003 | 0.078 | 0.078 | 0 | 0.07 | 0.08 | 0.09 | 0.48 | 0.07 | 0.14 |
| 2 | 173117510 | rs6717347 | -0.03 | -0.497 | -0.065 | -1.009 | -0.024 | 0.007 | 0.007 | 0.437 | 0.29 | 0.48 | 0.24 | 0.33 | 0.26 | 0.32 |
| 2 | 173118509 | rs6721348 | -0.342 | -0.518 | -0.266 | -0.804 | -0.005 | -0.012 | -0.012 | 0 | 0.88 | 0.77 | 0.93 | 0.88 | 0.91 | 0.84 |
| 2 | 173121662 | rs4561630 | 0.629 | -0.221 | -1.33 | -0.859 | -0.044 | 0.028 | 0.028 | 0.019 | 0.5 | 0.15 | 0.55 | 0.53 | 0.48 | 0.51 |
| 2 | 173124183 | rs7574002 | 0.247 | 0.273 | -1.124 | -0.596 | 0.001 | -0.014 | -0.014 | 0.358 | 0.64 | 0.5 | 0.64 | 0.34 | 0.62 | 0.46 |
| 2 | 173125108 | rs4531900 | -0.409 | 0.24 | -1.09 | -0.597 | -0.035 | 0.095 | 0.095 | 0.065 | 0.48 | 0.18 | 0.33 | 0.63 | 0.49 | 0.56 |
| 2 | 173125947 | rs6433346 | 1.132 | 0.407 | -1.179 | -1.477 | 0.009 | 0.026 | 0.026 | 0 | 0.36 | 0.26 | 0.42 | 0.02 | 0.33 | 0.19 |
| 2 | 173126490 | rs4289149 | 0.12 | 0.215 | -0.236 | -0.381 | 0.046 | -0.059 | -0.059 | 3.103 | 0.86 | 0.62 | 0.84 | 0.96 | 0.82 | 0.78 |
| 2 | 173127669 | rs7588418 | 0.285 | 0.244 | -1.096 | -0.3 | 0.001 | 0.016 | 0.016 | 0 | 0.62 | 0.33 | 0.57 | 0.25 | 0.62 | 0.44 |
| 2 | 173140874 | rs4972760 | 0.467 | -0.112 | -1.215 | -0.241 | 0.017 | 0.029 | 0.029 | 0 | 0.84 | 0.94 | 0.92 | 0.95 | 0.88 | 0.88 |
| 2 | 173141580 | rs4246617 | 0.364 | -1.044 | -1.224 | -0.812 | 0.005 | 0.003 | 0.003 | 0 | 0.45 | 0.56 | 0.6 | 0.18 | 0.62 | 0.35 |
| 2 | 173142480 | rs4349318 | 0.454 | -1.135 | -0.857 | -0.857 | -0.014 | -0.033 | -0.033 | 0.049 | 0.43 | 0.45 | 0.59 | 0.18 | 0.62 | 0.34 |
| 2 | 173143498 | rs4433964 | 0.258 | -1.167 | -0.744 | -0.837 | 0.057 | 0.045 | 0.045 | 0 | 0.39 | 0.28 | 0.31 | 0.77 | 0.26 | 0.53 |
| 2 | 173146356 | rs10195843 | -0.832 | -1.589 | -0.76 | -1.265 | -0.015 | -0.032 | -0.032 | 0 | 0.42 | 0.54 | 0.32 | 0.82 | 0.35 | 0.53 |
| 2 | 173150393 | rs4316911 | -0.453 | -1.811 | -0.164 | -1.537 | -0.012 | -0.024 | -0.024 | 0 | 0.46 | 0.6 | 0.34 | 0.88 | 0.28 | 0.55 |
| 2 | 173151384 | rs6720199 | 0.082 | -1.011 | -0.211 | -1.842 | -0.016 | 0.047 | 0.047 | 0 | 0.69 | 0.8 | 0.7 | 0.83 | 0.65 | 0.7 |
| 2 | 173152508 | rs7574709 | 1.79 | -1.143 | 0.095 | -1.904 | 0.005 | -0.008 | -0.008 | 0.03 | 0.22 | 0.23 | 0.12 | 0.02 | 0.11 | 0.14 |
| 2 | 173155718 | rs12471128 | 0.734 | -1.13 | 0.289 | -1.868 | -0.014 | 0.06 | 0.06 | 0 | 0.39 | 0.21 | 0.45 | 0.04 | 0.52 | 0.29 |
| 2 | 173159304 | rs10930541 | -0.388 | -1.137 | 0.204 | -1.696 | -0.006 | -0.033 | -0.033 | 0 | 0.61 | 0.55 | 0.65 | 0.82 | 0.67 | 0.68 |
| 2 | 173162998 | rs4438453 | -0.615 | -1.035 | 0.184 | -1.693 | -0.005 | -0.031 | -0.031 | 0 | 0.63 | 0.6 | 0.68 | 0.87 | 0.69 | 0.69 |
| 2 | 173165495 | rs6742057 | 1.78 | -1.026 | 0.269 | -1.696 | 0.035 | 0.038 | 0.038 | 0.081 | 0.21 | 0.15 | 0.12 | 0.03 | 0.11 | 0.14 |
| 2 | 173170965 | rs7605564 | 0.158 | -0.792 | 0.459 | -1.694 | 0.003 | 0.013 | 0.013 | 0 | 0.47 | 0.53 | 0.59 | 0.78 | 0.54 | 0.58 |
| 2 | 173173642 | rs12692999 | -0.016 | -0.821 | 0.483 | -1.708 | 0.018 | 0.096 | 0.096 | 0 | 0.49 | 0.65 | 0.6 | 0.78 | 0.54 | 0.58 |
| 2 | 173175211 | rs7594561 | -0.876 | -0.769 | 0.433 | -1.744 | 0.007 | -0.01 | -0.01 | 0 | 0.85 | 0.84 | 0.79 | 0.9 | 0.79 | 0.86 |
| 2 | 173178490 | rs4427989 | 0.674 | -1.218 | 0.773 | -2.27 | -0.013 | -0.068 | -0.068 | 0 | 0.71 | 0.47 | 0.69 | 0.84 | 0.75 | 0.8 |
| 2 | 173191687 | rs4246619 | 0.053 | -1.503 | 0.505 | -2.575 | 0.054 | 0.107 | 0.107 | 0 | 0.54 | 0.69 | 0.72 | 0.84 | 0.64 | 0.63 |
| 2 | 173195634 | rs6755520 | 0.77 | -1.449 | 0.499 | -2.542 | 0.004 | 0.016 | 0.016 | 0 | 0.44 | 0.36 | 0.6 | 0.77 | 0.57 | 0.57 |
| 2 | 173195660 | rs7591203 | 0.316 | -1.42 | 0.393 | -2.572 | 0.023 | 0.034 | 0.034 | 0 | 0.34 | 0.26 | 0.2 | 0.09 | 0.24 | 0.28 |
| 2 | 173198175 | rs7425380 | -0.351 | -1.259 | 0.378 | -2.359 | -0.001 | -0.014 | -0.014 | 0.241 | 0.82 | 0.8 | 0.82 | 0.9 | 0.82 | 0.85 |
| 2 | 173201204 | rs10209443 | 0.425 | -1.915 | 0.203 | -2.549 | 0.01 | 0.004 | 0.004 | 0 | 0.31 | 0.32 | 0.14 | 0.09 | 0.18 | 0.24 |
| 2 | 173201269 | rs6729717 | -0.134 | -1.635 | 0.326 | -2.533 | 0.001 | -0.039 | -0.039 | 0 | 0.8 | 0.68 | 0.82 | 0.86 | 0.82 | 0.83 |
| 2 | 173205214 | rs12614473 | 0.479 | -1.883 | -0.595 | -2.387 | 0.06 | 0.018 | 0.018 | 0.361 | 0.35 | 0.09 | 0.27 | 0.73 | 0.28 | 0.49 |
| 2 | 173205527 | rs10930550 | -0.288 | -1.643 | -0.182 | -2.254 | 0.002 | -0.027 | -0.027 | 0.028 | 0.81 | 0.73 | 0.83 | 0.9 | 0.82 | 0.81 |
| 2 | 173207781 | rs7605224 | -0.07 | -1.656 | -0.128 | -2.247 | -0.002 | -0.043 | -0.043 | 1.672 | 0.78 | 0.57 | 0.82 | 0.85 | 0.82 | 0.81 |
| 2 | 173214847 | rs4972807 | 0.287 | -1.953 | -0.422 | -2.275 | -0.013 | -0.046 | -0.046 | 0 | 0.63 | 0.55 | 0.77 | 0.81 | 0.72 | 0.76 |
| 2 | 173216405 | rs11688011 | 0.083 | -1.48 | -0.518 | -2.208 | -0.027 | -0.05 | -0.05 | 0 | 0.2 | 0.28 | 0.1 | 0.04 | 0.1 | 0.15 |
| 2 | 173217091 | rs4972809 | -0.174 | -1.171 | -0.497 | -1.895 | -0.014 | 0.044 | 0.044 | 0 | 0.86 | 0.75 | 0.79 | 0.25 | 0.88 | 0.62 |
| 2 | 173221247 | rs12998355 | 1.425 | 0.196 | -0.832 | -2.291 | -0.058 | 0.035 | 0.035 | 0 | 0.2 | 0.1 | 0.4 | 0.05 | 0.42 | 0.18 |
| 2 | 173227933 | rs6738142 | -0.842 | -0.295 | -0.558 | -2.619 | 0 | -0.03 | -0.03 | 0 | 0.37 | 0.43 | 0.22 | 0.07 | 0.31 | 0.21 |
| 2 | 173228629 | rs10930552 | 0.762 | 0.048 | -0.154 | -2.819 | 0.025 | -0.067 | -0.067 | 0 | 0.29 | 0.33 | 0.47 | 0.05 | 0.48 | 0.22 |
| 2 | 173229472 | rs10173487 | -0.306 | -0.187 | -0.174 | -2.965 | 0.013 | 0.013 | 0.013 | 0.007 | 0.46 | 0.55 | 0.57 | 0.11 | 0.59 | 0.29 |
| 2 | 173237175 | rs11687888 | 1.06 | -0.016 | -0.13 | -2.884 | -0.113 | 0.137 | 0.137 | 0 | 0.28 | 0.07 | 0.46 | 0.01 | 0.47 | 0.22 |

| | | | | | | | | | | | | | | | | |
|---|---|---|---|---|---|---|---|---|---|---|---|---|---|---|---|---|
| 2 | 173237498 | rs6725142 | -0.261 | -0.365 | -0.016 | -2.95 | -0.018 | -0.024 | -0.024 | 0 | 0.23 | 0.32 | 0.15 | 0.1 | 0.17 | 0.27 |
| 2 | 173238265 | rs10185752 | -0.638 | -0.468 | -0.31 | -3.039 | 0.031 | 0.036 | 0.036 | 0 | 0.5 | 0.56 | 0.61 | 0.16 | 0.66 | 0.42 |
| 2 | 173239722 | rs4283388 | -0.696 | 0.377 | -1.108 | -2.93 | -0.006 | 0.016 | 0.016 | 0.111 | 0.33 | 0.34 | 0.28 | 0.05 | 0.33 | 0.35 |
| 2 | 173240147 | rs6736756 | -1.13 | -0.363 | -1.465 | -2.2 | 0.052 | 0.061 | 0.061 | 1.522 | 0.67 | 0.92 | 0.69 | 0.17 | 0.75 | 0.56 |
| 2 | 173245780 | rs7557725 | -0.951 | -0.409 | -0.815 | -1.573 | 0.035 | 0.05 | 0.05 | 0.478 | 0.39 | 0.74 | 0.23 | 0.12 | 0.27 | 0.33 |
| 2 | 173249050 | rs7608333 | -1.038 | -0.361 | -1.228 | -1.52 | 0.08 | 0.114 | 0.114 | 1.451 | 0.4 | 0.86 | 0.24 | 0.15 | 0.27 | 0.33 |
| 2 | 173253372 | rs6727015 | -0.642 | 0.259 | -0.641 | -1.547 | 0.025 | -0.026 | -0.026 | 0 | 0.11 | 0.24 | 0.07 | 0.03 | 0.14 | 0.18 |
| 2 | 173255627 | rs2884228 | 0.146 | 0.335 | -0.348 | -1.5 | 0.034 | -0.002 | -0.002 | 0.014 | 0.08 | 0.19 | 0.1 | 0.06 | 0.11 | 0.16 |
| 2 | 173256383 | rs17286255 | 0.886 | 0.136 | -0.45 | -2.003 | -0.026 | 0.026 | 0.026 | 0 | 0.33 | 0.24 | 0.45 | 0.02 | 0.46 | 0.22 |
| 2 | 173262339 | rs12373564 | 0.069 | -0.455 | -0.47 | -1.78 | -0.032 | -0.052 | -0.052 | 0 | 0.68 | 0.62 | 0.66 | 0.09 | 0.78 | 0.53 |
| 2 | 173274750 | rs10166912 | -0.638 | -0.032 | -0.139 | -1.548 | 0.081 | -0.094 | -0.094 | 0.436 | 0.13 | 0.46 | 0.13 | 0 | 0.16 | 0.03 |
| 2 | 173275164 | rs4972828 | -1.117 | -0.565 | -1.278 | -1.825 | 0.015 | 0.107 | 0.107 | 0 | 0.79 | 0.94 | 0.89 | 0.97 | 0.85 | 0.87 |
| 2 | 173277783 | rs12621273 | -0.778 | -0.925 | -0.978 | -1.85 | -0.007 | -0.017 | -0.017 | 1.551 | 0.39 | 0.57 | 0.4 | 0.88 | 0.3 | 0.52 |
| 2 | 173279650 | rs17726737 | 0.855 | -0.657 | -1.78 | -1.88 | 0.022 | 0.089 | 0.089 | 0 | 0.22 | 0.06 | 0.08 | 0.01 | 0.17 | 0.18 |
| 2 | 173280280 | rs17726833 | 1.17 | 0.216 | -2.056 | -1.958 | -0.064 | 0.128 | 0.128 | 0.064 | 0.28 | 0.07 | 0.29 | 0.02 | 0.39 | 0.21 |
| 2 | 173287607 | rs11686909 | 0.551 | -0.436 | -2.191 | -1.918 | -0.048 | 0.205 | 0.205 | 0.202 | 0.38 | 0.09 | 0.44 | 0.05 | 0.57 | 0.27 |
| 2 | 173290395 | rs2141698 | -0.43 | -0.876 | -1.815 | -1.049 | 0.055 | 0.066 | 0.066 | 0 | 0.71 | 0.83 | 0.9 | 0.9 | 0.82 | 0.65 |
| 2 | 173294454 | rs6721624 | 0.373 | -0.35 | -1.619 | -0.79 | 0.002 | 0.135 | 0.135 | 0 | 0.46 | 0.8 | 0.52 | 0.25 | 0.55 | 0.31 |
| 2 | 173295297 | rs3108779 | -0.87 | -0.292 | -0.858 | -0.766 | 0 | -0.006 | -0.006 | 0.012 | 0.59 | 0.56 | 0.52 | 0.75 | 0.48 | 0.69 |
| 2 | 173298008 | rs2077921 | -1.228 | -0.163 | -0.639 | 0.126 | 0 | 0 | 0 | 0 | 0.26 | 0.27 | 0.39 | 0.59 | 0.24 | 0.43 |
| 2 | 173299861 | rs7567159 | -0.9 | -0.147 | -0.306 | 0.373 | -0.02 | 0.076 | 0.076 | 0 | 0.16 | 0.21 | 0.28 | 0.55 | 0.13 | 0.35 |
| 2 | 173300673 | rs13027811 | 1.412 | 0.462 | -0.457 | 0.2 | -0.001 | -0.002 | -0.002 | 0 | 0.06 | 0.04 | 0.08 | 0 | 0.12 | 0.03 |
| 2 | 173301379 | rs3934627 | -0.778 | 0.032 | -0.336 | 0.453 | 0.001 | 0.024 | 0.024 | 0 | 0.14 | 0.13 | 0.27 | 0.52 | 0.13 | 0.27 |
| 2 | 173317864 | rs6717877 | -0.055 | 0.126 | -0.127 | -0.025 | -0.008 | 0.043 | 0.043 | 0.205 | 0.17 | 0.07 | 0.13 | 0.03 | 0.18 | 0.16 |
| 2 | 173318571 | rs13421350 | -0.901 | -0.311 | 0.68 | -0.083 | -0.039 | 0.071 | 0.071 | 0 | 0.16 | 0.29 | 0.13 | 0.29 | 0.1 | 0.2 |
| 2 | 173320520 | rs6755175 | -0.787 | -0.349 | 0.746 | -0.153 | -0.05 | -0.045 | -0.045 | 0 | 0.14 | 0.37 | 0.08 | 0.03 | 0.07 | 0.09 |
| 2 | 173326436 | rs1076594 | -1.61 | -0.265 | 0.614 | -0.225 | -0.006 | -0.003 | -0.003 | 2.557 | 0.34 | 0.34 | 0.27 | 0.32 | 0.29 | 0.38 |
| 2 | 173327168 | rs10165340 | -0.229 | -0.337 | 0.575 | -0.339 | -0.051 | -0.033 | -0.033 | 0 | 0.13 | 0.22 | 0.06 | 0.03 | 0.06 | 0.09 |
| 2 | 173327273 | rs1076597 | -1.363 | -0.033 | 0.654 | -0.067 | -0.003 | 0.01 | 0.01 | 2.745 | 0.28 | 0.26 | 0.44 | 0.81 | 0.3 | 0.57 |
| 2 | 173333047 | rs3792256 | -0.241 | -0.071 | 0.571 | -0.12 | 0.002 | 0.044 | 0.044 | 0.137 | 0.18 | 0.1 | 0.12 | 0.04 | 0.16 | 0.17 |
| 2 | 173339808 | rs11895564 | -1.069 | -0.551 | 0.742 | -0.647 | -0.002 | -0.039 | -0.039 | 0.055 | 0.32 | 0.33 | 0.22 | 0.09 | 0.28 | 0.31 |
| 2 | 173342164 | rs6759557 | -1.566 | -0.614 | 0.839 | -0.456 | -0.006 | 0.005 | 0.005 | 0.597 | 0.37 | 0.44 | 0.45 | 0.6 | 0.34 | 0.51 |
| 2 | 173343823 | rs3828343 | -0.987 | -0.604 | 0.772 | -0.49 | -0.004 | -0.044 | -0.044 | 0 | 0.32 | 0.33 | 0.22 | 0.09 | 0.27 | 0.28 |
| 2 | 173350494 | rs2701270 | -0.193 | -0.561 | 0.932 | -0.49 | 0.007 | -0.078 | -0.078 | 0 | 0.16 | 0.48 | 0.12 | 0.06 | 0.12 | 0.12 |
| 2 | 173351199 | rs2701268 | -0.837 | -0.574 | 0.928 | -0.455 | -0.005 | 0.063 | 0.063 | 0.666 | 0.19 | 0.54 | 0.19 | 0.32 | 0.15 | 0.22 |
| 2 | 173352103 | rs2293649 | -0.755 | -0.534 | 0.888 | -0.436 | 0.013 | -0.038 | -0.038 | 1.364 | 0.25 | 0.2 | 0.36 | 0.54 | 0.22 | 0.46 |
| 2 | 173357401 | rs7564654 | -0.745 | -0.532 | 0.778 | -0.412 | 0.015 | -0.063 | -0.063 | 0.027 | 0.25 | 0.15 | 0.36 | 0.54 | 0.22 | 0.46 |
| 2 | 173369231 | rs17664 | -1.947 | -0.522 | 0.767 | -0.47 | 0.021 | 0.064 | 0.064 | 2.946 | 0.43 | 0.68 | 0.54 | 0.87 | 0.37 | 0.67 |
| 2 | 173378145 | rs6755100 | 0.309 | -0.464 | 0.409 | -0.316 | 0.003 | 0.314 | 0.314 | 0 | 0.05 | 0.15 | 0.22 | 0.51 | 0.06 | 0.23 |
| 2 | 173382300 | rs11688498 | 2.644 | -0.569 | -0.638 | -1.198 | -0.138 | 0.198 | 0.198 | 0.384 | 0.3 | 0.04 | 0.31 | 0.01 | 0.4 | 0.15 |
| 2 | 173396868 | rs13012170 | -1.834 | -0.544 | -0.275 | -0.915 | 0.016 | 0.004 | 0.004 | 0 | 0.88 | 0.97 | 0.96 | 0.9 | 0.92 | 0.93 |
| 2 | 173407137 | rs6433365 | -0.353 | -0.211 | 0.497 | -0.719 | -0.013 | -0.08 | -0.08 | 0 | 0.16 | 0.45 | 0.12 | 0.06 | 0.11 | 0.18 |
| 2 | 173415560 | rs12151618 | -1.27 | -0.215 | 0.218 | -0.656 | 0.005 | -0.001 | -0.001 | 0 | 0.3 | 0.69 | 0.3 | 0.33 | 0.2 | 0.39 |
| 2 | 173415896 | rs1920978 | 0.857 | -0.313 | 0.283 | -0.459 | 0.048 | 0.023 | 0.023 | 0 | 0.5 | 0.26 | 0.62 | 0.55 | 0.61 | 0.51 |
| 2 | 173430707 | rs10930562 | -1.467 | -0.064 | 1.519 | -0.193 | -0.043 | 0.014 | 0.014 | 0.911 | 0.22 | 0.45 | 0.21 | 0.24 | 0.15 | 0.24 |
| 2 | 173435757 | rs13394924 | 0.167 | -0.023 | 1.49 | -0.176 | 0.004 | 0.039 | 0.039 | 0.042 | 0.13 | 0.06 | 0.08 | 0.02 | 0.11 | 0.08 |
| 2 | 173461820 | rs11904158 | -1.639 | -0.017 | 1.595 | -0.187 | -0.032 | 0.004 | 0.004 | 1.291 | 0.23 | 0.55 | 0.23 | 0.24 | 0.16 | 0.24 |
| 2 | 173462230 | rs11904366 | -1.002 | -0.022 | 1.588 | -0.268 | -0.029 | -0.111 | -0.111 | 2.333 | 0.18 | 0.4 | 0.13 | 0.02 | 0.13 | 0.15 |
| 2 | 173475703 | rs11896032 | -0.843 | -0.023 | 1.643 | -0.269 | -0.02 | -0.081 | -0.081 | 0.011 | 0.17 | 0.28 | 0.11 | 0.02 | 0.13 | 0.19 |
| 2 | 173481200 | rs7566884 | -1.353 | 0.03 | 1.457 | -0.299 | -0.034 | -0.106 | -0.106 | 0.428 | 0.25 | 0.34 | 0.14 | 0.02 | 0.17 | 0.25 |
| 2 | 173495678 | rs2606067 | 1.626 | 0.192 | 1.352 | -0.232 | -0.018 | -0.013 | -0.013 | 0 | 0.7 | 0.6 | 0.74 | 0.74 | 0.77 | 0.66 |
| 2 | 173498212 | rs17753893 | 0.004 | 0.526 | 1.334 | -0.415 | 0.007 | -0.037 | -0.037 | 0 | 0.15 | 0.18 | 0.19 | 0.02 | 0.22 | 0.07 |
| 2 | 173500587 | rs9287957 | -0.531 | 0.353 | 1.385 | -0.305 | -0.002 | -0.053 | -0.053 | 0 | 0.16 | 0.21 | 0.11 | 0.02 | 0.14 | 0.1 |
| 2 | 173501069 | rs836628 | 0.884 | 0.386 | 1.386 | -0.26 | -0.02 | 0.026 | 0.026 | 0.113 | 0.86 | 0.75 | 0.85 | 0.77 | 0.91 | 0.76 |
| 2 | 173512756 | rs836616 | 0.653 | 0.44 | 1.036 | -0.238 | 0.001 | -0.019 | -0.019 | 0.18 | 0.76 | 0.9 | 0.78 | 0.74 | 0.8 | 0.67 |
| 2 | 173521724 | rs836610 | 1.137 | 0.482 | 1.053 | -0.081 | -0.015 | -0.012 | -0.012 | 0 | 0.65 | 0.42 | 0.58 | 0.24 | 0.74 | 0.49 |
| 2 | 173550622 | rs10190099 | -0.743 | 0.3 | 0.887 | -0.195 | -0.045 | -0.11 | -0.11 | 0 | 0.25 | 0.35 | 0.12 | 0.03 | 0.15 | 0.22 |
| 2 | 173557660 | rs10166107 | -0.688 | 0.298 | 0.878 | -0.196 | -0.029 | -0.078 | -0.078 | 0 | 0.24 | 0.29 | 0.11 | 0.03 | 0.15 | 0.22 |
| 2 | 173560072 | rs2060773 | -0.413 | 0.546 | 0.662 | -0.248 | 0.003 | -0.025 | -0.025 | 0 | 0.18 | 0.19 | 0.26 | 0.03 | 0.27 | 0.11 |
| 2 | 173568994 | rs12328591 | -0.697 | 0.838 | 0.386 | -0.612 | -0.041 | -0.1 | -0.1 | 0 | 0.24 | 0.34 | 0.12 | 0.03 | 0.15 | 0.23 |
| 2 | 173570130 | rs12618102 | -1.068 | 1.041 | 0.078 | -0.482 | -0.039 | -0.031 | -0.031 | 0.103 | 0.34 | 0.51 | 0.38 | 0.78 | 0.24 | 0.49 |
| 2 | 173574173 | rs13028638 | 0.053 | 2.119 | -0.269 | -0.002 | -0.07 | 0.061 | 0.061 | 0 | 0.1 | 0.01 | 0.21 | 0 | 0.21 | 0.06 |
| 2 | 173581583 | rs1451452 | -0.223 | 1.759 | 0.054 | 0.214 | -0.021 | 0.019 | 0.019 | 0 | 0.26 | 0.15 | 0.35 | 0.09 | 0.41 | 0.25 |
| 2 | 173582810 | rs1097473 | 0.195 | 1.726 | 0.236 | 0.215 | 0.039 | -0.056 | -0.056 | 0 | 0.72 | 0.64 | 0.62 | 0.91 | 0.59 | 0.75 |
| 2 | 173583779 | rs9989778 | -0.385 | 1.393 | -0.039 | -0.118 | -0.003 | -0.038 | -0.038 | 0 | 0.13 | 0.2 | 0.08 | 0.03 | 0.12 | 0.19 |
| 2 | 173585237 | rs836593 | 0.528 | 1.588 | -0.16 | 0.457 | 0.031 | -0.04 | -0.04 | 1.315 | 0.65 | 0.56 | 0.38 | 0.16 | 0.5 | 0.59 |
| 2 | 173592268 | rs1349499 | -0.915 | 1.239 | -0.146 | 1.587 | -0.005 | 0.18 | 0.18 | 0 | 0.05 | 0.09 | 0.14 | 0.51 | 0.04 | 0.1 |
| 2 | 173592606 | rs1349498 | -1.96 | 1.969 | -0.046 | 1.51 | 0.019 | 0.122 | 0.122 | 0 | 0.09 | 0.15 | 0.23 | 0.52 | 0.18 | 0.2 |
| 2 | 173608201 | rs10193942 | -1.527 | 1.901 | 0.419 | 0.604 | 0.039 | -0.053 | -0.053 | 0 | 0.16 | 0.28 | 0.2 | 0.04 | 0.25 | 0.3 |

| | | | | | | | | | | | | | | | | |
|---|---|---|---|---|---|---|---|---|---|---|---|---|---|---|---|---|
| 2 | 173617201 | rs13413998 | -1.365 | 1.857 | 0.551 | 0.572 | 0.012 | -0.041 | -0.041 | 0.005 | 0.11 | 0.18 | 0.1 | 0.02 | 0.13 | 0.16 |
| 2 | 173620465 | rs6433379 | 2.093 | 1.605 | 1.516 | 0.593 | 0.008 | -0.045 | -0.045 | 0 | 0.83 | 0.48 | 0.83 | 0.96 | 0.86 | 0.77 |
| 2 | 173622459 | rs12619176 | 1.402 | 1.604 | 1.622 | 0.594 | 0.069 | 0.126 | 0.126 | 0.309 | 0.95 | 0.61 | 0.93 | 0.98 | 0.99 | 0.93 |
| 2 | 173623248 | rs2081406 | -0.44 | 1.257 | 1.48 | 0.011 | 0 | -0.001 | -0.001 | 0 | 0.14 | 0.16 | 0.07 | 0.02 | 0.09 | 0.07 |
| 2 | 173623973 | rs10164414 | -1 | 1.27 | 1.504 | 0.183 | -0.002 | -0.003 | -0.003 | 0 | 0.11 | 0.09 | 0.09 | 0.02 | 0.13 | 0.17 |
| 2 | 173636245 | rs13007746 | 1.389 | 1.129 | 1.526 | 0.337 | 0.072 | 0.131 | 0.131 | 0.801 | 0.94 | 0.57 | 0.92 | 0.98 | 0.99 | 0.93 |
| 2 | 173654046 | rs6712008 | 0.796 | 1.022 | 1.11 | 1.17 | -0.015 | 0.084 | 0.084 | 0.064 | 0.84 | 0.75 | 0.78 | 0.44 | 0.92 | 0.69 |
| 2 | 173655524 | rs10203199 | 0.78 | 0.82 | 0.3 | -0.227 | 0.012 | 0.029 | 0.029 | 0 | 0.45 | 0.34 | 0.28 | 0.09 | 0.4 | 0.35 |
| 2 | 173664609 | rs2161994 | 0.567 | 1.155 | 0.342 | 0.154 | 0.004 | 0.026 | 0.026 | 0 | 0.24 | 0.32 | 0.34 | 0.44 | 0.29 | 0.39 |
| 2 | 173675359 | rs3769312 | 0.306 | 1.128 | 0.389 | 0.132 | 0.032 | 0.034 | 0.034 | 0 | 0.11 | 0.15 | 0.22 | 0.23 | 0.2 | 0.14 |
| 2 | 173679782 | rs3769309 | 0.303 | 0.824 | 0.12 | -0.258 | -0.01 | -0.006 | -0.006 | 0.838 | 0.58 | 0.33 | 0.5 | 0.49 | 0.56 | 0.36 |
| 2 | 173680085 | rs3820852 | 0.472 | 0.551 | -0.49 | -0.53 | 0.01 | -0.054 | -0.054 | 1.987 | 0.16 | 0.01 | 0.1 | 0.23 | 0.09 | 0.1 |
| 2 | 173686663 | rs2258180 | 0.137 | 0.61 | -0.415 | -0.463 | 0.028 | -0.004 | -0.004 | 2.337 | 0.37 | 0.5 | 0.51 | 0.59 | 0.49 | 0.54 |
| 2 | 173686798 | rs2258175 | 0.326 | 0.617 | -0.418 | -0.463 | 0.019 | 0.003 | 0.003 | 0 | 0.35 | 0.44 | 0.48 | 0.56 | 0.47 | 0.51 |
| 2 | 173694029 | rs6715990 | 1.854 | 0.477 | -0.707 | -0.597 | 0.008 | 0.14 | 0.14 | 0 | 0.29 | 0.1 | 0.17 | 0.03 | 0.27 | 0.12 |
| 2 | 173703518 | rs2553001 | -1.75 | 0.487 | -0.797 | -0.591 | 0.023 | 0.091 | 0.091 | 0 | 0.69 | 0.8 | 0.81 | 0.97 | 0.73 | 0.88 |
| 2 | 173703573 | rs7581330 | -0.03 | 0.489 | -0.83 | -0.509 | 0.007 | -0.019 | -0.019 | 0.124 | 0.13 | 0.13 | 0.14 | 0.05 | 0.13 | 0.18 |
| 2 | 173711641 | rs10175694 | 1.634 | 0.542 | -1.543 | -0.649 | 0.004 | 0.153 | 0.153 | 0 | 0.29 | 0.1 | 0.17 | 0.03 | 0.27 | 0.12 |
| 2 | 173717307 | rs2676511 | -0.853 | 0.411 | -1.618 | -0.677 | 0.007 | 0.05 | 0.05 | 0 | 0.6 | 0.7 | 0.67 | 0.84 | 0.63 | 0.69 |
| 2 | 173718391 | rs2113804 | -0.818 | 0.439 | -1.63 | -0.671 | 0.003 | 0.058 | 0.058 | 0 | 0.59 | 0.71 | 0.64 | 0.81 | 0.61 | 0.66 |
| 2 | 173720268 | rs956752 | -0.091 | 0.351 | -1.651 | -0.662 | 0.005 | -0.005 | -0.005 | 0.022 | 0.1 | 0.1 | 0.14 | 0.12 | 0.11 | 0.18 |
| 2 | 173727773 | rs3769289 | 0.219 | 0.481 | -1.949 | -0.547 | -0.056 | -0.082 | -0.082 | 0 | 0.16 | 0.01 | 0.14 | 0.25 | 0.18 | 0.12 |
| 2 | 173737895 | rs1991173 | 0.051 | -0.046 | -1.685 | -0.874 | -0.004 | 0.002 | 0.002 | 0 | 0.46 | 0.58 | 0.59 | 0.81 | 0.52 | 0.39 |
| 2 | 173744658 | rs17706710 | 1.048 | -0.082 | -1.733 | -0.958 | -0.023 | 0.055 | 0.055 | 0 | 0.1 | 0 | 0.03 | 0 | 0.07 | 0.07 |
| 2 | 173749682 | rs2676517 | -0.349 | -0.105 | -1.718 | -0.941 | 0.009 | 0.015 | 0.015 | 0 | 0.51 | 0.57 | 0.65 | 0.83 | 0.6 | 0.48 |
| 2 | 173754914 | rs2676520 | 0.126 | 0.129 | -1.444 | -0.63 | -0.007 | -0.055 | -0.055 | 0 | 0.72 | 0.61 | 0.79 | 0.85 | 0.8 | 0.52 |
| 2 | 173765808 | rs13030842 | -0.482 | 0.338 | -1.001 | 0.114 | 0 | 0 | 0 | 0.012 | 0.4 | 0.57 | 0.53 | 0.65 | 0.48 | 0.33 |
| 2 | 173779775 | rs2553011 | -0.501 | 0.76 | -1.32 | 0.241 | 0.006 | -0.005 | -0.005 | 0 | 0.38 | 0.54 | 0.52 | 0.65 | 0.48 | 0.33 |
| 2 | 173784658 | rs2113807 | -0.031 | 0.076 | -2.56 | -0.189 | 0.091 | 0.236 | 0.236 | 0.686 | 0.51 | 0.09 | 0.53 | 0.82 | 0.67 | 0.55 |
| 2 | 173784960 | rs2468392 | -1.075 | -1.059 | -1.594 | -0.056 | -0.039 | 0.003 | 0.003 | 1.089 | 0.29 | 0.55 | 0.16 | 0.28 | 0.19 | 0.41 |
| 2 | 173789160 | rs3769262 | -0.136 | -0.795 | -1.513 | -0.27 | 0.025 | 0.047 | 0.047 | 0 | 0.51 | 0.34 | 0.46 | 0.7 | 0.58 | 0.41 |
| 2 | 173796835 | rs3769254 | 0.198 | -1.167 | -1.204 | -0.312 | -0.006 | 0.027 | 0.027 | 0 | 0.45 | 0.2 | 0.45 | 0.35 | 0.42 | 0.46 |
| 2 | 173801439 | rs11680594 | 0.158 | -1.202 | -1.211 | -0.11 | -0.002 | -0.012 | -0.012 | 0 | 0.46 | 0.21 | 0.47 | 0.44 | 0.43 | 0.47 |
| 2 | 173809446 | rs3769252 | 0.256 | -1.201 | -1.184 | -0.229 | -0.013 | -0.039 | -0.039 | 0.449 | 0.44 | 0.18 | 0.45 | 0.47 | 0.42 | 0.47 |
| 2 | 173815967 | rs11695769 | 0.517 | -1.258 | -1.219 | -0.238 | -0.021 | -0.007 | -0.007 | 0 | 0.41 | 0.26 | 0.39 | 0.47 | 0.49 | 0.3 |
| 2 | 173820377 | rs745288 | 0.735 | -1.049 | -0.747 | -0.472 | -0.014 | -0.054 | -0.054 | 0 | 0.85 | 0.34 | 0.84 | 0.97 | 0.92 | 0.76 |
| 2 | 173824496 | rs4972521 | 0.671 | -1.031 | -0.999 | -0.371 | -0.042 | -0.093 | -0.093 | 0 | 0.35 | 0.08 | 0.39 | 0.45 | 0.34 | 0.4 |
| 2 | 173831934 | rs16861098 | 0.776 | -1.064 | -1.298 | -0.561 | 0.017 | 0.086 | 0.086 | 0 | 0.29 | 0.05 | 0.22 | 0.16 | 0.23 | 0.2 |
| 2 | 173834077 | rs750374 | 0.124 | -1.262 | -0.387 | -0.695 | -0.001 | -0.002 | -0.002 | 0 | 0.45 | 0.35 | 0.5 | 0.48 | 0.43 | 0.63 |
| 2 | 173835127 | rs3754757 | 0.744 | -1.13 | -0.881 | -0.65 | -0.045 | -0.095 | -0.095 | 0 | 0.34 | 0.04 | 0.38 | 0.45 | 0.33 | 0.4 |
| 2 | 173842735 | rs1867942 | 0.039 | -1.206 | 0.085 | -1.639 | 0.054 | 0.073 | 0.073 | 0.347 | 0.56 | 0.33 | 0.68 | 0.8 | 0.72 | 0.59 |
| 2 | 173846130 | rs16861119 | 0.435 | -0.946 | 0.58 | -1.476 | 0.03 | -0.016 | -0.016 | 0 | 0.06 | 0.17 | 0.02 | 0 | 0.04 | 0.18 |
| 2 | 173847492 | rs2592944 | 0.19 | -0.588 | 0.446 | -1.91 | -0.01 | 0.07 | 0.07 | 0 | 0.6 | 0.3 | 0.66 | 0.88 | 0.62 | 0.53 |
| 2 | 173849809 | rs2602216 | -0.83 | -0.753 | 0.508 | -1.983 | -0.026 | -0.048 | -0.048 | 0 | 0.66 | 0.6 | 0.66 | 0.91 | 0.78 | 0.81 |
| 2 | 173857654 | rs2602215 | 0.779 | -0.53 | 0.424 | -1.835 | -0.056 | 0.01 | 0.01 | 0 | 0.27 | 0.16 | 0.34 | 0.79 | 0.41 | 0.34 |
| 2 | 173861340 | rs2290374 | 0.629 | -0.524 | 0.495 | -1.85 | -0.021 | 0.01 | 0.01 | 0 | 0.29 | 0.24 | 0.35 | 0.79 | 0.4 | 0.35 |
| 2 | 173865284 | rs3754753 | 0.761 | -0.587 | 0.498 | -1.965 | -0.018 | 0.029 | 0.029 | 0 | 0.28 | 0.26 | 0.29 | 0.12 | 0.32 | 0.24 |
| 2 | 173866203 | rs2028934 | -0.183 | -0.755 | 0.271 | -1.974 | 0.026 | 0.014 | 0.014 | 0 | 0.56 | 0.46 | 0.64 | 0.91 | 0.72 | 0.59 |
| 2 | 173867808 | rs7604728 | -1.472 | -0.639 | 0.096 | -1.959 | -0.006 | -0.021 | -0.021 | 0.295 | 0.77 | 0.77 | 0.81 | 1 | 0.76 | 0.81 |
| 2 | 173868790 | rs12053389 | 0.353 | -0.727 | -0.532 | -1.78 | -0.065 | 0.079 | 0.079 | 0 | 0.31 | 0.18 | 0.45 | 0.9 | 0.48 | 0.39 |
| 2 | 173871359 | rs12473597 | -0.017 | -0.753 | -0.474 | -1.793 | -0.018 | 0.015 | 0.015 | 0 | 0.35 | 0.29 | 0.46 | 0.9 | 0.49 | 0.43 |
| 2 | 173875458 | rs3769232 | 0.095 | -0.59 | -0.867 | -1.693 | 0.071 | 0.094 | 0.094 | 0 | 0.4 | 0.14 | 0.43 | 0.21 | 0.3 | 0.23 |
| 2 | 173876920 | rs4972525 | -0.265 | 0.014 | -0.07 | -0.99 | 0.012 | -0.154 | -0.154 | 2.227 | 0.71 | 0.6 | 0.79 | 1 | 0.73 | 0.64 |
| 2 | 173877524 | rs1979186 | 0.114 | 0.012 | -1.212 | -0.815 | -0.009 | 0.081 | 0.081 | 0 | 0.62 | 0.25 | 0.66 | 0.88 | 0.65 | 0.55 |
| 2 | 173887019 | rs3754751 | 0.868 | -0.756 | -0.787 | -0.572 | -0.025 | -0.029 | -0.029 | 0 | 0.71 | 0.34 | 0.77 | 0.9 | 0.81 | 0.7 |
| 2 | 173887053 | rs3754750 | 0.72 | -0.874 | -0.779 | -0.652 | -0.051 | -0.037 | -0.037 | 0 | 0.75 | 0.37 | 0.82 | 0.9 | 0.86 | 0.72 |
| 2 | 173888638 | rs3769222 | 1.177 | -1.308 | -1.063 | -0.971 | -0.007 | -0.014 | -0.014 | 0 | 0.64 | 0.36 | 0.76 | 0.9 | 0.8 | 0.65 |
| 2 | 173888671 | rs16861191 | 0.591 | -1.034 | -1.078 | -0.888 | 0 | -0.01 | -0.01 | 0 | 0.09 | 0.08 | 0.07 | 0.04 | 0.08 | 0.06 |
| 2 | 173889333 | rs4972394 | 0.319 | -1.167 | -0.31 | -0.794 | -0.049 | -0.049 | -0.049 | 0 | 0.85 | 0.52 | 0.92 | 0.95 | 0.93 | 0.83 |
| 2 | 173895285 | rs2305462 | 0.911 | -1.134 | -0.839 | -0.504 | 0.001 | 0 | 0 | 0 | 0.77 | 0.25 | 0.67 | 0.83 | 0.85 | 0.71 |
| 2 | 173901863 | rs6751505 | -0.348 | -0.414 | -1.035 | -0.064 | 0.018 | 0.009 | 0.009 | 0 | 0.5 | 0.59 | 0.51 | 0.57 | 0.43 | 0.48 |
| 2 | 173903800 | rs3769216 | 0.434 | 0.019 | -1.042 | -0.204 | -0.032 | -0.015 | -0.015 | 0 | 0.3 | 0.2 | 0.4 | 0.31 | 0.44 | 0.44 |
| 2 | 173905497 | rs3769215 | 0.598 | -0.408 | -0.807 | 0.055 | 0.006 | 0.001 | 0.001 | 0 | 0.3 | 0.3 | 0.36 | 0.27 | 0.21 | 0.33 |
| 2 | 173909208 | rs895583 | -0.514 | -0.438 | -0.959 | 0.092 | 0 | -0.007 | -0.007 | 0.185 | 0.09 | 0.04 | 0.15 | 0.39 | 0.12 | 0.12 |
| 2 | 173910787 | rs3769213 | 0.57 | -0.582 | -0.899 | -0.077 | 0.019 | -0.034 | -0.034 | 1.933 | 0.29 | 0.24 | 0.29 | 0.61 | 0.22 | 0.32 |
| 2 | 173914775 | rs3769210 | -0.132 | -0.666 | -0.771 | -0.06 | 0.009 | 0.001 | 0.001 | 0.101 | 0.37 | 0.39 | 0.41 | 0.67 | 0.29 | 0.39 |
| 2 | 173915007 | rs4972527 | -0.528 | -0.638 | -0.494 | -0.282 | -0.001 | -0.046 | -0.046 | 0 | 0.66 | 0.72 | 0.83 | 0.97 | 0.73 | 0.83 |
| 2 | 173919485 | rs10497403 | 0.438 | -0.661 | -0.593 | 0.091 | 0.015 | -0.021 | -0.021 | 0 | 0.29 | 0.22 | 0.3 | 0.64 | 0.23 | 0.32 |
| 2 | 173923137 | rs6707813 | -0.442 | -0.948 | -0.434 | 0.085 | -0.005 | 0.005 | 0.005 | 0 | 0.69 | 0.68 | 0.48 | 0.62 | 0.76 | 0.55 |

| | | | | | | | | | | | | | | | | |
|---|---|---|---|---|---|---|---|---|---|---|---|---|---|---|---|---|
| 2 | 173932262 | rs7573673 | -0.225 | -0.709 | -0.636 | -0.769 | 0.02 | 0.135 | 0.135 | 0 | 0.5 | 0.72 | 0.5 | 0.12 | 0.42 | 0.29 |
| 2 | 173937983 | rs2358084 | 0.051 | -0.709 | -0.548 | -0.797 | 0.001 | -0.016 | -0.016 | 0 | 0.46 | 0.59 | 0.4 | 0.1 | 0.38 | 0.23 |
| 2 | 173943880 | rs6433395 | 0.77 | -0.372 | -0.826 | -0.615 | -0.045 | 0.002 | 0.002 | 0.185 | 0.24 | 0.12 | 0.27 | 0.22 | 0.39 | 0.37 |
| 2 | 173952189 | rs12987496 | 0.65 | -0.641 | -1.16 | -0.811 | -0.1 | 0.005 | 0.005 | 0 | 0.28 | 0.03 | 0.2 | 0.21 | 0.36 | 0.36 |
| 2 | 173965296 | rs3754747 | 1.465 | -0.724 | -1.155 | -0.779 | 0.005 | 0.043 | 0.043 | 0.474 | 0.23 | 0.05 | 0.22 | 0.12 | 0.17 | 0.31 |
| 2 | 173965366 | rs3754746 | 2.283 | -0.766 | -1.22 | -0.827 | -0.02 | -0.011 | -0.011 | 0.599 | 0.06 | 0 | 0.04 | 0.06 | 0.05 | 0.12 |
| 2 | 173968996 | rs6759847 | -1.079 | -0.713 | -0.903 | -0.747 | 0.021 | 0.07 | 0.07 | 0 | 0.71 | 0.88 | 0.61 | 0.86 | 0.78 | 0.62 |
| 2 | 174002629 | rs17760899 | -0.213 | -0.796 | -0.562 | -0.753 | 0.01 | 0.056 | 0.056 | 0.285 | 0.46 | 0.64 | 0.28 | 0.1 | 0.38 | 0.22 |
| 2 | 174007883 | rs10930577 | 0.813 | -1.052 | -0.223 | -0.943 | 0.036 | -0.056 | -0.056 | 0.627 | 0.24 | 0.37 | 0.34 | 0.09 | 0.28 | 0.08 |
| 2 | 174013499 | rs3769185 | 1.013 | -0.99 | -0.501 | -1.079 | -0.063 | 0.035 | 0.035 | 0.49 | 0.45 | 0.1 | 0.54 | 0.33 | 0.53 | 0.64 |
| 2 | 174015168 | rs7575189 | 0.725 | -1.008 | -0.488 | -1.046 | -0.021 | 0.07 | 0.07 | 0.877 | 0.48 | 0.1 | 0.48 | 0.32 | 0.57 | 0.69 |
| 2 | 174026324 | rs746179 | 1.39 | -1.823 | -0.204 | -1.003 | 0.004 | 0.005 | 0.005 | 0.014 | 0.26 | 0.25 | 0.37 | 0.13 | 0.19 | 0.35 |
| 2 | 174032545 | rs7599340 | -0.926 | -1.866 | -0.13 | -1.001 | -0.019 | -0.009 | -0.009 | 0 | 0.69 | 0.61 | 0.5 | 0.28 | 0.76 | 0.57 |
| 2 | 174034099 | rs17761169 | 1.343 | -1.623 | -0.448 | -1.053 | -0.174 | -0.041 | -0.041 | 0 | 0.24 | 0.02 | 0.3 | 0.22 | 0.44 | 0.35 |
| 2 | 174035039 | rs1816717 | 1.25 | -1.706 | -0.283 | -1.096 | 0.006 | -0.049 | -0.049 | 0 | 0.21 | 0.26 | 0.25 | 0.05 | 0.21 | 0.07 |
| 2 | 174039497 | rs16861304 | 0.119 | -1.758 | -0.252 | -1.072 | -0.071 | 0.077 | 0.077 | 0.068 | 0.24 | 0.39 | 0.18 | 0.58 | 0.14 | 0.18 |
| 2 | 174041358 | rs935007 | -1.385 | -1.617 | -0.245 | -1.002 | -0.003 | -0.007 | -0.007 | 0 | 0.73 | 0.69 | 0.74 | 0.85 | 0.8 | 0.6 |
| 2 | 174048371 | rs1837470 | 0.76 | -1.545 | -0.471 | -0.948 | 0.041 | 0.056 | 0.056 | 0.81 | 0.48 | 0.1 | 0.48 | 0.34 | 0.61 | 0.67 |
| 2 | 174074407 | rs2289396 | 0.159 | -2.11 | -0.022 | -0.551 | 0.051 | 0.054 | 0.054 | 0 | 0.55 | 0.32 | 0.56 | 0.3 | 0.68 | 0.52 |
| 2 | 174086492 | rs16861363 | 0.855 | -1.879 | 0.169 | -0.602 | -0.007 | -0.027 | -0.027 | 0.35 | 0.28 | 0.33 | 0.25 | 0.11 | 0.21 | 0.31 |
| 2 | 174086672 | rs11249 | 0.541 | -1.917 | 0.202 | -0.755 | -0.003 | -0.024 | -0.024 | 0 | 0.22 | 0.24 | 0.28 | 0.06 | 0.2 | 0.17 |
| 2 | 174087747 | rs2028382 | 1.422 | -2.001 | 0.154 | -0.623 | -0.103 | -0.002 | -0.002 | 0 | 0.31 | 0.09 | 0.28 | 0.24 | 0.47 | 0.35 |
| 2 | 174088266 | rs10497402 | 1.108 | -1.961 | 0.189 | -0.628 | 0.004 | 0.007 | 0.007 | 0.07 | 0.27 | 0.24 | 0.25 | 0.11 | 0.21 | 0.31 |
| 2 | 174089606 | rs10497401 | 0.739 | -1.798 | -0.059 | -0.39 | -0.007 | -0.042 | -0.042 | 0 | 0.19 | 0.24 | 0.21 | 0.05 | 0.15 | 0.06 |
| 2 | 174090542 | rs8446 | 0.865 | -1.808 | -0.023 | -0.366 | -0.004 | -0.001 | -0.001 | 0.474 | 0.59 | 0.43 | 0.54 | 0.34 | 0.68 | 0.66 |
| 2 | 174091088 | rs11887008 | 0.052 | -1.836 | 0.174 | -0.399 | -0.007 | -0.043 | -0.043 | 0.163 | 0.74 | 0.68 | 0.69 | 0.92 | 0.79 | 0.78 |
| 2 | 174091200 | rs4972399 | 0.352 | -1.822 | 0.122 | -0.39 | -0.002 | -0.023 | -0.023 | 0.038 | 0.22 | 0.24 | 0.28 | 0.06 | 0.2 | 0.16 |
| 2 | 174091964 | rs2119123 | 0.14 | -1.769 | 0.095 | -0.288 | -0.041 | 0.104 | 0.104 | 0 | 0.82 | 0.7 | 0.82 | 0.41 | 0.87 | 0.81 |
| 2 | 174093597 | rs4972538 | 1.072 | -1.669 | -0.004 | -0.212 | -0.01 | 0.268 | 0.268 | 0 | 0.94 | 0.83 | 0.89 | 0.41 | 0.96 | 0.85 |
| 2 | 174094145 | rs10182247 | -0.373 | -1.706 | -0.264 | -0.335 | -0.021 | -0.089 | -0.089 | 0 | 0.33 | 0.4 | 0.28 | 0.05 | 0.24 | 0.22 |
| 2 | 174097106 | rs16861406 | -0.257 | -1.415 | -0.396 | 0.003 | 0 | 0.104 | 0.104 | 0 | 0.06 | 0.06 | 0.16 | 0.59 | 0.07 | 0.08 |
| 2 | 174098201 | rs3769154 | -0.318 | -1.549 | -0.251 | -0.074 | -0.051 | 0.122 | 0.122 | 0 | 0.19 | 0.32 | 0.16 | 0.57 | 0.13 | 0.15 |
| 2 | 174103428 | rs3820831 | -0.678 | -1.135 | -0.985 | 0.226 | 0 | 0.078 | 0.078 | 0 | 0.08 | 0.06 | 0.16 | 0.57 | 0.09 | 0.19 |
| 2 | 174105564 | rs1583784 | -0.14 | -1.22 | -1.46 | -0.336 | 0.017 | 0.093 | 0.093 | 0.099 | 0.58 | 0.74 | 0.6 | 0.84 | 0.65 | 0.71 |
| 2 | 174107299 | rs4972539 | 0.582 | -0.949 | -1.812 | 0.008 | -0.034 | 0.026 | 0.026 | 0.331 | 0.24 | 0.02 | 0.3 | 0.13 | 0.22 | 0.24 |
| 2 | 174114111 | rs10497400 | -0.352 | -0.582 | -1.265 | 0.818 | -0.014 | 0.03 | 0.03 | 0 | 0.06 | 0.01 | 0.19 | 0.59 | 0.08 | 0.15 |
| 2 | 174114139 | rs10497399 | 0.016 | -1.172 | -0.833 | 0.465 | -0.074 | -0.112 | -0.112 | 0 | 0.19 | 0.44 | 0.16 | 0.04 | 0.08 | 0.23 |
| 2 | 174116129 | rs7591449 | -0.81 | -1.197 | -0.612 | 0.439 | -0.055 | 0.063 | 0.063 | 0 | 0.27 | 0.62 | 0.36 | 0.62 | 0.17 | 0.38 |
| 2 | 174117020 | rs6731322 | -0.098 | -1.049 | -0.538 | 0.637 | -0.066 | 0.164 | 0.164 | 0 | 0.21 | 0.45 | 0.25 | 0.59 | 0.11 | 0.25 |
| 2 | 174118046 | rs11686011 | 0.267 | -0.25 | -1.555 | 0.31 | -0.139 | 0.01 | 0.01 | 0 | 0.37 | 0.04 | 0.29 | 0.22 | 0.51 | 0.36 |
| 2 | 174123273 | rs820007 | -0.854 | -0.066 | -0.129 | 1.001 | 0.009 | 0.133 | 0.133 | 0 | 0.08 | 0.23 | 0.1 | 0.23 | 0.06 | 0.04 |
| 2 | 174127820 | rs4972540 | 0.064 | -0.252 | -0.825 | 0.501 | 0.01 | 0.017 | 0.017 | 0.072 | 0.12 | 0.04 | 0.2 | 0.05 | 0.08 | 0.17 |
| 2 | 174129804 | rs10497398 | 0.417 | -0.322 | -1.076 | 0.249 | -0.005 | 0.001 | 0.001 | 0 | 0.1 | 0.06 | 0.12 | 0.11 | 0.1 | 0.18 |
| 2 | 174132406 | rs10202480 | 0.489 | -0.396 | -0.665 | -0.023 | 0.01 | -0.035 | -0.035 | 0 | 0.07 | 0.19 | 0.04 | 0 | 0.03 | 0.01 |
| 2 | 174134529 | rs13031154 | 0.614 | 0.266 | -1.745 | -0.063 | -0.097 | -0.075 | -0.075 | 0.924 | 0.19 | 0.01 | 0.16 | 0.21 | 0.28 | 0.18 |
| 2 | 174139401 | rs4972542 | -1.088 | -0.328 | -0.84 | 0.374 | -0.002 | -0.001 | -0.001 | 3.777 | 0.34 | 0.36 | 0.39 | 0.69 | 0.25 | 0.42 |
| 2 | 174141670 | rs1442973 | -0.648 | -0.368 | -0.56 | 0.697 | -0.014 | 0.035 | 0.035 | 0 | 0.23 | 0.29 | 0.3 | 0.54 | 0.13 | 0.24 |
| 2 | 174145477 | rs820042 | -0.991 | 0.226 | -0.031 | 0.24 | 0.043 | -0.013 | -0.013 | 0 | 0.06 | 0.24 | 0.04 | 0 | 0.07 | 0.07 |
| 2 | 174149401 | rs6650777 | -0.023 | 0.051 | -0.89 | -0.091 | 0.002 | -0.001 | -0.001 | 0.007 | 0.26 | 0.25 | 0.25 | 0.08 | 0.22 | 0.23 |
| 2 | 174151250 | rs13025846 | 0.206 | 0.076 | -0.958 | 0.017 | 0.022 | 0.023 | 0.023 | 0 | 0.24 | 0.08 | 0.23 | 0.12 | 0.22 | 0.23 |
| 2 | 174155192 | rs868294 | -0.757 | 0.696 | -1.017 | 0.077 | 0.008 | -0.001 | -0.001 | 0 | 0.55 | 0.46 | 0.55 | 0.38 | 0.64 | 0.56 |
| 2 | 174157614 | rs1093977 | 0.645 | 0.735 | -1.039 | 0.096 | -0.002 | 0 | 0 | 0 | 0.43 | 0.5 | 0.45 | 0.62 | 0.36 | 0.44 |
| 2 | 174163804 | rs1838023 | -0.501 | 1.641 | -1.1 | 0.442 | -0.042 | -0.019 | -0.019 | 0 | 0.19 | 0.12 | 0.25 | 0.21 | 0.31 | 0.21 |
| 2 | 174182985 | rs6714014 | -0.928 | 0.717 | -0.826 | -0.133 | -0.001 | -0.037 | -0.037 | 0 | 0.18 | 0.21 | 0.09 | 0.01 | 0.1 | 0.12 |
| 2 | 174184206 | rs7595292 | -1.436 | 0.792 | -0.035 | -0.566 | -0.013 | -0.164 | -0.164 | 0 | 0.29 | 0.42 | 0.26 | 0.01 | 0.21 | 0.17 |
| 2 | 174206181 | rs10153883 | -1.069 | 0.312 | 0.914 | -0.712 | -0.012 | -0.064 | -0.064 | 0.172 | 0.31 | 0.44 | 0.2 | 0.11 | 0.23 | 0.1 |
| 2 | 174207268 | rs7562518 | 1.573 | 0.173 | 0.408 | -0.18 | -0.01 | -0.01 | -0.01 | 0 | 0.47 | 0.21 | 0.56 | 0.56 | 0.57 | 0.45 |
| 2 | 174207470 | rs7589172 | -0.198 | 0.132 | 0.38 | -0.279 | 0.016 | -0.041 | -0.041 | 5.751 | 0.88 | 0.78 | 0.81 | 0.99 | 0.85 | 0.87 |
| 2 | 174209822 | rs10204104 | 0.234 | 0.106 | 0.485 | -0.369 | 0.023 | -0.061 | -0.061 | 0 | 0.86 | 0.69 | 0.79 | 0.99 | 0.85 | 0.87 |
| 2 | 174212672 | rs1560834 | -0.449 | 0.169 | -0.34 | -0.413 | -0.041 | 0.106 | 0.106 | 0 | 0.3 | 0.07 | 0.17 | 0.13 | 0.29 | 0.17 |
| 2 | 174212910 | rs1550622 | -0.148 | 0.165 | -0.205 | -0.461 | 0.014 | -0.071 | -0.071 | 0.005 | 0.86 | 0.76 | 0.8 | 1 | 0.85 | 0.87 |
| 2 | 174214306 | rs752529 | -0.041 | 0.198 | -0.509 | -0.217 | -0.04 | 0.154 | 0.154 | 0 | 0.25 | 0.05 | 0.14 | 0 | 0.24 | 0.13 |
| 2 | 174221017 | rs4972554 | 0.289 | 0.051 | -0.393 | -0.103 | 0.004 | -0.029 | -0.029 | 0.179 | 0.85 | 0.81 | 0.82 | 0.98 | 0.85 | 0.88 |
| 2 | 174226867 | rs11691325 | 1.899 | -0.151 | -0.66 | 0.279 | -0.065 | -0.033 | -0.033 | 0 | 0.4 | 0.17 | 0.51 | 0.56 | 0.51 | 0.33 |
| 2 | 174227876 | rs767074 | -0.492 | -0.238 | -0.489 | 0.265 | -0.015 | -0.011 | -0.011 | 0 | 0.37 | 0.53 | 0.27 | 0.3 | 0.29 | 0.41 |
| 2 | 174229122 | rs11078 | 1.905 | -0.208 | -0.502 | 0.579 | -0.055 | -0.039 | -0.039 | 0.016 | 0.4 | 0.16 | 0.45 | 0.4 | 0.48 | 0.31 |
| 2 | 174230582 | rs6761787 | 0.1 | -0.203 | -0.413 | 0.422 | 0.012 | -0.048 | -0.048 | 0 | 0.85 | 0.77 | 0.82 | 0.97 | 0.85 | 0.88 |
| 2 | 174232845 | rs10193202 | -0.665 | -0.208 | -0.337 | 0.588 | -0.015 | -0.011 | -0.011 | 0 | 0.37 | 0.53 | 0.27 | 0.3 | 0.29 | 0.41 |
| 2 | 174233041 | rs1047885 | -0.601 | -0.221 | -0.345 | 0.582 | -0.014 | -0.01 | -0.01 | 0.971 | 0.36 | 0.53 | 0.27 | 0.3 | 0.29 | 0.41 |

| | | | | | | | | | | | | | | | | |
|---|---|---|---|---|---|---|---|---|---|---|---|---|---|---|---|---|
| 2 | 174243909 | rs10754981 | 0.274 | -0.262 | -0.478 | 0.667 | -0.006 | -0.002 | -0.002 | 0.179 | 0.66 | 0.43 | 0.74 | 0.71 | 0.75 | 0.65 |
| 2 | 174247919 | rs7591287 | 1.186 | 0.604 | -1.144 | 0.86 | -0.042 | 0.014 | 0.014 | 0.984 | 0.44 | 0.16 | 0.36 | 0.32 | 0.47 | 0.35 |
| 2 | 174251944 | rs7559958 | -0.424 | 1.116 | -0.571 | 0.839 | 0.004 | -0.031 | -0.031 | 0 | 0.35 | 0.36 | 0.29 | 0.22 | 0.37 | 0.32 |
| 2 | 174252625 | rs11679682 | 2.19 | 1.329 | -1.227 | 0.918 | 0.017 | 0.002 | 0.002 | 0 | 0.32 | 0.04 | 0.19 | 0.17 | 0.25 | 0.26 |
| 2 | 174252797 | rs16861601 | 1.044 | 1.374 | -1.195 | 0.833 | 0.006 | -0.076 | -0.076 | 0.022 | 0.23 | 0.3 | 0.18 | 0.09 | 0.23 | 0.12 |
| 2 | 174253310 | rs16861605 | 1.402 | 1.55 | -0.871 | 1.235 | 0.02 | 0.138 | 0.138 | 0 | 0.11 | 0.22 | 0.28 | 0.36 | 0.13 | 0.03 |
| 2 | 174253761 | rs6433403 | -1.079 | 1.155 | -1.212 | 1.151 | 0 | 0.001 | 0.001 | 0 | 0.47 | 0.47 | 0.34 | 0.43 | 0.47 | 0.59 |
| 2 | 174253799 | rs10930590 | 1.337 | 1.166 | -1.161 | 0.986 | 0.053 | -0.004 | -0.004 | 0.796 | 0.43 | 0.14 | 0.23 | 0.34 | 0.34 | 0.53 |
| 2 | 174253888 | rs10930591 | 0.592 | 1.264 | -0.993 | 1.384 | -0.007 | -0.008 | -0.008 | 0 | 0.1 | 0.06 | 0.1 | 0.14 | 0.14 | 0.13 |
| 2 | 174254430 | rs6717268 | 0.12 | 1.09 | -0.778 | 1.24 | 0.069 | 0.13 | 0.13 | 0 | 0.79 | 0.52 | 0.52 | 0.57 | 0.7 | 0.84 |
| 2 | 174259439 | rs11689493 | 0.884 | 1.06 | -1.275 | 0.914 | 0 | 0.011 | 0.011 | 0 | 0.41 | 0.16 | 0.29 | 0.33 | 0.39 | 0.32 |
| 2 | 174261612 | rs10200473 | 0.494 | 1.07 | -0.902 | 1.064 | 0.052 | 0.047 | 0.047 | 0 | 0.82 | 0.64 | 0.6 | 0.68 | 0.73 | 0.82 |
| 2 | 174267281 | rs1865236 | 0.384 | 0.208 | -1.616 | 0.514 | 0.003 | 0.012 | 0.012 | 0 | 0.61 | 0.41 | 0.6 | 0.8 | 0.65 | 0.52 |
| 2 | 174267348 | rs1865235 | 0.61 | -0.016 | -1.815 | 1.153 | 0.007 | -0.026 | -0.026 | 0 | 0.78 | 0.86 | 0.84 | 0.64 | 0.79 | 0.71 |
| 2 | 174271805 | rs6433406 | -0.483 | -0.484 | -1.831 | 0.882 | 0.013 | -0.007 | -0.007 | 0 | 0.4 | 0.55 | 0.37 | 0.55 | 0.29 | 0.27 |
| 2 | 174272761 | rs10432425 | 0.002 | -0.69 | -1.638 | 1.253 | 0 | -0.002 | -0.002 | 0 | 0.73 | 0.65 | 0.68 | 0.77 | 0.7 | 0.67 |
| 2 | 174275268 | rs16861616 | -0.588 | -0.455 | -1.664 | 1.61 | 0.047 | -0.078 | -0.078 | 0 | 0.14 | 0.03 | 0.18 | 0.38 | 0.07 | 0.16 |
| 2 | 174283773 | rs7567442 | -0.703 | -0.049 | -0.902 | 1.48 | -0.01 | 0.09 | 0.09 | 0.355 | 0.22 | 0.46 | 0.34 | 0.42 | 0.22 | 0.42 |
| 2 | 174286056 | rs6724653 | -0.649 | -0.297 | -1.568 | 1.344 | 0.024 | -0.064 | -0.064 | 0.224 | 0.29 | 0.14 | 0.33 | 0.34 | 0.26 | 0.28 |
| 2 | 174292382 | rs6433411 | -0.389 | -0.153 | -1.717 | 1.244 | -0.027 | -0.091 | -0.091 | 2.192 | 0.23 | 0.08 | 0.33 | 0.35 | 0.27 | 0.17 |
| 2 | 174297659 | rs10497394 | -0.325 | -0.142 | -1.567 | 1.296 | -0.017 | -0.068 | -0.068 | 0 | 0.22 | 0.1 | 0.33 | 0.36 | 0.28 | 0.17 |
| 2 | 174297735 | rs7573192 | -0.458 | -0.215 | -1.188 | 1.26 | -0.001 | -0.014 | -0.014 | 0 | 0.23 | 0.21 | 0.34 | 0.35 | 0.28 | 0.17 |
| 2 | 174299127 | rs13417028 | -1.466 | -0.255 | -0.678 | 1.429 | 0.009 | 0.035 | 0.035 | 0.153 | 0.41 | 0.69 | 0.53 | 0.78 | 0.39 | 0.61 |
| 2 | 174303726 | rs1435772 | -0.297 | -0.199 | -0.308 | 1.492 | -0.001 | 0.014 | 0.014 | 0 | 0.24 | 0.31 | 0.35 | 0.35 | 0.28 | 0.19 |
| 2 | 174306537 | rs16861690 | -0.566 | -0.075 | -0.095 | 1.328 | 0.023 | 0.231 | 0.231 | 1.35 | 0.11 | 0.36 | 0.14 | 0.39 | 0.07 | 0.26 |
| 2 | 174307600 | rs1865651 | -0.971 | 0.149 | -0.773 | 0.63 | 0.002 | 0.014 | 0.014 | 0 | 0.53 | 0.57 | 0.55 | 0.78 | 0.44 | 0.63 |
| 2 | 174307942 | rs930311 | -0.466 | 0.333 | -1.181 | -0.085 | -0.009 | 0.113 | 0.113 | 0 | 0.4 | 0.27 | 0.55 | 0.92 | 0.44 | 0.45 |
| 2 | 174314161 | rs4972566 | 0.798 | -0.946 | -0.498 | 0.401 | -0.084 | -0.004 | -0.004 | 0.639 | 0.71 | 0.44 | 0.76 | 0.78 | 0.89 | 0.58 |
| 2 | 174317510 | rs6433414 | -0.794 | -0.957 | 0.039 | 0.373 | -0.008 | 0.184 | 0.184 | 0 | 0.1 | 0.44 | 0.14 | 0.2 | 0.04 | 0.14 |
| 2 | 174317944 | rs7608717 | 0.563 | -1.194 | -0.289 | 0.144 | 0.043 | 0.048 | 0.048 | 0 | 0.82 | 0.95 | 0.91 | 0.98 | 0.93 | 0.75 |
| 2 | 174321693 | rs10183922 | 1.633 | -0.156 | -0.222 | 0.655 | -0.001 | -0.001 | -0.001 | 1.682 | 0.58 | 0.46 | 0.4 | 0.45 | 0.64 | 0.49 |
| 2 | 174321840 | rs10173092 | 1.322 | -0.189 | -0.053 | 0.73 | 0.004 | 0.01 | 0.01 | 0 | 0.66 | 0.49 | 0.5 | 0.63 | 0.68 | 0.63 |
| 2 | 174324905 | rs919868 | 0.082 | 0.131 | -0.342 | 0.999 | -0.071 | -0.069 | -0.069 | 0 | 0.11 | 0.01 | 0.36 | 0.39 | 0.26 | 0.11 |
| 2 | 174326790 | rs16861732 | -1.533 | -0.218 | 0.179 | 1.03 | -0.021 | 0.141 | 0.141 | 0 | 0.09 | 0.22 | 0.12 | 0.19 | 0.03 | 0.18 |
| 2 | 174327631 | rs13019811 | 1.481 | -0.821 | -0.838 | 0.316 | -0.091 | 0.12 | 0.12 | 0.133 | 0.4 | 0.04 | 0.3 | 0.2 | 0.46 | 0.3 |
| 2 | 174336177 | rs7349190 | 0.894 | -0.968 | -0.395 | 0.274 | 0.011 | 0.024 | 0.024 | 0 | 0.48 | 0.23 | 0.42 | 0.39 | 0.49 | 0.49 |
| 2 | 174336932 | rs12473093 | -0.395 | -1.179 | -0.128 | 0.112 | -0.078 | -0.07 | -0.07 | 0.162 | 0.36 | 0.11 | 0.47 | 0.42 | 0.43 | 0.37 |
| 2 | 174337978 | rs11688731 | 1.258 | -1.204 | -0.119 | 0.111 | 0.012 | 0.024 | 0.024 | 0 | 0.49 | 0.23 | 0.42 | 0.39 | 0.49 | 0.49 |
| 2 | 174338087 | rs1433681 | -0.053 | -1.157 | 0.087 | 0.117 | 0.006 | 0.05 | 0.05 | 1.683 | 0.13 | 0.48 | 0.09 | 0.19 | 0.07 | 0.15 |
| 2 | 174340647 | rs6722351 | 0.705 | -1.204 | -0.384 | 0.252 | -0.028 | -0.016 | -0.016 | 0 | 0.57 | 0.73 | 0.44 | 0.51 | 0.55 | 0.51 |
| 2 | 174344080 | rs12466967 | 1.409 | -1.139 | -0.211 | 0.393 | 0.042 | -0.044 | -0.044 | 0.254 | 0.76 | 0.15 | 0.78 | 0.62 | 0.9 | 0.67 |
| 2 | 174351233 | rs10930601 | 0.276 | -1.101 | 0.122 | -0.252 | 0.013 | 0.03 | 0.03 | 0 | 0.48 | 0.46 | 0.38 | 0.35 | 0.42 | 0.38 |
| 2 | 174353464 | rs1369648 | 1.296 | -1.178 | -0.01 | -0.312 | -0.029 | 0.099 | 0.099 | 0.037 | 0.36 | 0.08 | 0.28 | 0.23 | 0.35 | 0.31 |
| 2 | 174357864 | rs4972581 | 0.088 | -1.203 | 0.702 | -0.315 | 0.015 | 0.031 | 0.031 | 0 | 0.49 | 0.59 | 0.39 | 0.35 | 0.42 | 0.38 |
| 2 | 174358434 | rs16861826 | -0.703 | -1.173 | 0.362 | -0.327 | -0.015 | -0.033 | -0.033 | 0 | 0.11 | 0.35 | 0.08 | 0.04 | 0.06 | 0.07 |
| 2 | 174360696 | rs12617057 | 0.372 | -1.04 | 0.412 | -0.222 | 0.013 | 0.043 | 0.043 | 0 | 0.47 | 0.43 | 0.36 | 0.26 | 0.41 | 0.38 |
| 2 | 174377174 | rs7560077 | 2.676 | -0.944 | 1.342 | 0.347 | -0.056 | -0.001 | -0.001 | 0 | 0.86 | 0.29 | 0.89 | 0.88 | 0.97 | 0.94 |
| 2 | 174380427 | rs16861840 | -1.541 | -0.86 | 1.72 | 0.384 | -0.008 | 0.012 | 0.012 | 0 | 0.08 | 0.28 | 0.05 | 0.04 | 0.02 | 0.06 |
| 2 | 174401638 | rs11684559 | 0.578 | -0.584 | 1.171 | 0.235 | 0.022 | 0.015 | 0.015 | 0.842 | 0.77 | 0.67 | 0.51 | 0.35 | 0.68 | 0.64 |
| 2 | 174414897 | rs1526300 | -1.224 | -0.704 | 0.848 | 0.244 | -0.016 | -0.006 | -0.006 | 0 | 0.06 | 0.13 | 0.03 | 0.04 | 0.02 | 0.06 |
| 2 | 174421167 | rs6433428 | 0.689 | -0.782 | 0.782 | 0.211 | -0.014 | -0.008 | -0.008 | 0 | 0.73 | 0.7 | 0.48 | 0.36 | 0.66 | 0.62 |
| 2 | 174421675 | rs6433429 | -2.437 | -1.004 | 0.815 | 0.164 | -0.048 | 0.024 | 0.024 | 0.032 | 0.11 | 0.61 | 0.1 | 0.12 | 0.02 | 0.09 |
| 2 | 174432078 | rs501548 | 0.92 | -0.775 | 0.882 | 0.188 | 0.018 | 0.003 | 0.003 | 0.443 | 0.71 | 0.6 | 0.46 | 0.36 | 0.65 | 0.59 |
| 2 | 174437538 | rs10168874 | -0.934 | -0.737 | 0.602 | 0.199 | -0.007 | -0.003 | -0.003 | 0 | 0.27 | 0.28 | 0.53 | 0.64 | 0.35 | 0.41 |
| 2 | 174441493 | rs12328679 | -1.718 | -1.505 | 0.608 | 0.358 | -0.024 | -0.002 | -0.002 | 0 | 0.07 | 0.18 | 0.04 | 0.05 | 0.01 | 0.07 |
| 2 | 174450397 | rs7584596 | 1.305 | -1.515 | 0.633 | 1.089 | -0.056 | 0.234 | 0.234 | 0.24 | 0.84 | 0.51 | 0.81 | 0.59 | 0.93 | 0.77 |
| 2 | 174452807 | rs6724972 | -0.343 | -1.454 | 0.032 | 0.587 | 0.009 | 0.014 | 0.014 | 0 | 0.33 | 0.25 | 0.16 | 0.19 | 0.28 | 0.42 |
| 2 | 174454044 | rs6714145 | 0.769 | -1.558 | -0.247 | 0.619 | 0.075 | 0.08 | 0.08 | 0.012 | 0.56 | 0.28 | 0.74 | 0.68 | 0.72 | 0.53 |
| 2 | 174455978 | rs7599124 | 0.07 | -1.536 | -0.707 | 0.698 | -0.026 | -0.055 | -0.055 | 0.26 | 0.14 | 0.04 | 0.13 | 0.22 | 0.19 | 0.16 |
| 2 | 174459939 | rs4972590 | -0.379 | -1.349 | -0.239 | 0.479 | -0.061 | -0.057 | -0.057 | 0 | 0.28 | 0.41 | 0.11 | 0.22 | 0.13 | 0.41 |
| 2 | 174466249 | rs16861885 | 0.756 | -0.849 | -0.937 | 0.734 | -0.008 | -0.04 | -0.04 | 0.373 | 0.39 | 0.18 | 0.57 | 0.4 | 0.59 | 0.25 |
| 2 | 174472177 | rs2167624 | -0.854 | -0.539 | -0.493 | 0.626 | 0 | 0 | 0 | 0 | 0.18 | 0.15 | 0.17 | 0.15 | 0.12 | 0.27 |
| 2 | 174472685 | rs515309 | 0.533 | -0.287 | -0.412 | 0.579 | 0.014 | -0.009 | -0.009 | 0 | 0.11 | 0.19 | 0.14 | 0.07 | 0.14 | 0.14 |
| 2 | 174472810 | rs730912 | -0.726 | -0.542 | -0.78 | 0.526 | 0.007 | 0.007 | 0.007 | 0 | 0.17 | 0.09 | 0.16 | 0.16 | 0.12 | 0.27 |
| 2 | 174476200 | rs6731912 | -0.284 | -0.209 | -0.691 | 0.82 | -0.008 | -0.04 | -0.04 | 0 | 0.34 | 0.26 | 0.43 | 0.47 | 0.32 | 0.43 |
| 2 | 174478729 | rs2358235 | 0.694 | -0.477 | -1.471 | -0.214 | -0.061 | 0.05 | 0.05 | 0 | 0.37 | 0.18 | 0.32 | 0.32 | 0.47 | 0.37 |
| 2 | 174479004 | rs1465873 | 0.185 | -1.239 | -0.411 | -0.031 | -0.056 | -0.133 | -0.133 | 0 | 0.73 | 0.46 | 0.84 | 0.98 | 0.83 | 0.82 |
| 2 | 174480429 | rs731588 | 0.874 | -1.18 | -0.188 | 0.294 | -0.064 | -0.028 | -0.028 | 0.017 | 0.84 | 0.46 | 0.89 | 0.87 | 0.92 | 0.74 |
| 2 | 174480641 | rs755818 | 1.188 | -1.322 | -1.165 | 0.415 | -0.009 | 0.016 | 0.016 | 0 | 0.41 | 0.15 | 0.51 | 0.35 | 0.57 | 0.45 |

| | | | | | | | | | | | | | | | | |
|---|---|---|---|---|---|---|---|---|---|---|---|---|---|---|---|---|
| 2 | 174480695 | rs755817 | 0.741 | -1.539 | 0.359 | 0.997 | -0.054 | -0.017 | -0.017 | 0 | 0.84 | 0.55 | 0.88 | 0.85 | 0.93 | 0.74 |
| 2 | 174480881 | rs755816 | 1.198 | -1.251 | 0.659 | 1.309 | -0.026 | 0.097 | 0.097 | 1.293 | 0.76 | 0.47 | 0.63 | 0.41 | 0.85 | 0.63 |
| 2 | 174481149 | rs544938 | 1.855 | -1.328 | 0.974 | 1.482 | -0.02 | 0.032 | 0.032 | 0.492 | 0.86 | 0.39 | 0.79 | 0.85 | 0.92 | 0.84 |
| 2 | 174483240 | rs642108 | 1.053 | -1.684 | 0.871 | 1.632 | -0.018 | 0.012 | 0.012 | 0 | 0.7 | 0.42 | 0.77 | 0.45 | 0.8 | 0.78 |
| 2 | 174486590 | rs6722856 | -1.151 | -1.69 | 1.149 | 1.589 | -0.005 | 0.002 | 0.002 | 0 | 0.26 | 0.29 | 0.17 | 0.23 | 0.19 | 0.17 |
| 2 | 174494735 | rs631896 | -1.131 | -2.146 | 0.464 | 1.313 | 0.054 | 0.037 | 0.037 | 0.204 | 0.32 | 0.18 | 0.14 | 0.24 | 0.19 | 0.35 |
| 2 | 174495916 | rs10171980 | -0.183 | -2.119 | 0.439 | 1.323 | 0.048 | 0.003 | 0.003 | 0 | 0.2 | 0.07 | 0.09 | 0.2 | 0.1 | 0.3 |
| 2 | 174503303 | rs13396925 | -1.389 | -2.013 | 0.695 | 1.241 | -0.031 | -0.015 | -0.015 | 0 | 0.32 | 0.47 | 0.21 | 0.28 | 0.2 | 0.39 |
| 2 | 174504056 | rs13400085 | -0.697 | -2.016 | 0.624 | 1.244 | 0.057 | -0.016 | -0.016 | 0.93 | 0.25 | 0.12 | 0.16 | 0.28 | 0.13 | 0.34 |
| 2 | 174504924 | rs13028485 | -0.708 | -1.796 | 0.711 | 1.405 | 0 | 0.014 | 0.014 | 0 | 0.08 | 0.11 | 0.06 | 0.2 | 0.07 | 0.11 |
| 2 | 174505422 | rs2601029 | -1.645 | -1.801 | 0.711 | 1.402 | 0.028 | 0.085 | 0.085 | 0.13 | 0.16 | 0.42 | 0.12 | 0.2 | 0.14 | 0.16 |
| 2 | 174512937 | rs821147 | -2.216 | -1.875 | 0.627 | 1.377 | -0.003 | 0 | 0 | 0 | 0.46 | 0.55 | 0.3 | 0.48 | 0.32 | 0.52 |
| 2 | 174515825 | rs3936406 | 1.49 | -1.84 | 0.414 | 1.404 | 0.012 | -0.003 | -0.003 | 0 | 0.58 | 0.61 | 0.8 | 0.56 | 0.74 | 0.54 |
| 2 | 174519164 | rs6734550 | 1.036 | -1.83 | 0.455 | 1.234 | -0.009 | 0 | 0 | 0.254 | 0.64 | 0.6 | 0.82 | 0.38 | 0.82 | 0.61 |
| 2 | 174520048 | rs2358230 | -1.226 | -1.865 | 0.494 | 1.18 | 0.067 | -0.023 | -0.023 | 1.472 | 0.08 | 0.55 | 0.07 | 0 | 0.07 | 0.05 |
| 2 | 174521516 | rs2167625 | 2.159 | -1.82 | -0.416 | 1.206 | 0.058 | 0.32 | 0.32 | 0.946 | 0.49 | 0.05 | 0.63 | 0.14 | 0.66 | 0.44 |
| 2 | 174525929 | rs1809119 | 1.231 | -1.767 | -0.312 | 1.289 | 0.099 | -0.043 | -0.043 | 0 | 0.63 | 0.12 | 0.78 | 0.38 | 0.81 | 0.57 |
| 2 | 174529037 | rs526782 | -1.9 | -1.086 | -0.032 | 1.612 | 0.008 | 0.01 | 0.01 | 0 | 0.36 | 0.39 | 0.28 | 0.81 | 0.27 | 0.35 |
| 2 | 174530465 | rs9646736 | -1.241 | -1.059 | -0.485 | 1.592 | 0.019 | -0.023 | -0.023 | 1.705 | 0.28 | 0.28 | 0.15 | 0.57 | 0.16 | 0.26 |
| 2 | 174538452 | rs17312140 | -1.03 | -1 | -0.468 | 1.616 | -0.003 | 0.007 | 0.007 | 0 | 0.08 | 0.1 | 0.06 | 0.2 | 0.1 | 0.1 |
| 2 | 174539109 | rs2043920 | -0.648 | -0.999 | -0.752 | 1.644 | -0.01 | 0.014 | 0.014 | 0 | 0.22 | 0.29 | 0.13 | 0.57 | 0.11 | 0.24 |
| 2 | 174541769 | rs10930614 | -0.38 | -0.787 | -0.761 | 1.73 | -0.034 | -0.034 | -0.034 | 0 | 0.07 | 0.01 | 0.06 | 0.2 | 0.09 | 0.1 |
| 2 | 174544738 | rs2358229 | -0.35 | -0.776 | -0.747 | 1.735 | -0.03 | -0.03 | -0.03 | 0 | 0.07 | 0.01 | 0.06 | 0.2 | 0.09 | 0.1 |
| 2 | 174549552 | rs10166760 | -1.064 | -1.018 | -0.48 | 1.541 | 0.054 | -0.071 | -0.071 | 2.041 | 0.28 | 0.16 | 0.14 | 0.57 | 0.17 | 0.25 |
| 2 | 174550506 | rs10170039 | -1.478 | -1.064 | -0.532 | 1.472 | 0.022 | -0.023 | -0.023 | 1.117 | 0.33 | 0.32 | 0.2 | 0.57 | 0.25 | 0.31 |
| 2 | 174551768 | rs16861976 | -0.941 | -1.129 | -0.931 | 0.652 | 0.063 | 0.07 | 0.07 | 0.266 | 0.25 | 0.08 | 0.09 | 0.16 | 0.12 | 0.2 |
| 2 | 174553435 | rs752280 | -0.583 | -0.584 | -0.223 | 0.95 | -0.006 | -0.001 | -0.001 | 0 | 0.16 | 0.21 | 0.29 | 0.17 | 0.12 | 0.19 |
| 2 | 174554115 | rs12988452 | -1.572 | -0.334 | 0.79 | 1.602 | -0.006 | 0.201 | 0.201 | 0 | 0.18 | 0.43 | 0.15 | 0.46 | 0.13 | 0.13 |
| 2 | 174554558 | rs6713428 | -2.093 | -0.353 | 1.144 | 1.656 | -0.013 | 0.164 | 0.164 | 1.741 | 0.14 | 0.65 | 0.15 | 0.46 | 0.1 | 0.12 |
| 2 | 174560184 | rs10185554 | -1.676 | -0.658 | 1.306 | 1.512 | -0.068 | -0.004 | -0.004 | 0 | 0.12 | 0.27 | 0.06 | 0.12 | 0.03 | 0.04 |
| 2 | 174578522 | rs12329372 | -1.903 | -0.172 | 1.652 | 1.752 | 0.004 | 0.162 | 0.162 | 0 | 0.17 | 0.29 | 0.16 | 0.5 | 0.16 | 0.14 |
| 2 | 174590064 | rs3914403 | -2.199 | -0.181 | 1.518 | 1.75 | -0.011 | 0.179 | 0.179 | 0.392 | 0.19 | 0.54 | 0.19 | 0.5 | 0.15 | 0.14 |
| 2 | 174597656 | rs16823293 | -2.033 | 0.127 | 1.325 | 1.75 | 0.054 | 0.336 | 0.336 | 0.935 | 0.13 | 0.54 | 0.17 | 0.5 | 0.13 | 0.11 |
| 2 | 174604544 | rs7560771 | -0.07 | 0.533 | 0.008 | 0.443 | -0.015 | 0.022 | 0.022 | 0 | 0.69 | 0.71 | 0.69 | 0.9 | 0.6 | 0.8 |
| 2 | 174617984 | rs16823294 | -1.32 | 0.042 | 0.988 | 1.521 | -0.011 | 0.271 | 0.271 | 0.395 | 0.08 | 0.34 | 0.12 | 0.26 | 0.04 | 0.18 |
| 2 | 174622525 | rs3845743 | 0.433 | -0.463 | 0.125 | 1.004 | 0.006 | 0.009 | 0.009 | 0.02 | 0.35 | 0.31 | 0.17 | 0.16 | 0.27 | 0.19 |
| 2 | 174629287 | rs6433436 | -0.344 | 0.817 | -0.45 | 0.796 | -0.072 | 0.029 | 0.029 | 0.983 | 0.86 | 0.98 | 0.86 | 0.93 | 0.78 | 0.76 |
| 2 | 174654783 | rs16862084 | -0.936 | 0.468 | 0.073 | 1.084 | -0.025 | 0.097 | 0.097 | 0.309 | 0.19 | 0.36 | 0.32 | 0.34 | 0.15 | 0.31 |
| 2 | 174660974 | rs1828772 | -0.55 | 0.654 | -0.377 | 1.196 | -0.006 | 0.057 | 0.057 | 0 | 0.14 | 0.2 | 0.31 | 0.34 | 0.13 | 0.26 |
| 2 | 174666821 | rs16862092 | -0.302 | 0.796 | -0.598 | 1.192 | -0.001 | 0.021 | 0.021 | 0 | 0.13 | 0.14 | 0.31 | 0.29 | 0.15 | 0.25 |
| 2 | 174671104 | rs1877705 | 0.146 | 1.498 | -1.022 | 0.947 | -0.045 | 0.023 | 0.023 | 1.213 | 0.13 | 0.04 | 0.14 | 0.08 | 0.24 | 0.17 |
| 2 | 174680163 | rs907956 | 0.863 | 1.669 | -1.106 | 0.726 | -0.012 | 0.044 | 0.044 | 7.688 | 0.15 | 0.05 | 0.07 | 0.02 | 0.14 | 0.05 |
| 2 | 174680319 | rs16862115 | -0.715 | 1.778 | -0.798 | 1.49 | -0.021 | -0.03 | -0.03 | 0.073 | 0.14 | 0.09 | 0.36 | 0.3 | 0.21 | 0.27 |
| 2 | 174698705 | rs867458 | -0.131 | 1.432 | -0.567 | 1.654 | 0.006 | 0.065 | 0.065 | 0 | 0.08 | 0.11 | 0.13 | 0.33 | 0.12 | 0.1 |
| 2 | 174705545 | rs7602424 | 0.94 | 1.176 | -0.724 | 1.345 | -0.014 | -0.005 | -0.005 | 0.002 | 0.57 | 0.39 | 0.57 | 0.4 | 0.49 | 0.6 |
| 2 | 174705739 | rs17305716 | 0.007 | 1.608 | -0.922 | 1.598 | -0.018 | -0.013 | -0.013 | 0 | 0.1 | 0.04 | 0.11 | 0.18 | 0.16 | 0.12 |
| 2 | 174706535 | rs6433441 | -0.055 | 1.43 | -0.517 | 1.657 | 0.004 | 0.003 | 0.003 | 0 | 0.78 | 0.77 | 0.8 | 0.72 | 0.7 | 0.82 |
| 2 | 174707728 | rs16862130 | 0.086 | 1.507 | -0.546 | 1.575 | -0.018 | -0.018 | -0.018 | 2.055 | 0.09 | 0.04 | 0.11 | 0.23 | 0.16 | 0.12 |
| 2 | 174708199 | rs1024125 | 0.071 | 1.146 | -0.459 | 1.214 | 0.007 | -0.005 | -0.005 | 0 | 0.78 | 0.75 | 0.79 | 0.8 | 0.7 | 0.83 |
| 2 | 174709517 | rs4972412 | 0.342 | -0.272 | -1.01 | 0.897 | 0.056 | -0.014 | -0.014 | 0.1 | 0.32 | 0.17 | 0.28 | 0.29 | 0.21 | 0.28 |
| 2 | 174710364 | rs4972601 | 0.342 | -0.062 | -0.774 | 0.488 | 0.01 | 0.003 | 0.003 | 0 | 0.3 | 0.26 | 0.46 | 0.23 | 0.25 | 0.32 |
| 2 | 174712414 | rs962327 | 0.083 | -0.137 | -0.286 | 0.143 | -0.032 | -0.006 | -0.006 | 0 | 0.87 | 0.78 | 0.92 | 0.88 | 0.94 | 0.85 |
| 2 | 174714533 | rs17233751 | 0.111 | 0.382 | -0.885 | 0.193 | -0.055 | -0.074 | -0.074 | 0.074 | 0.12 | 0.01 | 0.1 | 0.22 | 0.14 | 0.12 |
| 2 | 174718059 | rs7591669 | 0.226 | 0.072 | -0.573 | -0.992 | -0.012 | 0.091 | 0.091 | 0 | 0.45 | 0.32 | 0.45 | 0.05 | 0.54 | 0.3 |
| 2 | 174721886 | rs4972603 | -0.038 | -0.106 | -0.013 | -1.107 | -0.001 | -0.025 | -0.025 | 0 | 0.51 | 0.61 | 0.63 | 0.88 | 0.45 | 0.63 |
| 2 | 174728683 | rs2063701 | 0.957 | -0.419 | 0.739 | -0.805 | -0.054 | -0.068 | -0.068 | 0 | 0.8 | 0.46 | 0.85 | 0.92 | 0.89 | 0.87 |
| 2 | 174731838 | rs1567852 | 0.884 | -0.407 | 0.983 | -0.817 | -0.045 | -0.071 | -0.071 | 0 | 0.81 | 0.63 | 0.76 | 0.92 | 0.87 | 0.85 |
| 2 | 174733318 | rs2137804 | -0.404 | -0.265 | 0.852 | -0.722 | -0.015 | -0.036 | -0.036 | 1.023 | 0.1 | 0.23 | 0.2 | 0.02 | 0.07 | 0.1 |
| 2 | 174738385 | rs11675772 | -0.878 | -0.536 | 0.528 | -0.727 | -0.016 | -0.003 | -0.003 | 2.387 | 0.66 | 0.65 | 0.7 | 0.33 | 0.79 | 0.44 |
| 2 | 174739890 | rs6716552 | 1.073 | 0.288 | 0.039 | -0.902 | -0.005 | -0.01 | -0.01 | 0 | 0.11 | 0.12 | 0.16 | 0 | 0.17 | 0.04 |
| 2 | 174745793 | rs10177396 | -0.333 | 0.518 | -0.349 | -1.193 | -0.053 | -0.019 | -0.019 | 0 | 0.27 | 0.15 | 0.31 | 0.32 | 0.35 | 0.24 |
| 2 | 174748349 | rs4972613 | 0.834 | 0.397 | 0.171 | -1.297 | 0.01 | -0.02 | -0.02 | 1.209 | 0.62 | 0.45 | 0.5 | 0.68 | 0.59 | 0.74 |
| 2 | 174761611 | rs6718188 | -0.416 | 0.338 | -0.174 | -1.316 | -0.033 | -0.001 | -0.001 | 0.022 | 0.29 | 0.2 | 0.31 | 0.3 | 0.35 | 0.24 |
| 2 | 174769344 | rs4972618 | -1.093 | 0.868 | 0.006 | -1.453 | -0.007 | -0.001 | -0.001 | 2.471 | 0.35 | 0.63 | 0.41 | 0.36 | 0.42 | 0.29 |
| 2 | 174775653 | rs6711060 | 1.095 | 0.695 | 0.211 | -1.615 | -0.007 | -0.035 | -0.035 | 0.673 | 0.1 | 0.15 | 0.08 | 0.02 | 0.06 | 0.1 |
| 2 | 174787496 | rs10173522 | -1.073 | 0.82 | 0.132 | -1.718 | -0.013 | 0.008 | 0.008 | 0 | 0.4 | 0.34 | 0.39 | 0.35 | 0.43 | 0.33 |
| 2 | 174792769 | rs9287968 | -0.193 | 0.894 | -0.21 | -1.634 | -0.044 | -0.032 | -0.032 | 0 | 0.3 | 0.19 | 0.32 | 0.34 | 0.37 | 0.24 |
| 2 | 174808899 | rs4325816 | 0.918 | 1.175 | -0.337 | -1.604 | 0 | -0.033 | -0.033 | 0 | 0.21 | 0.23 | 0.15 | 0.02 | 0.23 | 0.1 |
| 2 | 174820750 | rs1047640 | 1.481 | 1.336 | -0.758 | -1.472 | -0.044 | -0.034 | -0.034 | 0.325 | 0.08 | 0.01 | 0.09 | 0.34 | 0.12 | 0.11 |

| | | | | | | | | | | | | | | | | |
|---|---|---|---|---|---|---|---|---|---|---|---|---|---|---|---|---|
| 2 | 174831023 | rs1867858 | 1.366 | 0.984 | -0.621 | -1.479 | 0.019 | -0.038 | -0.038 | 0 | 0.35 | 0.14 | 0.29 | 0.6 | 0.32 | 0.48 |
| 2 | 174833833 | rs16866 | 0.223 | 1.013 | 0.119 | -1.367 | 0.009 | -0.097 | -0.097 | 0.781 | 0.22 | 0.33 | 0.15 | 0.02 | 0.23 | 0.1 |
| 2 | 174840107 | rs10497424 | -0.104 | 0.799 | 0.355 | -0.9 | -0.005 | 0.001 | 0.001 | 0 | 0.43 | 0.47 | 0.43 | 0.45 | 0.36 | 0.39 |
| 2 | 174844867 | rs6721460 | -0.18 | 0.865 | 0.402 | -0.955 | 0.006 | -0.028 | -0.028 | 0 | 0.13 | 0.16 | 0.18 | 0.03 | 0.14 | 0.15 |
| 2 | 174847693 | rs10497423 | -0.37 | 0.636 | 0.608 | -1.021 | 0.001 | 0 | 0 | 0 | 0.58 | 0.57 | 0.53 | 0.6 | 0.61 | 0.61 |
| 2 | 174849854 | rs16862245 | 0.259 | 0.804 | 0.383 | -0.844 | -0.005 | -0.013 | -0.013 | 0 | 0.11 | 0.16 | 0.09 | 0.04 | 0.07 | 0.11 |
| 2 | 174851673 | rs16862247 | -0.017 | 0.794 | 0.589 | -0.779 | -0.015 | 0.016 | 0.016 | 0 | 0.12 | 0.26 | 0.11 | 0.12 | 0.07 | 0.15 |
| 2 | 174852271 | rs7578920 | -0.067 | 0.792 | 0.594 | -0.78 | -0.016 | 0.015 | 0.015 | 0 | 0.12 | 0.26 | 0.11 | 0.12 | 0.07 | 0.15 |
| 2 | 174852785 | rs10497422 | 0.192 | 0.814 | 0.355 | -0.876 | -0.006 | -0.014 | -0.014 | 0 | 0.11 | 0.15 | 0.09 | 0.04 | 0.07 | 0.1 |
| 2 | 174854758 | rs4972626 | -0.639 | 0.756 | 0.203 | -0.831 | -0.006 | -0.015 | -0.015 | 0 | 0.55 | 0.7 | 0.61 | 0.47 | 0.49 | 0.53 |
| 2 | 174858646 | rs935398 | -0.418 | 1.019 | 0.624 | -1.565 | -0.002 | 0.005 | 0.005 | 0 | 0.55 | 0.55 | 0.46 | 0.25 | 0.5 | 0.55 |
| 2 | 174865555 | rs11890900 | -1.429 | 1.329 | 0.896 | -1.468 | 0.082 | -0.025 | -0.025 | 0 | 0.1 | 0.36 | 0.21 | 0.04 | 0.12 | 0.12 |
| 2 | 174865857 | rs2121362 | -0.609 | 1.348 | 0.814 | -1.569 | -0.002 | 0.131 | 0.131 | 0 | 0.58 | 0.85 | 0.76 | 0.8 | 0.63 | 0.6 |
| 2 | 174866777 | rs4423657 | -1.32 | 1.581 | 1.009 | -1.433 | 0.078 | -0.022 | -0.022 | 0 | 0.09 | 0.32 | 0.21 | 0.04 | 0.12 | 0.12 |
| 2 | 174874454 | rs2028889 | -0.251 | 1.69 | 0.584 | -1.461 | 0.003 | 0.027 | 0.027 | 0.126 | 0.54 | 0.33 | 0.59 | 0.78 | 0.61 | 0.56 |
| 2 | 174876391 | rs7590276 | 0.009 | 1.716 | 0.36 | -1.438 | 0.01 | 0.162 | 0.162 | 0.135 | 0.49 | 0.16 | 0.43 | 0.78 | 0.52 | 0.49 |
| 2 | 174879174 | rs7601227 | 0.211 | 1.966 | 0.145 | -1.424 | 0.009 | -0.018 | -0.018 | 0 | 0.31 | 0.29 | 0.44 | 0.78 | 0.37 | 0.45 |
| 2 | 174891414 | rs3813817 | 0.637 | 2.012 | 0.051 | -0.852 | -0.018 | 0.186 | 0.186 | 0.063 | 0.47 | 0.13 | 0.44 | 0.82 | 0.51 | 0.49 |
| 2 | 174893412 | rs4402725 | -0.436 | 1.617 | -0.048 | -0.886 | 0.017 | 0.051 | 0.051 | 0.032 | 0.46 | 0.37 | 0.36 | 0.14 | 0.36 | 0.38 |
| 2 | 174896455 | rs17308965 | 0.784 | 1.697 | -0.124 | -0.402 | -0.063 | 0.091 | 0.091 | 0 | 0.64 | 0.22 | 0.5 | 0.78 | 0.57 | 0.59 |
| 2 | 174902997 | rs17309203 | 1.103 | 1.666 | 0.225 | -0.27 | -0.033 | 0.028 | 0.028 | 0 | 0.66 | 0.27 | 0.5 | 0.81 | 0.6 | 0.6 |
| 2 | 174905977 | rs2121359 | -0.624 | 2.213 | -0.204 | -0.144 | -0.063 | -0.002 | -0.002 | 0 | 0.23 | 0.09 | 0.4 | 0.18 | 0.31 | 0.33 |
| 2 | 174909182 | rs17309392 | 0.817 | 1.748 | 0.029 | -0.109 | -0.03 | 0.016 | 0.016 | 0 | 0.7 | 0.77 | 0.54 | 0.82 | 0.6 | 0.6 |
| 2 | 174910616 | rs6433461 | -0.534 | 2.319 | 0.046 | 0.182 | 0.13 | -0.009 | -0.009 | 0 | 0.21 | 0.43 | 0.43 | 0.15 | 0.35 | 0.33 |
| 2 | 174919189 | rs6756828 | -0.778 | 1.547 | -0.754 | 0.389 | -0.011 | -0.022 | -0.022 | 0 | 0.23 | 0.15 | 0.33 | 0.69 | 0.27 | 0.45 |
| 2 | 174926587 | rs1562311 | 0.676 | 1.796 | 0.091 | -0.435 | 0.028 | -0.073 | -0.073 | 0 | 0.86 | 0.74 | 0.85 | 1 | 0.77 | 0.91 |
| 2 | 174928360 | rs10173820 | -0.042 | 0.351 | 0.333 | -1.176 | -0.063 | -0.085 | -0.085 | 0 | 0.17 | 0.31 | 0.07 | 0.01 | 0.07 | 0.12 |
| 2 | 174931752 | rs9287971 | -1.163 | 0.024 | -0.375 | -1.367 | 0.015 | 0.015 | 0.015 | 0.089 | 0.5 | 0.5 | 0.35 | 0.73 | 0.34 | 0.61 |
| 2 | 174931957 | rs2044471 | -0.325 | -0.172 | -0.502 | -1.539 | -0.015 | -0.089 | -0.089 | 0 | 0.3 | 0.36 | 0.18 | 0.01 | 0.27 | 0.18 |
| 2 | 174937278 | rs3739153 | -1.456 | 0.05 | -0.665 | -2.144 | 0.005 | -0.081 | -0.081 | 0 | 0.62 | 0.63 | 0.53 | 0.96 | 0.5 | 0.71 |
| 2 | 174939168 | rs6737629 | -0.968 | 0.099 | -0.716 | -2.142 | 0.014 | 0.122 | 0.122 | 0 | 0.46 | 0.3 | 0.45 | 0.95 | 0.43 | 0.59 |
| 2 | 174942311 | rs10930633 | -0.242 | -0.1 | -0.559 | -2.148 | 0.004 | 0.009 | 0.009 | 0 | 0.28 | 0.26 | 0.33 | 0.94 | 0.23 | 0.51 |
| 2 | 174943876 | rs3754812 | -0.242 | -0.1 | -0.559 | -2.148 | 0.004 | 0.009 | 0.009 | 0 | 0.28 | 0.26 | 0.33 | 0.94 | 0.23 | 0.51 |
| 2 | 174944105 | rs16862333 | -0.241 | -0.103 | -0.565 | -2.149 | 0.009 | 0.037 | 0.037 | 0 | 0.28 | 0.23 | 0.33 | 0.94 | 0.23 | 0.51 |
| 2 | 174944669 | rs16862337 | 0.676 | -0.145 | -0.591 | -2.115 | 0.035 | -0.049 | -0.049 | 0.476 | 0.16 | 0.1 | 0.15 | 0.71 | 0.08 | 0.4 |
| 2 | 174949358 | rs11694601 | -1.098 | -0.098 | -0.551 | -2.123 | 0.015 | 0.08 | 0.08 | 0 | 0.46 | 0.33 | 0.46 | 0.95 | 0.43 | 0.59 |
| 2 | 174983004 | rs16862397 | 1.258 | -0.259 | -0.361 | -2.118 | -0.026 | -0.045 | -0.045 | 0 | 0.16 | 0.2 | 0.08 | 0.01 | 0.09 | 0.12 |
| 2 | 174998583 | rs1595796 | -1.403 | 0.216 | -0.51 | -2.807 | 0.056 | -0.067 | -0.067 | 2.383 | 0.83 | 0.72 | 0.66 | 0.96 | 0.74 | 0.87 |
| 2 | 175000711 | rs10930641 | -0.651 | 0.049 | -0.537 | -2.838 | 0.019 | 0.12 | 0.12 | 0 | 0.46 | 0.33 | 0.44 | 0.95 | 0.43 | 0.59 |
| 2 | 175009066 | rs10497417 | 1.106 | 0.021 | -0.544 | -2.805 | 0.006 | 0.012 | 0.012 | 0 | 0.13 | 0.2 | 0.17 | 0.22 | 0.15 | 0.1 |
| 2 | 175025275 | rs16862442 | 1.277 | -0.066 | -0.609 | -2.846 | 0.038 | -0.053 | -0.053 | 0 | 0.16 | 0.08 | 0.15 | 0.71 | 0.08 | 0.4 |
| 2 | 175026805 | rs16862445 | 0.313 | -0.159 | -0.551 | -2.904 | -0.055 | -0.016 | -0.016 | 0 | 0.34 | 0.47 | 0.23 | 0.72 | 0.15 | 0.52 |
| 2 | 175031590 | rs10203886 | -1.096 | -0.125 | -0.586 | -2.937 | 0.011 | -0.109 | -0.109 | 0 | 0.62 | 0.59 | 0.52 | 0.96 | 0.5 | 0.71 |
| 2 | 175045794 | rs16862458 | 2.038 | -0.144 | -0.556 | -2.946 | -0.064 | -0.087 | -0.087 | 1.091 | 0.16 | 0.31 | 0.08 | 0.01 | 0.07 | 0.12 |
| 2 | 175054934 | rs10930647 | -0.302 | -0.146 | -0.581 | -2.952 | -0.018 | -0.086 | -0.086 | 0 | 0.44 | 0.55 | 0.39 | 0.94 | 0.3 | 0.62 |
| 2 | 175055150 | rs10930648 | 2.066 | -0.182 | -0.577 | -2.963 | -0.052 | -0.072 | -0.072 | 0.916 | 0.16 | 0.26 | 0.07 | 0.01 | 0.07 | 0.12 |
| 2 | 175059767 | rs4972434 | -0.989 | -0.072 | -0.518 | -2.89 | 0.011 | -0.112 | -0.112 | 0.022 | 0.62 | 0.58 | 0.52 | 0.96 | 0.5 | 0.71 |
| 2 | 175077961 | rs10180256 | -0.887 | -0.154 | -0.579 | -2.93 | 0.01 | -0.113 | -0.113 | 0.111 | 0.61 | 0.53 | 0.51 | 0.96 | 0.5 | 0.71 |
| 2 | 175079125 | rs12328930 | -0.03 | 0.005 | -0.689 | -2.704 | -0.004 | -0.027 | -0.027 | 0 | 0.4 | 0.4 | 0.34 | 0.23 | 0.42 | 0.28 |
| 2 | 175093978 | rs12999294 | 1.268 | 0.222 | -1.108 | -2.5 | -0.02 | -0.025 | -0.025 | 0.561 | 0.11 | 0.05 | 0.16 | 0.22 | 0.15 | 0.1 |
| 2 | 175106028 | rs13424932 | 1.546 | 0.328 | -1.056 | -2.583 | -0.045 | 0.091 | 0.091 | 0.994 | 0.21 | 0.06 | 0.14 | 0.01 | 0.23 | 0.16 |
| 2 | 175113512 | rs3820898 | -0.19 | 0.094 | -1.401 | -2.615 | 0.014 | 0.12 | 0.12 | 0.952 | 0.46 | 0.16 | 0.34 | 0.26 | 0.45 | 0.36 |
| 2 | 175127653 | rs4350713 | 0.381 | 0.408 | -0.901 | -2.321 | 0 | -0.003 | -0.003 | 3.672 | 0.52 | 0.51 | 0.38 | 0.29 | 0.43 | 0.4 |
| 2 | 175133282 | rs11687678 | 1.235 | 0.515 | -1.079 | -2.315 | -0.032 | 0.06 | 0.06 | 0.301 | 0.14 | 0.04 | 0.11 | 0 | 0.21 | 0.07 |
| 2 | 175143309 | rs6433472 | 0.083 | 0.485 | -0.662 | -1.983 | 0.009 | -0.013 | -0.013 | 0 | 0.67 | 0.57 | 0.49 | 0.28 | 0.63 | 0.48 |
| 2 | 175163818 | rs17320820 | 0.292 | 0.223 | -1.348 | -2.365 | -0.02 | 0.127 | 0.127 | 2.341 | 0.54 | 0.27 | 0.37 | 0.19 | 0.56 | 0.37 |
| 2 | 175170619 | rs10202742 | 0.817 | 0.107 | -1.414 | -2.806 | -0.025 | 0.087 | 0.087 | 2.422 | 0.33 | 0.14 | 0.21 | 0.02 | 0.38 | 0.14 |
| 2 | 175172197 | rs1597417 | 0.691 | 0.223 | -1.493 | -2.216 | -0.002 | -0.002 | -0.002 | 0 | 0.21 | 0.22 | 0.48 | 0.69 | 0.26 | 0.42 |
| 2 | 175175747 | rs13002825 | 0.711 | 0.187 | -1.577 | -2.122 | -0.054 | -0.058 | -0.058 | 1.867 | 0.2 | 0.04 | 0.46 | 0.69 | 0.25 | 0.43 |
| 2 | 175175807 | rs6736532 | 1.577 | 0.131 | -1.565 | -2.265 | -0.021 | 0.064 | 0.064 | 1.672 | 0.33 | 0.15 | 0.21 | 0.02 | 0.38 | 0.14 |
| 2 | 175176215 | rs17320933 | 0.619 | 0.064 | -1.262 | -1.917 | -0.029 | -0.03 | -0.03 | 0.647 | 0.19 | 0.1 | 0.47 | 0.7 | 0.26 | 0.44 |
| 2 | 175185780 | rs13035792 | -0.563 | -0.479 | -0.291 | -2.401 | -0.012 | -0.115 | -0.115 | 0 | 0.59 | 0.48 | 0.79 | 0.97 | 0.7 | 0.77 |
| 2 | 175192359 | rs2165213 | 0.156 | -0.768 | -0.094 | -2.418 | -0.005 | -0.012 | -0.012 | 3.072 | 0.43 | 0.39 | 0.19 | 0.21 | 0.27 | 0.3 |
| 2 | 175194550 | rs11891208 | -0.579 | -0.577 | 0.006 | -2.162 | -0.042 | 0.042 | 0.042 | 2.633 | 0.14 | 0.54 | 0.1 | 0.2 | 0.04 | 0.24 |
| 2 | 175205513 | rs7582977 | -0.368 | -0.493 | -0.26 | -2.015 | 0.019 | 0.021 | 0.021 | 3.166 | 0.14 | 0.09 | 0.04 | 0.02 | 0.08 | 0.19 |
| 2 | 175211288 | rs6735817 | -1.253 | -0.483 | 0.089 | -2.133 | -0.031 | -0.039 | -0.039 | 0 | 0.67 | 0.59 | 0.83 | 0.8 | 0.78 | 0.84 |
| 2 | 175220520 | rs6715304 | -0.321 | -0.348 | -0.396 | -2.063 | 0.075 | 0.131 | 0.131 | 0.183 | 0.53 | 0.22 | 0.75 | 0.79 | 0.68 | 0.75 |
| 2 | 175234490 | rs10930659 | -0.304 | -0.353 | -0.398 | -2.066 | 0.076 | 0.133 | 0.133 | 0.273 | 0.53 | 0.22 | 0.75 | 0.79 | 0.68 | 0.75 |
| 2 | 175239185 | rs17255455 | -0.157 | -0.161 | -0.415 | -2.082 | 0.039 | 0.157 | 0.157 | 0.195 | 0.5 | 0.22 | 0.71 | 0.79 | 0.57 | 0.74 |

| | | | | | | | | | | | | | | | | |
|---|---|---|---|---|---|---|---|---|---|---|---|---|---|---|---|---|
| 2 | 175243501 | rs2305292 | -0.473 | -0.436 | -0.13 | -1.901 | -0.07 | -0.052 | -0.052 | 0 | 0.2 | 0.48 | 0.1 | 0.1 | 0.11 | 0.27 |
| 2 | 175250289 | rs2122398 | -0.367 | -0.373 | -0.093 | -1.997 | 0.004 | 0.015 | 0.015 | 0.321 | 0.46 | 0.41 | 0.67 | 0.7 | 0.51 | 0.55 |
| 2 | 175263063 | rs10497410 | 1.299 | 0.229 | -0.755 | -2.052 | -0.066 | 0.061 | 0.061 | 0 | 0.13 | 0.01 | 0.16 | 0.01 | 0.21 | 0.08 |
| 2 | 175268334 | rs11695234 | 1.642 | 0.054 | -0.971 | -2.23 | 0.02 | 0.073 | 0.073 | 0 | 0.31 | 0.11 | 0.16 | 0.21 | 0.28 | 0.13 |
| 2 | 175275679 | rs7570614 | -0.284 | -0.012 | -0.839 | -2.39 | -0.003 | -0.02 | -0.02 | 0 | 0.29 | 0.22 | 0.22 | 0.01 | 0.32 | 0.2 |
| 2 | 175279049 | rs4972661 | 0.612 | 0.028 | -1.244 | -2.269 | -0.007 | 0.041 | 0.041 | 0 | 0.18 | 0.07 | 0.12 | 0.08 | 0.2 | 0.3 |
| 2 | 175287778 | rs2290158 | 1.373 | -0.153 | -1.39 | -1.923 | 0.005 | 0.068 | 0.068 | 0 | 0.31 | 0.06 | 0.15 | 0.21 | 0.28 | 0.12 |
| 2 | 175289314 | rs17255817 | -0.01 | -0.249 | -1.391 | -1.985 | 0.023 | 0.103 | 0.103 | 0 | 0.16 | 0.01 | 0.05 | 0 | 0.1 | 0.11 |
| 2 | 175292326 | rs7591631 | 0.099 | 0.233 | -0.9 | -1.821 | -0.016 | 0.053 | 0.053 | 0 | 0.36 | 0.75 | 0.33 | 0.09 | 0.42 | 0.39 |
| 2 | 175316379 | rs4972663 | -0.118 | 0.785 | -1.175 | -0.707 | -0.01 | -0.013 | -0.013 | 0.025 | 0.82 | 0.86 | 0.54 | 0.32 | 0.8 | 0.63 |
| 2 | 175329213 | rs1450043 | -0.68 | 0.601 | -1.259 | -0.882 | 0.001 | -0.002 | -0.002 | 0.218 | 0.13 | 0.15 | 0.04 | 0.09 | 0.09 | 0.12 |
| 2 | 175339504 | rs4972666 | -0.196 | 0.991 | -1.84 | 0.085 | -0.111 | -0.14 | -0.14 | 0 | 0.22 | 0.02 | 0.43 | 0.58 | 0.25 | 0.36 |
| 2 | 175348838 | rs10221679 | -0.559 | 1.284 | -1.662 | -0.371 | -0.01 | 0.035 | 0.035 | 0 | 0.18 | 0.11 | 0.1 | 0.05 | 0.16 | 0.19 |
| 2 | 175352590 | rs1077071 | 0.51 | 0.862 | -1.197 | -0.475 | -0.015 | 0.006 | 0.006 | 0 | 0.17 | 0.24 | 0.14 | 0.2 | 0.14 | 0.16 |
| 2 | 175353898 | rs7593976 | -1.279 | 0.785 | -0.919 | -0.325 | 0.002 | 0.002 | 0.002 | 0.234 | 0.62 | 0.6 | 0.72 | 0.73 | 0.63 | 0.66 |
| 2 | 175360010 | rs7563578 | 0.127 | 0.973 | -1.021 | -0.076 | -0.013 | -0.019 | -0.019 | 0 | 0.24 | 0.2 | 0.48 | 0.5 | 0.32 | 0.31 |
| 2 | 175365518 | rs7602123 | 0.684 | 0.473 | -0.745 | -0.455 | 0.003 | 0.002 | 0.002 | 0 | 0.36 | 0.32 | 0.22 | 0.28 | 0.3 | 0.36 |
| 2 | 175386861 | rs9287975 | -0.568 | 1.022 | 0.02 | -0.725 | 0.099 | 0.041 | 0.041 | 0 | 0.05 | 0.23 | 0.08 | 0.05 | 0.1 | 0.08 |
| 2 | 175393104 | rs13407318 | -0.029 | 1.23 | -0.391 | -1.419 | -0.013 | 0.096 | 0.096 | 0 | 0.3 | 0.14 | 0.28 | 0.06 | 0.33 | 0.24 |
| 2 | 175409011 | rs2033309 | -0.016 | 0.562 | -0.058 | -1.783 | -0.001 | -0.042 | -0.042 | 0 | 0.17 | 0.34 | 0.14 | 0.09 | 0.15 | 0.23 |
| 2 | 175409060 | rs2033310 | -1.204 | 0.372 | 0.06 | -1.925 | 0.006 | 0.042 | 0.042 | 0 | 0.49 | 0.54 | 0.42 | 0.15 | 0.48 | 0.47 |
| 2 | 175418960 | rs1010028 | 0.867 | 0.555 | 0.054 | -2.218 | -0.003 | 0.088 | 0.088 | 0 | 0.49 | 0.29 | 0.37 | 0.85 | 0.47 | 0.48 |
| 2 | 175424429 | rs7739 | -1.043 | 0.361 | 0.241 | -2.273 | -0.014 | -0.015 | -0.015 | 0 | 0.87 | 0.82 | 0.94 | 0.96 | 0.94 | 0.8 |
| 2 | 175425346 | rs3205060 | 0.602 | 0.509 | 0.625 | -2.203 | 0.015 | 0.216 | 0.216 | 0.389 | 0.51 | 0.19 | 0.56 | 0.85 | 0.53 | 0.5 |
| 2 | 175426750 | rs6732100 | 0.125 | 0.422 | 0.601 | -2.165 | 0.103 | 0.042 | 0.042 | 0 | 0.06 | 0.27 | 0.09 | 0.05 | 0.09 | 0.08 |
| 2 | 175432056 | rs2303891 | 2.02 | 0.146 | -0.032 | -2.178 | -0.063 | -0.048 | -0.048 | 0 | 0.23 | 0.01 | 0.1 | 0.26 | 0.25 | 0.14 |
| 2 | 175436940 | rs4972450 | -0.348 | -0.242 | 0.381 | -2.421 | -0.013 | -0.029 | -0.029 | 0 | 0.88 | 0.81 | 0.97 | 0.99 | 0.95 | 0.87 |
| 2 | 175438251 | rs6755710 | 0.304 | -0.21 | 0.525 | -2.404 | 0.005 | -0.074 | -0.074 | 0 | 0.85 | 0.53 | 0.89 | 0.96 | 0.87 | 0.8 |
| 2 | 175439536 | rs12624277 | 0.726 | -0.05 | 0.358 | -2.437 | 0.019 | 0.05 | 0.05 | 0 | 0.5 | 0.35 | 0.57 | 0.87 | 0.53 | 0.48 |
| 2 | 175447213 | rs6750611 | 0.317 | 0.117 | 0.162 | -2.255 | 0.077 | 0.016 | 0.016 | 0 | 0.05 | 0.21 | 0.08 | 0.03 | 0.08 | 0.07 |
| 2 | 175451014 | rs12994635 | 0.512 | 0.05 | -0.108 | -2.322 | 0.009 | 0.087 | 0.087 | 0 | 0.32 | 0.17 | 0.32 | 0.09 | 0.33 | 0.31 |
| 2 | 175452275 | rs2115872 | 0.409 | 0.109 | -0.241 | -2.294 | 0.014 | 0.069 | 0.069 | 0 | 0.33 | 0.22 | 0.32 | 0.1 | 0.34 | 0.31 |
| 2 | 175457155 | rs1430182 | 0.53 | 0.354 | -0.215 | -2.293 | -0.002 | 0.145 | 0.145 | 0.049 | 0.46 | 0.24 | 0.56 | 0.87 | 0.53 | 0.49 |
| 2 | 175463356 | rs13411041 | 0.088 | 0.377 | -0.511 | -2.308 | -0.003 | 0.091 | 0.091 | 0 | 0.35 | 0.22 | 0.25 | 0.1 | 0.37 | 0.3 |
| 2 | 175467769 | rs1991601 | -1.477 | -0.038 | -0.341 | -2.504 | 0.005 | 0.004 | 0.004 | 0 | 0.71 | 0.77 | 0.88 | 0.99 | 0.74 | 0.82 |
| 2 | 175471607 | rs4595937 | 0.343 | 0.125 | -0.773 | -2.596 | 0.022 | 0.089 | 0.089 | 0 | 0.35 | 0.21 | 0.23 | 0.07 | 0.31 | 0.29 |
| 2 | 175473587 | rs4972701 | 0.011 | 0.324 | -1.067 | -2.404 | -0.048 | 0.069 | 0.069 | 0.484 | 0.31 | 0.16 | 0.61 | 0.9 | 0.43 | 0.47 |
| 2 | 175487267 | rs7590328 | -1.206 | 0.059 | -1.264 | -2.397 | 0.013 | 0.114 | 0.114 | 0 | 0.67 | 0.85 | 0.87 | 0.97 | 0.72 | 0.72 |
| 2 | 175495311 | rs1991600 | 0.625 | 0.43 | -1.112 | -2.051 | -0.019 | -0.027 | -0.027 | 0 | 0.25 | 0.11 | 0.53 | 0.65 | 0.27 | 0.38 |
| 2 | 175513995 | rs12466643 | -1.355 | 0.388 | -1.286 | -2.123 | 0.008 | 0.101 | 0.101 | 0 | 0.68 | 0.83 | 0.86 | 0.96 | 0.71 | 0.73 |
| 2 | 175525782 | rs12470024 | 1.182 | 0.569 | -1.491 | -1.99 | -0.065 | -0.087 | -0.087 | 0 | 0.2 | 0.03 | 0.5 | 0.65 | 0.23 | 0.37 |
| 2 | 175534730 | rs4378760 | 0.024 | 0.582 | -0.602 | -1.852 | -0.001 | -0.001 | -0.001 | 0 | 0.59 | 0.62 | 0.67 | 0.69 | 0.53 | 0.65 |
| 2 | 175542474 | rs6716254 | 0.127 | 0.762 | -0.598 | -2.222 | -0.006 | -0.09 | -0.09 | 0 | 0.33 | 0.38 | 0.23 | 0.06 | 0.3 | 0.19 |
| 2 | 175550678 | rs4356633 | 0.187 | 0.976 | -0.481 | -1.438 | 0.035 | 0.03 | 0.03 | 0 | 0.91 | 0.86 | 0.91 | 0.79 | 0.85 | 0.9 |
| 2 | 175558804 | rs2358891 | 0.315 | 0.517 | -0.591 | -1.77 | -0.001 | 0 | 0 | 0 | 0.29 | 0.31 | 0.18 | 0.04 | 0.27 | 0.28 |
| 2 | 175562223 | rs6738929 | 0.445 | 0.556 | -0.832 | -1.573 | -0.004 | -0.098 | -0.098 | 0 | 0.3 | 0.58 | 0.19 | 0.05 | 0.32 | 0.24 |
| 2 | 175565083 | rs13391108 | 0.231 | 0.845 | -1.134 | -1.16 | -0.031 | 0.106 | 0.106 | 0 | 0.37 | 0.19 | 0.64 | 0.85 | 0.39 | 0.5 |
| 2 | 175565552 | rs12620150 | -0.086 | 0.875 | -1.289 | -0.998 | -0.019 | 0.157 | 0.157 | 0 | 0.46 | 0.18 | 0.69 | 0.86 | 0.47 | 0.56 |
| 2 | 175570795 | rs2646186 | -1.188 | 0.971 | -0.156 | -0.304 | 0.041 | 0.128 | 0.128 | 0.545 | 0.12 | 0.47 | 0.11 | 0.17 | 0.13 | 0.15 |
| 2 | 175575649 | rs12472540 | 0.518 | 0.642 | -0.504 | -0.381 | -0.023 | 0.054 | 0.054 | 0.521 | 0.87 | 0.45 | 0.91 | 0.87 | 0.94 | 0.89 |
| 2 | 175582267 | rs10497426 | -0.814 | 1.157 | -1.068 | -0.393 | -0.039 | 0.022 | 0.022 | 0 | 0.16 | 0.06 | 0.14 | 0.11 | 0.2 | 0.16 |
| 2 | 175583220 | rs17270421 | -0.55 | 1.068 | -0.927 | -0.362 | -0.016 | 0.04 | 0.04 | 0.013 | 0.12 | 0.05 | 0.05 | 0.01 | 0.12 | 0.07 |
| 2 | 175584157 | rs1463263 | 0.183 | 0.893 | -1.097 | -0.391 | -0.014 | 0.036 | 0.036 | 0 | 0.43 | 0.34 | 0.65 | 0.69 | 0.5 | 0.5 |
| 2 | 175590459 | rs1901500 | 0.698 | 0.917 | -0.188 | -0.446 | 0.034 | -0.06 | -0.06 | 0.11 | 0.88 | 0.73 | 0.89 | 0.98 | 0.85 | 0.96 |
| 2 | 175591073 | rs12611955 | 0.157 | 0.796 | -1.027 | -0.303 | -0.036 | -0.07 | -0.07 | 0 | 0.27 | 0.06 | 0.33 | 0.64 | 0.27 | 0.41 |
| 2 | 175594914 | rs2600698 | -1.059 | 0.357 | -0.067 | -0.63 | 0.004 | -0.061 | -0.061 | 0 | 0.22 | 0.54 | 0.15 | 0.08 | 0.2 | 0.08 |
| 2 | 175597601 | rs2646181 | -1.344 | 0.083 | 0.022 | -0.775 | -0.009 | -0.037 | -0.037 | 0 | 0.27 | 0.62 | 0.16 | 0.15 | 0.22 | 0.28 |
| 2 | 175598296 | rs2359319 | 0.314 | -0.424 | -0.236 | -0.868 | -0.007 | -0.02 | -0.02 | 0 | 0.35 | 0.1 | 0.38 | 0.66 | 0.33 | 0.47 |
| 2 | 175598518 | rs2646179 | -1.033 | -0.512 | -0.005 | -0.929 | 0.016 | 0.132 | 0.132 | 0 | 0.56 | 0.84 | 0.48 | 0.79 | 0.42 | 0.7 |
| 2 | 175599106 | rs7575145 | 0.349 | -0.426 | 0.417 | -0.854 | 0.01 | 0.003 | 0.003 | 0 | 0.35 | 0.19 | 0.39 | 0.66 | 0.34 | 0.47 |
| 2 | 175602014 | rs2123117 | 0.712 | -0.369 | 1.079 | -0.983 | 0.001 | -0.024 | -0.024 | 0 | 0.43 | 0.45 | 0.51 | 0.21 | 0.51 | 0.25 |
| 2 | 175621482 | rs2646165 | 0.557 | -0.482 | 0.791 | -0.914 | -0.043 | -0.089 | -0.089 | 2.074 | 0.78 | 0.39 | 0.9 | 0.94 | 0.87 | 0.75 |
| 2 | 175624597 | rs1376865 | 0.902 | -0.551 | 0.707 | -0.949 | 0.005 | -0.009 | -0.009 | 0 | 0.75 | 0.26 | 0.89 | 0.92 | 0.87 | 0.73 |
| 2 | 175629021 | rs7560774 | 0.997 | -0.262 | 0.13 | -0.954 | -0.082 | 0.093 | 0.093 | 0.019 | 0.4 | 0.06 | 0.49 | 0.22 | 0.52 | 0.25 |
| 2 | 175630553 | rs3755485 | 0.551 | -0.257 | 0.321 | -0.865 | 0 | 0.006 | 0.006 | 0 | 0.44 | 0.36 | 0.5 | 0.25 | 0.53 | 0.26 |
| 2 | 175636801 | rs935863 | 0.154 | -0.278 | 0.073 | -0.587 | -0.013 | -0.009 | -0.009 | 0 | 0.36 | 0.47 | 0.4 | 0.68 | 0.31 | 0.47 |
| 2 | 175641575 | rs2244340 | 0.541 | -0.104 | -0.627 | -0.411 | -0.025 | 0.067 | 0.067 | 0 | 0.4 | 0.17 | 0.46 | 0.28 | 0.43 | 0.26 |
| 2 | 175650669 | rs12693055 | 0.356 | -0.056 | -0.097 | 0.053 | -0.006 | -0.003 | -0.003 | 0.163 | 0.41 | 0.46 | 0.47 | 0.35 | 0.41 | 0.22 |
| 2 | 175656117 | rs7598080 | 0.484 | -0.398 | -0.1 | 0.391 | 0 | 0 | 0 | 0 | 0.73 | 0.71 | 0.81 | 0.74 | 0.82 | 0.57 |

| | | | | | | | | | | | | | | | | |
|---|---|---|---|---|---|---|---|---|---|---|---|---|---|---|---|---|
| 2 | 175666865 | rs12476097 | -0.063 | -0.156 | -0.794 | 0.578 | -0.112 | -0.131 | -0.131 | 1.706 | 0.33 | 0.03 | 0.21 | 0.46 | 0.37 | 0.49 |
| 2 | 175675092 | rs13034776 | -0.234 | -0.074 | -0.09 | 0.318 | -0.004 | -0.027 | -0.027 | 0 | 0.05 | 0.12 | 0.03 | 0 | 0.03 | 0.02 |
| 2 | 175687705 | rs10497430 | 1.252 | -0.384 | 0.564 | 0.224 | -0.033 | 0.054 | 0.054 | 0 | 0.42 | 0.1 | 0.64 | 0.3 | 0.44 | 0.23 |
| 2 | 175691945 | rs2646167 | 0.476 | -0.423 | 1.09 | 0.211 | 0.011 | 0.071 | 0.071 | 0 | 0.51 | 0.69 | 0.72 | 0.3 | 0.49 | 0.27 |
| 2 | 175716383 | rs7584204 | 1.125 | -0.368 | 1.229 | 0.175 | 0.006 | 0.024 | 0.024 | 0 | 0.4 | 0.27 | 0.53 | 0.3 | 0.39 | 0.24 |
| 2 | 175722369 | rs17270470 | 1.21 | -0.416 | 1.187 | 0.19 | -0.038 | 0.035 | 0.035 | 0 | 0.39 | 0.08 | 0.52 | 0.3 | 0.4 | 0.23 |
| 2 | 175737705 | rs2646166 | 0.494 | -0.42 | 1.272 | 0.18 | 0.013 | 0.041 | 0.041 | 0 | 0.48 | 0.62 | 0.59 | 0.3 | 0.44 | 0.27 |
| 2 | 175744588 | rs1378729 | 0.621 | -0.417 | 1.263 | 0.213 | 0.011 | 0.038 | 0.038 | 0 | 0.47 | 0.62 | 0.59 | 0.28 | 0.43 | 0.26 |
| 2 | 175775334 | rs6725402 | 0.563 | -0.432 | 1.226 | 0.225 | 0.022 | 0.076 | 0.076 | 0 | 0.47 | 0.68 | 0.59 | 0.28 | 0.43 | 0.26 |
| 2 | 175780603 | rs4324297 | -0.701 | -0.362 | 1.192 | 0.268 | 0.018 | -0.019 | -0.019 | 0 | 0.07 | 0.34 | 0.06 | 0 | 0.03 | 0.02 |
| 2 | 175876576 | rs17288308 | 1.129 | -0.134 | -0.028 | 0.402 | -0.016 | 0.027 | 0.027 | 0 | 0.37 | 0.13 | 0.52 | 0.29 | 0.39 | 0.22 |
| 2 | 175877586 | rs7567205 | 0.937 | -0.112 | -0.106 | 0.402 | 0.001 | 0.009 | 0.009 | 0 | 0.39 | 0.34 | 0.53 | 0.29 | 0.39 | 0.22 |
| 2 | 175890331 | rs6718911 | 0.768 | 0.029 | -1.211 | 0.594 | -0.013 | -0.014 | -0.014 | 0 | 0.19 | 0.15 | 0.16 | 0.35 | 0.27 | 0.28 |
| 2 | 175893279 | rs3856435 | 1.499 | -0.021 | -1.278 | 0.556 | -0.018 | 0.015 | 0.015 | 0 | 0.36 | 0.14 | 0.51 | 0.3 | 0.37 | 0.22 |
| 2 | 175893333 | rs1150186 | 0.015 | -0.012 | -1.068 | 0.633 | 0.024 | 0.01 | 0.01 | 0 | 0.75 | 0.81 | 0.87 | 0.81 | 0.83 | 0.67 |
| 2 | 175901341 | rs3886691 | -0.14 | 0.707 | -0.725 | 0.706 | -0.003 | -0.003 | -0.003 | 0.112 | 0.19 | 0.18 | 0.17 | 0.34 | 0.27 | 0.28 |
| 2 | 175904189 | rs1153719 | 0.102 | 0.175 | -0.741 | 0.183 | -0.006 | -0.047 | -0.047 | 0.002 | 0.85 | 0.77 | 0.83 | 0.98 | 0.88 | 0.94 |
| 2 | 175920757 | rs1153702 | -0.451 | 0.045 | -1.096 | 0.517 | -0.013 | 0.012 | 0.012 | 0 | 0.38 | 0.55 | 0.5 | 0.45 | 0.34 | 0.36 |
| 2 | 175922382 | rs7583431 | 0.048 | 0.413 | -1.447 | 0.42 | 0.005 | -0.016 | -0.016 | 0 | 0.39 | 0.42 | 0.22 | 0.32 | 0.42 | 0.3 |
| 2 | 175922786 | rs1153699 | -0.172 | -0.733 | -1.084 | 0.446 | -0.012 | 0.014 | 0.014 | 0 | 0.31 | 0.4 | 0.42 | 0.43 | 0.26 | 0.3 |
| 2 | 175944768 | rs212347 | -0.76 | -1.021 | -0.731 | 0.46 | -0.022 | -0.034 | -0.034 | 0 | 0.24 | 0.38 | 0.15 | 0.13 | 0.17 | 0.16 |
| 2 | 175962664 | rs2072538 | 0.538 | -1.118 | -0.801 | 0.413 | -0.001 | -0.034 | -0.034 | 4.1 | 0.07 | 0.01 | 0.05 | 0.13 | 0.07 | 0.07 |
| 2 | 175971267 | rs1205399 | -1.536 | -1.216 | -0.658 | 0.377 | -0.035 | -0.048 | -0.048 | 0.033 | 0.27 | 0.46 | 0.16 | 0.15 | 0.19 | 0.18 |
| 2 | 175979296 | rs212341 | 0.073 | -1.307 | -0.504 | 0.399 | 0.007 | 0.017 | 0.017 | 0 | 0.15 | 0.1 | 0.07 | 0 | 0.1 | 0.08 |
| 2 | 175980132 | rs1153677 | -0.123 | -1.31 | -0.429 | 0.398 | -0.001 | -0.032 | -0.032 | 0 | 0.16 | 0.16 | 0.08 | 0 | 0.1 | 0.08 |
| 2 | 175982957 | rs1153675 | 0.012 | -1.308 | -0.426 | 0.399 | -0.001 | -0.029 | -0.029 | 0 | 0.16 | 0.16 | 0.08 | 0 | 0.1 | 0.08 |
| 2 | 175986973 | rs1153672 | 0.073 | -1.308 | -0.432 | 0.399 | 0.002 | -0.013 | -0.013 | 0 | 0.15 | 0.12 | 0.08 | 0 | 0.1 | 0.08 |
| 2 | 176006027 | rs13388308 | -1.479 | -1.289 | -0.399 | 0.379 | -0.023 | -0.053 | -0.053 | 0 | 0.26 | 0.45 | 0.26 | 0.13 | 0.21 | 0.17 |
| 2 | 176015840 | rs10497434 | -0.112 | -1.332 | -0.408 | 0.397 | -0.008 | 0.046 | 0.046 | 0 | 0.08 | 0.24 | 0.07 | 0.13 | 0.08 | 0.08 |
| 2 | 176025463 | rs606959 | -0.116 | -1.347 | -0.451 | 0.367 | 0.002 | -0.019 | -0.019 | 0 | 0.84 | 0.87 | 0.92 | 1 | 0.9 | 0.92 |
| 2 | 176037482 | rs13006117 | -0.223 | -1.369 | -0.41 | 0.356 | 0.004 | -0.012 | -0.012 | 0 | 0.17 | 0.14 | 0.09 | 0.01 | 0.11 | 0.1 |
| 2 | 176046655 | rs268229 | 1.029 | -1.428 | -0.37 | 0.364 | -0.028 | -0.043 | -0.043 | 0 | 0.76 | 0.6 | 0.85 | 0.85 | 0.83 | 0.84 |
| 2 | 176047906 | rs1982235 | 1.303 | -2.121 | -1.036 | 0.471 | 0.079 | -0.1 | -0.1 | 2.873 | 0.49 | 0.04 | 0.49 | 0.51 | 0.65 | 0.65 |
| 2 | 176050714 | rs268237 | -0.263 | -1.422 | -0.426 | 0.657 | 0.015 | 0.087 | 0.087 | 0 | 0.92 | 0.72 | 0.94 | 0.86 | 0.94 | 0.93 |
| 2 | 176051503 | rs13030474 | -0.455 | -1.402 | -0.576 | 0.651 | 0.005 | 0.009 | 0.009 | 0 | 0.18 | 0.15 | 0.2 | 0.01 | 0.15 | 0.1 |
| 2 | 176068396 | rs268174 | 0.029 | -1.361 | -0.443 | 0.797 | -0.013 | 0.038 | 0.038 | 0 | 0.91 | 0.82 | 0.94 | 0.86 | 0.95 | 0.91 |
| 2 | 176070435 | rs10183675 | 0.066 | -1.285 | -0.514 | 0.836 | -0.006 | 0.009 | 0.009 | 0 | 0.08 | 0.1 | 0.05 | 0.13 | 0.05 | 0.07 |
| 2 | 176077055 | rs1433236 | 0.089 | -1.292 | -0.563 | 1.126 | 0.002 | 0.034 | 0.034 | 0.059 | 0.05 | 0.03 | 0.18 | 0.22 | 0.07 | 0.08 |
| 2 | 176084790 | rs374954 | -1.666 | -1.524 | 0.165 | 1.016 | -0.01 | 0.009 | 0.009 | 0 | 0.32 | 0.44 | 0.34 | 0.36 | 0.26 | 0.28 |
| 2 | 176089888 | rs7580618 | 0.883 | -1.458 | 0.13 | 1.121 | -0.002 | 0.024 | 0.024 | 0 | 0.8 | 0.76 | 0.8 | 0.79 | 0.81 | 0.8 |
| 2 | 176094819 | rs7591715 | 1.285 | -1.485 | 0.146 | 1.142 | -0.01 | 0.057 | 0.057 | 0 | 0.77 | 0.74 | 0.76 | 0.6 | 0.8 | 0.74 |
| 2 | 176098893 | rs268164 | -0.894 | -1.585 | 0.042 | 0.818 | -0.004 | 0.006 | 0.006 | 0 | 0.9 | 0.94 | 0.94 | 1 | 0.9 | 0.94 |
| 2 | 176100028 | rs268165 | 1.831 | -1.822 | 0.43 | 1.03 | -0.016 | 0.002 | 0.002 | 0 | 0.7 | 0.38 | 0.74 | 0.66 | 0.81 | 0.73 |
| 2 | 176101320 | rs6717770 | -0.579 | -1.821 | 0.471 | 1.029 | -0.033 | -0.04 | -0.04 | 0 | 0.08 | 0.3 | 0.03 | 0 | 0.03 | 0.04 |
| 2 | 176101662 | rs6721193 | -1.929 | -1.799 | 0.113 | 1.046 | -0.09 | 0.095 | 0.095 | 0 | 0.19 | 0.53 | 0.2 | 0.34 | 0.09 | 0.21 |
| 2 | 176102135 | rs268166 | -0.805 | -1.36 | -0.206 | 0.817 | -0.006 | 0.021 | 0.021 | 0 | 0.9 | 0.96 | 0.94 | 1 | 0.9 | 0.94 |
| 2 | 176124669 | rs7565266 | -1.415 | -1.468 | 0.109 | 1.118 | -0.02 | 0.057 | 0.057 | 0 | 0.25 | 0.43 | 0.27 | 0.38 | 0.2 | 0.21 |
| 2 | 176137592 | rs17198360 | 1.46 | -1.083 | -0.593 | 0.977 | -0.16 | -0.065 | -0.065 | 1.034 | 0.23 | 0.01 | 0.32 | 0.15 | 0.43 | 0.3 |
| 2 | 176146829 | rs16863112 | -0.567 | -1.619 | 0.262 | 1.244 | 0.005 | -0.025 | -0.025 | 0.456 | 0.1 | 0.05 | 0.17 | 0.37 | 0.1 | 0.13 |
| 2 | 176152722 | rs6743113 | -0.267 | -1.592 | 0.296 | 1.24 | -0.005 | -0.086 | -0.086 | 0.087 | 0.14 | 0.31 | 0.09 | 0.01 | 0.1 | 0.08 |
| 2 | 176160297 | rs1866925 | 1.805 | -1.685 | 0.389 | 1.225 | -0.028 | 0.012 | 0.012 | 0 | 0.7 | 0.45 | 0.72 | 0.62 | 0.78 | 0.78 |
| 2 | 176198759 | rs6757526 | -0.576 | -1.579 | 0.03 | 1.305 | 0 | 0.023 | 0.023 | 0 | 0.1 | 0.11 | 0.18 | 0.37 | 0.1 | 0.11 |
| 2 | 176207636 | rs6712279 | -2.008 | -1.705 | 0.047 | 1.319 | 0.007 | -0.007 | -0.007 | 2.24 | 0.33 | 0.66 | 0.29 | 0.5 | 0.22 | 0.34 |
| 2 | 176219945 | rs9973752 | -1.57 | -1.669 | 0.13 | 1.363 | -0.004 | 0.014 | 0.014 | 0 | 0.26 | 0.31 | 0.32 | 0.54 | 0.22 | 0.34 |
| 2 | 176236921 | rs4308106 | -1.981 | -1.764 | 0.326 | 1.333 | -0.014 | 0.025 | 0.025 | 0 | 0.29 | 0.58 | 0.35 | 0.53 | 0.22 | 0.36 |
| 2 | 176239405 | rs6748853 | -1.488 | -1.757 | 0.478 | 1.413 | -0.025 | 0.142 | 0.142 | 0 | 0.15 | 0.28 | 0.18 | 0.52 | 0.1 | 0.28 |
| 2 | 176243479 | rs7588119 | -1.101 | -1.638 | 0.603 | 1.186 | -0.007 | 0.078 | 0.078 | 0 | 0.06 | 0.21 | 0.02 | 0.14 | 0 | 0.15 |
| 2 | 176271521 | rs2461773 | -1.654 | -1.704 | 0.498 | 1.152 | -0.038 | 0.072 | 0.072 | 0 | 0.1 | 0.45 | 0.05 | 0.14 | 0.02 | 0.13 |
| 2 | 176289319 | rs2461751 | -0.738 | -2.303 | -0.009 | 1.028 | -0.006 | 0.005 | 0.005 | 0 | 0.28 | 0.32 | 0.24 | 0.56 | 0.18 | 0.23 |
| 2 | 176300657 | rs2461759 | -1.154 | -2.337 | 0.043 | 1.03 | -0.016 | 0.003 | 0.003 | 0 | 0.32 | 0.4 | 0.25 | 0.69 | 0.18 | 0.35 |
| 2 | 176304996 | rs1554531 | 0.137 | -2.319 | 0.025 | 1.041 | 0.023 | -0.055 | -0.055 | 0.378 | 0.22 | 0.15 | 0.22 | 0.54 | 0.16 | 0.19 |
| 2 | 176305624 | rs2461756 | -0.793 | -2.296 | -0.004 | 0.995 | -0.029 | 0.009 | 0.009 | 0 | 0.08 | 0.19 | 0.02 | 0.13 | 0.02 | 0.13 |
| 2 | 176312418 | rs1879953 | 0.186 | -2.286 | -0.064 | 1.004 | 0.029 | -0.085 | -0.085 | 0 | 0.22 | 0.1 | 0.21 | 0.54 | 0.16 | 0.18 |
| 2 | 176317989 | rs2437890 | -1.857 | -2.289 | -0.155 | 0.873 | 0.066 | 0.02 | 0.02 | 0 | 0.47 | 0.66 | 0.41 | 0.69 | 0.3 | 0.43 |
| 2 | 176322489 | rs2437896 | -1.857 | -2.289 | -0.148 | 0.869 | 0.064 | 0.017 | 0.017 | 0 | 0.47 | 0.66 | 0.41 | 0.68 | 0.3 | 0.43 |
| 2 | 176330366 | rs16863195 | -0.117 | -2.297 | -0.256 | 0.892 | 0.01 | 0.022 | 0.022 | 0 | 0.17 | 0.12 | 0.16 | 0.15 | 0.13 | 0.22 |
| 2 | 176333564 | rs16863205 | -0.207 | -2.351 | -0.159 | 0.903 | -0.062 | 0.126 | 0.126 | 0 | 0.25 | 0.46 | 0.23 | 0.53 | 0.15 | 0.21 |
| 2 | 176346170 | rs2177250 | -1.874 | -2.435 | -0.301 | 0.837 | 0.069 | 0.043 | 0.043 | 0 | 0.48 | 0.66 | 0.42 | 0.76 | 0.3 | 0.44 |
| 2 | 176358042 | rs6433539 | -0.051 | -2.535 | -0.151 | 0.933 | -0.022 | 0.004 | 0.004 | 0 | 0.18 | 0.31 | 0.19 | 0.23 | 0.13 | 0.2 |

| | | | | | | | | | | | | | | | | |
|---|---|---|---|---|---|---|---|---|---|---|---|---|---|---|---|---|
| 2 | 176368315 | rs7600040 | -0.785 | -2.502 | -0.24 | 0.963 | -0.003 | -0.004 | -0.004 | 2.786 | 0.3 | 0.34 | 0.22 | 0.52 | 0.17 | 0.24 |
| 2 | 176389860 | rs9646737 | 0.162 | -2.38 | -0.339 | 1.025 | 0.014 | -0.022 | -0.022 | 0 | 0.23 | 0.2 | 0.21 | 0.5 | 0.14 | 0.21 |
| 2 | 176416078 | rs7574494 | -1.334 | -2.335 | -0.458 | 0.693 | 0.024 | 0.013 | 0.013 | 0 | 0.44 | 0.59 | 0.41 | 0.76 | 0.3 | 0.43 |
| 2 | 176432399 | rs6433545 | -1.507 | -2.198 | -0.43 | 0.557 | -0.016 | 0.009 | 0.009 | 0 | 0.39 | 0.46 | 0.4 | 0.59 | 0.33 | 0.33 |
| 2 | 176435942 | rs10189341 | -1.345 | -2.259 | -0.291 | 0.535 | 0.043 | -0.036 | -0.036 | 0 | 0.39 | 0.21 | 0.39 | 0.58 | 0.32 | 0.34 |
| 2 | 176439109 | rs4972770 | -0.651 | -2.254 | -0.175 | 0.607 | -0.037 | -0.019 | -0.019 | 0.034 | 0.22 | 0.47 | 0.21 | 0.19 | 0.15 | 0.22 |
| 2 | 176439632 | rs7595293 | -0.638 | -2.253 | -0.174 | 0.607 | -0.037 | -0.018 | -0.018 | 0 | 0.21 | 0.47 | 0.21 | 0.19 | 0.15 | 0.22 |
| 2 | 176443757 | rs7600916 | -1.722 | -2.499 | -0.296 | 0.571 | 0.046 | 0.035 | 0.035 | 0 | 0.54 | 0.67 | 0.43 | 0.75 | 0.35 | 0.61 |
| 2 | 176445799 | rs12616030 | -0.535 | -2.224 | -0.119 | 0.63 | -0.037 | 0.04 | 0.04 | 0 | 0.33 | 0.46 | 0.36 | 0.59 | 0.25 | 0.29 |
| 2 | 176449962 | rs11693978 | -1.013 | -2.236 | -0.098 | 0.601 | 0.016 | -0.011 | -0.011 | 0.087 | 0.36 | 0.34 | 0.35 | 0.58 | 0.28 | 0.31 |
| 2 | 176451291 | rs4283390 | -0.973 | -2.298 | -0.122 | 0.65 | -0.022 | 0.017 | 0.017 | 0.357 | 0.1 | 0.16 | 0.03 | 0.19 | 0.04 | 0.14 |
| 2 | 176461268 | rs7572607 | -0.547 | -2.281 | -0.205 | 0.656 | 0.026 | -0.021 | -0.021 | 0.125 | 0.32 | 0.26 | 0.33 | 0.39 | 0.28 | 0.2 |
| 2 | 176471491 | rs4365432 | -1.29 | -2.262 | -0.274 | 0.662 | -0.011 | 0.003 | 0.003 | 0 | 0.39 | 0.58 | 0.37 | 0.58 | 0.28 | 0.31 |
| 2 | 176476136 | rs7582388 | -0.06 | -1.864 | -0.61 | 0.704 | 0.016 | -0.038 | -0.038 | 0 | 0.26 | 0.18 | 0.31 | 0.39 | 0.23 | 0.18 |
| 2 | 176483606 | rs7603261 | 0.493 | -1.872 | -0.327 | 0.475 | -0.053 | -0.03 | -0.03 | 0 | 0.17 | 0.31 | 0.08 | 0.17 | 0.08 | 0.29 |
| 2 | 176483881 | rs4972778 | -1.491 | -1.255 | -1.018 | 0.172 | 0.004 | -0.001 | -0.001 | 0 | 0.71 | 0.7 | 0.73 | 0.81 | 0.57 | 0.81 |
| 2 | 176485746 | rs12474376 | 0.633 | -1.673 | -1.483 | 0.061 | 0.005 | 0.005 | 0.005 | 0 | 0.24 | 0.09 | 0.32 | 0.23 | 0.21 | 0.47 |
| 2 | 176487261 | rs9287987 | -1.165 | -1.918 | -1.317 | -0.27 | 0.003 | 0.009 | 0.009 | 0 | 0.55 | 0.57 | 0.47 | 0.45 | 0.43 | 0.68 |
| 2 | 176488774 | rs11682476 | -0.182 | -1.926 | -1.197 | -0.283 | -0.038 | -0.07 | -0.07 | 0.114 | 0.32 | 0.39 | 0.15 | 0.21 | 0.22 | 0.21 |
| 2 | 176490148 | rs7588395 | 0.029 | -1.951 | -1.185 | -0.295 | -0.025 | -0.051 | -0.051 | 0 | 0.3 | 0.33 | 0.15 | 0.21 | 0.22 | 0.19 |
| 2 | 176494873 | rs4263082 | 0.935 | -1.823 | -1.134 | -0.274 | 0.01 | 0.004 | 0.004 | 0 | 0.24 | 0.09 | 0.3 | 0.23 | 0.2 | 0.45 |
| 2 | 176495726 | rs11897944 | 0.501 | -1.844 | -1.091 | -0.34 | -0.033 | -0.113 | -0.113 | 1.58 | 0.2 | 0.31 | 0.09 | 0.04 | 0.14 | 0.08 |
| 2 | 176505213 | rs9789760 | 0.287 | -1.919 | -1.149 | -0.217 | -0.006 | 0.001 | 0.001 | 0 | 0.61 | 0.48 | 0.61 | 0.57 | 0.68 | 0.66 |
| 2 | 176526784 | rs1446806 | -1.577 | -1.853 | -1.255 | -0.335 | -0.015 | 0 | 0 | 0 | 0.63 | 0.61 | 0.69 | 0.66 | 0.53 | 0.75 |
| 2 | 176528033 | rs17585824 | -0.326 | -1.975 | -1.492 | -0.357 | -0.001 | 0.146 | 0.146 | 0 | 0.4 | 0.04 | 0.34 | 0.19 | 0.32 | 0.45 |
| 2 | 176530770 | rs16863349 | 1.503 | -1.915 | -1.354 | -0.259 | 0.001 | -0.042 | -0.042 | 0 | 0.16 | 0.08 | 0.28 | 0.45 | 0.17 | 0.18 |
| 2 | 176543105 | rs10170575 | 0.839 | -1.967 | -1.386 | -0.413 | 0.031 | 0.034 | 0.034 | 0 | 0.17 | 0.1 | 0.06 | 0.02 | 0.11 | 0.04 |
| 2 | 176544182 | rs6738625 | 0.83 | -1.815 | -1.249 | -0.295 | -0.008 | 0.002 | 0.002 | 0 | 0.14 | 0.34 | 0.11 | 0.19 | 0.11 | 0.13 |
| 2 | 176545579 | rs7578330 | -1.737 | -1.484 | -1.678 | -0.288 | -0.001 | 0 | 0 | 0 | 0.72 | 0.69 | 0.75 | 0.84 | 0.6 | 0.84 |
| 2 | 176548510 | rs12479248 | 0.639 | -1.513 | -1.757 | -0.277 | -0.039 | 0.059 | 0.059 | 0 | 0.24 | 0.05 | 0.38 | 0.13 | 0.23 | 0.41 |
| 2 | 176549467 | rs7592980 | -0.48 | -1.478 | -1.345 | -0.181 | 0.032 | 0.022 | 0.022 | 0 | 0.48 | 0.56 | 0.36 | 0.7 | 0.37 | 0.43 |
| 2 | 176552107 | rs11891974 | 1.489 | -0.716 | -1.705 | -0.009 | 0.01 | 0.002 | 0.002 | 0 | 0.22 | 0.24 | 0.19 | 0.18 | 0.31 | 0.14 |
| 2 | 176552298 | rs17270679 | 1.172 | 0.007 | -1.901 | -0.076 | 0.04 | 0.03 | 0.03 | 0 | 0.16 | 0.27 | 0.12 | 0.13 | 0.22 | 0.11 |
| 2 | 176552756 | rs11683783 | -0.003 | -0.779 | -2.086 | -0.294 | 0.042 | 0.062 | 0.062 | 0.023 | 0.52 | 0.32 | 0.48 | 0.68 | 0.63 | 0.45 |
| 2 | 176580867 | rs11680722 | 0.514 | -0.09 | -1.708 | -0.04 | 0.003 | -0.002 | -0.002 | 0 | 0.27 | 0.27 | 0.2 | 0.16 | 0.34 | 0.22 |
| 2 | 176590041 | rs10209318 | -0.047 | -0.347 | -2.078 | -0.379 | -0.004 | 0.038 | 0.038 | 0 | 0.2 | 0.07 | 0.36 | 0.14 | 0.19 | 0.29 |
| 2 | 176596709 | rs4510185 | -0.15 | -0.239 | -2.048 | -0.481 | -0.002 | 0.033 | 0.033 | 0.244 | 0.23 | 0.1 | 0.34 | 0.14 | 0.22 | 0.38 |
| 2 | 176601784 | rs11689292 | 0.762 | -0.241 | -2.302 | -0.609 | 0.013 | 0.043 | 0.043 | 0 | 0.19 | 0.04 | 0.2 | 0.16 | 0.18 | 0.11 |
| 2 | 176604794 | rs12477983 | 0.961 | -0.145 | -2.24 | -0.554 | -0.026 | 0.003 | 0.003 | 0 | 0.13 | 0.04 | 0.25 | 0.08 | 0.15 | 0.19 |
| 2 | 176610873 | rs7579789 | -0.056 | -0.095 | -1.909 | -0.611 | 0.007 | 0.045 | 0.045 | 0.024 | 0.7 | 0.81 | 0.69 | 0.77 | 0.7 | 0.7 |
| 2 | 176614615 | rs1348843 | -0.975 | -0.192 | -1.799 | -0.758 | 0.023 | 0.031 | 0.031 | 0 | 0.78 | 0.85 | 0.91 | 0.98 | 0.87 | 0.86 |
| 2 | 176618964 | rs11685409 | 0.718 | -0.121 | -1.721 | -0.524 | 0.056 | 0.043 | 0.043 | 0 | 0.27 | 0.54 | 0.3 | 0.56 | 0.37 | 0.32 |
| 2 | 176621400 | rs13017316 | -0.102 | -0.215 | -1.789 | -0.602 | -0.003 | 0.017 | 0.017 | 0.012 | 0.52 | 0.41 | 0.64 | 0.44 | 0.53 | 0.54 |
| 2 | 176621926 | rs6752840 | -1.255 | -0.223 | -1.64 | -0.517 | 0.047 | 0.076 | 0.076 | 0 | 0.8 | 0.95 | 0.93 | 1 | 0.9 | 0.86 |
| 2 | 176631915 | rs6714708 | -1.255 | -0.223 | -1.64 | -0.517 | 0.047 | 0.076 | 0.076 | 0 | 0.8 | 0.95 | 0.93 | 1 | 0.9 | 0.86 |
| 2 | 176636201 | rs6746246 | -0.302 | 0.243 | -1.31 | -0.045 | 0.02 | 0.059 | 0.059 | 0 | 0.49 | 0.62 | 0.51 | 0.71 | 0.57 | 0.46 |
| 2 | 176641602 | rs10930709 | -0.236 | 0.548 | -0.806 | 0.541 | 0.077 | 0.066 | 0.066 | 0 | 0.3 | 0.5 | 0.3 | 0.48 | 0.46 | 0.3 |
| 2 | 176643561 | rs1446824 | -1 | -0.471 | -1.551 | -0.229 | 0.061 | 0.125 | 0.125 | 0 | 0.64 | 0.89 | 0.72 | 0.81 | 0.74 | 0.77 |
| 2 | 176646367 | rs2044492 | 0.774 | -0.577 | -1.352 | -0.35 | 0.002 | 0.001 | 0.001 | 0 | 0.27 | 0.24 | 0.38 | 0.15 | 0.22 | 0.41 |
| 2 | 176651117 | rs4356634 | 1.028 | -0.277 | -0.876 | 0.236 | 0.03 | -0.006 | -0.006 | 0 | 0.21 | 0.26 | 0.21 | 0.13 | 0.38 | 0.12 |
| 2 | 176652167 | rs10497444 | 1.019 | -0.506 | -1.417 | 0.183 | 0.003 | 0.005 | 0.005 | 0 | 0.06 | 0.03 | 0.03 | 0.02 | 0.04 | 0.03 |
| 2 | 176652998 | rs10197647 | -1.23 | -0.529 | -1.438 | 0.179 | 0.056 | 0.124 | 0.124 | 0 | 0.64 | 0.93 | 0.72 | 0.81 | 0.74 | 0.76 |
| 2 | 176654738 | rs7566934 | -0.705 | -0.557 | -1.401 | 0.134 | 0.081 | 0.172 | 0.172 | 0 | 0.58 | 0.88 | 0.7 | 0.8 | 0.7 | 0.73 |
| 2 | 176654894 | rs7567121 | -0.285 | -0.51 | -1.403 | -0.008 | 0.009 | 0.006 | 0.006 | 0 | 0.39 | 0.33 | 0.25 | 0.36 | 0.32 | 0.44 |
| 2 | 176657097 | rs726032 | 0.628 | -0.586 | -1.447 | -0.027 | 0.017 | 0.027 | 0.027 | 0 | 0.3 | 0.2 | 0.18 | 0.17 | 0.23 | 0.33 |
| 2 | 176657222 | rs726034 | -1.208 | -0.13 | -1.693 | 0.076 | 0.011 | -0.012 | -0.012 | 0.116 | 0.64 | 0.96 | 0.69 | 0.74 | 0.67 | 0.75 |
| 2 | 176660426 | rs1011143 | 0.167 | -0.232 | -1.398 | 0.116 | 0.078 | 0.055 | 0.055 | 0.699 | 0.29 | 0.52 | 0.49 | 0.42 | 0.45 | 0.38 |
| 2 | 176663264 | rs765578 | 0.079 | -0.281 | -1.325 | 0.082 | 0.045 | 0.083 | 0.083 | 0 | 0.36 | 0.16 | 0.2 | 0.19 | 0.25 | 0.36 |
| 2 | 176671391 | rs1446832 | 0.467 | -0.221 | -1.271 | 0.156 | -0.012 | -0.002 | -0.002 | 2.769 | 0.31 | 0.34 | 0.19 | 0.32 | 0.23 | 0.37 |
| 2 | 176672999 | rs12616125 | 0.671 | -0.219 | -1.353 | 0.17 | 0.019 | 0.027 | 0.027 | 0 | 0.29 | 0.2 | 0.17 | 0.17 | 0.21 | 0.33 |
| 2 | 176675393 | rs6754856 | -0.677 | -0.237 | -1.65 | 0.215 | 0.069 | -0.044 | -0.044 | 0 | 0.59 | 0.92 | 0.69 | 0.59 | 0.68 | 0.73 |
| 2 | 176676164 | rs4619574 | -0.704 | -0.233 | -1.648 | 0.215 | 0.067 | -0.045 | -0.045 | 0.088 | 0.59 | 0.92 | 0.69 | 0.59 | 0.68 | 0.73 |
| 2 | 176677084 | rs1446829 | -0.759 | -0.242 | -1.641 | 0.219 | 0.067 | 0.046 | 0.046 | 0.38 | 0.61 | 0.92 | 0.7 | 0.73 | 0.69 | 0.75 |
| 2 | 176680246 | rs11896469 | -0.472 | 0.727 | -1.771 | 0.217 | -0.021 | -0.036 | -0.036 | 0 | 0.56 | 0.93 | 0.65 | 0.59 | 0.59 | 0.72 |
| 2 | 176684549 | rs10201284 | -1.285 | 0.636 | -1.79 | 0.182 | 0.037 | -0.004 | -0.004 | 0 | 0.68 | 0.96 | 0.73 | 0.76 | 0.74 | 0.78 |
| 2 | 176687393 | rs2044491 | 0.457 | 0.761 | -1.341 | 0.282 | -0.008 | -0.014 | -0.014 | 0.054 | 0.3 | 0.31 | 0.19 | 0.18 | 0.22 | 0.37 |
| 2 | 176693074 | rs981599 | -0.387 | 1.28 | -0.771 | 0.839 | 0.084 | 0.078 | 0.078 | 0.103 | 0.25 | 0.49 | 0.46 | 0.39 | 0.37 | 0.35 |
| 2 | 176693188 | rs981600 | -0.035 | 1.368 | -0.618 | 0.067 | 0.095 | -0.093 | -0.093 | 1.866 | 0.2 | 0.45 | 0.4 | 0.08 | 0.31 | 0.1 |
| 2 | 176712492 | rs1446662 | -0.045 | 0.547 | -1.121 | -0.166 | -0.002 | -0.017 | -0.017 | 1.123 | 0.31 | 0.63 | 0.28 | 0.15 | 0.28 | 0.29 |

| | | | | | | | | | | | | | | | | |
|---|---|---|---|---|---|---|---|---|---|---|---|---|---|---|---|---|
| 2 | 176716469 | rs1012357 | -0.454 | 0.914 | -0.979 | -0.469 | 0.005 | -0.024 | -0.024 | 0 | 0.42 | 0.47 | 0.45 | 0.27 | 0.5 | 0.27 |
| 2 | 176716654 | rs10930713 | 0.807 | 0.993 | -1.542 | -0.545 | -0.031 | 0.021 | 0.021 | 0 | 0.13 | 0.04 | 0.1 | 0.1 | 0.17 | 0.15 |
| 2 | 176720147 | rs935406 | 0.634 | 1.014 | -1.315 | -0.559 | 0.006 | -0.009 | -0.009 | 0 | 0.19 | 0.2 | 0.19 | 0.13 | 0.23 | 0.21 |
| 2 | 176720802 | rs6747251 | 0.894 | 1.144 | -1.372 | -0.519 | -0.015 | 0.011 | 0.011 | 0 | 0.11 | 0.07 | 0.08 | 0.1 | 0.13 | 0.14 |
| 2 | 176721984 | rs1374369 | -0.394 | 0.847 | -1.476 | 0.067 | 0.001 | -0.001 | -0.001 | 0 | 0.57 | 0.53 | 0.39 | 0.59 | 0.5 | 0.67 |
| 2 | 176725040 | rs1838864 | -0.12 | 0.544 | -1.566 | -0.026 | -0.001 | 0.019 | 0.019 | 0 | 0.47 | 0.43 | 0.32 | 0.28 | 0.42 | 0.44 |
| 2 | 176728443 | rs12693083 | -0.552 | 0.15 | -1.477 | 0.056 | -0.001 | 0 | 0 | 0 | 0.53 | 0.49 | 0.34 | 0.6 | 0.41 | 0.63 |
| 2 | 176729944 | rs2166523 | 0.517 | -0.065 | -1.495 | 0.164 | -0.012 | 0.012 | 0.012 | 0 | 0.18 | 0.18 | 0.08 | 0.31 | 0.1 | 0.26 |
| 2 | 176730467 | rs2884887 | 1.45 | 0.36 | -1.206 | 0.386 | -0.013 | 0.108 | 0.108 | 0 | 0.1 | 0.15 | 0.07 | 0.31 | 0.08 | 0.22 |
| 2 | 176732191 | rs10196901 | 1.63 | 0.364 | -1.376 | 0.394 | -0.006 | 0.05 | 0.05 | 0 | 0.1 | 0.1 | 0.07 | 0.31 | 0.08 | 0.21 |
| 2 | 176734598 | rs4972489 | 0.396 | 0.726 | -1.518 | 0.286 | -0.059 | -0.091 | -0.091 | 0 | 0.15 | 0.01 | 0.34 | 0.26 | 0.22 | 0.12 |
| 2 | 176735728 | rs7579376 | 0.328 | 0.44 | -1.287 | 0.144 | -0.001 | 0 | 0 | 0 | 0.41 | 0.41 | 0.31 | 0.47 | 0.35 | 0.55 |
| 2 | 176736761 | rs12471824 | -0.161 | 0.554 | -1.169 | -0.574 | 0 | -0.009 | -0.009 | 0.077 | 0.45 | 0.48 | 0.59 | 0.21 | 0.56 | 0.38 |
| 2 | 176738540 | rs12693085 | -0.079 | -0.043 | -1.693 | -0.8 | 0.04 | 0.066 | 0.066 | 0 | 0.66 | 0.79 | 0.84 | 0.94 | 0.77 | 0.76 |
| 2 | 176741174 | rs4972491 | -0.942 | 0.13 | -1.811 | -0.458 | 0.02 | 0.058 | 0.058 | 1.234 | 0.78 | 0.94 | 0.86 | 0.91 | 0.82 | 0.81 |
| 2 | 176742753 | rs10930718 | -0.333 | -0.465 | -1.655 | -0.22 | 0.029 | 0.011 | 0.011 | 0 | 0.29 | 0.24 | 0.18 | 0.3 | 0.15 | 0.34 |
| 2 | 176744882 | rs921083 | 0.076 | 0.182 | -0.914 | 0.026 | 0.027 | 0.021 | 0.021 | 0.006 | 0.14 | 0.27 | 0.35 | 0.09 | 0.17 | 0.07 |
| 2 | 176745836 | rs1816922 | -1.251 | 0.246 | -1.127 | -0.3 | 0.009 | 0.009 | 0.009 | 0 | 0.5 | 0.55 | 0.57 | 0.4 | 0.41 | 0.37 |
| 2 | 176746065 | rs13397065 | 0.228 | 0.546 | -0.789 | -0.22 | 0.019 | 0.015 | 0.015 | 0.06 | 0.13 | 0.22 | 0.34 | 0.09 | 0.17 | 0.07 |
| 2 | 176747479 | rs11681949 | 0.572 | 0.424 | -1.002 | -0.43 | 0 | 0 | 0 | 0 | 0.1 | 0.07 | 0.04 | 0.01 | 0.08 | 0.07 |
| 2 | 176749778 | rs6433564 | -1.364 | 0.311 | -1.164 | -0.475 | -0.013 | 0.016 | 0.016 | 0.99 | 0.45 | 0.84 | 0.54 | 0.41 | 0.43 | 0.42 |
| 2 | 176750331 | rs1446666 | -1.008 | 0.473 | -0.956 | -0.292 | -0.002 | -0.012 | -0.012 | 0.931 | 0.36 | 0.8 | 0.51 | 0.4 | 0.35 | 0.43 |
| 2 | 176752831 | rs921084 | 0.893 | 1.391 | -1.284 | -1.014 | -0.06 | 0.023 | 0.023 | 0 | 0.13 | 0.02 | 0.23 | 0.11 | 0.22 | 0.07 |
| 2 | 176756644 | rs10197186 | -0.546 | 0.911 | -0.515 | -0.901 | -0.002 | -0.016 | -0.016 | 1.672 | 0.25 | 0.22 | 0.41 | 0.07 | 0.28 | 0.22 |
| 2 | 176756752 | rs6709287 | -0.231 | 0.607 | -0.277 | -0.391 | 0.038 | 0.009 | 0.009 | 0.444 | 0.06 | 0.35 | 0.03 | 0 | 0.02 | 0.07 |
| 2 | 176769442 | rs4972497 | -0.753 | 0.897 | -1.15 | -0.603 | -0.034 | 0.08 | 0.08 | 0 | 0.21 | 0.08 | 0.36 | 0.05 | 0.25 | 0.21 |
| 2 | 176769575 | rs7599364 | -1.05 | 0.334 | -0.611 | -0.629 | -0.015 | -0.016 | -0.016 | 0 | 0.08 | 0.17 | 0.02 | 0.04 | 0.04 | 0.07 |
| 2 | 176770136 | rs4972798 | -0.741 | 0.074 | -0.713 | -0.532 | 0.001 | -0.001 | -0.001 | 0 | 0.43 | 0.36 | 0.7 | 0.5 | 0.55 | 0.33 |
| 2 | 176771548 | rs10185549 | -0.463 | 0.179 | -0.419 | -0.646 | 0.121 | 0.104 | 0.104 | 0.014 | 0.75 | 0.49 | 0.69 | 0.64 | 0.53 | 0.79 |
| 2 | 176772269 | rs2884888 | 0.167 | -0.271 | -0.504 | -0.846 | -0.005 | 0 | 0 | 0 | 0.32 | 0.27 | 0.54 | 0.36 | 0.34 | 0.2 |
| 2 | 176780250 | rs2289969 | 1.144 | -0.226 | -0.573 | -0.921 | 0.007 | 0.014 | 0.014 | 0.149 | 0.21 | 0.22 | 0.3 | 0.38 | 0.26 | 0.11 |
| 2 | 176784138 | rs1020410 | 0.266 | -0.528 | -0.004 | -1.031 | 0.015 | 0.015 | 0.015 | 0.098 | 0.48 | 0.49 | 0.27 | 0.3 | 0.29 | 0.61 |
| 2 | 176784435 | rs17315753 | 1.316 | -0.341 | -0.021 | -1.03 | 0.092 | -0.028 | -0.028 | 0 | 0.05 | 0.08 | 0.1 | 0 | 0.19 | 0.03 |
| 2 | 176789816 | rs12477861 | 0.956 | -0.514 | -0.155 | -1.086 | -0.023 | -0.002 | -0.002 | 0 | 0.06 | 0.01 | 0.1 | 0.07 | 0.07 | 0.1 |
| 2 | 176791642 | rs12621733 | 0.997 | -0.514 | -0.155 | -1.086 | -0.023 | -0.002 | -0.002 | 0.049 | 0.06 | 0.01 | 0.1 | 0.07 | 0.07 | 0.1 |
| 2 | 176791759 | rs7571968 | -0.332 | -0.592 | -0.057 | -1.132 | -0.014 | 0.042 | 0.042 | 3.375 | 0.55 | 0.7 | 0.38 | 0.3 | 0.48 | 0.64 |
| 2 | 176792929 | rs12622749 | 0.945 | -0.644 | -0.121 | -1.104 | -0.023 | -0.002 | -0.002 | 0 | 0.06 | 0.01 | 0.1 | 0.06 | 0.07 | 0.1 |
| 2 | 176793016 | rs12622783 | 0.946 | -0.648 | -0.123 | -1.106 | -0.023 | -0.002 | -0.002 | 0 | 0.06 | 0.01 | 0.1 | 0.06 | 0.07 | 0.1 |
| 2 | 176794388 | rs2289966 | 2.596 | -1.368 | -0.555 | -1.482 | 0.072 | -0.071 | -0.071 | 0 | 0.19 | 0 | 0.05 | 0.22 | 0.06 | 0.19 |
| 2 | 176806653 | rs2166528 | 2.074 | -1.023 | -0.289 | -1.371 | -0.038 | 0.067 | 0.067 | 0 | 0.16 | 0.04 | 0.25 | 0.06 | 0.18 | 0.07 |
| 2 | 176809605 | rs4369827 | 0.225 | -1.027 | -0.149 | -1.378 | -0.014 | -0.012 | -0.012 | 0.205 | 0.45 | 0.53 | 0.26 | 0.3 | 0.26 | 0.52 |
| 2 | 176822227 | rs11691719 | 1.065 | -1.03 | -0.181 | -1.374 | 0.094 | 0.027 | 0.027 | 2.701 | 0.34 | 0.05 | 0.11 | 0.23 | 0.15 | 0.41 |
| 2 | 176838145 | rs9646792 | -1.207 | -0.988 | -0.097 | -1.362 | -0.026 | 0.015 | 0.015 | 1.265 | 0.64 | 0.77 | 0.54 | 0.31 | 0.52 | 0.69 |
| 2 | 176846592 | rs12621889 | 1.078 | -0.841 | -0.201 | -1.278 | 0.002 | -0.004 | -0.004 | 0 | 0.07 | 0.06 | 0.11 | 0.07 | 0.07 | 0.1 |
| 2 | 176848708 | rs819032 | -0.534 | -0.855 | -0.258 | -1.282 | -0.013 | 0.042 | 0.042 | 3.028 | 0.55 | 0.7 | 0.38 | 0.3 | 0.48 | 0.64 |
| 2 | 176855323 | rs6433567 | -0.822 | -0.755 | -0.572 | -1.181 | -0.003 | -0.001 | -0.001 | 0 | 0.6 | 0.54 | 0.84 | 0.77 | 0.78 | 0.57 |
| 2 | 176858163 | rs12475053 | 1.948 | -0.716 | -0.595 | -0.953 | -0.042 | 0.02 | 0.02 | 0.134 | 0.16 | 0.03 | 0.24 | 0.11 | 0.18 | 0.08 |
| 2 | 176874514 | rs3116565 | -0.243 | -0.639 | -0.897 | -1.064 | 0.004 | 0.013 | 0.013 | 2.173 | 0.52 | 0.4 | 0.73 | 0.77 | 0.59 | 0.46 |
| 2 | 176881892 | rs6715628 | 1.387 | -0.025 | -1.335 | -0.412 | -0.021 | -0.042 | -0.042 | 0 | 0.16 | 0.09 | 0.27 | 0.41 | 0.19 | 0.1 |
| 2 | 176888760 | rs711810 | 0.211 | -0.665 | -0.994 | -1.074 | -0.064 | -0.036 | -0.036 | 0 | 0.33 | 0.53 | 0.26 | 0.21 | 0.18 | 0.36 |
| 2 | 176889698 | rs11891328 | 0.459 | -0.306 | -1.318 | -1.129 | 0.097 | -0.037 | -0.037 | 0 | 0.07 | 0.14 | 0.11 | 0 | 0.19 | 0.03 |
| 2 | 176890664 | rs17223885 | 0.017 | -0.566 | -1.728 | -1.28 | 0.009 | 0.083 | 0.083 | 0.173 | 0.21 | 0.05 | 0.15 | 0.08 | 0.14 | 0.18 |
| 2 | 176892059 | rs16863504 | 1.543 | -0.64 | -1.717 | -1.25 | 0.061 | -0.037 | -0.037 | 0 | 0.2 | 0.02 | 0.19 | 0.21 | 0.1 | 0.31 |
| 2 | 176892928 | rs17223990 | 0.68 | 0.023 | -1.423 | -0.94 | 0 | 0.011 | 0.011 | 0 | 0.09 | 0.05 | 0.1 | 0.07 | 0.07 | 0.11 |
| 2 | 176900090 | rs974699 | 0.144 | -0.068 | -1.441 | -0.999 | -0.004 | -0.002 | -0.002 | 0 | 0.41 | 0.6 | 0.35 | 0.28 | 0.25 | 0.46 |
| 2 | 176901200 | rs847138 | -0.554 | 0.497 | -1.618 | -1.035 | -0.053 | 0.087 | 0.087 | 0.856 | 0.27 | 0.11 | 0.25 | 0.08 | 0.33 | 0.18 |
| 2 | 176914739 | rs847191 | -0.654 | 0.292 | -1.328 | -0.945 | -0.026 | 0.053 | 0.053 | 0 | 0.61 | 0.73 | 0.52 | 0.24 | 0.51 | 0.62 |
| 2 | 176914965 | rs847189 | -0.114 | 0.313 | -1.327 | -1.192 | 0.004 | -0.008 | -0.008 | 0 | 0.14 | 0.13 | 0.2 | 0.07 | 0.26 | 0.14 |
| 2 | 176916106 | rs847184 | -1.728 | 0.607 | -1.677 | -0.69 | -0.033 | -0.078 | -0.078 | 0 | 0.84 | 0.96 | 0.82 | 0.62 | 0.73 | 0.76 |
| 2 | 176918785 | rs16863530 | 0.956 | -0.092 | -1.296 | -1.034 | -0.066 | -0.072 | -0.072 | 0 | 0.33 | 0.52 | 0.25 | 0.16 | 0.18 | 0.36 |
| 2 | 176921784 | rs847175 | -0.543 | 0.301 | -1.442 | -0.805 | -0.051 | 0.106 | 0.106 | 0 | 0.61 | 0.8 | 0.51 | 0.21 | 0.49 | 0.61 |
| 2 | 176927667 | rs12474551 | 0.178 | 0.148 | -1.276 | -0.696 | 0 | -0.005 | -0.005 | 0 | 0.09 | 0.07 | 0.11 | 0.05 | 0.07 | 0.12 |
| 2 | 176928633 | rs7426104 | -0.383 | 0.366 | -1.315 | -0.437 | 0.027 | 0.022 | 0.022 | 0 | 0.57 | 0.85 | 0.42 | 0.51 | 0.37 | 0.64 |
| 2 | 176936834 | rs847126 | -0.211 | -0.305 | -1.137 | -0.968 | 0.158 | 0.186 | 0.186 | 0 | 0.53 | 0.81 | 0.34 | 0.19 | 0.22 | 0.55 |
| 2 | 176950926 | rs6752623 | -0.402 | -0.078 | -0.386 | -1.02 | -0.022 | -0.074 | -0.074 | 0.232 | 0.17 | 0.32 | 0.22 | 0.04 | 0.13 | 0.26 |
| 2 | 176951487 | rs12474905 | 1.043 | 0.339 | -0.848 | -1.04 | -0.049 | 0.036 | 0.036 | 0 | 0.16 | 0.04 | 0.31 | 0.1 | 0.2 | 0.07 |
| 2 | 176955545 | rs847200 | -0.369 | -0.131 | -0.132 | -0.934 | -0.037 | -0.081 | -0.081 | 0 | 0.15 | 0.34 | 0.21 | 0.04 | 0.1 | 0.23 |
| 2 | 176956510 | rs3754983 | -0.468 | -0.122 | -0.198 | -0.93 | -0.029 | -0.064 | -0.064 | 0 | 0.15 | 0.29 | 0.2 | 0.04 | 0.1 | 0.23 |
| 2 | 176962989 | rs847153 | -1.211 | -0.155 | -0.329 | -1.504 | 0 | -0.036 | -0.036 | 0.247 | 0.38 | 0.64 | 0.42 | 0.07 | 0.31 | 0.21 |

| | | | | | | | | | | | | | | | | |
|---|---|---|---|---|---|---|---|---|---|---|---|---|---|---|---|---|
| 2 | 176964304 | rs711812 | -1.211 | -0.155 | -0.329 | -1.504 | 0 | -0.036 | -0.036 | 0 | 0.38 | 0.64 | 0.42 | 0.07 | 0.31 | 0.21 |
| 2 | 176965086 | rs35817516 | 0.628 | -0.338 | -0.491 | -1.639 | -0.046 | 0.049 | 0.049 | 0 | 0.09 | 0.01 | 0.22 | 0.01 | 0.12 | 0.05 |
| 2 | 176966190 | rs711813 | -1.64 | -0.325 | -0.437 | -1.638 | 0.011 | 0.011 | 0.011 | 1.627 | 0.38 | 0.69 | 0.42 | 0.07 | 0.29 | 0.21 |
| 2 | 176968223 | rs847150 | -0.627 | -0.408 | -0.083 | -1.696 | 0.018 | -0.035 | -0.035 | 0 | 0.15 | 0.23 | 0.31 | 0.06 | 0.18 | 0.13 |
| 2 | 176969206 | rs741610 | -1.668 | -0.418 | -0.302 | -1.658 | 0.024 | 0.073 | 0.073 | 2.39 | 0.39 | 0.76 | 0.43 | 0.07 | 0.29 | 0.21 |
| 2 | 176970659 | rs847147 | -0.76 | -0.415 | -0.108 | -1.569 | -0.004 | -0.116 | -0.116 | 1.477 | 0.26 | 0.43 | 0.32 | 0.04 | 0.22 | 0.17 |
| 2 | 176974104 | rs863678 | -1.402 | -0.193 | -0.924 | -0.934 | -0.025 | -0.055 | -0.055 | 0 | 0.63 | 0.89 | 0.66 | 0.44 | 0.62 | 0.52 |
| 2 | 176974122 | rs6745764 | -0.626 | -0.53 | -0.199 | -0.742 | 0.005 | 0 | 0 | 1.282 | 0.21 | 0.34 | 0.22 | 0.23 | 0.2 | 0.14 |
| 2 | 176975973 | rs711815 | -1.089 | -0.624 | -0.596 | -0.683 | 0.022 | 0.031 | 0.031 | 4.131 | 0.37 | 0.73 | 0.33 | 0.25 | 0.23 | 0.17 |
| 2 | 176979396 | rs6709104 | -0.864 | -0.651 | -0.514 | -0.972 | -0.005 | -0.075 | -0.075 | 0 | 0.35 | 0.36 | 0.3 | 0.02 | 0.22 | 0.17 |
| 2 | 176981343 | rs7601234 | -0.906 | -0.595 | -0.547 | -0.953 | -0.002 | -0.047 | -0.047 | 0 | 0.36 | 0.37 | 0.31 | 0.05 | 0.24 | 0.17 |
| 2 | 176985696 | rs711819 | -1.135 | -0.65 | -1.204 | -0.217 | -0.057 | -0.074 | -0.074 | 0.055 | 0.86 | 0.97 | 0.84 | 0.4 | 0.76 | 0.85 |
| 2 | 176987220 | rs10445765 | 0.698 | -1.557 | -1.493 | -0.778 | 0.026 | -0.007 | -0.007 | 0 | 0.43 | 0.15 | 0.69 | 0.4 | 0.67 | 0.58 |
| 2 | 176988412 | rs698258 | 0.428 | -1.287 | -0.87 | -0.559 | -0.139 | -0.154 | -0.154 | 0.002 | 0.22 | 0.54 | 0.07 | 0 | 0.02 | 0.01 |
| 2 | 176991779 | rs2592394 | -0.308 | -1.1 | -1.003 | -0.46 | -0.016 | -0.032 | -0.032 | 0 | 0.75 | 0.83 | 0.78 | 0.42 | 0.71 | 0.75 |
| 2 | 176993129 | rs711825 | -0.301 | -1.1 | -0.99 | -0.469 | -0.015 | -0.031 | -0.031 | 0 | 0.75 | 0.83 | 0.78 | 0.44 | 0.71 | 0.77 |
| 2 | 176995262 | rs7580746 | 0.649 | -1.281 | -0.984 | -0.504 | -0.132 | -0.076 | -0.076 | 0 | 0.23 | 0.46 | 0.09 | 0.1 | 0.03 | 0.08 |
| 2 | 176999012 | rs2113563 | 0.803 | -1.069 | -1.416 | -0.422 | 0.082 | 0.077 | 0.077 | 1.442 | 0.49 | 0.12 | 0.68 | 0.34 | 0.68 | 0.68 |
| 2 | 177004115 | rs4078756 | 2.589 | -1.033 | -1.382 | 0.073 | 0.031 | -0.032 | -0.032 | 0.03 | 0.07 | 0.02 | 0.02 | 0.27 | 0.01 | 0.05 |
| 2 | 177005757 | rs1348807 | 0.131 | -0.88 | -1.244 | -0.045 | -0.012 | 0.008 | 0.008 | 0.271 | 0.61 | 0.48 | 0.74 | 0.62 | 0.68 | 0.78 |
| 2 | 177007664 | rs10188827 | -1.401 | -0.875 | -1.38 | -0.047 | -0.056 | -0.066 | -0.066 | 0 | 0.86 | 0.97 | 0.84 | 0.72 | 0.75 | 0.86 |
| 2 | 177008914 | rs6736786 | -0.868 | -0.971 | -1.616 | -0.554 | 0.032 | 0.09 | 0.09 | 0.94 | 0.53 | 0.84 | 0.32 | 0.38 | 0.39 | 0.31 |
| 2 | 177011655 | rs10196832 | 0.457 | -0.724 | -1.107 | -0.952 | -0.14 | -0.093 | -0.093 | 3.755 | 0.24 | 0.51 | 0.08 | 0.09 | 0.03 | 0.07 |
| 2 | 177015815 | rs13418078 | 0.532 | -0.693 | -1.084 | -0.946 | -0.138 | -0.095 | -0.095 | 0 | 0.24 | 0.47 | 0.09 | 0.09 | 0.04 | 0.07 |
| 2 | 177016012 | rs13418288 | 0.54 | -0.684 | -1.08 | -0.941 | -0.138 | -0.095 | -0.095 | 0.326 | 0.24 | 0.47 | 0.09 | 0.09 | 0.04 | 0.07 |
| 2 | 177018726 | rs4972504 | -0.838 | 0.916 | -0.653 | -0.272 | -0.064 | -0.061 | -0.061 | 1.043 | 0.85 | 0.96 | 0.8 | 0.75 | 0.69 | 0.86 |
| 2 | 177021401 | rs4972505 | 0.968 | 0.783 | -0.693 | -0.341 | -0.011 | -0.015 | -0.015 | 0 | 0.6 | 0.5 | 0.61 | 0.72 | 0.38 | 0.67 |
| 2 | 177028551 | rs7606110 | 0.202 | 0.876 | -0.81 | -0.487 | 0.003 | -0.038 | -0.038 | 0.387 | 0.2 | 0.17 | 0.18 | 0.02 | 0.31 | 0.18 |
| 2 | 177028767 | rs1542180 | 0.696 | 0.912 | -0.849 | -0.461 | 0.01 | 0.024 | 0.024 | 1.145 | 0.62 | 0.77 | 0.62 | 0.75 | 0.37 | 0.67 |
| 2 | 177032267 | rs2301301 | 0.004 | 0.861 | -0.7 | -0.498 | 0.018 | -0.065 | -0.065 | 1.732 | 0.21 | 0.21 | 0.18 | 0.02 | 0.31 | 0.18 |
| 2 | 177041370 | rs2305163 | 1.485 | 0.331 | -0.779 | -0.455 | -0.004 | 0.004 | 0.004 | 0.588 | 0.11 | 0.07 | 0.03 | 0.07 | 0.01 | 0.01 |
| 2 | 177042261 | rs13028842 | 0.294 | 1.513 | -0.459 | -0.309 | -0.023 | 0.018 | 0.018 | 0 | 0.17 | 0.08 | 0.15 | 0.02 | 0.27 | 0.15 |
| 2 | 177043971 | rs1374325 | -0.09 | 1.519 | -0.365 | -0.321 | -0.001 | -0.029 | -0.029 | 0.402 | 0.2 | 0.18 | 0.18 | 0.02 | 0.31 | 0.18 |
| 2 | 177047801 | rs1446575 | 0.915 | 1.493 | -0.251 | -0.27 | 0.008 | 0.023 | 0.023 | 1.036 | 0.62 | 0.75 | 0.62 | 0.75 | 0.37 | 0.68 |
| 2 | 177054411 | rs13390932 | -0.815 | 1.532 | -0.208 | -0.299 | 0.044 | -0.161 | -0.161 | 0 | 0.24 | 0.47 | 0.18 | 0.02 | 0.29 | 0.17 |
| 2 | 177058316 | rs1374326 | -0.571 | 1.683 | -0.461 | -0.194 | 0.064 | -0.14 | -0.14 | 0 | 0.21 | 0.45 | 0.18 | 0.02 | 0.29 | 0.16 |
| 2 | 177064823 | rs1026032 | -0.046 | 2.096 | -1.204 | 0.45 | -0.07 | -0.066 | -0.066 | 0 | 0.13 | 0.03 | 0.15 | 0.24 | 0.27 | 0.12 |
| 2 | 177065030 | rs1026031 | -0.437 | 2.112 | -1.08 | 0.412 | -0.013 | -0.012 | -0.012 | 0 | 0.15 | 0.12 | 0.16 | 0.25 | 0.29 | 0.13 |
| 2 | 177067780 | rs17395735 | -1.086 | 2.081 | -0.902 | 0.325 | 0.025 | 0.015 | 0.015 | 0 | 0.19 | 0.27 | 0.2 | 0.25 | 0.32 | 0.19 |
| 2 | 177068638 | rs17336374 | 0.142 | 1.787 | -1.511 | -0.122 | -0.052 | 0.115 | 0.115 | 0 | 0.26 | 0.06 | 0.16 | 0.06 | 0.31 | 0.18 |
| 2 | 177068722 | rs7593931 | -1.234 | 1.681 | -1.478 | -0.086 | 0.031 | 0.037 | 0.037 | 0 | 0.46 | 0.32 | 0.35 | 0.3 | 0.62 | 0.36 |
| 2 | 177070756 | rs1026028 | -0.371 | 1.454 | -1.262 | -0.3 | -0.016 | 0.068 | 0.068 | 0 | 0.29 | 0.19 | 0.19 | 0.06 | 0.34 | 0.21 |
| 2 | 177071181 | rs2166512 | -0.41 | 1.458 | -1.284 | -0.291 | -0.009 | 0.017 | 0.017 | 0 | 0.31 | 0.29 | 0.2 | 0.06 | 0.36 | 0.21 |
| 2 | 177075245 | rs13414698 | -0.552 | 1.783 | -0.695 | 0.302 | 0.049 | 0.013 | 0.013 | 0 | 0.3 | 0.42 | 0.28 | 0.28 | 0.46 | 0.2 |
| 2 | 177076890 | rs6714226 | -0.561 | 1.16 | -1.139 | 0.135 | 0 | 0.04 | 0.04 | 0 | 0.53 | 0.65 | 0.6 | 0.71 | 0.53 | 0.49 |
| 2 | 177077922 | rs1403994 | 1.057 | 0.832 | -1.152 | -0.278 | -0.004 | -0.093 | -0.093 | 0.674 | 0.19 | 0.29 | 0.13 | 0 | 0.18 | 0.05 |
| 2 | 177080965 | rs2140538 | -0.289 | 0.662 | -1.018 | -0.045 | 0.009 | -0.008 | -0.008 | 0 | 0.38 | 0.5 | 0.48 | 0.65 | 0.42 | 0.27 |
| 2 | 177088525 | rs4893873 | 0.124 | 0.752 | -1.113 | 0.225 | -0.018 | -0.013 | -0.013 | 0 | 0.3 | 0.23 | 0.37 | 0.62 | 0.41 | 0.22 |
| 2 | 177089140 | rs12151479 | 1.255 | 1.246 | -1.161 | 0.696 | -0.048 | -0.023 | -0.023 | 0 | 0.12 | 0.04 | 0.25 | 0.16 | 0.25 | 0.1 |
| 2 | 177096308 | rs3850167 | 0.094 | 0.85 | -1.33 | 0.647 | -0.002 | -0.004 | -0.004 | 0 | 0.53 | 0.49 | 0.48 | 0.33 | 0.44 | 0.52 |
| 2 | 177096752 | rs11885361 | 0.441 | 0.95 | -1.006 | 0.94 | 0.031 | 0.03 | 0.03 | 0 | 0.16 | 0.24 | 0.26 | 0.28 | 0.26 | 0.1 |
| 2 | 177111419 | rs1524082 | -1.272 | 0.026 | -1.385 | 0.333 | 0.029 | -0.024 | -0.024 | 0 | 0.66 | 0.9 | 0.64 | 0.69 | 0.71 | 0.5 |
| 2 | 177111819 | rs970797 | -1.144 | 0.297 | -0.845 | 0.449 | 0.005 | 0.011 | 0.011 | 0 | 0.49 | 0.6 | 0.54 | 0.69 | 0.57 | 0.47 |
| 2 | 177112082 | rs970796 | -0.507 | 0.779 | -0.68 | 0.411 | 0.042 | -0.008 | -0.008 | 0 | 0.32 | 0.49 | 0.43 | 0.35 | 0.4 | 0.35 |
| 2 | 177113353 | rs12471146 | 1.912 | 0.903 | -1.461 | 0.277 | 0.009 | 0.052 | 0.052 | 0 | 0.13 | 0.02 | 0.12 | 0.07 | 0.07 | 0.24 |
| 2 | 177115852 | rs13416955 | 0.036 | 1.222 | -1.027 | 0.688 | 0.052 | -0.005 | -0.005 | 0 | 0.15 | 0.22 | 0.27 | 0.14 | 0.28 | 0.08 |
| 2 | 177124224 | rs6740374 | -1.343 | 0.237 | -1.509 | 0.907 | 0.011 | -0.124 | -0.124 | 0 | 0.8 | 0.97 | 0.85 | 0.45 | 0.82 | 0.54 |
| 2 | 177134172 | rs2305161 | -0.085 | 0.574 | -0.569 | -0.676 | -0.002 | -0.013 | -0.013 | 0 | 0.25 | 0.32 | 0.29 | 0.12 | 0.2 | 0.3 |
| 2 | 177135473 | rs6720043 | -0.048 | 0.56 | -0.354 | -0.689 | -0.009 | -0.045 | -0.045 | 0 | 0.25 | 0.35 | 0.29 | 0.09 | 0.2 | 0.3 |
| 2 | 177137489 | rs9789537 | 0.241 | 0.569 | -0.394 | -0.678 | 0.003 | -0.008 | -0.008 | 0 | 0.22 | 0.24 | 0.24 | 0.07 | 0.13 | 0.28 |
| 2 | 177144029 | rs12151388 | 0.181 | 1.823 | -0.593 | -0.543 | -0.07 | -0.045 | -0.045 | 0 | 0.11 | 0.03 | 0.31 | 0.16 | 0.29 | 0.1 |
| 2 | 177146275 | rs6709357 | -0.083 | 1.33 | -0.247 | -0.754 | -0.007 | -0.037 | -0.037 | 0 | 0.24 | 0.35 | 0.29 | 0.12 | 0.21 | 0.31 |
| 2 | 177152343 | rs1403996 | -2.274 | 1.092 | -0.325 | -0.983 | 0.086 | -0.045 | -0.045 | 0 | 0.58 | 0.86 | 0.75 | 0.46 | 0.7 | 0.57 |
| 2 | 177152660 | rs17336785 | 1.022 | 1.116 | -0.339 | -0.878 | -0.013 | 0.07 | 0.07 | 0.095 | 0.17 | 0.03 | 0.23 | 0.07 | 0.12 | 0.28 |
| 2 | 177163078 | rs7597047 | -0.154 | 1.111 | 0.186 | -0.888 | -0.017 | -0.082 | -0.082 | 0.186 | 0.25 | 0.58 | 0.31 | 0.12 | 0.21 | 0.3 |
| 2 | 177170168 | rs13414530 | 0.667 | 0.99 | 0.215 | -0.878 | 0.003 | -0.001 | -0.001 | 0 | 0.19 | 0.19 | 0.24 | 0.07 | 0.12 | 0.28 |
| 2 | 177198249 | rs4346418 | -0.836 | 1.077 | 0.366 | -0.789 | -0.001 | 0 | 0 | 0 | 0.44 | 0.53 | 0.66 | 0.47 | 0.56 | 0.53 |
| 2 | 177198846 | rs2689951 | -0.808 | 1.081 | 0.343 | -0.782 | 0.002 | -0.004 | -0.004 | 0.003 | 0.44 | 0.47 | 0.65 | 0.47 | 0.56 | 0.53 |
| 2 | 177199125 | rs2689949 | -0.875 | 1.104 | -0.107 | -0.767 | -0.001 | 0 | 0 | 0 | 0.45 | 0.61 | 0.66 | 0.48 | 0.56 | 0.53 |

| | | | | | | | | | | | | | | | | |
|---|---|---|---|---|---|---|---|---|---|---|---|---|---|---|---|---|
| 2 | 177210511 | rs818361 | -1.259 | 0.203 | 0.258 | -0.599 | 0.074 | -0.021 | -0.021 | 0.126 | 0.56 | 0.78 | 0.58 | 0.46 | 0.68 | 0.57 |
| 2 | 177242425 | rs10930727 | 0.367 | -0.11 | 0.74 | -0.664 | -0.009 | -0.032 | -0.032 | 0 | 0.72 | 0.34 | 0.68 | 0.87 | 0.78 | 0.7 |
| 2 | 177250927 | rs10930728 | 0.044 | -0.09 | 0.66 | -0.736 | -0.018 | -0.103 | -0.103 | 0 | 0.74 | 0.52 | 0.69 | 0.9 | 0.78 | 0.7 |
| 2 | 177253799 | rs2102939 | 0.253 | -0.077 | 0.435 | -0.724 | -0.02 | -0.074 | -0.074 | 0 | 0.72 | 0.38 | 0.68 | 0.9 | 0.78 | 0.7 |
| 2 | 177260099 | rs7604433 | -0.49 | -0.241 | 0.308 | -0.476 | 0.017 | 0.048 | 0.048 | 0 | 0.41 | 0.35 | 0.51 | 0.18 | 0.62 | 0.26 |
| 2 | 177272297 | rs13024543 | -0.201 | 0.203 | 0.996 | 0.169 | -0.008 | 0.023 | 0.023 | 0 | 0.8 | 0.72 | 0.9 | 0.72 | 0.84 | 0.64 |
| 2 | 177278437 | rs7606424 | 0.402 | 0.314 | 0.329 | 0.224 | -0.001 | -0.022 | -0.022 | 0.079 | 0.17 | 0.07 | 0.08 | 0.27 | 0.16 | 0.36 |
| 2 | 177281955 | rs16863792 | 0.772 | -0.085 | 0.303 | 0.475 | 0.087 | -0.089 | -0.089 | 0.012 | 0.41 | 0.08 | 0.4 | 0.46 | 0.27 | 0.38 |
| 2 | 177292674 | rs16863814 | -1.567 | 0.183 | 1.014 | 0.786 | 0.028 | 0.438 | 0.438 | 4.575 | 0.07 | 0.5 | 0.06 | 0.31 | 0.01 | 0.08 |
| 2 | 177315153 | rs6722762 | 0.447 | 0.351 | 0.681 | 0.279 | 0.01 | 0.009 | 0.009 | 0 | 0.47 | 0.43 | 0.43 | 0.76 | 0.33 | 0.55 |
| 2 | 177318061 | rs6747963 | -0.335 | 0.483 | -0.051 | 0.42 | -0.004 | 0.027 | 0.027 | 0 | 0.66 | 0.79 | 0.61 | 0.77 | 0.66 | 0.6 |
| 2 | 177320069 | rs970859 | -0.258 | -0.65 | 0.213 | 0.994 | 0.049 | 0.07 | 0.07 | 0 | 0.49 | 0.63 | 0.45 | 0.77 | 0.37 | 0.54 |
| 2 | 177323409 | rs6734063 | 0.01 | -0.286 | -1.141 | 1.166 | 0.072 | 0.129 | 0.129 | 0 | 0.45 | 0.19 | 0.41 | 0.76 | 0.37 | 0.54 |
| 2 | 177329852 | rs12621021 | -1.286 | 0.154 | 0.223 | 1.762 | -0.017 | 0.184 | 0.184 | 0.003 | 0.06 | 0.21 | 0.02 | 0.22 | 0 | 0.06 |
| 2 | 177330638 | rs6718842 | -1.273 | -0.168 | 0.406 | 2.12 | -0.021 | 0.074 | 0.074 | 0.478 | 0.13 | 0.29 | 0.15 | 0.31 | 0.07 | 0.18 |
| 2 | 177330967 | rs1554406 | -1.764 | 0.868 | 1.454 | 0.936 | 0.142 | -0.044 | -0.044 | 5.392 | 0.1 | 0.55 | 0.13 | 0 | 0.16 | 0.02 |
| 2 | 177331138 | rs1554405 | 0.31 | 0.63 | 0.879 | -0.103 | 0.001 | -0.036 | -0.036 | 0.165 | 0.74 | 0.76 | 0.87 | 1 | 0.79 | 0.84 |
| 2 | 177331710 | rs1005932 | 0.476 | 0.931 | -0.123 | -0.002 | -0.074 | -0.054 | -0.054 | 3.225 | 0.24 | 0.1 | 0.54 | 0.64 | 0.34 | 0.32 |
| 2 | 177335291 | rs13419894 | 0.098 | 0.196 | -0.229 | -0.66 | -0.004 | 0.016 | 0.016 | 0.458 | 0.16 | 0.09 | 0.09 | 0.11 | 0.18 | 0.18 |
| 2 | 177338441 | rs13031685 | 0.205 | -0.799 | 0.12 | -1.533 | -0.055 | -0.087 | -0.087 | 0 | 0.68 | 0.44 | 0.83 | 0.9 | 0.82 | 0.72 |
| 2 | 177338662 | rs4894073 | 0.991 | -0.89 | 0.058 | -1.784 | -0.002 | -0.015 | -0.015 | 0 | 0.48 | 0.46 | 0.72 | 0.82 | 0.6 | 0.43 |
| 2 | 177340813 | rs3898818 | -0.191 | -0.979 | -0.363 | -1.982 | -0.005 | -0.031 | -0.031 | 1.339 | 0.1 | 0.17 | 0.06 | 0 | 0.08 | 0.07 |
| 2 | 177348025 | rs10201330 | 1.04 | -1.078 | -1.165 | -2.027 | -0.006 | 0.025 | 0.025 | 0 | 0.11 | 0.03 | 0.06 | 0.02 | 0.09 | 0.08 |
| 2 | 177348389 | rs10803899 | 1.783 | -0.884 | -1.403 | -1.949 | -0.017 | 0.061 | 0.061 | 0 | 0.4 | 0.21 | 0.65 | 0.77 | 0.52 | 0.4 |
| 2 | 177349591 | rs13403148 | 0.553 | -0.543 | -1.262 | -1.967 | -0.047 | 0.051 | 0.051 | 0 | 0.14 | 0.04 | 0.13 | 0 | 0.24 | 0.1 |
| 2 | 177351121 | rs6433594 | 1.068 | -0.419 | -1.094 | -1.771 | 0.017 | 0.152 | 0.152 | 0 | 0.47 | 0.23 | 0.65 | 0.85 | 0.54 | 0.56 |
| 2 | 177357019 | rs9749801 | 0.196 | -0.366 | -0.958 | -1.758 | 0.001 | -0.013 | -0.013 | 0.598 | 0.63 | 0.61 | 0.78 | 0.86 | 0.67 | 0.74 |
| 2 | 177357666 | rs11676114 | -0.134 | -0.153 | -0.865 | -1.75 | 0.001 | -0.061 | -0.061 | 0 | 0.71 | 0.62 | 0.8 | 0.92 | 0.68 | 0.78 |
| 2 | 177362096 | rs976973 | 0.368 | 0.303 | -1.256 | -1.455 | 0.024 | 0.088 | 0.088 | 0 | 0.48 | 0.31 | 0.71 | 0.9 | 0.58 | 0.6 |
| 2 | 177363159 | rs4893865 | 0.317 | 0.454 | -1.044 | -1.194 | 0.007 | 0.02 | 0.02 | 0.643 | 0.51 | 0.44 | 0.73 | 0.8 | 0.56 | 0.55 |
| 2 | 177365599 | rs2122013 | -0.452 | -0.15 | -1.035 | -1.908 | 0 | -0.01 | -0.01 | 0.876 | 0.41 | 0.37 | 0.63 | 0.92 | 0.43 | 0.52 |
| 2 | 177367427 | rs1375180 | 0.128 | -0.405 | -1.14 | -2.23 | -0.002 | -0.011 | -0.011 | 0 | 0.19 | 0.2 | 0.47 | 0.88 | 0.16 | 0.29 |
| 2 | 177367950 | rs6708551 | -0.08 | -1.064 | -1.491 | -2.988 | 0.023 | 0.063 | 0.063 | 0 | 0.58 | 0.38 | 0.4 | 0.09 | 0.66 | 0.52 |
| 2 | 177370519 | rs7568419 | 0.804 | -1.079 | -1.312 | -2.983 | -0.002 | -0.007 | -0.007 | 0 | 0.19 | 0.2 | 0.46 | 0.86 | 0.15 | 0.27 |
| 2 | 177370607 | rs7609112 | -1.634 | -1.401 | -0.715 | -3.089 | -0.028 | -0.081 | -0.081 | 0 | 0.8 | 0.7 | 0.89 | 0.97 | 0.85 | 0.82 |
| 2 | 177370744 | rs11679930 | 0.683 | -1.352 | -0.954 | -3.079 | -0.003 | 0.017 | 0.017 | 0 | 0.22 | 0.16 | 0.15 | 0.06 | 0.26 | 0.19 |
| 2 | 177376868 | rs10497446 | 0.896 | -1.678 | -1.082 | -3.166 | -0.042 | 0.184 | 0.184 | 0 | 0.4 | 0.17 | 0.29 | 0.08 | 0.52 | 0.34 |
| 2 | 177379051 | rs10204624 | 0.811 | -1.586 | -0.967 | -3.167 | 0.006 | 0.001 | 0.001 | 0 | 0.14 | 0.1 | 0.1 | 0 | 0.15 | 0.1 |
| 2 | 177384172 | rs2969348 | 0.516 | -1.812 | -0.789 | -3.298 | 0.019 | 0.035 | 0.035 | 0.274 | 0.6 | 0.34 | 0.4 | 0.11 | 0.66 | 0.58 |
| 2 | 177387653 | rs13387510 | 0.533 | -1.669 | -0.901 | -3.188 | 0.003 | 0.038 | 0.038 | 0 | 0.17 | 0.09 | 0.12 | 0.04 | 0.18 | 0.13 |
| 2 | 177389173 | rs1375175 | 0.862 | -1.527 | -1.09 | -3.317 | -0.059 | 0.083 | 0.083 | 0 | 0.33 | 0.18 | 0.26 | 0.05 | 0.47 | 0.3 |
| 2 | 177389867 | rs4243394 | 0.666 | -1.342 | -0.819 | -3.079 | -0.002 | -0.004 | -0.004 | 0 | 0.19 | 0.19 | 0.47 | 0.86 | 0.15 | 0.29 |
| 2 | 177397832 | rs4893878 | -0.628 | -1.167 | -0.497 | -2.978 | -0.027 | 0.019 | 0.019 | 0 | 0.69 | 0.71 | 0.77 | 0.94 | 0.6 | 0.74 |
| 2 | 177399532 | rs877738 | -0.72 | -1.337 | -0.094 | -2.659 | 0.01 | 0.033 | 0.033 | 0 | 0.63 | 0.75 | 0.46 | 0.17 | 0.65 | 0.59 |
| 2 | 177401312 | rs4894116 | 0.621 | -1.217 | -0.174 | -2.539 | -0.029 | 0.066 | 0.066 | 0 | 0.36 | 0.09 | 0.56 | 0.81 | 0.4 | 0.43 |
| 2 | 177401438 | rs1037222 | -0.291 | -1.199 | -0.088 | -2.228 | -0.008 | 0.028 | 0.028 | 0 | 0.94 | 0.87 | 0.91 | 0.91 | 0.97 | 0.9 |
| 2 | 177405454 | rs6744916 | 0.475 | -1.292 | -0.173 | -2.195 | -0.068 | 0.055 | 0.055 | 0.26 | 0.36 | 0.23 | 0.31 | 0.17 | 0.46 | 0.49 |
| 2 | 177409147 | rs6433596 | 0.64 | -1.266 | -0.475 | -2.143 | -0.013 | 0.025 | 0.025 | 0 | 0.36 | 0.13 | 0.56 | 0.77 | 0.39 | 0.44 |
| 2 | 177409908 | rs17341803 | 0.657 | -1.217 | 0.404 | -2.002 | 0.015 | 0.091 | 0.091 | 5.979 | 0.18 | 0.01 | 0.06 | 0.04 | 0.11 | 0.04 |
| 2 | 177412667 | rs6433597 | 0.177 | -1.087 | 0.534 | -1.967 | -0.063 | -0.114 | -0.114 | 0 | 0.73 | 0.5 | 0.89 | 0.94 | 0.86 | 0.94 |
| 2 | 177425416 | rs895837 | 1.008 | -0.254 | 0.128 | -1.447 | 0.01 | -0.044 | -0.044 | 0 | 0.9 | 0.79 | 0.85 | 0.98 | 0.91 | 0.76 |
| 2 | 177426526 | rs6744879 | 0.969 | 0.115 | 0.131 | -1.673 | 0.031 | -0.044 | -0.044 | 0 | 0.65 | 0.44 | 0.55 | 0.81 | 0.58 | 0.52 |
| 2 | 177447797 | rs12611670 | 0.65 | -0.369 | -0.816 | -2.559 | -0.032 | 0.268 | 0.268 | 1.735 | 0.49 | 0.1 | 0.69 | 0.86 | 0.55 | 0.7 |
| 2 | 177456444 | rs4893900 | 1.497 | -0.454 | -0.902 | -2.588 | -0.105 | 0.126 | 0.126 | 0 | 0.37 | 0.06 | 0.53 | 0.85 | 0.43 | 0.38 |
| 2 | 177460699 | rs2044642 | 0.823 | -0.398 | -0.465 | -2.658 | -0.013 | -0.066 | -0.066 | 0 | 0.25 | 0.23 | 0.09 | 0 | 0.14 | 0.12 |
| 2 | 177473322 | rs1448520 | 0.112 | 0.117 | -0.36 | -2.418 | 0.031 | -0.083 | -0.083 | 0 | 0.17 | 0.26 | 0.18 | 0.01 | 0.24 | 0.16 |
| 2 | 177480054 | rs2922309 | -0.854 | -0.291 | -0.86 | -2.358 | 0.004 | -0.049 | -0.049 | 0 | 0.13 | 0.2 | 0.16 | 0.02 | 0.13 | 0.3 |
| 2 | 177480820 | rs1839152 | 0.57 | -0.175 | -1.326 | -2.244 | -0.081 | 0.196 | 0.196 | 0 | 0.42 | 0.13 | 0.59 | 0.88 | 0.53 | 0.47 |
| 2 | 177483123 | rs2969363 | 0.054 | -0.929 | -0.71 | -1.869 | 0.069 | 0.143 | 0.143 | 0 | 0.49 | 0.26 | 0.78 | 0.81 | 0.62 | 0.46 |
| 2 | 177495608 | rs963649 | 0.043 | -0.868 | 0.287 | -1.827 | 0 | -0.018 | -0.018 | 0 | 0.91 | 0.87 | 0.96 | 1 | 0.91 | 0.93 |
| 2 | 177504593 | rs2029039 | 0.921 | -0.949 | 0.508 | -1.87 | 0.006 | -0.08 | -0.08 | 0 | 0.87 | 0.61 | 0.92 | 0.99 | 0.9 | 0.79 |
| 2 | 177519429 | rs6433602 | 0.914 | -1.201 | -0.045 | -2.136 | -0.088 | 0.041 | 0.041 | 0 | 0.33 | 0.1 | 0.69 | 0.81 | 0.55 | 0.31 |
| 2 | 177520106 | rs6740315 | -0.107 | -1.402 | 0.609 | -2.356 | 0.012 | 0.012 | 0.012 | 0 | 0.47 | 0.56 | 0.79 | 0.85 | 0.64 | 0.52 |
| 2 | 177522377 | rs4893911 | 1.032 | -1.396 | 0.522 | -2.354 | -0.049 | 0.069 | 0.069 | 0 | 0.33 | 0.14 | 0.72 | 0.85 | 0.55 | 0.31 |
| 2 | 177530860 | rs11694028 | 0.176 | -1.414 | 0.157 | -2.36 | -0.014 | 0.036 | 0.036 | 0 | 0.08 | 0.01 | 0.03 | 0 | 0.09 | 0.08 |
| 2 | 177531742 | rs17345806 | -0.128 | -1.54 | 0.064 | -2.444 | 0.012 | 0.015 | 0.015 | 0 | 0.08 | 0.04 | 0.04 | 0.01 | 0.05 | 0.24 |
| 2 | 177532212 | rs2710024 | 0.708 | -1.71 | -0.116 | -2.528 | 0.04 | 0.053 | 0.053 | 0 | 0.37 | 0.24 | 0.14 | 0.15 | 0.27 | 0.34 |
| 2 | 177532328 | rs1562611 | 1.084 | -1.436 | -0.156 | -2.369 | -0.075 | 0.076 | 0.076 | 0 | 0.33 | 0.1 | 0.71 | 0.85 | 0.55 | 0.31 |
| 2 | 177536629 | rs13019019 | 1.044 | -1.41 | -0.134 | -2.356 | -0.045 | 0.066 | 0.066 | 0 | 0.33 | 0.14 | 0.72 | 0.85 | 0.55 | 0.31 |

| | | | | | | | | | | | | | | | | |
|---|---|---|---|---|---|---|---|---|---|---|---|---|---|---|---|---|
| 2 | 177541263 | rs11679256 | -0.073 | -1.189 | -0.249 | -2.217 | 0.002 | -0.041 | -0.041 | 0 | 0.09 | 0.13 | 0.03 | 0 | 0.09 | 0.08 |
| 2 | 177542723 | rs4606966 | -0.215 | -1.185 | -0.758 | -2.133 | 0.043 | 0.061 | 0.061 | 0 | 0.54 | 0.64 | 0.82 | 0.85 | 0.69 | 0.63 |
| 2 | 177548513 | rs938278 | -1.178 | -1.093 | -0.72 | -2.197 | 0.021 | -0.012 | -0.012 | 0 | 0.32 | 0.73 | 0.16 | 0.01 | 0.19 | 0.37 |
| 2 | 177549497 | rs868179 | -0.869 | -1.213 | -0.563 | -2.156 | -0.023 | -0.067 | -0.067 | 0 | 0.25 | 0.74 | 0.12 | 0.01 | 0.1 | 0.33 |
| 2 | 177551886 | rs1554011 | 0.551 | -0.768 | -1.668 | -2.482 | -0.06 | 0.184 | 0.184 | 0.107 | 0.41 | 0.09 | 0.66 | 0.88 | 0.57 | 0.5 |
| 2 | 177556469 | rs17786781 | 0.61 | -0.725 | -1.773 | -2.731 | -0.044 | 0.047 | 0.047 | 0 | 0.2 | 0.1 | 0.15 | 0 | 0.3 | 0.12 |
| 2 | 177559549 | rs16864246 | 0.218 | -1.211 | -1.239 | -3.039 | 0.005 | -0.021 | -0.021 | 0.161 | 0.18 | 0.16 | 0.08 | 0 | 0.11 | 0.11 |
| 2 | 177561988 | rs1355849 | 1.743 | -0.682 | -1.714 | -3.038 | -0.054 | 0.035 | 0.035 | 1.47 | 0.33 | 0.06 | 0.44 | 0.82 | 0.32 | 0.32 |
| 2 | 177567562 | rs4407280 | 0.262 | -0.65 | -1.041 | -3.061 | 0.004 | 0.009 | 0.009 | 0 | 0.74 | 0.82 | 0.72 | 0.84 | 0.75 | 0.62 |
| 2 | 177567856 | rs6749847 | -0.203 | -0.679 | -0.843 | -3.016 | 0.013 | -0.017 | -0.017 | 0 | 0.68 | 0.63 | 0.62 | 0.84 | 0.56 | 0.58 |
| 2 | 177573878 | rs12623637 | 0.216 | -0.474 | -0.555 | -2.807 | 0.001 | 0.031 | 0.031 | 0 | 0.53 | 0.61 | 0.56 | 0.83 | 0.45 | 0.5 |
| 2 | 177576731 | rs16864287 | -0.743 | -0.109 | -0.073 | -2.738 | -0.001 | -0.029 | -0.029 | 0.19 | 0.33 | 0.72 | 0.21 | 0.02 | 0.34 | 0.24 |
| 2 | 177589065 | rs16864347 | -0.571 | -0.058 | -0.802 | -2.475 | -0.011 | -0.055 | -0.055 | 0 | 0.14 | 0.34 | 0.09 | 0.02 | 0.1 | 0.13 |
| 2 | 177595862 | rs713285 | -0.045 | 0.743 | -1.743 | -2.219 | -0.052 | 0.054 | 0.054 | 0 | 0.25 | 0.16 | 0.17 | 0 | 0.37 | 0.14 |
| 2 | 177611643 | rs2362870 | 0.438 | -0.173 | -1.747 | -1.992 | 0.041 | 0.074 | 0.074 | 1.737 | 0.72 | 0.16 | 0.82 | 0.92 | 0.8 | 0.75 |
| 2 | 177615308 | rs10164810 | -0.147 | -0.384 | -1.689 | -2.422 | 0.011 | 0.018 | 0.018 | 0 | 0.73 | 0.79 | 0.61 | 0.13 | 0.79 | 0.6 |
| 2 | 177616426 | rs13021051 | 0.588 | -1.183 | -1.219 | -3.079 | -0.011 | -0.049 | -0.049 | 0 | 0.55 | 0.55 | 0.53 | 0.13 | 0.71 | 0.49 |
| 2 | 177619231 | rs2885631 | 0.067 | -1.077 | -1.01 | -3.045 | -0.013 | -0.048 | -0.048 | 0.642 | 0.14 | 0.13 | 0.07 | 0.01 | 0.08 | 0.08 |
| 2 | 177622704 | rs4894196 | 0.113 | -0.064 | -1.184 | -3.522 | -0.055 | 0.049 | 0.049 | 0 | 0.28 | 0.21 | 0.39 | 0.08 | 0.42 | 0.27 |
| 2 | 177623339 | rs6714936 | -0.082 | -0.6 | -0.487 | -3.882 | -0.003 | -0.065 | -0.065 | 1.831 | 0.53 | 0.46 | 0.58 | 0.07 | 0.68 | 0.4 |
| 2 | 177623460 | rs17639578 | 1.31 | 0.233 | -0.876 | -4.065 | -0.043 | 0.064 | 0.064 | 0 | 0.12 | 0.01 | 0.06 | 0 | 0.14 | 0.06 |
| 2 | 177625920 | rs938275 | 1.877 | 0.6 | -0.456 | -4.157 | -0.016 | 0.026 | 0.026 | 0 | 0.22 | 0.11 | 0.16 | 0 | 0.27 | 0.12 |
| 2 | 177627904 | rs6705679 | 0.373 | 0.544 | -0.225 | -4.144 | -0.003 | -0.017 | -0.017 | 0 | 0.4 | 0.36 | 0.5 | 0.07 | 0.55 | 0.32 |
| 2 | 177628258 | rs6706063 | 0.459 | 0.091 | -0.32 | -4.006 | 0.077 | 0.116 | 0.116 | 0 | 0.39 | 0.23 | 0.35 | 0.87 | 0.25 | 0.53 |
| 2 | 177628917 | rs4894197 | 0.808 | 0.033 | -0.565 | -4.11 | 0.035 | 0.178 | 0.178 | 0 | 0.34 | 0.03 | 0.36 | 0.89 | 0.25 | 0.53 |
| 2 | 177629485 | rs17612243 | -0.195 | 0.203 | -0.286 | -4.23 | -0.009 | -0.01 | -0.01 | 0.117 | 0.22 | 0.21 | 0.33 | 0.04 | 0.28 | 0.26 |
| 2 | 177630608 | rs6758766 | 0.747 | 0.099 | -0.326 | -4.201 | 0.075 | 0.156 | 0.156 | 0 | 0.36 | 0.14 | 0.35 | 0.89 | 0.25 | 0.53 |
| 2 | 177633192 | rs6733830 | -0.893 | 0.12 | -0.093 | -4.249 | -0.013 | -0.008 | -0.008 | 0.389 | 0.62 | 0.37 | 0.74 | 1 | 0.57 | 0.84 |
| 2 | 177633313 | rs11889984 | 0.636 | -0.031 | -0.218 | -4.164 | 0.077 | 0.153 | 0.153 | 0 | 0.37 | 0.16 | 0.36 | 0.91 | 0.25 | 0.53 |
| 2 | 177640092 | rs12988851 | 1.76 | 0.017 | 0.759 | -4.353 | -0.009 | -0.004 | -0.004 | 1.729 | 0.23 | 0.19 | 0.16 | 0 | 0.28 | 0.12 |
| 2 | 177643866 | rs12470292 | -0.245 | -0.574 | 0.787 | -4.522 | 0.023 | 0.113 | 0.113 | 0 | 0.45 | 0.29 | 0.6 | 0.93 | 0.39 | 0.68 |
| 2 | 177643902 | rs10184742 | -1.689 | -0.676 | 1.373 | -4.461 | -0.042 | -0.132 | -0.132 | 0.388 | 0.18 | 0.46 | 0.19 | 0.03 | 0.12 | 0.16 |
| 2 | 177645122 | rs6740977 | -0.587 | -0.454 | 1.069 | -4.35 | 0.022 | -0.026 | -0.026 | 0 | 0.08 | 0.19 | 0.15 | 0.03 | 0.12 | 0.08 |
| 2 | 177646988 | rs1554824 | 0.842 | -0.625 | 0.68 | -4.586 | -0.024 | 0.111 | 0.111 | 1.384 | 0.31 | 0.1 | 0.48 | 0.89 | 0.32 | 0.56 |
| 2 | 177648762 | rs10177590 | 0.876 | -0.502 | 0.785 | -4.683 | -0.019 | 0.096 | 0.096 | 0.356 | 0.44 | 0.24 | 0.26 | 0 | 0.45 | 0.23 |
| 2 | 177649722 | rs7581696 | 0.851 | -0.498 | 0.793 | -4.682 | -0.016 | 0.084 | 0.084 | 0.296 | 0.44 | 0.25 | 0.26 | 0 | 0.45 | 0.23 |
| 2 | 177653757 | rs6433615 | -1.078 | -0.377 | 0.477 | -4.526 | 0.008 | -0.03 | -0.03 | 0 | 0.12 | 0.21 | 0.17 | 0.04 | 0.16 | 0.16 |
| 2 | 177669182 | rs840681 | 0.511 | -0.474 | 0.714 | -4.546 | -0.01 | 0.04 | 0.04 | 0.115 | 0.46 | 0.3 | 0.27 | 0 | 0.47 | 0.24 |
| 2 | 177677940 | rs1534472 | 0.635 | -0.413 | 1.178 | -4.342 | 0.048 | 0.001 | 0.001 | 2.703 | 0.93 | 0.67 | 0.94 | 0.97 | 0.93 | 0.96 |
| 2 | 177678844 | rs7577987 | -0.778 | -0.453 | 1.126 | -4.355 | -0.006 | -0.037 | -0.037 | 0 | 0.07 | 0.13 | 0.05 | 0.01 | 0.06 | 0.12 |
| 2 | 177680323 | rs13022517 | 1.811 | -0.279 | 0.609 | -4.608 | -0.013 | 0.012 | 0.012 | 0 | 0.29 | 0.22 | 0.21 | 0 | 0.35 | 0.11 |
| 2 | 177681162 | rs7592623 | 1.399 | -0.523 | 0.764 | -4.626 | -0.012 | -0.069 | -0.069 | 0.041 | 0.68 | 0.45 | 0.78 | 0.93 | 0.78 | 0.73 |
| 2 | 177683404 | rs13032587 | 1.936 | -0.553 | 0.455 | -4.741 | 0.053 | 0.165 | 0.165 | 0 | 0.55 | 0.25 | 0.77 | 0.92 | 0.68 | 0.65 |
| 2 | 177687497 | rs7588841 | 0.971 | -0.198 | 0.689 | -4.548 | -0.027 | -0.106 | -0.106 | 0 | 0.78 | 0.51 | 0.88 | 0.93 | 0.85 | 0.86 |
| 2 | 177691497 | rs10497460 | 0.106 | -0.216 | 0.04 | -4.496 | 0.008 | 0.023 | 0.023 | 0.851 | 0.37 | 0.29 | 0.51 | 0.95 | 0.34 | 0.69 |
| 2 | 177698447 | rs7583729 | -1.549 | 0.105 | 0.364 | -4.554 | 0.007 | -0.111 | -0.111 | 0.001 | 0.67 | 0.57 | 0.76 | 0.97 | 0.62 | 0.9 |
| 2 | 177704321 | rs17671252 | -0.73 | -0.319 | 0.225 | -4.45 | 0.007 | -0.019 | -0.019 | 1.429 | 0.27 | 0.21 | 0.15 | 0.01 | 0.24 | 0.19 |
| 2 | 177704758 | rs1513885 | -0.546 | -0.162 | 0.606 | -4.211 | -0.009 | 0.146 | 0.146 | 0.654 | 0.54 | 0.24 | 0.62 | 0.94 | 0.56 | 0.79 |
| 2 | 177705651 | rs1527269 | -1.794 | -0.076 | 0.411 | -4.115 | -0.006 | -0.017 | -0.017 | 0 | 0.7 | 0.68 | 0.78 | 0.97 | 0.64 | 0.91 |
| 2 | 177709967 | rs4894203 | 0.095 | -0.007 | 0.274 | -4.09 | 0.012 | -0.019 | -0.019 | 0 | 0.07 | 0.13 | 0.12 | 0.03 | 0.06 | 0.05 |
| 2 | 177710924 | rs1355845 | 0.286 | -0.098 | 0.106 | -4.242 | -0.036 | 0.224 | 0.224 | 1.519 | 0.39 | 0.09 | 0.52 | 0.94 | 0.4 | 0.7 |
| 2 | 177711791 | rs11679607 | -0.314 | -0.056 | 0.245 | -4.194 | -0.009 | -0.085 | -0.085 | 0.596 | 0.19 | 0.29 | 0.11 | 0.01 | 0.17 | 0.13 |
| 2 | 177713373 | rs6723196 | -0.305 | -0.043 | 0.314 | -4.164 | 0.002 | 0.019 | 0.019 | 1.555 | 0.48 | 0.36 | 0.65 | 0.98 | 0.46 | 0.78 |
| 2 | 177716131 | rs1513870 | -1.323 | 0.109 | 0.133 | -3.839 | 0.001 | -0.076 | -0.076 | 0.227 | 0.68 | 0.67 | 0.77 | 0.98 | 0.64 | 0.91 |
| 2 | 177720450 | rs1398967 | -0.528 | 0.253 | 0.041 | -3.972 | 0.002 | -0.111 | -0.111 | 0 | 0.57 | 0.6 | 0.64 | 0.99 | 0.52 | 0.84 |
| 2 | 177720581 | rs1398968 | 0.808 | -0.093 | -0.196 | -4.098 | -0.042 | 0.193 | 0.193 | 1.468 | 0.35 | 0.09 | 0.51 | 0.95 | 0.36 | 0.67 |
| 2 | 177720675 | rs1398969 | -1.549 | -0.059 | -0.041 | -3.89 | -0.013 | 0.027 | 0.027 | 0.452 | 0.68 | 0.77 | 0.78 | 0.98 | 0.63 | 0.86 |
| 2 | 177725934 | rs10930757 | -0.984 | -0.202 | 0.462 | -3.766 | 0.002 | -0.103 | -0.103 | 0.28 | 0.61 | 0.54 | 0.73 | 0.98 | 0.56 | 0.83 |
| 2 | 177742124 | rs9287993 | -1.349 | -0.277 | 0.54 | -3.766 | -0.011 | 0 | 0 | 1.641 | 0.64 | 0.72 | 0.76 | 0.98 | 0.56 | 0.83 |
| 2 | 177742445 | rs10497474 | -0.773 | -0.583 | 0.591 | -3.702 | 0.001 | -0.09 | -0.09 | 0 | 0.2 | 0.25 | 0.2 | 0.01 | 0.16 | 0.12 |
| 2 | 177744204 | rs6706678 | -0.796 | -0.58 | 0.593 | -3.701 | 0 | -0.09 | -0.09 | 0 | 0.2 | 0.25 | 0.2 | 0.01 | 0.16 | 0.12 |
| 2 | 177767048 | rs6743416 | -1.255 | -0.609 | 0.674 | -3.709 | -0.01 | -0.142 | -0.142 | 0.579 | 0.23 | 0.42 | 0.2 | 0.04 | 0.17 | 0.15 |
| 2 | 177768841 | rs6433620 | -0.182 | -0.614 | 0.693 | -3.722 | -0.004 | -0.041 | -0.041 | 0 | 0.14 | 0.13 | 0.07 | 0 | 0.11 | 0.1 |
| 2 | 177769236 | rs7586401 | -1.398 | -0.494 | 0.612 | -3.804 | -0.022 | 0.002 | 0.002 | 2.316 | 0.66 | 0.71 | 0.75 | 0.98 | 0.53 | 0.83 |
| 2 | 177770340 | rs6712128 | -1.754 | -0.505 | 0.604 | -3.817 | -0.001 | -0.043 | -0.043 | 0 | 0.28 | 0.65 | 0.24 | 0.03 | 0.19 | 0.16 |
| 2 | 177770364 | rs17672797 | -0.13 | -0.488 | 0.56 | -3.795 | 0.01 | 0.013 | 0.013 | 2.839 | 0.47 | 0.37 | 0.64 | 0.98 | 0.41 | 0.72 |
| 2 | 177770595 | rs17627058 | 0.609 | -0.464 | 0.377 | -3.769 | -0.027 | 0.265 | 0.265 | 2.339 | 0.38 | 0.06 | 0.51 | 0.95 | 0.35 | 0.67 |
| 2 | 177771124 | rs1358296 | -1.296 | -0.392 | 0.472 | -3.705 | 0.003 | -0.079 | -0.079 | 0.873 | 0.64 | 0.63 | 0.74 | 0.99 | 0.53 | 0.83 |
| 2 | 177787845 | rs7564166 | 0.555 | -0.463 | 0.345 | -3.719 | 0.015 | 0.162 | 0.162 | 2.208 | 0.38 | 0.15 | 0.51 | 0.94 | 0.35 | 0.67 |

| | | | | | | | | | | | | | | | | |
|---|---|---|---|---|---|---|---|---|---|---|---|---|---|---|---|---|
| 2 | 177796683 | rs16864736 | -0.992 | -0.424 | 0.509 | -3.724 | 0.008 | -0.14 | -0.14 | 0 | 0.6 | 0.44 | 0.62 | 0.99 | 0.48 | 0.81 |
| 2 | 177796823 | rs17679370 | -0.693 | -0.403 | 0.264 | -3.714 | -0.009 | 0.054 | 0.054 | 2.243 | 0.57 | 0.29 | 0.6 | 0.99 | 0.48 | 0.81 |
| 2 | 177799752 | rs7603168 | -1.295 | -0.442 | 0.036 | -3.647 | -0.009 | -0.149 | -0.149 | 0.164 | 0.23 | 0.52 | 0.21 | 0.01 | 0.17 | 0.27 |
| 2 | 177804965 | rs1370539 | 1.205 | -0.551 | -0.874 | -3.466 | 0.006 | 0.068 | 0.068 | 0 | 0.2 | 0.02 | 0.14 | 0.09 | 0.17 | 0.21 |
| 2 | 177811141 | rs10930771 | -0.442 | -0.127 | -0.234 | -3.16 | 0 | -0.067 | -0.067 | 0.689 | 0.57 | 0.44 | 0.7 | 0.96 | 0.51 | 0.67 |
| 2 | 177813333 | rs11691671 | -0.034 | 0.161 | -0.068 | -2.706 | 0.03 | -0.031 | -0.031 | 0 | 0.06 | 0.15 | 0.12 | 0 | 0.06 | 0.02 |
| 2 | 177819108 | rs11684592 | -1.304 | 0.136 | -0.196 | -2.789 | 0.016 | -0.051 | -0.051 | 0.328 | 0.13 | 0.22 | 0.15 | 0.04 | 0.12 | 0.16 |
| 2 | 177820169 | rs12476172 | -0.802 | 0.054 | -0.501 | -3.014 | -0.018 | 0.017 | 0.017 | 1.85 | 0.58 | 0.66 | 0.71 | 0.99 | 0.5 | 0.83 |
| 2 | 177822624 | rs12613021 | 0.766 | 0.027 | -1.11 | -3.018 | -0.04 | 0.266 | 0.266 | 2.192 | 0.37 | 0.03 | 0.51 | 0.95 | 0.35 | 0.55 |
| 2 | 177822887 | rs16864766 | 0.174 | 0.027 | -1.101 | -3.016 | 0.022 | -0.028 | -0.028 | 0 | 0.05 | 0.11 | 0.04 | 0 | 0.06 | 0.07 |
| 2 | 177825415 | rs745767 | 0.713 | 0.07 | -1.726 | -2.952 | -0.04 | 0.265 | 0.265 | 2.08 | 0.37 | 0.03 | 0.51 | 0.95 | 0.35 | 0.55 |
| 2 | 177834328 | rs11895189 | -0.45 | 0.341 | -1.233 | -2.629 | -0.028 | 0.063 | 0.063 | 2.168 | 0.58 | 0.71 | 0.71 | 0.99 | 0.5 | 0.83 |
| 2 | 177839636 | rs1113669 | 0.338 | 0.255 | -1.748 | -2.577 | 0.005 | 0.259 | 0.259 | 0.636 | 0.61 | 0.09 | 0.74 | 0.99 | 0.7 | 0.74 |
| 2 | 177852291 | rs10497468 | -0.616 | 0.097 | -0.448 | -2.374 | 0 | -0.024 | -0.024 | 0 | 0.05 | 0.17 | 0.11 | 0 | 0.03 | 0.02 |
| 2 | 177874316 | rs1881697 | 0.458 | 0.275 | -0.104 | -2.655 | -0.004 | 0.149 | 0.149 | 1.799 | 0.54 | 0.24 | 0.7 | 0.92 | 0.56 | 0.66 |
| 2 | 177884232 | rs9287995 | -0.683 | 0.253 | 0.641 | -2.63 | 0.004 | -0.075 | -0.075 | 0 | 0.42 | 0.4 | 0.36 | 0.07 | 0.44 | 0.29 |
| 2 | 177888234 | rs1997350 | 0.227 | 0.325 | 0.65 | -2.577 | -0.01 | 0.032 | 0.032 | 2.145 | 0.61 | 0.73 | 0.75 | 0.93 | 0.58 | 0.73 |
| 2 | 177888371 | rs7589434 | 0.285 | 0.317 | 0.716 | -2.402 | -0.002 | -0.013 | -0.013 | 0.06 | 0.59 | 0.6 | 0.73 | 0.9 | 0.58 | 0.73 |
| 2 | 177890113 | rs1104779 | 0.272 | 0.034 | -0.103 | -1.948 | 0.014 | 0.008 | 0.008 | 0 | 0.15 | 0.03 | 0.09 | 0.11 | 0.12 | 0.11 |
| 2 | 177891144 | rs6433636 | 0.214 | 0.176 | 0.092 | -1.833 | 0.003 | 0.005 | 0.005 | 0 | 0.14 | 0.12 | 0.08 | 0.09 | 0.1 | 0.11 |
| 2 | 177908411 | rs10497467 | 0.105 | 0.275 | -0.215 | -0.976 | -0.027 | -0.04 | -0.04 | 0.075 | 0.2 | 0.09 | 0.28 | 0.31 | 0.26 | 0.15 |
| 2 | 177909595 | rs1529093 | 1.915 | -0.297 | -0.155 | -0.751 | -0.02 | -0.028 | -0.028 | 0 | 0.45 | 0.2 | 0.55 | 0.51 | 0.54 | 0.55 |
| 2 | 177912203 | rs10198127 | -0.264 | -0.356 | 0.181 | -0.764 | -0.014 | -0.043 | -0.043 | 0 | 0.89 | 0.77 | 0.96 | 0.97 | 0.93 | 0.85 |
| 2 | 177914806 | rs1529098 | -0.639 | -0.346 | 0.424 | -0.751 | -0.002 | -0.004 | -0.004 | 0 | 0.35 | 0.32 | 0.37 | 0.4 | 0.36 | 0.26 |
| 2 | 177923700 | rs11898028 | 0.84 | -0.369 | 0.431 | -0.746 | -0.058 | -0.047 | -0.047 | 0.116 | 0.81 | 0.55 | 0.92 | 0.9 | 0.9 | 0.8 |
| 2 | 177924358 | rs17842284 | 0.696 | -0.273 | -0.134 | -0.781 | -0.013 | -0.024 | -0.024 | 2.903 | 0.17 | 0.1 | 0.27 | 0.31 | 0.23 | 0.14 |
| 2 | 177925893 | rs17683062 | 0.672 | -0.256 | -0.121 | -0.787 | 0.002 | 0.018 | 0.018 | 0.229 | 0.17 | 0.1 | 0.09 | 0.09 | 0.13 | 0.12 |
| 2 | 177928171 | rs4893924 | -0.431 | -0.311 | 0.093 | -0.682 | -0.001 | -0.005 | -0.005 | 0 | 0.9 | 0.91 | 0.97 | 0.97 | 0.93 | 0.85 |
| 2 | 177929006 | rs6705298 | -0.32 | -0.193 | -0.335 | -0.624 | -0.009 | -0.016 | -0.016 | 0 | 0.35 | 0.24 | 0.37 | 0.39 | 0.36 | 0.25 |
| 2 | 177929684 | rs17683237 | 1.088 | 1 | -1.582 | -0.534 | -0.042 | 0.043 | 0.043 | 0 | 0.08 | 0.02 | 0.12 | 0 | 0.24 | 0.07 |
| 2 | 177932011 | rs1534681 | 1.03 | -0.014 | -1.308 | -0.242 | 0.02 | -0.122 | -0.122 | 1.632 | 0.36 | 0.07 | 0.42 | 0.51 | 0.3 | 0.48 |
| 2 | 177936325 | rs10497466 | 0.816 | 0.354 | -1.271 | -0.24 | 0.034 | 0.08 | 0.08 | 0 | 0.47 | 0.2 | 0.53 | 0.74 | 0.39 | 0.65 |
| 2 | 177937881 | rs4894239 | 0.511 | 0.115 | -0.485 | -0.12 | 0.002 | -0.032 | -0.032 | 0.116 | 0.93 | 0.9 | 0.95 | 1 | 0.93 | 0.96 |
| 2 | 177943893 | rs4894241 | 1.059 | 0.367 | -0.095 | -0.196 | 0.001 | 0.006 | 0.006 | 0.015 | 0.6 | 0.67 | 0.72 | 0.73 | 0.58 | 0.74 |
| 2 | 177949579 | rs1554874 | 0.821 | 0.458 | -0.092 | -0.308 | -0.005 | -0.001 | -0.001 | 0 | 0.79 | 0.77 | 0.85 | 0.8 | 0.78 | 0.84 |
| 2 | 177952507 | rs10930778 | 1.45 | 0.317 | 0.818 | -0.375 | -0.025 | -0.012 | -0.012 | 0 | 0.75 | 0.34 | 0.82 | 0.8 | 0.79 | 0.84 |
| 2 | 177962141 | rs4894243 | 1.71 | 0.37 | 0.732 | -0.443 | -0.001 | 0.006 | 0.006 | 0 | 0.63 | 0.33 | 0.71 | 0.8 | 0.58 | 0.79 |
| 2 | 177972585 | rs2588874 | -0.082 | 0.311 | 0.681 | -0.367 | -0.008 | 0.004 | 0.004 | 0 | 0.85 | 0.84 | 0.9 | 0.8 | 0.86 | 0.87 |
| 2 | 177987024 | rs11688232 | -0.156 | 0.503 | 0.709 | -0.424 | 0.008 | -0.041 | -0.041 | 0 | 0.14 | 0.18 | 0.12 | 0 | 0.2 | 0.05 |
| 2 | 177995692 | rs2706134 | -0.94 | 0.418 | 0.58 | -0.471 | -0.013 | 0.003 | 0.003 | 0 | 0.19 | 0.15 | 0.16 | 0 | 0.26 | 0.09 |
| 2 | 178000177 | rs2588866 | 0.402 | 0.046 | 1.029 | -0.174 | -0.004 | -0.029 | -0.029 | 0 | 0.07 | 0.1 | 0.05 | 0 | 0.07 | 0.04 |
| 2 | 178000949 | rs12990744 | -0.012 | 0.166 | 0.559 | -0.216 | -0.041 | 0.013 | 0.013 | 0 | 0.09 | 0.01 | 0.08 | 0.04 | 0.12 | 0.05 |
| 2 | 178000985 | rs2588868 | 0.278 | -0.193 | 0.976 | 0.092 | -0.006 | 0.009 | 0.009 | 0 | 0.84 | 0.82 | 0.78 | 0.73 | 0.85 | 0.86 |
| 2 | 178011652 | rs2588876 | 0.652 | -0.264 | 1.118 | -0.05 | -0.016 | 0.012 | 0.012 | 0 | 0.86 | 0.81 | 0.93 | 0.8 | 0.91 | 0.86 |
| 2 | 178012446 | rs10197258 | 1.878 | -0.321 | 0.963 | -0.066 | -0.011 | -0.001 | -0.001 | 1.867 | 0.74 | 0.3 | 0.81 | 0.75 | 0.82 | 0.8 |
| 2 | 178015465 | rs2588877 | 0.294 | -0.122 | 0.543 | -0.152 | 0 | -0.034 | -0.034 | 0 | 0.07 | 0.15 | 0.06 | 0 | 0.08 | 0.04 |
| 2 | 178020447 | rs3109082 | 0.576 | -0.103 | 0.302 | -0.143 | 0 | -0.011 | -0.011 | 0.117 | 0.07 | 0.09 | 0.05 | 0 | 0.08 | 0.04 |
| 2 | 178023428 | rs16865043 | 0.26 | -0.352 | -0.015 | -0.125 | 0.008 | -0.022 | -0.022 | 0 | 0.09 | 0 | 0.04 | 0.07 | 0.06 | 0.11 |
| 2 | 178026558 | rs2706126 | 0.116 | -0.024 | 0.254 | -0.129 | 0.008 | -0.021 | -0.021 | 0 | 0.89 | 0.96 | 0.94 | 0.82 | 0.91 | 0.9 |
| 2 | 178032858 | rs2706121 | 0.573 | 0.048 | 0.264 | -0.127 | 0.011 | -0.015 | -0.015 | 0 | 0.87 | 0.95 | 0.93 | 0.81 | 0.9 | 0.86 |
| 2 | 178040695 | rs2706108 | 0.563 | 0.126 | 0.335 | -0.081 | 0.003 | -0.008 | -0.008 | 0.059 | 0.88 | 0.92 | 0.93 | 0.81 | 0.89 | 0.89 |
| 2 | 178040870 | rs2588873 | 1.778 | 0.085 | 0.905 | -0.297 | -0.011 | -0.006 | -0.006 | 0.946 | 0.74 | 0.34 | 0.81 | 0.76 | 0.79 | 0.84 |
| 2 | 178068159 | rs2251139 | 0.379 | 0.077 | 0.929 | -0.301 | 0.001 | -0.003 | -0.003 | 0 | 0.88 | 0.91 | 0.93 | 0.81 | 0.9 | 0.89 |
| 2 | 178073075 | rs2247763 | 1.774 | 0.09 | 0.834 | -0.303 | -0.01 | -0.004 | -0.004 | 0.643 | 0.75 | 0.35 | 0.81 | 0.77 | 0.79 | 0.84 |
| 2 | 178073166 | rs2247754 | 1.814 | 0.092 | 0.834 | -0.302 | -0.01 | -0.005 | -0.005 | 0 | 0.74 | 0.34 | 0.81 | 0.77 | 0.79 | 0.84 |
| 2 | 178087227 | rs13401026 | 0.185 | 0.128 | 1.598 | -0.306 | 0.041 | -0.056 | -0.056 | 0 | 0.1 | 0.35 | 0.1 | 0 | 0.11 | 0.03 |
| 2 | 178091822 | rs13035806 | 0.098 | 0.16 | 1.641 | -0.306 | 0.038 | -0.057 | -0.057 | 0 | 0.1 | 0.33 | 0.1 | 0 | 0.11 | 0.03 |
| 2 | 178092162 | rs2706110 | 1.794 | 0.123 | 1.519 | -0.305 | -0.007 | -0.003 | -0.003 | 0 | 0.75 | 0.31 | 0.8 | 0.77 | 0.79 | 0.84 |
| 2 | 178100425 | rs2001350 | 0.492 | 0.171 | 1.3 | -0.025 | -0.002 | 0.011 | 0.011 | 0 | 0.88 | 0.9 | 0.91 | 0.77 | 0.89 | 0.9 |
| 2 | 178103229 | rs6726395 | 1.688 | 0.74 | 0.838 | -0.262 | -0.008 | 0.004 | 0.004 | 0 | 0.6 | 0.36 | 0.66 | 0.65 | 0.51 | 0.77 |
| 2 | 178113484 | rs1962142 | -0.46 | 0.697 | 0.214 | -0.115 | -0.004 | -0.024 | -0.024 | 2.176 | 0.9 | 0.96 | 0.83 | 0.81 | 0.89 | 0.92 |
| 2 | 178114632 | rs10930781 | -0.045 | 0.711 | 0.217 | -0.173 | 0.002 | -0.022 | -0.022 | 0 | 0.88 | 0.94 | 0.82 | 0.77 | 0.88 | 0.89 |
| 2 | 178118152 | rs1806649 | 0.424 | 0.973 | 0.073 | -0.27 | -0.083 | 0.032 | 0.032 | 0.107 | 0.16 | 0.01 | 0.14 | 0.08 | 0.27 | 0.07 |
| 2 | 178118990 | rs13001694 | -0.637 | 0.625 | 0.567 | -0.238 | 0.005 | -0.025 | -0.025 | 0 | 0.29 | 0.33 | 0.23 | 0.11 | 0.38 | 0.13 |
| 2 | 178124787 | rs2364722 | 2.164 | 0.31 | -0.333 | -0.361 | -0.009 | -0.047 | -0.047 | 0.016 | 0.31 | 0.15 | 0.43 | 0.55 | 0.31 | 0.53 |
| 2 | 178134677 | rs6433657 | 0.701 | 0.367 | -0.575 | -0.569 | -0.061 | 0.005 | 0.005 | 0 | 0.6 | 0.81 | 0.6 | 0.68 | 0.49 | 0.72 |
| 2 | 178136077 | rs17647588 | -0.177 | 0.604 | -0.26 | -1.023 | -0.023 | 0.029 | 0.029 | 0 | 0.25 | 0.22 | 0.21 | 0.08 | 0.38 | 0.11 |
| 2 | 178146736 | rs2001349 | -0.57 | -0.139 | 0.077 | -0.647 | -0.015 | -0.075 | -0.075 | 0.01 | 0.26 | 0.48 | 0.17 | 0.09 | 0.22 | 0.17 |
| 2 | 178149364 | rs6433659 | 0.921 | -0.013 | 0.039 | -0.102 | 0.021 | 0.02 | 0.02 | 0 | 0.49 | 0.31 | 0.35 | 0.61 | 0.45 | 0.63 |

| | | | | | | | | | | | | | | | | |
|---|---|---|---|---|---|---|---|---|---|---|---|---|---|---|---|---|
| 2 | 178151626 | rs7559715 | -0.625 | -0.507 | 1.002 | -0.284 | 0.015 | -0.067 | -0.067 | 0 | 0.17 | 0.22 | 0.12 | 0.02 | 0.18 | 0.08 |
| 2 | 178158043 | rs12623239 | 1.649 | -0.995 | 1.419 | -0.374 | 0.012 | 0.048 | 0.048 | 4.583 | 0.7 | 0.24 | 0.79 | 0.91 | 0.78 | 0.82 |
| 2 | 178162586 | rs2444563 | 0.218 | -0.92 | 1.913 | -0.36 | -0.007 | -0.012 | -0.012 | 0.042 | 0.83 | 0.86 | 0.88 | 0.98 | 0.82 | 0.92 |
| 2 | 178171918 | rs13002088 | 1.133 | -1.014 | 2.339 | -0.365 | 0.023 | -0.112 | -0.112 | 0 | 0.77 | 0.52 | 0.82 | 0.93 | 0.8 | 0.86 |
| 2 | 178178466 | rs1453376 | 0.229 | -1.015 | 2.383 | -0.366 | -0.004 | -0.021 | -0.021 | 0 | 0.83 | 0.83 | 0.88 | 0.97 | 0.82 | 0.92 |
| 2 | 178179022 | rs6720934 | -1.559 | -1.017 | 2.372 | -0.308 | 0.048 | 0.072 | 0.072 | 0.021 | 0.06 | 0.33 | 0.06 | 0.05 | 0.02 | 0.06 |
| 2 | 178185645 | rs1869356 | -0.552 | -1.031 | 2.456 | -0.314 | -0.029 | -0.003 | -0.003 | 14.918 | 0.06 | 0.19 | 0.02 | 0.01 | 0.02 | 0.04 |
| 2 | 178193780 | rs10930785 | 0.297 | -0.93 | 2.65 | -0.378 | 0.006 | -0.043 | -0.043 | 0 | 0.83 | 0.8 | 0.88 | 0.98 | 0.82 | 0.92 |
| 2 | 178196885 | rs10179975 | 1.198 | -0.941 | 2.594 | -0.395 | 0.016 | -0.096 | -0.096 | 0 | 0.77 | 0.48 | 0.82 | 0.93 | 0.8 | 0.86 |
| 2 | 178225266 | rs1901826 | 1.976 | -1.504 | 2.733 | 0.08 | 0.001 | -0.001 | -0.001 | 2.591 | 0.65 | 0.54 | 0.78 | 0.68 | 0.68 | 0.7 |
| 2 | 178270734 | rs10497508 | -0.957 | -2.033 | 2.888 | 0.165 | 0.002 | -0.102 | -0.102 | 3.49 | 0.17 | 0.01 | 0.09 | 0.3 | 0.12 | 0.15 |
| 2 | 178285811 | rs4893823 | 2.088 | -2.063 | 3.181 | 0.151 | -0.022 | 0.009 | 0.009 | 0 | 0.65 | 0.52 | 0.79 | 0.64 | 0.75 | 0.71 |
| 2 | 178358281 | rs12469386 | 2.766 | -2.184 | 2.932 | 0.101 | 0.075 | -0.005 | -0.005 | 0.401 | 0.58 | 0.11 | 0.76 | 0.63 | 0.73 | 0.69 |
| 2 | 178387547 | rs12615367 | -1.441 | -2.133 | 2.883 | 0.151 | 0.005 | -0.006 | -0.006 | 3.173 | 0.19 | 0.15 | 0.09 | 0.31 | 0.11 | 0.21 |
| 2 | 178399810 | rs4893829 | 1.198 | -2.086 | 2.592 | 0.169 | 0.005 | -0.038 | -0.038 | 0.18 | 0.85 | 0.8 | 0.9 | 0.94 | 0.86 | 0.92 |
| 2 | 178401288 | rs7606235 | 1.961 | -2.084 | 2.594 | 0.179 | -0.001 | 0 | 0 | 0 | 0.69 | 0.7 | 0.82 | 0.63 | 0.77 | 0.77 |
| 2 | 178402786 | rs3213945 | -1.029 | -2.039 | 2.169 | 0.217 | 0.026 | -0.052 | -0.052 | 0.229 | 0.16 | 0.05 | 0.07 | 0.31 | 0.1 | 0.15 |
| 2 | 178417945 | rs12988366 | 0.967 | -1.974 | 2.518 | -0.054 | 0.005 | -0.046 | -0.046 | 0 | 0.86 | 0.83 | 0.91 | 0.97 | 0.87 | 0.89 |
| 2 | 178418575 | rs1348850 | 2.878 | -2.125 | 2.58 | -0.088 | 0.073 | -0.027 | -0.027 | 0.932 | 0.58 | 0.09 | 0.76 | 0.62 | 0.73 | 0.63 |
| 2 | 178419117 | rs6737342 | 1.577 | -2.067 | 3.003 | -0.13 | 0.016 | -0.091 | -0.091 | 0.561 | 0.83 | 0.6 | 0.9 | 0.94 | 0.87 | 0.84 |
| 2 | 178426793 | rs11889488 | 0.656 | -2.068 | 2.956 | -0.107 | 0.004 | -0.039 | -0.039 | 0 | 0.86 | 0.83 | 0.91 | 0.97 | 0.87 | 0.89 |
| 2 | 178430747 | rs4893963 | 2.144 | -2.083 | 3.101 | -0.106 | 0.001 | -0.016 | -0.016 | 0 | 0.73 | 0.26 | 0.83 | 0.92 | 0.83 | 0.81 |
| 2 | 178442592 | rs4893835 | 1.657 | -2.082 | 2.739 | -0.114 | -0.017 | -0.094 | -0.094 | 0 | 0.78 | 0.6 | 0.88 | 0.96 | 0.84 | 0.84 |
| 2 | 178445161 | rs11691740 | 1.641 | -2.081 | 2.739 | -0.101 | -0.017 | -0.09 | -0.09 | 0.004 | 0.78 | 0.6 | 0.87 | 0.96 | 0.84 | 0.84 |
| 2 | 178447253 | rs13404970 | -1.287 | -2.133 | 2.802 | -0.106 | -0.027 | 0.043 | 0.043 | 0 | 0.18 | 0.26 | 0.09 | 0.3 | 0.1 | 0.18 |
| 2 | 178457339 | rs12991272 | 2.069 | -2.108 | 2.61 | -0.1 | -0.029 | 0.011 | 0.011 | 0 | 0.65 | 0.49 | 0.81 | 0.66 | 0.75 | 0.69 |
| 2 | 178459669 | rs10930800 | 2.515 | -2.121 | 2.274 | -0.076 | 0.069 | 0.037 | 0.037 | 0.986 | 0.57 | 0.16 | 0.76 | 0.66 | 0.73 | 0.6 |
| 2 | 178467144 | rs12612857 | 2.122 | -2.08 | 2.507 | 0.008 | 0.016 | 0 | 0 | 0.224 | 0.63 | 0.29 | 0.79 | 0.64 | 0.77 | 0.71 |
| 2 | 178468493 | rs16865433 | -1.245 | -1.832 | 2.416 | 0.183 | -0.002 | 0.004 | 0.004 | 0 | 0.16 | 0.16 | 0.08 | 0.3 | 0.1 | 0.18 |
| 2 | 178470372 | rs11901206 | -2.057 | -1.761 | 2.699 | 0.06 | -0.027 | 0.059 | 0.059 | 0 | 0.12 | 0.36 | 0.06 | 0.18 | 0.06 | 0.1 |
| 2 | 178471407 | rs10201883 | 1.045 | -1.799 | 2.834 | -0.25 | 0.037 | -0.063 | -0.063 | 0.624 | 0.85 | 0.67 | 0.89 | 0.94 | 0.87 | 0.91 |
| 2 | 178474768 | rs763757 | -1.72 | -1.816 | 2.892 | -0.226 | -0.001 | 0.022 | 0.022 | 0 | 0.06 | 0.2 | 0.04 | 0.1 | 0.04 | 0.02 |
| 2 | 178475599 | rs3731802 | -2 | -1.82 | 2.752 | -0.229 | 0.001 | 0.028 | 0.028 | 0.061 | 0.07 | 0.27 | 0.05 | 0.1 | 0.04 | 0.02 |
| 2 | 178477114 | rs13031008 | 2.632 | -2.185 | 2.66 | -0.324 | 0.035 | 0.022 | 0.022 | 0.106 | 0.59 | 0.27 | 0.8 | 0.69 | 0.74 | 0.68 |
| 2 | 178479149 | rs16865446 | -1.578 | -2.19 | 2.614 | -0.325 | 0.067 | -0.022 | -0.022 | 0.008 | 0.19 | 0.06 | 0.04 | 0.19 | 0.09 | 0.18 |
| 2 | 178479248 | rs10497498 | -1.593 | -2.196 | 2.611 | -0.328 | 0.068 | -0.022 | -0.022 | 0 | 0.19 | 0.06 | 0.04 | 0.19 | 0.09 | 0.18 |
| 2 | 178479422 | rs3821003 | -1.58 | -2.184 | 2.682 | -0.322 | 0.053 | -0.015 | -0.015 | 0 | 0.19 | 0.07 | 0.05 | 0.19 | 0.09 | 0.18 |
| 2 | 178479661 | rs3731806 | -0.605 | -2.199 | 2.644 | -0.327 | 0.032 | -0.036 | -0.036 | 0 | 0.12 | 0.02 | 0.03 | 0.17 | 0.07 | 0.1 |
| 2 | 178480063 | rs3821004 | -1.707 | -2.199 | 2.644 | -0.327 | 0.067 | -0.022 | -0.022 | 0 | 0.19 | 0.06 | 0.04 | 0.19 | 0.09 | 0.18 |
| 2 | 178480405 | rs3821005 | -0.609 | -2.187 | 2.643 | -0.323 | 0.032 | -0.036 | -0.036 | 0 | 0.12 | 0.02 | 0.03 | 0.17 | 0.07 | 0.1 |
| 2 | 178481818 | rs10497497 | 0.329 | -2.191 | 2.804 | -0.329 | 0.019 | -0.039 | -0.039 | 0 | 0.88 | 0.84 | 0.92 | 0.98 | 0.88 | 0.92 |
| 2 | 178487729 | rs2293471 | 1.739 | -2.195 | 2.659 | -0.328 | 0.021 | -0.067 | -0.067 | 0.633 | 0.8 | 0.49 | 0.86 | 0.91 | 0.84 | 0.87 |
| 2 | 178488028 | rs2293473 | -0.613 | -2.184 | 2.493 | -0.318 | 0.032 | -0.036 | -0.036 | 0.02 | 0.12 | 0.02 | 0.03 | 0.17 | 0.07 | 0.1 |
| 2 | 178492722 | rs6730406 | 2.064 | -2.178 | 2.563 | -0.362 | -0.024 | -0.074 | -0.074 | 0.184 | 0.73 | 0.46 | 0.85 | 0.89 | 0.82 | 0.79 |
| 2 | 178493301 | rs13034886 | 2.35 | -2.181 | 2.375 | -0.39 | 0.016 | 0.032 | 0.032 | 3.951 | 0.7 | 0.21 | 0.82 | 0.86 | 0.81 | 0.79 |
| 2 | 178493713 | rs6705405 | 2.294 | -2.18 | 2.375 | -0.39 | 0.015 | 0.03 | 0.03 | 4.069 | 0.7 | 0.21 | 0.82 | 0.86 | 0.81 | 0.79 |
| 2 | 178493731 | rs6734528 | 2.771 | -2.191 | 2.363 | -0.389 | 0.07 | 0.05 | 0.05 | 0.476 | 0.58 | 0.2 | 0.79 | 0.7 | 0.74 | 0.69 |
| 2 | 178496874 | rs4893972 | 2.505 | -2.134 | 2.39 | -0.357 | 0.015 | 0.029 | 0.029 | 4.176 | 0.7 | 0.21 | 0.81 | 0.86 | 0.81 | 0.79 |
| 2 | 178498015 | rs10469714 | -2.789 | -2.274 | 2.506 | -0.372 | -0.071 | -0.038 | -0.038 | 0 | 0.18 | 0.59 | 0.11 | 0.11 | 0.07 | 0.14 |
| 2 | 178502353 | rs12471621 | 2.465 | -2.403 | 2.366 | -0.422 | 0.052 | 0.038 | 0.038 | 1.505 | 0.63 | 0.16 | 0.79 | 0.71 | 0.75 | 0.76 |
| 2 | 178511430 | rs2222712 | 0.377 | -2.415 | 2.654 | -0.422 | 0.009 | -0.042 | -0.042 | 0.024 | 0.88 | 0.82 | 0.92 | 0.97 | 0.89 | 0.93 |
| 2 | 178513894 | rs1527402 | 1.36 | -2.421 | 1.812 | -0.445 | 0.003 | -0.063 | -0.063 | 3.977 | 0.8 | 0.55 | 0.87 | 0.91 | 0.84 | 0.91 |
| 2 | 178514468 | rs13008857 | 2.461 | -2.409 | 1.802 | -0.434 | 0.055 | 0.04 | 0.04 | 1.465 | 0.62 | 0.16 | 0.79 | 0.71 | 0.75 | 0.77 |
| 2 | 178518154 | rs6746845 | -1.69 | -2.393 | 1.807 | -0.421 | -0.066 | 0.037 | 0.037 | 0 | 0.18 | 0.39 | 0.09 | 0.19 | 0.09 | 0.14 |
| 2 | 178522104 | rs1527401 | 2.927 | -2.396 | 1.645 | -0.419 | 0.106 | 0.067 | 0.067 | 1.762 | 0.55 | 0.1 | 0.77 | 0.7 | 0.72 | 0.69 |
| 2 | 178528703 | rs13025391 | 0.743 | -2.138 | 1.759 | -0.316 | 0.029 | -0.076 | -0.076 | 0 | 0.86 | 0.59 | 0.9 | 0.97 | 0.88 | 0.91 |
| 2 | 178531991 | rs7594290 | -0.826 | -2.146 | 1.699 | -0.316 | 0 | 0 | 0 | 0 | 0.07 | 0.07 | 0.04 | 0.08 | 0.06 | 0.06 |
| 2 | 178533715 | rs1358326 | 0.799 | -2.124 | 1.687 | -0.331 | 0.028 | -0.078 | -0.078 | 0 | 0.85 | 0.58 | 0.9 | 0.97 | 0.88 | 0.91 |
| 2 | 178535203 | rs7605757 | -1.954 | -2.148 | 1.707 | -0.337 | -0.043 | 0.016 | 0.016 | 0 | 0.22 | 0.33 | 0.11 | 0.26 | 0.14 | 0.15 |
| 2 | 178536783 | rs12612481 | -1.505 | -2.113 | 0.758 | -0.314 | 0.023 | -0.016 | -0.016 | 0 | 0.19 | 0.09 | 0.08 | 0.23 | 0.13 | 0.15 |
| 2 | 178537890 | rs3770011 | 1.592 | -1.974 | 0.302 | -0.236 | 0.03 | 0.024 | 0.024 | 0.639 | 0.63 | 0.25 | 0.79 | 0.71 | 0.74 | 0.77 |
| 2 | 178540293 | rs3770016 | 0.734 | -1.62 | 0.062 | -0.035 | -0.009 | -0.017 | -0.017 | 0.307 | 0.79 | 0.35 | 0.84 | 0.84 | 0.83 | 0.82 |
| 2 | 178544417 | rs3770019 | -1.472 | -1.632 | -0.013 | 0.076 | 0.012 | 0.004 | 0.004 | 0 | 0.23 | 0.18 | 0.06 | 0.18 | 0.1 | 0.15 |
| 2 | 178545736 | rs9677333 | -1.472 | -1.676 | -0.012 | 0.01 | 0.009 | 0.005 | 0.005 | 0 | 0.23 | 0.18 | 0.06 | 0.17 | 0.1 | 0.15 |
| 2 | 178551924 | rs17400121 | -0.954 | -1.528 | -0.341 | -0.068 | 0.029 | 0.053 | 0.053 | 0 | 0.13 | 0.03 | 0.03 | 0.01 | 0.06 | 0.06 |
| 2 | 178551958 | rs6433691 | -1.768 | -1.885 | 0.674 | 0.242 | -0.104 | 0.012 | 0.012 | 0 | 0.26 | 0.47 | 0.09 | 0.26 | 0.1 | 0.16 |
| 2 | 178552788 | rs13032794 | -1.466 | -1.867 | 0.903 | -0.47 | -0.046 | -0.101 | -0.101 | 1.198 | 0.2 | 0.37 | 0.07 | 0.01 | 0.11 | 0.1 |
| 2 | 178558967 | rs1438054 | 0.919 | -2.343 | 1.389 | -0.633 | -0.014 | -0.015 | -0.015 | 0 | 0.74 | 0.71 | 0.87 | 0.8 | 0.8 | 0.76 |
| 2 | 178561295 | rs16865645 | -1.026 | -2.422 | 1.348 | -0.673 | 0.03 | 0.03 | 0.03 | 0 | 0.22 | 0.12 | 0.09 | 0.14 | 0.17 | 0.18 |

| | | | | | | | | | | | | | | | | |
|---|---|---|---|---|---|---|---|---|---|---|---|---|---|---|---|---|
| 2 | 178563430 | rs1469973 | 1.533 | -2.658 | 0.937 | -1.134 | 0.074 | 0.099 | 0.099 | 0 | 0.61 | 0.24 | 0.8 | 0.82 | 0.73 | 0.7 |
| 2 | 178566437 | rs12989020 | -1.211 | -2.659 | 0.964 | -1.129 | 0.042 | 0.093 | 0.093 | 0.012 | 0.29 | 0.04 | 0.12 | 0.12 | 0.24 | 0.24 |
| 2 | 178576681 | rs13428806 | -1.68 | -2.684 | 1.565 | -1.173 | -0.048 | -0.066 | -0.066 | 0 | 0.34 | 0.54 | 0.17 | 0.16 | 0.23 | 0.24 |
| 2 | 178585514 | rs4893987 | 1.787 | -2.587 | 1.029 | -1.761 | 0.049 | 0.261 | 0.261 | 1.268 | 0.48 | 0.11 | 0.7 | 0.84 | 0.62 | 0.61 |
| 2 | 178596672 | rs10175364 | 1.225 | -2.454 | 1.331 | -1.763 | -0.005 | -0.03 | -0.03 | 0 | 0.56 | 0.55 | 0.77 | 0.84 | 0.67 | 0.66 |
| 2 | 178600616 | rs3770034 | 1.038 | -2.466 | 1.184 | -1.693 | 0.033 | 0.035 | 0.035 | 0 | 0.64 | 0.32 | 0.82 | 0.84 | 0.77 | 0.75 |
| 2 | 178607359 | rs1866209 | 0.933 | -2.468 | 1.135 | -1.694 | -0.031 | -0.045 | -0.045 | 1.142 | 0.65 | 0.45 | 0.83 | 0.84 | 0.77 | 0.75 |
| 2 | 178607998 | rs1370656 | 0.58 | -2.535 | 1.064 | -1.736 | -0.046 | -0.135 | -0.135 | 0 | 0.77 | 0.51 | 0.88 | 0.99 | 0.84 | 0.82 |
| 2 | 178609136 | rs2164857 | 1.123 | -2.479 | 0.661 | -1.708 | 0.06 | 0.07 | 0.07 | 0 | 0.64 | 0.26 | 0.82 | 0.84 | 0.77 | 0.75 |
| 2 | 178627664 | rs3754996 | -1.41 | -2.51 | -0.312 | -1.452 | 0.087 | 0.099 | 0.099 | 0.02 | 0.51 | 0.82 | 0.31 | 0.37 | 0.38 | 0.36 |
| 2 | 178631142 | rs6755503 | 0.688 | -2.524 | -0.223 | -1.464 | -0.002 | -0.019 | -0.019 | 1.193 | 0.14 | 0.15 | 0.1 | 0.01 | 0.13 | 0.1 |
| 2 | 178632479 | rs934798 | 0.933 | -2.535 | -0.259 | -1.392 | 0.014 | 0.015 | 0.015 | 0 | 0.12 | 0.06 | 0.05 | 0.09 | 0.11 | 0.1 |
| 2 | 178635553 | rs7605091 | -0.671 | -2.332 | -0.17 | -1.31 | -0.018 | -0.005 | -0.005 | 0 | 0.36 | 0.56 | 0.2 | 0.36 | 0.27 | 0.25 |
| 2 | 178640880 | rs1438040 | -1.894 | -1.101 | -0.695 | -1.04 | -0.043 | -0.027 | -0.027 | 0 | 0.7 | 0.83 | 0.44 | 0.4 | 0.62 | 0.59 |
| 2 | 178643119 | rs6718860 | -0.876 | -1.107 | -0.364 | -0.987 | -0.002 | 0.036 | 0.036 | 0 | 0.52 | 0.69 | 0.35 | 0.39 | 0.51 | 0.41 |
| 2 | 178679676 | rs1946814 | -0.641 | -1.547 | -0.63 | -0.927 | 0.069 | 0.035 | 0.035 | 0.359 | 0.45 | 0.74 | 0.28 | 0.38 | 0.34 | 0.32 |
| 2 | 178681534 | rs6433698 | -0.427 | -1.596 | -0.653 | -0.95 | 0.056 | 0.037 | 0.037 | 0.238 | 0.42 | 0.73 | 0.26 | 0.31 | 0.32 | 0.32 |
| 2 | 178682603 | rs6720891 | -0.427 | -1.596 | -0.653 | -0.95 | 0.056 | 0.037 | 0.037 | 0.238 | 0.42 | 0.73 | 0.26 | 0.31 | 0.32 | 0.32 |
| 2 | 178682655 | rs3821010 | -0.427 | -1.596 | -0.653 | -0.95 | 0.056 | 0.037 | 0.037 | 0.238 | 0.42 | 0.73 | 0.26 | 0.31 | 0.32 | 0.32 |
| 2 | 178683973 | rs1438060 | -0.427 | -1.596 | -0.653 | -0.95 | 0.056 | 0.037 | 0.037 | 0.238 | 0.42 | 0.73 | 0.26 | 0.31 | 0.32 | 0.32 |
| 2 | 178690786 | rs934803 | -0.122 | -1.373 | -0.768 | -1.041 | 0.017 | -0.046 | -0.046 | 0 | 0.88 | 0.76 | 0.9 | 0.99 | 0.84 | 0.87 |
| 2 | 178696741 | rs13007187 | -0.92 | -1.712 | -0.68 | -0.633 | 0.009 | 0.014 | 0.014 | 0 | 0.6 | 0.51 | 0.38 | 0.47 | 0.51 | 0.5 |
| 2 | 178697917 | rs753346 | 0.296 | -2.064 | -0.179 | -0.851 | -0.002 | -0.004 | -0.004 | 0.543 | 0.21 | 0.23 | 0.14 | 0.17 | 0.22 | 0.17 |
| 2 | 178703583 | rs1370661 | 1.527 | -2.399 | -0.429 | -0.96 | 0.056 | 0.094 | 0.094 | 0 | 0.53 | 0.17 | 0.73 | 0.71 | 0.62 | 0.55 |
| 2 | 178708156 | rs16865825 | -0.15 | -2.385 | -0.385 | -0.943 | -0.005 | -0.001 | -0.001 | 0 | 0.24 | 0.28 | 0.16 | 0.26 | 0.24 | 0.18 |
| 2 | 178710984 | rs1438051 | 1.302 | -2.286 | -0.225 | -0.894 | 0.04 | 0.061 | 0.061 | 0 | 0.54 | 0.24 | 0.74 | 0.71 | 0.63 | 0.55 |
| 2 | 178711358 | rs6758473 | 0.929 | -2.341 | -0.112 | -0.93 | 0.003 | 0.003 | 0.003 | 1.385 | 0.29 | 0.2 | 0.17 | 0.19 | 0.23 | 0.22 |
| 2 | 178712283 | rs7570981 | 0.033 | -2.447 | -0.067 | -0.981 | -0.035 | -0.012 | -0.012 | 0 | 0.25 | 0.35 | 0.1 | 0.18 | 0.12 | 0.2 |
| 2 | 178748294 | rs1405648 | -0.833 | -2.351 | -0.09 | -0.936 | 0.009 | 0.012 | 0.012 | 0 | 0.34 | 0.34 | 0.18 | 0.19 | 0.28 | 0.35 |
| 2 | 178768683 | rs17401468 | -0.982 | -2.149 | -0.093 | -0.838 | 0.019 | -0.003 | -0.003 | 0 | 0.11 | 0.24 | 0.07 | 0.1 | 0.12 | 0.16 |
| 2 | 178774704 | rs13419437 | -0.392 | -2.134 | -0.082 | -0.82 | 0.007 | 0.003 | 0.003 | 0 | 0.09 | 0.19 | 0.06 | 0.1 | 0.1 | 0.1 |
| 2 | 178781124 | rs3108476 | 2.257 | -2.197 | -0.231 | -0.857 | -0.089 | -0.101 | -0.101 | 0 | 0.25 | 0.09 | 0.56 | 0.52 | 0.39 | 0.33 |
| 2 | 178783330 | rs10208501 | -0.023 | -2.214 | -0.111 | -0.828 | 0.006 | 0.003 | 0.003 | 0 | 0.08 | 0.1 | 0.05 | 0.1 | 0.1 | 0.1 |
| 2 | 178800051 | rs2695737 | -0.96 | -2.365 | -0.072 | -0.953 | -0.035 | -0.031 | -0.031 | 0.087 | 0.69 | 0.58 | 0.8 | 0.81 | 0.78 | 0.78 |
| 2 | 178818102 | rs17401916 | 1.191 | -2.208 | -0.336 | -0.874 | 0.006 | -0.008 | -0.008 | 0 | 0.05 | 0.05 | 0.03 | 0.01 | 0.08 | 0.09 |
| 2 | 178818812 | rs7595479 | 0.008 | -2.05 | -0.384 | -0.945 | 0.009 | 0.006 | 0.006 | 0 | 0.25 | 0.24 | 0.09 | 0.17 | 0.14 | 0.26 |
| 2 | 178836333 | rs2573085 | -0.947 | -1.67 | -0.724 | -0.782 | 0.011 | -0.037 | -0.037 | 0 | 0.61 | 0.86 | 0.33 | 0.46 | 0.39 | 0.49 |
| 2 | 178845469 | rs2573083 | -1.303 | -1.648 | -0.746 | -0.794 | -0.04 | -0.089 | -0.089 | 0 | 0.64 | 0.91 | 0.35 | 0.47 | 0.43 | 0.52 |
| 2 | 178851177 | rs10497477 | 0.285 | -1.263 | -0.482 | -0.69 | 0.006 | -0.008 | -0.008 | 1.682 | 0.05 | 0.05 | 0.03 | 0.01 | 0.08 | 0.09 |
| 2 | 178856900 | rs13003047 | -0.084 | -1.227 | -0.778 | -0.758 | -0.065 | 0.056 | 0.056 | 0 | 0.13 | 0.01 | 0.1 | 0.01 | 0.21 | 0.19 |
| 2 | 178870030 | rs1020010 | 0.438 | -1.868 | -0.573 | -0.988 | 0.009 | 0.045 | 0.045 | 0 | 0.52 | 0.37 | 0.66 | 0.7 | 0.53 | 0.59 |
| 2 | 178877602 | rs1534290 | 2.156 | -2.089 | -0.863 | -1.055 | -0.16 | -0.162 | -0.162 | 2.675 | 0.27 | 0.02 | 0.56 | 0.53 | 0.39 | 0.38 |
| 2 | 178881409 | rs6726477 | 0.027 | -2.131 | -0.219 | -1.037 | 0.035 | 0.035 | 0.035 | 0 | 0.91 | 0.66 | 0.93 | 0.88 | 0.89 | 0.9 |
| 2 | 178884948 | rs6741976 | 1.053 | -2.019 | -0.113 | -1.07 | -0.007 | -0.009 | -0.009 | 0 | 0.08 | 0.1 | 0.03 | 0.02 | 0.03 | 0.07 |
| 2 | 178907633 | rs7608073 | 0.434 | -1.984 | -0.055 | -1.054 | -0.012 | -0.029 | -0.029 | 0 | 0.34 | 0.36 | 0.21 | 0.19 | 0.32 | 0.31 |
| 2 | 178912784 | rs16866032 | 1.326 | -1.872 | -0.209 | -0.895 | -0.013 | 0 | 0 | 0 | 0.14 | 0.18 | 0.06 | 0.15 | 0.09 | 0.08 |
| 2 | 178920669 | rs2009303 | -0.3 | -1.71 | -0.329 | -1.037 | 0.001 | 0.006 | 0.006 | 0 | 0.46 | 0.44 | 0.31 | 0.3 | 0.47 | 0.41 |
| 2 | 178938805 | rs920094 | -0.137 | -1.727 | -0.023 | -1.012 | -0.061 | -0.097 | -0.097 | 0 | 0.24 | 0.42 | 0.11 | 0.07 | 0.13 | 0.2 |
| 2 | 178939528 | rs11676719 | 0.921 | -1.564 | -0.424 | -1.32 | -0.001 | -0.005 | -0.005 | 0.075 | 0.09 | 0.05 | 0.06 | 0 | 0.12 | 0.07 |
| 2 | 178942302 | rs6710329 | 0.955 | -1.525 | -0.714 | -1.655 | 0.011 | 0.005 | 0.005 | 0 | 0.25 | 0.17 | 0.15 | 0.19 | 0.18 | 0.23 |
| 2 | 178942789 | rs1435572 | 1.205 | -1.566 | -0.732 | -1.449 | -0.032 | -0.007 | -0.007 | 0 | 0.44 | 0.22 | 0.64 | 0.59 | 0.48 | 0.53 |
| 2 | 178957158 | rs6433712 | -2.225 | -1.594 | -0.179 | -1.364 | -0.049 | 0.017 | 0.017 | 0 | 0.86 | 0.67 | 0.93 | 0.87 | 0.92 | 0.93 |
| 2 | 178958540 | rs1435571 | 0.715 | -1.476 | -0.457 | -1.372 | 0.013 | -0.001 | -0.001 | 0 | 0.27 | 0.2 | 0.16 | 0.28 | 0.2 | 0.23 |
| 2 | 178976514 | rs10176475 | 1.002 | -1.394 | -0.283 | -1.416 | -0.001 | 0 | 0 | 0.209 | 0.25 | 0.22 | 0.15 | 0.2 | 0.19 | 0.23 |
| 2 | 178979984 | rs13408924 | 1.03 | -0.194 | -0.624 | -1.541 | -0.003 | 0.006 | 0.006 | 0 | 0.29 | 0.23 | 0.27 | 0.2 | 0.26 | 0.26 |
| 2 | 178993213 | rs2366366 | 1.833 | 0.4 | -1.26 | -2.025 | -0.095 | -0.121 | -0.121 | 1.886 | 0.29 | 0.07 | 0.48 | 0.57 | 0.36 | 0.36 |
| 2 | 178993562 | rs2163647 | 1.708 | -0.124 | -1.129 | -1.995 | -0.013 | 0.013 | 0.013 | 0 | 0.27 | 0.17 | 0.26 | 0.2 | 0.26 | 0.25 |
| 2 | 179000291 | rs334062 | 1.116 | 0.065 | -1.12 | -1.885 | -0.074 | -0.016 | -0.016 | 0 | 0.36 | 0.19 | 0.55 | 0.66 | 0.48 | 0.46 |
| 2 | 179003272 | rs334066 | -1.128 | -0.023 | -0.701 | -1.893 | -0.044 | -0.048 | -0.048 | 0 | 0.77 | 0.58 | 0.86 | 0.87 | 0.79 | 0.86 |
| 2 | 179011166 | rs334074 | -2.125 | -0.13 | -0.555 | -1.737 | -0.031 | 0.033 | 0.033 | 0 | 0.86 | 0.65 | 0.9 | 0.87 | 0.88 | 0.95 |
| 2 | 179017996 | rs17638592 | -0.06 | -0.369 | -1.057 | -2.13 | 0.016 | 0.132 | 0.132 | 1.61 | 0.49 | 0.21 | 0.38 | 0.23 | 0.46 | 0.37 |
| 2 | 179021948 | rs11690846 | 2.127 | -0.457 | -1.446 | -2.22 | 0.002 | 0.002 | 0.002 | 0 | 0.11 | 0.07 | 0.06 | 0 | 0.09 | 0.02 |
| 2 | 179022610 | rs16866133 | -0.027 | -0.942 | -1.297 | -2.464 | 0.005 | -0.014 | -0.014 | 0 | 0.23 | 0.19 | 0.08 | 0.02 | 0.1 | 0.25 |
| 2 | 179022751 | rs10203979 | -0.611 | -0.742 | -1.317 | -2.501 | -0.023 | -0.065 | -0.065 | 0 | 0.3 | 0.37 | 0.17 | 0.12 | 0.2 | 0.3 |
| 2 | 179026301 | rs334092 | -1.021 | -0.752 | -1.354 | -2.54 | 0.001 | -0.013 | -0.013 | 0 | 0.66 | 0.65 | 0.68 | 0.79 | 0.64 | 0.81 |
| 2 | 179029309 | rs13013858 | 0.101 | -0.497 | -1.2 | -2.301 | 0 | -0.003 | -0.003 | 0 | 0.4 | 0.39 | 0.58 | 0.75 | 0.46 | 0.49 |
| 2 | 179034697 | rs438027 | 0.766 | -0.61 | -1.198 | -2.321 | -0.026 | -0.05 | -0.05 | 0 | 0.14 | 0.23 | 0.04 | 0 | 0.04 | 0.11 |
| 2 | 179037071 | rs334126 | -0.8 | -0.57 | -1.229 | -2.347 | 0 | 0.01 | 0.01 | 5.205 | 0.55 | 0.63 | 0.64 | 0.79 | 0.54 | 0.7 |
| 2 | 179042627 | rs11883528 | 2.407 | -0.352 | -1.244 | -2.387 | -0.003 | -0.003 | -0.003 | 0 | 0.25 | 0.26 | 0.46 | 0.64 | 0.29 | 0.32 |

| | | | | | | | | | | | | | | | | |
|---|---|---|---|---|---|---|---|---|---|---|---|---|---|---|---|---|
| 2 | 179043561 | rs12471750 | 1.754 | -0.546 | -1.161 | -2.355 | -0.018 | 0 | 0 | 1.421 | 0.3 | 0.26 | 0.53 | 0.74 | 0.4 | 0.37 |
| 2 | 179051952 | rs930191 | 0.031 | -0.596 | -1.161 | -2.343 | -0.001 | 0.003 | 0.003 | 2.145 | 0.43 | 0.5 | 0.57 | 0.74 | 0.42 | 0.48 |
| 2 | 179056079 | rs10930827 | 2.516 | -1.079 | -1.042 | -2.234 | -0.048 | -0.045 | -0.045 | 0 | 0.23 | 0.12 | 0.48 | 0.64 | 0.34 | 0.36 |
| 2 | 179060313 | rs334102 | -0.608 | -1.187 | -1.155 | -2.241 | -0.003 | 0.02 | 0.02 | 6.334 | 0.53 | 0.63 | 0.62 | 0.79 | 0.5 | 0.69 |
| 2 | 179066443 | rs334110 | -0.477 | -1.584 | 0.25 | -1.711 | -0.001 | -0.034 | -0.034 | 0 | 0.63 | 0.6 | 0.66 | 0.82 | 0.6 | 0.71 |
| 2 | 179071826 | rs7419749 | 1.748 | -1.307 | 0.46 | -1.23 | -0.176 | -0.054 | -0.054 | 0 | 0.34 | 0.03 | 0.56 | 0.66 | 0.5 | 0.37 |
| 2 | 179072181 | rs334113 | -0.842 | -1.212 | 0.694 | -1.19 | -0.025 | -0.052 | -0.052 | 0 | 0.22 | 0.43 | 0.12 | 0.15 | 0.15 | 0.26 |
| 2 | 179082987 | rs7581504 | 0.322 | -1.215 | 0.608 | -1.2 | -0.003 | -0.046 | -0.046 | 0 | 0.32 | 0.39 | 0.27 | 0.15 | 0.32 | 0.26 |
| 2 | 179086079 | rs334032 | -1.216 | -0.974 | 0.539 | -1.06 | -0.036 | -0.04 | -0.04 | 0.443 | 0.3 | 0.43 | 0.12 | 0.15 | 0.13 | 0.25 |
| 2 | 179097483 | rs334042 | 0.143 | -0.711 | 0.151 | -0.917 | -0.002 | -0.056 | -0.056 | 0 | 0.33 | 0.48 | 0.29 | 0.14 | 0.31 | 0.27 |
| 2 | 179097620 | rs13003146 | -0.566 | -0.701 | -0.091 | -0.953 | -0.036 | 0.076 | 0.076 | 0 | 0.2 | 0.03 | 0.13 | 0.08 | 0.21 | 0.26 |
| 2 | 179102232 | rs334044 | -0.609 | -0.866 | -0.149 | -0.963 | 0.013 | 0.1 | 0.1 | 2.799 | 0.44 | 0.93 | 0.34 | 0.26 | 0.35 | 0.36 |
| 2 | 179104988 | rs6433715 | -1.414 | -0.806 | 0.07 | -0.899 | -0.021 | -0.057 | -0.057 | 1.963 | 0.3 | 0.44 | 0.19 | 0.19 | 0.25 | 0.3 |
| 2 | 179127829 | rs919682 | -0.97 | -0.524 | 0.225 | -0.737 | -0.002 | -0.002 | -0.002 | 0 | 0.24 | 0.22 | 0.13 | 0.19 | 0.2 | 0.31 |
| 2 | 179129455 | rs9989738 | 1.098 | -0.899 | 0.346 | -0.59 | -0.132 | -0.044 | -0.044 | 0 | 0.36 | 0.04 | 0.52 | 0.65 | 0.44 | 0.39 |
| 2 | 179130831 | rs333989 | 0.897 | -0.86 | 0.802 | -0.555 | -0.045 | -0.075 | -0.075 | 0.073 | 0.72 | 0.57 | 0.83 | 0.85 | 0.79 | 0.69 |
| 2 | 179134960 | rs333985 | 0.742 | -0.72 | 0.788 | -0.458 | -0.063 | -0.022 | -0.022 | 0 | 0.82 | 0.66 | 0.92 | 0.85 | 0.92 | 0.8 |
| 2 | 179144028 | rs333993 | 0.42 | -0.494 | 0.667 | -0.394 | 0.006 | -0.011 | -0.011 | 0 | 0.34 | 0.37 | 0.28 | 0.23 | 0.35 | 0.35 |
| 2 | 179146883 | rs12471986 | -0.165 | -0.448 | 0.391 | -0.404 | -0.018 | -0.063 | -0.063 | 0 | 0.2 | 0.25 | 0.27 | 0.03 | 0.16 | 0.21 |
| 2 | 179156878 | rs1434080 | 1.028 | -0.47 | -0.084 | -0.236 | -0.064 | -0.029 | -0.029 | 1.294 | 0.35 | 0.14 | 0.4 | 0.65 | 0.44 | 0.38 |
| 2 | 179157946 | rs836882 | 0.208 | -0.513 | 0.017 | -0.267 | -0.072 | -0.034 | -0.034 | 0 | 0.85 | 0.67 | 0.93 | 0.93 | 0.95 | 0.82 |
| 2 | 179158325 | rs334620 | -0.234 | -0.504 | -0.076 | -0.246 | -0.001 | -0.005 | -0.005 | 0.018 | 0.49 | 0.53 | 0.53 | 0.28 | 0.51 | 0.45 |
| 2 | 179160218 | rs11680778 | 0.819 | -0.321 | -0.576 | -0.021 | -0.107 | -0.151 | -0.151 | 0 | 0.35 | 0.07 | 0.39 | 0.49 | 0.42 | 0.34 |
| 2 | 179166160 | rs13400738 | 0.84 | -0.14 | -0.45 | -0.021 | -0.096 | -0.151 | -0.151 | 0.555 | 0.36 | 0.08 | 0.39 | 0.48 | 0.42 | 0.36 |
| 2 | 179182695 | rs993598 | 0.297 | -0.1 | -0.062 | -0.127 | -0.013 | -0.006 | -0.006 | 1.041 | 0.47 | 0.23 | 0.44 | 0.52 | 0.51 | 0.41 |
| 2 | 179185051 | rs12478760 | 0.743 | -0.1 | 0.01 | -0.327 | 0.007 | 0.009 | 0.009 | 0 | 0.13 | 0.16 | 0.39 | 0.25 | 0.18 | 0.14 |
| 2 | 179186529 | rs7592114 | 0.584 | -0.171 | 0.145 | -0.345 | -0.001 | -0.016 | -0.016 | 0 | 0.31 | 0.39 | 0.25 | 0.22 | 0.29 | 0.3 |
| 2 | 179187786 | rs4133879 | -0.463 | -0.035 | 0.135 | -0.307 | 0.001 | -0.023 | -0.023 | 0.693 | 0.64 | 0.43 | 0.72 | 0.76 | 0.64 | 0.65 |
| 2 | 179191290 | rs12151447 | 0.117 | -0.197 | 0.316 | -0.075 | -0.023 | -0.026 | -0.026 | 0 | 0.13 | 0.07 | 0.38 | 0.24 | 0.18 | 0.15 |
| 2 | 179198946 | rs16866260 | 0.028 | -0.239 | 0.323 | -0.098 | 0.04 | 0.042 | 0.042 | 0 | 0.14 | 0.05 | 0.05 | 0.03 | 0.06 | 0.16 |
| 2 | 179201920 | rs12618595 | -0.051 | -0.231 | 0.323 | -0.083 | 0.032 | -0.038 | -0.038 | 0 | 0.14 | 0.05 | 0.18 | 0.28 | 0.07 | 0.18 |
| 2 | 179202472 | rs7585689 | 0.049 | -0.248 | 0.323 | -0.133 | 0.039 | 0.041 | 0.041 | 0 | 0.14 | 0.05 | 0.05 | 0.03 | 0.06 | 0.16 |
| 2 | 179203797 | rs1434087 | -0.739 | -0.221 | 0.284 | -0.127 | -0.002 | -0.008 | -0.008 | 0 | 0.9 | 0.87 | 0.94 | 0.97 | 0.9 | 0.95 |
| 2 | 179206754 | rs13402976 | -0.644 | 0.106 | 0.275 | 0.134 | 0.024 | -0.094 | -0.094 | 0 | 0.15 | 0.3 | 0.27 | 0 | 0.16 | 0.13 |
| 2 | 179208524 | rs4894013 | -0.224 | 0.264 | -0.062 | 0.212 | 0.021 | -0.074 | -0.074 | 0 | 0.13 | 0.24 | 0.26 | 0 | 0.16 | 0.12 |
| 2 | 179211925 | rs6725937 | -0.408 | 0.292 | -0.783 | 0.234 | 0.004 | 0 | 0 | 0 | 0.14 | 0.09 | 0.06 | 0.03 | 0.06 | 0.16 |
| 2 | 179212631 | rs12478837 | -0.101 | 0.295 | -0.781 | 0.231 | -0.011 | 0.013 | 0.013 | 0 | 0.13 | 0.08 | 0.24 | 0 | 0.16 | 0.13 |
| 2 | 179212806 | rs10497513 | -0.613 | 0.475 | -0.719 | 0.392 | 0.003 | 0.002 | 0.002 | 0.026 | 0.28 | 0.21 | 0.43 | 0.27 | 0.24 | 0.31 |
| 2 | 179215801 | rs1368907 | -0.565 | 0.32 | -1.052 | -0.411 | 0.011 | 0.014 | 0.014 | 0.086 | 0.27 | 0.17 | 0.3 | 0.03 | 0.22 | 0.29 |
| 2 | 179232560 | rs1434091 | 0.109 | 0.695 | -1.869 | 1.098 | -0.022 | -0.143 | -0.143 | 0 | 0.23 | 0.02 | 0.17 | 0.38 | 0.19 | 0.26 |
| 2 | 179236048 | rs1434094 | 0.003 | 1.034 | -1.451 | 0.232 | 0.003 | 0.071 | 0.071 | 0 | 0.52 | 0.63 | 0.53 | 0.88 | 0.5 | 0.58 |
| 2 | 179238507 | rs7595571 | -0.671 | 0.876 | -1.753 | -0.218 | -0.005 | -0.031 | -0.031 | 0.059 | 0.79 | 0.82 | 0.68 | 1 | 0.72 | 0.86 |
| 2 | 179240726 | rs12476608 | 0.236 | 0.779 | -1.795 | -0.259 | 0.016 | 0.049 | 0.049 | 0 | 0.25 | 0.15 | 0.14 | 0.12 | 0.22 | 0.28 |
| 2 | 179240924 | rs12468466 | 0.113 | 0.726 | -1.11 | -0.285 | -0.004 | -0.013 | -0.013 | 1.116 | 0.26 | 0.28 | 0.16 | 0.12 | 0.23 | 0.28 |
| 2 | 179241052 | rs12468511 | 0.207 | 0.76 | -1.469 | -0.268 | 0.017 | 0.05 | 0.05 | 0 | 0.26 | 0.15 | 0.14 | 0.12 | 0.22 | 0.28 |
| 2 | 179242526 | rs6706330 | 0.504 | 0.585 | -1.44 | -0.164 | -0.002 | 0.029 | 0.029 | 0 | 0.52 | 0.62 | 0.55 | 0.61 | 0.48 | 0.57 |
| 2 | 179243248 | rs6738749 | 0.212 | 0.738 | -1.282 | -0.058 | 0.006 | -0.006 | -0.006 | 0 | 0.66 | 0.63 | 0.59 | 0.72 | 0.55 | 0.75 |
| 2 | 179244560 | rs7565742 | -0.833 | 0.312 | -1.144 | -0.138 | -0.002 | -0.018 | -0.018 | 0 | 0.09 | 0.1 | 0.05 | 0 | 0.07 | 0.05 |
| 2 | 179244614 | rs1017397 | 0.169 | -0.021 | -1.318 | -0.266 | 0.027 | -0.013 | -0.013 | 0 | 0.38 | 0.29 | 0.6 | 0.62 | 0.35 | 0.5 |
| 2 | 179250840 | rs4404314 | 0.16 | -0.191 | -1.536 | -0.362 | 0.02 | -0.053 | -0.053 | 0.961 | 0.41 | 0.15 | 0.61 | 0.59 | 0.38 | 0.58 |
| 2 | 179251630 | rs10930830 | 1.316 | -0.654 | -1.694 | 0.213 | -0.053 | -0.118 | -0.118 | 0 | 0.25 | 0.03 | 0.39 | 0.41 | 0.27 | 0.23 |
| 2 | 179260382 | rs2304340 | -0.62 | -0.874 | -0.955 | -0.202 | 0.001 | -0.007 | -0.007 | 0.147 | 0.6 | 0.73 | 0.49 | 0.43 | 0.58 | 0.68 |
| 2 | 179263287 | rs890579 | -0.065 | -1.043 | -0.523 | -0.381 | 0 | 0 | 0 | 0 | 0.46 | 0.46 | 0.63 | 0.75 | 0.4 | 0.6 |
| 2 | 179269678 | rs2114581 | 0.359 | -1.025 | -0.587 | -0.375 | 0.03 | -0.029 | -0.029 | 2.978 | 0.42 | 0.2 | 0.61 | 0.6 | 0.38 | 0.58 |
| 2 | 179282337 | rs7589565 | 0.513 | -1.23 | -0.569 | 0.245 | 0.011 | -0.011 | -0.011 | 0 | 0.4 | 0.33 | 0.31 | 0.51 | 0.35 | 0.39 |
| 2 | 179284012 | rs4141635 | -2.885 | -1.271 | -0.181 | -0.174 | -0.001 | -0.021 | -0.021 | 0.262 | 0.81 | 0.84 | 0.86 | 1 | 0.82 | 0.96 |
| 2 | 179287834 | rs2114582 | -1.487 | -1.139 | -0.059 | 0.088 | -0.022 | 0.013 | 0.013 | 0 | 0.33 | 0.54 | 0.27 | 0.66 | 0.27 | 0.43 |
| 2 | 179288690 | rs4894020 | -0.984 | -1.117 | -0.025 | 0.124 | -0.035 | 0.074 | 0.074 | 0 | 0.26 | 0.48 | 0.23 | 0.66 | 0.21 | 0.37 |
| 2 | 179301884 | rs2059691 | -1.97 | -1.196 | -0.029 | 0.064 | 0 | -0.001 | -0.001 | 0.337 | 0.69 | 0.74 | 0.62 | 0.72 | 0.7 | 0.89 |
| 2 | 179319391 | rs6753709 | -1.075 | -1.132 | 0.117 | 0.406 | -0.034 | 0.077 | 0.077 | 0 | 0.26 | 0.48 | 0.23 | 0.65 | 0.21 | 0.37 |
| 2 | 179324025 | rs7600176 | -1.079 | -1.132 | 0.117 | 0.378 | -0.035 | 0.071 | 0.071 | 0 | 0.26 | 0.48 | 0.23 | 0.66 | 0.21 | 0.37 |
| 2 | 179333450 | rs7575643 | -0.446 | -1.069 | 0.206 | 0.535 | -0.007 | 0.026 | 0.026 | 0 | 0.21 | 0.21 | 0.2 | 0.63 | 0.18 | 0.36 |
| 2 | 179361881 | rs12052983 | -1.401 | -1.11 | -0.125 | 0.513 | -0.015 | 0.005 | 0.005 | 0 | 0.33 | 0.59 | 0.28 | 0.67 | 0.26 | 0.42 |
| 2 | 179366093 | rs2303537 | -1.588 | -1.011 | -0.608 | 0.544 | 0.001 | 0 | 0 | 0 | 0.37 | 0.65 | 0.43 | 0.7 | 0.3 | 0.62 |
| 2 | 179367560 | rs13026251 | 1.453 | -0.174 | -1.222 | 0.769 | -0.066 | 0.114 | 0.114 | 0 | 0.25 | 0.07 | 0.12 | 0.04 | 0.3 | 0.21 |
| 2 | 179368380 | rs16866352 | 0.726 | -0.144 | -0.544 | 1.607 | -0.026 | -0.005 | -0.005 | 0 | 0.07 | 0 | 0.13 | 0.48 | 0.1 | 0.19 |
| 2 | 179379190 | rs2288324 | 0.297 | -0.381 | -0.311 | 1.36 | -0.022 | 0.02 | 0.02 | 1.985 | 0.08 | 0.13 | 0.04 | 0.16 | 0.03 | 0.05 |
| 2 | 179388657 | rs13005601 | -0.253 | -0.475 | 0.06 | 1.564 | -0.001 | 0.064 | 0.064 | 0 | 0.13 | 0.18 | 0.16 | 0.44 | 0.14 | 0.26 |
| 2 | 179393111 | rs16866378 | 0.366 | -0.535 | 0.067 | 1.587 | -0.022 | 0.024 | 0.024 | 2.7 | 0.08 | 0.14 | 0.04 | 0.17 | 0.03 | 0.05 |

| | | | | | | | | | | | | | | | | |
|---|---|---|---|---|---|---|---|---|---|---|---|---|---|---|---|---|
| 2 | 179396354 | rs3829748 | -0.136 | -0.536 | 0.084 | 1.604 | 0 | 0.06 | 0.06 | 0.026 | 0.12 | 0.15 | 0.16 | 0.45 | 0.14 | 0.27 |
| 2 | 179398509 | rs3731752 | 0.695 | -0.556 | 0.098 | 1.583 | -0.004 | 0.012 | 0.012 | 1.017 | 0.07 | 0.08 | 0.04 | 0.17 | 0.03 | 0.05 |
| 2 | 179400895 | rs2278196 | 1.159 | -0.545 | 0.104 | 1.596 | -0.004 | 0.003 | 0.003 | 3.541 | 0.07 | 0.08 | 0.04 | 0.15 | 0.03 | 0.01 |
| 2 | 179413110 | rs3731748 | -0.137 | -0.609 | -0.032 | 1.483 | 0 | 0.062 | 0.062 | 0.018 | 0.12 | 0.15 | 0.16 | 0.45 | 0.14 | 0.28 |
| 2 | 179414318 | rs747122 | -1.038 | -0.767 | 0.718 | 1.407 | 0.006 | -0.015 | -0.015 | 0 | 0.08 | 0.27 | 0.18 | 0.02 | 0.05 | 0.2 |
| 2 | 179416556 | rs11897366 | -1.174 | -0.735 | 0.71 | 1.401 | 0.006 | -0.019 | -0.019 | 0.411 | 0.08 | 0.27 | 0.18 | 0.03 | 0.05 | 0.2 |
| 2 | 179430997 | rs3731746 | -1.302 | -0.607 | 0.849 | 1.579 | -0.028 | 0.082 | 0.082 | 0 | 0.21 | 0.34 | 0.21 | 0.63 | 0.17 | 0.32 |
| 2 | 179432185 | rs12463674 | 2.12 | -0.527 | 0.29 | 0.987 | -0.123 | 0.106 | 0.106 | 0 | 0.22 | 0.01 | 0.11 | 0.04 | 0.31 | 0.22 |
| 2 | 179435337 | rs6732060 | -0.751 | -0.493 | 0.405 | 0.998 | 0.005 | -0.019 | -0.019 | 0 | 0.07 | 0.24 | 0.18 | 0.03 | 0.05 | 0.2 |
| 2 | 179438866 | rs10164753 | 0.17 | -0.493 | 0.525 | 0.998 | -0.003 | -0.008 | -0.008 | 0 | 0.05 | 0.09 | 0.15 | 0.03 | 0.05 | 0.2 |
| 2 | 179440029 | rs12464787 | 2.102 | -0.542 | 0.797 | 0.972 | -0.059 | 0.082 | 0.082 | 0 | 0.23 | 0.08 | 0.12 | 0.04 | 0.31 | 0.21 |
| 2 | 179443540 | rs10497517 | -0.645 | -0.538 | 0.817 | 1.04 | 0.002 | -0.02 | -0.02 | 0 | 0.07 | 0.22 | 0.17 | 0.03 | 0.05 | 0.2 |
| 2 | 179444939 | rs2303838 | -1.194 | -0.649 | 0.761 | 1.051 | -0.03 | 0.069 | 0.069 | 0 | 0.22 | 0.34 | 0.21 | 0.63 | 0.18 | 0.34 |
| 2 | 179448911 | rs16866400 | -1.191 | -0.609 | 0.692 | 1.04 | -0.005 | 0.012 | 0.012 | 0 | 0.23 | 0.26 | 0.23 | 0.69 | 0.21 | 0.38 |
| 2 | 179451660 | rs1007025 | 0.171 | -0.635 | 0.814 | 1.087 | 0.002 | 0.122 | 0.122 | 0.018 | 0.13 | 0.22 | 0.17 | 0.46 | 0.14 | 0.28 |
| 2 | 179457147 | rs16866406 | 0.799 | -0.629 | 0.77 | 1.089 | -0.001 | 0.017 | 0.017 | 0.183 | 0.11 | 0.07 | 0.15 | 0.46 | 0.14 | 0.28 |
| 2 | 179461221 | rs4894030 | -1.956 | -0.727 | 0.772 | 0.263 | 0.013 | 0.002 | 0.002 | 0 | 0.69 | 0.75 | 0.66 | 0.77 | 0.75 | 0.89 |
| 2 | 179469386 | rs10497518 | -0.473 | -0.607 | 0.702 | 0.863 | -0.002 | 0.101 | 0.101 | 0 | 0.05 | 0.13 | 0.05 | 0.25 | 0.05 | 0.09 |
| 2 | 179490954 | rs16866418 | -0.421 | -0.645 | 0.739 | 0.774 | -0.026 | 0.082 | 0.082 | 0 | 0.22 | 0.32 | 0.21 | 0.6 | 0.18 | 0.34 |
| 2 | 179498042 | rs6723526 | 1.151 | -0.458 | 0.333 | 0.453 | -0.029 | 0.048 | 0.048 | 0.494 | 0.09 | 0.01 | 0.06 | 0.01 | 0.1 | 0.1 |
| 2 | 179528989 | rs2562848 | -1.812 | -0.281 | 0.437 | 0.898 | 0.001 | 0.047 | 0.047 | 0.659 | 0.12 | 0.5 | 0.23 | 0.13 | 0.08 | 0.27 |
| 2 | 179533609 | rs12998857 | 0.239 | -0.254 | 0.435 | 0.896 | 0.001 | -0.079 | -0.079 | 0.04 | 0.17 | 0.04 | 0.18 | 0.45 | 0.17 | 0.28 |
| 2 | 179539903 | rs2472751 | -1.61 | -0.268 | 0.288 | 0.899 | -0.021 | 0.062 | 0.062 | 0 | 0.28 | 0.57 | 0.4 | 0.58 | 0.23 | 0.52 |
| 2 | 179554305 | rs2244492 | 0.248 | -0.246 | 0.26 | 0.787 | 0.007 | -0.033 | -0.033 | 0 | 0.35 | 0.61 | 0.35 | 0.17 | 0.38 | 0.48 |
| 2 | 179558366 | rs2042995 | -1.693 | -0.283 | 0.593 | 1.47 | -0.016 | 0.047 | 0.047 | 0 | 0.29 | 0.59 | 0.42 | 0.57 | 0.25 | 0.52 |
| 2 | 179578704 | rs13398235 | -0.824 | -0.115 | 0.714 | 1.611 | -0.009 | -0.018 | -0.018 | 0 | 0.06 | 0.16 | 0.15 | 0.03 | 0.04 | 0.18 |
| 2 | 179578730 | rs2562839 | -1.787 | -0.325 | 0.97 | 1.754 | -0.022 | 0.051 | 0.051 | 0 | 0.25 | 0.46 | 0.38 | 0.66 | 0.22 | 0.47 |
| 2 | 179579093 | rs12693164 | -0.494 | -0.303 | 0.952 | 1.762 | -0.005 | 0.032 | 0.032 | 0 | 0.17 | 0.19 | 0.19 | 0.44 | 0.16 | 0.26 |
| 2 | 179580210 | rs10183237 | -0.658 | -0.291 | 0.942 | 1.752 | -0.009 | -0.018 | -0.018 | 0.363 | 0.06 | 0.16 | 0.15 | 0.03 | 0.04 | 0.18 |
| 2 | 179580583 | rs2562834 | -1.048 | -0.294 | 0.911 | 1.762 | -0.007 | 0.043 | 0.043 | 0 | 0.2 | 0.26 | 0.22 | 0.63 | 0.19 | 0.3 |
| 2 | 179582327 | rs13390491 | -0.427 | -0.263 | 0.896 | 1.776 | -0.004 | 0.034 | 0.034 | 0 | 0.17 | 0.19 | 0.18 | 0.44 | 0.16 | 0.25 |
| 2 | 179582537 | rs2627043 | -1.742 | -0.282 | 0.942 | 1.761 | -0.021 | 0.048 | 0.048 | 0 | 0.26 | 0.46 | 0.38 | 0.66 | 0.22 | 0.47 |
| 2 | 179583496 | rs16866465 | -0.32 | -0.242 | 0.789 | 1.772 | -0.002 | 0.03 | 0.03 | 0 | 0.16 | 0.17 | 0.18 | 0.44 | 0.16 | 0.25 |
| 2 | 179592724 | rs2627042 | -1.413 | -0.078 | 0.706 | 1.418 | -0.003 | 0.054 | 0.054 | 0 | 0.08 | 0.19 | 0.19 | 0.22 | 0.07 | 0.23 |
| 2 | 179615887 | rs922984 | 1.556 | -0.044 | 1.388 | 1.431 | 0.054 | 0.195 | 0.195 | 0.012 | 0.9 | 0.57 | 0.64 | 0.78 | 0.91 | 0.78 |
| 2 | 179620951 | rs7585334 | 1.041 | -0.029 | 1.34 | 1.44 | 0.01 | 0.063 | 0.063 | 0 | 0.92 | 0.83 | 0.67 | 0.78 | 0.91 | 0.78 |
| 2 | 179623758 | rs2291310 | 1.054 | -0.033 | 1.342 | 1.47 | 0.006 | 0.038 | 0.038 | 0 | 0.92 | 0.84 | 0.67 | 0.78 | 0.91 | 0.78 |
| 2 | 179629461 | rs2291311 | 1.054 | -0.033 | 1.345 | 1.469 | 0.008 | 0.047 | 0.047 | 0 | 0.92 | 0.83 | 0.67 | 0.78 | 0.91 | 0.78 |
| 2 | 179631214 | rs2291312 | -1.098 | -0.033 | 1.336 | 1.47 | -0.003 | 0.01 | 0.01 | 0 | 0.07 | 0.08 | 0.33 | 0.21 | 0.08 | 0.21 |
| 2 | 179632133 | rs4894044 | 0.348 | -0.288 | 1.572 | 1.677 | -0.028 | 0.106 | 0.106 | 0 | 0.83 | 0.71 | 0.84 | 0.54 | 0.86 | 0.77 |
| 2 | 179650701 | rs6715406 | 0.8 | -0.155 | 1.135 | 0.352 | -0.027 | 0.112 | 0.112 | 0.156 | 0.36 | 0.14 | 0.19 | 0.07 | 0.41 | 0.22 |
| 2 | 179650954 | rs6715901 | 0.387 | 0.15 | 1.044 | 0.307 | -0.048 | 0.116 | 0.116 | 0.51 | 0.4 | 0.15 | 0.39 | 0.13 | 0.51 | 0.27 |
| 2 | 179652047 | rs10497522 | -0.197 | 0.065 | 1.086 | 0.365 | -0.004 | 0.007 | 0.007 | 0 | 0.06 | 0.08 | 0.19 | 0.13 | 0.04 | 0.2 |
| 2 | 179652802 | rs10497523 | -0.634 | 0.078 | 1.195 | 0.439 | -0.001 | 0.136 | 0.136 | 0 | 0.53 | 0.73 | 0.69 | 0.86 | 0.56 | 0.68 |
| 2 | 179659912 | rs16866538 | -0.086 | 0.047 | 1.432 | 0.788 | 0.001 | 0.28 | 0.28 | 0.122 | 0.07 | 0.2 | 0.1 | 0.59 | 0.05 | 0.22 |
| 2 | 179680971 | rs12185628 | -0.009 | 0.13 | 0.129 | -0.127 | -0.007 | 0.015 | 0.015 | 3.119 | 0.21 | 0.14 | 0.15 | 0.15 | 0.22 | 0.29 |
| 2 | 179684839 | rs10497526 | 0.336 | 0.111 | 0.112 | -0.222 | -0.012 | 0.016 | 0.016 | 1.258 | 0.17 | 0.1 | 0.13 | 0.01 | 0.18 | 0.24 |
| 2 | 179690233 | rs6433735 | -0.044 | 0.058 | 0.188 | -1.134 | -0.008 | 0.042 | 0.042 | 0 | 0.55 | 0.63 | 0.68 | 0.86 | 0.54 | 0.64 |
| 2 | 179691670 | rs12469367 | 1.199 | -0.097 | 0.146 | -1.215 | -0.014 | 0.122 | 0.122 | 0.182 | 0.37 | 0.14 | 0.19 | 0.06 | 0.42 | 0.2 |
| 2 | 179693338 | rs7592323 | 1.363 | 0.564 | -0.022 | -1.34 | 0.024 | 0.09 | 0.09 | 0.789 | 0.33 | 0.14 | 0.15 | 0.06 | 0.31 | 0.17 |
| 2 | 179693775 | rs2366913 | 0.468 | 0.011 | 0.326 | -1.009 | 0.004 | 0.014 | 0.014 | 0 | 0.48 | 0.41 | 0.45 | 0.69 | 0.48 | 0.54 |
| 2 | 179695237 | rs11681211 | 1.598 | 0.22 | 0.231 | -0.905 | -0.012 | 0.074 | 0.074 | 0.276 | 0.23 | 0.11 | 0.15 | 0 | 0.22 | 0.07 |
| 2 | 179695728 | rs17453312 | 1.003 | 0.215 | 0.198 | -0.88 | 0 | 0.043 | 0.043 | 0 | 0.27 | 0.19 | 0.18 | 0.13 | 0.26 | 0.12 |
| 2 | 179698596 | rs10497527 | -0.235 | 0.149 | 0.226 | -0.787 | 0.02 | 0.003 | 0.003 | 0.036 | 0.08 | 0.12 | 0.22 | 0.07 | 0.1 | 0.09 |
| 2 | 179698945 | rs16866554 | 0.2 | -0.031 | 0.089 | -0.859 | -0.025 | 0.032 | 0.032 | 0 | 0.1 | 0.26 | 0.06 | 0.15 | 0.06 | 0.08 |
| 2 | 179710000 | rs17453396 | 0.314 | -0.015 | -0.267 | -0.912 | -0.045 | -0.022 | -0.022 | 0.104 | 0.08 | 0 | 0.19 | 0.07 | 0.09 | 0.06 |
| 2 | 179710706 | rs6433738 | -0.662 | -0.147 | -0.243 | -1.109 | 0 | -0.026 | -0.026 | 1.596 | 0.87 | 0.89 | 0.9 | 0.99 | 0.85 | 0.84 |
| 2 | 179710973 | rs726215 | 1.966 | -0.035 | -0.308 | -1.238 | -0.011 | 0.008 | 0.008 | 0.048 | 0.19 | 0.13 | 0.14 | 0 | 0.21 | 0.05 |
| 2 | 179713601 | rs4894054 | 0.379 | -0.069 | -0.284 | -1.154 | -0.002 | 0 | 0 | 0.436 | 0.81 | 0.83 | 0.71 | 0.92 | 0.77 | 0.76 |
| 2 | 179717217 | rs6711319 | 0.036 | -0.288 | -0.21 | -1.117 | -0.012 | -0.022 | -0.022 | 0.042 | 0.35 | 0.53 | 0.42 | 0.3 | 0.33 | 0.32 |
| 2 | 179729081 | rs6709708 | 1.904 | -0.415 | -0.597 | -1.284 | -0.077 | 0.195 | 0.195 | 0 | 0.36 | 0.06 | 0.18 | 0.06 | 0.38 | 0.23 |
| 2 | 179729341 | rs12465233 | -0.199 | -0.412 | -0.408 | -1.154 | 0.002 | -0.018 | -0.018 | 0 | 0.39 | 0.53 | 0.43 | 0.26 | 0.37 | 0.22 |
| 2 | 179732050 | rs10185678 | -0.363 | -0.667 | -0.278 | -1.424 | 0.003 | 0.005 | 0.005 | 0 | 0.4 | 0.59 | 0.63 | 0.77 | 0.33 | 0.55 |
| 2 | 179732845 | rs12988301 | 0.795 | -0.537 | -0.295 | -1.273 | -0.013 | 0.024 | 0.024 | 0 | 0.09 | 0.19 | 0.07 | 0.16 | 0.07 | 0.07 |
| 2 | 179737144 | rs12693170 | -0.337 | -0.544 | -0.367 | -1.786 | -0.006 | -0.091 | -0.091 | 0 | 0.66 | 0.43 | 0.73 | 0.98 | 0.68 | 0.79 |
| 2 | 179738269 | rs11693319 | 1.819 | -0.592 | -0.246 | -1.765 | -0.015 | -0.087 | -0.087 | 0 | 0.2 | 0.28 | 0.15 | 0.01 | 0.16 | 0.07 |
| 2 | 179739483 | rs13405116 | -0.462 | -0.551 | -0.4 | -1.717 | 0.02 | 0.026 | 0.026 | 0 | 0.4 | 0.62 | 0.49 | 0.86 | 0.33 | 0.56 |
| 2 | 179742232 | rs7565966 | -0.54 | -0.654 | -0.354 | -1.611 | 0.012 | 0.078 | 0.078 | 0 | 0.43 | 0.7 | 0.49 | 0.8 | 0.38 | 0.45 |
| 2 | 179753549 | rs1873164 | 0.423 | -0.551 | -0.301 | -1.247 | -0.01 | 0.08 | 0.08 | 0 | 0.8 | 0.63 | 0.8 | 0.3 | 0.79 | 0.64 |

| | | | | | | | | | | | | | | | | |
|---|---|---|---|---|---|---|---|---|---|---|---|---|---|---|---|---|
| 2 | 179758007 | rs1032282 | -0.692 | -0.445 | -0.536 | -1.374 | -0.015 | -0.088 | -0.088 | 1.749 | 0.85 | 0.61 | 0.87 | 0.98 | 0.86 | 0.89 |
| 2 | 179759434 | rs6747402 | 0.137 | -0.268 | -0.644 | -1.369 | -0.016 | 0.114 | 0.114 | 0 | 0.53 | 0.23 | 0.6 | 0.24 | 0.48 | 0.55 |
| 2 | 179759810 | rs6750760 | -0.49 | -0.079 | -0.645 | -1.143 | 0 | -0.009 | -0.009 | 0 | 0.66 | 0.55 | 0.69 | 0.27 | 0.6 | 0.62 |
| 2 | 179775893 | rs10179812 | -0.443 | -0.227 | -1.028 | -0.962 | -0.012 | -0.041 | -0.041 | 0 | 0.11 | 0.05 | 0.38 | 0.19 | 0.09 | 0.33 |
| 2 | 179793615 | rs10194023 | -0.568 | -0.203 | -0.988 | -1.249 | -0.003 | 0.008 | 0.008 | 0 | 0.59 | 0.59 | 0.72 | 0.78 | 0.59 | 0.68 |
| 2 | 179794447 | rs10171173 | 0.331 | -0.269 | -1.037 | -1.333 | -0.002 | -0.003 | -0.003 | 0 | 0.39 | 0.39 | 0.61 | 0.75 | 0.4 | 0.6 |
| 2 | 179801042 | rs6752606 | -1.186 | -0.227 | -0.985 | -1.332 | -0.003 | 0.018 | 0.018 | 0 | 0.67 | 0.72 | 0.81 | 0.94 | 0.67 | 0.78 |
| 2 | 179817206 | rs7561438 | -0.465 | -0.027 | -1.639 | -1.229 | 0.002 | -0.005 | -0.005 | 0 | 0.29 | 0.33 | 0.2 | 0.19 | 0.26 | 0.16 |
| 2 | 179837050 | rs6747725 | -0.165 | 0.028 | -1.549 | -0.969 | 0 | 0 | 0 | 0 | 0.43 | 0.56 | 0.64 | 0.75 | 0.41 | 0.62 |
| 2 | 179839555 | rs13019119 | 0.028 | 0.59 | -1.922 | -0.343 | -0.002 | 0 | 0 | 0 | 0.18 | 0.14 | 0.1 | 0.03 | 0.18 | 0.07 |
| 2 | 179839833 | rs10930846 | 0.237 | 0.718 | -1.9 | 0.413 | -0.026 | -0.057 | -0.057 | 0 | 0.23 | 0.06 | 0.34 | 0.7 | 0.27 | 0.33 |
| 2 | 179860406 | rs12992925 | -0.117 | 0.077 | -1.845 | 0.229 | -0.008 | 0.002 | 0.002 | 0.461 | 0.37 | 0.13 | 0.44 | 0.66 | 0.56 | 0.38 |
| 2 | 179860562 | rs13019215 | -0.407 | -0.432 | -1.179 | -0.083 | 0.033 | 0.031 | 0.031 | 0.371 | 0.46 | 0.31 | 0.69 | 0.88 | 0.65 | 0.48 |
| 2 | 179861233 | rs7564269 | -0.334 | -1.238 | -0.467 | -0.462 | -0.031 | -0.018 | -0.018 | 0 | 0.89 | 0.62 | 0.91 | 0.93 | 0.93 | 0.73 |
| 2 | 179866426 | rs17453612 | 0.829 | -1.992 | -0.785 | -0.937 | 0.067 | 0.078 | 0.078 | 0 | 0.29 | 0.19 | 0.15 | 0.05 | 0.17 | 0.22 |
| 2 | 179867426 | rs4893862 | 1.385 | -1.348 | -0.997 | -0.87 | -0.066 | -0.041 | -0.041 | 0 | 0.29 | 0.12 | 0.37 | 0.63 | 0.44 | 0.34 |
| 2 | 179871375 | rs4243393 | -0.119 | -2.126 | -0.648 | -0.892 | -0.056 | -0.054 | -0.054 | 0 | 0.65 | 0.43 | 0.79 | 0.8 | 0.78 | 0.55 |
| 2 | 179873140 | rs2886806 | 0.715 | -2.474 | -0.627 | -0.935 | 0.153 | 0.08 | 0.08 | 0 | 0.51 | 0.21 | 0.42 | 0.66 | 0.72 | 0.49 |
| 2 | 179873673 | rs6743556 | 0.491 | -2.674 | -0.076 | -1.047 | -0.048 | -0.021 | -0.021 | 0 | 0.76 | 0.56 | 0.68 | 0.8 | 0.83 | 0.68 |
| 2 | 179882262 | rs6433746 | -0.214 | -2.709 | -0.039 | -1.087 | -0.015 | -0.044 | -0.044 | 0 | 0.84 | 0.72 | 0.89 | 0.93 | 0.88 | 0.72 |
| 2 | 179884886 | rs12474600 | 1.462 | -2.837 | -0.046 | -1.132 | 0.02 | 0.006 | 0.006 | 0 | 0.18 | 0.13 | 0.23 | 0.14 | 0.1 | 0.17 |
| 2 | 179900720 | rs7581299 | 1.407 | -3.047 | -0.1 | -1.3 | 0.067 | 0.052 | 0.052 | 0 | 0.46 | 0.25 | 0.34 | 0.65 | 0.63 | 0.44 |
| 2 | 179900746 | rs4894075 | -0.667 | -3.035 | -0.067 | -1.294 | -0.024 | -0.015 | -0.015 | 0 | 0.19 | 0.28 | 0.3 | 0.13 | 0.16 | 0.12 |
| 2 | 179909467 | rs17362972 | 1.764 | -2.999 | -0.16 | -1.299 | 0.033 | -0.026 | -0.026 | 0 | 0.17 | 0.01 | 0.22 | 0.14 | 0.1 | 0.16 |
| 2 | 179915438 | rs340332 | 0.786 | -2.941 | -0.082 | -1.643 | -0.007 | -0.075 | -0.075 | 0.067 | 0.77 | 0.62 | 0.79 | 0.92 | 0.79 | 0.65 |
| 2 | 179923685 | rs6709090 | 1.375 | -3.05 | 0.164 | -1.565 | -0.014 | -0.036 | -0.036 | 0 | 0.18 | 0.23 | 0.09 | 0.02 | 0.1 | 0.16 |
| 2 | 179924203 | rs10497530 | 1.009 | -3.074 | 0.19 | -1.578 | -0.048 | -0.09 | -0.09 | 0.048 | 0.2 | 0.37 | 0.11 | 0.02 | 0.1 | 0.16 |
| 2 | 179932785 | rs340348 | 1.132 | -3.216 | 0.092 | -1.62 | 0.191 | 0.254 | 0.254 | 0.194 | 0.52 | 0.14 | 0.74 | 0.83 | 0.73 | 0.54 |
| 2 | 179933412 | rs340349 | -0.33 | -3.205 | 0.272 | -1.614 | -0.027 | -0.047 | -0.047 | 0 | 0.87 | 0.75 | 0.94 | 0.93 | 0.92 | 0.74 |
| 2 | 179934792 | rs340350 | 0.916 | -3.17 | 0.096 | -1.597 | -0.074 | -0.131 | -0.131 | 0.644 | 0.71 | 0.58 | 0.85 | 0.93 | 0.83 | 0.7 |
| 2 | 179935619 | rs168158 | 1.319 | -3.142 | -0.114 | -1.565 | 0.133 | 0.186 | 0.186 | 0.092 | 0.48 | 0.13 | 0.53 | 0.77 | 0.66 | 0.47 |
| 2 | 179939493 | rs2044480 | 1.691 | -2.965 | -0.258 | -1.636 | -0.029 | 0.031 | 0.031 | 0.122 | 0.06 | 0 | 0.04 | 0 | 0.09 | 0.01 |
| 2 | 179939876 | rs11897059 | 1.424 | -3.037 | -0.183 | -1.67 | 0.017 | 0.008 | 0.008 | 0 | 0.18 | 0.13 | 0.08 | 0.02 | 0.1 | 0.16 |
| 2 | 179939959 | rs340367 | 0.915 | -3.058 | -0.463 | -1.677 | 0.14 | 0.16 | 0.16 | 0.397 | 0.52 | 0.17 | 0.58 | 0.77 | 0.7 | 0.47 |
| 2 | 179966804 | rs1968618 | 1.154 | -2.773 | -0.377 | -1.747 | -0.001 | -0.012 | -0.012 | 0 | 0.68 | 0.33 | 0.68 | 0.8 | 0.66 | 0.83 |
| 2 | 179970037 | rs3731739 | 0.342 | -2.714 | -0.225 | -1.672 | 0.057 | 0.048 | 0.048 | 0.021 | 0.08 | 0.39 | 0.1 | 0.06 | 0.09 | 0.1 |
| 2 | 179977252 | rs2289994 | -0.327 | -2.759 | -0.49 | -1.752 | -0.036 | -0.074 | -0.074 | 0 | 0.22 | 0.29 | 0.11 | 0.06 | 0.17 | 0.29 |
| 2 | 179980070 | rs2044479 | -2.006 | -2.785 | -0.646 | -1.637 | -0.017 | 0.028 | 0.028 | 0.03 | 0.62 | 0.9 | 0.44 | 0.3 | 0.49 | 0.61 |
| 2 | 179983572 | rs6743405 | -1.358 | -2.759 | -0.507 | -1.574 | 0.111 | 0.169 | 0.169 | 0.061 | 0.52 | 0.84 | 0.32 | 0.18 | 0.36 | 0.55 |
| 2 | 179992164 | rs17363260 | 0.239 | -2.755 | -0.446 | -1.548 | -0.009 | -0.019 | -0.019 | 2.476 | 0.13 | 0.16 | 0.07 | 0.06 | 0.11 | 0.26 |
| 2 | 180030225 | rs6433764 | -1.388 | -2.669 | -0.701 | -1.516 | 0.109 | 0.171 | 0.171 | 0.089 | 0.53 | 0.86 | 0.33 | 0.19 | 0.36 | 0.54 |
| 2 | 180040551 | rs4894081 | 0.732 | -2.614 | -0.616 | -1.519 | -0.002 | -0.01 | -0.01 | 0.577 | 0.68 | 0.32 | 0.68 | 0.8 | 0.66 | 0.81 |
| 2 | 180045213 | rs873023 | -0.744 | -2.531 | -0.501 | -1.478 | 0 | -0.005 | -0.005 | 0 | 0.82 | 0.84 | 0.83 | 0.91 | 0.79 | 0.93 |
| 2 | 180051514 | rs10930856 | 0.245 | -2.66 | -0.515 | -1.541 | -0.011 | -0.021 | -0.021 | 0 | 0.13 | 0.16 | 0.07 | 0.06 | 0.11 | 0.26 |
| 2 | 180056341 | rs12693178 | 0.748 | -2.586 | -0.638 | -1.531 | 0.027 | 0.053 | 0.053 | 0 | 0.52 | 0.31 | 0.62 | 0.75 | 0.57 | 0.63 |
| 2 | 180088968 | rs10208593 | -0.398 | -2.518 | 0.006 | -1.309 | -0.037 | -0.071 | -0.071 | 0 | 0.21 | 0.29 | 0.13 | 0.06 | 0.16 | 0.25 |
| 2 | 180089983 | rs12987285 | -0.521 | -2.148 | -0.095 | -1.12 | -0.051 | -0.072 | -0.072 | 0 | 0.39 | 0.42 | 0.2 | 0.13 | 0.23 | 0.41 |
| 2 | 180096561 | rs10174774 | -0.943 | -1.846 | -0.8 | -0.911 | 0.12 | 0.17 | 0.17 | 0.043 | 0.52 | 0.84 | 0.32 | 0.19 | 0.36 | 0.57 |
| 2 | 180115065 | rs6707081 | -0.971 | -1.834 | -0.816 | -0.905 | 0.111 | 0.155 | 0.155 | 0.099 | 0.52 | 0.83 | 0.32 | 0.19 | 0.36 | 0.57 |
| 2 | 180137861 | rs10172996 | -0.933 | -0.952 | -0.118 | -0.313 | -0.02 | -0.022 | -0.022 | 0 | 0.19 | 0.55 | 0.17 | 0.11 | 0.16 | 0.3 |
| 2 | 180139783 | rs10179379 | -0.086 | -0.566 | -0.721 | -0.104 | 0 | -0.001 | -0.001 | 0 | 0.5 | 0.51 | 0.69 | 0.54 | 0.56 | 0.42 |
| 2 | 180140077 | rs13423876 | 1.408 | 0.583 | -1.358 | -0.604 | -0.021 | 0.064 | 0.064 | 0 | 0.19 | 0.06 | 0.13 | 0.07 | 0.19 | 0.1 |
| 2 | 180140151 | rs2087642 | -0.802 | 0.89 | -0.558 | -0.617 | 0.039 | 0.012 | 0.012 | 2.037 | 0.16 | 0.53 | 0.19 | 0.19 | 0.17 | 0.09 |
| 2 | 180141614 | rs6742175 | -0.109 | 0.713 | -1.045 | -0.933 | 0.002 | -0.002 | -0.002 | 0 | 0.72 | 0.72 | 0.48 | 0.36 | 0.64 | 0.49 |
| 2 | 180149795 | rs6749058 | -0.112 | 0.564 | -1.345 | -1.215 | -0.008 | -0.037 | -0.037 | 0.606 | 0.39 | 0.56 | 0.26 | 0.27 | 0.3 | 0.26 |
| 2 | 180150435 | rs1388182 | 0.349 | 0.475 | -0.908 | -1.163 | 0.065 | -0.013 | -0.013 | 0 | 0.08 | 0.37 | 0.05 | 0 | 0.05 | 0.02 |
| 2 | 180151555 | rs13425924 | 1.889 | 0.431 | -1.011 | -1.208 | -0.053 | 0.133 | 0.133 | 0 | 0.3 | 0.07 | 0.21 | 0.13 | 0.34 | 0.24 |
| 2 | 180153772 | rs16866687 | 0.113 | 0.446 | -0.91 | -1.208 | 0.08 | -0.001 | -0.001 | 0 | 0.07 | 0.39 | 0.06 | 0 | 0.05 | 0.02 |
| 2 | 180156048 | rs6752206 | -0.08 | 0.47 | -0.938 | -1.027 | -0.011 | 0.09 | 0.09 | 0.964 | 0.76 | 0.51 | 0.66 | 0.45 | 0.72 | 0.55 |
| 2 | 180157228 | rs7565599 | -0.267 | 0.45 | -0.943 | -1.122 | -0.003 | 0.035 | 0.035 | 0 | 0.44 | 0.28 | 0.44 | 0.37 | 0.39 | 0.31 |
| 2 | 180158041 | rs6433772 | 0.093 | 0.505 | -1.038 | -1.091 | -0.018 | 0.059 | 0.059 | 1.153 | 0.73 | 0.49 | 0.64 | 0.45 | 0.71 | 0.54 |
| 2 | 180158510 | rs6761161 | 1.968 | 0.544 | -1.585 | -0.999 | -0.026 | 0.111 | 0.111 | 0 | 0.3 | 0.14 | 0.21 | 0.09 | 0.33 | 0.24 |
| 2 | 180159285 | rs7559258 | 0.711 | 0.342 | -1.567 | -0.89 | -0.01 | -0.002 | -0.002 | 0 | 0.12 | 0.1 | 0.12 | 0.18 | 0.15 | 0.08 |
| 2 | 180159428 | rs16866690 | -0.742 | 0.272 | -1.788 | -0.751 | -0.009 | 0.038 | 0.038 | 0 | 0.47 | 0.66 | 0.47 | 0.37 | 0.41 | 0.32 |
| 2 | 180164474 | rs7558234 | 0.711 | 0.339 | -1.825 | -0.681 | -0.002 | 0.076 | 0.076 | 0 | 0.51 | 0.29 | 0.51 | 0.63 | 0.57 | 0.67 |
| 2 | 180168421 | rs6722892 | -0.338 | 0.259 | -1.263 | -0.352 | -0.001 | -0.037 | -0.037 | 0 | 0.26 | 0.54 | 0.2 | 0.2 | 0.2 | 0.14 |
| 2 | 180169014 | rs4894088 | 0.427 | 0.732 | -1.442 | 0.026 | -0.025 | -0.029 | -0.029 | 0.37 | 0.05 | 0 | 0.16 | 0.1 | 0.07 | 0.05 |
| 2 | 180169489 | rs1565521 | -0.225 | 0.804 | -1.465 | -0.095 | 0.009 | 0.018 | 0.018 | 1.831 | 0.79 | 0.76 | 0.67 | 0.35 | 0.69 | 0.53 |
| 2 | 180172781 | rs6735828 | 0.918 | 0.443 | -1.652 | -0.374 | 0 | -0.024 | -0.024 | 0 | 0.55 | 0.52 | 0.36 | 0.16 | 0.44 | 0.36 |

| | | | | | | | | | | | | | | | | |
|---|---|---|---|---|---|---|---|---|---|---|---|---|---|---|---|---|
| 2 | 180173051 | rs13002273 | -0.236 | 0.399 | -1.852 | -0.605 | 0.004 | -0.058 | -0.058 | 0.821 | 0.2 | 0.22 | 0.09 | 0 | 0.17 | 0.1 |
| 2 | 180173172 | rs12693185 | -0.554 | 0.596 | -1.526 | -0.126 | -0.007 | -0.008 | -0.008 | 0 | 0.8 | 0.82 | 0.67 | 0.38 | 0.67 | 0.55 |
| 2 | 180173318 | rs4365493 | 0.381 | 0.617 | -1.51 | -0.114 | 0.032 | 0.01 | 0.01 | 2.111 | 0.68 | 0.6 | 0.59 | 0.39 | 0.53 | 0.51 |
| 2 | 180173829 | rs13035089 | -0.101 | 0.462 | -1.061 | 0.128 | 0.019 | -0.066 | -0.066 | 0.421 | 0.14 | 0.22 | 0.09 | 0 | 0.15 | 0.05 |
| 2 | 180177128 | rs10185366 | 0.734 | 0.623 | -1.096 | 0.836 | 0.002 | 0.097 | 0.097 | 0.018 | 0.07 | 0.12 | 0.04 | 0.27 | 0.09 | 0.18 |
| 2 | 180177796 | rs10497537 | 0.825 | 0.524 | -1.088 | 0.843 | -0.005 | 0.102 | 0.102 | 0.003 | 0.07 | 0.12 | 0.04 | 0.27 | 0.08 | 0.17 |
| 2 | 180192356 | rs12619980 | 1.065 | 0.311 | -1.122 | 0.647 | 0.004 | -0.001 | -0.001 | 0 | 0.37 | 0.42 | 0.48 | 0.4 | 0.41 | 0.37 |
| 2 | 180192663 | rs7606501 | 0.631 | -0.421 | -1.61 | 1.107 | -0.002 | 0.086 | 0.086 | 0 | 0.29 | 0.13 | 0.17 | 0.08 | 0.27 | 0.25 |
| 2 | 180196356 | rs7591863 | 0.198 | -0.423 | -1.521 | 1.13 | 0.004 | 0.002 | 0.002 | 0 | 0.35 | 0.21 | 0.37 | 0.33 | 0.31 | 0.31 |
| 2 | 180198983 | rs6726305 | 0.228 | -0.1 | -1.237 | 0.634 | 0.013 | 0.026 | 0.026 | 0 | 0.48 | 0.61 | 0.55 | 0.39 | 0.55 | 0.42 |
| 2 | 180201455 | rs10930865 | -0.228 | 0.122 | -1.172 | 0.327 | -0.017 | 0.006 | 0.006 | 0 | 0.89 | 0.84 | 0.93 | 0.85 | 0.94 | 0.91 |
| 2 | 180202305 | rs7582128 | -0.053 | 0.618 | -1.254 | 0.017 | 0.026 | 0.096 | 0.096 | 0 | 0.5 | 0.33 | 0.59 | 0.71 | 0.45 | 0.59 |
| 2 | 180206856 | rs6751891 | 0.526 | 1.031 | -0.639 | 0.022 | 0.026 | -0.039 | -0.039 | 0 | 0.79 | 0.64 | 0.82 | 0.87 | 0.76 | 0.9 |
| 2 | 180209323 | rs7608378 | 0.418 | 1.089 | -0.785 | 0.057 | 0.008 | 0.006 | 0.006 | 0 | 0.06 | 0.15 | 0.07 | 0.06 | 0.05 | 0.05 |
| 2 | 180211052 | rs4894095 | 0.632 | 0.741 | -1.992 | 0.209 | -0.134 | -0.11 | -0.11 | 0.416 | 0.44 | 0.05 | 0.48 | 0.49 | 0.46 | 0.53 |
| 2 | 180215310 | rs6433781 | -0.318 | 0.838 | -1.21 | 0.286 | -0.001 | 0.008 | 0.008 | 0 | 0.9 | 0.82 | 0.92 | 0.88 | 0.91 | 0.91 |
| 2 | 180216013 | rs13021289 | 0.024 | 1.546 | -1.154 | 0.09 | 0.038 | -0.03 | -0.03 | 0 | 0.08 | 0.09 | 0.11 | 0 | 0.2 | 0.03 |
| 2 | 180216731 | rs13403289 | 0.773 | 1.108 | -1.533 | 0.186 | -0.023 | -0.032 | -0.032 | 0 | 0.6 | 0.17 | 0.55 | 0.56 | 0.6 | 0.6 |
| 2 | 180217735 | rs6759773 | 0.007 | 1.215 | -1.169 | 0.288 | -0.002 | -0.006 | -0.006 | 0 | 0.9 | 0.85 | 0.93 | 0.9 | 0.91 | 0.92 |
| 2 | 180220868 | rs7425027 | -0.058 | 1.328 | -1.648 | 0.392 | -0.066 | -0.047 | -0.047 | 0 | 0.26 | 0.1 | 0.54 | 0.61 | 0.31 | 0.35 |
| 2 | 180225657 | rs12328936 | -0.237 | 0.706 | -0.857 | -0.28 | -0.003 | -0.032 | -0.032 | 0 | 0.12 | 0.09 | 0.05 | 0 | 0.08 | 0.01 |
| 2 | 180233627 | rs7586173 | 0.803 | 1.308 | -1.503 | -0.974 | -0.012 | 0.01 | 0.01 | 0 | 0.68 | 0.73 | 0.76 | 0.95 | 0.62 | 0.85 |
| 2 | 180239030 | rs10175322 | 0.544 | 0.807 | -0.907 | -1.033 | 0.007 | -0.001 | -0.001 | 0 | 0.26 | 0.24 | 0.2 | 0.27 | 0.34 | 0.31 |
| 2 | 180240770 | rs13011836 | 1.334 | 1.277 | -1.404 | -1.21 | -0.071 | 0.093 | 0.093 | 0 | 0.16 | 0.01 | 0.11 | 0.04 | 0.23 | 0.18 |
| 2 | 180244477 | rs964327 | 0.937 | 1.304 | -1.414 | -1.285 | -0.062 | 0.077 | 0.077 | 0 | 0.18 | 0.06 | 0.16 | 0.01 | 0.29 | 0.12 |
| 2 | 180247946 | rs12471246 | 1.756 | 0.819 | -1.396 | -1.538 | 0.041 | 0.006 | 0.006 | 0 | 0.34 | 0.27 | 0.5 | 0.68 | 0.22 | 0.52 |
| 2 | 180258493 | rs1850523 | 1.47 | 1.135 | -1.555 | -1.327 | -0.014 | -0.006 | -0.006 | 0 | 0.41 | 0.47 | 0.57 | 0.68 | 0.28 | 0.54 |
| 2 | 180266506 | rs6713554 | 0.982 | 1.25 | -1.107 | -0.76 | 0.01 | -0.014 | -0.014 | 0.943 | 0.59 | 0.53 | 0.73 | 0.68 | 0.57 | 0.65 |
| 2 | 180269147 | rs2037241 | 0.374 | 1.543 | -1.588 | -0.417 | 0.006 | 0.029 | 0.029 | 0 | 0.09 | 0.09 | 0.07 | 0.23 | 0.1 | 0.13 |
| 2 | 180272486 | rs1022360 | -0.105 | 1.292 | -1.521 | -0.36 | -0.051 | -0.007 | -0.007 | 0 | 0.25 | 0.11 | 0.19 | 0.27 | 0.34 | 0.33 |
| 2 | 180287156 | rs3942366 | -0.309 | 0.656 | -1.622 | 0.135 | 0.011 | -0.038 | -0.038 | 0 | 0.34 | 0.17 | 0.41 | 0.52 | 0.27 | 0.27 |
| 2 | 180289739 | rs9808219 | 0.385 | 0.625 | -0.984 | 0.074 | 0.033 | 0.033 | 0.033 | 0 | 0.07 | 0.16 | 0.07 | 0.09 | 0.09 | 0.06 |
| 2 | 180292467 | rs995011 | 0.069 | 0.388 | -0.628 | 0.093 | 0.007 | 0.014 | 0.014 | 0.004 | 0.13 | 0.27 | 0.11 | 0.15 | 0.1 | 0.09 |
| 2 | 180293852 | rs16866726 | 0.831 | -0.043 | -1.054 | 0.805 | 0.015 | -0.025 | -0.025 | 1.318 | 0.09 | 0.02 | 0.26 | 0.39 | 0.06 | 0.18 |
| 2 | 180297960 | rs1554261 | -0.752 | 0.015 | -1.022 | 0.137 | 0.011 | -0.01 | -0.01 | 0 | 0.79 | 0.73 | 0.72 | 0.81 | 0.75 | 0.81 |
| 2 | 180298412 | rs10179194 | 0.788 | -0.622 | -1.357 | -0.13 | 0.019 | 0.014 | 0.014 | 2.172 | 0.28 | 0.21 | 0.11 | 0.22 | 0.17 | 0.35 |
| 2 | 180298424 | rs10497539 | 1.831 | -0.814 | -1.393 | -0.235 | 0 | -0.001 | -0.001 | 0.014 | 0.13 | 0.13 | 0.06 | 0.23 | 0.11 | 0.31 |
| 2 | 180303911 | rs12998067 | 0.132 | -0.649 | -1.37 | -0.151 | 0.001 | 0.001 | 0.001 | 0 | 0.52 | 0.52 | 0.62 | 0.58 | 0.58 | 0.46 |
| 2 | 180305196 | rs10930874 | 0.432 | -0.757 | -1.602 | -0.169 | 0.02 | 0.014 | 0.014 | 1.943 | 0.28 | 0.21 | 0.11 | 0.24 | 0.17 | 0.36 |
| 2 | 180305369 | rs34383794 | 1.439 | -0.061 | -1.988 | -0.275 | -0.105 | 0.067 | 0.067 | 0 | 0.2 | 0.03 | 0.19 | 0.03 | 0.32 | 0.1 |
| 2 | 180307783 | rs3749123 | 1.14 | -0.324 | -1.901 | 0.008 | 0.016 | -0.146 | -0.146 | 0.384 | 0.18 | 0.02 | 0.3 | 0.45 | 0.12 | 0.28 |
| 2 | 180311207 | rs2271760 | 1.791 | -0.446 | -1.858 | 0.229 | 0.011 | -0.013 | -0.013 | 0 | 0.12 | 0.08 | 0.2 | 0.33 | 0.11 | 0.27 |
| 2 | 180311444 | rs2271761 | 2.776 | -0.688 | -1.499 | 0.065 | 0.007 | 0.051 | 0.051 | 1.073 | 0.05 | 0.2 | 0.02 | 0.02 | 0.02 | 0.03 |
| 2 | 180313995 | rs6757845 | -1.288 | -0.645 | -1.6 | -0.275 | -0.033 | 0.037 | 0.037 | 0 | 0.73 | 0.8 | 0.64 | 0.92 | 0.59 | 0.84 |
| 2 | 180316070 | rs10192549 | 1.881 | -0.618 | -1.474 | -0.31 | 0.01 | -0.006 | -0.006 | 0 | 0.12 | 0.15 | 0.1 | 0.08 | 0.15 | 0.11 |
| 2 | 180316107 | rs16866735 | 1.903 | -0.723 | -1.581 | -0.293 | 0.008 | -0.103 | -0.103 | 0 | 0.16 | 0.02 | 0.16 | 0.34 | 0.12 | 0.26 |
| 2 | 180318028 | rs16866737 | 2.438 | -0.877 | -1.506 | -0.523 | 0.033 | 0.027 | 0.027 | 3.845 | 0.16 | 0.05 | 0.04 | 0 | 0.07 | 0.03 |
| 2 | 180318192 | rs9288032 | 0.626 | -0.661 | -1.358 | -0.423 | 0.041 | -0.009 | -0.009 | 0 | 0.25 | 0.42 | 0.2 | 0.23 | 0.28 | 0.25 |
| 2 | 180318474 | rs1517706 | -0.519 | -0.116 | -1.249 | -0.065 | -0.041 | 0.061 | 0.061 | 0 | 0.69 | 0.8 | 0.63 | 0.89 | 0.57 | 0.81 |
| 2 | 180324063 | rs1914498 | 0.798 | 0.427 | -0.862 | 0.456 | 0.04 | 0.018 | 0.018 | 0 | 0.5 | 0.31 | 0.36 | 0.43 | 0.4 | 0.42 |
| 2 | 180329583 | rs13024531 | 0.204 | 0.738 | -1.041 | 0.828 | 0.001 | 0.078 | 0.078 | 0.115 | 0.25 | 0.39 | 0.17 | 0.46 | 0.24 | 0.4 |
| 2 | 180333450 | rs1517708 | -1.058 | 1.612 | -1.29 | 0.991 | 0.039 | -0.019 | -0.019 | 0 | 0.88 | 0.8 | 0.86 | 0.93 | 0.76 | 0.93 |
| 2 | 180335135 | rs1581253 | -0.363 | 1.068 | -1.587 | 1.125 | 0.063 | -0.016 | -0.016 | 0.166 | 0.72 | 0.56 | 0.76 | 0.76 | 0.63 | 0.78 |
| 2 | 180335921 | rs1028179 | -0.351 | 0.762 | -2.135 | 1.514 | -0.043 | -0.11 | -0.11 | 0 | 0.21 | 0.02 | 0.33 | 0.36 | 0.19 | 0.29 |
| 2 | 180337809 | rs7588860 | -1.223 | 0.353 | -1.673 | 1.283 | -0.001 | 0.014 | 0.014 | 0 | 0.09 | 0.11 | 0.25 | 0.36 | 0.07 | 0.24 |
| 2 | 180337821 | rs12473469 | -1.154 | 0.63 | -2.104 | 0.342 | -0.053 | -0.013 | -0.013 | 0 | 0.17 | 0.05 | 0.16 | 0.18 | 0.22 | 0.21 |
| 2 | 180338995 | rs16866746 | 0.502 | -0.163 | -2.015 | 0.217 | 0.012 | 0 | 0 | 0 | 0.07 | 0.05 | 0.13 | 0.38 | 0.04 | 0.23 |
| 2 | 180342077 | rs12990747 | 0.844 | -0.094 | -2.047 | -0.508 | -0.01 | -0.014 | -0.014 | 0.038 | 0.2 | 0.14 | 0.48 | 0.5 | 0.24 | 0.31 |
| 2 | 180342763 | rs17803813 | 0.67 | -0.555 | -2.437 | -1.163 | 0.01 | 0.072 | 0.072 | 0.019 | 0.21 | 0.09 | 0.14 | 0.02 | 0.18 | 0.22 |
| 2 | 180342873 | rs6718486 | 0.974 | -0.777 | -2.011 | -1.575 | 0.002 | 0.005 | 0.005 | 0.058 | 0.25 | 0.28 | 0.17 | 0.42 | 0.28 | 0.27 |
| 2 | 180344210 | rs12052861 | -1.337 | -0.956 | -1.682 | -1.751 | -0.014 | -0.05 | -0.05 | 0 | 0.86 | 0.76 | 0.9 | 0.94 | 0.86 | 0.91 |
| 2 | 180352334 | rs17747468 | 0.639 | -0.787 | -1.681 | -1.669 | 0.008 | 0.047 | 0.047 | 0 | 0.18 | 0.09 | 0.08 | 0 | 0.16 | 0.1 |
| 2 | 180354642 | rs17747577 | -0.228 | -0.853 | -1.657 | -1.718 | 0.025 | 0.153 | 0.153 | 0 | 0.38 | 0.19 | 0.23 | 0.02 | 0.34 | 0.33 |
| 2 | 180355308 | rs11686958 | -0.179 | -0.846 | -1.152 | -1.24 | 0 | -0.001 | -0.001 | 0 | 0.6 | 0.43 | 0.72 | 0.52 | 0.57 | 0.61 |
| 2 | 180370044 | rs9288035 | 0.303 | -1.182 | -1.29 | -1.391 | -0.011 | -0.089 | -0.089 | 0 | 0.66 | 0.46 | 0.74 | 0.93 | 0.63 | 0.7 |
| 2 | 180370497 | rs7604698 | 0.138 | -1.382 | -0.663 | -0.824 | -0.063 | -0.114 | -0.114 | 0.095 | 0.67 | 0.52 | 0.82 | 0.93 | 0.77 | 0.72 |
| 2 | 180374441 | rs1400819 | 0.511 | -1.304 | -0.709 | -0.761 | -0.008 | 0.11 | 0.11 | 0.668 | 0.12 | 0.25 | 0.08 | 0.39 | 0.09 | 0.27 |
| 2 | 180374598 | rs1400820 | 1.433 | -1.27 | -0.789 | -0.872 | -0.053 | -0.017 | -0.017 | 0 | 0.36 | 0.2 | 0.61 | 0.53 | 0.42 | 0.39 |
| 2 | 180374761 | rs9288038 | 1.217 | -1.27 | -0.813 | -0.905 | -0.008 | -0.027 | -0.027 | 0 | 0.11 | 0.15 | 0.07 | 0.01 | 0.08 | 0.14 |

| | | | | | | | | | | | | | | | | |
|---|---|---|---|---|---|---|---|---|---|---|---|---|---|---|---|---|
| 2 | 180377479 | rs13003114 | -0.215 | -1.267 | -0.913 | -0.894 | -0.005 | -0.003 | -0.003 | 0 | 0.3 | 0.28 | 0.21 | 0.4 | 0.34 | 0.34 |
| 2 | 180378253 | rs10210118 | -0.634 | -1.193 | -0.615 | -0.682 | -0.046 | -0.07 | -0.07 | 1.491 | 0.77 | 0.71 | 0.9 | 0.94 | 0.85 | 0.86 |
| 2 | 180380818 | rs7593183 | 0.385 | -0.682 | -0.429 | -0.377 | 0.054 | 0.017 | 0.017 | 0.272 | 0.55 | 0.34 | 0.75 | 0.54 | 0.68 | 0.46 |
| 2 | 180383498 | rs4567894 | -0.181 | -0.528 | -0.545 | -0.274 | -0.043 | -0.081 | -0.081 | 0 | 0.67 | 0.65 | 0.83 | 0.92 | 0.77 | 0.72 |
| 2 | 180396869 | rs6715297 | 0.195 | 0.669 | -0.688 | 0.755 | 0.014 | 0.03 | 0.03 | 0.275 | 0.69 | 0.49 | 0.78 | 0.58 | 0.67 | 0.66 |
| 2 | 180400021 | rs10497541 | -0.41 | 0.498 | -1.155 | 1.273 | 0.001 | -0.039 | -0.039 | 0.043 | 0.11 | 0.03 | 0.12 | 0.23 | 0.13 | 0.11 |
| 2 | 180401233 | rs6738933 | -0.493 | 0.147 | -0.887 | 0.687 | -0.012 | -0.045 | -0.045 | 0 | 0.09 | 0.18 | 0.04 | 0 | 0.04 | 0.07 |
| 2 | 180401424 | rs6725869 | -0.318 | 0.141 | -0.862 | 0.684 | 0.007 | 0.005 | 0.005 | 0 | 0.09 | 0.08 | 0.03 | 0 | 0.04 | 0.07 |
| 2 | 180403128 | rs11884072 | -0.478 | -0.359 | -0.738 | -0.308 | -0.025 | -0.131 | -0.131 | 8.326 | 0.31 | 0.4 | 0.16 | 0.04 | 0.2 | 0.23 |
| 2 | 180404547 | rs4894108 | -0.044 | -0.238 | -0.349 | 0.146 | 0.003 | 0.062 | 0.062 | 0 | 0.8 | 0.68 | 0.79 | 0.42 | 0.79 | 0.62 |
| 2 | 180405781 | rs11888508 | -0.747 | 0.512 | -0.6 | -0.641 | -0.001 | -0.139 | -0.139 | 5.069 | 0.38 | 0.44 | 0.25 | 0.02 | 0.35 | 0.25 |
| 2 | 180408734 | rs13030456 | -0.484 | -0.138 | -1.294 | -1.072 | 0.014 | -0.01 | -0.01 | 0.055 | 0.35 | 0.33 | 0.19 | 0.03 | 0.3 | 0.18 |
| 2 | 180414913 | rs4530328 | 1.087 | -0.135 | -1.547 | -0.811 | 0.059 | 0.095 | 0.095 | 0.265 | 0.18 | 0.03 | 0.05 | 0.01 | 0.09 | 0.09 |
| 2 | 180426764 | rs16866778 | 1.695 | 0.232 | -1.428 | -0.356 | -0.028 | 0.001 | 0.001 | 0 | 0.06 | 0.01 | 0.06 | 0.32 | 0.08 | 0.23 |
| 2 | 180427409 | rs3112931 | 0.182 | -0.193 | -1.155 | -1.337 | 0.002 | -0.08 | -0.08 | 0.637 | 0.49 | 0.47 | 0.72 | 0.96 | 0.58 | 0.73 |
| 2 | 180435707 | rs1546603 | -1.091 | 0.254 | -0.916 | -0.974 | 0.001 | -0.016 | -0.016 | 0 | 0.91 | 0.89 | 0.9 | 0.99 | 0.84 | 0.97 |
| 2 | 180436657 | rs7594895 | 0.243 | -0.216 | -0.864 | -1.062 | -0.023 | -0.128 | -0.128 | 0 | 0.25 | 0.37 | 0.13 | 0.01 | 0.18 | 0.15 |
| 2 | 180439113 | rs13405069 | 0.572 | 0.218 | -0.938 | -0.988 | 0.018 | -0.036 | -0.036 | 0 | 0.19 | 0.24 | 0.24 | 0.09 | 0.24 | 0.13 |
| 2 | 180439410 | rs1356067 | 0.309 | 0.111 | -1.38 | -0.945 | 0.006 | 0.011 | 0.011 | 0 | 0.45 | 0.41 | 0.32 | 0.58 | 0.44 | 0.46 |
| 2 | 180439812 | rs1515306 | -0.139 | 0.116 | -1.277 | -1.24 | 0.001 | -0.024 | -0.024 | 0 | 0.64 | 0.59 | 0.6 | 0.72 | 0.62 | 0.74 |
| 2 | 180440437 | rs3112976 | -0.179 | -0.291 | -1.578 | -1.858 | 0.024 | 0.14 | 0.14 | 0.012 | 0.48 | 0.71 | 0.57 | 0.9 | 0.43 | 0.55 |
| 2 | 180440806 | rs17766255 | 1.918 | 0.146 | -1.861 | -1.854 | -0.089 | 0.065 | 0.065 | 0.612 | 0.18 | 0.01 | 0.21 | 0.07 | 0.24 | 0.29 |
| 2 | 180445308 | rs908287 | 1.808 | -0.089 | -1.428 | -1.615 | -0.022 | 0.104 | 0.104 | 0 | 0.05 | 0.13 | 0.15 | 0.23 | 0.01 | 0.05 |
| 2 | 180446934 | rs1515311 | 0.948 | -0.165 | -1.615 | -1.788 | 0.021 | 0.081 | 0.081 | 0 | 0.25 | 0.15 | 0.15 | 0.04 | 0.23 | 0.15 |
| 2 | 180449157 | rs3106713 | 1.074 | -0.114 | -1.46 | -1.769 | -0.007 | 0.011 | 0.011 | 0.17 | 0.22 | 0.29 | 0.24 | 0.64 | 0.19 | 0.35 |
| 2 | 180450382 | rs17818050 | -0.469 | -0.078 | -1.5 | -1.871 | 0.011 | 0.052 | 0.052 | 0 | 0.4 | 0.26 | 0.39 | 0.26 | 0.38 | 0.24 |
| 2 | 180452188 | rs2138196 | -2.095 | 0.299 | -1.281 | -1.747 | 0.009 | -0.024 | -0.024 | 0 | 0.82 | 0.8 | 0.87 | 0.91 | 0.78 | 0.88 |
| 2 | 180452731 | rs13028216 | 2.095 | 0.012 | -1.424 | -1.934 | -0.019 | -0.05 | -0.05 | 0 | 0.17 | 0.2 | 0.07 | 0.01 | 0.09 | 0.07 |
| 2 | 180453399 | rs11683314 | 1.117 | 0.196 | -1.519 | -1.979 | -0.098 | 0.108 | 0.108 | 2.488 | 0.25 | 0.04 | 0.28 | 0.07 | 0.34 | 0.29 |
| 2 | 180454264 | rs13012946 | 0.01 | 0.124 | -1.535 | -1.975 | 0.001 | 0.1 | 0.1 | 0.061 | 0.49 | 0.31 | 0.55 | 0.29 | 0.48 | 0.45 |
| 2 | 180455067 | rs13411180 | 1.254 | 0.079 | -1.457 | -1.908 | -0.009 | -0.001 | -0.001 | 0 | 0.24 | 0.26 | 0.27 | 0.23 | 0.14 | 0.16 |
| 2 | 180457484 | rs16866836 | 2.026 | 0.309 | -1.353 | -1.806 | 0.006 | -0.025 | -0.025 | 0 | 0.17 | 0.21 | 0.14 | 0.09 | 0.21 | 0.11 |
| 2 | 180460930 | rs1533385 | -1.214 | 0.295 | -1.366 | -1.829 | -0.005 | 0.028 | 0.028 | 0 | 0.79 | 0.68 | 0.77 | 0.4 | 0.82 | 0.63 |
| 2 | 180465299 | rs1515317 | -1.214 | 0.294 | -1.366 | -1.829 | -0.005 | 0.028 | 0.028 | 0 | 0.79 | 0.69 | 0.77 | 0.4 | 0.82 | 0.63 |
| 2 | 180477612 | rs2090219 | -0.645 | 0.448 | -1.073 | -1.918 | 0.02 | -0.008 | -0.008 | 0.099 | 0.81 | 0.66 | 0.78 | 0.84 | 0.82 | 0.82 |
| 2 | 180479625 | rs872158 | 0.739 | 0.349 | -1.213 | -1.899 | 0.027 | 0.059 | 0.059 | 0 | 0.41 | 0.23 | 0.45 | 0.3 | 0.35 | 0.41 |
| 2 | 180480167 | rs6745521 | 1.71 | 0.382 | -1.224 | -1.889 | -0.008 | 0.019 | 0.019 | 0 | 0.1 | 0.05 | 0.07 | 0.03 | 0.13 | 0.07 |
| 2 | 180484927 | rs4894115 | -0.181 | 0.267 | -1.149 | -1.926 | 0 | 0.024 | 0.024 | 0 | 0.54 | 0.42 | 0.53 | 0.33 | 0.48 | 0.48 |
| 2 | 180487430 | rs11904349 | -0.002 | 0.196 | -1.236 | -1.975 | -0.021 | 0.04 | 0.04 | 0.366 | 0.35 | 0.28 | 0.38 | 0.13 | 0.43 | 0.41 |
| 2 | 180488450 | rs16866864 | 0.98 | 0.218 | -1.134 | -1.951 | -0.009 | 0.037 | 0.037 | 0.23 | 0.16 | 0.5 | 0.19 | 0.16 | 0.09 | 0.17 |
| 2 | 180488818 | rs1515269 | -1.021 | 0.187 | -1.179 | -1.951 | -0.004 | -0.006 | -0.006 | 2.857 | 0.69 | 0.71 | 0.76 | 0.49 | 0.62 | 0.69 |
| 2 | 180489474 | rs984893 | 1.844 | 0.271 | -1.203 | -1.951 | -0.011 | 0.005 | 0.005 | 0 | 0.06 | 0.03 | 0.05 | 0 | 0.09 | 0.01 |
| 2 | 180489836 | rs16866865 | 0.538 | 0.27 | -1.244 | -2.009 | 0.033 | 0.042 | 0.042 | 0 | 0.33 | 0.15 | 0.3 | 0.27 | 0.27 | 0.2 |
| 2 | 180491521 | rs3106703 | 0.949 | 0.265 | -1.202 | -2.015 | -0.024 | 0.036 | 0.036 | 0 | 0.16 | 0.46 | 0.22 | 0.16 | 0.1 | 0.19 |
| 2 | 180491714 | rs3112962 | 0.302 | 0.272 | -1.193 | -1.999 | 0.01 | -0.046 | -0.046 | 0 | 0.35 | 0.49 | 0.45 | 0.22 | 0.35 | 0.49 |
| 2 | 180492282 | rs3112960 | -0.812 | 0.268 | -1.18 | -1.966 | -0.001 | -0.001 | -0.001 | 2.888 | 0.68 | 0.64 | 0.76 | 0.46 | 0.62 | 0.69 |
| 2 | 180492632 | rs1515274 | 1.858 | 0.215 | -1.162 | -1.933 | 0.023 | 0.002 | 0.002 | 0.25 | 0.22 | 0.11 | 0.23 | 0.24 | 0.14 | 0.14 |
| 2 | 180495199 | rs12476520 | 1.613 | 0.493 | -1.133 | -1.714 | 0 | 0.003 | 0.003 | 0 | 0.24 | 0.23 | 0.19 | 0.52 | 0.28 | 0.3 |
| 2 | 180501314 | rs3112945 | 1.687 | 0.417 | -0.955 | -1.669 | -0.032 | 0.057 | 0.057 | 0.021 | 0.12 | 0.35 | 0.16 | 0.16 | 0.06 | 0.15 |
| 2 | 180502356 | rs3112943 | -0.207 | 0.583 | -1.034 | -1.75 | -0.019 | 0.017 | 0.017 | 0 | 0.58 | 0.65 | 0.7 | 0.41 | 0.49 | 0.61 |
| 2 | 180503546 | rs3112940 | 0.352 | 1.478 | -0.761 | -1.058 | -0.026 | 0.016 | 0.016 | 0 | 0.19 | 0.38 | 0.21 | 0.17 | 0.15 | 0.14 |
| 2 | 180508524 | rs1515279 | 1.751 | 1.565 | -0.92 | -1.16 | 0.024 | 0.001 | 0.001 | 0 | 0.41 | 0.28 | 0.25 | 0.53 | 0.37 | 0.39 |
| 2 | 180509889 | rs10497543 | 0.111 | 1.702 | -1.069 | -1.123 | 0.001 | 0.005 | 0.005 | 0 | 0.13 | 0.13 | 0.07 | 0.07 | 0.13 | 0.07 |
| 2 | 180509950 | rs3106694 | -0.489 | 1.842 | -1.153 | -0.951 | -0.033 | 0.024 | 0.024 | 0.062 | 0.76 | 0.84 | 0.59 | 0.77 | 0.71 | 0.64 |
| 2 | 180510494 | rs954649 | 0.826 | 1.741 | -1.109 | -1.089 | -0.025 | 0.04 | 0.04 | 0.338 | 0.2 | 0.12 | 0.24 | 0.05 | 0.25 | 0.28 |
| 2 | 180512804 | rs10432487 | -1.093 | 1.534 | -0.691 | -0.867 | -0.005 | -0.012 | -0.012 | 0.435 | 0.28 | 0.59 | 0.31 | 0.24 | 0.25 | 0.14 |
| 2 | 180513046 | rs6751130 | 0.068 | 2.057 | -0.557 | -1.071 | 0.015 | -0.038 | -0.038 | 0.292 | 0.27 | 0.4 | 0.38 | 0.21 | 0.26 | 0.31 |
| 2 | 180515472 | rs7607628 | -0.184 | 2.078 | -0.627 | -1.166 | 0.005 | 0.001 | 0.001 | 0.294 | 0.19 | 0.4 | 0.21 | 0.15 | 0.13 | 0.08 |
| 2 | 180516544 | rs7558364 | -0.314 | 2.169 | -0.91 | -1.218 | -0.084 | 0.058 | 0.058 | 0 | 0.33 | 0.08 | 0.46 | 0.28 | 0.41 | 0.38 |
| 2 | 180518040 | rs956261 | 0.091 | 2.23 | -0.821 | -1.142 | -0.013 | -0.007 | -0.007 | 0 | 0.19 | 0.08 | 0.25 | 0.22 | 0.21 | 0.15 |
| 2 | 180518155 | rs1877941 | 1.788 | 2.129 | -0.672 | -1.135 | 0.034 | 0.02 | 0.02 | 0 | 0.43 | 0.23 | 0.25 | 0.57 | 0.42 | 0.53 |
| 2 | 180520630 | rs10497546 | 1.112 | 2.055 | -0.111 | -0.701 | -0.008 | 0.063 | 0.063 | 0.069 | 0.12 | 0.34 | 0.17 | 0.16 | 0.06 | 0.06 |
| 2 | 180521334 | rs1515281 | 1.2 | 2.102 | -0.135 | -0.705 | 0.024 | 0.072 | 0.072 | 0 | 0.05 | 0.15 | 0.17 | 0.15 | 0.04 | 0.08 |
| 2 | 180531008 | rs1877945 | 1.384 | 2.246 | -0.191 | -0.753 | -0.006 | -0.006 | -0.006 | 0 | 0.14 | 0.13 | 0.2 | 0.05 | 0.19 | 0.22 |
| 2 | 180531222 | rs17770642 | 0.76 | 2.231 | -0.201 | -0.761 | -0.017 | 0.017 | 0.017 | 0 | 0.06 | 0.02 | 0.04 | 0 | 0.07 | 0.01 |
| 2 | 180532277 | rs16866903 | 0.196 | 2.156 | -0.282 | -0.819 | -0.032 | -0.032 | -0.032 | 0.135 | 0.19 | 0.03 | 0.24 | 0.22 | 0.2 | 0.14 |
| 2 | 180533005 | rs7581866 | 0.035 | 2.152 | -0.25 | -0.818 | 0.003 | -0.004 | -0.004 | 0 | 0.18 | 0.36 | 0.21 | 0.16 | 0.13 | 0.07 |
| 2 | 180534096 | rs3106720 | -1.673 | 2.147 | -0.316 | -0.816 | 0.001 | 0.002 | 0.002 | 0 | 0.39 | 0.6 | 0.47 | 0.38 | 0.34 | 0.22 |
| 2 | 180535781 | rs1877944 | 0.01 | 2.015 | -0.304 | -0.809 | -0.035 | 0.084 | 0.084 | 0 | 0.28 | 0.14 | 0.29 | 0.13 | 0.35 | 0.29 |

| | | | | | | | | | | | | | | | | |
|---|---|---|---|---|---|---|---|---|---|---|---|---|---|---|---|---|
| 2 | 180538171 | rs6708429 | -0.31 | 1.754 | -0.251 | -0.695 | 0.004 | -0.008 | -0.008 | 0 | 0.19 | 0.42 | 0.21 | 0.16 | 0.13 | 0.08 |
| 2 | 180542062 | rs17822210 | 0.932 | 1.766 | -0.531 | -0.642 | -0.005 | 0.014 | 0.014 | 0 | 0.05 | 0.02 | 0.02 | 0 | 0.05 | 0.01 |
| 2 | 180542562 | rs2276572 | 1.224 | 1.725 | -0.215 | -0.643 | -0.005 | 0.05 | 0.05 | 0 | 0.1 | 0.27 | 0.16 | 0.15 | 0.06 | 0.06 |
| 2 | 180542687 | rs2276573 | 1.844 | 1.872 | -0.443 | -0.705 | 0.007 | 0.034 | 0.034 | 0 | 0.46 | 0.19 | 0.28 | 0.56 | 0.46 | 0.53 |
| 2 | 180546744 | rs10180972 | 0.343 | 1.95 | -0.324 | -0.669 | 0.003 | -0.044 | -0.044 | 0 | 0.21 | 0.22 | 0.23 | 0.05 | 0.23 | 0.22 |
| 2 | 180549443 | rs7590028 | -0.895 | 1.674 | -0.025 | -0.411 | 0.008 | -0.036 | -0.036 | 0 | 0.43 | 0.45 | 0.55 | 0.28 | 0.48 | 0.4 |
| 2 | 180550931 | rs11690551 | -0.275 | 1.731 | -0.751 | -0.418 | -0.013 | 0.048 | 0.048 | 0 | 0.35 | 0.23 | 0.31 | 0.13 | 0.39 | 0.29 |
| 2 | 180555884 | rs1022311 | -0.49 | 1.748 | -0.998 | -0.178 | 0.033 | 0.085 | 0.085 | 0 | 0.93 | 0.84 | 0.78 | 0.8 | 0.91 | 0.88 |
| 2 | 180557720 | rs1399689 | 0.124 | 1.557 | -1.713 | -0.236 | -0.023 | 0.066 | 0.066 | 0 | 0.28 | 0.15 | 0.29 | 0.13 | 0.35 | 0.28 |
| 2 | 180565422 | rs7557247 | 0.185 | 1.076 | -1.88 | 0.233 | 0.017 | 0.032 | 0.032 | 0 | 0.48 | 0.19 | 0.3 | 0.54 | 0.49 | 0.56 |
| 2 | 180566678 | rs16866933 | 0.032 | 1.215 | -1.259 | 1.167 | 0 | 0.114 | 0.114 | 0 | 0.06 | 0.05 | 0.06 | 0.36 | 0.1 | 0.24 |
| 2 | 180568073 | rs11885327 | -0.211 | 0.948 | -1.832 | 0.313 | 0.007 | -0.055 | -0.055 | 0.272 | 0.39 | 0.16 | 0.25 | 0.46 | 0.39 | 0.43 |
| 2 | 180569100 | rs12470217 | -0.314 | 1.451 | -1.401 | 0.713 | -0.001 | -0.011 | -0.011 | 0 | 0.12 | 0.1 | 0.16 | 0.24 | 0.11 | 0.09 |
| 2 | 180570920 | rs10803928 | -1.489 | 1.342 | -0.839 | 0.557 | 0.003 | 0.009 | 0.009 | 0 | 0.25 | 0.38 | 0.28 | 0.65 | 0.26 | 0.36 |
| 2 | 180571192 | rs16866934 | -0.794 | 1.451 | -0.576 | 0.663 | 0.02 | 0.203 | 0.203 | 0 | 0.12 | 0.28 | 0.12 | 0.45 | 0.15 | 0.28 |
| 2 | 180571917 | rs16866936 | -0.775 | 1.417 | -0.595 | 0.661 | 0.016 | 0.198 | 0.198 | 0 | 0.12 | 0.27 | 0.12 | 0.45 | 0.15 | 0.28 |
| 2 | 180572095 | rs17496658 | -0.134 | 1.46 | -0.732 | 0.688 | -0.009 | 0.027 | 0.027 | 0 | 0.1 | 0.1 | 0.09 | 0.44 | 0.14 | 0.25 |
| 2 | 180579145 | rs13407248 | -1.602 | 1.476 | -0.623 | 0.658 | 0.019 | 0.093 | 0.093 | 0 | 0.19 | 0.33 | 0.16 | 0.43 | 0.26 | 0.29 |
| 2 | 180582926 | rs17772191 | -0.908 | 1.515 | -0.86 | 0.401 | -0.019 | 0.016 | 0.016 | 0.525 | 0.1 | 0.06 | 0.07 | 0 | 0.16 | 0.08 |
| 2 | 180591421 | rs1964081 | -0.385 | 1.447 | -0.765 | 0.542 | 0.004 | 0.11 | 0.11 | 0 | 0.11 | 0.16 | 0.11 | 0.48 | 0.14 | 0.25 |
| 2 | 180593759 | rs17772551 | -0.924 | 1.512 | -0.9 | 0.205 | -0.035 | 0.042 | 0.042 | 0.7 | 0.1 | 0.02 | 0.07 | 0 | 0.16 | 0.08 |
| 2 | 180594128 | rs13405414 | -1.217 | 2.13 | -0.989 | -0.181 | -0.01 | 0.007 | 0.007 | 0.054 | 0.88 | 0.91 | 0.67 | 0.96 | 0.84 | 0.81 |
| 2 | 180595863 | rs12472888 | -0.786 | 2.534 | -0.925 | 0.094 | -0.003 | 0.019 | 0.019 | 0 | 0.15 | 0.13 | 0.11 | 0.45 | 0.24 | 0.29 |
| 2 | 180597262 | rs992334 | 0.65 | 2.287 | -0.652 | -0.375 | -0.007 | 0.037 | 0.037 | 0 | 0.2 | 0.27 | 0.14 | 0.31 | 0.21 | 0.19 |
| 2 | 180599126 | rs2367801 | -0.314 | 2.627 | -0.389 | 0.036 | -0.002 | -0.026 | -0.026 | 0.204 | 0.93 | 0.9 | 0.95 | 0.99 | 0.93 | 0.99 |
| 2 | 180604037 | rs17824301 | 0.292 | 2.719 | -0.426 | 0.057 | 0.019 | -0.014 | -0.014 | 0 | 0.05 | 0.08 | 0.05 | 0.01 | 0.1 | 0.08 |
| 2 | 180604436 | rs10497549 | 1.514 | 2.596 | -0.717 | -0.215 | -0.025 | 0.02 | 0.02 | 0 | 0.16 | 0.03 | 0.12 | 0.09 | 0.16 | 0.1 |
| 2 | 180613083 | rs7557892 | 0.317 | 2.271 | -0.479 | 0.019 | 0.034 | -0.021 | -0.021 | 0.074 | 0.05 | 0.09 | 0.05 | 0.01 | 0.1 | 0.08 |
| 2 | 180622833 | rs12053119 | 0.703 | 2.195 | -0.555 | 0.05 | -0.019 | -0.017 | -0.017 | 0.092 | 0.1 | 0.02 | 0.05 | 0.45 | 0.13 | 0.22 |
| 2 | 180625438 | rs11674376 | 0.894 | 2.266 | -0.389 | -0.031 | -0.005 | 0.005 | 0.005 | 0.006 | 0.27 | 0.27 | 0.32 | 0.13 | 0.27 | 0.26 |
| 2 | 180631925 | rs1486239 | -0.925 | 2.252 | -0.425 | -0.053 | 0.032 | 0.02 | 0.02 | 0 | 0.47 | 0.37 | 0.46 | 0.59 | 0.58 | 0.56 |
| 2 | 180635236 | rs7574345 | -0.675 | 2.288 | -0.475 | 0.023 | -0.019 | -0.027 | -0.027 | 0 | 0.2 | 0.15 | 0.14 | 0.46 | 0.31 | 0.31 |
| 2 | 180651001 | rs7589885 | 0.501 | 1.799 | -0.418 | -0.124 | 0.015 | 0.016 | 0.016 | 0 | 0.45 | 0.38 | 0.25 | 0.32 | 0.38 | 0.28 |
| 2 | 180659519 | rs723357 | -0.288 | 1.835 | -0.404 | -0.004 | -0.005 | -0.006 | -0.006 | 0 | 0.45 | 0.52 | 0.47 | 0.6 | 0.51 | 0.55 |
| 2 | 180659575 | rs10168253 | -0.102 | 1.419 | -0.812 | 0.164 | -0.008 | 0.001 | 0.001 | 0 | 0.4 | 0.34 | 0.4 | 0.58 | 0.41 | 0.49 |
| 2 | 180663034 | rs9288042 | 0.8 | 1.61 | -0.237 | 0.181 | 0.01 | -0.006 | -0.006 | 0 | 0.7 | 0.61 | 0.52 | 0.75 | 0.65 | 0.43 |
| 2 | 180664033 | rs357708 | -1.804 | 1.102 | 0.641 | 0.107 | 0.002 | -0.02 | -0.02 | 0 | 0.19 | 0.47 | 0.32 | 0.12 | 0.16 | 0.25 |
| 2 | 180665865 | rs7585592 | -1.087 | 1.309 | 0.576 | -0.268 | -0.002 | 0.002 | 0.002 | 0 | 0.26 | 0.29 | 0.52 | 0.16 | 0.32 | 0.33 |
| 2 | 180667929 | rs13410767 | -1.276 | 1.245 | 1.418 | -0.319 | 0.007 | -0.118 | -0.118 | 0 | 0.28 | 0.44 | 0.32 | 0.04 | 0.29 | 0.17 |
| 2 | 180668073 | rs13398994 | -1.291 | 1.251 | 1.42 | -0.316 | 0.006 | -0.118 | -0.118 | 0.837 | 0.29 | 0.44 | 0.32 | 0.04 | 0.29 | 0.21 |
| 2 | 180668301 | rs13399303 | -1.118 | 1.246 | 1.526 | -0.318 | 0.001 | -0.049 | -0.049 | 0.174 | 0.28 | 0.33 | 0.31 | 0.04 | 0.29 | 0.21 |
| 2 | 180668778 | rs357712 | -2.419 | 1.266 | 1.501 | -0.334 | -0.006 | 0.023 | 0.023 | 0 | 0.4 | 0.67 | 0.61 | 0.15 | 0.42 | 0.39 |
| 2 | 180675669 | rs1906012 | -0.834 | 1.218 | 1.561 | -0.354 | -0.002 | 0.002 | 0.002 | 0 | 0.23 | 0.25 | 0.52 | 0.14 | 0.3 | 0.29 |
| 2 | 180679735 | rs165372 | -0.34 | 0.896 | 1.788 | -0.171 | -0.022 | 0.04 | 0.04 | 0 | 0.92 | 0.83 | 0.96 | 0.83 | 0.96 | 0.87 |
| 2 | 180681868 | rs11680455 | 1.41 | 1.059 | 1.109 | 0.107 | -0.036 | -0.073 | -0.073 | 0.033 | 0.15 | 0.02 | 0.14 | 0.4 | 0.2 | 0.07 |
| 2 | 180684202 | rs13418648 | -0.359 | 0.868 | 2.109 | -0.012 | 0 | -0.015 | -0.015 | 0.01 | 0.19 | 0.19 | 0.29 | 0.03 | 0.26 | 0.19 |
| 2 | 180693147 | rs165380 | -2.411 | 0.796 | 2.163 | 0.029 | 0.001 | -0.012 | -0.012 | 0.41 | 0.4 | 0.6 | 0.62 | 0.16 | 0.42 | 0.39 |
| 2 | 180706328 | rs259833 | -2.446 | 0.736 | 2.14 | 0.032 | -0.007 | 0.028 | 0.028 | 0 | 0.4 | 0.67 | 0.61 | 0.16 | 0.42 | 0.4 |
| 2 | 180712985 | rs12328338 | -1.031 | 0.871 | 1.972 | 0.114 | 0.008 | -0.006 | -0.006 | 0 | 0.25 | 0.28 | 0.52 | 0.16 | 0.31 | 0.32 |
| 2 | 180715335 | rs10196283 | 0.428 | 1.001 | 1.683 | 0.03 | 0.003 | -0.028 | -0.028 | 0 | 0.13 | 0.12 | 0.21 | 0 | 0.16 | 0.16 |
| 2 | 180716475 | rs16867038 | -0.059 | 1.084 | 1.725 | 0.119 | 0.012 | -0.021 | -0.021 | 4.303 | 0.05 | 0.11 | 0.05 | 0 | 0.06 | 0.05 |
| 2 | 180718419 | rs6725456 | 2.085 | 0.956 | 1.513 | 0.124 | -0.015 | 0.07 | 0.07 | 0.145 | 0.53 | 0.16 | 0.35 | 0.66 | 0.54 | 0.48 |
| 2 | 180728060 | rs4894131 | -1.221 | 1.643 | 1.634 | 0.178 | -0.034 | 0.047 | 0.047 | 0 | 0.24 | 0.15 | 0.31 | 0.04 | 0.33 | 0.26 |
| 2 | 180736508 | rs259843 | -1.735 | 1.65 | 1.802 | 0.232 | 0.027 | -0.029 | -0.029 | 0 | 0.31 | 0.37 | 0.57 | 0.16 | 0.37 | 0.38 |
| 2 | 180738602 | rs11896563 | -1.074 | 1.746 | 1.738 | 0.138 | 0.02 | -0.03 | -0.03 | 0 | 0.2 | 0.25 | 0.28 | 0.1 | 0.26 | 0.2 |
| 2 | 180744430 | rs259839 | -1.096 | 1.326 | 1.638 | 0.344 | -0.011 | 0.005 | 0.005 | 0 | 0.12 | 0.06 | 0.08 | 0.01 | 0.19 | 0.08 |
| 2 | 180761005 | rs6758549 | -1.882 | 1.19 | 2.128 | 0.455 | 0.005 | 0.009 | 0.009 | 0 | 0.15 | 0.32 | 0.36 | 0.14 | 0.13 | 0.19 |
| 2 | 180763456 | rs13392155 | -1.994 | 1.105 | 2.372 | 0.388 | 0.047 | 0.037 | 0.037 | 2.291 | 0.08 | 0.49 | 0.07 | 0.02 | 0.03 | 0.01 |
| 2 | 180775704 | rs259822 | -2.362 | 1.108 | 1.869 | 0.373 | -0.004 | 0.04 | 0.04 | 0.193 | 0.33 | 0.72 | 0.48 | 0.16 | 0.33 | 0.27 |
| 2 | 180776022 | rs3912255 | -1.371 | 1.242 | 1.494 | 0.462 | 0.017 | -0.108 | -0.108 | 2.152 | 0.2 | 0.42 | 0.14 | 0.04 | 0.23 | 0.09 |
| 2 | 180777260 | rs259826 | -1.693 | 1.063 | 1.025 | 1.08 | 0.001 | 0.001 | 0.001 | 0.176 | 0.1 | 0.09 | 0.33 | 0.13 | 0.1 | 0.18 |
| 2 | 180783672 | rs260059 | 1.463 | 1.03 | 1.184 | 1.598 | 0.014 | 0.06 | 0.06 | 0 | 0.73 | 0.62 | 0.42 | 0.53 | 0.71 | 0.7 |
| 2 | 180793050 | rs17777566 | -0.781 | 1.112 | 1.277 | 1.89 | 0.014 | 0.045 | 0.045 | 0 | 0.09 | 0.13 | 0.37 | 0.23 | 0.12 | 0.14 |
| 2 | 180796377 | rs7558336 | 1.969 | 1.126 | 1.35 | 1.881 | 0.004 | 0.016 | 0.016 | 0 | 0.75 | 0.73 | 0.43 | 0.53 | 0.71 | 0.7 |
| 2 | 180807468 | rs3845699 | 1.799 | 1.119 | 1.489 | 1.886 | 0.014 | 0.011 | 0.011 | 0 | 0.83 | 0.77 | 0.52 | 0.76 | 0.77 | 0.75 |
| 2 | 180810202 | rs1046356 | -0.202 | 1.029 | 1.341 | 1.878 | 0.001 | 0.003 | 0.003 | 0 | 0.08 | 0.05 | 0.1 | 0.23 | 0.06 | 0.05 |
| 2 | 180825685 | rs12693214 | 0.526 | 1.069 | 1.061 | 1.871 | -0.009 | 0.003 | 0.003 | 0.005 | 0.93 | 0.95 | 0.9 | 0.99 | 0.9 | 0.9 |
| 2 | 180826451 | rs16867120 | -0.826 | 0.793 | 0.94 | 2.294 | -0.008 | 0.034 | 0.034 | 0 | 0.09 | 0.12 | 0.03 | 0.26 | 0.05 | 0.15 |
| 2 | 180830958 | rs6733426 | 0.959 | 0.88 | 0.798 | 2.106 | -0.003 | -0.011 | -0.011 | 0 | 0.92 | 0.91 | 0.89 | 0.99 | 0.89 | 0.9 |

| | | | | | | | | | | | | | | | | |
|---|---|---|---|---|---|---|---|---|---|---|---|---|---|---|---|---|
| 2 | 180846743 | rs3738943 | 1.377 | 0.877 | 0.801 | 2.14 | -0.002 | -0.006 | -0.006 | 0 | 0.85 | 0.9 | 0.8 | 0.76 | 0.84 | 0.84 |
| 2 | 180848488 | rs3845702 | 1.335 | 0.881 | 0.771 | 2.142 | -0.001 | -0.004 | -0.004 | 0 | 0.85 | 0.87 | 0.8 | 0.76 | 0.84 | 0.84 |
| 2 | 180864967 | rs10497555 | -1.502 | 0.895 | 0.694 | 2.15 | -0.004 | -0.011 | -0.011 | 0 | 0.15 | 0.08 | 0.2 | 0.25 | 0.17 | 0.16 |
| 2 | 180872820 | rs2672657 | -1.53 | 1.166 | 0.774 | 1.723 | 0.032 | 0.028 | 0.028 | 0 | 0.18 | 0.32 | 0.48 | 0.24 | 0.22 | 0.24 |
| 2 | 180874721 | rs2584201 | -1.513 | 1.141 | 0.514 | 1.852 | 0.004 | 0.087 | 0.087 | 0 | 0.22 | 0.36 | 0.5 | 0.44 | 0.23 | 0.35 |
| 2 | 180875602 | rs7558006 | -0.572 | 0.931 | 0.01 | 1.057 | 0.003 | 0.023 | 0.023 | 0 | 0.54 | 0.36 | 0.57 | 0.32 | 0.46 | 0.6 |
| 2 | 180877386 | rs262265 | 0.889 | 1.379 | 0.583 | 0.51 | 0.041 | -0.069 | -0.069 | 0.695 | 0.82 | 0.68 | 0.72 | 0.94 | 0.74 | 0.81 |
| 2 | 180889175 | rs3914461 | -0.358 | 1.206 | 0.463 | -0.079 | 0.001 | -0.057 | -0.057 | 0 | 0.34 | 0.3 | 0.29 | 0.01 | 0.33 | 0.19 |
| 2 | 180901804 | rs262256 | 0.4 | 1.316 | 0.742 | -0.175 | -0.005 | -0.113 | -0.113 | 2.766 | 0.62 | 0.63 | 0.69 | 0.99 | 0.64 | 0.75 |
| 2 | 180903932 | rs262254 | 1.271 | 1.147 | 0.933 | -0.15 | 0.011 | 0.153 | 0.153 | 0 | 0.51 | 0.3 | 0.53 | 0.91 | 0.51 | 0.63 |
| 2 | 180907132 | rs6727127 | -0.587 | 1.111 | 1.068 | -0.133 | 0.003 | 0.006 | 0.006 | 1.024 | 0.37 | 0.26 | 0.3 | 0.01 | 0.36 | 0.25 |
| 2 | 180910026 | rs17176528 | -0.046 | 1.098 | 1.012 | -0.133 | -0.02 | 0.024 | 0.024 | 0 | 0.1 | 0.05 | 0.16 | 0.01 | 0.17 | 0.09 |
| 2 | 180910880 | rs262274 | 0.001 | 1.024 | 0.946 | 0.286 | 0.018 | 0.318 | 0.318 | 0.088 | 0.1 | 0.32 | 0.27 | 0.42 | 0.08 | 0.2 |
| 2 | 180912075 | rs4894152 | 0.189 | 0.967 | 0.746 | 0.077 | 0.005 | 0.06 | 0.06 | 2.197 | 0.32 | 0.17 | 0.28 | 0.01 | 0.33 | 0.19 |
| 2 | 180914893 | rs7560818 | -0.366 | 0.954 | 0.722 | 0.082 | 0.004 | 0.011 | 0.011 | 0.976 | 0.37 | 0.26 | 0.31 | 0.01 | 0.36 | 0.25 |
| 2 | 180922263 | rs13009329 | -0.051 | 0.754 | 0.539 | 0.262 | 0.003 | -0.024 | -0.024 | 0 | 0.08 | 0.06 | 0.07 | 0.3 | 0.07 | 0.27 |
| 2 | 180925068 | rs6728772 | 1.377 | 0.912 | 0.744 | 0.137 | 0.018 | 0.002 | 0.002 | 1.418 | 0.23 | 0.14 | 0.12 | 0.01 | 0.16 | 0.1 |
| 2 | 180926576 | rs6761653 | 0.88 | 0.881 | 0.875 | 0.191 | -0.02 | 0.162 | 0.162 | 0 | 0.81 | 0.47 | 0.63 | 0.49 | 0.84 | 0.73 |
| 2 | 180934568 | rs16867162 | 1.281 | 0.932 | 0.855 | 0.145 | -0.002 | -0.043 | -0.043 | 2.069 | 0.24 | 0.21 | 0.12 | 0.01 | 0.15 | 0.1 |
| 2 | 180937368 | rs262283 | -1.333 | 1.004 | 0.956 | 0.223 | 0.004 | 0.092 | 0.092 | 0.066 | 0.22 | 0.36 | 0.41 | 0.5 | 0.22 | 0.35 |
| 2 | 180940802 | rs10497556 | 0.183 | 0.886 | 0.973 | 0.444 | 0.014 | 0.018 | 0.018 | 0.047 | 0.32 | 0.24 | 0.24 | 0.07 | 0.28 | 0.23 |
| 2 | 180945635 | rs262292 | -0.946 | 0.897 | 0.98 | 0.516 | 0 | 0 | 0 | 1.363 | 0.45 | 0.55 | 0.52 | 0.49 | 0.36 | 0.44 |
| 2 | 180946075 | rs4125973 | 1.354 | 0.896 | 0.738 | 0.476 | -0.099 | -0.031 | -0.031 | 0 | 0.43 | 0.09 | 0.3 | 0.43 | 0.47 | 0.43 |
| 2 | 180948331 | rs10203253 | 0.099 | 0.879 | 0.89 | 0.561 | 0.003 | 0.31 | 0.31 | 0.052 | 0.09 | 0.3 | 0.26 | 0.42 | 0.06 | 0.16 |
| 2 | 180954390 | rs1914695 | 0.469 | 0.92 | 0.794 | 0.502 | -0.001 | 0.001 | 0.001 | 0 | 0.08 | 0.06 | 0.11 | 0.06 | 0.1 | 0.12 |
| 2 | 180956889 | rs10180505 | 0.789 | 0.901 | 0.994 | 0.571 | 0.004 | 0.235 | 0.235 | 0 | 0.07 | 0.16 | 0.26 | 0.42 | 0.06 | 0.16 |
| 2 | 180958521 | rs1401102 | 1.193 | 0.881 | 1.047 | 0.44 | -0.018 | 0.015 | 0.015 | 0 | 0.44 | 0.29 | 0.31 | 0.43 | 0.48 | 0.43 |
| 2 | 180965039 | rs11679764 | 0.187 | 0.855 | 1.058 | 0.405 | -0.001 | 0 | 0 | 0.338 | 0.59 | 0.47 | 0.49 | 0.44 | 0.65 | 0.58 |
| 2 | 180971752 | rs10803930 | -1.236 | 0.84 | 1.034 | 0.419 | 0 | -0.003 | -0.003 | 0 | 0.53 | 0.49 | 0.67 | 0.57 | 0.52 | 0.57 |
| 2 | 180974882 | rs1878739 | -0.58 | 0.77 | 1.206 | 0.443 | 0.008 | 0.056 | 0.056 | 0.386 | 0.42 | 0.3 | 0.4 | 0.16 | 0.44 | 0.34 |
| 2 | 180976481 | rs16867190 | 1.822 | 1.118 | 0.84 | 0.387 | -0.059 | -0.06 | -0.06 | 0.173 | 0.37 | 0.09 | 0.23 | 0.42 | 0.37 | 0.35 |
| 2 | 180977109 | rs1473622 | 0.094 | 1.074 | 0.92 | 0.334 | 0.005 | 0.059 | 0.059 | 1.57 | 0.33 | 0.21 | 0.3 | 0.09 | 0.34 | 0.23 |
| 2 | 180981724 | rs4894157 | -0.294 | 1.103 | 0.818 | 0.381 | -0.037 | 0.064 | 0.064 | 0 | 0.13 | 0.02 | 0.17 | 0.01 | 0.18 | 0.15 |
| 2 | 180981960 | rs7562405 | 0.583 | 0.994 | 1.092 | 0.55 | 0.06 | 0.285 | 0.285 | 1.76 | 0.85 | 0.56 | 0.67 | 0.59 | 0.84 | 0.76 |
| 2 | 180982120 | rs13393435 | 0.781 | 1.018 | 0.482 | 0.508 | 0.018 | 0.038 | 0.038 | 0 | 0.52 | 0.28 | 0.42 | 0.43 | 0.55 | 0.5 |
| 2 | 180989224 | rs11684453 | -1.478 | 0.828 | 0.52 | 0.507 | 0.001 | 0.02 | 0.02 | 0 | 0.25 | 0.3 | 0.42 | 0.51 | 0.25 | 0.36 |
| 2 | 180991780 | rs1401108 | -1.186 | 0.914 | 0.982 | 0.332 | 0.032 | -0.035 | -0.035 | 0 | 0.15 | 0.27 | 0.16 | 0.06 | 0.19 | 0.2 |
| 2 | 180992501 | rs7566676 | -0.814 | 1.201 | 1.176 | 0.609 | 0.029 | -0.003 | -0.003 | 0 | 0.14 | 0.21 | 0.32 | 0.15 | 0.18 | 0.21 |
| 2 | 180992576 | rs1518434 | 2.213 | 1.13 | 1.296 | 0.603 | 0.004 | -0.006 | -0.006 | 0 | 0.7 | 0.3 | 0.49 | 0.79 | 0.63 | 0.59 |
| 2 | 180993295 | rs7570171 | -0.304 | 1.042 | 1.234 | 0.647 | 0.032 | -0.013 | -0.013 | 0.071 | 0.11 | 0.27 | 0.15 | 0.06 | 0.15 | 0.14 |
| 2 | 180995313 | rs10497560 | 1.229 | 0.719 | 0.422 | 0.375 | 0.031 | 0.062 | 0.062 | 1 | 0.2 | 0.05 | 0.09 | 0.07 | 0.15 | 0.14 |
| 2 | 180999625 | rs16867209 | -0.131 | 0.847 | 0.414 | 0.836 | 0 | -0.031 | -0.031 | 0 | 0.12 | 0.05 | 0.13 | 0.28 | 0.11 | 0.2 |
| 2 | 181002168 | rs10178913 | -1.245 | 0.876 | 0.697 | 0.416 | 0.039 | -0.013 | -0.013 | 0.602 | 0.19 | 0.37 | 0.35 | 0.18 | 0.22 | 0.24 |
| 2 | 181015276 | rs7560050 | 0.328 | 1.079 | 0.568 | 0.363 | -0.007 | 0.003 | 0.003 | 0 | 0.09 | 0.08 | 0.13 | 0.06 | 0.15 | 0.14 |
| 2 | 181015789 | rs7589542 | 1.562 | 1.001 | 0.721 | 0.329 | 0.048 | -0.028 | -0.028 | 0 | 0.7 | 0.45 | 0.5 | 0.76 | 0.62 | 0.63 |
| 2 | 181016133 | rs17180327 | 2.679 | 0.664 | -0.239 | 0.532 | -0.115 | 0.011 | 0.011 | 0.025 | 0.3 | 0.01 | 0.24 | 0.23 | 0.35 | 0.15 |
| 2 | 181020320 | rs16867220 | -0.195 | 0.661 | 0.303 | 0.906 | -0.001 | 0.028 | 0.028 | 0 | 0.12 | 0.1 | 0.14 | 0.31 | 0.13 | 0.23 |
| 2 | 181024926 | rs4893898 | -0.533 | 0.684 | 0.326 | 0.847 | 0.049 | 0.011 | 0.011 | 0.491 | 0.14 | 0.28 | 0.34 | 0.17 | 0.2 | 0.19 |
| 2 | 181025228 | rs17180572 | 1.247 | 0.707 | 0.217 | 0.816 | -0.008 | 0.001 | 0.001 | 0 | 0.07 | 0.05 | 0.09 | 0.06 | 0.07 | 0.12 |
| 2 | 181025248 | rs1401105 | -1.799 | 0.649 | 0.458 | 0.886 | 0.002 | 0.003 | 0.003 | 0 | 0.33 | 0.61 | 0.56 | 0.56 | 0.34 | 0.53 |
| 2 | 181027688 | rs6433833 | -1.838 | 0.636 | 0.47 | 0.879 | -0.001 | -0.001 | -0.001 | 0 | 0.33 | 0.62 | 0.56 | 0.55 | 0.34 | 0.53 |
| 2 | 181035163 | rs4413130 | 1.608 | 0.68 | 0.498 | 0.875 | -0.007 | -0.005 | -0.005 | 0.299 | 0.64 | 0.36 | 0.4 | 0.45 | 0.58 | 0.45 |
| 2 | 181036060 | rs2887173 | 0.855 | 0.455 | 0.405 | 0.984 | 0.011 | 0.096 | 0.096 | 0 | 0.79 | 0.62 | 0.74 | 0.62 | 0.78 | 0.65 |
| 2 | 181038491 | rs9288049 | 0.586 | 0.034 | 0.055 | 0.876 | 0.037 | 0.014 | 0.014 | 0.901 | 0.28 | 0.15 | 0.14 | 0.22 | 0.19 | 0.27 |
| 2 | 181041286 | rs16867225 | -0.386 | 0.862 | 0.806 | 1.442 | -0.024 | 0.178 | 0.178 | 0.317 | 0.11 | 0.28 | 0.13 | 0.32 | 0.06 | 0.21 |
| 2 | 181042370 | rs7597201 | 0.649 | 1.005 | 0.068 | 1.144 | -0.008 | 0.001 | 0.001 | 0 | 0.07 | 0.05 | 0.09 | 0.06 | 0.07 | 0.12 |
| 2 | 181044958 | rs2056790 | -0.453 | 1.065 | 0.224 | 1.532 | 0.001 | 0.012 | 0.012 | 0.137 | 0.12 | 0.1 | 0.17 | 0.26 | 0.18 | 0.24 |
| 2 | 181045656 | rs13026866 | 1.596 | 0.946 | 0.29 | 1.479 | -0.023 | -0.037 | -0.037 | 0 | 0.62 | 0.24 | 0.44 | 0.5 | 0.58 | 0.57 |
| 2 | 181046999 | rs13393322 | 1.567 | 0.96 | 0.451 | 1.105 | 0.009 | 0.016 | 0.016 | 0.005 | 0.49 | 0.42 | 0.6 | 0.45 | 0.58 | 0.31 |
| 2 | 181055597 | rs1529121 | 1.177 | 0.818 | 0.546 | 1.086 | -0.001 | -0.001 | -0.001 | 0 | 0.58 | 0.46 | 0.69 | 0.52 | 0.67 | 0.44 |
| 2 | 181059434 | rs10174149 | -0.002 | 0.745 | 0.635 | 1.11 | -0.031 | -0.075 | -0.075 | 0 | 0.73 | 0.56 | 0.86 | 0.87 | 0.84 | 0.65 |
| 2 | 181062674 | rs4516385 | 0.852 | 0.774 | 0.627 | 1.112 | 0.01 | -0.01 | -0.01 | 0 | 0.11 | 0.04 | 0.14 | 0.35 | 0.09 | 0.18 |
| 2 | 181062797 | rs1534692 | -0.827 | 0.712 | 0.696 | 0.617 | -0.034 | -0.117 | -0.117 | 0.296 | 0.8 | 0.66 | 0.9 | 1 | 0.9 | 0.88 |
| 2 | 181063623 | rs16867241 | 1.014 | 0.709 | 0.703 | 0.616 | -0.012 | 0.024 | 0.024 | 0 | 0.06 | 0.01 | 0.03 | 0 | 0.05 | 0.04 |
| 2 | 181066236 | rs10179768 | -0.943 | 0.707 | 0.704 | 0.625 | -0.034 | -0.116 | -0.116 | 0 | 0.81 | 0.66 | 0.9 | 1 | 0.9 | 0.88 |
| 2 | 181068775 | rs12329114 | -1.041 | 0.738 | 0.621 | 0.64 | -0.024 | -0.091 | -0.091 | 0 | 0.82 | 0.7 | 0.91 | 1 | 0.9 | 0.88 |
| 2 | 181071329 | rs7579494 | -0.843 | 0.85 | 0.768 | 0.687 | 0.038 | 0.05 | 0.05 | 0 | 0.26 | 0.37 | 0.51 | 0.58 | 0.38 | 0.38 |
| 2 | 181074667 | rs10191609 | 1.6 | 0.736 | 0.119 | 0.625 | 0.004 | 0.091 | 0.091 | 0 | 0.47 | 0.24 | 0.36 | 0.3 | 0.46 | 0.28 |
| 2 | 181076749 | rs13418455 | 2.256 | 0.821 | -0.06 | 0.637 | 0.006 | 0.068 | 0.068 | 0 | 0.36 | 0.15 | 0.26 | 0.23 | 0.37 | 0.14 |

| | | | | | | | | | | | | | | | | |
|---|---|---|---|---|---|---|---|---|---|---|---|---|---|---|---|---|
| 2 | 181085244 | rs10497563 | -0.274 | 1.474 | 0.004 | 0.473 | 0.001 | 0.045 | 0.045 | 0.106 | 0.27 | 0.1 | 0.18 | 0.17 | 0.2 | 0.39 |
| 2 | 181086493 | rs10497564 | -0.599 | 1.62 | 0.467 | 0.838 | 0.021 | 0.01 | 0.01 | 0.25 | 0.19 | 0.26 | 0.32 | 0.21 | 0.27 | 0.22 |
| 2 | 181091838 | rs2887175 | -1.264 | 1.341 | 0.547 | 0.701 | 0.011 | 0.003 | 0.003 | 0.125 | 0.27 | 0.37 | 0.41 | 0.27 | 0.34 | 0.35 |
| 2 | 181105861 | rs16867270 | -0.37 | 1.296 | 0.481 | 0.652 | 0.018 | 0.003 | 0.003 | 0.877 | 0.37 | 0.18 | 0.35 | 0.43 | 0.33 | 0.51 |
| 2 | 181106013 | rs2368105 | -1.525 | 1.415 | 0.554 | 0.631 | 0.005 | 0.003 | 0.003 | 0 | 0.55 | 0.45 | 0.65 | 0.63 | 0.59 | 0.74 |
| 2 | 181106772 | rs16867273 | 0.908 | 1.237 | 0.486 | 0.605 | -0.003 | -0.053 | -0.053 | 0 | 0.14 | 0.04 | 0.21 | 0.34 | 0.14 | 0.24 |
| 2 | 181106805 | rs1320386 | -2.445 | 1.266 | 0.884 | 0.593 | -0.002 | -0.001 | -0.001 | 0 | 0.64 | 0.67 | 0.76 | 0.69 | 0.68 | 0.86 |
| 2 | 181110770 | rs17189753 | 1.164 | 1.35 | 0.885 | 0.02 | -0.034 | -0.003 | -0.003 | 0.035 | 0.08 | 0.01 | 0.1 | 0.06 | 0.09 | 0.12 |
| 2 | 181113934 | rs6433839 | -0.538 | 1.33 | 0.931 | 0.01 | 0.014 | -0.021 | -0.021 | 0 | 0.9 | 0.76 | 0.89 | 0.94 | 0.91 | 0.88 |
| 2 | 181115711 | rs2368110 | 0.356 | 1.333 | 0.733 | 0.006 | -0.01 | 0.014 | 0.014 | 0 | 0.1 | 0.05 | 0.1 | 0.06 | 0.09 | 0.13 |
| 2 | 181115798 | rs4438449 | -1.048 | 1.366 | 0.772 | 0.014 | 0.014 | 0.004 | 0.004 | 0.602 | 0.27 | 0.37 | 0.4 | 0.26 | 0.34 | 0.36 |
| 2 | 181117047 | rs2887165 | 0.987 | 1.33 | 0.739 | -0.012 | 0 | 0.04 | 0.04 | 0.007 | 0.21 | 0.12 | 0.13 | 0.08 | 0.18 | 0.26 |
| 2 | 181119001 | rs2686145 | 0.34 | 1.415 | 0.771 | 0.081 | -0.001 | -0.001 | -0.001 | 0.676 | 0.1 | 0.15 | 0.1 | 0.06 | 0.09 | 0.12 |
| 2 | 181121917 | rs3936728 | -0.574 | 1.408 | 0.729 | 0.116 | 0.015 | 0.008 | 0.008 | 0.877 | 0.19 | 0.27 | 0.3 | 0.21 | 0.26 | 0.24 |
| 2 | 181122167 | rs1850680 | -0.087 | 1.474 | 0.7 | 0.127 | 0.006 | 0.004 | 0.004 | 2.245 | 0.17 | 0.2 | 0.29 | 0.21 | 0.25 | 0.24 |
| 2 | 181158949 | rs4893905 | 0.38 | 1.137 | 0.296 | 0.125 | 0.004 | -0.025 | -0.025 | 0 | 0.16 | 0.09 | 0.22 | 0.31 | 0.15 | 0.35 |
| 2 | 181169171 | rs16867289 | 0.785 | 1.122 | 0.258 | 0.128 | -0.003 | -0.025 | -0.025 | 0 | 0.14 | 0.04 | 0.22 | 0.23 | 0.14 | 0.22 |
| 2 | 181177756 | rs6755569 | -0.071 | 1.236 | 0.375 | 0.125 | 0.025 | 0.014 | 0.014 | 1.217 | 0.17 | 0.29 | 0.29 | 0.2 | 0.25 | 0.23 |
| 2 | 181184360 | rs1401262 | 1.375 | 1.147 | 0.089 | 0.081 | -0.015 | -0.031 | -0.031 | 0.158 | 0.59 | 0.18 | 0.44 | 0.49 | 0.59 | 0.42 |
| 2 | 181190720 | rs11686724 | 1.213 | 1.086 | -0.093 | -0.046 | -0.019 | 0.067 | 0.067 | 0 | 0.18 | 0.04 | 0.12 | 0 | 0.18 | 0.15 |
| 2 | 181206632 | rs6719837 | 0.142 | 1.2 | 0.713 | 0.816 | 0.003 | -0.007 | -0.007 | 0 | 0.15 | 0.12 | 0.22 | 0.22 | 0.14 | 0.24 |
| 2 | 181225083 | rs7589013 | 1.674 | 0.86 | 1.141 | 0.842 | -0.009 | 0.039 | 0.039 | 3.57 | 0.92 | 0.62 | 0.97 | 0.92 | 0.98 | 0.89 |
| 2 | 181242202 | rs16867303 | -0.193 | 1.113 | 0.903 | 0.824 | -0.053 | -0.006 | -0.006 | 0 | 0.24 | 0.1 | 0.29 | 0.27 | 0.28 | 0.39 |
| 2 | 181243761 | rs1829390 | 0.802 | 1.081 | 1.028 | 0.821 | 0.006 | -0.011 | -0.011 | 0 | 0.69 | 0.54 | 0.68 | 0.65 | 0.71 | 0.5 |
| 2 | 181249081 | rs12468410 | 0.473 | 1.068 | 0.678 | 0.82 | -0.011 | 0.002 | 0.002 | 0 | 0.08 | 0.04 | 0.11 | 0.07 | 0.1 | 0.13 |
| 2 | 181260626 | rs10439301 | 1.666 | 1.075 | 0.747 | 0.845 | -0.004 | 0.005 | 0.005 | 0.442 | 0.75 | 0.28 | 0.64 | 0.7 | 0.78 | 0.68 |
| 2 | 181264524 | rs13026665 | -0.417 | 1.177 | 1.111 | 0.944 | -0.029 | 0.004 | 0.004 | 0 | 0.24 | 0.13 | 0.3 | 0.27 | 0.28 | 0.39 |
| 2 | 181277132 | rs1823790 | 1.441 | 1.142 | 0.936 | 0.966 | -0.03 | -0.043 | -0.043 | 0 | 0.6 | 0.16 | 0.45 | 0.49 | 0.59 | 0.42 |
| 2 | 181279611 | rs10205905 | -0.999 | 1.258 | 1.011 | 1.149 | 0.038 | 0.184 | 0.184 | 0.45 | 0.12 | 0.53 | 0.23 | 0.24 | 0.11 | 0.19 |
| 2 | 181284675 | rs4894182 | -0.371 | 1.404 | 0.334 | 1.142 | -0.083 | -0.016 | -0.016 | 0 | 0.23 | 0.05 | 0.29 | 0.27 | 0.3 | 0.4 |
| 2 | 181290099 | rs1817147 | 1.406 | 0.651 | 1.218 | 1.251 | 0.023 | 0.103 | 0.103 | 2.673 | 0.87 | 0.42 | 0.75 | 0.84 | 0.88 | 0.91 |
| 2 | 181295340 | rs4894183 | -0.923 | 0.935 | 0.316 | 1.414 | 0.034 | 0.104 | 0.104 | 0 | 0.09 | 0.33 | 0.23 | 0.16 | 0.11 | 0.08 |
| 2 | 181314519 | rs7606036 | -0.397 | 0.821 | -0.408 | 0.979 | 0 | 0.008 | 0.008 | 0 | 0.49 | 0.59 | 0.46 | 0.43 | 0.5 | 0.67 |
| 2 | 181319803 | rs11886250 | -0.413 | 0.633 | 0.525 | 0.758 | -0.006 | 0.01 | 0.01 | 0.137 | 0.13 | 0.39 | 0.08 | 0.11 | 0.1 | 0.15 |
| 2 | 181333307 | rs12464199 | -0.319 | 0.794 | 0.631 | 0.975 | 0.006 | 0.042 | 0.042 | 0 | 0.16 | 0.23 | 0.38 | 0.34 | 0.19 | 0.3 |
| 2 | 181335172 | rs4894186 | -1.421 | 0.734 | 0.582 | 1.243 | -0.004 | 0.051 | 0.051 | 0 | 0.59 | 0.8 | 0.65 | 0.68 | 0.6 | 0.77 |
| 2 | 181335796 | rs717684 | -0.919 | 0.66 | 0.796 | 1.243 | 0.003 | 0.047 | 0.047 | 0 | 0.26 | 0.35 | 0.43 | 0.45 | 0.28 | 0.45 |
| 2 | 181339313 | rs1809134 | -1.002 | 0.84 | 0.664 | 1.059 | 0 | 0.002 | 0.002 | 0.121 | 0.27 | 0.27 | 0.44 | 0.45 | 0.33 | 0.5 |
| 2 | 181339431 | rs934882 | -1.017 | 0.843 | 0.665 | 1.06 | 0 | 0.002 | 0.002 | 0.151 | 0.27 | 0.27 | 0.44 | 0.45 | 0.34 | 0.5 |
| 2 | 181339971 | rs12693239 | -1.32 | 0.857 | 0.647 | 1.067 | 0.004 | 0.015 | 0.015 | 0.157 | 0.3 | 0.33 | 0.45 | 0.45 | 0.34 | 0.5 |
| 2 | 181345152 | rs980181 | -1.649 | 0.887 | 0.811 | 1.019 | -0.001 | -0.011 | -0.011 | 0 | 0.34 | 0.68 | 0.48 | 0.45 | 0.35 | 0.5 |
| 2 | 181351508 | rs1971135 | -0.937 | 0.954 | 0.953 | 0.984 | 0.004 | 0.022 | 0.022 | 0 | 0.28 | 0.36 | 0.45 | 0.45 | 0.29 | 0.48 |
| 2 | 181353800 | rs11889327 | 0.287 | 0.882 | 1.028 | 0.759 | 0.004 | 0.082 | 0.082 | 0 | 0.1 | 0.25 | 0.07 | 0.18 | 0.1 | 0.18 |
| 2 | 181361097 | rs16867319 | 0.349 | 0.879 | 0.998 | 0.757 | 0.001 | 0.074 | 0.074 | 0 | 0.1 | 0.25 | 0.07 | 0.19 | 0.1 | 0.18 |
| 2 | 181368236 | rs13025893 | -0.443 | 0.937 | 0.792 | 0.769 | 0.023 | 0.044 | 0.044 | 0.013 | 0.15 | 0.4 | 0.11 | 0.18 | 0.16 | 0.24 |
| 2 | 181369622 | rs1453062 | 1.028 | 0.891 | 0.485 | 0.65 | 0.001 | 0.014 | 0.014 | 0.059 | 0.22 | 0.18 | 0.13 | 0.05 | 0.2 | 0.13 |
| 2 | 181373822 | rs7600357 | -0.767 | 0.884 | 0.708 | 0.825 | 0.019 | 0.042 | 0.042 | 0.037 | 0.16 | 0.46 | 0.11 | 0.19 | 0.16 | 0.24 |
| 2 | 181378445 | rs719812 | -1.101 | 1.029 | 0.85 | 0.598 | 0.03 | -0.048 | -0.048 | 4.726 | 0.22 | 0.53 | 0.16 | 0.17 | 0.25 | 0.4 |
| 2 | 181381991 | rs714545 | -2.136 | 0.517 | 0.067 | 0.334 | -0.046 | -0.034 | -0.034 | 0 | 0.7 | 0.95 | 0.71 | 0.69 | 0.68 | 0.86 |
| 2 | 181387265 | rs721301 | -0.787 | 0.507 | 0.704 | 0.351 | -0.002 | 0.06 | 0.06 | 0 | 0.31 | 0.42 | 0.46 | 0.52 | 0.3 | 0.49 |
| 2 | 181388289 | rs7580616 | 0.509 | 0.529 | 0.88 | 0.293 | 0.071 | 0.008 | 0.008 | 0 | 0.05 | 0.3 | 0.04 | 0 | 0.05 | 0.05 |
| 2 | 181400836 | rs1349718 | 1.398 | 0.545 | 0.884 | 0.293 | 0 | 0 | 0 | 0 | 0.06 | 0.07 | 0.05 | 0.11 | 0.1 | 0.17 |
| 2 | 181403464 | rs10497567 | 0.527 | 0.554 | 1.045 | 0.275 | -0.008 | -0.051 | -0.051 | 0 | 0.11 | 0.21 | 0.07 | 0.06 | 0.09 | 0.19 |
| 2 | 181417009 | rs13025932 | -0.793 | 0.565 | 0.804 | 0.381 | 0.036 | 0.013 | 0.013 | 0.073 | 0.16 | 0.4 | 0.1 | 0.15 | 0.17 | 0.22 |
| 2 | 181433460 | rs1528437 | 1.168 | 0.398 | 0.487 | 0.151 | 0.005 | 0.055 | 0.055 | 0 | 0.21 | 0.09 | 0.13 | 0.05 | 0.19 | 0.13 |
| 2 | 181438352 | rs7575156 | -0.345 | 0.35 | 0.418 | 0.175 | 0.001 | -0.027 | -0.027 | 0 | 0.35 | 0.42 | 0.22 | 0.16 | 0.34 | 0.35 |
| 2 | 181440081 | rs1358520 | 0.179 | 0.308 | 0.639 | 0.2 | 0 | 0.02 | 0.02 | 0 | 0.19 | 0.18 | 0.38 | 0.46 | 0.2 | 0.3 |
| 2 | 181444035 | rs13028421 | 0.586 | 0.22 | 0.556 | 0.136 | -0.001 | 0.063 | 0.063 | 0 | 0.68 | 0.5 | 0.52 | 0.49 | 0.68 | 0.5 |
| 2 | 181448039 | rs13006569 | -1.168 | 0.199 | 0.213 | 0.12 | -0.039 | -0.011 | -0.011 | 0 | 0.55 | 0.84 | 0.62 | 0.56 | 0.55 | 0.68 |
| 2 | 181449266 | rs17198592 | 1.375 | 0.186 | 0.211 | 0.083 | -0.007 | 0.056 | 0.056 | 0 | 0.19 | 0.03 | 0.12 | 0.05 | 0.18 | 0.13 |
| 2 | 181457236 | rs10930926 | 0.233 | 0.158 | 0.453 | 0.279 | 0.004 | 0.014 | 0.014 | 0 | 0.3 | 0.23 | 0.2 | 0.16 | 0.29 | 0.34 |
| 2 | 181458934 | rs16867335 | -0.596 | 0.241 | 0.509 | 0.803 | -0.021 | -0.063 | -0.063 | 0 | 0.15 | 0.07 | 0.26 | 0.3 | 0.18 | 0.23 |
| 2 | 181463705 | rs4666665 | 0.358 | 0.255 | 0.7 | 0.807 | 0 | 0.017 | 0.017 | 0 | 0.15 | 0.18 | 0.19 | 0.27 | 0.13 | 0.23 |
| 2 | 181465853 | rs6739798 | -0.463 | 0.255 | 0.695 | 0.808 | -0.024 | -0.064 | -0.064 | 0 | 0.14 | 0.07 | 0.26 | 0.3 | 0.18 | 0.24 |
| 2 | 181467278 | rs11900625 | -0.733 | 0.246 | 0.887 | 0.796 | 0.043 | 0.003 | 0.003 | 0.576 | 0.22 | 0.48 | 0.17 | 0.23 | 0.26 | 0.41 |
| 2 | 181468010 | rs991055 | -0.841 | 0.238 | 0.875 | 0.794 | -0.005 | 0.036 | 0.036 | 0 | 0.27 | 0.49 | 0.28 | 0.38 | 0.24 | 0.43 |
| 2 | 181470028 | rs10164739 | -0.885 | 0.247 | 0.856 | 0.792 | 0.034 | 0.058 | 0.058 | 0 | 0.26 | 0.43 | 0.28 | 0.38 | 0.28 | 0.43 |
| 2 | 181470838 | rs1477499 | -2.024 | 0.238 | 0.542 | 0.794 | -0.021 | 0.07 | 0.07 | 0 | 0.46 | 0.78 | 0.56 | 0.68 | 0.47 | 0.71 |
| 2 | 181489592 | rs6746974 | -1.4 | 0.251 | 0.584 | 0.808 | 0.01 | 0.047 | 0.047 | 0.012 | 0.3 | 0.57 | 0.37 | 0.41 | 0.34 | 0.47 |

| | | | | | | | | | | | | | | | | |
|---|---|---|---|---|---|---|---|---|---|---|---|---|---|---|---|---|
| 2 | 181492024 | rs11896300 | -1.548 | 0.251 | 0.574 | 0.809 | -0.01 | -0.017 | -0.017 | 1.536 | 0.37 | 0.71 | 0.37 | 0.5 | 0.39 | 0.63 |
| 2 | 181492790 | rs13429179 | 0.314 | 0.271 | 0.369 | 0.846 | 0 | 0.017 | 0.017 | 0 | 0.15 | 0.17 | 0.19 | 0.27 | 0.13 | 0.23 |
| 2 | 181501242 | rs2368177 | -1.408 | 0.267 | 0.395 | 0.822 | 0 | 0.001 | 0.001 | 0 | 0.3 | 0.34 | 0.46 | 0.57 | 0.31 | 0.46 |
| 2 | 181513729 | rs918959 | -0.123 | 0.219 | 0.409 | 0.856 | 0.007 | -0.019 | -0.019 | 0 | 0.08 | 0.04 | 0.2 | 0.19 | 0.08 | 0.07 |
| 2 | 181516167 | rs2194691 | -0.155 | 0.187 | 0.526 | 0.738 | 0.025 | -0.039 | -0.039 | 0 | 0.16 | 0.49 | 0.11 | 0.11 | 0.16 | 0.25 |
| 2 | 181516693 | rs4667318 | -0.135 | 0.125 | 0.47 | 1.006 | 0 | 0.016 | 0.016 | 0 | 0.15 | 0.18 | 0.2 | 0.27 | 0.13 | 0.22 |
| 2 | 181516696 | rs16867346 | -0.054 | 0.106 | 0.339 | 0.992 | 0.002 | -0.002 | -0.002 | 0 | 0.07 | 0.04 | 0.2 | 0.19 | 0.08 | 0.08 |
| 2 | 181518008 | rs1528451 | -1.343 | 0.015 | -0.092 | 0.377 | 0.001 | 0.015 | 0.015 | 0.002 | 0.4 | 0.29 | 0.54 | 0.59 | 0.38 | 0.49 |
| 2 | 181521287 | rs12988791 | 0.16 | 0.13 | 0.341 | 0.594 | -0.001 | 0.002 | 0.002 | 0 | 0.33 | 0.34 | 0.22 | 0.19 | 0.34 | 0.37 |
| 2 | 181521345 | rs1406257 | -1.246 | 0.258 | 0.362 | 0.565 | -0.001 | -0.003 | -0.003 | 0 | 0.32 | 0.3 | 0.51 | 0.57 | 0.32 | 0.48 |
| 2 | 181522438 | rs11686983 | -0.103 | 0.259 | 0.375 | 0.696 | 0.001 | 0.004 | 0.004 | 0 | 0.18 | 0.22 | 0.24 | 0.24 | 0.15 | 0.23 |
| 2 | 181522783 | rs6727573 | -1.466 | 0.258 | 0.373 | 0.655 | 0 | 0.001 | 0.001 | 0 | 0.57 | 0.61 | 0.68 | 0.62 | 0.58 | 0.68 |
| 2 | 181522907 | rs6712389 | 0.898 | 0.292 | 0.305 | 0.593 | 0 | -0.037 | -0.037 | 0 | 0.25 | 0.33 | 0.17 | 0.05 | 0.25 | 0.2 |
| 2 | 181542055 | rs3845691 | -0.377 | 0.174 | 0.281 | 0.686 | 0 | -0.004 | -0.004 | 0.547 | 0.42 | 0.45 | 0.4 | 0.28 | 0.41 | 0.44 |
| 2 | 181550896 | rs1528436 | -0.941 | 0.156 | 0.281 | 0.696 | 0.004 | 0 | 0 | 0 | 0.5 | 0.47 | 0.45 | 0.42 | 0.5 | 0.6 |
| 2 | 181572460 | rs1528434 | 0.775 | 0.3 | 0.639 | 0.988 | -0.002 | 0.025 | 0.025 | 0.919 | 0.79 | 0.34 | 0.73 | 0.65 | 0.83 | 0.75 |
| 2 | 181574075 | rs16823635 | 0.028 | 0.372 | 0.075 | 1.045 | 0.005 | 0.003 | 0.003 | 0 | 0.07 | 0.04 | 0.21 | 0.2 | 0.08 | 0.08 |
| 2 | 181586333 | rs7556967 | -0.81 | 0.455 | 0.144 | 0.904 | 0.01 | 0.014 | 0.014 | 0 | 0.19 | 0.27 | 0.42 | 0.24 | 0.2 | 0.21 |
| 2 | 181586726 | rs13029379 | 0.992 | 0.515 | -0.043 | 1.024 | -0.022 | -0.04 | -0.04 | 0 | 0.07 | 0.02 | 0.12 | 0.14 | 0.09 | 0.17 |
| 2 | 181590915 | rs1025127 | -0.156 | 0.445 | 0.108 | 0.92 | 0.013 | 0.045 | 0.045 | 0 | 0.13 | 0.23 | 0.19 | 0.24 | 0.11 | 0.19 |
| 2 | 181593247 | rs13000621 | 0.032 | 0.341 | -0.343 | 0.694 | 0.001 | -0.011 | -0.011 | 0 | 0.54 | 0.61 | 0.5 | 0.54 | 0.53 | 0.54 |
| 2 | 181597561 | rs6728513 | 0.256 | 0.408 | -0.405 | 0.612 | 0.002 | 0.049 | 0.049 | 0 | 0.42 | 0.27 | 0.31 | 0.19 | 0.39 | 0.37 |
| 2 | 181602526 | rs1019612 | -1.503 | 0.318 | -0.42 | 0.646 | -0.003 | 0.013 | 0.013 | 0 | 0.69 | 0.74 | 0.69 | 0.75 | 0.65 | 0.76 |
| 2 | 181604341 | rs13398168 | -0.71 | 0.197 | 0.243 | 1.668 | 0.02 | 0.059 | 0.059 | 0.49 | 0.16 | 0.31 | 0.14 | 0.22 | 0.16 | 0.22 |
| 2 | 181609561 | rs4402718 | 0.942 | 0.297 | 0.137 | 1.048 | 0.001 | 0.072 | 0.072 | 0.274 | 0.74 | 0.56 | 0.62 | 0.46 | 0.74 | 0.65 |
| 2 | 181615426 | rs11893311 | -0.027 | 0.271 | -0.705 | 1.031 | -0.007 | 0.002 | 0.002 | 0 | 0.05 | 0.05 | 0.07 | 0 | 0.08 | 0.13 |
| 2 | 181625664 | rs4362518 | 1.208 | 0.195 | -0.775 | 1.083 | -0.002 | -0.003 | -0.003 | 0 | 0.65 | 0.35 | 0.5 | 0.31 | 0.71 | 0.5 |
| 2 | 181626522 | rs16867365 | 0.021 | 0.457 | -1.035 | 1.09 | -0.002 | -0.041 | -0.041 | 0 | 0.07 | 0 | 0.11 | 0.12 | 0.05 | 0.12 |
| 2 | 181628139 | rs10211365 | -0.113 | 0.617 | -0.781 | 0.94 | 0 | -0.002 | -0.002 | 0.494 | 0.42 | 0.39 | 0.55 | 0.51 | 0.45 | 0.34 |
| 2 | 181642212 | rs10930936 | 0.544 | 0.473 | 0.35 | 1.334 | -0.003 | 0.077 | 0.077 | 0.03 | 0.8 | 0.34 | 0.61 | 0.61 | 0.82 | 0.8 |
| 2 | 181647026 | rs10497572 | -0.211 | 0.312 | -0.216 | 1.195 | 0.019 | 0.014 | 0.014 | 0 | 0.45 | 0.28 | 0.28 | 0.39 | 0.34 | 0.48 |
| 2 | 181659051 | rs11686097 | -0.764 | 0.582 | 0.172 | 0.929 | 0.01 | 0.002 | 0.002 | 0 | 0.07 | 0.21 | 0.06 | 0.06 | 0.07 | 0.1 |
| 2 | 181664063 | rs12693257 | -1.061 | 0.232 | 0.353 | 0.946 | -0.011 | -0.007 | -0.007 | 0 | 0.36 | 0.54 | 0.49 | 0.34 | 0.3 | 0.41 |
| 2 | 181665771 | rs4550629 | -1.212 | 1.335 | 0.221 | 0.779 | -0.04 | 0.042 | 0.042 | 0 | 0.42 | 0.8 | 0.59 | 0.34 | 0.48 | 0.44 |
| 2 | 181678814 | rs10174914 | -1.701 | 0.931 | 1.05 | 1.402 | -0.022 | 0.179 | 0.179 | 0.819 | 0.18 | 0.59 | 0.4 | 0.39 | 0.12 | 0.27 |
| 2 | 181684892 | rs4563185 | 0.012 | 0.564 | 1.422 | 0.998 | -0.001 | -0.033 | -0.033 | 0 | 0.2 | 0.17 | 0.12 | 0.01 | 0.14 | 0.16 |
| 2 | 181693910 | rs6729229 | -1.677 | 0.731 | 1.777 | 1.426 | -0.025 | 0.106 | 0.106 | 1.514 | 0.12 | 0.51 | 0.09 | 0.16 | 0.08 | 0.13 |
| 2 | 181698136 | rs6758984 | -1.051 | 0.76 | 1.966 | 1.2 | -0.005 | -0.006 | -0.006 | 0 | 0.07 | 0.16 | 0.06 | 0.03 | 0.07 | 0.05 |
| 2 | 181716272 | rs12624291 | 2.491 | 1.147 | 1.931 | 1.199 | 0.051 | 0.035 | 0.035 | 1.27 | 0.82 | 0.32 | 0.52 | 0.75 | 0.72 | 0.88 |
| 2 | 181723942 | rs4128937 | 2.506 | 1.121 | 1.934 | 1.202 | 0.047 | 0.033 | 0.033 | 1.314 | 0.82 | 0.32 | 0.52 | 0.75 | 0.72 | 0.87 |
| 2 | 181733814 | rs12614968 | 2.149 | 1.141 | 1.808 | 1.025 | 0.097 | -0.059 | -0.059 | 0 | 0.86 | 0.38 | 0.82 | 0.97 | 0.76 | 0.94 |
| 2 | 181750912 | rs4614891 | 1.659 | 1.115 | 1.873 | 1.086 | 0.192 | 0.109 | 0.109 | 0 | 0.89 | 0.46 | 0.59 | 0.79 | 0.78 | 0.91 |
| 2 | 181757693 | rs13031974 | -1.034 | 0.627 | 1.44 | 1.148 | -0.001 | -0.002 | -0.002 | 0 | 0.07 | 0.17 | 0.07 | 0.04 | 0.08 | 0.07 |
| 2 | 181772262 | rs3922472 | -0.983 | 0.667 | 1.176 | 1.187 | 0.016 | 0.103 | 0.103 | 0 | 0.06 | 0.22 | 0.32 | 0.21 | 0.04 | 0.07 |
| 2 | 181784642 | rs13011269 | 1.187 | 0.147 | 0.074 | 1.43 | -0.006 | -0.075 | -0.075 | 0.02 | 0.12 | 0.01 | 0.14 | 0.23 | 0.12 | 0.06 |
| 2 | 181784667 | rs10048671 | 0.644 | 0.411 | 0.15 | 0.514 | 0.032 | -0.032 | -0.032 | 0.774 | 0.71 | 0.46 | 0.75 | 0.78 | 0.64 | 0.77 |
| 2 | 181785294 | rs10048799 | 0.568 | 0.244 | -0.768 | 0.134 | 0.012 | 0.038 | 0.038 | 0.06 | 0.48 | 0.31 | 0.56 | 0.6 | 0.44 | 0.51 |
| 2 | 181787645 | rs10209735 | -0.65 | -0.173 | -0.379 | -0.018 | -0.03 | -0.043 | -0.043 | 0.955 | 0.71 | 0.48 | 0.76 | 0.81 | 0.75 | 0.73 |
| 2 | 181787694 | rs10209813 | -0.098 | -0.21 | -0.507 | -0.119 | 0.014 | 0.019 | 0.019 | 0 | 0.62 | 0.32 | 0.68 | 0.78 | 0.67 | 0.64 |
| 2 | 181789840 | rs6433874 | 0.431 | 0.079 | -0.443 | -0.288 | -0.012 | -0.028 | -0.028 | 0.925 | 0.09 | 0.19 | 0.07 | 0.03 | 0.07 | 0.09 |
| 2 | 181793615 | rs4130586 | -1.006 | 0.018 | -0.725 | -0.399 | -0.008 | -0.051 | -0.051 | 0 | 0.71 | 0.68 | 0.82 | 0.93 | 0.74 | 0.76 |
| 2 | 181794257 | rs4638754 | 0.604 | 0.325 | -1.104 | -0.323 | -0.134 | 0.008 | 0.008 | 0 | 0.19 | 0.02 | 0.2 | 0.15 | 0.38 | 0.16 |
| 2 | 181802404 | rs4614892 | -0.137 | 0.008 | -0.806 | -0.151 | -0.007 | 0.02 | 0.02 | 0 | 0.28 | 0.39 | 0.45 | 0.37 | 0.25 | 0.28 |
| 2 | 181803345 | rs10184337 | 1.245 | -0.005 | -0.785 | -0.108 | -0.009 | 0.015 | 0.015 | 0.249 | 0.06 | 0.18 | 0.03 | 0.06 | 0.02 | 0.08 |
| 2 | 181805301 | rs10930945 | 0.186 | -0.114 | -1.441 | -0.312 | -0.036 | 0.008 | 0.008 | 0 | 0.42 | 0.15 | 0.42 | 0.59 | 0.51 | 0.51 |
| 2 | 181807231 | rs10173769 | -1.581 | 0.28 | -1.866 | -0.236 | 0.001 | -0.036 | -0.036 | 0 | 0.71 | 0.73 | 0.87 | 0.96 | 0.77 | 0.79 |
| 2 | 181807616 | rs6759936 | 0.234 | -0.055 | -1.294 | 0.448 | -0.002 | -0.016 | -0.016 | 0 | 0.22 | 0.17 | 0.42 | 0.33 | 0.24 | 0.2 |
| 2 | 181810948 | rs10192997 | -0.356 | 0.22 | -1.378 | -0.321 | 0.042 | 0.104 | 0.104 | 0 | 0.48 | 0.67 | 0.62 | 0.79 | 0.34 | 0.62 |
| 2 | 181811957 | rs12165234 | -0.159 | 0.248 | -1.319 | -0.159 | 0.024 | 0.049 | 0.049 | 0.004 | 0.46 | 0.64 | 0.57 | 0.72 | 0.32 | 0.57 |
| 2 | 181814381 | rs4233784 | -0.363 | 0.489 | -1.004 | -0.765 | 0.014 | -0.024 | -0.024 | 0 | 0.72 | 0.67 | 0.73 | 0.8 | 0.57 | 0.78 |
| 2 | 181815227 | rs4531896 | 1.591 | 0.557 | -1.83 | -0.47 | -0.076 | -0.121 | -0.121 | 0.019 | 0.15 | 0.01 | 0.35 | 0.46 | 0.17 | 0.25 |
| 2 | 181817351 | rs4618005 | 0.657 | -0.029 | -1.254 | -0.832 | -0.019 | -0.033 | -0.033 | 0 | 0.34 | 0.19 | 0.45 | 0.56 | 0.3 | 0.53 |
| 2 | 181823924 | rs13414852 | 0.687 | 0.021 | -1.152 | -0.658 | -0.098 | 0.059 | 0.059 | 0 | 0.24 | 0.05 | 0.25 | 0.16 | 0.39 | 0.16 |
| 2 | 181825225 | rs6717105 | 0.386 | -0.336 | -0.651 | -0.841 | -0.02 | 0.001 | 0.001 | 0 | 0.26 | 0.36 | 0.23 | 0.25 | 0.22 | 0.23 |
| 2 | 181825338 | rs4471850 | 0.482 | -0.49 | -0.843 | -0.698 | -0.018 | -0.006 | -0.006 | 0 | 0.43 | 0.21 | 0.47 | 0.56 | 0.36 | 0.55 |
| 2 | 181829296 | rs4638755 | 0.774 | -0.31 | -0.651 | -0.526 | -0.01 | -0.007 | -0.007 | 1.429 | 0.39 | 0.29 | 0.54 | 0.59 | 0.38 | 0.54 |
| 2 | 181834018 | rs4602187 | 0.333 | -0.25 | -0.424 | -0.598 | -0.004 | -0.005 | -0.005 | 0 | 0.58 | 0.46 | 0.72 | 0.68 | 0.67 | 0.72 |
| 2 | 181844316 | rs4595925 | -0.502 | -0.111 | -0.322 | -0.635 | 0.001 | 0.002 | 0.002 | 0.018 | 0.72 | 0.83 | 0.82 | 0.88 | 0.72 | 0.85 |
| 2 | 181862579 | rs4667005 | -0.474 | -0.089 | -0.352 | -0.611 | 0.004 | 0.026 | 0.026 | 0.047 | 0.72 | 0.85 | 0.82 | 0.87 | 0.72 | 0.85 |

| | | | | | | | | | | | | | | | | |
|---|---|---|---|---|---|---|---|---|---|---|---|---|---|---|---|---|
| 2 | 181863900 | rs1107786 | 1.025 | 0.079 | -0.346 | -0.57 | -0.028 | 0.02 | 0.02 | 0 | 0.19 | 0.12 | 0.18 | 0.09 | 0.29 | 0.17 |
| 2 | 181878770 | rs17668214 | 1.05 | 0.086 | -0.483 | -0.567 | -0.045 | 0.021 | 0.021 | 0 | 0.19 | 0.09 | 0.18 | 0.09 | 0.29 | 0.17 |
| 2 | 181903964 | rs6729509 | 0.14 | -0.128 | -0.12 | -0.654 | -0.01 | 0.078 | 0.078 | 0.56 | 0.54 | 0.73 | 0.64 | 0.8 | 0.44 | 0.69 |
| 2 | 181911489 | rs12477135 | 0.973 | -0.327 | 0.123 | -0.615 | -0.016 | -0.006 | -0.006 | 0 | 0.39 | 0.27 | 0.54 | 0.61 | 0.39 | 0.55 |
| 2 | 181913040 | rs13418851 | -0.256 | -0.295 | 0.164 | -0.572 | -0.009 | 0.099 | 0.099 | 0 | 0.06 | 0.19 | 0.07 | 0.2 | 0.03 | 0.11 |
| 2 | 181929264 | rs13406825 | -0.445 | -0.311 | 0.177 | -0.539 | -0.006 | 0.124 | 0.124 | 0 | 0.06 | 0.21 | 0.07 | 0.2 | 0.03 | 0.14 |
| 2 | 181952420 | rs11893940 | 0.911 | -0.276 | 0.059 | -0.56 | -0.011 | -0.006 | -0.006 | 0 | 0.39 | 0.31 | 0.55 | 0.6 | 0.39 | 0.55 |
| 2 | 181953354 | rs7349232 | 1.038 | -0.288 | -0.123 | -0.77 | -0.001 | -0.008 | -0.008 | 0 | 0.21 | 0.15 | 0.13 | 0.1 | 0.21 | 0.13 |
| 2 | 181967075 | rs2368196 | 0.132 | -0.364 | -0.049 | -0.697 | -0.004 | -0.004 | -0.004 | 0 | 0.67 | 0.46 | 0.73 | 0.7 | 0.66 | 0.69 |
| 2 | 181969931 | rs908733 | 0.997 | -0.139 | -0.195 | -1.001 | -0.002 | 0.025 | 0.025 | 0 | 0.41 | 0.31 | 0.55 | 0.7 | 0.38 | 0.6 |
| 2 | 181971800 | rs6706397 | -0.364 | -0.055 | -0.79 | -1.003 | -0.002 | -0.014 | -0.014 | 0 | 0.34 | 0.36 | 0.23 | 0.17 | 0.31 | 0.14 |
| 2 | 181980885 | rs1040228 | -0.942 | -0.042 | -0.776 | -1.053 | -0.011 | 0.02 | 0.02 | 0 | 0.77 | 0.81 | 0.81 | 0.9 | 0.7 | 0.82 |
| 2 | 181987553 | rs13020845 | 0.329 | -0.073 | -0.837 | -1.116 | 0.002 | -0.023 | -0.023 | 0 | 0.24 | 0.29 | 0.17 | 0.16 | 0.22 | 0.13 |
| 2 | 181988909 | rs7588217 | 0.992 | -0.084 | -0.88 | -1.119 | -0.001 | -0.013 | -0.013 | 0 | 0.43 | 0.49 | 0.59 | 0.73 | 0.39 | 0.68 |
| 2 | 181991389 | rs4077924 | -1.322 | -0.059 | -0.902 | -1.056 | -0.024 | 0.021 | 0.021 | 0 | 0.79 | 0.89 | 0.82 | 0.9 | 0.71 | 0.82 |
| 2 | 182018554 | rs7558428 | -0.356 | -0.178 | -0.715 | -0.917 | -0.002 | 0.022 | 0.022 | 0 | 0.31 | 0.23 | 0.21 | 0.11 | 0.31 | 0.13 |
| 2 | 182024283 | rs6738120 | 0.227 | -0.202 | -0.65 | -0.862 | -0.015 | -0.049 | -0.049 | 0 | 0.15 | 0.19 | 0.07 | 0 | 0.11 | 0.04 |
| 2 | 182030229 | rs10189776 | 1.063 | -0.366 | -0.765 | -0.865 | 0.008 | -0.018 | -0.018 | 0 | 0.42 | 0.47 | 0.58 | 0.68 | 0.44 | 0.58 |
| 2 | 182033294 | rs11693055 | 0.831 | -0.377 | -0.839 | -0.888 | -0.03 | 0.033 | 0.033 | 0 | 0.2 | 0.05 | 0.15 | 0.15 | 0.21 | 0.17 |
| 2 | 182034448 | rs6433897 | -0.92 | -0.431 | -0.808 | -0.908 | -0.002 | -0.005 | -0.005 | 1.339 | 0.77 | 0.71 | 0.8 | 0.82 | 0.75 | 0.79 |
| 2 | 182039477 | rs6735551 | 1.144 | -0.484 | -0.755 | -0.87 | 0.006 | -0.009 | -0.009 | 0.54 | 0.41 | 0.45 | 0.58 | 0.67 | 0.44 | 0.58 |
| 2 | 182042445 | rs4305233 | 0.062 | -0.405 | -1.214 | -0.873 | -0.005 | -0.015 | -0.015 | 0 | 0.64 | 0.58 | 0.75 | 0.82 | 0.66 | 0.75 |
| 2 | 182042947 | rs13031976 | -0.013 | -0.406 | -1.289 | -0.911 | 0 | 0 | 0 | 0 | 0.67 | 0.67 | 0.75 | 0.82 | 0.66 | 0.76 |
| 2 | 182045054 | rs4462758 | -1.454 | -0.296 | -1.268 | -0.815 | -0.005 | 0.01 | 0.01 | 0 | 0.82 | 0.89 | 0.84 | 0.88 | 0.76 | 0.81 |
| 2 | 182047921 | rs6433900 | -0.384 | -0.23 | -1.229 | -0.765 | 0.008 | 0.033 | 0.033 | 0 | 0.71 | 0.78 | 0.78 | 0.88 | 0.67 | 0.79 |
| 2 | 182062879 | rs12616042 | 0.566 | -0.086 | -1.319 | -0.68 | -0.003 | 0.009 | 0.009 | 0 | 0.46 | 0.55 | 0.6 | 0.7 | 0.44 | 0.66 |
| 2 | 182063698 | rs7599846 | 0.458 | 0.156 | -1.451 | -0.597 | -0.002 | -0.008 | -0.008 | 2.539 | 0.07 | 0.07 | 0.05 | 0.08 | 0.05 | 0.13 |
| 2 | 182068527 | rs13035276 | 1.013 | 0.166 | -1.417 | -0.703 | -0.04 | 0.028 | 0.028 | 0 | 0.17 | 0.04 | 0.15 | 0.13 | 0.21 | 0.09 |
| 2 | 182073998 | rs6713632 | 0.195 | 0.198 | -1.281 | -0.617 | -0.01 | 0.083 | 0.083 | 0.114 | 0.31 | 0.14 | 0.21 | 0.13 | 0.32 | 0.13 |
| 2 | 182083827 | rs6717590 | -0.341 | 0.209 | -0.81 | 0.04 | -0.005 | -0.008 | -0.008 | 0 | 0.77 | 0.73 | 0.82 | 0.83 | 0.76 | 0.79 |
| 2 | 182088349 | rs4233796 | -0.341 | 0.209 | -0.81 | 0.04 | -0.005 | -0.008 | -0.008 | 0.006 | 0.77 | 0.73 | 0.82 | 0.83 | 0.76 | 0.79 |
| 2 | 182111206 | rs16867384 | 0.633 | 1.051 | -1.715 | 0.816 | -0.085 | -0.14 | -0.14 | 0 | 0.17 | 0.01 | 0.4 | 0.51 | 0.23 | 0.3 |
| 2 | 182117625 | rs877226 | 0.843 | 0.542 | -1.451 | -0.12 | 0.003 | -0.055 | -0.055 | 0 | 0.3 | 0.32 | 0.21 | 0.1 | 0.3 | 0.21 |
| 2 | 182128798 | rs1899037 | -1.402 | 0.025 | -1.916 | 0.173 | -0.056 | -0.014 | -0.014 | 0 | 0.81 | 0.98 | 0.84 | 0.87 | 0.75 | 0.79 |
| 2 | 182132986 | rs1030964 | -1.293 | 0.001 | -1.939 | 0.141 | -0.054 | -0.011 | -0.011 | 0 | 0.81 | 0.98 | 0.83 | 0.87 | 0.75 | 0.79 |
| 2 | 182138557 | rs7566314 | -1.306 | 0.029 | -1.919 | 0.154 | -0.046 | -0.005 | -0.005 | 2.471 | 0.81 | 0.95 | 0.84 | 0.87 | 0.75 | 0.79 |
| 2 | 182139108 | rs12467604 | 0.014 | -0.394 | -1.682 | -0.136 | 0.005 | 0.001 | 0.001 | 0 | 0.61 | 0.62 | 0.66 | 0.63 | 0.57 | 0.71 |
| 2 | 182156402 | rs13393709 | -0.401 | -0.378 | -1.978 | -0.258 | 0.005 | 0.004 | 0.004 | 0 | 0.39 | 0.32 | 0.33 | 0.35 | 0.44 | 0.29 |
| 2 | 182160053 | rs13339833 | -0.293 | -0.353 | -2.014 | -0.237 | 0.003 | 0.007 | 0.007 | 0 | 0.41 | 0.26 | 0.31 | 0.38 | 0.45 | 0.41 |
| 2 | 182167100 | rs10206405 | -0.369 | -0.04 | -2.088 | 0.329 | 0.005 | 0.009 | 0.009 | 0 | 0.42 | 0.24 | 0.31 | 0.38 | 0.45 | 0.41 |
| 2 | 182167931 | rs16867401 | 0.61 | -0.101 | -2.018 | 0.314 | 0.019 | 0.055 | 0.055 | 0.377 | 0.17 | 0.08 | 0.08 | 0.03 | 0.13 | 0.16 |
| 2 | 182189405 | rs16867406 | 0.851 | -0.09 | -2.067 | 0.303 | 0.006 | 0.082 | 0.082 | 0.981 | 0.16 | 0.03 | 0.08 | 0.03 | 0.14 | 0.16 |
| 2 | 182190965 | rs1440046 | -0.792 | 0.011 | -2.218 | 0.405 | -0.043 | -0.057 | -0.057 | 0 | 0.78 | 0.94 | 0.63 | 0.72 | 0.78 | 0.71 |
| 2 | 182206080 | rs7557202 | -0.034 | 0.047 | -2.115 | 0.395 | -0.014 | -0.023 | -0.023 | 0 | 0.34 | 0.16 | 0.3 | 0.39 | 0.42 | 0.36 |
| 2 | 182206328 | rs6433908 | -0.034 | 0.047 | -2.115 | 0.395 | -0.014 | -0.023 | -0.023 | 1.308 | 0.34 | 0.16 | 0.3 | 0.39 | 0.42 | 0.36 |
| 2 | 182206802 | rs959382 | -0.044 | -0.002 | -2.116 | 0.394 | -0.011 | -0.022 | -0.022 | 2.21 | 0.34 | 0.16 | 0.3 | 0.39 | 0.42 | 0.36 |
| 2 | 182213375 | rs10497579 | -0.157 | -0.01 | -2.077 | 0.389 | -0.001 | -0.006 | -0.006 | 0 | 0.35 | 0.29 | 0.31 | 0.39 | 0.42 | 0.36 |
| 2 | 182220184 | rs10184153 | 0.182 | 0.226 | -1.869 | 0.264 | 0 | 0.006 | 0.006 | 0 | 0.25 | 0.3 | 0.24 | 0.38 | 0.3 | 0.22 |
| 2 | 182226778 | rs16867420 | 1.464 | 0.369 | -2.292 | 0.298 | -0.04 | -0.077 | -0.077 | 0 | 0.13 | 0.01 | 0.31 | 0.24 | 0.14 | 0.18 |
| 2 | 182228931 | rs6726597 | 1.217 | 0.634 | -1.822 | -0.076 | -0.057 | -0.026 | -0.026 | 0 | 0.16 | 0.01 | 0.13 | 0.12 | 0.23 | 0.18 |
| 2 | 182236208 | rs6705323 | 0.525 | 0.549 | -1.502 | -0.32 | 0.01 | -0.011 | -0.011 | 0 | 0.36 | 0.48 | 0.31 | 0.37 | 0.38 | 0.22 |
| 2 | 182236365 | rs6734093 | -0.32 | 0.502 | -1.501 | -0.336 | 0.003 | -0.003 | -0.003 | 0 | 0.58 | 0.56 | 0.46 | 0.48 | 0.64 | 0.43 |
| 2 | 182251813 | rs12616646 | -0.426 | 0.401 | -1.373 | -0.306 | 0 | -0.001 | -0.001 | 0.25 | 0.62 | 0.57 | 0.49 | 0.49 | 0.68 | 0.48 |
| 2 | 182253217 | rs10211229 | 0.693 | 0.299 | -1.443 | -1.34 | 0.014 | -0.107 | -0.107 | 0 | 0.36 | 0.45 | 0.27 | 0.11 | 0.35 | 0.21 |
| 2 | 182256688 | rs7577055 | 0.663 | 0.368 | -1.408 | -1.306 | 0.011 | -0.106 | -0.106 | 0 | 0.37 | 0.45 | 0.26 | 0.11 | 0.35 | 0.21 |
| 2 | 182258150 | rs12693275 | -0.136 | -0.196 | -0.959 | -0.768 | -0.012 | -0.027 | -0.027 | 0 | 0.6 | 0.51 | 0.42 | 0.22 | 0.65 | 0.48 |
| 2 | 182261869 | rs7602460 | 0.331 | -0.075 | -0.966 | -0.668 | 0.014 | 0.026 | 0.026 | 0 | 0.53 | 0.57 | 0.41 | 0.22 | 0.6 | 0.43 |
| 2 | 182263951 | rs831864 | 0.137 | -0.156 | -0.954 | -0.101 | 0.008 | -0.006 | -0.006 | 0 | 0.85 | 0.89 | 0.87 | 0.87 | 0.89 | 0.89 |
| 2 | 182273419 | rs16867426 | -0.241 | 0.062 | -0.605 | -0.764 | -0.006 | -0.023 | -0.023 | 0 | 0.41 | 0.49 | 0.5 | 0.84 | 0.29 | 0.56 |
| 2 | 182289461 | rs2035207 | -0.26 | 0.231 | -0.628 | -0.563 | -0.01 | -0.032 | -0.032 | 0.434 | 0.43 | 0.51 | 0.49 | 0.84 | 0.31 | 0.55 |
| 2 | 182293443 | rs165432 | 0.046 | 0.94 | -0.692 | 0.05 | -0.002 | -0.007 | -0.007 | 0 | 0.92 | 0.92 | 0.91 | 1 | 0.88 | 0.95 |
| 2 | 182305254 | rs155128 | -0.194 | 0.871 | -0.511 | -0.053 | 0.003 | -0.044 | -0.044 | 0 | 0.31 | 0.48 | 0.22 | 0.18 | 0.36 | 0.25 |
| 2 | 182305424 | rs16867428 | -0.395 | 1.258 | -0.778 | 0.912 | 0.012 | 0.044 | 0.044 | 0 | 0.06 | 0.07 | 0.08 | 0.24 | 0.11 | 0.13 |
| 2 | 182308352 | rs6740847 | -0.186 | 1.118 | -1.187 | 0.89 | 0.003 | 0.053 | 0.053 | 0 | 0.49 | 0.67 | 0.45 | 0.34 | 0.54 | 0.64 |
| 2 | 182311594 | rs13384671 | 0.294 | 0.642 | -1.732 | 0.927 | -0.017 | -0.059 | -0.059 | 1.801 | 0.36 | 0.2 | 0.42 | 0.48 | 0.34 | 0.21 |
| 2 | 182319301 | rs1449263 | 0.468 | 0.609 | -1.346 | 1.064 | 0 | 0.024 | 0.024 | 0 | 0.48 | 0.61 | 0.53 | 0.37 | 0.54 | 0.62 |
| 2 | 182325907 | rs3770138 | 0.814 | 0.519 | -1.641 | 1.22 | -0.018 | 0.031 | 0.031 | 0.06 | 0.27 | 0.04 | 0.12 | 0.21 | 0.21 | 0.14 |
| 2 | 182329647 | rs155149 | 0.409 | 0.575 | -1.366 | 1.244 | -0.007 | -0.017 | -0.017 | 0 | 0.83 | 0.83 | 0.7 | 0.58 | 0.79 | 0.8 |
| 2 | 182332961 | rs3770132 | 0.28 | 0.632 | -1.475 | 1.225 | -0.02 | -0.059 | -0.059 | 0.643 | 0.12 | 0.01 | 0.12 | 0.2 | 0.11 | 0.14 |

| | | | | | | | | | | | | | | | | |
|---|---|---|---|---|---|---|---|---|---|---|---|---|---|---|---|---|
| 2 | 182334873 | rs711801 | -1.053 | 0.635 | -1.28 | 1.167 | 0.003 | 0.023 | 0.023 | 0 | 0.11 | 0.19 | 0.21 | 0.22 | 0.13 | 0.1 |
| 2 | 182336474 | rs1449264 | 0.132 | 0.741 | -1.053 | 0.386 | 0.018 | -0.068 | -0.068 | 0 | 0.28 | 0.36 | 0.42 | 0.05 | 0.34 | 0.2 |
| 2 | 182336805 | rs1449265 | 0.504 | 0.524 | -1.126 | 0.585 | 0 | 0.001 | 0.001 | 0 | 0.54 | 0.4 | 0.31 | 0.59 | 0.49 | 0.63 |
| 2 | 182336884 | rs3770126 | 1.012 | 0.249 | -1.502 | 0.65 | -0.06 | -0.016 | -0.016 | 0 | 0.27 | 0.12 | 0.31 | 0.36 | 0.3 | 0.23 |
| 2 | 182337447 | rs3770125 | 1.217 | 0.304 | -1.363 | 0.745 | 0.007 | 0.024 | 0.024 | 0 | 0.06 | 0.11 | 0.19 | 0.23 | 0.1 | 0.07 |
| 2 | 182338027 | rs11694175 | 0.116 | -0.246 | -1.423 | 0.592 | 0.002 | -0.001 | -0.001 | 0.041 | 0.42 | 0.46 | 0.71 | 0.38 | 0.56 | 0.32 |
| 2 | 182341674 | rs155087 | -1.206 | -0.06 | -1.317 | 0.761 | -0.012 | 0.044 | 0.044 | 0 | 0.76 | 0.7 | 0.88 | 0.65 | 0.8 | 0.58 |
| 2 | 182348554 | rs17224699 | 1.814 | -0.036 | -1.426 | 0.658 | -0.02 | 0.056 | 0.056 | 0 | 0.21 | 0.07 | 0.13 | 0.14 | 0.2 | 0.15 |
| 2 | 182349409 | rs3755021 | 1.859 | -0.063 | -1.496 | 0.647 | -0.051 | 0.052 | 0.052 | 0 | 0.2 | 0.02 | 0.12 | 0.14 | 0.2 | 0.15 |
| 2 | 182351352 | rs155106 | -0.692 | -0.067 | -1.372 | 0.715 | -0.001 | 0 | 0 | 0 | 0.55 | 0.57 | 0.66 | 0.51 | 0.6 | 0.43 |
| 2 | 182352469 | rs3770120 | 1.099 | 0.261 | -1.459 | 0.191 | -0.061 | 0.102 | 0.102 | 0 | 0.26 | 0.06 | 0.18 | 0.13 | 0.29 | 0.16 |
| 2 | 182352568 | rs10172410 | -0.153 | 0.235 | -1.38 | 0.527 | 0.006 | 0.006 | 0.006 | 0 | 0.4 | 0.44 | 0.55 | 0.46 | 0.49 | 0.29 |
| 2 | 182352791 | rs155110 | 0.38 | 0.477 | -1.547 | 0.138 | -0.012 | 0.078 | 0.078 | 0 | 0.3 | 0.21 | 0.21 | 0.13 | 0.3 | 0.16 |
| 2 | 182359592 | rs16867434 | -0.101 | -0.209 | -0.609 | 0.343 | -0.026 | -0.018 | -0.018 | 0 | 0.14 | 0.22 | 0.19 | 0.05 | 0.1 | 0.03 |
| 2 | 182360182 | rs2305586 | 1.015 | -0.002 | -1.1 | 0.706 | -0.037 | -0.07 | -0.07 | 0 | 0.11 | 0.01 | 0.17 | 0.24 | 0.13 | 0.22 |
| 2 | 182366911 | rs7607758 | -0.207 | 0 | -0.9 | 0.48 | -0.001 | 0.004 | 0.004 | 0 | 0.46 | 0.57 | 0.5 | 0.29 | 0.55 | 0.26 |
| 2 | 182373981 | rs13032116 | -1.354 | 0.476 | -0.857 | 0.533 | -0.007 | 0.004 | 0.004 | 0 | 0.79 | 0.75 | 0.81 | 0.84 | 0.68 | 0.91 |
| 2 | 182379723 | rs12469449 | 1.305 | 0.475 | -0.853 | 0.532 | -0.033 | 0.019 | 0.019 | 0 | 0.2 | 0.16 | 0.19 | 0.16 | 0.34 | 0.09 |
| 2 | 182380705 | rs10490692 | -0.726 | 0.619 | -0.894 | 0.787 | -0.001 | -0.002 | -0.002 | 0 | 0.73 | 0.66 | 0.63 | 0.62 | 0.53 | 0.85 |
| 2 | 182382316 | rs3770111 | 1.493 | 0.621 | -0.914 | 0.831 | 0.001 | 0.021 | 0.021 | 0 | 0.05 | 0.05 | 0.17 | 0.22 | 0.13 | 0.06 |
| 2 | 182390953 | rs3770107 | -1.347 | 0.585 | -0.788 | 0.726 | 0.018 | -0.018 | -0.018 | 0.261 | 0.78 | 0.66 | 0.8 | 0.84 | 0.67 | 0.9 |
| 2 | 182394954 | rs6707704 | -0.773 | 0.585 | -0.719 | 0.769 | 0.03 | 0.012 | 0.012 | 0 | 0.74 | 0.62 | 0.62 | 0.62 | 0.54 | 0.84 |
| 2 | 182399097 | rs7562325 | -0.745 | 0.649 | -1.196 | 0.797 | 0.008 | 0.002 | 0.002 | 0 | 0.74 | 0.65 | 0.63 | 0.62 | 0.55 | 0.84 |
| 2 | 182399486 | rs7601843 | 0.567 | 0.674 | -0.955 | 0.943 | 0.031 | 0.07 | 0.07 | 0 | 0.07 | 0.17 | 0.18 | 0.22 | 0.13 | 0.06 |
| 2 | 182401116 | rs10490691 | 0.571 | 0.677 | -0.955 | 0.944 | 0.031 | 0.07 | 0.07 | 0.003 | 0.07 | 0.16 | 0.18 | 0.22 | 0.13 | 0.06 |
| 2 | 182415098 | rs971741 | 0.378 | 0.676 | -1.015 | 0.942 | 0.019 | 0.045 | 0.045 | 0 | 0.07 | 0.14 | 0.19 | 0.22 | 0.14 | 0.06 |
| 2 | 182418130 | rs10930971 | -0.198 | 0.431 | -1.041 | 0.855 | -0.026 | -0.027 | -0.027 | 0 | 0.26 | 0.39 | 0.3 | 0.14 | 0.22 | 0.17 |
| 2 | 182419986 | rs10490689 | -0.74 | 0.431 | -1.123 | 0.798 | 0.002 | 0.028 | 0.028 | 0 | 0.32 | 0.5 | 0.49 | 0.36 | 0.36 | 0.23 |
| 2 | 182439519 | rs12479005 | 0.984 | 0.668 | -1.661 | 0.956 | -0.052 | -0.07 | -0.07 | 0 | 0.13 | 0.01 | 0.18 | 0.23 | 0.16 | 0.22 |
| 2 | 182451737 | rs10490684 | 1.289 | 0.71 | -1.756 | 0.573 | -0.112 | 0.137 | 0.137 | 0.223 | 0.26 | 0.02 | 0.21 | 0.06 | 0.37 | 0.11 |
| 2 | 182454375 | rs1449253 | 0.249 | 0.452 | -1.382 | 0.58 | -0.025 | -0.003 | -0.003 | 0.197 | 0.16 | 0.55 | 0.23 | 0.11 | 0.08 | 0.12 |
| 2 | 182455994 | rs993648 | -0.748 | 0.561 | -1.37 | 0.864 | 0.002 | 0.01 | 0.01 | 3.664 | 0.23 | 0.77 | 0.44 | 0.38 | 0.25 | 0.18 |
| 2 | 182457803 | rs767463 | -0.748 | 0.561 | -1.37 | 0.864 | 0.002 | 0.01 | 0.01 | 3.664 | 0.23 | 0.77 | 0.44 | 0.38 | 0.25 | 0.18 |
| 2 | 182460381 | rs16867452 | 0.307 | 0.558 | -1.339 | 0.867 | -0.025 | -0.003 | -0.003 | 0 | 0.16 | 0.55 | 0.24 | 0.11 | 0.08 | 0.12 |
| 2 | 182460714 | rs10445770 | 0.403 | 0.56 | -1.845 | 0.836 | 0.06 | 0.032 | 0.032 | 0.076 | 0.51 | 0.17 | 0.34 | 0.56 | 0.38 | 0.68 |
| 2 | 182470425 | rs10930973 | -0.616 | 0.548 | -1.777 | 0.893 | 0.001 | 0.009 | 0.009 | 3.941 | 0.23 | 0.77 | 0.44 | 0.38 | 0.25 | 0.18 |
| 2 | 182485307 | rs16867461 | 1.064 | 0.574 | -1.701 | 0.85 | 0.051 | 0.071 | 0.071 | 0 | 0.06 | 0.14 | 0.19 | 0.23 | 0.16 | 0.06 |
| 2 | 182486386 | rs2053471 | 0.082 | 0.072 | -1.68 | 0.864 | -0.035 | -0.02 | -0.02 | 1.218 | 0.17 | 0.62 | 0.25 | 0.11 | 0.09 | 0.12 |
| 2 | 182487154 | rs12617685 | 0.802 | 0.057 | -2.128 | 1.011 | -0.051 | -0.097 | -0.097 | 0.812 | 0.19 | 0.01 | 0.22 | 0.31 | 0.21 | 0.24 |
| 2 | 182489754 | rs1596023 | 0.304 | 0.528 | -1.534 | 0.697 | 0.071 | 0.054 | 0.054 | 0.065 | 0.5 | 0.13 | 0.34 | 0.6 | 0.37 | 0.68 |
| 2 | 182504802 | rs7602326 | -0.663 | 0.922 | -1.08 | 0.455 | -0.041 | 0.072 | 0.072 | 0.411 | 0.72 | 0.83 | 0.76 | 0.92 | 0.61 | 0.86 |
| 2 | 182509184 | rs3910660 | 0.571 | 0.911 | -1.089 | 0.449 | -0.06 | 0.091 | 0.091 | 0.578 | 0.28 | 0.1 | 0.23 | 0.08 | 0.39 | 0.14 |
| 2 | 182510823 | rs3910661 | 0.306 | 0.979 | -0.805 | 0.651 | -0.036 | 0.05 | 0.05 | 0.541 | 0.61 | 0.19 | 0.54 | 0.77 | 0.45 | 0.79 |
| 2 | 182513160 | rs4018755 | 0.306 | 0.979 | -0.81 | 0.651 | -0.035 | 0.048 | 0.048 | 0.541 | 0.61 | 0.19 | 0.54 | 0.77 | 0.45 | 0.79 |
| 2 | 182515795 | rs3910664 | -0.076 | 0.983 | -0.829 | 0.648 | -0.014 | 0.06 | 0.06 | 0.687 | 0.65 | 0.21 | 0.6 | 0.9 | 0.58 | 0.85 |
| 2 | 182521578 | rs1473295 | 0.309 | 0.958 | -0.866 | 0.709 | -0.034 | 0.053 | 0.053 | 0.93 | 0.58 | 0.17 | 0.52 | 0.73 | 0.44 | 0.78 |
| 2 | 182530854 | rs4018756 | 0.301 | 0.97 | -0.86 | 0.717 | -0.035 | 0.047 | 0.047 | 0.863 | 0.59 | 0.17 | 0.52 | 0.72 | 0.45 | 0.78 |
| 2 | 182534947 | rs17198745 | 0.636 | 1.016 | -0.951 | 0.735 | -0.079 | 0.119 | 0.119 | 1.045 | 0.29 | 0.07 | 0.24 | 0.08 | 0.39 | 0.15 |
| 2 | 182543455 | rs1801262 | -0.747 | 0.977 | -0.594 | 0.693 | -0.011 | 0.024 | 0.024 | 1.039 | 0.69 | 0.73 | 0.75 | 0.92 | 0.61 | 0.85 |
| 2 | 182545218 | rs2583016 | -0.731 | 0.982 | -0.742 | 0.696 | -0.018 | 0.048 | 0.048 | 1.202 | 0.69 | 0.76 | 0.75 | 0.92 | 0.61 | 0.85 |
| 2 | 182548671 | rs7420169 | -1.686 | 0.492 | 0.219 | 1.371 | 0.097 | 0.167 | 0.167 | 4.568 | 0.11 | 0.55 | 0.22 | 0.2 | 0.17 | 0.08 |
| 2 | 182552583 | rs2950648 | -0.727 | 0.411 | 0.385 | 1.347 | 0.098 | 0.106 | 0.106 | 4.329 | 0.05 | 0.47 | 0.15 | 0.02 | 0.03 | 0.01 |
| 2 | 182553659 | rs16867470 | -0.654 | 0.542 | -0.674 | 1.487 | -0.012 | -0.01 | -0.01 | 0 | 0.08 | 0.06 | 0.08 | 0.21 | 0.16 | 0.06 |
| 2 | 182572502 | rs2583008 | 0.965 | 0.506 | -0.6 | 1.365 | 0.024 | 0.045 | 0.045 | 0 | 0.9 | 0.72 | 0.91 | 0.83 | 0.84 | 0.94 |
| 2 | 182580567 | rs10497585 | 0.036 | 0.134 | -1.11 | 1.525 | -0.095 | -0.163 | -0.163 | 3.021 | 0.22 | 0.02 | 0.38 | 0.47 | 0.31 | 0.28 |
| 2 | 182581492 | rs1441159 | 0.525 | -0.685 | -0.384 | 1.159 | -0.025 | 0.022 | 0.022 | 0.115 | 0.93 | 0.74 | 0.96 | 0.92 | 0.98 | 0.97 |
| 2 | 182603382 | rs10175898 | 0.369 | -0.651 | -0.443 | 1.176 | -0.024 | 0.021 | 0.021 | 0.061 | 0.94 | 0.76 | 0.96 | 0.92 | 0.98 | 0.97 |
| 2 | 182617191 | rs16867493 | 0.073 | -0.265 | -0.838 | 1.456 | -0.188 | -0.159 | -0.159 | 0 | 0.32 | 0.02 | 0.5 | 0.47 | 0.51 | 0.27 |
| 2 | 182639861 | rs6433931 | 0.222 | -0.819 | -0.254 | 1.216 | 0.011 | 0.013 | 0.013 | 0.261 | 0.52 | 0.63 | 0.41 | 0.37 | 0.4 | 0.48 |
| 2 | 182647932 | rs13428784 | -0.346 | -0.956 | -0.216 | 1.327 | -0.024 | 0.041 | 0.041 | 0.573 | 0.07 | 0.28 | 0.03 | 0.07 | 0.01 | 0.07 |
| 2 | 182651182 | rs10200243 | -0.923 | -1.044 | -0.246 | 1.556 | -0.029 | 0.156 | 0.156 | 0.015 | 0.11 | 0.26 | 0.11 | 0.39 | 0.07 | 0.22 |
| 2 | 182687039 | rs16867499 | 0.801 | -1.088 | -0.761 | 1.351 | -0.185 | -0.147 | -0.147 | 0 | 0.3 | 0.01 | 0.46 | 0.46 | 0.48 | 0.33 |
| 2 | 182690504 | rs950165 | -1.443 | -0.881 | -0.083 | 1.383 | -0.028 | 0.216 | 0.216 | 0 | 0.13 | 0.34 | 0.11 | 0.39 | 0.09 | 0.25 |
| 2 | 182692674 | rs937296 | -0.546 | -0.847 | -0.184 | 1.262 | -0.008 | 0.048 | 0.048 | 0 | 0.07 | 0.1 | 0.18 | 0.24 | 0.06 | 0.14 |
| 2 | 182696101 | rs16867500 | -1.573 | -0.897 | -0.14 | 1.276 | -0.031 | 0.128 | 0.128 | 0 | 0.14 | 0.35 | 0.23 | 0.34 | 0.1 | 0.25 |
| 2 | 182716212 | rs10176571 | -1.153 | -0.736 | 0.063 | 1.182 | -0.019 | 0.136 | 0.136 | 0.001 | 0.1 | 0.34 | 0.21 | 0.26 | 0.06 | 0.17 |
| 2 | 182733299 | rs13418310 | -1.017 | -0.728 | 0.115 | 0.711 | -0.014 | 0.063 | 0.063 | 0 | 0.1 | 0.31 | 0.21 | 0.13 | 0.05 | 0.16 |
| 2 | 182733535 | rs13393177 | -1.017 | -0.728 | 0.115 | 0.711 | -0.014 | 0.063 | 0.063 | 0 | 0.1 | 0.32 | 0.21 | 0.13 | 0.05 | 0.16 |
| 2 | 182743246 | rs7592216 | -0.253 | -0.744 | 0.261 | 0.488 | 0.035 | -0.078 | -0.078 | 0.411 | 0.87 | 0.67 | 0.91 | 0.99 | 0.89 | 0.88 |

| | | | | | | | | | | | | | | | | |
|---|---|---|---|---|---|---|---|---|---|---|---|---|---|---|---|---|
| 2 | 182749568 | rs1157595 | 0.34 | -1.026 | -0.637 | 0.593 | -0.12 | -0.071 | -0.071 | 0 | 0.39 | 0.09 | 0.35 | 0.68 | 0.54 | 0.4 |
| 2 | 182761117 | rs3815878 | -0.554 | -0.48 | -0.447 | 0.713 | 0.003 | 0.051 | 0.051 | 0.048 | 0.08 | 0.32 | 0.06 | 0.09 | 0.03 | 0.11 |
| 2 | 182776005 | rs6755017 | 0.265 | -0.298 | -0.498 | 0.985 | -0.003 | 0.011 | 0.011 | 0 | 0.06 | 0.07 | 0.06 | 0.11 | 0.03 | 0.11 |
| 2 | 182776851 | rs17587367 | 0.456 | -0.292 | -0.57 | 0.945 | -0.191 | -0.18 | -0.18 | 0.098 | 0.31 | 0.01 | 0.32 | 0.58 | 0.51 | 0.32 |
| 2 | 182806769 | rs7606321 | -0.73 | -0.168 | 0.976 | 1.04 | -0.005 | 0.007 | 0.007 | 0 | 0.06 | 0.11 | 0.04 | 0.09 | 0.03 | 0.11 |
| 2 | 182815579 | rs6433943 | -1.258 | -0.156 | 0.892 | 0.991 | -0.007 | 0.079 | 0.079 | 0.092 | 0.11 | 0.42 | 0.23 | 0.13 | 0.05 | 0.16 |
| 2 | 182838728 | rs12615866 | -1.171 | -0.087 | 0.325 | 0.881 | -0.002 | 0.077 | 0.077 | 0 | 0.11 | 0.49 | 0.23 | 0.11 | 0.06 | 0.16 |
| 2 | 182847922 | rs2840148 | -0.447 | 0.409 | -0.363 | 0.659 | 0.001 | -0.005 | -0.005 | 0.914 | 0.33 | 0.37 | 0.26 | 0.27 | 0.35 | 0.37 |
| 2 | 182852282 | rs1196291 | -0.612 | 0.584 | -0.849 | 0.659 | -0.022 | 0.055 | 0.055 | 0 | 0.58 | 0.84 | 0.65 | 0.7 | 0.51 | 0.56 |
| 2 | 182853855 | rs16867517 | 0.216 | 0.267 | -0.572 | 0.797 | -0.012 | -0.001 | -0.001 | 0 | 0.13 | 0.18 | 0.12 | 0.13 | 0.08 | 0.04 |
| 2 | 182855636 | rs11685642 | 0.471 | 0.15 | -0.152 | 0.793 | 0.001 | -0.005 | -0.005 | 0.228 | 0.24 | 0.28 | 0.17 | 0.17 | 0.22 | 0.31 |
| 2 | 182856074 | rs12621807 | -0.501 | 0.206 | -0.23 | 1.061 | 0.014 | -0.041 | -0.041 | 0.008 | 0.29 | 0.22 | 0.22 | 0.48 | 0.25 | 0.22 |
| 2 | 182859734 | rs1153744 | -1.169 | 0.029 | -0.403 | 0.311 | 0.034 | -0.106 | -0.106 | 0 | 0.81 | 0.72 | 0.84 | 1 | 0.75 | 0.85 |
| 2 | 182859928 | rs13424231 | 1.85 | -0.014 | -0.408 | 0.59 | -0.007 | 0.068 | 0.068 | 0 | 0.06 | 0.19 | 0.1 | 0.11 | 0.02 | 0.02 |
| 2 | 182863389 | rs1153738 | 1.101 | 0.03 | -0.426 | 0.605 | 0.013 | -0.024 | -0.024 | 0 | 0.25 | 0.17 | 0.45 | 0.34 | 0.25 | 0.33 |
| 2 | 182866563 | rs1356170 | 0.455 | 0.001 | -0.405 | 0.63 | -0.006 | 0.001 | 0.001 | 0.008 | 0.31 | 0.25 | 0.23 | 0.3 | 0.41 | 0.25 |
| 2 | 182869111 | rs7594388 | 0.323 | -0.141 | -0.346 | 0.767 | 0.016 | 0.004 | 0.004 | 0 | 0.32 | 0.33 | 0.25 | 0.38 | 0.43 | 0.28 |
| 2 | 182869208 | rs1153731 | 1.418 | -0.428 | -0.566 | 0.611 | 0.004 | -0.086 | -0.086 | 0.089 | 0.21 | 0.08 | 0.42 | 0.33 | 0.23 | 0.29 |
| 2 | 182878470 | rs1588928 | 0.633 | -0.483 | -0.081 | 0.574 | -0.002 | -0.045 | -0.045 | 0.3 | 0.33 | 0.44 | 0.21 | 0.17 | 0.28 | 0.33 |
| 2 | 182885196 | rs1196184 | -0.458 | -0.484 | 0.255 | 0.876 | -0.036 | 0.018 | 0.018 | 0 | 0.9 | 0.81 | 0.89 | 0.88 | 0.97 | 0.93 |
| 2 | 182887637 | rs12693291 | 0.537 | -0.61 | 0.28 | 0.774 | -0.02 | 0.018 | 0.018 | 0 | 0.65 | 0.44 | 0.44 | 0.56 | 0.73 | 0.63 |
| 2 | 182901257 | rs6732434 | 0.48 | -0.684 | 0.232 | 0.807 | -0.017 | 0.012 | 0.012 | 0 | 0.64 | 0.43 | 0.43 | 0.52 | 0.72 | 0.63 |
| 2 | 182915414 | rs1400009 | 0.899 | -0.389 | -0.105 | 1.204 | -0.009 | -0.041 | -0.041 | 0 | 0.19 | 0.04 | 0.21 | 0.21 | 0.2 | 0.27 |
| 2 | 182928012 | rs1196160 | -0.088 | -0.388 | -0.038 | 1.21 | -0.009 | 0.012 | 0.012 | 0 | 0.72 | 0.7 | 0.7 | 0.6 | 0.76 | 0.69 |
| 2 | 182938377 | rs4404233 | -0.282 | -0.165 | -0.193 | 1.122 | -0.001 | 0.003 | 0.003 | 0 | 0.38 | 0.39 | 0.51 | 0.51 | 0.61 | 0.55 |
| 2 | 182946485 | rs17648206 | 0.805 | 0.312 | -0.597 | 1.183 | -0.085 | -0.024 | -0.024 | 0 | 0.22 | 0.07 | 0.3 | 0.22 | 0.41 | 0.25 |
| 2 | 182959449 | rs1196146 | -0.66 | -0.307 | -0.305 | 1.091 | 0.087 | 0.062 | 0.062 | 0.434 | 0.5 | 0.78 | 0.73 | 0.56 | 0.66 | 0.61 |
| 2 | 182960414 | rs1107901 | 0.251 | -0.089 | 0.228 | 1.127 | 0.084 | 0.103 | 0.103 | 0.134 | 0.16 | 0.39 | 0.2 | 0.27 | 0.22 | 0.26 |
| 2 | 182962353 | rs9288078 | 0.342 | -0.101 | 0.31 | 1.163 | 0.101 | 0.089 | 0.089 | 0.189 | 0.14 | 0.48 | 0.21 | 0.23 | 0.2 | 0.26 |
| 2 | 182968268 | rs1400008 | -0.321 | 0.307 | 0.213 | 1.167 | 0 | -0.012 | -0.012 | 0 | 0.38 | 0.59 | 0.51 | 0.48 | 0.6 | 0.53 |
| 2 | 182976727 | rs16822596 | 1.163 | -0.314 | -0.242 | 0.958 | 0.017 | -0.074 | -0.074 | 0 | 0.15 | 0.02 | 0.06 | 0.29 | 0.11 | 0.12 |
| 2 | 182979076 | rs6705911 | 1.357 | -0.085 | -0.04 | 1.019 | -0.064 | -0.03 | -0.03 | 0 | 0.24 | 0.1 | 0.32 | 0.26 | 0.41 | 0.26 |
| 2 | 182981264 | rs1515894 | -0.758 | -0.144 | -0.059 | 0.933 | 0.07 | 0.064 | 0.064 | 0.366 | 0.48 | 0.76 | 0.72 | 0.58 | 0.64 | 0.64 |
| 2 | 182983202 | rs17457004 | 0.691 | 0.02 | 0.12 | 1.031 | 0.013 | 0.006 | 0.006 | 0 | 0.27 | 0.27 | 0.34 | 0.3 | 0.43 | 0.28 |
| 2 | 182983220 | rs13398100 | 0.971 | 0.13 | 0.551 | 0.945 | 0.09 | 0.091 | 0.091 | 0.01 | 0.13 | 0.32 | 0.19 | 0.22 | 0.19 | 0.27 |
| 2 | 182992305 | rs7593912 | 0.396 | 0.145 | 0.269 | 0.919 | 0.088 | 0.17 | 0.17 | 0.713 | 0.15 | 0.48 | 0.36 | 0.28 | 0.19 | 0.31 |
| 2 | 182994498 | rs4666813 | 0.247 | 0.142 | 0.216 | 0.903 | 0.071 | 0.142 | 0.142 | 1.242 | 0.16 | 0.53 | 0.37 | 0.28 | 0.19 | 0.31 |
| 2 | 182998122 | rs6711862 | -0.298 | -0.04 | -0.316 | 0.292 | 0.042 | 0.111 | 0.111 | 0 | 0.51 | 0.64 | 0.54 | 0.85 | 0.66 | 0.66 |
| 2 | 183003366 | rs1347081 | -0.867 | -0.168 | -0.356 | 0.142 | 0.024 | 0.144 | 0.144 | 0 | 0.59 | 0.78 | 0.75 | 0.89 | 0.69 | 0.73 |
| 2 | 183007250 | rs1999 | -0.35 | -0.158 | -0.35 | 0.148 | 0.037 | 0.118 | 0.118 | 0 | 0.53 | 0.67 | 0.73 | 0.89 | 0.67 | 0.68 |
| 2 | 183023392 | rs4666815 | -0.819 | 0.051 | -0.35 | 0.324 | 0.01 | 0.007 | 0.007 | 0 | 0.42 | 0.65 | 0.47 | 0.66 | 0.37 | 0.53 |
| 2 | 183023484 | rs1432523 | -0.247 | 0.02 | -0.322 | 0.342 | -0.004 | 0.003 | 0.003 | 0.105 | 0.36 | 0.53 | 0.44 | 0.66 | 0.34 | 0.47 |
| 2 | 183026612 | rs1897472 | -1.502 | 0.074 | -0.461 | 0.265 | 0.016 | 0.109 | 0.109 | 0 | 0.62 | 0.77 | 0.77 | 0.9 | 0.7 | 0.74 |
| 2 | 183031491 | rs13420620 | 0.939 | 0.076 | 0.282 | 0.716 | 0.036 | 0.132 | 0.132 | 0 | 0.11 | 0.23 | 0.31 | 0.32 | 0.16 | 0.27 |
| 2 | 183032495 | rs6721984 | 0.939 | 0.076 | 0.282 | 0.716 | 0.036 | 0.132 | 0.132 | 0 | 0.11 | 0.23 | 0.31 | 0.32 | 0.16 | 0.27 |
| 2 | 183037089 | rs3769811 | 0.735 | -0.019 | 0.232 | 0.614 | -0.004 | 0.003 | 0.003 | 0 | 0.22 | 0.22 | 0.12 | 0.36 | 0.17 | 0.2 |
| 2 | 183043334 | rs3769802 | 0.558 | -0.047 | 0.316 | 0.602 | -0.02 | 0.02 | 0.02 | 0 | 0.23 | 0.34 | 0.13 | 0.36 | 0.17 | 0.19 |
| 2 | 183044608 | rs12612546 | -1.602 | -0.078 | 0.15 | 0.449 | 0.024 | 0.098 | 0.098 | 0 | 0.62 | 0.75 | 0.77 | 0.92 | 0.71 | 0.74 |
| 2 | 183049580 | rs10206325 | -1.586 | -0.073 | 0.152 | 0.451 | 0.025 | 0.1 | 0.1 | 0 | 0.62 | 0.75 | 0.77 | 0.92 | 0.71 | 0.72 |
| 2 | 183059200 | rs3769789 | 0.915 | -0.048 | 0.274 | 0.535 | 0.01 | -0.016 | -0.016 | 0 | 0.22 | 0.14 | 0.11 | 0.36 | 0.17 | 0.18 |
| 2 | 183064460 | rs1344772 | -1.477 | 0.045 | 0.184 | 0.433 | 0.023 | 0.083 | 0.083 | 0 | 0.62 | 0.73 | 0.77 | 0.92 | 0.71 | 0.72 |
| 2 | 183065667 | rs17457303 | 1.431 | 0.22 | 0.252 | 0.562 | -0.003 | -0.003 | -0.003 | 0 | 0.18 | 0.16 | 0.29 | 0.24 | 0.34 | 0.21 |
| 2 | 183083378 | rs6728985 | -1.073 | 0.093 | 0.243 | 0.401 | 0.04 | 0.088 | 0.088 | 0 | 0.55 | 0.69 | 0.74 | 0.92 | 0.69 | 0.67 |
| 2 | 183088584 | rs3754929 | -0.913 | 0.693 | 0.471 | 0.892 | 0.02 | 0.011 | 0.011 | 0 | 0.79 | 0.74 | 0.69 | 0.76 | 0.65 | 0.78 |
| 2 | 183092726 | rs1432520 | -0.905 | 0.691 | 0.433 | 0.891 | -0.012 | -0.008 | -0.008 | 0.453 | 0.79 | 0.85 | 0.69 | 0.76 | 0.65 | 0.78 |
| 2 | 183101882 | rs10930995 | 0.636 | 0.602 | 0.778 | 1.102 | 0.016 | 0.075 | 0.075 | 0 | 0.11 | 0.2 | 0.3 | 0.32 | 0.16 | 0.26 |
| 2 | 183108485 | rs2568675 | 0.182 | 0.654 | 0.8 | 1.083 | 0.063 | 0.183 | 0.183 | 0 | 0.12 | 0.34 | 0.31 | 0.32 | 0.17 | 0.26 |
| 2 | 183120344 | rs2623423 | -1.092 | 0.404 | -0.123 | 0.81 | -0.037 | -0.022 | -0.022 | 0 | 0.78 | 0.92 | 0.7 | 0.78 | 0.65 | 0.78 |
| 2 | 183126026 | rs6759303 | 0.966 | 0.1 | -0.216 | 0.644 | -0.001 | 0.06 | 0.06 | 0.049 | 0.12 | 0.01 | 0.06 | 0 | 0.09 | 0.04 |
| 2 | 183151764 | rs12474371 | 0.606 | 0.577 | -0.349 | 1.282 | -0.073 | -0.066 | -0.066 | 0 | 0.12 | 0.01 | 0.15 | 0.18 | 0.21 | 0.2 |
| 2 | 183155786 | rs16823011 | 0.157 | 0.485 | 0.137 | 1.348 | 0.035 | 0.162 | 0.162 | 0 | 0.13 | 0.27 | 0.29 | 0.46 | 0.2 | 0.27 |
| 2 | 183162809 | rs833141 | -0.767 | 0.428 | -0.14 | 0.914 | -0.006 | 0.002 | 0.002 | 0 | 0.91 | 0.95 | 0.94 | 0.88 | 0.87 | 0.9 |
| 2 | 183163148 | rs833142 | 0.873 | -0.08 | -0.111 | 0.626 | 0.011 | 0.019 | 0.019 | 0.342 | 0.67 | 0.53 | 0.59 | 0.35 | 0.64 | 0.53 |
| 2 | 183167402 | rs2448980 | 0.165 | -0.531 | 0.095 | 0.321 | -0.031 | -0.046 | -0.046 | 0.183 | 0.91 | 0.77 | 0.98 | 1 | 0.98 | 0.94 |
| 2 | 183178206 | rs833119 | 0.433 | -0.907 | -0.959 | -0.13 | 0.026 | 0.071 | 0.071 | 0.661 | 0.36 | 0.15 | 0.31 | 0.19 | 0.29 | 0.24 |
| 2 | 183181858 | rs12470644 | 0.773 | -1.063 | -1.488 | -0.03 | -0.177 | -0.027 | -0.027 | 0 | 0.33 | 0.02 | 0.5 | 0.31 | 0.47 | 0.35 |
| 2 | 183182776 | rs833125 | 0.367 | -1.148 | -1.146 | 0.017 | 0.02 | -0.02 | -0.02 | 0 | 0.69 | 0.3 | 0.73 | 0.41 | 0.75 | 0.57 |
| 2 | 183183111 | rs11679657 | 0.838 | -0.96 | -0.673 | 0.102 | 0 | 0.035 | 0.035 | 0 | 0.18 | 0.21 | 0.15 | 0.5 | 0.22 | 0.34 |
| 2 | 183185164 | rs833126 | 0.736 | -0.892 | -0.75 | -0.203 | 0.007 | 0.011 | 0.011 | 1.129 | 0.35 | 0.36 | 0.24 | 0.1 | 0.28 | 0.22 |

| | | | | | | | | | | | | | | | | |
|---|---|---|---|---|---|---|---|---|---|---|---|---|---|---|---|---|
| 2 | 183195681 | rs6715692 | 0.297 | -0.498 | -0.989 | 0.076 | -0.008 | -0.062 | -0.062 | 0 | 0.2 | 0.11 | 0.22 | 0.59 | 0.24 | 0.37 |
| 2 | 183210271 | rs6736414 | 0.561 | -0.685 | -1.536 | -0.277 | 0.008 | 0.026 | 0.026 | 0.066 | 0.26 | 0.22 | 0.25 | 0.03 | 0.22 | 0.2 |
| 2 | 183213152 | rs17264887 | 0.453 | -0.189 | -2.089 | 0.082 | -0.156 | -0.035 | -0.035 | 0 | 0.4 | 0.03 | 0.28 | 0.38 | 0.46 | 0.34 |
| 2 | 183217916 | rs1564542 | 0.636 | 0.107 | -2.048 | 0.288 | -0.04 | -0.149 | -0.149 | 0.242 | 0.29 | 0.01 | 0.17 | 0.47 | 0.25 | 0.26 |
| 2 | 183218612 | rs833153 | 0.008 | -0.019 | -1.702 | 0.329 | -0.008 | 0.012 | 0.012 | 0 | 0.05 | 0.09 | 0.1 | 0.08 | 0.03 | 0.05 |
| 2 | 183227222 | rs11894978 | 0.135 | -0.199 | -0.404 | 0.419 | 0.057 | 0.287 | 0.287 | 0.037 | 0.05 | 0.33 | 0.04 | 0.23 | 0.02 | 0.07 |
| 2 | 183228308 | rs2623439 | -1.161 | -0.248 | -1.015 | -0.25 | -0.067 | 0.109 | 0.109 | 0 | 0.76 | 0.96 | 0.78 | 1 | 0.7 | 0.92 |
| 2 | 183238919 | rs10931008 | 0.4 | -0.267 | -0.702 | -0.268 | -0.002 | -0.006 | -0.006 | 0 | 0.25 | 0.27 | 0.43 | 0.52 | 0.31 | 0.48 |
| 2 | 183242337 | rs4643498 | -0.527 | -0.279 | -0.763 | -0.413 | -0.007 | -0.018 | -0.018 | 0 | 0.63 | 0.67 | 0.64 | 0.97 | 0.58 | 0.78 |
| 2 | 183243313 | rs17264929 | 1.081 | -0.024 | -1.145 | -0.253 | -0.033 | 0.043 | 0.043 | 0 | 0.08 | 0.02 | 0.07 | 0 | 0.11 | 0.06 |
| 2 | 183244353 | rs16823150 | 1.19 | -0.171 | -1.064 | -0.1 | -0.053 | -0.111 | -0.111 | 0 | 0.18 | 0.05 | 0.15 | 0.44 | 0.27 | 0.43 |
| 2 | 183256567 | rs2195611 | -0.003 | -0.297 | -0.586 | -0.445 | 0 | -0.018 | -0.018 | 0 | 0.57 | 0.45 | 0.54 | 0.89 | 0.55 | 0.73 |
| 2 | 183262128 | rs1430158 | 0.475 | -0.336 | -0.915 | -0.308 | -0.007 | -0.024 | -0.024 | 0 | 0.24 | 0.14 | 0.35 | 0.69 | 0.3 | 0.51 |
| 2 | 183263387 | rs10931010 | -0.175 | -0.47 | -1.191 | -0.506 | -0.004 | 0.043 | 0.043 | 0.02 | 0.14 | 0.06 | 0.1 | 0 | 0.12 | 0.08 |
| 2 | 183272945 | rs1430147 | 1.867 | -0.883 | -1.192 | -0.465 | -0.034 | -0.03 | -0.03 | 0 | 0.24 | 0.03 | 0.16 | 0.24 | 0.22 | 0.2 |
| 2 | 183278519 | rs1560370 | 0.956 | -0.947 | -1.234 | -0.276 | -0.092 | -0.126 | -0.126 | 0 | 0.24 | 0.06 | 0.21 | 0.43 | 0.37 | 0.48 |
| 2 | 183283295 | rs16823201 | -0.424 | -0.945 | -1.168 | -0.845 | 0.023 | 0.024 | 0.024 | 0 | 0.14 | 0.08 | 0.09 | 0.04 | 0.06 | 0.1 |
| 2 | 183284352 | rs2368253 | -0.366 | -0.672 | -0.55 | -0.731 | -0.01 | 0.045 | 0.045 | 0 | 0.65 | 0.77 | 0.65 | 0.93 | 0.64 | 0.78 |
| 2 | 183298160 | rs1946815 | 0.227 | -0.582 | -0.367 | -0.581 | 0.012 | 0.011 | 0.011 | 0 | 0.41 | 0.36 | 0.28 | 0.29 | 0.29 | 0.29 |
| 2 | 183300646 | rs10207132 | 0.802 | -0.588 | -0.67 | -0.574 | 0.036 | 0.036 | 0.036 | 0.037 | 0.34 | 0.1 | 0.19 | 0.25 | 0.25 | 0.24 |
| 2 | 183304603 | rs10209382 | -0.968 | -0.539 | -0.556 | -0.673 | -0.005 | -0.002 | -0.002 | 0 | 0.7 | 0.79 | 0.71 | 0.97 | 0.67 | 0.88 |
| 2 | 183306159 | rs7569537 | -0.435 | -0.535 | -0.534 | -0.484 | -0.001 | -0.018 | -0.018 | 0 | 0.63 | 0.66 | 0.63 | 0.91 | 0.64 | 0.82 |
| 2 | 183307061 | rs12468743 | 0.319 | -0.507 | -0.359 | -0.616 | -0.011 | -0.006 | -0.006 | 0 | 0.28 | 0.27 | 0.41 | 0.69 | 0.39 | 0.58 |
| 2 | 183316895 | rs4572561 | 0.492 | -0.569 | -0.385 | -0.763 | -0.007 | -0.018 | -0.018 | 0 | 0.38 | 0.4 | 0.2 | 0.17 | 0.24 | 0.23 |
| 2 | 183320846 | rs1125184 | 0.703 | -0.532 | -0.642 | -0.747 | 0.037 | 0.112 | 0.112 | 0.211 | 0.35 | 0.03 | 0.16 | 0.17 | 0.24 | 0.25 |
| 2 | 183321841 | rs12465662 | 0.703 | -0.532 | -0.642 | -0.733 | 0.042 | 0.102 | 0.102 | 0.211 | 0.35 | 0.03 | 0.32 | 0.18 | 0.23 | 0.25 |
| 2 | 183324227 | rs1367204 | 0.703 | -0.532 | -0.642 | -0.733 | 0.042 | 0.102 | 0.102 | 0.211 | 0.35 | 0.03 | 0.32 | 0.18 | 0.23 | 0.25 |
| 2 | 183343102 | rs10171703 | -0.632 | -0.484 | -0.96 | -0.858 | -0.002 | 0.002 | 0.002 | 0.292 | 0.39 | 0.39 | 0.36 | 0.78 | 0.46 | 0.62 |
| 2 | 183346071 | rs10169102 | -0.789 | -0.167 | -0.802 | -0.684 | 0.01 | -0.092 | -0.092 | 0 | 0.69 | 0.46 | 0.67 | 0.96 | 0.67 | 0.87 |
| 2 | 183347174 | rs4666836 | 0.305 | -0.231 | -0.807 | -0.691 | 0.076 | 0.097 | 0.097 | 0 | 0.44 | 0.25 | 0.41 | 0.22 | 0.27 | 0.31 |
| 2 | 183350379 | rs1897104 | 0.153 | 0.143 | -0.825 | -0.038 | -0.02 | 0 | 0 | 0.032 | 0.26 | 0.23 | 0.26 | 0.74 | 0.4 | 0.56 |
| 2 | 183354600 | rs1594305 | 0.451 | 0.208 | -0.802 | 0.066 | -0.008 | -0.008 | -0.008 | 0 | 0.23 | 0.21 | 0.48 | 0.61 | 0.37 | 0.46 |
| 2 | 183354887 | rs4666587 | 0.471 | -0.328 | -0.74 | -0.492 | 0 | -0.004 | -0.004 | 0.124 | 0.44 | 0.53 | 0.31 | 0.34 | 0.4 | 0.43 |
| 2 | 183354928 | rs4666844 | 1.059 | -0.418 | -0.578 | -0.565 | 0 | -0.007 | -0.007 | 0 | 0.37 | 0.39 | 0.27 | 0.21 | 0.36 | 0.33 |
| 2 | 183376081 | rs6433976 | -0.587 | -0.753 | -0.558 | -0.747 | -0.049 | -0.159 | -0.159 | 0.054 | 0.77 | 0.45 | 0.86 | 0.99 | 0.86 | 0.94 |
| 2 | 183378313 | rs2082136 | 0.259 | -0.684 | -0.628 | -0.643 | -0.013 | 0.034 | 0.034 | 0.215 | 0.54 | 0.25 | 0.37 | 0.4 | 0.49 | 0.49 |
| 2 | 183401673 | rs1430153 | 0.779 | -1.317 | -0.856 | -0.957 | 0.078 | 0.114 | 0.114 | 0.357 | 0.59 | 0.25 | 0.75 | 0.94 | 0.76 | 0.86 |
| 2 | 183408260 | rs2368281 | -0.555 | -1.368 | -0.637 | -0.983 | -0.003 | -0.049 | -0.049 | 0 | 0.74 | 0.77 | 0.88 | 0.99 | 0.84 | 0.9 |
| 2 | 183411197 | rs16823363 | 1.449 | -0.902 | -1.038 | -0.675 | -0.034 | -0.01 | -0.01 | 0 | 0.37 | 0.13 | 0.28 | 0.38 | 0.37 | 0.4 |
| 2 | 183416697 | rs6711313 | 0.777 | -0.951 | -1.025 | -0.686 | 0.083 | 0.123 | 0.123 | 0.32 | 0.59 | 0.23 | 0.75 | 0.94 | 0.76 | 0.86 |
| 2 | 183417141 | rs1978650 | 1.324 | -0.595 | -0.991 | -0.579 | -0.066 | -0.082 | -0.082 | 3.879 | 0.22 | 0.11 | 0.47 | 0.56 | 0.38 | 0.46 |
| 2 | 183423637 | rs1430156 | 0.234 | -0.848 | -0.549 | -0.763 | -0.027 | -0.096 | -0.096 | 0 | 0.72 | 0.37 | 0.85 | 0.99 | 0.84 | 0.9 |
| 2 | 183441026 | rs10931024 | -0.273 | -0.759 | -0.662 | -0.764 | -0.007 | -0.062 | -0.062 | 0 | 0.73 | 0.77 | 0.87 | 0.99 | 0.83 | 0.89 |
| 2 | 183448149 | rs17356575 | 0.915 | -0.538 | -0.755 | -0.576 | 0.005 | 0.009 | 0.009 | 0 | 0.08 | 0.04 | 0.1 | 0.05 | 0.07 | 0.03 |
| 2 | 183452578 | rs4146244 | 1.026 | -0.512 | -1.022 | -0.596 | -0.051 | -0.037 | -0.037 | 6.648 | 0.27 | 0.16 | 0.56 | 0.61 | 0.41 | 0.48 |
| 2 | 183454853 | rs17265482 | 1.576 | -0.825 | -1.112 | -0.836 | -0.021 | -0.001 | -0.001 | 0.321 | 0.33 | 0.13 | 0.23 | 0.33 | 0.33 | 0.33 |
| 2 | 183455377 | rs10460376 | 1.211 | -0.788 | -1.088 | -0.813 | -0.076 | -0.058 | -0.058 | 0.172 | 0.27 | 0.13 | 0.55 | 0.61 | 0.41 | 0.48 |
| 2 | 183457549 | rs1997240 | 1.2 | -0.787 | -1.094 | -0.812 | -0.053 | -0.04 | -0.04 | 0 | 0.27 | 0.16 | 0.56 | 0.61 | 0.41 | 0.48 |
| 2 | 183462762 | rs6742011 | 1.356 | -0.672 | -0.96 | -0.656 | -0.014 | 0 | 0 | 0.093 | 0.34 | 0.15 | 0.23 | 0.32 | 0.33 | 0.33 |
| 2 | 183463401 | rs12990399 | 0.501 | -0.84 | -1.136 | -0.79 | -0.01 | -0.062 | -0.062 | 0 | 0.19 | 0.23 | 0.1 | 0.02 | 0.14 | 0.09 |
| 2 | 183465070 | rs13429751 | -1.195 | -0.939 | -1.162 | -0.826 | 0.002 | -0.022 | -0.022 | 0 | 0.3 | 0.25 | 0.14 | 0.06 | 0.21 | 0.15 |
| 2 | 183467216 | rs13398082 | -1.202 | -0.751 | -1.22 | -0.868 | 0.005 | -0.015 | -0.015 | 0 | 0.31 | 0.25 | 0.15 | 0.06 | 0.24 | 0.15 |
| 2 | 183467456 | rs10497608 | -1.111 | -0.958 | -1.111 | -0.818 | -0.032 | -0.016 | -0.016 | 0 | 0.67 | 0.8 | 0.42 | 0.38 | 0.56 | 0.48 |
| 2 | 183467888 | rs10497610 | -0.308 | -0.951 | -1.058 | -0.815 | -0.012 | -0.049 | -0.049 | 0 | 0.23 | 0.24 | 0.1 | 0.03 | 0.14 | 0.1 |
| 2 | 183480546 | rs4490144 | -1.29 | -0.73 | -0.949 | -0.502 | 0.003 | 0.003 | 0.003 | 0 | 0.64 | 0.56 | 0.74 | 0.64 | 0.65 | 0.65 |
| 2 | 183487851 | rs12693319 | -0.661 | -0.716 | -0.94 | -0.439 | -0.001 | 0 | 0 | 0 | 0.56 | 0.42 | 0.68 | 0.59 | 0.57 | 0.61 |
| 2 | 183489249 | rs10193866 | -0.846 | -1.053 | -0.985 | -0.717 | -0.014 | -0.03 | -0.03 | 0 | 0.19 | 0.21 | 0.08 | 0.05 | 0.11 | 0.11 |
| 2 | 183494672 | rs10931027 | -0.039 | -0.915 | -0.969 | -0.412 | -0.005 | -0.012 | -0.012 | 1.158 | 0.45 | 0.32 | 0.66 | 0.54 | 0.54 | 0.53 |
| 2 | 183497242 | rs13009280 | -0.791 | -0.904 | -0.976 | -0.403 | -0.007 | 0.001 | 0.001 | 0.04 | 0.56 | 0.52 | 0.73 | 0.6 | 0.63 | 0.6 |
| 2 | 183498471 | rs7355517 | 0.772 | -0.985 | -1.069 | -0.492 | -0.056 | -0.079 | -0.079 | 0 | 0.31 | 0.17 | 0.58 | 0.52 | 0.44 | 0.49 |
| 2 | 183499313 | rs13023449 | 0.569 | -0.942 | -1.045 | -0.496 | -0.01 | -0.046 | -0.046 | 0 | 0.15 | 0.24 | 0.09 | 0.03 | 0.1 | 0.04 |
| 2 | 183502805 | rs7569593 | -0.948 | -1.006 | -1.087 | -0.506 | -0.007 | 0.002 | 0.002 | 0.079 | 0.57 | 0.53 | 0.73 | 0.6 | 0.64 | 0.59 |
| 2 | 183508696 | rs10497611 | 0.743 | -0.778 | -1.093 | -0.354 | -0.056 | -0.079 | -0.079 | 0 | 0.31 | 0.16 | 0.58 | 0.52 | 0.44 | 0.48 |
| 2 | 183512726 | rs10180264 | -1.394 | -1.815 | -1.112 | -0.354 | 0.008 | 0.02 | 0.02 | 0 | 0.65 | 0.56 | 0.75 | 0.6 | 0.64 | 0.68 |
| 2 | 183515517 | rs16823492 | -0.123 | -1.824 | -1.35 | -0.266 | -0.034 | -0.063 | -0.063 | 0.047 | 0.2 | 0.27 | 0.09 | 0.03 | 0.08 | 0.08 |
| 2 | 183519463 | rs12467753 | 0.548 | -1.621 | -1.518 | -0.154 | -0.103 | -0.128 | -0.128 | 0.365 | 0.33 | 0.07 | 0.59 | 0.51 | 0.46 | 0.52 |
| 2 | 183521984 | rs10169420 | -0.179 | -1.051 | -0.987 | -0.053 | -0.011 | 0.019 | 0.019 | 0 | 0.35 | 0.24 | 0.43 | 0.34 | 0.36 | 0.27 |
| 2 | 183523028 | rs2578007 | -0.663 | -1.028 | -0.847 | -0.032 | -0.009 | -0.005 | -0.005 | 0 | 0.75 | 0.76 | 0.62 | 0.67 | 0.69 | 0.71 |
| 2 | 183529352 | rs17356699 | 0.178 | -1.11 | -0.964 | -0.112 | 0.02 | 0.109 | 0.109 | 3.172 | 0.2 | 0.02 | 0.09 | 0.08 | 0.15 | 0.09 |

| | | | | | | | | | | | | | | | | |
|---|---|---|---|---|---|---|---|---|---|---|---|---|---|---|---|---|
| 2 | 183533806 | rs2577999 | -0.914 | -0.992 | -0.728 | 0.07 | 0.048 | 0.042 | 0.042 | 0.624 | 0.77 | 0.51 | 0.47 | 0.65 | 0.71 | 0.75 |
| 2 | 183535884 | rs4322805 | 0.988 | -1.053 | -0.725 | 0.156 | -0.003 | 0.008 | 0.008 | 0 | 0.47 | 0.27 | 0.33 | 0.5 | 0.51 | 0.58 |
| 2 | 183537399 | rs825662 | -0.202 | -0.436 | -0.132 | 0.881 | 0.047 | 0.041 | 0.041 | 0.732 | 0.76 | 0.5 | 0.47 | 0.65 | 0.71 | 0.74 |
| 2 | 183558687 | rs415994 | -0.662 | -0.064 | -0.83 | 0.786 | -0.026 | -0.033 | -0.033 | 0 | 0.75 | 0.8 | 0.65 | 0.54 | 0.65 | 0.74 |
| 2 | 183562648 | rs376699 | -0.65 | -0.1 | -0.892 | 0.814 | -0.033 | -0.039 | -0.039 | 0 | 0.75 | 0.81 | 0.65 | 0.54 | 0.65 | 0.74 |
| 2 | 183570131 | rs288268 | -0.722 | -0.104 | -0.89 | 0.817 | -0.044 | -0.061 | -0.061 | 0 | 0.76 | 0.84 | 0.53 | 0.51 | 0.65 | 0.75 |
| 2 | 183572496 | rs167013 | -0.735 | -0.102 | -0.89 | 0.809 | -0.031 | -0.037 | -0.037 | 0 | 0.76 | 0.82 | 0.65 | 0.54 | 0.65 | 0.76 |
| 2 | 183579698 | rs288264 | -0.372 | -0.212 | -0.821 | 0.597 | -0.001 | -0.001 | -0.001 | 5.657 | 0.72 | 0.71 | 0.52 | 0.51 | 0.64 | 0.7 |
| 2 | 183582170 | rs17356755 | 0.977 | -0.236 | -0.946 | 0.42 | -0.001 | 0.016 | 0.016 | 0 | 0.11 | 0.03 | 0.05 | 0 | 0.08 | 0.03 |
| 2 | 183584755 | rs6729801 | 1.215 | -0.269 | -0.296 | 0.984 | 0.012 | 0.072 | 0.072 | 0 | 0.52 | 0.09 | 0.38 | 0.3 | 0.56 | 0.52 |
| 2 | 183589615 | rs825668 | -0.282 | -0.258 | -0.28 | 0.961 | -0.049 | -0.076 | -0.076 | 0 | 0.72 | 0.83 | 0.53 | 0.51 | 0.64 | 0.67 |
| 2 | 183602821 | rs288259 | -0.338 | -0.251 | -0.276 | 0.964 | -0.05 | -0.076 | -0.076 | 1.376 | 0.73 | 0.83 | 0.53 | 0.51 | 0.64 | 0.68 |
| 2 | 183608627 | rs288254 | -0.338 | -0.251 | -0.276 | 0.964 | -0.05 | -0.076 | -0.076 | 0.126 | 0.73 | 0.83 | 0.53 | 0.51 | 0.64 | 0.68 |
| 2 | 183614670 | rs11678585 | 0.754 | -0.26 | -0.279 | 0.949 | -0.007 | -0.042 | -0.042 | 0 | 0.58 | 0.11 | 0.4 | 0.39 | 0.59 | 0.56 |
| 2 | 183617672 | rs12616928 | 0.636 | -0.252 | -0.281 | 0.951 | 0.01 | -0.028 | -0.028 | 0 | 0.05 | 0.01 | 0.02 | 0.09 | 0.03 | 0.03 |
| 2 | 183622547 | rs288334 | -0.524 | -0.248 | -0.265 | 0.945 | -0.051 | -0.072 | -0.072 | 0 | 0.74 | 0.83 | 0.53 | 0.51 | 0.65 | 0.7 |
| 2 | 183627504 | rs192618 | -0.207 | -0.237 | -0.265 | 0.95 | -0.003 | -0.004 | -0.004 | 0 | 0.72 | 0.71 | 0.52 | 0.51 | 0.64 | 0.68 |
| 2 | 183628964 | rs288295 | -0.612 | -0.345 | -0.269 | 0.951 | -0.053 | -0.074 | -0.074 | 0 | 0.75 | 0.83 | 0.53 | 0.51 | 0.65 | 0.72 |
| 2 | 183632923 | rs6731059 | 0.084 | -0.324 | -0.224 | 0.877 | -0.037 | -0.061 | -0.061 | 0 | 0.12 | 0.52 | 0.1 | 0.02 | 0.04 | 0.03 |
| 2 | 183642628 | rs288283 | -0.549 | -0.256 | -0.521 | 0.908 | -0.051 | -0.08 | -0.08 | 1.36 | 0.74 | 0.83 | 0.53 | 0.5 | 0.65 | 0.7 |
| 2 | 183642878 | rs13509 | -0.165 | -0.258 | -0.52 | 0.908 | -0.004 | -0.006 | -0.006 | 0 | 0.71 | 0.7 | 0.52 | 0.5 | 0.64 | 0.67 |
| 2 | 183650229 | rs288310 | -0.55 | -0.256 | -0.52 | 0.914 | -0.051 | -0.085 | -0.085 | 0 | 0.74 | 0.83 | 0.53 | 0.49 | 0.65 | 0.7 |
| 2 | 183653727 | rs288304 | -0.162 | -0.231 | -0.514 | 0.926 | -0.005 | -0.009 | -0.009 | 0 | 0.71 | 0.71 | 0.52 | 0.49 | 0.64 | 0.66 |
| 2 | 183657867 | rs288298 | -0.288 | -0.251 | -0.495 | 0.917 | -0.065 | 0.003 | 0.003 | 0.015 | 0.14 | 0.59 | 0.12 | 0.12 | 0.05 | 0.13 |
| 2 | 183665741 | rs10931040 | 1.064 | -0.19 | -0.628 | 0.924 | 0.005 | 0.084 | 0.084 | 0 | 0.52 | 0.11 | 0.38 | 0.29 | 0.54 | 0.52 |
| 2 | 183667233 | rs16823704 | -0.053 | -0.227 | -0.528 | 1.037 | -0.016 | 0.036 | 0.036 | 0 | 0.06 | 0.14 | 0.03 | 0.1 | 0.04 | 0.03 |
| 2 | 183679421 | rs288354 | 0.314 | 0 | -0.867 | 1.057 | -0.014 | -0.029 | -0.029 | 0 | 0.63 | 0.74 | 0.53 | 0.39 | 0.58 | 0.62 |
| 2 | 183685562 | rs10931041 | 0.63 | 0.411 | -0.523 | 1.116 | -0.001 | 0 | 0 | 0 | 0.6 | 0.44 | 0.47 | 0.35 | 0.58 | 0.58 |
| 2 | 183687283 | rs10497614 | -0.927 | -0.047 | 0.182 | 0.833 | -0.049 | 0.027 | 0.027 | 0 | 0.13 | 0.4 | 0.09 | 0.11 | 0.05 | 0.1 |
| 2 | 183689763 | rs12619566 | -1.119 | 0.756 | -0.455 | 0.925 | -0.004 | -0.003 | -0.003 | 0 | 0.08 | 0.07 | 0.06 | 0.14 | 0.11 | 0.21 |
| 2 | 183690254 | rs11902959 | -0.523 | 0.613 | -0.332 | 1.119 | -0.002 | 0.001 | 0.001 | 0 | 0.14 | 0.11 | 0.07 | 0.14 | 0.1 | 0.1 |
| 2 | 183690632 | rs17356894 | -0.989 | 0.603 | -0.492 | 1.057 | -0.003 | -0.003 | -0.003 | 0 | 0.08 | 0.06 | 0.07 | 0.17 | 0.11 | 0.26 |
| 2 | 183694177 | rs17356901 | 0.962 | 0.624 | -1.446 | 0.396 | -0.071 | 0.141 | 0.141 | 0.455 | 0.41 | 0.06 | 0.48 | 0.13 | 0.38 | 0.32 |
| 2 | 183694499 | rs7593100 | 0.37 | 0.524 | -0.564 | 0.656 | 0.001 | 0.047 | 0.047 | 0 | 0.1 | 0.24 | 0.08 | 0.14 | 0.09 | 0.07 |
| 2 | 183694845 | rs1530056 | 1.03 | 0.623 | -0.74 | 0.522 | -0.034 | 0.172 | 0.172 | 0.172 | 0.46 | 0.09 | 0.54 | 0.17 | 0.48 | 0.38 |
| 2 | 183695449 | rs288316 | -1.238 | 0.51 | 0.133 | 0.433 | -0.021 | -0.004 | -0.004 | 0 | 0.29 | 0.51 | 0.19 | 0.26 | 0.22 | 0.18 |
| 2 | 183698691 | rs16823799 | -0.817 | 0.217 | 0.249 | 0.194 | -0.01 | -0.067 | -0.067 | 0.035 | 0.12 | 0.32 | 0.09 | 0.01 | 0.09 | 0.06 |
| 2 | 183701588 | rs288324 | -0.558 | 0.481 | -0.431 | 0.179 | -0.008 | 0.002 | 0.002 | 0 | 0.47 | 0.66 | 0.39 | 0.59 | 0.49 | 0.59 |
| 2 | 183704583 | rs3768842 | 0.044 | 0.545 | -0.759 | 0.026 | -0.025 | -0.046 | -0.046 | 0 | 0.22 | 0.12 | 0.22 | 0.44 | 0.28 | 0.37 |
| 2 | 183706855 | rs2242070 | -0.424 | 0.357 | -0.744 | 0.006 | -0.009 | -0.053 | -0.053 | 0.025 | 0.27 | 0.16 | 0.24 | 0.54 | 0.28 | 0.54 |
| 2 | 183710951 | rs288244 | -0.442 | 0.484 | -0.454 | -0.34 | 0.008 | -0.029 | -0.029 | 0.099 | 0.92 | 0.85 | 0.9 | 0.98 | 0.95 | 0.94 |
| 2 | 183716807 | rs2118590 | 1.573 | 0.205 | 0.309 | -0.561 | -0.001 | -0.014 | -0.014 | 1.677 | 0.49 | 0.19 | 0.53 | 0.43 | 0.53 | 0.35 |
| 2 | 183721708 | rs10931045 | 1.49 | 0.25 | 0.335 | -0.542 | 0.006 | -0.011 | -0.011 | 6.53 | 0.5 | 0.19 | 0.53 | 0.43 | 0.53 | 0.39 |
| 2 | 183722966 | rs7602601 | -0.348 | 0.265 | 0.454 | -0.553 | -0.007 | -0.003 | -0.003 | 0.715 | 0.1 | 0.09 | 0.1 | 0 | 0.12 | 0.05 |
| 2 | 183723948 | rs6433993 | 0.051 | 0.192 | 0.482 | -0.575 | -0.012 | -0.018 | -0.018 | 0 | 0.69 | 0.77 | 0.68 | 0.53 | 0.67 | 0.55 |
| 2 | 183728290 | rs7607737 | -0.811 | 0.164 | 0.412 | -0.574 | 0.033 | -0.074 | -0.074 | 0.172 | 0.12 | 0.31 | 0.12 | 0 | 0.13 | 0.05 |
| 2 | 183729197 | rs12992123 | 1.743 | 0.249 | 0.384 | -0.562 | -0.005 | -0.01 | -0.01 | 0 | 0.5 | 0.17 | 0.53 | 0.43 | 0.54 | 0.39 |
| 2 | 183730527 | rs10206992 | 0.622 | 0.278 | 0.386 | -0.539 | -0.044 | -0.094 | -0.094 | 0.099 | 0.22 | 0.05 | 0.23 | 0.44 | 0.28 | 0.4 |
| 2 | 183732364 | rs12469777 | 1.179 | -0.176 | -0.068 | -0.951 | 0 | 0 | 0 | 0 | 0.47 | 0.48 | 0.56 | 0.43 | 0.54 | 0.37 |
| 2 | 183737017 | rs10931049 | 1.597 | -0.321 | -0.256 | -1.054 | -0.009 | -0.041 | -0.041 | 0 | 0.41 | 0.17 | 0.49 | 0.43 | 0.5 | 0.39 |
| 2 | 183737652 | rs10202627 | -1.251 | -0.308 | -0.393 | -0.996 | 0.008 | 0.001 | 0.001 | 0 | 0.5 | 0.3 | 0.42 | 0.57 | 0.46 | 0.55 |
| 2 | 183747442 | rs10181110 | 0.306 | -0.065 | -0.245 | -0.876 | -0.018 | -0.027 | -0.027 | 0 | 0.29 | 0.19 | 0.32 | 0.46 | 0.33 | 0.4 |
| 2 | 183752083 | rs10167133 | 0.018 | -0.102 | -0.185 | -0.785 | -0.02 | -0.001 | -0.001 | 0 | 0.94 | 0.78 | 0.97 | 0.98 | 1 | 0.96 |
| 2 | 183757561 | rs7558974 | -0.515 | 0.03 | -0.423 | -0.766 | -0.005 | -0.003 | -0.003 | 0 | 0.38 | 0.37 | 0.37 | 0.47 | 0.4 | 0.43 |
| 2 | 183760854 | rs7606391 | -0.521 | 0.034 | -0.427 | -0.74 | -0.006 | -0.004 | -0.004 | 0 | 0.38 | 0.37 | 0.38 | 0.47 | 0.41 | 0.43 |
| 2 | 183764930 | rs4666599 | 0.72 | 0.103 | 0.062 | -1.033 | 0.016 | -0.038 | -0.038 | 0.397 | 0.89 | 0.75 | 0.91 | 0.96 | 0.91 | 0.94 |
| 2 | 183765016 | rs4666600 | -0.279 | -0.099 | 0.136 | -1.171 | -0.029 | -0.069 | -0.069 | 0 | 0.21 | 0.46 | 0.16 | 0.13 | 0.11 | 0.18 |
| 2 | 183767134 | rs10164916 | 1.494 | 0.001 | -0.303 | -1.259 | 0.005 | 0.144 | 0.144 | 0.964 | 0.59 | 0.17 | 0.7 | 0.83 | 0.72 | 0.74 |
| 2 | 183785598 | rs4427983 | 1.35 | -0.419 | -0.385 | -1.519 | 0.029 | 0.106 | 0.106 | 1.407 | 0.59 | 0.21 | 0.73 | 0.86 | 0.73 | 0.74 |
| 2 | 183790072 | rs4666601 | -0.331 | -0.474 | -0.074 | -1.535 | -0.009 | -0.019 | -0.019 | 0 | 0.92 | 0.87 | 0.98 | 1 | 0.96 | 0.96 |
| 2 | 183793398 | rs9333280 | 1.409 | -0.473 | -0.081 | -1.596 | 0.002 | -0.024 | -0.024 | 0 | 0.69 | 0.28 | 0.77 | 0.97 | 0.79 | 0.87 |
| 2 | 183795633 | rs6718462 | -0.524 | -0.445 | 0.06 | -1.608 | 0.006 | -0.056 | -0.056 | 0 | 0.13 | 0.49 | 0.12 | 0.03 | 0.05 | 0.05 |
| 2 | 183803954 | rs10931052 | 1.49 | -0.448 | 0.018 | -1.619 | 0.002 | -0.028 | -0.028 | 0 | 0.7 | 0.28 | 0.77 | 0.97 | 0.79 | 0.87 |
| 2 | 183812553 | rs7566668 | 1.546 | -0.445 | 0.016 | -1.616 | 0.002 | -0.022 | -0.022 | 0 | 0.69 | 0.28 | 0.77 | 0.97 | 0.79 | 0.87 |
| 2 | 183820456 | rs4233770 | 0.35 | -0.468 | 0.044 | -1.611 | -0.023 | -0.034 | -0.034 | 0.18 | 0.08 | 0.16 | 0.02 | 0 | 0.04 | 0.04 |
| 2 | 183830649 | rs10177344 | 1.744 | -0.223 | -0.045 | -1.61 | 0.014 | 0.104 | 0.104 | 1.523 | 0.59 | 0.21 | 0.72 | 0.85 | 0.71 | 0.74 |
| 2 | 183832540 | rs6711655 | 0.16 | -0.15 | -0.124 | -1.583 | -0.027 | 0.025 | 0.025 | 0 | 0.1 | 0.04 | 0.1 | 0 | 0.14 | 0.04 |
| 2 | 183833514 | rs6758029 | 0.235 | -0.297 | 0.011 | -1.413 | 0 | -0.008 | -0.008 | 0 | 0.08 | 0.1 | 0.01 | 0 | 0.04 | 0.04 |
| 2 | 183859473 | rs16823914 | 1.457 | 0.073 | -0.584 | -1.045 | -0.042 | -0.062 | -0.062 | 0 | 0.11 | 0.01 | 0.09 | 0.19 | 0.13 | 0.14 |

| | | | | | | | | | | | | | | | | |
|---|---|---|---|---|---|---|---|---|---|---|---|---|---|---|---|---|
| 2 | 183893404 | rs10153554 | 0.465 | 0.025 | -0.181 | -1.142 | 0.007 | -0.022 | -0.022 | 0 | 0.06 | 0.12 | 0.04 | 0.02 | 0.05 | 0.02 |
| 2 | 183900274 | rs3791247 | 1.156 | 0.145 | -0.266 | -0.962 | -0.005 | -0.016 | -0.016 | 2.793 | 0.1 | 0 | 0.03 | 0.11 | 0.06 | 0.13 |
| 2 | 183900752 | rs3791246 | -0.212 | 0.303 | 0.393 | -0.922 | 0.006 | -0.035 | -0.035 | 0 | 0.11 | 0.25 | 0.1 | 0.04 | 0.05 | 0.05 |
| 2 | 183936915 | rs2675117 | 0.322 | 0.53 | 0.492 | -0.978 | 0.016 | -0.051 | -0.051 | 0 | 0.84 | 0.63 | 0.83 | 0.89 | 0.86 | 0.86 |
| 2 | 183939361 | rs16823967 | -0.416 | 0.557 | 0.443 | -0.977 | 0 | -0.004 | -0.004 | 0 | 0.14 | 0.14 | 0.15 | 0.11 | 0.13 | 0.14 |
| 2 | 183941444 | rs2675043 | -0.247 | 0.882 | 0.282 | -1.19 | 0.001 | -0.028 | -0.028 | 0 | 0.09 | 0.12 | 0.07 | 0.01 | 0.15 | 0.04 |
| 2 | 183952469 | rs12463411 | 1.918 | 1.319 | 0.473 | -1.419 | -0.027 | 0.07 | 0.07 | 0.045 | 0.66 | 0.18 | 0.66 | 0.88 | 0.63 | 0.74 |
| 2 | 183955300 | rs2705733 | -0.794 | 1.385 | 0.508 | -1.414 | 0.028 | -0.025 | -0.025 | 0 | 0.07 | 0.1 | 0.09 | 0.01 | 0.16 | 0.04 |
| 2 | 183958143 | rs17704684 | 0.969 | 1.393 | 0.426 | -1.416 | -0.025 | 0.026 | 0.026 | 0 | 0.05 | 0 | 0.06 | 0 | 0.07 | 0.02 |
| 2 | 183961668 | rs2368335 | 2.026 | 1.412 | 0.494 | -1.453 | -0.017 | 0.048 | 0.048 | 0 | 0.67 | 0.2 | 0.67 | 0.88 | 0.65 | 0.74 |
| 2 | 183963690 | rs2705730 | 0.639 | 1.152 | 0.792 | -1.558 | 0.011 | -0.102 | -0.102 | 2.08 | 0.82 | 0.66 | 0.87 | 0.99 | 0.84 | 0.92 |
| 2 | 183964169 | rs2944346 | -1.092 | 1.099 | 0.71 | -1.507 | 0.018 | -0.093 | -0.093 | 0 | 0.23 | 0.44 | 0.23 | 0.11 | 0.24 | 0.18 |
| 2 | 183978487 | rs2247840 | -1.547 | 1.097 | 0.543 | -1.388 | 0.006 | -0.067 | -0.067 | 1.53 | 0.3 | 0.33 | 0.26 | 0.12 | 0.32 | 0.23 |
| 2 | 183978549 | rs2675058 | -1.196 | 1.135 | 0.623 | -1.356 | 0.017 | -0.106 | -0.106 | 0 | 0.22 | 0.31 | 0.17 | 0.01 | 0.26 | 0.1 |
| 2 | 183979483 | rs6750185 | 0.049 | 1.093 | 0.609 | -1.349 | 0.005 | -0.066 | -0.066 | 0.844 | 0.11 | 0.25 | 0.06 | 0.01 | 0.07 | 0.06 |
| 2 | 183980395 | rs4465737 | -0.929 | 1.095 | 0.606 | -1.357 | -0.001 | -0.094 | -0.094 | 0 | 0.19 | 0.26 | 0.12 | 0.01 | 0.15 | 0.08 |
| 2 | 183988599 | rs2251775 | -0.727 | 1.015 | 0.822 | -1.381 | 0.051 | -0.093 | -0.093 | 0 | 0.15 | 0.42 | 0.15 | 0.01 | 0.19 | 0.06 |
| 2 | 183989863 | rs1473617 | 1.722 | 1.059 | 0.387 | -1.421 | 0 | -0.053 | -0.053 | 1.143 | 0.71 | 0.26 | 0.75 | 0.99 | 0.68 | 0.86 |
| 2 | 183991040 | rs2254866 | -1.638 | 1.789 | 0.359 | -1.123 | 0.004 | -0.016 | -0.016 | 0.022 | 0.28 | 0.72 | 0.23 | 0.12 | 0.32 | 0.18 |
| 2 | 183991266 | rs2254879 | -0.571 | 1.797 | -0.678 | -0.837 | -0.045 | -0.115 | -0.115 | 1.586 | 0.72 | 0.9 | 0.63 | 0.42 | 0.7 | 0.53 |
| 2 | 184000143 | rs2675045 | -1.823 | 1.936 | -0.578 | -0.744 | -0.026 | 0.038 | 0.038 | 0 | 0.39 | 0.71 | 0.35 | 0.24 | 0.43 | 0.37 |
| 2 | 184000361 | rs6705795 | 0.346 | 1.716 | -0.244 | -0.641 | -0.005 | 0.008 | 0.008 | 0 | 0.13 | 0.33 | 0.09 | 0.12 | 0.1 | 0.12 |
| 2 | 184001854 | rs2176232 | 1.762 | 1.732 | -0.528 | -0.578 | -0.012 | -0.018 | -0.018 | 0 | 0.09 | 0 | 0.09 | 0.12 | 0.06 | 0.17 |
| 2 | 184013706 | rs2675093 | 0.946 | 1.362 | -0.784 | -0.755 | 0.003 | -0.093 | -0.093 | 0 | 0.23 | 0.1 | 0.27 | 0.46 | 0.23 | 0.3 |
| 2 | 184016627 | rs3762477 | -0.068 | 1.659 | -0.488 | -0.858 | -0.003 | -0.029 | -0.029 | 0.325 | 0.21 | 0.33 | 0.14 | 0.12 | 0.18 | 0.15 |
| 2 | 184016791 | rs16824056 | 2.287 | 1.649 | -1.022 | -1.049 | -0.034 | -0.031 | -0.031 | 0 | 0.09 | 0.01 | 0.07 | 0.1 | 0.11 | 0.07 |
| 2 | 184022488 | rs17706149 | 1.106 | 1.347 | -0.642 | -1.04 | -0.015 | 0.005 | 0.005 | 0 | 0.12 | 0.04 | 0.12 | 0.12 | 0.11 | 0.2 |
| 2 | 184024190 | rs3748884 | 0.057 | 1.307 | -0.643 | -1.025 | 0.012 | -0.012 | -0.012 | 0 | 0.16 | 0.36 | 0.13 | 0.12 | 0.15 | 0.15 |
| 2 | 184024702 | rs6434006 | -0.432 | 1.321 | -0.463 | -1.029 | 0.189 | 0.018 | 0.018 | 1.135 | 0.94 | 0.66 | 0.92 | 0.99 | 0.86 | 0.96 |
| 2 | 184028531 | rs2887234 | 1.04 | 1.118 | -0.664 | -1.033 | -0.026 | 0.038 | 0.038 | 0 | 0.61 | 0.3 | 0.65 | 0.75 | 0.57 | 0.62 |
| 2 | 184066858 | rs6434019 | -0.38 | 1.007 | -0.548 | -1.1 | 0 | 0.007 | 0.007 | 0 | 0.68 | 0.6 | 0.63 | 0.42 | 0.65 | 0.58 |
| 2 | 184069487 | rs987607 | -0.017 | 1.157 | -0.409 | -1.056 | 0.015 | -0.019 | -0.019 | 0.009 | 0.22 | 0.25 | 0.22 | 0.13 | 0.28 | 0.24 |
| 2 | 184074201 | rs11691085 | 1.383 | 1.116 | -0.72 | -1.049 | 0.019 | 0.013 | 0.013 | 0 | 0.38 | 0.15 | 0.38 | 0.29 | 0.35 | 0.33 |
| 2 | 184076622 | rs11904393 | 0.454 | 1.287 | -0.409 | -0.811 | 0.04 | -0.008 | -0.008 | 0 | 0.13 | 0.22 | 0.14 | 0.13 | 0.19 | 0.19 |
| 2 | 184077430 | rs11894287 | -0.245 | 1.289 | -0.459 | -0.846 | 0.01 | -0.009 | -0.009 | 0.453 | 0.2 | 0.22 | 0.19 | 0.14 | 0.24 | 0.23 |
| 2 | 184077474 | rs6740639 | 0.601 | 0.982 | -0.428 | -0.98 | -0.002 | -0.002 | -0.002 | 0.045 | 0.54 | 0.51 | 0.47 | 0.4 | 0.45 | 0.43 |
| 2 | 184078964 | rs6748231 | 1.624 | 0.221 | -0.67 | -1.186 | -0.011 | -0.022 | -0.022 | 0 | 0.3 | 0.17 | 0.35 | 0.31 | 0.29 | 0.38 |
| 2 | 184079707 | rs12474605 | 0.643 | 0.659 | -0.936 | -0.981 | -0.059 | -0.157 | -0.157 | 0 | 0.23 | 0.02 | 0.33 | 0.58 | 0.26 | 0.35 |
| 2 | 184081074 | rs16824171 | 0.97 | 0.566 | -0.734 | -1.113 | -0.004 | -0.041 | -0.041 | 0 | 0.15 | 0.16 | 0.09 | 0.01 | 0.13 | 0.08 |
| 2 | 184085788 | rs17758247 | 0.182 | 0.566 | -0.765 | -0.937 | 0.009 | -0.05 | -0.05 | 0 | 0.16 | 0.17 | 0.13 | 0 | 0.24 | 0.05 |
| 2 | 184086289 | rs7576936 | -0.476 | 0.611 | -0.758 | -0.894 | -0.008 | -0.047 | -0.047 | 0 | 0.76 | 0.81 | 0.65 | 0.41 | 0.73 | 0.65 |
| 2 | 184095979 | rs1462531 | 0.585 | 0.444 | -0.668 | -0.88 | -0.003 | 0.075 | 0.075 | 0 | 0.13 | 0 | 0.07 | 0 | 0.09 | 0.03 |
| 2 | 184106930 | rs6739996 | 0.023 | 0.275 | -0.942 | -0.698 | 0.019 | -0.004 | -0.004 | 0 | 0.25 | 0.2 | 0.12 | 0 | 0.16 | 0.09 |
| 2 | 184125573 | rs898946 | -0.559 | 0.731 | -0.97 | -0.171 | 0 | -0.001 | -0.001 | 0 | 0.86 | 0.82 | 0.89 | 0.84 | 0.89 | 0.79 |
| 2 | 184130941 | rs2368419 | 0.083 | 1.099 | -1.124 | -0.169 | 0.005 | 0.001 | 0.001 | 0 | 0.16 | 0.16 | 0.16 | 0.16 | 0.25 | 0.24 |
| 2 | 184134742 | rs11684526 | 0.462 | 0.692 | -1.071 | -0.121 | 0 | 0.001 | 0.001 | 0 | 0.55 | 0.41 | 0.55 | 0.57 | 0.52 | 0.72 |
| 2 | 184137284 | rs12465639 | -0.089 | 1.133 | -1.359 | 0.625 | -0.001 | 0.004 | 0.004 | 0.343 | 0.19 | 0.19 | 0.29 | 0.43 | 0.32 | 0.2 |
| 2 | 184141374 | rs16824240 | -0.007 | 0.397 | -1.566 | 0.56 | -0.001 | -0.041 | -0.041 | 0.03 | 0.12 | 0.02 | 0.09 | 0.16 | 0.09 | 0.23 |
| 2 | 184142431 | rs10193946 | -0.398 | 0.403 | -1.286 | 0.317 | -0.005 | -0.005 | -0.005 | 0.064 | 0.43 | 0.52 | 0.42 | 0.37 | 0.37 | 0.28 |
| 2 | 184142975 | rs7573365 | 0.143 | 0.283 | -0.955 | -0.119 | -0.021 | -0.109 | -0.109 | 0 | 0.24 | 0.35 | 0.14 | 0 | 0.17 | 0.08 |
| 2 | 184143715 | rs10176594 | -0.734 | 0.462 | -1.312 | 0.204 | 0.011 | 0.007 | 0.007 | 0 | 0.61 | 0.56 | 0.56 | 0.56 | 0.53 | 0.52 |
| 2 | 184159036 | rs11683501 | 0.336 | 0.204 | -1.654 | 0.129 | -0.03 | -0.014 | -0.014 | 0 | 0.39 | 0.2 | 0.28 | 0.61 | 0.49 | 0.56 |
| 2 | 184164557 | rs11903398 | -0.271 | 0.002 | -0.609 | 0.231 | -0.021 | -0.017 | -0.017 | 0 | 0.13 | 0.59 | 0.1 | 0.08 | 0.06 | 0.09 |
| 2 | 184170353 | rs1972947 | 1.084 | 0.179 | -1.153 | 0.231 | -0.026 | -0.009 | -0.009 | 0 | 0.51 | 0.17 | 0.42 | 0.49 | 0.53 | 0.62 |
| 2 | 184172465 | rs555543 | 0.586 | 0.401 | -1.026 | 0.484 | -0.036 | -0.016 | -0.016 | 0 | 0.19 | 0.06 | 0.39 | 0.19 | 0.27 | 0.08 |
| 2 | 184181744 | rs7559989 | 0.008 | 0.145 | 0.172 | 0.461 | -0.001 | 0.071 | 0.071 | 0.404 | 0.13 | 0.36 | 0.09 | 0.21 | 0.08 | 0.13 |
| 2 | 184191329 | rs576301 | 0.468 | 0.048 | -0.242 | 0.334 | -0.03 | -0.015 | -0.015 | 0 | 0.18 | 0.05 | 0.36 | 0.17 | 0.24 | 0.08 |
| 2 | 184203775 | rs10172290 | -0.408 | -0.027 | 0.143 | 0.387 | -0.012 | -0.015 | -0.015 | 0 | 0.16 | 0.24 | 0.1 | 0.12 | 0.11 | 0.12 |
| 2 | 184211084 | rs7556778 | 0.257 | -0.01 | -0.109 | 0.286 | 0.003 | 0.002 | 0.002 | 0 | 0.4 | 0.39 | 0.29 | 0.59 | 0.45 | 0.52 |
| 2 | 184211228 | rs7556885 | 0.766 | -0.378 | 0.345 | 0.488 | -0.003 | -0.003 | -0.003 | 0 | 0.67 | 0.64 | 0.56 | 0.72 | 0.69 | 0.76 |
| 2 | 184213419 | rs2018674 | 0.115 | -0.22 | 0.313 | 0.647 | -0.043 | -0.059 | -0.059 | 0 | 0.17 | 0.04 | 0.35 | 0.21 | 0.2 | 0.14 |
| 2 | 184234352 | rs421636 | 0.731 | -0.431 | 1.56 | 0.902 | -0.007 | 0.101 | 0.101 | 0 | 0.78 | 0.38 | 0.56 | 0.49 | 0.78 | 0.77 |
| 2 | 184242657 | rs1837146 | 0.499 | -0.378 | 1.1 | 0.635 | -0.057 | -0.035 | -0.035 | 0 | 0.13 | 0.01 | 0.32 | 0.11 | 0.18 | 0.14 |
| 2 | 184272613 | rs2368388 | 0.315 | -0.411 | 1.108 | 0.619 | -0.012 | -0.002 | -0.002 | 0 | 0.14 | 0.12 | 0.33 | 0.1 | 0.18 | 0.14 |
| 2 | 184272825 | rs842583 | 0.753 | -0.447 | 1.098 | 0.709 | 0.032 | 0.427 | 0.427 | 1.029 | 0.94 | 0.46 | 0.88 | 0.63 | 0.97 | 0.95 |
| 2 | 184277964 | rs10497616 | 0.394 | -0.59 | 0.202 | 0.393 | 0.056 | 0.119 | 0.119 | 0 | 0.3 | 0.07 | 0.21 | 0.06 | 0.2 | 0.24 |
| 2 | 184306911 | rs6434026 | 0.831 | 0.033 | 1.109 | 1.096 | -0.008 | 0.089 | 0.089 | 0 | 0.77 | 0.4 | 0.57 | 0.51 | 0.78 | 0.77 |
| 2 | 184320036 | rs12693342 | 0.732 | 0.111 | 0.451 | 1.134 | -0.002 | 0.159 | 0.159 | 0 | 0.78 | 0.47 | 0.59 | 0.51 | 0.78 | 0.75 |
| 2 | 184325147 | rs17760522 | -0.371 | 0.481 | -0.865 | 0.558 | -0.01 | 0.008 | 0.008 | 0 | 0.17 | 0.11 | 0.32 | 0.1 | 0.19 | 0.17 |

| | | | | | | | | | | | | | | | | |
|---|---|---|---|---|---|---|---|---|---|---|---|---|---|---|---|---|
| 2 | 184339696 | rs10191108 | -0.486 | 0.427 | -0.668 | 0.718 | 0 | -0.004 | -0.004 | 4.215 | 0.31 | 0.32 | 0.52 | 0.58 | 0.27 | 0.38 |
| 2 | 184341553 | rs4233772 | -0.762 | 0.366 | -0.176 | 0.613 | 0.066 | 0.106 | 0.106 | 0.394 | 0.07 | 0.28 | 0.29 | 0.09 | 0.06 | 0.08 |
| 2 | 184353132 | rs2196690 | 0.885 | 0.814 | -0.825 | -0.207 | -0.024 | 0.014 | 0.014 | 0 | 0.21 | 0.12 | 0.23 | 0.02 | 0.33 | 0.19 |
| 2 | 184353567 | rs1370648 | 0.134 | 0.403 | -0.809 | 0.317 | -0.012 | -0.026 | -0.026 | 0 | 0.2 | 0.2 | 0.12 | 0.06 | 0.12 | 0.21 |
| 2 | 184353810 | rs13409979 | 0.119 | 0.402 | -0.81 | 0.317 | -0.012 | -0.025 | -0.025 | 0 | 0.2 | 0.2 | 0.12 | 0.06 | 0.12 | 0.21 |
| 2 | 184355309 | rs1438018 | -0.053 | 0.489 | -1.362 | 0.599 | -0.003 | -0.057 | -0.057 | 2.469 | 0.19 | 0.15 | 0.4 | 0.48 | 0.2 | 0.19 |
| 2 | 184355378 | rs10460317 | -0.508 | 0.348 | -0.997 | 0.427 | 0.001 | -0.031 | -0.031 | 0 | 0.54 | 0.53 | 0.62 | 0.83 | 0.49 | 0.74 |
| 2 | 184358402 | rs2368443 | -0.114 | 0.518 | -1.421 | 0.847 | -0.04 | -0.004 | -0.004 | 3.87 | 0.62 | 0.83 | 0.75 | 0.36 | 0.55 | 0.39 |
| 2 | 184360199 | rs1438019 | 0.125 | 0.646 | -0.588 | 0.6 | 0.04 | 0.03 | 0.03 | 0.05 | 0.92 | 0.75 | 0.84 | 0.92 | 0.92 | 0.93 |
| 2 | 184364553 | rs1898582 | 0.488 | 0.312 | -0.775 | 0.394 | 0.038 | 0.104 | 0.104 | 0.11 | 0.4 | 0.2 | 0.33 | 0.17 | 0.36 | 0.35 |
| 2 | 184371403 | rs9288091 | 0.223 | 0.451 | -0.139 | 0.544 | 0.003 | 0.006 | 0.006 | 0.683 | 0.52 | 0.52 | 0.43 | 0.65 | 0.47 | 0.63 |
| 2 | 184380033 | rs1406334 | -0.064 | 0.46 | -0.1 | 0.513 | 0.003 | 0.018 | 0.018 | 2.04 | 0.68 | 0.49 | 0.52 | 0.64 | 0.67 | 0.68 |
| 2 | 184382752 | rs12185705 | -0.571 | 0.451 | 0.052 | 0.414 | 0.005 | -0.04 | -0.04 | 0 | 0.9 | 0.87 | 0.84 | 1 | 0.89 | 0.91 |
| 2 | 184384952 | rs13413583 | 0.308 | 0.452 | -0.177 | 0.415 | -0.005 | -0.005 | -0.005 | 0 | 0.1 | 0.08 | 0.15 | 0 | 0.11 | 0.09 |
| 2 | 184385671 | rs6722136 | -0.904 | 0.47 | -0.026 | 0.628 | -0.004 | 0.076 | 0.076 | 1.487 | 0.79 | 0.6 | 0.66 | 0.66 | 0.77 | 0.77 |
| 2 | 184389762 | rs12623821 | 0.959 | 0.511 | -0.056 | 0.727 | 0.01 | 0.05 | 0.05 | 0 | 0.09 | 0.14 | 0.07 | 0.28 | 0.1 | 0.11 |
| 2 | 184390882 | rs16824710 | 1.211 | 0.327 | -0.104 | 0.393 | -0.005 | -0.034 | -0.034 | 0.678 | 0.16 | 0.09 | 0.29 | 0.35 | 0.2 | 0.15 |
| 2 | 184390974 | rs6750117 | -0.102 | 0.305 | -0.127 | 0.357 | 0.007 | -0.004 | -0.004 | 0 | 0.61 | 0.55 | 0.59 | 0.64 | 0.58 | 0.72 |
| 2 | 184391990 | rs6754252 | 0.57 | 0.345 | -0.523 | 0.231 | 0.004 | 0.006 | 0.006 | 0 | 0.45 | 0.47 | 0.42 | 0.27 | 0.42 | 0.53 |
| 2 | 184392834 | rs10198445 | 0.057 | 0.407 | -0.628 | 0.148 | 0.007 | -0.023 | -0.023 | 0 | 0.63 | 0.55 | 0.5 | 0.28 | 0.62 | 0.58 |
| 2 | 184401191 | rs13001300 | 1.482 | 0.522 | -1.148 | 0.108 | -0.032 | 0.037 | 0.037 | 0 | 0.17 | 0.07 | 0.09 | 0 | 0.21 | 0.05 |
| 2 | 184402831 | rs7584434 | 1.387 | 0.336 | -1.426 | 0.131 | -0.004 | -0.066 | -0.066 | 0 | 0.22 | 0.27 | 0.22 | 0 | 0.2 | 0.08 |
| 2 | 184407963 | rs2368446 | -0.437 | 0.481 | -1.81 | 1.093 | 0.001 | 0.01 | 0.01 | 0 | 0.84 | 0.86 | 0.8 | 0.65 | 0.83 | 0.81 |
| 2 | 184418780 | rs1528612 | 0.242 | 0.206 | -1.791 | 1.198 | 0.018 | 0.016 | 0.016 | 0.123 | 0.18 | 0.11 | 0.17 | 0.11 | 0.17 | 0.26 |
| 2 | 184440113 | rs16824771 | -0.175 | 0.725 | -1.224 | 1.517 | -0.006 | -0.035 | -0.035 | 3.14 | 0.16 | 0.09 | 0.25 | 0.39 | 0.19 | 0.17 |
| 2 | 184444370 | rs1950038 | 0.236 | 0.395 | -1.345 | 1.109 | 0.016 | 0.002 | 0.002 | 0.496 | 0.64 | 0.72 | 0.6 | 0.67 | 0.71 | 0.79 |
| 2 | 184446942 | rs1406333 | 0.296 | 0.885 | -2.129 | 1.223 | -0.059 | -0.054 | -0.054 | 0 | 0.3 | 0.08 | 0.34 | 0.36 | 0.35 | 0.41 |
| 2 | 184468466 | rs2368424 | 0.326 | 0.647 | -1.109 | 1.043 | -0.005 | -0.004 | -0.004 | 0 | 0.28 | 0.32 | 0.18 | 0.19 | 0.22 | 0.06 |
| 2 | 184484253 | rs6742026 | -0.317 | 0.555 | -0.744 | 0.967 | -0.005 | 0.016 | 0.016 | 0.073 | 0.39 | 0.49 | 0.34 | 0.46 | 0.34 | 0.25 |
| 2 | 184484622 | rs6742674 | 0.067 | 0.633 | -0.7 | 1.043 | -0.003 | -0.001 | -0.001 | 0 | 0.35 | 0.4 | 0.28 | 0.32 | 0.31 | 0.18 |
| 2 | 184495407 | rs6434039 | -0.911 | 0.389 | -0.79 | 0.777 | 0.004 | 0.005 | 0.005 | 0 | 0.57 | 0.54 | 0.46 | 0.49 | 0.53 | 0.46 |
| 2 | 184495461 | rs6434040 | -0.382 | 0.405 | -0.705 | 0.612 | -0.004 | 0.01 | 0.01 | 0 | 0.29 | 0.24 | 0.21 | 0.17 | 0.33 | 0.33 |
| 2 | 184499725 | rs12693351 | -0.542 | 0.344 | -0.304 | 0.581 | 0 | -0.027 | -0.027 | 0 | 0.31 | 0.34 | 0.22 | 0.17 | 0.33 | 0.33 |
| 2 | 184502493 | rs1463328 | -0.739 | 0.341 | -0.538 | 0.6 | -0.001 | 0.006 | 0.006 | 0 | 0.91 | 0.73 | 0.88 | 0.87 | 0.94 | 0.93 |
| 2 | 184516249 | rs1377760 | -0.811 | 0.345 | -0.493 | 0.593 | 0.004 | 0.008 | 0.008 | 0.02 | 0.91 | 0.75 | 0.89 | 0.87 | 0.95 | 0.94 |
| 2 | 184520495 | rs13023195 | 0.745 | 0.397 | -0.924 | 0.495 | -0.047 | 0.093 | 0.093 | 0 | 0.17 | 0.01 | 0.12 | 0.03 | 0.22 | 0.2 |
| 2 | 184524587 | rs6732860 | 1.284 | 0.978 | -1.125 | 0.798 | -0.073 | -0.032 | -0.032 | 0 | 0.23 | 0.03 | 0.14 | 0.25 | 0.24 | 0.33 |
| 2 | 184525209 | rs9678612 | 0.56 | 0.956 | -1.028 | 0.781 | -0.035 | 0.035 | 0.035 | 0 | 0.3 | 0.15 | 0.17 | 0.25 | 0.31 | 0.38 |
| 2 | 184538604 | rs7588905 | 1.102 | 0.863 | -1.156 | 0.909 | -0.101 | -0.107 | -0.107 | 0 | 0.25 | 0.02 | 0.17 | 0.36 | 0.28 | 0.36 |
| 2 | 184540102 | rs10803967 | -0.429 | 0.633 | -0.487 | 0.942 | 0.009 | 0.023 | 0.023 | 0 | 0.5 | 0.64 | 0.48 | 0.6 | 0.52 | 0.64 |
| 2 | 184541177 | rs1454034 | -0.586 | 0.629 | -0.322 | 0.957 | 0.001 | 0.001 | 0.001 | 0 | 0.5 | 0.54 | 0.47 | 0.56 | 0.52 | 0.64 |
| 2 | 184541981 | rs1454035 | 1.146 | 0.655 | -0.089 | 0.865 | 0.015 | 0.046 | 0.046 | 0 | 0.1 | 0.17 | 0.26 | 0.23 | 0.16 | 0.14 |
| 2 | 184544232 | rs1901977 | -0.481 | 0.435 | -0.054 | 0.681 | 0.006 | 0.018 | 0.018 | 0 | 0.53 | 0.66 | 0.48 | 0.62 | 0.52 | 0.67 |
| 2 | 184546456 | rs7594190 | -0.464 | 0.434 | 0.001 | 0.68 | 0 | 0 | 0 | 0.152 | 0.52 | 0.57 | 0.47 | 0.62 | 0.52 | 0.67 |
| 2 | 184551466 | rs13016089 | 0.838 | 0.114 | -0.342 | 0.36 | -0.002 | 0.08 | 0.08 | 0.731 | 0.22 | 0.04 | 0.12 | 0.05 | 0.17 | 0.05 |
| 2 | 184570407 | rs1551339 | -0.09 | 0.141 | -0.072 | 0.589 | 0.01 | 0.031 | 0.031 | 0.042 | 0.52 | 0.66 | 0.47 | 0.65 | 0.52 | 0.67 |
| 2 | 184573149 | rs4666934 | 0.998 | 0.411 | 0.295 | 0.748 | 0.012 | 0.04 | 0.04 | 0 | 0.1 | 0.15 | 0.24 | 0.23 | 0.16 | 0.14 |
| 2 | 184581717 | rs16824955 | 0.76 | 0.052 | -0.105 | 0.559 | -0.016 | -0.009 | -0.009 | 0 | 0.08 | 0 | 0.03 | 0.05 | 0.05 | 0.02 |
| 2 | 184587877 | rs11884398 | 0.12 | 0.105 | 0.191 | 0.595 | -0.013 | -0.025 | -0.025 | 0 | 0.14 | 0.25 | 0.05 | 0.05 | 0.11 | 0.14 |
| 2 | 184591219 | rs10177684 | 0.476 | 0.28 | 0.252 | 0.733 | 0.045 | 0.125 | 0.125 | 0 | 0.11 | 0.34 | 0.26 | 0.23 | 0.15 | 0.16 |
| 2 | 184604111 | rs6721530 | 1.003 | 0.06 | 0.405 | 0.208 | -0.017 | -0.068 | -0.068 | 0 | 0.12 | 0.27 | 0.05 | 0 | 0.06 | 0.06 |
| 2 | 184604185 | rs6750043 | 1.003 | 0.06 | 0.405 | 0.208 | -0.017 | -0.068 | -0.068 | 0.029 | 0.12 | 0.27 | 0.05 | 0 | 0.06 | 0.06 |
| 2 | 184613551 | rs10192234 | 0.881 | 0.046 | -0.279 | 0.158 | -0.074 | -0.021 | -0.021 | 0.006 | 0.34 | 0.12 | 0.37 | 0.67 | 0.45 | 0.49 |
| 2 | 184614354 | rs17715373 | 0.6 | -0.149 | -0.311 | -0.08 | 0.009 | 0.025 | 0.025 | 0.245 | 0.22 | 0.15 | 0.1 | 0.06 | 0.16 | 0.08 |
| 2 | 184619848 | rs12993478 | 0.702 | -0.135 | -0.511 | -0.073 | 0.008 | 0.054 | 0.054 | 0.571 | 0.22 | 0.06 | 0.1 | 0.06 | 0.16 | 0.08 |
| 2 | 184626750 | rs10173261 | 1.378 | 0.114 | -0.06 | 0.074 | -0.014 | -0.055 | -0.055 | 0.481 | 0.12 | 0.22 | 0.04 | 0 | 0.06 | 0.06 |
| 2 | 184632227 | rs1439771 | 0.833 | 0.154 | 0.108 | 0.086 | -0.03 | 0 | 0 | 0 | 0.35 | 0.24 | 0.4 | 0.67 | 0.45 | 0.49 |
| 2 | 184633519 | rs2119047 | -0.364 | 0.111 | 0.183 | 0.057 | 0.004 | 0.035 | 0.035 | 0 | 0.49 | 0.63 | 0.45 | 0.68 | 0.51 | 0.55 |
| 2 | 184635166 | rs16825054 | 0.946 | 0.233 | -0.278 | 0.324 | -0.068 | -0.057 | -0.057 | 0 | 0.32 | 0.14 | 0.38 | 0.63 | 0.43 | 0.39 |
| 2 | 184652758 | rs12477832 | 1.015 | 0.308 | -0.142 | 0.463 | -0.033 | -0.065 | -0.065 | 0 | 0.27 | 0.18 | 0.36 | 0.55 | 0.36 | 0.32 |
| 2 | 184663090 | rs7576053 | -0.325 | -0.001 | 0.491 | 0.263 | -0.002 | -0.005 | -0.005 | 0 | 0.46 | 0.54 | 0.44 | 0.64 | 0.47 | 0.51 |
| 2 | 184669864 | rs826151 | -1.38 | -0.183 | 0.377 | 0.011 | -0.018 | -0.011 | -0.011 | 0 | 0.7 | 0.79 | 0.52 | 0.67 | 0.61 | 0.58 |
| 2 | 184674424 | rs4666639 | 1.3 | -0.109 | 0.432 | 0.089 | -0.027 | -0.054 | -0.054 | 0 | 0.28 | 0.21 | 0.36 | 0.57 | 0.36 | 0.32 |
| 2 | 184680313 | rs10167905 | 2.167 | -0.162 | 0.383 | 0.053 | 0.02 | 0.051 | 0.051 | 0.084 | 0.14 | 0.05 | 0.05 | 0 | 0.08 | 0.09 |
| 2 | 184694169 | rs13431075 | 0.249 | -0.09 | 0.464 | 0.084 | 0 | 0.001 | 0.001 | 0 | 0.43 | 0.48 | 0.43 | 0.57 | 0.44 | 0.41 |
| 2 | 184701773 | rs3845692 | 1.201 | 0.116 | 0.031 | 0.263 | -0.029 | -0.056 | -0.056 | 0.028 | 0.27 | 0.19 | 0.36 | 0.57 | 0.36 | 0.32 |
| 2 | 184727745 | rs826165 | -0.005 | -0.213 | 0.293 | 0.018 | 0.003 | -0.009 | -0.009 | 0 | 0.58 | 0.45 | 0.84 | 0.9 | 0.75 | 0.74 |
| 2 | 184728413 | rs12475510 | 1.856 | 0.114 | -0.015 | 0.504 | 0.015 | 0.066 | 0.066 | 0 | 0.05 | 0.08 | 0.21 | 0.24 | 0.11 | 0.12 |
| 2 | 184732178 | rs826171 | -0.058 | -0.151 | 0.464 | 0.12 | -0.031 | -0.068 | -0.068 | 0.388 | 0.58 | 0.55 | 0.85 | 0.9 | 0.75 | 0.73 |

| | | | | | | | | | | | | | | | | |
|---|---|---|---|---|---|---|---|---|---|---|---|---|---|---|---|---|
| 2 | 184738091 | rs1682430 | -1.552 | -0.35 | 0.304 | 0.014 | 0.034 | 0.066 | 0.066 | 0.543 | 0.77 | 0.92 | 0.93 | 0.96 | 0.86 | 0.92 |
| 2 | 184738177 | rs1682431 | -1.175 | -0.379 | 0.605 | 0.059 | -0.005 | -0.005 | -0.005 | 0 | 0.75 | 0.74 | 0.91 | 0.88 | 0.84 | 0.85 |
| 2 | 184741522 | rs826185 | -0.536 | -0.394 | 0.573 | 0.047 | 0.023 | 0.026 | 0.026 | 0 | 0.61 | 0.68 | 0.86 | 0.88 | 0.75 | 0.76 |
| 2 | 184750212 | rs826115 | 0.997 | -0.102 | 0.178 | 0.272 | -0.014 | 0.004 | 0.004 | 0 | 0.31 | 0.25 | 0.48 | 0.29 | 0.37 | 0.44 |
| 2 | 184761417 | rs893736 | 0.959 | -0.142 | -0.017 | 0.257 | -0.029 | -0.041 | -0.041 | 0 | 0.3 | 0.18 | 0.35 | 0.59 | 0.36 | 0.31 |
| 2 | 184774274 | rs826136 | -1.873 | -0.232 | -0.125 | 0.108 | 0.019 | 0.058 | 0.058 | 0.177 | 0.79 | 0.93 | 0.92 | 0.96 | 0.84 | 0.91 |
| 2 | 184777132 | rs826131 | -1.748 | -0.25 | -0.122 | 0.099 | 0.02 | 0.051 | 0.051 | 0.037 | 0.78 | 0.91 | 0.92 | 0.96 | 0.84 | 0.91 |
| 2 | 184783736 | rs862236 | 0.848 | 0.202 | -0.001 | 0.36 | -0.018 | 0.005 | 0.005 | 0 | 0.31 | 0.25 | 0.48 | 0.29 | 0.38 | 0.44 |
| 2 | 184786881 | rs826127 | 0.558 | 0.208 | 0.042 | 0.367 | 0.015 | -0.01 | -0.01 | 0 | 0.32 | 0.43 | 0.49 | 0.29 | 0.37 | 0.44 |
| 2 | 184797826 | rs1817567 | 1.781 | 0.592 | -0.741 | 0.491 | -0.064 | 0.1 | 0.1 | 0 | 0.22 | 0.03 | 0.1 | 0.06 | 0.23 | 0.14 |
| 2 | 184800613 | rs12463971 | 0.366 | 0.226 | -0.94 | 0.581 | -0.05 | -0.03 | -0.03 | 0 | 0.38 | 0.2 | 0.4 | 0.59 | 0.48 | 0.33 |
| 2 | 184802609 | rs12466174 | 1.44 | 0.332 | -0.668 | 0.845 | 0.008 | 0.052 | 0.052 | 0 | 0.06 | 0.09 | 0.2 | 0.26 | 0.1 | 0.11 |
| 2 | 184818243 | rs10205814 | -1.129 | -0.251 | -0.621 | 0.525 | -0.074 | -0.14 | -0.14 | 0.132 | 0.71 | 0.56 | 0.89 | 0.96 | 0.85 | 0.84 |
| 2 | 184820021 | rs11883645 | 1.051 | 0.258 | -1.214 | 0.696 | 0 | -0.001 | -0.001 | 0.122 | 0.2 | 0.24 | 0.25 | 0.21 | 0.24 | 0.27 |
| 2 | 184826437 | rs12464511 | 1.398 | 0.735 | -1.465 | 0.879 | -0.006 | -0.001 | -0.001 | 0 | 0.08 | 0.09 | 0.21 | 0.26 | 0.12 | 0.1 |
| 2 | 184828432 | rs10497642 | 1.505 | 0.491 | -1.426 | 0.558 | -0.001 | -0.001 | -0.001 | 0 | 0.11 | 0.09 | 0.16 | 0.1 | 0.12 | 0.23 |
| 2 | 184849109 | rs4638756 | -0.082 | 0.008 | -1.303 | 0.577 | 0.071 | 0.065 | 0.065 | 0 | 0.49 | 0.27 | 0.63 | 0.68 | 0.59 | 0.54 |
| 2 | 184870226 | rs12693365 | 0.328 | 0.126 | -1.012 | 0.582 | 0 | -0.008 | -0.008 | 0 | 0.24 | 0.27 | 0.36 | 0.47 | 0.28 | 0.39 |
| 2 | 184881731 | rs12621228 | 1.57 | 0.484 | -1.397 | 0.502 | -0.054 | -0.067 | -0.067 | 0.647 | 0.11 | 0.01 | 0.16 | 0.25 | 0.14 | 0.07 |
| 2 | 184882169 | rs4505485 | 0.238 | 0.035 | -1.166 | 0.152 | 0.004 | -0.016 | -0.016 | 0 | 0.36 | 0.32 | 0.4 | 0.5 | 0.38 | 0.48 |
| 2 | 184887430 | rs4517938 | -1.047 | -0.441 | -1.163 | -0.22 | 0 | 0 | 0 | 0.259 | 0.76 | 0.85 | 0.88 | 0.91 | 0.84 | 0.92 |
| 2 | 184898361 | rs11677862 | 0.336 | -0.197 | -1.209 | -0.155 | -0.005 | 0 | 0 | 0 | 0.38 | 0.36 | 0.47 | 0.41 | 0.45 | 0.43 |
| 2 | 184909755 | rs10931115 | 0.019 | 0.072 | -1.28 | 0.17 | -0.022 | -0.063 | -0.063 | 0 | 0.25 | 0.13 | 0.37 | 0.6 | 0.34 | 0.37 |
| 2 | 184909956 | rs12465769 | 0.241 | -0.329 | -1.275 | -0.088 | 0.042 | 0.074 | 0.074 | 0 | 0.61 | 0.22 | 0.67 | 0.78 | 0.72 | 0.84 |
| 2 | 184933453 | rs6434064 | 2.016 | -0.103 | -1.235 | 0.036 | -0.02 | 0.07 | 0.07 | 0.373 | 0.12 | 0 | 0.05 | 0 | 0.1 | 0.07 |
| 2 | 184941469 | rs4427984 | 0.698 | 0.196 | -1.368 | 0.04 | -0.054 | 0.032 | 0.032 | 0 | 0.11 | 0.02 | 0.11 | 0.05 | 0.19 | 0.15 |
| 2 | 184941520 | rs10204944 | 0.518 | 0.122 | -1.208 | 0.227 | 0.024 | -0.043 | -0.043 | 0 | 0.26 | 0.15 | 0.22 | 0.46 | 0.21 | 0.31 |
| 2 | 184942934 | rs949972 | -0.773 | 0.266 | -0.969 | 0.12 | 0 | 0 | 0 | 0 | 0.53 | 0.44 | 0.38 | 0.6 | 0.51 | 0.55 |
| 2 | 184947327 | rs12614924 | 1.366 | -0.033 | -0.807 | 0.302 | -0.04 | -0.06 | -0.06 | 0 | 0.1 | 0 | 0.16 | 0.24 | 0.11 | 0.05 |
| 2 | 184954425 | rs12623132 | 1.011 | -0.509 | 0.327 | 0.535 | 0.023 | 0.041 | 0.041 | 0 | 0.49 | 0.16 | 0.6 | 0.4 | 0.54 | 0.54 |
| 2 | 184957166 | rs1983470 | -0.679 | -0.495 | 0.157 | 0.607 | -0.013 | 0.009 | 0.009 | 0 | 0.39 | 0.4 | 0.28 | 0.51 | 0.32 | 0.39 |
| 2 | 184966326 | rs1358093 | 0.866 | -0.246 | 0.191 | 0.6 | 0.009 | 0.031 | 0.031 | 0 | 0.48 | 0.34 | 0.61 | 0.37 | 0.51 | 0.51 |
| 2 | 184967629 | rs1526225 | 0.995 | -0.168 | 0.102 | 0.337 | -0.009 | 0.007 | 0.007 | 0 | 0.08 | 0.06 | 0.09 | 0.05 | 0.13 | 0.06 |
| 2 | 184971481 | rs1007465 | 0.599 | -0.371 | 0.077 | 0.461 | -0.004 | -0.06 | -0.06 | 0 | 0.16 | 0.08 | 0.17 | 0.44 | 0.19 | 0.29 |
| 2 | 184975795 | rs6751918 | -1.342 | -0.489 | 0.525 | 0.363 | 0.019 | 0.011 | 0.011 | 0 | 0.51 | 0.62 | 0.39 | 0.55 | 0.49 | 0.46 |
| 2 | 184976775 | rs17615865 | 0.151 | -0.687 | 0.137 | 0.592 | 0.024 | -0.031 | -0.031 | 0.822 | 0.16 | 0.07 | 0.12 | 0.3 | 0.09 | 0.21 |
| 2 | 184979740 | rs1405192 | 1.027 | -0.669 | 0.272 | 0.582 | 0 | -0.001 | -0.001 | 0 | 0.13 | 0.12 | 0.12 | 0.2 | 0.16 | 0.12 |
| 2 | 184987780 | rs1526195 | 0.138 | -0.714 | 0.335 | 0.543 | 0.007 | 0.008 | 0.008 | 0.021 | 0.24 | 0.15 | 0.17 | 0.2 | 0.26 | 0.19 |
| 2 | 184989114 | rs13011100 | -1.181 | -0.57 | 0.379 | 0.574 | 0.01 | -0.007 | -0.007 | 2.034 | 0.47 | 0.25 | 0.37 | 0.5 | 0.48 | 0.45 |
| 2 | 184994108 | rs13023858 | 1.078 | -0.563 | 0.379 | 0.578 | 0 | -0.001 | -0.001 | 0 | 0.13 | 0.12 | 0.11 | 0.2 | 0.15 | 0.12 |
| 2 | 185000058 | rs11680461 | -0.601 | -0.56 | 0.374 | 0.569 | -0.003 | 0.021 | 0.021 | 0.173 | 0.34 | 0.13 | 0.26 | 0.3 | 0.33 | 0.33 |
| 2 | 185021912 | rs1526208 | 0.169 | -0.567 | 0.809 | 0.547 | 0.011 | 0.011 | 0.011 | 0.057 | 0.71 | 0.21 | 0.78 | 0.8 | 0.75 | 0.81 |
| 2 | 185030292 | rs10497645 | 1.268 | -0.562 | 0.632 | 0.545 | 0.002 | -0.014 | -0.014 | 0 | 0.14 | 0.1 | 0.06 | 0 | 0.1 | 0.07 |
| 2 | 185031379 | rs16825540 | 1.132 | -0.552 | 0.549 | 0.567 | -0.001 | 0 | 0 | 0 | 0.12 | 0.12 | 0.11 | 0.2 | 0.15 | 0.13 |
| 2 | 185036013 | rs872194 | 0.839 | -0.606 | 0.532 | 0.541 | 0.037 | 0.02 | 0.02 | 0 | 0.58 | 0.16 | 0.69 | 0.65 | 0.67 | 0.66 |
| 2 | 185036752 | rs6434075 | -0.299 | -0.614 | 0.552 | 0.535 | -0.003 | -0.041 | -0.041 | 0.038 | 0.24 | 0.11 | 0.21 | 0.35 | 0.23 | 0.28 |
| 2 | 185037476 | rs1918001 | 0.851 | -0.614 | 0.551 | 0.544 | 0.037 | 0.017 | 0.017 | 0 | 0.58 | 0.16 | 0.69 | 0.64 | 0.67 | 0.66 |
| 2 | 185043362 | rs1997131 | -0.834 | -0.593 | 0.544 | 0.519 | -0.03 | -0.103 | -0.103 | 0 | 0.83 | 0.35 | 0.91 | 1 | 0.9 | 0.93 |
| 2 | 185052062 | rs7602025 | 0.63 | -0.556 | 0.396 | 0.579 | 0.026 | 0.015 | 0.015 | 0 | 0.59 | 0.24 | 0.7 | 0.65 | 0.67 | 0.66 |
| 2 | 185056639 | rs10931123 | 0.536 | -0.547 | 0.4 | 0.566 | 0.019 | 0.026 | 0.026 | 0 | 0.61 | 0.25 | 0.7 | 0.69 | 0.68 | 0.67 |
| 2 | 185067134 | rs12470103 | 1.293 | -0.526 | 0.215 | 0.59 | -0.048 | -0.052 | -0.052 | 1.923 | 0.46 | 0.04 | 0.58 | 0.45 | 0.52 | 0.53 |
| 2 | 185075781 | rs35416170 | 1.134 | -0.699 | 0.362 | 0.637 | -0.001 | 0.001 | 0.001 | 0 | 0.12 | 0.11 | 0.11 | 0.2 | 0.15 | 0.12 |
| 2 | 185088578 | rs1158848 | 0.828 | -0.806 | 0.9 | 0.562 | 0.001 | -0.003 | -0.003 | 1.061 | 0.61 | 0.33 | 0.64 | 0.45 | 0.62 | 0.6 |
| 2 | 185099836 | rs12621296 | 0.801 | -0.808 | 0.915 | 0.561 | 0.001 | -0.002 | -0.002 | 0 | 0.61 | 0.34 | 0.64 | 0.45 | 0.62 | 0.6 |
| 2 | 185100535 | rs7596901 | -0.144 | -0.822 | 1.06 | 0.569 | -0.003 | -0.02 | -0.02 | 0.318 | 0.77 | 0.88 | 0.79 | 0.69 | 0.77 | 0.74 |
| 2 | 185107332 | rs1455882 | 1.355 | -0.867 | 1.002 | 0.498 | -0.014 | -0.022 | -0.022 | 0 | 0.46 | 0.11 | 0.58 | 0.45 | 0.52 | 0.53 |
| 2 | 185111870 | rs985366 | 0.281 | -0.831 | 1.033 | 0.529 | -0.012 | 0.05 | 0.05 | 0 | 0.73 | 0.45 | 0.76 | 0.64 | 0.77 | 0.72 |
| 2 | 185157849 | rs13025307 | 1.04 | -0.807 | 0.933 | 0.496 | -0.004 | 0.016 | 0.016 | 0 | 0.13 | 0.17 | 0.12 | 0.2 | 0.15 | 0.13 |
| 2 | 185164544 | rs17582837 | 0.968 | -0.801 | 0.906 | 0.499 | -0.005 | 0.013 | 0.013 | 0 | 0.13 | 0.17 | 0.12 | 0.2 | 0.15 | 0.13 |
| 2 | 185164916 | rs4666970 | 0.173 | -0.808 | 0.592 | 0.521 | -0.003 | 0.009 | 0.009 | 0 | 0.74 | 0.73 | 0.78 | 0.64 | 0.77 | 0.72 |
| 2 | 185187144 | rs719736 | 0.894 | -0.321 | 0.692 | 0.801 | -0.099 | -0.075 | -0.075 | 0 | 0.45 | 0.03 | 0.57 | 0.45 | 0.51 | 0.53 |
| 2 | 185191288 | rs7588228 | 0.121 | -0.123 | 0.962 | 0.365 | -0.007 | -0.005 | -0.005 | 0 | 0.6 | 0.6 | 0.72 | 0.64 | 0.66 | 0.65 |
| 2 | 185196480 | rs2369343 | 0.006 | -0.121 | 0.648 | 0.366 | 0.007 | 0.003 | 0.003 | 1.91 | 0.62 | 0.83 | 0.74 | 0.64 | 0.67 | 0.65 |
| 2 | 185198135 | rs12465214 | 1.099 | -0.283 | 0.469 | 0.202 | -0.112 | -0.054 | -0.054 | 0 | 0.37 | 0.05 | 0.55 | 0.58 | 0.46 | 0.51 |
| 2 | 185198456 | rs7561599 | 0.117 | -0.172 | 0.651 | 0.442 | 0.007 | 0.005 | 0.005 | 1.232 | 0.61 | 0.75 | 0.74 | 0.64 | 0.67 | 0.65 |
| 2 | 185207504 | rs971784 | -0.96 | -0.162 | 0.374 | 0.18 | 0.02 | 0 | 0 | 1.546 | 0.74 | 0.84 | 0.86 | 0.98 | 0.87 | 0.88 |
| 2 | 185207659 | rs971782 | 0.697 | 0.023 | 0.314 | 0.229 | -0.009 | 0.006 | 0.006 | 0 | 0.09 | 0.06 | 0.09 | 0.07 | 0.15 | 0.07 |
| 2 | 185210610 | rs12328731 | 0.719 | -0.094 | 0.292 | 0.351 | 0.006 | -0.006 | -0.006 | 0.188 | 0.19 | 0.14 | 0.1 | 0.04 | 0.16 | 0.15 |
| 2 | 185212246 | rs2198715 | 0.719 | -0.091 | 0.292 | 0.351 | 0.007 | -0.006 | -0.006 | 0 | 0.19 | 0.14 | 0.1 | 0.04 | 0.16 | 0.15 |

| | | | | | | | | | | | | | | | | |
|---|---|---|---|---|---|---|---|---|---|---|---|---|---|---|---|---|
| 2 | 185217338 | rs13008055 | 1.043 | -0.133 | -0.123 | 0.329 | -0.153 | -0.12 | -0.12 | 1.326 | 0.38 | 0.03 | 0.59 | 0.52 | 0.51 | 0.45 |
| 2 | 185221268 | rs17501385 | 0.942 | -0.141 | -0.105 | 0.307 | 0.01 | 0.08 | 0.08 | 0 | 0.18 | 0.07 | 0.1 | 0.04 | 0.17 | 0.15 |
| 2 | 185221959 | rs10209231 | -0.405 | -0.22 | 0.142 | 0.302 | -0.015 | -0.09 | -0.09 | 2.792 | 0.64 | 0.57 | 0.76 | 0.91 | 0.73 | 0.81 |
| 2 | 185223403 | rs17429535 | 0.961 | -0.195 | -0.975 | 0.316 | 0.013 | 0.079 | 0.079 | 0 | 0.18 | 0.07 | 0.1 | 0.04 | 0.16 | 0.15 |
| 2 | 185227071 | rs1443022 | 0.564 | -0.187 | -0.956 | 0.292 | -0.009 | 0.203 | 0.203 | 0.218 | 0.44 | 0.2 | 0.63 | 0.87 | 0.56 | 0.66 |
| 2 | 185228736 | rs12466190 | 1.285 | 0.092 | -1.136 | 0.446 | -0.041 | -0.05 | -0.05 | 0.882 | 0.1 | 0.01 | 0.32 | 0.27 | 0.18 | 0.18 |
| 2 | 185229307 | rs4666976 | 0.854 | -0.516 | -0.604 | 0.03 | -0.009 | 0.203 | 0.203 | 0.218 | 0.44 | 0.2 | 0.63 | 0.87 | 0.56 | 0.65 |
| 2 | 185235861 | rs1372949 | 0.958 | -0.561 | -0.618 | 0.014 | 0.043 | 0.048 | 0.048 | 0.573 | 0.24 | 0.11 | 0.13 | 0.02 | 0.13 | 0.13 |
| 2 | 185242856 | rs13028278 | 0.709 | -0.388 | -0.748 | 0.1 | 0.013 | 0.079 | 0.079 | 0 | 0.18 | 0.07 | 0.09 | 0.04 | 0.16 | 0.15 |
| 2 | 185250355 | rs12478223 | 1.103 | -0.3 | -0.737 | 0.202 | -0.149 | -0.12 | -0.12 | 0 | 0.38 | 0.03 | 0.59 | 0.52 | 0.51 | 0.45 |
| 2 | 185256424 | rs1348341 | 0.228 | -0.371 | -0.672 | 0.226 | 0.009 | -0.017 | -0.017 | 0 | 0.24 | 0.26 | 0.14 | 0.39 | 0.22 | 0.36 |
| 2 | 185258714 | rs10931132 | -0.843 | -0.226 | -1.327 | 0.161 | 0.014 | -0.01 | -0.01 | 0 | 0.74 | 0.83 | 0.86 | 0.98 | 0.87 | 0.87 |
| 2 | 185277167 | rs4363994 | 0.565 | 0.249 | -1.3 | 0.619 | -0.131 | -0.118 | -0.118 | 0.171 | 0.38 | 0.04 | 0.6 | 0.51 | 0.51 | 0.45 |
| 2 | 185277576 | rs1443033 | 0.104 | 0.28 | -0.438 | 0.28 | -0.003 | -0.012 | -0.012 | 0.329 | 0.62 | 0.66 | 0.76 | 0.83 | 0.73 | 0.79 |
| 2 | 185279087 | rs10204602 | 0.143 | 0.332 | -0.811 | 0.319 | 0.006 | 0.009 | 0.009 | 2.65 | 0.61 | 0.66 | 0.8 | 0.81 | 0.72 | 0.77 |
| 2 | 185280417 | rs10931138 | -0.797 | 1.113 | -1.242 | 0.776 | -0.043 | -0.022 | -0.022 | 0 | 0.28 | 0.12 | 0.47 | 0.63 | 0.36 | 0.34 |
| 2 | 185282179 | rs2034055 | 0.206 | 1.271 | -0.63 | 0.701 | 0.068 | 0.06 | 0.06 | 0 | 0.94 | 0.78 | 0.94 | 0.9 | 0.9 | 0.82 |
| 2 | 185283979 | rs1596274 | -0.728 | 1.143 | -1.324 | 1.434 | -0.049 | -0.114 | -0.114 | 0.035 | 0.16 | 0.01 | 0.09 | 0.36 | 0.18 | 0.19 |
| 2 | 185287387 | rs1443029 | -0.962 | 1.104 | -1.517 | 1.368 | -0.036 | -0.063 | -0.063 | 0 | 0.16 | 0.06 | 0.13 | 0.37 | 0.24 | 0.32 |
| 2 | 185289177 | rs10931139 | -0.585 | 1.19 | -1.711 | 1.323 | -0.074 | -0.144 | -0.144 | 0.042 | 0.18 | 0.01 | 0.35 | 0.56 | 0.25 | 0.25 |
| 2 | 185300193 | rs10188196 | -0.158 | 0.506 | -0.444 | 0.605 | 0.006 | 0.004 | 0.004 | 0 | 0.06 | 0.31 | 0.05 | 0 | 0.01 | 0 |
| 2 | 185318914 | rs1443032 | -1.47 | 0.125 | -0.281 | 0.203 | -0.006 | 0.113 | 0.113 | 0 | 0.48 | 0.36 | 0.57 | 0.93 | 0.54 | 0.59 |
| 2 | 185327680 | rs6751688 | -1.383 | 0.07 | 0.009 | -0.069 | 0.001 | -0.017 | -0.017 | 0 | 0.51 | 0.55 | 0.57 | 0.94 | 0.54 | 0.62 |
| 2 | 185349505 | rs1348342 | 0.246 | 0.346 | 0.539 | 0.046 | 0.087 | -0.003 | -0.003 | 0.214 | 0.89 | 0.81 | 0.88 | 0.85 | 0.79 | 0.88 |
| 2 | 185350711 | rs1372952 | 1.753 | 0.496 | -0.137 | -0.991 | -0.022 | 0.045 | 0.045 | 0 | 0.22 | 0.09 | 0.17 | 0.05 | 0.26 | 0.15 |
| 2 | 185350744 | rs17616975 | 1.005 | 0.263 | -0.128 | -0.396 | -0.001 | -0.032 | -0.032 | 0 | 0.1 | 0.03 | 0.05 | 0.23 | 0.1 | 0.27 |
| 2 | 185352219 | rs1427149 | -0.535 | 0.223 | 0.089 | -0.492 | 0.003 | -0.001 | -0.001 | 0 | 0.41 | 0.51 | 0.47 | 0.8 | 0.34 | 0.54 |
| 2 | 185353473 | rs1014024 | 1.364 | 0.245 | -0.183 | -0.493 | -0.017 | 0.181 | 0.181 | 0.056 | 0.46 | 0.17 | 0.4 | 0.05 | 0.45 | 0.34 |
| 2 | 185354045 | rs10497652 | 0.946 | 0.998 | -0.301 | -0.119 | -0.025 | 0.023 | 0.023 | 0 | 0.06 | 0.01 | 0.06 | 0 | 0.13 | 0.02 |
| 2 | 185357149 | rs12471708 | -0.545 | 0.889 | -0.261 | -0.095 | 0.092 | -0.002 | -0.002 | 0 | 0.89 | 0.8 | 0.88 | 0.85 | 0.79 | 0.88 |
| 2 | 185362149 | rs12467551 | 0.23 | 0.463 | -0.686 | -0.05 | -0.002 | -0.103 | -0.103 | 0 | 0.23 | 0.11 | 0.37 | 0.55 | 0.22 | 0.23 |
| 2 | 185383451 | rs12620876 | -0.07 | 0.407 | -1.122 | -0.118 | -0.033 | -0.138 | -0.138 | 3.851 | 0.29 | 0.07 | 0.35 | 0.56 | 0.27 | 0.28 |
| 2 | 185390094 | rs4666659 | 0.496 | 0.586 | -0.558 | -0.174 | 0.012 | 0.001 | 0.001 | 1.142 | 0.13 | 0.15 | 0.13 | 0.18 | 0.18 | 0.11 |
| 2 | 185401265 | rs17430153 | 0.735 | 0.082 | -0.396 | -0.487 | -0.001 | -0.002 | -0.002 | 0 | 0.25 | 0.23 | 0.17 | 0.17 | 0.25 | 0.37 |
| 2 | 185405580 | rs2033137 | 0.251 | 0.06 | -0.117 | -0.327 | -0.005 | -0.003 | -0.003 | 0 | 0.61 | 0.52 | 0.58 | 0.58 | 0.63 | 0.52 |
| 2 | 185410563 | rs7558116 | 1.543 | 0.08 | -0.568 | -0.679 | -0.046 | 0.14 | 0.14 | 0 | 0.29 | 0.09 | 0.19 | 0.04 | 0.3 | 0.16 |
| 2 | 185410800 | rs6434087 | 1.612 | 0.054 | -0.579 | -0.684 | -0.053 | 0.144 | 0.144 | 0 | 0.29 | 0.07 | 0.19 | 0.04 | 0.3 | 0.16 |
| 2 | 185421477 | rs4389295 | 0.858 | -0.052 | -0.212 | -0.658 | -0.006 | -0.013 | -0.013 | 0 | 0.22 | 0.19 | 0.35 | 0.54 | 0.31 | 0.29 |
| 2 | 185428946 | rs10803976 | -0.867 | -0.11 | -0.022 | -0.67 | -0.001 | 0.038 | 0.038 | 0.139 | 0.84 | 0.7 | 0.73 | 0.75 | 0.88 | 0.88 |
| 2 | 185430182 | rs10803977 | 0.711 | -0.171 | -0.296 | -0.638 | -0.002 | 0.019 | 0.019 | 0 | 0.57 | 0.4 | 0.36 | 0.21 | 0.56 | 0.54 |
| 2 | 185434554 | rs12999003 | 0.774 | -0.17 | -0.349 | -0.637 | -0.004 | 0.072 | 0.072 | 0 | 0.56 | 0.26 | 0.35 | 0.21 | 0.55 | 0.54 |
| 2 | 185436262 | rs17430279 | 1.147 | -0.166 | -0.118 | -0.447 | 0.002 | 0.004 | 0.004 | 0.121 | 0.24 | 0.19 | 0.15 | 0.17 | 0.23 | 0.37 |
| 2 | 185444506 | rs10198713 | 0.286 | 0.013 | 0.219 | -0.201 | -0.013 | 0.026 | 0.026 | 0.112 | 0.79 | 0.54 | 0.71 | 0.75 | 0.86 | 0.83 |
| 2 | 185454461 | rs1021041 | 0.225 | 0.023 | 0.133 | -0.201 | -0.012 | 0.009 | 0.009 | 0.05 | 0.8 | 0.63 | 0.72 | 0.75 | 0.86 | 0.83 |
| 2 | 185460295 | rs11888068 | 0.51 | 0.294 | -0.477 | -0.002 | -0.013 | -0.03 | -0.03 | 0 | 0.22 | 0.16 | 0.3 | 0.49 | 0.31 | 0.27 |
| 2 | 185462041 | rs10497655 | 0.626 | 0.289 | -0.658 | -0.009 | -0.069 | -0.125 | -0.125 | 0 | 0.21 | 0.05 | 0.31 | 0.5 | 0.32 | 0.27 |
| 2 | 185467661 | rs12693382 | 1.504 | 0.401 | -1.066 | -0.551 | -0.006 | 0.052 | 0.052 | 0 | 0.17 | 0.1 | 0.12 | 0.02 | 0.16 | 0.09 |
| 2 | 185478036 | rs12693384 | 0.373 | 0.179 | -0.828 | -0.662 | -0.003 | -0.039 | -0.039 | 1.19 | 0.43 | 0.48 | 0.29 | 0.14 | 0.41 | 0.39 |
| 2 | 185480633 | rs1427150 | 1.609 | 0.218 | -0.742 | -0.504 | -0.001 | 0.009 | 0.009 | 0 | 0.24 | 0.16 | 0.16 | 0.12 | 0.26 | 0.3 |
| 2 | 185489430 | rs17617468 | 1.537 | 0.075 | -0.867 | -0.266 | -0.012 | -0.058 | -0.058 | 0 | 0.12 | 0.02 | 0.26 | 0.29 | 0.11 | 0.15 |
| 2 | 185526003 | rs899846 | -1.018 | -0.011 | -0.806 | -0.181 | 0.01 | -0.048 | -0.048 | 0 | 0.84 | 0.91 | 0.72 | 0.65 | 0.89 | 0.76 |
| 2 | 185530491 | rs899847 | 0.32 | 0.01 | -0.842 | -0.475 | -0.003 | 0.084 | 0.084 | 0 | 0.58 | 0.69 | 0.38 | 0.16 | 0.58 | 0.48 |
| 2 | 185533580 | rs7597593 | 0.298 | -0.278 | -0.82 | -0.525 | 0.033 | 0.02 | 0.02 | 0 | 0.51 | 0.32 | 0.58 | 0.57 | 0.61 | 0.49 |
| 2 | 185535807 | rs17508767 | 2.835 | -0.179 | -1.041 | -0.601 | -0.021 | 0.026 | 0.026 | 0 | 0.08 | 0 | 0.03 | 0.02 | 0.07 | 0.05 |
| 2 | 185538930 | rs6728764 | 0.284 | -0.409 | -0.933 | -0.788 | 0.015 | 0.06 | 0.06 | 0 | 0.42 | 0.62 | 0.28 | 0.14 | 0.38 | 0.34 |
| 2 | 185559043 | rs4308093 | 0.264 | -0.436 | -0.454 | -0.773 | -0.021 | -0.125 | -0.125 | 1.881 | 0.32 | 0.51 | 0.19 | 0.03 | 0.25 | 0.15 |
| 2 | 185564834 | rs7585738 | 1.54 | -0.441 | -0.412 | -0.746 | 0.007 | 0.024 | 0.024 | 0 | 0.21 | 0.18 | 0.13 | 0.02 | 0.18 | 0.1 |
| 2 | 185579987 | rs1480476 | 1.657 | -0.428 | -0.488 | -0.74 | 0.004 | 0.047 | 0.047 | 0 | 0.19 | 0.14 | 0.11 | 0.02 | 0.17 | 0.1 |
| 2 | 185616128 | rs13384546 | 1.625 | -0.464 | -0.375 | -0.757 | 0.005 | 0.027 | 0.027 | 0 | 0.19 | 0.17 | 0.11 | 0.02 | 0.17 | 0.1 |
| 2 | 185626426 | rs2369595 | -1.868 | -0.67 | -0.351 | -0.855 | 0.003 | 0.007 | 0.007 | 0 | 0.77 | 0.8 | 0.83 | 0.85 | 0.8 | 0.68 |
| 2 | 185636177 | rs6759491 | -0.001 | -0.682 | -0.47 | -0.874 | 0.016 | 0.031 | 0.031 | 0.501 | 0.34 | 0.72 | 0.2 | 0.04 | 0.25 | 0.15 |
| 2 | 185638875 | rs17617913 | 2.428 | -0.68 | -0.619 | -0.3 | -0.017 | -0.057 | -0.057 | 0 | 0.11 | 0.01 | 0.2 | 0.32 | 0.13 | 0.12 |
| 2 | 185647985 | rs17584494 | 1.992 | -0.405 | -0.988 | -0.039 | -0.058 | -0.058 | -0.058 | 0 | 0.11 | 0.04 | 0.21 | 0.28 | 0.19 | 0.16 |
| 2 | 185655092 | rs6716235 | 0.241 | -0.581 | -0.915 | -0.178 | 0.017 | 0.032 | 0.032 | 0.46 | 0.34 | 0.72 | 0.2 | 0.03 | 0.25 | 0.15 |
| 2 | 185669127 | rs6751335 | 1.959 | -0.471 | -1.072 | -0.158 | 0.005 | 0.028 | 0.028 | 0 | 0.2 | 0.16 | 0.11 | 0.02 | 0.17 | 0.1 |
| 2 | 185719174 | rs12693396 | 0.351 | -0.453 | -1.102 | -0.151 | 0.005 | -0.013 | -0.013 | 0 | 0.33 | 0.66 | 0.2 | 0.03 | 0.25 | 0.15 |
| 2 | 185735599 | rs7588907 | 0.148 | -0.626 | -1.197 | -0.261 | 0.018 | 0.015 | 0.015 | 1 | 0.37 | 0.66 | 0.2 | 0.03 | 0.25 | 0.15 |
| 2 | 185769921 | rs10497662 | 3.09 | -0.375 | -1.464 | 0.169 | -0.021 | -0.036 | -0.036 | 2.368 | 0.08 | 0 | 0.03 | 0.1 | 0.07 | 0.06 |
| 2 | 185771745 | rs4666994 | -1.614 | -0.189 | -1.454 | -0.027 | -0.001 | -0.004 | -0.004 | 0 | 0.73 | 0.71 | 0.8 | 0.85 | 0.75 | 0.62 |

| | | | | | | | | | | | | | | | | |
|---|---|---|---|---|---|---|---|---|---|---|---|---|---|---|---|---|
| 2 | 185778428 | rs1344706 | 1.13 | 0.121 | -1.488 | 0.44 | -0.119 | -0.15 | -0.15 | 0 | 0.25 | 0.04 | 0.33 | 0.52 | 0.38 | 0.35 |
| 2 | 185783141 | rs1583048 | 1.858 | 0.207 | -1.34 | 0.398 | -0.056 | -0.06 | -0.06 | 0 | 0.09 | 0.02 | 0.2 | 0.32 | 0.17 | 0.12 |
| 2 | 185785610 | rs11901504 | -0.583 | -0.536 | -0.946 | -0.169 | 0.008 | 0.032 | 0.032 | 0.041 | 0.54 | 0.56 | 0.69 | 0.84 | 0.58 | 0.52 |
| 2 | 185796571 | rs6751736 | -1.048 | -1.045 | -1.018 | -0.228 | -0.001 | -0.008 | -0.008 | 0 | 0.65 | 0.61 | 0.75 | 0.85 | 0.69 | 0.64 |
| 2 | 185797408 | rs17509873 | 2.921 | -1.089 | -1.12 | -0.297 | -0.007 | 0.025 | 0.025 | 0 | 0.08 | 0.02 | 0.03 | 0.02 | 0.07 | 0.05 |
| 2 | 185797687 | rs10931156 | 0.491 | -1.223 | -1.067 | -0.361 | 0.049 | 0.211 | 0.211 | 1.432 | 0.51 | 0.14 | 0.66 | 0.84 | 0.62 | 0.59 |
| 2 | 185800905 | rs12476147 | -0.328 | -1.187 | -0.928 | -0.353 | 0 | -0.01 | -0.01 | 0 | 0.46 | 0.44 | 0.69 | 0.84 | 0.62 | 0.59 |
| 2 | 185802363 | rs12477430 | -1.049 | -1.09 | -1.009 | -0.281 | -0.001 | -0.01 | -0.01 | 0.009 | 0.65 | 0.62 | 0.75 | 0.87 | 0.69 | 0.64 |
| 2 | 185811940 | rs4380187 | 1.209 | -0.866 | -1.49 | 0.078 | -0.164 | -0.099 | -0.099 | 0.583 | 0.3 | 0.05 | 0.37 | 0.44 | 0.45 | 0.38 |
| 2 | 185823466 | rs6434107 | 0.519 | -1.097 | -1.027 | -0.251 | -0.048 | -0.1 | -0.1 | 0 | 0.19 | 0.37 | 0.09 | 0.03 | 0.08 | 0.12 |
| 2 | 185829672 | rs6709418 | 0.559 | -1.388 | -1.286 | -0.376 | 0.042 | 0.081 | 0.081 | 0.204 | 0.6 | 0.19 | 0.71 | 0.8 | 0.71 | 0.61 |
| 2 | 185835898 | rs6741522 | 0.821 | -1.461 | -1.295 | -0.448 | 0.061 | 0.075 | 0.075 | 1.814 | 0.27 | 0.16 | 0.12 | 0.03 | 0.14 | 0.15 |
| 2 | 185846132 | rs1366841 | 0.06 | -1.387 | -1.177 | -0.414 | -0.054 | -0.127 | -0.127 | 0 | 0.37 | 0.49 | 0.18 | 0.05 | 0.19 | 0.2 |
| 2 | 185865740 | rs12477914 | 2.274 | -0.792 | -1.584 | 0.281 | -0.015 | -0.068 | -0.068 | 0.53 | 0.12 | 0.01 | 0.25 | 0.35 | 0.14 | 0.14 |
| 2 | 185905031 | rs263765 | 1.872 | -0.677 | -1.152 | -0.187 | 0.01 | 0.022 | 0.022 | 0.153 | 0.05 | 0.11 | 0.09 | 0.11 | 0.08 | 0.14 |
| 2 | 185921692 | rs263771 | 1.053 | -0.789 | -1.281 | -0.305 | -0.018 | -0.015 | -0.015 | 0 | 0.19 | 0.11 | 0.18 | 0.18 | 0.21 | 0.26 |
| 2 | 185926550 | rs10202378 | 1.443 | -1.023 | -0.85 | -0.48 | -0.049 | -0.067 | -0.067 | 0.029 | 0.12 | 0.21 | 0.04 | 0.01 | 0.03 | 0.06 |
| 2 | 185930659 | rs965472 | 0.998 | -0.442 | -1.288 | -0.245 | 0.023 | 0.036 | 0.036 | 1.219 | 0.18 | 0.12 | 0.08 | 0.02 | 0.12 | 0.09 |
| 2 | 185932256 | rs6752032 | 0.247 | -0.355 | -1.101 | -0.376 | 0.007 | 0.011 | 0.011 | 0 | 0.26 | 0.23 | 0.13 | 0.07 | 0.19 | 0.19 |
| 2 | 185932315 | rs263767 | -0.48 | -0.252 | -1.056 | -0.315 | -0.022 | -0.027 | -0.027 | 0 | 0.62 | 0.58 | 0.79 | 0.77 | 0.75 | 0.64 |
| 2 | 185935405 | rs17584997 | 2.228 | 0.243 | -1.565 | 0.322 | -0.045 | -0.054 | -0.054 | 4.092 | 0.1 | 0.01 | 0.26 | 0.31 | 0.16 | 0.1 |
| 2 | 185937040 | rs1483247 | 1.755 | -0.118 | -1.706 | -0.134 | -0.021 | 0.015 | 0.015 | 0 | 0.13 | 0.01 | 0.08 | 0.06 | 0.15 | 0.12 |
| 2 | 185952716 | rs1021147 | -0.435 | -0.641 | -1.44 | -0.511 | -0.033 | -0.111 | -0.111 | 0 | 0.59 | 0.51 | 0.83 | 0.95 | 0.81 | 0.78 |
| 2 | 185969194 | rs10204328 | 0.248 | -0.302 | -1.766 | -0.308 | -0.019 | 0.062 | 0.062 | 0 | 0.4 | 0.21 | 0.66 | 0.77 | 0.63 | 0.52 |
| 2 | 185989325 | rs7604162 | -2.384 | -0.403 | -1.78 | -0.377 | 0.003 | -0.024 | -0.024 | 0 | 0.94 | 0.92 | 0.96 | 1 | 0.95 | 0.97 |
| 2 | 185992567 | rs4667014 | 0.471 | -0.452 | -2.241 | -0.403 | -0.045 | 0.056 | 0.056 | 0 | 0.38 | 0.13 | 0.64 | 0.77 | 0.63 | 0.52 |
| 2 | 185992761 | rs12991795 | 1.254 | -0.382 | -2.132 | -0.405 | 0.03 | 0.056 | 0.056 | 0.889 | 0.2 | 0.12 | 0.09 | 0.03 | 0.12 | 0.11 |
| 2 | 186043667 | rs4666673 | 1.492 | -0.167 | -2.06 | -0.041 | -0.035 | -0.053 | -0.053 | 0.005 | 0.16 | 0.08 | 0.31 | 0.33 | 0.22 | 0.15 |
| 2 | 186044627 | rs1483242 | 0.479 | -0.295 | -2.414 | -0.144 | -0.065 | -0.042 | -0.042 | 0 | 0.42 | 0.06 | 0.37 | 0.45 | 0.45 | 0.49 |
| 2 | 186050380 | rs1483246 | 0.039 | -0.188 | -2.127 | -0.088 | -0.055 | -0.027 | -0.027 | 0 | 0.61 | 0.23 | 0.48 | 0.48 | 0.58 | 0.6 |
| 2 | 186061623 | rs2291274 | 1.485 | -0.146 | -2.035 | -0.227 | -0.027 | -0.041 | -0.041 | 0 | 0.17 | 0.09 | 0.31 | 0.33 | 0.22 | 0.15 |
| 2 | 186062357 | rs13018827 | 0.243 | -0.32 | -2.266 | -0.517 | 0.023 | 0.167 | 0.167 | 0 | 0.47 | 0.22 | 0.38 | 0.25 | 0.44 | 0.36 |
| 2 | 186063153 | rs17762216 | 1.34 | 0.268 | -1.453 | 0.029 | -0.015 | 0.022 | 0.022 | 0 | 0.14 | 0.01 | 0.07 | 0.06 | 0.12 | 0.12 |
| 2 | 186067813 | rs17762453 | 1.683 | 0.557 | -1.35 | 0.106 | -0.066 | -0.042 | -0.042 | 0 | 0.11 | 0.02 | 0.22 | 0.2 | 0.27 | 0.16 |
| 2 | 186084222 | rs6747787 | 0.724 | -0.222 | -1.373 | 0.095 | 0.028 | 0.013 | 0.013 | 0.469 | 0.31 | 0.1 | 0.19 | 0.25 | 0.25 | 0.33 |
| 2 | 186088401 | rs11683056 | 0.439 | 0.377 | -0.908 | 0.502 | 0.014 | -0.005 | -0.005 | 0 | 0.51 | 0.3 | 0.36 | 0.47 | 0.46 | 0.6 |
| 2 | 186107423 | rs7590299 | 0.245 | 0.453 | -1.338 | 0.538 | -0.106 | 0.055 | 0.055 | 0 | 0.33 | 0.05 | 0.3 | 0.23 | 0.36 | 0.27 |
| 2 | 186111022 | rs4667021 | -0.443 | 0.205 | -1.283 | 0.561 | 0.01 | -0.032 | -0.032 | 0.21 | 0.47 | 0.26 | 0.61 | 0.53 | 0.56 | 0.39 |
| 2 | 186120083 | rs13392489 | -0.438 | 0.216 | -1.235 | 0.566 | 0.013 | -0.021 | -0.021 | 0 | 0.46 | 0.3 | 0.62 | 0.53 | 0.56 | 0.39 |
| 2 | 186164824 | rs2059349 | 1.304 | 0.332 | -1.404 | 0.609 | 0.002 | 0.02 | 0.02 | 0.315 | 0.13 | 0.19 | 0.3 | 0.3 | 0.18 | 0.12 |
| 2 | 186165639 | rs4600609 | 0.271 | 0.316 | -1.469 | 0.544 | -0.106 | 0.062 | 0.062 | 0 | 0.32 | 0.06 | 0.31 | 0.23 | 0.39 | 0.27 |
| 2 | 186172142 | rs7557829 | 0.24 | 0.316 | -1.469 | 0.528 | -0.106 | 0.059 | 0.059 | 0 | 0.32 | 0.05 | 0.31 | 0.23 | 0.39 | 0.28 |
| 2 | 186173063 | rs10197781 | -0.719 | 0.254 | -1.205 | 0.604 | 0.101 | 0.207 | 0.207 | 3.454 | 0.91 | 0.5 | 0.87 | 0.84 | 0.9 | 0.85 |
| 2 | 186178814 | rs1424532 | 0.54 | 0.205 | -1.303 | 0.311 | 0.035 | 0.041 | 0.041 | 0.008 | 0.46 | 0.33 | 0.28 | 0.3 | 0.35 | 0.46 |
| 2 | 186195271 | rs1100829 | 0.113 | 0.769 | -0.975 | 0.675 | 0.054 | 0.178 | 0.178 | 0.061 | 0.19 | 0.41 | 0.39 | 0.44 | 0.26 | 0.25 |
| 2 | 186206092 | rs850882 | -0.328 | 0.769 | -1.3 | 0.473 | 0.009 | 0.01 | 0.01 | 0 | 0.42 | 0.53 | 0.53 | 0.46 | 0.49 | 0.41 |
| 2 | 186208756 | rs1159985 | 1.496 | 0.773 | -1.618 | 0.511 | -0.049 | -0.062 | -0.062 | 0 | 0.13 | 0.02 | 0.21 | 0.24 | 0.2 | 0.16 |
| 2 | 186209004 | rs1364471 | 0.547 | 0.773 | -1.567 | 0.428 | 0.03 | 0.037 | 0.037 | 0 | 0.44 | 0.29 | 0.24 | 0.3 | 0.31 | 0.44 |
| 2 | 186209495 | rs17765227 | 0.793 | 0.996 | -1.373 | 0.714 | -0.043 | 0.115 | 0.115 | 0.86 | 0.22 | 0.04 | 0.13 | 0.01 | 0.23 | 0.16 |
| 2 | 186218867 | rs850906 | -0.133 | 0.798 | -0.272 | 1.891 | 0.062 | 0.167 | 0.167 | 0.128 | 0.08 | 0.34 | 0.11 | 0.15 | 0.09 | 0.12 |
| 2 | 186220642 | rs850908 | -0.085 | 0.7 | -0.424 | 1.543 | -0.019 | -0.012 | -0.012 | 2.47 | 0.73 | 0.67 | 0.83 | 0.74 | 0.8 | 0.63 |
| 2 | 186223503 | rs850914 | -1.336 | 1.206 | -0.3 | 1.977 | 0.047 | 0.171 | 0.171 | 0.1 | 0.2 | 0.44 | 0.4 | 0.44 | 0.26 | 0.25 |
| 2 | 186240504 | rs9288114 | -0.467 | 0.983 | -1.308 | 1.147 | -0.008 | -0.011 | -0.011 | 0 | 0.12 | 0.08 | 0.29 | 0.19 | 0.16 | 0.12 |
| 2 | 186245653 | rs17403192 | -0.775 | 0.332 | -1.407 | 0.708 | 0.016 | 0.019 | 0.019 | 0 | 0.14 | 0.06 | 0.06 | 0.09 | 0.11 | 0.17 |
| 2 | 186247637 | rs10931170 | 0.595 | 0.194 | -0.527 | 0.608 | 0.03 | 0.036 | 0.036 | 0 | 0.94 | 0.73 | 0.87 | 0.94 | 0.95 | 0.97 |
| 2 | 186249405 | rs12997946 | -0.423 | -0.835 | -1.166 | 0.924 | 0.049 | -0.061 | -0.061 | 0 | 0.25 | 0.12 | 0.15 | 0.37 | 0.15 | 0.31 |
| 2 | 186249631 | rs16823795 | -0.423 | -0.835 | -1.079 | 1.113 | 0.046 | -0.066 | -0.066 | 0 | 0.25 | 0.13 | 0.14 | 0.42 | 0.15 | 0.31 |
| 2 | 186260603 | rs17331432 | 0.623 | -0.212 | -1.001 | 0.543 | -0.012 | -0.002 | -0.002 | 0 | 0.11 | 0.09 | 0.11 | 0.08 | 0.21 | 0.18 |
| 2 | 186265033 | rs6711108 | -0.357 | -0.389 | -1.088 | 0.438 | -0.019 | 0.016 | 0.016 | 0 | 0.16 | 0.11 | 0.14 | 0.09 | 0.24 | 0.22 |
| 2 | 186267921 | rs13006333 | 0.733 | -0.419 | -1.004 | 0.059 | -0.014 | -0.013 | -0.013 | 0 | 0.4 | 0.24 | 0.43 | 0.47 | 0.42 | 0.32 |
| 2 | 186272059 | rs16826817 | -0.611 | -0.597 | -1.001 | 0.014 | 0.008 | -0.008 | -0.008 | 0.206 | 0.37 | 0.31 | 0.27 | 0.39 | 0.37 | 0.5 |
| 2 | 186272255 | rs12622713 | 1.075 | -0.293 | -1.518 | -0.413 | -0.058 | -0.149 | -0.149 | 0.356 | 0.28 | 0.02 | 0.28 | 0.43 | 0.27 | 0.18 |
| 2 | 186275756 | rs1424537 | -0.399 | -0.176 | -1.389 | -0.257 | 0.008 | -0.018 | -0.018 | 0 | 0.37 | 0.31 | 0.27 | 0.44 | 0.37 | 0.5 |
| 2 | 186281516 | rs2194509 | -0.397 | -0.223 | -1.363 | -0.313 | -0.009 | -0.004 | -0.004 | 0 | 0.56 | 0.34 | 0.46 | 0.55 | 0.57 | 0.68 |
| 2 | 186283241 | rs10177733 | 0.291 | -0.418 | -1.474 | -0.273 | -0.017 | -0.041 | -0.041 | 0 | 0.38 | 0.24 | 0.37 | 0.5 | 0.44 | 0.59 |
| 2 | 186291818 | rs17403547 | 0.303 | -0.407 | -1.483 | -0.307 | 0.022 | 0.042 | 0.042 | 0 | 0.19 | 0.1 | 0.1 | 0.05 | 0.13 | 0.09 |
| 2 | 186297074 | rs2113533 | 0.708 | -0.473 | -1.45 | -0.281 | -0.016 | -0.051 | -0.051 | 1.628 | 0.17 | 0.1 | 0.16 | 0.39 | 0.24 | 0.42 |
| 2 | 186300310 | rs6731674 | 1.032 | -0.383 | -1.483 | -0.326 | 0.009 | 0.026 | 0.026 | 0 | 0.14 | 0.07 | 0.08 | 0.05 | 0.12 | 0.06 |
| 2 | 186303487 | rs767036 | 0.739 | -0.362 | -1.472 | -0.297 | -0.016 | -0.045 | -0.045 | 0 | 0.17 | 0.1 | 0.16 | 0.34 | 0.25 | 0.41 |

| | | | | | | | | | | | | | | | | |
|---|---|---|---|---|---|---|---|---|---|---|---|---|---|---|---|---|
| 2 | 186304202 | rs2113530 | 0.524 | -0.369 | -1.432 | -0.3 | -0.008 | -0.032 | -0.032 | 0 | 0.18 | 0.13 | 0.17 | 0.34 | 0.25 | 0.41 |
| 2 | 186309738 | rs10202955 | 1.984 | -0.495 | -1.435 | -0.498 | -0.029 | 0.038 | 0.038 | 0 | 0.18 | 0.03 | 0.21 | 0.11 | 0.19 | 0.18 |
| 2 | 186338274 | rs2081326 | 0.566 | -0.513 | -1.385 | -0.416 | -0.018 | 0.088 | 0.088 | 0 | 0.34 | 0.09 | 0.33 | 0.16 | 0.33 | 0.24 |
| 2 | 186345148 | rs11675192 | 0.81 | -0.575 | -1.403 | -0.449 | -0.002 | 0.001 | 0.001 | 0 | 0.09 | 0.08 | 0.08 | 0.05 | 0.11 | 0.05 |
| 2 | 186387129 | rs10931178 | 0.672 | -0.44 | -1.079 | -0.414 | 0.001 | -0.001 | -0.001 | 0 | 0.37 | 0.32 | 0.35 | 0.39 | 0.37 | 0.29 |
| 2 | 186410095 | rs12232985 | 1.764 | -0.46 | -1.074 | -0.391 | -0.002 | -0.035 | -0.035 | 0 | 0.07 | 0.01 | 0.06 | 0.33 | 0.05 | 0.23 |
| 2 | 186421690 | rs4667044 | 0.544 | -0.498 | -1.052 | -0.445 | 0 | -0.004 | -0.004 | 0 | 0.61 | 0.59 | 0.64 | 0.5 | 0.59 | 0.49 |
| 2 | 186434458 | rs4667046 | 1.755 | -0.538 | -1.376 | -0.749 | -0.046 | -0.107 | -0.107 | 0 | 0.16 | 0.02 | 0.23 | 0.38 | 0.17 | 0.09 |
| 2 | 186442089 | rs7593451 | 2.148 | -0.473 | -1.183 | -0.548 | -0.014 | 0.041 | 0.041 | 0 | 0.21 | 0.09 | 0.26 | 0.11 | 0.22 | 0.19 |
| 2 | 186444443 | rs6753011 | 0.573 | -0.478 | -1.126 | -0.806 | -0.005 | 0.011 | 0.011 | 0 | 0.59 | 0.44 | 0.62 | 0.49 | 0.59 | 0.48 |
| 2 | 186445393 | rs7604780 | -0.066 | -0.386 | -1.15 | -0.79 | 0.002 | 0.026 | 0.026 | 0.023 | 0.75 | 0.67 | 0.72 | 0.54 | 0.7 | 0.54 |
| 2 | 186458903 | rs9679452 | -0.812 | -0.365 | -1.121 | -0.777 | 0.004 | 0.034 | 0.034 | 0.251 | 0.82 | 0.76 | 0.75 | 0.55 | 0.74 | 0.58 |
| 2 | 186466450 | rs888425 | -0.661 | -0.356 | -1.145 | -0.776 | 0.012 | 0.082 | 0.082 | 0.398 | 0.81 | 0.68 | 0.74 | 0.55 | 0.74 | 0.58 |
| 2 | 186475590 | rs2111710 | -0.061 | -0.334 | -1.066 | -0.96 | -0.001 | -0.002 | -0.002 | 0 | 0.24 | 0.24 | 0.22 | 0.44 | 0.29 | 0.46 |
| 2 | 186504175 | rs759686 | -0.122 | -0.319 | -1.043 | -0.962 | -0.004 | -0.02 | -0.02 | 0 | 0.75 | 0.73 | 0.7 | 0.5 | 0.7 | 0.54 |
| 2 | 186509076 | rs763330 | 0.327 | -0.332 | -1.058 | -1.055 | -0.002 | 0.009 | 0.009 | 0 | 0.39 | 0.5 | 0.38 | 0.46 | 0.36 | 0.29 |
| 2 | 186521491 | rs2160846 | 1.941 | -0.318 | -1.322 | -1.082 | -0.004 | 0.01 | 0.01 | 0 | 0.22 | 0.15 | 0.27 | 0.04 | 0.23 | 0.2 |
| 2 | 186526579 | rs2193615 | 0.357 | -0.326 | -1.259 | -1.077 | -0.003 | 0.011 | 0.011 | 0 | 0.6 | 0.46 | 0.62 | 0.5 | 0.58 | 0.49 |
| 2 | 186541545 | rs9917247 | 0.403 | -0.38 | -1.264 | -1.059 | -0.001 | -0.003 | -0.003 | 0 | 0.38 | 0.31 | 0.36 | 0.46 | 0.36 | 0.29 |
| 2 | 186556464 | rs16827022 | 1.081 | -0.333 | -1.256 | -1.08 | 0.007 | 0.005 | 0.005 | 0 | 0.14 | 0.1 | 0.06 | 0 | 0.12 | 0.05 |
| 2 | 186577266 | rs6434140 | 0.355 | -0.353 | -1.258 | -1.072 | 0.005 | 0.004 | 0.004 | 0.03 | 0.42 | 0.54 | 0.48 | 0.47 | 0.47 | 0.61 |
| 2 | 186583961 | rs4334457 | 0.543 | -0.329 | -1.394 | -0.948 | 0.018 | 0.094 | 0.094 | 0.375 | 0.19 | 0.36 | 0.21 | 0.43 | 0.25 | 0.42 |
| 2 | 186591557 | rs10202023 | 0.678 | -0.274 | -1.449 | -0.744 | 0.007 | 0.013 | 0.013 | 0 | 0.15 | 0.1 | 0.07 | 0.05 | 0.12 | 0.05 |
| 2 | 186617835 | rs13030170 | 0.77 | -0.255 | -1.598 | -0.825 | 0.005 | 0.007 | 0.007 | 0.021 | 0.15 | 0.13 | 0.07 | 0.06 | 0.11 | 0.05 |
| 2 | 186623388 | rs7592227 | 0.688 | -0.381 | -1.599 | -0.979 | 0.008 | -0.058 | -0.058 | 0.023 | 0.4 | 0.52 | 0.41 | 0.11 | 0.46 | 0.39 |
| 2 | 186646161 | rs7566901 | 0.391 | -0.297 | -1.414 | -0.924 | -0.001 | 0.002 | 0.002 | 0.007 | 0.49 | 0.56 | 0.49 | 0.47 | 0.51 | 0.65 |
| 2 | 186670979 | rs4666689 | -0.909 | -0.158 | -1.136 | -0.776 | -0.002 | -0.001 | -0.001 | 0 | 0.85 | 0.8 | 0.85 | 0.88 | 0.88 | 0.94 |
| 2 | 186671912 | rs1862066 | 0.124 | -0.158 | -1.136 | -0.776 | -0.001 | 0.002 | 0.002 | 0.007 | 0.49 | 0.56 | 0.49 | 0.47 | 0.51 | 0.65 |
| 2 | 186672897 | rs4666690 | 0.472 | -0.157 | -1.286 | -0.775 | -0.005 | -0.007 | -0.007 | 0 | 0.36 | 0.25 | 0.36 | 0.41 | 0.37 | 0.29 |
| 2 | 186678877 | rs17230036 | 0.724 | -0.112 | -1.054 | -0.523 | -0.008 | 0.038 | 0.038 | 0.009 | 0.14 | 0 | 0.05 | 0.06 | 0.11 | 0.05 |
| 2 | 186689559 | rs767042 | -0.02 | 0.074 | -0.819 | -0.388 | -0.006 | 0.001 | 0.001 | 0.006 | 0.47 | 0.44 | 0.48 | 0.47 | 0.51 | 0.65 |
| 2 | 186694044 | rs16827202 | 0.404 | 0.073 | -1.365 | -0.223 | -0.029 | -0.089 | -0.089 | 0.82 | 0.25 | 0.12 | 0.22 | 0.43 | 0.29 | 0.47 |
| 2 | 186722205 | rs12464986 | 0.104 | 0.079 | -0.955 | 0.188 | 0.002 | -0.008 | -0.008 | 3.752 | 0.37 | 0.24 | 0.33 | 0.41 | 0.34 | 0.29 |
| 2 | 186726208 | rs4343438 | 0.013 | 0.113 | -1.072 | 0.027 | 0 | -0.002 | -0.002 | 0.015 | 0.38 | 0.35 | 0.42 | 0.47 | 0.35 | 0.29 |
| 2 | 186736211 | rs13031528 | 0.295 | 0.144 | -1.204 | -0.033 | 0.006 | 0.024 | 0.024 | 0.067 | 0.12 | 0.07 | 0.05 | 0.06 | 0.1 | 0.05 |
| 2 | 186756348 | rs10931206 | -0.428 | -0.004 | -1.172 | -0.146 | 0.006 | 0.005 | 0.005 | 0 | 0.51 | 0.42 | 0.47 | 0.53 | 0.45 | 0.34 |
| 2 | 186770538 | rs12475245 | 0.109 | -0.09 | -0.786 | -0.15 | 0.005 | -0.007 | -0.007 | 0.548 | 0.39 | 0.23 | 0.33 | 0.41 | 0.35 | 0.3 |
| 2 | 186782880 | rs12616454 | -0.527 | -0.086 | -0.786 | -0.174 | 0.003 | 0.003 | 0.003 | 0 | 0.52 | 0.42 | 0.47 | 0.53 | 0.45 | 0.35 |
| 2 | 186801246 | rs1487354 | -0.506 | -0.143 | -0.369 | -0.744 | -0.005 | -0.069 | -0.069 | 0 | 0.68 | 0.6 | 0.76 | 0.91 | 0.7 | 0.77 |
| 2 | 186805925 | rs4666694 | 1.855 | -0.285 | -0.887 | -1.048 | -0.039 | -0.121 | -0.121 | 0 | 0.16 | 0.02 | 0.22 | 0.41 | 0.17 | 0.08 |
| 2 | 186807268 | rs6760904 | 1.645 | -0.188 | -0.484 | -0.747 | 0 | -0.022 | -0.022 | 0 | 0.21 | 0.18 | 0.18 | 0.03 | 0.22 | 0.18 |
| 2 | 186809595 | rs10196671 | 0.438 | -0.144 | -0.641 | -0.77 | 0 | 0.003 | 0.003 | 0 | 0.27 | 0.3 | 0.43 | 0.49 | 0.35 | 0.47 |
| 2 | 186813272 | rs2101414 | 0.609 | -0.148 | -0.616 | -0.752 | -0.039 | -0.071 | -0.071 | 0 | 0.26 | 0.13 | 0.33 | 0.43 | 0.34 | 0.46 |
| 2 | 186844579 | rs10182124 | 0.299 | -0.149 | -0.672 | -0.792 | -0.003 | -0.008 | -0.008 | 0.281 | 0.27 | 0.29 | 0.41 | 0.49 | 0.34 | 0.46 |
| 2 | 186849059 | rs7560510 | -0.382 | -0.157 | -0.725 | -0.71 | 0.025 | -0.035 | -0.035 | 0 | 0.56 | 0.26 | 0.54 | 0.52 | 0.64 | 0.68 |
| 2 | 186857379 | rs16827374 | 1.251 | -0.023 | -0.897 | -0.606 | -0.012 | -0.048 | -0.048 | 0.094 | 0.1 | 0.04 | 0.18 | 0.36 | 0.1 | 0.25 |
| 2 | 186860130 | rs4564739 | 1.633 | -0.009 | -0.679 | -0.662 | -0.022 | 0.072 | 0.072 | 0 | 0.2 | 0.07 | 0.18 | 0.03 | 0.22 | 0.2 |
| 2 | 186861063 | rs12618471 | 0.093 | 0.032 | -0.81 | -0.659 | -0.037 | -0.07 | -0.07 | 0 | 0.36 | 0.16 | 0.36 | 0.49 | 0.42 | 0.48 |
| 2 | 186863405 | rs4666697 | 0.065 | 0.069 | -0.692 | -0.453 | 0.004 | -0.006 | -0.006 | 0 | 0.4 | 0.28 | 0.34 | 0.42 | 0.35 | 0.31 |
| 2 | 186880577 | rs16827418 | -0.337 | 0.222 | -1.52 | -0.209 | -0.023 | -0.067 | -0.067 | 0.005 | 0.4 | 0.19 | 0.37 | 0.51 | 0.43 | 0.48 |
| 2 | 186880719 | rs7574377 | -0.763 | -0.073 | -0.967 | -0.633 | -0.007 | 0.051 | 0.051 | 0 | 0.68 | 0.53 | 0.78 | 0.57 | 0.72 | 0.77 |
| 2 | 186882149 | rs964744 | -0.1 | -0.09 | -0.979 | -1.059 | 0.005 | 0.018 | 0.018 | 0 | 0.76 | 0.74 | 0.84 | 0.94 | 0.79 | 0.75 |
| 2 | 186896757 | rs875162 | 0.713 | -0.474 | -0.87 | -1.073 | -0.003 | 0.005 | 0.005 | 0.019 | 0.2 | 0.24 | 0.23 | 0.42 | 0.14 | 0.19 |
| 2 | 186897066 | rs6434158 | 0.202 | -0.519 | -0.946 | -1.147 | 0.011 | -0.007 | -0.007 | 0 | 0.63 | 0.6 | 0.65 | 0.52 | 0.71 | 0.64 |
| 2 | 186897323 | rs901613 | -1.159 | -0.515 | -0.96 | -1.132 | 0.006 | 0.016 | 0.016 | 0 | 0.83 | 0.84 | 0.88 | 0.94 | 0.85 | 0.83 |
| 2 | 186905790 | rs1038587 | -0.897 | -0.512 | -0.88 | -1.13 | 0 | -0.002 | -0.002 | 0 | 0.82 | 0.79 | 0.87 | 0.94 | 0.84 | 0.83 |
| 2 | 186915437 | rs6434159 | 0.693 | -0.452 | -0.83 | -1.129 | 0.004 | 0 | 0 | 0 | 0.56 | 0.5 | 0.6 | 0.52 | 0.63 | 0.55 |
| 2 | 186971595 | rs13028175 | 0.705 | -0.405 | -0.907 | -1.077 | 0.006 | -0.018 | -0.018 | 0 | 0.19 | 0.18 | 0.24 | 0.42 | 0.16 | 0.2 |
| 2 | 186997541 | rs12373744 | 0.005 | -0.413 | -0.887 | -1.091 | 0.005 | -0.009 | -0.009 | 2.395 | 0.26 | 0.25 | 0.27 | 0.42 | 0.21 | 0.29 |
| 2 | 187010476 | rs10931227 | 0.02 | -0.414 | -0.879 | -1.07 | 0.008 | -0.017 | -0.017 | 0.34 | 0.26 | 0.22 | 0.26 | 0.42 | 0.21 | 0.29 |
| 2 | 187028772 | rs13419562 | 0.101 | -0.451 | -0.924 | -1.062 | -0.002 | 0.006 | 0.006 | 0.068 | 0.21 | 0.25 | 0.26 | 0.42 | 0.18 | 0.23 |
| 2 | 187040643 | rs4284795 | 0.902 | -0.454 | -0.926 | -1.064 | 0.001 | 0.003 | 0.003 | 0.199 | 0.14 | 0.14 | 0.23 | 0.42 | 0.13 | 0.14 |
| 2 | 187040829 | rs16827526 | 0.818 | -0.532 | -0.983 | -0.993 | 0.013 | 0.015 | 0.015 | 0 | 0.18 | 0.11 | 0.12 | 0.1 | 0.15 | 0.17 |
| 2 | 187041149 | rs13425057 | 0.061 | -0.323 | -0.845 | -0.876 | -0.002 | 0.006 | 0.006 | 0.068 | 0.21 | 0.25 | 0.26 | 0.42 | 0.18 | 0.23 |
| 2 | 187051713 | rs10175904 | 1.056 | -0.292 | -0.892 | -0.932 | 0 | -0.011 | -0.011 | 0 | 0.06 | 0.08 | 0.03 | 0 | 0.06 | 0.09 |
| 2 | 187063564 | rs10208050 | 1.013 | -0.293 | -0.892 | -0.932 | -0.001 | -0.02 | -0.02 | 0.002 | 0.06 | 0.1 | 0.03 | 0 | 0.06 | 0.09 |
| 2 | 187081525 | rs16827624 | 0.377 | -0.734 | -0.987 | -1.199 | 0.03 | 0.037 | 0.037 | 0 | 0.23 | 0.16 | 0.17 | 0.11 | 0.17 | 0.21 |
| 2 | 187094545 | rs13414850 | 2.016 | -0.083 | -0.592 | -1.413 | 0.018 | -0.072 | -0.072 | 2.76 | 0.2 | 0.26 | 0.21 | 0.04 | 0.26 | 0.21 |
| 2 | 187103445 | rs3107406 | -0.893 | -0.036 | -0.644 | -1.206 | -0.006 | 0.001 | 0.001 | 1.264 | 0.44 | 0.55 | 0.44 | 0.53 | 0.36 | 0.46 |

| | | | | | | | | | | | | | | | | |
|---|---|---|---|---|---|---|---|---|---|---|---|---|---|---|---|---|
| 2 | 187152664 | rs6743356 | -1.858 | 0.314 | -1.138 | -1.096 | -0.102 | -0.172 | -0.172 | 2.175 | 0.79 | 0.98 | 0.72 | 0.58 | 0.72 | 0.79 |
| 2 | 187210847 | rs1448860 | -0.025 | -0.043 | -0.607 | -1.156 | 0.005 | -0.006 | -0.006 | 0 | 0.63 | 0.6 | 0.59 | 0.47 | 0.7 | 0.64 |
| 2 | 187213730 | rs10497668 | 0.664 | -0.179 | -0.742 | -1.221 | 0.026 | 0.033 | 0.033 | 0 | 0.23 | 0.16 | 0.17 | 0.11 | 0.17 | 0.21 |
| 2 | 187240797 | rs16827790 | 1.994 | -0.111 | -0.672 | -0.892 | 0.024 | -0.036 | -0.036 | 0 | 0.11 | 0.06 | 0.11 | 0.42 | 0.07 | 0.13 |
| 2 | 187282986 | rs12614800 | 0.75 | -0.058 | -0.614 | -1.092 | -0.016 | -0.04 | -0.04 | 3.916 | 0.36 | 0.2 | 0.35 | 0.43 | 0.38 | 0.34 |
| 2 | 187297391 | rs10804005 | -0.683 | -0.123 | -0.514 | -1.173 | 0 | -0.004 | -0.004 | 0 | 0.74 | 0.71 | 0.71 | 0.89 | 0.77 | 0.76 |
| 2 | 187344191 | rs7562659 | -0.142 | -0.165 | -0.45 | -1.169 | 0.007 | -0.009 | -0.009 | 0 | 0.63 | 0.61 | 0.6 | 0.47 | 0.7 | 0.64 |
| 2 | 187349063 | rs12473076 | 0.799 | -0.145 | -0.507 | -1.195 | -0.017 | -0.04 | -0.04 | 0.394 | 0.36 | 0.19 | 0.35 | 0.42 | 0.38 | 0.34 |
| 2 | 187350241 | rs1451874 | 0.792 | -0.147 | -0.508 | -1.196 | -0.017 | -0.04 | -0.04 | 0.465 | 0.36 | 0.19 | 0.35 | 0.42 | 0.38 | 0.34 |
| 2 | 187356004 | rs6709401 | 1.722 | 0.04 | -0.611 | -1.391 | 0.015 | -0.067 | -0.067 | 0.336 | 0.21 | 0.26 | 0.22 | 0.05 | 0.26 | 0.21 |
| 2 | 187368733 | rs10164423 | 1.222 | 0.024 | -0.791 | -1.302 | -0.001 | 0.001 | 0.001 | 0 | 0.06 | 0.06 | 0.02 | 0 | 0.06 | 0.09 |
| 2 | 187377481 | rs4144784 | 0.427 | 0.041 | -0.921 | -1.338 | -0.017 | 0.058 | 0.058 | 0.154 | 0.25 | 0.08 | 0.27 | 0.11 | 0.24 | 0.22 |
| 2 | 187380976 | rs13400819 | 1.038 | 0.35 | -0.649 | -1.161 | -0.005 | 0.001 | 0.001 | 0 | 0.06 | 0.06 | 0.04 | 0 | 0.08 | 0.09 |
| 2 | 187388234 | rs16827994 | 1.39 | 0.378 | -0.425 | -1.244 | 0.019 | -0.074 | -0.074 | 0 | 0.2 | 0.28 | 0.21 | 0.04 | 0.26 | 0.21 |
| 2 | 187393729 | rs13397032 | 0.058 | 0.103 | -0.367 | -1.112 | 0.001 | -0.002 | -0.002 | 1.782 | 0.38 | 0.57 | 0.38 | 0.41 | 0.4 | 0.43 |
| 2 | 187398463 | rs2083678 | 0.016 | -0.073 | -1.103 | -2.052 | 0.011 | 0.072 | 0.072 | 0 | 0.28 | 0.12 | 0.27 | 0.11 | 0.25 | 0.24 |
| 2 | 187410817 | rs6717629 | -0.179 | -0.032 | -1.127 | -2.066 | 0.022 | 0.061 | 0.061 | 0 | 0.72 | 0.83 | 0.74 | 0.89 | 0.77 | 0.76 |
| 2 | 187428110 | rs4667106 | 0.161 | -0.186 | -1.119 | -2.121 | 0.002 | -0.002 | -0.002 | 11.064 | 0.68 | 0.65 | 0.71 | 0.89 | 0.75 | 0.75 |
| 2 | 187434444 | rs10194631 | 0.036 | -0.119 | -0.979 | -2.353 | -0.015 | -0.046 | -0.046 | 1.646 | 0.69 | 0.56 | 0.71 | 0.89 | 0.75 | 0.75 |
| 2 | 187434619 | rs10209563 | -0.15 | -0.12 | -0.983 | -2.354 | 0.013 | 0.049 | 0.049 | 0 | 0.71 | 0.82 | 0.73 | 0.89 | 0.75 | 0.75 |
| 2 | 187441142 | rs11690463 | 0.01 | -0.087 | -1.261 | -2.338 | 0.009 | 0.077 | 0.077 | 0 | 0.28 | 0.11 | 0.26 | 0.11 | 0.25 | 0.25 |
| 2 | 187456121 | rs3768777 | 0.271 | -0.139 | -1.105 | -2.238 | -0.002 | -0.01 | -0.01 | 0 | 0.6 | 0.53 | 0.61 | 0.8 | 0.66 | 0.63 |
| 2 | 187462581 | rs4233788 | -0.173 | -0.175 | -1.144 | -2.276 | 0.024 | 0.061 | 0.061 | 0.313 | 0.67 | 0.78 | 0.73 | 0.84 | 0.73 | 0.72 |
| 2 | 187470311 | rs3911239 | 0.557 | -0.194 | -1.051 | -2.273 | -0.017 | -0.044 | -0.044 | 0 | 0.58 | 0.46 | 0.59 | 0.8 | 0.66 | 0.63 |
| 2 | 187475104 | rs2370693 | 0.101 | -0.156 | -1.26 | -2.116 | 0.001 | -0.004 | -0.004 | 0.282 | 0.65 | 0.65 | 0.73 | 0.84 | 0.73 | 0.72 |
| 2 | 187475357 | rs3911084 | 0.032 | -0.156 | -1.299 | -2.115 | -0.014 | -0.036 | -0.036 | 0 | 0.64 | 0.54 | 0.71 | 0.84 | 0.73 | 0.72 |
| 2 | 187481835 | rs3768781 | 1.227 | -0.076 | -1.724 | -2.284 | -0.104 | -0.077 | -0.077 | 0.107 | 0.22 | 0.02 | 0.31 | 0.73 | 0.31 | 0.27 |
| 2 | 187487269 | rs6741369 | -0.44 | -0.361 | -0.85 | -1.853 | 0.025 | 0.069 | 0.069 | 0.371 | 0.33 | 0.18 | 0.26 | 0.16 | 0.26 | 0.27 |
| 2 | 187489647 | rs12620429 | 1.064 | -0.411 | -1.075 | -1.906 | -0.049 | 0.074 | 0.074 | 0 | 0.15 | 0 | 0.16 | 0.05 | 0.19 | 0.12 |
| 2 | 187493841 | rs2197138 | 0.2 | -0.202 | -0.898 | -1.778 | -0.011 | -0.031 | -0.031 | 0 | 0.65 | 0.55 | 0.72 | 0.84 | 0.74 | 0.73 |
| 2 | 187494427 | rs7596996 | 0.185 | -0.089 | -1.433 | -1.719 | 0.013 | 0.025 | 0.025 | 0.123 | 0.66 | 0.67 | 0.72 | 0.84 | 0.73 | 0.72 |
| 2 | 187498107 | rs9333289 | 0.616 | 0.529 | -1.725 | -1.532 | -0.074 | 0.079 | 0.079 | 0.022 | 0.32 | 0.01 | 0.22 | 0.15 | 0.3 | 0.29 |
| 2 | 187503226 | rs4667108 | 0.094 | 0.811 | -1.046 | -1.971 | 0.001 | -0.041 | -0.041 | 0 | 0.39 | 0.49 | 0.5 | 0.74 | 0.41 | 0.34 |
| 2 | 187505486 | rs3816375 | 0.121 | 0.793 | -0.966 | -1.98 | 0.001 | -0.026 | -0.026 | 0 | 0.39 | 0.44 | 0.5 | 0.74 | 0.41 | 0.34 |
| 2 | 187518260 | rs3768787 | 1.031 | 0.782 | -0.816 | -1.993 | -0.001 | -0.001 | -0.001 | 0.143 | 0.17 | 0.15 | 0.17 | 0.08 | 0.19 | 0.2 |
| 2 | 187521260 | rs3738919 | -0.015 | 0.829 | -0.997 | -2.137 | 0.025 | 0.128 | 0.128 | 0 | 0.37 | 0.16 | 0.24 | 0.08 | 0.33 | 0.34 |
| 2 | 187534183 | rs2595389 | -0.447 | 0.84 | -0.962 | -2.176 | 0.008 | 0.162 | 0.162 | 0 | 0.43 | 0.21 | 0.29 | 0.08 | 0.42 | 0.44 |
| 2 | 187536155 | rs12475476 | 1.276 | 0.956 | -0.292 | -2.205 | 0.025 | 0.023 | 0.023 | 0 | 0.17 | 0.25 | 0.31 | 0.72 | 0.26 | 0.18 |
| 2 | 187537847 | rs12151442 | 1.077 | 0.932 | 0.201 | -2.079 | 0.01 | 0.099 | 0.099 | 0 | 0.37 | 0.13 | 0.27 | 0.17 | 0.31 | 0.38 |
| 2 | 187539721 | rs16828163 | 1.348 | 0.884 | 0.087 | -2.19 | 0 | -0.009 | -0.009 | 0.435 | 0.17 | 0.16 | 0.17 | 0.08 | 0.19 | 0.17 |
| 2 | 187541318 | rs12620821 | 0.076 | 0.872 | 0.542 | -2.191 | 0.002 | -0.023 | -0.023 | 0.667 | 0.54 | 0.45 | 0.45 | 0.25 | 0.5 | 0.55 |
| 2 | 187546583 | rs11891192 | 0.642 | 1.239 | 0.396 | -2.281 | -0.019 | 0.019 | 0.019 | 0 | 0.23 | 0.16 | 0.3 | 0.08 | 0.36 | 0.23 |
| 2 | 187553817 | rs2121975 | -0.069 | 0.975 | 0.619 | -2.233 | -0.001 | -0.003 | -0.003 | 0.73 | 0.58 | 0.37 | 0.47 | 0.25 | 0.56 | 0.59 |
| 2 | 187557294 | rs3886170 | 0.936 | 1.003 | 0.597 | -2.213 | -0.004 | 0.011 | 0.011 | 0 | 0.21 | 0.15 | 0.19 | 0.08 | 0.25 | 0.21 |
| 2 | 187560212 | rs11678136 | 0.947 | 1.012 | 0.602 | -2.204 | -0.004 | 0.011 | 0.011 | 0 | 0.21 | 0.15 | 0.18 | 0.08 | 0.24 | 0.21 |
| 2 | 187560308 | rs11678190 | 1.139 | 1.003 | 0.653 | -2.213 | 0.013 | 0.064 | 0.064 | 0 | 0.37 | 0.19 | 0.27 | 0.17 | 0.31 | 0.37 |
| 2 | 187561806 | rs2595375 | -1.205 | 1.192 | 0.789 | -2.227 | 0 | -0.013 | -0.013 | 0.969 | 0.61 | 0.61 | 0.6 | 0.8 | 0.63 | 0.62 |
| 2 | 187562355 | rs11695399 | 0.81 | 1.425 | 0.835 | -2.06 | -0.004 | -0.018 | -0.018 | 0.061 | 0.2 | 0.19 | 0.16 | 0.08 | 0.16 | 0.23 |
| 2 | 187573478 | rs1072901 | 0.698 | 1.422 | 0.744 | -2.085 | -0.012 | 0.012 | 0.012 | 0 | 0.31 | 0.21 | 0.36 | 0.72 | 0.34 | 0.27 |
| 2 | 187577810 | rs17334436 | 2.097 | 1.361 | 0.708 | -2.145 | -0.006 | 0.031 | 0.031 | 0 | 0.07 | 0.01 | 0.06 | 0 | 0.07 | 0.03 |
| 2 | 187580694 | rs17406346 | 0.695 | 1.426 | 0.857 | -2.015 | -0.012 | 0.012 | 0.012 | 0 | 0.31 | 0.2 | 0.36 | 0.72 | 0.34 | 0.27 |
| 2 | 187602523 | rs930657 | -0.206 | 1.48 | 0.701 | -1.984 | -0.004 | 0.01 | 0.01 | 0 | 0.62 | 0.31 | 0.44 | 0.26 | 0.56 | 0.63 |
| 2 | 187609216 | rs13030356 | 0.944 | 1.463 | 0.526 | -1.988 | -0.071 | 0.137 | 0.137 | 0.07 | 0.4 | 0.07 | 0.28 | 0.18 | 0.37 | 0.4 |
| 2 | 187626338 | rs13026081 | 1.388 | 1.332 | 0.343 | -2.175 | -0.063 | 0.246 | 0.246 | 0.139 | 0.36 | 0.01 | 0.23 | 0.05 | 0.32 | 0.29 |
| 2 | 187633716 | rs2595384 | 0.202 | 1.383 | 0.746 | -1.945 | 0 | -0.01 | -0.01 | 0.095 | 0.34 | 0.39 | 0.37 | 0.74 | 0.34 | 0.27 |
| 2 | 187638337 | rs7421739 | 0.009 | 1.229 | 0.79 | -1.908 | -0.002 | -0.005 | -0.005 | 0 | 0.6 | 0.36 | 0.41 | 0.21 | 0.51 | 0.61 |
| 2 | 187643406 | rs2370708 | -1.824 | 1.386 | 1.988 | -1.277 | 0.071 | 0.054 | 0.054 | 1.255 | 0.08 | 0.49 | 0.16 | 0.05 | 0.06 | 0.07 |
| 2 | 187644547 | rs3114942 | 0.962 | 1.712 | 1.964 | -1.417 | 0.011 | -0.049 | -0.049 | 0 | 0.68 | 0.34 | 0.65 | 0.86 | 0.62 | 0.58 |
| 2 | 187664331 | rs2887833 | 0.204 | 1.718 | 1.808 | -1.411 | 0.011 | -0.034 | -0.034 | 0 | 0.75 | 0.68 | 0.73 | 0.89 | 0.68 | 0.65 |
| 2 | 187672415 | rs6728163 | -1.304 | 1.701 | 1.619 | -1.397 | 0.066 | 0.019 | 0.019 | 0.595 | 0.07 | 0.4 | 0.08 | 0.03 | 0.05 | 0.07 |
| 2 | 187683048 | rs7572652 | -1.339 | 1.68 | 1.542 | -1.425 | -0.01 | 0.028 | 0.028 | 0.824 | 0.33 | 0.71 | 0.35 | 0.13 | 0.37 | 0.42 |
| 2 | 187685195 | rs10931256 | 0.174 | 1.677 | 0.776 | -1.29 | -0.004 | 0.012 | 0.012 | 0 | 0.21 | 0.15 | 0.21 | 0.1 | 0.24 | 0.25 |
| 2 | 187686062 | rs11891963 | 0.036 | 1.373 | 0.744 | -1.25 | 0.001 | 0.006 | 0.006 | 0 | 0.22 | 0.17 | 0.23 | 0.12 | 0.21 | 0.25 |
| 2 | 187692305 | rs12616053 | -0.176 | 1.28 | 0.568 | -1.216 | -0.014 | 0.046 | 0.046 | 0 | 0.26 | 0.1 | 0.25 | 0.11 | 0.27 | 0.3 |
| 2 | 187695570 | rs10179296 | -1.274 | 1.293 | 0.75 | -1.253 | -0.017 | 0.098 | 0.098 | 0.541 | 0.36 | 0.74 | 0.36 | 0.13 | 0.38 | 0.42 |
| 2 | 187697883 | rs10164815 | -0.974 | 1.252 | 1.127 | -1.118 | 0.087 | -0.001 | -0.001 | 0.137 | 0.08 | 0.42 | 0.11 | 0.02 | 0.11 | 0.11 |
| 2 | 187698707 | rs6434197 | -0.974 | 1.252 | 1.127 | -1.118 | 0.087 | -0.001 | -0.001 | 0.024 | 0.08 | 0.42 | 0.11 | 0.02 | 0.11 | 0.11 |
| 2 | 187698751 | rs7559533 | -0.069 | 1.309 | 1.058 | -1.244 | -0.014 | 0.045 | 0.045 | 0 | 0.26 | 0.1 | 0.25 | 0.11 | 0.27 | 0.3 |
| 2 | 187701549 | rs13414169 | -0.556 | 1.305 | 0.92 | -1.217 | 0.092 | 0.009 | 0.009 | 0.52 | 0.07 | 0.35 | 0.1 | 0.02 | 0.11 | 0.11 |

| | | | | | | | | | | | | | | | | |
|---|---|---|---|---|---|---|---|---|---|---|---|---|---|---|---|---|
| 2 | 187704866 | rs3114949 | 0.867 | 1.258 | 0.973 | -1.239 | -0.011 | 0.047 | 0.047 | 0.119 | 0.65 | 0.33 | 0.64 | 0.86 | 0.62 | 0.59 |
| 2 | 187707440 | rs10201512 | -0.357 | 1.31 | 0.748 | -1.199 | 0.07 | 0.02 | 0.02 | 0.096 | 0.06 | 0.31 | 0.08 | 0.02 | 0.08 | 0.1 |
| 2 | 187710756 | rs2084753 | -0.493 | 1.318 | 0.745 | -1.199 | 0.076 | 0.024 | 0.024 | 0.456 | 0.07 | 0.35 | 0.08 | 0.02 | 0.08 | 0.1 |
| 2 | 187711022 | rs3114950 | 0.795 | 1.288 | 0.697 | -1.229 | -0.025 | 0.115 | 0.115 | 0.571 | 0.64 | 0.21 | 0.63 | 0.87 | 0.62 | 0.59 |
| 2 | 187713396 | rs3100021 | 0.393 | 1.302 | 0.728 | -1.22 | 0.011 | -0.086 | -0.086 | 0 | 0.7 | 0.57 | 0.71 | 0.89 | 0.7 | 0.69 |
| 2 | 187714262 | rs1451556 | -0.611 | 1.456 | 0.74 | -0.759 | 0.072 | 0.017 | 0.017 | 0.226 | 0.06 | 0.32 | 0.08 | 0.02 | 0.08 | 0.1 |
| 2 | 187718869 | rs3114922 | -0.005 | 1.917 | -0.311 | -0.337 | -0.053 | -0.061 | -0.061 | 0 | 0.18 | 0.09 | 0.2 | 0.66 | 0.3 | 0.21 |
| 2 | 187723119 | rs1376625 | 0.348 | 1.715 | -0.397 | -0.835 | -0.024 | 0.036 | 0.036 | 0 | 0.23 | 0.12 | 0.27 | 0.13 | 0.29 | 0.3 |
| 2 | 187735749 | rs3114927 | 0.317 | 1.411 | -0.09 | -0.767 | -0.061 | -0.063 | -0.063 | 0 | 0.17 | 0.08 | 0.21 | 0.67 | 0.3 | 0.19 |
| 2 | 187737712 | rs3114929 | 0.244 | 1.4 | -0.095 | -0.773 | -0.061 | -0.063 | -0.063 | 0 | 0.17 | 0.08 | 0.21 | 0.67 | 0.31 | 0.19 |
| 2 | 187747487 | rs718195 | 0.356 | 1.773 | -0.139 | -0.483 | -0.107 | 0.005 | 0.005 | 0 | 0.23 | 0.04 | 0.3 | 0.18 | 0.37 | 0.26 |
| 2 | 187754997 | rs17700953 | 0.628 | 0.972 | 0.096 | -0.534 | -0.08 | 0.145 | 0.145 | 0 | 0.42 | 0.09 | 0.36 | 0.18 | 0.45 | 0.35 |
| 2 | 187775780 | rs1451548 | 0.199 | 0.66 | 0.675 | -0.399 | 0.014 | 0.014 | 0.014 | 0 | 0.37 | 0.34 | 0.41 | 0.22 | 0.24 | 0.46 |
| 2 | 187778830 | rs2084750 | -0.788 | 0.574 | 0.755 | -0.398 | -0.005 | -0.002 | -0.002 | 0 | 0.67 | 0.64 | 0.64 | 0.82 | 0.82 | 0.68 |
| 2 | 187801925 | rs10931264 | 0.316 | 1.506 | -1.078 | 0.679 | -0.07 | -0.078 | -0.078 | 0.3 | 0.17 | 0.08 | 0.21 | 0.6 | 0.34 | 0.2 |
| 2 | 187802572 | rs1562893 | 0.619 | 1.339 | -1.291 | 0.345 | -0.118 | 0.141 | 0.141 | 0 | 0.42 | 0.04 | 0.34 | 0.18 | 0.42 | 0.34 |
| 2 | 187808330 | rs11686527 | -0.446 | 1.611 | -1.095 | 1.101 | -0.133 | -0.145 | -0.145 | 0 | 0.24 | 0.04 | 0.31 | 0.54 | 0.39 | 0.28 |
| 2 | 187813262 | rs6721015 | 0.052 | 1.411 | -0.305 | 1.257 | 0.03 | 0.066 | 0.066 | 0.38 | 0.11 | 0.13 | 0.19 | 0.36 | 0.26 | 0.15 |
| 2 | 187843646 | rs4667136 | 0.109 | 1.408 | -0.328 | 1.233 | 0.018 | 0.089 | 0.089 | 0 | 0.09 | 0.1 | 0.18 | 0.4 | 0.23 | 0.16 |
| 2 | 187846104 | rs12621194 | 1.124 | 0.824 | 0.142 | 1.044 | -0.035 | 0.047 | 0.047 | 0.917 | 0.82 | 0.4 | 0.67 | 0.74 | 0.85 | 0.74 |
| 2 | 187866742 | rs12614343 | 0.408 | 0.898 | 0.18 | 1.091 | 0.054 | 0.171 | 0.171 | 0 | 0.09 | 0.16 | 0.19 | 0.41 | 0.22 | 0.11 |
| 2 | 187867340 | rs34938912 | 0.758 | 0.876 | 0.073 | 1.067 | -0.049 | -0.019 | -0.019 | 0 | 0.08 | 0.01 | 0.18 | 0.4 | 0.22 | 0.11 |
| 2 | 187880937 | rs2370736 | 0.888 | 0.157 | -0.44 | 0.598 | 0.053 | 0.044 | 0.044 | 0.008 | 0.23 | 0.09 | 0.11 | 0.15 | 0.1 | 0.18 |
| 2 | 187882931 | rs17754256 | 1.244 | 0.201 | -1.151 | 0.157 | -0.151 | -0.027 | -0.027 | 0 | 0.25 | 0.03 | 0.25 | 0.19 | 0.39 | 0.31 |
| 2 | 187942514 | rs12693461 | -0.071 | -0.006 | -0.038 | 0.162 | -0.002 | -0.004 | -0.004 | 0 | 0.88 | 0.86 | 0.83 | 0.91 | 0.89 | 0.77 |
| 2 | 187976717 | rs9646745 | 1.095 | 0.111 | 0.009 | 0.286 | 0.021 | 0.097 | 0.097 | 0 | 0.08 | 0.1 | 0.19 | 0.41 | 0.22 | 0.11 |
| 2 | 188004669 | rs1487400 | -0.916 | 0.986 | 0.665 | 0.613 | 0.039 | 0.139 | 0.139 | 0 | 0.92 | 0.86 | 0.81 | 0.59 | 0.78 | 0.89 |
| 2 | 188087887 | rs11683016 | -0.022 | 0.844 | 0.835 | 0.4 | -0.042 | 0.066 | 0.066 | 0 | 0.65 | 0.81 | 0.48 | 0.23 | 0.45 | 0.57 |
| 2 | 188103476 | rs16828714 | 0.846 | 0.76 | 1.015 | 0.365 | 0.018 | 0.022 | 0.022 | 0 | 0.53 | 0.38 | 0.4 | 0.31 | 0.34 | 0.36 |
| 2 | 188116484 | rs6434216 | -0.974 | 0.615 | 1.192 | 0.417 | -0.01 | -0.024 | -0.024 | 0 | 0.09 | 0.4 | 0.12 | 0.01 | 0.03 | 0.05 |
| 2 | 188131502 | rs9646806 | 0.524 | 0.604 | 1.104 | 0.419 | -0.003 | -0.007 | -0.007 | 0 | 0.93 | 0.47 | 0.94 | 1 | 0.99 | 1 |
| 2 | 188143677 | rs1918901 | -0.19 | 0.685 | 0.511 | 0.462 | -0.078 | -0.036 | -0.036 | 0.46 | 0.34 | 0.12 | 0.47 | 0.61 | 0.52 | 0.43 |
| 2 | 188156578 | rs1706252 | -0.848 | 0.638 | 0.498 | 0.366 | 0.038 | 0.178 | 0.178 | 0 | 0.42 | 0.17 | 0.67 | 0.86 | 0.66 | 0.66 |
| 2 | 188162128 | rs17705798 | 0.937 | 0.828 | 0.459 | 0.503 | -0.096 | -0.015 | -0.015 | 0 | 0.23 | 0.05 | 0.2 | 0.18 | 0.3 | 0.29 |
| 2 | 188162612 | rs17705846 | 0.445 | 0.707 | 0.685 | 0.534 | -0.015 | -0.005 | -0.005 | 2.295 | 0.06 | 0.02 | 0.06 | 0.07 | 0.1 | 0.2 |
| 2 | 188163025 | rs2048487 | -0.503 | 0.655 | 1.069 | 0.494 | 0.022 | 0.018 | 0.018 | 2.168 | 0.07 | 0.4 | 0.05 | 0 | 0.01 | 0 |
| 2 | 188186079 | rs840584 | -0.726 | 0.721 | 0.678 | 0.526 | 0.038 | 0.178 | 0.178 | 0 | 0.42 | 0.18 | 0.67 | 0.86 | 0.66 | 0.66 |
| 2 | 188196469 | rs840616 | -0.773 | 0.71 | 0.879 | 0.542 | 0.028 | 0.08 | 0.08 | 0 | 0.43 | 0.26 | 0.67 | 0.86 | 0.65 | 0.66 |
| 2 | 188207245 | rs1528233 | -0.874 | 0.707 | 0.961 | 0.572 | 0.014 | 0.023 | 0.023 | 0 | 0.44 | 0.34 | 0.69 | 0.86 | 0.66 | 0.66 |
| 2 | 188210214 | rs696092 | -0.874 | 0.703 | 0.959 | 0.574 | 0.014 | 0.023 | 0.023 | 0 | 0.44 | 0.34 | 0.69 | 0.85 | 0.66 | 0.66 |
| 2 | 188211112 | rs3771095 | 0.429 | 0.735 | 0.908 | 0.565 | -0.015 | -0.005 | -0.005 | 0 | 0.06 | 0.02 | 0.06 | 0.07 | 0.1 | 0.2 |
| 2 | 188219052 | rs858750 | 0.701 | 0.726 | 0.941 | 0.615 | -0.005 | -0.008 | -0.008 | 0 | 0.91 | 0.91 | 0.8 | 0.75 | 0.86 | 0.77 |
| 2 | 188228516 | rs696574 | 1.614 | 0.702 | 1.056 | 0.616 | -0.008 | 0.047 | 0.047 | 0 | 0.86 | 0.71 | 0.77 | 0.74 | 0.85 | 0.76 |
| 2 | 188230146 | rs3771089 | -0.073 | 0.736 | 1.008 | 0.607 | -0.026 | -0.007 | -0.007 | 0 | 0.34 | 0.21 | 0.47 | 0.6 | 0.52 | 0.43 |
| 2 | 188238334 | rs17366895 | -0.043 | 0.782 | 1.023 | 0.576 | 0.011 | 0.001 | 0.001 | 0 | 0.06 | 0.1 | 0.07 | 0.09 | 0.1 | 0.19 |
| 2 | 188243633 | rs3771084 | -0.685 | 0.799 | 1.049 | 0.605 | 0 | 0.01 | 0.01 | 0 | 0.42 | 0.37 | 0.63 | 0.76 | 0.58 | 0.48 |
| 2 | 188258904 | rs1157699 | 0.599 | 0.927 | 0.567 | 0.619 | -0.06 | 0.014 | 0.014 | 0 | 0.26 | 0.09 | 0.29 | 0.19 | 0.32 | 0.33 |
| 2 | 188291105 | rs7601254 | -0.497 | 0.518 | 0.65 | 0.433 | -0.001 | 0.016 | 0.016 | 0 | 0.42 | 0.32 | 0.51 | 0.65 | 0.55 | 0.55 |
| 2 | 188297133 | rs12104503 | 0.549 | 0.528 | 0.723 | 0.414 | 0.015 | 0.018 | 0.018 | 0 | 0.53 | 0.62 | 0.33 | 0.17 | 0.36 | 0.41 |
| 2 | 188326763 | rs12693471 | 1.231 | 0.13 | 0.726 | -0.028 | 0.024 | 0.081 | 0.081 | 0.004 | 0.39 | 0.26 | 0.26 | 0.14 | 0.3 | 0.28 |
| 2 | 188332692 | rs8176592 | 1.134 | 0.308 | 1.095 | 0.061 | -0.004 | -0.031 | -0.031 | 0.842 | 0.41 | 0.44 | 0.28 | 0.14 | 0.3 | 0.28 |
| 2 | 188340396 | rs8176546 | 1.17 | 0.199 | 1.02 | 0.032 | 0.009 | 0.016 | 0.016 | 0.049 | 0.39 | 0.37 | 0.26 | 0.14 | 0.3 | 0.28 |
| 2 | 188341047 | rs8176541 | 1.366 | 0.216 | 0.704 | 0.04 | 0.02 | 0.13 | 0.13 | 0.366 | 0.37 | 0.18 | 0.24 | 0.14 | 0.3 | 0.28 |
| 2 | 188355835 | rs8176490 | 0.636 | 0.06 | 0.779 | 0.304 | -0.013 | -0.009 | -0.009 | 0 | 0.12 | 0.12 | 0.04 | 0.07 | 0.06 | 0.11 |
| 2 | 188368791 | rs2192824 | 0.41 | 0.683 | 0.207 | 0.513 | -0.113 | 0.059 | 0.059 | 0 | 0.33 | 0.1 | 0.35 | 0.18 | 0.39 | 0.34 |
| 2 | 188370232 | rs3755248 | 0.425 | 0.407 | 0.291 | 0.37 | 0.076 | 0.118 | 0.118 | 0 | 0.51 | 0.71 | 0.29 | 0.22 | 0.31 | 0.39 |
| 2 | 188388311 | rs12613071 | -0.527 | 0.867 | 0.109 | 0.837 | -0.025 | -0.026 | -0.026 | 0 | 0.11 | 0.09 | 0.18 | 0.3 | 0.25 | 0.15 |
| 2 | 188390819 | rs2192825 | -0.661 | 0.87 | 0.187 | 0.398 | -0.058 | 0.081 | 0.081 | 0 | 0.67 | 0.86 | 0.69 | 0.84 | 0.61 | 0.64 |
| 2 | 188391908 | rs16829088 | 1.433 | 0.35 | 0.767 | 0.42 | 0.002 | 0.003 | 0.003 | 0.136 | 0.31 | 0.27 | 0.19 | 0.22 | 0.24 | 0.26 |
| 2 | 188395700 | rs16829090 | 0.075 | 0.327 | 0.914 | 0.66 | 0.007 | 0.089 | 0.089 | 0 | 0.06 | 0.08 | 0.12 | 0.31 | 0.14 | 0.12 |
| 2 | 188404283 | rs13427829 | 0.202 | 0.244 | 1.214 | 0.602 | -0.043 | 0.046 | 0.046 | 0 | 0.19 | 0.33 | 0.21 | 0.31 | 0.09 | 0.22 |
| 2 | 188410351 | rs10179730 | -0.106 | 0.301 | 1.183 | 0.639 | 0.019 | 0.12 | 0.12 | 0 | 0.94 | 0.88 | 0.87 | 0.69 | 0.86 | 0.88 |
| 2 | 188419590 | rs10153820 | 0.163 | -0.052 | 1.293 | -0.034 | -0.033 | 0.033 | 0.033 | 0 | 0.2 | 0.3 | 0.22 | 0.31 | 0.11 | 0.22 |
| 2 | 188435643 | rs6730608 | 0.274 | 0.102 | 1.077 | -0.132 | -0.044 | 0.093 | 0.093 | 0 | 0.35 | 0.14 | 0.34 | 0.15 | 0.42 | 0.34 |
| 2 | 188447756 | rs10187989 | 0.261 | 0.103 | 1.063 | -0.161 | -0.034 | 0.077 | 0.077 | 0 | 0.35 | 0.19 | 0.34 | 0.15 | 0.42 | 0.34 |
| 2 | 188450136 | rs11902917 | 0.286 | 0.095 | 1.075 | -0.086 | -0.007 | 0.007 | 0.007 | 0 | 0.19 | 0.17 | 0.21 | 0.31 | 0.11 | 0.25 |
| 2 | 188459720 | rs11904140 | -0.668 | 0.668 | 0.741 | 0.264 | 0.055 | 0.005 | 0.005 | 0.46 | 0.06 | 0.34 | 0.1 | 0.01 | 0.05 | 0.02 |
| 2 | 188464253 | rs6705455 | -0.346 | 0.719 | 0.328 | 0.399 | 0.003 | 0 | 0 | 1.555 | 0.56 | 0.46 | 0.57 | 0.46 | 0.53 | 0.6 |
| 2 | 188482933 | rs7561970 | -0.961 | 0.819 | 0.523 | 0.6 | -0.003 | -0.001 | -0.001 | 0.057 | 0.16 | 0.12 | 0.19 | 0.21 | 0.19 | 0.32 |

| | | | | | | | | | | | | | | | | |
|---|---|---|---|---|---|---|---|---|---|---|---|---|---|---|---|---|
| 2 | 188491863 | rs1356868 | 1.686 | 0.537 | 1.555 | 0.704 | -0.017 | -0.011 | -0.011 | 0 | 0.79 | 0.45 | 0.74 | 0.76 | 0.82 | 0.66 |
| 2 | 188508011 | rs1464339 | 1.686 | 0.537 | 1.556 | 0.684 | -0.017 | -0.011 | -0.011 | 0.591 | 0.79 | 0.45 | 0.74 | 0.77 | 0.82 | 0.66 |
| 2 | 188509060 | rs10198468 | 0.481 | 0.538 | 1.281 | 0.699 | 0 | -0.009 | -0.009 | 0 | 0.88 | 0.92 | 0.88 | 0.78 | 0.89 | 0.71 |
| 2 | 188521234 | rs10203300 | -1.789 | 0.796 | 1.654 | 0.708 | 0.033 | -0.031 | -0.031 | 1.111 | 0.09 | 0.28 | 0.13 | 0.01 | 0.1 | 0.06 |
| 2 | 188530578 | rs7580316 | -1.373 | 0.563 | 1.528 | 0.953 | 0.005 | 0.023 | 0.023 | 0 | 0.14 | 0.38 | 0.16 | 0.18 | 0.12 | 0.24 |
| 2 | 188535099 | rs13392433 | -1.463 | 0.554 | 1.361 | 0.95 | -0.006 | 0.032 | 0.032 | 0.023 | 0.16 | 0.41 | 0.16 | 0.22 | 0.12 | 0.33 |
| 2 | 188545602 | rs4432415 | 1.329 | 1.089 | 0.546 | 0.703 | -0.016 | -0.002 | -0.002 | 0 | 0.66 | 0.3 | 0.62 | 0.66 | 0.61 | 0.58 |
| 2 | 188550378 | rs10497686 | -0.817 | 0.91 | 0.274 | 0.812 | -0.035 | -0.051 | -0.051 | 0 | 0.12 | 0 | 0.12 | 0.21 | 0.12 | 0.29 |
| 2 | 188553074 | rs16829228 | -0.72 | 0.916 | 0.682 | 0.808 | 0 | 0.002 | 0.002 | 0 | 0.11 | 0.13 | 0.13 | 0.17 | 0.12 | 0.23 |
| 2 | 188573244 | rs13388743 | -1.073 | 0.915 | 1.374 | 0.674 | -0.003 | -0.002 | -0.002 | 0.339 | 0.17 | 0.21 | 0.16 | 0.2 | 0.15 | 0.32 |
| 2 | 188573740 | rs2090751 | -1.913 | 0.872 | 1.662 | 0.622 | -0.019 | -0.018 | -0.018 | 1.886 | 0.25 | 0.58 | 0.2 | 0.21 | 0.17 | 0.35 |
| 2 | 188583791 | rs9288146 | -1.885 | 0.872 | 1.666 | 0.622 | -0.02 | -0.02 | -0.02 | 0.019 | 0.25 | 0.54 | 0.19 | 0.21 | 0.17 | 0.35 |
| 2 | 188585156 | rs11675886 | 1.864 | 0.388 | 0.17 | 0.614 | -0.058 | -0.033 | -0.033 | 0 | 0.46 | 0.02 | 0.3 | 0.37 | 0.41 | 0.35 |
| 2 | 188611039 | rs1520477 | -1.031 | 0.268 | 0.804 | -0.107 | -0.004 | -0.014 | -0.014 | 0 | 0.26 | 0.24 | 0.19 | 0.17 | 0.19 | 0.38 |
| 2 | 188626214 | rs4667193 | 1.504 | 0.147 | 0.543 | -0.173 | 0.058 | 0.103 | 0.103 | 0.521 | 0.62 | 0.15 | 0.74 | 0.79 | 0.76 | 0.52 |
| 2 | 188640654 | rs2139631 | 0.29 | 0.179 | 1.394 | 0.016 | -0.03 | -0.05 | -0.05 | 0 | 0.85 | 0.66 | 0.92 | 0.95 | 0.93 | 0.89 |
| 2 | 188646078 | rs10931308 | -0.345 | 0.157 | 1.259 | 0.017 | -0.024 | -0.042 | -0.042 | 0 | 0.89 | 0.74 | 0.95 | 0.97 | 0.95 | 0.93 |
| 2 | 188653232 | rs4667194 | 0.963 | 0.385 | 0.975 | -0.156 | -0.01 | 0.008 | 0.008 | 0.358 | 0.64 | 0.36 | 0.65 | 0.83 | 0.55 | 0.49 |
| 2 | 188656082 | rs13027479 | 1.188 | 0.465 | 0.621 | -0.101 | -0.023 | 0.045 | 0.045 | 0.113 | 0.62 | 0.29 | 0.64 | 0.81 | 0.53 | 0.55 |
| 2 | 188683655 | rs7371285 | 1.16 | 0.502 | 0.658 | -0.213 | -0.004 | -0.008 | -0.008 | 0.453 | 0.64 | 0.38 | 0.67 | 0.89 | 0.62 | 0.62 |
| 2 | 188684507 | rs4471853 | -1.264 | 0.501 | 0.932 | -0.238 | -0.03 | 0.058 | 0.058 | 0.07 | 0.3 | 0.18 | 0.27 | 0.08 | 0.36 | 0.37 |
| 2 | 188706144 | rs7580146 | -1.087 | 0.627 | 1.281 | 0.011 | -0.014 | 0.011 | 0.011 | 0 | 0.17 | 0.11 | 0.18 | 0.17 | 0.2 | 0.29 |
| 2 | 188721717 | rs10931316 | 1.168 | 0.614 | 1.551 | 0.078 | 0.033 | 0.006 | 0.006 | 0.133 | 0.83 | 0.73 | 0.77 | 0.8 | 0.77 | 0.7 |
| 2 | 188724638 | rs16829324 | -1.717 | 0.589 | 1.583 | 0.093 | 0.033 | 0.005 | 0.005 | 0.563 | 0.18 | 0.28 | 0.23 | 0.2 | 0.23 | 0.3 |
| 2 | 188734097 | rs12476617 | 1.977 | 0.725 | 1.503 | 0.864 | 0.021 | -0.014 | -0.014 | 0 | 0.75 | 0.53 | 0.66 | 0.76 | 0.74 | 0.66 |
| 2 | 188744807 | rs12474851 | 2.018 | 0.644 | 1.173 | 0.801 | 0.014 | -0.018 | -0.018 | 0 | 0.74 | 0.53 | 0.66 | 0.76 | 0.74 | 0.66 |
| 2 | 188755657 | rs12619322 | 1.773 | 0.6 | 1.062 | 0.924 | 0.014 | 0.016 | 0.016 | 0.546 | 0.77 | 0.6 | 0.68 | 0.69 | 0.76 | 0.68 |
| 2 | 188755699 | rs12998150 | 2.448 | 0.254 | 0.687 | 1.046 | -0.017 | 0.114 | 0.114 | 0.717 | 0.46 | 0.19 | 0.32 | 0.23 | 0.45 | 0.31 |
| 2 | 188759951 | rs6721831 | -0.089 | 0.587 | 0.488 | 0.972 | -0.046 | -0.029 | -0.029 | 0.25 | 0.1 | 0.02 | 0.17 | 0.17 | 0.21 | 0.28 |
| 2 | 188767101 | rs12468071 | -0.328 | 0.543 | 0.493 | 1.033 | -0.061 | -0.045 | -0.045 | 0 | 0.14 | 0.02 | 0.27 | 0.22 | 0.24 | 0.32 |
| 2 | 188773871 | rs17326435 | 0.074 | 0.43 | 0.544 | 0.867 | -0.035 | 0.071 | 0.071 | 0 | 0.13 | 0.01 | 0.08 | 0 | 0.16 | 0.09 |
| 2 | 188780717 | rs9646808 | -0.596 | 0.399 | 0.914 | 0.851 | -0.004 | -0.068 | -0.068 | 1.257 | 0.18 | 0.22 | 0.1 | 0 | 0.16 | 0.09 |
| 2 | 188782216 | rs6732052 | -1.534 | 0.38 | 1.103 | 1.077 | -0.003 | -0.002 | -0.002 | 0 | 0.3 | 0.33 | 0.47 | 0.68 | 0.37 | 0.6 |
| 2 | 188789727 | rs35429436 | -0.082 | 0.278 | 1.056 | 1.2 | -0.006 | 0.088 | 0.088 | 0 | 0.11 | 0.2 | 0.17 | 0.42 | 0.12 | 0.25 |
| 2 | 188796294 | rs10181512 | -2.172 | 0.343 | 0.931 | 1.076 | 0 | 0.035 | 0.035 | 0.105 | 0.51 | 0.66 | 0.58 | 0.68 | 0.54 | 0.72 |
| 2 | 188798760 | rs1317323 | -1.519 | 0.53 | 0.697 | 0.952 | -0.004 | 0.003 | 0.003 | 0.015 | 0.32 | 0.28 | 0.29 | 0.22 | 0.39 | 0.4 |
| 2 | 188804659 | rs17327157 | -0.136 | 0.329 | -0.165 | 0.24 | -0.019 | 0.012 | 0.012 | 0 | 0.2 | 0.03 | 0.14 | 0.14 | 0.17 | 0.33 |
| 2 | 188806662 | rs10497691 | -0.181 | -0.003 | 0.133 | 0.503 | 0.002 | 0.017 | 0.017 | 0 | 0.13 | 0.14 | 0.15 | 0.43 | 0.09 | 0.18 |
| 2 | 188808468 | rs4299304 | 1.263 | -0.149 | 0.531 | 0.326 | 0.003 | -0.005 | -0.005 | 0 | 0.7 | 0.27 | 0.73 | 0.53 | 0.73 | 0.73 |
| 2 | 188811935 | rs7581351 | -0.834 | -0.153 | 1.278 | 0.376 | -0.026 | 0.205 | 0.205 | 0.023 | 0.16 | 0.39 | 0.27 | 0.45 | 0.12 | 0.21 |
| 2 | 188814444 | rs10170218 | -1.145 | -0.006 | 0.698 | 0.348 | -0.004 | -0.063 | -0.063 | 0 | 0.23 | 0.18 | 0.24 | 0.44 | 0.27 | 0.27 |
| 2 | 188829691 | rs11901865 | 0.142 | 0.119 | 0.464 | 0.137 | 0.002 | -0.002 | -0.002 | 0.082 | 0.18 | 0.04 | 0.11 | 0.16 | 0.15 | 0.32 |
| 2 | 188838592 | rs12999094 | 0.193 | 0.059 | 0.934 | 0.17 | -0.007 | -0.033 | -0.033 | 0.024 | 0.75 | 0.62 | 0.82 | 0.84 | 0.81 | 0.67 |
| 2 | 188842652 | rs17262129 | 2.018 | 0.085 | 0.344 | 0.153 | -0.02 | 0.007 | 0.007 | 0 | 0.47 | 0.14 | 0.48 | 0.38 | 0.53 | 0.39 |
| 2 | 188842803 | rs6713963 | 0.378 | 0.236 | 0.296 | 0.202 | -0.015 | 0.022 | 0.022 | 0 | 0.11 | 0.07 | 0.07 | 0 | 0.17 | 0.07 |
| 2 | 188857787 | rs1439563 | -0.279 | 0.112 | 0.393 | 0.508 | -0.001 | 0.013 | 0.013 | 0 | 0.14 | 0.15 | 0.17 | 0.39 | 0.1 | 0.3 |
| 2 | 188859657 | rs7579998 | 0.189 | 0.088 | 0.586 | 0.353 | 0 | 0.001 | 0.001 | 0 | 0.75 | 0.78 | 0.83 | 0.82 | 0.81 | 0.74 |
| 2 | 188860470 | rs4667205 | 1.703 | 0.236 | 0.778 | 0.15 | -0.032 | 0.013 | 0.013 | 0 | 0.46 | 0.17 | 0.48 | 0.38 | 0.52 | 0.36 |
| 2 | 188865131 | rs10181019 | -1.131 | 0.497 | 1.443 | -0.024 | 0.055 | -0.057 | -0.057 | 0 | 0.14 | 0.47 | 0.18 | 0.05 | 0.18 | 0.1 |
| 2 | 188869904 | rs1439560 | 0.983 | 0.462 | 1.424 | -0.04 | 0.043 | -0.07 | -0.07 | 0 | 0.85 | 0.44 | 0.81 | 0.95 | 0.82 | 0.9 |
| 2 | 188874902 | rs1946875 | 0.654 | 0.497 | 1.435 | -0.023 | 0.047 | -0.05 | -0.05 | 0 | 0.86 | 0.6 | 0.83 | 0.95 | 0.82 | 0.9 |
| 2 | 188875158 | rs1400079 | 0.806 | 0.495 | 1.443 | -0.024 | 0.049 | -0.056 | -0.056 | 0 | 0.86 | 0.54 | 0.82 | 0.95 | 0.82 | 0.9 |
| 2 | 188879205 | rs2165150 | 0.607 | 0.544 | 0.945 | -0.391 | 0.025 | -0.125 | -0.125 | 0 | 0.76 | 0.48 | 0.76 | 0.93 | 0.72 | 0.87 |
| 2 | 188888319 | rs10198982 | -0.242 | 0.57 | 0.585 | -0.32 | 0.008 | -0.061 | -0.061 | 0 | 0.19 | 0.28 | 0.12 | 0.03 | 0.24 | 0.1 |
| 2 | 188916374 | rs893746 | -0.769 | 0.72 | 0.248 | 0.212 | -0.003 | -0.006 | -0.006 | 0 | 0.17 | 0.17 | 0.24 | 0.33 | 0.24 | 0.25 |
| 2 | 188923773 | rs1579690 | 0.234 | 0.43 | 0.437 | -0.45 | -0.016 | -0.083 | -0.083 | 0 | 0.71 | 0.63 | 0.84 | 0.94 | 0.75 | 0.69 |
| 2 | 188924330 | rs2043945 | 0.147 | 0.42 | 0.555 | -0.454 | 0 | -0.026 | -0.026 | 0 | 0.72 | 0.76 | 0.85 | 0.94 | 0.75 | 0.69 |
| 2 | 188942940 | rs1371523 | 0.599 | 0.523 | 0.541 | -0.577 | -0.005 | -0.07 | -0.07 | 0.022 | 0.66 | 0.61 | 0.8 | 0.95 | 0.7 | 0.68 |
| 2 | 188944218 | rs1439877 | 0.065 | 0.516 | 0.604 | -0.577 | 0.009 | 0.012 | 0.012 | 0.33 | 0.72 | 0.79 | 0.86 | 0.95 | 0.75 | 0.69 |
| 2 | 188947841 | rs7582658 | 1.312 | 0.478 | 0.889 | -0.466 | 0.006 | 0.026 | 0.026 | 0.265 | 0.55 | 0.37 | 0.59 | 0.61 | 0.53 | 0.49 |
| 2 | 188949715 | rs16829731 | 1.568 | 0.277 | 0.821 | -0.326 | 0.004 | -0.006 | -0.006 | 0.015 | 0.11 | 0.11 | 0.06 | 0.18 | 0.06 | 0.14 |
| 2 | 188953427 | rs7569881 | 1.306 | 0.422 | 0.608 | -0.495 | 0.004 | 0.046 | 0.046 | 0.011 | 0.51 | 0.37 | 0.55 | 0.61 | 0.49 | 0.48 |
| 2 | 188957640 | rs1439912 | 0.363 | 0.323 | 0.724 | -0.315 | -0.005 | -0.07 | -0.07 | 0 | 0.66 | 0.61 | 0.8 | 0.95 | 0.7 | 0.68 |
| 2 | 188961179 | rs11680615 | 0.112 | 0.308 | 0.625 | -0.317 | -0.015 | -0.03 | -0.03 | 0.003 | 0.27 | 0.01 | 0.13 | 0.22 | 0.23 | 0.32 |
| 2 | 188961699 | rs12997059 | 1.968 | 0.416 | 0.622 | -0.646 | -0.019 | -0.048 | -0.048 | 0 | 0.37 | 0.2 | 0.38 | 0.39 | 0.35 | 0.3 |
| 2 | 188962426 | rs11886097 | 0.372 | 0.352 | 0.802 | -1.19 | 0.037 | -0.024 | -0.024 | 0 | 0.06 | 0.19 | 0.06 | 0 | 0.05 | 0.01 |
| 2 | 188963148 | rs11893148 | -0.531 | 0.32 | 0.623 | -1.155 | 0 | 0.111 | 0.111 | 1.346 | 0.34 | 0.14 | 0.22 | 0.1 | 0.38 | 0.36 |
| 2 | 188965391 | rs1866542 | 0.754 | 0.328 | 0.514 | -1.122 | -0.003 | -0.067 | -0.067 | 0.206 | 0.6 | 0.53 | 0.7 | 0.9 | 0.57 | 0.64 |
| 2 | 188984445 | rs6707652 | -0.256 | 0.552 | 0.999 | -0.742 | 0.054 | -0.022 | -0.022 | 1.469 | 0.13 | 0.44 | 0.15 | 0.05 | 0.18 | 0.05 |

| | | | | | | | | | | | | | | | |
|---|---|---|---|---|---|---|---|---|---|---|---|---|---|---|---|
| 2 | 188986175 | rs12992671 | 1.095 | 0.554 | 0.797 | -0.743 | -0.009 | 0.004 | 0.004 | 0 | 0.09 | 0.05 | 0.1 | 0.05 | 0.15 | 0.05 |
| 2 | 188996350 | rs971268 | 0.249 | 0.532 | 0.799 | -0.754 | 0.045 | -0.032 | -0.032 | 3.051 | 0.86 | 0.46 | 0.84 | 0.95 | 0.82 | 0.95 |
| 2 | 189001642 | rs12469006 | 0.748 | 0.478 | 0.606 | -0.87 | 0.014 | 0.069 | 0.069 | 0 | 0.49 | 0.33 | 0.71 | 0.9 | 0.54 | 0.56 |
| 2 | 189016215 | rs2119094 | 1.048 | 0.413 | 1.025 | -0.514 | 0.012 | -0.043 | -0.043 | 0 | 0.91 | 0.58 | 0.95 | 1 | 0.96 | 0.97 |
| 2 | 189022609 | rs997045 | 0.921 | 0.426 | 0.965 | -0.536 | 0.002 | -0.072 | -0.072 | 0.405 | 0.82 | 0.5 | 0.85 | 0.95 | 0.81 | 0.91 |
| 2 | 189025084 | rs2882449 | 0.932 | 0.428 | 0.797 | -0.497 | -0.007 | 0.004 | 0.004 | 0 | 0.09 | 0.06 | 0.11 | 0.04 | 0.15 | 0.06 |
| 2 | 189029385 | rs1439903 | -0.544 | 0.425 | 0.968 | -0.5 | 0.037 | -0.009 | -0.009 | 2.959 | 0.89 | 0.65 | 0.86 | 0.96 | 0.83 | 0.94 |
| 2 | 189058585 | rs10931334 | -0.706 | 0.37 | 0.483 | -0.434 | -0.01 | -0.047 | -0.047 | 0.118 | 0.16 | 0.33 | 0.13 | 0.04 | 0.16 | 0.09 |
| 2 | 189065718 | rs7582899 | -0.836 | 0.215 | 0.5 | -0.384 | -0.018 | -0.03 | -0.03 | 0 | 0.08 | 0.28 | 0.02 | 0 | 0.01 | 0.03 |
| 2 | 189073346 | rs1439887 | -0.275 | 0.325 | -0.273 | -0.731 | -0.001 | 0.099 | 0.099 | 0 | 0.7 | 0.86 | 0.84 | 0.96 | 0.73 | 0.69 |
| 2 | 189080087 | rs1595763 | 0.001 | 0.13 | -0.217 | -0.162 | -0.022 | 0.031 | 0.031 | 1.008 | 0.88 | 0.83 | 0.96 | 0.78 | 0.95 | 0.77 |
| 2 | 189088249 | rs893745 | 1.047 | 0.198 | 0.194 | -0.299 | 0 | 0.007 | 0.007 | 0 | 0.46 | 0.38 | 0.69 | 0.7 | 0.52 | 0.41 |
| 2 | 189091513 | rs13416775 | -0.525 | 0.172 | 0.376 | -0.289 | -0.041 | 0.08 | 0.08 | 1.51 | 0.13 | 0.32 | 0.05 | 0.22 | 0.05 | 0.23 |
| 2 | 189106576 | rs11692963 | -0.281 | 0.188 | 0.274 | -0.39 | -0.014 | 0.159 | 0.159 | 0 | 0.41 | 0.21 | 0.26 | 0.08 | 0.43 | 0.37 |
| 2 | 189112348 | rs4132426 | 1.294 | 0.486 | 0.504 | 0.132 | -0.008 | -0.027 | -0.027 | 0 | 0.41 | 0.24 | 0.52 | 0.57 | 0.38 | 0.34 |
| 2 | 189122229 | rs7582137 | -0.608 | 0.345 | 0.362 | -0.119 | -0.017 | 0.027 | 0.027 | 0.542 | 0.23 | 0.4 | 0.15 | 0.25 | 0.21 | 0.3 |
| 2 | 189139485 | rs11901438 | 0.411 | 0.538 | 0.341 | -0.133 | 0.004 | 0.09 | 0.09 | 0 | 0.3 | 0.21 | 0.15 | 0.04 | 0.27 | 0.31 |
| 2 | 189154515 | rs10469735 | 1.101 | 0.087 | 0.57 | 0.027 | 0.019 | -0.012 | -0.012 | 0 | 0.1 | 0.21 | 0.13 | 0.04 | 0.16 | 0.07 |
| 2 | 189154925 | rs9288153 | 1.101 | 0.087 | 0.567 | 0.027 | 0.017 | -0.012 | -0.012 | 0 | 0.1 | 0.21 | 0.13 | 0.04 | 0.16 | 0.07 |
| 2 | 189156963 | rs2004888 | 1.101 | 0.087 | 0.567 | 0.027 | 0.017 | -0.012 | -0.012 | 0 | 0.1 | 0.21 | 0.13 | 0.04 | 0.16 | 0.07 |
| 2 | 189158620 | rs12693491 | 1.179 | 0.087 | 0.555 | 0.027 | 0.014 | -0.011 | -0.011 | 1.628 | 0.1 | 0.19 | 0.13 | 0.04 | 0.16 | 0.07 |
| 2 | 189174910 | rs10193506 | 0.18 | 0.272 | 0.582 | -0.135 | -0.006 | 0.049 | 0.049 | 0 | 0.59 | 0.67 | 0.72 | 0.92 | 0.53 | 0.63 |
| 2 | 189181182 | rs7559554 | 0.644 | 0.206 | 0.604 | -0.003 | -0.01 | 0.123 | 0.123 | 0.364 | 0.09 | 0.23 | 0.04 | 0.21 | 0.04 | 0.2 |
| 2 | 189186150 | rs12988577 | 1.525 | 0.026 | 0.416 | 0.32 | -0.091 | 0.008 | 0.008 | 0.862 | 0.35 | 0.06 | 0.35 | 0.39 | 0.39 | 0.28 |
| 2 | 189187396 | rs12693496 | -0.622 | 0.273 | 0.389 | 0.135 | -0.008 | 0.126 | 0.126 | 0 | 0.69 | 0.87 | 0.85 | 0.96 | 0.7 | 0.69 |
| 2 | 189189808 | rs12151502 | 0.822 | 0.27 | 0.409 | 0.169 | -0.01 | 0.126 | 0.126 | 0.364 | 0.09 | 0.23 | 0.04 | 0.21 | 0.04 | 0.2 |
| 2 | 189192535 | rs13008931 | 0.692 | 0.291 | 0.448 | 0.029 | 0.002 | 0.038 | 0.038 | 0 | 0.49 | 0.35 | 0.68 | 0.71 | 0.49 | 0.43 |
| 2 | 189193142 | rs12105671 | 0.813 | 0.282 | 0.389 | -0.003 | 0.017 | 0.143 | 0.143 | 0 | 0.27 | 0.06 | 0.08 | 0.04 | 0.2 | 0.28 |
| 2 | 189194029 | rs7574262 | -0.641 | 0.277 | 0.533 | -0.057 | -0.005 | -0.008 | -0.008 | 0 | 0.46 | 0.53 | 0.25 | 0.28 | 0.4 | 0.54 |
| 2 | 189195208 | rs4433962 | 0.362 | 0.358 | 0.427 | 0.032 | -0.001 | 0.007 | 0.007 | 0.271 | 0.17 | 0.19 | 0.15 | 0.25 | 0.18 | 0.26 |
| 2 | 189269721 | rs4340469 | -0.322 | 0.302 | 0.403 | 0.079 | -0.008 | 0.072 | 0.072 | 0.248 | 0.21 | 0.58 | 0.18 | 0.29 | 0.2 | 0.27 |
| 2 | 189298316 | rs3890574 | 0.059 | 0.329 | 0.353 | 0.072 | 0 | 0.039 | 0.039 | 0.191 | 0.18 | 0.31 | 0.16 | 0.29 | 0.2 | 0.26 |
| 2 | 189308284 | rs10931359 | 0.607 | 0.269 | 0.192 | 0.042 | 0.002 | 0.033 | 0.033 | 0 | 0.49 | 0.37 | 0.66 | 0.68 | 0.49 | 0.42 |
| 2 | 189363100 | rs7424417 | 1.223 | 0.36 | 0.277 | -0.559 | -0.02 | 0.099 | 0.099 | 0 | 0.2 | 0.01 | 0.09 | 0.04 | 0.18 | 0.19 |
| 2 | 189397071 | rs11685321 | 0.209 | 0.4 | 0.297 | -0.624 | 0.002 | -0.072 | -0.072 | 0.629 | 0.62 | 0.44 | 0.68 | 0.92 | 0.57 | 0.72 |
| 2 | 189414364 | rs13034731 | 1.69 | 0.046 | -0.581 | -0.655 | -0.156 | -0.117 | -0.117 | 0 | 0.28 | 0.02 | 0.36 | 0.42 | 0.37 | 0.32 |
| 2 | 189441521 | rs1354903 | 0.461 | 0.622 | 0.089 | -1.001 | -0.045 | 0.094 | 0.094 | 0 | 0.17 | 0.05 | 0.13 | 0 | 0.23 | 0.09 |
| 2 | 189446471 | rs1354905 | 1.09 | 0.606 | 0.417 | -0.977 | 0.006 | 0.05 | 0.05 | 0 | 0.51 | 0.39 | 0.53 | 0.83 | 0.44 | 0.64 |
| 2 | 189456969 | rs12463744 | 0.729 | 0.472 | 1.306 | -0.183 | 0.139 | 0.029 | 0.029 | 0.874 | 0.92 | 0.61 | 0.85 | 0.96 | 0.89 | 0.97 |
| 2 | 189460553 | rs8273 | 1.048 | 0.461 | 1.307 | -0.177 | 0.129 | 0.017 | 0.017 | 1.433 | 0.91 | 0.54 | 0.85 | 0.96 | 0.89 | 0.97 |
| 2 | 189501952 | rs1507558 | 1.058 | 0.5 | 1.301 | -0.35 | 0.134 | 0.01 | 0.01 | 1.637 | 0.91 | 0.54 | 0.85 | 0.96 | 0.89 | 0.97 |
| 2 | 189519890 | rs4666763 | 0.586 | 0.488 | 1.304 | -0.422 | -0.004 | -0.069 | -0.069 | 0 | 0.71 | 0.42 | 0.76 | 0.92 | 0.72 | 0.79 |
| 2 | 189526890 | rs11889220 | 0.971 | 0.62 | 1.399 | -0.292 | 0.148 | 0.023 | 0.023 | 1.572 | 0.92 | 0.54 | 0.85 | 0.96 | 0.89 | 0.98 |
| 2 | 189527809 | rs1395867 | -0.203 | 0.631 | 1.096 | -0.295 | 0.005 | -0.017 | -0.017 | 0.153 | 0.94 | 0.93 | 0.94 | 1 | 0.91 | 0.98 |
| 2 | 189553843 | rs1092261 | -0.2 | 0.635 | 1.146 | -0.293 | 0.024 | -0.03 | -0.03 | 0 | 0.94 | 0.9 | 0.93 | 1 | 0.91 | 0.98 |
| 2 | 189564693 | rs10173369 | 0.496 | 0.63 | 1.219 | -0.297 | 0.056 | -0.022 | -0.022 | 0 | 0.93 | 0.78 | 0.87 | 0.99 | 0.9 | 0.97 |
| 2 | 189569061 | rs1395866 | 0.479 | 0.563 | 1.223 | -0.396 | -0.002 | -0.073 | -0.073 | 0.548 | 0.72 | 0.45 | 0.77 | 0.92 | 0.72 | 0.79 |
| 2 | 189572285 | rs13018057 | 0.319 | 0.622 | 1.097 | -0.184 | 0.122 | 0.02 | 0.02 | 0 | 0.93 | 0.68 | 0.86 | 0.99 | 0.9 | 0.98 |
| 2 | 189579463 | rs1066770 | -0.085 | 0.631 | 0.609 | -0.186 | 0.035 | -0.031 | -0.031 | 0.471 | 0.94 | 0.85 | 0.93 | 1 | 0.91 | 0.98 |
| 2 | 189584537 | rs1995702 | 0.595 | 0.488 | 0.644 | -0.406 | -0.001 | -0.038 | -0.038 | 0 | 0.66 | 0.36 | 0.74 | 0.92 | 0.68 | 0.77 |
| 2 | 189591884 | rs1916961 | -0.084 | 0.453 | 0.802 | -0.442 | 0.001 | -0.039 | -0.039 | 0 | 0.9 | 0.85 | 0.92 | 1 | 0.91 | 0.98 |
| 2 | 189597457 | rs12693514 | 0.542 | 0.427 | 1.137 | -0.462 | -0.001 | -0.114 | -0.114 | 0 | 0.68 | 0.52 | 0.81 | 0.93 | 0.69 | 0.77 |
| 2 | 189602559 | rs4667244 | 0.404 | 0.405 | 1.296 | -0.473 | -0.003 | 0.023 | 0.023 | 0 | 0.7 | 0.77 | 0.82 | 0.93 | 0.7 | 0.77 |
| 2 | 189614748 | rs6434295 | 0.481 | 0.404 | 1.41 | -0.471 | 0 | -0.061 | -0.061 | 0 | 0.69 | 0.64 | 0.81 | 0.93 | 0.69 | 0.77 |
| 2 | 189622513 | rs13026165 | 0.567 | 0.393 | 1.395 | -0.374 | 0 | -0.112 | -0.112 | 0.046 | 0.68 | 0.52 | 0.81 | 0.93 | 0.69 | 0.77 |
| 2 | 189627031 | rs17709220 | 0.367 | 0.373 | 1.26 | -0.378 | -0.01 | 0.086 | 0.086 | 0 | 0.2 | 0.04 | 0.1 | 0.04 | 0.19 | 0.2 |
| 2 | 189635696 | rs7594142 | 0.548 | 0.355 | 1.385 | -0.401 | 0 | -0.048 | -0.048 | 0.195 | 0.69 | 0.64 | 0.76 | 0.92 | 0.69 | 0.77 |
| 2 | 189643316 | rs11901929 | 0.795 | 0.337 | 1.296 | -0.186 | 0.002 | 0.017 | 0.017 | 0 | 0.64 | 0.33 | 0.64 | 0.82 | 0.65 | 0.7 |
| 2 | 189654868 | rs781288 | -0.185 | 0.409 | 0.97 | -0.225 | 0.002 | -0.044 | -0.044 | 0 | 0.09 | 0.15 | 0.08 | 0 | 0.09 | 0.02 |
| 2 | 189682074 | rs16830657 | -0.342 | 0.406 | 0.948 | -0.227 | 0.001 | -0.05 | -0.05 | 3.062 | 0.1 | 0.19 | 0.08 | 0 | 0.09 | 0.02 |
| 2 | 189699756 | rs10172458 | -0.754 | 0.427 | 0.971 | -0.226 | 0.035 | -0.056 | -0.056 | 1.144 | 0.12 | 0.34 | 0.09 | 0.04 | 0.11 | 0.03 |
| 2 | 189700120 | rs4667249 | 0.57 | 0.362 | 0.841 | -0.294 | -0.001 | -0.099 | -0.099 | 0.072 | 0.68 | 0.53 | 0.76 | 0.91 | 0.69 | 0.77 |
| 2 | 189711790 | rs12475677 | 0.66 | 0.209 | 0.511 | -0.377 | -0.006 | -0.07 | -0.07 | 0.587 | 0.67 | 0.41 | 0.75 | 0.95 | 0.71 | 0.79 |
| 2 | 189723976 | rs1201151 | 0.079 | 0.299 | 0.537 | -0.341 | -0.001 | -0.034 | -0.034 | 0.602 | 0.09 | 0.14 | 0.08 | 0 | 0.07 | 0.01 |
| 2 | 189743056 | rs1878204 | 0.398 | 0.262 | -0.963 | -0.165 | 0.001 | -0.008 | -0.008 | 0 | 0.37 | 0.36 | 0.42 | 0.44 | 0.36 | 0.37 |
| 2 | 189748646 | rs925825 | 0.617 | 0.479 | -1.569 | 0.214 | -0.073 | -0.111 | -0.111 | 0.119 | 0.3 | 0.05 | 0.32 | 0.51 | 0.35 | 0.42 |
| 2 | 189750863 | rs997277 | -0.544 | -0.217 | -1.637 | -0.503 | 0.029 | 0.111 | 0.111 | 0 | 0.38 | 0.14 | 0.41 | 0.17 | 0.35 | 0.28 |
| 2 | 189753823 | rs1914030 | 0.207 | -0.103 | -1.344 | -0.698 | 0.002 | -0.028 | -0.028 | 0 | 0.09 | 0.06 | 0.13 | 0 | 0.12 | 0.02 |
| 2 | 189757531 | rs2067991 | 0.422 | -0.145 | -1.557 | -0.731 | -0.001 | -0.002 | -0.002 | 0 | 0.17 | 0.13 | 0.14 | 0.17 | 0.16 | 0.16 |

| | | | | | | | | | | | | | | | | |
|---|---|---|---|---|---|---|---|---|---|---|---|---|---|---|---|---|
| 2 | 189760814 | rs2090357 | -0.109 | -0.575 | -1.159 | -0.872 | -0.001 | -0.005 | -0.005 | 0 | 0.78 | 0.71 | 0.83 | 0.8 | 0.79 | 0.82 |
| 2 | 189761153 | rs2708204 | 0.631 | -0.745 | -1.291 | -0.607 | -0.002 | 0.013 | 0.013 | 0 | 0.08 | 0.09 | 0.03 | 0.17 | 0.05 | 0.05 |
| 2 | 189762024 | rs1516445 | -0.564 | -0.786 | -1.167 | -0.681 | -0.004 | -0.003 | -0.003 | 0.665 | 0.81 | 0.7 | 0.84 | 0.82 | 0.84 | 0.84 |
| 2 | 189762841 | rs1996312 | -0.715 | -0.773 | -1.293 | -0.674 | 0 | 0 | 0 | 0.038 | 0.82 | 0.8 | 0.85 | 0.82 | 0.84 | 0.84 |
| 2 | 189766881 | rs4666767 | 0.277 | -0.781 | -1.305 | -0.713 | 0 | -0.003 | -0.003 | 1.502 | 0.73 | 0.74 | 0.72 | 0.82 | 0.72 | 0.82 |
| 2 | 189777898 | rs12474463 | 0.841 | -1.768 | -1.255 | -0.986 | 0.012 | -0.011 | -0.011 | 0 | 0.19 | 0.17 | 0.14 | 0.25 | 0.08 | 0.09 |
| 2 | 189782047 | rs11686968 | 0.428 | -1.721 | -1.228 | -0.982 | -0.007 | 0.007 | 0.007 | 0.236 | 0.13 | 0.05 | 0.13 | 0.03 | 0.16 | 0.04 |
| 2 | 189782825 | rs16830823 | -0.003 | -1.909 | -1.247 | -0.891 | 0.023 | -0.008 | -0.008 | 0 | 0.24 | 0.19 | 0.18 | 0.24 | 0.13 | 0.15 |
| 2 | 189784930 | rs17357652 | -0.875 | -2.018 | -1.678 | -0.953 | 0.099 | 0.053 | 0.053 | 0 | 0.47 | 0.2 | 0.33 | 0.31 | 0.31 | 0.33 |
| 2 | 189796272 | rs16830840 | -0.26 | -1.595 | -1.4 | -0.637 | -0.004 | 0.024 | 0.024 | 0 | 0.11 | 0.15 | 0.15 | 0.25 | 0.09 | 0.12 |
| 2 | 189800385 | rs17241561 | -0.684 | -1.648 | -1.262 | -0.67 | 0.019 | -0.006 | -0.006 | 0 | 0.35 | 0.25 | 0.31 | 0.31 | 0.28 | 0.31 |
| 2 | 189802107 | rs17241582 | 0.631 | -1.482 | -1.496 | -0.837 | -0.02 | 0.069 | 0.069 | 0.238 | 0.22 | 0.01 | 0.16 | 0.07 | 0.19 | 0.19 |
| 2 | 189821391 | rs17292158 | 1.518 | -1.608 | -1.157 | -0.798 | 0 | 0.066 | 0.066 | 1.626 | 0.13 | 0.04 | 0.04 | 0 | 0.08 | 0.03 |
| 2 | 189823392 | rs12693523 | 1.163 | -2.077 | -0.992 | -0.975 | 0.001 | -0.001 | -0.001 | 1.952 | 0.38 | 0.39 | 0.52 | 0.65 | 0.56 | 0.62 |
| 2 | 189829688 | rs1122869 | 0.534 | -1.855 | -0.447 | -0.627 | 0.001 | -0.002 | -0.002 | 0 | 0.56 | 0.63 | 0.67 | 0.79 | 0.67 | 0.72 |
| 2 | 189833911 | rs6434304 | 0.662 | -1.849 | -0.614 | -0.59 | 0.006 | 0.004 | 0.004 | 0 | 0.54 | 0.44 | 0.54 | 0.66 | 0.67 | 0.69 |
| 2 | 189837995 | rs17358566 | 0.292 | -1.155 | -0.84 | -0.887 | -0.001 | -0.015 | -0.015 | 0 | 0.06 | 0.08 | 0.05 | 0 | 0.07 | 0.06 |
| 2 | 189844000 | rs1516454 | 0.402 | -1.424 | -1.059 | -1.022 | -0.007 | 0.044 | 0.044 | 1.052 | 0.1 | 0.02 | 0.14 | 0 | 0.12 | 0.02 |
| 2 | 189847300 | rs17292943 | -0.124 | -1.899 | -1.096 | -1.246 | 0.033 | 0.038 | 0.038 | 0.329 | 0.35 | 0.18 | 0.25 | 0.23 | 0.27 | 0.3 |
| 2 | 189849773 | rs3106796 | 2.146 | -1.905 | -1.44 | -1.321 | -0.136 | -0.132 | -0.132 | 0 | 0.23 | 0.04 | 0.46 | 0.61 | 0.42 | 0.46 |
| 2 | 189855166 | rs13306275 | 0.39 | -2.025 | -0.979 | -1.352 | 0.004 | 0.02 | 0.02 | 2.965 | 0.14 | 0.02 | 0.15 | 0.07 | 0.13 | 0.04 |
| 2 | 189857993 | rs12693525 | -0.347 | -2.025 | -0.866 | -1.37 | 0.017 | 0.008 | 0.008 | 2.611 | 0.42 | 0.63 | 0.31 | 0.33 | 0.29 | 0.37 |
| 2 | 189862097 | rs7579903 | 1.547 | -2.026 | -0.988 | -1.389 | -0.003 | -0.068 | -0.068 | 0 | 0.2 | 0.23 | 0.08 | 0 | 0.16 | 0.13 |
| 2 | 189864080 | rs1800255 | 0.199 | -1.777 | -0.85 | -1.182 | 0.032 | 0.043 | 0.043 | 3.746 | 0.33 | 0.11 | 0.25 | 0.23 | 0.26 | 0.29 |
| 2 | 189864582 | rs1801184 | 0.115 | -1.484 | -0.612 | -0.961 | 0.006 | 0.014 | 0.014 | 6.468 | 0.35 | 0.28 | 0.23 | 0.07 | 0.29 | 0.17 |
| 2 | 189874873 | rs10204508 | 0.202 | -1.583 | -0.152 | -0.602 | 0.017 | 0.018 | 0.018 | 0.293 | 0.36 | 0.31 | 0.14 | 0.07 | 0.24 | 0.22 |
| 2 | 189878769 | rs7596514 | -0.137 | -1.497 | -0.86 | -0.571 | 0.015 | 0.015 | 0.015 | 0 | 0.31 | 0.2 | 0.24 | 0.24 | 0.25 | 0.32 |
| 2 | 189879843 | rs3134646 | 1.417 | -1.411 | -1.622 | -0.499 | -0.161 | -0.144 | -0.144 | 0 | 0.29 | 0.04 | 0.57 | 0.59 | 0.51 | 0.44 |
| 2 | 189880810 | rs4667257 | 0.647 | -1.159 | -0.374 | -0.019 | -0.028 | -0.028 | -0.028 | 0.961 | 0.69 | 0.55 | 0.74 | 0.76 | 0.75 | 0.68 |
| 2 | 189881677 | rs7421775 | -0.691 | -0.935 | -1.077 | 0.196 | 0.026 | -0.014 | -0.014 | 0 | 0.24 | 0.07 | 0.23 | 0.27 | 0.2 | 0.27 |
| 2 | 189881888 | rs7419983 | -0.612 | -0.953 | -0.327 | 0.327 | -0.003 | 0 | 0 | 0 | 0.35 | 0.38 | 0.27 | 0.32 | 0.25 | 0.26 |
| 2 | 189882685 | rs7579312 | 1.564 | -0.832 | 0.813 | 0.285 | -0.064 | -0.012 | -0.012 | 0 | 0.84 | 0.58 | 0.87 | 0.88 | 0.95 | 0.84 |
| 2 | 189883166 | rs6434309 | 2.095 | -0.906 | 1.322 | 0.339 | -0.049 | -0.009 | -0.009 | 0 | 0.8 | 0.31 | 0.72 | 0.82 | 0.91 | 0.82 |
| 2 | 189883644 | rs7424137 | 2.206 | -0.709 | 0.756 | 0.164 | 0.027 | 0.035 | 0.035 | 0 | 0.6 | 0.05 | 0.6 | 0.7 | 0.76 | 0.66 |
| 2 | 189896170 | rs6717231 | 1.315 | -1.136 | 1.344 | 0.231 | -0.021 | 0.022 | 0.022 | 0 | 0.9 | 0.81 | 0.79 | 0.82 | 0.96 | 0.84 |
| 2 | 189897394 | rs11186 | -0.697 | -1.111 | 1.45 | 0.208 | 0.004 | 0.024 | 0.024 | 0 | 0.18 | 0.07 | 0.11 | 0.11 | 0.12 | 0.16 |
| 2 | 189897631 | rs13914 | -0.544 | -1.116 | 1.506 | 0.222 | -0.012 | 0.003 | 0.003 | 0 | 0.06 | 0.1 | 0.04 | 0.08 | 0.02 | 0.05 |
| 2 | 189902190 | rs7424790 | 2.329 | -1.132 | 1.488 | 0.211 | 0.079 | 0.039 | 0.039 | 0 | 0.62 | 0.09 | 0.61 | 0.7 | 0.79 | 0.66 |
| 2 | 189904203 | rs10208525 | -0.685 | -1.071 | 1.274 | 0.241 | 0.004 | 0.023 | 0.023 | 0 | 0.17 | 0.07 | 0.11 | 0.11 | 0.11 | 0.16 |
| 2 | 189904233 | rs10197596 | -0.658 | -1.071 | 1.274 | 0.276 | 0.003 | 0.019 | 0.019 | 0 | 0.17 | 0.07 | 0.11 | 0.12 | 0.11 | 0.16 |
| 2 | 189906743 | rs10194682 | 2.232 | -1.116 | 1.32 | 0.219 | -0.04 | -0.022 | -0.022 | 0 | 0.63 | 0.48 | 0.64 | 0.68 | 0.79 | 0.66 |
| 2 | 189907676 | rs13392501 | -0.622 | -1.189 | 1.241 | 0.166 | -0.003 | 0.043 | 0.043 | 0 | 0.17 | 0.03 | 0.1 | 0.11 | 0.12 | 0.16 |
| 2 | 189913748 | rs13383261 | -1.455 | -1.038 | 1.379 | 0.278 | 0.03 | 0.059 | 0.059 | 0 | 0.27 | 0.17 | 0.14 | 0.11 | 0.17 | 0.18 |
| 2 | 189974794 | rs4128538 | 1.394 | -1.025 | 1.329 | 0.297 | -0.03 | 0.089 | 0.089 | 0 | 0.89 | 0.64 | 0.76 | 0.78 | 0.96 | 0.84 |
| 2 | 189975790 | rs7557945 | 1.686 | -0.927 | 0.997 | 0.31 | -0.036 | -0.021 | -0.021 | 0 | 0.62 | 0.5 | 0.62 | 0.69 | 0.79 | 0.66 |
| 2 | 189981212 | rs13024858 | 1.85 | -0.791 | 0.914 | 0.317 | -0.042 | -0.013 | -0.013 | 0.034 | 0.8 | 0.34 | 0.78 | 0.84 | 0.91 | 0.93 |
| 2 | 190012746 | rs17270966 | -0.869 | -0.016 | 0.765 | 0.313 | -0.074 | 0.077 | 0.077 | 0 | 0.21 | 0.04 | 0.18 | 0.06 | 0.25 | 0.19 |
| 2 | 190018099 | rs1356168 | 1.973 | -0.061 | 0.995 | 0.291 | 0 | -0.005 | -0.005 | 2.054 | 0.63 | 0.35 | 0.61 | 0.77 | 0.66 | 0.71 |
| 2 | 190058461 | rs12999125 | 1.992 | -0.06 | 1.072 | 0.256 | -0.037 | 0.042 | 0.042 | 0.048 | 0.82 | 0.42 | 0.69 | 0.78 | 0.87 | 0.81 |
| 2 | 190073366 | rs1878085 | 2.124 | -0.05 | 1.071 | 0.231 | 0 | 0.01 | 0.01 | 0 | 0.62 | 0.32 | 0.55 | 0.7 | 0.65 | 0.61 |
| 2 | 190080363 | rs10184895 | -1.227 | -0.075 | 1.692 | 0.223 | -0.002 | 0.154 | 0.154 | 0.25 | 0.1 | 0.38 | 0.21 | 0.22 | 0.06 | 0.1 |
| 2 | 190096685 | rs7609495 | 2.191 | -0.061 | 1.706 | 0.218 | 0 | 0.002 | 0.002 | 0 | 0.62 | 0.33 | 0.53 | 0.7 | 0.67 | 0.61 |
| 2 | 190097106 | rs1515875 | 0.779 | -0.04 | 1.62 | 0.202 | -0.022 | 0.048 | 0.048 | 0 | 0.79 | 0.85 | 0.87 | 0.95 | 0.8 | 0.8 |
| 2 | 190111412 | rs1515867 | 1.277 | -0.163 | 1.65 | 0.216 | -0.008 | -0.031 | -0.031 | 0 | 0.94 | 0.84 | 0.95 | 1 | 0.96 | 0.95 |
| 2 | 190115961 | rs10204942 | 1.605 | -0.188 | 1.869 | 0.257 | -0.005 | -0.035 | -0.035 | 0 | 0.93 | 0.78 | 0.95 | 1 | 0.96 | 0.95 |
| 2 | 190130511 | rs4667274 | 2.286 | -0.227 | 1.895 | 0.203 | 0.005 | 0.056 | 0.056 | 0 | 0.59 | 0.1 | 0.52 | 0.7 | 0.66 | 0.61 |
| 2 | 190137632 | rs12473931 | -0.659 | -0.248 | 1.801 | 0.229 | 0.002 | 0.002 | 0.002 | 0.139 | 0.07 | 0.09 | 0.14 | 0.04 | 0.04 | 0.03 |
| 2 | 190146586 | rs10207860 | -2.404 | -0.244 | 1.641 | 0.228 | -0.003 | -0.043 | -0.043 | 0 | 0.09 | 0.26 | 0.12 | 0.01 | 0.08 | 0.09 |
| 2 | 190150205 | rs16831319 | -2.492 | -0.252 | 1.886 | 0.242 | -0.017 | 0.07 | 0.07 | 0.402 | 0.09 | 0.26 | 0.12 | 0.18 | 0.06 | 0.13 |
| 2 | 190153035 | rs7576165 | 2.236 | -0.199 | 1.368 | 0.207 | 0.003 | 0.009 | 0.009 | 0 | 0.61 | 0.3 | 0.54 | 0.7 | 0.66 | 0.61 |
| 2 | 190160414 | rs4667276 | 2.371 | -0.201 | 1.399 | 0.2 | 0.01 | 0.026 | 0.026 | 0 | 0.6 | 0.25 | 0.53 | 0.7 | 0.66 | 0.59 |
| 2 | 190171008 | rs10188246 | 0.819 | -0.193 | 1.422 | 0.204 | -0.009 | 0.148 | 0.148 | 0.068 | 0.9 | 0.6 | 0.78 | 0.76 | 0.94 | 0.89 |
| 2 | 190174516 | rs1996576 | -1.613 | -0.172 | 1.396 | 0.18 | -0.005 | 0.028 | 0.028 | 0 | 0.26 | 0.18 | 0.17 | 0.06 | 0.25 | 0.26 |
| 2 | 190192867 | rs17271015 | -0.856 | -0.165 | 1.302 | 0.186 | -0.074 | 0.067 | 0.067 | 0 | 0.19 | 0.01 | 0.12 | 0.06 | 0.21 | 0.21 |
| 2 | 190215527 | rs10177513 | -2.044 | -0.169 | 1.353 | 0.182 | 0 | -0.03 | -0.03 | 0 | 0.29 | 0.33 | 0.24 | 0.06 | 0.28 | 0.3 |
| 2 | 190310982 | rs1520854 | 1.148 | -0.128 | 1.128 | 0.217 | -0.031 | 0.074 | 0.074 | 0.018 | 0.85 | 0.54 | 0.73 | 0.76 | 0.9 | 0.83 |
| 2 | 190316198 | rs17271043 | -0.814 | -0.127 | 1.116 | 0.333 | -0.025 | 0.037 | 0.037 | 0 | 0.2 | 0.1 | 0.12 | 0.08 | 0.21 | 0.21 |
| 2 | 190323530 | rs1157910 | 2.125 | -0.128 | 1.123 | 0.334 | 0.009 | 0.013 | 0.013 | 0 | 0.61 | 0.27 | 0.54 | 0.67 | 0.66 | 0.58 |
| 2 | 190345166 | rs10497702 | -1.103 | -0.071 | 0.876 | 0.278 | 0.007 | -0.004 | -0.004 | 0 | 0.14 | 0.16 | 0.11 | 0.08 | 0.17 | 0.16 |

| | | | | | | | | | | | | | | | | |
|---|---|---|---|---|---|---|---|---|---|---|---|---|---|---|---|---|
| 2 | 190350153 | rs11686908 | -0.962 | -0.092 | 0.105 | 0.175 | -0.076 | -0.005 | -0.005 | 0 | 0.15 | 0.01 | 0.12 | 0.08 | 0.2 | 0.17 |
| 2 | 190350282 | rs745742 | 1.934 | -0.355 | 0.533 | 0.165 | -0.049 | -0.008 | -0.008 | 0.224 | 0.76 | 0.38 | 0.7 | 0.77 | 0.86 | 0.78 |
| 2 | 190352703 | rs12623394 | 2.054 | -0.353 | 0.499 | 0.165 | -0.028 | -0.009 | -0.009 | 0 | 0.73 | 0.32 | 0.64 | 0.76 | 0.83 | 0.74 |
| 2 | 190356197 | rs1866605 | 1.492 | -0.183 | 0.416 | 0.387 | -0.044 | 0.044 | 0.044 | 0.519 | 0.83 | 0.39 | 0.72 | 0.77 | 0.9 | 0.83 |
| 2 | 190357442 | rs10198663 | 1.449 | -0.158 | 0.429 | 0.399 | -0.042 | 0.046 | 0.046 | 0.522 | 0.83 | 0.38 | 0.72 | 0.77 | 0.9 | 0.83 |
| 2 | 190367057 | rs11674773 | -1.381 | 0.656 | -0.631 | -0.164 | -0.125 | 0.149 | 0.149 | 0 | 0.26 | 0.03 | 0.19 | 0.05 | 0.34 | 0.18 |
| 2 | 190376493 | rs11680808 | -0.899 | 1.105 | -0.53 | -0.156 | -0.1 | 0.047 | 0.047 | 0.122 | 0.16 | 0.01 | 0.15 | 0.06 | 0.27 | 0.13 |
| 2 | 190382243 | rs12622601 | 1.943 | 0.897 | -0.195 | -0.255 | -0.02 | 0.016 | 0.016 | 0 | 0.71 | 0.73 | 0.77 | 0.91 | 0.6 | 0.69 |
| 2 | 190389803 | rs17271134 | -0.883 | 1.012 | -0.278 | -0.052 | -0.047 | 0.041 | 0.041 | 0.067 | 0.16 | 0.07 | 0.15 | 0.05 | 0.26 | 0.13 |
| 2 | 190412377 | rs11685197 | -1.156 | 0.861 | -0.07 | 0.039 | -0.005 | 0.005 | 0.005 | 0 | 0.14 | 0.11 | 0.14 | 0.05 | 0.27 | 0.11 |
| 2 | 190425474 | rs11539983 | -1.56 | 0.429 | 0.915 | 0.21 | 0.021 | -0.016 | -0.016 | 0 | 0.07 | 0.27 | 0.03 | 0 | 0.03 | 0.02 |
| 2 | 190428193 | rs7596205 | -1.087 | 0.279 | 0.699 | 0.016 | -0.002 | 0.066 | 0.066 | 0.436 | 0.12 | 0.01 | 0.04 | 0.02 | 0.11 | 0.18 |
| 2 | 190431875 | rs4667287 | 1.604 | 0.165 | 1.817 | -0.037 | -0.021 | -0.029 | -0.029 | 0 | 0.85 | 0.72 | 0.86 | 0.88 | 0.89 | 0.81 |
| 2 | 190431970 | rs13404407 | 0.985 | -0.349 | 1.192 | -0.065 | 0.007 | -0.009 | -0.009 | 0 | 0.05 | 0.04 | 0 | 0.07 | 0.01 | 0.01 |
| 2 | 190433289 | rs16831659 | -0.883 | -0.223 | 1.192 | -0.111 | -0.003 | 0.065 | 0.065 | 0.409 | 0.12 | 0.01 | 0.04 | 0.02 | 0.11 | 0.18 |
| 2 | 190440548 | rs1439812 | -0.857 | 0.068 | 1.209 | 0.064 | -0.008 | 0.007 | 0.007 | 0 | 0.14 | 0.19 | 0.15 | 0.05 | 0.27 | 0.13 |
| 2 | 190444199 | rs1123110 | 1.825 | -0.066 | 0.895 | -0.062 | -0.002 | 0.047 | 0.047 | 1.219 | 0.59 | 0.29 | 0.67 | 0.8 | 0.62 | 0.63 |
| 2 | 190444392 | rs1123109 | -0.287 | 0.242 | 0.575 | 0.039 | -0.07 | 0.043 | 0.043 | 0 | 0.12 | 0.01 | 0.11 | 0.04 | 0.23 | 0.12 |
| 2 | 190447811 | rs17198983 | 2.254 | -0.134 | 0.433 | 0.329 | -0.028 | 0.007 | 0.007 | 0.654 | 0.56 | 0.11 | 0.64 | 0.66 | 0.59 | 0.54 |
| 2 | 190448965 | rs2352267 | -2.063 | -0.21 | 1.086 | 0.331 | 0.017 | -0.019 | -0.019 | 0 | 0.34 | 0.51 | 0.26 | 0.21 | 0.4 | 0.34 |
| 2 | 190452586 | rs6718798 | -2.039 | -0.267 | 1.092 | 0.222 | 0.014 | -0.036 | -0.036 | 0.2 | 0.34 | 0.52 | 0.27 | 0.17 | 0.39 | 0.34 |
| 2 | 190452748 | rs6706281 | 1.55 | -0.251 | 1.179 | 0.214 | 0.014 | 0.008 | 0.008 | 0 | 0.77 | 0.6 | 0.75 | 0.77 | 0.74 | 0.71 |
| 2 | 190459468 | rs1437883 | 1.14 | -0.189 | 1.406 | 0.082 | 0.014 | -0.005 | -0.005 | 5.515 | 0.82 | 0.75 | 0.76 | 0.86 | 0.74 | 0.71 |
| 2 | 190463032 | rs1370593 | 2.211 | -0.208 | 1.303 | 0.059 | -0.019 | 0.06 | 0.06 | 1.111 | 0.61 | 0.18 | 0.64 | 0.77 | 0.6 | 0.55 |
| 2 | 190466451 | rs17271183 | -0.35 | -0.191 | 1.263 | 0.067 | -0.081 | 0.006 | 0.006 | 0 | 0.13 | 0.01 | 0.15 | 0.06 | 0.25 | 0.18 |
| 2 | 190468920 | rs1347708 | 1.892 | -0.167 | 0.96 | 0.086 | -0.007 | 0.024 | 0.024 | 0.661 | 0.61 | 0.29 | 0.65 | 0.77 | 0.6 | 0.56 |
| 2 | 190472176 | rs10188692 | -2.022 | -0.412 | 0.959 | -0.211 | -0.009 | -0.046 | -0.046 | 0 | 0.29 | 0.55 | 0.22 | 0.14 | 0.27 | 0.2 |
| 2 | 190474499 | rs13415781 | -0.609 | -0.419 | 1.031 | -0.265 | -0.009 | -0.025 | -0.025 | 0 | 0.1 | 0.21 | 0.06 | 0.02 | 0.06 | 0.06 |
| 2 | 190482320 | rs11901582 | -0.773 | -0.252 | 0.662 | -0.344 | 0.018 | -0.066 | -0.066 | 0 | 0.22 | 0.34 | 0.17 | 0.04 | 0.27 | 0.12 |
| 2 | 190486244 | rs13025114 | 2.079 | -0.392 | 0.593 | -0.092 | -0.008 | 0.026 | 0.026 | 0.835 | 0.59 | 0.06 | 0.71 | 0.71 | 0.65 | 0.74 |
| 2 | 190487810 | rs1437889 | 1.61 | -0.979 | 1.009 | 0.26 | -0.056 | 0.044 | 0.044 | 3.552 | 0.81 | 0.41 | 0.88 | 0.75 | 0.92 | 0.86 |
| 2 | 190492014 | rs10497705 | 0.243 | -0.955 | -0.024 | -0.352 | -0.031 | -0.013 | -0.013 | 0 | 0.29 | 0.18 | 0.56 | 0.72 | 0.36 | 0.6 |
| 2 | 190493757 | rs6434353 | 1.588 | -1.668 | 1.667 | 0.336 | 0.088 | 0.126 | 0.126 | 0.33 | 0.95 | 0.63 | 0.93 | 0.96 | 0.94 | 0.91 |
| 2 | 190501523 | rs6434355 | 1.436 | -2.355 | 1.422 | -0.048 | 0.06 | 0.013 | 0.013 | 2.649 | 0.6 | 0.29 | 0.81 | 0.71 | 0.74 | 0.77 |
| 2 | 190502035 | rs12613410 | -0.686 | -2.526 | 1.783 | -0.016 | -0.021 | 0.035 | 0.035 | 1.158 | 0.12 | 0.15 | 0.03 | 0.18 | 0.07 | 0.11 |
| 2 | 190503961 | rs4667294 | 0.553 | -2.657 | 1.957 | -0.462 | -0.041 | -0.013 | -0.013 | 0.637 | 0.9 | 0.6 | 0.96 | 0.97 | 0.99 | 0.93 |
| 2 | 190521054 | rs7568449 | 1.896 | -2.847 | 2.247 | -0.446 | 0.019 | 0.016 | 0.016 | 1.517 | 0.59 | 0.34 | 0.8 | 0.74 | 0.75 | 0.72 |
| 2 | 190528996 | rs4667297 | 1.811 | -2.57 | 2.288 | -0.443 | 0.019 | 0.017 | 0.017 | 3.027 | 0.6 | 0.33 | 0.75 | 0.74 | 0.73 | 0.76 |
| 2 | 190529231 | rs12693551 | 1.811 | -2.571 | 2.322 | -0.443 | 0.01 | 0.01 | 0.01 | 3.253 | 0.6 | 0.35 | 0.75 | 0.74 | 0.73 | 0.76 |
| 2 | 190531615 | rs3828267 | -0.764 | -2.611 | 2.434 | -0.459 | -0.028 | 0.033 | 0.033 | 2.2 | 0.13 | 0.35 | 0.11 | 0.12 | 0.04 | 0.05 |
| 2 | 190536283 | rs893784 | 2.019 | -2.597 | 2.315 | -0.454 | 0.1 | 0.083 | 0.083 | 3.606 | 0.59 | 0.16 | 0.73 | 0.74 | 0.73 | 0.76 |
| 2 | 190542186 | rs10931430 | 1.94 | -2.539 | 2.324 | -0.41 | 0.091 | 0.072 | 0.072 | 2.532 | 0.6 | 0.17 | 0.73 | 0.74 | 0.73 | 0.76 |
| 2 | 190546300 | rs11679767 | 2.056 | -2.556 | 2.335 | -0.417 | 0.095 | 0.084 | 0.084 | 3.048 | 0.6 | 0.13 | 0.73 | 0.74 | 0.73 | 0.76 |
| 2 | 190562708 | rs1437891 | 1.936 | -2.684 | 2.349 | -0.419 | 0.089 | 0.071 | 0.071 | 2.532 | 0.6 | 0.18 | 0.73 | 0.74 | 0.73 | 0.76 |
| 2 | 190583187 | rs7557519 | -1.098 | -2.7 | 2.486 | -0.388 | -0.018 | -0.016 | -0.016 | 0.629 | 0.26 | 0.33 | 0.24 | 0.22 | 0.2 | 0.12 |
| 2 | 190607153 | rs12053254 | -0.149 | -2.722 | 2.591 | -0.399 | 0.009 | 0.065 | 0.065 | 0.006 | 0.06 | 0.13 | 0.12 | 0.12 | 0.04 | 0.04 |
| 2 | 190612339 | rs1440006 | 0.838 | -2.73 | 2.583 | -0.404 | 0.01 | 0.009 | 0.009 | 0 | 0.88 | 0.85 | 0.96 | 0.95 | 0.93 | 0.87 |
| 2 | 190617700 | rs16831960 | -0.854 | -2.73 | 2.58 | -0.404 | 0.035 | 0.023 | 0.023 | 0 | 0.11 | 0.09 | 0.03 | 0.05 | 0.07 | 0.13 |
| 2 | 190635143 | rs1055091 | 2.128 | -2.736 | 2.628 | -0.394 | 0.086 | 0.065 | 0.065 | 2.567 | 0.6 | 0.17 | 0.7 | 0.73 | 0.73 | 0.76 |
| 2 | 190636014 | rs2289404 | -1.162 | -2.724 | 2.593 | -0.389 | -0.017 | -0.014 | -0.014 | 0.858 | 0.26 | 0.33 | 0.24 | 0.22 | 0.2 | 0.12 |
| 2 | 190639915 | rs3791767 | -1.187 | -2.723 | 2.597 | -0.388 | -0.013 | -0.01 | -0.01 | 0 | 0.26 | 0.29 | 0.24 | 0.22 | 0.2 | 0.17 |
| 2 | 190647919 | rs2352709 | 1.128 | -2.626 | 2.529 | -0.343 | -0.047 | -0.03 | -0.03 | 0 | 0.87 | 0.63 | 0.95 | 0.95 | 0.93 | 0.87 |
| 2 | 190648805 | rs5742926 | 0.383 | -2.583 | 2.062 | -0.326 | -0.01 | 0.03 | 0.03 | 0 | 0.14 | 0.03 | 0.11 | 0.09 | 0.15 | 0.07 |
| 2 | 190653750 | rs5742967 | -0.155 | -2.531 | 1.972 | -0.291 | 0.01 | 0.067 | 0.067 | 0.015 | 0.05 | 0.13 | 0.11 | 0.12 | 0.04 | 0.04 |
| 2 | 190684782 | rs1233276 | -1.376 | -2.543 | 1.648 | -0.399 | 0 | -0.003 | -0.003 | 0 | 0.27 | 0.3 | 0.15 | 0.14 | 0.22 | 0.19 |
| 2 | 190696924 | rs5743072 | -0.757 | -2.502 | 1.324 | -0.335 | -0.022 | -0.047 | -0.047 | 0.07 | 0.23 | 0.28 | 0.14 | 0.12 | 0.18 | 0.16 |
| 2 | 190707034 | rs1233258 | -1.885 | -2.754 | 0.757 | -0.232 | 0.144 | 0.074 | 0.074 | 1.734 | 0.5 | 0.78 | 0.3 | 0.4 | 0.28 | 0.33 |
| 2 | 190712137 | rs13404927 | -0.922 | -2.219 | 1.834 | 0.007 | -0.048 | -0.046 | -0.046 | 0 | 0.1 | 0.32 | 0.03 | 0.02 | 0.03 | 0.07 |
| 2 | 190714621 | rs256575 | -0.275 | -2.287 | 2.041 | -0.004 | -0.038 | -0.049 | -0.049 | 0 | 0.1 | 0.23 | 0.03 | 0.02 | 0.03 | 0.07 |
| 2 | 190717628 | rs5743130 | -0.782 | -2.291 | 2.143 | -0.03 | -0.02 | -0.018 | -0.018 | 0 | 0.09 | 0.14 | 0.01 | 0.02 | 0.03 | 0.07 |
| 2 | 190724141 | rs733318 | -0.719 | -2.33 | 2.233 | -0.001 | -0.005 | 0.085 | 0.085 | 0.755 | 0.07 | 0.2 | 0.13 | 0.18 | 0.04 | 0.04 |
| 2 | 190731832 | rs256563 | -2.513 | -2.338 | 2.221 | -0.005 | 0.03 | 0.011 | 0.011 | 3.276 | 0.27 | 0.77 | 0.2 | 0.22 | 0.1 | 0.19 |
| 2 | 190733398 | rs5743175 | -0.738 | -2.327 | 2.207 | -0.003 | -0.009 | -0.007 | -0.007 | 0 | 0.09 | 0.11 | 0.01 | 0.02 | 0.03 | 0.07 |
| 2 | 190744738 | rs256550 | -2.459 | -2.332 | 2.15 | -0.001 | 0.042 | 0.014 | 0.014 | 3.629 | 0.28 | 0.79 | 0.19 | 0.22 | 0.1 | 0.19 |
| 2 | 190751163 | rs256546 | -2.516 | -2.457 | 1.982 | -0.096 | 0.137 | 0.115 | 0.115 | 2.702 | 0.46 | 0.89 | 0.35 | 0.29 | 0.29 | 0.27 |
| 2 | 190751439 | rs256545 | -0.094 | -2.43 | 2.12 | -0.078 | -0.035 | -0.04 | -0.04 | 0 | 0.1 | 0.15 | 0.03 | 0.02 | 0.03 | 0.08 |
| 2 | 190758688 | rs16832158 | -0.774 | -2.42 | 1.857 | -0.071 | -0.005 | 0.088 | 0.088 | 0.907 | 0.07 | 0.24 | 0.14 | 0.17 | 0.04 | 0.04 |
| 2 | 190775484 | rs4381805 | -0.027 | -2.361 | 1.413 | -0.14 | -0.04 | -0.051 | -0.051 | 0 | 0.89 | 0.77 | 0.97 | 0.98 | 0.97 | 0.92 |
| 2 | 190780698 | rs12466395 | 1.342 | -2.227 | 0.326 | -0.076 | 0.046 | 0.054 | 0.054 | 0.15 | 0.61 | 0.3 | 0.78 | 0.88 | 0.76 | 0.78 |

| | | | | | | | | | | | | | | | | |
|---|---|---|---|---|---|---|---|---|---|---|---|---|---|---|---|---|
| 2 | 190794653 | rs7603201 | 1.376 | -2.116 | 0.122 | 0.325 | 0.03 | 0.011 | 0.011 | 3.368 | 0.73 | 0.23 | 0.8 | 0.78 | 0.9 | 0.81 |
| 2 | 190798570 | rs11694671 | 1.458 | -1.781 | 0.184 | 0.713 | 0.08 | 0.016 | 0.016 | 1.322 | 0.64 | 0.15 | 0.76 | 0.71 | 0.78 | 0.77 |
| 2 | 190802120 | rs11901349 | 1.014 | -1.739 | 0.606 | 0.515 | -0.001 | -0.008 | -0.008 | 0.011 | 0.9 | 0.91 | 0.96 | 0.97 | 0.93 | 0.96 |
| 2 | 190805662 | rs7571613 | -0.86 | -1.784 | 1.58 | 0.473 | -0.01 | -0.04 | -0.04 | 0.003 | 0.21 | 0.28 | 0.13 | 0.12 | 0.17 | 0.17 |
| 2 | 190813716 | rs2168059 | 0.434 | -1.75 | 1.493 | 0.494 | -0.004 | -0.011 | -0.011 | 0 | 0.92 | 0.92 | 0.97 | 0.97 | 0.94 | 0.96 |
| 2 | 190827023 | rs2053163 | 2.334 | -1.852 | 1.919 | 1.254 | 0.05 | 0.03 | 0.03 | 0.812 | 0.59 | 0.05 | 0.59 | 0.31 | 0.7 | 0.7 |
| 2 | 190835473 | rs6749643 | -1.423 | -1.697 | 2.108 | 0.329 | -0.04 | -0.101 | -0.101 | 0 | 0.25 | 0.43 | 0.15 | 0.11 | 0.18 | 0.17 |
| 2 | 190883360 | rs10931440 | 1.894 | -1.624 | 1.91 | 0.423 | 0.088 | 0.023 | 0.023 | 0.905 | 0.64 | 0.16 | 0.76 | 0.69 | 0.78 | 0.78 |
| 2 | 190889261 | rs16832254 | -1.972 | -1.64 | 1.941 | 0.228 | -0.037 | -0.081 | -0.081 | 0.311 | 0.33 | 0.51 | 0.17 | 0.14 | 0.21 | 0.21 |
| 2 | 190895573 | rs1369319 | 1.767 | -1.632 | 1.707 | 0.241 | 0.069 | 0.121 | 0.121 | 1.394 | 0.64 | 0.16 | 0.75 | 0.83 | 0.77 | 0.79 |
| 2 | 190898218 | rs1011584 | -1.438 | -1.688 | 2.149 | 0.137 | 0.037 | 0.027 | 0.027 | 0.855 | 0.06 | 0.44 | 0.03 | 0 | 0 | 0 |
| 2 | 190900963 | rs13010250 | 1.88 | -1.673 | 1.726 | 0.109 | 0.06 | 0.153 | 0.153 | 1.528 | 0.63 | 0.13 | 0.73 | 0.85 | 0.75 | 0.79 |
| 2 | 190916072 | rs17732729 | -0.391 | -1.548 | 1.779 | 0.145 | -0.015 | 0.063 | 0.063 | 0 | 0.18 | 0.06 | 0.12 | 0.09 | 0.18 | 0.16 |
| 2 | 190921491 | rs16832285 | -0.649 | -1.401 | 1.77 | 0.222 | 0.006 | -0.017 | -0.017 | 0 | 0.05 | 0.22 | 0.05 | 0 | 0.03 | 0 |
| 2 | 190931436 | rs4853626 | -1.42 | -1.291 | 0.321 | 0.302 | 0.046 | 0.044 | 0.044 | 0.653 | 0.34 | 0.84 | 0.26 | 0.25 | 0.22 | 0.18 |
| 2 | 190934874 | rs13420500 | -1.305 | -1.199 | 0.193 | 0.328 | 0 | 0 | 0 | 0.149 | 0.33 | 0.71 | 0.25 | 0.25 | 0.22 | 0.17 |
| 2 | 190944165 | rs785247 | -1.124 | -1.35 | 0.393 | -0.29 | -0.071 | -0.115 | -0.115 | 0 | 0.23 | 0.44 | 0.09 | 0.04 | 0.08 | 0.04 |
| 2 | 190946324 | rs4853633 | -0.243 | -2.259 | 0.174 | -0.797 | -0.024 | -0.012 | -0.012 | 0.055 | 0.69 | 0.65 | 0.78 | 0.76 | 0.78 | 0.87 |
| 2 | 190947260 | rs785242 | 0.83 | -1.511 | 0.249 | -0.69 | -0.019 | -0.017 | -0.017 | 0 | 0.37 | 0.28 | 0.53 | 0.57 | 0.55 | 0.74 |
| 2 | 190948115 | rs4853637 | 0.515 | -1.159 | 0.086 | -0.863 | 0.024 | -0.014 | -0.014 | 0 | 0.89 | 0.63 | 0.88 | 0.94 | 0.9 | 0.93 |
| 2 | 190948685 | rs4853638 | 1.53 | -1.18 | -0.265 | -0.756 | -0.101 | -0.114 | -0.114 | 0.053 | 0.28 | 0.1 | 0.45 | 0.56 | 0.46 | 0.7 |
| 2 | 190950202 | rs6737521 | -0.363 | -1.333 | -0.035 | -1.508 | -0.003 | 0.011 | 0.011 | 0 | 0.82 | 0.87 | 0.79 | 0.9 | 0.83 | 0.85 |
| 2 | 190952343 | rs785237 | 0.916 | -1.429 | 0.593 | -1.609 | -0.04 | 0.013 | 0.013 | 1.638 | 0.38 | 0.22 | 0.48 | 0.67 | 0.5 | 0.71 |
| 2 | 190954001 | rs13021283 | 1.701 | -1.615 | 0.369 | -1.618 | -0.13 | -0.05 | -0.05 | 0.637 | 0.29 | 0.06 | 0.45 | 0.68 | 0.46 | 0.69 |
| 2 | 190959523 | rs13393016 | -1.013 | -1.643 | 0.576 | -1.68 | -0.005 | -0.016 | -0.016 | 0 | 0.31 | 0.65 | 0.31 | 0.17 | 0.21 | 0.19 |
| 2 | 190968863 | rs11674449 | 1.695 | -1.636 | 0.403 | -1.682 | -0.116 | -0.039 | -0.039 | 0.112 | 0.31 | 0.09 | 0.46 | 0.67 | 0.47 | 0.69 |
| 2 | 190972168 | rs2115553 | -0.312 | -1.804 | 0.7 | -1.673 | -0.015 | -0.052 | -0.052 | 0 | 0.12 | 0.22 | 0.06 | 0.01 | 0.04 | 0.01 |
| 2 | 190973233 | rs785232 | 0.395 | -1.831 | 0.725 | -1.643 | 0.025 | 0.046 | 0.046 | 0 | 0.35 | 0.2 | 0.2 | 0.15 | 0.31 | 0.1 |
| 2 | 190980437 | rs4853659 | -0.273 | -1.636 | -0.212 | -1.904 | 0.008 | 0.021 | 0.021 | 0.528 | 0.25 | 0.14 | 0.26 | 0.16 | 0.18 | 0.18 |
| 2 | 190980751 | rs4853660 | -0.797 | -1.622 | -0.304 | -1.924 | 0.027 | 0.049 | 0.049 | 3.951 | 0.47 | 0.37 | 0.42 | 0.22 | 0.37 | 0.22 |
| 2 | 190993716 | rs6750727 | 0.584 | -1.328 | -0.601 | -1.688 | 0.002 | 0.001 | 0.001 | 0.307 | 0.1 | 0.07 | 0.08 | 0.06 | 0.07 | 0.05 |
| 2 | 191002267 | rs4531957 | -0.543 | -1.659 | -0.665 | -1.909 | 0.007 | 0.019 | 0.019 | 1.364 | 0.33 | 0.28 | 0.4 | 0.18 | 0.25 | 0.15 |
| 2 | 191008125 | rs2562817 | 0.258 | -1.834 | -0.156 | -1.67 | 0.01 | -0.011 | -0.011 | 0.761 | 0.93 | 0.67 | 0.95 | 1 | 0.98 | 1 |
| 2 | 191018607 | rs11901445 | -0.119 | -1.888 | -0.717 | -1.938 | 0.021 | 0.101 | 0.101 | 0 | 0.34 | 0.19 | 0.25 | 0.1 | 0.31 | 0.27 |
| 2 | 191020013 | rs785260 | -1.688 | -1.879 | -0.605 | -1.95 | -0.001 | 0 | 0 | 0 | 0.67 | 0.68 | 0.64 | 0.29 | 0.53 | 0.43 |
| 2 | 191020292 | rs785259 | -0.815 | -1.984 | -0.711 | -2.041 | 0.011 | 0.036 | 0.036 | 0.08 | 0.52 | 0.53 | 0.51 | 0.19 | 0.42 | 0.35 |
| 2 | 191020850 | rs6738692 | -0.007 | -2.001 | -0.532 | -2.04 | -0.024 | -0.048 | -0.048 | 0 | 0.31 | 0.4 | 0.39 | 0.19 | 0.24 | 0.15 |
| 2 | 191021281 | rs6710331 | 1.112 | -1.851 | -0.717 | -1.951 | 0.013 | 0.003 | 0.003 | 0 | 0.15 | 0.14 | 0.14 | 0.1 | 0.11 | 0.06 |
| 2 | 191023961 | rs4853680 | 1.254 | -1.19 | -1.171 | -1.547 | -0.123 | -0.026 | -0.026 | 0.456 | 0.31 | 0.1 | 0.34 | 0.71 | 0.47 | 0.58 |
| 2 | 191026633 | rs10804032 | -0.186 | -1.053 | -0.829 | -1.488 | -0.026 | -0.152 | -0.152 | 0 | 0.76 | 0.51 | 0.85 | 0.99 | 0.81 | 0.79 |
| 2 | 191029604 | rs785248 | -0.417 | -0.39 | -0.879 | -0.81 | -0.008 | -0.018 | -0.018 | 0.649 | 0.41 | 0.45 | 0.48 | 0.22 | 0.3 | 0.19 |
| 2 | 191039370 | rs698601 | -0.465 | -0.401 | -1.163 | -0.419 | 0.018 | 0.101 | 0.101 | 0 | 0.23 | 0.06 | 0.11 | 0 | 0.18 | 0.23 |
| 2 | 191039604 | rs785209 | 0.393 | 0.054 | -0.284 | 0.507 | 0.003 | 0.019 | 0.019 | 0.145 | 0.88 | 0.83 | 0.74 | 0.8 | 0.84 | 0.86 |
| 2 | 191045969 | rs6720084 | -1.134 | -0.219 | 0.443 | 0.7 | -0.046 | 0.016 | 0.016 | 0 | 0.14 | 0.32 | 0.05 | 0.15 | 0.03 | 0.09 |
| 2 | 191046105 | rs785213 | -1.109 | -0.216 | 0.444 | 0.701 | -0.045 | 0.018 | 0.018 | 0 | 0.14 | 0.32 | 0.05 | 0.15 | 0.03 | 0.09 |
| 2 | 191052117 | rs698607 | -1.173 | 0.004 | 0.629 | 0.75 | -0.018 | 0.043 | 0.043 | 0 | 0.14 | 0.37 | 0.06 | 0.15 | 0.06 | 0.15 |
| 2 | 191053129 | rs16832404 | -0.821 | 0.259 | 0.262 | 1.269 | 0.008 | 0.05 | 0.05 | 0 | 0.09 | 0.13 | 0.14 | 0.25 | 0.1 | 0.09 |
| 2 | 191060269 | rs6755406 | -1.255 | -0.49 | 0.878 | 0.916 | -0.053 | 0.043 | 0.043 | 0 | 0.14 | 0.47 | 0.07 | 0.15 | 0.03 | 0.1 |
| 2 | 191060568 | rs6755967 | -1.929 | -0.474 | 0.837 | 1.293 | -0.027 | 0.076 | 0.076 | 0.262 | 0.19 | 0.59 | 0.12 | 0.33 | 0.11 | 0.19 |
| 2 | 191062031 | rs11894667 | -1.271 | -0.764 | 0.67 | 1.049 | -0.016 | 0.017 | 0.017 | 0 | 0.35 | 0.39 | 0.26 | 0.43 | 0.24 | 0.23 |
| 2 | 191065595 | rs2271270 | -1.358 | -0.833 | 0.996 | 1.015 | -0.027 | 0.022 | 0.022 | 0.001 | 0.36 | 0.54 | 0.28 | 0.44 | 0.24 | 0.24 |
| 2 | 191066970 | rs3749016 | -1.22 | -0.761 | 1.144 | 1.017 | -0.032 | 0.037 | 0.037 | 0 | 0.35 | 0.5 | 0.27 | 0.45 | 0.24 | 0.24 |
| 2 | 191078163 | rs3791788 | -1.204 | -0.769 | 1.548 | 1.04 | 0.009 | -0.01 | -0.01 | 0 | 0.35 | 0.33 | 0.26 | 0.43 | 0.24 | 0.24 |
| 2 | 191113253 | rs7590991 | 0.931 | -0.411 | 0.282 | 0.76 | -0.14 | -0.004 | -0.004 | 0 | 0.27 | 0.03 | 0.21 | 0.24 | 0.43 | 0.5 |
| 2 | 191114229 | rs2244076 | -0.623 | -0.368 | 0.563 | 1 | -0.033 | 0.028 | 0.028 | 0.011 | 0.36 | 0.45 | 0.26 | 0.43 | 0.24 | 0.25 |
| 2 | 191143591 | rs2582734 | -0.568 | -0.433 | 0.587 | 1.051 | 0 | -0.002 | -0.002 | 0 | 0.93 | 0.96 | 0.99 | 0.85 | 0.98 | 0.93 |
| 2 | 191148558 | rs2664253 | 0.897 | -0.37 | 0.33 | 0.88 | -0.002 | -0.037 | -0.037 | 0.026 | 0.55 | 0.49 | 0.35 | 0.16 | 0.6 | 0.6 |
| 2 | 191160345 | rs291454 | -0.82 | -0.367 | 0.25 | 0.886 | 0.008 | -0.009 | -0.009 | 0 | 0.93 | 0.97 | 0.99 | 0.85 | 0.98 | 0.93 |
| 2 | 191161622 | rs1058180 | 0.125 | -0.376 | 0.304 | 0.899 | -0.012 | 0.009 | 0.009 | 0.052 | 0.65 | 0.63 | 0.75 | 0.57 | 0.76 | 0.75 |
| 2 | 191166159 | rs12467835 | 0.618 | -0.333 | 0.215 | 0.93 | -0.043 | -0.023 | -0.023 | 0 | 0.09 | 0.03 | 0.38 | 0.42 | 0.16 | 0.15 |
| 2 | 191166611 | rs1595997 | -0.799 | -0.361 | 0.23 | 0.861 | 0.008 | -0.009 | -0.009 | 0.052 | 0.93 | 0.97 | 0.99 | 0.85 | 0.98 | 0.93 |
| 2 | 191181481 | rs291458 | 0.718 | -0.375 | 0.28 | 0.817 | 0 | -0.005 | -0.005 | 0 | 0.57 | 0.59 | 0.37 | 0.24 | 0.6 | 0.6 |
| 2 | 191185725 | rs1372053 | 0.639 | -0.321 | 0.167 | 1.155 | -0.027 | -0.02 | -0.02 | 0 | 0.08 | 0.04 | 0.39 | 0.31 | 0.16 | 0.15 |
| 2 | 191186951 | rs291467 | 0.826 | -0.68 | 0.535 | 0.902 | 0.053 | 0.011 | 0.011 | 0 | 0.05 | 0.27 | 0.05 | 0 | 0.03 | 0.02 |
| 2 | 191190180 | rs291471 | -0.306 | -0.68 | 0.12 | 1.009 | -0.007 | 0.016 | 0.016 | 0.723 | 0.93 | 0.92 | 0.98 | 0.8 | 0.97 | 0.92 |
| 2 | 191207447 | rs2067385 | 1.199 | -0.836 | -0.12 | 0.688 | 0.022 | 0.049 | 0.049 | 0.594 | 0.09 | 0 | 0.04 | 0 | 0.04 | 0.02 |
| 2 | 191213664 | rs1882892 | 0.582 | -0.704 | 0.053 | 0.814 | 0.062 | -0.042 | -0.042 | 0 | 0.58 | 0.2 | 0.66 | 0.51 | 0.7 | 0.72 |
| 2 | 191219614 | rs3791815 | -0.045 | -0.748 | 0.399 | 0.789 | 0.012 | -0.023 | -0.023 | 0 | 0.31 | 0.24 | 0.23 | 0.49 | 0.23 | 0.25 |
| 2 | 191219850 | rs7601609 | -0.335 | -0.747 | 0.594 | 0.765 | 0.034 | -0.037 | -0.037 | 0 | 0.09 | 0.41 | 0.08 | 0 | 0.07 | 0.03 |

| | | | | | | | | | | | | | | | | |
|---|---|---|---|---|---|---|---|---|---|---|---|---|---|---|---|---|
| 2 | 191224981 | rs4656 | -0.928 | -0.759 | 0.534 | 0.777 | 0.042 | -0.023 | -0.023 | 0.132 | 0.41 | 0.72 | 0.33 | 0.49 | 0.3 | 0.28 |
| 2 | 191226742 | rs2067416 | -0.937 | -0.759 | 0.521 | 0.777 | 0.051 | -0.031 | -0.031 | 0 | 0.41 | 0.77 | 0.34 | 0.49 | 0.3 | 0.28 |
| 2 | 191227073 | rs2067419 | 0.248 | -0.769 | 0.528 | 0.939 | 0.024 | -0.058 | -0.058 | 0.076 | 0.28 | 0.13 | 0.21 | 0.39 | 0.23 | 0.18 |
| 2 | 191227734 | rs972690 | -0.86 | -0.738 | 0.459 | 0.81 | 0.022 | -0.013 | -0.013 | 0 | 0.41 | 0.65 | 0.32 | 0.49 | 0.3 | 0.28 |
| 2 | 191231503 | rs10931450 | -0.05 | -0.684 | 0.232 | 0.992 | 0.015 | -0.028 | -0.028 | 0 | 0.31 | 0.22 | 0.22 | 0.46 | 0.23 | 0.25 |
| 2 | 191234190 | rs1108939 | 0.684 | -0.247 | 0.121 | 1.193 | 0 | 0.015 | 0.015 | 0 | 0.08 | 0.06 | 0.39 | 0.29 | 0.15 | 0.11 |
| 2 | 191235610 | rs7592352 | -0.262 | -0.466 | 0.495 | 0.695 | -0.016 | -0.04 | -0.04 | 0 | 0.07 | 0.18 | 0.03 | 0 | 0.02 | 0.02 |
| 2 | 191235901 | rs1882891 | -1.275 | -0.037 | -0.209 | 0.374 | 0 | -0.008 | -0.008 | 0 | 0.84 | 0.8 | 0.92 | 0.91 | 0.87 | 0.93 |
| 2 | 191237072 | rs2736619 | -0.801 | -0.061 | -0.235 | 0.361 | -0.001 | -0.006 | -0.006 | 0 | 0.82 | 0.8 | 0.92 | 0.89 | 0.86 | 0.86 |
| 2 | 191241186 | rs1866852 | -0.465 | -0.237 | -0.796 | 0.473 | -0.039 | 0.035 | 0.035 | 0 | 0.59 | 0.79 | 0.68 | 0.64 | 0.48 | 0.46 |
| 2 | 191245071 | rs10804035 | -0.357 | -0.501 | -1.266 | 0.161 | -0.006 | 0.028 | 0.028 | 0 | 0.57 | 0.63 | 0.72 | 0.74 | 0.53 | 0.48 |
| 2 | 191246782 | rs13390311 | 1.554 | -0.745 | -1.312 | 0.035 | -0.011 | -0.004 | -0.004 | 0 | 0.22 | 0.23 | 0.1 | 0.14 | 0.14 | 0.09 |
| 2 | 191247547 | rs4853698 | -0.229 | -0.901 | -1.258 | -0.134 | -0.005 | 0.01 | 0.01 | 0 | 0.6 | 0.61 | 0.72 | 0.74 | 0.55 | 0.47 |
| 2 | 191250446 | rs1545212 | -1.591 | -0.677 | -1.262 | -0.039 | -0.045 | 0.02 | 0.02 | 0 | 0.85 | 0.9 | 0.84 | 0.88 | 0.68 | 0.63 |
| 2 | 191255883 | rs11892526 | -0.519 | -0.662 | -1.378 | 0.249 | -0.004 | -0.023 | -0.023 | 0 | 0.84 | 0.87 | 0.55 | 0.65 | 0.84 | 0.72 |
| 2 | 191261942 | rs11695439 | 0.665 | -0.688 | -0.995 | 0.119 | 0.014 | 0.024 | 0.024 | 0.07 | 0.36 | 0.29 | 0.15 | 0.13 | 0.25 | 0.17 |
| 2 | 191264390 | rs4853700 | -0.736 | -0.715 | -1.025 | 0.139 | -0.011 | 0.03 | 0.03 | 0 | 0.55 | 0.68 | 0.67 | 0.62 | 0.49 | 0.42 |
| 2 | 191272150 | rs894072 | -2.278 | -0.717 | -0.974 | 0.069 | -0.01 | -0.005 | -0.005 | 0.089 | 0.84 | 0.94 | 0.93 | 0.87 | 0.85 | 0.91 |
| 2 | 191278341 | rs6725814 | -0.517 | -0.381 | -0.753 | 0.219 | 0 | -0.017 | -0.017 | 0.451 | 0.66 | 0.65 | 0.77 | 0.87 | 0.6 | 0.55 |
| 2 | 191288071 | rs2354877 | 0.72 | -0.734 | -0.925 | 0.097 | -0.014 | -0.002 | -0.002 | 0.095 | 0.32 | 0.42 | 0.19 | 0.26 | 0.28 | 0.12 |
| 2 | 191299996 | rs4853707 | -2.107 | -0.739 | -0.833 | -0.217 | -0.026 | -0.073 | -0.073 | 0.021 | 0.81 | 0.75 | 0.93 | 1 | 0.91 | 0.92 |
| 2 | 191301368 | rs9646748 | -0.079 | -0.71 | -0.92 | -0.145 | 0.005 | 0.017 | 0.017 | 0 | 0.49 | 0.43 | 0.64 | 0.62 | 0.45 | 0.39 |
| 2 | 191305647 | rs7564829 | -0.004 | -0.669 | -0.854 | -0.132 | 0.002 | 0.006 | 0.006 | 0.017 | 0.48 | 0.46 | 0.64 | 0.62 | 0.45 | 0.35 |
| 2 | 191306726 | rs4261750 | -0.862 | -0.673 | -0.892 | -0.176 | -0.024 | -0.024 | -0.024 | 0 | 0.64 | 0.57 | 0.81 | 0.74 | 0.76 | 0.74 |
| 2 | 191307408 | rs4129081 | 0.402 | -0.674 | -0.931 | -0.124 | 0.006 | 0.104 | 0.104 | 0 | 0.24 | 0.41 | 0.54 | 0.48 | 0.27 | 0.22 |
| 2 | 191307562 | rs4853708 | -0.427 | -0.689 | -0.936 | -0.133 | 0.004 | -0.001 | -0.001 | 0 | 0.4 | 0.46 | 0.71 | 0.6 | 0.57 | 0.61 |
| 2 | 191312906 | rs4638823 | 1.984 | -0.611 | -1.155 | -0.118 | -0.063 | 0.024 | 0.024 | 0 | 0.16 | 0.05 | 0.17 | 0.12 | 0.31 | 0.39 |
| 2 | 191318776 | rs13409841 | -0.048 | -0.746 | -0.953 | -0.115 | 0.006 | 0.029 | 0.029 | 0.256 | 0.48 | 0.41 | 0.63 | 0.65 | 0.45 | 0.34 |
| 2 | 191323138 | rs4283453 | 2.155 | -0.715 | -1.009 | -0.15 | 0.003 | 0.034 | 0.034 | 0 | 0.24 | 0.06 | 0.1 | 0.14 | 0.18 | 0.13 |
| 2 | 191330837 | rs4853712 | -0.943 | -0.778 | -0.889 | -0.206 | -0.014 | -0.038 | -0.038 | 0 | 0.62 | 0.56 | 0.83 | 0.86 | 0.74 | 0.79 |
| 2 | 191333728 | rs4340549 | -0.753 | -0.782 | -0.945 | -0.168 | -0.008 | -0.005 | -0.005 | 3.2 | 0.64 | 0.42 | 0.79 | 0.72 | 0.77 | 0.74 |
| 2 | 191343763 | rs12613807 | 0.045 | -0.733 | -0.999 | -0.179 | 0.007 | 0.041 | 0.041 | 0.344 | 0.48 | 0.35 | 0.62 | 0.63 | 0.46 | 0.35 |
| 2 | 191354491 | rs3811609 | 0.528 | -0.714 | -0.824 | -0.155 | 0.002 | 0.04 | 0.04 | 0.033 | 0.23 | 0.29 | 0.53 | 0.49 | 0.27 | 0.21 |
| 2 | 191366579 | rs7721 | 2.019 | -0.692 | -0.859 | -0.17 | -0.068 | 0.043 | 0.043 | 0 | 0.16 | 0.05 | 0.17 | 0.1 | 0.31 | 0.38 |
| 2 | 191366707 | rs1128723 | 2.276 | -0.74 | -0.818 | -0.188 | 0.002 | -0.002 | -0.002 | 0 | 0.22 | 0.18 | 0.13 | 0.27 | 0.16 | 0.19 |
| 2 | 191369852 | rs7564107 | 2.279 | -0.76 | -0.835 | -0.201 | -0.005 | 0.033 | 0.033 | 0 | 0.24 | 0.06 | 0.09 | 0.14 | 0.2 | 0.14 |
| 2 | 191370268 | rs11685221 | 2.135 | -0.752 | -0.826 | -0.199 | -0.083 | 0.041 | 0.041 | 0 | 0.16 | 0.02 | 0.17 | 0.1 | 0.31 | 0.38 |
| 2 | 191371041 | rs13392968 | 0.966 | -0.79 | -0.779 | -0.18 | 0.04 | 0.014 | 0.014 | 0 | 0.46 | 0.24 | 0.22 | 0.41 | 0.36 | 0.34 |
| 2 | 191379268 | rs4586658 | -0.509 | -0.648 | -0.885 | -0.091 | -0.007 | -0.006 | -0.006 | 0 | 0.39 | 0.33 | 0.7 | 0.58 | 0.58 | 0.59 |
| 2 | 191386158 | rs7584794 | -0.509 | -0.648 | -0.885 | -0.091 | -0.007 | -0.006 | -0.006 | 0 | 0.39 | 0.33 | 0.7 | 0.58 | 0.58 | 0.59 |
| 2 | 191390126 | rs10931458 | -0.972 | -0.681 | -0.903 | -0.096 | -0.018 | -0.012 | -0.012 | 0 | 0.64 | 0.5 | 0.8 | 0.72 | 0.77 | 0.73 |
| 2 | 191410588 | rs13417310 | 1.905 | -0.48 | -1.077 | -0.104 | 0.012 | 0.032 | 0.032 | 0 | 0.25 | 0.17 | 0.1 | 0.14 | 0.2 | 0.14 |
| 2 | 191424949 | rs10179262 | 1.918 | -0.478 | -1.128 | -0.103 | 0.011 | 0.039 | 0.039 | 0 | 0.25 | 0.13 | 0.1 | 0.14 | 0.2 | 0.14 |
| 2 | 191438741 | rs7603849 | 0.736 | -0.508 | -1.003 | -0.01 | -0.035 | 0.057 | 0.057 | 0 | 0.41 | 0.23 | 0.27 | 0.29 | 0.51 | 0.52 |
| 2 | 191448249 | rs3934784 | 1.909 | -0.435 | -1.035 | -0.001 | -0.079 | 0.037 | 0.037 | 0 | 0.16 | 0.02 | 0.17 | 0.11 | 0.31 | 0.38 |
| 2 | 191466532 | rs6724607 | -0.951 | -0.494 | -1.018 | -0.012 | -0.032 | 0.05 | 0.05 | 0 | 0.6 | 0.76 | 0.73 | 0.71 | 0.49 | 0.48 |
| 2 | 191467149 | rs12621493 | 0.962 | -0.599 | -0.714 | 0.322 | 0 | 0.01 | 0.01 | 0.141 | 0.2 | 0.2 | 0.51 | 0.45 | 0.26 | 0.24 |
| 2 | 191468706 | rs13028201 | 0.445 | -0.587 | -0.706 | 0.329 | -0.002 | -0.006 | -0.006 | 0.83 | 0.23 | 0.2 | 0.52 | 0.45 | 0.27 | 0.25 |
| 2 | 191469731 | rs10737936 | -1.486 | -0.808 | -0.542 | 0.436 | -0.002 | -0.003 | -0.003 | 0.178 | 0.74 | 0.68 | 0.89 | 0.82 | 0.8 | 0.86 |
| 2 | 191472880 | rs12476782 | 0.967 | -0.592 | 0.044 | 0.671 | 0.002 | 0.002 | 0.002 | 0 | 0.21 | 0.16 | 0.16 | 0.2 | 0.21 | 0.16 |
| 2 | 191493166 | rs887695 | -0.091 | -0.489 | -0.826 | 0.743 | 0.024 | -0.013 | -0.013 | 0 | 0.35 | 0.75 | 0.23 | 0.39 | 0.26 | 0.17 |
| 2 | 191504064 | rs1558473 | 1.277 | 0.115 | -0.692 | 0.772 | 0 | 0 | 0 | 1.432 | 0.19 | 0.16 | 0.15 | 0.21 | 0.24 | 0.16 |
| 2 | 191504093 | rs12991526 | 0.059 | -0.448 | -0.512 | 0.603 | 0.014 | 0.013 | 0.013 | 0 | 0.46 | 0.45 | 0.28 | 0.27 | 0.33 | 0.26 |
| 2 | 191506773 | rs1558471 | 0.707 | -0.216 | -0.421 | 0.739 | -0.002 | 0.032 | 0.032 | 0 | 0.17 | 0.23 | 0.38 | 0.35 | 0.15 | 0.22 |
| 2 | 191520845 | rs2293765 | 0.069 | -0.797 | -0.393 | 0.215 | -0.017 | 0.059 | 0.059 | 0 | 0.54 | 0.22 | 0.38 | 0.37 | 0.56 | 0.55 |
| 2 | 191528162 | rs958770 | -1.164 | -0.899 | -0.456 | 0.184 | 0.012 | 0.012 | 0.012 | 0 | 0.71 | 0.76 | 0.79 | 0.82 | 0.75 | 0.86 |
| 2 | 191532039 | rs1978273 | -0.532 | -0.918 | -0.468 | 0.185 | -0.012 | 0.083 | 0.083 | 0 | 0.54 | 0.74 | 0.63 | 0.69 | 0.45 | 0.49 |
| 2 | 191535576 | rs2286896 | 0.932 | -0.631 | -0.612 | 0.423 | -0.001 | 0.062 | 0.062 | 0 | 0.17 | 0.26 | 0.39 | 0.41 | 0.15 | 0.21 |
| 2 | 191538562 | rs10931468 | 0.809 | -0.719 | -0.658 | 0.182 | -0.003 | 0.065 | 0.065 | 0 | 0.17 | 0.27 | 0.27 | 0.38 | 0.15 | 0.2 |
| 2 | 191545821 | rs12618075 | 2.394 | -0.685 | -0.514 | 0.153 | 0.001 | 0.003 | 0.003 | 0 | 0.19 | 0.18 | 0.15 | 0.27 | 0.25 | 0.19 |
| 2 | 191552398 | rs6716985 | 2.168 | -0.656 | -0.669 | -0.082 | -0.112 | 0.051 | 0.051 | 0.365 | 0.17 | 0.02 | 0.16 | 0.11 | 0.3 | 0.37 |
| 2 | 191553931 | rs4853516 | 0.303 | -0.788 | -0.224 | 0.163 | 0.019 | -0.017 | -0.017 | 0.054 | 0.35 | 0.45 | 0.56 | 0.67 | 0.4 | 0.39 |
| 2 | 191575308 | rs9288176 | 1.454 | -0.578 | -0.34 | -0.109 | 0.002 | -0.053 | -0.053 | 0 | 0.19 | 0.21 | 0.14 | 0 | 0.21 | 0.03 |
| 2 | 191585275 | rs6434412 | 1.008 | -0.792 | -0.246 | 0.018 | -0.002 | 0.001 | 0.001 | 0 | 0.31 | 0.36 | 0.2 | 0.29 | 0.29 | 0.25 |
| 2 | 191591310 | rs10176654 | 1.527 | -0.502 | -0.461 | -0.002 | -0.001 | -0.027 | -0.027 | 0 | 0.19 | 0.16 | 0.14 | 0 | 0.21 | 0.03 |
| 2 | 191592247 | rs4366932 | 0.495 | -0.65 | -0.431 | 0.118 | 0 | 0.004 | 0.004 | 0 | 0.33 | 0.32 | 0.21 | 0.45 | 0.38 | 0.29 |
| 2 | 191598603 | rs1263128 | -0.354 | -0.728 | -0.328 | 0.098 | -0.047 | -0.034 | -0.034 | 0.121 | 0.68 | 0.6 | 0.88 | 0.8 | 0.82 | 0.83 |
| 2 | 191599327 | rs1263126 | -0.14 | -0.744 | -0.44 | 0.146 | 0.01 | 0.034 | 0.034 | 0 | 0.52 | 0.41 | 0.65 | 0.64 | 0.54 | 0.44 |
| 2 | 191611464 | rs1263119 | 0.523 | -0.746 | -0.474 | 0.26 | -0.025 | 0.004 | 0.004 | 0 | 0.39 | 0.28 | 0.29 | 0.33 | 0.46 | 0.23 |

| | | | | | | | | | | | | | | | | |
|---|---|---|---|---|---|---|---|---|---|---|---|---|---|---|---|---|
| 2 | 191624519 | rs1476896 | 2.728 | -0.674 | -0.556 | 0.242 | 0.011 | -0.004 | -0.004 | 0 | 0.17 | 0.16 | 0.23 | 0.15 | 0.28 | 0.38 |
| 2 | 191634597 | rs6705471 | 0.614 | -0.565 | -0.663 | 0.396 | 0.004 | -0.01 | -0.01 | 0 | 0.13 | 0.08 | 0.36 | 0.35 | 0.08 | 0.22 |
| 2 | 191634958 | rs13412750 | 2.418 | -0.543 | -0.77 | 0.43 | -0.058 | -0.078 | -0.078 | 0 | 0.18 | 0.05 | 0.14 | 0.3 | 0.25 | 0.18 |
| 2 | 191649639 | rs7601608 | -0.109 | -0.553 | -0.558 | 0.299 | 0.002 | 0.005 | 0.005 | 0 | 0.49 | 0.57 | 0.34 | 0.36 | 0.47 | 0.56 |
| 2 | 191668301 | rs13423600 | 2.1 | -0.839 | -0.622 | 0.591 | -0.008 | -0.01 | -0.01 | 0 | 0.18 | 0.15 | 0.14 | 0.29 | 0.24 | 0.14 |
| 2 | 191668839 | rs1013895 | 2.642 | -0.759 | -0.765 | 0.439 | -0.049 | 0.023 | 0.023 | 0.129 | 0.15 | 0.05 | 0.21 | 0.13 | 0.27 | 0.38 |
| 2 | 191677664 | rs6719798 | 2.196 | -0.857 | -0.624 | 0.476 | -0.008 | -0.011 | -0.011 | 5.872 | 0.19 | 0.15 | 0.14 | 0.29 | 0.24 | 0.15 |
| 2 | 191683388 | rs7596437 | -0.341 | -1.224 | -0.355 | 0.269 | -0.018 | 0.001 | 0.001 | 0 | 0.44 | 0.5 | 0.43 | 0.55 | 0.26 | 0.42 |
| 2 | 191684231 | rs7573809 | 0.177 | -1.222 | -0.503 | 0.27 | 0.058 | -0.006 | -0.006 | 0 | 0.55 | 0.31 | 0.55 | 0.45 | 0.74 | 0.58 |
| 2 | 191691482 | rs13033350 | 2.255 | -1.14 | -0.683 | 0.248 | -0.054 | -0.004 | -0.004 | 0.86 | 0.16 | 0.05 | 0.26 | 0.17 | 0.27 | 0.4 |
| 2 | 191691968 | rs6434420 | 0.795 | -1.162 | -0.718 | 0.254 | -0.058 | 0.012 | 0.012 | 0 | 0.36 | 0.17 | 0.28 | 0.29 | 0.46 | 0.18 |
| 2 | 191701184 | rs1860173 | -0.376 | -1.908 | -0.422 | 0.088 | 0.01 | 0 | 0 | 0 | 0.42 | 0.39 | 0.42 | 0.42 | 0.26 | 0.41 |
| 2 | 191712570 | rs11898764 | 1.507 | -2.085 | -0.116 | 0.287 | 0.001 | 0.017 | 0.017 | 0 | 0.19 | 0.17 | 0.14 | 0.41 | 0.24 | 0.2 |
| 2 | 191716012 | rs1882396 | -0.67 | -2.515 | -0.361 | 0.002 | 0.058 | 0.099 | 0.099 | 0 | 0.48 | 0.77 | 0.58 | 0.69 | 0.34 | 0.5 |
| 2 | 191728044 | rs11893307 | 1.387 | -2.333 | -0.171 | 0.402 | -0.005 | -0.006 | -0.006 | 0.014 | 0.19 | 0.16 | 0.14 | 0.42 | 0.24 | 0.16 |
| 2 | 191736873 | rs2204859 | -0.707 | -2.472 | -0.141 | 0.29 | 0.058 | 0.084 | 0.084 | 0 | 0.45 | 0.75 | 0.56 | 0.67 | 0.32 | 0.35 |
| 2 | 191749035 | rs3771316 | 0.001 | -2.315 | 0.435 | 0.083 | 0.002 | -0.002 | -0.002 | 0 | 0.17 | 0.1 | 0.38 | 0.18 | 0.08 | 0.17 |
| 2 | 191752327 | rs1921915 | -0.934 | -2.457 | 0.074 | -0.021 | 0.01 | -0.004 | -0.004 | 0.588 | 0.87 | 0.88 | 0.96 | 0.85 | 0.93 | 0.88 |
| 2 | 191765206 | rs3771313 | 1.197 | -2.486 | 0.152 | -0.047 | 0.051 | 0.175 | 0.175 | 0 | 0.48 | 0.22 | 0.4 | 0.23 | 0.6 | 0.55 |
| 2 | 191781169 | rs2355570 | 1.222 | -2.426 | 0.329 | 0.001 | -0.003 | 0.002 | 0.002 | 0 | 0.19 | 0.17 | 0.14 | 0.44 | 0.25 | 0.17 |
| 2 | 191800905 | rs3771308 | 1.277 | -2.408 | 0.328 | 0 | -0.003 | 0.005 | 0.005 | 0 | 0.19 | 0.17 | 0.14 | 0.44 | 0.25 | 0.16 |
| 2 | 191815866 | rs1517354 | -0.091 | -2.449 | 0.536 | 0.03 | -0.067 | 0.004 | 0.004 | 0.134 | 0.84 | 0.53 | 0.93 | 0.85 | 0.93 | 0.88 |
| 2 | 191825769 | rs883844 | 0.328 | -2.155 | 0.734 | 0.122 | -0.002 | -0.003 | -0.003 | 0 | 0.56 | 0.5 | 0.76 | 0.36 | 0.63 | 0.62 |
| 2 | 191828686 | rs10437 | 0.223 | -2.097 | 0.672 | 0.193 | -0.022 | 0.088 | 0.088 | 0 | 0.08 | 0.13 | 0.38 | 0.2 | 0.04 | 0.11 |
| 2 | 191830805 | rs17748980 | 2.028 | -2.145 | 0.592 | 0.027 | -0.031 | 0.041 | 0.041 | 0 | 0.31 | 0.21 | 0.26 | 0.14 | 0.44 | 0.39 |
| 2 | 191835596 | rs3771300 | 1.06 | -2.106 | 0.466 | 0.036 | 0.008 | -0.01 | -0.01 | 0 | 0.4 | 0.34 | 0.64 | 0.34 | 0.48 | 0.5 |
| 2 | 191837634 | rs16824035 | 0.188 | -1.865 | 0.478 | 0.114 | 0 | -0.013 | -0.013 | 0 | 0.18 | 0.27 | 0.15 | 0.08 | 0.16 | 0.16 |
| 2 | 191837841 | rs11894425 | -0.519 | -1.955 | 0.501 | 0.116 | 0.005 | 0.007 | 0.007 | 0 | 0.38 | 0.46 | 0.29 | 0.52 | 0.41 | 0.33 |
| 2 | 191838514 | rs13395505 | -0.51 | -1.953 | 0.493 | 0.117 | 0.005 | 0.007 | 0.007 | 0 | 0.38 | 0.46 | 0.29 | 0.52 | 0.41 | 0.33 |
| 2 | 191841759 | rs2066804 | 0.992 | -1.936 | 0.061 | 0.119 | -0.005 | -0.008 | -0.008 | 0 | 0.2 | 0.14 | 0.14 | 0.44 | 0.25 | 0.17 |
| 2 | 191843445 | rs13010343 | 0.885 | -1.734 | -0.032 | -0.189 | -0.008 | 0.025 | 0.025 | 0.047 | 0.13 | 0.08 | 0.08 | 0 | 0.14 | 0.02 |
| 2 | 191850566 | rs2280233 | 1.698 | -1.759 | 0.008 | -0.158 | -0.04 | 0.051 | 0.051 | 0 | 0.32 | 0.21 | 0.27 | 0.15 | 0.45 | 0.44 |
| 2 | 191850766 | rs2280232 | 0.358 | -2.067 | 0.158 | -0.356 | 0.016 | 0.026 | 0.026 | 0 | 0.63 | 0.68 | 0.68 | 0.83 | 0.75 | 0.72 |
| 2 | 191851897 | rs2066795 | -0.241 | -2.357 | 0.597 | 0.093 | -0.055 | -0.029 | -0.029 | 0.345 | 0.2 | 0.33 | 0.22 | 0.15 | 0.13 | 0.13 |
| 2 | 191854874 | rs11887698 | -1.101 | -2.43 | 0.544 | 0.106 | -0.035 | -0.004 | -0.004 | 0 | 0.28 | 0.57 | 0.44 | 0.32 | 0.16 | 0.21 |
| 2 | 191855521 | rs7562024 | 0.856 | -2.347 | 0.027 | 0.046 | -0.002 | -0.039 | -0.039 | 1.384 | 0.47 | 0.48 | 0.37 | 0.14 | 0.57 | 0.47 |
| 2 | 191858636 | rs12693590 | 0.924 | -1.65 | 0.135 | 0.628 | -0.016 | -0.055 | -0.055 | 0 | 0.11 | 0.14 | 0.06 | 0 | 0.08 | 0.01 |
| 2 | 191858830 | rs34997637 | 0.702 | -1.494 | -0.263 | 0.791 | -0.031 | -0.082 | -0.082 | 1.894 | 0.21 | 0.09 | 0.16 | 0.44 | 0.26 | 0.16 |
| 2 | 191858990 | rs11904548 | -0.173 | -1.085 | -0.135 | 0.723 | -0.019 | 0.035 | 0.035 | 0 | 0.08 | 0.13 | 0.16 | 0.16 | 0.05 | 0.12 |
| 2 | 191869163 | rs2030171 | -0.348 | -0.854 | -0.676 | 0.732 | -0.001 | 0 | 0 | 0.107 | 0.42 | 0.4 | 0.53 | 0.6 | 0.36 | 0.2 |
| 2 | 191874667 | rs2066802 | -0.953 | -0.851 | -0.876 | 1.033 | 0.014 | -0.016 | -0.016 | 0.356 | 0.13 | 0.07 | 0.05 | 0.23 | 0.07 | 0.15 |
| 2 | 191875901 | rs13029532 | -0.004 | -1.473 | -0.249 | 0.695 | -0.001 | 0 | 0 | 0 | 0.86 | 0.87 | 0.88 | 0.89 | 0.92 | 0.84 |
| 2 | 191882263 | rs17817966 | 0.934 | -1.832 | -0.91 | 0.344 | -0.054 | 0.007 | 0.007 | 0.253 | 0.38 | 0.11 | 0.44 | 0.37 | 0.56 | 0.6 |
| 2 | 191889582 | rs6740131 | -0.85 | -2.482 | -0.891 | 0.283 | 0.034 | 0.026 | 0.026 | 0.336 | 0.35 | 0.25 | 0.35 | 0.26 | 0.22 | 0.24 |
| 2 | 191891763 | rs3024918 | -0.695 | -2.449 | -0.889 | 0.272 | 0.041 | 0.036 | 0.036 | 0.089 | 0.33 | 0.25 | 0.34 | 0.2 | 0.18 | 0.24 |
| 2 | 191893087 | rs3024912 | 0.464 | -2.336 | -0.754 | 0.338 | 0.024 | 0.124 | 0.124 | 0 | 0.26 | 0.46 | 0.2 | 0.44 | 0.26 | 0.16 |
| 2 | 191894141 | rs3024908 | 0.332 | -2.414 | -0.946 | 0.286 | -0.003 | -0.019 | -0.019 | 0.674 | 0.14 | 0.06 | 0.12 | 0.16 | 0.11 | 0.16 |
| 2 | 191896054 | rs3024935 | 1.233 | -2.719 | -0.906 | 0.175 | 0.016 | 0.068 | 0.068 | 0.002 | 0.12 | 0 | 0.04 | 0 | 0.08 | 0.01 |
| 2 | 191896716 | rs3024896 | 1.007 | -2.713 | -0.623 | 0.367 | -0.02 | -0.078 | -0.078 | 0.02 | 0.2 | 0.08 | 0.15 | 0.33 | 0.21 | 0.15 |
| 2 | 191897540 | rs925847 | -0.476 | -2.836 | -0.126 | 0.321 | -0.012 | 0.013 | 0.013 | 0 | 0.37 | 0.43 | 0.29 | 0.44 | 0.28 | 0.31 |
| 2 | 191900449 | rs3024886 | 0.441 | -2.526 | 0.217 | 0.538 | -0.007 | 0.045 | 0.045 | 0 | 0.22 | 0.29 | 0.46 | 0.41 | 0.2 | 0.2 |
| 2 | 191902758 | rs3821236 | 0.447 | -2.529 | 0.215 | 0.54 | -0.007 | 0.046 | 0.046 | 0 | 0.22 | 0.29 | 0.46 | 0.41 | 0.2 | 0.2 |
| 2 | 191913799 | rs16833215 | -0.879 | -2.423 | 0.252 | 0.404 | -0.015 | 0.021 | 0.021 | 0 | 0.35 | 0.42 | 0.53 | 0.42 | 0.31 | 0.21 |
| 2 | 191917317 | rs4853540 | 1.242 | -2.908 | 0.141 | 0.003 | 0.064 | 0.083 | 0.083 | 0.05 | 0.63 | 0.21 | 0.78 | 0.87 | 0.78 | 0.81 |
| 2 | 191918031 | rs11686127 | 0.824 | -2.955 | 0.252 | -0.017 | 0 | 0 | 0 | 0.72 | 0.1 | 0.11 | 0.06 | 0.1 | 0.05 | 0.1 |
| 2 | 191922841 | rs3024866 | 0.993 | -3.112 | 0.156 | 0.131 | 0.064 | -0.076 | -0.076 | 0 | 0.61 | 0.14 | 0.52 | 0.52 | 0.74 | 0.68 |
| 2 | 191931464 | rs1517352 | 2.06 | -2.975 | -0.122 | 0.195 | -0.044 | -0.141 | -0.141 | 0 | 0.41 | 0.04 | 0.43 | 0.52 | 0.6 | 0.62 |
| 2 | 191935804 | rs10168266 | 0.416 | -2.832 | 0.075 | 0.294 | 0.002 | -0.02 | -0.02 | 0 | 0.21 | 0.14 | 0.35 | 0.33 | 0.18 | 0.18 |
| 2 | 191939187 | rs2459611 | 0.19 | -3.389 | 0.284 | 0.171 | -0.024 | -0.104 | -0.104 | 0 | 0.8 | 0.36 | 0.88 | 1 | 0.87 | 0.99 |
| 2 | 191940451 | rs7601754 | 1.533 | -3.349 | 0.193 | 0.184 | 0.046 | 0.069 | 0.069 | 1.121 | 0.67 | 0.15 | 0.79 | 0.86 | 0.82 | 0.88 |
| 2 | 191943742 | rs11889341 | 0.051 | -3.153 | 0.262 | 0.364 | 0.011 | -0.037 | -0.037 | 0.64 | 0.26 | 0.1 | 0.36 | 0.34 | 0.23 | 0.24 |
| 2 | 191953389 | rs12990918 | 0.174 | -3.106 | 0.679 | 0.273 | -0.016 | -0.09 | -0.09 | 0.429 | 0.14 | 0.37 | 0.09 | 0 | 0.1 | 0.01 |
| 2 | 191954852 | rs10931481 | 1.423 | -2.563 | 0.382 | 1.065 | 0.002 | -0.002 | -0.002 | 0.121 | 0.59 | 0.64 | 0.53 | 0.51 | 0.69 | 0.64 |
| 2 | 191964633 | rs7574865 | 0.333 | -1.947 | 0.447 | 1.16 | 0.014 | -0.037 | -0.037 | 0.035 | 0.73 | 0.88 | 0.64 | 0.65 | 0.77 | 0.7 |
| 2 | 191968420 | rs13407419 | -0.478 | -1.92 | 0.443 | 1.176 | -0.001 | -0.001 | -0.001 | 0.648 | 0.11 | 0.11 | 0.08 | 0.1 | 0.08 | 0.09 |
| 2 | 191970120 | rs7582694 | 0.644 | -1.572 | 0.375 | 1.195 | 0.015 | -0.026 | -0.026 | 0.239 | 0.73 | 0.82 | 0.63 | 0.65 | 0.77 | 0.7 |
| 2 | 191973563 | rs6752770 | -1.265 | -1.381 | 0.894 | 1.118 | -0.013 | -0.019 | -0.019 | 0.713 | 0.3 | 0.41 | 0.29 | 0.22 | 0.28 | 0.23 |
| 2 | 191979359 | rs4073699 | 1.114 | -1.158 | 1.029 | 1.107 | 0.027 | 0.047 | 0.047 | 0.156 | 0.54 | 0.28 | 0.69 | 0.71 | 0.65 | 0.77 |
| 2 | 191989356 | rs6738544 | 0.911 | -0.922 | 0.906 | 1.239 | -0.007 | -0.023 | -0.023 | 2.135 | 0.58 | 0.46 | 0.74 | 0.74 | 0.66 | 0.81 |

| | | | | | | | | | | | | | | | | |
|---|---|---|---|---|---|---|---|---|---|---|---|---|---|---|---|---|
| 2 | 191991459 | rs4555370 | 0.79 | -0.93 | 1.06 | 1.235 | 0 | 0 | 0 | 0.276 | 0.59 | 0.59 | 0.75 | 0.74 | 0.66 | 0.81 |
| 2 | 191993626 | rs6705162 | -1.114 | -0.949 | 1.137 | 1.225 | 0.002 | 0.052 | 0.052 | 0.028 | 0.39 | 0.21 | 0.23 | 0.26 | 0.34 | 0.19 |
| 2 | 191996518 | rs1551443 | 0.905 | -0.949 | 1.114 | 1.224 | -0.001 | -0.003 | -0.003 | 0 | 0.58 | 0.56 | 0.73 | 0.71 | 0.66 | 0.78 |
| 2 | 192001538 | rs6744380 | 1.299 | -0.865 | 0.503 | 1.021 | 0.003 | 0.002 | 0.002 | 0 | 0.52 | 0.46 | 0.68 | 0.45 | 0.65 | 0.64 |
| 2 | 192002538 | rs2356350 | 1.001 | -0.597 | -0.279 | 1.074 | 0.016 | -0.007 | -0.007 | 0 | 0.55 | 0.67 | 0.7 | 0.48 | 0.63 | 0.67 |
| 2 | 192008749 | rs10189819 | -0.982 | -0.498 | -0.145 | 1.019 | -0.015 | -0.038 | -0.038 | 0 | 0.27 | 0.34 | 0.2 | 0.1 | 0.17 | 0.18 |
| 2 | 192009455 | rs11685878 | 1.284 | -0.229 | -0.816 | 1.312 | 0.031 | 0.006 | 0.006 | 0.872 | 0.52 | 0.3 | 0.64 | 0.48 | 0.6 | 0.64 |
| 2 | 192009517 | rs12991409 | -0.457 | -0.35 | -0.911 | 1.273 | -0.001 | -0.001 | -0.001 | 1.157 | 0.1 | 0.11 | 0.05 | 0.18 | 0.06 | 0.06 |
| 2 | 192009652 | rs4853546 | 0.576 | -0.29 | -0.936 | 1.337 | 0 | -0.002 | -0.002 | 0 | 0.62 | 0.38 | 0.68 | 0.6 | 0.67 | 0.76 |
| 2 | 192010189 | rs1031509 | 0.394 | -0.302 | -0.704 | 1.526 | -0.002 | 0.007 | 0.007 | 7.876 | 0.68 | 0.39 | 0.72 | 0.59 | 0.73 | 0.78 |
| 2 | 192013921 | rs10497711 | -0.466 | -0.212 | -0.712 | 1.424 | 0.007 | -0.008 | -0.008 | 0 | 0.1 | 0.09 | 0.05 | 0.17 | 0.06 | 0.06 |
| 2 | 192015072 | rs7572482 | 1.114 | -0.215 | -0.691 | 1.425 | 0.019 | 0.009 | 0.009 | 3.037 | 0.58 | 0.3 | 0.67 | 0.39 | 0.66 | 0.69 |
| 2 | 192020729 | rs13022931 | -0.591 | -0.216 | -0.717 | 1.457 | 0.006 | -0.008 | -0.008 | 0 | 0.1 | 0.08 | 0.05 | 0.17 | 0.06 | 0.06 |
| 2 | 192024904 | rs13001658 | 0.6 | 0.318 | -0.508 | 1.371 | 0 | 0.019 | 0.019 | 8.981 | 0.68 | 0.41 | 0.71 | 0.56 | 0.7 | 0.74 |
| 2 | 192031227 | rs4853550 | 1.043 | -0.009 | -0.341 | 1.615 | -0.014 | 0.039 | 0.039 | 0.523 | 0.77 | 0.38 | 0.78 | 0.41 | 0.8 | 0.76 |
| 2 | 192031263 | rs4853551 | 1.31 | -0.172 | 0.9 | 1.835 | 0 | 0.035 | 0.035 | 0.016 | 0.88 | 0.61 | 0.92 | 0.88 | 0.93 | 0.93 |
| 2 | 192038624 | rs6434450 | -2.327 | -0.302 | 0.22 | 1.818 | 0.017 | -0.048 | -0.048 | 0 | 0.34 | 0.87 | 0.25 | 0.42 | 0.25 | 0.24 |
| 2 | 192038738 | rs7595886 | -1.87 | -0.354 | 1.024 | 1.956 | -0.012 | 0.038 | 0.038 | 0.266 | 0.22 | 0.61 | 0.15 | 0.37 | 0.13 | 0.21 |
| 2 | 192045160 | rs7606940 | -2.864 | -0.449 | 1.108 | 2.27 | 0.011 | -0.024 | -0.024 | 0 | 0.33 | 0.73 | 0.24 | 0.46 | 0.25 | 0.24 |
| 2 | 192046201 | rs7584366 | -2.38 | -0.503 | 1.082 | 2.151 | -0.028 | -0.108 | -0.108 | 0 | 0.19 | 0.64 | 0.12 | 0 | 0.12 | 0.03 |
| 2 | 192047465 | rs4145348 | -1.613 | -0.43 | 0.835 | 2.276 | -0.013 | -0.04 | -0.04 | 0 | 0.13 | 0.05 | 0.11 | 0.41 | 0.12 | 0.2 |
| 2 | 192055486 | rs17585427 | -2.466 | -0.809 | 1.754 | 2.869 | -0.028 | 0.131 | 0.131 | 0 | 0.18 | 0.34 | 0.14 | 0.55 | 0.12 | 0.2 |
| 2 | 192060780 | rs6709982 | 1.347 | -1.571 | 1.801 | 2.261 | -0.019 | 0.019 | 0.019 | 0 | 0.7 | 0.64 | 0.82 | 0.6 | 0.8 | 0.7 |
| 2 | 192062401 | rs6756937 | -0.636 | -2.539 | 1.071 | 1.253 | 0.007 | -0.027 | -0.027 | 0.136 | 0.23 | 0.24 | 0.14 | 0 | 0.15 | 0.06 |
| 2 | 192072088 | rs4853555 | 1.763 | -2.524 | 0.85 | 1.301 | 0.056 | 0.031 | 0.031 | 0.365 | 0.48 | 0.28 | 0.66 | 0.6 | 0.62 | 0.66 |
| 2 | 192085141 | rs6434455 | 1.183 | -2.387 | 0.28 | 0.829 | 0.063 | 0.022 | 0.022 | 0 | 0.57 | 0.85 | 0.74 | 0.65 | 0.68 | 0.69 |
| 2 | 192086682 | rs13015187 | -0.44 | -2.044 | -0.488 | 0.307 | -0.041 | -0.017 | -0.017 | 3.917 | 0.35 | 0.06 | 0.27 | 0.31 | 0.34 | 0.34 |
| 2 | 192086890 | rs6728759 | 0.63 | -1.457 | 1.123 | 0.421 | 0.015 | 0.022 | 0.022 | 0.171 | 0.46 | 0.37 | 0.6 | 0.66 | 0.61 | 0.72 |
| 2 | 192091308 | rs17439247 | 0.868 | -1.11 | 0.674 | 0.428 | -0.099 | -0.018 | -0.018 | 0 | 0.39 | 0.09 | 0.53 | 0.64 | 0.56 | 0.55 |
| 2 | 192101498 | rs10445783 | 0.654 | -1.162 | 0.916 | 0.64 | 0.028 | 0.034 | 0.034 | 0.377 | 0.48 | 0.09 | 0.6 | 0.63 | 0.66 | 0.56 |
| 2 | 192109558 | rs6704784 | 0.615 | -1.172 | 0.909 | 0.645 | 0.038 | 0.035 | 0.035 | 0.477 | 0.48 | 0.09 | 0.6 | 0.63 | 0.67 | 0.56 |
| 2 | 192118047 | rs1823913 | 0.592 | -1.164 | 0.935 | 0.609 | 0.05 | 0.074 | 0.074 | 0.373 | 0.48 | 0.1 | 0.6 | 0.65 | 0.67 | 0.57 |
| 2 | 192123804 | rs4853462 | 0.671 | -1.176 | 1.015 | 0.602 | 0.048 | 0.048 | 0.048 | 0.568 | 0.47 | 0.1 | 0.61 | 0.63 | 0.67 | 0.56 |
| 2 | 192144038 | rs10497713 | -0.128 | 0.138 | 2.146 | 0.746 | 0 | 0.018 | 0.018 | 0 | 0.56 | 0.37 | 0.39 | 0.33 | 0.42 | 0.54 |
| 2 | 192153214 | rs17346475 | -0.065 | 0.128 | 2.021 | 0.741 | -0.006 | 0.072 | 0.072 | 0 | 0.54 | 0.18 | 0.38 | 0.33 | 0.41 | 0.54 |
| 2 | 192160875 | rs4853575 | 0.19 | 0.173 | 2.119 | 0.766 | -0.005 | 0.009 | 0.009 | 0 | 0.41 | 0.33 | 0.59 | 0.66 | 0.58 | 0.47 |
| 2 | 192170980 | rs10931497 | 0.164 | 0.177 | 2.096 | 0.768 | -0.005 | 0.01 | 0.01 | 0 | 0.42 | 0.33 | 0.59 | 0.65 | 0.58 | 0.47 |
| 2 | 192190800 | rs4613316 | -0.248 | 0.427 | 2.192 | 0.868 | 0 | 0.029 | 0.029 | 0.009 | 0.55 | 0.7 | 0.41 | 0.34 | 0.4 | 0.53 |
| 2 | 192197929 | rs6722183 | -0.225 | 0.435 | 2.18 | 0.872 | 0.001 | 0.018 | 0.018 | 0.033 | 0.55 | 0.67 | 0.4 | 0.33 | 0.41 | 0.53 |
| 2 | 192204885 | rs11892998 | 0.026 | -0.062 | 2.27 | 0.869 | 0.019 | 0.028 | 0.028 | 0 | 0.5 | 0.63 | 0.4 | 0.33 | 0.37 | 0.53 |
| 2 | 192224782 | rs2138914 | 0.304 | 0.015 | 2.183 | 0.944 | 0.025 | 0.087 | 0.087 | 0 | 0.47 | 0.19 | 0.37 | 0.32 | 0.4 | 0.52 |
| 2 | 192229592 | rs13431098 | 0.04 | 0.203 | 2.83 | 1.426 | 0 | 0.003 | 0.003 | 0 | 0.59 | 0.45 | 0.6 | 0.35 | 0.58 | 0.6 |
| 2 | 192230877 | rs12619838 | -2.797 | -0.018 | 3.936 | 1.952 | 0.004 | 0.06 | 0.06 | 0.155 | 0.08 | 0.33 | 0.12 | 0.06 | 0.01 | 0.06 |
| 2 | 192231585 | rs6741861 | -3.286 | -0.024 | 4.041 | 1.948 | -0.019 | 0.031 | 0.031 | 1.769 | 0.1 | 0.57 | 0.15 | 0.06 | 0.01 | 0.06 |
| 2 | 192250568 | rs13026888 | 3.143 | 0.013 | 4.072 | 1.95 | -0.002 | 0.007 | 0.007 | 2.376 | 0.9 | 0.38 | 0.84 | 0.94 | 0.96 | 0.94 |
| 2 | 192272933 | rs2271768 | -3.395 | 0.001 | 4.002 | 2.071 | 0.016 | 0.027 | 0.027 | 1.9 | 0.11 | 0.55 | 0.15 | 0.06 | 0.04 | 0.06 |
| 2 | 192276192 | rs4344879 | 3.175 | 0.002 | 3.875 | 2.072 | -0.014 | -0.006 | -0.006 | 0.743 | 0.89 | 0.35 | 0.84 | 0.94 | 0.96 | 0.94 |
| 2 | 192289938 | rs12623842 | 3.36 | -0.007 | 3.758 | 2.067 | -0.02 | -0.015 | -0.015 | 0 | 0.87 | 0.26 | 0.84 | 0.94 | 0.96 | 0.94 |
| 2 | 192316038 | rs7560131 | 2.165 | -0.172 | 3.186 | 1.923 | 0.022 | -0.016 | -0.016 | 0 | 0.92 | 0.71 | 0.96 | 1 | 0.98 | 0.96 |
| 2 | 192322509 | rs4853597 | 2.56 | -0.184 | 2.837 | 2.001 | -0.024 | 0.027 | 0.027 | 4.644 | 0.86 | 0.24 | 0.81 | 0.78 | 0.97 | 0.93 |
| 2 | 192327964 | rs7571217 | -2.016 | 0.244 | 1.203 | 2.167 | 0 | 0.273 | 0.273 | 1.118 | 0.11 | 0.49 | 0.18 | 0.21 | 0.04 | 0.05 |
| 2 | 192335294 | rs7604359 | -1.571 | 0.457 | 0.861 | 1.768 | 0.048 | 0.271 | 0.271 | 2.372 | 0.07 | 0.42 | 0.07 | 0.22 | 0.03 | 0.02 |
| 2 | 192339531 | rs16833762 | -1.033 | 0.896 | 0.294 | 1.277 | 0.007 | -0.012 | -0.012 | 0 | 0.05 | 0.07 | 0.04 | 0.01 | 0.09 | 0.05 |
| 2 | 192339802 | rs4502362 | -1.848 | 0.91 | 0.376 | 1.292 | 0.002 | -0.001 | -0.001 | 0 | 0.06 | 0.11 | 0.07 | 0.05 | 0.07 | 0.05 |
| 2 | 192340847 | rs9750499 | -0.24 | 0.613 | 0.252 | 1.151 | -0.008 | -0.004 | -0.004 | 1.363 | 0.4 | 0.44 | 0.4 | 0.43 | 0.33 | 0.4 |
| 2 | 192340920 | rs16833765 | -0.102 | 0.407 | 0.315 | 1.011 | -0.001 | -0.009 | -0.009 | 0 | 0.32 | 0.34 | 0.25 | 0.15 | 0.26 | 0.26 |
| 2 | 192341466 | rs9751523 | 0.481 | 0.833 | -0.358 | 0.64 | -0.036 | 0.024 | 0.024 | 0.008 | 0.19 | 0.07 | 0.13 | 0.15 | 0.2 | 0.19 |
| 2 | 192341763 | rs9750812 | -0.76 | 0.573 | -0.212 | 0.866 | 0.002 | -0.002 | -0.002 | 0 | 0.6 | 0.52 | 0.44 | 0.48 | 0.54 | 0.64 |
| 2 | 192345577 | rs4525637 | -0.328 | 0.013 | 0.13 | 0.876 | -0.019 | 0.004 | 0.004 | 0 | 0.32 | 0.35 | 0.49 | 0.64 | 0.25 | 0.49 |
| 2 | 192349702 | rs16833768 | 0.312 | -0.094 | 0.1 | 0.778 | 0.001 | -0.002 | -0.002 | 0 | 0.27 | 0.21 | 0.43 | 0.5 | 0.23 | 0.32 |
| 2 | 192350205 | rs1517848 | 0.258 | -0.145 | 0.026 | 0.724 | -0.029 | -0.01 | -0.01 | 4.766 | 0.63 | 0.53 | 0.49 | 0.31 | 0.72 | 0.49 |
| 2 | 192358445 | rs5015847 | -0.142 | 0 | 0.037 | 0.772 | -0.045 | 0.077 | 0.077 | 0 | 0.73 | 0.54 | 0.61 | 0.5 | 0.83 | 0.79 |
| 2 | 192364713 | rs12466214 | 0.081 | 0.196 | -0.93 | 0.35 | -0.012 | 0.064 | 0.064 | 0 | 0.55 | 0.18 | 0.49 | 0.74 | 0.52 | 0.7 |
| 2 | 192366298 | rs9653314 | 0.288 | 0.419 | -0.207 | 0.172 | -0.003 | -0.003 | -0.003 | 1.134 | 0.32 | 0.45 | 0.38 | 0.23 | 0.3 | 0.26 |
| 2 | 192369816 | rs10177310 | -1.041 | 0.341 | -0.191 | 0.128 | 0.025 | 0.202 | 0.202 | 1.574 | 0.07 | 0.38 | 0.19 | 0.17 | 0.05 | 0.11 |
| 2 | 192377327 | rs10183577 | -0.244 | 0.436 | -0.105 | -0.406 | -0.001 | -0.014 | -0.014 | 0 | 0.63 | 0.67 | 0.71 | 0.89 | 0.6 | 0.78 |
| 2 | 192377470 | rs17416887 | 1.105 | 0.281 | -0.202 | -0.413 | -0.036 | 0.091 | 0.091 | 0.415 | 0.24 | 0.07 | 0.19 | 0.06 | 0.25 | 0.19 |
| 2 | 192381221 | rs16833795 | -0.086 | 0.308 | -0.36 | -0.173 | -0.016 | -0.027 | -0.027 | 0 | 0.06 | 0.01 | 0.04 | 0.15 | 0.07 | 0.2 |
| 2 | 192381471 | rs13411587 | 0.354 | 0.303 | -0.205 | -0.312 | -0.025 | 0.015 | 0.015 | 0 | 0.08 | 0.03 | 0.08 | 0.05 | 0.15 | 0.04 |

| | | | | | | | | | | | | | | | | |
|---|---|---|---|---|---|---|---|---|---|---|---|---|---|---|---|---|
| 2 | 192381934 | rs2356656 | 0.278 | 0.258 | 0.196 | -0.335 | 0.087 | -0.026 | -0.026 | 0 | 0.89 | 0.69 | 0.9 | 0.95 | 0.85 | 0.96 |
| 2 | 192383966 | rs2356653 | 0.579 | -0.104 | -0.036 | -0.661 | -0.001 | -0.034 | -0.034 | 0.014 | 0.51 | 0.55 | 0.59 | 0.88 | 0.49 | 0.68 |
| 2 | 192389932 | rs10208726 | 0.653 | 0.182 | -0.168 | -0.41 | -0.016 | 0.008 | 0.008 | 0 | 0.08 | 0.03 | 0.08 | 0 | 0.14 | 0.02 |
| 2 | 192391171 | rs4853610 | 1.513 | -0.552 | -0.139 | -0.509 | -0.015 | 0.06 | 0.06 | 0 | 0.23 | 0.13 | 0.2 | 0.05 | 0.26 | 0.2 |
| 2 | 192391282 | rs4853611 | -0.231 | -0.455 | -0.024 | -0.605 | -0.007 | 0.122 | 0.122 | 0 | 0.45 | 0.28 | 0.37 | 0.13 | 0.47 | 0.31 |
| 2 | 192395165 | rs883405 | 0.698 | -0.848 | 0.283 | 0.221 | -0.005 | 0.003 | 0.003 | 0 | 0.43 | 0.36 | 0.42 | 0.44 | 0.46 | 0.34 |
| 2 | 192397191 | rs2203748 | -0.117 | -0.756 | 0.144 | 0.312 | 0.007 | 0.01 | 0.01 | 0 | 0.12 | 0.1 | 0.21 | 0.51 | 0.08 | 0.34 |
| 2 | 192403359 | rs11690795 | 0.76 | -0.899 | 0.168 | 0.273 | -0.037 | 0.074 | 0.074 | 0 | 0.42 | 0.21 | 0.4 | 0.3 | 0.48 | 0.37 |
| 2 | 192405450 | rs1517843 | 0.86 | -0.764 | -0.031 | 0.408 | -0.03 | 0.043 | 0.043 | 0 | 0.37 | 0.22 | 0.28 | 0.28 | 0.41 | 0.33 |
| 2 | 192408469 | rs7561808 | -0.288 | -0.525 | -0.472 | 1.2 | 0.021 | -0.01 | -0.01 | 0 | 0.1 | 0.07 | 0.17 | 0.55 | 0.05 | 0.31 |
| 2 | 192417071 | rs4476296 | -0.395 | -0.443 | -0.5 | 1.362 | 0.025 | -0.063 | -0.063 | 0 | 0.14 | 0.08 | 0.18 | 0.52 | 0.1 | 0.23 |
| 2 | 192417747 | rs4586589 | -0.424 | -0.568 | -0.126 | 0.905 | -0.003 | -0.006 | -0.006 | 1.452 | 0.46 | 0.47 | 0.54 | 0.66 | 0.42 | 0.61 |
| 2 | 192421268 | rs10178390 | -0.04 | -0.559 | -0.492 | 1.026 | 0.024 | -0.059 | -0.059 | 0.284 | 0.14 | 0.08 | 0.19 | 0.55 | 0.1 | 0.24 |
| 2 | 192422875 | rs7604362 | 0.76 | -0.606 | -0.702 | 0.713 | -0.064 | 0.03 | 0.03 | 0 | 0.35 | 0.13 | 0.24 | 0.27 | 0.38 | 0.3 |
| 2 | 192423103 | rs6434482 | -0.11 | -0.162 | -0.108 | 1.296 | -0.025 | 0.284 | 0.284 | 0.311 | 0.88 | 0.67 | 0.8 | 0.51 | 0.93 | 0.83 |
| 2 | 192428701 | rs13425331 | -0.204 | 0.062 | -0.682 | 1.172 | -0.031 | 0.126 | 0.126 | 0.072 | 0.38 | 0.17 | 0.43 | 0.12 | 0.44 | 0.35 |
| 2 | 192429183 | rs7574064 | 0.752 | 0.442 | 0.005 | 1.727 | -0.015 | 0.036 | 0.036 | 0 | 0.23 | 0.14 | 0.28 | 0.11 | 0.26 | 0.28 |
| 2 | 192435264 | rs4640333 | -0.694 | 1.001 | 0.411 | 1.611 | -0.001 | 0.004 | 0.004 | 0 | 0.62 | 0.45 | 0.54 | 0.5 | 0.64 | 0.63 |
| 2 | 192436267 | rs4853614 | 0.307 | 1.067 | 0.211 | 1.322 | -0.023 | 0.016 | 0.016 | 0 | 0.31 | 0.23 | 0.3 | 0.31 | 0.36 | 0.26 |
| 2 | 192442429 | rs6713885 | -0.024 | 0.835 | 0.031 | 1.226 | 0.018 | 0.05 | 0.05 | 0 | 0.31 | 0.18 | 0.22 | 0.15 | 0.28 | 0.35 |
| 2 | 192450588 | rs13013800 | 0.08 | 0.948 | -0.568 | 1.003 | 0.001 | -0.032 | -0.032 | 0 | 0.3 | 0.17 | 0.24 | 0.4 | 0.27 | 0.25 |
| 2 | 192456857 | rs4853619 | 0.355 | 1.37 | -0.179 | 1.049 | -0.005 | -0.005 | -0.005 | 0 | 0.67 | 0.68 | 0.67 | 0.6 | 0.6 | 0.59 |
| 2 | 192465598 | rs6706693 | -1.425 | 1.429 | 0.364 | 0.676 | 0.01 | -0.008 | -0.008 | 0.524 | 0.25 | 0.34 | 0.28 | 0.2 | 0.32 | 0.28 |
| 2 | 192469816 | rs4134381 | -1.222 | 1.454 | 0.441 | 0.662 | -0.001 | 0.001 | 0.001 | 0.655 | 0.16 | 0.19 | 0.19 | 0.2 | 0.15 | 0.13 |
| 2 | 192470385 | rs4318338 | 1.89 | 1.327 | 0.988 | 0.579 | 0.018 | -0.016 | -0.016 | 0 | 0.74 | 0.6 | 0.7 | 0.8 | 0.7 | 0.69 |
| 2 | 192473543 | rs4853629 | -1.538 | 1.144 | 1.721 | 0.553 | -0.005 | 0.133 | 0.133 | 2.17 | 0.09 | 0.34 | 0.17 | 0.2 | 0.06 | 0.11 |
| 2 | 192474316 | rs11683305 | -2.171 | 1.448 | 1.542 | 0.452 | -0.02 | 0 | 0 | 0 | 0.2 | 0.18 | 0.16 | 0 | 0.26 | 0.21 |
| 2 | 192479007 | rs3935827 | 2.465 | 1.399 | 1.749 | 0.461 | 0.044 | -0.033 | -0.033 | 0 | 0.73 | 0.52 | 0.68 | 0.81 | 0.68 | 0.75 |
| 2 | 192479674 | rs4401164 | 1.188 | 1.423 | 1.789 | 0.461 | -0.001 | 0.095 | 0.095 | 0 | 0.92 | 0.81 | 0.84 | 0.81 | 0.93 | 0.89 |
| 2 | 192479789 | rs4438431 | -1.964 | 1.445 | 1.574 | 0.464 | -0.031 | 0.021 | 0.021 | 0 | 0.18 | 0.14 | 0.16 | 0 | 0.25 | 0.2 |
| 2 | 192490640 | rs11899661 | 2.376 | 1.435 | 1.709 | 0.472 | 0.041 | -0.03 | -0.03 | 0.888 | 0.73 | 0.53 | 0.68 | 0.81 | 0.68 | 0.74 |
| 2 | 192491453 | rs6434494 | 0.822 | 1.38 | 1.146 | 0.683 | -0.005 | 0.026 | 0.026 | 0 | 0.93 | 0.91 | 0.86 | 0.81 | 0.94 | 0.89 |
| 2 | 192506225 | rs6434497 | 1.768 | 1.3 | 1.177 | 0.687 | 0.005 | -0.003 | -0.003 | 0 | 0.78 | 0.77 | 0.7 | 0.81 | 0.71 | 0.76 |
| 2 | 192510265 | rs5008133 | 1.802 | 1.205 | 1.293 | 0.657 | 0.018 | -0.014 | -0.014 | 0 | 0.77 | 0.58 | 0.69 | 0.81 | 0.72 | 0.76 |
| 2 | 192514750 | rs13006848 | -1.254 | 1.355 | 1.172 | 0.514 | -0.016 | 0.007 | 0.007 | 0 | 0.09 | 0.06 | 0.11 | 0 | 0.19 | 0.06 |
| 2 | 192515992 | rs10931512 | 1.614 | 0.696 | 1.876 | 1.036 | -0.034 | 0.082 | 0.082 | 0 | 0.83 | 0.63 | 0.71 | 0.73 | 0.88 | 0.74 |
| 2 | 192519227 | rs12617890 | -1.005 | 0.644 | 1.953 | 1.042 | -0.008 | 0.004 | 0.004 | 0 | 0.07 | 0.13 | 0.04 | 0.07 | 0.05 | 0.04 |
| 2 | 192521641 | rs4467214 | 1.274 | 0.576 | 2.22 | 1.359 | 0.028 | 0.106 | 0.106 | 0 | 0.8 | 0.56 | 0.64 | 0.63 | 0.76 | 0.73 |
| 2 | 192525536 | rs12472343 | 2.305 | 0.373 | 1.758 | 1.214 | -0.001 | -0.091 | -0.091 | 0.831 | 0.67 | 0.08 | 0.57 | 0.55 | 0.7 | 0.68 |
| 2 | 192527407 | rs7579713 | -0.947 | 0.367 | 2.152 | 1.051 | -0.016 | -0.002 | -0.002 | 0.004 | 0.08 | 0.2 | 0.04 | 0.07 | 0.05 | 0.04 |
| 2 | 192529395 | rs13019682 | -0.947 | 0.367 | 2.152 | 1.051 | -0.016 | -0.002 | -0.002 | 0 | 0.08 | 0.2 | 0.04 | 0.07 | 0.05 | 0.04 |
| 2 | 192530721 | rs13021612 | -0.947 | 0.433 | 2.152 | 1.051 | -0.013 | -0.002 | -0.002 | 0 | 0.08 | 0.2 | 0.04 | 0.07 | 0.05 | 0.04 |
| 2 | 192532017 | rs4592792 | -2.103 | 0.382 | 2.603 | 1.081 | -0.018 | -0.007 | -0.007 | 0 | 0.16 | 0.39 | 0.16 | 0.17 | 0.1 | 0.07 |
| 2 | 192560825 | rs10192358 | 2.014 | 0.478 | 2.688 | 2.264 | 0.017 | 0.174 | 0.174 | 0 | 0.81 | 0.53 | 0.68 | 0.53 | 0.77 | 0.7 |
| 2 | 192561207 | rs12996017 | -0.137 | 0.42 | 2.484 | 2.264 | 0.001 | -0.001 | -0.001 | 0 | 0.07 | 0.06 | 0.03 | 0.11 | 0.05 | 0.03 |
| 2 | 192572805 | rs6730901 | -0.616 | 0.972 | 1.987 | 1.806 | -0.026 | 0.032 | 0.032 | 0 | 0.12 | 0.06 | 0.17 | 0.07 | 0.17 | 0.08 |
| 2 | 192582719 | rs11682444 | -1.253 | 1.131 | 2.413 | 2.185 | 0.006 | 0.018 | 0.018 | 0 | 0.09 | 0.11 | 0.17 | 0.23 | 0.15 | 0.1 |
| 2 | 192586602 | rs7596512 | -2.502 | 1.089 | 2.711 | 1.969 | 0.03 | -0.051 | -0.051 | 0 | 0.29 | 0.48 | 0.33 | 0.2 | 0.36 | 0.17 |
| 2 | 192588465 | rs6434503 | -1.872 | 0.86 | 2.891 | 2.006 | -0.02 | 0.148 | 0.148 | 1.232 | 0.11 | 0.47 | 0.17 | 0.22 | 0.06 | 0.2 |
| 2 | 192594975 | rs10201681 | 0.579 | 0.731 | 1.402 | 1.095 | 0 | 0.001 | 0.001 | 0 | 0.55 | 0.46 | 0.64 | 0.69 | 0.55 | 0.65 |
| 2 | 192598953 | rs4853632 | -0.331 | 0.142 | 0.893 | 0.767 | -0.061 | 0.002 | 0.002 | 0 | 0.24 | 0.46 | 0.18 | 0.24 | 0.16 | 0.24 |
| 2 | 192601140 | rs6434504 | 1.271 | 0.601 | 0.752 | 0.738 | 0.012 | 0.048 | 0.048 | 0 | 0.52 | 0.32 | 0.62 | 0.65 | 0.53 | 0.63 |
| 2 | 192604268 | rs6720388 | 2.103 | 0.679 | -0.074 | 0.67 | -0.025 | -0.011 | -0.011 | 0 | 0.37 | 0.14 | 0.47 | 0.64 | 0.34 | 0.54 |
| 2 | 192605627 | rs10208071 | -0.092 | 0.71 | 0.181 | 0.583 | -0.031 | 0.052 | 0.052 | 0 | 0.73 | 0.82 | 0.79 | 0.89 | 0.66 | 0.85 |
| 2 | 192614107 | rs6739080 | -0.169 | 0.344 | 0.61 | 1.019 | -0.038 | 0.019 | 0.019 | 0 | 0.22 | 0.61 | 0.18 | 0.29 | 0.13 | 0.21 |
| 2 | 192615315 | rs12466549 | 1.929 | 0.828 | 0.112 | 0.841 | 0.013 | -0.058 | -0.058 | 0.771 | 0.47 | 0.14 | 0.52 | 0.56 | 0.39 | 0.59 |
| 2 | 192616165 | rs6718527 | -0.727 | 0.056 | 1.166 | 1.284 | -0.056 | 0.118 | 0.118 | 0 | 0.24 | 0.42 | 0.17 | 0.41 | 0.15 | 0.25 |
| 2 | 192618369 | rs6434508 | -0.016 | 0.145 | 0.778 | 1.399 | -0.021 | 0.054 | 0.054 | 0 | 0.81 | 0.77 | 0.84 | 0.58 | 0.87 | 0.79 |
| 2 | 192620452 | rs6706341 | -1.596 | -0.018 | 0.912 | 0.664 | -0.017 | -0.071 | -0.071 | 0 | 0.12 | 0.36 | 0.11 | 0.01 | 0.09 | 0.06 |
| 2 | 192623122 | rs7368902 | 0.23 | -0.114 | 1.282 | 1.241 | -0.026 | 0.121 | 0.121 | 0 | 0.16 | 0.21 | 0.28 | 0.49 | 0.09 | 0.23 |
| 2 | 192636163 | rs10173668 | 0.334 | 0.193 | 0.667 | 0.623 | -0.078 | 0.069 | 0.069 | 0 | 0.75 | 0.91 | 0.8 | 0.92 | 0.65 | 0.85 |
| 2 | 192636199 | rs4527158 | 2.025 | 0.169 | 0.572 | 0.57 | 0.004 | 0.016 | 0.016 | 0.029 | 0.52 | 0.32 | 0.42 | 0.47 | 0.49 | 0.61 |
| 2 | 192640624 | rs7593737 | 1.467 | 0.395 | 0.564 | 0.425 | -0.012 | -0.009 | -0.009 | 1.618 | 0.58 | 0.37 | 0.5 | 0.51 | 0.56 | 0.62 |
| 2 | 192643022 | rs11890239 | 0.815 | -0.074 | 0.348 | 0.502 | -0.074 | 0.112 | 0.112 | 0 | 0.74 | 0.55 | 0.65 | 0.51 | 0.86 | 0.69 |
| 2 | 192653312 | rs4325700 | 0.261 | 0.126 | 0.233 | 0.622 | -0.014 | 0.062 | 0.062 | 0 | 0.17 | 0.19 | 0.24 | 0.51 | 0.1 | 0.28 |
| 2 | 192658185 | rs4483984 | 2.077 | -0.037 | -0.572 | 0.287 | -0.081 | 0.043 | 0.043 | 0.012 | 0.31 | 0.02 | 0.26 | 0.17 | 0.34 | 0.34 |
| 2 | 192666465 | rs6708269 | 0.028 | 0.707 | 0.837 | 0.652 | -0.011 | 0.008 | 0.008 | 0 | 0.78 | 0.86 | 0.75 | 0.89 | 0.71 | 0.86 |
| 2 | 192683003 | rs12693614 | 1.139 | 0.817 | 0.413 | 0.623 | 0.003 | 0.005 | 0.005 | 0.029 | 0.61 | 0.5 | 0.59 | 0.49 | 0.62 | 0.64 |
| 2 | 192689978 | rs4349283 | 1.104 | 0.894 | 0.338 | 0.661 | -0.001 | -0.001 | -0.001 | 0 | 0.62 | 0.46 | 0.58 | 0.49 | 0.62 | 0.64 |

| | | | | | | | | | | | | | | | | |
|---|---|---|---|---|---|---|---|---|---|---|---|---|---|---|---|---|
| 2 | 192694720 | rs7608278 | -1.07 | 0.076 | 0.212 | 1.323 | -0.008 | 0.021 | 0.021 | 0 | 0.25 | 0.32 | 0.35 | 0.58 | 0.19 | 0.31 |
| 2 | 192695107 | rs7569546 | -0.908 | 0.191 | -0.155 | 0.723 | -0.003 | 0.023 | 0.023 | 0 | 0.24 | 0.3 | 0.24 | 0.48 | 0.19 | 0.25 |
| 2 | 192697988 | rs35896030 | -1.15 | 0.464 | -0.196 | 0.268 | -0.021 | -0.05 | -0.05 | 0 | 0.12 | 0.04 | 0.11 | 0.24 | 0.12 | 0.16 |
| 2 | 192704219 | rs4853646 | -1.956 | 0.284 | -0.121 | 0.044 | 0.015 | -0.029 | -0.029 | 0 | 0.22 | 0.1 | 0.15 | 0.28 | 0.16 | 0.25 |
| 2 | 192714781 | rs4241281 | 0.555 | 0.489 | 0.373 | -0.95 | 0.054 | -0.156 | -0.156 | 0 | 0.71 | 0.56 | 0.77 | 1 | 0.64 | 0.86 |
| 2 | 192716848 | rs6434514 | 1.32 | 0.123 | 0.675 | -0.896 | -0.03 | 0.082 | 0.082 | 0 | 0.58 | 0.28 | 0.65 | 0.77 | 0.59 | 0.7 |
| 2 | 192720975 | rs10174723 | 0.127 | 0.298 | 0.869 | -1.088 | 0.051 | -0.148 | -0.148 | 0.044 | 0.75 | 0.58 | 0.8 | 1 | 0.7 | 0.91 |
| 2 | 192743811 | rs7568281 | 2.005 | -0.523 | 0.49 | -0.253 | 0.027 | 0.042 | 0.042 | 1.407 | 0.71 | 0.25 | 0.79 | 0.86 | 0.84 | 0.78 |
| 2 | 192752136 | rs4853647 | -1.101 | 0.167 | 0.458 | 0.346 | -0.013 | -0.026 | -0.026 | 0 | 0.1 | 0.05 | 0.1 | 0.26 | 0.1 | 0.15 |
| 2 | 192770914 | rs13426894 | -1.819 | -0.438 | 1.086 | -0.472 | -0.049 | -0.11 | -0.11 | 0.044 | 0.15 | 0.49 | 0.07 | 0 | 0.04 | 0.06 |
| 2 | 192772592 | rs6710083 | -1.917 | -0.737 | 0.537 | -0.553 | 0.019 | 0.014 | 0.014 | 1.864 | 0.2 | 0.8 | 0.11 | 0.04 | 0.05 | 0.12 |
| 2 | 192776190 | rs7575939 | -1.884 | -0.879 | 0.943 | -0.751 | -0.077 | -0.146 | -0.146 | 0.711 | 0.23 | 0.47 | 0.11 | 0 | 0.09 | 0.12 |
| 2 | 192797427 | rs13413304 | -1.944 | -0.477 | 0.871 | -0.757 | -0.01 | -0.042 | -0.042 | 0 | 0.25 | 0.31 | 0.26 | 0.06 | 0.2 | 0.2 |
| 2 | 192797804 | rs7422307 | -1.606 | -0.44 | 0.609 | -0.711 | 0.018 | 0.05 | 0.05 | 0.059 | 0.15 | 0.08 | 0.16 | 0.01 | 0.12 | 0.15 |
| 2 | 192806206 | rs4630705 | -1.009 | -0.265 | 0.41 | -0.151 | -0.022 | -0.055 | -0.055 | 0 | 0.1 | 0.01 | 0.16 | 0.2 | 0.11 | 0.13 |
| 2 | 192808024 | rs7583281 | 1.277 | 0.138 | 1.077 | -0.723 | -0.006 | 0.061 | 0.061 | 0.226 | 0.68 | 0.78 | 0.73 | 0.91 | 0.67 | 0.78 |
| 2 | 192810074 | rs13001482 | -1.39 | 0.167 | 1.052 | -0.717 | -0.025 | 0.116 | 0.116 | 0.154 | 0.3 | 0.14 | 0.26 | 0.07 | 0.33 | 0.22 |
| 2 | 192813453 | rs10187928 | -1.091 | 0.217 | 1.785 | -0.658 | 0.005 | -0.039 | -0.039 | 0 | 0.24 | 0.23 | 0.24 | 0.07 | 0.27 | 0.14 |
| 2 | 192822383 | rs6747698 | 0.895 | 0.114 | 2.137 | -0.624 | 0.005 | -0.064 | -0.064 | 0 | 0.77 | 0.71 | 0.77 | 0.94 | 0.78 | 0.86 |
| 2 | 192826028 | rs6434520 | 1.45 | 0.011 | 2.46 | -0.646 | -0.01 | -0.063 | -0.063 | 0 | 0.7 | 0.45 | 0.74 | 0.92 | 0.78 | 0.81 |
| 2 | 192843949 | rs6758825 | 1.047 | 0.263 | 1.557 | -0.522 | 0.009 | -0.062 | -0.062 | 0 | 0.77 | 0.68 | 0.77 | 0.92 | 0.78 | 0.84 |
| 2 | 192846499 | rs7600875 | -0.854 | 0.386 | 1.602 | -0.483 | -0.01 | 0.015 | 0.015 | 0 | 0.19 | 0.13 | 0.2 | 0.08 | 0.22 | 0.15 |
| 2 | 192867728 | rs4474831 | 1.115 | 0.041 | 1.388 | 0.613 | -0.027 | 0.2 | 0.2 | 1.128 | 0.85 | 0.48 | 0.71 | 0.57 | 0.87 | 0.8 |
| 2 | 192874825 | rs12612937 | 1.42 | 0.733 | 0.656 | -0.155 | -0.061 | -0.017 | -0.017 | 0 | 0.59 | 0.1 | 0.51 | 0.53 | 0.58 | 0.61 |
| 2 | 192879000 | rs7558259 | 1.433 | 0.832 | 0.53 | -0.505 | -0.012 | 0.045 | 0.045 | 1.823 | 0.52 | 0.18 | 0.48 | 0.56 | 0.5 | 0.55 |
| 2 | 192898315 | rs12465925 | -1.068 | 0.731 | 1.335 | -0.192 | -0.004 | 0.08 | 0.08 | 0 | 0.13 | 0.2 | 0.23 | 0.38 | 0.13 | 0.21 |
| 2 | 192905094 | rs4241288 | -0.796 | 0.714 | 1.313 | -0.07 | -0.001 | 0.026 | 0.026 | 0 | 0.08 | 0.07 | 0.1 | 0.32 | 0.09 | 0.16 |
| 2 | 192918492 | rs6434533 | 1.651 | 0.612 | 2.26 | -0.119 | 0.012 | 0.224 | 0.224 | 0.459 | 0.88 | 0.58 | 0.86 | 0.67 | 0.91 | 0.84 |
| 2 | 192923303 | rs7585423 | -2 | 0.417 | 1.976 | -0.084 | 0.002 | -0.034 | -0.034 | 0 | 0.13 | 0.07 | 0.22 | 0.37 | 0.13 | 0.19 |
| 2 | 192924987 | rs10185068 | 0.26 | 0.616 | 2.193 | -0.388 | 0.03 | -0.075 | -0.075 | 1.888 | 0.73 | 0.56 | 0.78 | 0.94 | 0.65 | 0.81 |
| 2 | 192932791 | rs10931519 | 1.407 | 0.494 | 2.211 | -0.396 | -0.037 | 0.019 | 0.019 | 0 | 0.6 | 0.25 | 0.64 | 0.58 | 0.56 | 0.6 |
| 2 | 192942966 | rs2884024 | -1.441 | 0.467 | 2.62 | -0.392 | 0.009 | 0.04 | 0.04 | 0.037 | 0.07 | 0.24 | 0.14 | 0.07 | 0.03 | 0.03 |
| 2 | 192943798 | rs2356758 | 1.914 | 0.547 | 2.082 | -0.393 | -0.013 | 0.015 | 0.015 | 0.144 | 0.56 | 0.22 | 0.53 | 0.55 | 0.53 | 0.57 |
| 2 | 192949003 | rs13001304 | -0.15 | 0.911 | 1.57 | -0.475 | -0.017 | 0.014 | 0.014 | 0 | 0.23 | 0.19 | 0.16 | 0 | 0.29 | 0.14 |
| 2 | 192949895 | rs13393294 | -1.36 | 0.809 | 1.549 | -0.392 | 0 | -0.041 | -0.041 | 0 | 0.11 | 0.05 | 0.11 | 0.36 | 0.11 | 0.16 |
| 2 | 192953602 | rs7567063 | -1.687 | 0.82 | 1.513 | -0.39 | -0.043 | 0.001 | 0.001 | 0 | 0.39 | 0.28 | 0.4 | 0.44 | 0.46 | 0.4 |
| 2 | 192954320 | rs7556774 | 1.997 | 0.838 | 0.869 | -0.389 | -0.028 | 0 | 0 | 0.325 | 0.52 | 0.1 | 0.51 | 0.51 | 0.51 | 0.57 |
| 2 | 192960408 | rs4419188 | 0.623 | 0.417 | 0.998 | -0.634 | 0.003 | -0.053 | -0.053 | 0 | 0.69 | 0.59 | 0.68 | 0.96 | 0.67 | 0.79 |
| 2 | 192963676 | rs16834182 | -1.831 | -0.334 | 0.794 | -0.042 | -0.003 | -0.03 | -0.03 | 0 | 0.08 | 0.04 | 0.03 | 0.2 | 0.07 | 0.2 |
| 2 | 192965939 | rs16834198 | -1.491 | -0.325 | 0.739 | -0.038 | -0.004 | -0.017 | -0.017 | 0 | 0.07 | 0.04 | 0.03 | 0.18 | 0.07 | 0.2 |
| 2 | 192969237 | rs10166928 | -1.245 | -0.996 | 0.629 | 1.005 | -0.037 | 0.175 | 0.175 | 0 | 0.19 | 0.41 | 0.23 | 0.51 | 0.12 | 0.18 |
| 2 | 192975204 | rs13413029 | -1.676 | -0.873 | 0.009 | 0.981 | 0.015 | -0.024 | -0.024 | 0 | 0.25 | 0.72 | 0.25 | 0.61 | 0.14 | 0.31 |
| 2 | 192981038 | rs6434535 | 0.184 | -0.688 | -0.398 | 0.238 | 0.031 | 0.055 | 0.055 | 0.211 | 0.51 | 0.36 | 0.48 | 0.27 | 0.45 | 0.49 |
| 2 | 192986033 | rs10164776 | 0.66 | -0.806 | -0.953 | -0.166 | -0.08 | 0.114 | 0.114 | 0.339 | 0.34 | 0.13 | 0.24 | 0 | 0.44 | 0.15 |
| 2 | 192989148 | rs13008224 | -0.225 | -0.884 | -0.321 | 0.196 | 0.065 | 0.18 | 0.18 | 0 | 0.08 | 0.34 | 0.17 | 0.26 | 0.07 | 0.22 |
| 2 | 192989689 | rs17367154 | 0.725 | -0.885 | -0.479 | 0.175 | -0.068 | 0.093 | 0.093 | 0.256 | 0.34 | 0.16 | 0.24 | 0 | 0.44 | 0.15 |
| 2 | 192991640 | rs6434538 | -0.11 | -0.86 | -0.682 | 0.251 | 0.005 | 0.094 | 0.094 | 0 | 0.56 | 0.26 | 0.57 | 0.71 | 0.49 | 0.63 |
| 2 | 193000979 | rs4853661 | 0.416 | -0.426 | -0.862 | 0.391 | -0.123 | -0.07 | -0.07 | 2.795 | 0.26 | 0.02 | 0.36 | 0.23 | 0.39 | 0.34 |
| 2 | 193001863 | rs10166176 | -0.11 | -0.426 | -0.557 | 0.421 | 0.04 | 0.086 | 0.086 | 0.006 | 0.08 | 0.27 | 0.16 | 0.2 | 0.07 | 0.16 |
| 2 | 193008931 | rs4853491 | -0.009 | -0.602 | -1.224 | 0.31 | 0.038 | -0.128 | -0.128 | 0 | 0.17 | 0.01 | 0.13 | 0.53 | 0.08 | 0.29 |
| 2 | 193011395 | rs17433560 | 0.507 | -0.564 | -1.218 | 0.258 | -0.05 | -0.018 | -0.018 | 6.975 | 0.25 | 0.07 | 0.34 | 0.21 | 0.38 | 0.35 |
| 2 | 193013528 | rs2356757 | -0.667 | -0.482 | -1.102 | 0.235 | 0 | 0.147 | 0.147 | 0 | 0.42 | 0.16 | 0.49 | 0.74 | 0.45 | 0.65 |
| 2 | 193023767 | rs4392190 | 0.114 | -0.663 | -0.72 | 0.125 | -0.07 | 0.02 | 0.02 | 1.713 | 0.74 | 0.6 | 0.7 | 0.25 | 0.86 | 0.58 |
| 2 | 193028594 | rs11678001 | -0.205 | 0.607 | -0.962 | 0.639 | -0.003 | -0.001 | -0.001 | 0 | 0.23 | 0.23 | 0.34 | 0.22 | 0.32 | 0.35 |
| 2 | 193030524 | rs4368280 | -0.098 | 0.561 | -0.984 | 0.588 | 0.009 | 0.009 | 0.009 | 0 | 0.52 | 0.47 | 0.53 | 0.58 | 0.56 | 0.65 |
| 2 | 193031408 | rs10210380 | -0.046 | 0.729 | -0.82 | 0.483 | 0 | 0.011 | 0.011 | 0.105 | 0.51 | 0.5 | 0.57 | 0.57 | 0.52 | 0.49 |
| 2 | 193032744 | rs4853492 | -0.568 | 0.789 | -0.602 | 0.044 | -0.001 | 0.001 | 0.001 | 0 | 0.66 | 0.73 | 0.69 | 0.69 | 0.69 | 0.69 |
| 2 | 193035023 | rs2356955 | 0.444 | 0.787 | -0.55 | 0.038 | 0.002 | 0.029 | 0.029 | 0.201 | 0.5 | 0.46 | 0.57 | 0.63 | 0.52 | 0.51 |
| 2 | 193036678 | rs2356956 | -0.157 | 0.749 | -0.389 | -0.014 | -0.003 | -0.013 | -0.013 | 0.237 | 0.61 | 0.61 | 0.66 | 0.69 | 0.66 | 0.66 |
| 2 | 193037634 | rs17368777 | 1.447 | 0.786 | -0.705 | -0.235 | -0.065 | -0.081 | -0.081 | 0.037 | 0.2 | 0.01 | 0.22 | 0.31 | 0.2 | 0.22 |
| 2 | 193038770 | rs12463739 | 0.55 | 0.536 | -0.663 | 0.013 | 0.004 | 0.019 | 0.019 | 0 | 0.4 | 0.37 | 0.43 | 0.33 | 0.37 | 0.31 |
| 2 | 193041060 | rs2356960 | -0.368 | -0.299 | -0.463 | 0.168 | -0.009 | -0.109 | -0.109 | 0 | 0.84 | 0.62 | 0.91 | 1 | 0.88 | 0.91 |
| 2 | 193046220 | rs2356961 | 1.156 | -0.322 | -0.47 | 0.065 | -0.077 | -0.001 | -0.001 | 0.642 | 0.29 | 0.09 | 0.38 | 0.24 | 0.36 | 0.2 |
| 2 | 193046999 | rs10931531 | 0.837 | -1.158 | 0.072 | -0.116 | 0.002 | 0.012 | 0.012 | 0 | 0.11 | 0.03 | 0.05 | 0 | 0.09 | 0.09 |
| 2 | 193050042 | rs4267464 | -0.894 | -1.154 | 0.319 | -0.114 | 0 | -0.058 | -0.058 | 0 | 0.89 | 0.81 | 0.93 | 1 | 0.91 | 0.91 |
| 2 | 193051320 | rs10497726 | 0.493 | -1.334 | 0.264 | -0.164 | 0.025 | -0.001 | -0.001 | 0 | 0.33 | 0.23 | 0.26 | 0.36 | 0.23 | 0.33 |
| 2 | 193052353 | rs1113932 | 0.867 | -1.319 | 0.191 | -0.186 | -0.001 | -0.022 | -0.022 | 0.026 | 0.11 | 0.06 | 0.06 | 0 | 0.09 | 0.09 |
| 2 | 193053820 | rs4458153 | 0.181 | -1.318 | 0.053 | -0.166 | -0.009 | -0.009 | -0.009 | 0 | 0.55 | 0.53 | 0.67 | 0.63 | 0.68 | 0.58 |
| 2 | 193056737 | rs3768702 | 0.201 | -1.225 | 0.165 | -0.054 | 0.047 | 0.014 | 0.014 | 0 | 0.37 | 0.2 | 0.28 | 0.37 | 0.25 | 0.37 |

| | | | | | | | | | | | | | | | | |
|---|---|---|---|---|---|---|---|---|---|---|---|---|---|---|---|---|
| 2 | 193057128 | rs3768703 | -0.011 | -0.878 | 0.221 | 0.249 | 0.003 | 0.005 | 0.005 | 0 | 0.48 | 0.51 | 0.61 | 0.63 | 0.6 | 0.53 |
| 2 | 193071438 | rs2658997 | -0.442 | -1.277 | 1.232 | 1.008 | -0.035 | 0.04 | 0.04 | 0 | 0.29 | 0.43 | 0.27 | 0.38 | 0.2 | 0.19 |
| 2 | 193081247 | rs2252769 | 0.227 | -1.287 | 1.425 | 0.99 | -0.031 | 0.033 | 0.033 | 0 | 0.71 | 0.6 | 0.74 | 0.62 | 0.8 | 0.81 |
| 2 | 193085020 | rs2592276 | 0.27 | -1.305 | 1.839 | 1.024 | -0.029 | 0.034 | 0.034 | 0 | 0.71 | 0.59 | 0.72 | 0.59 | 0.8 | 0.81 |
| 2 | 193089321 | rs16834303 | -0.201 | -1.32 | 1.941 | 0.911 | 0.018 | -0.03 | -0.03 | 0.112 | 0.27 | 0.14 | 0.24 | 0.38 | 0.2 | 0.19 |
| 2 | 193096199 | rs1823109 | 0.288 | -1.506 | 1.953 | 0.908 | -0.047 | 0.032 | 0.032 | 0 | 0.71 | 0.55 | 0.72 | 0.62 | 0.82 | 0.81 |
| 2 | 193140029 | rs2581225 | -0.261 | -1.525 | 1.249 | 0.773 | 0.029 | -0.093 | -0.093 | 0 | 0.76 | 0.92 | 0.73 | 0.53 | 0.83 | 0.81 |
| 2 | 193140700 | rs1838821 | 0.853 | -2.052 | 1.138 | 0.371 | 0.004 | 0 | 0 | 0 | 0.54 | 0.55 | 0.62 | 0.54 | 0.72 | 0.73 |
| 2 | 193143171 | rs1596882 | 0.78 | -1.488 | 2.12 | 0.569 | -0.068 | -0.068 | -0.068 | 0 | 0.75 | 0.42 | 0.86 | 0.91 | 0.89 | 0.9 |
| 2 | 193153552 | rs11899841 | -0.486 | -1.412 | 1.416 | 0.619 | 0.049 | 0.042 | 0.042 | 0 | 0.2 | 0.1 | 0.09 | 0.09 | 0.11 | 0.09 |
| 2 | 193178320 | rs10931539 | 0.634 | -1.415 | 1.586 | 0.617 | -0.071 | -0.065 | -0.065 | 0 | 0.77 | 0.56 | 0.87 | 0.91 | 0.89 | 0.9 |
| 2 | 193203397 | rs12993533 | 0.858 | -1.494 | 0.802 | 0.603 | -0.047 | -0.04 | -0.04 | 0 | 0.75 | 0.37 | 0.86 | 0.91 | 0.89 | 0.9 |
| 2 | 193206075 | rs13014116 | 0.05 | -0.94 | -0.508 | 0.914 | -0.046 | -0.073 | -0.073 | 0.002 | 0.12 | 0.03 | 0.18 | 0.33 | 0.22 | 0.37 |
| 2 | 193218969 | rs1843238 | -0.743 | -1.338 | 0.042 | 0.146 | 0.049 | 0.042 | 0.042 | 0 | 0.2 | 0.11 | 0.09 | 0.09 | 0.1 | 0.1 |
| 2 | 193232090 | rs10931542 | 0.823 | -0.547 | 0.646 | 0.363 | -0.013 | -0.061 | -0.061 | 0 | 0.89 | 0.7 | 0.93 | 1 | 0.93 | 0.91 |
| 2 | 193232686 | rs1446397 | 1.252 | -0.667 | 1.076 | 0.883 | -0.017 | 0.079 | 0.079 | 0.33 | 0.87 | 0.58 | 0.92 | 0.86 | 0.93 | 0.89 |
| 2 | 193233396 | rs12993326 | 0.848 | -1.595 | 0.702 | 0.164 | 0.048 | 0.045 | 0.045 | 0.542 | 0.63 | 0.27 | 0.8 | 0.8 | 0.79 | 0.74 |
| 2 | 193238289 | rs1550850 | 1.036 | -1.104 | 1.291 | 0.389 | -0.015 | -0.085 | -0.085 | 0 | 0.86 | 0.69 | 0.91 | 1 | 0.89 | 0.94 |
| 2 | 193255360 | rs1351905 | 1.236 | -1.23 | 1.598 | 0.327 | -0.029 | -0.113 | -0.113 | 2.373 | 0.84 | 0.55 | 0.9 | 0.99 | 0.89 | 0.91 |
| 2 | 193261854 | rs2357135 | 1.472 | -1.274 | 1.797 | 0.552 | -0.041 | -0.004 | -0.004 | 0 | 0.83 | 0.43 | 0.85 | 0.88 | 0.9 | 0.9 |
| 2 | 193291047 | rs10931545 | 1.44 | -1.726 | 1.642 | 0.46 | -0.062 | -0.028 | -0.028 | 0 | 0.77 | 0.4 | 0.79 | 0.86 | 0.88 | 0.87 |
| 2 | 193303647 | rs7558031 | 1.226 | -1.676 | 1.616 | 0.725 | -0.061 | 0.012 | 0.012 | 0 | 0.8 | 0.59 | 0.72 | 0.83 | 0.88 | 0.89 |
| 2 | 193312496 | rs1484884 | 1.294 | -1.672 | 1.186 | 0.727 | -0.059 | -0.004 | -0.004 | 0.943 | 0.78 | 0.42 | 0.71 | 0.81 | 0.88 | 0.88 |
| 2 | 193326547 | rs16834614 | -0.148 | -0.514 | -0.146 | 1.308 | -0.046 | -0.065 | -0.065 | 0 | 0.16 | 0.09 | 0.35 | 0.51 | 0.3 | 0.33 |
| 2 | 193327381 | rs1484887 | 0.746 | -0.597 | -0.143 | 1.138 | -0.004 | 0 | 0 | 0 | 0.65 | 0.67 | 0.49 | 0.32 | 0.57 | 0.53 |
| 2 | 193331208 | rs6741441 | -0.244 | -0.268 | -0.401 | 0.997 | 0.004 | 0.003 | 0.003 | 0.999 | 0.33 | 0.29 | 0.26 | 0.21 | 0.28 | 0.22 |
| 2 | 193333428 | rs12990943 | 0.412 | 0.213 | -0.065 | 1.336 | 0.013 | 0.022 | 0.022 | 0.124 | 0.5 | 0.39 | 0.49 | 0.36 | 0.57 | 0.31 |
| 2 | 193335578 | rs10173564 | 0.349 | 0.068 | -0.045 | 1.302 | 0.007 | 0.002 | 0.002 | 1.232 | 0.31 | 0.27 | 0.21 | 0.2 | 0.19 | 0.22 |
| 2 | 193337095 | rs1601324 | -0.26 | 0.149 | 0.194 | 1.386 | -0.011 | -0.014 | -0.014 | 0 | 0.2 | 0.18 | 0.31 | 0.44 | 0.3 | 0.46 |
| 2 | 193339122 | rs13431198 | 0.329 | 0.062 | 0.189 | 1.441 | -0.008 | -0.003 | -0.003 | 0 | 0.32 | 0.31 | 0.21 | 0.17 | 0.17 | 0.2 |
| 2 | 193344064 | rs2357149 | -0.191 | 0.167 | 0.125 | 1.597 | -0.014 | -0.024 | -0.024 | 0 | 0.18 | 0.11 | 0.29 | 0.34 | 0.25 | 0.37 |
| 2 | 193345479 | rs6713179 | -0.028 | -0.011 | 0.461 | 1.413 | -0.02 | -0.012 | -0.012 | 0 | 0.34 | 0.39 | 0.25 | 0.27 | 0.21 | 0.28 |
| 2 | 193359852 | rs4850494 | 0.423 | -0.25 | 0.042 | 1.298 | 0.05 | 0.038 | 0.038 | 2.779 | 0.31 | 0.17 | 0.2 | 0.15 | 0.18 | 0.16 |
| 2 | 193360531 | rs1994245 | 0.75 | 0.588 | 0.326 | 1.581 | -0.001 | 0.003 | 0.003 | 0.495 | 0.45 | 0.44 | 0.45 | 0.32 | 0.48 | 0.39 |
| 2 | 193366488 | rs10176810 | 0.036 | 0.7 | 0.25 | 1.647 | 0.007 | 0.021 | 0.021 | 0 | 0.8 | 0.75 | 0.71 | 0.57 | 0.74 | 0.63 |
| 2 | 193366879 | rs13017965 | -0.304 | 0.321 | 0.235 | 1.24 | -0.01 | -0.011 | -0.011 | 0.656 | 0.65 | 0.59 | 0.73 | 0.75 | 0.75 | 0.74 |
| 2 | 193368001 | rs1484900 | -1.517 | 1.067 | 0.155 | 1.408 | 0.03 | 0.008 | 0.008 | 0 | 0.06 | 0.13 | 0.05 | 0.11 | 0.08 | 0.1 |
| 2 | 193383203 | rs17442261 | -0.836 | 1.765 | -0.11 | 1.546 | -0.004 | -0.004 | -0.004 | 0.061 | 0.33 | 0.29 | 0.56 | 0.48 | 0.45 | 0.41 |
| 2 | 193388876 | rs6734271 | 0.678 | 1.709 | 0.255 | 1.515 | 0.03 | 0.031 | 0.031 | 0 | 0.66 | 0.57 | 0.43 | 0.5 | 0.55 | 0.59 |
| 2 | 193401723 | rs7425787 | -0.782 | 1.878 | 0.349 | 1.51 | -0.028 | -0.033 | -0.033 | 0.016 | 0.32 | 0.2 | 0.55 | 0.48 | 0.43 | 0.41 |
| 2 | 193420670 | rs12989619 | -0.808 | 1.852 | 0.432 | 1.497 | -0.031 | -0.029 | -0.029 | 0 | 0.32 | 0.21 | 0.56 | 0.51 | 0.43 | 0.42 |
| 2 | 193441245 | rs9941538 | -0.782 | 1.953 | 0.425 | 1.508 | -0.022 | -0.024 | -0.024 | 1.785 | 0.32 | 0.22 | 0.55 | 0.51 | 0.42 | 0.42 |
| 2 | 193445606 | rs4850546 | 0.962 | 1.639 | 0.273 | 1.198 | 0.001 | 0 | 0 | 0 | 0.6 | 0.58 | 0.41 | 0.47 | 0.52 | 0.51 |
| 2 | 193480135 | rs968538 | 1.184 | 1.401 | 0.935 | 1.138 | 0.004 | -0.011 | -0.011 | 0 | 0.55 | 0.3 | 0.35 | 0.49 | 0.49 | 0.5 |
| 2 | 193507209 | rs12615009 | -0.248 | 1.358 | 0.721 | 1.065 | -0.068 | -0.073 | -0.073 | 0.296 | 0.3 | 0.14 | 0.55 | 0.49 | 0.46 | 0.41 |
| 2 | 193559014 | rs1439664 | 0.876 | 1.359 | 0.728 | 1.054 | -0.011 | -0.008 | -0.008 | 0 | 0.61 | 0.71 | 0.4 | 0.48 | 0.49 | 0.52 |
| 2 | 193584810 | rs1898920 | 1.421 | 1.196 | 0.924 | 0.98 | 0.036 | 0.006 | 0.006 | 0.601 | 0.53 | 0.22 | 0.34 | 0.48 | 0.46 | 0.51 |
| 2 | 193598053 | rs13035689 | -0.004 | 1.253 | 0.464 | 0.992 | 0.019 | 0.032 | 0.032 | 0.581 | 0.07 | 0.02 | 0.03 | 0 | 0.05 | 0.07 |
| 2 | 193629044 | rs11686149 | 0.784 | 1.454 | 0.572 | 1.191 | -0.007 | -0.009 | -0.009 | 0.4 | 0.61 | 0.67 | 0.44 | 0.5 | 0.51 | 0.53 |
| 2 | 193671303 | rs1439659 | 0.805 | 1.384 | 0.418 | 1.287 | -0.02 | -0.019 | -0.019 | 0.431 | 0.61 | 0.75 | 0.45 | 0.47 | 0.5 | 0.55 |
| 2 | 193679049 | rs10931571 | 0.86 | 1.383 | 0.529 | 1.287 | -0.001 | -0.001 | -0.001 | 0 | 0.6 | 0.63 | 0.44 | 0.47 | 0.5 | 0.55 |
| 2 | 193706320 | rs13003012 | 0 | 1.606 | 0.153 | 1.652 | -0.136 | -0.107 | -0.107 | 0 | 0.26 | 0.05 | 0.49 | 0.4 | 0.41 | 0.32 |
| 2 | 193712184 | rs4850793 | 0.533 | 1.577 | -0.184 | 1.175 | -0.038 | -0.038 | -0.038 | 0.259 | 0.61 | 0.79 | 0.45 | 0.46 | 0.49 | 0.54 |
| 2 | 193724767 | rs10165191 | 0.484 | 1.575 | -0.185 | 1.174 | -0.039 | -0.039 | -0.039 | 0 | 0.62 | 0.79 | 0.46 | 0.46 | 0.49 | 0.54 |
| 2 | 193728069 | rs6758874 | 0.916 | 1.629 | -0.411 | 1.099 | 0.037 | -0.001 | -0.001 | 0 | 0.53 | 0.67 | 0.4 | 0.46 | 0.38 | 0.47 |
| 2 | 193742078 | rs2218495 | 0.215 | 0.892 | -0.787 | 0.833 | 0.005 | -0.02 | -0.02 | 0 | 0.83 | 0.92 | 0.89 | 0.73 | 0.87 | 0.83 |
| 2 | 193742317 | rs2034219 | -1.517 | 1.678 | -0.944 | 0.808 | -0.013 | 0.002 | 0.002 | 0 | 0.25 | 0.19 | 0.18 | 0.24 | 0.28 | 0.27 |
| 2 | 193765033 | rs9288211 | 1.11 | 0.97 | -0.6 | 0.606 | -0.026 | 0.028 | 0.028 | 0 | 0.74 | 0.44 | 0.76 | 0.62 | 0.8 | 0.66 |
| 2 | 193811641 | rs12478299 | 2.025 | 0.736 | -1.177 | 0.145 | -0.02 | 0.177 | 0.177 | 3.876 | 0.32 | 0.03 | 0.23 | 0.04 | 0.29 | 0.26 |
| 2 | 193825169 | rs6434583 | -0.216 | 0.926 | -1.143 | 0.191 | -0.036 | -0.005 | -0.005 | 0 | 0.39 | 0.26 | 0.55 | 0.58 | 0.48 | 0.38 |
| 2 | 193829017 | rs920978 | 0.341 | 0.742 | -1.126 | 0.272 | -0.004 | -0.028 | -0.028 | 0 | 0.11 | 0.03 | 0.05 | 0.15 | 0.12 | 0.14 |
| 2 | 193858558 | rs4547545 | -0.266 | 0.868 | -1.094 | 0.312 | -0.003 | 0 | 0 | 0 | 0.4 | 0.37 | 0.57 | 0.58 | 0.51 | 0.38 |
| 2 | 193878916 | rs10172318 | -0.197 | 0.885 | -1.325 | 0.352 | -0.011 | -0.002 | -0.002 | 0 | 0.4 | 0.28 | 0.57 | 0.58 | 0.52 | 0.38 |
| 2 | 193911926 | rs1947571 | -0.159 | 0.862 | -1.357 | 0.33 | -0.073 | -0.048 | -0.048 | 1.793 | 0.38 | 0.11 | 0.55 | 0.58 | 0.51 | 0.38 |
| 2 | 193933194 | rs10206411 | -0.298 | 0.944 | -1.257 | 0.283 | -0.003 | 0 | 0 | 0 | 0.41 | 0.37 | 0.57 | 0.58 | 0.51 | 0.38 |
| 2 | 193954458 | rs2357537 | 2.199 | 0.862 | -1.381 | 0.163 | 0.029 | 0.151 | 0.151 | 2.894 | 0.31 | 0.04 | 0.22 | 0.05 | 0.24 | 0.24 |
| 2 | 193995395 | rs2357533 | -0.508 | 0.757 | -1.294 | 0.494 | -0.002 | 0.001 | 0.001 | 0 | 0.42 | 0.37 | 0.6 | 0.58 | 0.53 | 0.39 |
| 2 | 194002782 | rs12619354 | 0.831 | 0.903 | -1.519 | 0.708 | -0.151 | -0.141 | -0.141 | 1.274 | 0.25 | 0.02 | 0.48 | 0.42 | 0.36 | 0.29 |
| 2 | 194003301 | rs6753418 | -0.405 | 0.841 | -1.304 | 0.407 | -0.003 | 0.001 | 0.001 | 0 | 0.42 | 0.37 | 0.6 | 0.58 | 0.53 | 0.39 |

| | | | | | | | | | | | | | | | | |
|---|---|---|---|---|---|---|---|---|---|---|---|---|---|---|---|---|
| 2 | 194017714 | rs1902745 | 1.711 | 0.458 | -1.509 | -0.201 | 0.037 | 0.118 | 0.118 | 1.491 | 0.3 | 0.05 | 0.17 | 0.06 | 0.24 | 0.22 |
| 2 | 194068734 | rs1125878 | 0.166 | 0.949 | -1.033 | 0.093 | -0.05 | 0.078 | 0.078 | 0 | 0.13 | 0.01 | 0.09 | 0 | 0.17 | 0.11 |
| 2 | 194098094 | rs2357539 | 0.123 | 0.774 | -0.602 | 0.084 | -0.037 | 0.073 | 0.073 | 0 | 0.12 | 0.01 | 0.08 | 0 | 0.15 | 0.1 |
| 2 | 194100270 | rs1997458 | 0.035 | 0.719 | -0.051 | 0.339 | 0 | 0.056 | 0.056 | 0.625 | 0.82 | 0.53 | 0.82 | 0.68 | 0.82 | 0.73 |
| 2 | 194142585 | rs2083295 | 0.575 | 0.646 | -0.014 | 0.625 | -0.005 | 0.013 | 0.013 | 0.016 | 0.64 | 0.47 | 0.69 | 0.54 | 0.62 | 0.55 |
| 2 | 194148040 | rs16835141 | 0.872 | 0.797 | -0.194 | 0.305 | 0.002 | 0.011 | 0.011 | 1.314 | 0.59 | 0.26 | 0.66 | 0.62 | 0.55 | 0.5 |
| 2 | 194189578 | rs6434599 | 0.755 | 0.823 | -0.235 | 0.313 | 0.002 | -0.006 | -0.006 | 0 | 0.61 | 0.44 | 0.68 | 0.62 | 0.55 | 0.53 |
| 2 | 194189678 | rs7589368 | 0.851 | 0.833 | -0.5 | 0.319 | 0.001 | 0.009 | 0.009 | 1.622 | 0.59 | 0.27 | 0.67 | 0.62 | 0.54 | 0.49 |
| 2 | 194191457 | rs16835154 | 1.08 | 0.806 | -0.606 | 0.305 | -0.014 | 0.021 | 0.021 | 2.137 | 0.57 | 0.08 | 0.64 | 0.62 | 0.54 | 0.48 |
| 2 | 194206882 | rs10192841 | 0.743 | 0.833 | -0.354 | 0.326 | 0 | -0.002 | -0.002 | 0 | 0.59 | 0.36 | 0.67 | 0.62 | 0.55 | 0.53 |
| 2 | 194217082 | rs7578712 | -0.269 | 1.121 | -0.541 | 0.439 | 0.038 | 0.049 | 0.049 | 0 | 0.28 | 0.36 | 0.53 | 0.57 | 0.33 | 0.28 |
| 2 | 194228442 | rs11692549 | -0.381 | 1.166 | -0.479 | 0.5 | -0.056 | -0.069 | -0.069 | 0 | 0.7 | 0.9 | 0.49 | 0.44 | 0.68 | 0.72 |
| 2 | 194243822 | rs4420715 | -0.352 | 1.226 | -0.521 | 0.541 | -0.088 | -0.103 | -0.103 | 0.134 | 0.69 | 0.95 | 0.51 | 0.43 | 0.68 | 0.7 |
| 2 | 194247107 | rs2357645 | 0.023 | 1.241 | -0.481 | 0.536 | -0.008 | 0.001 | 0.001 | 0 | 0.66 | 0.74 | 0.48 | 0.35 | 0.65 | 0.64 |
| 2 | 194250514 | rs16835181 | 0.386 | 1.229 | -0.353 | 0.568 | 0.013 | -0.031 | -0.031 | 0.044 | 0.61 | 0.56 | 0.41 | 0.23 | 0.58 | 0.56 |
| 2 | 194275048 | rs2718443 | 0.203 | 1.236 | -0.37 | 0.487 | -0.093 | -0.103 | -0.103 | 0 | 0.31 | 0.06 | 0.49 | 0.57 | 0.34 | 0.3 |
| 2 | 194296294 | rs2245901 | -0.424 | 1.249 | -0.403 | 0.439 | 0.011 | -0.011 | -0.011 | 0.247 | 0.39 | 0.43 | 0.58 | 0.65 | 0.41 | 0.39 |
| 2 | 194305624 | rs1371638 | 0.164 | 1.276 | -0.436 | 0.485 | 0.001 | -0.001 | -0.001 | 0 | 0.34 | 0.31 | 0.53 | 0.64 | 0.35 | 0.36 |
| 2 | 194341313 | rs2256763 | -0.284 | 1.333 | -0.39 | 0.422 | 0.004 | 0.019 | 0.019 | 0 | 0.4 | 0.49 | 0.54 | 0.51 | 0.4 | 0.46 |
| 2 | 194344883 | rs1440057 | 0.686 | 1.336 | -0.343 | 0.429 | -0.033 | -0.055 | -0.055 | 0 | 0.31 | 0.14 | 0.45 | 0.42 | 0.32 | 0.37 |
| 2 | 194353714 | rs6713417 | -0.113 | 1.433 | -0.322 | 0.358 | 0.005 | 0.017 | 0.017 | 0 | 0.39 | 0.49 | 0.55 | 0.52 | 0.4 | 0.49 |
| 2 | 194413474 | rs2719161 | -1.204 | 1.643 | -0.177 | 0.524 | 0.074 | 0.069 | 0.069 | 0.867 | 0.07 | 0.27 | 0.09 | 0.08 | 0.1 | 0.08 |
| 2 | 194437889 | rs2719164 | 0.193 | 1.344 | -0.552 | 0.379 | 0.001 | 0.006 | 0.006 | 0 | 0.41 | 0.43 | 0.55 | 0.52 | 0.41 | 0.51 |
| 2 | 194439091 | rs10209930 | 0.836 | 1.35 | -0.512 | 0.385 | -0.007 | -0.022 | -0.022 | 0 | 0.35 | 0.2 | 0.48 | 0.44 | 0.32 | 0.44 |
| 2 | 194448287 | rs10931601 | -0.287 | 1.347 | -0.499 | 0.394 | 0.028 | 0.048 | 0.048 | 0.02 | 0.95 | 0.85 | 0.94 | 0.92 | 0.92 | 0.93 |
| 2 | 194449488 | rs11890112 | 0.608 | 1.371 | -0.848 | 0.331 | -0.016 | -0.021 | -0.021 | 0 | 0.38 | 0.18 | 0.5 | 0.44 | 0.35 | 0.47 |
| 2 | 194479605 | rs16835263 | 0.745 | 1.442 | -0.595 | 0.514 | -0.005 | 0.013 | 0.013 | 0.109 | 0.07 | 0.08 | 0.1 | 0.22 | 0.12 | 0.13 |
| 2 | 194481544 | rs10497743 | -0.458 | 1.375 | -0.625 | 0.613 | 0.002 | 0 | 0 | 0.258 | 0.61 | 0.61 | 0.46 | 0.4 | 0.6 | 0.49 |
| 2 | 194482136 | rs767853 | 0.896 | 1.307 | -0.641 | 0.621 | -0.019 | -0.069 | -0.069 | 0 | 0.32 | 0.16 | 0.47 | 0.51 | 0.31 | 0.43 |
| 2 | 194500272 | rs995929 | 0 | 1.31 | -0.625 | 0.352 | 0.024 | 0.039 | 0.039 | 0.141 | 0.94 | 0.87 | 0.94 | 0.92 | 0.91 | 0.93 |
| 2 | 194518477 | rs6757627 | -0.576 | 1.26 | -0.6 | 0.027 | -0.002 | -0.008 | -0.008 | 1.111 | 0.63 | 0.69 | 0.48 | 0.49 | 0.61 | 0.5 |
| 2 | 194538059 | rs1351086 | 1.003 | 1.267 | -0.616 | 0.041 | -0.018 | -0.038 | -0.038 | 0 | 0.32 | 0.17 | 0.46 | 0.42 | 0.31 | 0.42 |
| 2 | 194543505 | rs11885434 | 0.925 | 1.261 | -0.618 | 0.06 | -0.02 | -0.052 | -0.052 | 0 | 0.32 | 0.17 | 0.46 | 0.46 | 0.31 | 0.43 |
| 2 | 194557065 | rs1478129 | 1.502 | 1.1 | -0.858 | -0.31 | -0.015 | 0.127 | 0.127 | 0.155 | 0.26 | 0.02 | 0.15 | 0.05 | 0.22 | 0.16 |
| 2 | 194563864 | rs4605388 | 0.167 | 1.389 | -0.129 | 0.277 | 0.017 | 0.033 | 0.033 | 0.005 | 0.95 | 0.9 | 0.94 | 0.91 | 0.92 | 0.93 |
| 2 | 194566960 | rs11685549 | 0.566 | 1.426 | -0.193 | 0.231 | -0.03 | 0.044 | 0.044 | 0 | 0.09 | 0.01 | 0.07 | 0 | 0.13 | 0.06 |
| 2 | 194639832 | rs11695748 | -0.042 | 1.299 | 0.269 | 0.601 | 0.003 | 0.012 | 0.012 | 0 | 0.61 | 0.6 | 0.47 | 0.49 | 0.6 | 0.51 |
| 2 | 194731362 | rs13385986 | 2.52 | 1.494 | 1.386 | 0.364 | 0.103 | 0.068 | 0.068 | 1.055 | 0.91 | 0.66 | 0.91 | 0.87 | 0.86 | 0.93 |
| 2 | 194748016 | rs6705791 | 1.265 | 1.494 | 1.391 | 0.364 | 0.045 | 0.077 | 0.077 | 0.595 | 0.94 | 0.78 | 0.94 | 0.87 | 0.91 | 0.94 |
| 2 | 194749557 | rs13431912 | 2.206 | 1.475 | 1.239 | 0.366 | 0.101 | 0.066 | 0.066 | 1.357 | 0.91 | 0.64 | 0.91 | 0.87 | 0.86 | 0.93 |
| 2 | 194805093 | rs6750903 | -1.409 | 1.07 | 0.703 | 0.397 | 0.035 | 0.048 | 0.048 | 0.366 | 0.07 | 0.24 | 0.08 | 0.12 | 0.09 | 0.07 |
| 2 | 194849951 | rs6434642 | 1.102 | 1.011 | 0.914 | 0.364 | 0.042 | 0.062 | 0.062 | 0.704 | 0.93 | 0.65 | 0.93 | 0.9 | 0.93 | 0.94 |
| 2 | 194888500 | rs7600163 | 0.049 | 0.916 | 0.665 | 0.121 | -0.006 | 0.005 | 0.005 | 0 | 0.43 | 0.27 | 0.53 | 0.63 | 0.4 | 0.46 |
| 2 | 194911328 | rs6723021 | 0.27 | 0.586 | 0.021 | -0.096 | 0 | -0.016 | -0.016 | 0 | 0.1 | 0.09 | 0.05 | 0 | 0.14 | 0.12 |
| 2 | 194972526 | rs1607345 | 0.39 | 0.668 | -0.031 | -0.038 | 0.001 | -0.033 | -0.033 | 0 | 0.82 | 0.63 | 0.86 | 0.9 | 0.76 | 0.82 |
| 2 | 194990422 | rs12476410 | 1.139 | 0.623 | -0.534 | -0.137 | -0.037 | 0.019 | 0.019 | 0.006 | 0.33 | 0.07 | 0.25 | 0.19 | 0.35 | 0.25 |
| 2 | 194990534 | rs2358030 | 0.67 | 0.514 | -0.532 | -0.228 | -0.042 | 0.029 | 0.029 | 0 | 0.34 | 0.1 | 0.44 | 0.75 | 0.28 | 0.46 |
| 2 | 194994028 | rs12052724 | 1.031 | 0.676 | -0.748 | -0.238 | -0.096 | -0.09 | -0.09 | 0 | 0.3 | 0.04 | 0.4 | 0.68 | 0.27 | 0.4 |
| 2 | 195039470 | rs6750206 | -0.316 | 0.862 | -0.462 | -0.303 | -0.003 | -0.002 | -0.002 | 0 | 0.14 | 0.1 | 0.11 | 0.05 | 0.23 | 0.16 |
| 2 | 195134095 | rs13003177 | -0.273 | 0.305 | -0.275 | -0.174 | 0.004 | 0.005 | 0.005 | 0 | 0.14 | 0.1 | 0.07 | 0 | 0.13 | 0.05 |
| 2 | 195151349 | rs4578878 | 0.699 | 0.38 | -0.546 | -0.289 | -0.049 | 0.07 | 0.07 | 0.016 | 0.28 | 0.03 | 0.21 | 0.12 | 0.3 | 0.25 |
| 2 | 195160638 | rs4255977 | 1.858 | 0.644 | 1.149 | 0.438 | 0.086 | 0.049 | 0.049 | 0 | 0.93 | 0.69 | 0.82 | 0.93 | 0.91 | 0.89 |
| 2 | 195180067 | rs6736325 | 1.627 | 0.664 | 1.17 | 0.748 | 0.119 | 0.207 | 0.207 | 0 | 0.94 | 0.75 | 0.74 | 0.85 | 0.9 | 0.88 |
| 2 | 195205040 | rs1376696 | 1.696 | 0.66 | 1.109 | 0.859 | 0.11 | 0.223 | 0.223 | 0.016 | 0.94 | 0.78 | 0.73 | 0.81 | 0.89 | 0.89 |
| 2 | 195266232 | rs896441 | 1.326 | 0.451 | 0.943 | 0.689 | -0.005 | -0.002 | -0.002 | 0 | 0.65 | 0.33 | 0.69 | 0.71 | 0.73 | 0.73 |
| 2 | 195291870 | rs11688042 | -1.282 | 0.521 | -0.434 | 0.134 | 0.016 | 0.051 | 0.051 | 0 | 0.19 | 0.1 | 0.11 | 0.02 | 0.15 | 0.11 |
| 2 | 195296472 | rs2122952 | -1.316 | 0.222 | -0.03 | 0.343 | -0.008 | -0.024 | -0.024 | 0 | 0.19 | 0.22 | 0.2 | 0.08 | 0.14 | 0.14 |
| 2 | 195300730 | rs2060798 | 1.867 | 0.249 | 1.279 | 0.392 | 0.128 | 0.052 | 0.052 | 1.648 | 0.91 | 0.46 | 0.86 | 0.94 | 0.91 | 0.85 |
| 2 | 195309148 | rs6731269 | 1.671 | 0.224 | 1.763 | 0.395 | 0.156 | 0.101 | 0.101 | 0 | 0.95 | 0.58 | 0.89 | 0.94 | 0.94 | 0.88 |
| 2 | 195344783 | rs2360090 | 1.766 | -0.474 | 1.576 | 0.027 | 0.026 | 0.049 | 0.049 | 0 | 0.71 | 0.18 | 0.72 | 0.91 | 0.78 | 0.74 |
| 2 | 195352886 | rs7421656 | 1.618 | -0.055 | 1.245 | 0.75 | 0.007 | -0.006 | -0.006 | 0 | 0.63 | 0.27 | 0.62 | 0.7 | 0.68 | 0.67 |
| 2 | 195362314 | rs2360101 | 1.672 | -0.048 | 1.225 | 0.725 | 0.008 | -0.01 | -0.01 | 0 | 0.65 | 0.24 | 0.68 | 0.7 | 0.69 | 0.68 |
| 2 | 195366084 | rs13383716 | -1.805 | -0.771 | 0.933 | 0.558 | 0.01 | 0.001 | 0.001 | 0 | 0.38 | 0.62 | 0.26 | 0.29 | 0.24 | 0.22 |
| 2 | 195373284 | rs12465821 | -0.899 | -0.991 | 0.828 | 0.44 | -0.04 | -0.01 | -0.01 | 0.339 | 0.27 | 0.54 | 0.26 | 0.29 | 0.21 | 0.23 |
| 2 | 195377437 | rs16831761 | -2.032 | -0.649 | 0.794 | 0.654 | 0.087 | 0.014 | 0.014 | 0 | 0.07 | 0.2 | 0.19 | 0.04 | 0.13 | 0.13 |
| 2 | 195379647 | rs17513288 | -0.823 | -1.158 | 0.444 | 0.773 | -0.022 | -0.031 | -0.031 | 0 | 0.13 | 0.04 | 0.18 | 0.18 | 0.18 | 0.21 |
| 2 | 195381344 | rs11885263 | -1.923 | -1.197 | 1.275 | 0.918 | 0.007 | 0.043 | 0.043 | 0.053 | 0.2 | 0.4 | 0.23 | 0.24 | 0.24 | 0.33 |
| 2 | 195382519 | rs11892547 | -1.322 | -1.195 | 1.238 | 0.915 | -0.001 | 0.001 | 0.001 | 0 | 0.17 | 0.21 | 0.21 | 0.18 | 0.22 | 0.22 |
| 2 | 195386011 | rs1528003 | 2.363 | -1.672 | 0.783 | 0.932 | 0.026 | -0.047 | -0.047 | 0 | 0.47 | 0.1 | 0.44 | 0.52 | 0.56 | 0.35 |

| | | | | | | | | | | | | | | | | |
|---|---|---|---|---|---|---|---|---|---|---|---|---|---|---|---|---|
| 2 | 195414927 | rs11673827 | -2.149 | -1.081 | 1.646 | 1.341 | -0.001 | -0.005 | -0.005 | 0 | 0.22 | 0.36 | 0.37 | 0.2 | 0.26 | 0.38 |
| 2 | 195417908 | rs10198324 | -2.906 | -1.144 | 1.601 | 1.351 | 0.06 | 0.234 | 0.234 | 5.676 | 0.06 | 0.44 | 0.06 | 0.17 | 0.02 | 0.13 |
| 2 | 195441018 | rs12475012 | -2.41 | -1.111 | 1.303 | 1.223 | 0.004 | 0.029 | 0.029 | 0 | 0.05 | 0.13 | 0.15 | 0.07 | 0.03 | 0.16 |
| 2 | 195448392 | rs17626258 | -0.01 | -1.13 | 1.165 | 0.704 | -0.008 | -0.02 | -0.02 | 0.58 | 0.07 | 0.01 | 0.09 | 0.13 | 0.06 | 0.08 |
| 2 | 195471298 | rs13009328 | -0.193 | -1.113 | 1.218 | 0.658 | -0.01 | 0.009 | 0.009 | 0 | 0.05 | 0.02 | 0.07 | 0.05 | 0.1 | 0.13 |
| 2 | 195476042 | rs12993717 | -1.271 | -1.155 | 1.248 | 0.722 | -0.011 | -0.005 | -0.005 | 0.283 | 0.17 | 0.07 | 0.35 | 0.16 | 0.19 | 0.36 |
| 2 | 195484228 | rs11679788 | -0.677 | -1.158 | 1.249 | 0.697 | 0 | -0.009 | -0.009 | 1.634 | 0.13 | 0.06 | 0.21 | 0.14 | 0.16 | 0.2 |
| 2 | 195501868 | rs12693735 | -1.969 | -1.185 | 1.435 | 0.69 | -0.013 | 0.001 | 0.001 | 0.174 | 0.22 | 0.36 | 0.42 | 0.26 | 0.23 | 0.38 |
| 2 | 195514148 | rs6727455 | 1.937 | -1.14 | 1.546 | 0.735 | -0.025 | 0.01 | 0.01 | 0.649 | 0.77 | 0.47 | 0.57 | 0.74 | 0.78 | 0.67 |
| 2 | 195519360 | rs13013910 | 2.195 | -1.12 | 1.571 | 0.739 | -0.01 | -0.002 | -0.002 | 0 | 0.75 | 0.34 | 0.54 | 0.73 | 0.75 | 0.66 |
| 2 | 195526300 | rs17368354 | -0.415 | -0.883 | 0.252 | 0.823 | -0.007 | -0.021 | -0.021 | 0.753 | 0.11 | 0.05 | 0.21 | 0.15 | 0.15 | 0.16 |
| 2 | 195541397 | rs13033486 | -0.546 | -0.848 | 0.231 | 0.843 | -0.007 | -0.028 | -0.028 | 0 | 0.13 | 0.04 | 0.22 | 0.15 | 0.15 | 0.17 |
| 2 | 195552313 | rs16927 | -1.202 | -1.208 | 0.887 | 0.658 | -0.006 | -0.002 | -0.002 | 0 | 0.28 | 0.3 | 0.17 | 0.2 | 0.22 | 0.29 |
| 2 | 195552753 | rs801392 | 1.736 | -1.187 | 0.131 | 0.617 | 0 | -0.031 | -0.031 | 0.284 | 0.93 | 0.86 | 0.94 | 0.98 | 0.92 | 0.87 |
| 2 | 195556215 | rs13025498 | 1.678 | -1.205 | 0.078 | 0.66 | 0.057 | 0.015 | 0.015 | 0 | 0.62 | 0.14 | 0.74 | 0.68 | 0.7 | 0.56 |
| 2 | 195564847 | rs699315 | 1.225 | -1.451 | 0.267 | 0.695 | -0.006 | -0.024 | -0.024 | 0 | 0.92 | 0.88 | 0.94 | 1 | 0.94 | 0.98 |
| 2 | 195579293 | rs801301 | -1.892 | -1.521 | 0.332 | 0.822 | 0.029 | 0.011 | 0.011 | 0 | 0.32 | 0.73 | 0.2 | 0.23 | 0.2 | 0.38 |
| 2 | 195583436 | rs1439937 | 1.955 | -1.548 | 0.294 | 0.775 | 0.041 | 0.023 | 0.023 | 0 | 0.65 | 0.25 | 0.77 | 0.77 | 0.77 | 0.61 |
| 2 | 195588645 | rs801295 | -1.862 | -1.629 | 0.364 | 0.74 | -0.014 | -0.002 | -0.002 | 1.819 | 0.27 | 0.67 | 0.2 | 0.21 | 0.18 | 0.29 |
| 2 | 195596411 | rs801289 | 1.074 | -1.509 | 0.426 | 0.794 | -0.003 | -0.025 | -0.025 | 0 | 0.92 | 0.98 | 0.92 | 0.87 | 0.91 | 0.89 |
| 2 | 195643802 | rs801340 | -2.284 | -1.631 | 0.478 | 0.967 | -0.026 | -0.013 | -0.013 | 0 | 0.23 | 0.72 | 0.15 | 0.1 | 0.12 | 0.28 |
| 2 | 195651706 | rs12618745 | -1.12 | -1.574 | 0.976 | 1.003 | -0.005 | -0.026 | -0.026 | 0.264 | 0.08 | 0.02 | 0.07 | 0.13 | 0.08 | 0.1 |
| 2 | 195663691 | rs2215453 | -1.84 | -1.563 | 1.074 | 0.996 | -0.033 | -0.021 | -0.021 | 0 | 0.18 | 0.68 | 0.12 | 0.1 | 0.1 | 0.24 |
| 2 | 195698234 | rs1860602 | 2.597 | -1.593 | 1.016 | 1.01 | 0.083 | -0.039 | -0.039 | 0.032 | 0.66 | 0.12 | 0.73 | 0.62 | 0.77 | 0.6 |
| 2 | 195731002 | rs17587873 | 2.456 | -1.557 | 0.966 | 1 | 0.089 | -0.04 | -0.04 | 0.642 | 0.65 | 0.11 | 0.73 | 0.64 | 0.76 | 0.6 |
| 2 | 195746156 | rs7563950 | 2.493 | -1.215 | 1.217 | 0.954 | 0 | 0 | 0 | 0.02 | 0.73 | 0.28 | 0.81 | 0.79 | 0.81 | 0.7 |
| 2 | 195751653 | rs1377652 | 2.304 | -1.248 | 1.284 | 0.952 | -0.038 | -0.018 | -0.018 | 0 | 0.84 | 0.31 | 0.9 | 0.91 | 0.95 | 0.82 |
| 2 | 195753257 | rs16830373 | -2.472 | -1.183 | 1.543 | 0.967 | 0.019 | -0.01 | -0.01 | 0 | 0.06 | 0.24 | 0.03 | 0 | 0.02 | 0.04 |
| 2 | 195764850 | rs1453808 | 2.631 | -1.242 | 1.996 | 1.022 | -0.003 | -0.02 | -0.02 | 0 | 0.84 | 0.54 | 0.88 | 0.88 | 0.86 | 0.83 |
| 2 | 195771824 | rs11888197 | -2.113 | -1.164 | 1.971 | 1.088 | -0.001 | 0.168 | 0.168 | 0 | 0.11 | 0.39 | 0.11 | 0.24 | 0.05 | 0.16 |
| 2 | 195779715 | rs1349803 | 1.958 | -1.067 | 1.834 | 0.805 | -0.021 | -0.049 | -0.049 | 0 | 0.91 | 0.72 | 0.94 | 1 | 0.94 | 0.92 |
| 2 | 195785764 | rs4850568 | 2.609 | -1.062 | 1.996 | 0.863 | -0.003 | -0.004 | -0.004 | 1.628 | 0.84 | 0.64 | 0.88 | 0.85 | 0.85 | 0.81 |
| 2 | 195786904 | rs4850569 | -2.452 | -1.336 | 2.04 | 1.049 | -0.012 | 0.037 | 0.037 | 0.212 | 0.18 | 0.29 | 0.28 | 0.31 | 0.12 | 0.21 |
| 2 | 195791620 | rs12611555 | 3.086 | -1.414 | 2.319 | 1.033 | 0.044 | -0.028 | -0.028 | 0 | 0.72 | 0.2 | 0.65 | 0.67 | 0.83 | 0.69 |
| 2 | 195808307 | rs17626732 | -1.363 | -1.397 | 2.23 | 1.058 | -0.004 | -0.008 | -0.008 | 0 | 0.07 | 0.04 | 0.05 | 0.13 | 0.08 | 0.1 |
| 2 | 195809618 | rs10184058 | -2.097 | -1.406 | 2.303 | 1.037 | -0.008 | -0.033 | -0.033 | 0.111 | 0.09 | 0.39 | 0.07 | 0 | 0.05 | 0.08 |
| 2 | 195815416 | rs4334512 | -1.594 | -1.411 | 2.228 | 1.051 | -0.027 | 0.034 | 0.034 | 0.051 | 0.12 | 0.25 | 0.23 | 0.18 | 0.04 | 0.12 |
| 2 | 195816892 | rs10931664 | -2.202 | -1.342 | 2.041 | 1.088 | -0.037 | 0.037 | 0.037 | 0.63 | 0.23 | 0.43 | 0.31 | 0.31 | 0.15 | 0.29 |
| 2 | 195828622 | rs891522 | 2.191 | -1.34 | 1.073 | 1.078 | 0.061 | -0.057 | -0.057 | 0 | 0.73 | 0.13 | 0.65 | 0.67 | 0.85 | 0.7 |
| 2 | 195843011 | rs1377656 | 1.868 | -1.284 | 0.971 | 0.994 | 0.078 | -0.058 | -0.058 | 0 | 0.72 | 0.11 | 0.78 | 0.64 | 0.83 | 0.71 |
| 2 | 195846449 | rs4144840 | -1.714 | -1.154 | 1.527 | 1.017 | -0.009 | 0.021 | 0.021 | 0 | 0.18 | 0.18 | 0.13 | 0.32 | 0.16 | 0.26 |
| 2 | 195861792 | rs2217471 | 0.304 | -0.579 | 0.311 | 0.538 | 0.003 | -0.002 | -0.002 | 0.016 | 0.58 | 0.35 | 0.69 | 0.62 | 0.59 | 0.67 |
| 2 | 195864023 | rs1430692 | 0.523 | -0.634 | 1.106 | 0.688 | 0.025 | -0.031 | -0.031 | 0 | 0.86 | 0.73 | 0.9 | 0.96 | 0.85 | 0.91 |
| 2 | 195864232 | rs6727001 | 1.605 | -0.638 | 0.977 | 0.835 | -0.001 | -0.014 | -0.014 | 0 | 0.7 | 0.6 | 0.8 | 0.8 | 0.7 | 0.71 |
| 2 | 195868972 | rs11681386 | -1.406 | -0.528 | 1.219 | 1.004 | 0.002 | -0.006 | -0.006 | 0 | 0.16 | 0.13 | 0.12 | 0.2 | 0.16 | 0.23 |
| 2 | 195870770 | rs1835438 | 2.312 | -0.451 | 1.358 | 0.968 | -0.011 | -0.05 | -0.05 | 0 | 0.62 | 0.16 | 0.69 | 0.5 | 0.64 | 0.55 |
| 2 | 195871343 | rs16832040 | -1.32 | -0.437 | 1.356 | 0.975 | 0.001 | -0.007 | -0.007 | 0 | 0.16 | 0.13 | 0.12 | 0.2 | 0.18 | 0.23 |
| 2 | 195871819 | rs10172965 | -1.243 | -0.43 | 1.405 | 0.974 | 0.009 | 0.059 | 0.059 | 4.359 | 0.2 | 0.61 | 0.18 | 0.3 | 0.18 | 0.22 |
| 2 | 195872980 | rs16832060 | -1.39 | -0.506 | 1.5 | 0.957 | -0.003 | 0.006 | 0.006 | 0 | 0.15 | 0.17 | 0.1 | 0.18 | 0.14 | 0.28 |
| 2 | 195874296 | rs7581004 | -1.757 | -0.504 | 1.534 | 0.947 | 0.008 | 0.005 | 0.005 | 0 | 0.27 | 0.36 | 0.2 | 0.22 | 0.3 | 0.37 |
| 2 | 195881635 | rs10200877 | -0.911 | -0.571 | 1.638 | 1.028 | 0.075 | 0.182 | 0.182 | 3.436 | 0.06 | 0.35 | 0.08 | 0.14 | 0.03 | 0.06 |
| 2 | 195885106 | rs16832336 | -0.793 | -0.685 | 1.589 | 0.849 | 0.014 | -0.054 | -0.054 | 0 | 0.09 | 0.29 | 0.07 | 0 | 0.05 | 0.04 |
| 2 | 195887318 | rs13024059 | 1.157 | -0.926 | 0.591 | 0.771 | -0.025 | 0.012 | 0.012 | 0 | 0.64 | 0.58 | 0.59 | 0.45 | 0.69 | 0.44 |
| 2 | 195889744 | rs6731550 | 2.455 | -0.936 | 0.524 | 0.731 | -0.085 | 0.016 | 0.016 | 0.584 | 0.43 | 0.1 | 0.44 | 0.42 | 0.51 | 0.33 |
| 2 | 195890398 | rs718645 | -0.847 | -0.939 | 0.522 | 0.736 | -0.003 | 0.021 | 0.021 | 0 | 0.15 | 0.17 | 0.1 | 0.28 | 0.13 | 0.29 |
| 2 | 195892736 | rs12693755 | 0.465 | -0.889 | 0.533 | 0.754 | 0.011 | -0.103 | -0.103 | 0.125 | 0.77 | 0.45 | 0.85 | 0.97 | 0.82 | 0.89 |
| 2 | 195896262 | rs1430676 | -1.701 | -1.018 | 0.678 | 1.205 | 0.025 | -0.041 | -0.041 | 0 | 0.33 | 0.17 | 0.34 | 0.39 | 0.3 | 0.47 |
| 2 | 195897450 | rs17823179 | 0.111 | -0.723 | 0.867 | 0.784 | 0.011 | 0.029 | 0.029 | 0 | 0.15 | 0.04 | 0.22 | 0.09 | 0.12 | 0.21 |
| 2 | 195913337 | rs1155301 | -0.511 | -0.724 | 0.866 | 0.813 | 0.017 | 0.014 | 0.014 | 0.018 | 0.18 | 0.04 | 0.25 | 0.13 | 0.15 | 0.23 |
| 2 | 195913361 | rs17772818 | 0.17 | -0.7 | 0.78 | 0.815 | -0.013 | 0.006 | 0.006 | 0 | 0.14 | 0.01 | 0.22 | 0.09 | 0.12 | 0.21 |
| 2 | 195922433 | rs1367424 | 0.455 | -0.659 | 0.862 | 0.833 | 0.008 | 0.001 | 0.001 | 0 | 0.9 | 0.78 | 0.92 | 0.91 | 0.9 | 0.88 |
| 2 | 195928055 | rs12478243 | 1.119 | -0.859 | 0.555 | 0.711 | -0.06 | 0.008 | 0.008 | 0 | 0.72 | 0.57 | 0.66 | 0.72 | 0.83 | 0.64 |
| 2 | 195954734 | rs777717 | 1.325 | -0.488 | 0.582 | 0.892 | -0.044 | 0.032 | 0.032 | 0 | 0.83 | 0.66 | 0.87 | 0.81 | 0.93 | 0.86 |
| 2 | 195955184 | rs777715 | 1.586 | -0.262 | 0.672 | 1.23 | 0.026 | 0.198 | 0.198 | 0.678 | 0.89 | 0.58 | 0.86 | 0.77 | 0.9 | 0.86 |
| 2 | 195958665 | rs11684338 | -1.235 | -0.224 | -0.179 | 1.236 | -0.003 | 0.009 | 0.009 | 0 | 0.11 | 0.15 | 0.16 | 0.21 | 0.09 | 0.12 |
| 2 | 195958753 | rs11695739 | -1.235 | -0.224 | -0.179 | 1.236 | -0.003 | 0.009 | 0.009 | 0 | 0.11 | 0.15 | 0.16 | 0.21 | 0.09 | 0.12 |
| 2 | 195967027 | rs1849068 | 0.899 | -0.106 | -0.917 | 0.838 | 0.022 | 0.095 | 0.095 | 0.726 | 0.52 | 0.3 | 0.52 | 0.28 | 0.57 | 0.36 |
| 2 | 195967073 | rs1849067 | 0.757 | -0.629 | -0.996 | 0.785 | 0.005 | 0.004 | 0.004 | 0 | 0.67 | 0.31 | 0.6 | 0.73 | 0.75 | 0.59 |
| 2 | 195968050 | rs4850358 | -0.416 | -0.96 | -0.917 | 0.982 | 0.021 | -0.003 | -0.003 | 0 | 0.23 | 0.12 | 0.22 | 0.24 | 0.19 | 0.21 |

| | | | | | | | | | | | | | | | | |
|---|---|---|---|---|---|---|---|---|---|---|---|---|---|---|---|---|
| 2 | 195968887 | rs17824823 | -0.348 | -0.89 | -0.97 | 1.687 | -0.005 | 0.002 | 0.002 | 0 | 0.1 | 0 | 0.05 | 0.06 | 0.07 | 0.05 |
| 2 | 195974363 | rs1599755 | -0.308 | -0.867 | 0.316 | 1.506 | -0.017 | 0 | 0 | 0 | 0.09 | 0.25 | 0.07 | 0.06 | 0.05 | 0.05 |
| 2 | 195978752 | rs2667146 | 0.342 | -0.679 | -0.082 | 1.082 | 0.02 | 0.025 | 0.025 | 0 | 0.5 | 0.63 | 0.56 | 0.56 | 0.55 | 0.61 |
| 2 | 195986610 | rs2566717 | -1.211 | -0.64 | 0.668 | 1.725 | -0.029 | 0.062 | 0.062 | 0.105 | 0.12 | 0.32 | 0.12 | 0.15 | 0.07 | 0.11 |
| 2 | 195993306 | rs777734 | -1.714 | -0.625 | 0.702 | 1.939 | 0.046 | 0.188 | 0.188 | 0.096 | 0.05 | 0.3 | 0.09 | 0.11 | 0.04 | 0.11 |
| 2 | 195993600 | rs12473173 | -0.888 | -0.676 | 0.678 | 1.777 | 0.005 | -0.004 | -0.004 | 0 | 0.11 | 0.13 | 0.11 | 0.12 | 0.07 | 0.09 |
| 2 | 196000458 | rs1355433 | -1.001 | -0.623 | 0.671 | 2.116 | 0.013 | -0.049 | -0.049 | 0.038 | 0.14 | 0.08 | 0.27 | 0.31 | 0.14 | 0.18 |
| 2 | 196002614 | rs2217469 | 2.15 | -0.838 | 1.206 | 1.949 | -0.027 | 0.035 | 0.035 | 1.155 | 0.73 | 0.33 | 0.59 | 0.6 | 0.79 | 0.73 |
| 2 | 196044448 | rs13422281 | -1.537 | -1.037 | 1.245 | 2.201 | 0.016 | -0.043 | -0.043 | 1.014 | 0.18 | 0.11 | 0.31 | 0.31 | 0.16 | 0.21 |
| 2 | 196053038 | rs12990705 | 1.728 | -1.076 | 1.54 | 2.388 | -0.03 | 0.052 | 0.052 | 2.397 | 0.77 | 0.3 | 0.66 | 0.61 | 0.81 | 0.75 |
| 2 | 196061096 | rs1511258 | 1.409 | -1.075 | 1.933 | 2.484 | -0.045 | 0.18 | 0.18 | 0.459 | 0.8 | 0.52 | 0.67 | 0.57 | 0.83 | 0.79 |
| 2 | 196067184 | rs2171816 | 2.489 | -1.092 | 1.939 | 2.382 | 0.006 | -0.008 | -0.008 | 0 | 0.7 | 0.25 | 0.61 | 0.62 | 0.76 | 0.74 |
| 2 | 196084383 | rs985978 | -0.701 | -1.384 | 1.849 | 2.294 | -0.01 | -0.016 | -0.016 | 1.646 | 0.18 | 0.2 | 0.23 | 0.12 | 0.17 | 0.15 |
| 2 | 196087150 | rs13018723 | -2.991 | -1.248 | 2.365 | 2.497 | 0.012 | -0.05 | -0.05 | 0 | 0.11 | 0.46 | 0.09 | 0.02 | 0.07 | 0.05 |
| 2 | 196102559 | rs1021335 | 2.28 | -1.254 | 2.406 | 2.587 | -0.021 | 0.031 | 0.031 | 1.647 | 0.73 | 0.28 | 0.62 | 0.56 | 0.8 | 0.74 |
| 2 | 196116079 | rs6755097 | -2.249 | -1.244 | 2.39 | 2.596 | 0.004 | 0.016 | 0.016 | 0.143 | 0.08 | 0.28 | 0.07 | 0.08 | 0.06 | 0.07 |
| 2 | 196134798 | rs1489771 | 2.182 | -1.26 | 2.201 | 2.767 | -0.021 | 0.03 | 0.03 | 1.75 | 0.73 | 0.28 | 0.63 | 0.56 | 0.8 | 0.74 |
| 2 | 196146395 | rs7584938 | -2.378 | -1.002 | 1.801 | 2.694 | 0 | 0.045 | 0.045 | 0.862 | 0.08 | 0.31 | 0.13 | 0.1 | 0.06 | 0.06 |
| 2 | 196167335 | rs777289 | 1.871 | -0.987 | 1.532 | 2.928 | -0.014 | 0.11 | 0.11 | 0 | 0.94 | 0.8 | 0.93 | 0.82 | 0.95 | 0.93 |
| 2 | 196186077 | rs1524039 | 1.862 | -0.981 | 1.452 | 2.828 | -0.015 | 0.095 | 0.095 | 0 | 0.94 | 0.82 | 0.93 | 0.83 | 0.96 | 0.94 |
| 2 | 196196134 | rs1357640 | 1.895 | -1.154 | 1.58 | 2.707 | -0.062 | 0.11 | 0.11 | 0.763 | 0.76 | 0.42 | 0.58 | 0.52 | 0.8 | 0.72 |
| 2 | 196206373 | rs12618482 | -1.053 | -1.088 | 1.289 | 2.772 | -0.027 | 0.104 | 0.104 | 0 | 0.13 | 0.22 | 0.25 | 0.37 | 0.1 | 0.2 |
| 2 | 196256958 | rs1524040 | -0.558 | -1.209 | 0.756 | 2.873 | -0.034 | -0.048 | -0.048 | 0 | 0.14 | 0.01 | 0.14 | 0.17 | 0.15 | 0.21 |
| 2 | 196260290 | rs16840070 | 1.867 | -1.866 | 0.937 | 2.633 | 0.03 | -0.033 | -0.033 | 0.098 | 0.62 | 0.17 | 0.49 | 0.5 | 0.67 | 0.64 |
| 2 | 196260762 | rs16840081 | -0.893 | -1.824 | 1.184 | 2.702 | -0.008 | 0.013 | 0.013 | 0.054 | 0.14 | 0.16 | 0.26 | 0.28 | 0.11 | 0.2 |
| 2 | 196261366 | rs16840084 | -0.954 | -1.821 | 1.236 | 2.703 | -0.032 | 0.055 | 0.055 | 0.034 | 0.14 | 0.25 | 0.27 | 0.27 | 0.11 | 0.19 |
| 2 | 196272301 | rs17780773 | 2.628 | -2.012 | 0.776 | 2.834 | 0.034 | 0.275 | 0.275 | 1.027 | 0.49 | 0.07 | 0.3 | 0.18 | 0.59 | 0.45 |
| 2 | 196273054 | rs16840306 | -0.634 | -1.873 | 1.111 | 2.823 | -0.031 | 0.01 | 0.01 | 0 | 0.13 | 0.27 | 0.15 | 0.11 | 0.08 | 0.17 |
| 2 | 196274566 | rs10176269 | -1.814 | -1.897 | 1.226 | 2.695 | -0.007 | 0.018 | 0.018 | 0.096 | 0.3 | 0.52 | 0.35 | 0.29 | 0.28 | 0.3 |
| 2 | 196275148 | rs7593614 | -0.281 | -1.785 | 0.195 | 2.15 | 0.012 | -0.024 | -0.024 | 0.194 | 0.15 | 0.2 | 0.18 | 0.06 | 0.18 | 0.1 |
| 2 | 196277506 | rs10497759 | 1.336 | -1.618 | -0.466 | 2.173 | 0.029 | -0.042 | -0.042 | 0 | 0.59 | 0.13 | 0.46 | 0.48 | 0.64 | 0.66 |
| 2 | 196288136 | rs4505549 | 1.727 | -1.588 | -0.638 | 1.885 | 0.041 | 0.094 | 0.094 | 0.247 | 0.51 | 0.07 | 0.45 | 0.34 | 0.63 | 0.61 |
| 2 | 196289817 | rs10176418 | 1.613 | -1.868 | 0.566 | 2.184 | -0.061 | 0.096 | 0.096 | 0.097 | 0.77 | 0.45 | 0.59 | 0.6 | 0.86 | 0.66 |
| 2 | 196294622 | rs6706746 | -0.008 | -1.883 | 0.585 | 2.22 | 0.02 | -0.018 | -0.018 | 0 | 0.07 | 0.02 | 0.12 | 0.17 | 0.03 | 0.16 |
| 2 | 196300251 | rs6731123 | 0.048 | -1.839 | 0.98 | 2.239 | -0.017 | 0.015 | 0.015 | 0 | 0.92 | 0.88 | 0.87 | 0.83 | 0.97 | 0.85 |
| 2 | 196304657 | rs12614118 | 2.195 | -2.262 | 0.887 | 2.383 | -0.065 | 0.032 | 0.032 | 0 | 0.68 | 0.44 | 0.47 | 0.49 | 0.83 | 0.61 |
| 2 | 196305450 | rs10931697 | 1.24 | -2.27 | 1.262 | 2.393 | -0.016 | 0.142 | 0.142 | 0 | 0.86 | 0.56 | 0.71 | 0.76 | 0.89 | 0.84 |
| 2 | 196309463 | rs10497760 | -1.191 | -2.284 | 1.515 | 2.374 | 0.003 | 0.138 | 0.138 | 0 | 0.06 | 0.15 | 0.23 | 0.24 | 0.07 | 0.16 |
| 2 | 196309536 | rs1542865 | 2.368 | -2.296 | 1.504 | 2.344 | -0.065 | 0.034 | 0.034 | 0 | 0.68 | 0.45 | 0.47 | 0.48 | 0.83 | 0.61 |
| 2 | 196316701 | rs16841191 | -1.323 | -2.329 | 1.536 | 2.329 | -0.018 | 0.061 | 0.061 | 0 | 0.15 | 0.17 | 0.36 | 0.34 | 0.11 | 0.22 |
| 2 | 196316861 | rs10208012 | -0.591 | -2.297 | 1.837 | 2.122 | 0.009 | -0.058 | -0.058 | 0 | 0.17 | 0.21 | 0.14 | 0.04 | 0.22 | 0.1 |
| 2 | 196319300 | rs7559362 | 3.316 | -2.298 | 1.693 | 2.232 | -0.053 | 0.147 | 0.147 | 0.369 | 0.45 | 0.03 | 0.29 | 0.26 | 0.56 | 0.48 |
| 2 | 196326146 | rs12693770 | -0.268 | -2.364 | 2.321 | 2.35 | 0.001 | -0.011 | -0.011 | 0 | 0.12 | 0.1 | 0.12 | 0.04 | 0.19 | 0.08 |
| 2 | 196326799 | rs10497775 | 0.23 | -2.358 | 2.255 | 2.385 | 0.012 | -0.011 | -0.011 | 0 | 0.07 | 0.03 | 0.11 | 0.17 | 0.03 | 0.16 |
| 2 | 196328390 | rs1392658 | 3.147 | -2.372 | 2.361 | 2.423 | 0.109 | 0.02 | 0.02 | 0 | 0.51 | 0.17 | 0.32 | 0.44 | 0.61 | 0.51 |
| 2 | 196335282 | rs6434780 | -2.222 | -2.341 | 2.222 | 2.449 | -0.018 | 0.026 | 0.026 | 1.086 | 0.34 | 0.52 | 0.51 | 0.38 | 0.33 | 0.29 |
| 2 | 196351189 | rs4355137 | 2.967 | -2.296 | 2.227 | 2.426 | 0.093 | 0.024 | 0.024 | 0 | 0.51 | 0.25 | 0.32 | 0.44 | 0.61 | 0.51 |
| 2 | 196360157 | rs2889103 | -1.867 | -2.294 | 2.118 | 2.427 | -0.028 | 0.003 | 0.003 | 0 | 0.23 | 0.25 | 0.27 | 0.28 | 0.09 | 0.26 |
| 2 | 196365890 | rs4591358 | -1.647 | -2.247 | 2.148 | 2.398 | 0.002 | 0.016 | 0.016 | 0.31 | 0.25 | 0.4 | 0.39 | 0.28 | 0.3 | 0.24 |
| 2 | 196370612 | rs6745113 | -2.954 | -2.268 | 1.853 | 2.383 | 0.038 | 0.007 | 0.007 | 0 | 0.47 | 0.64 | 0.65 | 0.53 | 0.38 | 0.52 |
| 2 | 196373911 | rs13393013 | 0.176 | -2.347 | 1.432 | 2.366 | 0 | -0.001 | -0.001 | 0 | 0.07 | 0.07 | 0.12 | 0.18 | 0.03 | 0.16 |
| 2 | 196374552 | rs13008940 | -1.106 | -2.409 | 1.69 | 2.406 | 0.007 | -0.051 | -0.051 | 1.033 | 0.2 | 0.23 | 0.16 | 0.04 | 0.23 | 0.17 |
| 2 | 196375520 | rs17177643 | 2.83 | -2.375 | 1.635 | 2.481 | 0.108 | 0.22 | 0.22 | 0.209 | 0.49 | 0.08 | 0.32 | 0.24 | 0.62 | 0.44 |
| 2 | 196400096 | rs7579393 | -2.452 | -2.125 | 1.825 | 2.099 | -0.021 | -0.003 | -0.003 | 0 | 0.37 | 0.39 | 0.39 | 0.34 | 0.28 | 0.44 |
| 2 | 196400949 | rs13415067 | 0.067 | -2.04 | 1.646 | 2.084 | -0.003 | 0.002 | 0.002 | 0 | 0.07 | 0.05 | 0.12 | 0.19 | 0.03 | 0.16 |
| 2 | 196414587 | rs10200021 | -1.506 | -1.972 | 1.591 | 2.08 | -0.009 | 0.011 | 0.011 | 0 | 0.23 | 0.38 | 0.25 | 0.19 | 0.22 | 0.25 |
| 2 | 196436054 | rs6746966 | 2.241 | -1.897 | 0.764 | 2.09 | 0.095 | 0.007 | 0.007 | 0 | 0.55 | 0.2 | 0.35 | 0.42 | 0.67 | 0.51 |
| 2 | 196438150 | rs4850368 | -0.369 | -1.585 | 0.868 | 2.007 | 0.003 | -0.004 | -0.004 | 0 | 0.07 | 0.06 | 0.09 | 0.18 | 0.03 | 0.14 |
| 2 | 196442199 | rs10497761 | -1.286 | -1.384 | 1.181 | 1.631 | -0.018 | 0.002 | 0.002 | 0 | 0.26 | 0.5 | 0.28 | 0.2 | 0.21 | 0.22 |
| 2 | 196442418 | rs7566549 | 1.642 | -1.367 | 0.833 | 1.706 | 0.039 | -0.022 | -0.022 | 3.093 | 0.64 | 0.24 | 0.5 | 0.65 | 0.73 | 0.66 |
| 2 | 196444229 | rs16837470 | -0.575 | -0.594 | 0.651 | 1.441 | -0.001 | 0.121 | 0.121 | 0 | 0.06 | 0.2 | 0.2 | 0.16 | 0.05 | 0.08 |
| 2 | 196446217 | rs6759802 | -1.088 | -0.35 | 0.038 | 0.991 | 0.005 | -0.025 | -0.025 | 0 | 0.17 | 0.18 | 0.12 | 0.05 | 0.18 | 0.18 |
| 2 | 196448961 | rs6758272 | -1.39 | -0.662 | 0.053 | 1.939 | -0.004 | 0.063 | 0.063 | 0.238 | 0.14 | 0.19 | 0.3 | 0.45 | 0.1 | 0.24 |
| 2 | 196452537 | rs6434788 | -0.498 | -2.021 | 0.578 | 0.766 | -0.044 | -0.086 | -0.086 | 4.013 | 0.3 | 0.36 | 0.19 | 0.07 | 0.18 | 0.16 |
| 2 | 196454192 | rs16837576 | -1.18 | -2.069 | 0.542 | 1.246 | -0.021 | -0.023 | -0.023 | 0 | 0.37 | 0.38 | 0.27 | 0.19 | 0.24 | 0.22 |
| 2 | 196455295 | rs17179105 | -0.25 | -2.155 | 0.927 | 1.174 | 0.059 | 0.088 | 0.088 | 0 | 0.27 | 0.09 | 0.12 | 0 | 0.18 | 0.12 |
| 2 | 196457566 | rs16837590 | 0.331 | -2.049 | 1.189 | 1.242 | 0.001 | -0.002 | -0.002 | 0 | 0.08 | 0.08 | 0.09 | 0.12 | 0.06 | 0.06 |
| 2 | 196461111 | rs10497762 | -0.886 | -2.015 | 0.723 | 0.925 | -0.001 | 0.056 | 0.056 | 0 | 0.18 | 0.24 | 0.41 | 0.39 | 0.17 | 0.29 |
| 2 | 196461451 | rs12473922 | 0.37 | -1.931 | 0.665 | 0.884 | 0.001 | 0.004 | 0.004 | 0 | 0.11 | 0.09 | 0.19 | 0.06 | 0.11 | 0.07 |

| | | | | | | | | | | | | | | | | |
|---|---|---|---|---|---|---|---|---|---|---|---|---|---|---|---|---|
| 2 | 196465136 | rs17794270 | 0.608 | -2.039 | 0.527 | 0.959 | 0 | -0.014 | -0.014 | 5.4 | 0.09 | 0.01 | 0.1 | 0.09 | 0.07 | 0.06 |
| 2 | 196467695 | rs17794360 | 0.09 | -2.529 | 0.188 | 0.571 | 0.074 | 0.128 | 0.128 | 0 | 0.3 | 0.09 | 0.13 | 0.01 | 0.21 | 0.17 |
| 2 | 196468431 | rs10931700 | 1.498 | -2.589 | 0.346 | 0.646 | -0.021 | 0.086 | 0.086 | 0 | 0.71 | 0.48 | 0.47 | 0.52 | 0.76 | 0.65 |
| 2 | 196469154 | rs4850632 | 2.08 | -2.574 | 0.37 | 0.656 | -0.039 | -0.055 | -0.055 | 2.136 | 0.38 | 0.13 | 0.26 | 0.45 | 0.54 | 0.43 |
| 2 | 196470225 | rs17794503 | 0.06 | -2.737 | 0.358 | 0.573 | 0.073 | 0.189 | 0.189 | 0 | 0.3 | 0.08 | 0.13 | 0.01 | 0.21 | 0.17 |
| 2 | 196472286 | rs13410347 | -0.527 | -2.687 | 0.496 | 0.662 | 0.01 | 0.158 | 0.158 | 0.007 | 0.08 | 0.17 | 0.22 | 0.33 | 0.06 | 0.22 |
| 2 | 196478618 | rs10497763 | -0.009 | -2.699 | 0.411 | 0.382 | 0.07 | 0.11 | 0.11 | 0 | 0.3 | 0.15 | 0.14 | 0.01 | 0.21 | 0.18 |
| 2 | 196484192 | rs1554168 | -1.328 | -3.207 | 0.296 | 0.774 | -0.034 | 0.007 | 0.007 | 2.846 | 0.31 | 0.38 | 0.42 | 0.64 | 0.17 | 0.3 |
| 2 | 196493768 | rs12471439 | -1.226 | -3.18 | 0.183 | 0.954 | 0.022 | -0.02 | -0.02 | 0.271 | 0.29 | 0.2 | 0.37 | 0.57 | 0.16 | 0.3 |
| 2 | 196506802 | rs6734593 | -0.744 | -3.151 | 0.127 | 0.923 | 0.014 | -0.027 | -0.027 | 0.015 | 0.24 | 0.12 | 0.31 | 0.31 | 0.15 | 0.29 |
| 2 | 196513007 | rs12996251 | 2.062 | -3.341 | 0.215 | 0.539 | -0.007 | 0.016 | 0.016 | 0.197 | 0.36 | 0.22 | 0.38 | 0.34 | 0.54 | 0.42 |
| 2 | 196517509 | rs11682521 | 1.049 | -3.775 | 0.437 | 0.317 | -0.016 | -0.01 | -0.01 | 0 | 0.55 | 0.49 | 0.49 | 0.34 | 0.7 | 0.51 |
| 2 | 196527469 | rs1898771 | 1.933 | -3.827 | 0.808 | 0.32 | 0.002 | 0.001 | 0.001 | 0.227 | 0.39 | 0.38 | 0.44 | 0.41 | 0.61 | 0.45 |
| 2 | 196528101 | rs11889699 | 2.537 | -3.89 | 0.813 | 0.31 | -0.012 | 0.008 | 0.008 | 0.522 | 0.35 | 0.11 | 0.38 | 0.34 | 0.57 | 0.43 |
| 2 | 196542002 | rs10178320 | 0.77 | -3.895 | 0.799 | 0.307 | 0.056 | 0.02 | 0.02 | 0 | 0.58 | 0.74 | 0.55 | 0.41 | 0.73 | 0.55 |
| 2 | 196550734 | rs17180544 | 0.678 | -3.817 | 0.666 | 0.203 | 0.047 | 0.125 | 0.125 | 0.11 | 0.21 | 0.01 | 0.11 | 0.01 | 0.16 | 0.11 |
| 2 | 196561132 | rs10194124 | 0.535 | -3.737 | 0.724 | 0.381 | 0.033 | -0.008 | -0.008 | 0 | 0.62 | 0.74 | 0.62 | 0.67 | 0.73 | 0.57 |
| 2 | 196571804 | rs10165594 | 1.972 | -3.735 | 0.744 | 0.313 | 0.002 | 0.001 | 0.001 | 0.193 | 0.39 | 0.38 | 0.44 | 0.41 | 0.6 | 0.45 |
| 2 | 196572611 | rs7565283 | 1.869 | -3.745 | 0.737 | 0.303 | -0.017 | -0.048 | -0.048 | 0.973 | 0.08 | 0.15 | 0.03 | 0 | 0.04 | 0.01 |
| 2 | 196618135 | rs898934 | 0.59 | -3.705 | 0.648 | 0.299 | 0.021 | -0.003 | -0.003 | 0 | 0.62 | 0.7 | 0.61 | 0.66 | 0.73 | 0.54 |
| 2 | 196622252 | rs1026628 | 1.98 | -3.527 | 0.587 | 0.251 | 0.003 | 0.003 | 0.003 | 0.461 | 0.38 | 0.37 | 0.45 | 0.41 | 0.6 | 0.47 |
| 2 | 196636986 | rs16839847 | -0.107 | -3.328 | 0.571 | 0.218 | -0.035 | -0.103 | -0.103 | 0.077 | 0.19 | 0.28 | 0.1 | 0 | 0.13 | 0.08 |
| 2 | 196659034 | rs17442764 | 0.672 | -3.022 | 0.27 | 0.598 | -0.01 | -0.004 | -0.004 | 0.736 | 0.07 | 0.03 | 0.18 | 0.21 | 0.06 | 0.21 |
| 2 | 196661774 | rs2280451 | 0.696 | -3.001 | 0.211 | 0.62 | -0.039 | -0.03 | -0.03 | 0 | 0.07 | 0.01 | 0.21 | 0.25 | 0.09 | 0.22 |
| 2 | 196664879 | rs12613279 | 1.306 | -2.749 | -0.019 | 0.423 | -0.042 | -0.033 | -0.033 | 0 | 0.08 | 0.01 | 0.08 | 0.12 | 0.14 | 0.1 |
| 2 | 196678777 | rs13035759 | 1.589 | -3.609 | 0.602 | 0.144 | 0.002 | 0.002 | 0.002 | 0 | 0.44 | 0.38 | 0.43 | 0.33 | 0.61 | 0.46 |
| 2 | 196682630 | rs975417 | 0.567 | -3.699 | 0.532 | 0.068 | 0.04 | -0.012 | -0.012 | 0 | 0.62 | 0.77 | 0.62 | 0.61 | 0.72 | 0.57 |
| 2 | 196694902 | rs12471921 | 0.053 | -3.673 | 0.483 | 0.046 | 0.007 | -0.035 | -0.035 | 0.22 | 0.13 | 0.07 | 0.23 | 0.32 | 0.08 | 0.29 |
| 2 | 196698255 | rs7582441 | 1.003 | -3.619 | 0.71 | -0.09 | -0.023 | -0.012 | -0.012 | 1.094 | 0.56 | 0.51 | 0.52 | 0.31 | 0.72 | 0.55 |
| 2 | 196702403 | rs6434802 | 0.849 | -3.657 | 0.605 | -0.077 | 0.046 | 0.035 | 0.035 | 0 | 0.57 | 0.67 | 0.52 | 0.29 | 0.71 | 0.54 |
| 2 | 196703316 | rs10198943 | 1.559 | -3.687 | 0.603 | -0.088 | -0.009 | -0.011 | -0.011 | 2.13 | 0.46 | 0.46 | 0.45 | 0.3 | 0.62 | 0.46 |
| 2 | 196720705 | rs16841018 | -0.028 | -3.976 | 1.189 | -0.526 | -0.029 | -0.079 | -0.079 | 0.692 | 0.13 | 0.29 | 0.08 | 0 | 0.09 | 0.07 |
| 2 | 196749544 | rs341947 | -1.07 | -4.117 | 1.283 | -0.649 | -0.026 | -0.057 | -0.057 | 0.009 | 0.24 | 0.68 | 0.17 | 0.07 | 0.16 | 0.1 |
| 2 | 196765294 | rs382397 | 1.527 | -4.098 | 1.32 | -0.638 | -0.017 | -0.045 | -0.045 | 0 | 0.08 | 0.13 | 0.04 | 0 | 0.04 | 0.01 |
| 2 | 196771780 | rs6710775 | -0.949 | -4.123 | 1.369 | -0.652 | -0.047 | -0.091 | -0.091 | 0.462 | 0.23 | 0.6 | 0.17 | 0.07 | 0.16 | 0.1 |
| 2 | 196775588 | rs16841955 | 0.622 | -4.109 | 1.368 | -0.646 | -0.005 | -0.029 | -0.029 | 1.643 | 0.11 | 0.13 | 0.06 | 0 | 0.08 | 0.07 |
| 2 | 196791276 | rs1489802 | 2.172 | -4.364 | 1.194 | -0.63 | 0.116 | -0.075 | -0.075 | 0 | 0.41 | 0.09 | 0.58 | 0.56 | 0.65 | 0.68 |
| 2 | 196791792 | rs7570606 | -0.735 | -4.376 | 1.37 | -0.624 | -0.061 | -0.119 | -0.119 | 0 | 0.22 | 0.48 | 0.16 | 0.07 | 0.16 | 0.1 |
| 2 | 196825402 | rs6719500 | -0.313 | -4.352 | 1.165 | -0.591 | -0.034 | -0.063 | -0.063 | 0.072 | 0.2 | 0.3 | 0.12 | 0.07 | 0.14 | 0.09 |
| 2 | 196851911 | rs10931715 | 1.738 | -4.474 | 0.941 | -0.597 | 0.176 | 0.065 | 0.065 | 0 | 0.48 | 0.17 | 0.63 | 0.64 | 0.66 | 0.76 |
| 2 | 196852494 | rs1519612 | -1.333 | -4.536 | 0.999 | -0.617 | -0.019 | 0.002 | 0.002 | 1.662 | 0.29 | 0.67 | 0.24 | 0.35 | 0.18 | 0.12 |
| 2 | 196856041 | rs17297290 | 0.422 | -4.432 | 0.902 | -0.588 | 0.067 | 0.13 | 0.13 | 0.129 | 0.22 | 0.01 | 0.11 | 0.01 | 0.16 | 0.12 |
| 2 | 196858665 | rs10189449 | -1.893 | -4.592 | 1.032 | -0.649 | 0.059 | -0.005 | -0.005 | 0 | 0.45 | 0.69 | 0.31 | 0.44 | 0.27 | 0.28 |
| 2 | 196865910 | rs7557335 | 1.394 | -4.164 | 1.423 | -0.557 | -0.012 | -0.044 | -0.044 | 0 | 0.09 | 0.14 | 0.03 | 0 | 0.05 | 0.01 |
| 2 | 196866420 | rs6434811 | 0.924 | -4.192 | 1.233 | -0.547 | 0.009 | -0.013 | -0.013 | 2.637 | 0.79 | 0.19 | 0.79 | 0.65 | 0.9 | 0.94 |
| 2 | 196880762 | rs6751734 | -0.805 | -4.157 | 1.023 | -0.54 | -0.072 | 0.023 | 0.023 | 8.975 | 0.18 | 0.6 | 0.1 | 0.21 | 0.07 | 0.04 |
| 2 | 196894770 | rs6414150 | 0.529 | -4.348 | 0.884 | -0.818 | 0.065 | 0.066 | 0.066 | 0 | 0.62 | 0.75 | 0.78 | 0.91 | 0.74 | 0.73 |
| 2 | 196912067 | rs4850381 | 0.595 | -4.221 | 0.94 | -0.765 | 0.087 | 0.094 | 0.094 | 0 | 0.58 | 0.75 | 0.77 | 0.9 | 0.72 | 0.72 |
| 2 | 196914283 | rs6434817 | 1.132 | -4.226 | 1.044 | -0.73 | 0.041 | 0.008 | 0.008 | 0 | 0.53 | 0.56 | 0.69 | 0.62 | 0.72 | 0.7 |
| 2 | 196924449 | rs10207336 | -0.181 | -4.027 | 1.013 | -0.714 | -0.025 | -0.051 | -0.051 | 0 | 0.14 | 0.66 | 0.13 | 0.07 | 0.11 | 0.03 |
| 2 | 196942213 | rs4622750 | 2.006 | -4.105 | 0.494 | -0.682 | 0.074 | -0.075 | -0.075 | 0.016 | 0.4 | 0.11 | 0.58 | 0.54 | 0.61 | 0.66 |
| 2 | 196945385 | rs16844650 | -0.714 | -4.249 | 0.707 | -0.76 | 0.001 | -0.001 | -0.001 | 3.751 | 0.37 | 0.34 | 0.2 | 0.09 | 0.23 | 0.23 |
| 2 | 196949416 | rs4850382 | -0.423 | -4.262 | 0.741 | -0.77 | -0.002 | 0 | 0 | 0 | 0.19 | 0.19 | 0.17 | 0.29 | 0.15 | 0.11 |
| 2 | 196952010 | rs6712905 | -1.447 | -4.262 | 0.727 | -0.768 | 0.074 | 0 | 0 | 0 | 0.46 | 0.67 | 0.34 | 0.44 | 0.26 | 0.28 |
| 2 | 196984757 | rs7575006 | 0.522 | -4.114 | 0.648 | -0.721 | -0.005 | -0.012 | -0.012 | 0 | 0.14 | 0.17 | 0.1 | 0.08 | 0.13 | 0.08 |
| 2 | 196985112 | rs7587769 | -1.876 | -4.207 | 0.674 | -0.766 | 0.075 | 0.096 | 0.096 | 0 | 0.54 | 0.68 | 0.33 | 0.17 | 0.39 | 0.32 |
| 2 | 196987326 | rs6733543 | 1.669 | -4.215 | 0.615 | -0.743 | 0.006 | 0.001 | 0.001 | 0.985 | 0.46 | 0.58 | 0.67 | 0.62 | 0.64 | 0.68 |
| 2 | 196995578 | rs6434828 | 0.412 | -4.205 | 0.616 | -0.75 | -0.025 | -0.03 | -0.03 | 0 | 0.62 | 0.6 | 0.8 | 0.91 | 0.77 | 0.77 |
| 2 | 196996654 | rs12994633 | 2.02 | -4.101 | 0.356 | -0.67 | 0.075 | -0.063 | -0.063 | 0.017 | 0.4 | 0.11 | 0.59 | 0.55 | 0.61 | 0.65 |
| 2 | 197021086 | rs6712631 | 1.363 | -4.046 | 0.478 | -0.65 | 0.03 | 0.031 | 0.031 | 0 | 0.48 | 0.43 | 0.7 | 0.9 | 0.62 | 0.68 |
| 2 | 197021276 | rs10931726 | 1.984 | -3.952 | 0.061 | -0.627 | 0.069 | -0.063 | -0.063 | 0 | 0.4 | 0.1 | 0.59 | 0.55 | 0.61 | 0.66 |
| 2 | 197037730 | rs3811630 | -0.832 | -3.717 | 0.05 | -0.641 | 0.014 | 0.007 | 0.007 | 0 | 0.37 | 0.34 | 0.2 | 0.07 | 0.23 | 0.18 |
| 2 | 197045077 | rs1378624 | 0.002 | -3.578 | 0.101 | -0.469 | -0.008 | -0.012 | -0.012 | 0.088 | 0.31 | 0.36 | 0.32 | 0.13 | 0.25 | 0.22 |
| 2 | 197045688 | rs13003408 | 0.817 | -3.622 | -0.604 | -0.688 | 0.121 | -0.051 | -0.051 | 0 | 0.58 | 0.15 | 0.65 | 0.55 | 0.72 | 0.73 |
| 2 | 197048768 | rs11693514 | 1.938 | -3.618 | -0.69 | -0.824 | 0.028 | -0.101 | -0.101 | 0.002 | 0.38 | 0.08 | 0.55 | 0.54 | 0.58 | 0.66 |
| 2 | 197049786 | rs12052865 | -0.842 | -3.501 | -0.444 | -0.798 | -0.06 | 0.003 | 0.003 | 0 | 0.28 | 0.37 | 0.19 | 0.3 | 0.14 | 0.16 |
| 2 | 197051708 | rs10931727 | -0.695 | -3.509 | -0.407 | -0.795 | -0.038 | 0 | 0 | 0 | 0.27 | 0.3 | 0.16 | 0.29 | 0.12 | 0.15 |
| 2 | 197053774 | rs3214010 | -0.755 | -3.407 | -0.863 | -0.779 | -0.007 | -0.001 | -0.001 | 0 | 0.28 | 0.26 | 0.2 | 0.3 | 0.16 | 0.11 |
| 2 | 197056124 | rs17302390 | 0.568 | -3.231 | -1.003 | -0.821 | 0.035 | 0.073 | 0.073 | 0.26 | 0.17 | 0.03 | 0.1 | 0 | 0.14 | 0.08 |

| | | | | | | | | | | | | | | | | |
|---|---|---|---|---|---|---|---|---|---|---|---|---|---|---|---|---|
| 2 | 197059471 | rs2889153 | -0.295 | -3.047 | -0.953 | -0.735 | -0.05 | -0.142 | -0.142 | 0 | 0.69 | 0.46 | 0.84 | 0.99 | 0.78 | 0.81 |
| 2 | 197063250 | rs16846841 | 0.041 | -2.929 | -1.166 | -0.546 | -0.028 | 0.026 | 0.026 | 1.154 | 0.1 | 0.22 | 0.1 | 0.16 | 0.05 | 0.05 |
| 2 | 197065179 | rs3748875 | 1.572 | -2.821 | -1.778 | -0.513 | -0.017 | -0.142 | -0.142 | 3.878 | 0.35 | 0.04 | 0.56 | 0.54 | 0.58 | 0.66 |
| 2 | 197065889 | rs3795921 | 0.097 | -1.9 | -0.769 | -0.745 | -0.044 | -0.133 | -0.133 | 0.013 | 0.72 | 0.6 | 0.84 | 0.99 | 0.78 | 0.81 |
| 2 | 197065964 | rs3820908 | 0.199 | -1.129 | -0.591 | -0.561 | -0.021 | -0.032 | -0.032 | 0 | 0.63 | 0.57 | 0.77 | 0.83 | 0.7 | 0.77 |
| 2 | 197068127 | rs17375180 | -0.787 | -0.178 | -0.586 | -0.466 | 0 | -0.027 | -0.027 | 0 | 0.27 | 0.24 | 0.23 | 0.01 | 0.3 | 0.12 |
| 2 | 197070382 | rs10206935 | -0.871 | 0.039 | 0.134 | -0.005 | 0.022 | -0.016 | -0.016 | 0 | 0.07 | 0.2 | 0.11 | 0.04 | 0.09 | 0.07 |
| 2 | 197073204 | rs10186734 | -1.975 | -0.582 | 0.941 | 0.185 | -0.032 | 0.018 | 0.018 | 0.547 | 0.19 | 0.56 | 0.15 | 0.2 | 0.13 | 0.08 |
| 2 | 197075021 | rs10804070 | -0.509 | -0.48 | 0.613 | 0.456 | 0 | 0.009 | 0.009 | 0.709 | 0.26 | 0.26 | 0.32 | 0.52 | 0.27 | 0.34 |
| 2 | 197078179 | rs7606889 | -0.958 | -0.656 | 1.45 | 0.303 | -0.042 | -0.073 | -0.073 | 0 | 0.12 | 0.37 | 0.05 | 0 | 0.05 | 0.02 |
| 2 | 197078453 | rs6434831 | -1.818 | -0.635 | 1.773 | 0.68 | 0.007 | -0.011 | -0.011 | 0.268 | 0.38 | 0.68 | 0.36 | 0.48 | 0.33 | 0.36 |
| 2 | 197080331 | rs9288258 | -1.524 | -0.667 | 2.338 | 0.636 | -0.056 | -0.088 | -0.088 | 4.821 | 0.13 | 0.59 | 0.08 | 0.01 | 0.05 | 0.02 |
| 2 | 197081215 | rs6707519 | -1.877 | -0.671 | 2.038 | 0.74 | 0.023 | 0.015 | 0.015 | 0 | 0.38 | 0.77 | 0.38 | 0.64 | 0.33 | 0.37 |
| 2 | 197081843 | rs1869793 | -0.383 | -0.632 | 2.043 | 0.717 | -0.024 | -0.045 | -0.045 | 0.696 | 0.09 | 0.15 | 0.03 | 0 | 0.05 | 0.02 |
| 2 | 197084089 | rs10205756 | -1.549 | -0.652 | 2.165 | 0.733 | -0.012 | 0.037 | 0.037 | 0 | 0.35 | 0.48 | 0.35 | 0.46 | 0.33 | 0.35 |
| 2 | 197085823 | rs2290512 | 0.024 | -0.567 | 1.735 | 1.108 | -0.012 | -0.039 | -0.039 | 0 | 0.17 | 0.04 | 0.24 | 0.22 | 0.2 | 0.11 |
| 2 | 197092013 | rs13431857 | -1.104 | -1.074 | 1.675 | 0.629 | -0.049 | 0.011 | 0.011 | 0 | 0.18 | 0.26 | 0.13 | 0.2 | 0.08 | 0.14 |
| 2 | 197093005 | rs3748874 | -1.121 | -1.081 | 1.846 | 0.626 | -0.052 | 0.012 | 0.012 | 0 | 0.18 | 0.27 | 0.13 | 0.2 | 0.08 | 0.14 |
| 2 | 197098024 | rs10183339 | -0.985 | -1.044 | 1.834 | 0.62 | -0.043 | 0.018 | 0.018 | 0 | 0.17 | 0.25 | 0.13 | 0.2 | 0.09 | 0.14 |
| 2 | 197098638 | rs6740409 | -1.172 | -1.061 | 1.8 | 0.625 | -0.079 | 0.046 | 0.046 | 0 | 0.17 | 0.42 | 0.15 | 0.2 | 0.08 | 0.14 |
| 2 | 197110184 | rs12622742 | -0.198 | -0.969 | 1.299 | 0.662 | -0.079 | -0.158 | -0.158 | 0 | 0.23 | 0.03 | 0.23 | 0.44 | 0.27 | 0.32 |
| 2 | 197110296 | rs7575508 | 1.364 | -1.128 | 1.386 | 0.406 | -0.01 | -0.007 | -0.007 | 0.464 | 0.58 | 0.58 | 0.62 | 0.37 | 0.65 | 0.54 |
| 2 | 197117081 | rs7593952 | -1.223 | -1.243 | 0.941 | 0.306 | -0.092 | -0.106 | -0.106 | 0.005 | 0.16 | 0.39 | 0.06 | 0.02 | 0.04 | 0.11 |
| 2 | 197118022 | rs1869796 | 1.579 | -1.109 | 0.538 | 0.382 | 0.072 | 0.001 | 0.001 | 0 | 0.57 | 0.17 | 0.6 | 0.4 | 0.69 | 0.55 |
| 2 | 197118312 | rs6749366 | 0.904 | -0.741 | 0.761 | 0.612 | -0.005 | 0.015 | 0.015 | 0.145 | 0.7 | 0.44 | 0.65 | 0.41 | 0.72 | 0.66 |
| 2 | 197122493 | rs6731238 | 1.242 | -0.887 | -0.191 | 0.22 | 0.097 | 0.119 | 0.119 | 0.245 | 0.51 | 0.14 | 0.55 | 0.33 | 0.61 | 0.49 |
| 2 | 197123050 | rs17231279 | -1.271 | -0.686 | 0.162 | 0.309 | 0.042 | -0.019 | -0.019 | 0 | 0.45 | 0.71 | 0.35 | 0.52 | 0.37 | 0.49 |
| 2 | 197123318 | rs13422185 | -0.992 | -0.894 | 0.236 | 0.369 | -0.034 | 0.062 | 0.062 | 0.428 | 0.18 | 0.64 | 0.15 | 0.29 | 0.09 | 0.39 |
| 2 | 197127249 | rs17231419 | 0.049 | -0.797 | 0.022 | 0.388 | -0.012 | 0.046 | 0.046 | 0.089 | 0.27 | 0.07 | 0.2 | 0.23 | 0.27 | 0.1 |
| 2 | 197127830 | rs9288260 | -0.859 | -0.547 | 0.259 | 0.582 | -0.037 | 0.119 | 0.119 | 0.7 | 0.16 | 0.56 | 0.15 | 0.29 | 0.08 | 0.38 |
| 2 | 197129767 | rs10189788 | -0.921 | -0.526 | 0.254 | 0.576 | -0.027 | 0.111 | 0.111 | 0.719 | 0.17 | 0.57 | 0.16 | 0.29 | 0.1 | 0.39 |
| 2 | 197134790 | rs6746308 | 1.025 | 0.19 | -0.65 | -0.115 | -0.023 | 0.028 | 0.028 | 0 | 0.51 | 0.13 | 0.57 | 0.57 | 0.52 | 0.7 |
| 2 | 197137698 | rs11893319 | -0.618 | -0.528 | -0.144 | -0.667 | -0.025 | -0.038 | -0.038 | 0 | 0.3 | 0.4 | 0.23 | 0.26 | 0.26 | 0.15 |
| 2 | 197141271 | rs4145361 | 0.546 | -0.821 | -0.23 | -1.091 | -0.011 | -0.076 | -0.076 | 0.008 | 0.7 | 0.44 | 0.81 | 0.97 | 0.81 | 0.8 |
| 2 | 197152881 | rs10804072 | 0.293 | -0.973 | 0.604 | -1.186 | -0.013 | -0.027 | -0.027 | 0 | 0.93 | 0.81 | 0.97 | 0.97 | 0.99 | 0.92 |
| 2 | 197153259 | rs4850695 | 1.281 | -1.753 | 0.456 | -1.598 | 0.088 | 0.156 | 0.156 | 0.264 | 0.6 | 0.19 | 0.77 | 0.94 | 0.79 | 0.67 |
| 2 | 197153880 | rs16849064 | -0.877 | -1.78 | 0.764 | -1.612 | -0.045 | -0.156 | -0.156 | 7.272 | 0.31 | 0.47 | 0.18 | 0.02 | 0.19 | 0.23 |
| 2 | 197154052 | rs1902271 | 1.264 | -1.773 | 0.733 | -1.608 | 0.061 | 0.094 | 0.094 | 0.132 | 0.61 | 0.24 | 0.78 | 0.96 | 0.8 | 0.69 |
| 2 | 197163582 | rs17829296 | 0.065 | -1.739 | 1.182 | -1.631 | 0.003 | 0.066 | 0.066 | 0.005 | 0.21 | 0.08 | 0.13 | 0 | 0.19 | 0.12 |
| 2 | 197164952 | rs13394918 | -0.981 | -1.921 | 1.266 | -1.887 | 0.003 | -0.002 | -0.002 | 0 | 0.33 | 0.32 | 0.21 | 0.18 | 0.27 | 0.23 |
| 2 | 197169190 | rs2124384 | -0.031 | -1.92 | 1.278 | -1.887 | -0.006 | -0.058 | -0.058 | 0.409 | 0.23 | 0.28 | 0.16 | 0.01 | 0.2 | 0.14 |
| 2 | 197172538 | rs2305566 | 0.298 | -1.908 | 1.589 | -1.87 | -0.002 | 0 | 0 | 0 | 0.1 | 0.04 | 0.06 | 0.16 | 0.07 | 0.1 |
| 2 | 197172816 | rs2305567 | 0.49 | -1.675 | 1.68 | -1.906 | -0.019 | -0.148 | -0.148 | 1.99 | 0.69 | 0.53 | 0.8 | 0.96 | 0.76 | 0.78 |
| 2 | 197174387 | rs6713102 | 1.03 | -1.632 | 1.629 | -1.86 | -0.009 | -0.063 | -0.063 | 0 | 0.59 | 0.49 | 0.74 | 0.8 | 0.69 | 0.69 |
| 2 | 197175982 | rs7558972 | 1.523 | -1.563 | 1.324 | -1.815 | 0.062 | 0.262 | 0.262 | 0 | 0.5 | 0.15 | 0.69 | 0.78 | 0.67 | 0.57 |
| 2 | 197184321 | rs1531111 | 0.673 | -2.086 | 1.981 | -2.042 | 0.03 | 0.011 | 0.011 | 0 | 0.64 | 0.39 | 0.82 | 0.97 | 0.83 | 0.77 |
| 2 | 197190821 | rs6725009 | 1.234 | -2.057 | 2.324 | -2.067 | 0.105 | 0.181 | 0.181 | 0 | 0.57 | 0.21 | 0.76 | 0.95 | 0.79 | 0.71 |
| 2 | 197195099 | rs16849716 | -0.771 | -2.028 | 2.497 | -2.035 | 0.022 | -0.001 | -0.001 | 0 | 0.33 | 0.28 | 0.13 | 0.01 | 0.16 | 0.22 |
| 2 | 197196727 | rs10166118 | 0.691 | -2.068 | 2.423 | -2.055 | 0.007 | -0.027 | -0.027 | 0 | 0.63 | 0.43 | 0.82 | 0.97 | 0.83 | 0.77 |
| 2 | 197200191 | rs1014255 | 0.691 | -2.045 | 2.094 | -2.044 | -0.025 | -0.078 | -0.078 | 0 | 0.63 | 0.44 | 0.83 | 0.97 | 0.83 | 0.77 |
| 2 | 197208366 | rs6757981 | -0.183 | -2.274 | 1.711 | -2.086 | -0.01 | -0.034 | -0.034 | 0.015 | 0.09 | 0.23 | 0.07 | 0.06 | 0.05 | 0.07 |
| 2 | 197217035 | rs17830265 | 0.832 | -2.267 | 1.506 | -2.083 | -0.011 | 0.027 | 0.027 | 2.05 | 0.07 | 0 | 0.04 | 0.05 | 0.04 | 0.06 |
| 2 | 197219340 | rs16850199 | 0.426 | -2.268 | 1.589 | -2.084 | -0.001 | -0.032 | -0.032 | 0 | 0.08 | 0.19 | 0.05 | 0.05 | 0.04 | 0.06 |
| 2 | 197232054 | rs12327933 | -1.484 | -2.386 | 1.387 | -2.202 | 0.096 | 0.12 | 0.12 | 0.188 | 0.44 | 0.71 | 0.23 | 0.07 | 0.2 | 0.28 |
| 2 | 197236215 | rs6749624 | -0.562 | -2.564 | 1.453 | -2.256 | -0.058 | -0.12 | -0.12 | 0 | 0.22 | 0.55 | 0.11 | 0.01 | 0.1 | 0.19 |
| 2 | 197247672 | rs11899924 | -1.371 | -2.723 | 1.268 | -2.482 | -0.034 | -0.077 | -0.077 | 0 | 0.34 | 0.51 | 0.19 | 0.18 | 0.2 | 0.26 |
| 2 | 197251253 | rs11902313 | 0.549 | -2.813 | 1.362 | -2.421 | -0.009 | -0.03 | -0.03 | 0 | 0.08 | 0.15 | 0.05 | 0.04 | 0.03 | 0.08 |
| 2 | 197252086 | rs4398290 | 1.157 | -2.865 | 1.408 | -2.454 | -0.018 | -0.035 | -0.035 | 0 | 0.67 | 0.39 | 0.81 | 0.82 | 0.84 | 0.75 |
| 2 | 197254889 | rs4414697 | -0.146 | -2.76 | 1.044 | -2.384 | -0.018 | -0.036 | -0.036 | 0 | 0.89 | 0.8 | 0.93 | 0.95 | 0.96 | 0.9 |
| 2 | 197254937 | rs13427710 | -0.12 | -2.786 | 1.084 | -2.48 | -0.054 | -0.12 | -0.12 | 0.051 | 0.22 | 0.4 | 0.11 | 0.01 | 0.1 | 0.2 |
| 2 | 197255486 | rs6735220 | 0.789 | -2.7 | 0.787 | -2.432 | -0.024 | -0.069 | -0.069 | 0 | 0.67 | 0.41 | 0.83 | 0.93 | 0.86 | 0.7 |
| 2 | 197256269 | rs16850723 | 1.043 | -2.537 | -0.122 | -1.386 | -0.02 | -0.075 | -0.075 | 0 | 0.11 | 0.01 | 0.21 | 0.34 | 0.12 | 0.16 |
| 2 | 197262413 | rs6713635 | 1.356 | -2.505 | 0.319 | -1.964 | 0.098 | 0.188 | 0.188 | 0 | 0.47 | 0.21 | 0.73 | 0.8 | 0.73 | 0.54 |
| 2 | 197270594 | rs1358504 | 1.149 | -2.069 | 0.539 | -2.002 | 0.072 | 0.154 | 0.154 | 0 | 0.53 | 0.24 | 0.73 | 0.89 | 0.71 | 0.53 |
| 2 | 197273364 | rs6434849 | 0.726 | -2.144 | 0.601 | -2.091 | 0.043 | 0.055 | 0.055 | 0 | 0.59 | 0.28 | 0.77 | 0.93 | 0.75 | 0.65 |
| 2 | 197273377 | rs1528400 | 0.291 | -2.306 | 0.679 | -2.158 | -0.047 | -0.065 | -0.065 | 2.282 | 0.89 | 0.76 | 0.96 | 0.99 | 0.97 | 0.94 |
| 2 | 197274355 | rs12471412 | -0.275 | -2.311 | 0.732 | -2.16 | -0.025 | -0.04 | -0.04 | 0.321 | 0.91 | 0.86 | 0.97 | 1 | 0.97 | 0.95 |
| 2 | 197292383 | rs6434859 | 0.893 | -2.314 | 0.721 | -2.175 | 0.021 | -0.002 | -0.002 | 0 | 0.59 | 0.33 | 0.78 | 0.93 | 0.75 | 0.65 |
| 2 | 197295756 | rs1406218 | 1.38 | -2.277 | 0.659 | -2.138 | 0.057 | 0.096 | 0.096 | 0 | 0.53 | 0.29 | 0.74 | 0.89 | 0.71 | 0.51 |

| | | | | | | | | | | | | | | | | |
|---|---|---|---|---|---|---|---|---|---|---|---|---|---|---|---|---|
| 2 | 197298222 | rs2204641 | -0.196 | -2.025 | 0.286 | -1.891 | -0.017 | -0.03 | -0.03 | 0 | 0.91 | 0.88 | 0.97 | 0.99 | 0.97 | 0.95 |
| 2 | 197298232 | rs2204640 | -0.753 | -1.909 | 0.128 | -1.834 | -0.003 | -0.013 | -0.013 | 0 | 0.93 | 0.95 | 0.98 | 1 | 0.97 | 0.96 |
| 2 | 197300379 | rs6745575 | -0.805 | -1.427 | 0.178 | -1.525 | -0.004 | -0.041 | -0.041 | 0.078 | 0.3 | 0.31 | 0.18 | 0.07 | 0.22 | 0.3 |
| 2 | 197309433 | rs12693795 | -0.997 | -0.914 | -0.403 | -1.182 | -0.009 | -0.06 | -0.06 | 0.895 | 0.39 | 0.59 | 0.2 | 0.1 | 0.25 | 0.41 |
| 2 | 197309807 | rs10497786 | -1.272 | -1.268 | -0.11 | -0.257 | 0.002 | -0.004 | -0.004 | 0 | 0.17 | 0.15 | 0.38 | 0.35 | 0.15 | 0.26 |
| 2 | 197311265 | rs1528398 | -1.893 | -1.627 | 0.059 | -0.924 | 0.07 | 0.106 | 0.106 | 0 | 0.42 | 0.78 | 0.27 | 0.11 | 0.25 | 0.31 |
| 2 | 197315114 | rs6721157 | -0.546 | -1.816 | 0.777 | -1.044 | -0.032 | -0.032 | -0.032 | 0 | 0.17 | 0.28 | 0.11 | 0.08 | 0.1 | 0.1 |
| 2 | 197318655 | rs10469750 | -2.12 | -1.891 | 1.167 | -1.093 | -0.064 | -0.09 | -0.09 | 0.073 | 0.29 | 0.51 | 0.19 | 0.11 | 0.17 | 0.29 |
| 2 | 197320088 | rs13402395 | -0.547 | -1.833 | 1.062 | -1.054 | -0.031 | -0.031 | -0.031 | 0 | 0.17 | 0.28 | 0.11 | 0.08 | 0.1 | 0.1 |
| 2 | 197321747 | rs1358402 | -1.909 | -1.849 | 1.263 | -1.077 | -0.017 | -0.05 | -0.05 | 0 | 0.28 | 0.38 | 0.18 | 0.09 | 0.18 | 0.13 |
| 2 | 197327735 | rs9677260 | -2.716 | -2.075 | 1.524 | -1.15 | -0.026 | -0.093 | -0.093 | 0.051 | 0.3 | 0.41 | 0.18 | 0.03 | 0.17 | 0.25 |
| 2 | 197332633 | rs16851952 | -1.156 | -1.851 | 1.366 | -1.04 | 0 | -0.008 | -0.008 | 0 | 0.11 | 0.11 | 0.07 | 0.02 | 0.07 | 0.19 |
| 2 | 197351151 | rs17244073 | -0.573 | -1.826 | 1.286 | -1.047 | -0.005 | 0.001 | 0.001 | 0 | 0.08 | 0.07 | 0.08 | 0 | 0.09 | 0.03 |
| 2 | 197354190 | rs2375856 | -2.818 | -1.978 | 1.721 | -1.1 | 0.009 | -0.007 | -0.007 | 0 | 0.34 | 0.55 | 0.25 | 0.08 | 0.21 | 0.23 |
| 2 | 197360596 | rs3902848 | -1.805 | -1.954 | 1.71 | -1.083 | 0.008 | 0.016 | 0.016 | 0.06 | 0.19 | 0.15 | 0.15 | 0.06 | 0.15 | 0.11 |
| 2 | 197404714 | rs4341955 | 2.676 | -2.186 | 1.469 | -1.231 | 0.114 | 0.266 | 0.266 | 0.733 | 0.58 | 0.03 | 0.72 | 0.92 | 0.77 | 0.71 |
| 2 | 197414906 | rs6719725 | -2.066 | -2.105 | 1.84 | -1.188 | -0.018 | -0.052 | -0.052 | 0 | 0.24 | 0.29 | 0.18 | 0.06 | 0.17 | 0.12 |
| 2 | 197417132 | rs1534382 | -1.854 | -2.109 | 1.873 | -1.189 | 0.027 | 0.063 | 0.063 | 1.08 | 0.23 | 0.11 | 0.16 | 0.05 | 0.18 | 0.11 |
| 2 | 197433521 | rs2222728 | -0.719 | -2.028 | 2.036 | -1.176 | -0.019 | -0.031 | -0.031 | 0 | 0.1 | 0.16 | 0.05 | 0.02 | 0.05 | 0.17 |
| 2 | 197457241 | rs7577213 | 1.616 | -1.961 | 1.785 | -1.143 | 0.007 | -0.004 | -0.004 | 0 | 0.78 | 0.81 | 0.88 | 0.96 | 0.83 | 0.89 |
| 2 | 197470002 | rs6434862 | 2.863 | -1.881 | 1.956 | -1.2 | 0.061 | 0.104 | 0.104 | 0.039 | 0.62 | 0.19 | 0.77 | 0.94 | 0.75 | 0.71 |
| 2 | 197470459 | rs16855046 | -0.621 | -1.833 | 1.864 | -1.074 | -0.011 | -0.016 | -0.016 | 0 | 0.09 | 0.14 | 0.05 | 0.03 | 0.06 | 0.16 |
| 2 | 197475599 | rs12994106 | 0.549 | -1.707 | 1.634 | -1.062 | 0.039 | 0.052 | 0.052 | 0.157 | 0.87 | 0.97 | 0.97 | 0.97 | 0.93 | 0.92 |
| 2 | 197482207 | rs4241191 | 1.623 | -1.705 | 1.757 | -1.069 | 0.012 | 0.035 | 0.035 | 0.044 | 0.78 | 0.86 | 0.9 | 0.95 | 0.83 | 0.89 |
| 2 | 197484762 | rs7581407 | 1.773 | -1.709 | 1.992 | -1.067 | -0.026 | -0.093 | -0.093 | 0 | 0.78 | 0.65 | 0.87 | 0.95 | 0.83 | 0.9 |
| 2 | 197490846 | rs10497792 | -1.061 | -1.629 | 1.864 | -1.053 | 0.008 | -0.037 | -0.037 | 0 | 0.09 | 0.15 | 0.08 | 0.01 | 0.11 | 0.03 |
| 2 | 197491974 | rs10497791 | -1.458 | -1.635 | 1.955 | -1.055 | 0.024 | -0.046 | -0.046 | 0.001 | 0.1 | 0.24 | 0.1 | 0.03 | 0.11 | 0.03 |
| 2 | 197500771 | rs7575378 | 1.534 | -1.609 | 2.282 | -1.182 | -0.032 | -0.074 | -0.074 | 0.566 | 0.74 | 0.64 | 0.87 | 0.95 | 0.83 | 0.73 |
| 2 | 197504764 | rs16855610 | -0.598 | -1.609 | 2.257 | -1.16 | -0.019 | -0.032 | -0.032 | 0 | 0.1 | 0.21 | 0.06 | 0.02 | 0.06 | 0.15 |
| 2 | 197504826 | rs6434873 | 2.664 | -1.983 | 2.061 | -1.436 | 0.067 | 0.151 | 0.151 | 0 | 0.58 | 0.19 | 0.74 | 0.93 | 0.7 | 0.7 |
| 2 | 197509306 | rs1533660 | 1.728 | -1.947 | 1.923 | -1.413 | 0.013 | 0.027 | 0.027 | 0 | 0.71 | 0.78 | 0.85 | 0.97 | 0.76 | 0.85 |
| 2 | 197522966 | rs10197202 | 1.875 | -1.974 | 1.897 | -0.907 | 0.017 | 0.028 | 0.028 | 0 | 0.7 | 0.78 | 0.85 | 0.83 | 0.75 | 0.84 |
| 2 | 197534692 | rs7590083 | 0.51 | -1.972 | 1.832 | -0.875 | 0.021 | -0.007 | -0.007 | 0 | 0.84 | 0.92 | 0.94 | 0.85 | 0.88 | 0.88 |
| 2 | 197541296 | rs17271281 | -0.644 | -1.936 | 1.809 | -1.128 | -0.003 | 0.004 | 0.004 | 0 | 0.13 | 0.1 | 0.09 | 0.01 | 0.13 | 0.03 |
| 2 | 197583379 | rs6751627 | -0.448 | -1.924 | 1.939 | -1.089 | -0.018 | -0.024 | -0.024 | 0 | 0.1 | 0.19 | 0.06 | 0.03 | 0.06 | 0.15 |
| 2 | 197605400 | rs2577583 | 2.404 | -1.932 | 1.981 | -1.104 | -0.013 | -0.059 | -0.059 | 0 | 0.6 | 0.45 | 0.79 | 0.92 | 0.71 | 0.71 |
| 2 | 197613153 | rs17383482 | -1.775 | -1.932 | 1.937 | -1.037 | 0.018 | 0.066 | 0.066 | 0 | 0.29 | 0.18 | 0.26 | 0.06 | 0.24 | 0.15 |
| 2 | 197617106 | rs16857304 | -0.454 | -1.909 | 1.818 | -1.019 | -0.015 | -0.016 | -0.016 | 0 | 0.1 | 0.19 | 0.06 | 0.05 | 0.06 | 0.15 |
| 2 | 197623781 | rs6434883 | -2.46 | -1.879 | 1.675 | -1.006 | -0.01 | -0.048 | -0.048 | 0.633 | 0.4 | 0.55 | 0.35 | 0.11 | 0.31 | 0.3 |
| 2 | 197626234 | rs6434885 | 2.524 | -1.779 | 1.487 | -1.088 | 0.087 | 0.234 | 0.234 | 0.064 | 0.55 | 0.11 | 0.73 | 0.92 | 0.68 | 0.69 |
| 2 | 197626466 | rs2289381 | 1.958 | -1.838 | 1.434 | -0.688 | 0.013 | -0.008 | -0.008 | 0.242 | 0.58 | 0.41 | 0.48 | 0.52 | 0.65 | 0.51 |
| 2 | 197628215 | rs1574424 | -0.8 | -1.665 | 1.407 | -1.416 | 0.02 | 0.04 | 0.04 | 0.045 | 0.16 | 0.07 | 0.05 | 0.02 | 0.11 | 0.14 |
| 2 | 197633664 | rs17442969 | -0.772 | -1.62 | 1.441 | -1.424 | -0.011 | -0.025 | -0.025 | 0 | 0.12 | 0.2 | 0.07 | 0.03 | 0.08 | 0.16 |
| 2 | 197669189 | rs2289380 | 3.027 | -1.403 | 1.359 | -1.473 | 0.05 | 0.3 | 0.3 | 0 | 0.53 | 0.06 | 0.72 | 0.92 | 0.65 | 0.65 |
| 2 | 197675680 | rs1992170 | -0.907 | -1.177 | 0.913 | -1.366 | 0.007 | 0.026 | 0.026 | 0.018 | 0.17 | 0.1 | 0.07 | 0.02 | 0.13 | 0.15 |
| 2 | 197676017 | rs11900782 | -0.794 | -1.166 | 0.924 | -1.35 | -0.009 | 0.006 | 0.006 | 0.324 | 0.12 | 0.08 | 0.09 | 0 | 0.14 | 0.04 |
| 2 | 197681043 | rs16858489 | -2.135 | -1.196 | 1.073 | -1.304 | -0.054 | -0.11 | -0.11 | 0.471 | 0.3 | 0.5 | 0.15 | 0.08 | 0.19 | 0.31 |
| 2 | 197683558 | rs16858525 | -0.86 | -1.197 | 1.088 | -1.304 | -0.012 | -0.028 | -0.028 | 0 | 0.12 | 0.22 | 0.07 | 0.03 | 0.08 | 0.16 |
| 2 | 197684441 | rs16858528 | -0.949 | -1.187 | 1.109 | -1.308 | 0.007 | 0.027 | 0.027 | 0.014 | 0.17 | 0.1 | 0.07 | 0.02 | 0.13 | 0.15 |
| 2 | 197701704 | rs7602892 | -0.846 | -1.243 | 1.047 | -1.308 | -0.011 | -0.025 | -0.025 | 0 | 0.12 | 0.2 | 0.08 | 0.03 | 0.08 | 0.16 |
| 2 | 197703076 | rs3795920 | -0.846 | -1.243 | 1.047 | -1.308 | -0.011 | -0.025 | -0.025 | 0 | 0.12 | 0.2 | 0.08 | 0.03 | 0.08 | 0.16 |
| 2 | 197715078 | rs1866186 | -2.204 | -1.247 | 1.09 | -1.311 | -0.054 | -0.11 | -0.11 | 0 | 0.3 | 0.5 | 0.15 | 0.08 | 0.19 | 0.31 |
| 2 | 197756153 | rs332297 | -2.367 | -1.247 | 1.091 | -1.311 | -0.053 | -0.109 | -0.109 | 0 | 0.3 | 0.5 | 0.15 | 0.08 | 0.19 | 0.31 |
| 2 | 197764931 | rs7569332 | -0.988 | -1.126 | 0.945 | -1.242 | -0.016 | -0.026 | -0.026 | 0 | 0.12 | 0.25 | 0.08 | 0.05 | 0.08 | 0.17 |
| 2 | 197797061 | rs11693566 | 1.352 | -0.923 | 1.175 | -1.192 | 0.008 | -0.07 | -0.07 | 0 | 0.85 | 0.75 | 0.91 | 0.97 | 0.87 | 0.96 |
| 2 | 197797224 | rs10931765 | 2.092 | -0.945 | 1.142 | -1.092 | -0.019 | -0.062 | -0.062 | 0 | 0.69 | 0.47 | 0.76 | 0.85 | 0.75 | 0.82 |
| 2 | 197799124 | rs6749960 | -1.109 | -0.809 | 0.447 | -1.054 | 0.01 | 0.01 | 0.01 | 0 | 0.17 | 0.1 | 0.14 | 0.12 | 0.13 | 0.14 |
| 2 | 197804986 | rs2376102 | 0.904 | -0.687 | 0.304 | -1.02 | -0.001 | -0.054 | -0.054 | 0 | 0.87 | 0.84 | 0.92 | 1 | 0.88 | 0.96 |
| 2 | 197816284 | rs2376187 | -1.196 | -0.649 | -0.198 | -0.904 | 0.01 | 0.003 | 0.003 | 0 | 0.17 | 0.11 | 0.16 | 0.13 | 0.13 | 0.15 |
| 2 | 197823816 | rs6714155 | -0.907 | -0.628 | -0.22 | -0.802 | 0.01 | -0.017 | -0.017 | 0 | 0.12 | 0.07 | 0.13 | 0.14 | 0.07 | 0.14 |
| 2 | 197824902 | rs10497798 | -1.064 | -0.429 | -0.118 | -0.859 | -0.003 | -0.004 | -0.004 | 0.158 | 0.12 | 0.07 | 0.07 | 0 | 0.12 | 0.04 |
| 2 | 197828599 | rs16859921 | -1.1 | -0.458 | -0.188 | -0.871 | -0.006 | 0.005 | 0.005 | 0.008 | 0.12 | 0.07 | 0.08 | 0 | 0.12 | 0.04 |
| 2 | 197828651 | rs6434895 | -1.765 | -0.61 | -0.333 | -1.342 | 0.013 | 0.092 | 0.092 | 0 | 0.39 | 0.24 | 0.2 | 0.16 | 0.3 | 0.36 |
| 2 | 197831878 | rs1437871 | -0.682 | -0.384 | 0.237 | -0.925 | 0 | -0.017 | -0.017 | 0 | 0.08 | 0.09 | 0.04 | 0 | 0.09 | 0.1 |
| 2 | 197833037 | rs13404405 | -1.551 | -0.437 | 0.693 | -0.951 | 0.004 | -0.052 | -0.052 | 0 | 0.1 | 0.23 | 0.06 | 0 | 0.09 | 0.1 |
| 2 | 197834551 | rs6734285 | -2.213 | -0.523 | 0.866 | -1.137 | -0.007 | -0.046 | -0.046 | 0 | 0.28 | 0.44 | 0.21 | 0.14 | 0.25 | 0.23 |
| 2 | 197836360 | rs7591162 | 2.637 | -0.691 | 1.137 | -1.196 | 0.031 | 0.109 | 0.109 | 0 | 0.59 | 0.19 | 0.72 | 0.83 | 0.69 | 0.72 |
| 2 | 197845449 | rs6735887 | 2.629 | -0.695 | 1.154 | -1.176 | 0.034 | 0.098 | 0.098 | 0.036 | 0.61 | 0.18 | 0.73 | 0.83 | 0.72 | 0.74 |
| 2 | 197848176 | rs5016260 | 0.617 | -0.674 | 1.377 | -1.032 | -0.009 | -0.077 | -0.077 | 0.522 | 0.88 | 0.67 | 0.93 | 1 | 0.94 | 0.96 |

| | | | | | | | | | | | | | | | | |
|---|---|---|---|---|---|---|---|---|---|---|---|---|---|---|---|---|
| 2 | 197850176 | rs16860099 | -1.183 | -0.542 | 1.118 | -0.864 | -0.002 | 0.001 | 0.001 | 0 | 0.16 | 0.23 | 0.13 | 0.17 | 0.14 | 0.12 |
| 2 | 197850652 | rs11888210 | -1.532 | -0.524 | 0.851 | -0.854 | -0.015 | 0.003 | 0.003 | 0.1 | 0.19 | 0.4 | 0.15 | 0.17 | 0.14 | 0.12 |
| 2 | 197857070 | rs10931768 | 1.887 | -0.11 | -0.287 | -1.22 | -0.035 | 0.152 | 0.152 | 0 | 0.48 | 0.15 | 0.59 | 0.79 | 0.56 | 0.68 |
| 2 | 197858129 | rs1036542 | 1.589 | -0.087 | 0.07 | -1.08 | 0.015 | 0.029 | 0.029 | 0 | 0.69 | 0.22 | 0.82 | 0.96 | 0.79 | 0.86 |
| 2 | 197860491 | rs16860250 | -1.38 | -0.117 | 1.033 | -1.128 | -0.001 | -0.031 | -0.031 | 0 | 0.12 | 0.14 | 0.05 | 0 | 0.1 | 0.06 |
| 2 | 197862903 | rs2376209 | 1.029 | -0.164 | 1.079 | -1.128 | -0.014 | -0.096 | -0.096 | 0.11 | 0.77 | 0.37 | 0.85 | 0.97 | 0.85 | 0.87 |
| 2 | 197870808 | rs2376211 | 2.469 | -0.111 | 0.951 | -1.128 | 0.009 | 0.109 | 0.109 | 0 | 0.56 | 0.19 | 0.66 | 0.81 | 0.65 | 0.75 |
| 2 | 197882236 | rs10497802 | -1.735 | -0.389 | 1.147 | -1.062 | 0 | 0.006 | 0.006 | 0 | 0.24 | 0.21 | 0.21 | 0.16 | 0.2 | 0.15 |
| 2 | 197889022 | rs13015419 | 0.406 | -0.377 | 0.942 | -1.059 | -0.016 | -0.095 | -0.095 | 1.557 | 0.83 | 0.57 | 0.88 | 0.97 | 0.89 | 0.92 |
| 2 | 197903464 | rs6714359 | -0.082 | -0.328 | 0.371 | -1.043 | 0 | -0.025 | -0.025 | 0 | 0.85 | 0.87 | 0.91 | 1 | 0.89 | 0.93 |
| 2 | 197904195 | rs16860949 | -2.042 | -0.188 | 0.512 | -0.794 | -0.009 | -0.013 | -0.013 | 0 | 0.27 | 0.64 | 0.25 | 0.19 | 0.2 | 0.16 |
| 2 | 197907294 | rs4850749 | -0.962 | -0.115 | -0.004 | -0.833 | -0.023 | -0.011 | -0.011 | 0 | 0.14 | 0.03 | 0.12 | 0.12 | 0.12 | 0.14 |
| 2 | 197908675 | rs13019481 | 1.968 | -0.4 | 0.426 | -1.109 | -0.003 | -0.02 | -0.02 | 0 | 0.63 | 0.41 | 0.73 | 0.86 | 0.72 | 0.77 |
| 2 | 197910432 | rs6434900 | 2.355 | -0.522 | 0.434 | -0.583 | 0.032 | 0.046 | 0.046 | 0 | 0.58 | 0.18 | 0.65 | 0.67 | 0.68 | 0.74 |
| 2 | 197920169 | rs4850410 | 1.914 | -0.62 | 0.602 | -0.969 | -0.021 | -0.045 | -0.045 | 0.949 | 0.69 | 0.44 | 0.81 | 0.87 | 0.8 | 0.79 |
| 2 | 197942576 | rs12478675 | -0.517 | -0.642 | 0.489 | -0.985 | -0.047 | -0.031 | -0.031 | 5.123 | 0.16 | 0.02 | 0.1 | 0.13 | 0.16 | 0.12 |
| 2 | 197942832 | rs4850415 | 2.327 | -0.589 | 0.559 | -0.943 | 0.038 | 0.102 | 0.102 | 0 | 0.59 | 0.18 | 0.75 | 0.85 | 0.73 | 0.76 |
| 2 | 197950058 | rs893401 | 2.27 | -0.571 | 0.565 | -0.934 | 0.028 | 0.07 | 0.07 | 0 | 0.6 | 0.23 | 0.75 | 0.85 | 0.73 | 0.76 |
| 2 | 197976089 | rs4395261 | 2.362 | -0.786 | 0.612 | -1.362 | 0.055 | 0.117 | 0.117 | 0 | 0.59 | 0.17 | 0.78 | 0.96 | 0.75 | 0.76 |
| 2 | 197978625 | rs2697269 | 2.22 | -0.84 | 1.034 | -1.484 | -0.056 | -0.101 | -0.101 | 0 | 0.82 | 0.36 | 0.89 | 0.99 | 0.93 | 0.94 |
| 2 | 197986048 | rs2697270 | 0.608 | -0.855 | 1.706 | -1.491 | -0.022 | -0.048 | -0.048 | 0 | 0.91 | 0.78 | 0.95 | 0.99 | 0.96 | 0.96 |
| 2 | 197990509 | rs2579394 | 2.817 | -0.879 | 1.695 | -1.515 | 0.028 | 0.024 | 0.024 | 0 | 0.65 | 0.24 | 0.81 | 0.96 | 0.8 | 0.79 |
| 2 | 197990527 | rs6434907 | -2.014 | -0.873 | 1.708 | -1.502 | 0.005 | -0.021 | -0.021 | 0 | 0.09 | 0.42 | 0.06 | 0 | 0.03 | 0.03 |
| 2 | 197998115 | rs2579390 | 0.455 | -0.821 | 1.273 | -1.462 | -0.022 | -0.048 | -0.048 | 0 | 0.91 | 0.81 | 0.96 | 1 | 0.97 | 0.97 |
| 2 | 198006719 | rs6728379 | -0.723 | -0.816 | 1.095 | -1.466 | -0.001 | 0.036 | 0.036 | 0 | 0.16 | 0.09 | 0.08 | 0.03 | 0.13 | 0.15 |
| 2 | 198011904 | rs2249470 | 1.895 | -0.791 | 1.158 | -1.507 | -0.045 | -0.079 | -0.079 | 0 | 0.87 | 0.61 | 0.92 | 0.99 | 0.96 | 0.96 |
| 2 | 198018181 | rs2579384 | 1.614 | -0.742 | 1.202 | -1.463 | 0 | -0.003 | -0.003 | 0.177 | 0.94 | 0.65 | 0.96 | 1 | 1 | 1 |
| 2 | 198026894 | rs1534540 | 2.356 | -0.791 | 1.118 | -1.508 | -0.037 | -0.14 | -0.14 | 0.003 | 0.67 | 0.43 | 0.83 | 0.96 | 0.81 | 0.78 |
| 2 | 198030920 | rs7594879 | -1.234 | -0.509 | 0.317 | -1.371 | 0.005 | 0.073 | 0.073 | 0 | 0.22 | 0.1 | 0.1 | 0.03 | 0.16 | 0.19 |
| 2 | 198037641 | rs2697246 | -1.525 | -0.805 | 0.85 | -1.224 | -0.051 | -0.077 | -0.077 | 0 | 0.21 | 0.41 | 0.18 | 0.09 | 0.14 | 0.19 |
| 2 | 198044985 | rs11901946 | -1.665 | -0.769 | 0.459 | -1.485 | 0.027 | 0.072 | 0.072 | 0 | 0.38 | 0.81 | 0.26 | 0.13 | 0.25 | 0.29 |
| 2 | 198048140 | rs2697232 | 1.599 | -0.766 | 0.477 | -1.518 | 0.046 | 0.182 | 0.182 | 0 | 0.57 | 0.11 | 0.71 | 0.87 | 0.7 | 0.69 |
| 2 | 198049461 | rs17388610 | 0.009 | -0.751 | 0.508 | -1.544 | -0.039 | 0.064 | 0.064 | 0 | 0.19 | 0.06 | 0.11 | 0.09 | 0.18 | 0.16 |
| 2 | 198065054 | rs6434916 | 0.862 | -0.427 | 0.84 | -1.34 | -0.009 | -0.081 | -0.081 | 0 | 0.62 | 0.5 | 0.78 | 0.91 | 0.72 | 0.72 |
| 2 | 198073045 | rs2712902 | 1.33 | -0.443 | 0.92 | -1.187 | 0.022 | -0.014 | -0.014 | 0 | 0.93 | 0.72 | 0.95 | 1 | 0.95 | 0.98 |
| 2 | 198076664 | rs2697303 | -0.913 | -0.211 | 0.496 | -0.993 | -0.017 | -0.058 | -0.058 | 0 | 0.36 | 0.51 | 0.23 | 0.2 | 0.27 | 0.3 |
| 2 | 198077950 | rs9283513 | 0.464 | 0.019 | -0.412 | -1.075 | 0.03 | 0.153 | 0.153 | 0 | 0.5 | 0.25 | 0.66 | 0.77 | 0.59 | 0.55 |
| 2 | 198081584 | rs2697298 | -0.412 | -0.036 | -0.385 | -1.024 | -0.041 | -0.028 | -0.028 | 0 | 0.72 | 0.79 | 0.5 | 0.36 | 0.55 | 0.48 |
| 2 | 198089911 | rs16863365 | 0.996 | -0.064 | -0.527 | -1.131 | 0 | -0.016 | -0.016 | 0 | 0.13 | 0.12 | 0.07 | 0 | 0.12 | 0.08 |
| 2 | 198090919 | rs2697237 | 1.054 | -0.139 | -0.577 | -1.177 | 0 | -0.006 | -0.006 | 1.038 | 0.27 | 0.29 | 0.16 | 0.17 | 0.22 | 0.22 |
| 2 | 198096952 | rs7559692 | 1.081 | -0.162 | -0.489 | -1.315 | 0 | 0 | 0 | 0.58 | 0.31 | 0.23 | 0.26 | 0.21 | 0.22 | 0.18 |
| 2 | 198097758 | rs11902332 | 0.43 | 0.053 | -0.815 | -1.264 | -0.007 | -0.087 | -0.087 | 0 | 0.15 | 0.32 | 0.09 | 0 | 0.12 | 0.08 |
| 2 | 198108977 | rs2712885 | 0.83 | 0.18 | -1.739 | -1.17 | 0 | -0.019 | -0.019 | 0 | 0.27 | 0.29 | 0.14 | 0.13 | 0.22 | 0.22 |
| 2 | 198120197 | rs6434921 | 0.689 | 0.538 | -1.843 | -1.007 | -0.148 | -0.125 | -0.125 | 0.412 | 0.24 | 0.03 | 0.43 | 0.64 | 0.4 | 0.45 |
| 2 | 198129370 | rs7580108 | 0.714 | 0.435 | -1.734 | -0.911 | 0.004 | 0.004 | 0.004 | 0 | 0.14 | 0.11 | 0.07 | 0 | 0.12 | 0.09 |
| 2 | 198132601 | rs1318141 | -0.764 | 0.313 | -1.843 | -0.769 | 0.001 | -0.01 | -0.01 | 0.201 | 0.69 | 0.69 | 0.84 | 0.86 | 0.77 | 0.7 |
| 2 | 198135412 | rs2712867 | 0.155 | 0.838 | -2.067 | 0.113 | 0.01 | 0.003 | 0.003 | 0.564 | 0.25 | 0.14 | 0.14 | 0.17 | 0.21 | 0.08 |
| 2 | 198142210 | rs11693873 | -0.419 | 1.665 | -2.246 | 0.485 | -0.135 | -0.095 | -0.095 | 2.741 | 0.25 | 0.03 | 0.56 | 0.71 | 0.4 | 0.56 |
| 2 | 198143591 | rs10931777 | 0.04 | 1.441 | -1.734 | 0.243 | 0 | 0.004 | 0.004 | 0 | 0.64 | 0.71 | 0.73 | 0.77 | 0.66 | 0.84 |
| 2 | 198144002 | rs1429411 | -0.504 | 2.409 | -2.186 | 0.518 | -0.125 | -0.123 | -0.123 | 4.55 | 0.19 | 0.03 | 0.53 | 0.61 | 0.37 | 0.33 |
| 2 | 198148570 | rs6715936 | 0.002 | 2.221 | -2.232 | 0.078 | 0.002 | 0 | 0 | 0 | 0.27 | 0.12 | 0.21 | 0.2 | 0.26 | 0.14 |
| 2 | 198148767 | rs12619296 | -1.131 | 2.502 | -1.298 | 0.36 | 0.003 | 0.05 | 0.05 | 0 | 0.46 | 0.37 | 0.62 | 0.7 | 0.56 | 0.49 |
| 2 | 198162003 | rs7593063 | -0.19 | 1.911 | -0.4 | 0.104 | 0.01 | 0.002 | 0.002 | 0.237 | 0.08 | 0.39 | 0.05 | 0.03 | 0.02 | 0.04 |
| 2 | 198166825 | rs10931779 | -1.28 | 1.86 | -0.418 | 0.01 | -0.004 | -0.011 | -0.011 | 0 | 0.31 | 0.72 | 0.48 | 0.52 | 0.33 | 0.4 |
| 2 | 198176766 | rs10804081 | -0.585 | 1.795 | -0.177 | 0.034 | -0.024 | -0.05 | -0.05 | 0 | 0.11 | 0.38 | 0.06 | 0.03 | 0.04 | 0.05 |
| 2 | 198182334 | rs6738721 | -1.478 | 1.725 | -0.502 | -0.005 | -0.027 | 0.111 | 0.111 | 0 | 0.56 | 0.89 | 0.67 | 0.74 | 0.58 | 0.53 |
| 2 | 198198852 | rs7578368 | 0.754 | 1.825 | -0.016 | 0.26 | 0.003 | -0.031 | -0.031 | 0 | 0.2 | 0.04 | 0.18 | 0.2 | 0.21 | 0.07 |
| 2 | 198232639 | rs788008 | 0.338 | 1.87 | 0.094 | 0.29 | -0.008 | -0.024 | -0.024 | 0 | 0.22 | 0.08 | 0.19 | 0.22 | 0.24 | 0.12 |
| 2 | 198263146 | rs16865262 | 0.358 | 1.835 | 0.198 | 0.43 | -0.013 | -0.053 | -0.053 | 0 | 0.22 | 0.05 | 0.19 | 0.25 | 0.25 | 0.09 |
| 2 | 198265350 | rs12621129 | 0.21 | 1.87 | 0.157 | 0.715 | -0.066 | -0.069 | -0.069 | 0 | 0.13 | 0.03 | 0.36 | 0.48 | 0.27 | 0.24 |
| 2 | 198274929 | rs12151767 | 0.846 | 1.746 | 1.307 | 0.64 | -0.011 | 0.012 | 0.012 | 0 | 0.6 | 0.36 | 0.41 | 0.29 | 0.51 | 0.62 |
| 2 | 198283305 | rs788023 | -0.443 | 1.746 | 1.431 | 0.64 | -0.013 | -0.002 | -0.002 | 0 | 0.87 | 0.89 | 0.63 | 0.52 | 0.73 | 0.76 |
| 2 | 198297156 | rs16865650 | 0.624 | 1.754 | 1.425 | 0.548 | 0.004 | -0.038 | -0.038 | 0 | 0.21 | 0.05 | 0.17 | 0.23 | 0.21 | 0.1 |
| 2 | 198308260 | rs3849362 | 0.875 | 1.726 | 1.473 | 0.569 | -0.014 | 0.022 | 0.022 | 0 | 0.6 | 0.35 | 0.41 | 0.24 | 0.51 | 0.58 |
| 2 | 198327314 | rs3754822 | 0.606 | 1.699 | 1.534 | 0.562 | 0.005 | -0.036 | -0.036 | 0 | 0.21 | 0.05 | 0.17 | 0.23 | 0.22 | 0.1 |
| 2 | 198349536 | rs6745660 | 0.854 | 1.675 | 1.762 | 0.556 | -0.01 | 0.016 | 0.016 | 0 | 0.6 | 0.37 | 0.42 | 0.24 | 0.51 | 0.57 |
| 2 | 198352293 | rs788016 | 0.888 | 1.704 | 1.942 | 0.551 | -0.02 | 0.035 | 0.035 | 0 | 0.59 | 0.31 | 0.41 | 0.24 | 0.51 | 0.57 |
| 2 | 198354414 | rs2565163 | -0.314 | 1.682 | 1.951 | 0.555 | -0.017 | -0.006 | -0.006 | 0.002 | 0.86 | 0.89 | 0.63 | 0.48 | 0.73 | 0.69 |
| 2 | 198355153 | rs11682567 | 0.397 | 1.679 | 2.04 | 0.554 | 0.014 | -0.021 | -0.021 | 0.014 | 0.23 | 0.1 | 0.17 | 0.23 | 0.21 | 0.1 |

| | | | | | | | | | | | | | | | | |
|---|---|---|---|---|---|---|---|---|---|---|---|---|---|---|---|---|
| 2 | 198360886 | rs2340690 | 0.613 | 1.679 | 2.068 | 0.554 | 0.008 | -0.036 | -0.036 | 0 | 0.21 | 0.05 | 0.17 | 0.23 | 0.21 | 0.1 |
| 2 | 198362524 | rs17730989 | 0.903 | 1.679 | 2.056 | 0.554 | -0.021 | 0.036 | 0.036 | 0 | 0.59 | 0.31 | 0.41 | 0.24 | 0.51 | 0.57 |
| 2 | 198369370 | rs6434930 | -0.213 | 1.691 | 2.381 | 0.562 | 0.037 | 0.081 | 0.081 | 0 | 0.86 | 0.79 | 0.63 | 0.48 | 0.73 | 0.69 |
| 2 | 198371687 | rs13386066 | 0 | 1.689 | 2.385 | 0.585 | -0.026 | 0.004 | 0.004 | 0 | 0.25 | 0.37 | 0.21 | 0.22 | 0.21 | 0.1 |
| 2 | 198378546 | rs7590697 | 0.806 | 1.692 | 2.308 | 0.584 | 0.004 | -0.04 | -0.04 | 0.263 | 0.2 | 0.04 | 0.17 | 0.21 | 0.2 | 0.07 |
| 2 | 198407177 | rs3792158 | 0.857 | 1.69 | 2.327 | 0.559 | -0.024 | 0.035 | 0.035 | 0 | 0.6 | 0.31 | 0.41 | 0.25 | 0.51 | 0.57 |
| 2 | 198434319 | rs11891750 | 0.852 | 1.633 | 2.295 | 0.531 | -0.01 | 0.015 | 0.015 | 0 | 0.59 | 0.37 | 0.41 | 0.25 | 0.51 | 0.57 |
| 2 | 198461941 | rs10198804 | 0.048 | 1.593 | 2.303 | 0.488 | -0.024 | 0.001 | 0.001 | 0 | 0.25 | 0.35 | 0.21 | 0.21 | 0.21 | 0.1 |
| 2 | 198463090 | rs10178946 | -0.001 | 1.756 | 2.285 | 0.676 | 0.04 | 0.081 | 0.081 | 0 | 0.85 | 0.75 | 0.62 | 0.49 | 0.73 | 0.66 |
| 2 | 198477620 | rs4389332 | 1.124 | 1.631 | 2.424 | 0.636 | -0.02 | 0.028 | 0.028 | 0 | 0.6 | 0.32 | 0.41 | 0.24 | 0.51 | 0.57 |
| 2 | 198480072 | rs6719729 | 0.248 | 1.628 | 2.165 | 0.706 | 0.051 | 0.104 | 0.104 | 0 | 0.84 | 0.71 | 0.62 | 0.49 | 0.73 | 0.67 |
| 2 | 198497512 | rs970552 | 0.936 | 1.434 | 2.073 | 0.446 | 0 | -0.003 | -0.003 | 0.432 | 0.61 | 0.6 | 0.42 | 0.22 | 0.52 | 0.52 |
| 2 | 198508951 | rs10497806 | 0.198 | 1.482 | 2.091 | 0.552 | -0.022 | -0.027 | -0.027 | 0 | 0.83 | 0.87 | 0.62 | 0.46 | 0.74 | 0.64 |
| 2 | 198511086 | rs16866970 | 1.089 | 1.461 | 2.382 | 0.542 | 0.002 | -0.006 | -0.006 | 0.573 | 0.6 | 0.59 | 0.41 | 0.22 | 0.52 | 0.52 |
| 2 | 198511917 | rs10166328 | 0.176 | 1.46 | 2.372 | 0.542 | -0.022 | -0.026 | -0.026 | 0 | 0.83 | 0.87 | 0.62 | 0.46 | 0.74 | 0.64 |
| 2 | 198513930 | rs10196961 | 1.05 | 1.46 | 2.369 | 0.541 | 0.011 | -0.028 | -0.028 | 0 | 0.6 | 0.51 | 0.4 | 0.22 | 0.52 | 0.52 |
| 2 | 198517592 | rs7603283 | 0.362 | 1.461 | 2.332 | 0.548 | 0.006 | -0.011 | -0.011 | 0 | 0.21 | 0.15 | 0.18 | 0.24 | 0.22 | 0.1 |
| 2 | 198520615 | rs10931787 | 0.374 | 1.477 | 2.333 | 0.553 | 0.018 | 0.054 | 0.054 | 0 | 0.82 | 0.76 | 0.61 | 0.46 | 0.74 | 0.64 |
| 2 | 198533369 | rs6719832 | 0.374 | 1.499 | 2.545 | 0.563 | 0.018 | 0.054 | 0.054 | 0 | 0.82 | 0.76 | 0.61 | 0.46 | 0.74 | 0.64 |
| 2 | 198572461 | rs7592344 | 0.11 | 1.449 | 2.609 | 0.683 | -0.009 | 0.01 | 0.01 | 0 | 0.22 | 0.28 | 0.19 | 0.26 | 0.22 | 0.12 |
| 2 | 198573612 | rs10931791 | -0.432 | 1.428 | 2.576 | 0.665 | -0.035 | -0.046 | -0.046 | 0 | 0.18 | 0.1 | 0.38 | 0.51 | 0.27 | 0.35 |
| 2 | 198574523 | rs6751690 | 0.464 | 1.485 | 2.601 | 0.64 | 0.007 | -0.034 | -0.034 | 1.409 | 0.2 | 0.08 | 0.18 | 0.26 | 0.21 | 0.1 |
| 2 | 198576142 | rs4850808 | -1.303 | 1.433 | 2.608 | 0.631 | -0.001 | -0.002 | -0.002 | 0.024 | 0.4 | 0.41 | 0.58 | 0.77 | 0.48 | 0.48 |
| 2 | 198582601 | rs13029495 | -1.303 | 1.433 | 2.608 | 0.631 | -0.001 | -0.002 | -0.002 | 0 | 0.4 | 0.41 | 0.58 | 0.77 | 0.48 | 0.48 |
| 2 | 198584209 | rs16823148 | 0.008 | 1.388 | 2.604 | 0.608 | -0.015 | 0.015 | 0.015 | 0 | 0.23 | 0.31 | 0.2 | 0.26 | 0.21 | 0.12 |
| 2 | 198590886 | rs700640 | 1.086 | 1.425 | 2.593 | 0.573 | 0 | -0.004 | -0.004 | 0 | 0.6 | 0.59 | 0.42 | 0.23 | 0.52 | 0.52 |
| 2 | 198614612 | rs1065953 | 1.046 | 1.44 | 2.582 | 0.564 | -0.007 | 0.01 | 0.01 | 0 | 0.61 | 0.64 | 0.42 | 0.23 | 0.52 | 0.52 |
| 2 | 198631714 | rs700651 | 0.192 | 1.443 | 2.574 | 0.616 | -0.06 | -0.081 | -0.081 | 0 | 0.83 | 0.94 | 0.62 | 0.49 | 0.73 | 0.64 |
| 2 | 198643631 | rs700655 | 1.125 | 1.42 | 2.72 | 0.592 | -0.004 | 0.007 | 0.007 | 0 | 0.59 | 0.41 | 0.4 | 0.23 | 0.52 | 0.52 |
| 2 | 198706916 | rs700679 | 0.885 | 1.501 | 2.287 | 0.618 | -0.016 | 0.02 | 0.02 | 0 | 0.62 | 0.69 | 0.43 | 0.23 | 0.52 | 0.53 |
| 2 | 198713231 | rs6707521 | 0.456 | 1.498 | 2.246 | 0.609 | 0.005 | -0.024 | -0.024 | 0.189 | 0.19 | 0.12 | 0.18 | 0.26 | 0.21 | 0.11 |
| 2 | 198805930 | rs10207433 | 0.982 | 1.495 | 2.337 | 0.593 | -0.005 | 0.006 | 0.006 | 0 | 0.61 | 0.65 | 0.42 | 0.23 | 0.52 | 0.54 |
| 2 | 198809975 | rs1518364 | 1.111 | 1.491 | 2.357 | 0.591 | 0.009 | -0.019 | -0.019 | 0 | 0.6 | 0.55 | 0.41 | 0.23 | 0.53 | 0.51 |
| 2 | 198819824 | rs4410288 | 1.056 | 1.515 | 2.362 | 0.594 | -0.001 | -0.001 | -0.001 | 0 | 0.61 | 0.59 | 0.42 | 0.23 | 0.52 | 0.51 |
| 2 | 198823828 | rs1401096 | 0.334 | 1.567 | 2.217 | 0.669 | -0.041 | -0.058 | -0.058 | 0.274 | 0.82 | 0.9 | 0.61 | 0.49 | 0.74 | 0.65 |
| 2 | 198875464 | rs1595825 | 0.031 | 1.577 | 2.285 | 0.658 | -0.006 | 0.04 | 0.04 | 0 | 0.2 | 0.31 | 0.2 | 0.26 | 0.21 | 0.14 |
| 2 | 198881668 | rs1016883 | 0.173 | 1.547 | 2.274 | 0.649 | -0.006 | 0.021 | 0.021 | 0 | 0.19 | 0.23 | 0.18 | 0.26 | 0.2 | 0.13 |
| 2 | 198891799 | rs1595823 | 1.093 | 1.544 | 2.348 | 0.639 | 0.016 | -0.027 | -0.027 | 0.043 | 0.62 | 0.5 | 0.42 | 0.22 | 0.55 | 0.55 |
| 2 | 198898265 | rs6715070 | 0.496 | 1.565 | 2.172 | 0.657 | 0.017 | 0.045 | 0.045 | 0 | 0.81 | 0.74 | 0.61 | 0.48 | 0.74 | 0.68 |
| 2 | 198901395 | rs1147169 | 0.696 | 1.769 | 2.338 | 0.75 | 0.081 | 0.159 | 0.159 | 0.039 | 0.83 | 0.7 | 0.61 | 0.47 | 0.75 | 0.68 |
| 2 | 198904914 | rs2196176 | 0.436 | 1.784 | 2.118 | 0.757 | 0.06 | 0.12 | 0.12 | 0 | 0.85 | 0.73 | 0.61 | 0.48 | 0.75 | 0.71 |
| 2 | 198906354 | rs1440086 | 0.382 | 1.81 | 1.341 | 0.797 | 0.001 | -0.009 | -0.009 | 0 | 0.17 | 0.14 | 0.17 | 0.25 | 0.2 | 0.11 |
| 2 | 198925980 | rs1866665 | 0.472 | 1.77 | 1.013 | 0.783 | -0.001 | 0.009 | 0.009 | 0 | 0.16 | 0.18 | 0.17 | 0.25 | 0.19 | 0.09 |
| 2 | 198929716 | rs6434954 | 0.535 | 1.898 | 0.692 | 0.804 | 0.017 | 0.039 | 0.039 | 0 | 0.83 | 0.81 | 0.62 | 0.47 | 0.74 | 0.66 |
| 2 | 198946551 | rs6434955 | 0.083 | 1.634 | 0.131 | 0.81 | -0.006 | 0.089 | 0.089 | 0 | 0.18 | 0.36 | 0.19 | 0.25 | 0.19 | 0.12 |
| 2 | 198950029 | rs2228135 | 0.009 | 1.642 | 0.113 | 0.809 | -0.007 | 0.084 | 0.084 | 0 | 0.18 | 0.36 | 0.19 | 0.25 | 0.19 | 0.12 |
| 2 | 198952637 | rs2033570 | 0.559 | 2.098 | -1.08 | 0.882 | -0.072 | -0.078 | -0.078 | 0 | 0.84 | 0.94 | 0.62 | 0.46 | 0.71 | 0.69 |
| 2 | 198959603 | rs3755333 | -0.43 | 0.989 | -1.762 | -0.014 | -0.017 | 0.109 | 0.109 | 0 | 0.43 | 0.88 | 0.32 | 0.2 | 0.38 | 0.25 |
| 2 | 198962243 | rs7595412 | -0.312 | 1.169 | -1.82 | -0.132 | -0.015 | 0.011 | 0.011 | 0 | 0.93 | 0.95 | 0.93 | 1 | 0.9 | 0.96 |
| 2 | 198962295 | rs10497815 | 0.626 | 0.75 | -1.995 | -0.711 | 0.006 | -0.005 | -0.005 | 0.057 | 0.12 | 0.08 | 0.05 | 0 | 0.08 | 0.16 |
| 2 | 198962831 | rs3771362 | 0.964 | 0.408 | -1.89 | -0.898 | -0.008 | 0.003 | 0.003 | 0 | 0.27 | 0.09 | 0.18 | 0.2 | 0.27 | 0.19 |
| 2 | 198965495 | rs2880366 | -0.066 | 0.411 | -0.391 | -0.913 | 0.003 | -0.03 | -0.03 | 0.444 | 0.61 | 0.46 | 0.72 | 0.8 | 0.63 | 0.76 |
| 2 | 198966257 | rs3732231 | -0.348 | 0.415 | -0.112 | -0.911 | -0.02 | 0.032 | 0.032 | 0 | 0.66 | 0.77 | 0.75 | 0.8 | 0.63 | 0.78 |
| 2 | 198969583 | rs1304358 | 0.787 | 0.431 | 0.214 | -0.941 | -0.01 | 0.123 | 0.123 | 0 | 0.48 | 0.23 | 0.66 | 0.8 | 0.55 | 0.6 |
| 2 | 198977987 | rs11681576 | 0.013 | 0.383 | -0.289 | -1.095 | -0.021 | -0.045 | -0.045 | 0.649 | 0.41 | 0.22 | 0.55 | 0.56 | 0.45 | 0.64 |
| 2 | 198978661 | rs1808850 | -0.809 | 0.452 | 0.383 | -1.087 | 0.049 | -0.044 | -0.044 | 0 | 0.92 | 0.8 | 0.9 | 1 | 0.9 | 0.96 |
| 2 | 198988387 | rs10173738 | 0.338 | 0.343 | 0.585 | -1.139 | -0.028 | -0.153 | -0.153 | 0 | 0.2 | 0.39 | 0.12 | 0 | 0.14 | 0.17 |
| 2 | 198996212 | rs11884188 | 1.004 | 0.292 | 0.402 | -1.151 | 0.025 | -0.019 | -0.019 | 0.128 | 0.46 | 0.25 | 0.31 | 0.43 | 0.42 | 0.3 |
| 2 | 199008506 | rs11674075 | 1.184 | 0.413 | 0.363 | -1.162 | -0.001 | -0.033 | -0.033 | 0 | 0.52 | 0.15 | 0.32 | 0.43 | 0.48 | 0.34 |
| 2 | 199023800 | rs714644 | 0.745 | 0.333 | 0.425 | -1.193 | 0.008 | -0.005 | -0.005 | 0 | 0.18 | 0.15 | 0.09 | 0 | 0.12 | 0.16 |
| 2 | 199027024 | rs992603 | -1.641 | 0.398 | 0.433 | -1.198 | -0.006 | -0.001 | -0.001 | 0 | 0.94 | 0.94 | 0.93 | 1 | 0.9 | 0.96 |
| 2 | 199032498 | rs1434288 | -0.216 | 0.33 | 0.47 | -1.149 | -0.003 | -0.01 | -0.01 | 0.107 | 0.76 | 0.76 | 0.83 | 1 | 0.78 | 0.81 |
| 2 | 199043041 | rs10210770 | -0.23 | 0.388 | 0.491 | -1.156 | -0.004 | -0.029 | -0.029 | 0 | 0.75 | 0.73 | 0.82 | 1 | 0.75 | 0.8 |
| 2 | 199047626 | rs9789480 | 1.084 | 0.517 | 0.058 | -1.081 | -0.051 | -0.087 | -0.087 | 0 | 0.17 | 0.06 | 0.46 | 0.56 | 0.27 | 0.44 |
| 2 | 199049108 | rs6434957 | 0.531 | -0.007 | 0.214 | -0.82 | 0.017 | -0.033 | -0.033 | 0 | 0.51 | 0.15 | 0.32 | 0.43 | 0.47 | 0.34 |
| 2 | 199056513 | rs6742863 | 0.794 | -0.395 | 0.426 | -1.242 | -0.024 | -0.042 | -0.042 | 0 | 0.27 | 0.38 | 0.16 | 0.12 | 0.21 | 0.16 |
| 2 | 199070749 | rs1456549 | 0.855 | -0.518 | 0.685 | -1.316 | -0.001 | -0.003 | -0.003 | 0.467 | 0.27 | 0.27 | 0.15 | 0.12 | 0.22 | 0.16 |
| 2 | 199082114 | rs10199534 | 0.695 | -0.557 | 0.59 | -1.399 | 0.055 | 0.115 | 0.115 | 0 | 0.4 | 0.19 | 0.15 | 0.12 | 0.3 | 0.34 |
| 2 | 199084258 | rs11682314 | 0.474 | -0.337 | 0.456 | -1.604 | -0.03 | 0.017 | 0.017 | 0.785 | 0.28 | 0.16 | 0.56 | 0.82 | 0.4 | 0.55 |

| | | | | | | | | | | | | | | | | |
|---|---|---|---|---|---|---|---|---|---|---|---|---|---|---|---|---|
| 2 | 199085661 | rs4850833 | -0.042 | -0.26 | 0.69 | -1.565 | 0.056 | -0.088 | -0.088 | 0 | 0.87 | 0.55 | 0.87 | 1 | 0.88 | 0.96 |
| 2 | 199086265 | rs1350208 | -0.117 | -0.199 | 0.287 | -1.48 | 0.009 | 0.159 | 0.159 | 0 | 0.45 | 0.19 | 0.67 | 0.88 | 0.59 | 0.61 |
| 2 | 199089918 | rs17799440 | 0.765 | -0.541 | 0.48 | -1.241 | 0.008 | -0.002 | -0.002 | 0.022 | 0.08 | 0.12 | 0.08 | 0.05 | 0.14 | 0.05 |
| 2 | 199094168 | rs10181932 | 0.316 | -0.728 | 0.8 | -1.466 | 0.039 | 0.069 | 0.069 | 0 | 0.48 | 0.64 | 0.26 | 0.14 | 0.33 | 0.34 |
| 2 | 199095862 | rs9808268 | 0.387 | -0.745 | 0.642 | -1.474 | 0.05 | 0.097 | 0.097 | 0 | 0.48 | 0.64 | 0.26 | 0.14 | 0.33 | 0.34 |
| 2 | 199095875 | rs9807926 | 0.764 | -0.672 | 0.068 | -1.438 | 0.043 | 0.148 | 0.148 | 0 | 0.43 | 0.17 | 0.17 | 0.14 | 0.33 | 0.34 |
| 2 | 199109322 | rs1456562 | 0.759 | -0.203 | 0.565 | -1.383 | -0.011 | -0.076 | -0.076 | 0.463 | 0.85 | 0.34 | 0.82 | 1 | 0.87 | 0.96 |
| 2 | 199117457 | rs6761616 | -0.588 | -0.162 | 0.932 | -1.363 | 0.026 | -0.086 | -0.086 | 0 | 0.13 | 0.39 | 0.11 | 0 | 0.13 | 0.04 |
| 2 | 199118338 | rs10931806 | -0.313 | -0.091 | -0.164 | -1.261 | -0.074 | 0.051 | 0.051 | 1.308 | 0.73 | 0.91 | 0.54 | 0.15 | 0.64 | 0.44 |
| 2 | 199122020 | rs4850836 | 0.645 | -0.114 | -0.176 | -0.69 | -0.142 | -0.146 | -0.146 | 0 | 0.21 | 0.02 | 0.32 | 0.37 | 0.37 | 0.24 |
| 2 | 199122447 | rs1456556 | -0.178 | -0.504 | 1.034 | -1.544 | 0.026 | -0.033 | -0.033 | 0 | 0.09 | 0.41 | 0.12 | 0 | 0.05 | 0.02 |
| 2 | 199123438 | rs10178814 | -0.193 | -0.755 | 1.009 | -1.723 | -0.001 | -0.004 | -0.004 | 2.068 | 0.1 | 0.13 | 0.06 | 0.05 | 0.12 | 0.05 |
| 2 | 199133114 | rs1456554 | 0.653 | -0.584 | 1.269 | -1.589 | 0.001 | -0.038 | -0.038 | 0 | 0.07 | 0.19 | 0.09 | 0 | 0.05 | 0.02 |
| 2 | 199137857 | rs1379020 | -0.846 | -0.65 | 1.365 | -1.621 | -0.012 | -0.068 | -0.068 | 1.418 | 0.11 | 0.51 | 0.13 | 0 | 0.05 | 0.02 |
| 2 | 199144404 | rs7591713 | 0.15 | -0.486 | 0.362 | -1.53 | -0.001 | -0.038 | -0.038 | 0 | 0.07 | 0.21 | 0.04 | 0 | 0.04 | 0.02 |
| 2 | 199156137 | rs1379021 | 0.373 | -0.569 | -0.112 | -1.69 | -0.118 | 0.127 | 0.127 | 4.349 | 0.32 | 0.11 | 0.62 | 0.88 | 0.48 | 0.61 |
| 2 | 199158911 | rs758063 | 0.076 | -0.742 | 0.086 | -1.669 | -0.053 | 0.167 | 0.167 | 0.971 | 0.39 | 0.16 | 0.66 | 0.88 | 0.55 | 0.65 |
| 2 | 199160257 | rs7578662 | -0.205 | -0.733 | 0.31 | -1.608 | 0.004 | -0.011 | -0.011 | 0 | 0.07 | 0.21 | 0.05 | 0.05 | 0.09 | 0.06 |
| 2 | 199164126 | rs2190650 | 0.203 | -0.752 | -0.272 | -1.676 | -0.061 | 0.15 | 0.15 | 1.035 | 0.37 | 0.16 | 0.65 | 0.88 | 0.54 | 0.65 |
| 2 | 199166382 | rs13387754 | 0.27 | -0.931 | -0.346 | -1.893 | 0.02 | 0.031 | 0.031 | 0.745 | 0.43 | 0.36 | 0.17 | 0.08 | 0.33 | 0.21 |
| 2 | 199169096 | rs13398206 | 0.393 | -0.775 | -0.547 | -1.805 | 0.018 | 0.069 | 0.069 | 0.566 | 0.39 | 0.29 | 0.16 | 0.09 | 0.33 | 0.2 |
| 2 | 199170703 | rs4998599 | -0.325 | -0.381 | -0.291 | -1.041 | 0.004 | 0.017 | 0.017 | 0 | 0.87 | 0.93 | 0.93 | 0.96 | 0.87 | 0.86 |
| 2 | 199176477 | rs2529672 | -0.733 | -0.349 | 0.254 | -0.588 | -0.004 | 0.064 | 0.064 | 0.721 | 0.2 | 0.46 | 0.27 | 0.24 | 0.23 | 0.2 |
| 2 | 199176823 | rs7575825 | 0.3 | -0.698 | 0.061 | -1.785 | 0.005 | -0.001 | -0.001 | 0.086 | 0.27 | 0.24 | 0.11 | 0.07 | 0.19 | 0.15 |
| 2 | 199182704 | rs997467 | 0.452 | -0.544 | 0.175 | -2.111 | 0.001 | 0.261 | 0.261 | 0.611 | 0.47 | 0.14 | 0.59 | 0.9 | 0.58 | 0.78 |
| 2 | 199183461 | rs2529664 | -0.947 | 0.01 | 0.751 | -1.861 | 0.044 | -0.054 | -0.054 | 0.001 | 0.11 | 0.36 | 0.16 | 0.02 | 0.13 | 0.06 |
| 2 | 199185059 | rs2727783 | -0.398 | 0.049 | 0.684 | -1.843 | 0.015 | -0.031 | -0.031 | 0 | 0.08 | 0.13 | 0.14 | 0.02 | 0.13 | 0.06 |
| 2 | 199206396 | rs2727765 | -0.315 | -0.031 | 0.243 | -1.823 | 0.005 | 0.023 | 0.023 | 0.235 | 0.65 | 0.73 | 0.79 | 0.92 | 0.72 | 0.84 |
| 2 | 199228515 | rs7576530 | -0.47 | 0.168 | 0.078 | -1.725 | -0.005 | 0.003 | 0.003 | 0 | 0.08 | 0.08 | 0.09 | 0.02 | 0.13 | 0.06 |
| 2 | 199231431 | rs2467041 | -0.42 | 0.14 | 0.252 | -1.92 | -0.007 | -0.08 | -0.08 | 4.423 | 0.41 | 0.5 | 0.37 | 0.11 | 0.36 | 0.2 |
| 2 | 199236870 | rs12477557 | 0.22 | 0.143 | 0.132 | -1.784 | -0.01 | -0.058 | -0.058 | 0.764 | 0.33 | 0.37 | 0.28 | 0.08 | 0.22 | 0.14 |
| 2 | 199254349 | rs2215023 | 0.843 | 0.555 | 0.28 | -1.259 | 0.041 | -0.092 | -0.092 | 0.284 | 0.84 | 0.64 | 0.85 | 0.96 | 0.8 | 0.94 |
| 2 | 199282688 | rs13012598 | -0.482 | 0.112 | 0.063 | -1.338 | -0.011 | -0.066 | -0.066 | 0.021 | 0.14 | 0.25 | 0.12 | 0.02 | 0.12 | 0.04 |
| 2 | 199299078 | rs11688831 | 0.494 | 0.751 | -1.152 | -1.119 | -0.144 | -0.104 | -0.104 | 0 | 0.24 | 0.02 | 0.37 | 0.68 | 0.38 | 0.5 |
| 2 | 199305825 | rs4241196 | 0.891 | 0.226 | -0.115 | -1.374 | -0.045 | -0.127 | -0.127 | 0 | 0.8 | 0.43 | 0.79 | 0.97 | 0.86 | 0.95 |
| 2 | 199312684 | rs12693845 | -1.295 | 1.139 | -0.634 | -0.976 | -0.034 | 0.025 | 0.025 | 0 | 0.27 | 0.15 | 0.3 | 0.21 | 0.36 | 0.2 |
| 2 | 199317135 | rs4499421 | 0.346 | 0.881 | 0.011 | -1.343 | 0.009 | -0.054 | -0.054 | 0 | 0.63 | 0.45 | 0.44 | 0.17 | 0.55 | 0.62 |
| 2 | 199321325 | rs7587814 | 0.291 | 0.724 | -0.301 | -1.569 | -0.01 | 0.052 | 0.052 | 0 | 0.12 | 0.03 | 0.08 | 0 | 0.14 | 0.12 |
| 2 | 199323526 | rs4502401 | 0.532 | 0.702 | 0.039 | -1.686 | -0.012 | 0.013 | 0.013 | 0 | 0.25 | 0.19 | 0.45 | 0.79 | 0.35 | 0.28 |
| 2 | 199330674 | rs6757018 | 1.014 | 0.574 | 0.484 | -1.712 | -0.015 | -0.12 | -0.12 | 0 | 0.74 | 0.61 | 0.79 | 0.95 | 0.75 | 0.81 |
| 2 | 199339346 | rs4315518 | 1.082 | 0.504 | 0.476 | -1.729 | -0.021 | -0.088 | -0.088 | 0 | 0.85 | 0.42 | 0.85 | 0.95 | 0.89 | 0.93 |
| 2 | 199346565 | rs10183458 | -0.063 | 0.618 | -0.343 | -1.664 | -0.005 | -0.04 | -0.04 | 0.345 | 0.9 | 0.82 | 0.9 | 0.96 | 0.9 | 0.93 |
| 2 | 199348965 | rs4408724 | 0.559 | 0.606 | -0.338 | -1.649 | -0.006 | -0.071 | -0.071 | 0 | 0.75 | 0.68 | 0.8 | 0.95 | 0.76 | 0.81 |
| 2 | 199352368 | rs4413156 | 0.626 | 0.77 | -1.394 | -1.264 | -0.1 | 0.211 | 0.211 | 0.463 | 0.44 | 0.06 | 0.31 | 0.16 | 0.4 | 0.52 |
| 2 | 199356751 | rs10201699 | -1.263 | 1.325 | -0.164 | -0.749 | 0.025 | -0.041 | -0.041 | 0 | 0.12 | 0.18 | 0.12 | 0.05 | 0.16 | 0.09 |
| 2 | 199368722 | rs10804083 | 1.277 | 1.08 | 0.257 | -0.837 | 0.008 | -0.046 | -0.046 | 0 | 0.82 | 0.33 | 0.81 | 0.92 | 0.82 | 0.92 |
| 2 | 199369099 | rs6434973 | 1.012 | 1.034 | 1.324 | -0.86 | 0.028 | -0.039 | -0.039 | 0.208 | 0.85 | 0.7 | 0.84 | 0.92 | 0.83 | 0.92 |
| 2 | 199375335 | rs4355104 | 0.24 | 0.618 | 0.779 | -1.245 | -0.023 | 0.105 | 0.105 | 0 | 0.37 | 0.1 | 0.54 | 0.76 | 0.44 | 0.41 |
| 2 | 199451092 | rs7594192 | 0.611 | -0.119 | 1.637 | -1.588 | -0.003 | -0.062 | -0.062 | 1.014 | 0.7 | 0.31 | 0.78 | 0.95 | 0.78 | 0.86 |
| 2 | 199453491 | rs7602048 | -0.596 | -0.346 | 1.557 | -1.668 | 0.03 | 0.022 | 0.022 | 0.12 | 0.06 | 0.31 | 0.06 | 0.02 | 0.04 | 0.02 |
| 2 | 199465260 | rs10931818 | 0.941 | -0.362 | 1.071 | -1.753 | 0.024 | 0.224 | 0.224 | 1.751 | 0.64 | 0.1 | 0.74 | 0.94 | 0.74 | 0.84 |
| 2 | 199475972 | rs10931819 | 1.173 | -0.33 | 1.206 | -0.796 | -0.028 | 0.049 | 0.049 | 1.363 | 0.86 | 0.32 | 0.85 | 0.74 | 0.94 | 0.89 |
| 2 | 199476070 | rs16830242 | -0.479 | -0.293 | 1.219 | -0.778 | -0.003 | 0.177 | 0.177 | 0 | 0.1 | 0.27 | 0.11 | 0.26 | 0.06 | 0.11 |
| 2 | 199479235 | rs983733 | 1.147 | -0.478 | 0.732 | -0.865 | 0.013 | 0.031 | 0.031 | 0.639 | 0.62 | 0.15 | 0.69 | 0.69 | 0.71 | 0.8 |
| 2 | 199490528 | rs16830272 | -0.164 | -0.41 | 0.885 | -0.728 | -0.007 | 0.105 | 0.105 | 0 | 0.09 | 0.16 | 0.08 | 0.25 | 0.05 | 0.1 |
| 2 | 199494728 | rs16830276 | -0.441 | -0.414 | 0.946 | -0.73 | -0.007 | 0.136 | 0.136 | 0 | 0.1 | 0.21 | 0.09 | 0.25 | 0.06 | 0.1 |
| 2 | 199495674 | rs12477961 | -1.138 | -0.141 | 0.363 | -0.646 | -0.001 | -0.004 | -0.004 | 0.015 | 0.68 | 0.69 | 0.57 | 0.49 | 0.57 | 0.76 |
| 2 | 199496171 | rs17814334 | 0.838 | 0.15 | 0.099 | -0.613 | -0.044 | 0.037 | 0.037 | 0 | 0.32 | 0.1 | 0.28 | 0.2 | 0.29 | 0.56 |
| 2 | 199520404 | rs7562655 | 0.269 | -0.435 | 0.513 | -0.113 | 0.023 | 0.051 | 0.051 | 0 | 0.45 | 0.22 | 0.44 | 0.38 | 0.48 | 0.34 |
| 2 | 199523122 | rs17229285 | -0.096 | -0.421 | 0.484 | 0.087 | -0.014 | 0.152 | 0.152 | 0 | 0.49 | 0.09 | 0.42 | 0.27 | 0.5 | 0.64 |
| 2 | 199533654 | rs4266009 | 0.678 | -0.384 | 0.623 | 0.09 | 0.01 | 0.018 | 0.018 | 0 | 0.41 | 0.29 | 0.44 | 0.39 | 0.47 | 0.27 |
| 2 | 199557466 | rs12475616 | -0.274 | -0.369 | 0.875 | 0.214 | 0.032 | 0.337 | 0.337 | 0 | 0.93 | 0.71 | 0.88 | 0.67 | 0.96 | 0.89 |
| 2 | 199557725 | rs17826222 | 0.598 | -0.337 | 0.853 | -0.091 | 0.013 | 0.012 | 0.012 | 0 | 0.39 | 0.28 | 0.43 | 0.39 | 0.45 | 0.27 |
| 2 | 199558179 | rs17826270 | 0.724 | -0.319 | 0.728 | -0.09 | 0.007 | 0.004 | 0.004 | 0 | 0.38 | 0.17 | 0.41 | 0.39 | 0.45 | 0.27 |
| 2 | 199568260 | rs1996464 | 0.214 | -0.103 | 0.463 | -0.119 | -0.049 | 0.07 | 0.07 | 0 | 0.17 | 0.05 | 0.17 | 0.04 | 0.23 | 0.1 |
| 2 | 199571750 | rs1356485 | 1.027 | -0.126 | 0.261 | -0.199 | -0.044 | 0.006 | 0.006 | 0 | 0.4 | 0.11 | 0.44 | 0.37 | 0.51 | 0.26 |
| 2 | 199572540 | rs4674510 | -0.106 | -0.108 | 0.393 | -0.194 | 0.023 | -0.079 | -0.079 | 0 | 0.2 | 0.29 | 0.19 | 0.04 | 0.23 | 0.1 |
| 2 | 199576145 | rs6434983 | -0.472 | -0.142 | 0.451 | -0.165 | -0.008 | -0.001 | -0.001 | 0 | 0.62 | 0.53 | 0.65 | 0.42 | 0.74 | 0.36 |
| 2 | 199577125 | rs1518099 | -0.485 | -0.149 | 0.537 | -0.073 | -0.001 | 0 | 0 | 0.486 | 0.62 | 0.61 | 0.66 | 0.43 | 0.74 | 0.36 |

| | | | | | | | | | | | | | | | | |
|---|---|---|---|---|---|---|---|---|---|---|---|---|---|---|---|---|
| 2 | 199581722 | rs10439328 | 0.968 | -0.189 | 0.278 | 0.141 | -0.038 | -0.098 | -0.098 | 0 | 0.3 | 0.07 | 0.27 | 0.38 | 0.23 | 0.53 |
| 2 | 199582660 | rs4673212 | -0.393 | -0.063 | 0.416 | -0.256 | 0.02 | 0.011 | 0.011 | 0.014 | 0.59 | 0.21 | 0.61 | 0.41 | 0.73 | 0.36 |
| 2 | 199583657 | rs13013140 | 0.058 | 0.05 | 0.455 | -0.281 | 0.006 | -0.037 | -0.037 | 0.777 | 0.18 | 0.15 | 0.18 | 0.04 | 0.22 | 0.1 |
| 2 | 199590635 | rs7600760 | 0.681 | 0.226 | 0.086 | -0.341 | -0.004 | -0.007 | -0.007 | 0 | 0.13 | 0.12 | 0.15 | 0.03 | 0.18 | 0.08 |
| 2 | 199592900 | rs6742486 | -0.572 | 0.116 | 0.141 | -0.32 | 0.009 | 0.012 | 0.012 | 0 | 0.59 | 0.69 | 0.61 | 0.41 | 0.72 | 0.35 |
| 2 | 199594244 | rs1607331 | -0.338 | 0.117 | 0.226 | -0.252 | 0.007 | 0.026 | 0.026 | 0 | 0.56 | 0.7 | 0.59 | 0.41 | 0.68 | 0.34 |
| 2 | 199601925 | rs11893063 | 0.604 | 0.259 | -1.379 | -0.119 | -0.041 | -0.011 | -0.011 | 0 | 0.39 | 0.08 | 0.39 | 0.38 | 0.48 | 0.25 |
| 2 | 199606419 | rs4675429 | -0.275 | 0.774 | -0.691 | -0.116 | -0.02 | 0.023 | 0.023 | 0 | 0.15 | 0.09 | 0.16 | 0.04 | 0.2 | 0.08 |
| 2 | 199620941 | rs6705243 | -0.064 | 0.887 | -0.696 | -0.132 | -0.062 | 0.055 | 0.055 | 0 | 0.16 | 0.07 | 0.17 | 0.04 | 0.27 | 0.12 |
| 2 | 199623598 | rs1533534 | 0.232 | 0.879 | -0.69 | -0.141 | -0.049 | 0.039 | 0.039 | 0 | 0.14 | 0.07 | 0.17 | 0.04 | 0.26 | 0.11 |
| 2 | 199623902 | rs2048968 | -0.392 | 0.266 | -1.015 | -0.019 | 0.024 | 0.015 | 0.015 | 0 | 0.49 | 0.38 | 0.58 | 0.42 | 0.63 | 0.46 |
| 2 | 199627907 | rs10202112 | -0.628 | 0.233 | -1.633 | -0.137 | -0.004 | -0.001 | -0.001 | 0 | 0.58 | 0.57 | 0.65 | 0.44 | 0.73 | 0.49 |
| 2 | 199632946 | rs16830436 | -0.101 | 0.319 | -1.721 | -0.107 | 0.014 | 0.01 | 0.01 | 0 | 0.52 | 0.59 | 0.6 | 0.45 | 0.63 | 0.42 |
| 2 | 199633768 | rs1518081 | 0.624 | 0.476 | -3.101 | -0.138 | -0.055 | 0.043 | 0.043 | 0 | 0.16 | 0.07 | 0.18 | 0.08 | 0.27 | 0.08 |
| 2 | 199635641 | rs16830444 | 1.236 | 0.351 | -3.132 | -0.153 | -0.012 | -0.047 | -0.047 | 0 | 0.3 | 0.12 | 0.39 | 0.37 | 0.36 | 0.33 |
| 2 | 199638824 | rs1996466 | 0.058 | 0.289 | -3.09 | -0.17 | 0.02 | 0.01 | 0.01 | 0 | 0.47 | 0.12 | 0.36 | 0.56 | 0.37 | 0.57 |
| 2 | 199639129 | rs17829022 | 0.326 | 0.767 | -3.295 | 0.076 | -0.034 | -0.008 | -0.008 | 0 | 0.4 | 0.12 | 0.46 | 0.38 | 0.47 | 0.37 |
| 2 | 199640420 | rs11680034 | 0.143 | 1.519 | -2.399 | 0.339 | -0.047 | 0.037 | 0.037 | 0 | 0.15 | 0.07 | 0.17 | 0.08 | 0.26 | 0.08 |
| 2 | 199646763 | rs13025012 | -0.341 | 0.349 | -2.034 | 0.944 | -0.011 | -0.088 | -0.088 | 0.11 | 0.24 | 0.01 | 0.17 | 0.29 | 0.19 | 0.3 |
| 2 | 199660637 | rs1878662 | -0.726 | 0.853 | -1.318 | 1.117 | -0.002 | -0.008 | -0.008 | 0 | 0.18 | 0.15 | 0.21 | 0.29 | 0.15 | 0.47 |
| 2 | 199666111 | rs13035389 | 0.032 | 0.92 | -1.037 | 0.729 | 0.016 | 0.016 | 0.016 | 0 | 0.37 | 0.48 | 0.5 | 0.55 | 0.42 | 0.3 |
| 2 | 199671852 | rs10497823 | 0.224 | 0.752 | -0.637 | 0.757 | 0.013 | 0.054 | 0.054 | 0 | 0.16 | 0.22 | 0.2 | 0.28 | 0.13 | 0.45 |
| 2 | 199676607 | rs17829897 | 0.373 | 0.223 | -0.443 | 0.054 | 0.055 | 0.11 | 0.11 | 0 | 0.44 | 0.18 | 0.26 | 0.16 | 0.37 | 0.25 |
| 2 | 199679110 | rs10189905 | -0.19 | 0.313 | 0.331 | 0.211 | 0.001 | -0.032 | -0.032 | 0 | 0.13 | 0.24 | 0.1 | 0.09 | 0.09 | 0.08 |
| 2 | 199696908 | rs1533536 | -0.292 | -0.809 | 0.126 | -0.43 | 0.043 | 0.046 | 0.046 | 0 | 0.53 | 0.58 | 0.76 | 0.83 | 0.7 | 0.79 |
| 2 | 199697068 | rs1533537 | -0.533 | -1.597 | -0.121 | -0.794 | 0.071 | 0.136 | 0.136 | 0 | 0.63 | 0.83 | 0.83 | 0.94 | 0.74 | 0.83 |
| 2 | 199697571 | rs2693278 | 0.321 | -1.687 | 0.359 | -0.839 | -0.007 | -0.015 | -0.015 | 1.183 | 0.51 | 0.45 | 0.75 | 0.83 | 0.7 | 0.79 |
| 2 | 199698669 | rs1608945 | 0.255 | -0.638 | 0.353 | -1.291 | -0.008 | 0.005 | 0.005 | 0.185 | 0.33 | 0.26 | 0.55 | 0.83 | 0.43 | 0.69 |
| 2 | 199703618 | rs16830580 | 1.47 | -0.741 | 0.574 | -0.9 | -0.035 | 0 | 0 | 2.49 | 0.1 | 0.19 | 0.06 | 0.1 | 0.03 | 0.04 |
| 2 | 199753674 | rs12614240 | 0.51 | -0.693 | 0.332 | -1.076 | -0.171 | -0.015 | -0.015 | 0 | 0.29 | 0.06 | 0.52 | 0.78 | 0.43 | 0.45 |
| 2 | 199765094 | rs7596739 | 1.251 | -0.937 | 0.831 | -1.067 | -0.055 | 0.007 | 0.007 | 0.872 | 0.11 | 0.25 | 0.08 | 0.11 | 0.03 | 0.04 |
| 2 | 199772591 | rs1456520 | -0.413 | -1.169 | 0.812 | -1.205 | -0.098 | 0.02 | 0.02 | 0.456 | 0.84 | 0.53 | 0.9 | 0.84 | 0.95 | 0.72 |
| 2 | 199780624 | rs1379013 | 1.35 | -1.718 | -0.033 | -1.964 | -0.007 | 0.118 | 0.118 | 0.185 | 0.18 | 0.01 | 0.06 | 0.01 | 0.15 | 0.16 |
| 2 | 199782944 | rs7571513 | 1.968 | -1.638 | 0.059 | -2.008 | -0.032 | 0.031 | 0.031 | 0.353 | 0.17 | 0.09 | 0.19 | 0 | 0.27 | 0.1 |
| 2 | 199795231 | rs1456547 | -0.855 | -1.748 | 0.176 | -2.049 | -0.004 | -0.053 | -0.053 | 0 | 0.7 | 0.45 | 0.73 | 0.89 | 0.7 | 0.86 |
| 2 | 199801528 | rs720527 | 0.281 | -1.36 | -0.066 | -1.745 | -0.059 | 0.06 | 0.06 | 0.937 | 0.44 | 0.15 | 0.56 | 0.73 | 0.47 | 0.54 |
| 2 | 199808873 | rs12693871 | -0.031 | -1.292 | -0.026 | -1.788 | -0.023 | 0.183 | 0.183 | 1.363 | 0.48 | 0.17 | 0.58 | 0.83 | 0.5 | 0.79 |
| 2 | 199809716 | rs966744 | 1.47 | -1.207 | -0.279 | -0.431 | -0.127 | -0.149 | -0.149 | 0 | 0.23 | 0.02 | 0.41 | 0.44 | 0.31 | 0.34 |
| 2 | 199827930 | rs4533423 | 1.328 | -1.273 | -0.062 | -0.615 | -0.007 | 0.051 | 0.051 | 0 | 0.2 | 0.1 | 0.13 | 0.06 | 0.2 | 0.07 |
| 2 | 199838991 | rs2091340 | -1.419 | -1.091 | 0.5 | -0.339 | -0.078 | 0.016 | 0.016 | 5.946 | 0.87 | 0.57 | 0.92 | 0.89 | 0.97 | 0.96 |
| 2 | 199887785 | rs3916066 | 1.642 | -0.441 | -0.082 | -0.213 | -0.004 | -0.017 | -0.017 | 0 | 0.18 | 0.14 | 0.19 | 0 | 0.28 | 0.1 |
| 2 | 199889554 | rs2881064 | 1.29 | -0.211 | -0.16 | 0.735 | -0.051 | -0.107 | -0.107 | 4.496 | 0.14 | 0.01 | 0.34 | 0.52 | 0.14 | 0.22 |
| 2 | 200007302 | rs3963519 | 0.018 | -0.445 | -0.251 | 0.359 | 0.018 | 0.099 | 0.099 | 2.187 | 0.49 | 0.12 | 0.5 | 0.33 | 0.56 | 0.44 |
| 2 | 200008894 | rs1376584 | -0.142 | -0.484 | 0.184 | 0.339 | 0.008 | 0.018 | 0.018 | 1.804 | 0.51 | 0.42 | 0.52 | 0.32 | 0.56 | 0.44 |
| 2 | 200010475 | rs16831047 | 1.485 | -0.353 | -0.093 | 0.233 | -0.059 | -0.07 | -0.07 | 0 | 0.13 | 0.01 | 0.22 | 0.24 | 0.16 | 0.21 |
| 2 | 200015094 | rs1451473 | 0.115 | -0.563 | 0.167 | 0.313 | 0.025 | 0.059 | 0.059 | 2.206 | 0.49 | 0.32 | 0.52 | 0.32 | 0.56 | 0.44 |
| 2 | 200031387 | rs16831114 | -0.09 | -0.594 | -0.006 | 0.293 | 0.034 | 0.079 | 0.079 | 2.041 | 0.5 | 0.25 | 0.5 | 0.32 | 0.56 | 0.46 |
| 2 | 200036945 | rs4675343 | -0.062 | -0.61 | -0.075 | 0.303 | 0.036 | 0.083 | 0.083 | 2.12 | 0.49 | 0.23 | 0.5 | 0.33 | 0.56 | 0.46 |
| 2 | 200053196 | rs10497830 | 0.952 | -1.006 | -0.656 | 0.395 | -0.021 | -0.031 | -0.031 | 0.007 | 0.22 | 0.12 | 0.28 | 0.32 | 0.24 | 0.13 |
| 2 | 200056608 | rs13012540 | 1.861 | -1.236 | -0.866 | 0.201 | -0.007 | 0.115 | 0.115 | 0.178 | 0.19 | 0.03 | 0.09 | 0.01 | 0.17 | 0.14 |
| 2 | 200060715 | rs1982411 | -1.299 | -1.125 | 0.35 | 0.215 | -0.003 | 0.062 | 0.062 | 0 | 0.89 | 0.83 | 0.83 | 0.68 | 0.93 | 0.95 |
| 2 | 200068160 | rs1376591 | -0.649 | -1.177 | 0.262 | 0.248 | 0.026 | 0.071 | 0.071 | 2.276 | 0.5 | 0.32 | 0.48 | 0.68 | 0.45 | 0.55 |
| 2 | 200073190 | rs12616812 | -0.704 | -1.172 | 0.241 | 0.338 | 0.023 | 0.08 | 0.08 | 2.938 | 0.49 | 0.26 | 0.48 | 0.68 | 0.46 | 0.56 |
| 2 | 200125150 | rs1374360 | 0.113 | -1.037 | 0.099 | 0.607 | -0.009 | -0.025 | -0.025 | 0.111 | 0.25 | 0.11 | 0.28 | 0.33 | 0.25 | 0.14 |
| 2 | 200126137 | rs4449074 | 0.599 | -1.252 | -0.223 | 0.471 | -0.034 | -0.02 | -0.02 | 0.503 | 0.38 | 0.17 | 0.47 | 0.38 | 0.52 | 0.43 |
| 2 | 200134004 | rs12105015 | 1.824 | -0.758 | -0.076 | 0.55 | 0.02 | 0.02 | 0.02 | 0 | 0.1 | 0.16 | 0.09 | 0.14 | 0.12 | 0.22 |
| 2 | 200137608 | rs2881208 | 0.743 | -1.192 | -0.45 | 0.436 | -0.012 | -0.037 | -0.037 | 0 | 0.56 | 0.13 | 0.56 | 0.4 | 0.63 | 0.61 |
| 2 | 200142591 | rs994185 | -1.058 | -0.934 | 0.414 | 0.605 | 0.021 | 0.006 | 0.006 | 0 | 0.87 | 0.48 | 0.89 | 0.87 | 0.89 | 0.77 |
| 2 | 200142649 | rs10497831 | 1.474 | -0.922 | 0.498 | 0.612 | 0.014 | 0.005 | 0.005 | 0 | 0.11 | 0.19 | 0.09 | 0.12 | 0.11 | 0.22 |
| 2 | 200152198 | rs17196295 | 2.193 | -0.152 | -0.001 | 0.154 | -0.094 | 0.11 | 0.11 | 0.643 | 0.17 | 0.02 | 0.18 | 0 | 0.29 | 0.08 |
| 2 | 200154449 | rs7592248 | -0.811 | -0.558 | 0.368 | 0.478 | 0.004 | 0.025 | 0.025 | 0.115 | 0.85 | 0.41 | 0.83 | 0.83 | 0.89 | 0.76 |
| 2 | 200157595 | rs1446636 | -1.243 | -0.548 | 0.485 | 0.483 | 0.037 | 0.066 | 0.066 | 0 | 0.87 | 0.5 | 0.85 | 0.83 | 0.89 | 0.77 |
| 2 | 200162425 | rs895526 | 0.338 | -0.565 | 0.236 | 0.41 | -0.011 | 0.11 | 0.11 | 0 | 0.76 | 0.47 | 0.72 | 0.54 | 0.83 | 0.86 |
| 2 | 200167236 | rs2009622 | 0.313 | -0.377 | 0.111 | 0.494 | 0.005 | -0.016 | -0.016 | 0.214 | 0.64 | 0.24 | 0.57 | 0.36 | 0.72 | 0.49 |
| 2 | 200170530 | rs16831252 | 0.191 | -0.096 | 0.046 | 0.765 | 0 | 0.037 | 0.037 | 0 | 0.13 | 0.14 | 0.23 | 0.47 | 0.13 | 0.09 |
| 2 | 200189323 | rs4673313 | 0.254 | -0.103 | 0.073 | 0.762 | 0.021 | 0.25 | 0.25 | 0.048 | 0.84 | 0.46 | 0.73 | 0.53 | 0.87 | 0.91 |
| 2 | 200214273 | rs3828186 | 1.016 | -0.038 | -0.157 | 0.802 | -0.02 | -0.027 | -0.027 | 0 | 0.1 | 0.03 | 0.09 | 0.15 | 0.12 | 0.02 |
| 2 | 200243869 | rs1446646 | 1.68 | 0.228 | -0.269 | 0.469 | -0.102 | 0.107 | 0.107 | 0.531 | 0.18 | 0.03 | 0.2 | 0 | 0.33 | 0.1 |
| 2 | 200273376 | rs4673339 | 0.865 | 0.01 | -0.843 | 0.639 | -0.151 | -0.016 | -0.016 | 0.156 | 0.27 | 0.03 | 0.35 | 0.21 | 0.41 | 0.16 |

| | | | | | | | | | | | | | | | | |
|---|---|---|---|---|---|---|---|---|---|---|---|---|---|---|---|---|
| 2 | 200296044 | rs6726276 | -0.122 | -0.224 | -0.212 | 0.927 | 0.003 | 0.06 | 0.06 | 0 | 0.14 | 0.21 | 0.25 | 0.45 | 0.13 | 0.08 |
| 2 | 200311014 | rs7579004 | 0.663 | -0.388 | -0.602 | 0.676 | 0.049 | 0.038 | 0.038 | 0.161 | 0.53 | 0.16 | 0.54 | 0.35 | 0.68 | 0.45 |
| 2 | 200313323 | rs6752494 | -0.348 | -0.079 | 0.186 | 0.817 | -0.007 | 0.015 | 0.015 | 2.119 | 0.24 | 0.28 | 0.27 | 0.45 | 0.14 | 0.13 |
| 2 | 200332082 | rs6714133 | -0.02 | -0.099 | 0.166 | 0.489 | 0.013 | -0.01 | -0.01 | 0 | 0.21 | 0.15 | 0.07 | 0.3 | 0.11 | 0.12 |
| 2 | 200341637 | rs11684919 | 1.233 | 1.175 | 0.292 | 0.095 | -0.066 | 0.069 | 0.069 | 0 | 0.12 | 0.02 | 0.1 | 0 | 0.19 | 0.06 |
| 2 | 200371741 | rs7577740 | -1.637 | 0.045 | 0.926 | 0.229 | 0.041 | 0.036 | 0.036 | 8.754 | 0.06 | 0.52 | 0.04 | 0.01 | 0 | 0.01 |
| 2 | 200381686 | rs6731539 | -1.225 | 0.062 | 0.918 | 0.238 | 0.052 | 0.049 | 0.049 | 6.151 | 0.05 | 0.45 | 0.03 | 0.01 | 0 | 0.01 |
| 2 | 200391076 | rs6704623 | -0.659 | -0.083 | 0.456 | 0.353 | -0.012 | 0.011 | 0.011 | 0.36 | 0.27 | 0.69 | 0.3 | 0.47 | 0.17 | 0.13 |
| 2 | 200440820 | rs17528188 | 0.245 | 1.087 | -0.258 | 0.535 | -0.107 | -0.081 | -0.081 | 0 | 0.18 | 0.05 | 0.38 | 0.28 | 0.38 | 0.16 |
| 2 | 200452591 | rs10182928 | 0.806 | 1.027 | -0.415 | 0.215 | -0.016 | 0.005 | 0.005 | 0 | 0.27 | 0.12 | 0.16 | 0.24 | 0.27 | 0.55 |
| 2 | 200460345 | rs1551085 | 0.475 | 0.639 | -0.572 | -0.327 | -0.006 | -0.121 | -0.121 | 0.244 | 0.2 | 0.32 | 0.12 | 0 | 0.17 | 0.04 |
| 2 | 200473658 | rs12693889 | -1.169 | 0.043 | -0.613 | -0.279 | 0.09 | -0.101 | -0.101 | 0 | 0.67 | 0.93 | 0.68 | 0.53 | 0.81 | 0.74 |
| 2 | 200481331 | rs13012419 | -0.505 | 1.046 | 0.435 | 0.672 | -0.025 | -0.121 | -0.121 | 0.039 | 0.2 | 0.32 | 0.12 | 0 | 0.15 | 0.04 |
| 2 | 200498970 | rs1376132 | -0.329 | 1.864 | -1.065 | 0.047 | -0.089 | 0.008 | 0.008 | 0 | 0.13 | 0.02 | 0.22 | 0.1 | 0.34 | 0.11 |
| 2 | 200518102 | rs2346110 | 0.306 | 0.284 | -0.186 | -0.168 | -0.003 | -0.101 | -0.101 | 1.32 | 0.78 | 0.69 | 0.85 | 0.99 | 0.79 | 0.96 |
| 2 | 200523384 | rs6711885 | 0.389 | -0.152 | -0.515 | 0.012 | -0.019 | -0.021 | -0.021 | 0 | 0.33 | 0.25 | 0.38 | 0.49 | 0.45 | 0.61 |
| 2 | 200527705 | rs17590164 | 0.42 | -0.257 | -0.283 | 0.022 | 0.055 | -0.005 | -0.005 | 0.127 | 0.46 | 0.34 | 0.47 | 0.5 | 0.34 | 0.35 |
| 2 | 200528519 | rs9677698 | 0.519 | -0.821 | -0.619 | -0.241 | -0.007 | -0.032 | -0.032 | 0 | 0.15 | 0.13 | 0.37 | 0.46 | 0.18 | 0.17 |
| 2 | 200549085 | rs12620740 | 0.009 | -1.306 | -0.114 | -0.205 | 0.089 | 0.098 | 0.098 | 0 | 0.48 | 0.24 | 0.52 | 0.7 | 0.63 | 0.73 |
| 2 | 200559028 | rs4673422 | -0.713 | -1.313 | -0.188 | -0.262 | -0.003 | -0.063 | -0.063 | 0 | 0.83 | 0.8 | 0.88 | 1 | 0.82 | 0.97 |
| 2 | 200567136 | rs10211241 | 0.004 | -1.198 | -0.299 | -0.171 | 0.1 | 0.112 | 0.112 | 0 | 0.48 | 0.22 | 0.52 | 0.7 | 0.62 | 0.73 |
| 2 | 200576280 | rs6741826 | 0.159 | -1.348 | -0.124 | -0.205 | -0.007 | -0.001 | -0.001 | 0 | 0.75 | 0.74 | 0.64 | 0.74 | 0.75 | 0.92 |
| 2 | 200578867 | rs1357582 | -0.038 | -2.052 | -0.439 | -0.626 | 0.15 | 0.104 | 0.104 | 0.052 | 0.54 | 0.2 | 0.59 | 0.7 | 0.73 | 0.76 |
| 2 | 200579502 | rs4673435 | -0.051 | -2.076 | -0.451 | -0.638 | 0.148 | 0.103 | 0.103 | 0.082 | 0.54 | 0.2 | 0.59 | 0.7 | 0.73 | 0.76 |
| 2 | 200580494 | rs4493243 | 0.761 | -2.402 | -0.232 | -0.042 | -0.054 | 0.143 | 0.143 | 0.024 | 0.81 | 0.49 | 0.68 | 0.56 | 0.89 | 0.92 |
| 2 | 200592510 | rs12052622 | 0.287 | -2.582 | -0.36 | 0.024 | 0.044 | 0.015 | 0.015 | 0 | 0.37 | 0.26 | 0.32 | 0.3 | 0.18 | 0.23 |
| 2 | 200597218 | rs7576880 | 0.193 | -2.663 | -0.024 | -0.021 | -0.021 | -0.003 | -0.003 | 0 | 0.38 | 0.38 | 0.33 | 0.3 | 0.17 | 0.33 |
| 2 | 200606148 | rs10931874 | 0.003 | -2.56 | 0.125 | -0.069 | -0.031 | -0.094 | -0.094 | 0 | 0.86 | 0.43 | 0.89 | 0.99 | 0.92 | 0.98 |
| 2 | 200607333 | rs11695675 | 0.137 | -2.535 | -0.084 | -0.122 | 0.168 | 0.122 | 0.122 | 0 | 0.5 | 0.18 | 0.55 | 0.7 | 0.71 | 0.63 |
| 2 | 200612911 | rs11685147 | 0.777 | -2.476 | -0.033 | -0.062 | 0.01 | -0.006 | -0.006 | 0.092 | 0.79 | 0.2 | 0.66 | 0.75 | 0.89 | 0.93 |
| 2 | 200613782 | rs12693898 | 0.424 | -2.453 | -0.035 | -0.047 | -0.037 | -0.003 | -0.003 | 0 | 0.82 | 0.43 | 0.88 | 0.85 | 0.91 | 0.97 |
| 2 | 200615496 | rs11887310 | -0.445 | -2.404 | 0.054 | -0.076 | -0.017 | -0.093 | -0.093 | 1.036 | 0.87 | 0.64 | 0.9 | 0.99 | 0.92 | 0.98 |
| 2 | 200621420 | rs4530349 | -0.031 | -2.23 | -0.663 | -0.172 | 0.108 | 0.118 | 0.118 | 0 | 0.49 | 0.24 | 0.52 | 0.73 | 0.69 | 0.61 |
| 2 | 200625581 | rs2346586 | 0.157 | -2.455 | -1.04 | 0.428 | 0.077 | -0.035 | -0.035 | 0 | 0.44 | 0.13 | 0.49 | 0.6 | 0.66 | 0.58 |
| 2 | 200628118 | rs4673480 | 1.416 | -0.929 | -1.12 | 0.867 | -0.051 | -0.053 | -0.053 | 0.092 | 0.1 | 0.01 | 0.07 | 0.19 | 0.17 | 0.1 |
| 2 | 200634917 | rs7572473 | 0.775 | -0.998 | -1.172 | 0.579 | 0.04 | 0.018 | 0.018 | 2.573 | 0.68 | 0.12 | 0.57 | 0.78 | 0.76 | 0.86 |
| 2 | 200638509 | rs12615435 | 0.383 | -0.817 | -1.273 | 0.959 | 0.002 | -0.015 | -0.015 | 0.036 | 0.07 | 0.04 | 0.08 | 0.2 | 0.09 | 0.03 |
| 2 | 200648286 | rs12473716 | -0.936 | -1.311 | -0.744 | 0.677 | -0.011 | -0.038 | -0.038 | 0 | 0.54 | 0.48 | 0.76 | 0.83 | 0.68 | 0.64 |
| 2 | 200656557 | rs17529497 | 1.137 | -0.424 | -1.698 | 0.963 | -0.11 | -0.069 | -0.069 | 6.379 | 0.23 | 0.02 | 0.24 | 0.29 | 0.32 | 0.34 |
| 2 | 200666787 | rs10931878 | -0.238 | -0.802 | -1.679 | 1.265 | -0.02 | -0.041 | -0.041 | 0 | 0.39 | 0.14 | 0.43 | 0.59 | 0.55 | 0.53 |
| 2 | 200676926 | rs7605378 | 0.689 | -0.881 | -1.145 | 1.621 | 0.007 | 0.038 | 0.038 | 0 | 0.67 | 0.57 | 0.58 | 0.56 | 0.68 | 0.79 |
| 2 | 200677625 | rs2030653 | -1.007 | -0.849 | -1.228 | 1.777 | 0.014 | 0.021 | 0.021 | 0.029 | 0.86 | 0.86 | 0.91 | 0.82 | 0.81 | 0.9 |
| 2 | 200696446 | rs10203122 | 0.268 | -0.904 | -1.232 | 1.679 | 0.003 | -0.009 | -0.009 | 0 | 0.17 | 0.18 | 0.33 | 0.28 | 0.13 | 0.12 |
| 2 | 200700685 | rs2577850 | 0.612 | 0.005 | -1.799 | 1.446 | 0.019 | -0.009 | -0.009 | 0 | 0.43 | 0.35 | 0.45 | 0.51 | 0.38 | 0.48 |
| 2 | 200701788 | rs10497841 | 0.298 | 0.412 | -2.269 | 1.398 | -0.02 | -0.076 | -0.076 | 0 | 0.31 | 0.14 | 0.51 | 0.49 | 0.38 | 0.45 |
| 2 | 200702885 | rs1000507 | 0.84 | 0.454 | -2.38 | 1.478 | -0.067 | -0.075 | -0.075 | 1.073 | 0.24 | 0.01 | 0.31 | 0.33 | 0.29 | 0.34 |
| 2 | 200710706 | rs769960 | -0.749 | 0.136 | -1.933 | 1.162 | -0.013 | 0.014 | 0.014 | 0.399 | 0.72 | 0.85 | 0.82 | 0.83 | 0.66 | 0.86 |
| 2 | 200711563 | rs4673509 | 0.576 | -0.09 | -1.716 | 1.359 | -0.001 | 0.017 | 0.017 | 0.005 | 0.06 | 0.06 | 0.18 | 0.16 | 0.06 | 0.06 |
| 2 | 200726221 | rs17529746 | 2.36 | -0.337 | -2.54 | 0.509 | -0.098 | 0.083 | 0.083 | 0 | 0.24 | 0.07 | 0.16 | 0.11 | 0.31 | 0.33 |
| 2 | 200741720 | rs11694369 | 2.337 | -0.309 | -2.018 | 1.078 | -0.062 | -0.043 | -0.043 | 0 | 0.1 | 0.01 | 0.07 | 0.1 | 0.17 | 0.06 |
| 2 | 200767512 | rs1658807 | -2.019 | -0.402 | -1.868 | 1.138 | 0.027 | 0.038 | 0.038 | 0.013 | 0.88 | 0.86 | 0.91 | 0.82 | 0.82 | 0.93 |
| 2 | 200769539 | rs1704186 | -2.044 | -0.402 | -1.868 | 1.128 | 0.027 | 0.037 | 0.037 | 0.018 | 0.88 | 0.86 | 0.91 | 0.82 | 0.82 | 0.93 |
| 2 | 200778866 | rs166849 | 0.161 | -0.442 | -1.658 | 1.116 | -0.004 | -0.001 | -0.001 | 0 | 0.64 | 0.73 | 0.69 | 0.65 | 0.61 | 0.72 |
| 2 | 200784238 | rs281787 | -0.072 | -0.437 | -1.652 | 1.098 | -0.002 | 0.003 | 0.003 | 0.036 | 0.67 | 0.74 | 0.79 | 0.82 | 0.65 | 0.86 |
| 2 | 200792881 | rs971232 | -2.046 | -0.435 | -1.664 | 1.233 | 0.027 | 0.036 | 0.036 | 0.013 | 0.88 | 0.86 | 0.91 | 0.82 | 0.82 | 0.93 |
| 2 | 200793205 | rs10202253 | 0.963 | -0.523 | -1.467 | 1.308 | 0 | -0.008 | -0.008 | 0 | 0.57 | 0.54 | 0.38 | 0.39 | 0.54 | 0.64 |
| 2 | 200793784 | rs16832051 | 1.355 | -0.525 | -1.46 | 1.305 | 0.016 | 0.051 | 0.051 | 0 | 0.2 | 0.09 | 0.12 | 0 | 0.18 | 0.07 |
| 2 | 200795571 | rs6435036 | 1.023 | -0.442 | -1.489 | 1.349 | 0 | -0.011 | -0.011 | 0 | 0.57 | 0.52 | 0.38 | 0.39 | 0.55 | 0.65 |
| 2 | 200819812 | rs281765 | -0.115 | -0.354 | -1.476 | 1.261 | -0.004 | 0.122 | 0.122 | 0 | 0.07 | 0.17 | 0.31 | 0.26 | 0.06 | 0.07 |
| 2 | 200820505 | rs281766 | -2.116 | -0.246 | -1.383 | 1.331 | 0.013 | 0.021 | 0.021 | 0.042 | 0.88 | 0.89 | 0.92 | 0.8 | 0.82 | 0.93 |
| 2 | 200823281 | rs281767 | -0.334 | -0.251 | -1.389 | 1.256 | -0.006 | 0.008 | 0.008 | 0.109 | 0.68 | 0.77 | 0.8 | 0.8 | 0.64 | 0.86 |
| 2 | 200829993 | rs10178177 | -2.126 | -0.205 | -1.357 | 1.263 | 0.014 | 0.025 | 0.025 | 0.108 | 0.88 | 0.89 | 0.92 | 0.8 | 0.82 | 0.93 |
| 2 | 200848896 | rs4673627 | 0.668 | -0.281 | -1.322 | 1.165 | 0 | -0.001 | -0.001 | 0 | 0.57 | 0.59 | 0.46 | 0.49 | 0.55 | 0.76 |
| 2 | 200850378 | rs281792 | 0.051 | -0.144 | -1.235 | 1.212 | -0.006 | 0.076 | 0.076 | 0.012 | 0.07 | 0.13 | 0.32 | 0.2 | 0.06 | 0.07 |
| 2 | 200853425 | rs17445724 | 1.351 | -0.248 | -1.49 | 1.01 | -0.009 | 0.003 | 0.003 | 0 | 0.17 | 0.01 | 0.13 | 0.11 | 0.15 | 0.28 |
| 2 | 200872831 | rs2136585 | 1.133 | -0.084 | -0.778 | 1.241 | -0.001 | -0.005 | -0.005 | 0.074 | 0.53 | 0.53 | 0.37 | 0.4 | 0.54 | 0.65 |
| 2 | 200884246 | rs7608932 | 1.092 | -0.086 | -0.781 | 1.238 | -0.001 | -0.006 | -0.006 | 0 | 0.54 | 0.53 | 0.37 | 0.4 | 0.54 | 0.65 |
| 2 | 200901481 | rs10175678 | -0.001 | -0.117 | -0.697 | 1.47 | -0.007 | 0.057 | 0.057 | 0.013 | 0.07 | 0.1 | 0.32 | 0.2 | 0.06 | 0.07 |
| 2 | 200902672 | rs203766 | 0.896 | -0.02 | -0.644 | 1.512 | 0.03 | 0.046 | 0.046 | 0 | 0.08 | 0.13 | 0.09 | 0.19 | 0.15 | 0.07 |

| | | | | | | | | | | | | | | | | |
|---|---|---|---|---|---|---|---|---|---|---|---|---|---|---|---|---|
| 2 | 200904011 | rs7604950 | 0.463 | -0.15 | -0.804 | 1.32 | 0.032 | 0.072 | 0.072 | 0 | 0.26 | 0.09 | 0.12 | 0.1 | 0.21 | 0.11 |
| 2 | 200913244 | rs3115414 | 0.866 | -0.162 | -0.788 | 1.313 | -0.004 | 0.101 | 0.101 | 4.758 | 0.09 | 0.22 | 0.31 | 0.16 | 0.06 | 0.04 |
| 2 | 200915783 | rs3115413 | -0.833 | -0.132 | -0.779 | 1.297 | -0.009 | -0.003 | -0.003 | 0 | 0.35 | 0.35 | 0.44 | 0.26 | 0.27 | 0.14 |
| 2 | 200946126 | rs10497847 | 0.391 | -0.087 | -1.236 | 1.239 | 0.018 | 0.046 | 0.046 | 0 | 0.23 | 0.14 | 0.19 | 0.11 | 0.19 | 0.17 |
| 2 | 200946393 | rs2347286 | 0.093 | -0.146 | -1.132 | 1.317 | -0.031 | 0.075 | 0.075 | 7.324 | 0.16 | 0.23 | 0.35 | 0.3 | 0.12 | 0.11 |
| 2 | 200951540 | rs3094758 | -1.714 | 0.002 | -1.141 | 1.352 | -0.001 | -0.009 | -0.009 | 0 | 0.46 | 0.5 | 0.68 | 0.6 | 0.46 | 0.35 |
| 2 | 200957953 | rs10182513 | 0.19 | 0.066 | -1.146 | 1.444 | -0.002 | 0.029 | 0.029 | 0.075 | 0.06 | 0.06 | 0.31 | 0.19 | 0.06 | 0.07 |
| 2 | 200975897 | rs13026730 | -0.345 | 0.021 | -1.16 | 1.325 | 0.022 | 0.02 | 0.02 | 0 | 0.3 | 0.23 | 0.23 | 0.25 | 0.24 | 0.22 |
| 2 | 200999256 | rs12613687 | 0.545 | 0.155 | -1.307 | 1.439 | -0.029 | -0.039 | -0.039 | 0 | 0.13 | 0.07 | 0.15 | 0.36 | 0.19 | 0.1 |
| 2 | 201003837 | rs1106399 | 0.235 | 0.108 | -1.334 | 1.435 | -0.057 | -0.057 | -0.057 | 0.014 | 0.09 | 0.02 | 0.16 | 0.24 | 0.18 | 0.11 |
| 2 | 201009778 | rs1919366 | 1.573 | -0.17 | -1.133 | 1.499 | 0 | 0 | 0 | 0 | 0.54 | 0.49 | 0.32 | 0.34 | 0.54 | 0.67 |
| 2 | 201010789 | rs1919367 | 0.246 | -0.271 | -1.054 | 1.444 | -0.018 | -0.021 | -0.021 | 0 | 0.71 | 0.68 | 0.57 | 0.84 | 0.76 | 0.9 |
| 2 | 201012142 | rs13020689 | 1.593 | -0.395 | -0.769 | 1.768 | -0.02 | 0.021 | 0.021 | 0.036 | 0.07 | 0.15 | 0.02 | 0.15 | 0.03 | 0.03 |
| 2 | 201021954 | rs1569175 | -1.068 | -0.269 | -0.745 | 1.431 | 0 | 0.124 | 0.124 | 5.998 | 0.91 | 0.79 | 0.69 | 0.84 | 0.94 | 0.96 |
| 2 | 201044937 | rs1115187 | 0.979 | -0.089 | -1.029 | 1.068 | -0.001 | -0.041 | -0.041 | 5.959 | 0.17 | 0.16 | 0.09 | 0 | 0.15 | 0.05 |
| 2 | 201051150 | rs11675980 | -0.198 | 0.168 | -0.989 | 1.54 | -0.031 | -0.051 | -0.051 | 0 | 0.14 | 0.07 | 0.17 | 0.38 | 0.21 | 0.13 |
| 2 | 201059371 | rs6722092 | 0.028 | 0.174 | -0.959 | 1.497 | 0.04 | 0.114 | 0.114 | 0 | 0.85 | 0.71 | 0.82 | 0.63 | 0.8 | 0.87 |
| 2 | 201072852 | rs4673782 | 0.151 | -0.001 | -1.741 | 1.607 | 0.002 | -0.003 | -0.003 | 0 | 0.14 | 0.16 | 0.13 | 0.3 | 0.08 | 0.15 |
| 2 | 201073133 | rs1534599 | 1.343 | -0.288 | -1.971 | 1.465 | 0.002 | 0.025 | 0.025 | 0 | 0.55 | 0.6 | 0.35 | 0.33 | 0.57 | 0.69 |
| 2 | 201073161 | rs4673785 | 1.685 | -0.194 | -1.95 | 1.565 | -0.012 | 0.019 | 0.019 | 1.618 | 0.06 | 0.12 | 0.1 | 0.12 | 0.04 | 0.09 |
| 2 | 201075483 | rs1653297 | 0.536 | -0.144 | -1.96 | 1.589 | 0.01 | 0.068 | 0.068 | 0.017 | 0.8 | 0.7 | 0.73 | 0.51 | 0.76 | 0.77 |
| 2 | 201076401 | rs1653301 | 0.467 | -0.132 | -2.237 | 1.321 | -0.005 | -0.008 | -0.008 | 1.83 | 0.66 | 0.67 | 0.67 | 0.43 | 0.61 | 0.48 |
| 2 | 201082457 | rs4673802 | -0.489 | -0.271 | -2.273 | 1.329 | 0.035 | -0.063 | -0.063 | 1.122 | 0.32 | 0.15 | 0.48 | 0.57 | 0.27 | 0.48 |
| 2 | 201084274 | rs16842086 | 1.095 | -0.226 | -2.22 | 1.099 | 0.013 | 0.027 | 0.027 | 0 | 0.19 | 0.11 | 0.09 | 0 | 0.16 | 0.04 |
| 2 | 201086663 | rs1729413 | 0.821 | -0.21 | -2.218 | 1.253 | 0.001 | 0.013 | 0.013 | 0 | 0.48 | 0.58 | 0.57 | 0.41 | 0.46 | 0.43 |
| 2 | 201088171 | rs1369843 | -0.794 | 0.287 | -1.795 | 1.615 | 0.052 | 0.107 | 0.107 | 0 | 0.89 | 0.77 | 0.74 | 0.69 | 0.79 | 0.81 |
| 2 | 201096881 | rs2078279 | 0.172 | 0.203 | -1.871 | 1.574 | 0.023 | 0.011 | 0.011 | 0 | 0.26 | 0.15 | 0.37 | 0.17 | 0.19 | 0.08 |
| 2 | 201102905 | rs296797 | 0.451 | 0.134 | -1.839 | 1.253 | -0.001 | -0.001 | -0.001 | 0 | 0.63 | 0.61 | 0.37 | 0.53 | 0.58 | 0.73 |
| 2 | 201111241 | rs12614633 | 0.366 | -0.319 | -2.272 | 1.077 | 0.053 | -0.001 | -0.001 | 0 | 0.28 | 0.11 | 0.11 | 0.27 | 0.2 | 0.33 |
| 2 | 201120228 | rs17446700 | 0.612 | -0.316 | -2.321 | 1.067 | 0.031 | -0.013 | -0.013 | 0 | 0.26 | 0.03 | 0.1 | 0.22 | 0.19 | 0.31 |
| 2 | 201135222 | rs13006859 | 0.305 | -0.275 | -2.34 | 1.083 | 0.052 | -0.018 | -0.018 | 0.005 | 0.3 | 0.04 | 0.11 | 0.26 | 0.21 | 0.36 |
| 2 | 201137782 | rs17630248 | -0.331 | -0.182 | -2.511 | 1.203 | 0.043 | -0.05 | -0.05 | 0 | 0.43 | 0.15 | 0.43 | 0.42 | 0.36 | 0.41 |
| 2 | 201143242 | rs10931895 | 0.462 | 0.085 | -2.246 | 1.38 | 0.033 | 0.083 | 0.083 | 0 | 0.12 | 0.19 | 0.25 | 0.39 | 0.21 | 0.23 |
| 2 | 201144004 | rs295118 | 0.15 | 0.054 | -2.173 | 1.36 | 0.032 | 0.034 | 0.034 | 0 | 0.75 | 0.64 | 0.42 | 0.44 | 0.62 | 0.72 |
| 2 | 201146831 | rs13003304 | 0.578 | 0.072 | -2.262 | 1.601 | 0.072 | 0.002 | 0.002 | 0.687 | 0.29 | 0.03 | 0.1 | 0.22 | 0.19 | 0.33 |
| 2 | 201148951 | rs11691757 | -0.22 | 0.194 | -2.109 | 1.689 | 0.034 | 0.067 | 0.067 | 0.628 | 0.15 | 0.22 | 0.51 | 0.56 | 0.25 | 0.27 |
| 2 | 201158122 | rs12612420 | 0.592 | -0.19 | -2.367 | 1.385 | 0.068 | -0.005 | -0.005 | 0 | 0.29 | 0.03 | 0.1 | 0.22 | 0.18 | 0.33 |
| 2 | 201160699 | rs295140 | 0.282 | -0.049 | -2.243 | 1.284 | -0.002 | 0.086 | 0.086 | 0.256 | 0.57 | 0.69 | 0.4 | 0.27 | 0.58 | 0.42 |
| 2 | 201163783 | rs4673905 | 0.231 | -0.048 | -2.33 | 1.444 | -0.041 | -0.153 | -0.153 | 3.945 | 0.33 | 0.05 | 0.23 | 0.44 | 0.31 | 0.45 |
| 2 | 201168993 | rs4673912 | -0.444 | 0.032 | -2.311 | 1.092 | 0.018 | 0.11 | 0.11 | 0.51 | 0.43 | 0.17 | 0.56 | 0.71 | 0.38 | 0.56 |
| 2 | 201195602 | rs295114 | 1.263 | -0.525 | -2.297 | 0.633 | 0.018 | -0.038 | -0.038 | 0 | 0.34 | 0.52 | 0.3 | 0.23 | 0.42 | 0.39 |
| 2 | 201200818 | rs6435049 | 0.076 | -0.653 | -3.012 | 0.314 | 0.058 | 0.112 | 0.112 | 1 | 0.65 | 0.18 | 0.77 | 0.81 | 0.76 | 0.91 |
| 2 | 201201190 | rs995521 | -0.351 | -0.734 | -2.987 | 0.295 | 0 | -0.002 | -0.002 | 0 | 0.75 | 0.29 | 0.84 | 0.86 | 0.89 | 0.95 |
| 2 | 201203110 | rs883999 | 1.607 | -0.776 | -3.057 | 0.427 | 0.033 | 0.061 | 0.061 | 0 | 0.23 | 0.02 | 0.09 | 0.09 | 0.14 | 0.23 |
| 2 | 201208692 | rs985256 | -0.452 | -0.723 | -2.968 | 0.419 | 0.014 | 0.042 | 0.042 | 2.035 | 0.75 | 0.18 | 0.79 | 0.81 | 0.78 | 0.9 |
| 2 | 201211605 | rs3769466 | 1.131 | -0.653 | -3.119 | 0.601 | 0.052 | -0.076 | -0.076 | 0 | 0.31 | 0.02 | 0.19 | 0.33 | 0.2 | 0.36 |
| 2 | 201214071 | rs1900706 | -0.54 | -0.145 | -2.608 | 0.42 | -0.023 | 0.041 | 0.041 | 2.222 | 0.66 | 0.27 | 0.7 | 0.77 | 0.6 | 0.6 |
| 2 | 201226248 | rs1550293 | -0.092 | -0.172 | -2.062 | 0.663 | -0.008 | 0.028 | 0.028 | 2.788 | 0.81 | 0.29 | 0.62 | 0.71 | 0.88 | 0.92 |
| 2 | 201229473 | rs295129 | -0.215 | -0.253 | -2.387 | 0.634 | -0.006 | 0.06 | 0.06 | 0.966 | 0.46 | 0.68 | 0.38 | 0.3 | 0.56 | 0.47 |
| 2 | 201239957 | rs3754798 | 1.625 | -0.09 | -2.491 | 0.608 | -0.108 | -0.035 | -0.035 | 0.231 | 0.22 | 0.01 | 0.19 | 0.16 | 0.34 | 0.4 |
| 2 | 201240305 | rs12990174 | 0.272 | -0.158 | -2.464 | 0.603 | 0.032 | -0.015 | -0.015 | 0 | 0.49 | 0.18 | 0.37 | 0.54 | 0.41 | 0.52 |
| 2 | 201241967 | rs295120 | 0.677 | -0.11 | -2.162 | 0.549 | -0.014 | -0.01 | -0.01 | 0 | 0.14 | 0.6 | 0.12 | 0.14 | 0.08 | 0.05 |
| 2 | 201246198 | rs3754794 | 1.358 | -0.335 | -2.77 | 0.472 | 0.035 | 0.059 | 0.059 | 0 | 0.25 | 0.01 | 0.09 | 0.09 | 0.15 | 0.22 |
| 2 | 201247479 | rs7598349 | -0.185 | 0.283 | -1.787 | 0.768 | -0.007 | -0.006 | -0.006 | 0 | 0.62 | 0.36 | 0.46 | 0.56 | 0.55 | 0.52 |
| 2 | 201247491 | rs842828 | 0.946 | 0.134 | -0.926 | 0.606 | 0.01 | 0.043 | 0.043 | 1.451 | 0.11 | 0.37 | 0.1 | 0.19 | 0.08 | 0.09 |
| 2 | 201252355 | rs3769450 | 0.443 | 0.091 | -1.42 | 0.646 | 0.064 | 0.031 | 0.031 | 0 | 0.48 | 0.17 | 0.53 | 0.4 | 0.38 | 0.42 |
| 2 | 201253769 | rs3739118 | 1.051 | 0.621 | -1.196 | 0.613 | -0.04 | -0.003 | -0.003 | 0.348 | 0.2 | 0.06 | 0.16 | 0.14 | 0.28 | 0.33 |
| 2 | 201253956 | rs842823 | 0.515 | 0.528 | -0.556 | 0.548 | -0.005 | 0.004 | 0.004 | 0.414 | 0.24 | 0.18 | 0.19 | 0.15 | 0.31 | 0.34 |
| 2 | 201254171 | rs4674015 | -0.24 | -0.178 | -0.276 | 0.352 | 0.001 | 0 | 0 | 0.117 | 0.71 | 0.43 | 0.75 | 0.7 | 0.72 | 0.83 |
| 2 | 201256574 | rs7585698 | 0.253 | -0.293 | -0.097 | 0.182 | 0.04 | 0.067 | 0.067 | 0.164 | 0.08 | 0.33 | 0.09 | 0.19 | 0.07 | 0.08 |
| 2 | 201257203 | rs733365 | 0.191 | -0.402 | -0.203 | 0.133 | 0.031 | 0.007 | 0.007 | 0.812 | 0.52 | 0.3 | 0.41 | 0.55 | 0.42 | 0.51 |
| 2 | 201270538 | rs4674036 | 0.027 | -0.456 | -0.389 | 0.136 | 0.011 | 0.019 | 0.019 | 2.917 | 0.46 | 0.38 | 0.54 | 0.26 | 0.57 | 0.46 |
| 2 | 201285773 | rs13428321 | 2.18 | -0.398 | -0.519 | 0.052 | 0.004 | -0.011 | -0.011 | 0.009 | 0.15 | 0.12 | 0.07 | 0 | 0.13 | 0.02 |
| 2 | 201286358 | rs17447424 | 0.498 | -0.207 | -0.675 | 0.205 | -0.017 | -0.005 | -0.005 | 0 | 0.3 | 0.24 | 0.45 | 0.25 | 0.41 | 0.43 |
| 2 | 201290536 | rs13417802 | 0.018 | -0.812 | -0.711 | 0.1 | -0.036 | -0.021 | -0.021 | 0 | 0.38 | 0.5 | 0.27 | 0.31 | 0.31 | 0.26 |
| 2 | 201296480 | rs1866148 | -0.992 | -0.307 | -0.017 | 0.344 | 0.027 | 0.097 | 0.097 | 0.693 | 0.94 | 0.75 | 0.95 | 0.8 | 0.97 | 0.93 |
| 2 | 201300132 | rs2043766 | 0.02 | -0.413 | -0.266 | 0.273 | 0 | 0 | 0 | 0 | 0.51 | 0.52 | 0.42 | 0.58 | 0.43 | 0.49 |
| 2 | 201303010 | rs2028201 | 0.782 | -0.564 | -0.385 | 0.192 | 0.004 | -0.024 | -0.024 | 0 | 0.37 | 0.16 | 0.31 | 0.4 | 0.38 | 0.43 |
| 2 | 201308117 | rs1561296 | 0.423 | -0.459 | -0.563 | 0.274 | -0.036 | 0.055 | 0.055 | 5.457 | 0.42 | 0.15 | 0.52 | 0.22 | 0.54 | 0.44 |

| | | | | | | | | | | | | | | | | |
|---|---|---|---|---|---|---|---|---|---|---|---|---|---|---|---|---|
| 2 | 201309547 | rs3769446 | 0.057 | -0.452 | -0.461 | 0.252 | -0.001 | -0.001 | -0.001 | 0.014 | 0.48 | 0.46 | 0.34 | 0.4 | 0.38 | 0.46 |
| 2 | 201312329 | rs920249 | 0.901 | -0.771 | -0.261 | 0.167 | 0.003 | -0.004 | -0.004 | 0 | 0.78 | 0.19 | 0.8 | 0.63 | 0.88 | 0.87 |
| 2 | 201312663 | rs6722649 | -0.594 | -0.711 | -0.27 | -0.183 | -0.008 | -0.045 | -0.045 | 0 | 0.78 | 0.59 | 0.85 | 0.81 | 0.8 | 0.9 |
| 2 | 201313305 | rs1370558 | 0.051 | -0.615 | -0.129 | -0.096 | -0.036 | -0.014 | -0.014 | 0 | 0.85 | 0.68 | 0.92 | 0.82 | 0.94 | 0.91 |
| 2 | 201313988 | rs13004353 | 0.825 | 0.139 | -0.736 | 0.168 | -0.047 | -0.067 | -0.067 | 0.857 | 0.38 | 0.17 | 0.46 | 0.52 | 0.43 | 0.42 |
| 2 | 201314274 | rs17447598 | 0.587 | 0.41 | -0.575 | 0.238 | -0.041 | -0.004 | -0.004 | 0 | 0.23 | 0.12 | 0.27 | 0.17 | 0.33 | 0.41 |
| 2 | 201318373 | rs3769441 | 0.683 | 0.094 | -0.715 | -0.127 | 0.038 | 0.123 | 0.123 | 0.081 | 0.29 | 0.07 | 0.22 | 0.03 | 0.22 | 0.13 |
| 2 | 201319021 | rs3769439 | 0.887 | 0.087 | -0.686 | 0.017 | -0.053 | -0.048 | -0.048 | 1.516 | 0.4 | 0.17 | 0.5 | 0.58 | 0.45 | 0.53 |
| 2 | 201320531 | rs1318721 | -0.351 | 0.096 | -0.51 | 0.052 | -0.007 | 0.069 | 0.069 | 0.213 | 0.51 | 0.25 | 0.4 | 0.25 | 0.51 | 0.41 |
| 2 | 201325033 | rs17532176 | 1.251 | 0.073 | -0.632 | 0.018 | 0.039 | 0.099 | 0.099 | 0.486 | 0.26 | 0.06 | 0.12 | 0 | 0.18 | 0.1 |
| 2 | 201327631 | rs6730788 | -0.76 | 0.184 | -0.397 | 0.241 | 0.041 | 0.1 | 0.1 | 0 | 0.94 | 0.75 | 0.94 | 0.82 | 0.96 | 0.95 |
| 2 | 201329124 | rs6435050 | 0.785 | 0.232 | -1.42 | 0.2 | -0.036 | -0.032 | -0.032 | 2.843 | 0.41 | 0.23 | 0.51 | 0.58 | 0.45 | 0.53 |
| 2 | 201329213 | rs6435051 | 0.76 | -0.042 | -0.556 | 0.485 | -0.017 | 0.107 | 0.107 | 2 | 0.87 | 0.4 | 0.83 | 0.7 | 0.91 | 0.93 |
| 2 | 201333627 | rs11885025 | 0.411 | -0.189 | -1.385 | -0.066 | 0.049 | 0.096 | 0.096 | 4.5 | 0.31 | 0.08 | 0.18 | 0.1 | 0.25 | 0.13 |
| 2 | 201337667 | rs6723687 | -0.049 | -0.308 | -0.538 | -0.097 | 0.013 | -0.003 | -0.003 | 0 | 0.06 | 0.24 | 0.03 | 0.04 | 0.03 | 0.02 |
| 2 | 201339557 | rs12470550 | 1.114 | -0.285 | -1.519 | 0.083 | -0.034 | -0.056 | -0.056 | 0 | 0.21 | 0.01 | 0.24 | 0.24 | 0.21 | 0.25 |
| 2 | 201340734 | rs3769432 | -0.225 | 0.008 | -1.094 | 0.04 | 0 | 0.012 | 0.012 | 1.012 | 0.1 | 0.17 | 0.19 | 0.12 | 0.12 | 0.08 |
| 2 | 201344160 | rs733055 | 1.301 | 0.638 | -1.384 | -0.322 | -0.009 | 0.167 | 0.167 | 5.569 | 0.31 | 0.05 | 0.19 | 0 | 0.31 | 0.12 |
| 2 | 201344491 | rs2043770 | -0.065 | 1.111 | -0.573 | 0.472 | 0.012 | 0.053 | 0.053 | 1.866 | 0.71 | 0.5 | 0.79 | 0.64 | 0.73 | 0.62 |
| 2 | 201344732 | rs2043769 | -0.342 | 1.347 | -0.471 | 0.546 | 0.05 | 0.178 | 0.178 | 1.195 | 0.81 | 0.52 | 0.83 | 0.64 | 0.82 | 0.69 |
| 2 | 201346859 | rs6719002 | 0.198 | 1.393 | -0.283 | 0.638 | 0.04 | 0.131 | 0.131 | 0.463 | 0.8 | 0.6 | 0.83 | 0.66 | 0.81 | 0.7 |
| 2 | 201349037 | rs11690787 | 0.552 | 1.366 | -0.523 | 0.63 | -0.031 | -0.047 | -0.047 | 0.708 | 0.13 | 0.06 | 0.11 | 0.34 | 0.16 | 0.28 |
| 2 | 201349631 | rs13019534 | 1.241 | 1.524 | -0.624 | 0.628 | -0.046 | -0.056 | -0.056 | 1.93 | 0.64 | 0.17 | 0.58 | 0.53 | 0.59 | 0.61 |
| 2 | 201350954 | rs9288311 | -0.757 | 1.82 | -0.688 | 0.762 | 0.011 | 0.022 | 0.022 | 0 | 0.15 | 0.27 | 0.24 | 0.25 | 0.18 | 0.09 |
| 2 | 201352021 | rs17447933 | 0.561 | 1.848 | -0.867 | 0.587 | -0.007 | -0.007 | -0.007 | 0 | 0.11 | 0.11 | 0.2 | 0.11 | 0.15 | 0.07 |
| 2 | 201353083 | rs10497863 | 0.271 | 1.615 | -0.749 | 0.415 | -0.005 | 0.001 | 0.001 | 0 | 0.07 | 0.08 | 0.05 | 0.04 | 0.11 | 0.01 |
| 2 | 201356309 | rs12233018 | 0.07 | 1.367 | -0.648 | 0.753 | 0.004 | 0.058 | 0.058 | 0 | 0.11 | 0.12 | 0.1 | 0.34 | 0.12 | 0.27 |
| 2 | 201356552 | rs10931901 | -0.102 | 0.954 | -0.779 | 0.391 | 0 | 0.012 | 0.012 | 0 | 0.5 | 0.47 | 0.55 | 0.6 | 0.52 | 0.44 |
| 2 | 201357235 | rs713396 | -0.374 | 0.443 | -1.084 | 0.561 | 0.009 | -0.001 | -0.001 | 0 | 0.19 | 0.11 | 0.08 | 0.13 | 0.13 | 0.29 |
| 2 | 201357624 | rs713397 | -0.924 | 0.016 | -1.431 | 0.562 | 0.011 | -0.037 | -0.037 | 0 | 0.19 | 0.06 | 0.16 | 0.22 | 0.15 | 0.29 |
| 2 | 201358597 | rs7591915 | -0.927 | 0.151 | -0.776 | 0.043 | 0.007 | 0.011 | 0.011 | 0.157 | 0.52 | 0.57 | 0.53 | 0.25 | 0.57 | 0.62 |
| 2 | 201358614 | rs11896250 | 0.195 | 0.193 | -0.605 | -0.133 | -0.013 | -0.008 | -0.008 | 0 | 0.23 | 0.33 | 0.34 | 0.17 | 0.2 | 0.37 |
| 2 | 201359381 | rs6718287 | 1.328 | 0.033 | -0.611 | 0.022 | 0.039 | 0.008 | 0.008 | 0 | 0.44 | 0.16 | 0.41 | 0.6 | 0.41 | 0.35 |
| 2 | 201367356 | rs10084263 | 0.171 | 0.116 | -0.484 | 0.019 | 0.042 | -0.043 | -0.043 | 1.496 | 0.67 | 0.51 | 0.76 | 0.78 | 0.61 | 0.73 |
| 2 | 201368368 | rs7596859 | 2.061 | 0.504 | -0.468 | -0.518 | 0.002 | -0.032 | -0.032 | 0.036 | 0.14 | 0.15 | 0.07 | 0 | 0.15 | 0.09 |
| 2 | 201371768 | rs3739116 | -0.659 | 0.414 | -0.314 | -0.518 | -0.002 | -0.001 | -0.001 | 0 | 0.78 | 0.81 | 0.68 | 0.82 | 0.8 | 0.63 |
| 2 | 201381937 | rs1347691 | -1.938 | 0.141 | -0.44 | -0.517 | 0.014 | -0.069 | -0.069 | 0 | 0.81 | 0.92 | 0.76 | 0.56 | 0.83 | 0.77 |
| 2 | 201391491 | rs999576 | 2.136 | 0.144 | -0.47 | -0.586 | 0.011 | 0.019 | 0.019 | 0 | 0.27 | 0.16 | 0.2 | 0.29 | 0.25 | 0.16 |
| 2 | 201395161 | rs842941 | -1.872 | 0.132 | -0.32 | -0.589 | -0.016 | 0.067 | 0.067 | 0 | 0.79 | 0.76 | 0.73 | 0.56 | 0.83 | 0.77 |
| 2 | 201397345 | rs17636455 | 2.014 | 0.218 | -0.401 | -0.473 | 0.011 | 0.019 | 0.019 | 0 | 0.27 | 0.16 | 0.2 | 0.29 | 0.25 | 0.16 |
| 2 | 201403980 | rs1015501 | 2.04 | 0.224 | -0.387 | -0.463 | 0.01 | 0.018 | 0.018 | 0 | 0.27 | 0.16 | 0.2 | 0.29 | 0.25 | 0.16 |
| 2 | 201404694 | rs842938 | 0.15 | 0.248 | -0.349 | -0.481 | 0.006 | 0.006 | 0.006 | 0 | 0.49 | 0.46 | 0.54 | 0.48 | 0.46 | 0.52 |
| 2 | 201404947 | rs842937 | 0.472 | 0.37 | -0.261 | -0.405 | -0.011 | -0.004 | -0.004 | 0 | 0.22 | 0.3 | 0.34 | 0.18 | 0.2 | 0.36 |
| 2 | 201412611 | rs12053340 | 0.711 | 0.571 | -0.447 | -0.317 | -0.032 | 0.02 | 0.02 | 0 | 0.14 | 0.07 | 0.11 | 0.08 | 0.22 | 0.15 |
| 2 | 201434903 | rs2881746 | 0.711 | 0.612 | -0.513 | -0.296 | -0.032 | 0.02 | 0.02 | 0 | 0.14 | 0.07 | 0.11 | 0.08 | 0.22 | 0.15 |
| 2 | 201437048 | rs17532665 | 2.526 | 0.65 | -0.578 | -0.491 | -0.035 | 0.046 | 0.046 | 0 | 0.12 | 0.05 | 0.07 | 0 | 0.15 | 0.08 |
| 2 | 201437214 | rs17532679 | 0.781 | 0.618 | -0.535 | -0.416 | -0.032 | 0.02 | 0.02 | 0 | 0.14 | 0.07 | 0.11 | 0.08 | 0.22 | 0.15 |
| 2 | 201440299 | rs3764778 | 1.944 | 0.366 | -0.653 | -0.478 | 0.011 | 0.019 | 0.019 | 0 | 0.27 | 0.16 | 0.2 | 0.29 | 0.25 | 0.16 |
| 2 | 201444441 | rs13026134 | -0.944 | 0.287 | -0.627 | -0.439 | 0 | 0 | 0 | 0 | 0.79 | 0.83 | 0.67 | 0.82 | 0.8 | 0.64 |
| 2 | 201445152 | rs4674194 | -0.402 | 0.292 | -0.615 | -0.434 | 0.005 | 0.004 | 0.004 | 0 | 0.51 | 0.54 | 0.46 | 0.52 | 0.54 | 0.48 |
| 2 | 201451006 | rs4470327 | -0.857 | 0.425 | -0.706 | -0.371 | 0 | 0 | 0 | 0 | 0.79 | 0.83 | 0.67 | 0.82 | 0.8 | 0.64 |
| 2 | 201453948 | rs11689822 | 0.644 | 0.622 | -0.792 | -0.298 | -0.05 | 0.024 | 0.024 | 0 | 0.14 | 0.05 | 0.1 | 0.08 | 0.22 | 0.15 |
| 2 | 201456245 | rs1527944 | 0.53 | 0.62 | -0.805 | -0.299 | -0.044 | 0.026 | 0.026 | 0 | 0.14 | 0.07 | 0.09 | 0.08 | 0.22 | 0.15 |
| 2 | 201456390 | rs17532798 | 2.018 | 0.649 | -0.662 | -0.529 | 0.006 | -0.039 | -0.039 | 2.204 | 0.14 | 0.16 | 0.08 | 0 | 0.15 | 0.09 |
| 2 | 201457610 | rs1527947 | 0.622 | 0.646 | -0.627 | -0.522 | -0.028 | 0.021 | 0.021 | 0 | 0.14 | 0.12 | 0.09 | 0.08 | 0.22 | 0.15 |
| 2 | 201458460 | rs1918720 | -0.734 | 0.424 | -0.451 | -0.578 | -0.011 | -0.004 | -0.004 | 0 | 0.78 | 0.7 | 0.66 | 0.82 | 0.8 | 0.64 |
| 2 | 201463018 | rs17448443 | 0.948 | 0.447 | -0.466 | -0.576 | -0.047 | 0.024 | 0.024 | 0 | 0.14 | 0.06 | 0.09 | 0.08 | 0.22 | 0.15 |
| 2 | 201463510 | rs17532861 | 2.129 | 0.441 | -0.458 | -0.584 | 0.005 | -0.039 | -0.039 | 0.986 | 0.14 | 0.17 | 0.07 | 0 | 0.15 | 0.09 |
| 2 | 201465291 | rs17532875 | 2.176 | 0.449 | -0.454 | -0.581 | 0.006 | -0.039 | -0.039 | 0 | 0.14 | 0.16 | 0.07 | 0 | 0.15 | 0.09 |
| 2 | 201465718 | rs2463492 | 0.24 | 0.439 | -0.49 | -0.582 | 0.002 | -0.004 | -0.004 | 0.474 | 0.64 | 0.64 | 0.57 | 0.74 | 0.58 | 0.49 |
| 2 | 201466335 | rs4337457 | -1.343 | 0.44 | -0.43 | -0.591 | -0.012 | -0.015 | -0.015 | 0 | 0.83 | 0.81 | 0.78 | 0.91 | 0.88 | 0.69 |
| 2 | 201466732 | rs6742109 | 0.345 | 0.442 | -0.456 | -0.587 | -0.001 | 0 | 0 | 0 | 0.64 | 0.64 | 0.57 | 0.74 | 0.58 | 0.49 |
| 2 | 201467231 | rs2293524 | 2.215 | 0.26 | -0.518 | -0.569 | 0.012 | 0.018 | 0.018 | 0 | 0.27 | 0.17 | 0.22 | 0.3 | 0.25 | 0.16 |
| 2 | 201469149 | rs2293528 | 0.66 | 0.256 | -0.534 | -0.555 | -0.004 | 0.029 | 0.029 | 0 | 0.36 | 0.42 | 0.35 | 0.44 | 0.32 | 0.33 |
| 2 | 201469948 | rs17532973 | 2.661 | 0.294 | -0.557 | -0.572 | -0.035 | 0.049 | 0.049 | 0 | 0.13 | 0.05 | 0.07 | 0 | 0.15 | 0.1 |
| 2 | 201473674 | rs2470902 | -0.663 | 0.292 | -0.509 | -0.532 | -0.01 | -0.003 | -0.003 | 0 | 0.78 | 0.7 | 0.66 | 0.82 | 0.8 | 0.64 |
| 2 | 201474436 | rs10497866 | -0.638 | 0.295 | -0.528 | -0.53 | -0.01 | -0.003 | -0.003 | 0 | 0.78 | 0.7 | 0.66 | 0.82 | 0.8 | 0.64 |
| 2 | 201478264 | rs2465665 | 0.261 | 0.324 | -0.554 | -0.533 | 0.002 | -0.004 | -0.004 | 0.095 | 0.64 | 0.64 | 0.57 | 0.74 | 0.58 | 0.49 |
| 2 | 201481512 | rs2002957 | 2.761 | 0.248 | -0.564 | -0.58 | -0.061 | 0.071 | 0.071 | 0 | 0.12 | 0.01 | 0.06 | 0 | 0.15 | 0.08 |

| | | | | | | | | | | | | | | | | |
|---|---|---|---|---|---|---|---|---|---|---|---|---|---|---|---|---|
| 2 | 201482866 | rs2470899 | -2.287 | 0.214 | -0.489 | -0.596 | 0.004 | -0.037 | -0.037 | 0 | 0.86 | 0.84 | 0.93 | 1 | 0.85 | 0.91 |
| 2 | 201483925 | rs16833889 | -0.258 | 0.247 | -0.46 | -0.594 | 0.003 | -0.022 | -0.022 | 0 | 0.32 | 0.35 | 0.32 | 0.17 | 0.34 | 0.46 |
| 2 | 201485649 | rs4672866 | 1.783 | 0.202 | -0.47 | -0.574 | -0.006 | -0.012 | -0.012 | 1.072 | 0.32 | 0.22 | 0.34 | 0.39 | 0.34 | 0.22 |
| 2 | 201485830 | rs6761375 | 1.244 | 0.204 | -0.382 | -0.567 | -0.008 | -0.009 | -0.009 | 0 | 0.17 | 0.18 | 0.22 | 0.09 | 0.13 | 0.31 |
| 2 | 201487116 | rs4428003 | -0.73 | 0.249 | -0.421 | -0.393 | -0.004 | 0.033 | 0.033 | 0 | 0.64 | 0.57 | 0.66 | 0.56 | 0.68 | 0.67 |
| 2 | 201488166 | rs4614918 | -0.73 | 0.249 | -0.421 | -0.393 | -0.004 | 0.033 | 0.033 | 0 | 0.64 | 0.57 | 0.66 | 0.56 | 0.68 | 0.67 |
| 2 | 201488413 | rs4035120 | -2.397 | 0.246 | -0.489 | -0.398 | 0.006 | -0.039 | -0.039 | 0 | 0.86 | 0.84 | 0.93 | 1 | 0.85 | 0.92 |
| 2 | 201488758 | rs2348021 | -0.744 | 0.28 | -0.594 | -0.327 | -0.005 | 0.032 | 0.032 | 0 | 0.63 | 0.52 | 0.7 | 0.56 | 0.68 | 0.67 |
| 2 | 201489465 | rs16833921 | -0.011 | 0.026 | -0.318 | -0.056 | 0.002 | -0.021 | -0.021 | 0 | 0.31 | 0.35 | 0.32 | 0.17 | 0.34 | 0.46 |
| 2 | 201489836 | rs6720189 | -0.718 | 0.353 | -0.774 | 0.028 | -0.018 | 0.042 | 0.042 | 0 | 0.54 | 0.68 | 0.59 | 0.61 | 0.51 | 0.7 |
| 2 | 201491450 | rs1986414 | -0.635 | -0.333 | -0.009 | -0.033 | -0.029 | -0.036 | -0.036 | 0 | 0.89 | 0.69 | 0.95 | 0.96 | 0.97 | 0.96 |
| 2 | 201491560 | rs1986415 | 0.618 | -1.059 | -0.791 | 0.128 | 0.037 | -0.156 | -0.156 | 3.613 | 0.2 | 0.02 | 0.17 | 0.45 | 0.12 | 0.26 |
| 2 | 201496111 | rs2254576 | -0.958 | -0.751 | -0.031 | -0.405 | -0.023 | 0.055 | 0.055 | 0.551 | 0.59 | 0.76 | 0.52 | 0.79 | 0.53 | 0.47 |
| 2 | 201500183 | rs2256977 | -0.765 | -0.745 | 0.053 | -0.426 | -0.004 | 0.007 | 0.007 | 0.434 | 0.57 | 0.61 | 0.51 | 0.8 | 0.54 | 0.48 |
| 2 | 201500631 | rs2465661 | 0.137 | -0.518 | 0.012 | -0.504 | 0.015 | 0.019 | 0.019 | 1.578 | 0.49 | 0.55 | 0.43 | 0.75 | 0.37 | 0.41 |
| 2 | 201501431 | rs17533238 | 0.879 | -0.505 | 0.405 | 0.038 | -0.01 | 0.009 | 0.009 | 0 | 0.08 | 0.06 | 0.08 | 0.04 | 0.16 | 0.07 |
| 2 | 201502213 | rs6741421 | 1.617 | -0.629 | 0.558 | 0.01 | -0.033 | -0.019 | -0.019 | 0.308 | 0.07 | 0.18 | 0.03 | 0.02 | 0.01 | 0.01 |
| 2 | 201502395 | rs10172572 | 1.009 | -0.725 | 0.443 | 0.122 | 0.031 | -0.046 | -0.046 | 1.224 | 0.19 | 0.11 | 0.18 | 0.44 | 0.11 | 0.23 |
| 2 | 201503176 | rs2465663 | 0.264 | -0.616 | 0.47 | 0.057 | -0.002 | -0.036 | -0.036 | 4.071 | 0.27 | 0.18 | 0.26 | 0.49 | 0.27 | 0.3 |
| 2 | 201505269 | rs7587871 | -1.426 | -0.574 | 0.28 | -0.076 | -0.003 | -0.01 | -0.01 | 0.298 | 0.4 | 0.59 | 0.38 | 0.79 | 0.32 | 0.43 |
| 2 | 201505953 | rs11887334 | -0.056 | -0.321 | -0.255 | 0.017 | -0.001 | 0.003 | 0.003 | 0 | 0.29 | 0.31 | 0.27 | 0.5 | 0.28 | 0.31 |
| 2 | 201514892 | rs13404701 | 0.432 | 0.319 | -0.518 | -0.278 | 0.03 | -0.021 | -0.021 | 0.265 | 0.08 | 0.11 | 0.1 | 0.01 | 0.2 | 0.03 |
| 2 | 201515305 | rs2348023 | 0.601 | 0.332 | -0.744 | -0.272 | -0.045 | 0.038 | 0.038 | 0 | 0.08 | 0.03 | 0.1 | 0.01 | 0.2 | 0.03 |
| 2 | 201516685 | rs2348024 | 0.836 | -0.398 | -0.899 | -0.316 | 0.053 | 0.058 | 0.058 | 0.018 | 0.51 | 0.29 | 0.53 | 0.33 | 0.6 | 0.48 |
| 2 | 201520598 | rs2540069 | 0.086 | -0.426 | -0.388 | -0.362 | -0.008 | -0.013 | -0.013 | 0 | 0.6 | 0.5 | 0.62 | 0.27 | 0.66 | 0.56 |
| 2 | 201521472 | rs2287011 | 0.252 | 0.023 | -0.842 | -0.694 | -0.006 | -0.019 | -0.019 | 0 | 0.21 | 0.11 | 0.37 | 0.25 | 0.23 | 0.32 |
| 2 | 201522755 | rs2348025 | -0.396 | 0.158 | -0.289 | -0.692 | -0.001 | -0.009 | -0.009 | 2.077 | 0.41 | 0.61 | 0.5 | 0.8 | 0.4 | 0.49 |
| 2 | 201526746 | rs2881811 | 1.111 | 0.524 | -0.539 | -1.04 | -0.021 | 0.051 | 0.051 | 0 | 0.28 | 0.13 | 0.17 | 0.01 | 0.33 | 0.17 |
| 2 | 201527239 | rs2241080 | 0.226 | -1.581 | -0.308 | -1.323 | -0.004 | 0.012 | 0.012 | 1.455 | 0.27 | 0.32 | 0.36 | 0.55 | 0.24 | 0.34 |
| 2 | 201529900 | rs6435060 | -0.694 | -1.567 | -0.481 | -1.52 | -0.001 | -0.01 | -0.01 | 0.418 | 0.41 | 0.61 | 0.5 | 0.81 | 0.4 | 0.49 |
| 2 | 201531844 | rs4674311 | -0.329 | -1.861 | 0.203 | -1.344 | 0 | -0.012 | -0.012 | 0 | 0.55 | 0.6 | 0.6 | 0.83 | 0.54 | 0.52 |
| 2 | 201535075 | rs2540066 | 0.219 | -1.996 | 0.537 | -1.069 | -0.001 | 0.018 | 0.018 | 0 | 0.46 | 0.37 | 0.32 | 0.35 | 0.48 | 0.45 |
| 2 | 201536463 | rs2715906 | -0.029 | -2.028 | 0.694 | -1.078 | 0 | 0.004 | 0.004 | 0 | 0.5 | 0.5 | 0.48 | 0.43 | 0.55 | 0.55 |
| 2 | 201538081 | rs2540064 | 0.396 | -2.04 | 0.7 | -1.087 | 0.001 | 0.052 | 0.052 | 0 | 0.44 | 0.31 | 0.29 | 0.13 | 0.44 | 0.33 |
| 2 | 201539206 | rs7563682 | 0.532 | -2.188 | 0.763 | -1.211 | -0.013 | -0.002 | -0.002 | 0.017 | 0.37 | 0.29 | 0.6 | 0.63 | 0.43 | 0.49 |
| 2 | 201540533 | rs11684227 | 0.104 | -1.9 | 0.96 | -1.055 | 0 | 0.014 | 0.014 | 0 | 0.45 | 0.43 | 0.64 | 0.63 | 0.46 | 0.52 |
| 2 | 201540988 | rs16834021 | -0.662 | -1.93 | 0.971 | -1.095 | 0.011 | -0.04 | -0.04 | 0.008 | 0.08 | 0.17 | 0.04 | 0 | 0.06 | 0.02 |
| 2 | 201542014 | rs16834034 | -0.943 | -1.856 | 1.032 | -1.059 | 0.022 | -0.045 | -0.045 | 0 | 0.09 | 0.29 | 0.05 | 0 | 0.06 | 0.02 |
| 2 | 201543496 | rs6729738 | -0.587 | -1.839 | 0.782 | -1.042 | 0 | 0.024 | 0.024 | 0 | 0.54 | 0.64 | 0.69 | 0.86 | 0.56 | 0.64 |
| 2 | 201552876 | rs12473437 | 0.359 | -1.889 | 0.635 | -1 | 0.002 | -0.013 | -0.013 | 0.095 | 0.4 | 0.21 | 0.46 | 0.57 | 0.38 | 0.41 |
| 2 | 201553438 | rs4674325 | 0.48 | -1.854 | 0.636 | -0.978 | 0.004 | -0.008 | -0.008 | 0 | 0.41 | 0.3 | 0.46 | 0.57 | 0.38 | 0.42 |
| 2 | 201558699 | rs10515940 | 0.033 | -1.769 | 0.414 | -1.605 | 0.023 | -0.028 | -0.028 | 0 | 0.07 | 0.11 | 0.07 | 0 | 0.12 | 0.08 |
| 2 | 201558766 | rs2540061 | 0 | -2.141 | 0.389 | -1.471 | 0.004 | 0.039 | 0.039 | 0 | 0.47 | 0.34 | 0.29 | 0.15 | 0.43 | 0.35 |
| 2 | 201560740 | rs2247016 | 1.102 | -2.938 | 1.198 | -0.768 | -0.063 | -0.092 | -0.092 | 0 | 0.62 | 0.53 | 0.49 | 0.09 | 0.76 | 0.44 |
| 2 | 201561779 | rs2540058 | -0.554 | -2.042 | 1.392 | 0.016 | -0.001 | -0.002 | -0.002 | 0 | 0.19 | 0.13 | 0.43 | 0.7 | 0.14 | 0.29 |
| 2 | 201562344 | rs13428164 | -0.743 | -2.068 | 1.491 | -0.051 | -0.015 | 0.02 | 0.02 | 0 | 0.14 | 0.38 | 0.07 | 0.21 | 0.1 | 0.23 |
| 2 | 201562553 | rs6735028 | 1.823 | -2.099 | 0.59 | -0.086 | 0.091 | 0.133 | 0.133 | 0 | 0.56 | 0.24 | 0.46 | 0.08 | 0.71 | 0.4 |
| 2 | 201567968 | rs2540053 | 1.063 | -1.605 | 0.065 | 0.192 | -0.046 | -0.004 | -0.004 | 0 | 0.77 | 0.65 | 0.92 | 0.74 | 0.89 | 0.76 |
| 2 | 201568999 | rs2540052 | 0.169 | -0.872 | -0.208 | 0.615 | -0.014 | 0.012 | 0.012 | 0 | 0.89 | 0.83 | 0.96 | 0.78 | 0.93 | 0.83 |
| 2 | 201571143 | rs2540051 | -1.241 | -0.379 | -0.599 | 1.053 | 0.006 | -0.005 | -0.005 | 0.443 | 0.21 | 0.15 | 0.46 | 0.7 | 0.19 | 0.38 |
| 2 | 201575480 | rs2715902 | -1.2 | -0.558 | -0.42 | 1.303 | -0.026 | 0.041 | 0.041 | 0 | 0.18 | 0.21 | 0.43 | 0.63 | 0.13 | 0.26 |
| 2 | 201575564 | rs2540050 | -1.458 | -0.646 | -0.351 | 1.423 | -0.039 | 0.062 | 0.062 | 0 | 0.17 | 0.21 | 0.32 | 0.58 | 0.09 | 0.22 |
| 2 | 201582193 | rs2348114 | 0.954 | -1.051 | 0.509 | 0.809 | -0.033 | -0.063 | -0.063 | 0 | 0.87 | 0.49 | 0.89 | 0.98 | 0.93 | 0.97 |
| 2 | 201585916 | rs4674353 | -0.587 | -1.03 | -0.007 | 1.494 | -0.001 | -0.04 | -0.04 | 0 | 0.12 | 0.05 | 0.23 | 0.35 | 0.1 | 0.3 |
| 2 | 201588986 | rs4672912 | 1.246 | -0.587 | -0.037 | 1.629 | 0.004 | 0.003 | 0.003 | 0.715 | 0.76 | 0.2 | 0.8 | 0.82 | 0.82 | 0.8 |
| 2 | 201596926 | rs4416197 | 1.432 | -1.075 | -0.589 | 1.075 | 0.022 | 0.044 | 0.044 | 0 | 0.67 | 0.16 | 0.75 | 0.82 | 0.75 | 0.77 |
| 2 | 201597760 | rs6729962 | 0.235 | -0.996 | -0.55 | 1.058 | 0.002 | 0.037 | 0.037 | 0 | 0.09 | 0.05 | 0.06 | 0 | 0.07 | 0.03 |
| 2 | 201604436 | rs951183 | 1.093 | -1.578 | -1.089 | 0.632 | 0.026 | -0.005 | -0.005 | 1.668 | 0.56 | 0.27 | 0.55 | 0.49 | 0.62 | 0.4 |
| 2 | 201606722 | rs11684485 | 1.043 | -1.335 | -0.783 | 0.722 | 0.008 | -0.006 | -0.006 | 0 | 0.57 | 0.36 | 0.56 | 0.49 | 0.62 | 0.42 |
| 2 | 201607818 | rs10432515 | 0.752 | -1.367 | -0.781 | 0.715 | -0.029 | 0.013 | 0.013 | 0 | 0.09 | 0.03 | 0.06 | 0.04 | 0.11 | 0.17 |
| 2 | 201610584 | rs2348121 | 0.589 | -1.449 | -0.208 | 0.69 | -0.01 | 0.009 | 0.009 | 0.172 | 0.66 | 0.44 | 0.63 | 0.53 | 0.73 | 0.58 |
| 2 | 201616145 | rs2348124 | 0.315 | -1.33 | -1.067 | 0.346 | 0.011 | 0.026 | 0.026 | 0 | 0.23 | 0.15 | 0.14 | 0.13 | 0.18 | 0.14 |
| 2 | 201616442 | rs1002414 | -0.633 | -1.446 | 0.104 | 0.28 | -0.006 | -0.036 | -0.036 | 0 | 0.35 | 0.4 | 0.22 | 0.17 | 0.29 | 0.31 |
| 2 | 201627550 | rs11693950 | 1.258 | -1.132 | 0.128 | 0.455 | 0.013 | 0.019 | 0.019 | 0 | 0.35 | 0.44 | 0.42 | 0.37 | 0.44 | 0.28 |
| 2 | 201627921 | rs11695319 | 0.131 | -1.411 | 0.182 | 0.272 | -0.006 | 0.004 | 0.004 | 0.156 | 0.58 | 0.57 | 0.54 | 0.44 | 0.67 | 0.54 |
| 2 | 201632988 | rs10185328 | 0.112 | -1.448 | 0.193 | 0.275 | -0.009 | 0.006 | 0.006 | 0.057 | 0.59 | 0.56 | 0.54 | 0.46 | 0.67 | 0.55 |
| 2 | 201634161 | rs2348128 | -0.511 | -1.576 | 0.383 | 0.257 | -0.019 | 0.026 | 0.026 | 0 | 0.86 | 0.68 | 0.9 | 0.83 | 0.9 | 0.92 |
| 2 | 201638647 | rs12622452 | 1.627 | -1.4 | 0.226 | 0.242 | 0.003 | 0.007 | 0.007 | 0 | 0.34 | 0.36 | 0.42 | 0.38 | 0.44 | 0.27 |
| 2 | 201643484 | rs10515941 | 0.095 | -1.15 | 0.033 | 0.217 | 0.003 | -0.002 | -0.002 | 0.064 | 0.13 | 0.12 | 0.06 | 0.04 | 0.13 | 0.1 |

| | | | | | | | | | | | | | | | | |
|---|---|---|---|---|---|---|---|---|---|---|---|---|---|---|---|---|
| 2 | 201644232 | rs2348129 | -0.364 | -1.142 | 0.028 | 0.222 | -0.006 | 0.036 | 0.036 | 0 | 0.22 | 0.15 | 0.12 | 0.08 | 0.24 | 0.27 |
| 2 | 201656457 | rs3752657 | -0.183 | -1.215 | 0.772 | 0.35 | 0.009 | 0.002 | 0.002 | 0.228 | 0.72 | 0.8 | 0.78 | 0.66 | 0.8 | 0.62 |
| 2 | 201664304 | rs4674449 | 0.519 | -1.331 | 0.825 | 0.25 | 0 | 0 | 0 | 2.105 | 0.62 | 0.38 | 0.7 | 0.63 | 0.7 | 0.52 |
| 2 | 201664399 | rs4672951 | 1.028 | -1.266 | 0.777 | 1.114 | -0.001 | -0.006 | -0.006 | 0 | 0.53 | 0.34 | 0.63 | 0.44 | 0.55 | 0.34 |
| 2 | 201676441 | rs2270282 | 0.984 | -1.687 | 0.122 | 0.815 | -0.004 | 0.045 | 0.045 | 0 | 0.45 | 0.34 | 0.49 | 0.25 | 0.55 | 0.3 |
| 2 | 201677791 | rs2270280 | 0.208 | -1.686 | 0.187 | 0.833 | -0.025 | 0.024 | 0.024 | 0 | 0.63 | 0.45 | 0.58 | 0.44 | 0.72 | 0.48 |
| 2 | 201682395 | rs11685853 | 1.014 | -1.486 | 0.028 | 0.808 | -0.007 | 0.043 | 0.043 | 0 | 0.45 | 0.34 | 0.49 | 0.25 | 0.54 | 0.3 |
| 2 | 201686122 | rs3815501 | -0.508 | -1.488 | 0.251 | 0.796 | 0 | -0.001 | -0.001 | 0 | 0.72 | 0.77 | 0.65 | 0.47 | 0.83 | 0.58 |
| 2 | 201689742 | rs12995928 | 0.244 | -1.423 | -0.182 | 0.832 | -0.001 | -0.001 | -0.001 | 0 | 0.62 | 0.38 | 0.57 | 0.44 | 0.72 | 0.48 |
| 2 | 201692776 | rs12477038 | 1.179 | -1.093 | -0.742 | 1.026 | -0.017 | -0.086 | -0.086 | 0.013 | 0.13 | 0.01 | 0.25 | 0.32 | 0.11 | 0.16 |
| 2 | 201724360 | rs12465952 | 0.364 | -1.102 | -0.594 | 1.005 | -0.01 | -0.03 | -0.03 | 0 | 0.19 | 0.1 | 0.26 | 0.3 | 0.19 | 0.2 |
| 2 | 201736166 | rs7562391 | 0.28 | -1.275 | 0.135 | 0.429 | 0.06 | -0.078 | -0.078 | 1.944 | 0.12 | 0.45 | 0.13 | 0.01 | 0.14 | 0.04 |
| 2 | 201736734 | rs7606251 | -1.116 | -1.342 | 0.065 | 0.138 | -0.002 | -0.057 | -0.057 | 3.177 | 0.23 | 0.72 | 0.17 | 0.01 | 0.24 | 0.15 |
| 2 | 201739327 | rs4035021 | 0.152 | 0.362 | -0.714 | 0.25 | -0.082 | -0.065 | -0.065 | 0 | 0.33 | 0.08 | 0.41 | 0.58 | 0.36 | 0.33 |
| 2 | 201760937 | rs13384245 | 0.901 | 0.526 | 0.676 | 0.227 | 0.031 | -0.068 | -0.068 | 0 | 0.12 | 0.23 | 0.13 | 0.01 | 0.19 | 0.06 |
| 2 | 201762147 | rs10204787 | 0.853 | 0.524 | 0.675 | 0.226 | 0.029 | -0.068 | -0.068 | 0 | 0.13 | 0.24 | 0.13 | 0.01 | 0.19 | 0.06 |
| 2 | 201773545 | rs11894842 | -0.764 | 0.52 | 0.766 | 0.207 | 0.008 | -0.077 | -0.077 | 0.541 | 0.2 | 0.68 | 0.18 | 0.01 | 0.25 | 0.14 |
| 2 | 201776988 | rs4381763 | 0.013 | 0.522 | 0.759 | 0.208 | 0.019 | -0.09 | -0.09 | 1.134 | 0.16 | 0.64 | 0.16 | 0.01 | 0.21 | 0.07 |
| 2 | 201942480 | rs10194347 | 0.826 | 0.53 | 0.771 | 0.274 | 0.026 | -0.067 | -0.067 | 0.056 | 0.13 | 0.24 | 0.13 | 0.01 | 0.19 | 0.06 |
| 2 | 201971024 | rs13410282 | -0.639 | 0.51 | 0.46 | 0.058 | 0.015 | -0.097 | -0.097 | 0 | 0.2 | 0.65 | 0.18 | 0.01 | 0.25 | 0.13 |
| 2 | 201988238 | rs7583529 | -0.369 | 0.51 | 0.508 | 0.069 | 0.026 | -0.1 | -0.1 | 0 | 0.18 | 0.59 | 0.17 | 0.01 | 0.22 | 0.1 |
| 2 | 201992407 | rs6728771 | -1.092 | 0.441 | 0.433 | 0.03 | 0.003 | -0.07 | -0.07 | 0.205 | 0.21 | 0.69 | 0.17 | 0.01 | 0.22 | 0.1 |
| 2 | 201995779 | rs12620435 | 0.547 | 0.443 | 0.564 | 0.02 | 0.007 | -0.03 | -0.03 | 0 | 0.08 | 0.12 | 0.07 | 0 | 0.12 | 0.07 |
| 2 | 201999845 | rs16836710 | -0.457 | 0.401 | 0.626 | 0.028 | 0.042 | -0.13 | -0.13 | 0 | 0.18 | 0.43 | 0.15 | 0.01 | 0.21 | 0.1 |
| 2 | 202006096 | rs10190751 | -0.793 | 0.321 | 0.585 | 0.049 | 0.024 | -0.13 | -0.13 | 0 | 0.2 | 0.46 | 0.15 | 0.05 | 0.21 | 0.11 |
| 2 | 202016245 | rs12105811 | 0.289 | 0.332 | 0.649 | 0.068 | 0.048 | -0.069 | -0.069 | 0.058 | 0.13 | 0.34 | 0.13 | 0.05 | 0.17 | 0.05 |
| 2 | 202025131 | rs2518142 | -1.38 | -0.223 | -0.01 | -0.246 | 0.012 | 0.011 | 0.011 | 0 | 0.79 | 0.83 | 0.89 | 0.86 | 0.93 | 0.7 |
| 2 | 202025621 | rs1594 | 0.278 | -0.068 | -0.332 | -0.24 | 0.015 | -0.036 | -0.036 | 0 | 0.38 | 0.46 | 0.4 | 0.26 | 0.51 | 0.34 |
| 2 | 202036478 | rs7558475 | -2.318 | -0.252 | -0.247 | -0.29 | -0.008 | -0.003 | -0.003 | 0.146 | 0.84 | 0.85 | 0.9 | 0.86 | 0.95 | 0.7 |
| 2 | 202044716 | rs11900753 | 0.094 | 0.104 | 0.01 | -0.258 | 0.033 | -0.022 | -0.022 | 0 | 0.08 | 0.37 | 0.05 | 0 | 0.04 | 0.06 |
| 2 | 202047833 | rs16836789 | 0.094 | 0.104 | 0.01 | -0.258 | 0.033 | -0.022 | -0.022 | 0 | 0.08 | 0.37 | 0.05 | 0 | 0.04 | 0.06 |
| 2 | 202050677 | rs3900115 | 0.117 | 0.076 | -0.178 | -0.318 | 0.018 | -0.052 | -0.052 | 0.301 | 0.38 | 0.49 | 0.4 | 0.22 | 0.51 | 0.32 |
| 2 | 202052754 | rs16836813 | 1.055 | 0.074 | -0.286 | -0.29 | -0.004 | -0.029 | -0.029 | 0 | 0.07 | 0.12 | 0.02 | 0 | 0.04 | 0.06 |
| 2 | 202055312 | rs12613347 | 0.937 | 0.11 | -0.682 | -0.19 | -0.052 | -0.107 | -0.107 | 0 | 0.23 | 0.04 | 0.16 | 0.33 | 0.25 | 0.33 |
| 2 | 202060820 | rs3731714 | 2.24 | 0.397 | -0.042 | -0.445 | -0.08 | -0.032 | -0.032 | 0.114 | 0.18 | 0.01 | 0.25 | 0.18 | 0.29 | 0.22 |
| 2 | 202074528 | rs7576306 | 1.022 | -0.057 | 0.029 | -0.292 | -0.04 | 0.041 | 0.041 | 0 | 0.3 | 0.15 | 0.37 | 0.21 | 0.47 | 0.26 |
| 2 | 202077613 | rs10188598 | 1.015 | 0.004 | 0.02 | -0.308 | -0.045 | 0.043 | 0.043 | 0 | 0.3 | 0.15 | 0.38 | 0.21 | 0.49 | 0.26 |
| 2 | 202093395 | rs12693932 | -0.219 | -0.029 | -0.014 | -0.26 | -0.017 | 0.053 | 0.053 | 0.014 | 0.4 | 0.72 | 0.44 | 0.22 | 0.52 | 0.3 |
| 2 | 202096271 | rs6743518 | 1.088 | 0.173 | -0.305 | -0.154 | 0.003 | -0.019 | -0.019 | 0.112 | 0.05 | 0.1 | 0.02 | 0.01 | 0.04 | 0.06 |
| 2 | 202102685 | rs3769827 | 0.729 | 0.206 | -0.405 | -0.081 | -0.001 | -0.001 | -0.001 | 0 | 0.31 | 0.31 | 0.39 | 0.22 | 0.48 | 0.22 |
| 2 | 202111380 | rs3769825 | 0.268 | 0.037 | -0.488 | -0.166 | -0.005 | 0.002 | 0.002 | 0 | 0.36 | 0.35 | 0.44 | 0.2 | 0.53 | 0.25 |
| 2 | 202119272 | rs3754935 | 0.687 | -1.004 | -0.198 | -0.16 | -0.069 | 0.015 | 0.015 | 0 | 0.2 | 0.27 | 0.06 | 0.22 | 0.07 | 0.3 |
| 2 | 202119767 | rs13402616 | -1.158 | -1.121 | -0.038 | -0.362 | -0.004 | -0.063 | -0.063 | 0 | 0.11 | 0.4 | 0.06 | 0.01 | 0.08 | 0.24 |
| 2 | 202119789 | rs10931934 | 0.988 | -0.93 | -0.461 | -0.221 | -0.047 | -0.057 | -0.057 | 0 | 0.37 | 0.13 | 0.5 | 0.48 | 0.57 | 0.32 |
| 2 | 202122995 | rs3769823 | -0.487 | -1.043 | -0.108 | -0.253 | 0.001 | 0.001 | 0.001 | 0 | 0.62 | 0.59 | 0.59 | 0.7 | 0.69 | 0.67 |
| 2 | 202131587 | rs2293554 | -0.212 | -1.172 | -0.001 | -0.179 | -0.111 | -0.001 | -0.001 | 0 | 0.23 | 0.45 | 0.09 | 0.2 | 0.08 | 0.47 |
| 2 | 202149696 | rs1045487 | 0.609 | -1.266 | 0.04 | -0.182 | -0.083 | 0.017 | 0.017 | 0 | 0.18 | 0.26 | 0.05 | 0.19 | 0.04 | 0.27 |
| 2 | 202151781 | rs1045494 | 0.625 | -1.266 | 0.04 | -0.18 | -0.083 | 0.019 | 0.019 | 0 | 0.18 | 0.26 | 0.05 | 0.19 | 0.04 | 0.28 |
| 2 | 202153078 | rs1035142 | -0.255 | -1.198 | -0.256 | -0.341 | 0.001 | 0.004 | 0.004 | 0 | 0.58 | 0.38 | 0.47 | 0.68 | 0.61 | 0.66 |
| 2 | 202153225 | rs700635 | -1.366 | -1.214 | -0.375 | -0.39 | -0.002 | 0.004 | 0.004 | 0 | 0.69 | 0.81 | 0.57 | 0.7 | 0.72 | 0.86 |
| 2 | 202153252 | rs700636 | 0.89 | -1.133 | -0.535 | -0.363 | -0.058 | -0.098 | -0.098 | 0.34 | 0.4 | 0.12 | 0.42 | 0.49 | 0.57 | 0.37 |
| 2 | 202154200 | rs17468277 | 1.736 | -0.835 | -0.676 | -0.512 | -0.018 | 0.02 | 0.02 | 0 | 0.07 | 0.04 | 0.07 | 0 | 0.12 | 0.02 |
| 2 | 202161636 | rs13009885 | 1.076 | -0.883 | -0.624 | -0.42 | -0.127 | -0.134 | -0.134 | 9.143 | 0.32 | 0.07 | 0.34 | 0.49 | 0.45 | 0.34 |
| 2 | 202173812 | rs16837131 | 0.224 | -0.392 | -0.063 | -0.258 | 0.014 | -0.042 | -0.042 | 0 | 0.08 | 0.23 | 0.08 | 0.01 | 0.1 | 0.07 |
| 2 | 202196063 | rs9288318 | -0.552 | -0.31 | -0.113 | -0.176 | -0.023 | 0.05 | 0.05 | 0 | 0.61 | 0.78 | 0.49 | 0.71 | 0.61 | 0.84 |
| 2 | 202205197 | rs10185177 | -0.257 | -0.616 | 0.522 | -0.099 | -0.066 | 0.068 | 0.068 | 0 | 0.19 | 0.51 | 0.12 | 0.25 | 0.1 | 0.27 |
| 2 | 202207993 | rs7577057 | 0.723 | -0.583 | 0.456 | -0.129 | -0.018 | -0.012 | -0.012 | 0 | 0.43 | 0.27 | 0.37 | 0.45 | 0.5 | 0.57 |
| 2 | 202213382 | rs6710974 | 0.393 | -0.568 | 0.481 | -0.182 | 0.001 | -0.002 | -0.002 | 0 | 0.47 | 0.43 | 0.45 | 0.48 | 0.58 | 0.62 |
| 2 | 202215819 | rs3817579 | 1.583 | -0.652 | 0.442 | -0.138 | -0.053 | 0.063 | 0.063 | 0 | 0.11 | 0.14 | 0.03 | 0.25 | 0.02 | 0.21 |
| 2 | 202241594 | rs12052257 | -0.289 | -0.48 | 0.546 | -0.102 | -0.001 | 0.016 | 0.016 | 0 | 0.58 | 0.54 | 0.53 | 0.74 | 0.61 | 0.83 |
| 2 | 202241907 | rs10202963 | 0.325 | -0.505 | 0.45 | -0.232 | 0 | -0.001 | -0.001 | 0 | 0.47 | 0.44 | 0.5 | 0.5 | 0.58 | 0.63 |
| 2 | 202245244 | rs3795966 | 0.43 | -0.155 | 0.773 | 0.114 | 0.005 | -0.008 | -0.008 | 0.503 | 0.59 | 0.38 | 0.57 | 0.5 | 0.66 | 0.71 |
| 2 | 202252539 | rs2244438 | 0.402 | -0.186 | 0.815 | 0.116 | -0.021 | 0.013 | 0.013 | 0 | 0.28 | 0.21 | 0.39 | 0.24 | 0.31 | 0.09 |
| 2 | 202264156 | rs13022344 | 0.504 | -0.191 | 0.813 | 0.092 | 0.009 | -0.012 | -0.012 | 0 | 0.58 | 0.35 | 0.56 | 0.49 | 0.66 | 0.7 |
| 2 | 202271347 | rs2540334 | 1.228 | -0.098 | 0.732 | 0.061 | -0.003 | -0.008 | -0.008 | 0.105 | 0.21 | 0.15 | 0.31 | 0.25 | 0.18 | 0.05 |
| 2 | 202284867 | rs10497867 | 0.732 | -0.194 | 0.912 | 0.108 | 0.021 | -0.029 | -0.029 | 0 | 0.55 | 0.23 | 0.51 | 0.49 | 0.61 | 0.69 |
| 2 | 202285100 | rs4507084 | -0.536 | -0.205 | 0.931 | 0.105 | -0.007 | 0.005 | 0.005 | 0 | 0.71 | 0.67 | 0.6 | 0.75 | 0.69 | 0.91 |
| 2 | 202305145 | rs7603584 | 0.473 | -0.175 | 1.024 | 0.096 | -0.002 | 0.003 | 0.003 | 0.105 | 0.58 | 0.44 | 0.56 | 0.49 | 0.64 | 0.66 |
| 2 | 202315207 | rs6717641 | 0.598 | -0.175 | 0.982 | 0.096 | 0.005 | -0.012 | -0.012 | 1.839 | 0.57 | 0.35 | 0.55 | 0.49 | 0.64 | 0.65 |

| | | | | | | | | | | | | | | | |
|---|---|---|---|---|---|---|---|---|---|---|---|---|---|---|---|
| 2 | 202315759 | rs2540449 | 0.448 | -0.169 | 0.97 | 0.097 | -0.021 | 0.01 | 0.01 | 0 | 0.28 | 0.21 | 0.39 | 0.25 | 0.32 | 0.09 |
| 2 | 202323922 | rs10931948 | -0.375 | -0.246 | 0.645 | 0.143 | -0.01 | 0.007 | 0.007 | 0 | 0.7 | 0.67 | 0.59 | 0.74 | 0.66 | 0.87 |
| 2 | 202337236 | rs12991600 | -0.049 | 0.046 | 0.025 | -0.072 | -0.002 | 0.01 | 0.01 | 0 | 0.6 | 0.53 | 0.56 | 0.49 | 0.64 | 0.65 |
| 2 | 202348303 | rs2080326 | -0.56 | 0.166 | -0.551 | -0.025 | 0.011 | -0.016 | -0.016 | 0 | 0.46 | 0.72 | 0.5 | 0.5 | 0.41 | 0.35 |
| 2 | 202352867 | rs2540436 | -0.486 | 0.184 | -0.555 | -0.056 | 0.008 | -0.021 | -0.021 | 0 | 0.45 | 0.72 | 0.5 | 0.51 | 0.41 | 0.35 |
| 2 | 202383316 | rs11694706 | -0.088 | 0.567 | -0.929 | -0.806 | -0.018 | 0.07 | 0.07 | 0.023 | 0.59 | 0.29 | 0.64 | 0.96 | 0.59 | 0.65 |
| 2 | 202383408 | rs11694829 | -0.236 | 0.742 | -1.008 | -0.926 | -0.002 | -0.006 | -0.006 | 0 | 0.68 | 0.3 | 0.75 | 0.96 | 0.69 | 0.72 |
| 2 | 202384125 | rs2098354 | -0.75 | 0.752 | -0.445 | -0.998 | 0.012 | -0.152 | -0.152 | 0.451 | 0.75 | 0.57 | 0.8 | 0.97 | 0.74 | 0.73 |
| 2 | 202395424 | rs1914257 | -0.093 | 0.749 | -0.496 | -0.999 | -0.001 | -0.024 | -0.024 | 0 | 0.69 | 0.33 | 0.74 | 0.96 | 0.68 | 0.72 |
| 2 | 202402532 | rs10931953 | -0.615 | 0.725 | -0.43 | -0.903 | 0.004 | -0.114 | -0.114 | 0 | 0.74 | 0.52 | 0.79 | 0.95 | 0.74 | 0.73 |
| 2 | 202415228 | rs7567196 | -0.49 | 0.657 | -0.764 | -0.232 | 0.001 | 0.042 | 0.042 | 0 | 0.43 | 0.32 | 0.56 | 0.76 | 0.46 | 0.57 |
| 2 | 202428465 | rs4675195 | -0.878 | 0.679 | -0.743 | -0.256 | 0.005 | 0.044 | 0.044 | 0 | 0.48 | 0.41 | 0.61 | 0.76 | 0.51 | 0.58 |
| 2 | 202435352 | rs6754666 | 1.062 | 0.334 | -0.934 | 0.128 | -0.008 | -0.033 | -0.033 | 0 | 0.22 | 0.16 | 0.47 | 0.54 | 0.3 | 0.26 |
| 2 | 202437990 | rs12693942 | -0.588 | 0.431 | -0.822 | -0.042 | -0.01 | 0.047 | 0.047 | 0 | 0.45 | 0.32 | 0.57 | 0.76 | 0.48 | 0.57 |
| 2 | 202451042 | rs11690916 | -0.124 | 0.392 | -0.359 | 0.111 | 0.013 | -0.007 | -0.007 | 0 | 0.94 | 0.7 | 0.94 | 0.97 | 0.95 | 0.99 |
| 2 | 202451984 | rs2160241 | -0.025 | 0.363 | -0.145 | 0.101 | 0.038 | 0.018 | 0.018 | 0.037 | 0.93 | 0.59 | 0.94 | 0.97 | 0.95 | 0.99 |
| 2 | 202491995 | rs2540450 | 0.222 | 0.168 | -1.009 | -0.04 | 0.002 | -0.053 | -0.053 | 0 | 0.31 | 0.16 | 0.42 | 0.65 | 0.33 | 0.33 |
| 2 | 202493504 | rs1914261 | 1.113 | 0.296 | -0.818 | -0.088 | -0.008 | 0.02 | 0.02 | 0 | 0.45 | 0.3 | 0.36 | 0.24 | 0.45 | 0.42 |
| 2 | 202497586 | rs1208082 | -0.816 | 0.459 | -1.437 | 0.707 | -0.013 | -0.029 | -0.029 | 0 | 0.75 | 0.85 | 0.59 | 0.64 | 0.67 | 0.74 |
| 2 | 202501448 | rs1208081 | -0.174 | 0.522 | -1.304 | 0.39 | -0.001 | -0.013 | -0.013 | 0.116 | 0.65 | 0.7 | 0.55 | 0.35 | 0.62 | 0.65 |
| 2 | 202503225 | rs2676325 | 0.008 | 0.417 | -1.036 | 0.363 | 0.01 | 0.076 | 0.076 | 0 | 0.5 | 0.68 | 0.44 | 0.25 | 0.45 | 0.41 |
| 2 | 202509606 | rs2110738 | 0.095 | 0.585 | -0.367 | 0.34 | -0.002 | 0.009 | 0.009 | 0.049 | 0.55 | 0.59 | 0.47 | 0.68 | 0.48 | 0.45 |
| 2 | 202509814 | rs3754932 | 0.33 | 0.47 | -0.051 | 0.256 | -0.001 | 0.013 | 0.013 | 0 | 0.52 | 0.43 | 0.46 | 0.68 | 0.5 | 0.46 |
| 2 | 202511579 | rs10184654 | -0.864 | 0.364 | 0.271 | 0.961 | 0.008 | 0.374 | 0.374 | 0 | 0.07 | 0.28 | 0.03 | 0.4 | 0.03 | 0.1 |
| 2 | 202512449 | rs11894115 | -0.715 | 0.411 | 0.313 | 0.985 | 0.033 | 0.444 | 0.444 | 0 | 0.06 | 0.27 | 0.02 | 0.4 | 0.03 | 0.11 |
| 2 | 202513765 | rs4675211 | -0.083 | 0.401 | 0.299 | 0.94 | -0.009 | 0.028 | 0.028 | 0 | 0.56 | 0.66 | 0.48 | 0.67 | 0.5 | 0.46 |
| 2 | 202514201 | rs10194353 | -0.814 | 0.377 | 0.165 | 1.108 | 0.033 | 0.444 | 0.444 | 0 | 0.06 | 0.28 | 0.02 | 0.4 | 0.03 | 0.11 |
| 2 | 202521426 | rs888011 | -0.318 | 0.675 | 0.094 | 0.508 | 0.009 | 0.025 | 0.025 | 0.028 | 0.63 | 0.7 | 0.61 | 0.69 | 0.62 | 0.52 |
| 2 | 202521824 | rs11890273 | -0.655 | 0.848 | 0.217 | 0.44 | 0.058 | -0.072 | -0.072 | 0.204 | 0.11 | 0.31 | 0.14 | 0.01 | 0.12 | 0.08 |
| 2 | 202523347 | rs2597900 | 0.153 | 0.704 | -0.146 | 0.322 | 0.002 | 0.013 | 0.013 | 0 | 0.49 | 0.46 | 0.49 | 0.57 | 0.42 | 0.48 |
| 2 | 202523547 | rs7602825 | 0.605 | 0.039 | 0.111 | 0.144 | -0.002 | -0.094 | -0.094 | 0 | 0.81 | 0.53 | 0.87 | 0.95 | 0.83 | 0.94 |
| 2 | 202525551 | rs7584026 | -0.615 | -0.101 | -0.406 | 0.401 | 0.024 | 0.086 | 0.086 | 0 | 0.24 | 0.44 | 0.21 | 0.36 | 0.24 | 0.21 |
| 2 | 202527467 | rs10931967 | 1.173 | 0.041 | -1.333 | 0.29 | -0.146 | 0.025 | 0.025 | 0.047 | 0.42 | 0.06 | 0.45 | 0.31 | 0.5 | 0.42 |
| 2 | 202529514 | rs17468821 | 0.689 | 0.025 | -1.335 | 0.192 | 0.035 | 0.04 | 0.04 | 0.121 | 0.19 | 0.07 | 0.26 | 0.1 | 0.15 | 0.31 |
| 2 | 202531547 | rs7559992 | 1.203 | 0.514 | -1.042 | 0.308 | -0.001 | -0.005 | -0.005 | 0.018 | 0.1 | 0.06 | 0.05 | 0.21 | 0.1 | 0.13 |
| 2 | 202531624 | rs16837959 | 0.258 | 0.583 | -0.939 | 0.139 | 0.031 | -0.065 | -0.065 | 0.501 | 0.13 | 0.25 | 0.17 | 0.02 | 0.16 | 0.06 |
| 2 | 202540338 | rs9646756 | 0.98 | 0.347 | -1.55 | 0.349 | -0.154 | -0.036 | -0.036 | 5.595 | 0.35 | 0.02 | 0.29 | 0.29 | 0.41 | 0.37 |
| 2 | 202540420 | rs6731576 | -0.166 | 0.119 | -0.952 | 0.614 | -0.012 | 0.009 | 0.009 | 0.583 | 0.71 | 0.72 | 0.61 | 0.62 | 0.79 | 0.6 |
| 2 | 202541707 | rs13420006 | -0.538 | 0.695 | -0.923 | 0.432 | -0.001 | 0.001 | 0.001 | 0 | 0.36 | 0.37 | 0.3 | 0.32 | 0.4 | 0.23 |
| 2 | 202542836 | rs1914267 | -0.219 | 0.87 | -1.229 | 0.504 | 0.002 | 0.007 | 0.007 | 0.054 | 0.52 | 0.57 | 0.58 | 0.57 | 0.45 | 0.6 |
| 2 | 202543510 | rs4675217 | 0.486 | 0.97 | -1.828 | 0.24 | -0.055 | 0 | 0 | 0.678 | 0.43 | 0.03 | 0.51 | 0.31 | 0.42 | 0.27 |
| 2 | 202547964 | rs13402805 | -0.716 | 1.011 | -0.138 | 0.346 | 0.022 | 0.022 | 0.022 | 0 | 0.16 | 0.31 | 0.13 | 0.25 | 0.19 | 0.28 |
| 2 | 202554092 | rs13008898 | 0.552 | 1.051 | 0.435 | 0.066 | 0.067 | -0.097 | -0.097 | 0 | 0.86 | 0.67 | 0.88 | 0.99 | 0.85 | 0.86 |
| 2 | 202555132 | rs2287050 | -0.97 | 0.296 | 0.522 | 0.226 | -0.008 | 0 | 0 | 0.224 | 0.3 | 0.54 | 0.15 | 0.31 | 0.22 | 0.24 |
| 2 | 202558415 | rs7590550 | -0.385 | 0.258 | 0.609 | 0.669 | 0.002 | -0.002 | -0.002 | 0 | 0.19 | 0.2 | 0.14 | 0.39 | 0.21 | 0.42 |
| 2 | 202559855 | rs1468794 | -0.745 | 0.332 | 0.764 | 0.447 | 0.028 | -0.027 | -0.027 | 0.067 | 0.94 | 0.82 | 0.92 | 1 | 0.94 | 0.98 |
| 2 | 202560012 | rs1468793 | -1.4 | -0.09 | 1.067 | 0.428 | -0.012 | -0.001 | -0.001 | 0 | 0.33 | 0.41 | 0.23 | 0.65 | 0.27 | 0.54 |
| 2 | 202560099 | rs1468792 | 0.451 | -0.126 | 1.136 | 0.431 | -0.026 | 0.063 | 0.063 | 5.75 | 0.82 | 0.65 | 0.88 | 0.74 | 0.9 | 0.85 |
| 2 | 202561571 | rs888012 | 1.288 | -0.132 | 1.03 | 0.384 | -0.009 | -0.005 | -0.005 | 0 | 0.63 | 0.44 | 0.74 | 0.35 | 0.69 | 0.43 |
| 2 | 202566764 | rs3731707 | -1.11 | 0.109 | 1.07 | 0.388 | -0.005 | 0.003 | 0.003 | 0.13 | 0.29 | 0.32 | 0.16 | 0.63 | 0.25 | 0.55 |
| 2 | 202575821 | rs3219168 | -0.224 | 0.126 | 1.109 | 0.371 | 0.023 | -0.019 | -0.019 | 0 | 0.92 | 0.81 | 0.89 | 0.97 | 0.92 | 0.97 |
| 2 | 202578325 | rs6707021 | -0.224 | 0.126 | 1.109 | 0.371 | 0.023 | -0.019 | -0.019 | 0 | 0.92 | 0.8 | 0.89 | 0.97 | 0.92 | 0.97 |
| 2 | 202579648 | rs13384229 | -0.032 | 0.108 | 1.113 | 0.415 | -0.014 | 0.076 | 0.076 | 0 | 0.18 | 0.29 | 0.12 | 0.39 | 0.19 | 0.4 |
| 2 | 202580370 | rs3219167 | 0.735 | 0.162 | 0.8 | 0.438 | -0.003 | 0.021 | 0.021 | 4.278 | 0.81 | 0.77 | 0.85 | 0.74 | 0.85 | 0.84 |
| 2 | 202598113 | rs2276615 | 1.932 | -0.081 | 0.227 | -0.035 | 0.022 | 0.081 | 0.081 | 0 | 0.51 | 0.1 | 0.65 | 0.35 | 0.59 | 0.34 |
| 2 | 202600117 | rs2192762 | 1.029 | -0.126 | 0.899 | -0.064 | -0.012 | -0.003 | -0.003 | 5.96 | 0.63 | 0.45 | 0.79 | 0.38 | 0.7 | 0.4 |
| 2 | 202607126 | rs1981725 | 1.562 | -0.109 | 0.862 | -0.068 | 0.038 | 0.037 | 0.037 | 0 | 0.57 | 0.22 | 0.73 | 0.37 | 0.65 | 0.38 |
| 2 | 202608877 | rs4673207 | 1.274 | -0.128 | 0.922 | -0.076 | 0.048 | 0.022 | 0.022 | 0 | 0.59 | 0.24 | 0.75 | 0.37 | 0.7 | 0.39 |
| 2 | 202625615 | rs3219156 | -0.041 | -0.019 | 1.371 | -0.02 | -0.005 | -0.067 | -0.067 | 0.131 | 0.88 | 0.76 | 0.93 | 1 | 0.9 | 0.95 |
| 2 | 202632938 | rs4675226 | 1.236 | -0.064 | 1.633 | 0.072 | 0.007 | -0.016 | -0.016 | 0.038 | 0.67 | 0.27 | 0.74 | 0.58 | 0.7 | 0.53 |
| 2 | 202633582 | rs3219153 | 0.454 | -0.046 | 1.572 | 0.093 | -0.002 | 0.031 | 0.031 | 0 | 0.17 | 0.21 | 0.11 | 0.4 | 0.18 | 0.39 |
| 2 | 202635604 | rs7561592 | 1.976 | -0.079 | 1.502 | 0.069 | -0.008 | 0.052 | 0.052 | 0 | 0.55 | 0.11 | 0.66 | 0.34 | 0.6 | 0.35 |
| 2 | 202636911 | rs970595 | 2.185 | 0.22 | 1.214 | -0.032 | -0.095 | 0.061 | 0.061 | 0.457 | 0.5 | 0.03 | 0.52 | 0.34 | 0.5 | 0.32 |
| 2 | 202639198 | rs193607 | 0.557 | -0.402 | 1.777 | 0.201 | -0.009 | 0.147 | 0.147 | 0 | 0.78 | 0.47 | 0.81 | 0.58 | 0.8 | 0.58 |
| 2 | 202652583 | rs10178881 | 0.071 | -0.299 | 1.052 | 0.219 | -0.002 | 0.034 | 0.034 | 0 | 0.16 | 0.2 | 0.11 | 0.41 | 0.18 | 0.4 |
| 2 | 202656963 | rs7571046 | -0.172 | -0.28 | 1.012 | 0.248 | 0 | -0.004 | -0.004 | 0.392 | 0.22 | 0.22 | 0.14 | 0.4 | 0.24 | 0.42 |
| 2 | 202657818 | rs9784022 | -0.404 | -0.295 | 1.009 | 0.273 | 0.001 | 0.049 | 0.049 | 0 | 0.25 | 0.36 | 0.17 | 0.4 | 0.26 | 0.44 |
| 2 | 202665059 | rs9973862 | -0.965 | -0.277 | 1.126 | 0.153 | 0.007 | -0.044 | -0.044 | 0.097 | 0.09 | 0.29 | 0.07 | 0.01 | 0.06 | 0.04 |
| 2 | 202665227 | rs2052249 | -1.1 | -0.447 | 1.408 | 0.182 | -0.006 | 0.045 | 0.045 | 0.239 | 0.31 | 0.52 | 0.23 | 0.42 | 0.29 | 0.49 |

| | | | | | | | | | | | | | | | | |
|---|---|---|---|---|---|---|---|---|---|---|---|---|---|---|---|---|
| 2 | 202673058 | rs6742520 | 1.642 | -0.937 | 1.274 | 0.138 | 0.067 | 0.031 | 0.031 | 0 | 0.53 | 0.16 | 0.7 | 0.34 | 0.64 | 0.37 |
| 2 | 202674612 | rs2052247 | -1.077 | -0.438 | 1.695 | 0.151 | -0.001 | 0.023 | 0.023 | 0.096 | 0.3 | 0.61 | 0.23 | 0.4 | 0.32 | 0.47 |
| 2 | 202679085 | rs16838219 | -0.468 | -0.441 | 1.336 | 0.173 | -0.001 | -0.002 | -0.002 | 0 | 0.25 | 0.24 | 0.17 | 0.42 | 0.27 | 0.45 |
| 2 | 202682229 | rs12053321 | -0.198 | -0.422 | 1.23 | 0.188 | -0.003 | -0.011 | -0.011 | 0.375 | 0.23 | 0.23 | 0.16 | 0.4 | 0.26 | 0.45 |
| 2 | 202687014 | rs7585095 | 1.092 | -0.466 | 1.338 | 0.146 | -0.009 | -0.021 | -0.021 | 0 | 0.93 | 0.75 | 0.97 | 1 | 0.96 | 0.98 |
| 2 | 202688293 | rs4675229 | 0.862 | -0.463 | 1.26 | 0.148 | -0.013 | -0.024 | -0.024 | 0.097 | 0.94 | 0.77 | 0.97 | 1 | 0.96 | 0.98 |
| 2 | 202688907 | rs11680161 | 0.635 | -0.397 | 1.189 | 0.196 | 0 | 0.042 | 0.042 | 0.056 | 0.7 | 0.47 | 0.78 | 0.58 | 0.68 | 0.53 |
| 2 | 202699201 | rs193599 | 1.136 | -1.05 | 0.315 | -0.419 | -0.004 | -0.035 | -0.035 | 2.252 | 0.16 | 0.16 | 0.07 | 0 | 0.09 | 0.03 |
| 2 | 202703593 | rs2110646 | 0.075 | -1.062 | 0.252 | -0.152 | -0.001 | 0.01 | 0.01 | 0 | 0.2 | 0.22 | 0.15 | 0.38 | 0.2 | 0.45 |
| 2 | 202704137 | rs4675231 | 0.395 | -0.896 | 0.107 | -0.13 | -0.018 | -0.009 | -0.009 | 0 | 0.58 | 0.48 | 0.75 | 0.62 | 0.65 | 0.49 |
| 2 | 202713771 | rs6707595 | -0.122 | -0.805 | 0.193 | 0.11 | -0.011 | 0.062 | 0.062 | 0 | 0.23 | 0.53 | 0.18 | 0.36 | 0.2 | 0.44 |
| 2 | 202721424 | rs2882324 | -0.849 | -0.151 | -0.354 | -0.092 | 0.003 | 0.108 | 0.108 | 0.319 | 0.41 | 0.81 | 0.31 | 0.23 | 0.38 | 0.25 |
| 2 | 202722785 | rs13425147 | -1.017 | 0.078 | 0.19 | -0.413 | 0.098 | -0.029 | -0.029 | 0.299 | 0.09 | 0.39 | 0.1 | 0.01 | 0.11 | 0.02 |
| 2 | 202725173 | rs16838312 | 1.755 | -0.069 | -0.965 | -0.359 | -0.124 | -0.148 | -0.148 | 0 | 0.24 | 0.01 | 0.54 | 0.33 | 0.38 | 0.26 |
| 2 | 202727036 | rs16838316 | 1.235 | -0.181 | -0.113 | -0.576 | 0.003 | -0.012 | -0.012 | 0.702 | 0.06 | 0.08 | 0.05 | 0.01 | 0.07 | 0.07 |
| 2 | 202728891 | rs11889379 | -0.402 | -0.239 | -0.095 | -0.611 | -0.023 | -0.133 | -0.133 | 1.76 | 0.31 | 0.44 | 0.17 | 0.01 | 0.22 | 0.16 |
| 2 | 202732028 | rs16838323 | -0.94 | 0.338 | 0.555 | 0.342 | 0.072 | -0.094 | -0.094 | 3.951 | 0.15 | 0.52 | 0.16 | 0.01 | 0.19 | 0.09 |
| 2 | 202739991 | rs6435114 | -0.248 | -0.145 | 0.039 | 0.998 | -0.015 | 0.039 | 0.039 | 0.015 | 0.26 | 0.37 | 0.15 | 0.49 | 0.21 | 0.26 |
| 2 | 202741064 | rs2072504 | 1.257 | -0.277 | 0.006 | 1.094 | 0.041 | 0.076 | 0.076 | 0.662 | 0.56 | 0.15 | 0.61 | 0.23 | 0.64 | 0.58 |
| 2 | 202742182 | rs6752497 | 0.631 | -0.161 | 0.356 | 0.756 | 0.01 | -0.022 | -0.022 | 0 | 0.85 | 0.57 | 0.86 | 0.89 | 0.87 | 0.85 |
| 2 | 202745638 | rs2192879 | 1.517 | -0.231 | 0.362 | 0.766 | -0.004 | -0.008 | -0.008 | 0 | 0.86 | 0.59 | 0.87 | 0.87 | 0.91 | 0.92 |
| 2 | 202747624 | rs13425892 | -0.035 | -0.098 | 0.263 | 0.892 | 0.033 | 0.085 | 0.085 | 0.001 | 0.05 | 0.24 | 0.07 | 0.11 | 0.03 | 0.06 |
| 2 | 202759326 | rs12472067 | 0.362 | 0.082 | -0.088 | 1.09 | -0.001 | -0.04 | -0.04 | 0.884 | 0.64 | 0.2 | 0.71 | 0.37 | 0.64 | 0.61 |
| 2 | 202763587 | rs3918349 | -0.996 | 0.307 | 0.185 | 1.027 | 0.003 | 0.045 | 0.045 | 0 | 0.46 | 0.71 | 0.37 | 0.75 | 0.45 | 0.43 |
| 2 | 202767721 | rs7604971 | -0.459 | 0.283 | 0.277 | 1.027 | 0.001 | 0.007 | 0.007 | 0 | 0.37 | 0.44 | 0.31 | 0.64 | 0.43 | 0.35 |
| 2 | 202771029 | rs16838426 | -0.21 | 0.309 | 0.324 | 1.201 | 0.012 | 0.189 | 0.189 | 2.126 | 0.11 | 0.36 | 0.14 | 0.3 | 0.04 | 0.1 |
| 2 | 202775055 | rs10931978 | -0.315 | 0.339 | 0.052 | 1.142 | -0.002 | -0.003 | -0.003 | 0 | 0.35 | 0.37 | 0.25 | 0.46 | 0.43 | 0.34 |
| 2 | 202793266 | rs6722501 | -0.199 | 0.082 | 0.727 | 1.11 | -0.009 | 0.044 | 0.044 | 3.625 | 0.1 | 0.37 | 0.09 | 0.12 | 0.02 | 0.08 |
| 2 | 202793520 | rs7573217 | 0.761 | 0.117 | -0.119 | 0.982 | -0.026 | -0.025 | -0.025 | 3.415 | 0.1 | 0.02 | 0.13 | 0.09 | 0.12 | 0.23 |
| 2 | 202795537 | rs4140880 | 1.435 | 0.122 | 0.287 | 0.954 | -0.002 | -0.009 | -0.009 | 0 | 0.78 | 0.26 | 0.82 | 0.87 | 0.83 | 0.9 |
| 2 | 202798759 | rs16838452 | -0.943 | -0.07 | 0.089 | 0.869 | 0.041 | -0.053 | -0.053 | 1.376 | 0.1 | 0.29 | 0.08 | 0.01 | 0.13 | 0.02 |
| 2 | 202799297 | rs7595229 | 1.021 | -0.011 | -1.132 | 1.044 | 0.017 | -0.021 | -0.021 | 0 | 0.77 | 0.19 | 0.77 | 0.67 | 0.82 | 0.89 |
| 2 | 202806638 | rs6725131 | 0.161 | 0.723 | -1.165 | 0.653 | -0.001 | -0.001 | -0.001 | 0 | 0.31 | 0.28 | 0.41 | 0.26 | 0.34 | 0.39 |
| 2 | 202807196 | rs17385675 | -0.962 | 0.549 | -1.195 | 0.53 | 0.006 | 0.051 | 0.051 | 2.669 | 0.11 | 0.02 | 0.05 | 0 | 0.08 | 0.02 |
| 2 | 202815530 | rs7562987 | 0.693 | 0.719 | -0.82 | 0.707 | 0.007 | -0.039 | -0.039 | 0 | 0.72 | 0.52 | 0.78 | 0.79 | 0.75 | 0.65 |
| 2 | 202828666 | rs10210937 | -0.475 | 0.517 | -1.336 | 0.489 | -0.004 | -0.024 | -0.024 | 0 | 0.32 | 0.63 | 0.36 | 0.11 | 0.27 | 0.3 |
| 2 | 202832130 | rs10931982 | 0.367 | 0.801 | -1.678 | 0.503 | -0.001 | 0.027 | 0.027 | 0 | 0.79 | 0.87 | 0.84 | 0.88 | 0.8 | 0.85 |
| 2 | 202833838 | rs1985266 | -0.807 | 0.889 | -2.279 | 0.447 | -0.054 | -0.013 | -0.013 | 0 | 0.24 | 0.02 | 0.18 | 0.13 | 0.25 | 0.22 |
| 2 | 202834128 | rs1123917 | -0.478 | 1.568 | -2.602 | 0.723 | -0.065 | -0.084 | -0.084 | 0 | 0.23 | 0.09 | 0.38 | 0.59 | 0.31 | 0.32 |
| 2 | 202835148 | rs10804128 | -0.969 | 0.751 | -1.83 | 0.101 | 0.015 | 0.079 | 0.079 | 0 | 0.5 | 0.66 | 0.62 | 0.78 | 0.55 | 0.55 |
| 2 | 202836995 | rs13004821 | -1.028 | 0.594 | -0.834 | 0.01 | -0.01 | -0.012 | -0.012 | 0 | 0.1 | 0.21 | 0.21 | 0.09 | 0.1 | 0.22 |
| 2 | 202837158 | rs17385875 | -0.287 | 0.727 | -0.751 | 0.217 | 0.001 | 0.004 | 0.004 | 0 | 0.06 | 0.07 | 0.07 | 0.09 | 0.07 | 0.12 |
| 2 | 202837219 | rs16838552 | -1.042 | 0.735 | -0.752 | 0.179 | -0.009 | -0.012 | -0.012 | 0 | 0.11 | 0.2 | 0.22 | 0.09 | 0.11 | 0.22 |
| 2 | 202838353 | rs967693 | -0.434 | 0.987 | -1.082 | -0.068 | -0.045 | 0.086 | 0.086 | 0 | 0.42 | 0.22 | 0.36 | 0.21 | 0.47 | 0.3 |
| 2 | 202839644 | rs17385923 | 0.146 | 1.053 | -1.246 | -0.047 | -0.081 | 0.113 | 0.113 | 0 | 0.34 | 0.03 | 0.18 | 0.1 | 0.36 | 0.17 |
| 2 | 202839768 | rs12998820 | -0.766 | 0.885 | -0.86 | 0.139 | 0.008 | 0.069 | 0.069 | 0 | 0.52 | 0.29 | 0.49 | 0.3 | 0.58 | 0.52 |
| 2 | 202845843 | rs6722215 | -0.642 | 1.158 | -0.8 | 0.17 | 0.001 | 0.026 | 0.026 | 0 | 0.49 | 0.48 | 0.42 | 0.25 | 0.53 | 0.37 |
| 2 | 202851115 | rs7571877 | 0.595 | 1.115 | -1.84 | -0.153 | -0.06 | 0.134 | 0.134 | 0.053 | 0.31 | 0.02 | 0.2 | 0.08 | 0.27 | 0.33 |
| 2 | 202859438 | rs4673221 | -0.66 | 1.368 | -0.899 | 0.754 | 0.004 | 0.02 | 0.02 | 0.063 | 0.15 | 0.17 | 0.29 | 0.3 | 0.19 | 0.22 |
| 2 | 202861219 | rs6740467 | -0.482 | 1.437 | -0.294 | 1.318 | 0.046 | 0.312 | 0.312 | 0 | 0.08 | 0.24 | 0.17 | 0.42 | 0.06 | 0.17 |
| 2 | 202864487 | rs4675246 | -0.024 | 2.009 | -0.899 | 0.742 | -0.032 | 0.023 | 0.023 | 0 | 0.06 | 0.01 | 0.2 | 0 | 0.19 | 0.06 |
| 2 | 202867453 | rs10173592 | 0.45 | 1.663 | -1.283 | 0.444 | -0.011 | 0.036 | 0.036 | 0.346 | 0.23 | 0.13 | 0.21 | 0.11 | 0.24 | 0.14 |
| 2 | 202874850 | rs13412938 | 0.333 | 1.514 | -1.298 | 0.366 | -0.008 | 0.049 | 0.049 | 0.647 | 0.24 | 0.13 | 0.21 | 0.11 | 0.24 | 0.15 |
| 2 | 202875696 | rs1477032 | -0.55 | 0.74 | -1.074 | 0.711 | -0.026 | 0.009 | 0.009 | 0 | 0.72 | 0.42 | 0.81 | 0.71 | 0.82 | 0.73 |
| 2 | 202882110 | rs13035955 | 0.05 | 1.021 | -1.097 | 0.548 | 0.008 | -0.011 | -0.011 | 1.354 | 0.08 | 0.12 | 0.22 | 0 | 0.21 | 0.07 |
| 2 | 202883828 | rs11681902 | 0.367 | 1.028 | -1.155 | 0.92 | 0.01 | 0.082 | 0.082 | 0 | 0.08 | 0.14 | 0.18 | 0.28 | 0.09 | 0.13 |
| 2 | 202884393 | rs1477031 | -0.12 | 0.826 | -1.258 | 0.778 | 0.008 | 0.007 | 0.007 | 0 | 0.64 | 0.31 | 0.61 | 0.72 | 0.62 | 0.66 |
| 2 | 202901279 | rs13034579 | 0.185 | 1.114 | -0.793 | 0.485 | 0.025 | -0.048 | -0.048 | 0 | 0.15 | 0.24 | 0.27 | 0 | 0.29 | 0.08 |
| 2 | 202901910 | rs4673222 | 0.596 | 0.975 | -0.863 | 0.675 | 0 | 0.055 | 0.055 | 0 | 0.09 | 0.12 | 0.18 | 0.3 | 0.08 | 0.14 |
| 2 | 202911239 | rs13017513 | -0.076 | 0.871 | -1.168 | 0.527 | -0.019 | 0.051 | 0.051 | 0.359 | 0.41 | 0.3 | 0.42 | 0.27 | 0.53 | 0.47 |
| 2 | 202911548 | rs16838664 | 1.344 | 0.859 | -1.253 | 0.52 | -0.059 | -0.054 | -0.054 | 0 | 0.25 | 0.02 | 0.15 | 0.26 | 0.24 | 0.38 |
| 2 | 202912115 | rs6725357 | 0.547 | 0.896 | -1.194 | 0.509 | 0.019 | -0.041 | -0.041 | 0 | 0.15 | 0.24 | 0.27 | 0 | 0.29 | 0.08 |
| 2 | 202912549 | rs6725902 | 1.448 | 0.719 | -0.39 | 0.427 | -0.008 | -0.039 | -0.039 | 0.464 | 0.27 | 0.38 | 0.21 | 0.14 | 0.22 | 0.14 |
| 2 | 202912688 | rs7592801 | -0.675 | 0.66 | -0.398 | 0.46 | -0.013 | 0.012 | 0.012 | 0 | 0.45 | 0.4 | 0.62 | 0.59 | 0.54 | 0.48 |
| 2 | 202914491 | rs4675250 | -0.638 | 0.788 | -0.29 | 0.449 | 0.011 | 0.004 | 0.004 | 0 | 0.24 | 0.35 | 0.45 | 0.31 | 0.37 | 0.22 |
| 2 | 202918708 | rs4675252 | 0.416 | 0.691 | 0.002 | 0.568 | 0.003 | 0.126 | 0.126 | 0 | 0.11 | 0.26 | 0.19 | 0.28 | 0.08 | 0.13 |
| 2 | 202922665 | rs7596320 | -0.057 | 0.593 | -0.171 | 0.608 | -0.021 | 0.021 | 0.021 | 0 | 0.65 | 0.48 | 0.6 | 0.5 | 0.76 | 0.61 |
| 2 | 202925675 | rs1402873 | 1.097 | 0.922 | -0.834 | 0.507 | -0.08 | 0.066 | 0.066 | 0 | 0.12 | 0.01 | 0.25 | 0 | 0.28 | 0.08 |
| 2 | 202928940 | rs7566288 | 1.373 | 0.532 | -0.695 | 0.594 | 0.004 | 0 | 0 | 0 | 0.26 | 0.21 | 0.2 | 0.21 | 0.24 | 0.14 |

| | | | | | | | | | | | | | | | | |
|---|---|---|---|---|---|---|---|---|---|---|---|---|---|---|---|---|
| 2 | 202937284 | rs871949 | 1.231 | 0.388 | -0.846 | 0.591 | 0.024 | -0.01 | -0.01 | 2.186 | 0.21 | 0.06 | 0.19 | 0.21 | 0.16 | 0.26 |
| 2 | 202940099 | rs1521886 | -0.169 | 0.388 | -0.448 | 0.551 | 0.029 | 0.013 | 0.013 | 1.636 | 0.5 | 0.6 | 0.64 | 0.51 | 0.61 | 0.37 |
| 2 | 202941310 | rs939745 | -1.246 | 0.561 | -0.546 | 0.529 | 0.003 | 0.002 | 0.002 | 1.2 | 0.74 | 0.84 | 0.83 | 0.72 | 0.77 | 0.63 |
| 2 | 202950603 | rs10170697 | 0.849 | 0.36 | -0.747 | 0.378 | 0.027 | -0.046 | -0.046 | 0 | 0.38 | 0.18 | 0.31 | 0.41 | 0.32 | 0.31 |
| 2 | 202951113 | rs13389388 | 0.849 | 0.36 | -0.747 | 0.378 | 0.027 | -0.046 | -0.046 | 0 | 0.38 | 0.18 | 0.31 | 0.41 | 0.32 | 0.31 |
| 2 | 202951712 | rs1521880 | -0.789 | 0.352 | -0.605 | 0.342 | 0 | 0 | 0 | 0 | 0.36 | 0.61 | 0.52 | 0.31 | 0.45 | 0.32 |
| 2 | 202953227 | rs1521879 | -1.53 | 0.336 | -0.567 | 0.299 | -0.001 | 0 | 0 | 1.145 | 0.75 | 0.79 | 0.83 | 0.72 | 0.78 | 0.63 |
| 2 | 202953787 | rs16838805 | 0.099 | 0.296 | -0.164 | 0.171 | -0.036 | -0.077 | -0.077 | 1.805 | 0.14 | 0.5 | 0.13 | 0.04 | 0.1 | 0.1 |
| 2 | 202954511 | rs13022378 | -0.38 | 0.511 | -0.103 | 0.13 | 0.046 | -0.132 | -0.132 | 0.324 | 0.25 | 0.42 | 0.34 | 0.04 | 0.37 | 0.18 |
| 2 | 202955820 | rs1568534 | 0.79 | 0.304 | -0.772 | -0.024 | 0.023 | -0.012 | -0.012 | 4.338 | 0.51 | 0.23 | 0.38 | 0.47 | 0.44 | 0.56 |
| 2 | 202956470 | rs11690765 | 0.798 | 0.25 | -0.751 | 0.323 | -0.006 | -0.065 | -0.065 | 0 | 0.22 | 0.13 | 0.26 | 0.49 | 0.19 | 0.27 |
| 2 | 202961665 | rs2177083 | -0.265 | -0.264 | -0.933 | -0.107 | -0.001 | -0.002 | -0.002 | 0.384 | 0.41 | 0.57 | 0.45 | 0.26 | 0.5 | 0.49 |
| 2 | 202964026 | rs10931991 | -0.563 | -0.341 | -1.233 | 0.118 | 0.014 | 0.017 | 0.017 | 0 | 0.65 | 0.71 | 0.77 | 0.74 | 0.71 | 0.62 |
| 2 | 202964528 | rs2350722 | -0.301 | -0.365 | -1.217 | 0.003 | -0.005 | 0.058 | 0.058 | 0.018 | 0.48 | 0.59 | 0.48 | 0.24 | 0.52 | 0.5 |
| 2 | 202969687 | rs6746162 | -1.191 | -0.211 | -1.142 | 0.22 | -0.003 | -0.012 | -0.012 | 0 | 0.78 | 0.81 | 0.69 | 0.47 | 0.78 | 0.85 |
| 2 | 202974099 | rs12611699 | 0.647 | -0.496 | -0.605 | -0.072 | -0.012 | -0.009 | -0.009 | 0.302 | 0.33 | 0.47 | 0.24 | 0.27 | 0.28 | 0.29 |
| 2 | 202980253 | rs2882485 | -0.096 | -0.457 | -0.595 | -0.09 | -0.02 | 0.07 | 0.07 | 0 | 0.44 | 0.21 | 0.43 | 0.2 | 0.5 | 0.57 |
| 2 | 202980521 | rs6708948 | 0.538 | -0.405 | -0.939 | -0.292 | -0.098 | 0.05 | 0.05 | 0 | 0.28 | 0.03 | 0.32 | 0.13 | 0.36 | 0.34 |
| 2 | 202980887 | rs6748072 | -0.425 | -0.376 | -0.644 | -0.213 | 0.002 | -0.029 | -0.029 | 0 | 0.46 | 0.43 | 0.46 | 0.2 | 0.5 | 0.57 |
| 2 | 202986369 | rs16838864 | 0.955 | -0.18 | -0.44 | 0.048 | -0.001 | 0.05 | 0.05 | 0 | 0.06 | 0.07 | 0.16 | 0.32 | 0.06 | 0.06 |
| 2 | 202990187 | rs4675261 | -0.638 | -0.078 | -0.374 | -0.867 | 0.007 | -0.002 | -0.002 | 0 | 0.62 | 0.73 | 0.71 | 0.66 | 0.66 | 0.66 |
| 2 | 203001990 | rs34829484 | 1.295 | -0.046 | -0.282 | -0.965 | -0.006 | -0.043 | -0.043 | 0.761 | 0.16 | 0.17 | 0.1 | 0.09 | 0.13 | 0.25 |
| 2 | 203004983 | rs11690066 | -0.722 | -0.098 | -0.468 | -0.839 | 0.006 | -0.002 | -0.002 | 0 | 0.63 | 0.77 | 0.72 | 0.67 | 0.66 | 0.67 |
| 2 | 203009333 | rs2049193 | 0.055 | 0.074 | -0.406 | -0.887 | 0.018 | -0.12 | -0.12 | 0 | 0.28 | 0.41 | 0.32 | 0.09 | 0.32 | 0.33 |
| 2 | 203021290 | rs2177086 | 0.539 | 0.13 | -0.764 | -0.776 | 0 | -0.005 | -0.005 | 0 | 0.37 | 0.42 | 0.42 | 0.58 | 0.37 | 0.34 |
| 2 | 203022536 | rs6435130 | -0.651 | 0.045 | -1.028 | -0.621 | -0.01 | -0.038 | -0.038 | 0 | 0.65 | 0.93 | 0.73 | 0.66 | 0.69 | 0.66 |
| 2 | 203047431 | rs10174937 | 0.368 | 0.041 | -0.786 | -0.703 | 0.018 | 0.013 | 0.013 | 0 | 0.36 | 0.24 | 0.26 | 0.31 | 0.29 | 0.36 |
| 2 | 203056987 | rs6714212 | -0.231 | 0.18 | -0.458 | -0.864 | 0.003 | 0.012 | 0.012 | 0 | 0.46 | 0.53 | 0.44 | 0.32 | 0.41 | 0.46 |
| 2 | 203060299 | rs4560092 | 0.076 | 0.139 | -0.312 | -0.884 | -0.012 | 0.044 | 0.044 | 0 | 0.42 | 0.12 | 0.41 | 0.31 | 0.4 | 0.43 |
| 2 | 203062078 | rs12988414 | 0.076 | 0.139 | -0.317 | -0.924 | -0.016 | 0.047 | 0.047 | 0 | 0.42 | 0.11 | 0.4 | 0.31 | 0.4 | 0.43 |
| 2 | 203062103 | rs13015146 | -0.198 | 0.142 | -0.293 | -0.923 | 0.005 | 0.017 | 0.017 | 0 | 0.46 | 0.53 | 0.44 | 0.32 | 0.4 | 0.46 |
| 2 | 203063848 | rs13001933 | 0.901 | 0.206 | -0.394 | -0.932 | -0.002 | -0.003 | -0.003 | 0 | 0.08 | 0.04 | 0.17 | 0 | 0.12 | 0.09 |
| 2 | 203069878 | rs3181153 | -1.422 | 0.054 | -0.41 | -1.077 | -0.001 | 0.016 | 0.016 | 0 | 0.85 | 0.92 | 0.85 | 0.88 | 0.86 | 0.74 |
| 2 | 203098559 | rs6709162 | 0.437 | 0.01 | -0.459 | -1.103 | 0.025 | 0.024 | 0.024 | 0 | 0.4 | 0.65 | 0.28 | 0.31 | 0.29 | 0.36 |
| 2 | 203102345 | rs6761234 | 0.142 | 0.057 | -0.561 | -1.085 | -0.009 | 0.011 | 0.011 | 0 | 0.45 | 0.22 | 0.56 | 0.57 | 0.58 | 0.38 |
| 2 | 203107868 | rs4675274 | -1.46 | 0.059 | -0.787 | -1.09 | 0.002 | 0.018 | 0.018 | 0 | 0.84 | 0.93 | 0.85 | 0.88 | 0.86 | 0.74 |
| 2 | 203126453 | rs6758247 | -0.866 | 0.052 | -0.826 | -1.15 | -0.022 | -0.007 | -0.007 | 0 | 0.6 | 0.69 | 0.59 | 0.42 | 0.54 | 0.69 |
| 2 | 203126559 | rs6758561 | 0.989 | 0.006 | -0.814 | -1.197 | -0.004 | -0.002 | -0.002 | 0 | 0.34 | 0.06 | 0.23 | 0.29 | 0.28 | 0.34 |
| 2 | 203134839 | rs13392645 | 0.28 | 0.029 | -0.521 | -1.191 | -0.019 | -0.033 | -0.033 | 0 | 0.19 | 0.61 | 0.19 | 0.12 | 0.14 | 0.26 |
| 2 | 203152355 | rs7572505 | -0.435 | 0.055 | -1.184 | -1.195 | -0.001 | -0.019 | -0.019 | 0.107 | 0.26 | 0.63 | 0.36 | 0.13 | 0.25 | 0.35 |
| 2 | 203155936 | rs3731700 | 1.32 | 0.042 | -1.233 | -1.191 | -0.031 | -0.006 | -0.006 | 0 | 0.14 | 0.01 | 0.14 | 0.12 | 0.14 | 0.26 |
| 2 | 203165534 | rs6753156 | -0.365 | 0.104 | -1.177 | -1.15 | -0.001 | -0.02 | -0.02 | 0 | 0.26 | 0.6 | 0.36 | 0.13 | 0.26 | 0.35 |
| 2 | 203171672 | rs10084427 | -0.343 | 0.111 | -1.21 | -1.147 | -0.001 | -0.017 | -0.017 | 0 | 0.26 | 0.63 | 0.36 | 0.13 | 0.26 | 0.35 |
| 2 | 203194256 | rs6435143 | -0.759 | 0.336 | -1.516 | -1.201 | -0.016 | 0.104 | 0.104 | 0 | 0.56 | 0.83 | 0.65 | 0.73 | 0.58 | 0.63 |
| 2 | 203203865 | rs4516412 | 0.18 | 0.569 | -1.658 | -1.219 | -0.044 | 0.115 | 0.115 | 0 | 0.28 | 0.07 | 0.3 | 0.05 | 0.33 | 0.17 |
| 2 | 203204567 | rs12473579 | 0.18 | 0.569 | -1.658 | -1.219 | -0.044 | 0.115 | 0.115 | 0 | 0.28 | 0.07 | 0.3 | 0.05 | 0.33 | 0.17 |
| 2 | 203222966 | rs16824599 | 1.092 | 0.41 | -1.769 | -1.092 | -0.027 | -0.005 | -0.005 | 0 | 0.14 | 0 | 0.15 | 0.12 | 0.14 | 0.26 |
| 2 | 203223295 | rs6715945 | 0.256 | 0.519 | -1.819 | -1.287 | -0.021 | 0.037 | 0.037 | 0 | 0.14 | 0.06 | 0.22 | 0.01 | 0.19 | 0.14 |
| 2 | 203246411 | rs4303700 | 0.812 | 0.486 | -2.039 | -1.452 | -0.012 | 0.084 | 0.084 | 0 | 0.21 | 0.05 | 0.13 | 0.05 | 0.22 | 0.08 |
| 2 | 203255485 | rs13426118 | 0.78 | 0.308 | -0.712 | -1.607 | -0.005 | -0.043 | -0.043 | 0 | 0.18 | 0.45 | 0.18 | 0.12 | 0.14 | 0.27 |
| 2 | 203260701 | rs13025713 | 1.34 | 0.348 | -0.759 | -1.615 | -0.037 | 0.032 | 0.032 | 0 | 0.07 | 0.01 | 0.17 | 0 | 0.12 | 0.09 |
| 2 | 203273559 | rs7583801 | -1.16 | 0.224 | -0.489 | -1.7 | 0.004 | 0.092 | 0.092 | 0.054 | 0.48 | 0.65 | 0.5 | 0.18 | 0.48 | 0.44 |
| 2 | 203286422 | rs7586383 | 0.727 | 0.229 | -0.431 | -1.665 | 0 | -0.01 | -0.01 | 0 | 0.08 | 0.11 | 0.18 | 0 | 0.11 | 0.09 |
| 2 | 203313638 | rs7589059 | -0.167 | 0.206 | -0.418 | -1.675 | 0.004 | -0.072 | -0.072 | 0.859 | 0.26 | 0.56 | 0.36 | 0.11 | 0.26 | 0.36 |
| 2 | 203315612 | rs6747756 | -0.167 | 0.206 | -0.418 | -1.677 | 0.004 | -0.061 | -0.061 | 1.503 | 0.26 | 0.55 | 0.36 | 0.13 | 0.26 | 0.36 |
| 2 | 203352937 | rs6751210 | -1.212 | 0.128 | -0.753 | -1.704 | 0.004 | 0.095 | 0.095 | 0.025 | 0.48 | 0.65 | 0.49 | 0.2 | 0.48 | 0.48 |
| 2 | 203358214 | rs12621870 | 1.452 | 0.125 | -1.119 | -1.771 | -0.023 | -0.066 | -0.066 | 0.287 | 0.21 | 0.11 | 0.24 | 0.54 | 0.25 | 0.23 |
| 2 | 203372389 | rs7605442 | 1.861 | 0.142 | -1.091 | -1.846 | -0.001 | -0.017 | -0.017 | 0 | 0.08 | 0.05 | 0.06 | 0.01 | 0.08 | 0.05 |
| 2 | 203388253 | rs12472248 | 1.437 | 0.212 | -0.987 | -1.686 | -0.029 | 0.032 | 0.032 | 0 | 0.07 | 0.01 | 0.17 | 0 | 0.12 | 0.09 |
| 2 | 203421199 | rs1061157 | 1.624 | 0.437 | -0.882 | -1.584 | -0.023 | 0.027 | 0.027 | 0 | 0.14 | 0 | 0.11 | 0.1 | 0.14 | 0.27 |
| 2 | 203425475 | rs6435156 | -0.141 | 0.431 | -0.853 | -1.587 | 0.002 | -0.095 | -0.095 | 0.002 | 0.26 | 0.53 | 0.32 | 0.11 | 0.26 | 0.36 |
| 2 | 203426668 | rs17199249 | 0.761 | 0.561 | -1.203 | -1.466 | 0.003 | -0.016 | -0.016 | 0 | 0.09 | 0.11 | 0.18 | 0 | 0.12 | 0.09 |
| 2 | 203430456 | rs1048829 | 0.416 | 0.509 | -1.325 | -1.555 | 0.006 | 0.094 | 0.094 | 0.056 | 0.52 | 0.36 | 0.54 | 0.84 | 0.53 | 0.56 |
| 2 | 203430667 | rs7600694 | 0.389 | 0.648 | -1.474 | -1.04 | -0.002 | -0.006 | -0.006 | 0 | 0.09 | 0.1 | 0.18 | 0 | 0.11 | 0.09 |
| 2 | 203431804 | rs3731696 | 1.083 | 0.577 | -1.528 | -0.804 | -0.033 | -0.005 | -0.005 | 0 | 0.14 | 0 | 0.14 | 0.13 | 0.14 | 0.27 |
| 2 | 203465357 | rs4675285 | 1.36 | 0.763 | -1.285 | -0.353 | -0.03 | -0.032 | -0.032 | 0 | 0.09 | 0.01 | 0.17 | 0.52 | 0.11 | 0.13 |
| 2 | 203469588 | rs13432396 | 0.186 | 0.762 | -1.357 | -1.239 | -0.023 | 0.094 | 0.094 | 0 | 0.58 | 0.71 | 0.37 | 0.22 | 0.45 | 0.46 |
| 2 | 203469960 | rs13035933 | -0.182 | 0.79 | -1 | -1.054 | -0.001 | -0.001 | -0.001 | 0.479 | 0.67 | 0.64 | 0.55 | 0.27 | 0.62 | 0.57 |
| 2 | 203488449 | rs10189685 | 0.912 | 0.642 | -0.559 | -1.326 | 0.007 | -0.089 | -0.089 | 0.016 | 0.27 | 0.34 | 0.2 | 0.05 | 0.3 | 0.1 |

| | | | | | | | | | | | | | | | | |
|---|---|---|---|---|---|---|---|---|---|---|---|---|---|---|---|---|
| 2 | 203497786 | rs7581601 | 0.977 | 0.667 | -1.038 | -1.323 | -0.037 | 0.105 | 0.105 | 0 | 0.27 | 0.07 | 0.19 | 0.05 | 0.32 | 0.1 |
| 2 | 203499512 | rs11692610 | 0.688 | 0.154 | -1.209 | -1.509 | 0.049 | 0.091 | 0.091 | 0.465 | 0.34 | 0.17 | 0.22 | 0.2 | 0.23 | 0.44 |
| 2 | 203511571 | rs12466268 | 0.926 | 0.16 | -0.622 | -1.536 | 0.027 | 0.084 | 0.084 | 0 | 0.32 | 0.15 | 0.23 | 0.2 | 0.24 | 0.45 |
| 2 | 203520272 | rs7571824 | -1.647 | 0.24 | -1.071 | -1.457 | 0.002 | -0.012 | -0.012 | 0 | 0.83 | 0.89 | 0.81 | 0.76 | 0.83 | 0.78 |
| 2 | 203527979 | rs7569317 | 0.107 | 0.283 | -1.174 | -1.548 | 0.011 | 0.111 | 0.111 | 0 | 0.51 | 0.3 | 0.59 | 0.71 | 0.51 | 0.68 |
| 2 | 203532304 | rs11694172 | 1.484 | 0.239 | -1.639 | -1.569 | -0.012 | 0.076 | 0.076 | 0 | 0.31 | 0.02 | 0.21 | 0.2 | 0.22 | 0.44 |
| 2 | 203547106 | rs7568438 | 1.548 | 0.402 | -1.48 | -1.472 | -0.035 | 0.119 | 0.119 | 0 | 0.27 | 0.07 | 0.17 | 0.06 | 0.31 | 0.11 |
| 2 | 203553836 | rs4675287 | 1.701 | 0.561 | -1.494 | -1.463 | -0.015 | -0.018 | -0.018 | 0.504 | 0.12 | 0.11 | 0.2 | 0.51 | 0.15 | 0.13 |
| 2 | 203555828 | rs10755019 | 1.815 | 0.522 | -1.538 | -1.474 | -0.008 | -0.036 | -0.036 | 0 | 0.16 | 0.07 | 0.19 | 0.22 | 0.17 | 0.22 |
| 2 | 203557151 | rs1039968 | 1.418 | 0.505 | -1.403 | -1.768 | -0.029 | 0.11 | 0.11 | 0 | 0.27 | 0.08 | 0.17 | 0.06 | 0.31 | 0.11 |
| 2 | 203566078 | rs12474453 | 1.017 | 0.823 | -0.912 | -1.227 | -0.031 | 0.034 | 0.034 | 0 | 0.08 | 0.04 | 0.16 | 0 | 0.14 | 0.1 |
| 2 | 203602207 | rs7602496 | 0.171 | 0.594 | -0.784 | -1.223 | -0.04 | 0.068 | 0.068 | 0 | 0.63 | 0.74 | 0.44 | 0.25 | 0.54 | 0.54 |
| 2 | 203606754 | rs7581854 | 0.25 | 0.6 | -0.653 | -0.958 | 0.005 | -0.013 | -0.013 | 0 | 0.63 | 0.39 | 0.43 | 0.32 | 0.56 | 0.54 |
| 2 | 203628757 | rs11685819 | 0.395 | 0.609 | -0.718 | -1.003 | -0.009 | 0.02 | 0.02 | 0.215 | 0.61 | 0.35 | 0.42 | 0.32 | 0.55 | 0.54 |
| 2 | 203631359 | rs11680951 | 0.392 | 0.602 | -0.722 | -1.007 | -0.009 | 0.02 | 0.02 | 0.336 | 0.61 | 0.35 | 0.42 | 0.32 | 0.55 | 0.54 |
| 2 | 203632510 | rs11544338 | 0.327 | 0.607 | -0.718 | -1.004 | -0.005 | 0.008 | 0.008 | 0.278 | 0.61 | 0.37 | 0.43 | 0.33 | 0.56 | 0.54 |
| 2 | 203635605 | rs6435166 | 0.431 | 1.498 | -0.94 | -0.898 | -0.016 | -0.022 | -0.022 | 0 | 0.79 | 0.79 | 0.64 | 0.39 | 0.7 | 0.62 |
| 2 | 203636012 | rs6713203 | 1.529 | 1.248 | -1.105 | -1.39 | 0.015 | 0.032 | 0.032 | 0 | 0.14 | 0.05 | 0.07 | 0.01 | 0.1 | 0.06 |
| 2 | 203640713 | rs2036927 | 0.846 | 2.05 | -0.511 | -1.038 | 0.037 | 0.005 | 0.005 | 0 | 0.8 | 0.66 | 0.75 | 0.84 | 0.75 | 0.84 |
| 2 | 203665782 | rs13429334 | 0.661 | 1.979 | -0.362 | -0.968 | 0.068 | 0.132 | 0.132 | 0.141 | 0.91 | 0.7 | 0.82 | 0.85 | 0.88 | 0.86 |
| 2 | 203689744 | rs4675295 | 1.094 | 2.038 | -0.521 | -1.03 | -0.018 | -0.046 | -0.046 | 0 | 0.09 | 0.01 | 0.18 | 0.51 | 0.13 | 0.13 |
| 2 | 203690387 | rs1344959 | 0.482 | 1.966 | -0.439 | -1.015 | 0.056 | 0.067 | 0.067 | 1.052 | 0.93 | 0.7 | 0.89 | 0.93 | 0.95 | 0.95 |
| 2 | 203691170 | rs12470845 | -0.536 | 2.215 | -0.342 | -1.034 | 0.018 | 0.054 | 0.054 | 0 | 0.18 | 0.28 | 0.35 | 0.51 | 0.26 | 0.22 |
| 2 | 203696607 | rs4675297 | -0.84 | 2.237 | -0.347 | -1.036 | -0.001 | -0.001 | -0.001 | 0.255 | 0.21 | 0.23 | 0.43 | 0.59 | 0.32 | 0.31 |
| 2 | 203700897 | rs10048682 | -0.419 | 2.214 | -0.355 | -1.057 | -0.004 | 0.002 | 0.002 | 0 | 0.18 | 0.18 | 0.34 | 0.51 | 0.25 | 0.22 |
| 2 | 203703612 | rs7579055 | 0.007 | 2.207 | -0.277 | -1.107 | -0.005 | -0.006 | -0.006 | 0 | 0.09 | 0.05 | 0.15 | 0 | 0.11 | 0.07 |
| 2 | 203708307 | rs934287 | 0.441 | 2.298 | -0.258 | -1.161 | 0.024 | -0.039 | -0.039 | 0 | 0.82 | 0.68 | 0.81 | 0.91 | 0.83 | 0.93 |
| 2 | 203730854 | rs4673239 | -0.787 | 2.358 | -0.309 | -1.131 | 0 | 0.001 | 0.001 | 0 | 0.2 | 0.2 | 0.41 | 0.58 | 0.32 | 0.31 |
| 2 | 203740672 | rs3845802 | 2.016 | 2.422 | -0.098 | -1.022 | -0.027 | 0.052 | 0.052 | 0.608 | 0.6 | 0.27 | 0.38 | 0.33 | 0.5 | 0.62 |
| 2 | 203750049 | rs10189499 | -1.078 | 2.561 | -0.22 | -1.062 | -0.006 | -0.005 | -0.005 | 0.397 | 0.23 | 0.24 | 0.43 | 0.58 | 0.36 | 0.32 |
| 2 | 203753072 | rs7582720 | 1.123 | 2.307 | -0.229 | -1.027 | -0.037 | 0.047 | 0.047 | 0 | 0.11 | 0.02 | 0.07 | 0.02 | 0.13 | 0.01 |
| 2 | 203765756 | rs35212307 | 1.185 | 2.307 | -0.229 | -1.027 | -0.037 | 0.046 | 0.046 | 0 | 0.1 | 0.01 | 0.07 | 0.02 | 0.13 | 0.01 |
| 2 | 203775711 | rs34048914 | 0.978 | 2.147 | -0.117 | -0.918 | -0.003 | 0.031 | 0.031 | 0 | 0.11 | 0.1 | 0.19 | 0.51 | 0.17 | 0.16 |
| 2 | 203805939 | rs10932004 | -0.692 | 2.107 | -0.043 | -0.923 | 0.01 | 0.012 | 0.012 | 0 | 0.2 | 0.25 | 0.4 | 0.59 | 0.29 | 0.31 |
| 2 | 203845301 | rs10205697 | -0.184 | 1.912 | -0.062 | -0.902 | -0.008 | -0.006 | -0.006 | 0 | 0.17 | 0.16 | 0.33 | 0.51 | 0.26 | 0.22 |
| 2 | 203845646 | rs10197426 | 0.461 | 1.861 | 0.001 | -0.922 | 0.052 | 0.058 | 0.058 | 0.488 | 0.93 | 0.72 | 0.89 | 0.93 | 0.95 | 0.94 |
| 2 | 203856457 | rs1864466 | 1.116 | 1.911 | -0.021 | -0.972 | 0.04 | 0.023 | 0.023 | 0.358 | 0.91 | 0.72 | 0.87 | 0.93 | 0.93 | 0.94 |
| 2 | 203880834 | rs4675310 | -1.149 | 1.866 | -0.017 | -0.918 | -0.017 | 0.03 | 0.03 | 0 | 0.89 | 0.95 | 0.92 | 0.98 | 0.87 | 0.98 |
| 2 | 203888858 | rs10187018 | -0.534 | 1.887 | 0.095 | -0.902 | 0.015 | 0.024 | 0.024 | 0.103 | 0.2 | 0.28 | 0.39 | 0.58 | 0.27 | 0.31 |
| 2 | 203905175 | rs6733394 | 0.431 | 1.822 | 0.119 | -0.886 | 0.051 | 0.057 | 0.057 | 0.465 | 0.93 | 0.72 | 0.89 | 0.93 | 0.95 | 0.94 |
| 2 | 204005877 | rs4297863 | 0.056 | 1.68 | -0.041 | -0.881 | 0.06 | 0.089 | 0.089 | 0.063 | 0.94 | 0.74 | 0.94 | 0.93 | 0.96 | 0.95 |
| 2 | 204025388 | rs10172342 | -0.478 | 1.77 | -0.115 | -0.888 | 0.008 | 0.029 | 0.029 | 0.016 | 0.19 | 0.23 | 0.34 | 0.51 | 0.26 | 0.22 |
| 2 | 204048840 | rs10932008 | 0.754 | 1.606 | -0.006 | -1.011 | 0.025 | -0.041 | -0.041 | 0 | 0.82 | 0.47 | 0.83 | 0.9 | 0.81 | 0.93 |
| 2 | 204051009 | rs10182274 | -0.637 | 1.62 | 0.065 | -0.949 | 0.004 | 0.009 | 0.009 | 0 | 0.21 | 0.24 | 0.41 | 0.58 | 0.28 | 0.31 |
| 2 | 204062061 | rs4675323 | 0.321 | 1.484 | 0.109 | -0.892 | 0.093 | 0.138 | 0.138 | 0.236 | 0.94 | 0.63 | 0.94 | 0.92 | 0.96 | 0.95 |
| 2 | 204101328 | rs6742900 | 0.461 | 1.532 | 0.136 | -0.933 | 0.084 | 0.116 | 0.116 | 0.202 | 0.94 | 0.66 | 0.94 | 0.93 | 0.96 | 0.95 |
| 2 | 204105996 | rs7584129 | 0.012 | 1.531 | 0.198 | -0.932 | 0.042 | 0.072 | 0.072 | 0 | 0.95 | 0.79 | 0.94 | 0.93 | 0.96 | 0.95 |
| 2 | 204108732 | rs6745954 | 0.314 | 1.52 | 0.197 | -0.938 | 0.034 | 0.064 | 0.064 | 0.254 | 0.94 | 0.78 | 0.94 | 0.93 | 0.96 | 0.95 |
| 2 | 204111443 | rs6435176 | 0.67 | 1.541 | 0.197 | -0.959 | 0.075 | 0.106 | 0.106 | 0.12 | 0.94 | 0.66 | 0.94 | 0.93 | 0.96 | 0.95 |
| 2 | 204114244 | rs4673251 | 0.49 | 1.611 | 0.217 | -0.939 | 0.09 | 0.116 | 0.116 | 0.202 | 0.94 | 0.66 | 0.94 | 0.93 | 0.96 | 0.95 |
| 2 | 204116690 | rs2043449 | 0.49 | 1.588 | 0.217 | -0.939 | 0.084 | 0.116 | 0.116 | 0.202 | 0.94 | 0.66 | 0.94 | 0.93 | 0.96 | 0.95 |
| 2 | 204116887 | rs2351771 | 0.49 | 1.588 | 0.217 | -0.929 | 0.084 | 0.122 | 0.122 | 0.202 | 0.94 | 0.66 | 0.94 | 0.93 | 0.96 | 0.95 |
| 2 | 204126463 | rs10174606 | -0.064 | 1.587 | 0.219 | -0.962 | 0.046 | 0.072 | 0.072 | 0 | 0.95 | 0.8 | 0.94 | 0.93 | 0.96 | 0.95 |
| 2 | 204130718 | rs6713579 | 0.048 | 1.588 | 0.219 | -0.962 | 0.084 | 0.116 | 0.116 | 0.202 | 0.94 | 0.66 | 0.94 | 0.93 | 0.96 | 0.95 |
| 2 | 204144460 | rs11690137 | 1.498 | 1.529 | -0.341 | -1.217 | 0.002 | 0.041 | 0.041 | 0 | 0.16 | 0.05 | 0.11 | 0.01 | 0.19 | 0.09 |
| 2 | 204147222 | rs12470569 | -1.673 | 2.313 | 0.672 | -0.288 | -0.004 | 0 | 0 | 0.203 | 0.31 | 0.33 | 0.47 | 0.61 | 0.4 | 0.32 |
| 2 | 204148987 | rs7608286 | 0.673 | 2.496 | 0.931 | -0.106 | 0.084 | 0.116 | 0.116 | 0.202 | 0.94 | 0.66 | 0.94 | 0.93 | 0.96 | 0.95 |
| 2 | 204153646 | rs4675328 | 0.207 | 2.497 | 0.931 | -0.105 | 0.111 | 0.145 | 0.145 | 0.041 | 0.95 | 0.66 | 0.94 | 0.93 | 0.96 | 0.95 |
| 2 | 204165171 | rs11888559 | 1.797 | 2.481 | 0.941 | -0.089 | 0.008 | 0.118 | 0.118 | 1.812 | 0.91 | 0.43 | 0.87 | 0.84 | 0.95 | 0.87 |
| 2 | 204169264 | rs6435181 | 0.19 | 2.501 | 0.761 | -0.074 | 0.109 | 0.143 | 0.143 | 0.041 | 0.95 | 0.67 | 0.94 | 0.93 | 0.96 | 0.95 |
| 2 | 204169274 | rs6435182 | 0.241 | 2.499 | 0.76 | -0.075 | 0.106 | 0.14 | 0.14 | 0.102 | 0.95 | 0.67 | 0.94 | 0.93 | 0.96 | 0.95 |
| 2 | 204181332 | rs6732652 | 0.597 | 2.606 | 0.673 | -0.055 | 0.056 | 0.197 | 0.197 | 0.22 | 0.85 | 0.52 | 0.75 | 0.43 | 0.83 | 0.8 |
| 2 | 204190907 | rs4675332 | -0.874 | 2.713 | 0.587 | -0.134 | -0.027 | 0.028 | 0.028 | 0 | 0.17 | 0.11 | 0.23 | 0.02 | 0.25 | 0.09 |
| 2 | 204192201 | rs11682759 | 0.64 | 2.659 | 0.636 | -0.143 | 0.037 | 0.187 | 0.187 | 0.456 | 0.84 | 0.52 | 0.75 | 0.43 | 0.84 | 0.8 |
| 2 | 204226331 | rs12466364 | -1.022 | 2.678 | 0.137 | -0.093 | -0.023 | 0.02 | 0.02 | 0 | 0.18 | 0.12 | 0.24 | 0.02 | 0.26 | 0.09 |
| 2 | 204238402 | rs10172421 | 0.297 | 2.549 | 0.13 | -0.057 | -0.009 | 0.002 | 0.002 | 0 | 0.07 | 0.05 | 0.15 | 0 | 0.11 | 0.07 |
| 2 | 204292517 | rs2469954 | 0.991 | 2.288 | 0.102 | 0.156 | -0.021 | -0.019 | -0.019 | 0 | 0.72 | 0.74 | 0.57 | 0.49 | 0.6 | 0.76 |
| 2 | 204301296 | rs2250522 | 0.955 | 2.312 | 0.012 | 0.144 | -0.018 | -0.018 | -0.018 | 0 | 0.72 | 0.74 | 0.57 | 0.5 | 0.61 | 0.76 |
| 2 | 204309325 | rs2246849 | 1.006 | 2.23 | -0.043 | 0.016 | -0.024 | 0.007 | 0.007 | 0.009 | 0.67 | 0.71 | 0.52 | 0.4 | 0.59 | 0.67 |

| | | | | | | | | | | | | | | | | |
|---|---|---|---|---|---|---|---|---|---|---|---|---|---|---|---|---|
| 2 | 204315394 | rs11679740 | 1.515 | 2.09 | 0.159 | -0.075 | -0.069 | 0.096 | 0.096 | 0.981 | 0.59 | 0.13 | 0.44 | 0.33 | 0.55 | 0.62 |
| 2 | 204322298 | rs2469953 | 0.959 | 2.084 | 0.151 | -0.102 | -0.023 | 0.006 | 0.006 | 0.007 | 0.67 | 0.73 | 0.52 | 0.41 | 0.59 | 0.68 |
| 2 | 204360201 | rs2305417 | 1.605 | 2.034 | 0.652 | -0.12 | -0.078 | 0.109 | 0.109 | 2.533 | 0.58 | 0.09 | 0.43 | 0.33 | 0.56 | 0.63 |
| 2 | 204366776 | rs16839858 | 0.716 | 2.324 | 0.579 | -0.05 | -0.044 | -0.054 | -0.054 | 0.169 | 0.1 | 0.02 | 0.2 | 0.49 | 0.17 | 0.13 |
| 2 | 204399335 | rs2469961 | -1.07 | 1.97 | 0.515 | -0.018 | 0.061 | 0.091 | 0.091 | 1.545 | 0.07 | 0.49 | 0.07 | 0.07 | 0.03 | 0.05 |
| 2 | 204402793 | rs10176600 | 0.077 | 2.107 | 0.465 | -0.026 | -0.01 | -0.008 | -0.008 | 0 | 0.1 | 0.05 | 0.11 | 0.1 | 0.14 | 0.1 |
| 2 | 204403952 | rs2245613 | -0.571 | 2.074 | 0.524 | -0.027 | 0.093 | 0.125 | 0.125 | 0.159 | 0.05 | 0.34 | 0.06 | 0.07 | 0.03 | 0.05 |
| 2 | 204425613 | rs2443796 | -0.256 | 1.989 | -0.159 | 0.079 | -0.007 | -0.014 | -0.014 | 0 | 0.77 | 0.74 | 0.64 | 0.51 | 0.71 | 0.79 |
| 2 | 204427340 | rs10174652 | -0.056 | 2.059 | -0.249 | 0.078 | -0.012 | -0.008 | -0.008 | 0 | 0.11 | 0.05 | 0.09 | 0.1 | 0.13 | 0.12 |
| 2 | 204432604 | rs6730171 | -0.011 | 2.053 | -0.287 | 0.082 | 0.002 | 0.012 | 0.012 | 0 | 0.24 | 0.33 | 0.37 | 0.49 | 0.29 | 0.21 |
| 2 | 204436759 | rs7557216 | -1.691 | 2.151 | -0.532 | -0.081 | 0.095 | 0.115 | 0.115 | 0 | 0.53 | 0.81 | 0.66 | 0.65 | 0.65 | 0.5 |
| 2 | 204440428 | rs12989194 | -1.443 | 2.165 | 0.035 | -0.081 | 0.029 | -0.027 | -0.027 | 0 | 0.29 | 0.53 | 0.29 | 0.16 | 0.37 | 0.29 |
| 2 | 204440491 | rs13013366 | -0.53 | 1.781 | 0.191 | -0.122 | 0.092 | -0.075 | -0.075 | 0 | 0.17 | 0.42 | 0.19 | 0.06 | 0.23 | 0.17 |
| 2 | 204449103 | rs13422031 | -0.281 | 1.279 | -0.168 | 0.079 | -0.002 | -0.01 | -0.01 | 0 | 0.12 | 0.14 | 0.1 | 0.09 | 0.09 | 0.27 |
| 2 | 204454414 | rs17419055 | -0.297 | 1.825 | -0.345 | -0.221 | -0.021 | 0.024 | 0.024 | 0 | 0.11 | 0.08 | 0.11 | 0.01 | 0.14 | 0.03 |
| 2 | 204459206 | rs4675350 | -1.17 | 1.791 | 0.054 | -0.234 | 0.033 | 0.079 | 0.079 | 0 | 0.55 | 0.76 | 0.64 | 0.75 | 0.58 | 0.56 |
| 2 | 204465119 | rs1879878 | -1.187 | 1.581 | 0.318 | -0.252 | 0.027 | 0.057 | 0.057 | 0 | 0.53 | 0.64 | 0.62 | 0.76 | 0.57 | 0.54 |
| 2 | 204466812 | rs4675351 | 0.284 | 1.471 | 0.428 | -0.215 | 0.038 | -0.009 | -0.009 | 0 | 0.22 | 0.34 | 0.31 | 0.19 | 0.29 | 0.14 |
| 2 | 204471473 | rs4675354 | -0.925 | 1.457 | 0.558 | -0.142 | -0.001 | 0.006 | 0.006 | 0 | 0.33 | 0.45 | 0.31 | 0.57 | 0.28 | 0.4 |
| 2 | 204477552 | rs7583807 | 0.765 | 1.4 | 0.294 | -0.171 | -0.007 | 0.003 | 0.003 | 0 | 0.66 | 0.46 | 0.68 | 0.41 | 0.73 | 0.61 |
| 2 | 204512294 | rs7370984 | 0.588 | 1.354 | 0.222 | -0.094 | -0.001 | 0 | 0 | 0.005 | 0.59 | 0.61 | 0.46 | 0.68 | 0.48 | 0.57 |
| 2 | 204513610 | rs3925039 | 0.507 | 1.543 | 0.015 | -0.011 | -0.012 | 0.015 | 0.015 | 0 | 0.13 | 0.07 | 0.15 | 0.08 | 0.18 | 0.05 |
| 2 | 204525230 | rs16839991 | -0.221 | 0.934 | 0.112 | -0.24 | 0.004 | 0.113 | 0.113 | 0 | 0.19 | 0.32 | 0.24 | 0.48 | 0.19 | 0.19 |
| 2 | 204529949 | rs3115961 | -0.415 | 0.45 | -0.04 | -0.154 | -0.008 | 0.07 | 0.07 | 0 | 0.24 | 0.39 | 0.25 | 0.43 | 0.23 | 0.19 |
| 2 | 204537170 | rs3115968 | -0.564 | 0.008 | 0.1 | -0.22 | -0.049 | 0.113 | 0.113 | 0.136 | 0.19 | 0.35 | 0.14 | 0.48 | 0.1 | 0.11 |
| 2 | 204539973 | rs4675357 | -1.97 | 0.399 | 0.009 | -0.33 | 0.009 | -0.026 | -0.026 | 0 | 0.95 | 0.94 | 0.95 | 1 | 0.94 | 0.98 |
| 2 | 204545178 | rs16840016 | 0.853 | 0.217 | 0.031 | -0.416 | -0.009 | -0.012 | -0.012 | 0.01 | 0.14 | 0.19 | 0.14 | 0.11 | 0.12 | 0.29 |
| 2 | 204570000 | rs1879877 | -0.714 | 0.428 | 0.219 | -0.064 | -0.004 | 0.005 | 0.005 | 0 | 0.28 | 0.33 | 0.28 | 0.56 | 0.25 | 0.14 |
| 2 | 204570092 | rs3181096 | 1.639 | 0.444 | 0.136 | -0.339 | 0.008 | 0.041 | 0.041 | 0.191 | 0.4 | 0.3 | 0.31 | 0.28 | 0.36 | 0.64 |
| 2 | 204570139 | rs3181097 | -0.191 | 0.44 | 0.247 | -0.113 | 0.013 | 0.375 | 0.375 | 0.733 | 0.06 | 0.22 | 0.12 | 0.46 | 0.03 | 0.09 |
| 2 | 204575951 | rs1181388 | 0.601 | 0.435 | 0.259 | -0.282 | -0.024 | 0.07 | 0.07 | 0.191 | 0.79 | 0.68 | 0.8 | 0.48 | 0.86 | 0.9 |
| 2 | 204577905 | rs10932017 | 0.985 | 0.577 | 0.015 | -0.453 | -0.004 | 0.016 | 0.016 | 0.463 | 0.56 | 0.38 | 0.47 | 0.35 | 0.55 | 0.69 |
| 2 | 204579600 | rs7425641 | -0.351 | 0.02 | -0.091 | 0.025 | -0.002 | -0.021 | -0.021 | 0 | 0.91 | 0.88 | 0.92 | 0.99 | 0.92 | 0.97 |
| 2 | 204583163 | rs10490573 | 0.253 | 0.031 | -0.276 | 0.028 | -0.016 | 0.051 | 0.051 | 0 | 0.16 | 0.07 | 0.17 | 0.06 | 0.2 | 0.04 |
| 2 | 204587269 | rs10515944 | 0.003 | -0.072 | -0.219 | 0.057 | 0 | 0.011 | 0.011 | 0 | 0.17 | 0.13 | 0.17 | 0.06 | 0.19 | 0.05 |
| 2 | 204590071 | rs4675363 | 0.582 | -0.111 | -0.318 | 0.019 | 0.004 | 0.041 | 0.041 | 0 | 0.76 | 0.83 | 0.75 | 0.93 | 0.74 | 0.93 |
| 2 | 204593862 | rs3181108 | -0.711 | -0.081 | -0.316 | 0.022 | -0.003 | -0.019 | -0.019 | 0.034 | 0.92 | 0.89 | 0.92 | 0.99 | 0.92 | 0.97 |
| 2 | 204594512 | rs3116496 | -0.41 | -0.035 | -0.382 | 0.117 | -0.003 | 0.067 | 0.067 | 0.229 | 0.22 | 0.04 | 0.11 | 0.09 | 0.21 | 0.05 |
| 2 | 204633156 | rs10932020 | 1.067 | -0.123 | 0.202 | 0.778 | -0.006 | 0.014 | 0.014 | 0 | 0.67 | 0.6 | 0.75 | 0.5 | 0.72 | 0.87 |
| 2 | 204634767 | rs231390 | 1.324 | -0.591 | -0.059 | 0.558 | -0.006 | -0.002 | -0.002 | 0 | 0.54 | 0.52 | 0.64 | 0.35 | 0.65 | 0.67 |
| 2 | 204639764 | rs4673268 | -0.372 | -0.189 | 0.518 | 1.076 | 0.01 | 0.193 | 0.193 | 0 | 0.1 | 0.23 | 0.13 | 0.4 | 0.08 | 0.08 |
| 2 | 204643194 | rs4675369 | -0.575 | -0.176 | 0.495 | 1.078 | -0.017 | 0.107 | 0.107 | 0 | 0.16 | 0.28 | 0.14 | 0.44 | 0.1 | 0.09 |
| 2 | 204648363 | rs17187469 | -0.02 | -0.201 | 0.116 | 1.104 | 0.002 | 0.023 | 0.023 | 0 | 0.17 | 0.1 | 0.1 | 0.07 | 0.18 | 0.05 |
| 2 | 204661197 | rs10932022 | 0.754 | 0.206 | -0.528 | 1.082 | -0.044 | -0.032 | -0.032 | 0 | 0.07 | 0.01 | 0.3 | 0.21 | 0.2 | 0.16 |
| 2 | 204676238 | rs7591187 | -1.273 | -0.959 | 0.061 | 0.958 | -0.01 | 0.011 | 0.011 | 0.109 | 0.32 | 0.58 | 0.25 | 0.51 | 0.22 | 0.13 |
| 2 | 204684835 | rs1018361 | -0.79 | -0.565 | -0.052 | 1.086 | -0.007 | 0.022 | 0.022 | 0 | 0.24 | 0.28 | 0.18 | 0.51 | 0.16 | 0.13 |
| 2 | 204688432 | rs231747 | 0.229 | -0.694 | 0.321 | 1.106 | -0.031 | 0.211 | 0.211 | 0 | 0.86 | 0.59 | 0.87 | 0.6 | 0.94 | 0.9 |
| 2 | 204694263 | rs13030124 | 1.317 | -0.597 | 0.156 | 0.4 | 0.004 | 0.125 | 0.125 | 0 | 0.51 | 0.28 | 0.39 | 0.27 | 0.45 | 0.62 |
| 2 | 204694351 | rs11571308 | 0.229 | -0.604 | 0.388 | 0.416 | -0.003 | -0.002 | -0.002 | 0 | 0.1 | 0.12 | 0.09 | 0.08 | 0.11 | 0.05 |
| 2 | 204694611 | rs3116513 | -0.759 | -0.621 | 0.575 | 0.374 | 0.017 | -0.003 | -0.003 | 0 | 0.31 | 0.42 | 0.47 | 0.65 | 0.38 | 0.3 |
| 2 | 204694805 | rs3115969 | -1.393 | -0.599 | 0.461 | 0.328 | -0.002 | -0.002 | -0.002 | 0 | 0.41 | 0.61 | 0.56 | 0.73 | 0.48 | 0.36 |
| 2 | 204711588 | rs11903660 | 0.915 | -0.603 | 0.348 | 0.367 | -0.002 | -0.025 | -0.025 | 4.864 | 0.07 | 0.07 | 0.05 | 0 | 0.06 | 0.02 |
| 2 | 204718729 | rs2162610 | 0.654 | -0.672 | 0.558 | 0.446 | -0.006 | -0.01 | -0.01 | 0.999 | 0.81 | 0.78 | 0.85 | 0.9 | 0.81 | 0.93 |
| 2 | 204721132 | rs11571291 | 1.879 | -0.804 | 0.571 | 0.336 | 0.009 | 0.11 | 0.11 | 0 | 0.5 | 0.31 | 0.37 | 0.27 | 0.45 | 0.63 |
| 2 | 204730901 | rs11571315 | 0.754 | -0.796 | 0.49 | 0.348 | 0.021 | -0.005 | -0.005 | 0.969 | 0.67 | 0.47 | 0.51 | 0.36 | 0.61 | 0.68 |
| 2 | 204730944 | rs733618 | -0.258 | -0.78 | 0.498 | 0.644 | -0.012 | 0.055 | 0.055 | 0.004 | 0.11 | 0.14 | 0.08 | 0.39 | 0.06 | 0.13 |
| 2 | 204732714 | rs231775 | -0.602 | -0.836 | -0.142 | 0.087 | 0.009 | 0.001 | 0.001 | 0 | 0.3 | 0.39 | 0.46 | 0.64 | 0.36 | 0.31 |
| 2 | 204738919 | rs3087243 | 1.438 | -0.786 | -0.086 | 0.094 | -0.017 | 0.165 | 0.165 | 0 | 0.52 | 0.18 | 0.38 | 0.26 | 0.47 | 0.63 |
| 2 | 204740866 | rs231726 | 0.032 | -0.737 | 0.027 | 0.308 | 0.039 | 0.038 | 0.038 | 0 | 0.23 | 0.38 | 0.43 | 0.59 | 0.32 | 0.27 |
| 2 | 204746767 | rs11571300 | 1.019 | -0.743 | -0.75 | 0.22 | -0.04 | -0.004 | -0.004 | 0.144 | 0.11 | 0 | 0.13 | 0.15 | 0.15 | 0.18 |
| 2 | 204756905 | rs6748358 | 1.09 | -0.639 | -0.581 | 0.301 | -0.031 | 0.107 | 0.107 | 0 | 0.57 | 0.21 | 0.43 | 0.31 | 0.51 | 0.71 |
| 2 | 204762327 | rs10497873 | 0.47 | -1.176 | -0.98 | -0.037 | 0.024 | 0.062 | 0.062 | 0 | 0.26 | 0.08 | 0.19 | 0.2 | 0.21 | 0.27 |
| 2 | 204763044 | rs10197319 | 0.514 | -1.141 | -0.941 | -0.029 | 0.005 | 0.046 | 0.046 | 0 | 0.51 | 0.42 | 0.36 | 0.21 | 0.49 | 0.52 |
| 2 | 204765194 | rs7607760 | 0.563 | -1.125 | -1.096 | -0.009 | 0.011 | 0.099 | 0.099 | 0 | 0.51 | 0.33 | 0.36 | 0.21 | 0.49 | 0.52 |
| 2 | 204773679 | rs11571311 | 1.709 | -0.946 | -1.471 | 0.029 | -0.041 | 0.119 | 0.119 | 0 | 0.3 | 0.1 | 0.23 | 0.11 | 0.3 | 0.44 |
| 2 | 204788156 | rs2033171 | -1.356 | -0.8 | -0.661 | 0.738 | -0.017 | 0.074 | 0.074 | 0 | 0.42 | 0.72 | 0.56 | 0.67 | 0.47 | 0.28 |
| 2 | 204794952 | rs3096866 | -0.168 | -0.716 | -0.387 | 0.893 | 0.061 | 0.066 | 0.066 | 0 | 0.23 | 0.4 | 0.44 | 0.56 | 0.31 | 0.21 |
| 2 | 204797790 | rs4325730 | 1.301 | -2.279 | -0.12 | 0.782 | 0.046 | -0.029 | -0.029 | 0 | 0.65 | 0.27 | 0.78 | 0.6 | 0.78 | 0.82 |
| 2 | 204799697 | rs4452124 | 0.291 | -2.308 | -0.079 | 0.7 | -0.042 | 0.022 | 0.022 | 2.38 | 0.79 | 0.35 | 0.87 | 0.72 | 0.92 | 0.88 |

| | | | | | | | | | | | | | | | | |
|---|---|---|---|---|---|---|---|---|---|---|---|---|---|---|---|---|
| 2 | 204800845 | rs11883722 | -0.991 | -2.127 | -0.626 | 0.793 | -0.062 | 0.044 | 0.044 | 0 | 0.31 | 0.51 | 0.2 | 0.39 | 0.19 | 0.18 |
| 2 | 204801980 | rs4335928 | -0.035 | -1.615 | -1.644 | 0.447 | 0.004 | -0.034 | -0.034 | 0 | 0.12 | 0.01 | 0.08 | 0.11 | 0.11 | 0.05 |
| 2 | 204812993 | rs10932034 | 0.318 | -1.324 | -1.595 | 0.53 | 0.137 | 0.064 | 0.064 | 0.411 | 0.57 | 0.24 | 0.77 | 0.71 | 0.78 | 0.82 |
| 2 | 204816382 | rs4355090 | -0.52 | -0.684 | -1.25 | 0.747 | 0.007 | -0.007 | -0.007 | 0.612 | 0.38 | 0.33 | 0.35 | 0.49 | 0.33 | 0.24 |
| 2 | 204816575 | rs4521021 | -0.634 | 0.832 | -0.21 | 0.192 | 0.177 | -0.059 | -0.059 | 0 | 0.1 | 0.41 | 0.19 | 0 | 0.22 | 0.06 |
| 2 | 204817281 | rs10166287 | -0.134 | -0.186 | -0.814 | -0.244 | 0.017 | 0.019 | 0.019 | 0 | 0.22 | 0.18 | 0.16 | 0.18 | 0.13 | 0.32 |
| 2 | 204820058 | rs11571312 | -0.261 | -0.363 | -0.992 | -0.368 | 0.056 | 0.083 | 0.083 | 0.043 | 0.23 | 0.08 | 0.15 | 0.16 | 0.13 | 0.29 |
| 2 | 204824283 | rs10172036 | -0.99 | -0.054 | -1.33 | -0.351 | -0.005 | -0.002 | -0.002 | 0.223 | 0.6 | 0.48 | 0.54 | 0.45 | 0.62 | 0.49 |
| 2 | 204824324 | rs10183087 | -0.343 | 0.415 | -1.369 | -0.424 | -0.011 | -0.058 | -0.058 | 0 | 0.32 | 0.42 | 0.26 | 0.21 | 0.24 | 0.37 |
| 2 | 204825286 | rs4404254 | -0.36 | 0.4 | -1.37 | -0.432 | -0.011 | -0.056 | -0.056 | 0 | 0.32 | 0.42 | 0.26 | 0.21 | 0.24 | 0.37 |
| 2 | 204825346 | rs10932037 | 0.507 | 0.401 | -1.339 | -0.442 | -0.007 | 0.005 | 0.005 | 0 | 0.06 | 0.05 | 0.08 | 0.04 | 0.1 | 0.05 |
| 2 | 204825584 | rs10932038 | 0.507 | 0.401 | -1.339 | -0.442 | -0.007 | 0.005 | 0.005 | 0 | 0.06 | 0.05 | 0.08 | 0.04 | 0.1 | 0.05 |
| 2 | 204825729 | rs1559931 | -0.357 | 0.403 | -1.316 | -0.437 | -0.009 | -0.048 | -0.048 | 0 | 0.32 | 0.41 | 0.26 | 0.21 | 0.24 | 0.37 |
| 2 | 204832438 | rs1896286 | 0.37 | -0.132 | -1.575 | -0.1 | 0.008 | 0.053 | 0.053 | 0.235 | 0.62 | 0.17 | 0.48 | 0.63 | 0.64 | 0.46 |
| 2 | 204833769 | rs3116534 | 1.226 | 0.086 | -1.407 | 0.091 | -0.032 | -0.106 | -0.106 | 0.02 | 0.31 | 0.06 | 0.27 | 0.33 | 0.31 | 0.32 |
| 2 | 204834613 | rs3116536 | -0.089 | 0.253 | -0.679 | 0.231 | 0.039 | 0.014 | 0.014 | 0 | 0.83 | 0.42 | 0.65 | 0.79 | 0.78 | 0.79 |
| 2 | 204842468 | rs4675391 | 0.228 | -0.257 | -1.201 | 0.072 | 0.009 | 0.007 | 0.007 | 0.139 | 0.41 | 0.38 | 0.31 | 0.4 | 0.32 | 0.38 |
| 2 | 204846228 | rs12613960 | -0.399 | -0.082 | -0.572 | 0.144 | 0.048 | -0.008 | -0.008 | 0 | 0.76 | 0.59 | 0.69 | 0.84 | 0.69 | 0.8 |
| 2 | 204848818 | rs13003194 | -0.996 | -0.323 | -0.144 | 0.181 | -0.02 | 0.019 | 0.019 | 0 | 0.84 | 0.77 | 0.88 | 0.85 | 0.89 | 0.82 |
| 2 | 204853697 | rs10932040 | 1.877 | -0.521 | -0.747 | 0.029 | 0.019 | 0.024 | 0.024 | 0.496 | 0.26 | 0.08 | 0.13 | 0.19 | 0.21 | 0.09 |
| 2 | 204858820 | rs6754077 | -0.058 | -0.751 | -0.678 | 0.032 | 0.067 | 0.034 | 0.034 | 0 | 0.45 | 0.29 | 0.28 | 0.38 | 0.32 | 0.28 |
| 2 | 204865802 | rs13425717 | -0.101 | -0.665 | -0.725 | -0.008 | 0.024 | 0.014 | 0.014 | 0.023 | 0.46 | 0.39 | 0.3 | 0.38 | 0.32 | 0.28 |
| 2 | 204872034 | rs3096735 | -0.195 | -0.695 | -0.842 | -0.003 | 0.058 | 0.056 | 0.056 | 0 | 0.5 | 0.25 | 0.33 | 0.37 | 0.35 | 0.32 |
| 2 | 204875234 | rs4675397 | -0.421 | -0.934 | -0.71 | -0.09 | -0.037 | -0.004 | -0.004 | 0 | 0.2 | 0.3 | 0.17 | 0.2 | 0.11 | 0.19 |
| 2 | 204884340 | rs3096734 | 1.508 | -0.976 | -0.337 | 0.279 | -0.018 | 0.022 | 0.022 | 0 | 0.17 | 0.26 | 0.19 | 0.19 | 0.11 | 0.18 |
| 2 | 204886384 | rs2353319 | -1.143 | -1.086 | -0.589 | 0.397 | -0.001 | -0.003 | -0.003 | 0 | 0.6 | 0.61 | 0.5 | 0.57 | 0.44 | 0.61 |
| 2 | 204892565 | rs6717395 | 1.251 | -0.949 | -0.667 | 0.444 | -0.01 | 0.054 | 0.054 | 0 | 0.09 | 0.16 | 0.13 | 0.2 | 0.06 | 0.16 |
| 2 | 204916206 | rs11688848 | 2.113 | -1.202 | -1.339 | 0.324 | -0.031 | 0.003 | 0.003 | 0 | 0.2 | 0.03 | 0.12 | 0.14 | 0.2 | 0.06 |
| 2 | 204921552 | rs1344638 | -3.362 | -0.415 | -1.119 | 0.564 | -0.062 | -0.074 | -0.074 | 0 | 0.83 | 0.95 | 0.82 | 0.74 | 0.74 | 0.83 |
| 2 | 204924187 | rs7575195 | 0.889 | -0.73 | -0.508 | 0.803 | -0.021 | -0.008 | -0.008 | 0 | 0.11 | 0.32 | 0.11 | 0.04 | 0.05 | 0.03 |
| 2 | 204930722 | rs1863691 | 1.786 | -0.723 | -1.182 | 0.791 | -0.02 | 0.084 | 0.084 | 0.087 | 0.29 | 0.03 | 0.14 | 0.12 | 0.24 | 0.1 |
| 2 | 204933578 | rs1896284 | 0.836 | -0.664 | -1.053 | 0.835 | -0.003 | -0.017 | -0.017 | 0.393 | 0.12 | 0.16 | 0.05 | 0.02 | 0.08 | 0.08 |
| 2 | 204957945 | rs4675420 | -1.41 | -0.65 | -1.083 | 0.448 | -0.007 | -0.015 | -0.015 | 0 | 0.61 | 0.68 | 0.51 | 0.55 | 0.43 | 0.61 |
| 2 | 204967728 | rs4673282 | 1.073 | -0.463 | -1.368 | 0.444 | 0.003 | -0.005 | -0.005 | 0.551 | 0.08 | 0.05 | 0.05 | 0.18 | 0.05 | 0.16 |
| 2 | 204969138 | rs11685977 | 1.363 | -0.347 | -1.484 | 0.118 | -0.016 | 0.081 | 0.081 | 0.167 | 0.29 | 0.04 | 0.14 | 0.13 | 0.22 | 0.09 |
| 2 | 204977614 | rs13017295 | 1.688 | 0.175 | -1.4 | 0.074 | -0.012 | 0.012 | 0.012 | 0 | 0.08 | 0.06 | 0.13 | 0 | 0.25 | 0.08 |
| 2 | 204979611 | rs12478622 | 1.502 | -0.155 | -1.424 | 0.3 | 0.007 | -0.016 | -0.016 | 0.494 | 0.1 | 0.08 | 0.16 | 0.23 | 0.07 | 0.26 |
| 2 | 204979663 | rs13430376 | 1.258 | -0.339 | -1.461 | 0.074 | 0.006 | -0.005 | -0.005 | 0 | 0.09 | 0.07 | 0.03 | 0 | 0.05 | 0.03 |
| 2 | 204981173 | rs12619716 | 1.541 | -0.356 | -1.774 | 0.052 | -0.018 | 0.076 | 0.076 | 0.159 | 0.29 | 0.03 | 0.14 | 0.13 | 0.21 | 0.09 |
| 2 | 204986334 | rs11673985 | 2.446 | 0.189 | -1.483 | 0.234 | 0.01 | 0.001 | 0.001 | 0 | 0.06 | 0.05 | 0.12 | 0 | 0.2 | 0.05 |
| 2 | 204989546 | rs1344637 | -0.441 | -0.463 | -1.252 | 0.516 | 0.022 | 0.005 | 0.005 | 0 | 0.53 | 0.62 | 0.45 | 0.47 | 0.38 | 0.44 |
| 2 | 204991426 | rs6726769 | 0.41 | -0.722 | -1.549 | 0.631 | 0.01 | -0.031 | -0.031 | 0.799 | 0.13 | 0.05 | 0.18 | 0.24 | 0.08 | 0.3 |
| 2 | 204999473 | rs933984 | 0.549 | -0.618 | -1.158 | 0.471 | -0.005 | 0.017 | 0.017 | 0 | 0.08 | 0.11 | 0.07 | 0.12 | 0.07 | 0.16 |
| 2 | 204999862 | rs6747093 | 0.053 | -0.599 | -1.364 | 0.201 | -0.001 | -0.006 | -0.006 | 0 | 0.29 | 0.3 | 0.32 | 0.13 | 0.32 | 0.35 |
| 2 | 205000176 | rs6747746 | 1.394 | -0.731 | -1.483 | 0.18 | 0.003 | -0.014 | -0.014 | 0 | 0.14 | 0.02 | 0.15 | 0.13 | 0.09 | 0.23 |
| 2 | 205002784 | rs6738139 | 0.304 | -1.027 | -1.73 | 0.244 | 0.017 | -0.011 | -0.011 | 0.515 | 0.37 | 0.31 | 0.21 | 0.43 | 0.26 | 0.31 |
| 2 | 205004122 | rs10932047 | 1.869 | -0.86 | -2.072 | 0.364 | 0.039 | -0.062 | -0.062 | 0 | 0.23 | 0.05 | 0.12 | 0.33 | 0.17 | 0.08 |
| 2 | 205008562 | rs219221 | 0.844 | -0.827 | -2.011 | 0.009 | 0.004 | 0 | 0 | 0 | 0.33 | 0.35 | 0.41 | 0.38 | 0.39 | 0.33 |
| 2 | 205009691 | rs10932048 | 1.615 | -0.71 | -2.549 | 0.185 | 0.033 | -0.082 | -0.082 | 0.158 | 0.23 | 0.05 | 0.18 | 0.37 | 0.18 | 0.08 |
| 2 | 205011169 | rs219224 | -1.257 | -0.582 | -2.496 | 0.159 | -0.006 | -0.017 | -0.017 | 0 | 0.86 | 0.98 | 0.85 | 0.87 | 0.91 | 0.77 |
| 2 | 205014049 | rs219226 | -0.635 | -0.553 | -2.413 | 0.127 | 0.064 | -0.056 | -0.056 | 0 | 0.63 | 0.79 | 0.68 | 0.52 | 0.76 | 0.69 |
| 2 | 205016921 | rs17765750 | 1.599 | 0.188 | -2.625 | 0.255 | -0.089 | -0.04 | -0.04 | 0 | 0.17 | 0.01 | 0.18 | 0.17 | 0.28 | 0.13 |
| 2 | 205024829 | rs11892294 | 0.338 | -0.066 | -1.74 | 0.472 | 0.011 | -0.019 | -0.019 | 0.105 | 0.23 | 0.19 | 0.18 | 0.36 | 0.2 | 0.18 |
| 2 | 205027279 | rs2042580 | -0.018 | -0.183 | -1.818 | 0.013 | 0.011 | 0.025 | 0.025 | 1.237 | 0.8 | 0.89 | 0.85 | 0.88 | 0.85 | 0.64 |
| 2 | 205030241 | rs16840855 | 1.112 | -0.162 | -1.796 | 0.163 | 0.044 | -0.072 | -0.072 | 0.207 | 0.17 | 0.05 | 0.14 | 0.37 | 0.12 | 0.13 |
| 2 | 205031069 | rs6730424 | -0.205 | 1.253 | -1.036 | -0.245 | -0.007 | -0.016 | -0.016 | 0 | 0.19 | 0.16 | 0.16 | 0 | 0.25 | 0.15 |
| 2 | 205032057 | rs4419220 | 0.754 | 0.595 | -0.755 | 0.651 | -0.016 | -0.041 | -0.041 | 0 | 0.8 | 0.97 | 0.82 | 0.77 | 0.83 | 0.59 |
| 2 | 205041492 | rs7579680 | 0.831 | 0.585 | -0.33 | 0.427 | -0.01 | 0.018 | 0.018 | 0.096 | 0.8 | 0.9 | 0.82 | 0.87 | 0.84 | 0.68 |
| 2 | 205043409 | rs12617227 | 1.492 | 0.534 | 0.3 | 0.462 | -0.011 | -0.006 | -0.006 | 0.153 | 0.61 | 0.52 | 0.62 | 0.63 | 0.71 | 0.5 |
| 2 | 205043436 | rs10932049 | 0.482 | 0.604 | -0.123 | 0.467 | -0.014 | -0.009 | -0.009 | 0 | 0.83 | 0.97 | 0.83 | 0.87 | 0.88 | 0.73 |
| 2 | 205053197 | rs16840880 | -0.338 | 0.714 | 0.274 | 0.703 | -0.006 | 0.007 | 0.007 | 0.16 | 0.15 | 0.23 | 0.19 | 0.2 | 0.12 | 0.13 |
| 2 | 205054041 | rs7607984 | 1.526 | 0.836 | 0.536 | 0.647 | 0.001 | -0.003 | -0.003 | 0 | 0.65 | 0.47 | 0.62 | 0.66 | 0.71 | 0.6 |
| 2 | 205054059 | rs2194766 | 0.693 | 1.004 | 0.398 | 0.716 | 0.02 | -0.01 | -0.01 | 0 | 0.81 | 0.7 | 0.81 | 0.86 | 0.83 | 0.72 |
| 2 | 205057607 | rs13413530 | -0.697 | 0.579 | 0.255 | 0.699 | 0.019 | 0.041 | 0.041 | 0.224 | 0.16 | 0.34 | 0.19 | 0.23 | 0.14 | 0.14 |
| 2 | 205057668 | rs13401647 | -0.625 | 0.516 | 0.269 | 0.606 | 0.01 | 0.033 | 0.033 | 0.214 | 0.15 | 0.35 | 0.19 | 0.2 | 0.13 | 0.13 |
| 2 | 205062633 | rs6714929 | -0.895 | 0.272 | -0.2 | 0.45 | -0.048 | -0.03 | -0.03 | 0 | 0.25 | 0.46 | 0.13 | 0.18 | 0.17 | 0.28 |
| 2 | 205067509 | rs35321733 | -1.171 | 1.233 | -0.83 | 0.388 | -0.019 | 0.019 | 0.019 | 0 | 0.16 | 0.09 | 0.15 | 0.07 | 0.2 | 0.15 |
| 2 | 205072783 | rs10208393 | -0.793 | 0.634 | -0.566 | 0.826 | -0.009 | 0.016 | 0.016 | 0 | 0.3 | 0.39 | 0.33 | 0.45 | 0.27 | 0.3 |
| 2 | 205080949 | rs7596432 | -0.67 | 0.843 | -1.195 | 0.319 | -0.025 | -0.076 | -0.076 | 0.087 | 0.65 | 0.85 | 0.67 | 0.53 | 0.64 | 0.42 |

| | | | | | | | | | | | | | | | | |
|---|---|---|---|---|---|---|---|---|---|---|---|---|---|---|---|---|
| 2 | 205084439 | rs11692619 | -0.044 | 0.921 | -1.6 | -0.135 | -0.031 | -0.059 | -0.059 | 0.115 | 0.26 | 0.14 | 0.35 | 0.44 | 0.35 | 0.25 |
| 2 | 205090645 | rs12473084 | 1.053 | 0.949 | -1.971 | -0.183 | 0.017 | 0.059 | 0.059 | 0 | 0.15 | 0.06 | 0.06 | 0 | 0.11 | 0.04 |
| 2 | 205091301 | rs6710075 | -0.864 | 0.981 | -0.882 | -0.299 | -0.003 | 0.001 | 0.001 | 0 | 0.45 | 0.57 | 0.54 | 0.44 | 0.48 | 0.28 |
| 2 | 205092319 | rs7591705 | -1.204 | 1.359 | -0.189 | -0.515 | 0.003 | 0.004 | 0.004 | 0 | 0.56 | 0.53 | 0.5 | 0.54 | 0.51 | 0.41 |
| 2 | 205093597 | rs6718710 | -1.322 | 1.337 | -0.366 | -0.487 | -0.019 | -0.035 | -0.035 | 0 | 0.6 | 0.75 | 0.61 | 0.54 | 0.56 | 0.4 |
| 2 | 205099183 | rs2288218 | -1.762 | 1.309 | -0.434 | -0.369 | -0.021 | -0.072 | -0.072 | 1.276 | 0.66 | 0.85 | 0.67 | 0.57 | 0.63 | 0.42 |
| 2 | 205105619 | rs6745801 | -0.095 | 1.103 | 0.2 | -0.582 | -0.02 | -0.145 | -0.145 | 5.095 | 0.26 | 0.4 | 0.15 | 0 | 0.18 | 0.06 |
| 2 | 205106093 | rs4675431 | 0.107 | 0.908 | -0.249 | -0.202 | -0.004 | 0.004 | 0.004 | 0 | 0.41 | 0.53 | 0.57 | 0.61 | 0.38 | 0.58 |
| 2 | 205110646 | rs1559639 | -0.636 | 0.915 | -0.369 | -0.403 | 0.003 | 0.091 | 0.091 | 0 | 0.46 | 0.37 | 0.31 | 0.22 | 0.48 | 0.25 |
| 2 | 205111927 | rs16840921 | 1.411 | 0.768 | -0.901 | 0.089 | -0.055 | -0.085 | -0.085 | 0 | 0.35 | 0.07 | 0.38 | 0.33 | 0.37 | 0.59 |
| 2 | 205112368 | rs6710279 | 0.062 | 0.954 | -0.9 | -0.035 | 0.029 | 0.164 | 0.164 | 0 | 0.4 | 0.25 | 0.2 | 0.12 | 0.34 | 0.21 |
| 2 | 205113409 | rs13417432 | 0.25 | 1.064 | -0.828 | 0.099 | 0.02 | 0.015 | 0.015 | 0 | 0.06 | 0.12 | 0.1 | 0.09 | 0.14 | 0.04 |
| 2 | 205114083 | rs13407072 | 0.425 | 0.575 | -0.557 | -0.113 | 0.002 | -0.051 | -0.051 | 0.017 | 0.11 | 0.3 | 0.08 | 0.01 | 0.09 | 0.06 |
| 2 | 205114593 | rs7581837 | 1.012 | 0.494 | -0.881 | -0.237 | -0.045 | 0.12 | 0.12 | 0 | 0.27 | 0.08 | 0.16 | 0.1 | 0.31 | 0.16 |
| 2 | 205115622 | rs6707500 | -0.078 | 0.456 | -0.513 | -0.069 | -0.003 | 0.057 | 0.057 | 0.778 | 0.72 | 0.63 | 0.66 | 0.46 | 0.77 | 0.81 |
| 2 | 205116002 | rs6726035 | 0.557 | 0.444 | -0.554 | -0.092 | 0.002 | 0.025 | 0.025 | 0.461 | 0.67 | 0.57 | 0.6 | 0.45 | 0.69 | 0.76 |
| 2 | 205116824 | rs6714893 | 0.091 | 0.376 | -0.54 | -0.013 | -0.004 | 0.036 | 0.036 | 0.42 | 0.72 | 0.42 | 0.64 | 0.46 | 0.77 | 0.81 |
| 2 | 205118871 | rs13424874 | 0.309 | 0.742 | -0.823 | 0.141 | -0.002 | 0.008 | 0.008 | 0 | 0.51 | 0.81 | 0.52 | 0.36 | 0.49 | 0.66 |
| 2 | 205120223 | rs11677768 | -1.827 | 0.828 | -0.833 | 0.33 | -0.055 | -0.106 | -0.106 | 0 | 0.86 | 0.99 | 0.76 | 0.54 | 0.83 | 0.84 |
| 2 | 205121598 | rs759876 | 1.123 | 0.748 | -0.619 | 0.294 | -0.012 | 0.011 | 0.011 | 0.872 | 0.22 | 0.46 | 0.34 | 0.19 | 0.19 | 0.38 |
| 2 | 205121831 | rs9288348 | 0.494 | 0.72 | -0.731 | 0.219 | -0.001 | -0.001 | -0.001 | 0.03 | 0.33 | 0.39 | 0.23 | 0.26 | 0.32 | 0.31 |
| 2 | 205122018 | rs4673291 | -0.501 | 0.603 | -1.038 | 0.25 | -0.076 | -0.039 | -0.039 | 0 | 0.55 | 0.85 | 0.57 | 0.45 | 0.51 | 0.69 |
| 2 | 205124890 | rs4675438 | 1.088 | 0.466 | -0.365 | 0.596 | -0.024 | 0.014 | 0.014 | 0.647 | 0.21 | 0.4 | 0.32 | 0.19 | 0.18 | 0.38 |
| 2 | 205125606 | rs12466870 | 0.601 | 0.528 | -0.729 | 0.578 | -0.009 | 0.026 | 0.026 | 0.727 | 0.23 | 0.34 | 0.36 | 0.27 | 0.21 | 0.43 |
| 2 | 205129576 | rs759881 | -0.724 | 0.682 | -0.762 | 0.684 | -0.01 | -0.054 | -0.054 | 0 | 0.83 | 0.91 | 0.74 | 0.5 | 0.83 | 0.83 |
| 2 | 205132626 | rs6733573 | 1.095 | 0.884 | -0.611 | 0.592 | 0.003 | 0.026 | 0.026 | 0 | 0.24 | 0.37 | 0.29 | 0.25 | 0.24 | 0.27 |
| 2 | 205140805 | rs723021 | -0.093 | 0.88 | -0.989 | 0.277 | -0.006 | 0.065 | 0.065 | 0.028 | 0.54 | 0.32 | 0.41 | 0.26 | 0.56 | 0.53 |
| 2 | 205145765 | rs726833 | -0.499 | 1.237 | -0.729 | 0.451 | -0.028 | -0.023 | -0.023 | 0.32 | 0.59 | 0.74 | 0.57 | 0.47 | 0.57 | 0.62 |
| 2 | 205147331 | rs10198084 | 0.113 | 1.002 | -0.664 | -0.152 | 0.017 | 0.01 | 0.01 | 0 | 0.43 | 0.65 | 0.28 | 0.37 | 0.32 | 0.48 |
| 2 | 205151228 | rs4675449 | -0.299 | 1.084 | -0.412 | -0.243 | 0.046 | 0.064 | 0.064 | 0 | 0.5 | 0.81 | 0.38 | 0.37 | 0.43 | 0.5 |
| 2 | 205153119 | rs918967 | -0.39 | 1.12 | -0.351 | -0.203 | 0.031 | 0.037 | 0.037 | 0 | 0.09 | 0.21 | 0.14 | 0.12 | 0.14 | 0.03 |
| 2 | 205153827 | rs16841113 | 1.407 | 1.171 | -0.477 | -0.157 | 0 | 0 | 0 | 0.145 | 0.07 | 0.08 | 0.12 | 0.09 | 0.08 | 0.04 |
| 2 | 205154872 | rs10171678 | -1.284 | 1.197 | -0.537 | -0.194 | -0.066 | -0.048 | -0.048 | 0 | 0.62 | 0.89 | 0.53 | 0.46 | 0.57 | 0.54 |
| 2 | 205161476 | rs10514647 | 0.37 | 1.313 | -0.051 | -0.192 | 0.024 | -0.029 | -0.029 | 0 | 0.06 | 0.12 | 0.05 | 0 | 0.08 | 0.01 |
| 2 | 205167348 | rs10205524 | -0.589 | 1.568 | -0.115 | -0.287 | -0.006 | 0.019 | 0.019 | 0 | 0.54 | 0.69 | 0.44 | 0.37 | 0.53 | 0.49 |
| 2 | 205169149 | rs1981913 | 0.092 | 1.501 | 0.049 | -0.365 | -0.006 | -0.007 | -0.007 | 0 | 0.45 | 0.49 | 0.31 | 0.37 | 0.4 | 0.46 |
| 2 | 205169372 | rs4080156 | -1.547 | 1.396 | -0.031 | -0.31 | -0.02 | -0.056 | -0.056 | 0 | 0.66 | 0.9 | 0.6 | 0.46 | 0.67 | 0.55 |
| 2 | 205172592 | rs6747433 | 0.773 | 1.323 | -0.085 | -0.325 | 0.022 | 0.026 | 0.026 | 0 | 0.36 | 0.26 | 0.22 | 0.24 | 0.29 | 0.44 |
| 2 | 205177507 | rs2353849 | 0.446 | 1.132 | -0.032 | -0.348 | 0.018 | 0.004 | 0.004 | 0 | 0.39 | 0.35 | 0.25 | 0.37 | 0.32 | 0.45 |
| 2 | 205177698 | rs7603542 | 0.112 | 1.235 | -0.101 | -0.118 | 0.03 | -0.031 | -0.031 | 0 | 0.08 | 0.21 | 0.1 | 0 | 0.1 | 0.02 |
| 2 | 205186323 | rs7587879 | -1.168 | 1.337 | -0.498 | 0.055 | -0.061 | -0.106 | -0.106 | 0.146 | 0.67 | 0.93 | 0.6 | 0.45 | 0.67 | 0.55 |
| 2 | 205189419 | rs4673296 | -1.148 | 1.267 | -0.28 | 0.188 | -0.013 | -0.125 | -0.125 | 0 | 0.67 | 0.93 | 0.63 | 0.47 | 0.72 | 0.54 |
| 2 | 205192151 | rs6741142 | -1.211 | 1.261 | -0.275 | 0.172 | -0.013 | -0.133 | -0.133 | 0.003 | 0.67 | 0.94 | 0.63 | 0.47 | 0.72 | 0.54 |
| 2 | 205194890 | rs6724499 | -0.163 | 1.136 | 0.084 | 0.04 | -0.001 | 0.001 | 0.001 | 0 | 0.47 | 0.55 | 0.43 | 0.34 | 0.48 | 0.32 |
| 2 | 205204850 | rs6743373 | 0.052 | 1.205 | -0.258 | 0.003 | -0.005 | 0.009 | 0.009 | 0 | 0.42 | 0.64 | 0.41 | 0.26 | 0.49 | 0.27 |
| 2 | 205206832 | rs7599906 | 0.237 | 1.218 | -0.835 | 0.003 | 0.01 | -0.02 | -0.02 | 0 | 0.4 | 0.51 | 0.4 | 0.25 | 0.49 | 0.26 |
| 2 | 205209435 | rs13035748 | 0.597 | 1.261 | -0.98 | 0.008 | -0.031 | -0.001 | -0.001 | 0 | 0.11 | 0.04 | 0.17 | 0.09 | 0.16 | 0.07 |
| 2 | 205209449 | rs6760579 | 0.125 | 1.166 | -0.966 | -0.069 | 0.031 | -0.049 | -0.049 | 0.021 | 0.09 | 0.22 | 0.15 | 0 | 0.15 | 0.02 |
| 2 | 205209652 | rs6760812 | 0.784 | 0.963 | -1.007 | 0.042 | -0.015 | 0.018 | 0.018 | 0 | 0.32 | 0.22 | 0.25 | 0.25 | 0.34 | 0.23 |
| 2 | 205212629 | rs7599719 | -0.797 | 0.627 | -0.84 | 0.337 | -0.028 | 0.014 | 0.014 | 0 | 0.26 | 0.42 | 0.23 | 0.24 | 0.22 | 0.28 |
| 2 | 205219203 | rs6750362 | 0.108 | 0.357 | -0.788 | 0.467 | -0.047 | 0.055 | 0.055 | 0.019 | 0.15 | 0.39 | 0.07 | 0.17 | 0.08 | 0.21 |
| 2 | 205220924 | rs6708278 | -0.78 | 0.709 | -1.855 | 0.439 | -0.022 | -0.113 | -0.113 | 0 | 0.67 | 0.92 | 0.64 | 0.48 | 0.71 | 0.53 |
| 2 | 205222127 | rs6435231 | -0.127 | 0.755 | -1.387 | 0.165 | 0.001 | 0.013 | 0.013 | 0 | 0.35 | 0.34 | 0.27 | 0.25 | 0.32 | 0.23 |
| 2 | 205225459 | rs6435232 | -0.354 | 0.863 | -1.994 | 0.104 | -0.048 | 0.123 | 0.123 | 0.287 | 0.47 | 0.85 | 0.34 | 0.3 | 0.51 | 0.34 |
| 2 | 205228510 | rs4075322 | 0.104 | 0.77 | -1.575 | 1.058 | -0.003 | 0.099 | 0.099 | 0 | 0.9 | 0.89 | 0.77 | 0.58 | 0.89 | 0.74 |
| 2 | 205230974 | rs4539778 | -0.026 | 0.768 | -1.593 | 1.118 | -0.007 | -0.032 | -0.032 | 0.388 | 0.91 | 0.98 | 0.77 | 0.53 | 0.9 | 0.75 |
| 2 | 205231284 | rs7577929 | 0.281 | 0.741 | -1.564 | 1.104 | -0.002 | 0.077 | 0.077 | 0 | 0.9 | 0.9 | 0.76 | 0.52 | 0.89 | 0.74 |
| 2 | 205232320 | rs7581860 | -0.743 | 0.916 | -0.783 | 0.938 | 0.02 | 0.112 | 0.112 | 0.274 | 0.08 | 0.26 | 0.1 | 0.22 | 0.07 | 0.06 |
| 2 | 205239583 | rs11684723 | 0.41 | 0.715 | -1.364 | -0.007 | -0.059 | 0.067 | 0.067 | 0.184 | 0.27 | 0.14 | 0.37 | 0.1 | 0.37 | 0.25 |
| 2 | 205244699 | rs6713530 | 0.935 | 0.391 | -0.991 | 0.287 | -0.001 | 0.042 | 0.042 | 0.204 | 0.81 | 0.63 | 0.66 | 0.31 | 0.83 | 0.69 |
| 2 | 205245469 | rs4132730 | -0.017 | -0.128 | -1.149 | -0.19 | -0.005 | -0.003 | -0.003 | 0.356 | 0.3 | 0.33 | 0.17 | 0.36 | 0.21 | 0.21 |
| 2 | 205253568 | rs6712212 | 0.243 | -0.282 | -0.943 | -0.474 | -0.002 | -0.029 | -0.029 | 0 | 0.12 | 0.12 | 0.1 | 0 | 0.11 | 0.03 |
| 2 | 205255807 | rs10932062 | 1.268 | -0.032 | -1.379 | -0.699 | -0.032 | 0.003 | 0.003 | 0 | 0.09 | 0 | 0.14 | 0.05 | 0.11 | 0.08 |
| 2 | 205256040 | rs6708542 | -1.283 | -0.887 | -1.258 | -0.481 | -0.001 | 0.001 | 0.001 | 1.883 | 0.33 | 0.63 | 0.4 | 0.41 | 0.25 | 0.33 |
| 2 | 205258852 | rs6748817 | -1.898 | -1.173 | -1.2 | -0.399 | -0.018 | 0.08 | 0.08 | 0 | 0.74 | 0.99 | 0.84 | 0.86 | 0.81 | 0.84 |
| 2 | 205260409 | rs13394087 | 0.006 | -1.241 | -0.072 | 0.003 | 0.001 | 0.052 | 0.052 | 0 | 0.15 | 0.13 | 0.27 | 0.4 | 0.15 | 0.31 |
| 2 | 205270870 | rs4347813 | 0.383 | -1.366 | -0.098 | -0.104 | 0.008 | -0.015 | -0.015 | 0 | 0.72 | 0.76 | 0.67 | 0.47 | 0.74 | 0.52 |
| 2 | 205274728 | rs6435238 | 1.26 | -1.324 | 0.034 | -0.089 | 0.014 | 0.001 | 0.001 | 0.156 | 0.56 | 0.43 | 0.57 | 0.46 | 0.64 | 0.48 |
| 2 | 205287435 | rs4414662 | -0.122 | -1.292 | 0.177 | -0.038 | -0.024 | -0.033 | -0.033 | 0 | 0.26 | 0.29 | 0.16 | 0.12 | 0.22 | 0.19 |

| | | | | | | | | | | | | | | | | |
|---|---|---|---|---|---|---|---|---|---|---|---|---|---|---|---|---|
| 2 | 205293339 | rs12620515 | 0.385 | -1.144 | -0.023 | 0.187 | -0.001 | 0.169 | 0.169 | 0 | 0.09 | 0.16 | 0.2 | 0.36 | 0.07 | 0.28 |
| 2 | 205307882 | rs7602656 | 0.523 | -1.249 | 0.196 | -0.192 | -0.033 | -0.068 | -0.068 | 0 | 0.69 | 0.46 | 0.8 | 0.87 | 0.73 | 0.78 |
| 2 | 205316761 | rs1374708 | -0.478 | -0.733 | 0.248 | 0.211 | -0.008 | -0.015 | -0.015 | 0 | 0.84 | 0.73 | 0.9 | 0.87 | 0.84 | 0.81 |
| 2 | 205317811 | rs1156250 | 0.684 | -0.85 | 0.112 | 0.314 | -0.004 | -0.101 | -0.101 | 0 | 0.13 | 0.01 | 0.24 | 0.42 | 0.11 | 0.32 |
| 2 | 205320570 | rs1447489 | 0.014 | -0.632 | 1.045 | -0.175 | 0.088 | 0.017 | 0.017 | 2.668 | 0.95 | 0.75 | 0.96 | 1 | 0.96 | 0.98 |
| 2 | 205321371 | rs1447488 | -0.429 | -0.661 | 0.725 | -0.141 | -0.048 | -0.009 | -0.009 | 0 | 0.86 | 0.7 | 0.91 | 0.88 | 0.89 | 0.83 |
| 2 | 205346667 | rs12613305 | -0.524 | -0.858 | 0.387 | -0.468 | 0.014 | -0.061 | -0.061 | 0.078 | 0.62 | 0.55 | 0.61 | 0.92 | 0.54 | 0.72 |
| 2 | 205347379 | rs12373835 | -0.318 | -0.927 | 0.733 | -0.142 | 0.026 | 0.116 | 0.116 | 0 | 0.5 | 0.31 | 0.54 | 0.8 | 0.42 | 0.55 |
| 2 | 205348724 | rs12373801 | -0.253 | -0.864 | 0.558 | -0.117 | 0.009 | 0.03 | 0.03 | 0 | 0.51 | 0.41 | 0.55 | 0.79 | 0.42 | 0.54 |
| 2 | 205350649 | rs4673300 | 0.064 | -0.75 | -0.31 | 0.074 | 0.01 | 0.121 | 0.121 | 0 | 0.43 | 0.2 | 0.49 | 0.78 | 0.35 | 0.53 |
| 2 | 205363596 | rs6743255 | 1.178 | -1.267 | 0.879 | 1.342 | -0.031 | 0.1 | 0.1 | 1.809 | 0.77 | 0.44 | 0.88 | 0.54 | 0.88 | 0.83 |
| 2 | 205365299 | rs10932068 | -0.718 | -1.281 | 0.735 | 1.243 | -0.001 | -0.016 | -0.016 | 0 | 0.15 | 0.24 | 0.14 | 0.09 | 0.12 | 0.06 |
| 2 | 205366339 | rs4556959 | 0.212 | -1.214 | 1.161 | 1.26 | 0.012 | -0.021 | -0.021 | 0 | 0.94 | 0.78 | 0.96 | 1 | 0.96 | 0.99 |
| 2 | 205374700 | rs10932071 | 1.599 | -1.256 | 1.359 | 1.615 | 0.019 | 0.228 | 0.228 | 2.161 | 0.89 | 0.54 | 0.92 | 0.71 | 0.93 | 0.87 |
| 2 | 205378570 | rs9288354 | 1.239 | -1.368 | 1.445 | 1.334 | -0.017 | -0.122 | -0.122 | 0.21 | 0.84 | 0.52 | 0.92 | 1 | 0.92 | 0.98 |
| 2 | 205379823 | rs12104992 | 0.254 | -1.315 | 1.308 | 1.506 | -0.002 | -0.041 | -0.041 | 0 | 0.08 | 0.13 | 0.03 | 0 | 0.04 | 0.01 |
| 2 | 205380227 | rs12105014 | 0.254 | -1.315 | 1.308 | 1.52 | -0.002 | -0.037 | -0.037 | 0 | 0.08 | 0.13 | 0.03 | 0 | 0.04 | 0.01 |
| 2 | 205385386 | rs7569476 | 1.881 | -1.773 | 0.996 | 1.531 | -0.01 | -0.01 | -0.01 | 0.507 | 0.65 | 0.42 | 0.81 | 0.82 | 0.82 | 0.79 |
| 2 | 205385454 | rs7581907 | 1.069 | -1.765 | 1.046 | 1.555 | -0.059 | -0.016 | -0.016 | 0 | 0.76 | 0.6 | 0.85 | 0.82 | 0.88 | 0.8 |
| 2 | 205386242 | rs6735824 | 1.728 | -1.752 | 0.512 | 1.687 | 0.012 | 0.006 | 0.006 | 0 | 0.6 | 0.4 | 0.79 | 0.33 | 0.8 | 0.65 |
| 2 | 205388729 | rs1447498 | 1.685 | -1.649 | 0.483 | 1.328 | 0.013 | -0.009 | -0.009 | 0 | 0.61 | 0.4 | 0.79 | 0.5 | 0.79 | 0.67 |
| 2 | 205390559 | rs2166733 | -0.849 | -1.358 | 0.003 | 1.468 | 0.03 | -0.029 | -0.029 | 0 | 0.19 | 0.04 | 0.11 | 0.18 | 0.12 | 0.19 |
| 2 | 205391934 | rs4675467 | 0.938 | -1.358 | 0.417 | 1.127 | 0.007 | -0.008 | -0.008 | 0 | 0.62 | 0.63 | 0.8 | 0.49 | 0.75 | 0.67 |
| 2 | 205393780 | rs989629 | 1.414 | -1.149 | 1.388 | 1.013 | -0.02 | -0.015 | -0.015 | 1.45 | 0.87 | 0.53 | 0.88 | 0.93 | 0.92 | 0.87 |
| 2 | 205399643 | rs6435243 | -1.902 | -1.1 | 1.765 | 1.451 | 0.002 | 0.087 | 0.087 | 1.404 | 0.19 | 0.56 | 0.14 | 0.28 | 0.14 | 0.23 |
| 2 | 205434906 | rs4673304 | 1.847 | -1.182 | 2.506 | 1.552 | 0.011 | 0 | 0 | 0.362 | 0.79 | 0.48 | 0.81 | 0.8 | 0.83 | 0.68 |
| 2 | 205441717 | rs1374713 | 0.382 | -1.176 | 2.582 | 1.557 | 0.02 | 0.024 | 0.024 | 0 | 0.92 | 0.73 | 0.89 | 0.93 | 0.92 | 0.9 |
| 2 | 205467232 | rs1861766 | 2.137 | -1.225 | 2.101 | 1.604 | -0.001 | 0.001 | 0.001 | 1.582 | 0.73 | 0.29 | 0.77 | 0.63 | 0.79 | 0.66 |
| 2 | 205471376 | rs10174747 | 1.87 | -1.246 | 2.104 | 1.52 | -0.006 | -0.014 | -0.014 | 0 | 0.75 | 0.39 | 0.78 | 0.8 | 0.8 | 0.65 |
| 2 | 205475415 | rs9283519 | 1.944 | -1.286 | 2.118 | 1.536 | -0.006 | -0.025 | -0.025 | 0.621 | 0.75 | 0.38 | 0.78 | 0.82 | 0.79 | 0.67 |
| 2 | 205478454 | rs12474806 | 1.96 | -1.26 | 2.066 | 1.546 | -0.007 | -0.008 | -0.008 | 0.819 | 0.74 | 0.39 | 0.79 | 0.77 | 0.79 | 0.64 |
| 2 | 205480867 | rs12621708 | 2.468 | -1.502 | 1.533 | 1.239 | 0.05 | -0.054 | -0.054 | 0 | 0.6 | 0.15 | 0.71 | 0.55 | 0.71 | 0.45 |
| 2 | 205487689 | rs2193449 | 1.756 | -1.09 | 1.786 | 1.141 | 0.007 | 0.002 | 0.002 | 0.993 | 0.76 | 0.48 | 0.8 | 0.78 | 0.79 | 0.65 |
| 2 | 205525273 | rs11691772 | 2.231 | -1.061 | 1.798 | 1.039 | 0.003 | -0.001 | -0.001 | 0 | 0.73 | 0.24 | 0.76 | 0.74 | 0.82 | 0.82 |
| 2 | 205536648 | rs7593604 | -2.476 | -1.018 | 2.153 | 1.076 | -0.006 | 0 | 0 | 1.507 | 0.25 | 0.71 | 0.24 | 0.25 | 0.18 | 0.18 |
| 2 | 205538672 | rs17595154 | 0.028 | -1.03 | 2.478 | 1.089 | -0.028 | 0.035 | 0.035 | 0 | 0.08 | 0.11 | 0.02 | 0.15 | 0.02 | 0.01 |
| 2 | 205539092 | rs1016338 | 2.029 | -1.041 | 2.813 | 1.085 | 0.004 | 0.076 | 0.076 | 0.366 | 0.81 | 0.46 | 0.78 | 0.73 | 0.84 | 0.82 |
| 2 | 205542008 | rs7577059 | -2.806 | -1.039 | 2.803 | 1.082 | -0.014 | 0.005 | 0.005 | 1.021 | 0.25 | 0.68 | 0.24 | 0.25 | 0.18 | 0.18 |
| 2 | 205561950 | rs10208062 | -1.266 | -1.052 | 2.808 | 1.082 | -0.052 | 0.091 | 0.091 | 1.075 | 0.11 | 0.34 | 0.05 | 0.15 | 0.02 | 0.01 |
| 2 | 205572718 | rs2160690 | 1.764 | -0.882 | 2.567 | 0.957 | 0.024 | -0.033 | -0.033 | 0 | 0.83 | 0.53 | 0.8 | 0.89 | 0.84 | 0.83 |
| 2 | 205581824 | rs6435250 | 1.478 | -0.83 | 2.429 | 0.983 | 0.053 | -0.01 | -0.01 | 0 | 0.85 | 0.51 | 0.8 | 0.89 | 0.84 | 0.83 |
| 2 | 205587422 | rs16836537 | -0.925 | -0.786 | 2.009 | 0.944 | 0.03 | -0.018 | -0.018 | 0.082 | 0.11 | 0.25 | 0.15 | 0.08 | 0.15 | 0.15 |
| 2 | 205596994 | rs980388 | 1.131 | -0.783 | 2.008 | 0.903 | 0.029 | -0.016 | -0.016 | 0.009 | 0.86 | 0.73 | 0.81 | 0.89 | 0.81 | 0.83 |
| 2 | 205606092 | rs12613874 | 2.208 | -1.174 | 1.462 | 0.691 | 0.042 | 0.038 | 0.038 | 1.016 | 0.6 | 0.18 | 0.66 | 0.69 | 0.69 | 0.7 |
| 2 | 205608382 | rs10180248 | -2.362 | -1.155 | 1.667 | 0.667 | 0.003 | -0.012 | -0.012 | 0 | 0.32 | 0.68 | 0.25 | 0.12 | 0.22 | 0.24 |
| 2 | 205618711 | rs2098462 | -2.415 | -1.107 | 1.641 | 0.69 | 0.005 | 0 | 0 | 0 | 0.33 | 0.7 | 0.25 | 0.14 | 0.23 | 0.25 |
| 2 | 205620551 | rs16836575 | -1.128 | -1.274 | 1.662 | 0.686 | -0.001 | -0.01 | -0.01 | 0 | 0.08 | 0.09 | 0.04 | 0.04 | 0.04 | 0.08 |
| 2 | 205623306 | rs1861764 | 2.407 | -1.268 | 1.56 | 0.892 | 0.057 | 0.005 | 0.005 | 1.196 | 0.66 | 0.16 | 0.74 | 0.69 | 0.77 | 0.77 |
| 2 | 205623839 | rs16836582 | -0.222 | -1.288 | 1.334 | 1.09 | -0.008 | -0.042 | -0.042 | 0 | 0.07 | 0 | 0.04 | 0.16 | 0.04 | 0.18 |
| 2 | 205624129 | rs6715511 | -2.649 | -1.1 | 1.625 | 0.828 | 0.041 | 0.009 | 0.009 | 0.521 | 0.33 | 0.79 | 0.25 | 0.31 | 0.23 | 0.22 |
| 2 | 205626400 | rs10490273 | 1.063 | -1.065 | 1.742 | 0.86 | -0.014 | 0.05 | 0.05 | 0.057 | 0.06 | 0.1 | 0.03 | 0.17 | 0.02 | 0.03 |
| 2 | 205626483 | rs16836589 | 1.44 | -1.061 | 1.663 | 0.862 | 0.005 | 0.01 | 0.01 | 0 | 0.05 | 0.03 | 0.02 | 0.17 | 0.02 | 0.03 |
| 2 | 205631567 | rs9653327 | 0.666 | -1.064 | 1.961 | 0.834 | 0.023 | -0.002 | -0.002 | 0 | 0.87 | 0.75 | 0.82 | 0.87 | 0.83 | 0.87 |
| 2 | 205636870 | rs12467431 | 2.584 | -1.074 | 1.757 | 0.818 | 0.049 | 0 | 0 | 1.034 | 0.65 | 0.16 | 0.7 | 0.66 | 0.75 | 0.75 |
| 2 | 205639108 | rs2878473 | 1.087 | -0.967 | 2.202 | 0.683 | 0.032 | -0.01 | -0.01 | 0 | 0.84 | 0.63 | 0.79 | 0.87 | 0.82 | 0.87 |
| 2 | 205639924 | rs7579849 | -1.29 | -0.973 | 2.18 | 0.655 | -0.026 | -0.033 | -0.033 | 0 | 0.08 | 0.26 | 0.04 | 0.04 | 0.02 | 0.07 |
| 2 | 205640829 | rs12104689 | -1.655 | -0.753 | 2.008 | 0.753 | 0.012 | -0.046 | -0.046 | 0 | 0.18 | 0.46 | 0.2 | 0.1 | 0.17 | 0.12 |
| 2 | 205645815 | rs975894 | 0.987 | -0.738 | 1.88 | 0.891 | 0.043 | 0.036 | 0.036 | 0.014 | 0.95 | 0.78 | 0.92 | 0.9 | 0.95 | 0.92 |
| 2 | 205646827 | rs10192244 | 0.929 | -0.749 | 2.047 | 0.884 | 0.072 | 0.059 | 0.059 | 0.127 | 0.95 | 0.71 | 0.94 | 0.93 | 0.95 | 0.92 |
| 2 | 205646988 | rs10204710 | 0.829 | -0.777 | 1.906 | 0.87 | 0.044 | 0.033 | 0.033 | 0.143 | 0.95 | 0.77 | 0.95 | 0.93 | 0.95 | 0.92 |
| 2 | 205658016 | rs6435252 | -2.208 | -0.75 | 2.05 | 0.975 | -0.009 | 0.029 | 0.029 | 2.331 | 0.16 | 0.66 | 0.14 | 0.23 | 0.11 | 0.1 |
| 2 | 205658682 | rs6730405 | -1.879 | -0.667 | 1.994 | 0.953 | 0.068 | 0.021 | 0.021 | 0 | 0.08 | 0.31 | 0.12 | 0.1 | 0.09 | 0.1 |
| 2 | 205685674 | rs6760087 | -2.744 | -0.807 | 1.386 | 0.895 | 0.008 | -0.003 | -0.003 | 0.139 | 0.27 | 0.75 | 0.19 | 0.31 | 0.15 | 0.22 |
| 2 | 205688132 | rs12614659 | -0.96 | -0.691 | 1.054 | 1.035 | -0.002 | 0.064 | 0.064 | 0 | 0.12 | 0.22 | 0.11 | 0.26 | 0.08 | 0.11 |
| 2 | 205695976 | rs10203626 | -0.588 | -0.641 | 0.464 | 0.854 | -0.008 | -0.003 | -0.003 | 0 | 0.05 | 0.07 | 0.01 | 0.05 | 0.02 | 0.09 |
| 2 | 205697101 | rs17450916 | 0.424 | -0.514 | -0.493 | 0.894 | -0.014 | -0.032 | -0.032 | 0 | 0.15 | 0.03 | 0.1 | 0.21 | 0.15 | 0.12 |
| 2 | 205700834 | rs6735981 | -1.37 | -0.871 | -0.435 | 0.79 | 0.033 | -0.004 | -0.004 | 0 | 0.23 | 0.8 | 0.19 | 0.24 | 0.11 | 0.09 |
| 2 | 205703928 | rs13408553 | -0.146 | -0.925 | 0.317 | 0.798 | -0.011 | -0.022 | -0.022 | 0 | 0.11 | 0.27 | 0.08 | 0.06 | 0.08 | 0.05 |
| 2 | 205710514 | rs7605954 | 0.045 | -0.934 | 0.381 | 0.785 | -0.015 | -0.035 | -0.035 | 0 | 0.1 | 0.26 | 0.05 | 0.03 | 0.07 | 0.05 |

| | | | | | | | | | | | | | | | | |
|---|---|---|---|---|---|---|---|---|---|---|---|---|---|---|---|---|
| 2 | 205714902 | rs10184471 | -1.772 | -0.956 | 0.64 | 0.795 | -0.011 | -0.001 | -0.001 | 0 | 0.21 | 0.74 | 0.16 | 0.21 | 0.1 | 0.08 |
| 2 | 205715245 | rs10469651 | -1.077 | -0.91 | 0.557 | 0.858 | -0.002 | 0.185 | 0.185 | 1.192 | 0.06 | 0.3 | 0.05 | 0.19 | 0.01 | 0.07 |
| 2 | 205728234 | rs4675474 | 0.664 | -0.57 | -0.397 | 1.019 | -0.06 | -0.067 | -0.067 | 0.026 | 0.16 | 0.01 | 0.28 | 0.2 | 0.16 | 0.36 |
| 2 | 205733847 | rs6733159 | -0.841 | -0.278 | -0.351 | 0.841 | 0.01 | -0.037 | -0.037 | 0 | 0.24 | 0.15 | 0.17 | 0.4 | 0.21 | 0.16 |
| 2 | 205759917 | rs4429453 | -0.739 | 0.151 | -0.701 | 0.134 | -0.003 | 0.001 | 0.001 | 0.136 | 0.32 | 0.4 | 0.27 | 0.31 | 0.29 | 0.15 |
| 2 | 205763480 | rs13031723 | 0.922 | 0.239 | -1.137 | 0.132 | -0.024 | -0.048 | -0.048 | 0 | 0.16 | 0.05 | 0.16 | 0.24 | 0.19 | 0.09 |
| 2 | 205764917 | rs11896131 | 1.289 | 0.128 | -1.077 | -0.403 | 0.004 | 0.003 | 0.003 | 0.09 | 0.09 | 0.08 | 0.06 | 0.03 | 0.07 | 0 |
| 2 | 205767001 | rs10804145 | -0.805 | 0.089 | -1.596 | -0.683 | -0.031 | -0.028 | -0.028 | 0 | 0.66 | 0.85 | 0.74 | 0.64 | 0.62 | 0.72 |
| 2 | 205777605 | rs10490275 | 0.502 | 0.446 | -1.296 | -1.712 | -0.051 | 0.079 | 0.079 | 0.527 | 0.19 | 0.06 | 0.18 | 0.01 | 0.25 | 0.08 |
| 2 | 205778583 | rs10207414 | 0.498 | 0.429 | -1.003 | -1.72 | -0.018 | 0.014 | 0.014 | 0.152 | 0.19 | 0.12 | 0.18 | 0.01 | 0.25 | 0.07 |
| 2 | 205783813 | rs7560192 | -0.116 | 0.306 | -0.777 | -1.809 | -0.009 | -0.002 | -0.002 | 0.272 | 0.27 | 0.21 | 0.22 | 0.01 | 0.31 | 0.08 |
| 2 | 205783828 | rs7572649 | -0.935 | 0.353 | -0.718 | -1.89 | -0.004 | 0.019 | 0.019 | 0.996 | 0.36 | 0.69 | 0.29 | 0.06 | 0.35 | 0.13 |
| 2 | 205790839 | rs1510775 | -1.018 | 0.563 | -1.263 | -2.073 | 0.022 | 0.131 | 0.131 | 7.848 | 0.54 | 0.74 | 0.59 | 0.05 | 0.58 | 0.18 |
| 2 | 205794074 | rs2216852 | -0.929 | 0.672 | -1.16 | -1.999 | 0.005 | 0.138 | 0.138 | 3.116 | 0.48 | 0.71 | 0.56 | 0.05 | 0.52 | 0.17 |
| 2 | 205794531 | rs13002022 | 0.443 | 1.087 | -1.227 | -1.358 | -0.051 | 0.066 | 0.066 | 0.409 | 0.17 | 0.06 | 0.17 | 0 | 0.24 | 0.07 |
| 2 | 205799241 | rs13421467 | 0.375 | 0.76 | -0.522 | -0.962 | 0.006 | -0.012 | -0.012 | 0 | 0.07 | 0.21 | 0.04 | 0.02 | 0.05 | 0.08 |
| 2 | 205800837 | rs236825 | 0.018 | 0.875 | -1.543 | -1.017 | -0.084 | 0.012 | 0.012 | 0 | 0.7 | 0.94 | 0.79 | 0.77 | 0.65 | 0.85 |
| 2 | 205802814 | rs6727754 | -0.633 | 0.7 | -1.525 | -1.844 | -0.007 | 0.15 | 0.15 | 2.452 | 0.48 | 0.7 | 0.32 | 0.05 | 0.49 | 0.16 |
| 2 | 205806424 | rs236822 | 1.266 | 0.372 | -1.315 | -1.841 | 0.025 | 0.004 | 0.004 | 0.789 | 0.33 | 0.18 | 0.53 | 0.71 | 0.29 | 0.68 |
| 2 | 205808353 | rs13384038 | 0.569 | 0.52 | -0.631 | -2.05 | -0.006 | -0.107 | -0.107 | 0 | 0.28 | 0.43 | 0.17 | 0.02 | 0.25 | 0.1 |
| 2 | 205809041 | rs6435262 | 0.957 | 0.468 | -0.63 | -2.084 | 0.001 | -0.015 | -0.015 | 0 | 0.24 | 0.26 | 0.13 | 0.02 | 0.22 | 0.09 |
| 2 | 205809480 | rs6435264 | 0.897 | 0.477 | -0.55 | -2.095 | -0.002 | -0.05 | -0.05 | 0.038 | 0.24 | 0.31 | 0.13 | 0.02 | 0.22 | 0.09 |
| 2 | 205811482 | rs236831 | -0.958 | 0.725 | -0.65 | -2.204 | -0.052 | -0.006 | -0.006 | 0 | 0.75 | 0.95 | 0.82 | 0.78 | 0.71 | 0.83 |
| 2 | 205811844 | rs7605665 | 0.426 | 0.756 | -0.514 | -2.236 | 0.006 | -0.005 | -0.005 | 0.593 | 0.29 | 0.29 | 0.16 | 0.02 | 0.26 | 0.11 |
| 2 | 205812418 | rs7592342 | -0.323 | 0.79 | 0.084 | -2.246 | 0.002 | -0.077 | -0.077 | 0 | 0.36 | 0.59 | 0.24 | 0.05 | 0.35 | 0.15 |
| 2 | 205815073 | rs10490277 | 0.237 | 0.561 | 0.07 | -2.142 | -0.017 | -0.106 | -0.106 | 1.612 | 0.2 | 0.36 | 0.1 | 0.02 | 0.15 | 0.07 |
| 2 | 205815842 | rs6435265 | -1.368 | 0.507 | -0.224 | -2.355 | 0 | -0.056 | -0.056 | 0 | 0.39 | 0.58 | 0.24 | 0.05 | 0.39 | 0.15 |
| 2 | 205818211 | rs236830 | 0.1 | 0.264 | -0.845 | -2.382 | 0.015 | 0.135 | 0.135 | 0.142 | 0.51 | 0.71 | 0.68 | 0.76 | 0.49 | 0.75 |
| 2 | 205818746 | rs17535946 | 1.22 | 0.208 | -0.992 | -2.316 | -0.028 | 0.009 | 0.009 | 2.525 | 0.08 | 0.02 | 0.09 | 0.03 | 0.15 | 0.09 |
| 2 | 205818886 | rs236829 | -0.026 | 0.226 | -0.92 | -2.339 | 0 | 0.077 | 0.077 | 0.952 | 0.53 | 0.63 | 0.7 | 0.76 | 0.56 | 0.78 |
| 2 | 205823152 | rs17535981 | 0.855 | 0.166 | -1.023 | -1.855 | -0.043 | -0.003 | -0.003 | 0.379 | 0.08 | 0.01 | 0.09 | 0.03 | 0.15 | 0.09 |
| 2 | 205829991 | rs236843 | -0.772 | 0.142 | -1.16 | -2.01 | -0.078 | 0.071 | 0.071 | 0.825 | 0.61 | 0.96 | 0.75 | 0.77 | 0.62 | 0.85 |
| 2 | 205833521 | rs236841 | 1.594 | -0.044 | -1.255 | -1.763 | 0.024 | -0.055 | -0.055 | 0 | 0.27 | 0.17 | 0.23 | 0.6 | 0.22 | 0.53 |
| 2 | 205834111 | rs236838 | -1.048 | 0.048 | -1.284 | -1.739 | -0.053 | 0.081 | 0.081 | 1.01 | 0.62 | 0.92 | 0.75 | 0.79 | 0.61 | 0.85 |
| 2 | 205843303 | rs1355339 | -1.752 | 0.018 | -0.978 | -1.667 | -0.013 | 0.032 | 0.032 | 0.096 | 0.77 | 0.72 | 0.78 | 0.39 | 0.81 | 0.49 |
| 2 | 205844259 | rs13017914 | -0.802 | 0.127 | -0.873 | -1.626 | -0.005 | 0.006 | 0.006 | 0.969 | 0.63 | 0.55 | 0.66 | 0.38 | 0.65 | 0.43 |
| 2 | 205847150 | rs2063407 | 0.433 | 0.552 | -1.442 | -1.33 | -0.016 | 0.002 | 0.002 | 0 | 0.21 | 0.22 | 0.39 | 0.16 | 0.26 | 0.24 |
| 2 | 205847411 | rs1510777 | 0.287 | -0.136 | -1.013 | -1.132 | 0.047 | 0.039 | 0.039 | 0.69 | 0.5 | 0.39 | 0.62 | 0.37 | 0.57 | 0.36 |
| 2 | 205864532 | rs2335705 | -0.122 | -0.128 | -0.629 | -1.075 | -0.008 | -0.046 | -0.046 | 0 | 0.88 | 0.8 | 0.9 | 0.99 | 0.86 | 0.93 |
| 2 | 205866960 | rs11695066 | 0.339 | -0.053 | -0.543 | -1.394 | 0.001 | -0.029 | -0.029 | 0.701 | 0.57 | 0.48 | 0.44 | 0.82 | 0.53 | 0.71 |
| 2 | 205883547 | rs11896055 | -0.114 | -0.157 | -0.6 | -1.272 | 0.006 | -0.032 | -0.032 | 0 | 0.32 | 0.37 | 0.46 | 0.13 | 0.33 | 0.2 |
| 2 | 205889602 | rs6741267 | 0.009 | -0.4 | -0.764 | -0.705 | -0.01 | -0.017 | -0.017 | 0 | 0.4 | 0.3 | 0.59 | 0.5 | 0.51 | 0.34 |
| 2 | 205894489 | rs1397481 | 2.114 | -0.777 | -0.904 | -0.598 | -0.009 | -0.034 | -0.034 | 2.618 | 0.24 | 0.13 | 0.16 | 0.4 | 0.28 | 0.51 |
| 2 | 205898117 | rs1397482 | 0.47 | -0.919 | -0.468 | -0.544 | 0.001 | 0 | 0 | 0 | 0.43 | 0.39 | 0.26 | 0.42 | 0.4 | 0.54 |
| 2 | 205912210 | rs7563474 | 0.317 | -0.99 | -0.026 | -0.358 | 0.002 | 0.013 | 0.013 | 0 | 0.7 | 0.71 | 0.51 | 0.47 | 0.72 | 0.64 |
| 2 | 205912403 | rs1510765 | 0.699 | -1.022 | 0.059 | -0.411 | 0.005 | 0.032 | 0.032 | 0 | 0.65 | 0.53 | 0.39 | 0.48 | 0.64 | 0.65 |
| 2 | 205917744 | rs6736058 | 0.317 | -0.98 | -0.213 | -0.437 | 0.003 | 0.044 | 0.044 | 0 | 0.7 | 0.53 | 0.5 | 0.47 | 0.73 | 0.64 |
| 2 | 205924277 | rs6754606 | 0.213 | -0.387 | -0.562 | -0.273 | 0.012 | 0.017 | 0.017 | 0 | 0.13 | 0.08 | 0.14 | 0.04 | 0.12 | 0.08 |
| 2 | 205942560 | rs4675490 | -0.94 | -0.45 | -0.434 | -0.23 | 0.006 | 0.009 | 0.009 | 0 | 0.31 | 0.33 | 0.61 | 0.56 | 0.39 | 0.42 |
| 2 | 205943592 | rs985164 | -1.23 | -0.569 | -0.972 | -0.365 | -0.016 | -0.056 | -0.056 | 0 | 0.84 | 0.78 | 0.94 | 1 | 0.92 | 0.98 |
| 2 | 205949173 | rs17283572 | 1.176 | -0.223 | -1.341 | 0.028 | -0.061 | -0.065 | -0.065 | 0.883 | 0.12 | 0.02 | 0.15 | 0.25 | 0.19 | 0.09 |
| 2 | 205951603 | rs7559302 | 0.293 | -0.296 | -1.302 | -0.031 | -0.089 | -0.113 | -0.113 | 0 | 0.2 | 0.02 | 0.28 | 0.38 | 0.32 | 0.19 |
| 2 | 205951940 | rs16836906 | 1.189 | -0.302 | -0.65 | 0.082 | 0.027 | 0.072 | 0.072 | 0 | 0.05 | 0.14 | 0.31 | 0.11 | 0.05 | 0.16 |
| 2 | 205952511 | rs10187682 | 1.457 | -0.621 | -0.828 | -0.285 | 0.009 | 0.032 | 0.032 | 0.053 | 0.11 | 0.05 | 0.05 | 0 | 0.08 | 0.02 |
| 2 | 205954308 | rs10194091 | -0.71 | -0.556 | -0.767 | -0.129 | 0 | -0.001 | -0.001 | 0.162 | 0.17 | 0.1 | 0.4 | 0.22 | 0.24 | 0.18 |
| 2 | 205955471 | rs1434159 | -1.442 | -1.072 | -0.585 | -0.444 | -0.003 | 0.001 | 0.001 | 0 | 0.71 | 0.69 | 0.86 | 0.61 | 0.78 | 0.58 |
| 2 | 205957557 | rs4675494 | 0.142 | -1.004 | -0.786 | -0.682 | -0.033 | -0.047 | -0.047 | 0 | 0.39 | 0.22 | 0.55 | 0.42 | 0.39 | 0.34 |
| 2 | 205967315 | rs13429972 | 0.031 | -0.825 | -0.474 | -0.559 | -0.005 | -0.001 | -0.001 | 0.007 | 0.14 | 0.11 | 0.13 | 0.16 | 0.17 | 0.16 |
| 2 | 205969220 | rs2289025 | 0.553 | -1.188 | -0.368 | -0.798 | -0.009 | -0.043 | -0.043 | 3.334 | 0.58 | 0.52 | 0.72 | 0.81 | 0.59 | 0.75 |
| 2 | 205971820 | rs13000345 | 0.856 | -1.041 | -0.265 | -0.734 | -0.015 | 0.098 | 0.098 | 0.584 | 0.25 | 0.12 | 0.12 | 0.03 | 0.22 | 0.1 |
| 2 | 205975381 | rs12477364 | 2.381 | -1.187 | -0.06 | -0.666 | 0.002 | -0.029 | -0.029 | 0.935 | 0.23 | 0.11 | 0.12 | 0.37 | 0.21 | 0.37 |
| 2 | 205977739 | rs13385430 | 0.697 | -1.157 | -0.084 | -0.667 | -0.005 | 0.005 | 0.005 | 0 | 0.11 | 0.1 | 0.13 | 0.15 | 0.16 | 0.13 |
| 2 | 205990030 | rs2276677 | 0.25 | -1.171 | -0.03 | -0.7 | -0.009 | -0.043 | -0.043 | 0 | 0.61 | 0.61 | 0.74 | 0.82 | 0.61 | 0.77 |
| 2 | 205990353 | rs2276676 | 1.742 | -1.023 | -0.269 | -0.662 | -0.049 | -0.063 | -0.063 | 0.008 | 0.21 | 0.11 | 0.49 | 0.42 | 0.3 | 0.33 |
| 2 | 205995406 | rs16837000 | 2.059 | -1.07 | -0.125 | -0.548 | 0.005 | -0.019 | -0.019 | 1.044 | 0.25 | 0.16 | 0.16 | 0.39 | 0.22 | 0.4 |
| 2 | 205997457 | rs919936 | -0.51 | -1.108 | -0.171 | -0.649 | 0.022 | 0.119 | 0.119 | 0.941 | 0.41 | 0.17 | 0.26 | 0.15 | 0.42 | 0.21 |
| 2 | 205999801 | rs16837012 | 1.728 | -1.043 | -0.055 | -0.588 | -0.007 | 0.012 | 0.012 | 0 | 0.27 | 0.31 | 0.18 | 0.4 | 0.23 | 0.42 |
| 2 | 206011030 | rs34045998 | 1.674 | -1.15 | -0.166 | -0.564 | -0.04 | 0.078 | 0.078 | 0.802 | 0.18 | 0.01 | 0.08 | 0.03 | 0.17 | 0.07 |
| 2 | 206016832 | rs1100393 | 1.246 | -1.008 | 0.022 | -0.481 | 0.026 | 0.04 | 0.04 | 0.006 | 0.26 | 0.29 | 0.53 | 0.43 | 0.34 | 0.36 |

| | | | | | | | | | | | | | | | | |
|---|---|---|---|---|---|---|---|---|---|---|---|---|---|---|---|---|
| 2 | 206023893 | rs6435271 | -0.699 | -1.512 | 0.145 | -0.388 | -0.016 | 0.011 | 0.011 | 0 | 0.65 | 0.49 | 0.79 | 0.57 | 0.73 | 0.55 |
| 2 | 206023947 | rs849121 | -0.634 | -1.497 | 0.049 | -0.297 | 0.024 | -0.001 | -0.001 | 0 | 0.62 | 0.8 | 0.8 | 0.59 | 0.69 | 0.56 |
| 2 | 206025813 | rs849116 | -1.04 | -1.225 | 0.286 | -0.125 | -0.013 | 0.028 | 0.028 | 0 | 0.7 | 0.55 | 0.8 | 0.6 | 0.74 | 0.59 |
| 2 | 206026296 | rs849115 | -1.547 | -1.074 | -0.54 | -0.079 | 0.009 | -0.004 | -0.004 | 0 | 0.74 | 0.82 | 0.83 | 0.67 | 0.77 | 0.59 |
| 2 | 206030543 | rs10202338 | -0.691 | -0.852 | -0.402 | -0.399 | -0.009 | -0.033 | -0.033 | 1.804 | 0.43 | 0.45 | 0.31 | 0.17 | 0.42 | 0.22 |
| 2 | 206031122 | rs17188399 | 0.868 | -0.655 | -0.529 | -0.346 | -0.007 | 0.04 | 0.04 | 0.339 | 0.19 | 0.09 | 0.12 | 0.05 | 0.19 | 0.1 |
| 2 | 206033781 | rs13417480 | 0.631 | -0.661 | -0.623 | -0.339 | 0.018 | 0.025 | 0.025 | 0 | 0.19 | 0.1 | 0.17 | 0.12 | 0.22 | 0.13 |
| 2 | 206035066 | rs1468997 | -0.274 | -0.438 | -0.89 | -0.215 | -0.001 | -0.01 | -0.01 | 0.341 | 0.4 | 0.4 | 0.32 | 0.18 | 0.42 | 0.23 |
| 2 | 206036918 | rs736320 | -0.58 | -0.251 | -1.251 | 0.309 | 0.009 | -0.016 | -0.016 | 0 | 0.71 | 0.81 | 0.7 | 0.61 | 0.77 | 0.63 |
| 2 | 206046172 | rs7558785 | -0.203 | 0.252 | -1.212 | -0.205 | 0 | -0.005 | -0.005 | 0 | 0.19 | 0.21 | 0.14 | 0.07 | 0.17 | 0.1 |
| 2 | 206048814 | rs10170188 | 0.548 | -0.185 | -1.694 | -0.651 | -0.005 | 0.108 | 0.108 | 0.014 | 0.27 | 0.1 | 0.14 | 0.02 | 0.26 | 0.29 |
| 2 | 206051516 | rs11892047 | -0.136 | -0.185 | -1.268 | -0.361 | -0.004 | 0.012 | 0.012 | 0 | 0.11 | 0.19 | 0.09 | 0.18 | 0.14 | 0.14 |
| 2 | 206057391 | rs12468546 | 0.753 | -0.417 | -1.729 | -0.741 | -0.018 | 0.139 | 0.139 | 0.088 | 0.3 | 0.09 | 0.17 | 0.02 | 0.29 | 0.25 |
| 2 | 206057649 | rs10932098 | 0.436 | -0.845 | -1.928 | -0.294 | 0.026 | -0.041 | -0.041 | 1.474 | 0.15 | 0.1 | 0.06 | 0.26 | 0.09 | 0.06 |
| 2 | 206058350 | rs10932099 | -1.042 | -1.035 | -1.238 | -0.42 | -0.002 | -0.014 | -0.014 | 0.356 | 0.55 | 0.53 | 0.34 | 0.28 | 0.5 | 0.4 |
| 2 | 206059325 | rs969516 | -1.325 | -0.994 | -1.365 | -0.461 | -0.007 | 0.114 | 0.114 | 0.108 | 0.7 | 0.86 | 0.83 | 0.98 | 0.71 | 0.74 |
| 2 | 206060511 | rs1008644 | 0.896 | -1.143 | -1.176 | -0.44 | -0.001 | -0.001 | -0.001 | 0.001 | 0.46 | 0.52 | 0.67 | 0.63 | 0.51 | 0.58 |
| 2 | 206062549 | rs1921788 | -0.835 | -1.013 | -1.534 | 0.056 | -0.04 | 0.025 | 0.025 | 0.017 | 0.62 | 0.92 | 0.73 | 0.76 | 0.66 | 0.65 |
| 2 | 206063298 | rs11681930 | -0.85 | -1.511 | -1.255 | -0.703 | -0.026 | 0.031 | 0.031 | 0 | 0.22 | 0.34 | 0.16 | 0.34 | 0.22 | 0.19 |
| 2 | 206075239 | rs11901441 | -0.028 | -1.504 | -1.22 | -0.805 | -0.002 | -0.021 | -0.021 | 0.019 | 0.11 | 0.28 | 0.12 | 0.08 | 0.13 | 0.13 |
| 2 | 206076974 | rs11904275 | -0.028 | -1.504 | -1.22 | -0.805 | -0.002 | -0.021 | -0.021 | 0 | 0.11 | 0.28 | 0.12 | 0.08 | 0.13 | 0.13 |
| 2 | 206077638 | rs1921782 | 0.252 | -1.951 | -1.348 | -0.944 | 0.023 | 0.003 | 0.003 | 0 | 0.56 | 0.83 | 0.8 | 0.66 | 0.62 | 0.7 |
| 2 | 206081546 | rs1921779 | -0.689 | -1.863 | -1.333 | -0.978 | -0.026 | 0.148 | 0.148 | 0.029 | 0.7 | 0.93 | 0.86 | 0.95 | 0.72 | 0.76 |
| 2 | 206081610 | rs1921778 | -0.771 | -1.862 | -1.333 | -0.978 | -0.025 | 0.15 | 0.15 | 0.018 | 0.7 | 0.93 | 0.85 | 0.95 | 0.72 | 0.76 |
| 2 | 206085473 | rs6750663 | -0.768 | -2.297 | -1.295 | -0.973 | 0.018 | 0.136 | 0.136 | 0.017 | 0.7 | 0.89 | 0.85 | 0.98 | 0.75 | 0.77 |
| 2 | 206087119 | rs1921797 | 1.206 | -2.401 | -1.203 | -0.979 | -0.002 | -0.001 | -0.001 | 0.314 | 0.46 | 0.51 | 0.68 | 0.58 | 0.57 | 0.58 |
| 2 | 206088211 | rs10182802 | -1.483 | -2.232 | -0.916 | -0.709 | -0.038 | 0.065 | 0.065 | 0 | 0.24 | 0.38 | 0.17 | 0.39 | 0.18 | 0.19 |
| 2 | 206090248 | rs10165609 | -0.226 | -2.143 | -1.415 | -0.686 | 0.015 | -0.042 | -0.042 | 0 | 0.13 | 0.08 | 0.06 | 0.32 | 0.1 | 0.07 |
| 2 | 206096536 | rs11888554 | -0.664 | -2.121 | -1.344 | -0.724 | -0.009 | 0.014 | 0.014 | 0 | 0.2 | 0.26 | 0.15 | 0.32 | 0.16 | 0.16 |
| 2 | 206105258 | rs1012337 | 1.415 | -1.879 | -1.305 | -1.283 | -0.062 | -0.046 | -0.046 | 0 | 0.31 | 0.16 | 0.57 | 0.56 | 0.42 | 0.51 |
| 2 | 206115788 | rs11682188 | -1.143 | -2.677 | -0.389 | -1.254 | -0.007 | -0.02 | -0.02 | 0 | 0.13 | 0.23 | 0.11 | 0.12 | 0.14 | 0.15 |
| 2 | 206118484 | rs10490297 | -0.426 | -2.863 | -0.415 | -1.272 | -0.017 | -0.026 | -0.026 | 0 | 0.11 | 0.18 | 0.09 | 0.08 | 0.09 | 0.12 |
| 2 | 206125313 | rs11892551 | -1.279 | -3.177 | -0.368 | -1.368 | -0.024 | -0.002 | -0.002 | 2.102 | 0.25 | 0.31 | 0.15 | 0.28 | 0.16 | 0.16 |
| 2 | 206127438 | rs10202153 | -1.646 | -3.177 | -0.372 | -1.359 | -0.017 | 0 | 0 | 0 | 0.28 | 0.34 | 0.18 | 0.33 | 0.21 | 0.2 |
| 2 | 206131317 | rs2336413 | 0.93 | -3.482 | -0.762 | -1.616 | 0.066 | 0.08 | 0.08 | 0 | 0.45 | 0.28 | 0.71 | 0.7 | 0.63 | 0.58 |
| 2 | 206132248 | rs1540364 | 0.785 | -3.734 | -0.034 | -1.135 | 0.005 | -0.006 | -0.006 | 0.579 | 0.83 | 0.85 | 0.85 | 0.76 | 0.86 | 0.93 |
| 2 | 206135584 | rs1540360 | 0.04 | -3.658 | -0.133 | -1.059 | 0.021 | -0.028 | -0.028 | 0 | 0.12 | 0.04 | 0.05 | 0.2 | 0.07 | 0.05 |
| 2 | 206139804 | rs6435279 | 1.24 | -3.908 | 0.245 | -1.245 | 0.098 | 0.064 | 0.064 | 0 | 0.51 | 0.28 | 0.71 | 0.59 | 0.63 | 0.56 |
| 2 | 206139895 | rs1882381 | 0.712 | -4.043 | -0.073 | -1.375 | 0.039 | 0.113 | 0.113 | 0 | 0.2 | 0.02 | 0.08 | 0.02 | 0.13 | 0.22 |
| 2 | 206140982 | rs16837138 | -0.051 | -3.957 | 0.268 | -1.354 | 0.005 | 0.007 | 0.007 | 0 | 0.1 | 0.05 | 0.08 | 0.1 | 0.1 | 0.13 |
| 2 | 206144833 | rs7606468 | 0.661 | -3.78 | 1.072 | -1.239 | -0.055 | -0.016 | -0.016 | 0 | 0.64 | 0.52 | 0.83 | 0.67 | 0.77 | 0.71 |
| 2 | 206150594 | rs1207421 | -0.024 | -2.748 | 1.02 | -1.026 | 0 | -0.025 | -0.025 | 1.403 | 0.9 | 0.93 | 0.95 | 0.72 | 0.9 | 0.93 |
| 2 | 206155107 | rs1207425 | -1.394 | -3.229 | 1.484 | -1.42 | -0.012 | -0.082 | -0.082 | 0.264 | 0.37 | 0.53 | 0.19 | 0 | 0.22 | 0.09 |
| 2 | 206158317 | rs2704638 | 1.225 | -2.529 | 2.282 | -0.648 | -0.005 | 0.045 | 0.045 | 0 | 0.85 | 0.74 | 0.91 | 0.7 | 0.85 | 0.92 |
| 2 | 206159496 | rs1921799 | 0.467 | -2.57 | 2.566 | -0.673 | -0.001 | -0.023 | -0.023 | 0.056 | 0.12 | 0.08 | 0.07 | 0 | 0.1 | 0.06 |
| 2 | 206160453 | rs4673317 | 1.298 | -2.456 | 2.348 | -0.609 | -0.003 | 0.043 | 0.043 | 0.028 | 0.84 | 0.71 | 0.9 | 0.7 | 0.84 | 0.92 |
| 2 | 206182051 | rs4119081 | 1.906 | -2.592 | 2.475 | -0.765 | 0.013 | 0.005 | 0.005 | 0 | 0.61 | 0.44 | 0.82 | 0.68 | 0.72 | 0.8 |
| 2 | 206183690 | rs1207440 | -0.021 | -2.592 | 2.351 | -0.765 | 0 | -0.026 | -0.026 | 0 | 0.9 | 0.96 | 0.96 | 0.71 | 0.91 | 0.93 |
| 2 | 206184330 | rs1207439 | -1.946 | -2.603 | 2.409 | -0.855 | 0.004 | -0.029 | -0.029 | 0.095 | 0.32 | 0.67 | 0.17 | 0.03 | 0.23 | 0.14 |
| 2 | 206185567 | rs1207438 | 1.367 | -2.262 | 2.566 | -0.681 | -0.006 | 0.047 | 0.047 | 0 | 0.84 | 0.74 | 0.92 | 0.72 | 0.85 | 0.91 |
| 2 | 206188122 | rs849189 | 0.791 | -2.261 | 2.523 | -0.679 | -0.007 | 0.062 | 0.062 | 0 | 0.88 | 0.82 | 0.94 | 0.72 | 0.9 | 0.93 |
| 2 | 206195233 | rs2541155 | -1.585 | -2.29 | 2.505 | -0.701 | -0.031 | -0.079 | -0.079 | 0 | 0.27 | 0.58 | 0.12 | 0.03 | 0.18 | 0.12 |
| 2 | 206198319 | rs3845806 | 2.351 | -2.28 | 2.108 | -0.694 | 0.091 | 0.045 | 0.045 | 0.611 | 0.57 | 0.12 | 0.78 | 0.69 | 0.67 | 0.79 |
| 2 | 206198786 | rs849221 | 0.157 | -2.274 | 2.126 | -0.681 | 0.002 | -0.018 | -0.018 | 0 | 0.9 | 0.94 | 0.95 | 0.72 | 0.9 | 0.93 |
| 2 | 206203291 | rs12476682 | 2.023 | -2.263 | 2.064 | -0.669 | 0.095 | 0.031 | 0.031 | 0.224 | 0.6 | 0.19 | 0.82 | 0.69 | 0.72 | 0.8 |
| 2 | 206208171 | rs4505506 | 1.906 | -2.254 | 1.962 | -0.666 | 0.093 | 0.026 | 0.026 | 0.31 | 0.6 | 0.16 | 0.81 | 0.69 | 0.72 | 0.8 |
| 2 | 206211404 | rs849252 | -0.414 | -2.327 | 2.425 | -0.687 | -0.019 | -0.07 | -0.07 | 0.015 | 0.17 | 0.22 | 0.09 | 0 | 0.12 | 0.06 |
| 2 | 206223884 | rs10490292 | 2.045 | -2.297 | 1.98 | -0.659 | 0.087 | 0.024 | 0.024 | 0.243 | 0.6 | 0.17 | 0.81 | 0.68 | 0.71 | 0.8 |
| 2 | 206232244 | rs1990667 | 0.744 | -2.209 | 1.901 | -0.736 | 0.003 | -0.034 | -0.034 | 0 | 0.87 | 0.92 | 0.94 | 0.71 | 0.87 | 0.88 |
| 2 | 206236189 | rs849205 | -1.788 | -2.006 | 2.503 | -1.569 | -0.049 | -0.163 | -0.163 | 0.455 | 0.27 | 0.51 | 0.14 | 0 | 0.2 | 0.08 |
| 2 | 206236440 | rs849203 | -0.587 | -1.806 | 1.768 | -1.286 | 0.01 | 0.006 | 0.006 | 0 | 0.16 | 0.13 | 0.08 | 0 | 0.15 | 0.06 |
| 2 | 206256232 | rs7600287 | 1.749 | -1.681 | 2.073 | -1.233 | -0.023 | -0.095 | -0.095 | 1.461 | 0.67 | 0.44 | 0.83 | 0.96 | 0.72 | 0.85 |
| 2 | 206257794 | rs849235 | 0.84 | -1.721 | 1.869 | -1.179 | -0.022 | -0.042 | -0.042 | 0.485 | 0.87 | 0.72 | 0.94 | 0.97 | 0.92 | 0.94 |
| 2 | 206259137 | rs698906 | 1.515 | -1.737 | 1.26 | -1.176 | -0.017 | -0.033 | -0.033 | 2.084 | 0.78 | 0.34 | 0.89 | 0.96 | 0.86 | 0.89 |
| 2 | 206262570 | rs849227 | -1.435 | -1.27 | 1.159 | -0.789 | -0.012 | -0.057 | -0.057 | 0.015 | 0.23 | 0.67 | 0.16 | 0.01 | 0.15 | 0.09 |
| 2 | 206266711 | rs2111079 | 1.478 | -0.952 | 1.525 | -0.295 | -0.046 | -0.06 | -0.06 | 0 | 0.88 | 0.52 | 0.95 | 0.99 | 0.96 | 0.94 |
| 2 | 206284251 | rs849259 | 1.723 | -0.949 | 0.864 | -0.293 | -0.064 | -0.075 | -0.075 | 0.425 | 0.85 | 0.42 | 0.93 | 0.98 | 0.95 | 0.94 |
| 2 | 206295119 | rs849243 | 1.608 | -0.615 | 0.741 | -0.264 | -0.004 | -0.038 | -0.038 | 2.468 | 0.92 | 0.81 | 0.96 | 0.99 | 0.94 | 0.97 |
| 2 | 206306315 | rs6435284 | -2.218 | -1.218 | 0.887 | -0.656 | -0.014 | -0.043 | -0.043 | 0.174 | 0.21 | 0.68 | 0.12 | 0.02 | 0.12 | 0.07 |

| | | | | | | | | | | | | | | | | |
|---|---|---|---|---|---|---|---|---|---|---|---|---|---|---|---|---|
| 2 | 206345419 | rs6714542 | -0.133 | -1.248 | 0.538 | -0.466 | -0.012 | -0.007 | -0.007 | 0 | 0.41 | 0.53 | 0.26 | 0.37 | 0.31 | 0.44 |
| 2 | 206346002 | rs6760289 | -0.467 | -1.595 | 0.7 | -0.458 | -0.011 | -0.015 | -0.015 | 0 | 0.1 | 0.14 | 0.07 | 0.07 | 0.07 | 0.07 |
| 2 | 206346565 | rs4675530 | 0.378 | -1.169 | -0.039 | 0.235 | 0.019 | 0.022 | 0.022 | 0 | 0.44 | 0.18 | 0.64 | 0.38 | 0.57 | 0.37 |
| 2 | 206363651 | rs4675531 | 0.99 | -0.456 | 0.722 | 0.303 | 0.008 | 0.075 | 0.075 | 0 | 0.93 | 0.73 | 0.97 | 0.88 | 0.97 | 0.86 |
| 2 | 206363729 | rs4675532 | -0.237 | -0.204 | 0.429 | 0.152 | 0.004 | 0.005 | 0.005 | 0 | 0.34 | 0.36 | 0.57 | 0.47 | 0.41 | 0.47 |
| 2 | 206370333 | rs994044 | 0.445 | 0.039 | 0.678 | 0.297 | -0.114 | -0.109 | -0.109 | 0.872 | 0.22 | 0.02 | 0.49 | 0.39 | 0.32 | 0.38 |
| 2 | 206371240 | rs4346365 | -1.021 | 0.002 | 0.98 | 0.325 | 0.001 | 0.001 | 0.001 | 0.014 | 0.37 | 0.37 | 0.58 | 0.58 | 0.43 | 0.62 |
| 2 | 206374024 | rs12474620 | -0.686 | -0.018 | 0.971 | 0.35 | 0.001 | 0.002 | 0.002 | 0.078 | 0.32 | 0.33 | 0.56 | 0.47 | 0.41 | 0.47 |
| 2 | 206377815 | rs7576084 | 1.165 | -0.091 | 1.181 | 0.285 | -0.009 | -0.017 | -0.017 | 0 | 0.58 | 0.32 | 0.4 | 0.42 | 0.57 | 0.39 |
| 2 | 206391545 | rs992159 | 1.973 | -0.147 | 1.481 | 0.165 | -0.033 | -0.018 | -0.018 | 0 | 0.87 | 0.27 | 0.91 | 0.89 | 0.93 | 0.85 |
| 2 | 206399128 | rs6724376 | 0.901 | 0.002 | 1.479 | 0.138 | -0.008 | -0.014 | -0.014 | 0 | 0.63 | 0.34 | 0.41 | 0.43 | 0.59 | 0.56 |
| 2 | 206425066 | rs2192933 | 1.122 | 0.171 | 1.377 | 0.387 | 0.016 | 0.036 | 0.036 | 0 | 0.66 | 0.49 | 0.48 | 0.5 | 0.6 | 0.69 |
| 2 | 206428228 | rs3919545 | -0.119 | 0.271 | 0.934 | 0.536 | -0.045 | -0.055 | -0.055 | 0.264 | 0.2 | 0.07 | 0.4 | 0.38 | 0.3 | 0.2 |
| 2 | 206432563 | rs7566612 | 0.884 | 0.498 | 1.202 | 0.518 | 0.006 | 0.017 | 0.017 | 0.054 | 0.69 | 0.42 | 0.69 | 0.56 | 0.65 | 0.73 |
| 2 | 206432930 | rs16837439 | -0.637 | 0.353 | 1.559 | 0.823 | -0.009 | 0.042 | 0.042 | 0.467 | 0.12 | 0.34 | 0.08 | 0.13 | 0.07 | 0.1 |
| 2 | 206435709 | rs7420812 | -0.431 | 0.712 | 0.774 | 1.085 | -0.008 | -0.014 | -0.014 | 0 | 0.14 | 0.07 | 0.12 | 0.29 | 0.16 | 0.15 |
| 2 | 206437137 | rs2041832 | 1.047 | 0.591 | 0.162 | 1.27 | -0.004 | -0.004 | -0.004 | 0.975 | 0.31 | 0.22 | 0.28 | 0.23 | 0.34 | 0.28 |
| 2 | 206438036 | rs12694029 | 1.511 | 0.388 | 0.027 | 1.099 | -0.021 | 0.026 | 0.026 | 0.412 | 0.54 | 0.23 | 0.72 | 0.57 | 0.55 | 0.62 |
| 2 | 206443038 | rs9288370 | 0.125 | 0.73 | 0.301 | 1.245 | 0.005 | 0.031 | 0.031 | 0 | 0.13 | 0.16 | 0.12 | 0.31 | 0.16 | 0.15 |
| 2 | 206443651 | rs6747305 | 1.324 | 0.643 | 0.39 | 1.177 | 0.001 | 0.027 | 0.027 | 0 | 0.34 | 0.23 | 0.28 | 0.18 | 0.34 | 0.29 |
| 2 | 206446775 | rs10932118 | -0.307 | 0.606 | 1.106 | 1.169 | 0.001 | 0.002 | 0.002 | 0 | 0.52 | 0.44 | 0.58 | 0.52 | 0.52 | 0.58 |
| 2 | 206449649 | rs17596643 | 1.681 | 0.569 | 1.119 | 1.075 | 0.014 | -0.051 | -0.051 | 0 | 0.25 | 0.16 | 0.49 | 0.48 | 0.21 | 0.33 |
| 2 | 206452025 | rs2058712 | 0.877 | 0.605 | 1.966 | 0.831 | 0.009 | 0.024 | 0.024 | 0.104 | 0.36 | 0.25 | 0.29 | 0.18 | 0.34 | 0.29 |
| 2 | 206454215 | rs12618000 | -1.061 | 0.59 | 1.83 | 0.887 | 0.008 | 0.031 | 0.031 | 0.074 | 0.63 | 0.72 | 0.71 | 0.85 | 0.65 | 0.71 |
| 2 | 206455402 | rs11675753 | -0.742 | 0.588 | 1.629 | 0.857 | -0.007 | -0.013 | -0.013 | 0 | 0.59 | 0.49 | 0.5 | 0.82 | 0.65 | 0.71 |
| 2 | 206460083 | rs17682318 | 1.159 | 0.635 | 1.348 | 0.069 | 0.022 | 0.092 | 0.092 | 0.926 | 0.32 | 0.17 | 0.23 | 0 | 0.26 | 0.07 |
| 2 | 206461610 | rs6435293 | -0.868 | 0.708 | 1.551 | 0.152 | -0.002 | 0.014 | 0.014 | 0.096 | 0.59 | 0.59 | 0.4 | 0.26 | 0.57 | 0.35 |
| 2 | 206464465 | rs1079943 | -0.879 | 0.776 | 1.227 | -0.048 | -0.003 | 0.017 | 0.017 | 0 | 0.59 | 0.66 | 0.43 | 0.18 | 0.56 | 0.35 |
| 2 | 206480054 | rs13387277 | -0.531 | 0.915 | 0.963 | 0.092 | 0.037 | -0.035 | -0.035 | 1.238 | 0.24 | 0.44 | 0.15 | 0.18 | 0.28 | 0.26 |
| 2 | 206481573 | rs4673336 | -0.852 | 0.511 | 0.597 | -0.176 | 0 | -0.07 | -0.07 | 2.129 | 0.65 | 0.7 | 0.77 | 1 | 0.73 | 0.93 |
| 2 | 206484652 | rs11675303 | 0.697 | 0.788 | 0.745 | 0.965 | -0.03 | 0.049 | 0.049 | 0 | 0.09 | 0.01 | 0.05 | 0 | 0.11 | 0.03 |
| 2 | 206485381 | rs11675671 | 0.179 | 0.786 | 0.798 | 0.979 | -0.003 | 0.007 | 0.007 | 0 | 0.11 | 0.06 | 0.06 | 0.07 | 0.12 | 0.13 |
| 2 | 206490951 | rs6435297 | 0.493 | 0.656 | 0.963 | 1.319 | 0.035 | 0.243 | 0.243 | 0 | 0.89 | 0.59 | 0.74 | 0.64 | 0.9 | 0.62 |
| 2 | 206495714 | rs6739327 | -0.138 | 0.311 | -0.041 | 0.41 | 0 | -0.015 | -0.015 | 0 | 0.88 | 0.89 | 0.94 | 1 | 0.89 | 0.92 |
| 2 | 206496702 | rs918058 | -0.084 | 0.191 | -0.282 | 0.524 | -0.027 | -0.031 | -0.031 | 0 | 0.36 | 0.29 | 0.29 | 0.46 | 0.5 | 0.46 |
| 2 | 206496749 | rs1861441 | -0.317 | -0.073 | -0.069 | 0.47 | -0.033 | -0.04 | -0.04 | 0 | 0.23 | 0.36 | 0.13 | 0.13 | 0.18 | 0.2 |
| 2 | 206500903 | rs2358495 | -0.607 | 0.245 | -0.538 | 0.468 | 0.01 | 0.011 | 0.011 | 0 | 0.56 | 0.61 | 0.59 | 0.6 | 0.67 | 0.68 |
| 2 | 206505305 | rs759455 | 0.244 | 0.386 | -1.117 | 0.472 | -0.003 | 0.139 | 0.139 | 0.54 | 0.48 | 0.17 | 0.43 | 0.31 | 0.46 | 0.35 |
| 2 | 206508387 | rs849506 | -1.443 | 0.095 | -0.065 | 0.953 | -0.027 | 0.088 | 0.088 | 2.306 | 0.08 | 0.27 | 0.04 | 0.16 | 0.02 | 0.11 |
| 2 | 206511008 | rs6435300 | -0.189 | 0.461 | -0.662 | 0.925 | -0.015 | 0.001 | 0.001 | 0 | 0.59 | 0.68 | 0.52 | 0.57 | 0.53 | 0.59 |
| 2 | 206518838 | rs849511 | -0.22 | 0.987 | -0.972 | 0.978 | -0.001 | -0.002 | -0.002 | 0 | 0.45 | 0.3 | 0.48 | 0.55 | 0.48 | 0.6 |
| 2 | 206528291 | rs7569933 | 0.67 | 0.573 | -1.156 | 0.67 | -0.005 | -0.035 | -0.035 | 0 | 0.37 | 0.17 | 0.36 | 0.41 | 0.37 | 0.32 |
| 2 | 206536329 | rs2160327 | 0.15 | 0.557 | -1.169 | 0.58 | -0.019 | 0.009 | 0.009 | 0.954 | 0.39 | 0.15 | 0.46 | 0.37 | 0.45 | 0.48 |
| 2 | 206544665 | rs6711044 | 0.128 | 0.52 | -1.348 | 0.738 | -0.024 | -0.004 | -0.004 | 0 | 0.45 | 0.16 | 0.47 | 0.42 | 0.47 | 0.28 |
| 2 | 206545465 | rs2192884 | 0.368 | 0.538 | -0.693 | 0.361 | 0.007 | -0.05 | -0.05 | 0.4 | 0.84 | 0.73 | 0.87 | 0.95 | 0.88 | 0.94 |
| 2 | 206546150 | rs3806577 | 0.423 | 0.172 | -1.024 | 0.572 | -0.071 | -0.085 | -0.085 | 0 | 0.4 | 0.1 | 0.42 | 0.45 | 0.46 | 0.59 |
| 2 | 206547545 | rs10090 | -0.268 | -0.248 | 0.023 | 0.404 | -0.001 | 0 | 0 | 3.538 | 0.54 | 0.42 | 0.5 | 0.54 | 0.49 | 0.44 |
| 2 | 206551978 | rs861078 | 1.409 | -0.006 | -0.252 | -0.175 | -0.002 | 0.013 | 0.013 | 1.344 | 0.65 | 0.37 | 0.65 | 0.9 | 0.68 | 0.84 |
| 2 | 206554544 | rs698909 | 0.859 | 0.089 | 0.082 | -0.329 | 0.011 | -0.084 | -0.084 | 0 | 0.72 | 0.67 | 0.7 | 0.96 | 0.69 | 0.87 |
| 2 | 206560613 | rs950219 | -0.958 | -0.331 | 0.225 | -0.514 | 0.003 | -0.11 | -0.11 | 0 | 0.32 | 0.38 | 0.32 | 0.04 | 0.3 | 0.13 |
| 2 | 206561427 | rs3771052 | -0.636 | -0.343 | 0.063 | -0.524 | -0.025 | 0.073 | 0.073 | 0 | 0.28 | 0.14 | 0.3 | 0.04 | 0.31 | 0.13 |
| 2 | 206563258 | rs849556 | 1.057 | -0.356 | 0.255 | -0.339 | -0.003 | -0.051 | -0.051 | 0 | 0.65 | 0.45 | 0.72 | 0.9 | 0.71 | 0.82 |
| 2 | 206565500 | rs3771051 | -0.02 | -0.255 | 0.11 | -0.372 | -0.023 | 0.048 | 0.048 | 0 | 0.18 | 0.08 | 0.14 | 0.04 | 0.22 | 0.1 |
| 2 | 206569334 | rs849552 | 1.164 | -0.431 | 0.402 | -0.072 | 0.012 | 0.039 | 0.039 | 2.126 | 0.92 | 0.65 | 0.91 | 0.94 | 0.98 | 0.95 |
| 2 | 206571135 | rs3771048 | 0.284 | -0.415 | -0.755 | 0.422 | -0.022 | -0.088 | -0.088 | 0.147 | 0.15 | 0.05 | 0.22 | 0.44 | 0.18 | 0.3 |
| 2 | 206577411 | rs3771044 | -0.294 | -0.742 | -0.542 | -0.029 | -0.032 | -0.02 | -0.02 | 0.068 | 0.19 | 0.35 | 0.19 | 0.15 | 0.11 | 0.13 |
| 2 | 206586412 | rs3771038 | 0.049 | -0.522 | -1.08 | 0.134 | -0.009 | 0.004 | 0.004 | 0 | 0.14 | 0.18 | 0.09 | 0.15 | 0.08 | 0.11 |
| 2 | 206589121 | rs861079 | 0.631 | -0.086 | -2.099 | -0.119 | -0.136 | 0.134 | 0.134 | 3.224 | 0.43 | 0.06 | 0.47 | 0.8 | 0.53 | 0.74 |
| 2 | 206592525 | rs849527 | 0.368 | -0.071 | -2.099 | -0.125 | -0.098 | 0.187 | 0.187 | 1.237 | 0.47 | 0.08 | 0.55 | 0.81 | 0.56 | 0.73 |
| 2 | 206592695 | rs849526 | 0.383 | -0.061 | -2.092 | -0.12 | -0.1 | 0.184 | 0.184 | 1.196 | 0.47 | 0.08 | 0.55 | 0.81 | 0.56 | 0.74 |
| 2 | 206595810 | rs3771033 | 1.204 | 0.107 | -1.956 | 0.307 | -0.085 | -0.085 | -0.085 | 0 | 0.28 | 0.09 | 0.37 | 0.34 | 0.35 | 0.47 |
| 2 | 206595936 | rs849523 | -0.768 | -0.178 | -1.048 | 0.705 | -0.012 | 0.028 | 0.028 | 0.033 | 0.28 | 0.29 | 0.26 | 0.59 | 0.24 | 0.38 |
| 2 | 206602710 | rs849582 | -0.665 | -0.292 | -0.974 | 0.891 | -0.02 | 0.082 | 0.082 | 0.02 | 0.13 | 0.2 | 0.12 | 0.42 | 0.1 | 0.26 |
| 2 | 206605180 | rs849575 | -0.696 | -0.276 | -1.089 | 0.863 | -0.011 | 0.046 | 0.046 | 0 | 0.16 | 0.19 | 0.15 | 0.45 | 0.14 | 0.29 |
| 2 | 206609988 | rs3771021 | 0.234 | -0.197 | -1.855 | 0.52 | -0.066 | -0.012 | -0.012 | 0 | 0.38 | 0.14 | 0.45 | 0.34 | 0.45 | 0.29 |
| 2 | 206610051 | rs849565 | -1.458 | -0.66 | -1.065 | 0.711 | 0.003 | -0.005 | -0.005 | 0 | 0.28 | 0.67 | 0.29 | 0.46 | 0.2 | 0.31 |
| 2 | 206610502 | rs849563 | -0.597 | -0.4 | -1.439 | 1.083 | -0.018 | 0.053 | 0.053 | 0 | 0.16 | 0.21 | 0.22 | 0.46 | 0.12 | 0.28 |
| 2 | 206612585 | rs1996412 | 0.004 | -0.126 | -2.472 | 0.721 | -0.14 | -0.178 | -0.178 | 0.053 | 0.35 | 0.04 | 0.42 | 0.51 | 0.5 | 0.32 |
| 2 | 206612714 | rs12472412 | 0.323 | -0.064 | -2.209 | 0.831 | -0.035 | -0.039 | -0.039 | 0 | 0.17 | 0 | 0.18 | 0.16 | 0.14 | 0.28 |

| | | | | | | | | | | | | | | | | |
|---|---|---|---|---|---|---|---|---|---|---|---|---|---|---|---|---|
| 2 | 206614091 | rs13026243 | 0.261 | -0.118 | -1.594 | 0.939 | -0.004 | -0.01 | -0.01 | 0 | 0.69 | 0.37 | 0.61 | 0.53 | 0.66 | 0.39 |
| 2 | 206614160 | rs849560 | 1.068 | -0.609 | -0.459 | 0.835 | -0.038 | 0.07 | 0.07 | 0 | 0.77 | 0.54 | 0.81 | 0.69 | 0.82 | 0.71 |
| 2 | 206616011 | rs3771014 | -0.057 | -0.409 | -0.647 | 0.519 | 0.011 | 0.02 | 0.02 | 0 | 0.07 | 0.02 | 0.02 | 0.01 | 0.05 | 0.03 |
| 2 | 206622576 | rs10180293 | -1.684 | -0.531 | -0.071 | 0.288 | -0.023 | -0.052 | -0.052 | 0 | 0.2 | 0.41 | 0.14 | 0.12 | 0.16 | 0.16 |
| 2 | 206627473 | rs872943 | -1.188 | -0.409 | -0.17 | 0.506 | -0.02 | -0.011 | -0.011 | 0.003 | 0.14 | 0.28 | 0.11 | 0.12 | 0.1 | 0.16 |
| 2 | 206630887 | rs3771004 | 0.277 | -0.008 | -1.26 | 0.707 | 0.011 | -0.01 | -0.01 | 0.07 | 0.3 | 0.22 | 0.21 | 0.36 | 0.28 | 0.17 |
| 2 | 206631254 | rs16837637 | -0.228 | 0.233 | -1.659 | 0.451 | 0.007 | 0.041 | 0.041 | 1.24 | 0.53 | 0.25 | 0.64 | 0.64 | 0.61 | 0.43 |
| 2 | 206633843 | rs3771003 | -0.283 | 0.021 | -0.968 | 0.543 | -0.002 | 0.022 | 0.022 | 0 | 0.31 | 0.39 | 0.31 | 0.51 | 0.32 | 0.29 |
| 2 | 206634869 | rs16837641 | -0.557 | -0.306 | -1.36 | 0.44 | 0.019 | -0.039 | -0.039 | 0 | 0.37 | 0.17 | 0.38 | 0.57 | 0.34 | 0.32 |
| 2 | 206636123 | rs2241153 | 0.034 | 0.115 | -1.379 | 0.224 | -0.039 | -0.006 | -0.006 | 0 | 0.28 | 0.18 | 0.38 | 0.32 | 0.4 | 0.18 |
| 2 | 206639831 | rs3755233 | 0.664 | -0.432 | -0.563 | -0.247 | -0.021 | 0.027 | 0.027 | 0 | 0.16 | 0.33 | 0.17 | 0.21 | 0.11 | 0.08 |
| 2 | 206640640 | rs13027631 | -0.377 | -0.548 | -0.692 | -0.371 | -0.047 | -0.027 | -0.027 | 0 | 0.24 | 0.49 | 0.22 | 0.21 | 0.14 | 0.11 |
| 2 | 206642220 | rs3732088 | 0.975 | -0.436 | -0.878 | -0.512 | 0 | -0.007 | -0.007 | 0 | 0.12 | 0.13 | 0.19 | 0.02 | 0.11 | 0.04 |
| 2 | 206642337 | rs2160328 | 0.73 | -0.606 | -0.779 | -0.194 | -0.003 | -0.003 | -0.003 | 0 | 0.27 | 0.23 | 0.37 | 0.34 | 0.4 | 0.19 |
| 2 | 206645166 | rs3771000 | 0.856 | -0.504 | -1.526 | -0.228 | -0.001 | -0.006 | -0.006 | 0.324 | 0.26 | 0.29 | 0.19 | 0.16 | 0.18 | 0.4 |
| 2 | 206649743 | rs3770996 | 0.262 | -0.593 | -1.597 | -0.217 | -0.004 | -0.003 | -0.003 | 0 | 0.43 | 0.19 | 0.38 | 0.45 | 0.43 | 0.69 |
| 2 | 206652963 | rs3755232 | 0.28 | -0.227 | -1.475 | -0.199 | 0.008 | 0.025 | 0.025 | 0 | 0.33 | 0.32 | 0.24 | 0.16 | 0.26 | 0.43 |
| 2 | 206654123 | rs1990708 | -0.25 | -0.185 | -0.764 | 0.172 | -0.02 | 0.006 | 0.006 | 0 | 0.19 | 0.31 | 0.25 | 0.22 | 0.13 | 0.09 |
| 2 | 206662343 | rs9706 | 0.228 | 0.26 | -1.375 | 0.41 | -0.005 | 0.024 | 0.024 | 0 | 0.44 | 0.23 | 0.4 | 0.42 | 0.44 | 0.66 |
| 2 | 206665680 | rs13005956 | 0.397 | -0.212 | -0.781 | 0.323 | -0.004 | -0.003 | -0.003 | 1.022 | 0.69 | 0.39 | 0.61 | 0.76 | 0.78 | 0.89 |
| 2 | 206667368 | rs7590905 | -0.496 | -0.122 | -0.309 | -0.172 | -0.007 | -0.118 | -0.118 | 1.439 | 0.21 | 0.36 | 0.28 | 0 | 0.18 | 0.1 |
| 2 | 206667756 | rs10932127 | 0.33 | 0.253 | -0.962 | 0.484 | -0.064 | -0.059 | -0.059 | 0 | 0.4 | 0.23 | 0.46 | 0.5 | 0.48 | 0.64 |
| 2 | 206669214 | rs2041915 | 1.616 | 0.119 | 0.051 | 0.921 | -0.038 | 0.171 | 0.171 | 3.32 | 0.86 | 0.38 | 0.91 | 0.64 | 0.93 | 0.94 |
| 2 | 206673842 | rs2058803 | 1.105 | 0.667 | 0.042 | 0.053 | 0.004 | -0.04 | -0.04 | 0.344 | 0.78 | 0.37 | 0.83 | 0.87 | 0.79 | 0.62 |
| 2 | 206690019 | rs10182735 | -0.723 | 0.751 | 0.492 | -0.05 | 0.024 | -0.04 | -0.04 | 0 | 0.08 | 0.29 | 0.08 | 0.02 | 0.08 | 0.04 |
| 2 | 206699721 | rs10932131 | -0.117 | 0.549 | 0.337 | -0.722 | 0.001 | -0.059 | -0.059 | 0 | 0.5 | 0.52 | 0.44 | 0.06 | 0.42 | 0.36 |
| 2 | 206714121 | rs867851 | 0.287 | 0.798 | 0.383 | -0.215 | 0.013 | 0.014 | 0.014 | 0.608 | 0.46 | 0.15 | 0.54 | 0.55 | 0.58 | 0.6 |
| 2 | 206720318 | rs13017212 | -0.849 | 0.925 | 0.512 | -0.595 | 0.015 | -0.017 | -0.017 | 0.008 | 0.17 | 0.28 | 0.14 | 0.09 | 0.2 | 0.31 |
| 2 | 206723882 | rs12470444 | 0.276 | 0.859 | 0.433 | -0.47 | -0.016 | -0.029 | -0.029 | 0 | 0.45 | 0.25 | 0.51 | 0.48 | 0.52 | 0.61 |
| 2 | 206725525 | rs7583160 | 0.093 | 0.961 | 0.461 | -0.265 | 0.056 | 0.47 | 0.47 | 0.015 | 0.95 | 0.78 | 0.92 | 0.51 | 0.93 | 0.93 |
| 2 | 206733483 | rs16837745 | 0.601 | 0.928 | -0.351 | -0.274 | -0.003 | -0.02 | -0.02 | 0.174 | 0.38 | 0.2 | 0.43 | 0.39 | 0.38 | 0.3 |
| 2 | 206734416 | rs7596827 | -0.358 | 0.262 | -0.552 | -0.219 | 0 | -0.003 | -0.003 | 0 | 0.48 | 0.39 | 0.47 | 0.52 | 0.48 | 0.39 |
| 2 | 206736522 | rs6435309 | 0.232 | -0.479 | -0.536 | -1.246 | -0.025 | -0.105 | -0.105 | 0.013 | 0.66 | 0.45 | 0.87 | 0.99 | 0.8 | 0.74 |
| 2 | 206738529 | rs11895885 | 0.692 | -0.232 | -0.886 | -1.301 | 0 | -0.015 | -0.015 | 0 | 0.44 | 0.46 | 0.68 | 0.83 | 0.57 | 0.5 |
| 2 | 206742710 | rs4675557 | 0.719 | -0.244 | -0.818 | -1.309 | 0 | -0.032 | -0.032 | 0 | 0.46 | 0.51 | 0.68 | 0.84 | 0.56 | 0.55 |
| 2 | 206743798 | rs1861619 | -0.001 | -0.428 | -0.737 | -1.426 | -0.037 | -0.132 | -0.132 | 2.093 | 0.64 | 0.48 | 0.84 | 0.99 | 0.76 | 0.75 |
| 2 | 206748062 | rs10932134 | -0.749 | 0.209 | -0.246 | -0.795 | -0.002 | 0.012 | 0.012 | 0 | 0.37 | 0.64 | 0.26 | 0.17 | 0.35 | 0.42 |
| 2 | 206754920 | rs13396172 | -0.534 | -0.105 | 0.28 | -0.338 | -0.021 | -0.029 | -0.029 | 0 | 0.06 | 0.13 | 0.01 | 0 | 0.01 | 0.03 |
| 2 | 206763713 | rs7577743 | -0.501 | -0.112 | 0.237 | -0.341 | -0.018 | -0.026 | -0.026 | 0 | 0.06 | 0.09 | 0.01 | 0 | 0.01 | 0.02 |
| 2 | 206771423 | rs12694035 | -1.058 | 0.41 | 0.357 | -0.213 | -0.005 | 0.025 | 0.025 | 0 | 0.38 | 0.3 | 0.46 | 0.82 | 0.44 | 0.52 |
| 2 | 206787601 | rs4675563 | -0.293 | 0.632 | -0.203 | 0.59 | -0.014 | 0.042 | 0.042 | 0 | 0.25 | 0.27 | 0.28 | 0.53 | 0.21 | 0.23 |
| 2 | 206787733 | rs4675564 | -1.316 | 0.87 | -0.29 | 0.503 | -0.002 | -0.003 | -0.003 | 0.012 | 0.39 | 0.35 | 0.45 | 0.82 | 0.46 | 0.48 |
| 2 | 206792547 | rs6748524 | -0.916 | 1.342 | -0.748 | 0.815 | -0.006 | -0.005 | -0.005 | 0 | 0.15 | 0.19 | 0.16 | 0.27 | 0.25 | 0.24 |
| 2 | 206801824 | rs4675570 | 0.543 | 0.807 | -0.801 | 1 | -0.001 | 0.031 | 0.031 | 0 | 0.77 | 0.71 | 0.71 | 0.51 | 0.8 | 0.7 |
| 2 | 206806002 | rs4673340 | 0.511 | 0.759 | -1.082 | 0.34 | 0 | 0.003 | 0.003 | 0.133 | 0.52 | 0.43 | 0.61 | 0.42 | 0.52 | 0.42 |
| 2 | 206806379 | rs2080566 | 0.175 | 0.542 | -1.237 | 0.256 | -0.009 | -0.046 | -0.046 | 0 | 0.59 | 0.43 | 0.8 | 0.84 | 0.65 | 0.68 |
| 2 | 206808059 | rs7583781 | -0.713 | 0.727 | -1.649 | -0.412 | 0.006 | 0.075 | 0.075 | 0 | 0.32 | 0.17 | 0.17 | 0.1 | 0.32 | 0.22 |
| 2 | 206809159 | rs12694036 | -0.599 | 0.869 | -1.623 | -0.438 | 0.035 | -0.059 | -0.059 | 0.243 | 0.8 | 0.56 | 0.83 | 0.88 | 0.78 | 0.84 |
| 2 | 206810935 | rs7562683 | -1.18 | 0.243 | -2.465 | -0.663 | -0.008 | -0.008 | -0.008 | 1.805 | 0.48 | 0.28 | 0.43 | 0.51 | 0.53 | 0.55 |
| 2 | 206812263 | rs13429746 | -0.183 | 0.998 | -2.032 | -0.223 | 0.005 | -0.021 | -0.021 | 0 | 0.15 | 0.16 | 0.14 | 0.04 | 0.22 | 0.1 |
| 2 | 206813228 | rs4675571 | -0.007 | 1.036 | -2.088 | -0.149 | -0.014 | -0.109 | -0.109 | 0 | 0.21 | 0.1 | 0.36 | 0.44 | 0.2 | 0.33 |
| 2 | 206814829 | rs6712220 | -0.606 | 1.23 | -1.516 | -0.409 | 0.022 | -0.021 | -0.021 | 0 | 0.19 | 0.2 | 0.26 | 0.09 | 0.4 | 0.23 |
| 2 | 206822533 | rs12618329 | -0.509 | 0.874 | -1.482 | -0.025 | -0.003 | -0.005 | -0.005 | 0.407 | 0.19 | 0.18 | 0.22 | 0.17 | 0.22 | 0.16 |
| 2 | 206822771 | rs10179924 | -0.374 | 1.056 | -1.745 | 0.267 | 0.009 | 0.044 | 0.044 | 0 | 0.52 | 0.42 | 0.72 | 0.7 | 0.55 | 0.69 |
| 2 | 206824322 | rs7592024 | -0.262 | 0.671 | -1.012 | -0.199 | -0.009 | -0.065 | -0.065 | 0 | 0.13 | 0.21 | 0.26 | 0.01 | 0.13 | 0.06 |
| 2 | 206824722 | rs13032812 | 0.326 | 1.063 | -1.385 | -0.537 | -0.033 | 0.051 | 0.051 | 0 | 0.12 | 0.04 | 0.09 | 0 | 0.18 | 0.05 |
| 2 | 206825955 | rs6735654 | 1.321 | 0.417 | -1.2 | -0.739 | 0 | 0.004 | 0.004 | 0.17 | 0.21 | 0.16 | 0.12 | 0.2 | 0.18 | 0.19 |
| 2 | 206829006 | rs1861720 | -1.02 | 0.231 | -1.146 | -0.818 | -0.005 | -0.015 | -0.015 | 0 | 0.74 | 0.6 | 0.73 | 0.82 | 0.74 | 0.82 |
| 2 | 206830070 | rs7600189 | 0.233 | 0.247 | -1.165 | -0.817 | 0.001 | 0.006 | 0.006 | 0.555 | 0.53 | 0.44 | 0.61 | 0.62 | 0.56 | 0.61 |
| 2 | 206833189 | rs6757928 | -0.859 | 0.21 | -1.467 | -0.914 | -0.013 | 0.066 | 0.066 | 0 | 0.46 | 0.33 | 0.45 | 0.22 | 0.48 | 0.31 |
| 2 | 206836612 | rs7603514 | 0.964 | 0.155 | -1.471 | -0.782 | 0.002 | 0.012 | 0.012 | 0 | 0.25 | 0.19 | 0.13 | 0.19 | 0.2 | 0.2 |
| 2 | 206843819 | rs12992648 | 0.569 | 0.058 | -1.832 | -0.38 | -0.066 | -0.022 | -0.022 | 0 | 0.18 | 0.03 | 0.18 | 0.18 | 0.26 | 0.11 |
| 2 | 206847296 | rs12618490 | -0.032 | -0.512 | -1.786 | -0.493 | 0.024 | -0.005 | -0.005 | 0 | 0.64 | 0.25 | 0.48 | 0.64 | 0.64 | 0.73 |
| 2 | 206862015 | rs10192718 | -0.31 | -0.565 | -1.876 | -0.6 | -0.021 | 0.061 | 0.061 | 0 | 0.27 | 0.16 | 0.39 | 0.14 | 0.35 | 0.14 |
| 2 | 206863874 | rs818026 | -0.393 | -0.716 | -1.662 | -0.722 | -0.009 | 0.003 | 0.003 | 0.002 | 0.68 | 0.43 | 0.72 | 0.65 | 0.71 | 0.75 |
| 2 | 206867762 | rs17309042 | 1.854 | -0.72 | -1.705 | -0.754 | -0.004 | 0.027 | 0.027 | 0.848 | 0.05 | 0.01 | 0.22 | 0 | 0.07 | 0.03 |
| 2 | 206870784 | rs818024 | -1.145 | -0.836 | -1.009 | -0.734 | -0.01 | -0.023 | -0.023 | 0.834 | 0.77 | 0.67 | 0.78 | 0.87 | 0.76 | 0.87 |
| 2 | 206874339 | rs817991 | 1.481 | -0.727 | -1.364 | -0.712 | -0.033 | -0.031 | -0.031 | 0.132 | 0.42 | 0.2 | 0.32 | 0.5 | 0.35 | 0.6 |
| 2 | 206885193 | rs817999 | -0.431 | -0.646 | -1.212 | -0.634 | -0.012 | 0.016 | 0.016 | 0.011 | 0.71 | 0.45 | 0.73 | 0.64 | 0.7 | 0.75 |

| | | | | | | | | | | | | | | | | |
|---|---|---|---|---|---|---|---|---|---|---|---|---|---|---|---|---|
| 2 | 206886765 | rs2969766 | -0.337 | -0.649 | -1.235 | -0.636 | -0.002 | 0.001 | 0.001 | 0.076 | 0.71 | 0.34 | 0.72 | 0.64 | 0.7 | 0.75 |
| 2 | 206889814 | rs6759124 | 1.095 | -0.639 | -1.272 | -0.635 | -0.02 | -0.02 | -0.02 | 0 | 0.44 | 0.23 | 0.33 | 0.5 | 0.34 | 0.6 |
| 2 | 206911228 | rs2909111 | 1.109 | -0.648 | -1.276 | -0.444 | -0.021 | -0.032 | -0.032 | 0.075 | 0.44 | 0.24 | 0.32 | 0.48 | 0.34 | 0.6 |
| 2 | 206921451 | rs818009 | -1.097 | -0.563 | -1.261 | -0.59 | -0.014 | 0.017 | 0.017 | 0.302 | 0.77 | 0.55 | 0.77 | 0.75 | 0.75 | 0.87 |
| 2 | 206927850 | rs17286674 | -0.347 | -0.509 | -1.332 | -0.615 | -0.056 | 0.07 | 0.07 | 0 | 0.26 | 0.07 | 0.38 | 0.14 | 0.35 | 0.14 |
| 2 | 206948124 | rs4675587 | 0.341 | -0.598 | -0.923 | -0.43 | -0.016 | 0.013 | 0.013 | 0.065 | 0.26 | 0.46 | 0.25 | 0.26 | 0.27 | 0.18 |
| 2 | 206959970 | rs752919 | -0.305 | -0.776 | -1.419 | -0.692 | 0.051 | 0.124 | 0.124 | 0 | 0.51 | 0.73 | 0.58 | 0.26 | 0.48 | 0.3 |
| 2 | 206982546 | rs12694040 | -0.404 | -0.753 | -1.343 | -0.678 | 0.04 | 0.09 | 0.09 | 0 | 0.49 | 0.65 | 0.58 | 0.26 | 0.48 | 0.31 |
| 2 | 206986628 | rs1564163 | -0.468 | -0.755 | -1.384 | -0.679 | 0.059 | 0.151 | 0.151 | 0.007 | 0.5 | 0.77 | 0.58 | 0.26 | 0.48 | 0.31 |
| 2 | 206987015 | rs1053517 | -0.274 | -0.744 | -1.384 | -0.679 | 0.063 | 0.152 | 0.152 | 0 | 0.5 | 0.77 | 0.58 | 0.26 | 0.48 | 0.31 |
| 2 | 207025444 | rs11689362 | -0.532 | -0.78 | -1.402 | -0.681 | 0.046 | 0.096 | 0.096 | 0 | 0.5 | 0.65 | 0.56 | 0.26 | 0.48 | 0.32 |
| 2 | 207032341 | rs7584715 | 0.273 | -0.876 | -1.323 | -0.732 | -0.011 | -0.019 | -0.019 | 0 | 0.38 | 0.52 | 0.29 | 0.21 | 0.34 | 0.21 |
| 2 | 207041053 | rs3732083 | -0.459 | -0.802 | -1.371 | -0.667 | 0.045 | 0.09 | 0.09 | 0 | 0.5 | 0.65 | 0.57 | 0.26 | 0.48 | 0.32 |
| 2 | 207041441 | rs3732082 | 0.208 | -0.777 | -1.248 | -0.62 | -0.018 | -0.029 | -0.029 | 0 | 0.39 | 0.45 | 0.31 | 0.26 | 0.37 | 0.29 |
| 2 | 207041933 | rs3755227 | 1.749 | -0.795 | -1.513 | -0.236 | -0.007 | 0.027 | 0.027 | 0 | 0.18 | 0 | 0.1 | 0.07 | 0.13 | 0.16 |
| 2 | 207048940 | rs12694045 | 0.624 | -0.42 | -0.953 | 0.173 | -0.017 | -0.027 | -0.027 | 0 | 0.39 | 0.49 | 0.31 | 0.26 | 0.37 | 0.29 |
| 2 | 207052179 | rs17222648 | 0.503 | -0.458 | -1.024 | 0.42 | 0 | -0.013 | -0.013 | 0.65 | 0.09 | 0.07 | 0.26 | 0 | 0.11 | 0.04 |
| 2 | 207053416 | rs16838023 | 0.218 | 0.136 | -1.25 | 0.867 | -0.001 | 0.01 | 0.01 | 0 | 0.2 | 0.19 | 0.21 | 0.46 | 0.2 | 0.36 |
| 2 | 207058522 | rs12478085 | -0.94 | -0.315 | -1.035 | 0.667 | -0.011 | 0.013 | 0.013 | 0 | 0.17 | 0.19 | 0.09 | 0.28 | 0.08 | 0.14 |
| 2 | 207058820 | rs17288678 | 0.673 | -0.319 | -1.119 | 0.547 | 0.015 | 0.027 | 0.027 | 0.037 | 0.11 | 0.03 | 0.24 | 0 | 0.1 | 0.05 |
| 2 | 207063142 | rs17222948 | 0.673 | -0.319 | -1.119 | 0.547 | 0.015 | 0.027 | 0.027 | 0.037 | 0.11 | 0.03 | 0.24 | 0 | 0.1 | 0.05 |
| 2 | 207063825 | rs2010572 | 0.058 | -0.336 | -1.116 | 0.601 | 0.002 | -0.004 | -0.004 | 0 | 0.11 | 0.08 | 0.04 | 0.16 | 0.06 | 0.08 |
| 2 | 207067503 | rs16838070 | -0.905 | -0.32 | -1.027 | 0.616 | -0.012 | -0.002 | -0.002 | 2.794 | 0.27 | 0.29 | 0.34 | 0.29 | 0.18 | 0.19 |
| 2 | 207069563 | rs6715398 | 1.092 | -0.125 | -1.111 | 0.767 | -0.018 | -0.019 | -0.019 | 0 | 0.42 | 0.23 | 0.41 | 0.44 | 0.52 | 0.46 |
| 2 | 207082016 | rs2058709 | -0.722 | -0.035 | -0.773 | 0.774 | 0.001 | -0.006 | -0.006 | 0 | 0.92 | 0.95 | 0.97 | 0.87 | 0.95 | 0.92 |
| 2 | 207083630 | rs12468934 | 0.983 | -0.107 | -0.735 | 0.768 | -0.004 | -0.018 | -0.018 | 1.038 | 0.46 | 0.27 | 0.47 | 0.47 | 0.56 | 0.46 |
| 2 | 207091609 | rs6435336 | -0.751 | 0.131 | -0.482 | 0.901 | -0.005 | -0.002 | -0.002 | 0 | 0.39 | 0.63 | 0.36 | 0.55 | 0.41 | 0.48 |
| 2 | 207096550 | rs6718087 | 0.428 | 0.069 | -0.317 | 0.861 | 0.031 | 0.001 | 0.001 | 0 | 0.26 | 0.38 | 0.18 | 0.23 | 0.27 | 0.35 |
| 2 | 207097363 | rs750504 | 0.096 | 0.169 | -0.258 | 0.97 | -0.003 | 0.012 | 0.012 | 0 | 0.09 | 0.07 | 0.16 | 0.27 | 0.12 | 0.12 |
| 2 | 207097533 | rs750505 | -0.555 | 0.111 | -0.359 | 0.897 | 0.027 | 0.021 | 0.021 | 0 | 0.36 | 0.45 | 0.34 | 0.51 | 0.4 | 0.48 |
| 2 | 207097741 | rs888084 | 0.651 | 0.06 | -0.588 | 0.973 | -0.034 | -0.04 | -0.04 | 0.163 | 0.07 | 0.01 | 0.13 | 0.16 | 0.11 | 0.09 |
| 2 | 207101446 | rs726165 | -1.37 | -0.031 | -0.635 | 0.823 | 0.022 | -0.02 | -0.02 | 0 | 0.53 | 0.86 | 0.62 | 0.51 | 0.51 | 0.55 |
| 2 | 207105300 | rs1024809 | 0.524 | -0.039 | -0.544 | 0.793 | 0.041 | 0.002 | 0.002 | 0 | 0.26 | 0.4 | 0.19 | 0.23 | 0.28 | 0.35 |
| 2 | 207116449 | rs1362467 | -1.337 | -0.035 | -1.225 | 0.742 | 0.019 | -0.04 | -0.04 | 0.021 | 0.54 | 0.89 | 0.62 | 0.51 | 0.52 | 0.57 |
| 2 | 207121782 | rs2058710 | -0.697 | 0.316 | -1.25 | 0.848 | -0.029 | -0.003 | -0.003 | 0 | 0.22 | 0.08 | 0.37 | 0.2 | 0.25 | 0.16 |
| 2 | 207124756 | rs1861437 | -1.292 | 0.14 | -1.069 | 0.84 | 0.044 | 0.023 | 0.023 | 0 | 0.49 | 0.34 | 0.57 | 0.38 | 0.52 | 0.53 |
| 2 | 207138479 | rs6758398 | -1.275 | 0.199 | -0.945 | 0.747 | 0.021 | 0.018 | 0.018 | 0 | 0.48 | 0.41 | 0.58 | 0.37 | 0.52 | 0.52 |
| 2 | 207145650 | rs13000165 | -0.976 | 0.01 | -0.977 | 0.657 | 0.07 | 0.05 | 0.05 | 0.093 | 0.5 | 0.31 | 0.6 | 0.36 | 0.56 | 0.53 |
| 2 | 207145906 | rs13000757 | -0.315 | 0.091 | -1.014 | 0.613 | -0.066 | -0.03 | -0.03 | 0 | 0.36 | 0.15 | 0.34 | 0.36 | 0.45 | 0.47 |
| 2 | 207168163 | rs723606 | -0.589 | 0.113 | -0.866 | 0.634 | -0.025 | 0.058 | 0.058 | 0 | 0.42 | 0.3 | 0.49 | 0.2 | 0.43 | 0.42 |
| 2 | 207169731 | rs10932150 | -0.865 | 0.109 | -0.847 | 0.63 | 0.003 | -0.02 | -0.02 | 0.068 | 0.44 | 0.48 | 0.51 | 0.2 | 0.43 | 0.42 |
| 2 | 207172627 | rs7582864 | -1.032 | 0.087 | -0.789 | 0.648 | 0.056 | 0.041 | 0.041 | 0.031 | 0.5 | 0.33 | 0.61 | 0.36 | 0.56 | 0.53 |
| 2 | 207174316 | rs3732084 | -1.193 | 0.092 | -1.018 | 0.661 | 0.038 | 0.028 | 0.028 | 0 | 0.52 | 0.38 | 0.62 | 0.36 | 0.57 | 0.53 |
| 2 | 207178422 | rs1448902 | -1.915 | -0.246 | -1.334 | 0.751 | -0.014 | 0.001 | 0.001 | 0 | 0.63 | 0.58 | 0.69 | 0.68 | 0.64 | 0.65 |
| 2 | 207178872 | rs4673350 | -1.007 | 0.346 | -1.492 | 0.484 | 0.061 | 0.027 | 0.027 | 0.96 | 0.47 | 0.14 | 0.59 | 0.36 | 0.56 | 0.52 |
| 2 | 207197155 | rs12989476 | 0.427 | 0.396 | -1.367 | 0.247 | -0.077 | 0.019 | 0.019 | 0 | 0.28 | 0.11 | 0.22 | 0.2 | 0.31 | 0.35 |
| 2 | 207227020 | rs4675602 | 0.249 | 0.464 | -1.402 | 0.416 | -0.044 | -0.036 | -0.036 | 0 | 0.08 | 0.01 | 0.12 | 0.13 | 0.14 | 0.11 |
| 2 | 207229528 | rs10196764 | 0.64 | 0.679 | -1.498 | 0.309 | -0.027 | 0.002 | 0.002 | 0.002 | 0.23 | 0.15 | 0.16 | 0.2 | 0.24 | 0.34 |
| 2 | 207229821 | rs1031293 | -0.534 | 0.671 | -1.485 | 0.296 | -0.048 | -0.008 | -0.008 | 0 | 0.33 | 0.23 | 0.29 | 0.33 | 0.38 | 0.45 |
| 2 | 207248234 | rs7423990 | 0.665 | 0.583 | -1.414 | 0.362 | -0.026 | 0.003 | 0.003 | 0.004 | 0.22 | 0.16 | 0.17 | 0.2 | 0.24 | 0.33 |
| 2 | 207252744 | rs10804151 | -0.843 | 0.175 | -1.022 | 0.483 | 0.005 | 0.004 | 0.004 | 0 | 0.55 | 0.47 | 0.65 | 0.33 | 0.58 | 0.52 |
| 2 | 207267616 | rs4675609 | -0.81 | 0.407 | -1.162 | 0.538 | -0.004 | 0.001 | 0.001 | 0 | 0.56 | 0.49 | 0.67 | 0.47 | 0.58 | 0.53 |
| 2 | 207270798 | rs4271750 | 0.153 | 0.855 | -1.33 | 0.195 | -0.042 | -0.036 | -0.036 | 0.011 | 0.07 | 0.01 | 0.12 | 0.13 | 0.12 | 0.1 |
| 2 | 207270954 | rs4675612 | -0.991 | 0.608 | -1.268 | 0 | -0.005 | -0.015 | -0.015 | 0 | 0.6 | 0.59 | 0.67 | 0.31 | 0.59 | 0.5 |
| 2 | 207285883 | rs11903628 | -1.045 | 0.585 | -1.283 | -0.134 | -0.003 | 0.01 | 0.01 | 0 | 0.29 | 0.27 | 0.47 | 0.14 | 0.31 | 0.18 |
| 2 | 207287770 | rs13021402 | 0.018 | 0.55 | -1.496 | -0.172 | -0.012 | 0.069 | 0.069 | 0 | 0.16 | 0.04 | 0.3 | 0 | 0.17 | 0.06 |
| 2 | 207302819 | rs10469652 | -0.582 | 0.312 | -1.679 | -0.269 | 0.032 | 0.111 | 0.111 | 0.83 | 0.48 | 0.3 | 0.62 | 0.23 | 0.41 | 0.42 |
| 2 | 207305654 | rs1562660 | 0.204 | 0.501 | -1.565 | -0.29 | -0.014 | 0.014 | 0.014 | 0.053 | 0.14 | 0.09 | 0.33 | 0.05 | 0.2 | 0.12 |
| 2 | 207307823 | rs10932151 | 0.164 | 0.45 | -1.476 | -0.409 | 0.006 | 0.05 | 0.05 | 0 | 0.47 | 0.37 | 0.31 | 0.25 | 0.39 | 0.41 |
| 2 | 207311915 | rs1448905 | 0.269 | 0.364 | -1.162 | -0.469 | -0.002 | 0.06 | 0.06 | 0 | 0.45 | 0.32 | 0.3 | 0.25 | 0.39 | 0.4 |
| 2 | 207317934 | rs2300977 | 0.519 | 0.517 | -1.389 | -0.834 | 0 | 0 | 0 | 0 | 0.31 | 0.33 | 0.26 | 0.26 | 0.3 | 0.33 |
| 2 | 207322291 | rs16838253 | 2.013 | 0.364 | -1.814 | -0.687 | -0.003 | -0.02 | -0.02 | 0 | 0.13 | 0.01 | 0.07 | 0.14 | 0.15 | 0.1 |
| 2 | 207326256 | rs6757230 | -1.588 | 0.513 | -0.82 | -0.25 | -0.008 | 0.03 | 0.03 | 0 | 0.79 | 0.77 | 0.86 | 0.54 | 0.82 | 0.64 |
| 2 | 207326937 | rs1035583 | -0.443 | 0.592 | -0.744 | -0.245 | 0 | 0 | 0 | 1.428 | 0.62 | 0.65 | 0.65 | 0.31 | 0.6 | 0.52 |
| 2 | 207330823 | rs2113378 | 2.217 | 0.272 | -1.059 | -0.245 | -0.048 | -0.02 | -0.02 | 0 | 0.13 | 0.03 | 0.08 | 0.13 | 0.18 | 0.12 |
| 2 | 207335853 | rs2300973 | 0.878 | 0.278 | -0.968 | -0.316 | 0.004 | 0.002 | 0.002 | 0 | 0.3 | 0.22 | 0.24 | 0.26 | 0.3 | 0.32 |
| 2 | 207344021 | rs2300971 | 1.414 | 0.505 | -0.835 | -0.257 | 0.036 | 0.001 | 0.001 | 0 | 0.15 | 0.34 | 0.25 | 0.08 | 0.17 | 0.04 |
| 2 | 207351621 | rs2300970 | 0.428 | 0.593 | -0.847 | -0.265 | 0.071 | 0.02 | 0.02 | 0 | 0.19 | 0.42 | 0.47 | 0.14 | 0.27 | 0.13 |
| 2 | 207351674 | rs2113379 | -0.796 | 0.576 | -0.834 | -0.294 | 0 | -0.01 | -0.01 | 0.441 | 0.68 | 0.74 | 0.76 | 0.54 | 0.69 | 0.61 |

| | | | | | | | | | | | | | | | | |
|---|---|---|---|---|---|---|---|---|---|---|---|---|---|---|---|---|
| 2 | 207355354 | rs2269183 | -1.665 | 0.577 | -0.917 | -0.241 | -0.001 | -0.02 | -0.02 | 0.795 | 0.8 | 0.86 | 0.86 | 0.54 | 0.78 | 0.62 |
| 2 | 207356705 | rs3845807 | 0.004 | 0.248 | -1.138 | -0.388 | -0.002 | 0.002 | 0.002 | 0 | 0.55 | 0.52 | 0.56 | 0.46 | 0.52 | 0.54 |
| 2 | 207369450 | rs1921668 | 1.013 | 0.495 | -1.011 | -0.168 | 0 | 0.003 | 0.003 | 0 | 0.31 | 0.25 | 0.26 | 0.26 | 0.31 | 0.33 |
| 2 | 207373828 | rs1025077 | -0.472 | 0.414 | -0.792 | -0.181 | -0.015 | 0.022 | 0.022 | 0 | 0.62 | 0.57 | 0.76 | 0.4 | 0.63 | 0.53 |
| 2 | 207386623 | rs17211421 | 1.922 | 0.417 | -1.068 | -0.301 | -0.019 | 0.082 | 0.082 | 0 | 0.14 | 0.01 | 0.04 | 0 | 0.06 | 0.07 |
| 2 | 207387703 | rs733237 | -1.285 | 0.407 | -0.943 | -0.072 | -0.001 | -0.004 | -0.004 | 0 | 0.77 | 0.81 | 0.85 | 0.62 | 0.77 | 0.65 |
| 2 | 207388536 | rs1921666 | 1.037 | 0.383 | -0.82 | -0.104 | -0.001 | -0.001 | -0.001 | 0 | 0.32 | 0.27 | 0.41 | 0.31 | 0.37 | 0.36 |
| 2 | 207393424 | rs13007621 | 2.804 | 0.416 | -0.931 | -0.131 | -0.016 | 0.049 | 0.049 | 0.599 | 0.1 | 0.03 | 0.09 | 0 | 0.09 | 0.02 |
| 2 | 207396228 | rs759845 | -0.193 | 0.377 | -0.806 | -0.079 | -0.012 | -0.015 | -0.015 | 0.547 | 0.59 | 0.68 | 0.59 | 0.51 | 0.62 | 0.49 |
| 2 | 207418899 | rs3770977 | 2.104 | 0.311 | -0.99 | 0.011 | 0.006 | -0.034 | -0.034 | 0 | 0.13 | 0.01 | 0.07 | 0.2 | 0.16 | 0.12 |
| 2 | 207425094 | rs4675615 | 1.225 | 0.452 | -0.814 | -0.014 | 0 | 0.01 | 0.01 | 0 | 0.29 | 0.2 | 0.27 | 0.23 | 0.3 | 0.33 |
| 2 | 207437125 | rs6736588 | 0.26 | 0.761 | -0.482 | 0.192 | 0.004 | 0 | 0 | 0.039 | 0.6 | 0.52 | 0.41 | 0.46 | 0.53 | 0.53 |
| 2 | 207459077 | rs10497883 | -0.708 | 1.098 | -0.524 | 0.818 | 0.005 | 0.071 | 0.071 | 0.063 | 0.79 | 0.71 | 0.85 | 0.54 | 0.8 | 0.59 |
| 2 | 207465830 | rs6435342 | 0.815 | 0.867 | -0.778 | 0.562 | 0.002 | -0.005 | -0.005 | 0 | 0.45 | 0.35 | 0.5 | 0.54 | 0.55 | 0.46 |
| 2 | 207470884 | rs4085933 | 1.036 | 0.876 | -0.958 | 0.603 | 0.007 | 0 | 0 | 0 | 0.42 | 0.25 | 0.35 | 0.46 | 0.48 | 0.44 |
| 2 | 207480608 | rs3770983 | -0.245 | 0.876 | -1.078 | 0.395 | -0.006 | 0.002 | 0.002 | 0.265 | 0.69 | 0.85 | 0.76 | 0.71 | 0.72 | 0.62 |
| 2 | 207486889 | rs10186957 | -0.904 | 0.853 | -1.18 | 0.406 | -0.002 | -0.03 | -0.03 | 0 | 0.81 | 0.94 | 0.87 | 0.74 | 0.84 | 0.64 |
| 2 | 207488021 | rs17730135 | 1.051 | 0.706 | -1.146 | 0.333 | 0.009 | -0.014 | -0.014 | 0 | 0.42 | 0.27 | 0.34 | 0.49 | 0.48 | 0.43 |
| 2 | 207492886 | rs13004168 | 1.701 | 1.03 | -1.217 | 0.643 | -0.014 | 0.048 | 0.048 | 0.168 | 0.11 | 0.05 | 0.11 | 0 | 0.11 | 0.02 |
| 2 | 207496427 | rs2163035 | 0.161 | 1.001 | -0.866 | 0.957 | 0.028 | 0.06 | 0.06 | 1.701 | 0.78 | 0.66 | 0.66 | 0.59 | 0.75 | 0.56 |
| 2 | 207503668 | rs11895936 | -0.28 | 0.604 | -1.341 | 0.58 | -0.002 | 0.002 | 0.002 | 0.014 | 0.42 | 0.45 | 0.51 | 0.44 | 0.45 | 0.47 |
| 2 | 207503991 | rs10932158 | -1.244 | 0.465 | -1.7 | 0.3 | 0.003 | 0.019 | 0.019 | 4.808 | 0.52 | 0.7 | 0.74 | 0.54 | 0.51 | 0.5 |
| 2 | 207508741 | rs4673360 | -0.37 | 0.227 | -1.536 | 0.099 | -0.012 | 0.019 | 0.019 | 0.631 | 0.54 | 0.18 | 0.48 | 0.46 | 0.53 | 0.51 |
| 2 | 207512319 | rs3845808 | -1.658 | 0.431 | -1.347 | -0.004 | 0.021 | 0.02 | 0.02 | 0 | 0.73 | 0.71 | 0.67 | 0.65 | 0.66 | 0.68 |
| 2 | 207518810 | rs13395560 | 0.126 | 0.313 | -1.088 | -0.622 | -0.015 | 0.023 | 0.023 | 1.299 | 0.22 | 0.15 | 0.31 | 0 | 0.24 | 0.07 |
| 2 | 207527840 | rs2115591 | -1.608 | 0.145 | -1.058 | -0.61 | -0.003 | -0.018 | -0.018 | 0 | 0.4 | 0.64 | 0.49 | 0.17 | 0.37 | 0.24 |
| 2 | 207533025 | rs7582155 | -0.781 | 0.109 | -0.831 | -0.717 | 0.008 | 0.011 | 0.011 | 0 | 0.29 | 0.29 | 0.27 | 0.17 | 0.27 | 0.19 |
| 2 | 207533219 | rs13382730 | 1.063 | 0.11 | -0.791 | -0.712 | 0 | -0.017 | -0.017 | 2.996 | 0.14 | 0.11 | 0.11 | 0 | 0.14 | 0.02 |
| 2 | 207533256 | rs13382742 | -1.758 | 0.102 | -0.826 | -0.726 | -0.002 | -0.014 | -0.014 | 0 | 0.41 | 0.65 | 0.49 | 0.17 | 0.38 | 0.26 |
| 2 | 207535453 | rs2099131 | -1.314 | 0.024 | -0.411 | -0.394 | -0.014 | -0.025 | -0.025 | 0.028 | 0.28 | 0.64 | 0.39 | 0.17 | 0.24 | 0.24 |
| 2 | 207536319 | rs16838459 | -0.033 | 0.273 | -0.46 | -0.424 | 0.004 | -0.003 | -0.003 | 0 | 0.18 | 0.18 | 0.18 | 0.17 | 0.19 | 0.2 |
| 2 | 207537673 | rs1012525 | 0.921 | -0.216 | -0.374 | -0.374 | 0.004 | -0.06 | -0.06 | 0 | 0.11 | 0.01 | 0.14 | 0.17 | 0.11 | 0.18 |
| 2 | 207543023 | rs13425179 | -1.86 | -0.251 | -0.303 | -0.425 | -0.006 | -0.006 | -0.006 | 0 | 0.33 | 0.64 | 0.44 | 0.22 | 0.32 | 0.26 |
| 2 | 207545346 | rs4675623 | -1.324 | -0.806 | 0.139 | -0.085 | -0.001 | 0.001 | 0.001 | 2.927 | 0.23 | 0.28 | 0.19 | 0.21 | 0.17 | 0.21 |
| 2 | 207554267 | rs10490746 | 0.366 | -0.614 | 0.008 | -0.128 | 0.012 | -0.045 | -0.045 | 0 | 0.09 | 0.36 | 0.04 | 0 | 0.03 | 0.04 |
| 2 | 207560889 | rs16838595 | 0.508 | -0.819 | -1.106 | 0.155 | 0.01 | -0.008 | -0.008 | 1.905 | 0.22 | 0.01 | 0.1 | 0.21 | 0.19 | 0.19 |
| 2 | 207563641 | rs12993580 | -0.197 | -0.73 | -1.086 | 0.129 | 0.004 | 0.056 | 0.056 | 0.09 | 0.28 | 0.03 | 0.14 | 0.21 | 0.29 | 0.21 |
| 2 | 207564094 | rs1864265 | -1.105 | -0.809 | -1.066 | 0.228 | 0.046 | -0.041 | -0.041 | 0.299 | 0.38 | 0.19 | 0.26 | 0.43 | 0.33 | 0.36 |
| 2 | 207564996 | rs16838601 | -0.497 | -0.895 | -1.009 | 0.288 | -0.004 | 0.021 | 0.021 | 0.868 | 0.1 | 0.14 | 0.12 | 0.22 | 0.04 | 0.15 |
| 2 | 207567542 | rs16838621 | -0.934 | -0.854 | -1.021 | 0.186 | 0.034 | -0.014 | -0.014 | 0.518 | 0.38 | 0.3 | 0.27 | 0.42 | 0.31 | 0.33 |
| 2 | 207571816 | rs4675625 | -0.298 | -0.776 | -1.186 | 0.148 | -0.001 | -0.007 | -0.007 | 0 | 0.17 | 0.19 | 0.18 | 0.21 | 0.17 | 0.18 |
| 2 | 207572396 | rs10490747 | -0.952 | -0.776 | -1.059 | 0.135 | 0.022 | -0.002 | -0.002 | 0 | 0.42 | 0.34 | 0.28 | 0.44 | 0.34 | 0.38 |
| 2 | 207574944 | rs10490749 | 0.044 | -0.809 | -0.757 | -0.693 | 0.01 | 0.014 | 0.014 | 0 | 0.23 | 0.15 | 0.12 | 0.23 | 0.2 | 0.17 |
| 2 | 207578440 | rs11887929 | -0.88 | -0.961 | -0.459 | -1.298 | 0.01 | 0.015 | 0.015 | 0 | 0.55 | 0.66 | 0.57 | 0.95 | 0.46 | 0.78 |
| 2 | 207581835 | rs16838677 | 0.504 | -1.034 | -0.227 | -0.819 | 0.001 | 0.049 | 0.049 | 0 | 0.09 | 0.08 | 0.25 | 0.5 | 0.06 | 0.26 |
| 2 | 207582852 | rs16838691 | 1.511 | -1.106 | -0.382 | -0.882 | -0.005 | -0.058 | -0.058 | 0.26 | 0.1 | 0.01 | 0.07 | 0.2 | 0.12 | 0.2 |
| 2 | 207583215 | rs1001110 | -1.536 | -1.196 | -0.382 | -1.173 | -0.054 | 0.171 | 0.171 | 0 | 0.6 | 0.86 | 0.62 | 0.96 | 0.52 | 0.81 |
| 2 | 207583406 | rs1001111 | -1.339 | -1.408 | -0.182 | -0.953 | 0.029 | 0.138 | 0.138 | 0 | 0.54 | 0.79 | 0.55 | 0.96 | 0.44 | 0.78 |
| 2 | 207587317 | rs10490542 | -0.533 | -1.354 | 0.704 | -0.48 | -0.021 | 0.025 | 0.025 | 0 | 0.22 | 0.34 | 0.17 | 0.3 | 0.2 | 0.36 |
| 2 | 207594276 | rs13419306 | -0.609 | -1.331 | 0.544 | -0.469 | -0.027 | 0.032 | 0.032 | 0 | 0.22 | 0.38 | 0.18 | 0.3 | 0.2 | 0.36 |
| 2 | 207599165 | rs4624363 | -1.14 | -1.451 | 0.465 | -0.484 | -0.069 | 0.144 | 0.144 | 0.937 | 0.2 | 0.58 | 0.33 | 0.53 | 0.1 | 0.34 |
| 2 | 207599663 | rs12473690 | 0.073 | -1.554 | 0.324 | -0.572 | 0.028 | -0.033 | -0.033 | 0 | 0.17 | 0.08 | 0.11 | 0.22 | 0.17 | 0.22 |
| 2 | 207602424 | rs2193884 | -1.554 | -1.287 | 0.701 | -0.35 | -0.075 | 0.094 | 0.094 | 0.082 | 0.25 | 0.57 | 0.36 | 0.53 | 0.14 | 0.38 |
| 2 | 207602890 | rs2287633 | -0.978 | -1.311 | 0.865 | -0.362 | -0.003 | 0.007 | 0.007 | 0 | 0.2 | 0.21 | 0.34 | 0.53 | 0.14 | 0.38 |
| 2 | 207603221 | rs2287632 | -0.091 | -1.298 | 0.823 | -0.339 | -0.01 | -0.011 | -0.011 | 0 | 0.09 | 0.19 | 0.08 | 0.06 | 0.07 | 0.08 |
| 2 | 207603234 | rs2287631 | -1.461 | -1.264 | 0.715 | -0.344 | -0.064 | 0.074 | 0.074 | 0.147 | 0.25 | 0.6 | 0.36 | 0.53 | 0.14 | 0.38 |
| 2 | 207605882 | rs10173707 | -1.438 | -1.25 | 0.722 | -0.337 | -0.064 | 0.076 | 0.076 | 0.186 | 0.25 | 0.6 | 0.36 | 0.54 | 0.14 | 0.38 |
| 2 | 207611230 | rs2302922 | -1.406 | -1.196 | 0.596 | -0.327 | -0.051 | 0.063 | 0.063 | 0.229 | 0.25 | 0.55 | 0.37 | 0.59 | 0.15 | 0.43 |
| 2 | 207611257 | rs2302921 | -0.477 | -1.122 | 0.531 | -0.204 | -0.022 | 0.07 | 0.07 | 0 | 0.15 | 0.22 | 0.27 | 0.48 | 0.07 | 0.32 |
| 2 | 207613980 | rs13427620 | 0.05 | -0.655 | 0.087 | -1.07 | -0.023 | 0.065 | 0.065 | 0 | 0.59 | 0.74 | 0.53 | 0.3 | 0.54 | 0.42 |
| 2 | 207616325 | rs10932164 | -0.335 | -0.872 | 0.198 | -1.18 | -0.004 | -0.001 | -0.001 | 0 | 0.25 | 0.31 | 0.35 | 0.7 | 0.18 | 0.51 |
| 2 | 207618271 | rs13004504 | 0.895 | -0.812 | 0.435 | -1.244 | 0.004 | 0.119 | 0.119 | 0 | 0.49 | 0.28 | 0.46 | 0.24 | 0.5 | 0.34 |
| 2 | 207630093 | rs16838839 | 0.719 | -0.774 | 0.769 | -1.067 | 0.015 | -0.001 | -0.001 | 0 | 0.08 | 0.26 | 0.05 | 0.05 | 0.06 | 0.08 |
| 2 | 207632952 | rs2287630 | -1.206 | -1.036 | 0.352 | -1.523 | -0.015 | 0.152 | 0.152 | 0 | 0.58 | 0.81 | 0.58 | 0.97 | 0.46 | 0.78 |
| 2 | 207633002 | rs3815686 | -0.005 | -1.089 | 0.144 | -1.549 | 0.027 | 0.009 | 0.009 | 0 | 0.27 | 0.21 | 0.38 | 0.71 | 0.2 | 0.51 |
| 2 | 207634236 | rs2277912 | 1.694 | -1.114 | 0.103 | -1.568 | -0.057 | 0.007 | 0.007 | 0 | 0.18 | 0.01 | 0.09 | 0.15 | 0.18 | 0.18 |
| 2 | 207649350 | rs888621 | -0.353 | -1.001 | 0.361 | -1.342 | 0.022 | 0.01 | 0.01 | 0 | 0.28 | 0.26 | 0.38 | 0.71 | 0.2 | 0.51 |
| 2 | 207659531 | rs1001805 | 0.816 | -0.969 | 0.414 | -1.195 | 0.032 | 0.019 | 0.019 | 0 | 0.08 | 0.3 | 0.05 | 0.05 | 0.06 | 0.06 |
| 2 | 207678216 | rs12694062 | 0.505 | -1.002 | 0.354 | -1.222 | -0.02 | -0.032 | -0.032 | 0 | 0.1 | 0.19 | 0.06 | 0.05 | 0.07 | 0.08 |

| | | | | | | | | | | | | | | | | |
|---|---|---|---|---|---|---|---|---|---|---|---|---|---|---|---|---|
| 2 | 207678245 | rs2059022 | 0.118 | -1.109 | 0.062 | -1.275 | 0.033 | 0.002 | 0.002 | 0 | 0.24 | 0.18 | 0.35 | 0.71 | 0.17 | 0.48 |
| 2 | 207686299 | rs7596719 | -1.12 | -1.054 | -0.161 | -1.295 | 0.026 | 0.029 | 0.029 | 0.136 | 0.38 | 0.68 | 0.46 | 0.82 | 0.26 | 0.6 |
| 2 | 207689161 | rs7593959 | 0.625 | -0.972 | -0.12 | -1.227 | -0.008 | -0.005 | -0.005 | 0 | 0.11 | 0.26 | 0.1 | 0.11 | 0.09 | 0.12 |
| 2 | 207692740 | rs1862090 | 1.663 | -0.877 | -0.395 | -1.409 | -0.016 | 0.037 | 0.037 | 0.042 | 0.24 | 0.16 | 0.32 | 0.03 | 0.3 | 0.16 |
| 2 | 207701151 | rs6435354 | 1.45 | -0.793 | 0.113 | -0.803 | -0.039 | 0.143 | 0.143 | 0 | 0.44 | 0.2 | 0.41 | 0.18 | 0.47 | 0.33 |
| 2 | 207705755 | rs13003233 | 0.849 | -0.462 | -0.005 | -0.328 | 0 | -0.036 | -0.036 | 0.008 | 0.58 | 0.51 | 0.51 | 0.29 | 0.56 | 0.45 |
| 2 | 207707077 | rs6435356 | -0.916 | -0.279 | -0.492 | -0.234 | -0.052 | 0.062 | 0.062 | 0 | 0.82 | 0.95 | 0.87 | 1 | 0.73 | 0.93 |
| 2 | 207711974 | rs7581388 | 0.688 | -0.026 | -0.976 | 0.252 | -0.005 | 0.018 | 0.018 | 0 | 0.1 | 0.11 | 0.07 | 0.2 | 0.09 | 0.17 |
| 2 | 207717444 | rs6737274 | 0.199 | 0.401 | -1.114 | 0.341 | -0.003 | -0.011 | -0.011 | 0 | 0.66 | 0.71 | 0.57 | 0.49 | 0.65 | 0.6 |
| 2 | 207723592 | rs16838934 | -0.577 | 0.194 | -1.263 | 0.555 | 0.019 | -0.026 | -0.026 | 0 | 0.14 | 0.08 | 0.06 | 0.19 | 0.11 | 0.04 |
| 2 | 207723962 | rs7565773 | -0.217 | 0.782 | -0.73 | -0.414 | 0.054 | -0.124 | -0.124 | 0 | 0.24 | 0.48 | 0.13 | 0.04 | 0.28 | 0.17 |
| 2 | 207725616 | rs10192834 | -0.525 | 0.64 | -0.528 | -0.437 | -0.008 | -0.006 | -0.006 | 1.452 | 0.36 | 0.41 | 0.27 | 0.31 | 0.33 | 0.24 |
| 2 | 207726297 | rs2042135 | -0.99 | 1.011 | -1.054 | -0.394 | -0.003 | 0.015 | 0.015 | 0 | 0.48 | 0.78 | 0.35 | 0.56 | 0.48 | 0.42 |
| 2 | 207733111 | rs4673364 | 1.031 | 1.147 | -0.928 | -0.314 | -0.011 | -0.014 | -0.014 | 0 | 0.09 | 0.07 | 0.07 | 0.21 | 0.15 | 0.18 |
| 2 | 207733167 | rs10515948 | 0.443 | 0.921 | -0.634 | -0.461 | 0 | 0.01 | 0.01 | 0 | 0.2 | 0.16 | 0.16 | 0.14 | 0.21 | 0.18 |
| 2 | 207733838 | rs6705071 | 0.987 | 0.843 | -0.616 | -0.444 | 0.001 | 0.004 | 0.004 | 0 | 0.16 | 0.13 | 0.12 | 0.13 | 0.16 | 0.11 |
| 2 | 207735837 | rs6435358 | 0.095 | 0.72 | -0.5 | -0.847 | 0.018 | -0.085 | -0.085 | 0 | 0.31 | 0.32 | 0.19 | 0.04 | 0.35 | 0.26 |
| 2 | 207737913 | rs10515949 | 0.698 | 0.589 | -0.125 | -0.656 | 0.034 | -0.049 | -0.049 | 1.786 | 0.22 | 0.08 | 0.14 | 0.31 | 0.18 | 0.12 |
| 2 | 207738671 | rs10515950 | 0.601 | 0.57 | -0.103 | -0.662 | 0.037 | -0.05 | -0.05 | 0 | 0.22 | 0.08 | 0.14 | 0.31 | 0.18 | 0.12 |
| 2 | 207738878 | rs12469482 | 0.389 | 0.552 | -0.052 | -0.671 | 0.022 | -0.018 | -0.018 | 0 | 0.24 | 0.19 | 0.17 | 0.31 | 0.19 | 0.12 |
| 2 | 207740730 | rs1031432 | -1.333 | 0.867 | 0.357 | -0.436 | 0.011 | -0.005 | -0.005 | 0 | 0.28 | 0.6 | 0.23 | 0.26 | 0.34 | 0.39 |
| 2 | 207742630 | rs918343 | -1.272 | 0.807 | 0.368 | -0.474 | -0.005 | 0.007 | 0.007 | 0 | 0.68 | 0.64 | 0.46 | 0.55 | 0.72 | 0.57 |
| 2 | 207743908 | rs11692974 | -0.568 | 0.829 | 0.46 | -0.491 | 0.029 | -0.055 | -0.055 | 0 | 0.13 | 0.29 | 0.11 | 0.04 | 0.14 | 0.17 |
| 2 | 207744508 | rs4675641 | -0.365 | 0.828 | 0.449 | -0.483 | -0.005 | 0.005 | 0.005 | 0 | 0.57 | 0.51 | 0.36 | 0.51 | 0.58 | 0.4 |
| 2 | 207748122 | rs11897520 | 0.604 | 0.728 | 0.444 | -0.494 | -0.002 | 0.041 | 0.041 | 0 | 0.42 | 0.26 | 0.25 | 0.3 | 0.39 | 0.18 |
| 2 | 207749505 | rs10187986 | 1.603 | 0.79 | 0.388 | -0.473 | -0.031 | 0.053 | 0.053 | 0 | 0.28 | 0.09 | 0.13 | 0.16 | 0.28 | 0.1 |
| 2 | 207749855 | rs6728532 | -0.504 | 0.842 | 0.449 | -0.474 | -0.004 | 0.037 | 0.037 | 0 | 0.56 | 0.71 | 0.37 | 0.33 | 0.53 | 0.35 |
| 2 | 207750902 | rs1031434 | 0.634 | 0.774 | 0.416 | -0.466 | -0.01 | 0.046 | 0.046 | 0 | 0.41 | 0.19 | 0.25 | 0.3 | 0.39 | 0.18 |
| 2 | 207755367 | rs1377390 | 0.522 | 0.763 | 0.435 | -0.48 | 0.087 | 0.056 | 0.056 | 1.166 | 0.05 | 0.29 | 0.04 | 0.04 | 0.03 | 0.07 |
| 2 | 207768601 | rs7607350 | -1.389 | 0.739 | 0.128 | -0.433 | 0.008 | -0.031 | -0.031 | 0.874 | 0.7 | 0.77 | 0.46 | 0.54 | 0.72 | 0.56 |
| 2 | 207783156 | rs10170650 | 1.432 | 0.855 | 0.043 | -0.442 | -0.03 | 0.06 | 0.06 | 0 | 0.27 | 0.09 | 0.13 | 0.15 | 0.28 | 0.1 |
| 2 | 207789394 | rs6759178 | -0.624 | 0.804 | 0.045 | -0.301 | 0.001 | -0.01 | -0.01 | 0 | 0.6 | 0.61 | 0.38 | 0.55 | 0.59 | 0.48 |
| 2 | 207789418 | rs12464342 | 0.649 | 0.899 | -0.303 | -0.282 | -0.044 | 0.028 | 0.028 | 0.002 | 0.4 | 0.08 | 0.23 | 0.3 | 0.37 | 0.19 |
| 2 | 207790337 | rs6705661 | 0.101 | 0.957 | -0.156 | -0.245 | -0.01 | -0.012 | -0.012 | 0 | 0.15 | 0.08 | 0.1 | 0.21 | 0.19 | 0.23 |
| 2 | 207798923 | rs6720430 | -0.654 | 0.869 | -0.108 | -0.256 | 0.001 | -0.015 | -0.015 | 0.08 | 0.61 | 0.38 | 0.38 | 0.52 | 0.63 | 0.45 |
| 2 | 207800908 | rs4675643 | -0.165 | 0.904 | -0.272 | -0.257 | -0.011 | -0.005 | -0.005 | 0.754 | 0.18 | 0.17 | 0.13 | 0.22 | 0.26 | 0.24 |
| 2 | 207802116 | rs10932172 | -1.093 | 0.819 | 0.035 | -0.242 | -0.009 | 0.025 | 0.025 | 0 | 0.67 | 0.56 | 0.42 | 0.58 | 0.67 | 0.54 |
| 2 | 207803548 | rs7605998 | -0.006 | 0.521 | -0.258 | -0.15 | 0.008 | 0.028 | 0.028 | 0 | 0.46 | 0.35 | 0.27 | 0.33 | 0.39 | 0.25 |
| 2 | 207806431 | rs2359858 | 0.965 | 0.354 | -0.844 | -0.031 | -0.051 | 0.051 | 0.051 | 0 | 0.28 | 0.06 | 0.14 | 0.14 | 0.27 | 0.11 |
| 2 | 207813700 | rs993024 | -1.081 | 0.559 | -0.4 | 0.21 | -0.008 | -0.015 | -0.015 | 0.188 | 0.74 | 0.8 | 0.5 | 0.56 | 0.7 | 0.51 |
| 2 | 207816620 | rs1349735 | 0.097 | 0.469 | -0.371 | 0.065 | 0.009 | 0.014 | 0.014 | 0.256 | 0.49 | 0.4 | 0.26 | 0.41 | 0.45 | 0.4 |
| 2 | 207820276 | rs13420911 | 0.146 | 0.475 | -0.369 | 0.067 | 0.009 | 0.014 | 0.014 | 0.464 | 0.48 | 0.38 | 0.26 | 0.41 | 0.46 | 0.37 |
| 2 | 207820319 | rs6719240 | 0.665 | 0.376 | -0.234 | 0.058 | 0.003 | 0.055 | 0.055 | 0.869 | 0.06 | 0.2 | 0.05 | 0.09 | 0.02 | 0.08 |
| 2 | 207824384 | rs11903403 | 0.001 | 0.385 | -0.252 | 0.066 | 0.001 | 0.002 | 0.002 | 0.17 | 0.5 | 0.55 | 0.28 | 0.41 | 0.46 | 0.4 |
| 2 | 207824650 | rs1453169 | 0.175 | 0.394 | -0.354 | 0.073 | 0.007 | 0.01 | 0.01 | 0.518 | 0.48 | 0.39 | 0.26 | 0.41 | 0.46 | 0.39 |
| 2 | 207830363 | rs7561116 | 1.278 | 0.414 | -0.603 | 0.34 | 0.001 | 0.076 | 0.076 | 0.04 | 0.33 | 0.11 | 0.14 | 0.17 | 0.28 | 0.17 |
| 2 | 207831069 | rs17536565 | 0.472 | 0.873 | -0.478 | 0.488 | -0.003 | -0.001 | -0.001 | 0 | 0.48 | 0.47 | 0.27 | 0.49 | 0.41 | 0.44 |
| 2 | 207831954 | rs17458475 | 0.799 | 0.721 | -0.419 | 0.489 | 0.011 | 0.019 | 0.019 | 0 | 0.39 | 0.3 | 0.19 | 0.26 | 0.3 | 0.25 |
| 2 | 207833983 | rs7582305 | 0.848 | 0.994 | -0.701 | 0.582 | 0.005 | 0.006 | 0.006 | 0 | 0.41 | 0.37 | 0.2 | 0.26 | 0.3 | 0.25 |
| 2 | 207835223 | rs13385857 | -0.252 | 1.254 | -0.476 | 0.864 | 0.015 | -0.031 | -0.031 | 0 | 0.07 | 0.26 | 0.05 | 0 | 0.06 | 0.03 |
| 2 | 207836111 | rs4468795 | 0.119 | 1.371 | -1.29 | 1.234 | -0.008 | -0.012 | -0.012 | 0 | 0.76 | 0.81 | 0.51 | 0.58 | 0.71 | 0.56 |
| 2 | 207843712 | rs1006389 | -0.634 | 1.016 | -1.423 | 1.036 | -0.027 | -0.046 | -0.046 | 0 | 0.09 | 0.01 | 0.07 | 0.1 | 0.16 | 0.04 |
| 2 | 207849856 | rs7563853 | -1.039 | 0.353 | -1.103 | 1.169 | -0.001 | -0.001 | -0.001 | 0.007 | 0.36 | 0.36 | 0.45 | 0.64 | 0.31 | 0.39 |
| 2 | 207851374 | rs11695049 | -0.096 | 0.175 | -1.009 | 1.09 | 0.019 | -0.096 | -0.096 | 0 | 0.23 | 0.06 | 0.35 | 0.54 | 0.15 | 0.35 |
| 2 | 207853432 | rs7568351 | -1.311 | 0.063 | -0.791 | 0.989 | 0.014 | -0.004 | -0.004 | 0.243 | 0.34 | 0.24 | 0.44 | 0.64 | 0.31 | 0.39 |
| 2 | 207858148 | rs1453170 | 0.971 | -0.094 | -0.51 | 0.341 | 0.009 | 0.036 | 0.036 | 0.398 | 0.5 | 0.43 | 0.44 | 0.21 | 0.56 | 0.55 |
| 2 | 207858919 | rs10188585 | 0.99 | -0.138 | -0.48 | 0.319 | 0.01 | 0.04 | 0.04 | 0.49 | 0.49 | 0.41 | 0.44 | 0.21 | 0.56 | 0.55 |
| 2 | 207859422 | rs7564361 | -0.505 | -0.491 | -0.668 | 0.08 | -0.002 | -0.005 | -0.005 | 0 | 0.76 | 0.73 | 0.83 | 0.84 | 0.8 | 0.84 |
| 2 | 207859734 | rs11686465 | 0.566 | -0.511 | -0.702 | 0.111 | -0.017 | -0.012 | -0.012 | 0 | 0.63 | 0.47 | 0.73 | 0.74 | 0.64 | 0.81 |
| 2 | 207861861 | rs935912 | 0.522 | -0.511 | -0.625 | 0.111 | -0.012 | -0.009 | -0.009 | 0 | 0.63 | 0.51 | 0.74 | 0.74 | 0.64 | 0.81 |
| 2 | 207864250 | rs2253368 | 0.24 | -0.531 | -0.626 | 0.1 | -0.008 | -0.015 | -0.015 | 0 | 0.24 | 0.32 | 0.17 | 0.16 | 0.2 | 0.16 |
| 2 | 207872075 | rs2080711 | -1.008 | -0.55 | -0.164 | 0.092 | -0.004 | -0.003 | -0.003 | 0 | 0.38 | 0.58 | 0.27 | 0.26 | 0.36 | 0.19 |
| 2 | 207874372 | rs1377389 | 0.207 | -0.532 | -0.26 | 0.104 | -0.009 | -0.016 | -0.016 | 0 | 0.24 | 0.32 | 0.17 | 0.16 | 0.2 | 0.16 |
| 2 | 207890135 | rs6732038 | 0.209 | -0.18 | -0.517 | -0.568 | -0.001 | 0.153 | 0.153 | 0 | 0.49 | 0.23 | 0.44 | 0.17 | 0.55 | 0.31 |
| 2 | 207894593 | rs11898699 | -0.158 | -0.179 | -0.276 | -0.554 | 0.002 | -0.011 | -0.011 | 1.16 | 0.74 | 0.82 | 0.57 | 0.37 | 0.72 | 0.67 |
| 2 | 207897877 | rs1990810 | -1.778 | -0.182 | -0.369 | -0.541 | -0.001 | -0.029 | -0.029 | 0.053 | 0.87 | 0.94 | 0.67 | 0.47 | 0.88 | 0.71 |
| 2 | 207902848 | rs1990809 | 0.893 | -0.151 | -0.361 | -0.618 | -0.024 | 0.112 | 0.112 | 0 | 0.38 | 0.21 | 0.35 | 0.07 | 0.39 | 0.28 |
| 2 | 207915477 | rs11902322 | 0.128 | -0.034 | -0.22 | -0.489 | 0 | -0.002 | -0.002 | 6.586 | 0.34 | 0.4 | 0.2 | 0.3 | 0.33 | 0.37 |
| 2 | 207915849 | rs16839086 | 0.137 | -0.04 | -0.223 | -0.497 | 0.001 | -0.003 | -0.003 | 0 | 0.34 | 0.4 | 0.2 | 0.3 | 0.33 | 0.37 |

| | | | | | | | | | | | | | | | | |
|---|---|---|---|---|---|---|---|---|---|---|---|---|---|---|---|---|
| 2 | 207929757 | rs1124215 | -0.964 | 0.144 | -1.073 | -0.433 | 0.006 | -0.08 | -0.08 | 0.908 | 0.8 | 0.94 | 0.66 | 0.51 | 0.84 | 0.64 |
| 2 | 207931865 | rs17281469 | 1.505 | 0.226 | -0.95 | -0.377 | -0.001 | 0.002 | 0.002 | 0.06 | 0.08 | 0.06 | 0.09 | 0.15 | 0.13 | 0.03 |
| 2 | 207934200 | rs1263632 | 0.618 | -0.027 | -1.009 | -0.679 | -0.005 | 0.056 | 0.056 | 0 | 0.51 | 0.71 | 0.39 | 0.35 | 0.49 | 0.49 |
| 2 | 207934267 | rs1263631 | -1.235 | 0.055 | -0.983 | -0.634 | 0.012 | -0.046 | -0.046 | 0 | 0.79 | 0.9 | 0.66 | 0.51 | 0.84 | 0.64 |
| 2 | 207936541 | rs6739008 | 0.597 | 0.131 | -0.838 | -0.576 | -0.037 | 0.065 | 0.065 | 0 | 0.2 | 0.13 | 0.18 | 0.02 | 0.23 | 0.12 |
| 2 | 207941850 | rs1263599 | 0.537 | 0.01 | -0.3 | -0.537 | -0.006 | 0.025 | 0.025 | 0.821 | 0.22 | 0.3 | 0.1 | 0.3 | 0.18 | 0.31 |
| 2 | 207947086 | rs2287505 | 0.43 | -0.162 | -0.383 | -0.575 | 0 | 0.022 | 0.022 | 1.799 | 0.22 | 0.33 | 0.21 | 0.19 | 0.27 | 0.17 |
| 2 | 207947476 | rs2287504 | 0.529 | -0.261 | -0.281 | -0.488 | -0.011 | 0.12 | 0.12 | 1.034 | 0.12 | 0.33 | 0.1 | 0.18 | 0.14 | 0.09 |
| 2 | 207954639 | rs1263619 | -0.396 | -0.046 | -0.719 | -0.725 | -0.015 | -0.055 | -0.055 | 0 | 0.65 | 0.83 | 0.51 | 0.49 | 0.68 | 0.52 |
| 2 | 207956108 | rs1263618 | -0.309 | -0.21 | -0.545 | -0.911 | 0.001 | -0.003 | -0.003 | 0 | 0.34 | 0.66 | 0.21 | 0.49 | 0.33 | 0.37 |
| 2 | 207957351 | rs7582048 | 1.322 | -0.153 | -0.591 | -1.007 | -0.012 | 0.057 | 0.057 | 0 | 0.29 | 0.16 | 0.27 | 0 | 0.31 | 0.13 |
| 2 | 207960996 | rs13012163 | 0.093 | -0.114 | -0.699 | -1.192 | 0.01 | 0.013 | 0.013 | 0 | 0.52 | 0.38 | 0.49 | 0.03 | 0.58 | 0.31 |
| 2 | 207962456 | rs6759901 | 0.093 | -0.114 | -0.708 | -1.192 | 0.01 | 0.008 | 0.008 | 0 | 0.52 | 0.38 | 0.49 | 0.03 | 0.58 | 0.31 |
| 2 | 207965189 | rs12464132 | 0.329 | -0.062 | -0.39 | -1.159 | 0.002 | 0.027 | 0.027 | 0.151 | 0.51 | 0.46 | 0.47 | 0.17 | 0.54 | 0.24 |
| 2 | 207965794 | rs1263611 | -0.193 | 0.019 | -0.428 | -1.016 | 0.025 | 0.064 | 0.064 | 0.006 | 0.32 | 0.53 | 0.25 | 0.48 | 0.36 | 0.36 |
| 2 | 207973097 | rs1269416 | -1.424 | -0.004 | -0.779 | -1.021 | 0.011 | -0.082 | -0.082 | 0 | 0.78 | 0.92 | 0.66 | 0.51 | 0.84 | 0.62 |
| 2 | 207974110 | rs12472554 | 1.378 | -0.005 | -0.889 | -1.043 | -0.016 | 0.105 | 0.105 | 0 | 0.29 | 0.1 | 0.27 | 0 | 0.31 | 0.12 |
| 2 | 207974818 | rs2244923 | -1.03 | -0.039 | -0.656 | -0.679 | 0.001 | 0.001 | 0.001 | 0 | 0.77 | 0.8 | 0.66 | 0.51 | 0.84 | 0.62 |
| 2 | 207975828 | rs1881935 | -0.431 | 0.148 | -0.635 | -0.504 | 0.011 | -0.017 | -0.017 | 0 | 0.61 | 0.67 | 0.52 | 0.49 | 0.67 | 0.49 |
| 2 | 207979050 | rs2244399 | 0.099 | 0.488 | -0.433 | 0.02 | 0.006 | 0.037 | 0.037 | 0.233 | 0.16 | 0.23 | 0.07 | 0.34 | 0.16 | 0.27 |
| 2 | 207983196 | rs1263621 | -0.349 | 0.568 | -0.649 | 0.029 | 0 | 0.001 | 0.001 | 0 | 0.53 | 0.44 | 0.4 | 0.39 | 0.43 | 0.53 |
| 2 | 207984215 | rs6706279 | -0.206 | 0.052 | -0.2 | -0.245 | 0.019 | 0.013 | 0.013 | 0 | 0.6 | 0.3 | 0.72 | 0.66 | 0.65 | 0.63 |
| 2 | 207986297 | rs2269077 | 0.305 | -0.11 | 0.086 | -0.317 | -0.007 | 0 | 0 | 0 | 0.35 | 0.49 | 0.27 | 0.33 | 0.34 | 0.3 |
| 2 | 207986713 | rs2543596 | 0.405 | -0.105 | 0.056 | -0.314 | -0.002 | 0.001 | 0.001 | 0 | 0.34 | 0.34 | 0.26 | 0.33 | 0.34 | 0.3 |
| 2 | 207990793 | rs2300941 | 0.129 | -0.114 | -0.064 | -0.3 | 0.016 | 0.004 | 0.004 | 0 | 0.57 | 0.36 | 0.71 | 0.63 | 0.66 | 0.57 |
| 2 | 207992155 | rs16839275 | 1.359 | 0.324 | -0.879 | -0.433 | -0.07 | -0.117 | -0.117 | 0.185 | 0.22 | 0.09 | 0.26 | 0.51 | 0.31 | 0.35 |
| 2 | 207998800 | rs2284934 | -0.913 | -0.166 | -0.495 | -0.532 | -0.012 | -0.006 | -0.006 | 0 | 0.63 | 0.53 | 0.76 | 0.66 | 0.67 | 0.69 |
| 2 | 207999571 | rs7594919 | 0.192 | 0.08 | -0.491 | -0.648 | -0.007 | -0.031 | -0.031 | 0 | 0.37 | 0.36 | 0.35 | 0.13 | 0.29 | 0.26 |
| 2 | 208001268 | rs764487 | 1.037 | 0.107 | -0.831 | -0.697 | -0.067 | -0.063 | -0.063 | 0 | 0.27 | 0.15 | 0.41 | 0.53 | 0.4 | 0.41 |
| 2 | 208006413 | rs10181757 | 1.444 | -0.193 | -0.925 | -0.895 | 0 | 0.037 | 0.037 | 0 | 0.21 | 0.09 | 0.13 | 0.09 | 0.24 | 0.1 |
| 2 | 208007503 | rs1406791 | -0.27 | -0.325 | -0.713 | -0.979 | -0.019 | -0.061 | -0.061 | 2.086 | 0.37 | 0.46 | 0.35 | 0.16 | 0.31 | 0.23 |
| 2 | 208009536 | rs929695 | 1.494 | -0.233 | -0.788 | -0.787 | -0.005 | 0.053 | 0.053 | 0 | 0.2 | 0.07 | 0.13 | 0.08 | 0.23 | 0.1 |
| 2 | 208009919 | rs3791994 | -0.353 | -0.264 | -0.646 | -0.496 | -0.012 | -0.001 | -0.001 | 0 | 0.61 | 0.54 | 0.61 | 0.61 | 0.67 | 0.59 |
| 2 | 208013555 | rs12466923 | -0.94 | -0.262 | -0.71 | -0.608 | -0.034 | 0.026 | 0.026 | 0.049 | 0.74 | 0.56 | 0.83 | 0.67 | 0.79 | 0.75 |
| 2 | 208020941 | rs7603172 | 0.414 | -0.17 | -1.548 | -0.999 | -0.019 | 0.142 | 0.142 | 0 | 0.3 | 0.05 | 0.18 | 0.1 | 0.32 | 0.26 |
| 2 | 208025054 | rs9288381 | 0.395 | -0.306 | -1.11 | -1.066 | 0 | 0.101 | 0.101 | 0 | 0.3 | 0.08 | 0.19 | 0.1 | 0.32 | 0.26 |
| 2 | 208027775 | rs2241262 | 1.606 | -0.061 | -1.817 | -1.075 | -0.097 | -0.155 | -0.155 | 0 | 0.19 | 0.01 | 0.24 | 0.48 | 0.25 | 0.31 |
| 2 | 208029014 | rs3791997 | -0.632 | -1.247 | -1.288 | -1.41 | -0.014 | 0.05 | 0.05 | 0.313 | 0.56 | 0.59 | 0.52 | 0.28 | 0.57 | 0.4 |
| 2 | 208033676 | rs1920048 | -0.284 | -1.303 | -1.308 | -1.364 | 0.025 | 0.07 | 0.07 | 0 | 0.59 | 0.34 | 0.73 | 0.88 | 0.64 | 0.7 |
| 2 | 208043412 | rs11888248 | 1.488 | -1.203 | -1.591 | -1.172 | -0.022 | -0.033 | -0.033 | 0 | 0.16 | 0.03 | 0.25 | 0.17 | 0.18 | 0.1 |
| 2 | 208046557 | rs7600286 | -0.121 | -1.228 | -1.381 | -1.354 | 0.01 | 0.019 | 0.019 | 0 | 0.39 | 0.6 | 0.27 | 0.13 | 0.37 | 0.3 |
| 2 | 208049581 | rs2111592 | 0.626 | -1.274 | -1.372 | -1.399 | -0.008 | 0.117 | 0.117 | 0 | 0.3 | 0.07 | 0.19 | 0.12 | 0.32 | 0.27 |
| 2 | 208050149 | rs1263686 | 0.438 | -1.313 | -0.553 | -1.411 | 0 | -0.018 | -0.018 | 0.141 | 0.32 | 0.24 | 0.2 | 0.13 | 0.33 | 0.27 |
| 2 | 208055723 | rs1263674 | 0.43 | -1.388 | -0.303 | -1.448 | 0 | -0.018 | -0.018 | 2.413 | 0.32 | 0.24 | 0.2 | 0.12 | 0.33 | 0.27 |
| 2 | 208055947 | rs1263673 | 1.529 | -1.396 | -0.638 | -1.452 | 0.037 | 0.05 | 0.05 | 0.093 | 0.26 | 0.35 | 0.42 | 0.47 | 0.38 | 0.4 |
| 2 | 208059640 | rs2360675 | 1.265 | -1.821 | -0.394 | -1.449 | -0.03 | -0.012 | -0.012 | 0 | 0.39 | 0.17 | 0.52 | 0.56 | 0.54 | 0.41 |
| 2 | 208068379 | rs1920049 | 0.372 | -1.736 | -0.312 | -1.801 | 0.054 | 0.135 | 0.135 | 0 | 0.55 | 0.29 | 0.58 | 0.8 | 0.62 | 0.62 |
| 2 | 208070211 | rs10932182 | 0.119 | -1.65 | -0.232 | -1.823 | 0.026 | 0.07 | 0.07 | 0 | 0.59 | 0.34 | 0.73 | 0.88 | 0.64 | 0.68 |
| 2 | 208079942 | rs2216375 | 0.006 | -1.529 | -0.382 | -1.756 | -0.005 | -0.041 | -0.041 | 0 | 0.34 | 0.33 | 0.21 | 0.12 | 0.33 | 0.28 |
| 2 | 208086714 | rs2111598 | 0.356 | -1.695 | -0.605 | -1.632 | -0.045 | -0.043 | -0.043 | 0 | 0.78 | 0.4 | 0.87 | 0.92 | 0.91 | 0.86 |
| 2 | 208087139 | rs741475 | 0.788 | -1.604 | -0.853 | -1.607 | 0.043 | 0.146 | 0.146 | 0 | 0.48 | 0.25 | 0.54 | 0.73 | 0.57 | 0.51 |
| 2 | 208088537 | rs960379 | 0.366 | -1.707 | -0.638 | -1.686 | 0.025 | 0.075 | 0.075 | 0 | 0.53 | 0.7 | 0.72 | 0.81 | 0.6 | 0.54 |
| 2 | 208088987 | rs11695409 | 0.09 | -1.591 | -0.611 | -1.549 | 0.009 | 0.031 | 0.031 | 0 | 0.6 | 0.73 | 0.76 | 0.8 | 0.63 | 0.61 |
| 2 | 208089904 | rs2216374 | 0.846 | -1.596 | -0.591 | -1.45 | 0.052 | 0.097 | 0.097 | 0 | 0.54 | 0.34 | 0.57 | 0.77 | 0.64 | 0.61 |
| 2 | 208090279 | rs17285604 | -0.69 | -1.37 | -0.344 | -1.41 | 0 | 0.003 | 0.003 | 0 | 0.38 | 0.3 | 0.31 | 0.29 | 0.42 | 0.36 |
| 2 | 208091112 | rs717973 | 1.443 | -1.502 | -0.465 | -1.253 | -0.016 | 0.018 | 0.018 | 0 | 0.39 | 0.32 | 0.46 | 0.66 | 0.54 | 0.52 |
| 2 | 208096604 | rs6435367 | -1.662 | -1.055 | 0.278 | -0.615 | 0.054 | 0.074 | 0.074 | 0 | 0.45 | 0.75 | 0.33 | 0.28 | 0.35 | 0.36 |
| 2 | 208097209 | rs12329153 | -0.058 | -0.855 | 0.534 | -0.497 | -0.037 | 0.011 | 0.011 | 0 | 0.16 | 0.21 | 0.11 | 0.16 | 0.06 | 0.12 |
| 2 | 208103052 | rs6435369 | -0.924 | -0.675 | 0.673 | -0.484 | 0.005 | -0.052 | -0.052 | 0.055 | 0.1 | 0.53 | 0.07 | 0 | 0.06 | 0.04 |
| 2 | 208106829 | rs716630 | 1.009 | -0.568 | 0.285 | -0.501 | -0.001 | -0.004 | -0.004 | 0.432 | 0.1 | 0.07 | 0.07 | 0 | 0.12 | 0.04 |
| 2 | 208112070 | rs12469545 | 1.687 | -0.648 | -0.204 | -0.539 | 0.038 | 0.063 | 0.063 | 0 | 0.56 | 0.28 | 0.6 | 0.71 | 0.66 | 0.57 |
| 2 | 208112602 | rs6739693 | 0.242 | -0.635 | -0.13 | -0.539 | 0.005 | -0.003 | -0.003 | 1.009 | 0.17 | 0.17 | 0.19 | 0.17 | 0.1 | 0.12 |
| 2 | 208114630 | rs1035263 | 1.131 | -0.429 | -0.177 | -0.554 | -0.002 | -0.004 | -0.004 | 2.525 | 0.09 | 0.06 | 0.07 | 0 | 0.13 | 0.03 |
| 2 | 208115178 | rs4675654 | 1.043 | -0.2 | -0.127 | -0.355 | 0.039 | 0.083 | 0.083 | 0 | 0.55 | 0.27 | 0.6 | 0.73 | 0.65 | 0.57 |
| 2 | 208116942 | rs12694083 | 0.407 | -0.163 | -0.07 | -0.289 | 0.051 | 0.025 | 0.025 | 0 | 0.89 | 0.61 | 0.9 | 0.89 | 0.91 | 0.77 |
| 2 | 208118301 | rs12621900 | 0.296 | -0.171 | -1.119 | -0.238 | -0.008 | -0.017 | -0.017 | 0 | 0.62 | 0.4 | 0.67 | 0.84 | 0.74 | 0.77 |
| 2 | 208119043 | rs2072570 | -0.031 | 0.649 | -1.682 | 0.374 | 0.006 | 0.04 | 0.04 | 0 | 0.5 | 0.21 | 0.43 | 0.56 | 0.55 | 0.56 |
| 2 | 208119704 | rs7591154 | -0.78 | 1.252 | -0.931 | 0.247 | 0.029 | -0.017 | -0.017 | 0 | 0.22 | 0.26 | 0.21 | 0.22 | 0.22 | 0.28 |
| 2 | 208120208 | rs12466346 | 0.734 | 0.448 | -2.124 | -0.385 | -0.027 | 0.276 | 0.276 | 0.37 | 0.51 | 0.08 | 0.4 | 0.19 | 0.62 | 0.51 |

| | | | | | | | | | | | | | | | | |
|---|---|---|---|---|---|---|---|---|---|---|---|---|---|---|---|---|
| 2 | 208126655 | rs4675657 | -0.085 | 0.567 | -1.568 | 0.167 | -0.007 | 0.001 | 0.001 | 0 | 0.65 | 0.58 | 0.75 | 0.71 | 0.7 | 0.82 |
| 2 | 208129918 | rs6732962 | -0.313 | 0.6 | -1.793 | -0.405 | -0.008 | 0.011 | 0.011 | 0 | 0.52 | 0.39 | 0.35 | 0.53 | 0.54 | 0.58 |
| 2 | 208133793 | rs7577873 | -0.062 | 0.488 | -1.54 | -0.403 | 0.037 | 0.032 | 0.032 | 0 | 0.15 | 0.48 | 0.17 | 0.17 | 0.12 | 0.16 |
| 2 | 208145220 | rs4675658 | 1.308 | 0.491 | -0.986 | -0.414 | 0.027 | 0.027 | 0.027 | 0.152 | 0.07 | 0.12 | 0.11 | 0.15 | 0.1 | 0.14 |
| 2 | 208148401 | rs7592640 | 0.4 | 0.369 | -1.055 | -0.49 | 0.012 | -0.006 | -0.006 | 0 | 0.33 | 0.33 | 0.28 | 0.3 | 0.27 | 0.16 |
| 2 | 208150499 | rs918193 | 0.846 | 0.342 | -0.763 | -0.376 | -0.052 | -0.012 | -0.012 | 1.211 | 0.33 | 0.11 | 0.45 | 0.32 | 0.37 | 0.28 |
| 2 | 208161729 | rs11674656 | -0.237 | 0.431 | -0.339 | -0.382 | 0.045 | -0.025 | -0.025 | 0.548 | 0.21 | 0.47 | 0.25 | 0.15 | 0.23 | 0.21 |
| 2 | 208161788 | rs11674696 | -1.052 | 0.078 | -0.511 | -0.591 | 0.001 | 0.083 | 0.083 | 1.205 | 0.52 | 0.84 | 0.42 | 0.37 | 0.53 | 0.6 |
| 2 | 208165513 | rs11883547 | 0.526 | -0.034 | -0.48 | -0.655 | -0.011 | 0.003 | 0.003 | 0 | 0.18 | 0.11 | 0.21 | 0.15 | 0.22 | 0.19 |
| 2 | 208169592 | rs16839580 | 0.391 | 0.083 | -0.105 | -0.644 | -0.004 | 0.019 | 0.019 | 0 | 0.28 | 0.22 | 0.16 | 0.22 | 0.3 | 0.39 |
| 2 | 208173545 | rs16839586 | 1.384 | 0.107 | -0.276 | -0.675 | -0.015 | 0.012 | 0.012 | 0 | 0.2 | 0.12 | 0.12 | 0.21 | 0.23 | 0.35 |
| 2 | 208176487 | rs2194374 | -1.37 | -0.017 | -0.472 | -0.494 | 0.009 | 0.024 | 0.024 | 1.029 | 0.56 | 0.71 | 0.48 | 0.7 | 0.57 | 0.81 |
| 2 | 208176734 | rs763617 | -0.17 | -0.173 | -0.113 | -0.321 | 0.047 | -0.031 | -0.031 | 1.036 | 0.22 | 0.47 | 0.26 | 0.15 | 0.24 | 0.21 |
| 2 | 208179396 | rs6435370 | -0.478 | -0.918 | -0.116 | -0.001 | -0.061 | 0.037 | 0.037 | 0 | 0.78 | 0.65 | 0.69 | 0.37 | 0.89 | 0.67 |
| 2 | 208191797 | rs7604355 | -0.456 | -0.513 | -0.174 | 0.039 | -0.037 | 0.021 | 0.021 | 0.088 | 0.39 | 0.55 | 0.37 | 0.51 | 0.29 | 0.46 |
| 2 | 208194639 | rs16839639 | -0.462 | -0.411 | -0.554 | 0.07 | 0.007 | -0.006 | -0.006 | 0 | 0.39 | 0.41 | 0.38 | 0.51 | 0.31 | 0.46 |
| 2 | 208196247 | rs16839653 | 1.586 | -0.325 | -1.062 | 0.262 | 0.03 | -0.045 | -0.045 | 1.091 | 0.15 | 0.05 | 0.23 | 0.29 | 0.1 | 0.14 |
| 2 | 208207317 | rs6710342 | 0.832 | 0.051 | -1.005 | -0.001 | 0.002 | 0.005 | 0.005 | 0 | 0.19 | 0.09 | 0.1 | 0.23 | 0.2 | 0.29 |
| 2 | 208207603 | rs6754615 | 0.467 | 0.061 | -0.989 | -0.014 | 0.001 | 0.016 | 0.016 | 0.108 | 0.22 | 0.12 | 0.12 | 0.23 | 0.22 | 0.32 |
| 2 | 208220191 | rs17202235 | 0.505 | 0.019 | -1.13 | 0.116 | 0.059 | 0.035 | 0.035 | 0 | 0.35 | 0.16 | 0.2 | 0.3 | 0.3 | 0.33 |
| 2 | 208221168 | rs2113383 | -1.21 | 0.64 | -0.62 | -0.125 | 0.027 | -0.026 | -0.026 | 0 | 0.7 | 0.56 | 0.68 | 0.74 | 0.68 | 0.69 |
| 2 | 208222261 | rs2161767 | -0.337 | 0.648 | -0.406 | -0.003 | 0.007 | 0.014 | 0.014 | 0 | 0.34 | 0.35 | 0.48 | 0.44 | 0.37 | 0.36 |
| 2 | 208229097 | rs11677973 | 0.464 | 0.835 | -0.781 | 0.039 | -0.03 | 0.005 | 0.005 | 0 | 0.15 | 0.11 | 0.14 | 0.13 | 0.18 | 0.18 |
| 2 | 208230143 | rs10804160 | -0.958 | 0.875 | -0.455 | -0.178 | -0.001 | 0 | 0 | 0 | 0.62 | 0.62 | 0.59 | 0.43 | 0.63 | 0.56 |
| 2 | 208231478 | rs17808718 | 1.481 | 0.676 | -0.688 | -0.369 | 0.031 | 0.004 | 0.004 | 0 | 0.28 | 0.16 | 0.19 | 0.3 | 0.26 | 0.26 |
| 2 | 208231640 | rs4675666 | 1.35 | 0.723 | -0.683 | -0.255 | -0.003 | 0.018 | 0.018 | 0 | 0.17 | 0.15 | 0.25 | 0.31 | 0.12 | 0.16 |
| 2 | 208236850 | rs7561038 | -2.246 | 1.02 | -0.911 | -0.397 | -0.01 | -0.005 | -0.005 | 0 | 0.81 | 0.85 | 0.84 | 0.74 | 0.79 | 0.72 |
| 2 | 208253030 | rs4675672 | -0.332 | 0.748 | -0.834 | -0.515 | -0.005 | 0.055 | 0.055 | 0 | 0.56 | 0.67 | 0.57 | 0.73 | 0.52 | 0.58 |
| 2 | 208254232 | rs1862884 | -1.829 | 0.912 | -0.949 | -0.473 | -0.002 | 0.031 | 0.031 | 0 | 0.89 | 0.85 | 0.85 | 0.73 | 0.88 | 0.75 |
| 2 | 208255518 | rs17203016 | 1.696 | 1.029 | -1.126 | -0.294 | -0.062 | -0.052 | -0.052 | 0 | 0.1 | 0.01 | 0.1 | 0.14 | 0.19 | 0.11 |
| 2 | 208263279 | rs11692326 | 1.031 | 0.903 | -1.017 | -0.538 | -0.038 | 0.022 | 0.022 | 0 | 0.16 | 0.11 | 0.15 | 0.09 | 0.22 | 0.2 |
| 2 | 208267610 | rs6719582 | -1.56 | 0.885 | -1.052 | -0.594 | -0.016 | 0.011 | 0.011 | 0 | 0.73 | 0.83 | 0.74 | 0.73 | 0.68 | 0.7 |
| 2 | 208272732 | rs6721035 | 1.827 | -0.136 | -1.279 | -0.176 | 0.047 | -0.068 | -0.068 | 0 | 0.25 | 0.09 | 0.16 | 0.41 | 0.21 | 0.26 |
| 2 | 208289161 | rs11892543 | 1.644 | 0.361 | -0.478 | 0.027 | -0.034 | 0.165 | 0.165 | 0 | 0.12 | 0.31 | 0.25 | 0.22 | 0.08 | 0.12 |
| 2 | 208292744 | rs11684288 | 0.665 | 0.818 | -1.12 | -0.129 | -0.003 | 0.012 | 0.012 | 0 | 0.14 | 0.11 | 0.13 | 0.08 | 0.13 | 0.18 |
| 2 | 208297345 | rs13421357 | 0.664 | 0.807 | -1.121 | -0.127 | -0.006 | 0.014 | 0.014 | 0 | 0.14 | 0.11 | 0.13 | 0.08 | 0.14 | 0.18 |
| 2 | 208309509 | rs17809990 | 1.855 | 0.984 | -0.939 | 0.025 | -0.053 | 0.093 | 0.093 | 0.016 | 0.16 | 0.01 | 0.1 | 0 | 0.19 | 0.04 |
| 2 | 208326488 | rs1030290 | 1.407 | 0.574 | -0.171 | 0.658 | -0.028 | 0.133 | 0.133 | 0 | 0.11 | 0.21 | 0.25 | 0.22 | 0.08 | 0.13 |
| 2 | 208334096 | rs6725292 | -0.26 | 0.922 | 0.023 | 0.296 | 0 | 0.009 | 0.009 | 0 | 0.78 | 0.57 | 0.59 | 0.85 | 0.81 | 0.76 |
| 2 | 208334341 | rs17810085 | 0.411 | 1.133 | -0.668 | -0.221 | -0.011 | 0.077 | 0.077 | 0.012 | 0.47 | 0.22 | 0.37 | 0.76 | 0.51 | 0.55 |
| 2 | 208334716 | rs17810115 | 2.428 | 0.967 | -0.936 | -0.45 | -0.056 | 0.084 | 0.084 | 0 | 0.14 | 0.01 | 0.11 | 0 | 0.18 | 0.04 |
| 2 | 208335510 | rs4142221 | 1.124 | 1 | -1.304 | -0.282 | 0.005 | -0.164 | -0.164 | 0 | 0.32 | 0.02 | 0.2 | 0.5 | 0.28 | 0.25 |
| 2 | 208336308 | rs10165827 | 1.896 | 1.168 | -0.554 | -0.774 | 0.027 | -0.052 | -0.052 | 0.991 | 0.1 | 0.15 | 0.14 | 0 | 0.11 | 0.1 |
| 2 | 208338451 | rs6741811 | 0.585 | 0.969 | -1.29 | -0.921 | 0.018 | 0.004 | 0.004 | 0 | 0.34 | 0.13 | 0.24 | 0.76 | 0.33 | 0.48 |
| 2 | 208345380 | rs722761 | 0.168 | 1.133 | -0.648 | -0.722 | -0.011 | 0.016 | 0.016 | 0 | 0.61 | 0.32 | 0.47 | 0.75 | 0.6 | 0.54 |
| 2 | 208349258 | rs981901 | 0.318 | 1.134 | -0.857 | -0.696 | 0.001 | 0.006 | 0.006 | 0 | 0.28 | 0.39 | 0.39 | 0.25 | 0.3 | 0.36 |
| 2 | 208350039 | rs7340418 | 0.318 | 1.134 | -0.857 | -0.696 | 0.001 | 0.006 | 0.006 | 0 | 0.28 | 0.39 | 0.39 | 0.25 | 0.3 | 0.36 |
| 2 | 208363062 | rs2042484 | 0.148 | 0.117 | -1.34 | -0.541 | 0.019 | 0.037 | 0.037 | 0 | 0.46 | 0.32 | 0.34 | 0.75 | 0.43 | 0.51 |
| 2 | 208369213 | rs13010104 | 1.51 | 0.498 | -1.477 | -0.384 | -0.048 | 0.076 | 0.076 | 0 | 0.13 | 0.01 | 0.11 | 0 | 0.17 | 0.04 |
| 2 | 208372996 | rs2360969 | 0.696 | 0.056 | -1.712 | -0.479 | -0.177 | 0.061 | 0.061 | 0.598 | 0.26 | 0.03 | 0.39 | 0.12 | 0.43 | 0.33 |
| 2 | 208386024 | rs2709373 | 1.754 | -0.06 | -1.426 | -0.732 | -0.001 | -0.03 | -0.03 | 0 | 0.15 | 0.14 | 0.12 | 0 | 0.17 | 0.05 |
| 2 | 208387685 | rs10932200 | 0.447 | -0.242 | -0.872 | -0.216 | -0.041 | -0.026 | -0.026 | 0 | 0.32 | 0.24 | 0.44 | 0.37 | 0.42 | 0.55 |
| 2 | 208390388 | rs2709376 | 0.009 | -0.692 | -0.295 | -0.457 | 0.021 | -0.024 | -0.024 | 0 | 0.92 | 0.66 | 0.94 | 1 | 0.95 | 0.98 |
| 2 | 208391978 | rs2253206 | 0.002 | -0.75 | -0.201 | -0.485 | -0.016 | 0 | 0 | 0 | 0.59 | 0.33 | 0.5 | 0.63 | 0.53 | 0.43 |
| 2 | 208393998 | rs2551638 | 0.227 | -0.959 | 0.25 | -0.271 | 0.004 | -0.004 | -0.004 | 0.273 | 0.93 | 0.68 | 0.97 | 1 | 0.99 | 0.98 |
| 2 | 208426257 | rs10932201 | 0.462 | -0.864 | 0.157 | -0.371 | 0.026 | 0.041 | 0.041 | 0 | 0.5 | 0.13 | 0.45 | 0.63 | 0.48 | 0.43 |
| 2 | 208444028 | rs2254137 | -0.353 | -0.863 | 0.284 | -0.478 | 0.005 | 0.007 | 0.007 | 0 | 0.67 | 0.36 | 0.59 | 0.63 | 0.65 | 0.49 |
| 2 | 208454151 | rs2709393 | -0.287 | -0.627 | 0.481 | -0.558 | 0.05 | 0.01 | 0.01 | 2.545 | 0.94 | 0.66 | 0.95 | 1 | 0.96 | 0.99 |
| 2 | 208465778 | rs2551928 | -1.262 | -0.661 | 0.289 | -0.595 | 0.002 | -0.07 | -0.07 | 0 | 0.81 | 0.78 | 0.87 | 1 | 0.82 | 0.94 |
| 2 | 208467150 | rs1045780 | -2.069 | -0.623 | 0.059 | -0.576 | -0.001 | -0.03 | -0.03 | 0 | 0.85 | 0.86 | 0.88 | 1 | 0.83 | 0.95 |
| 2 | 208469184 | rs2256941 | -2.06 | -0.618 | 0.062 | -0.573 | -0.001 | -0.03 | -0.03 | 0.024 | 0.85 | 0.86 | 0.88 | 1 | 0.83 | 0.95 |
| 2 | 208477479 | rs2709417 | -1.085 | -0.622 | 0.098 | -0.575 | 0.017 | -0.035 | -0.035 | 0 | 0.09 | 0.44 | 0.06 | 0 | 0.05 | 0.02 |
| 2 | 208477852 | rs2551949 | -2.048 | -0.587 | 0.115 | -0.556 | -0.001 | -0.03 | -0.03 | 0.024 | 0.85 | 0.86 | 0.88 | 1 | 0.83 | 0.95 |
| 2 | 208480505 | rs2551946 | -0.722 | -0.718 | -0.085 | -0.467 | 0.005 | -0.073 | -0.073 | 0.918 | 0.15 | 0.44 | 0.1 | 0.03 | 0.12 | 0.05 |
| 2 | 208489101 | rs4234080 | -0.037 | -1 | -0.469 | -0.369 | -0.041 | -0.028 | -0.028 | 0 | 0.27 | 0.39 | 0.27 | 0.14 | 0.19 | 0.17 |
| 2 | 208494234 | rs2952768 | 0.846 | -0.992 | -0.633 | -0.182 | -0.011 | -0.021 | -0.021 | 0.1 | 0.3 | 0.29 | 0.41 | 0.4 | 0.35 | 0.51 |
| 2 | 208497965 | rs2464975 | -2.227 | -1.145 | -0.624 | -0.261 | 0.009 | -0.055 | -0.055 | 0 | 0.85 | 0.8 | 0.87 | 1 | 0.83 | 0.95 |
| 2 | 208501730 | rs7591784 | -0.387 | -1.178 | -0.661 | -0.25 | 0.001 | -0.001 | -0.001 | 0 | 0.54 | 0.51 | 0.46 | 0.59 | 0.47 | 0.44 |
| 2 | 208505879 | rs7594560 | 0.497 | -1.29 | -0.636 | -0.265 | -0.038 | -0.018 | -0.018 | 0 | 0.25 | 0.31 | 0.23 | 0.13 | 0.16 | 0.18 |

| | | | | | | | | | | | | | | | | |
|---|---|---|---|---|---|---|---|---|---|---|---|---|---|---|---|---|
| 2 | 208506371 | rs2709390 | 0.543 | -0.957 | -0.436 | 0.101 | -0.019 | -0.032 | -0.032 | 0 | 0.3 | 0.26 | 0.41 | 0.4 | 0.36 | 0.51 |
| 2 | 208507807 | rs4675690 | 0.055 | -1.018 | -0.486 | 0.145 | 0.002 | -0.001 | -0.001 | 0.096 | 0.54 | 0.51 | 0.46 | 0.59 | 0.47 | 0.44 |
| 2 | 208520043 | rs2709363 | -0.25 | -0.695 | -0.578 | -0.412 | -0.001 | 0.005 | 0.005 | 0 | 0.38 | 0.4 | 0.41 | 0.68 | 0.38 | 0.55 |
| 2 | 208526488 | rs9646834 | 1.135 | -0.603 | -1.323 | -0.443 | -0.036 | -0.037 | -0.037 | 0.683 | 0.12 | 0.18 | 0.03 | 0.08 | 0.03 | 0.16 |
| 2 | 208531122 | rs12477677 | 0.791 | 0.369 | -1.177 | 0.119 | -0.104 | -0.165 | -0.165 | 0 | 0.22 | 0.01 | 0.32 | 0.46 | 0.28 | 0.33 |
| 2 | 208531553 | rs1055228 | 0.905 | 0.563 | -1.039 | 0.163 | 0.02 | 0.061 | 0.061 | 0 | 0.44 | 0.29 | 0.44 | 0.32 | 0.43 | 0.37 |
| 2 | 208531683 | rs1055229 | -0.047 | 1.222 | -0.633 | 0.675 | 0.047 | 0.043 | 0.043 | 0.084 | 0.21 | 0.42 | 0.15 | 0.22 | 0.2 | 0.24 |
| 2 | 208536536 | rs2551968 | -0.336 | 0.85 | -0.977 | 0.11 | -0.003 | 0.003 | 0.003 | 0.416 | 0.4 | 0.38 | 0.44 | 0.69 | 0.46 | 0.6 |
| 2 | 208544280 | rs6722248 | -0.128 | 0.692 | -1.23 | -0.469 | -0.006 | -0.001 | -0.001 | 0 | 0.18 | 0.12 | 0.16 | 0 | 0.22 | 0.05 |
| 2 | 208544582 | rs2432726 | -0.563 | 0.121 | -1.485 | -0.585 | 0.005 | 0.038 | 0.038 | 0 | 0.51 | 0.39 | 0.56 | 0.67 | 0.55 | 0.56 |
| 2 | 208548378 | rs906848 | -0.272 | -0.23 | -1.387 | -0.802 | 0.062 | 0.066 | 0.066 | 0 | 0.49 | 0.71 | 0.46 | 0.67 | 0.41 | 0.53 |
| 2 | 208555031 | rs2663867 | -0.089 | 0.076 | -1.576 | -0.294 | -0.044 | -0.006 | -0.006 | 0 | 0.3 | 0.13 | 0.48 | 0.33 | 0.41 | 0.32 |
| 2 | 208560343 | rs17458775 | 0.112 | -0.02 | -1.844 | -0.408 | -0.134 | -0.046 | -0.046 | 0 | 0.28 | 0.02 | 0.49 | 0.28 | 0.39 | 0.19 |
| 2 | 208566316 | rs4673374 | 0.147 | -0.542 | -1.035 | 0.145 | -0.116 | 0.009 | 0.009 | 6.676 | 0.3 | 0.07 | 0.48 | 0.26 | 0.5 | 0.39 |
| 2 | 208566869 | rs2663887 | -0.787 | -0.809 | -0.452 | 0.206 | -0.064 | 0.169 | 0.169 | 0 | 0.17 | 0.38 | 0.08 | 0.4 | 0.08 | 0.35 |
| 2 | 208605594 | rs2663879 | -0.482 | -0.863 | -0.367 | 0.241 | -0.049 | 0.196 | 0.196 | 0 | 0.14 | 0.3 | 0.06 | 0.4 | 0.07 | 0.34 |
| 2 | 208606083 | rs11683791 | 0.967 | -0.925 | -0.072 | 0.196 | 0.028 | 0.072 | 0.072 | 6.112 | 0.48 | 0.23 | 0.43 | 0.34 | 0.42 | 0.25 |
| 2 | 208610801 | rs9636269 | 0.142 | -0.455 | -0.499 | 0.422 | -0.104 | 0.02 | 0.02 | 6.665 | 0.31 | 0.1 | 0.49 | 0.26 | 0.51 | 0.4 |
| 2 | 208623154 | rs6755821 | -1.184 | -0.356 | 0.29 | 0.976 | -0.065 | 0.205 | 0.205 | 0 | 0.16 | 0.37 | 0.06 | 0.41 | 0.07 | 0.34 |
| 2 | 208639876 | rs7567627 | -1.352 | 0.333 | 0.29 | 1.389 | 0 | 0.463 | 0.463 | 0 | 0.09 | 0.36 | 0.05 | 0.41 | 0.05 | 0.28 |
| 2 | 208641174 | rs6708488 | 0.352 | 0.523 | -0.257 | 1.19 | -0.011 | 0.187 | 0.187 | 0 | 0.84 | 0.66 | 0.88 | 0.62 | 0.85 | 0.78 |
| 2 | 208641908 | rs4675713 | -0.491 | 1.52 | -1.469 | 0.676 | -0.102 | 0.006 | 0.006 | 2.424 | 0.26 | 0.11 | 0.45 | 0.28 | 0.46 | 0.32 |
| 2 | 208643043 | rs6716597 | -0.385 | 1.615 | -1.523 | 0.644 | -0.116 | 0.003 | 0.003 | 1.656 | 0.23 | 0.07 | 0.4 | 0.24 | 0.42 | 0.3 |
| 2 | 208649423 | rs10188753 | -0.332 | 1.124 | -1.348 | 0.263 | -0.006 | 0.067 | 0.067 | 0 | 0.54 | 0.68 | 0.49 | 0.73 | 0.46 | 0.51 |
| 2 | 208651612 | rs2136448 | 0.05 | 1.146 | -1.639 | 0.396 | -0.063 | -0.015 | -0.015 | 0 | 0.36 | 0.17 | 0.49 | 0.34 | 0.41 | 0.3 |
| 2 | 208664252 | rs16840032 | -0.449 | 0.542 | -0.381 | 0.051 | -0.007 | -0.069 | -0.069 | 0 | 0.11 | 0.35 | 0.09 | 0 | 0.09 | 0.05 |
| 2 | 208686410 | rs12619380 | -0.641 | 0.457 | -0.246 | 0.563 | -0.006 | 0.052 | 0.052 | 0 | 0.25 | 0.4 | 0.2 | 0.39 | 0.24 | 0.28 |
| 2 | 208690281 | rs2306433 | -0.896 | 0.128 | 0.031 | 0.402 | -0.019 | 0.054 | 0.054 | 0 | 0.29 | 0.53 | 0.22 | 0.39 | 0.24 | 0.28 |
| 2 | 208691531 | rs2021858 | 0.188 | -0.004 | -0.812 | 0.072 | -0.001 | -0.071 | -0.071 | 2.603 | 0.2 | 0.1 | 0.32 | 0.54 | 0.24 | 0.33 |
| 2 | 208715314 | rs6435396 | 0.179 | 0.007 | -0.654 | -0.943 | -0.001 | -0.002 | -0.002 | 0 | 0.61 | 0.55 | 0.46 | 0.35 | 0.56 | 0.45 |
| 2 | 208716745 | rs10192712 | 0.054 | -0.028 | -0.667 | -0.976 | -0.001 | -0.001 | -0.001 | 0 | 0.65 | 0.6 | 0.61 | 0.37 | 0.68 | 0.49 |
| 2 | 208717783 | rs17537169 | 1.1 | 0.076 | -1.728 | -1.308 | -0.142 | 0.158 | 0.158 | 0 | 0.3 | 0.02 | 0.33 | 0.1 | 0.34 | 0.26 |
| 2 | 208729540 | rs12622006 | -0.298 | 0.034 | -0.984 | -0.962 | -0.009 | -0.009 | -0.009 | 0.581 | 0.27 | 0.13 | 0.23 | 0.27 | 0.29 | 0.16 |
| 2 | 208735106 | rs10932212 | -0.147 | 0.063 | -0.992 | -0.939 | -0.006 | -0.008 | -0.008 | 0 | 0.25 | 0.16 | 0.23 | 0.27 | 0.29 | 0.17 |
| 2 | 208742259 | rs13432421 | -0.14 | 0.071 | -1.059 | -0.933 | -0.016 | -0.016 | -0.016 | 0 | 0.25 | 0.12 | 0.23 | 0.27 | 0.29 | 0.16 |
| 2 | 208748931 | rs13419848 | -0.306 | -0.232 | -1.006 | -1.069 | 0.008 | -0.003 | -0.003 | 0.574 | 0.34 | 0.18 | 0.22 | 0.34 | 0.29 | 0.22 |
| 2 | 208761235 | rs10497888 | -0.207 | -0.143 | -0.665 | -0.899 | 0 | -0.006 | -0.006 | 0.232 | 0.28 | 0.19 | 0.2 | 0.27 | 0.26 | 0.14 |
| 2 | 208768367 | rs2621470 | 1.274 | -0.104 | -0.949 | -1.118 | -0.02 | 0.111 | 0.111 | 0 | 0.34 | 0.17 | 0.26 | 0.11 | 0.34 | 0.26 |
| 2 | 208775856 | rs2621472 | -0.266 | -0.125 | -0.81 | -1.044 | -0.006 | -0.017 | -0.017 | 0.215 | 0.67 | 0.78 | 0.51 | 0.39 | 0.62 | 0.41 |
| 2 | 208795331 | rs2046078 | 0.966 | 0.21 | -0.897 | -0.694 | -0.02 | 0.083 | 0.083 | 0 | 0.32 | 0.17 | 0.27 | 0.17 | 0.34 | 0.27 |
| 2 | 208796078 | rs1871671 | -0.52 | 0.116 | -0.936 | -0.62 | 0.012 | 0.014 | 0.014 | 3.107 | 0.71 | 0.71 | 0.5 | 0.41 | 0.61 | 0.43 |
| 2 | 208809499 | rs12998109 | 0.27 | 0.766 | -2.35 | -0.228 | -0.045 | -0.036 | -0.036 | 0 | 0.27 | 0.03 | 0.17 | 0.24 | 0.25 | 0.13 |
| 2 | 208822120 | rs11691125 | 0.154 | 0.501 | -1.347 | 0.396 | -0.002 | -0.003 | -0.003 | 0.308 | 0.6 | 0.32 | 0.46 | 0.41 | 0.59 | 0.4 |
| 2 | 208829527 | rs6435401 | 0.248 | 0.007 | -0.798 | 0.371 | 0.01 | 0.006 | 0.006 | 0 | 0.51 | 0.43 | 0.56 | 0.4 | 0.67 | 0.4 |
| 2 | 208834227 | rs12694096 | -0.77 | -0.598 | -1.098 | 0.78 | 0.001 | -0.085 | -0.085 | 0 | 0.88 | 1 | 0.8 | 0.61 | 0.93 | 0.68 |
| 2 | 208834477 | rs11679492 | 0.685 | -0.416 | -1.248 | 0.424 | -0.011 | -0.086 | -0.086 | 0 | 0.45 | 0.05 | 0.51 | 0.47 | 0.62 | 0.42 |
| 2 | 208842394 | rs2044971 | 0.149 | -0.976 | -0.41 | 0.115 | -0.03 | 0.012 | 0.012 | 0 | 0.61 | 0.46 | 0.65 | 0.47 | 0.75 | 0.43 |
| 2 | 208848865 | rs2123294 | -0.18 | -0.996 | -0.387 | 0.106 | -0.021 | 0.013 | 0.013 | 0 | 0.64 | 0.56 | 0.66 | 0.47 | 0.75 | 0.43 |
| 2 | 208853299 | rs10497890 | 1.39 | -0.357 | -1.21 | -0.067 | -0.123 | -0.078 | -0.078 | 3.544 | 0.21 | 0.01 | 0.37 | 0.28 | 0.34 | 0.13 |
| 2 | 208859893 | rs930745 | 1.376 | -0.752 | -0.928 | -0.082 | -0.01 | 0.017 | 0.017 | 2.456 | 0.12 | 0.09 | 0.12 | 0 | 0.12 | 0.02 |
| 2 | 208874694 | rs11676872 | -0.156 | -0.837 | -0.844 | -0.084 | 0.074 | -0.031 | -0.031 | 0 | 0.59 | 0.24 | 0.63 | 0.48 | 0.75 | 0.42 |
| 2 | 208885654 | rs6740362 | 0.689 | -0.752 | -0.832 | -0.088 | 0 | -0.049 | -0.049 | 0 | 0.15 | 0.2 | 0.12 | 0 | 0.12 | 0.02 |
| 2 | 208890373 | rs1037416 | 0.578 | -0.754 | -0.38 | -0.056 | 0.025 | -0.003 | -0.003 | 0 | 0.47 | 0.28 | 0.53 | 0.48 | 0.63 | 0.4 |
| 2 | 208899477 | rs9789678 | -0.028 | -0.739 | -0.248 | 0.224 | 0.025 | -0.01 | -0.01 | 0 | 0.61 | 0.32 | 0.65 | 0.52 | 0.74 | 0.43 |
| 2 | 208906654 | rs2084870 | -0.27 | -0.804 | 0.026 | 0.148 | -0.006 | 0 | 0 | 0.01 | 0.39 | 0.63 | 0.35 | 0.45 | 0.26 | 0.57 |
| 2 | 208909308 | rs4675722 | 0.04 | -0.809 | 0.325 | 0.133 | 0.003 | -0.003 | -0.003 | 0.768 | 0.36 | 0.39 | 0.34 | 0.46 | 0.26 | 0.56 |
| 2 | 208909332 | rs4675723 | -0.151 | -0.781 | -0.133 | 0.147 | -0.026 | 0.008 | 0.008 | 0 | 0.37 | 0.55 | 0.34 | 0.46 | 0.27 | 0.56 |
| 2 | 208916427 | rs10497894 | 0.034 | -0.716 | -0.527 | 0.175 | 0.059 | -0.031 | -0.031 | 0.107 | 0.63 | 0.21 | 0.63 | 0.54 | 0.74 | 0.44 |
| 2 | 208925719 | rs17538082 | 1.365 | -0.462 | -0.871 | -0.075 | -0.051 | -0.074 | -0.074 | 0.256 | 0.23 | 0.04 | 0.12 | 0.26 | 0.27 | 0.26 |
| 2 | 208926542 | rs12464255 | 1.251 | -0.367 | -0.705 | -0.13 | -0.003 | -0.001 | -0.001 | 0.048 | 0.13 | 0.12 | 0.12 | 0 | 0.12 | 0.02 |
| 2 | 208927892 | rs11683477 | -0.057 | -0.642 | -1.102 | -0.282 | 0.026 | 0.088 | 0.088 | 0 | 0.48 | 0.23 | 0.71 | 0.74 | 0.59 | 0.7 |
| 2 | 208938866 | rs4234085 | -0.82 | -0.332 | -0.808 | 0.06 | 0 | -0.001 | -0.001 | 0 | 0.69 | 0.37 | 0.84 | 0.74 | 0.72 | 0.72 |
| 2 | 208952782 | rs1993173 | 0.313 | 0.266 | -1.656 | 0.603 | -0.084 | -0.042 | -0.042 | 0 | 0.28 | 0.08 | 0.45 | 0.31 | 0.38 | 0.19 |
| 2 | 208957750 | rs896637 | -0.301 | 0.149 | -1.349 | 0.465 | 0.005 | 0.03 | 0.03 | 0 | 0.57 | 0.32 | 0.74 | 0.72 | 0.63 | 0.72 |
| 2 | 208961424 | rs6713275 | 0.382 | 0.094 | -1.544 | 0.4 | 0.002 | 0 | 0 | 0 | 0.19 | 0.12 | 0.13 | 0.17 | 0.21 | 0.22 |
| 2 | 208966158 | rs6733487 | -0.459 | 0.523 | -1.775 | 0.419 | -0.009 | -0.007 | -0.007 | 1.557 | 0.44 | 0.3 | 0.45 | 0.48 | 0.56 | 0.52 |
| 2 | 208966609 | rs2363456 | -0.442 | 0.856 | -1.551 | 0.317 | -0.006 | -0.005 | -0.005 | 0 | 0.43 | 0.33 | 0.36 | 0.49 | 0.55 | 0.52 |
| 2 | 208969865 | rs2084872 | 0.698 | 0.437 | -0.772 | 0.12 | -0.018 | -0.024 | -0.024 | 1.082 | 0.22 | 0.41 | 0.14 | 0.15 | 0.21 | 0.23 |
| 2 | 208971007 | rs10181061 | -0.44 | 0.527 | -0.541 | 0.053 | -0.031 | -0.012 | -0.012 | 0 | 0.39 | 0.18 | 0.51 | 0.37 | 0.53 | 0.34 |

| | | | | | | | | | | | | | | | | |
|---|---|---|---|---|---|---|---|---|---|---|---|---|---|---|---|---|
| 2 | 208974411 | rs6744366 | 0.427 | 0.599 | -0.474 | 0.092 | -0.025 | -0.017 | -0.017 | 0 | 0.24 | 0.15 | 0.33 | 0.33 | 0.35 | 0.3 |
| 2 | 208976843 | rs10200650 | -0.193 | 0.29 | -0.128 | 0.205 | -0.054 | -0.028 | -0.028 | 0.566 | 0.33 | 0.18 | 0.48 | 0.37 | 0.48 | 0.29 |
| 2 | 208976955 | rs10932241 | 0.881 | -0.176 | -0.1 | 0.106 | 0.02 | -0.005 | -0.005 | 0 | 0.46 | 0.39 | 0.42 | 0.54 | 0.4 | 0.58 |
| 2 | 208977797 | rs10497893 | 1.748 | -0.342 | -0.303 | 0.003 | -0.014 | 0.081 | 0.081 | 0.005 | 0.32 | 0.06 | 0.22 | 0.15 | 0.36 | 0.34 |
| 2 | 208979528 | rs7355261 | 0.187 | -0.299 | -0.079 | 0.041 | -0.032 | -0.016 | -0.016 | 0.718 | 0.33 | 0.27 | 0.48 | 0.37 | 0.47 | 0.29 |
| 2 | 208985903 | rs966932 | 0.235 | -0.362 | 0.251 | 0 | 0.006 | 0.067 | 0.067 | 0 | 0.54 | 0.74 | 0.33 | 0.25 | 0.48 | 0.46 |
| 2 | 208986049 | rs6435415 | 0.259 | -0.365 | 0.249 | -0.001 | 0.006 | 0.068 | 0.068 | 0 | 0.54 | 0.74 | 0.33 | 0.25 | 0.48 | 0.46 |
| 2 | 208986385 | rs2305429 | -1.524 | -0.458 | 0.125 | 0.05 | -0.003 | 0.011 | 0.011 | 0.157 | 0.87 | 0.91 | 0.8 | 0.61 | 0.95 | 0.75 |
| 2 | 208986637 | rs2305430 | -0.589 | -0.503 | 0.416 | 0.001 | -0.006 | 0.011 | 0.011 | 0.772 | 0.82 | 0.83 | 0.78 | 0.61 | 0.92 | 0.73 |
| 2 | 208989708 | rs12328374 | 0.778 | -0.44 | 0.72 | -0.278 | 0.004 | 0.007 | 0.007 | 0 | 0.47 | 0.42 | 0.29 | 0.25 | 0.44 | 0.49 |
| 2 | 208990345 | rs2601495 | 1.439 | -0.424 | 0.599 | -0.306 | 0.008 | 0.047 | 0.047 | 0 | 0.34 | 0.25 | 0.22 | 0.15 | 0.37 | 0.38 |
| 2 | 208994697 | rs2242071 | -0.167 | -0.298 | 0.811 | -0.138 | -0.002 | 0.004 | 0.004 | 0 | 0.67 | 0.69 | 0.56 | 0.58 | 0.75 | 0.69 |
| 2 | 208994849 | rs13392731 | 0.077 | -0.316 | 0.918 | -0.05 | 0.002 | 0 | 0 | 0 | 0.58 | 0.62 | 0.53 | 0.47 | 0.73 | 0.55 |
| 2 | 208995098 | rs2601488 | -0.973 | -0.107 | 1.208 | 0.016 | 0.005 | 0 | 0 | 0 | 0.49 | 0.55 | 0.55 | 0.47 | 0.54 | 0.43 |
| 2 | 208996214 | rs12463771 | 1.433 | 0.123 | 1.141 | 0.092 | -0.003 | 0.004 | 0.004 | 1.777 | 0.16 | 0.15 | 0.22 | 0.04 | 0.17 | 0.06 |
| 2 | 209000573 | rs4449112 | -0.496 | -0.021 | 1.214 | -0.354 | -0.004 | 0.095 | 0.095 | 0.046 | 0.75 | 0.59 | 0.58 | 0.59 | 0.79 | 0.75 |
| 2 | 209000738 | rs2854712 | 1.289 | -0.083 | 1.114 | -0.43 | 0.032 | 0.064 | 0.064 | 0 | 0.46 | 0.27 | 0.26 | 0.27 | 0.43 | 0.49 |
| 2 | 209002036 | rs971493 | -0.02 | -0.294 | 0.952 | -0.656 | -0.039 | -0.004 | -0.004 | 0 | 0.27 | 0.2 | 0.31 | 0.32 | 0.36 | 0.23 |
| 2 | 209007559 | rs796287 | 0.069 | 0.019 | 1.054 | -0.557 | 0 | 0.022 | 0.022 | 0 | 0.66 | 0.62 | 0.54 | 0.6 | 0.73 | 0.71 |
| 2 | 209008605 | rs2362791 | 0.883 | 0.056 | 0.712 | -0.66 | 0.026 | 0.092 | 0.092 | 0 | 0.48 | 0.27 | 0.27 | 0.27 | 0.48 | 0.49 |
| 2 | 209011014 | rs6750890 | 1.618 | 0.234 | 0.503 | -0.638 | -0.006 | 0.004 | 0.004 | 0.489 | 0.15 | 0.08 | 0.21 | 0 | 0.15 | 0.06 |
| 2 | 209011127 | rs796283 | -1.218 | 0.006 | 0.675 | -0.676 | -0.006 | 0.012 | 0.012 | 0.184 | 0.57 | 0.68 | 0.58 | 0.42 | 0.53 | 0.5 |
| 2 | 209012367 | rs2362790 | 0.456 | 0.181 | 0.653 | -0.677 | 0.018 | -0.057 | -0.057 | 2.915 | 0.63 | 0.53 | 0.47 | 0.17 | 0.59 | 0.4 |
| 2 | 209013401 | rs3754821 | 1.318 | -0.556 | 1.158 | -0.372 | 0.034 | 0.037 | 0.037 | 0.314 | 0.58 | 0.15 | 0.64 | 0.37 | 0.71 | 0.49 |
| 2 | 209013586 | rs3754819 | -0.161 | -0.57 | 1.418 | -0.376 | -0.003 | -0.016 | -0.016 | 0.208 | 0.26 | 0.27 | 0.11 | 0.09 | 0.17 | 0.24 |
| 2 | 209014071 | rs2592158 | -0.495 | -0.584 | 1.601 | -0.311 | 0.034 | 0.297 | 0.297 | 0.038 | 0.9 | 0.71 | 0.79 | 0.59 | 0.93 | 0.83 |
| 2 | 209015976 | rs796266 | -0.155 | -0.572 | 1.269 | -0.307 | -0.001 | 0.049 | 0.049 | 0 | 0.06 | 0.15 | 0.02 | 0.13 | 0.05 | 0.1 |
| 2 | 209020181 | rs6758738 | 1.711 | -0.616 | 1.009 | -0.328 | -0.075 | 0.044 | 0.044 | 0.571 | 0.44 | 0.11 | 0.43 | 0.37 | 0.57 | 0.43 |
| 2 | 209021301 | rs796272 | -0.946 | -0.579 | 1.089 | -0.277 | 0 | 0.078 | 0.078 | 0 | 0.92 | 0.92 | 0.8 | 0.59 | 0.93 | 0.82 |
| 2 | 209022909 | rs6720791 | 1.195 | -0.473 | 0.896 | -0.753 | 0.025 | 0.023 | 0.023 | 0.245 | 0.58 | 0.31 | 0.65 | 0.37 | 0.71 | 0.49 |
| 2 | 209024565 | rs2601474 | -1.119 | -0.874 | 0.724 | -0.422 | -0.005 | 0.02 | 0.02 | 0 | 0.36 | 0.53 | 0.46 | 0.51 | 0.26 | 0.28 |
| 2 | 209025027 | rs6729051 | -0.211 | -0.848 | 0.81 | -0.419 | 0 | -0.044 | -0.044 | 0.414 | 0.26 | 0.27 | 0.25 | 0 | 0.19 | 0.09 |
| 2 | 209025923 | rs2441351 | -0.147 | -0.306 | 0.435 | 0.118 | 0.014 | 0.065 | 0.065 | 0 | 0.24 | 0.36 | 0.43 | 0.51 | 0.22 | 0.24 |
| 2 | 209028191 | rs796280 | 0.147 | -0.865 | -0.345 | 0.191 | 0 | -0.015 | -0.015 | 0 | 0.22 | 0.18 | 0.41 | 0.53 | 0.18 | 0.24 |
| 2 | 209033557 | rs7589531 | 0.424 | -0.807 | 0.223 | -0.361 | -0.04 | -0.012 | -0.012 | 0 | 0.84 | 0.81 | 0.94 | 0.86 | 0.93 | 0.82 |
| 2 | 209034801 | rs1348751 | 0.418 | -0.784 | 0.212 | -0.348 | -0.041 | -0.012 | -0.012 | 0 | 0.84 | 0.81 | 0.94 | 0.86 | 0.93 | 0.82 |
| 2 | 209036712 | rs10804166 | 0.447 | -0.784 | 0.06 | -0.347 | -0.045 | -0.01 | -0.01 | 0 | 0.84 | 0.79 | 0.94 | 0.85 | 0.93 | 0.82 |
| 2 | 209042963 | rs6738940 | -0.812 | -0.519 | -1.157 | -0.418 | 0.039 | 0.064 | 0.064 | 0 | 0.25 | 0.05 | 0.11 | 0.14 | 0.17 | 0.25 |
| 2 | 209046100 | rs6751157 | -0.907 | -0.201 | -0.558 | -0.535 | 0.003 | 0.003 | 0.003 | 0 | 0.14 | 0.11 | 0.09 | 0.03 | 0.12 | 0.23 |
| 2 | 209048136 | rs12995271 | -1.641 | -0.13 | 0.211 | -0.947 | 0.001 | -0.118 | -0.118 | 0.007 | 0.28 | 0.41 | 0.22 | 0.03 | 0.25 | 0.31 |
| 2 | 209050447 | rs6710543 | -0.849 | -0.195 | -0.075 | -0.91 | -0.004 | 0.013 | 0.013 | 0 | 0.16 | 0.11 | 0.12 | 0.03 | 0.18 | 0.24 |
| 2 | 209068854 | rs6714981 | 0.41 | -0.306 | 1.11 | -0.841 | -0.004 | -0.079 | -0.079 | 0.432 | 0.87 | 0.54 | 0.92 | 1 | 0.93 | 0.93 |
| 2 | 209069303 | rs12373582 | -1.478 | -0.163 | 0.668 | -0.862 | 0.007 | -0.015 | -0.015 | 0 | 0.09 | 0.11 | 0.09 | 0.03 | 0.15 | 0.2 |
| 2 | 209074635 | rs13387151 | -1.894 | -0.373 | 0.89 | -0.967 | -0.002 | -0.043 | -0.043 | 1.02 | 0.24 | 0.72 | 0.18 | 0.03 | 0.21 | 0.26 |
| 2 | 209087335 | rs4675743 | 2.368 | -0.529 | 1.077 | -0.898 | -0.015 | -0.052 | -0.052 | 0 | 0.86 | 0.26 | 0.9 | 1 | 0.91 | 0.92 |
| 2 | 209093069 | rs6734788 | 2.21 | -0.489 | 1.457 | -0.896 | 0.014 | -0.076 | -0.076 | 1.048 | 0.86 | 0.54 | 0.88 | 0.96 | 0.87 | 0.87 |
| 2 | 209099697 | rs12105636 | -0.921 | -0.906 | 0.396 | -0.121 | 0.016 | -0.063 | -0.063 | 0 | 0.2 | 0.1 | 0.13 | 0.54 | 0.15 | 0.24 |
| 2 | 209100914 | rs16840781 | -1.041 | -0.897 | 0.504 | -0.058 | 0.011 | -0.035 | -0.035 | 0 | 0.2 | 0.12 | 0.13 | 0.53 | 0.15 | 0.21 |
| 2 | 209112551 | rs6435435 | 0.946 | -0.907 | 1.013 | -0.015 | 0.01 | -0.026 | -0.026 | 0 | 0.94 | 0.85 | 0.95 | 0.99 | 0.95 | 0.96 |
| 2 | 209116753 | rs3769521 | 0.014 | -1.314 | 0.12 | -0.5 | -0.013 | 0.015 | 0.015 | 1.877 | 0.16 | 0.01 | 0.18 | 0.07 | 0.15 | 0.26 |
| 2 | 209117317 | rs1437410 | 1.555 | -1.312 | 0.008 | -1.156 | -0.054 | 0.031 | 0.031 | 1.22 | 0.4 | 0.2 | 0.47 | 0.35 | 0.45 | 0.33 |
| 2 | 209120258 | rs12478635 | 0.28 | -1.262 | -0.179 | -1.094 | -0.034 | -0.156 | -0.156 | 0 | 0.21 | 0.03 | 0.18 | 0.48 | 0.22 | 0.25 |
| 2 | 209130114 | rs11895188 | -0.56 | -1.184 | -0.064 | -1.131 | -0.002 | -0.007 | -0.007 | 0 | 0.08 | 0.08 | 0.09 | 0.01 | 0.1 | 0.07 |
| 2 | 209135339 | rs6746926 | 1.821 | -1.118 | -0.196 | -1.287 | -0.126 | -0.029 | -0.029 | 0 | 0.39 | 0.07 | 0.47 | 0.38 | 0.45 | 0.33 |
| 2 | 209149617 | rs11678092 | -0.795 | -1.203 | 1.379 | -1.427 | -0.006 | -0.013 | -0.013 | 0 | 0.3 | 0.34 | 0.2 | 0.12 | 0.25 | 0.43 |
| 2 | 209162775 | rs4675751 | 1.586 | -0.988 | 1.817 | -1.194 | 0.007 | -0.042 | -0.042 | 0 | 0.85 | 0.58 | 0.86 | 0.93 | 0.87 | 0.83 |
| 2 | 209179732 | rs6736599 | 2.049 | -1.008 | 1.967 | -1.204 | -0.014 | -0.054 | -0.054 | 0 | 0.82 | 0.4 | 0.85 | 0.93 | 0.87 | 0.83 |
| 2 | 209185355 | rs10932260 | 2.013 | -1.014 | 2.049 | -1.206 | -0.012 | -0.057 | -0.057 | 0 | 0.82 | 0.44 | 0.85 | 0.93 | 0.87 | 0.83 |
| 2 | 209190632 | rs999890 | -1.052 | -0.977 | 1.82 | -1.188 | 0.001 | -0.024 | -0.024 | 0 | 0.11 | 0.15 | 0.11 | 0.01 | 0.13 | 0.08 |
| 2 | 209190649 | rs999891 | -1.052 | -0.977 | 1.82 | -1.188 | 0.001 | -0.024 | -0.024 | 2.572 | 0.11 | 0.15 | 0.11 | 0.01 | 0.13 | 0.08 |
| 2 | 209233517 | rs35217101 | -1.019 | -1.22 | 1.49 | -1.149 | -0.008 | -0.049 | -0.049 | 0 | 0.11 | 0.21 | 0.1 | 0.01 | 0.1 | 0.07 |
| 2 | 209235962 | rs12694110 | 1.187 | -1.333 | 1.277 | -1.181 | 0.038 | 0.061 | 0.061 | 0 | 0.51 | 0.28 | 0.58 | 0.42 | 0.57 | 0.42 |
| 2 | 209236372 | rs35444703 | -0.738 | -1.374 | 1.541 | -1.18 | -0.001 | -0.012 | -0.012 | 0 | 0.42 | 0.23 | 0.37 | 0.52 | 0.43 | 0.48 |
| 2 | 209237839 | rs6728530 | 1.197 | -1.368 | 1.502 | -1.204 | 0.036 | 0.057 | 0.057 | 0 | 0.51 | 0.28 | 0.59 | 0.42 | 0.57 | 0.43 |
| 2 | 209239742 | rs4675764 | 1.213 | -1.369 | 1.483 | -1.204 | 0.037 | 0.06 | 0.06 | 0 | 0.51 | 0.27 | 0.58 | 0.41 | 0.56 | 0.43 |
| 2 | 209239790 | rs7586020 | -0.8 | -1.334 | 1.522 | -1.19 | 0.004 | -0.001 | -0.001 | 0 | 0.43 | 0.32 | 0.37 | 0.53 | 0.43 | 0.48 |
| 2 | 209264186 | rs11897777 | -1.092 | -1.258 | 1.529 | -1.115 | -0.01 | -0.052 | -0.052 | 0 | 0.11 | 0.21 | 0.11 | 0.01 | 0.1 | 0.07 |
| 2 | 209280931 | rs11902124 | -0.483 | -1.267 | 1.512 | -1.118 | 0 | -0.008 | -0.008 | 0.444 | 0.08 | 0.08 | 0.08 | 0.01 | 0.07 | 0.06 |
| 2 | 209286233 | rs12617462 | 0.819 | -1.342 | 1.326 | -1.129 | -0.016 | -0.036 | -0.036 | 0 | 0.1 | 0.02 | 0.1 | 0.46 | 0.1 | 0.22 |

| | | | | | | | | | | | | | | | | |
|---|---|---|---|---|---|---|---|---|---|---|---|---|---|---|---|---|
| 2 | 209293977 | rs13016936 | 0.435 | -1.499 | 0.862 | -1.432 | 0 | 0.008 | 0.008 | 0 | 0.26 | 0.18 | 0.21 | 0.07 | 0.25 | 0.25 |
| 2 | 209296943 | rs10497900 | 2.311 | -1.666 | 0.669 | -1.454 | -0.141 | -0.068 | -0.068 | 0.084 | 0.34 | 0.07 | 0.47 | 0.41 | 0.44 | 0.35 |
| 2 | 209298030 | rs1370369 | 0.506 | -1.654 | 0.738 | -1.489 | 0.002 | 0.069 | 0.069 | 0 | 0.25 | 0.09 | 0.2 | 0.07 | 0.24 | 0.25 |
| 2 | 209315573 | rs1370371 | 0.74 | -1.583 | 0.776 | -1.391 | 0.059 | 0.16 | 0.16 | 0.61 | 0.57 | 0.22 | 0.65 | 0.87 | 0.66 | 0.6 |
| 2 | 209332455 | rs2576810 | -0.265 | -1.119 | 1.023 | -0.986 | -0.047 | -0.15 | -0.15 | 0 | 0.79 | 0.97 | 0.83 | 0.54 | 0.77 | 0.75 |
| 2 | 209334978 | rs879707 | 1.405 | -1.112 | 1.493 | -1.023 | -0.007 | 0.01 | 0.01 | 0 | 0.65 | 0.38 | 0.72 | 0.48 | 0.7 | 0.62 |
| 2 | 209342094 | rs897083 | 0.042 | -1.133 | 1.396 | -1.018 | -0.006 | -0.088 | -0.088 | 0 | 0.77 | 0.86 | 0.81 | 0.54 | 0.78 | 0.75 |
| 2 | 209350128 | rs3820902 | 1.403 | -1.149 | 1.496 | -1.028 | 0.011 | -0.012 | -0.012 | 3.971 | 0.63 | 0.29 | 0.71 | 0.48 | 0.7 | 0.56 |
| 2 | 209350308 | rs3820901 | 1.529 | -1.156 | 1.5 | -1.031 | 0.021 | -0.017 | -0.017 | 0 | 0.62 | 0.27 | 0.7 | 0.48 | 0.7 | 0.56 |
| 2 | 209357870 | rs6717082 | -0.942 | -1.144 | 1.569 | -1.03 | 0.012 | -0.036 | -0.036 | 0 | 0.09 | 0.35 | 0.09 | 0.01 | 0.06 | 0.04 |
| 2 | 209358027 | rs3820900 | -0.067 | -1.137 | 1.287 | -0.987 | -0.008 | -0.085 | -0.085 | 2.786 | 0.77 | 0.86 | 0.81 | 0.54 | 0.78 | 0.7 |
| 2 | 209358951 | rs1057390 | -1.428 | -1.141 | 1.328 | -0.988 | -0.039 | -0.03 | -0.03 | 0.867 | 0.13 | 0.56 | 0.1 | 0.06 | 0.07 | 0.13 |
| 2 | 209359062 | rs1057392 | -0.051 | -1.048 | 1.054 | -0.941 | -0.009 | -0.085 | -0.085 | 0 | 0.77 | 0.87 | 0.81 | 0.54 | 0.78 | 0.7 |
| 2 | 209378138 | rs6755377 | 1.002 | -0.992 | 0.573 | -0.653 | 0.001 | -0.001 | -0.001 | 5.202 | 0.65 | 0.35 | 0.68 | 0.47 | 0.72 | 0.62 |
| 2 | 209412884 | rs10932263 | 1.507 | -0.721 | 0.112 | -0.759 | -0.105 | -0.083 | -0.083 | 0 | 0.33 | 0.06 | 0.42 | 0.42 | 0.4 | 0.39 |
| 2 | 209414011 | rs7605626 | -1.943 | -0.839 | 1.415 | -0.583 | -0.038 | -0.034 | -0.034 | 0.088 | 0.15 | 0.69 | 0.11 | 0.06 | 0.07 | 0.13 |
| 2 | 209414499 | rs7567228 | -0.578 | -0.741 | 1.544 | -0.546 | 0.029 | 0.003 | 0.003 | 0 | 0.06 | 0.24 | 0.05 | 0.01 | 0.04 | 0.03 |
| 2 | 209436386 | rs10497902 | -1.522 | -0.753 | 0.324 | -0.641 | -0.01 | -0.039 | -0.039 | 0 | 0.27 | 0.67 | 0.16 | 0.06 | 0.18 | 0.23 |
| 2 | 209465672 | rs10932265 | -0.385 | -0.511 | 0.454 | -0.633 | 0.001 | 0.007 | 0.007 | 0.971 | 0.14 | 0.1 | 0.06 | 0 | 0.11 | 0.11 |
| 2 | 209467044 | rs7580611 | 0.526 | -0.656 | 0.861 | 0.218 | -0.038 | 0.181 | 0.181 | 0.102 | 0.8 | 0.53 | 0.84 | 0.5 | 0.87 | 0.71 |
| 2 | 209470028 | rs10190156 | -0.14 | -0.976 | 0.688 | 0.021 | -0.014 | -0.05 | -0.05 | 0 | 0.11 | 0.15 | 0.08 | 0 | 0.09 | 0.03 |
| 2 | 209470820 | rs10198932 | 0.978 | -0.962 | 0.707 | 0.044 | -0.057 | 0.09 | 0.09 | 0.091 | 0.73 | 0.54 | 0.8 | 0.49 | 0.81 | 0.68 |
| 2 | 209472551 | rs13018313 | -0.423 | -0.998 | 0.518 | -0.399 | -0.033 | -0.092 | -0.092 | 0 | 0.18 | 0.27 | 0.11 | 0 | 0.12 | 0.05 |
| 2 | 209472582 | rs12694115 | 0.965 | -0.704 | -0.527 | -0.474 | -0.005 | -0.002 | -0.002 | 0 | 0.43 | 0.26 | 0.53 | 0.46 | 0.55 | 0.45 |
| 2 | 209472769 | rs6733352 | -0.186 | 0.086 | -0.85 | -0.147 | -0.099 | -0.09 | -0.09 | 0 | 0.64 | 0.9 | 0.61 | 0.47 | 0.53 | 0.55 |
| 2 | 209474223 | rs1377908 | -0.674 | 0.371 | -0.26 | -0.043 | -0.023 | -0.02 | -0.02 | 0 | 0.52 | 0.73 | 0.44 | 0.54 | 0.48 | 0.53 |
| 2 | 209485037 | rs6732025 | -0.086 | 0.211 | 0.04 | -0.425 | -0.001 | -0.024 | -0.024 | 0 | 0.37 | 0.56 | 0.29 | 0.08 | 0.25 | 0.32 |
| 2 | 209496949 | rs13394305 | -0.301 | 0.245 | -0.348 | -0.421 | 0.057 | 0.098 | 0.098 | 0.053 | 0.4 | 0.69 | 0.31 | 0.08 | 0.27 | 0.32 |
| 2 | 209503800 | rs1921738 | 0.301 | 0.252 | -0.523 | -0.416 | 0.036 | 0.117 | 0.117 | 0 | 0.33 | 0.12 | 0.27 | 0.05 | 0.26 | 0.24 |
| 2 | 209509097 | rs1429889 | -0.671 | 0.475 | -0.379 | 0.112 | 0.002 | 0.056 | 0.056 | 0 | 0.52 | 0.67 | 0.43 | 0.65 | 0.45 | 0.53 |
| 2 | 209510934 | rs1864352 | 0.596 | 0.831 | -0.763 | 0.055 | -0.001 | -0.002 | -0.002 | 0 | 0.35 | 0.37 | 0.47 | 0.43 | 0.37 | 0.43 |
| 2 | 209511062 | rs2099159 | -1.288 | 0.935 | -1.597 | 0.152 | -0.008 | -0.025 | -0.025 | 0 | 0.89 | 0.96 | 0.92 | 0.53 | 0.9 | 0.79 |
| 2 | 209513915 | rs1517347 | 0.82 | 1.307 | -1.898 | -0.125 | -0.096 | -0.087 | -0.087 | 0 | 0.22 | 0.01 | 0.22 | 0.31 | 0.27 | 0.3 |
| 2 | 209516321 | rs13020347 | -0.138 | 0.825 | -1.078 | -0.602 | 0.026 | 0.164 | 0.164 | 0 | 0.46 | 0.22 | 0.36 | 0.07 | 0.44 | 0.3 |
| 2 | 209517915 | rs2048920 | 0.407 | 0.753 | -1.509 | -0.112 | -0.037 | 0.037 | 0.037 | 0 | 0.2 | 0.01 | 0.22 | 0.08 | 0.21 | 0.24 |
| 2 | 209520458 | rs1568167 | -0.06 | 0.799 | -1.031 | -0.008 | 0.001 | 0.001 | 0.001 | 0 | 0.41 | 0.37 | 0.41 | 0.4 | 0.39 | 0.51 |
| 2 | 209532165 | rs4293512 | -0.194 | 0.952 | -0.451 | -0.003 | -0.002 | -0.007 | -0.007 | 0 | 0.42 | 0.48 | 0.41 | 0.33 | 0.38 | 0.51 |
| 2 | 209534957 | rs9288396 | 0.552 | 0.721 | -0.825 | -0.312 | 0.01 | 0.033 | 0.033 | 0.41 | 0.22 | 0.35 | 0.34 | 0.34 | 0.25 | 0.39 |
| 2 | 209535670 | rs1517344 | -0.483 | 0.776 | -0.422 | -0.039 | -0.036 | 0.063 | 0.063 | 0 | 0.5 | 0.88 | 0.5 | 0.37 | 0.47 | 0.51 |
| 2 | 209555119 | rs2090465 | 0.069 | 0.773 | -0.165 | -0.031 | 0.002 | 0.002 | 0.002 | 0 | 0.41 | 0.38 | 0.42 | 0.37 | 0.39 | 0.49 |
| 2 | 209557324 | rs1921735 | -0.497 | 0.651 | 0.025 | 0.04 | -0.006 | 0.048 | 0.048 | 0 | 0.49 | 0.79 | 0.49 | 0.37 | 0.46 | 0.5 |
| 2 | 209572263 | rs1914406 | 0.018 | 0.69 | -0.219 | 0.019 | -0.006 | 0.014 | 0.014 | 0 | 0.43 | 0.38 | 0.44 | 0.31 | 0.45 | 0.49 |
| 2 | 209587434 | rs6435472 | -0.286 | 0.845 | -1.467 | 0.19 | 0.024 | -0.002 | -0.002 | 0 | 0.48 | 0.84 | 0.46 | 0.45 | 0.4 | 0.5 |
| 2 | 209600093 | rs7604543 | -0.642 | 1.375 | -1.399 | 0.67 | -0.063 | -0.094 | -0.094 | 0 | 0.81 | 0.97 | 0.75 | 0.62 | 0.77 | 0.81 |
| 2 | 209603868 | rs9808141 | 0.013 | 1.153 | -1.474 | 0.294 | -0.053 | -0.065 | -0.065 | 0 | 0.59 | 0.86 | 0.55 | 0.53 | 0.55 | 0.5 |
| 2 | 209641274 | rs982404 | -0.584 | 1.276 | -1.169 | 0.281 | -0.026 | -0.062 | -0.062 | 1.193 | 0.79 | 0.89 | 0.64 | 0.47 | 0.74 | 0.8 |
| 2 | 209655509 | rs7580077 | 0.59 | 0.847 | -0.006 | -0.19 | 0.007 | 0.005 | 0.005 | 0 | 0.41 | 0.34 | 0.44 | 0.4 | 0.38 | 0.45 |
| 2 | 209659222 | rs6732651 | 0.058 | 0.97 | -0.478 | -0.268 | 0 | -0.06 | -0.06 | 0 | 0.36 | 0.47 | 0.31 | 0.09 | 0.37 | 0.35 |
| 2 | 209676037 | rs1980188 | -0.143 | 0.855 | -0.576 | -0.15 | 0.02 | 0.038 | 0.038 | 0 | 0.73 | 0.62 | 0.7 | 0.49 | 0.66 | 0.76 |
| 2 | 209693319 | rs6435477 | -0.176 | 0.601 | -0.968 | -0.262 | -0.002 | -0.051 | -0.051 | 0 | 0.72 | 0.83 | 0.73 | 0.47 | 0.74 | 0.77 |
| 2 | 209720180 | rs4459678 | 1.044 | 0.5 | -1.334 | -0.479 | -0.027 | 0.087 | 0.087 | 0.575 | 0.22 | 0.03 | 0.19 | 0.06 | 0.24 | 0.31 |
| 2 | 209722605 | rs9288401 | 0.297 | 0.692 | -0.356 | 0.136 | 0.002 | 0.006 | 0.006 | 0 | 0.67 | 0.6 | 0.7 | 0.4 | 0.71 | 0.71 |
| 2 | 209723212 | rs4261669 | -0.041 | 0.602 | -0.961 | 0.245 | -0.002 | -0.04 | -0.04 | 0 | 0.72 | 0.79 | 0.73 | 0.47 | 0.74 | 0.77 |
| 2 | 209736883 | rs12467529 | 0.679 | 0.891 | -1.276 | 0.44 | -0.04 | -0.065 | -0.065 | 1.864 | 0.14 | 0.04 | 0.3 | 0.34 | 0.2 | 0.3 |
| 2 | 209748709 | rs6435493 | -0.52 | 0.528 | -0.882 | 0.691 | -0.029 | -0.081 | -0.081 | 0.011 | 0.79 | 0.89 | 0.77 | 0.47 | 0.77 | 0.79 |
| 2 | 209794478 | rs10192840 | -0.228 | 0.242 | 0.36 | 0.678 | 0.035 | 0.085 | 0.085 | 0 | 0.24 | 0.51 | 0.27 | 0.43 | 0.28 | 0.24 |
| 2 | 209813987 | rs10497916 | 0.194 | 0.213 | 0.967 | 0.734 | 0.054 | 0.068 | 0.068 | 0 | 0.21 | 0.35 | 0.21 | 0.36 | 0.28 | 0.22 |
| 2 | 209814683 | rs870908 | 1.042 | -0.361 | 0.477 | 0.228 | 0.026 | 0.03 | 0.03 | 0 | 0.14 | 0.08 | 0.1 | 0 | 0.09 | 0.03 |
| 2 | 209849299 | rs1402996 | -0.347 | 0.005 | 0.987 | 0.597 | 0.025 | 0.071 | 0.071 | 0 | 0.26 | 0.53 | 0.27 | 0.44 | 0.29 | 0.26 |
| 2 | 209873988 | rs2364820 | -0.378 | 0.105 | 0.256 | 0.486 | -0.011 | -0.01 | -0.01 | 0 | 0.42 | 0.24 | 0.43 | 0.57 | 0.52 | 0.37 |
| 2 | 209875890 | rs13025406 | -0.805 | 0.088 | 0.319 | 0.476 | 0.004 | 0.038 | 0.038 | 0 | 0.47 | 0.76 | 0.47 | 0.63 | 0.52 | 0.41 |
| 2 | 209878841 | rs13387460 | 0.599 | 0.029 | 0.238 | 0.415 | 0.004 | -0.014 | -0.014 | 3.579 | 0.24 | 0.44 | 0.27 | 0.2 | 0.24 | 0.16 |
| 2 | 209886096 | rs7601750 | -0.86 | 0.077 | 0.172 | 0.428 | 0.008 | 0.045 | 0.045 | 0 | 0.47 | 0.76 | 0.52 | 0.64 | 0.53 | 0.41 |
| 2 | 209899788 | rs1996808 | -1.545 | 0.052 | -0.234 | 0.254 | -0.05 | -0.101 | -0.101 | 1.694 | 0.85 | 0.99 | 0.77 | 0.68 | 0.82 | 0.72 |
| 2 | 209900229 | rs7608394 | 0.35 | 0.057 | -0.188 | 0.278 | 0.019 | 0.045 | 0.045 | 0 | 0.24 | 0.31 | 0.25 | 0.43 | 0.29 | 0.26 |
| 2 | 209910168 | rs4673438 | 1.138 | -0.289 | -0.556 | -0.816 | -0.011 | -0.001 | -0.001 | 0 | 0.21 | 0.12 | 0.24 | 0.21 | 0.24 | 0.15 |
| 2 | 209912119 | rs16842309 | 0.608 | -0.086 | -1.116 | -0.875 | 0.035 | 0.127 | 0.127 | 0.025 | 0.36 | 0.17 | 0.25 | 0.04 | 0.29 | 0.31 |
| 2 | 209922229 | rs16842335 | -0.324 | 0.028 | -0.222 | -0.01 | -0.026 | -0.027 | -0.027 | 2.585 | 0.06 | 0.16 | 0.02 | 0 | 0.02 | 0.04 |
| 2 | 209929978 | rs10211095 | -0.447 | 0.234 | -0.918 | 0.097 | 0.019 | 0.013 | 0.013 | 4.288 | 0.31 | 0.77 | 0.3 | 0.27 | 0.25 | 0.22 |

| | | | | | | | | | | | | | | | | |
|---|---|---|---|---|---|---|---|---|---|---|---|---|---|---|---|---|
| 2 | 209942078 | rs16842384 | -0.271 | 0.837 | -0.405 | -0.414 | 0.024 | -0.102 | -0.102 | 0 | 0.25 | 0.4 | 0.33 | 0.05 | 0.28 | 0.38 |
| 2 | 209949958 | rs3845635 | 0.276 | 0.323 | -0.651 | -0.827 | 0.007 | 0.013 | 0.013 | 0 | 0.33 | 0.3 | 0.42 | 0.77 | 0.32 | 0.39 |
| 2 | 209950811 | rs1561530 | 0.276 | 0.323 | -0.609 | -0.827 | 0.007 | 0.013 | 0.013 | 0 | 0.33 | 0.3 | 0.42 | 0.77 | 0.32 | 0.39 |
| 2 | 209963585 | rs966284 | 0.867 | 0.538 | -0.34 | -0.973 | 0.018 | -0.046 | -0.046 | 0.217 | 0.21 | 0.27 | 0.32 | 0.05 | 0.28 | 0.31 |
| 2 | 209963889 | rs1158709 | -0.816 | 0.384 | 0.023 | -1.041 | -0.01 | 0.135 | 0.135 | 0 | 0.58 | 0.84 | 0.77 | 0.82 | 0.62 | 0.78 |
| 2 | 209965827 | rs13422892 | 1.06 | 0.508 | -0.076 | -1.339 | 0.005 | -0.035 | -0.035 | 0 | 0.1 | 0.09 | 0.1 | 0 | 0.13 | 0.07 |
| 2 | 209966941 | rs2364826 | 0.57 | 0.501 | -0.049 | -1.35 | -0.02 | -0.038 | -0.038 | 0 | 0.34 | 0.48 | 0.44 | 0.77 | 0.32 | 0.4 |
| 2 | 209992353 | rs13428628 | 1.177 | 0.518 | -0.267 | -1.294 | -0.021 | 0.032 | 0.032 | 0 | 0.09 | 0.03 | 0.1 | 0 | 0.13 | 0.07 |
| 2 | 210000798 | rs2886188 | -0.795 | 0.478 | -0.231 | -1.294 | 0.006 | 0.011 | 0.011 | 0 | 0.64 | 0.68 | 0.84 | 0.82 | 0.74 | 0.78 |
| 2 | 210009863 | rs10202412 | -0.197 | 0.263 | -0.505 | -0.694 | 0.001 | -0.003 | -0.003 | 0 | 0.44 | 0.57 | 0.48 | 0.38 | 0.46 | 0.53 |
| 2 | 210011962 | rs991724 | 0.294 | 0.252 | -0.428 | -0.741 | 0 | -0.012 | -0.012 | 0 | 0.28 | 0.18 | 0.24 | 0.33 | 0.3 | 0.2 |
| 2 | 210012699 | rs12999691 | 1.148 | 0.244 | -0.41 | -0.716 | 0.006 | -0.014 | -0.014 | 0 | 0.19 | 0.15 | 0.15 | 0.33 | 0.17 | 0.15 |
| 2 | 210013527 | rs6740556 | -1.339 | 0.293 | -0.307 | -0.802 | 0.009 | 0.028 | 0.028 | 0 | 0.67 | 0.74 | 0.85 | 0.82 | 0.76 | 0.84 |
| 2 | 210013794 | rs10932285 | 0.917 | 0.416 | -0.175 | -0.676 | 0.04 | -0.083 | -0.083 | 0 | 0.21 | 0.34 | 0.32 | 0.05 | 0.29 | 0.33 |
| 2 | 210024301 | rs16842496 | 0.903 | 0.416 | -0.175 | -0.676 | 0.039 | -0.083 | -0.083 | 0 | 0.21 | 0.34 | 0.32 | 0.05 | 0.29 | 0.31 |
| 2 | 210024938 | rs7587994 | 0.132 | 0.154 | -0.662 | -0.634 | 0 | -0.028 | -0.028 | 0 | 0.35 | 0.19 | 0.3 | 0.4 | 0.37 | 0.26 |
| 2 | 210044418 | rs13418760 | 0.588 | 0.162 | -0.643 | -0.653 | 0.018 | -0.024 | -0.024 | 0.054 | 0.29 | 0.23 | 0.2 | 0.41 | 0.22 | 0.2 |
| 2 | 210044484 | rs10198749 | -1.594 | 0.05 | -0.59 | -0.748 | 0.027 | 0.039 | 0.039 | 0 | 0.74 | 0.85 | 0.9 | 0.84 | 0.85 | 0.89 |
| 2 | 210044774 | rs6756445 | 0.104 | 0.117 | -0.644 | -0.751 | 0.01 | -0.011 | -0.011 | 1.558 | 0.37 | 0.26 | 0.29 | 0.41 | 0.35 | 0.27 |
| 2 | 210051578 | rs12694157 | 0.737 | 0.109 | -0.643 | -0.747 | 0.019 | -0.024 | -0.024 | 0.104 | 0.29 | 0.23 | 0.2 | 0.41 | 0.22 | 0.21 |
| 2 | 210053121 | rs1530133 | -0.248 | 0.058 | -0.43 | -0.79 | -0.005 | -0.002 | -0.002 | 1.437 | 0.42 | 0.44 | 0.32 | 0.41 | 0.37 | 0.35 |
| 2 | 210053781 | rs12464060 | 1.708 | 0.149 | -0.519 | -0.774 | -0.016 | -0.041 | -0.041 | 1.238 | 0.12 | 0.06 | 0.25 | 0.38 | 0.17 | 0.22 |
| 2 | 210058068 | rs17738962 | 1.575 | -0.181 | -0.448 | -0.665 | -0.025 | 0.033 | 0.033 | 0 | 0.08 | 0.03 | 0.09 | 0 | 0.13 | 0.08 |
| 2 | 210065783 | rs17677155 | 0.083 | -0.168 | -0.394 | -0.668 | 0.005 | 0.003 | 0.003 | 0 | 0.38 | 0.55 | 0.62 | 0.43 | 0.52 | 0.61 |
| 2 | 210073419 | rs7570897 | -1.436 | -0.168 | -0.567 | -0.666 | 0.019 | 0.026 | 0.026 | 0 | 0.74 | 0.82 | 0.9 | 0.84 | 0.85 | 0.89 |
| 2 | 210077614 | rs2028355 | -0.472 | -0.133 | -0.505 | -0.606 | -0.001 | -0.015 | -0.015 | 0.299 | 0.62 | 0.63 | 0.77 | 0.84 | 0.7 | 0.75 |
| 2 | 210083966 | rs10932287 | 0.558 | -0.155 | -0.463 | -0.597 | -0.005 | -0.004 | -0.004 | 0 | 0.32 | 0.31 | 0.59 | 0.42 | 0.49 | 0.53 |
| 2 | 210102616 | rs1446603 | 0.383 | -0.138 | -0.59 | -0.434 | -0.005 | 0.004 | 0.004 | 0.227 | 0.29 | 0.32 | 0.18 | 0.42 | 0.21 | 0.24 |
| 2 | 210118415 | rs1446609 | -0.029 | 0.104 | -0.543 | -0.447 | 0.028 | 0.018 | 0.018 | 0.115 | 0.37 | 0.44 | 0.62 | 0.42 | 0.52 | 0.6 |
| 2 | 210123479 | rs12478877 | -0.076 | -0.008 | -0.565 | -0.559 | 0 | -0.001 | -0.001 | 0 | 0.41 | 0.39 | 0.69 | 0.42 | 0.62 | 0.59 |
| 2 | 210147107 | rs4673461 | 1.003 | -0.054 | -1.02 | -0.533 | -0.014 | -0.072 | -0.072 | 0 | 0.18 | 0.08 | 0.3 | 0.36 | 0.24 | 0.23 |
| 2 | 210153220 | rs12614235 | -1.076 | 0.044 | -0.842 | -0.556 | 0 | 0.006 | 0.006 | 1.356 | 0.77 | 0.76 | 0.82 | 0.59 | 0.8 | 0.77 |
| 2 | 210155558 | rs6746221 | -0.427 | -0.065 | -0.758 | -0.652 | 0.021 | 0.003 | 0.003 | 0.473 | 0.52 | 0.64 | 0.72 | 0.42 | 0.66 | 0.67 |
| 2 | 210178242 | rs1473329 | 0.475 | -0.03 | -0.801 | -0.636 | -0.083 | -0.077 | -0.077 | 0.211 | 0.37 | 0.14 | 0.57 | 0.43 | 0.51 | 0.53 |
| 2 | 210187488 | rs7576085 | -0.002 | -0.152 | -0.857 | -0.448 | 0.001 | 0 | 0 | 0 | 0.42 | 0.46 | 0.62 | 0.4 | 0.53 | 0.6 |
| 2 | 210192750 | rs4673469 | -1.062 | -0.07 | -0.549 | -0.194 | -0.009 | -0.054 | -0.054 | 0 | 0.78 | 0.89 | 0.83 | 0.57 | 0.8 | 0.77 |
| 2 | 210195717 | rs10183434 | 0.524 | -0.415 | -0.807 | -0.465 | 0.002 | 0 | 0 | 0.843 | 0.53 | 0.48 | 0.65 | 0.41 | 0.6 | 0.64 |
| 2 | 210204470 | rs1020407 | -1.266 | -0.41 | -0.817 | -0.464 | -0.009 | -0.054 | -0.054 | 0 | 0.78 | 0.89 | 0.83 | 0.57 | 0.8 | 0.77 |
| 2 | 210208746 | rs10439339 | 0.515 | -0.416 | -0.81 | -0.483 | 0.002 | 0 | 0 | 0.196 | 0.53 | 0.48 | 0.65 | 0.4 | 0.6 | 0.64 |
| 2 | 210215272 | rs12611509 | 1.179 | -0.388 | -0.673 | -0.448 | 0.006 | 0 | 0 | 0 | 0.15 | 0.14 | 0.07 | 0.16 | 0.08 | 0.06 |
| 2 | 210219690 | rs1470498 | 0.279 | -0.363 | -0.783 | -0.44 | -0.019 | -0.046 | -0.046 | 0 | 0.25 | 0.4 | 0.17 | 0.16 | 0.2 | 0.14 |
| 2 | 210228698 | rs11681397 | 0.873 | -0.389 | -0.967 | -0.437 | -0.042 | -0.034 | -0.034 | 0 | 0.46 | 0.1 | 0.56 | 0.37 | 0.57 | 0.57 |
| 2 | 210228910 | rs995565 | -1.043 | -0.508 | -1.119 | -0.403 | 0.015 | -0.001 | -0.001 | 1.613 | 0.77 | 0.89 | 0.81 | 0.72 | 0.85 | 0.84 |
| 2 | 210229386 | rs16842843 | -0.181 | -0.146 | -0.167 | -0.184 | 0.031 | -0.017 | -0.017 | 0.084 | 0.07 | 0.38 | 0.06 | 0 | 0.06 | 0.09 |
| 2 | 210232466 | rs975719 | -0.415 | -0.195 | -0.283 | -0.311 | 0.009 | 0.001 | 0.001 | 0.166 | 0.33 | 0.35 | 0.35 | 0.35 | 0.41 | 0.42 |
| 2 | 210234855 | rs4417658 | -1.543 | -0.213 | -0.77 | -0.313 | -0.002 | 0.09 | 0.09 | 1.606 | 0.5 | 0.87 | 0.49 | 0.35 | 0.56 | 0.57 |
| 2 | 210241321 | rs12623783 | -0.334 | -0.513 | -0.263 | -0.504 | -0.05 | -0.015 | -0.015 | 0 | 0.22 | 0.6 | 0.12 | 0.17 | 0.12 | 0.16 |
| 2 | 210245922 | rs7606091 | 1.166 | -0.379 | -0.486 | -0.449 | 0.003 | -0.044 | -0.044 | 2.938 | 0.09 | 0.16 | 0.09 | 0 | 0.1 | 0.06 |
| 2 | 210269845 | rs6435526 | -1.326 | -0.38 | -1.017 | -0.458 | -0.016 | 0.215 | 0.215 | 1.037 | 0.48 | 0.85 | 0.44 | 0.18 | 0.48 | 0.51 |
| 2 | 210287968 | rs12614974 | 0.455 | -0.351 | -0.397 | 0.383 | 0.038 | -0.043 | -0.043 | 0 | 0.13 | 0.03 | 0.04 | 0.17 | 0.06 | 0.06 |
| 2 | 210330576 | rs13027724 | -0.134 | -0.002 | -0.082 | 0.387 | -0.003 | -0.024 | -0.024 | 0.024 | 0.05 | 0.1 | 0.03 | 0 | 0.05 | 0.11 |
| 2 | 210330754 | rs10737940 | 0.354 | 0.003 | 0.123 | 0.545 | 0.003 | 0.012 | 0.012 | 0.084 | 0.68 | 0.57 | 0.61 | 0.62 | 0.66 | 0.6 |
| 2 | 210335716 | rs10208979 | 0.747 | -0.026 | 0.241 | 0.523 | -0.001 | 0 | 0 | 0 | 0.63 | 0.35 | 0.58 | 0.62 | 0.61 | 0.5 |
| 2 | 210346084 | rs6728378 | 0.503 | -0.019 | 0.187 | 0.527 | 0 | 0.006 | 0.006 | 0.283 | 0.67 | 0.45 | 0.6 | 0.62 | 0.66 | 0.61 |
| 2 | 210366089 | rs17237719 | 0.07 | -0.16 | 0.105 | 0.687 | -0.003 | -0.023 | -0.023 | 0 | 0.05 | 0.09 | 0.03 | 0 | 0.05 | 0.11 |
| 2 | 210374052 | rs288068 | -0.226 | -0.292 | 0.141 | 0.609 | -0.063 | 0.123 | 0.123 | 0 | 0.83 | 0.63 | 0.8 | 0.63 | 0.92 | 0.84 |
| 2 | 210382709 | rs16843022 | -0.39 | -0.255 | 0.059 | 0.286 | -0.068 | 0.007 | 0.007 | 0.85 | 0.21 | 0.45 | 0.15 | 0.19 | 0.13 | 0.23 |
| 2 | 210391837 | rs2365661 | 0.214 | 0.241 | -0.415 | 0.547 | -0.063 | 0.018 | 0.018 | 0.652 | 0.16 | 0.05 | 0.18 | 0.07 | 0.27 | 0.27 |
| 2 | 210392711 | rs288057 | 0.421 | -0.74 | 0.18 | 0.79 | -0.085 | 0.023 | 0.023 | 1.363 | 0.77 | 0.58 | 0.86 | 0.77 | 0.9 | 0.79 |
| 2 | 210404224 | rs288094 | -1.165 | -1.204 | -0.211 | 0.387 | 0.134 | 0.044 | 0.044 | 4.379 | 0.32 | 0.89 | 0.17 | 0.2 | 0.13 | 0.38 |
| 2 | 210408416 | rs288071 | -0.385 | -1.172 | 0.632 | 0.405 | -0.045 | 0.011 | 0.011 | 2.097 | 0.19 | 0.65 | 0.14 | 0.2 | 0.11 | 0.15 |
| 2 | 210410333 | rs7593381 | 0.056 | -1.172 | 0.894 | 0.413 | -0.075 | 0.059 | 0.059 | 0.627 | 0.17 | 0.47 | 0.12 | 0.2 | 0.08 | 0.14 |
| 2 | 210441906 | rs13419929 | -0.314 | -1.189 | 1.107 | 0.429 | -0.06 | 0.025 | 0.025 | 1.166 | 0.19 | 0.52 | 0.14 | 0.2 | 0.11 | 0.15 |
| 2 | 210454751 | rs17733579 | 0.732 | -0.811 | 0.302 | 0.497 | -0.078 | -0.103 | -0.103 | 0.584 | 0.2 | 0.04 | 0.28 | 0.43 | 0.3 | 0.31 |
| 2 | 210484445 | rs2663644 | 0.044 | -1.488 | 0.617 | 0.251 | -0.066 | -0.078 | -0.078 | 0 | 0.13 | 0.28 | 0.02 | 0.01 | 0.02 | 0.26 |
| 2 | 210499568 | rs2710475 | -0.623 | -1.56 | 0.461 | 0.304 | -0.007 | 0.005 | 0.005 | 3.236 | 0.3 | 0.69 | 0.15 | 0.21 | 0.09 | 0.4 |
| 2 | 210508524 | rs2663652 | -2.444 | -1.144 | -0.064 | 0.194 | -0.083 | -0.085 | -0.085 | 0.002 | 0.72 | 0.98 | 0.63 | 0.67 | 0.67 | 0.82 |
| 2 | 210509617 | rs16843326 | -0.72 | -1.904 | 1.133 | 0.27 | -0.106 | -0.026 | -0.026 | 0.242 | 0.29 | 0.51 | 0.13 | 0.21 | 0.1 | 0.42 |
| 2 | 210517915 | rs2271252 | -0.134 | -1.259 | 1.595 | 0.512 | -0.02 | 0.035 | 0.035 | 2.366 | 0.17 | 0.56 | 0.1 | 0.2 | 0.11 | 0.15 |

| | | | | | | | | | | | | | | | | |
|---|---|---|---|---|---|---|---|---|---|---|---|---|---|---|---|---|
| 2 | 210519462 | rs2710482 | -0.3 | -1.289 | 1.619 | 0.498 | -0.023 | 0.016 | 0.016 | 3.228 | 0.18 | 0.64 | 0.11 | 0.2 | 0.11 | 0.14 |
| 2 | 210545085 | rs3820748 | -0.026 | -1.222 | 1.713 | 0.531 | -0.01 | 0.065 | 0.065 | 2.341 | 0.16 | 0.46 | 0.09 | 0.2 | 0.11 | 0.15 |
| 2 | 210549474 | rs6709925 | -0.119 | -1.222 | 1.683 | 0.531 | -0.018 | 0.043 | 0.043 | 2.586 | 0.17 | 0.53 | 0.1 | 0.2 | 0.11 | 0.15 |
| 2 | 210552162 | rs3768815 | -0.137 | -1.223 | 1.673 | 0.531 | -0.019 | 0.038 | 0.038 | 2.878 | 0.17 | 0.56 | 0.1 | 0.2 | 0.11 | 0.15 |
| 2 | 210555776 | rs3768812 | 1.218 | -1.181 | 1.546 | 0.661 | 0.029 | -0.005 | -0.005 | 0 | 0.58 | 0.22 | 0.68 | 0.61 | 0.6 | 0.31 |
| 2 | 210557380 | rs6720659 | 0.403 | -1.078 | 2.077 | 0.875 | -0.016 | 0.06 | 0.06 | 0.47 | 0.13 | 0.27 | 0.07 | 0.2 | 0.08 | 0.14 |
| 2 | 210557542 | rs741007 | -0.382 | -1.079 | 2.1 | 0.874 | -0.02 | 0.03 | 0.03 | 2.901 | 0.17 | 0.59 | 0.1 | 0.2 | 0.11 | 0.15 |
| 2 | 210558162 | rs741006 | 0.36 | -1.103 | 1.962 | 0.903 | -0.016 | 0.06 | 0.06 | 0.47 | 0.13 | 0.27 | 0.07 | 0.2 | 0.08 | 0.14 |
| 2 | 210559960 | rs2239672 | 0.638 | -1.105 | 1.986 | 0.903 | -0.011 | 0.026 | 0.026 | 0.265 | 0.12 | 0.16 | 0.06 | 0.2 | 0.08 | 0.14 |
| 2 | 210580399 | rs6742070 | -0.131 | -1.119 | 1.914 | 0.891 | -0.02 | -0.028 | -0.028 | 0 | 0.06 | 0.1 | 0.01 | 0.01 | 0.01 | 0.25 |
| 2 | 210594768 | rs7594601 | -0.378 | -0.797 | 2.014 | 1.073 | -0.013 | 0.087 | 0.087 | 0.567 | 0.87 | 0.63 | 0.92 | 0.8 | 0.92 | 0.86 |
| 2 | 210595324 | rs16843614 | 0.276 | -0.796 | 1.92 | 1.073 | -0.017 | 0.042 | 0.042 | 0.399 | 0.13 | 0.23 | 0.07 | 0.2 | 0.08 | 0.14 |
| 2 | 210600134 | rs11692345 | 0.276 | -0.796 | 1.92 | 1.073 | -0.017 | 0.042 | 0.042 | 0.399 | 0.13 | 0.23 | 0.07 | 0.2 | 0.07 | 0.14 |
| 2 | 210601209 | rs2215625 | -0.051 | -0.793 | 1.448 | 1.081 | -0.025 | 0.085 | 0.085 | 1.346 | 0.85 | 0.52 | 0.92 | 0.79 | 0.93 | 0.86 |
| 2 | 210618203 | rs12622598 | 0.376 | -0.665 | 1.22 | 1.085 | -0.003 | 0.13 | 0.13 | 0.668 | 0.13 | 0.31 | 0.07 | 0.22 | 0.07 | 0.16 |
| 2 | 210642687 | rs2191915 | 0.92 | -0.655 | 1.072 | 0.949 | -0.024 | -0.031 | -0.031 | 0 | 0.63 | 0.23 | 0.61 | 0.6 | 0.6 | 0.49 |
| 2 | 210644422 | rs11692608 | 0.451 | -0.586 | 1.036 | 0.931 | -0.004 | 0.128 | 0.128 | 0.71 | 0.13 | 0.32 | 0.07 | 0.22 | 0.08 | 0.16 |
| 2 | 210669245 | rs2370833 | -1.033 | -0.57 | 0.599 | 0.678 | -0.001 | -0.001 | -0.001 | 0 | 0.36 | 0.62 | 0.38 | 0.37 | 0.4 | 0.48 |
| 2 | 210672638 | rs10932310 | 0.723 | -0.56 | 0.2 | 0.683 | -0.007 | -0.005 | -0.005 | 0 | 0.64 | 0.32 | 0.62 | 0.63 | 0.6 | 0.51 |
| 2 | 210673714 | rs2286851 | 0.811 | -0.523 | 0.218 | 0.701 | -0.014 | -0.011 | -0.011 | 0.821 | 0.62 | 0.23 | 0.62 | 0.63 | 0.61 | 0.52 |
| 2 | 210686883 | rs7569393 | 0.048 | -0.966 | 0.783 | 1.006 | -0.031 | 0.102 | 0.102 | 4.326 | 0.85 | 0.6 | 0.93 | 0.79 | 0.93 | 0.81 |
| 2 | 210704200 | rs9288414 | -0.897 | -0.98 | 0.141 | -0.037 | -0.011 | -0.052 | -0.052 | 1.83 | 0.3 | 0.28 | 0.11 | 0.01 | 0.12 | 0.28 |
| 2 | 210714162 | rs2191906 | -0.937 | -0.815 | 0.223 | -0.063 | -0.03 | -0.094 | -0.094 | 0 | 0.32 | 0.35 | 0.12 | 0.01 | 0.15 | 0.3 |
| 2 | 210733445 | rs10490026 | -0.773 | -0.203 | 0.732 | 0.336 | -0.038 | -0.107 | -0.107 | 0 | 0.17 | 0.39 | 0.1 | 0.01 | 0.11 | 0.27 |
| 2 | 210770527 | rs11688226 | -0.648 | -0.2 | 0.425 | 0.338 | 0.027 | 0.041 | 0.041 | 0 | 0.16 | 0.12 | 0.08 | 0 | 0.11 | 0.29 |
| 2 | 210799863 | rs10200921 | -0.579 | -0.16 | 0.809 | 0.812 | -0.003 | 0.006 | 0.006 | 0 | 0.13 | 0.16 | 0.19 | 0.26 | 0.14 | 0.12 |
| 2 | 210801263 | rs2215669 | -1.413 | 0.333 | 1.102 | 1.127 | 0.037 | 0.052 | 0.052 | 0.587 | 0.19 | 0.58 | 0.26 | 0.22 | 0.21 | 0.34 |
| 2 | 210808905 | rs7584375 | -1.812 | 0.339 | 0.969 | 0.883 | 0.002 | 0.016 | 0.016 | 0 | 0.34 | 0.63 | 0.45 | 0.49 | 0.36 | 0.47 |
| 2 | 210810599 | rs10178327 | -1.381 | 0.234 | 1.054 | 0.572 | 0.015 | -0.012 | -0.012 | 0.341 | 0.07 | 0.33 | 0.05 | 0 | 0.02 | 0.16 |
| 2 | 210849283 | rs10804178 | -0.538 | 1.029 | 0.682 | 1.026 | 0.05 | 0.062 | 0.062 | 0 | 0.32 | 0.51 | 0.42 | 0.47 | 0.49 | 0.43 |
| 2 | 210851634 | rs1861072 | 0.929 | 0.59 | 0.129 | 0.732 | -0.036 | -0.073 | -0.073 | 0.018 | 0.56 | 0.07 | 0.5 | 0.48 | 0.45 | 0.5 |
| 2 | 210859726 | rs2539861 | 0.082 | 0.039 | -0.24 | 0.819 | -0.001 | -0.05 | -0.05 | 0 | 0.08 | 0 | 0.08 | 0.18 | 0.09 | 0.07 |
| 2 | 210879917 | rs13006132 | 0.47 | 0.249 | 0.005 | 0.892 | 0.012 | 0.029 | 0.029 | 0.011 | 0.09 | 0.01 | 0.05 | 0 | 0.08 | 0.07 |
| 2 | 210880893 | rs7570090 | -0.12 | 0.675 | 0.829 | 0.867 | 0.004 | 0.003 | 0.003 | 0 | 0.29 | 0.34 | 0.35 | 0.51 | 0.44 | 0.39 |
| 2 | 210885022 | rs7585265 | 0.29 | 1.541 | 0.935 | 1.377 | -0.052 | -0.019 | -0.019 | 0 | 0.19 | 0.09 | 0.26 | 0.2 | 0.31 | 0.28 |
| 2 | 210889961 | rs12989455 | 1.087 | 1.468 | 0.886 | 1.331 | 0.005 | 0.039 | 0.039 | 0.164 | 0.07 | 0 | 0.05 | 0 | 0.07 | 0.04 |
| 2 | 210898710 | rs2041688 | 1.486 | 1.451 | 0.969 | 1.324 | -0.075 | -0.071 | -0.071 | 0 | 0.57 | 0.09 | 0.55 | 0.48 | 0.48 | 0.5 |
| 2 | 210907710 | rs2539871 | 1.668 | 1.809 | 0.952 | 1.038 | -0.095 | -0.12 | -0.12 | 6.364 | 0.58 | 0.03 | 0.61 | 0.48 | 0.6 | 0.45 |
| 2 | 210941838 | rs13021055 | 0.477 | 1.942 | 1.21 | 1.056 | -0.001 | -0.016 | -0.016 | 0 | 0.1 | 0.11 | 0.07 | 0 | 0.08 | 0.07 |
| 2 | 210980437 | rs6736257 | 0.356 | 2.056 | 1.748 | 1.186 | 0.014 | 0.134 | 0.134 | 0 | 0.1 | 0.27 | 0.06 | 0.3 | 0.11 | 0.1 |
| 2 | 211000909 | rs2302539 | -1.488 | 1.997 | 1.739 | 1.158 | 0.006 | 0.01 | 0.01 | 0.002 | 0.36 | 0.6 | 0.35 | 0.52 | 0.39 | 0.42 |
| 2 | 211007521 | rs12613036 | 0.1 | 1.565 | 1.563 | 1.269 | -0.05 | -0.022 | -0.022 | 1.804 | 0.2 | 0.09 | 0.26 | 0.23 | 0.32 | 0.28 |
| 2 | 211042400 | rs2098279 | 0.5 | 0.948 | 0.8 | 0.688 | -0.098 | -0.083 | -0.083 | 0 | 0.16 | 0.01 | 0.34 | 0.29 | 0.24 | 0.27 |
| 2 | 211045482 | rs1396828 | 1.43 | 1.294 | 1.144 | 0.86 | -0.085 | -0.076 | -0.076 | 0.236 | 0.57 | 0.08 | 0.55 | 0.47 | 0.49 | 0.53 |
| 2 | 211056657 | rs7601356 | 0.328 | 1.335 | 1.194 | 0.851 | -0.054 | -0.022 | -0.022 | 0 | 0.2 | 0.09 | 0.27 | 0.23 | 0.32 | 0.29 |
| 2 | 211060050 | rs2286963 | 0.356 | 1.376 | 1.201 | 0.851 | -0.052 | -0.004 | -0.004 | 0 | 0.2 | 0.09 | 0.22 | 0.19 | 0.31 | 0.29 |
| 2 | 211069011 | rs1509568 | 0.279 | 1.36 | 1.348 | 0.784 | -0.032 | -0.086 | -0.086 | 0 | 0.86 | 0.51 | 0.9 | 0.99 | 0.92 | 0.89 |
| 2 | 211073548 | rs1509571 | -0.537 | 1.356 | 1.302 | 0.78 | -0.016 | -0.061 | -0.061 | 0 | 0.89 | 0.75 | 0.92 | 1 | 0.92 | 0.93 |
| 2 | 211074909 | rs3764913 | 0.367 | 1.423 | 1.098 | 0.863 | -0.053 | -0.005 | -0.005 | 2.353 | 0.2 | 0.09 | 0.22 | 0.19 | 0.32 | 0.29 |
| 2 | 211112436 | rs12476559 | 1.019 | 0.804 | 0.525 | 0.757 | -0.079 | -0.117 | -0.117 | 0.055 | 0.56 | 0.08 | 0.59 | 0.53 | 0.48 | 0.53 |
| 2 | 211132415 | rs1509569 | -0.085 | 0.832 | 0.803 | 0.819 | 0.017 | 0.015 | 0.015 | 0 | 0.29 | 0.35 | 0.28 | 0.49 | 0.43 | 0.35 |
| 2 | 211138809 | rs10932321 | 0.761 | 0.678 | 0.469 | 0.785 | 0.007 | -0.002 | -0.002 | 0.001 | 0.56 | 0.36 | 0.62 | 0.49 | 0.47 | 0.57 |
| 2 | 211151774 | rs2048277 | 0.006 | 0.568 | 0.896 | 0.676 | -0.023 | -0.06 | -0.06 | 0 | 0.12 | 0.24 | 0.07 | 0.01 | 0.09 | 0.13 |
| 2 | 211151969 | rs6734950 | 0.009 | 0.455 | 1 | 0.72 | -0.001 | -0.002 | -0.002 | 0.158 | 0.64 | 0.62 | 0.59 | 0.47 | 0.5 | 0.52 |
| 2 | 211154128 | rs11889646 | 0.221 | 0.439 | 0.942 | 0.726 | 0.001 | 0.005 | 0.005 | 0.167 | 0.63 | 0.41 | 0.57 | 0.47 | 0.5 | 0.52 |
| 2 | 211164884 | rs4673510 | 0.197 | 0.369 | 0.549 | 0.754 | -0.058 | -0.059 | -0.059 | 0.17 | 0.1 | 0.02 | 0.33 | 0.31 | 0.21 | 0.25 |
| 2 | 211173237 | rs7557043 | 0.105 | 0.723 | 0.999 | 0.689 | -0.018 | -0.062 | -0.062 | 0.282 | 0.11 | 0.22 | 0.08 | 0 | 0.09 | 0.08 |
| 2 | 211175043 | rs1074158 | 1.224 | 0.719 | 0.368 | 0.727 | 0.099 | 0.011 | 0.011 | 0.441 | 0.51 | 0.15 | 0.49 | 0.46 | 0.41 | 0.36 |
| 2 | 211175921 | rs11686587 | 0.073 | 0.381 | 0.36 | 0.627 | -0.016 | -0.061 | -0.061 | 2.26 | 0.11 | 0.21 | 0.08 | 0 | 0.09 | 0.08 |
| 2 | 211186894 | rs10932323 | 0.905 | 0.853 | 0.301 | 0.937 | 0.04 | 0.013 | 0.013 | 1.405 | 0.51 | 0.39 | 0.44 | 0.45 | 0.34 | 0.32 |
| 2 | 211187769 | rs10932324 | 1.305 | 0.777 | -0.171 | 0.892 | 0.123 | -0.023 | -0.023 | 0 | 0.45 | 0.06 | 0.41 | 0.44 | 0.32 | 0.24 |
| 2 | 211189875 | rs1396829 | 0.895 | 0.826 | -0.101 | 0.926 | 0.086 | 0.027 | 0.027 | 1.875 | 0.49 | 0.31 | 0.43 | 0.44 | 0.34 | 0.32 |
| 2 | 211208213 | rs12477545 | 1.172 | 1.088 | -0.373 | 0.936 | 0.116 | 0.04 | 0.04 | 2.417 | 0.46 | 0.14 | 0.43 | 0.41 | 0.33 | 0.25 |
| 2 | 211210554 | rs12471867 | 1.156 | 1.056 | -0.345 | 0.937 | 0.104 | 0.043 | 0.043 | 2.335 | 0.47 | 0.16 | 0.43 | 0.42 | 0.33 | 0.25 |
| 2 | 211232032 | rs10932328 | 0.258 | 1.544 | -0.09 | 1.242 | -0.004 | 0.019 | 0.019 | 0 | 0.74 | 0.51 | 0.77 | 0.66 | 0.75 | 0.72 |
| 2 | 211235485 | rs1551610 | -0.17 | 1.63 | -0.308 | 1.26 | -0.012 | -0.005 | -0.005 | 0.462 | 0.27 | 0.22 | 0.33 | 0.24 | 0.42 | 0.47 |
| 2 | 211237918 | rs1379836 | 1.011 | 1.295 | -0.765 | 1.123 | 0.1 | 0.06 | 0.06 | 2.5 | 0.49 | 0.19 | 0.45 | 0.42 | 0.36 | 0.27 |
| 2 | 211241608 | rs4673519 | 1.011 | 1.295 | -0.777 | 1.08 | 0.101 | 0.081 | 0.081 | 2.5 | 0.49 | 0.19 | 0.45 | 0.4 | 0.37 | 0.27 |
| 2 | 211258344 | rs1458281 | -0.505 | 1.861 | -0.421 | 1.566 | -0.075 | -0.018 | -0.018 | 0 | 0.28 | 0.16 | 0.32 | 0.24 | 0.42 | 0.46 |

| | | | | | | | | | | | | | | | | |
|---|---|---|---|---|---|---|---|---|---|---|---|---|---|---|---|---|
| 2 | 211287749 | rs3856338 | -0.529 | 1.93 | -0.519 | 1.362 | -0.058 | -0.013 | -0.013 | 0 | 0.28 | 0.2 | 0.32 | 0.25 | 0.42 | 0.46 |
| 2 | 211295019 | rs17770211 | -0.196 | 2.066 | -0.543 | 1.342 | -0.081 | -0.033 | -0.033 | 0.368 | 0.25 | 0.1 | 0.28 | 0.24 | 0.38 | 0.45 |
| 2 | 211296239 | rs7893 | -1.024 | 0.923 | 0.007 | 1.199 | -0.037 | 0.164 | 0.164 | 1.258 | 0.19 | 0.48 | 0.38 | 0.63 | 0.13 | 0.26 |
| 2 | 211297355 | rs1044710 | -0.997 | 0.927 | 0.005 | 1.197 | -0.039 | 0.167 | 0.167 | 1.282 | 0.19 | 0.48 | 0.38 | 0.63 | 0.12 | 0.25 |
| 2 | 211302685 | rs13420747 | -0.427 | 0.923 | -0.266 | 0.841 | -0.008 | -0.021 | -0.021 | 0.545 | 0.1 | 0.07 | 0.15 | 0.24 | 0.12 | 0.14 |
| 2 | 211317210 | rs3755187 | -1.153 | 0.788 | 0.028 | 0.892 | -0.023 | 0.044 | 0.044 | 0.846 | 0.13 | 0.25 | 0.14 | 0.22 | 0.09 | 0.06 |
| 2 | 211328663 | rs3770693 | -0.843 | 0.781 | 0.025 | 0.884 | -0.015 | 0.059 | 0.059 | 1.513 | 0.11 | 0.24 | 0.14 | 0.22 | 0.09 | 0.06 |
| 2 | 211334708 | rs3732055 | -0.436 | 0.963 | -0.035 | 0.795 | -0.059 | -0.025 | -0.025 | 0 | 0.31 | 0.25 | 0.54 | 0.63 | 0.45 | 0.59 |
| 2 | 211335599 | rs3856342 | 1.37 | 0.479 | -0.456 | 0.352 | 0.072 | 0.039 | 0.039 | 0.631 | 0.46 | 0.23 | 0.32 | 0.37 | 0.37 | 0.27 |
| 2 | 211338832 | rs3770685 | -0.635 | 0.504 | -0.226 | 0.283 | 0 | 0 | 0 | 0 | 0.36 | 0.36 | 0.61 | 0.63 | 0.56 | 0.64 |
| 2 | 211341349 | rs10932334 | -0.435 | 0.479 | -0.362 | 0.32 | -0.062 | -0.023 | -0.023 | 0 | 0.33 | 0.21 | 0.55 | 0.63 | 0.49 | 0.64 |
| 2 | 211357218 | rs759688 | 0.544 | 0.319 | -0.087 | 0.393 | 0.047 | 0.246 | 0.246 | 0 | 0.93 | 0.66 | 0.86 | 0.74 | 0.92 | 0.92 |
| 2 | 211376416 | rs918233 | -1.229 | 0.261 | -0.096 | 0.236 | 0.002 | 0.002 | 0.002 | 1.703 | 0.5 | 0.49 | 0.67 | 0.6 | 0.65 | 0.71 |
| 2 | 211387772 | rs10188695 | -0.012 | 1 | -0.227 | 0.355 | -0.022 | 0.008 | 0.008 | 0.007 | 0.19 | 0.3 | 0.31 | 0.21 | 0.15 | 0.26 |
| 2 | 211396680 | rs981024 | 1.223 | 1.005 | -0.129 | 0.351 | 0.003 | 0.005 | 0.005 | 0 | 0.55 | 0.38 | 0.48 | 0.35 | 0.42 | 0.46 |
| 2 | 211404035 | rs2250976 | 0.638 | 1.008 | -0.511 | 0.338 | 0.003 | 0.007 | 0.007 | 0.253 | 0.6 | 0.61 | 0.51 | 0.35 | 0.42 | 0.49 |
| 2 | 211420207 | rs2887913 | 1.215 | 0.965 | -0.42 | 0.464 | -0.009 | -0.012 | -0.012 | 0 | 0.55 | 0.39 | 0.49 | 0.35 | 0.41 | 0.46 |
| 2 | 211426974 | rs2287603 | 0.85 | 1.592 | -1.344 | 0.572 | -0.073 | -0.079 | -0.079 | 0 | 0.11 | 0.01 | 0.19 | 0.38 | 0.22 | 0.28 |
| 2 | 211428354 | rs10804185 | 2.087 | -0.213 | -1.179 | 0.147 | 0.064 | 0.129 | 0.129 | 0.533 | 0.35 | 0.07 | 0.17 | 0.14 | 0.26 | 0.2 |
| 2 | 211436065 | rs6714124 | 0.813 | 0.339 | -0.285 | 0.755 | 0 | -0.002 | -0.002 | 0.425 | 0.65 | 0.59 | 0.37 | 0.29 | 0.49 | 0.56 |
| 2 | 211446121 | rs10165443 | 0.498 | 0.285 | -0.342 | 0.73 | -0.033 | -0.035 | -0.035 | 0 | 0.68 | 0.72 | 0.49 | 0.56 | 0.55 | 0.62 |
| 2 | 211448659 | rs4399666 | 0.26 | 0.272 | -0.354 | 0.724 | -0.009 | -0.009 | -0.009 | 0 | 0.72 | 0.75 | 0.5 | 0.56 | 0.55 | 0.63 |
| 2 | 211457261 | rs10184633 | 0.573 | 0.274 | -0.331 | 0.711 | -0.008 | -0.009 | -0.009 | 0 | 0.68 | 0.67 | 0.49 | 0.56 | 0.55 | 0.62 |
| 2 | 211465780 | rs2371000 | 0.025 | 0.288 | -0.327 | 0.711 | -0.041 | -0.045 | -0.045 | 0.313 | 0.73 | 0.82 | 0.51 | 0.56 | 0.57 | 0.63 |
| 2 | 211467548 | rs2371001 | 0.82 | 0.221 | -0.33 | 0.671 | 0.006 | -0.012 | -0.012 | 0 | 0.63 | 0.56 | 0.37 | 0.29 | 0.49 | 0.55 |
| 2 | 211469123 | rs6734696 | 0.044 | 0.394 | -0.838 | 0.798 | -0.042 | -0.046 | -0.046 | 0 | 0.74 | 0.83 | 0.51 | 0.57 | 0.56 | 0.63 |
| 2 | 211470493 | rs13010668 | -0.605 | 0.41 | -0.797 | 0.641 | -0.001 | 0.048 | 0.048 | 0 | 0.17 | 0.31 | 0.19 | 0.27 | 0.17 | 0.19 |
| 2 | 211471071 | rs2371002 | 0.087 | 0.463 | -1.209 | 0.592 | -0.108 | -0.123 | -0.123 | 1.188 | 0.71 | 0.9 | 0.49 | 0.52 | 0.57 | 0.64 |
| 2 | 211486368 | rs3821135 | -0.437 | 0.367 | -0.313 | 1.129 | 0.02 | 0.129 | 0.129 | 0.019 | 0.08 | 0.16 | 0.16 | 0.29 | 0.12 | 0.16 |
| 2 | 211503556 | rs2302909 | -1.073 | 0.102 | -0.452 | 0.384 | -0.035 | -0.024 | -0.024 | 0 | 0.17 | 0.66 | 0.11 | 0.14 | 0.08 | 0.16 |
| 2 | 211508287 | rs4356603 | -1.048 | 0.057 | -0.751 | 0.425 | -0.017 | -0.008 | -0.008 | 2.14 | 0.18 | 0.72 | 0.11 | 0.14 | 0.07 | 0.16 |
| 2 | 211514492 | rs2371011 | 0.19 | 1.074 | -1.389 | 0.833 | -0.105 | -0.116 | -0.116 | 0 | 0.18 | 0.03 | 0.43 | 0.43 | 0.31 | 0.35 |
| 2 | 211516625 | rs11686101 | -0.269 | 0.218 | -1.466 | 0.525 | 0.01 | 0.011 | 0.011 | 0 | 0.17 | 0.16 | 0.15 | 0.09 | 0.13 | 0.16 |
| 2 | 211517423 | rs7599931 | -1.12 | 0.306 | -0.869 | 0.314 | -0.004 | -0.026 | -0.026 | 0 | 0.27 | 0.41 | 0.29 | 0.2 | 0.28 | 0.29 |
| 2 | 211523163 | rs2287598 | 0.133 | -0.243 | -0.787 | 0.149 | -0.03 | 0.101 | 0.101 | 0.035 | 0.77 | 0.56 | 0.8 | 0.62 | 0.83 | 0.8 |
| 2 | 211523457 | rs6435580 | 1.853 | -0.206 | -1.572 | 0.027 | 0.011 | 0.169 | 0.169 | 0 | 0.37 | 0.05 | 0.19 | 0.1 | 0.31 | 0.27 |
| 2 | 211525764 | rs2270476 | 0.654 | -0.048 | -1.031 | 0.446 | -0.004 | 0.018 | 0.018 | 0 | 0.13 | 0.12 | 0.1 | 0.27 | 0.06 | 0.09 |
| 2 | 211529850 | rs12997383 | -0.442 | 0.633 | -1.106 | 0.379 | -0.003 | -0.004 | -0.004 | 0 | 0.17 | 0.21 | 0.14 | 0.09 | 0.14 | 0.16 |
| 2 | 211532703 | rs2287596 | -0.076 | 1.237 | -1.236 | 0.285 | -0.127 | -0.122 | -0.122 | 0.003 | 0.2 | 0.03 | 0.36 | 0.43 | 0.34 | 0.36 |
| 2 | 211534457 | rs16844773 | 0.219 | 1.115 | -1.182 | 0.218 | -0.001 | 0.002 | 0.002 | 0 | 0.07 | 0.07 | 0.07 | 0.04 | 0.09 | 0.1 |
| 2 | 211536846 | rs4672587 | -0.274 | 1.266 | -0.973 | 0.413 | -0.016 | -0.016 | -0.016 | 0.064 | 0.21 | 0.19 | 0.42 | 0.44 | 0.34 | 0.37 |
| 2 | 211541064 | rs6748782 | 0.218 | 0.194 | -1.086 | -0.595 | 0.019 | 0.014 | 0.014 | 0 | 0.13 | 0.07 | 0.06 | 0 | 0.09 | 0.06 |
| 2 | 211543147 | rs7684 | -1.039 | 0.596 | -0.742 | -1.582 | 0 | -0.1 | -0.1 | 0 | 0.4 | 0.34 | 0.34 | 0.01 | 0.41 | 0.25 |
| 2 | 211547400 | rs13386028 | -1.234 | 0.99 | -0.39 | -1.173 | 0 | 0.005 | 0.005 | 0 | 0.39 | 0.35 | 0.45 | 0.65 | 0.42 | 0.35 |
| 2 | 211547469 | rs13386056 | 0.205 | 0.969 | -0.33 | -1.197 | -0.002 | -0.027 | -0.027 | 0 | 0.19 | 0.17 | 0.18 | 0.01 | 0.21 | 0.18 |
| 2 | 211553159 | rs4672588 | 0.76 | 1.044 | -0.343 | -1.16 | 0.005 | 0.072 | 0.072 | 0 | 0.05 | 0.07 | 0.18 | 0.64 | 0.09 | 0.11 |
| 2 | 211556975 | rs10490320 | -0.998 | 0.929 | -0.315 | -1.241 | -0.001 | 0.004 | 0.004 | 0.145 | 0.35 | 0.33 | 0.4 | 0.65 | 0.37 | 0.34 |
| 2 | 211559526 | rs2371023 | -0.425 | 0.947 | -0.328 | -1.23 | 0.002 | 0.001 | 0.001 | 0.68 | 0.25 | 0.28 | 0.36 | 0.65 | 0.31 | 0.32 |
| 2 | 211560510 | rs13427306 | 1.275 | 0.448 | -0.674 | -1.918 | -0.01 | 0.029 | 0.029 | 0.177 | 0.27 | 0.17 | 0.2 | 0.21 | 0.26 | 0.36 |
| 2 | 211565692 | rs4673546 | 1.488 | 0.53 | -1.318 | -2.289 | -0.054 | -0.036 | -0.036 | 0.138 | 0.11 | 0.01 | 0.25 | 0.14 | 0.17 | 0.19 |
| 2 | 211569722 | rs2371030 | -1.142 | 0.399 | -1.087 | -2.427 | 0.002 | 0.074 | 0.074 | 0.449 | 0.58 | 0.31 | 0.7 | 0.85 | 0.64 | 0.61 |
| 2 | 211570448 | rs1016396 | -0.177 | 0.322 | -1.173 | -2.456 | -0.004 | 0.094 | 0.094 | 0 | 0.43 | 0.24 | 0.42 | 0.71 | 0.45 | 0.38 |
| 2 | 211570616 | rs12999612 | 0.203 | 0.309 | -1.175 | -2.48 | 0.01 | 0.129 | 0.129 | 0 | 0.37 | 0.18 | 0.24 | 0.01 | 0.36 | 0.28 |
| 2 | 211576458 | rs2111713 | -0.833 | 0.349 | -1.127 | -2.454 | 0.012 | 0.132 | 0.132 | 0.657 | 0.56 | 0.29 | 0.67 | 0.85 | 0.63 | 0.6 |
| 2 | 211583066 | rs7583500 | -0.231 | 0.361 | -1.211 | -2.414 | -0.009 | 0.105 | 0.105 | 0 | 0.44 | 0.22 | 0.42 | 0.71 | 0.45 | 0.39 |
| 2 | 211585271 | rs1861897 | 1.478 | 0.355 | -1.161 | -2.363 | -0.006 | 0.016 | 0.016 | 0 | 0.06 | 0.05 | 0.18 | 0.69 | 0.09 | 0.11 |
| 2 | 211593520 | rs1024898 | -1.163 | 0.315 | -1.111 | -2.389 | 0.012 | 0.085 | 0.085 | 0.94 | 0.59 | 0.31 | 0.72 | 0.85 | 0.67 | 0.62 |
| 2 | 211600311 | rs10932355 | 0.576 | 0.425 | -1.068 | -2.157 | -0.007 | -0.001 | -0.001 | 0 | 0.14 | 0.08 | 0.28 | 0.14 | 0.19 | 0.22 |
| 2 | 211608379 | rs4673553 | -0.579 | 0.39 | -1.169 | -2.316 | -0.032 | 0.147 | 0.147 | 0 | 0.4 | 0.17 | 0.55 | 0.84 | 0.46 | 0.44 |
| 2 | 211614638 | rs10206976 | 1.382 | 0.503 | -1.12 | -2.191 | -0.02 | -0.01 | -0.01 | 0 | 0.11 | 0.08 | 0.25 | 0.14 | 0.17 | 0.19 |
| 2 | 211617598 | rs2287412 | 1.531 | 0.466 | -0.637 | -2.416 | -0.004 | -0.002 | -0.002 | 0 | 0.29 | 0.28 | 0.2 | 0.15 | 0.27 | 0.36 |
| 2 | 211619881 | rs10490327 | 1.88 | 0.069 | -0.894 | -2.519 | -0.033 | 0.064 | 0.064 | 0 | 0.13 | 0 | 0.06 | 0.01 | 0.15 | 0.1 |
| 2 | 211623562 | rs2111716 | -0.841 | 0.291 | -0.267 | -2.343 | 0.007 | -0.032 | -0.032 | 0 | 0.57 | 0.49 | 0.53 | 0.71 | 0.55 | 0.46 |
| 2 | 211628123 | rs2887930 | 1.277 | 0.33 | -0.193 | -2.049 | 0.008 | 0.065 | 0.065 | 0 | 0.06 | 0.09 | 0.19 | 0.69 | 0.1 | 0.11 |
| 2 | 211628462 | rs2371074 | 0.113 | 0.112 | 0.101 | -2.238 | -0.019 | 0.021 | 0.021 | 0 | 0.22 | 0.37 | 0.3 | 0.69 | 0.21 | 0.18 |
| 2 | 211629715 | rs6736693 | 0.361 | 0.083 | 0.102 | -2.237 | -0.034 | 0.03 | 0.03 | 0 | 0.2 | 0.37 | 0.27 | 0.69 | 0.18 | 0.14 |
| 2 | 211641003 | rs764637 | -2.075 | 0.05 | -0.41 | -2.232 | -0.001 | -0.02 | -0.02 | 0 | 0.89 | 0.9 | 0.95 | 1 | 0.92 | 0.98 |
| 2 | 211644461 | rs12997044 | 1.586 | 0.11 | -0.506 | -2.482 | 0.014 | -0.075 | -0.075 | 0 | 0.28 | 0.4 | 0.22 | 0.15 | 0.26 | 0.34 |
| 2 | 211644647 | rs10172053 | 1.267 | 0.343 | -0.475 | -2.243 | -0.012 | -0.005 | -0.005 | 0 | 0.12 | 0.09 | 0.25 | 0.14 | 0.17 | 0.2 |

| | | | | | | | | | | | | | | | | |
|---|---|---|---|---|---|---|---|---|---|---|---|---|---|---|---|---|
| 2 | 211647733 | rs10932361 | -0.083 | 0.127 | -0.428 | -2.068 | 0.012 | 0.107 | 0.107 | 0 | 0.88 | 0.73 | 0.75 | 0.33 | 0.86 | 0.85 |
| 2 | 211647954 | rs11687290 | -0.114 | 0.014 | -0.529 | -2.182 | 0.013 | 0.07 | 0.07 | 0.962 | 0.3 | 0.18 | 0.22 | 0.01 | 0.28 | 0.21 |
| 2 | 211648697 | rs10932362 | 0.569 | 0.092 | -0.78 | -2.108 | -0.047 | 0.127 | 0.127 | 2.728 | 0.19 | 0.01 | 0.17 | 0.01 | 0.21 | 0.19 |
| 2 | 211648806 | rs4672596 | 0.135 | 0.188 | -0.501 | -1.839 | -0.002 | -0.006 | -0.006 | 0 | 0.58 | 0.56 | 0.53 | 0.32 | 0.58 | 0.64 |
| 2 | 211655965 | rs6758081 | 0.405 | 0.045 | -0.773 | -2.093 | 0.008 | 0.019 | 0.019 | 0 | 0.31 | 0.26 | 0.32 | 0.77 | 0.21 | 0.47 |
| 2 | 211657226 | rs10195020 | -0.333 | 0.465 | -0.468 | -2.503 | 0.007 | -0.089 | -0.089 | 0 | 0.59 | 0.58 | 0.38 | 0.08 | 0.59 | 0.34 |
| 2 | 211657851 | rs10490328 | 0.944 | 0.257 | -0.823 | -2.886 | -0.022 | 0.023 | 0.023 | 1.498 | 0.18 | 0.09 | 0.19 | 0.07 | 0.29 | 0.14 |
| 2 | 211661897 | rs4673558 | 1.251 | 0.28 | -0.84 | -2.859 | -0.077 | -0.036 | -0.036 | 4.567 | 0.12 | 0.01 | 0.3 | 0.15 | 0.23 | 0.22 |
| 2 | 211661935 | rs13021675 | 0.691 | -0.232 | -0.56 | -3.123 | 0.031 | 0.051 | 0.051 | 0.909 | 0.28 | 0.17 | 0.11 | 0.01 | 0.21 | 0.15 |
| 2 | 211664333 | rs10164524 | 0.057 | -0.202 | -0.698 | -2.908 | -0.009 | 0.042 | 0.042 | 0 | 0.28 | 0.14 | 0.22 | 0.07 | 0.35 | 0.17 |
| 2 | 211672261 | rs986293 | 0.362 | 0.007 | -0.825 | -2.956 | -0.025 | 0.039 | 0.039 | 0 | 0.2 | 0.1 | 0.2 | 0.07 | 0.32 | 0.12 |
| 2 | 211679181 | rs13031255 | 0.677 | 0.134 | -0.851 | -2.902 | -0.034 | 0.018 | 0.018 | 0.02 | 0.05 | 0.01 | 0.11 | 0.05 | 0.16 | 0.05 |
| 2 | 211681243 | rs12478788 | 0.953 | 0.145 | -1.122 | -3.426 | -0.112 | 0.006 | 0.006 | 0 | 0.21 | 0.02 | 0.25 | 0.13 | 0.31 | 0.27 |
| 2 | 211682425 | rs2719968 | 1.151 | -0.905 | -1.451 | -3.496 | 0.021 | -0.01 | -0.01 | 0 | 0.21 | 0.07 | 0.27 | 0.74 | 0.12 | 0.3 |
| 2 | 211682987 | rs12464940 | -0.448 | -0.995 | -1.054 | -3.555 | -0.027 | -0.025 | -0.025 | 0 | 0.65 | 0.57 | 0.65 | 0.26 | 0.79 | 0.67 |
| 2 | 211685063 | rs1158374 | 1.504 | -1.1 | -1.456 | -3.489 | 0.017 | 0.08 | 0.08 | 2.425 | 0.21 | 0.07 | 0.1 | 0.06 | 0.12 | 0.19 |
| 2 | 211690173 | rs11686803 | 1.469 | -0.668 | -1.037 | -3.28 | 0.001 | 0.002 | 0.002 | 0 | 0.05 | 0.04 | 0.1 | 0.05 | 0.15 | 0.05 |
| 2 | 211695915 | rs12694212 | -0.662 | -1.115 | -1.084 | -3.374 | 0.007 | 0.002 | 0.002 | 0 | 0.74 | 0.76 | 0.69 | 0.28 | 0.84 | 0.69 |
| 2 | 211696395 | rs2542927 | -1.742 | -1.378 | -1.113 | -3.526 | 0.008 | -0.013 | -0.013 | 0 | 0.81 | 0.83 | 0.93 | 1 | 0.91 | 0.98 |
| 2 | 211697118 | rs2542925 | 1.244 | -0.934 | -1.032 | -3.273 | 0.024 | -0.018 | -0.018 | 0.318 | 0.21 | 0.07 | 0.27 | 0.72 | 0.11 | 0.3 |
| 2 | 211699011 | rs7608489 | 0.263 | -0.243 | -0.755 | -3.053 | 0.053 | -0.042 | -0.042 | 0 | 0.31 | 0.5 | 0.41 | 0.21 | 0.49 | 0.37 |
| 2 | 211701811 | rs13019409 | 1.132 | -0.244 | -1.515 | -2.928 | -0.128 | 0.001 | 0.001 | 0 | 0.23 | 0.02 | 0.26 | 0.16 | 0.34 | 0.3 |
| 2 | 211703869 | rs7580498 | 0.311 | -0.057 | -0.181 | -2.301 | 0.024 | -0.074 | -0.074 | 0.035 | 0.22 | 0.28 | 0.28 | 0.06 | 0.36 | 0.2 |
| 2 | 211711552 | rs2719965 | 0.593 | -0.591 | 0.237 | -1.748 | -0.021 | 0.266 | 0.266 | 0.53 | 0.09 | 0.37 | 0.23 | 0.65 | 0.02 | 0.08 |
| 2 | 211717695 | rs12995607 | 0.865 | -0.329 | -0.277 | -1.927 | -0.031 | 0.026 | 0.026 | 0 | 0.06 | 0.02 | 0.09 | 0.05 | 0.11 | 0.07 |
| 2 | 211724958 | rs7593111 | 0.912 | -1.043 | -0.459 | -2.384 | 0.026 | 0.016 | 0.016 | 0.207 | 0.18 | 0.1 | 0.08 | 0 | 0.11 | 0.03 |
| 2 | 211730557 | rs4130275 | 0.632 | -1.29 | -0.376 | -2.799 | -0.013 | -0.048 | -0.048 | 0 | 0.28 | 0.3 | 0.14 | 0.07 | 0.16 | 0.19 |
| 2 | 211738452 | rs6733934 | 1.132 | -1.023 | -0.466 | -2.811 | -0.002 | 0 | 0 | 0 | 0.06 | 0.05 | 0.06 | 0 | 0.09 | 0.02 |
| 2 | 211740016 | rs6726552 | 0.553 | -1.144 | -0.13 | -2.466 | 0.031 | 0.081 | 0.081 | 0.04 | 0.53 | 0.32 | 0.49 | 0.24 | 0.61 | 0.64 |
| 2 | 211741194 | rs16845044 | 0.46 | -1.215 | 0.167 | -2.481 | -0.016 | -0.091 | -0.091 | 0 | 0.23 | 0.29 | 0.15 | 0.05 | 0.18 | 0.07 |
| 2 | 211741314 | rs2068914 | 2.018 | -1.231 | 0.059 | -2.468 | -0.008 | -0.025 | -0.025 | 0 | 0.14 | 0.13 | 0.04 | 0 | 0.04 | 0.01 |
| 2 | 211744633 | rs6759308 | 1.14 | -1.257 | -0.114 | -2.605 | 0.021 | 0.14 | 0.14 | 4.017 | 0.45 | 0.17 | 0.47 | 0.24 | 0.6 | 0.61 |
| 2 | 211750920 | rs4673564 | -1.388 | -1.184 | 0.292 | -2.419 | -0.009 | 0.01 | 0.01 | 0 | 0.81 | 0.68 | 0.69 | 0.3 | 0.86 | 0.74 |
| 2 | 211751804 | rs17792034 | 0.798 | -1.084 | -0.102 | -2.38 | 0.008 | 0.005 | 0.005 | 0 | 0.12 | 0.07 | 0.06 | 0 | 0.1 | 0.03 |
| 2 | 211765267 | rs4520971 | -0.307 | -1.21 | 0.11 | -2.411 | -0.034 | -0.022 | -0.022 | 0 | 0.7 | 0.48 | 0.63 | 0.3 | 0.78 | 0.71 |
| 2 | 211778272 | rs4505460 | -1.427 | -1.127 | 0.175 | -2.36 | -0.009 | 0.018 | 0.018 | 0 | 0.81 | 0.65 | 0.69 | 0.3 | 0.86 | 0.74 |
| 2 | 211785292 | rs4605308 | -0.434 | -1.101 | -0.557 | -2.96 | -0.001 | -0.094 | -0.094 | 0.152 | 0.44 | 0.42 | 0.29 | 0.01 | 0.38 | 0.16 |
| 2 | 211787329 | rs3924144 | 1.584 | -1.172 | -0.593 | -3.002 | 0.006 | -0.069 | -0.069 | 1.042 | 0.18 | 0.22 | 0.19 | 0.01 | 0.26 | 0.12 |
| 2 | 211795919 | rs4461202 | 1.263 | -1.1 | -0.892 | -3.157 | -0.095 | 0.069 | 0.069 | 0 | 0.31 | 0.1 | 0.33 | 0.23 | 0.45 | 0.48 |
| 2 | 211797987 | rs17740443 | 0.85 | -1.123 | -0.895 | -3.172 | 0.061 | 0.062 | 0.062 | 1.836 | 0.25 | 0.12 | 0.09 | 0 | 0.13 | 0.06 |
| 2 | 211807729 | rs13007741 | 1.611 | -0.696 | -0.802 | -2.951 | -0.032 | 0.032 | 0.032 | 0 | 0.07 | 0.02 | 0.11 | 0.05 | 0.13 | 0.06 |
| 2 | 211808629 | rs6746197 | 0.612 | -0.74 | -0.703 | -2.938 | 0.007 | -0.037 | -0.037 | 0 | 0.24 | 0.25 | 0.26 | 0.08 | 0.29 | 0.2 |
| 2 | 211813132 | rs13006521 | -1.636 | -0.776 | -0.661 | -2.821 | -0.01 | 0.02 | 0.02 | 0 | 0.82 | 0.69 | 0.7 | 0.3 | 0.86 | 0.76 |
| 2 | 211829497 | rs186471 | 0.557 | -1.308 | -0.707 | -2.806 | -0.042 | 0.01 | 0.01 | 0.05 | 0.17 | 0.34 | 0.29 | 0.76 | 0.06 | 0.32 |
| 2 | 211840292 | rs17740874 | 2.191 | -1.217 | -1.088 | -2.908 | -0.005 | -0.015 | -0.015 | 0 | 0.14 | 0.11 | 0.16 | 0.01 | 0.22 | 0.1 |
| 2 | 211840828 | rs12991650 | 1.453 | -1.087 | -1.218 | -3.25 | -0.014 | 0.003 | 0.003 | 0 | 0.14 | 0.11 | 0.2 | 0.05 | 0.25 | 0.08 |
| 2 | 211853395 | rs389035 | 1.179 | -1.152 | -1.131 | -3.261 | -0.018 | -0.001 | -0.001 | 0 | 0.26 | 0.55 | 0.39 | 0.69 | 0.24 | 0.34 |
| 2 | 211856725 | rs651797 | -0.369 | -1.084 | -1.092 | -3.183 | 0.024 | 0.172 | 0.172 | 0.008 | 0.49 | 0.74 | 0.67 | 0.82 | 0.55 | 0.52 |
| 2 | 211859726 | rs1657863 | 0.217 | -1.077 | -1.045 | -3.172 | 0 | -0.005 | -0.005 | 0 | 0.42 | 0.55 | 0.59 | 0.76 | 0.51 | 0.43 |
| 2 | 211863745 | rs3097415 | -1.535 | -1.745 | -1.03 | -2.821 | -0.002 | -0.02 | -0.02 | 0.053 | 0.85 | 0.87 | 0.96 | 1 | 0.96 | 0.99 |
| 2 | 211864403 | rs1652311 | 0.157 | -1.277 | -0.941 | -2.597 | 0.002 | 0.006 | 0.006 | 0.68 | 0.35 | 0.62 | 0.47 | 0.77 | 0.3 | 0.44 |
| 2 | 211871631 | rs7567073 | 0.173 | -0.354 | -1.46 | -2.096 | -0.042 | 0.094 | 0.094 | 0.058 | 0.36 | 0.15 | 0.42 | 0.22 | 0.55 | 0.43 |
| 2 | 211872193 | rs13402304 | -0.364 | -0.522 | -0.744 | -1.603 | -0.008 | -0.04 | -0.04 | 0.169 | 0.08 | 0.15 | 0.06 | 0 | 0.08 | 0.05 |
| 2 | 211873461 | rs11684145 | 0.255 | -0.317 | -1.558 | -1.687 | -0.043 | 0.06 | 0.06 | 0.062 | 0.3 | 0.08 | 0.29 | 0.22 | 0.34 | 0.42 |
| 2 | 211878255 | rs167648 | 0.359 | -0.694 | -0.898 | -1.91 | 0.002 | -0.001 | -0.001 | 0 | 0.55 | 0.39 | 0.62 | 0.77 | 0.55 | 0.49 |
| 2 | 211881975 | rs9288423 | 1.23 | -1.037 | -1.039 | -2.222 | 0.008 | -0.016 | -0.016 | 0 | 0.18 | 0.14 | 0.06 | 0 | 0.09 | 0.03 |
| 2 | 211893388 | rs319719 | -0.572 | -0.589 | -0.615 | -1.968 | 0.003 | -0.047 | -0.047 | 1.098 | 0.68 | 0.71 | 0.87 | 1 | 0.83 | 0.85 |
| 2 | 211897179 | rs10804187 | 0.322 | -0.482 | -1.342 | -1.916 | -0.003 | -0.004 | -0.004 | 0 | 0.38 | 0.34 | 0.59 | 0.79 | 0.48 | 0.45 |
| 2 | 211897927 | rs11692975 | 0.769 | -0.422 | -1.431 | -1.892 | -0.05 | 0.053 | 0.053 | 0.155 | 0.28 | 0.07 | 0.26 | 0.21 | 0.34 | 0.37 |
| 2 | 211900425 | rs192927 | 0.459 | -0.401 | -1.309 | -1.764 | -0.033 | 0.045 | 0.045 | 0.471 | 0.36 | 0.21 | 0.58 | 0.77 | 0.48 | 0.45 |
| 2 | 211903193 | rs191180 | 0.362 | -0.252 | -1.322 | -1.622 | 0.014 | -0.029 | -0.029 | 0 | 0.36 | 0.4 | 0.6 | 0.81 | 0.48 | 0.57 |
| 2 | 211904482 | rs7577965 | -0.236 | 0.566 | -1.658 | -0.638 | -0.015 | 0.055 | 0.055 | 0 | 0.32 | 0.17 | 0.28 | 0.21 | 0.36 | 0.27 |
| 2 | 211927188 | rs13016652 | -0.284 | 0.73 | -1.752 | -0.5 | -0.018 | 0.084 | 0.084 | 0 | 0.39 | 0.16 | 0.36 | 0.77 | 0.41 | 0.5 |
| 2 | 211935990 | rs12464316 | -0.15 | 0.987 | -1.604 | -0.513 | -0.03 | 0.083 | 0.083 | 0 | 0.37 | 0.11 | 0.34 | 0.78 | 0.4 | 0.49 |
| 2 | 211937214 | rs6739390 | 0.169 | 0.918 | -1.608 | -1.295 | 0.005 | 0.092 | 0.092 | 0 | 0.35 | 0.2 | 0.2 | 0.07 | 0.28 | 0.2 |
| 2 | 211946057 | rs6435599 | 0.293 | 0.875 | -1.61 | -1.317 | 0.008 | 0.106 | 0.106 | 0 | 0.33 | 0.16 | 0.14 | 0.07 | 0.25 | 0.19 |
| 2 | 211951445 | rs11682145 | 0.629 | 1.205 | -1.846 | -0.247 | -0.077 | -0.024 | -0.024 | 1.741 | 0.34 | 0.02 | 0.26 | 0.3 | 0.34 | 0.45 |
| 2 | 211953919 | rs10497942 | 0.643 | 1.141 | -2.116 | 0.305 | -0.042 | -0.076 | -0.076 | 0 | 0.11 | 0 | 0.1 | 0.3 | 0.14 | 0.05 |
| 2 | 211954880 | rs10932366 | 0.492 | 1.094 | -2.028 | 0.084 | -0.066 | -0.007 | -0.007 | 0.351 | 0.38 | 0.07 | 0.35 | 0.64 | 0.43 | 0.48 |

| | | | | | | | | | | | | | | | | |
|---|---|---|---|---|---|---|---|---|---|---|---|---|---|---|---|---|
| 2 | 211960740 | rs13005623 | 0.128 | 0.918 | -1.096 | 0.109 | 0.005 | 0.062 | 0.062 | 0 | 0.47 | 0.34 | 0.41 | 0.8 | 0.47 | 0.51 |
| 2 | 211966616 | rs1469926 | 0.085 | 0.777 | -0.328 | 0.034 | 0 | -0.004 | -0.004 | 0 | 0.5 | 0.47 | 0.48 | 0.68 | 0.5 | 0.52 |
| 2 | 211969078 | rs319729 | -0.77 | 0.742 | -0.301 | -0.038 | 0.004 | 0.028 | 0.028 | 0 | 0.73 | 0.8 | 0.84 | 0.96 | 0.81 | 0.83 |
| 2 | 211969338 | rs16845301 | 0.865 | 0.728 | -0.375 | -0.08 | 0.004 | 0.009 | 0.009 | 0 | 0.21 | 0.16 | 0.14 | 0.04 | 0.15 | 0.16 |
| 2 | 211972676 | rs182657 | 0.887 | 0.749 | -0.334 | -0.043 | 0.02 | 0.019 | 0.019 | 0.166 | 0.23 | 0.34 | 0.41 | 0.27 | 0.32 | 0.31 |
| 2 | 211973600 | rs319724 | -0.379 | 0.728 | -0.539 | 0.053 | -0.011 | 0.08 | 0.08 | 0 | 0.48 | 0.8 | 0.61 | 0.34 | 0.47 | 0.49 |
| 2 | 211982126 | rs10208692 | -0.02 | 0.646 | 0.238 | -0.233 | 0.003 | -0.028 | -0.028 | 0 | 0.59 | 0.5 | 0.46 | 0.72 | 0.59 | 0.58 |
| 2 | 211990188 | rs1370619 | 0.044 | 0.566 | 0.318 | -0.051 | 0 | 0.041 | 0.041 | 0.273 | 0.19 | 0.28 | 0.17 | 0.34 | 0.19 | 0.07 |
| 2 | 211990354 | rs17799315 | 1.197 | 0.464 | 0.154 | -0.137 | -0.045 | -0.043 | -0.043 | 0 | 0.37 | 0.07 | 0.27 | 0.37 | 0.39 | 0.51 |
| 2 | 212005919 | rs10166020 | 1.131 | 0.437 | 0.24 | -0.172 | -0.017 | -0.024 | -0.024 | 0.973 | 0.37 | 0.14 | 0.27 | 0.37 | 0.39 | 0.51 |
| 2 | 212010493 | rs11901164 | -0.784 | 0.509 | 0.529 | -0.179 | 0.004 | 0.034 | 0.034 | 0.319 | 0.28 | 0.65 | 0.48 | 0.46 | 0.32 | 0.21 |
| 2 | 212021755 | rs6435604 | -0.022 | 0.418 | 0.776 | -0.254 | 0.096 | 0.112 | 0.112 | 0 | 0.08 | 0.39 | 0.31 | 0.12 | 0.13 | 0.11 |
| 2 | 212026295 | rs13392643 | 0.713 | 0.388 | 0.68 | -0.371 | -0.036 | 0.156 | 0.156 | 0 | 0.49 | 0.13 | 0.38 | 0.7 | 0.54 | 0.57 |
| 2 | 212030850 | rs6713384 | 0.177 | 0.41 | 0.758 | -0.355 | -0.001 | 0.011 | 0.011 | 0 | 0.57 | 0.4 | 0.42 | 0.71 | 0.59 | 0.58 |
| 2 | 212035162 | rs4673585 | -0.425 | 0.404 | 0.763 | -0.381 | 0.007 | 0.085 | 0.085 | 0 | 0.65 | 0.81 | 0.76 | 0.83 | 0.72 | 0.69 |
| 2 | 212036651 | rs10432525 | -0.56 | 0.379 | 0.75 | -0.393 | 0.01 | 0.09 | 0.09 | 0 | 0.64 | 0.81 | 0.74 | 0.83 | 0.72 | 0.67 |
| 2 | 212039636 | rs4673587 | 0.064 | 0.458 | 0.722 | -0.379 | 0 | -0.009 | -0.009 | 0 | 0.58 | 0.4 | 0.59 | 0.82 | 0.63 | 0.64 |
| 2 | 212045199 | rs4356605 | 0.001 | 0.507 | 0.767 | -0.263 | 0.007 | 0.028 | 0.028 | 0 | 0.91 | 0.79 | 0.75 | 0.89 | 0.92 | 0.91 |
| 2 | 212047246 | rs4673595 | 0.169 | 0.384 | 0.766 | -0.285 | 0.005 | -0.059 | -0.059 | 0 | 0.2 | 0.12 | 0.31 | 0.45 | 0.21 | 0.14 |
| 2 | 212051150 | rs1437927 | 0.117 | 0.11 | 0.509 | -0.675 | -0.008 | 0.029 | 0.029 | 0 | 0.67 | 0.31 | 0.64 | 0.84 | 0.66 | 0.74 |
| 2 | 212053326 | rs13035922 | 0.935 | 0.112 | 0.394 | -0.66 | -0.027 | 0.06 | 0.06 | 0 | 0.22 | 0.07 | 0.18 | 0.15 | 0.22 | 0.22 |
| 2 | 212065767 | rs1437914 | -1.056 | 0.105 | 0.407 | -0.642 | -0.02 | 0.051 | 0.051 | 0 | 0.77 | 0.91 | 0.79 | 0.85 | 0.78 | 0.78 |
| 2 | 212067654 | rs17738493 | 0.026 | 0.099 | 0.614 | -0.623 | 0.068 | -0.033 | -0.033 | 0 | 0.08 | 0.28 | 0.16 | 0.01 | 0.11 | 0.04 |
| 2 | 212068615 | rs3910518 | 0.249 | 0 | 0.283 | -0.641 | 0.03 | -0.075 | -0.075 | 0 | 0.68 | 0.82 | 0.65 | 0.54 | 0.73 | 0.75 |
| 2 | 212076352 | rs7609319 | -1.523 | 0.086 | 0.239 | -0.61 | -0.007 | 0.006 | 0.006 | 0 | 0.88 | 0.95 | 0.88 | 0.87 | 0.87 | 0.87 |
| 2 | 212077276 | rs12478428 | 0.476 | 0.044 | 0.285 | -0.625 | -0.003 | -0.032 | -0.032 | 0 | 0.29 | 0.17 | 0.27 | 0.37 | 0.28 | 0.15 |
| 2 | 212078938 | rs4672608 | 0.364 | 0.084 | 0.454 | -0.635 | 0.006 | -0.029 | -0.029 | 0.491 | 0.78 | 0.36 | 0.67 | 0.86 | 0.75 | 0.83 |
| 2 | 212079450 | rs10497937 | 1.858 | 0.11 | 0.285 | -0.575 | 0 | -0.001 | -0.001 | 0 | 0.1 | 0.04 | 0.1 | 0.13 | 0.1 | 0.11 |
| 2 | 212097711 | rs1465828 | -1.301 | 0.135 | 0.631 | -0.581 | 0.03 | -0.005 | -0.005 | 0 | 0.87 | 0.74 | 0.8 | 0.86 | 0.83 | 0.85 |
| 2 | 212105386 | rs2371172 | -1.08 | 0.004 | 0.648 | -0.606 | -0.003 | 0.001 | 0.001 | 0 | 0.52 | 0.68 | 0.62 | 0.51 | 0.48 | 0.37 |
| 2 | 212106926 | rs750576 | -0.378 | -0.11 | 0.596 | -0.675 | 0.019 | -0.053 | -0.053 | 0 | 0.4 | 0.21 | 0.44 | 0.5 | 0.34 | 0.31 |
| 2 | 212110840 | rs1437919 | -0.554 | -0.15 | 0.238 | -0.639 | -0.005 | -0.002 | -0.002 | 0 | 0.27 | 0.3 | 0.35 | 0.25 | 0.21 | 0.27 |
| 2 | 212112490 | rs1437917 | -1.366 | 0.038 | 0.09 | -0.519 | 0.015 | 0.001 | 0.001 | 0.188 | 0.61 | 0.52 | 0.63 | 0.62 | 0.53 | 0.44 |
| 2 | 212126069 | rs16845743 | 1.418 | -0.063 | -0.54 | -0.497 | -0.016 | 0.006 | 0.006 | 0 | 0.13 | 0.04 | 0.15 | 0.13 | 0.13 | 0.12 |
| 2 | 212137665 | rs17691926 | 1.527 | -0.046 | -0.743 | -0.549 | -0.128 | -0.075 | -0.075 | 0 | 0.33 | 0.04 | 0.27 | 0.35 | 0.43 | 0.51 |
| 2 | 212137884 | rs10171524 | 1.201 | -0.1 | -0.364 | -0.545 | 0.013 | -0.001 | -0.001 | 0 | 0.37 | 0.38 | 0.38 | 0.36 | 0.49 | 0.54 |
| 2 | 212139537 | rs4442936 | 0.237 | -0.207 | -0.466 | -0.613 | 0.015 | 0.023 | 0.023 | 0.632 | 0.69 | 0.75 | 0.62 | 0.75 | 0.74 | 0.7 |
| 2 | 212141680 | rs10497933 | 1.005 | -0.135 | -0.566 | -0.615 | 0.004 | -0.06 | -0.06 | 3.647 | 0.25 | 0.12 | 0.2 | 0.38 | 0.21 | 0.14 |
| 2 | 212151252 | rs12618997 | 1.081 | -0.163 | -0.411 | -0.603 | 0.001 | -0.004 | -0.004 | 0 | 0.15 | 0.19 | 0.2 | 0.13 | 0.14 | 0.13 |
| 2 | 212157104 | rs4632271 | -0.233 | -0.164 | -0.629 | -0.6 | 0.02 | 0.01 | 0.01 | 1.257 | 0.83 | 0.89 | 0.8 | 0.88 | 0.89 | 0.84 |
| 2 | 212159787 | rs10209471 | 0.736 | -0.178 | -0.572 | -0.61 | -0.031 | -0.041 | -0.041 | 0 | 0.64 | 0.48 | 0.58 | 0.75 | 0.73 | 0.69 |
| 2 | 212165522 | rs13007747 | -0.536 | -0.14 | -0.689 | -0.548 | -0.004 | 0.001 | 0.001 | 0 | 0.47 | 0.55 | 0.42 | 0.52 | 0.39 | 0.3 |
| 2 | 212167904 | rs1595063 | -1.141 | -0.126 | -0.706 | -0.549 | 0 | 0.004 | 0.004 | 0 | 0.95 | 0.97 | 0.82 | 0.89 | 0.92 | 0.9 |
| 2 | 212169952 | rs6728697 | -0.112 | -0.088 | -0.715 | -0.543 | -0.011 | -0.021 | -0.021 | 0.057 | 0.81 | 0.66 | 0.69 | 0.86 | 0.82 | 0.81 |
| 2 | 212172094 | rs10190934 | 0.23 | 0.111 | -1.298 | -0.421 | -0.01 | -0.044 | -0.044 | 0 | 0.19 | 0.41 | 0.22 | 0.13 | 0.15 | 0.15 |
| 2 | 212181355 | rs1992027 | 1.144 | 0.733 | -2.486 | -0.158 | -0.027 | 0.027 | 0.027 | 0 | 0.18 | 0.1 | 0.23 | 0.13 | 0.25 | 0.14 |
| 2 | 212184018 | rs1836726 | 0.564 | 0.707 | -2.527 | 0.196 | -0.01 | -0.093 | -0.093 | 0 | 0.26 | 0.09 | 0.2 | 0.38 | 0.24 | 0.17 |
| 2 | 212191113 | rs6707202 | 0.53 | 0.855 | -1.979 | 0.484 | -0.059 | -0.091 | -0.091 | 0.012 | 0.29 | 0.07 | 0.19 | 0.4 | 0.33 | 0.5 |
| 2 | 212195610 | rs11694182 | 0.329 | 0.402 | -2.049 | 0.399 | -0.022 | 0.005 | 0.005 | 0 | 0.28 | 0.06 | 0.27 | 0.23 | 0.24 | 0.48 |
| 2 | 212196284 | rs10804193 | 0.475 | 0.223 | -1.517 | 0.323 | -0.003 | 0.004 | 0.004 | 0.155 | 0.31 | 0.45 | 0.38 | 0.31 | 0.3 | 0.19 |
| 2 | 212196304 | rs1836733 | -1.43 | 0.008 | -1.384 | 0.304 | -0.021 | -0.022 | -0.022 | 0 | 0.74 | 0.81 | 0.69 | 0.63 | 0.64 | 0.77 |
| 2 | 212199994 | rs1595068 | 0.247 | -0.067 | -1.297 | 0.394 | 0.004 | -0.009 | -0.009 | 0 | 0.41 | 0.11 | 0.3 | 0.39 | 0.34 | 0.57 |
| 2 | 212200881 | rs7562566 | -0.818 | -0.211 | -0.358 | 0.279 | 0.012 | 0.011 | 0.011 | 0 | 0.69 | 0.58 | 0.67 | 0.63 | 0.63 | 0.75 |
| 2 | 212202491 | rs7566827 | -1.576 | -0.228 | -0.568 | 0.261 | -0.027 | -0.027 | -0.027 | 0.028 | 0.76 | 0.81 | 0.7 | 0.63 | 0.65 | 0.75 |
| 2 | 212203600 | rs16845921 | 0.053 | -0.387 | -0.478 | 0.132 | -0.001 | -0.001 | -0.001 | 0 | 0.15 | 0.12 | 0.03 | 0.09 | 0.1 | 0.08 |
| 2 | 212214642 | rs1836744 | -0.421 | -0.237 | -0.179 | 0.259 | 0.007 | -0.004 | -0.004 | 0 | 0.66 | 0.39 | 0.63 | 0.67 | 0.6 | 0.73 |
| 2 | 212221824 | rs1595067 | 0.98 | -0.217 | -0.622 | 0.233 | -0.019 | -0.029 | -0.029 | 0 | 0.32 | 0.1 | 0.32 | 0.35 | 0.29 | 0.5 |
| 2 | 212221870 | rs2033643 | 0.628 | -0.347 | -0.64 | 0.201 | 0.008 | -0.028 | -0.028 | 0.256 | 0.28 | 0.09 | 0.3 | 0.29 | 0.26 | 0.17 |
| 2 | 212226177 | rs6712652 | 1.171 | -0.315 | -0.527 | 0.239 | -0.005 | -0.008 | -0.008 | 0 | 0.29 | 0.09 | 0.28 | 0.28 | 0.24 | 0.5 |
| 2 | 212228438 | rs10427172 | -0.977 | -0.383 | 0.126 | 0.028 | -0.025 | -0.022 | -0.022 | 0.482 | 0.14 | 0.35 | 0.05 | 0.09 | 0.09 | 0.11 |
| 2 | 212236141 | rs1579599 | -0.048 | -0.397 | 0.288 | 0.072 | 0.003 | -0.001 | -0.001 | 0 | 0.43 | 0.59 | 0.38 | 0.44 | 0.34 | 0.59 |
| 2 | 212238541 | rs1971802 | 0.468 | -0.403 | 0.107 | 0.081 | -0.013 | -0.005 | -0.005 | 0 | 0.38 | 0.12 | 0.31 | 0.35 | 0.33 | 0.57 |
| 2 | 212238982 | rs1821659 | 0.485 | -0.444 | 0.138 | 0.054 | 0.012 | -0.006 | -0.006 | 0.009 | 0.29 | 0.22 | 0.29 | 0.28 | 0.27 | 0.17 |
| 2 | 212243011 | rs16845990 | -0.085 | -0.448 | 0.152 | 0.068 | -0.006 | 0 | 0 | 0 | 0.44 | 0.46 | 0.37 | 0.44 | 0.36 | 0.61 |
| 2 | 212244403 | rs10932374 | 0.754 | -0.552 | -0.377 | -0.083 | -0.006 | -0.023 | -0.023 | 0 | 0.28 | 0.13 | 0.28 | 0.34 | 0.26 | 0.49 |
| 2 | 212244657 | rs12467225 | 0.6 | -0.578 | -0.415 | -0.114 | -0.01 | -0.021 | -0.021 | 1.116 | 0.3 | 0.14 | 0.29 | 0.34 | 0.29 | 0.51 |
| 2 | 212244952 | rs1836724 | -1.013 | -0.61 | -0.286 | -0.147 | 0.007 | -0.012 | -0.012 | 0 | 0.67 | 0.4 | 0.64 | 0.72 | 0.6 | 0.75 |
| 2 | 212250195 | rs16846013 | 1.643 | -0.339 | -0.544 | 0.017 | -0.014 | -0.037 | -0.037 | 0 | 0.21 | 0.09 | 0.28 | 0.34 | 0.2 | 0.43 |
| 2 | 212251864 | rs3748962 | 0.53 | -0.477 | -0.521 | 0.057 | -0.011 | -0.029 | -0.029 | 0 | 0.33 | 0.08 | 0.3 | 0.29 | 0.3 | 0.22 |

| | | | | | | | | | | | | | | | | |
|---|---|---|---|---|---|---|---|---|---|---|---|---|---|---|---|---|
| 2 | 212261968 | rs2076818 | -0.194 | -0.389 | 0.069 | 0.091 | 0.023 | 0.048 | 0.048 | 0 | 0.93 | 0.73 | 0.92 | 0.9 | 0.93 | 0.93 |
| 2 | 212264560 | rs3791691 | 0.82 | -0.367 | -0.104 | 0.135 | 0.002 | -0.029 | -0.029 | 0 | 0.28 | 0.15 | 0.28 | 0.34 | 0.24 | 0.48 |
| 2 | 212264598 | rs3791692 | -1.032 | -0.679 | -0.082 | 0.025 | 0.015 | -0.029 | -0.029 | 0 | 0.63 | 0.54 | 0.62 | 0.71 | 0.56 | 0.75 |
| 2 | 212264818 | rs7585000 | -0.226 | -0.464 | -0.36 | 0.041 | 0.008 | -0.018 | -0.018 | 0 | 0.29 | 0.27 | 0.32 | 0.35 | 0.28 | 0.17 |
| 2 | 212267373 | rs12992515 | 0.116 | -0.465 | -0.347 | 0.072 | 0.007 | -0.018 | -0.018 | 0 | 0.28 | 0.24 | 0.32 | 0.37 | 0.26 | 0.17 |
| 2 | 212278218 | rs16846100 | 0.891 | -0.399 | -0.543 | 0.041 | -0.007 | -0.036 | -0.036 | 0.018 | 0.32 | 0.09 | 0.29 | 0.35 | 0.26 | 0.54 |
| 2 | 212281381 | rs4673616 | 0.743 | -0.334 | -0.511 | 0.025 | -0.012 | -0.025 | -0.025 | 0.726 | 0.33 | 0.11 | 0.31 | 0.35 | 0.28 | 0.55 |
| 2 | 212283146 | rs13030304 | 1.131 | -0.382 | -0.563 | 0.016 | -0.028 | 0.048 | 0.048 | 0.21 | 0.14 | 0.05 | 0.23 | 0.04 | 0.19 | 0.07 |
| 2 | 212284621 | rs3791704 | 0.47 | -0.353 | -0.464 | 0.111 | -0.023 | -0.031 | -0.031 | 0 | 0.35 | 0.1 | 0.32 | 0.36 | 0.31 | 0.56 |
| 2 | 212285785 | rs16846127 | 0.23 | -0.37 | -0.436 | 0.121 | 0.007 | -0.01 | -0.01 | 0 | 0.36 | 0.15 | 0.32 | 0.36 | 0.3 | 0.57 |
| 2 | 212292699 | rs1836717 | 0.216 | -0.364 | -0.591 | 0.078 | -0.01 | -0.016 | -0.016 | 0 | 0.36 | 0.11 | 0.32 | 0.36 | 0.32 | 0.57 |
| 2 | 212297838 | rs16846161 | -0.63 | -0.387 | -0.297 | 0.229 | 0.003 | 0.019 | 0.019 | 0.612 | 0.07 | 0.12 | 0.11 | 0.17 | 0.09 | 0.12 |
| 2 | 212300055 | rs6742399 | 0.188 | -0.568 | -0.043 | 0.133 | 0.013 | -0.017 | -0.017 | 0 | 0.57 | 0.28 | 0.55 | 0.44 | 0.64 | 0.35 |
| 2 | 212302127 | rs2033647 | 0.232 | -0.555 | -0.2 | 0.102 | -0.002 | -0.009 | -0.009 | 0 | 0.36 | 0.23 | 0.32 | 0.37 | 0.31 | 0.56 |
| 2 | 212303073 | rs10165449 | 0.604 | -0.554 | -0.139 | -0.048 | 0 | -0.019 | -0.019 | 0.018 | 0.38 | 0.38 | 0.41 | 0.15 | 0.41 | 0.19 |
| 2 | 212303506 | rs13395652 | 1.619 | -0.645 | -1.077 | -0.125 | -0.016 | 0.025 | 0.025 | 0 | 0.27 | 0.23 | 0.37 | 0.15 | 0.34 | 0.13 |
| 2 | 212304565 | rs16825005 | 1.131 | -0.605 | -1.271 | 0.105 | -0.016 | -0.063 | -0.063 | 0 | 0.31 | 0.09 | 0.14 | 0.4 | 0.27 | 0.51 |
| 2 | 212304841 | rs16825008 | 0.739 | -0.589 | -1.274 | 0.102 | -0.026 | -0.064 | -0.064 | 0 | 0.33 | 0.09 | 0.31 | 0.41 | 0.31 | 0.54 |
| 2 | 212305314 | rs9808028 | 0.614 | -0.512 | -1.275 | -0.283 | -0.035 | 0.047 | 0.047 | 1.348 | 0.36 | 0.15 | 0.46 | 0.27 | 0.45 | 0.27 |
| 2 | 212307077 | rs1836715 | -0.496 | -0.637 | -1.29 | -0.087 | -0.009 | -0.035 | -0.035 | 1.097 | 0.79 | 0.86 | 0.72 | 0.72 | 0.77 | 0.83 |
| 2 | 212308833 | rs7605095 | -0.697 | -0.793 | -1.369 | -0.286 | 0.01 | 0.014 | 0.014 | 0 | 0.28 | 0.22 | 0.16 | 0.15 | 0.21 | 0.26 |
| 2 | 212310843 | rs4673618 | 0.901 | -0.246 | -1.308 | -0.06 | -0.062 | 0.117 | 0.117 | 1.66 | 0.36 | 0.06 | 0.4 | 0.17 | 0.42 | 0.18 |
| 2 | 212322255 | rs16846210 | 1.066 | -0.294 | -0.72 | 0.49 | 0.01 | -0.032 | -0.032 | 0 | 0.2 | 0.15 | 0.15 | 0.38 | 0.15 | 0.37 |
| 2 | 212325228 | rs4597456 | -0.656 | 0.001 | -0.818 | 0.257 | -0.012 | 0.036 | 0.036 | 0.079 | 0.56 | 0.74 | 0.49 | 0.7 | 0.47 | 0.72 |
| 2 | 212327353 | rs10804195 | 0.565 | -0.331 | -0.839 | 0.312 | 0.005 | -0.008 | -0.008 | 0.01 | 0.65 | 0.31 | 0.69 | 0.65 | 0.7 | 0.71 |
| 2 | 212330564 | rs4569408 | 0.108 | -0.11 | -0.544 | 0.346 | 0 | 0.064 | 0.064 | 0.365 | 0.79 | 0.39 | 0.73 | 0.71 | 0.81 | 0.84 |
| 2 | 212336730 | rs13383927 | 0.512 | 0.228 | -1.263 | 0.004 | -0.095 | 0.041 | 0.041 | 0 | 0.3 | 0.05 | 0.38 | 0.17 | 0.34 | 0.13 |
| 2 | 212350088 | rs4672614 | 0.802 | -0.105 | 0.211 | 0.347 | -0.009 | 0.101 | 0.101 | 0 | 0.82 | 0.65 | 0.76 | 0.7 | 0.82 | 0.86 |
| 2 | 212356992 | rs4672615 | -1.319 | 0.203 | 0.432 | 0.147 | 0.005 | 0.077 | 0.077 | 0 | 0.28 | 0.48 | 0.37 | 0.38 | 0.26 | 0.27 |
| 2 | 212360399 | rs13028369 | -0.003 | 0.209 | 0.375 | 0.131 | -0.009 | -0.046 | -0.046 | 0 | 0.14 | 0.09 | 0.23 | 0.38 | 0.15 | 0.19 |
| 2 | 212361074 | rs4278873 | -1.019 | -0.192 | 0.5 | -0.206 | -0.04 | -0.12 | -0.12 | 0.469 | 0.28 | 0.49 | 0.1 | 0.03 | 0.17 | 0.2 |
| 2 | 212361511 | rs4321319 | 0.224 | -0.252 | 0.576 | -0.112 | -0.036 | -0.043 | -0.043 | 0.569 | 0.17 | 0.25 | 0.06 | 0.03 | 0.07 | 0.11 |
| 2 | 212362263 | rs10932377 | -0.953 | 0.185 | 0.469 | 0.109 | 0.035 | -0.009 | -0.009 | 0 | 0.44 | 0.7 | 0.43 | 0.4 | 0.33 | 0.38 |
| 2 | 212367912 | rs6435629 | -0.709 | 0.245 | 0.361 | 0.176 | 0.001 | -0.001 | -0.001 | 0 | 0.42 | 0.57 | 0.41 | 0.41 | 0.32 | 0.37 |
| 2 | 212375386 | rs6725395 | -0.624 | 0.487 | -0.473 | 0.247 | 0.01 | 0.018 | 0.018 | 0 | 0.49 | 0.4 | 0.46 | 0.38 | 0.53 | 0.41 |
| 2 | 212378242 | rs16846352 | -0.903 | 0.348 | -0.234 | 0.122 | 0 | -0.067 | -0.067 | 0 | 0.11 | 0.59 | 0.05 | 0 | 0.08 | 0.06 |
| 2 | 212378524 | rs6435632 | 0.134 | 0.189 | -1.296 | 0.507 | 0.001 | -0.068 | -0.068 | 0.009 | 0.3 | 0.18 | 0.48 | 0.5 | 0.31 | 0.17 |
| 2 | 212378991 | rs10200942 | -0.334 | 0.001 | -1.129 | 0.077 | 0.019 | 0.05 | 0.05 | 0 | 0.38 | 0.21 | 0.25 | 0.23 | 0.31 | 0.37 |
| 2 | 212379238 | rs13408475 | -0.771 | -0.224 | -0.533 | -0.289 | -0.049 | -0.107 | -0.107 | 0 | 0.28 | 0.56 | 0.14 | 0.09 | 0.16 | 0.18 |
| 2 | 212382646 | rs10196354 | -0.236 | 0.023 | -0.218 | 0.318 | -0.006 | 0.008 | 0.008 | 0.042 | 0.27 | 0.57 | 0.36 | 0.34 | 0.25 | 0.09 |
| 2 | 212384231 | rs6740117 | 0.123 | -0.142 | -0.513 | -0.254 | 0.003 | 0.029 | 0.029 | 0 | 0.61 | 0.67 | 0.51 | 0.28 | 0.61 | 0.38 |
| 2 | 212390350 | rs10932380 | -0.865 | 0.163 | -0.491 | -0.381 | 0.004 | 0.028 | 0.028 | 0 | 0.62 | 0.69 | 0.65 | 0.79 | 0.63 | 0.68 |
| 2 | 212395971 | rs10932382 | 0.651 | 0.148 | -1.243 | 0.19 | -0.042 | -0.089 | -0.089 | 1.583 | 0.15 | 0.03 | 0.31 | 0.47 | 0.22 | 0.29 |
| 2 | 212403387 | rs17406681 | 1.15 | -0.053 | -1.124 | -0.476 | -0.081 | 0.027 | 0.027 | 0 | 0.18 | 0.04 | 0.26 | 0.11 | 0.25 | 0.05 |
| 2 | 212406789 | rs6747637 | -0.362 | -0.357 | -0.689 | -0.674 | -0.004 | -0.02 | -0.02 | 0 | 0.45 | 0.53 | 0.52 | 0.72 | 0.43 | 0.65 |
| 2 | 212407077 | rs9288433 | -1.01 | -0.082 | -0.461 | -0.453 | -0.007 | -0.043 | -0.043 | 0.007 | 0.68 | 0.63 | 0.81 | 0.83 | 0.69 | 0.7 |
| 2 | 212409676 | rs10932383 | -0.237 | 0.141 | -0.899 | -0.168 | 0.016 | 0.038 | 0.038 | 0 | 0.42 | 0.22 | 0.48 | 0.65 | 0.36 | 0.58 |
| 2 | 212409848 | rs10932384 | -0.283 | -0.098 | -0.636 | -0.184 | 0.006 | 0.063 | 0.063 | 0.112 | 0.59 | 0.79 | 0.68 | 0.73 | 0.58 | 0.59 |
| 2 | 212411658 | rs4435392 | -0.78 | -0.631 | 0.141 | 0.087 | -0.03 | -0.056 | -0.056 | 0.087 | 0.19 | 0.29 | 0.06 | 0.04 | 0.13 | 0.18 |
| 2 | 212412738 | rs11895509 | -0.048 | -0.322 | -0.27 | 0.793 | -0.023 | 0.029 | 0.029 | 0 | 0.93 | 0.99 | 0.79 | 0.51 | 0.89 | 0.71 |
| 2 | 212414621 | rs12470850 | -0.793 | -0.807 | 0.573 | -0.125 | -0.044 | -0.092 | -0.092 | 1.033 | 0.2 | 0.38 | 0.08 | 0.04 | 0.15 | 0.18 |
| 2 | 212415948 | rs12475861 | -1.513 | -0.877 | 0.816 | -0.207 | -0.015 | -0.037 | -0.037 | 0 | 0.25 | 0.66 | 0.13 | 0.06 | 0.16 | 0.25 |
| 2 | 212416029 | rs13421680 | -2.058 | -1.12 | 0.474 | -0.316 | 0.043 | 0.044 | 0.044 | 0.122 | 0.39 | 0.7 | 0.24 | 0.19 | 0.24 | 0.31 |
| 2 | 212418110 | rs10497945 | -1.974 | -1.2 | 1.063 | -0.987 | -0.013 | -0.055 | -0.055 | 0 | 0.37 | 0.61 | 0.13 | 0.06 | 0.2 | 0.28 |
| 2 | 212419760 | rs17413177 | 0.159 | -0.566 | 1.478 | -0.684 | 0.003 | -0.002 | -0.002 | 0 | 0.1 | 0.08 | 0.04 | 0.01 | 0.06 | 0.04 |
| 2 | 212423302 | rs7576673 | -1.412 | -0.644 | 1.732 | -0.726 | 0.01 | 0.009 | 0.009 | 0 | 0.24 | 0.17 | 0.08 | 0.02 | 0.17 | 0.15 |
| 2 | 212428192 | rs6733265 | -0.866 | -0.591 | 2.099 | -0.727 | -0.006 | -0.085 | -0.085 | 0.031 | 0.14 | 0.37 | 0.07 | 0.01 | 0.09 | 0.04 |
| 2 | 212428439 | rs7595473 | -1.78 | -0.577 | 1.797 | -0.716 | -0.007 | -0.047 | -0.047 | 0 | 0.3 | 0.64 | 0.13 | 0.06 | 0.2 | 0.2 |
| 2 | 212434037 | rs6735769 | -1.731 | -0.584 | 1.63 | -0.692 | -0.007 | -0.04 | -0.04 | 0.063 | 0.31 | 0.64 | 0.14 | 0.06 | 0.2 | 0.2 |
| 2 | 212435213 | rs12622730 | 1.521 | -0.593 | 1.443 | -0.58 | -0.038 | -0.063 | -0.063 | 0 | 0.7 | 0.47 | 0.88 | 0.88 | 0.83 | 0.8 |
| 2 | 212437161 | rs17335169 | 0.25 | -0.565 | 0.916 | -0.652 | 0 | 0.038 | 0.038 | 1.295 | 0.1 | 0.03 | 0.04 | 0 | 0.08 | 0.05 |
| 2 | 212440041 | rs10198754 | 1.495 | -0.413 | 1.566 | -0.421 | -0.038 | -0.056 | -0.056 | 2.101 | 0.78 | 0.44 | 0.91 | 0.9 | 0.89 | 0.84 |
| 2 | 212440500 | rs1009142 | 0.62 | -0.423 | 1.34 | -0.414 | 0.016 | 0.015 | 0.015 | 0 | 0.85 | 0.97 | 0.93 | 0.88 | 0.9 | 0.83 |
| 2 | 212453901 | rs7561282 | 1.485 | -0.359 | 1.817 | -0.414 | -0.031 | -0.054 | -0.054 | 0.865 | 0.79 | 0.47 | 0.88 | 0.88 | 0.88 | 0.83 |
| 2 | 212458790 | rs10203022 | 1.602 | -0.424 | 1.793 | -0.292 | -0.04 | -0.026 | -0.026 | 0 | 0.79 | 0.39 | 0.87 | 0.86 | 0.91 | 0.84 |
| 2 | 212464224 | rs1965462 | 0.686 | -0.502 | 1.685 | -0.285 | 0.024 | 0.011 | 0.011 | 0 | 0.85 | 0.88 | 0.93 | 0.88 | 0.93 | 0.84 |
| 2 | 212465328 | rs7556879 | 1.365 | -0.494 | 1.758 | -0.284 | -0.041 | -0.023 | -0.023 | 0 | 0.81 | 0.59 | 0.88 | 0.86 | 0.9 | 0.85 |
| 2 | 212471642 | rs7559679 | 0.801 | -0.467 | 1.517 | -0.149 | -0.018 | -0.001 | -0.001 | 1.807 | 0.83 | 0.77 | 0.91 | 0.83 | 0.92 | 0.89 |
| 2 | 212475771 | rs10177211 | 0.696 | -0.347 | 1.127 | -0.027 | -0.038 | 0.056 | 0.056 | 0.558 | 0.83 | 0.43 | 0.87 | 0.77 | 0.92 | 0.88 |

| | | | | | | | | | | | | | | | | |
|---|---|---|---|---|---|---|---|---|---|---|---|---|---|---|---|---|
| 2 | 212486350 | rs10469656 | 0.786 | -0.138 | 0.546 | -0.135 | -0.046 | 0.028 | 0.028 | 1.89 | 0.8 | 0.46 | 0.89 | 0.79 | 0.91 | 0.9 |
| 2 | 212519028 | rs6730119 | 0.172 | 0.899 | -0.486 | -0.545 | -0.005 | 0.003 | 0.003 | 0 | 0.64 | 0.65 | 0.54 | 0.13 | 0.57 | 0.41 |
| 2 | 212521325 | rs7567959 | 0.681 | 0.917 | -0.281 | -0.282 | 0.009 | -0.003 | -0.003 | 1.668 | 0.7 | 0.71 | 0.62 | 0.29 | 0.57 | 0.55 |
| 2 | 212527729 | rs1851185 | 0.391 | 0.979 | 0.846 | -0.4 | 0.02 | 0.029 | 0.029 | 0 | 0.78 | 0.62 | 0.74 | 0.34 | 0.78 | 0.64 |
| 2 | 212529506 | rs4672621 | 1.125 | 0.767 | 0.786 | -0.498 | 0.013 | -0.017 | -0.017 | 0 | 0.65 | 0.48 | 0.58 | 0.3 | 0.58 | 0.5 |
| 2 | 212530466 | rs6725181 | -0.719 | 0.746 | 0.737 | -0.829 | -0.024 | -0.024 | -0.024 | 0 | 0.15 | 0.41 | 0.23 | 0.09 | 0.09 | 0.14 |
| 2 | 212534520 | rs12476100 | -0.708 | 0.871 | 0.347 | -0.89 | -0.033 | 0.07 | 0.07 | 0 | 0.3 | 0.16 | 0.39 | 0.88 | 0.33 | 0.51 |
| 2 | 212538415 | rs13019783 | 0.574 | 0.787 | 0.307 | -0.952 | 0.003 | 0.003 | 0.003 | 0 | 0.1 | 0.06 | 0.18 | 0.09 | 0.1 | 0.13 |
| 2 | 212541377 | rs10207288 | 0.411 | 1.058 | 0.427 | -0.995 | -0.005 | 0.038 | 0.038 | 0 | 0.7 | 0.79 | 0.62 | 0.1 | 0.7 | 0.49 |
| 2 | 212542748 | rs7565327 | 1.147 | 1.184 | 0.454 | -1.018 | -0.001 | -0.07 | -0.07 | 3.243 | 0.54 | 0.56 | 0.37 | 0.01 | 0.45 | 0.32 |
| 2 | 212555085 | rs2371276 | 1.113 | 1.188 | -0.003 | -0.641 | -0.001 | 0.019 | 0.019 | 0 | 0.54 | 0.48 | 0.41 | 0.14 | 0.52 | 0.37 |
| 2 | 212558540 | rs7425448 | -0.799 | 1.467 | -0.062 | -0.32 | 0.014 | -0.014 | -0.014 | 0 | 0.08 | 0.11 | 0.08 | 0.03 | 0.17 | 0.05 |
| 2 | 212564895 | rs6435659 | -0.888 | 1.444 | -0.326 | -0.006 | -0.008 | 0.005 | 0.005 | 0 | 0.35 | 0.33 | 0.5 | 0.35 | 0.44 | 0.47 |
| 2 | 212571939 | rs6435660 | 0.442 | 1.065 | -0.554 | -0.417 | 0.009 | 0.033 | 0.033 | 1.734 | 0.55 | 0.44 | 0.38 | 0.09 | 0.6 | 0.31 |
| 2 | 212577231 | rs6745717 | -1.273 | 1.261 | -0.331 | -0.365 | 0.026 | -0.039 | -0.039 | 0 | 0.13 | 0.25 | 0.09 | 0.04 | 0.16 | 0.1 |
| 2 | 212578978 | rs13032114 | -0.903 | 1.238 | -1.157 | -0.19 | -0.038 | -0.064 | -0.064 | 0.012 | 0.6 | 0.85 | 0.66 | 0.55 | 0.57 | 0.71 |
| 2 | 212579897 | rs12162287 | -0.862 | 1.342 | -1.828 | -0.498 | -0.039 | -0.124 | -0.124 | 0.08 | 0.75 | 0.96 | 0.79 | 0.55 | 0.73 | 0.74 |
| 2 | 212581902 | rs6729362 | 0.531 | 1.059 | -1.788 | -0.494 | 0.001 | -0.005 | -0.005 | 0 | 0.13 | 0.11 | 0.13 | 0.19 | 0.08 | 0.26 |
| 2 | 212582246 | rs6758063 | -0.367 | 0.721 | -1.451 | -0.698 | 0 | 0.002 | 0.002 | 0.504 | 0.67 | 0.64 | 0.71 | 0.46 | 0.64 | 0.64 |
| 2 | 212584780 | rs7604206 | -0.697 | 0.525 | -1.423 | -0.687 | -0.003 | 0.009 | 0.009 | 0 | 0.7 | 0.65 | 0.73 | 0.46 | 0.71 | 0.66 |
| 2 | 212585422 | rs7591137 | -0.411 | 0.409 | -1.773 | -0.995 | -0.007 | -0.019 | -0.019 | 0 | 0.33 | 0.22 | 0.42 | 0.44 | 0.29 | 0.53 |
| 2 | 212587321 | rs13002712 | -1.427 | 0.378 | -1.998 | -0.909 | -0.11 | -0.112 | -0.112 | 0.448 | 0.64 | 0.95 | 0.65 | 0.49 | 0.55 | 0.74 |
| 2 | 212588627 | rs7421820 | -2.226 | 0.525 | -1.926 | -0.459 | -0.031 | -0.057 | -0.057 | 0 | 0.86 | 0.96 | 0.82 | 0.56 | 0.8 | 0.87 |
| 2 | 212591641 | rs6435665 | 0.822 | 0.199 | -1.85 | -0.691 | -0.006 | 0.017 | 0.017 | 0 | 0.17 | 0.18 | 0.14 | 0.28 | 0.11 | 0.37 |
| 2 | 212596915 | rs13026255 | 1.428 | -0.158 | -1.911 | -0.976 | -0.031 | 0.091 | 0.091 | 0.201 | 0.29 | 0.12 | 0.28 | 0 | 0.39 | 0.11 |
| 2 | 212597041 | rs6711487 | -1.774 | -0.193 | -1.681 | -1.063 | -0.037 | -0.063 | -0.063 | 0.398 | 0.69 | 0.86 | 0.6 | 0.53 | 0.61 | 0.81 |
| 2 | 212599229 | rs13015253 | -0.43 | -0.746 | -1.62 | -1.091 | 0.027 | -0.021 | -0.021 | 0 | 0.34 | 0.7 | 0.3 | 0.48 | 0.13 | 0.51 |
| 2 | 212604921 | rs6722322 | -0.165 | -0.292 | -1.254 | -0.981 | 0.007 | 0.003 | 0.003 | 0 | 0.39 | 0.38 | 0.28 | 0.28 | 0.34 | 0.47 |
| 2 | 212615516 | rs6757068 | -0.088 | -0.132 | -0.903 | -0.725 | -0.001 | 0.007 | 0.007 | 0.384 | 0.55 | 0.43 | 0.43 | 0.29 | 0.52 | 0.51 |
| 2 | 212626917 | rs7588792 | -2.532 | 0.086 | -1.139 | -0.331 | -0.008 | -0.01 | -0.01 | 0 | 0.84 | 0.9 | 0.73 | 0.6 | 0.8 | 0.84 |
| 2 | 212627259 | rs10932398 | 0.486 | -0.05 | -1.08 | -0.665 | 0.011 | 0.052 | 0.052 | 0.128 | 0.52 | 0.29 | 0.42 | 0.29 | 0.5 | 0.51 |
| 2 | 212629579 | rs7570613 | -2.083 | 0.119 | -1.098 | -0.637 | -0.026 | -0.035 | -0.035 | 0 | 0.83 | 0.91 | 0.7 | 0.54 | 0.73 | 0.83 |
| 2 | 212642288 | rs13008313 | -0.541 | -0.34 | -1.224 | -0.997 | 0.005 | 0.036 | 0.036 | 0.475 | 0.44 | 0.24 | 0.27 | 0.32 | 0.37 | 0.6 |
| 2 | 212642374 | rs13008370 | -0.548 | -0.337 | -1.212 | -0.97 | 0.005 | 0.029 | 0.029 | 0.536 | 0.44 | 0.24 | 0.27 | 0.33 | 0.37 | 0.6 |
| 2 | 212645885 | rs7423708 | 1.348 | -0.406 | -0.834 | -1.2 | 0.002 | 0.007 | 0.007 | 0 | 0.31 | 0.24 | 0.4 | 0.19 | 0.34 | 0.21 |
| 2 | 212650162 | rs6756725 | 0.935 | -0.027 | -0.895 | -1.135 | -0.042 | 0.038 | 0.038 | 1.118 | 0.16 | 0.11 | 0.12 | 0.06 | 0.23 | 0.16 |
| 2 | 212650985 | rs7588431 | -0.321 | -0.196 | -0.623 | -1.203 | 0 | -0.031 | -0.031 | 2.549 | 0.6 | 0.57 | 0.55 | 0.25 | 0.59 | 0.43 |
| 2 | 212655885 | rs6435670 | -0.372 | -0.309 | -0.275 | -0.721 | 0.005 | -0.026 | -0.026 | 0 | 0.24 | 0.32 | 0.15 | 0.07 | 0.25 | 0.18 |
| 2 | 212658297 | rs4673633 | -0.087 | -0.406 | -0.149 | -0.198 | -0.005 | 0.005 | 0.005 | 0.021 | 0.22 | 0.23 | 0.16 | 0.27 | 0.16 | 0.4 |
| 2 | 212670425 | rs4672626 | -0.428 | -0.268 | -0.223 | -0.594 | -0.01 | -0.013 | -0.013 | 0 | 0.87 | 0.83 | 0.95 | 0.95 | 0.9 | 0.89 |
| 2 | 212683946 | rs1371203 | 0.621 | -0.1 | -0.443 | -0.639 | 0 | 0 | 0 | 1.512 | 0.76 | 0.73 | 0.73 | 0.71 | 0.79 | 0.76 |
| 2 | 212688964 | rs12475812 | 0.171 | -0.024 | -0.416 | -0.594 | 0.002 | 0.002 | 0.002 | 0 | 0.85 | 0.8 | 0.78 | 0.71 | 0.86 | 0.78 |
| 2 | 212695168 | rs16846896 | 0.361 | -0.025 | -0.308 | -0.565 | -0.018 | 0.052 | 0.052 | 0 | 0.08 | 0.25 | 0.17 | 0.19 | 0.05 | 0.1 |
| 2 | 212697542 | rs16846917 | 0.231 | -0.052 | -0.72 | -0.916 | 0.007 | 0.001 | 0.001 | 0 | 0.28 | 0.23 | 0.19 | 0.26 | 0.25 | 0.35 |
| 2 | 212698718 | rs714393 | 1.081 | -0.079 | -0.689 | -1.029 | -0.029 | -0.029 | -0.029 | 0 | 0.39 | 0.19 | 0.51 | 0.41 | 0.48 | 0.32 |
| 2 | 212703221 | rs10208623 | 1.188 | -0.11 | -0.942 | -1.047 | -0.121 | -0.129 | -0.129 | 0.067 | 0.38 | 0.04 | 0.5 | 0.42 | 0.48 | 0.32 |
| 2 | 212703601 | rs1020126 | -0.826 | -0.021 | -0.733 | -0.984 | 0.003 | 0.003 | 0.003 | 0 | 0.83 | 0.74 | 0.79 | 0.72 | 0.86 | 0.82 |
| 2 | 212709028 | rs1439235 | 1.113 | 0.063 | -1.172 | -0.821 | -0.005 | -0.025 | -0.025 | 0.066 | 0.18 | 0.1 | 0.15 | 0.26 | 0.18 | 0.33 |
| 2 | 212712835 | rs13407198 | 1.751 | 0.132 | -1.259 | -0.804 | -0.07 | -0.156 | -0.156 | 0 | 0.27 | 0.03 | 0.38 | 0.41 | 0.33 | 0.24 |
| 2 | 212719203 | rs13028597 | 0.916 | 0.079 | -1.106 | -0.794 | 0 | -0.001 | -0.001 | 0 | 0.21 | 0.22 | 0.15 | 0.26 | 0.18 | 0.33 |
| 2 | 212722915 | rs1025753 | 1.784 | 0.013 | -1.251 | -0.801 | -0.071 | -0.144 | -0.144 | 0 | 0.27 | 0.05 | 0.38 | 0.4 | 0.33 | 0.24 |
| 2 | 212722988 | rs16847102 | 0.927 | -0.031 | -1.072 | -0.906 | -0.007 | -0.042 | -0.042 | 0 | 0.15 | 0.27 | 0.07 | 0.04 | 0.13 | 0.12 |
| 2 | 212723026 | rs984773 | 0.594 | 0.091 | -1.065 | -0.685 | -0.003 | -0.007 | -0.007 | 0 | 0.22 | 0.21 | 0.15 | 0.26 | 0.19 | 0.33 |
| 2 | 212723459 | rs1025752 | 0.436 | 0.319 | -0.954 | -0.478 | -0.001 | 0 | 0 | 0 | 0.43 | 0.38 | 0.46 | 0.5 | 0.46 | 0.38 |
| 2 | 212736394 | rs16847152 | -0.015 | 0.336 | -1.179 | -0.373 | 0.002 | 0 | 0 | 0 | 0.61 | 0.48 | 0.64 | 0.54 | 0.64 | 0.49 |
| 2 | 212737233 | rs12470608 | 0.374 | 0.67 | -1.69 | -0.345 | -0.004 | 0.038 | 0.038 | 0 | 0.14 | 0.07 | 0.12 | 0.05 | 0.17 | 0.08 |
| 2 | 212744186 | rs2068401 | -0.164 | 0.701 | -1.595 | -0.33 | -0.022 | 0.037 | 0.037 | 0 | 0.57 | 0.69 | 0.36 | 0.27 | 0.53 | 0.58 |
| 2 | 212752193 | rs6727114 | 0.472 | 0.636 | -2.079 | 0.008 | -0.073 | -0.121 | -0.121 | 0.836 | 0.28 | 0.03 | 0.41 | 0.37 | 0.32 | 0.19 |
| 2 | 212764825 | rs16847170 | 0.315 | 0.227 | -1.503 | -0.73 | -0.002 | -0.026 | -0.026 | 5.201 | 0.11 | 0.14 | 0.06 | 0 | 0.09 | 0.03 |
| 2 | 212765359 | rs16847171 | 0.296 | 0.235 | -1.435 | -0.73 | -0.007 | -0.044 | -0.044 | 0.908 | 0.11 | 0.16 | 0.06 | 0 | 0.09 | 0.03 |
| 2 | 212768779 | rs2043888 | 0.543 | 0.58 | -1.754 | -0.47 | -0.036 | 0.09 | 0.09 | 0.411 | 0.15 | 0.01 | 0.13 | 0 | 0.18 | 0.04 |
| 2 | 212771670 | rs13391359 | -0.814 | 0.464 | -1.074 | 0.002 | -0.004 | -0.009 | -0.009 | 0.325 | 0.15 | 0.14 | 0.18 | 0.14 | 0.14 | 0.32 |
| 2 | 212781914 | rs1347889 | -0.246 | 0.394 | -1.365 | -0.021 | -0.005 | 0.036 | 0.036 | 0 | 0.48 | 0.69 | 0.63 | 0.64 | 0.49 | 0.61 |
| 2 | 212783175 | rs6435681 | 0.188 | -0.667 | -1.617 | -0.756 | 0.042 | 0.058 | 0.058 | 0 | 0.72 | 0.81 | 0.79 | 0.91 | 0.78 | 0.8 |
| 2 | 212791249 | rs16847207 | 0.39 | -0.438 | -0.708 | -0.687 | 0.026 | -0.004 | -0.004 | 0 | 0.1 | 0.29 | 0.12 | 0.11 | 0.08 | 0.05 |
| 2 | 212793198 | rs7598440 | 0.54 | -0.735 | -1.619 | -0.606 | -0.101 | -0.135 | -0.135 | 0.145 | 0.29 | 0.11 | 0.38 | 0.52 | 0.53 | 0.33 |
| 2 | 212796981 | rs10180494 | 0.711 | -1.094 | -1.588 | -0.668 | -0.087 | -0.133 | -0.133 | 0 | 0.29 | 0.12 | 0.39 | 0.54 | 0.55 | 0.33 |
| 2 | 212802765 | rs839530 | 1.004 | -1.073 | -0.937 | -0.57 | 0.036 | 0.012 | 0.012 | 0 | 0.09 | 0.27 | 0.13 | 0.12 | 0.11 | 0.05 |
| 2 | 212808107 | rs839533 | -0.779 | -2.16 | -1.164 | -1.094 | 0.11 | -0.047 | -0.047 | 0 | 0.66 | 0.86 | 0.71 | 0.66 | 0.84 | 0.55 |

| | | | | | | | | | | | | | | | | |
|---|---|---|---|---|---|---|---|---|---|---|---|---|---|---|---|---|
| 2 | 212816089 | rs839523 | -1.162 | -1.541 | -0.522 | -0.992 | 0.043 | 0.01 | 0.01 | 0 | 0.45 | 0.64 | 0.46 | 0.37 | 0.3 | 0.49 |
| 2 | 212817010 | rs839521 | -0.945 | -1.633 | -0.205 | -1.05 | -0.005 | 0 | 0 | 0.397 | 0.41 | 0.52 | 0.43 | 0.37 | 0.27 | 0.44 |
| 2 | 212820953 | rs13428633 | 1.104 | -1.673 | 0.225 | -1.087 | 0.037 | -0.005 | -0.005 | 0 | 0.06 | 0.25 | 0.03 | 0 | 0.03 | 0.02 |
| 2 | 212824999 | rs12616932 | -0.61 | -1.683 | -0.22 | -1.097 | 0.031 | -0.034 | -0.034 | 0 | 0.34 | 0.17 | 0.39 | 0.38 | 0.24 | 0.44 |
| 2 | 212828793 | rs6744796 | -0.944 | -1.627 | -0.211 | -1.056 | -0.003 | 0 | 0 | 0 | 0.38 | 0.34 | 0.43 | 0.38 | 0.27 | 0.47 |
| 2 | 212832434 | rs1402708 | -1.26 | -1.541 | -0.205 | -1.046 | 0.029 | -0.012 | -0.012 | 0 | 0.93 | 0.72 | 0.97 | 1 | 0.97 | 0.98 |
| 2 | 212839046 | rs707284 | 1.114 | -1.395 | -1.273 | -0.953 | 0.056 | 0.037 | 0.037 | 0 | 0.51 | 0.36 | 0.49 | 0.63 | 0.67 | 0.49 |
| 2 | 212841868 | rs2102996 | 0.544 | -0.869 | -1.529 | -0.026 | 0.047 | -0.089 | -0.089 | 0 | 0.14 | 0.01 | 0.19 | 0.25 | 0.05 | 0.3 |
| 2 | 212845292 | rs12473131 | -0.396 | -0.859 | -1.484 | -0.062 | 0.018 | -0.038 | -0.038 | 0 | 0.23 | 0.13 | 0.33 | 0.36 | 0.17 | 0.35 |
| 2 | 212845994 | rs12613571 | -0.103 | -0.956 | -1.666 | -0.118 | 0.059 | -0.102 | -0.102 | 0 | 0.19 | 0.01 | 0.2 | 0.27 | 0.07 | 0.35 |
| 2 | 212846898 | rs7563378 | 0.075 | -0.413 | -0.975 | 0.067 | 0.008 | 0.148 | 0.148 | 0.988 | 0.82 | 0.55 | 0.76 | 0.41 | 0.82 | 0.74 |
| 2 | 212847371 | rs9288445 | 0.671 | -1.018 | -1.002 | -0.6 | 0.001 | -0.004 | -0.004 | 0.017 | 0.73 | 0.23 | 0.72 | 0.4 | 0.84 | 0.75 |
| 2 | 212852895 | rs2860059 | 0.892 | -0.961 | -0.939 | -1.01 | 0.011 | -0.009 | -0.009 | 0 | 0.69 | 0.22 | 0.72 | 0.38 | 0.81 | 0.69 |
| 2 | 212856539 | rs12987596 | -0.17 | -1.235 | -1.691 | -1.579 | 0 | 0.001 | 0.001 | 0.025 | 0.42 | 0.39 | 0.51 | 0.35 | 0.39 | 0.46 |
| 2 | 212861363 | rs10176432 | -0.637 | -1.258 | -1.539 | -1.447 | -0.062 | 0.183 | 0.183 | 0.073 | 0.82 | 0.53 | 0.8 | 0.4 | 0.91 | 0.74 |
| 2 | 212864338 | rs10174063 | 0.987 | -1.454 | -1.828 | -1.68 | 0.03 | 0.025 | 0.025 | 1.708 | 0.56 | 0.18 | 0.58 | 0.32 | 0.64 | 0.53 |
| 2 | 212866473 | rs13020474 | 1.357 | -1.51 | -1.951 | -1.736 | -0.016 | -0.116 | -0.116 | 0 | 0.19 | 0.01 | 0.29 | 0.31 | 0.15 | 0.3 |
| 2 | 212867291 | rs7423124 | 1.243 | -1.385 | -2.177 | -1.978 | 0.041 | 0.042 | 0.042 | 0.188 | 0.51 | 0.18 | 0.54 | 0.33 | 0.61 | 0.46 |
| 2 | 212868624 | rs1879637 | 1.052 | -1.411 | -1.835 | -2.144 | 0.002 | -0.157 | -0.157 | 1.196 | 0.67 | 0.46 | 0.75 | 0.94 | 0.71 | 0.68 |
| 2 | 212871592 | rs960824 | 1.068 | -1.47 | -1.954 | -2.101 | 0.011 | 0.014 | 0.014 | 0.66 | 0.66 | 0.24 | 0.74 | 0.93 | 0.75 | 0.73 |
| 2 | 212882190 | rs1473636 | -0.099 | -1.479 | -1.976 | -2.076 | -0.009 | -0.028 | -0.028 | 0.07 | 0.8 | 0.22 | 0.86 | 0.93 | 0.9 | 0.81 |
| 2 | 212883383 | rs1473638 | 0.071 | -1.35 | -1.751 | -2.058 | 0.005 | -0.107 | -0.107 | 0 | 0.79 | 0.49 | 0.82 | 0.93 | 0.81 | 0.81 |
| 2 | 212891401 | rs2177035 | 0.061 | -1.316 | -1.642 | -2.056 | 0.034 | -0.112 | -0.112 | 0 | 0.83 | 0.58 | 0.86 | 1 | 0.83 | 0.86 |
| 2 | 212895366 | rs12694259 | 0.958 | -1.529 | -1.912 | -2.041 | 0.047 | 0.063 | 0.063 | 2.276 | 0.53 | 0.2 | 0.53 | 0.66 | 0.64 | 0.45 |
| 2 | 212896904 | rs12464239 | 1.456 | -1.61 | -1.943 | -2.078 | 0.012 | -0.097 | -0.097 | 0 | 0.19 | 0.01 | 0.26 | 0.27 | 0.12 | 0.33 |
| 2 | 212902339 | rs1546717 | 0.484 | -1.302 | -1.474 | -2.427 | 0.017 | 0.012 | 0.012 | 9.362 | 0.76 | 0.18 | 0.85 | 0.94 | 0.88 | 0.8 |
| 2 | 212913183 | rs10445735 | 0.611 | -1.339 | -1.468 | -2.46 | 0.018 | -0.083 | -0.083 | 0.063 | 0.13 | 0.35 | 0.09 | 0 | 0.12 | 0.05 |
| 2 | 212914726 | rs1464443 | 1.205 | -1.367 | -1.489 | -2.5 | -0.008 | 0.161 | 0.161 | 6.058 | 0.67 | 0.1 | 0.74 | 0.92 | 0.7 | 0.77 |
| 2 | 212916157 | rs6712126 | -0.293 | -1.389 | -1.428 | -2.415 | -0.035 | -0.11 | -0.11 | 0 | 0.23 | 0.43 | 0.15 | 0.08 | 0.16 | 0.18 |
| 2 | 212916758 | rs939645 | -0.363 | -1.385 | -1.405 | -2.397 | -0.028 | -0.112 | -0.112 | 1.234 | 0.24 | 0.43 | 0.17 | 0.08 | 0.18 | 0.18 |
| 2 | 212927789 | rs17343843 | 1.161 | -1.343 | -1.555 | -2.377 | -0.009 | 0.05 | 0.05 | 0 | 0.09 | 0.01 | 0.05 | 0 | 0.05 | 0.09 |
| 2 | 212929113 | rs10197270 | 1.119 | -1.409 | -1.548 | -2.459 | 0.045 | 0.119 | 0.119 | 3.329 | 0.63 | 0.16 | 0.74 | 0.92 | 0.76 | 0.71 |
| 2 | 212931980 | rs12694262 | -0.389 | -1.442 | -1.398 | -2.44 | 0.017 | -0.07 | -0.07 | 0 | 0.88 | 0.57 | 0.9 | 1 | 0.93 | 0.93 |
| 2 | 212936891 | rs12373751 | 1.15 | -1.452 | -1.446 | -2.523 | 0.053 | 0.117 | 0.117 | 3.028 | 0.63 | 0.16 | 0.74 | 0.93 | 0.77 | 0.71 |
| 2 | 212938165 | rs1521545 | 0.081 | -1.44 | -1.345 | -2.444 | -0.028 | -0.123 | -0.123 | 0 | 0.83 | 0.56 | 0.9 | 1 | 0.92 | 0.88 |
| 2 | 212945959 | rs10497958 | 0.975 | -1.537 | -1.66 | -2.633 | -0.044 | 0.053 | 0.053 | 0 | 0.14 | 0.02 | 0.13 | 0.08 | 0.18 | 0.16 |
| 2 | 212948287 | rs1357127 | -0.786 | -1.503 | -1.441 | -2.615 | -0.009 | -0.081 | -0.081 | 0 | 0.87 | 0.63 | 0.92 | 1 | 0.95 | 0.89 |
| 2 | 212958457 | rs1521550 | 1.377 | -1.511 | -1.633 | -2.634 | 0.011 | 0.199 | 0.199 | 0.471 | 0.56 | 0.17 | 0.68 | 0.93 | 0.64 | 0.65 |
| 2 | 212961992 | rs7564926 | 0.482 | -1.994 | -1.341 | -2.599 | -0.01 | -0.159 | -0.159 | 0 | 0.76 | 0.5 | 0.84 | 1 | 0.83 | 0.82 |
| 2 | 212965531 | rs2371438 | 1.13 | -2.038 | -1.487 | -2.39 | 0.044 | 0.021 | 0.021 | 0 | 0.14 | 0.06 | 0.09 | 0.06 | 0.06 | 0.27 |
| 2 | 212969560 | rs10189975 | 1.438 | -1.383 | -1.12 | -2.577 | 0.075 | -0.064 | -0.064 | 0 | 0.25 | 0.33 | 0.25 | 0.1 | 0.45 | 0.21 |
| 2 | 212973223 | rs17344252 | 1.348 | -1.94 | -1.354 | -2.424 | 0.061 | 0.004 | 0.004 | 0 | 0.13 | 0 | 0.08 | 0.06 | 0.05 | 0.25 |
| 2 | 212974828 | rs17416172 | 1.025 | -2.024 | -1.085 | -2.79 | -0.027 | 0.033 | 0.033 | 0 | 0.34 | 0.25 | 0.31 | 0.08 | 0.53 | 0.28 |
| 2 | 212975064 | rs6435689 | -0.759 | -2.408 | -1.342 | -3.034 | -0.05 | 0.124 | 0.124 | 0.312 | 0.67 | 0.92 | 0.51 | 0.14 | 0.66 | 0.67 |
| 2 | 212975239 | rs6435690 | 0.443 | -2.336 | -1.148 | -2.989 | 0.011 | 0.11 | 0.11 | 0 | 0.48 | 0.28 | 0.39 | 0.14 | 0.58 | 0.53 |
| 2 | 212975953 | rs2371444 | 1.738 | -2.369 | -1.097 | -3.002 | -0.064 | 0.065 | 0.065 | 0 | 0.29 | 0.15 | 0.28 | 0.08 | 0.49 | 0.24 |
| 2 | 212984455 | rs7426206 | -0.286 | -2.081 | -0.466 | -2.837 | 0.003 | 0.052 | 0.052 | 0.091 | 0.63 | 0.77 | 0.47 | 0.13 | 0.65 | 0.61 |
| 2 | 212987580 | rs16847741 | 0.357 | -2.081 | -0.345 | -2.816 | -0.006 | -0.04 | -0.04 | 0 | 0.09 | 0.29 | 0.03 | 0.01 | 0.04 | 0.04 |
| 2 | 212989308 | rs6738304 | 0.643 | -2.131 | -0.41 | -2.863 | 0.007 | 0.029 | 0.029 | 0 | 0.51 | 0.44 | 0.39 | 0.12 | 0.57 | 0.52 |
| 2 | 212990868 | rs7560730 | -1.354 | -2.201 | -0.949 | -2.857 | -0.034 | -0.047 | -0.047 | 7.084 | 0.86 | 0.98 | 0.66 | 0.2 | 0.84 | 0.76 |
| 2 | 212997024 | rs11680307 | 0.605 | -2.193 | -0.735 | -3.022 | 0.002 | -0.012 | -0.012 | 0.936 | 0.55 | 0.48 | 0.45 | 0.13 | 0.65 | 0.52 |
| 2 | 212999007 | rs2888051 | 1.642 | -2.209 | -0.888 | -3.024 | -0.107 | 0.191 | 0.191 | 1.494 | 0.4 | 0.07 | 0.36 | 0.13 | 0.54 | 0.47 |
| 2 | 213003179 | rs12694263 | 1.889 | -2.199 | -0.889 | -3.008 | -0.004 | 0.064 | 0.064 | 0 | 0.14 | 0.02 | 0.12 | 0.01 | 0.15 | 0.1 |
| 2 | 213006392 | rs1568519 | 1.125 | -2.015 | -0.426 | -2.761 | 0.033 | -0.041 | -0.041 | 0.269 | 0.08 | 0.29 | 0.07 | 0 | 0.06 | 0.09 |
| 2 | 213008886 | rs6435692 | -0.055 | -1.96 | -1.01 | -2.74 | 0.006 | 0.057 | 0.057 | 0 | 0.62 | 0.77 | 0.48 | 0.12 | 0.64 | 0.59 |
| 2 | 213009392 | rs6732730 | 0.959 | -1.716 | -1.847 | -2.491 | -0.009 | -0.042 | -0.042 | 0 | 0.07 | 0.2 | 0.03 | 0 | 0.03 | 0.04 |
| 2 | 213017462 | rs1357142 | -0.621 | -1.513 | -2.938 | -2.324 | -0.034 | 0.01 | 0.01 | 0.117 | 0.72 | 0.96 | 0.55 | 0.22 | 0.68 | 0.67 |
| 2 | 213017607 | rs6715314 | 0.318 | -0.824 | -2.434 | -1.949 | -0.096 | 0.218 | 0.218 | 2.165 | 0.42 | 0.08 | 0.36 | 0.12 | 0.53 | 0.47 |
| 2 | 213028059 | rs16847823 | -0.04 | -1.283 | -1.704 | -1.347 | -0.018 | -0.007 | -0.007 | 0 | 0.23 | 0.23 | 0.13 | 0.17 | 0.12 | 0.34 |
| 2 | 213033301 | rs1402766 | 0.711 | -1.189 | -2.176 | -1.663 | 0.03 | 0.073 | 0.073 | 0 | 0.56 | 0.68 | 0.49 | 0.09 | 0.66 | 0.42 |
| 2 | 213033533 | rs17416620 | 0.689 | -0.673 | -2.817 | -1.432 | -0.026 | 0.024 | 0.024 | 0 | 0.08 | 0.04 | 0.09 | 0 | 0.13 | 0.06 |
| 2 | 213040127 | rs1521647 | -0.938 | -0.584 | -2.771 | -1.171 | -0.017 | -0.068 | -0.068 | 0.279 | 0.9 | 0.99 | 0.71 | 0.26 | 0.91 | 0.8 |
| 2 | 213040157 | rs17344783 | -0.272 | -0.961 | -2.803 | -1.451 | 0.069 | 0.118 | 0.118 | 0.267 | 0.39 | 0.85 | 0.26 | 0.17 | 0.27 | 0.38 |
| 2 | 213040853 | rs10200506 | 0.917 | -0.639 | -2.667 | -1.911 | 0.004 | -0.002 | -0.002 | 3.826 | 0.16 | 0.1 | 0.14 | 0.01 | 0.17 | 0.08 |
| 2 | 213040865 | rs7591480 | 0.392 | -0.777 | -1.994 | -1.561 | 0.039 | 0.081 | 0.081 | 0.019 | 0.57 | 0.27 | 0.47 | 0.09 | 0.68 | 0.5 |
| 2 | 213042489 | rs3942465 | -0.51 | -0.306 | -1.936 | -0.974 | -0.027 | -0.002 | -0.002 | 0 | 0.79 | 0.98 | 0.61 | 0.17 | 0.78 | 0.7 |
| 2 | 213043071 | rs10497961 | -0.245 | -0.604 | -0.915 | -1.058 | -0.013 | -0.045 | -0.045 | 0 | 0.1 | 0.12 | 0.04 | 0.01 | 0.06 | 0.08 |
| 2 | 213050781 | rs1357139 | 0.511 | -0.47 | -0.438 | -1.418 | 0.037 | -0.039 | -0.039 | 0 | 0.72 | 0.47 | 0.74 | 0.82 | 0.64 | 0.73 |
| 2 | 213053239 | rs6755027 | 1.159 | -1.197 | -0.469 | -1.75 | 0.003 | 0.165 | 0.165 | 0 | 0.5 | 0.12 | 0.54 | 0.79 | 0.52 | 0.42 |

| | | | | | | | | | | | | | | | | |
|---|---|---|---|---|---|---|---|---|---|---|---|---|---|---|---|---|
| 2 | 213054058 | rs13017883 | 0.445 | -1.21 | 0.028 | -1.909 | -0.008 | -0.058 | -0.058 | 0.611 | 0.11 | 0.21 | 0.04 | 0.01 | 0.08 | 0.13 |
| 2 | 213061726 | rs6435699 | 0.024 | -1.338 | 0.209 | -1.96 | 0.026 | 0.031 | 0.031 | 0 | 0.18 | 0.15 | 0.18 | 0.02 | 0.12 | 0.3 |
| 2 | 213067749 | rs6707285 | -1.924 | -1.246 | 0.134 | -2.084 | -0.001 | 0.233 | 0.233 | 0 | 0.53 | 0.86 | 0.4 | 0.05 | 0.5 | 0.6 |
| 2 | 213073599 | rs10932426 | 0.743 | -1.261 | 0.225 | -2.044 | -0.011 | 0.004 | 0.004 | 0 | 0.79 | 0.74 | 0.78 | 0.83 | 0.72 | 0.85 |
| 2 | 213073735 | rs6710350 | -0.2 | -1.477 | 0.068 | -2.178 | 0.033 | 0.082 | 0.082 | 0 | 0.2 | 0.05 | 0.18 | 0.04 | 0.14 | 0.39 |
| 2 | 213074560 | rs12475565 | -0.874 | -1.483 | 0.33 | -2.184 | 0.033 | 0.039 | 0.039 | 0 | 0.29 | 0.18 | 0.2 | 0.04 | 0.18 | 0.46 |
| 2 | 213076113 | rs17417113 | 0.489 | -1.363 | 0.111 | -2.125 | -0.034 | 0.028 | 0.028 | 0 | 0.1 | 0.09 | 0.1 | 0.01 | 0.23 | 0.07 |
| 2 | 213083282 | rs10048673 | 2.633 | -1.434 | 0.29 | -2.143 | -0.071 | 0.098 | 0.098 | 0 | 0.4 | 0.08 | 0.5 | 0.79 | 0.45 | 0.32 |
| 2 | 213089095 | rs939224 | 2.283 | -1.477 | 0.661 | -2.143 | -0.001 | -0.001 | -0.001 | 0 | 0.44 | 0.48 | 0.54 | 0.78 | 0.46 | 0.32 |
| 2 | 213090009 | rs4580336 | 1.676 | -1.474 | 0.735 | -2.09 | 0.023 | 0.027 | 0.027 | 0 | 0.54 | 0.57 | 0.64 | 0.78 | 0.69 | 0.39 |
| 2 | 213107619 | rs10207020 | 1.199 | -1.514 | 0.73 | -2.092 | 0.034 | 0.033 | 0.033 | 0 | 0.6 | 0.68 | 0.68 | 0.79 | 0.76 | 0.46 |
| 2 | 213110424 | rs1949652 | 2.567 | -1.453 | 0.842 | -2.071 | -0.026 | 0.061 | 0.061 | 0.064 | 0.43 | 0.23 | 0.48 | 0.76 | 0.47 | 0.32 |
| 2 | 213110812 | rs16848048 | 0.787 | -1.264 | 0.745 | -2.027 | 0 | 0.016 | 0.016 | 0 | 0.06 | 0.04 | 0.04 | 0 | 0.07 | 0.06 |
| 2 | 213111238 | rs12151481 | -0.948 | -1.277 | 0.831 | -2.039 | 0.015 | -0.04 | -0.04 | 1.05 | 0.14 | 0.27 | 0.13 | 0.01 | 0.25 | 0.1 |
| 2 | 213113079 | rs10497966 | 0.495 | -1.265 | 0.791 | -2.025 | -0.021 | 0.015 | 0.015 | 0 | 0.08 | 0.09 | 0.09 | 0.01 | 0.22 | 0.07 |
| 2 | 213117008 | rs12466094 | -0.261 | -1.418 | 0.783 | -2.045 | 0.037 | 0.058 | 0.058 | 0 | 0.2 | 0.05 | 0.16 | 0.06 | 0.13 | 0.39 |
| 2 | 213117976 | rs1521636 | -1.522 | -1.121 | 1.001 | -1.977 | 0.062 | -0.127 | -0.127 | 0 | 0.21 | 0.45 | 0.17 | 0.01 | 0.33 | 0.17 |
| 2 | 213119367 | rs7572366 | -0.178 | -1.102 | 0.895 | -1.964 | 0.005 | -0.03 | -0.03 | 0.479 | 0.06 | 0.18 | 0.03 | 0 | 0.04 | 0.04 |
| 2 | 213119487 | rs7586610 | 0.209 | -1.1 | 0.818 | -1.963 | 0.004 | -0.023 | -0.023 | 0 | 0.05 | 0.13 | 0.02 | 0 | 0.04 | 0.03 |
| 2 | 213120523 | rs16848108 | -0.682 | -1.056 | 0.823 | -1.915 | 0.003 | -0.027 | -0.027 | 0 | 0.14 | 0.16 | 0.13 | 0.01 | 0.29 | 0.13 |
| 2 | 213129211 | rs1521662 | 1.201 | -1.091 | 0.432 | -1.831 | 0.053 | 0.036 | 0.036 | 0.011 | 0.62 | 0.75 | 0.67 | 0.75 | 0.72 | 0.47 |
| 2 | 213130571 | rs6745249 | 1.94 | -1.1 | 0.418 | -1.833 | 0.013 | 0.067 | 0.067 | 0 | 0.49 | 0.3 | 0.55 | 0.74 | 0.51 | 0.39 |
| 2 | 213136747 | rs2135161 | -1.396 | -1.076 | 0.024 | -1.922 | -0.007 | -0.052 | -0.052 | 0 | 0.17 | 0.2 | 0.07 | 0 | 0.12 | 0.09 |
| 2 | 213137150 | rs931817 | -0.479 | -0.869 | -0.14 | -1.779 | -0.002 | -0.033 | -0.033 | 0.001 | 0.08 | 0.11 | 0.02 | 0 | 0.04 | 0.05 |
| 2 | 213146827 | rs979042 | 0.899 | -0.786 | -0.425 | -1.728 | -0.013 | 0.017 | 0.017 | 0 | 0.71 | 0.75 | 0.75 | 0.81 | 0.66 | 0.65 |
| 2 | 213156498 | rs17346713 | -1.492 | -1.207 | -0.656 | -1.886 | 0.042 | 0.155 | 0.155 | 0 | 0.3 | 0.07 | 0.17 | 0.03 | 0.21 | 0.37 |
| 2 | 213162300 | rs6711391 | 0.668 | -1.473 | -0.595 | -2.062 | 0 | 0 | 0 | 0 | 0.82 | 0.26 | 0.83 | 0.89 | 0.9 | 0.81 |
| 2 | 213164193 | rs10932432 | 2.048 | -1.488 | -0.658 | -2.107 | 0.134 | 0.207 | 0.207 | 0.462 | 0.54 | 0.1 | 0.67 | 0.85 | 0.72 | 0.47 |
| 2 | 213164792 | rs17418814 | -0.818 | -1.469 | -0.627 | -2.105 | 0.043 | 0.067 | 0.067 | 0 | 0.24 | 0.12 | 0.15 | 0.03 | 0.17 | 0.32 |
| 2 | 213173172 | rs16848312 | -0.831 | -1.481 | -0.571 | -2.116 | 0.045 | 0.049 | 0.049 | 0 | 0.21 | 0.12 | 0.14 | 0.03 | 0.13 | 0.24 |
| 2 | 213183696 | rs1910866 | 1.025 | -1.44 | -0.281 | -1.674 | -0.01 | -0.002 | -0.002 | 0 | 0.76 | 0.29 | 0.74 | 0.78 | 0.85 | 0.71 |
| 2 | 213187368 | rs10164827 | -1.056 | -1.611 | 0.388 | -1.798 | -0.031 | -0.138 | -0.138 | 0 | 0.19 | 0.48 | 0.1 | 0.01 | 0.1 | 0.23 |
| 2 | 213188733 | rs905882 | 0.565 | -1.226 | 0.653 | -1.554 | 0.009 | 0.162 | 0.162 | 0.742 | 0.93 | 0.78 | 0.83 | 0.8 | 0.94 | 0.91 |
| 2 | 213189366 | rs905879 | 1.781 | -1.389 | 0.528 | -1.69 | -0.053 | -0.016 | -0.016 | 0.715 | 0.67 | 0.56 | 0.67 | 0.76 | 0.77 | 0.62 |
| 2 | 213189994 | rs4146051 | 0.333 | -1.236 | 0.583 | -1.634 | 0.009 | 0.16 | 0.16 | 0.593 | 0.93 | 0.8 | 0.83 | 0.79 | 0.94 | 0.92 |
| 2 | 213190493 | rs7601515 | 1.741 | -1.227 | 0.575 | -1.629 | -0.048 | -0.011 | -0.011 | 0 | 0.69 | 0.58 | 0.67 | 0.77 | 0.77 | 0.63 |
| 2 | 213191389 | rs1505366 | 0.205 | -1.19 | 0.599 | -1.639 | 0.017 | 0.174 | 0.174 | 0.744 | 0.93 | 0.79 | 0.83 | 0.8 | 0.95 | 0.92 |
| 2 | 213192198 | rs12473610 | -1.148 | -1.145 | 0.166 | -1.599 | 0.011 | -0.001 | -0.001 | 0 | 0.23 | 0.18 | 0.15 | 0.02 | 0.17 | 0.27 |
| 2 | 213192268 | rs7562782 | 1.424 | -1.124 | 0.163 | -1.603 | 0.017 | 0.005 | 0.005 | 0.013 | 0.72 | 0.79 | 0.69 | 0.78 | 0.78 | 0.66 |
| 2 | 213194322 | rs16848380 | -0.174 | -0.858 | 0.319 | -1.953 | -0.001 | -0.026 | -0.026 | 0.082 | 0.1 | 0.15 | 0.05 | 0 | 0.08 | 0.12 |
| 2 | 213200363 | rs6743763 | -1.564 | -0.704 | 0.493 | -2.027 | -0.006 | -0.036 | -0.036 | 0.48 | 0.14 | 0.19 | 0.15 | 0.03 | 0.13 | 0.15 |
| 2 | 213200920 | rs1394782 | 1.961 | -0.913 | 0.986 | -2.063 | 0.006 | 0.009 | 0.009 | 0.416 | 0.64 | 0.35 | 0.66 | 0.8 | 0.71 | 0.53 |
| 2 | 213201725 | rs1394783 | -0.75 | -0.427 | 0.652 | -2.421 | -0.004 | -0.024 | -0.024 | 0 | 0.23 | 0.21 | 0.13 | 0.02 | 0.29 | 0.17 |
| 2 | 213203057 | rs1505353 | 3.127 | -0.416 | 0.527 | -2.42 | -0.057 | 0.148 | 0.148 | 0 | 0.41 | 0.07 | 0.52 | 0.77 | 0.42 | 0.35 |
| 2 | 213205047 | rs4673661 | 1.406 | -0.428 | 0.982 | -2.399 | -0.004 | -0.021 | -0.021 | 0 | 0.7 | 0.64 | 0.69 | 0.8 | 0.73 | 0.54 |
| 2 | 213206488 | rs13395316 | -0.617 | -0.404 | 1.002 | -2.402 | 0.018 | -0.069 | -0.069 | 0 | 0.21 | 0.28 | 0.14 | 0.02 | 0.29 | 0.16 |
| 2 | 213210229 | rs1505357 | 1.851 | -0.416 | 0.882 | -2.393 | -0.002 | 0 | 0 | 0 | 0.57 | 0.43 | 0.68 | 0.79 | 0.56 | 0.5 |
| 2 | 213211349 | rs1394788 | 0.528 | -0.366 | 0.511 | -2.35 | -0.012 | -0.021 | -0.021 | 0 | 0.78 | 0.67 | 0.81 | 0.82 | 0.84 | 0.67 |
| 2 | 213211407 | rs1910861 | 2.849 | -0.307 | 0.383 | -2.336 | -0.053 | 0.159 | 0.159 | 0.224 | 0.43 | 0.07 | 0.52 | 0.77 | 0.43 | 0.36 |
| 2 | 213216431 | rs10192514 | -0.346 | -0.027 | 0.664 | -2.321 | -0.004 | -0.025 | -0.025 | 0 | 0.17 | 0.17 | 0.08 | 0.09 | 0.13 | 0.23 |
| 2 | 213219641 | rs1948877 | 1.545 | -0.023 | 0.713 | -2.33 | -0.006 | -0.046 | -0.046 | 0.043 | 0.7 | 0.54 | 0.68 | 0.79 | 0.72 | 0.54 |
| 2 | 213221111 | rs13018908 | -0.589 | -0.143 | 0.562 | -2.345 | -0.011 | -0.04 | -0.04 | 0 | 0.17 | 0.25 | 0.08 | 0.1 | 0.13 | 0.24 |
| 2 | 213222446 | rs17347530 | -0.756 | 0.033 | 0.032 | -2.407 | -0.061 | 0.038 | 0.038 | 0 | 0.09 | 0.01 | 0.06 | 0.02 | 0.18 | 0.14 |
| 2 | 213223044 | rs13006832 | -1.749 | -0.167 | 0.303 | -2.422 | -0.018 | 0.045 | 0.045 | 0 | 0.28 | 0.2 | 0.22 | 0.09 | 0.33 | 0.35 |
| 2 | 213224540 | rs13014726 | -0.56 | 0.251 | 0.247 | -2.433 | -0.026 | 0.06 | 0.06 | 0 | 0.18 | 0.07 | 0.12 | 0.04 | 0.21 | 0.25 |
| 2 | 213226396 | rs12478244 | -0.787 | 0.237 | 0.258 | -2.428 | -0.026 | 0.002 | 0.002 | 0 | 0.09 | 0.02 | 0.1 | 0.05 | 0.12 | 0.09 |
| 2 | 213226544 | rs11892564 | -1.756 | -0.136 | 0.384 | -2.751 | 0.009 | -0.001 | -0.001 | 0 | 0.29 | 0.73 | 0.24 | 0.07 | 0.19 | 0.26 |
| 2 | 213227254 | rs16848458 | -0.859 | -0.185 | 0.836 | -2.775 | -0.03 | -0.115 | -0.115 | 0.964 | 0.17 | 0.33 | 0.08 | 0 | 0.1 | 0.09 |
| 2 | 213233631 | rs6734836 | -3.282 | -0.144 | 0.786 | -2.803 | -0.036 | 0.15 | 0.15 | 0 | 0.56 | 0.84 | 0.47 | 0.16 | 0.52 | 0.57 |
| 2 | 213233739 | rs4511675 | -2.613 | -0.163 | 0.831 | -2.809 | 0.003 | -0.001 | -0.001 | 0 | 0.36 | 0.64 | 0.34 | 0.12 | 0.31 | 0.32 |
| 2 | 213236480 | rs6435700 | -1.104 | -0.146 | 0.667 | -2.761 | -0.007 | -0.025 | -0.025 | 0 | 0.12 | 0.39 | 0.15 | 0.07 | 0.09 | 0.15 |
| 2 | 213240767 | rs13034566 | -1.967 | -0.104 | 0.685 | -2.748 | -0.006 | -0.149 | -0.149 | 0 | 0.34 | 0.43 | 0.2 | 0.04 | 0.31 | 0.35 |
| 2 | 213249244 | rs13015805 | -0.54 | 0.038 | 0.354 | -2.571 | -0.017 | 0.06 | 0.06 | 0 | 0.18 | 0.08 | 0.11 | 0.04 | 0.2 | 0.26 |
| 2 | 213251503 | rs13019524 | -1.706 | 0.036 | 0.416 | -2.558 | -0.02 | 0.084 | 0.084 | 0 | 0.3 | 0.18 | 0.23 | 0.1 | 0.32 | 0.39 |
| 2 | 213275304 | rs10188926 | -1.276 | 0.294 | 0.595 | -2.432 | 0.005 | -0.011 | -0.011 | 0 | 0.09 | 0.17 | 0.1 | 0.04 | 0.11 | 0.16 |
| 2 | 213277649 | rs4673664 | -2.83 | 0.312 | 0.454 | -2.866 | -0.009 | 0.088 | 0.088 | 0 | 0.58 | 0.79 | 0.46 | 0.14 | 0.55 | 0.6 |
| 2 | 213279029 | rs6757140 | -1.833 | 0.259 | 0.665 | -2.95 | 0.001 | -0.072 | -0.072 | 0.013 | 0.41 | 0.38 | 0.24 | 0.01 | 0.37 | 0.27 |
| 2 | 213282779 | rs13417523 | -0.879 | 0.269 | 0.691 | -2.933 | 0.025 | -0.017 | -0.017 | 0.335 | 0.09 | 0.22 | 0.1 | 0.04 | 0.11 | 0.16 |
| 2 | 213283190 | rs1394780 | -1.916 | 0.214 | 0.58 | -3.018 | -0.004 | -0.006 | -0.006 | 0 | 0.24 | 0.23 | 0.18 | 0.04 | 0.24 | 0.25 |

| | | | | | | | | | | | | | | | | |
|---|---|---|---|---|---|---|---|---|---|---|---|---|---|---|---|---|
| 2 | 213283512 | rs6435701 | -3.121 | 0.209 | 0.6 | -3.025 | -0.016 | 0.146 | 0.146 | 0 | 0.57 | 0.84 | 0.47 | 0.14 | 0.55 | 0.59 |
| 2 | 213284021 | rs13383863 | -1.2 | 0.319 | 0.591 | -2.833 | 0.026 | -0.019 | -0.019 | 0.706 | 0.09 | 0.22 | 0.1 | 0.04 | 0.11 | 0.16 |
| 2 | 213291199 | rs10166403 | -1.528 | 0.347 | 0.593 | -2.813 | 0.022 | -0.002 | -0.002 | 0 | 0.1 | 0.21 | 0.12 | 0.09 | 0.14 | 0.22 |
| 2 | 213292130 | rs16848535 | -0.886 | 0.232 | 0.51 | -2.77 | 0.004 | 0.027 | 0.027 | 0.344 | 0.06 | 0.25 | 0.1 | 0.04 | 0.04 | 0.11 |
| 2 | 213294557 | rs7569142 | -2.339 | 0.182 | 0.273 | -2.798 | -0.001 | 0.044 | 0.044 | 0 | 0.39 | 0.26 | 0.26 | 0.06 | 0.37 | 0.32 |
| 2 | 213297810 | rs1394791 | -0.936 | 0.178 | -0.091 | -2.653 | -0.015 | 0.109 | 0.109 | 0 | 0.22 | 0.07 | 0.13 | 0.01 | 0.22 | 0.16 |
| 2 | 213303947 | rs1505376 | -2.469 | 0.088 | 0.028 | -2.795 | 0.012 | 0.123 | 0.123 | 0.049 | 0.49 | 0.72 | 0.33 | 0.1 | 0.41 | 0.36 |
| 2 | 213304043 | rs1505377 | -0.849 | 0.117 | -0.135 | -2.818 | -0.021 | 0.128 | 0.128 | 0 | 0.25 | 0.05 | 0.13 | 0 | 0.23 | 0.15 |
| 2 | 213316586 | rs17259208 | -1.322 | 0.391 | -0.412 | -3.028 | -0.044 | 0.16 | 0.16 | 3.129 | 0.3 | 0.05 | 0.19 | 0.05 | 0.31 | 0.24 |
| 2 | 213316826 | rs2204853 | -0.208 | 0.401 | 0.031 | -3.057 | 0.012 | -0.052 | -0.052 | 0 | 0.13 | 0.25 | 0.12 | 0 | 0.13 | 0.08 |
| 2 | 213316881 | rs972489 | 0.272 | 0.397 | 0.043 | -3.042 | -0.017 | 0.017 | 0.017 | 0 | 0.11 | 0.08 | 0.11 | 0 | 0.13 | 0.08 |
| 2 | 213317329 | rs6711080 | -2.571 | 0.339 | 0.47 | -3.077 | 0 | -0.008 | -0.008 | 0.8 | 0.47 | 0.5 | 0.37 | 0.1 | 0.43 | 0.36 |
| 2 | 213317794 | rs11689233 | -0.194 | 0.404 | 0.622 | -2.894 | 0.006 | -0.038 | -0.038 | 0 | 0.12 | 0.2 | 0.12 | 0 | 0.13 | 0.08 |
| 2 | 213318641 | rs974968 | -2.714 | 0.354 | 0.711 | -2.869 | 0.007 | 0.131 | 0.131 | 0.088 | 0.51 | 0.74 | 0.39 | 0.13 | 0.44 | 0.4 |
| 2 | 213319545 | rs7570124 | -0.944 | 0.355 | 0.627 | -2.852 | -0.051 | -0.012 | -0.012 | 0.008 | 0.09 | 0.25 | 0.08 | 0.04 | 0.01 | 0.07 |
| 2 | 213328749 | rs2170529 | -2.337 | 0.388 | 0.595 | -2.861 | -0.001 | -0.037 | -0.037 | 14.07 | 0.43 | 0.5 | 0.35 | 0.1 | 0.39 | 0.34 |
| 2 | 213334523 | rs10168850 | -1.605 | 0.426 | 0.483 | -2.848 | -0.057 | 0.128 | 0.128 | 0 | 0.34 | 0.15 | 0.27 | 0.05 | 0.39 | 0.27 |
| 2 | 213336689 | rs17325821 | -0.481 | 0.456 | 0.392 | -2.861 | -0.043 | 0.093 | 0.093 | 0 | 0.22 | 0.03 | 0.15 | 0.05 | 0.25 | 0.19 |
| 2 | 213339192 | rs1384289 | -0.105 | 0.416 | 0.912 | -2.753 | 0.013 | -0.005 | -0.005 | 0.375 | 0.07 | 0.31 | 0.03 | 0 | 0.02 | 0.02 |
| 2 | 213341614 | rs985995 | -2.484 | 0.371 | 0.861 | -2.792 | -0.009 | -0.03 | -0.03 | 0 | 0.28 | 0.69 | 0.26 | 0.09 | 0.18 | 0.22 |
| 2 | 213347104 | rs1482378 | -1.818 | 0.632 | 0.946 | -2.709 | -0.039 | 0.014 | 0.014 | 0 | 0.12 | 0.52 | 0.14 | 0.09 | 0.06 | 0.17 |
| 2 | 213351973 | rs7568763 | -0.028 | 0.815 | 1.027 | -2.612 | 0.019 | -0.044 | -0.044 | 0 | 0.1 | 0.16 | 0.12 | 0 | 0.13 | 0.07 |
| 2 | 213354898 | rs13423636 | 0.034 | 0.813 | 1.111 | -2.613 | 0.01 | -0.033 | -0.033 | 0 | 0.1 | 0.11 | 0.12 | 0 | 0.12 | 0.07 |
| 2 | 213385817 | rs4404217 | -1.629 | 1.214 | 0.366 | -2.726 | 0.031 | -0.051 | -0.051 | 0 | 0.21 | 0.32 | 0.21 | 0.05 | 0.29 | 0.26 |
| 2 | 213386112 | rs6435707 | -3.035 | 1.124 | 0.216 | -2.789 | -0.021 | 0.129 | 0.129 | 0 | 0.6 | 0.91 | 0.48 | 0.17 | 0.58 | 0.59 |
| 2 | 213386267 | rs7571561 | -1.581 | 1.105 | 0.293 | -2.981 | 0.03 | -0.05 | -0.05 | 0 | 0.21 | 0.32 | 0.21 | 0.05 | 0.29 | 0.26 |
| 2 | 213387389 | rs1026882 | -2.667 | 0.998 | -0.059 | -2.919 | -0.027 | 0.225 | 0.225 | 0 | 0.58 | 0.9 | 0.46 | 0.1 | 0.55 | 0.49 |
| 2 | 213394605 | rs1351593 | -1.5 | 1.447 | 0.696 | -2.432 | 0.023 | -0.029 | -0.029 | 1.612 | 0.12 | 0.25 | 0.1 | 0.05 | 0.17 | 0.19 |
| 2 | 213404389 | rs7355673 | -2.028 | 1.225 | 0.413 | -2.411 | -0.037 | 0.131 | 0.131 | 0.404 | 0.41 | 0.77 | 0.3 | 0.07 | 0.42 | 0.29 |
| 2 | 213413231 | rs7599312 | -1.609 | 1.693 | 0.79 | -1.857 | 0.059 | -0.087 | -0.087 | 0.107 | 0.2 | 0.42 | 0.2 | 0.03 | 0.27 | 0.12 |
| 2 | 213421764 | rs10191896 | -1.865 | 1.948 | 0.873 | -1.691 | 0.046 | -0.057 | -0.057 | 0.202 | 0.29 | 0.54 | 0.39 | 0.1 | 0.37 | 0.19 |
| 2 | 213422621 | rs17262541 | -0.112 | 1.561 | 0.896 | -1.717 | 0 | -0.007 | -0.007 | 0 | 0.08 | 0.1 | 0.04 | 0.02 | 0.08 | 0.06 |
| 2 | 213432392 | rs7578094 | -2.471 | 1.615 | 1.302 | -1.848 | 0.012 | -0.042 | -0.042 | 0 | 0.33 | 0.67 | 0.41 | 0.09 | 0.37 | 0.24 |
| 2 | 213456994 | rs17263937 | -1.673 | 1.322 | 1.168 | -1.782 | -0.018 | 0.023 | 0.023 | 0 | 0.19 | 0.1 | 0.31 | 0.08 | 0.24 | 0.17 |
| 2 | 213463406 | rs1002618 | 0.142 | 1.252 | 0.628 | -2.055 | -0.03 | 0.045 | 0.045 | 0 | 0.17 | 0.01 | 0.15 | 0.07 | 0.18 | 0.18 |
| 2 | 213469506 | rs11688976 | -2.564 | 1.136 | 1.645 | -1.753 | 0.001 | 0.035 | 0.035 | 0 | 0.33 | 0.77 | 0.39 | 0.11 | 0.31 | 0.22 |
| 2 | 213473115 | rs7606703 | -1.156 | 1.041 | 1.873 | -1.776 | 0.038 | -0.072 | -0.072 | 0.051 | 0.13 | 0.21 | 0.13 | 0 | 0.19 | 0.08 |
| 2 | 213480637 | rs4672649 | -2.995 | 0.945 | 1.85 | -1.735 | -0.02 | 0.043 | 0.043 | 0.473 | 0.13 | 0.52 | 0.23 | 0.09 | 0.07 | 0.09 |
| 2 | 213484752 | rs2054615 | -2.455 | 0.943 | 1.84 | -1.743 | -0.016 | 0.06 | 0.06 | 0.411 | 0.33 | 0.83 | 0.41 | 0.12 | 0.35 | 0.23 |
| 2 | 213485715 | rs11692809 | -1.82 | 1.034 | 1.835 | -1.755 | 0.025 | -0.041 | -0.041 | 0 | 0.24 | 0.54 | 0.34 | 0.12 | 0.29 | 0.17 |
| 2 | 213486423 | rs6704686 | -1.228 | 1.033 | 1.827 | -1.748 | 0.028 | -0.049 | -0.049 | 0 | 0.09 | 0.29 | 0.06 | 0 | 0.06 | 0.07 |
| 2 | 213488021 | rs6712252 | -2.545 | 1.036 | 1.737 | -1.741 | -0.007 | 0.042 | 0.042 | 0 | 0.34 | 0.74 | 0.39 | 0.12 | 0.35 | 0.23 |
| 2 | 213490883 | rs2128324 | -2.492 | 1.157 | 1.65 | -1.683 | -0.03 | 0.088 | 0.088 | 0.598 | 0.33 | 0.88 | 0.41 | 0.12 | 0.35 | 0.23 |
| 2 | 213504183 | rs13397735 | -1.643 | 1.428 | 1.655 | -1.592 | 0.026 | -0.063 | -0.063 | 0 | 0.24 | 0.3 | 0.2 | 0.02 | 0.34 | 0.16 |
| 2 | 213510285 | rs16848991 | -1.068 | 1.331 | 1.621 | -1.483 | -0.011 | 0.009 | 0.009 | 0 | 0.06 | 0.08 | 0.19 | 0.1 | 0.03 | 0.11 |
| 2 | 213521462 | rs2371618 | -0.857 | 1.357 | 1.698 | -1.48 | -0.011 | 0.013 | 0.013 | 0 | 0.06 | 0.08 | 0.18 | 0.1 | 0.03 | 0.11 |
| 2 | 213526121 | rs13425327 | -1.833 | 1.418 | 1.726 | -1.469 | 0.06 | -0.068 | -0.068 | 0 | 0.12 | 0.3 | 0.08 | 0 | 0.12 | 0.09 |
| 2 | 213530069 | rs2371710 | -0.75 | 1.436 | 1.52 | -1.465 | 0 | -0.001 | -0.001 | 0 | 0.19 | 0.21 | 0.31 | 0.12 | 0.25 | 0.19 |
| 2 | 213535235 | rs16849057 | -1.349 | 1.296 | 1.533 | -1.583 | 0.024 | -0.063 | -0.063 | 0.164 | 0.11 | 0.23 | 0.05 | 0 | 0.08 | 0.07 |
| 2 | 213535830 | rs6742049 | -0.88 | 1.298 | 1.481 | -1.582 | 0.008 | -0.035 | -0.035 | 0 | 0.09 | 0.1 | 0.05 | 0 | 0.08 | 0.07 |
| 2 | 213542950 | rs13422625 | -2.114 | 1.216 | 1.143 | -1.587 | -0.012 | 0.058 | 0.058 | 0.579 | 0.35 | 0.77 | 0.39 | 0.09 | 0.37 | 0.28 |
| 2 | 213544609 | rs6435720 | -1.37 | 1.247 | 1.039 | -1.578 | 0.032 | -0.091 | -0.091 | 0 | 0.14 | 0.39 | 0.07 | 0 | 0.11 | 0.08 |
| 2 | 213545335 | rs17331933 | -0.73 | 1.049 | 0.105 | -1.47 | 0 | -0.002 | -0.002 | 0 | 0.19 | 0.2 | 0.3 | 0.08 | 0.26 | 0.2 |
| 2 | 213551094 | rs6435721 | -1.311 | 1.06 | -1.337 | -1.372 | -0.017 | 0.087 | 0.087 | 0.655 | 0.36 | 0.83 | 0.39 | 0.09 | 0.37 | 0.3 |
| 2 | 213555564 | rs11674822 | -1.247 | 0.738 | -1.28 | -1.103 | 0.007 | 0.093 | 0.093 | 0 | 0.26 | 0.1 | 0.15 | 0.06 | 0.23 | 0.3 |
| 2 | 213558906 | rs17272806 | -1.38 | 1.191 | -1.429 | -0.971 | -0.036 | 0.032 | 0.032 | 0.466 | 0.15 | 0.03 | 0.12 | 0.11 | 0.15 | 0.2 |
| 2 | 213559229 | rs7580519 | -1.666 | 1.156 | -0.962 | -1.039 | 0 | -0.007 | -0.007 | 0.99 | 0.17 | 0.19 | 0.1 | 0.07 | 0.17 | 0.25 |
| 2 | 213561598 | rs6731231 | -2.158 | 0.723 | -1.096 | -0.671 | -0.02 | 0.13 | 0.13 | 0.642 | 0.49 | 0.82 | 0.4 | 0.39 | 0.48 | 0.58 |
| 2 | 213561811 | rs13390015 | -0.761 | 1.011 | -0.682 | -0.596 | 0 | -0.001 | -0.001 | 0.483 | 0.1 | 0.09 | 0.06 | 0.12 | 0.1 | 0.19 |
| 2 | 213563072 | rs7575556 | -1.196 | 0.771 | -0.042 | -0.748 | -0.005 | -0.058 | -0.058 | 1.403 | 0.29 | 0.49 | 0.28 | 0.21 | 0.27 | 0.22 |
| 2 | 213572539 | rs10188254 | -1.497 | 0.954 | 0.461 | -0.412 | 0.032 | 0.047 | 0.047 | 0.598 | 0.22 | 0.43 | 0.21 | 0.32 | 0.22 | 0.31 |
| 2 | 213575237 | rs16849118 | -0.345 | 0.927 | 0.808 | -0.37 | 0 | 0.014 | 0.014 | 0 | 0.12 | 0.16 | 0.13 | 0.2 | 0.13 | 0.13 |
| 2 | 213575506 | rs7587255 | -2.015 | 0.752 | 0.708 | 0.421 | 0.004 | 0.007 | 0.007 | 0.022 | 0.41 | 0.55 | 0.35 | 0.57 | 0.41 | 0.51 |
| 2 | 213576272 | rs16849123 | -0.037 | 0.557 | 0.518 | 0.479 | -0.005 | -0.013 | -0.013 | 0 | 0.05 | 0.08 | 0.01 | 0.01 | 0.05 | 0.05 |
| 2 | 213577479 | rs17332777 | -1.42 | 0.679 | 0.463 | 0.569 | 0.006 | -0.037 | -0.037 | 0 | 0.1 | 0.11 | 0.07 | 0.07 | 0.09 | 0.15 |
| 2 | 213578777 | rs13393014 | -1.725 | 0.449 | 0.522 | 0.822 | -0.002 | 0.023 | 0.023 | 0 | 0.23 | 0.42 | 0.15 | 0.27 | 0.17 | 0.27 |
| 2 | 213580723 | rs6435723 | 1.311 | 0.029 | 0.196 | 1.786 | 0.013 | -0.029 | -0.029 | 0 | 0.84 | 0.89 | 0.6 | 0.44 | 0.9 | 0.68 |
| 2 | 213582322 | rs16849142 | -0.697 | 0.078 | 0.279 | 1.724 | 0.004 | 0.02 | 0.02 | 0 | 0.09 | 0.13 | 0.07 | 0.19 | 0.09 | 0.07 |
| 2 | 213582942 | rs12619430 | -0.224 | 0.042 | 0.223 | 2.409 | -0.007 | -0.009 | -0.009 | 0 | 0.05 | 0.02 | 0.12 | 0.21 | 0.07 | 0.07 |

| | | | | | | | | | | | | | | | | |
|---|---|---|---|---|---|---|---|---|---|---|---|---|---|---|---|---|
| 2 | 213584688 | rs10932444 | 1.623 | 0.007 | 0.635 | 2.335 | -0.01 | -0.04 | -0.04 | 0 | 0.72 | 0.57 | 0.76 | 0.81 | 0.77 | 0.73 |
| 2 | 213585180 | rs7369041 | 0.112 | 0.118 | 0.547 | 2.234 | -0.012 | -0.013 | -0.013 | 0 | 0.07 | 0.02 | 0.07 | 0.2 | 0.08 | 0.06 |
| 2 | 213591785 | rs2371621 | 0.091 | 0.1 | 0.644 | 2.231 | 0.002 | -0.003 | -0.003 | 0.373 | 0.91 | 0.94 | 0.9 | 0.88 | 0.97 | 0.89 |
| 2 | 213593970 | rs7565158 | 2.05 | 0.011 | 1.044 | 2.259 | 0.004 | -0.014 | -0.014 | 0 | 0.62 | 0.3 | 0.5 | 0.44 | 0.69 | 0.51 |
| 2 | 213595110 | rs16849174 | -0.68 | -0.31 | 1.674 | 2.215 | -0.001 | 0.093 | 0.093 | 0 | 0.08 | 0.14 | 0.14 | 0.24 | 0.09 | 0.08 |
| 2 | 213597773 | rs6758369 | -1.802 | -0.368 | 2.367 | 2.179 | -0.024 | 0.083 | 0.083 | 1.591 | 0.18 | 0.43 | 0.2 | 0.27 | 0.17 | 0.19 |
| 2 | 213598952 | rs16849192 | -0.31 | -0.354 | 2.382 | 2.113 | -0.01 | -0.032 | -0.032 | 0 | 0.09 | 0.14 | 0.04 | 0.03 | 0.08 | 0.11 |
| 2 | 213602048 | rs6725004 | -2.112 | -0.334 | 2.422 | 2.107 | 0.013 | 0.006 | 0.006 | 1.775 | 0.19 | 0.41 | 0.31 | 0.26 | 0.18 | 0.29 |
| 2 | 213611221 | rs16849209 | -1.48 | -0.32 | 2.217 | 1.948 | 0.025 | 0.039 | 0.039 | 0 | 0.17 | 0.27 | 0.2 | 0.24 | 0.22 | 0.13 |
| 2 | 213617274 | rs10469770 | -1.451 | -0.324 | 2.16 | 1.962 | 0.007 | 0.015 | 0.015 | 0 | 0.17 | 0.2 | 0.21 | 0.25 | 0.22 | 0.13 |
| 2 | 213635052 | rs6435726 | 0.347 | -0.425 | 2.111 | 2.025 | 0.004 | 0.005 | 0.005 | 0 | 0.9 | 0.94 | 0.72 | 0.67 | 0.97 | 0.75 |
| 2 | 213635698 | rs10497982 | -0.642 | -0.426 | 2.316 | 1.925 | 0.001 | -0.049 | -0.049 | 0 | 0.1 | 0.26 | 0.06 | 0.03 | 0.1 | 0.13 |
| 2 | 213640577 | rs4375811 | -0.815 | -0.462 | 2.332 | 1.927 | -0.006 | -0.048 | -0.048 | 0 | 0.11 | 0.41 | 0.06 | 0.03 | 0.08 | 0.11 |
| 2 | 213646473 | rs12986480 | 2.478 | -0.551 | 2.296 | 2.01 | 0.024 | -0.003 | -0.003 | 0 | 0.6 | 0.29 | 0.45 | 0.36 | 0.68 | 0.46 |
| 2 | 213649587 | rs6710703 | 1.931 | -0.547 | 2.106 | 2.038 | 0 | 0.008 | 0.008 | 0.856 | 0.72 | 0.5 | 0.75 | 0.69 | 0.71 | 0.75 |
| 2 | 213657426 | rs11693864 | 2.394 | -0.529 | 1.946 | 1.953 | 0.04 | -0.015 | -0.015 | 0 | 0.59 | 0.15 | 0.45 | 0.36 | 0.68 | 0.46 |
| 2 | 213662939 | rs6708555 | -1.33 | -0.139 | 1.798 | 1.516 | 0.003 | 0.003 | 0.003 | 0 | 0.16 | 0.16 | 0.15 | 0.2 | 0.21 | 0.14 |
| 2 | 213665065 | rs7594160 | 0.583 | -0.286 | 1.542 | 1.131 | -0.008 | -0.042 | -0.042 | 0.024 | 0.9 | 0.74 | 0.96 | 0.97 | 0.92 | 0.9 |
| 2 | 213676322 | rs9288457 | -1.578 | -0.553 | 0.879 | 1.412 | -0.028 | 0.033 | 0.033 | 3.175 | 0.27 | 0.61 | 0.42 | 0.47 | 0.15 | 0.4 |
| 2 | 213681716 | rs13015944 | 1.854 | -0.542 | 1.254 | 1.389 | 0.032 | -0.062 | -0.062 | 0 | 0.65 | 0.09 | 0.51 | 0.57 | 0.74 | 0.56 |
| 2 | 213685014 | rs3923957 | -0.223 | -0.553 | 1.319 | 1.311 | 0.003 | -0.034 | -0.034 | 0 | 0.08 | 0.21 | 0.03 | 0.04 | 0.07 | 0.09 |
| 2 | 213687346 | rs6435735 | -1.897 | -0.803 | 0.541 | 0.538 | 0.044 | 0.001 | 0.001 | 0 | 0.48 | 0.83 | 0.63 | 0.46 | 0.41 | 0.48 |
| 2 | 213689498 | rs4260187 | -1.222 | -0.991 | 0.494 | 0.659 | -0.021 | 0.023 | 0.023 | 1.078 | 0.27 | 0.3 | 0.37 | 0.38 | 0.18 | 0.41 |
| 2 | 213690991 | rs4673694 | -0.091 | -0.783 | 0.292 | 0.638 | -0.005 | -0.036 | -0.036 | 0 | 0.09 | 0.15 | 0.06 | 0.03 | 0.1 | 0.1 |
| 2 | 213695702 | rs4673696 | 0.924 | -0.765 | 0.044 | 0.595 | 0.056 | -0.022 | -0.022 | 0.529 | 0.59 | 0.2 | 0.46 | 0.54 | 0.7 | 0.56 |
| 2 | 213707759 | rs4131363 | 0.755 | -0.893 | 0.179 | 0.506 | 0.026 | 0.008 | 0.008 | 1.296 | 0.71 | 0.21 | 0.65 | 0.71 | 0.78 | 0.73 |
| 2 | 213718301 | rs6709858 | -0.224 | -0.774 | -0.01 | -0.04 | -0.008 | 0.072 | 0.072 | 0 | 0.16 | 0.05 | 0.1 | 0.03 | 0.15 | 0.13 |
| 2 | 213722554 | rs4480940 | 1.596 | -0.439 | 0.003 | -0.23 | -0.001 | -0.001 | -0.001 | 0 | 0.31 | 0.29 | 0.33 | 0.38 | 0.36 | 0.45 |
| 2 | 213728381 | rs4132518 | 0.528 | -0.584 | 0.104 | -0.372 | -0.027 | -0.138 | -0.138 | 0 | 0.66 | 0.42 | 0.77 | 0.97 | 0.73 | 0.81 |
| 2 | 213737261 | rs7606842 | 0.498 | -0.594 | 0.273 | -0.375 | -0.03 | -0.148 | -0.148 | 0 | 0.67 | 0.52 | 0.79 | 0.97 | 0.72 | 0.82 |
| 2 | 213738972 | rs7600221 | 0.157 | -0.514 | -0.037 | -0.294 | -0.008 | -0.004 | -0.004 | 0 | 0.36 | 0.24 | 0.45 | 0.6 | 0.37 | 0.36 |
| 2 | 213759117 | rs1482581 | 1.256 | -0.297 | 0.063 | 0.103 | 0.004 | 0.005 | 0.005 | 0 | 0.32 | 0.34 | 0.33 | 0.37 | 0.35 | 0.45 |
| 2 | 213760193 | rs7581865 | 0.771 | -0.337 | -0.136 | 0.054 | -0.011 | -0.041 | -0.041 | 1.194 | 0.14 | 0.16 | 0.11 | 0 | 0.12 | 0.05 |
| 2 | 213761541 | rs1038323 | 0.48 | 0.003 | -0.458 | 0.333 | -0.001 | 0.006 | 0.006 | 0 | 0.47 | 0.55 | 0.43 | 0.38 | 0.48 | 0.51 |
| 2 | 213765108 | rs6715589 | -0.579 | 0.61 | -1.004 | 0.481 | -0.006 | -0.004 | -0.004 | 0 | 0.23 | 0.2 | 0.29 | 0.3 | 0.37 | 0.19 |
| 2 | 213765361 | rs7566816 | 1.092 | 0.388 | -1.079 | 0.248 | 0.005 | 0.004 | 0.004 | 0 | 0.06 | 0.02 | 0.03 | 0 | 0.04 | 0.01 |
| 2 | 213765929 | rs13388768 | -0.107 | 0.751 | -1.192 | 0.164 | 0.02 | -0.047 | -0.047 | 0 | 0.09 | 0.17 | 0.05 | 0 | 0.09 | 0.08 |
| 2 | 213766025 | rs12151552 | -0.505 | 0.435 | -1.085 | 0.108 | -0.028 | -0.032 | -0.032 | 0 | 0.2 | 0.09 | 0.26 | 0.31 | 0.3 | 0.11 |
| 2 | 213768573 | rs1471641 | 0.126 | -0.268 | -1.694 | 0.13 | -0.015 | -0.002 | -0.002 | 0 | 0.71 | 0.32 | 0.69 | 0.72 | 0.82 | 0.78 |
| 2 | 213769574 | rs12473314 | 1.583 | -0.194 | -2.056 | 0.058 | -0.046 | -0.054 | -0.054 | 0 | 0.41 | 0.09 | 0.38 | 0.42 | 0.42 | 0.58 |
| 2 | 213776349 | rs900369 | 1.438 | 0.283 | -2.392 | -0.007 | -0.164 | -0.12 | -0.12 | 0.206 | 0.34 | 0.03 | 0.37 | 0.4 | 0.47 | 0.48 |
| 2 | 213777980 | rs6761763 | 0.255 | 0.271 | -1.617 | 0.001 | 0.027 | -0.062 | -0.062 | 0 | 0.11 | 0.29 | 0.06 | 0 | 0.09 | 0.11 |
| 2 | 213788877 | rs7563759 | -0.961 | 0.108 | -1.59 | 0.025 | 0.005 | -0.002 | -0.002 | 0 | 0.47 | 0.52 | 0.35 | 0.15 | 0.33 | 0.39 |
| 2 | 213793411 | rs16849449 | 1.531 | -0.021 | -1.663 | 0.18 | 0.001 | -0.018 | -0.018 | 0 | 0.08 | 0.03 | 0.11 | 0.12 | 0.04 | 0.12 |
| 2 | 213794754 | rs11893132 | 0.911 | 0.139 | -1.324 | 0.119 | 0.004 | 0.033 | 0.033 | 0 | 0.05 | 0.06 | 0.13 | 0.18 | 0.06 | 0.04 |
| 2 | 213797880 | rs4672667 | -1.053 | 0.162 | -1.409 | -0.159 | -0.008 | 0.004 | 0.004 | 0 | 0.57 | 0.59 | 0.59 | 0.6 | 0.47 | 0.49 |
| 2 | 213809326 | rs17275294 | -0.048 | 0.032 | -1.396 | -0.607 | -0.017 | 0.047 | 0.047 | 0 | 0.16 | 0.08 | 0.06 | 0 | 0.15 | 0.17 |
| 2 | 213816631 | rs975381 | -1.293 | -0.334 | -1.008 | -0.849 | -0.006 | 0.004 | 0.004 | 2.459 | 0.63 | 0.64 | 0.68 | 0.66 | 0.56 | 0.54 |
| 2 | 213818578 | rs7560454 | 1.19 | -0.727 | -1.061 | -0.973 | -0.001 | 0.01 | 0.01 | 0 | 0.2 | 0.17 | 0.37 | 0.57 | 0.25 | 0.14 |
| 2 | 213820871 | rs4673710 | 1.295 | -0.842 | -1.37 | -1.304 | -0.021 | 0.048 | 0.048 | 0.85 | 0.14 | 0.01 | 0.1 | 0.03 | 0.13 | 0.12 |
| 2 | 213824045 | rs12619285 | 0.776 | -0.882 | -1.021 | -1.295 | -0.001 | 0.048 | 0.048 | 0 | 0.25 | 0.35 | 0.47 | 0.63 | 0.25 | 0.21 |
| 2 | 213824781 | rs1871944 | 1.048 | -0.886 | -1.338 | -1.267 | -0.017 | 0.049 | 0.049 | 0 | 0.16 | 0.08 | 0.06 | 0 | 0.15 | 0.17 |
| 2 | 213825327 | rs13006331 | 2.141 | -0.758 | -1.306 | -1.502 | 0.007 | -0.008 | -0.008 | 0 | 0.21 | 0.17 | 0.14 | 0.33 | 0.18 | 0.39 |
| 2 | 213829682 | rs4673711 | -0.175 | -0.645 | -1.61 | -1.378 | -0.029 | 0.209 | 0.209 | 0 | 0.36 | 0.09 | 0.23 | 0.03 | 0.31 | 0.32 |
| 2 | 213835124 | rs1482577 | 0.483 | -0.555 | -0.907 | -0.928 | 0 | 0 | 0 | 0 | 0.32 | 0.31 | 0.46 | 0.64 | 0.31 | 0.23 |
| 2 | 213838959 | rs1384393 | -1.547 | -0.658 | -0.86 | -1.164 | -0.001 | 0.003 | 0.003 | 0 | 0.92 | 0.95 | 0.9 | 0.98 | 0.93 | 0.89 |
| 2 | 213839773 | rs13023481 | -0.113 | -1.012 | -0.728 | -1.442 | 0.008 | 0.012 | 0.012 | 0 | 0.61 | 0.62 | 0.7 | 0.66 | 0.69 | 0.37 |
| 2 | 213842454 | rs16849504 | 1.58 | -0.975 | -0.677 | -1.419 | -0.001 | 0.016 | 0.016 | 0 | 0.08 | 0.05 | 0.11 | 0.01 | 0.08 | 0.11 |
| 2 | 213843147 | rs7601145 | -0.625 | -1.136 | -0.474 | -1.523 | -0.042 | -0.015 | -0.015 | 0 | 0.7 | 0.5 | 0.73 | 0.66 | 0.77 | 0.5 |
| 2 | 213845835 | rs17276404 | 1.028 | -0.903 | -0.469 | -1.697 | -0.007 | 0.016 | 0.016 | 0.009 | 0.35 | 0.19 | 0.5 | 0.66 | 0.39 | 0.28 |
| 2 | 213849767 | rs10497988 | -0.317 | -1.022 | -0.261 | -1.56 | 0.014 | 0.021 | 0.021 | 0 | 0.42 | 0.4 | 0.3 | 0.34 | 0.31 | 0.64 |
| 2 | 213853203 | rs2371788 | -1.703 | -1.018 | -0.247 | -1.37 | -0.031 | 0.024 | 0.024 | 0.075 | 0.8 | 0.72 | 0.84 | 0.67 | 0.85 | 0.6 |
| 2 | 213855871 | rs7576850 | -0.363 | -0.996 | -0.26 | -1.521 | -0.001 | -0.016 | -0.016 | 0.006 | 0.61 | 0.56 | 0.67 | 0.67 | 0.59 | 0.52 |
| 2 | 213856166 | rs7577413 | 0.213 | -1.023 | -0.313 | -1.571 | -0.001 | 0.007 | 0.007 | 2.498 | 0.51 | 0.54 | 0.64 | 0.66 | 0.51 | 0.4 |
| 2 | 213857830 | rs17355277 | 1.157 | -1.002 | -0.536 | -1.363 | -0.011 | -0.004 | -0.004 | 0.194 | 0.2 | 0.15 | 0.18 | 0 | 0.26 | 0.09 |
| 2 | 213858069 | rs6734978 | 0.114 | -1.021 | -0.499 | -1.187 | 0.015 | 0.025 | 0.025 | 2.902 | 0.57 | 0.36 | 0.64 | 0.63 | 0.58 | 0.52 |
| 2 | 213860159 | rs17355424 | 2.525 | -0.739 | -0.737 | -1.164 | -0.076 | 0.067 | 0.067 | 0.015 | 0.11 | 0.01 | 0.14 | 0 | 0.24 | 0.03 |
| 2 | 213864027 | rs12617251 | 0.179 | -0.945 | -0.63 | -1.177 | 0.009 | 0.047 | 0.047 | 3.933 | 0.57 | 0.21 | 0.62 | 0.64 | 0.57 | 0.52 |
| 2 | 213869354 | rs17277107 | 1.468 | -0.814 | -0.415 | -1.156 | 0.013 | -0.046 | -0.046 | 0.235 | 0.17 | 0.16 | 0.17 | 0 | 0.26 | 0.08 |

| | | | | | | | | | | | | | | | | |
|---|---|---|---|---|---|---|---|---|---|---|---|---|---|---|---|---|
| 2 | 213870678 | rs7607184 | -0.152 | -0.912 | -0.314 | -1.168 | -0.003 | 0.018 | 0.018 | 4.325 | 0.62 | 0.4 | 0.65 | 0.65 | 0.58 | 0.55 |
| 2 | 213871709 | rs3768792 | -1.395 | -0.884 | -0.215 | -1.524 | -0.047 | 0.033 | 0.033 | 0.235 | 0.82 | 0.58 | 0.86 | 0.79 | 0.86 | 0.68 |
| 2 | 213877864 | rs9808132 | -1.438 | -0.886 | -0.219 | -1.548 | -0.045 | 0.013 | 0.013 | 0.198 | 0.83 | 0.58 | 0.86 | 0.8 | 0.86 | 0.7 |
| 2 | 213879274 | rs1384397 | 1.209 | -0.84 | -0.29 | -1.576 | 0.008 | -0.04 | -0.04 | 0 | 0.18 | 0.18 | 0.17 | 0.01 | 0.25 | 0.07 |
| 2 | 213879407 | rs1871946 | 0.969 | -1.064 | -0.744 | -1.631 | -0.104 | 0.021 | 0.021 | 0.89 | 0.39 | 0.04 | 0.51 | 0.68 | 0.43 | 0.31 |
| 2 | 213892890 | rs10178238 | 0.011 | -0.919 | -0.536 | -1.523 | -0.004 | 0.053 | 0.053 | 3.6 | 0.62 | 0.35 | 0.67 | 0.79 | 0.58 | 0.58 |
| 2 | 213897308 | rs10932459 | -1.498 | -1.361 | -0.344 | -1.329 | -0.065 | 0.039 | 0.039 | 0.196 | 0.84 | 0.63 | 0.88 | 0.78 | 0.92 | 0.67 |
| 2 | 213903517 | rs13023118 | 1.667 | -1.371 | -0.42 | -1.386 | 0.004 | -0.002 | -0.002 | 0 | 0.14 | 0.11 | 0.1 | 0.21 | 0.08 | 0.33 |
| 2 | 213909647 | rs7585103 | 0.194 | -1.339 | -0.765 | -1.522 | 0.002 | -0.018 | -0.018 | 0 | 0.31 | 0.37 | 0.29 | 0.23 | 0.35 | 0.4 |
| 2 | 213911042 | rs1482579 | 0.247 | -1.257 | -0.825 | -1.639 | 0.009 | 0.044 | 0.044 | 2.732 | 0.57 | 0.39 | 0.65 | 0.75 | 0.58 | 0.55 |
| 2 | 213911121 | rs12612034 | 0.438 | -1.188 | -1.204 | -1.062 | -0.144 | -0.01 | -0.01 | 0 | 0.39 | 0.03 | 0.52 | 0.65 | 0.48 | 0.29 |
| 2 | 213923915 | rs11885466 | 0.496 | -0.871 | -0.082 | -0.359 | -0.059 | 0.011 | 0.011 | 0.177 | 0.16 | 0.33 | 0.12 | 0.21 | 0.08 | 0.32 |
| 2 | 213929224 | rs13008255 | -0.935 | -0.079 | -0.167 | -0.189 | 0.02 | -0.024 | -0.024 | 0 | 0.08 | 0.17 | 0.05 | 0.05 | 0.07 | 0.06 |
| 2 | 213929286 | rs10186029 | -0.158 | -0.379 | -0.451 | -0.665 | -0.003 | 0.045 | 0.045 | 0 | 0.43 | 0.28 | 0.47 | 0.76 | 0.45 | 0.36 |
| 2 | 213935900 | rs10203838 | 0.242 | -1.248 | -0.701 | -1.17 | 0.002 | 0.045 | 0.045 | 0 | 0.46 | 0.62 | 0.28 | 0.02 | 0.42 | 0.26 |
| 2 | 213937653 | rs16849639 | -0.067 | -1.599 | -0.543 | -1.296 | -0.083 | -0.046 | -0.046 | 4.077 | 0.13 | 0.39 | 0.03 | 0.02 | 0.02 | 0.03 |
| 2 | 213943634 | rs1530239 | -1.355 | -1.114 | -1.663 | -1.032 | 0.008 | -0.012 | -0.012 | 0 | 0.88 | 0.91 | 0.92 | 0.81 | 0.91 | 0.64 |
| 2 | 213965322 | rs4673725 | -0.195 | -1.214 | -1.902 | -1.2 | 0.008 | -0.008 | -0.008 | 0.581 | 0.8 | 0.9 | 0.9 | 0.8 | 0.84 | 0.58 |
| 2 | 213968775 | rs10497991 | 0.867 | -1.343 | -2.053 | -1.083 | -0.135 | -0.054 | -0.054 | 0 | 0.33 | 0.03 | 0.58 | 0.71 | 0.43 | 0.29 |
| 2 | 213976427 | rs10207421 | 0.891 | -1.161 | -2.104 | -0.465 | -0.112 | -0.174 | -0.174 | 0 | 0.32 | 0.02 | 0.41 | 0.54 | 0.39 | 0.25 |
| 2 | 213978117 | rs7565643 | 0.668 | -1.252 | -1.779 | -0.679 | -0.04 | -0.007 | -0.007 | 0 | 0.08 | 0.25 | 0.02 | 0.02 | 0.01 | 0.02 |
| 2 | 213978526 | rs10179508 | 0.596 | -1.194 | -1.962 | -0.669 | -0.138 | -0.034 | -0.034 | 0 | 0.36 | 0.03 | 0.6 | 0.71 | 0.46 | 0.31 |
| 2 | 213980181 | rs10196879 | 0.404 | -0.977 | -1.659 | -0.94 | 0.058 | 0.095 | 0.095 | 4.91 | 0.07 | 0.46 | 0.06 | 0.06 | 0.04 | 0.08 |
| 2 | 213980453 | rs6435757 | 1.34 | -0.799 | -2.984 | -0.909 | -0.121 | 0.217 | 0.217 | 0.002 | 0.31 | 0.03 | 0.23 | 0 | 0.36 | 0.18 |
| 2 | 213986695 | rs920525 | 0.517 | -0.867 | -2.945 | -0.856 | 0.053 | 0.05 | 0.05 | 1.942 | 0.77 | 0.14 | 0.89 | 0.92 | 0.9 | 0.8 |
| 2 | 213987806 | rs4672674 | -1.463 | -0.718 | -2.975 | -0.76 | 0.001 | -0.01 | -0.01 | 1.179 | 0.94 | 0.91 | 0.97 | 0.99 | 0.95 | 0.91 |
| 2 | 214001948 | rs4672676 | 0.011 | -0.618 | -3.311 | -0.73 | -0.135 | 0.252 | 0.252 | 1.075 | 0.46 | 0.05 | 0.65 | 0.91 | 0.54 | 0.62 |
| 2 | 214006439 | rs16849846 | 0.67 | 0.328 | -2.856 | -0.078 | -0.087 | -0.116 | -0.116 | 0 | 0.16 | 0.01 | 0.26 | 0.38 | 0.25 | 0.15 |
| 2 | 214012508 | rs6709554 | 0.501 | -0.064 | -2.44 | -0.729 | -0.096 | 0.193 | 0.193 | 0 | 0.3 | 0.05 | 0.24 | 0 | 0.35 | 0.18 |
| 2 | 214015305 | rs2197025 | 0.052 | -0.102 | -2.007 | 0 | -0.071 | -0.062 | -0.062 | 0 | 0.35 | 0.15 | 0.39 | 0.48 | 0.45 | 0.29 |
| 2 | 214015745 | rs3768788 | 0.431 | 0.052 | -2.115 | -0.081 | -0.064 | -0.097 | -0.097 | 0 | 0.3 | 0.11 | 0.48 | 0.47 | 0.37 | 0.19 |
| 2 | 214023052 | rs12471471 | -0.173 | -0.485 | -1.94 | -0.784 | 0.001 | 0.102 | 0.102 | 0 | 0.51 | 0.28 | 0.69 | 0.94 | 0.53 | 0.61 |
| 2 | 214026818 | rs11889674 | 0.814 | -0.201 | -1.936 | -0.385 | -0.125 | -0.149 | -0.149 | 0.483 | 0.28 | 0.02 | 0.46 | 0.42 | 0.36 | 0.19 |
| 2 | 214036796 | rs12997801 | 0.444 | -0.43 | -1.561 | -0.391 | -0.002 | 0.007 | 0.007 | 0 | 0.22 | 0.16 | 0.21 | 0.52 | 0.17 | 0.42 |
| 2 | 214037703 | rs1441172 | 0.447 | 0.118 | -1.265 | -1.086 | -0.089 | 0.177 | 0.177 | 0 | 0.3 | 0.08 | 0.24 | 0.01 | 0.38 | 0.21 |
| 2 | 214038302 | rs10932472 | 0.057 | -0.072 | -0.619 | -0.942 | 0.004 | -0.056 | -0.056 | 0 | 0.86 | 0.56 | 0.94 | 0.96 | 0.91 | 0.85 |
| 2 | 214039733 | rs4567882 | 0.045 | 0.082 | -0.714 | -0.749 | -0.002 | -0.021 | -0.021 | 0.073 | 0.58 | 0.37 | 0.72 | 0.95 | 0.6 | 0.66 |
| 2 | 214047628 | rs935093 | -0.65 | -0.176 | 0.204 | -0.279 | -0.01 | -0.068 | -0.068 | 0 | 0.13 | 0.31 | 0.1 | 0.02 | 0.11 | 0.09 |
| 2 | 214049210 | rs7582450 | -0.529 | -0.526 | 0.047 | -0.057 | -0.001 | -0.001 | -0.001 | 0 | 0.43 | 0.6 | 0.39 | 0.43 | 0.37 | 0.26 |
| 2 | 214061022 | rs16849921 | 0.754 | -0.589 | -0.026 | -0.09 | 0.007 | -0.043 | -0.043 | 1.305 | 0.28 | 0.12 | 0.28 | 0.37 | 0.26 | 0.17 |
| 2 | 214067583 | rs16849941 | -0.65 | -0.846 | 0.867 | -0.29 | -0.059 | -0.004 | -0.004 | 0 | 0.1 | 0.21 | 0.02 | 0.08 | 0.01 | 0.03 |
| 2 | 214080362 | rs6435758 | -0.915 | -0.275 | 0.88 | -0.197 | -0.013 | -0.097 | -0.097 | 0.021 | 0.15 | 0.49 | 0.12 | 0.02 | 0.11 | 0.09 |
| 2 | 214094200 | rs6435760 | -0.14 | -0.081 | 0.176 | -0.068 | 0.008 | -0.052 | -0.052 | 0 | 0.09 | 0.25 | 0.08 | 0 | 0.08 | 0.02 |
| 2 | 214101248 | rs17361996 | -0.485 | 0.493 | -0.331 | -0.014 | -0.051 | 0.008 | 0.008 | 0.021 | 0.08 | 0.01 | 0.12 | 0.05 | 0.14 | 0.16 |
| 2 | 214113474 | rs13001761 | -1.007 | -0.401 | 0.116 | 0.125 | -0.065 | -0.021 | -0.021 | 0.684 | 0.2 | 0.4 | 0.13 | 0.15 | 0.09 | 0.06 |
| 2 | 214115067 | rs16850034 | -1.52 | -0.387 | -0.745 | -0.002 | -0.021 | -0.045 | -0.045 | 0 | 0.23 | 0.09 | 0.17 | 0.24 | 0.19 | 0.36 |
| 2 | 214115898 | rs10181294 | -0.567 | -0.091 | -1.566 | -1.017 | -0.099 | 0.111 | 0.111 | 0 | 0.39 | 0.09 | 0.31 | 0.17 | 0.45 | 0.42 |
| 2 | 214117303 | rs10185045 | 0.659 | -0.196 | -1.612 | -1.109 | -0.04 | 0.122 | 0.122 | 0 | 0.26 | 0.07 | 0.2 | 0 | 0.28 | 0.14 |
| 2 | 214135710 | rs16850094 | 0.969 | -0.361 | -1.775 | -0.569 | -0.035 | -0.115 | -0.115 | 0.344 | 0.15 | 0.01 | 0.28 | 0.48 | 0.18 | 0.39 |
| 2 | 214138425 | rs1219973 | 0.546 | -0.38 | -0.831 | -0.899 | 0.009 | -0.161 | -0.161 | 0 | 0.29 | 0.45 | 0.23 | 0.01 | 0.27 | 0.12 |
| 2 | 214148299 | rs10164670 | 1.549 | -0.417 | -0.944 | -0.911 | -0.094 | 0.138 | 0.138 | 0.274 | 0.22 | 0.02 | 0.18 | 0 | 0.26 | 0.08 |
| 2 | 214153984 | rs2708175 | 0.266 | -0.643 | -0.76 | -1.032 | 0 | -0.001 | -0.001 | 0 | 0.65 | 0.65 | 0.54 | 0.3 | 0.61 | 0.31 |
| 2 | 214156385 | rs6723575 | 1.636 | -0.636 | -0.952 | -0.878 | -0.005 | -0.013 | -0.013 | 0 | 0.31 | 0.12 | 0.3 | 0.29 | 0.33 | 0.18 |
| 2 | 214163007 | rs12472656 | 1.543 | -0.598 | -0.982 | -0.896 | -0.015 | -0.015 | -0.015 | 0.837 | 0.32 | 0.12 | 0.3 | 0.3 | 0.34 | 0.19 |
| 2 | 214171778 | rs10203794 | 0.857 | -0.705 | -0.75 | -1.078 | 0.009 | -0.116 | -0.116 | 0.239 | 0.28 | 0.37 | 0.22 | 0.01 | 0.27 | 0.11 |
| 2 | 214174298 | rs11678617 | 1.341 | -0.728 | -0.761 | -1.089 | -0.046 | 0.096 | 0.096 | 0.01 | 0.24 | 0.06 | 0.19 | 0.01 | 0.27 | 0.11 |
| 2 | 214214297 | rs6435770 | 1.765 | -0.786 | -0.82 | -1.109 | -0.012 | -0.018 | -0.018 | 0.65 | 0.32 | 0.08 | 0.3 | 0.3 | 0.33 | 0.19 |
| 2 | 214217706 | rs10804214 | -0.613 | -0.788 | -0.639 | -1.072 | 0 | -0.001 | -0.001 | 0 | 0.35 | 0.35 | 0.47 | 0.7 | 0.39 | 0.69 |
| 2 | 214296544 | rs2134602 | -1.09 | -0.964 | -0.679 | -1.068 | -0.055 | -0.099 | -0.099 | 0.134 | 0.13 | 0.46 | 0.07 | 0 | 0.04 | 0.06 |
| 2 | 214307024 | rs1604692 | 0.784 | -0.896 | -0.654 | -0.978 | -0.069 | -0.062 | -0.062 | 0 | 0.12 | 0.01 | 0.16 | 0.19 | 0.18 | 0.28 |
| 2 | 214354100 | rs7563592 | 0.393 | -0.992 | -0.611 | -0.988 | 0.03 | 0.048 | 0.048 | 1.077 | 0.51 | 0.62 | 0.37 | 0.28 | 0.39 | 0.3 |
| 2 | 214385908 | rs1813708 | -0.43 | -0.867 | -0.071 | -0.95 | -0.011 | -0.036 | -0.036 | 0 | 0.92 | 0.77 | 0.98 | 1 | 0.98 | 0.97 |
| 2 | 214394505 | rs2113473 | 1.8 | -0.871 | -0.495 | -0.927 | -0.022 | -0.027 | -0.027 | 3.375 | 0.32 | 0.08 | 0.3 | 0.3 | 0.34 | 0.23 |
| 2 | 214469961 | rs4673748 | -0.371 | -0.551 | -0.346 | -0.835 | 0.023 | 0.001 | 0.001 | 0 | 0.19 | 0.3 | 0.2 | 0.17 | 0.2 | 0.28 |
| 2 | 214493601 | rs1554177 | 0.735 | -0.803 | -0.711 | -1.001 | -0.005 | -0.005 | -0.005 | 0.038 | 0.45 | 0.53 | 0.36 | 0.33 | 0.38 | 0.29 |
| 2 | 214513147 | rs6435780 | 1.07 | -0.841 | -0.661 | -1.026 | 0.021 | 0.014 | 0.014 | 0.01 | 0.4 | 0.22 | 0.33 | 0.33 | 0.36 | 0.25 |
| 2 | 214527841 | rs1515915 | 1.075 | -0.856 | -0.741 | -1.036 | -0.16 | -0.181 | -0.181 | 0 | 0.31 | 0.02 | 0.44 | 0.49 | 0.42 | 0.42 |
| 2 | 214529976 | rs13024335 | 0.784 | -0.777 | -0.349 | -0.92 | 0.012 | 0.007 | 0.007 | 1.838 | 0.39 | 0.34 | 0.33 | 0.33 | 0.34 | 0.23 |
| 2 | 214556527 | rs4132244 | 1.182 | -0.782 | -0.397 | -0.807 | 0.001 | 0.006 | 0.006 | 0 | 0.34 | 0.25 | 0.32 | 0.29 | 0.35 | 0.24 |

| | | | | | | | | | | | | | | | | |
|---|---|---|---|---|---|---|---|---|---|---|---|---|---|---|---|---|
| 2 | 214561068 | rs13020757 | 1.414 | -0.692 | -0.596 | -0.844 | -0.026 | -0.047 | -0.047 | 0 | 0.32 | 0.07 | 0.29 | 0.33 | 0.34 | 0.24 |
| 2 | 214563795 | rs7592552 | 0.275 | -0.539 | -0.414 | -0.697 | 0.009 | 0.014 | 0.014 | 0.039 | 0.47 | 0.62 | 0.36 | 0.34 | 0.4 | 0.29 |
| 2 | 214572679 | rs6758514 | 0.227 | -0.515 | 0.051 | -0.451 | -0.006 | -0.008 | -0.008 | 0 | 0.05 | 0.28 | 0.02 | 0 | 0 | 0.01 |
| 2 | 214580309 | rs727503 | -0.323 | -0.361 | -0.378 | -0.496 | 0.001 | 0.002 | 0.002 | 0.071 | 0.63 | 0.62 | 0.55 | 0.47 | 0.61 | 0.55 |
| 2 | 214584354 | rs2162505 | 1.49 | -0.127 | -1.273 | -0.419 | -0.04 | 0.006 | 0.006 | 0.275 | 0.24 | 0.1 | 0.22 | 0.26 | 0.31 | 0.17 |
| 2 | 214594071 | rs16850783 | 0.572 | -0.261 | -1.443 | -0.193 | 0.01 | -0.03 | -0.03 | 0 | 0.19 | 0.13 | 0.3 | 0.5 | 0.16 | 0.39 |
| 2 | 214594856 | rs9283526 | 0.098 | -0.279 | -1.369 | -0.244 | -0.006 | 0.025 | 0.025 | 0 | 0.23 | 0.22 | 0.32 | 0.52 | 0.21 | 0.41 |
| 2 | 214600292 | rs724954 | 0.366 | -0.247 | -1.792 | -0.162 | -0.034 | -0.004 | -0.004 | 0.589 | 0.35 | 0.14 | 0.45 | 0.59 | 0.4 | 0.46 |
| 2 | 214610452 | rs13007000 | 0.074 | -0.057 | -1.873 | -0.039 | -0.014 | 0.038 | 0.038 | 1.29 | 0.48 | 0.18 | 0.48 | 0.42 | 0.54 | 0.47 |
| 2 | 214628482 | rs17754969 | 0.151 | 0.023 | -1.797 | -0.437 | -0.011 | 0.105 | 0.105 | 0 | 0.25 | 0.1 | 0.23 | 0.05 | 0.23 | 0.28 |
| 2 | 214639412 | rs6739421 | 0.488 | -0.014 | -2.015 | -0.32 | -0.063 | 0.058 | 0.058 | 1.214 | 0.33 | 0.14 | 0.25 | 0.28 | 0.34 | 0.15 |
| 2 | 214640685 | rs4672684 | 0.726 | -0.421 | -1.776 | -0.213 | -0.032 | -0.01 | -0.01 | 0 | 0.35 | 0.11 | 0.47 | 0.62 | 0.39 | 0.52 |
| 2 | 214662575 | rs6720110 | 0.983 | -0.243 | -1.808 | 0.007 | -0.044 | -0.012 | -0.012 | 0.064 | 0.3 | 0.13 | 0.32 | 0.32 | 0.3 | 0.13 |
| 2 | 214667730 | rs2556296 | 0.843 | -0.319 | -0.88 | -0.01 | -0.001 | 0.002 | 0.002 | 2.295 | 0.31 | 0.32 | 0.35 | 0.28 | 0.31 | 0.13 |
| 2 | 214673201 | rs10166032 | -0.773 | -0.433 | -0.262 | -0.068 | 0.003 | 0.02 | 0.02 | 2.89 | 0.65 | 0.3 | 0.67 | 0.73 | 0.69 | 0.87 |
| 2 | 214677478 | rs1834774 | 0.883 | -0.56 | -0.416 | -0.131 | -0.064 | -0.094 | -0.094 | 0 | 0.35 | 0.07 | 0.39 | 0.56 | 0.38 | 0.5 |
| 2 | 214683214 | rs1582839 | 0.826 | -0.354 | -0.364 | -0.1 | -0.007 | 0.097 | 0.097 | 0 | 0.22 | 0.08 | 0.23 | 0.04 | 0.24 | 0.32 |
| 2 | 214698536 | rs2556323 | 0.872 | -0.45 | -0.107 | -0.081 | -0.002 | 0.013 | 0.013 | 3.464 | 0.32 | 0.3 | 0.31 | 0.29 | 0.31 | 0.13 |
| 2 | 214702812 | rs2372257 | 0.318 | -0.41 | -0.072 | -0.18 | -0.046 | 0.072 | 0.072 | 0.14 | 0.26 | 0.06 | 0.25 | 0.09 | 0.3 | 0.36 |
| 2 | 214703618 | rs4673763 | 0.114 | -0.419 | 0.349 | -0.179 | -0.005 | -0.017 | -0.017 | 0.089 | 0.69 | 0.39 | 0.7 | 0.88 | 0.69 | 0.64 |
| 2 | 214717579 | rs10498015 | 0.583 | -0.361 | 0.505 | 0.112 | -0.002 | -0.021 | -0.021 | 0 | 0.38 | 0.25 | 0.4 | 0.5 | 0.37 | 0.49 |
| 2 | 214717896 | rs13388769 | 0.58 | -0.348 | 0.433 | 0.119 | -0.005 | -0.036 | -0.036 | 0.193 | 0.38 | 0.21 | 0.4 | 0.5 | 0.37 | 0.5 |
| 2 | 214727221 | rs2042791 | 0.53 | -0.391 | 0.417 | 0.1 | -0.005 | -0.037 | -0.037 | 0 | 0.38 | 0.21 | 0.4 | 0.5 | 0.37 | 0.51 |
| 2 | 214729674 | rs7572850 | 0.284 | -0.377 | 0.381 | 0.079 | -0.022 | 0.075 | 0.075 | 0.186 | 0.27 | 0.08 | 0.25 | 0.09 | 0.3 | 0.37 |
| 2 | 214740120 | rs13416548 | 0.692 | -0.375 | 0.445 | 0.109 | -0.007 | -0.035 | -0.035 | 4.72 | 0.37 | 0.21 | 0.4 | 0.48 | 0.37 | 0.5 |
| 2 | 214756500 | rs10191167 | -0.733 | -0.349 | 0.261 | 0.123 | 0 | 0.001 | 0.001 | 4.439 | 0.66 | 0.3 | 0.65 | 0.61 | 0.67 | 0.87 |
| 2 | 214776240 | rs16851079 | 0.291 | -0.323 | 0.138 | -0.008 | -0.023 | 0.059 | 0.059 | 0.362 | 0.27 | 0.09 | 0.25 | 0.11 | 0.29 | 0.36 |
| 2 | 214785551 | rs1588453 | 0.3 | -0.285 | 0.484 | -0.009 | -0.009 | 0.007 | 0.007 | 3.311 | 0.89 | 0.51 | 0.95 | 0.96 | 0.95 | 0.97 |
| 2 | 214794743 | rs12623569 | 1.359 | -0.182 | 0.163 | 0.227 | -0.001 | -0.001 | -0.001 | 0 | 0.26 | 0.27 | 0.31 | 0.41 | 0.28 | 0.11 |
| 2 | 214802586 | rs1027939 | -1.57 | -0.109 | -0.126 | 0.252 | -0.014 | -0.038 | -0.038 | 0 | 0.75 | 0.85 | 0.7 | 0.58 | 0.71 | 0.87 |
| 2 | 214806075 | rs953133 | -0.232 | -0.209 | -0.084 | 0.177 | 0.001 | -0.018 | -0.018 | 0 | 0.6 | 0.26 | 0.59 | 0.47 | 0.62 | 0.79 |
| 2 | 214810651 | rs11675655 | 0.761 | -0.163 | -0.261 | 0.122 | -0.015 | -0.02 | -0.02 | 0.038 | 0.37 | 0.19 | 0.37 | 0.43 | 0.37 | 0.5 |
| 2 | 214829765 | rs1877050 | -1.631 | -0.127 | -0.279 | 0.121 | -0.003 | -0.008 | -0.008 | 0.495 | 0.74 | 0.78 | 0.69 | 0.58 | 0.71 | 0.87 |
| 2 | 214832161 | rs10167662 | -0.107 | -0.011 | -0.318 | 0.29 | -0.002 | -0.01 | -0.01 | 0 | 0.45 | 0.37 | 0.43 | 0.47 | 0.42 | 0.53 |
| 2 | 214838513 | rs9789347 | 0.917 | 0.079 | -0.831 | 0.51 | -0.027 | -0.068 | -0.068 | 0 | 0.26 | 0.05 | 0.29 | 0.4 | 0.24 | 0.46 |
| 2 | 214842289 | rs6735162 | -0.974 | -0.137 | 0.24 | 1.001 | -0.009 | 0.052 | 0.052 | 0.058 | 0.17 | 0.59 | 0.19 | 0.19 | 0.15 | 0.19 |
| 2 | 214848645 | rs9283527 | -0.913 | -0.282 | 0.006 | 0.684 | 0.033 | 0.015 | 0.015 | 0 | 0.52 | 0.62 | 0.43 | 0.55 | 0.38 | 0.69 |
| 2 | 214859650 | rs1510547 | -1.139 | -0.183 | -0.193 | 0.573 | 0.02 | -0.038 | -0.038 | 0 | 0.55 | 0.77 | 0.48 | 0.5 | 0.43 | 0.8 |
| 2 | 214865555 | rs7578505 | 0.296 | 0.086 | -0.139 | 0.625 | 0.058 | 0.043 | 0.043 | 0 | 0.1 | 0.29 | 0.06 | 0.08 | 0.1 | 0.08 |
| 2 | 214867770 | rs10210989 | -1.161 | -0.207 | -0.273 | 0.598 | 0.019 | -0.013 | -0.013 | 0.133 | 0.54 | 0.62 | 0.47 | 0.5 | 0.43 | 0.8 |
| 2 | 214871591 | rs1510548 | 0.106 | -0.316 | -0.661 | 0.434 | 0.078 | -0.024 | -0.024 | 0.009 | 0.36 | 0.1 | 0.29 | 0.39 | 0.25 | 0.61 |
| 2 | 214880748 | rs1033070 | -0.214 | -0.03 | -0.544 | 0.486 | 0.013 | 0.007 | 0.007 | 0 | 0.43 | 0.33 | 0.43 | 0.43 | 0.39 | 0.73 |
| 2 | 214884987 | rs4673807 | -0.195 | -0.04 | -0.535 | 0.494 | 0.013 | 0.006 | 0.006 | 0 | 0.43 | 0.34 | 0.43 | 0.43 | 0.38 | 0.73 |
| 2 | 214885653 | rs2055867 | -0.808 | 0.091 | -0.407 | 0.69 | -0.008 | 0.105 | 0.105 | 0 | 0.12 | 0.29 | 0.09 | 0.27 | 0.08 | 0.09 |
| 2 | 214887022 | rs11675561 | 0.106 | 0.368 | -1.181 | 0.074 | -0.048 | 0.027 | 0.027 | 1.983 | 0.42 | 0.2 | 0.43 | 0.34 | 0.5 | 0.18 |
| 2 | 214888568 | rs13002799 | 0.277 | 0.571 | -0.462 | 0.439 | 0.002 | 0.044 | 0.044 | 0 | 0.08 | 0.1 | 0.08 | 0.22 | 0.09 | 0.07 |
| 2 | 214890634 | rs1510551 | 0.431 | -0.036 | -0.031 | 0.088 | -0.008 | 0.002 | 0.002 | 0 | 0.61 | 0.48 | 0.74 | 0.5 | 0.68 | 0.69 |
| 2 | 214891902 | rs2193786 | 1.18 | 0.464 | -0.067 | -0.177 | -0.007 | 0 | 0 | 0 | 0.33 | 0.27 | 0.25 | 0.04 | 0.32 | 0.28 |
| 2 | 214895245 | rs6756845 | 0.091 | 0.175 | -0.224 | -0.289 | -0.042 | -0.055 | -0.055 | 0 | 0.34 | 0.16 | 0.52 | 0.47 | 0.42 | 0.52 |
| 2 | 214897080 | rs11901290 | 0.226 | 0.037 | 0.05 | -0.235 | -0.003 | 0.057 | 0.057 | 0 | 0.12 | 0.19 | 0.1 | 0.22 | 0.11 | 0.09 |
| 2 | 214908936 | rs17207263 | 1.709 | 0.099 | -0.082 | -0.399 | -0.003 | -0.036 | -0.036 | 0 | 0.29 | 0.25 | 0.18 | 0.01 | 0.28 | 0.12 |
| 2 | 214916394 | rs1397354 | 0.373 | 0.032 | 0.145 | -0.311 | -0.059 | -0.057 | -0.057 | 1.877 | 0.32 | 0.19 | 0.52 | 0.5 | 0.44 | 0.53 |
| 2 | 214917099 | rs17207529 | 0.173 | 0.109 | 0.148 | -0.29 | 0.007 | 0.03 | 0.03 | 0 | 0.14 | 0.17 | 0.24 | 0.29 | 0.2 | 0.17 |
| 2 | 214917402 | rs1510552 | -1.342 | 0.049 | 0.215 | -0.332 | -0.017 | 0.016 | 0.016 | 0 | 0.81 | 0.66 | 0.86 | 0.81 | 0.84 | 0.91 |
| 2 | 214919665 | rs2372221 | -0.453 | 0.22 | 0.229 | -0.232 | -0.039 | -0.046 | -0.046 | 0 | 0.38 | 0.21 | 0.58 | 0.59 | 0.46 | 0.69 |
| 2 | 214928336 | rs11897357 | 0.366 | 0.231 | 0.258 | -0.243 | -0.002 | 0.014 | 0.014 | 0 | 0.12 | 0.12 | 0.09 | 0.21 | 0.11 | 0.09 |
| 2 | 214929313 | rs11689583 | 1.975 | 0.231 | 0.224 | -0.333 | -0.029 | 0.136 | 0.136 | 0 | 0.28 | 0.08 | 0.17 | 0 | 0.25 | 0.13 |
| 2 | 214933509 | rs11687649 | -0.72 | 0.073 | 0.356 | -0.318 | -0.001 | 0.002 | 0.002 | 0 | 0.72 | 0.66 | 0.6 | 0.28 | 0.71 | 0.55 |
| 2 | 214936393 | rs17227489 | 1.25 | 0.243 | 0.438 | -0.48 | -0.001 | 0.038 | 0.038 | 0 | 0.35 | 0.23 | 0.26 | 0.03 | 0.32 | 0.32 |
| 2 | 214940734 | rs6715771 | 1.426 | 0.244 | 0.445 | -0.483 | -0.002 | -0.036 | -0.036 | 0 | 0.19 | 0.2 | 0.1 | 0.01 | 0.15 | 0.07 |
| 2 | 214940909 | rs6715997 | -0.856 | 0.267 | 0.372 | -0.404 | -0.002 | -0.011 | -0.011 | 0.022 | 0.72 | 0.76 | 0.65 | 0.49 | 0.72 | 0.61 |
| 2 | 214941899 | rs1030143 | -0.125 | 0.287 | 0.358 | -0.366 | -0.001 | -0.009 | -0.009 | 0 | 0.12 | 0.13 | 0.19 | 0.03 | 0.18 | 0.15 |
| 2 | 214943848 | rs7590446 | -0.01 | 0.026 | 0.582 | -0.248 | -0.007 | 0.012 | 0.012 | 0 | 0.68 | 0.41 | 0.77 | 0.54 | 0.73 | 0.82 |
| 2 | 214947117 | rs16851422 | -0.272 | 0.108 | 0.469 | -0.234 | -0.003 | 0.029 | 0.029 | 0 | 0.14 | 0.14 | 0.15 | 0.3 | 0.14 | 0.23 |
| 2 | 214956523 | rs2080776 | -0.027 | 0.05 | 0.532 | -0.342 | 0 | -0.036 | -0.036 | 0 | 0.6 | 0.63 | 0.51 | 0.07 | 0.58 | 0.46 |
| 2 | 214958515 | rs11688016 | 0.528 | 0.066 | 0.635 | -0.379 | -0.002 | 0.01 | 0.01 | 0 | 0.52 | 0.45 | 0.34 | 0.04 | 0.46 | 0.36 |
| 2 | 214959639 | rs7601427 | 0.467 | 0.076 | 0.608 | -0.314 | 0.025 | -0.026 | -0.026 | 0.044 | 0.09 | 0.18 | 0.17 | 0.03 | 0.13 | 0.1 |
| 2 | 214960114 | rs13014300 | 1.257 | 0.044 | 0.595 | -0.298 | 0.004 | 0 | 0 | 0 | 0.2 | 0.17 | 0.13 | 0.16 | 0.17 | 0.09 |
| 2 | 214966578 | rs13387383 | -0.072 | 0.043 | 0.593 | -0.309 | -0.002 | -0.014 | -0.014 | 0 | 0.31 | 0.35 | 0.32 | 0.19 | 0.32 | 0.24 |

| | | | | | | | | | | | | | | | | |
|---|---|---|---|---|---|---|---|---|---|---|---|---|---|---|---|---|
| 2 | 214967092 | rs2193788 | -0.286 | 0.078 | 0.591 | -1.016 | 0.001 | 0.037 | 0.037 | 0 | 0.33 | 0.26 | 0.33 | 0.07 | 0.33 | 0.39 |
| 2 | 214968118 | rs7561816 | -1.02 | 0.207 | 0.637 | -0.858 | -0.001 | -0.007 | -0.007 | 0 | 0.7 | 0.7 | 0.58 | 0.34 | 0.67 | 0.55 |
| 2 | 214968212 | rs10514629 | 0.632 | 0.282 | 0.587 | -0.877 | 0.038 | -0.021 | -0.021 | 1.045 | 0.08 | 0.19 | 0.18 | 0.03 | 0.13 | 0.1 |
| 2 | 214969001 | rs4541202 | -0.278 | 0.298 | 0.282 | -0.884 | 0 | -0.032 | -0.032 | 0 | 0.6 | 0.63 | 0.51 | 0.07 | 0.59 | 0.46 |
| 2 | 214969898 | rs13387629 | 0.484 | 0.573 | 0.56 | -0.138 | 0 | -0.001 | -0.001 | 0 | 0.07 | 0.07 | 0.17 | 0.03 | 0.12 | 0.1 |
| 2 | 214974352 | rs11685260 | 0.421 | 0.137 | 0.515 | 0.112 | -0.018 | 0.089 | 0.089 | 0 | 0.29 | 0.15 | 0.18 | 0.03 | 0.27 | 0.14 |
| 2 | 214990437 | rs11900320 | 0.078 | 0.433 | -0.985 | 0.23 | -0.001 | 0.085 | 0.085 | 0 | 0.09 | 0.12 | 0.07 | 0.26 | 0.09 | 0.08 |
| 2 | 214993997 | rs2161022 | -0.783 | 0.432 | -1.063 | 0.224 | -0.006 | 0.075 | 0.075 | 0 | 0.12 | 0.18 | 0.13 | 0.27 | 0.11 | 0.21 |
| 2 | 214994793 | rs1014450 | -0.197 | 0.633 | -0.923 | 0.502 | -0.005 | 0.011 | 0.011 | 0 | 0.38 | 0.27 | 0.26 | 0.3 | 0.36 | 0.21 |
| 2 | 215019834 | rs6755912 | -0.789 | 0.541 | 0.062 | 0.789 | 0.022 | 0.03 | 0.03 | 0 | 0.19 | 0.43 | 0.25 | 0.18 | 0.17 | 0.31 |
| 2 | 215024392 | rs16851554 | -0.126 | 0.262 | -0.986 | 0.884 | -0.029 | -0.074 | -0.074 | 0 | 0.21 | 0.01 | 0.26 | 0.28 | 0.14 | 0.13 |
| 2 | 215032580 | rs11895074 | 0.297 | -0.199 | -0.723 | 0.505 | -0.005 | -0.09 | -0.09 | 3.217 | 0.84 | 0.58 | 0.9 | 0.96 | 0.9 | 0.96 |
| 2 | 215041348 | rs10460418 | 0.368 | -0.142 | -1.481 | 0.687 | -0.045 | -0.001 | -0.001 | 3.22 | 0.28 | 0.12 | 0.22 | 0.26 | 0.32 | 0.34 |
| 2 | 215045596 | rs7581545 | 0.819 | -0.029 | -1.509 | 0.655 | 0.001 | 0.002 | 0.002 | 0 | 0.48 | 0.53 | 0.57 | 0.61 | 0.46 | 0.4 |
| 2 | 215048212 | rs10190431 | 1.499 | -0.013 | -1.499 | 0.858 | 0.002 | 0.002 | 0.002 | 0 | 0.37 | 0.52 | 0.37 | 0.37 | 0.44 | 0.31 |
| 2 | 215048355 | rs10490497 | 1.139 | -0.123 | -1.518 | 1.544 | 0.036 | 0.333 | 0.333 | 2.298 | 0.05 | 0.37 | 0.07 | 0.2 | 0.01 | 0.01 |
| 2 | 215050520 | rs1912196 | 0.716 | 0.017 | -1.508 | 1.08 | -0.04 | -0.007 | -0.007 | 0 | 0.24 | 0.06 | 0.14 | 0.22 | 0.25 | 0.25 |
| 2 | 215052524 | rs11887857 | -0.137 | -0.042 | -0.537 | 1.742 | -0.002 | -0.026 | -0.026 | 0 | 0.86 | 0.93 | 0.85 | 0.8 | 0.88 | 0.75 |
| 2 | 215054694 | rs1912199 | -0.436 | -0.043 | -0.503 | 1.513 | -0.047 | 0.118 | 0.118 | 1.56 | 0.2 | 0.37 | 0.3 | 0.44 | 0.08 | 0.12 |
| 2 | 215067753 | rs6435818 | 0.122 | 0.673 | -0.857 | 1.04 | -0.006 | -0.003 | -0.003 | 0.478 | 0.86 | 0.93 | 0.85 | 0.87 | 0.87 | 0.77 |
| 2 | 215077380 | rs13007666 | -1.011 | 0.73 | -0.175 | 1.116 | 0.01 | 0 | 0 | 0 | 0.22 | 0.35 | 0.23 | 0.21 | 0.21 | 0.18 |
| 2 | 215081910 | rs16851611 | -1.349 | 0.439 | 0.24 | 1.54 | 0.003 | 0.074 | 0.074 | 0 | 0.2 | 0.35 | 0.26 | 0.34 | 0.18 | 0.15 |
| 2 | 215090921 | rs1912177 | 1.321 | -0.033 | 0.085 | 1.509 | 0.014 | -0.033 | -0.033 | 0.017 | 0.59 | 0.16 | 0.55 | 0.42 | 0.69 | 0.72 |
| 2 | 215104440 | rs16851644 | 0.04 | -0.034 | 0.211 | 1.653 | 0.007 | 0.016 | 0.016 | 0 | 0.1 | 0.17 | 0.12 | 0.18 | 0.1 | 0.16 |
| 2 | 215107556 | rs2052440 | -1.312 | -0.097 | 0.255 | 1.606 | 0 | -0.004 | -0.004 | 0 | 0.34 | 0.31 | 0.4 | 0.51 | 0.29 | 0.27 |
| 2 | 215108398 | rs16825126 | 1.684 | -0.094 | 0.39 | 1.732 | 0.032 | -0.004 | -0.004 | 0.136 | 0.59 | 0.2 | 0.51 | 0.41 | 0.7 | 0.72 |
| 2 | 215121752 | rs6741126 | 0.159 | -0.057 | 0.387 | 1.731 | -0.017 | -0.022 | -0.022 | 0 | 0.09 | 0.06 | 0.11 | 0.17 | 0.1 | 0.15 |
| 2 | 215124975 | rs1912183 | 1.177 | -0.167 | 0.778 | 1.648 | -0.006 | 0.032 | 0.032 | 1.717 | 0.65 | 0.53 | 0.55 | 0.48 | 0.71 | 0.73 |
| 2 | 215135270 | rs1912209 | 1.949 | -0.131 | 0.155 | 1.42 | -0.045 | 0.102 | 0.102 | 4.053 | 0.46 | 0.11 | 0.38 | 0.29 | 0.55 | 0.58 |
| 2 | 215147973 | rs10439348 | -1.149 | -0.076 | 0.337 | 1.483 | -0.002 | 0.007 | 0.007 | 1.925 | 0.36 | 0.38 | 0.46 | 0.52 | 0.3 | 0.28 |
| 2 | 215148115 | rs10439273 | 0.023 | -0.03 | -0.233 | 1.388 | -0.042 | -0.001 | -0.001 | 0 | 0.22 | 0.01 | 0.26 | 0.18 | 0.2 | 0.13 |
| 2 | 215153267 | rs13394429 | -0.593 | 0.323 | 0.329 | 1.423 | 0.028 | 0.017 | 0.017 | 0 | 0.14 | 0.32 | 0.18 | 0.13 | 0.17 | 0.12 |
| 2 | 215157046 | rs1912195 | -0.353 | -0.035 | 0.015 | 1.531 | -0.023 | -0.067 | -0.067 | 0 | 0.21 | 0.08 | 0.31 | 0.36 | 0.18 | 0.25 |
| 2 | 215159153 | rs2052442 | -1.276 | -0.064 | 0.05 | 1.366 | -0.003 | 0.026 | 0.026 | 1.069 | 0.35 | 0.43 | 0.43 | 0.53 | 0.3 | 0.29 |
| 2 | 215168395 | rs7559483 | -0.998 | 0.023 | 0.312 | 1.511 | -0.036 | 0.019 | 0.019 | 0.436 | 0.22 | 0.5 | 0.22 | 0.21 | 0.16 | 0.13 |
| 2 | 215179277 | rs6731110 | -0.267 | 0.316 | 0.124 | 1.267 | 0.03 | 0.062 | 0.062 | 0 | 0.11 | 0.28 | 0.16 | 0.19 | 0.1 | 0.16 |
| 2 | 215188619 | rs16851771 | 0.521 | 0.234 | 0.11 | 1.17 | 0.004 | -0.008 | -0.008 | 0 | 0.08 | 0.21 | 0.08 | 0.03 | 0.06 | 0.01 |
| 2 | 215188809 | rs16851773 | 1.833 | 0.223 | -0.56 | 1.32 | -0.02 | 0.01 | 0.01 | 2.297 | 0.57 | 0.04 | 0.43 | 0.31 | 0.67 | 0.66 |
| 2 | 215189950 | rs16851779 | 0.115 | 0.083 | -0.183 | 1.289 | 0.013 | -0.003 | -0.003 | 0.502 | 0.09 | 0.34 | 0.1 | 0.03 | 0.06 | 0.01 |
| 2 | 215192040 | rs1912184 | -0.482 | -0.008 | -1.246 | 1.294 | -0.008 | -0.032 | -0.032 | 0.539 | 0.23 | 0.11 | 0.25 | 0.34 | 0.18 | 0.21 |
| 2 | 215194668 | rs1912185 | -0.761 | -0.022 | -1.225 | 1.28 | 0.001 | 0.008 | 0.008 | 0 | 0.91 | 0.87 | 0.85 | 0.81 | 0.9 | 0.86 |
| 2 | 215195125 | rs2372254 | 0.291 | -0.012 | -1.167 | 1.206 | 0.001 | -0.032 | -0.032 | 0 | 0.76 | 0.86 | 0.74 | 0.65 | 0.81 | 0.79 |
| 2 | 215196213 | rs6728177 | -1.293 | -0.035 | -1.085 | 1.236 | 0.027 | -0.045 | -0.045 | 0 | 0.24 | 0.8 | 0.33 | 0.36 | 0.16 | 0.14 |
| 2 | 215215856 | rs6737326 | -0.909 | 0.066 | -0.993 | 1.286 | -0.01 | 0.027 | 0.027 | 1.459 | 0.2 | 0.69 | 0.31 | 0.36 | 0.15 | 0.13 |
| 2 | 215218657 | rs1912190 | 1.403 | 0.175 | -1.613 | 1.26 | 0.02 | 0.08 | 0.08 | 4.106 | 0.55 | 0.08 | 0.42 | 0.28 | 0.67 | 0.66 |
| 2 | 215234693 | rs7423982 | -0.038 | 0.328 | -0.99 | 1.448 | -0.001 | 0.063 | 0.063 | 0 | 0.92 | 0.89 | 0.8 | 0.66 | 0.92 | 0.88 |
| 2 | 215239943 | rs6749380 | 0.089 | 0.136 | -1.746 | 0.707 | -0.006 | 0.078 | 0.078 | 0.071 | 0.64 | 0.89 | 0.71 | 0.76 | 0.68 | 0.77 |
| 2 | 215243552 | rs6734329 | -0.296 | 0.199 | -1.837 | 0.811 | 0.007 | -0.111 | -0.111 | 0 | 0.82 | 0.95 | 0.74 | 0.56 | 0.84 | 0.82 |
| 2 | 215243598 | rs6705674 | 0.167 | 0.201 | -2.592 | 0.273 | 0.004 | -0.129 | -0.129 | 2.542 | 0.69 | 0.94 | 0.67 | 0.55 | 0.72 | 0.78 |
| 2 | 215243743 | rs10804226 | 2.213 | 0.334 | -2.688 | 0.352 | -0.093 | -0.076 | -0.076 | 0 | 0.25 | 0.04 | 0.32 | 0.32 | 0.33 | 0.51 |
| 2 | 215243815 | rs4673821 | 0.82 | 0.027 | -1.549 | 0.193 | -0.001 | -0.001 | -0.001 | 0 | 0.45 | 0.58 | 0.58 | 0.54 | 0.53 | 0.73 |
| 2 | 215244282 | rs4673822 | 0.014 | 0.3 | -0.935 | 0.91 | 0.021 | -0.061 | -0.061 | 0 | 0.82 | 0.92 | 0.79 | 0.57 | 0.85 | 0.82 |
| 2 | 215249581 | rs4673823 | 0.963 | 0.478 | -0.146 | 0.591 | -0.009 | -0.007 | -0.007 | 0 | 0.71 | 0.41 | 0.69 | 0.74 | 0.77 | 0.78 |
| 2 | 215251226 | rs6731626 | -0.123 | 0.529 | 0.506 | 0.583 | -0.017 | -0.101 | -0.101 | 0 | 0.16 | 0.41 | 0.1 | 0.01 | 0.13 | 0.04 |
| 2 | 215254537 | rs6743939 | -1.266 | 1.212 | 0.173 | 0.604 | -0.106 | -0.107 | -0.107 | 0 | 0.33 | 0.11 | 0.42 | 0.46 | 0.44 | 0.32 |
| 2 | 215260644 | rs9636337 | 2.392 | 1.14 | 0.189 | 0.515 | -0.007 | 0.105 | 0.105 | 0.72 | 0.48 | 0.06 | 0.32 | 0.28 | 0.42 | 0.51 |
| 2 | 215264632 | rs6705720 | -0.456 | 1.13 | 0.843 | 0.496 | -0.024 | -0.101 | -0.101 | 3.214 | 0.18 | 0.53 | 0.12 | 0.01 | 0.15 | 0.05 |
| 2 | 215275609 | rs4380236 | -1.831 | 1.296 | 0.789 | 0.52 | -0.008 | 0.018 | 0.018 | 0.251 | 0.31 | 0.81 | 0.35 | 0.26 | 0.3 | 0.22 |
| 2 | 215275625 | rs4585026 | 0.51 | 1.27 | 0.799 | 0.526 | 0.003 | 0.028 | 0.028 | 0 | 0.1 | 0.16 | 0.19 | 0.25 | 0.1 | 0.17 |
| 2 | 215279604 | rs3924563 | -0.504 | 1.094 | 0.867 | 0.491 | -0.029 | -0.089 | -0.089 | 0 | 0.19 | 0.58 | 0.11 | 0.01 | 0.14 | 0.05 |
| 2 | 215283259 | rs4264565 | -0.073 | 1.12 | 0.645 | 0.558 | -0.07 | -0.12 | -0.12 | 0 | 0.2 | 0.07 | 0.3 | 0.45 | 0.26 | 0.26 |
| 2 | 215284439 | rs6728732 | -0.476 | 1.019 | 0.882 | 0.423 | 0.046 | 0.083 | 0.083 | 0.379 | 0.14 | 0.45 | 0.24 | 0.24 | 0.15 | 0.17 |
| 2 | 215295727 | rs9646849 | -2.38 | 1.129 | 0.733 | 0.428 | 0.015 | 0.113 | 0.113 | 0.374 | 0.51 | 0.93 | 0.66 | 0.71 | 0.57 | 0.49 |
| 2 | 215325694 | rs7563874 | -0.175 | 0.944 | 1.101 | 0.47 | -0.014 | -0.084 | -0.084 | 0.702 | 0.16 | 0.31 | 0.1 | 0.01 | 0.15 | 0.05 |
| 2 | 215335207 | rs7564279 | 0.67 | 0.921 | 0.777 | 0.458 | -0.029 | -0.087 | -0.087 | 4.33 | 0.8 | 0.41 | 0.88 | 0.99 | 0.85 | 0.95 |
| 2 | 215339569 | rs13031969 | -0.634 | 1.052 | 0.411 | 0.672 | -0.035 | -0.095 | -0.095 | 0.306 | 0.2 | 0.05 | 0.33 | 0.3 | 0.23 | 0.22 |
| 2 | 215342676 | rs4673833 | 0.612 | 1.028 | 0.477 | 0.534 | -0.036 | -0.047 | -0.047 | 0.746 | 0.1 | 0.02 | 0.23 | 0.24 | 0.15 | 0.17 |
| 2 | 215345289 | rs12614846 | -0.379 | 0.991 | 0.547 | 0.504 | -0.003 | -0.003 | -0.003 | 0 | 0.17 | 0.17 | 0.28 | 0.25 | 0.23 | 0.2 |
| 2 | 215349689 | rs6760280 | 2.174 | 0.939 | 0.419 | 0.462 | -0.016 | -0.008 | -0.008 | 0 | 0.52 | 0.09 | 0.4 | 0.52 | 0.45 | 0.6 |

| | | | | | | | | | | | | | | | | |
|---|---|---|---|---|---|---|---|---|---|---|---|---|---|---|---|---|
| 2 | 215355340 | rs4673839 | 0.496 | 0.949 | 0.521 | 0.489 | 0.013 | 0.042 | 0.042 | 0 | 0.11 | 0.18 | 0.28 | 0.26 | 0.15 | 0.17 |
| 2 | 215355718 | rs3935486 | -1.14 | 0.94 | 0.676 | 0.479 | -0.044 | 0.051 | 0.051 | 0 | 0.31 | 0.16 | 0.28 | 0.23 | 0.39 | 0.23 |
| 2 | 215358560 | rs4577276 | -0.767 | 0.895 | 1.229 | 0.479 | 0.022 | 0.024 | 0.024 | 0 | 0.19 | 0.32 | 0.36 | 0.26 | 0.25 | 0.2 |
| 2 | 215361240 | rs4378808 | -0.287 | 0.889 | 0.937 | 0.577 | 0.011 | -0.003 | -0.003 | 0 | 0.19 | 0.21 | 0.21 | 0.22 | 0.27 | 0.17 |
| 2 | 215365050 | rs4362557 | 0.151 | 0.911 | 0.499 | 0.685 | -0.01 | 0.011 | 0.011 | 0 | 0.07 | 0.04 | 0.26 | 0.3 | 0.09 | 0.17 |
| 2 | 215381996 | rs4673850 | 0.224 | 0.613 | 0.976 | 0.457 | 0.039 | 0.068 | 0.068 | 0.194 | 0.12 | 0.21 | 0.29 | 0.26 | 0.17 | 0.18 |
| 2 | 215384251 | rs4673851 | -2.019 | 0.443 | 1.005 | 0.212 | 0 | -0.002 | -0.002 | 0 | 0.47 | 0.47 | 0.69 | 0.71 | 0.55 | 0.51 |
| 2 | 215384466 | rs13013351 | 2.081 | 0.393 | 0.542 | 0.187 | -0.041 | 0.132 | 0.132 | 0.156 | 0.47 | 0.1 | 0.28 | 0.28 | 0.45 | 0.49 |
| 2 | 215390981 | rs10932542 | -0.319 | 0.504 | 0.652 | 0.35 | -0.009 | -0.031 | -0.031 | 0.042 | 0.22 | 0.2 | 0.28 | 0.46 | 0.27 | 0.27 |
| 2 | 215394416 | rs12467438 | -1.259 | 0.463 | 0.771 | 0.302 | -0.01 | 0.005 | 0.005 | 0 | 0.34 | 0.32 | 0.57 | 0.71 | 0.44 | 0.46 |
| 2 | 215400721 | rs4672712 | -1.654 | 0.33 | 1.03 | 0.326 | 0.001 | -0.007 | -0.007 | 0 | 0.44 | 0.43 | 0.61 | 0.7 | 0.51 | 0.47 |
| 2 | 215403685 | rs12612233 | 0.455 | 0.423 | 0.586 | 0.364 | -0.039 | -0.04 | -0.04 | 0 | 0.16 | 0.05 | 0.32 | 0.26 | 0.23 | 0.21 |
| 2 | 215404421 | rs13386380 | -0.441 | 0.559 | 0.79 | 0.503 | 0.013 | 0.068 | 0.068 | 0 | 0.21 | 0.3 | 0.28 | 0.47 | 0.26 | 0.25 |
| 2 | 215405482 | rs7563747 | -0.335 | 0.587 | -0.071 | 0.515 | -0.042 | -0.101 | -0.101 | 0 | 0.2 | 0.09 | 0.27 | 0.46 | 0.26 | 0.25 |
| 2 | 215406125 | rs11904475 | -1.114 | 0.285 | 0.077 | -0.331 | -0.002 | 0.105 | 0.105 | 0 | 0.48 | 0.7 | 0.65 | 0.71 | 0.54 | 0.49 |
| 2 | 215407021 | rs9653261 | -1.036 | -0.091 | 0.948 | -0.61 | -0.023 | -0.042 | -0.042 | 0 | 0.32 | 0.45 | 0.27 | 0.18 | 0.25 | 0.18 |
| 2 | 215409170 | rs10932543 | -1.324 | 0.023 | 0.986 | -0.683 | 0.012 | 0.054 | 0.054 | 0 | 0.35 | 0.72 | 0.2 | 0.18 | 0.3 | 0.19 |
| 2 | 215409277 | rs11678130 | 0.126 | -0.386 | 1.14 | -0.644 | 0.012 | 0.01 | 0.01 | 0 | 0.13 | 0.07 | 0.07 | 0 | 0.08 | 0.04 |
| 2 | 215411621 | rs7566531 | 0.743 | -0.377 | 1.801 | -0.652 | -0.033 | -0.065 | -0.065 | 2.276 | 0.69 | 0.56 | 0.85 | 0.83 | 0.78 | 0.83 |
| 2 | 215414537 | rs13428970 | -0.997 | -0.328 | 1.811 | -0.662 | -0.006 | -0.018 | -0.018 | 0 | 0.3 | 0.31 | 0.15 | 0.17 | 0.22 | 0.17 |
| 2 | 215437176 | rs6720010 | -0.888 | -0.279 | 1.512 | -0.632 | -0.015 | -0.033 | -0.033 | 0 | 0.3 | 0.59 | 0.23 | 0.19 | 0.22 | 0.17 |
| 2 | 215447373 | rs6734403 | -0.226 | -0.294 | 1.06 | -0.468 | -0.007 | -0.004 | -0.004 | 0 | 0.21 | 0.22 | 0.14 | 0.21 | 0.14 | 0.14 |
| 2 | 215447953 | rs7425093 | 0.229 | -0.101 | 1.457 | -0.491 | -0.017 | -0.022 | -0.022 | 0 | 0.78 | 0.62 | 0.85 | 0.8 | 0.84 | 0.86 |
| 2 | 215461986 | rs13404189 | 0.397 | -0.562 | 1.004 | -0.998 | 0.023 | 0.032 | 0.032 | 0 | 0.21 | 0.12 | 0.11 | 0 | 0.13 | 0.06 |
| 2 | 215472318 | rs16852229 | 0.255 | -0.248 | 1.72 | -0.598 | -0.039 | -0.029 | -0.029 | 0 | 0.78 | 0.41 | 0.77 | 0.8 | 0.85 | 0.86 |
| 2 | 215476931 | rs10932551 | 0.114 | -0.245 | 1.796 | -0.535 | -0.028 | -0.011 | -0.011 | 0.106 | 0.78 | 0.65 | 0.78 | 0.78 | 0.85 | 0.86 |
| 2 | 215478550 | rs1119776 | -0.097 | -0.259 | 1.771 | -0.542 | 0.001 | 0 | 0 | 0.507 | 0.8 | 0.8 | 0.79 | 0.78 | 0.85 | 0.86 |
| 2 | 215485519 | rs1400863 | 0.25 | -0.287 | 1.45 | -0.544 | -0.042 | -0.016 | -0.016 | 0 | 0.78 | 0.45 | 0.76 | 0.79 | 0.85 | 0.86 |
| 2 | 215489130 | rs2222182 | 0.185 | -0.284 | 1.445 | -0.542 | -0.038 | -0.013 | -0.013 | 0 | 0.78 | 0.61 | 0.78 | 0.79 | 0.85 | 0.86 |
| 2 | 215501876 | rs7603357 | -0.084 | -0.262 | 0.694 | -0.515 | 0.006 | 0.001 | 0.001 | 0.486 | 0.19 | 0.17 | 0.2 | 0.21 | 0.14 | 0.14 |
| 2 | 215510824 | rs1356438 | -0.066 | -0.272 | 0.561 | -0.393 | 0.025 | -0.022 | -0.022 | 0 | 0.19 | 0.14 | 0.2 | 0.25 | 0.14 | 0.14 |
| 2 | 215513430 | rs4672722 | 0.04 | -0.095 | 0.199 | -0.546 | 0.018 | 0.042 | 0.042 | 0 | 0.49 | 0.23 | 0.54 | 0.56 | 0.44 | 0.47 |
| 2 | 215514980 | rs13031916 | 1.263 | 0.377 | 0.054 | -0.491 | -0.109 | -0.016 | -0.016 | 0.031 | 0.23 | 0.04 | 0.19 | 0.23 | 0.37 | 0.38 |
| 2 | 215515382 | rs13009173 | 1.213 | 0.396 | 0.254 | -0.446 | -0.026 | 0 | 0 | 1.138 | 0.23 | 0.19 | 0.26 | 0.24 | 0.38 | 0.39 |
| 2 | 215519819 | rs1878607 | 0.976 | -0.022 | 0.94 | -0.07 | -0.018 | 0 | 0 | 0.183 | 0.78 | 0.28 | 0.76 | 0.74 | 0.88 | 0.93 |
| 2 | 215529096 | rs10932556 | -0.237 | 0.112 | 0.67 | 0.023 | -0.029 | 0.046 | 0.046 | 1.222 | 0.13 | 0.25 | 0.17 | 0.26 | 0.07 | 0.06 |
| 2 | 215536329 | rs6714965 | -0.202 | 0.095 | 0.305 | 0.022 | -0.029 | 0.032 | 0.032 | 0.858 | 0.13 | 0.22 | 0.18 | 0.26 | 0.07 | 0.06 |
| 2 | 215544960 | rs16852485 | -0.052 | 0.459 | -0.721 | -0.032 | -0.113 | -0.137 | -0.137 | 0.022 | 0.19 | 0.03 | 0.47 | 0.5 | 0.32 | 0.36 |
| 2 | 215546304 | rs7579026 | 0.652 | -0.515 | 0.208 | -0.232 | -0.076 | 0 | 0 | 2.214 | 0.8 | 0.52 | 0.92 | 0.78 | 0.89 | 0.95 |
| 2 | 215562577 | rs10427252 | 0.939 | -0.278 | -0.67 | -0.294 | 0.051 | 0.061 | 0.061 | 0.892 | 0.62 | 0.77 | 0.83 | 0.76 | 0.78 | 0.85 |
| 2 | 215564101 | rs280626 | 0.144 | -0.34 | -0.983 | -0.348 | -0.015 | 0.157 | 0.157 | 0 | 0.77 | 0.98 | 0.89 | 1 | 0.85 | 0.94 |
| 2 | 215565107 | rs10932562 | 0.96 | -0.29 | -0.516 | -0.244 | -0.02 | -0.022 | -0.022 | 0 | 0.61 | 0.53 | 0.82 | 0.75 | 0.78 | 0.85 |
| 2 | 215567903 | rs10932563 | 0.74 | 0.338 | -0.468 | -0.024 | -0.006 | -0.008 | -0.008 | 0 | 0.75 | 0.59 | 0.84 | 0.76 | 0.8 | 0.87 |
| 2 | 215570362 | rs13010891 | 0.711 | 0.021 | -0.459 | -0.001 | -0.026 | -0.012 | -0.012 | 2.423 | 0.73 | 0.61 | 0.89 | 0.76 | 0.86 | 0.88 |
| 2 | 215574029 | rs6720448 | -0.898 | 0.991 | -0.723 | 0.429 | -0.006 | -0.001 | -0.001 | 0.326 | 0.33 | 0.3 | 0.36 | 0.65 | 0.42 | 0.25 |
| 2 | 215574857 | rs7586962 | 0.75 | 1.063 | -0.778 | 0.332 | 0.011 | 0.03 | 0.03 | 0 | 0.52 | 0.4 | 0.5 | 0.65 | 0.55 | 0.54 |
| 2 | 215575104 | rs13405209 | 0.323 | 1.245 | -0.409 | 0.273 | 0 | 0.008 | 0.008 | 0.31 | 0.63 | 0.46 | 0.63 | 0.6 | 0.66 | 0.48 |
| 2 | 215575422 | rs16852556 | -0.027 | 1.082 | -0.214 | 0.012 | -0.01 | -0.043 | -0.043 | 0.065 | 0.08 | 0.16 | 0.04 | 0 | 0.05 | 0.02 |
| 2 | 215576391 | rs10165920 | -0.752 | 1.434 | -0.545 | -0.089 | -0.026 | 0.03 | 0.03 | 0 | 0.12 | 0.06 | 0.15 | 0 | 0.2 | 0.1 |
| 2 | 215589182 | rs2372334 | 1.359 | 1.077 | 0.035 | -0.009 | 0.013 | -0.051 | -0.051 | 0 | 0.74 | 0.58 | 0.72 | 0.9 | 0.7 | 0.84 |
| 2 | 215590582 | rs280621 | 0.459 | 1.013 | -0.014 | -0.032 | 0.005 | 0.006 | 0.006 | 0 | 0.86 | 0.88 | 0.9 | 0.92 | 0.9 | 0.92 |
| 2 | 215592306 | rs7585356 | 0.896 | 1.019 | -0.714 | -0.175 | -0.022 | -0.027 | -0.027 | 0.131 | 0.3 | 0.14 | 0.27 | 0.31 | 0.32 | 0.4 |
| 2 | 215607896 | rs1838802 | -0.174 | 0.869 | -0.079 | -0.361 | 0 | 0 | 0 | 0 | 0.59 | 0.55 | 0.53 | 0.61 | 0.47 | 0.46 |
| 2 | 215619921 | rs10498020 | 0.955 | 0.038 | -0.338 | -0.274 | 0.001 | 0.003 | 0.003 | 0 | 0.18 | 0.14 | 0.18 | 0.31 | 0.15 | 0.14 |
| 2 | 215622330 | rs10179855 | 0.099 | -0.069 | -0.118 | -0.579 | -0.007 | -0.009 | -0.009 | 0 | 0.12 | 0.17 | 0.1 | 0.07 | 0.08 | 0.11 |
| 2 | 215622955 | rs1972958 | 0.433 | 0.138 | -0.078 | -0.666 | -0.001 | -0.001 | -0.001 | 0 | 0.42 | 0.48 | 0.37 | 0.39 | 0.44 | 0.5 |
| 2 | 215624679 | rs10176759 | 0.617 | 0.146 | -0.165 | -0.663 | -0.001 | -0.001 | -0.001 | 0 | 0.41 | 0.4 | 0.37 | 0.39 | 0.44 | 0.5 |
| 2 | 215629322 | rs7584646 | 0.266 | 0.14 | -0.436 | -0.786 | -0.041 | -0.003 | -0.003 | 0.026 | 0.41 | 0.19 | 0.48 | 0.61 | 0.47 | 0.45 |
| 2 | 215632192 | rs5031011 | 1.123 | 0.002 | -0.066 | -0.852 | 0 | 0 | 0 | 0 | 0.36 | 0.33 | 0.3 | 0.34 | 0.37 | 0.4 |
| 2 | 215633463 | rs4234006 | 0.692 | -0.498 | -0.057 | -0.86 | -0.025 | -0.089 | -0.089 | 0 | 0.76 | 0.55 | 0.76 | 0.95 | 0.82 | 0.84 |
| 2 | 215633624 | rs11685215 | 1.305 | -0.543 | -0.516 | -0.735 | 0.03 | -0.08 | -0.08 | 0 | 0.15 | 0.02 | 0.16 | 0.42 | 0.1 | 0.15 |
| 2 | 215634993 | rs3820727 | 1.884 | -0.423 | -0.496 | -0.828 | -0.084 | -0.03 | -0.03 | 0 | 0.3 | 0.15 | 0.36 | 0.34 | 0.5 | 0.44 |
| 2 | 215635273 | rs16852689 | -0.322 | -0.433 | -0.232 | -0.691 | -0.002 | -0.031 | -0.031 | 0 | 0.14 | 0.18 | 0.15 | 0 | 0.12 | 0.06 |
| 2 | 215642236 | rs3768710 | 0.735 | -0.516 | -0.276 | -0.589 | 0.033 | -0.08 | -0.08 | 0 | 0.17 | 0.03 | 0.17 | 0.43 | 0.11 | 0.15 |
| 2 | 215643483 | rs4672729 | -0.277 | -0.563 | 0.072 | -0.71 | 0.004 | 0.002 | 0.002 | 0 | 0.6 | 0.63 | 0.6 | 0.35 | 0.7 | 0.53 |
| 2 | 215643500 | rs17487792 | 1.874 | -0.4 | 0.161 | -0.811 | -0.051 | 0.018 | 0.018 | 0.029 | 0.19 | 0.02 | 0.18 | 0.18 | 0.18 | 0.24 |
| 2 | 215645046 | rs3768708 | -0.875 | -0.382 | 0.335 | -0.8 | -0.006 | -0.031 | -0.031 | 0 | 0.73 | 0.57 | 0.75 | 0.77 | 0.8 | 0.67 |
| 2 | 215650580 | rs16852755 | -0.452 | -0.353 | 0.299 | -0.747 | -0.007 | -0.054 | -0.054 | 0.058 | 0.12 | 0.19 | 0.11 | 0 | 0.08 | 0.05 |
| 2 | 215661710 | rs10498023 | 0.673 | -0.321 | 0.501 | -0.643 | -0.011 | 0.017 | 0.017 | 0 | 0.06 | 0.15 | 0.04 | 0.05 | 0.02 | 0.08 |

| | | | | | | | | | | | | | | | | |
|---|---|---|---|---|---|---|---|---|---|---|---|---|---|---|---|---|
| 2 | 215664391 | rs7591615 | -1.438 | -0.284 | 0.469 | -0.875 | -0.006 | -0.074 | -0.074 | 2.357 | 0.23 | 0.36 | 0.25 | 0.05 | 0.22 | 0.17 |
| 2 | 215666904 | rs6712055 | 0.775 | -0.293 | 0.268 | -0.75 | -0.002 | -0.035 | -0.035 | 0 | 0.3 | 0.15 | 0.25 | 0.61 | 0.29 | 0.39 |
| 2 | 215672527 | rs16852804 | 0.149 | -0.29 | 0.552 | -0.754 | -0.015 | -0.046 | -0.046 | 0 | 0.09 | 0.14 | 0.1 | 0 | 0.05 | 0.05 |
| 2 | 215672546 | rs6435862 | -0.998 | -0.41 | 0.483 | -0.925 | 0 | -0.012 | -0.012 | 0 | 0.73 | 0.72 | 0.84 | 0.83 | 0.75 | 0.73 |
| 2 | 215673664 | rs12477063 | 0.829 | -0.37 | 0.286 | -0.991 | 0.026 | 0.142 | 0.142 | 0 | 0.5 | 0.29 | 0.64 | 0.78 | 0.63 | 0.59 |
| 2 | 215677308 | rs6744811 | 1.964 | -0.475 | 0.153 | -0.916 | -0.025 | -0.024 | -0.024 | 0.011 | 0.31 | 0.19 | 0.4 | 0.33 | 0.39 | 0.4 |
| 2 | 215679686 | rs10190149 | 0.563 | -0.433 | 0.15 | -0.846 | -0.009 | -0.046 | -0.046 | 0 | 0.19 | 0.08 | 0.25 | 0.46 | 0.24 | 0.19 |
| 2 | 215681580 | rs6726763 | 1.258 | -0.321 | 0.201 | -0.833 | -0.003 | 0.008 | 0.008 | 0.063 | 0.24 | 0.16 | 0.16 | 0.2 | 0.21 | 0.34 |
| 2 | 215716169 | rs2166458 | 1.722 | -0.314 | 0.058 | -0.726 | -0.019 | -0.02 | -0.02 | 0 | 0.32 | 0.19 | 0.39 | 0.34 | 0.39 | 0.4 |
| 2 | 215748105 | rs13401954 | 0.155 | -0.31 | 0.466 | -0.637 | 0.001 | 0.024 | 0.024 | 0 | 0.22 | 0.23 | 0.26 | 0.42 | 0.25 | 0.19 |
| 2 | 215755678 | rs737453 | -0.285 | -0.333 | 0.311 | -0.636 | 0.003 | 0.06 | 0.06 | 0 | 0.25 | 0.31 | 0.34 | 0.42 | 0.28 | 0.23 |
| 2 | 215758159 | rs7560809 | 0.562 | -0.428 | 0.152 | -0.752 | 0.004 | 0.048 | 0.048 | 0.168 | 0.32 | 0.15 | 0.19 | 0.26 | 0.26 | 0.34 |
| 2 | 215758510 | rs11899149 | -0.953 | -0.525 | 0.623 | -0.791 | 0.005 | -0.008 | -0.008 | 0 | 0.6 | 0.53 | 0.55 | 0.68 | 0.55 | 0.58 |
| 2 | 215761713 | rs10932578 | -0.766 | -0.595 | 0.118 | -0.888 | 0 | -0.006 | -0.006 | 0 | 0.58 | 0.39 | 0.57 | 0.69 | 0.56 | 0.58 |
| 2 | 215766291 | rs943293 | 1.701 | -0.517 | -0.798 | -0.647 | -0.123 | -0.106 | -0.106 | 0 | 0.25 | 0.01 | 0.35 | 0.31 | 0.37 | 0.38 |
| 2 | 215778534 | rs4672731 | -0.18 | -0.459 | -0.516 | -0.324 | -0.03 | -0.04 | -0.04 | 0.098 | 0.55 | 0.9 | 0.51 | 0.44 | 0.45 | 0.52 |
| 2 | 215780104 | rs4672732 | 1.055 | -0.295 | -0.504 | -0.448 | -0.012 | 0.013 | 0.013 | 0 | 0.36 | 0.19 | 0.36 | 0.3 | 0.39 | 0.43 |
| 2 | 215780583 | rs6756170 | 0.431 | -0.266 | -0.404 | -0.463 | 0 | 0.033 | 0.033 | 0 | 0.43 | 0.29 | 0.46 | 0.3 | 0.44 | 0.47 |
| 2 | 215792343 | rs10932580 | 0.072 | -0.262 | -0.947 | -0.472 | 0.009 | 0.108 | 0.108 | 0 | 0.5 | 0.22 | 0.49 | 0.31 | 0.47 | 0.48 |
| 2 | 215794078 | rs4672734 | 0.456 | -0.127 | -0.992 | -0.344 | -0.022 | 0.055 | 0.055 | 0 | 0.42 | 0.13 | 0.46 | 0.31 | 0.44 | 0.47 |
| 2 | 215795913 | rs12614872 | 0.879 | -0.015 | -1.197 | -0.262 | -0.078 | -0.02 | -0.02 | 1.09 | 0.36 | 0.09 | 0.4 | 0.3 | 0.43 | 0.44 |
| 2 | 215798408 | rs7578759 | 0.26 | -0.277 | -0.678 | -0.297 | 0.023 | 0.026 | 0.026 | 0 | 0.44 | 0.74 | 0.45 | 0.35 | 0.39 | 0.52 |
| 2 | 215806919 | rs2225064 | 0.904 | -0.286 | 0.175 | -0.315 | -0.001 | 0.005 | 0.005 | 0 | 0.36 | 0.33 | 0.38 | 0.29 | 0.38 | 0.46 |
| 2 | 215819853 | rs6760738 | -0.742 | -0.421 | 0.259 | -0.47 | -0.05 | -0.036 | -0.036 | 0 | 0.15 | 0.48 | 0.17 | 0.09 | 0.07 | 0.12 |
| 2 | 215820013 | rs10498027 | 0.681 | -0.568 | -0.047 | -0.524 | 0.028 | 0.084 | 0.084 | 0 | 0.45 | 0.3 | 0.39 | 0.23 | 0.4 | 0.46 |
| 2 | 215821825 | rs16852994 | -0.623 | -0.498 | -0.08 | -0.451 | -0.044 | -0.034 | -0.034 | 0.473 | 0.15 | 0.42 | 0.16 | 0.08 | 0.07 | 0.12 |
| 2 | 215823334 | rs956132 | 1.069 | -0.38 | -0.442 | -0.598 | -0.003 | 0.058 | 0.058 | 0.047 | 0.37 | 0.11 | 0.34 | 0.24 | 0.37 | 0.45 |
| 2 | 215832746 | rs13024862 | 0.354 | -0.331 | -0.122 | -0.534 | 0.02 | 0.043 | 0.043 | 0 | 0.5 | 0.55 | 0.5 | 0.26 | 0.43 | 0.49 |
| 2 | 215839570 | rs16853022 | -0.038 | -0.327 | -0.141 | -0.332 | -0.018 | -0.003 | -0.003 | 1.085 | 0.08 | 0.33 | 0.04 | 0.04 | 0.01 | 0.08 |
| 2 | 215840414 | rs17493319 | -0.151 | -0.152 | -0.367 | -0.32 | 0.002 | -0.005 | -0.005 | 0 | 0.6 | 0.43 | 0.42 | 0.3 | 0.5 | 0.57 |
| 2 | 215843954 | rs7579031 | 0.27 | -0.112 | -0.663 | -0.316 | 0.011 | 0.019 | 0.019 | 0 | 0.5 | 0.46 | 0.44 | 0.28 | 0.6 | 0.5 |
| 2 | 215855780 | rs4533467 | 0.932 | 0.102 | -1.173 | -0.434 | -0.137 | 0.038 | 0.038 | 0.675 | 0.45 | 0.03 | 0.33 | 0.29 | 0.5 | 0.46 |
| 2 | 215858828 | rs1831024 | 0.928 | -0.391 | -0.597 | -0.645 | -0.011 | -0.033 | -0.033 | 0 | 0.73 | 0.3 | 0.72 | 0.96 | 0.91 | 0.84 |
| 2 | 215864312 | rs4673928 | -0.783 | -0.108 | -0.677 | -0.505 | -0.016 | -0.039 | -0.039 | 0 | 0.22 | 0.25 | 0.23 | 0.04 | 0.09 | 0.15 |
| 2 | 215865575 | rs10498030 | 0.898 | 0.466 | -1.015 | 0.32 | -0.085 | -0.113 | -0.113 | 0 | 0.17 | 0.02 | 0.19 | 0.34 | 0.28 | 0.2 |
| 2 | 215889265 | rs4673930 | -0.146 | 0.205 | -0.762 | -0.054 | -0.01 | -0.035 | -0.035 | 0 | 0.12 | 0.16 | 0.19 | 0 | 0.05 | 0.04 |
| 2 | 215893773 | rs7575071 | -0.336 | 0.299 | -0.88 | -0.121 | -0.041 | 0 | 0 | 0.732 | 0.35 | 0.21 | 0.4 | 0.66 | 0.44 | 0.36 |
| 2 | 215896093 | rs4672743 | -0.198 | 0.312 | -0.46 | -0.086 | 0.009 | -0.002 | -0.002 | 0 | 0.11 | 0.12 | 0.19 | 0 | 0.04 | 0.05 |
| 2 | 215901774 | rs17501837 | -0.708 | 0.953 | -0.149 | 0.677 | 0.01 | 0.046 | 0.046 | 0 | 0.13 | 0.14 | 0.26 | 0.41 | 0.21 | 0.14 |
| 2 | 215910724 | rs11890512 | -1.22 | 0.685 | 1.186 | 0.491 | 0.065 | 0.045 | 0.045 | 0.508 | 0.05 | 0.56 | 0.06 | 0 | 0.01 | 0.03 |
| 2 | 215914734 | rs16853241 | 0.66 | 0.65 | 0.301 | 0.069 | -0.025 | 0.032 | 0.032 | 0 | 0.42 | 0.11 | 0.41 | 0.29 | 0.39 | 0.53 |
| 2 | 215914847 | rs12468887 | -1.051 | 0.668 | 0.737 | 0.11 | 0.028 | 0.105 | 0.105 | 0 | 0.18 | 0.67 | 0.3 | 0.4 | 0.21 | 0.14 |
| 2 | 215917789 | rs17430358 | -0.237 | 0.664 | 0.773 | 0.1 | 0.016 | 0.063 | 0.063 | 0 | 0.14 | 0.18 | 0.26 | 0.41 | 0.21 | 0.14 |
| 2 | 215919061 | rs10206806 | -0.701 | 0.667 | 0.783 | 0.102 | 0.053 | 0.176 | 0.176 | 0 | 0.16 | 0.42 | 0.28 | 0.42 | 0.21 | 0.14 |
| 2 | 215922612 | rs2030628 | -0.797 | 0.618 | 0.142 | 0.058 | 0.003 | 0.044 | 0.044 | 0.096 | 0.57 | 0.82 | 0.57 | 0.71 | 0.61 | 0.46 |
| 2 | 215923761 | rs2030627 | 0.506 | 0.636 | 0.124 | 0.063 | 0.005 | 0.032 | 0.032 | 0.379 | 0.39 | 0.28 | 0.27 | 0.3 | 0.4 | 0.29 |
| 2 | 215927329 | rs2888330 | 0.162 | 0.6 | -0.14 | 0.103 | 0.02 | 0.043 | 0.043 | 0 | 0.48 | 0.27 | 0.47 | 0.34 | 0.42 | 0.62 |
| 2 | 215936199 | rs16853324 | 0.76 | -0.042 | -0.568 | 0.242 | 0 | 0.05 | 0.05 | 0 | 0.08 | 0.09 | 0.16 | 0.36 | 0.06 | 0.07 |
| 2 | 215937865 | rs6744831 | 0.522 | 0.046 | -0.984 | 0.165 | -0.062 | 0.044 | 0.044 | 0.323 | 0.44 | 0.11 | 0.48 | 0.3 | 0.51 | 0.57 |
| 2 | 215938189 | rs1403826 | 0.424 | 0.132 | -0.762 | 0.114 | -0.002 | 0.034 | 0.034 | 0.804 | 0.38 | 0.21 | 0.26 | 0.29 | 0.39 | 0.29 |
| 2 | 215940242 | rs6711179 | -0.988 | 0.097 | -0.701 | -0.289 | -0.002 | 0.009 | 0.009 | 0.021 | 0.71 | 0.79 | 0.8 | 0.74 | 0.68 | 0.8 |
| 2 | 215941620 | rs6731654 | -1 | 0.104 | -0.703 | -0.286 | -0.001 | 0.004 | 0.004 | 0.026 | 0.71 | 0.77 | 0.79 | 0.74 | 0.68 | 0.8 |
| 2 | 215947252 | rs1880130 | -0.056 | 0.241 | -1.174 | -0.325 | 0.02 | 0.035 | 0.035 | 0 | 0.56 | 0.16 | 0.57 | 0.33 | 0.61 | 0.69 |
| 2 | 215964050 | rs2970958 | 0.206 | 0.185 | -0.451 | -0.961 | 0.001 | 0.021 | 0.021 | 1.446 | 0.5 | 0.41 | 0.49 | 0.22 | 0.54 | 0.38 |
| 2 | 215968929 | rs2948974 | -1.329 | -0.227 | 0.079 | -1.177 | 0.001 | -0.042 | -0.042 | 0 | 0.24 | 0.73 | 0.25 | 0 | 0.13 | 0.17 |
| 2 | 215980432 | rs10221973 | -1.203 | 0.727 | 0.357 | -0.935 | -0.026 | 0.043 | 0.043 | 0 | 0.16 | 0.11 | 0.21 | 0.02 | 0.17 | 0.17 |
| 2 | 216003934 | rs2948983 | -1.228 | 0.746 | 0.941 | -0.929 | 0.031 | -0.079 | -0.079 | 0 | 0.14 | 0.28 | 0.21 | 0 | 0.15 | 0.15 |
| 2 | 216010489 | rs1004302 | -2.166 | 0.593 | 0.727 | -1.006 | -0.002 | 0.004 | 0.004 | 3.516 | 0.23 | 0.74 | 0.27 | 0.02 | 0.18 | 0.2 |
| 2 | 216023682 | rs7419736 | 0.253 | 0.295 | 0.186 | -0.57 | 0.027 | -0.086 | -0.086 | 0 | 0.17 | 0.01 | 0.11 | 0.28 | 0.08 | 0.38 |
| 2 | 216039339 | rs4563222 | 1.734 | 1.175 | 1.314 | -0.999 | 0.071 | -0.142 | -0.142 | 0 | 0.79 | 0.47 | 0.8 | 0.98 | 0.78 | 0.88 |
| 2 | 216062462 | rs10460420 | 1.351 | 0.799 | 1.418 | -0.815 | 0.129 | 0.092 | 0.092 | 0.965 | 0.95 | 0.55 | 0.95 | 0.98 | 0.97 | 0.98 |
| 2 | 216067031 | rs4673954 | 0.938 | 0.617 | 1.272 | -0.88 | 0.037 | -0.107 | -0.107 | 0 | 0.85 | 0.58 | 0.84 | 1 | 0.87 | 0.93 |
| 2 | 216080128 | rs12474265 | 0.912 | 0.858 | 0.621 | -1.115 | -0.008 | -0.003 | -0.003 | 0 | 0.68 | 0.31 | 0.67 | 1 | 0.65 | 0.87 |
| 2 | 216102969 | rs4531931 | 0.759 | 0.921 | 0.132 | -1.057 | 0.021 | -0.179 | -0.179 | 0 | 0.69 | 0.52 | 0.68 | 1 | 0.65 | 0.87 |
| 2 | 216105295 | rs4476347 | 0.403 | 0.825 | 0.145 | -0.877 | 0.016 | -0.148 | -0.148 | 0 | 0.78 | 0.54 | 0.8 | 1 | 0.75 | 0.94 |
| 2 | 216106637 | rs4673965 | 0.261 | 0.539 | -0.766 | -0.312 | 0.02 | -0.008 | -0.008 | 0 | 0.5 | 0.23 | 0.49 | 0.48 | 0.6 | 0.39 |
| 2 | 216117488 | rs11683778 | -0.987 | 0.444 | -0.042 | -0.215 | -0.012 | -0.051 | -0.051 | 0 | 0.09 | 0.27 | 0.02 | 0 | 0.04 | 0.04 |
| 2 | 216129158 | rs4673981 | 0.049 | 0.635 | -0.838 | 0.135 | -0.016 | -0.006 | -0.006 | 0.445 | 0.63 | 0.28 | 0.53 | 0.55 | 0.58 | 0.52 |
| 2 | 216130934 | rs7565010 | 0.022 | 0.729 | -0.528 | 0.141 | 0.004 | 0.004 | 0.004 | 0.416 | 0.67 | 0.46 | 0.55 | 0.56 | 0.63 | 0.54 |

| | | | | | | | | | | | | | | | | |
|---|---|---|---|---|---|---|---|---|---|---|---|---|---|---|---|---|
| 2 | 216137072 | rs11687225 | 0.28 | 0.704 | -1.275 | 0.665 | -0.02 | 0.005 | 0.005 | 1.46 | 0.59 | 0.27 | 0.46 | 0.36 | 0.56 | 0.43 |
| 2 | 216137364 | rs1808119 | 0.228 | 0.427 | -1.633 | 1.015 | 0.018 | -0.07 | -0.07 | 0.01 | 0.25 | 0.06 | 0.2 | 0.32 | 0.2 | 0.2 |
| 2 | 216137978 | rs16853680 | -0.933 | 0.707 | -0.797 | 1.184 | 0.009 | 0.179 | 0.179 | 0.242 | 0.05 | 0.15 | 0.09 | 0.24 | 0.05 | 0.12 |
| 2 | 216139373 | rs1404774 | -0.389 | 0.786 | -1.459 | 0.504 | -0.055 | -0.003 | -0.003 | 0 | 0.15 | 0.01 | 0.19 | 0.13 | 0.21 | 0.32 |
| 2 | 216141237 | rs10498034 | -0.349 | 0.249 | -0.671 | 0.685 | -0.03 | 0.01 | 0.01 | 0 | 0.23 | 0.54 | 0.15 | 0.22 | 0.16 | 0.24 |
| 2 | 216141357 | rs2372535 | 0.224 | 0.402 | -1.172 | 0.99 | -0.015 | -0.045 | -0.045 | 3.486 | 0.14 | 0.06 | 0.21 | 0.46 | 0.15 | 0.13 |
| 2 | 216143939 | rs1464864 | 0.416 | 0.174 | -0.518 | 0.416 | 0.001 | 0.04 | 0.04 | 0 | 0.71 | 0.65 | 0.65 | 0.54 | 0.72 | 0.78 |
| 2 | 216149093 | rs9789612 | -1.183 | -0.073 | -0.046 | 0.217 | 0.023 | 0.08 | 0.08 | 0 | 0.5 | 0.69 | 0.48 | 0.68 | 0.43 | 0.46 |
| 2 | 216155860 | rs10197653 | 1.05 | 0.46 | -0.739 | -0.412 | -0.037 | 0.031 | 0.031 | 0 | 0.18 | 0.08 | 0.19 | 0 | 0.31 | 0.05 |
| 2 | 216156435 | rs16853697 | -0.231 | 0.246 | -0.541 | -0.037 | -0.001 | 0.003 | 0.003 | 0 | 0.29 | 0.32 | 0.25 | 0.41 | 0.25 | 0.2 |
| 2 | 216181828 | rs11892429 | 0.847 | 0.531 | -0.185 | -0.624 | -0.069 | 0.066 | 0.066 | 0 | 0.16 | 0.06 | 0.19 | 0 | 0.32 | 0.05 |
| 2 | 216185825 | rs10197559 | 0.999 | 0.534 | -0.208 | -0.622 | -0.083 | 0.08 | 0.08 | 0 | 0.16 | 0.05 | 0.18 | 0 | 0.32 | 0.05 |
| 2 | 216195281 | rs1983462 | 0.067 | 0.192 | 0.175 | -0.787 | 0.051 | -0.137 | -0.137 | 0.009 | 0.23 | 0.41 | 0.21 | 0 | 0.32 | 0.1 |
| 2 | 216197971 | rs12995526 | 0.381 | -0.049 | 0.081 | -0.719 | -0.021 | 0.052 | 0.052 | 0.134 | 0.57 | 0.26 | 0.55 | 0.77 | 0.51 | 0.66 |
| 2 | 216198049 | rs10179873 | 0.86 | 0.026 | 0.266 | -0.703 | -0.024 | 0.009 | 0.009 | 0 | 0.19 | 0.14 | 0.2 | 0 | 0.32 | 0.05 |
| 2 | 216203296 | rs16853823 | 0.177 | -0.086 | -0.222 | -0.639 | 0 | -0.013 | -0.013 | 0 | 0.24 | 0.19 | 0.24 | 0.48 | 0.2 | 0.18 |
| 2 | 216207225 | rs7586969 | 0.187 | -0.277 | -0.169 | -0.904 | -0.011 | -0.019 | -0.019 | 0.226 | 0.67 | 0.33 | 0.69 | 1 | 0.61 | 0.84 |
| 2 | 216210236 | rs16853834 | 1.45 | -0.326 | -0.484 | -0.884 | -0.006 | -0.083 | -0.083 | 1.718 | 0.18 | 0.01 | 0.21 | 0.23 | 0.16 | 0.24 |
| 2 | 216210299 | rs1404772 | -0.998 | -0.214 | -0.196 | -1.764 | -0.002 | -0.026 | -0.026 | 0.154 | 0.91 | 0.93 | 0.93 | 1 | 0.94 | 0.94 |
| 2 | 216212089 | rs4673991 | 1.281 | -0.208 | -0.489 | -1.765 | 0.003 | 0.048 | 0.048 | 0.092 | 0.34 | 0.1 | 0.31 | 0.29 | 0.31 | 0.49 |
| 2 | 216217639 | rs7604984 | -0.681 | -0.196 | -0.481 | -1.755 | -0.002 | -0.009 | -0.009 | 0.527 | 0.55 | 0.56 | 0.6 | 0.7 | 0.62 | 0.45 |
| 2 | 216218717 | rs6706415 | 0.836 | -0.123 | -0.926 | -1.718 | 0.042 | -0.095 | -0.095 | 0.008 | 0.21 | 0.31 | 0.21 | 0 | 0.33 | 0.1 |
| 2 | 216229692 | rs11651 | 0.908 | -0.038 | -1.761 | -1.615 | 0.004 | 0.051 | 0.051 | 0.177 | 0.35 | 0.07 | 0.31 | 0.3 | 0.31 | 0.51 |
| 2 | 216232978 | rs2289200 | -0.05 | 0.104 | -1.289 | -1.026 | -0.02 | 0.023 | 0.023 | 0 | 0.32 | 0.08 | 0.37 | 0.7 | 0.3 | 0.35 |
| 2 | 216240584 | rs6707530 | -0.19 | 0.347 | -0.23 | -1.392 | -0.019 | -0.046 | -0.046 | 0 | 0.33 | 0.38 | 0.43 | 0.77 | 0.25 | 0.26 |
| 2 | 216243650 | rs2304573 | -0.003 | 0.307 | 0.035 | -1.413 | 0.006 | 0.015 | 0.015 | 0 | 0.31 | 0.23 | 0.41 | 0.77 | 0.25 | 0.26 |
| 2 | 216246080 | rs7608342 | 1.038 | 0.22 | -0.906 | -2.215 | -0.068 | 0.069 | 0.069 | 0.16 | 0.31 | 0.05 | 0.35 | 0.23 | 0.4 | 0.54 |
| 2 | 216249587 | rs13652 | -0.734 | 0.073 | -0.493 | -2.487 | 0.011 | -0.017 | -0.017 | 0 | 0.77 | 0.78 | 0.89 | 1 | 0.88 | 0.83 |
| 2 | 216253454 | rs7588661 | 0.014 | 0.28 | 0.025 | -2.309 | 0.005 | 0.009 | 0.009 | 0 | 0.45 | 0.38 | 0.47 | 0.19 | 0.62 | 0.57 |
| 2 | 216256707 | rs10804242 | -1.054 | 0.357 | -0.114 | -2.211 | -0.017 | 0.051 | 0.051 | 0.02 | 0.68 | 0.84 | 0.64 | 0.83 | 0.6 | 0.48 |
| 2 | 216256981 | rs11883812 | 0.26 | -0.169 | 0.054 | -2.367 | -0.012 | -0.054 | -0.054 | 0.167 | 0.22 | 0.25 | 0.09 | 0 | 0.08 | 0.13 |
| 2 | 216258251 | rs1968510 | -0.314 | -0.144 | 0.04 | -2.367 | -0.002 | -0.034 | -0.034 | 0 | 0.83 | 0.74 | 0.91 | 1 | 0.9 | 0.85 |
| 2 | 216260298 | rs1561302 | 0.896 | 0.297 | 0.386 | -2.619 | 0.067 | -0.071 | -0.071 | 0 | 0.14 | 0.29 | 0.14 | 0 | 0.25 | 0.04 |
| 2 | 216263366 | rs1525351 | -1 | 0.038 | 0.312 | -2.931 | -0.002 | -0.102 | -0.102 | 0 | 0.59 | 0.54 | 0.66 | 0.93 | 0.58 | 0.42 |
| 2 | 216266736 | rs6744921 | 0.996 | 0.013 | 0.277 | -2.931 | 0 | -0.054 | -0.054 | 0 | 0.37 | 0.32 | 0.31 | 0.07 | 0.38 | 0.53 |
| 2 | 216268902 | rs7575234 | 0.717 | 0.045 | -0.234 | -2.764 | 0.009 | 0.066 | 0.066 | 0 | 0.3 | 0.14 | 0.47 | 0.93 | 0.27 | 0.27 |
| 2 | 216276991 | rs13004592 | 0.803 | -0.06 | -0.344 | -2.802 | -0.014 | 0.154 | 0.154 | 0 | 0.39 | 0.16 | 0.3 | 0.07 | 0.4 | 0.53 |
| 2 | 216277672 | rs7588830 | 0.783 | 0.078 | -0.201 | -2.841 | 0.02 | 0.08 | 0.08 | 0 | 0.31 | 0.17 | 0.47 | 0.93 | 0.26 | 0.27 |
| 2 | 216279273 | rs17516906 | 1.245 | 0.019 | -0.341 | -2.855 | 0.009 | 0.05 | 0.05 | 0.139 | 0.09 | 0.01 | 0.04 | 0 | 0.06 | 0.1 |
| 2 | 216282481 | rs724617 | 1.194 | -0.161 | 0.222 | -3.07 | -0.05 | 0.182 | 0.182 | 0 | 0.37 | 0.12 | 0.29 | 0.07 | 0.41 | 0.53 |
| 2 | 216282993 | rs1437799 | -0.28 | -0.134 | 0.441 | -3.051 | 0.005 | -0.09 | -0.09 | 0 | 0.27 | 0.3 | 0.19 | 0 | 0.3 | 0.17 |
| 2 | 216284774 | rs17517509 | 0.825 | -0.075 | 0.401 | -2.871 | 0.01 | 0.013 | 0.013 | 0 | 0.31 | 0.24 | 0.48 | 0.92 | 0.26 | 0.25 |
| 2 | 216287497 | rs16854041 | -0.279 | 0.261 | 0.144 | -3.196 | -0.001 | -0.073 | -0.073 | 0 | 0.29 | 0.35 | 0.17 | 0 | 0.27 | 0.27 |
| 2 | 216289343 | rs1250250 | 1.133 | 0.131 | 0.021 | -3.021 | 0.002 | 0.002 | 0.002 | 0 | 0.28 | 0.26 | 0.48 | 0.89 | 0.26 | 0.25 |
| 2 | 216294663 | rs1250238 | 1.349 | 0.098 | -0.132 | -3.09 | 0.002 | 0.046 | 0.046 | 0 | 0.26 | 0.16 | 0.47 | 0.92 | 0.26 | 0.24 |
| 2 | 216298084 | rs2289202 | 1.268 | 0.187 | 0.132 | -3.233 | 0.053 | -0.087 | -0.087 | 0 | 0.16 | 0.25 | 0.14 | 0 | 0.24 | 0.04 |
| 2 | 216299281 | rs1250246 | -0.109 | 0.221 | 0.244 | -3.244 | -0.004 | -0.033 | -0.033 | 0.398 | 0.43 | 0.53 | 0.62 | 0.93 | 0.5 | 0.28 |
| 2 | 216299550 | rs2577290 | 1.148 | 0.224 | 0.092 | -3.128 | -0.001 | -0.037 | -0.037 | 0.27 | 0.27 | 0.27 | 0.48 | 0.92 | 0.26 | 0.24 |
| 2 | 216300185 | rs1250258 | -1.793 | -0.037 | -0.391 | -3.198 | 0.011 | 0.118 | 0.118 | 0.053 | 0.73 | 0.9 | 0.79 | 0.92 | 0.77 | 0.55 |
| 2 | 216307103 | rs1250233 | 0.79 | 0.427 | -0.052 | -3.028 | 0 | -0.012 | -0.012 | 0.086 | 0.25 | 0.26 | 0.46 | 0.92 | 0.27 | 0.22 |
| 2 | 216313591 | rs1250252 | -0.794 | 0.457 | -0.169 | -3.339 | 0.004 | 0.027 | 0.027 | 0.529 | 0.48 | 0.64 | 0.63 | 0.94 | 0.55 | 0.44 |
| 2 | 216316487 | rs1250255 | -1.599 | 0.324 | -0.48 | -3.456 | 0.1 | 0.256 | 0.256 | 0.115 | 0.63 | 0.97 | 0.78 | 0.94 | 0.8 | 0.48 |
| 2 | 216317657 | rs17459063 | 0.019 | 0.474 | 0.096 | -3.136 | 0.021 | -0.095 | -0.095 | 0 | 0.21 | 0.3 | 0.17 | 0 | 0.29 | 0.2 |
| 2 | 216320050 | rs1250220 | -0.66 | 0.378 | -0.162 | -2.963 | 0 | -0.004 | -0.004 | 0.053 | 0.46 | 0.43 | 0.62 | 0.94 | 0.55 | 0.43 |
| 2 | 216323133 | rs16854092 | 0.071 | 0.644 | 0.113 | -1.358 | 0.017 | 0.082 | 0.082 | 0 | 0.16 | 0.34 | 0.33 | 0.28 | 0.14 | 0.16 |
| 2 | 216325274 | rs1250224 | -0.309 | 1.024 | 0.064 | -0.968 | 0.024 | -0.02 | -0.02 | 1.092 | 0.21 | 0.32 | 0.47 | 0.8 | 0.24 | 0.23 |
| 2 | 216325768 | rs10498039 | 0.582 | 1.175 | -0.193 | -1.291 | -0.035 | 0.047 | 0.047 | 0 | 0.16 | 0.06 | 0.14 | 0 | 0.27 | 0.12 |
| 2 | 216328307 | rs13017714 | 1.411 | 1.261 | -0.119 | -1.119 | 0.001 | -0.009 | -0.009 | 0 | 0.05 | 0.05 | 0.06 | 0 | 0.13 | 0.08 |
| 2 | 216328360 | rs10179068 | 1.823 | 1.007 | 0.203 | -1.176 | 0.018 | 0.086 | 0.086 | 0 | 0.52 | 0.36 | 0.33 | 0.19 | 0.41 | 0.58 |
| 2 | 216329797 | rs1250226 | 0.079 | 1.114 | 0.267 | -1.085 | 0.023 | -0.019 | -0.019 | 0 | 0.22 | 0.32 | 0.47 | 0.8 | 0.24 | 0.22 |
| 2 | 216331654 | rs12694364 | 0.702 | 0.888 | 0.182 | -0.948 | -0.004 | -0.001 | -0.001 | 1.474 | 0.6 | 0.41 | 0.38 | 0.2 | 0.49 | 0.67 |
| 2 | 216335950 | rs1272628 | -0.386 | 1.041 | 0.363 | -0.727 | 0.044 | -0.12 | -0.12 | 0 | 0.79 | 0.6 | 0.82 | 1 | 0.7 | 0.93 |
| 2 | 216343401 | rs1437790 | 0.113 | 0.902 | 0.354 | -0.017 | 0.018 | 0.14 | 0.14 | 0 | 0.12 | 0.19 | 0.24 | 0.49 | 0.15 | 0.11 |
| 2 | 216345776 | rs10192231 | -0.834 | 0.86 | 0.353 | -0.068 | 0.007 | 0.181 | 0.181 | 0 | 0.19 | 0.41 | 0.28 | 0.49 | 0.18 | 0.16 |
| 2 | 216346048 | rs6730873 | 0.839 | 0.813 | -0.339 | -0.072 | -0.059 | -0.077 | -0.077 | 0 | 0.22 | 0.04 | 0.32 | 0.32 | 0.26 | 0.19 |
| 2 | 216356666 | rs10174972 | 1.309 | 0.506 | -0.123 | -0.217 | 0.061 | 0.053 | 0.053 | 0 | 0.33 | 0.22 | 0.2 | 0.21 | 0.18 | 0.54 |
| 2 | 216358592 | rs2372565 | -0.791 | 0.78 | 0.267 | -0.276 | 0.028 | -0.084 | -0.084 | 0 | 0.81 | 0.74 | 0.84 | 1 | 0.71 | 0.93 |
| 2 | 216363390 | rs13384892 | 0.148 | 0.34 | -0.102 | 0.005 | 0.012 | 0.014 | 0.014 | 0 | 0.66 | 0.42 | 0.52 | 0.52 | 0.56 | 0.78 |
| 2 | 216364035 | rs1437793 | 0.178 | 0.35 | -0.222 | -0.011 | -0.025 | 0.002 | 0.002 | 0 | 0.32 | 0.15 | 0.31 | 0.32 | 0.38 | 0.23 |

| | | | | | | | | | | | | | | | | |
|---|---|---|---|---|---|---|---|---|---|---|---|---|---|---|---|---|
| 2 | 216367956 | rs17460876 | 0.335 | 0.199 | -0.227 | -0.438 | -0.007 | -0.028 | -0.028 | 0 | 0.09 | 0.12 | 0.06 | 0 | 0.07 | 0.05 |
| 2 | 216369179 | rs13392737 | 1.535 | 0.136 | -0.178 | -0.372 | 0.029 | 0.024 | 0.024 | 0 | 0.3 | 0.21 | 0.18 | 0.18 | 0.14 | 0.53 |
| 2 | 216370187 | rs6728285 | 0.353 | 0.302 | -0.45 | 0.028 | -0.104 | -0.084 | -0.084 | 0 | 0.31 | 0.04 | 0.4 | 0.34 | 0.37 | 0.23 |
| 2 | 216377226 | rs1898537 | -1 | 0.598 | 0.116 | 0.093 | 0.073 | 0.13 | 0.13 | 0 | 0.78 | 0.66 | 0.71 | 0.51 | 0.64 | 0.83 |
| 2 | 216385015 | rs1866413 | -0.536 | 0.729 | -0.218 | 0.263 | 0.045 | 0.078 | 0.078 | 0 | 0.69 | 0.57 | 0.64 | 0.51 | 0.58 | 0.77 |
| 2 | 216386016 | rs7424885 | 1.208 | 0.718 | -0.71 | 0.125 | -0.065 | -0.002 | -0.002 | 0 | 0.17 | 0.02 | 0.18 | 0.11 | 0.22 | 0.07 |
| 2 | 216394251 | rs11673781 | -0.223 | 0.623 | 0.094 | 0.173 | 0.021 | 0.03 | 0.03 | 0 | 0.68 | 0.5 | 0.63 | 0.55 | 0.56 | 0.77 |
| 2 | 216401599 | rs13030883 | 0.75 | 0.526 | -0.295 | 0.169 | -0.091 | -0.098 | -0.098 | 0 | 0.31 | 0.04 | 0.37 | 0.36 | 0.36 | 0.23 |
| 2 | 216403098 | rs10932618 | -0.47 | 0.542 | -0.174 | 0.178 | 0.01 | 0.012 | 0.012 | 0 | 0.69 | 0.67 | 0.64 | 0.56 | 0.56 | 0.77 |
| 2 | 216409359 | rs11887728 | 0.386 | 0.399 | -0.198 | 0.065 | -0.007 | -0.01 | -0.01 | 0.119 | 0.37 | 0.56 | 0.27 | 0.19 | 0.21 | 0.53 |
| 2 | 216411370 | rs16854271 | 0.812 | 0.458 | -0.368 | 0.086 | -0.089 | -0.093 | -0.093 | 0.471 | 0.31 | 0.04 | 0.36 | 0.36 | 0.35 | 0.23 |
| 2 | 216412887 | rs11885509 | -0.427 | 0.49 | -0.199 | 0.095 | 0.039 | 0.046 | 0.046 | 0 | 0.68 | 0.55 | 0.63 | 0.55 | 0.56 | 0.76 |
| 2 | 216413141 | rs11885618 | 0.704 | 0.35 | -0.273 | 0.086 | 0.027 | 0.034 | 0.034 | 0 | 0.33 | 0.23 | 0.25 | 0.19 | 0.2 | 0.53 |
| 2 | 216417880 | rs12478512 | 1.045 | 0.26 | -0.428 | 0.05 | -0.017 | 0.014 | 0.014 | 0 | 0.3 | 0.01 | 0.21 | 0.19 | 0.2 | 0.54 |
| 2 | 216421048 | rs2118773 | 0.571 | 0.405 | -0.271 | 0.078 | 0.003 | -0.004 | -0.004 | 0.269 | 0.33 | 0.21 | 0.36 | 0.36 | 0.33 | 0.23 |
| 2 | 216421067 | rs10932619 | 0.498 | 1.29 | 0.621 | 0.341 | -0.027 | -0.025 | -0.025 | 7.932 | 0.65 | 0.25 | 0.6 | 0.56 | 0.54 | 0.78 |
| 2 | 216426979 | rs6715587 | 0.404 | 1.139 | 0.088 | 0.417 | 0.021 | 0.017 | 0.017 | 0 | 0.63 | 0.5 | 0.54 | 0.54 | 0.53 | 0.78 |
| 2 | 216429439 | rs4672772 | 0.388 | 1.146 | 0.083 | 0.405 | 0.026 | 0.02 | 0.02 | 0 | 0.63 | 0.52 | 0.54 | 0.53 | 0.53 | 0.78 |
| 2 | 216436519 | rs13034471 | 0.371 | 1.143 | -0.014 | 0.404 | 0.011 | 0.008 | 0.008 | 0 | 0.63 | 0.48 | 0.53 | 0.53 | 0.52 | 0.78 |
| 2 | 216439570 | rs4672773 | 0.581 | 1.069 | -1.005 | 0.395 | -0.047 | -0.038 | -0.038 | 0 | 0.59 | 0.23 | 0.49 | 0.53 | 0.5 | 0.76 |
| 2 | 216449620 | rs2083247 | 0.545 | 0.79 | -0.794 | 0.324 | 0.003 | -0.003 | -0.003 | 1.782 | 0.78 | 0.28 | 0.69 | 0.7 | 0.84 | 0.79 |
| 2 | 216449837 | rs2083246 | 0.177 | 0.404 | -0.766 | -0.193 | -0.002 | -0.004 | -0.004 | 0 | 0.4 | 0.59 | 0.46 | 0.65 | 0.34 | 0.56 |
| 2 | 216449970 | rs1898722 | -0.555 | 0.898 | -0.442 | 0.119 | 0.012 | -0.017 | -0.017 | 0 | 0.3 | 0.32 | 0.32 | 0.3 | 0.33 | 0.26 |
| 2 | 216450871 | rs7604503 | 0.054 | 0.491 | -0.456 | 0.332 | -0.002 | 0.002 | 0.002 | 0 | 0.43 | 0.55 | 0.27 | 0.42 | 0.42 | 0.34 |
| 2 | 216450969 | rs10190384 | 0.236 | 0.016 | -0.347 | 0.118 | -0.001 | -0.015 | -0.015 | 0.005 | 0.43 | 0.34 | 0.36 | 0.44 | 0.59 | 0.34 |
| 2 | 216452339 | rs1898721 | 0.432 | -0.388 | -0.183 | 0.29 | 0 | 0.005 | 0.005 | 0 | 0.38 | 0.25 | 0.49 | 0.33 | 0.36 | 0.45 |
| 2 | 216452392 | rs10181605 | 0.018 | -0.592 | 0.105 | 0.427 | 0.023 | -0.026 | -0.026 | 0 | 0.1 | 0.06 | 0.13 | 0.29 | 0.03 | 0.17 |
| 2 | 216454245 | rs7589782 | -1.061 | -0.813 | -0.004 | 0.247 | -0.098 | -0.018 | -0.018 | 0.366 | 0.25 | 0.54 | 0.18 | 0.25 | 0.07 | 0.26 |
| 2 | 216458974 | rs4672775 | 0.316 | -0.468 | -0.56 | 0.339 | 0.005 | -0.026 | -0.026 | 0 | 0.28 | 0.18 | 0.32 | 0.32 | 0.25 | 0.44 |
| 2 | 216465787 | rs1898716 | 1.065 | -0.506 | -0.025 | 0.235 | -0.033 | -0.01 | -0.01 | 0 | 0.38 | 0.3 | 0.34 | 0.42 | 0.55 | 0.23 |
| 2 | 216476874 | rs2947 | -0.522 | -0.746 | 0.088 | 0.177 | -0.021 | -0.006 | -0.006 | 0.22 | 0.23 | 0.37 | 0.29 | 0.25 | 0.12 | 0.2 |
| 2 | 216477172 | rs11891673 | -0.567 | -0.526 | -0.041 | 0.079 | 0.005 | 0.011 | 0.011 | 0 | 0.44 | 0.6 | 0.42 | 0.31 | 0.35 | 0.54 |
| 2 | 216484582 | rs2678273 | 0.415 | -0.696 | 0.375 | -0.586 | -0.023 | -0.163 | -0.163 | 0 | 0.77 | 0.52 | 0.85 | 1 | 0.84 | 0.9 |
| 2 | 216484751 | rs16854526 | -0.322 | -1.604 | 0.223 | -0.372 | -0.058 | 0.08 | 0.08 | 0 | 0.27 | 0.45 | 0.28 | 0.42 | 0.15 | 0.54 |
| 2 | 216484768 | rs2579005 | 1.125 | -1.411 | 0.124 | -0.3 | 0.036 | 0.013 | 0.013 | 0.027 | 0.52 | 0.41 | 0.61 | 0.63 | 0.68 | 0.37 |
| 2 | 216485049 | rs2680185 | 1.415 | -1.103 | -0.674 | -0.753 | -0.009 | -0.013 | -0.013 | 0.616 | 0.27 | 0.26 | 0.44 | 0.5 | 0.44 | 0.3 |
| 2 | 216486285 | rs1519018 | -0.877 | -2.247 | -0.443 | -0.688 | -0.033 | 0.017 | 0.017 | 0 | 0.36 | 0.49 | 0.33 | 0.43 | 0.19 | 0.55 |
| 2 | 216486659 | rs7597588 | -0.036 | -2.463 | -0.326 | -0.596 | -0.035 | 0.013 | 0.013 | 0 | 0.18 | 0.34 | 0.21 | 0.24 | 0.06 | 0.18 |
| 2 | 216486673 | rs1519016 | -1.072 | -2.402 | -0.291 | -0.559 | -0.033 | 0.017 | 0.017 | 0 | 0.36 | 0.49 | 0.33 | 0.43 | 0.19 | 0.55 |
| 2 | 216489509 | rs16854563 | 0.68 | -2.262 | -0.508 | -0.486 | 0.001 | 0 | 0 | 0 | 0.13 | 0.13 | 0.1 | 0.18 | 0.09 | 0.33 |
| 2 | 216489767 | rs2678272 | 0.68 | -2.262 | -0.508 | -0.486 | 0.001 | 0 | 0 | 0 | 0.13 | 0.13 | 0.1 | 0.18 | 0.09 | 0.34 |
| 2 | 216492242 | rs6435916 | 1.77 | -2.242 | -0.613 | -0.618 | -0.022 | -0.027 | -0.027 | 0 | 0.4 | 0.2 | 0.53 | 0.61 | 0.62 | 0.34 |
| 2 | 216498833 | rs1250043 | -1.632 | -2.264 | -0.545 | -0.502 | 0.006 | -0.001 | -0.001 | 0.463 | 0.45 | 0.6 | 0.36 | 0.43 | 0.22 | 0.62 |
| 2 | 216504834 | rs16854581 | 0.542 | -2.142 | -0.716 | -0.662 | 0.005 | -0.005 | -0.005 | 0.682 | 0.13 | 0.12 | 0.16 | 0.19 | 0.07 | 0.34 |
| 2 | 216505585 | rs6726101 | 0.624 | -2.133 | -0.799 | -0.837 | 0.003 | 0.02 | 0.02 | 0 | 0.15 | 0.08 | 0.06 | 0 | 0.08 | 0.08 |
| 2 | 216506362 | rs13431127 | 2.117 | -2.186 | -0.971 | -0.863 | -0.097 | -0.133 | -0.133 | 0.025 | 0.37 | 0.09 | 0.46 | 0.53 | 0.58 | 0.35 |
| 2 | 216507249 | rs1250034 | -0.91 | -1.852 | -0.261 | -0.952 | 0.012 | -0.073 | -0.073 | 0 | 0.86 | 0.79 | 0.89 | 1 | 0.84 | 0.96 |
| 2 | 216507445 | rs1250036 | -0.343 | -1.864 | -0.084 | -0.631 | -0.041 | 0.024 | 0.024 | 0 | 0.15 | 0.29 | 0.17 | 0.2 | 0.07 | 0.2 |
| 2 | 216515088 | rs6744383 | 0.762 | -1.904 | -0.245 | -0.634 | -0.015 | 0.002 | 0.002 | 0 | 0.62 | 0.43 | 0.59 | 0.59 | 0.8 | 0.55 |
| 2 | 216517743 | rs17525056 | 0.879 | -1.883 | -0.533 | -0.684 | -0.024 | 0.06 | 0.06 | 0.127 | 0.2 | 0.07 | 0.11 | 0 | 0.19 | 0.09 |
| 2 | 216521207 | rs949951 | 1.09 | -1.58 | -0.376 | -0.472 | -0.014 | -0.05 | -0.05 | 0.869 | 0.42 | 0.1 | 0.47 | 0.59 | 0.6 | 0.46 |
| 2 | 216524456 | rs1316689 | 0.236 | -1.642 | 0.009 | -0.393 | -0.018 | 0.065 | 0.065 | 0 | 0.08 | 0.24 | 0.19 | 0.19 | 0.04 | 0.07 |
| 2 | 216524861 | rs2678289 | -0.035 | -1.617 | 0.053 | -0.408 | 0.01 | 0.013 | 0.013 | 0 | 0.59 | 0.63 | 0.72 | 0.83 | 0.71 | 0.62 |
| 2 | 216527422 | rs1250074 | 0.049 | -1.295 | -0.272 | -0.196 | 0.006 | 0.001 | 0.001 | 0 | 0.51 | 0.5 | 0.43 | 0.57 | 0.63 | 0.5 |
| 2 | 216532931 | rs1250079 | 0.725 | -1.058 | -1.237 | -0.257 | -0.089 | -0.105 | -0.105 | 0 | 0.38 | 0.09 | 0.36 | 0.59 | 0.56 | 0.41 |
| 2 | 216534235 | rs869140 | -0.033 | -1.048 | -0.854 | -0.474 | 0.065 | 0.139 | 0.139 | 0 | 0.54 | 0.24 | 0.56 | 0.8 | 0.69 | 0.73 |
| 2 | 216544927 | rs1250055 | 0.04 | -0.359 | -1.281 | -0.099 | 0.019 | 0.017 | 0.017 | 0 | 0.54 | 0.26 | 0.46 | 0.62 | 0.64 | 0.47 |
| 2 | 216548145 | rs16854686 | -1.009 | -0.106 | -0.86 | 0.632 | 0.004 | 0.033 | 0.033 | 0 | 0.08 | 0.1 | 0.28 | 0.36 | 0.06 | 0.34 |
| 2 | 216550255 | rs6744457 | -0.053 | 0.416 | -0.8 | -0.467 | 0 | -0.045 | -0.045 | 0 | 0.21 | 0.22 | 0.16 | 0 | 0.24 | 0.07 |
| 2 | 216559610 | rs11677106 | -0.506 | 0.835 | -0.597 | 0.203 | 0.021 | 0.005 | 0.005 | 0 | 0.23 | 0.27 | 0.43 | 0.29 | 0.35 | 0.32 |
| 2 | 216564272 | rs6716449 | -0.713 | -0.019 | -0.262 | 0.361 | -0.009 | 0.088 | 0.088 | 0 | 0.06 | 0.09 | 0.24 | 0.25 | 0.05 | 0.22 |
| 2 | 216576315 | rs615857 | -0.029 | -0.944 | -0.991 | -0.372 | -0.008 | -0.003 | -0.003 | 0 | 0.67 | 0.39 | 0.76 | 0.3 | 0.8 | 0.66 |
| 2 | 216577703 | rs10498043 | 0.775 | -1.252 | -1.247 | -0.297 | 0.109 | 0.247 | 0.247 | 1.006 | 0.53 | 0.21 | 0.5 | 0.05 | 0.71 | 0.38 |
| 2 | 216581110 | rs3107233 | -1.733 | -1.588 | -0.433 | -0.361 | -0.067 | 0.102 | 0.102 | 0 | 0.19 | 0.42 | 0.31 | 0.34 | 0.13 | 0.33 |
| 2 | 216587597 | rs672324 | -1.478 | -1.611 | -0.266 | -0.374 | -0.035 | 0.111 | 0.111 | 0 | 0.15 | 0.35 | 0.29 | 0.3 | 0.11 | 0.28 |
| 2 | 216603052 | rs10932625 | -0.103 | -1.634 | -0.37 | -0.435 | -0.028 | -0.018 | -0.018 | 0 | 0.69 | 0.56 | 0.79 | 0.26 | 0.81 | 0.59 |
| 2 | 216604360 | rs4672779 | 0.755 | -1.762 | -0.451 | -0.533 | 0.057 | 0.057 | 0.057 | 4.587 | 0.57 | 0.26 | 0.74 | 0.26 | 0.75 | 0.54 |
| 2 | 216611057 | rs4672781 | -1.698 | -1.64 | -0.551 | -0.385 | -0.071 | 0.078 | 0.078 | 3.758 | 0.23 | 0.47 | 0.36 | 0.36 | 0.14 | 0.33 |
| 2 | 216614926 | rs6758001 | 0.043 | -1.582 | -0.6 | -0.389 | -0.021 | 0.012 | 0.012 | 0.054 | 0.06 | 0.15 | 0.04 | 0.06 | 0.03 | 0.07 |

| | | | | | | | | | | | | | | | | |
|---|---|---|---|---|---|---|---|---|---|---|---|---|---|---|---|---|
| 2 | 216618960 | rs6758995 | 0.032 | -1.838 | -1.026 | -0.62 | -0.009 | -0.004 | -0.004 | 0 | 0.65 | 0.61 | 0.65 | 0.29 | 0.76 | 0.51 |
| 2 | 216624563 | rs4255961 | -1.451 | -1.147 | -0.338 | 0.085 | -0.057 | 0.09 | 0.09 | 3.733 | 0.23 | 0.51 | 0.34 | 0.42 | 0.15 | 0.34 |
| 2 | 216625794 | rs1357368 | -1.081 | -1.028 | -0.562 | -0.811 | -0.006 | -0.084 | -0.084 | 0.38 | 0.37 | 0.36 | 0.46 | 1 | 0.24 | 0.7 |
| 2 | 216629374 | rs940020 | -0.787 | -0.41 | -0.6 | -0.687 | 0.008 | 0.001 | 0.001 | 0 | 0.56 | 0.56 | 0.74 | 0.91 | 0.57 | 0.69 |
| 2 | 216631366 | rs4674024 | -0.48 | 0.252 | -0.28 | 0.756 | 0 | 0 | 0 | 0 | 0.36 | 0.35 | 0.57 | 0.44 | 0.42 | 0.39 |
| 2 | 216659531 | rs2222438 | -0.971 | 0.221 | -0.495 | 1.712 | -0.043 | -0.019 | -0.019 | 0 | 0.08 | 0 | 0.26 | 0.51 | 0.14 | 0.19 |
| 2 | 216659581 | rs2222437 | -1.582 | 0.153 | 0.166 | 1.663 | 0.019 | 0.144 | 0.144 | 0 | 0.1 | 0.19 | 0.27 | 0.51 | 0.15 | 0.19 |
| 2 | 216661413 | rs1107123 | -2.314 | -0.073 | 0.179 | 1.401 | 0.007 | 0.065 | 0.065 | 0 | 0.18 | 0.25 | 0.33 | 0.51 | 0.21 | 0.29 |
| 2 | 216676912 | rs6435922 | 1.237 | -0.79 | 0.455 | 1.223 | 0.03 | 0.041 | 0.041 | 0 | 0.53 | 0.33 | 0.4 | 0.31 | 0.6 | 0.54 |
| 2 | 216683448 | rs11682295 | -0.099 | -0.91 | 0.398 | 1.616 | -0.017 | 0.025 | 0.025 | 0 | 0.08 | 0.07 | 0.27 | 0.42 | 0.15 | 0.12 |
| 2 | 216694061 | rs4674035 | -0.915 | -1.333 | 0.744 | 1.499 | -0.01 | 0.028 | 0.028 | 0 | 0.25 | 0.33 | 0.39 | 0.39 | 0.23 | 0.19 |
| 2 | 216694165 | rs12470373 | -1.884 | -1.429 | 0.563 | 1.586 | 0.027 | 0.017 | 0.017 | 0 | 0.42 | 0.64 | 0.55 | 0.69 | 0.34 | 0.46 |
| 2 | 216697115 | rs12473821 | -1.324 | -1.452 | 0.214 | 1.51 | -0.046 | -0.013 | -0.013 | 0.061 | 0.33 | 0.41 | 0.28 | 0.29 | 0.18 | 0.35 |
| 2 | 216703958 | rs4467263 | 0.352 | -1.163 | -0.098 | 1.562 | -0.002 | -0.032 | -0.032 | 0.06 | 0.1 | 0.01 | 0.11 | 0.18 | 0.07 | 0.2 |
| 2 | 216705979 | rs12477965 | -0.173 | -1.139 | -0.072 | 1.597 | 0.008 | -0.025 | -0.025 | 0.496 | 0.14 | 0.03 | 0.14 | 0.19 | 0.1 | 0.2 |
| 2 | 216725663 | rs4422154 | 0.026 | -1 | -0.033 | 1.806 | -0.036 | -0.024 | -0.024 | 0.022 | 0.08 | 0.03 | 0.11 | 0.07 | 0.16 | 0.07 |
| 2 | 216725945 | rs4453695 | -0.937 | -1.165 | 0.068 | 1.819 | -0.025 | 0.057 | 0.057 | 0 | 0.19 | 0.28 | 0.33 | 0.63 | 0.11 | 0.37 |
| 2 | 216734070 | rs10432532 | -1.688 | -1.412 | -0.28 | 1.546 | 0.047 | 0.061 | 0.061 | 0 | 0.45 | 0.67 | 0.59 | 0.71 | 0.37 | 0.51 |
| 2 | 216742443 | rs4561658 | 0.134 | -1.47 | -0.049 | 1.761 | -0.047 | 0.101 | 0.101 | 0 | 0.17 | 0.29 | 0.3 | 0.38 | 0.09 | 0.19 |
| 2 | 216746811 | rs12694376 | 1.817 | -1.641 | -1.566 | 1.112 | -0.145 | 0.081 | 0.081 | 0 | 0.37 | 0.05 | 0.35 | 0.21 | 0.55 | 0.23 |
| 2 | 216754805 | rs4311059 | -1.515 | -1.418 | -0.982 | 0.945 | 0.04 | 0.026 | 0.026 | 0 | 0.44 | 0.67 | 0.46 | 0.36 | 0.36 | 0.39 |
| 2 | 216755489 | rs11673908 | -0.531 | -0.56 | -1.23 | 1.164 | -0.047 | -0.039 | -0.039 | 0 | 0.11 | 0.02 | 0.1 | 0.11 | 0.16 | 0.09 |
| 2 | 216758382 | rs4577277 | 0.497 | -1.17 | -0.908 | 1.274 | 0.001 | -0.002 | -0.002 | 0 | 0.8 | 0.84 | 0.79 | 0.73 | 0.85 | 0.7 |
| 2 | 216758854 | rs4257398 | 0.286 | -1.395 | -0.934 | 1.002 | -0.01 | 0.005 | 0.005 | 0 | 0.87 | 0.87 | 0.91 | 0.87 | 0.93 | 0.89 |
| 2 | 216763506 | rs13418044 | -0.016 | -0.952 | -1.042 | 0.55 | -0.009 | 0.03 | 0.03 | 0 | 0.19 | 0.12 | 0.1 | 0 | 0.15 | 0.12 |
| 2 | 216767198 | rs7562243 | 0.206 | -1.028 | -0.765 | 0.587 | -0.006 | -0.019 | -0.019 | 0 | 0.75 | 0.71 | 0.85 | 0.88 | 0.83 | 0.83 |
| 2 | 216768578 | rs13410394 | 0.735 | -1.673 | -0.91 | 0.284 | 0.043 | -0.027 | -0.027 | 0 | 0.58 | 0.3 | 0.69 | 0.45 | 0.71 | 0.51 |
| 2 | 216770735 | rs6704859 | 0.366 | -0.48 | -0.92 | 0.305 | -0.001 | -0.009 | -0.009 | 0 | 0.52 | 0.52 | 0.35 | 0.22 | 0.47 | 0.46 |
| 2 | 216770807 | rs11891588 | 0.563 | 0.277 | -0.802 | 0.789 | -0.007 | -0.006 | -0.006 | 0 | 0.22 | 0.21 | 0.44 | 0.36 | 0.34 | 0.2 |
| 2 | 216778244 | rs4365463 | -0.084 | 0.095 | -0.777 | 0.156 | -0.017 | 0.002 | 0.002 | 0 | 0.29 | 0.26 | 0.34 | 0.25 | 0.44 | 0.32 |
| 2 | 216779667 | rs10932631 | 0.168 | -0.313 | -1.171 | -0.147 | -0.04 | 0.072 | 0.072 | 2.39 | 0.42 | 0.17 | 0.38 | 0.26 | 0.54 | 0.34 |
| 2 | 216783409 | rs7582760 | -0.544 | -0.821 | -0.88 | -0.356 | -0.007 | 0.047 | 0.047 | 0.505 | 0.63 | 0.75 | 0.69 | 0.79 | 0.64 | 0.65 |
| 2 | 216783829 | rs11683175 | 0.755 | -0.712 | -0.892 | -0.441 | -0.018 | 0.015 | 0.015 | 0.231 | 0.35 | 0.35 | 0.29 | 0.07 | 0.45 | 0.26 |
| 2 | 216784966 | rs13407338 | 0.763 | -0.72 | -0.896 | -0.439 | -0.016 | 0.012 | 0.012 | 0.234 | 0.35 | 0.35 | 0.29 | 0.07 | 0.45 | 0.26 |
| 2 | 216794024 | rs4674039 | -0.15 | -0.684 | -0.991 | -0.352 | -0.019 | 0.019 | 0.019 | 0 | 0.28 | 0.4 | 0.41 | 0.73 | 0.28 | 0.38 |
| 2 | 216795091 | rs10209954 | -0.104 | -0.824 | -1.841 | -0.306 | 0.028 | 0.061 | 0.061 | 0.018 | 0.65 | 0.94 | 0.73 | 0.8 | 0.74 | 0.66 |
| 2 | 216800372 | rs12995228 | 0.699 | -0.177 | -1.063 | -0.076 | -0.002 | 0.001 | 0.001 | 0 | 0.41 | 0.38 | 0.39 | 0.3 | 0.4 | 0.28 |
| 2 | 216802161 | rs4614933 | -0.13 | -0.197 | -1.089 | -0.075 | 0 | 0 | 0 | 0 | 0.74 | 0.75 | 0.65 | 0.67 | 0.65 | 0.61 |
| 2 | 216806264 | rs6740786 | 1.459 | -0.222 | -1.372 | -0.208 | -0.012 | 0.025 | 0.025 | 0 | 0.05 | 0 | 0.05 | 0 | 0.05 | 0.01 |
| 2 | 216807502 | rs4258792 | -0.871 | -0.046 | -1.372 | -0.149 | -0.042 | 0.007 | 0.007 | 0 | 0.85 | 0.94 | 0.74 | 0.88 | 0.75 | 0.74 |
| 2 | 216807726 | rs8911 | 0.142 | -0.273 | -1.175 | -0.432 | 0.002 | 0.028 | 0.028 | 0.039 | 0.23 | 0.15 | 0.13 | 0 | 0.19 | 0.17 |
| 2 | 216807914 | rs13553 | 0.553 | 0.035 | -0.781 | -0.079 | -0.002 | -0.005 | -0.005 | 0 | 0.6 | 0.46 | 0.52 | 0.31 | 0.53 | 0.43 |
| 2 | 216811707 | rs1344695 | 0.613 | -0.35 | -0.729 | 0.079 | -0.004 | -0.002 | -0.002 | 0 | 0.6 | 0.41 | 0.53 | 0.3 | 0.58 | 0.43 |
| 2 | 216817646 | rs2372674 | -0.067 | -0.036 | -0.734 | -0.057 | 0.005 | 0.005 | 0.005 | 0 | 0.75 | 0.7 | 0.66 | 0.69 | 0.66 | 0.61 |
| 2 | 216817663 | rs6706891 | 1.154 | -0.489 | -1.048 | -0.185 | -0.035 | 0.05 | 0.05 | 0 | 0.34 | 0.17 | 0.29 | 0.15 | 0.39 | 0.24 |
| 2 | 216817984 | rs1124134 | 0.494 | -0.156 | -0.792 | -0.306 | 0.021 | -0.008 | -0.008 | 0 | 0.68 | 0.57 | 0.62 | 0.69 | 0.56 | 0.58 |
| 2 | 216818577 | rs2115568 | -0.779 | -0.915 | -1.234 | -0.739 | 0.01 | -0.018 | -0.018 | 0.649 | 0.21 | 0.17 | 0.07 | 0 | 0.08 | 0.13 |
| 2 | 216818705 | rs3770523 | -0.358 | -0.278 | -1.265 | -0.261 | 0.006 | 0.031 | 0.031 | 0 | 0.25 | 0.16 | 0.25 | 0.12 | 0.24 | 0.17 |
| 2 | 216820354 | rs10211101 | 0.225 | -0.184 | -0.605 | 0.314 | 0.002 | 0.008 | 0.008 | 0.324 | 0.64 | 0.69 | 0.5 | 0.31 | 0.58 | 0.41 |
| 2 | 216822197 | rs3755152 | 0.832 | -0.968 | -0.085 | 0.932 | -0.035 | 0.01 | 0.01 | 0.92 | 0.77 | 0.3 | 0.82 | 0.68 | 0.89 | 0.75 |
| 2 | 216825842 | rs2372675 | 0.059 | -0.453 | -0.136 | 0.62 | -0.026 | 0.064 | 0.064 | 0 | 0.27 | 0.14 | 0.16 | 0.12 | 0.27 | 0.29 |
| 2 | 216826136 | rs4622721 | -0.345 | -0.642 | -0.132 | 0.754 | 0.016 | -0.005 | -0.005 | 0 | 0.84 | 0.87 | 0.89 | 0.8 | 0.9 | 0.79 |
| 2 | 216826606 | rs3770530 | 1.053 | -0.694 | 0.115 | 0.892 | 0.027 | 0.039 | 0.039 | 0 | 0.5 | 0.32 | 0.65 | 0.57 | 0.59 | 0.46 |
| 2 | 216833017 | rs12694381 | -0.325 | -0.538 | -0.313 | 1.098 | 0.032 | -0.019 | -0.019 | 0 | 0.85 | 0.92 | 0.91 | 0.79 | 0.91 | 0.77 |
| 2 | 216838699 | rs3770535 | 1.229 | -0.363 | -0.241 | 0.541 | -0.009 | -0.005 | -0.005 | 0 | 0.42 | 0.31 | 0.49 | 0.45 | 0.45 | 0.32 |
| 2 | 216838820 | rs3770536 | 0.494 | -0.421 | -0.194 | 0.445 | -0.004 | -0.031 | -0.031 | 0 | 0.56 | 0.47 | 0.74 | 0.81 | 0.63 | 0.49 |
| 2 | 216838931 | rs11692400 | -0.557 | -0.817 | -0.136 | 0.711 | -0.028 | -0.048 | -0.048 | 0 | 0.85 | 0.77 | 0.93 | 0.96 | 0.92 | 0.79 |
| 2 | 216843865 | rs10932644 | -0.125 | -0.265 | -0.487 | 1.332 | -0.003 | 0.066 | 0.066 | 0 | 0.1 | 0.11 | 0.17 | 0.4 | 0.09 | 0.15 |
| 2 | 216847565 | rs13014394 | 0.067 | -0.144 | -0.569 | 1.256 | -0.001 | 0 | 0 | 0 | 0.22 | 0.21 | 0.15 | 0.14 | 0.21 | 0.25 |
| 2 | 216850582 | rs3770545 | -0.863 | 0.03 | -0.543 | 1.517 | 0.001 | -0.01 | -0.01 | 0.496 | 0.16 | 0.1 | 0.23 | 0.42 | 0.19 | 0.18 |
| 2 | 216860059 | rs3770546 | -0.958 | -0.058 | -0.088 | 1.008 | 0.002 | 0.011 | 0.011 | 0 | 0.12 | 0.27 | 0.12 | 0.15 | 0.1 | 0.11 |
| 2 | 216861460 | rs13014738 | -0.663 | -0.442 | -0.48 | 0.691 | 0.023 | 0.032 | 0.032 | 0 | 0.49 | 0.57 | 0.53 | 0.34 | 0.58 | 0.46 |
| 2 | 216862444 | rs3770550 | 0.121 | -0.789 | -0.52 | 1.176 | 0 | 0.071 | 0.071 | 0 | 0.09 | 0.12 | 0.2 | 0.32 | 0.09 | 0.17 |
| 2 | 216862467 | rs3770551 | 0.44 | -0.927 | -0.532 | 0.985 | -0.017 | 0.027 | 0.027 | 0 | 0.35 | 0.23 | 0.45 | 0.26 | 0.52 | 0.4 |
| 2 | 216862623 | rs11676895 | -0.671 | -0.889 | -0.841 | 0.841 | 0.041 | 0.022 | 0.022 | 0 | 0.58 | 0.67 | 0.72 | 0.66 | 0.67 | 0.64 |
| 2 | 216864594 | rs3755155 | -0.417 | -0.602 | -1.166 | 0.58 | 0.021 | 0.014 | 0.014 | 0 | 0.63 | 0.7 | 0.75 | 0.63 | 0.73 | 0.66 |
| 2 | 216864687 | rs3770557 | -0.031 | -0.329 | -1.361 | 0.441 | -0.007 | 0.002 | 0.002 | 0 | 0.42 | 0.61 | 0.6 | 0.52 | 0.45 | 0.48 |
| 2 | 216866439 | rs3770560 | -0.65 | -0.114 | -1.347 | 0.586 | 0.018 | -0.009 | -0.009 | 0 | 0.17 | 0.04 | 0.17 | 0.23 | 0.17 | 0.2 |
| 2 | 216871480 | rs3770564 | 0.966 | 0.606 | -0.834 | 0.328 | 0.011 | -0.036 | -0.036 | 0 | 0.88 | 0.84 | 0.87 | 0.96 | 0.83 | 0.85 |

| | | | | | | | | | | | | | | | | |
|---|---|---|---|---|---|---|---|---|---|---|---|---|---|---|---|---|
| 2 | 216874573 | rs2372682 | -0.723 | 1.69 | -1.005 | 0.173 | -0.014 | 0.006 | 0.006 | 0 | 0.15 | 0.12 | 0.15 | 0.1 | 0.23 | 0.21 |
| 2 | 216876363 | rs17393335 | -0.723 | 1.718 | -1.005 | 0.173 | -0.014 | 0.006 | 0.006 | 0 | 0.15 | 0.12 | 0.15 | 0.1 | 0.23 | 0.21 |
| 2 | 216876828 | rs3770565 | -0.269 | 1.927 | -1.862 | 0.842 | -0.04 | -0.114 | -0.114 | 0 | 0.2 | 0.04 | 0.37 | 0.39 | 0.28 | 0.12 |
| 2 | 216877338 | rs764932 | 0.016 | 1.919 | -1.876 | 1.264 | -0.054 | -0.053 | -0.053 | 0 | 0.17 | 0.01 | 0.29 | 0.2 | 0.25 | 0.1 |
| 2 | 216880767 | rs3755156 | -0.604 | 1.951 | -1.011 | 1.606 | 0.025 | 0.09 | 0.09 | 0 | 0.1 | 0.23 | 0.13 | 0.19 | 0.1 | 0.22 |
| 2 | 216885707 | rs7564348 | -0.103 | 1.198 | -0.567 | 0.875 | -0.007 | 0.007 | 0.007 | 0.139 | 0.34 | 0.35 | 0.47 | 0.39 | 0.33 | 0.17 |
| 2 | 216885852 | rs6435930 | -1.766 | 1.405 | -0.143 | 0.757 | 0.04 | 0.043 | 0.043 | 0 | 0.61 | 0.78 | 0.79 | 0.69 | 0.71 | 0.61 |
| 2 | 216887593 | rs12104693 | -0.739 | 1.378 | -0.109 | 0.76 | 0.079 | 0.044 | 0.044 | 0 | 0.25 | 0.51 | 0.29 | 0.28 | 0.33 | 0.42 |
| 2 | 216890504 | rs934153 | 0.067 | 1.389 | -0.212 | 0.763 | 0.004 | -0.007 | -0.007 | 0.411 | 0.34 | 0.27 | 0.46 | 0.4 | 0.33 | 0.17 |
| 2 | 216905728 | rs10180583 | -0.063 | 1.403 | -0.177 | 0.768 | 0.02 | 0.084 | 0.084 | 0 | 0.1 | 0.23 | 0.12 | 0.19 | 0.1 | 0.23 |
| 2 | 216905867 | rs12468812 | -0.876 | 1.297 | -0.208 | 0.776 | 0.006 | -0.006 | -0.006 | 0.287 | 0.48 | 0.32 | 0.62 | 0.51 | 0.57 | 0.36 |
| 2 | 216908679 | rs2303842 | 0.599 | 1.409 | -0.176 | 0.949 | -0.063 | 0.002 | 0.002 | 0 | 0.13 | 0.04 | 0.15 | 0.12 | 0.23 | 0.19 |
| 2 | 216923679 | rs1429148 | -0.429 | 0.735 | 0.231 | 0.998 | -0.004 | 0.097 | 0.097 | 0 | 0.1 | 0.25 | 0.12 | 0.19 | 0.05 | 0.23 |
| 2 | 216927447 | rs10168653 | -0.411 | 0.732 | 0.198 | 0.997 | -0.003 | 0.101 | 0.101 | 0 | 0.1 | 0.25 | 0.12 | 0.19 | 0.05 | 0.2 |
| 2 | 216928870 | rs16855357 | -0.733 | 0.746 | 0.129 | 1.002 | 0.048 | 0.16 | 0.16 | 0 | 0.12 | 0.44 | 0.16 | 0.19 | 0.1 | 0.25 |
| 2 | 216933792 | rs705648 | -1.002 | 0.274 | -0.814 | 0.52 | 0.061 | 0.056 | 0.056 | 0.048 | 0.61 | 0.82 | 0.79 | 0.71 | 0.75 | 0.63 |
| 2 | 216943032 | rs7587968 | -0.079 | 0.254 | -1.15 | 0.422 | -0.008 | -0.017 | -0.017 | 0 | 0.61 | 0.74 | 0.54 | 0.39 | 0.63 | 0.66 |
| 2 | 216947351 | rs1078103 | 0.396 | 0.288 | -1.51 | 0.033 | -0.002 | -0.004 | -0.004 | 0.064 | 0.25 | 0.26 | 0.49 | 0.68 | 0.3 | 0.19 |
| 2 | 216955335 | rs828920 | -0.916 | -0.112 | -2.197 | -0.816 | 0.029 | -0.019 | -0.019 | 0 | 0.79 | 0.99 | 0.94 | 0.86 | 0.9 | 0.69 |
| 2 | 216956668 | rs828922 | 1.253 | 0.324 | -1.964 | -0.944 | -0.061 | 0.078 | 0.078 | 0 | 0.21 | 0.07 | 0.17 | 0.07 | 0.3 | 0.2 |
| 2 | 216958412 | rs6721207 | 1.598 | 0.138 | -0.915 | -0.787 | -0.005 | -0.017 | -0.017 | 0 | 0.21 | 0.23 | 0.19 | 0.11 | 0.21 | 0.25 |
| 2 | 216972732 | rs828907 | 0.119 | 0.119 | -0.827 | -0.861 | -0.013 | 0.16 | 0.16 | 0.019 | 0.51 | 0.19 | 0.26 | 0.2 | 0.44 | 0.56 |
| 2 | 216977028 | rs828910 | -0.217 | 0.143 | -1.165 | -0.826 | -0.001 | -0.017 | -0.017 | 0.33 | 0.54 | 0.51 | 0.33 | 0.2 | 0.49 | 0.56 |
| 2 | 216993611 | rs828704 | -1.569 | 0.338 | -1.442 | -0.752 | 0.002 | 0.018 | 0.018 | 0 | 0.8 | 0.86 | 0.82 | 0.89 | 0.79 | 0.77 |
| 2 | 216998661 | rs207878 | 0.056 | 0.191 | -1.835 | -0.819 | -0.072 | 0.2 | 0.2 | 0.014 | 0.49 | 0.07 | 0.28 | 0.22 | 0.49 | 0.56 |
| 2 | 217003152 | rs2303400 | 0.272 | 0.276 | -1.541 | -0.786 | -0.016 | 0.015 | 0.015 | 0 | 0.37 | 0.22 | 0.55 | 0.68 | 0.43 | 0.32 |
| 2 | 217010821 | rs16855458 | 1.839 | 0.086 | -1.234 | -0.503 | -0.067 | -0.042 | -0.042 | 0 | 0.2 | 0.01 | 0.2 | 0.19 | 0.23 | 0.17 |
| 2 | 217012092 | rs2160981 | 0.394 | 0.172 | -0.993 | -0.549 | -0.005 | 0.012 | 0.012 | 0.045 | 0.36 | 0.22 | 0.54 | 0.69 | 0.39 | 0.32 |
| 2 | 217012901 | rs207906 | -1.079 | -0.528 | -0.473 | -0.574 | -0.001 | -0.007 | -0.007 | 0 | 0.88 | 0.88 | 0.9 | 0.93 | 0.87 | 0.83 |
| 2 | 217018256 | rs16855489 | 0.525 | -0.734 | -0.886 | -0.722 | 0.015 | 0.034 | 0.034 | 0.047 | 0.6 | 0.28 | 0.5 | 0.27 | 0.65 | 0.51 |
| 2 | 217018422 | rs207910 | -0.052 | -0.996 | 0.228 | -1.091 | -0.023 | -0.04 | -0.04 | 4.761 | 0.84 | 0.74 | 0.85 | 0.93 | 0.87 | 0.83 |
| 2 | 217022813 | rs1364726 | 0.206 | -0.892 | 0.169 | -0.944 | -0.009 | -0.023 | -0.023 | 2.222 | 0.13 | 0.2 | 0.1 | 0.07 | 0.11 | 0.18 |
| 2 | 217027560 | rs207916 | -0.712 | -0.893 | -0.003 | -1.059 | -0.017 | 0.045 | 0.045 | 0 | 0.61 | 0.75 | 0.4 | 0.28 | 0.55 | 0.68 |
| 2 | 217031328 | rs3821107 | -0.492 | -1.113 | 0.13 | -1.259 | -0.026 | -0.013 | -0.013 | 0.01 | 0.31 | 0.5 | 0.18 | 0.23 | 0.23 | 0.48 |
| 2 | 217031509 | rs207922 | 1.632 | -1.045 | -0.204 | -1.306 | -0.094 | 0.158 | 0.158 | 0.06 | 0.28 | 0.05 | 0.21 | 0.06 | 0.33 | 0.2 |
| 2 | 217041881 | rs207938 | -0.847 | -1.072 | -0.159 | -1.164 | -0.036 | 0.028 | 0.028 | 0 | 0.61 | 0.83 | 0.41 | 0.32 | 0.56 | 0.69 |
| 2 | 217046443 | rs207946 | 0.281 | -1.108 | 0.08 | -1.434 | 0.006 | -0.072 | -0.072 | 0 | 0.42 | 0.52 | 0.33 | 0.13 | 0.44 | 0.38 |
| 2 | 217048565 | rs6729441 | 0.597 | -1.214 | 0.284 | -1.305 | -0.005 | -0.023 | -0.023 | 0 | 0.13 | 0.22 | 0.1 | 0.07 | 0.11 | 0.17 |
| 2 | 217057937 | rs7583902 | 0.15 | -1.227 | 0.055 | -1.312 | -0.012 | -0.047 | -0.047 | 0 | 0.15 | 0.47 | 0.12 | 0.07 | 0.11 | 0.17 |
| 2 | 217061720 | rs9288518 | 0.64 | -1.154 | -0.28 | -1.203 | -0.007 | 0.037 | 0.037 | 0 | 0.38 | 0.19 | 0.59 | 0.72 | 0.4 | 0.32 |
| 2 | 217064902 | rs2032765 | 0.449 | -1.194 | 0.218 | -0.743 | 0.007 | -0.002 | -0.002 | 0 | 0.17 | 0.14 | 0.06 | 0.15 | 0.11 | 0.3 |
| 2 | 217070248 | rs1051677 | 0.449 | -1.194 | 0.218 | -0.746 | 0.007 | -0.001 | -0.001 | 0 | 0.17 | 0.14 | 0.06 | 0.15 | 0.11 | 0.3 |
| 2 | 217070652 | rs6941 | 0.464 | -1.193 | 0.219 | -0.745 | 0.007 | -0.002 | -0.002 | 0.004 | 0.17 | 0.14 | 0.06 | 0.15 | 0.11 | 0.3 |
| 2 | 217075168 | rs207887 | -1.223 | -1.167 | 0.203 | -0.597 | -0.003 | 0.031 | 0.031 | 0 | 0.84 | 0.84 | 0.69 | 0.47 | 0.88 | 0.85 |
| 2 | 217103149 | rs207840 | -1.26 | -1.205 | 0.203 | -0.596 | -0.001 | 0.022 | 0.022 | 0 | 0.84 | 0.86 | 0.69 | 0.47 | 0.88 | 0.85 |
| 2 | 217105695 | rs207844 | 1.048 | -1.316 | 0.415 | -0.969 | 0.035 | -0.124 | -0.124 | 0 | 0.32 | 0.52 | 0.25 | 0.07 | 0.37 | 0.2 |
| 2 | 217105707 | rs7597209 | -0.408 | -1.304 | 0.259 | -0.949 | 0.006 | 0.023 | 0.023 | 0 | 0.52 | 0.35 | 0.45 | 0.4 | 0.51 | 0.65 |
| 2 | 217110611 | rs206364 | 0.761 | -1.585 | 0.223 | -1.026 | 0.083 | 0.002 | 0.002 | 0.942 | 0.58 | 0.22 | 0.55 | 0.4 | 0.73 | 0.57 |
| 2 | 217120038 | rs206375 | -0.732 | -1.384 | 0.542 | -0.892 | -0.04 | 0.121 | 0.121 | 1.04 | 0.8 | 0.42 | 0.66 | 0.47 | 0.88 | 0.86 |
| 2 | 217122402 | rs206374 | -0.684 | -1.423 | 0.521 | -0.885 | -0.039 | 0.094 | 0.094 | 1.664 | 0.8 | 0.39 | 0.67 | 0.47 | 0.88 | 0.86 |
| 2 | 217125314 | rs721774 | 1.801 | -1.467 | 0.401 | -1.093 | -0.071 | -0.015 | -0.015 | 0.16 | 0.23 | 0.04 | 0.29 | 0.19 | 0.3 | 0.19 |
| 2 | 217127309 | rs207873 | 0.862 | -1.452 | 0.553 | -0.969 | 0.078 | -0.007 | -0.007 | 0 | 0.59 | 0.22 | 0.56 | 0.43 | 0.75 | 0.59 |
| 2 | 217137889 | rs1370701 | 0.184 | -1.323 | 0.818 | -1.02 | 0.017 | 0.044 | 0.044 | 0 | 0.2 | 0.12 | 0.09 | 0.07 | 0.12 | 0.23 |
| 2 | 217143359 | rs207808 | 1.303 | -1.149 | 0.457 | -0.972 | 0.036 | -0.104 | -0.104 | 0.315 | 0.29 | 0.42 | 0.24 | 0.07 | 0.35 | 0.19 |
| 2 | 217144907 | rs207811 | 1.268 | -1.121 | 0.194 | -0.957 | 0.046 | -0.127 | -0.127 | 0 | 0.3 | 0.48 | 0.25 | 0.07 | 0.36 | 0.17 |
| 2 | 217145087 | rs207813 | 0.071 | -1.273 | 0.03 | -1.066 | -0.025 | 0.178 | 0.178 | 0.154 | 0.44 | 0.78 | 0.32 | 0.07 | 0.42 | 0.29 |
| 2 | 217146409 | rs2544062 | 0.522 | -1.145 | 0.441 | -0.693 | -0.031 | -0.074 | -0.074 | 0 | 0.14 | 0.29 | 0.07 | 0 | 0.06 | 0.12 |
| 2 | 217146914 | rs207818 | -1.169 | -1.047 | -0.178 | -0.441 | 0.001 | 0.018 | 0.018 | 0 | 0.84 | 0.89 | 0.71 | 0.51 | 0.89 | 0.85 |
| 2 | 217151143 | rs6746802 | 0.091 | -0.404 | 0.118 | -0.365 | -0.054 | 0.009 | 0.009 | 0 | 0.31 | 0.15 | 0.37 | 0.26 | 0.43 | 0.33 |
| 2 | 217151315 | rs2544066 | 0.871 | -0.492 | 0.422 | -0.398 | 0 | 0.002 | 0.002 | 0 | 0.48 | 0.38 | 0.51 | 0.57 | 0.45 | 0.34 |
| 2 | 217153043 | rs2160980 | 1.137 | -0.491 | 0.231 | -0.372 | -0.043 | -0.028 | -0.028 | 0 | 0.46 | 0.12 | 0.49 | 0.57 | 0.45 | 0.33 |
| 2 | 217153197 | rs440866 | 0.607 | -0.494 | 0.299 | -0.399 | -0.003 | 0.033 | 0.033 | 0 | 0.53 | 0.31 | 0.54 | 0.74 | 0.51 | 0.56 |
| 2 | 217159344 | rs2193747 | -0.685 | -0.203 | 0.057 | -0.206 | -0.032 | -0.004 | -0.004 | 0.12 | 0.18 | 0.33 | 0.09 | 0.17 | 0.12 | 0.28 |
| 2 | 217159587 | rs12467128 | 0.762 | -0.075 | -0.15 | -0.236 | -0.099 | -0.03 | -0.03 | 0 | 0.25 | 0.04 | 0.29 | 0.2 | 0.35 | 0.2 |
| 2 | 217160363 | rs6741495 | -0.06 | -0.073 | 0.021 | -0.189 | 0.031 | 0.15 | 0.15 | 0.264 | 0.06 | 0.23 | 0.05 | 0.16 | 0.06 | 0.22 |
| 2 | 217171643 | rs10498047 | -0.494 | -0.085 | 0.091 | -0.237 | 0.018 | -0.015 | -0.015 | 0 | 0.14 | 0.47 | 0.12 | 0.08 | 0.1 | 0.21 |
| 2 | 217172384 | rs16855891 | 0.251 | 0.03 | -0.103 | -0.099 | 0.006 | 0.087 | 0.087 | 0.161 | 0.05 | 0.15 | 0.06 | 0.16 | 0.06 | 0.22 |
| 2 | 217173897 | rs6739597 | -1.045 | 0.069 | -0.169 | -0.154 | 0 | -0.007 | -0.007 | 0 | 0.44 | 0.48 | 0.41 | 0.37 | 0.48 | 0.49 |
| 2 | 217184715 | rs16855901 | -1.144 | -0.033 | 0.448 | 0.045 | -0.038 | -0.022 | -0.022 | 0 | 0.18 | 0.44 | 0.12 | 0.13 | 0.12 | 0.3 |

| | | | | | | | | | | | | | | | | |
|---|---|---|---|---|---|---|---|---|---|---|---|---|---|---|---|---|
| 2 | 217186171 | rs6435938 | -1.036 | 0.148 | 0.389 | 0.248 | 0.015 | 0.08 | 0.08 | 1.3 | 0.09 | 0.34 | 0.1 | 0.15 | 0.08 | 0.28 |
| 2 | 217186741 | rs6728666 | -0.732 | 0.353 | -0.5 | -0.232 | -0.009 | 0.091 | 0.091 | 0.023 | 0.57 | 0.77 | 0.68 | 0.79 | 0.52 | 0.66 |
| 2 | 217194911 | rs12622637 | 0.82 | 0.342 | -0.829 | -0.007 | -0.115 | -0.168 | -0.168 | 0 | 0.26 | 0.02 | 0.47 | 0.56 | 0.29 | 0.25 |
| 2 | 217199624 | rs955816 | -0.917 | 0.269 | -0.782 | -0.164 | 0.01 | 0.131 | 0.131 | 0.004 | 0.51 | 0.77 | 0.64 | 0.78 | 0.42 | 0.63 |
| 2 | 217203920 | rs4674076 | -0.947 | 0.132 | -0.761 | -0.121 | -0.015 | 0.126 | 0.126 | 0 | 0.51 | 0.76 | 0.64 | 0.78 | 0.46 | 0.64 |
| 2 | 217205588 | rs4674077 | 0.062 | 0.026 | -0.261 | -0.165 | -0.033 | 0.003 | 0.003 | 0 | 0.36 | 0.12 | 0.55 | 0.7 | 0.4 | 0.5 |
| 2 | 217208571 | rs6756590 | -1.047 | 0.017 | -0.09 | -0.168 | 0.003 | 0.041 | 0.041 | 0.007 | 0.53 | 0.66 | 0.64 | 0.78 | 0.45 | 0.64 |
| 2 | 217209138 | rs2030493 | -0.305 | -0.029 | -0.371 | -0.088 | 0.003 | 0.018 | 0.018 | 0 | 0.42 | 0.28 | 0.55 | 0.65 | 0.38 | 0.39 |
| 2 | 217211217 | rs12467575 | 0.715 | 0 | -1.089 | -0.053 | 0.019 | 0.002 | 0.002 | 0 | 0.11 | 0.06 | 0.05 | 0.08 | 0.05 | 0.14 |
| 2 | 217216651 | rs7561922 | 0.466 | 0.272 | -1.014 | 0.218 | -0.06 | -0.125 | -0.125 | 0 | 0.29 | 0.06 | 0.49 | 0.56 | 0.33 | 0.26 |
| 2 | 217220258 | rs784553 | -1.106 | 0.436 | -1.534 | -0.403 | -0.08 | 0.21 | 0.21 | 0 | 0.65 | 0.96 | 0.74 | 0.96 | 0.65 | 0.7 |
| 2 | 217220670 | rs10191931 | 0.259 | 0.366 | -0.542 | 0.473 | 0.008 | 0.037 | 0.037 | 0.014 | 0.28 | 0.33 | 0.41 | 0.54 | 0.3 | 0.2 |
| 2 | 217228194 | rs17550937 | -0.209 | 0.811 | -1.578 | 0.255 | -0.021 | -0.028 | -0.028 | 0.505 | 0.09 | 0.01 | 0.06 | 0.14 | 0.12 | 0.3 |
| 2 | 217233740 | rs1510839 | -0.458 | 0.801 | -1.274 | 0.048 | 0.003 | 0.094 | 0.094 | 0.216 | 0.46 | 0.77 | 0.51 | 0.79 | 0.44 | 0.64 |
| 2 | 217244944 | rs4672798 | -0.28 | 0.881 | -1.061 | 0.422 | 0.002 | 0.01 | 0.01 | 0 | 0.51 | 0.59 | 0.53 | 0.43 | 0.45 | 0.6 |
| 2 | 217245058 | rs784554 | -0.376 | 0.295 | -0.445 | 0.106 | -0.021 | -0.038 | -0.038 | 2.543 | 0.32 | 0.51 | 0.24 | 0.22 | 0.24 | 0.39 |
| 2 | 217247182 | rs1589493 | -1.155 | 0.313 | -0.011 | 0.231 | 0.001 | -0.002 | -0.002 | 0 | 0.66 | 0.65 | 0.55 | 0.76 | 0.63 | 0.7 |
| 2 | 217251859 | rs9636338 | 0.027 | 0.184 | -0.468 | 0.496 | -0.009 | -0.068 | -0.068 | 0 | 0.15 | 0.05 | 0.2 | 0.32 | 0.16 | 0.1 |
| 2 | 217252413 | rs1111341 | -0.54 | 0.151 | -0.252 | 0.46 | -0.001 | 0.028 | 0.028 | 0.418 | 0.53 | 0.72 | 0.36 | 0.43 | 0.47 | 0.59 |
| 2 | 217252700 | rs925382 | -1.845 | 0.073 | -0.284 | 0.409 | -0.076 | -0.008 | -0.008 | 0 | 0.7 | 0.91 | 0.57 | 0.75 | 0.64 | 0.7 |
| 2 | 217253277 | rs10199099 | 0.458 | -0.228 | -0.148 | 0.219 | -0.058 | -0.002 | -0.002 | 0 | 0.23 | 0.07 | 0.19 | 0.24 | 0.34 | 0.3 |
| 2 | 217258982 | rs784541 | 0.172 | -0.684 | 0.65 | 0.158 | 0.012 | -0.007 | -0.007 | 0 | 0.06 | 0.2 | 0.07 | 0.02 | 0.04 | 0.04 |
| 2 | 217261557 | rs2666288 | -1.299 | -0.77 | 0.638 | 0.211 | 0.099 | 0.002 | 0.002 | 0.248 | 0.47 | 0.82 | 0.46 | 0.54 | 0.3 | 0.4 |
| 2 | 217263742 | rs784551 | -1.166 | -0.445 | 0.609 | 0.124 | -0.011 | 0.031 | 0.031 | 0.266 | 0.53 | 0.73 | 0.37 | 0.43 | 0.5 | 0.59 |
| 2 | 217265357 | rs784548 | -1.473 | -0.421 | 0.635 | 0.121 | -0.01 | 0.006 | 0.006 | 1.309 | 0.58 | 0.76 | 0.4 | 0.43 | 0.55 | 0.63 |
| 2 | 217269954 | rs4674084 | 0.137 | -0.165 | 0.645 | 0.261 | -0.001 | 0 | 0 | 0 | 0.24 | 0.18 | 0.21 | 0.24 | 0.36 | 0.3 |
| 2 | 217278023 | rs3806573 | 0.456 | -0.19 | 0.466 | 0.385 | 0.007 | -0.017 | -0.017 | 0 | 0.15 | 0.19 | 0.23 | 0.32 | 0.12 | 0.09 |
| 2 | 217278730 | rs1510841 | 0.358 | -0.08 | 0.264 | 0.323 | -0.143 | -0.04 | -0.04 | 0 | 0.23 | 0.01 | 0.19 | 0.25 | 0.35 | 0.29 |
| 2 | 217289413 | rs11886888 | 0.284 | -0.223 | 0.762 | 0.389 | -0.001 | 0.036 | 0.036 | 0.014 | 0.15 | 0.29 | 0.29 | 0.33 | 0.17 | 0.14 |
| 2 | 217292714 | rs16856088 | -0.008 | -0.422 | 0.784 | 0.179 | -0.004 | 0.004 | 0.004 | 0 | 0.15 | 0.17 | 0.05 | 0.18 | 0.11 | 0.28 |
| 2 | 217297418 | rs2066521 | 1.476 | -0.552 | 0.975 | 0.158 | -0.026 | 0.028 | 0.028 | 0 | 0.07 | 0.09 | 0.01 | 0.17 | 0.01 | 0.06 |
| 2 | 217298205 | rs11686241 | 0.334 | -0.563 | 1.043 | 0.147 | -0.003 | 0.062 | 0.062 | 0.113 | 0.16 | 0.32 | 0.3 | 0.34 | 0.17 | 0.16 |
| 2 | 217306698 | rs13411858 | 0.22 | -0.572 | 1.125 | 0.141 | 0 | 0.093 | 0.093 | 0.198 | 0.16 | 0.36 | 0.31 | 0.34 | 0.17 | 0.16 |
| 2 | 217309111 | rs284544 | -2.785 | -0.608 | 0.647 | 0.03 | -0.028 | 0.009 | 0.009 | 0.268 | 0.7 | 0.91 | 0.7 | 0.78 | 0.68 | 0.73 |
| 2 | 217311609 | rs2738284 | 0.651 | -0.65 | 0.605 | 0.028 | -0.144 | -0.025 | -0.025 | 0 | 0.24 | 0.01 | 0.27 | 0.24 | 0.36 | 0.28 |
| 2 | 217311630 | rs284550 | -1.165 | -0.735 | 0.925 | 0.092 | 0.091 | -0.055 | -0.055 | 0.21 | 0.36 | 0.8 | 0.37 | 0.51 | 0.21 | 0.22 |
| 2 | 217313044 | rs284552 | 0.619 | -0.73 | 1.102 | -0.097 | -0.035 | 0.133 | 0.133 | 0 | 0.09 | 0.29 | 0.03 | 0.17 | 0.01 | 0.06 |
| 2 | 217315787 | rs2066527 | 0.024 | -0.703 | 0.894 | -0.108 | -0.021 | 0.028 | 0.028 | 0 | 0.09 | 0.04 | 0.05 | 0.01 | 0.1 | 0.22 |
| 2 | 217326879 | rs11888959 | -1.718 | -0.675 | 0.838 | -0.02 | 0.075 | -0.008 | -0.008 | 0.651 | 0.45 | 0.86 | 0.42 | 0.55 | 0.32 | 0.45 |
| 2 | 217329184 | rs2129119 | -0.281 | -0.619 | 0.97 | 0.001 | -0.048 | 0.058 | 0.058 | 0.456 | 0.28 | 0.52 | 0.35 | 0.37 | 0.2 | 0.16 |
| 2 | 217340816 | rs184026 | -1.31 | -0.562 | 0.975 | 0.012 | -0.002 | 0 | 0 | 0 | 0.38 | 0.65 | 0.42 | 0.37 | 0.32 | 0.39 |
| 2 | 217342357 | rs2280042 | 1.376 | -0.513 | 0.722 | 0.095 | -0.008 | 0.007 | 0.007 | 0 | 0.07 | 0.07 | 0.01 | 0.17 | 0.01 | 0.06 |
| 2 | 217350717 | rs6743777 | -1.231 | -0.571 | 0.969 | 0.044 | -0.004 | 0.012 | 0.012 | 0.775 | 0.32 | 0.46 | 0.38 | 0.38 | 0.29 | 0.43 |
| 2 | 217358023 | rs16856243 | -0.711 | -0.545 | 0.801 | 0.018 | -0.007 | 0.011 | 0.011 | 0 | 0.17 | 0.27 | 0.07 | 0.18 | 0.12 | 0.29 |
| 2 | 217363049 | rs3755140 | 1.445 | -0.549 | 0.587 | 0.037 | -0.024 | 0.04 | 0.04 | 0 | 0.07 | 0.08 | 0.02 | 0.19 | 0.01 | 0.06 |
| 2 | 217363061 | rs284574 | 0.481 | -0.443 | 0.486 | -0.034 | 0.001 | 0 | 0 | 0.057 | 0.59 | 0.53 | 0.44 | 0.42 | 0.48 | 0.58 |
| 2 | 217364539 | rs284572 | -0.242 | -0.459 | 0.482 | 0.019 | -0.011 | 0.142 | 0.142 | 0.077 | 0.81 | 0.55 | 0.67 | 0.61 | 0.82 | 0.82 |
| 2 | 217367286 | rs16856272 | 2.048 | -0.507 | -0.127 | -0.012 | -0.041 | -0.058 | -0.058 | 0 | 0.11 | 0.16 | 0.05 | 0 | 0.04 | 0.01 |
| 2 | 217368839 | rs284541 | 2.303 | -0.383 | -0.953 | 0.05 | -0.017 | 0.041 | 0.041 | 0.195 | 0.31 | 0.13 | 0.31 | 0.2 | 0.33 | 0.28 |
| 2 | 217369186 | rs998102 | 0.352 | 0.084 | -0.81 | 0.288 | -0.13 | -0.013 | -0.013 | 0 | 0.22 | 0.02 | 0.23 | 0.22 | 0.34 | 0.26 |
| 2 | 217398581 | rs12612079 | 0.735 | -0.144 | -0.437 | 0.034 | 0.005 | -0.018 | -0.018 | 0 | 0.11 | 0.07 | 0.03 | 0.19 | 0.06 | 0.1 |
| 2 | 217405203 | rs4550645 | -0.005 | -0.201 | -0.801 | -0.117 | -0.099 | -0.086 | -0.086 | 0.064 | 0.62 | 0.9 | 0.46 | 0.44 | 0.48 | 0.57 |
| 2 | 217408239 | rs4672803 | 1.076 | -0.099 | -0.606 | -0.291 | 0.02 | 0.076 | 0.076 | 3.551 | 0.39 | 0.17 | 0.33 | 0.2 | 0.35 | 0.29 |
| 2 | 217411246 | rs922987 | 0.558 | 0.408 | -0.051 | -0.033 | 0 | -0.003 | -0.003 | 0 | 0.59 | 0.4 | 0.43 | 0.26 | 0.49 | 0.47 |
| 2 | 217412087 | rs2012243 | 0.91 | 0.545 | 0.114 | -0.022 | -0.004 | -0.029 | -0.029 | 1.376 | 0.39 | 0.45 | 0.35 | 0.21 | 0.39 | 0.37 |
| 2 | 217414502 | rs10197526 | -0.064 | 0.216 | -0.141 | -0.487 | 0.043 | 0.063 | 0.063 | 0 | 0.34 | 0.19 | 0.15 | 0.06 | 0.2 | 0.32 |
| 2 | 217424109 | rs12463731 | 0.708 | 1.368 | 0.692 | 0.571 | 0.052 | -0.035 | -0.035 | 0.004 | 0.79 | 0.69 | 0.76 | 0.88 | 0.65 | 0.82 |
| 2 | 217428055 | rs4674093 | 0.766 | 1.416 | 0.696 | 0.585 | 0.043 | -0.035 | -0.035 | 0 | 0.8 | 0.69 | 0.77 | 0.88 | 0.68 | 0.83 |
| 2 | 217428582 | rs9917209 | -0.985 | 0.513 | -0.014 | 0.736 | -0.019 | 0.021 | 0.021 | 0.064 | 0.07 | 0.15 | 0.03 | 0.14 | 0.02 | 0.16 |
| 2 | 217446590 | rs10170284 | -1.107 | 0.849 | 0.097 | 0.55 | 0.012 | -0.017 | -0.017 | 0.054 | 0.2 | 0.28 | 0.21 | 0.14 | 0.25 | 0.16 |
| 2 | 217453875 | rs11674057 | -0.693 | 0.785 | -0.315 | 0.586 | -0.027 | 0.006 | 0.006 | 0 | 0.18 | 0.06 | 0.19 | 0.16 | 0.25 | 0.17 |
| 2 | 217453939 | rs17561224 | 1.305 | 0.855 | -0.233 | 0.904 | -0.011 | 0.011 | 0.011 | 0.065 | 0.61 | 0.28 | 0.47 | 0.39 | 0.59 | 0.54 |
| 2 | 217456567 | rs940794 | -1.243 | 0.908 | 0.312 | 1.074 | -0.001 | -0.013 | -0.013 | 0 | 0.33 | 0.62 | 0.45 | 0.68 | 0.36 | 0.47 |
| 2 | 217457187 | rs4234019 | -0.014 | 1.042 | 0.633 | 1.136 | 0.063 | 0.027 | 0.027 | 0.054 | 0.91 | 0.7 | 0.89 | 0.94 | 0.92 | 0.86 |
| 2 | 217463442 | rs7590938 | 0.94 | 0.653 | -0.186 | 0.701 | 0.005 | 0.028 | 0.028 | 0.228 | 0.43 | 0.39 | 0.4 | 0.25 | 0.45 | 0.38 |
| 2 | 217469833 | rs10932668 | -0.931 | 0.514 | -0.019 | 1.323 | -0.01 | 0.169 | 0.169 | 0 | 0.22 | 0.52 | 0.29 | 0.46 | 0.18 | 0.29 |
| 2 | 217475199 | rs16856415 | -0.137 | 0.872 | -0.734 | 1.33 | 0.005 | 0.122 | 0.122 | 0 | 0.1 | 0.18 | 0.25 | 0.33 | 0.11 | 0.17 |
| 2 | 217477217 | rs11693848 | -0.297 | 1.117 | -1.361 | 0.286 | -0.087 | 0.001 | 0.001 | 0 | 0.21 | 0.02 | 0.21 | 0.16 | 0.26 | 0.2 |
| 2 | 217480229 | rs10932669 | -0.323 | 0.668 | -0.66 | 0.653 | 0 | 0.054 | 0.054 | 0 | 0.13 | 0.23 | 0.24 | 0.31 | 0.14 | 0.24 |

| | | | | | | | | | | | | | | | | |
|---|---|---|---|---|---|---|---|---|---|---|---|---|---|---|---|---|
| 2 | 217480424 | rs4674100 | -0.087 | 0.563 | -0.898 | 0.33 | 0.002 | -0.007 | -0.007 | 0 | 0.29 | 0.25 | 0.35 | 0.37 | 0.27 | 0.24 |
| 2 | 217482379 | rs12694392 | -0.481 | 1.024 | -0.931 | 0.286 | 0 | 0.005 | 0.005 | 0.809 | 0.18 | 0.17 | 0.28 | 0.31 | 0.22 | 0.27 |
| 2 | 217484157 | rs10165302 | -0.507 | 0.89 | -0.946 | 0.205 | 0.001 | -0.003 | -0.003 | 1.149 | 0.19 | 0.14 | 0.27 | 0.33 | 0.21 | 0.24 |
| 2 | 217484229 | rs17496232 | 0.257 | 1.116 | -1.67 | -0.271 | -0.09 | 0.015 | 0.015 | 0.043 | 0.28 | 0.05 | 0.21 | 0.21 | 0.33 | 0.17 |
| 2 | 217485550 | rs10203969 | 0.305 | 1.088 | -0.817 | 0.258 | -0.016 | 0.001 | 0.001 | 0.015 | 0.32 | 0.08 | 0.22 | 0.28 | 0.32 | 0.31 |
| 2 | 217493755 | rs1525608 | -0.306 | 1.19 | -0.516 | 0.343 | 0 | 0.005 | 0.005 | 0 | 0.61 | 0.57 | 0.56 | 0.44 | 0.58 | 0.62 |
| 2 | 217500819 | rs9341105 | -0.596 | 1.042 | -0.584 | 0.236 | 0.012 | 0.044 | 0.044 | 0 | 0.36 | 0.25 | 0.2 | 0.03 | 0.28 | 0.26 |
| 2 | 217504112 | rs2372848 | 1.036 | 1.903 | 0.021 | 1.465 | 0.051 | -0.048 | -0.048 | 2.031 | 0.9 | 0.7 | 0.93 | 0.99 | 0.9 | 0.89 |
| 2 | 217521648 | rs9341191 | -1.036 | 1.218 | -1.057 | 1.312 | -0.027 | -0.02 | -0.02 | 0 | 0.06 | 0 | 0.05 | 0.11 | 0.07 | 0.14 |
| 2 | 217525243 | rs9341201 | -1.147 | 1.119 | -0.673 | 1.095 | 0.006 | -0.027 | -0.027 | 0.003 | 0.05 | 0.13 | 0.02 | 0 | 0.04 | 0.05 |
| 2 | 217532631 | rs11888965 | -1.998 | 0.9 | 0.437 | 1.896 | 0.015 | 0.03 | 0.03 | 2.591 | 0.08 | 0.34 | 0.07 | 0.07 | 0.03 | 0.08 |
| 2 | 217534244 | rs2067041 | -2.322 | 0.848 | 0.665 | 1.876 | 0.04 | 0.235 | 0.235 | 1.542 | 0.09 | 0.43 | 0.19 | 0.25 | 0.07 | 0.13 |
| 2 | 217536354 | rs4674107 | 0.062 | 0.543 | -0.774 | 1.559 | -0.037 | -0.072 | -0.072 | 1.103 | 0.33 | 0.02 | 0.26 | 0.36 | 0.3 | 0.22 |
| 2 | 217538816 | rs2241199 | -1.662 | 0.6 | 0.067 | 1.43 | -0.005 | 0.001 | 0.001 | 0.691 | 0.06 | 0.14 | 0.06 | 0.07 | 0.04 | 0.08 |
| 2 | 217539576 | rs11575213 | -1.405 | 0.689 | 0.09 | 1.474 | 0.021 | 0.036 | 0.036 | 2.342 | 0.07 | 0.33 | 0.07 | 0.07 | 0.03 | 0.08 |
| 2 | 217550529 | rs7420849 | -0.68 | 0.851 | -1.003 | 0.834 | -0.004 | 0.131 | 0.131 | 0 | 0.52 | 0.76 | 0.66 | 0.82 | 0.55 | 0.48 |
| 2 | 217554213 | rs2241193 | 0.77 | 0.618 | -0.722 | 1.43 | -0.006 | 0.016 | 0.016 | 0 | 0.85 | 0.77 | 0.72 | 0.74 | 0.85 | 0.82 |
| 2 | 217558735 | rs3770472 | -0.887 | 1.01 | -1.175 | 1.226 | -0.039 | 0.021 | 0.021 | 0 | 0.16 | 0.04 | 0.18 | 0.13 | 0.24 | 0.16 |
| 2 | 217561467 | rs1978346 | -1.417 | 1.063 | -1.369 | 1.516 | 0.012 | -0.03 | -0.03 | 0.557 | 0.45 | 0.77 | 0.58 | 0.5 | 0.37 | 0.37 |
| 2 | 217562362 | rs1978345 | -1.424 | 1.05 | -1.354 | 1.654 | 0.013 | -0.009 | -0.009 | 0 | 0.45 | 0.79 | 0.58 | 0.56 | 0.37 | 0.37 |
| 2 | 217563445 | rs6727330 | -1.335 | 1.053 | -1.041 | 1.789 | 0.007 | -0.018 | -0.018 | 0 | 0.43 | 0.72 | 0.55 | 0.54 | 0.36 | 0.38 |
| 2 | 217567718 | rs888186 | -0.008 | 1.281 | -1.078 | 1.931 | 0 | 0.004 | 0.004 | 0 | 0.9 | 0.89 | 0.88 | 0.87 | 0.92 | 0.8 |
| 2 | 217575726 | rs6757568 | -0.665 | 1.123 | -0.467 | 2.234 | -0.019 | 0.069 | 0.069 | 1.203 | 0.2 | 0.59 | 0.35 | 0.34 | 0.16 | 0.26 |
| 2 | 217576692 | rs17773051 | 1.971 | 1.582 | -0.807 | 2.381 | -0.038 | 0.061 | 0.061 | 0.476 | 0.57 | 0.13 | 0.41 | 0.36 | 0.59 | 0.47 |
| 2 | 217583263 | rs16856553 | -0.706 | 1.708 | -0.253 | 2.424 | -0.005 | 0.108 | 0.108 | 0 | 0.07 | 0.1 | 0.15 | 0.3 | 0.06 | 0.14 |
| 2 | 217585034 | rs759545 | -0.786 | 1.711 | -0.36 | 2.434 | 0.001 | 0.043 | 0.043 | 0 | 0.07 | 0.07 | 0.15 | 0.31 | 0.06 | 0.14 |
| 2 | 217585982 | rs2372866 | -0.111 | 2.003 | 0.045 | 2.365 | 0.007 | -0.009 | -0.009 | 0 | 0.06 | 0.06 | 0.14 | 0.03 | 0.09 | 0.12 |
| 2 | 217604130 | rs888188 | 1.183 | 2.166 | 0.101 | 2.484 | 0.009 | 0.055 | 0.055 | 0.003 | 0.85 | 0.81 | 0.69 | 0.55 | 0.78 | 0.69 |
| 2 | 217605925 | rs1861631 | 1.28 | 2.037 | 0.219 | 1.938 | 0.022 | 0 | 0 | 0 | 0.82 | 0.76 | 0.69 | 0.83 | 0.74 | 0.7 |
| 2 | 217610649 | rs10932676 | -1.837 | 1.485 | 0.836 | 2.082 | -0.017 | 0.004 | 0.004 | 0 | 0.2 | 0.32 | 0.12 | 0.19 | 0.1 | 0.12 |
| 2 | 217618048 | rs7590178 | -1.652 | 1.845 | 0.815 | 2.383 | 0.005 | -0.013 | -0.013 | 0.137 | 0.17 | 0.09 | 0.18 | 0.2 | 0.14 | 0.15 |
| 2 | 217619036 | rs1548942 | -0.058 | 2.329 | 0.751 | 2.595 | -0.015 | -0.023 | -0.023 | 0 | 0.05 | 0.01 | 0.14 | 0.17 | 0.15 | 0.1 |
| 2 | 217625331 | rs11691253 | -1.514 | 2.129 | 1.56 | 2.574 | 0.004 | -0.006 | -0.006 | 0.221 | 0.17 | 0.12 | 0.23 | 0.22 | 0.13 | 0.15 |
| 2 | 217625523 | rs13015993 | 2.567 | 2.142 | 1.538 | 2.581 | 0.034 | 0.076 | 0.076 | 0 | 0.72 | 0.48 | 0.6 | 0.53 | 0.72 | 0.74 |
| 2 | 217628087 | rs6435953 | 1.698 | 2.202 | 1.289 | 2.393 | 0.076 | 0.08 | 0.08 | 0 | 0.9 | 0.69 | 0.83 | 0.75 | 0.85 | 0.89 |
| 2 | 217632422 | rs2712180 | 1.758 | 2.231 | 1.007 | 2.383 | 0.02 | 0.027 | 0.027 | 0 | 0.9 | 0.84 | 0.86 | 0.74 | 0.84 | 0.86 |
| 2 | 217633814 | rs2541408 | 1.948 | 2.199 | 1.18 | 2.393 | 0.035 | 0.049 | 0.049 | 0 | 0.89 | 0.75 | 0.85 | 0.74 | 0.84 | 0.86 |
| 2 | 217635146 | rs2193230 | 2.015 | 2.261 | 1.428 | 2.37 | 0.062 | 0.069 | 0.069 | 0 | 0.89 | 0.71 | 0.84 | 0.74 | 0.84 | 0.86 |
| 2 | 217636105 | rs11688933 | -1.185 | 2.772 | 1.495 | 2.375 | -0.001 | -0.011 | -0.011 | 0 | 0.17 | 0.11 | 0.25 | 0.2 | 0.17 | 0.18 |
| 2 | 217636736 | rs2541404 | 2.239 | 3.029 | 1.988 | 2.686 | 0.08 | 0.081 | 0.081 | 0 | 0.78 | 0.51 | 0.72 | 0.56 | 0.7 | 0.58 |
| 2 | 217637885 | rs2541403 | 2.93 | 3.017 | 2.073 | 2.643 | -0.014 | -0.006 | -0.006 | 0 | 0.6 | 0.34 | 0.47 | 0.36 | 0.53 | 0.4 |
| 2 | 217637960 | rs6708249 | -0.784 | 3.054 | 2.073 | 2.429 | 0.004 | 0.001 | 0.001 | 0 | 0.11 | 0.14 | 0.12 | 0.18 | 0.14 | 0.27 |
| 2 | 217640818 | rs7421283 | -2.298 | 3.089 | 2.173 | 2.47 | 0.064 | 0.047 | 0.047 | 0.1 | 0.2 | 0.35 | 0.27 | 0.36 | 0.31 | 0.41 |
| 2 | 217642427 | rs13010690 | -1.018 | 3.038 | 2.065 | 2.485 | 0 | 0.004 | 0.004 | 0 | 0.17 | 0.11 | 0.19 | 0.09 | 0.16 | 0.18 |
| 2 | 217643055 | rs11890102 | -0.715 | 3.042 | 2.084 | 2.518 | 0.006 | 0.002 | 0.002 | 0 | 0.11 | 0.13 | 0.11 | 0.19 | 0.15 | 0.26 |
| 2 | 217644852 | rs9989901 | -3.084 | 3.247 | 2.011 | 2.404 | 0 | 0.001 | 0.001 | 0 | 0.42 | 0.52 | 0.46 | 0.51 | 0.5 | 0.62 |
| 2 | 217646555 | rs10189682 | -2.262 | 3.155 | 2.004 | 2.458 | 0.098 | 0.089 | 0.089 | 0.206 | 0.2 | 0.47 | 0.25 | 0.42 | 0.28 | 0.39 |
| 2 | 217660346 | rs2712163 | 2.562 | 3.135 | 1.983 | 2.423 | -0.009 | -0.01 | -0.01 | 0.441 | 0.6 | 0.35 | 0.6 | 0.49 | 0.51 | 0.4 |
| 2 | 217661530 | rs2024489 | 2.503 | 3.133 | 2.072 | 2.422 | 0.01 | 0.011 | 0.011 | 0 | 0.61 | 0.53 | 0.61 | 0.49 | 0.51 | 0.4 |
| 2 | 217661788 | rs2712162 | 1.861 | 3.202 | 2.01 | 2.406 | 0.007 | -0.002 | -0.002 | 0 | 0.72 | 0.67 | 0.76 | 0.73 | 0.69 | 0.54 |
| 2 | 217669595 | rs2541385 | 2.378 | 3.225 | 1.886 | 2.475 | 0.005 | 0.005 | 0.005 | 0 | 0.63 | 0.42 | 0.63 | 0.5 | 0.57 | 0.45 |
| 2 | 217680944 | rs1015785 | 1.778 | 3.359 | 2.106 | 2.488 | 0.026 | 0.046 | 0.046 | 0 | 0.85 | 0.57 | 0.87 | 0.76 | 0.83 | 0.69 |
| 2 | 217681961 | rs2014998 | -1.719 | 3.413 | 2.102 | 2.563 | 0.008 | -0.026 | -0.026 | 0 | 0.22 | 0.3 | 0.19 | 0.07 | 0.26 | 0.24 |
| 2 | 217682453 | rs10204070 | -1.07 | 3.157 | 1.848 | 3.154 | -0.001 | 0.089 | 0.089 | 0 | 0.1 | 0.16 | 0.26 | 0.45 | 0.11 | 0.33 |
| 2 | 217682779 | rs2712184 | 2.042 | 2.892 | 1.765 | 2.152 | 0.019 | 0.013 | 0.013 | 0 | 0.64 | 0.44 | 0.63 | 0.53 | 0.57 | 0.46 |
| 2 | 217703091 | rs1179714 | 1.52 | 3.386 | 1.308 | 1.958 | 0.076 | 0.025 | 0.025 | 0 | 0.77 | 0.71 | 0.7 | 0.66 | 0.59 | 0.59 |
| 2 | 217714948 | rs1179704 | 1.007 | 3.328 | 0.79 | 1.625 | 0.036 | 0.015 | 0.015 | 0 | 0.89 | 0.85 | 0.85 | 0.75 | 0.71 | 0.68 |
| 2 | 217720468 | rs10178069 | 0.628 | 1.627 | 0.252 | 0.228 | -0.001 | -0.041 | -0.041 | 3.655 | 0.22 | 0.21 | 0.1 | 0 | 0.17 | 0.13 |
| 2 | 217725044 | rs1179735 | -0.27 | 1.034 | 0.158 | 0.014 | -0.013 | -0.009 | -0.009 | 0.203 | 0.73 | 0.76 | 0.7 | 0.57 | 0.61 | 0.61 |
| 2 | 217728801 | rs1468860 | 1.556 | 0.163 | -0.608 | -0.699 | 0.033 | -0.101 | -0.101 | 0.024 | 0.24 | 0.08 | 0.39 | 0.5 | 0.17 | 0.31 |
| 2 | 217730567 | rs1179727 | 1.052 | 0.167 | -0.11 | -0.794 | -0.004 | 0.007 | 0.007 | 0 | 0.24 | 0.22 | 0.19 | 0.06 | 0.26 | 0.16 |
| 2 | 217731308 | rs1179726 | -0.478 | 0.067 | -0.009 | -0.851 | 0 | -0.003 | -0.003 | 0 | 0.56 | 0.56 | 0.63 | 0.56 | 0.46 | 0.58 |
| 2 | 217733190 | rs1179723 | -0.244 | 0.077 | 0.097 | -0.858 | 0.033 | 0.01 | 0.01 | 0.673 | 0.66 | 0.52 | 0.66 | 0.57 | 0.57 | 0.5 |
| 2 | 217733397 | rs1922006 | 0.886 | 0.078 | -0.044 | -0.824 | -0.005 | 0.002 | 0.002 | 0 | 0.3 | 0.31 | 0.43 | 0.51 | 0.19 | 0.42 |
| 2 | 217733731 | rs1179722 | 0.758 | 0.51 | -0.114 | -0.46 | -0.005 | 0.007 | 0.007 | 0 | 0.24 | 0.23 | 0.2 | 0.06 | 0.27 | 0.17 |
| 2 | 217736131 | rs2241191 | -0.204 | -0.168 | 0.109 | -0.177 | 0.02 | 0.012 | 0.012 | 0 | 0.48 | 0.57 | 0.5 | 0.56 | 0.34 | 0.45 |
| 2 | 217746145 | rs1179718 | 0.455 | 0.001 | -0.335 | -0.062 | 0.024 | 0.008 | 0.008 | 0 | 0.41 | 0.29 | 0.49 | 0.62 | 0.32 | 0.33 |
| 2 | 217747908 | rs10206884 | -0.191 | 0.37 | -0.385 | -0.343 | -0.001 | 0.006 | 0.006 | 0 | 0.47 | 0.45 | 0.49 | 0.43 | 0.53 | 0.67 |
| 2 | 217755282 | rs1922000 | 0.088 | 0.096 | 0.15 | -0.446 | -0.004 | 0.032 | 0.032 | 0 | 0.49 | 0.34 | 0.44 | 0.39 | 0.49 | 0.65 |

| | | | | | | | | | | | | | | | | |
|---|---|---|---|---|---|---|---|---|---|---|---|---|---|---|---|---|
| 2 | 217761884 | rs1179692 | 1.127 | -0.012 | 0.504 | -0.512 | 0.014 | -0.035 | -0.035 | 0.427 | 0.15 | 0.25 | 0.13 | 0.07 | 0.17 | 0.03 |
| 2 | 217762183 | rs1179691 | -0.183 | -0.239 | 0.63 | -0.409 | 0 | -0.049 | -0.049 | 0 | 0.56 | 0.8 | 0.56 | 0.51 | 0.57 | 0.69 |
| 2 | 217768442 | rs10932680 | 0.324 | -0.198 | 0.801 | -0.433 | -0.003 | 0.025 | 0.025 | 0 | 0.49 | 0.31 | 0.45 | 0.38 | 0.49 | 0.65 |
| 2 | 217775937 | rs7589256 | -0.119 | -0.193 | 0.569 | -0.426 | 0.009 | -0.005 | -0.005 | 0 | 0.56 | 0.71 | 0.57 | 0.47 | 0.58 | 0.66 |
| 2 | 217799675 | rs2542213 | -1.344 | 0.329 | 1.03 | 0.005 | 0.094 | 0.122 | 0.122 | 2.918 | 0.08 | 0.55 | 0.14 | 0.15 | 0.09 | 0.03 |
| 2 | 217803756 | rs2705592 | -0.244 | 0.108 | 0.237 | -0.07 | -0.017 | 0.104 | 0.104 | 0 | 0.48 | 0.81 | 0.55 | 0.72 | 0.47 | 0.34 |
| 2 | 217808729 | rs754752 | 0.575 | 0.225 | -0.202 | -0.222 | -0.04 | 0.019 | 0.019 | 0 | 0.32 | 0.12 | 0.38 | 0.26 | 0.36 | 0.48 |
| 2 | 217810502 | rs6714809 | 0.239 | 0.205 | 0.287 | -0.148 | 0.006 | 0.01 | 0.01 | 0 | 0.36 | 0.42 | 0.46 | 0.47 | 0.41 | 0.49 |
| 2 | 217811464 | rs2542209 | -0.834 | 0.078 | -0.269 | -0.266 | 0.016 | -0.023 | -0.023 | 0 | 0.54 | 0.84 | 0.61 | 0.49 | 0.58 | 0.59 |
| 2 | 217826219 | rs17776232 | 0.592 | -0.002 | -0.363 | -0.255 | -0.01 | 0.01 | 0.01 | 0 | 0.14 | 0.11 | 0.11 | 0.02 | 0.17 | 0.1 |
| 2 | 217827849 | rs2372928 | 1.022 | -0.082 | -0.765 | -0.186 | -0.077 | -0.129 | -0.129 | 0 | 0.38 | 0.02 | 0.28 | 0.56 | 0.35 | 0.32 |
| 2 | 217831965 | rs2705579 | -0.744 | 0.022 | -0.96 | -0.09 | -0.051 | -0.027 | -0.027 | 0.483 | 0.68 | 0.87 | 0.59 | 0.65 | 0.61 | 0.52 |
| 2 | 217835147 | rs1882425 | -0.777 | -0.005 | -1.07 | -0.153 | -0.061 | -0.006 | -0.006 | 3.976 | 0.68 | 0.89 | 0.6 | 0.73 | 0.62 | 0.57 |
| 2 | 217837588 | rs13420604 | 0.531 | -0.144 | -0.974 | -0.03 | 0.033 | -0.054 | -0.054 | 0 | 0.42 | 0.17 | 0.39 | 0.5 | 0.37 | 0.46 |
| 2 | 217839801 | rs10207131 | 0.92 | 0.152 | -1.149 | -0.267 | 0.027 | -0.063 | -0.063 | 0 | 0.39 | 0.08 | 0.24 | 0.58 | 0.33 | 0.36 |
| 2 | 217840620 | rs2888447 | -0.188 | 0.379 | -0.181 | -0.621 | -0.01 | 0.026 | 0.026 | 0 | 0.16 | 0.08 | 0.08 | 0 | 0.15 | 0.16 |
| 2 | 217849509 | rs1882423 | -0.394 | 0.32 | 0.115 | -0.54 | -0.002 | -0.015 | -0.015 | 0 | 0.08 | 0.26 | 0.04 | 0.03 | 0.03 | 0.1 |
| 2 | 217850417 | rs1882421 | -1.723 | 0.444 | -0.371 | -0.679 | -0.011 | 0.065 | 0.065 | 0.836 | 0.29 | 0.77 | 0.19 | 0.05 | 0.27 | 0.27 |
| 2 | 217851509 | rs1882420 | -1.803 | 0.232 | 0.113 | -0.088 | -0.004 | -0.006 | -0.006 | 1.239 | 0.17 | 0.77 | 0.28 | 0.14 | 0.14 | 0.13 |
| 2 | 217855088 | rs2141817 | -0.748 | 0.133 | 0.436 | -0.112 | 0.009 | -0.004 | -0.004 | 0 | 0.08 | 0.31 | 0.05 | 0.03 | 0.03 | 0.1 |
| 2 | 217856142 | rs2705594 | -0.14 | 0.251 | -0.22 | -0.479 | -0.001 | -0.011 | -0.011 | 0 | 0.48 | 0.52 | 0.31 | 0.21 | 0.43 | 0.42 |
| 2 | 217856192 | rs2542197 | -0.661 | 0.583 | -0.844 | -1.434 | -0.01 | 0.033 | 0.033 | 0 | 0.4 | 0.72 | 0.53 | 0.76 | 0.37 | 0.55 |
| 2 | 217856966 | rs1882419 | -0.361 | 0.57 | -0.583 | -1.579 | 0.001 | 0.011 | 0.011 | 0 | 0.32 | 0.32 | 0.55 | 0.84 | 0.33 | 0.38 |
| 2 | 217857450 | rs2252488 | -0.979 | 1.209 | -0.599 | -1.663 | -0.067 | 0.081 | 0.081 | 0 | 0.78 | 0.94 | 0.8 | 0.95 | 0.69 | 0.73 |
| 2 | 217861242 | rs16856807 | 0.013 | 1.079 | -0.115 | -1.848 | 0.019 | -0.017 | -0.017 | 0 | 0.08 | 0.18 | 0.14 | 0.03 | 0.13 | 0.02 |
| 2 | 217862696 | rs10195963 | -0.787 | 0.966 | 0.107 | -1.877 | -0.031 | 0.047 | 0.047 | 0 | 0.76 | 0.86 | 0.78 | 0.96 | 0.66 | 0.72 |
| 2 | 217869406 | rs10169454 | -0.611 | 0.679 | 0.602 | -1.861 | -0.006 | 0.025 | 0.025 | 0.112 | 0.06 | 0.28 | 0.02 | 0.03 | 0 | 0.02 |
| 2 | 217874896 | rs6733648 | -1.086 | 0.744 | 0.434 | -1.899 | -0.073 | 0.085 | 0.085 | 0 | 0.77 | 0.94 | 0.79 | 0.95 | 0.68 | 0.72 |
| 2 | 217874931 | rs13389571 | -1.685 | 0.818 | 0.71 | -1.796 | 0.014 | 0.003 | 0.003 | 1.885 | 0.1 | 0.53 | 0.07 | 0.04 | 0.03 | 0.05 |
| 2 | 217884314 | rs10171745 | -1.18 | 0.769 | 0.271 | -1.809 | -0.003 | -0.012 | -0.012 | 0 | 0.1 | 0.39 | 0.05 | 0.03 | 0.04 | 0.02 |
| 2 | 217885659 | rs10177578 | -0.958 | 0.817 | -0.286 | -1.888 | -0.035 | 0.053 | 0.053 | 0 | 0.75 | 0.85 | 0.79 | 0.93 | 0.66 | 0.73 |
| 2 | 217894024 | rs17833945 | 0.142 | 0.517 | -0.518 | -2.177 | 0.004 | 0.011 | 0.011 | 0 | 0.23 | 0.22 | 0.11 | 0.02 | 0.18 | 0.2 |
| 2 | 217900280 | rs6735174 | 0.5 | 0.548 | -0.637 | -2.157 | -0.005 | 0.045 | 0.045 | 0 | 0.2 | 0.13 | 0.09 | 0.02 | 0.18 | 0.2 |
| 2 | 217900296 | rs6721811 | -1.274 | 0.608 | -0.661 | -2.187 | -0.036 | 0.057 | 0.057 | 0 | 0.76 | 0.86 | 0.78 | 0.95 | 0.67 | 0.73 |
| 2 | 217903776 | rs2372943 | 1.993 | 0.46 | -0.637 | -2.294 | 0.032 | 0.057 | 0.057 | 0.007 | 0.29 | 0.13 | 0.15 | 0.08 | 0.19 | 0.17 |
| 2 | 217905832 | rs13387042 | 0.684 | 0.452 | -0.561 | -2.232 | -0.006 | 0.124 | 0.124 | 0 | 0.43 | 0.22 | 0.64 | 0.9 | 0.48 | 0.52 |
| 2 | 217908985 | rs4621152 | -0.144 | 0.456 | -0.307 | -2.085 | 0.012 | -0.085 | -0.085 | 0.591 | 0.65 | 0.49 | 0.73 | 0.92 | 0.64 | 0.71 |
| 2 | 217910910 | rs6723013 | -1.286 | 0.763 | -0.636 | -2.032 | -0.058 | 0.087 | 0.087 | 0 | 0.76 | 0.92 | 0.79 | 0.96 | 0.67 | 0.76 |
| 2 | 217915414 | rs12694403 | 0.701 | 0.363 | -0.078 | -1.496 | -0.019 | -0.053 | -0.053 | 0 | 0.32 | 0.36 | 0.19 | 0.08 | 0.24 | 0.18 |
| 2 | 217930669 | rs16856888 | 0.127 | 0.319 | -0.307 | -1.331 | -0.003 | -0.014 | -0.014 | 0 | 0.45 | 0.44 | 0.26 | 0.18 | 0.34 | 0.32 |
| 2 | 217935431 | rs6734010 | -0.691 | 0.483 | -1.214 | -1.634 | -0.059 | 0.164 | 0.164 | 0.129 | 0.59 | 0.8 | 0.7 | 0.92 | 0.52 | 0.73 |
| 2 | 217938410 | rs13011326 | 0.84 | 0.358 | -0.971 | -1.444 | -0.017 | -0.054 | -0.054 | 0.026 | 0.32 | 0.31 | 0.19 | 0.08 | 0.22 | 0.17 |
| 2 | 217938519 | rs4674132 | -0.091 | 0.81 | -1.663 | -1.025 | -0.039 | 0.16 | 0.16 | 0.581 | 0.61 | 0.81 | 0.74 | 0.92 | 0.59 | 0.72 |
| 2 | 217944627 | rs7579306 | -0.139 | 0.249 | -1.064 | -0.406 | -0.017 | -0.039 | -0.039 | 0 | 0.43 | 0.47 | 0.32 | 0.11 | 0.29 | 0.34 |
| 2 | 217947644 | rs12694405 | -0.499 | 0.65 | -1.068 | 0.526 | -0.022 | -0.011 | -0.011 | 0.127 | 0.63 | 0.76 | 0.61 | 0.57 | 0.52 | 0.6 |
| 2 | 217950825 | rs6435962 | 0.789 | 0.189 | -0.239 | 0.856 | 0.008 | 0.049 | 0.049 | 0 | 0.71 | 0.62 | 0.66 | 0.47 | 0.69 | 0.69 |
| 2 | 217951726 | rs6739040 | 0.148 | -0.395 | -0.278 | -0.011 | -0.045 | -0.076 | -0.076 | 0 | 0.15 | 0.27 | 0.1 | 0.03 | 0.06 | 0.19 |
| 2 | 217954247 | rs12694406 | 0.46 | -0.191 | -0.713 | 0.505 | -0.029 | -0.075 | -0.075 | 0 | 0.76 | 0.93 | 0.75 | 0.43 | 0.73 | 0.72 |
| 2 | 217963545 | rs16856932 | 0.006 | -0.344 | -0.297 | 0.03 | 0.001 | -0.005 | -0.005 | 0 | 0.19 | 0.16 | 0.1 | 0.06 | 0.19 | 0.15 |
| 2 | 217971141 | rs2372972 | 0.285 | -0.114 | 0.127 | 0.315 | 0.069 | 0.257 | 0.257 | 0 | 0.82 | 0.53 | 0.74 | 0.54 | 0.79 | 0.87 |
| 2 | 217971848 | rs13414870 | -0.517 | -0.256 | -0.388 | 0.338 | 0.005 | 0.015 | 0.015 | 0 | 0.25 | 0.19 | 0.14 | 0.12 | 0.25 | 0.28 |
| 2 | 217973448 | rs2068218 | -0.491 | -0.242 | -0.112 | 0.379 | 0.013 | -0.064 | -0.064 | 2.73 | 0.22 | 0.63 | 0.18 | 0.06 | 0.25 | 0.14 |
| 2 | 217974444 | rs13015994 | 1.429 | -0.764 | -1.009 | 0.337 | -0.14 | -0.096 | -0.096 | 0 | 0.37 | 0.04 | 0.48 | 0.4 | 0.42 | 0.38 |
| 2 | 217976591 | rs6723845 | -0.143 | -1.179 | 0.399 | 0.527 | -0.001 | -0.034 | -0.034 | 0.137 | 0.15 | 0.19 | 0.09 | 0.06 | 0.16 | 0.09 |
| 2 | 217977296 | rs10188411 | -1.06 | -1.017 | 0.186 | 0.595 | -0.047 | -0.094 | -0.094 | 0 | 0.29 | 0.42 | 0.15 | 0.12 | 0.21 | 0.3 |
| 2 | 217980433 | rs12466929 | -1.237 | -0.634 | 0.135 | 1.48 | -0.032 | 0.076 | 0.076 | 0 | 0.2 | 0.31 | 0.23 | 0.54 | 0.12 | 0.24 |
| 2 | 217986622 | rs10179787 | -1.501 | -0.492 | 0.498 | 1.462 | 0.001 | 0.299 | 0.299 | 0.415 | 0.11 | 0.45 | 0.18 | 0.28 | 0.04 | 0.19 |
| 2 | 217988422 | rs6735740 | -2.032 | -0.658 | 0.069 | 1.396 | -0.015 | 0.033 | 0.033 | 3.283 | 0.14 | 0.78 | 0.21 | 0.28 | 0.04 | 0.15 |
| 2 | 217995074 | rs6742032 | -0.979 | -0.501 | -0.154 | 1.513 | -0.017 | 0.158 | 0.158 | 0 | 0.07 | 0.19 | 0.03 | 0.25 | 0.03 | 0.07 |
| 2 | 218016330 | rs12623324 | -0.264 | -0.352 | -1.623 | 0.268 | -0.015 | 0.11 | 0.11 | 0 | 0.62 | 0.97 | 0.76 | 0.78 | 0.68 | 0.55 |
| 2 | 218023000 | rs1864444 | -0.979 | -0.128 | -0.882 | 0.245 | 0.006 | 0.028 | 0.028 | 0 | 0.25 | 0.18 | 0.3 | 0.13 | 0.2 | 0.38 |
| 2 | 218023013 | rs11674430 | 0.894 | 0.204 | -0.16 | 0.201 | 0.011 | -0.06 | -0.06 | 0 | 0.72 | 0.62 | 0.64 | 0.86 | 0.72 | 0.6 |
| 2 | 218026864 | rs10498055 | -0.825 | -0.102 | -0.109 | 0.806 | -0.004 | 0.027 | 0.027 | 0 | 0.19 | 0.25 | 0.38 | 0.43 | 0.22 | 0.2 |
| 2 | 218033277 | rs1864440 | -0.631 | -0.177 | -0.13 | 0.785 | -0.005 | 0.023 | 0.023 | 0 | 0.14 | 0.17 | 0.25 | 0.42 | 0.14 | 0.15 |
| 2 | 218042959 | rs2033306 | 0.184 | -0.374 | -0.874 | 0.785 | -0.025 | -0.104 | -0.104 | 0 | 0.15 | 0.01 | 0.08 | 0.24 | 0.1 | 0.07 |
| 2 | 218052964 | rs12615123 | -1.471 | -0.551 | 0.154 | 0.64 | -0.02 | 0.03 | 0.03 | 0.007 | 0.24 | 0.71 | 0.42 | 0.42 | 0.2 | 0.35 |
| 2 | 218058184 | rs10804254 | -2.098 | -0.808 | -0.27 | 0.378 | 0.042 | 0.217 | 0.217 | 0 | 0.48 | 0.92 | 0.6 | 0.77 | 0.38 | 0.57 |
| 2 | 218061696 | rs11691144 | -0.213 | -0.748 | -0.024 | 0.551 | -0.018 | 0.06 | 0.06 | 0 | 0.15 | 0.22 | 0.24 | 0.41 | 0.14 | 0.15 |
| 2 | 218069071 | rs2195610 | -0.686 | -0.936 | 0.592 | 0.49 | -0.03 | 0.091 | 0.091 | 0 | 0.18 | 0.43 | 0.25 | 0.39 | 0.15 | 0.14 |

| | | | | | | | | | | | | | | | | |
|---|---|---|---|---|---|---|---|---|---|---|---|---|---|---|---|---|
| 2 | 218081443 | rs12477810 | -1.315 | -1.139 | 0.772 | 0.44 | -0.046 | 0.015 | 0.015 | 0 | 0.26 | 0.58 | 0.31 | 0.66 | 0.19 | 0.27 |
| 2 | 218084589 | rs17653058 | -0.612 | -1.132 | 0.581 | 0.462 | 0.015 | -0.028 | -0.028 | 1.156 | 0.21 | 0.11 | 0.27 | 0.66 | 0.18 | 0.19 |
| 2 | 218088388 | rs3856477 | -1.244 | -1.133 | 0.72 | 0.445 | -0.042 | 0.022 | 0.022 | 0 | 0.25 | 0.47 | 0.3 | 0.67 | 0.19 | 0.27 |
| 2 | 218090609 | rs2553038 | -0.635 | -1.297 | 0.534 | 0.199 | -0.014 | -0.097 | -0.097 | 0 | 0.22 | 0.45 | 0.21 | 0.1 | 0.18 | 0.19 |
| 2 | 218093880 | rs6435966 | -0.927 | -1.441 | 0.468 | 0.104 | -0.032 | -0.108 | -0.108 | 0 | 0.25 | 0.5 | 0.21 | 0.1 | 0.18 | 0.28 |
| 2 | 218097716 | rs2194748 | 2.147 | -1.532 | -0.197 | -0.03 | 0.032 | 0.194 | 0.194 | 0.086 | 0.47 | 0.08 | 0.47 | 0.22 | 0.6 | 0.43 |
| 2 | 218108789 | rs1609871 | 1.059 | -1.571 | 0.285 | 0.287 | -0.007 | 0.006 | 0.006 | 0.004 | 0.64 | 0.36 | 0.72 | 0.62 | 0.77 | 0.7 |
| 2 | 218114695 | rs10932707 | -1.774 | -1.904 | 0.561 | 0.356 | 0.072 | 0.107 | 0.107 | 0 | 0.51 | 0.65 | 0.36 | 0.77 | 0.32 | 0.55 |
| 2 | 218121776 | rs4674138 | -0.233 | -1.838 | 0.426 | 0.475 | -0.013 | 0.001 | 0.001 | 0.561 | 0.26 | 0.28 | 0.21 | 0.35 | 0.18 | 0.14 |
| 2 | 218122764 | rs11684828 | -1.808 | -1.74 | 0.533 | 0.437 | 0.005 | 0.004 | 0.004 | 0.613 | 0.47 | 0.47 | 0.32 | 0.73 | 0.31 | 0.37 |
| 2 | 218139904 | rs2888479 | -0.693 | -1.767 | 0.276 | 0.493 | -0.018 | 0.064 | 0.064 | 0 | 0.18 | 0.19 | 0.11 | 0.38 | 0.12 | 0.18 |
| 2 | 218146599 | rs12694412 | 1.224 | -1.539 | -0.445 | -0.024 | 0.115 | 0.173 | 0.173 | 0.047 | 0.5 | 0.2 | 0.64 | 0.25 | 0.69 | 0.55 |
| 2 | 218153274 | rs1863187 | -0.776 | -0.718 | -0.683 | 0.392 | 0.018 | 0.01 | 0.01 | 0.766 | 0.35 | 0.22 | 0.27 | 0.65 | 0.26 | 0.2 |
| 2 | 218154769 | rs17796839 | -0.623 | -0.002 | -0.808 | 0.387 | -0.021 | -0.032 | -0.032 | 0.76 | 0.4 | 0.28 | 0.32 | 0.54 | 0.44 | 0.27 |
| 2 | 218159072 | rs17797102 | -0.154 | 0.135 | -0.725 | 0.217 | 0.01 | 0.044 | 0.044 | 0 | 0.15 | 0.09 | 0.11 | 0.03 | 0.13 | 0.13 |
| 2 | 218159244 | rs6752033 | 0.847 | -0.009 | 0.083 | 0.549 | -0.016 | 0.009 | 0.009 | 0 | 0.8 | 0.65 | 0.8 | 0.74 | 0.83 | 0.78 |
| 2 | 218164218 | rs7599287 | 0.198 | -0.256 | -0.118 | 0.454 | -0.013 | -0.068 | -0.068 | 1.62 | 0.28 | 0.16 | 0.21 | 0.5 | 0.28 | 0.15 |
| 2 | 218167851 | rs4674141 | -0.362 | 0.049 | 0.891 | 0.604 | 0.007 | -0.009 | -0.009 | 0.132 | 0.55 | 0.59 | 0.69 | 0.47 | 0.63 | 0.66 |
| 2 | 218175073 | rs16857280 | -2.232 | 0.107 | 1.933 | 0.73 | 0.007 | 0.067 | 0.067 | 0 | 0.07 | 0.17 | 0.13 | 0.23 | 0.09 | 0.16 |
| 2 | 218175206 | rs4141838 | 1.126 | 0.099 | 2.461 | 0.65 | -0.025 | -0.087 | -0.087 | 0 | 0.81 | 0.54 | 0.82 | 0.97 | 0.86 | 0.75 |
| 2 | 218175472 | rs4141835 | 1.584 | 0.234 | 1.917 | 0.723 | -0.007 | -0.001 | -0.001 | 1.396 | 0.77 | 0.63 | 0.72 | 0.74 | 0.77 | 0.66 |
| 2 | 218177933 | rs16857299 | -1.999 | 0.334 | 1.492 | 0.775 | 0.016 | 0.069 | 0.069 | 0 | 0.06 | 0.18 | 0.08 | 0.24 | 0.08 | 0.07 |
| 2 | 218188700 | rs10804257 | -0.263 | 0.385 | 1.331 | 0.682 | -0.011 | 0.074 | 0.074 | 0 | 0.13 | 0.02 | 0.09 | 0 | 0.12 | 0.08 |
| 2 | 218208991 | rs16857347 | -2.22 | 0.66 | 1.781 | 0.875 | 0.002 | 0.013 | 0.013 | 0.224 | 0.05 | 0.19 | 0.12 | 0.07 | 0.05 | 0.29 |
| 2 | 218217618 | rs899283 | 2.25 | 0.691 | 1.774 | 0.943 | 0.074 | 0.149 | 0.149 | 0.686 | 0.9 | 0.48 | 0.79 | 0.8 | 0.9 | 0.76 |
| 2 | 218219166 | rs6435978 | -0.963 | 0.794 | 1.311 | 0.819 | 0.016 | -0.023 | -0.023 | 0 | 0.11 | 0.23 | 0.18 | 0.05 | 0.15 | 0.22 |
| 2 | 218224382 | rs16857377 | -1.829 | 0.809 | 1.306 | 0.956 | 0.062 | 0.123 | 0.123 | 0 | 0.06 | 0.19 | 0.17 | 0.18 | 0.1 | 0.26 |
| 2 | 218228349 | rs7561131 | -1.834 | 0.725 | 1.266 | 0.989 | 0.038 | 0.065 | 0.065 | 0.264 | 0.18 | 0.55 | 0.3 | 0.25 | 0.21 | 0.29 |
| 2 | 218231649 | rs6435979 | -1.526 | 0.966 | 1.03 | 1.293 | 0.011 | 0.05 | 0.05 | 0.73 | 0.27 | 0.59 | 0.33 | 0.44 | 0.25 | 0.47 |
| 2 | 218232676 | rs11680648 | -1.306 | 1.064 | 0.993 | 1.324 | 0.018 | 0.079 | 0.079 | 0 | 0.08 | 0.2 | 0.09 | 0.2 | 0.08 | 0.09 |
| 2 | 218236786 | rs6724073 | -0.135 | 0.853 | 0.384 | 1.346 | -0.005 | 0.046 | 0.046 | 0 | 0.25 | 0.31 | 0.39 | 0.51 | 0.27 | 0.43 |
| 2 | 218238881 | rs12620884 | 1.216 | 0.737 | -0.649 | 0.715 | -0.107 | 0.026 | 0.026 | 0.816 | 0.41 | 0.07 | 0.35 | 0.27 | 0.5 | 0.12 |
| 2 | 218239426 | rs714862 | -0.104 | 0.379 | -0.394 | 0.641 | 0.002 | -0.005 | -0.005 | 0 | 0.27 | 0.23 | 0.3 | 0.48 | 0.15 | 0.25 |
| 2 | 218248681 | rs6716223 | 0.66 | 0.358 | -0.448 | 0.44 | -0.037 | 0.024 | 0.024 | 0.866 | 0.09 | 0 | 0.09 | 0.03 | 0.09 | 0.18 |
| 2 | 218257643 | rs6435981 | 0.915 | 0.271 | 0.103 | 0.492 | 0.031 | 0.056 | 0.056 | 0 | 0.92 | 0.82 | 0.83 | 0.85 | 0.88 | 0.82 |
| 2 | 218259783 | rs10932715 | -0.42 | 0.284 | 0.329 | 0.653 | 0.012 | -0.03 | -0.03 | 0 | 0.23 | 0.18 | 0.29 | 0.56 | 0.18 | 0.44 |
| 2 | 218262593 | rs899280 | 1.2 | 0.278 | 0.486 | 0.575 | 0.025 | 0.051 | 0.051 | 0 | 0.91 | 0.78 | 0.83 | 0.85 | 0.88 | 0.82 |
| 2 | 218267899 | rs1351162 | 1.073 | 0.377 | 0.522 | 0.589 | 0.041 | 0.067 | 0.067 | 0.094 | 0.92 | 0.77 | 0.83 | 0.85 | 0.88 | 0.82 |
| 2 | 218269504 | rs6712901 | 1.047 | 0.377 | 0.522 | 0.588 | 0.043 | 0.066 | 0.066 | 0.07 | 0.92 | 0.77 | 0.83 | 0.85 | 0.88 | 0.82 |
| 2 | 218271719 | rs6759952 | 1.805 | 0.226 | 1.179 | 0.419 | 0.005 | 0.068 | 0.068 | 1.76 | 0.53 | 0.35 | 0.4 | 0.23 | 0.56 | 0.15 |
| 2 | 218275370 | rs4674167 | 1.217 | 0.216 | 1.552 | 0.404 | -0.011 | -0.018 | -0.018 | 0.204 | 0.63 | 0.57 | 0.51 | 0.26 | 0.66 | 0.34 |
| 2 | 218277119 | rs12478389 | 0.787 | 0.233 | 1.518 | 0.408 | 0.029 | -0.034 | -0.034 | 0 | 0.1 | 0.24 | 0.11 | 0.03 | 0.1 | 0.19 |
| 2 | 218280268 | rs1118150 | 0.827 | 0.191 | 1.463 | 0.444 | -0.005 | -0.003 | -0.003 | 0.641 | 0.68 | 0.61 | 0.54 | 0.27 | 0.7 | 0.38 |
| 2 | 218283439 | rs13432615 | -1.744 | 0.182 | 1.483 | 0.437 | -0.008 | -0.011 | -0.011 | 0 | 0.38 | 0.43 | 0.48 | 0.74 | 0.34 | 0.67 |
| 2 | 218284524 | rs1478576 | 1.389 | 0.18 | 1.683 | 0.452 | -0.007 | -0.01 | -0.01 | 0 | 0.62 | 0.46 | 0.51 | 0.26 | 0.66 | 0.33 |
| 2 | 218292652 | rs1478581 | -0.407 | 0.189 | 1.354 | 0.533 | 0.017 | -0.032 | -0.032 | 0 | 0.24 | 0.18 | 0.3 | 0.58 | 0.18 | 0.45 |
| 2 | 218293616 | rs10514638 | -0.602 | 0.229 | 1.422 | 0.549 | 0.024 | 0.041 | 0.041 | 0 | 0.07 | 0.13 | 0.16 | 0.15 | 0.12 | 0.18 |
| 2 | 218295406 | rs13388294 | -0.866 | 0.008 | 1.298 | 0.53 | 0.006 | -0.051 | -0.051 | 0 | 0.33 | 0.18 | 0.34 | 0.58 | 0.29 | 0.57 |
| 2 | 218296003 | rs1382435 | 1.661 | -0.071 | 1.421 | 0.523 | 0.039 | 0.048 | 0.048 | 2.661 | 0.54 | 0.26 | 0.54 | 0.31 | 0.65 | 0.18 |
| 2 | 218300530 | rs16857630 | 0.571 | -0.183 | 1.236 | 0.699 | 0.011 | -0.07 | -0.07 | 0 | 0.15 | 0.04 | 0.16 | 0.52 | 0.13 | 0.31 |
| 2 | 218304106 | rs2618148 | -1.697 | -0.024 | 1.537 | 0.593 | 0.01 | -0.006 | -0.006 | 0.341 | 0.38 | 0.71 | 0.45 | 0.62 | 0.31 | 0.71 |
| 2 | 218305064 | rs6715218 | -1.331 | -0.001 | 1.454 | 0.562 | -0.001 | 0 | 0 | 0 | 0.34 | 0.35 | 0.42 | 0.65 | 0.3 | 0.7 |
| 2 | 218312398 | rs17806990 | 0.02 | 0.186 | 1.168 | 0.401 | -0.002 | -0.008 | -0.008 | 0 | 0.13 | 0.11 | 0.07 | 0.02 | 0.15 | 0.21 |
| 2 | 218320757 | rs1564241 | -0.279 | -0.174 | 1.664 | 0.828 | -0.011 | 0.21 | 0.21 | 0 | 0.17 | 0.36 | 0.26 | 0.54 | 0.13 | 0.28 |
| 2 | 218326801 | rs2618139 | -2.458 | -0.907 | 1.235 | 0.396 | 0.057 | 0.137 | 0.137 | 1.758 | 0.5 | 0.82 | 0.54 | 0.72 | 0.36 | 0.81 |
| 2 | 218329039 | rs970833 | 2.003 | -0.969 | 0.031 | -0.644 | -0.001 | 0 | 0 | 0 | 0.21 | 0.2 | 0.14 | 0.2 | 0.18 | 0.1 |
| 2 | 218339504 | rs2568153 | -0.229 | -1.273 | -0.14 | -0.688 | -0.001 | -0.001 | -0.001 | 0 | 0.32 | 0.28 | 0.43 | 0.56 | 0.28 | 0.48 |
| 2 | 218352602 | rs4555323 | 1.068 | -1.267 | -0.194 | -0.708 | -0.007 | 0.008 | 0.008 | 1.253 | 0.14 | 0.08 | 0.16 | 0.03 | 0.15 | 0.23 |
| 2 | 218354130 | rs874840 | -0.353 | -1.308 | -0.093 | -0.71 | -0.001 | 0.011 | 0.011 | 0.087 | 0.32 | 0.32 | 0.44 | 0.56 | 0.29 | 0.49 |
| 2 | 218356528 | rs10490765 | 0.913 | -1.113 | -0.292 | -0.783 | -0.012 | 0.019 | 0.019 | 0.917 | 0.14 | 0.08 | 0.16 | 0.03 | 0.15 | 0.23 |
| 2 | 218356758 | rs6725886 | 0.595 | -1.14 | -0.136 | -0.797 | 0.023 | -0.056 | -0.056 | 0.307 | 0.16 | 0.28 | 0.18 | 0.03 | 0.15 | 0.23 |
| 2 | 218358312 | rs12472818 | -1.31 | -1.338 | -0.682 | -0.907 | -0.081 | 0.105 | 0.105 | 0.022 | 0.68 | 0.85 | 0.68 | 0.87 | 0.52 | 0.83 |
| 2 | 218360362 | rs2373067 | -1.434 | -1.806 | -0.723 | -0.968 | -0.016 | 0.182 | 0.182 | 3.961 | 0.55 | 0.79 | 0.62 | 0.85 | 0.44 | 0.79 |
| 2 | 218363032 | rs6728963 | 0.503 | -2.241 | -0.55 | -1.088 | -0.069 | -0.06 | -0.06 | 0 | 0.24 | 0.51 | 0.15 | 0.16 | 0.09 | 0.24 |
| 2 | 218365337 | rs10198000 | -0.339 | -2.051 | -0.411 | -0.939 | -0.006 | -0.025 | -0.025 | 0.041 | 0.42 | 0.53 | 0.32 | 0.2 | 0.29 | 0.52 |
| 2 | 218365548 | rs4372880 | 1.616 | -2.1 | -0.345 | -0.82 | -0.007 | 0.04 | 0.04 | 0 | 0.18 | 0.22 | 0.27 | 0.53 | 0.13 | 0.28 |
| 2 | 218366794 | rs10191778 | -0.311 | -2.096 | -0.375 | -0.885 | -0.007 | -0.029 | -0.029 | 0 | 0.42 | 0.53 | 0.31 | 0.21 | 0.29 | 0.52 |
| 2 | 218374066 | rs2373072 | 0.163 | -2.04 | -0.254 | -0.76 | -0.015 | -0.017 | -0.017 | 0 | 0.31 | 0.42 | 0.37 | 0.72 | 0.25 | 0.39 |
| 2 | 218390014 | rs13008340 | -0.415 | -1.513 | -0.194 | -0.468 | 0 | -0.003 | -0.003 | 0 | 0.55 | 0.58 | 0.53 | 0.65 | 0.43 | 0.67 |

| | | | | | | | | | | | | | | | | |
|---|---|---|---|---|---|---|---|---|---|---|---|---|---|---|---|---|
| 2 | 218392555 | rs2373078 | 0.783 | -1.317 | -0.663 | -0.213 | -0.01 | 0.011 | 0.011 | 0 | 0.13 | 0.19 | 0.08 | 0.18 | 0.1 | 0.13 |
| 2 | 218394329 | rs7582879 | -0.893 | -1.353 | -0.707 | -0.166 | -0.012 | 0.013 | 0.013 | 0 | 0.6 | 0.63 | 0.52 | 0.7 | 0.43 | 0.79 |
| 2 | 218395145 | rs1002353 | -1.545 | -2.166 | 0.129 | 0.214 | -0.047 | 0.018 | 0.018 | 0 | 0.38 | 0.55 | 0.33 | 0.45 | 0.23 | 0.38 |
| 2 | 218398137 | rs2373079 | -0.828 | -2.417 | 0.171 | 0.069 | 0.048 | 0.015 | 0.015 | 0 | 0.27 | 0.17 | 0.14 | 0.2 | 0.14 | 0.31 |
| 2 | 218404073 | rs3796025 | -1.726 | -2.233 | 0.542 | -0.11 | -0.02 | -0.038 | -0.038 | 0 | 0.38 | 0.48 | 0.23 | 0.21 | 0.28 | 0.46 |
| 2 | 218404208 | rs17203333 | 0.682 | -2.223 | 0.419 | -0.111 | -0.052 | 0.049 | 0.049 | 0 | 0.09 | 0 | 0.07 | 0.01 | 0.14 | 0.13 |
| 2 | 218413982 | rs4674189 | -0.782 | -2.202 | 0.388 | -0.1 | 0.045 | 0.016 | 0.016 | 0 | 0.27 | 0.18 | 0.14 | 0.2 | 0.15 | 0.33 |
| 2 | 218416197 | rs13018792 | -2.091 | -2.242 | 0.426 | 0.063 | 0.07 | -0.009 | -0.009 | 0 | 0.47 | 0.76 | 0.38 | 0.43 | 0.34 | 0.48 |
| 2 | 218423237 | rs13007483 | -1.385 | -2.322 | 0.615 | -0.026 | -0.024 | -0.081 | -0.081 | 0.374 | 0.38 | 0.5 | 0.23 | 0.15 | 0.28 | 0.46 |
| 2 | 218423580 | rs12185655 | -1.98 | -2.414 | 0.217 | 0.088 | 0.041 | 0.005 | 0.005 | 0 | 0.53 | 0.75 | 0.4 | 0.43 | 0.37 | 0.56 |
| 2 | 218433164 | rs7596630 | -2.393 | -2.496 | 0.867 | 0.069 | 0.071 | 0.062 | 0.062 | 0 | 0.49 | 0.74 | 0.38 | 0.3 | 0.34 | 0.48 |
| 2 | 218433524 | rs6705979 | -0.938 | -2.516 | 1.409 | -0.126 | 0.031 | 0.037 | 0.037 | 0 | 0.27 | 0.2 | 0.15 | 0.07 | 0.14 | 0.2 |
| 2 | 218437453 | rs4672834 | -2.006 | -2.525 | 1.529 | -0.157 | -0.005 | -0.074 | -0.074 | 0.243 | 0.39 | 0.57 | 0.24 | 0.03 | 0.28 | 0.25 |
| 2 | 218438843 | rs6714500 | -0.632 | -2.5 | 1.553 | -0.124 | -0.033 | -0.043 | -0.043 | 0 | 0.26 | 0.34 | 0.15 | 0.07 | 0.14 | 0.16 |
| 2 | 218446232 | rs4399701 | -2.618 | -2.525 | 1.189 | -0.071 | 0.079 | 0.083 | 0.083 | 0.138 | 0.53 | 0.78 | 0.38 | 0.32 | 0.34 | 0.48 |
| 2 | 218450174 | rs7582926 | -2.071 | -1.823 | 1.631 | -0.145 | -0.007 | -0.031 | -0.031 | 0 | 0.4 | 0.6 | 0.28 | 0.11 | 0.28 | 0.29 |
| 2 | 218451834 | rs6719545 | -2.071 | -1.823 | 1.634 | -0.165 | -0.008 | -0.065 | -0.065 | 0 | 0.4 | 0.6 | 0.28 | 0.06 | 0.28 | 0.29 |
| 2 | 218458029 | rs11686077 | -0.626 | -1.563 | 1.406 | 0.017 | -0.044 | 0.063 | 0.063 | 0 | 0.11 | 0.01 | 0.09 | 0.01 | 0.14 | 0.12 |
| 2 | 218461792 | rs1897103 | -1.9 | -1.717 | 2.064 | 0.358 | -0.07 | -0.066 | -0.066 | 0 | 0.32 | 0.54 | 0.18 | 0.13 | 0.15 | 0.34 |
| 2 | 218462340 | rs2568179 | -2.183 | -1.722 | 2.052 | 0.22 | -0.036 | -0.069 | -0.069 | 1.608 | 0.35 | 0.56 | 0.19 | 0.11 | 0.19 | 0.38 |
| 2 | 218476953 | rs2254682 | -2.463 | -1.579 | 2.07 | 0.163 | 0.048 | -0.021 | -0.021 | 0 | 0.4 | 0.68 | 0.39 | 0.44 | 0.24 | 0.45 |
| 2 | 218481552 | rs1814354 | -1.83 | -1.404 | 2.49 | -1.002 | 0.018 | 0.015 | 0.015 | 0.095 | 0.06 | 0.34 | 0.02 | 0.02 | 0.01 | 0.08 |
| 2 | 218481987 | rs2568189 | -1.798 | -1.526 | 2.415 | -1.073 | -0.03 | -0.092 | -0.092 | 0 | 0.32 | 0.61 | 0.18 | 0.05 | 0.16 | 0.31 |
| 2 | 218505964 | rs12694420 | 3.208 | -0.493 | 2.845 | -0.447 | -0.055 | -0.063 | -0.063 | 0 | 0.89 | 0.4 | 0.93 | 0.98 | 0.97 | 0.79 |
| 2 | 218512598 | rs6733585 | 2.369 | -0.461 | 2.779 | -0.448 | -0.012 | -0.064 | -0.064 | 0 | 0.87 | 0.72 | 0.95 | 0.97 | 0.92 | 0.7 |
| 2 | 218530865 | rs1367194 | 1.503 | -0.343 | 2.408 | -0.433 | 0.026 | -0.042 | -0.042 | 1.565 | 0.9 | 0.76 | 0.95 | 0.99 | 0.92 | 0.8 |
| 2 | 218531716 | rs16858076 | -2.371 | -0.399 | 1.885 | -0.339 | -0.004 | -0.017 | -0.017 | 0 | 0.07 | 0.34 | 0.04 | 0.02 | 0.02 | 0.09 |
| 2 | 218570310 | rs10199911 | 2.137 | -0.355 | 1.575 | -0.375 | -0.004 | -0.082 | -0.082 | 0.075 | 0.86 | 0.45 | 0.92 | 0.99 | 0.9 | 0.76 |
| 2 | 218578035 | rs934240 | 1.283 | -0.095 | 1.605 | -0.304 | 0.077 | -0.046 | -0.046 | 0.071 | 0.89 | 0.55 | 0.94 | 0.99 | 0.9 | 0.76 |
| 2 | 218580550 | rs10168801 | 1.742 | -0.476 | 1.239 | -0.272 | 0.048 | 0.037 | 0.037 | 0 | 0.61 | 0.33 | 0.69 | 0.77 | 0.78 | 0.63 |
| 2 | 218588959 | rs744564 | 1.153 | -0.156 | 1.107 | -0.362 | 0.051 | -0.059 | -0.059 | 0 | 0.88 | 0.57 | 0.94 | 0.99 | 0.91 | 0.73 |
| 2 | 218615634 | rs17600246 | -0.858 | -0.67 | -0.155 | -0.098 | 0.056 | 0.019 | 0.019 | 0.906 | 0.23 | 0.1 | 0.18 | 0.15 | 0.12 | 0.1 |
| 2 | 218617530 | rs7574419 | 1.4 | -0.425 | 0.564 | -0.277 | -0.05 | -0.101 | -0.101 | 0 | 0.78 | 0.53 | 0.9 | 0.94 | 0.86 | 0.68 |
| 2 | 218618574 | rs12466441 | 1.759 | 0.293 | 0.615 | -0.558 | 0.008 | 0.072 | 0.072 | 0 | 0.51 | 0.38 | 0.55 | 0.82 | 0.47 | 0.53 |
| 2 | 218618906 | rs7607467 | 0.092 | 0.128 | 0.708 | -0.572 | -0.003 | -0.024 | -0.024 | 0.613 | 0.94 | 0.91 | 0.97 | 1 | 0.94 | 0.95 |
| 2 | 218620552 | rs2052928 | 1.175 | -0.06 | 1.049 | -0.701 | -0.021 | -0.044 | -0.044 | 0 | 0.74 | 0.67 | 0.9 | 0.97 | 0.85 | 0.85 |
| 2 | 218625581 | rs1814355 | -0.072 | 0.025 | 0.934 | -0.678 | -0.003 | -0.024 | -0.024 | 0.109 | 0.94 | 0.91 | 0.97 | 1 | 0.95 | 0.95 |
| 2 | 218625787 | rs7560625 | -1.24 | 0.083 | 1.109 | -0.654 | 0.067 | -0.035 | -0.035 | 0 | 0.25 | 0.39 | 0.37 | 0.15 | 0.4 | 0.34 |
| 2 | 218634019 | rs1583109 | -1.005 | 0.138 | 1.208 | -0.672 | 0.01 | 0.063 | 0.063 | 0.424 | 0.11 | 0.34 | 0.14 | 0.14 | 0.06 | 0.11 |
| 2 | 218634639 | rs11692882 | -1.481 | 0.131 | 1.191 | -0.226 | 0.065 | 0.017 | 0.017 | 0.025 | 0.25 | 0.37 | 0.41 | 0.23 | 0.39 | 0.34 |
| 2 | 218634992 | rs2099182 | 2.401 | 0.115 | 1.161 | -0.23 | 0.001 | 0.024 | 0.024 | 0 | 0.52 | 0.4 | 0.51 | 0.74 | 0.47 | 0.53 |
| 2 | 218635840 | rs7604895 | 1.171 | 0.119 | 1.091 | -0.244 | 0.024 | -0.017 | -0.017 | 1.147 | 0.77 | 0.71 | 0.73 | 0.94 | 0.59 | 0.71 |
| 2 | 218636444 | rs1835335 | 2.364 | 0.128 | 1 | -0.233 | 0.001 | 0.027 | 0.027 | 0 | 0.52 | 0.4 | 0.56 | 0.77 | 0.47 | 0.53 |
| 2 | 218637207 | rs1346769 | 0.783 | 0.605 | 0.392 | -0.22 | 0.041 | -0.02 | -0.02 | 0 | 0.81 | 0.69 | 0.77 | 0.93 | 0.69 | 0.74 |
| 2 | 218638176 | rs2373146 | -0.208 | 0.298 | -0.068 | -0.214 | -0.035 | 0.011 | 0.011 | 0 | 0.16 | 0.1 | 0.26 | 0.07 | 0.33 | 0.25 |
| 2 | 218650612 | rs16825222 | -0.588 | 0.163 | -0.113 | -0.27 | 0.044 | 0.049 | 0.049 | 0.325 | 0.19 | 0.09 | 0.04 | 0.03 | 0.1 | 0.1 |
| 2 | 218653590 | rs16858224 | -0.555 | 0.152 | -0.131 | -0.295 | 0.061 | 0.071 | 0.071 | 0.361 | 0.19 | 0.06 | 0.04 | 0.03 | 0.09 | 0.1 |
| 2 | 218654088 | rs2571460 | 2.008 | 0.188 | -0.038 | -0.098 | 0 | 0.001 | 0.001 | 0 | 0.53 | 0.48 | 0.52 | 0.68 | 0.47 | 0.53 |
| 2 | 218657728 | rs11684943 | -0.371 | 0.325 | -0.211 | -0.027 | -0.003 | 0.005 | 0.005 | 0 | 0.09 | 0.07 | 0.06 | 0 | 0.1 | 0.07 |
| 2 | 218658984 | rs2571457 | 1.312 | 0.245 | -0.359 | 0.004 | -0.018 | 0.005 | 0.005 | 0 | 0.61 | 0.76 | 0.57 | 0.69 | 0.58 | 0.58 |
| 2 | 218659070 | rs4674213 | -0.942 | 0.064 | -0.42 | 0.2 | 0.027 | -0.013 | -0.013 | 0 | 0.23 | 0.14 | 0.2 | 0.27 | 0.1 | 0.2 |
| 2 | 218663979 | rs768921 | -0.097 | 0.64 | -0.187 | 0.304 | 0 | 0.03 | 0.03 | 2.08 | 0.06 | 0.09 | 0.13 | 0.18 | 0.05 | 0.09 |
| 2 | 218664633 | rs3747 | 0.534 | 0.61 | -0.396 | 0.301 | -0.007 | 0.023 | 0.023 | 0 | 0.55 | 0.57 | 0.57 | 0.73 | 0.5 | 0.55 |
| 2 | 218667836 | rs3088214 | 0.232 | 0.82 | -0.307 | 0.757 | 0.013 | 0.036 | 0.036 | 0 | 0.66 | 0.56 | 0.58 | 0.5 | 0.58 | 0.48 |
| 2 | 218668732 | rs1063281 | 0.216 | 0.786 | 0.081 | 1.049 | 0.011 | 0.027 | 0.027 | 0 | 0.66 | 0.6 | 0.58 | 0.61 | 0.59 | 0.49 |
| 2 | 218674484 | rs1035671 | 0.258 | -0.478 | -0.027 | 0.691 | -0.045 | -0.007 | -0.007 | 0 | 0.21 | 0.24 | 0.22 | 0.16 | 0.06 | 0.13 |
| 2 | 218674697 | rs918949 | -0.423 | -0.484 | 0.261 | 0.722 | -0.027 | -0.052 | -0.052 | 0.355 | 0.67 | 0.81 | 0.59 | 0.51 | 0.59 | 0.49 |
| 2 | 218676890 | rs1991161 | 0.233 | -0.839 | 0.191 | 0.661 | 0.003 | -0.005 | -0.005 | 0 | 0.55 | 0.68 | 0.56 | 0.6 | 0.55 | 0.47 |
| 2 | 218679599 | rs2059408 | 0.463 | 0.002 | -0.47 | 0.512 | -0.063 | -0.075 | -0.075 | 0.261 | 0.12 | 0.03 | 0.15 | 0.43 | 0.19 | 0.13 |
| 2 | 218680570 | rs2161969 | -0.234 | -0.115 | -0.524 | 0.18 | -0.036 | 0.021 | 0.021 | 1.487 | 0.55 | 0.26 | 0.49 | 0.46 | 0.46 | 0.35 |
| 2 | 218681340 | rs4674216 | 0.264 | 0.008 | -0.474 | 0.463 | 0 | 0.065 | 0.065 | 0.831 | 0.4 | 0.18 | 0.38 | 0.25 | 0.36 | 0.4 |
| 2 | 218682771 | rs3796026 | 0.143 | 0.165 | -0.398 | 0.411 | -0.003 | 0.089 | 0.089 | 0.806 | 0.43 | 0.24 | 0.41 | 0.25 | 0.41 | 0.41 |
| 2 | 218683154 | rs2571445 | -1.622 | 0.09 | -0.361 | 0.482 | -0.034 | -0.05 | -0.05 | 0.047 | 0.7 | 0.84 | 0.62 | 0.62 | 0.59 | 0.56 |
| 2 | 218688596 | rs2552527 | -0.237 | 0.146 | -0.148 | 0.428 | 0.013 | 0.054 | 0.054 | 0 | 0.48 | 0.66 | 0.46 | 0.25 | 0.41 | 0.43 |
| 2 | 218690487 | rs11897815 | 0.832 | 0.031 | -0.181 | 0.381 | 0.012 | 0.013 | 0.013 | 1.397 | 0.36 | 0.32 | 0.38 | 0.23 | 0.27 | 0.3 |
| 2 | 218693550 | rs2571438 | 0.648 | -0.082 | -0.188 | 0.49 | -0.011 | -0.007 | -0.007 | 0 | 0.36 | 0.37 | 0.3 | 0.27 | 0.26 | 0.31 |
| 2 | 218694153 | rs6729308 | 0.329 | 0.185 | -0.436 | 0.346 | 0.015 | 0.02 | 0.02 | 0.05 | 0.34 | 0.45 | 0.5 | 0.44 | 0.47 | 0.48 |
| 2 | 218694296 | rs6729330 | 0.368 | 0.19 | -0.463 | 0.417 | 0.013 | 0.005 | 0.005 | 0.027 | 0.33 | 0.44 | 0.49 | 0.35 | 0.47 | 0.48 |
| 2 | 218694551 | rs16858298 | 1.434 | 0.301 | -0.822 | 0.38 | 0.041 | -0.005 | -0.005 | 0.05 | 0.32 | 0.14 | 0.29 | 0.27 | 0.23 | 0.3 |

| | | | | | | | | | | | | | | | | |
|---|---|---|---|---|---|---|---|---|---|---|---|---|---|---|---|---|
| 2 | 218695102 | rs3796028 | 0.785 | 0.447 | -0.618 | 0.656 | 0.007 | 0.022 | 0.022 | 0.016 | 0.25 | 0.32 | 0.43 | 0.43 | 0.36 | 0.37 |
| 2 | 218695686 | rs2571440 | -0.204 | 0.64 | -0.869 | 0.667 | -0.054 | -0.071 | -0.071 | 0 | 0.84 | 0.97 | 0.86 | 0.72 | 0.82 | 0.8 |
| 2 | 218698323 | rs2552531 | -0.585 | 0.535 | 0.043 | 0.445 | 0.1 | 0.024 | 0.024 | 0 | 0.05 | 0.24 | 0.04 | 0.01 | 0.07 | 0.06 |
| 2 | 218709145 | rs929936 | -0.122 | 0.513 | -0.783 | 0.549 | 0.048 | 0.054 | 0.054 | 0.209 | 0.45 | 0.71 | 0.32 | 0.34 | 0.35 | 0.44 |
| 2 | 218711738 | rs987338 | -0.203 | 0.735 | -0.493 | 0.548 | 0.019 | 0.032 | 0.032 | 0.054 | 0.45 | 0.62 | 0.32 | 0.34 | 0.37 | 0.44 |
| 2 | 218712306 | rs1004814 | -0.181 | 0.68 | -0.495 | 0.543 | 0.026 | 0.029 | 0.029 | 0.041 | 0.44 | 0.62 | 0.31 | 0.34 | 0.35 | 0.45 |
| 2 | 218712606 | rs1364641 | -0.181 | 0.68 | -0.495 | 0.543 | 0.026 | 0.029 | 0.029 | 0.041 | 0.44 | 0.62 | 0.32 | 0.34 | 0.35 | 0.45 |
| 2 | 218712726 | rs1364642 | -0.168 | 0.674 | -0.498 | 0.54 | 0.025 | 0.028 | 0.028 | 0.022 | 0.44 | 0.62 | 0.32 | 0.35 | 0.35 | 0.45 |
| 2 | 218713282 | rs3796033 | -0.113 | 0.622 | -0.208 | 0.631 | 0.014 | 0.025 | 0.025 | 0.084 | 0.43 | 0.6 | 0.31 | 0.29 | 0.35 | 0.43 |
| 2 | 218714686 | rs3791978 | -0.288 | 0.715 | -0.523 | 0.685 | 0.01 | 0.021 | 0.021 | 0 | 0.51 | 0.62 | 0.44 | 0.42 | 0.52 | 0.49 |
| 2 | 218715071 | rs3791975 | 1.692 | -0.088 | -0.746 | 1.045 | -0.003 | 0.003 | 0.003 | 1.617 | 0.21 | 0.18 | 0.26 | 0.32 | 0.12 | 0.26 |
| 2 | 218720671 | rs3791962 | 0.543 | -0.193 | -0.401 | 0.723 | -0.007 | -0.019 | -0.019 | 0.009 | 0.36 | 0.38 | 0.25 | 0.19 | 0.28 | 0.38 |
| 2 | 218721834 | rs3791961 | 0.564 | -0.531 | -0.582 | 0.834 | 0.047 | -0.016 | -0.016 | 1.223 | 0.54 | 0.71 | 0.48 | 0.49 | 0.34 | 0.6 |
| 2 | 218723084 | rs3791955 | -0.088 | -0.129 | -0.167 | 1.048 | 0.056 | 0.035 | 0.035 | 0 | 0.13 | 0.33 | 0.15 | 0.21 | 0.18 | 0.06 |
| 2 | 218731915 | rs1863132 | 0.375 | -1.35 | -0.575 | 0.661 | -0.008 | 0.001 | 0.001 | 0 | 0.35 | 0.67 | 0.31 | 0.33 | 0.14 | 0.37 |
| 2 | 218732851 | rs17790748 | 0.034 | -1.022 | -0.763 | 0.63 | -0.014 | -0.003 | -0.003 | 0.3 | 0.34 | 0.18 | 0.29 | 0.36 | 0.35 | 0.36 |
| 2 | 218736178 | rs2042541 | 0 | -1.01 | -0.654 | 0.65 | -0.005 | 0 | 0 | 0 | 0.34 | 0.23 | 0.3 | 0.35 | 0.35 | 0.36 |
| 2 | 218736292 | rs2042542 | -0.047 | -1.025 | -0.391 | 0.642 | 0.002 | -0.002 | -0.002 | 0 | 0.35 | 0.38 | 0.31 | 0.36 | 0.35 | 0.36 |
| 2 | 218744803 | rs11898107 | 1.764 | -1.703 | -0.208 | 0.63 | -0.011 | -0.002 | -0.002 | 0 | 0.18 | 0.16 | 0.2 | 0.29 | 0.06 | 0.2 |
| 2 | 218744886 | rs890049 | 0.388 | -1.364 | -0.23 | 0.546 | -0.006 | -0.001 | -0.001 | 0 | 0.36 | 0.35 | 0.34 | 0.32 | 0.44 | 0.32 |
| 2 | 218745685 | rs10199795 | 0.172 | -2.472 | -0.188 | 0.604 | -0.002 | 0 | 0 | 0 | 0.44 | 0.58 | 0.54 | 0.53 | 0.61 | 0.37 |
| 2 | 218745862 | rs7421896 | 0.75 | -2.721 | 0.188 | 1.135 | -0.005 | -0.001 | -0.001 | 0 | 0.21 | 0.19 | 0.09 | 0.25 | 0.07 | 0.27 |
| 2 | 218749379 | rs3791940 | 0.634 | -2.514 | -0.204 | 1.151 | -0.093 | 0.036 | 0.036 | 0.063 | 0.22 | 0.37 | 0.11 | 0.31 | 0.1 | 0.29 |
| 2 | 218750817 | rs10211238 | 0.634 | -2.514 | -0.277 | 1.158 | -0.099 | 0.043 | 0.043 | 0.063 | 0.22 | 0.39 | 0.11 | 0.31 | 0.11 | 0.29 |
| 2 | 218753636 | rs3791938 | -0.777 | -2.35 | -1.314 | 1.089 | 0.098 | 0.011 | 0.011 | 0.508 | 0.49 | 0.23 | 0.34 | 0.54 | 0.33 | 0.58 |
| 2 | 218754326 | rs10174633 | -0.201 | -2.287 | -1.314 | 1.07 | 0.041 | -0.034 | -0.034 | 0 | 0.26 | 0.19 | 0.12 | 0.46 | 0.12 | 0.33 |
| 2 | 218756117 | rs975441 | -0.2 | -2.182 | -1.259 | 0.861 | 0.044 | -0.023 | -0.023 | 0.154 | 0.27 | 0.17 | 0.12 | 0.42 | 0.12 | 0.34 |
| 2 | 218768422 | rs3791936 | -0.693 | -1.853 | -1.226 | 0.465 | 0.044 | 0.007 | 0.007 | 0.862 | 0.52 | 0.65 | 0.47 | 0.43 | 0.36 | 0.54 |
| 2 | 218768581 | rs3791935 | 0.025 | -1.397 | -1.309 | 0.247 | 0.057 | -0.025 | -0.025 | 1.039 | 0.22 | 0.03 | 0.06 | 0.23 | 0.12 | 0.29 |
| 2 | 218776372 | rs6755375 | -0.565 | -1.031 | -0.651 | -0.075 | 0.017 | 0.022 | 0.022 | 2.408 | 0.44 | 0.41 | 0.43 | 0.08 | 0.35 | 0.33 |
| 2 | 218782054 | rs890051 | -0.532 | -0.815 | -0.635 | 0.571 | 0.041 | 0.029 | 0.029 | 0.266 | 0.46 | 0.65 | 0.42 | 0.34 | 0.35 | 0.45 |
| 2 | 218784225 | rs3828280 | -1.535 | -0.668 | 0.644 | 1.171 | -0.039 | 0.137 | 0.137 | 1.02 | 0.15 | 0.5 | 0.17 | 0.27 | 0.07 | 0.15 |
| 2 | 218789397 | rs4672859 | -2.048 | -0.366 | -0.1 | 0.929 | 0.008 | -0.006 | -0.006 | 0.437 | 0.32 | 0.85 | 0.3 | 0.3 | 0.25 | 0.34 |
| 2 | 218793983 | rs3791911 | -1.809 | 0.142 | 0.672 | 0.797 | 0.011 | -0.043 | -0.043 | 1.413 | 0.14 | 0.19 | 0.1 | 0.02 | 0.19 | 0.07 |
| 2 | 218799554 | rs1365964 | -1.985 | 0.14 | 0.827 | 1.051 | 0.014 | 0.011 | 0.011 | 0.131 | 0.17 | 0.21 | 0.13 | 0.25 | 0.2 | 0.21 |
| 2 | 218803329 | rs3791906 | -2.479 | -0.33 | 0.957 | 1.308 | 0.001 | -0.003 | -0.003 | 0 | 0.24 | 0.74 | 0.33 | 0.32 | 0.18 | 0.24 |
| 2 | 218804984 | rs3791904 | -2.157 | -0.315 | 1.619 | 1.334 | -0.016 | 0.089 | 0.089 | 0.018 | 0.14 | 0.36 | 0.14 | 0.25 | 0.09 | 0.16 |
| 2 | 218810908 | rs6753621 | -2.804 | -0.336 | 1.669 | 1.394 | 0.008 | -0.008 | -0.008 | 0.65 | 0.31 | 0.76 | 0.3 | 0.37 | 0.23 | 0.33 |
| 2 | 218816078 | rs744753 | -2.39 | -0.291 | 1.934 | 1.196 | -0.034 | -0.012 | -0.012 | 0.024 | 0.14 | 0.51 | 0.1 | 0.1 | 0.07 | 0.16 |
| 2 | 218821586 | rs3791890 | -2.18 | -0.266 | 1.658 | 1.504 | -0.009 | 0.119 | 0.119 | 2.056 | 0.12 | 0.32 | 0.11 | 0.26 | 0.09 | 0.15 |
| 2 | 218822874 | rs7573770 | -1.121 | -0.072 | 1.398 | 1.453 | 0.018 | -0.047 | -0.047 | 0 | 0.09 | 0.17 | 0.07 | 0 | 0.12 | 0.05 |
| 2 | 218824450 | rs16858496 | -1.238 | -0.211 | 1.472 | 1.438 | 0.006 | 0.029 | 0.029 | 1.762 | 0.08 | 0.29 | 0.04 | 0.06 | 0.04 | 0.12 |
| 2 | 218825490 | rs16858505 | -2.215 | -0.24 | 1.352 | 1.396 | 0.03 | -0.052 | -0.052 | 0 | 0.21 | 0.49 | 0.23 | 0.11 | 0.22 | 0.22 |
| 2 | 218826077 | rs3791886 | -2.224 | -0.24 | 1.352 | 1.396 | 0.03 | -0.052 | -0.052 | 0 | 0.22 | 0.49 | 0.23 | 0.11 | 0.22 | 0.21 |
| 2 | 218827913 | rs11694996 | -1.862 | -0.595 | 0.546 | 1.544 | -0.06 | 0.079 | 0.079 | 3.199 | 0.19 | 0.59 | 0.12 | 0.34 | 0.08 | 0.24 |
| 2 | 218832951 | rs12619872 | -0.455 | -0.702 | -0.691 | 1.216 | 0.049 | -0.075 | -0.075 | 0 | 0.33 | 0.13 | 0.39 | 0.49 | 0.28 | 0.37 |
| 2 | 218838347 | rs11674313 | -0.485 | -0.808 | -0.469 | 1.108 | 0.04 | -0.077 | -0.077 | 0 | 0.34 | 0.13 | 0.39 | 0.49 | 0.28 | 0.37 |
| 2 | 218843998 | rs16858524 | -0.471 | -0.517 | 0.143 | 0.474 | -0.02 | -0.117 | -0.117 | 0 | 0.2 | 0.43 | 0.12 | 0.02 | 0.13 | 0.29 |
| 2 | 218853391 | rs934039 | -0.413 | -0.68 | -0.237 | 0.607 | 0.034 | -0.023 | -0.023 | 0 | 0.38 | 0.24 | 0.4 | 0.52 | 0.29 | 0.52 |
| 2 | 218862241 | rs1001098 | 0.022 | -0.825 | -0.293 | 0.855 | -0.002 | 0 | 0 | 0.344 | 0.28 | 0.26 | 0.4 | 0.43 | 0.18 | 0.3 |
| 2 | 218864319 | rs6712327 | -0.889 | -0.616 | -0.277 | 0.61 | 0.005 | 0 | 0 | 0 | 0.51 | 0.59 | 0.55 | 0.48 | 0.4 | 0.61 |
| 2 | 218869388 | rs934036 | 0.943 | -0.397 | -0.357 | 0.435 | 0.006 | -0.039 | -0.039 | 0 | 0.74 | 0.45 | 0.79 | 0.75 | 0.75 | 0.74 |
| 2 | 218876567 | rs1427675 | 1.171 | 0.11 | -0.58 | 0.251 | 0.018 | -0.037 | -0.037 | 0.242 | 0.7 | 0.43 | 0.77 | 0.73 | 0.68 | 0.74 |
| 2 | 218877398 | rs10932737 | 0.884 | -0.605 | -1.037 | 0.379 | 0.029 | 0.063 | 0.063 | 0.781 | 0.48 | 0.13 | 0.48 | 0.3 | 0.61 | 0.37 |
| 2 | 218878318 | rs10932738 | 0.604 | -0.19 | -0.5 | 0.014 | 0.036 | -0.074 | -0.074 | 0 | 0.78 | 0.59 | 0.83 | 0.93 | 0.75 | 0.8 |
| 2 | 218879547 | rs6736691 | 1.142 | 0.162 | -0.955 | 0.119 | -0.012 | 0.069 | 0.069 | 0 | 0.32 | 0.19 | 0.28 | 0.14 | 0.33 | 0.11 |
| 2 | 218880952 | rs876961 | 0.723 | 0.673 | -0.18 | 0.75 | 0.001 | 0.014 | 0.014 | 0 | 0.62 | 0.55 | 0.63 | 0.57 | 0.62 | 0.5 |
| 2 | 218883186 | rs7580997 | 0.64 | 0.668 | 0.322 | 0.707 | 0.009 | -0.03 | -0.03 | 0 | 0.79 | 0.56 | 0.83 | 0.87 | 0.77 | 0.82 |
| 2 | 218892066 | rs6746015 | -1.101 | 0.202 | 0.121 | 1.111 | 0.013 | -0.011 | -0.011 | 0.253 | 0.25 | 0.74 | 0.22 | 0.32 | 0.14 | 0.41 |
| 2 | 218894127 | rs1427667 | -0.357 | 0.24 | -0.01 | 0.875 | -0.023 | -0.042 | -0.042 | 0 | 0.89 | 0.99 | 0.93 | 0.87 | 0.88 | 0.89 |
| 2 | 218899066 | rs10175708 | -1.115 | 0.279 | -0.061 | 0.742 | -0.004 | 0.049 | 0.049 | 0.031 | 0.44 | 0.67 | 0.55 | 0.74 | 0.42 | 0.72 |
| 2 | 218917004 | rs12989035 | 1.616 | 0.343 | -0.384 | 0.724 | -0.034 | 0.193 | 0.193 | 0.049 | 0.43 | 0.16 | 0.36 | 0.13 | 0.45 | 0.16 |
| 2 | 218922941 | rs13397894 | 0.754 | 0.847 | -0.347 | 1.352 | -0.098 | -0.114 | -0.114 | 0.311 | 0.2 | 0.04 | 0.34 | 0.4 | 0.29 | 0.33 |
| 2 | 218923606 | rs3181381 | -1.287 | 0.664 | -0.203 | 1.263 | -0.012 | 0.027 | 0.027 | 0 | 0.39 | 0.3 | 0.53 | 0.74 | 0.42 | 0.66 |
| 2 | 218928644 | rs3181211 | 0.113 | 1.214 | -0.693 | 1.208 | -0.088 | -0.093 | -0.093 | 0 | 0.21 | 0.06 | 0.35 | 0.39 | 0.31 | 0.35 |
| 2 | 218931549 | rs12471723 | 0.183 | 1.275 | -0.563 | 1.219 | -0.018 | -0.019 | -0.019 | 0 | 0.21 | 0.16 | 0.36 | 0.39 | 0.31 | 0.35 |
| 2 | 218939266 | rs4674245 | 0.149 | 1.304 | -1.084 | 1.141 | -0.09 | -0.077 | -0.077 | 1.24 | 0.21 | 0.06 | 0.35 | 0.35 | 0.31 | 0.35 |
| 2 | 218942848 | rs6436025 | -1.2 | 0.503 | -0.552 | 0.854 | -0.004 | 0.002 | 0.002 | 0 | 0.1 | 0.1 | 0.19 | 0.1 | 0.07 | 0.19 |
| 2 | 218945674 | rs6737563 | -0.403 | 1.034 | -1.169 | 0.979 | -0.176 | -0.139 | -0.139 | 0 | 0.74 | 0.96 | 0.6 | 0.37 | 0.51 | 0.59 |

| | | | | | | | | | | | | | | | | |
|---|---|---|---|---|---|---|---|---|---|---|---|---|---|---|---|---|
| 2 | 218955809 | rs10804264 | 0.455 | 0.556 | -0.051 | 0.735 | 0.008 | 0 | 0 | 0 | 0.63 | 0.59 | 0.49 | 0.32 | 0.46 | 0.49 |
| 2 | 218966315 | rs7426289 | -0.029 | 0.414 | 0.071 | 0.647 | -0.071 | -0.036 | -0.036 | 0 | 0.69 | 0.83 | 0.55 | 0.34 | 0.5 | 0.54 |
| 2 | 218988774 | rs4674257 | -0.39 | 0.491 | 0.277 | 0.693 | -0.077 | -0.051 | -0.051 | 0.04 | 0.73 | 0.86 | 0.55 | 0.34 | 0.51 | 0.6 |
| 2 | 218991005 | rs4674259 | -0.38 | 0.491 | 0.437 | 0.692 | -0.077 | -0.051 | -0.051 | 0 | 0.73 | 0.85 | 0.55 | 0.34 | 0.51 | 0.59 |
| 2 | 218997580 | rs6723449 | -0.392 | 0.49 | 0.44 | 0.692 | -0.114 | -0.079 | -0.079 | 0 | 0.73 | 0.91 | 0.56 | 0.34 | 0.51 | 0.59 |
| 2 | 219007060 | rs11677534 | -0.301 | 0.547 | 0.749 | 0.723 | -0.033 | -0.021 | -0.021 | 0 | 0.71 | 0.78 | 0.54 | 0.35 | 0.5 | 0.57 |
| 2 | 219010146 | rs11676348 | 0.314 | 0.524 | 0.727 | 0.727 | -0.001 | 0 | 0 | 0 | 0.63 | 0.66 | 0.5 | 0.32 | 0.46 | 0.48 |
| 2 | 219021385 | rs4672875 | -1.417 | -0.039 | 1.262 | 0.551 | 0.041 | 0.144 | 0.144 | 0 | 0.07 | 0.38 | 0.16 | 0.08 | 0.03 | 0.17 |
| 2 | 219037597 | rs10175470 | 1.685 | -0.221 | 0.78 | 0.505 | -0.027 | 0.014 | 0.014 | 0 | 0.88 | 0.44 | 0.81 | 0.9 | 0.93 | 0.79 |
| 2 | 219051314 | rs10171839 | 0.22 | -0.215 | -0.391 | 0.331 | -0.068 | -0.045 | -0.045 | 0 | 0.38 | 0.1 | 0.46 | 0.61 | 0.52 | 0.57 |
| 2 | 219058743 | rs6752254 | 0.223 | -0.659 | 0.694 | 0.268 | 0.001 | 0 | 0 | 0.98 | 0.41 | 0.4 | 0.5 | 0.56 | 0.52 | 0.61 |
| 2 | 219066518 | rs6718220 | 0.371 | -1.535 | 0.55 | -0.05 | 0.025 | 0.14 | 0.14 | 1.706 | 0.43 | 0.2 | 0.3 | 0.19 | 0.4 | 0.23 |
| 2 | 219066980 | rs10206984 | 1.064 | -1.587 | 0.591 | -0.045 | -0.057 | -0.057 | -0.057 | 0 | 0.36 | 0.13 | 0.47 | 0.57 | 0.52 | 0.6 |
| 2 | 219068115 | rs11695623 | 1.24 | -1.582 | 0.655 | -0.035 | -0.033 | 0.007 | 0.007 | 0.355 | 0.8 | 0.35 | 0.77 | 0.77 | 0.92 | 0.83 |
| 2 | 219069595 | rs4674272 | 0.215 | -1.648 | 1.471 | -0.069 | 0.008 | 0.004 | 0.004 | 1.982 | 0.93 | 0.64 | 0.97 | 1 | 1 | 0.99 |
| 2 | 219074398 | rs4674273 | -0.06 | -1.636 | 1.464 | -0.062 | 0.014 | -0.038 | -0.038 | 0 | 0.11 | 0.03 | 0.18 | 0.23 | 0.08 | 0.16 |
| 2 | 219076403 | rs4672880 | -0.827 | -1.635 | 1.447 | -0.062 | -0.024 | 0.09 | 0.09 | 0 | 0.13 | 0.29 | 0.2 | 0.23 | 0.08 | 0.17 |
| 2 | 219080978 | rs6720449 | 0.434 | -1.64 | 1.333 | -0.08 | 0.025 | 0.14 | 0.14 | 1.706 | 0.43 | 0.2 | 0.3 | 0.19 | 0.4 | 0.23 |
| 2 | 219082119 | rs2271541 | 1.066 | -1.532 | 1.356 | -0.155 | -0.082 | -0.082 | -0.082 | 0.241 | 0.36 | 0.09 | 0.47 | 0.58 | 0.52 | 0.6 |
| 2 | 219103580 | rs10169718 | 1.372 | -1.884 | 1.038 | -0.469 | -0.082 | -0.082 | -0.082 | 0 | 0.36 | 0.1 | 0.47 | 0.58 | 0.52 | 0.6 |
| 2 | 219109882 | rs17462354 | 1.985 | -2.849 | 1.137 | -0.615 | 0.03 | 0.057 | 0.057 | 0 | 0.1 | 0 | 0.02 | 0 | 0.07 | 0.05 |
| 2 | 219113538 | rs907982 | -0.165 | -2.454 | 1.631 | -0.364 | 0.001 | -0.002 | -0.002 | 0.313 | 0.95 | 0.72 | 0.98 | 1 | 1 | 1 |
| 2 | 219120904 | rs1473900 | -0.369 | -2.454 | 1.665 | -0.364 | 0.002 | 0 | 0 | 0.839 | 0.95 | 0.77 | 0.99 | 1 | 1 | 1 |
| 2 | 219122090 | rs10932766 | 1.276 | -2.465 | 1.527 | -0.377 | -0.041 | -0.103 | -0.103 | 0 | 0.37 | 0.08 | 0.5 | 0.58 | 0.59 | 0.61 |
| 2 | 219123769 | rs10203642 | -0.128 | -2.456 | 1.621 | -0.401 | 0.013 | 0.011 | 0.011 | 0.227 | 0.95 | 0.71 | 0.98 | 1 | 1 | 1 |
| 2 | 219124222 | rs3731859 | 1.268 | -2.451 | 1.377 | -0.377 | -0.026 | -0.083 | -0.083 | 0 | 0.37 | 0.1 | 0.5 | 0.57 | 0.59 | 0.61 |
| 2 | 219125809 | rs11554825 | 1.278 | -2.461 | 1.372 | -0.382 | -0.026 | -0.083 | -0.083 | 0 | 0.37 | 0.1 | 0.5 | 0.57 | 0.59 | 0.61 |
| 2 | 219128506 | rs2292549 | 1.551 | -2.342 | 1.444 | -0.534 | -0.037 | -0.063 | -0.063 | 0 | 0.87 | 0.31 | 0.91 | 0.99 | 0.95 | 0.98 |
| 2 | 219154781 | rs992157 | 1.031 | -2.117 | 1.088 | -0.553 | 0.003 | -0.025 | -0.025 | 0 | 0.38 | 0.17 | 0.52 | 0.62 | 0.58 | 0.62 |
| 2 | 219156979 | rs3731860 | -0.414 | -2.098 | 1.213 | -0.431 | 0.099 | 0.025 | 0.025 | 0.516 | 0.51 | 0.14 | 0.39 | 0.37 | 0.37 | 0.37 |
| 2 | 219160864 | rs7587220 | -0.534 | -2.112 | 1.464 | -0.502 | 0.088 | 0.025 | 0.025 | 0.326 | 0.52 | 0.18 | 0.4 | 0.38 | 0.37 | 0.36 |
| 2 | 219162836 | rs4996257 | 0.82 | -2.082 | 1.589 | -0.478 | 0.011 | 0.001 | 0.001 | 0.413 | 0.42 | 0.33 | 0.56 | 0.62 | 0.63 | 0.63 |
| 2 | 219170525 | rs1017698 | 0.82 | -2.082 | 1.551 | -0.478 | 0.007 | 0.001 | 0.001 | 0.413 | 0.42 | 0.35 | 0.56 | 0.62 | 0.63 | 0.64 |
| 2 | 219171191 | rs2014615 | 0.551 | -2.036 | 1.005 | -0.46 | 0.064 | 0.017 | 0.017 | 0.15 | 0.46 | 0.72 | 0.6 | 0.62 | 0.63 | 0.63 |
| 2 | 219184872 | rs4552182 | 0.592 | -1.97 | 1.046 | -0.584 | -0.003 | -0.001 | -0.001 | 0 | 0.46 | 0.47 | 0.65 | 0.65 | 0.66 | 0.67 |
| 2 | 219187603 | rs10193189 | 0.66 | -1.969 | 0.748 | -0.584 | 0.002 | 0.001 | 0.001 | 0.098 | 0.46 | 0.45 | 0.71 | 0.64 | 0.66 | 0.63 |
| 2 | 219191569 | rs10932774 | -0.593 | -1.928 | -0.111 | -0.561 | 0.088 | 0.043 | 0.043 | 0.338 | 0.49 | 0.19 | 0.33 | 0.35 | 0.34 | 0.33 |
| 2 | 219195799 | rs10932775 | 0.889 | -1.674 | 0.1 | -0.527 | -0.009 | -0.014 | -0.014 | 0.086 | 0.38 | 0.32 | 0.54 | 0.59 | 0.51 | 0.64 |
| 2 | 219202484 | rs12471773 | 0.298 | -1.692 | 0.258 | -1.039 | 0.005 | 0.016 | 0.016 | 0.659 | 0.23 | 0.18 | 0.23 | 0.1 | 0.2 | 0.08 |
| 2 | 219209796 | rs921970 | -1.317 | -1.888 | 0.445 | -1.15 | 0.01 | 0.026 | 0.026 | 2.366 | 0.39 | 0.34 | 0.35 | 0.14 | 0.37 | 0.16 |
| 2 | 219210781 | rs9076 | 1.896 | -1.883 | 0.088 | -1.239 | -0.005 | -0.013 | -0.013 | 0.268 | 0.43 | 0.14 | 0.57 | 0.65 | 0.55 | 0.57 |
| 2 | 219221846 | rs4324314 | 0.563 | -1.099 | 0.319 | -1.197 | -0.032 | 0.012 | 0.012 | 1.216 | 0.2 | 0.04 | 0.24 | 0.1 | 0.21 | 0.06 |
| 2 | 219229641 | rs11696005 | -0.16 | -1.084 | 0.591 | -1.233 | 0.006 | 0.013 | 0.013 | 0.439 | 0.26 | 0.24 | 0.28 | 0.12 | 0.21 | 0.09 |
| 2 | 219229762 | rs6745037 | 0.182 | -1.079 | 0.3 | -1.255 | 0.003 | 0.024 | 0.024 | 0 | 0.1 | 0.05 | 0.05 | 0.03 | 0.11 | 0.07 |
| 2 | 219229842 | rs6745172 | 1.903 | -1.217 | 0.641 | -1.264 | 0.036 | 0.038 | 0.038 | 0.227 | 0.47 | 0.3 | 0.61 | 0.71 | 0.61 | 0.65 |
| 2 | 219233452 | rs7586267 | -0.056 | -0.466 | 0.784 | -0.969 | -0.025 | 0.053 | 0.053 | 0.222 | 0.26 | 0.09 | 0.28 | 0.1 | 0.29 | 0.06 |
| 2 | 219233864 | rs7576929 | -0.405 | -0.428 | 1.456 | -0.937 | -0.001 | -0.021 | -0.021 | 0 | 0.06 | 0.21 | 0.03 | 0 | 0.02 | 0.04 |
| 2 | 219234705 | rs11695562 | -1.3 | -0.25 | 1.027 | -0.83 | -0.002 | -0.004 | -0.004 | 0 | 0.39 | 0.41 | 0.36 | 0.22 | 0.34 | 0.24 |
| 2 | 219242862 | rs4674301 | -0.607 | 0.168 | 0.645 | -0.683 | -0.017 | 0.029 | 0.029 | 0.842 | 0.26 | 0.16 | 0.3 | 0.14 | 0.31 | 0.15 |
| 2 | 219249013 | rs2276631 | -0.179 | 0.115 | 0.703 | -0.681 | 0 | 0.02 | 0.02 | 0.001 | 0.22 | 0.15 | 0.27 | 0.1 | 0.25 | 0.11 |
| 2 | 219252208 | rs3731863 | -1.18 | -0.069 | 0.699 | -0.65 | -0.007 | -0.001 | -0.001 | 1.869 | 0.11 | 0.16 | 0.06 | 0.11 | 0.05 | 0.11 |
| 2 | 219252377 | rs2290708 | -0.253 | -0.028 | 0.58 | -0.691 | 0.007 | 0.046 | 0.046 | 2.122 | 0.24 | 0.12 | 0.18 | 0.08 | 0.25 | 0.13 |
| 2 | 219255423 | rs2695343 | 1.386 | -0.236 | 0.605 | -0.624 | 0.037 | 0.045 | 0.045 | 0.054 | 0.56 | 0.22 | 0.59 | 0.7 | 0.64 | 0.66 |
| 2 | 219259732 | rs17235409 | -0.907 | 0.09 | 0.68 | -0.474 | -0.007 | 0.003 | 0.003 | 0 | 0.06 | 0.06 | 0.08 | 0.14 | 0.01 | 0.07 |
| 2 | 219259844 | rs1059823 | -1.767 | 0.462 | 0.627 | -0.549 | -0.011 | 0.022 | 0.022 | 0.868 | 0.42 | 0.84 | 0.43 | 0.28 | 0.39 | 0.35 |
| 2 | 219260651 | rs13062 | -1.113 | 0.524 | 1.034 | -0.588 | -0.001 | -0.008 | -0.008 | 0 | 0.33 | 0.39 | 0.36 | 0.24 | 0.34 | 0.27 |
| 2 | 219261176 | rs2279014 | -1.142 | 0.549 | 1.021 | -0.586 | -0.002 | -0.007 | -0.007 | 0 | 0.33 | 0.4 | 0.37 | 0.24 | 0.34 | 0.26 |
| 2 | 219264107 | rs2739049 | 1.226 | 0.785 | 0.522 | -0.496 | -0.003 | 0.011 | 0.011 | 0 | 0.63 | 0.29 | 0.61 | 0.68 | 0.62 | 0.61 |
| 2 | 219266204 | rs2252235 | 1.184 | 0.846 | 0.56 | -0.572 | -0.006 | 0.017 | 0.017 | 0 | 0.63 | 0.3 | 0.61 | 0.7 | 0.63 | 0.61 |
| 2 | 219266423 | rs2227251 | -0.132 | 0.856 | -0.04 | -0.599 | -0.048 | 0.048 | 0.048 | 0.211 | 0.2 | 0.07 | 0.23 | 0.07 | 0.28 | 0.24 |
| 2 | 219267781 | rs2227255 | 1.258 | 0.743 | -0.106 | -0.577 | -0.078 | 0.001 | 0.001 | 0.268 | 0.6 | 0.03 | 0.58 | 0.69 | 0.62 | 0.58 |
| 2 | 219283097 | rs16858987 | -1.607 | 0.075 | 1.427 | -0.603 | 0.029 | 0.026 | 0.026 | 11.146 | 0.05 | 0.3 | 0.03 | 0 | 0 | 0 |
| 2 | 219287276 | rs10189479 | -1.465 | 0.225 | 1.708 | -0.925 | -0.002 | -0.006 | -0.006 | 0 | 0.4 | 0.59 | 0.4 | 0.25 | 0.41 | 0.34 |
| 2 | 219287370 | rs10165754 | -0.211 | 0.24 | 1.643 | -0.923 | -0.03 | 0.058 | 0.058 | 0 | 0.25 | 0.09 | 0.31 | 0.11 | 0.33 | 0.21 |
| 2 | 219293326 | rs4674305 | 1.286 | 0.272 | 2.202 | -0.925 | -0.027 | 0.093 | 0.093 | 1.108 | 0.58 | 0.07 | 0.59 | 0.77 | 0.59 | 0.67 |
| 2 | 219295249 | rs3821031 | -0.59 | 0.266 | 2.413 | -0.985 | -0.004 | 0.013 | 0.013 | 0 | 0.31 | 0.27 | 0.35 | 0.1 | 0.38 | 0.26 |
| 2 | 219310483 | rs4674308 | 0.313 | 0.263 | 3.035 | -1.01 | -0.007 | 0.035 | 0.035 | 0 | 0.69 | 0.75 | 0.65 | 0.9 | 0.62 | 0.73 |
| 2 | 219338789 | rs523396 | 0.712 | 0.26 | 3.223 | -0.977 | 0 | -0.007 | -0.007 | 0 | 0.66 | 0.62 | 0.63 | 0.78 | 0.59 | 0.68 |
| 2 | 219349752 | rs492400 | 0.998 | 0.251 | 3.411 | -0.982 | -0.001 | -0.009 | -0.009 | 0.64 | 0.61 | 0.43 | 0.61 | 0.78 | 0.59 | 0.68 |

| | | | | | | | | | | | | | | | | |
|---|---|---|---|---|---|---|---|---|---|---|---|---|---|---|---|---|
| 2 | 219353195 | rs10932782 | 0.271 | 0.259 | 3.382 | -0.982 | -0.007 | 0.033 | 0.033 | 0 | 0.69 | 0.74 | 0.65 | 0.9 | 0.62 | 0.73 |
| 2 | 219432995 | rs500422 | 1.523 | 0.211 | 3.019 | -1.001 | -0.041 | 0.1 | 0.1 | 0.898 | 0.56 | 0.03 | 0.57 | 0.78 | 0.58 | 0.6 |
| 2 | 219472325 | rs611203 | 0.715 | 0.446 | 3.393 | -0.846 | 0.002 | -0.013 | -0.013 | 0.446 | 0.66 | 0.58 | 0.64 | 0.78 | 0.6 | 0.68 |
| 2 | 219472734 | rs3731867 | -0.386 | 0.433 | 3.046 | -0.853 | -0.106 | 0.092 | 0.092 | 0 | 0.28 | 0.02 | 0.33 | 0.1 | 0.38 | 0.27 |
| 2 | 219492924 | rs612874 | 1.292 | 0.419 | 3.062 | -0.881 | -0.033 | 0.085 | 0.085 | 0.386 | 0.59 | 0.07 | 0.59 | 0.78 | 0.59 | 0.68 |
| 2 | 219497928 | rs12989189 | -0.929 | 0.496 | 2.988 | -1.303 | -0.001 | 0 | 0 | 0 | 0.13 | 0.15 | 0.07 | 0.12 | 0.13 | 0.09 |
| 2 | 219498759 | rs7574429 | 0.402 | 0.344 | 3.451 | -0.987 | -0.025 | 0.091 | 0.091 | 0 | 0.7 | 0.81 | 0.66 | 0.94 | 0.62 | 0.74 |
| 2 | 219501133 | rs2278530 | 1.138 | 0.337 | 3.726 | -0.991 | 0 | 0.006 | 0.006 | 0 | 0.62 | 0.4 | 0.61 | 0.82 | 0.6 | 0.67 |
| 2 | 219501328 | rs1055816 | 1.354 | 0.337 | 3.681 | -0.99 | -0.033 | 0.129 | 0.129 | 0.386 | 0.59 | 0.07 | 0.59 | 0.82 | 0.59 | 0.68 |
| 2 | 219508988 | rs3770214 | 0.402 | 0.343 | 3.696 | -0.988 | -0.025 | 0.091 | 0.091 | 0 | 0.7 | 0.81 | 0.66 | 0.94 | 0.62 | 0.74 |
| 2 | 219511368 | rs4674321 | 0.719 | 0.343 | 3.701 | -0.988 | 0.003 | -0.018 | -0.018 | 0.073 | 0.67 | 0.58 | 0.64 | 0.82 | 0.59 | 0.68 |
| 2 | 219528995 | rs13022721 | -2.865 | 0.356 | 3.47 | -0.751 | 0.001 | 0.127 | 0.127 | 3.477 | 0.09 | 0.38 | 0.05 | 0.12 | 0.03 | 0.06 |
| 2 | 219529838 | rs2303563 | -2.499 | 0.359 | 3.319 | -0.749 | 0.022 | 0.154 | 0.154 | 1.86 | 0.08 | 0.37 | 0.05 | 0.12 | 0.03 | 0.06 |
| 2 | 219533378 | rs16859170 | -2.499 | 0.359 | 3.319 | -0.749 | 0.022 | 0.154 | 0.154 | 1.86 | 0.08 | 0.37 | 0.05 | 0.12 | 0.03 | 0.06 |
| 2 | 219536127 | rs13009016 | -2.499 | 0.359 | 3.327 | -0.749 | 0.021 | 0.152 | 0.152 | 1.86 | 0.08 | 0.37 | 0.05 | 0.12 | 0.03 | 0.06 |
| 2 | 219541056 | rs2556388 | -1.314 | 0.366 | 3.074 | -1.01 | -0.024 | 0.213 | 0.213 | 0.256 | 0.44 | 0.93 | 0.43 | 0.14 | 0.43 | 0.33 |
| 2 | 219543433 | rs2710251 | -1.168 | 0.359 | 3.169 | -1.001 | 0.011 | 0.165 | 0.165 | 0 | 0.42 | 0.83 | 0.42 | 0.14 | 0.43 | 0.33 |
| 2 | 219544388 | rs1863703 | -2.422 | 0.359 | 3.329 | -1.017 | -0.031 | 0.021 | 0.021 | 1.101 | 0.12 | 0.42 | 0.07 | 0.09 | 0.05 | 0.06 |
| 2 | 219555262 | rs1344642 | -1.145 | 0.36 | 3.259 | -1.044 | 0.013 | 0.076 | 0.076 | 0 | 0.42 | 0.71 | 0.41 | 0.14 | 0.42 | 0.33 |
| 2 | 219562675 | rs1863704 | 0.206 | 0.389 | 2.886 | -1.03 | -0.106 | 0.139 | 0.139 | 0 | 0.28 | 0.02 | 0.33 | 0.06 | 0.38 | 0.27 |
| 2 | 219563602 | rs12993599 | -0.418 | 0.372 | 3.09 | -0.884 | -0.003 | 0.028 | 0.028 | 0 | 0.05 | 0.13 | 0.04 | 0.09 | 0.04 | 0.06 |
| 2 | 219602819 | rs3731876 | -1.1 | 0.561 | 2.935 | -0.928 | -0.034 | 0.203 | 0.203 | 0.214 | 0.43 | 0.86 | 0.44 | 0.14 | 0.47 | 0.33 |
| 2 | 219609762 | rs2272192 | -0.386 | 0.551 | 2.875 | -0.935 | 0 | -0.001 | -0.001 | 0 | 0.05 | 0.03 | 0.03 | 0.08 | 0.05 | 0.06 |
| 2 | 219610433 | rs586194 | -1.125 | 0.553 | 2.811 | -0.932 | -0.055 | 0.212 | 0.212 | 0.339 | 0.43 | 0.92 | 0.44 | 0.14 | 0.47 | 0.33 |
| 2 | 219623504 | rs12613545 | -0.981 | 0.573 | 2.886 | -0.922 | -0.001 | 0.039 | 0.039 | 0 | 0.42 | 0.63 | 0.42 | 0.14 | 0.47 | 0.33 |
| 2 | 219640125 | rs6436086 | -0.974 | 0.581 | 2.89 | -0.918 | -0.001 | 0.039 | 0.039 | 0 | 0.42 | 0.63 | 0.42 | 0.14 | 0.47 | 0.33 |
| 2 | 219642601 | rs7603709 | -0.926 | 0.581 | 2.907 | -0.918 | -0.001 | 0.022 | 0.022 | 0 | 0.42 | 0.6 | 0.42 | 0.14 | 0.47 | 0.33 |
| 2 | 219647455 | rs4646534 | -0.672 | 0.581 | 2.816 | -0.909 | -0.008 | 0.017 | 0.017 | 0 | 0.06 | 0.13 | 0.05 | 0.08 | 0.05 | 0.06 |
| 2 | 219647487 | rs4674338 | 0.037 | 0.573 | 2.904 | -0.924 | -0.013 | 0.018 | 0.018 | 0 | 0.31 | 0.24 | 0.33 | 0.06 | 0.41 | 0.27 |
| 2 | 219650616 | rs933994 | -1.031 | 0.584 | 2.943 | -0.912 | -0.004 | 0.106 | 0.106 | 0.037 | 0.43 | 0.72 | 0.41 | 0.14 | 0.47 | 0.33 |
| 2 | 219651349 | rs1996992 | -0.418 | 0.628 | 2.667 | -0.883 | -0.004 | 0.023 | 0.023 | 0 | 0.06 | 0.13 | 0.04 | 0.08 | 0.05 | 0.06 |
| 2 | 219651995 | rs17470271 | -0.043 | 0.63 | 2.153 | -0.868 | -0.018 | 0.041 | 0.041 | 0 | 0.31 | 0.22 | 0.34 | 0.06 | 0.42 | 0.27 |
| 2 | 219658170 | rs12987009 | -0.078 | 0.512 | 2.161 | -0.861 | -0.02 | 0.045 | 0.045 | 0 | 0.32 | 0.22 | 0.35 | 0.06 | 0.43 | 0.28 |
| 2 | 219664519 | rs7566656 | -0.38 | 0.359 | 1.812 | -0.632 | 0.005 | -0.057 | -0.057 | 0 | 0.42 | 0.46 | 0.4 | 0.16 | 0.51 | 0.36 |
| 2 | 219664663 | rs4646535 | -0.31 | 0.357 | 1.866 | -0.633 | -0.035 | 0.113 | 0.113 | 0 | 0.39 | 0.16 | 0.38 | 0.16 | 0.51 | 0.36 |
| 2 | 219664977 | rs12694443 | -0.291 | 0.464 | 0.939 | -0.574 | 0.005 | -0.057 | -0.057 | 1.026 | 0.42 | 0.46 | 0.4 | 0.16 | 0.51 | 0.36 |
| 2 | 219666509 | rs13033409 | -0.29 | 0.465 | 0.939 | -0.574 | 0.005 | -0.057 | -0.057 | 0.681 | 0.42 | 0.46 | 0.4 | 0.16 | 0.51 | 0.36 |
| 2 | 219668813 | rs6436089 | -0.585 | 0.488 | 0.835 | -0.549 | -0.003 | 0.052 | 0.052 | 0.116 | 0.44 | 0.63 | 0.42 | 0.16 | 0.51 | 0.36 |
| 2 | 219671598 | rs4674345 | -0.313 | 0.307 | 0.795 | -0.519 | 0.003 | -0.041 | -0.041 | 0 | 0.41 | 0.41 | 0.4 | 0.16 | 0.51 | 0.36 |
| 2 | 219675067 | rs6709815 | -0.312 | 0.265 | 0.731 | -0.45 | 0.004 | -0.041 | -0.041 | 0 | 0.41 | 0.41 | 0.4 | 0.16 | 0.52 | 0.35 |
| 2 | 219681601 | rs647952 | -0.979 | -0.504 | 1.371 | 0.173 | -0.04 | -0.06 | -0.06 | 0 | 0.17 | 0.37 | 0.11 | 0.05 | 0.08 | 0.1 |
| 2 | 219682257 | rs645163 | -0.652 | -1.114 | 1.058 | 0.991 | 0.008 | -0.011 | -0.011 | 0.109 | 0.21 | 0.2 | 0.19 | 0.36 | 0.13 | 0.17 |
| 2 | 219692133 | rs6706738 | -1.4 | -1.108 | 2.246 | 0.807 | -0.008 | -0.011 | -0.011 | 0.176 | 0.07 | 0.65 | 0.05 | 0.01 | 0 | 0.01 |
| 2 | 219703298 | rs730947 | -0.585 | -0.946 | 1.389 | 1.171 | -0.037 | -0.054 | -0.054 | 0 | 0.09 | 0.01 | 0.15 | 0.23 | 0.11 | 0.09 |
| 2 | 219720349 | rs751135 | -1.978 | -0.929 | 1.3 | 1.618 | 0.024 | -0.075 | -0.075 | 0.126 | 0.31 | 0.84 | 0.28 | 0.51 | 0.2 | 0.3 |
| 2 | 219721190 | rs11695967 | -1.36 | -0.877 | 1.778 | 1.665 | 0 | 0.014 | 0.014 | 0 | 0.19 | 0.17 | 0.18 | 0.49 | 0.16 | 0.16 |
| 2 | 219722664 | rs690877 | -0.481 | -1.203 | 0.763 | 1.643 | -0.014 | -0.012 | -0.012 | 0.488 | 0.7 | 0.98 | 0.77 | 0.72 | 0.76 | 0.59 |
| 2 | 219733973 | rs12053317 | -2.505 | -1.097 | 1.725 | 2.452 | 0.039 | -0.059 | -0.059 | 0 | 0.27 | 0.87 | 0.27 | 0.38 | 0.17 | 0.41 |
| 2 | 219743874 | rs3806557 | -2.497 | -1.028 | 2.533 | 2.58 | -0.019 | 0.096 | 0.096 | 0 | 0.24 | 0.59 | 0.24 | 0.43 | 0.2 | 0.36 |
| 2 | 219746561 | rs10177996 | -2.756 | -0.923 | 2.565 | 2.175 | 0.016 | -0.004 | -0.004 | 0 | 0.27 | 0.82 | 0.26 | 0.26 | 0.19 | 0.34 |
| 2 | 219750746 | rs2385199 | -2.257 | -0.918 | 2.88 | 2.18 | -0.013 | 0.025 | 0.025 | 0 | 0.23 | 0.44 | 0.22 | 0.24 | 0.18 | 0.34 |
| 2 | 219766672 | rs10932789 | -1.602 | -0.804 | 3.044 | 2.365 | 0.014 | -0.04 | -0.04 | 0 | 0.16 | 0.09 | 0.18 | 0.31 | 0.12 | 0.24 |
| 2 | 219774700 | rs6723140 | -2.594 | -0.758 | 2.94 | 2.398 | 0.009 | -0.013 | -0.013 | 0.215 | 0.24 | 0.83 | 0.3 | 0.35 | 0.17 | 0.25 |
| 2 | 219820444 | rs6436101 | -1.929 | -0.655 | 2.344 | 2.447 | 0.005 | 0.048 | 0.048 | 0 | 0.18 | 0.52 | 0.2 | 0.16 | 0.16 | 0.25 |
| 2 | 219821659 | rs6436102 | -1.892 | -0.573 | 2.066 | 2.388 | -0.034 | 0.107 | 0.107 | 0 | 0.18 | 0.49 | 0.23 | 0.3 | 0.12 | 0.19 |
| 2 | 219829206 | rs359980 | -2.788 | -0.199 | 1.495 | 1.891 | -0.002 | 0.062 | 0.062 | 0 | 0.09 | 0.17 | 0.08 | 0.15 | 0.06 | 0.26 |
| 2 | 219835093 | rs452400 | -3.142 | -0.204 | 1.513 | 1.888 | 0.002 | 0.083 | 0.083 | 0 | 0.1 | 0.21 | 0.08 | 0.15 | 0.06 | 0.26 |
| 2 | 219838802 | rs168612 | -1.851 | -0.387 | 1.5 | 1.851 | -0.017 | 0.098 | 0.098 | 0 | 0.06 | 0.16 | 0.04 | 0.14 | 0.02 | 0.18 |
| 2 | 219840891 | rs359965 | -1.951 | -0.388 | 1.499 | 1.85 | -0.019 | 0.092 | 0.092 | 0 | 0.06 | 0.17 | 0.03 | 0.13 | 0.02 | 0.18 |
| 2 | 219842203 | rs359964 | -2.569 | -0.397 | 1.516 | 1.846 | -0.005 | 0.16 | 0.16 | 1.216 | 0.08 | 0.34 | 0.05 | 0.16 | 0.02 | 0.19 |
| 2 | 219846749 | rs860573 | -2.05 | -0.308 | 1.043 | 1.796 | -0.015 | 0.102 | 0.102 | 0.487 | 0.07 | 0.27 | 0.04 | 0.14 | 0.02 | 0.18 |
| 2 | 219863589 | rs398926 | -1.642 | 0.376 | 0.684 | 2.289 | 0.021 | 0.146 | 0.146 | 0 | 0.13 | 0.17 | 0.18 | 0.52 | 0.16 | 0.17 |
| 2 | 219878756 | rs9967806 | -2.174 | -0.423 | 1.102 | 1.709 | -0.069 | -0.026 | -0.026 | 0 | 0.14 | 0.62 | 0.08 | 0.08 | 0.04 | 0.12 |
| 2 | 219896981 | rs10932793 | -2.492 | -0.411 | 3.292 | 1.701 | -0.008 | -0.016 | -0.016 | 0 | 0.17 | 0.44 | 0.14 | 0.16 | 0.12 | 0.15 |
| 2 | 219903258 | rs6736922 | 0.068 | -0.88 | 1.821 | 1.18 | -0.022 | 0.008 | 0.008 | 0.128 | 0.63 | 0.55 | 0.62 | 0.43 | 0.76 | 0.62 |
| 2 | 219905247 | rs3736403 | -1.262 | -0.899 | 1.932 | 1.168 | 0.003 | 0.007 | 0.007 | 0 | 0.09 | 0.13 | 0.09 | 0.16 | 0.09 | 0.09 |
| 2 | 219909780 | rs10445821 | -1.657 | -0.923 | 2.073 | 1.184 | 0.048 | 0.088 | 0.088 | 0 | 0.1 | 0.4 | 0.12 | 0.16 | 0.09 | 0.09 |
| 2 | 219910164 | rs10445823 | -2.056 | -0.952 | 2.099 | 1.168 | 0.039 | 0.081 | 0.081 | 0 | 0.11 | 0.49 | 0.13 | 0.16 | 0.09 | 0.09 |
| 2 | 219915871 | rs7600987 | -1.433 | -0.954 | 2.055 | 1.167 | 0.008 | 0.03 | 0.03 | 0 | 0.09 | 0.19 | 0.1 | 0.16 | 0.08 | 0.09 |

| | | | | | | | | | | | | | | | | |
|---|---|---|---|---|---|---|---|---|---|---|---|---|---|---|---|---|
| 2 | 219920037 | rs394452 | -0.586 | -0.913 | 1.853 | 1.099 | -0.031 | 0.123 | 0.123 | 0.163 | 0.73 | 0.99 | 0.85 | 0.92 | 0.81 | 0.83 |
| 2 | 219920412 | rs3731881 | 1.173 | -0.903 | 1.951 | 1.17 | 0.005 | 0.007 | 0.007 | 0 | 0.54 | 0.44 | 0.53 | 0.28 | 0.67 | 0.53 |
| 2 | 219920565 | rs3731878 | -1.406 | -0.925 | 1.954 | 1.173 | -0.001 | 0.002 | 0.002 | 0 | 0.09 | 0.11 | 0.1 | 0.16 | 0.09 | 0.09 |
| 2 | 219930481 | rs17655003 | -1.421 | -0.922 | 1.956 | 1.196 | -0.001 | 0.003 | 0.003 | 0 | 0.09 | 0.11 | 0.09 | 0.16 | 0.09 | 0.05 |
| 2 | 219934348 | rs1052483 | -1.671 | -0.921 | 1.922 | 1.196 | 0.001 | 0.024 | 0.024 | 0 | 0.1 | 0.21 | 0.11 | 0.16 | 0.09 | 0.05 |
| 2 | 219938270 | rs4547519 | 1.848 | -0.328 | 0.873 | 0.444 | -0.122 | 0.025 | 0.025 | 0 | 0.23 | 0.01 | 0.17 | 0.1 | 0.36 | 0.11 |
| 2 | 219941063 | rs897477 | -0.363 | -0.661 | 1.245 | 0.682 | 0.008 | -0.007 | -0.007 | 0 | 0.65 | 0.7 | 0.65 | 0.44 | 0.76 | 0.62 |
| 2 | 219941586 | rs883045 | 0.711 | -0.66 | 1.239 | 0.682 | 0.017 | 0.023 | 0.023 | 0 | 0.54 | 0.42 | 0.53 | 0.28 | 0.68 | 0.57 |
| 2 | 219955093 | rs897476 | 0.77 | -0.681 | 1.214 | 0.633 | 0.029 | 0.041 | 0.041 | 0 | 0.54 | 0.39 | 0.53 | 0.28 | 0.68 | 0.57 |
| 2 | 219972984 | rs12104660 | -1.289 | -0.603 | 1.138 | 0.648 | 0.009 | 0.029 | 0.029 | 0 | 0.11 | 0.25 | 0.11 | 0.15 | 0.09 | 0.05 |
| 2 | 220006583 | rs1402508 | 0.87 | -0.543 | 0.781 | 0.555 | 0.048 | 0.095 | 0.095 | 0 | 0.52 | 0.28 | 0.5 | 0.28 | 0.64 | 0.55 |
| 2 | 220011092 | rs2293081 | -1.498 | -0.51 | 0.675 | 0.572 | 0.005 | 0.007 | 0.007 | 0 | 0.09 | 0.12 | 0.12 | 0.17 | 0.13 | 0.06 |
| 2 | 220019484 | rs7572601 | -0.16 | -0.519 | 0.553 | 0.573 | -0.015 | 0.005 | 0.005 | 0.005 | 0.62 | 0.65 | 0.64 | 0.44 | 0.76 | 0.6 |
| 2 | 220026369 | rs6713887 | 0.733 | -0.516 | 0.165 | 0.502 | 0.037 | 0.096 | 0.096 | 0.042 | 0.54 | 0.2 | 0.49 | 0.28 | 0.63 | 0.55 |
| 2 | 220033956 | rs6436121 | 1.349 | -0.465 | 0.004 | 0.137 | -0.002 | 0.055 | 0.055 | 0.62 | 0.6 | 0.24 | 0.42 | 0.23 | 0.65 | 0.58 |
| 2 | 220035118 | rs13029742 | -0.563 | -0.453 | -0.009 | 0.196 | -0.077 | 0.022 | 0.022 | 0 | 0.26 | 0.06 | 0.34 | 0.23 | 0.3 | 0.24 |
| 2 | 220036390 | rs6436122 | -0.037 | -0.338 | 0.157 | 0.284 | 0.005 | 0.343 | 0.343 | 0.135 | 0.92 | 0.72 | 0.8 | 0.47 | 0.95 | 0.83 |
| 2 | 220037097 | rs11236 | 1.039 | -0.226 | -0.726 | 0.334 | 0 | 0.078 | 0.078 | 0.328 | 0.57 | 0.24 | 0.41 | 0.23 | 0.62 | 0.57 |
| 2 | 220037444 | rs17655123 | -0.348 | -0.223 | -0.737 | 0.328 | -0.026 | -0.024 | -0.024 | 0 | 0.07 | 0.01 | 0.12 | 0.16 | 0.13 | 0.05 |
| 2 | 220037666 | rs1127102 | 0.993 | -0.25 | -0.717 | 0.324 | -0.002 | 0.078 | 0.078 | 0.328 | 0.57 | 0.24 | 0.4 | 0.23 | 0.62 | 0.57 |
| 2 | 220037756 | rs1043160 | 0.943 | -0.256 | -0.711 | 0.321 | -0.002 | 0.066 | 0.066 | 0.299 | 0.58 | 0.27 | 0.41 | 0.23 | 0.62 | 0.58 |
| 2 | 220040330 | rs1127101 | 0.978 | -0.239 | -0.812 | 0.329 | -0.002 | 0.078 | 0.078 | 0.328 | 0.57 | 0.24 | 0.4 | 0.23 | 0.62 | 0.57 |
| 2 | 220041091 | rs6436123 | 0.996 | -0.239 | -0.811 | 0.33 | -0.001 | 0.078 | 0.078 | 0.326 | 0.57 | 0.24 | 0.4 | 0.23 | 0.62 | 0.57 |
| 2 | 220042675 | rs10170244 | 0.996 | -0.239 | -0.811 | 0.33 | -0.001 | 0.078 | 0.078 | 0.326 | 0.57 | 0.24 | 0.4 | 0.23 | 0.62 | 0.57 |
| 2 | 220046840 | rs3210652 | -0.382 | -0.039 | -0.713 | 0.38 | -0.023 | -0.023 | -0.023 | 0.6 | 0.06 | 0.01 | 0.12 | 0.17 | 0.14 | 0.06 |
| 2 | 220046975 | rs3731900 | 0.637 | -0.223 | -0.592 | 0.003 | -0.006 | 0.051 | 0.051 | 0.149 | 0.61 | 0.24 | 0.4 | 0.24 | 0.64 | 0.57 |
| 2 | 220070301 | rs10190834 | 0.757 | -0.791 | -0.24 | -0.014 | -0.051 | -0.028 | -0.028 | 0 | 0.78 | 0.41 | 0.84 | 0.86 | 0.91 | 0.8 |
| 2 | 220072852 | rs4674367 | 1.56 | -0.734 | -0.708 | 0.166 | 0.059 | 0.096 | 0.096 | 0 | 0.52 | 0.23 | 0.43 | 0.25 | 0.72 | 0.4 |
| 2 | 220083279 | rs1109866 | 0.347 | -0.426 | -0.055 | 0.134 | -0.052 | -0.03 | -0.03 | 0 | 0.78 | 0.41 | 0.84 | 0.86 | 0.91 | 0.8 |
| 2 | 220084938 | rs10799 | 1.882 | -0.169 | -0.722 | -0.446 | -0.093 | 0.103 | 0.103 | 0 | 0.16 | 0.01 | 0.13 | 0 | 0.32 | 0.03 |
| 2 | 220086280 | rs9989854 | 0.199 | -1.574 | -0.019 | 0.183 | -0.053 | -0.03 | -0.03 | 0.017 | 0.78 | 0.42 | 0.84 | 0.86 | 0.91 | 0.8 |
| 2 | 220087441 | rs2276635 | 0.331 | -1.667 | 0.107 | 0.128 | 0.018 | 0.059 | 0.059 | 0.708 | 0.15 | 0.03 | 0.11 | 0.05 | 0.07 | 0.18 |
| 2 | 220087796 | rs2385397 | 0.243 | -1.667 | 0.431 | 0.128 | 0.011 | 0.014 | 0.014 | 0.506 | 0.16 | 0.13 | 0.12 | 0.05 | 0.07 | 0.18 |
| 2 | 220088248 | rs908196 | 0.795 | -1.668 | 0.469 | 0.129 | -0.051 | -0.03 | -0.03 | 0 | 0.78 | 0.41 | 0.84 | 0.86 | 0.91 | 0.8 |
| 2 | 220093423 | rs11689596 | 0.795 | -1.675 | 0.856 | 0.19 | -0.051 | -0.03 | -0.03 | 0 | 0.78 | 0.4 | 0.84 | 0.86 | 0.91 | 0.8 |
| 2 | 220108423 | rs35328001 | 0.318 | -1.748 | 0.865 | 0.251 | 0 | 0.006 | 0.006 | 2.312 | 0.24 | 0.16 | 0.15 | 0.07 | 0.18 | 0.32 |
| 2 | 220113384 | rs8447 | 1.673 | -1.761 | 0.695 | 0.313 | 0.049 | 0.085 | 0.085 | 0 | 0.54 | 0.23 | 0.43 | 0.25 | 0.72 | 0.41 |
| 2 | 220117458 | rs1024131 | -0.569 | -1.779 | 1.244 | 0.72 | -0.004 | -0.002 | -0.002 | 0 | 0.18 | 0.23 | 0.13 | 0.13 | 0.09 | 0.19 |
| 2 | 220135208 | rs2891733 | 0.897 | -1.834 | 1.244 | 0.73 | -0.048 | -0.025 | -0.025 | 0 | 0.79 | 0.45 | 0.85 | 0.87 | 0.9 | 0.81 |
| 2 | 220146651 | rs3731897 | -0.606 | -1.975 | 1.187 | 0.508 | 0.007 | -0.009 | -0.009 | 0 | 0.24 | 0.2 | 0.15 | 0.06 | 0.15 | 0.21 |
| 2 | 220147166 | rs3770224 | -1.544 | -2.033 | 1.069 | 0.737 | 0.026 | 0.005 | 0.005 | 0.028 | 0.34 | 0.75 | 0.48 | 0.69 | 0.19 | 0.4 |
| 2 | 220151858 | rs6436131 | -2.337 | -2.178 | 0.767 | 0.565 | 0.087 | 0.099 | 0.099 | 0.089 | 0.45 | 0.91 | 0.57 | 0.74 | 0.29 | 0.52 |
| 2 | 220152359 | rs7596956 | -1.32 | -2.118 | 0.929 | 0.591 | -0.021 | -0.003 | -0.003 | 1.583 | 0.33 | 0.59 | 0.46 | 0.7 | 0.19 | 0.4 |
| 2 | 220155834 | rs1983298 | -0.105 | -1.923 | 0.494 | 0.353 | -0.015 | 0.031 | 0.031 | 0 | 0.1 | 0.04 | 0.08 | 0.04 | 0.1 | 0.12 |
| 2 | 220157418 | rs4485556 | -0.981 | -1.995 | 0.83 | 0.541 | -0.012 | -0.011 | -0.011 | 0 | 0.28 | 0.31 | 0.25 | 0.17 | 0.18 | 0.32 |
| 2 | 220160077 | rs11887718 | -1.25 | -2.119 | 1.07 | 0.442 | -0.039 | -0.043 | -0.043 | 0 | 0.28 | 0.42 | 0.19 | 0.19 | 0.17 | 0.28 |
| 2 | 220160230 | rs4674374 | -1.412 | -2.364 | 1.317 | 0.928 | -0.011 | 0.003 | 0.003 | 0 | 0.29 | 0.66 | 0.47 | 0.66 | 0.16 | 0.37 |
| 2 | 220160644 | rs6436132 | -1.586 | -2.405 | 1.126 | 0.833 | -0.014 | 0.013 | 0.013 | 0 | 0.33 | 0.62 | 0.53 | 0.62 | 0.23 | 0.38 |
| 2 | 220165763 | rs2292606 | 0.067 | -2.147 | 0.692 | 0.148 | -0.014 | 0.032 | 0.032 | 0 | 0.1 | 0.02 | 0.08 | 0.04 | 0.1 | 0.13 |
| 2 | 220168607 | rs2271593 | -0.531 | -2.262 | 0.943 | 0.064 | -0.029 | -0.094 | -0.094 | 0.22 | 0.25 | 0.35 | 0.2 | 0.07 | 0.16 | 0.24 |
| 2 | 220172576 | rs7591986 | -1.14 | -2.198 | 1.055 | 0.166 | -0.014 | 0.01 | 0.01 | 0.53 | 0.16 | 0.42 | 0.12 | 0.13 | 0.11 | 0.2 |
| 2 | 220191861 | rs6751714 | -2.087 | -2.347 | 0.698 | 0.117 | 0.071 | 0.162 | 0.162 | 0 | 0.43 | 0.94 | 0.33 | 0.18 | 0.28 | 0.44 |
| 2 | 220200183 | rs13027600 | -1.248 | -1.627 | 1.484 | 0.482 | -0.004 | 0.02 | 0.02 | 0 | 0.14 | 0.37 | 0.11 | 0.12 | 0.12 | 0.19 |
| 2 | 220200242 | rs6436140 | -1.376 | -1.623 | 1.19 | 0.484 | -0.021 | 0.003 | 0.003 | 0.684 | 0.16 | 0.56 | 0.12 | 0.12 | 0.12 | 0.19 |
| 2 | 220212719 | rs6751428 | 0.442 | -1.877 | 0.821 | 0.186 | -0.039 | -0.12 | -0.12 | 2.542 | 0.81 | 0.51 | 0.88 | 1 | 0.91 | 0.85 |
| 2 | 220218007 | rs6728330 | 1.123 | -2.121 | 0.38 | 0.651 | 0.084 | -0.092 | -0.092 | 0.094 | 0.62 | 0.15 | 0.58 | 0.55 | 0.79 | 0.68 |
| 2 | 220226393 | rs6711142 | 0.249 | -2.04 | 0.606 | 0.362 | -0.04 | -0.118 | -0.118 | 0 | 0.81 | 0.53 | 0.88 | 0.99 | 0.91 | 0.85 |
| 2 | 220228371 | rs10177476 | 1.065 | -1.443 | 0.339 | 0.754 | -0.009 | -0.029 | -0.029 | 1.435 | 0.44 | 0.28 | 0.55 | 0.58 | 0.55 | 0.49 |
| 2 | 220238496 | rs4672926 | -0.009 | -1.245 | 1.013 | 0.693 | -0.031 | 0.003 | 0.003 | 0 | 0.82 | 0.71 | 0.83 | 0.82 | 0.89 | 0.84 |
| 2 | 220239322 | rs3821040 | 0.353 | -1.292 | 1.188 | 0.709 | -0.064 | 0.032 | 0.032 | 0 | 0.8 | 0.49 | 0.81 | 0.74 | 0.88 | 0.83 |
| 2 | 220239534 | rs3731904 | 0.318 | -1.288 | 1.187 | 0.711 | -0.051 | 0.03 | 0.03 | 0 | 0.8 | 0.6 | 0.82 | 0.74 | 0.89 | 0.83 |
| 2 | 220241274 | rs6708740 | -0.152 | -1.287 | 1.225 | 0.711 | -0.031 | 0.023 | 0.023 | 0 | 0.83 | 0.72 | 0.84 | 0.74 | 0.89 | 0.84 |
| 2 | 220241550 | rs6709228 | -0.403 | -1.202 | 1.262 | 0.92 | -0.006 | -0.014 | -0.014 | 0.327 | 0.39 | 0.4 | 0.36 | 0.27 | 0.34 | 0.25 |
| 2 | 220243295 | rs6717249 | -0.228 | -0.894 | 1.706 | 0.436 | -0.045 | 0.103 | 0.103 | 1.297 | 0.85 | 0.58 | 0.86 | 0.72 | 0.92 | 0.9 |
| 2 | 220250322 | rs878437 | 1.927 | -0.42 | 1.807 | -0.146 | 0.034 | 0.079 | 0.079 | 4.017 | 0.95 | 0.61 | 0.97 | 0.92 | 0.99 | 0.98 |
| 2 | 220267512 | rs1512693 | 1.71 | -0.413 | 1.217 | -0.135 | 0.062 | 0.139 | 0.139 | 3.008 | 0.94 | 0.51 | 0.96 | 0.91 | 0.99 | 0.98 |
| 2 | 220282513 | rs2854885 | 1.698 | -0.39 | 1.164 | -0.481 | 0.069 | 0.059 | 0.059 | 1.826 | 0.94 | 0.47 | 0.95 | 1 | 0.99 | 0.98 |
| 2 | 220285309 | rs1058261 | -0.546 | -0.437 | 0.763 | -0.632 | -0.006 | -0.053 | -0.053 | 0 | 0.4 | 0.48 | 0.34 | 0.18 | 0.36 | 0.28 |
| 2 | 220286142 | rs1058284 | -0.543 | -0.434 | 0.752 | -0.63 | -0.006 | -0.051 | -0.051 | 0.147 | 0.4 | 0.46 | 0.34 | 0.18 | 0.36 | 0.28 |

| | | | | | | | | | | | | | | | | |
|---|---|---|---|---|---|---|---|---|---|---|---|---|---|---|---|---|
| 2 | 220293242 | rs12469789 | 0.657 | -0.334 | -0.225 | -0.597 | 0.014 | 0.215 | 0.215 | 0.919 | 0.54 | 0.06 | 0.61 | 0.82 | 0.63 | 0.7 |
| 2 | 220297038 | rs7564856 | 0.622 | -0.478 | -0.186 | -0.511 | 0.044 | 0.194 | 0.194 | 1.181 | 0.54 | 0.07 | 0.61 | 0.8 | 0.64 | 0.74 |
| 2 | 220297176 | rs6436151 | -1.819 | 0.04 | 1.255 | 0.099 | 0.06 | 0.03 | 0.03 | 0.07 | 0.05 | 0.37 | 0.05 | 0 | 0.02 | 0.02 |
| 2 | 220299272 | rs907684 | 1.685 | 0.067 | 0.683 | 0.093 | 0.034 | -0.016 | -0.016 | 0.409 | 0.92 | 0.64 | 0.94 | 1 | 0.97 | 0.96 |
| 2 | 220301208 | rs11896268 | -1.612 | 0.105 | 0.498 | 0.115 | 0.046 | 0.018 | 0.018 | 1.952 | 0.05 | 0.28 | 0.05 | 0 | 0.02 | 0.05 |
| 2 | 220311039 | rs748515 | 0.066 | 0.558 | -0.455 | -0.256 | -0.014 | 0.035 | 0.035 | 0 | 0.36 | 0.28 | 0.36 | 0.25 | 0.41 | 0.45 |
| 2 | 220315970 | rs10932806 | 0.168 | 0.468 | -0.896 | -0.07 | -0.028 | 0.115 | 0.115 | 0.334 | 0.5 | 0.14 | 0.49 | 0.67 | 0.51 | 0.46 |
| 2 | 220334821 | rs6724274 | -0.286 | 0.353 | -0.691 | -0.299 | 0.004 | 0.078 | 0.078 | 0 | 0.54 | 0.32 | 0.46 | 0.27 | 0.55 | 0.67 |
| 2 | 220337196 | rs1039897 | 0.781 | 0.956 | -0.355 | -0.286 | -0.008 | 0.051 | 0.051 | 0.263 | 0.61 | 0.16 | 0.57 | 0.78 | 0.65 | 0.49 |
| 2 | 220338065 | rs4674402 | -0.745 | 0.877 | 0.412 | -0.28 | 0.003 | -0.038 | -0.038 | 0 | 0.36 | 0.43 | 0.37 | 0.16 | 0.34 | 0.5 |
| 2 | 220340540 | rs6745082 | 0.821 | 0.892 | 0.312 | -0.279 | -0.001 | 0.051 | 0.051 | 0.263 | 0.61 | 0.16 | 0.57 | 0.78 | 0.65 | 0.49 |
| 2 | 220346814 | rs3731908 | 0.609 | 0.738 | 0.536 | -0.303 | 0.025 | 0.034 | 0.034 | 0 | 0.66 | 0.26 | 0.66 | 0.8 | 0.73 | 0.55 |
| 2 | 220350502 | rs10167337 | 0.871 | 0.86 | 0.62 | -0.301 | 0.016 | 0.044 | 0.044 | 0.172 | 0.61 | 0.22 | 0.57 | 0.77 | 0.67 | 0.5 |
| 2 | 220351379 | rs6726806 | -0.437 | 0.866 | 0.84 | -0.279 | 0.001 | 0.004 | 0.004 | 0 | 0.3 | 0.29 | 0.27 | 0.15 | 0.27 | 0.45 |
| 2 | 220355529 | rs12464085 | -1.113 | 0.821 | 1.174 | -0.304 | -0.003 | -0.013 | -0.013 | 0 | 0.38 | 0.57 | 0.39 | 0.17 | 0.33 | 0.49 |
| 2 | 220356520 | rs12473286 | -0.312 | 0.823 | 1.065 | -0.305 | 0.002 | 0.008 | 0.008 | 0 | 0.3 | 0.27 | 0.27 | 0.15 | 0.27 | 0.45 |
| 2 | 220358198 | rs1050816 | -1.045 | 0.829 | 1.132 | -0.298 | -0.003 | -0.009 | -0.009 | 0 | 0.37 | 0.55 | 0.42 | 0.23 | 0.33 | 0.5 |
| 2 | 220360683 | rs4674406 | -1.12 | 0.828 | 1.09 | -0.327 | -0.003 | -0.006 | -0.006 | 0.006 | 0.38 | 0.55 | 0.42 | 0.23 | 0.35 | 0.5 |
| 2 | 220363488 | rs2280039 | -1.18 | 0.816 | 1.049 | -0.355 | 0.002 | 0.005 | 0.005 | 0 | 0.39 | 0.6 | 0.42 | 0.23 | 0.35 | 0.5 |
| 2 | 220372633 | rs16860002 | -0.986 | 0.867 | 0.719 | -0.309 | 0.068 | 0.109 | 0.109 | 0.641 | 0.07 | 0.29 | 0.14 | 0.09 | 0.07 | 0.05 |
| 2 | 220376150 | rs3755062 | 0.691 | 0.886 | 0.496 | -0.291 | 0.007 | 0.047 | 0.047 | 0 | 0.59 | 0.19 | 0.56 | 0.76 | 0.64 | 0.49 |
| 2 | 220384126 | rs746233 | -0.895 | 0.734 | 0.514 | -0.294 | 0.001 | 0.007 | 0.007 | 0 | 0.57 | 0.64 | 0.56 | 0.3 | 0.55 | 0.6 |
| 2 | 220385220 | rs3755064 | 0.725 | 0.662 | 0.254 | -0.155 | -0.057 | 0.036 | 0.036 | 0.18 | 0.47 | 0.1 | 0.45 | 0.7 | 0.58 | 0.4 |
| 2 | 220389729 | rs6704759 | -0.209 | 0.759 | 1.181 | 0.18 | -0.004 | 0.038 | 0.038 | 0 | 0.09 | 0.21 | 0.14 | 0.13 | 0.06 | 0.09 |
| 2 | 220399951 | rs2276643 | -0.213 | 1.09 | 1.291 | 0.337 | 0.027 | 0.011 | 0.011 | 0 | 0.2 | 0.47 | 0.27 | 0.15 | 0.2 | 0.15 |
| 2 | 220401128 | rs3770236 | -0.133 | 1.082 | 1.454 | 0.339 | 0.015 | 0.005 | 0.005 | 0 | 0.19 | 0.31 | 0.26 | 0.15 | 0.2 | 0.14 |
| 2 | 220404648 | rs1043833 | -0.272 | 1.084 | 1.106 | 0.687 | -0.027 | 0.007 | 0.007 | 0 | 0.33 | 0.19 | 0.48 | 0.27 | 0.45 | 0.22 |
| 2 | 220413134 | rs7597875 | 0.326 | 1.064 | 1.051 | 0.771 | -0.009 | -0.006 | -0.006 | 0 | 0.63 | 0.33 | 0.45 | 0.62 | 0.54 | 0.78 |
| 2 | 220427395 | rs3183099 | 0.383 | 0.988 | 0.127 | 0.646 | -0.021 | -0.015 | -0.015 | 0 | 0.19 | 0.08 | 0.26 | 0.21 | 0.24 | 0.09 |
| 2 | 220435034 | rs10180675 | 0.015 | 0.656 | 0.351 | 0.647 | -0.026 | 0.009 | 0.009 | 0.04 | 0.85 | 0.55 | 0.82 | 0.9 | 0.91 | 0.82 |
| 2 | 220439102 | rs7588807 | 0.237 | 0.908 | -0.115 | 0.666 | 0.002 | 0.003 | 0.003 | 0 | 0.5 | 0.42 | 0.4 | 0.57 | 0.52 | 0.45 |
| 2 | 220442490 | rs2059693 | -0.124 | 0.863 | -0.396 | 0.11 | -0.009 | -0.011 | -0.011 | 0 | 0.33 | 0.4 | 0.34 | 0.24 | 0.28 | 0.44 |
| 2 | 220443516 | rs6729914 | -0.36 | 0.734 | -0.336 | 0.407 | -0.001 | 0.003 | 0.003 | 0 | 0.5 | 0.54 | 0.54 | 0.3 | 0.49 | 0.59 |
| 2 | 220462640 | rs681747 | 1.1 | 0.338 | 1.069 | 0.298 | 0.031 | 0.016 | 0.016 | 4.149 | 0.93 | 0.44 | 0.94 | 0.99 | 0.97 | 0.98 |
| 2 | 220466199 | rs620698 | 1.326 | 0.291 | 1.329 | 0.275 | -0.003 | -0.016 | -0.016 | 1.2 | 0.93 | 0.34 | 0.93 | 0.99 | 0.97 | 0.98 |
| 2 | 220473355 | rs673951 | -0.071 | 1.227 | 0.307 | 0.384 | -0.131 | -0.074 | -0.074 | 0.047 | 0.22 | 0.03 | 0.39 | 0.68 | 0.36 | 0.41 |
| 2 | 220476443 | rs627530 | 0.694 | 0.82 | 0.846 | 0.157 | 0.045 | 0.029 | 0.029 | 4.466 | 0.93 | 0.47 | 0.94 | 0.99 | 0.97 | 0.98 |
| 2 | 220505265 | rs684428 | -0.008 | -0.316 | 0.009 | -0.532 | -0.042 | -0.042 | -0.042 | 0.033 | 0.68 | 0.43 | 0.81 | 0.81 | 0.82 | 0.71 |
| 2 | 220506522 | rs2305054 | 0.059 | -0.293 | 0.397 | -0.555 | 0.027 | 0.028 | 0.028 | 0.04 | 0.3 | 0.26 | 0.17 | 0.18 | 0.18 | 0.28 |
| 2 | 220510817 | rs12694470 | -0.198 | -0.2 | 0.554 | -0.924 | -0.001 | -0.041 | -0.041 | 0.016 | 0.17 | 0.34 | 0.11 | 0.06 | 0.16 | 0.11 |
| 2 | 220521602 | rs628917 | -0.062 | -0.145 | 0.722 | -1.023 | -0.024 | -0.035 | -0.035 | 0 | 0.37 | 0.5 | 0.24 | 0.26 | 0.33 | 0.37 |
| 2 | 220529636 | rs520102 | -0.292 | -0.312 | -0.438 | -1.1 | -0.02 | 0.063 | 0.063 | 0.165 | 0.62 | 0.93 | 0.76 | 0.72 | 0.64 | 0.53 |
| 2 | 220534468 | rs7569125 | 0.214 | 0.164 | -0.58 | -1.469 | -0.022 | 0.075 | 0.075 | 0 | 0.22 | 0.09 | 0.25 | 0.03 | 0.27 | 0.14 |
| 2 | 220535148 | rs4674417 | -0.329 | -0.279 | -0.36 | -1.684 | 0.043 | 0.073 | 0.073 | 0.492 | 0.53 | 0.73 | 0.67 | 0.65 | 0.61 | 0.73 |
| 2 | 220535733 | rs890086 | -0.965 | -0.203 | -0.26 | -1.588 | 0.013 | 0.016 | 0.016 | 0 | 0.64 | 0.73 | 0.71 | 0.65 | 0.7 | 0.74 |
| 2 | 220535982 | rs7576278 | 0.833 | 0.122 | -0.132 | -1.499 | -0.063 | 0.028 | 0.028 | 0 | 0.12 | 0.01 | 0.19 | 0.03 | 0.19 | 0.11 |
| 2 | 220545506 | rs7571961 | -0.476 | -0.197 | 0.66 | -1.427 | 0.022 | 0.001 | 0.001 | 0 | 0.05 | 0.23 | 0.03 | 0 | 0.02 | 0.07 |
| 2 | 220551797 | rs10932821 | 0.782 | -0.361 | 0.064 | -1.895 | 0.04 | 0.105 | 0.105 | 0 | 0.28 | 0.12 | 0.12 | 0.04 | 0.17 | 0.2 |
| 2 | 220554225 | rs6736062 | 1.149 | -0.413 | 0.051 | -1.907 | 0.031 | 0.092 | 0.092 | 0 | 0.25 | 0.09 | 0.11 | 0.04 | 0.16 | 0.14 |
| 2 | 220554811 | rs1318125 | 1.089 | -0.412 | 0.051 | -1.907 | 0.033 | 0.095 | 0.095 | 0 | 0.26 | 0.09 | 0.11 | 0.04 | 0.16 | 0.14 |
| 2 | 220556378 | rs639259 | -1.436 | -0.109 | -0.062 | -1.493 | -0.065 | -0.045 | -0.045 | 0.065 | 0.77 | 0.92 | 0.62 | 0.31 | 0.68 | 0.61 |
| 2 | 220564035 | rs7602880 | 0.579 | -0.052 | 0.338 | -1.506 | -0.003 | -0.005 | -0.005 | 0 | 0.3 | 0.32 | 0.25 | 0.23 | 0.3 | 0.28 |
| 2 | 220566547 | rs6721542 | 0.464 | -0.045 | 0.463 | -1.522 | 0.019 | -0.068 | -0.068 | 0.754 | 0.21 | 0.34 | 0.26 | 0.08 | 0.24 | 0.2 |
| 2 | 220569845 | rs1544859 | -0.494 | -0.051 | 0.208 | -1.524 | 0 | 0.029 | 0.029 | 0 | 0.49 | 0.61 | 0.49 | 0.27 | 0.52 | 0.45 |
| 2 | 220573045 | rs514898 | 1.025 | -0.354 | 0.143 | -1.499 | -0.011 | -0.013 | -0.013 | 1.649 | 0.26 | 0.34 | 0.23 | 0.19 | 0.28 | 0.2 |
| 2 | 220574103 | rs6745199 | 0.435 | -0.313 | -0.181 | -1.489 | 0.027 | -0.089 | -0.089 | 0 | 0.22 | 0.39 | 0.26 | 0.08 | 0.23 | 0.2 |
| 2 | 220575527 | rs539166 | -1.496 | -0.35 | -0.232 | -1.532 | -0.013 | -0.007 | -0.007 | 1.682 | 0.76 | 0.82 | 0.62 | 0.31 | 0.68 | 0.61 |
| 2 | 220577351 | rs7605568 | 1.244 | -0.282 | -1.134 | -1.489 | 0.006 | 0.009 | 0.009 | 0 | 0.24 | 0.14 | 0.22 | 0.19 | 0.27 | 0.21 |
| 2 | 220580548 | rs678134 | -0.406 | -0.292 | -1.14 | -1.496 | -0.005 | 0.106 | 0.106 | 0 | 0.48 | 0.73 | 0.5 | 0.27 | 0.51 | 0.4 |
| 2 | 220580671 | rs611328 | -0.017 | 0.198 | -1.292 | -1.194 | 0.011 | -0.081 | -0.081 | 0 | 0.24 | 0.59 | 0.28 | 0.08 | 0.24 | 0.2 |
| 2 | 220590633 | rs7569439 | -0.066 | 0.545 | -2.287 | -0.888 | -0.03 | 0.054 | 0.054 | 0 | 0.62 | 0.79 | 0.41 | 0.27 | 0.48 | 0.48 |
| 2 | 220593514 | rs10498067 | -0.217 | 0.294 | -1.023 | -0.263 | -0.01 | 0.134 | 0.134 | 0 | 0.39 | 0.18 | 0.32 | 0.08 | 0.36 | 0.27 |
| 2 | 220599533 | rs6733609 | -1.198 | 0.235 | 0.21 | 1.221 | 0.024 | 0.18 | 0.18 | 0.937 | 0.15 | 0.47 | 0.29 | 0.34 | 0.13 | 0.08 |
| 2 | 220607058 | rs4674423 | -0.527 | -0.712 | 0.167 | 1.365 | -0.058 | 0.046 | 0.046 | 0 | 0.27 | 0.5 | 0.25 | 0.4 | 0.15 | 0.25 |
| 2 | 220608798 | rs16860247 | -1.016 | -1.053 | 1.158 | 0.963 | 0.011 | -0.05 | -0.05 | 0 | 0.11 | 0.41 | 0.06 | 0.01 | 0.07 | 0.03 |
| 2 | 220609811 | rs6711869 | -0.427 | -1.035 | 0.518 | 0.981 | -0.013 | -0.033 | -0.033 | 0 | 0.8 | 0.89 | 0.71 | 0.75 | 0.81 | 0.83 |
| 2 | 220621838 | rs1477511 | 0.774 | -1.168 | 0.696 | 1.247 | 0.039 | 0.025 | 0.025 | 0.058 | 0.22 | 0.12 | 0.18 | 0.18 | 0.13 | 0.18 |
| 2 | 220624215 | rs777 | 1.581 | -1.181 | 0.724 | 1.422 | 0.005 | 0.071 | 0.071 | 0.008 | 0.43 | 0.19 | 0.41 | 0.31 | 0.59 | 0.56 |
| 2 | 220624390 | rs1035681 | 1.434 | -1.18 | 0.818 | 1.448 | 0.023 | -0.013 | -0.013 | 0 | 0.46 | 0.22 | 0.47 | 0.55 | 0.6 | 0.62 |

| | | | | | | | | | | | | | | | | |
|---|---|---|---|---|---|---|---|---|---|---|---|---|---|---|---|---|
| 2 | 220625929 | rs1364697 | 1.716 | -1.472 | 0.556 | 1.262 | -0.102 | -0.003 | -0.003 | 0.038 | 0.39 | 0.03 | 0.4 | 0.31 | 0.58 | 0.56 |
| 2 | 220627017 | rs1346012 | 0.524 | -1.478 | 0.719 | 1.22 | -0.004 | 0.003 | 0.003 | 0 | 0.7 | 0.63 | 0.65 | 0.71 | 0.73 | 0.81 |
| 2 | 220629468 | rs7583114 | -0.552 | -1.417 | 0.59 | 1.234 | -0.015 | -0.043 | -0.043 | 0.052 | 0.8 | 0.89 | 0.77 | 0.73 | 0.8 | 0.83 |
| 2 | 220631109 | rs1346010 | -0.53 | -1.404 | 0.674 | 1.249 | -0.006 | -0.028 | -0.028 | 0 | 0.79 | 0.83 | 0.76 | 0.74 | 0.8 | 0.83 |
| 2 | 220631353 | rs4674430 | 0.851 | -1.53 | 0.81 | 1.251 | -0.003 | -0.004 | -0.004 | 0 | 0.52 | 0.46 | 0.52 | 0.34 | 0.66 | 0.58 |
| 2 | 220632013 | rs4674431 | -0.128 | -1.453 | 0.747 | 1.222 | 0 | -0.04 | -0.04 | 0 | 0.77 | 0.79 | 0.7 | 0.49 | 0.79 | 0.76 |
| 2 | 220634291 | rs4674434 | -0.667 | -1.445 | 0.706 | 1.271 | -0.016 | -0.051 | -0.051 | 0.057 | 0.8 | 0.88 | 0.76 | 0.71 | 0.8 | 0.83 |
| 2 | 220635070 | rs12476246 | 1.55 | -1.453 | 0.742 | 1.212 | -0.048 | 0.023 | 0.023 | 0.184 | 0.4 | 0.11 | 0.46 | 0.32 | 0.58 | 0.54 |
| 2 | 220638012 | rs1541725 | 1.626 | -1.049 | 0.885 | 0.992 | -0.022 | 0.011 | 0.011 | 0 | 0.38 | 0.24 | 0.44 | 0.33 | 0.5 | 0.51 |
| 2 | 220639696 | rs4674436 | 0.981 | -1.391 | 0.878 | 0.96 | 0.042 | 0.036 | 0.036 | 0.033 | 0.23 | 0.14 | 0.16 | 0.16 | 0.13 | 0.19 |
| 2 | 220643119 | rs4674438 | 1.168 | -1.332 | 0.919 | 0.979 | 0.001 | -0.003 | -0.003 | 0 | 0.6 | 0.38 | 0.62 | 0.49 | 0.65 | 0.71 |
| 2 | 220645599 | rs6720347 | -0.022 | -1.285 | 0.887 | 0.943 | -0.009 | -0.019 | -0.019 | 0 | 0.68 | 0.68 | 0.76 | 0.79 | 0.75 | 0.77 |
| 2 | 220656956 | rs12473089 | -0.067 | -1.217 | 0.845 | 0.998 | -0.001 | 0.002 | 0.002 | 0 | 0.23 | 0.24 | 0.26 | 0.3 | 0.18 | 0.14 |
| 2 | 220658503 | rs4674441 | 0.689 | -1.061 | 0.65 | 0.986 | -0.016 | -0.008 | -0.008 | 0 | 0.6 | 0.44 | 0.68 | 0.61 | 0.69 | 0.66 |
| 2 | 220663412 | rs982459 | 1.684 | -0.971 | 0.523 | 0.927 | -0.019 | 0.012 | 0.012 | 0 | 0.39 | 0.25 | 0.43 | 0.33 | 0.51 | 0.52 |
| 2 | 220663477 | rs982458 | -0.59 | -0.929 | 0.296 | 0.79 | 0.013 | 0.04 | 0.04 | 2.598 | 0.74 | 0.82 | 0.79 | 0.85 | 0.79 | 0.78 |
| 2 | 220665739 | rs17575566 | 0.052 | -0.853 | 0.282 | 0.808 | -0.017 | 0.04 | 0.04 | 0.439 | 0.08 | 0.03 | 0.06 | 0 | 0.09 | 0.04 |
| 2 | 220666208 | rs4672945 | -0.781 | -0.846 | 0.3 | 0.749 | 0 | 0 | 0 | 0 | 0.51 | 0.51 | 0.46 | 0.46 | 0.35 | 0.34 |
| 2 | 220666777 | rs13429827 | -1.508 | -0.84 | 0.36 | 0.797 | -0.001 | 0.001 | 0.001 | 0 | 0.6 | 0.62 | 0.54 | 0.66 | 0.45 | 0.46 |
| 2 | 220667933 | rs719325 | 1.707 | -0.832 | 0.372 | 0.802 | -0.009 | 0.001 | 0.001 | 0 | 0.36 | 0.31 | 0.42 | 0.34 | 0.48 | 0.49 |
| 2 | 220677486 | rs2891829 | -0.616 | -1.033 | 0.456 | 1.031 | 0.02 | -0.005 | -0.005 | 0 | 0.44 | 0.4 | 0.42 | 0.46 | 0.29 | 0.32 |
| 2 | 220677869 | rs12615743 | 0.963 | -1.039 | 0.433 | 1.021 | 0.068 | 0.008 | 0.008 | 0.073 | 0.5 | 0.26 | 0.52 | 0.54 | 0.64 | 0.63 |
| 2 | 220687059 | rs1016201 | 1.404 | -1.038 | -0.502 | 0.622 | -0.079 | -0.013 | -0.013 | 0 | 0.42 | 0.03 | 0.46 | 0.33 | 0.57 | 0.53 |
| 2 | 220690131 | rs16860351 | -0.285 | -1.72 | -0.07 | 1.192 | 0.03 | 0.007 | 0.007 | 4.348 | 0.31 | 0.21 | 0.2 | 0.28 | 0.18 | 0.21 |
| 2 | 220691829 | rs10175587 | 0.109 | -1.408 | -0.156 | 1.07 | 0.028 | 0.01 | 0.01 | 4.564 | 0.2 | 0.11 | 0.15 | 0.17 | 0.11 | 0.17 |
| 2 | 220692981 | rs13016645 | -0.765 | -0.67 | 0.294 | 1.193 | -0.021 | 0.129 | 0.129 | 0 | 0.13 | 0.26 | 0.25 | 0.38 | 0.11 | 0.2 |
| 2 | 220696155 | rs16860370 | -1.237 | -0.717 | 0.339 | 0.172 | -0.033 | -0.087 | -0.087 | 0 | 0.15 | 0.36 | 0.1 | 0 | 0.08 | 0.02 |
| 2 | 220696298 | rs6719949 | -0.301 | -0.715 | 0.023 | 0.199 | 0.009 | 0 | 0 | 0 | 0.46 | 0.57 | 0.38 | 0.53 | 0.36 | 0.41 |
| 2 | 220697602 | rs11688142 | -0.457 | -0.948 | 0.564 | -0.284 | -0.007 | 0 | 0 | 1.291 | 0.08 | 0.05 | 0.07 | 0 | 0.1 | 0.02 |
| 2 | 220698125 | rs10498070 | -1.964 | -1.037 | 0.262 | -0.052 | 0.001 | -0.01 | -0.01 | 0 | 0.28 | 0.76 | 0.36 | 0.48 | 0.26 | 0.28 |
| 2 | 220702824 | rs10932829 | -1.567 | -1.54 | -0.481 | -0.368 | 0.124 | -0.06 | -0.06 | 0.365 | 0.48 | 0.88 | 0.41 | 0.47 | 0.31 | 0.3 |
| 2 | 220703292 | rs13000695 | -0.527 | -1.296 | -0.385 | -0.222 | -0.004 | 0.069 | 0.069 | 0 | 0.14 | 0.22 | 0.19 | 0.47 | 0.12 | 0.22 |
| 2 | 220707363 | rs1425033 | -0.127 | -1.142 | -0.699 | -0.569 | -0.025 | 0.052 | 0.052 | 0.761 | 0.09 | 0 | 0.06 | 0 | 0.1 | 0.02 |
| 2 | 220709039 | rs13398816 | -0.131 | -1.267 | -0.357 | -0.474 | -0.097 | -0.154 | -0.154 | 1.354 | 0.29 | 0.54 | 0.13 | 0.01 | 0.11 | 0.07 |
| 2 | 220712045 | rs7590480 | -0.524 | -0.504 | -1.069 | 0.093 | 0.085 | 0.012 | 0.012 | 1.967 | 0.51 | 0.9 | 0.44 | 0.58 | 0.37 | 0.38 |
| 2 | 220714502 | rs7557937 | 0.618 | 0.227 | 0.179 | -0.419 | 0.045 | -0.07 | -0.07 | 0 | 0.76 | 0.69 | 0.71 | 0.9 | 0.67 | 0.8 |
| 2 | 220719283 | rs2664220 | -0.641 | -1.529 | 0.313 | -0.508 | -0.077 | -0.06 | -0.06 | 0 | 0.34 | 0.44 | 0.12 | 0.18 | 0.18 | 0.2 |
| 2 | 220722698 | rs10498071 | -0.561 | -1.715 | 0.682 | -0.553 | -0.007 | 0.018 | 0.018 | 0.641 | 0.09 | 0.13 | 0.05 | 0.14 | 0.08 | 0.07 |
| 2 | 220723823 | rs2385624 | -0.155 | -1.752 | 0.917 | -0.609 | -0.036 | -0.076 | -0.076 | 0 | 0.76 | 0.74 | 0.95 | 1 | 0.9 | 0.86 |
| 2 | 220726247 | rs12694485 | 0.872 | -1.824 | 1.094 | -0.396 | -0.055 | -0.023 | -0.023 | 0.912 | 0.66 | 0.49 | 0.88 | 0.75 | 0.81 | 0.79 |
| 2 | 220729620 | rs10178122 | 0.513 | -1.682 | 0.979 | -0.845 | -0.027 | -0.143 | -0.143 | 0 | 0.64 | 0.61 | 0.85 | 1 | 0.71 | 0.81 |
| 2 | 220733584 | rs12694490 | 1.106 | -1.58 | 0.988 | -0.738 | 0.002 | 0 | 0 | 2.878 | 0.57 | 0.49 | 0.82 | 0.77 | 0.68 | 0.74 |
| 2 | 220735707 | rs6751315 | 0.111 | -1.62 | 0.988 | -0.769 | -0.02 | -0.098 | -0.098 | 0.021 | 0.71 | 0.72 | 0.91 | 1 | 0.77 | 0.82 |
| 2 | 220738491 | rs7595755 | 0.227 | -1.58 | 0.673 | -0.746 | -0.04 | -0.102 | -0.102 | 1.44 | 0.67 | 0.61 | 0.82 | 0.96 | 0.76 | 0.79 |
| 2 | 220740379 | rs6740059 | 0.966 | -1.345 | 0.59 | -0.601 | 0.008 | 0.008 | 0.008 | 4.665 | 0.53 | 0.49 | 0.75 | 0.71 | 0.62 | 0.71 |
| 2 | 220752532 | rs4142173 | 0.518 | -1.307 | 0.899 | -0.443 | -0.03 | -0.023 | -0.023 | 0.008 | 0.59 | 0.54 | 0.77 | 0.74 | 0.72 | 0.72 |
| 2 | 220753783 | rs11687676 | 0.874 | -1.393 | 0.83 | -0.44 | 0.001 | 0.001 | 0.001 | 0.375 | 0.55 | 0.48 | 0.73 | 0.72 | 0.66 | 0.72 |
| 2 | 220768879 | rs2253615 | -0.317 | -1.165 | 0.007 | -0.261 | 0.068 | 0.024 | 0.024 | 0 | 0.73 | 0.87 | 0.93 | 0.76 | 0.86 | 0.78 |
| 2 | 220771575 | rs9288556 | -0.28 | -1.306 | -0.295 | 0.114 | -0.014 | 0.05 | 0.05 | 0.02 | 0.29 | 0.39 | 0.41 | 0.47 | 0.28 | 0.43 |
| 2 | 220771707 | rs2891835 | -0.06 | -0.798 | -0.434 | 0.123 | 0.04 | 0.021 | 0.021 | 0 | 0.7 | 0.81 | 0.9 | 0.73 | 0.81 | 0.78 |
| 2 | 220773422 | rs17288469 | -1.125 | 0.339 | -0.161 | 0.464 | 0.008 | -0.019 | -0.019 | 0 | 0.1 | 0.2 | 0.1 | 0.03 | 0.14 | 0.06 |
| 2 | 220779861 | rs4433991 | -1.177 | -0.015 | -0.094 | 1.585 | -0.017 | 0.216 | 0.216 | 0.015 | 0.05 | 0.2 | 0.12 | 0.37 | 0.01 | 0.22 |
| 2 | 220781006 | rs750764 | -0.305 | 0.249 | -0.574 | 1.204 | 0.003 | 0 | 0 | 0 | 0.57 | 0.48 | 0.46 | 0.48 | 0.49 | 0.59 |
| 2 | 220783322 | rs4672960 | 0.303 | -0.223 | -1.325 | 0.659 | -0.004 | 0.058 | 0.058 | 3.073 | 0.32 | 0.19 | 0.25 | 0.09 | 0.32 | 0.19 |
| 2 | 220784986 | rs13006717 | -0.054 | -0.3 | -1.368 | 0.744 | 0.014 | -0.021 | -0.021 | 0 | 0.25 | 0.22 | 0.22 | 0.48 | 0.18 | 0.41 |
| 2 | 220786974 | rs11692519 | 0.091 | -0.327 | -1.443 | 0.75 | 0.021 | -0.029 | -0.029 | 0.721 | 0.23 | 0.18 | 0.12 | 0.41 | 0.17 | 0.37 |
| 2 | 220789370 | rs731412 | 1.204 | -0.157 | -1.93 | 0.457 | -0.149 | -0.144 | -0.144 | 2.193 | 0.32 | 0.03 | 0.33 | 0.39 | 0.41 | 0.31 |
| 2 | 220790637 | rs1007684 | -0.574 | -0.093 | -1.803 | 0.478 | -0.02 | -0.016 | -0.016 | 0.746 | 0.55 | 0.28 | 0.48 | 0.57 | 0.5 | 0.6 |
| 2 | 220791581 | rs11693842 | 0.639 | 0.104 | -1.917 | 0.215 | -0.02 | 0.035 | 0.035 | 2.417 | 0.22 | 0.08 | 0.23 | 0.09 | 0.24 | 0.11 |
| 2 | 220792262 | rs17288923 | -0.002 | 1.052 | -2.029 | 0.339 | -0.023 | 0.017 | 0.017 | 2.28 | 0.25 | 0.14 | 0.21 | 0.15 | 0.28 | 0.15 |
| 2 | 220792320 | rs7607421 | -1.147 | 0.737 | -1.259 | 0.326 | -0.005 | 0.006 | 0.006 | 0.143 | 0.65 | 0.58 | 0.7 | 0.62 | 0.64 | 0.69 |
| 2 | 220792739 | rs2385826 | -0.427 | -0.029 | -0.916 | 0.442 | -0.043 | 0.019 | 0.019 | 0 | 0.39 | 0.48 | 0.43 | 0.49 | 0.22 | 0.49 |
| 2 | 220794320 | rs11692459 | 0.935 | -0.164 | -1.689 | 0.475 | -0.162 | -0.031 | -0.031 | 0 | 0.36 | 0.04 | 0.35 | 0.31 | 0.48 | 0.33 |
| 2 | 220803438 | rs2385828 | -0.104 | -0.702 | -1.322 | 0.496 | 0.001 | 0.002 | 0.002 | 0 | 0.38 | 0.38 | 0.48 | 0.25 | 0.36 | 0.27 |
| 2 | 220803938 | rs2385836 | 0.185 | -0.708 | -1.65 | 0.518 | 0.002 | 0.037 | 0.037 | 0 | 0.34 | 0.18 | 0.4 | 0.18 | 0.34 | 0.24 |
| 2 | 220812228 | rs6725376 | 1.339 | -0.683 | -1.768 | 0.832 | -0.162 | -0.138 | -0.138 | 0 | 0.36 | 0.04 | 0.35 | 0.41 | 0.49 | 0.37 |
| 2 | 220813461 | rs4674482 | 1.207 | -0.744 | -1.904 | 0.876 | 0.057 | 0.012 | 0.012 | 0 | 0.22 | 0.07 | 0.09 | 0.13 | 0.12 | 0.1 |
| 2 | 220817726 | rs6706564 | 0.043 | -0.209 | -1.973 | 0.881 | -0.003 | -0.017 | -0.017 | 0 | 0.23 | 0.2 | 0.26 | 0.38 | 0.24 | 0.34 |
| 2 | 220828626 | rs10185419 | 1.611 | -0.222 | -2.161 | 0.847 | -0.096 | -0.148 | -0.148 | 1.413 | 0.31 | 0.04 | 0.19 | 0.41 | 0.35 | 0.32 |

| | | | | | | | | | | | | | | | | |
|---|---|---|---|---|---|---|---|---|---|---|---|---|---|---|---|---|
| 2 | 220829165 | rs13005658 | 0.22 | -0.306 | -2.143 | 0.826 | -0.002 | -0.003 | -0.003 | 0 | 0.22 | 0.2 | 0.25 | 0.29 | 0.24 | 0.33 |
| 2 | 220829849 | rs10199944 | 0.465 | -0.385 | -1.97 | 0.738 | 0.031 | -0.032 | -0.032 | 0 | 0.45 | 0.23 | 0.42 | 0.48 | 0.58 | 0.46 |
| 2 | 220833222 | rs10221578 | 0.43 | -0.414 | -2.202 | 0.737 | 0.032 | 0.004 | 0.004 | 0.534 | 0.46 | 0.2 | 0.5 | 0.4 | 0.59 | 0.47 |
| 2 | 220839453 | rs2385844 | 0.561 | -0.298 | -2.195 | 0.866 | 0.003 | -0.03 | -0.03 | 0.453 | 0.43 | 0.17 | 0.49 | 0.39 | 0.57 | 0.47 |
| 2 | 220841109 | rs34497856 | 0.105 | -0.261 | -2.309 | 0.887 | -0.002 | -0.008 | -0.008 | 0 | 0.23 | 0.2 | 0.27 | 0.36 | 0.24 | 0.34 |
| 2 | 220845194 | rs750894 | 0.943 | 0.186 | -2.384 | 0.971 | -0.011 | 0.006 | 0.006 | 0 | 0.12 | 0.09 | 0.22 | 0.04 | 0.22 | 0.14 |
| 2 | 220847180 | rs11689678 | 1.032 | -0.277 | -2.471 | 1.074 | -0.098 | -0.111 | -0.111 | 0 | 0.3 | 0.04 | 0.18 | 0.34 | 0.33 | 0.32 |
| 2 | 220851174 | rs13414052 | 1.018 | -0.3 | -2.469 | 0.765 | -0.094 | -0.056 | -0.056 | 0.859 | 0.3 | 0.04 | 0.18 | 0.26 | 0.33 | 0.32 |
| 2 | 220853614 | rs6714200 | -0.259 | -0.07 | -2.071 | 0.941 | -0.02 | -0.008 | -0.008 | 0 | 0.37 | 0.29 | 0.42 | 0.46 | 0.46 | 0.51 |
| 2 | 220854992 | rs16860653 | 0.441 | 0.936 | -1.95 | 1.063 | -0.094 | -0.07 | -0.07 | 0 | 0.21 | 0.03 | 0.17 | 0.25 | 0.31 | 0.28 |
| 2 | 220861593 | rs995842 | -0.532 | 0.512 | -1.64 | 0.654 | 0.016 | 0.072 | 0.072 | 0 | 0.48 | 0.26 | 0.56 | 0.62 | 0.57 | 0.68 |
| 2 | 220861711 | rs11680001 | -0.393 | 0.047 | -1.376 | 1.238 | -0.004 | 0.02 | 0.02 | 3.26 | 0.16 | 0.16 | 0.11 | 0.54 | 0.06 | 0.29 |
| 2 | 220863024 | rs6717409 | 0.27 | 0.344 | -1.074 | 0.417 | 0.021 | 0.003 | 0.003 | 0.005 | 0.59 | 0.26 | 0.43 | 0.41 | 0.68 | 0.56 |
| 2 | 220863380 | rs7595011 | -0.666 | 0.99 | -0.324 | 0.262 | 0.001 | 0.002 | 0.002 | 0.453 | 0.39 | 0.65 | 0.47 | 0.81 | 0.33 | 0.44 |
| 2 | 220864654 | rs10198756 | 0.441 | 0.937 | 0.377 | 0.275 | -0.025 | 0.041 | 0.041 | 1.828 | 0.26 | 0.38 | 0.26 | 0.6 | 0.17 | 0.32 |
| 2 | 220873544 | rs2664226 | -0.956 | 1.059 | 0.897 | 0.148 | 0.04 | -0.03 | -0.03 | 0 | 0.94 | 0.88 | 0.78 | 1 | 0.87 | 0.94 |
| 2 | 220874148 | rs1963367 | 0.355 | 1.038 | 0.823 | 0.214 | 0.066 | 0.079 | 0.079 | 0.228 | 0.07 | 0.18 | 0.19 | 0.2 | 0.14 | 0.05 |
| 2 | 220877302 | rs10210670 | -0.636 | 0.992 | 0.805 | 0.229 | -0.003 | -0.03 | -0.03 | 0 | 0.37 | 0.58 | 0.46 | 0.82 | 0.33 | 0.41 |
| 2 | 220878921 | rs2385848 | 0.507 | 1.047 | 0.428 | 0.258 | 0.053 | 0.063 | 0.063 | 0.064 | 0.06 | 0.15 | 0.14 | 0.19 | 0.14 | 0.05 |
| 2 | 220879384 | rs7608770 | 0.455 | 0.765 | 0.253 | 0.221 | 0.026 | -0.049 | -0.049 | 0 | 0.26 | 0.12 | 0.24 | 0.62 | 0.17 | 0.33 |
| 2 | 220883373 | rs6756097 | 0.198 | 0.431 | 0.595 | -0.356 | -0.009 | 0.024 | 0.024 | 0 | 0.57 | 0.34 | 0.34 | 0.13 | 0.52 | 0.56 |
| 2 | 220889178 | rs7573142 | 0.609 | 0.157 | -0.162 | -0.792 | 0.028 | 0.039 | 0.039 | 0 | 0.35 | 0.23 | 0.16 | 0.07 | 0.24 | 0.24 |
| 2 | 220893404 | rs1862987 | 0.999 | 0.22 | -0.567 | -0.772 | -0.091 | 0.071 | 0.071 | 0 | 0.22 | 0.01 | 0.15 | 0.06 | 0.29 | 0.33 |
| 2 | 220899751 | rs6745046 | -0.072 | -0.236 | 0.438 | -0.454 | -0.021 | -0.053 | -0.053 | 0 | 0.37 | 0.56 | 0.25 | 0.07 | 0.24 | 0.24 |
| 2 | 220915008 | rs992654 | -0.103 | 0.268 | -0.176 | -0.033 | 0.002 | 0.014 | 0.014 | 0.159 | 0.6 | 0.64 | 0.61 | 0.85 | 0.63 | 0.72 |
| 2 | 220917579 | rs7424109 | -0.585 | 0.478 | 0.205 | 0.828 | 0.009 | 0.061 | 0.061 | 0 | 0.29 | 0.49 | 0.31 | 0.4 | 0.38 | 0.4 |
| 2 | 220918879 | rs2216843 | 0.482 | 0.234 | -0.266 | 0.859 | 0.003 | -0.017 | -0.017 | 1.773 | 0.4 | 0.13 | 0.39 | 0.44 | 0.35 | 0.43 |
| 2 | 220921565 | rs1559520 | -0.467 | 0.012 | 0.259 | 0.752 | 0 | 0.009 | 0.009 | 0 | 0.66 | 0.7 | 0.69 | 0.74 | 0.71 | 0.79 |
| 2 | 220921709 | rs17305387 | 0.121 | 0.185 | 0.502 | 0.929 | 0.013 | 0.045 | 0.045 | 0 | 0.28 | 0.49 | 0.31 | 0.34 | 0.38 | 0.37 |
| 2 | 220923635 | rs2971613 | 0.33 | 0.145 | 0.659 | 0.864 | -0.004 | -0.003 | -0.003 | 0.264 | 0.43 | 0.42 | 0.41 | 0.44 | 0.35 | 0.42 |
| 2 | 220926429 | rs4674493 | 0.39 | 0.124 | 0.907 | 0.779 | 0.008 | 0.011 | 0.011 | 0.062 | 0.43 | 0.33 | 0.37 | 0.32 | 0.33 | 0.39 |
| 2 | 220927290 | rs744499 | -0.686 | 0.114 | 0.872 | 0.786 | 0.002 | -0.005 | -0.005 | 0.223 | 0.69 | 0.62 | 0.69 | 0.75 | 0.71 | 0.78 |
| 2 | 220927449 | rs2385852 | 0.603 | 0.248 | 0.622 | 1.009 | -0.009 | -0.02 | -0.02 | 0 | 0.21 | 0.18 | 0.24 | 0.33 | 0.32 | 0.33 |
| 2 | 220931577 | rs6728853 | -0.721 | 0.268 | 0.471 | 0.776 | -0.008 | 0.003 | 0.003 | 0 | 0.71 | 0.76 | 0.69 | 0.76 | 0.71 | 0.78 |
| 2 | 220975474 | rs7606731 | -0.444 | 0.308 | 0.41 | 0.811 | -0.002 | -0.007 | -0.007 | 0 | 0.65 | 0.58 | 0.63 | 0.69 | 0.69 | 0.65 |
| 2 | 220978325 | rs13383613 | 0.45 | 0.271 | -0.222 | 0.849 | 0.02 | 0.003 | 0.003 | 0 | 0.46 | 0.16 | 0.39 | 0.42 | 0.36 | 0.46 |
| 2 | 220981041 | rs3935261 | 0.663 | 0.355 | -0.044 | 0.934 | 0 | -0.01 | -0.01 | 0.08 | 0.43 | 0.24 | 0.42 | 0.45 | 0.38 | 0.46 |
| 2 | 220992143 | rs13001386 | -0.066 | 0.582 | -0.212 | 0.885 | -0.046 | 0.007 | 0.007 | 0 | 0.58 | 0.18 | 0.54 | 0.61 | 0.57 | 0.63 |
| 2 | 220997227 | rs4073777 | 0.198 | 0.779 | 0.071 | 0.845 | -0.003 | -0.014 | -0.014 | 0 | 0.39 | 0.47 | 0.3 | 0.31 | 0.41 | 0.31 |
| 2 | 220999334 | rs4480974 | 1.849 | 0.753 | -0.678 | 0.812 | -0.041 | -0.065 | -0.065 | 0.041 | 0.17 | 0.02 | 0.18 | 0.24 | 0.17 | 0.18 |
| 2 | 220999979 | rs12694510 | -1.092 | 0.755 | -0.377 | 0.815 | 0.015 | -0.007 | -0.007 | 0 | 0.93 | 0.77 | 0.83 | 0.95 | 0.95 | 0.92 |
| 2 | 221000018 | rs12614221 | 0.292 | 0.767 | -0.839 | 0.866 | 0.009 | 0.002 | 0.002 | 0 | 0.42 | 0.27 | 0.38 | 0.43 | 0.34 | 0.46 |
| 2 | 221003602 | rs12052391 | 0.743 | 0.752 | -0.895 | 1.051 | -0.029 | -0.041 | -0.041 | 0 | 0.12 | 0.07 | 0.15 | 0.21 | 0.19 | 0.16 |
| 2 | 221013438 | rs6754489 | 0.439 | 0.778 | -0.873 | 1.064 | 0.001 | 0.003 | 0.003 | 0 | 0.14 | 0.18 | 0.16 | 0.2 | 0.2 | 0.16 |
| 2 | 221016421 | rs10193131 | 0.313 | 0.429 | -1.337 | 0.505 | 0.003 | 0.051 | 0.051 | 0 | 0.42 | 0.24 | 0.32 | 0.29 | 0.43 | 0.41 |
| 2 | 221016706 | rs11886925 | -0.226 | 0.044 | -1.43 | 0.773 | 0.035 | -0.016 | -0.016 | 0.375 | 0.16 | 0.07 | 0.09 | 0.17 | 0.07 | 0.17 |
| 2 | 221019458 | rs10932835 | -0.118 | 0.423 | -1.387 | 0.48 | -0.01 | 0.028 | 0.028 | 0 | 0.43 | 0.3 | 0.56 | 0.67 | 0.52 | 0.44 |
| 2 | 221024539 | rs6748896 | 0.293 | -0.766 | -0.588 | 0.289 | -0.025 | -0.015 | -0.015 | 0 | 0.66 | 0.54 | 0.77 | 0.76 | 0.8 | 0.65 |
| 2 | 221025443 | rs10205602 | -1.267 | -0.383 | -0.667 | -0.433 | 0.044 | -0.136 | -0.136 | 0.229 | 0.73 | 0.48 | 0.65 | 1 | 0.63 | 0.83 |
| 2 | 221031021 | rs12613666 | 0.345 | -1.06 | -1.068 | 0.239 | 0.002 | -0.002 | -0.002 | 0 | 0.15 | 0.17 | 0.14 | 0.28 | 0.09 | 0.22 |
| 2 | 221035322 | rs4459724 | 0.345 | -1.06 | -1.068 | 0.273 | 0.002 | -0.003 | -0.003 | 0 | 0.15 | 0.16 | 0.14 | 0.25 | 0.09 | 0.21 |
| 2 | 221035778 | rs11679411 | 0.795 | -0.915 | -1.122 | 0.403 | -0.008 | -0.069 | -0.069 | 0 | 0.26 | 0.11 | 0.27 | 0.46 | 0.29 | 0.29 |
| 2 | 221044785 | rs10932837 | 0.372 | -0.94 | -0.892 | 0.185 | -0.026 | 0.042 | 0.042 | 0 | 0.47 | 0.18 | 0.44 | 0.64 | 0.49 | 0.55 |
| 2 | 221047792 | rs6711809 | -1.704 | -1.145 | -0.451 | -0.046 | 0.046 | -0.128 | -0.128 | 0 | 0.75 | 0.61 | 0.64 | 1 | 0.62 | 0.83 |
| 2 | 221051296 | rs4132708 | -0.537 | -1.196 | -0.316 | 0.044 | 0.002 | -0.023 | -0.023 | 0 | 0.63 | 0.43 | 0.57 | 0.87 | 0.57 | 0.77 |
| 2 | 221053277 | rs4583451 | -0.776 | -1.194 | -0.4 | 0.253 | -0.054 | 0.038 | 0.038 | 0 | 0.28 | 0.41 | 0.2 | 0.37 | 0.13 | 0.28 |
| 2 | 221063851 | rs6718891 | -0.508 | -0.942 | -0.604 | 0.034 | -0.004 | -0.005 | -0.005 | 0.02 | 0.62 | 0.39 | 0.57 | 0.9 | 0.57 | 0.77 |
| 2 | 221067691 | rs11688377 | 0.411 | -1.048 | -0.774 | 0.208 | -0.026 | 0.039 | 0.039 | 0 | 0.47 | 0.21 | 0.44 | 0.64 | 0.49 | 0.55 |
| 2 | 221074372 | rs4674511 | -0.119 | -1.298 | -1.226 | 0.435 | 0.036 | 0.033 | 0.033 | 0 | 0.49 | 0.18 | 0.39 | 0.61 | 0.4 | 0.58 |
| 2 | 221074922 | rs4234030 | -1.032 | -1.31 | -0.961 | 0.403 | 0 | -0.024 | -0.024 | 0 | 0.63 | 0.42 | 0.57 | 0.89 | 0.57 | 0.77 |
| 2 | 221080165 | rs17656266 | 1.212 | -1.321 | -1.009 | 0.513 | 0.013 | -0.065 | -0.065 | 0 | 0.13 | 0.03 | 0.12 | 0.25 | 0.08 | 0.22 |
| 2 | 221081421 | rs17610130 | 1.116 | -1.273 | -0.992 | 0.544 | -0.053 | -0.078 | -0.078 | 0 | 0.12 | 0.01 | 0.17 | 0.28 | 0.17 | 0.19 |
| 2 | 221081739 | rs10207046 | 0.088 | -1.416 | -0.586 | 0.448 | -0.017 | 0.013 | 0.013 | 1.269 | 0.11 | 0.37 | 0.08 | 0.1 | 0.05 | 0.06 |
| 2 | 221093338 | rs4597515 | 0.802 | -1.34 | -0.895 | 0.415 | -0.019 | -0.007 | -0.007 | 0 | 0.33 | 0.1 | 0.25 | 0.34 | 0.32 | 0.36 |
| 2 | 221103834 | rs17656376 | 0.299 | -1.19 | -0.696 | 0.42 | -0.033 | 0.066 | 0.066 | 0 | 0.17 | 0.27 | 0.14 | 0.27 | 0.09 | 0.23 |
| 2 | 221105606 | rs17610269 | 0.299 | -1.19 | -0.69 | 0.42 | -0.032 | 0.062 | 0.062 | 0 | 0.17 | 0.27 | 0.14 | 0.27 | 0.09 | 0.23 |
| 2 | 221110281 | rs6705130 | 1.744 | -1.115 | -0.671 | 0.006 | 0.027 | -0.099 | -0.099 | 0 | 0.25 | 0.35 | 0.36 | 0 | 0.38 | 0.18 |
| 2 | 221110435 | rs1551385 | -0.069 | -1.009 | -0.926 | 0.395 | 0.019 | 0.071 | 0.071 | 0 | 0.52 | 0.68 | 0.56 | 0.28 | 0.58 | 0.46 |
| 2 | 221110780 | rs1993313 | 0.561 | -1.067 | -1.034 | 0.351 | 0 | -0.006 | -0.006 | 0.098 | 0.44 | 0.58 | 0.5 | 0.26 | 0.47 | 0.41 |

| | | | | | | | | | | | | | | | | |
|---|---|---|---|---|---|---|---|---|---|---|---|---|---|---|---|---|
| 2 | 221111295 | rs10932846 | 1.325 | -0.768 | -1.688 | 0.439 | -0.03 | -0.068 | -0.068 | 0 | 0.17 | 0.05 | 0.14 | 0.29 | 0.18 | 0.24 |
| 2 | 221112306 | rs17472512 | 0.077 | -0.671 | -1.272 | 0.25 | 0.002 | -0.013 | -0.013 | 3.114 | 0.1 | 0.13 | 0.07 | 0.05 | 0.12 | 0.04 |
| 2 | 221125602 | rs17472561 | -0.179 | -0.703 | -1.286 | 0.237 | -0.001 | -0.014 | -0.014 | 2.687 | 0.11 | 0.13 | 0.07 | 0.05 | 0.12 | 0.05 |
| 2 | 221134279 | rs10498074 | 0.926 | -0.989 | -1.61 | 0.117 | -0.005 | 0.02 | 0.02 | 0 | 0.16 | 0 | 0.11 | 0.12 | 0.13 | 0.08 |
| 2 | 221136622 | rs9917205 | -0.047 | -1.09 | -1.562 | 0.251 | 0 | -0.002 | -0.002 | 0.873 | 0.22 | 0.15 | 0.19 | 0.24 | 0.21 | 0.26 |
| 2 | 221139131 | rs6436196 | 0.847 | -1.055 | -1.464 | 0.276 | 0.02 | -0.01 | -0.01 | 0 | 0.33 | 0.61 | 0.48 | 0.23 | 0.43 | 0.32 |
| 2 | 221139537 | rs17656530 | 1.687 | -0.824 | -1.787 | 0.64 | -0.019 | -0.021 | -0.021 | 0 | 0.06 | 0 | 0.1 | 0.22 | 0.06 | 0.14 |
| 2 | 221144573 | rs17472652 | 0.319 | -1.058 | -1.929 | 0.366 | -0.025 | -0.027 | -0.027 | 0 | 0.2 | 0.04 | 0.19 | 0.23 | 0.21 | 0.26 |
| 2 | 221144862 | rs6711764 | 0.007 | -1.56 | -2.319 | 0.102 | 0.039 | -0.087 | -0.087 | 0.036 | 0.46 | 0.13 | 0.31 | 0.53 | 0.35 | 0.45 |
| 2 | 221146465 | rs10804287 | 0.926 | -1.773 | -2.487 | 0.079 | -0.003 | -0.089 | -0.089 | 0.11 | 0.27 | 0.09 | 0.2 | 0.41 | 0.23 | 0.35 |
| 2 | 221149605 | rs10932849 | 1.363 | -1.976 | -2.591 | -0.009 | 0.041 | -0.004 | -0.004 | 0.019 | 0.11 | 0.04 | 0.02 | 0.14 | 0.04 | 0.09 |
| 2 | 221150805 | rs17610467 | 1.092 | -1.985 | -2.668 | -0.036 | -0.053 | -0.119 | -0.119 | 0 | 0.23 | 0.06 | 0.28 | 0.51 | 0.25 | 0.39 |
| 2 | 221157107 | rs1454558 | 1.101 | -2.098 | -1.886 | -0.282 | 0.004 | -0.043 | -0.043 | 0 | 0.42 | 0.5 | 0.48 | 0.12 | 0.5 | 0.27 |
| 2 | 221158425 | rs11680840 | -0.203 | -2.185 | -1.801 | -0.29 | 0.011 | 0 | 0 | 0 | 0.32 | 0.25 | 0.22 | 0.38 | 0.24 | 0.33 |
| 2 | 221160422 | rs10186311 | 1.421 | -2.196 | -1.715 | -0.339 | 0.021 | 0.026 | 0.026 | 0 | 0.18 | 0.06 | 0.1 | 0.12 | 0.13 | 0.09 |
| 2 | 221160610 | rs1378023 | -0.119 | -2.153 | -1.649 | -0.329 | 0.003 | -0.001 | -0.001 | 0.066 | 0.31 | 0.28 | 0.22 | 0.38 | 0.25 | 0.34 |
| 2 | 221161465 | rs1454561 | -2.094 | -2.064 | -1.585 | -0.374 | 0.031 | -0.119 | -0.119 | 0 | 0.74 | 0.4 | 0.6 | 1 | 0.63 | 0.83 |
| 2 | 221166925 | rs4674516 | -0.504 | -2.082 | -1.539 | -0.323 | -0.005 | -0.009 | -0.009 | 0 | 0.76 | 0.62 | 0.78 | 0.77 | 0.78 | 0.75 |
| 2 | 221170656 | rs11888018 | 2.309 | -2.098 | -1.173 | -0.364 | 0.076 | -0.159 | -0.159 | 0.314 | 0.24 | 0.42 | 0.36 | 0 | 0.38 | 0.17 |
| 2 | 221175234 | rs7590177 | 0.926 | -2.081 | -1.22 | -0.321 | -0.027 | -0.016 | -0.016 | 0 | 0.2 | 0.04 | 0.19 | 0.22 | 0.22 | 0.25 |
| 2 | 221180112 | rs11676335 | 0.816 | -1.948 | -1.131 | -0.252 | -0.003 | -0.009 | -0.009 | 0 | 0.42 | 0.55 | 0.48 | 0.12 | 0.5 | 0.27 |
| 2 | 221186275 | rs897144 | -0.046 | -2.09 | -1.232 | -0.228 | 0.026 | 0.024 | 0.024 | 0.109 | 0.6 | 0.67 | 0.66 | 0.35 | 0.71 | 0.51 |
| 2 | 221187667 | rs724158 | -0.507 | -2.102 | -1.254 | -0.062 | 0.028 | 0.043 | 0.043 | 1.545 | 0.48 | 0.35 | 0.37 | 0.55 | 0.39 | 0.5 |
| 2 | 221189253 | rs4672976 | 0.613 | -2.034 | -1.257 | 0.002 | -0.013 | -0.11 | -0.11 | 0 | 0.3 | 0.14 | 0.25 | 0.43 | 0.26 | 0.41 |
| 2 | 221191009 | rs1454550 | -1.503 | -2.023 | -1.234 | -0.006 | 0.001 | 0.006 | 0.006 | 0 | 0.87 | 0.87 | 0.86 | 0.73 | 0.93 | 0.85 |
| 2 | 221197071 | rs12623779 | 1.087 | -1.718 | -0.896 | -0.582 | 0.006 | -0.027 | -0.027 | 0 | 0.31 | 0.24 | 0.28 | 0.47 | 0.26 | 0.41 |
| 2 | 221201418 | rs10171102 | 0.962 | -1.785 | -0.584 | -0.648 | -0.001 | -0.006 | -0.006 | 2.015 | 0.18 | 0.15 | 0.09 | 0.08 | 0.13 | 0.09 |
| 2 | 221201659 | rs10208333 | -0.384 | -1.794 | -0.576 | -0.699 | 0.002 | 0.009 | 0.009 | 0 | 0.35 | 0.27 | 0.28 | 0.3 | 0.34 | 0.33 |
| 2 | 221201901 | rs12478433 | -0.884 | -1.823 | -0.593 | -0.729 | -0.001 | -0.005 | -0.005 | 0 | 0.64 | 0.35 | 0.54 | 0.77 | 0.6 | 0.73 |
| 2 | 221203877 | rs17657213 | 0.384 | -1.866 | -0.659 | -0.753 | 0.038 | 0.056 | 0.056 | 0.063 | 0.4 | 0.23 | 0.34 | 0.69 | 0.29 | 0.5 |
| 2 | 221204273 | rs13386574 | -0.677 | -1.6 | -0.365 | -0.759 | 0.012 | -0.05 | -0.05 | 0 | 0.66 | 0.47 | 0.55 | 0.77 | 0.6 | 0.73 |
| 2 | 221205862 | rs1454540 | -1.018 | -1.569 | -0.835 | -0.647 | -0.018 | 0.007 | 0.007 | 0 | 0.89 | 0.84 | 0.92 | 0.77 | 0.98 | 0.91 |
| 2 | 221210328 | rs10188578 | 0.232 | -1.502 | -0.869 | -0.739 | -0.015 | 0.062 | 0.062 | 0 | 0.42 | 0.64 | 0.47 | 0.08 | 0.48 | 0.27 |
| 2 | 221211949 | rs6747324 | 0.252 | -1.352 | -1.123 | -0.645 | -0.041 | 0.15 | 0.15 | 0 | 0.47 | 0.21 | 0.45 | 0.69 | 0.49 | 0.64 |
| 2 | 221216221 | rs17611441 | 0.432 | -1.18 | -1.026 | -0.501 | 0.01 | 0.016 | 0.016 | 0 | 0.18 | 0.12 | 0.14 | 0.14 | 0.14 | 0.09 |
| 2 | 221218910 | rs12386135 | 1.042 | -1.112 | -1.067 | -0.465 | -0.001 | -0.001 | -0.001 | 0 | 0.11 | 0.11 | 0.1 | 0.13 | 0.09 | 0.04 |
| 2 | 221220210 | rs2068848 | 0.034 | -1.198 | -1.321 | -0.573 | 0.031 | 0.107 | 0.107 | 0.613 | 0.52 | 0.17 | 0.35 | 0.66 | 0.39 | 0.57 |
| 2 | 221223201 | rs11694802 | -1.004 | -1.092 | -0.898 | -0.634 | 0.01 | -0.038 | -0.038 | 0 | 0.67 | 0.44 | 0.55 | 0.86 | 0.57 | 0.79 |
| 2 | 221229840 | rs6741459 | 1.085 | -1.258 | -1.201 | -0.624 | -0.061 | -0.115 | -0.115 | 0 | 0.33 | 0.08 | 0.28 | 0.5 | 0.3 | 0.42 |
| 2 | 221229922 | rs1454535 | -0.85 | -0.804 | -0.633 | -0.56 | 0.027 | -0.071 | -0.071 | 0.017 | 0.67 | 0.48 | 0.55 | 0.86 | 0.57 | 0.79 |
| 2 | 221231339 | rs997701 | -0.7 | -0.765 | -1.183 | -0.531 | -0.015 | 0.025 | 0.025 | 0.054 | 0.66 | 0.34 | 0.53 | 0.86 | 0.57 | 0.77 |
| 2 | 221231486 | rs1961966 | 0.235 | -0.45 | -0.893 | -0.163 | -0.004 | -0.004 | -0.004 | 0 | 0.14 | 0.11 | 0.18 | 0.19 | 0.18 | 0.21 |
| 2 | 221232625 | rs4143572 | 0.355 | -0.488 | -0.988 | -0.261 | 0.032 | 0.016 | 0.016 | 0 | 0.39 | 0.19 | 0.43 | 0.58 | 0.3 | 0.52 |
| 2 | 221234146 | rs1454532 | 0.044 | -0.596 | 0 | 0.032 | -0.027 | -0.046 | -0.046 | 0 | 0.75 | 0.65 | 0.83 | 0.83 | 0.79 | 0.86 |
| 2 | 221239894 | rs13012805 | 0.05 | -0.641 | 0.296 | -0.018 | -0.023 | 0.023 | 0.023 | 0 | 0.15 | 0.35 | 0.15 | 0.17 | 0.11 | 0.19 |
| 2 | 221241860 | rs4674529 | 0.98 | -0.605 | -0.069 | 0.024 | -0.018 | -0.012 | -0.012 | 0 | 0.59 | 0.5 | 0.68 | 0.65 | 0.67 | 0.67 |
| 2 | 221246428 | rs34216516 | 0.207 | -0.039 | -0.455 | 0.331 | 0.003 | -0.007 | -0.007 | 0.522 | 0.13 | 0.11 | 0.13 | 0.17 | 0.11 | 0.18 |
| 2 | 221254709 | rs4674531 | -0.054 | -0.029 | -0.406 | 0.349 | 0.007 | -0.008 | -0.008 | 4.926 | 0.14 | 0.11 | 0.14 | 0.17 | 0.12 | 0.18 |
| 2 | 221259296 | rs4674532 | -0.925 | 0.279 | -0.296 | 0.597 | -0.007 | -0.008 | -0.008 | 0 | 0.73 | 0.74 | 0.66 | 0.6 | 0.61 | 0.75 |
| 2 | 221260058 | rs6721684 | -0.859 | 0.133 | -0.141 | 0.488 | -0.015 | -0.02 | -0.02 | 0 | 0.71 | 0.76 | 0.65 | 0.54 | 0.59 | 0.76 |
| 2 | 221271128 | rs4553816 | 0.331 | -0.309 | -0.322 | 0.266 | 0.001 | 0.046 | 0.046 | 0 | 0.4 | 0.22 | 0.45 | 0.29 | 0.35 | 0.59 |
| 2 | 221281392 | rs6742567 | 1.018 | -0.352 | -0.623 | 0.187 | 0.007 | 0.098 | 0.098 | 0.213 | 0.3 | 0.1 | 0.15 | 0.15 | 0.25 | 0.41 |
| 2 | 221289722 | rs4629147 | 0.561 | -0.324 | 0.067 | 0.214 | 0.01 | 0.02 | 0.02 | 0 | 0.13 | 0.23 | 0.34 | 0.13 | 0.12 | 0.18 |
| 2 | 221290227 | rs2068114 | 0.369 | -0.376 | 0.181 | 0.249 | 0.014 | 0.004 | 0.004 | 0 | 0.6 | 0.35 | 0.51 | 0.61 | 0.68 | 0.68 |
| 2 | 221296802 | rs7578324 | 1.077 | -0.555 | -0.126 | -0.019 | 0.012 | 0.081 | 0.081 | 0 | 0.29 | 0.14 | 0.15 | 0.14 | 0.24 | 0.4 |
| 2 | 221297322 | rs4674538 | -0.426 | -0.449 | -0.047 | 0.072 | -0.001 | 0 | 0 | 0 | 0.74 | 0.71 | 0.86 | 0.74 | 0.8 | 0.86 |
| 2 | 221300772 | rs4549077 | 0.223 | 0.101 | -0.483 | 0.34 | 0.021 | 0.079 | 0.079 | 0.803 | 0.53 | 0.17 | 0.25 | 0.39 | 0.43 | 0.55 |
| 2 | 221312040 | rs4535036 | 0.131 | 0.265 | 0.234 | 0.621 | 0.005 | 0.013 | 0.013 | 0.055 | 0.82 | 0.57 | 0.62 | 0.83 | 0.83 | 0.83 |
| 2 | 221312732 | rs4362550 | -0.909 | 0.346 | -0.545 | 0.906 | -0.083 | -0.06 | -0.06 | 0 | 0.7 | 0.91 | 0.63 | 0.61 | 0.58 | 0.7 |
| 2 | 221313310 | rs4539786 | -0.683 | 0.338 | 0.368 | 0.878 | 0.015 | -0.003 | -0.003 | 0.621 | 0.19 | 0.35 | 0.39 | 0.16 | 0.23 | 0.18 |
| 2 | 221315436 | rs10932864 | -0.067 | -0.389 | -0.12 | 0.892 | -0.013 | 0.005 | 0.005 | 0 | 0.15 | 0.19 | 0.07 | 0.19 | 0.1 | 0.1 |
| 2 | 221320545 | rs13424036 | 0.106 | -0.03 | -0.963 | 0.822 | -0.054 | 0.002 | 0.002 | 0 | 0.14 | 0.02 | 0.26 | 0.13 | 0.15 | 0.09 |
| 2 | 221323389 | rs4592842 | -0.29 | -0.238 | -0.883 | 0.921 | -0.013 | 0.016 | 0.016 | 0 | 0.18 | 0.24 | 0.15 | 0.3 | 0.14 | 0.12 |
| 2 | 221326622 | rs10208119 | -1.071 | -0.226 | -1.128 | 1.215 | -0.005 | -0.069 | -0.069 | 0 | 0.31 | 0.16 | 0.46 | 0.51 | 0.28 | 0.22 |
| 2 | 221326873 | rs7592124 | -1.034 | -0.221 | -1.099 | 0.717 | -0.005 | -0.031 | -0.031 | 0.512 | 0.31 | 0.16 | 0.39 | 0.43 | 0.28 | 0.22 |
| 2 | 221327084 | rs4444513 | -1.055 | -0.219 | -1.167 | 0.718 | -0.005 | -0.032 | -0.032 | 0.48 | 0.31 | 0.16 | 0.39 | 0.43 | 0.28 | 0.22 |
| 2 | 221329785 | rs12613270 | -0.219 | -0.268 | -1.129 | 0.691 | 0.008 | -0.011 | -0.011 | 0 | 0.18 | 0.13 | 0.13 | 0.3 | 0.13 | 0.12 |
| 2 | 221331107 | rs4452127 | 0.885 | -0.431 | -1.073 | 0.416 | -0.003 | -0.019 | -0.019 | 0 | 0.76 | 0.3 | 0.66 | 0.86 | 0.82 | 0.87 |
| 2 | 221339243 | rs4611634 | 0.715 | -0.353 | -0.987 | 0.444 | -0.001 | -0.03 | -0.03 | 0 | 0.77 | 0.35 | 0.65 | 0.86 | 0.79 | 0.86 |

| | | | | | | | | | | | | | | | | |
|---|---|---|---|---|---|---|---|---|---|---|---|---|---|---|---|---|
| 2 | 221345473 | rs7588046 | 1.356 | -0.178 | -1.155 | 0.413 | 0.007 | 0.013 | 0.013 | 0 | 0.58 | 0.24 | 0.48 | 0.6 | 0.61 | 0.71 |
| 2 | 221348373 | rs2708138 | -0.016 | -0.109 | -1.327 | 0.444 | 0 | -0.042 | -0.042 | 0 | 0.18 | 0.05 | 0.16 | 0.26 | 0.17 | 0.16 |
| 2 | 221354872 | rs2068384 | 0.152 | -0.093 | -1.346 | 0.448 | -0.052 | -0.003 | -0.003 | 0 | 0.15 | 0.02 | 0.26 | 0.14 | 0.18 | 0.1 |
| 2 | 221356166 | rs2708129 | -0.148 | -0.067 | -1.294 | 0.472 | 0.004 | -0.023 | -0.023 | 0.339 | 0.19 | 0.09 | 0.17 | 0.26 | 0.18 | 0.16 |
| 2 | 221361301 | rs9789718 | 1.102 | 0.111 | -1.68 | 0.527 | 0.014 | 0.026 | 0.026 | 0 | 0.56 | 0.24 | 0.48 | 0.61 | 0.61 | 0.67 |
| 2 | 221362838 | rs13010568 | 1.032 | -0.061 | -1.638 | 0.555 | 0.019 | 0.024 | 0.024 | 0 | 0.57 | 0.24 | 0.5 | 0.61 | 0.63 | 0.67 |
| 2 | 221364518 | rs1922772 | 0.694 | -0.57 | -1.124 | 0.714 | 0.003 | 0.001 | 0.001 | 0 | 0.72 | 0.3 | 0.79 | 0.75 | 0.82 | 0.78 |
| 2 | 221367957 | rs2602288 | -0.277 | -0.797 | -1.703 | 0.809 | 0.04 | -0.134 | -0.134 | 0 | 0.23 | 0.02 | 0.14 | 0.5 | 0.11 | 0.43 |
| 2 | 221368321 | rs13020102 | 0.663 | -1.351 | -2.48 | 0.204 | -0.012 | 0.074 | 0.074 | 0 | 0.21 | 0.01 | 0.16 | 0.04 | 0.18 | 0.18 |
| 2 | 221392046 | rs10190940 | -0.213 | -1.094 | -1.013 | 0.011 | 0.071 | -0.006 | -0.006 | 0.167 | 0.07 | 0.42 | 0.07 | 0 | 0.07 | 0.03 |
| 2 | 221394310 | rs4234035 | 1.15 | -1.541 | -0.675 | -0.136 | 0.027 | -0.009 | -0.009 | 0 | 0.45 | 0.28 | 0.59 | 0.44 | 0.61 | 0.37 |
| 2 | 221400478 | rs1955401 | -0.237 | -1.267 | -0.748 | 0.02 | -0.013 | 0.124 | 0.124 | 1.084 | 0.78 | 0.51 | 0.79 | 0.49 | 0.83 | 0.65 |
| 2 | 221412389 | rs4674541 | -1.272 | -1.222 | -0.762 | 0.037 | 0.014 | -0.032 | -0.032 | 0 | 0.85 | 0.92 | 0.86 | 0.49 | 0.9 | 0.68 |
| 2 | 221418204 | rs13415553 | -0.556 | -1.144 | -0.696 | -0.141 | -0.008 | -0.101 | -0.101 | 0.048 | 0.15 | 0.49 | 0.09 | 0.01 | 0.09 | 0.13 |
| 2 | 221419951 | rs13422266 | -0.408 | -1.135 | -0.861 | -0.119 | -0.009 | -0.086 | -0.086 | 1.271 | 0.14 | 0.43 | 0.09 | 0.01 | 0.08 | 0.14 |
| 2 | 221421266 | rs16861480 | 0.193 | -1.144 | -0.861 | -0.119 | -0.048 | -0.07 | -0.07 | 0 | 0.12 | 0.41 | 0.04 | 0 | 0.02 | 0.12 |
| 2 | 221423114 | rs17474341 | 1.302 | -0.169 | -1.581 | -0.114 | -0.127 | -0.054 | -0.054 | 0.647 | 0.24 | 0.01 | 0.23 | 0.21 | 0.3 | 0.21 |
| 2 | 221423839 | rs2392716 | 0.252 | -0.795 | -0.824 | -0.126 | 0 | 0.036 | 0.036 | 0 | 0.25 | 0.15 | 0.22 | 0.05 | 0.26 | 0.18 |
| 2 | 221435466 | rs1620682 | -0.727 | -0.367 | -0.395 | 0.714 | -0.042 | 0.053 | 0.053 | 3.56 | 0.31 | 0.52 | 0.23 | 0.49 | 0.19 | 0.47 |
| 2 | 221442745 | rs1712234 | -0.441 | -0.796 | -1.045 | 0.875 | -0.001 | -0.003 | -0.003 | 0 | 0.26 | 0.29 | 0.17 | 0.48 | 0.11 | 0.46 |
| 2 | 221447084 | rs1796432 | -0.477 | -0.042 | -1.404 | 0.519 | 0.01 | 0.048 | 0.048 | 0.007 | 0.4 | 0.24 | 0.46 | 0.29 | 0.38 | 0.49 |
| 2 | 221448538 | rs2708121 | -0.912 | 0.014 | -1.105 | 0.876 | 0.005 | -0.018 | -0.018 | 0 | 0.38 | 0.69 | 0.34 | 0.5 | 0.36 | 0.56 |
| 2 | 221450147 | rs2708123 | 0.121 | 0.015 | -0.567 | 1.116 | -0.007 | 0.037 | 0.037 | 0.645 | 0.17 | 0.2 | 0.17 | 0.35 | 0.14 | 0.37 |
| 2 | 221452342 | rs938730 | 1.496 | -0.464 | -0.51 | 1.069 | 0.014 | 0.011 | 0.011 | 0.151 | 0.52 | 0.42 | 0.46 | 0.39 | 0.56 | 0.35 |
| 2 | 221454938 | rs938731 | 0.475 | -0.567 | -0.565 | 1.071 | -0.047 | -0.054 | -0.054 | 0 | 0.72 | 0.96 | 0.74 | 0.72 | 0.67 | 0.74 |
| 2 | 221455163 | rs938732 | -1.224 | -0.51 | -0.529 | 1.081 | 0.014 | -0.039 | -0.039 | 0 | 0.88 | 0.99 | 0.87 | 0.86 | 0.9 | 0.91 |
| 2 | 221471181 | rs1850492 | 1.109 | -0.378 | -0.409 | 0.908 | 0.003 | 0.004 | 0.004 | 0 | 0.54 | 0.64 | 0.51 | 0.38 | 0.54 | 0.35 |
| 2 | 221479073 | rs1517442 | 0.114 | -0.39 | -0.541 | 1.008 | -0.042 | -0.086 | -0.086 | 0 | 0.72 | 0.95 | 0.68 | 0.61 | 0.67 | 0.74 |
| 2 | 221483430 | rs9288562 | 0.776 | -0.438 | -0.287 | 0.992 | 0.002 | -0.007 | -0.007 | 0 | 0.14 | 0.07 | 0.15 | 0.23 | 0.11 | 0.34 |
| 2 | 221489559 | rs10190825 | 1 | -0.378 | -0.264 | 0.983 | 0.014 | 0.003 | 0.003 | 0 | 0.56 | 0.76 | 0.58 | 0.4 | 0.56 | 0.38 |
| 2 | 221490435 | rs6719602 | 1.134 | -0.401 | -0.208 | 0.983 | 0 | 0 | 0 | 0.069 | 0.54 | 0.57 | 0.56 | 0.39 | 0.54 | 0.35 |
| 2 | 221506737 | rs10187249 | 0.472 | -0.45 | -0.302 | 1.015 | -0.008 | 0.016 | 0.016 | 0.686 | 0.17 | 0.18 | 0.11 | 0.24 | 0.13 | 0.37 |
| 2 | 221509069 | rs1533503 | 1.257 | -0.303 | -0.638 | 0.981 | 0.002 | 0.005 | 0.005 | 0 | 0.52 | 0.46 | 0.53 | 0.36 | 0.52 | 0.34 |
| 2 | 221516241 | rs1400697 | 0.122 | -0.272 | -0.656 | 0.926 | -0.043 | -0.054 | -0.054 | 0 | 0.72 | 0.94 | 0.68 | 0.64 | 0.66 | 0.74 |
| 2 | 221529878 | rs11896343 | 0.771 | -0.097 | -0.379 | 0.764 | 0.001 | 0.003 | 0.003 | 0.027 | 0.52 | 0.47 | 0.53 | 0.38 | 0.52 | 0.35 |
| 2 | 221540944 | rs10932880 | -0.155 | 0.413 | -0.422 | 0.809 | -0.002 | 0 | 0 | 0 | 0.85 | 0.87 | 0.9 | 0.85 | 0.78 | 0.83 |
| 2 | 221541332 | rs17612835 | -0.977 | 0.581 | 0.033 | 1.13 | 0.019 | 0.034 | 0.034 | 0 | 0.14 | 0.22 | 0.35 | 0.35 | 0.19 | 0.4 |
| 2 | 221542433 | rs4674544 | -0.772 | -0.081 | 0.588 | -0.043 | -0.001 | -0.014 | -0.014 | 0 | 0.44 | 0.52 | 0.51 | 0.16 | 0.42 | 0.29 |
| 2 | 221549153 | rs7600191 | 0.077 | -0.343 | 0.854 | -0.045 | 0.016 | 0.026 | 0.026 | 0.081 | 0.28 | 0.16 | 0.27 | 0.16 | 0.21 | 0.23 |
| 2 | 221550136 | rs12694529 | -0.5 | -0.367 | 0.942 | -0.075 | 0 | -0.032 | -0.032 | 0 | 0.14 | 0.13 | 0.15 | 0 | 0.17 | 0.04 |
| 2 | 221553342 | rs11687245 | -0.507 | -0.245 | 1.119 | 0.103 | 0.004 | -0.035 | -0.035 | 0 | 0.07 | 0.11 | 0.1 | 0 | 0.07 | 0.03 |
| 2 | 221556078 | rs7606753 | -1.237 | -0.308 | 1.085 | 0.283 | 0.028 | 0.054 | 0.054 | 0 | 0.49 | 0.72 | 0.55 | 0.34 | 0.41 | 0.35 |
| 2 | 221557049 | rs12477600 | -0.566 | -0.308 | 1.116 | 0.282 | -0.008 | -0.045 | -0.045 | 0 | 0.43 | 0.49 | 0.42 | 0.15 | 0.37 | 0.27 |
| 2 | 221559762 | rs1517443 | -1.011 | -0.288 | 1.082 | 0.272 | 0.037 | 0.078 | 0.078 | 0 | 0.5 | 0.77 | 0.61 | 0.34 | 0.42 | 0.36 |
| 2 | 221567918 | rs16861659 | 0.852 | -0.266 | 0.678 | 0.232 | 0.001 | -0.013 | -0.013 | 0 | 0.09 | 0.07 | 0.03 | 0 | 0.07 | 0.05 |
| 2 | 221576851 | rs12694534 | -0.921 | -0.149 | 0.763 | 0.491 | 0.042 | 0.146 | 0.146 | 0 | 0.46 | 0.77 | 0.51 | 0.14 | 0.37 | 0.27 |
| 2 | 221578398 | rs6705773 | -0.949 | -0.097 | 0.893 | 0.516 | 0.01 | 0.025 | 0.025 | 0 | 0.45 | 0.64 | 0.5 | 0.15 | 0.39 | 0.27 |
| 2 | 221597490 | rs7589343 | -0.985 | -0.289 | 0.897 | 0.798 | 0.024 | 0.004 | 0.004 | 0 | 0.45 | 0.71 | 0.55 | 0.39 | 0.37 | 0.35 |
| 2 | 221607036 | rs6743505 | -0.518 | -0.261 | 0.734 | 0.413 | -0.007 | -0.027 | -0.027 | 0.941 | 0.41 | 0.47 | 0.39 | 0.14 | 0.33 | 0.26 |
| 2 | 221632617 | rs10183570 | -1.051 | -0.229 | -0.242 | 0.426 | 0.037 | 0.04 | 0.04 | 0 | 0.45 | 0.76 | 0.52 | 0.34 | 0.37 | 0.35 |
| 2 | 221643640 | rs7591149 | -1.828 | 0.074 | 0.501 | 1.058 | 0.006 | 0.074 | 0.074 | 0 | 0.13 | 0.37 | 0.2 | 0.18 | 0.1 | 0.12 |
| 2 | 221644928 | rs1509634 | -1.017 | 0.007 | -0.14 | 0.602 | -0.001 | -0.003 | -0.003 | 0 | 0.28 | 0.32 | 0.23 | 0.15 | 0.25 | 0.23 |
| 2 | 221654925 | rs7562441 | -0.107 | 0.503 | -0.61 | 1.001 | -0.017 | -0.028 | -0.028 | 1.602 | 0.8 | 0.84 | 0.75 | 0.61 | 0.69 | 0.64 |
| 2 | 221660614 | rs2893611 | 1.483 | 0.326 | -0.774 | 0.543 | 0.001 | 0.027 | 0.027 | 0.263 | 0.52 | 0.43 | 0.47 | 0.27 | 0.49 | 0.37 |
| 2 | 221662586 | rs11692804 | -0.696 | 0.243 | -0.858 | 0.549 | -0.014 | -0.012 | -0.012 | 0 | 0.27 | 0.34 | 0.22 | 0.17 | 0.24 | 0.21 |
| 2 | 221669822 | rs2136495 | -0.677 | 0.352 | -1.315 | 0.595 | -0.008 | 0.005 | 0.005 | 0.337 | 0.28 | 0.36 | 0.25 | 0.32 | 0.25 | 0.25 |
| 2 | 221671043 | rs1980001 | -0.975 | 0.405 | -1.297 | 0.702 | -0.076 | -0.111 | -0.111 | 0 | 0.86 | 0.99 | 0.78 | 0.59 | 0.75 | 0.65 |
| 2 | 221676683 | rs12694546 | -0.97 | 0.423 | -1.483 | 0.684 | -0.081 | -0.119 | -0.119 | 1.045 | 0.85 | 0.99 | 0.78 | 0.59 | 0.75 | 0.65 |
| 2 | 221678075 | rs10205272 | 1.321 | 0.556 | -1.419 | 0.645 | -0.001 | -0.011 | -0.011 | 0 | 0.47 | 0.46 | 0.48 | 0.28 | 0.44 | 0.37 |
| 2 | 221679980 | rs1396865 | -1.32 | 0.163 | -1.654 | 0.675 | 0.004 | 0 | 0 | 0 | 0.33 | 0.33 | 0.27 | 0.31 | 0.31 | 0.26 |
| 2 | 221685029 | rs6753144 | -0.686 | 0.454 | -2.28 | 0.447 | -0.031 | -0.014 | -0.014 | 1.085 | 0.21 | 0.01 | 0.15 | 0.13 | 0.21 | 0.17 |
| 2 | 221689038 | rs4674556 | 0.008 | 0.659 | -2.472 | 0.148 | -0.065 | 0.127 | 0.127 | 0 | 0.43 | 0.1 | 0.52 | 0.26 | 0.51 | 0.32 |
| 2 | 221691587 | rs10804292 | 0.076 | 0.56 | -2.541 | 0.26 | -0.024 | 0.02 | 0.02 | 0 | 0.41 | 0.16 | 0.49 | 0.36 | 0.44 | 0.44 |
| 2 | 221697522 | rs7600490 | -0.176 | 0.154 | -1.918 | 0.582 | 0.004 | 0.014 | 0.014 | 4.679 | 0.78 | 0.71 | 0.74 | 0.72 | 0.71 | 0.82 |
| 2 | 221703746 | rs1463991 | 0.551 | 0.09 | -1.513 | 0.435 | -0.002 | 0.016 | 0.016 | 0 | 0.43 | 0.32 | 0.52 | 0.37 | 0.43 | 0.43 |
| 2 | 221704514 | rs1463990 | -0.829 | 0.037 | -1.54 | 0.334 | -0.024 | -0.031 | -0.031 | 0 | 0.87 | 0.94 | 0.75 | 0.71 | 0.75 | 0.87 |
| 2 | 221705657 | rs16861825 | 0.617 | -0.178 | -1.672 | 0.425 | -0.078 | -0.039 | -0.039 | 0 | 0.31 | 0.02 | 0.17 | 0.29 | 0.27 | 0.35 |
| 2 | 221709623 | rs1467060 | 0.277 | -0.47 | -1.065 | 0.212 | -0.008 | 0.02 | 0.02 | 0 | 0.41 | 0.23 | 0.51 | 0.36 | 0.43 | 0.43 |
| 2 | 221714058 | rs8179787 | 0.423 | -0.198 | -0.516 | 0.027 | 0 | -0.003 | -0.003 | 0.186 | 0.3 | 0.26 | 0.29 | 0.19 | 0.34 | 0.28 |

| | | | | | | | | | | | | | | | | |
|---|---|---|---|---|---|---|---|---|---|---|---|---|---|---|---|---|
| 2 | 221722531 | rs631486 | -0.091 | -0.144 | -0.239 | 0.043 | 0.014 | 0.063 | 0.063 | 0.858 | 0.61 | 0.25 | 0.65 | 0.76 | 0.64 | 0.71 |
| 2 | 221724838 | rs2444838 | 1.005 | -0.039 | 0.731 | 0.508 | 0.085 | 0.146 | 0.146 | 1.063 | 0.95 | 0.54 | 0.95 | 0.94 | 1 | 0.93 |
| 2 | 221731290 | rs1869565 | -1.032 | -0.148 | 0.209 | 0.361 | -0.005 | -0.004 | -0.004 | 0 | 0.3 | 0.67 | 0.16 | 0.26 | 0.17 | 0.23 |
| 2 | 221743989 | rs679975 | 0.944 | 0.336 | -0.177 | -0.186 | -0.035 | 0.081 | 0.081 | 0 | 0.31 | 0.14 | 0.35 | 0.09 | 0.39 | 0.24 |
| 2 | 221745571 | rs10498097 | 0.548 | 0.282 | 0.316 | 0.155 | -0.003 | 0.016 | 0.016 | 0 | 0.16 | 0.31 | 0.27 | 0.19 | 0.16 | 0.14 |
| 2 | 221745652 | rs664024 | 1.389 | 0.243 | 0.438 | 0.104 | -0.021 | 0.061 | 0.061 | 0 | 0.32 | 0.17 | 0.36 | 0.09 | 0.39 | 0.24 |
| 2 | 221750516 | rs634813 | -1.017 | 0.195 | 0.947 | 0.213 | -0.001 | 0.003 | 0.003 | 0 | 0.73 | 0.74 | 0.77 | 0.52 | 0.72 | 0.6 |
| 2 | 221761576 | rs6713495 | 0.289 | 0.16 | 0.768 | 0.123 | 0.004 | -0.016 | -0.016 | 0 | 0.23 | 0.06 | 0.11 | 0.24 | 0.15 | 0.19 |
| 2 | 221762952 | rs830747 | 0.139 | 0.079 | 1.015 | 0.066 | 0 | -0.001 | -0.001 | 0 | 0.74 | 0.68 | 0.63 | 0.62 | 0.73 | 0.77 |
| 2 | 221768934 | rs597256 | 0.782 | 0.112 | 1.175 | -0.277 | -0.002 | 0.021 | 0.021 | 0.158 | 0.45 | 0.35 | 0.41 | 0.15 | 0.45 | 0.38 |
| 2 | 221768959 | rs605008 | -1.25 | 0.179 | 0.882 | -0.137 | -0.007 | -0.068 | -0.068 | 0 | 0.75 | 0.9 | 0.76 | 0.43 | 0.74 | 0.65 |
| 2 | 221770276 | rs2457408 | -0.177 | 0.215 | 0.883 | -0.359 | 0 | -0.002 | -0.002 | 0 | 0.62 | 0.62 | 0.68 | 0.34 | 0.62 | 0.52 |
| 2 | 221772886 | rs16861897 | -0.65 | 0.385 | 0.738 | -0.153 | 0.001 | -0.001 | -0.001 | 1.505 | 0.14 | 0.12 | 0.04 | 0.06 | 0.08 | 0.14 |
| 2 | 221773296 | rs17339993 | -0.37 | 0.399 | 0.555 | -0.149 | 0.006 | 0.035 | 0.035 | 0 | 0.13 | 0.03 | 0.04 | 0.06 | 0.08 | 0.14 |
| 2 | 221790557 | rs1075304 | -0.924 | 0.439 | 1.076 | 0.021 | -0.001 | -0.007 | -0.007 | 0 | 0.27 | 0.33 | 0.25 | 0.23 | 0.27 | 0.22 |
| 2 | 221798062 | rs1378006 | 1.369 | 0.333 | 0.347 | -0.032 | -0.004 | -0.012 | -0.012 | 0.236 | 0.61 | 0.36 | 0.71 | 0.73 | 0.7 | 0.69 |
| 2 | 221800201 | rs12053297 | -0.917 | 0.264 | -0.556 | -0.056 | -0.039 | 0.033 | 0.033 | 0 | 0.27 | 0.09 | 0.26 | 0.22 | 0.28 | 0.26 |
| 2 | 221802941 | rs617222 | -1.339 | 0.569 | -0.128 | 0.005 | 0.001 | -0.003 | -0.003 | 0 | 0.31 | 0.71 | 0.18 | 0.1 | 0.23 | 0.22 |
| 2 | 221803552 | rs604293 | -1.14 | 0.572 | -0.042 | 0.109 | -0.003 | -0.033 | -0.033 | 0 | 0.27 | 0.69 | 0.16 | 0.1 | 0.21 | 0.19 |
| 2 | 221810486 | rs12465335 | -1.205 | 0.362 | -0.25 | 0.654 | 0.007 | -0.005 | -0.005 | 0 | 0.09 | 0.07 | 0.11 | 0.11 | 0.06 | 0.19 |
| 2 | 221812691 | rs6436229 | 0.399 | 0.057 | 0.409 | 0.803 | -0.022 | 0.018 | 0.018 | 1.536 | 0.76 | 0.57 | 0.73 | 0.72 | 0.81 | 0.7 |
| 2 | 221819192 | rs10932896 | -0.213 | 0 | 0.704 | 0.791 | 0 | -0.002 | -0.002 | 0 | 0.19 | 0.16 | 0.24 | 0.26 | 0.18 | 0.2 |
| 2 | 221826031 | rs1550068 | 1.614 | -0.443 | 0.229 | 0.64 | -0.034 | -0.037 | -0.037 | 0.127 | 0.45 | 0.15 | 0.49 | 0.48 | 0.53 | 0.39 |
| 2 | 221835326 | rs1542355 | 0.094 | -0.487 | 0.904 | 1.326 | -0.017 | 0.077 | 0.077 | 0.721 | 0.23 | 0.52 | 0.27 | 0.4 | 0.19 | 0.2 |
| 2 | 221835642 | rs6733642 | -0.517 | -0.552 | 0.715 | 1.162 | 0.015 | 0.001 | 0.001 | 0 | 0.21 | 0.12 | 0.17 | 0.21 | 0.15 | 0.23 |
| 2 | 221838139 | rs7607818 | -0.279 | -0.588 | 0.479 | 1.215 | 0.012 | -0.027 | -0.027 | 0 | 0.13 | 0.01 | 0.11 | 0.14 | 0.09 | 0.16 |
| 2 | 221844215 | rs6755709 | 2.089 | -0.458 | 0.449 | 1.112 | -0.058 | -0.016 | -0.016 | 0 | 0.44 | 0.12 | 0.48 | 0.37 | 0.49 | 0.38 |
| 2 | 221847734 | rs7579067 | 0.137 | -0.654 | 0.859 | 0.891 | -0.005 | -0.033 | -0.033 | 0 | 0.85 | 0.73 | 0.9 | 0.93 | 0.86 | 0.85 |
| 2 | 221852295 | rs2034747 | -0.454 | -0.679 | 0.416 | 1.65 | 0.02 | -0.097 | -0.097 | 0 | 0.14 | 0.02 | 0.18 | 0.39 | 0.08 | 0.18 |
| 2 | 221860170 | rs1369337 | 2.151 | -0.687 | 0.279 | 1.743 | -0.045 | 0.038 | 0.038 | 1.361 | 0.43 | 0.12 | 0.44 | 0.32 | 0.51 | 0.37 |
| 2 | 221860558 | rs12476074 | -0.4 | -0.765 | 0.884 | 1.652 | 0.008 | -0.023 | -0.023 | 0 | 0.17 | 0.26 | 0.12 | 0.12 | 0.18 | 0.16 |
| 2 | 221862562 | rs897124 | 2.286 | -0.773 | 0.837 | 1.82 | -0.05 | 0.12 | 0.12 | 1.313 | 0.42 | 0.11 | 0.44 | 0.21 | 0.51 | 0.37 |
| 2 | 221864808 | rs10498092 | -0.197 | -0.896 | 0.884 | 1.724 | 0.007 | -0.021 | -0.021 | 0 | 0.17 | 0.25 | 0.11 | 0.13 | 0.17 | 0.14 |
| 2 | 221868801 | rs6756080 | 2.385 | -1.012 | 0.608 | 1.707 | -0.051 | 0.067 | 0.067 | 0.537 | 0.4 | 0.11 | 0.46 | 0.29 | 0.52 | 0.37 |
| 2 | 221876452 | rs1157763 | -0.422 | -1.06 | 0.913 | 1.776 | -0.024 | -0.04 | -0.04 | 0.153 | 0.76 | 0.55 | 0.77 | 0.82 | 0.79 | 0.78 |
| 2 | 221880457 | rs16862036 | -0.869 | -1.109 | 0.961 | 1.72 | -0.005 | 0.01 | 0.01 | 0 | 0.21 | 0.23 | 0.22 | 0.4 | 0.11 | 0.27 |
| 2 | 221890618 | rs12465212 | 0.66 | -0.956 | 0.985 | 1.678 | -0.02 | -0.01 | -0.01 | 0.021 | 0.6 | 0.48 | 0.57 | 0.38 | 0.72 | 0.55 |
| 2 | 221899444 | rs10498091 | -0.665 | -0.794 | 1.164 | 1.409 | 0.003 | -0.019 | -0.019 | 0.015 | 0.17 | 0.27 | 0.12 | 0.13 | 0.16 | 0.15 |
| 2 | 221902399 | rs7596207 | 1.172 | -1.047 | 1.009 | 1.457 | 0.036 | 0.041 | 0.041 | 0.893 | 0.55 | 0.29 | 0.59 | 0.61 | 0.64 | 0.57 |
| 2 | 221903690 | rs2392819 | -0.375 | -2.045 | 1.258 | 1.38 | 0.004 | 0.002 | 0.002 | 0 | 0.73 | 0.78 | 0.73 | 0.75 | 0.81 | 0.75 |
| 2 | 221905185 | rs12471062 | -0.474 | -2.005 | 1.249 | 1.404 | -0.003 | -0.015 | -0.015 | 0 | 0.17 | 0.27 | 0.12 | 0.13 | 0.16 | 0.14 |
| 2 | 221906696 | rs7587424 | 1.359 | -1.988 | 0.929 | 1.38 | 0.067 | 0.07 | 0.07 | 0.43 | 0.51 | 0.22 | 0.59 | 0.61 | 0.64 | 0.54 |
| 2 | 221920025 | rs1025586 | -1.397 | -2.028 | 1.054 | 1.399 | 0.039 | 0.027 | 0.027 | 0.192 | 0.43 | 0.68 | 0.36 | 0.38 | 0.31 | 0.45 |
| 2 | 221920200 | rs11677430 | -0.87 | -1.715 | 1.423 | 1.235 | -0.008 | 0 | 0 | 0 | 0.15 | 0.44 | 0.09 | 0.14 | 0.12 | 0.18 |
| 2 | 221924429 | rs11686456 | -1.32 | -1.731 | 1.021 | 1.191 | -0.019 | -0.037 | -0.037 | 0 | 0.2 | 0.41 | 0.11 | 0.14 | 0.14 | 0.21 |
| 2 | 221925929 | rs1344814 | 1.651 | -2.475 | 0.062 | 1.158 | 0.088 | -0.028 | -0.028 | 4.045 | 0.6 | 0.15 | 0.71 | 0.46 | 0.75 | 0.58 |
| 2 | 221926405 | rs1435179 | 1.13 | -2.707 | 1.273 | 1.343 | -0.067 | 0.075 | 0.075 | 0.049 | 0.74 | 0.48 | 0.79 | 0.48 | 0.87 | 0.69 |
| 2 | 221927852 | rs9808155 | 0.209 | -2.902 | 1.329 | 1.096 | -0.002 | 0.004 | 0.004 | 0 | 0.23 | 0.28 | 0.18 | 0.34 | 0.18 | 0.25 |
| 2 | 221935421 | rs6717445 | -0.222 | -2.939 | 1.577 | 1.04 | -0.005 | -0.025 | -0.025 | 0 | 0.05 | 0.14 | 0.03 | 0 | 0.01 | 0.08 |
| 2 | 221938126 | rs1991966 | -1.166 | -3.055 | 1.686 | 1.2 | -0.048 | 0.109 | 0.109 | 0 | 0.18 | 0.29 | 0.22 | 0.37 | 0.07 | 0.21 |
| 2 | 221938210 | rs1991967 | -0.118 | -3.049 | 1.673 | 1.213 | -0.019 | 0.075 | 0.075 | 0 | 0.13 | 0.16 | 0.2 | 0.37 | 0.06 | 0.12 |
| 2 | 221940361 | rs16862137 | -1.254 | -3.059 | 1.645 | 1.202 | -0.076 | 0.12 | 0.12 | 0 | 0.22 | 0.55 | 0.25 | 0.37 | 0.07 | 0.25 |
| 2 | 221948730 | rs7567692 | 1.019 | -2.84 | 0.423 | 0.7 | 0.073 | 0.104 | 0.104 | 0.13 | 0.58 | 0.16 | 0.57 | 0.25 | 0.75 | 0.51 |
| 2 | 221953533 | rs2392820 | 0.175 | -2.568 | 0.496 | 0.061 | 0 | -0.001 | -0.001 | 0 | 0.28 | 0.32 | 0.25 | 0.28 | 0.28 | 0.2 |
| 2 | 221954500 | rs10498101 | 0.52 | -2.353 | 0.164 | -0.319 | 0.007 | 0.022 | 0.022 | 0.356 | 0.1 | 0.02 | 0.05 | 0 | 0.06 | 0.04 |
| 2 | 221965126 | rs6739121 | -0.408 | -2.396 | 0.823 | -0.124 | -0.051 | 0.008 | 0.008 | 0.178 | 0.18 | 0.41 | 0.13 | 0.16 | 0.08 | 0.14 |
| 2 | 221967358 | rs12694558 | 1.44 | -2.529 | 0.348 | -0.225 | -0.024 | -0.043 | -0.043 | 0 | 0.4 | 0.12 | 0.56 | 0.56 | 0.58 | 0.53 |
| 2 | 221967525 | rs6705296 | -0.487 | -2.382 | 0.584 | -0.278 | -0.013 | -0.006 | -0.006 | 0.135 | 0.21 | 0.24 | 0.16 | 0.17 | 0.13 | 0.25 |
| 2 | 221968989 | rs7564113 | 0.455 | -2.308 | 0.222 | -0.524 | 0 | 0 | 0 | 2.145 | 0.53 | 0.55 | 0.6 | 0.6 | 0.62 | 0.56 |
| 2 | 221971645 | rs1369327 | 1.679 | -2.143 | -0.368 | -0.594 | -0.027 | -0.011 | -0.011 | 0.807 | 0.36 | 0.22 | 0.54 | 0.59 | 0.55 | 0.49 |
| 2 | 221974518 | rs7563641 | 1.256 | -2.047 | -0.233 | -0.571 | 0 | 0.001 | 0.001 | 0 | 0.42 | 0.36 | 0.55 | 0.6 | 0.59 | 0.53 |
| 2 | 221977638 | rs11883646 | 1.264 | -1.811 | -0.244 | -0.311 | -0.054 | -0.056 | -0.056 | 0 | 0.21 | 0.01 | 0.21 | 0.26 | 0.21 | 0.17 |
| 2 | 221987040 | rs7604236 | -0.398 | -1.271 | 0.206 | -0.071 | -0.008 | -0.041 | -0.041 | 0.891 | 0.11 | 0.15 | 0.05 | 0 | 0.07 | 0.07 |
| 2 | 221996939 | rs4527188 | 0.125 | -1.304 | 0.146 | -0.062 | 0.007 | 0.004 | 0.004 | 0.138 | 0.52 | 0.4 | 0.6 | 0.59 | 0.67 | 0.56 |
| 2 | 222000041 | rs6730381 | 0.752 | -1.022 | 0.075 | 0.138 | -0.003 | 0.002 | 0.002 | 0.171 | 0.41 | 0.25 | 0.56 | 0.59 | 0.61 | 0.49 |
| 2 | 222007110 | rs6708587 | -0.661 | -0.544 | 0.594 | 0.391 | -0.012 | 0.014 | 0.014 | 0 | 0.13 | 0.24 | 0.13 | 0.14 | 0.09 | 0.16 |
| 2 | 222013644 | rs4536641 | 0.869 | -0.177 | 0.182 | 0.351 | -0.012 | -0.043 | -0.043 | 0.751 | 0.92 | 0.85 | 0.97 | 1 | 0.95 | 0.94 |
| 2 | 222019583 | rs4672995 | 0.865 | -0.229 | 0.664 | 0.7 | 0.005 | 0.122 | 0.122 | 0 | 0.91 | 0.54 | 0.91 | 0.86 | 0.94 | 0.95 |
| 2 | 222026796 | rs17346918 | 0.36 | -0.363 | -0.075 | 0.125 | 0.004 | 0.098 | 0.098 | 0 | 0.41 | 0.16 | 0.3 | 0.24 | 0.37 | 0.3 |

| | | | | | | | | | | | | | | | | |
|---|---|---|---|---|---|---|---|---|---|---|---|---|---|---|---|---|
| 2 | 222027144 | rs13012474 | 0.193 | -0.2 | -0.568 | 0.353 | -0.009 | -0.021 | -0.021 | 0 | 0.08 | 0 | 0.05 | 0.06 | 0.06 | 0.14 |
| 2 | 222027666 | rs2019083 | 0.156 | -0.357 | -0.346 | 0.23 | -0.045 | 0.011 | 0.011 | 0 | 0.43 | 0.08 | 0.52 | 0.62 | 0.55 | 0.57 |
| 2 | 222027894 | rs16862205 | -0.807 | -0.056 | 0.492 | 0.569 | 0.015 | 0.123 | 0.123 | 0 | 0.09 | 0.41 | 0.09 | 0.13 | 0.06 | 0.05 |
| 2 | 222035912 | rs1864803 | 0.453 | 0.375 | -0.594 | 0.364 | 0.044 | 0.005 | 0.005 | 0 | 0.47 | 0.16 | 0.4 | 0.62 | 0.39 | 0.55 |
| 2 | 222041066 | rs7603664 | 0.089 | 1.28 | 0.665 | 0.15 | -0.001 | -0.003 | -0.003 | 0.005 | 0.55 | 0.5 | 0.51 | 0.64 | 0.54 | 0.7 |
| 2 | 222044974 | rs10084181 | 0.419 | 1.284 | 0.521 | 0.175 | 0.041 | 0.023 | 0.023 | 0 | 0.5 | 0.16 | 0.42 | 0.61 | 0.43 | 0.57 |
| 2 | 222054233 | rs6714776 | -0.002 | 1.508 | 0.801 | 0.083 | -0.001 | -0.001 | -0.001 | 1.114 | 0.29 | 0.29 | 0.36 | 0.21 | 0.25 | 0.32 |
| 2 | 222058767 | rs1583236 | 0.23 | 1.553 | 0.766 | 0.05 | 0.012 | -0.022 | -0.022 | 0 | 0.11 | 0.15 | 0.1 | 0.1 | 0.15 | 0.06 |
| 2 | 222065248 | rs16862242 | 0.011 | 1.262 | 0.771 | 0.101 | 0.002 | 0.003 | 0.003 | 0 | 0.29 | 0.22 | 0.35 | 0.21 | 0.26 | 0.32 |
| 2 | 222067150 | rs12477147 | 0.264 | 1.357 | 0.677 | 0.146 | 0.001 | 0 | 0 | 0 | 0.27 | 0.29 | 0.34 | 0.18 | 0.23 | 0.2 |
| 2 | 222068646 | rs7559847 | 1.477 | 1.68 | 0.921 | 0.224 | 0.091 | -0.048 | -0.048 | 0 | 0.75 | 0.53 | 0.76 | 0.82 | 0.63 | 0.86 |
| 2 | 222068994 | rs2116540 | 0.622 | 0.775 | 0.424 | 0.512 | -0.007 | 0.016 | 0.016 | 0 | 0.91 | 0.74 | 0.93 | 0.93 | 0.94 | 0.96 |
| 2 | 222073239 | rs10932906 | -0.083 | 0.783 | -0.49 | 0.552 | -0.001 | 0.02 | 0.02 | 0.91 | 0.59 | 0.72 | 0.58 | 0.62 | 0.44 | 0.5 |
| 2 | 222074201 | rs10174463 | 0.009 | 0.679 | -0.531 | 0.67 | -0.022 | 0.068 | 0.068 | 0.328 | 0.37 | 0.19 | 0.38 | 0.21 | 0.54 | 0.39 |
| 2 | 222078384 | rs10498102 | 0.254 | 0.849 | 0.193 | 1.114 | -0.164 | 0.057 | 0.057 | 0.361 | 0.33 | 0.05 | 0.33 | 0.21 | 0.46 | 0.38 |
| 2 | 222081456 | rs12476870 | -0.161 | 0.123 | 0.247 | 1.106 | -0.013 | 0.011 | 0.011 | 0.029 | 0.27 | 0.3 | 0.33 | 0.34 | 0.18 | 0.31 |
| 2 | 222088183 | rs16862321 | 0.942 | -0.19 | -0.292 | 1.079 | 0.045 | 0.037 | 0.037 | 4.972 | 0.31 | 0.07 | 0.22 | 0.21 | 0.26 | 0.19 |
| 2 | 222094603 | rs1469623 | 1.049 | -0.228 | -0.339 | 1.04 | 0.02 | 0.109 | 0.109 | 5.275 | 0.29 | 0.04 | 0.2 | 0.08 | 0.25 | 0.15 |
| 2 | 222105363 | rs13029409 | 0.013 | 0.466 | -0.315 | 1.024 | -0.043 | -0.057 | -0.057 | 0 | 0.28 | 0.16 | 0.4 | 0.41 | 0.33 | 0.24 |
| 2 | 222110858 | rs7580353 | 0.684 | 0.331 | 0.212 | 1.023 | 0.014 | -0.004 | -0.004 | 0.804 | 0.36 | 0.42 | 0.35 | 0.34 | 0.42 | 0.38 |
| 2 | 222110876 | rs4508589 | 0.635 | 0.209 | 0.088 | 0.876 | -0.036 | -0.045 | -0.045 | 3.689 | 0.18 | 0.13 | 0.34 | 0.41 | 0.28 | 0.14 |
| 2 | 222111919 | rs1946596 | -1.133 | 0.173 | 0.222 | 1.397 | -0.015 | 0.001 | 0.001 | 4.66 | 0.64 | 0.61 | 0.67 | 0.65 | 0.75 | 0.62 |
| 2 | 222119829 | rs1432254 | 0.578 | 0.235 | 0.306 | 1.098 | -0.003 | 0.002 | 0.002 | 0.431 | 0.38 | 0.64 | 0.37 | 0.35 | 0.42 | 0.38 |
| 2 | 222130384 | rs1002827 | -0.608 | 0.277 | -0.09 | 1.093 | -0.05 | -0.063 | -0.063 | 0 | 0.3 | 0.15 | 0.4 | 0.41 | 0.33 | 0.24 |
| 2 | 222132699 | rs13014378 | -0.602 | 0.331 | -0.664 | 1.116 | -0.054 | -0.07 | -0.07 | 0 | 0.3 | 0.14 | 0.4 | 0.41 | 0.33 | 0.24 |
| 2 | 222150408 | rs6436247 | -0.989 | 0.464 | -0.41 | 1.066 | 0.002 | 0.008 | 0.008 | 3.284 | 0.64 | 0.77 | 0.72 | 0.73 | 0.64 | 0.62 |
| 2 | 222150590 | rs1991736 | -0.235 | 0.359 | -1.147 | 0.995 | 0.003 | 0.075 | 0.075 | 0 | 0.53 | 0.19 | 0.58 | 0.67 | 0.55 | 0.42 |
| 2 | 222150749 | rs4359642 | 0.771 | 0.18 | -1.577 | 0.845 | 0.021 | 0.048 | 0.048 | 0 | 0.1 | 0.04 | 0.06 | 0 | 0.04 | 0.13 |
| 2 | 222151274 | rs12994243 | 1.014 | 0.012 | -1.694 | 0.944 | -0.059 | -0.032 | -0.032 | 0.589 | 0.31 | 0.02 | 0.24 | 0.27 | 0.3 | 0.36 |
| 2 | 222152611 | rs17351792 | 1.323 | 0.129 | -1.731 | 0.829 | -0.047 | 0.03 | 0.03 | 1.387 | 0.27 | 0.02 | 0.22 | 0.19 | 0.26 | 0.18 |
| 2 | 222153932 | rs982757 | 0.738 | 0.21 | -0.675 | 0.921 | -0.009 | -0.01 | -0.01 | 3.573 | 0.62 | 0.38 | 0.5 | 0.54 | 0.58 | 0.6 |
| 2 | 222156270 | rs4672997 | -0.585 | 0.284 | -1.409 | 0.986 | -0.014 | -0.02 | -0.02 | 0.038 | 0.32 | 0.22 | 0.42 | 0.41 | 0.34 | 0.27 |
| 2 | 222167434 | rs4674579 | -0.278 | -0.155 | -1.982 | 0.704 | -0.014 | -0.097 | -0.097 | 0.35 | 0.69 | 0.97 | 0.61 | 0.58 | 0.77 | 0.58 |
| 2 | 222172363 | rs1432251 | -0.548 | -0.28 | -2.434 | 0.283 | -0.047 | -0.055 | -0.055 | 0 | 0.47 | 0.96 | 0.47 | 0.51 | 0.58 | 0.45 |
| 2 | 222174820 | rs1821178 | 0.489 | -0.038 | -2.37 | 0.163 | 0.014 | 0.023 | 0.023 | 0 | 0.11 | 0.04 | 0.06 | 0 | 0.08 | 0.08 |
| 2 | 222175679 | rs10203169 | 0.001 | 0.447 | -1.119 | 0.643 | 0.026 | 0.022 | 0.022 | 0 | 0.26 | 0.36 | 0.29 | 0.32 | 0.44 | 0.25 |
| 2 | 222184640 | rs1430242 | -0.007 | 0.375 | -0.751 | 0.299 | 0.026 | 0.022 | 0.022 | 0 | 0.26 | 0.36 | 0.29 | 0.33 | 0.43 | 0.25 |
| 2 | 222192743 | rs12623432 | -0.301 | 0.523 | -0.447 | 0.313 | 0.08 | 0.082 | 0.082 | 0.678 | 0.27 | 0.48 | 0.29 | 0.37 | 0.41 | 0.27 |
| 2 | 222194001 | rs7608336 | -0.225 | 0.11 | -0.995 | -0.18 | -0.02 | 0.002 | 0.002 | 0 | 0.61 | 0.89 | 0.52 | 0.31 | 0.62 | 0.49 |
| 2 | 222198941 | rs1430237 | -0.194 | 0.081 | -0.499 | -0.216 | -0.001 | -0.002 | -0.002 | 0 | 0.6 | 0.61 | 0.5 | 0.26 | 0.62 | 0.49 |
| 2 | 222199604 | rs919505 | -0.179 | 0.076 | -0.432 | -0.218 | 0 | 0.01 | 0.01 | 0.02 | 0.6 | 0.69 | 0.51 | 0.26 | 0.62 | 0.49 |
| 2 | 222202862 | rs1430230 | -0.158 | 0.077 | -0.477 | -0.218 | -0.003 | -0.01 | -0.01 | 0.282 | 0.59 | 0.57 | 0.49 | 0.26 | 0.62 | 0.49 |
| 2 | 222203245 | rs6436252 | -0.148 | 0.133 | -0.772 | -0.192 | 0 | 0.002 | 0.002 | 0.01 | 0.6 | 0.66 | 0.51 | 0.26 | 0.62 | 0.49 |
| 2 | 222207613 | rs12622686 | -0.055 | 0.16 | -1.466 | -0.318 | 0.026 | 0.013 | 0.013 | 0 | 0.49 | 0.12 | 0.49 | 0.59 | 0.43 | 0.69 |
| 2 | 222209188 | rs10432449 | 0.399 | 0.385 | -0.624 | -0.57 | 0 | -0.001 | -0.001 | 1.249 | 0.3 | 0.35 | 0.33 | 0.2 | 0.33 | 0.22 |
| 2 | 222210212 | rs1430209 | 0.14 | 0.016 | -0.625 | -1.121 | -0.001 | -0.006 | -0.006 | 0 | 0.55 | 0.54 | 0.42 | 0.2 | 0.46 | 0.37 |
| 2 | 222211083 | rs1025369 | 0.744 | 0.092 | -0.76 | -1.046 | -0.034 | -0.114 | -0.114 | 0 | 0.22 | 0.03 | 0.34 | 0.61 | 0.21 | 0.47 |
| 2 | 222213043 | rs6737417 | 0.021 | -0.097 | -0.133 | -1.065 | -0.001 | 0.066 | 0.066 | 0 | 0.56 | 0.76 | 0.43 | 0.2 | 0.46 | 0.37 |
| 2 | 222213174 | rs11687096 | 0.082 | -0.099 | -0.109 | -1.066 | -0.001 | -0.008 | -0.008 | 0 | 0.54 | 0.53 | 0.41 | 0.2 | 0.46 | 0.37 |
| 2 | 222216234 | rs1864463 | 0.963 | 0.317 | 0.123 | -0.827 | 0.032 | 0.025 | 0.025 | 0 | 0.25 | 0.14 | 0.11 | 0 | 0.15 | 0.16 |
| 2 | 222219604 | rs10168570 | 0.45 | 0.074 | 0.167 | -0.781 | 0.001 | -0.001 | -0.001 | 0 | 0.3 | 0.39 | 0.31 | 0.21 | 0.32 | 0.27 |
| 2 | 222223711 | rs10211371 | 0.367 | 0.048 | -0.091 | -0.977 | 0.015 | -0.012 | -0.012 | 0 | 0.3 | 0.51 | 0.35 | 0.2 | 0.33 | 0.26 |
| 2 | 222225453 | rs13411626 | 0.849 | 0.083 | -1.083 | -1.051 | -0.027 | -0.068 | -0.068 | 0.107 | 0.15 | 0.03 | 0.31 | 0.62 | 0.18 | 0.37 |
| 2 | 222228975 | rs12998099 | -0.097 | -0.173 | -0.942 | -1.157 | -0.008 | 0.094 | 0.094 | 0 | 0.55 | 0.81 | 0.47 | 0.2 | 0.47 | 0.44 |
| 2 | 222229386 | rs16862590 | 1.204 | -0.213 | -1.203 | -1.047 | 0.052 | 0.06 | 0.06 | 0 | 0.23 | 0.1 | 0.11 | 0 | 0.14 | 0.17 |
| 2 | 222241514 | rs11689951 | 0.649 | 0.511 | -1.499 | -0.882 | -0.068 | -0.061 | -0.061 | 0 | 0.25 | 0.04 | 0.28 | 0.22 | 0.27 | 0.29 |
| 2 | 222250513 | rs2195614 | 0.089 | 0.345 | -1.02 | -1.096 | 0.036 | 0.09 | 0.09 | 0 | 0.52 | 0.28 | 0.42 | 0.22 | 0.43 | 0.48 |
| 2 | 222251031 | rs6714968 | -0.798 | 0.6 | -1.015 | -1.105 | -0.019 | 0.04 | 0.04 | 1.528 | 0.71 | 0.85 | 0.78 | 0.84 | 0.67 | 0.85 |
| 2 | 222259245 | rs2217431 | 0.024 | 0.181 | -0.174 | -1.018 | 0.005 | 0.009 | 0.009 | 0 | 0.53 | 0.56 | 0.43 | 0.22 | 0.43 | 0.48 |
| 2 | 222261724 | rs10196742 | 0.722 | 0.109 | -0.264 | -0.97 | 0 | 0.006 | 0.006 | 0 | 0.23 | 0.23 | 0.36 | 0.57 | 0.22 | 0.33 |
| 2 | 222264036 | rs6745962 | -0.193 | 0.015 | -0.249 | -1.093 | 0.019 | 0.036 | 0.036 | 0 | 0.52 | 0.61 | 0.38 | 0.22 | 0.42 | 0.47 |
| 2 | 222276477 | rs2052941 | 0.386 | -0.039 | -0.265 | -1.059 | -0.008 | -0.01 | -0.01 | 1.094 | 0.28 | 0.15 | 0.26 | 0.22 | 0.28 | 0.29 |
| 2 | 222277888 | rs2069212 | -0.48 | 0.158 | -0.383 | -1.16 | -0.005 | 0.023 | 0.023 | 0 | 0.57 | 0.62 | 0.42 | 0.22 | 0.47 | 0.46 |
| 2 | 222280501 | rs12996275 | 0.543 | -0.445 | -0.515 | -0.995 | -0.009 | -0.015 | -0.015 | 0 | 0.25 | 0.11 | 0.25 | 0.22 | 0.27 | 0.24 |
| 2 | 222282959 | rs8508 | -0.533 | -0.433 | -0.757 | -1.01 | 0.021 | 0.049 | 0.049 | 0 | 0.55 | 0.65 | 0.39 | 0.22 | 0.42 | 0.44 |
| 2 | 222283281 | rs3087584 | -0.544 | -0.441 | -0.761 | -1.014 | 0.02 | 0.048 | 0.048 | 0 | 0.55 | 0.65 | 0.39 | 0.22 | 0.42 | 0.44 |
| 2 | 222283504 | rs957466 | -0.533 | -0.425 | -0.753 | -1.006 | 0.021 | 0.049 | 0.049 | 0 | 0.55 | 0.65 | 0.39 | 0.22 | 0.42 | 0.44 |
| 2 | 222283514 | rs957465 | -0.533 | -0.425 | -0.753 | -1.006 | 0.021 | 0.049 | 0.049 | 0 | 0.55 | 0.65 | 0.39 | 0.22 | 0.42 | 0.44 |
| 2 | 222284277 | rs17370971 | 0.945 | -0.561 | -0.779 | -1.06 | 0.069 | 0.099 | 0.099 | 0 | 0.26 | 0.1 | 0.11 | 0 | 0.15 | 0.21 |

| | | | | | | | | | | | | | | | | |
|---|---|---|---|---|---|---|---|---|---|---|---|---|---|---|---|---|
| 2 | 222284646 | rs3770208 | -0.583 | -0.465 | -0.743 | -0.986 | 0.023 | 0.045 | 0.045 | 0 | 0.55 | 0.64 | 0.39 | 0.26 | 0.42 | 0.44 |
| 2 | 222285699 | rs16862630 | 0.253 | -0.261 | -1.153 | -0.787 | 0.035 | 0.026 | 0.026 | 0 | 0.28 | 0.23 | 0.13 | 0.04 | 0.15 | 0.21 |
| 2 | 222290027 | rs10196918 | 0.055 | 0.413 | -1.459 | -0.19 | -0.038 | -0.044 | -0.044 | 0 | 0.25 | 0.08 | 0.25 | 0.26 | 0.28 | 0.3 |
| 2 | 222290902 | rs2303901 | -0.309 | 0.588 | -1.175 | -0.478 | -0.031 | 0.213 | 0.213 | 0.107 | 0.52 | 0.84 | 0.31 | 0.15 | 0.48 | 0.39 |
| 2 | 222294268 | rs10211091 | -0.414 | 0.359 | -0.426 | -0.434 | -0.031 | -0.143 | -0.143 | 1.815 | 0.74 | 0.54 | 0.83 | 1 | 0.8 | 0.85 |
| 2 | 222295760 | rs3770205 | 0.441 | 0.309 | -0.897 | -0.34 | 0.007 | 0.032 | 0.032 | 0 | 0.46 | 0.25 | 0.55 | 0.6 | 0.45 | 0.55 |
| 2 | 222296320 | rs3770204 | 0.389 | 0.373 | -1.568 | -0.309 | -0.01 | 0.042 | 0.042 | 0 | 0.45 | 0.12 | 0.54 | 0.61 | 0.45 | 0.55 |
| 2 | 222297107 | rs4340507 | 0.077 | 0.01 | -0.869 | -0.301 | 0.009 | -0.017 | -0.017 | 0 | 0.2 | 0.17 | 0.11 | 0 | 0.11 | 0.13 |
| 2 | 222299999 | rs3770199 | 0.208 | 0.468 | -0.783 | -0.224 | 0.001 | -0.006 | -0.006 | 0 | 0.71 | 0.68 | 0.77 | 0.95 | 0.73 | 0.77 |
| 2 | 222300116 | rs3770197 | -0.835 | 0.076 | -0.592 | -0.215 | -0.004 | -0.016 | -0.016 | 0.125 | 0.38 | 0.42 | 0.25 | 0.17 | 0.3 | 0.38 |
| 2 | 222301357 | rs3213844 | 0.525 | -0.071 | -0.531 | 0.197 | 0.02 | -0.006 | -0.006 | 0 | 0.61 | 0.29 | 0.57 | 0.6 | 0.7 | 0.68 |
| 2 | 222301713 | rs2303900 | 1.059 | 0.582 | -1.249 | -0.015 | 0.011 | -0.009 | -0.009 | 0 | 0.19 | 0.23 | 0.15 | 0.13 | 0.24 | 0.1 |
| 2 | 222307310 | rs2288629 | -1.002 | 0.343 | -0.338 | -0.163 | -0.006 | -0.054 | -0.054 | 0 | 0.75 | 0.73 | 0.82 | 0.98 | 0.78 | 0.9 |
| 2 | 222307534 | rs2288628 | 1.049 | 0.362 | 0.251 | -0.118 | 0.035 | -0.025 | -0.025 | 0 | 0.19 | 0.3 | 0.15 | 0.13 | 0.24 | 0.1 |
| 2 | 222308844 | rs16862666 | 0.58 | 0.319 | 0.439 | 0.03 | 0.046 | 0.023 | 0.023 | 0 | 0.22 | 0.55 | 0.25 | 0.22 | 0.25 | 0.18 |
| 2 | 222314482 | rs3770178 | -0.722 | 0.255 | 0.13 | 0.011 | -0.024 | -0.045 | -0.045 | 0 | 0.73 | 0.55 | 0.72 | 0.89 | 0.77 | 0.82 |
| 2 | 222314806 | rs13392880 | 0.322 | 0.229 | 0.186 | 0.152 | -0.006 | 0.101 | 0.101 | 0 | 0.11 | 0.24 | 0.16 | 0.32 | 0.06 | 0.18 |
| 2 | 222315958 | rs12476016 | 1.474 | 0.194 | -0.744 | 0.099 | -0.03 | 0.017 | 0.017 | 0 | 0.14 | 0.02 | 0.1 | 0.07 | 0.16 | 0.17 |
| 2 | 222316575 | rs2052942 | -0.351 | 0.161 | -0.518 | 0.186 | -0.015 | 0.01 | 0.01 | 0 | 0.63 | 0.52 | 0.59 | 0.58 | 0.71 | 0.65 |
| 2 | 222317545 | rs3770176 | 1.59 | 0.279 | -1.434 | 0.198 | -0.083 | 0.025 | 0.025 | 0.116 | 0.17 | 0.03 | 0.12 | 0.11 | 0.24 | 0.1 |
| 2 | 222318513 | rs4674599 | 0.606 | 0.236 | -1.394 | 0.215 | -0.091 | -0.122 | -0.122 | 0 | 0.41 | 0.05 | 0.42 | 0.46 | 0.47 | 0.54 |
| 2 | 222325139 | rs919503 | 1.017 | 0.688 | -1.226 | 0.006 | -0.098 | 0.022 | 0.022 | 0.068 | 0.19 | 0.02 | 0.15 | 0.13 | 0.27 | 0.08 |
| 2 | 222330122 | rs16862708 | 0.977 | 0.441 | -1.501 | 0.327 | -0.008 | -0.039 | -0.039 | 0 | 0.17 | 0.05 | 0.27 | 0.24 | 0.2 | 0.21 |
| 2 | 222337626 | rs975194 | -0.909 | 0.099 | -0.876 | 0.378 | -0.01 | -0.025 | -0.025 | 0 | 0.68 | 0.69 | 0.81 | 0.86 | 0.75 | 0.71 |
| 2 | 222340239 | rs12463452 | 0.068 | 0.141 | -0.755 | 0.434 | 0.002 | -0.001 | -0.001 | 0.009 | 0.49 | 0.54 | 0.46 | 0.53 | 0.54 | 0.39 |
| 2 | 222344196 | rs3770146 | 0.261 | 0.377 | -0.492 | 0.58 | -0.002 | -0.003 | -0.003 | 0.174 | 0.37 | 0.4 | 0.38 | 0.54 | 0.4 | 0.37 |
| 2 | 222352629 | rs10498113 | 0.201 | 0.349 | -0.41 | 0.587 | 0.004 | 0.009 | 0.009 | 0.01 | 0.38 | 0.45 | 0.39 | 0.54 | 0.4 | 0.4 |
| 2 | 222357235 | rs6436266 | -1.087 | 0.372 | -0.835 | 0.32 | -0.039 | 0.147 | 0.147 | 0 | 0.59 | 0.89 | 0.75 | 0.86 | 0.59 | 0.69 |
| 2 | 222358909 | rs10498114 | 1.569 | 0.534 | -0.677 | 0.476 | -0.041 | -0.026 | -0.026 | 0 | 0.09 | 0.01 | 0.08 | 0.13 | 0.13 | 0.07 |
| 2 | 222366631 | rs13408240 | 0.192 | 0.372 | -0.184 | 0.452 | -0.018 | 0.037 | 0.037 | 1.638 | 0.27 | 0.33 | 0.29 | 0.38 | 0.23 | 0.4 |
| 2 | 222369870 | rs7575433 | -0.792 | 0.671 | -0.645 | 0.231 | -0.016 | 0.071 | 0.071 | 0 | 0.66 | 0.83 | 0.76 | 1 | 0.63 | 0.87 |
| 2 | 222377409 | rs10498115 | 2.268 | 0.87 | -1.012 | 0.362 | -0.038 | -0.031 | -0.031 | 0.265 | 0.06 | 0.01 | 0.1 | 0.13 | 0.13 | 0.07 |
| 2 | 222378256 | rs2710505 | -0.945 | 0.749 | -1.077 | 0.232 | -0.044 | 0.167 | 0.167 | 0 | 0.66 | 0.91 | 0.77 | 1 | 0.63 | 0.87 |
| 2 | 222378994 | rs9288570 | -0.252 | 0.516 | -1.029 | 0.308 | -0.005 | 0.002 | 0.002 | 0 | 0.28 | 0.66 | 0.31 | 0.36 | 0.21 | 0.36 |
| 2 | 222379489 | rs2710506 | 1.201 | 0.512 | -1.232 | 0.41 | -0.097 | -0.096 | -0.096 | 0.018 | 0.36 | 0.1 | 0.36 | 0.56 | 0.41 | 0.42 |
| 2 | 222380888 | rs2710508 | -0.48 | 0.498 | -1.19 | 0.467 | -0.041 | 0.115 | 0.115 | 0 | 0.58 | 0.83 | 0.59 | 0.76 | 0.48 | 0.72 |
| 2 | 222383983 | rs2710510 | -1.656 | 0.727 | -1.309 | 0.695 | -0.019 | -0.101 | -0.101 | 0 | 0.71 | 0.92 | 0.74 | 0.59 | 0.72 | 0.64 |
| 2 | 222393635 | rs4613266 | -1.194 | 1.031 | -0.624 | 1.059 | -0.011 | 0.047 | 0.047 | 0 | 0.52 | 0.71 | 0.55 | 0.37 | 0.54 | 0.39 |
| 2 | 222394305 | rs4302206 | -1.885 | 1.065 | -0.672 | 1.085 | 0.037 | -0.042 | -0.042 | 0 | 0.61 | 0.85 | 0.71 | 0.59 | 0.69 | 0.54 |
| 2 | 222400223 | rs2250204 | -1.916 | 1.033 | -0.695 | 1.089 | 0.025 | -0.051 | -0.051 | 0 | 0.61 | 0.86 | 0.71 | 0.59 | 0.69 | 0.53 |
| 2 | 222401086 | rs16862811 | 1.461 | 1.15 | -0.714 | 1.133 | -0.021 | -0.019 | -0.019 | 0 | 0.07 | 0.03 | 0.07 | 0.14 | 0.14 | 0.06 |
| 2 | 222408278 | rs10191992 | 0.441 | 0.861 | -0.264 | 0.974 | 0.009 | 0.035 | 0.035 | 0 | 0.08 | 0.21 | 0.06 | 0.07 | 0.05 | 0.13 |
| 2 | 222410331 | rs2680856 | -1.325 | 0.905 | -0.943 | 1.065 | -0.022 | -0.001 | -0.001 | 0 | 0.54 | 0.77 | 0.63 | 0.44 | 0.54 | 0.47 |
| 2 | 222410399 | rs6716153 | 0.312 | 0.508 | -0.467 | 1.453 | -0.004 | -0.045 | -0.045 | 1.143 | 0.08 | 0 | 0.17 | 0.27 | 0.1 | 0.11 |
| 2 | 222421725 | rs10932919 | -0.548 | 0.586 | 0.432 | 1.479 | 0.02 | 0.239 | 0.239 | 0 | 0.07 | 0.24 | 0.27 | 0.37 | 0.08 | 0.17 |
| 2 | 222423389 | rs10498118 | -1.095 | 0.955 | 0.215 | 1.426 | 0.052 | 0.145 | 0.145 | 0 | 0.18 | 0.38 | 0.39 | 0.5 | 0.27 | 0.26 |
| 2 | 222424544 | rs10498119 | 0.611 | 0.99 | 0.058 | 1.352 | -0.049 | -0.02 | -0.02 | 4.25 | 0.1 | 0.04 | 0.11 | 0.13 | 0.19 | 0.08 |
| 2 | 222427871 | rs2056290 | -2.033 | 0.719 | 0.024 | 1.001 | -0.007 | 0.029 | 0.029 | 0 | 0.54 | 0.78 | 0.61 | 0.63 | 0.52 | 0.45 |
| 2 | 222430983 | rs3770143 | 0.84 | 0.729 | 0.347 | 1.075 | -0.023 | -0.023 | -0.023 | 0.073 | 0.05 | 0.02 | 0.07 | 0.13 | 0.15 | 0.08 |
| 2 | 222432320 | rs7340471 | -0.333 | 0.258 | 0.666 | 0.774 | 0.004 | -0.026 | -0.026 | 0.112 | 0.27 | 0.18 | 0.15 | 0 | 0.24 | 0.09 |
| 2 | 222438968 | rs2248489 | 1.324 | -0.048 | 1.008 | 1.32 | -0.003 | -0.001 | -0.001 | 0.029 | 0.56 | 0.45 | 0.52 | 0.42 | 0.62 | 0.61 |
| 2 | 222448817 | rs2710489 | 1.772 | 0.186 | 1.03 | 1.391 | 0.046 | 0.033 | 0.033 | 0 | 0.49 | 0.28 | 0.44 | 0.42 | 0.58 | 0.57 |
| 2 | 222450720 | rs2946559 | -0.378 | 0.208 | 0.906 | 1.383 | 0 | 0 | 0 | 0 | 0.89 | 0.88 | 0.91 | 0.89 | 0.9 | 0.88 |
| 2 | 222464939 | rs2710530 | -0.564 | 0.23 | 0.744 | 1.434 | 0.001 | -0.005 | -0.005 | 0 | 0.9 | 0.91 | 0.93 | 0.84 | 0.91 | 0.88 |
| 2 | 222465766 | rs2680835 | 1.891 | 0.189 | 0.683 | 1.355 | 0.03 | 0.064 | 0.064 | 0.389 | 0.47 | 0.15 | 0.48 | 0.37 | 0.58 | 0.57 |
| 2 | 222469594 | rs2710526 | 1.335 | 0.523 | 1.235 | 1.581 | -0.027 | 0.163 | 0.163 | 0 | 0.81 | 0.59 | 0.66 | 0.52 | 0.84 | 0.72 |
| 2 | 222471516 | rs2680837 | 0.094 | 0.643 | 0.784 | 1.478 | -0.006 | 0.001 | 0.001 | 0 | 0.88 | 0.87 | 0.92 | 0.81 | 0.93 | 0.89 |
| 2 | 222473631 | rs2710521 | 1.946 | 0.9 | 1.076 | 1.467 | -0.003 | 0.001 | 0.001 | 0.141 | 0.52 | 0.44 | 0.48 | 0.52 | 0.51 | 0.64 |
| 2 | 222474346 | rs2710519 | 0.01 | 1.178 | 1.326 | 1.657 | 0.009 | -0.006 | -0.006 | 0 | 0.87 | 0.9 | 0.88 | 0.82 | 0.91 | 0.87 |
| 2 | 222478526 | rs13403047 | -0.317 | 1.296 | 1.404 | 1.361 | -0.006 | -0.016 | -0.016 | 0 | 0.17 | 0.12 | 0.12 | 0 | 0.19 | 0.08 |
| 2 | 222479220 | rs17300372 | -0.207 | 1.296 | 1.391 | 1.362 | -0.02 | 0.022 | 0.022 | 0 | 0.16 | 0.07 | 0.12 | 0 | 0.19 | 0.08 |
| 2 | 222479303 | rs10804299 | 0.601 | 1.563 | 1.486 | 1.557 | -0.009 | -0.026 | -0.026 | 0 | 0.79 | 0.87 | 0.77 | 0.63 | 0.75 | 0.77 |
| 2 | 222482917 | rs872217 | 0.825 | 1.565 | 1.752 | 1.558 | 0.012 | 0.055 | 0.055 | 0 | 0.77 | 0.69 | 0.74 | 0.63 | 0.74 | 0.77 |
| 2 | 222491734 | rs6748185 | -0.121 | 1.513 | 1.027 | 1.526 | -0.021 | 0.021 | 0.021 | 0 | 0.16 | 0.07 | 0.12 | 0 | 0.19 | 0.08 |
| 2 | 222493429 | rs2176694 | -0.881 | 1.293 | 1.014 | 1.744 | 0.004 | 0.136 | 0.136 | 0 | 0.05 | 0.2 | 0.19 | 0.2 | 0.06 | 0.08 |
| 2 | 222493572 | rs2139351 | -1.716 | 1.265 | 1.06 | 1.734 | -0.049 | 0.008 | 0.008 | 0 | 0.46 | 0.79 | 0.58 | 0.57 | 0.5 | 0.39 |
| 2 | 222512569 | rs2037327 | -1.332 | 0.642 | 0.857 | 2.286 | -0.003 | 0.002 | 0.002 | 0 | 0.17 | 0.16 | 0.18 | 0.28 | 0.13 | 0.16 |
| 2 | 222514104 | rs16862912 | -0.84 | 0.682 | 0.878 | 2.306 | -0.007 | 0.013 | 0.013 | 0 | 0.15 | 0.16 | 0.18 | 0.27 | 0.13 | 0.16 |
| 2 | 222519213 | rs16862915 | -1.542 | 0.751 | 0.757 | 2.277 | -0.014 | 0.152 | 0.152 | 0 | 0.24 | 0.45 | 0.41 | 0.48 | 0.21 | 0.26 |

| | | | | | | | | | | | | | | | | |
|---|---|---|---|---|---|---|---|---|---|---|---|---|---|---|---|---|
| 2 | 222519992 | rs7609471 | -1.028 | 0.492 | -0.38 | 1.646 | 0.033 | -0.007 | -0.007 | 0.547 | 0.26 | 0.14 | 0.2 | 0.28 | 0.17 | 0.2 |
| 2 | 222522255 | rs825282 | -1.231 | 1.029 | 0.323 | 2.071 | -0.01 | 0.054 | 0.054 | 0 | 0.21 | 0.28 | 0.28 | 0.43 | 0.2 | 0.18 |
| 2 | 222532318 | rs11682490 | -1.46 | 1.244 | 1.526 | 1.877 | -0.014 | 0.001 | 0.001 | 0 | 0.08 | 0.18 | 0.06 | 0.05 | 0.06 | 0.13 |
| 2 | 222533936 | rs13420950 | -2.07 | 1.25 | 1.74 | 1.536 | -0.021 | -0.064 | -0.064 | 0.152 | 0.18 | 0.32 | 0.13 | 0.05 | 0.13 | 0.2 |
| 2 | 222538149 | rs17374455 | -0.797 | 1.27 | 1.661 | 1.545 | 0.002 | 0.001 | 0.001 | 0.371 | 0.1 | 0.07 | 0.06 | 0 | 0.08 | 0.08 |
| 2 | 222540194 | rs6742877 | -1.229 | 1.232 | 1.712 | 1.545 | -0.014 | 0.002 | 0.002 | 0 | 0.08 | 0.21 | 0.07 | 0.05 | 0.05 | 0.12 |
| 2 | 222542378 | rs10209807 | -2.492 | 1.758 | 1.624 | 2.593 | -0.028 | -0.06 | -0.06 | 0 | 0.34 | 0.82 | 0.48 | 0.49 | 0.32 | 0.44 |
| 2 | 222547283 | rs12053343 | -1.468 | 1.943 | 1.886 | 2.776 | -0.007 | 0.069 | 0.069 | 0 | 0.07 | 0.06 | 0.27 | 0.41 | 0.11 | 0.14 |
| 2 | 222550973 | rs1519486 | -1.416 | 1.881 | 2.084 | 2.512 | 0.056 | 0.008 | 0.008 | 0 | 0.11 | 0.25 | 0.24 | 0.12 | 0.17 | 0.15 |
| 2 | 222553382 | rs825288 | 1.93 | 1.389 | 2.18 | 2.89 | 0.003 | 0.103 | 0.103 | 0 | 0.92 | 0.89 | 0.77 | 0.44 | 0.92 | 0.92 |
| 2 | 222562778 | rs825277 | 1.498 | 0.999 | 2.977 | 2.473 | 0.038 | 0.253 | 0.253 | 0.09 | 0.95 | 0.63 | 0.86 | 0.84 | 0.98 | 0.97 |
| 2 | 222572601 | rs825269 | 1.464 | 1.001 | 3.051 | 2.469 | 0.016 | 0.216 | 0.216 | 1.385 | 0.94 | 0.67 | 0.86 | 0.84 | 0.98 | 0.98 |
| 2 | 222585070 | rs825262 | 2.218 | 0.909 | 3.596 | 2.423 | -0.009 | 0.205 | 0.205 | 4.403 | 0.91 | 0.47 | 0.84 | 0.84 | 0.98 | 0.97 |
| 2 | 222624677 | rs1364386 | 2.098 | 1.081 | 1.76 | 2.45 | 0.011 | -0.13 | -0.13 | 0 | 0.67 | 0.09 | 0.58 | 0.59 | 0.72 | 0.72 |
| 2 | 222636828 | rs11893149 | -1.339 | 1.025 | 2.207 | 2.449 | -0.027 | 0.003 | 0.003 | 0.019 | 0.09 | 0.15 | 0.09 | 0.07 | 0.04 | 0.15 |
| 2 | 222653136 | rs1424339 | 1.47 | 1.003 | 1.475 | 2.529 | -0.073 | 0.094 | 0.094 | 1.412 | 0.84 | 0.39 | 0.74 | 0.78 | 0.94 | 0.81 |
| 2 | 222660766 | rs16863084 | -1.273 | 1.675 | 0.37 | 2.462 | -0.046 | -0.044 | -0.044 | 3.665 | 0.07 | 0 | 0.16 | 0.15 | 0.15 | 0.05 |
| 2 | 222681528 | rs10210217 | -1.606 | 1.467 | 0.463 | 1.821 | 0 | 0.001 | 0.001 | 2.86 | 0.21 | 0.74 | 0.25 | 0.16 | 0.18 | 0.17 |
| 2 | 222682936 | rs2551201 | -0.961 | 1.477 | -0.885 | 1.64 | -0.011 | 0.026 | 0.026 | 0 | 0.54 | 0.86 | 0.57 | 0.58 | 0.55 | 0.5 |
| 2 | 222683948 | rs1424341 | -0.735 | 1.885 | 0.358 | 1.747 | 0.006 | -0.003 | -0.003 | 0 | 0.48 | 0.58 | 0.41 | 0.46 | 0.42 | 0.47 |
| 2 | 222685565 | rs2161765 | -0.409 | 2.135 | 0.646 | 1.499 | 0.037 | -0.025 | -0.025 | 0 | 0.37 | 0.25 | 0.31 | 0.43 | 0.24 | 0.28 |
| 2 | 222689249 | rs16863122 | -0.869 | 2.303 | 0.769 | 1.246 | 0.107 | -0.015 | -0.015 | 0 | 0.1 | 0.36 | 0.1 | 0.03 | 0.15 | 0.18 |
| 2 | 222692046 | rs1955086 | -1.269 | 1.901 | 1.232 | 1.474 | 0.019 | -0.016 | -0.016 | 0 | 0.23 | 0.45 | 0.22 | 0.17 | 0.25 | 0.3 |
| 2 | 222695048 | rs13405578 | -1.356 | 1.865 | 0.985 | 1.477 | 0.016 | 0.102 | 0.102 | 0 | 0.13 | 0.42 | 0.16 | 0.13 | 0.09 | 0.03 |
| 2 | 222713878 | rs6706361 | 0.301 | 1.829 | 1.4 | 1.351 | 0.039 | -0.031 | -0.031 | 0 | 0.13 | 0.04 | 0.09 | 0.13 | 0.04 | 0.1 |
| 2 | 222716083 | rs4099413 | 0.011 | 1.645 | 0.775 | 1.433 | -0.031 | -0.014 | -0.014 | 0 | 0.19 | 0.08 | 0.21 | 0.24 | 0.23 | 0.24 |
| 2 | 222717778 | rs4674616 | -2.451 | 1.765 | 0.9 | 1.481 | -0.013 | -0.019 | -0.019 | 0 | 0.49 | 0.78 | 0.5 | 0.44 | 0.47 | 0.37 |
| 2 | 222721065 | rs10932926 | 0.625 | 1.71 | 0.85 | 1.405 | 0.037 | -0.031 | -0.031 | 0 | 0.14 | 0.04 | 0.09 | 0.13 | 0.04 | 0.1 |
| 2 | 222729505 | rs4084367 | -1.011 | 1.747 | 0.92 | 1.391 | 0.011 | -0.014 | -0.014 | 0 | 0.34 | 0.25 | 0.31 | 0.37 | 0.27 | 0.33 |
| 2 | 222733842 | rs10932929 | -0.934 | 1.749 | 1.013 | 1.407 | 0.009 | 0.085 | 0.085 | 0 | 0.13 | 0.39 | 0.16 | 0.13 | 0.09 | 0.02 |
| 2 | 222737511 | rs4085398 | -0.832 | 1.745 | 0.873 | 1.408 | -0.001 | -0.047 | -0.047 | 0.424 | 0.33 | 0.13 | 0.3 | 0.37 | 0.27 | 0.33 |
| 2 | 222739011 | rs4099418 | -2.249 | 1.808 | 0.84 | 1.428 | -0.024 | 0.018 | 0.018 | 0.15 | 0.5 | 0.85 | 0.54 | 0.5 | 0.47 | 0.37 |
| 2 | 222755728 | rs17312759 | -0.096 | 1.946 | 1.124 | 1.361 | -0.008 | 0.001 | 0.001 | 0 | 0.21 | 0.16 | 0.22 | 0.23 | 0.23 | 0.3 |
| 2 | 222763536 | rs10172187 | -1.284 | 1.652 | 1.199 | 1.275 | 0.045 | 0.098 | 0.098 | 0 | 0.11 | 0.4 | 0.16 | 0.12 | 0.1 | 0.02 |
| 2 | 222782424 | rs16825445 | -1.506 | 1.689 | 1.327 | 1.258 | 0.049 | 0.096 | 0.096 | 0 | 0.11 | 0.38 | 0.15 | 0.11 | 0.1 | 0.02 |
| 2 | 222783386 | rs16863167 | -1.197 | 1.694 | 1.362 | 1.376 | 0.06 | 0.151 | 0.151 | 0 | 0.09 | 0.3 | 0.14 | 0.15 | 0.1 | 0.02 |
| 2 | 222785645 | rs824918 | 0.028 | 1.621 | 1.162 | 1.218 | -0.016 | -0.003 | -0.003 | 0 | 0.79 | 0.73 | 0.83 | 0.8 | 0.85 | 0.72 |
| 2 | 222786528 | rs824921 | 1.799 | 2.132 | 1.031 | 1.139 | -0.145 | -0.082 | -0.082 | 0 | 0.6 | 0.05 | 0.55 | 0.63 | 0.53 | 0.5 |
| 2 | 222789037 | rs824922 | 0.081 | 1.829 | 1.067 | 1.125 | -0.015 | -0.003 | -0.003 | 0.615 | 0.78 | 0.73 | 0.84 | 0.79 | 0.85 | 0.72 |
| 2 | 222794169 | rs13021449 | -2.008 | 2.141 | 1.176 | 1.122 | 0.105 | 0.074 | 0.074 | 0 | 0.13 | 0.5 | 0.2 | 0.12 | 0.19 | 0.04 |
| 2 | 222800141 | rs10932930 | -1.906 | 2.159 | 1.115 | 1.135 | 0.12 | 0.092 | 0.092 | 0 | 0.12 | 0.46 | 0.19 | 0.12 | 0.19 | 0.04 |
| 2 | 222818891 | rs13022647 | -0.5 | 2.383 | 1.04 | 0.951 | 0.127 | 0.008 | 0.008 | 0.018 | 0.07 | 0.21 | 0.12 | 0.05 | 0.18 | 0.15 |
| 2 | 222821561 | rs824939 | -0.241 | 2.168 | 1.122 | 0.946 | -0.018 | 0 | 0 | 0.155 | 0.82 | 0.8 | 0.89 | 0.79 | 0.87 | 0.71 |
| 2 | 222827492 | rs13424672 | -0.236 | 2.281 | 0.791 | 0.874 | -0.002 | 0.014 | 0.014 | 0.019 | 0.18 | 0.21 | 0.22 | 0.23 | 0.15 | 0.11 |
| 2 | 222835057 | rs889838 | -1.764 | 2.567 | 0.759 | 0.855 | 0.033 | 0.016 | 0.016 | 0.185 | 0.29 | 0.69 | 0.46 | 0.29 | 0.45 | 0.3 |
| 2 | 222836455 | rs7600373 | -0.012 | 2.565 | 0.554 | 0.865 | -0.001 | 0.017 | 0.017 | 0.379 | 0.17 | 0.2 | 0.22 | 0.23 | 0.15 | 0.11 |
| 2 | 222840137 | rs7588270 | -1.002 | 2.581 | 0.25 | 0.782 | 0.207 | -0.012 | -0.012 | 0 | 0.11 | 0.32 | 0.22 | 0.06 | 0.3 | 0.19 |
| 2 | 222845062 | rs889835 | -2.093 | 2.153 | -0.845 | 0.72 | 0.06 | -0.087 | -0.087 | 0 | 0.55 | 0.97 | 0.67 | 0.48 | 0.69 | 0.55 |
| 2 | 222856726 | rs10203019 | -0.927 | 2.158 | 0.166 | 0.828 | 0.015 | 0.041 | 0.041 | 0.001 | 0.22 | 0.39 | 0.34 | 0.23 | 0.26 | 0.15 |
| 2 | 222857113 | rs1834134 | -1.446 | 2.025 | -0.192 | 0.911 | 0.025 | 0.023 | 0.023 | 2.047 | 0.25 | 0.71 | 0.39 | 0.29 | 0.35 | 0.29 |
| 2 | 222858354 | rs6748217 | 0.072 | 1.539 | -0.724 | 0.488 | 0.012 | -0.013 | -0.013 | 0.27 | 0.89 | 0.84 | 0.89 | 0.95 | 0.85 | 0.89 |
| 2 | 222859340 | rs13418105 | -0.734 | 0.593 | -1.109 | 0.751 | 0.011 | -0.009 | -0.009 | 0 | 0.37 | 0.15 | 0.23 | 0.39 | 0.34 | 0.4 |
| 2 | 222861275 | rs6738997 | -0.724 | 0.242 | -0.222 | 0.716 | -0.027 | 0.001 | 0.001 | 0 | 0.24 | 0.41 | 0.25 | 0.24 | 0.18 | 0.33 |
| 2 | 222870106 | rs6436285 | 0.39 | 0.154 | -0.1 | 0.594 | -0.013 | -0.026 | -0.026 | 1.387 | 0.93 | 0.8 | 0.95 | 0.97 | 0.97 | 0.99 |
| 2 | 222873925 | rs16825454 | -0.254 | 0.184 | 0.197 | 1.25 | 0.013 | 0.099 | 0.099 | 0 | 0.08 | 0.15 | 0.22 | 0.17 | 0.13 | 0.13 |
| 2 | 222881439 | rs6748403 | -1.579 | -0.054 | 0.549 | 1.43 | -0.008 | 0.157 | 0.157 | 0.406 | 0.22 | 0.44 | 0.4 | 0.45 | 0.22 | 0.33 |
| 2 | 222881518 | rs1559473 | 0.486 | -0.188 | 0.079 | 0.655 | 0 | -0.008 | -0.008 | 0 | 0.27 | 0.23 | 0.19 | 0.12 | 0.27 | 0.14 |
| 2 | 222881776 | rs11678073 | -0.393 | -0.147 | 0.075 | 0.658 | 0.015 | 0.26 | 0.26 | 0 | 0.14 | 0.35 | 0.28 | 0.45 | 0.16 | 0.19 |
| 2 | 222882027 | rs1025082 | 1.271 | -0.354 | -0.36 | 0.492 | -0.099 | 0.019 | 0.019 | 0.56 | 0.44 | 0.09 | 0.35 | 0.39 | 0.48 | 0.52 |
| 2 | 222883459 | rs7592247 | 0.004 | -0.426 | 0.12 | 0.265 | -0.001 | -0.028 | -0.028 | 0 | 0.66 | 0.61 | 0.71 | 0.92 | 0.66 | 0.85 |
| 2 | 222890875 | rs6738615 | 0.055 | -0.534 | -0.135 | 0.164 | 0.003 | 0.008 | 0.008 | 0 | 0.09 | 0.04 | 0.03 | 0 | 0.06 | 0.05 |
| 2 | 222895598 | rs7603435 | 0.451 | -0.679 | 0.188 | 0.575 | 0.001 | -0.021 | -0.021 | 0 | 0.24 | 0.26 | 0.25 | 0.1 | 0.29 | 0.13 |
| 2 | 222895715 | rs4673010 | 0.931 | -0.788 | 0.386 | 0.521 | -0.001 | -0.04 | -0.04 | 0 | 0.55 | 0.41 | 0.56 | 0.87 | 0.57 | 0.7 |
| 2 | 222898526 | rs12477596 | 1.285 | -0.735 | 0.581 | 0.639 | 0.055 | 0.202 | 0.202 | 0 | 0.06 | 0.17 | 0.19 | 0.19 | 0.07 | 0.09 |
| 2 | 222903188 | rs4673011 | 0.828 | -0.743 | 0.47 | 0.613 | -0.004 | -0.004 | -0.004 | 0 | 0.56 | 0.42 | 0.58 | 0.6 | 0.57 | 0.64 |
| 2 | 222921656 | rs348974 | 1.445 | -0.904 | -0.176 | 0.45 | -0.077 | -0.007 | -0.007 | 0 | 0.45 | 0.11 | 0.34 | 0.42 | 0.47 | 0.55 |
| 2 | 222922524 | rs1007119 | 0.446 | -1.015 | 0.086 | -0.352 | -0.001 | -0.005 | -0.005 | 0.008 | 0.24 | 0.24 | 0.21 | 0.09 | 0.27 | 0.13 |
| 2 | 222923470 | rs16863338 | 0.003 | -0.85 | 0.01 | -0.138 | 0.019 | 0.073 | 0.073 | 0 | 0.08 | 0.17 | 0.2 | 0.16 | 0.1 | 0.09 |
| 2 | 222925667 | rs348978 | 0.461 | -0.943 | -0.418 | -0.241 | 0.002 | 0.011 | 0.011 | 0 | 0.53 | 0.49 | 0.52 | 0.72 | 0.55 | 0.73 |

| | | | | | | | | | | | | | | | | |
|---|---|---|---|---|---|---|---|---|---|---|---|---|---|---|---|---|
| 2 | 222926680 | rs348980 | 0.854 | -0.777 | -1.408 | -0.142 | -0.059 | 0.12 | 0.12 | 0.099 | 0.46 | 0.11 | 0.38 | 0.72 | 0.5 | 0.61 |
| 2 | 222932490 | rs348984 | 0.553 | -0.667 | -1.557 | -0.258 | -0.143 | 0.013 | 0.013 | 1.828 | 0.37 | 0.05 | 0.32 | 0.36 | 0.49 | 0.35 |
| 2 | 222936264 | rs10167714 | 0.565 | -0.65 | -1.198 | -0.309 | 0.001 | -0.011 | -0.011 | 0 | 0.07 | 0.06 | 0.03 | 0 | 0.04 | 0.01 |
| 2 | 222936992 | rs13389057 | 0.036 | -0.696 | -1.064 | -0.331 | -0.017 | -0.04 | -0.04 | 0 | 0.09 | 0.16 | 0.04 | 0 | 0.04 | 0.01 |
| 2 | 222941085 | rs2113399 | -0.329 | -0.742 | -0.654 | -0.354 | -0.026 | -0.057 | -0.057 | 0 | 0.1 | 0.28 | 0.06 | 0 | 0.04 | 0.01 |
| 2 | 222943945 | rs348970 | 0.163 | -0.917 | -0.848 | -0.359 | -0.016 | 0.062 | 0.062 | 0 | 0.45 | 0.28 | 0.37 | 0.34 | 0.53 | 0.39 |
| 2 | 222946417 | rs348989 | -0.107 | -1.071 | -0.186 | -0.438 | 0.001 | 0.048 | 0.048 | 0 | 0.49 | 0.62 | 0.4 | 0.34 | 0.53 | 0.42 |
| 2 | 222946649 | rs348990 | 0.396 | -1.058 | -0.231 | -0.431 | -0.05 | 0.078 | 0.078 | 0.222 | 0.43 | 0.14 | 0.35 | 0.34 | 0.53 | 0.39 |
| 2 | 222955922 | rs349011 | -0.415 | -1.171 | -0.532 | -0.456 | -0.009 | 0.05 | 0.05 | 0 | 0.67 | 0.78 | 0.64 | 0.82 | 0.69 | 0.6 |
| 2 | 222961756 | rs717760 | 0.64 | -1.173 | -0.572 | -0.417 | -0.005 | 0.025 | 0.025 | 0 | 0.19 | 0.17 | 0.24 | 0.47 | 0.15 | 0.18 |
| 2 | 222965299 | rs349020 | 0.744 | -1.193 | -0.533 | -0.429 | -0.054 | 0.078 | 0.078 | 0.359 | 0.43 | 0.15 | 0.35 | 0.34 | 0.54 | 0.39 |
| 2 | 222965927 | rs349022 | -0.21 | -0.924 | -1.564 | -0.392 | 0.002 | 0.009 | 0.009 | 0 | 0.62 | 0.32 | 0.59 | 0.82 | 0.68 | 0.56 |
| 2 | 222970051 | rs12104556 | 0.534 | -0.335 | -1.547 | -0.255 | -0.003 | 0.003 | 0.003 | 0 | 0.39 | 0.65 | 0.28 | 0.17 | 0.41 | 0.32 |
| 2 | 222971274 | rs6728176 | 0.08 | -0.007 | -1.88 | 0.003 | -0.009 | 0.117 | 0.117 | 0 | 0.48 | 0.73 | 0.49 | 0.18 | 0.47 | 0.49 |
| 2 | 222976013 | rs4673012 | -0.859 | 0.126 | -2.344 | 0.151 | -0.061 | -0.002 | -0.002 | 0 | 0.69 | 0.93 | 0.74 | 0.66 | 0.63 | 0.66 |
| 2 | 222984939 | rs4674635 | -0.029 | -0.164 | -1.469 | 0.943 | 0.008 | -0.027 | -0.027 | 0.193 | 0.19 | 0.11 | 0.23 | 0.48 | 0.15 | 0.18 |
| 2 | 222986114 | rs10932941 | -0.812 | -0.314 | -1.399 | 0.684 | 0.007 | -0.004 | -0.004 | 0.139 | 0.31 | 0.23 | 0.46 | 0.62 | 0.21 | 0.35 |
| 2 | 222986215 | rs16863407 | -0.068 | -0.243 | -1.493 | 0.94 | 0.007 | -0.023 | -0.023 | 0 | 0.2 | 0.11 | 0.27 | 0.6 | 0.16 | 0.21 |
| 2 | 222986395 | rs6436298 | -0.348 | -0.131 | -2.005 | 0.852 | -0.016 | -0.122 | -0.122 | 0.211 | 0.74 | 0.92 | 0.79 | 0.51 | 0.76 | 0.69 |
| 2 | 222988705 | rs16863412 | -0.492 | -0.559 | -1.014 | 0.335 | -0.016 | 0.003 | 0.003 | 0 | 0.25 | 0.44 | 0.25 | 0.29 | 0.2 | 0.34 |
| 2 | 222989917 | rs13420000 | -0.117 | -0.962 | -2.039 | -0.357 | -0.001 | 0 | 0 | 1.323 | 0.5 | 0.55 | 0.6 | 0.38 | 0.68 | 0.56 |
| 2 | 222992139 | rs10932942 | -1.173 | -1.173 | -1.137 | -0.656 | 0.013 | -0.042 | -0.042 | 0 | 0.92 | 0.92 | 0.93 | 1 | 0.92 | 0.9 |
| 2 | 222993315 | rs2395845 | 1.247 | -0.951 | -1.344 | -0.839 | -0.02 | 0.029 | 0.029 | 0.355 | 0.06 | 0.01 | 0.17 | 0.01 | 0.08 | 0.14 |
| 2 | 222993733 | rs12694574 | 0.721 | -1.775 | -1.265 | -1.294 | 0.028 | 0.066 | 0.066 | 0 | 0.27 | 0.12 | 0.16 | 0.07 | 0.2 | 0.14 |
| 2 | 222994084 | rs17399261 | 1.975 | -1.821 | -1.28 | -1.328 | -0.12 | 0.053 | 0.053 | 0 | 0.28 | 0.07 | 0.23 | 0.15 | 0.44 | 0.2 |
| 2 | 222994221 | rs6749789 | 1.484 | -1.885 | -1.066 | -1.363 | -0.008 | -0.082 | -0.082 | 1.271 | 0.19 | 0.26 | 0.13 | 0 | 0.15 | 0.02 |
| 2 | 222995155 | rs7601880 | -0.256 | -1.889 | -1.057 | -1.356 | -0.036 | 0.067 | 0.067 | 0 | 0.77 | 0.68 | 0.76 | 0.46 | 0.88 | 0.7 |
| 2 | 222995369 | rs10182674 | -0.449 | -1.891 | -1.062 | -1.357 | -0.004 | 0.01 | 0.01 | 0 | 0.78 | 0.82 | 0.77 | 0.46 | 0.88 | 0.7 |
| 2 | 222997104 | rs2894450 | 0.805 | -1.879 | -1.061 | -1.456 | -0.006 | -0.015 | -0.015 | 0 | 0.44 | 0.55 | 0.44 | 0.34 | 0.59 | 0.41 |
| 2 | 222999840 | rs1370923 | 1.363 | -1.922 | -0.523 | -1.446 | 0.004 | 0.004 | 0.004 | 0 | 0.14 | 0.19 | 0.17 | 0.22 | 0.15 | 0.22 |
| 2 | 223002705 | rs13385121 | 1.846 | -1.966 | -0.294 | -1.481 | -0.002 | 0.043 | 0.043 | 0 | 0.17 | 0.06 | 0.11 | 0 | 0.15 | 0.02 |
| 2 | 223005125 | rs1370920 | -0.397 | -1.952 | -0.247 | -1.47 | 0.02 | 0.049 | 0.049 | 0.584 | 0.44 | 0.69 | 0.37 | 0.29 | 0.36 | 0.36 |
| 2 | 223008570 | rs1370919 | 0.659 | -1.991 | -0.315 | -1.382 | 0 | 0 | 0 | 0 | 0.5 | 0.54 | 0.6 | 0.35 | 0.68 | 0.56 |
| 2 | 223021856 | rs11683100 | -0.005 | -1.63 | -0.099 | -1.199 | -0.041 | 0.07 | 0.07 | 0 | 0.77 | 0.61 | 0.75 | 0.42 | 0.88 | 0.7 |
| 2 | 223031155 | rs7565380 | -1.282 | -1.32 | -0.523 | -1.328 | 0.012 | -0.041 | -0.041 | 0.401 | 0.92 | 0.92 | 0.93 | 1 | 0.92 | 0.9 |
| 2 | 223040476 | rs11885004 | 1.257 | -1.319 | -0.636 | -1.414 | 0.002 | -0.005 | -0.005 | 0.184 | 0.18 | 0.12 | 0.12 | 0 | 0.15 | 0.02 |
| 2 | 223044442 | rs6710838 | 0.596 | -1.483 | -0.536 | -1.48 | -0.003 | -0.04 | -0.04 | 0 | 0.47 | 0.49 | 0.38 | 0.12 | 0.59 | 0.22 |
| 2 | 223051502 | rs10169326 | 0.89 | -1.335 | -0.536 | -1.55 | -0.009 | -0.091 | -0.091 | 0 | 0.21 | 0.32 | 0.15 | 0 | 0.17 | 0.02 |
| 2 | 223051837 | rs6736007 | 0.335 | -1.558 | -0.85 | -1.5 | -0.005 | -0.002 | -0.002 | 0 | 0.57 | 0.46 | 0.62 | 0.41 | 0.68 | 0.61 |
| 2 | 223059951 | rs1438607 | 1.942 | -1.489 | -1.754 | -1.564 | -0.112 | 0.035 | 0.035 | 0 | 0.25 | 0.07 | 0.22 | 0.11 | 0.42 | 0.17 |
| 2 | 223063546 | rs7593694 | 1.609 | -1.511 | -1.603 | -1.486 | 0.016 | 0.01 | 0.01 | 0 | 0.06 | 0.04 | 0.01 | 0.03 | 0.01 | 0.08 |
| 2 | 223064768 | rs12620338 | 0.984 | -1.475 | -1.63 | -1.51 | -0.018 | -0.042 | -0.042 | 5.604 | 0.23 | 0.02 | 0.16 | 0.28 | 0.18 | 0.29 |
| 2 | 223082331 | rs1978859 | 0.286 | -1.561 | -1.647 | -1.561 | -0.016 | 0.105 | 0.105 | 1.171 | 0.41 | 0.08 | 0.28 | 0.28 | 0.35 | 0.31 |
| 2 | 223086976 | rs6741337 | 0.048 | -1.328 | -1.412 | -1.317 | 0.017 | -0.017 | -0.017 | 0 | 0.14 | 0.12 | 0.17 | 0.59 | 0.07 | 0.25 |
| 2 | 223090779 | rs1013262 | 1.423 | -0.707 | -1.396 | -1.214 | -0.097 | 0.05 | 0.05 | 0 | 0.26 | 0.09 | 0.23 | 0.11 | 0.42 | 0.17 |
| 2 | 223095968 | rs1549773 | 0.995 | -0.597 | -1.194 | -1.194 | -0.001 | 0.025 | 0.025 | 0.174 | 0.18 | 0.11 | 0.12 | 0 | 0.17 | 0.02 |
| 2 | 223100523 | rs2118666 | 0.409 | -0.131 | -0.779 | -0.82 | -0.011 | 0.006 | 0.006 | 0 | 0.34 | 0.35 | 0.4 | 0.66 | 0.48 | 0.33 |
| 2 | 223101131 | rs6436301 | -0.522 | 0.148 | -0.675 | -0.893 | 0.01 | -0.041 | -0.041 | 0 | 0.91 | 0.89 | 0.93 | 1 | 0.9 | 0.88 |
| 2 | 223102757 | rs1367409 | 0.191 | 0.361 | -0.755 | -1.091 | 0 | -0.025 | -0.025 | 0 | 0.4 | 0.43 | 0.5 | 0.11 | 0.44 | 0.31 |
| 2 | 223102937 | rs13410020 | 0.79 | 0.133 | -1.053 | -1.135 | -0.014 | 0.029 | 0.029 | 0 | 0.14 | 0.05 | 0.11 | 0 | 0.17 | 0.02 |
| 2 | 223104980 | rs10498134 | -0.519 | -0.062 | -1.016 | -1.469 | -0.001 | 0 | 0 | 0 | 0.37 | 0.4 | 0.59 | 0.8 | 0.38 | 0.6 |
| 2 | 223105652 | rs1367410 | -0.95 | -0.316 | -0.727 | -1.648 | 0.001 | 0.056 | 0.056 | 0 | 0.5 | 0.64 | 0.71 | 0.8 | 0.51 | 0.61 |
| 2 | 223106018 | rs1367413 | 0.838 | -0.354 | -0.909 | -0.977 | 0.003 | -0.01 | -0.01 | 0 | 0.13 | 0.08 | 0.09 | 0.27 | 0.11 | 0.31 |
| 2 | 223106322 | rs6436302 | -1.029 | -0.187 | -0.823 | -0.789 | 0.001 | 0.029 | 0.029 | 0 | 0.5 | 0.58 | 0.7 | 0.8 | 0.51 | 0.61 |
| 2 | 223106526 | rs12053273 | 0.419 | -0.004 | -0.83 | -0.634 | 0.014 | 0.085 | 0.085 | 0.092 | 0.16 | 0.28 | 0.25 | 0.5 | 0.2 | 0.16 |
| 2 | 223107780 | rs1835432 | -1.368 | -0.106 | -1.281 | -0.978 | -0.073 | 0.055 | 0.055 | 0 | 0.7 | 0.9 | 0.81 | 0.87 | 0.62 | 0.8 |
| 2 | 223108399 | rs6436304 | -0.339 | -0.042 | -0.983 | -0.741 | -0.001 | 0 | 0 | 0 | 0.45 | 0.57 | 0.46 | 0.77 | 0.44 | 0.5 |
| 2 | 223109002 | rs1367415 | -0.045 | 0.031 | -0.819 | -0.769 | -0.002 | -0.007 | -0.007 | 0 | 0.23 | 0.25 | 0.1 | 0.11 | 0.15 | 0.16 |
| 2 | 223109247 | rs11695486 | 0.329 | 0.509 | -1.007 | -1.095 | -0.004 | 0.011 | 0.011 | 2.051 | 0.6 | 0.32 | 0.46 | 0.77 | 0.54 | 0.61 |
| 2 | 223109307 | rs12328631 | 0.767 | -0.101 | -0.77 | -1.426 | -0.048 | -0.083 | -0.083 | 0 | 0.16 | 0.26 | 0.11 | 0 | 0.07 | 0.02 |
| 2 | 223114678 | rs1430657 | 1.52 | 0.553 | -1.218 | -1.409 | -0.021 | 0.058 | 0.058 | 0 | 0.16 | 0.06 | 0.13 | 0 | 0.18 | 0.02 |
| 2 | 223115115 | rs17403188 | 0.475 | 1.103 | -0.957 | -0.876 | -0.001 | 0 | 0 | 0.519 | 0.07 | 0.07 | 0.07 | 0.05 | 0.12 | 0.14 |
| 2 | 223120910 | rs1430659 | -0.38 | 1.117 | -1.266 | -0.26 | 0.002 | -0.059 | -0.059 | 0 | 0.14 | 0.04 | 0.19 | 0.29 | 0.15 | 0.19 |
| 2 | 223122565 | rs7589708 | -0.354 | 1.126 | -1.28 | -0.262 | 0.001 | -0.063 | -0.063 | 0 | 0.14 | 0.04 | 0.19 | 0.3 | 0.14 | 0.19 |
| 2 | 223124093 | rs4674639 | 0.819 | 1.401 | -1.127 | 0.067 | 0.013 | -0.051 | -0.051 | 0.049 | 0.19 | 0.09 | 0.38 | 0.39 | 0.18 | 0.31 |
| 2 | 223126967 | rs12617763 | 0.162 | 1.668 | -0.829 | 0.179 | -0.001 | 0.009 | 0.009 | 0 | 0.1 | 0.07 | 0.16 | 0.3 | 0.11 | 0.16 |
| 2 | 223131625 | rs10932949 | -0.539 | 1.895 | -0.401 | -0.403 | 0.051 | -0.057 | -0.057 | 0.075 | 0.88 | 0.73 | 0.88 | 0.99 | 0.79 | 0.94 |
| 2 | 223135983 | rs17344688 | 0.156 | 1.615 | -1.123 | -0.277 | -0.008 | 0.011 | 0.011 | 0 | 0.1 | 0.06 | 0.06 | 0.07 | 0.13 | 0.17 |
| 2 | 223137001 | rs12995399 | 0.616 | 1.38 | -1.297 | -0.072 | 0.008 | -0.03 | -0.03 | 0.013 | 0.2 | 0.11 | 0.4 | 0.37 | 0.2 | 0.33 |

| | | | | | | | | | | | | | | | | |
|---|---|---|---|---|---|---|---|---|---|---|---|---|---|---|---|---|
| 2 | 223148292 | rs930140 | 0.068 | 1.571 | -1.27 | 0.011 | -0.022 | 0.024 | 0.024 | 2.377 | 0.56 | 0.23 | 0.42 | 0.56 | 0.47 | 0.43 |
| 2 | 223153374 | rs10174636 | 0.394 | 1.199 | -1.423 | -0.499 | 0.002 | 0.009 | 0.009 | 0 | 0.23 | 0.19 | 0.17 | 0 | 0.21 | 0.05 |
| 2 | 223156587 | rs7600206 | 0.573 | 0.761 | -1.633 | -0.107 | 0.034 | -0.034 | -0.034 | 0 | 0.23 | 0.12 | 0.33 | 0.36 | 0.18 | 0.38 |
| 2 | 223158039 | rs2276630 | 0.616 | 0.886 | -1.83 | 0.301 | 0.02 | -0.094 | -0.094 | 6.183 | 0.23 | 0.01 | 0.05 | 0.28 | 0.13 | 0.26 |
| 2 | 223161606 | rs13405641 | 0.51 | 1.537 | -1.099 | 0.12 | -0.008 | 0 | 0 | 0 | 0.15 | 0.11 | 0.14 | 0 | 0.18 | 0.03 |
| 2 | 223164518 | rs16863657 | -0.131 | 1.531 | -0.177 | 0.984 | 0.056 | 0.167 | 0.167 | 0 | 0.08 | 0.22 | 0.18 | 0.25 | 0.15 | 0.11 |
| 2 | 223167710 | rs6436310 | -0.233 | 0.933 | -0.517 | 0.593 | 0.025 | 0.001 | 0.001 | 1.277 | 0.64 | 0.57 | 0.39 | 0.34 | 0.52 | 0.43 |
| 2 | 223167873 | rs7557746 | 0.205 | 0.58 | -0.764 | 0.106 | 0.041 | 0.068 | 0.068 | 0 | 0.47 | 0.37 | 0.25 | 0.29 | 0.34 | 0.39 |
| 2 | 223172227 | rs10202724 | 0.011 | -0.311 | -0.875 | 0.4 | 0.001 | 0.023 | 0.023 | 0.79 | 0.67 | 0.62 | 0.75 | 0.57 | 0.65 | 0.67 |
| 2 | 223177624 | rs1864556 | 1.5 | -0.288 | -1.163 | 0.242 | 0.038 | -0.045 | -0.045 | 0 | 0.3 | 0.14 | 0.45 | 0.38 | 0.25 | 0.31 |
| 2 | 223203549 | rs13422755 | 0.44 | -0.56 | -0.681 | 0.306 | -0.001 | 0.012 | 0.012 | 0 | 0.3 | 0.22 | 0.19 | 0.19 | 0.28 | 0.27 |
| 2 | 223217087 | rs6755099 | -0.542 | -0.456 | -0.56 | 0.433 | 0.047 | 0.062 | 0.062 | 0.095 | 0.86 | 0.82 | 0.84 | 0.7 | 0.76 | 0.79 |
| 2 | 223226794 | rs1430665 | 1.406 | -0.629 | -0.777 | 0.389 | -0.015 | 0.027 | 0.027 | 1.237 | 0.46 | 0.17 | 0.59 | 0.38 | 0.46 | 0.42 |
| 2 | 223234004 | rs16863671 | 1.345 | -0.665 | -0.699 | 0.362 | 0.004 | 0.003 | 0.003 | 0.074 | 0.16 | 0.12 | 0.08 | 0.14 | 0.1 | 0.14 |
| 2 | 223235543 | rs1430666 | -0.496 | -0.541 | -0.741 | 0.408 | 0.065 | 0.085 | 0.085 | 0.014 | 0.86 | 0.77 | 0.83 | 0.7 | 0.76 | 0.79 |
| 2 | 223251464 | rs4673017 | 1.23 | -0.488 | -1.025 | 0.418 | -0.003 | 0.033 | 0.033 | 1.058 | 0.46 | 0.23 | 0.6 | 0.37 | 0.47 | 0.42 |
| 2 | 223271022 | rs724150 | 1.138 | -0.283 | -1.062 | 0.435 | 0.029 | 0.037 | 0.037 | 0 | 0.48 | 0.33 | 0.56 | 0.39 | 0.43 | 0.32 |
| 2 | 223273313 | rs6436315 | 0.944 | 0.093 | -0.795 | 0.638 | -0.008 | -0.002 | -0.002 | 0 | 0.2 | 0.15 | 0.13 | 0.18 | 0.2 | 0.21 |
| 2 | 223281386 | rs7565920 | 0.958 | 0.12 | -0.834 | 0.56 | -0.009 | -0.009 | -0.009 | 0 | 0.61 | 0.36 | 0.67 | 0.49 | 0.53 | 0.43 |
| 2 | 223288369 | rs6758392 | -0.429 | 0.567 | -1.257 | 0.777 | 0.111 | 0.098 | 0.098 | 0 | 0.87 | 0.71 | 0.83 | 0.73 | 0.74 | 0.65 |
| 2 | 223295475 | rs10202965 | 0.716 | 0.529 | -1.323 | 0.745 | -0.014 | -0.007 | -0.007 | 0 | 0.22 | 0.15 | 0.14 | 0.19 | 0.22 | 0.17 |
| 2 | 223298246 | rs7556867 | 0.227 | 0.786 | -0.961 | 0.912 | 0.095 | 0.098 | 0.098 | 0 | 0.87 | 0.73 | 0.84 | 0.72 | 0.76 | 0.72 |
| 2 | 223326724 | rs10932956 | -0.229 | -0.346 | -1.916 | 0.18 | 0.015 | 0.071 | 0.071 | 0 | 0.29 | 0.06 | 0.22 | 0.13 | 0.28 | 0.24 |
| 2 | 223329125 | rs6723696 | -0.562 | 0.12 | -0.951 | 0.599 | -0.048 | -0.007 | -0.007 | 2.644 | 0.56 | 0.8 | 0.45 | 0.57 | 0.51 | 0.55 |
| 2 | 223329929 | rs6712500 | -0.267 | 0.349 | -0.946 | 0.772 | 0.004 | 0.013 | 0.013 | 0 | 0.07 | 0.1 | 0.06 | 0.13 | 0.09 | 0.1 |
| 2 | 223331920 | rs7604173 | -0.306 | 0.215 | -0.545 | 0.616 | 0.002 | 0.001 | 0.001 | 0 | 0.45 | 0.37 | 0.35 | 0.4 | 0.42 | 0.45 |
| 2 | 223338071 | rs13021671 | -0.335 | 0.047 | -0.627 | 0.548 | 0.018 | -0.011 | -0.011 | 0 | 0.41 | 0.37 | 0.37 | 0.43 | 0.35 | 0.42 |
| 2 | 223339582 | rs735678 | -1.41 | 0.621 | 0.229 | 0.474 | 0.022 | -0.024 | -0.024 | 1.336 | 0.3 | 0.56 | 0.44 | 0.22 | 0.35 | 0.24 |
| 2 | 223343855 | rs4674656 | -1.114 | 0.33 | -0.146 | 0.474 | -0.002 | 0.005 | 0.005 | 0.921 | 0.26 | 0.5 | 0.38 | 0.2 | 0.24 | 0.3 |
| 2 | 223348008 | rs6722708 | -0.379 | 0.66 | 0.022 | 1.194 | 0.031 | 0.071 | 0.071 | 0 | 0.07 | 0.2 | 0.05 | 0.14 | 0.04 | 0.08 |
| 2 | 223349309 | rs10176846 | -1.708 | 0.599 | -0.304 | 1.04 | -0.004 | 0.077 | 0.077 | 0 | 0.27 | 0.44 | 0.42 | 0.38 | 0.23 | 0.36 |
| 2 | 223349415 | rs10176933 | -0.887 | 0.599 | -0.294 | 1.055 | 0.023 | 0.128 | 0.128 | 0 | 0.18 | 0.33 | 0.36 | 0.36 | 0.16 | 0.31 |
| 2 | 223350001 | rs10203329 | -1.005 | 0.718 | -0.471 | 1.028 | -0.002 | 0.067 | 0.067 | 0 | 0.28 | 0.45 | 0.42 | 0.38 | 0.24 | 0.37 |
| 2 | 223351343 | rs13397034 | -0.997 | 0.749 | -0.455 | 1.038 | -0.002 | 0.065 | 0.065 | 0.003 | 0.28 | 0.43 | 0.39 | 0.37 | 0.24 | 0.38 |
| 2 | 223352259 | rs2033694 | 0.67 | 1.227 | -1.084 | 0.8 | -0.052 | -0.019 | -0.019 | 0 | 0.83 | 0.96 | 0.8 | 0.85 | 0.76 | 0.75 |
| 2 | 223355051 | rs6757951 | 0.185 | 1.362 | -1.451 | 0.737 | -0.058 | -0.047 | -0.047 | 0 | 0.58 | 0.21 | 0.65 | 0.5 | 0.53 | 0.62 |
| 2 | 223359365 | rs6745812 | -0.139 | 1.333 | -0.323 | 0.208 | 0.036 | -0.118 | -0.118 | 0 | 0.74 | 0.59 | 0.78 | 0.99 | 0.68 | 0.85 |
| 2 | 223359573 | rs10169007 | 0.016 | 1.323 | -0.158 | 0.168 | 0.035 | -0.052 | -0.052 | 0 | 0.91 | 0.68 | 0.92 | 1 | 0.93 | 0.99 |
| 2 | 223361479 | rs17562982 | 1.256 | 1.584 | -0.229 | 0.231 | 0.048 | 0.132 | 0.132 | 0 | 0.49 | 0.29 | 0.53 | 0.2 | 0.4 | 0.5 |
| 2 | 223366131 | rs17563113 | -0.893 | 1.513 | -0.366 | 0.198 | -0.112 | 0.072 | 0.072 | 0 | 0.39 | 0.07 | 0.37 | 0.79 | 0.55 | 0.44 |
| 2 | 223366832 | rs10804308 | 0.481 | 1.59 | 0.698 | 0.332 | 0.005 | -0.033 | -0.033 | 0 | 0.89 | 0.35 | 0.87 | 0.98 | 0.93 | 0.96 |
| 2 | 223368935 | rs7559017 | -0.289 | 1.618 | 0.893 | 0.342 | 0.009 | -0.038 | -0.038 | 3.14 | 0.92 | 0.76 | 0.93 | 1 | 0.93 | 0.99 |
| 2 | 223371747 | rs16863807 | 0.68 | 1.646 | 0.662 | 0.591 | -0.038 | -0.051 | -0.051 | 0 | 0.1 | 0 | 0.09 | 0.46 | 0.14 | 0.2 |
| 2 | 223384515 | rs6740937 | -0.677 | 1.329 | 0.644 | 0.182 | -0.016 | 0.044 | 0.044 | 0.57 | 0.41 | 0.3 | 0.55 | 0.81 | 0.49 | 0.46 |
| 2 | 223386425 | rs3820782 | -0.55 | 1.245 | 0.97 | 0.151 | 0.039 | 0.041 | 0.041 | 0.121 | 0.05 | 0.28 | 0.03 | 0.01 | 0.02 | 0.07 |
| 2 | 223400520 | rs10195337 | -0.601 | 1.245 | 1.231 | 0.134 | 0.033 | -0.029 | -0.029 | 0 | 0.94 | 0.79 | 0.93 | 1 | 0.94 | 0.99 |
| 2 | 223412143 | rs13382464 | -0.004 | 1.376 | 0.991 | 0.184 | -0.061 | -0.02 | -0.02 | 0 | 0.31 | 0.08 | 0.28 | 0.29 | 0.37 | 0.25 |
| 2 | 223412813 | rs4592839 | 1.381 | 1.329 | 1.198 | 0.172 | 0.016 | 0.086 | 0.086 | 2.313 | 0.47 | 0.3 | 0.35 | 0.14 | 0.42 | 0.54 |
| 2 | 223427854 | rs1347630 | -0.325 | 1.491 | 1.352 | 0.16 | 0.047 | -0.02 | -0.02 | 0.661 | 0.93 | 0.8 | 0.87 | 0.97 | 0.88 | 0.93 |
| 2 | 223435651 | rs17564655 | 0.086 | 1.526 | 1.273 | 0.165 | -0.043 | -0.008 | -0.008 | 0 | 0.3 | 0.12 | 0.27 | 0.29 | 0.37 | 0.22 |
| 2 | 223436607 | rs7185 | 0.253 | 1.521 | 1.544 | 0.14 | 0.1 | -0.013 | -0.013 | 0 | 0.91 | 0.62 | 0.86 | 0.97 | 0.88 | 0.93 |
| 2 | 223457193 | rs16863916 | 1.479 | 1.447 | 1.167 | 0.097 | 0.032 | 0.101 | 0.101 | 0.963 | 0.44 | 0.3 | 0.29 | 0.11 | 0.36 | 0.47 |
| 2 | 223477160 | rs2028953 | 1.679 | 1.417 | 1.037 | 0.096 | 0.042 | 0.168 | 0.168 | 1.423 | 0.42 | 0.16 | 0.25 | 0.1 | 0.34 | 0.44 |
| 2 | 223488778 | rs1542198 | -1.104 | 1.433 | 0.791 | 0.204 | -0.032 | -0.059 | -0.059 | 0 | 0.9 | 0.99 | 0.74 | 0.45 | 0.86 | 0.79 |
| 2 | 223491922 | rs12151627 | 0.125 | 1.454 | 0.826 | 0.196 | -0.108 | -0.06 | -0.06 | 0.908 | 0.3 | 0.04 | 0.26 | 0.3 | 0.36 | 0.22 |
| 2 | 223510646 | rs8179677 | 0.041 | 1.436 | 0.863 | 0.202 | -0.063 | -0.021 | -0.021 | 0.001 | 0.31 | 0.1 | 0.27 | 0.29 | 0.37 | 0.22 |
| 2 | 223520797 | rs12467164 | -0.846 | 1.387 | 1.567 | 0.774 | -0.014 | -0.007 | -0.007 | 0 | 0.06 | 0.2 | 0.05 | 0.02 | 0.01 | 0.02 |
| 2 | 223538538 | rs6436328 | -0.376 | 1.544 | 0.717 | 0.805 | 0.013 | -0.012 | -0.012 | 0 | 0.07 | 0.13 | 0.11 | 0.03 | 0.12 | 0.07 |
| 2 | 223542231 | rs1447108 | 0.955 | 1.36 | 0.357 | 0.839 | 0.005 | 0.045 | 0.045 | 0.821 | 0.72 | 0.4 | 0.55 | 0.41 | 0.71 | 0.66 |
| 2 | 223546047 | rs12611436 | -0.117 | 1.462 | -0.11 | 0.864 | -0.009 | -0.007 | -0.007 | 0.476 | 0.11 | 0.07 | 0.13 | 0.3 | 0.17 | 0.06 |
| 2 | 223547137 | rs6716022 | -1.87 | 1.397 | -0.053 | 0.605 | -0.01 | -0.014 | -0.014 | 0 | 0.32 | 0.78 | 0.33 | 0.38 | 0.33 | 0.21 |
| 2 | 223550985 | rs6710583 | -1.436 | 1.185 | 0.25 | 0.906 | 0.009 | 0.081 | 0.081 | 0.263 | 0.18 | 0.26 | 0.18 | 0.34 | 0.21 | 0.08 |
| 2 | 223551147 | rs6739216 | -0.267 | 1.482 | -0.45 | 1.063 | -0.005 | 0.006 | 0.006 | 0 | 0.11 | 0.07 | 0.14 | 0.32 | 0.17 | 0.06 |
| 2 | 223558890 | rs7592038 | -0.177 | 1.05 | -0.742 | -0.221 | 0.002 | 0.016 | 0.016 | 0.038 | 0.43 | 0.61 | 0.38 | 0.21 | 0.34 | 0.31 |
| 2 | 223563574 | rs7587974 | 0 | 1.004 | -0.774 | -0.942 | 0 | 0.011 | 0.011 | 0 | 0.31 | 0.28 | 0.26 | 0.13 | 0.26 | 0.18 |
| 2 | 223563644 | rs7601761 | -0.874 | 0.813 | -0.157 | -1.263 | -0.001 | -0.023 | -0.023 | 1.952 | 0.47 | 0.53 | 0.36 | 0.18 | 0.39 | 0.25 |
| 2 | 223565624 | rs6436329 | 0.117 | 0.933 | -0.366 | -1.561 | -0.009 | 0.024 | 0.024 | 0 | 0.33 | 0.28 | 0.31 | 0.11 | 0.35 | 0.2 |
| 2 | 223565830 | rs6749783 | -0.995 | 0.957 | -0.403 | -1.547 | -0.001 | -0.022 | -0.022 | 0.117 | 0.52 | 0.54 | 0.41 | 0.15 | 0.46 | 0.27 |
| 2 | 223567016 | rs951997 | -0.853 | 0.823 | -0.703 | -1.984 | 0 | -0.006 | -0.006 | 0 | 0.6 | 0.38 | 0.61 | 0.17 | 0.57 | 0.37 |

| | | | | | | | | | | | | | | | | |
|---|---|---|---|---|---|---|---|---|---|---|---|---|---|---|---|---|
| 2 | 223568242 | rs4480975 | 0.795 | 0.982 | -0.823 | -1.754 | -0.042 | 0.032 | 0.032 | 0.292 | 0.13 | 0.01 | 0.09 | 0.04 | 0.12 | 0.07 |
| 2 | 223574708 | rs10173147 | 0.335 | 1.032 | -0.657 | -1.757 | -0.002 | 0.001 | 0.001 | 0 | 0.14 | 0.13 | 0.11 | 0.03 | 0.13 | 0.07 |
| 2 | 223577268 | rs16864047 | -1.008 | 0.766 | 0.225 | -1.516 | 0.063 | 0.043 | 0.043 | 0 | 0.06 | 0.29 | 0.04 | 0.02 | 0.04 | 0.02 |
| 2 | 223579883 | rs2084288 | 0.751 | 0.785 | -0.435 | -1.483 | -0.017 | 0.032 | 0.032 | 0 | 0.09 | 0 | 0.22 | 0.03 | 0.1 | 0.09 |
| 2 | 223588671 | rs6436333 | -1.287 | 0.789 | -0.801 | -1.682 | -0.074 | -0.051 | -0.051 | 0.1 | 0.78 | 0.98 | 0.7 | 0.2 | 0.69 | 0.53 |
| 2 | 223593164 | rs7601126 | -0.435 | 0.685 | -0.624 | -1.767 | -0.036 | 0.028 | 0.028 | 0.408 | 0.71 | 0.87 | 0.67 | 0.21 | 0.61 | 0.39 |
| 2 | 223594372 | rs7564567 | 0.233 | 1.083 | 0.246 | -1.402 | 0.007 | -0.01 | -0.01 | 0 | 0.87 | 0.81 | 0.87 | 0.91 | 0.87 | 0.8 |
| 2 | 223594490 | rs895466 | 0.584 | 1.078 | 0.567 | -1.428 | 0.037 | -0.03 | -0.03 | 0.002 | 0.84 | 0.6 | 0.83 | 0.91 | 0.83 | 0.78 |
| 2 | 223610075 | rs10173577 | -0.152 | 1.293 | 0.056 | -1.286 | -0.001 | 0 | 0 | 1.304 | 0.88 | 0.89 | 0.92 | 0.9 | 0.87 | 0.79 |
| 2 | 223610799 | rs959327 | 0.976 | 1.195 | -0.089 | -1.959 | 0.017 | 0.054 | 0.054 | 0 | 0.47 | 0.39 | 0.46 | 0.81 | 0.49 | 0.61 |
| 2 | 223613962 | rs4674687 | -0.221 | 1.296 | -0.196 | -1.934 | -0.011 | -0.002 | -0.002 | 0 | 0.87 | 0.88 | 0.92 | 0.87 | 0.81 | 0.78 |
| 2 | 223615362 | rs17779311 | 0.498 | 1.071 | -0.396 | -2.215 | -0.121 | 0.128 | 0.128 | 0.991 | 0.26 | 0.03 | 0.2 | 0.04 | 0.31 | 0.13 |
| 2 | 223618017 | rs7601953 | 0.388 | 1.069 | -0.27 | -2.232 | -0.084 | 0.109 | 0.109 | 1.225 | 0.27 | 0.09 | 0.21 | 0.04 | 0.32 | 0.12 |
| 2 | 223620151 | rs7595587 | 1.437 | 1.133 | -0.081 | -2.351 | 0.003 | -0.007 | -0.007 | 0 | 0.46 | 0.44 | 0.48 | 0.85 | 0.43 | 0.63 |
| 2 | 223621273 | rs895468 | -0.68 | 1.485 | -0.52 | -2.182 | -0.021 | 0.012 | 0.012 | 0.239 | 0.88 | 0.94 | 0.94 | 0.91 | 0.84 | 0.8 |
| 2 | 223621511 | rs870998 | 0.117 | 1.526 | -0.505 | -2.167 | -0.074 | 0.096 | 0.096 | 1.537 | 0.29 | 0.15 | 0.23 | 0.04 | 0.35 | 0.13 |
| 2 | 223625518 | rs10176458 | 0.411 | 1.528 | -0.632 | -2.145 | -0.089 | 0.11 | 0.11 | 1.94 | 0.26 | 0.07 | 0.2 | 0.04 | 0.31 | 0.12 |
| 2 | 223636135 | rs4674690 | -0.414 | 1.467 | -0.53 | -1.976 | -0.008 | 0.004 | 0.004 | 0.211 | 0.87 | 0.9 | 0.93 | 0.91 | 0.84 | 0.8 |
| 2 | 223648291 | rs17779768 | 0.432 | 1.561 | -0.809 | -2.126 | -0.051 | 0.08 | 0.08 | 0 | 0.23 | 0.07 | 0.18 | 0.03 | 0.27 | 0.11 |
| 2 | 223653986 | rs6749903 | 1.355 | 1.171 | -0.282 | -2.074 | 0.018 | 0.03 | 0.03 | 0.009 | 0.46 | 0.42 | 0.49 | 0.85 | 0.44 | 0.61 |
| 2 | 223654070 | rs2395926 | 0.202 | 1.422 | -0.204 | -1.556 | 0.001 | -0.002 | -0.002 | 0 | 0.86 | 0.86 | 0.93 | 0.91 | 0.84 | 0.8 |
| 2 | 223657553 | rs10206340 | -1.143 | 1.298 | 0.245 | -1.665 | 0.04 | -0.076 | -0.076 | 1.909 | 0.15 | 0.49 | 0.09 | 0.03 | 0.16 | 0.19 |
| 2 | 223661303 | rs9653359 | -1.037 | 1.472 | 0.167 | -1.737 | 0.032 | -0.081 | -0.081 | 0 | 0.16 | 0.42 | 0.27 | 0.03 | 0.18 | 0.2 |
| 2 | 223663875 | rs6436344 | 0.138 | 1.932 | -0.975 | -2.152 | -0.098 | 0.12 | 0.12 | 2.826 | 0.24 | 0.03 | 0.17 | 0.03 | 0.27 | 0.11 |
| 2 | 223664488 | rs13408246 | 0.202 | 1.614 | -0.483 | -2.013 | -0.003 | 0 | 0 | 0 | 0.11 | 0.09 | 0.23 | 0.03 | 0.14 | 0.17 |
| 2 | 223665111 | rs4673036 | 0.656 | 0.974 | -0.569 | -2.314 | -0.02 | 0.097 | 0.097 | 0 | 0.56 | 0.73 | 0.75 | 0.82 | 0.55 | 0.72 |
| 2 | 223667882 | rs1867802 | -0.317 | 0.691 | -0.105 | -1.913 | -0.004 | -0.044 | -0.044 | 0 | 0.14 | 0.17 | 0.08 | 0.02 | 0.12 | 0.06 |
| 2 | 223669074 | rs6760807 | -0.114 | 0.978 | -0.487 | -2.169 | -0.022 | 0.047 | 0.047 | 0 | 0.15 | 0.03 | 0.07 | 0.04 | 0.13 | 0.07 |
| 2 | 223674322 | rs6436346 | -0.575 | 0.689 | -0.129 | -1.611 | -0.03 | -0.04 | -0.04 | 0 | 0.7 | 0.84 | 0.59 | 0.59 | 0.68 | 0.71 |
| 2 | 223674479 | rs6718169 | -0.48 | 0.692 | -0.081 | -1.973 | -0.001 | -0.021 | -0.021 | 0 | 0.19 | 0.19 | 0.11 | 0.04 | 0.16 | 0.07 |
| 2 | 223682484 | rs7565176 | 1.726 | 0.611 | 0.558 | -1.655 | 0.001 | 0.001 | 0.001 | 0 | 0.37 | 0.33 | 0.41 | 0.65 | 0.44 | 0.58 |
| 2 | 223693230 | rs6751670 | 1.208 | 0.62 | 0.581 | -1.639 | 0.004 | 0.008 | 0.008 | 0 | 0.42 | 0.64 | 0.46 | 0.65 | 0.47 | 0.58 |
| 2 | 223698442 | rs10197778 | 0.327 | 0.571 | 0.369 | -1.701 | -0.034 | 0.066 | 0.066 | 0.099 | 0.55 | 0.81 | 0.52 | 0.7 | 0.57 | 0.66 |
| 2 | 223700432 | rs12994390 | 1.302 | 0.671 | 0.342 | -1.596 | -0.004 | -0.015 | -0.015 | 0.083 | 0.41 | 0.47 | 0.44 | 0.66 | 0.47 | 0.6 |
| 2 | 223709176 | rs41343745 | 0.544 | 0.682 | -0.017 | -1.681 | 0.008 | 0.024 | 0.024 | 0 | 0.13 | 0.08 | 0.05 | 0.03 | 0.1 | 0.06 |
| 2 | 223709493 | rs11885806 | 0.183 | 0.601 | 0.242 | -1.67 | 0.001 | 0.031 | 0.031 | 0 | 0.54 | 0.68 | 0.5 | 0.68 | 0.54 | 0.66 |
| 2 | 223715899 | rs2053717 | 1.857 | 0.603 | 0.074 | -1.754 | 0.005 | -0.012 | -0.012 | 0 | 0.33 | 0.23 | 0.38 | 0.65 | 0.37 | 0.58 |
| 2 | 223724690 | rs7606898 | -0.714 | 0.257 | 0.157 | -1.996 | -0.002 | 0.015 | 0.015 | 0 | 0.47 | 0.56 | 0.52 | 0.21 | 0.45 | 0.29 |
| 2 | 223727213 | rs10439277 | -0.143 | 0.252 | 0.174 | -1.92 | -0.006 | 0.036 | 0.036 | 0 | 0.43 | 0.31 | 0.47 | 0.2 | 0.42 | 0.28 |
| 2 | 223732283 | rs10469668 | -2.119 | 0.138 | 0.081 | -1.9 | -0.024 | 0.038 | 0.038 | 0 | 0.68 | 0.85 | 0.63 | 0.19 | 0.61 | 0.36 |
| 2 | 223743187 | rs12105595 | 1.101 | -0.08 | 0.274 | -1.843 | 0.02 | -0.016 | -0.016 | 0 | 0.05 | 0.21 | 0.05 | 0 | 0.06 | 0.02 |
| 2 | 223747117 | rs795892 | 1.878 | -0.064 | 0.058 | -1.831 | -0.016 | 0.01 | 0.01 | 0 | 0.33 | 0.15 | 0.37 | 0.76 | 0.39 | 0.63 |
| 2 | 223748542 | rs795882 | 1.055 | 0.19 | -0.184 | -1.698 | 0.001 | 0.007 | 0.007 | 0 | 0.42 | 0.61 | 0.46 | 0.76 | 0.45 | 0.66 |
| 2 | 223749562 | rs705727 | -1.131 | 0.293 | -0.295 | -1.708 | -0.017 | 0.01 | 0.01 | 0 | 0.93 | 0.97 | 0.94 | 0.98 | 0.89 | 0.98 |
| 2 | 223758070 | rs795887 | -0.35 | 0.225 | -0.202 | -1.625 | -0.027 | 0.08 | 0.08 | 0 | 0.81 | 0.94 | 0.89 | 0.97 | 0.79 | 0.92 |
| 2 | 223765491 | rs2119160 | -0.068 | 0.2 | -0.156 | -1.586 | -0.003 | 0.019 | 0.019 | 0 | 0.38 | 0.3 | 0.47 | 0.21 | 0.34 | 0.27 |
| 2 | 223766830 | rs6722420 | -0.205 | 0.255 | -0.174 | -1.638 | -0.007 | 0.041 | 0.041 | 0 | 0.8 | 0.88 | 0.88 | 0.97 | 0.79 | 0.92 |
| 2 | 223771053 | rs6717500 | 1.403 | 0.172 | -0.347 | -1.69 | -0.061 | -0.044 | -0.044 | 0.009 | 0.15 | 0.01 | 0.14 | 0.13 | 0.14 | 0.17 |
| 2 | 223785740 | rs6756323 | 0.845 | -0.064 | 0.549 | -1.293 | -0.004 | -0.019 | -0.019 | 0 | 0.37 | 0.46 | 0.4 | 0.76 | 0.39 | 0.63 |
| 2 | 223793149 | rs1485 | 1.028 | -0.061 | 0.091 | -1.293 | -0.001 | 0.012 | 0.012 | 0 | 0.34 | 0.27 | 0.37 | 0.76 | 0.38 | 0.63 |
| 2 | 223798501 | rs6756107 | 0.513 | 0.043 | -0.525 | -1.234 | 0 | -0.023 | -0.023 | 0.352 | 0.75 | 0.75 | 0.84 | 0.97 | 0.73 | 0.9 |
| 2 | 223807257 | rs1046108 | 0.281 | 0.171 | -1.237 | -0.974 | -0.004 | 0.02 | 0.02 | 0 | 0.38 | 0.3 | 0.43 | 0.21 | 0.34 | 0.27 |
| 2 | 223808401 | rs10755042 | 1.004 | 0.081 | -0.859 | -0.426 | -0.014 | 0.062 | 0.062 | 0 | 0.81 | 0.92 | 0.88 | 0.97 | 0.79 | 0.92 |
| 2 | 223810266 | rs6436364 | 0.89 | 0.084 | -0.86 | -0.42 | -0.007 | 0.045 | 0.045 | 0 | 0.8 | 0.9 | 0.87 | 0.97 | 0.79 | 0.92 |
| 2 | 223820025 | rs7607175 | -1.081 | -0.618 | -0.663 | -0.296 | 0.01 | 0.008 | 0.008 | 0 | 0.15 | 0.09 | 0.04 | 0.04 | 0.06 | 0.08 |
| 2 | 223823141 | rs6735297 | -0.762 | -0.918 | -0.877 | -0.492 | 0.019 | 0.062 | 0.062 | 1.024 | 0.33 | 0.16 | 0.38 | 0.13 | 0.32 | 0.21 |
| 2 | 223824537 | rs16864264 | 0.057 | -0.964 | -0.918 | -0.55 | 0.003 | 0.012 | 0.012 | 3.13 | 0.21 | 0.13 | 0.25 | 0.1 | 0.2 | 0.09 |
| 2 | 223832789 | rs16864271 | -0.768 | -0.983 | -0.326 | -0.515 | 0.005 | 0.023 | 0.023 | 0.02 | 0.12 | 0.06 | 0.14 | 0.05 | 0.13 | 0.11 |
| 2 | 223836467 | rs10168780 | 0.714 | -0.979 | -0.403 | -0.721 | 0.022 | 0.023 | 0.023 | 0 | 0.83 | 0.94 | 0.94 | 1 | 0.91 | 0.94 |
| 2 | 223838477 | rs6717855 | -0.939 | -0.953 | -0.375 | -0.713 | 0.008 | -0.021 | -0.021 | 0 | 0.2 | 0.16 | 0.09 | 0 | 0.12 | 0.08 |
| 2 | 223840963 | rs9808129 | 1.717 | -0.998 | -0.236 | -0.682 | 0.028 | 0.05 | 0.05 | 0 | 0.7 | 0.83 | 0.8 | 0.94 | 0.79 | 0.84 |
| 2 | 223845942 | rs7584993 | 1.712 | -1.076 | 0.217 | -0.158 | 0.049 | 0.053 | 0.053 | 0.578 | 0.49 | 0.38 | 0.59 | 0.7 | 0.66 | 0.75 |
| 2 | 223852353 | rs1440063 | 2.187 | -1.02 | 0.224 | -0.104 | 0.01 | 0.014 | 0.014 | 1.589 | 0.45 | 0.38 | 0.56 | 0.61 | 0.56 | 0.7 |
| 2 | 223872815 | rs1371648 | -1.219 | -1.052 | 0.216 | -0.308 | -0.021 | -0.08 | -0.08 | 0.473 | 0.34 | 0.38 | 0.17 | 0.03 | 0.2 | 0.12 |
| 2 | 223873878 | rs16864337 | -0.624 | -0.65 | -0.109 | 0.293 | 0.004 | -0.006 | -0.006 | 0 | 0.12 | 0.06 | 0.09 | 0.26 | 0.1 | 0.05 |
| 2 | 223875639 | rs11693836 | 0.035 | -0.853 | 0.229 | 0.052 | -0.017 | -0.051 | -0.051 | 0 | 0.17 | 0.2 | 0.09 | 0.01 | 0.09 | 0.06 |
| 2 | 223883460 | rs7565240 | 1.453 | -0.865 | 0.54 | 0.013 | -0.018 | -0.068 | -0.068 | 0 | 0.66 | 0.58 | 0.79 | 0.91 | 0.75 | 0.82 |
| 2 | 223884438 | rs13426685 | -1.059 | -0.827 | 0.493 | -0.013 | -0.008 | -0.055 | -0.055 | 0 | 0.13 | 0.21 | 0.06 | 0.02 | 0.1 | 0.07 |
| 2 | 223889682 | rs16864374 | -0.866 | -0.985 | 0.384 | 0.117 | 0.02 | -0.043 | -0.043 | 0.117 | 0.14 | 0.06 | 0.11 | 0.27 | 0.11 | 0.06 |

| | | | | | | | | | | | | | | | | |
|---|---|---|---|---|---|---|---|---|---|---|---|---|---|---|---|---|
| 2 | 223890168 | rs747869 | 0.412 | -1 | 0.513 | 0.105 | 0 | -0.024 | -0.024 | 0.359 | 0.16 | 0.13 | 0.07 | 0.01 | 0.09 | 0.06 |
| 2 | 223890360 | rs2395945 | 1.81 | -1.003 | 0.682 | 0.091 | -0.017 | 0 | 0 | 0 | 0.68 | 0.58 | 0.78 | 0.7 | 0.74 | 0.86 |
| 2 | 223891324 | rs1440073 | 1.81 | -1.003 | 0.682 | 0.091 | -0.017 | 0 | 0 | 0 | 0.68 | 0.58 | 0.78 | 0.7 | 0.74 | 0.86 |
| 2 | 223892964 | rs895772 | 2.211 | -1.011 | 0.674 | 0.087 | 0.016 | 0.015 | 0.015 | 0.065 | 0.52 | 0.43 | 0.7 | 0.7 | 0.65 | 0.8 |
| 2 | 223897721 | rs1440076 | 0.433 | -0.789 | 0.588 | 0.217 | 0.033 | -0.017 | -0.017 | 0 | 0.94 | 0.75 | 0.94 | 0.99 | 0.95 | 0.99 |
| 2 | 223897744 | rs17794731 | -1.688 | -0.823 | 0.414 | 0.264 | 0.017 | -0.014 | -0.014 | 0 | 0.26 | 0.17 | 0.16 | 0.27 | 0.21 | 0.13 |
| 2 | 223910943 | rs1899039 | 2.002 | -0.963 | 0.614 | 0.429 | 0.026 | 0.016 | 0.016 | 1.918 | 0.51 | 0.41 | 0.72 | 0.67 | 0.65 | 0.77 |
| 2 | 223911503 | rs4673048 | 1.005 | -0.955 | 0.624 | 0.406 | 0.016 | 0.008 | 0.008 | 4.793 | 0.68 | 0.75 | 0.84 | 0.79 | 0.8 | 0.91 |
| 2 | 223917983 | rs12621643 | 1.658 | -0.588 | 0.671 | 0.541 | -0.02 | -0.011 | -0.011 | 0 | 0.58 | 0.48 | 0.75 | 0.69 | 0.71 | 0.81 |
| 2 | 223918363 | rs3795884 | 0.015 | -0.797 | 0.27 | 0.506 | 0.051 | 0.045 | 0.045 | 3.015 | 0.81 | 0.92 | 0.96 | 0.99 | 0.93 | 0.93 |
| 2 | 223921771 | rs2017069 | 0.031 | -0.819 | 0.253 | 0.5 | 0.08 | 0.091 | 0.091 | 2.697 | 0.81 | 0.97 | 0.96 | 0.99 | 0.93 | 0.91 |
| 2 | 223930506 | rs9283536 | 1.761 | -0.755 | 0.373 | 0.585 | -0.005 | -0.001 | -0.001 | 0 | 0.59 | 0.55 | 0.76 | 0.7 | 0.74 | 0.86 |
| 2 | 223932598 | rs2119158 | 1.017 | -0.758 | 0.287 | 0.443 | 0.049 | 0.07 | 0.07 | 0.718 | 0.72 | 0.86 | 0.87 | 0.97 | 0.83 | 0.89 |
| 2 | 223937559 | rs960246 | 0.484 | -0.759 | 0.247 | 0.39 | 0.056 | 0.072 | 0.072 | 1.848 | 0.77 | 0.91 | 0.9 | 0.97 | 0.88 | 0.92 |
| 2 | 223948795 | rs7608931 | -0.064 | -0.574 | 0.218 | 0.27 | 0.072 | 0.077 | 0.077 | 2.592 | 0.81 | 0.96 | 0.96 | 0.99 | 0.93 | 0.94 |
| 2 | 223950293 | rs11898211 | -0.953 | -0.543 | 0.245 | 0.28 | -0.001 | -0.015 | -0.015 | 0 | 0.07 | 0.14 | 0.05 | 0 | 0.08 | 0.03 |
| 2 | 223954609 | rs7594650 | 1.325 | -0.508 | 0.325 | 0.272 | -0.039 | -0.12 | -0.12 | 0.361 | 0.67 | 0.54 | 0.83 | 0.97 | 0.8 | 0.89 |
| 2 | 223959874 | rs7577495 | -1.108 | -0.388 | 0.299 | 0.271 | -0.003 | 0.037 | 0.037 | 0 | 0.14 | 0.23 | 0.13 | 0.25 | 0.14 | 0.07 |
| 2 | 223960235 | rs2197779 | 1.446 | -0.434 | 0.28 | 0.275 | -0.012 | -0.009 | -0.009 | 0 | 0.58 | 0.53 | 0.76 | 0.74 | 0.71 | 0.83 |
| 2 | 223964797 | rs6436374 | 1.218 | -0.232 | -0.037 | 0.221 | 0.01 | 0.025 | 0.025 | 1.217 | 0.58 | 0.65 | 0.65 | 0.75 | 0.65 | 0.79 |
| 2 | 223974443 | rs978917 | 1.494 | -0.351 | -0.001 | 0.297 | -0.002 | -0.001 | -0.001 | 0 | 0.53 | 0.51 | 0.7 | 0.6 | 0.59 | 0.73 |
| 2 | 223986045 | rs1448315 | 0.554 | -0.442 | -0.207 | -0.095 | -0.013 | -0.027 | -0.027 | 0 | 0.79 | 0.8 | 0.94 | 0.96 | 0.93 | 0.91 |
| 2 | 224002355 | rs2034374 | 0.486 | -0.329 | -0.833 | -0.04 | 0.062 | 0.071 | 0.071 | 3.208 | 0.81 | 0.94 | 0.96 | 0.97 | 0.93 | 0.91 |
| 2 | 224006732 | rs6737886 | -1.187 | 1.088 | 0.046 | 0.122 | 0.092 | -0.017 | -0.017 | 0 | 0.15 | 0.38 | 0.13 | 0.12 | 0.23 | 0.15 |
| 2 | 224007819 | rs13003917 | -0.496 | 0.267 | -0.328 | 0.414 | -0.042 | 0.085 | 0.085 | 0 | 0.24 | 0.41 | 0.35 | 0.41 | 0.19 | 0.23 |
| 2 | 224009758 | rs2166877 | 1.715 | 0.374 | -1.07 | -0.384 | -0.007 | -0.055 | -0.055 | 2.772 | 0.44 | 0.13 | 0.45 | 0.47 | 0.42 | 0.52 |
| 2 | 224015791 | rs10181857 | 0.661 | 0.364 | -1.075 | -0.335 | -0.018 | -0.004 | -0.004 | 0.361 | 0.59 | 0.51 | 0.59 | 0.6 | 0.66 | 0.68 |
| 2 | 224018770 | rs16864505 | 0.117 | 0.396 | -0.832 | -0.294 | -0.002 | -0.002 | -0.002 | 0.018 | 0.05 | 0.05 | 0.02 | 0.05 | 0.03 | 0.02 |
| 2 | 224023217 | rs11899732 | -0.105 | 0.664 | -0.61 | -0.399 | 0.04 | 0.063 | 0.063 | 1.867 | 0.08 | 0.32 | 0.04 | 0.12 | 0.07 | 0.09 |
| 2 | 224023296 | rs895767 | 0.21 | 0.561 | -0.195 | -0.419 | 0.004 | -0.005 | -0.005 | 0 | 0.17 | 0.12 | 0.29 | 0.35 | 0.16 | 0.21 |
| 2 | 224025078 | rs6739367 | 0.486 | 0.716 | -0.19 | -0.532 | 0.004 | -0.007 | -0.007 | 0.541 | 0.07 | 0.08 | 0.09 | 0 | 0.16 | 0.07 |
| 2 | 224028510 | rs10204910 | 0.488 | 0.575 | -0.233 | -0.549 | -0.007 | 0.042 | 0.042 | 0.409 | 0.16 | 0.06 | 0.06 | 0.01 | 0.14 | 0.09 |
| 2 | 224031666 | rs11883605 | -0.325 | 0.784 | 0.9 | -0.364 | 0.058 | -0.055 | -0.055 | 0 | 0.09 | 0.31 | 0.12 | 0 | 0.15 | 0.07 |
| 2 | 224036595 | rs16864539 | -0.213 | 0.643 | 0.987 | -0.178 | 0 | 0.006 | 0.006 | 0 | 0.05 | 0.06 | 0.05 | 0.1 | 0.09 | 0.02 |
| 2 | 224038175 | rs10172933 | -1.344 | 0.605 | 0.729 | -0.869 | 0.001 | 0.002 | 0.002 | 0 | 0.4 | 0.33 | 0.25 | 0.21 | 0.43 | 0.44 |
| 2 | 224038765 | rs10175867 | -0.835 | 0.537 | 0.534 | -0.989 | 0 | 0.003 | 0.003 | 0 | 0.26 | 0.22 | 0.16 | 0.19 | 0.29 | 0.36 |
| 2 | 224044250 | rs1597167 | -2.603 | 0.513 | 0.404 | -0.742 | -0.011 | -0.04 | -0.04 | 0 | 0.57 | 0.88 | 0.57 | 0.49 | 0.62 | 0.63 |
| 2 | 224046645 | rs17190891 | 0.703 | 0.541 | 0.37 | -0.795 | -0.017 | 0.069 | 0.069 | 0 | 0.12 | 0.01 | 0.09 | 0.01 | 0.14 | 0.08 |
| 2 | 224047812 | rs1448303 | -2.268 | 0.532 | 0.584 | -0.774 | 0.014 | 0.07 | 0.07 | 0 | 0.5 | 0.68 | 0.38 | 0.26 | 0.58 | 0.51 |
| 2 | 224069989 | rs4234047 | -2.252 | 0.46 | 0.832 | -0.644 | 0.012 | 0.004 | 0.004 | 0 | 0.47 | 0.54 | 0.29 | 0.5 | 0.46 | 0.56 |
| 2 | 224071098 | rs4674761 | -0.704 | 0.433 | 0.883 | -0.654 | -0.005 | 0.001 | 0.001 | 0 | 0.21 | 0.19 | 0.11 | 0.25 | 0.17 | 0.2 |
| 2 | 224081654 | rs10804318 | -0.878 | 0.511 | 0.937 | -0.558 | 0 | 0.001 | 0.001 | 0 | 0.25 | 0.22 | 0.15 | 0.21 | 0.28 | 0.36 |
| 2 | 224087707 | rs4673053 | 1.602 | 0.469 | 0.851 | -0.486 | -0.004 | 0.009 | 0.009 | 0 | 0.66 | 0.46 | 0.79 | 0.51 | 0.68 | 0.52 |
| 2 | 224089192 | rs12623221 | 0.002 | 0.459 | 0.833 | -0.426 | -0.009 | 0.073 | 0.073 | 0 | 0.94 | 0.9 | 0.96 | 0.77 | 0.96 | 0.87 |
| 2 | 224093101 | rs2203733 | 2.039 | 0.712 | 0.575 | -0.485 | 0.002 | -0.004 | -0.004 | 0 | 0.47 | 0.41 | 0.68 | 0.46 | 0.5 | 0.4 |
| 2 | 224096286 | rs10197080 | -0.369 | 1.079 | 0.832 | -0.503 | 0.005 | -0.034 | -0.034 | 0 | 0.09 | 0.12 | 0.06 | 0.01 | 0.11 | 0.16 |
| 2 | 224099427 | rs11887070 | -0.377 | 1.247 | 0.993 | -0.456 | 0.137 | 0.006 | 0.006 | 2.801 | 0.05 | 0.33 | 0.08 | 0 | 0.1 | 0.05 |
| 2 | 224101021 | rs10933001 | -0.118 | 1.173 | 0.609 | -0.568 | -0.039 | 0.081 | 0.081 | 0 | 0.19 | 0.01 | 0.12 | 0.07 | 0.19 | 0.12 |
| 2 | 224102083 | rs10933002 | -0.338 | 1.082 | 0.656 | -0.446 | 0.007 | -0.022 | -0.022 | 2.346 | 0.06 | 0.02 | 0.03 | 0.17 | 0.03 | 0.09 |
| 2 | 224111323 | rs2395967 | 0.882 | 1.16 | 0.862 | -0.459 | 0.042 | 0.173 | 0.173 | 0.724 | 0.88 | 0.43 | 0.87 | 0.71 | 0.89 | 0.82 |
| 2 | 224112539 | rs4674771 | 2.175 | 1.468 | 0.21 | -0.502 | -0.005 | 0.042 | 0.042 | 0.384 | 0.52 | 0.15 | 0.64 | 0.55 | 0.48 | 0.37 |
| 2 | 224115686 | rs12694603 | -0.913 | 1.47 | 0.262 | -0.494 | -0.004 | 0.003 | 0.003 | 0.222 | 0.14 | 0.14 | 0.1 | 0.09 | 0.22 | 0.34 |
| 2 | 224128542 | rs4674774 | 1.627 | 1.386 | -0.475 | -0.407 | 0.001 | 0.041 | 0.041 | 0.314 | 0.52 | 0.15 | 0.64 | 0.55 | 0.47 | 0.37 |
| 2 | 224135966 | rs4674775 | -1.173 | 1.89 | -0.124 | -0.222 | 0.044 | 0.009 | 0.009 | 0 | 0.14 | 0.21 | 0.14 | 0.17 | 0.25 | 0.23 |
| 2 | 224140247 | rs12468161 | 0.254 | 2.057 | -0.724 | -0.466 | -0.087 | 0.085 | 0.085 | 0 | 0.21 | 0.01 | 0.14 | 0.06 | 0.25 | 0.16 |
| 2 | 224140630 | rs12479209 | 0.265 | 2.056 | -0.725 | -0.466 | -0.087 | 0.084 | 0.084 | 0 | 0.21 | 0.01 | 0.14 | 0.06 | 0.25 | 0.16 |
| 2 | 224145451 | rs1517657 | -0.61 | 2.346 | -0.55 | -0.41 | 0.015 | 0 | 0 | 0.069 | 0.12 | 0.13 | 0.14 | 0.12 | 0.27 | 0.22 |
| 2 | 224146067 | rs1914484 | 1.886 | 2.211 | -0.626 | -0.332 | 0.075 | -0.004 | -0.004 | 0 | 0.48 | 0.16 | 0.6 | 0.53 | 0.37 | 0.43 |
| 2 | 224149783 | rs1607248 | -0.661 | 2.342 | -0.462 | -0.231 | 0.013 | 0 | 0 | 1.633 | 0.12 | 0.13 | 0.13 | 0.12 | 0.27 | 0.22 |
| 2 | 224151518 | rs13424509 | -0.87 | 1.538 | -0.043 | -0.377 | 0.001 | 0.012 | 0.012 | 1.264 | 0.11 | 0.24 | 0.07 | 0.13 | 0.12 | 0.09 |
| 2 | 224175831 | rs1517628 | 0.525 | 1.349 | -0.387 | -1.101 | 0 | -0.029 | -0.029 | 0.181 | 0.63 | 0.38 | 0.74 | 0.9 | 0.6 | 0.71 |
| 2 | 224179428 | rs2894500 | -0.587 | 0.999 | 0.225 | -1.141 | 0.006 | -0.114 | -0.114 | 4.931 | 0.16 | 0.42 | 0.11 | 0.02 | 0.16 | 0.11 |
| 2 | 224180509 | rs1517630 | 0.719 | 0.774 | -0.507 | -1.189 | -0.001 | 0.004 | 0.004 | 0.026 | 0.18 | 0.14 | 0.13 | 0.1 | 0.17 | 0.2 |
| 2 | 224186245 | rs10445738 | -1.588 | 0.26 | -1.271 | -0.803 | 0.024 | -0.131 | -0.131 | 0 | 0.7 | 0.94 | 0.52 | 0.43 | 0.78 | 0.65 |
| 2 | 224193250 | rs1400795 | 0.944 | 0.838 | -0.761 | -0.675 | -0.006 | -0.005 | -0.005 | 0.083 | 0.24 | 0.19 | 0.17 | 0.29 | 0.25 | 0.24 |
| 2 | 224204815 | rs12151816 | 0.09 | 0.687 | -0.433 | -0.975 | -0.003 | 0.015 | 0.015 | 0 | 0.29 | 0.24 | 0.21 | 0.11 | 0.3 | 0.28 |
| 2 | 224204870 | rs10184043 | -1.793 | 0.467 | -0.872 | -1.079 | 0.022 | -0.137 | -0.137 | 0 | 0.69 | 0.94 | 0.51 | 0.43 | 0.76 | 0.59 |
| 2 | 224211363 | rs7580582 | 0.355 | 0.755 | -0.274 | -1.303 | 0.01 | -0.024 | -0.024 | 2.49 | 0.12 | 0.17 | 0.09 | 0.03 | 0.18 | 0.08 |
| 2 | 224226065 | rs2395981 | 1.663 | 0.775 | -0.346 | -1.381 | -0.023 | 0.012 | 0.012 | 0 | 0.14 | 0.03 | 0.12 | 0.08 | 0.17 | 0.16 |

| | | | | | | | | | | | | | | | | |
|---|---|---|---|---|---|---|---|---|---|---|---|---|---|---|---|---|
| 2 | 224226772 | rs1400797 | -1.351 | 0.622 | -0.082 | -1.346 | 0.032 | 0.024 | 0.024 | 0 | 0.63 | 0.72 | 0.51 | 0.24 | 0.76 | 0.5 |
| 2 | 224227452 | rs1517641 | -0.731 | 0.732 | -0.062 | -1.346 | 0.007 | -0.037 | -0.037 | 0 | 0.42 | 0.53 | 0.31 | 0.15 | 0.51 | 0.35 |
| 2 | 224232297 | rs990296 | 0.675 | 0.742 | -0.064 | -1.359 | -0.008 | 0.033 | 0.033 | 0 | 0.28 | 0.18 | 0.2 | 0.1 | 0.29 | 0.27 |
| 2 | 224232527 | rs1250924 | -0.502 | 0.539 | -0.401 | -1.214 | -0.003 | 0 | 0 | 0 | 0.43 | 0.59 | 0.32 | 0.15 | 0.51 | 0.35 |
| 2 | 224235559 | rs1250927 | -0.891 | 0.253 | -0.552 | -0.961 | 0.032 | 0.024 | 0.024 | 0 | 0.63 | 0.72 | 0.51 | 0.24 | 0.76 | 0.5 |
| 2 | 224239650 | rs997671 | -0.376 | 0.188 | -0.311 | -0.73 | -0.008 | -0.018 | -0.018 | 0 | 0.34 | 0.38 | 0.36 | 0.16 | 0.33 | 0.31 |
| 2 | 224244085 | rs12466239 | 0.937 | 0.331 | -0.487 | -0.9 | -0.03 | 0.047 | 0.047 | 0 | 0.29 | 0.16 | 0.25 | 0.11 | 0.32 | 0.18 |
| 2 | 224244450 | rs1250937 | 0.54 | 0.332 | -0.628 | -0.829 | -0.002 | -0.005 | -0.005 | 0.37 | 0.25 | 0.24 | 0.2 | 0.11 | 0.22 | 0.28 |
| 2 | 224244634 | rs1250939 | -0.178 | 0.182 | -0.729 | -0.014 | -0.02 | 0.045 | 0.045 | 0 | 0.92 | 0.85 | 0.97 | 0.83 | 0.97 | 0.9 |
| 2 | 224257311 | rs2395982 | 0.431 | 0.562 | -1.225 | 0.081 | -0.003 | -0.036 | -0.036 | 0 | 0.28 | 0.23 | 0.36 | 0.54 | 0.32 | 0.39 |
| 2 | 224261774 | rs1578934 | -0.772 | 0.542 | -0.964 | 0.108 | -0.001 | -0.001 | -0.001 | 0 | 0.07 | 0.05 | 0.15 | 0.05 | 0.08 | 0.03 |
| 2 | 224267984 | rs2395983 | -0.169 | 0.572 | -1.451 | 0.163 | -0.032 | 0.063 | 0.063 | 0 | 0.59 | 0.84 | 0.63 | 0.69 | 0.56 | 0.61 |
| 2 | 224269949 | rs1921647 | 1.157 | 0.427 | -1.607 | 0.48 | -0.029 | -0.029 | -0.029 | 0 | 0.28 | 0.16 | 0.27 | 0.34 | 0.3 | 0.28 |
| 2 | 224275952 | rs16864757 | 0.203 | 0.164 | -0.418 | 0.479 | -0.006 | 0.051 | 0.051 | 0 | 0.22 | 0.3 | 0.23 | 0.32 | 0.21 | 0.31 |
| 2 | 224277827 | rs1921635 | -0.601 | -0.027 | -0.283 | 0.535 | -0.019 | 0.057 | 0.057 | 0.57 | 0.3 | 0.47 | 0.45 | 0.53 | 0.27 | 0.52 |
| 2 | 224282988 | rs7600591 | -1.131 | -0.055 | -0.633 | 0.297 | -0.1 | 0.161 | 0.161 | 0 | 0.66 | 0.98 | 0.78 | 0.9 | 0.62 | 0.82 |
| 2 | 224300080 | rs16864786 | -0.386 | 0.026 | -0.394 | 0.451 | -0.01 | 0.035 | 0.035 | 0 | 0.28 | 0.36 | 0.32 | 0.37 | 0.26 | 0.34 |
| 2 | 224300897 | rs16864788 | 1.577 | -0.014 | -0.139 | 0.666 | 0.025 | 0.089 | 0.089 | 0 | 0.07 | 0.13 | 0.16 | 0.19 | 0.1 | 0.22 |
| 2 | 224301025 | rs1882354 | -0.632 | 0.02 | 0.01 | 0.42 | -0.016 | 0.041 | 0.041 | 0 | 0.29 | 0.48 | 0.34 | 0.36 | 0.26 | 0.34 |
| 2 | 224306142 | rs1317691 | 1.438 | 0.064 | -0.053 | 0.405 | -0.038 | -0.025 | -0.025 | 0 | 0.3 | 0.16 | 0.28 | 0.35 | 0.33 | 0.28 |
| 2 | 224308931 | rs2395986 | -1.161 | 0.047 | 0.123 | 0.412 | -0.04 | 0.149 | 0.149 | 0 | 0.66 | 0.9 | 0.79 | 0.9 | 0.61 | 0.8 |
| 2 | 224314224 | rs7563838 | 0.582 | 0 | 0.171 | 0.413 | -0.004 | 0.053 | 0.053 | 0 | 0.21 | 0.27 | 0.23 | 0.35 | 0.21 | 0.32 |
| 2 | 224318539 | rs934995 | 1.057 | 0.017 | 0.088 | 0.41 | -0.03 | -0.005 | -0.005 | 0 | 0.34 | 0.21 | 0.3 | 0.34 | 0.35 | 0.31 |
| 2 | 224321540 | rs1950123 | -0.422 | 0.095 | 0.123 | 0.286 | -0.013 | 0.125 | 0.125 | 0 | 0.6 | 0.81 | 0.69 | 0.88 | 0.56 | 0.79 |
| 2 | 224323672 | rs4674783 | 0.538 | 0.128 | 0.146 | 0.355 | 0 | 0 | 0 | 0.011 | 0.06 | 0.06 | 0.09 | 0.02 | 0.05 | 0.02 |
| 2 | 224344833 | rs7601486 | -0.937 | 0.363 | -0.272 | -0.073 | -0.009 | 0.085 | 0.085 | 0 | 0.65 | 0.83 | 0.78 | 0.9 | 0.61 | 0.8 |
| 2 | 224360395 | rs10173406 | -0.238 | 0.482 | -0.215 | 0.333 | 0 | 0 | 0 | 0 | 0.44 | 0.49 | 0.54 | 0.54 | 0.41 | 0.51 |
| 2 | 224363430 | rs6726892 | 0.568 | 0.432 | -0.504 | 0.238 | -0.006 | -0.011 | -0.011 | 0 | 0.34 | 0.29 | 0.44 | 0.54 | 0.37 | 0.51 |
| 2 | 224364255 | rs6730755 | 0.076 | 0.401 | -0.737 | 0.358 | -0.001 | 0 | 0 | 0 | 0.47 | 0.58 | 0.57 | 0.55 | 0.44 | 0.56 |
| 2 | 224365038 | rs6721111 | 0.064 | 0.285 | -1.321 | -0.233 | -0.007 | 0.125 | 0.125 | 0 | 0.6 | 0.83 | 0.77 | 0.85 | 0.59 | 0.81 |
| 2 | 224365778 | rs1439933 | 0.255 | -0.852 | -1.493 | -0.562 | 0.002 | 0.121 | 0.121 | 0 | 0.68 | 0.91 | 0.84 | 0.87 | 0.72 | 0.83 |
| 2 | 224365894 | rs1439932 | 0.359 | -1.277 | -1.048 | -0.741 | 0.003 | 0.006 | 0.006 | 0 | 0.67 | 0.75 | 0.63 | 0.79 | 0.73 | 0.64 |
| 2 | 224366779 | rs721357 | 1.326 | -1.326 | -0.958 | -0.601 | 0.011 | 0.02 | 0.02 | 0 | 0.51 | 0.62 | 0.56 | 0.64 | 0.63 | 0.56 |
| 2 | 224366956 | rs721358 | 0.566 | -1.735 | -0.974 | -0.707 | -0.013 | -0.054 | -0.054 | 0 | 0.1 | 0 | 0.23 | 0.22 | 0.06 | 0.24 |
| 2 | 224373759 | rs714943 | 0.799 | -2.064 | -0.848 | -0.705 | -0.014 | 0.013 | 0.013 | 0.092 | 0.25 | 0.39 | 0.2 | 0.31 | 0.21 | 0.14 |
| 2 | 224380120 | rs1439930 | 0.905 | -2.035 | -0.751 | -0.684 | 0.01 | 0.024 | 0.024 | 0.281 | 0.28 | 0.15 | 0.19 | 0.23 | 0.24 | 0.15 |
| 2 | 224388481 | rs6728104 | -1.003 | -1.987 | -0.33 | -0.573 | 0.008 | 0.004 | 0.004 | 0 | 0.27 | 0.23 | 0.3 | 0.25 | 0.13 | 0.29 |
| 2 | 224390501 | rs2396006 | 0.023 | -2.002 | -0.289 | -0.586 | 0.024 | 0.032 | 0.032 | 0.545 | 0.15 | 0.09 | 0.05 | 0.03 | 0.07 | 0.05 |
| 2 | 224394506 | rs893766 | 0.487 | -1.707 | -0.193 | -0.398 | -0.008 | 0.015 | 0.015 | 0 | 0.12 | 0.13 | 0.24 | 0.22 | 0.06 | 0.24 |
| 2 | 224397461 | rs7565690 | 1.128 | -1.619 | -0.327 | -0.386 | 0.006 | 0.01 | 0.01 | 0 | 0.12 | 0.19 | 0.1 | 0.17 | 0.14 | 0.12 |
| 2 | 224397999 | rs1439929 | 0.514 | -1.419 | -0.373 | -0.238 | 0.006 | -0.013 | -0.013 | 0.015 | 0.11 | 0.08 | 0.24 | 0.22 | 0.06 | 0.24 |
| 2 | 224408490 | rs11675944 | -1.8 | -1.334 | -0.272 | -0.306 | 0.019 | 0.016 | 0.016 | 0.375 | 0.73 | 0.74 | 0.63 | 0.65 | 0.57 | 0.57 |
| 2 | 224408554 | rs13387403 | 0.776 | -1.471 | -0.326 | -0.189 | -0.005 | 0.033 | 0.033 | 0 | 0.11 | 0.12 | 0.24 | 0.22 | 0.07 | 0.24 |
| 2 | 224410192 | rs16864923 | 0.19 | -1.634 | -0.692 | -0.395 | 0.042 | 0.086 | 0.086 | 0 | 0.23 | 0.06 | 0.1 | 0.1 | 0.16 | 0.12 |
| 2 | 224413227 | rs11885052 | 0.048 | -1.421 | -0.455 | -0.413 | 0.005 | 0.03 | 0.03 | 0 | 0.34 | 0.29 | 0.25 | 0.23 | 0.31 | 0.17 |
| 2 | 224413393 | rs1371548 | -1.227 | -1.784 | -0.296 | -0.813 | -0.01 | -0.005 | -0.005 | 1.042 | 0.4 | 0.42 | 0.37 | 0.33 | 0.24 | 0.37 |
| 2 | 224418252 | rs1371549 | -0.661 | -1.884 | -0.488 | -0.702 | 0.014 | 0.168 | 0.168 | 0 | 0.35 | 0.06 | 0.18 | 0.07 | 0.29 | 0.18 |
| 2 | 224439486 | rs10498152 | -0.056 | -1.833 | -0.671 | -0.679 | -0.059 | 0.151 | 0.151 | 0 | 0.27 | 0.02 | 0.16 | 0.07 | 0.27 | 0.16 |
| 2 | 224453952 | rs959100 | 0.211 | -1.916 | 0.353 | 0.044 | -0.028 | 0.068 | 0.068 | 0 | 0.18 | 0.07 | 0.16 | 0.08 | 0.22 | 0.13 |
| 2 | 224459683 | rs16864968 | -0.426 | -1.941 | 0.169 | 0.121 | -0.056 | 0.038 | 0.038 | 1.933 | 0.12 | 0.4 | 0.05 | 0.11 | 0.02 | 0.04 |
| 2 | 224465827 | rs2894511 | -0.104 | -1.665 | 0.205 | 0.713 | 0 | 0.003 | 0.003 | 0 | 0.89 | 0.91 | 0.77 | 0.78 | 0.95 | 0.74 |
| 2 | 224469380 | rs3754632 | -0.989 | -0.95 | 0.399 | 0.189 | -0.03 | -0.051 | -0.051 | 0 | 0.31 | 0.48 | 0.21 | 0.19 | 0.24 | 0.17 |
| 2 | 224473767 | rs13402129 | -1.881 | -1.042 | 0.983 | 0.505 | -0.046 | -0.015 | -0.015 | 0 | 0.14 | 0.5 | 0.1 | 0.11 | 0.08 | 0.06 |
| 2 | 224474315 | rs16864994 | -2.538 | -1.239 | 1.692 | 0.595 | -0.022 | 0.016 | 0.016 | 2.243 | 0.08 | 0.28 | 0.03 | 0.06 | 0.01 | 0.04 |
| 2 | 224476078 | rs10177693 | -2.741 | -1.419 | 1.814 | 0.62 | -0.084 | 0.014 | 0.014 | 4.089 | 0.17 | 0.44 | 0.07 | 0.15 | 0.06 | 0.07 |
| 2 | 224483652 | rs7585110 | -3.154 | -1.463 | 2.068 | 0.606 | -0.004 | 0.034 | 0.034 | 2.359 | 0.08 | 0.35 | 0.03 | 0.06 | 0.01 | 0.04 |
| 2 | 224498382 | rs1014492 | -1.764 | -1.557 | 2.6 | 0.631 | -0.036 | 0.011 | 0.011 | 2.029 | 0.1 | 0.23 | 0.05 | 0.1 | 0.04 | 0.03 |
| 2 | 224498568 | rs1014491 | -3.141 | -1.591 | 2.777 | 0.741 | -0.092 | 0.04 | 0.04 | 5.714 | 0.18 | 0.46 | 0.07 | 0.19 | 0.06 | 0.08 |
| 2 | 224508025 | rs13385650 | -3.014 | -1.605 | 2.832 | 0.733 | -0.066 | 0.053 | 0.053 | 4.303 | 0.15 | 0.43 | 0.06 | 0.15 | 0.05 | 0.06 |
| 2 | 224512860 | rs7557395 | -3.421 | -1.6 | 2.598 | 0.732 | -0.09 | 0.024 | 0.024 | 6.521 | 0.19 | 0.53 | 0.07 | 0.2 | 0.06 | 0.08 |
| 2 | 224522028 | rs6436419 | -2.769 | -1.369 | 2.583 | 0.856 | 0.012 | 0.163 | 0.163 | 3.09 | 0.08 | 0.47 | 0.06 | 0.14 | 0.04 | 0.04 |
| 2 | 224531444 | rs7585813 | -2.877 | -1.398 | 2.44 | 0.94 | -0.067 | 0.086 | 0.086 | 6.225 | 0.15 | 0.56 | 0.08 | 0.19 | 0.06 | 0.08 |
| 2 | 224544818 | rs4673070 | -0.153 | -1.398 | 1.498 | 0.775 | 0.006 | -0.001 | -0.001 | 0 | 0.61 | 0.7 | 0.52 | 0.36 | 0.71 | 0.67 |
| 2 | 224555134 | rs7590719 | -0.86 | -1.281 | 1.598 | 0.912 | -0.006 | 0.037 | 0.037 | 0.155 | 0.05 | 0.1 | 0.04 | 0.11 | 0.04 | 0.04 |
| 2 | 224562229 | rs7570601 | -1.913 | -1.273 | 1.946 | 0.956 | 0.029 | 0.133 | 0.133 | 1.086 | 0.08 | 0.47 | 0.05 | 0.06 | 0.02 | 0.02 |
| 2 | 224562525 | rs4674808 | 0.371 | -1.51 | 1.254 | 0.734 | 0.014 | 0.031 | 0.031 | 0 | 0.56 | 0.68 | 0.52 | 0.26 | 0.69 | 0.61 |
| 2 | 224563288 | rs11888087 | 0.154 | -1.401 | 0.788 | 1.422 | 0.008 | -0.003 | -0.003 | 0 | 0.89 | 0.95 | 0.77 | 0.58 | 0.94 | 0.8 |
| 2 | 224568089 | rs12992947 | 0.269 | -1.302 | 1.033 | 1.397 | -0.014 | 0.257 | 0.257 | 0.038 | 0.87 | 0.59 | 0.81 | 0.6 | 0.94 | 0.81 |
| 2 | 224569899 | rs1866246 | 0.306 | -1.241 | 0.89 | 1.41 | -0.019 | 0.214 | 0.214 | 0.597 | 0.87 | 0.62 | 0.81 | 0.59 | 0.94 | 0.81 |

| | | | | | | | | | | | | | | | | |
|---|---|---|---|---|---|---|---|---|---|---|---|---|---|---|---|---|
| 2 | 224570349 | rs893451 | 0.536 | -1.26 | -0.168 | 1.404 | -0.037 | 0.114 | 0.114 | 2.095 | 0.84 | 0.38 | 0.79 | 0.59 | 0.93 | 0.81 |
| 2 | 224576402 | rs10176854 | 0.508 | -1.019 | -1.282 | 0.944 | 0.069 | 0.051 | 0.051 | 0.184 | 0.52 | 0.11 | 0.55 | 0.34 | 0.7 | 0.62 |
| 2 | 224581443 | rs7589996 | -0.499 | -0.024 | -1.036 | 0.901 | 0.005 | 0 | 0 | 0 | 0.46 | 0.6 | 0.42 | 0.44 | 0.41 | 0.4 |
| 2 | 224584078 | rs10185440 | 0.18 | 0.529 | -1.447 | 0.847 | -0.034 | 0.013 | 0.013 | 0 | 0.61 | 0.8 | 0.62 | 0.67 | 0.53 | 0.77 |
| 2 | 224584355 | rs12474832 | 0.18 | 0.529 | -1.447 | 0.847 | -0.034 | 0.013 | 0.013 | 0 | 0.61 | 0.8 | 0.62 | 0.67 | 0.53 | 0.77 |
| 2 | 224589375 | rs4673073 | -0.235 | 0.959 | -1.431 | 0.679 | -0.003 | -0.016 | -0.016 | 1.277 | 0.72 | 0.84 | 0.74 | 0.67 | 0.74 | 0.83 |
| 2 | 224594801 | rs7607927 | -0.325 | 0.79 | -1.428 | 0.697 | -0.002 | -0.023 | -0.023 | 0 | 0.73 | 0.85 | 0.74 | 0.68 | 0.74 | 0.84 |
| 2 | 224596619 | rs10498153 | 0.903 | 1.185 | -2.11 | 0.496 | -0.069 | 0.033 | 0.033 | 0 | 0.2 | 0.04 | 0.2 | 0.12 | 0.25 | 0.13 |
| 2 | 224596756 | rs11902634 | 1.37 | 0.782 | -1.54 | 0.616 | 0.01 | -0.007 | -0.007 | 0 | 0.34 | 0.33 | 0.37 | 0.36 | 0.28 | 0.5 |
| 2 | 224599175 | rs655783 | -0.054 | 0.836 | -1.614 | 0.546 | 0.01 | 0.002 | 0.002 | 0 | 0.53 | 0.77 | 0.53 | 0.56 | 0.49 | 0.71 |
| 2 | 224600826 | rs684002 | 0.587 | 0.844 | -1.372 | 0.817 | -0.036 | 0.045 | 0.045 | 0 | 0.1 | 0.02 | 0.09 | 0 | 0.18 | 0.03 |
| 2 | 224605340 | rs755018 | 1.065 | 0.684 | -1.456 | 1.029 | -0.023 | -0.004 | -0.004 | 0 | 0.55 | 0.31 | 0.49 | 0.4 | 0.52 | 0.63 |
| 2 | 224610232 | rs4673075 | -0.137 | 0.719 | -1.371 | 0.545 | 0 | 0.155 | 0.155 | 0 | 0.06 | 0.24 | 0.14 | 0.19 | 0.03 | 0.04 |
| 2 | 224619161 | rs3099468 | -0.811 | 0.946 | -2.048 | 0.247 | -0.046 | 0.06 | 0.06 | 0 | 0.89 | 0.98 | 0.91 | 1 | 0.8 | 0.96 |
| 2 | 224620222 | rs12151380 | 0.709 | 0.82 | -2.152 | 0.184 | -0.076 | 0.116 | 0.116 | 0 | 0.19 | 0.02 | 0.17 | 0.02 | 0.2 | 0.12 |
| 2 | 224627775 | rs4674815 | 0.035 | 0.458 | -1.977 | 0.375 | -0.088 | 0.094 | 0.094 | 0 | 0.67 | 0.87 | 0.71 | 0.85 | 0.55 | 0.83 |
| 2 | 224631953 | rs10193515 | 0.506 | 0.688 | -1.991 | 0.384 | -0.01 | 0.123 | 0.123 | 0.16 | 0.47 | 0.19 | 0.35 | 0.24 | 0.46 | 0.58 |
| 2 | 224647136 | rs581815 | 0.088 | 0.411 | -1.353 | 0.874 | -0.017 | 0.002 | 0.002 | 0 | 0.43 | 0.3 | 0.48 | 0.43 | 0.55 | 0.52 |
| 2 | 224658733 | rs7604002 | -0.516 | 0.326 | -1.388 | 1.232 | 0.012 | -0.034 | -0.034 | 0.679 | 0.1 | 0.04 | 0.2 | 0.27 | 0.07 | 0.23 |
| 2 | 224662848 | rs7599342 | -0.408 | 0.281 | -0.927 | 0.968 | -0.003 | 0.002 | 0.002 | 0 | 0.57 | 0.62 | 0.56 | 0.63 | 0.5 | 0.52 |
| 2 | 224665694 | rs7583875 | 0.116 | -0.275 | -0.112 | 1.147 | -0.001 | -0.006 | -0.006 | 0.499 | 0.43 | 0.44 | 0.35 | 0.29 | 0.42 | 0.31 |
| 2 | 224669429 | rs10933018 | 0.271 | -0.372 | -0.155 | 1.041 | 0 | -0.035 | -0.035 | 5.813 | 0.39 | 0.34 | 0.25 | 0.07 | 0.38 | 0.27 |
| 2 | 224673015 | rs12467861 | 0.275 | -0.529 | 0.284 | 0.963 | 0 | -0.05 | -0.05 | 0 | 0.4 | 0.38 | 0.26 | 0.07 | 0.38 | 0.27 |
| 2 | 224674868 | rs17243807 | 0.291 | -0.655 | 0.258 | 0.864 | -0.001 | -0.007 | -0.007 | 0 | 0.38 | 0.3 | 0.27 | 0.07 | 0.37 | 0.26 |
| 2 | 224679435 | rs7587148 | -0.591 | -0.829 | 0.619 | 1.189 | -0.001 | 0 | 0 | 0 | 0.53 | 0.54 | 0.55 | 0.44 | 0.48 | 0.54 |
| 2 | 224693416 | rs12468846 | -0.444 | -0.815 | 0.572 | 1.224 | 0 | 0.024 | 0.024 | 0 | 0.13 | 0.15 | 0.31 | 0.38 | 0.11 | 0.27 |
| 2 | 224695232 | rs4673077 | -0.372 | -0.791 | 0.684 | 1.228 | 0 | 0 | 0 | 0.333 | 0.51 | 0.45 | 0.58 | 0.45 | 0.48 | 0.54 |
| 2 | 224696267 | rs7558186 | -0.372 | -0.741 | 0.684 | 1.228 | 0 | 0 | 0 | 3.397 | 0.51 | 0.45 | 0.58 | 0.45 | 0.48 | 0.54 |
| 2 | 224722446 | rs6436445 | 0.017 | -0.634 | 0.542 | 1.022 | -0.013 | 0.007 | 0.007 | 0.073 | 0.18 | 0.12 | 0.25 | 0.14 | 0.24 | 0.26 |
| 2 | 224723032 | rs7599348 | -0.367 | -0.973 | 0.579 | 0.979 | 0.005 | 0.068 | 0.068 | 0 | 0.48 | 0.33 | 0.42 | 0.19 | 0.47 | 0.46 |
| 2 | 224724493 | rs7564664 | -0.594 | -1.033 | 0.64 | 1.007 | 0 | -0.001 | -0.001 | 0 | 0.52 | 0.46 | 0.53 | 0.46 | 0.49 | 0.54 |
| 2 | 224738957 | rs633956 | -0.929 | -1.495 | 0.446 | 1.193 | -0.043 | 0.028 | 0.028 | 0.188 | 0.88 | 0.74 | 0.94 | 0.83 | 0.94 | 0.98 |
| 2 | 224740705 | rs666438 | 0.285 | -1.547 | 0.484 | 1.275 | -0.072 | 0.182 | 0.182 | 0.491 | 0.83 | 0.6 | 0.83 | 0.58 | 0.93 | 0.9 |
| 2 | 224741642 | rs10498156 | -0.416 | -1.517 | 0.264 | 1.275 | -0.016 | 0.141 | 0.141 | 0 | 0.05 | 0.14 | 0.11 | 0.25 | 0.01 | 0.07 |
| 2 | 224743373 | rs571111 | -0.016 | -1.097 | 0.108 | 1.104 | 0.001 | 0.035 | 0.035 | 0 | 0.49 | 0.37 | 0.43 | 0.19 | 0.53 | 0.48 |
| 2 | 224754537 | rs632138 | -1.015 | -1.02 | 0.031 | 1.241 | -0.001 | -0.002 | -0.002 | 0 | 0.89 | 0.88 | 0.95 | 0.83 | 0.94 | 0.98 |
| 2 | 224763816 | rs530065 | -0.983 | -1.016 | 0.033 | 1.243 | 0 | -0.003 | -0.003 | 0.047 | 0.89 | 0.88 | 0.96 | 0.83 | 0.94 | 0.98 |
| 2 | 224765894 | rs604990 | -0.983 | -1.016 | 0.033 | 1.243 | 0 | -0.003 | -0.003 | 0 | 0.89 | 0.88 | 0.96 | 0.83 | 0.94 | 0.98 |
| 2 | 224773548 | rs548470 | -0.158 | -0.992 | 0.011 | 0.987 | -0.026 | 0.067 | 0.067 | 0.104 | 0.85 | 0.78 | 0.85 | 0.58 | 0.93 | 0.92 |
| 2 | 224787773 | rs509818 | 1.864 | -0.516 | -0.609 | 0.885 | -0.093 | -0.079 | -0.079 | 0 | 0.22 | 0.02 | 0.32 | 0.28 | 0.27 | 0.35 |
| 2 | 224792256 | rs6746036 | -0.14 | -1.251 | -0.13 | 0.743 | -0.064 | 0.108 | 0.108 | 0 | 0.81 | 0.69 | 0.84 | 0.58 | 0.91 | 0.88 |
| 2 | 224795038 | rs10933025 | 0.221 | -1.314 | -0.032 | 0.71 | -0.093 | 0.124 | 0.124 | 0.375 | 0.78 | 0.44 | 0.83 | 0.58 | 0.91 | 0.87 |
| 2 | 224800016 | rs6721089 | 0.075 | -1.169 | -0.254 | 0.491 | -0.018 | 0.058 | 0.058 | 0 | 0.44 | 0.28 | 0.41 | 0.26 | 0.51 | 0.48 |
| 2 | 224808313 | rs1530085 | 1.3 | -1.076 | -0.539 | 0.316 | -0.086 | -0.024 | -0.024 | 0.673 | 0.31 | 0.11 | 0.4 | 0.32 | 0.41 | 0.39 |
| 2 | 224814186 | rs6734908 | 0.59 | -1.159 | -0.414 | 0.255 | 0.004 | -0.013 | -0.013 | 0 | 0.42 | 0.51 | 0.43 | 0.31 | 0.43 | 0.42 |
| 2 | 224819470 | rs12465843 | 1.139 | -1.012 | -1.192 | -0.061 | -0.035 | 0.006 | 0.006 | 0 | 0.18 | 0.01 | 0.18 | 0.11 | 0.16 | 0.17 |
| 2 | 224820329 | rs10209883 | 0.544 | -0.766 | -0.775 | -0.031 | -0.032 | 0.034 | 0.034 | 0 | 0.13 | 0.08 | 0.17 | 0.01 | 0.26 | 0.11 |
| 2 | 224835369 | rs1025734 | 0.801 | -1.092 | -1.336 | 0.219 | -0.122 | 0.027 | 0.027 | 0.15 | 0.41 | 0.08 | 0.4 | 0.3 | 0.47 | 0.48 |
| 2 | 224836562 | rs16865366 | 0.702 | -1.714 | -0.423 | 0.084 | -0.038 | -0.056 | -0.056 | 0 | 0.1 | 0.24 | 0.04 | 0 | 0.03 | 0.01 |
| 2 | 224837750 | rs12694625 | -1.027 | -1.951 | -0.504 | 0.426 | -0.013 | -0.031 | -0.031 | 0.072 | 0.57 | 0.75 | 0.56 | 0.47 | 0.47 | 0.53 |
| 2 | 224839696 | rs6754561 | -0.913 | -2.051 | -0.239 | 0.354 | 0 | 0.007 | 0.007 | 0 | 0.34 | 0.6 | 0.44 | 0.4 | 0.35 | 0.36 |
| 2 | 224846015 | rs16865390 | -0.082 | -2.94 | 0.46 | 0.996 | -0.063 | 0.065 | 0.065 | 0.333 | 0.14 | 0.37 | 0.23 | 0.16 | 0.04 | 0.04 |
| 2 | 224847707 | rs975278 | -0.118 | -3.176 | 0.358 | 1.007 | -0.003 | -0.003 | -0.003 | 0 | 0.68 | 0.71 | 0.82 | 0.43 | 0.83 | 0.64 |
| 2 | 224847806 | rs7605945 | 0.36 | -2.809 | -0.229 | 0.883 | 0.075 | 0.047 | 0.047 | 0.379 | 0.55 | 0.33 | 0.6 | 0.28 | 0.79 | 0.6 |
| 2 | 224854068 | rs7608941 | 0.318 | -2.083 | -0.256 | 0.866 | 0.021 | 0.003 | 0.003 | 1.829 | 0.53 | 0.51 | 0.61 | 0.48 | 0.75 | 0.71 |
| 2 | 224857266 | rs10191694 | 0.203 | -1.641 | -0.366 | 1.081 | -0.024 | 0.035 | 0.035 | 0 | 0.64 | 0.51 | 0.74 | 0.42 | 0.68 | 0.56 |
| 2 | 224858093 | rs16865421 | 0.168 | -1.518 | -0.377 | 1.074 | -0.005 | 0.06 | 0.06 | 0 | 0.07 | 0.13 | 0.07 | 0.22 | 0.07 | 0.21 |
| 2 | 224858750 | rs6738983 | 0.407 | -1.651 | -0.849 | 0.674 | 0.142 | 0.081 | 0.081 | 0.256 | 0.47 | 0.21 | 0.52 | 0.28 | 0.67 | 0.56 |
| 2 | 224860216 | rs13396978 | -0.851 | -1.413 | 0.075 | 0.562 | -0.018 | -0.037 | -0.037 | 0 | 0.33 | 0.63 | 0.23 | 0.13 | 0.2 | 0.16 |
| 2 | 224866576 | rs12436 | -1.215 | -1.499 | 0.667 | 1.027 | -0.039 | 0.058 | 0.058 | 0 | 0.17 | 0.35 | 0.18 | 0.24 | 0.11 | 0.24 |
| 2 | 224870700 | rs6436453 | 1.565 | -1.022 | 1.303 | 1.456 | -0.008 | 0.076 | 0.076 | 0 | 0.74 | 0.52 | 0.7 | 0.57 | 0.75 | 0.74 |
| 2 | 224873690 | rs7590948 | 1.994 | -1.074 | 1.128 | 1.502 | -0.027 | -0.052 | -0.052 | 1.007 | 0.46 | 0.09 | 0.51 | 0.42 | 0.55 | 0.57 |
| 2 | 224876707 | rs17196253 | -0.127 | -1.097 | 1.363 | 1.488 | -0.017 | -0.027 | -0.027 | 0 | 0.27 | 0.36 | 0.19 | 0.16 | 0.2 | 0.16 |
| 2 | 224877511 | rs7602990 | 1.114 | -0.977 | 1.702 | 1.579 | -0.027 | 0.07 | 0.07 | 0.057 | 0.93 | 0.78 | 0.96 | 0.79 | 0.98 | 0.93 |
| 2 | 224877561 | rs7563931 | 1.404 | -0.973 | 1.777 | 1.591 | -0.049 | 0.162 | 0.162 | 0.173 | 0.83 | 0.59 | 0.8 | 0.58 | 0.9 | 0.77 |
| 2 | 224878301 | rs1438828 | -0.17 | -0.897 | 1.65 | 1.569 | -0.007 | 0.075 | 0.075 | 0 | 0.1 | 0.17 | 0.16 | 0.21 | 0.09 | 0.16 |
| 2 | 224880336 | rs6436459 | -1.182 | -0.88 | 1.651 | 1.558 | 0.011 | 0.009 | 0.009 | 0 | 0.18 | 0.22 | 0.25 | 0.21 | 0.24 | 0.19 |
| 2 | 224887340 | rs4368321 | -0.254 | -0.904 | 1.609 | 1.284 | -0.019 | -0.031 | -0.031 | 0 | 0.26 | 0.38 | 0.18 | 0.16 | 0.19 | 0.16 |
| 2 | 224889383 | rs13008520 | -0.2 | -1.045 | 1.294 | 1.414 | 0.011 | -0.027 | -0.027 | 0.049 | 0.09 | 0.11 | 0.1 | 0 | 0.15 | 0.03 |

| | | | | | | | | | | | | | | | | |
|---|---|---|---|---|---|---|---|---|---|---|---|---|---|---|---|---|
| 2 | 224892945 | rs920251 | 0.795 | -0.783 | 1.331 | 1.404 | -0.001 | -0.001 | -0.001 | 0.245 | 0.63 | 0.59 | 0.73 | 0.6 | 0.66 | 0.72 |
| 2 | 224893429 | rs6719480 | -0.303 | -0.799 | 1.361 | 1.384 | 0.018 | 0.039 | 0.039 | 0 | 0.27 | 0.19 | 0.16 | 0.17 | 0.19 | 0.16 |
| 2 | 224893921 | rs12476888 | 1.983 | -0.908 | 1.319 | 1.704 | -0.018 | 0.093 | 0.093 | 0 | 0.47 | 0.12 | 0.52 | 0.3 | 0.56 | 0.54 |
| 2 | 224902897 | rs861442 | 0.83 | -0.849 | 1.36 | 1.603 | -0.002 | -0.003 | -0.003 | 0 | 0.63 | 0.55 | 0.72 | 0.61 | 0.66 | 0.72 |
| 2 | 224914800 | rs282258 | 1.848 | -0.871 | 0.625 | 1.529 | 0.024 | 0.123 | 0.123 | 0 | 0.51 | 0.17 | 0.54 | 0.28 | 0.57 | 0.53 |
| 2 | 224920176 | rs282268 | -2.016 | -0.915 | 0.634 | 1.583 | 0.036 | -0.044 | -0.044 | 0.505 | 0.35 | 0.81 | 0.35 | 0.53 | 0.27 | 0.33 |
| 2 | 224920925 | rs3957256 | -0.23 | -0.927 | 0.809 | 1.771 | -0.011 | 0.034 | 0.034 | 0 | 0.11 | 0.17 | 0.16 | 0.24 | 0.1 | 0.16 |
| 2 | 224924960 | rs2894520 | -2.126 | -0.857 | 0.34 | 1.526 | 0.014 | 0.028 | 0.028 | 0 | 0.49 | 0.79 | 0.45 | 0.44 | 0.45 | 0.37 |
| 2 | 224925121 | rs1530018 | -0.487 | -0.856 | 0.44 | 1.526 | 0.014 | 0.039 | 0.039 | 0 | 0.27 | 0.19 | 0.16 | 0.19 | 0.21 | 0.16 |
| 2 | 224925417 | rs4673085 | -0.452 | -0.713 | 0.144 | 1.654 | -0.005 | 0.013 | 0.013 | 1.403 | 0.21 | 0.56 | 0.19 | 0.21 | 0.18 | 0.11 |
| 2 | 224926358 | rs6731577 | 0.303 | -0.819 | 0.05 | 0.991 | -0.005 | -0.017 | -0.017 | 2.734 | 0.09 | 0.19 | 0.14 | 0.04 | 0.08 | 0.1 |
| 2 | 224929412 | rs2577284 | -1.302 | -0.684 | -0.697 | 0.902 | 0.033 | -0.056 | -0.056 | 0 | 0.34 | 0.8 | 0.34 | 0.49 | 0.26 | 0.33 |
| 2 | 224929659 | rs12694628 | -0.957 | -0.507 | -1.155 | 0.711 | 0 | -0.008 | -0.008 | 0 | 0.35 | 0.39 | 0.26 | 0.2 | 0.34 | 0.2 |
| 2 | 224931733 | rs282249 | -0.949 | -0.471 | -1.696 | 1.253 | 0.036 | -0.121 | -0.121 | 0 | 0.34 | 0.89 | 0.34 | 0.52 | 0.26 | 0.31 |
| 2 | 224932899 | rs2396040 | 0.679 | 0.061 | -1.13 | 0.995 | -0.012 | -0.004 | -0.004 | 0 | 0.95 | 0.98 | 0.97 | 0.92 | 0.91 | 0.87 |
| 2 | 224933339 | rs2118410 | -0.498 | -0.379 | -0.763 | 0.876 | -0.018 | -0.014 | -0.014 | 0 | 0.36 | 0.52 | 0.3 | 0.3 | 0.29 | 0.37 |
| 2 | 224936872 | rs893359 | 0.238 | -0.677 | -0.586 | 1.366 | -0.006 | 0.048 | 0.048 | 0.207 | 0.88 | 0.88 | 0.9 | 0.68 | 0.91 | 0.83 |
| 2 | 224941842 | rs2123714 | 1.35 | -1.439 | -0.008 | 0.982 | -0.005 | 0.006 | 0.006 | 3.414 | 0.66 | 0.41 | 0.73 | 0.51 | 0.76 | 0.7 |
| 2 | 224942253 | rs3997614 | 0.059 | -1.414 | 0.011 | 1.001 | -0.01 | 0.066 | 0.066 | 1.65 | 0.88 | 0.84 | 0.9 | 0.68 | 0.91 | 0.83 |
| 2 | 224943685 | rs6740660 | -0.486 | -1.218 | 0.584 | 1.005 | -0.004 | -0.008 | -0.008 | 0.071 | 0.19 | 0.22 | 0.15 | 0.17 | 0.15 | 0.12 |
| 2 | 224945517 | rs16865540 | -0.584 | -1.37 | 1.092 | 1.067 | -0.009 | 0.051 | 0.051 | 0 | 0.21 | 0.35 | 0.22 | 0.33 | 0.17 | 0.22 |
| 2 | 224959877 | rs2167939 | 0.166 | -1.642 | 1.049 | 1.058 | -0.033 | -0.004 | -0.004 | 0 | 0.73 | 0.43 | 0.76 | 0.71 | 0.81 | 0.67 |
| 2 | 224962122 | rs13016085 | 1.504 | -1.658 | 0.272 | 1.085 | 0.053 | -0.022 | -0.022 | 0 | 0.52 | 0.09 | 0.56 | 0.38 | 0.64 | 0.45 |
| 2 | 224962961 | rs16865577 | 0.237 | -1.387 | 0.362 | 1.027 | -0.005 | 0.005 | 0.005 | 0 | 0.19 | 0.01 | 0.17 | 0.16 | 0.15 | 0.27 |
| 2 | 224966354 | rs6745242 | -0.079 | -1.413 | 0.388 | 1.042 | 0.009 | -0.012 | -0.012 | 0 | 0.78 | 0.88 | 0.81 | 0.71 | 0.81 | 0.67 |
| 2 | 224968744 | rs1453669 | 0.205 | -1.467 | 0.207 | 0.875 | 0.025 | 0.025 | 0.025 | 0 | 0.19 | 0.08 | 0.21 | 0.16 | 0.14 | 0.26 |
| 2 | 224976957 | rs6436465 | 1.146 | -1.314 | 0.135 | 0.723 | 0.036 | -0.016 | -0.016 | 0.089 | 0.46 | 0.24 | 0.47 | 0.43 | 0.59 | 0.44 |
| 2 | 224988072 | rs6436467 | 0.134 | -1.457 | 0.313 | 0.738 | 0 | 0.001 | 0.001 | 0 | 0.88 | 0.91 | 0.89 | 0.71 | 0.85 | 0.81 |
| 2 | 224993943 | rs7586193 | -0.398 | -1.545 | 0.489 | 0.679 | 0.045 | 0.06 | 0.06 | 0.012 | 0.36 | 0.19 | 0.38 | 0.28 | 0.26 | 0.37 |
| 2 | 225006135 | rs4257392 | 0.558 | -1.539 | 0.844 | 0.687 | 0.001 | 0.083 | 0.083 | 0 | 0.87 | 0.73 | 0.88 | 0.71 | 0.85 | 0.81 |
| 2 | 225006599 | rs2123717 | 0.348 | -1.604 | 0.387 | 0.477 | 0.066 | 0.069 | 0.069 | 0 | 0.26 | 0.1 | 0.18 | 0.15 | 0.15 | 0.29 |
| 2 | 225007484 | rs6711302 | -0.831 | -1.556 | 0.445 | 0.504 | -0.012 | -0.017 | -0.017 | 0 | 0.4 | 0.48 | 0.41 | 0.28 | 0.26 | 0.38 |
| 2 | 225008442 | rs7605202 | 0.09 | -1.489 | -0.17 | 0.641 | -0.021 | -0.014 | -0.014 | 0.617 | 0.9 | 0.99 | 0.94 | 0.9 | 0.87 | 0.88 |
| 2 | 225020251 | rs6727070 | 0.108 | -1.689 | -0.22 | 0.602 | 0.061 | 0.056 | 0.056 | 0 | 0.26 | 0.1 | 0.18 | 0.15 | 0.14 | 0.25 |
| 2 | 225022528 | rs11689046 | 0.193 | -1.425 | -0.336 | 0.61 | -0.022 | -0.014 | -0.014 | 0 | 0.9 | 0.98 | 0.94 | 0.89 | 0.87 | 0.88 |
| 2 | 225022657 | rs11690142 | 1.123 | -1.661 | 0.045 | 0.669 | 0.065 | 0.006 | 0.006 | 0 | 0.55 | 0.25 | 0.68 | 0.55 | 0.7 | 0.55 |
| 2 | 225023616 | rs7569163 | 1.288 | -1.964 | -0.122 | 0.428 | 0.084 | 0.04 | 0.04 | 0.017 | 0.5 | 0.25 | 0.66 | 0.56 | 0.69 | 0.52 |
| 2 | 225025120 | rs1453672 | 1.626 | -1.42 | 0.532 | 0.76 | -0.031 | 0.031 | 0.031 | 0.871 | 0.78 | 0.4 | 0.85 | 0.72 | 0.83 | 0.77 |
| 2 | 225025431 | rs4673090 | 0.906 | -1.172 | 0.053 | 0.55 | -0.003 | 0.014 | 0.014 | 0 | 0.91 | 0.91 | 0.96 | 0.99 | 0.91 | 0.97 |
| 2 | 225028919 | rs6756793 | -1.2 | -0.758 | -0.321 | 0.733 | 0.015 | 0.037 | 0.037 | 0 | 0.45 | 0.8 | 0.46 | 0.46 | 0.42 | 0.5 |
| 2 | 225030129 | rs6761041 | -1.116 | -0.813 | 0.426 | 1.233 | 0.002 | -0.005 | -0.005 | 0.03 | 0.43 | 0.62 | 0.49 | 0.59 | 0.42 | 0.5 |
| 2 | 225045264 | rs2396058 | -0.789 | -0.715 | 1.206 | 1.389 | 0.015 | 0.133 | 0.133 | 0 | 0.08 | 0.31 | 0.09 | 0.23 | 0.06 | 0.16 |
| 2 | 225047744 | rs2629046 | -1.364 | -0.883 | 1.129 | 1.246 | 0.013 | -0.011 | -0.011 | 0 | 0.45 | 0.7 | 0.49 | 0.6 | 0.42 | 0.47 |
| 2 | 225073276 | rs1563127 | -1.341 | -0.918 | 1.251 | 1.297 | 0.006 | -0.002 | -0.002 | 0 | 0.45 | 0.65 | 0.52 | 0.61 | 0.42 | 0.48 |
| 2 | 225076480 | rs978888 | -1.665 | -0.191 | 0.846 | 1.435 | -0.048 | 0.033 | 0.033 | 0 | 0.65 | 0.89 | 0.67 | 0.82 | 0.56 | 0.69 |
| 2 | 225078491 | rs2629043 | -1.429 | 0.045 | 1.652 | 1.888 | 0.009 | -0.017 | -0.017 | 0 | 0.41 | 0.74 | 0.5 | 0.61 | 0.38 | 0.46 |
| 2 | 225087829 | rs6717510 | 0.652 | 0.049 | 1.789 | 1.931 | 0 | 0 | 0 | 1.234 | 0.67 | 0.66 | 0.59 | 0.69 | 0.68 | 0.72 |
| 2 | 225088153 | rs6732868 | 0.068 | 0.324 | 1.97 | 2.183 | -0.013 | 0.009 | 0.009 | 0 | 0.77 | 0.71 | 0.68 | 0.69 | 0.8 | 0.76 |
| 2 | 225111488 | rs7556741 | -1.015 | 0.39 | 1.884 | 2.268 | 0.009 | 0.153 | 0.153 | 0 | 0.08 | 0.27 | 0.08 | 0.3 | 0.06 | 0.18 |
| 2 | 225112637 | rs974685 | -0.124 | 0.437 | 1.624 | 2.122 | 0 | -0.001 | -0.001 | 0 | 0.78 | 0.81 | 0.69 | 0.69 | 0.8 | 0.76 |
| 2 | 225133282 | rs6706618 | -0.277 | 0.333 | 1.432 | 2.152 | 0.01 | -0.015 | -0.015 | 0 | 0.8 | 0.87 | 0.8 | 0.7 | 0.82 | 0.78 |
| 2 | 225134322 | rs6436470 | 0.896 | 0.525 | 1.673 | 2.211 | -0.016 | 0.04 | 0.04 | 0 | 0.72 | 0.39 | 0.6 | 0.37 | 0.75 | 0.61 |
| 2 | 225135115 | rs10933051 | 0.022 | 0.547 | 1.371 | 2.248 | -0.017 | 0.027 | 0.027 | 0 | 0.78 | 0.74 | 0.69 | 0.68 | 0.82 | 0.74 |
| 2 | 225136413 | rs2287527 | -0.941 | 0.533 | 1.361 | 2.292 | 0.002 | 0.034 | 0.034 | 0 | 0.07 | 0.12 | 0.06 | 0.26 | 0.07 | 0.16 |
| 2 | 225136607 | rs2287526 | -0.002 | 0.526 | 1.393 | 2.305 | -0.018 | 0.028 | 0.028 | 0 | 0.78 | 0.74 | 0.69 | 0.68 | 0.83 | 0.74 |
| 2 | 225136760 | rs2287524 | 0.869 | 0.534 | 1.283 | 2.104 | -0.021 | 0.053 | 0.053 | 0 | 0.71 | 0.61 | 0.63 | 0.42 | 0.76 | 0.59 |
| 2 | 225140153 | rs1035316 | -0.261 | 0.908 | 0.985 | 2.325 | 0 | 0.005 | 0.005 | 0 | 0.22 | 0.21 | 0.37 | 0.45 | 0.2 | 0.25 |
| 2 | 225151655 | rs1008415 | -0.929 | 0.952 | 1.123 | 2.522 | 0.001 | -0.012 | -0.012 | 0.311 | 0.27 | 0.26 | 0.39 | 0.56 | 0.26 | 0.35 |
| 2 | 225155065 | rs17793362 | -0.976 | 1.042 | 1.123 | 2.537 | 0.006 | 0.05 | 0.05 | 0 | 0.07 | 0.14 | 0.06 | 0.26 | 0.07 | 0.1 |
| 2 | 225156620 | rs4674875 | -2.259 | 0.988 | 1.649 | 2.456 | 0.063 | 0.236 | 0.236 | 0 | 0.16 | 0.59 | 0.22 | 0.46 | 0.21 | 0.15 |
| 2 | 225159450 | rs17178608 | 0.581 | 1.328 | 0.834 | 2.001 | -0.027 | 0.034 | 0.034 | 0.033 | 0.07 | 0.01 | 0.07 | 0 | 0.11 | 0.03 |
| 2 | 225173446 | rs1035312 | -1.48 | 1.175 | 1.513 | 2.209 | 0 | 0.001 | 0.001 | 0.944 | 0.32 | 0.37 | 0.39 | 0.55 | 0.4 | 0.41 |
| 2 | 225177666 | rs35160719 | -0.707 | 1.247 | 1.066 | 2.2 | 0 | -0.064 | -0.064 | 0.016 | 0.25 | 0.16 | 0.29 | 0.54 | 0.26 | 0.36 |
| 2 | 225178583 | rs6755111 | -1.43 | 1.188 | 1.089 | 2.201 | -0.023 | -0.044 | -0.044 | 0.502 | 0.32 | 0.25 | 0.38 | 0.54 | 0.39 | 0.41 |
| 2 | 225182419 | rs2098533 | -1.659 | 1.196 | 1.068 | 2.147 | -0.003 | -0.006 | -0.006 | 0 | 0.34 | 0.34 | 0.4 | 0.54 | 0.42 | 0.44 |
| 2 | 225186069 | rs6708010 | -0.904 | 0.585 | 1.049 | 2.344 | 0.005 | -0.016 | -0.016 | 0 | 0.25 | 0.21 | 0.23 | 0.54 | 0.24 | 0.34 |
| 2 | 225189016 | rs16865781 | -0.757 | 0.541 | 1.227 | 2.533 | 0.006 | 0.064 | 0.064 | 0 | 0.05 | 0.12 | 0.05 | 0.26 | 0.05 | 0.1 |
| 2 | 225189305 | rs16865783 | -1.822 | 0.47 | 1.23 | 2.574 | 0 | 0.077 | 0.077 | 0 | 0.07 | 0.15 | 0.05 | 0.31 | 0.05 | 0.1 |
| 2 | 225189782 | rs10432452 | -1.59 | 0.555 | 1.487 | 2.808 | -0.01 | 0.022 | 0.022 | 0 | 0.24 | 0.38 | 0.24 | 0.72 | 0.22 | 0.3 |

| | | | | | | | | | | | | | | | | |
|---|---|---|---|---|---|---|---|---|---|---|---|---|---|---|---|---|
| 2 | 225192649 | rs16865794 | -0.425 | 0.568 | 1.343 | 2.749 | -0.008 | 0.06 | 0.06 | 0 | 0.18 | 0.23 | 0.19 | 0.42 | 0.17 | 0.2 |
| 2 | 225207978 | rs2287520 | -1.251 | 0.553 | 2.027 | 2.844 | 0.023 | 0.318 | 0.318 | 0 | 0.06 | 0.22 | 0.16 | 0.45 | 0.05 | 0.1 |
| 2 | 225209245 | rs714056 | -1.667 | 0.609 | 2.464 | 2.815 | 0.003 | 0.034 | 0.034 | 0 | 0.24 | 0.44 | 0.33 | 0.7 | 0.25 | 0.31 |
| 2 | 225214052 | rs12623539 | -0.852 | 0.735 | 1.826 | 2.896 | 0.004 | 0.189 | 0.189 | 0 | 0.05 | 0.11 | 0.14 | 0.46 | 0.05 | 0.1 |
| 2 | 225222900 | rs2042086 | -1.31 | 0.462 | 1.782 | 2.492 | -0.008 | 0.074 | 0.074 | 0 | 0.24 | 0.37 | 0.29 | 0.43 | 0.24 | 0.23 |
| 2 | 225224042 | rs16865861 | -0.319 | 0.457 | 1.478 | 2.495 | 0.011 | -0.011 | -0.011 | 0.204 | 0.18 | 0.09 | 0.15 | 0.24 | 0.17 | 0.2 |
| 2 | 225229888 | rs16865873 | -0.569 | 0.452 | 1.563 | 2.162 | -0.007 | 0.003 | 0.003 | 0 | 0.2 | 0.19 | 0.16 | 0.28 | 0.17 | 0.23 |
| 2 | 225244888 | rs3738954 | -0.371 | 0.474 | 1.576 | 2.166 | 0.009 | -0.009 | -0.009 | 0 | 0.18 | 0.1 | 0.16 | 0.24 | 0.17 | 0.25 |
| 2 | 225244923 | rs3738953 | -0.75 | 0.477 | 1.501 | 2.16 | -0.02 | 0.015 | 0.015 | 0 | 0.2 | 0.25 | 0.17 | 0.28 | 0.17 | 0.25 |
| 2 | 225258573 | rs12992690 | -0.053 | 0.492 | 0.709 | 2.048 | -0.011 | -0.053 | -0.053 | 1.346 | 0.17 | 0.01 | 0.13 | 0.21 | 0.15 | 0.2 |
| 2 | 225259356 | rs10170836 | -1.867 | 0.74 | 0.328 | 1.714 | 0.017 | -0.04 | -0.04 | 0 | 0.42 | 0.83 | 0.37 | 0.41 | 0.36 | 0.46 |
| 2 | 225260545 | rs16865942 | -0.173 | 0.646 | 1.429 | 1.425 | 0.009 | -0.006 | -0.006 | 0 | 0.17 | 0.26 | 0.09 | 0.1 | 0.17 | 0.23 |
| 2 | 225265645 | rs13010329 | 0.145 | 0.793 | 1.315 | 1.655 | 0.013 | -0.031 | -0.031 | 0.265 | 0.18 | 0.07 | 0.24 | 0.26 | 0.17 | 0.24 |
| 2 | 225266667 | rs3845837 | -0.172 | 0.909 | 1.455 | 1.409 | 0.018 | -0.045 | -0.045 | 0 | 0.11 | 0.22 | 0.08 | 0 | 0.16 | 0.15 |
| 2 | 225267000 | rs4674892 | -1.079 | 0.932 | 1.514 | 1.438 | -0.03 | 0.088 | 0.088 | 0 | 0.28 | 0.42 | 0.29 | 0.37 | 0.24 | 0.35 |
| 2 | 225276261 | rs2052409 | -0.937 | 0.981 | 1.141 | 1.053 | 0.028 | -0.081 | -0.081 | 0 | 0.15 | 0.35 | 0.11 | 0 | 0.18 | 0.18 |
| 2 | 225279474 | rs7605583 | -0.891 | 0.982 | 1.058 | 1.071 | 0.033 | -0.068 | -0.068 | 0 | 0.14 | 0.31 | 0.1 | 0 | 0.19 | 0.21 |
| 2 | 225281602 | rs918287 | -2.012 | 0.845 | 0.481 | 1.221 | -0.011 | -0.052 | -0.052 | 0 | 0.45 | 0.87 | 0.42 | 0.37 | 0.42 | 0.54 |
| 2 | 225290792 | rs1916762 | 0.126 | 0.85 | 0.669 | 1.375 | 0.013 | -0.029 | -0.029 | 1.883 | 0.22 | 0.07 | 0.26 | 0.25 | 0.18 | 0.27 |
| 2 | 225292912 | rs4674898 | -1.912 | 0.859 | 0.864 | 1.266 | 0.002 | -0.019 | -0.019 | 0.544 | 0.42 | 0.75 | 0.41 | 0.37 | 0.42 | 0.52 |
| 2 | 225298112 | rs6734283 | -0.575 | 0.736 | 1.157 | 1.149 | 0.046 | 0.052 | 0.052 | 0 | 0.16 | 0.53 | 0.1 | 0.12 | 0.18 | 0.16 |
| 2 | 225304551 | rs7587019 | 0.571 | 0.677 | 0.588 | 1.153 | -0.021 | -0.082 | -0.082 | 0 | 0.19 | 0.01 | 0.25 | 0.25 | 0.18 | 0.26 |
| 2 | 225305589 | rs1721282 | 2.357 | 0.734 | 0.076 | 1.16 | -0.021 | 0.08 | 0.08 | 0 | 0.38 | 0.14 | 0.33 | 0.19 | 0.39 | 0.28 |
| 2 | 225305832 | rs1642922 | -0.688 | 0.738 | -0.051 | 1.237 | -0.022 | -0.089 | -0.089 | 0 | 0.84 | 0.96 | 0.74 | 0.56 | 0.85 | 0.85 |
| 2 | 225310582 | rs2462140 | 2.424 | 0.743 | -0.056 | 0.852 | -0.01 | 0.06 | 0.06 | 0 | 0.36 | 0.18 | 0.31 | 0.19 | 0.35 | 0.28 |
| 2 | 225318971 | rs12611811 | -0.251 | 0.713 | 0.5 | 0.963 | -0.003 | 0.041 | 0.041 | 0 | 0.2 | 0.25 | 0.29 | 0.55 | 0.18 | 0.17 |
| 2 | 225322828 | rs7564678 | 0.382 | 0.704 | 0.73 | 0.94 | -0.011 | 0.004 | 0.004 | 0 | 0.25 | 0.33 | 0.28 | 0.24 | 0.23 | 0.32 |
| 2 | 225324391 | rs1568751 | -1.355 | 0.709 | 0.741 | 0.929 | 0.006 | 0.023 | 0.023 | 0 | 0.36 | 0.53 | 0.39 | 0.57 | 0.4 | 0.37 |
| 2 | 225326693 | rs6436490 | -2.452 | 0.691 | 0.39 | 0.961 | -0.015 | 0.064 | 0.064 | 0 | 0.64 | 0.84 | 0.69 | 0.81 | 0.65 | 0.73 |
| 2 | 225330266 | rs7605197 | -2.455 | 0.882 | 0.322 | 0.962 | -0.008 | 0.062 | 0.062 | 0 | 0.64 | 0.87 | 0.7 | 0.8 | 0.65 | 0.73 |
| 2 | 225335290 | rs2396092 | 0.455 | 0.626 | 0.775 | 0.942 | -0.016 | 0.021 | 0.021 | 0 | 0.19 | 0.25 | 0.23 | 0.24 | 0.13 | 0.31 |
| 2 | 225335595 | rs4674908 | -1.731 | 0.752 | 0.603 | 0.789 | 0 | 0.004 | 0.004 | 0 | 0.5 | 0.46 | 0.62 | 0.66 | 0.48 | 0.56 |
| 2 | 225336309 | rs3768899 | 0.745 | 0.578 | 0.605 | 0.75 | -0.016 | 0.021 | 0.021 | 0 | 0.18 | 0.25 | 0.23 | 0.24 | 0.13 | 0.31 |
| 2 | 225336531 | rs10498160 | 0.762 | 0.578 | 0.605 | 0.766 | -0.016 | 0.023 | 0.023 | 0 | 0.18 | 0.25 | 0.23 | 0.24 | 0.13 | 0.31 |
| 2 | 225336889 | rs10498161 | 0.745 | 0.578 | 0.605 | 0.766 | -0.016 | 0.022 | 0.022 | 0 | 0.18 | 0.25 | 0.23 | 0.24 | 0.13 | 0.31 |
| 2 | 225337188 | rs13016316 | 1.71 | 0.537 | 0.566 | 0.731 | 0.012 | -0.081 | -0.081 | 1.999 | 0.14 | 0.01 | 0.19 | 0.24 | 0.08 | 0.23 |
| 2 | 225338022 | rs3768897 | 0.626 | 0.547 | 0.571 | 0.735 | -0.016 | 0.022 | 0.022 | 0.003 | 0.19 | 0.25 | 0.23 | 0.24 | 0.13 | 0.31 |
| 2 | 225346646 | rs2070127 | 0.512 | 0.528 | 0.561 | 0.726 | -0.017 | 0.017 | 0.017 | 0 | 0.19 | 0.25 | 0.23 | 0.24 | 0.13 | 0.31 |
| 2 | 225347713 | rs2047134 | -0.574 | 0.555 | 0.726 | 0.866 | -0.014 | -0.005 | -0.005 | 4.482 | 0.31 | 0.16 | 0.37 | 0.41 | 0.33 | 0.26 |
| 2 | 225362478 | rs3738952 | 1.86 | 0.555 | 0.725 | 0.843 | 0.009 | -0.079 | -0.079 | 0.074 | 0.13 | 0.01 | 0.19 | 0.24 | 0.08 | 0.23 |
| 2 | 225365635 | rs4674916 | -0.645 | 0.58 | 0.7 | 0.928 | -0.007 | 0.007 | 0.007 | 0 | 0.32 | 0.2 | 0.35 | 0.37 | 0.33 | 0.26 |
| 2 | 225393122 | rs11688390 | 0.739 | 0.58 | 0.7 | 0.928 | -0.016 | 0.021 | 0.021 | 0.003 | 0.19 | 0.25 | 0.22 | 0.24 | 0.13 | 0.31 |
| 2 | 225397170 | rs4674919 | -1.752 | 0.647 | 0.61 | 0.895 | 0.001 | 0.001 | 0.001 | 0 | 0.51 | 0.46 | 0.58 | 0.6 | 0.47 | 0.56 |
| 2 | 225398684 | rs6751391 | 0.3 | 0.675 | 0.584 | 0.875 | 0.042 | 0.059 | 0.059 | 0 | 0.18 | 0.45 | 0.13 | 0.19 | 0.2 | 0.19 |
| 2 | 225399864 | rs17480230 | -0.518 | 0.694 | -0.017 | 0.821 | -0.037 | -0.008 | -0.008 | 0 | 0.31 | 0.1 | 0.35 | 0.36 | 0.33 | 0.26 |
| 2 | 225411429 | rs10197090 | 0.913 | 0.78 | -0.004 | 0.737 | 0.001 | -0.025 | -0.025 | 0 | 0.15 | 0.17 | 0.09 | 0.01 | 0.2 | 0.18 |
| 2 | 225443901 | rs6749402 | -2.833 | 0.706 | -0.676 | 0.896 | -0.036 | 0.034 | 0.034 | 0 | 0.69 | 0.92 | 0.72 | 0.8 | 0.67 | 0.75 |
| 2 | 225462004 | rs11682897 | 0.538 | 0.861 | -0.485 | 0.9 | 0.003 | -0.029 | -0.029 | 0 | 0.18 | 0.11 | 0.22 | 0.25 | 0.15 | 0.31 |
| 2 | 225469413 | rs6736188 | -0.732 | 1.077 | -0.459 | 0.604 | -0.002 | -0.009 | -0.009 | 0 | 0.21 | 0.15 | 0.3 | 0.35 | 0.23 | 0.22 |
| 2 | 225473263 | rs7580488 | -1.138 | 1.281 | -0.265 | 0.515 | -0.01 | -0.008 | -0.008 | 0 | 0.3 | 0.31 | 0.35 | 0.45 | 0.37 | 0.31 |
| 2 | 225473789 | rs6749153 | 0.563 | 0.918 | -0.261 | 0.217 | 0.003 | -0.097 | -0.097 | 0 | 0.19 | 0.31 | 0.12 | 0.02 | 0.17 | 0.13 |
| 2 | 225478370 | rs16866077 | -0.01 | 0.833 | -0.432 | 0.53 | -0.015 | -0.047 | -0.047 | 0 | 0.21 | 0.12 | 0.24 | 0.42 | 0.2 | 0.32 |
| 2 | 225479503 | rs17480503 | 0.67 | 0.808 | -0.26 | 0.296 | -0.007 | -0.09 | -0.09 | 1.015 | 0.2 | 0.31 | 0.12 | 0.02 | 0.18 | 0.13 |
| 2 | 225484736 | rs1908252 | -1.683 | 0.815 | -0.268 | 0.253 | 0 | -0.005 | -0.005 | 0 | 0.47 | 0.49 | 0.42 | 0.41 | 0.49 | 0.47 |
| 2 | 225491143 | rs6731785 | -2.763 | 0.518 | -0.89 | 0.204 | -0.03 | 0.066 | 0.066 | 0 | 0.67 | 0.87 | 0.66 | 0.84 | 0.63 | 0.72 |
| 2 | 225495599 | rs16866084 | 1.752 | 0.477 | -1.078 | 0.269 | 0.014 | -0.078 | -0.078 | 0 | 0.14 | 0.03 | 0.2 | 0.37 | 0.11 | 0.19 |
| 2 | 225503101 | rs1565074 | 1.752 | 0.477 | -1.134 | 0.269 | 0.001 | -0.105 | -0.105 | 2.813 | 0.14 | 0.01 | 0.2 | 0.37 | 0.11 | 0.2 |
| 2 | 225505092 | rs16866089 | 1.493 | 0.722 | -0.967 | 0.208 | -0.05 | -0.071 | -0.071 | 0 | 0.11 | 0.01 | 0.21 | 0.3 | 0.15 | 0.23 |
| 2 | 225511361 | rs2047136 | -2.564 | 0.635 | -0.958 | 0.105 | -0.073 | 0.074 | 0.074 | 0 | 0.68 | 0.95 | 0.67 | 0.84 | 0.63 | 0.72 |
| 2 | 225511657 | rs16866096 | 0.176 | 0.574 | -0.766 | 0.129 | -0.014 | -0.039 | -0.039 | 0 | 0.21 | 0.12 | 0.24 | 0.4 | 0.2 | 0.32 |
| 2 | 225534192 | rs11884368 | 0.142 | 0.499 | -0.588 | 0.051 | 0.001 | -0.003 | -0.003 | 0.192 | 0.35 | 0.32 | 0.34 | 0.44 | 0.3 | 0.34 |
| 2 | 225548072 | rs12623277 | -2.009 | 0.604 | -0.733 | -0.019 | -0.024 | 0.079 | 0.079 | 0 | 0.65 | 0.81 | 0.65 | 0.83 | 0.62 | 0.68 |
| 2 | 225553128 | rs4624372 | -2.343 | 0.609 | -0.992 | -0.016 | -0.067 | 0.081 | 0.081 | 0 | 0.69 | 0.92 | 0.67 | 0.83 | 0.64 | 0.73 |
| 2 | 225561035 | rs6717108 | -0.603 | 0.618 | -0.727 | -0.054 | 0.005 | -0.006 | -0.006 | 0 | 0.44 | 0.38 | 0.44 | 0.44 | 0.43 | 0.37 |
| 2 | 225563591 | rs16866119 | 0.657 | 0.754 | -0.904 | -0.005 | -0.015 | -0.02 | -0.02 | 0 | 0.15 | 0.11 | 0.18 | 0.3 | 0.17 | 0.28 |
| 2 | 225566563 | rs10173549 | 0.765 | 0.752 | -0.986 | -0.005 | -0.032 | -0.041 | -0.041 | 0.019 | 0.14 | 0.07 | 0.17 | 0.3 | 0.17 | 0.28 |
| 2 | 225574650 | rs10197382 | -2.283 | 0.762 | -1.004 | -0.181 | -0.049 | 0.052 | 0.052 | 0.023 | 0.65 | 0.91 | 0.65 | 0.74 | 0.62 | 0.69 |
| 2 | 225579262 | rs10198809 | 0.753 | 0.407 | -0.932 | -0.045 | -0.035 | -0.048 | -0.048 | 0 | 0.14 | 0.06 | 0.17 | 0.31 | 0.17 | 0.28 |
| 2 | 225587479 | rs7585688 | -0.091 | 0.413 | -0.931 | -0.208 | -0.015 | 0.005 | 0.005 | 0.311 | 0.36 | 0.39 | 0.28 | 0.36 | 0.26 | 0.35 |

| | | | | | | | | | | | | | | | | |
|---|---|---|---|---|---|---|---|---|---|---|---|---|---|---|---|---|
| 2 | 225588620 | rs13384654 | 0.391 | 0.978 | -1.174 | 0.146 | -0.001 | 0.008 | 0.008 | 1.359 | 0.16 | 0.16 | 0.24 | 0.37 | 0.18 | 0.28 |
| 2 | 225618741 | rs10173714 | -1.811 | 0.715 | -1.62 | -0.209 | -0.093 | 0.134 | 0.134 | 2.092 | 0.56 | 0.91 | 0.58 | 0.75 | 0.51 | 0.62 |
| 2 | 225630435 | rs1045653 | 1.085 | 0.752 | -1.466 | 0.018 | -0.016 | -0.036 | -0.036 | 0.016 | 0.14 | 0.08 | 0.23 | 0.36 | 0.16 | 0.25 |
| 2 | 225634693 | rs1565073 | -1.211 | 0.737 | -1.241 | -1.626 | -0.054 | 0.092 | 0.092 | 0 | 0.58 | 0.86 | 0.59 | 0.73 | 0.5 | 0.66 |
| 2 | 225639220 | rs6436508 | -0.154 | 0.249 | -1.048 | -1.645 | -0.004 | 0.001 | 0.001 | 0.072 | 0.71 | 0.73 | 0.56 | 0.26 | 0.71 | 0.52 |
| 2 | 225644800 | rs3768872 | -0.919 | -0.258 | -1.116 | -1.64 | -0.012 | 0.088 | 0.088 | 0.023 | 0.56 | 0.71 | 0.54 | 0.8 | 0.42 | 0.7 |
| 2 | 225659217 | rs16866183 | 0.744 | -0.355 | -0.817 | -1.459 | 0.002 | -0.017 | -0.017 | 1.427 | 0.23 | 0.15 | 0.4 | 0.74 | 0.2 | 0.48 |
| 2 | 225669996 | rs4674940 | 1.301 | -0.362 | -0.45 | -1.588 | -0.025 | -0.06 | -0.06 | 2.268 | 0.11 | 0.24 | 0.04 | 0 | 0.04 | 0.04 |
| 2 | 225672460 | rs12328236 | 0.016 | -0.427 | -0.416 | -1.656 | -0.029 | -0.108 | -0.108 | 0.227 | 0.16 | 0.43 | 0.07 | 0.02 | 0.11 | 0.08 |
| 2 | 225684166 | rs7572725 | -0.568 | -0.386 | -0.734 | -1.635 | -0.011 | -0.059 | -0.059 | 0.741 | 0.28 | 0.66 | 0.12 | 0.02 | 0.15 | 0.12 |
| 2 | 225688041 | rs13405571 | -1.061 | 0.263 | -0.448 | -1.378 | -0.003 | -0.074 | -0.074 | 0.143 | 0.25 | 0.66 | 0.15 | 0.02 | 0.23 | 0.1 |
| 2 | 225692190 | rs7556687 | -1.178 | 0.242 | -0.523 | -1.453 | -0.002 | -0.066 | -0.066 | 0 | 0.27 | 0.66 | 0.15 | 0.02 | 0.25 | 0.11 |
| 2 | 225696401 | rs1821409 | 1.178 | 0.196 | -0.846 | -1.509 | -0.007 | -0.016 | -0.016 | 0 | 0.3 | 0.07 | 0.4 | 0.74 | 0.23 | 0.54 |
| 2 | 225697363 | rs11693393 | -0.343 | -0.129 | -0.141 | -1.091 | -0.016 | 0.016 | 0.016 | 0 | 0.11 | 0.06 | 0.07 | 0 | 0.14 | 0.03 |
| 2 | 225709926 | rs7577271 | -0.449 | -0.162 | 0.102 | -1.059 | -0.033 | -0.113 | -0.113 | 0 | 0.17 | 0.48 | 0.08 | 0.02 | 0.11 | 0.08 |
| 2 | 225719693 | rs16866236 | -0.245 | -0.055 | -0.03 | -0.928 | -0.036 | 0.048 | 0.048 | 0 | 0.1 | 0.02 | 0.06 | 0 | 0.14 | 0.03 |
| 2 | 225721125 | rs7605551 | -0.4 | -0.143 | 0.188 | -1.023 | -0.032 | -0.092 | -0.092 | 0.153 | 0.14 | 0.48 | 0.06 | 0.02 | 0.09 | 0.07 |
| 2 | 225727431 | rs7589474 | -0.441 | -0.146 | 0.178 | -1.025 | -0.035 | -0.095 | -0.095 | 1.372 | 0.15 | 0.48 | 0.05 | 0.03 | 0.09 | 0.07 |
| 2 | 225727663 | rs6436515 | -0.813 | -0.054 | 0.225 | -1.059 | -0.007 | -0.029 | -0.029 | 0 | 0.2 | 0.69 | 0.12 | 0.06 | 0.15 | 0.17 |
| 2 | 225730189 | rs7559750 | -1.795 | -0.096 | 0.167 | -1.08 | -0.007 | 0.082 | 0.082 | 0.054 | 0.31 | 0.8 | 0.19 | 0.06 | 0.28 | 0.19 |
| 2 | 225736752 | rs7592442 | 0.771 | -0.101 | 0.871 | -0.877 | -0.004 | -0.029 | -0.029 | 0.037 | 0.08 | 0.21 | 0.02 | 0.02 | 0.06 | 0.05 |
| 2 | 225737546 | rs7607230 | -0.03 | -0.066 | 0.469 | -0.858 | -0.019 | -0.069 | -0.069 | 0.045 | 0.13 | 0.32 | 0.03 | 0.02 | 0.09 | 0.07 |
| 2 | 225750753 | rs10197805 | -0.89 | -0.037 | 0.108 | -0.84 | -0.008 | -0.066 | -0.066 | 0 | 0.19 | 0.53 | 0.09 | 0.06 | 0.14 | 0.17 |
| 2 | 225759978 | rs16866294 | 0.904 | -0.094 | -0.246 | -0.918 | -0.01 | -0.021 | -0.021 | 0 | 0.16 | 0.04 | 0.22 | 0.19 | 0.18 | 0.17 |
| 2 | 225760431 | rs16866296 | -0.587 | 0.403 | 0.384 | -0.565 | -0.033 | -0.091 | -0.091 | 0 | 0.14 | 0.48 | 0.06 | 0.02 | 0.09 | 0.07 |
| 2 | 225771834 | rs11677647 | 0.263 | 0.566 | -0.282 | -0.242 | 0.027 | -0.008 | -0.008 | 0 | 0.49 | 0.21 | 0.53 | 0.54 | 0.43 | 0.43 |
| 2 | 225773085 | rs17199382 | 0.204 | 0.642 | -0.389 | -0.384 | -0.018 | 0.035 | 0.035 | 0 | 0.16 | 0.09 | 0.1 | 0.05 | 0.18 | 0.16 |
| 2 | 225774390 | rs6731100 | 0.809 | 1.015 | 0.27 | 0.027 | 0.022 | 0.033 | 0.033 | 1.191 | 0.45 | 0.32 | 0.32 | 0.38 | 0.34 | 0.36 |
| 2 | 225774612 | rs13024433 | 1.364 | 1.024 | 0.249 | 0.018 | 0.016 | -0.013 | -0.013 | 0 | 0.38 | 0.11 | 0.26 | 0.38 | 0.28 | 0.35 |
| 2 | 225775245 | rs11688439 | 0.28 | 0.99 | 0.274 | -0.133 | -0.021 | -0.002 | -0.002 | 0 | 0.21 | 0.08 | 0.27 | 0.22 | 0.28 | 0.24 |
| 2 | 225786241 | rs13016606 | -0.202 | 1.05 | 0.198 | -0.116 | 0.002 | 0.013 | 0.013 | 0 | 0.66 | 0.42 | 0.59 | 0.54 | 0.61 | 0.6 |
| 2 | 225792835 | rs7558808 | -0.245 | 1.056 | 0.221 | -0.113 | 0.003 | 0.015 | 0.015 | 0 | 0.66 | 0.41 | 0.59 | 0.54 | 0.61 | 0.59 |
| 2 | 225795103 | rs17271567 | 0.93 | 1.079 | 0.048 | -0.1 | 0.002 | 0.066 | 0.066 | 0.553 | 0.43 | 0.15 | 0.29 | 0.33 | 0.34 | 0.35 |
| 2 | 225809036 | rs6750683 | -0.238 | 1.233 | 0.323 | -0.04 | 0.013 | 0.054 | 0.054 | 0 | 0.67 | 0.53 | 0.61 | 0.54 | 0.62 | 0.59 |
| 2 | 225810786 | rs1072581 | -0.301 | 1.237 | -0.055 | -0.038 | 0.005 | 0.022 | 0.022 | 0 | 0.67 | 0.62 | 0.62 | 0.54 | 0.62 | 0.59 |
| 2 | 225812560 | rs869836 | -1.075 | 1.245 | -1.263 | -0.042 | -0.069 | -0.125 | -0.125 | 0.01 | 0.75 | 0.93 | 0.64 | 0.54 | 0.64 | 0.63 |
| 2 | 225818564 | rs11686538 | 0.934 | 1.481 | -1.404 | 0.213 | 0.036 | 0.034 | 0.034 | 0.783 | 0.39 | 0.11 | 0.25 | 0.33 | 0.27 | 0.34 |
| 2 | 225820260 | rs4674952 | -0.586 | 1.345 | -1.337 | 0.254 | 0.001 | 0.011 | 0.011 | 0 | 0.66 | 0.42 | 0.59 | 0.54 | 0.62 | 0.59 |
| 2 | 225824421 | rs12999154 | -0.062 | 1.066 | -1.77 | 0.532 | 0.01 | 0.065 | 0.065 | 0.805 | 0.43 | 0.21 | 0.31 | 0.33 | 0.34 | 0.35 |
| 2 | 225834004 | rs35573107 | -1.442 | 1.468 | -0.886 | 0.868 | 0.007 | 0.004 | 0.004 | 0 | 0.11 | 0.16 | 0.09 | 0.12 | 0.14 | 0.05 |
| 2 | 225839321 | rs6436519 | -1.185 | 1.967 | -1.203 | 1.029 | -0.024 | -0.012 | -0.012 | 0 | 0.18 | 0.15 | 0.39 | 0.17 | 0.29 | 0.26 |
| 2 | 225840782 | rs13411031 | -0.6 | 1.811 | -1.736 | 0.609 | -0.009 | 0.039 | 0.039 | 0 | 0.41 | 0.7 | 0.3 | 0.2 | 0.45 | 0.31 |
| 2 | 225844421 | rs12329131 | -0.668 | 1.787 | -2.498 | 0.966 | -0.065 | -0.145 | -0.145 | 0 | 0.8 | 0.99 | 0.71 | 0.59 | 0.77 | 0.7 |
| 2 | 225848579 | rs375205 | -0.72 | 1.857 | -1.757 | 0.876 | -0.019 | -0.072 | -0.072 | 0.559 | 0.79 | 0.9 | 0.7 | 0.59 | 0.77 | 0.69 |
| 2 | 225850905 | rs377571 | -0.647 | 1.83 | -1.956 | 0.031 | 0.001 | -0.001 | -0.001 | 0 | 0.61 | 0.62 | 0.57 | 0.52 | 0.62 | 0.61 |
| 2 | 225850923 | rs9989765 | 0.01 | 1.861 | -1.876 | 0.026 | -0.029 | -0.022 | -0.022 | 0 | 0.06 | 0.01 | 0.04 | 0.06 | 0.06 | 0.08 |
| 2 | 225851064 | rs16866402 | 0.567 | 1.845 | -1.972 | -0.206 | 0.003 | 0.006 | 0.006 | 1.197 | 0.14 | 0.1 | 0.1 | 0.07 | 0.11 | 0.03 |
| 2 | 225854120 | rs921281 | -0.347 | 1.902 | -1.775 | -0.14 | -0.006 | -0.054 | -0.054 | 0.54 | 0.41 | 0.55 | 0.29 | 0.08 | 0.42 | 0.31 |
| 2 | 225856191 | rs281525 | 0.257 | 1.58 | -1.125 | -0.243 | 0.005 | 0.012 | 0.012 | 0 | 0.5 | 0.43 | 0.42 | 0.66 | 0.38 | 0.56 |
| 2 | 225856744 | rs6730803 | -0.216 | 1.343 | -1.14 | -0.42 | 0.006 | 0.002 | 0.002 | 0 | 0.09 | 0.07 | 0.03 | 0 | 0.04 | 0.05 |
| 2 | 225856755 | rs6759015 | -0.405 | 1.504 | -0.629 | -0.389 | 0.002 | 0 | 0 | 0.115 | 0.71 | 0.77 | 0.43 | 0.32 | 0.61 | 0.59 |
| 2 | 225857707 | rs11897700 | -0.99 | 1.195 | 0.83 | -0.382 | -0.01 | -0.112 | -0.112 | 0 | 0.18 | 0.48 | 0.1 | 0.01 | 0.1 | 0.08 |
| 2 | 225858116 | rs13416139 | -1.265 | 1.063 | 0.956 | -0.425 | -0.009 | -0.024 | -0.024 | 0 | 0.22 | 0.74 | 0.14 | 0.11 | 0.12 | 0.17 |
| 2 | 225861869 | rs386549 | -1.139 | 1.109 | 0.978 | -0.304 | 0.01 | 0.014 | 0.014 | 0 | 0.89 | 0.89 | 0.74 | 0.76 | 0.79 | 0.84 |
| 2 | 225866470 | rs281529 | 1.295 | 0.996 | 0.842 | -0.327 | -0.014 | 0.007 | 0.007 | 1.677 | 0.67 | 0.15 | 0.61 | 0.65 | 0.68 | 0.67 |
| 2 | 225867366 | rs281527 | -1.078 | 1.114 | 0.511 | -0.169 | -0.011 | -0.012 | -0.012 | 0 | 0.9 | 0.94 | 0.74 | 0.76 | 0.79 | 0.84 |
| 2 | 225875158 | rs590233 | -1.006 | 1.16 | 0.455 | -0.368 | 0.007 | 0.003 | 0.003 | 0 | 0.9 | 0.92 | 0.76 | 0.83 | 0.79 | 0.85 |
| 2 | 225880074 | rs375154 | 0.847 | 0.776 | 0.636 | -0.399 | 0.008 | 0.015 | 0.015 | 0 | 0.72 | 0.36 | 0.54 | 0.59 | 0.55 | 0.67 |
| 2 | 225884211 | rs204249 | -0.19 | 0.847 | 0.441 | -0.416 | 0.076 | 0.078 | 0.078 | 0 | 0.86 | 0.57 | 0.72 | 0.79 | 0.79 | 0.85 |
| 2 | 225896014 | rs10176032 | 0.812 | 0.821 | 0.249 | -0.431 | -0.056 | 0.032 | 0.032 | 0 | 0.11 | 0.01 | 0.14 | 0.02 | 0.22 | 0.16 |
| 2 | 225896524 | rs441098 | -0.108 | 0.642 | 0.56 | -0.321 | 0.083 | 0.089 | 0.089 | 0 | 0.86 | 0.54 | 0.72 | 0.79 | 0.79 | 0.85 |
| 2 | 225900695 | rs602291 | -1.041 | 0.745 | 0.159 | -0.281 | 0.022 | 0.023 | 0.023 | 0 | 0.9 | 0.9 | 0.76 | 0.79 | 0.79 | 0.85 |
| 2 | 225919929 | rs380873 | 1.022 | 0.337 | -0.357 | -0.533 | -0.07 | -0.152 | -0.152 | 0.56 | 0.31 | 0.02 | 0.38 | 0.56 | 0.31 | 0.34 |
| 2 | 225921137 | rs4674964 | -1.136 | 0.592 | 0.642 | 0.107 | -0.001 | -0.012 | -0.012 | 0 | 0.31 | 0.65 | 0.26 | 0.2 | 0.32 | 0.27 |
| 2 | 225929498 | rs448375 | -0.163 | 0.552 | 1.008 | 0.738 | 0.02 | 0.017 | 0.017 | 0 | 0.86 | 0.83 | 0.78 | 0.79 | 0.78 | 0.84 |
| 2 | 225938996 | rs388208 | 0.512 | 0.609 | 0.319 | 0.573 | 0.029 | 0.084 | 0.084 | 0 | 0.79 | 0.51 | 0.72 | 0.63 | 0.73 | 0.8 |
| 2 | 225946531 | rs11677682 | -1.115 | 0.176 | -0.258 | -0.261 | -0.006 | -0.01 | -0.01 | 0 | 0.41 | 0.5 | 0.39 | 0.34 | 0.37 | 0.3 |
| 2 | 225952959 | rs6728535 | -0.922 | -0.22 | -0.09 | -0.475 | -0.006 | -0.001 | -0.001 | 0 | 0.27 | 0.29 | 0.21 | 0.26 | 0.14 | 0.14 |
| 2 | 225960921 | rs4673132 | 0.661 | -0.296 | -0.709 | -0.829 | 0.007 | 0.068 | 0.068 | 0 | 0.16 | 0.06 | 0.1 | 0.02 | 0.17 | 0.19 |

| | | | | | | | | | | | | | | | | |
|---|---|---|---|---|---|---|---|---|---|---|---|---|---|---|---|---|
| 2 | 225974964 | rs7568255 | -1.742 | -0.198 | -0.444 | -0.773 | -0.074 | 0.007 | 0.007 | 0 | 0.58 | 0.86 | 0.52 | 0.37 | 0.54 | 0.49 |
| 2 | 225977951 | rs16866521 | 0.16 | 0.065 | -0.074 | -0.312 | -0.005 | 0.136 | 0.136 | 0 | 0.06 | 0.13 | 0.14 | 0.26 | 0.04 | 0.06 |
| 2 | 225979968 | rs10498181 | 0.195 | 0.097 | -0.096 | -0.406 | 0.014 | -0.025 | -0.025 | 0 | 0.17 | 0.24 | 0.18 | 0.09 | 0.23 | 0.16 |
| 2 | 225985821 | rs1449511 | 0.012 | 0.016 | -0.187 | -0.131 | -0.005 | 0.141 | 0.141 | 0 | 0.06 | 0.13 | 0.14 | 0.26 | 0.04 | 0.07 |
| 2 | 225986779 | rs6757717 | -0.751 | -0.036 | 0.016 | -0.619 | 0.004 | -0.065 | -0.065 | 0.009 | 0.34 | 0.43 | 0.3 | 0.11 | 0.42 | 0.35 |
| 2 | 225987420 | rs12104926 | -0.966 | -0.008 | 0.147 | -0.617 | -0.007 | 0 | 0 | 1.658 | 0.35 | 0.43 | 0.34 | 0.35 | 0.31 | 0.27 |
| 2 | 225989070 | rs6436522 | -0.617 | 0.083 | -0.329 | -1.45 | 0.04 | 0.11 | 0.11 | 0 | 0.46 | 0.72 | 0.29 | 0.1 | 0.35 | 0.4 |
| 2 | 225995545 | rs13415546 | -0.213 | 0.245 | -0.01 | -1.606 | -0.01 | 0.031 | 0.031 | 0 | 0.38 | 0.3 | 0.29 | 0.1 | 0.42 | 0.4 |
| 2 | 226004087 | rs1562675 | -0.819 | -0.146 | -0.146 | -1.97 | -0.003 | -0.048 | -0.048 | 0.199 | 0.54 | 0.48 | 0.45 | 0.14 | 0.54 | 0.53 |
| 2 | 226013860 | rs6717227 | 0.441 | -0.149 | 0.078 | -1.976 | 0.001 | -0.064 | -0.064 | 2.584 | 0.19 | 0.31 | 0.2 | 0.04 | 0.17 | 0.16 |
| 2 | 226016494 | rs726553 | 0.223 | -0.161 | 0.05 | -2.003 | -0.027 | 0.109 | 0.109 | 0 | 0.36 | 0.15 | 0.25 | 0.09 | 0.37 | 0.37 |
| 2 | 226023294 | rs7582304 | -1.217 | -0.263 | 0.218 | -2.017 | 0.001 | 0.039 | 0.039 | 0 | 0.6 | 0.7 | 0.49 | 0.14 | 0.57 | 0.54 |
| 2 | 226025235 | rs1037246 | -1.283 | -0.26 | 0.268 | -2.016 | 0 | -0.028 | -0.028 | 0 | 0.61 | 0.6 | 0.48 | 0.14 | 0.56 | 0.54 |
| 2 | 226030132 | rs728654 | 0.396 | -0.125 | 0.36 | -1.886 | -0.001 | -0.015 | -0.015 | 2.331 | 0.19 | 0.2 | 0.19 | 0.04 | 0.17 | 0.16 |
| 2 | 226035360 | rs2165927 | -0.715 | -0.205 | 0.286 | -1.967 | 0.004 | 0.001 | 0.001 | 0 | 0.52 | 0.57 | 0.46 | 0.13 | 0.52 | 0.52 |
| 2 | 226041219 | rs7605435 | -0.025 | -0.179 | 0.169 | -1.773 | -0.012 | -0.018 | -0.018 | 0 | 0.07 | 0.1 | 0.03 | 0 | 0.02 | 0.01 |
| 2 | 226058673 | rs13403401 | 0.652 | -0.161 | -0.015 | -1.535 | -0.002 | -0.01 | -0.01 | 0 | 0.14 | 0.14 | 0.1 | 0.05 | 0.12 | 0.15 |
| 2 | 226062278 | rs7577796 | -0.056 | -0.07 | -0.191 | -1.52 | 0.003 | 0.001 | 0.001 | 0 | 0.24 | 0.21 | 0.21 | 0.04 | 0.2 | 0.17 |
| 2 | 226065765 | rs6436533 | -0.736 | -0.142 | 0.087 | -1.437 | 0.004 | -0.01 | -0.01 | 0 | 0.41 | 0.6 | 0.34 | 0.1 | 0.34 | 0.3 |
| 2 | 226071280 | rs1075705 | 1.16 | -0.002 | -0.244 | -1.452 | -0.018 | 0.037 | 0.037 | 0 | 0.17 | 0.1 | 0.12 | 0.04 | 0.18 | 0.23 |
| 2 | 226074083 | rs1443663 | 0.402 | 0.02 | -0.315 | -1.138 | -0.028 | -0.003 | -0.003 | 0.703 | 0.23 | 0.15 | 0.24 | 0.22 | 0.27 | 0.27 |
| 2 | 226082156 | rs6736537 | -1.184 | 0.065 | -0.16 | -1.158 | -0.006 | 0.062 | 0.062 | 0 | 0.57 | 0.74 | 0.49 | 0.3 | 0.52 | 0.5 |
| 2 | 226092297 | rs6727407 | -0.834 | 0.026 | 0.109 | -0.755 | -0.029 | -0.064 | -0.064 | 0 | 0.12 | 0.28 | 0.09 | 0 | 0.06 | 0.05 |
| 2 | 226097873 | rs4674977 | -0.193 | 0.216 | 0.187 | -0.725 | -0.002 | 0.002 | 0.002 | 0 | 0.28 | 0.25 | 0.28 | 0.25 | 0.28 | 0.24 |
| 2 | 226105129 | rs1373213 | -0.111 | 0.227 | 0.079 | -0.72 | -0.005 | 0.007 | 0.007 | 0 | 0.27 | 0.19 | 0.28 | 0.25 | 0.28 | 0.24 |
| 2 | 226112612 | rs4674980 | -0.782 | 0.369 | -0.206 | -0.586 | -0.005 | -0.037 | -0.037 | 0 | 0.38 | 0.54 | 0.38 | 0.26 | 0.35 | 0.29 |
| 2 | 226115446 | rs7571208 | 0 | 0.644 | 0.148 | 0.373 | -0.001 | 0.017 | 0.017 | 0 | 0.46 | 0.62 | 0.47 | 0.69 | 0.44 | 0.61 |
| 2 | 226116965 | rs6728230 | -0.409 | 0.536 | 0.231 | -0.162 | -0.016 | -0.051 | -0.051 | 1.299 | 0.1 | 0.26 | 0.08 | 0 | 0.09 | 0.03 |
| 2 | 226118444 | rs16866546 | 0.329 | 0.657 | 0.09 | 0.097 | 0.004 | -0.012 | -0.012 | 0.488 | 0.35 | 0.38 | 0.39 | 0.74 | 0.35 | 0.57 |
| 2 | 226140372 | rs13018232 | 0.655 | 0.718 | -0.453 | 0.112 | -0.039 | -0.036 | -0.036 | 0 | 0.31 | 0.14 | 0.3 | 0.66 | 0.32 | 0.55 |
| 2 | 226150336 | rs935158 | -0.761 | 0.816 | -0.694 | -0.22 | -0.017 | 0.018 | 0.018 | 0 | 0.83 | 0.89 | 0.7 | 0.89 | 0.76 | 0.9 |
| 2 | 226154132 | rs6743368 | -0.538 | 0.96 | -0.916 | -0.478 | -0.064 | 0.075 | 0.075 | 0 | 0.59 | 0.83 | 0.48 | 0.26 | 0.53 | 0.42 |
| 2 | 226157800 | rs2044236 | 0.474 | 1.038 | -1.173 | -0.725 | -0.042 | 0.038 | 0.038 | 0 | 0.14 | 0.05 | 0.18 | 0.07 | 0.16 | 0.15 |
| 2 | 226168931 | rs13419540 | 0.447 | 0.871 | -0.337 | -0.779 | -0.006 | -0.014 | -0.014 | 0 | 0.4 | 0.56 | 0.25 | 0.19 | 0.34 | 0.26 |
| 2 | 226174087 | rs6753175 | -1.815 | 0.909 | -0.452 | -0.651 | -0.048 | -0.091 | -0.091 | 0 | 0.74 | 0.93 | 0.75 | 0.38 | 0.73 | 0.52 |
| 2 | 226174116 | rs10498183 | 0.354 | 0.404 | 0.057 | -0.643 | -0.005 | 0.06 | 0.06 | 0 | 0.09 | 0.3 | 0.08 | 0.11 | 0.07 | 0.02 |
| 2 | 226174464 | rs6711329 | 1.16 | 0.452 | -0.041 | -0.652 | -0.013 | 0.01 | 0.01 | 0 | 0.15 | 0.11 | 0.27 | 0.12 | 0.19 | 0.1 |
| 2 | 226178411 | rs17409709 | -1.135 | 0.46 | -0.112 | -0.657 | -0.102 | 0.016 | 0.016 | 0 | 0.6 | 0.89 | 0.56 | 0.31 | 0.57 | 0.36 |
| 2 | 226183385 | rs6436549 | 0.659 | 0.425 | -0.072 | -0.629 | 0.005 | 0.001 | 0.001 | 0 | 0.31 | 0.26 | 0.17 | 0.08 | 0.27 | 0.22 |
| 2 | 226183487 | rs6436551 | -0.747 | 0.477 | -0.543 | -0.578 | -0.008 | 0.022 | 0.022 | 0 | 0.56 | 0.67 | 0.51 | 0.31 | 0.53 | 0.35 |
| 2 | 226216097 | rs1517481 | -0.918 | 0.531 | -0.772 | -0.343 | -0.028 | 0.018 | 0.018 | 0 | 0.55 | 0.79 | 0.53 | 0.38 | 0.53 | 0.35 |
| 2 | 226219320 | rs4347829 | 0.836 | 0.637 | -1.128 | -0.575 | -0.037 | 0.042 | 0.042 | 0 | 0.28 | 0.01 | 0.14 | 0.15 | 0.26 | 0.22 |
| 2 | 226220594 | rs10498184 | 0.331 | 0.657 | -0.465 | -0.439 | 0.005 | -0.021 | -0.021 | 0 | 0.18 | 0.28 | 0.29 | 0.12 | 0.2 | 0.11 |
| 2 | 226237910 | rs1517484 | -0.601 | 0.627 | -0.822 | -0.745 | 0.001 | -0.03 | -0.03 | 2.98 | 0.45 | 0.52 | 0.34 | 0.22 | 0.47 | 0.3 |
| 2 | 226238479 | rs17481876 | 1.395 | 0.547 | -1.113 | -0.77 | -0.003 | 0.026 | 0.026 | 0 | 0.18 | 0 | 0.06 | 0.14 | 0.13 | 0.19 |
| 2 | 226239709 | rs1517485 | 0.932 | 0.682 | -0.974 | -0.796 | 0.05 | -0.042 | -0.042 | 1.032 | 0.17 | 0.29 | 0.2 | 0.08 | 0.25 | 0.1 |
| 2 | 226240872 | rs932181 | 1.148 | 0.604 | -1.633 | -0.549 | 0.019 | 0.031 | 0.031 | 0 | 0.19 | 0.1 | 0.07 | 0.14 | 0.13 | 0.19 |
| 2 | 226242049 | rs16866572 | -0.004 | 0.805 | -1.555 | 0.126 | -0.002 | -0.001 | -0.001 | 0.004 | 0.17 | 0.27 | 0.11 | 0.17 | 0.17 | 0.07 |
| 2 | 226244129 | rs9973944 | -0.162 | 0.868 | -2.24 | 0.52 | -0.091 | -0.044 | -0.044 | 0.143 | 0.63 | 0.95 | 0.46 | 0.39 | 0.6 | 0.37 |
| 2 | 226251703 | rs10498185 | -0.269 | 0.826 | -1.762 | 1.2 | -0.031 | -0.098 | -0.098 | 0.012 | 0.82 | 0.93 | 0.68 | 0.51 | 0.78 | 0.55 |
| 2 | 226254785 | rs1517494 | -0.714 | 0.184 | -0.652 | 0.774 | -0.031 | 0.016 | 0.016 | 0 | 0.12 | 0.38 | 0.07 | 0.12 | 0.05 | 0.1 |
| 2 | 226272295 | rs9288607 | -0.458 | 0.332 | -0.638 | 1.358 | 0 | 0.014 | 0.014 | 0 | 0.83 | 0.81 | 0.67 | 0.59 | 0.82 | 0.58 |
| 2 | 226273490 | rs2048935 | -0.715 | 0.322 | -0.695 | 1.373 | -0.02 | -0.076 | -0.076 | 0.17 | 0.84 | 0.95 | 0.78 | 0.59 | 0.82 | 0.58 |
| 2 | 226273689 | rs2048936 | -0.718 | 0.32 | -0.696 | 1.391 | -0.022 | -0.079 | -0.079 | 0.17 | 0.84 | 0.95 | 0.78 | 0.59 | 0.82 | 0.58 |
| 2 | 226275520 | rs1517505 | -0.408 | 0.28 | -0.439 | 1.372 | 0 | 0.019 | 0.019 | 0 | 0.83 | 0.79 | 0.76 | 0.59 | 0.82 | 0.58 |
| 2 | 226278417 | rs4402753 | 1.046 | -0.676 | -0.204 | 0.928 | -0.008 | 0.005 | 0.005 | 0 | 0.63 | 0.37 | 0.68 | 0.47 | 0.74 | 0.46 |
| 2 | 226299309 | rs6753020 | 0.691 | -0.822 | -0.432 | 0.836 | -0.02 | 0.03 | 0.03 | 0 | 0.68 | 0.55 | 0.69 | 0.57 | 0.76 | 0.52 |
| 2 | 226308019 | rs1517499 | -1.385 | -0.791 | -0.558 | 0.839 | -0.034 | -0.107 | -0.107 | 0 | 0.85 | 0.98 | 0.78 | 0.59 | 0.83 | 0.57 |
| 2 | 226317818 | rs13393228 | 1.231 | -0.294 | -1.488 | -0.184 | -0.097 | -0.004 | -0.004 | 0 | 0.29 | 0.02 | 0.37 | 0.23 | 0.36 | 0.3 |
| 2 | 226319552 | rs7597786 | -0.473 | -0.397 | -1.082 | 0.075 | -0.047 | -0.102 | -0.102 | 0.008 | 0.16 | 0.35 | 0.07 | 0.01 | 0.06 | 0.03 |
| 2 | 226336889 | rs17410253 | 0.953 | 0.387 | -2.205 | -0.083 | -0.149 | 0.04 | 0.04 | 0 | 0.36 | 0.02 | 0.43 | 0.23 | 0.44 | 0.28 |
| 2 | 226353626 | rs13433050 | -0.632 | 0.394 | -0.703 | 0.434 | -0.015 | 0.039 | 0.039 | 0.18 | 0.19 | 0.53 | 0.19 | 0.23 | 0.12 | 0.06 |
| 2 | 226356518 | rs7597924 | -0.779 | 0.32 | -0.598 | 0.397 | -0.02 | 0.023 | 0.023 | 0.863 | 0.19 | 0.58 | 0.2 | 0.24 | 0.12 | 0.06 |
| 2 | 226357281 | rs9288608 | -0.449 | 0.308 | 0.02 | 0.391 | -0.007 | 0.038 | 0.038 | 0 | 0.17 | 0.35 | 0.17 | 0.24 | 0.12 | 0.06 |
| 2 | 226358003 | rs1431089 | -1.298 | 0.489 | -0.435 | 0.658 | -0.03 | -0.043 | -0.043 | 0.577 | 0.92 | 0.99 | 0.89 | 0.85 | 0.9 | 0.93 |
| 2 | 226358289 | rs1897229 | -0.963 | 0.375 | -0.642 | 0.486 | 0.015 | 0.042 | 0.042 | 0 | 0.46 | 0.82 | 0.49 | 0.66 | 0.41 | 0.64 |
| 2 | 226359147 | rs6747620 | -0.35 | 0.863 | -0.101 | 0.504 | -0.008 | 0.055 | 0.055 | 0.061 | 0.18 | 0.46 | 0.19 | 0.24 | 0.12 | 0.06 |
| 2 | 226366322 | rs16866626 | 0.043 | 1.332 | 0.431 | 0.589 | 0.009 | -0.015 | -0.015 | 0 | 0.34 | 0.21 | 0.32 | 0.42 | 0.31 | 0.56 |
| 2 | 226375085 | rs1431080 | 0.152 | 1.431 | 0.827 | 0.467 | 0 | 0.002 | 0.002 | 0.098 | 0.21 | 0.27 | 0.24 | 0.24 | 0.19 | 0.06 |

| | | | | | | | | | | | | | | | | |
|---|---|---|---|---|---|---|---|---|---|---|---|---|---|---|---|---|
| 2 | 226380926 | rs1157324 | -1.307 | 1.359 | 0.831 | 0.402 | -0.046 | 0.002 | 0.002 | 0 | 0.58 | 0.81 | 0.59 | 0.65 | 0.51 | 0.61 |
| 2 | 226394889 | rs10804326 | 0.18 | 1.361 | 1.094 | 0.484 | -0.009 | 0.01 | 0.01 | 0 | 0.35 | 0.39 | 0.33 | 0.42 | 0.32 | 0.55 |
| 2 | 226396737 | rs13392113 | 0.179 | 1.366 | 1.139 | 0.48 | -0.001 | -0.001 | -0.001 | 0 | 0.21 | 0.22 | 0.23 | 0.23 | 0.19 | 0.06 |
| 2 | 226397028 | rs7606323 | -1.294 | 1.357 | 0.934 | 0.485 | -0.051 | 0.001 | 0.001 | 0 | 0.58 | 0.81 | 0.58 | 0.65 | 0.51 | 0.61 |
| 2 | 226413483 | rs7586576 | 0.669 | 1.404 | 0.963 | 0.566 | -0.002 | -0.003 | -0.003 | 0 | 0.72 | 0.26 | 0.69 | 0.63 | 0.75 | 0.87 |
| 2 | 226417918 | rs1367587 | 1.448 | 1.224 | 0.63 | 0.576 | -0.094 | 0.022 | 0.022 | 0 | 0.36 | 0.04 | 0.36 | 0.21 | 0.42 | 0.3 |
| 2 | 226423124 | rs11887399 | 0.029 | 0.964 | 1.173 | 0.553 | -0.032 | 0.011 | 0.011 | 0 | 0.19 | 0.36 | 0.17 | 0.23 | 0.12 | 0.04 |
| 2 | 226428226 | rs16866649 | 0.273 | 0.97 | 0.759 | 0.559 | -0.016 | 0.003 | 0.003 | 0.01 | 0.18 | 0.23 | 0.16 | 0.23 | 0.11 | 0.04 |
| 2 | 226444488 | rs17482442 | 0.949 | 1.101 | 0.121 | 0.622 | 0 | 0.017 | 0.017 | 0 | 0.16 | 0.07 | 0.06 | 0.08 | 0.13 | 0.13 |
| 2 | 226447735 | rs3828256 | 0.83 | 1.083 | 0.207 | 0.61 | 0.023 | -0.009 | -0.009 | 0 | 0.58 | 0.19 | 0.71 | 0.53 | 0.69 | 0.71 |
| 2 | 226456461 | rs1993323 | 0.912 | 1.43 | 0.225 | 0.663 | -0.006 | 0.001 | 0.001 | 0 | 0.57 | 0.3 | 0.68 | 0.53 | 0.58 | 0.69 |
| 2 | 226460340 | rs7585186 | -0.853 | 1.433 | 0.325 | 0.645 | -0.001 | -0.005 | -0.005 | 0.853 | 0.38 | 0.39 | 0.27 | 0.32 | 0.37 | 0.18 |
| 2 | 226465409 | rs715678 | 0.481 | 1.415 | 0.282 | 0.633 | 0.017 | 0.036 | 0.036 | 0 | 0.73 | 0.42 | 0.75 | 0.61 | 0.7 | 0.8 |
| 2 | 226467954 | rs12619063 | 2.503 | 1.363 | -0.309 | 0.56 | -0.022 | 0.036 | 0.036 | 0.019 | 0.26 | 0.03 | 0.25 | 0.14 | 0.25 | 0.26 |
| 2 | 226482672 | rs1431086 | 1.184 | 1.261 | -0.215 | 0.605 | 0 | 0.028 | 0.028 | 0 | 0.16 | 0.05 | 0.06 | 0.09 | 0.12 | 0.12 |
| 2 | 226482872 | rs12477972 | 0.221 | 1.306 | -0.186 | 0.463 | -0.05 | -0.025 | -0.025 | 0.103 | 0.2 | 0.05 | 0.18 | 0.24 | 0.24 | 0.06 |
| 2 | 226493536 | rs13428386 | 0.133 | 1.056 | 0.098 | 0.549 | 0.008 | 0.005 | 0.005 | 0 | 0.2 | 0.34 | 0.2 | 0.24 | 0.22 | 0.06 |
| 2 | 226501136 | rs936070 | 0.77 | 1.153 | 0.13 | 0.697 | -0.008 | -0.021 | -0.021 | 0 | 0.59 | 0.29 | 0.69 | 0.42 | 0.6 | 0.63 |
| 2 | 226502137 | rs6436574 | 0.736 | 1.16 | 0.009 | 0.701 | -0.002 | -0.003 | -0.003 | 0 | 0.6 | 0.38 | 0.7 | 0.42 | 0.6 | 0.63 |
| 2 | 226515698 | rs7595617 | 2.153 | 1.185 | -0.111 | 0.648 | 0.001 | 0.03 | 0.03 | 0.005 | 0.25 | 0.05 | 0.21 | 0.14 | 0.22 | 0.26 |
| 2 | 226516346 | rs11884382 | 0.086 | 1.549 | 0.619 | 0.761 | 0.078 | -0.051 | -0.051 | 0 | 0.09 | 0.32 | 0.09 | 0.01 | 0.17 | 0.03 |
| 2 | 226517182 | rs4627553 | -0.024 | 1.553 | 0.624 | 0.775 | 0.015 | -0.061 | -0.061 | 0.074 | 0.26 | 0.07 | 0.17 | 0.42 | 0.18 | 0.23 |
| 2 | 226517237 | rs4641920 | -0.001 | 1.564 | 0.629 | 0.78 | 0.011 | -0.062 | -0.062 | 0 | 0.26 | 0.07 | 0.17 | 0.42 | 0.18 | 0.23 |
| 2 | 226517551 | rs6723679 | -0.257 | 1.813 | 1.193 | 0.458 | 0.078 | -0.051 | -0.051 | 0 | 0.09 | 0.32 | 0.09 | 0.01 | 0.17 | 0.03 |
| 2 | 226529024 | rs16866695 | 1.12 | 1.79 | 1.077 | 0.476 | 0.016 | 0.044 | 0.044 | 0.009 | 0.75 | 0.38 | 0.76 | 0.61 | 0.73 | 0.83 |
| 2 | 226538647 | rs6736992 | 1.088 | 1.791 | 0.949 | 0.476 | 0.004 | 0.017 | 0.017 | 0.037 | 0.75 | 0.31 | 0.75 | 0.61 | 0.72 | 0.83 |
| 2 | 226559224 | rs962426 | -0.79 | 1.839 | 0.435 | 0.494 | -0.094 | -0.006 | -0.006 | 0 | 0.74 | 0.98 | 0.57 | 0.84 | 0.66 | 0.66 |
| 2 | 226563259 | rs16866701 | 0.672 | 1.529 | 1.422 | 1.211 | 0.004 | -0.004 | -0.004 | 0 | 0.1 | 0.09 | 0.11 | 0.24 | 0.06 | 0.03 |
| 2 | 226583962 | rs6737960 | -0.806 | 1.55 | 1.403 | 1.155 | 0.079 | -0.056 | -0.056 | 0 | 0.1 | 0.32 | 0.09 | 0.01 | 0.17 | 0.03 |
| 2 | 226591712 | rs1122817 | -1.527 | 1.539 | 0.963 | 1.202 | 0.021 | 0.057 | 0.057 | 0 | 0.25 | 0.56 | 0.24 | 0.4 | 0.28 | 0.17 |
| 2 | 226618383 | rs10172113 | 1.262 | 1.179 | 0.058 | 0.979 | 0.002 | -0.011 | -0.011 | 0 | 0.73 | 0.26 | 0.73 | 0.56 | 0.74 | 0.84 |
| 2 | 226629084 | rs7558381 | 1.301 | 1.18 | 0.014 | 0.988 | -0.004 | -0.01 | -0.01 | 0 | 0.72 | 0.25 | 0.65 | 0.56 | 0.66 | 0.82 |
| 2 | 226629841 | rs11687544 | 1.238 | 1.181 | 0.037 | 1.022 | -0.004 | -0.007 | -0.007 | 0.782 | 0.73 | 0.26 | 0.64 | 0.57 | 0.66 | 0.84 |
| 2 | 226631165 | rs16866708 | -0.079 | 1.341 | -0.287 | 0.951 | 0.138 | 0.001 | 0.001 | 0.031 | 0.07 | 0.33 | 0.09 | 0 | 0.15 | 0.02 |
| 2 | 226632146 | rs6754709 | 0.148 | 0.635 | -1.666 | 1.204 | -0.082 | 0.061 | 0.061 | 0 | 0.3 | 0.01 | 0.36 | 0.13 | 0.28 | 0.31 |
| 2 | 226636274 | rs974412 | 0.261 | 0.315 | -0.015 | 1.928 | -0.005 | -0.002 | -0.002 | 0 | 0.92 | 0.91 | 0.88 | 0.8 | 0.86 | 0.88 |
| 2 | 226655383 | rs12988196 | -0.544 | 0.313 | -0.074 | 1.883 | -0.015 | 0.022 | 0.022 | 1.478 | 0.11 | 0.16 | 0.14 | 0.21 | 0.07 | 0.02 |
| 2 | 226659586 | rs12621711 | 0.694 | 0.773 | -0.431 | 1.493 | -0.013 | -0.015 | -0.015 | 0 | 0.91 | 0.92 | 0.88 | 0.82 | 0.86 | 0.89 |
| 2 | 226672288 | rs9288611 | 0.102 | -0.406 | 0.016 | 1.291 | 0.005 | 0.002 | 0.002 | 0 | 0.55 | 0.46 | 0.69 | 0.43 | 0.68 | 0.57 |
| 2 | 226683986 | rs11903376 | -1.698 | -0.346 | 0.329 | 1.792 | -0.034 | 0.046 | 0.046 | 4.9 | 0.09 | 0.82 | 0.15 | 0.23 | 0.01 | 0 |
| 2 | 226697772 | rs873024 | -1.204 | -0.41 | 0.32 | 1.703 | -0.007 | 0.006 | 0.006 | 0.081 | 0.33 | 0.62 | 0.34 | 0.55 | 0.27 | 0.43 |
| 2 | 226714032 | rs12995191 | -1.189 | -0.608 | 1.173 | 1.971 | -0.017 | 0.052 | 0.052 | 0 | 0.23 | 0.67 | 0.16 | 0.45 | 0.15 | 0.31 |
| 2 | 226730588 | rs1518307 | -0.92 | -0.618 | 1.589 | 1.966 | -0.021 | 0.121 | 0.121 | 0 | 0.22 | 0.42 | 0.14 | 0.45 | 0.15 | 0.31 |
| 2 | 226735227 | rs35969878 | 0.35 | -1.202 | 0.986 | 1.579 | 0.026 | 0.039 | 0.039 | 0 | 0.17 | 0.07 | 0.07 | 0.09 | 0.12 | 0.29 |
| 2 | 226743260 | rs1190118 | -2.032 | -0.845 | 0.818 | 1.479 | -0.02 | 0.047 | 0.047 | 0 | 0.56 | 0.89 | 0.45 | 0.65 | 0.5 | 0.74 |
| 2 | 226747064 | rs1190116 | -1.957 | -0.519 | 1.491 | 2.05 | 0.028 | -0.023 | -0.023 | 0.037 | 0.34 | 0.8 | 0.35 | 0.57 | 0.27 | 0.43 |
| 2 | 226749827 | rs11883457 | -0.602 | -0.529 | 2.08 | 2.333 | -0.008 | 0.009 | 0.009 | 0 | 0.18 | 0.27 | 0.09 | 0.18 | 0.14 | 0.3 |
| 2 | 226750426 | rs12987704 | -0.791 | -0.526 | 1.965 | 2.334 | -0.023 | 0.016 | 0.016 | 0 | 0.21 | 0.46 | 0.12 | 0.18 | 0.15 | 0.3 |
| 2 | 226754597 | rs12694684 | 0.036 | -0.543 | 1.92 | 2.324 | -0.003 | -0.024 | -0.024 | 0.501 | 0.9 | 0.97 | 0.81 | 0.88 | 0.88 | 0.87 |
| 2 | 226774322 | rs10451554 | -1.081 | -0.457 | 2.112 | 2.27 | -0.031 | 0.005 | 0.005 | 0 | 0.22 | 0.57 | 0.12 | 0.18 | 0.15 | 0.3 |
| 2 | 226783634 | rs1878696 | -1.499 | -0.46 | 1.887 | 2.308 | 0.008 | -0.016 | -0.016 | 0 | 0.24 | 0.77 | 0.17 | 0.46 | 0.15 | 0.31 |
| 2 | 226783683 | rs1878697 | -0.937 | -0.379 | 2.07 | 1.708 | -0.011 | 0.006 | 0.006 | 0 | 0.2 | 0.36 | 0.11 | 0.19 | 0.15 | 0.31 |
| 2 | 226790973 | rs4675018 | -1.137 | -0.374 | 1.506 | 1.71 | -0.015 | -0.002 | -0.002 | 0.086 | 0.22 | 0.68 | 0.13 | 0.19 | 0.15 | 0.31 |
| 2 | 226795228 | rs13031468 | -1.258 | -0.354 | 1.279 | 1.722 | -0.011 | -0.003 | -0.003 | 0 | 0.23 | 0.71 | 0.14 | 0.19 | 0.15 | 0.31 |
| 2 | 226805730 | rs13387874 | 1.544 | 0.078 | 1.069 | 2.067 | -0.001 | -0.009 | -0.009 | 0 | 0.75 | 0.71 | 0.7 | 0.58 | 0.73 | 0.61 |
| 2 | 226807953 | rs6732525 | 0.433 | 0.088 | 0.949 | 2.177 | 0.005 | -0.011 | -0.011 | 0.587 | 0.9 | 0.94 | 0.81 | 0.87 | 0.89 | 0.89 |
| 2 | 226808378 | rs12986824 | -1.165 | 0.104 | 0.959 | 1.985 | -0.005 | -0.008 | -0.008 | 0.796 | 0.14 | 0.17 | 0.1 | 0.29 | 0.17 | 0.28 |
| 2 | 226814991 | rs4673151 | -1.571 | 0.015 | 1.328 | 1.8 | 0.02 | 0.119 | 0.119 | 0.043 | 0.16 | 0.45 | 0.13 | 0.28 | 0.15 | 0.23 |
| 2 | 226827127 | rs10168662 | -1.947 | -0.01 | 1.138 | 1.785 | -0.003 | 0.068 | 0.068 | 0.596 | 0.21 | 0.51 | 0.16 | 0.34 | 0.19 | 0.37 |
| 2 | 226845937 | rs6737505 | -1.857 | 0.043 | 1.145 | 1.79 | 0.004 | 0.074 | 0.074 | 0.776 | 0.2 | 0.51 | 0.16 | 0.33 | 0.19 | 0.37 |
| 2 | 226857323 | rs12694692 | -1.36 | 0.171 | 0.61 | 1.827 | 0.001 | 0.015 | 0.015 | 0 | 0.14 | 0.18 | 0.11 | 0.27 | 0.15 | 0.23 |
| 2 | 226863834 | rs12474644 | 0.471 | 0.288 | 0.816 | 1.826 | 0.006 | 0.034 | 0.034 | 0 | 0.91 | 0.87 | 0.78 | 0.82 | 0.9 | 0.89 |
| 2 | 226865961 | rs908719 | 0.819 | 0.228 | 1.231 | 1.783 | 0.016 | 0.064 | 0.064 | 0 | 0.9 | 0.74 | 0.77 | 0.82 | 0.9 | 0.89 |
| 2 | 226866680 | rs13428728 | -1.485 | 0.073 | 1.786 | 1.763 | 0.008 | 0.014 | 0.014 | 0.041 | 0.08 | 0.27 | 0.05 | 0.05 | 0.03 | 0.17 |
| 2 | 226872183 | rs7584507 | 2.183 | 0.037 | 1.989 | 1.745 | -0.013 | 0.031 | 0.031 | 0 | 0.81 | 0.39 | 0.71 | 0.76 | 0.87 | 0.71 |
| 2 | 226890028 | rs2396274 | 0.646 | 0.047 | 1.968 | 1.765 | 0.012 | 0.06 | 0.06 | 0 | 0.9 | 0.8 | 0.77 | 0.81 | 0.9 | 0.89 |
| 2 | 226890057 | rs715832 | -2.021 | 0.043 | 2.013 | 1.78 | 0.012 | 0.018 | 0.018 | 0.054 | 0.09 | 0.38 | 0.06 | 0.05 | 0.03 | 0.17 |
| 2 | 226890684 | rs12469813 | 1.907 | 0.06 | 1.71 | 1.815 | -0.005 | 0.069 | 0.069 | 0 | 0.83 | 0.5 | 0.71 | 0.76 | 0.87 | 0.72 |
| 2 | 226894981 | rs908720 | 0.512 | 0.232 | 1.107 | 1.814 | 0.004 | 0.022 | 0.022 | 0 | 0.91 | 0.89 | 0.77 | 0.81 | 0.89 | 0.88 |

| | | | | | | | | | | | | | | | | |
|---|---|---|---|---|---|---|---|---|---|---|---|---|---|---|---|---|
| 2 | 226899712 | rs1518297 | 0.946 | -0.235 | 0.599 | 1.533 | -0.02 | -0.063 | -0.063 | 1.059 | 0.84 | 0.99 | 0.76 | 0.8 | 0.87 | 0.81 |
| 2 | 226902562 | rs13018599 | 1.229 | -0.504 | 0.186 | 1.476 | 0.041 | -0.017 | -0.017 | 0 | 0.56 | 0.19 | 0.58 | 0.48 | 0.62 | 0.66 |
| 2 | 226907209 | rs925902 | -0.72 | -0.351 | -0.49 | 0.976 | -0.025 | 0.07 | 0.07 | 0 | 0.51 | 0.84 | 0.48 | 0.68 | 0.5 | 0.56 |
| 2 | 226915702 | rs4503980 | 0.237 | -0.486 | -0.266 | 1.063 | -0.003 | -0.007 | -0.007 | 0 | 0.24 | 0.19 | 0.31 | 0.37 | 0.26 | 0.41 |
| 2 | 226917839 | rs10191186 | 0.608 | -0.551 | 0.289 | 1.024 | -0.015 | 0.02 | 0.02 | 0 | 0.25 | 0.37 | 0.14 | 0.3 | 0.24 | 0.16 |
| 2 | 226923667 | rs4675037 | -0.6 | -0.64 | 0.983 | 1.062 | -0.002 | 0.008 | 0.008 | 0 | 0.47 | 0.43 | 0.45 | 0.66 | 0.49 | 0.57 |
| 2 | 226935229 | rs2204173 | 0.994 | -0.637 | 0.956 | 1.045 | 0.001 | -0.037 | -0.037 | 0 | 0.22 | 0.12 | 0.12 | 0.31 | 0.24 | 0.16 |
| 2 | 226940969 | rs16866816 | 1.397 | -0.656 | 0.996 | 1.006 | -0.007 | -0.011 | -0.011 | 0 | 0.1 | 0.14 | 0.1 | 0.08 | 0.07 | 0.06 |
| 2 | 226946435 | rs4675041 | -0.469 | -0.639 | 0.904 | 0.983 | 0.005 | 0.018 | 0.018 | 0 | 0.25 | 0.31 | 0.32 | 0.37 | 0.26 | 0.41 |
| 2 | 226951412 | rs4675044 | -1.211 | -0.678 | 0.389 | 0.725 | -0.02 | 0.061 | 0.061 | 0 | 0.53 | 0.74 | 0.48 | 0.68 | 0.52 | 0.58 |
| 2 | 226957178 | rs4675047 | -1.393 | -0.679 | 0.352 | 0.844 | -0.038 | 0.096 | 0.096 | 0 | 0.53 | 0.82 | 0.49 | 0.72 | 0.52 | 0.58 |
| 2 | 226960697 | rs6436598 | -1.202 | -0.677 | 0.358 | 0.805 | -0.014 | 0.054 | 0.054 | 0 | 0.53 | 0.74 | 0.48 | 0.68 | 0.52 | 0.58 |
| 2 | 226961688 | rs12611824 | -0.9 | -0.647 | 0.368 | 0.817 | -0.003 | 0.007 | 0.007 | 0 | 0.3 | 0.58 | 0.35 | 0.38 | 0.28 | 0.41 |
| 2 | 226989328 | rs1403356 | 1.003 | -0.629 | 0.141 | 0.87 | 0.003 | -0.02 | -0.02 | 0 | 0.23 | 0.16 | 0.13 | 0.31 | 0.24 | 0.16 |
| 2 | 226997454 | rs1522812 | 0.403 | -0.507 | -0.582 | 0.298 | -0.081 | 0.122 | 0.122 | 1.148 | 0.62 | 0.94 | 0.46 | 0.2 | 0.56 | 0.41 |
| 2 | 227005713 | rs10178651 | 0.447 | -0.513 | -0.614 | 0.24 | -0.078 | 0.134 | 0.134 | 1.144 | 0.61 | 0.94 | 0.42 | 0.2 | 0.56 | 0.4 |
| 2 | 227009627 | rs2948556 | -0.715 | -0.81 | 0.542 | 0.174 | -0.009 | 0.016 | 0.016 | 0 | 0.45 | 0.63 | 0.45 | 0.33 | 0.54 | 0.48 |
| 2 | 227010224 | rs2140237 | -0.222 | -0.985 | 0.061 | 0.039 | -0.015 | -0.016 | -0.016 | 0 | 0.29 | 0.14 | 0.28 | 0.29 | 0.33 | 0.36 |
| 2 | 227018472 | rs2972132 | -1.24 | -1.13 | 0.106 | -0.077 | -0.031 | 0.075 | 0.075 | 0 | 0.49 | 0.84 | 0.47 | 0.33 | 0.54 | 0.47 |
| 2 | 227019461 | rs10181515 | 0.908 | -0.99 | 0.193 | -0.08 | 0.006 | -0.068 | -0.068 | 0.888 | 0.19 | 0.39 | 0.19 | 0.05 | 0.23 | 0.14 |
| 2 | 227019577 | rs2948567 | -0.361 | -0.999 | -0.12 | 0.173 | -0.014 | -0.018 | -0.018 | 0 | 0.28 | 0.16 | 0.28 | 0.31 | 0.33 | 0.36 |
| 2 | 227020494 | rs1522802 | -0.402 | -0.932 | -0.757 | 0.177 | -0.032 | -0.07 | -0.07 | 0 | 0.83 | 0.96 | 0.88 | 0.71 | 0.8 | 0.72 |
| 2 | 227020521 | rs1522803 | 0.175 | -1.046 | -1.193 | -0.169 | -0.114 | -0.08 | -0.08 | 0.616 | 0.57 | 0.93 | 0.65 | 0.54 | 0.45 | 0.56 |
| 2 | 227022036 | rs13405357 | 0.496 | -1.076 | -0.435 | -0.001 | -0.017 | 0.03 | 0.03 | 0 | 0.31 | 0.18 | 0.26 | 0.2 | 0.35 | 0.16 |
| 2 | 227034499 | rs6713510 | -1.006 | -0.985 | 0.183 | -0.185 | -0.008 | 0.025 | 0.025 | 0 | 0.54 | 0.72 | 0.58 | 0.59 | 0.48 | 0.43 |
| 2 | 227041406 | rs10221889 | 0.902 | -1.067 | 0.258 | -0.333 | 0.001 | -0.07 | -0.07 | 0.94 | 0.16 | 0.11 | 0.3 | 0.45 | 0.14 | 0.23 |
| 2 | 227041527 | rs10221655 | 0.861 | -1.076 | 0.254 | -0.337 | 0.002 | -0.072 | -0.072 | 0 | 0.17 | 0.11 | 0.31 | 0.45 | 0.14 | 0.23 |
| 2 | 227044268 | rs7563595 | 0.238 | -1.078 | 0.646 | -0.338 | -0.027 | 0.197 | 0.197 | 0 | 0.2 | 0.45 | 0.33 | 0.44 | 0.14 | 0.23 |
| 2 | 227054881 | rs2943633 | 0.77 | -0.978 | 0.541 | -0.397 | -0.004 | 0.014 | 0.014 | 0 | 0.46 | 0.29 | 0.43 | 0.46 | 0.52 | 0.57 |
| 2 | 227060657 | rs16866883 | -0.26 | -0.917 | 0.818 | -0.377 | 0.029 | 0.002 | 0.002 | 0.298 | 0.07 | 0.4 | 0.04 | 0 | 0.01 | 0.04 |
| 2 | 227093745 | rs2943641 | -0.4 | -0.806 | 0.402 | -0.41 | 0.004 | 0.06 | 0.06 | 0 | 0.64 | 0.69 | 0.77 | 0.93 | 0.63 | 0.77 |
| 2 | 227098387 | rs1515114 | 0.657 | -0.815 | 0.497 | -0.35 | -0.005 | 0.001 | 0.001 | 0 | 0.44 | 0.36 | 0.61 | 0.49 | 0.48 | 0.52 |
| 2 | 227099180 | rs2943645 | -0.285 | -0.713 | 0.539 | -0.672 | 0.003 | 0.011 | 0.011 | 0 | 0.63 | 0.62 | 0.76 | 0.93 | 0.62 | 0.77 |
| 2 | 227116365 | rs2972143 | -0.347 | -0.492 | 0.313 | -0.546 | -0.001 | 0.008 | 0.008 | 0 | 0.66 | 0.69 | 0.77 | 0.93 | 0.63 | 0.75 |
| 2 | 227122216 | rs1515110 | -0.646 | -0.124 | 0.425 | -0.448 | 0.012 | -0.089 | -0.089 | 0.832 | 0.69 | 0.59 | 0.77 | 0.93 | 0.62 | 0.77 |
| 2 | 227123439 | rs2943657 | -0.226 | -0.187 | 0.867 | -0.487 | -0.003 | 0.036 | 0.036 | 0 | 0.66 | 0.26 | 0.75 | 0.93 | 0.62 | 0.77 |
| 2 | 227123584 | rs2943658 | -0.86 | -0.262 | 1.208 | -0.531 | -0.006 | 0.004 | 0.004 | 0 | 0.72 | 0.7 | 0.79 | 0.93 | 0.63 | 0.81 |
| 2 | 227125254 | rs2943659 | -0.38 | -0.251 | 1.138 | -0.526 | 0.003 | -0.072 | -0.072 | 0.457 | 0.67 | 0.39 | 0.76 | 0.93 | 0.62 | 0.77 |
| 2 | 227132974 | rs11675231 | 0.671 | -0.301 | 0.771 | -0.496 | -0.006 | -0.036 | -0.036 | 0 | 0.16 | 0.07 | 0.17 | 0.34 | 0.16 | 0.28 |
| 2 | 227134516 | rs2713538 | 0.429 | -0.277 | 0.792 | -0.514 | -0.003 | 0.031 | 0.031 | 0 | 0.52 | 0.33 | 0.59 | 0.6 | 0.46 | 0.49 |
| 2 | 227137125 | rs2713539 | -1.221 | -0.298 | 0.898 | -0.522 | 0.002 | -0.024 | -0.024 | 0.481 | 0.72 | 0.66 | 0.79 | 0.93 | 0.63 | 0.81 |
| 2 | 227142666 | rs1849878 | 0.554 | -0.286 | 0.874 | -0.51 | -0.019 | 0.06 | 0.06 | 0 | 0.51 | 0.2 | 0.55 | 0.6 | 0.46 | 0.49 |
| 2 | 227144141 | rs1515103 | 1.31 | -0.407 | 0.61 | -0.595 | -0.024 | -0.134 | -0.134 | 1.796 | 0.17 | 0.02 | 0.16 | 0.47 | 0.15 | 0.27 |
| 2 | 227148722 | rs4510219 | 0.574 | -0.291 | 1.022 | -0.511 | -0.002 | -0.009 | -0.009 | 2.385 | 0.16 | 0.12 | 0.18 | 0.34 | 0.17 | 0.28 |
| 2 | 227149804 | rs2713545 | 0.437 | -0.3 | 1.065 | -0.529 | -0.005 | 0.043 | 0.043 | 0 | 0.51 | 0.27 | 0.58 | 0.6 | 0.45 | 0.49 |
| 2 | 227151291 | rs6733028 | -0.6 | -0.159 | 1.278 | 0.002 | -0.015 | 0.047 | 0.047 | 0 | 0.21 | 0.52 | 0.22 | 0.35 | 0.18 | 0.32 |
| 2 | 227180398 | rs13026427 | 0.477 | 0.394 | -0.553 | 0.3 | -0.015 | -0.038 | -0.038 | 0.684 | 0.13 | 0.04 | 0.19 | 0.34 | 0.16 | 0.27 |
| 2 | 227184631 | rs10933137 | 0.168 | 0.264 | -0.54 | 0.157 | 0.019 | 0.013 | 0.013 | 0 | 0.43 | 0.32 | 0.47 | 0.63 | 0.35 | 0.34 |
| 2 | 227185764 | rs2396306 | -0.13 | 0.023 | -0.187 | 0.379 | -0.019 | 0.336 | 0.336 | 0 | 0.12 | 0.36 | 0.05 | 0.44 | 0.06 | 0.16 |
| 2 | 227190317 | rs2894593 | -0.867 | 0.443 | -1.099 | 0.002 | -0.036 | 0.064 | 0.064 | 0 | 0.66 | 0.82 | 0.63 | 0.86 | 0.55 | 0.64 |
| 2 | 227197598 | rs10167282 | 0.162 | 0.748 | -1.17 | 0.119 | -0.002 | 0.001 | 0.001 | 0 | 0.2 | 0.2 | 0.12 | 0.21 | 0.23 | 0.29 |
| 2 | 227199263 | rs10195178 | 0.08 | 0.673 | -1.078 | 0.144 | 0.001 | 0.068 | 0.068 | 0.753 | 0.5 | 0.32 | 0.48 | 0.39 | 0.47 | 0.45 |
| 2 | 227203200 | rs2396328 | 0.569 | 1.221 | -1.186 | -0.209 | -0.005 | 0.103 | 0.103 | 0 | 0.52 | 0.38 | 0.51 | 0.19 | 0.55 | 0.39 |
| 2 | 227204004 | rs2396327 | 0.261 | 1.201 | -0.327 | -0.076 | -0.002 | -0.036 | -0.036 | 0 | 0.61 | 0.59 | 0.65 | 0.81 | 0.62 | 0.55 |
| 2 | 227205102 | rs13005747 | 1.743 | 1.25 | -0.749 | -0.466 | 0.033 | 0.094 | 0.094 | 0 | 0.35 | 0.16 | 0.36 | 0.18 | 0.26 | 0.27 |
| 2 | 227207379 | rs7587413 | 0.398 | 1.26 | -0.351 | -0.555 | -0.003 | -0.006 | -0.006 | 0.403 | 0.25 | 0.25 | 0.25 | 0.15 | 0.22 | 0.21 |
| 2 | 227209828 | rs10933140 | -0.1 | 1.234 | -0.301 | -0.656 | -0.006 | 0.003 | 0.003 | 0 | 0.26 | 0.25 | 0.23 | 0.7 | 0.24 | 0.26 |
| 2 | 227209845 | rs12615325 | 0.732 | 1.258 | -0.277 | -0.616 | 0.041 | 0.24 | 0.24 | 0.007 | 0.08 | 0.2 | 0.14 | 0.62 | 0.07 | 0.16 |
| 2 | 227210398 | rs2894597 | -0.686 | 1.249 | -0.269 | -0.643 | -0.012 | -0.039 | -0.039 | 0 | 0.34 | 0.44 | 0.38 | 0.77 | 0.29 | 0.37 |
| 2 | 227221412 | rs16866988 | 1.209 | 1.04 | -0.25 | -0.543 | 0.018 | 0.315 | 0.315 | 0 | 0.06 | 0.2 | 0.11 | 0.62 | 0.04 | 0.16 |
| 2 | 227224367 | rs7570745 | 0.967 | 1.161 | -0.197 | -0.815 | 0.007 | 0.086 | 0.086 | 0 | 0.55 | 0.37 | 0.53 | 0.19 | 0.58 | 0.4 |
| 2 | 227237862 | rs2894595 | 0.672 | 1.204 | -0.122 | -1.032 | 0.02 | -0.064 | -0.064 | 0 | 0.17 | 0.24 | 0.15 | 0 | 0.29 | 0.12 |
| 2 | 227238030 | rs4675077 | -1.301 | 1.151 | -0.195 | -1.019 | -0.033 | 0.051 | 0.051 | 0 | 0.87 | 0.97 | 0.9 | 0.96 | 0.84 | 0.76 |
| 2 | 227241474 | rs7602912 | -0.313 | 1.144 | -0.099 | -1.052 | -0.015 | -0.024 | -0.024 | 0 | 0.31 | 0.34 | 0.36 | 0.78 | 0.26 | 0.37 |
| 2 | 227243909 | rs7574670 | 0.257 | 1.116 | -0.093 | -1.044 | 0.013 | 0.034 | 0.034 | 0 | 0.24 | 0.13 | 0.24 | 0.15 | 0.22 | 0.2 |
| 2 | 227244381 | rs10933143 | -0.198 | 1.084 | -0.185 | -1.228 | -0.015 | -0.024 | -0.024 | 0 | 0.31 | 0.34 | 0.36 | 0.78 | 0.26 | 0.37 |
| 2 | 227246379 | rs2396332 | 1.795 | 1.086 | -0.199 | -1.191 | 0.021 | 0.108 | 0.108 | 0 | 0.38 | 0.21 | 0.37 | 0.18 | 0.28 | 0.28 |
| 2 | 227247737 | rs7355577 | 0.491 | 1.1 | -0.119 | -1.275 | 0.04 | -0.079 | -0.079 | 0 | 0.16 | 0.27 | 0.14 | 0 | 0.28 | 0.12 |
| 2 | 227249236 | rs13398058 | 0.904 | 0.874 | -0.184 | -1.274 | 0.004 | 0.009 | 0.009 | 0 | 0.54 | 0.44 | 0.52 | 0.17 | 0.59 | 0.41 |

| | | | | | | | | | | | | | | | | |
|---|---|---|---|---|---|---|---|---|---|---|---|---|---|---|---|---|
| 2 | 227249312 | rs2894601 | -0.369 | 0.731 | -0.11 | -1.171 | -0.02 | -0.028 | -0.028 | 0 | 0.31 | 0.35 | 0.38 | 0.8 | 0.27 | 0.38 |
| 2 | 227254695 | rs13020901 | 0.089 | 1.396 | -0.083 | -1.228 | 0.014 | -0.064 | -0.064 | 0 | 0.22 | 0.24 | 0.18 | 0.01 | 0.34 | 0.3 |
| 2 | 227259267 | rs2894604 | -0.056 | 1.228 | -0.101 | -1.3 | -0.023 | 0.001 | 0.001 | 0 | 0.24 | 0.34 | 0.28 | 0.76 | 0.21 | 0.25 |
| 2 | 227263582 | rs7592897 | 1.85 | 1.185 | -0.22 | -1.511 | 0.017 | 0.142 | 0.142 | 0 | 0.43 | 0.19 | 0.34 | 0.16 | 0.34 | 0.32 |
| 2 | 227266027 | rs12185715 | 0.794 | 1.194 | -0.216 | -1.503 | -0.014 | -0.051 | -0.051 | 0 | 0.65 | 0.53 | 0.53 | 0.18 | 0.63 | 0.61 |
| 2 | 227282999 | rs2001303 | 1.991 | 1.187 | -0.408 | -1.486 | -0.007 | 0.161 | 0.161 | 0 | 0.41 | 0.07 | 0.33 | 0.16 | 0.3 | 0.31 |
| 2 | 227286027 | rs4395242 | -0.708 | 1.226 | -0.322 | -1.438 | 0.005 | -0.005 | -0.005 | 0 | 0.91 | 0.86 | 0.81 | 0.94 | 0.88 | 0.87 |
| 2 | 227291415 | rs11686139 | 1.624 | 1.618 | -0.491 | -1.49 | -0.055 | 0.057 | 0.057 | 0 | 0.11 | 0.03 | 0.1 | 0.01 | 0.19 | 0.22 |
| 2 | 227292162 | rs4600636 | 0.76 | 1.512 | -0.192 | -1.495 | 0.027 | -0.09 | -0.09 | 0 | 0.64 | 0.45 | 0.6 | 0.92 | 0.51 | 0.56 |
| 2 | 227293781 | rs6704649 | 1.172 | 1.322 | 0.181 | -1.331 | -0.011 | 0.135 | 0.135 | 0 | 0.07 | 0.15 | 0.11 | 0.66 | 0.04 | 0.18 |
| 2 | 227294482 | rs4129068 | 1.019 | 1.37 | -0.607 | -1.37 | -0.007 | 0.008 | 0.008 | 0 | 0.15 | 0.02 | 0.15 | 0.1 | 0.17 | 0.07 |
| 2 | 227301483 | rs12469727 | 1.289 | 1.45 | -0.314 | -1.278 | 0.005 | -0.047 | -0.047 | 0.057 | 0.11 | 0.17 | 0.08 | 0 | 0.15 | 0.08 |
| 2 | 227328729 | rs4522589 | 1.09 | 1.269 | -0.338 | -1.251 | 0.029 | -0.062 | -0.062 | 0.14 | 0.1 | 0.25 | 0.08 | 0 | 0.14 | 0.08 |
| 2 | 227337822 | rs1500989 | 1.709 | 1.156 | -1.26 | -1.19 | 0.002 | 0.148 | 0.148 | 0 | 0.41 | 0.1 | 0.33 | 0.18 | 0.29 | 0.32 |
| 2 | 227345365 | rs1392841 | 0.645 | 0.111 | -0.941 | -1.022 | -0.017 | -0.003 | -0.003 | 0 | 0.13 | 0.02 | 0.15 | 0.1 | 0.18 | 0.05 |
| 2 | 227346034 | rs931725 | 0.615 | -0.007 | -0.437 | -0.905 | 0.006 | -0.016 | -0.016 | 0 | 0.87 | 0.8 | 0.77 | 0.93 | 0.83 | 0.86 |
| 2 | 227351770 | rs1013243 | 0.906 | -0.462 | -0.42 | -0.623 | 0.004 | 0.042 | 0.042 | 0 | 0.06 | 0.04 | 0.09 | 0.65 | 0.03 | 0.19 |
| 2 | 227353523 | rs10498206 | -0.605 | -0.781 | -0.398 | -1.004 | 0 | -0.002 | -0.002 | 0 | 0.2 | 0.16 | 0.21 | 0.13 | 0.25 | 0.2 |
| 2 | 227353550 | rs13392647 | -1.117 | -0.784 | -0.399 | -1.053 | -0.001 | 0.002 | 0.002 | 0 | 0.26 | 0.2 | 0.3 | 0.78 | 0.28 | 0.39 |
| 2 | 227355975 | rs10498208 | -2.353 | -1.665 | -0.975 | -1.685 | -0.042 | 0.169 | 0.169 | 0 | 0.6 | 0.87 | 0.57 | 0.88 | 0.52 | 0.54 |
| 2 | 227358764 | rs13387692 | 1.676 | -1.997 | -1.122 | -1.758 | -0.005 | 0.078 | 0.078 | 0 | 0.14 | 0.02 | 0.13 | 0.01 | 0.15 | 0.13 |
| 2 | 227359131 | rs1846861 | -1.668 | -2.11 | -0.71 | -1.424 | 0.018 | 0.025 | 0.025 | 0 | 0.41 | 0.61 | 0.42 | 0.28 | 0.42 | 0.34 |
| 2 | 227360364 | rs16867044 | 1.335 | -1.985 | -0.979 | -1.433 | -0.001 | -0.021 | -0.021 | 0.744 | 0.1 | 0.04 | 0.13 | 0.2 | 0.11 | 0.12 |
| 2 | 227360446 | rs11903929 | -0.107 | -1.969 | -0.646 | -1.302 | -0.023 | 0.027 | 0.027 | 0.021 | 0.2 | 0.28 | 0.19 | 0.66 | 0.09 | 0.21 |
| 2 | 227371725 | rs11674718 | 0.482 | -1.709 | -0.89 | -1.207 | 0.018 | 0.022 | 0.022 | 0 | 0.23 | 0.15 | 0.18 | 0.04 | 0.17 | 0.14 |
| 2 | 227372171 | rs10166909 | 0.218 | -1.29 | -1.089 | -1.033 | 0.006 | 0.01 | 0.01 | 0 | 0.2 | 0.23 | 0.22 | 0.2 | 0.25 | 0.2 |
| 2 | 227373795 | rs1109991 | -1.373 | -1.666 | -1.453 | -1.145 | -0.036 | 0.146 | 0.146 | 0 | 0.59 | 0.86 | 0.56 | 0.83 | 0.52 | 0.53 |
| 2 | 227380359 | rs1909892 | -1.414 | -1.468 | -1.592 | -1.168 | -0.059 | 0.18 | 0.18 | 0 | 0.59 | 0.86 | 0.59 | 0.83 | 0.53 | 0.5 |
| 2 | 227393876 | rs1501002 | -1.658 | -1.714 | -0.707 | -1.487 | -0.15 | 0.028 | 0.028 | 0 | 0.66 | 0.92 | 0.48 | 0.23 | 0.58 | 0.37 |
| 2 | 227395509 | rs7599150 | -1.577 | -1.646 | -0.522 | -1.432 | -0.014 | 0.186 | 0.186 | 0.358 | 0.48 | 0.91 | 0.44 | 0.19 | 0.47 | 0.26 |
| 2 | 227410349 | rs2102101 | 0.243 | -1.892 | 0.102 | -1.416 | -0.043 | 0.053 | 0.053 | 0 | 0.1 | 0.04 | 0.08 | 0 | 0.18 | 0.11 |
| 2 | 227413223 | rs1875336 | -1.653 | -2.004 | 0.171 | -1.499 | 0.042 | 0.128 | 0.128 | 0.412 | 0.43 | 0.83 | 0.38 | 0.19 | 0.39 | 0.24 |
| 2 | 227430988 | rs6710574 | 0.411 | -2.117 | 0.429 | -1.542 | -0.002 | 0.004 | 0.004 | 0 | 0.12 | 0.15 | 0.16 | 0.1 | 0.14 | 0.12 |
| 2 | 227441399 | rs876496 | -2.052 | -2.12 | 0.364 | -1.549 | 0.046 | 0.136 | 0.136 | 0.873 | 0.44 | 0.88 | 0.39 | 0.2 | 0.39 | 0.24 |
| 2 | 227443549 | rs1273793 | -1.232 | -2.065 | 0.701 | -1.48 | 0.001 | 0.001 | 0.001 | 0.953 | 0.29 | 0.29 | 0.26 | 0.19 | 0.2 | 0.13 |
| 2 | 227447917 | rs16867099 | 0.913 | -1.642 | -0.198 | -1.167 | -0.004 | -0.008 | -0.008 | 0 | 0.08 | 0.07 | 0.03 | 0.01 | 0.02 | 0.01 |
| 2 | 227450527 | rs1273788 | 0.283 | -1.566 | -0.88 | -1.112 | 0 | -0.008 | -0.008 | 1.036 | 0.11 | 0.08 | 0.15 | 0.11 | 0.13 | 0.11 |
| 2 | 227455230 | rs2894606 | -1.899 | -1.61 | -1.035 | -1.154 | -0.006 | 0.079 | 0.079 | 0 | 0.39 | 0.9 | 0.38 | 0.19 | 0.38 | 0.23 |
| 2 | 227462877 | rs1273827 | 0.195 | -1.328 | -0.722 | -0.99 | 0 | -0.004 | -0.004 | 0 | 0.11 | 0.07 | 0.15 | 0.11 | 0.13 | 0.12 |
| 2 | 227464813 | rs1392844 | -1.256 | -1.445 | -1.053 | -1.056 | -0.005 | 0.119 | 0.119 | 0 | 0.38 | 0.9 | 0.34 | 0.12 | 0.35 | 0.23 |
| 2 | 227475163 | rs1273811 | 0.208 | -1.615 | 0.129 | -0.523 | 0.002 | 0.019 | 0.019 | 0.11 | 0.78 | 0.7 | 0.79 | 0.72 | 0.72 | 0.8 |
| 2 | 227486133 | rs2396384 | 1.606 | -2.155 | 0.413 | -0.899 | 0.013 | 0.027 | 0.027 | 0 | 0.43 | 0.13 | 0.59 | 0.67 | 0.57 | 0.71 |
| 2 | 227486156 | rs1273821 | -0.316 | -2.008 | 0.678 | -0.815 | 0.005 | 0.018 | 0.018 | 0.043 | 0.8 | 0.67 | 0.8 | 0.76 | 0.76 | 0.83 |
| 2 | 227506614 | rs16822391 | -0.142 | -1.753 | 0.697 | -0.828 | -0.037 | -0.022 | -0.022 | 0 | 0.3 | 0.48 | 0.19 | 0.23 | 0.24 | 0.16 |
| 2 | 227518448 | rs1882580 | -1.011 | -1.697 | 0.165 | -0.773 | -0.005 | -0.028 | -0.028 | 2.72 | 0.95 | 1 | 0.95 | 0.91 | 0.95 | 0.99 |
| 2 | 227521410 | rs9288615 | -0.235 | -1.711 | 0.195 | -0.78 | -0.017 | -0.017 | -0.017 | 0 | 0.33 | 0.59 | 0.23 | 0.24 | 0.28 | 0.18 |
| 2 | 227523774 | rs2707537 | 1.178 | -1.712 | -0.097 | -0.799 | 0.018 | 0.036 | 0.036 | 0 | 0.44 | 0.25 | 0.63 | 0.68 | 0.57 | 0.71 |
| 2 | 227526430 | rs1962277 | 0.718 | -1.602 | -0.704 | -0.74 | 0.023 | 0.035 | 0.035 | 0 | 0.16 | 0.07 | 0.08 | 0 | 0.1 | 0.09 |
| 2 | 227529244 | rs2222890 | -1.488 | -1.509 | -0.816 | -0.637 | -0.026 | 0.074 | 0.074 | 0 | 0.6 | 0.83 | 0.68 | 0.79 | 0.49 | 0.48 |
| 2 | 227530721 | rs2709423 | 0.539 | -1.584 | -0.643 | -0.264 | -0.005 | 0.119 | 0.119 | 0 | 0.09 | 0.13 | 0.4 | 0.56 | 0.11 | 0.18 |
| 2 | 227530857 | rs1922529 | -0.227 | -1.246 | -0.829 | -1.271 | 0.004 | -0.006 | -0.006 | 0 | 0.6 | 0.64 | 0.39 | 0.12 | 0.66 | 0.71 |
| 2 | 227530935 | rs11681198 | -0.652 | -0.482 | -0.907 | -0.472 | 0.018 | 0.023 | 0.023 | 0 | 0.16 | 0.11 | 0.08 | 0 | 0.1 | 0.09 |
| 2 | 227547136 | rs11675961 | -0.69 | -0.262 | -0.683 | -0.178 | 0.001 | -0.011 | -0.011 | 0 | 0.08 | 0.1 | 0.04 | 0 | 0.06 | 0.01 |
| 2 | 227548310 | rs1316328 | -0.753 | -0.643 | -0.33 | -0.141 | -0.02 | 0.01 | 0.01 | 0 | 0.12 | 0.24 | 0.08 | 0.09 | 0.12 | 0.09 |
| 2 | 227548504 | rs16822488 | -0.79 | -0.655 | -0.279 | -0.147 | -0.021 | 0.01 | 0.01 | 0 | 0.12 | 0.25 | 0.08 | 0.09 | 0.12 | 0.09 |
| 2 | 227550091 | rs6730045 | -1.912 | -1.062 | -0.673 | -0.34 | 0.074 | 0.057 | 0.057 | 0.328 | 0.42 | 0.85 | 0.3 | 0.28 | 0.37 | 0.18 |
| 2 | 227553794 | rs7574538 | -1.1 | -0.798 | -0.694 | -0.629 | -0.039 | -0.067 | -0.067 | 0 | 0.29 | 0.42 | 0.18 | 0.09 | 0.26 | 0.1 |
| 2 | 227555875 | rs10427370 | -0.117 | -0.763 | -0.66 | -0.607 | -0.024 | -0.127 | -0.127 | 0 | 0.18 | 0.34 | 0.11 | 0 | 0.14 | 0.03 |
| 2 | 227556383 | rs10933153 | -1.679 | -0.426 | -1.669 | -0.419 | 0.038 | 0.11 | 0.11 | 0.489 | 0.5 | 0.81 | 0.47 | 0.28 | 0.49 | 0.21 |
| 2 | 227558737 | rs17201100 | -0.943 | 0.084 | -1.576 | 0.138 | 0.009 | 0.003 | 0.003 | 0 | 0.16 | 0.13 | 0.1 | 0.09 | 0.17 | 0.07 |
| 2 | 227559317 | rs12185555 | -1.311 | 0.681 | -2.099 | 0.293 | -0.141 | -0.038 | -0.038 | 0.013 | 0.29 | 0.03 | 0.32 | 0.25 | 0.35 | 0.17 |
| 2 | 227559414 | rs12185556 | -1.116 | 0.658 | -1.529 | 0.756 | -0.064 | -0.057 | -0.057 | 0 | 0.14 | 0.02 | 0.23 | 0.22 | 0.18 | 0.07 |
| 2 | 227560786 | rs13019681 | -1.135 | -0.063 | -0.884 | 1.02 | -0.002 | -0.003 | -0.003 | 0.508 | 0.33 | 0.56 | 0.54 | 0.66 | 0.36 | 0.31 |
| 2 | 227561738 | rs12185691 | -0.964 | 0.078 | -1.135 | 1.23 | -0.004 | -0.008 | -0.008 | 0 | 0.17 | 0.18 | 0.27 | 0.28 | 0.21 | 0.11 |
| 2 | 227564025 | rs1922531 | 0.128 | -0.068 | -1.249 | 1.076 | -0.034 | -0.053 | -0.053 | 0 | 0.7 | 0.81 | 0.54 | 0.54 | 0.62 | 0.76 |
| 2 | 227566750 | rs1882575 | -1.096 | -1.221 | -0.643 | 0.682 | -0.007 | -0.003 | -0.003 | 0 | 0.23 | 0.23 | 0.19 | 0.17 | 0.1 | 0.18 |
| 2 | 227574221 | rs1317063 | -0.739 | -1.323 | -0.663 | 0.632 | 0.001 | 0 | 0 | 0 | 0.16 | 0.14 | 0.17 | 0.17 | 0.08 | 0.13 |
| 2 | 227579207 | rs2178704 | -1.217 | -1.262 | -0.512 | 0.663 | -0.059 | -0.04 | -0.04 | 0 | 0.25 | 0.37 | 0.23 | 0.18 | 0.12 | 0.19 |
| 2 | 227586606 | rs11683087 | 0.81 | -1.488 | -1.403 | 0.85 | 0.097 | 0.02 | 0.02 | 0.402 | 0.63 | 0.12 | 0.62 | 0.7 | 0.8 | 0.71 |

| | | | | | | | | | | | | | | | | |
|---|---|---|---|---|---|---|---|---|---|---|---|---|---|---|---|---|
| 2 | 227587688 | rs16822553 | -0.728 | -1.68 | -0.881 | 1.134 | -0.026 | 0.01 | 0.01 | 0 | 0.13 | 0.17 | 0.16 | 0.19 | 0.05 | 0.12 |
| 2 | 227589780 | rs2251692 | 1.263 | -2.261 | -1.622 | 1.489 | 0.149 | 0.06 | 0.06 | 1.276 | 0.52 | 0.04 | 0.52 | 0.33 | 0.74 | 0.57 |
| 2 | 227595233 | rs2894610 | -0.642 | -1.48 | -0.674 | 1.467 | 0.017 | -0.015 | -0.015 | 0 | 0.12 | 0.1 | 0.15 | 0.19 | 0.04 | 0.12 |
| 2 | 227599533 | rs16822570 | -0.529 | -1.459 | -0.795 | 1.477 | 0.043 | -0.04 | -0.04 | 0 | 0.12 | 0.05 | 0.15 | 0.19 | 0.04 | 0.12 |
| 2 | 227603355 | rs9288616 | -0.529 | -1.459 | -0.794 | 1.477 | 0.039 | -0.034 | -0.034 | 0.206 | 0.12 | 0.07 | 0.15 | 0.19 | 0.04 | 0.12 |
| 2 | 227614005 | rs10205233 | -0.87 | -1.475 | -0.747 | 1.407 | -0.002 | -0.002 | -0.002 | 4.064 | 0.12 | 0.14 | 0.16 | 0.19 | 0.04 | 0.12 |
| 2 | 227646959 | rs16822633 | -0.389 | -1.404 | -0.618 | 1.322 | 0.034 | -0.043 | -0.043 | 0.546 | 0.11 | 0 | 0.14 | 0.18 | 0.04 | 0.12 |
| 2 | 227647907 | rs16822640 | -1.1 | -1.492 | -0.612 | 1.197 | 0.028 | -0.005 | -0.005 | 0 | 0.16 | 0.14 | 0.17 | 0.23 | 0.07 | 0.24 |
| 2 | 227653948 | rs1560251 | -1.366 | -1.513 | -0.623 | 1.173 | -0.015 | 0.001 | 0.001 | 0 | 0.19 | 0.25 | 0.24 | 0.26 | 0.08 | 0.26 |
| 2 | 227660544 | rs1801278 | -1.032 | -0.687 | -0.482 | 0.814 | 0.018 | -0.018 | -0.018 | 0 | 0.05 | 0.06 | 0.03 | 0.02 | 0.09 | 0.05 |
| 2 | 227662753 | rs2234931 | -1.032 | -0.687 | -0.482 | 0.814 | 0.018 | -0.018 | -0.018 | 3.021 | 0.05 | 0.06 | 0.03 | 0.02 | 0.09 | 0.05 |
| 2 | 227666857 | rs6725330 | -1.131 | -1.36 | -0.348 | 0.544 | -0.009 | -0.007 | -0.007 | 0 | 0.2 | 0.2 | 0.12 | 0.09 | 0.12 | 0.26 |
| 2 | 227666992 | rs6725556 | -1.035 | -1.317 | 0.022 | 0.479 | -0.021 | -0.054 | -0.054 | 0 | 0.12 | 0.22 | 0.06 | 0.01 | 0.06 | 0.15 |
| 2 | 227675755 | rs16822696 | -0.272 | -1.621 | -0.173 | 0.354 | 0.006 | 0.004 | 0.004 | 0 | 0.13 | 0.04 | 0.06 | 0.15 | 0.1 | 0.08 |
| 2 | 227676743 | rs957797 | -1.676 | -1.526 | -0.102 | 0.717 | 0.001 | -0.002 | -0.002 | 0 | 0.24 | 0.74 | 0.23 | 0.31 | 0.17 | 0.25 |
| 2 | 227686907 | rs10933155 | 1.935 | -1.978 | -0.749 | 1.375 | 0.057 | 0.056 | 0.056 | 0 | 0.57 | 0.09 | 0.53 | 0.26 | 0.69 | 0.55 |
| 2 | 227699918 | rs10192769 | -1.982 | -1.988 | 0.471 | 1.351 | 0.031 | -0.026 | -0.026 | 0 | 0.33 | 0.9 | 0.27 | 0.32 | 0.21 | 0.37 |
| 2 | 227705016 | rs13386125 | -1.051 | -1.889 | 1.025 | 1.414 | 0.037 | 0.154 | 0.154 | 0.101 | 0.1 | 0.63 | 0.16 | 0.17 | 0.07 | 0.12 |
| 2 | 227721460 | rs16822770 | -0.743 | -1.884 | 1.134 | 1.472 | 0.083 | 0.115 | 0.115 | 0 | 0.08 | 0.4 | 0.14 | 0.1 | 0.07 | 0.12 |
| 2 | 227757333 | rs13427360 | -0.97 | -1.894 | 1.184 | 1.436 | 0.081 | 0.227 | 0.227 | 0 | 0.09 | 0.47 | 0.15 | 0.17 | 0.07 | 0.11 |
| 2 | 227769794 | rs4675117 | 1.552 | -1.409 | -0.033 | 0.526 | -0.146 | -0.062 | -0.062 | 0 | 0.29 | 0.04 | 0.41 | 0.24 | 0.38 | 0.47 |
| 2 | 227770592 | rs6723774 | 0.667 | -1.429 | 0.03 | 0.486 | -0.012 | -0.013 | -0.013 | 1.23 | 0.37 | 0.71 | 0.48 | 0.36 | 0.4 | 0.57 |
| 2 | 227770844 | rs17416354 | -0.101 | -1.453 | 0.008 | 0.546 | 0.022 | 0.063 | 0.063 | 0 | 0.5 | 0.78 | 0.6 | 0.57 | 0.51 | 0.68 |
| 2 | 227774635 | rs4675119 | 0.87 | -1.422 | 0.098 | 0.536 | 0.013 | 0.017 | 0.017 | 0 | 0.35 | 0.45 | 0.47 | 0.41 | 0.4 | 0.53 |
| 2 | 227791703 | rs7582251 | 0.746 | -1.441 | 0.242 | 0.534 | 0.022 | 0.021 | 0.021 | 0 | 0.38 | 0.55 | 0.47 | 0.41 | 0.42 | 0.54 |
| 2 | 227802354 | rs4675124 | -0.005 | -1.463 | 0.238 | 0.546 | 0.02 | 0.042 | 0.042 | 0 | 0.52 | 0.78 | 0.54 | 0.6 | 0.52 | 0.7 |
| 2 | 227856552 | rs6728239 | 0.772 | -1.376 | 0.346 | 0.549 | 0.005 | 0.005 | 0.005 | 0 | 0.38 | 0.44 | 0.47 | 0.41 | 0.42 | 0.59 |
| 2 | 227859693 | rs13398103 | 1.405 | -1.386 | 0.791 | 0.562 | -0.084 | -0.019 | -0.019 | 0.002 | 0.31 | 0.1 | 0.41 | 0.29 | 0.4 | 0.49 |
| 2 | 227860671 | rs10933164 | 0.813 | -1.304 | 0.431 | 0.283 | -0.053 | 0.204 | 0.204 | 0 | 0.3 | 0.02 | 0.16 | 0.01 | 0.28 | 0.14 |
| 2 | 227872182 | rs2228557 | 0.908 | -1.262 | 0.629 | 0.378 | 0.001 | 0.001 | 0.001 | 0 | 0.4 | 0.64 | 0.47 | 0.4 | 0.42 | 0.59 |
| 2 | 227872617 | rs9288618 | 1.008 | -1.262 | 0.568 | 0.378 | -0.021 | -0.02 | -0.02 | 1.136 | 0.4 | 0.73 | 0.48 | 0.4 | 0.42 | 0.59 |
| 2 | 227880511 | rs12475134 | -0.675 | -1.173 | 0.587 | 0.444 | -0.032 | -0.07 | -0.07 | 0.308 | 0.66 | 0.95 | 0.65 | 0.6 | 0.64 | 0.8 |
| 2 | 227886773 | rs3752895 | 0.598 | -1.168 | 0.906 | 0.432 | -0.009 | -0.009 | -0.009 | 0 | 0.44 | 0.69 | 0.48 | 0.41 | 0.43 | 0.56 |
| 2 | 227898434 | rs2276593 | 0.956 | -1.1 | 1.365 | 0.424 | -0.01 | -0.01 | -0.01 | 0 | 0.42 | 0.39 | 0.47 | 0.4 | 0.44 | 0.57 |
| 2 | 227908665 | rs4675140 | 0.433 | -0.49 | 1.308 | 0.058 | 0.009 | 0.002 | 0.002 | 0 | 0.52 | 0.31 | 0.49 | 0.48 | 0.49 | 0.64 |
| 2 | 227914975 | rs2251223 | -0.729 | -0.509 | 1.245 | 0.036 | -0.019 | -0.079 | -0.079 | 1.179 | 0.64 | 0.9 | 0.62 | 0.49 | 0.6 | 0.75 |
| 2 | 227915924 | rs2272205 | 1.292 | -0.512 | 1.153 | 0.044 | 0.004 | -0.008 | -0.008 | 0 | 0.12 | 0.04 | 0.06 | 0.19 | 0.1 | 0.15 |
| 2 | 227921926 | rs4296416 | -0.569 | -0.512 | 1.422 | 0.043 | -0.001 | 0 | 0 | 0 | 0.63 | 0.68 | 0.67 | 0.49 | 0.59 | 0.75 |
| 2 | 227922321 | rs3769641 | 1.372 | -0.391 | 1.011 | 0.089 | 0.001 | -0.007 | -0.007 | 1.001 | 0.11 | 0.04 | 0.06 | 0.19 | 0.1 | 0.13 |
| 2 | 227922470 | rs2141828 | 1.221 | -0.37 | 0.956 | 0.03 | -0.107 | -0.028 | -0.028 | 0 | 0.36 | 0.07 | 0.45 | 0.29 | 0.38 | 0.44 |
| 2 | 227925991 | rs12151732 | 0.018 | -0.277 | 0.951 | 0.078 | 0.036 | 0.018 | 0.018 | 0 | 0.06 | 0.27 | 0.16 | 0.06 | 0.05 | 0.12 |
| 2 | 227935910 | rs13411764 | 1.173 | -0.38 | 0.46 | 0.05 | -0.025 | 0.024 | 0.024 | 0 | 0.34 | 0.15 | 0.33 | 0.23 | 0.34 | 0.38 |
| 2 | 227937611 | rs1917124 | 1.237 | -0.365 | 0.131 | 0.049 | -0.109 | -0.033 | -0.033 | 0 | 0.34 | 0.02 | 0.31 | 0.23 | 0.34 | 0.38 |
| 2 | 227943480 | rs7574757 | -0.231 | -0.527 | 0.825 | 0.061 | 0.003 | -0.071 | -0.071 | 0.015 | 0.11 | 0.28 | 0.1 | 0.01 | 0.1 | 0.13 |
| 2 | 227956067 | rs13382950 | 0.34 | -0.277 | 0.248 | 0.411 | 0.013 | 0.036 | 0.036 | 0 | 0.5 | 0.25 | 0.51 | 0.47 | 0.5 | 0.57 |
| 2 | 227958469 | rs7558081 | 0.148 | -0.057 | -0.237 | 0.502 | 0 | 0 | 0 | 0 | 0.49 | 0.47 | 0.63 | 0.57 | 0.5 | 0.57 |
| 2 | 227959973 | rs12989872 | -0.239 | 0.237 | -1.014 | 0.387 | 0.016 | 0.03 | 0.03 | 2.078 | 0.67 | 0.83 | 0.82 | 0.76 | 0.72 | 0.82 |
| 2 | 227962226 | rs6758296 | -1.068 | 0.52 | -0.263 | 0.584 | 0.001 | -0.019 | -0.019 | 0.35 | 0.08 | 0.22 | 0.14 | 0.05 | 0.05 | 0.13 |
| 2 | 227966148 | rs16823202 | -0.214 | 0.654 | -1.104 | 0.766 | -0.028 | -0.005 | -0.005 | 0 | 0.59 | 0.83 | 0.51 | 0.39 | 0.64 | 0.44 |
| 2 | 227967105 | rs1574392 | -0.303 | 0.422 | -0.373 | 0.981 | 0.007 | 0.007 | 0.007 | 0 | 0.64 | 0.57 | 0.57 | 0.62 | 0.69 | 0.57 |
| 2 | 227970914 | rs10179820 | -0.025 | 0.408 | -0.072 | 0.97 | -0.005 | -0.01 | -0.01 | 0.028 | 0.6 | 0.37 | 0.5 | 0.53 | 0.65 | 0.51 |
| 2 | 227973193 | rs10174459 | 0.046 | 0.149 | -0.12 | 1.046 | -0.034 | 0.075 | 0.075 | 0 | 0.13 | 0.21 | 0.12 | 0.32 | 0.06 | 0.18 |
| 2 | 227977893 | rs6743614 | -0.792 | 0.636 | -1.002 | 0.842 | -0.106 | -0.166 | -0.166 | 0 | 0.73 | 0.98 | 0.6 | 0.56 | 0.73 | 0.64 |
| 2 | 227978032 | rs11686140 | 0.545 | 0.489 | -0.046 | 0.368 | 0 | 0.052 | 0.052 | 0 | 0.53 | 0.35 | 0.38 | 0.21 | 0.59 | 0.42 |
| 2 | 227978479 | rs11675161 | 0.329 | 0.438 | 0.514 | 0.333 | 0.001 | 0.005 | 0.005 | 0 | 0.54 | 0.57 | 0.43 | 0.21 | 0.63 | 0.43 |
| 2 | 227979775 | rs1922020 | -0.716 | 0.346 | 0.073 | -0.098 | 0.038 | 0.06 | 0.06 | 0 | 0.5 | 0.63 | 0.57 | 0.75 | 0.34 | 0.55 |
| 2 | 227990914 | rs6753676 | 0.956 | 0.459 | -0.438 | -0.351 | -0.017 | 0.016 | 0.016 | 0 | 0.28 | 0.18 | 0.31 | 0.21 | 0.42 | 0.31 |
| 2 | 227993937 | rs6708440 | 0.736 | 0.472 | -0.452 | -0.356 | 0 | -0.001 | -0.001 | 0 | 0.3 | 0.27 | 0.32 | 0.21 | 0.42 | 0.31 |
| 2 | 227997971 | rs6436656 | 1.008 | 0.502 | -0.646 | -0.341 | -0.018 | 0.014 | 0.014 | 0 | 0.28 | 0.18 | 0.31 | 0.21 | 0.41 | 0.31 |
| 2 | 228001180 | rs2396453 | -0.345 | 0.587 | -1.069 | -0.338 | 0.008 | 0.057 | 0.057 | 0 | 0.48 | 0.8 | 0.5 | 0.38 | 0.41 | 0.42 |
| 2 | 228005906 | rs7558432 | 0.008 | 0.555 | -0.694 | -0.09 | -0.013 | 0.071 | 0.071 | 0 | 0.42 | 0.69 | 0.4 | 0.22 | 0.5 | 0.33 |
| 2 | 228010257 | rs10498217 | 0.628 | 0.576 | -0.799 | -0.031 | 0 | 0 | 0 | 0 | 0.05 | 0.07 | 0.07 | 0.06 | 0.07 | 0.03 |
| 2 | 228013535 | rs12694713 | 0.37 | 0.733 | -1.711 | -0.073 | -0.002 | 0.016 | 0.016 | 0 | 0.37 | 0.67 | 0.37 | 0.28 | 0.43 | 0.32 |
| 2 | 228014602 | rs12468501 | 1.102 | 0.546 | -2.483 | 0.252 | -0.06 | -0.095 | -0.095 | 0.06 | 0.23 | 0.03 | 0.19 | 0.31 | 0.19 | 0.31 |
| 2 | 228014767 | rs12476572 | 0.289 | 0.773 | -1.502 | 0.329 | -0.024 | -0.079 | -0.079 | 0 | 0.69 | 0.91 | 0.76 | 0.59 | 0.7 | 0.67 |
| 2 | 228020491 | rs2078635 | 0.612 | 1.131 | -1.104 | 0.168 | -0.045 | -0.002 | -0.002 | 0 | 0.59 | 0.22 | 0.65 | 0.6 | 0.57 | 0.58 |
| 2 | 228025756 | rs1922021 | -0.995 | 0.567 | -0.978 | 0.256 | -0.011 | -0.007 | -0.007 | 0.336 | 0.39 | 0.77 | 0.33 | 0.4 | 0.37 | 0.42 |
| 2 | 228028119 | rs12613226 | -0.323 | 1.208 | -0.999 | 0.67 | -0.003 | -0.003 | -0.003 | 0.007 | 0.26 | 0.25 | 0.16 | 0.31 | 0.3 | 0.38 |
| 2 | 228030362 | rs1922022 | -0.354 | 1.214 | -0.984 | 0.657 | -0.002 | -0.002 | -0.002 | 0 | 0.27 | 0.25 | 0.16 | 0.31 | 0.31 | 0.38 |

| | | | | | | | | | | | | | | | | |
|---|---|---|---|---|---|---|---|---|---|---|---|---|---|---|---|---|
| 2 | 228044253 | rs16823288 | -0.681 | 0.773 | -0.971 | 0.167 | 0 | -0.003 | -0.003 | 0 | 0.17 | 0.19 | 0.33 | 0.07 | 0.18 | 0.11 |
| 2 | 228045315 | rs2396464 | -0.681 | 0.773 | -0.988 | 0.167 | 0 | -0.001 | -0.001 | 0 | 0.17 | 0.19 | 0.33 | 0.07 | 0.18 | 0.11 |
| 2 | 228052600 | rs2204862 | -0.469 | 0.586 | -0.95 | 0.057 | -0.001 | -0.031 | -0.031 | 0 | 0.15 | 0.18 | 0.23 | 0 | 0.15 | 0.08 |
| 2 | 228056670 | rs6436661 | 0.372 | 0.726 | -0.828 | 0.451 | -0.001 | 0.002 | 0.002 | 0 | 0.84 | 0.73 | 0.92 | 0.85 | 0.89 | 0.9 |
| 2 | 228057792 | rs4325745 | -0.196 | 0.791 | -1.073 | 0.145 | 0.002 | 0.012 | 0.012 | 0.016 | 0.38 | 0.59 | 0.4 | 0.33 | 0.41 | 0.47 |
| 2 | 228058913 | rs13429913 | -0.11 | 0.897 | -1.661 | 0.122 | -0.023 | -0.048 | -0.048 | 0 | 0.39 | 0.16 | 0.45 | 0.44 | 0.4 | 0.36 |
| 2 | 228059546 | rs6737679 | 0.224 | 1.074 | -1.769 | 0.126 | -0.021 | -0.086 | -0.086 | 0.763 | 0.36 | 0.09 | 0.27 | 0.44 | 0.35 | 0.32 |
| 2 | 228060507 | rs6741679 | -0.071 | 1.042 | -0.948 | 0.328 | -0.002 | 0.003 | 0.003 | 0 | 0.64 | 0.62 | 0.47 | 0.67 | 0.64 | 0.65 |
| 2 | 228061398 | rs10198349 | 0.951 | 0.987 | -0.32 | 0.499 | 0.034 | 0.121 | 0.121 | 0 | 0.92 | 0.73 | 0.78 | 0.77 | 0.91 | 0.76 |
| 2 | 228062729 | rs10498215 | 0.328 | 0.832 | -0.779 | 0.096 | 0.001 | 0.011 | 0.011 | 0 | 0.06 | 0.03 | 0.04 | 0 | 0.07 | 0.06 |
| 2 | 228062790 | rs1950135 | 0.438 | 0.855 | -1.2 | -0.062 | 0.016 | 0.044 | 0.044 | 0.022 | 0.49 | 0.63 | 0.52 | 0.75 | 0.37 | 0.61 |
| 2 | 228063071 | rs13425557 | 0.025 | 1.406 | -0.832 | -0.053 | -0.006 | -0.003 | -0.003 | 0.103 | 0.07 | 0.05 | 0.2 | 0 | 0.14 | 0.08 |
| 2 | 228064180 | rs10498214 | -1.04 | 1.571 | -0.955 | -0.146 | 0.025 | -0.043 | -0.043 | 0 | 0.16 | 0.18 | 0.27 | 0.03 | 0.28 | 0.16 |
| 2 | 228064828 | rs17367956 | -0.108 | 1.199 | -0.836 | 0.079 | 0.031 | -0.029 | -0.029 | 0 | 0.09 | 0.15 | 0.08 | 0.03 | 0.15 | 0.07 |
| 2 | 228065095 | rs6739977 | -0.108 | 1.199 | -0.836 | 0.065 | 0.031 | -0.032 | -0.032 | 0 | 0.09 | 0.15 | 0.08 | 0.03 | 0.14 | 0.07 |
| 2 | 228078194 | rs10933170 | 0.186 | 0.017 | -1.55 | 0.301 | 0.128 | 0.05 | 0.05 | 0 | 0.51 | 0.26 | 0.64 | 0.32 | 0.7 | 0.47 |
| 2 | 228078708 | rs4430946 | 0.331 | -0.45 | -1.036 | -0.229 | -0.006 | -0.05 | -0.05 | 0 | 0.15 | 0.2 | 0.19 | 0 | 0.14 | 0.11 |
| 2 | 228084164 | rs6436664 | 0.997 | -0.698 | -1.577 | -0.389 | -0.006 | 0.1 | 0.1 | 0 | 0.38 | 0.25 | 0.46 | 0.13 | 0.57 | 0.3 |
| 2 | 228085517 | rs11694554 | -1.304 | -1.432 | -0.897 | -0.053 | -0.023 | 0.014 | 0.014 | 0.198 | 0.81 | 0.76 | 0.93 | 0.8 | 0.87 | 0.87 |
| 2 | 228086920 | rs11884770 | 0.2 | -1.633 | -0.884 | -0.075 | -0.021 | -0.02 | -0.02 | 0 | 0.6 | 0.49 | 0.72 | 0.78 | 0.71 | 0.73 |
| 2 | 228087478 | rs2178631 | 1.075 | -1.673 | -0.873 | 0.299 | 0.047 | 0.056 | 0.056 | 0 | 0.46 | 0.34 | 0.5 | 0.37 | 0.6 | 0.52 |
| 2 | 228090768 | rs13021572 | 2.158 | -0.819 | -1.174 | 0.057 | -0.047 | 0.009 | 0.009 | 0.077 | 0.19 | 0.09 | 0.26 | 0.12 | 0.27 | 0.2 |
| 2 | 228091661 | rs13028299 | -0.075 | -1.024 | -1.061 | -0.02 | -0.002 | -0.002 | -0.002 | 3.588 | 0.47 | 0.52 | 0.54 | 0.5 | 0.45 | 0.45 |
| 2 | 228092334 | rs6728149 | -0.853 | -1.025 | -0.69 | 0.097 | -0.041 | 0.052 | 0.052 | 0 | 0.73 | 0.66 | 0.89 | 0.55 | 0.84 | 0.65 |
| 2 | 228092406 | rs6756898 | -0.335 | -1.157 | -0.767 | 0.139 | -0.04 | 0.033 | 0.033 | 0 | 0.66 | 0.59 | 0.83 | 0.51 | 0.75 | 0.75 |
| 2 | 228093667 | rs11884740 | 1.6 | -0.765 | -0.999 | -0.005 | -0.01 | 0.004 | 0.004 | 0 | 0.19 | 0.1 | 0.22 | 0.13 | 0.22 | 0.15 |
| 2 | 228094994 | rs13000521 | 1.023 | -0.822 | -1.012 | -0.342 | 0.001 | -0.019 | -0.019 | 0.381 | 0.16 | 0.08 | 0.17 | 0.27 | 0.13 | 0.37 |
| 2 | 228097707 | rs10169660 | 1.172 | -0.797 | -0.9 | -0.473 | 0.005 | -0.046 | -0.046 | 0.012 | 0.27 | 0.34 | 0.26 | 0.05 | 0.35 | 0.14 |
| 2 | 228098933 | rs7577732 | 1.135 | -0.777 | -0.912 | -0.463 | 0.003 | -0.042 | -0.042 | 0 | 0.27 | 0.34 | 0.26 | 0.05 | 0.35 | 0.14 |
| 2 | 228100150 | rs10187805 | -0.357 | -0.919 | -0.778 | -0.151 | -0.014 | 0.001 | 0.001 | 0.136 | 0.63 | 0.66 | 0.74 | 0.55 | 0.76 | 0.52 |
| 2 | 228102752 | rs1882435 | 0.335 | -0.897 | -0.496 | 0.24 | -0.002 | -0.017 | -0.017 | 2.243 | 0.29 | 0.25 | 0.29 | 0.42 | 0.32 | 0.26 |
| 2 | 228103702 | rs4675157 | -0.532 | -1.331 | -0.117 | 0.171 | -0.034 | -0.015 | -0.015 | 0 | 0.79 | 0.78 | 0.91 | 0.84 | 0.89 | 0.88 |
| 2 | 228109957 | rs6750355 | 0.052 | -1.082 | -0.253 | -0.033 | -0.08 | -0.097 | -0.097 | 0 | 0.67 | 0.52 | 0.88 | 0.9 | 0.84 | 0.85 |
| 2 | 228111435 | rs10178458 | 0.02 | -1.1 | -0.528 | -0.02 | -0.056 | -0.068 | -0.068 | 1.132 | 0.68 | 0.6 | 0.88 | 0.9 | 0.84 | 0.85 |
| 2 | 228113175 | rs6436669 | 0.02 | -1.1 | -0.528 | -0.02 | -0.056 | -0.068 | -0.068 | 0 | 0.68 | 0.6 | 0.88 | 0.9 | 0.84 | 0.85 |
| 2 | 228121147 | rs10933172 | -0.084 | 0.03 | -0.547 | 0.575 | 0.002 | 0 | 0 | 0 | 0.06 | 0.05 | 0.11 | 0.06 | 0.12 | 0.08 |
| 2 | 228123661 | rs10167041 | 0.487 | 0.152 | -0.701 | 0.608 | -0.01 | 0.009 | 0.009 | 0 | 0.56 | 0.65 | 0.5 | 0.57 | 0.5 | 0.66 |
| 2 | 228124702 | rs4332918 | 0.495 | 0.191 | -0.709 | 0.6 | -0.011 | 0.01 | 0.01 | 0 | 0.56 | 0.65 | 0.5 | 0.57 | 0.5 | 0.66 |
| 2 | 228126494 | rs6436670 | -0.708 | 0.176 | -0.846 | 0.4 | -0.035 | -0.018 | -0.018 | 0 | 0.75 | 0.85 | 0.64 | 0.63 | 0.66 | 0.79 |
| 2 | 228127905 | rs10169757 | 1.478 | -0.333 | -1.367 | 0.764 | -0.099 | -0.078 | -0.078 | 1.021 | 0.36 | 0.09 | 0.41 | 0.45 | 0.44 | 0.53 |
| 2 | 228130088 | rs4603754 | 0.736 | -0.531 | -0.978 | 1.023 | 0.047 | 0.056 | 0.056 | 0.366 | 0.47 | 0.21 | 0.57 | 0.65 | 0.64 | 0.65 |
| 2 | 228131169 | rs11677877 | 1.644 | -0.626 | -0.409 | 1.277 | 0.005 | -0.007 | -0.007 | 2.199 | 0.1 | 0.07 | 0.18 | 0.16 | 0.07 | 0.12 |
| 2 | 228131904 | rs7582825 | 1.156 | -0.609 | 0.289 | 1.165 | -0.038 | 0.009 | 0.009 | 0 | 0.1 | 0.34 | 0.03 | 0.05 | 0.03 | 0.05 |
| 2 | 228133512 | rs10933175 | -0.17 | -0.561 | 0.319 | 1.407 | 0.036 | 0.006 | 0.006 | 0.495 | 0.62 | 0.23 | 0.7 | 0.61 | 0.73 | 0.76 |
| 2 | 228133809 | rs4643535 | -0.726 | -0.637 | 0.581 | 1.353 | -0.009 | -0.009 | -0.009 | 0 | 0.65 | 0.81 | 0.57 | 0.58 | 0.66 | 0.72 |
| 2 | 228140460 | rs12468719 | -0.284 | -0.598 | 1.259 | 1.375 | -0.014 | 0.1 | 0.1 | 0 | 0.85 | 0.64 | 0.83 | 0.72 | 0.87 | 0.85 |
| 2 | 228143087 | rs4390760 | 0.842 | -0.586 | 1.002 | 1.327 | 0.001 | -0.001 | -0.001 | 0 | 0.09 | 0.1 | 0.12 | 0.06 | 0.15 | 0.08 |
| 2 | 228144706 | rs6729152 | -1.726 | -0.575 | 0.919 | 1.298 | 0.009 | -0.011 | -0.011 | 0 | 0.9 | 0.96 | 0.89 | 0.86 | 0.93 | 0.89 |
| 2 | 228156446 | rs9646881 | 1.809 | -0.577 | 0.984 | 1.289 | -0.049 | 0.037 | 0.037 | 2.029 | 0.36 | 0.13 | 0.25 | 0.23 | 0.37 | 0.44 |
| 2 | 228159641 | rs4675163 | 1.013 | -0.595 | 1.015 | 1.332 | -0.006 | -0.008 | -0.008 | 0 | 0.09 | 0.06 | 0.12 | 0.09 | 0.15 | 0.08 |
| 2 | 228167014 | rs4619602 | 1.845 | -0.602 | 0.971 | 1.327 | -0.072 | 0.024 | 0.024 | 1.684 | 0.36 | 0.09 | 0.25 | 0.24 | 0.37 | 0.44 |
| 2 | 228167544 | rs4271760 | 0.339 | -0.558 | 1.353 | 1.32 | -0.066 | 0.07 | 0.07 | 0.054 | 0.82 | 0.54 | 0.85 | 0.74 | 0.9 | 0.82 |
| 2 | 228177479 | rs7587228 | -0.324 | -0.492 | 1.194 | 1.287 | -0.033 | -0.006 | -0.006 | 0.421 | 0.84 | 0.77 | 0.87 | 0.87 | 0.92 | 0.84 |
| 2 | 228177567 | rs1134745 | 0.162 | -0.276 | 1.053 | 1.196 | 0.032 | 0.017 | 0.017 | 0 | 0.12 | 0.21 | 0.17 | 0.1 | 0.19 | 0.1 |
| 2 | 228179038 | rs4470338 | 0.162 | -0.276 | 1.008 | 1.196 | 0.027 | 0.014 | 0.014 | 0 | 0.12 | 0.2 | 0.17 | 0.1 | 0.2 | 0.1 |
| 2 | 228189493 | rs7559435 | 0.855 | -0.295 | 0.929 | 1.239 | 0.018 | 0.079 | 0.079 | 1.551 | 0.82 | 0.51 | 0.82 | 0.78 | 0.79 | 0.85 |
| 2 | 228191202 | rs6722598 | 0.303 | -0.286 | 0.865 | 1.192 | 0.009 | 0.07 | 0.07 | 2.083 | 0.91 | 0.64 | 0.92 | 0.87 | 0.93 | 0.92 |
| 2 | 228194481 | rs3211098 | 1.394 | -0.334 | 0.373 | 1.31 | -0.06 | 0.06 | 0.06 | 1.467 | 0.39 | 0.1 | 0.25 | 0.25 | 0.37 | 0.45 |
| 2 | 228215993 | rs11692634 | 1.248 | -0.391 | 0.482 | 1.435 | -0.001 | -0.001 | -0.001 | 0 | 0.09 | 0.08 | 0.11 | 0.1 | 0.07 | 0.1 |
| 2 | 228220769 | rs7560053 | -0.697 | -0.429 | 0.425 | 1.488 | -0.006 | -0.013 | -0.013 | 0 | 0.66 | 0.77 | 0.55 | 0.57 | 0.66 | 0.71 |
| 2 | 228224400 | rs4234071 | 0.579 | -0.187 | 1.108 | 1.516 | -0.001 | 0.049 | 0.049 | 0.718 | 0.92 | 0.72 | 0.93 | 0.88 | 0.93 | 0.92 |
| 2 | 228229617 | rs7577008 | -1.844 | -0.14 | 1.122 | 1.543 | -0.003 | 0.006 | 0.006 | 0.71 | 0.27 | 0.68 | 0.3 | 0.32 | 0.29 | 0.25 |
| 2 | 228230049 | rs7594089 | -0.68 | 0.079 | 0.756 | 1.579 | -0.018 | 0.079 | 0.079 | 0.804 | 0.07 | 0.27 | 0.03 | 0.13 | 0.02 | 0.06 |
| 2 | 228233553 | rs10933181 | -0.211 | 0.458 | -0.06 | 1.47 | 0.017 | 0.022 | 0.022 | 0 | 0.08 | 0.13 | 0.1 | 0.09 | 0.15 | 0.07 |
| 2 | 228240766 | rs12694728 | 0.49 | 0.263 | -0.382 | 1.227 | -0.005 | -0.007 | -0.007 | 0 | 0.15 | 0.16 | 0.21 | 0.28 | 0.19 | 0.13 |
| 2 | 228243905 | rs7574414 | 0.95 | 0.18 | -0.497 | 1.436 | -0.056 | -0.06 | -0.06 | 0 | 0.12 | 0.01 | 0.08 | 0.19 | 0.15 | 0.12 |
| 2 | 228244107 | rs11678432 | 0.594 | 0.174 | -0.698 | 1.131 | -0.018 | -0.009 | -0.009 | 0 | 0.07 | 0.03 | 0.15 | 0.07 | 0.09 | 0.05 |
| 2 | 228246529 | rs4234072 | -0.576 | -0.148 | -0.714 | 0.991 | -0.001 | -0.004 | -0.004 | 0 | 0.86 | 0.86 | 0.86 | 0.91 | 0.91 | 0.9 |
| 2 | 228248642 | rs11686056 | 0.268 | -0.088 | -0.541 | 1.076 | -0.005 | -0.021 | -0.021 | 1.529 | 0.11 | 0.06 | 0.13 | 0.16 | 0.12 | 0.19 |

| | | | | | | | | | | | | | | | | |
|---|---|---|---|---|---|---|---|---|---|---|---|---|---|---|---|---|
| 2 | 228254272 | rs4075148 | 0.497 | -0.637 | 0.44 | 0.943 | 0.004 | 0.039 | 0.039 | 0 | 0.18 | 0.22 | 0.12 | 0.33 | 0.18 | 0.18 |
| 2 | 228258288 | rs4485560 | -0.461 | -1.072 | 0.236 | 0.6 | -0.012 | -0.004 | -0.004 | 0 | 0.37 | 0.45 | 0.25 | 0.4 | 0.26 | 0.25 |
| 2 | 228258857 | rs6436685 | 0.444 | -0.962 | 0.103 | 0.612 | 0.014 | 0.003 | 0.003 | 0 | 0.57 | 0.64 | 0.61 | 0.52 | 0.64 | 0.69 |
| 2 | 228264258 | rs7557146 | 1.139 | -0.131 | 0.349 | 0.978 | -0.018 | 0.009 | 0.009 | 0 | 0.5 | 0.16 | 0.44 | 0.41 | 0.54 | 0.63 |
| 2 | 228267108 | rs7582023 | 0.352 | -0.273 | 0.694 | 0.332 | -0.02 | -0.039 | -0.039 | 0 | 0.73 | 0.41 | 0.7 | 0.83 | 0.81 | 0.85 |
| 2 | 228268085 | rs6749521 | 0.37 | 0.186 | 0.667 | 0.412 | 0.037 | 0.05 | 0.05 | 0 | 0.9 | 0.67 | 0.74 | 0.84 | 0.86 | 0.92 |
| 2 | 228272364 | rs12993552 | 0.863 | 0.349 | -0.323 | 0.142 | -0.032 | -0.024 | -0.024 | 0 | 0.06 | 0 | 0.15 | 0.09 | 0.1 | 0.04 |
| 2 | 228277430 | rs10184663 | 0.539 | 0.316 | -0.285 | 0.091 | -0.004 | 0.004 | 0.004 | 0 | 0.68 | 0.73 | 0.67 | 0.71 | 0.66 | 0.75 |
| 2 | 228279610 | rs11693555 | -1.118 | 0.479 | -0.616 | 0.056 | 0.001 | -0.002 | -0.002 | 0 | 0.14 | 0.17 | 0.22 | 0.11 | 0.16 | 0.11 |
| 2 | 228288185 | rs4673194 | 0.012 | 0.496 | -0.894 | 0.024 | -0.001 | 0.054 | 0.054 | 0 | 0.53 | 0.29 | 0.42 | 0.28 | 0.54 | 0.51 |
| 2 | 228288664 | rs11693569 | 0.302 | 0.759 | -0.713 | 0.099 | -0.032 | -0.026 | -0.026 | 0 | 0.06 | 0 | 0.15 | 0.09 | 0.1 | 0.04 |
| 2 | 228293381 | rs4470337 | 1.033 | 0.275 | -0.352 | -0.145 | 0.001 | 0.005 | 0.005 | 0 | 0.5 | 0.5 | 0.61 | 0.67 | 0.58 | 0.65 |
| 2 | 228296144 | rs11688520 | -0.523 | 0.013 | -0.317 | -0.202 | 0 | 0.001 | 0.001 | 0 | 0.09 | 0.11 | 0.06 | 0.04 | 0.06 | 0.07 |
| 2 | 228315672 | rs6708202 | 0.408 | 0.06 | -0.253 | -0.206 | 0.005 | 0.007 | 0.007 | 0 | 0.61 | 0.71 | 0.69 | 0.71 | 0.7 | 0.75 |
| 2 | 228336315 | rs13382412 | 0.135 | -0.006 | -0.021 | -0.225 | -0.013 | 0.006 | 0.006 | 0 | 0.05 | 0.11 | 0.03 | 0.03 | 0.02 | 0.06 |
| 2 | 228343348 | rs10170294 | 0.135 | -0.006 | -0.021 | -0.225 | -0.013 | 0.006 | 0.006 | 0 | 0.05 | 0.11 | 0.03 | 0.03 | 0.02 | 0.06 |
| 2 | 228358043 | rs7566800 | 0.538 | -0.24 | 0.057 | -0.359 | 0.003 | -0.012 | -0.012 | 0 | 0.16 | 0.11 | 0.03 | 0 | 0.02 | 0.07 |
| 2 | 228363432 | rs4496327 | 1.183 | -0.164 | 0.118 | -0.334 | -0.001 | -0.001 | -0.001 | 0 | 0.54 | 0.54 | 0.63 | 0.65 | 0.61 | 0.66 |
| 2 | 228372956 | rs10182259 | 0.234 | -0.162 | 0.099 | -0.332 | -0.013 | 0.006 | 0.006 | 0 | 0.05 | 0.11 | 0.03 | 0.03 | 0.02 | 0.06 |
| 2 | 228376038 | rs4508591 | 0.656 | -0.155 | 0.085 | -0.332 | 0.002 | 0.02 | 0.02 | 0 | 0.59 | 0.68 | 0.66 | 0.69 | 0.64 | 0.72 |
| 2 | 228399480 | rs7588620 | -0.888 | 0.007 | -0.002 | -0.246 | 0 | 0 | 0 | 7.899 | 0.83 | 0.86 | 0.86 | 0.79 | 0.81 | 0.86 |
| 2 | 228414974 | rs4635528 | 0.88 | 0.098 | -0.342 | -0.161 | -0.028 | -0.029 | -0.029 | 0 | 0.05 | 0 | 0.15 | 0.11 | 0.1 | 0.04 |
| 2 | 228419373 | rs13408988 | 0.084 | 0.009 | -0.306 | -0.209 | -0.013 | 0.006 | 0.006 | 0 | 0.05 | 0.11 | 0.03 | 0.03 | 0.02 | 0.06 |
| 2 | 228421789 | rs13400297 | 0.084 | 0.009 | -0.341 | -0.209 | -0.013 | 0.003 | 0.003 | 0 | 0.05 | 0.11 | 0.03 | 0.03 | 0.02 | 0.06 |
| 2 | 228438238 | rs11676303 | 0.686 | 0.133 | -0.234 | -0.303 | -0.012 | -0.041 | -0.041 | 0 | 0.59 | 0.48 | 0.78 | 0.79 | 0.7 | 0.77 |
| 2 | 228438549 | rs4129709 | 1.036 | 0.264 | -0.257 | -0.246 | 0.002 | 0.008 | 0.008 | 0 | 0.54 | 0.46 | 0.64 | 0.7 | 0.6 | 0.72 |
| 2 | 228446321 | rs10933193 | 0.108 | 0.973 | -0.64 | 0.12 | -0.001 | 0.004 | 0.004 | 0 | 0.78 | 0.8 | 0.84 | 0.79 | 0.77 | 0.86 |
| 2 | 228458917 | rs7608226 | -0.826 | 1.187 | -0.834 | 0.188 | -0.024 | 0.024 | 0.024 | 0 | 0.08 | 0.08 | 0.08 | 0 | 0.14 | 0.07 |
| 2 | 228460244 | rs11694508 | -0.524 | 1.395 | -0.987 | 0.149 | -0.04 | 0.01 | 0.01 | 0.011 | 0.06 | 0 | 0.13 | 0.03 | 0.13 | 0.09 |
| 2 | 228461771 | rs1467180 | -1.495 | 0.958 | -0.465 | -0.089 | -0.013 | -0.032 | -0.032 | 0 | 0.07 | 0.17 | 0.02 | 0 | 0.03 | 0.02 |
| 2 | 228463310 | rs6755969 | -0.778 | 1.703 | 0.103 | 0.066 | 0.029 | 0.009 | 0.009 | 0 | 0.26 | 0.46 | 0.27 | 0.27 | 0.34 | 0.12 |
| 2 | 228465561 | rs6722082 | 1.249 | 1.547 | 0.84 | -0.001 | -0.023 | -0.032 | -0.032 | 0.085 | 0.91 | 0.61 | 0.97 | 1 | 0.97 | 0.98 |
| 2 | 228465580 | rs10207659 | 0.396 | 1.937 | 0.753 | 0.066 | -0.055 | -0.06 | -0.06 | 0 | 0.14 | 0.01 | 0.15 | 0.22 | 0.2 | 0.08 |
| 2 | 228465748 | rs10175370 | -0.62 | 1.315 | -0.093 | 0.06 | -0.014 | -0.003 | -0.003 | 0 | 0.58 | 0.44 | 0.69 | 0.6 | 0.69 | 0.64 |
| 2 | 228465987 | rs6725452 | 0.652 | 1.459 | -0.211 | 0.174 | 0.004 | 0.002 | 0.002 | 0 | 0.18 | 0.12 | 0.24 | 0.27 | 0.31 | 0.11 |
| 2 | 228474106 | rs16823699 | 1.303 | 0.727 | -0.57 | 0.384 | 0.002 | 0.017 | 0.017 | 0 | 0.1 | 0.08 | 0.1 | 0.29 | 0.04 | 0.1 |
| 2 | 228475472 | rs2176189 | -1.433 | -0.218 | -0.97 | -0.304 | -0.019 | -0.044 | -0.044 | 0.613 | 0.84 | 0.87 | 0.95 | 1 | 0.94 | 0.9 |
| 2 | 228475552 | rs2176188 | 1.503 | -0.01 | -1.234 | -0.332 | -0.007 | 0.081 | 0.081 | 0 | 0.25 | 0.12 | 0.25 | 0.12 | 0.24 | 0.27 |
| 2 | 228476140 | rs2176186 | 0.73 | 0.006 | -1.11 | -0.299 | 0.023 | -0.038 | -0.038 | 0 | 0.34 | 0.23 | 0.34 | 0.41 | 0.28 | 0.36 |
| 2 | 228477310 | rs7583413 | -1.228 | 0.22 | -0.545 | -0.062 | 0.01 | 0.022 | 0.022 | 0 | 0.82 | 0.79 | 0.73 | 0.72 | 0.8 | 0.66 |
| 2 | 228481316 | rs16823719 | 1.237 | -0.226 | -0.553 | 0.418 | -0.005 | 0.031 | 0.031 | 0 | 0.1 | 0.11 | 0.1 | 0.26 | 0.04 | 0.1 |
| 2 | 228481419 | rs11889798 | 0.557 | -0.236 | -0.347 | 0.403 | -0.016 | 0.088 | 0.088 | 0 | 0.12 | 0.24 | 0.16 | 0.33 | 0.06 | 0.13 |
| 2 | 228483491 | rs16823728 | 0.78 | -0.604 | 0.086 | -0.05 | -0.056 | -0.09 | -0.09 | 3.761 | 0.16 | 0.25 | 0.06 | 0 | 0.07 | 0.08 |
| 2 | 228487798 | rs10933198 | 0.066 | -0.743 | -0.371 | -0.118 | -0.013 | -0.04 | -0.04 | 0 | 0.33 | 0.48 | 0.42 | 0.22 | 0.28 | 0.41 |
| 2 | 228488044 | rs7577918 | -0.635 | -0.749 | -0.38 | 0.006 | 0.027 | -0.009 | -0.009 | 1.103 | 0.44 | 0.7 | 0.42 | 0.49 | 0.3 | 0.51 |
| 2 | 228489184 | rs13425632 | 0.536 | -0.537 | -0.648 | -0.138 | 0.033 | 0.007 | 0.007 | 0.01 | 0.09 | 0.12 | 0.12 | 0.07 | 0.16 | 0.13 |
| 2 | 228489694 | rs12616684 | 1.065 | -0.993 | -0.77 | -0.386 | -0.032 | -0.006 | -0.006 | 0 | 0.27 | 0.13 | 0.38 | 0.3 | 0.43 | 0.27 |
| 2 | 228492640 | rs7573692 | 0.73 | -0.959 | -0.788 | -0.363 | -0.023 | -0.041 | -0.041 | 0 | 0.32 | 0.47 | 0.31 | 0.21 | 0.26 | 0.41 |
| 2 | 228494424 | rs4602219 | 0.707 | -0.961 | -0.81 | -0.363 | -0.023 | -0.041 | -0.041 | 0 | 0.32 | 0.47 | 0.31 | 0.22 | 0.26 | 0.41 |
| 2 | 228495796 | rs10193698 | 0.903 | -0.96 | -0.83 | -0.361 | -0.006 | -0.001 | -0.001 | 0 | 0.29 | 0.22 | 0.38 | 0.3 | 0.43 | 0.27 |
| 2 | 228498857 | rs6722305 | -1.27 | -0.941 | -0.947 | -0.366 | 0.017 | -0.012 | -0.012 | 0 | 0.75 | 0.79 | 0.81 | 0.71 | 0.86 | 0.87 |
| 2 | 228501121 | rs12996329 | 1.089 | -0.862 | -1.204 | -0.325 | -0.016 | -0.016 | -0.016 | 0 | 0.12 | 0.06 | 0.12 | 0.2 | 0.17 | 0.19 |
| 2 | 228501594 | rs17370706 | 1.754 | -0.978 | -1.077 | -0.085 | -0.01 | -0.005 | -0.005 | 0 | 0.18 | 0.13 | 0.25 | 0.21 | 0.19 | 0.33 |
| 2 | 228506427 | rs7593751 | -0.545 | -1.017 | -0.931 | -0.115 | 0.02 | -0.005 | -0.005 | 0.203 | 0.55 | 0.54 | 0.6 | 0.51 | 0.67 | 0.52 |
| 2 | 228506941 | rs10804335 | -0.018 | -0.748 | -1.1 | 0.007 | 0.003 | -0.006 | -0.006 | 0 | 0.41 | 0.31 | 0.54 | 0.51 | 0.6 | 0.44 |
| 2 | 228507831 | rs4973161 | 0.291 | -0.739 | -1.116 | 0.039 | 0.016 | -0.002 | -0.002 | 0 | 0.46 | 0.38 | 0.58 | 0.51 | 0.63 | 0.59 |
| 2 | 228509458 | rs997363 | -0.206 | -0.98 | -1.664 | -0.213 | 0.032 | 0.026 | 0.026 | 0 | 0.54 | 0.79 | 0.55 | 0.59 | 0.59 | 0.45 |
| 2 | 228513217 | rs6436722 | -0.173 | -0.417 | -1.095 | -0.027 | 0.004 | 0.025 | 0.025 | 0 | 0.46 | 0.68 | 0.53 | 0.63 | 0.45 | 0.61 |
| 2 | 228516984 | rs10204702 | 0.584 | -0.334 | -1.149 | -0.106 | 0.001 | -0.001 | -0.001 | 0 | 0.35 | 0.3 | 0.35 | 0.35 | 0.41 | 0.28 |
| 2 | 228517147 | rs10204884 | 1.776 | -0.107 | -1.561 | -0.044 | -0.001 | -0.011 | -0.011 | 0 | 0.19 | 0.16 | 0.21 | 0.25 | 0.17 | 0.29 |
| 2 | 228518649 | rs2048813 | 1.328 | 0.032 | -1.633 | -0.172 | 0.007 | 0.091 | 0.091 | 4.584 | 0.18 | 0.03 | 0.12 | 0.03 | 0.15 | 0.11 |
| 2 | 228519378 | rs10188499 | 1.439 | 0.284 | -1.327 | 0.133 | 0.001 | 0 | 0 | 0 | 0.26 | 0.26 | 0.35 | 0.33 | 0.33 | 0.21 |
| 2 | 228520709 | rs7609459 | 0.926 | -0.052 | -1.339 | -0.318 | 0.01 | -0.006 | -0.006 | 0 | 0.41 | 0.49 | 0.54 | 0.62 | 0.48 | 0.37 |
| 2 | 228521287 | rs7557243 | 0.492 | -0.153 | -1.239 | -0.424 | -0.006 | -0.001 | -0.001 | 0 | 0.27 | 0.37 | 0.15 | 0.25 | 0.22 | 0.36 |
| 2 | 228521978 | rs6436723 | -0.422 | -0.056 | -1.14 | -0.587 | 0.027 | 0.043 | 0.043 | 0.018 | 0.79 | 0.91 | 0.84 | 0.97 | 0.84 | 0.86 |
| 2 | 228522954 | rs4972892 | 0.266 | 0.079 | -1.256 | -0.569 | 0.014 | 0.133 | 0.133 | 0.198 | 0.68 | 0.9 | 0.79 | 0.95 | 0.7 | 0.74 |
| 2 | 228523140 | rs12469987 | 0.479 | 0.47 | -0.235 | 0.132 | 0.035 | -0.01 | -0.01 | 0 | 0.88 | 0.76 | 0.85 | 0.89 | 0.85 | 0.85 |
| 2 | 228524110 | rs6730730 | -0.53 | -0.243 | -0.094 | 0.392 | -0.062 | 0.017 | 0.017 | 0.904 | 0.24 | 0.52 | 0.17 | 0.27 | 0.11 | 0.16 |
| 2 | 228524310 | rs4321359 | 0.261 | 0.064 | -0.216 | -0.137 | -0.017 | -0.025 | -0.025 | 0 | 0.67 | 0.49 | 0.76 | 0.67 | 0.7 | 0.65 |

| | | | | | | | | | | | | | | | | |
|---|---|---|---|---|---|---|---|---|---|---|---|---|---|---|---|---|
| 2 | 228524407 | rs7568490 | -0.42 | 0.16 | -0.385 | -0.484 | -0.001 | -0.004 | -0.004 | 0 | 0.47 | 0.52 | 0.49 | 0.31 | 0.49 | 0.45 |
| 2 | 228526100 | rs2396465 | -0.717 | -0.02 | -0.091 | -0.487 | 0.019 | 0.042 | 0.042 | 0 | 0.07 | 0.18 | 0.13 | 0.11 | 0.11 | 0.08 |
| 2 | 228526762 | rs2048816 | 0.231 | 0.086 | -0.488 | -1.093 | -0.069 | 0.088 | 0.088 | 0.062 | 0.31 | 0.17 | 0.2 | 0.04 | 0.42 | 0.17 |
| 2 | 228527799 | rs17371490 | -0.721 | -0.432 | 0.029 | -0.514 | -0.016 | 0.015 | 0.015 | 0 | 0.16 | 0.25 | 0.22 | 0.23 | 0.15 | 0.15 |
| 2 | 228532825 | rs13009733 | -0.45 | -0.607 | 0.004 | -0.953 | -0.009 | -0.03 | -0.03 | 0 | 0.58 | 0.49 | 0.62 | 0.23 | 0.66 | 0.38 |
| 2 | 228535921 | rs7579047 | -0.849 | -0.723 | 0.429 | -0.982 | 0.069 | 0.053 | 0.053 | 3.596 | 0.06 | 0.46 | 0.04 | 0 | 0.01 | 0 |
| 2 | 228537398 | rs13383658 | 0.338 | -0.716 | 0.238 | -1.018 | 0.01 | 0.023 | 0.023 | 0 | 0.42 | 0.32 | 0.36 | 0.77 | 0.34 | 0.63 |
| 2 | 228542429 | rs11679813 | 0.404 | -0.648 | -0.127 | -1.205 | -0.068 | 0.095 | 0.095 | 0 | 0.33 | 0.14 | 0.24 | 0 | 0.49 | 0.2 |
| 2 | 228547823 | rs7585777 | -0.574 | -0.575 | -0.222 | -1.112 | 0.033 | 0.094 | 0.094 | 0 | 0.49 | 0.33 | 0.58 | 0.18 | 0.59 | 0.34 |
| 2 | 228565656 | rs11694828 | 0.01 | 0.305 | -0.694 | -0.905 | -0.016 | -0.023 | -0.023 | 0 | 0.35 | 0.27 | 0.25 | 0 | 0.48 | 0.22 |
| 2 | 228567098 | rs6713116 | -0.579 | -0.201 | -0.695 | -0.192 | -0.001 | 0 | 0 | 0 | 0.17 | 0.19 | 0.26 | 0.19 | 0.13 | 0.15 |
| 2 | 228567907 | rs932134 | -0.302 | -0.447 | -0.695 | -0.143 | 0.001 | -0.001 | -0.001 | 0 | 0.38 | 0.39 | 0.23 | 0.63 | 0.31 | 0.57 |
| 2 | 228567967 | rs7567984 | -0.43 | -0.418 | -0.624 | -0.192 | -0.003 | -0.003 | -0.003 | 0 | 0.3 | 0.3 | 0.21 | 0.26 | 0.3 | 0.22 |
| 2 | 228568652 | rs11682956 | 0.133 | -0.499 | -0.233 | -0.455 | -0.021 | -0.062 | -0.062 | 0 | 0.77 | 0.64 | 0.75 | 0.92 | 0.82 | 0.62 |
| 2 | 228570875 | rs7585761 | 0.237 | -0.771 | 0.014 | -0.618 | -0.038 | -0.067 | -0.067 | 0 | 0.21 | 0.35 | 0.17 | 0.05 | 0.12 | 0.13 |
| 2 | 228571475 | rs6708238 | -0.02 | -0.569 | 0.202 | -0.359 | -0.018 | -0.022 | -0.022 | 0 | 0.3 | 0.34 | 0.33 | 0.79 | 0.24 | 0.32 |
| 2 | 228577809 | rs7585481 | 0.003 | -0.666 | -0.415 | -0.375 | 0.013 | 0.015 | 0.015 | 0 | 0.67 | 0.29 | 0.62 | 0.21 | 0.76 | 0.68 |
| 2 | 228584416 | rs17438244 | 0.441 | -0.873 | -1.052 | -0.742 | -0.007 | 0.061 | 0.061 | 0 | 0.22 | 0.02 | 0.22 | 0.08 | 0.18 | 0.35 |
| 2 | 228584898 | rs11691652 | 1.001 | -0.568 | -0.988 | -0.436 | 0.007 | 0.002 | 0.002 | 0 | 0.24 | 0.23 | 0.22 | 0.24 | 0.33 | 0.23 |
| 2 | 228586517 | rs2396492 | 0.646 | -0.411 | -1.479 | -0.523 | 0.012 | 0.058 | 0.058 | 0 | 0.22 | 0.12 | 0.17 | 0.01 | 0.23 | 0.11 |
| 2 | 228589715 | rs4973237 | -0.602 | -0.338 | -1.409 | -0.52 | 0.015 | 0.019 | 0.019 | 0 | 0.73 | 0.82 | 0.85 | 0.91 | 0.82 | 0.64 |
| 2 | 228589902 | rs13002078 | 0.461 | -0.186 | -1.638 | -0.314 | 0.008 | 0.079 | 0.079 | 0 | 0.21 | 0.07 | 0.15 | 0 | 0.2 | 0.12 |
| 2 | 228590027 | rs4973239 | -0.225 | 0.005 | -1.132 | 0.867 | -0.035 | -0.118 | -0.118 | 0.017 | 0.3 | 0.05 | 0.47 | 0.63 | 0.31 | 0.34 |
| 2 | 228595308 | rs1995629 | -0.93 | 0.305 | 0.169 | 1.325 | 0.052 | 0.151 | 0.151 | 0 | 0.14 | 0.34 | 0.26 | 0.41 | 0.21 | 0.2 |
| 2 | 228600612 | rs4973257 | 0.542 | 0.143 | -0.075 | 1.342 | 0 | 0.011 | 0.011 | 0 | 0.82 | 0.82 | 0.75 | 0.52 | 0.76 | 0.77 |
| 2 | 228601654 | rs4973261 | 1.057 | 0.008 | -0.071 | 1.305 | -0.001 | 0.014 | 0.014 | 0 | 0.76 | 0.71 | 0.69 | 0.45 | 0.74 | 0.71 |
| 2 | 228605191 | rs16823876 | -0.412 | 0.244 | -0.274 | 1.365 | -0.026 | -0.053 | -0.053 | 0 | 0.16 | 0.07 | 0.24 | 0.41 | 0.23 | 0.21 |
| 2 | 228607356 | rs11673884 | -1.076 | -0.167 | -0.218 | 1.142 | 0.023 | 0.043 | 0.043 | 0 | 0.48 | 0.68 | 0.57 | 0.71 | 0.47 | 0.4 |
| 2 | 228608458 | rs13411616 | 0.86 | -0.165 | -0.184 | 1.076 | 0.033 | 0.075 | 0.075 | 0 | 0.52 | 0.3 | 0.42 | 0.26 | 0.52 | 0.57 |
| 2 | 228608762 | rs12465514 | -0.299 | -0.44 | 0.762 | 0.955 | 0.001 | -0.012 | -0.012 | 0 | 0.28 | 0.26 | 0.23 | 0.07 | 0.24 | 0.15 |
| 2 | 228612016 | rs4972941 | -1.019 | -0.107 | 0.673 | 0.233 | -0.026 | -0.11 | -0.11 | 3.904 | 0.2 | 0.5 | 0.17 | 0.04 | 0.16 | 0.07 |
| 2 | 228615752 | rs7595272 | 0.479 | -0.101 | 0.473 | 0.497 | -0.001 | 0 | 0 | 0 | 0.92 | 0.89 | 0.89 | 0.91 | 0.9 | 0.91 |
| 2 | 228622128 | rs1506563 | 0.744 | 0.388 | 0.555 | 0.172 | 0.005 | -0.016 | -0.016 | 1.538 | 0.77 | 0.69 | 0.8 | 0.85 | 0.78 | 0.85 |
| 2 | 228637113 | rs4458204 | -0.343 | 0.525 | 0.358 | -0.172 | -0.008 | -0.052 | -0.052 | 0 | 0.18 | 0.3 | 0.09 | 0.03 | 0.13 | 0.25 |
| 2 | 228638562 | rs12994381 | 0.182 | 0.913 | 0.429 | 0.469 | 0.049 | 0.059 | 0.059 | 0 | 0.09 | 0.13 | 0.08 | 0.21 | 0.13 | 0.11 |
| 2 | 228638856 | rs11886482 | -1.113 | 0.584 | 1.019 | 0.669 | -0.015 | 0.044 | 0.044 | 0.104 | 0.24 | 0.59 | 0.29 | 0.31 | 0.17 | 0.32 |
| 2 | 228639557 | rs10184090 | 0.998 | 0.468 | 0.779 | 0.471 | -0.013 | 0.01 | 0.01 | 0 | 0.89 | 0.82 | 0.94 | 0.85 | 0.91 | 0.91 |
| 2 | 228646129 | rs2396494 | -1.213 | 0.446 | 0.946 | 0.173 | -0.036 | -0.048 | -0.048 | 1.512 | 0.27 | 0.48 | 0.14 | 0.16 | 0.15 | 0.36 |
| 2 | 228647253 | rs11681222 | 0.977 | 0.169 | 1.112 | 0.28 | -0.031 | -0.026 | -0.026 | 0.038 | 0.72 | 0.47 | 0.65 | 0.78 | 0.79 | 0.65 |
| 2 | 228651252 | rs12614570 | 1.119 | 0.03 | 0.05 | -0.089 | 0 | 0.045 | 0.045 | 0 | 0.28 | 0.13 | 0.23 | 0.12 | 0.28 | 0.2 |
| 2 | 228652347 | rs2048054 | 1.214 | 0.051 | -0.228 | -0.383 | 0.012 | 0.008 | 0.008 | 0.15 | 0.55 | 0.21 | 0.52 | 0.61 | 0.61 | 0.44 |
| 2 | 228652593 | rs16823913 | 1.893 | 0.048 | -0.277 | -0.384 | -0.093 | -0.16 | -0.16 | 3.74 | 0.25 | 0.03 | 0.29 | 0.48 | 0.32 | 0.24 |
| 2 | 228652949 | rs7340134 | -0.549 | -0.003 | 0.287 | -0.222 | 0.01 | 0.05 | 0.05 | 0 | 0.13 | 0.21 | 0.26 | 0.34 | 0.19 | 0.17 |
| 2 | 228653164 | rs13024444 | 1.596 | 0.022 | 0.192 | -0.217 | 0.014 | 0.009 | 0.009 | 0.233 | 0.55 | 0.18 | 0.52 | 0.61 | 0.61 | 0.44 |
| 2 | 228653239 | rs17246885 | -0.239 | 0.032 | -0.156 | -0.103 | 0.012 | 0.036 | 0.036 | 0 | 0.2 | 0.13 | 0.13 | 0.04 | 0.16 | 0.3 |
| 2 | 228654932 | rs7604823 | -0.388 | 0.378 | -0.161 | 0.254 | 0.011 | 0.024 | 0.024 | 0 | 0.11 | 0.21 | 0.08 | 0.17 | 0.14 | 0.12 |
| 2 | 228667258 | rs2037596 | -0.536 | -0.265 | -0.547 | -0.215 | -0.002 | 0.001 | 0.001 | 0.037 | 0.17 | 0.14 | 0.28 | 0.19 | 0.18 | 0.32 |
| 2 | 228669735 | rs1827924 | -0.12 | -0.609 | -0.374 | -0.372 | -0.021 | -0.022 | -0.022 | 0.96 | 0.41 | 0.47 | 0.32 | 0.28 | 0.31 | 0.32 |
| 2 | 228671423 | rs11694155 | -0.225 | -0.465 | 0.024 | -0.418 | 0 | -0.106 | -0.106 | 0 | 0.18 | 0.36 | 0.13 | 0 | 0.13 | 0.08 |
| 2 | 228672579 | rs13034664 | 0.689 | -0.69 | -0.477 | -0.342 | 0.029 | 0.041 | 0.041 | 0.556 | 0.61 | 0.27 | 0.62 | 0.79 | 0.74 | 0.66 |
| 2 | 228677842 | rs6749704 | 0.424 | -0.721 | -0.787 | 0.091 | -0.003 | -0.004 | -0.004 | 1.398 | 0.28 | 0.14 | 0.25 | 0.29 | 0.23 | 0.26 |
| 2 | 228680580 | rs2396506 | 0.146 | -0.659 | -0.868 | -0.187 | -0.017 | -0.001 | -0.001 | 0 | 0.9 | 0.86 | 0.95 | 0.9 | 0.95 | 0.84 |
| 2 | 228685262 | rs940339 | -0.151 | 0.009 | -1.341 | -0.212 | -0.056 | 0.092 | 0.092 | 0 | 0.39 | 0.19 | 0.5 | 0.23 | 0.52 | 0.37 |
| 2 | 228691605 | rs950817 | 0.766 | -0.28 | -0.373 | 0.178 | -0.043 | -0.028 | -0.028 | 0.005 | 0.84 | 0.76 | 0.92 | 0.91 | 0.92 | 0.9 |
| 2 | 228696351 | rs6736469 | 0.538 | 0.468 | -0.815 | 0.505 | -0.048 | 0.042 | 0.042 | 1.379 | 0.43 | 0.2 | 0.57 | 0.39 | 0.44 | 0.44 |
| 2 | 228697684 | rs17248116 | 0.556 | 0.461 | -0.819 | 0.502 | -0.048 | 0.04 | 0.04 | 1.343 | 0.43 | 0.19 | 0.57 | 0.39 | 0.44 | 0.44 |
| 2 | 228713241 | rs2056566 | -0.382 | 0.642 | -0.019 | 0.202 | 0.032 | 0.129 | 0.129 | 0 | 0.82 | 0.61 | 0.78 | 0.45 | 0.77 | 0.72 |
| 2 | 228720202 | rs6436742 | 1.036 | 0.584 | -0.053 | 0.517 | -0.062 | -0.016 | -0.016 | 0.306 | 0.44 | 0.18 | 0.57 | 0.45 | 0.45 | 0.46 |
| 2 | 228723220 | rs13400894 | -0.193 | 0.655 | 0.321 | 0.601 | 0.039 | 0.102 | 0.102 | 0.038 | 0.36 | 0.17 | 0.16 | 0.07 | 0.3 | 0.26 |
| 2 | 228724368 | rs2396499 | -0.371 | 0.663 | 0.343 | 0.619 | 0.037 | 0.208 | 0.208 | 0 | 0.82 | 0.59 | 0.79 | 0.53 | 0.77 | 0.72 |
| 2 | 228724726 | rs940341 | -0.426 | 0.659 | 0.333 | 0.617 | 0.032 | 0.181 | 0.181 | 0 | 0.82 | 0.62 | 0.78 | 0.53 | 0.77 | 0.72 |
| 2 | 228725864 | rs2396500 | 1.193 | 0.654 | 0.38 | 0.623 | -0.045 | -0.005 | -0.005 | 0.094 | 0.43 | 0.22 | 0.58 | 0.44 | 0.45 | 0.45 |
| 2 | 228735267 | rs6737993 | -1.161 | 0.896 | 0.741 | 1.043 | 0.015 | 0.065 | 0.065 | 0 | 0.9 | 0.79 | 0.89 | 0.81 | 0.88 | 0.88 |
| 2 | 228735732 | rs10498223 | 0.949 | 0.734 | 0.696 | 1.029 | 0.001 | 0 | 0 | 0 | 0.51 | 0.49 | 0.65 | 0.47 | 0.49 | 0.5 |
| 2 | 228749278 | rs13405511 | 0.581 | 0.721 | 0.806 | 1.099 | 0.002 | 0.033 | 0.033 | 0 | 0.09 | 0.11 | 0.09 | 0.25 | 0.11 | 0.15 |
| 2 | 228750239 | rs4973445 | 0.386 | 0.661 | 0.711 | 0.982 | 0.004 | 0.012 | 0.012 | 0 | 0.59 | 0.5 | 0.66 | 0.53 | 0.53 | 0.57 |
| 2 | 228753452 | rs1524025 | -1.254 | 0.661 | 0.718 | 0.981 | 0.018 | 0.065 | 0.065 | 0.004 | 0.9 | 0.78 | 0.89 | 0.8 | 0.88 | 0.88 |
| 2 | 228753972 | rs1721357 | 0.877 | 0.66 | 0.716 | 0.977 | 0.002 | -0.002 | -0.002 | 2.256 | 0.23 | 0.16 | 0.14 | 0.01 | 0.24 | 0.16 |
| 2 | 228754056 | rs6736707 | 1.201 | 0.645 | 0.567 | 0.989 | -0.04 | -0.016 | -0.016 | 0.221 | 0.48 | 0.16 | 0.6 | 0.46 | 0.46 | 0.5 |

| | | | | | | | | | | | | | | | | |
|---|---|---|---|---|---|---|---|---|---|---|---|---|---|---|---|---|
| 2 | 228754349 | rs16824126 | -0.129 | 0.619 | 0.59 | 0.918 | -0.016 | -0.001 | -0.001 | 0 | 0.1 | 0.34 | 0.06 | 0.07 | 0.07 | 0.07 |
| 2 | 228759554 | rs7566347 | -0.165 | 0.639 | 0.457 | 0.924 | -0.001 | -0.031 | -0.031 | 0 | 0.68 | 0.61 | 0.75 | 0.8 | 0.64 | 0.71 |
| 2 | 228760336 | rs16824173 | 0.038 | 0.666 | 0.209 | 0.964 | 0 | 0 | 0 | 0 | 0.09 | 0.08 | 0.05 | 0.07 | 0.07 | 0.07 |
| 2 | 228767631 | rs2946931 | -0.809 | 0.664 | -0.087 | 0.896 | 0.007 | 0.016 | 0.016 | 0.405 | 0.43 | 0.34 | 0.29 | 0.33 | 0.43 | 0.4 |
| 2 | 228773287 | rs11680252 | 0.566 | 1.258 | -0.235 | 0.828 | -0.007 | -0.006 | -0.006 | 0.012 | 0.14 | 0.13 | 0.15 | 0.25 | 0.25 | 0.17 |
| 2 | 228776938 | rs3748863 | 1.402 | 1.239 | -0.183 | 0.857 | 0.013 | -0.009 | -0.009 | 0.003 | 0.41 | 0.33 | 0.52 | 0.46 | 0.32 | 0.46 |
| 2 | 228778565 | rs17251109 | 0.88 | 1.382 | -0.914 | 0.938 | -0.081 | -0.08 | -0.08 | 0 | 0.12 | 0 | 0.14 | 0.25 | 0.24 | 0.17 |
| 2 | 228782976 | rs1721355 | -0.971 | 1.34 | -0.876 | 0.951 | 0.005 | 0.03 | 0.03 | 0.161 | 0.48 | 0.36 | 0.36 | 0.34 | 0.57 | 0.41 |
| 2 | 228797628 | rs4973503 | 1.237 | 1.143 | -1.278 | 0.987 | -0.036 | -0.059 | -0.059 | 0.727 | 0.1 | 0.01 | 0.07 | 0.25 | 0.09 | 0.26 |
| 2 | 228798514 | rs1403970 | 0.15 | 1.289 | -0.712 | 1.022 | 0.074 | 0.116 | 0.116 | 0.224 | 0.76 | 0.48 | 0.72 | 0.55 | 0.64 | 0.7 |
| 2 | 228801982 | rs6737663 | 0.18 | 1.165 | -1.406 | 1.344 | 0.01 | -0.001 | -0.001 | 0.072 | 0.54 | 0.31 | 0.58 | 0.55 | 0.4 | 0.54 |
| 2 | 228804459 | rs7568614 | 0.461 | 1.079 | -1.437 | 1.337 | 0.04 | 0.001 | 0.001 | 0 | 0.48 | 0.31 | 0.57 | 0.55 | 0.38 | 0.51 |
| 2 | 228804641 | rs2894645 | 0.716 | 1.114 | -1.385 | 1.349 | 0.027 | -0.005 | -0.005 | 0 | 0.45 | 0.33 | 0.53 | 0.5 | 0.35 | 0.47 |
| 2 | 228805894 | rs2103106 | 0.793 | 1.429 | -1.479 | 1.59 | 0.037 | 0.035 | 0.035 | 0.504 | 0.45 | 0.27 | 0.53 | 0.4 | 0.36 | 0.46 |
| 2 | 228808187 | rs17265317 | 0.147 | 1.722 | -1.624 | 1.805 | -0.087 | -0.095 | -0.095 | 0.174 | 0.13 | 0.01 | 0.14 | 0.32 | 0.27 | 0.22 |
| 2 | 228808696 | rs1916804 | 1.064 | 1.513 | -1.551 | 1.733 | 0.031 | 0.036 | 0.036 | 1.179 | 0.42 | 0.26 | 0.51 | 0.35 | 0.32 | 0.4 |
| 2 | 228808890 | rs6436743 | -0.196 | 1.605 | -1.259 | 1.088 | -0.004 | -0.013 | -0.013 | 0 | 0.66 | 0.45 | 0.71 | 0.75 | 0.65 | 0.72 |
| 2 | 228809625 | rs13414787 | -1.006 | 1.986 | -1.172 | 1.403 | -0.03 | -0.007 | -0.007 | 0 | 0.17 | 0.1 | 0.16 | 0.15 | 0.26 | 0.24 |
| 2 | 228812013 | rs10196736 | -0.965 | 2.333 | -0.691 | 1.613 | 0.018 | -0.014 | -0.014 | 0 | 0.17 | 0.22 | 0.19 | 0.1 | 0.25 | 0.24 |
| 2 | 228812298 | rs1524028 | 0.581 | 2.047 | -0.156 | 1.888 | 0.015 | 0.243 | 0.243 | 0 | 0.94 | 0.78 | 0.77 | 0.64 | 0.91 | 0.93 |
| 2 | 228812878 | rs1524029 | 0.561 | 1.317 | -0.468 | 0.849 | 0.02 | 0.098 | 0.098 | 2 | 0.53 | 0.32 | 0.59 | 0.78 | 0.56 | 0.67 |
| 2 | 228812961 | rs7582143 | -0.706 | 1.009 | 0.136 | 0.615 | -0.005 | -0.062 | -0.062 | 0.066 | 0.38 | 0.5 | 0.34 | 0.15 | 0.35 | 0.3 |
| 2 | 228815975 | rs6744449 | -0.753 | 0.965 | -0.199 | 0.603 | 0.016 | -0.03 | -0.03 | 0 | 0.24 | 0.34 | 0.23 | 0.18 | 0.31 | 0.24 |
| 2 | 228819272 | rs1715826 | 1.048 | 0.965 | -0.695 | 0.594 | -0.007 | 0.005 | 0.005 | 1.168 | 0.48 | 0.18 | 0.37 | 0.45 | 0.47 | 0.6 |
| 2 | 228825101 | rs1715832 | 1.442 | 1.071 | -1.548 | 0.53 | -0.024 | -0.022 | -0.022 | 0 | 0.29 | 0.03 | 0.21 | 0.23 | 0.24 | 0.43 |
| 2 | 228838054 | rs7583252 | -0.485 | 0.939 | -0.945 | 0.56 | 0.015 | -0.02 | -0.02 | 0 | 0.32 | 0.38 | 0.3 | 0.31 | 0.43 | 0.27 |
| 2 | 228838093 | rs16824273 | 1.116 | 0.878 | -0.844 | 0.54 | -0.006 | -0.008 | -0.008 | 0 | 0.08 | 0.14 | 0.06 | 0.07 | 0.1 | 0.04 |
| 2 | 228840360 | rs11688249 | 0.26 | 1.189 | -0.908 | 0.757 | -0.1 | -0.016 | -0.016 | 0.067 | 0.23 | 0.06 | 0.22 | 0.23 | 0.37 | 0.21 |
| 2 | 228843948 | rs4973044 | -0.8 | 1.171 | -0.863 | 1.022 | 0.021 | -0.001 | -0.001 | 0 | 0.35 | 0.39 | 0.5 | 0.67 | 0.53 | 0.3 |
| 2 | 228851619 | rs13394238 | 0.26 | 1.135 | -0.869 | 1.046 | -0.075 | -0.006 | -0.006 | 0 | 0.22 | 0.09 | 0.22 | 0.24 | 0.37 | 0.21 |
| 2 | 228857159 | rs1343845 | -0.818 | 1.164 | -0.871 | 0.999 | 0.021 | -0.002 | -0.002 | 0.227 | 0.35 | 0.4 | 0.5 | 0.67 | 0.53 | 0.32 |
| 2 | 228864662 | rs11680053 | 0.484 | 1.147 | -0.869 | 1.001 | 0.019 | -0.002 | -0.002 | 0 | 0.64 | 0.61 | 0.5 | 0.33 | 0.47 | 0.68 |
| 2 | 228874280 | rs6743957 | 0.227 | 1.175 | -0.847 | 1.025 | 0.028 | 0.178 | 0.178 | 0 | 0.13 | 0.2 | 0.29 | 0.4 | 0.22 | 0.1 |
| 2 | 228883029 | rs3811515 | 1.775 | 1.059 | -1.269 | 0.776 | -0.048 | -0.006 | -0.006 | 0 | 0.3 | 0.03 | 0.25 | 0.22 | 0.24 | 0.44 |
| 2 | 228883721 | rs3811514 | 0.436 | 1.343 | -0.931 | 0.926 | -0.046 | 0.002 | 0.002 | 0 | 0.65 | 0.77 | 0.66 | 0.62 | 0.49 | 0.63 |
| 2 | 228888652 | rs11678867 | -0.282 | 1.389 | -0.937 | 0.916 | -0.029 | 0.008 | 0.008 | 0.145 | 0.14 | 0.12 | 0.15 | 0.11 | 0.21 | 0.2 |
| 2 | 228903932 | rs4464263 | -0.827 | 1.389 | -0.938 | 0.953 | -0.072 | -0.015 | -0.015 | 0 | 0.8 | 0.94 | 0.81 | 0.79 | 0.7 | 0.83 |
| 2 | 228907788 | rs6730930 | 1.21 | 1.138 | -0.855 | 0.943 | -0.005 | -0.015 | -0.015 | 0 | 0.52 | 0.49 | 0.38 | 0.22 | 0.33 | 0.52 |
| 2 | 228907927 | rs6436750 | 0.003 | 1.195 | -0.903 | 0.922 | -0.064 | -0.004 | -0.004 | 0.26 | 0.7 | 0.78 | 0.68 | 0.65 | 0.53 | 0.72 |
| 2 | 228912194 | rs4294998 | -0.732 | 1.256 | -0.869 | 0.94 | -0.048 | 0.002 | 0.002 | 0 | 0.35 | 0.23 | 0.34 | 0.37 | 0.51 | 0.37 |
| 2 | 228912778 | rs13430892 | 1.176 | 1.211 | -0.935 | 0.857 | -0.004 | -0.013 | -0.013 | 0.731 | 0.53 | 0.5 | 0.38 | 0.22 | 0.33 | 0.53 |
| 2 | 228916117 | rs12694769 | -1.484 | 1.214 | -0.934 | 0.826 | -0.004 | -0.013 | -0.013 | 0.196 | 0.48 | 0.51 | 0.62 | 0.77 | 0.67 | 0.46 |
| 2 | 228916212 | rs10196885 | -1.484 | 1.214 | -0.934 | 0.826 | -0.004 | -0.013 | -0.013 | 0 | 0.48 | 0.51 | 0.62 | 0.77 | 0.67 | 0.46 |
| 2 | 228919347 | rs10172089 | 1.247 | 1.166 | -0.959 | 0.802 | -0.002 | -0.01 | -0.01 | 0 | 0.52 | 0.49 | 0.38 | 0.23 | 0.33 | 0.53 |
| 2 | 228920753 | rs11694195 | -0.743 | 1.172 | -0.894 | 0.904 | 0.01 | 0.108 | 0.108 | 0 | 0.19 | 0.32 | 0.34 | 0.38 | 0.26 | 0.23 |
| 2 | 228923951 | rs13388871 | 0.057 | 1.168 | -0.727 | 0.9 | 0.003 | -0.002 | -0.002 | 0.045 | 0.06 | 0.14 | 0.04 | 0.04 | 0.04 | 0.1 |
| 2 | 228925597 | rs6757751 | 0.331 | 1.168 | -0.727 | 0.9 | 0 | 0.001 | 0.001 | 0 | 0.05 | 0.08 | 0.04 | 0.04 | 0.04 | 0.09 |
| 2 | 228925600 | rs4591331 | 0.146 | 1.171 | -0.734 | 0.902 | 0.006 | 0.078 | 0.078 | 0 | 0.14 | 0.18 | 0.3 | 0.34 | 0.22 | 0.12 |
| 2 | 228925898 | rs6760875 | -0.003 | 1.123 | -0.824 | 0.846 | -0.053 | -0.005 | -0.005 | 0 | 0.7 | 0.8 | 0.72 | 0.66 | 0.59 | 0.74 |
| 2 | 228928690 | rs12466849 | -1.024 | 0.829 | -0.771 | 1.085 | 0.009 | 0.122 | 0.122 | 0 | 0.2 | 0.33 | 0.34 | 0.43 | 0.26 | 0.23 |
| 2 | 228943517 | rs4384783 | 0.37 | 0.611 | -0.921 | 1.008 | -0.002 | 0.037 | 0.037 | 0.165 | 0.12 | 0.11 | 0.33 | 0.32 | 0.16 | 0.08 |
| 2 | 228944282 | rs13409615 | 0.107 | 0.198 | -1.201 | 0.722 | 0.047 | 0.108 | 0.108 | 0 | 0.22 | 0.09 | 0.13 | 0.02 | 0.12 | 0.16 |
| 2 | 228950588 | rs4641927 | -0.507 | 0.393 | -1.057 | 0.858 | 0.039 | -0.012 | -0.012 | 0 | 0.33 | 0.19 | 0.46 | 0.34 | 0.29 | 0.25 |
| 2 | 228950780 | rs4530360 | -0.259 | 0.879 | -1.059 | 0.895 | -0.008 | -0.01 | -0.01 | 0 | 0.77 | 0.83 | 0.8 | 0.7 | 0.71 | 0.8 |
| 2 | 228968195 | rs12479183 | -0.499 | 0.413 | -0.936 | 0.779 | 0.003 | -0.002 | -0.002 | 0 | 0.29 | 0.24 | 0.44 | 0.32 | 0.25 | 0.15 |
| 2 | 228971078 | rs12994876 | -1.001 | 0.811 | -1.255 | 0.727 | -0.014 | -0.023 | -0.023 | 1.421 | 0.12 | 0.03 | 0.12 | 0.19 | 0.14 | 0.14 |
| 2 | 228973660 | rs7561798 | -1.072 | 0.803 | -1.589 | 0.539 | 0.008 | 0.041 | 0.041 | 0.463 | 0.45 | 0.31 | 0.64 | 0.68 | 0.48 | 0.4 |
| 2 | 228977228 | rs4455149 | -0.406 | 0.748 | -1.543 | 0.538 | 0.014 | -0.035 | -0.035 | 0 | 0.35 | 0.3 | 0.52 | 0.49 | 0.34 | 0.27 |
| 2 | 228991140 | rs7562755 | 0.499 | 0.098 | -2.074 | 0.033 | 0.016 | 0.067 | 0.067 | 0 | 0.15 | 0.02 | 0.07 | 0.02 | 0.13 | 0.06 |
| 2 | 228999691 | rs6436755 | -0.277 | 0.986 | -1.993 | 0.103 | -0.016 | -0.039 | -0.039 | 0.473 | 0.38 | 0.29 | 0.56 | 0.49 | 0.44 | 0.31 |
| 2 | 229014719 | rs4438470 | -0.302 | 0.723 | -1.134 | 0.195 | -0.035 | -0.045 | -0.045 | 1.252 | 0.1 | 0.01 | 0.12 | 0.19 | 0.14 | 0.12 |
| 2 | 229021192 | rs6739688 | -0.72 | 0.649 | -0.472 | 0.012 | 0.02 | -0.009 | -0.009 | 0 | 0.2 | 0.37 | 0.24 | 0.2 | 0.25 | 0.17 |
| 2 | 229028291 | rs10171038 | 0.583 | 0.586 | -0.59 | -0.055 | -0.007 | 0.006 | 0.006 | 0.748 | 0.07 | 0.07 | 0.07 | 0 | 0.09 | 0.04 |
| 2 | 229029615 | rs4327223 | 0.617 | -0.087 | -0.61 | 0.08 | 0.011 | -0.006 | -0.006 | 0 | 0.17 | 0.13 | 0.12 | 0.18 | 0.12 | 0.16 |
| 2 | 229031147 | rs4641922 | 1.747 | -0.495 | -0.365 | -0.153 | -0.091 | 0.019 | 0.019 | 2.047 | 0.4 | 0.12 | 0.29 | 0.31 | 0.44 | 0.56 |
| 2 | 229037643 | rs13418687 | 1.056 | -0.497 | -0.435 | -0.141 | 0.013 | -0.007 | -0.007 | 0 | 0.17 | 0.11 | 0.11 | 0.18 | 0.12 | 0.14 |
| 2 | 229037885 | rs4973620 | -1.761 | -0.545 | -0.353 | -0.162 | -0.02 | 0.032 | 0.032 | 0.97 | 0.58 | 0.7 | 0.7 | 0.69 | 0.56 | 0.44 |
| 2 | 229038803 | rs7570716 | -1.069 | -0.412 | -0.302 | -0.14 | 0 | 0.001 | 0.001 | 0.23 | 0.4 | 0.56 | 0.59 | 0.51 | 0.45 | 0.28 |
| 2 | 229047536 | rs4277511 | -0.955 | -0.319 | -0.615 | -0.336 | -0.019 | 0.05 | 0.05 | 0.311 | 0.55 | 0.66 | 0.68 | 0.68 | 0.52 | 0.42 |

| | | | | | | | | | | | | | | | | |
|---|---|---|---|---|---|---|---|---|---|---|---|---|---|---|---|---|
| 2 | 229056924 | rs13398361 | -0.552 | 0.522 | -0.757 | -0.269 | -0.007 | 0.008 | 0.008 | 0 | 0.13 | 0.14 | 0.14 | 0.08 | 0.18 | 0.15 |
| 2 | 229077527 | rs4428009 | -0.557 | 0.354 | -0.704 | -0.267 | -0.043 | 0.103 | 0.103 | 0.226 | 0.5 | 0.12 | 0.64 | 0.68 | 0.52 | 0.42 |
| 2 | 229082758 | rs4419223 | -0.057 | 0.174 | -0.401 | -0.152 | -0.017 | 0.031 | 0.031 | 0 | 0.33 | 0.49 | 0.46 | 0.5 | 0.3 | 0.25 |
| 2 | 229101189 | rs12623683 | -1.966 | 0.57 | -0.6 | -0.23 | -0.005 | -0.002 | -0.002 | 0 | 0.71 | 0.83 | 0.82 | 0.75 | 0.74 | 0.57 |
| 2 | 229106949 | rs10804339 | -1.078 | 0.584 | -0.473 | -0.249 | 0.001 | -0.003 | -0.003 | 0 | 0.61 | 0.53 | 0.67 | 0.7 | 0.59 | 0.47 |
| 2 | 229107336 | rs6753687 | -1.132 | 0.592 | -0.591 | -0.245 | -0.017 | 0.018 | 0.018 | 0.603 | 0.62 | 0.66 | 0.69 | 0.7 | 0.59 | 0.47 |
| 2 | 229133568 | rs1528223 | 1.055 | 0.528 | -0.697 | -0.153 | 0.005 | -0.008 | -0.008 | 0 | 0.45 | 0.28 | 0.31 | 0.41 | 0.4 | 0.58 |
| 2 | 229133647 | rs1406144 | -0.268 | 0.546 | -0.52 | -0.221 | 0.002 | -0.055 | -0.055 | 0 | 0.44 | 0.15 | 0.53 | 0.53 | 0.45 | 0.32 |
| 2 | 229135267 | rs1528224 | -2.108 | 0.558 | -0.578 | -0.24 | 0.001 | 0.005 | 0.005 | 0.513 | 0.7 | 0.81 | 0.82 | 0.75 | 0.74 | 0.57 |
| 2 | 229135813 | rs9636276 | 1.95 | 0.555 | -0.623 | -0.23 | -0.018 | -0.039 | -0.039 | 0 | 0.13 | 0.04 | 0.12 | 0.16 | 0.14 | 0.15 |
| 2 | 229137908 | rs2091516 | -1.276 | 0.589 | -0.366 | 0.049 | 0.004 | -0.001 | -0.001 | 0 | 0.54 | 0.57 | 0.67 | 0.59 | 0.6 | 0.42 |
| 2 | 229152490 | rs2091515 | -0.382 | 0.615 | -0.688 | 0.069 | 0.002 | -0.055 | -0.055 | 0 | 0.44 | 0.15 | 0.53 | 0.53 | 0.45 | 0.32 |
| 2 | 229155830 | rs10498227 | 0.621 | 0.944 | -0.459 | 0.011 | -0.003 | 0.004 | 0.004 | 0 | 0.07 | 0.09 | 0.12 | 0.05 | 0.15 | 0.1 |
| 2 | 229168274 | rs11689427 | 0.104 | 0.674 | -0.622 | -0.162 | -0.006 | 0.031 | 0.031 | 0.022 | 0.28 | 0.2 | 0.2 | 0.17 | 0.28 | 0.2 |
| 2 | 229168389 | rs2091513 | 0.434 | 0.782 | -0.542 | -0.123 | 0.01 | -0.01 | -0.01 | 0 | 0.08 | 0.13 | 0.12 | 0.05 | 0.14 | 0.09 |
| 2 | 229173032 | rs10184282 | 1.136 | 0.554 | -0.598 | 0.007 | -0.008 | -0.009 | -0.009 | 0 | 0.17 | 0.1 | 0.17 | 0.16 | 0.2 | 0.16 |
| 2 | 229180510 | rs7559216 | -0.326 | 0.548 | -0.636 | -0.057 | -0.01 | 0.021 | 0.021 | 0 | 0.33 | 0.3 | 0.25 | 0.22 | 0.34 | 0.25 |
| 2 | 229181417 | rs6747061 | -0.33 | 0.541 | -0.614 | -0.056 | -0.007 | 0.01 | 0.01 | 0.029 | 0.33 | 0.31 | 0.25 | 0.22 | 0.34 | 0.25 |
| 2 | 229186057 | rs4973079 | 0.254 | 0.457 | -0.46 | 0.423 | 0.002 | 0.007 | 0.007 | 0.054 | 0.56 | 0.51 | 0.37 | 0.31 | 0.43 | 0.62 |
| 2 | 229188012 | rs12052730 | 0.549 | -0.141 | -0.426 | 0.483 | -0.091 | 0.133 | 0.133 | 0.382 | 0.24 | 0.43 | 0.35 | 0.47 | 0.15 | 0.14 |
| 2 | 229201683 | rs12479264 | 0.71 | -0.072 | -0.475 | 0.325 | -0.069 | -0.057 | -0.057 | 0 | 0.23 | 0.34 | 0.13 | 0.15 | 0.14 | 0.12 |
| 2 | 229207194 | rs13021389 | -0.401 | 0.698 | -0.63 | 0.479 | 0.008 | 0.009 | 0.009 | 0.304 | 0.42 | 0.39 | 0.42 | 0.37 | 0.58 | 0.37 |
| 2 | 229211957 | rs17235156 | -0.068 | 0.89 | -0.901 | 0.521 | -0.052 | 0.003 | 0.003 | 0.461 | 0.36 | 0.21 | 0.34 | 0.32 | 0.51 | 0.31 |
| 2 | 229231976 | rs16824763 | -0.228 | 0.961 | -0.049 | 1.136 | 0.027 | 0.155 | 0.155 | 0 | 0.07 | 0.17 | 0.27 | 0.35 | 0.08 | 0.09 |
| 2 | 229232050 | rs7595353 | 0.585 | 0.499 | -0.511 | 0.826 | 0.011 | 0.024 | 0.024 | 2.978 | 0.33 | 0.21 | 0.23 | 0.18 | 0.26 | 0.47 |
| 2 | 229232445 | rs7608239 | -0.47 | 0.427 | -0.149 | 1.203 | 0.002 | -0.005 | -0.005 | 2.324 | 0.65 | 0.68 | 0.69 | 0.46 | 0.67 | 0.44 |
| 2 | 229233118 | rs4973083 | 0.105 | -0.61 | 0.136 | 1.048 | 0 | 0.049 | 0.049 | 0 | 0.18 | 0.26 | 0.42 | 0.54 | 0.18 | 0.14 |
| 2 | 229240145 | rs2551985 | -0.59 | -0.897 | -0.261 | 0.524 | 0.016 | 0.108 | 0.108 | 0.143 | 0.72 | 0.9 | 0.74 | 0.94 | 0.72 | 0.74 |
| 2 | 229240608 | rs6722551 | 1.796 | -0.892 | 0.064 | 0.498 | 0 | 0 | 0 | 1.751 | 0.3 | 0.25 | 0.17 | 0.21 | 0.31 | 0.41 |
| 2 | 229241233 | rs1528219 | 0.528 | -0.601 | 0.253 | 0.661 | -0.005 | -0.002 | -0.002 | 0.365 | 0.18 | 0.12 | 0.1 | 0.17 | 0.2 | 0.11 |
| 2 | 229241777 | rs16824774 | 0.184 | -0.604 | 0.331 | 0.684 | 0.005 | 0.058 | 0.058 | 0 | 0.19 | 0.29 | 0.43 | 0.56 | 0.2 | 0.2 |
| 2 | 229241978 | rs1155462 | 0.643 | -0.685 | 0.375 | 0.621 | 0.001 | 0.021 | 0.021 | 0.535 | 0.47 | 0.46 | 0.59 | 0.76 | 0.5 | 0.58 |
| 2 | 229242118 | rs17239335 | -1 | -0.773 | 0.093 | 0.165 | 0.001 | -0.003 | -0.003 | 0.058 | 0.39 | 0.59 | 0.56 | 0.74 | 0.42 | 0.34 |
| 2 | 229242746 | rs929925 | -0.246 | -0.87 | 0.456 | -0.292 | -0.029 | -0.112 | -0.112 | 0.985 | 0.65 | 0.57 | 0.69 | 0.94 | 0.7 | 0.69 |
| 2 | 229247589 | rs2860071 | 0.795 | -1.403 | 0.273 | 0.181 | 0.02 | 0.011 | 0.011 | 0.4 | 0.62 | 0.35 | 0.58 | 0.31 | 0.72 | 0.78 |
| 2 | 229252925 | rs800958 | -0.052 | -1.494 | 0.562 | 0.264 | -0.072 | 0.124 | 0.124 | 0 | 0.8 | 0.63 | 0.69 | 0.43 | 0.89 | 0.9 |
| 2 | 229269825 | rs1469181 | 0.369 | -1.429 | 0.31 | -0.023 | -0.023 | 0.068 | 0.068 | 0 | 0.37 | 0.26 | 0.35 | 0.17 | 0.44 | 0.44 |
| 2 | 229283211 | rs2194545 | -0.614 | -1.496 | 0.363 | -0.002 | -0.014 | 0.038 | 0.038 | 0 | 0.63 | 0.69 | 0.64 | 0.82 | 0.54 | 0.53 |
| 2 | 229293255 | rs7566610 | -0.51 | -1.544 | 0.537 | 0.17 | -0.023 | 0.085 | 0.085 | 1.322 | 0.27 | 0.44 | 0.37 | 0.49 | 0.22 | 0.14 |
| 2 | 229296965 | rs2194549 | 2.163 | -1.535 | 0.423 | 0.134 | -0.042 | 0.048 | 0.048 | 0.841 | 0.25 | 0.1 | 0.22 | 0.14 | 0.28 | 0.34 |
| 2 | 229300887 | rs6726383 | -1.266 | -1.557 | 0.5 | 0.179 | -0.002 | 0 | 0 | 0 | 0.37 | 0.59 | 0.42 | 0.68 | 0.26 | 0.2 |
| 2 | 229302742 | rs12611880 | 0.381 | -0.833 | 0.004 | -0.056 | -0.055 | -0.001 | -0.001 | 0.082 | 0.15 | 0.02 | 0.09 | 0.12 | 0.17 | 0.09 |
| 2 | 229307813 | rs6712246 | 0.084 | -0.818 | 0.13 | -0.048 | 0.003 | -0.012 | -0.012 | 0 | 0.17 | 0.21 | 0.11 | 0.13 | 0.17 | 0.1 |
| 2 | 229324709 | rs7573585 | 0.787 | -0.69 | 0.08 | 0.285 | -0.019 | 0.013 | 0.013 | 0 | 0.1 | 0.11 | 0.04 | 0.19 | 0.04 | 0.06 |
| 2 | 229325525 | rs6752389 | -0.124 | -0.697 | 0.074 | 0.247 | 0 | -0.001 | -0.001 | 0 | 0.53 | 0.46 | 0.38 | 0.46 | 0.49 | 0.5 |
| 2 | 229326626 | rs10490601 | -0.019 | -0.03 | -0.052 | 0.128 | -0.008 | 0.024 | 0.024 | 0 | 0.43 | 0.36 | 0.34 | 0.26 | 0.45 | 0.44 |
| 2 | 229330321 | rs7592831 | -0.396 | 0.534 | -0.477 | 0.211 | -0.12 | 0.057 | 0.057 | 0.18 | 0.25 | 0.03 | 0.14 | 0.11 | 0.3 | 0.19 |
| 2 | 229331340 | rs13402716 | -0.614 | 0.493 | 0.027 | 0.191 | -0.009 | 0.008 | 0.008 | 0.559 | 0.27 | 0.25 | 0.15 | 0.11 | 0.3 | 0.19 |
| 2 | 229332427 | rs2059389 | 1.179 | 0.358 | 0.021 | -0.782 | -0.003 | 0.005 | 0.005 | 0.038 | 0.29 | 0.27 | 0.22 | 0.16 | 0.3 | 0.32 |
| 2 | 229332491 | rs2059390 | -0.438 | 0.506 | -0.769 | -0.601 | 0.047 | 0.038 | 0.038 | 1.068 | 0.58 | 0.73 | 0.39 | 0.28 | 0.61 | 0.51 |
| 2 | 229333848 | rs6738261 | -0.408 | 0.516 | -0.762 | -0.597 | 0.044 | 0.036 | 0.036 | 0.844 | 0.57 | 0.7 | 0.39 | 0.27 | 0.61 | 0.51 |
| 2 | 229339187 | rs10933256 | 0.638 | 1.247 | -1.238 | -0.48 | -0.033 | 0.045 | 0.045 | 0 | 0.68 | 0.78 | 0.69 | 0.91 | 0.58 | 0.72 |
| 2 | 229343602 | rs16824970 | 0.614 | 1.417 | -1.78 | -0.468 | -0.04 | -0.012 | -0.012 | 0.675 | 0.13 | 0.02 | 0.05 | 0.09 | 0.11 | 0.18 |
| 2 | 229344752 | rs10188715 | -0.697 | 0.566 | -1.686 | -0.398 | 0.021 | 0.028 | 0.028 | 0 | 0.87 | 0.91 | 0.82 | 1 | 0.87 | 0.93 |
| 2 | 229362382 | rs13033979 | 0.344 | 1.215 | -1.271 | 0.167 | 0.001 | -0.013 | -0.013 | 0 | 0.32 | 0.59 | 0.25 | 0.21 | 0.31 | 0.32 |
| 2 | 229365795 | rs10498226 | -0.844 | 2.063 | -1.324 | 0.507 | -0.122 | 0.007 | 0.007 | 0.105 | 0.18 | 0.02 | 0.12 | 0.13 | 0.28 | 0.21 |
| 2 | 229371034 | rs17255781 | -1.588 | 1.261 | -0.808 | 0.616 | -0.02 | 0.027 | 0.027 | 0 | 0.07 | 0.05 | 0.07 | 0 | 0.12 | 0.03 |
| 2 | 229380688 | rs17236584 | -2.02 | 1.289 | 0.984 | 0.45 | 0.003 | -0.006 | -0.006 | 5.892 | 0.18 | 0.14 | 0.16 | 0.09 | 0.26 | 0.13 |
| 2 | 229382616 | rs7597603 | -1.717 | 1.293 | 0.989 | 0.437 | -0.019 | 0.03 | 0.03 | 0 | 0.08 | 0.02 | 0.1 | 0 | 0.13 | 0.04 |
| 2 | 229389990 | rs1435859 | -2.432 | 1.067 | 2.045 | 1.194 | 0.022 | 0.04 | 0.04 | 0.127 | 0.23 | 0.53 | 0.19 | 0.33 | 0.28 | 0.18 |
| 2 | 229417323 | rs2396569 | -2.469 | 1.017 | 2.135 | 1.199 | 0.014 | 0.035 | 0.035 | 0.051 | 0.24 | 0.55 | 0.19 | 0.33 | 0.28 | 0.17 |
| 2 | 229446145 | rs6744195 | -0.927 | 0.821 | 3.001 | 1.248 | 0.005 | 0.125 | 0.125 | 0 | 0.08 | 0.22 | 0.07 | 0.2 | 0.06 | 0.13 |
| 2 | 229446995 | rs6748070 | 2.284 | 0.844 | 2.994 | 0.962 | 0.002 | -0.043 | -0.043 | 0 | 0.76 | 0.35 | 0.8 | 0.92 | 0.73 | 0.84 |
| 2 | 229480848 | rs11884202 | -2.008 | 0.858 | 2.548 | 1.099 | -0.033 | 0.03 | 0.03 | 0 | 0.19 | 0.07 | 0.16 | 0.1 | 0.26 | 0.15 |
| 2 | 229481486 | rs6743558 | -2.603 | 0.841 | 2.759 | 1.092 | 0.022 | -0.049 | -0.049 | 0 | 0.25 | 0.54 | 0.23 | 0.14 | 0.29 | 0.17 |
| 2 | 229493854 | rs10199039 | -2.143 | 0.91 | 2.226 | 1.1 | 0.006 | -0.02 | -0.02 | 1.323 | 0.2 | 0.29 | 0.19 | 0.11 | 0.26 | 0.16 |
| 2 | 229502619 | rs7604908 | -2.261 | 0.967 | 2.402 | 1.154 | 0.042 | 0.029 | 0.029 | 0 | 0.17 | 0.41 | 0.22 | 0.17 | 0.22 | 0.2 |
| 2 | 229504065 | rs10498231 | -0.948 | 1.011 | 2.32 | 1.359 | -0.02 | -0.03 | -0.03 | 0 | 0.07 | 0.01 | 0.04 | 0.12 | 0.09 | 0.1 |
| 2 | 229504348 | rs1865684 | -2.208 | 1.009 | 2.342 | 1.358 | 0.042 | 0.029 | 0.029 | 0 | 0.17 | 0.41 | 0.22 | 0.17 | 0.22 | 0.2 |

| | | | | | | | | | | | | | | | | |
|---|---|---|---|---|---|---|---|---|---|---|---|---|---|---|---|---|
| 2 | 229510488 | rs3943747 | -2.767 | 1.001 | 2.431 | 1.354 | 0.029 | 0.036 | 0.036 | 0 | 0.25 | 0.46 | 0.26 | 0.3 | 0.31 | 0.29 |
| 2 | 229510929 | rs1435867 | -0.817 | 0.99 | 2.64 | 1.369 | 0.001 | -0.001 | -0.001 | 0 | 0.07 | 0.06 | 0.06 | 0.13 | 0.06 | 0.15 |
| 2 | 229520063 | rs11885362 | -2.176 | 0.997 | 2.652 | 1.361 | 0.098 | 0.004 | 0.004 | 0.976 | 0.11 | 0.39 | 0.16 | 0.04 | 0.17 | 0.05 |
| 2 | 229529113 | rs6737010 | -2.049 | 0.996 | 2.629 | 1.361 | 0.108 | 0.015 | 0.015 | 0.442 | 0.1 | 0.34 | 0.15 | 0.04 | 0.17 | 0.05 |
| 2 | 229533099 | rs11883439 | -1.768 | 0.993 | 2.845 | 1.359 | 0.071 | 0.005 | 0.005 | 0 | 0.08 | 0.24 | 0.14 | 0.04 | 0.17 | 0.05 |
| 2 | 229533885 | rs1881204 | -0.788 | 0.988 | 2.793 | 1.396 | 0.001 | 0 | 0 | 0 | 0.07 | 0.06 | 0.06 | 0.13 | 0.06 | 0.14 |
| 2 | 229545235 | rs6754248 | -2.914 | 1.053 | 2.966 | 1.379 | 0.054 | 0.008 | 0.008 | 0.483 | 0.19 | 0.36 | 0.21 | 0.17 | 0.26 | 0.15 |
| 2 | 229550456 | rs11890623 | -2.089 | 1.054 | 2.976 | 1.375 | 0.062 | -0.005 | -0.005 | 0 | 0.1 | 0.27 | 0.15 | 0.04 | 0.17 | 0.05 |
| 2 | 229555490 | rs16825257 | -2.243 | 1.046 | 2.798 | 1.383 | 0 | 0.002 | 0.002 | 0 | 0.14 | 0.12 | 0.11 | 0.25 | 0.15 | 0.22 |
| 2 | 229562892 | rs4973111 | -2.745 | 1.018 | 2.679 | 1.398 | 0.034 | 0.035 | 0.035 | 0 | 0.25 | 0.43 | 0.27 | 0.3 | 0.32 | 0.29 |
| 2 | 229572846 | rs10203187 | -2.932 | 0.894 | 2.558 | 1.367 | 0.011 | 0.015 | 0.015 | 0.007 | 0.27 | 0.51 | 0.28 | 0.3 | 0.32 | 0.29 |
| 2 | 229587738 | rs10177317 | -3.018 | 0.837 | 2.283 | 1.379 | 0 | 0 | 0 | 0.088 | 0.28 | 0.65 | 0.29 | 0.31 | 0.32 | 0.29 |
| 2 | 229590473 | rs10184640 | -2.42 | 0.907 | 2.259 | 1.271 | 0.015 | -0.003 | -0.003 | 0 | 0.21 | 0.62 | 0.23 | 0.18 | 0.26 | 0.15 |
| 2 | 229595305 | rs1609248 | -1.134 | 0.681 | 1.174 | 0.964 | -0.004 | 0.009 | 0.009 | 0 | 0.36 | 0.74 | 0.32 | 0.25 | 0.38 | 0.26 |
| 2 | 229600563 | rs1534458 | -1.706 | 1.254 | 0.995 | 1.202 | -0.062 | -0.061 | -0.061 | 0 | 0.45 | 0.89 | 0.37 | 0.42 | 0.47 | 0.36 |
| 2 | 229608271 | rs6722400 | -1.784 | 1.276 | 1.324 | 1.201 | -0.061 | -0.06 | -0.06 | 0 | 0.46 | 0.89 | 0.37 | 0.43 | 0.47 | 0.37 |
| 2 | 229609201 | rs6726331 | -1.425 | 1.282 | 2.04 | 1.045 | 0.076 | -0.073 | -0.073 | 0 | 0.14 | 0.37 | 0.15 | 0.02 | 0.21 | 0.06 |
| 2 | 229611155 | rs13390723 | -0.272 | 1.233 | 1.844 | 1.045 | -0.018 | 0.015 | 0.015 | 1.383 | 0.19 | 0.27 | 0.12 | 0.23 | 0.13 | 0.19 |
| 2 | 229616236 | rs6719964 | -1.678 | 1.214 | 1.876 | 1.038 | 0.017 | 0.023 | 0.023 | 0.016 | 0.12 | 0.18 | 0.09 | 0.18 | 0.13 | 0.11 |
| 2 | 229617535 | rs10451593 | -0.694 | 1.195 | 1.663 | 1.056 | -0.006 | 0.005 | 0.005 | 0 | 0.05 | 0.04 | 0.03 | 0.18 | 0.06 | 0.09 |
| 2 | 229631893 | rs2222720 | -0.244 | 1.187 | 1.685 | 1.05 | -0.034 | 0.035 | 0.035 | 1.417 | 0.19 | 0.32 | 0.12 | 0.23 | 0.13 | 0.2 |
| 2 | 229639826 | rs12053225 | -1.142 | 1.208 | 1.82 | 1.049 | 0.004 | 0.019 | 0.019 | 0.013 | 0.08 | 0.1 | 0.05 | 0.18 | 0.09 | 0.11 |
| 2 | 229641638 | rs7572812 | -1.199 | 1.177 | 1.879 | 1.032 | -0.017 | 0.064 | 0.064 | 0 | 0.26 | 0.38 | 0.17 | 0.41 | 0.22 | 0.3 |
| 2 | 229642422 | rs11679180 | -2.267 | 1.133 | 1.524 | 0.709 | -0.035 | -0.009 | -0.009 | 0 | 0.19 | 0.08 | 0.14 | 0.19 | 0.23 | 0.16 |
| 2 | 229652463 | rs1527669 | -1.025 | 1.065 | 1.667 | 0.684 | -0.016 | 0.051 | 0.051 | 0 | 0.26 | 0.35 | 0.17 | 0.37 | 0.22 | 0.27 |
| 2 | 229656721 | rs6707063 | -0.5 | 1.091 | 1.771 | 0.57 | 0.028 | -0.04 | -0.04 | 0 | 0.08 | 0.2 | 0.1 | 0 | 0.15 | 0.03 |
| 2 | 229663969 | rs776975 | 1.928 | 0.971 | 1.846 | 0.552 | 0.041 | -0.094 | -0.094 | 1.62 | 0.82 | 0.62 | 0.79 | 1 | 0.74 | 0.92 |
| 2 | 229664290 | rs776977 | 1.445 | 0.987 | 1.825 | 0.555 | 0.015 | -0.04 | -0.04 | 0.498 | 0.86 | 0.77 | 0.82 | 1 | 0.77 | 0.93 |
| 2 | 229669531 | rs16825387 | -1.351 | 0.912 | 1.708 | 0.661 | -0.027 | 0.048 | 0.048 | 0 | 0.3 | 0.51 | 0.21 | 0.38 | 0.25 | 0.31 |
| 2 | 229722957 | rs16825447 | -0.116 | 1.164 | 1.65 | 0.576 | 0.002 | -0.011 | -0.011 | 0.018 | 0.06 | 0.07 | 0.1 | 0 | 0.14 | 0.02 |
| 2 | 229726713 | rs13002075 | 1.391 | 1.099 | 1.768 | 0.49 | 0.045 | -0.081 | -0.081 | 0 | 0.84 | 0.69 | 0.79 | 1 | 0.76 | 0.87 |
| 2 | 229729917 | rs800736 | -2.877 | 0.879 | 1.551 | 0.41 | 0.016 | -0.03 | -0.03 | 0 | 0.31 | 0.61 | 0.33 | 0.14 | 0.37 | 0.24 |
| 2 | 229743009 | rs7578485 | -0.381 | 1.702 | 1.047 | 0.18 | -0.002 | 0.016 | 0.016 | 0 | 0.23 | 0.28 | 0.15 | 0.27 | 0.21 | 0.23 |
| 2 | 229745063 | rs2695689 | 0.545 | 1.747 | 0.997 | 0.163 | -0.034 | 0.03 | 0.03 | 0 | 0.07 | 0 | 0.09 | 0.01 | 0.08 | 0.07 |
| 2 | 229748368 | rs13428414 | -2.035 | 1.684 | 1.028 | 0.05 | -0.001 | -0.026 | -0.026 | 0 | 0.21 | 0.37 | 0.2 | 0.14 | 0.22 | 0.11 |
| 2 | 229750263 | rs800748 | -1.531 | 1.785 | 1.113 | 0.102 | 0.025 | -0.011 | -0.011 | 0.325 | 0.34 | 0.44 | 0.28 | 0.27 | 0.35 | 0.35 |
| 2 | 229753798 | rs9288639 | -1.869 | 1.699 | 1.012 | 0.102 | 0.063 | -0.034 | -0.034 | 0.045 | 0.09 | 0.39 | 0.08 | 0 | 0.07 | 0.06 |
| 2 | 229757034 | rs800726 | -2.843 | 1.681 | 0.623 | 0.069 | 0.037 | 0 | 0 | 0.003 | 0.52 | 0.82 | 0.47 | 0.41 | 0.57 | 0.47 |
| 2 | 229758040 | rs800727 | -0.4 | 1.522 | 0.643 | -0.032 | -0.013 | 0.005 | 0.005 | 0 | 0.13 | 0.11 | 0.18 | 0.01 | 0.21 | 0.1 |
| 2 | 229759328 | rs800729 | -2.307 | 1.559 | 0.532 | 0.027 | -0.03 | 0.066 | 0.066 | 0.008 | 0.43 | 0.76 | 0.39 | 0.27 | 0.46 | 0.36 |
| 2 | 229768415 | rs7593771 | -0.679 | 1.274 | 0.556 | 0.23 | -0.005 | 0.038 | 0.038 | 0.012 | 0.24 | 0.38 | 0.17 | 0.28 | 0.21 | 0.23 |
| 2 | 229778669 | rs7600415 | -1.064 | 0.787 | -1.57 | -0.08 | -0.039 | -0.122 | -0.122 | 0 | 0.63 | 0.95 | 0.48 | 0.6 | 0.61 | 0.46 |
| 2 | 229780970 | rs2396595 | -0.455 | 1.032 | -0.898 | -0.013 | 0.003 | 0.005 | 0.005 | 0 | 0.48 | 0.62 | 0.39 | 0.6 | 0.5 | 0.41 |
| 2 | 229783318 | rs10193759 | 0.275 | 1.351 | -0.713 | -0.224 | 0.026 | -0.057 | -0.057 | 0 | 0.13 | 0.23 | 0.16 | 0.01 | 0.14 | 0.15 |
| 2 | 229789967 | rs13387642 | 0.176 | 1.341 | -0.74 | -0.239 | 0.045 | -0.077 | -0.077 | 0 | 0.13 | 0.31 | 0.17 | 0.01 | 0.14 | 0.15 |
| 2 | 229797694 | rs10180563 | 0.301 | 1.358 | -0.844 | -0.164 | 0.005 | 0.003 | 0.003 | 0 | 0.1 | 0.22 | 0.06 | 0.09 | 0.09 | 0.08 |
| 2 | 229813769 | rs9808412 | 1.297 | 1.389 | -0.856 | -0.121 | -0.053 | 0.049 | 0.049 | 0 | 0.1 | 0.01 | 0.14 | 0.01 | 0.12 | 0.13 |
| 2 | 229822600 | rs6705266 | 0.918 | 0.894 | -0.582 | 0.307 | -0.014 | -0.047 | -0.047 | 0 | 0.25 | 0.16 | 0.17 | 0.46 | 0.29 | 0.19 |
| 2 | 229833153 | rs10203445 | 0.151 | 1.028 | -0.247 | 0.414 | -0.008 | -0.025 | -0.025 | 0.762 | 0.16 | 0.29 | 0.08 | 0.09 | 0.11 | 0.09 |
| 2 | 229841602 | rs10490036 | 0.63 | 1.471 | -0.434 | 0.803 | -0.052 | -0.054 | -0.054 | 0 | 0.11 | 0.04 | 0.18 | 0.28 | 0.22 | 0.27 |
| 2 | 229871093 | rs2160643 | 0.662 | 0.649 | -1.14 | 0.959 | 0.001 | -0.049 | -0.049 | 2.734 | 0.22 | 0.15 | 0.14 | 0.5 | 0.22 | 0.17 |
| 2 | 229872265 | rs3903945 | -0.845 | 0.625 | -1.407 | 0.703 | -0.028 | -0.049 | -0.049 | 0 | 0.75 | 0.95 | 0.51 | 0.69 | 0.71 | 0.57 |
| 2 | 229874279 | rs1544688 | -1.064 | 0.113 | -0.784 | 0.354 | 0.046 | 0.046 | 0.046 | 0 | 0.49 | 0.75 | 0.51 | 0.58 | 0.37 | 0.29 |
| 2 | 229876728 | rs2072556 | 0.762 | -0.068 | -0.922 | 0.427 | 0.019 | -0.061 | -0.061 | 0.21 | 0.18 | 0.03 | 0.16 | 0.36 | 0.11 | 0.14 |
| 2 | 229883422 | rs10211472 | -0.535 | -0.071 | -0.111 | 0.317 | 0 | 0.002 | 0.002 | 0 | 0.36 | 0.32 | 0.31 | 0.5 | 0.3 | 0.19 |
| 2 | 229883497 | rs7563172 | 0.891 | -0.048 | -0.07 | 0.336 | 0.014 | -0.006 | -0.006 | 0 | 0.09 | 0.09 | 0.19 | 0.1 | 0.08 | 0.06 |
| 2 | 229889356 | rs9646887 | 1.012 | 0.2 | -0.116 | 0.294 | -0.008 | 0.002 | 0.002 | 1.18 | 0.05 | 0.04 | 0.07 | 0 | 0.12 | 0.07 |
| 2 | 229889483 | rs3771286 | -0.766 | 0.133 | 0.122 | 0.455 | 0 | 0.001 | 0.001 | 0 | 0.44 | 0.51 | 0.38 | 0.51 | 0.42 | 0.27 |
| 2 | 229893149 | rs7594377 | -0.152 | 0.35 | -0.072 | 0.521 | -0.035 | 0 | 0 | 0 | 0.13 | 0.32 | 0.22 | 0.1 | 0.11 | 0.07 |
| 2 | 229897032 | rs10933273 | -0.559 | 1.05 | -0.763 | 0.6 | 0.004 | 0 | 0 | 0 | 0.16 | 0.15 | 0.11 | 0.13 | 0.17 | 0.07 |
| 2 | 229906252 | rs2109818 | -0.447 | 0.773 | -1.059 | 0.475 | -0.01 | 0.071 | 0.071 | 0 | 0.55 | 0.77 | 0.65 | 0.64 | 0.52 | 0.44 |
| 2 | 229908245 | rs2215598 | -1.102 | 0.434 | -0.039 | 0.495 | -0.037 | -0.059 | -0.059 | 0.486 | 0.15 | 0.36 | 0.07 | 0.04 | 0.06 | 0.16 |
| 2 | 229909492 | rs13010266 | -0.312 | 1.45 | -0.756 | 1.065 | -0.051 | -0.068 | -0.068 | 3.818 | 0.22 | 0.07 | 0.33 | 0.47 | 0.36 | 0.15 |
| 2 | 229917808 | rs13034774 | -0.623 | 1.23 | -0.682 | 0.986 | -0.061 | -0.066 | -0.066 | 1.817 | 0.27 | 0.11 | 0.37 | 0.51 | 0.39 | 0.27 |
| 2 | 229922254 | rs31278 | 1.186 | 0.411 | 0.509 | 0.611 | -0.018 | -0.046 | -0.046 | 0.678 | 0.86 | 0.56 | 0.93 | 0.97 | 0.94 | 0.9 |
| 2 | 229927173 | rs12105612 | -1.773 | 0.333 | 1.269 | 1.131 | 0.032 | 0.212 | 0.212 | 4.37 | 0.08 | 0.51 | 0.26 | 0.13 | 0.04 | 0.02 |
| 2 | 229929352 | rs7563963 | -1.006 | 0.341 | 1.136 | 1.12 | -0.003 | -0.007 | -0.007 | 0 | 0.08 | 0.12 | 0.05 | 0.03 | 0.06 | 0.1 |
| 2 | 229940635 | rs1860762 | 0.13 | 1.342 | 0.463 | 1.523 | -0.07 | -0.091 | -0.091 | 0 | 0.85 | 0.97 | 0.77 | 0.45 | 0.76 | 0.67 |
| 2 | 229942970 | rs2191835 | 1.54 | 0.83 | 0.869 | 1.2 | 0.017 | 0.015 | 0.015 | 0 | 0.67 | 0.52 | 0.67 | 0.39 | 0.66 | 0.52 |

| | | | | | | | | | | | | | | | | |
|---|---|---|---|---|---|---|---|---|---|---|---|---|---|---|---|---|
| 2 | 229945688 | rs13020025 | -0.178 | 1.038 | 0.546 | 1.335 | -0.096 | -0.123 | -0.123 | 0 | 0.15 | 0.02 | 0.23 | 0.55 | 0.24 | 0.32 |
| 2 | 229951719 | rs2014401 | -2.096 | 0.556 | 1.66 | 1.402 | -0.013 | 0.097 | 0.097 | 0 | 0.15 | 0.51 | 0.31 | 0.16 | 0.09 | 0.15 |
| 2 | 229953526 | rs16825679 | -1.168 | 1.006 | 1.743 | 1.179 | 0.092 | 0.17 | 0.17 | 0 | 0.1 | 0.32 | 0.3 | 0.28 | 0.17 | 0.12 |
| 2 | 229956318 | rs990672 | -2.151 | 1.006 | 1.529 | 1.152 | -0.002 | -0.01 | -0.01 | 0 | 0.22 | 0.74 | 0.39 | 0.32 | 0.23 | 0.24 |
| 2 | 229957003 | rs2109820 | 0.238 | 1.1 | 1.261 | 1.043 | 0.078 | 0.05 | 0.05 | 0 | 0.92 | 0.77 | 0.92 | 0.83 | 0.86 | 0.9 |
| 2 | 229963404 | rs17676036 | -0.772 | 0.296 | 0.433 | 1.072 | 0.006 | -0.005 | -0.005 | 0 | 0.11 | 0.1 | 0.06 | 0.18 | 0.07 | 0.12 |
| 2 | 229964444 | rs4972886 | 0.761 | -0.076 | 0.708 | 1.227 | -0.033 | 0.014 | 0.014 | 0 | 0.87 | 0.85 | 0.95 | 0.82 | 0.94 | 0.81 |
| 2 | 229969798 | rs6436839 | -0.682 | 0.605 | 0.884 | 1.8 | 0.017 | 0.047 | 0.047 | 0 | 0.23 | 0.38 | 0.31 | 0.38 | 0.26 | 0.34 |
| 2 | 229973786 | rs3845826 | -1.208 | 0.793 | 0.668 | 1.629 | 0.02 | -0.004 | -0.004 | 0.708 | 0.32 | 0.38 | 0.6 | 0.72 | 0.41 | 0.45 |
| 2 | 229974922 | rs2765936 | 0.324 | 0.397 | 0.552 | 1.607 | -0.034 | 0.012 | 0.012 | 0 | 0.87 | 0.85 | 0.95 | 0.83 | 0.94 | 0.81 |
| 2 | 229975243 | rs17614019 | -1.114 | 0.594 | 0.546 | 1.825 | -0.002 | 0.033 | 0.033 | 0 | 0.31 | 0.36 | 0.34 | 0.53 | 0.31 | 0.46 |
| 2 | 229978470 | rs10451556 | -1.21 | 0.307 | 0.44 | 1.906 | -0.018 | 0.032 | 0.032 | 0 | 0.11 | 0.16 | 0.08 | 0.22 | 0.07 | 0.25 |
| 2 | 229981613 | rs10490031 | 0.192 | 1.31 | -0.097 | 2.331 | -0.065 | -0.062 | -0.062 | 0.229 | 0.11 | 0.02 | 0.41 | 0.38 | 0.21 | 0.11 |
| 2 | 229987952 | rs13018035 | -0.148 | 0.552 | -0.631 | 1.176 | -0.017 | -0.023 | -0.023 | 0 | 0.62 | 0.54 | 0.76 | 0.73 | 0.66 | 0.64 |
| 2 | 229992221 | rs2159724 | 0.614 | 0.495 | -0.366 | 1.099 | 0.005 | 0 | 0 | 0 | 0.52 | 0.53 | 0.68 | 0.5 | 0.59 | 0.41 |
| 2 | 229995411 | rs6436842 | -1.342 | 0.012 | -0.551 | 0.726 | 0.012 | -0.061 | -0.061 | 0.327 | 0.85 | 0.8 | 0.83 | 1 | 0.76 | 0.92 |
| 2 | 229997075 | rs6745202 | 1.331 | 0.035 | -0.611 | 0.805 | 0.043 | 0.01 | 0.01 | 0 | 0.5 | 0.36 | 0.67 | 0.5 | 0.57 | 0.4 |
| 2 | 229999532 | rs17676147 | 0.114 | -0.058 | -0.801 | 0.919 | 0.015 | -0.045 | -0.045 | 0.507 | 0.1 | 0.01 | 0.06 | 0.21 | 0.07 | 0.24 |
| 2 | 230002606 | rs4973160 | -1.583 | 0.226 | -0.637 | 0.777 | 0.008 | 0.043 | 0.043 | 0 | 0.55 | 0.39 | 0.71 | 0.87 | 0.61 | 0.69 |
| 2 | 230006859 | rs2023499 | -0.94 | -0.151 | -0.597 | 0.852 | 0.019 | -0.02 | -0.02 | 0 | 0.3 | 0.25 | 0.14 | 0.49 | 0.18 | 0.51 |
| 2 | 230014441 | rs1345281 | -1.322 | -0.012 | -0.541 | 0.833 | -0.005 | -0.001 | -0.001 | 0 | 0.47 | 0.49 | 0.31 | 0.51 | 0.44 | 0.62 |
| 2 | 230015954 | rs17676208 | 0.043 | -0.034 | -1.073 | 1.011 | 0.016 | -0.04 | -0.04 | 0 | 0.14 | 0.09 | 0.05 | 0.31 | 0.09 | 0.29 |
| 2 | 230020507 | rs17614185 | 0.152 | -0.001 | -1.049 | 1.04 | 0.011 | -0.028 | -0.028 | 0 | 0.13 | 0.08 | 0.05 | 0.26 | 0.09 | 0.28 |
| 2 | 230021166 | rs2396615 | -0.883 | 0.203 | -0.852 | 0.864 | -0.001 | -0.002 | -0.002 | 0 | 0.28 | 0.31 | 0.49 | 0.69 | 0.26 | 0.41 |
| 2 | 230025610 | rs4973166 | -2.016 | 0.093 | -0.961 | 0.354 | -0.002 | -0.036 | -0.036 | 0 | 0.6 | 0.49 | 0.75 | 0.88 | 0.63 | 0.75 |
| 2 | 230029360 | rs1419956 | -0.55 | 0.066 | -1.025 | 0.334 | 0.006 | 0.03 | 0.03 | 0 | 0.33 | 0.28 | 0.27 | 0.18 | 0.36 | 0.34 |
| 2 | 230030405 | rs4973171 | -0.861 | 0.11 | -1.182 | 0.521 | -0.058 | -0.059 | -0.059 | 0.015 | 0.41 | 0.21 | 0.65 | 0.52 | 0.51 | 0.48 |
| 2 | 230031082 | rs10171051 | 1.549 | 0.088 | -1.295 | 0.438 | -0.008 | 0.007 | 0.007 | 0 | 0.16 | 0.15 | 0.18 | 0 | 0.25 | 0.07 |
| 2 | 230032129 | rs10804349 | -0.026 | 0.051 | -1.21 | 0.444 | 0 | 0.003 | 0.003 | 0 | 0.18 | 0.15 | 0.08 | 0.34 | 0.11 | 0.29 |
| 2 | 230033560 | rs1419958 | 1.705 | 0.194 | -1.375 | 0.416 | -0.032 | -0.023 | -0.023 | 0.881 | 0.09 | 0.04 | 0.39 | 0.34 | 0.15 | 0.1 |
| 2 | 230033996 | rs1419959 | 0.822 | -0.323 | -1.215 | 0.593 | -0.06 | -0.05 | -0.05 | 0 | 0.25 | 0.16 | 0.57 | 0.36 | 0.4 | 0.17 |
| 2 | 230034082 | rs12477036 | 0.151 | -0.358 | -1.18 | 0.664 | -0.005 | -0.037 | -0.037 | 1.483 | 0.15 | 0.01 | 0.08 | 0.18 | 0.11 | 0.25 |
| 2 | 230039041 | rs1362256 | 0.386 | -0.187 | -1.139 | 0.571 | -0.104 | -0.036 | -0.036 | 0.097 | 0.3 | 0.13 | 0.59 | 0.32 | 0.44 | 0.26 |
| 2 | 230039817 | rs13013873 | 1.12 | -0.371 | -0.871 | -0.023 | -0.003 | -0.046 | -0.046 | 0 | 0.39 | 0.4 | 0.25 | 0.11 | 0.35 | 0.26 |
| 2 | 230042560 | rs1362260 | -0.616 | -0.327 | -1.392 | 0.199 | 0.002 | -0.003 | -0.003 | 0.064 | 0.59 | 0.42 | 0.46 | 0.67 | 0.51 | 0.71 |
| 2 | 230046668 | rs2109819 | 0.675 | -0.393 | -1.556 | 0.597 | 0.005 | 0.014 | 0.014 | 0 | 0.11 | 0.09 | 0.14 | 0.4 | 0.08 | 0.2 |
| 2 | 230048263 | rs7588090 | -0.27 | -0.424 | -1.605 | 0.515 | 0.009 | 0.002 | 0.002 | 0 | 0.86 | 0.95 | 0.92 | 0.85 | 0.89 | 0.73 |
| 2 | 230056151 | rs7585821 | 0.67 | -0.378 | -1.564 | 0.578 | 0.005 | 0.012 | 0.012 | 0 | 0.12 | 0.09 | 0.15 | 0.41 | 0.08 | 0.2 |
| 2 | 230058860 | rs1362238 | -1.16 | -0.24 | -1.573 | -0.008 | -0.002 | 0.015 | 0.015 | 0 | 0.73 | 0.73 | 0.81 | 0.89 | 0.72 | 0.77 |
| 2 | 230059595 | rs10184838 | 0.455 | -0.161 | -1.473 | 0.119 | -0.001 | 0.003 | 0.003 | 0.102 | 0.46 | 0.52 | 0.58 | 0.33 | 0.53 | 0.3 |
| 2 | 230065011 | rs6736122 | 0.268 | 0.406 | -1.652 | -0.164 | -0.009 | 0.055 | 0.055 | 0.025 | 0.3 | 0.19 | 0.2 | 0.11 | 0.31 | 0.27 |
| 2 | 230073155 | rs16825867 | -0.257 | 0.076 | -2.02 | -0.347 | 0.051 | -0.001 | -0.001 | 0 | 0.4 | 0.17 | 0.35 | 0.66 | 0.29 | 0.46 |
| 2 | 230074837 | rs17254849 | -0.555 | 0.412 | -2.405 | -0.718 | 0.022 | 0.106 | 0.106 | 0.433 | 0.52 | 0.24 | 0.57 | 0.76 | 0.47 | 0.65 |
| 2 | 230076347 | rs1362240 | 0.793 | 1.261 | -2.815 | -1.019 | -0.055 | -0.063 | -0.063 | 0.053 | 0.1 | 0.01 | 0.2 | 0.2 | 0.18 | 0.16 |
| 2 | 230081043 | rs883731 | 0.756 | 0.708 | -2.956 | -1.444 | 0.03 | 0.072 | 0.072 | 2.062 | 0.51 | 0.11 | 0.39 | 0.69 | 0.39 | 0.47 |
| 2 | 230092137 | rs6738963 | -0.068 | 0.438 | -1.539 | -1.364 | -0.002 | -0.012 | -0.012 | 0 | 0.45 | 0.46 | 0.4 | 0.24 | 0.53 | 0.35 |
| 2 | 230094437 | rs10933281 | -0.185 | 0.461 | -1.912 | -1.353 | 0.012 | 0.067 | 0.067 | 0.06 | 0.54 | 0.3 | 0.58 | 0.77 | 0.47 | 0.65 |
| 2 | 230096093 | rs1226981 | 0.792 | 0.458 | -2.029 | -1.369 | 0.055 | -0.006 | -0.006 | 0 | 0.38 | 0.13 | 0.34 | 0.68 | 0.28 | 0.44 |
| 2 | 230100791 | rs1226988 | 0.063 | 0.406 | -1.654 | -1.355 | -0.008 | -0.029 | -0.029 | 0.258 | 0.83 | 0.74 | 0.76 | 0.93 | 0.8 | 0.78 |
| 2 | 230103371 | rs6747101 | 0.107 | 0.563 | -1.571 | -1.61 | 0.005 | -0.025 | -0.025 | 0 | 0.42 | 0.51 | 0.38 | 0.23 | 0.48 | 0.33 |
| 2 | 230103927 | rs1226993 | 0.772 | 0.354 | -1.769 | -1.755 | 0.039 | 0.012 | 0.012 | 0 | 0.4 | 0.12 | 0.49 | 0.7 | 0.35 | 0.45 |
| 2 | 230109580 | rs1226997 | -0.406 | 0.394 | -1.693 | -1.75 | -0.001 | 0.001 | 0.001 | 0 | 0.57 | 0.41 | 0.6 | 0.77 | 0.51 | 0.67 |
| 2 | 230116035 | rs10498235 | 0.416 | 0.445 | -1.708 | -1.775 | 0 | 0.013 | 0.013 | 0.377 | 0.15 | 0.08 | 0.11 | 0.07 | 0.17 | 0.21 |
| 2 | 230117059 | rs6739881 | 0.71 | 0.909 | -2.043 | -2.093 | -0.051 | 0.092 | 0.092 | 0 | 0.28 | 0.09 | 0.18 | 0.08 | 0.34 | 0.28 |
| 2 | 230119278 | rs1227003 | -1.473 | 0.156 | -2.059 | -2.151 | 0.018 | 0.025 | 0.025 | 0 | 0.87 | 0.94 | 0.95 | 0.99 | 0.91 | 0.92 |
| 2 | 230122247 | rs6724020 | 1.043 | 0.186 | -1.975 | -2.181 | 0.016 | -0.028 | -0.028 | 0 | 0.34 | 0.51 | 0.33 | 0.23 | 0.34 | 0.28 |
| 2 | 230122385 | rs1227005 | 0.502 | 0.476 | -1.96 | -1.811 | 0.005 | -0.016 | -0.016 | 0 | 0.32 | 0.13 | 0.45 | 0.7 | 0.3 | 0.43 |
| 2 | 230129493 | rs4321351 | 0.118 | 0.545 | -1.559 | -1.799 | -0.022 | 0.022 | 0.022 | 0.475 | 0.19 | 0.09 | 0.25 | 0.1 | 0.28 | 0.24 |
| 2 | 230131445 | rs6754229 | 0.079 | 0.258 | -1.319 | -1.725 | 0.005 | 0 | 0 | 0 | 0.19 | 0.17 | 0.13 | 0 | 0.16 | 0.07 |
| 2 | 230146591 | rs7579884 | 0.456 | 0.139 | -1.001 | -1.489 | -0.01 | -0.043 | -0.043 | 0.899 | 0.13 | 0.2 | 0.06 | 0 | 0.07 | 0.05 |
| 2 | 230152783 | rs1226940 | -0.063 | 0.048 | -0.579 | -1.721 | 0.006 | 0.021 | 0.021 | 0 | 0.25 | 0.17 | 0.14 | 0.07 | 0.23 | 0.19 |
| 2 | 230166647 | rs4973184 | 1.001 | 0.188 | -0.091 | -1.685 | -0.015 | 0.039 | 0.039 | 0.015 | 0.42 | 0.32 | 0.57 | 0.84 | 0.49 | 0.61 |
| 2 | 230177257 | rs13389321 | -0.144 | -0.069 | 0.545 | -1.28 | -0.009 | -0.008 | -0.008 | 0 | 0.12 | 0.47 | 0.07 | 0.07 | 0.06 | 0.02 |
| 2 | 230179536 | rs13383306 | 0.758 | -0.008 | 0.176 | -1.226 | -0.006 | -0.002 | -0.002 | 0 | 0.09 | 0.21 | 0.06 | 0.07 | 0.06 | 0.02 |
| 2 | 230197262 | rs6436859 | -0.572 | 0.221 | -0.521 | -0.986 | -0.051 | 0.009 | 0.009 | 0.002 | 0.83 | 0.96 | 0.78 | 0.91 | 0.78 | 0.86 |
| 2 | 230198136 | rs12621121 | 1.511 | -0.148 | -0.538 | -1.32 | -0.047 | 0.093 | 0.093 | 0 | 0.41 | 0.16 | 0.52 | 0.81 | 0.48 | 0.61 |
| 2 | 230202430 | rs6741898 | 0.316 | -0.604 | -0.591 | -1.243 | -0.041 | 0.158 | 0.158 | 0.272 | 0.59 | 0.86 | 0.61 | 0.88 | 0.54 | 0.64 |
| 2 | 230203242 | rs13394622 | 0.49 | -0.599 | -0.354 | -1.211 | -0.003 | 0.065 | 0.065 | 0.239 | 0.57 | 0.72 | 0.6 | 0.8 | 0.54 | 0.64 |
| 2 | 230208956 | rs4973189 | 0.3 | -0.529 | 0.084 | -0.905 | -0.05 | 0.096 | 0.096 | 2.034 | 0.68 | 0.87 | 0.77 | 0.91 | 0.62 | 0.69 |

| | | | | | | | | | | | | | | | | |
|---|---|---|---|---|---|---|---|---|---|---|---|---|---|---|---|---|
| 2 | 230210521 | rs6717477 | 1.84 | -0.811 | 0.133 | -0.438 | -0.018 | 0.035 | 0.035 | 0.677 | 0.41 | 0.16 | 0.4 | 0.39 | 0.44 | 0.38 |
| 2 | 230211114 | rs6756789 | 0.803 | -0.807 | 0.361 | -0.431 | -0.017 | -0.063 | -0.063 | 1.055 | 0.12 | 0.22 | 0.08 | 0 | 0.08 | 0.03 |
| 2 | 230211434 | rs4973190 | -0.289 | -0.834 | 0.378 | -0.44 | -0.008 | 0.049 | 0.049 | 8.127 | 0.19 | 0.25 | 0.32 | 0.52 | 0.16 | 0.3 |
| 2 | 230214932 | rs9288645 | 1.478 | -0.447 | 0.702 | 0.095 | -0.001 | -0.001 | -0.001 | 0 | 0.54 | 0.53 | 0.48 | 0.39 | 0.52 | 0.42 |
| 2 | 230216155 | rs888283 | -1.024 | -0.259 | 0.085 | -0.048 | -0.066 | 0.048 | 0.048 | 0.178 | 0.85 | 0.99 | 0.89 | 0.95 | 0.81 | 0.86 |
| 2 | 230220266 | rs1003817 | -0.295 | -0.307 | 0.175 | -0.005 | -0.026 | 0.064 | 0.064 | 2.828 | 0.77 | 0.91 | 0.84 | 0.94 | 0.75 | 0.79 |
| 2 | 230224469 | rs6436860 | 0.532 | -0.411 | 0.229 | 0.004 | -0.035 | 0.088 | 0.088 | 1.875 | 0.67 | 0.85 | 0.78 | 0.91 | 0.63 | 0.7 |
| 2 | 230234526 | rs2396644 | -0.323 | -0.329 | 0.152 | 0.037 | -0.024 | 0.077 | 0.077 | 1.196 | 0.75 | 0.91 | 0.83 | 0.94 | 0.76 | 0.8 |
| 2 | 230237477 | rs4973194 | 0.285 | -0.423 | 0.327 | 0.132 | -0.008 | 0.019 | 0.019 | 1.246 | 0.64 | 0.74 | 0.78 | 0.91 | 0.63 | 0.67 |
| 2 | 230240232 | rs933602 | 0.338 | -0.228 | 0.569 | 0.641 | -0.007 | 0.021 | 0.021 | 5.8 | 0.08 | 0.03 | 0.25 | 0.53 | 0.08 | 0.29 |
| 2 | 230253849 | rs207667 | 0.161 | -0.248 | 0.348 | 0.544 | -0.005 | 0.001 | 0.001 | 2.347 | 0.67 | 0.74 | 0.8 | 0.91 | 0.64 | 0.72 |
| 2 | 230256586 | rs207671 | 0.742 | -0.167 | 0.01 | 0.538 | -0.002 | 0.003 | 0.003 | 1.553 | 0.56 | 0.45 | 0.52 | 0.39 | 0.56 | 0.43 |
| 2 | 230270559 | rs10164537 | -0.792 | -0.273 | -0.151 | 1.212 | 0.002 | 0.01 | 0.01 | 0.904 | 0.13 | 0.09 | 0.27 | 0.49 | 0.11 | 0.31 |
| 2 | 230270746 | rs10184352 | -1.075 | -0.292 | -0.157 | 1.215 | 0.002 | 0 | 0 | 0 | 0.14 | 0.12 | 0.28 | 0.5 | 0.13 | 0.29 |
| 2 | 230270946 | rs973322 | 0.89 | -0.611 | 0.132 | 0.911 | -0.013 | 0.006 | 0.006 | 0 | 0.68 | 0.41 | 0.83 | 0.65 | 0.77 | 0.63 |
| 2 | 230271297 | rs10933295 | -0.31 | -0.49 | 0.028 | 0.87 | 0 | -0.01 | -0.01 | 0 | 0.17 | 0.15 | 0.3 | 0.64 | 0.16 | 0.27 |
| 2 | 230272648 | rs6712266 | 0.491 | -0.447 | 0.004 | 0.457 | 0.009 | 0.072 | 0.072 | 0 | 0.37 | 0.21 | 0.21 | 0.09 | 0.33 | 0.35 |
| 2 | 230273403 | rs12694811 | 0.313 | -0.341 | -0.637 | 0.528 | 0.011 | 0.044 | 0.044 | 0 | 0.39 | 0.27 | 0.22 | 0.09 | 0.35 | 0.35 |
| 2 | 230276998 | rs6731881 | -0.071 | 0.143 | -1.691 | 0.985 | -0.001 | -0.02 | -0.02 | 0 | 0.83 | 0.88 | 0.8 | 0.48 | 0.84 | 0.72 |
| 2 | 230277144 | rs6732014 | 0.517 | 0.097 | -1.669 | 0.377 | -0.052 | 0.058 | 0.058 | 0.78 | 0.4 | 0.2 | 0.44 | 0.29 | 0.52 | 0.37 |
| 2 | 230278731 | rs6739440 | 0.792 | 0.255 | -1.106 | 1.522 | -0.001 | -0.019 | -0.019 | 0 | 0.83 | 0.88 | 0.69 | 0.36 | 0.84 | 0.72 |
| 2 | 230283954 | rs6738575 | -0.485 | -0.137 | -1.06 | 1.243 | 0.002 | -0.004 | -0.004 | 0 | 0.1 | 0.09 | 0.04 | 0.15 | 0.04 | 0.15 |
| 2 | 230284815 | rs2396663 | -0.162 | 0.045 | -1.842 | 0.65 | -0.047 | 0.068 | 0.068 | 0 | 0.19 | 0.02 | 0.12 | 0.06 | 0.17 | 0.23 |
| 2 | 230285770 | rs10192937 | 0.07 | -0.424 | -1.907 | 0.876 | -0.011 | 0.022 | 0.022 | 0.924 | 0.27 | 0.65 | 0.36 | 0.46 | 0.22 | 0.23 |
| 2 | 230288990 | rs7580533 | 0.41 | -0.271 | -2.099 | 0.501 | 0.066 | -0.034 | -0.034 | 0.24 | 0.61 | 0.11 | 0.58 | 0.39 | 0.72 | 0.6 |
| 2 | 230297454 | rs12466167 | -0.594 | -0.271 | -1.3 | 0.359 | 0.015 | 0.047 | 0.047 | 0.052 | 0.47 | 0.3 | 0.47 | 0.71 | 0.39 | 0.6 |
| 2 | 230308906 | rs12694814 | 0.407 | -0.247 | -0.751 | 0.435 | 0.008 | 0.023 | 0.023 | 1.782 | 0.53 | 0.39 | 0.52 | 0.31 | 0.61 | 0.41 |
| 2 | 230308989 | rs12694815 | 0.703 | -0.274 | -1.203 | 0.41 | -0.042 | 0.068 | 0.068 | 5.141 | 0.46 | 0.12 | 0.47 | 0.31 | 0.56 | 0.4 |
| 2 | 230315858 | rs12611671 | 0.637 | 0.261 | -1.335 | 0.676 | -0.032 | -0.037 | -0.037 | 0 | 0.2 | 0.12 | 0.27 | 0.34 | 0.29 | 0.16 |
| 2 | 230319228 | rs1966469 | 0.345 | 0.001 | -0.775 | 0.819 | 0.001 | 0 | 0 | 0 | 0.58 | 0.45 | 0.58 | 0.44 | 0.55 | 0.6 |
| 2 | 230322544 | rs7572143 | -1.054 | 0.105 | -0.642 | 0.844 | -0.011 | -0.013 | -0.013 | 0 | 0.8 | 0.81 | 0.73 | 0.66 | 0.71 | 0.84 |
| 2 | 230325712 | rs16826047 | 0.977 | -0.025 | -1.16 | 0.649 | -0.008 | -0.007 | -0.007 | 0 | 0.32 | 0.34 | 0.34 | 0.25 | 0.25 | 0.4 |
| 2 | 230326068 | rs11675728 | -0.472 | 0.168 | -1.523 | 0.656 | -0.025 | -0.007 | -0.007 | 0 | 0.43 | 0.19 | 0.34 | 0.4 | 0.42 | 0.44 |
| 2 | 230329343 | rs17483944 | -0.063 | 0.226 | -1.172 | 0.483 | -0.025 | 0.033 | 0.033 | 0 | 0.06 | 0 | 0.03 | 0 | 0.07 | 0.07 |
| 2 | 230337831 | rs4973206 | 0.242 | -0.262 | -0.892 | 0.718 | -0.018 | 0.016 | 0.016 | 0.253 | 0.33 | 0.38 | 0.34 | 0.41 | 0.24 | 0.45 |
| 2 | 230343104 | rs12464136 | 0.424 | -0.477 | -0.727 | 0.528 | -0.002 | 0.026 | 0.026 | 0.075 | 0.31 | 0.23 | 0.38 | 0.16 | 0.29 | 0.32 |
| 2 | 230344697 | rs1013437 | 0.578 | -0.479 | -0.697 | 0.534 | -0.004 | 0.026 | 0.026 | 0 | 0.29 | 0.2 | 0.36 | 0.16 | 0.27 | 0.31 |
| 2 | 230348025 | rs6721342 | 0.763 | -0.597 | -1.111 | 0.341 | -0.003 | 0.018 | 0.018 | 0.065 | 0.15 | 0.06 | 0.11 | 0.1 | 0.14 | 0.22 |
| 2 | 230348440 | rs17620481 | -0.097 | -0.846 | -1.309 | 0.466 | 0.016 | -0.038 | -0.038 | 0.179 | 0.19 | 0.15 | 0.11 | 0.38 | 0.16 | 0.28 |
| 2 | 230349801 | rs2193906 | 1.245 | -0.789 | -1.283 | 0.469 | 0.001 | 0.029 | 0.029 | 0.092 | 0.3 | 0.22 | 0.37 | 0.15 | 0.28 | 0.3 |
| 2 | 230350848 | rs977041 | -0.328 | -0.895 | -1.103 | 0.446 | 0.005 | 0.009 | 0.009 | 0 | 0.52 | 0.48 | 0.52 | 0.56 | 0.47 | 0.6 |
| 2 | 230351558 | rs12620995 | 2.136 | -0.987 | -1.458 | 0.257 | 0.016 | -0.028 | -0.028 | 0 | 0.08 | 0.02 | 0.13 | 0.27 | 0.05 | 0.07 |
| 2 | 230360087 | rs7597072 | 0.092 | -0.837 | -0.7 | -0.014 | 0.005 | 0.059 | 0.059 | 0 | 0.51 | 0.59 | 0.51 | 0.26 | 0.44 | 0.54 |
| 2 | 230360787 | rs12052301 | 1.204 | -0.805 | -1.289 | 0.043 | -0.007 | 0.019 | 0.019 | 0.084 | 0.15 | 0.03 | 0.11 | 0.1 | 0.14 | 0.22 |
| 2 | 230363126 | rs10187046 | 0.423 | -0.788 | -1.203 | 0.251 | 0.001 | -0.03 | -0.03 | 0 | 0.16 | 0.09 | 0.11 | 0.22 | 0.18 | 0.17 |
| 2 | 230363167 | rs13007913 | 1.154 | -0.47 | -1.073 | 0.231 | -0.001 | -0.001 | -0.001 | 0 | 0.29 | 0.28 | 0.36 | 0.15 | 0.27 | 0.3 |
| 2 | 230368047 | rs759791 | -0.349 | -0.172 | -1.472 | 0.371 | -0.092 | -0.096 | -0.096 | 0.24 | 0.66 | 0.9 | 0.6 | 0.48 | 0.56 | 0.67 |
| 2 | 230368876 | rs973440 | -0.126 | -0.072 | -1.073 | 0.245 | -0.003 | -0.005 | -0.005 | 0 | 0.42 | 0.42 | 0.44 | 0.32 | 0.36 | 0.44 |
| 2 | 230370114 | rs10197959 | -0.035 | 0.294 | -1.521 | 0.108 | -0.006 | 0.045 | 0.045 | 0 | 0.37 | 0.24 | 0.39 | 0.25 | 0.43 | 0.38 |
| 2 | 230372314 | rs13033846 | 0.515 | 0.482 | -1.901 | 0.193 | -0.045 | 0.045 | 0.045 | 0 | 0.25 | 0.07 | 0.31 | 0.17 | 0.31 | 0.28 |
| 2 | 230373798 | rs10193281 | 0.55 | 0.447 | -1.645 | 0.426 | 0.004 | -0.014 | -0.014 | 0.011 | 0.23 | 0.16 | 0.19 | 0.26 | 0.17 | 0.31 |
| 2 | 230376066 | rs6755403 | 0.488 | 0.841 | -1.261 | 0.291 | 0.002 | -0.003 | -0.003 | 0.3 | 0.18 | 0.2 | 0.18 | 0.14 | 0.23 | 0.23 |
| 2 | 230378541 | rs6735195 | -0.366 | 0.57 | -1.215 | 0.582 | -0.008 | 0.021 | 0.021 | 0 | 0.12 | 0.27 | 0.1 | 0.13 | 0.12 | 0.12 |
| 2 | 230379379 | rs4973212 | 0.235 | 0.899 | -1.565 | -0.058 | -0.048 | 0.08 | 0.08 | 0 | 0.31 | 0.11 | 0.27 | 0.16 | 0.36 | 0.33 |
| 2 | 230379524 | rs13010630 | -0.306 | 0.779 | -1.482 | 0.438 | -0.008 | 0.033 | 0.033 | 0 | 0.72 | 0.63 | 0.68 | 0.5 | 0.76 | 0.79 |
| 2 | 230383826 | rs17581255 | -0.178 | 1.303 | -2.04 | -0.123 | -0.069 | 0.07 | 0.07 | 0 | 0.29 | 0.11 | 0.27 | 0.16 | 0.37 | 0.36 |
| 2 | 230384365 | rs11695055 | -0.113 | 0.543 | -2.279 | -0.408 | -0.037 | 0.215 | 0.215 | 0.822 | 0.45 | 0.13 | 0.4 | 0.13 | 0.41 | 0.4 |
| 2 | 230386043 | rs958110 | 0.544 | 0.509 | -0.995 | 0.474 | -0.004 | -0.006 | -0.006 | 0.001 | 0.23 | 0.18 | 0.33 | 0.33 | 0.3 | 0.16 |
| 2 | 230396934 | rs7589154 | -0.119 | -0.369 | -0.305 | 0.194 | -0.069 | -0.034 | -0.034 | 0 | 0.23 | 0.47 | 0.19 | 0.13 | 0.09 | 0.12 |
| 2 | 230398622 | rs13013785 | 1.238 | -0.625 | -0.482 | 0.184 | -0.019 | -0.017 | -0.017 | 0.797 | 0.3 | 0.07 | 0.29 | 0.37 | 0.3 | 0.39 |
| 2 | 230401082 | rs7584721 | -0.316 | -0.644 | 0.233 | 0.132 | -0.092 | -0.032 | -0.032 | 0 | 0.23 | 0.48 | 0.18 | 0.13 | 0.08 | 0.13 |
| 2 | 230418426 | rs1477111 | -0.702 | 0.222 | 0.442 | 0.118 | 0.01 | -0.031 | -0.031 | 0 | 0.87 | 0.81 | 0.85 | 0.94 | 0.77 | 0.78 |
| 2 | 230451886 | rs7604611 | -0.018 | -0.181 | -0.334 | 0.22 | -0.053 | -0.033 | -0.033 | 0 | 0.73 | 0.61 | 0.81 | 0.83 | 0.88 | 0.84 |
| 2 | 230453614 | rs17581619 | 0.554 | 0.669 | -1.245 | 0.759 | -0.073 | -0.056 | -0.056 | 0 | 0.22 | 0.11 | 0.32 | 0.28 | 0.33 | 0.24 |
| 2 | 230459651 | rs6728781 | -0.499 | -0.173 | -0.709 | 0.465 | 0.009 | 0.047 | 0.047 | 0 | 0.07 | 0.24 | 0.11 | 0.12 | 0.03 | 0.03 |
| 2 | 230461591 | rs10193150 | -0.034 | -0.344 | -0.894 | 0.538 | 0.043 | 0.021 | 0.021 | 0 | 0.36 | 0.17 | 0.33 | 0.37 | 0.32 | 0.38 |
| 2 | 230462275 | rs1861616 | -0.239 | -0.23 | -0.741 | 0.45 | 0.031 | 0.003 | 0.003 | 0 | 0.42 | 0.3 | 0.43 | 0.49 | 0.35 | 0.42 |
| 2 | 230468480 | rs10933305 | 0.087 | -0.187 | -1.347 | 0.469 | 0.057 | 0.009 | 0.009 | 1.165 | 0.56 | 0.75 | 0.58 | 0.51 | 0.64 | 0.56 |
| 2 | 230480924 | rs11898749 | -0.807 | 0.067 | -0.902 | 0.462 | -0.005 | -0.001 | -0.001 | 0 | 0.19 | 0.11 | 0.13 | 0.24 | 0.19 | 0.28 |

| | | | | | | | | | | | | | | | | |
|---|---|---|---|---|---|---|---|---|---|---|---|---|---|---|---|---|
| 2 | 230481535 | rs10490186 | 0.906 | -0.473 | -1.145 | -0.081 | 0.052 | 0.073 | 0.073 | 0.436 | 0.53 | 0.19 | 0.7 | 0.56 | 0.64 | 0.56 |
| 2 | 230485415 | rs6727393 | 0.366 | -1.04 | 0.058 | -0.23 | 0.011 | -0.1 | -0.1 | 0.352 | 0.17 | 0.38 | 0.15 | 0.01 | 0.19 | 0.06 |
| 2 | 230485450 | rs12986692 | 1.056 | -1.156 | 0.162 | -0.501 | 0.002 | -0.025 | -0.025 | 0 | 0.66 | 0.38 | 0.68 | 0.97 | 0.7 | 0.84 |
| 2 | 230488467 | rs13393112 | 0.503 | -0.992 | 0.843 | -0.37 | 0.002 | -0.068 | -0.068 | 0 | 0.17 | 0.29 | 0.14 | 0.01 | 0.19 | 0.06 |
| 2 | 230492411 | rs1861617 | 0.977 | -0.758 | 0.924 | 0.183 | 0.009 | -0.052 | -0.052 | 0 | 0.89 | 0.78 | 0.86 | 0.98 | 0.91 | 0.93 |
| 2 | 230492607 | rs1861618 | 1.043 | -0.683 | 0.823 | 0.336 | -0.003 | 0.016 | 0.016 | 0.267 | 0.76 | 0.7 | 0.77 | 0.54 | 0.8 | 0.61 |
| 2 | 230494182 | rs877860 | -0.539 | -0.416 | 0.754 | 0.753 | -0.008 | 0.067 | 0.067 | 0.253 | 0.13 | 0.23 | 0.08 | 0.44 | 0.08 | 0.29 |
| 2 | 230505033 | rs6436883 | -1.072 | -0.676 | 0.058 | 0.731 | 0.032 | -0.023 | -0.023 | 0 | 0.52 | 0.82 | 0.43 | 0.5 | 0.4 | 0.54 |
| 2 | 230506712 | rs7605496 | -0.407 | -1.062 | 0.523 | 0.671 | -0.036 | -0.028 | -0.028 | 0.017 | 0.31 | 0.5 | 0.26 | 0.26 | 0.23 | 0.35 |
| 2 | 230510230 | rs6713110 | -0.215 | -1.039 | 0.716 | 0.734 | -0.018 | 0.034 | 0.034 | 1 | 0.22 | 0.56 | 0.19 | 0.25 | 0.16 | 0.2 |
| 2 | 230518233 | rs6752370 | -0.087 | -1.058 | 0.843 | 0.561 | -0.02 | 0.006 | 0.006 | 0.413 | 0.23 | 0.39 | 0.21 | 0.26 | 0.17 | 0.3 |
| 2 | 230520613 | rs1544673 | 0.798 | -0.917 | 1.056 | 0.496 | -0.043 | -0.014 | -0.014 | 0 | 0.87 | 0.66 | 0.83 | 0.93 | 0.95 | 0.87 |
| 2 | 230522128 | rs9646889 | 0.918 | -0.747 | 0.341 | 0.539 | -0.095 | -0.046 | -0.046 | 0 | 0.42 | 0.09 | 0.37 | 0.38 | 0.54 | 0.39 |
| 2 | 230522398 | rs1861612 | 0.311 | -0.84 | 0.712 | 0.51 | 0 | 0 | 0 | 0.027 | 0.49 | 0.51 | 0.5 | 0.47 | 0.55 | 0.51 |
| 2 | 230524304 | rs12052464 | 0.782 | -0.725 | 0.435 | 0.486 | -0.063 | -0.039 | -0.039 | 0 | 0.44 | 0.13 | 0.38 | 0.44 | 0.55 | 0.4 |
| 2 | 230526908 | rs7587835 | 0.54 | -0.744 | 0.496 | 0.457 | 0.018 | 0.103 | 0.103 | 7.725 | 0.93 | 0.55 | 0.86 | 0.9 | 0.99 | 0.92 |
| 2 | 230527801 | rs17678034 | 0.025 | -0.579 | -0.528 | 0.442 | -0.002 | -0.007 | -0.007 | 0.164 | 0.39 | 0.28 | 0.43 | 0.46 | 0.38 | 0.42 |
| 2 | 230532819 | rs6756438 | -1.211 | -0.538 | 0.385 | 0.329 | -0.068 | -0.055 | -0.055 | 0 | 0.18 | 0.54 | 0.18 | 0.07 | 0.07 | 0.18 |
| 2 | 230545220 | rs2193218 | 0.46 | -0.006 | 0.561 | 0.826 | 0.003 | 0.105 | 0.105 | 0 | 0.74 | 0.51 | 0.63 | 0.63 | 0.77 | 0.64 |
| 2 | 230545356 | rs2216249 | 0.048 | 0.136 | -0.086 | 0.464 | 0 | 0.01 | 0.01 | 2.226 | 0.22 | 0.21 | 0.22 | 0.37 | 0.22 | 0.32 |
| 2 | 230545657 | rs7558924 | -1.139 | 0.136 | 0.447 | 0.657 | -0.001 | 0.021 | 0.021 | 1.828 | 0.38 | 0.52 | 0.41 | 0.43 | 0.37 | 0.41 |
| 2 | 230546078 | rs7574501 | -0.162 | 0.117 | 0.773 | 0.657 | -0.006 | 0.045 | 0.045 | 4.046 | 0.26 | 0.37 | 0.23 | 0.37 | 0.23 | 0.31 |
| 2 | 230548739 | rs2396697 | -0.732 | 0.065 | 1.377 | 0.336 | -0.002 | -0.017 | -0.017 | 0 | 0.34 | 0.4 | 0.42 | 0.67 | 0.32 | 0.58 |
| 2 | 230553339 | rs208789 | 1.432 | -0.013 | 1.166 | 0.283 | -0.027 | 0.239 | 0.239 | 0.801 | 0.51 | 0.17 | 0.4 | 0.26 | 0.55 | 0.33 |
| 2 | 230554795 | rs208792 | -0.221 | -0.051 | 1.439 | 0.326 | 0 | -0.006 | -0.006 | 1.453 | 0.86 | 0.79 | 0.82 | 0.94 | 0.86 | 0.9 |
| 2 | 230563118 | rs1323595 | 0.24 | -0.108 | 1.383 | 0.299 | -0.002 | 0.003 | 0.003 | 0 | 0.13 | 0.13 | 0.17 | 0.06 | 0.14 | 0.1 |
| 2 | 230571420 | rs6761377 | -0.653 | 0.388 | 1.255 | 0.153 | 0.005 | -0.012 | -0.012 | 0 | 0.4 | 0.44 | 0.49 | 0.64 | 0.41 | 0.6 |
| 2 | 230603619 | rs13392809 | 0.402 | 0.705 | 0.999 | -0.08 | 0.003 | 0.048 | 0.048 | 1.401 | 0.55 | 0.75 | 0.48 | 0.36 | 0.5 | 0.39 |
| 2 | 230609149 | rs6707272 | 1.143 | 0.707 | 0.554 | -0.257 | 0.011 | 0.012 | 0.012 | 0.125 | 0.39 | 0.38 | 0.24 | 0.26 | 0.3 | 0.3 |
| 2 | 230612451 | rs6756038 | 1.25 | 0.696 | 0.499 | -0.263 | 0.047 | 0.073 | 0.073 | 0.081 | 0.38 | 0.23 | 0.23 | 0.26 | 0.3 | 0.3 |
| 2 | 230623048 | rs6436891 | 1.17 | 0.63 | 0.496 | -0.252 | 0.033 | 0.081 | 0.081 | 0.088 | 0.39 | 0.23 | 0.27 | 0.26 | 0.36 | 0.3 |
| 2 | 230626528 | rs9808110 | 0.274 | 0.603 | 0.594 | -0.295 | 0 | 0.003 | 0.003 | 0.457 | 0.55 | 0.59 | 0.47 | 0.37 | 0.51 | 0.38 |
| 2 | 230632268 | rs6687 | 0.199 | 0.63 | 0.613 | -0.141 | -0.002 | 0.003 | 0.003 | 0 | 0.15 | 0.21 | 0.19 | 0.11 | 0.15 | 0.08 |
| 2 | 230641763 | rs7598508 | 1.081 | 0.508 | -0.04 | -0.347 | 0.029 | 0.057 | 0.057 | 0.149 | 0.39 | 0.28 | 0.27 | 0.26 | 0.35 | 0.3 |
| 2 | 230641870 | rs7584686 | 1.584 | -0.123 | -0.616 | -0.794 | -0.049 | 0.059 | 0.059 | 0.006 | 0.22 | 0.02 | 0.17 | 0.14 | 0.24 | 0.15 |
| 2 | 230704946 | rs533241 | -0.468 | 0.318 | 0.146 | -0.773 | -0.006 | -0.003 | -0.003 | 0 | 0.82 | 0.67 | 0.79 | 0.9 | 0.83 | 0.92 |
| 2 | 230712174 | rs2053461 | 0.749 | 0.319 | 0.124 | -0.773 | -0.003 | -0.006 | -0.006 | 0 | 0.15 | 0.12 | 0.2 | 0.1 | 0.16 | 0.08 |
| 2 | 230729421 | rs10498242 | 0.789 | 0.3 | 0.245 | -0.784 | -0.001 | -0.005 | -0.005 | 0 | 0.15 | 0.12 | 0.19 | 0.1 | 0.16 | 0.08 |
| 2 | 230731397 | rs9288651 | 0.842 | 0.212 | 0.43 | -0.536 | -0.028 | -0.049 | -0.049 | 0 | 0.25 | 0.13 | 0.36 | 0.39 | 0.31 | 0.34 |
| 2 | 230752964 | rs546530 | -0.101 | -0.033 | 0.778 | -0.813 | -0.007 | 0.008 | 0.008 | 0.597 | 0.39 | 0.3 | 0.52 | 0.64 | 0.5 | 0.62 |
| 2 | 230763690 | rs581150 | -0.239 | -0.028 | 0.785 | -0.81 | 0.007 | -0.015 | -0.015 | 0 | 0.4 | 0.43 | 0.53 | 0.64 | 0.49 | 0.62 |
| 2 | 230787061 | rs12993164 | 1.014 | -0.169 | 0.409 | -0.835 | 0.013 | 0.136 | 0.136 | 0.126 | 0.4 | 0.03 | 0.25 | 0.25 | 0.34 | 0.3 |
| 2 | 230822296 | rs560304 | -0.749 | -0.083 | 0.529 | -0.812 | -0.01 | -0.029 | -0.029 | 0 | 0.55 | 0.51 | 0.71 | 0.75 | 0.66 | 0.7 |
| 2 | 230829594 | rs519707 | -0.304 | 0.141 | 0.155 | -0.232 | -0.006 | -0.006 | -0.006 | 0 | 0.71 | 0.67 | 0.73 | 0.8 | 0.75 | 0.72 |
| 2 | 230853000 | rs11679182 | 0.39 | 0.165 | -0.383 | -0.216 | 0.008 | 0.013 | 0.013 | 2.077 | 0.27 | 0.12 | 0.25 | 0.2 | 0.24 | 0.28 |
| 2 | 230873123 | rs720041 | 0.93 | 0.286 | -1.278 | -0.213 | 0.056 | 0.081 | 0.081 | 0 | 0.37 | 0.17 | 0.27 | 0.28 | 0.31 | 0.29 |
| 2 | 230875311 | rs11675143 | 0.548 | 0.329 | -1.101 | -0.233 | -0.037 | -0.039 | -0.039 | 0 | 0.22 | 0.03 | 0.23 | 0.2 | 0.23 | 0.27 |
| 2 | 230896678 | rs13407773 | -0.036 | 0.474 | -1.057 | -0.283 | 0.006 | -0.058 | -0.058 | 0 | 0.27 | 0.35 | 0.3 | 0.09 | 0.3 | 0.11 |
| 2 | 230899765 | rs7355333 | 0.371 | 0.334 | -1.754 | -0.24 | 0.01 | -0.01 | -0.01 | 0 | 0.22 | 0.09 | 0.21 | 0.24 | 0.19 | 0.25 |
| 2 | 230901647 | rs13432082 | -0.287 | 0.53 | -1.256 | -0.229 | 0 | -0.036 | -0.036 | 0 | 0.28 | 0.53 | 0.32 | 0.1 | 0.3 | 0.11 |
| 2 | 230903875 | rs4973245 | 0.018 | 0.569 | -2.197 | 0.13 | -0.028 | -0.007 | -0.007 | 0 | 0.39 | 0.06 | 0.46 | 0.38 | 0.39 | 0.38 |
| 2 | 230907854 | rs12478048 | -0.443 | 1.155 | -1.9 | -0.045 | -0.016 | 0.013 | 0.013 | 0 | 0.16 | 0.1 | 0.18 | 0.07 | 0.24 | 0.07 |
| 2 | 230911791 | rs4973246 | -0.618 | 0.951 | -1.497 | -0.067 | 0.002 | 0.017 | 0.017 | 0 | 0.62 | 0.68 | 0.62 | 0.68 | 0.65 | 0.56 |
| 2 | 230915321 | rs12990532 | -0.567 | 0.718 | -1.159 | -0.08 | -0.034 | -0.086 | -0.086 | 0.113 | 0.87 | 0.68 | 0.91 | 0.99 | 0.92 | 0.92 |
| 2 | 230919116 | rs4375854 | 0.554 | 0.646 | -0.988 | 0.085 | -0.013 | 0.073 | 0.073 | 0.89 | 0.69 | 0.47 | 0.66 | 0.51 | 0.72 | 0.56 |
| 2 | 230920379 | rs4973252 | 0.241 | 0.629 | -0.849 | 0.111 | -0.001 | 0.004 | 0.004 | 0 | 0.73 | 0.63 | 0.68 | 0.51 | 0.75 | 0.57 |
| 2 | 230929448 | rs11897719 | 0.755 | 0.806 | -0.816 | -0.177 | 0.015 | 0.043 | 0.043 | 0.469 | 0.11 | 0.04 | 0.08 | 0 | 0.08 | 0.07 |
| 2 | 230935754 | rs12475755 | -0.088 | 1.166 | -0.604 | -0.012 | -0.021 | 0.015 | 0.015 | 0 | 0.17 | 0.1 | 0.19 | 0.08 | 0.27 | 0.09 |
| 2 | 230943451 | rs4613264 | 0.273 | 0.316 | -0.444 | 0.065 | -0.005 | -0.019 | -0.019 | 0 | 0.57 | 0.5 | 0.45 | 0.7 | 0.43 | 0.6 |
| 2 | 230944429 | rs4973263 | -0.696 | 0.065 | -0.128 | 0.786 | -0.013 | 0.112 | 0.112 | 0 | 0.19 | 0.29 | 0.18 | 0.5 | 0.15 | 0.29 |
| 2 | 230945993 | rs10169375 | -0.114 | 0.042 | -0.093 | 0.879 | -0.008 | 0.072 | 0.072 | 0 | 0.13 | 0.12 | 0.12 | 0.48 | 0.09 | 0.27 |
| 2 | 230947503 | rs7599478 | -1.302 | 0.168 | 0.268 | 0.716 | -0.004 | 0.021 | 0.021 | 0 | 0.3 | 0.38 | 0.26 | 0.39 | 0.3 | 0.32 |
| 2 | 230954621 | rs10181349 | -0.437 | 0.26 | 0.322 | 0.246 | 0.015 | -0.095 | -0.095 | 0 | 0.8 | 0.6 | 0.79 | 0.94 | 0.78 | 0.86 |
| 2 | 230959540 | rs4972937 | -0.688 | 0.26 | 0.673 | 0.257 | 0.005 | -0.049 | -0.049 | 0.012 | 0.82 | 0.79 | 0.81 | 0.93 | 0.78 | 0.86 |
| 2 | 230992756 | rs6743476 | -0.843 | 0.172 | 0.665 | 0.254 | -0.002 | 0.031 | 0.031 | 0 | 0.29 | 0.45 | 0.27 | 0.38 | 0.3 | 0.32 |
| 2 | 230993221 | rs6743904 | 0.983 | 0.247 | 0.682 | 0.224 | -0.018 | -0.05 | -0.05 | 0 | 0.11 | 0.04 | 0.13 | 0.37 | 0.15 | 0.28 |
| 2 | 230995326 | rs6436907 | -0.79 | 0.254 | 0.699 | 0.207 | 0.005 | -0.049 | -0.049 | 0.64 | 0.82 | 0.78 | 0.81 | 0.93 | 0.78 | 0.86 |
| 2 | 230995447 | rs6436908 | 1.539 | 0.226 | 0.574 | 0.209 | -0.021 | -0.017 | -0.017 | 0 | 0.52 | 0.19 | 0.53 | 0.55 | 0.48 | 0.54 |
| 2 | 230996889 | rs1549572 | -1.089 | 0.251 | 0.29 | -0.002 | -0.004 | -0.021 | -0.021 | 0.19 | 0.3 | 0.4 | 0.23 | 0.21 | 0.28 | 0.31 |

| | | | | | | | | | | | | | | | | |
|---|---|---|---|---|---|---|---|---|---|---|---|---|---|---|---|---|
| 2 | 230997403 | rs12694840 | 1.031 | 0.168 | 0.323 | 0.108 | -0.001 | 0.003 | 0.003 | 0.764 | 0.57 | 0.55 | 0.57 | 0.71 | 0.51 | 0.62 |
| 2 | 231002347 | rs1549567 | 0.151 | 0.192 | 0.375 | 0.038 | 0.002 | -0.041 | -0.041 | 0.89 | 0.69 | 0.63 | 0.68 | 0.92 | 0.63 | 0.88 |
| 2 | 231005473 | rs13005048 | 1.059 | 0.542 | -0.12 | 0.242 | -0.045 | -0.04 | -0.04 | 0 | 0.14 | 0.07 | 0.33 | 0.31 | 0.21 | 0.24 |
| 2 | 231008417 | rs16826700 | 1.032 | 0.312 | -0.091 | 0.117 | -0.009 | -0.032 | -0.032 | 0 | 0.11 | 0.05 | 0.11 | 0.21 | 0.12 | 0.26 |
| 2 | 231009554 | rs1365773 | -0.49 | 0.022 | 0.348 | -0.227 | 0.003 | -0.026 | -0.026 | 0 | 0.79 | 0.7 | 0.75 | 0.92 | 0.74 | 0.9 |
| 2 | 231014505 | rs4973275 | -0.453 | -0.011 | 0.458 | -0.215 | 0.005 | -0.046 | -0.046 | 0 | 0.79 | 0.64 | 0.74 | 0.92 | 0.75 | 0.91 |
| 2 | 231021064 | rs6735555 | 0.158 | -0.097 | 0.357 | -0.246 | -0.009 | -0.053 | -0.053 | 0.062 | 0.74 | 0.39 | 0.72 | 0.9 | 0.75 | 0.81 |
| 2 | 231024518 | rs4972939 | -0.147 | -0.074 | 0.124 | -0.447 | 0.002 | -0.043 | -0.043 | 0 | 0.74 | 0.7 | 0.71 | 0.92 | 0.68 | 0.87 |
| 2 | 231024807 | rs16825905 | 1.478 | -0.178 | -0.656 | -0.176 | 0.006 | -0.061 | -0.061 | 1.073 | 0.27 | 0.13 | 0.15 | 0.33 | 0.2 | 0.28 |
| 2 | 231025024 | rs17326933 | 1.198 | -0.129 | -0.693 | -0.231 | 0 | -0.028 | -0.028 | 0.289 | 0.09 | 0.1 | 0.06 | 0 | 0.1 | 0.03 |
| 2 | 231026074 | rs6727261 | 0.81 | -0.089 | -0.79 | -0.128 | 0.008 | -0.011 | -0.011 | 0 | 0.37 | 0.25 | 0.22 | 0.34 | 0.3 | 0.3 |
| 2 | 231027162 | rs890580 | -0.229 | 0.03 | -0.635 | -0.511 | -0.007 | -0.04 | -0.04 | 0.054 | 0.75 | 0.67 | 0.73 | 0.87 | 0.7 | 0.78 |
| 2 | 231029373 | rs6726336 | 0.278 | 0.127 | -1.422 | -0.519 | -0.012 | 0.112 | 0.112 | 0 | 0.57 | 0.18 | 0.54 | 0.75 | 0.5 | 0.66 |
| 2 | 231029409 | rs17327304 | 0.611 | 0.302 | -1.415 | -0.567 | -0.027 | 0.061 | 0.061 | 0 | 0.1 | 0.02 | 0.06 | 0 | 0.12 | 0.03 |
| 2 | 231029693 | rs10173517 | -0.553 | -0.064 | -1.152 | -0.527 | -0.005 | -0.012 | -0.012 | 0 | 0.65 | 0.62 | 0.79 | 0.8 | 0.76 | 0.62 |
| 2 | 231032372 | rs10210254 | 0.117 | -0.25 | -0.757 | -0.433 | 0.018 | 0.032 | 0.032 | 0 | 0.72 | 0.57 | 0.58 | 0.41 | 0.7 | 0.62 |
| 2 | 231033516 | rs722555 | 0.467 | -0.599 | -1.122 | -0.777 | 0 | -0.005 | -0.005 | 0 | 0.6 | 0.54 | 0.47 | 0.41 | 0.59 | 0.57 |
| 2 | 231034524 | rs6436915 | 0.456 | -0.724 | -1.214 | -1.116 | -0.017 | 0.047 | 0.047 | 0 | 0.44 | 0.32 | 0.3 | 0.3 | 0.44 | 0.47 |
| 2 | 231034867 | rs10208770 | 1.679 | -0.778 | -1.303 | -1.377 | -0.01 | -0.012 | -0.012 | 0 | 0.21 | 0.25 | 0.1 | 0.12 | 0.1 | 0.21 |
| 2 | 231035575 | rs9784019 | 0.713 | -0.824 | -1.086 | -1.14 | 0.001 | -0.016 | -0.016 | 1.034 | 0.13 | 0.12 | 0.11 | 0 | 0.11 | 0.05 |
| 2 | 231037833 | rs10498244 | 1.532 | -0.576 | -1.394 | -1.392 | -0.018 | 0.043 | 0.043 | 0 | 0.3 | 0.19 | 0.19 | 0.16 | 0.29 | 0.3 |
| 2 | 231038160 | rs6436917 | -1.085 | -0.301 | -0.882 | -1.069 | 0.021 | 0.064 | 0.064 | 0 | 0.84 | 0.73 | 0.64 | 0.44 | 0.8 | 0.68 |
| 2 | 231039578 | rs10210465 | 1.43 | -0.206 | -1.018 | -0.984 | 0.004 | 0.001 | 0.001 | 0 | 0.12 | 0.15 | 0.11 | 0.14 | 0.14 | 0.13 |
| 2 | 231040633 | rs6749579 | 1.351 | -0.296 | -1.019 | -1.078 | 0.029 | 0.031 | 0.031 | 0 | 0.18 | 0.08 | 0.17 | 0.1 | 0.17 | 0.11 |
| 2 | 231042802 | rs2241525 | 1.199 | -0.208 | -1.019 | -1.052 | 0.004 | -0.002 | -0.002 | 0 | 0.12 | 0.15 | 0.11 | 0.14 | 0.14 | 0.11 |
| 2 | 231043270 | rs957682 | 0.102 | -0.157 | -1.188 | -1.053 | 0.008 | 0.143 | 0.143 | 0 | 0.49 | 0.3 | 0.31 | 0.16 | 0.41 | 0.4 |
| 2 | 231044255 | rs2396718 | 0.604 | -0.198 | -1.209 | -0.662 | -0.014 | 0.029 | 0.029 | 0.134 | 0.09 | 0.03 | 0.06 | 0.03 | 0.1 | 0.08 |
| 2 | 231044367 | rs11678451 | 0.265 | 0.025 | -1.034 | -0.713 | 0.001 | -0.058 | -0.058 | 0 | 0.45 | 0.51 | 0.29 | 0.13 | 0.45 | 0.41 |
| 2 | 231044464 | rs919178 | 0.544 | 0.024 | -1.503 | -0.12 | 0 | 0.002 | 0.002 | 0 | 0.36 | 0.62 | 0.42 | 0.36 | 0.4 | 0.28 |
| 2 | 231046438 | rs12615975 | 0.791 | 0.077 | -1.789 | 0.328 | 0.021 | -0.087 | -0.087 | 3.403 | 0.15 | 0.02 | 0.04 | 0.21 | 0.09 | 0.29 |
| 2 | 231047645 | rs10755048 | 0.561 | 0.214 | -1.712 | -0.345 | 0.026 | 0 | 0 | 0.489 | 0.71 | 0.82 | 0.83 | 0.78 | 0.78 | 0.65 |
| 2 | 231050569 | rs2114591 | 0.698 | 0.21 | -1.543 | -0.464 | -0.002 | -0.001 | -0.001 | 0 | 0.4 | 0.45 | 0.61 | 0.67 | 0.45 | 0.34 |
| 2 | 231050715 | rs3948464 | 0.242 | 0.321 | -1.834 | -0.223 | -0.001 | -0.021 | -0.021 | 0.461 | 0.87 | 0.85 | 0.87 | 1 | 0.87 | 0.94 |
| 2 | 231064469 | rs1124534 | -0.165 | 0.038 | -1.531 | 0.02 | 0 | -0.002 | -0.002 | 0 | 0.22 | 0.24 | 0.07 | 0.21 | 0.1 | 0.31 |
| 2 | 231077868 | rs7601176 | 0.379 | 1.21 | -1.61 | -0.362 | -0.056 | 0.093 | 0.093 | 0 | 0.71 | 0.87 | 0.66 | 0.91 | 0.54 | 0.82 |
| 2 | 231081384 | rs7580900 | -0.742 | 1.161 | -1.173 | -0.025 | -0.006 | -0.005 | -0.005 | 0 | 0.38 | 0.65 | 0.42 | 0.4 | 0.47 | 0.47 |
| 2 | 231081405 | rs7580912 | -0.359 | 1.279 | -0.461 | -0.036 | 0.042 | 0.048 | 0.048 | 0 | 0.24 | 0.37 | 0.32 | 0.32 | 0.34 | 0.4 |
| 2 | 231083111 | rs7573954 | -0.424 | 1.249 | -0.191 | -0.003 | 0.044 | 0.051 | 0.051 | 0 | 0.25 | 0.4 | 0.31 | 0.31 | 0.34 | 0.4 |
| 2 | 231083171 | rs6738805 | 0.964 | 0.798 | -0.81 | -0.854 | -0.001 | 0.062 | 0.062 | 0 | 0.2 | 0.07 | 0.09 | 0 | 0.13 | 0.13 |
| 2 | 231088666 | rs17263269 | 1.925 | 0.369 | -1.077 | -1.349 | -0.007 | 0.091 | 0.091 | 0 | 0.19 | 0.04 | 0.09 | 0.04 | 0.16 | 0.22 |
| 2 | 231089943 | rs10209615 | 1.142 | 0.351 | -1.109 | -1.408 | 0 | 0.076 | 0.076 | 0 | 0.23 | 0.08 | 0.11 | 0 | 0.16 | 0.14 |
| 2 | 231099204 | rs11687292 | 1.129 | 0.096 | -1.024 | -1.441 | -0.002 | -0.039 | -0.039 | 0.022 | 0.25 | 0.3 | 0.11 | 0.04 | 0.17 | 0.22 |
| 2 | 231103185 | rs2278197 | -0.78 | 0.058 | -0.983 | -1.395 | 0.014 | -0.014 | -0.014 | 3.431 | 0.6 | 0.72 | 0.52 | 0.41 | 0.63 | 0.6 |
| 2 | 231104479 | rs9288660 | -0.534 | 0.058 | -0.98 | -1.395 | 0.014 | -0.006 | -0.006 | 0 | 0.57 | 0.64 | 0.51 | 0.41 | 0.62 | 0.6 |
| 2 | 231107829 | rs12477362 | -0.097 | 0.067 | -1.107 | -1.42 | 0.001 | 0.026 | 0.026 | 0.293 | 0.46 | 0.65 | 0.43 | 0.31 | 0.5 | 0.53 |
| 2 | 231109268 | rs16826932 | 1.188 | 0.211 | -0.646 | -1.389 | -0.003 | 0.002 | 0.002 | 0 | 0.08 | 0.16 | 0.05 | 0.1 | 0.05 | 0.21 |
| 2 | 231110582 | rs28445040 | 1.334 | 0.267 | -0.64 | -1.442 | -0.001 | 0.033 | 0.033 | 0 | 0.22 | 0.11 | 0.11 | 0 | 0.17 | 0.15 |
| 2 | 231111422 | rs10498245 | -0.016 | 0.256 | -0.547 | -1.444 | 0.007 | 0.011 | 0.011 | 0 | 0.38 | 0.33 | 0.23 | 0.1 | 0.3 | 0.22 |
| 2 | 231115603 | rs9989792 | 0.003 | 0.284 | -0.614 | -1.444 | 0.008 | 0.019 | 0.019 | 0 | 0.38 | 0.31 | 0.23 | 0.1 | 0.3 | 0.22 |
| 2 | 231116874 | rs7423615 | 1.356 | 0.279 | -0.737 | -1.433 | -0.004 | 0.048 | 0.048 | 0 | 0.22 | 0.09 | 0.11 | 0 | 0.17 | 0.15 |
| 2 | 231118186 | rs3769840 | 0.999 | 0.573 | -0.707 | -1.201 | 0.039 | 0.042 | 0.042 | 0 | 0.13 | 0.16 | 0.24 | 0.21 | 0.27 | 0.17 |
| 2 | 231119264 | rs3769844 | 0.73 | 0.153 | -0.946 | -1.311 | -0.004 | -0.005 | -0.005 | 0 | 0.16 | 0.18 | 0.11 | 0.1 | 0.13 | 0.07 |
| 2 | 231130887 | rs2162480 | -0.211 | 0.647 | -0.903 | -1.037 | 0.001 | -0.002 | -0.002 | 0 | 0.49 | 0.55 | 0.31 | 0.21 | 0.41 | 0.44 |
| 2 | 231148009 | rs11689231 | 1.277 | 0.765 | -1.588 | -0.852 | -0.028 | 0.118 | 0.118 | 0.154 | 0.22 | 0.03 | 0.15 | 0.05 | 0.23 | 0.27 |
| 2 | 231160521 | rs1010491 | 0.25 | 0.925 | -0.922 | -0.301 | 0.002 | 0.005 | 0.005 | 0 | 0.08 | 0.16 | 0.05 | 0.1 | 0.05 | 0.21 |
| 2 | 231166371 | rs6436926 | -0.679 | 0.909 | -1.142 | 0.269 | -0.009 | -0.036 | -0.036 | 0.052 | 0.85 | 0.93 | 0.62 | 0.52 | 0.8 | 0.83 |
| 2 | 231172642 | rs2894708 | -0.538 | 0.941 | -1.157 | 0.341 | -0.011 | -0.043 | -0.043 | 0 | 0.84 | 0.93 | 0.61 | 0.48 | 0.79 | 0.83 |
| 2 | 231177822 | rs10498247 | 0.226 | 0.923 | -0.766 | 0.157 | 0.013 | -0.003 | -0.003 | 0 | 0.08 | 0.17 | 0.06 | 0.08 | 0.05 | 0.21 |
| 2 | 231177904 | rs933957 | 0.455 | 0.852 | -1.771 | -0.737 | -0.06 | 0.163 | 0.163 | 0 | 0.27 | 0.05 | 0.11 | 0 | 0.27 | 0.17 |
| 2 | 231178876 | rs730010 | 0.131 | 0.64 | -1.221 | -1 | 0.002 | -0.014 | -0.014 | 0 | 0.68 | 0.58 | 0.42 | 0.26 | 0.63 | 0.7 |
| 2 | 231179460 | rs933956 | -0.933 | 0.664 | -1.251 | -0.858 | -0.01 | -0.055 | -0.055 | 1.309 | 0.81 | 0.9 | 0.59 | 0.53 | 0.76 | 0.82 |
| 2 | 231180915 | rs4973310 | 0.635 | 0.25 | -1.213 | -1.187 | -0.003 | -0.005 | -0.005 | 0 | 0.2 | 0.46 | 0.18 | 0.19 | 0.16 | 0.3 |
| 2 | 231181082 | rs2396744 | -1.019 | 0.24 | -1.278 | -1.195 | -0.036 | -0.066 | -0.066 | 0 | 0.61 | 0.88 | 0.45 | 0.45 | 0.53 | 0.57 |
| 2 | 231182687 | rs9288664 | -2.309 | 0.277 | -1.368 | -1.316 | -0.04 | 0.031 | 0.031 | 0 | 0.93 | 0.99 | 0.9 | 1 | 0.84 | 0.98 |
| 2 | 231183321 | rs720977 | -0.261 | 0.197 | -1.128 | -1.351 | 0.008 | 0.069 | 0.069 | 0 | 0.48 | 0.64 | 0.29 | 0.19 | 0.4 | 0.46 |
| 2 | 231183786 | rs11684950 | 0.258 | 0.004 | -0.989 | -1.068 | -0.007 | -0.031 | -0.031 | 0.736 | 0.44 | 0.48 | 0.31 | 0.26 | 0.39 | 0.54 |
| 2 | 231189047 | rs933961 | -0.273 | -0.021 | -0.776 | -0.682 | 0 | 0 | 0 | 0 | 0.29 | 0.3 | 0.44 | 0.57 | 0.21 | 0.24 |
| 2 | 231190446 | rs17330963 | 0.274 | -0.604 | -1.431 | -1.237 | 0.013 | 0.013 | 0.013 | 0.01 | 0.53 | 0.32 | 0.58 | 0.65 | 0.44 | 0.5 |
| 2 | 231196132 | rs16827136 | 1.652 | -0.473 | -0.989 | -1.06 | 0.016 | 0.126 | 0.126 | 0 | 0.07 | 0.14 | 0.14 | 0.26 | 0.1 | 0.1 |

| | | | | | | | | | | | | | | | | |
|---|---|---|---|---|---|---|---|---|---|---|---|---|---|---|---|---|
| 2 | 231199033 | rs11896138 | 1.003 | -0.599 | -0.97 | -1.069 | -0.006 | -0.006 | -0.006 | 0 | 0.17 | 0.22 | 0.14 | 0.11 | 0.13 | 0.09 |
| 2 | 231213614 | rs7573405 | 1.372 | -0.636 | -0.728 | -1.076 | -0.008 | 0.053 | 0.053 | 0 | 0.12 | 0.13 | 0.31 | 0.46 | 0.09 | 0.16 |
| 2 | 231213706 | rs10206500 | 0.893 | -0.731 | -0.417 | -1.176 | -0.001 | 0.007 | 0.007 | 0 | 0.08 | 0.23 | 0.05 | 0.09 | 0.04 | 0.24 |
| 2 | 231221928 | rs4973316 | 0.362 | -0.721 | -0.46 | -1.169 | -0.004 | 0.02 | 0.02 | 0 | 0.2 | 0.27 | 0.45 | 0.72 | 0.19 | 0.25 |
| 2 | 231222340 | rs16827148 | 1.11 | -0.294 | -0.884 | -0.959 | 0.003 | 0.027 | 0.027 | 0.004 | 0.13 | 0.14 | 0.15 | 0.26 | 0.16 | 0.11 |
| 2 | 231222655 | rs10498250 | 1.047 | -0.578 | -0.706 | -0.805 | 0.006 | 0.007 | 0.007 | 1.715 | 0.16 | 0.13 | 0.13 | 0.1 | 0.13 | 0.09 |
| 2 | 231226488 | rs11891724 | -0.019 | -0.627 | -0.923 | -0.793 | -0.051 | 0.13 | 0.13 | 0 | 0.23 | 0.52 | 0.38 | 0.56 | 0.13 | 0.39 |
| 2 | 231227439 | rs7579078 | -0.893 | -0.623 | -0.947 | -0.818 | 0 | -0.001 | -0.001 | 0 | 0.38 | 0.62 | 0.5 | 0.66 | 0.26 | 0.48 |
| 2 | 231229636 | rs11886574 | -0.072 | -0.485 | -0.662 | -0.59 | -0.046 | 0.171 | 0.171 | 0 | 0.21 | 0.47 | 0.37 | 0.56 | 0.13 | 0.39 |
| 2 | 231234020 | rs11895690 | 1.037 | -0.164 | -1.174 | -2.076 | 0.003 | 0.057 | 0.057 | 0 | 0.31 | 0.19 | 0.18 | 0.08 | 0.3 | 0.27 |
| 2 | 231235886 | rs11689260 | 0.214 | -0.226 | -0.57 | -2.037 | 0.036 | 0.04 | 0.04 | 0.034 | 0.56 | 0.32 | 0.5 | 0.27 | 0.66 | 0.39 |
| 2 | 231236505 | rs12474823 | 1.587 | 0.097 | -0.507 | -1.902 | 0.025 | 0.099 | 0.099 | 5.5 | 0.24 | 0.01 | 0.14 | 0.05 | 0.16 | 0.17 |
| 2 | 231236963 | rs12694847 | 0.16 | 0.397 | -0.11 | -1.822 | 0.019 | 0.039 | 0.039 | 0.163 | 0.48 | 0.42 | 0.45 | 0.23 | 0.59 | 0.3 |
| 2 | 231241276 | rs1816208 | 0.978 | 0.746 | -0.164 | -1.803 | -0.012 | -0.002 | -0.002 | 0 | 0.17 | 0.16 | 0.22 | 0.16 | 0.27 | 0.08 |
| 2 | 231243335 | rs12105723 | 0.323 | 0.71 | 0.168 | -1.749 | -0.004 | 0.009 | 0.009 | 0 | 0.6 | 0.59 | 0.39 | 0.22 | 0.48 | 0.72 |
| 2 | 231245110 | rs1820550 | -0.767 | 0.612 | 0.079 | -1.628 | -0.001 | 0.006 | 0.006 | 0 | 0.77 | 0.75 | 0.61 | 0.38 | 0.75 | 0.8 |
| 2 | 231250415 | rs3754939 | 1.238 | 0.416 | 0.046 | -2.284 | 0.017 | 0.014 | 0.014 | 0.173 | 0.28 | 0.2 | 0.16 | 0.17 | 0.16 | 0.51 |
| 2 | 231250943 | rs6728027 | -0.262 | 0.679 | 0.071 | -2.334 | 0 | -0.001 | -0.001 | 0 | 0.47 | 0.55 | 0.39 | 0.26 | 0.43 | 0.58 |
| 2 | 231252048 | rs12994857 | 1.017 | 0.831 | -0.062 | -2.31 | -0.035 | 0.024 | 0.024 | 0 | 0.17 | 0.12 | 0.21 | 0.08 | 0.27 | 0.07 |
| 2 | 231254570 | rs12694859 | 0.908 | 0.787 | -0.06 | -2.311 | -0.04 | 0.029 | 0.029 | 0 | 0.18 | 0.12 | 0.21 | 0.08 | 0.28 | 0.08 |
| 2 | 231256812 | rs7559665 | 1.111 | 0.771 | -0.013 | -2.379 | -0.005 | 0.023 | 0.023 | 0.013 | 0.29 | 0.2 | 0.22 | 0.07 | 0.32 | 0.22 |
| 2 | 231258150 | rs7590429 | 0.066 | 0.724 | 0.115 | -2.407 | 0 | 0.001 | 0.001 | 0.29 | 0.47 | 0.48 | 0.39 | 0.21 | 0.44 | 0.53 |
| 2 | 231258450 | rs6436932 | -0.644 | 0.746 | 0.125 | -2.373 | -0.001 | 0.011 | 0.011 | 0 | 0.76 | 0.7 | 0.62 | 0.28 | 0.76 | 0.75 |
| 2 | 231267925 | rs11692506 | 0.564 | 1.469 | 0.226 | -2.463 | -0.004 | 0.016 | 0.016 | 0 | 0.3 | 0.21 | 0.23 | 0.07 | 0.32 | 0.23 |
| 2 | 231268324 | rs17273079 | 0.564 | 1.469 | 0.226 | -2.463 | -0.004 | 0.016 | 0.016 | 0 | 0.3 | 0.21 | 0.23 | 0.07 | 0.32 | 0.23 |
| 2 | 231270213 | rs12694862 | 0.442 | 1.464 | 0.318 | -2.446 | -0.011 | 0.009 | 0.009 | 0 | 0.18 | 0.17 | 0.21 | 0.08 | 0.28 | 0.07 |
| 2 | 231271572 | rs11898466 | -0.352 | 1.457 | 0.212 | -2.406 | -0.003 | 0.011 | 0.011 | 0.366 | 0.44 | 0.41 | 0.36 | 0.15 | 0.51 | 0.22 |
| 2 | 231277567 | rs10182438 | -1.254 | 1.719 | -0.033 | -2.36 | -0.009 | 0.044 | 0.044 | 0.247 | 0.34 | 0.74 | 0.33 | 0.12 | 0.32 | 0.31 |
| 2 | 231278861 | rs3754940 | -0.223 | 1.91 | 0.02 | -2.318 | -0.019 | 0.021 | 0.021 | 0 | 0.36 | 0.3 | 0.34 | 0.08 | 0.48 | 0.31 |
| 2 | 231278929 | rs3754941 | 0.737 | 1.898 | -0.141 | -2.289 | -0.078 | 0.135 | 0.135 | 0 | 0.27 | 0.08 | 0.23 | 0.07 | 0.37 | 0.17 |
| 2 | 231279221 | rs12694864 | 1.534 | 2.021 | -0.067 | -2.276 | -0.011 | 0.005 | 0.005 | 0 | 0.1 | 0.1 | 0.11 | 0 | 0.17 | 0.04 |
| 2 | 231279616 | rs3754944 | -0.932 | 1.952 | 0.083 | -2.301 | -0.004 | -0.029 | -0.029 | 0.577 | 0.46 | 0.53 | 0.48 | 0.18 | 0.59 | 0.36 |
| 2 | 231279792 | rs13398034 | -0.903 | 1.993 | 0.202 | -1.962 | 0.045 | -0.03 | -0.03 | 0 | 0.19 | 0.41 | 0.25 | 0.11 | 0.22 | 0.19 |
| 2 | 231279802 | rs13411586 | -0.903 | 1.993 | 0.191 | -1.962 | 0.047 | -0.03 | -0.03 | 0 | 0.19 | 0.41 | 0.25 | 0.11 | 0.22 | 0.19 |
| 2 | 231282791 | rs1427283 | 0.421 | 2.038 | -0.412 | -1.768 | 0.002 | 0 | 0 | 0 | 0.08 | 0.08 | 0.13 | 0.09 | 0.11 | 0.04 |
| 2 | 231283711 | rs9678342 | -0.894 | 1.824 | -0.263 | -1.977 | -0.002 | 0.022 | 0.022 | 0.03 | 0.42 | 0.64 | 0.42 | 0.12 | 0.5 | 0.33 |
| 2 | 231284556 | rs12694866 | 1.159 | 1.964 | -0.462 | -1.804 | 0.027 | -0.035 | -0.035 | 0 | 0.1 | 0.16 | 0.11 | 0 | 0.19 | 0.04 |
| 2 | 231285570 | rs10202501 | -0.008 | 2.029 | -1.022 | -1.798 | 0.017 | -0.04 | -0.04 | 0 | 0.17 | 0.19 | 0.21 | 0.01 | 0.28 | 0.18 |
| 2 | 231309217 | rs17273810 | 0.47 | 1.459 | -0.839 | -1.605 | 0.018 | -0.038 | -0.038 | 4.431 | 0.08 | 0.17 | 0.11 | 0.01 | 0.1 | 0.14 |
| 2 | 231310620 | rs999888 | 0.79 | 1.28 | -1.724 | -1.628 | -0.071 | 0.106 | 0.106 | 0 | 0.26 | 0.1 | 0.23 | 0.06 | 0.38 | 0.18 |
| 2 | 231316105 | rs7582700 | -0.094 | 0.375 | -1.309 | -0.95 | 0.043 | 0.059 | 0.059 | 0.744 | 0.61 | 0.76 | 0.56 | 0.18 | 0.71 | 0.49 |
| 2 | 231328728 | rs2306276 | -0.283 | 0.946 | -1.58 | -1 | -0.029 | 0.183 | 0.183 | 0.97 | 0.36 | 0.12 | 0.18 | 0.06 | 0.32 | 0.31 |
| 2 | 231331799 | rs1649869 | -0.164 | 1.103 | -0.99 | -0.396 | 0.026 | 0.102 | 0.102 | 0 | 0.8 | 0.62 | 0.65 | 0.49 | 0.76 | 0.85 |
| 2 | 231331896 | rs1050224 | -0.164 | 1.103 | -0.99 | -0.396 | 0.026 | 0.102 | 0.102 | 0 | 0.8 | 0.61 | 0.65 | 0.48 | 0.76 | 0.85 |
| 2 | 231334613 | rs13413034 | 1.515 | 1.2 | -1.508 | -0.682 | -0.029 | 0.047 | 0.047 | 0 | 0.08 | 0.01 | 0.05 | 0 | 0.11 | 0.02 |
| 2 | 231335466 | rs1649883 | 0.365 | 1.166 | -1.439 | -0.626 | -0.046 | 0.196 | 0.196 | 1 | 0.35 | 0.08 | 0.17 | 0.06 | 0.32 | 0.31 |
| 2 | 231337608 | rs1678168 | 0.365 | 1.166 | -1.439 | -0.612 | -0.046 | 0.193 | 0.193 | 1 | 0.35 | 0.08 | 0.17 | 0.07 | 0.32 | 0.31 |
| 2 | 231339731 | rs4539783 | 0.867 | 0.856 | -0.454 | -0.188 | 0.008 | 0.004 | 0.004 | 0.031 | 0.31 | 0.22 | 0.31 | 0.25 | 0.29 | 0.47 |
| 2 | 231345991 | rs1678198 | 0.25 | 1.082 | 0.03 | 0.127 | 0.001 | 0.002 | 0.002 | 1.435 | 0.51 | 0.42 | 0.51 | 0.49 | 0.48 | 0.67 |
| 2 | 231346602 | rs9678598 | -0.432 | 1.294 | 1.092 | 0.61 | 0.062 | 0.101 | 0.101 | 0 | 0.12 | 0.44 | 0.15 | 0.15 | 0.14 | 0.14 |
| 2 | 231355960 | rs4973326 | 1.075 | 0.958 | 0.646 | 0.5 | -0.014 | 0.032 | 0.032 | 0.755 | 0.54 | 0.12 | 0.44 | 0.35 | 0.47 | 0.59 |
| 2 | 231357373 | rs6436943 | -0.45 | 1.239 | 1.346 | 0.477 | -0.011 | -0.137 | -0.137 | 0.001 | 0.22 | 0.45 | 0.13 | 0.03 | 0.17 | 0.11 |
| 2 | 231365088 | rs1649921 | -0.645 | 1.42 | 1.239 | 0.808 | 0.015 | 0.094 | 0.094 | 0 | 0.93 | 0.86 | 0.73 | 0.61 | 0.87 | 0.87 |
| 2 | 231366562 | rs6751402 | -0.397 | 1.362 | 1.283 | 0.655 | -0.002 | -0.03 | -0.03 | 0.563 | 0.2 | 0.2 | 0.11 | 0.03 | 0.15 | 0.1 |
| 2 | 231368897 | rs6705605 | -0.375 | 1.308 | 1.421 | 0.686 | -0.007 | -0.045 | -0.045 | 0.141 | 0.19 | 0.24 | 0.11 | 0.04 | 0.14 | 0.1 |
| 2 | 231375974 | rs16827319 | 0.26 | 1.259 | 1.125 | 0.653 | 0.001 | -0.004 | -0.004 | 0 | 0.13 | 0.12 | 0.07 | 0.04 | 0.09 | 0.09 |
| 2 | 231376320 | rs6436944 | -0.714 | 1.292 | 0.681 | 0.786 | 0.003 | 0.065 | 0.065 | 0 | 0.93 | 0.89 | 0.73 | 0.59 | 0.87 | 0.87 |
| 2 | 231385231 | rs890669 | 1.901 | 1.304 | 0.645 | 0.743 | -0.026 | 0.031 | 0.031 | 0.807 | 0.56 | 0.16 | 0.45 | 0.34 | 0.49 | 0.59 |
| 2 | 231406070 | rs17275036 | 0.763 | 1.515 | 0.705 | 0.395 | -0.023 | 0.039 | 0.039 | 11.397 | 0.11 | 0.01 | 0.06 | 0.04 | 0.08 | 0.09 |
| 2 | 231406529 | rs6728423 | -0.356 | 1.285 | 0.945 | 0.265 | -0.023 | -0.139 | -0.139 | 0 | 0.24 | 0.46 | 0.13 | 0.04 | 0.18 | 0.11 |
| 2 | 231406680 | rs836237 | -0.293 | 1.312 | 0.046 | 0.53 | 0.049 | 0.289 | 0.289 | 0 | 0.91 | 0.73 | 0.69 | 0.46 | 0.87 | 0.87 |
| 2 | 231407519 | rs6717269 | -0.449 | 1.329 | 0.054 | 0.538 | 0.056 | 0.308 | 0.308 | 0 | 0.92 | 0.74 | 0.69 | 0.47 | 0.87 | 0.87 |
| 2 | 231407663 | rs9288668 | 1.316 | 1.207 | -0.964 | 0.505 | 0.027 | 0.194 | 0.194 | 0 | 0.51 | 0.16 | 0.4 | 0.22 | 0.46 | 0.57 |
| 2 | 231411801 | rs1678188 | -0.571 | 1.469 | -0.612 | 0.796 | -0.057 | -0.059 | -0.059 | 0 | 0.14 | 0.05 | 0.15 | 0.22 | 0.23 | 0.18 |
| 2 | 231413681 | rs1678192 | 0.072 | 0.951 | -0.499 | 0.77 | -0.017 | -0.041 | -0.041 | 0.38 | 0.09 | 0.03 | 0.1 | 0.19 | 0.1 | 0.19 |
| 2 | 231414688 | rs1612907 | -1.068 | 0.816 | -0.375 | 0.643 | 0.018 | 0.011 | 0.011 | 0 | 0.5 | 0.64 | 0.55 | 0.55 | 0.59 | 0.53 |
| 2 | 231415013 | rs1649911 | 0.705 | 0.863 | -0.375 | 0.696 | -0.008 | -0.022 | -0.022 | 1.566 | 0.09 | 0.05 | 0.11 | 0.19 | 0.1 | 0.19 |
| 2 | 231415189 | rs6756289 | -0.575 | 0.834 | 0.018 | 0.554 | 0.011 | -0.014 | -0.014 | 0 | 0.41 | 0.49 | 0.44 | 0.36 | 0.49 | 0.34 |
| 2 | 231420401 | rs17275498 | 0.106 | 0.751 | 0.091 | 0.156 | 0.013 | 0.039 | 0.039 | 5.768 | 0.16 | 0.07 | 0.08 | 0.01 | 0.12 | 0.09 |

| | | | | | | | | | | | | | | | | |
|---|---|---|---|---|---|---|---|---|---|---|---|---|---|---|---|---|
| 2 | 231421301 | rs11695625 | -0.013 | 0.834 | 0.267 | 0.265 | 0.003 | 0.001 | 0.001 | 0.361 | 0.25 | 0.28 | 0.33 | 0.33 | 0.35 | 0.25 |
| 2 | 231421683 | rs1678155 | 0.485 | 0.584 | 0.226 | 0.452 | 0.004 | 0.055 | 0.055 | 0 | 0.1 | 0.18 | 0.12 | 0.2 | 0.1 | 0.19 |
| 2 | 231422554 | rs1678156 | -0.991 | 0.657 | 0.566 | 0.325 | 0.013 | 0.014 | 0.014 | 0.648 | 0.39 | 0.49 | 0.45 | 0.51 | 0.45 | 0.43 |
| 2 | 231422963 | rs11682065 | -0.058 | 0.708 | 0.128 | 0.373 | -0.005 | -0.002 | -0.002 | 0.418 | 0.28 | 0.28 | 0.33 | 0.32 | 0.34 | 0.24 |
| 2 | 231427877 | rs1649917 | -0.668 | 0.445 | -0.059 | 0.15 | -0.012 | -0.005 | -0.005 | 0 | 0.27 | 0.53 | 0.21 | 0.2 | 0.22 | 0.28 |
| 2 | 231429220 | rs2059755 | -1.578 | 0.421 | -0.062 | 0.149 | -0.024 | -0.035 | -0.035 | 0.409 | 0.55 | 0.84 | 0.55 | 0.52 | 0.56 | 0.51 |
| 2 | 231430266 | rs7561418 | -0.212 | 0.241 | 0.293 | -0.246 | 0.002 | 0 | 0 | 0.578 | 0.29 | 0.31 | 0.33 | 0.32 | 0.34 | 0.24 |
| 2 | 231433237 | rs13405308 | -0.031 | 0.273 | 0.158 | -0.241 | -0.032 | -0.095 | -0.095 | 0 | 0.17 | 0.5 | 0.1 | 0.02 | 0.11 | 0.08 |
| 2 | 231434272 | rs16827412 | -0.558 | 0.695 | 0.06 | 0.147 | 0.004 | 0 | 0 | 0 | 0.3 | 0.35 | 0.34 | 0.32 | 0.34 | 0.23 |
| 2 | 231437079 | rs2396763 | -0.566 | 0.697 | 0.061 | 0.149 | 0.003 | 0 | 0 | 0.177 | 0.3 | 0.35 | 0.34 | 0.32 | 0.34 | 0.23 |
| 2 | 231440375 | rs16827417 | -0.314 | 0.708 | -0.104 | -0.346 | 0.006 | -0.107 | -0.107 | 0 | 0.24 | 0.36 | 0.15 | 0.02 | 0.25 | 0.09 |
| 2 | 231440555 | rs2288356 | -0.45 | 0.557 | -1.069 | -0.344 | -0.013 | 0.107 | 0.107 | 0 | 0.5 | 0.19 | 0.45 | 0.34 | 0.53 | 0.3 |
| 2 | 231442496 | rs7562389 | -0.097 | 1.061 | -0.794 | -0.147 | -0.003 | -0.001 | -0.001 | 0 | 0.67 | 0.61 | 0.59 | 0.61 | 0.56 | 0.59 |
| 2 | 231442893 | rs11674658 | 0.126 | 0.472 | -0.864 | -0.696 | 0.004 | 0.006 | 0.006 | 0 | 0.35 | 0.4 | 0.43 | 0.57 | 0.36 | 0.5 |
| 2 | 231443428 | rs10498252 | 1.263 | 0.259 | -0.63 | -0.604 | 0.001 | 0.006 | 0.006 | 0 | 0.19 | 0.17 | 0.18 | 0.29 | 0.23 | 0.31 |
| 2 | 231443567 | rs2042839 | 1.319 | 0.158 | -0.722 | -0.656 | 0.046 | 0.15 | 0.15 | 0.552 | 0.3 | 0.06 | 0.15 | 0.04 | 0.2 | 0.08 |
| 2 | 231443924 | rs2241536 | 0.417 | 0.268 | -0.576 | -0.47 | -0.006 | 0.035 | 0.035 | 0 | 0.16 | 0.23 | 0.24 | 0.28 | 0.13 | 0.19 |
| 2 | 231444896 | rs11694266 | 0.417 | 0.268 | -0.596 | -0.47 | -0.006 | 0.033 | 0.033 | 0 | 0.16 | 0.22 | 0.24 | 0.28 | 0.13 | 0.19 |
| 2 | 231447717 | rs10933344 | -0.379 | 0.232 | -0.713 | -0.501 | 0.003 | 0.008 | 0.008 | 0 | 0.36 | 0.46 | 0.44 | 0.56 | 0.36 | 0.5 |
| 2 | 231451851 | rs6742549 | -0.548 | 0.365 | -0.869 | -1.111 | -0.017 | -0.015 | -0.015 | 0 | 0.66 | 0.75 | 0.47 | 0.36 | 0.61 | 0.44 |
| 2 | 231454050 | rs13391467 | -0.354 | -0.04 | -0.669 | -0.904 | 0.011 | 0.019 | 0.019 | 1.579 | 0.54 | 0.64 | 0.38 | 0.34 | 0.43 | 0.39 |
| 2 | 231454354 | rs13005078 | 0.172 | 0.519 | -0.551 | -0.295 | 0.006 | 0.049 | 0.049 | 0 | 0.22 | 0.3 | 0.21 | 0.3 | 0.23 | 0.3 |
| 2 | 231457433 | rs6735285 | -0.293 | 0.888 | -0.434 | -0.12 | 0.016 | 0.15 | 0.15 | 0 | 0.14 | 0.33 | 0.14 | 0.28 | 0.15 | 0.24 |
| 2 | 231457618 | rs6735336 | -0.384 | 1.061 | -0.718 | -0.369 | -0.002 | 0.026 | 0.026 | 0.024 | 0.51 | 0.34 | 0.34 | 0.32 | 0.53 | 0.43 |
| 2 | 231457819 | rs6436951 | -0.196 | 0.907 | -0.685 | -0.048 | 0.008 | 0.009 | 0.009 | 1.275 | 0.41 | 0.26 | 0.31 | 0.32 | 0.39 | 0.33 |
| 2 | 231477223 | rs4972964 | -0.64 | 0.981 | -0.537 | -0.053 | -0.023 | -0.034 | -0.034 | 0 | 0.54 | 0.78 | 0.49 | 0.54 | 0.53 | 0.45 |
| 2 | 231477414 | rs10167683 | 0.239 | 1.103 | -0.758 | -0.052 | -0.051 | 0.077 | 0.077 | 0 | 0.15 | 0.02 | 0.1 | 0 | 0.17 | 0.08 |
| 2 | 231477415 | rs4972965 | -0.437 | 0.786 | 0.452 | 0.342 | -0.013 | 0.263 | 0.263 | 0.071 | 0.11 | 0.36 | 0.23 | 0.45 | 0.09 | 0.18 |
| 2 | 231493060 | rs11681644 | -0.401 | 0.924 | 0.206 | -0.117 | 0.036 | 0.021 | 0.021 | 0.221 | 0.49 | 0.28 | 0.48 | 0.57 | 0.59 | 0.45 |
| 2 | 231493293 | rs10169182 | -1.558 | 1.175 | 1.399 | 0.665 | -0.007 | 0.066 | 0.066 | 0.886 | 0.25 | 0.66 | 0.33 | 0.49 | 0.26 | 0.32 |
| 2 | 231508389 | rs7584576 | -1.714 | 1.179 | 1.421 | 0.586 | 0.001 | 0.008 | 0.008 | 0 | 0.32 | 0.69 | 0.4 | 0.52 | 0.38 | 0.33 |
| 2 | 231515321 | rs7590845 | -1.955 | 0.832 | 1.984 | -0.124 | 0.037 | 0.024 | 0.024 | 2.096 | 0.06 | 0.43 | 0.04 | 0 | 0.01 | 0 |
| 2 | 231515970 | rs13394080 | -1.476 | 0.863 | 1.754 | -0.091 | 0.01 | -0.084 | -0.084 | 0.082 | 0.2 | 0.56 | 0.13 | 0.07 | 0.19 | 0.13 |
| 2 | 231520821 | rs11903702 | -1.596 | 0.815 | 1.682 | -0.104 | -0.002 | -0.098 | -0.098 | 0 | 0.21 | 0.58 | 0.14 | 0.07 | 0.19 | 0.13 |
| 2 | 231521950 | rs7609123 | -1.416 | 0.847 | 1.548 | 0.043 | 0.007 | -0.063 | -0.063 | 0 | 0.19 | 0.67 | 0.12 | 0.08 | 0.17 | 0.15 |
| 2 | 231527626 | rs11892913 | -1.538 | 0.549 | 1.442 | 0.323 | 0.079 | 0.064 | 0.064 | 4.048 | 0.05 | 0.44 | 0.04 | 0 | 0.02 | 0.01 |
| 2 | 231536037 | rs1401063 | -0.057 | 0.806 | -0.215 | 0.363 | -0.021 | 0.012 | 0.012 | 0.196 | 0.6 | 0.72 | 0.61 | 0.56 | 0.55 | 0.44 |
| 2 | 231539173 | rs13021124 | -0.099 | -0.14 | -0.036 | -0.647 | -0.018 | -0.082 | -0.082 | 0.046 | 0.27 | 0.33 | 0.17 | 0 | 0.22 | 0.14 |
| 2 | 231539694 | rs16827499 | 0.001 | 0.366 | -0.44 | -0.502 | -0.016 | -0.014 | -0.014 | 0.367 | 0.09 | 0.06 | 0.09 | 0.15 | 0.1 | 0.24 |
| 2 | 231542102 | rs6709168 | -0.899 | 0.834 | -0.51 | -0.198 | -0.041 | -0.032 | -0.032 | 0.127 | 0.63 | 0.83 | 0.57 | 0.6 | 0.6 | 0.59 |
| 2 | 231543033 | rs6726604 | 0.452 | 0.567 | -0.43 | -0.33 | -0.034 | -0.118 | -0.118 | 0.255 | 0.28 | 0.38 | 0.17 | 0 | 0.22 | 0.13 |
| 2 | 231544263 | rs11901268 | 0.837 | 0.929 | -0.815 | 0.256 | -0.021 | -0.018 | -0.018 | 0 | 0.23 | 0.12 | 0.2 | 0.23 | 0.27 | 0.18 |
| 2 | 231546204 | rs13431868 | 0.339 | 0.508 | -0.771 | -0.249 | 0.019 | -0.006 | -0.006 | 0 | 0.31 | 0.2 | 0.17 | 0 | 0.25 | 0.13 |
| 2 | 231546750 | rs16827514 | 0.113 | 0.907 | -0.72 | 0.198 | -0.003 | 0.009 | 0.009 | 0.693 | 0.1 | 0.12 | 0.16 | 0.34 | 0.15 | 0.3 |
| 2 | 231547476 | rs13384784 | 0.221 | 0.47 | -0.762 | -0.37 | 0.004 | -0.037 | -0.037 | 2.129 | 0.31 | 0.24 | 0.17 | 0 | 0.22 | 0.13 |
| 2 | 231548319 | rs6746999 | -0.274 | 0.455 | -1.01 | -0.277 | 0.003 | 0.007 | 0.007 | 0.84 | 0.4 | 0.32 | 0.26 | 0.15 | 0.37 | 0.21 |
| 2 | 231552660 | rs6713780 | 0.054 | 0.719 | -2.342 | 0.003 | -0.044 | -0.019 | -0.019 | 0.789 | 0.4 | 0.15 | 0.42 | 0.39 | 0.39 | 0.55 |
| 2 | 231553094 | rs12995904 | 0.58 | 0.777 | -2.347 | 0.111 | -0.056 | -0.022 | -0.022 | 0.469 | 0.3 | 0.06 | 0.32 | 0.34 | 0.33 | 0.49 |
| 2 | 231555684 | rs938906 | -0.718 | 0.762 | -1.812 | 0.141 | 0.014 | 0.08 | 0.08 | 0 | 0.81 | 0.72 | 0.68 | 0.51 | 0.75 | 0.73 |
| 2 | 231556188 | rs4973343 | -0.219 | 0.403 | -2.148 | -0.288 | -0.003 | -0.011 | -0.011 | 0 | 0.57 | 0.64 | 0.56 | 0.51 | 0.55 | 0.62 |
| 2 | 231556273 | rs6729885 | 1.466 | 0.43 | -2.625 | -0.445 | 0.019 | 0.103 | 0.103 | 0 | 0.21 | 0.06 | 0.11 | 0.01 | 0.17 | 0.08 |
| 2 | 231556663 | rs4972969 | 0.413 | 0.909 | -2.161 | 0.016 | -0.037 | -0.023 | -0.023 | 8.43 | 0.3 | 0.11 | 0.31 | 0.35 | 0.32 | 0.52 |
| 2 | 231561178 | rs17762734 | 0.056 | 1.022 | -2.18 | 0.057 | -0.051 | -0.018 | -0.018 | 0 | 0.34 | 0.13 | 0.35 | 0.39 | 0.39 | 0.52 |
| 2 | 231569204 | rs6436969 | -0.273 | 0.817 | -2.442 | -0.035 | -0.036 | -0.11 | -0.11 | 1.196 | 0.68 | 0.89 | 0.61 | 0.55 | 0.67 | 0.64 |
| 2 | 231577363 | rs2466154 | -1.486 | 1.176 | -2.654 | -0.118 | -0.047 | -0.134 | -0.134 | 0.123 | 0.82 | 0.98 | 0.71 | 0.54 | 0.79 | 0.6 |
| 2 | 231579273 | rs2459925 | 0.647 | 1.75 | -2.88 | -0.332 | -0.106 | -0.124 | -0.124 | 0 | 0.27 | 0.07 | 0.4 | 0.43 | 0.36 | 0.33 |
| 2 | 231582196 | rs10933355 | 0.275 | 1.136 | -1.856 | -0.663 | -0.03 | -0.045 | -0.045 | 0.891 | 0.13 | 0.38 | 0.09 | 0.07 | 0.08 | 0.07 |
| 2 | 231582797 | rs6436971 | -0.235 | 0.889 | -1.962 | -0.797 | -0.024 | -0.043 | -0.043 | 0 | 0.19 | 0.74 | 0.13 | 0.05 | 0.09 | 0.07 |
| 2 | 231606083 | rs7561687 | 2.006 | 0.692 | -2.218 | -0.988 | -0.006 | 0.009 | 0.009 | 0.338 | 0.14 | 0.05 | 0.08 | 0.08 | 0.15 | 0.05 |
| 2 | 231623555 | rs10168275 | 2.012 | 0.391 | -2.29 | -0.887 | 0.012 | 0.047 | 0.047 | 0 | 0.19 | 0.06 | 0.09 | 0.04 | 0.16 | 0.12 |
| 2 | 231628878 | rs13420598 | -0.355 | 0.468 | -2.048 | -0.703 | 0.038 | 0.056 | 0.056 | 0.786 | 0.32 | 0.78 | 0.21 | 0.14 | 0.24 | 0.12 |
| 2 | 231629360 | rs2720162 | -1.313 | 0.482 | -2.07 | -0.697 | -0.017 | -0.054 | -0.054 | 0 | 0.63 | 0.92 | 0.62 | 0.57 | 0.63 | 0.47 |
| 2 | 231631689 | rs1464205 | 0.666 | 0.506 | -2.07 | -0.684 | -0.115 | -0.109 | -0.109 | 0 | 0.31 | 0.09 | 0.41 | 0.43 | 0.39 | 0.35 |
| 2 | 231642471 | rs6750664 | -0.909 | 0.023 | -1.135 | -1.007 | 0.035 | 0.051 | 0.051 | 0.737 | 0.32 | 0.78 | 0.21 | 0.14 | 0.24 | 0.12 |
| 2 | 231672572 | rs3792069 | 0.985 | 0.731 | -1.172 | -0.675 | -0.044 | -0.013 | -0.013 | 1.237 | 0.13 | 0.01 | 0.08 | 0.08 | 0.17 | 0.05 |
| 2 | 231683849 | rs7596575 | -0.474 | 0.419 | -0.342 | -0.418 | -0.016 | 0.096 | 0.096 | 0 | 0.22 | 0.46 | 0.25 | 0.33 | 0.21 | 0.22 |
| 2 | 231685198 | rs726379 | -0.369 | 0.581 | -0.03 | -1.218 | -0.002 | -0.008 | -0.008 | 0.42 | 0.29 | 0.3 | 0.16 | 0.16 | 0.23 | 0.12 |
| 2 | 231687306 | rs6760721 | -0.412 | 0.691 | -0.413 | -1.418 | -0.002 | 0.136 | 0.136 | 0 | 0.41 | 0.16 | 0.23 | 0.12 | 0.38 | 0.21 |
| 2 | 231690981 | rs10933358 | -0.424 | 0.696 | 0.304 | -1.021 | -0.012 | -0.092 | -0.092 | 0 | 0.3 | 0.4 | 0.18 | 0.08 | 0.27 | 0.12 |

| | | | | | | | | | | | | | | | |
|---|---|---|---|---|---|---|---|---|---|---|---|---|---|---|---|
| 2 | 231692685 | rs881109 | -0.384 | 0.762 | 0.047 | -0.97 | -0.014 | -0.101 | -0.101 | 0.048 | 0.3 | 0.4 | 0.18 | 0.08 | 0.27 | 0.12 |
| 2 | 231704107 | rs7579281 | -0.682 | 1.052 | 0.58 | -0.453 | 0.033 | 0.06 | 0.06 | 0.766 | 0.12 | 0.36 | 0.17 | 0.12 | 0.1 | 0.13 |
| 2 | 231707159 | rs13390515 | -0.969 | 1.461 | 0.574 | -0.784 | 0.024 | -0.004 | -0.004 | 0.003 | 0.14 | 0.18 | 0.19 | 0.12 | 0.2 | 0.2 |
| 2 | 231707741 | rs9288670 | -1.578 | 1.447 | 0.585 | -1.111 | 0.014 | -0.037 | -0.037 | 5.419 | 0.22 | 0.32 | 0.2 | 0.11 | 0.28 | 0.24 |
| 2 | 231709953 | rs10182071 | -1.59 | 1.533 | 0.293 | -1.17 | 0.021 | -0.057 | -0.057 | 0 | 0.24 | 0.5 | 0.2 | 0.09 | 0.25 | 0.24 |
| 2 | 231711860 | rs6726497 | -0.579 | 1.658 | -0.395 | -1.093 | -0.008 | 0.007 | 0.007 | 0 | 0.12 | 0.1 | 0.11 | 0.03 | 0.17 | 0.14 |
| 2 | 231720362 | rs2466150 | 0.407 | 1.414 | -0.865 | -1.474 | -0.007 | 0.047 | 0.047 | 0 | 0.77 | 0.86 | 0.83 | 0.92 | 0.75 | 0.87 |
| 2 | 231722128 | rs2466149 | 0.936 | 1.334 | -0.49 | -1.623 | -0.001 | -0.025 | -0.025 | 0 | 0.66 | 0.68 | 0.76 | 0.87 | 0.68 | 0.73 |
| 2 | 231725311 | rs6714365 | 1.633 | 1.571 | -0.98 | -2.031 | 0.02 | 0.186 | 0.186 | 0 | 0.46 | 0.25 | 0.59 | 0.85 | 0.44 | 0.56 |
| 2 | 231726018 | rs4355094 | 0.889 | 1.701 | -0.603 | -1.984 | -0.018 | 0.023 | 0.023 | 0 | 0.69 | 0.77 | 0.72 | 0.89 | 0.58 | 0.74 |
| 2 | 231727399 | rs1438145 | 1.341 | 1.315 | -0.521 | -2.071 | 0.004 | -0.072 | -0.072 | 0 | 0.69 | 0.43 | 0.76 | 0.88 | 0.69 | 0.74 |
| 2 | 231736610 | rs2164869 | -0.938 | 1.39 | -0.723 | -1.848 | -0.018 | 0.026 | 0.026 | 0.221 | 0.14 | 0.08 | 0.12 | 0.08 | 0.2 | 0.14 |
| 2 | 231737798 | rs4972973 | 1.602 | 0.95 | -1.052 | -1.994 | 0.005 | 0.151 | 0.151 | 0.138 | 0.54 | 0.23 | 0.53 | 0.81 | 0.5 | 0.57 |
| 2 | 231738038 | rs3111750 | 0.142 | 0.956 | -1.127 | -2.005 | -0.023 | -0.024 | -0.024 | 0.108 | 0.06 | 0.01 | 0.11 | 0.07 | 0.07 | 0.11 |
| 2 | 231738269 | rs2289233 | 1.411 | 0.753 | -1.015 | -1.919 | 0.002 | 0.163 | 0.163 | 0.168 | 0.54 | 0.23 | 0.53 | 0.81 | 0.5 | 0.57 |
| 2 | 231746780 | rs6726593 | -2.161 | -0.392 | -0.973 | -1.41 | 0.067 | 0.204 | 0.204 | 0 | 0.5 | 0.92 | 0.33 | 0.26 | 0.37 | 0.47 |
| 2 | 231747426 | rs934820 | -1.784 | -0.629 | 0.068 | -1.591 | -0.004 | -0.026 | -0.026 | 0.877 | 0.35 | 0.57 | 0.28 | 0.21 | 0.31 | 0.33 |
| 2 | 231750356 | rs6724157 | -2.299 | -0.681 | 0.301 | -1.53 | 0.026 | 0.028 | 0.028 | 0 | 0.31 | 0.72 | 0.2 | 0.12 | 0.19 | 0.23 |
| 2 | 231756585 | rs7577585 | -1.769 | -0.873 | 0.97 | -1.508 | -0.049 | -0.083 | -0.083 | 0 | 0.14 | 0.32 | 0.07 | 0.03 | 0.06 | 0.11 |
| 2 | 231757332 | rs6731224 | -1.04 | -0.794 | 1.249 | -1.545 | -0.005 | 0.005 | 0.005 | 0.001 | 0.15 | 0.11 | 0.11 | 0.04 | 0.17 | 0.08 |
| 2 | 231757359 | rs6759606 | -1.723 | -0.821 | 1.307 | -1.366 | -0.003 | -0.062 | -0.062 | 0 | 0.24 | 0.49 | 0.17 | 0.16 | 0.22 | 0.22 |
| 2 | 231757630 | rs10182844 | -1.944 | -0.7 | 1.083 | -1.075 | -0.042 | -0.04 | -0.04 | 0 | 0.21 | 0.34 | 0.12 | 0.11 | 0.11 | 0.2 |
| 2 | 231764443 | rs16827643 | -2.41 | -0.51 | 0.97 | -0.982 | 0.025 | 0.04 | 0.04 | 0 | 0.41 | 0.7 | 0.28 | 0.25 | 0.32 | 0.41 |
| 2 | 231772436 | rs1992188 | -2.667 | -0.518 | 0.568 | -0.945 | 0.064 | 0.148 | 0.148 | 0.02 | 0.43 | 0.93 | 0.31 | 0.27 | 0.33 | 0.44 |
| 2 | 231774377 | rs10201627 | -1.498 | -0.604 | 1.378 | -0.942 | 0.018 | 0.081 | 0.081 | 0 | 0.06 | 0.28 | 0.02 | 0.15 | 0.01 | 0.16 |
| 2 | 231774844 | rs2396777 | -2.358 | -0.664 | 1.332 | -0.994 | -0.033 | -0.016 | -0.016 | 0 | 0.27 | 0.68 | 0.17 | 0.23 | 0.15 | 0.34 |
| 2 | 231775144 | rs1992187 | -0.01 | -0.641 | 0.637 | -0.982 | -0.008 | -0.024 | -0.024 | 0 | 0.06 | 0 | 0.06 | 0.07 | 0.06 | 0.09 |
| 2 | 231775297 | rs1992186 | -2.828 | -0.386 | 0.652 | -0.996 | 0.032 | 0.14 | 0.14 | 0.015 | 0.44 | 0.92 | 0.32 | 0.27 | 0.35 | 0.44 |
| 2 | 231778776 | rs6747728 | -2.333 | -0.546 | 1.06 | -0.841 | -0.011 | -0.012 | -0.012 | 0.105 | 0.32 | 0.66 | 0.21 | 0.24 | 0.22 | 0.39 |
| 2 | 231784291 | rs2918006 | -1.363 | -0.444 | 0.727 | -0.458 | 0.02 | 0.017 | 0.017 | 0.028 | 0.16 | 0.08 | 0.12 | 0.09 | 0.14 | 0.16 |
| 2 | 231785026 | rs16827668 | -1.433 | -0.451 | 1.046 | 0.54 | -0.003 | 0.052 | 0.052 | 0 | 0.19 | 0.33 | 0.2 | 0.29 | 0.16 | 0.12 |
| 2 | 231790097 | rs3111780 | -2.858 | -0.481 | 0.318 | 0.384 | 0.049 | -0.074 | -0.074 | 0 | 0.46 | 0.93 | 0.41 | 0.51 | 0.37 | 0.48 |
| 2 | 231790350 | rs4973351 | -1.942 | -0.327 | 1.107 | 0.568 | -0.02 | 0.063 | 0.063 | 0.05 | 0.26 | 0.52 | 0.25 | 0.42 | 0.22 | 0.3 |
| 2 | 231793015 | rs3111779 | -1.289 | -0.093 | 0.416 | 0.333 | 0.008 | 0.007 | 0.007 | 0 | 0.13 | 0.06 | 0.11 | 0.07 | 0.11 | 0.12 |
| 2 | 231797549 | rs6750171 | -1.381 | 0.119 | 0.421 | 0.521 | -0.017 | 0.055 | 0.055 | 0 | 0.19 | 0.09 | 0.14 | 0.04 | 0.24 | 0.12 |
| 2 | 231799552 | rs1866233 | -1.788 | 0.147 | 0.415 | 0.764 | 0.007 | 0.041 | 0.041 | 0.188 | 0.32 | 0.79 | 0.27 | 0.19 | 0.3 | 0.31 |
| 2 | 231800199 | rs13407893 | -1.788 | 0.147 | 0.415 | 0.764 | 0.007 | 0.041 | 0.041 | 0.188 | 0.32 | 0.79 | 0.27 | 0.19 | 0.3 | 0.31 |
| 2 | 231808291 | rs4973353 | -1.907 | 0.194 | 0.124 | 0.932 | 0.012 | -0.072 | -0.072 | 0.019 | 0.34 | 0.83 | 0.36 | 0.54 | 0.3 | 0.42 |
| 2 | 231814203 | rs7569545 | -1.019 | 0.624 | 0.44 | 0.4 | 0.002 | -0.02 | -0.02 | 0 | 0.19 | 0.22 | 0.13 | 0.06 | 0.25 | 0.25 |
| 2 | 231825834 | rs3106075 | -0.769 | 0.709 | 1.077 | 0.905 | 0.012 | -0.052 | -0.052 | 0 | 0.2 | 0.29 | 0.16 | 0.04 | 0.27 | 0.15 |
| 2 | 231830047 | rs3111770 | -1.518 | 0.587 | 1.498 | 0.914 | 0.015 | 0.114 | 0.114 | 0 | 0.26 | 0.49 | 0.29 | 0.49 | 0.3 | 0.4 |
| 2 | 231833839 | rs1876820 | -0.149 | 0.287 | 0.738 | -0.44 | -0.018 | 0.071 | 0.071 | 1.19 | 0.45 | 0.3 | 0.32 | 0.04 | 0.49 | 0.46 |
| 2 | 231838134 | rs3106301 | -0.555 | 0.599 | -0.208 | -0.32 | 0.039 | 0.102 | 0.102 | 3.2 | 0.5 | 0.66 | 0.44 | 0.22 | 0.56 | 0.51 |
| 2 | 231839240 | rs1848728 | 0.05 | 0.524 | -0.91 | -0.671 | -0.021 | 0.106 | 0.106 | 0.468 | 0.41 | 0.26 | 0.29 | 0.04 | 0.43 | 0.35 |
| 2 | 231840796 | rs12463788 | -0.06 | 0.515 | -0.999 | -0.624 | -0.007 | -0.005 | -0.005 | 0 | 0.45 | 0.43 | 0.32 | 0.02 | 0.45 | 0.38 |
| 2 | 231842338 | rs4973358 | -0.442 | -0.096 | -1.464 | 0.199 | 0.02 | 0.101 | 0.101 | 0.504 | 0.22 | 0.02 | 0.17 | 0.05 | 0.18 | 0.19 |
| 2 | 231845762 | rs3115376 | -0.53 | -0.281 | -1.129 | 0.815 | 0.012 | 0.03 | 0.03 | 0.261 | 0.27 | 0.21 | 0.22 | 0.14 | 0.21 | 0.15 |
| 2 | 231849914 | rs7598647 | 0.024 | -0.839 | -0.105 | 0.999 | -0.014 | 0.055 | 0.055 | 0.169 | 0.06 | 0.18 | 0.11 | 0.07 | 0.01 | 0.04 |
| 2 | 231852394 | rs1060780 | -0.411 | -0.555 | -0.422 | 1.045 | -0.006 | 0.001 | 0.001 | 0.88 | 0.48 | 0.44 | 0.42 | 0.53 | 0.52 | 0.48 |
| 2 | 231857076 | rs10933363 | -0.267 | -1.022 | -0.31 | 0.467 | -0.004 | 0.061 | 0.061 | 0 | 0.53 | 0.71 | 0.49 | 0.81 | 0.57 | 0.66 |
| 2 | 231859544 | rs6736980 | -1.125 | -1.171 | 0.368 | 0.661 | -0.052 | -0.056 | -0.056 | 3.235 | 0.19 | 0.45 | 0.1 | 0.12 | 0.1 | 0.23 |
| 2 | 231864289 | rs2136474 | -0.928 | -0.671 | 0.284 | 0.866 | -0.007 | 0.006 | 0.006 | 1.7 | 0.39 | 0.47 | 0.36 | 0.47 | 0.41 | 0.5 |
| 2 | 231872525 | rs13413140 | -0.289 | -0.832 | 0.562 | 0.782 | 0.01 | 0.019 | 0.019 | 0.301 | 0.28 | 0.32 | 0.32 | 0.46 | 0.39 | 0.46 |
| 2 | 231874602 | rs11691545 | -0.02 | -0.894 | 0.608 | 0.862 | 0.011 | 0.002 | 0.002 | 0 | 0.27 | 0.31 | 0.25 | 0.31 | 0.37 | 0.25 |
| 2 | 231875286 | rs12694884 | -0.625 | -1.196 | 0.153 | 0.552 | 0.018 | 0.05 | 0.05 | 0 | 0.6 | 0.76 | 0.7 | 0.79 | 0.7 | 0.71 |
| 2 | 231875433 | rs13398605 | 0.187 | -1.158 | -0.154 | 0.509 | -0.01 | 0.007 | 0.007 | 0 | 0.32 | 0.24 | 0.27 | 0.31 | 0.41 | 0.26 |
| 2 | 231877207 | rs2175149 | -0.536 | -1.148 | -0.014 | 0.622 | 0.012 | 0 | 0 | 0.108 | 0.45 | 0.64 | 0.5 | 0.48 | 0.34 | 0.51 |
| 2 | 231878834 | rs4972979 | 0.22 | -0.524 | -0.489 | 0.678 | -0.001 | -0.003 | -0.003 | 0 | 0.31 | 0.36 | 0.37 | 0.37 | 0.3 | 0.26 |
| 2 | 231882097 | rs4973369 | 0.406 | -0.482 | -0.53 | 0.576 | 0 | 0 | 0 | 0 | 0.28 | 0.33 | 0.34 | 0.23 | 0.27 | 0.23 |
| 2 | 231884504 | rs11686658 | 0.364 | -0.192 | -1.177 | 0.105 | -0.001 | -0.012 | -0.012 | 0.335 | 0.22 | 0.21 | 0.22 | 0.05 | 0.28 | 0.19 |
| 2 | 231886444 | rs2276623 | 0.271 | 0.687 | -1.528 | 0.717 | -0.026 | -0.028 | -0.028 | 0 | 0.3 | 0.27 | 0.38 | 0.38 | 0.36 | 0.29 |
| 2 | 231891530 | rs6436988 | 0.247 | 0.723 | -1.587 | 0.718 | -0.021 | -0.022 | -0.022 | 0 | 0.31 | 0.27 | 0.38 | 0.38 | 0.36 | 0.29 |
| 2 | 231896979 | rs906881 | -0.575 | 0.057 | -2.139 | 0.396 | -0.021 | 0.016 | 0.016 | 0.072 | 0.72 | 0.99 | 0.76 | 0.78 | 0.74 | 0.78 |
| 2 | 231913056 | rs3731783 | 0.163 | -0.315 | -1.08 | 0.509 | -0.027 | 0.001 | 0.001 | 0.249 | 0.22 | 0.43 | 0.19 | 0.18 | 0.21 | 0.16 |
| 2 | 231914845 | rs6748332 | -0.557 | -0.859 | -0.782 | 0.798 | 0.005 | -0.005 | -0.005 | 0 | 0.33 | 0.72 | 0.26 | 0.41 | 0.26 | 0.23 |
| 2 | 231918067 | rs7608781 | 0.485 | -0.798 | -0.971 | 0.939 | 0.005 | -0.006 | -0.006 | 0.488 | 0.64 | 0.31 | 0.66 | 0.5 | 0.66 | 0.52 |
| 2 | 231921182 | rs16827740 | -0.608 | -0.883 | -0.974 | 0.879 | 0.056 | -0.056 | -0.056 | 0 | 0.33 | 0.85 | 0.33 | 0.5 | 0.27 | 0.44 |
| 2 | 231940960 | rs1472959 | -1.214 | -1.269 | -1.711 | 0.498 | 0.076 | 0.03 | 0.03 | 0.436 | 0.52 | 0.88 | 0.42 | 0.62 | 0.43 | 0.52 |
| 2 | 231942492 | rs6436999 | -0.862 | -1.361 | -1.454 | 0.479 | 0.107 | -0.009 | -0.009 | 0.823 | 0.45 | 0.88 | 0.38 | 0.6 | 0.34 | 0.48 |

| | | | | | | | | | | | | | | | | |
|---|---|---|---|---|---|---|---|---|---|---|---|---|---|---|---|---|
| 2 | 231949946 | rs2303357 | -0.972 | -1.36 | -1.288 | 0.496 | 0.075 | 0.023 | 0.023 | 1.231 | 0.46 | 0.75 | 0.38 | 0.61 | 0.37 | 0.48 |
| 2 | 231968920 | rs13394402 | 1.3 | -1.499 | -1.204 | 0.488 | -0.001 | -0.004 | -0.004 | 0 | 0.14 | 0.15 | 0.14 | 0.07 | 0.2 | 0.15 |
| 2 | 231969554 | rs10199752 | -1.26 | -1.529 | -1.164 | 0.483 | 0.118 | 0.026 | 0.026 | 1.217 | 0.48 | 0.88 | 0.4 | 0.62 | 0.37 | 0.48 |
| 2 | 231972175 | rs17619588 | 1.086 | -1.548 | -1.071 | 0.552 | -0.049 | 0.09 | 0.09 | 5.919 | 0.13 | 0.42 | 0.08 | 0.22 | 0.04 | 0.06 |
| 2 | 231976460 | rs17619600 | 0.964 | -1.551 | -1.106 | 0.553 | 0.021 | -0.034 | -0.034 | 0.411 | 0.16 | 0.02 | 0.13 | 0.2 | 0.09 | 0.25 |
| 2 | 231980019 | rs10194776 | -1.4 | -1.468 | -1.076 | 0.517 | 0.088 | 0.03 | 0.03 | 0.333 | 0.51 | 0.88 | 0.42 | 0.61 | 0.42 | 0.51 |
| 2 | 231980777 | rs16827801 | -0.709 | -1.482 | -1.049 | 0.548 | -0.01 | 0.001 | 0.001 | 0.77 | 0.43 | 0.6 | 0.34 | 0.5 | 0.33 | 0.46 |
| 2 | 231981992 | rs4973377 | -2.056 | -1.381 | -1.326 | 0.267 | -0.009 | 0.012 | 0.012 | 0 | 0.9 | 0.98 | 0.89 | 1 | 0.86 | 0.91 |
| 2 | 231982826 | rs1549339 | -0.939 | -1.432 | -1.004 | 0.583 | 0.03 | 0.009 | 0.009 | 1.022 | 0.46 | 0.65 | 0.37 | 0.61 | 0.37 | 0.48 |
| 2 | 231990371 | rs10187149 | -0.729 | -1.432 | -1.01 | 0.569 | -0.01 | 0 | 0 | 1.165 | 0.43 | 0.6 | 0.35 | 0.49 | 0.33 | 0.46 |
| 2 | 232043850 | rs2887283 | 0.953 | -1.473 | -1.05 | 0.5 | -0.012 | -0.036 | -0.036 | 0 | 0.16 | 0.25 | 0.17 | 0.07 | 0.2 | 0.16 |
| 2 | 232047155 | rs16827840 | -0.397 | -1.327 | -1.069 | 0.693 | -0.049 | 0.037 | 0.037 | 5.886 | 0.32 | 0.48 | 0.24 | 0.55 | 0.17 | 0.33 |
| 2 | 232054576 | rs7606037 | -1.11 | -1.421 | -1.383 | 0.702 | 0.127 | 0.05 | 0.05 | 0.341 | 0.52 | 0.9 | 0.43 | 0.63 | 0.4 | 0.48 |
| 2 | 232062351 | rs7601741 | 0.216 | -1.487 | -0.808 | 0.633 | -0.02 | -0.052 | -0.052 | 0 | 0.24 | 0.48 | 0.23 | 0.11 | 0.29 | 0.18 |
| 2 | 232066651 | rs753855 | -1.569 | -1.41 | -1.15 | 0.738 | -0.02 | 0.024 | 0.024 | 0.426 | 0.58 | 0.92 | 0.49 | 0.67 | 0.5 | 0.53 |
| 2 | 232069121 | rs17586589 | 1.256 | -1.548 | -0.885 | 0.923 | -0.038 | 0.113 | 0.113 | 7.767 | 0.11 | 0.3 | 0.04 | 0.29 | 0.04 | 0.06 |
| 2 | 232071682 | rs17586603 | -0.151 | -1.567 | -0.89 | 0.913 | -0.042 | 0.035 | 0.035 | 8.138 | 0.28 | 0.35 | 0.19 | 0.55 | 0.12 | 0.31 |
| 2 | 232077075 | rs5018532 | 0.131 | -1.446 | -1.02 | 0.971 | -0.022 | -0.039 | -0.039 | 0 | 0.24 | 0.49 | 0.24 | 0.14 | 0.29 | 0.18 |
| 2 | 232077670 | rs16827863 | 0.131 | -1.446 | -1.02 | 0.971 | -0.022 | -0.039 | -0.039 | 0.753 | 0.24 | 0.49 | 0.24 | 0.14 | 0.29 | 0.18 |
| 2 | 232091479 | rs3731776 | 1.691 | -1.354 | -0.951 | 1.014 | 0.036 | 0.32 | 0.32 | 1.902 | 0.06 | 0.34 | 0.03 | 0.29 | 0.02 | 0.02 |
| 2 | 232093297 | rs1729077 | 1.002 | -1.094 | -1.215 | 1.252 | -0.001 | 0.001 | 0.001 | 0 | 0.11 | 0.09 | 0.16 | 0.27 | 0.09 | 0.05 |
| 2 | 232107363 | rs12612219 | 1.002 | -1.609 | -1.041 | 0.817 | 0 | 0.001 | 0.001 | 0.605 | 0.18 | 0.18 | 0.16 | 0.1 | 0.24 | 0.16 |
| 2 | 232108246 | rs7562116 | 0.64 | -1.619 | -1.071 | 0.863 | -0.004 | -0.008 | -0.008 | 0 | 0.22 | 0.3 | 0.21 | 0.15 | 0.28 | 0.18 |
| 2 | 232109816 | rs6704770 | -0.53 | -1.577 | -1.627 | 0.815 | 0.041 | -0.031 | -0.031 | 0.605 | 0.34 | 0.83 | 0.26 | 0.4 | 0.32 | 0.24 |
| 2 | 232118813 | rs12621567 | -0.48 | -1.557 | -1.688 | 0.837 | 0.053 | -0.022 | -0.022 | 2.648 | 0.35 | 0.88 | 0.27 | 0.38 | 0.32 | 0.25 |
| 2 | 232122576 | rs11897387 | -1.261 | -1.639 | -1.775 | 0.745 | 0.113 | 0.069 | 0.069 | 0.809 | 0.51 | 0.93 | 0.42 | 0.66 | 0.4 | 0.5 |
| 2 | 232126290 | rs9677476 | -0.523 | -1.541 | -1.696 | 0.587 | 0.038 | -0.025 | -0.025 | 0.033 | 0.35 | 0.82 | 0.27 | 0.4 | 0.32 | 0.25 |
| 2 | 232141729 | rs2290275 | 0.163 | -1.127 | -1.657 | 0.762 | 0.066 | -0.022 | -0.022 | 0 | 0.21 | 0.05 | 0.16 | 0.22 | 0.09 | 0.28 |
| 2 | 232145852 | rs1729084 | 1.345 | -0.238 | -1.563 | 0.778 | -0.083 | -0.027 | -0.027 | 0.039 | 0.22 | 0.04 | 0.34 | 0.21 | 0.28 | 0.2 |
| 2 | 232158246 | rs2053921 | -0.626 | -0.819 | -1.544 | 0.809 | 0.107 | 0.037 | 0.037 | 0.867 | 0.47 | 0.93 | 0.41 | 0.65 | 0.36 | 0.48 |
| 2 | 232158885 | rs3731779 | 0.026 | -0.922 | -1.309 | 0.894 | 0.027 | -0.004 | -0.004 | 1.434 | 0.37 | 0.75 | 0.27 | 0.39 | 0.34 | 0.2 |
| 2 | 232159212 | rs745962 | -1.663 | -0.882 | -1.433 | 0.843 | -0.056 | 0.035 | 0.035 | 0 | 0.76 | 0.99 | 0.82 | 0.86 | 0.76 | 0.73 |
| 2 | 232159243 | rs3731781 | 0.181 | -1.152 | -0.201 | 1.27 | -0.005 | -0.033 | -0.033 | 0 | 0.29 | 0.36 | 0.21 | 0.09 | 0.32 | 0.16 |
| 2 | 232160291 | rs16827912 | 0.022 | -1.067 | -0.529 | 1.32 | 0.031 | 0.343 | 0.343 | 1.98 | 0.08 | 0.39 | 0.06 | 0.3 | 0.03 | 0.03 |
| 2 | 232171195 | rs6710104 | -1.213 | -1.042 | -0.694 | 0.926 | 0.108 | 0.038 | 0.038 | 0.846 | 0.47 | 0.92 | 0.4 | 0.64 | 0.36 | 0.48 |
| 2 | 232171968 | rs11894854 | 0.685 | -1.08 | -0.657 | 1.011 | 0.061 | 0.294 | 0.294 | 2 | 0.06 | 0.36 | 0.04 | 0.25 | 0.02 | 0.02 |
| 2 | 232176541 | rs7580717 | 0.108 | -0.897 | -1.937 | 0.107 | 0.001 | 0.008 | 0.008 | 0.364 | 0.3 | 0.24 | 0.22 | 0.09 | 0.36 | 0.17 |
| 2 | 232183591 | rs1868929 | -0.232 | -0.906 | -1.823 | 0.3 | -0.062 | 0.09 | 0.09 | 7.214 | 0.19 | 0.68 | 0.2 | 0.55 | 0.05 | 0.32 |
| 2 | 232187155 | rs6753112 | -0.411 | -0.071 | -2.484 | 0.257 | 0.081 | 0.024 | 0.024 | 0.499 | 0.48 | 0.93 | 0.43 | 0.61 | 0.41 | 0.48 |
| 2 | 232190925 | rs4973382 | -0.078 | 0.331 | -1.675 | -0.153 | 0.004 | 0.06 | 0.06 | 0.019 | 0.57 | 0.73 | 0.72 | 0.81 | 0.61 | 0.67 |
| 2 | 232192305 | rs1669070 | -0.484 | -0.112 | -1.353 | -0.008 | 0.035 | 0.042 | 0.042 | 0 | 0.46 | 0.25 | 0.37 | 0.3 | 0.41 | 0.4 |
| 2 | 232193217 | rs1669071 | -0.45 | -0.187 | -1.217 | -0.222 | 0.002 | 0.009 | 0.009 | 0 | 0.33 | 0.27 | 0.34 | 0.24 | 0.36 | 0.3 |
| 2 | 232196564 | rs1669083 | -0.272 | -0.227 | -1.19 | -0.248 | 0.007 | 0.016 | 0.016 | 0 | 0.32 | 0.21 | 0.31 | 0.22 | 0.32 | 0.28 |
| 2 | 232196597 | rs1669084 | -0.492 | -0.201 | -1.18 | -0.248 | 0.001 | 0.013 | 0.013 | 0 | 0.33 | 0.28 | 0.34 | 0.22 | 0.36 | 0.3 |
| 2 | 232198327 | rs6719593 | -1.53 | -0.517 | -0.397 | -0.439 | -0.026 | -0.057 | -0.057 | 0 | 0.11 | 0.73 | 0.07 | 0 | 0.04 | 0.01 |
| 2 | 232206025 | rs1729089 | -1.322 | -0.428 | -0.774 | -0.307 | -0.14 | 0.092 | 0.092 | 0 | 0.6 | 0.95 | 0.52 | 0.83 | 0.53 | 0.62 |
| 2 | 232216224 | rs7597595 | -1.193 | -0.506 | 0.131 | -0.05 | 0 | -0.008 | -0.008 | 0 | 0.07 | 0.32 | 0.04 | 0 | 0.01 | 0 |
| 2 | 232216533 | rs7598124 | -1.093 | -0.499 | 0.094 | -0.062 | -0.003 | -0.015 | -0.015 | 0.067 | 0.07 | 0.3 | 0.04 | 0 | 0.01 | 0 |
| 2 | 232236627 | rs6758989 | -0.79 | -0.589 | -0.056 | 0.259 | 0.013 | -0.013 | -0.013 | 0 | 0.35 | 0.69 | 0.24 | 0.56 | 0.23 | 0.36 |
| 2 | 232250398 | rs7594399 | -0.989 | 0.098 | 0.195 | 0.335 | 0.002 | -0.038 | -0.038 | 0 | 0.08 | 0.2 | 0.04 | 0 | 0.06 | 0.02 |
| 2 | 232263127 | rs2290130 | -0.115 | 0.352 | -0.234 | 0.478 | -0.002 | 0 | 0 | 0 | 0.24 | 0.16 | 0.24 | 0.24 | 0.28 | 0.28 |
| 2 | 232263973 | rs10445830 | -0.069 | 0.365 | -0.392 | 0.483 | -0.006 | -0.001 | -0.001 | 0 | 0.24 | 0.14 | 0.24 | 0.23 | 0.28 | 0.28 |
| 2 | 232271284 | rs13030174 | -0.043 | 0.362 | -0.463 | 0.486 | -0.009 | -0.006 | -0.006 | 0 | 0.24 | 0.1 | 0.23 | 0.23 | 0.29 | 0.29 |
| 2 | 232275879 | rs7592642 | -0.942 | 0.678 | 0.092 | 0.939 | -0.007 | -0.008 | -0.008 | 0 | 0.33 | 0.32 | 0.35 | 0.42 | 0.42 | 0.39 |
| 2 | 232276482 | rs11890076 | -0.884 | 0.741 | 0.209 | 0.757 | 0.054 | 0.057 | 0.057 | 0 | 0.1 | 0.22 | 0.11 | 0.2 | 0.14 | 0.12 |
| 2 | 232282909 | rs6750502 | -1.51 | 0.003 | -0.221 | 0.081 | -0.043 | 0.184 | 0.184 | 0 | 0.57 | 0.84 | 0.57 | 0.9 | 0.57 | 0.69 |
| 2 | 232286789 | rs16828053 | -0.036 | -0.11 | 0.535 | 0.74 | -0.053 | 0.008 | 0.008 | 0.092 | 0.24 | 0.39 | 0.2 | 0.26 | 0.15 | 0.3 |
| 2 | 232289244 | rs2369271 | -0.454 | 0.259 | 0.87 | 0.93 | 0.002 | 0.004 | 0.004 | 0 | 0.29 | 0.32 | 0.28 | 0.63 | 0.36 | 0.36 |
| 2 | 232290640 | rs12614599 | 0.28 | 0.352 | 0.462 | 1.031 | -0.004 | -0.026 | -0.026 | 0 | 0.23 | 0.16 | 0.24 | 0.64 | 0.28 | 0.34 |
| 2 | 232294380 | rs10186875 | -0.72 | 0.334 | 0.853 | 1.055 | -0.023 | -0.002 | -0.002 | 0 | 0.27 | 0.52 | 0.25 | 0.25 | 0.23 | 0.3 |
| 2 | 232301670 | rs7563453 | -1.121 | 1.134 | 0.911 | 1.502 | 0.085 | 0.262 | 0.262 | 0.37 | 0.18 | 0.48 | 0.2 | 0.45 | 0.21 | 0.26 |
| 2 | 232304779 | rs4973401 | -1.275 | 0.836 | 0.28 | 0.79 | 0.019 | -0.003 | -0.003 | 0 | 0.33 | 0.48 | 0.45 | 0.69 | 0.36 | 0.55 |
| 2 | 232316617 | rs10180468 | 0.085 | 0.584 | -0.613 | 0.029 | 0.002 | 0.008 | 0.008 | 0 | 0.29 | 0.22 | 0.22 | 0.06 | 0.28 | 0.2 |
| 2 | 232321956 | rs7598759 | -0.684 | 0.632 | -0.458 | 0.637 | 0.001 | -0.005 | -0.005 | 0 | 0.45 | 0.58 | 0.47 | 0.49 | 0.42 | 0.44 |
| 2 | 232322829 | rs6712148 | 0.511 | 0.612 | -0.487 | 0.443 | 0.001 | 0.013 | 0.013 | 1.231 | 0.27 | 0.2 | 0.21 | 0.06 | 0.26 | 0.15 |
| 2 | 232324780 | rs2442322 | -0.769 | 0.697 | -1.234 | 0.324 | -0.039 | -0.028 | -0.028 | 0.739 | 0.82 | 0.89 | 0.68 | 0.75 | 0.72 | 0.89 |
| 2 | 232325692 | rs12996154 | 0.256 | 0.449 | -1.044 | 0.583 | -0.035 | 0.1 | 0.1 | 1.229 | 0.33 | 0.13 | 0.24 | 0.13 | 0.33 | 0.2 |
| 2 | 232330209 | rs3806550 | 0.459 | 0.607 | -1.013 | 0.855 | 0.001 | 0.01 | 0.01 | 0 | 0.1 | 0.05 | 0.22 | 0.35 | 0.1 | 0.25 |
| 2 | 232331734 | rs4973410 | -0.689 | 0.512 | -0.896 | 0.123 | -0.002 | 0.028 | 0.028 | 0 | 0.54 | 0.59 | 0.53 | 0.73 | 0.49 | 0.63 |

| | | | | | | | | | | | | | | | | |
|---|---|---|---|---|---|---|---|---|---|---|---|---|---|---|---|---|
| 2 | 232343110 | rs4972995 | -0.649 | 0.539 | -0.601 | 0.327 | 0.005 | 0.012 | 0.012 | 0 | 0.31 | 0.48 | 0.39 | 0.67 | 0.32 | 0.5 |
| 2 | 232354247 | rs4973422 | 0.036 | 0.39 | -0.711 | 0.903 | 0 | 0.013 | 0.013 | 0 | 0.12 | 0.08 | 0.31 | 0.41 | 0.12 | 0.4 |
| 2 | 232368905 | rs7597348 | -1.328 | 0.151 | -0.57 | 0.702 | 0.002 | -0.002 | -0.002 | 0 | 0.37 | 0.33 | 0.48 | 0.67 | 0.38 | 0.67 |
| 2 | 232369533 | rs1584460 | 0.95 | 0.105 | -0.631 | 0.703 | -0.006 | -0.017 | -0.017 | 0 | 0.1 | 0.06 | 0.07 | 0.23 | 0.11 | 0.11 |
| 2 | 232373296 | rs7580705 | 1.539 | 0.186 | -1.074 | 0.099 | 0.003 | 0.092 | 0.092 | 8.712 | 0.24 | 0.07 | 0.14 | 0.08 | 0.21 | 0.08 |
| 2 | 232378231 | rs6750795 | -0.951 | 0.295 | -0.9 | 0.14 | -0.004 | 0.005 | 0.005 | 1.144 | 0.6 | 0.4 | 0.62 | 0.74 | 0.58 | 0.72 |
| 2 | 232379712 | rs12464121 | 0.621 | 0.357 | -1.062 | 0.094 | 0.011 | -0.048 | -0.048 | 0 | 0.34 | 0.13 | 0.41 | 0.48 | 0.28 | 0.45 |
| 2 | 232381245 | rs6754952 | -1.16 | 0.037 | -0.592 | -0.043 | 0.018 | -0.024 | -0.024 | 0.658 | 0.63 | 0.48 | 0.64 | 0.75 | 0.6 | 0.74 |
| 2 | 232381298 | rs11685196 | -0.51 | -0.038 | -0.515 | -0.062 | 0.009 | -0.006 | -0.006 | 0 | 0.52 | 0.36 | 0.55 | 0.53 | 0.48 | 0.63 |
| 2 | 232386454 | rs13413289 | 1.262 | -0.032 | -0.534 | -0.062 | 0.015 | 0.025 | 0.025 | 0 | 0.14 | 0.08 | 0.09 | 0.02 | 0.13 | 0.13 |
| 2 | 232387063 | rs10172595 | -0.479 | -0.148 | -0.591 | -0.12 | 0.013 | -0.005 | -0.005 | 0 | 0.51 | 0.34 | 0.55 | 0.53 | 0.48 | 0.65 |
| 2 | 232389301 | rs4973442 | 0.805 | -0.146 | -0.576 | -0.062 | 0.005 | -0.045 | -0.045 | 0 | 0.33 | 0.17 | 0.42 | 0.51 | 0.28 | 0.48 |
| 2 | 232394519 | rs10933376 | -0.65 | 0.42 | -0.424 | -0.206 | 0.057 | 0.075 | 0.075 | 0 | 0.87 | 0.58 | 0.8 | 0.81 | 0.87 | 0.93 |
| 2 | 232400839 | rs12992677 | 1.375 | 0.554 | -0.674 | -0.147 | -0.002 | -0.001 | -0.001 | 0 | 0.11 | 0.11 | 0.09 | 0.22 | 0.12 | 0.11 |
| 2 | 232401468 | rs4973444 | 1.106 | 0.6 | -0.322 | -0.141 | 0.019 | 0.032 | 0.032 | 0.001 | 0.1 | 0.16 | 0.17 | 0.19 | 0.13 | 0.07 |
| 2 | 232404469 | rs1797397 | 0.081 | 0.295 | -0.975 | -0.079 | 0.003 | -0.002 | -0.002 | 0.031 | 0.24 | 0.22 | 0.17 | 0.24 | 0.23 | 0.22 |
| 2 | 232406479 | rs1667309 | 1.359 | 0.296 | -1.023 | -0.177 | 0.006 | 0.007 | 0.007 | 0 | 0.13 | 0.11 | 0.07 | 0.02 | 0.11 | 0.11 |
| 2 | 232419666 | rs10165533 | -0.112 | 0.32 | -1.14 | -0.173 | 0.009 | 0.011 | 0.011 | 0 | 0.23 | 0.27 | 0.43 | 0.71 | 0.24 | 0.49 |
| 2 | 232423757 | rs1797393 | -0.529 | 0.313 | -1.402 | -0.384 | 0.019 | 0.008 | 0.008 | 0 | 0.54 | 0.67 | 0.55 | 0.42 | 0.6 | 0.43 |
| 2 | 232424046 | rs1797394 | -0.529 | 0.313 | -1.395 | -0.384 | 0.013 | 0.006 | 0.006 | 0 | 0.54 | 0.65 | 0.55 | 0.42 | 0.6 | 0.43 |
| 2 | 232433761 | rs1434519 | 1.436 | 0.274 | -1.902 | -0.519 | -0.089 | 0.169 | 0.169 | 0.488 | 0.25 | 0.02 | 0.17 | 0 | 0.3 | 0.22 |
| 2 | 232434661 | rs10460338 | -1.507 | 0.024 | -1.716 | -0.34 | -0.008 | 0.001 | 0.001 | 0 | 0.86 | 0.85 | 0.91 | 0.9 | 0.88 | 0.89 |
| 2 | 232438139 | rs12992206 | -0.638 | 0.356 | -1.777 | -0.549 | 0.005 | 0.004 | 0.004 | 0 | 0.51 | 0.54 | 0.51 | 0.43 | 0.54 | 0.39 |
| 2 | 232438403 | rs17199600 | -0.037 | 0.341 | -2.661 | -0.366 | -0.093 | 0.053 | 0.053 | 0.035 | 0.36 | 0.07 | 0.32 | 0.24 | 0.41 | 0.32 |
| 2 | 232455654 | rs1075708 | 0.924 | 0.238 | -2.553 | -0.444 | 0.001 | 0.086 | 0.086 | 0.472 | 0.36 | 0.13 | 0.27 | 0.19 | 0.33 | 0.19 |
| 2 | 232457335 | rs13028440 | 0.695 | 0.483 | -2.227 | -0.007 | -0.033 | -0.061 | -0.061 | 1.825 | 0.11 | 0.02 | 0.15 | 0.24 | 0.13 | 0.12 |
| 2 | 232461115 | rs4973009 | -0.693 | 0.332 | -1.697 | -0.512 | 0.003 | -0.018 | -0.018 | 0.927 | 0.86 | 0.89 | 0.92 | 1 | 0.88 | 0.89 |
| 2 | 232464308 | rs13384421 | 0.43 | 0.52 | -1.845 | -0.342 | -0.045 | -0.008 | -0.008 | 0.342 | 0.43 | 0.2 | 0.39 | 0.47 | 0.47 | 0.31 |
| 2 | 232465521 | rs10933379 | 0.056 | 0.567 | -1.538 | -0.533 | 0.005 | 0.019 | 0.019 | 0 | 0.46 | 0.6 | 0.48 | 0.57 | 0.39 | 0.57 |
| 2 | 232467731 | rs2060413 | -0.923 | 0.222 | -1.294 | -0.329 | 0.002 | -0.019 | -0.019 | 0.751 | 0.87 | 0.9 | 0.95 | 1 | 0.91 | 0.9 |
| 2 | 232472141 | rs10175445 | 0.275 | 0.528 | -1.593 | -0.185 | -0.011 | 0.006 | 0.006 | 0 | 0.4 | 0.27 | 0.42 | 0.42 | 0.43 | 0.31 |
| 2 | 232485833 | rs1991942 | 0.747 | 0.12 | -0.45 | -0.329 | -0.038 | -0.034 | -0.034 | 0 | 0.76 | 0.4 | 0.83 | 0.85 | 0.82 | 0.66 |
| 2 | 232495138 | rs6437030 | -0.814 | 0.779 | -0.518 | -0.27 | 0.02 | 0.008 | 0.008 | 0.212 | 0.11 | 0.14 | 0.15 | 0.17 | 0.17 | 0.26 |
| 2 | 232500132 | rs10204481 | 1.076 | 0.737 | -0.683 | -0.57 | -0.002 | 0.011 | 0.011 | 6.252 | 0.55 | 0.55 | 0.44 | 0.71 | 0.48 | 0.33 |
| 2 | 232517754 | rs6710658 | 1.072 | 0.247 | -0.668 | -0.331 | -0.007 | -0.011 | -0.011 | 0 | 0.61 | 0.57 | 0.53 | 0.7 | 0.66 | 0.44 |
| 2 | 232533884 | rs4973469 | 0.724 | 0.149 | -0.923 | -0.186 | -0.001 | -0.001 | -0.001 | 0 | 0.67 | 0.61 | 0.66 | 0.75 | 0.73 | 0.53 |
| 2 | 232540702 | rs4530357 | -0.449 | 0.558 | -1.099 | 0.19 | -0.002 | -0.001 | -0.001 | 0 | 0.46 | 0.45 | 0.38 | 0.49 | 0.49 | 0.34 |
| 2 | 232544906 | rs3856531 | -0.042 | 0.693 | -1.626 | -0.163 | -0.03 | 0.067 | 0.067 | 0.761 | 0.27 | 0.1 | 0.26 | 0.15 | 0.31 | 0.19 |
| 2 | 232550626 | rs4973473 | -0.189 | 0.166 | -1.703 | -0.771 | -0.024 | 0.118 | 0.118 | 0.043 | 0.42 | 0.21 | 0.58 | 0.86 | 0.52 | 0.6 |
| 2 | 232553362 | rs4973475 | -0.136 | 0.099 | -1.731 | -0.797 | -0.037 | 0.19 | 0.19 | 0.128 | 0.43 | 0.13 | 0.59 | 0.86 | 0.55 | 0.6 |
| 2 | 232558842 | rs6713758 | 0.224 | 0.008 | -1.321 | -0.509 | -0.01 | -0.007 | -0.007 | 0.27 | 0.15 | 0.07 | 0.19 | 0.17 | 0.21 | 0.15 |
| 2 | 232559206 | rs7604437 | -0.817 | -0.461 | -0.583 | -0.689 | -0.05 | -0.06 | -0.06 | 0 | 0.11 | 0.25 | 0.04 | 0 | 0.01 | 0.04 |
| 2 | 232560984 | rs10166046 | -0.785 | -0.393 | -0.14 | -0.659 | -0.008 | -0.009 | -0.009 | 0 | 0.22 | 0.66 | 0.22 | 0.18 | 0.25 | 0.17 |
| 2 | 232571214 | rs838441 | -1.819 | -0.823 | -0.323 | -0.985 | 0.041 | 0.077 | 0.077 | 2.034 | 0.38 | 0.82 | 0.29 | 0.2 | 0.31 | 0.3 |
| 2 | 232572011 | rs838440 | -0.677 | -0.85 | -0.227 | -1 | 0.008 | 0.004 | 0.004 | 0 | 0.17 | 0.14 | 0.07 | 0.03 | 0.08 | 0.15 |
| 2 | 232572381 | rs6737601 | -0.313 | -0.829 | -0.082 | -0.969 | -0.007 | -0.013 | -0.013 | 0 | 0.21 | 0.61 | 0.21 | 0.17 | 0.23 | 0.16 |
| 2 | 232576431 | rs2070936 | 0.94 | -0.566 | -0.227 | -0.368 | 0.003 | 0.001 | 0.001 | 0.29 | 0.45 | 0.57 | 0.33 | 0.39 | 0.28 | 0.48 |
| 2 | 232580258 | rs838437 | -1.165 | -0.185 | 0.209 | 0.074 | -0.021 | 0.088 | 0.088 | 0 | 0.19 | 0.32 | 0.26 | 0.34 | 0.1 | 0.17 |
| 2 | 232583513 | rs838436 | -0.713 | -0.112 | 0.412 | -0.844 | -0.017 | -0.048 | -0.048 | 0 | 0.12 | 0.29 | 0.06 | 0.03 | 0.07 | 0.09 |
| 2 | 232584807 | rs11695646 | -0.343 | 0.141 | 0.822 | -0.848 | 0.023 | -0.004 | -0.004 | 0 | 0.2 | 0.37 | 0.21 | 0.17 | 0.26 | 0.16 |
| 2 | 232597042 | rs838425 | 0.023 | -0.012 | 0.73 | -1.532 | 0.002 | -0.003 | -0.003 | 0.007 | 0.05 | 0.12 | 0.06 | 0.03 | 0.06 | 0.07 |
| 2 | 232607456 | rs838448 | -1.189 | -0.08 | 1.231 | -1.735 | -0.011 | -0.072 | -0.072 | 0.108 | 0.29 | 0.46 | 0.27 | 0.11 | 0.32 | 0.25 |
| 2 | 232608294 | rs2570991 | -1.171 | -0.072 | 1.236 | -1.731 | -0.01 | -0.071 | -0.071 | 0 | 0.29 | 0.46 | 0.27 | 0.11 | 0.32 | 0.25 |
| 2 | 232609805 | rs17199642 | 0.371 | -0.039 | 0.906 | -1.696 | -0.021 | 0.033 | 0.033 | 0 | 0.17 | 0.05 | 0.19 | 0.08 | 0.26 | 0.15 |
| 2 | 232630459 | rs17271824 | 0.371 | -0.039 | 0.906 | -1.725 | -0.021 | 0.036 | 0.036 | 0 | 0.17 | 0.05 | 0.19 | 0.08 | 0.26 | 0.15 |
| 2 | 232647754 | rs13424754 | -1.043 | -0.313 | 1.418 | -1.9 | 0 | 0.02 | 0.02 | 0.879 | 0.28 | 0.3 | 0.26 | 0.11 | 0.35 | 0.27 |
| 2 | 232661068 | rs13408416 | -0.244 | -0.321 | 1.425 | -1.908 | -0.006 | -0.007 | -0.007 | 2.664 | 0.07 | 0.1 | 0.01 | 0 | 0.01 | 0.05 |
| 2 | 232667302 | rs10185448 | -0.284 | -0.329 | 1.432 | -1.912 | -0.009 | -0.009 | -0.009 | 0.019 | 0.07 | 0.1 | 0.01 | 0 | 0.01 | 0.05 |
| 2 | 232672932 | rs1246014 | 1.814 | -0.173 | 0.975 | -2.237 | -0.013 | 0.181 | 0.181 | 0.874 | 0.41 | 0.04 | 0.46 | 0.86 | 0.31 | 0.45 |
| 2 | 232684296 | rs6705872 | -0.149 | -0.741 | 1.569 | -1.886 | 0.038 | 0.024 | 0.024 | 0 | 0.05 | 0.27 | 0.02 | 0 | 0.01 | 0.02 |
| 2 | 232717857 | rs920400 | -1.811 | -0.883 | 0.608 | -1.911 | 0.049 | 0.15 | 0.15 | 0.347 | 0.4 | 0.82 | 0.32 | 0.11 | 0.36 | 0.29 |
| 2 | 232718088 | rs11890626 | 0.677 | -0.887 | 0.299 | -1.917 | -0.004 | 0.015 | 0.015 | 0 | 0.16 | 0.09 | 0.2 | 0.08 | 0.26 | 0.14 |
| 2 | 232720739 | rs10195385 | -1.662 | -0.798 | -0.116 | -1.857 | 0.041 | 0.196 | 0.196 | 0.515 | 0.41 | 0.88 | 0.33 | 0.11 | 0.38 | 0.3 |
| 2 | 232722314 | rs13394372 | -0.414 | -0.726 | 0.063 | -1.739 | -0.012 | -0.013 | -0.013 | 0.152 | 0.07 | 0.1 | 0.01 | 0 | 0.01 | 0.05 |
| 2 | 232726300 | rs1866599 | -0.968 | -0.627 | -0.035 | -1.762 | 0.045 | 0.119 | 0.119 | 0.199 | 0.35 | 0.78 | 0.27 | 0.08 | 0.33 | 0.22 |
| 2 | 232728437 | rs2067564 | -0.437 | -0.162 | 0.33 | -1.542 | 0.033 | -0.033 | -0.033 | 0 | 0.33 | 0.51 | 0.32 | 0.18 | 0.45 | 0.33 |
| 2 | 232730151 | rs4973492 | 0.91 | 0.065 | -0.328 | -1.504 | -0.116 | 0.028 | 0.028 | 0.56 | 0.21 | 0.03 | 0.23 | 0.1 | 0.34 | 0.18 |
| 2 | 232744209 | rs10179086 | 0.009 | -0.134 | 0.211 | -1.362 | -0.03 | -0.09 | -0.09 | 1.671 | 0.24 | 0.48 | 0.22 | 0.08 | 0.27 | 0.2 |
| 2 | 232745518 | rs4973022 | -1.099 | -0.153 | -0.886 | -1.326 | 0.006 | 0.037 | 0.037 | 3.799 | 0.62 | 0.9 | 0.55 | 0.25 | 0.74 | 0.56 |

| | | | | | | | | | | | | | | | | |
|---|---|---|---|---|---|---|---|---|---|---|---|---|---|---|---|---|
| 2 | 232747892 | rs736326 | -0.315 | 0.006 | -0.146 | -1.106 | -0.018 | -0.098 | -0.098 | 0 | 0.25 | 0.48 | 0.24 | 0.07 | 0.29 | 0.2 |
| 2 | 232749836 | rs2580850 | -0.626 | 0.192 | -0.924 | -1.037 | -0.011 | 0.071 | 0.071 | 0.074 | 0.6 | 0.8 | 0.53 | 0.25 | 0.67 | 0.48 |
| 2 | 232756528 | rs2679180 | -0.703 | 0.151 | -0.851 | -1.035 | 0.006 | 0.064 | 0.064 | 0.038 | 0.6 | 0.75 | 0.52 | 0.25 | 0.69 | 0.48 |
| 2 | 232772215 | rs16828513 | -0.479 | -0.344 | -0.511 | -0.812 | -0.019 | -0.052 | -0.052 | 1.552 | 0.26 | 0.41 | 0.29 | 0.1 | 0.28 | 0.21 |
| 2 | 232779223 | rs2679184 | -0.854 | -0.53 | -1.154 | -0.923 | -0.014 | -0.02 | -0.02 | 0 | 0.36 | 0.54 | 0.31 | 0.17 | 0.28 | 0.24 |
| 2 | 232788325 | rs3107179 | 0.271 | -0.235 | -1.542 | -0.503 | 0.039 | 0.171 | 0.171 | 0 | 0.48 | 0.14 | 0.37 | 0.2 | 0.58 | 0.4 |
| 2 | 232791948 | rs5262 | -0.538 | 0.318 | -0.343 | -0.198 | -0.01 | -0.024 | -0.024 | 0 | 0.21 | 0.51 | 0.28 | 0.18 | 0.2 | 0.22 |
| 2 | 232796002 | rs893740 | 0.466 | 1.022 | -0.643 | -0.674 | 0.001 | -0.017 | -0.017 | 0.031 | 0.75 | 0.71 | 0.74 | 0.87 | 0.67 | 0.81 |
| 2 | 232796610 | rs749052 | 0.139 | 0.28 | -0.779 | -1.319 | 0.011 | 0.003 | 0.003 | 0 | 0.17 | 0.13 | 0.05 | 0.01 | 0.07 | 0.09 |
| 2 | 232797150 | rs12464230 | 0.092 | 0.063 | -0.123 | -1.667 | 0.001 | -0.045 | -0.045 | 0 | 0.09 | 0.21 | 0.14 | 0.03 | 0.07 | 0.15 |
| 2 | 232797966 | rs2580816 | -0.925 | 0.176 | 0.279 | -0.727 | -0.02 | 0.019 | 0.019 | 0.057 | 0.31 | 0.39 | 0.46 | 0.51 | 0.21 | 0.32 |
| 2 | 232798527 | rs2679177 | -0.352 | -0.157 | -0.282 | -1.321 | 0.002 | -0.041 | -0.041 | 0.131 | 0.61 | 0.53 | 0.7 | 0.8 | 0.6 | 0.73 |
| 2 | 232798723 | rs2580817 | 0.774 | -0.213 | -0.377 | -1.094 | 0.004 | 0.01 | 0.01 | 0 | 0.17 | 0.11 | 0.07 | 0.01 | 0.11 | 0.08 |
| 2 | 232803452 | rs7588259 | 0.099 | 0.328 | -0.791 | -1.689 | -0.019 | 0.028 | 0.028 | 0.297 | 0.57 | 0.73 | 0.51 | 0.46 | 0.58 | 0.77 |
| 2 | 232804155 | rs2580821 | 1.508 | 0.546 | -0.698 | -1.575 | 0.022 | 0.066 | 0.066 | 0 | 0.16 | 0.04 | 0.06 | 0.01 | 0.09 | 0.07 |
| 2 | 232804522 | rs10185460 | -0.523 | 0.549 | -0.67 | -1.544 | 0.008 | 0.006 | 0.006 | 0 | 0.76 | 0.71 | 0.57 | 0.46 | 0.67 | 0.84 |
| 2 | 232809066 | rs16825969 | 1.861 | -0.067 | -0.644 | -1.326 | -0.004 | -0.041 | -0.041 | 0 | 0.19 | 0.04 | 0.13 | 0.3 | 0.16 | 0.37 |
| 2 | 232816237 | rs2679163 | -0.625 | -0.043 | -0.724 | -1.352 | 0.007 | 0.156 | 0.156 | 0.002 | 0.53 | 0.74 | 0.65 | 0.79 | 0.39 | 0.72 |
| 2 | 232820171 | rs1000837 | -0.286 | -0.035 | -0.514 | -1.366 | 0.023 | 0.058 | 0.058 | 0 | 0.5 | 0.61 | 0.41 | 0.33 | 0.37 | 0.63 |
| 2 | 232838890 | rs12988522 | 1.914 | 0.025 | -0.762 | -1.258 | -0.035 | -0.07 | -0.07 | 0.476 | 0.19 | 0 | 0.13 | 0.29 | 0.17 | 0.35 |
| 2 | 232843189 | rs13029658 | 0.679 | -0.108 | -0.746 | -1.383 | 0.015 | 0.033 | 0.033 | 0 | 0.36 | 0.03 | 0.19 | 0.29 | 0.26 | 0.42 |
| 2 | 232857909 | rs3116154 | -1.726 | -0.126 | -0.84 | -1.421 | 0.004 | 0.084 | 0.084 | 0 | 0.85 | 0.99 | 0.94 | 1 | 0.91 | 0.93 |
| 2 | 232860077 | rs3116162 | -1.484 | -0.113 | -0.828 | -1.414 | 0.027 | 0.073 | 0.073 | 0 | 0.83 | 0.98 | 0.94 | 1 | 0.91 | 0.93 |
| 2 | 232863822 | rs3100598 | -1.014 | -0.002 | -0.715 | -1.312 | -0.02 | 0.023 | 0.023 | 0 | 0.64 | 0.97 | 0.81 | 0.71 | 0.72 | 0.57 |
| 2 | 232870178 | rs6745958 | -0.225 | 0.026 | -0.16 | -1.268 | 0.002 | -0.067 | -0.067 | 0 | 0.13 | 0.55 | 0.17 | 0.03 | 0.1 | 0.18 |
| 2 | 232879671 | rs723044 | 0.107 | -0.103 | 0.154 | -1.043 | -0.001 | -0.022 | -0.022 | 0 | 0.06 | 0.24 | 0.06 | 0 | 0.02 | 0.06 |
| 2 | 232880712 | rs11885823 | 0.792 | -0.072 | 0.239 | -1.044 | 0.016 | -0.026 | -0.026 | 0 | 0.08 | 0.21 | 0.12 | 0.03 | 0.08 | 0.12 |
| 2 | 232891737 | rs3116230 | -1.115 | -0.173 | -0.044 | -1.161 | 0.01 | 0.036 | 0.036 | 0 | 0.64 | 0.95 | 0.81 | 0.71 | 0.74 | 0.57 |
| 2 | 232907292 | rs3100612 | 0.672 | -0.229 | 0.116 | -1.116 | -0.049 | 0.185 | 0.185 | 0 | 0.43 | 0.14 | 0.37 | 0.19 | 0.59 | 0.28 |
| 2 | 232911825 | rs11901253 | 1.03 | -0.207 | 0.182 | -1.123 | 0.014 | -0.026 | -0.026 | 0.071 | 0.08 | 0.2 | 0.11 | 0.03 | 0.08 | 0.12 |
| 2 | 232915543 | rs3116237 | -1.126 | -0.217 | 0.183 | -1.126 | 0.005 | 0.045 | 0.045 | 0 | 0.63 | 0.92 | 0.8 | 0.7 | 0.72 | 0.57 |
| 2 | 232920430 | rs3116240 | 0.233 | -0.208 | 0.298 | -1.122 | 0.017 | 0.067 | 0.067 | 0 | 0.47 | 0.35 | 0.62 | 0.67 | 0.62 | 0.39 |
| 2 | 232929252 | rs12466111 | 1.052 | -0.127 | 0.364 | -1.071 | 0.008 | -0.026 | -0.026 | 0.586 | 0.08 | 0.16 | 0.11 | 0.03 | 0.08 | 0.12 |
| 2 | 232950374 | rs3100585 | -0.561 | -0.161 | 0.34 | -1.065 | -0.001 | -0.004 | -0.004 | 0 | 0.58 | 0.56 | 0.74 | 0.7 | 0.7 | 0.51 |
| 2 | 232954048 | rs3116155 | -1.136 | -0.163 | 0.061 | -1.051 | 0.022 | 0.035 | 0.035 | 0 | 0.64 | 0.94 | 0.81 | 0.71 | 0.74 | 0.58 |
| 2 | 232970387 | rs3100593 | -1.066 | -0.185 | 0.008 | -1.03 | 0.008 | 0.01 | 0.01 | 0 | 0.66 | 0.96 | 0.81 | 0.7 | 0.75 | 0.58 |
| 2 | 232989831 | rs3116168 | -0.769 | -0.121 | 0.214 | -1.079 | 0.022 | 0.074 | 0.074 | 0.005 | 0.62 | 0.8 | 0.57 | 0.24 | 0.71 | 0.47 |
| 2 | 232990738 | rs3103264 | 0.478 | -0.14 | 0.226 | -1.133 | 0.004 | 0.197 | 0.197 | 0 | 0.47 | 0.18 | 0.38 | 0.2 | 0.61 | 0.28 |
| 2 | 233005775 | rs3103257 | -0.985 | -0.151 | 0.28 | -1.034 | 0.016 | 0.019 | 0.019 | 0 | 0.64 | 0.85 | 0.79 | 0.69 | 0.72 | 0.56 |
| 2 | 233031057 | rs4973034 | 2.228 | -0.378 | -0.074 | -1.169 | -0.02 | -0.089 | -0.089 | 4.893 | 0.19 | 0 | 0.12 | 0.3 | 0.15 | 0.35 |
| 2 | 233034472 | rs3103295 | 0.339 | -0.338 | 0.042 | -1.199 | 0.037 | 0.077 | 0.077 | 0 | 0.5 | 0.3 | 0.62 | 0.66 | 0.64 | 0.36 |
| 2 | 233056621 | rs10498260 | 1.892 | -0.249 | -0.321 | -1.256 | -0.019 | -0.074 | -0.074 | 4.351 | 0.22 | 0.02 | 0.2 | 0.3 | 0.19 | 0.43 |
| 2 | 233067382 | rs3103286 | -0.855 | 0.256 | -0.506 | -1.368 | -0.036 | 0.131 | 0.131 | 0 | 0.67 | 0.89 | 0.74 | 0.83 | 0.6 | 0.75 |
| 2 | 233076420 | rs3116203 | 0.415 | -0.016 | 0.044 | -1.202 | 0.01 | 0.009 | 0.009 | 0 | 0.4 | 0.41 | 0.25 | 0.31 | 0.26 | 0.52 |
| 2 | 233094365 | rs6437055 | -0.842 | 0.298 | -0.391 | -1.363 | -0.075 | 0.215 | 0.215 | 0 | 0.57 | 0.86 | 0.72 | 0.85 | 0.46 | 0.71 |
| 2 | 233097786 | rs2087229 | 2.213 | 0.204 | -0.406 | -0.893 | -0.043 | -0.113 | -0.113 | 1.123 | 0.19 | 0.03 | 0.13 | 0.39 | 0.17 | 0.39 |
| 2 | 233098655 | rs3116217 | 2.323 | 0.213 | -0.406 | -0.892 | -0.016 | -0.11 | -0.11 | 0 | 0.19 | 0.03 | 0.13 | 0.39 | 0.16 | 0.38 |
| 2 | 233101304 | rs1872920 | 0.853 | 0.115 | -0.381 | -0.952 | -0.037 | -0.074 | -0.074 | 0 | 0.34 | 0.07 | 0.21 | 0.4 | 0.29 | 0.48 |
| 2 | 233108857 | rs882098 | -0.993 | 0.078 | 0.151 | -0.831 | -0.023 | 0.36 | 0.36 | 0.805 | 0.12 | 0.5 | 0.28 | 0.39 | 0.05 | 0.12 |
| 2 | 233120758 | rs2633266 | 1.581 | 0.156 | -0.041 | -0.922 | 0.005 | 0.031 | 0.031 | 0 | 0.15 | 0.05 | 0.07 | 0.01 | 0.11 | 0.09 |
| 2 | 233122412 | rs2853362 | -0.881 | 0.15 | -0.084 | -1.006 | -0.02 | 0.244 | 0.244 | 0 | 0.54 | 0.85 | 0.62 | 0.85 | 0.39 | 0.67 |
| 2 | 233122885 | rs2633265 | -0.402 | 0.187 | 0.409 | -0.921 | 0.029 | 0.025 | 0.025 | 0 | 0.49 | 0.61 | 0.36 | 0.4 | 0.34 | 0.58 |
| 2 | 233124685 | rs2633264 | 0.54 | 0.191 | 0.281 | -0.875 | 0 | -0.058 | -0.058 | 0 | 0.22 | 0.22 | 0.18 | 0.04 | 0.18 | 0.14 |
| 2 | 233131825 | rs2633261 | 0.729 | 0.194 | 0.399 | -0.871 | -0.001 | -0.036 | -0.036 | 0 | 0.19 | 0.16 | 0.1 | 0.01 | 0.14 | 0.1 |
| 2 | 233132810 | rs2633260 | 0.054 | 0.182 | 0.297 | -0.875 | -0.018 | -0.089 | -0.089 | 0 | 0.25 | 0.29 | 0.19 | 0.03 | 0.17 | 0.13 |
| 2 | 233137318 | rs2633254 | -1.253 | 0.249 | -0.159 | -1.052 | -0.053 | 0.128 | 0.128 | 0.791 | 0.7 | 0.93 | 0.73 | 0.87 | 0.63 | 0.69 |
| 2 | 233145683 | rs11686328 | 0.564 | -0.098 | 0.681 | -0.435 | 0.052 | -0.005 | -0.005 | 0 | 0.08 | 0.23 | 0.11 | 0.03 | 0.07 | 0.05 |
| 2 | 233149600 | rs13020779 | -0.809 | -0.191 | 0.707 | -0.394 | -0.005 | 0.021 | 0.021 | 0 | 0.3 | 0.54 | 0.46 | 0.46 | 0.23 | 0.22 |
| 2 | 233161818 | rs4973517 | 0.594 | -0.122 | 0.284 | -0.403 | -0.018 | -0.06 | -0.06 | 1.412 | 0.33 | 0.04 | 0.2 | 0.38 | 0.27 | 0.48 |
| 2 | 233166406 | rs4973518 | -1.431 | 0.008 | 1.264 | 0.035 | 0.004 | 0.262 | 0.262 | 0.114 | 0.15 | 0.45 | 0.38 | 0.48 | 0.11 | 0.13 |
| 2 | 233169114 | rs11903728 | 0.189 | 0.006 | 1.346 | -0.002 | 0.008 | -0.026 | -0.026 | 0 | 0.08 | 0.16 | 0.11 | 0.03 | 0.06 | 0.04 |
| 2 | 233171039 | rs6437059 | -0.755 | 0.011 | 1.28 | 0.011 | 0.037 | -0.004 | -0.004 | 4.188 | 0.1 | 0.37 | 0.17 | 0.05 | 0.08 | 0.07 |
| 2 | 233180919 | rs871411 | -0.954 | -0.006 | 1.221 | 0.143 | -0.005 | 0.08 | 0.08 | 0 | 0.27 | 0.44 | 0.44 | 0.45 | 0.22 | 0.22 |
| 2 | 233193002 | rs6713534 | -0.229 | 0.088 | 1.362 | 0.179 | 0.039 | -0.019 | -0.019 | 0 | 0.08 | 0.25 | 0.12 | 0.03 | 0.07 | 0.05 |
| 2 | 233201328 | rs3811578 | -1.406 | -0.018 | 1.306 | 0.662 | -0.003 | 0.145 | 0.145 | 0.026 | 0.1 | 0.17 | 0.26 | 0.45 | 0.09 | 0.07 |
| 2 | 233204359 | rs11896101 | -0.694 | 0.178 | 1.115 | -0.025 | 0.032 | -0.029 | -0.029 | 0.038 | 0.1 | 0.29 | 0.12 | 0.03 | 0.07 | 0.05 |
| 2 | 233208988 | rs6756905 | -0.828 | 0.173 | 0.988 | -0.027 | 0.038 | -0.033 | -0.033 | 0.197 | 0.11 | 0.37 | 0.13 | 0.03 | 0.07 | 0.05 |
| 2 | 233211861 | rs6747223 | -0.512 | 0.16 | 0.287 | -0.306 | 0.004 | -0.025 | -0.025 | 0.475 | 0.3 | 0.25 | 0.12 | 0 | 0.21 | 0.21 |
| 2 | 233215403 | rs883013 | -0.491 | 0.171 | 0.255 | 0.315 | -0.002 | 0.024 | 0.024 | 0 | 0.25 | 0.28 | 0.37 | 0.49 | 0.17 | 0.21 |

| | | | | | | | | | | | | | | | | |
|---|---|---|---|---|---|---|---|---|---|---|---|---|---|---|---|---|
| 2 | 233218894 | rs7583232 | -0.131 | 0.361 | 0.471 | 0.223 | -0.006 | -0.081 | -0.081 | 0 | 0.17 | 0.25 | 0.07 | 0.01 | 0.13 | 0.14 |
| 2 | 233227833 | rs2741302 | 0.442 | 0.846 | 0.218 | 0.717 | 0.032 | 0.171 | 0.171 | 0.137 | 0.92 | 0.83 | 0.76 | 0.49 | 0.86 | 0.86 |
| 2 | 233232064 | rs1850593 | 0.726 | 0.583 | 0.147 | 0.409 | 0.002 | 0.013 | 0.013 | 0.505 | 0.69 | 0.37 | 0.66 | 0.53 | 0.75 | 0.73 |
| 2 | 233235331 | rs2741309 | -0.299 | 0.459 | -0.492 | -0.029 | -0.01 | -0.029 | -0.029 | 0.037 | 0.79 | 0.83 | 0.78 | 0.6 | 0.72 | 0.77 |
| 2 | 233236078 | rs2741310 | 0.26 | 0.338 | -0.657 | -0.311 | -0.012 | 0.026 | 0.026 | 1.155 | 0.32 | 0.39 | 0.5 | 0.56 | 0.32 | 0.48 |
| 2 | 233242964 | rs2014683 | -0.015 | 0.394 | -1.111 | -0.46 | 0.041 | 0.044 | 0.044 | 0.819 | 0.5 | 0.3 | 0.58 | 0.61 | 0.45 | 0.53 |
| 2 | 233250683 | rs3748971 | 1.252 | 0.24 | -0.909 | -0.073 | -0.021 | -0.008 | -0.008 | 0 | 0.11 | 0.01 | 0.04 | 0.1 | 0.1 | 0.08 |
| 2 | 233264910 | rs2139026 | 0.309 | 0.415 | -0.545 | 0.013 | -0.003 | 0.008 | 0.008 | 0 | 0.73 | 0.34 | 0.75 | 0.68 | 0.81 | 0.74 |
| 2 | 233271287 | rs11678251 | 0.344 | 0.786 | -0.739 | 0.044 | -0.003 | 0.021 | 0.021 | 0 | 0.05 | 0.04 | 0.06 | 0.22 | 0.08 | 0.07 |
| 2 | 233280239 | rs2741321 | 0.426 | 0.796 | -0.749 | 0.012 | -0.01 | 0.006 | 0.006 | 0 | 0.05 | 0.03 | 0.06 | 0.2 | 0.08 | 0.07 |
| 2 | 233282536 | rs790051 | 0.45 | 0.425 | -0.592 | -0.448 | 0.008 | -0.027 | -0.027 | 0 | 0.31 | 0.29 | 0.14 | 0.04 | 0.22 | 0.27 |
| 2 | 233286473 | rs2853383 | 0.64 | 0.576 | -0.656 | -0.291 | -0.01 | 0.006 | 0.006 | 0 | 0.05 | 0.03 | 0.06 | 0.2 | 0.08 | 0.07 |
| 2 | 233290980 | rs2853386 | 0.026 | 0.692 | -0.654 | -0.579 | 0.005 | 0.021 | 0.021 | 0 | 0.44 | 0.59 | 0.53 | 0.79 | 0.39 | 0.51 |
| 2 | 233300816 | rs12996863 | 1.31 | 0.975 | -1.296 | -0.561 | -0.105 | 0.078 | 0.078 | 0 | 0.22 | 0.04 | 0.2 | 0.1 | 0.33 | 0.2 |
| 2 | 233303133 | rs790025 | 0.49 | 0.584 | -1.176 | -0.516 | 0.053 | 0.14 | 0.14 | 0.005 | 0.3 | 0.11 | 0.13 | 0.04 | 0.22 | 0.25 |
| 2 | 233309066 | rs2678505 | -0.111 | 1.466 | -0.506 | -0.024 | 0.013 | 0.052 | 0.052 | 0 | 0.06 | 0.08 | 0.06 | 0.2 | 0.08 | 0.07 |
| 2 | 233309548 | rs790040 | 0.043 | 0.997 | -0.382 | -0.114 | 0.026 | 0.001 | 0.001 | 0 | 0.67 | 0.58 | 0.59 | 0.63 | 0.53 | 0.69 |
| 2 | 233310149 | rs790044 | 0.051 | 1.016 | -0.593 | -0.104 | 0.025 | 0 | 0 | 0 | 0.67 | 0.58 | 0.59 | 0.63 | 0.53 | 0.69 |
| 2 | 233320885 | rs3762519 | -0.271 | 0.573 | -1.14 | -0.044 | 0.001 | 0.001 | 0.001 | 0.028 | 0.12 | 0.1 | 0.08 | 0.08 | 0.11 | 0.08 |
| 2 | 233327208 | rs4521030 | 0.204 | 0.79 | -1.517 | -0.097 | -0.019 | -0.01 | -0.01 | 0 | 0.44 | 0.26 | 0.53 | 0.46 | 0.49 | 0.48 |
| 2 | 233328630 | rs2678484 | -0.16 | 0.746 | -1.33 | -0.079 | 0 | 0.015 | 0.015 | 0.006 | 0.14 | 0.18 | 0.09 | 0.24 | 0.13 | 0.15 |
| 2 | 233331507 | rs2741271 | -0.556 | 0.821 | -1.206 | -0.028 | -0.009 | -0.016 | -0.016 | 3.092 | 0.59 | 0.46 | 0.64 | 0.7 | 0.65 | 0.64 |
| 2 | 233340715 | rs12616972 | -0.317 | 0.496 | -1.398 | 0.128 | 0.012 | -0.034 | -0.034 | 0 | 0.36 | 0.22 | 0.5 | 0.5 | 0.39 | 0.4 |
| 2 | 233346851 | rs1190430 | -0.032 | 0.12 | -1.855 | -0.389 | -0.006 | 0.017 | 0.017 | 0 | 0.6 | 0.27 | 0.52 | 0.34 | 0.62 | 0.45 |
| 2 | 233346865 | rs909431 | 0.748 | 0.612 | -1.78 | 0.189 | -0.006 | 0.01 | 0.01 | 0.36 | 0.15 | 0.12 | 0.34 | 0.45 | 0.21 | 0.28 |
| 2 | 233348647 | rs2241755 | -0.264 | -0.057 | -1.519 | -0.405 | 0.009 | 0.034 | 0.034 | 0 | 0.18 | 0.01 | 0.13 | 0.07 | 0.18 | 0.12 |
| 2 | 233360483 | rs7571362 | -0.098 | -0.013 | -1.356 | -0.486 | 0.015 | 0.038 | 0.038 | 0 | 0.17 | 0.09 | 0.1 | 0.1 | 0.11 | 0.22 |
| 2 | 233360507 | rs2697806 | 0.73 | 0.123 | -0.568 | -0.376 | -0.028 | -0.08 | -0.08 | 0 | 0.73 | 0.51 | 0.83 | 0.92 | 0.75 | 0.78 |
| 2 | 233362267 | rs1190462 | -0.515 | 0.177 | -1.572 | -0.147 | 0.014 | 0.054 | 0.054 | 0 | 0.22 | 0.05 | 0.15 | 0.08 | 0.23 | 0.22 |
| 2 | 233375784 | rs2117770 | 0.341 | 0.505 | -0.727 | -0.708 | 0.002 | -0.078 | -0.078 | 2.811 | 0.68 | 0.66 | 0.79 | 0.94 | 0.67 | 0.78 |
| 2 | 233381606 | rs17360319 | 0.422 | 0.581 | -1.114 | -0.659 | -0.019 | 0.056 | 0.056 | 0 | 0.1 | 0 | 0.07 | 0.01 | 0.09 | 0.08 |
| 2 | 233383416 | rs2697800 | 0.506 | 0.869 | -1.261 | -0.286 | -0.006 | -0.012 | -0.012 | 0 | 0.16 | 0.15 | 0.09 | 0.26 | 0.2 | 0.13 |
| 2 | 233387635 | rs733603 | -0.6 | 0.76 | -0.821 | -0.423 | 0.002 | -0.021 | -0.021 | 0 | 0.83 | 0.83 | 0.85 | 0.94 | 0.79 | 0.89 |
| 2 | 233388311 | rs733602 | 1.203 | 0.662 | -1.51 | -0.148 | -0.042 | -0.088 | -0.088 | 0 | 0.22 | 0.09 | 0.44 | 0.36 | 0.28 | 0.35 |
| 2 | 233390937 | rs2245601 | -0.027 | 0.618 | -1.082 | -0.218 | -0.015 | 0.011 | 0.011 | 0 | 0.39 | 0.35 | 0.55 | 0.65 | 0.45 | 0.48 |
| 2 | 233400074 | rs2767 | 0.459 | 0.357 | -1.16 | -0.232 | -0.005 | -0.019 | -0.019 | 0 | 0.3 | 0.27 | 0.56 | 0.56 | 0.35 | 0.43 |
| 2 | 233406998 | rs1881492 | -0.707 | 0.434 | -1.1 | -0.306 | 0.001 | -0.01 | -0.01 | 0 | 0.81 | 0.79 | 0.83 | 0.87 | 0.78 | 0.84 |
| 2 | 233407120 | rs2853462 | 0.738 | 0.555 | -1.46 | -0.13 | -0.026 | -0.03 | -0.03 | 0.717 | 0.16 | 0.11 | 0.11 | 0.29 | 0.21 | 0.14 |
| 2 | 233410294 | rs2099489 | 0.627 | -0.027 | -1.416 | -0.422 | 0.024 | 0.035 | 0.035 | 0 | 0.25 | 0.11 | 0.13 | 0.12 | 0.19 | 0.22 |
| 2 | 233411532 | rs11688448 | 0.627 | 0.013 | -1.382 | -0.419 | 0.024 | 0.035 | 0.035 | 0 | 0.25 | 0.11 | 0.13 | 0.12 | 0.19 | 0.22 |
| 2 | 233412820 | rs4973539 | 0.627 | 0.013 | -1.382 | -0.419 | 0.024 | 0.035 | 0.035 | 0 | 0.25 | 0.11 | 0.13 | 0.12 | 0.19 | 0.22 |
| 2 | 233414721 | rs11683262 | 0.634 | 0.01 | -1.383 | -0.412 | 0.024 | 0.033 | 0.033 | 0 | 0.25 | 0.1 | 0.13 | 0.12 | 0.19 | 0.22 |
| 2 | 233417552 | rs1190452 | -0.204 | 0.532 | -0.976 | -0.025 | 0.02 | 0.047 | 0.047 | 0 | 0.85 | 0.79 | 0.82 | 0.59 | 0.73 | 0.82 |
| 2 | 233418920 | rs1190449 | -0.035 | 0.118 | -1.158 | -0.164 | -0.002 | 0.039 | 0.039 | 0.046 | 0.56 | 0.73 | 0.77 | 0.59 | 0.57 | 0.61 |
| 2 | 233426526 | rs1656402 | 0.425 | 0.49 | -1.339 | -0.072 | -0.008 | -0.004 | -0.004 | 0.53 | 0.3 | 0.41 | 0.42 | 0.28 | 0.27 | 0.24 |
| 2 | 233438786 | rs2293338 | 0.055 | 1.607 | -1.435 | 0.766 | -0.036 | -0.036 | -0.036 | 0.796 | 0.12 | 0.07 | 0.12 | 0.33 | 0.26 | 0.14 |
| 2 | 233440924 | rs7571154 | -0.322 | 1.552 | -1.478 | 0.621 | -0.002 | 0.007 | 0.007 | 0 | 0.14 | 0.15 | 0.14 | 0.37 | 0.27 | 0.19 |
| 2 | 233446568 | rs17296210 | 0.112 | 1.012 | -1.559 | 0.398 | -0.09 | -0.117 | -0.117 | 0.094 | 0.21 | 0.01 | 0.31 | 0.31 | 0.3 | 0.37 |
| 2 | 233470843 | rs7571133 | 0.182 | 0.909 | -0.962 | -0.007 | -0.033 | 0.074 | 0.074 | 0 | 0.6 | 0.75 | 0.46 | 0.23 | 0.48 | 0.35 |
| 2 | 233485696 | rs7607256 | 0.23 | 0.947 | -0.693 | 0.155 | -0.002 | 0.034 | 0.034 | 0 | 0.58 | 0.68 | 0.56 | 0.65 | 0.6 | 0.33 |
| 2 | 233490451 | rs2344369 | -0.769 | 0.982 | 0.413 | 0.825 | 0.054 | 0.208 | 0.208 | 0 | 0.11 | 0.23 | 0.16 | 0.56 | 0.19 | 0.11 |
| 2 | 233496975 | rs11902420 | -1.103 | 0.778 | 0.594 | 0.607 | -0.017 | 0.138 | 0.138 | 1.659 | 0.12 | 0.42 | 0.26 | 0.22 | 0.06 | 0.13 |
| 2 | 233498669 | rs2276557 | -0.707 | 1.085 | -0.166 | 0.7 | 0.017 | 0.072 | 0.072 | 0 | 0.1 | 0.13 | 0.16 | 0.63 | 0.19 | 0.11 |
| 2 | 233508677 | rs4072149 | 0.5 | 0.905 | -0.26 | 0.103 | 0 | -0.001 | -0.001 | 0.481 | 0.62 | 0.61 | 0.49 | 0.31 | 0.54 | 0.56 |
| 2 | 233510622 | rs10193825 | 0.934 | 0.693 | -1.038 | -0.262 | -0.014 | 0.191 | 0.191 | 0.198 | 0.48 | 0.15 | 0.38 | 0.23 | 0.45 | 0.48 |
| 2 | 233520379 | rs13396021 | 0.672 | 0.766 | -0.847 | -0.162 | -0.017 | 0.087 | 0.087 | 0.092 | 0.56 | 0.26 | 0.43 | 0.23 | 0.51 | 0.59 |
| 2 | 233520942 | rs12694911 | 0.661 | 0.774 | -0.842 | -0.157 | -0.018 | 0.086 | 0.086 | 0.076 | 0.56 | 0.26 | 0.43 | 0.23 | 0.51 | 0.59 |
| 2 | 233526097 | rs13424032 | 0.081 | 0.947 | -1.143 | 0.255 | -0.023 | -0.047 | -0.047 | 0.263 | 0.16 | 0.05 | 0.25 | 0.23 | 0.19 | 0.19 |
| 2 | 233530570 | rs867838 | -0.47 | 0.937 | -0.816 | -0.242 | 0.002 | -0.014 | -0.014 | 0 | 0.38 | 0.64 | 0.55 | 0.72 | 0.41 | 0.5 |
| 2 | 233534676 | rs10933405 | 0.298 | 0.952 | -1.448 | 0.4 | -0.037 | -0.089 | -0.089 | 0.824 | 0.26 | 0.14 | 0.45 | 0.49 | 0.33 | 0.37 |
| 2 | 233537125 | rs11550699 | 0.243 | 0.922 | -1.442 | 0.406 | -0.038 | -0.087 | -0.087 | 0.512 | 0.27 | 0.15 | 0.46 | 0.49 | 0.35 | 0.38 |
| 2 | 233537477 | rs13009166 | -0.841 | 0.869 | -1.636 | -0.641 | 0.007 | -0.015 | -0.015 | 0 | 0.59 | 0.65 | 0.71 | 0.93 | 0.65 | 0.75 |
| 2 | 233537714 | rs17297626 | 0.24 | 0.974 | -1.779 | -0.414 | -0.023 | -0.02 | -0.02 | 0.159 | 0.23 | 0.04 | 0.14 | 0.21 | 0.22 | 0.21 |
| 2 | 233546985 | rs7286 | -0.547 | 0.661 | -1.318 | -0.669 | 0.004 | -0.022 | -0.022 | 0 | 0.67 | 0.76 | 0.75 | 0.95 | 0.72 | 0.9 |
| 2 | 233552244 | rs7565128 | -0.128 | 0.583 | -2.367 | -0.618 | -0.092 | 0.165 | 0.165 | 0 | 0.59 | 0.95 | 0.65 | 0.85 | 0.56 | 0.7 |
| 2 | 233567342 | rs11689851 | -0.385 | 0.604 | -2.346 | -0.872 | -0.052 | 0.032 | 0.032 | 0.012 | 0.78 | 0.9 | 0.64 | 0.88 | 0.62 | 0.91 |
| 2 | 233568875 | rs6747479 | 0.816 | 0.338 | -2.371 | -1.08 | -0.005 | 0.011 | 0.011 | 0 | 0.2 | 0.13 | 0.14 | 0.13 | 0.24 | 0.08 |
| 2 | 233591551 | rs12052869 | 0.71 | 0.222 | -1.974 | -1.096 | -0.017 | 0.028 | 0.028 | 0 | 0.2 | 0.1 | 0.13 | 0.11 | 0.24 | 0.08 |
| 2 | 233597196 | rs10205801 | -0.281 | -0.304 | -0.995 | -0.33 | 0.021 | 0.017 | 0.017 | 0 | 0.64 | 0.5 | 0.5 | 0.48 | 0.52 | 0.72 |

| | | | | | | | | | | | | | | | | |
|---|---|---|---|---|---|---|---|---|---|---|---|---|---|---|---|---|
| 2 | 233599904 | rs11555646 | 0.731 | -0.411 | -0.63 | -0.357 | 0.042 | 0.051 | 0.051 | 0 | 0.43 | 0.24 | 0.31 | 0.35 | 0.29 | 0.64 |
| 2 | 233602028 | rs6738386 | 0.639 | -0.42 | -0.556 | -0.333 | 0.026 | 0.03 | 0.03 | 0 | 0.44 | 0.32 | 0.31 | 0.35 | 0.29 | 0.64 |
| 2 | 233612557 | rs3817311 | 0.751 | -0.426 | -0.573 | -0.349 | 0.041 | 0.05 | 0.05 | 0 | 0.43 | 0.24 | 0.31 | 0.35 | 0.29 | 0.64 |
| 2 | 233628937 | rs12463466 | 1.837 | -0.569 | -0.863 | -0.44 | -0.025 | -0.013 | -0.013 | 0.944 | 0.09 | 0.02 | 0.11 | 0.11 | 0.08 | 0.28 |
| 2 | 233632766 | rs8192539 | 0.894 | -0.683 | -0.836 | -0.577 | -0.017 | 0.028 | 0.028 | 0 | 0.16 | 0.07 | 0.16 | 0.15 | 0.15 | 0.33 |
| 2 | 233633460 | rs1801251 | 0.249 | -0.589 | -0.823 | -0.554 | 0.039 | 0.05 | 0.05 | 0 | 0.43 | 0.24 | 0.31 | 0.35 | 0.29 | 0.64 |
| 2 | 233636450 | rs1973675 | -0.984 | -0.624 | -0.224 | 0.13 | -0.017 | -0.02 | -0.02 | 0 | 0.69 | 0.77 | 0.56 | 0.65 | 0.59 | 0.81 |
| 2 | 233639748 | rs9288676 | 1.626 | -0.592 | -0.247 | 0.174 | -0.049 | 0.03 | 0.03 | 0 | 0.19 | 0.03 | 0.13 | 0.11 | 0.24 | 0.08 |
| 2 | 233659553 | rs2289912 | 1.57 | -0.619 | -0.279 | 0.203 | 0.029 | -0.028 | -0.028 | 0 | 0.07 | 0.01 | 0.06 | 0.23 | 0.03 | 0.1 |
| 2 | 233684097 | rs10207739 | 2.005 | -0.666 | -0.153 | 0.043 | -0.043 | 0.007 | 0.007 | 0.384 | 0.15 | 0.03 | 0.1 | 0.11 | 0.2 | 0.06 |
| 2 | 233697200 | rs6751801 | 0.427 | -0.691 | -0.053 | 0.052 | 0.027 | 0.036 | 0.036 | 0.046 | 0.25 | 0.46 | 0.26 | 0.29 | 0.3 | 0.17 |
| 2 | 233699415 | rs10211596 | -0.015 | -0.706 | -0.067 | -0.003 | 0.005 | 0 | 0 | 0.098 | 0.48 | 0.43 | 0.62 | 0.48 | 0.62 | 0.25 |
| 2 | 233708806 | rs3816334 | -0.584 | -0.672 | -0.092 | 0.081 | 0.039 | 0.055 | 0.055 | 0 | 0.57 | 0.76 | 0.69 | 0.65 | 0.71 | 0.36 |
| 2 | 233715487 | rs13024330 | 1.627 | -0.654 | -0.08 | 0.081 | -0.052 | 0.039 | 0.039 | 0.137 | 0.23 | 0.1 | 0.36 | 0.12 | 0.38 | 0.09 |
| 2 | 233723182 | rs1284 | -1.574 | -0.646 | -0.091 | -0.045 | 0.038 | 0.017 | 0.017 | 0.148 | 0.73 | 0.87 | 0.86 | 0.76 | 0.86 | 0.68 |
| 2 | 233733144 | rs709933 | -0.534 | -0.373 | -0.045 | 0.039 | 0.03 | 0.057 | 0.057 | 0 | 0.56 | 0.76 | 0.65 | 0.64 | 0.63 | 0.36 |
| 2 | 233736349 | rs2592115 | -0.577 | -0.336 | 0.014 | 0.057 | 0.026 | 0.045 | 0.045 | 0 | 0.55 | 0.69 | 0.64 | 0.64 | 0.63 | 0.36 |
| 2 | 233743363 | rs778370 | 0.684 | -0.238 | -0.02 | 0.088 | -0.01 | -0.02 | -0.02 | 0 | 0.3 | 0.23 | 0.39 | 0.35 | 0.33 | 0.19 |
| 2 | 233744021 | rs709937 | 0.844 | -0.233 | -0.004 | 0.091 | -0.03 | -0.051 | -0.051 | 0 | 0.29 | 0.15 | 0.38 | 0.35 | 0.32 | 0.19 |
| 2 | 233750667 | rs11694000 | 0.588 | -0.202 | 0.043 | 0.109 | 0.028 | 0.027 | 0.027 | 0.031 | 0.24 | 0.38 | 0.25 | 0.28 | 0.31 | 0.17 |
| 2 | 233757679 | rs748002 | 0.626 | -0.103 | -0.107 | 0.191 | -0.006 | -0.012 | -0.012 | 0 | 0.29 | 0.28 | 0.4 | 0.35 | 0.33 | 0.18 |
| 2 | 233758884 | rs1554218 | 1.392 | -0.126 | -0.029 | 0.284 | -0.015 | -0.005 | -0.005 | 0 | 0.09 | 0.04 | 0.1 | 0.12 | 0.08 | 0.26 |
| 2 | 233759155 | rs2166440 | 0.417 | -0.088 | 0.243 | 0.317 | 0.034 | 0.083 | 0.083 | 0 | 0.13 | 0.3 | 0.15 | 0.28 | 0.13 | 0.36 |
| 2 | 233763362 | rs12477794 | 1.319 | -0.07 | -0.217 | 0.188 | -0.001 | 0 | 0 | 1.028 | 0.1 | 0.12 | 0.12 | 0.11 | 0.08 | 0.27 |
| 2 | 233774183 | rs809017 | -0.589 | -0.299 | -0.102 | 0.155 | -0.013 | -0.018 | -0.018 | 0 | 0.62 | 0.61 | 0.74 | 0.75 | 0.7 | 0.62 |
| 2 | 233780186 | rs778362 | -1.295 | -0.362 | -0.507 | 0.304 | 0.034 | 0.052 | 0.052 | 0.97 | 0.75 | 0.93 | 0.88 | 0.8 | 0.8 | 0.7 |
| 2 | 233780244 | rs778361 | -0.446 | -0.447 | -0.198 | 0.503 | 0 | -0.001 | -0.001 | 0 | 0.64 | 0.66 | 0.77 | 0.4 | 0.7 | 0.55 |
| 2 | 233784667 | rs778357 | -1.358 | -0.464 | -0.687 | 0.443 | 0.032 | 0.053 | 0.053 | 0.948 | 0.74 | 0.93 | 0.89 | 0.8 | 0.79 | 0.7 |
| 2 | 233784718 | rs778356 | -0.626 | -0.401 | -0.626 | 0.566 | 0.069 | -0.059 | -0.059 | 0.551 | 0.67 | 0.86 | 0.82 | 0.56 | 0.76 | 0.62 |
| 2 | 233790447 | rs2004203 | 0.195 | 0.021 | -0.44 | 0.337 | 0.044 | 0.014 | 0.014 | 2.967 | 0.33 | 0.43 | 0.48 | 0.34 | 0.44 | 0.24 |
| 2 | 233793517 | rs11685503 | 0.874 | 0.098 | -0.675 | 0.335 | -0.001 | -0.001 | -0.001 | 0 | 0.12 | 0.14 | 0.2 | 0.16 | 0.08 | 0.3 |
| 2 | 233799400 | rs6437075 | 0.26 | 0.163 | -0.682 | 0.431 | 0.023 | 0.04 | 0.04 | 0.011 | 0.09 | 0.16 | 0.09 | 0.16 | 0.13 | 0.1 |
| 2 | 233803391 | rs778351 | -0.168 | -0.001 | -0.815 | 0.276 | 0.007 | 0.004 | 0.004 | 0 | 0.63 | 0.68 | 0.73 | 0.62 | 0.67 | 0.61 |
| 2 | 233812756 | rs13004808 | -0.015 | 0.31 | -1.573 | -0.242 | -0.008 | 0.117 | 0.117 | 0.453 | 0.38 | 0.16 | 0.35 | 0.13 | 0.39 | 0.39 |
| 2 | 233818331 | rs6707071 | 0.02 | 0.901 | -1.397 | 0.053 | -0.01 | -0.006 | -0.006 | 0 | 0.4 | 0.29 | 0.34 | 0.39 | 0.46 | 0.32 |
| 2 | 233822436 | rs11903971 | 0.855 | 0.694 | -0.687 | -0.126 | 0.044 | -0.01 | -0.01 | 0 | 0.07 | 0.26 | 0.02 | 0.02 | 0.04 | 0.02 |
| 2 | 233824116 | rs2138580 | -0.674 | 0.626 | -1.592 | -0.221 | 0.018 | 0.016 | 0.016 | 0 | 0.49 | 0.77 | 0.54 | 0.49 | 0.54 | 0.4 |
| 2 | 233824203 | rs2176308 | -1.595 | 0.591 | -1.724 | -0.184 | 0.016 | 0.049 | 0.049 | 1.113 | 0.69 | 0.96 | 0.72 | 0.81 | 0.72 | 0.65 |
| 2 | 233826626 | rs1467188 | -1.576 | 0.594 | -1.723 | -0.191 | 0.017 | 0.054 | 0.054 | 1.176 | 0.68 | 0.95 | 0.71 | 0.82 | 0.72 | 0.65 |
| 2 | 233827266 | rs13397982 | -0.658 | 0.65 | -1.765 | -0.334 | -0.027 | 0.02 | 0.02 | 0 | 0.5 | 0.88 | 0.39 | 0.41 | 0.52 | 0.34 |
| 2 | 233830950 | rs11678490 | 1.139 | 0.704 | -1.492 | -0.194 | -0.001 | 0.001 | 0.001 | 1.839 | 0.19 | 0.18 | 0.13 | 0.33 | 0.18 | 0.25 |
| 2 | 233830994 | rs11684260 | -0.065 | 0.778 | -1.024 | -0.28 | -0.001 | 0.033 | 0.033 | 0.599 | 0.29 | 0.59 | 0.38 | 0.44 | 0.24 | 0.33 |
| 2 | 233831820 | rs6720335 | 0.814 | 0.958 | -1.19 | -0.829 | -0.056 | 0.078 | 0.078 | 0.696 | 0.19 | 0.07 | 0.16 | 0.04 | 0.3 | 0.07 |
| 2 | 233837896 | rs10933420 | -0.253 | 0.757 | -0.417 | -0.857 | 0.043 | -0.061 | -0.061 | 0 | 0.32 | 0.51 | 0.33 | 0.13 | 0.45 | 0.2 |
| 2 | 233838028 | rs10933421 | -0.296 | 0.739 | -0.218 | -0.866 | 0.05 | -0.072 | -0.072 | 0 | 0.32 | 0.57 | 0.34 | 0.13 | 0.45 | 0.2 |
| 2 | 233839052 | rs3817321 | 0.465 | 0.262 | -0.517 | -0.792 | 0.002 | -0.007 | -0.007 | 0 | 0.3 | 0.18 | 0.43 | 0.71 | 0.26 | 0.31 |
| 2 | 233839174 | rs6437079 | -0.118 | -0.159 | -0.723 | -1.262 | 0.028 | 0.147 | 0.147 | 0 | 0.52 | 0.76 | 0.47 | 0.15 | 0.59 | 0.37 |
| 2 | 233839239 | rs6748986 | 0.54 | 0.147 | -0.072 | -0.76 | 0.064 | 0.051 | 0.051 | 0.218 | 0.08 | 0.33 | 0.15 | 0.08 | 0.1 | 0.11 |
| 2 | 233843826 | rs10165217 | -0.712 | -0.055 | -0.912 | -0.735 | 0.003 | -0.004 | -0.004 | 0 | 0.76 | 0.93 | 0.86 | 0.75 | 0.81 | 0.61 |
| 2 | 233845755 | rs4973591 | 0.691 | -0.063 | -1.078 | -1.031 | -0.003 | -0.012 | -0.012 | 0 | 0.33 | 0.18 | 0.39 | 0.63 | 0.31 | 0.32 |
| 2 | 233854397 | rs11885626 | -1.553 | 0.125 | -0.327 | -1.026 | 0.005 | 0.064 | 0.064 | 0.287 | 0.84 | 0.97 | 0.9 | 0.99 | 0.84 | 0.86 |
| 2 | 233858636 | rs4973062 | -0.306 | 0.225 | -0.263 | -1.219 | -0.003 | -0.014 | -0.014 | 0 | 0.67 | 0.71 | 0.71 | 0.92 | 0.57 | 0.79 |
| 2 | 233862990 | rs6760560 | -1.698 | 0.259 | -0.412 | -1.223 | -0.001 | 0.064 | 0.064 | 0.287 | 0.84 | 0.97 | 0.9 | 0.99 | 0.84 | 0.86 |
| 2 | 233863591 | rs1400348 | -1.382 | 0.257 | -0.256 | -1.229 | -0.007 | -0.067 | -0.067 | 0 | 0.83 | 0.82 | 0.9 | 0.99 | 0.84 | 0.86 |
| 2 | 233864457 | rs12470285 | 0.034 | 0.521 | -0.412 | -0.808 | -0.008 | -0.035 | -0.035 | 0 | 0.14 | 0.09 | 0.17 | 0.28 | 0.13 | 0.33 |
| 2 | 233864497 | rs13404304 | 1.156 | 0.553 | 0.115 | -0.842 | 0.058 | -0.059 | -0.059 | 0 | 0.15 | 0.27 | 0.19 | 0.05 | 0.27 | 0.07 |
| 2 | 233868678 | rs6727341 | 0.75 | -0.227 | -0.019 | -0.301 | -0.001 | 0.019 | 0.019 | 0 | 0.08 | 0.09 | 0.05 | 0.23 | 0.05 | 0.22 |
| 2 | 233871467 | rs10933423 | -0.698 | -0.373 | 0.452 | -0.709 | 0.034 | -0.019 | -0.019 | 0 | 0.14 | 0.51 | 0.18 | 0.07 | 0.16 | 0.15 |
| 2 | 233872049 | rs10169752 | -0.42 | -0.683 | 0.136 | -1.204 | -0.001 | -0.017 | -0.017 | 0.755 | 0.59 | 0.61 | 0.51 | 0.14 | 0.65 | 0.48 |
| 2 | 233872100 | rs10193545 | 0.588 | -0.649 | -0.354 | -1.177 | -0.037 | 0.246 | 0.246 | 9.63 | 0.45 | 0.12 | 0.33 | 0.05 | 0.49 | 0.33 |
| 2 | 233874399 | rs13383577 | 0.908 | -1.019 | 0.142 | -1.067 | -0.005 | 0.065 | 0.065 | 1.07 | 0.47 | 0.35 | 0.35 | 0.07 | 0.51 | 0.25 |
| 2 | 233875716 | rs3811589 | -0.861 | -1.397 | 0.664 | -0.971 | -0.04 | -0.036 | -0.036 | 0 | 0.16 | 0.37 | 0.16 | 0.08 | 0.1 | 0.2 |
| 2 | 233875975 | rs3811588 | 0.483 | -1.128 | 0.518 | -0.533 | -0.002 | 0.006 | 0.006 | 0 | 0.14 | 0.12 | 0.12 | 0.23 | 0.11 | 0.26 |
| 2 | 233877065 | rs6725244 | 0.974 | -0.981 | 0.464 | -0.916 | -0.007 | 0.047 | 0.047 | 0.701 | 0.47 | 0.32 | 0.3 | 0.04 | 0.5 | 0.25 |
| 2 | 233882493 | rs13405304 | 0.464 | -0.604 | 0.598 | -0.016 | -0.003 | 0.202 | 0.202 | 0 | 0.1 | 0.26 | 0.1 | 0.31 | 0.06 | 0.19 |
| 2 | 233885426 | rs11683286 | -0.273 | -0.299 | -0.025 | 0.212 | 0 | 0.001 | 0.001 | 0 | 0.75 | 0.72 | 0.67 | 0.38 | 0.73 | 0.65 |
| 2 | 233894496 | rs13386477 | 0.052 | -0.848 | 0.007 | 0.758 | 0 | 0 | 0 | 0.303 | 0.77 | 0.76 | 0.72 | 0.61 | 0.79 | 0.77 |
| 2 | 233896498 | rs2233375 | -0.889 | -0.565 | 0.641 | 0.315 | 0.018 | -0.034 | -0.034 | 0 | 0.09 | 0.29 | 0.24 | 0.01 | 0.08 | 0.13 |
| 2 | 233897101 | rs2233381 | 0.763 | -1.005 | 0.103 | -0.126 | -0.045 | 0.182 | 0.182 | 0.563 | 0.41 | 0.17 | 0.31 | 0.03 | 0.52 | 0.22 |

| | | | | | | | | | | | | | | | | |
|---|---|---|---|---|---|---|---|---|---|---|---|---|---|---|---|---|
| 2 | 233898653 | rs11683026 | -0.349 | -1.098 | 0.266 | -0.091 | -0.014 | 0.016 | 0.016 | 0 | 0.12 | 0.27 | 0.26 | 0.13 | 0.08 | 0.19 |
| 2 | 233899126 | rs2233391 | 0.812 | -1.035 | -0.144 | -0.132 | -0.052 | 0.175 | 0.175 | 0.502 | 0.41 | 0.16 | 0.3 | 0.03 | 0.5 | 0.21 |
| 2 | 233905490 | rs4374365 | -0.108 | -1.04 | 0.275 | 0.055 | 0 | -0.008 | -0.008 | 0 | 0.55 | 0.57 | 0.58 | 0.17 | 0.6 | 0.4 |
| 2 | 233912755 | rs12694922 | 0.015 | -0.805 | 0.476 | 0.401 | -0.006 | 0.018 | 0.018 | 0 | 0.45 | 0.36 | 0.3 | 0.33 | 0.47 | 0.43 |
| 2 | 233924009 | rs7558755 | 0.51 | -0.793 | 0.522 | 0.395 | -0.008 | 0.079 | 0.079 | 0 | 0.86 | 0.71 | 0.73 | 0.69 | 0.88 | 0.7 |
| 2 | 233926201 | rs4973063 | 0.249 | -0.884 | 0.588 | 0.274 | 0.015 | -0.004 | -0.004 | 0 | 0.45 | 0.2 | 0.46 | 0.6 | 0.44 | 0.39 |
| 2 | 233928776 | rs4439944 | 0.405 | -0.808 | 0.796 | -0.264 | -0.006 | -0.022 | -0.022 | 0 | 0.39 | 0.35 | 0.27 | 0.04 | 0.44 | 0.21 |
| 2 | 233929510 | rs4257390 | -0.358 | -0.829 | 0.803 | -0.19 | 0 | -0.057 | -0.057 | 0 | 0.56 | 0.56 | 0.53 | 0.94 | 0.5 | 0.74 |
| 2 | 233929545 | rs4073363 | -0.298 | -0.801 | 0.626 | -0.148 | 0 | 0.005 | 0.005 | 0 | 0.54 | 0.45 | 0.51 | 0.9 | 0.49 | 0.69 |
| 2 | 233934575 | rs4973599 | 0.298 | -0.795 | 0.709 | 0.279 | -0.006 | 0.003 | 0.003 | 0 | 0.08 | 0.06 | 0.1 | 0 | 0.07 | 0.06 |
| 2 | 233939615 | rs4586623 | -1.034 | -1.625 | 0.997 | 0.461 | -0.09 | 0.044 | 0.044 | 0 | 0.29 | 0.45 | 0.16 | 0.38 | 0.1 | 0.45 |
| 2 | 233943293 | rs6708049 | 0.29 | -1.321 | 1.186 | 0.538 | -0.007 | 0.107 | 0.107 | 0 | 0.88 | 0.74 | 0.79 | 0.71 | 0.91 | 0.67 |
| 2 | 233943999 | rs13419542 | 0.198 | -1.499 | 1.067 | -0.04 | -0.026 | -0.046 | -0.046 | 0 | 0.14 | 0.18 | 0.04 | 0 | 0.05 | 0.05 |
| 2 | 233945565 | rs7425956 | 1.042 | -1.138 | 0.332 | 0.195 | -0.013 | 0.143 | 0.143 | 0 | 0.44 | 0.14 | 0.38 | 0.18 | 0.58 | 0.26 |
| 2 | 233949255 | rs13387054 | 0.467 | -0.923 | 0.565 | 0.501 | 0.037 | 0.004 | 0.004 | 0 | 0.31 | 0.2 | 0.2 | 0.38 | 0.21 | 0.28 |
| 2 | 233950265 | rs4356648 | -0.51 | -0.642 | 0.682 | 0.413 | 0.056 | 0.05 | 0.05 | 0 | 0.46 | 0.62 | 0.33 | 0.76 | 0.27 | 0.57 |
| 2 | 233950732 | rs7608422 | 0.424 | -0.642 | 0.378 | 0.445 | 0.019 | -0.003 | -0.003 | 0 | 0.33 | 0.26 | 0.21 | 0.42 | 0.21 | 0.33 |
| 2 | 233953940 | rs7566856 | -0.635 | -0.312 | -0.575 | 0.342 | 0.099 | 0.161 | 0.161 | 0 | 0.51 | 0.82 | 0.46 | 0.79 | 0.34 | 0.62 |
| 2 | 233955723 | rs4335931 | -1.188 | -0.438 | -0.579 | 0.34 | -0.043 | 0.103 | 0.103 | 0 | 0.58 | 0.86 | 0.63 | 0.79 | 0.44 | 0.74 |
| 2 | 233957372 | rs10933428 | -0.613 | -0.492 | -0.541 | 0.362 | 0.132 | 0.145 | 0.145 | 0 | 0.49 | 0.79 | 0.35 | 0.78 | 0.3 | 0.58 |
| 2 | 233962515 | rs6437089 | 1.629 | 0.456 | 0.317 | 0.989 | 0.014 | 0.028 | 0.028 | 0.07 | 0.78 | 0.71 | 0.6 | 0.64 | 0.78 | 0.6 |
| 2 | 233962551 | rs6437090 | 0.598 | 0.771 | -0.224 | -0.05 | -0.039 | 0.036 | 0.036 | 0.014 | 0.89 | 0.92 | 0.71 | 0.99 | 0.82 | 0.84 |
| 2 | 233968153 | rs9288684 | -1.45 | -0.334 | 0.325 | 0.27 | -0.022 | -0.003 | -0.003 | 0.022 | 0.21 | 0.67 | 0.12 | 0.16 | 0.1 | 0.13 |
| 2 | 233969443 | rs9752615 | -1.308 | -0.251 | 1.052 | 0.39 | 0.019 | 0.167 | 0.167 | 0 | 0.07 | 0.25 | 0.04 | 0.16 | 0.04 | 0.08 |
| 2 | 233969889 | rs9750891 | -1.932 | -0.342 | 1.3 | 0.279 | -0.037 | 0.066 | 0.066 | 0 | 0.13 | 0.41 | 0.07 | 0.15 | 0.05 | 0.11 |
| 2 | 233977318 | rs7570061 | -1.064 | -0.369 | 0.813 | -0.166 | -0.032 | -0.056 | -0.056 | 0.283 | 0.29 | 0.45 | 0.11 | 0.15 | 0.16 | 0.2 |
| 2 | 233982841 | rs13432536 | -1.877 | -0.327 | 1.517 | 0.115 | 0.046 | 0.087 | 0.087 | 0 | 0.07 | 0.33 | 0.04 | 0.12 | 0.04 | 0.07 |
| 2 | 233987114 | rs9288685 | 0.199 | -0.192 | 0.872 | -0.235 | -0.001 | -0.013 | -0.013 | 0.583 | 0.41 | 0.5 | 0.38 | 0.34 | 0.53 | 0.35 |
| 2 | 233987722 | rs10193128 | 0.458 | -0.284 | 1 | -0.349 | 0.001 | 0.015 | 0.015 | 0 | 0.35 | 0.26 | 0.35 | 0.34 | 0.48 | 0.34 |
| 2 | 233999081 | rs11693862 | -0.194 | -0.378 | 0.946 | 0.203 | 0.001 | -0.003 | -0.003 | 0 | 0.56 | 0.55 | 0.39 | 0.5 | 0.46 | 0.47 |
| 2 | 234003227 | rs7421653 | 0.805 | -0.453 | 0.572 | 0.428 | -0.015 | 0.11 | 0.11 | 0 | 0.93 | 0.82 | 0.75 | 0.69 | 0.97 | 0.76 |
| 2 | 234072738 | rs11673739 | 0.468 | -0.371 | -0.112 | 0.487 | -0.008 | -0.024 | -0.024 | 0 | 0.25 | 0.2 | 0.21 | 0.42 | 0.25 | 0.29 |
| 2 | 234073243 | rs6715810 | 0.993 | -0.901 | -0.504 | 0.186 | 0.01 | -0.02 | -0.02 | 0 | 0.23 | 0.11 | 0.35 | 0.26 | 0.18 | 0.35 |
| 2 | 234075691 | rs7570320 | 1.692 | -0.745 | -0.62 | -0.065 | -0.114 | 0.169 | 0.169 | 0 | 0.26 | 0.05 | 0.31 | 0.02 | 0.43 | 0.24 |
| 2 | 234078406 | rs7569837 | 0.237 | -0.618 | -0.4 | 0.164 | 0.008 | -0.037 | -0.037 | 7.074 | 0.32 | 0.24 | 0.4 | 0.47 | 0.24 | 0.42 |
| 2 | 234081187 | rs13408647 | 0.59 | -0.361 | -0.528 | 0.538 | -0.004 | -0.003 | -0.003 | 0 | 0.2 | 0.19 | 0.18 | 0.4 | 0.21 | 0.27 |
| 2 | 234086584 | rs6720896 | 0.324 | -0.232 | -0.601 | 0.429 | -0.009 | -0.013 | -0.013 | 0.276 | 0.23 | 0.2 | 0.22 | 0.42 | 0.29 | 0.29 |
| 2 | 234089949 | rs4395246 | -0.16 | -0.647 | -0.49 | 0.035 | 0.003 | -0.02 | -0.02 | 0 | 0.17 | 0.1 | 0.05 | 0 | 0.07 | 0.06 |
| 2 | 234092392 | rs4436949 | -0.543 | -0.006 | -0.18 | 0.596 | 0.019 | 0.062 | 0.062 | 0 | 0.24 | 0.38 | 0.44 | 0.6 | 0.23 | 0.44 |
| 2 | 234093035 | rs6740918 | -0.21 | 0.026 | -0.116 | 1.195 | 0.003 | 0.065 | 0.065 | 0 | 0.2 | 0.28 | 0.2 | 0.31 | 0.2 | 0.25 |
| 2 | 234093805 | rs7584458 | -0.285 | -0.123 | -1.26 | 0.538 | 0.037 | -0.02 | -0.02 | 0.495 | 0.62 | 0.81 | 0.59 | 0.55 | 0.71 | 0.7 |
| 2 | 234098802 | rs3890760 | -0.405 | 0.636 | -0.917 | 0.308 | -0.004 | -0.004 | -0.004 | 0 | 0.34 | 0.35 | 0.37 | 0.39 | 0.43 | 0.47 |
| 2 | 234102070 | rs10203185 | -0.04 | 0.676 | -0.901 | 0.332 | 0.001 | 0.001 | 0.001 | 0.354 | 0.3 | 0.33 | 0.36 | 0.39 | 0.41 | 0.45 |
| 2 | 234102738 | rs11681058 | -0.033 | 0.594 | -0.887 | 0.251 | 0 | 0.001 | 0.001 | 0.351 | 0.29 | 0.3 | 0.34 | 0.39 | 0.4 | 0.44 |
| 2 | 234107070 | rs7569827 | -0.582 | 0.317 | -1.222 | -0.503 | 0.022 | 0.112 | 0.112 | 0.757 | 0.61 | 0.85 | 0.76 | 0.8 | 0.65 | 0.77 |
| 2 | 234110954 | rs4468807 | 0.274 | 1.16 | -0.435 | -0.405 | 0.018 | -0.032 | -0.032 | 0 | 0.15 | 0.25 | 0.17 | 0.04 | 0.24 | 0.22 |
| 2 | 234113301 | rs9247 | 0.378 | 1.091 | -0.591 | -0.068 | -0.004 | -0.011 | -0.011 | 0 | 0.16 | 0.19 | 0.23 | 0.35 | 0.2 | 0.24 |
| 2 | 234113545 | rs1135727 | 0.033 | 0.78 | -0.216 | -0.544 | -0.009 | 0.023 | 0.023 | 0 | 0.57 | 0.65 | 0.66 | 0.79 | 0.5 | 0.69 |
| 2 | 234114837 | rs4663784 | 0.697 | 0.275 | -0.059 | -0.571 | 0.001 | -0.034 | -0.034 | 0 | 0.23 | 0.16 | 0.17 | 0 | 0.25 | 0.08 |
| 2 | 234115337 | rs3792117 | 0.604 | -0.237 | 0.247 | -0.304 | 0 | 0.008 | 0.008 | 0 | 0.07 | 0.04 | 0.03 | 0.17 | 0.05 | 0.05 |
| 2 | 234115629 | rs1057258 | -0.621 | -0.459 | 0.514 | -0.545 | -0.012 | 0.138 | 0.138 | 0.427 | 0.21 | 0.57 | 0.26 | 0.41 | 0.19 | 0.32 |
| 2 | 234115739 | rs14243 | -0.294 | -0.116 | -0.618 | -0.507 | -0.001 | -0.031 | -0.031 | 0 | 0.21 | 0.24 | 0.24 | 0.37 | 0.22 | 0.33 |
| 2 | 234118583 | rs4663834 | -0.617 | -0.293 | -0.706 | -0.837 | 0.011 | 0.025 | 0.025 | 0.03 | 0.67 | 0.78 | 0.85 | 0.95 | 0.75 | 0.79 |
| 2 | 234119635 | rs6431586 | -0.762 | -0.158 | -0.425 | 0.206 | -0.014 | 0.025 | 0.025 | 0 | 0.21 | 0.2 | 0.12 | 0.35 | 0.12 | 0.19 |
| 2 | 234123010 | rs7580869 | 0.598 | 0.133 | -0.636 | -0.607 | 0.012 | 0.02 | 0.02 | 0 | 0.84 | 0.81 | 0.82 | 0.96 | 0.88 | 0.86 |
| 2 | 234130678 | rs13025632 | -1.261 | 0.762 | -0.138 | -0.547 | 0.009 | -0.036 | -0.036 | 0 | 0.18 | 0.23 | 0.22 | 0.04 | 0.21 | 0.1 |
| 2 | 234135138 | rs2341565 | -1.372 | 0.543 | 0.171 | -0.673 | -0.006 | -0.095 | -0.095 | 0.117 | 0.22 | 0.57 | 0.18 | 0.04 | 0.16 | 0.08 |
| 2 | 234136796 | rs4663337 | -1.145 | 0.728 | 0.346 | -0.443 | 0.047 | -0.092 | -0.092 | 0 | 0.16 | 0.46 | 0.25 | 0.02 | 0.19 | 0.09 |
| 2 | 234139467 | rs10211468 | -0.699 | 0.889 | -0.743 | -0.53 | -0.022 | 0.043 | 0.043 | 4.631 | 0.39 | 0.3 | 0.58 | 0.84 | 0.46 | 0.7 |
| 2 | 234143244 | rs7563345 | -0.36 | 0.606 | -0.842 | 0.087 | 0.059 | -0.008 | -0.008 | 0 | 0.55 | 0.65 | 0.46 | 0.44 | 0.64 | 0.63 |
| 2 | 234145971 | rs13412102 | -0.164 | 1.016 | -0.317 | 0.267 | 0.022 | 0.016 | 0.016 | 1.12 | 0.52 | 0.51 | 0.38 | 0.36 | 0.6 | 0.57 |
| 2 | 234162415 | rs6431655 | -0.885 | 0.615 | 1 | 0.277 | 0.012 | -0.013 | -0.013 | 0 | 0.64 | 0.75 | 0.52 | 0.49 | 0.69 | 0.8 |
| 2 | 234168951 | rs6758317 | -1.919 | 1.045 | 1.462 | 0.079 | 0.043 | -0.016 | -0.016 | 0.053 | 0.15 | 0.38 | 0.2 | 0.13 | 0.17 | 0.29 |
| 2 | 234179690 | rs3792111 | 0.175 | 0.946 | 1.814 | 0.031 | 0.003 | 0.045 | 0.045 | 0.622 | 0.48 | 0.21 | 0.31 | 0.32 | 0.54 | 0.49 |
| 2 | 234180410 | rs3828309 | 0.091 | 0.946 | 1.786 | 0.031 | 0.011 | 0.048 | 0.048 | 0.487 | 0.48 | 0.22 | 0.31 | 0.32 | 0.54 | 0.49 |
| 2 | 234181848 | rs2289474 | 0.139 | 0.947 | 1.786 | 0.032 | 0.01 | 0.041 | 0.041 | 0.611 | 0.48 | 0.31 | 0.32 | 0.32 | 0.54 | 0.49 |
| 2 | 234183368 | rs2241880 | 0.054 | 0.947 | 1.786 | 0.032 | 0.014 | 0.046 | 0.046 | 0.49 | 0.49 | 0.31 | 0.32 | 0.32 | 0.54 | 0.5 |
| 2 | 234183468 | rs2241879 | 0.138 | 0.948 | 1.786 | 0.032 | 0.01 | 0.041 | 0.041 | 0.611 | 0.48 | 0.31 | 0.32 | 0.32 | 0.54 | 0.49 |
| 2 | 234186734 | rs2241876 | -1.154 | 0.971 | 1.764 | 0.035 | -0.006 | 0.004 | 0.004 | 2.203 | 0.64 | 0.69 | 0.52 | 0.46 | 0.71 | 0.78 |

| | | | | | | | | | | | | | | | | |
|---|---|---|---|---|---|---|---|---|---|---|---|---|---|---|---|---|
| 2 | 234189359 | rs7600743 | -0.401 | 0.954 | 1.701 | 0.032 | 0 | 0 | 0 | 0 | 0.06 | 0.09 | 0.05 | 0.06 | 0.06 | 0.18 |
| 2 | 234189406 | rs7585217 | -0.941 | 0.95 | 1.717 | 0.023 | -0.001 | -0.005 | -0.005 | 0 | 0.07 | 0.1 | 0.05 | 0.05 | 0.06 | 0.16 |
| 2 | 234190740 | rs3792106 | -0.461 | 0.954 | 1.73 | 0.026 | 0.002 | 0.002 | 0.002 | 0.932 | 0.56 | 0.5 | 0.39 | 0.37 | 0.6 | 0.65 |
| 2 | 234201657 | rs11679064 | -0.478 | 0.982 | 1.619 | 0.062 | 0 | -0.002 | -0.002 | 0 | 0.05 | 0.08 | 0.05 | 0.03 | 0.06 | 0.14 |
| 2 | 234203547 | rs1045095 | -1.238 | 1.066 | 1.927 | 0.008 | 0.031 | -0.033 | -0.033 | 0 | 0.63 | 0.81 | 0.5 | 0.49 | 0.69 | 0.8 |
| 2 | 234203597 | rs1045100 | 0.134 | 1.078 | 1.983 | -0.003 | -0.002 | 0.041 | 0.041 | 0.497 | 0.47 | 0.23 | 0.29 | 0.32 | 0.51 | 0.49 |
| 2 | 234203876 | rs10803619 | -1.021 | 1.076 | 1.939 | -0.003 | 0 | -0.003 | -0.003 | 0 | 0.07 | 0.09 | 0.06 | 0.05 | 0.07 | 0.16 |
| 2 | 234204113 | rs6861 | -1.134 | 1.075 | 1.952 | 0.003 | 0.011 | -0.011 | -0.011 | 0 | 0.63 | 0.74 | 0.5 | 0.46 | 0.69 | 0.78 |
| 2 | 234206474 | rs10187822 | 0.1 | 1.031 | 1.8 | -0.002 | -0.063 | 0.012 | 0.012 | 0.882 | 0.47 | 0.08 | 0.28 | 0.32 | 0.51 | 0.49 |
| 2 | 234208258 | rs13411660 | 0.116 | 1.099 | 1.822 | -0.001 | -0.005 | 0.037 | 0.037 | 0.516 | 0.47 | 0.23 | 0.29 | 0.33 | 0.51 | 0.49 |
| 2 | 234210972 | rs6759896 | 0.106 | 1.098 | 1.579 | -0.218 | 0.039 | -0.035 | -0.035 | 0 | 0.46 | 0.21 | 0.37 | 0.44 | 0.57 | 0.64 |
| 2 | 234214947 | rs7588509 | -1.649 | 1.062 | 1.781 | -0.242 | 0.018 | -0.079 | -0.079 | 0 | 0.14 | 0.45 | 0.09 | 0.03 | 0.1 | 0.06 |
| 2 | 234221687 | rs3792105 | -1.864 | 1.109 | 1.734 | -0.251 | 0.008 | -0.09 | -0.09 | 2.28 | 0.15 | 0.47 | 0.1 | 0.03 | 0.1 | 0.06 |
| 2 | 234229320 | rs7565275 | -0.5 | 1.12 | 1.353 | -0.229 | 0.005 | -0.016 | -0.016 | 0.255 | 0.05 | 0.11 | 0.04 | 0.03 | 0.05 | 0.11 |
| 2 | 234237765 | rs894100 | -1.333 | 1.04 | 1.242 | 0.014 | 0.067 | -0.068 | -0.068 | 0 | 0.65 | 0.86 | 0.59 | 0.52 | 0.72 | 0.77 |
| 2 | 234242347 | rs1000141 | 0.267 | 1.268 | 0.996 | 0.014 | -0.086 | -0.092 | -0.092 | 0 | 0.42 | 0.07 | 0.32 | 0.41 | 0.54 | 0.55 |
| 2 | 234244395 | rs3828307 | -0.719 | 1.321 | 1.102 | 0.087 | -0.001 | -0.013 | -0.013 | 0 | 0.06 | 0.12 | 0.07 | 0.04 | 0.05 | 0.13 |
| 2 | 234247627 | rs2241874 | -1.25 | 1.118 | 1.007 | 0.479 | 0.043 | -0.073 | -0.073 | 0 | 0.61 | 0.81 | 0.46 | 0.45 | 0.68 | 0.72 |
| 2 | 234251322 | rs3792091 | -0.562 | 1.907 | 1.087 | 0.793 | 0.002 | 0.025 | 0.025 | 0.048 | 0.51 | 0.55 | 0.31 | 0.28 | 0.55 | 0.56 |
| 2 | 234255547 | rs1046974 | 0.307 | 2.207 | 0.499 | 0.754 | -0.141 | -0.162 | -0.162 | 7.547 | 0.3 | 0.05 | 0.25 | 0.4 | 0.43 | 0.45 |
| 2 | 234259470 | rs4663540 | -1.909 | 1.914 | 0.542 | 0.64 | 0.016 | 0.047 | 0.047 | 1.328 | 0.59 | 0.89 | 0.64 | 0.61 | 0.64 | 0.75 |
| 2 | 234266842 | rs13035837 | 0.859 | 2.382 | 0.292 | 0.665 | 0.02 | -0.076 | -0.076 | 0 | 0.2 | 0.04 | 0.07 | 0.23 | 0.12 | 0.34 |
| 2 | 234273242 | rs838705 | 1.216 | 2.525 | 0.929 | 0.634 | 0.038 | 0.012 | 0.012 | 0 | 0.74 | 0.62 | 0.56 | 0.73 | 0.59 | 0.77 |
| 2 | 234282210 | rs6722763 | -1.617 | 2.52 | 1.205 | 0.784 | -0.02 | -0.049 | -0.049 | 0 | 0.23 | 0.47 | 0.16 | 0.13 | 0.18 | 0.25 |
| 2 | 234283288 | rs7580412 | -1.643 | 2.517 | 1.203 | 0.783 | -0.021 | -0.048 | -0.048 | 0 | 0.24 | 0.47 | 0.16 | 0.13 | 0.18 | 0.25 |
| 2 | 234284744 | rs838709 | -0.14 | 2.655 | 1.024 | 0.837 | -0.017 | -0.023 | -0.023 | 0 | 0.9 | 0.9 | 0.61 | 0.75 | 0.67 | 0.85 |
| 2 | 234287221 | rs6726046 | -1.715 | 2.57 | 1.02 | 0.815 | 0.085 | 0.03 | 0.03 | 0 | 0.46 | 0.77 | 0.24 | 0.36 | 0.3 | 0.58 |
| 2 | 234292684 | rs838715 | 2.227 | 2.396 | 1.027 | 0.839 | 0.04 | 0.063 | 0.063 | 0.25 | 0.43 | 0.15 | 0.37 | 0.36 | 0.37 | 0.25 |
| 2 | 234296650 | rs838718 | 1.448 | 2.466 | 1.347 | 0.831 | 0 | -0.003 | -0.003 | 0 | 0.53 | 0.5 | 0.48 | 0.4 | 0.46 | 0.42 |
| 2 | 234299919 | rs706848 | 0.234 | 2.479 | 1.017 | 0.856 | -0.089 | -0.082 | -0.082 | 5.192 | 0.69 | 0.83 | 0.54 | 0.51 | 0.56 | 0.51 |
| 2 | 234337940 | rs7572257 | -0.677 | 2.254 | 1.183 | 0.816 | -0.037 | -0.01 | -0.01 | 0 | 0.16 | 0.27 | 0.06 | 0.14 | 0.1 | 0.1 |
| 2 | 234346660 | rs2242102 | -2.479 | 2.335 | 0.706 | 0.816 | -0.047 | -0.057 | -0.057 | 0.615 | 0.4 | 0.83 | 0.6 | 0.45 | 0.54 | 0.41 |
| 2 | 234357513 | rs13016839 | 1.219 | 2.097 | 0.577 | 0.868 | -0.005 | -0.065 | -0.065 | 0.741 | 0.17 | 0 | 0.04 | 0.19 | 0.12 | 0.33 |
| 2 | 234358286 | rs2242100 | -2.302 | 2.275 | 0.608 | 0.77 | 0.06 | 0.038 | 0.038 | 0.203 | 0.58 | 0.9 | 0.65 | 0.63 | 0.68 | 0.69 |
| 2 | 234364426 | rs2034761 | -1.189 | 2.561 | 1.034 | 0.767 | 0.001 | 0.001 | 0.001 | 0.907 | 0.39 | 0.41 | 0.48 | 0.51 | 0.39 | 0.6 |
| 2 | 234373537 | rs2971855 | -2.035 | 2.662 | 0.816 | 0.712 | -0.005 | 0.013 | 0.013 | 0 | 0.3 | 0.69 | 0.27 | 0.18 | 0.35 | 0.27 |
| 2 | 234378902 | rs1053895 | -2.392 | 2.352 | 0.605 | 0.75 | 0.015 | 0.034 | 0.034 | 0.481 | 0.59 | 0.92 | 0.64 | 0.62 | 0.64 | 0.71 |
| 2 | 234380666 | rs2026 | -0.766 | 2.296 | 0.81 | 0.627 | -0.002 | -0.02 | -0.02 | 0 | 0.16 | 0.19 | 0.16 | 0.06 | 0.13 | 0.16 |
| 2 | 234380988 | rs2971869 | -0.576 | 2.353 | 1.056 | 0.676 | -0.009 | 0.01 | 0.01 | 0 | 0.17 | 0.1 | 0.1 | 0.11 | 0.23 | 0.11 |
| 2 | 234381817 | rs10175828 | 1.031 | 2.307 | 0.986 | 0.729 | -0.002 | 0 | 0 | 0 | 0.18 | 0.18 | 0.05 | 0.18 | 0.1 | 0.33 |
| 2 | 234386134 | rs2971863 | -0.86 | 2.424 | 1.091 | 0.743 | 0.026 | -0.036 | -0.036 | 1.583 | 0.2 | 0.51 | 0.15 | 0.14 | 0.25 | 0.12 |
| 2 | 234386803 | rs745155 | -1.118 | 2.379 | 1.102 | 0.739 | -0.007 | -0.009 | -0.009 | 0 | 0.37 | 0.5 | 0.2 | 0.32 | 0.35 | 0.38 |
| 2 | 234390469 | rs3796092 | -0.06 | 2.486 | 1.089 | 0.691 | 0.03 | -0.03 | -0.03 | 0 | 0.09 | 0.21 | 0.11 | 0.05 | 0.12 | 0.15 |
| 2 | 234394487 | rs1048603 | -2.034 | 2.489 | 1.112 | 0.698 | -0.019 | 0.036 | 0.036 | 0 | 0.3 | 0.75 | 0.27 | 0.19 | 0.36 | 0.27 |
| 2 | 234394569 | rs2603547 | -0.841 | 2.502 | 1.01 | 0.708 | 0.026 | -0.033 | -0.033 | 0 | 0.2 | 0.51 | 0.15 | 0.15 | 0.25 | 0.12 |
| 2 | 234404913 | rs768811 | -2.537 | 2.614 | 0.896 | 0.781 | 0.039 | 0.034 | 0.034 | 0.17 | 0.58 | 0.91 | 0.65 | 0.64 | 0.65 | 0.63 |
| 2 | 234422029 | rs838552 | -1.26 | 2.873 | 1.544 | 0.768 | 0.036 | -0.033 | -0.033 | 0 | 0.19 | 0.43 | 0.15 | 0.15 | 0.25 | 0.12 |
| 2 | 234422291 | rs838554 | -2.461 | 2.881 | 1.041 | 0.771 | -0.013 | 0.024 | 0.024 | 0 | 0.3 | 0.73 | 0.27 | 0.2 | 0.36 | 0.27 |
| 2 | 234432017 | rs838543 | -1.246 | 2.72 | 0.546 | 0.995 | 0.006 | -0.01 | -0.01 | 0 | 0.19 | 0.23 | 0.13 | 0.15 | 0.24 | 0.12 |
| 2 | 234449316 | rs2167884 | 1.031 | 2.72 | 0.546 | 0.995 | 0.006 | -0.01 | -0.01 | 0 | 0.81 | 0.77 | 0.87 | 0.85 | 0.76 | 0.88 |
| 2 | 234453888 | rs4047199 | 1.018 | 2.723 | 0.521 | 0.769 | 0.006 | -0.037 | -0.037 | 0.021 | 0.82 | 0.78 | 0.87 | 0.95 | 0.76 | 0.89 |
| 2 | 234466843 | rs12472244 | -0.839 | 2.144 | 0.026 | 1.137 | -0.007 | 0.003 | 0.003 | 0 | 0.11 | 0.12 | 0.33 | 0.28 | 0.19 | 0.1 |
| 2 | 234482610 | rs13387539 | 0.167 | 1.249 | 0.146 | 0.383 | -0.023 | -0.035 | -0.035 | 0 | 0.07 | 0.13 | 0.02 | 0 | 0.03 | 0.02 |
| 2 | 234493048 | rs4129945 | -0.459 | 1.225 | 0.233 | 0.46 | -0.029 | -0.002 | -0.002 | 0 | 0.08 | 0.19 | 0.02 | 0.08 | 0.03 | 0.03 |
| 2 | 234494513 | rs13430633 | -0.407 | 1.224 | -0.044 | 0.46 | -0.014 | -0.003 | -0.003 | 0 | 0.08 | 0.09 | 0.01 | 0.08 | 0.03 | 0.03 |
| 2 | 234495742 | rs17864661 | -0.78 | 0.952 | 0.074 | 0.137 | -0.008 | -0.043 | -0.043 | 0 | 0.14 | 0.19 | 0.14 | 0.07 | 0.12 | 0.16 |
| 2 | 234496534 | rs17864662 | -0.524 | 1.365 | -0.346 | 0.372 | 0 | -0.003 | -0.003 | 0 | 0.09 | 0.1 | 0.03 | 0.08 | 0.07 | 0.03 |
| 2 | 234504098 | rs1597944 | 0.903 | 1.039 | -1.246 | 0.339 | -0.036 | -0.016 | -0.016 | 0 | 0.56 | 0.16 | 0.46 | 0.54 | 0.5 | 0.67 |
| 2 | 234505285 | rs1115381 | 0.414 | 1.182 | -0.214 | 0.293 | 0.005 | 0.033 | 0.033 | 0 | 0.75 | 0.65 | 0.64 | 0.68 | 0.71 | 0.78 |
| 2 | 234506808 | rs17868299 | 1.314 | 1.247 | 0.09 | 0.222 | 0.019 | -0.026 | -0.026 | 0.901 | 0.06 | 0.14 | 0.04 | 0 | 0.05 | 0.01 |
| 2 | 234508692 | rs6728792 | -0.147 | 1.245 | 0.756 | 0.161 | 0.006 | -0.023 | -0.023 | 0 | 0.17 | 0.29 | 0.16 | 0.13 | 0.21 | 0.1 |
| 2 | 234508963 | rs2741012 | 0.787 | 0.943 | 0.024 | -0.82 | 0.001 | 0.063 | 0.063 | 0.075 | 0.29 | 0.1 | 0.18 | 0.03 | 0.29 | 0.38 |
| 2 | 234512976 | rs2269352 | 1.678 | 0.777 | -0.22 | -0.899 | 0.022 | -0.091 | -0.091 | 0 | 0.34 | 0.1 | 0.3 | 0.51 | 0.25 | 0.3 |
| 2 | 234516714 | rs2741023 | 0.411 | 0.84 | -0.246 | -0.956 | -0.042 | 0.207 | 0.207 | 0 | 0.33 | 0.04 | 0.21 | 0.03 | 0.34 | 0.41 |
| 2 | 234518011 | rs2741027 | 0.898 | 0.84 | -0.226 | -0.956 | -0.027 | 0.161 | 0.161 | 0.041 | 0.28 | 0.04 | 0.17 | 0.03 | 0.28 | 0.4 |
| 2 | 234518422 | rs17864668 | 0.449 | 0.961 | 0.073 | -0.936 | 0.009 | -0.002 | -0.002 | 0 | 0.09 | 0.17 | 0.07 | 0.11 | 0.16 | 0.05 |
| 2 | 234530160 | rs2741029 | 0.924 | 1.009 | -0.107 | -1.053 | -0.027 | 0.161 | 0.161 | 0.041 | 0.28 | 0.04 | 0.17 | 0.03 | 0.28 | 0.4 |
| 2 | 234536637 | rs1377460 | -0.605 | 0.969 | 0.059 | -0.99 | -0.003 | -0.012 | -0.012 | 0 | 0.81 | 0.82 | 0.69 | 0.67 | 0.77 | 0.79 |
| 2 | 234537447 | rs13394704 | 1.78 | 0.932 | -0.007 | -0.985 | -0.03 | -0.19 | -0.19 | 1.398 | 0.34 | 0.02 | 0.3 | 0.5 | 0.25 | 0.28 |

| | | | | | | | | | | | | | | | | |
|---|---|---|---|---|---|---|---|---|---|---|---|---|---|---|---|---|
| 2 | 234545861 | rs17854828 | 0.489 | 0.968 | 0.268 | -0.948 | 0.018 | -0.002 | -0.002 | 0 | 0.08 | 0.15 | 0.07 | 0.11 | 0.16 | 0.05 |
| 2 | 234555809 | rs2741035 | -0.57 | 0.919 | 0.265 | -0.946 | -0.002 | -0.01 | -0.01 | 0.274 | 0.81 | 0.76 | 0.69 | 0.67 | 0.77 | 0.78 |
| 2 | 234556742 | rs12995772 | 2.366 | 0.839 | -0.07 | -1.257 | -0.012 | 0.052 | 0.052 | 0 | 0.09 | 0 | 0.05 | 0 | 0.06 | 0.03 |
| 2 | 234560591 | rs1817154 | -0.518 | 0.735 | 0.334 | -1.163 | 0 | 0.004 | 0.004 | 0 | 0.8 | 0.74 | 0.69 | 0.67 | 0.77 | 0.78 |
| 2 | 234561953 | rs2602373 | 0.957 | 0.726 | 0.289 | -1.215 | -0.025 | 0.158 | 0.158 | 0.041 | 0.28 | 0.04 | 0.17 | 0.03 | 0.28 | 0.4 |
| 2 | 234565283 | rs11892031 | 0.796 | 0.728 | 0.662 | -1.202 | 0.022 | -0.034 | -0.034 | 0 | 0.08 | 0.19 | 0.13 | 0.03 | 0.07 | 0.05 |
| 2 | 234565438 | rs1453322 | 1.693 | 0.724 | 0.579 | -1.203 | 0.022 | -0.089 | -0.089 | 0 | 0.34 | 0.1 | 0.3 | 0.51 | 0.25 | 0.28 |
| 2 | 234566418 | rs7571337 | 0.348 | 0.734 | 0.625 | -1.203 | -0.001 | 0.009 | 0.009 | 0 | 0.51 | 0.7 | 0.51 | 0.54 | 0.48 | 0.38 |
| 2 | 234567051 | rs7608713 | -0.721 | 0.788 | 0.839 | -1.175 | 0.079 | -0.047 | -0.047 | 0.424 | 0.16 | 0.51 | 0.21 | 0.13 | 0.23 | 0.1 |
| 2 | 234567916 | rs1112310 | 1.644 | 0.731 | 0.691 | -1.18 | 0.031 | -0.07 | -0.07 | 0 | 0.35 | 0.15 | 0.31 | 0.51 | 0.26 | 0.29 |
| 2 | 234568964 | rs2602374 | 0.952 | 0.858 | 0.546 | -1.229 | -0.017 | 0.163 | 0.163 | 0.997 | 0.28 | 0.04 | 0.18 | 0.03 | 0.27 | 0.39 |
| 2 | 234569137 | rs1113193 | -0.765 | 0.63 | 0.807 | -1.127 | 0.096 | -0.042 | -0.042 | 0.038 | 0.17 | 0.46 | 0.26 | 0.14 | 0.25 | 0.13 |
| 2 | 234572783 | rs6760588 | 1.418 | 0.544 | 0.574 | -1.091 | 0.008 | -0.01 | -0.01 | 0 | 0.36 | 0.28 | 0.25 | 0.51 | 0.25 | 0.27 |
| 2 | 234573896 | rs17863773 | 0.077 | 0.61 | 0.517 | -0.981 | -0.01 | 0.006 | 0.006 | 0 | 0.1 | 0.12 | 0.14 | 0.1 | 0.18 | 0.11 |
| 2 | 234579363 | rs17864684 | 0.143 | 0.621 | 0.522 | -0.909 | -0.009 | -0.001 | -0.001 | 0.923 | 0.1 | 0.12 | 0.14 | 0.15 | 0.18 | 0.1 |
| 2 | 234580140 | rs2741045 | 1.001 | 0.457 | 0.38 | -1.088 | -0.052 | 0.165 | 0.165 | 0.841 | 0.27 | 0.03 | 0.17 | 0.03 | 0.27 | 0.34 |
| 2 | 234581834 | rs2741049 | -1.487 | 0.341 | 0.76 | -0.977 | -0.001 | 0.002 | 0.002 | 0.053 | 0.6 | 0.62 | 0.51 | 0.44 | 0.6 | 0.69 |
| 2 | 234581920 | rs7349250 | 0.772 | 0.311 | 1.016 | -0.972 | -0.022 | 0.068 | 0.068 | 0 | 0.1 | 0.18 | 0.05 | 0.17 | 0.04 | 0.07 |
| 2 | 234587639 | rs12053462 | -0.253 | 0.573 | 1.028 | -0.826 | -0.043 | -0.013 | -0.013 | 0 | 0.14 | 0.09 | 0.15 | 0.21 | 0.22 | 0.22 |
| 2 | 234588045 | rs7595138 | -1.326 | 0.56 | 1.092 | -0.84 | -0.002 | 0.002 | 0.002 | 0.152 | 0.6 | 0.61 | 0.51 | 0.44 | 0.6 | 0.69 |
| 2 | 234589190 | rs10197460 | 0.114 | 0.548 | 1.122 | -0.847 | 0.013 | 0.049 | 0.049 | 0 | 0.42 | 0.25 | 0.27 | 0.2 | 0.36 | 0.43 |
| 2 | 234589504 | rs10199293 | -1.394 | 0.549 | 1.305 | -0.846 | 0 | -0.009 | -0.009 | 0.025 | 0.61 | 0.74 | 0.51 | 0.44 | 0.6 | 0.69 |
| 2 | 234589616 | rs12472689 | -0.105 | 0.561 | 1.193 | -0.831 | -0.042 | -0.014 | -0.014 | 0.294 | 0.13 | 0.08 | 0.15 | 0.21 | 0.22 | 0.22 |
| 2 | 234592635 | rs4433994 | -1.452 | 0.592 | 1.308 | -0.819 | -0.029 | -0.056 | -0.056 | 1.149 | 0.62 | 0.9 | 0.52 | 0.44 | 0.61 | 0.69 |
| 2 | 234592816 | rs6724485 | -0.196 | 0.593 | 1.363 | -0.823 | 0.007 | 0.021 | 0.021 | 0 | 0.45 | 0.36 | 0.34 | 0.22 | 0.38 | 0.48 |
| 2 | 234593041 | rs4347832 | -0.196 | 0.593 | 1.363 | -0.823 | 0.007 | 0.021 | 0.021 | 0 | 0.45 | 0.36 | 0.34 | 0.22 | 0.38 | 0.48 |
| 2 | 234593117 | rs4261716 | -0.196 | 0.593 | 1.363 | -0.823 | 0.007 | 0.021 | 0.021 | 0 | 0.45 | 0.36 | 0.34 | 0.22 | 0.38 | 0.48 |
| 2 | 234593374 | rs4583459 | -1.286 | 0.593 | 1.379 | -0.822 | -0.003 | 0.005 | 0.005 | 0.007 | 0.6 | 0.56 | 0.5 | 0.44 | 0.6 | 0.69 |
| 2 | 234594083 | rs4553819 | -0.196 | 0.593 | 1.379 | -0.823 | 0.007 | 0.021 | 0.021 | 0 | 0.45 | 0.36 | 0.34 | 0.22 | 0.38 | 0.48 |
| 2 | 234595615 | rs6753320 | -0.092 | 0.593 | 1.379 | -0.823 | 0.005 | 0.016 | 0.016 | 0 | 0.45 | 0.38 | 0.35 | 0.22 | 0.38 | 0.48 |
| 2 | 234595747 | rs6736508 | -0.092 | 0.593 | 1.379 | -0.823 | 0.005 | 0.016 | 0.016 | 0 | 0.45 | 0.38 | 0.34 | 0.22 | 0.38 | 0.48 |
| 2 | 234595817 | rs6753569 | -0.092 | 0.593 | 1.379 | -0.823 | 0.005 | 0.016 | 0.016 | 0 | 0.45 | 0.38 | 0.34 | 0.22 | 0.38 | 0.48 |
| 2 | 234595950 | rs6736743 | -0.092 | 0.593 | 1.379 | -0.823 | 0.005 | 0.016 | 0.016 | 0 | 0.45 | 0.38 | 0.34 | 0.22 | 0.38 | 0.48 |
| 2 | 234596214 | rs6754100 | -1.442 | 0.593 | 1.424 | -0.824 | -0.022 | -0.047 | -0.047 | 2.668 | 0.61 | 0.89 | 0.52 | 0.44 | 0.61 | 0.7 |
| 2 | 234596836 | rs10168155 | -0.124 | 0.595 | 1.3 | -0.823 | 0.005 | 0.017 | 0.017 | 0 | 0.45 | 0.38 | 0.34 | 0.22 | 0.38 | 0.48 |
| 2 | 234597087 | rs10168416 | 0.474 | 0.613 | 1.316 | -0.826 | 0.012 | 0.025 | 0.025 | 0.014 | 0.37 | 0.23 | 0.22 | 0.2 | 0.31 | 0.42 |
| 2 | 234597483 | rs11680450 | -0.114 | 0.617 | 1.24 | -0.823 | 0.005 | 0.016 | 0.016 | 0 | 0.45 | 0.38 | 0.34 | 0.22 | 0.38 | 0.48 |
| 2 | 234597825 | rs10179094 | 0.118 | 0.617 | 1.24 | -0.824 | 0.012 | 0.048 | 0.048 | 0 | 0.42 | 0.25 | 0.26 | 0.2 | 0.36 | 0.43 |
| 2 | 234598991 | rs7563561 | -0.113 | 0.617 | 1.235 | -0.828 | 0.005 | 0.016 | 0.016 | 0 | 0.45 | 0.38 | 0.35 | 0.22 | 0.38 | 0.48 |
| 2 | 234599941 | rs12623271 | -0.128 | 0.617 | 1.252 | -0.827 | 0.005 | 0.016 | 0.016 | 0 | 0.55 | 0.38 | 0.34 | 0.22 | 0.38 | 0.48 |
| 2 | 234600116 | rs12474980 | -0.094 | 0.602 | 1.253 | -0.821 | -0.042 | -0.019 | -0.019 | 0 | 0.13 | 0.08 | 0.15 | 0.22 | 0.22 | 0.22 |
| 2 | 234600999 | rs13015720 | -0.128 | 0.597 | 1.228 | -0.837 | 0.005 | 0.016 | 0.016 | 0 | 0.45 | 0.38 | 0.34 | 0.22 | 0.38 | 0.48 |
| 2 | 234601669 | rs6759892 | -0.128 | 0.597 | 1.244 | -0.837 | 0.005 | 0.014 | 0.014 | 0 | 0.45 | 0.38 | 0.34 | 0.22 | 0.38 | 0.48 |
| 2 | 234601965 | rs1105880 | 0.239 | 0.596 | 1.239 | -0.826 | 0 | 0 | 0 | 0 | 0.4 | 0.37 | 0.3 | 0.22 | 0.33 | 0.47 |
| 2 | 234602191 | rs2070959 | 0.447 | 0.595 | 1.241 | -0.826 | 0.011 | 0.024 | 0.024 | 0.014 | 0.37 | 0.24 | 0.22 | 0.2 | 0.31 | 0.42 |
| 2 | 234602202 | rs1105879 | 0.169 | 0.595 | 1.248 | -0.827 | 0.007 | 0.014 | 0.014 | 0.079 | 0.4 | 0.3 | 0.3 | 0.22 | 0.33 | 0.47 |
| 2 | 234602906 | rs7592624 | -0.84 | 0.588 | 0.98 | -0.725 | 0 | -0.001 | -0.001 | 0 | 0.56 | 0.6 | 0.47 | 0.44 | 0.55 | 0.68 |
| 2 | 234603439 | rs6761246 | -1.237 | 0.6 | 0.609 | -0.677 | 0 | -0.012 | -0.012 | 0 | 0.61 | 0.75 | 0.51 | 0.44 | 0.6 | 0.69 |
| 2 | 234607394 | rs12988520 | -0.354 | 0.662 | 0.644 | -0.967 | 0.006 | 0.015 | 0.015 | 1.37 | 0.5 | 0.43 | 0.48 | 0.29 | 0.54 | 0.63 |
| 2 | 234610912 | rs17862866 | -1.306 | -0.266 | 0.72 | -0.657 | -0.064 | 0.068 | 0.068 | 0.512 | 0.6 | 0.89 | 0.53 | 0.35 | 0.45 | 0.66 |
| 2 | 234611094 | rs17863787 | 0.403 | -0.37 | 0.83 | -0.76 | 0.006 | -0.01 | -0.01 | 0 | 0.37 | 0.28 | 0.3 | 0.07 | 0.29 | 0.39 |
| 2 | 234615400 | rs6725478 | -0.214 | -0.434 | 0.712 | -0.614 | 0.022 | 0.056 | 0.056 | 0 | 0.45 | 0.68 | 0.36 | 0.07 | 0.34 | 0.4 |
| 2 | 234616357 | rs4663325 | -0.737 | -0.443 | 0.709 | -0.597 | -0.013 | 0.016 | 0.016 | 0.045 | 0.14 | 0.2 | 0.17 | 0.28 | 0.11 | 0.27 |
| 2 | 234616571 | rs4663326 | 0.006 | -0.446 | 0.627 | -0.594 | 0.004 | -0.027 | -0.027 | 0.509 | 0.12 | 0.04 | 0.1 | 0.23 | 0.1 | 0.22 |
| 2 | 234617236 | rs7572563 | -1.345 | -0.446 | 0.623 | -0.602 | -0.063 | 0.07 | 0.07 | 0.324 | 0.59 | 0.89 | 0.53 | 0.35 | 0.45 | 0.66 |
| 2 | 234617407 | rs7583278 | -0.199 | -0.449 | 0.701 | -0.604 | 0.025 | 0.073 | 0.073 | 0.011 | 0.45 | 0.69 | 0.36 | 0.07 | 0.34 | 0.4 |
| 2 | 234617632 | rs4663327 | -0.878 | -0.365 | 0.77 | -0.521 | -0.015 | 0.019 | 0.019 | 0.019 | 0.15 | 0.22 | 0.17 | 0.28 | 0.11 | 0.27 |
| 2 | 234621157 | rs2013018 | -0.221 | -0.31 | 0.578 | -0.505 | 0.004 | -0.007 | -0.007 | 0 | 0.12 | 0.08 | 0.1 | 0.23 | 0.1 | 0.22 |
| 2 | 234621404 | rs4556969 | -0.221 | -0.31 | 0.577 | -0.506 | 0.004 | -0.007 | -0.007 | 0.062 | 0.12 | 0.08 | 0.1 | 0.23 | 0.1 | 0.22 |
| 2 | 234621825 | rs3755321 | -0.221 | -0.323 | 0.609 | -0.515 | 0.004 | -0.007 | -0.007 | 0.046 | 0.12 | 0.08 | 0.1 | 0.23 | 0.1 | 0.22 |
| 2 | 234622742 | rs17862870 | -0.175 | -0.313 | 0.615 | -0.51 | 0.004 | -0.007 | -0.007 | 0 | 0.12 | 0.08 | 0.1 | 0.23 | 0.1 | 0.22 |
| 2 | 234623117 | rs4663945 | -0.029 | -0.314 | 0.616 | -0.511 | 0.003 | -0.027 | -0.027 | 0.284 | 0.11 | 0.05 | 0.1 | 0.23 | 0.1 | 0.22 |
| 2 | 234624817 | rs17868334 | -0.17 | -0.313 | 0.612 | -0.51 | 0.004 | -0.007 | -0.007 | 0 | 0.12 | 0.08 | 0.1 | 0.23 | 0.1 | 0.22 |
| 2 | 234625126 | rs17863792 | -0.17 | -0.313 | 0.612 | -0.51 | 0.004 | -0.007 | -0.007 | 0.844 | 0.12 | 0.08 | 0.1 | 0.23 | 0.1 | 0.22 |
| 2 | 234625297 | rs6744284 | 0.185 | -0.372 | 0.613 | -0.547 | 0.013 | 0.024 | 0.024 | 0.09 | 0.39 | 0.66 | 0.38 | 0.11 | 0.27 | 0.43 |
| 2 | 234625622 | rs1875263 | -0.109 | -0.358 | 0.588 | -0.531 | 0.024 | 0.078 | 0.078 | 0 | 0.44 | 0.7 | 0.42 | 0.11 | 0.32 | 0.44 |
| 2 | 234626199 | rs3806591 | -0.28 | 0.114 | 0.688 | -0.294 | 0.003 | -0.006 | -0.006 | 0 | 0.12 | 0.08 | 0.1 | 0.23 | 0.1 | 0.22 |
| 2 | 234627010 | rs3732221 | -0.28 | 0.114 | 0.654 | -0.294 | 0.004 | -0.007 | -0.007 | 1.148 | 0.12 | 0.08 | 0.1 | 0.23 | 0.1 | 0.22 |
| 2 | 234627304 | rs3732218 | -0.299 | 0.115 | 0.655 | -0.294 | 0.004 | -0.007 | -0.007 | 0.006 | 0.12 | 0.08 | 0.1 | 0.23 | 0.1 | 0.22 |

| | | | | | | | | | | | | | | | | |
|---|---|---|---|---|---|---|---|---|---|---|---|---|---|---|---|---|
| 2 | 234628529 | rs871514 | -1.178 | -0.179 | 0.655 | -0.307 | -0.012 | 0.104 | 0.104 | 0.383 | 0.55 | 0.83 | 0.52 | 0.36 | 0.41 | 0.65 |
| 2 | 234628647 | rs904856 | 0.019 | -0.158 | 0.662 | -0.272 | 0.004 | -0.006 | -0.006 | 0.017 | 0.11 | 0.07 | 0.09 | 0.23 | 0.07 | 0.2 |
| 2 | 234629239 | rs12468543 | 0.019 | -0.158 | 0.662 | -0.272 | 0.004 | -0.006 | -0.006 | 0 | 0.11 | 0.07 | 0.09 | 0.23 | 0.07 | 0.2 |
| 2 | 234630900 | rs12479208 | 0.019 | -0.158 | 0.662 | -0.272 | 0.004 | -0.006 | -0.006 | 0 | 0.11 | 0.07 | 0.09 | 0.23 | 0.07 | 0.2 |
| 2 | 234635467 | rs4294999 | -1.038 | -0.162 | 0.598 | -0.29 | -0.014 | 0.104 | 0.104 | 0.408 | 0.55 | 0.83 | 0.52 | 0.36 | 0.42 | 0.65 |
| 2 | 234637192 | rs2008595 | -1.097 | -0.169 | 0.599 | -0.293 | -0.03 | 0.102 | 0.102 | 0.705 | 0.56 | 0.89 | 0.52 | 0.36 | 0.42 | 0.64 |
| 2 | 234637569 | rs3806597 | -1.069 | -0.171 | 0.618 | -0.294 | -0.025 | 0.104 | 0.104 | 0.795 | 0.55 | 0.88 | 0.52 | 0.36 | 0.41 | 0.64 |
| 2 | 234647478 | rs6431628 | -1.096 | -0.147 | 0.742 | -0.297 | -0.032 | 0.102 | 0.102 | 0.626 | 0.56 | 0.89 | 0.53 | 0.36 | 0.42 | 0.64 |
| 2 | 234648860 | rs2018985 | -0.414 | -0.207 | 0.937 | -0.342 | 0.031 | 0.149 | 0.149 | 0 | 0.45 | 0.75 | 0.43 | 0.13 | 0.35 | 0.44 |
| 2 | 234649302 | rs17862875 | 0.43 | -0.291 | 1.325 | -0.383 | -0.001 | -0.016 | -0.016 | 0.072 | 0.34 | 0.31 | 0.36 | 0.12 | 0.27 | 0.42 |
| 2 | 234650193 | rs4663963 | -1.126 | -0.283 | 1.212 | -0.376 | -0.025 | 0.104 | 0.104 | 0.469 | 0.55 | 0.86 | 0.52 | 0.36 | 0.42 | 0.64 |
| 2 | 234650236 | rs4663964 | 0.203 | -0.26 | 1.214 | -0.376 | 0.009 | -0.028 | -0.028 | 0 | 0.1 | 0.04 | 0.09 | 0.23 | 0.07 | 0.2 |
| 2 | 234650604 | rs4663965 | -1.165 | -0.292 | 1.217 | -0.375 | -0.027 | 0.101 | 0.101 | 0.635 | 0.56 | 0.87 | 0.53 | 0.36 | 0.42 | 0.64 |
| 2 | 234655144 | rs4663968 | 0.128 | -0.263 | 1.227 | -0.363 | 0 | 0 | 0 | 0 | 0.1 | 0.1 | 0.09 | 0.23 | 0.07 | 0.2 |
| 2 | 234655303 | rs4663333 | -1.157 | -0.331 | 1.206 | -0.355 | -0.029 | 0.097 | 0.097 | 0.585 | 0.56 | 0.89 | 0.53 | 0.36 | 0.42 | 0.65 |
| 2 | 234657983 | rs10179091 | -1.057 | -0.378 | 1.376 | -0.331 | 0.007 | 0.043 | 0.043 | 0.395 | 0.54 | 0.64 | 0.51 | 0.36 | 0.41 | 0.64 |
| 2 | 234658116 | rs7604115 | -0.084 | -0.393 | 1.404 | -0.337 | -0.007 | -0.072 | -0.072 | 0 | 0.39 | 0.46 | 0.4 | 0.13 | 0.32 | 0.43 |
| 2 | 234658250 | rs7556676 | -0.698 | -0.381 | 1.38 | -0.331 | 0.013 | 0.039 | 0.039 | 0 | 0.5 | 0.59 | 0.5 | 0.35 | 0.39 | 0.63 |
| 2 | 234665498 | rs11568318 | 1.543 | -0.338 | 0.931 | -0.31 | 0 | -0.001 | -0.001 | 0 | 0.05 | 0.06 | 0.04 | 0 | 0.05 | 0.01 |
| 2 | 234665659 | rs4124874 | -1.212 | -0.363 | 0.877 | -0.321 | -0.036 | 0.091 | 0.091 | 0.242 | 0.56 | 0.9 | 0.53 | 0.36 | 0.42 | 0.63 |
| 2 | 234666461 | rs4399719 | -1.164 | -0.31 | 0.896 | -0.295 | -0.029 | 0.096 | 0.096 | 0.436 | 0.56 | 0.89 | 0.53 | 0.36 | 0.42 | 0.63 |
| 2 | 234666581 | rs12052787 | 0.148 | -0.29 | 0.796 | -0.268 | 0.012 | -0.028 | -0.028 | 0 | 0.1 | 0.04 | 0.09 | 0.22 | 0.07 | 0.18 |
| 2 | 234667582 | rs3755319 | -1.193 | -0.288 | 0.907 | -0.282 | 0.003 | 0.069 | 0.069 | 0.507 | 0.55 | 0.76 | 0.52 | 0.36 | 0.41 | 0.63 |
| 2 | 234668570 | rs887829 | 0.054 | -0.133 | 0.731 | -0.237 | -0.012 | -0.075 | -0.075 | 0 | 0.37 | 0.49 | 0.38 | 0.13 | 0.3 | 0.44 |
| 2 | 234672639 | rs6742078 | 0.085 | -0.106 | 0.753 | -0.835 | -0.014 | -0.085 | -0.085 | 1.748 | 0.37 | 0.47 | 0.38 | 0.13 | 0.3 | 0.43 |
| 2 | 234672663 | rs3771342 | -0.518 | -0.113 | 0.761 | -0.839 | 0.006 | -0.001 | -0.001 | 0.41 | 0.16 | 0.14 | 0.13 | 0.22 | 0.11 | 0.19 |
| 2 | 234672722 | rs4148324 | 0.061 | -0.115 | 0.727 | -0.84 | -0.012 | -0.073 | -0.073 | 0 | 0.37 | 0.49 | 0.38 | 0.13 | 0.3 | 0.44 |
| 2 | 234673239 | rs3771341 | 0.346 | -0.118 | 0.647 | -0.843 | -0.009 | -0.077 | -0.077 | 1.531 | 0.34 | 0.44 | 0.37 | 0.13 | 0.27 | 0.42 |
| 2 | 234673309 | rs4148325 | 0.064 | -0.112 | 0.649 | -0.84 | -0.012 | -0.075 | -0.075 | 0.483 | 0.37 | 0.49 | 0.38 | 0.13 | 0.3 | 0.43 |
| 2 | 234673462 | rs4148326 | -1.071 | -0.114 | 0.651 | -0.841 | 0.007 | 0.047 | 0.047 | 0.314 | 0.53 | 0.65 | 0.51 | 0.36 | 0.41 | 0.63 |
| 2 | 234673516 | rs17868342 | -0.606 | -0.189 | 0.483 | -0.796 | 0.004 | -0.001 | -0.001 | 1.427 | 0.16 | 0.16 | 0.13 | 0.22 | 0.12 | 0.19 |
| 2 | 234673588 | rs12479045 | 0.047 | -0.19 | 0.338 | -0.698 | 0.012 | -0.028 | -0.028 | 0 | 0.1 | 0.04 | 0.09 | 0.22 | 0.07 | 0.18 |
| 2 | 234674476 | rs929596 | 0.197 | -0.013 | 0.507 | -0.699 | -0.021 | -0.091 | -0.091 | 0 | 0.33 | 0.45 | 0.35 | 0.13 | 0.25 | 0.41 |
| 2 | 234677659 | rs4148328 | 0.722 | 0.344 | -0.536 | -0.392 | -0.035 | 0.014 | 0.014 | 3.19 | 0.37 | 0.15 | 0.35 | 0.57 | 0.4 | 0.25 |
| 2 | 234679384 | rs11563251 | -0.567 | -0.105 | 0.149 | -0.404 | -0.035 | -0.063 | -0.063 | 0 | 0.19 | 0.42 | 0.1 | 0.12 | 0.12 | 0.18 |
| 2 | 234681416 | rs10929303 | 0.181 | 0.072 | 0.176 | -0.47 | -0.001 | -0.056 | -0.056 | 1.478 | 0.78 | 0.58 | 0.78 | 0.87 | 0.77 | 0.83 |
| 2 | 234681544 | rs1042640 | -0.045 | 0.101 | 0.296 | -0.491 | 0 | 0.002 | 0.002 | 0 | 0.81 | 0.8 | 0.81 | 0.87 | 0.8 | 0.83 |
| 2 | 234682062 | rs4148329 | 1.348 | -0.019 | 0.127 | -0.719 | 0.012 | 0.052 | 0.052 | 0 | 0.56 | 0.27 | 0.52 | 0.63 | 0.57 | 0.61 |
| 2 | 234682119 | rs6717546 | 0.909 | -0.001 | 0.335 | -0.689 | -0.01 | -0.038 | -0.038 | 0 | 0.62 | 0.47 | 0.56 | 0.63 | 0.62 | 0.62 |
| 2 | 234683038 | rs1500482 | -0.291 | 0.001 | 0.264 | -0.686 | 0 | 0.002 | 0.002 | 0 | 0.81 | 0.8 | 0.8 | 0.87 | 0.8 | 0.83 |
| 2 | 234683350 | rs11563250 | 0.188 | -0.014 | 0.211 | -0.691 | -0.018 | -0.009 | -0.009 | 0 | 0.15 | 0.01 | 0.21 | 0.23 | 0.14 | 0.21 |
| 2 | 234683763 | rs6719561 | 1.855 | 0.036 | -0.093 | -0.831 | -0.074 | -0.062 | -0.062 | 0 | 0.3 | 0.06 | 0.27 | 0.51 | 0.34 | 0.22 |
| 2 | 234683892 | rs4663972 | -0.265 | 0.027 | 0 | -0.83 | -0.001 | -0.029 | -0.029 | 0 | 0.79 | 0.64 | 0.78 | 0.86 | 0.78 | 0.82 |
| 2 | 234685264 | rs10199882 | 0.341 | 0.117 | -0.324 | -0.777 | 0.01 | 0.012 | 0.012 | 0 | 0.16 | 0.17 | 0.22 | 0.23 | 0.14 | 0.21 |
| 2 | 234685527 | rs7586006 | 0.341 | 0.115 | -0.323 | -0.777 | 0.01 | 0.012 | 0.012 | 0 | 0.16 | 0.17 | 0.22 | 0.23 | 0.14 | 0.21 |
| 2 | 234687583 | rs10209214 | 1.795 | 0.066 | -0.422 | -0.759 | -0.071 | -0.059 | -0.059 | 0.17 | 0.3 | 0.07 | 0.27 | 0.51 | 0.35 | 0.22 |
| 2 | 234688036 | rs6431631 | -0.524 | 0.081 | -0.476 | -0.783 | 0 | 0.007 | 0.007 | 0 | 0.81 | 0.84 | 0.8 | 0.87 | 0.8 | 0.83 |
| 2 | 234689090 | rs4663335 | 0.274 | 0.092 | -0.625 | -0.776 | 0.004 | 0.017 | 0.017 | 0 | 0.17 | 0.13 | 0.22 | 0.23 | 0.16 | 0.21 |
| 2 | 234691401 | rs10199512 | 0.487 | 0.092 | -0.562 | -0.78 | 0 | 0.001 | 0.001 | 0 | 0.64 | 0.59 | 0.58 | 0.64 | 0.64 | 0.61 |
| 2 | 234692617 | rs11690786 | 0.13 | 0.232 | -0.659 | -0.733 | -0.001 | -0.012 | -0.012 | 0.056 | 0.33 | 0.31 | 0.29 | 0.12 | 0.3 | 0.39 |
| 2 | 234694177 | rs7596785 | -0.197 | 0.151 | -0.729 | -0.561 | -0.001 | -0.008 | -0.008 | 0 | 0.81 | 0.77 | 0.8 | 0.87 | 0.8 | 0.83 |
| 2 | 234698790 | rs2361502 | -0.07 | 0.24 | -0.968 | -0.36 | -0.008 | -0.023 | -0.023 | 0.006 | 0.31 | 0.32 | 0.32 | 0.13 | 0.25 | 0.39 |
| 2 | 234700385 | rs10207194 | 0.362 | 0.26 | -2.001 | 0.246 | -0.03 | 0.05 | 0.05 | 0.031 | 0.5 | 0.13 | 0.6 | 0.69 | 0.46 | 0.42 |
| 2 | 234701822 | rs6755193 | 0.57 | 0.578 | -2.219 | 0.517 | -0.051 | 0.029 | 0.029 | 0 | 0.36 | 0.05 | 0.37 | 0.35 | 0.34 | 0.35 |
| 2 | 234702459 | rs1500481 | -0.268 | 0.463 | -1.714 | 0.865 | -0.014 | 0.008 | 0.008 | 0 | 0.61 | 0.7 | 0.65 | 0.69 | 0.56 | 0.52 |
| 2 | 234702905 | rs10169532 | 0.589 | 0.363 | -1.289 | 1.183 | 0.015 | 0.046 | 0.046 | 0 | 0.42 | 0.21 | 0.43 | 0.39 | 0.41 | 0.42 |
| 2 | 234703136 | rs6431634 | -0.071 | 0.779 | -1.024 | 1.184 | -0.018 | 0.008 | 0.008 | 0 | 0.62 | 0.73 | 0.66 | 0.69 | 0.57 | 0.53 |
| 2 | 234703667 | rs11890704 | 0.34 | 0.74 | -1.025 | 1.289 | -0.005 | 0.04 | 0.04 | 0 | 0.06 | 0.1 | 0.05 | 0.13 | 0.08 | 0.08 |
| 2 | 234704495 | rs11563248 | 0.087 | 0.909 | -1.159 | 0.842 | -0.02 | 0.018 | 0.018 | 0 | 0.6 | 0.67 | 0.64 | 0.69 | 0.55 | 0.52 |
| 2 | 234704690 | rs759173 | 0.02 | 1.376 | -0.958 | 0.535 | 0.004 | -0.027 | -0.027 | 0 | 0.87 | 0.85 | 0.9 | 0.99 | 0.82 | 0.9 |
| 2 | 234705389 | rs741160 | -0.748 | 1.123 | -0.634 | 0.982 | 0.007 | 0.025 | 0.025 | 0 | 0.19 | 0.31 | 0.12 | 0.26 | 0.18 | 0.2 |
| 2 | 234707921 | rs9808308 | -0.353 | 0.776 | -0.756 | 0.831 | 0.001 | 0.008 | 0.008 | 0 | 0.47 | 0.57 | 0.4 | 0.37 | 0.49 | 0.39 |
| 2 | 234708152 | rs9808231 | -0.441 | -0.311 | -0.89 | 0.969 | -0.01 | 0.004 | 0.004 | 0 | 0.75 | 0.75 | 0.68 | 0.57 | 0.87 | 0.87 |
| 2 | 234708428 | rs13013882 | -0.16 | -0.318 | -0.198 | 1.241 | -0.046 | 0.023 | 0.023 | 0 | 0.7 | 0.6 | 0.55 | 0.38 | 0.82 | 0.8 |
| 2 | 234708629 | rs13018934 | 0.103 | 0.27 | -0.6 | 0.844 | 0.009 | -0.03 | -0.03 | 0 | 0.59 | 0.8 | 0.54 | 0.53 | 0.62 | 0.63 |
| 2 | 234709540 | rs1500474 | -0.357 | 0.067 | -0.027 | 1.06 | -0.014 | 0.013 | 0.013 | 0.515 | 0.23 | 0.65 | 0.33 | 0.34 | 0.21 | 0.15 |
| 2 | 234710666 | rs10929305 | -0.211 | 0.007 | 0.148 | 1.065 | -0.016 | 0.04 | 0.04 | 0 | 0.23 | 0.55 | 0.33 | 0.36 | 0.21 | 0.15 |
| 2 | 234712921 | rs10175949 | -1.717 | -0.22 | -0.307 | 0.986 | 0.013 | 0.03 | 0.03 | 0 | 0.82 | 0.96 | 0.79 | 0.9 | 0.87 | 0.88 |
| 2 | 234713429 | rs6716168 | 0.473 | -0.35 | -0.159 | 1.309 | 0.042 | 0.001 | 0.001 | 0 | 0.58 | 0.27 | 0.44 | 0.35 | 0.65 | 0.72 |

| | | | | | | | | | | | | | | | | |
|---|---|---|---|---|---|---|---|---|---|---|---|---|---|---|---|---|
| 2 | 234713706 | rs4430949 | 0.081 | -0.295 | -0.09 | 1.321 | -0.015 | 0.033 | 0.033 | 0 | 0.23 | 0.57 | 0.33 | 0.35 | 0.21 | 0.15 |
| 2 | 234716827 | rs10203123 | -0.34 | -0.253 | -0.08 | 1.103 | 0.001 | 0.009 | 0.009 | 0 | 0.65 | 0.62 | 0.52 | 0.5 | 0.65 | 0.66 |
| 2 | 234717669 | rs10803663 | -0.359 | -0.262 | -0.267 | 1.058 | -0.011 | 0.007 | 0.007 | 0 | 0.66 | 0.78 | 0.64 | 0.69 | 0.66 | 0.68 |
| 2 | 234718294 | rs7585768 | 0.761 | -0.586 | -0.065 | 0.879 | 0.001 | 0.002 | 0.002 | 0 | 0.44 | 0.46 | 0.4 | 0.42 | 0.47 | 0.57 |
| 2 | 234718524 | rs11674078 | 0.915 | -0.553 | -0.685 | 0.833 | -0.017 | -0.008 | -0.008 | 0.018 | 0.19 | 0.12 | 0.15 | 0.21 | 0.21 | 0.2 |
| 2 | 234718864 | rs17863814 | 1.837 | -0.492 | -1.096 | 0.699 | -0.012 | -0.009 | -0.009 | 0 | 0.22 | 0.07 | 0.11 | 0.2 | 0.21 | 0.11 |
| 2 | 234720734 | rs988344 | -1.733 | -0.695 | -0.959 | 0.672 | 0.019 | 0.026 | 0.026 | 0 | 0.82 | 0.91 | 0.78 | 0.89 | 0.87 | 0.88 |
| 2 | 234720967 | rs6735725 | -0.385 | -0.603 | -0.983 | 0.821 | -0.013 | -0.011 | -0.011 | 0 | 0.64 | 0.79 | 0.64 | 0.64 | 0.66 | 0.69 |
| 2 | 234721444 | rs6431635 | -0.207 | -0.659 | -1.088 | 0.599 | -0.017 | 0.059 | 0.059 | 0.146 | 0.66 | 0.75 | 0.62 | 0.89 | 0.66 | 0.64 |
| 2 | 234721774 | rs7569824 | -0.677 | -1.071 | -0.915 | 0.926 | -0.003 | 0.046 | 0.046 | 0 | 0.34 | 0.55 | 0.48 | 0.53 | 0.33 | 0.28 |
| 2 | 234722065 | rs10929308 | 0.979 | -1.137 | -0.995 | 0.806 | 0.003 | 0.129 | 0.129 | 1.012 | 0.49 | 0.24 | 0.31 | 0.24 | 0.53 | 0.6 |
| 2 | 234724225 | rs13001408 | 1.231 | -0.817 | -0.835 | 1.096 | -0.043 | 0.153 | 0.153 | 2.491 | 0.47 | 0.11 | 0.33 | 0.25 | 0.51 | 0.61 |
| 2 | 234725502 | rs726016 | -1.052 | -0.757 | -0.592 | 1.092 | 0 | -0.013 | -0.013 | 0 | 0.82 | 0.79 | 0.76 | 0.89 | 0.85 | 0.85 |
| 2 | 234726058 | rs13015369 | 0.791 | -0.831 | -0.327 | 1.108 | -0.001 | -0.005 | -0.005 | 0.011 | 0.22 | 0.31 | 0.28 | 0.16 | 0.25 | 0.19 |
| 2 | 234726080 | rs715210 | -0.502 | -0.846 | -0.253 | 1.13 | -0.009 | 0.031 | 0.031 | 1.167 | 0.33 | 0.54 | 0.41 | 0.43 | 0.33 | 0.23 |
| 2 | 234727986 | rs6754859 | 1.318 | -0.85 | -0.202 | 0.999 | 0.025 | 0.007 | 0.007 | 1.043 | 0.66 | 0.22 | 0.52 | 0.31 | 0.74 | 0.77 |
| 2 | 234728078 | rs1500480 | -0.867 | -0.688 | -0.476 | 0.79 | -0.001 | -0.013 | -0.013 | 0.132 | 0.82 | 0.79 | 0.75 | 0.89 | 0.84 | 0.85 |
| 2 | 234728911 | rs11563256 | 0.503 | -0.463 | -1.163 | 0.832 | -0.008 | 0.069 | 0.069 | 1.485 | 0.11 | 0.24 | 0.13 | 0.27 | 0.08 | 0.04 |
| 2 | 234730807 | rs6734083 | 0.998 | -0.295 | -2.101 | 0.42 | -0.002 | -0.001 | -0.001 | 0 | 0.09 | 0.14 | 0.07 | 0.09 | 0.08 | 0.02 |
| 2 | 234731883 | rs7603146 | -1.9 | -0.193 | -2.486 | 0.217 | -0.001 | -0.095 | -0.095 | 0.533 | 0.71 | 0.95 | 0.77 | 0.6 | 0.73 | 0.83 |
| 2 | 234732311 | rs17868361 | 0.966 | -0.223 | -1.693 | 0.729 | -0.021 | 0.01 | 0.01 | 0 | 0.15 | 0.06 | 0.2 | 0.09 | 0.16 | 0.14 |
| 2 | 234732816 | rs7606893 | -1.262 | -0.024 | -1.713 | 0.651 | -0.046 | -0.028 | -0.028 | 0 | 0.52 | 0.92 | 0.57 | 0.55 | 0.48 | 0.36 |
| 2 | 234733064 | rs7557532 | -0.09 | 0.175 | -0.55 | 1.078 | -0.002 | -0.001 | -0.001 | 0 | 0.09 | 0.14 | 0.07 | 0.09 | 0.08 | 0.02 |
| 2 | 234733303 | rs7607494 | -0.09 | 0.175 | -0.55 | 1.078 | -0.002 | -0.001 | -0.001 | 0 | 0.09 | 0.14 | 0.07 | 0.09 | 0.08 | 0.02 |
| 2 | 234737372 | rs7564261 | -0.467 | 0.189 | -0.449 | 1.161 | -0.004 | 0.035 | 0.035 | 0.64 | 0.23 | 0.37 | 0.27 | 0.35 | 0.2 | 0.1 |
| 2 | 234738175 | rs502059 | -0.949 | 0.063 | -1.269 | 0.796 | -0.109 | -0.112 | -0.112 | 0.877 | 0.66 | 0.92 | 0.51 | 0.52 | 0.54 | 0.67 |
| 2 | 234739340 | rs2361506 | -0.044 | -0.705 | -0.117 | 0.505 | 0.034 | 0.012 | 0.012 | 0 | 0.48 | 0.16 | 0.37 | 0.4 | 0.42 | 0.62 |
| 2 | 234740136 | rs879665 | -0.176 | -1.02 | 0.18 | 0.386 | 0.006 | 0 | 0 | 0 | 0.23 | 0.26 | 0.14 | 0.28 | 0.14 | 0.11 |
| 2 | 234740157 | rs879664 | 0.533 | -1 | 0.133 | 0.391 | -0.007 | -0.009 | -0.009 | 0.516 | 0.1 | 0.21 | 0.07 | 0.09 | 0.07 | 0.02 |
| 2 | 234740490 | rs10185631 | 1.38 | -0.918 | 0.307 | 0.403 | 0.02 | -0.008 | -0.008 | 0 | 0.13 | 0.08 | 0.08 | 0.19 | 0.08 | 0.08 |
| 2 | 234740758 | rs213547 | -1.27 | -0.953 | 0.358 | 0.44 | 0.001 | 0.002 | 0.002 | 0 | 0.62 | 0.62 | 0.45 | 0.51 | 0.5 | 0.65 |
| 2 | 234741808 | rs2270856 | 1.414 | -0.985 | 0.161 | 0.457 | 0.022 | -0.008 | -0.008 | 0 | 0.13 | 0.08 | 0.07 | 0.19 | 0.06 | 0.08 |
| 2 | 234744924 | rs13024275 | 0.655 | -0.861 | 0.179 | 0.404 | 0.003 | 0.003 | 0.003 | 0 | 0.09 | 0.11 | 0.07 | 0.09 | 0.07 | 0.02 |
| 2 | 234745389 | rs213551 | 0.325 | -0.882 | 0.081 | 0.348 | -0.024 | 0.043 | 0.043 | 0 | 0.37 | 0.13 | 0.31 | 0.23 | 0.37 | 0.55 |
| 2 | 234747267 | rs213554 | -0.581 | -0.886 | 0.326 | 0.743 | -0.001 | 0 | 0 | 0 | 0.47 | 0.51 | 0.41 | 0.48 | 0.43 | 0.59 |
| 2 | 234749151 | rs213555 | -1.392 | -0.944 | -0.338 | 0.73 | -0.101 | -0.018 | -0.018 | 0 | 0.64 | 0.91 | 0.52 | 0.71 | 0.51 | 0.68 |
| 2 | 234749354 | rs12582 | -0.185 | -0.997 | -0.264 | 0.817 | -0.001 | 0.002 | 0.002 | 0 | 0.14 | 0.2 | 0.12 | 0.28 | 0.09 | 0.07 |
| 2 | 234750831 | rs3806589 | -0.369 | -0.799 | -0.366 | 0.883 | -0.039 | 0.025 | 0.025 | 0 | 0.66 | 0.85 | 0.67 | 0.76 | 0.61 | 0.61 |
| 2 | 234752638 | rs7583713 | -0.279 | -0.804 | -0.424 | 0.82 | -0.062 | 0.113 | 0.113 | 0 | 0.66 | 0.89 | 0.67 | 0.87 | 0.61 | 0.6 |
| 2 | 234755575 | rs556691 | 0.07 | -0.741 | -0.179 | 0.771 | 0.004 | 0.026 | 0.026 | 0.285 | 0.72 | 0.38 | 0.65 | 0.54 | 0.69 | 0.46 |
| 2 | 234758500 | rs529963 | -0.208 | -0.972 | -0.246 | 0.642 | -0.002 | -0.001 | -0.001 | 0 | 0.27 | 0.28 | 0.14 | 0.14 | 0.17 | 0.12 |
| 2 | 234759800 | rs13384669 | -0.191 | -0.955 | -0.239 | 0.649 | -0.003 | -0.002 | -0.002 | 0 | 0.27 | 0.28 | 0.14 | 0.15 | 0.18 | 0.12 |
| 2 | 234763500 | rs506632 | 0.172 | -0.918 | -0.407 | 0.579 | 0.05 | 0.094 | 0.094 | 0.159 | 0.25 | 0.07 | 0.12 | 0.06 | 0.16 | 0.11 |
| 2 | 234767759 | rs658607 | 0.173 | -0.911 | -0.406 | 0.58 | 0.05 | 0.094 | 0.094 | 0.161 | 0.25 | 0.07 | 0.12 | 0.06 | 0.16 | 0.11 |
| 2 | 234768067 | rs579327 | 0.173 | -0.911 | -0.406 | 0.58 | 0.05 | 0.094 | 0.094 | 0.161 | 0.25 | 0.07 | 0.12 | 0.06 | 0.16 | 0.11 |
| 2 | 234768730 | rs2173746 | 0.173 | -0.92 | -0.172 | 0.575 | 0.05 | 0.094 | 0.094 | 0.161 | 0.25 | 0.07 | 0.12 | 0.06 | 0.16 | 0.11 |
| 2 | 234771057 | rs596509 | 0.173 | -0.913 | -0.179 | 0.591 | 0.05 | 0.094 | 0.094 | 0.161 | 0.25 | 0.07 | 0.12 | 0.06 | 0.16 | 0.11 |
| 2 | 234773700 | rs6431643 | 0.341 | -0.84 | -0.196 | 0.645 | 0 | 0 | 0 | 0 | 0.61 | 0.42 | 0.61 | 0.73 | 0.6 | 0.46 |
| 2 | 234775675 | rs213544 | 1.278 | -0.832 | -0.347 | 0.608 | 0.004 | 0.04 | 0.04 | 0.021 | 0.12 | 0.02 | 0.04 | 0.05 | 0.08 | 0.06 |
| 2 | 234776751 | rs968402 | 1.274 | -0.845 | -0.363 | 0.705 | -0.038 | 0.004 | 0.004 | 0.006 | 0.45 | 0.11 | 0.51 | 0.37 | 0.52 | 0.38 |
| 2 | 234790111 | rs28900731 | 1.079 | -0.926 | 0.133 | 0.668 | -0.06 | -0.097 | -0.097 | 1.91 | 0.14 | 0.3 | 0.08 | 0 | 0.07 | 0.03 |
| 2 | 234791098 | rs2361515 | -0.727 | -0.909 | -0.448 | 0.706 | -0.047 | -0.028 | -0.028 | 1.287 | 0.77 | 0.88 | 0.71 | 0.73 | 0.69 | 0.5 |
| 2 | 234796385 | rs11563065 | 1.021 | -0.705 | -0.581 | 0.844 | 0.006 | -0.004 | -0.004 | 0 | 0.13 | 0.09 | 0.05 | 0.17 | 0.08 | 0.07 |
| 2 | 234799176 | rs719154 | 0.457 | -0.683 | -0.507 | 0.935 | 0.023 | -0.026 | -0.026 | 0 | 0.57 | 0.19 | 0.56 | 0.54 | 0.6 | 0.44 |
| 2 | 234803432 | rs4663979 | -0.505 | -0.679 | -0.476 | 1.111 | 0.024 | 0.012 | 0.012 | 0 | 0.34 | 0.8 | 0.38 | 0.23 | 0.31 | 0.3 |
| 2 | 234804250 | rs6745386 | -1.195 | -0.742 | -0.572 | 1.085 | 0.066 | 0.018 | 0.018 | 0.006 | 0.47 | 0.9 | 0.43 | 0.63 | 0.37 | 0.54 |
| 2 | 234811911 | rs10929317 | -0.054 | -0.195 | 0.254 | 1.807 | 0.003 | -0.036 | -0.036 | 0 | 0.14 | 0.06 | 0.06 | 0.28 | 0.09 | 0.24 |
| 2 | 234819199 | rs10519356 | -0.983 | -0.209 | 0.398 | 1.745 | 0.017 | 0.094 | 0.094 | 2.965 | 0.08 | 0.55 | 0.17 | 0.1 | 0.07 | 0.22 |
| 2 | 234829120 | rs17862921 | 0.399 | -0.163 | -0.329 | 1.597 | -0.011 | 0.045 | 0.045 | 0 | 0.12 | 0.03 | 0.06 | 0.05 | 0.09 | 0.05 |
| 2 | 234837266 | rs735552 | -0.816 | -0.078 | -0.045 | 1.824 | 0.043 | 0.074 | 0.074 | 2.325 | 0.08 | 0.5 | 0.19 | 0.09 | 0.09 | 0.22 |
| 2 | 234837737 | rs758277 | -1.125 | 0.108 | -0.671 | 1.535 | -0.041 | 0.072 | 0.072 | 2.841 | 0.57 | 0.94 | 0.67 | 0.75 | 0.57 | 0.68 |
| 2 | 234843494 | rs10490012 | -1.258 | 0.126 | -0.408 | 1.471 | -0.013 | 0.048 | 0.048 | 1.61 | 0.23 | 0.62 | 0.35 | 0.41 | 0.18 | 0.31 |
| 2 | 234844945 | rs12473889 | 0.365 | 0.232 | -0.737 | 1.46 | -0.003 | 0.027 | 0.027 | 0 | 0.12 | 0.03 | 0.12 | 0.07 | 0.08 | 0.06 |
| 2 | 234848903 | rs7577157 | -1.018 | 0.505 | -0.06 | 1.875 | 0.038 | 0.372 | 0.372 | 1.316 | 0.08 | 0.34 | 0.22 | 0.34 | 0.09 | 0.26 |
| 2 | 234857018 | rs12466401 | -0.491 | 0.512 | -0.041 | 1.802 | 0.059 | 0.227 | 0.227 | 0.838 | 0.07 | 0.3 | 0.18 | 0.21 | 0.09 | 0.23 |
| 2 | 234868385 | rs10929320 | 0.069 | 1.002 | -2.481 | 1.2 | -0.144 | 0.106 | 0.106 | 0 | 0.37 | 0.03 | 0.34 | 0.18 | 0.41 | 0.19 |
| 2 | 234875433 | rs12185625 | -0.397 | 1.573 | -1.171 | 1.398 | -0.015 | 0.007 | 0.007 | 0 | 0.11 | 0.07 | 0.11 | 0.09 | 0.14 | 0.08 |
| 2 | 234876037 | rs13401339 | 0.241 | 0.998 | -1.218 | 1.649 | -0.016 | 0.016 | 0.016 | 0.005 | 0.57 | 0.35 | 0.47 | 0.59 | 0.56 | 0.53 |
| 2 | 234883944 | rs13414162 | 0.322 | 0.211 | -0.884 | 1.875 | -0.01 | 0.007 | 0.007 | 0.558 | 0.7 | 0.59 | 0.74 | 0.41 | 0.8 | 0.62 |

| | | | | | | | | | | | | | | | | |
|---|---|---|---|---|---|---|---|---|---|---|---|---|---|---|---|---|
| 2 | 234885030 | rs11685673 | 0.301 | 1.152 | -1.366 | 1.058 | -0.042 | 0.043 | 0.043 | 0 | 0.15 | 0.04 | 0.19 | 0.04 | 0.24 | 0.12 |
| 2 | 234885853 | rs10803667 | -0.324 | 0.629 | -0.848 | 1.378 | -0.007 | 0.015 | 0.015 | 0.055 | 0.84 | 0.75 | 0.86 | 0.78 | 0.88 | 0.79 |
| 2 | 234888161 | rs10207672 | 0.443 | 0.511 | -0.812 | 0.589 | -0.001 | -0.074 | -0.074 | 2.185 | 0.38 | 0.37 | 0.21 | 0.11 | 0.38 | 0.16 |
| 2 | 234889177 | rs4663992 | 0.465 | 0.398 | -0.851 | 0.49 | -0.012 | 0.047 | 0.047 | 0 | 0.28 | 0.2 | 0.33 | 0.07 | 0.3 | 0.17 |
| 2 | 234890391 | rs6708995 | 0.205 | 0.215 | -0.813 | 0.987 | 0 | -0.001 | -0.001 | 0 | 0.54 | 0.48 | 0.37 | 0.51 | 0.53 | 0.31 |
| 2 | 234892127 | rs4663995 | -0.64 | 0.076 | -0.943 | 1.238 | -0.003 | 0.032 | 0.032 | 0.676 | 0.83 | 0.74 | 0.7 | 0.59 | 0.84 | 0.49 |
| 2 | 234894893 | rs1016062 | 0.405 | 0.16 | -1.059 | 1.208 | -0.002 | -0.015 | -0.015 | 0 | 0.17 | 0.13 | 0.16 | 0.4 | 0.15 | 0.18 |
| 2 | 234897344 | rs12692252 | -0.923 | 0.131 | -1.282 | 1.136 | 0 | -0.018 | -0.018 | 0 | 0.84 | 0.85 | 0.73 | 0.76 | 0.82 | 0.69 |
| 2 | 234897718 | rs17864768 | 0.99 | 0.019 | -1.422 | 1.341 | -0.033 | -0.06 | -0.06 | 4.328 | 0.1 | 0.01 | 0.07 | 0.22 | 0.1 | 0.1 |
| 2 | 234899273 | rs11695247 | 1.756 | 0.014 | -1.103 | 0.441 | -0.061 | -0.004 | -0.004 | 0 | 0.16 | 0.01 | 0.13 | 0.12 | 0.21 | 0.1 |
| 2 | 234900513 | rs2362294 | -0.038 | -0.016 | -0.839 | 0.61 | -0.003 | 0.01 | 0.01 | 0 | 0.42 | 0.43 | 0.46 | 0.5 | 0.41 | 0.5 |
| 2 | 234901170 | rs11563056 | 0.91 | -0.03 | -1.208 | 0.256 | -0.011 | 0.032 | 0.032 | 0.466 | 0.34 | 0.26 | 0.2 | 0.12 | 0.37 | 0.15 |
| 2 | 234901424 | rs28948673 | 0.725 | 0.113 | -1.362 | 0.548 | -0.002 | 0.001 | 0.001 | 0.438 | 0.09 | 0.12 | 0.08 | 0.14 | 0.05 | 0.05 |
| 2 | 234905078 | rs11563208 | 0.125 | 0.654 | -1.697 | 0.315 | -0.008 | 0.056 | 0.056 | 0.049 | 0.24 | 0.08 | 0.32 | 0.08 | 0.25 | 0.13 |
| 2 | 234906526 | rs2362295 | -0.01 | 0.653 | -1.33 | 0.699 | -0.001 | -0.004 | -0.004 | 0 | 0.77 | 0.86 | 0.7 | 0.78 | 0.77 | 0.68 |
| 2 | 234908496 | rs6723922 | -0.206 | 0.498 | -1.381 | 0.674 | 0.008 | -0.01 | -0.01 | 0 | 0.4 | 0.73 | 0.52 | 0.57 | 0.41 | 0.49 |
| 2 | 234909799 | rs6746331 | -0.052 | 0.165 | -1.561 | 0.029 | 0.001 | -0.004 | -0.004 | 1 | 0.26 | 0.23 | 0.26 | 0.08 | 0.21 | 0.13 |
| 2 | 234910928 | rs11682848 | 1.196 | 0.03 | -2.002 | 0.662 | -0.022 | -0.088 | -0.088 | 0 | 0.14 | 0.02 | 0.15 | 0.36 | 0.19 | 0.2 |
| 2 | 234912215 | rs7560562 | -1.408 | -0.284 | -2.064 | 0.328 | -0.053 | 0.042 | 0.042 | 0 | 0.7 | 0.82 | 0.78 | 0.79 | 0.64 | 0.62 |
| 2 | 234912661 | rs6721761 | -1.391 | -0.283 | -1.075 | 0.502 | 0.02 | -0.012 | -0.012 | 0 | 0.7 | 0.65 | 0.7 | 0.73 | 0.65 | 0.62 |
| 2 | 234913033 | rs6712962 | -0.762 | -0.429 | -1.106 | 0.492 | 0.002 | -0.004 | -0.004 | 0.007 | 0.58 | 0.57 | 0.56 | 0.63 | 0.57 | 0.53 |
| 2 | 234917377 | rs11563204 | 0.707 | -0.451 | -1.114 | 0.476 | 0.006 | 0.01 | 0.01 | 0.045 | 0.26 | 0.19 | 0.26 | 0.08 | 0.21 | 0.14 |
| 2 | 234917732 | rs10490018 | 0.701 | -0.449 | -1.098 | 0.523 | 0.006 | -0.101 | -0.101 | 0 | 0.28 | 0.11 | 0.23 | 0.49 | 0.33 | 0.31 |
| 2 | 234919118 | rs17864777 | 1.892 | -0.631 | -1.094 | 0.431 | 0.014 | -0.005 | -0.005 | 0 | 0.1 | 0.04 | 0.12 | 0.11 | 0.06 | 0.08 |
| 2 | 234919314 | rs17865678 | -0.341 | -0.582 | -0.809 | 0.325 | -0.004 | -0.009 | -0.009 | 0 | 0.39 | 0.4 | 0.4 | 0.19 | 0.28 | 0.26 |
| 2 | 234919588 | rs17865679 | 1.585 | -0.353 | -0.601 | 0.157 | 0.006 | -0.001 | -0.001 | 0 | 0.11 | 0.08 | 0.12 | 0.11 | 0.06 | 0.08 |
| 2 | 234923124 | rs3732214 | 1.086 | -0.332 | -0.525 | 0.169 | 0.005 | 0.061 | 0.061 | 0.379 | 0.13 | 0.32 | 0.22 | 0.18 | 0.09 | 0.09 |
| 2 | 234926750 | rs17865681 | 0.945 | -0.147 | -0.676 | 0.063 | -0.011 | 0.019 | 0.019 | 0 | 0.05 | 0.01 | 0.02 | 0 | 0.04 | 0.02 |
| 2 | 234928612 | rs1118259 | -1.399 | -0.126 | -1.058 | 0.242 | -0.044 | 0.04 | 0.04 | 0 | 0.73 | 0.87 | 0.8 | 0.81 | 0.68 | 0.65 |
| 2 | 234929686 | rs17866592 | 1.045 | -0.288 | -1.776 | 0.131 | 0 | 0.002 | 0.002 | 3.624 | 0.14 | 0.11 | 0.22 | 0.05 | 0.16 | 0.09 |
| 2 | 234930311 | rs17866593 | -0.58 | -0.292 | -1.803 | -0.089 | 0.022 | 0.01 | 0.01 | 0.032 | 0.33 | 0.76 | 0.52 | 0.26 | 0.28 | 0.26 |
| 2 | 234931434 | rs599294 | 0.062 | -0.057 | -1.849 | 0.015 | 0.017 | -0.078 | -0.078 | 0 | 0.4 | 0.1 | 0.26 | 0.55 | 0.37 | 0.4 |
| 2 | 234932257 | rs585300 | -0.646 | 0.236 | -1.467 | 0.381 | -0.006 | 0.006 | 0.006 | 0.383 | 0.6 | 0.68 | 0.59 | 0.63 | 0.6 | 0.56 |
| 2 | 234933559 | rs676959 | -0.385 | -0.092 | -2.705 | 0.285 | -0.011 | 0.002 | 0.002 | 0 | 0.54 | 0.22 | 0.5 | 0.57 | 0.53 | 0.48 |
| 2 | 234936429 | rs11682121 | 0.373 | 0.013 | -2.451 | 0.285 | 0.017 | 0.002 | 0.002 | 0 | 0.14 | 0.08 | 0.08 | 0.12 | 0.12 | 0.12 |
| 2 | 234938264 | rs680841 | 0.294 | 0.427 | -2.969 | -0.15 | -0.14 | 0.08 | 0.08 | 2.063 | 0.26 | 0.03 | 0.41 | 0.09 | 0.36 | 0.34 |
| 2 | 234938888 | rs1395835 | 0.501 | -0.121 | -3.305 | 0.08 | -0.044 | -0.043 | -0.043 | 0 | 0.51 | 0.06 | 0.3 | 0.58 | 0.45 | 0.52 |
| 2 | 234939481 | rs11680332 | -0.948 | -0.054 | -2.816 | -0.148 | 0.014 | -0.017 | -0.017 | 0 | 0.76 | 0.74 | 0.47 | 0.9 | 0.64 | 0.67 |
| 2 | 234940575 | rs17866606 | 0.557 | 0.037 | -3.225 | -0.081 | 0.008 | 0.026 | 0.026 | 0 | 0.15 | 0.11 | 0.07 | 0 | 0.14 | 0.03 |
| 2 | 234941290 | rs4616477 | 0.147 | 0.212 | -3.127 | 0.537 | -0.005 | -0.129 | -0.129 | 0.026 | 0.32 | 0.02 | 0.23 | 0.5 | 0.3 | 0.39 |
| 2 | 234942536 | rs11684379 | -0.555 | 0.703 | -2.167 | 0.335 | -0.03 | 0.037 | 0.037 | 0.356 | 0.17 | 0.03 | 0.07 | 0.04 | 0.17 | 0.12 |
| 2 | 234943592 | rs10190663 | -0.518 | 0.526 | -1.771 | 0.498 | 0.017 | 0.021 | 0.021 | 0 | 0.54 | 0.7 | 0.51 | 0.66 | 0.41 | 0.64 |
| 2 | 234944028 | rs624460 | 0.56 | 0.373 | -1.169 | 0.383 | -0.004 | -0.015 | -0.015 | 0 | 0.55 | 0.52 | 0.53 | 0.29 | 0.63 | 0.61 |
| 2 | 234944043 | rs17864794 | -0.426 | 0.254 | -0.849 | 0.133 | -0.001 | -0.076 | -0.076 | 0.711 | 0.12 | 0.31 | 0.06 | 0 | 0.06 | 0.03 |
| 2 | 234945105 | rs4664004 | -0.341 | 0.025 | -1.275 | 0.718 | 0.05 | -0.012 | -0.012 | 0.551 | 0.33 | 0.11 | 0.33 | 0.3 | 0.24 | 0.51 |
| 2 | 234949004 | rs620654 | 0.628 | 0.113 | -1.137 | 1.031 | 0.006 | -0.002 | -0.002 | 0 | 0.57 | 0.67 | 0.59 | 0.55 | 0.65 | 0.47 |
| 2 | 234954002 | rs7585488 | -1.472 | -0.056 | -0.51 | 0.865 | 0.042 | 0.033 | 0.033 | 0 | 0.48 | 0.62 | 0.4 | 0.29 | 0.28 | 0.52 |
| 2 | 234954619 | rs634613 | 0.117 | -0.065 | -0.548 | 0.869 | 0.028 | 0.006 | 0.006 | 0.829 | 0.64 | 0.89 | 0.65 | 0.71 | 0.7 | 0.48 |
| 2 | 234957128 | rs2215375 | 1.027 | 0.334 | -0.215 | 1.009 | 0.01 | 0.029 | 0.029 | 0 | 0.05 | 0.09 | 0.15 | 0.12 | 0.09 | 0.05 |
| 2 | 234960021 | rs11563197 | -1.129 | 0.352 | -0.006 | 0.96 | -0.013 | -0.001 | -0.001 | 2.198 | 0.15 | 0.58 | 0.1 | 0.15 | 0.08 | 0.04 |
| 2 | 234961435 | rs1354895 | -0.841 | 0.203 | 0.017 | 1.087 | 0.015 | -0.003 | -0.003 | 0 | 0.41 | 0.43 | 0.39 | 0.45 | 0.28 | 0.51 |
| 2 | 234962085 | rs10171587 | -0.841 | 0.203 | 0.017 | 1.087 | 0.015 | -0.003 | -0.003 | 2.442 | 0.41 | 0.43 | 0.39 | 0.45 | 0.28 | 0.51 |
| 2 | 234962989 | rs617970 | -0.316 | 0.265 | -0.082 | 1.124 | -0.042 | -0.054 | -0.054 | 0 | 0.9 | 0.98 | 0.89 | 0.66 | 0.87 | 0.92 |
| 2 | 234963083 | rs633847 | 2.274 | 0.281 | 0.039 | 0.944 | -0.032 | 0.088 | 0.088 | 0 | 0.36 | 0.18 | 0.31 | 0.1 | 0.47 | 0.35 |
| 2 | 234965870 | rs13395783 | -0.041 | 0.092 | -0.037 | 1.321 | 0.068 | -0.07 | -0.07 | 0.111 | 0.32 | 0.07 | 0.32 | 0.38 | 0.21 | 0.47 |
| 2 | 234966238 | rs13396164 | -0.761 | 0.106 | -0.061 | 1.314 | 0.008 | -0.006 | -0.006 | 0 | 0.39 | 0.42 | 0.37 | 0.53 | 0.25 | 0.49 |
| 2 | 234968051 | rs2239536 | -0.738 | 0.11 | -0.059 | 1.29 | -0.027 | 0 | 0 | 0 | 0.39 | 0.5 | 0.37 | 0.37 | 0.25 | 0.48 |
| 2 | 234969639 | rs10210816 | -0.379 | 0.148 | 0.012 | 1.302 | 0.018 | -0.007 | -0.007 | 0.026 | 0.35 | 0.29 | 0.35 | 0.37 | 0.25 | 0.5 |
| 2 | 234970505 | rs9636246 | -1.318 | 0.14 | 0.012 | 1.24 | 0.036 | -0.004 | -0.004 | 0.148 | 0.47 | 0.66 | 0.43 | 0.53 | 0.31 | 0.52 |
| 2 | 234973949 | rs12692253 | -0.663 | 0.835 | -0.069 | 1.19 | -0.011 | 0.016 | 0.016 | 0.678 | 0.41 | 0.26 | 0.57 | 0.64 | 0.42 | 0.56 |
| 2 | 234978363 | rs250977 | 2.202 | 0.797 | 0.293 | 0.862 | 0.002 | 0.01 | 0.01 | 0 | 0.36 | 0.26 | 0.27 | 0.1 | 0.41 | 0.34 |
| 2 | 234978658 | rs6716659 | 0.791 | 0.005 | 0.4 | 1.149 | 0.023 | -0.022 | -0.022 | 0 | 0.05 | 0.16 | 0.03 | 0 | 0.03 | 0.01 |
| 2 | 234979872 | rs7557035 | -0.145 | 0.012 | 0.391 | 1.196 | 0.004 | 0.014 | 0.014 | 0 | 0.31 | 0.21 | 0.33 | 0.21 | 0.28 | 0.44 |
| 2 | 234980170 | rs11563027 | 0.552 | -0.022 | 0.602 | 1.072 | 0.021 | -0.023 | -0.023 | 0 | 0.05 | 0.16 | 0.03 | 0 | 0.03 | 0.01 |
| 2 | 234984811 | rs6719734 | 0.321 | -0.029 | 0.615 | 1.068 | 0.04 | -0.012 | -0.012 | 1.427 | 0.06 | 0.24 | 0.03 | 0 | 0.03 | 0.01 |
| 2 | 234990709 | rs250963 | 1.887 | -0.18 | 0.219 | 0.953 | -0.013 | 0.123 | 0.123 | 0 | 0.44 | 0.18 | 0.43 | 0.22 | 0.51 | 0.4 |
| 2 | 234990787 | rs10803670 | -1.052 | -0.242 | 0.611 | 1.233 | 0.022 | 0.212 | 0.212 | 0.244 | 0.11 | 0.53 | 0.13 | 0.32 | 0.06 | 0.06 |
| 2 | 234990853 | rs10929333 | -1.169 | -0.289 | 0.658 | 1.225 | -0.018 | 0.011 | 0.011 | 1.019 | 0.13 | 0.58 | 0.07 | 0.16 | 0.06 | 0.02 |
| 2 | 234991336 | rs250962 | 0.78 | -0.27 | 0.771 | 1.092 | -0.001 | 0 | 0 | 0 | 0.59 | 0.57 | 0.56 | 0.38 | 0.59 | 0.47 |

| | | | | | | | | | | | | | | | | |
|---|---|---|---|---|---|---|---|---|---|---|---|---|---|---|---|---|
| 2 | 234991650 | rs250960 | 0.149 | -0.222 | 0.403 | 1.287 | -0.044 | -0.057 | -0.057 | 0 | 0.88 | 0.96 | 0.87 | 0.71 | 0.84 | 0.91 |
| 2 | 234992912 | rs250958 | 1.87 | -0.296 | 0.413 | 1.293 | -0.001 | 0.054 | 0.054 | 0 | 0.47 | 0.23 | 0.48 | 0.38 | 0.53 | 0.46 |
| 2 | 234993308 | rs7581053 | -0.045 | -0.295 | 0.133 | 1.297 | -0.021 | -0.044 | -0.044 | 0 | 0.27 | 0.05 | 0.29 | 0.32 | 0.25 | 0.42 |
| 2 | 234993563 | rs250956 | 1.188 | -0.224 | 0.152 | 1.212 | 0.014 | 0.027 | 0.027 | 0 | 0.77 | 0.71 | 0.81 | 0.71 | 0.78 | 0.89 |
| 2 | 234997218 | rs250944 | 2.033 | -0.289 | -0.291 | 1.079 | -0.115 | 0.103 | 0.103 | 0 | 0.41 | 0.03 | 0.41 | 0.22 | 0.51 | 0.4 |
| 2 | 235000529 | rs250939 | 1.571 | -0.055 | 0.299 | 1.152 | -0.027 | 0.053 | 0.053 | 0 | 0.53 | 0.17 | 0.45 | 0.37 | 0.57 | 0.47 |
| 2 | 235001537 | rs16850379 | -0.303 | 0.019 | 0.582 | 1.254 | -0.001 | -0.001 | -0.001 | 0 | 0.25 | 0.23 | 0.37 | 0.33 | 0.25 | 0.36 |
| 2 | 235005739 | rs250936 | 0.845 | -0.1 | 0.449 | 1.354 | 0.019 | 0.078 | 0.078 | 0 | 0.81 | 0.66 | 0.83 | 0.71 | 0.83 | 0.83 |
| 2 | 235006892 | rs2271153 | 0.215 | -0.527 | 0.085 | 1.331 | 0.022 | -0.027 | -0.027 | 0.69 | 0.19 | 0.06 | 0.3 | 0.27 | 0.15 | 0.18 |
| 2 | 235007491 | rs250933 | 0.487 | -0.31 | 0.053 | 1.461 | -0.001 | 0.013 | 0.013 | 0 | 0.88 | 0.89 | 0.87 | 0.71 | 0.8 | 0.75 |
| 2 | 235008881 | rs250928 | 1.603 | -0.439 | 0.113 | 1.038 | -0.041 | 0.02 | 0.02 | 0 | 0.44 | 0.13 | 0.41 | 0.37 | 0.54 | 0.39 |
| 2 | 235009790 | rs250927 | 0.812 | -0.286 | 0.044 | 1.145 | -0.005 | 0.006 | 0.006 | 0 | 0.57 | 0.62 | 0.41 | 0.39 | 0.58 | 0.41 |
| 2 | 235011203 | rs13384101 | -0.769 | -0.247 | 0.442 | 1.238 | -0.021 | 0.049 | 0.049 | 0 | 0.24 | 0.17 | 0.12 | 0.08 | 0.23 | 0.2 |
| 2 | 235016487 | rs669466 | -1.236 | -1.198 | 1.131 | 1.794 | 0.03 | -0.019 | -0.019 | 0 | 0.32 | 0.74 | 0.43 | 0.66 | 0.23 | 0.33 |
| 2 | 235017927 | rs250916 | -0.006 | -1.23 | 0.306 | 2.018 | 0.057 | -0.072 | -0.072 | 1.646 | 0.2 | 0.05 | 0.24 | 0.36 | 0.15 | 0.16 |
| 2 | 235020980 | rs6731887 | -0.991 | -1.185 | 0.423 | 1.971 | 0.004 | 0.039 | 0.039 | 0.233 | 0.08 | 0.15 | 0.17 | 0.27 | 0.07 | 0.18 |
| 2 | 235034610 | rs250900 | -0.032 | -1.249 | -0.008 | 1.998 | 0.053 | -0.057 | -0.057 | 0 | 0.2 | 0.07 | 0.24 | 0.35 | 0.15 | 0.16 |
| 2 | 235034766 | rs250899 | -1.213 | -1.113 | -0.082 | 1.554 | 0.054 | -0.032 | -0.032 | 0 | 0.35 | 0.76 | 0.45 | 0.63 | 0.24 | 0.34 |
| 2 | 235039555 | rs1860700 | -1.687 | -0.055 | 1.147 | 1.737 | 0.032 | 0.112 | 0.112 | 0 | 0.15 | 0.54 | 0.27 | 0.34 | 0.15 | 0.34 |
| 2 | 235043481 | rs12617590 | -1.285 | -0.092 | 1.271 | 1.635 | -0.02 | 0.019 | 0.019 | 0 | 0.25 | 0.45 | 0.25 | 0.32 | 0.16 | 0.17 |
| 2 | 235047989 | rs7586829 | -0.541 | -0.195 | 1.371 | 1.532 | -0.009 | -0.044 | -0.044 | 1.329 | 0.14 | 0.17 | 0.05 | 0 | 0.06 | 0.1 |
| 2 | 235051708 | rs12621510 | -1.578 | -0.147 | 1.107 | 1.727 | -0.022 | 0.047 | 0.047 | 0.018 | 0.25 | 0.53 | 0.2 | 0.34 | 0.17 | 0.17 |
| 2 | 235058350 | rs16850533 | -1.613 | -0.183 | 1.16 | 1.709 | -0.02 | 0.031 | 0.031 | 0.032 | 0.26 | 0.58 | 0.21 | 0.34 | 0.17 | 0.18 |
| 2 | 235071568 | rs6711646 | -1.672 | -0.198 | 0.912 | 1.525 | -0.022 | -0.014 | -0.014 | 0.658 | 0.27 | 0.56 | 0.22 | 0.23 | 0.19 | 0.18 |
| 2 | 235072203 | rs6431233 | -2.026 | -0.194 | 0.42 | 1.659 | 0.012 | -0.047 | -0.047 | 0 | 0.35 | 0.84 | 0.42 | 0.63 | 0.27 | 0.32 |
| 2 | 235073458 | rs10929012 | -0.677 | -0.336 | 0.488 | 1.09 | 0.02 | 0.087 | 0.087 | 0 | 0.07 | 0.17 | 0.12 | 0.2 | 0.08 | 0.06 |
| 2 | 235073684 | rs10929014 | -1.137 | -0.331 | 0.009 | 1.084 | -0.029 | -0.028 | -0.028 | 0 | 0.24 | 0.51 | 0.19 | 0.2 | 0.16 | 0.16 |
| 2 | 235078194 | rs11884879 | -1.057 | -0.688 | -1.329 | 1.182 | 0.054 | -0.115 | -0.115 | 2.648 | 0.29 | 0.93 | 0.34 | 0.47 | 0.18 | 0.33 |
| 2 | 235079680 | rs6431235 | -0.065 | -0.413 | -1.717 | -0.059 | 0.03 | 0.108 | 0.108 | 3.948 | 0.42 | 0.67 | 0.3 | 0.14 | 0.36 | 0.29 |
| 2 | 235079857 | rs6723008 | -0.304 | -0.038 | -1.486 | -0.199 | 0 | -0.004 | -0.004 | 7.405 | 0.52 | 0.5 | 0.39 | 0.74 | 0.43 | 0.51 |
| 2 | 235082853 | rs7584695 | -0.572 | 0.838 | -1.143 | -0.099 | -0.002 | 0.006 | 0.006 | 0 | 0.44 | 0.55 | 0.5 | 0.75 | 0.5 | 0.33 |
| 2 | 235099369 | rs973368 | -0.062 | 0.957 | -0.706 | 0.442 | 0.003 | -0.009 | -0.009 | 0 | 0.35 | 0.28 | 0.33 | 0.55 | 0.34 | 0.26 |
| 2 | 235103289 | rs6758970 | -0.378 | 1.279 | -0.546 | 0.407 | 0.077 | 0.012 | 0.012 | 0 | 0.08 | 0.18 | 0.16 | 0.11 | 0.16 | 0.06 |
| 2 | 235110179 | rs6708323 | 0.015 | 1.005 | -0.718 | 0.341 | -0.001 | 0.001 | 0.001 | 0 | 0.33 | 0.26 | 0.26 | 0.62 | 0.27 | 0.23 |
| 2 | 235111962 | rs7567269 | -0.082 | 1.206 | -1.177 | 0.155 | 0 | 0 | 0 | 0.091 | 0.06 | 0.05 | 0.09 | 0.05 | 0.06 | 0.15 |
| 2 | 235124067 | rs17585471 | -0.549 | 1.339 | -1.269 | 0.118 | -0.052 | -0.014 | -0.014 | 0.004 | 0.1 | 0.03 | 0.19 | 0.1 | 0.23 | 0.07 |
| 2 | 235127466 | rs1860699 | -1.095 | 0.763 | -0.862 | -0.044 | 0.034 | 0.157 | 0.157 | 0 | 0.53 | 0.73 | 0.56 | 0.86 | 0.57 | 0.45 |
| 2 | 235130693 | rs10929015 | 1.003 | 0.632 | -1.142 | -0.178 | -0.046 | 0.232 | 0.232 | 0 | 0.43 | 0.15 | 0.46 | 0.09 | 0.5 | 0.49 |
| 2 | 235131943 | rs1878973 | -0.633 | 0.123 | -0.482 | 0.346 | -0.042 | 0.116 | 0.116 | 0.138 | 0.19 | 0.41 | 0.16 | 0.64 | 0.11 | 0.21 |
| 2 | 235132912 | rs1356660 | -0.567 | 0.446 | -0.997 | 0.464 | -0.022 | 0.054 | 0.054 | 0.015 | 0.74 | 0.86 | 0.75 | 0.88 | 0.71 | 0.85 |
| 2 | 235144787 | rs4663116 | -0.354 | 0.546 | -1.155 | 0.506 | 0.004 | 0.031 | 0.031 | 0 | 0.41 | 0.25 | 0.38 | 0.67 | 0.38 | 0.35 |
| 2 | 235161448 | rs7571897 | -0.478 | 0.606 | -1.568 | 0.711 | -0.019 | 0 | 0 | 0 | 0.57 | 0.82 | 0.53 | 0.4 | 0.54 | 0.54 |
| 2 | 235166726 | rs1519186 | 0.088 | 0.385 | -0.604 | 0.964 | -0.001 | 0.152 | 0.152 | 0 | 0.16 | 0.25 | 0.12 | 0.36 | 0.15 | 0.17 |
| 2 | 235166761 | rs1878971 | 0.444 | 0.355 | -1.388 | 0.286 | -0.025 | -0.074 | -0.074 | 0 | 0.29 | 0.11 | 0.23 | 0.44 | 0.29 | 0.28 |
| 2 | 235170697 | rs1464268 | 0.361 | 0.031 | -1.283 | -0.367 | 0.012 | -0.015 | -0.015 | 0 | 0.22 | 0.25 | 0.33 | 0.14 | 0.31 | 0.21 |
| 2 | 235172026 | rs6431242 | -0.848 | -0.416 | -1.064 | -0.603 | 0.006 | 0.132 | 0.132 | 0 | 0.51 | 0.78 | 0.6 | 0.28 | 0.48 | 0.39 |
| 2 | 235172880 | rs4663367 | -0.408 | -0.643 | -0.711 | -0.795 | 0.019 | -0.068 | -0.068 | 0 | 0.34 | 0.47 | 0.44 | 0.18 | 0.4 | 0.24 |
| 2 | 235181704 | rs4663369 | -1.643 | -0.587 | -1.154 | -0.615 | -0.095 | 0.087 | 0.087 | 0 | 0.63 | 0.95 | 0.76 | 0.78 | 0.6 | 0.58 |
| 2 | 235188422 | rs1401506 | -1.65 | -0.611 | -1.128 | -0.628 | -0.08 | 0.092 | 0.092 | 0 | 0.63 | 0.93 | 0.77 | 0.78 | 0.6 | 0.6 |
| 2 | 235190483 | rs6431245 | -0.734 | -0.722 | -0.581 | -0.63 | 0 | -0.01 | -0.01 | 0 | 0.46 | 0.48 | 0.6 | 0.68 | 0.52 | 0.44 |
| 2 | 235192719 | rs2049026 | -0.457 | -0.788 | -0.486 | -0.81 | -0.023 | -0.027 | -0.027 | 0 | 0.34 | 0.42 | 0.36 | 0.73 | 0.29 | 0.36 |
| 2 | 235193276 | rs7587925 | -0.447 | -0.592 | -0.409 | -0.797 | 0.006 | -0.114 | -0.114 | 0.043 | 0.33 | 0.56 | 0.4 | 0.05 | 0.34 | 0.24 |
| 2 | 235193436 | rs11886348 | 0.212 | -0.885 | -0.39 | -0.615 | -0.048 | -0.067 | -0.067 | 0 | 0.14 | 0.37 | 0.12 | 0.04 | 0.05 | 0.07 |
| 2 | 235194246 | rs4258786 | 1.579 | -1.107 | -0.673 | -0.575 | 0.004 | -0.019 | -0.019 | 0 | 0.18 | 0.12 | 0.17 | 0.51 | 0.15 | 0.18 |
| 2 | 235195230 | rs6717293 | -1.226 | -1.137 | -0.395 | -0.639 | -0.055 | -0.02 | -0.02 | 0 | 0.33 | 0.57 | 0.31 | 0.27 | 0.19 | 0.25 |
| 2 | 235195326 | rs11894176 | 0.437 | -1.152 | -0.38 | -0.648 | -0.042 | -0.054 | -0.054 | 0.018 | 0.13 | 0.28 | 0.12 | 0.04 | 0.05 | 0.07 |
| 2 | 235195409 | rs6707182 | -0.573 | -0.72 | -0.703 | -0.389 | -0.016 | 0.007 | 0.007 | 0.224 | 0.19 | 0.28 | 0.19 | 0.23 | 0.14 | 0.18 |
| 2 | 235199587 | rs1464264 | -1.039 | -0.449 | -1.108 | -0.34 | -0.029 | 0.07 | 0.07 | 0 | 0.61 | 0.75 | 0.66 | 0.75 | 0.58 | 0.64 |
| 2 | 235201218 | rs1519187 | -0.699 | -0.647 | -1.112 | -0.51 | 0.052 | 0.031 | 0.031 | 0.042 | 0.46 | 0.75 | 0.47 | 0.64 | 0.34 | 0.37 |
| 2 | 235201649 | rs4663126 | -1.854 | -0.536 | -1.24 | -0.564 | -0.037 | 0.011 | 0.011 | 0 | 0.67 | 0.82 | 0.76 | 0.7 | 0.6 | 0.6 |
| 2 | 235204421 | rs12467783 | -0.125 | -0.684 | -0.93 | -0.519 | 0.015 | -0.037 | -0.037 | 0 | 0.38 | 0.26 | 0.41 | 0.5 | 0.32 | 0.34 |
| 2 | 235209416 | rs2176652 | 1.085 | -0.658 | -1.279 | -0.971 | -0.066 | 0.083 | 0.083 | 0 | 0.15 | 0.03 | 0.25 | 0.01 | 0.23 | 0.15 |
| 2 | 235209795 | rs4663379 | 0.91 | -0.598 | -1.346 | -0.996 | -0.03 | 0.016 | 0.016 | 0 | 0.1 | 0.02 | 0.13 | 0.07 | 0.11 | 0.06 |
| 2 | 235209980 | rs7584309 | -1.723 | -0.542 | -1.298 | -0.929 | -0.035 | 0.009 | 0.009 | 0 | 0.68 | 0.82 | 0.75 | 0.72 | 0.59 | 0.6 |
| 2 | 235210293 | rs7575512 | -0.257 | -0.853 | -0.728 | -0.867 | -0.061 | 0.011 | 0.011 | 0.18 | 0.13 | 0.36 | 0.06 | 0.14 | 0.05 | 0.11 |
| 2 | 235212056 | rs6733065 | -0.861 | -0.624 | -1.197 | -0.904 | 0.027 | 0.071 | 0.071 | 0 | 0.53 | 0.64 | 0.53 | 0.72 | 0.42 | 0.47 |
| 2 | 235212137 | rs6715295 | -0.191 | -0.531 | -1.233 | -0.85 | 0.005 | -0.001 | -0.001 | 0 | 0.4 | 0.34 | 0.44 | 0.58 | 0.31 | 0.34 |
| 2 | 235215133 | rs11695811 | -0.275 | -0.473 | -0.953 | -0.602 | -0.003 | 0.014 | 0.014 | 0.103 | 0.52 | 0.37 | 0.59 | 0.59 | 0.48 | 0.48 |
| 2 | 235216549 | rs4663382 | 0.01 | -0.397 | -0.941 | -0.8 | 0 | -0.034 | -0.034 | 0 | 0.35 | 0.34 | 0.42 | 0.1 | 0.35 | 0.29 |

| | | | | | | | | | | | | | | | | |
|---|---|---|---|---|---|---|---|---|---|---|---|---|---|---|---|---|
| 2 | 235216752 | rs4663384 | -0.968 | 0.068 | -1.464 | -0.442 | -0.037 | 0.008 | 0.008 | 0 | 0.68 | 0.82 | 0.75 | 0.73 | 0.59 | 0.6 |
| 2 | 235219536 | rs10209413 | 0.495 | 0.01 | -1.664 | -0.557 | -0.07 | 0.027 | 0.027 | 0 | 0.33 | 0.09 | 0.28 | 0.23 | 0.4 | 0.43 |
| 2 | 235229127 | rs2885958 | 0.604 | 0.211 | -0.875 | -0.735 | 0.004 | 0.044 | 0.044 | 0 | 0.52 | 0.35 | 0.55 | 0.73 | 0.56 | 0.61 |
| 2 | 235230310 | rs4663388 | 0.178 | 0.324 | -1.152 | -0.557 | -0.003 | 0.033 | 0.033 | 0.396 | 0.05 | 0.08 | 0.07 | 0.16 | 0.03 | 0.04 |
| 2 | 235235092 | rs1714950 | -0.343 | 0.339 | -1.496 | -0.912 | -0.024 | 0.065 | 0.065 | 0 | 0.35 | 0.25 | 0.32 | 0.12 | 0.38 | 0.3 |
| 2 | 235235475 | rs1703557 | -0.965 | 0.136 | -2.028 | -0.559 | -0.093 | 0.09 | 0.09 | 0.019 | 0.52 | 0.97 | 0.48 | 0.32 | 0.45 | 0.41 |
| 2 | 235241961 | rs772783 | -0.411 | 0.738 | -0.805 | -0.081 | 0.017 | -0.016 | -0.016 | 0.17 | 0.25 | 0.31 | 0.25 | 0.16 | 0.31 | 0.22 |
| 2 | 235242143 | rs772781 | -0.748 | 0.673 | -0.88 | 0.163 | 0.018 | 0.001 | 0.001 | 2.021 | 0.34 | 0.49 | 0.29 | 0.3 | 0.38 | 0.31 |
| 2 | 235247521 | rs772789 | -0.986 | 0.254 | -1.838 | -0.298 | -0.002 | 0.08 | 0.08 | 5.075 | 0.42 | 0.9 | 0.43 | 0.29 | 0.36 | 0.45 |
| 2 | 235247664 | rs13023712 | -0.762 | -0.17 | -0.706 | -0.435 | -0.033 | -0.016 | -0.016 | 3.794 | 0.14 | 0.4 | 0.19 | 0.13 | 0.1 | 0.17 |
| 2 | 235248562 | rs772788 | -0.576 | 0.028 | -1.936 | -0.947 | -0.087 | 0.116 | 0.116 | 0 | 0.67 | 0.94 | 0.67 | 0.81 | 0.59 | 0.54 |
| 2 | 235255122 | rs7594935 | -1.275 | -0.922 | -1.04 | -1.022 | 0.015 | -0.007 | -0.007 | 0 | 0.25 | 0.77 | 0.13 | 0.04 | 0.09 | 0.23 |
| 2 | 235258102 | rs10221751 | -1.192 | -0.869 | -0.7 | -0.981 | -0.054 | -0.086 | -0.086 | 0 | 0.23 | 0.6 | 0.11 | 0.04 | 0.09 | 0.23 |
| 2 | 235258599 | rs6722406 | -0.778 | -0.386 | -0.22 | -0.705 | -0.035 | -0.069 | -0.069 | 0 | 0.12 | 0.51 | 0.08 | 0 | 0.05 | 0.16 |
| 2 | 235271063 | rs1703568 | 0.463 | -0.024 | -0.585 | -1.104 | -0.001 | 0.014 | 0.014 | 0 | 0.58 | 0.4 | 0.65 | 0.83 | 0.55 | 0.42 |
| 2 | 235271926 | rs2885897 | 0.317 | -0.115 | -0.205 | -1.15 | 0.005 | -0.054 | -0.054 | 0.032 | 0.6 | 0.56 | 0.65 | 0.83 | 0.55 | 0.43 |
| 2 | 235272133 | rs4613271 | 0.283 | 0.055 | 0.29 | -1.353 | 0 | -0.004 | -0.004 | 0 | 0.68 | 0.73 | 0.74 | 0.84 | 0.64 | 0.55 |
| 2 | 235284258 | rs7589325 | -1.137 | -0.161 | 0.454 | -1.455 | -0.019 | -0.081 | -0.081 | 0 | 0.21 | 0.39 | 0.15 | 0.04 | 0.15 | 0.31 |
| 2 | 235284355 | rs7572928 | -0.171 | -0.191 | 0.385 | -1.45 | -0.022 | -0.046 | -0.046 | 0 | 0.09 | 0.22 | 0.05 | 0.01 | 0.04 | 0.13 |
| 2 | 235284391 | rs7572941 | 0.857 | -0.195 | 0.541 | -1.45 | -0.007 | -0.021 | -0.021 | 0 | 0.05 | 0.07 | 0.02 | 0 | 0.02 | 0.05 |
| 2 | 235303181 | rs10754938 | 0.711 | -0.118 | 0.391 | -1.662 | 0.002 | -0.025 | -0.025 | 0 | 0.7 | 0.72 | 0.76 | 0.84 | 0.66 | 0.61 |
| 2 | 235304659 | rs4663398 | 2.005 | -0.202 | 0.281 | -2.103 | 0.001 | 0.057 | 0.057 | 0.486 | 0.52 | 0.4 | 0.57 | 0.8 | 0.48 | 0.44 |
| 2 | 235305076 | rs9784089 | 0.058 | -0.187 | 0.298 | -2.082 | 0.003 | -0.047 | -0.047 | 0 | 0.11 | 0.29 | 0.1 | 0.02 | 0.09 | 0.05 |
| 2 | 235306754 | rs13007476 | 1.474 | -0.288 | 0.106 | -2.108 | 0 | 0.057 | 0.057 | 0.075 | 0.57 | 0.34 | 0.66 | 0.83 | 0.56 | 0.56 |
| 2 | 235308118 | rs11692463 | 0.493 | -0.226 | 0.096 | -2.075 | -0.027 | 0.008 | 0.008 | 0 | 0.1 | 0.03 | 0.1 | 0.03 | 0.12 | 0.16 |
| 2 | 235308752 | rs2136261 | 2.309 | -0.771 | -0.4 | -1.304 | -0.005 | -0.015 | -0.015 | 0 | 0.37 | 0.29 | 0.52 | 0.43 | 0.41 | 0.29 |
| 2 | 235312149 | rs4663404 | 0.774 | -0.567 | -0.638 | -1.274 | 0 | -0.006 | -0.006 | 0.217 | 0.6 | 0.7 | 0.7 | 0.46 | 0.61 | 0.52 |
| 2 | 235313619 | rs6721410 | 2.397 | -0.68 | -0.414 | -1.33 | -0.023 | -0.047 | -0.047 | 3.154 | 0.34 | 0.19 | 0.51 | 0.42 | 0.4 | 0.27 |
| 2 | 235321635 | rs10165420 | 0.92 | -0.497 | -0.395 | -1.193 | 0.012 | -0.058 | -0.058 | 0.177 | 0.69 | 0.53 | 0.77 | 0.82 | 0.64 | 0.68 |
| 2 | 235326552 | rs6713036 | -0.712 | 0.091 | -0.884 | -0.811 | -0.093 | 0.003 | 0.003 | 0 | 0.84 | 0.98 | 0.82 | 0.84 | 0.71 | 0.8 |
| 2 | 235332731 | rs1472931 | 0.228 | 0.117 | -0.804 | -0.683 | -0.01 | -0.068 | -0.068 | 0.45 | 0.78 | 0.87 | 0.84 | 0.58 | 0.78 | 0.79 |
| 2 | 235332854 | rs3115355 | -0.29 | 0.226 | -0.747 | -0.78 | -0.015 | 0.018 | 0.018 | 0.696 | 0.85 | 0.87 | 0.85 | 0.96 | 0.77 | 0.89 |
| 2 | 235334750 | rs4074423 | 1.875 | 0.169 | -0.725 | -0.365 | -0.001 | -0.013 | -0.013 | 0 | 0.07 | 0 | 0.08 | 0.36 | 0.03 | 0.14 |
| 2 | 235335455 | rs3115356 | -0.33 | 0.418 | -0.774 | -1.471 | -0.096 | 0.013 | 0.013 | 0.073 | 0.83 | 0.98 | 0.79 | 0.86 | 0.67 | 0.86 |
| 2 | 235335472 | rs11691744 | 0.35 | 0.245 | -0.251 | -1.682 | -0.009 | -0.034 | -0.034 | 0 | 0.2 | 0.23 | 0.15 | 0.06 | 0.18 | 0.3 |
| 2 | 235336166 | rs1509084 | 0.437 | 0.364 | 0.033 | -1.767 | -0.005 | 0.001 | 0.001 | 0 | 0.74 | 0.68 | 0.73 | 0.85 | 0.61 | 0.79 |
| 2 | 235338531 | rs11686781 | -0.024 | -0.618 | 0.348 | -1.505 | -0.01 | -0.038 | -0.038 | 1.633 | 0.21 | 0.24 | 0.15 | 0.06 | 0.17 | 0.3 |
| 2 | 235339940 | rs9646711 | -0.913 | -0.511 | 0.495 | -1.511 | -0.004 | -0.049 | -0.049 | 2.161 | 0.33 | 0.34 | 0.21 | 0.04 | 0.27 | 0.38 |
| 2 | 235350736 | rs6743014 | 0.517 | -0.835 | 0.503 | -1.189 | -0.077 | -0.133 | -0.133 | 0 | 0.22 | 0.52 | 0.09 | 0.01 | 0.09 | 0.2 |
| 2 | 235352188 | rs4663414 | 0.127 | -0.434 | -0.096 | -1.312 | -0.034 | 0.035 | 0.035 | 0 | 0.77 | 0.85 | 0.78 | 0.91 | 0.64 | 0.8 |
| 2 | 235357879 | rs4663138 | -0.174 | -0.282 | -0.352 | -1.237 | -0.024 | 0.025 | 0.025 | 0 | 0.79 | 0.78 | 0.82 | 0.92 | 0.68 | 0.83 |
| 2 | 235363663 | rs4663416 | 0.398 | -0.074 | -0.388 | -1.102 | 0.014 | 0.108 | 0.108 | 0 | 0.51 | 0.26 | 0.68 | 0.92 | 0.55 | 0.58 |
| 2 | 235369660 | rs6742249 | -0.241 | -0.099 | -0.095 | 0.029 | 0.036 | 0.027 | 0.027 | 0 | 0.47 | 0.35 | 0.35 | 0.25 | 0.27 | 0.52 |
| 2 | 235370826 | rs11883650 | -0.451 | -0.121 | -0.002 | 0.018 | 0.01 | 0.009 | 0.009 | 0 | 0.49 | 0.49 | 0.37 | 0.25 | 0.27 | 0.54 |
| 2 | 235375390 | rs11687376 | -0.994 | 0.738 | 0.731 | 0.241 | -0.003 | -0.086 | -0.086 | 0.006 | 0.21 | 0.47 | 0.22 | 0.03 | 0.17 | 0.35 |
| 2 | 235375794 | rs6737220 | 0.127 | 0.738 | 0.883 | 0.227 | 0.008 | 0.004 | 0.004 | 0.053 | 0.07 | 0.17 | 0.11 | 0.03 | 0.07 | 0.17 |
| 2 | 235378731 | rs6736336 | 0.073 | 1.348 | 0.376 | 0.69 | -0.047 | -0.04 | -0.04 | 0.02 | 0.78 | 0.9 | 0.66 | 0.52 | 0.58 | 0.67 |
| 2 | 235383488 | rs6735084 | -0.272 | 1.361 | 0.351 | 0.702 | 0.002 | -0.055 | -0.055 | 1.251 | 0.19 | 0.33 | 0.22 | 0.03 | 0.17 | 0.33 |
| 2 | 235386650 | rs11679226 | 1.638 | 1.271 | 0.383 | 0.746 | 0.019 | -0.008 | -0.008 | 0 | 0.53 | 0.33 | 0.43 | 0.43 | 0.41 | 0.3 |
| 2 | 235390014 | rs6723554 | -0.328 | 1.684 | -0.029 | 0.809 | -0.052 | 0.009 | 0.009 | 0 | 0.83 | 0.95 | 0.83 | 0.88 | 0.68 | 0.72 |
| 2 | 235392893 | rs11900995 | -0.721 | 0.919 | 0.448 | 0.986 | 0.006 | 0.082 | 0.082 | 0 | 0.09 | 0.2 | 0.18 | 0.37 | 0.1 | 0.15 |
| 2 | 235395795 | rs1546456 | -0.993 | 0.89 | -0.019 | 0.378 | -0.002 | -0.048 | -0.048 | 0 | 0.19 | 0.32 | 0.21 | 0.04 | 0.16 | 0.27 |
| 2 | 235401020 | rs3115360 | -0.889 | 1.325 | -0.31 | 0.865 | -0.009 | -0.006 | -0.006 | 0 | 0.39 | 0.62 | 0.45 | 0.56 | 0.46 | 0.42 |
| 2 | 235402656 | rs10194289 | -0.232 | 1.719 | 0.865 | 1.426 | 0.077 | 0.114 | 0.114 | 0.005 | 0.18 | 0.4 | 0.21 | 0.38 | 0.24 | 0.22 |
| 2 | 235408036 | rs4663142 | 0.957 | 1.743 | 0.366 | 1.414 | -0.012 | -0.019 | -0.019 | 0 | 0.69 | 0.35 | 0.64 | 0.37 | 0.63 | 0.65 |
| 2 | 235411142 | rs4663146 | 0.201 | 1.634 | 0.147 | 1.507 | 0.015 | 0.123 | 0.123 | 0 | 0.84 | 0.48 | 0.82 | 0.76 | 0.86 | 0.86 |
| 2 | 235412208 | rs3935670 | 1.098 | 1.447 | -1.698 | 0.972 | 0.074 | 0.021 | 0.021 | 3.477 | 0.43 | 0.09 | 0.4 | 0.33 | 0.29 | 0.5 |
| 2 | 235419415 | rs6431266 | -0.22 | 2.036 | -1.326 | 1.203 | -0.058 | -0.055 | -0.055 | 0.561 | 0.17 | 0.05 | 0.21 | 0.3 | 0.26 | 0.25 |
| 2 | 235421603 | rs6745638 | -0.716 | 1.63 | -1.138 | 1.283 | -0.018 | -0.006 | -0.006 | 2.752 | 0.25 | 0.22 | 0.26 | 0.39 | 0.33 | 0.19 |
| 2 | 235425433 | rs10177805 | 0.467 | 0.785 | -1.078 | 1.843 | 0.003 | 0.04 | 0.04 | 0 | 0.9 | 0.91 | 0.96 | 0.56 | 0.93 | 0.89 |
| 2 | 235426711 | rs6431267 | 0.839 | 0.589 | -1.092 | 1.721 | 0.008 | 0.003 | 0.003 | 0 | 0.87 | 0.9 | 0.94 | 0.56 | 0.93 | 0.88 |
| 2 | 235430724 | rs10803620 | 1.253 | 0.593 | -1.237 | 1.648 | 0.013 | -0.037 | -0.037 | 0.539 | 0.83 | 0.88 | 0.9 | 0.59 | 0.88 | 0.89 |
| 2 | 235435440 | rs12472396 | 0.005 | 0.931 | -1.809 | 0.983 | -0.037 | 0.036 | 0.036 | 0 | 0.19 | 0.05 | 0.34 | 0.11 | 0.21 | 0.39 |
| 2 | 235442524 | rs940950 | 0.609 | 0.134 | -1.547 | 0.913 | 0.004 | -0.01 | -0.01 | 0 | 0.63 | 0.73 | 0.58 | 0.47 | 0.71 | 0.48 |
| 2 | 235442600 | rs940951 | 1.374 | 0.129 | -1.876 | 0.271 | -0.006 | 0.014 | 0.014 | 0 | 0.12 | 0.07 | 0.12 | 0 | 0.15 | 0.1 |
| 2 | 235444616 | rs10193604 | 0.86 | -0.026 | -1.443 | 0.369 | 0.003 | -0.006 | -0.006 | 0.02 | 0.44 | 0.55 | 0.43 | 0.4 | 0.49 | 0.38 |
| 2 | 235446253 | rs7592582 | 0.87 | -0.04 | -1.09 | 0.363 | -0.004 | 0.004 | 0.004 | 0 | 0.44 | 0.45 | 0.42 | 0.4 | 0.49 | 0.38 |
| 2 | 235452615 | rs1880745 | -0.254 | -0.35 | -1.213 | 0.66 | 0.053 | -0.09 | -0.09 | 0.248 | 0.82 | 0.95 | 0.93 | 0.6 | 0.92 | 0.87 |
| 2 | 235456684 | rs1880746 | -0.201 | -0.465 | -0.907 | 0.638 | 0.01 | -0.011 | -0.011 | 0.3 | 0.83 | 0.89 | 0.95 | 0.6 | 0.92 | 0.88 |

| | | | | | | | | | | | | | | | | |
|---|---|---|---|---|---|---|---|---|---|---|---|---|---|---|---|---|
| 2 | 235458125 | rs1880747 | -0.048 | -0.433 | -0.839 | 0.257 | 0.029 | 0.036 | 0.036 | 0 | 0.57 | 0.71 | 0.71 | 0.31 | 0.62 | 0.68 |
| 2 | 235464033 | rs13006968 | 0.829 | -0.398 | -0.882 | 0.254 | 0.001 | 0.003 | 0.003 | 0.402 | 0.28 | 0.2 | 0.23 | 0.3 | 0.3 | 0.2 |
| 2 | 235464046 | rs11685523 | -0.223 | -0.404 | -0.866 | 0.262 | 0.034 | 0.044 | 0.044 | 0 | 0.59 | 0.76 | 0.73 | 0.3 | 0.63 | 0.68 |
| 2 | 235469549 | rs12469734 | 0.399 | -0.112 | -0.098 | 0.213 | 0.001 | 0.001 | 0.001 | 0 | 0.55 | 0.43 | 0.55 | 0.44 | 0.63 | 0.55 |
| 2 | 235475892 | rs940942 | 0.745 | -0.052 | 0.081 | 0.148 | 0.003 | 0.037 | 0.037 | 0 | 0.5 | 0.4 | 0.45 | 0.29 | 0.57 | 0.36 |
| 2 | 235477886 | rs2386499 | -0.72 | 0.285 | 0.123 | 0.32 | 0.011 | 0.027 | 0.027 | 0 | 0.09 | 0.19 | 0.13 | 0.13 | 0.08 | 0.22 |
| 2 | 235479343 | rs1405129 | -0.133 | 0.526 | 0.035 | 0.298 | 0.016 | 0.001 | 0.001 | 0 | 0.27 | 0.36 | 0.47 | 0.31 | 0.34 | 0.55 |
| 2 | 235479516 | rs4663435 | 0.455 | 0.547 | -0.14 | 0.132 | -0.049 | 0.009 | 0.009 | 0 | 0.22 | 0.14 | 0.45 | 0.18 | 0.32 | 0.29 |
| 2 | 235479565 | rs4663436 | 0.855 | 0.403 | -0.354 | 0.12 | 0.015 | 0.129 | 0.129 | 0 | 0.43 | 0.23 | 0.36 | 0.18 | 0.43 | 0.23 |
| 2 | 235479716 | rs4663437 | -0.079 | 0.264 | 0.366 | 0.402 | 0.04 | 0 | 0 | 0 | 0.28 | 0.5 | 0.45 | 0.26 | 0.29 | 0.35 |
| 2 | 235480193 | rs6731361 | -0.561 | 0.285 | 0.19 | 0.485 | -0.011 | 0.007 | 0.007 | 1.814 | 0.29 | 0.37 | 0.17 | 0.61 | 0.18 | 0.37 |
| 2 | 235480729 | rs9287578 | -0.811 | 0.43 | -0.641 | 0.166 | 0.021 | 0.277 | 0.277 | 0.757 | 0.5 | 0.82 | 0.59 | 0.9 | 0.4 | 0.68 |
| 2 | 235484375 | rs9653208 | -0.068 | 0.914 | -0.15 | 0.899 | 0.011 | 0.038 | 0.038 | 0 | 0.27 | 0.41 | 0.44 | 0.44 | 0.27 | 0.56 |
| 2 | 235486158 | rs11677392 | 0.358 | 0.441 | -0.907 | 0.544 | 0.021 | 0.01 | 0.01 | 4.667 | 0.43 | 0.21 | 0.43 | 0.45 | 0.4 | 0.19 |
| 2 | 235486991 | rs13011210 | 1.111 | 0.37 | -1.004 | 0.076 | -0.074 | 0.046 | 0.046 | 0.011 | 0.2 | 0.01 | 0.16 | 0.08 | 0.28 | 0.08 |
| 2 | 235488853 | rs11678113 | 0.407 | 0.333 | -0.691 | -0.013 | -0.008 | 0.025 | 0.025 | 0 | 0.26 | 0.17 | 0.21 | 0.11 | 0.35 | 0.1 |
| 2 | 235490801 | rs1526107 | -0.108 | 0.339 | -0.597 | 0.263 | -0.002 | 0.022 | 0.022 | 0 | 0.47 | 0.37 | 0.43 | 0.54 | 0.51 | 0.41 |
| 2 | 235491138 | rs7576760 | -0.419 | 0.437 | -0.356 | 0.181 | 0.002 | 0.015 | 0.015 | 0 | 0.5 | 0.47 | 0.43 | 0.53 | 0.5 | 0.58 |
| 2 | 235493171 | rs6749297 | -0.188 | 0.384 | -0.096 | 0.23 | 0.002 | 0.004 | 0.004 | 0 | 0.49 | 0.52 | 0.42 | 0.51 | 0.48 | 0.4 |
| 2 | 235495729 | rs13035090 | 0.992 | 0.451 | -0.643 | 0.017 | -0.013 | 0.055 | 0.055 | 0 | 0.27 | 0.11 | 0.19 | 0.1 | 0.33 | 0.08 |
| 2 | 235495918 | rs12052580 | 0.899 | 0.318 | -0.71 | 0.519 | -0.028 | -0.107 | -0.107 | 0 | 0.2 | 0.02 | 0.21 | 0.38 | 0.19 | 0.33 |
| 2 | 235501001 | rs1405134 | -0.794 | 0.412 | -0.745 | -0.451 | -0.009 | 0.012 | 0.012 | 1.791 | 0.81 | 0.81 | 0.8 | 0.99 | 0.73 | 0.84 |
| 2 | 235501122 | rs716085 | 1.099 | 0.429 | -0.474 | -0.177 | -0.003 | -0.028 | -0.028 | 0 | 0.21 | 0.13 | 0.27 | 0.54 | 0.19 | 0.35 |
| 2 | 235501150 | rs7594340 | 1.227 | 0.448 | -0.672 | -0.167 | -0.038 | -0.125 | -0.125 | 0.083 | 0.2 | 0.02 | 0.27 | 0.54 | 0.19 | 0.35 |
| 2 | 235501912 | rs6431272 | 0.054 | 0.344 | 0.06 | -0.116 | 0.001 | -0.001 | -0.001 | 3.216 | 0.58 | 0.51 | 0.52 | 0.45 | 0.53 | 0.49 |
| 2 | 235502345 | rs940947 | -0.14 | 0.226 | -0.216 | -0.394 | -0.041 | 0.037 | 0.037 | 2.399 | 0.27 | 0.51 | 0.31 | 0.36 | 0.14 | 0.36 |
| 2 | 235502923 | rs7574244 | -0.993 | 0.579 | -0.627 | -0.849 | 0.004 | -0.027 | -0.027 | 0 | 0.89 | 0.86 | 0.88 | 1 | 0.84 | 0.9 |
| 2 | 235503852 | rs4663445 | 0.091 | 0.488 | -0.883 | -1.17 | -0.024 | 0.01 | 0.01 | 0.247 | 0.66 | 0.67 | 0.71 | 0.95 | 0.56 | 0.59 |
| 2 | 235504056 | rs4663447 | -0.243 | -0.145 | -1.016 | -0.775 | -0.011 | -0.039 | -0.039 | 0.962 | 0.4 | 0.38 | 0.51 | 0.85 | 0.26 | 0.53 |
| 2 | 235505182 | rs11899787 | 0.152 | -0.176 | -1.302 | -0.121 | 0.013 | 0.003 | 0.003 | 0 | 0.68 | 0.78 | 0.64 | 0.71 | 0.77 | 0.72 |
| 2 | 235505863 | rs7573392 | -0.161 | 0.09 | -1.526 | 0.055 | 0.001 | -0.005 | -0.005 | 0 | 0.73 | 0.74 | 0.64 | 0.53 | 0.8 | 0.79 |
| 2 | 235508990 | rs11901926 | -0.517 | -0.008 | -1.843 | -1.07 | 0.014 | 0.03 | 0.03 | 0 | 0.35 | 0.74 | 0.25 | 0.06 | 0.35 | 0.2 |
| 2 | 235509738 | rs10190832 | -0.804 | 0.03 | -1.846 | -1.084 | 0.018 | 0.064 | 0.064 | 0 | 0.4 | 0.74 | 0.29 | 0.06 | 0.41 | 0.24 |
| 2 | 235512068 | rs13426909 | -0.601 | 0.228 | -1.506 | -0.751 | 0.002 | -0.009 | -0.009 | 0 | 0.3 | 0.73 | 0.23 | 0.07 | 0.33 | 0.23 |
| 2 | 235522554 | rs1880742 | -0.423 | 0.243 | -1.734 | -0.419 | -0.031 | 0.113 | 0.113 | 0 | 0.56 | 0.8 | 0.57 | 0.75 | 0.51 | 0.51 |
| 2 | 235525599 | rs2386501 | -0.163 | -0.706 | -0.925 | 0.188 | -0.047 | 0.081 | 0.081 | 0 | 0.22 | 0.36 | 0.25 | 0.57 | 0.09 | 0.31 |
| 2 | 235532167 | rs11681954 | 0.004 | -0.184 | -1.056 | 0.035 | -0.007 | -0.007 | -0.007 | 0.007 | 0.11 | 0.07 | 0.12 | 0.09 | 0.13 | 0.05 |
| 2 | 235538902 | rs4663449 | 0.272 | -0.533 | -0.784 | -0.391 | 0.005 | 0.008 | 0.008 | 0 | 0.47 | 0.46 | 0.52 | 0.41 | 0.6 | 0.47 |
| 2 | 235539546 | rs6431278 | -0.975 | -0.896 | -0.733 | -0.685 | 0.017 | -0.059 | -0.059 | 0.936 | 0.77 | 0.72 | 0.8 | 0.97 | 0.72 | 0.81 |
| 2 | 235542430 | rs13402575 | 0.453 | -0.887 | -0.707 | -0.729 | 0 | -0.001 | -0.001 | 0 | 0.1 | 0.09 | 0.12 | 0.11 | 0.13 | 0.02 |
| 2 | 235549882 | rs1526117 | 1.485 | -0.992 | -0.918 | -1.1 | -0.168 | -0.012 | -0.012 | 0 | 0.33 | 0.02 | 0.38 | 0.29 | 0.48 | 0.44 |
| 2 | 235550191 | rs10206814 | -0.129 | -1.104 | -0.378 | -1.059 | 0.009 | -0.057 | -0.057 | 0 | 0.72 | 0.63 | 0.79 | 0.9 | 0.71 | 0.69 |
| 2 | 235570608 | rs884063 | -0.255 | -1.15 | -0.145 | -1.022 | 0.004 | -0.037 | -0.037 | 0 | 0.69 | 0.38 | 0.78 | 0.87 | 0.7 | 0.77 |
| 2 | 235570734 | rs884062 | 0.817 | -1.726 | -0.858 | -1.262 | -0.077 | 0.127 | 0.127 | 2.655 | 0.45 | 0.08 | 0.47 | 0.27 | 0.53 | 0.51 |
| 2 | 235575490 | rs13011120 | 0.465 | -1.771 | -0.593 | -1.293 | -0.004 | 0.114 | 0.114 | 1.287 | 0.51 | 0.21 | 0.48 | 0.3 | 0.56 | 0.58 |
| 2 | 235576220 | rs2386495 | 0.428 | -1.821 | -0.578 | -1.259 | -0.022 | 0.157 | 0.157 | 2.213 | 0.5 | 0.15 | 0.48 | 0.27 | 0.53 | 0.56 |
| 2 | 235576903 | rs7588146 | -0.123 | -1.921 | -0.531 | -1.382 | 0.02 | 0.067 | 0.067 | 0.197 | 0.6 | 0.25 | 0.73 | 0.9 | 0.69 | 0.74 |
| 2 | 235578565 | rs12475976 | 0.308 | -1.828 | -0.218 | -1.097 | -0.004 | -0.058 | -0.058 | 1.436 | 0.18 | 0.07 | 0.28 | 0.6 | 0.17 | 0.19 |
| 2 | 235590167 | rs13411060 | -0.213 | -1.762 | -0.054 | -1.072 | -0.002 | 0.013 | 0.013 | 0.009 | 0.23 | 0.21 | 0.28 | 0.6 | 0.2 | 0.25 |
| 2 | 235594600 | rs12466858 | -0.332 | -1.853 | -0.205 | -1.229 | 0.011 | -0.013 | -0.013 | 0 | 0.31 | 0.19 | 0.29 | 0.6 | 0.23 | 0.22 |
| 2 | 235599903 | rs11677044 | -0.62 | -1.949 | -0.016 | -1.574 | 0.005 | 0.01 | 0.01 | 0.339 | 0.65 | 0.72 | 0.77 | 0.91 | 0.72 | 0.81 |
| 2 | 235600627 | rs4663474 | 0.757 | -1.982 | -0.015 | -1.585 | 0.013 | -0.017 | -0.017 | 0 | 0.39 | 0.53 | 0.54 | 0.31 | 0.51 | 0.59 |
| 2 | 235603435 | rs1358069 | -0.62 | -1.858 | -0.127 | -1.744 | -0.002 | -0.027 | -0.027 | 0 | 0.64 | 0.67 | 0.76 | 0.91 | 0.71 | 0.81 |
| 2 | 235604411 | rs4663479 | 1.456 | -1.917 | -0.262 | -1.716 | -0.108 | 0.019 | 0.019 | 7.233 | 0.3 | 0.15 | 0.46 | 0.26 | 0.5 | 0.47 |
| 2 | 235604871 | rs13410622 | -0.242 | -1.844 | -0.206 | -1.596 | 0.008 | -0.005 | -0.005 | 0 | 0.3 | 0.2 | 0.25 | 0.65 | 0.24 | 0.25 |
| 2 | 235607892 | rs12478541 | -0.821 | -1.773 | -0.139 | -1.57 | -0.027 | -0.007 | -0.007 | 1.11 | 0.38 | 0.45 | 0.35 | 0.64 | 0.29 | 0.36 |
| 2 | 235609613 | rs4233612 | -0.66 | -1.698 | -0.118 | -1.636 | -0.013 | -0.076 | -0.076 | 0 | 0.67 | 0.54 | 0.75 | 0.9 | 0.74 | 0.81 |
| 2 | 235609674 | rs7606512 | 0.574 | -1.664 | 0.046 | -1.404 | -0.044 | -0.001 | -0.001 | 0 | 0.11 | 0.32 | 0.08 | 0.07 | 0.03 | 0.02 |
| 2 | 235611137 | rs4663483 | 0.723 | -1.595 | -0.494 | -1.365 | -0.023 | 0.019 | 0.019 | 8.175 | 0.35 | 0.31 | 0.45 | 0.28 | 0.47 | 0.51 |
| 2 | 235617634 | rs13387193 | 0.039 | -1.519 | -0.833 | -1.374 | -0.019 | 0.086 | 0.086 | 1.837 | 0.47 | 0.2 | 0.38 | 0.66 | 0.48 | 0.33 |
| 2 | 235621868 | rs1441616 | 0.172 | -1.243 | -0.692 | -1.373 | -0.001 | -0.024 | -0.024 | 0.663 | 0.34 | 0.33 | 0.31 | 0.11 | 0.3 | 0.26 |
| 2 | 235624385 | rs4271722 | -1.237 | -0.902 | -1.032 | -1.111 | 0.003 | -0.001 | -0.001 | 0 | 0.73 | 0.7 | 0.61 | 0.69 | 0.63 | 0.51 |
| 2 | 235624835 | rs13390788 | 0.232 | -1.199 | -1.495 | -0.972 | 0.025 | -0.072 | -0.072 | 0 | 0.22 | 0.06 | 0.15 | 0.6 | 0.2 | 0.17 |
| 2 | 235625587 | rs7607157 | -0.296 | -0.803 | -1.327 | -1.169 | -0.035 | -0.096 | -0.096 | 0 | 0.21 | 0.37 | 0.39 | 0.06 | 0.17 | 0.19 |
| 2 | 235626160 | rs4663163 | -0.962 | -0.292 | -1.837 | -0.751 | -0.144 | -0.022 | -0.022 | 0.04 | 0.72 | 0.97 | 0.8 | 0.73 | 0.61 | 0.53 |
| 2 | 235626227 | rs4663164 | -0.149 | -0.101 | -1.002 | -0.911 | -0.001 | -0.064 | -0.064 | 0 | 0.42 | 0.5 | 0.52 | 0.09 | 0.38 | 0.29 |
| 2 | 235628051 | rs2218249 | -0.595 | 0.063 | -1.216 | 0.158 | 0.013 | -0.025 | -0.025 | 0 | 0.39 | 0.68 | 0.27 | 0.45 | 0.31 | 0.47 |
| 2 | 235632999 | rs2380262 | 0.043 | -0.101 | -1.749 | -0.007 | 0.035 | 0 | 0 | 0.612 | 0.35 | 0.21 | 0.48 | 0.38 | 0.3 | 0.35 |
| 2 | 235635006 | rs6730767 | 0.191 | -0.304 | -0.786 | 0.219 | 0.031 | 0.112 | 0.112 | 0 | 0.06 | 0.22 | 0.07 | 0.1 | 0.06 | 0.12 |

| | | | | | | | | | | | | | | | | |
|---|---|---|---|---|---|---|---|---|---|---|---|---|---|---|---|---|
| 2 | 235638698 | rs6718805 | -0.614 | -0.327 | -0.779 | 0.128 | 0.009 | 0.019 | 0.019 | 0 | 0.08 | 0.24 | 0.08 | 0.05 | 0.07 | 0.14 |
| 2 | 235639080 | rs7586477 | 1.083 | -0.433 | -0.629 | -0.113 | -0.007 | 0.027 | 0.027 | 0.157 | 0.35 | 0.29 | 0.23 | 0.2 | 0.37 | 0.16 |
| 2 | 235641180 | rs2380261 | 0.683 | -0.382 | -0.699 | -0.11 | 0.035 | -0.021 | -0.021 | 0 | 0.32 | 0.17 | 0.46 | 0.4 | 0.28 | 0.34 |
| 2 | 235642969 | rs4474836 | -1.259 | -0.358 | -0.627 | -0.064 | 0.013 | 0.018 | 0.018 | 0 | 0.79 | 0.77 | 0.79 | 0.65 | 0.73 | 0.66 |
| 2 | 235646122 | rs11676712 | -0.271 | -0.326 | -0.516 | -0.38 | -0.002 | -0.008 | -0.008 | 0 | 0.44 | 0.44 | 0.32 | 0.25 | 0.44 | 0.3 |
| 2 | 235656278 | rs7557724 | -1.217 | -0.366 | 0.088 | -0.139 | 0.017 | 0.028 | 0.028 | 0 | 0.09 | 0.31 | 0.08 | 0.05 | 0.07 | 0.12 |
| 2 | 235659902 | rs7588886 | -1.137 | -0.236 | -0.605 | -0.11 | -0.004 | -0.021 | -0.021 | 0 | 0.05 | 0.16 | 0.06 | 0.01 | 0.06 | 0.01 |
| 2 | 235667715 | rs4380175 | -0.348 | -0.188 | -1.061 | -0.432 | -0.002 | 0.018 | 0.018 | 0.745 | 0.56 | 0.61 | 0.36 | 0.23 | 0.48 | 0.29 |
| 2 | 235668810 | rs4561611 | 0.83 | -0.361 | -1.92 | -0.296 | -0.178 | -0.086 | -0.086 | 1.051 | 0.34 | 0.03 | 0.5 | 0.3 | 0.41 | 0.46 |
| 2 | 235668847 | rs10201406 | 0.028 | -0.685 | -1.495 | -0.435 | -0.032 | -0.071 | -0.071 | 0.01 | 0.12 | 0.58 | 0.08 | 0 | 0.04 | 0.05 |
| 2 | 235668886 | rs4663492 | -0.487 | -1.289 | -1.874 | -0.643 | -0.005 | -0.002 | -0.002 | 0.801 | 0.28 | 0.61 | 0.19 | 0.53 | 0.12 | 0.27 |
| 2 | 235670433 | rs12613932 | -0.858 | -1.374 | -1.788 | -0.744 | 0.071 | -0.054 | -0.054 | 0 | 0.34 | 0.75 | 0.24 | 0.54 | 0.18 | 0.29 |
| 2 | 235672254 | rs11674223 | 1.38 | -1.351 | -1.905 | -0.832 | -0.085 | -0.006 | -0.006 | 0.227 | 0.37 | 0.16 | 0.52 | 0.3 | 0.43 | 0.46 |
| 2 | 235673205 | rs7563078 | -0.967 | -1.023 | -1.96 | -0.694 | -0.029 | 0.041 | 0.041 | 0.037 | 0.73 | 0.93 | 0.8 | 0.83 | 0.69 | 0.78 |
| 2 | 235673464 | rs4614878 | -0.068 | -1.331 | -1.813 | -0.665 | -0.08 | 0.121 | 0.121 | 0 | 0.25 | 0.47 | 0.18 | 0.53 | 0.14 | 0.23 |
| 2 | 235673539 | rs7602550 | 1.03 | -1.318 | -1.812 | -0.665 | 0.008 | -0.034 | -0.034 | 0 | 0.08 | 0.31 | 0.06 | 0 | 0.04 | 0.07 |
| 2 | 235673638 | rs4233613 | -0.667 | -1.262 | -1.845 | -0.612 | -0.07 | 0.061 | 0.061 | 0.008 | 0.71 | 0.91 | 0.76 | 0.83 | 0.61 | 0.76 |
| 2 | 235674274 | rs1992430 | -0.382 | -1.176 | -1.34 | -0.21 | -0.076 | 0.115 | 0.115 | 0 | 0.25 | 0.45 | 0.18 | 0.53 | 0.14 | 0.24 |
| 2 | 235677678 | rs7578453 | -0.023 | -0.899 | -1.079 | -0.348 | 0.012 | -0.027 | -0.027 | 0 | 0.09 | 0.41 | 0.06 | 0.01 | 0.06 | 0.02 |
| 2 | 235677775 | rs7590552 | -0.738 | -0.969 | -1.323 | -0.421 | -0.022 | -0.061 | -0.061 | 0.303 | 0.17 | 0.71 | 0.11 | 0.01 | 0.09 | 0.05 |
| 2 | 235680447 | rs13008181 | -0.48 | -0.961 | -1.99 | -0.461 | -0.038 | 0.05 | 0.05 | 0 | 0.49 | 0.1 | 0.37 | 0.71 | 0.52 | 0.5 |
| 2 | 235682144 | rs7574103 | 0.423 | -1.095 | -2.259 | -0.403 | -0.135 | 0.015 | 0.015 | 0 | 0.33 | 0.05 | 0.25 | 0.22 | 0.47 | 0.28 |
| 2 | 235683121 | rs13403639 | -0.358 | -1.175 | -2.065 | -0.464 | -0.02 | -0.054 | -0.054 | 6.303 | 0.17 | 0.7 | 0.11 | 0.01 | 0.09 | 0.05 |
| 2 | 235690982 | rs1000016 | 1.433 | -1.03 | -2.334 | -0.331 | 0.035 | -0.127 | -0.127 | 0.876 | 0.16 | 0.01 | 0.12 | 0.5 | 0.08 | 0.24 |
| 2 | 235691089 | rs1000017 | 0.15 | -0.641 | -2.031 | -0.583 | -0.069 | 0.052 | 0.052 | 0 | 0.33 | 0.16 | 0.26 | 0.21 | 0.47 | 0.28 |
| 2 | 235691219 | rs10208234 | 0.233 | -0.656 | -1.226 | -0.184 | 0.021 | 0.002 | 0.002 | 0.056 | 0.08 | 0.34 | 0.06 | 0.02 | 0.06 | 0.02 |
| 2 | 235695228 | rs7605821 | -1.011 | -0.319 | -2.465 | -0.153 | -0.049 | 0.063 | 0.063 | 0 | 0.62 | 0.88 | 0.44 | 0.8 | 0.5 | 0.5 |
| 2 | 235697004 | rs12988431 | 0.787 | -0.388 | -2.46 | 0.245 | -0.015 | -0.08 | -0.08 | 0 | 0.15 | 0.02 | 0.13 | 0.25 | 0.15 | 0.15 |
| 2 | 235701101 | rs6757665 | -0.413 | -0.583 | -2.112 | 0.02 | -0.001 | -0.001 | -0.001 | 0 | 0.24 | 0.69 | 0.19 | 0.24 | 0.2 | 0.17 |
| 2 | 235704072 | rs4663507 | 0.567 | -0.909 | -2.792 | 0.056 | 0.043 | -0.121 | -0.121 | 0.064 | 0.24 | 0.06 | 0.16 | 0.55 | 0.17 | 0.28 |
| 2 | 235704447 | rs6431302 | -1.689 | -0.036 | -2.802 | -0.356 | -0.15 | -0.013 | -0.013 | 0 | 0.71 | 0.98 | 0.52 | 0.81 | 0.68 | 0.52 |
| 2 | 235707172 | rs11692799 | 0.102 | -0.533 | -1.879 | -0.214 | -0.039 | 0.071 | 0.071 | 0 | 0.17 | 0.07 | 0.12 | 0.03 | 0.23 | 0.08 |
| 2 | 235708742 | rs4233615 | 1.578 | -0.857 | -1.247 | 0.177 | -0.013 | -0.002 | -0.002 | 0.029 | 0.43 | 0.29 | 0.56 | 0.38 | 0.44 | 0.56 |
| 2 | 235709572 | rs4487050 | 0.617 | -1.154 | -0.398 | 0.052 | 0.001 | -0.046 | -0.046 | 1.823 | 0.75 | 0.76 | 0.87 | 1 | 0.8 | 0.9 |
| 2 | 235709805 | rs4500898 | 1.831 | -1.323 | -0.407 | 0.205 | 0.004 | 0.006 | 0.006 | 1.809 | 0.49 | 0.28 | 0.6 | 0.42 | 0.5 | 0.62 |
| 2 | 235710513 | rs13033621 | -0.301 | -1.264 | -0.21 | -0.028 | -0.027 | 0.063 | 0.063 | 0 | 0.23 | 0.15 | 0.16 | 0.04 | 0.29 | 0.11 |
| 2 | 235717794 | rs4663509 | 0.389 | -1.17 | 0.181 | 0.3 | -0.001 | 0.011 | 0.011 | 0 | 0.15 | 0.15 | 0.14 | 0.45 | 0.15 | 0.24 |
| 2 | 235719875 | rs7570140 | 0.308 | -1.283 | 0.443 | 0.205 | -0.016 | -0.072 | -0.072 | 0.453 | 0.77 | 0.7 | 0.88 | 1 | 0.84 | 0.96 |
| 2 | 235721740 | rs1441612 | 2.135 | -1.157 | 0.913 | 0.278 | -0.008 | 0.004 | 0.004 | 2.127 | 0.51 | 0.22 | 0.62 | 0.51 | 0.5 | 0.65 |
| 2 | 235724801 | rs1441611 | 2.057 | -1.009 | 0.988 | 0.267 | 0.006 | 0.005 | 0.005 | 1.813 | 0.5 | 0.39 | 0.6 | 0.51 | 0.43 | 0.65 |
| 2 | 235730405 | rs11681769 | 0.415 | -0.762 | 1.014 | 0.2 | -0.025 | 0.014 | 0.014 | 0 | 0.08 | 0.06 | 0.1 | 0.03 | 0.19 | 0.04 |
| 2 | 235733155 | rs10929060 | 1.379 | -0.801 | 1.199 | 0.157 | 0.019 | -0.055 | -0.055 | 0.044 | 0.79 | 0.7 | 0.83 | 0.97 | 0.69 | 0.89 |
| 2 | 235735930 | rs1427696 | 0.823 | -1.138 | 1.022 | 0.298 | -0.002 | -0.028 | -0.028 | 1.332 | 0.88 | 0.86 | 0.93 | 1 | 0.88 | 0.97 |
| 2 | 235744236 | rs894184 | 1.209 | -1.515 | 0.779 | 0.553 | -0.031 | -0.006 | -0.006 | 0.035 | 0.61 | 0.51 | 0.59 | 0.6 | 0.7 | 0.74 |
| 2 | 235744397 | rs10929063 | -1.117 | -0.743 | 0.707 | 0.773 | 0.001 | -0.002 | -0.002 | 0.946 | 0.09 | 0.08 | 0.04 | 0.22 | 0.07 | 0.09 |
| 2 | 235745098 | rs972514 | 0.931 | -0.785 | 1.045 | 0.808 | 0.004 | 0.006 | 0.006 | 0 | 0.88 | 0.82 | 0.92 | 0.78 | 0.86 | 0.93 |
| 2 | 235753876 | rs6750155 | -1.105 | -0.675 | 1.134 | 0.969 | -0.001 | -0.005 | -0.005 | 0 | 0.32 | 0.32 | 0.41 | 0.57 | 0.34 | 0.28 |
| 2 | 235760592 | rs1946949 | 1.713 | -0.621 | 1.148 | 1.04 | -0.018 | 0.002 | 0.002 | 0 | 0.65 | 0.56 | 0.59 | 0.29 | 0.67 | 0.71 |
| 2 | 235766037 | rs1946950 | 1.987 | -0.634 | 0.844 | 1.135 | -0.012 | 0.071 | 0.071 | 0.26 | 0.56 | 0.27 | 0.5 | 0.2 | 0.52 | 0.66 |
| 2 | 235769182 | rs7585293 | -0.695 | -1.04 | 0.556 | 1.435 | 0.011 | -0.026 | -0.026 | 0.001 | 0.19 | 0.12 | 0.29 | 0.46 | 0.13 | 0.21 |
| 2 | 235775041 | rs1529413 | -1.013 | -0.888 | 0.462 | 1.407 | -0.036 | 0.068 | 0.068 | 1.006 | 0.24 | 0.31 | 0.31 | 0.47 | 0.14 | 0.21 |
| 2 | 235776166 | rs2033994 | 0.239 | -0.637 | 0.686 | 1.635 | 0.032 | 0.054 | 0.054 | 0 | 0.92 | 0.81 | 0.93 | 0.8 | 0.9 | 0.93 |
| 2 | 235779418 | rs6744853 | 0.242 | -0.629 | 0.441 | 1.639 | 0.046 | 0.072 | 0.072 | 0 | 0.92 | 0.75 | 0.93 | 0.8 | 0.9 | 0.93 |
| 2 | 235780667 | rs13394889 | -0.541 | -0.699 | 0.32 | 1.539 | -0.054 | 0.071 | 0.071 | 0.017 | 0.16 | 0.23 | 0.25 | 0.34 | 0.07 | 0.12 |
| 2 | 235784645 | rs4663515 | 0.479 | -0.505 | -0.328 | 0.679 | -0.036 | 0.058 | 0.058 | 0 | 0.75 | 0.5 | 0.67 | 0.32 | 0.82 | 0.8 |
| 2 | 235794359 | rs10202685 | 0.211 | 0.051 | -1.105 | 0.549 | 0.031 | 0.024 | 0.024 | 0 | 0.51 | 0.63 | 0.53 | 0.31 | 0.58 | 0.59 |
| 2 | 235795095 | rs4663517 | 0.195 | 0.386 | -1.365 | 0.417 | -0.009 | 0.052 | 0.052 | 0 | 0.55 | 0.73 | 0.71 | 0.7 | 0.49 | 0.52 |
| 2 | 235795528 | rs722161 | -0.591 | 0.326 | -0.94 | 0.581 | -0.002 | -0.012 | -0.012 | 0.109 | 0.26 | 0.31 | 0.41 | 0.2 | 0.23 | 0.19 |
| 2 | 235805656 | rs890657 | -0.035 | -0.309 | -1.575 | 0.628 | 0.057 | 0.012 | 0.012 | 6.653 | 0.69 | 0.85 | 0.77 | 0.69 | 0.75 | 0.8 |
| 2 | 235806038 | rs9789458 | -0.672 | 1.079 | -0.866 | 1.228 | 0.05 | 0.03 | 0.03 | 0 | 0.13 | 0.18 | 0.3 | 0.26 | 0.21 | 0.19 |
| 2 | 235806200 | rs6722544 | -0.268 | 0.027 | -0.337 | 1.083 | -0.023 | 0.019 | 0.019 | 0 | 0.29 | 0.41 | 0.3 | 0.53 | 0.24 | 0.4 |
| 2 | 235806512 | rs11685757 | -0.211 | -0.039 | -0.378 | 1.718 | -0.009 | 0.18 | 0.18 | 0 | 0.15 | 0.38 | 0.18 | 0.37 | 0.16 | 0.29 |
| 2 | 235806859 | rs10929070 | -1.173 | -0.14 | -0.432 | 1.625 | -0.018 | 0.04 | 0.04 | 0 | 0.24 | 0.41 | 0.37 | 0.57 | 0.22 | 0.38 |
| 2 | 235808211 | rs6431311 | -1.918 | -0.103 | -0.44 | 1.332 | -0.008 | -0.031 | -0.031 | 0.028 | 0.4 | 0.55 | 0.63 | 0.77 | 0.35 | 0.54 |
| 2 | 235809515 | rs11684741 | -1.105 | -0.128 | -0.233 | 1.313 | -0.006 | 0.02 | 0.02 | 0 | 0.28 | 0.43 | 0.43 | 0.66 | 0.27 | 0.47 |
| 2 | 235811887 | rs10929071 | -1.064 | -0.245 | 0.527 | 1.304 | -0.012 | 0.037 | 0.037 | 0.175 | 0.27 | 0.45 | 0.41 | 0.68 | 0.24 | 0.47 |
| 2 | 235812756 | rs12692157 | -1.518 | -0.421 | 0.666 | 1.109 | 0.001 | -0.059 | -0.059 | 0 | 0.22 | 0.48 | 0.27 | 0.09 | 0.18 | 0.11 |
| 2 | 235819508 | rs13424401 | -1.206 | -0.973 | 1.117 | 1.177 | -0.056 | -0.048 | -0.048 | 0 | 0.18 | 0.45 | 0.21 | 0.06 | 0.08 | 0.05 |
| 2 | 235821065 | rs6750091 | -1.303 | -0.986 | 1.211 | 1.188 | -0.051 | -0.031 | -0.031 | 0 | 0.18 | 0.42 | 0.21 | 0.07 | 0.08 | 0.07 |

| | | | | | | | | | | | | | | | | |
|---|---|---|---|---|---|---|---|---|---|---|---|---|---|---|---|---|
| 2 | 235825527 | rs12477521 | 0.152 | -0.973 | 1.158 | 1.294 | -0.008 | -0.012 | -0.012 | 0.192 | 0.11 | 0.09 | 0.24 | 0.19 | 0.1 | 0.17 |
| 2 | 235834973 | rs4663523 | -1.755 | -1 | 1.777 | 1.277 | -0.022 | -0.006 | -0.006 | 0 | 0.3 | 0.48 | 0.45 | 0.26 | 0.19 | 0.23 |
| 2 | 235840020 | rs6431324 | -1.874 | -0.998 | 1.54 | 1.274 | -0.011 | -0.005 | -0.005 | 1.552 | 0.32 | 0.65 | 0.48 | 0.27 | 0.19 | 0.24 |
| 2 | 235850039 | rs10203538 | -0.567 | -0.839 | 1.071 | 1.308 | 0.003 | 0.001 | 0.001 | 1.098 | 0.12 | 0.13 | 0.16 | 0.08 | 0.07 | 0.08 |
| 2 | 235857114 | rs2042831 | 0.633 | -0.947 | 0.882 | 1.441 | 0.022 | -0.064 | -0.064 | 0 | 0.86 | 0.94 | 0.89 | 0.61 | 0.89 | 0.72 |
| 2 | 235857310 | rs2379187 | 1.58 | -0.993 | 1.39 | 1.337 | 0.001 | 0 | 0 | 4.194 | 0.58 | 0.43 | 0.44 | 0.29 | 0.72 | 0.45 |
| 2 | 235859740 | rs12476919 | -0.743 | -0.931 | 1.121 | 1.393 | -0.002 | 0.002 | 0.002 | 0 | 0.25 | 0.23 | 0.43 | 0.3 | 0.18 | 0.27 |
| 2 | 235860196 | rs13019776 | 1.365 | -0.861 | 0.905 | 1.311 | 0.012 | 0.015 | 0.015 | 3.92 | 0.58 | 0.61 | 0.45 | 0.29 | 0.72 | 0.44 |
| 2 | 235867385 | rs4429425 | 1.257 | -0.67 | 0.843 | 1.679 | -0.001 | 0.011 | 0.011 | 3.726 | 0.82 | 0.79 | 0.79 | 0.58 | 0.87 | 0.69 |
| 2 | 235879707 | rs6754925 | -1.69 | -0.848 | 1.521 | 1.508 | -0.069 | 0.04 | 0.04 | 0.422 | 0.22 | 0.51 | 0.36 | 0.69 | 0.12 | 0.35 |
| 2 | 235890317 | rs11684843 | -0.368 | -0.664 | 1.037 | 1.474 | -0.005 | -0.01 | -0.01 | 0 | 0.05 | 0.02 | 0.04 | 0.06 | 0.06 | 0.02 |
| 2 | 235890728 | rs6720830 | -1.846 | -0.503 | 0.848 | 1.789 | 0.001 | -0.003 | -0.003 | 0 | 0.31 | 0.34 | 0.35 | 0.58 | 0.21 | 0.32 |
| 2 | 235891645 | rs10803629 | -1.956 | -0.502 | 0.938 | 1.808 | -0.003 | 0.005 | 0.005 | 0.452 | 0.3 | 0.33 | 0.22 | 0.58 | 0.21 | 0.31 |
| 2 | 235893205 | rs4663533 | -0.509 | -0.383 | 0.687 | 1.764 | -0.007 | 0.016 | 0.016 | 0.131 | 0.14 | 0.11 | 0.06 | 0.04 | 0.12 | 0.1 |
| 2 | 235894236 | rs4663176 | -1.058 | -0.393 | 0.844 | 2.08 | -0.012 | 0.134 | 0.134 | 0 | 0.07 | 0.08 | 0.11 | 0.48 | 0.04 | 0.21 |
| 2 | 235897062 | rs10176321 | -1.852 | 0.58 | 0.976 | 1.862 | 0.027 | 0.12 | 0.12 | 0 | 0.17 | 0.29 | 0.19 | 0.47 | 0.24 | 0.27 |
| 2 | 235913226 | rs6725062 | -1.834 | 0.549 | 1.413 | 1.684 | -0.019 | 0.032 | 0.032 | 0 | 0.21 | 0.4 | 0.43 | 0.69 | 0.14 | 0.35 |
| 2 | 235915286 | rs4663178 | -1.672 | 0.522 | 0.986 | 1.714 | 0.003 | -0.015 | -0.015 | 0 | 0.17 | 0.1 | 0.41 | 0.69 | 0.13 | 0.35 |
| 2 | 235921753 | rs10194249 | -2.325 | -0.006 | 0.987 | 1.822 | 0.002 | -0.002 | -0.002 | 0 | 0.25 | 0.73 | 0.39 | 0.3 | 0.14 | 0.2 |
| 2 | 235922791 | rs6743498 | -2.554 | -0.097 | 0.937 | 1.782 | 0.025 | -0.014 | -0.014 | 0 | 0.27 | 0.76 | 0.39 | 0.32 | 0.14 | 0.23 |
| 2 | 235926285 | rs13402968 | -1.568 | 0.02 | 1.477 | 1.686 | -0.018 | -0.043 | -0.043 | 0 | 0.1 | 0.4 | 0.05 | 0.01 | 0.02 | 0.02 |
| 2 | 235932581 | rs12471688 | -1.793 | 0.422 | 0.523 | 2.005 | -0.002 | 0.025 | 0.025 | 0 | 0.14 | 0.13 | 0.2 | 0.42 | 0.18 | 0.25 |
| 2 | 235941537 | rs10190447 | -2.23 | 0.685 | 0.634 | 1.53 | -0.003 | 0.001 | 0.001 | 0.092 | 0.35 | 0.65 | 0.47 | 0.31 | 0.38 | 0.36 |
| 2 | 235942799 | rs11692659 | -1.701 | 0.355 | 0.377 | 1.896 | -0.032 | -0.057 | -0.057 | 0 | 0.09 | 0 | 0.28 | 0.28 | 0.09 | 0.17 |
| 2 | 235949877 | rs3731644 | -0.53 | -0.358 | 1.103 | 1.987 | -0.031 | 0.204 | 0.204 | 0 | 0.16 | 0.4 | 0.41 | 0.41 | 0.1 | 0.39 |
| 2 | 235949920 | rs3731645 | -0.514 | -0.357 | 1.103 | 1.988 | -0.03 | 0.206 | 0.206 | 0 | 0.16 | 0.4 | 0.34 | 0.41 | 0.09 | 0.39 |
| 2 | 235950187 | rs1469375 | -2.119 | -0.827 | -0.026 | 1.046 | -0.054 | 0.199 | 0.199 | 0.339 | 0.6 | 0.86 | 0.64 | 0.91 | 0.45 | 0.68 |
| 2 | 235950284 | rs3731648 | -0.044 | -0.85 | 0.214 | 1.405 | -0.023 | 0.105 | 0.105 | 0 | 0.16 | 0.29 | 0.33 | 0.41 | 0.09 | 0.39 |
| 2 | 235953565 | rs6732663 | -0.703 | -0.768 | -0.097 | 1.194 | -0.01 | 0.02 | 0.02 | 0 | 0.27 | 0.38 | 0.32 | 0.57 | 0.21 | 0.21 |
| 2 | 235953839 | rs4663538 | -0.115 | -0.406 | -0.513 | 1.333 | 0.005 | -0.008 | -0.008 | 0 | 0.19 | 0.15 | 0.22 | 0.24 | 0.13 | 0.38 |
| 2 | 235957657 | rs1559986 | -1.254 | -0.625 | -0.731 | 1.038 | 0.002 | -0.004 | -0.004 | 0 | 0.4 | 0.31 | 0.32 | 0.56 | 0.37 | 0.29 |
| 2 | 235960254 | rs10203697 | -0.167 | -0.796 | -0.416 | 1.059 | -0.014 | -0.014 | -0.014 | 0 | 0.15 | 0.26 | 0.11 | 0.12 | 0.12 | 0.15 |
| 2 | 235960824 | rs4505466 | -1.403 | -1.091 | -0.72 | 1.185 | 0.033 | 0.029 | 0.029 | 0 | 0.51 | 0.6 | 0.32 | 0.63 | 0.32 | 0.58 |
| 2 | 235961111 | rs6754815 | -0.55 | -0.628 | -1.498 | 1.461 | 0.004 | -0.07 | -0.07 | 0.171 | 0.24 | 0.15 | 0.19 | 0.52 | 0.16 | 0.36 |
| 2 | 235962210 | rs10177808 | -0.028 | -0.333 | -1.338 | 1.066 | -0.012 | -0.021 | -0.021 | 0.927 | 0.12 | 0.15 | 0.08 | 0.08 | 0.1 | 0.11 |
| 2 | 235965087 | rs4318343 | 0.107 | -0.051 | -1.938 | 1.525 | 0.007 | -0.06 | -0.06 | 0 | 0.16 | 0.07 | 0.18 | 0.34 | 0.1 | 0.29 |
| 2 | 235965403 | rs4334446 | -0.706 | 0.479 | -1.81 | 0.864 | -0.039 | 0.062 | 0.062 | 0 | 0.57 | 0.85 | 0.53 | 0.72 | 0.43 | 0.62 |
| 2 | 235969515 | rs4528714 | -0.597 | 0.773 | -1.195 | 1.245 | -0.006 | -0.013 | -0.013 | 0 | 0.17 | 0.15 | 0.11 | 0.26 | 0.17 | 0.1 |
| 2 | 235970536 | rs6756314 | -1.022 | 0.276 | -0.609 | 0.725 | -0.019 | -0.019 | -0.019 | 0.377 | 0.19 | 0.65 | 0.11 | 0.17 | 0.11 | 0.13 |
| 2 | 235970791 | rs6756573 | -1.036 | 0.251 | -0.692 | 0.73 | -0.005 | -0.005 | -0.005 | 0 | 0.19 | 0.73 | 0.12 | 0.17 | 0.11 | 0.13 |
| 2 | 235971971 | rs13404220 | -1.13 | 0.562 | -0.376 | 0.374 | 0.008 | -0.038 | -0.038 | 0.139 | 0.19 | 0.25 | 0.2 | 0.06 | 0.19 | 0.22 |
| 2 | 235972328 | rs10177149 | -0.638 | 0.58 | -0.712 | 0.465 | -0.001 | -0.001 | -0.001 | 0 | 0.39 | 0.6 | 0.5 | 0.62 | 0.35 | 0.3 |
| 2 | 235973681 | rs12468832 | -0.13 | 1.025 | -0.984 | 0.584 | -0.019 | -0.016 | -0.016 | 0 | 0.7 | 0.78 | 0.63 | 0.6 | 0.58 | 0.66 |
| 2 | 235976565 | rs7573787 | 0.18 | 1.234 | -1.178 | 1.041 | -0.069 | -0.074 | -0.074 | 0 | 0.7 | 0.89 | 0.65 | 0.54 | 0.56 | 0.64 |
| 2 | 235976967 | rs4413110 | 0.811 | 0.758 | -1.086 | 0.459 | 0.012 | 0.092 | 0.092 | 0 | 0.41 | 0.17 | 0.28 | 0.17 | 0.34 | 0.37 |
| 2 | 235978207 | rs10205183 | -1.338 | 0.985 | -1.526 | 0.459 | -0.048 | -0.029 | -0.029 | 0 | 0.81 | 0.92 | 0.78 | 0.73 | 0.71 | 0.77 |
| 2 | 235979171 | rs4663541 | 0.452 | 0.796 | -0.722 | 0.424 | 0.001 | 0 | 0 | 0 | 0.47 | 0.46 | 0.34 | 0.22 | 0.39 | 0.4 |
| 2 | 235984919 | rs11684611 | -0.359 | 0.72 | -0.634 | 0.813 | -0.01 | -0.013 | -0.013 | 0 | 0.12 | 0.06 | 0.14 | 0.16 | 0.16 | 0.15 |
| 2 | 235986028 | rs6738858 | -1.323 | 0.477 | -0.085 | 0.821 | -0.002 | 0.042 | 0.042 | 0.015 | 0.22 | 0.47 | 0.21 | 0.28 | 0.2 | 0.25 |
| 2 | 235991454 | rs10929094 | -0.8 | 0.677 | -0.356 | 0.723 | -0.011 | -0.003 | -0.003 | 0 | 0.76 | 0.86 | 0.78 | 0.68 | 0.69 | 0.73 |
| 2 | 235991748 | rs13413269 | 0.971 | 0.658 | -0.159 | 0.745 | 0.001 | 0.008 | 0.008 | 0.032 | 0.05 | 0.09 | 0.02 | 0.07 | 0.03 | 0.07 |
| 2 | 235991856 | rs4663543 | 0.689 | 0.471 | -0.286 | 0.604 | -0.001 | 0 | 0 | 0 | 0.36 | 0.43 | 0.49 | 0.38 | 0.31 | 0.26 |
| 2 | 235993234 | rs10172863 | -0.849 | 0.478 | -0.294 | 0.556 | -0.005 | -0.003 | -0.003 | 0 | 0.71 | 0.77 | 0.76 | 0.6 | 0.65 | 0.66 |
| 2 | 235993933 | rs11883732 | 0.93 | 0.501 | 0.076 | 0.485 | 0.004 | -0.023 | -0.023 | 0.044 | 0.28 | 0.25 | 0.24 | 0.12 | 0.29 | 0.31 |
| 2 | 235997858 | rs7560067 | 0.495 | 0.535 | 0.123 | 0.492 | 0.06 | 0.077 | 0.077 | 0 | 0.06 | 0.31 | 0.02 | 0.08 | 0.03 | 0.03 |
| 2 | 236001347 | rs7561280 | 0.205 | 0.478 | 0.116 | 0.581 | 0.002 | -0.024 | -0.024 | 0.441 | 0.37 | 0.51 | 0.28 | 0.26 | 0.34 | 0.43 |
| 2 | 236003413 | rs7568837 | -0.285 | 0.555 | -0.152 | 0.616 | -0.001 | 0.009 | 0.009 | 0.222 | 0.42 | 0.57 | 0.3 | 0.34 | 0.38 | 0.49 |
| 2 | 236006914 | rs10190780 | -0.231 | 0.591 | -0.407 | 0.608 | 0.001 | -0.011 | -0.011 | 0 | 0.41 | 0.46 | 0.3 | 0.33 | 0.38 | 0.46 |
| 2 | 236008035 | rs6710014 | 0.19 | 0.595 | -0.538 | 0.535 | 0.001 | -0.012 | -0.012 | 0 | 0.37 | 0.36 | 0.28 | 0.26 | 0.34 | 0.39 |
| 2 | 236011222 | rs4277473 | -0.22 | 0.634 | -0.642 | 0.501 | -0.001 | 0.007 | 0.007 | 0.475 | 0.67 | 0.63 | 0.71 | 0.68 | 0.63 | 0.69 |
| 2 | 236014138 | rs4663545 | -0.191 | 0.273 | -1.025 | 0.351 | -0.006 | -0.007 | -0.007 | 0 | 0.43 | 0.42 | 0.39 | 0.28 | 0.39 | 0.26 |
| 2 | 236015535 | rs4233619 | 0.839 | 0.462 | -1.12 | 0.373 | 0.005 | 0.037 | 0.037 | 0 | 0.16 | 0.21 | 0.27 | 0.25 | 0.22 | 0.15 |
| 2 | 236018682 | rs6750398 | 0.176 | 0.351 | -1.658 | -0.189 | -0.015 | -0.061 | -0.061 | 0 | 0.34 | 0.45 | 0.31 | 0.17 | 0.22 | 0.38 |
| 2 | 236021615 | rs11692903 | 0.977 | 0.098 | -1.918 | 0.079 | 0.009 | 0.039 | 0.039 | 0.433 | 0.08 | 0.01 | 0.05 | 0.01 | 0.06 | 0.07 |
| 2 | 236023370 | rs4340455 | -0.989 | 0.644 | -1.86 | 0.043 | -0.021 | 0.046 | 0.046 | 0 | 0.68 | 0.82 | 0.67 | 0.85 | 0.71 | 0.63 |
| 2 | 236030089 | rs11680191 | 0.918 | 0.507 | -1.988 | 0.729 | -0.065 | -0.104 | -0.104 | 0 | 0.19 | 0.03 | 0.18 | 0.38 | 0.28 | 0.33 |
| 2 | 236033167 | rs11684900 | 1.44 | 0.475 | -1.933 | 0.768 | -0.041 | -0.07 | -0.07 | 0.399 | 0.16 | 0.03 | 0.18 | 0.31 | 0.22 | 0.29 |
| 2 | 236034085 | rs4381740 | 1.277 | 0.538 | -1.373 | 1 | 0.006 | 0.061 | 0.061 | 0 | 0.15 | 0.33 | 0.28 | 0.19 | 0.17 | 0.11 |
| 2 | 236038689 | rs6741246 | -0.139 | 0.246 | -1.292 | 0.423 | 0.001 | -0.004 | -0.004 | 2.023 | 0.47 | 0.5 | 0.52 | 0.52 | 0.53 | 0.31 |

| | | | | | | | | | | | | | | | | |
|---|---|---|---|---|---|---|---|---|---|---|---|---|---|---|---|---|
| 2 | 236038867 | rs4507050 | -1.077 | 0.165 | -1.325 | 0.407 | -0.013 | 0.041 | 0.041 | 0 | 0.7 | 0.81 | 0.69 | 0.84 | 0.73 | 0.66 |
| 2 | 236039869 | rs3922490 | -0.312 | -0.14 | -0.958 | 0.907 | -0.005 | 0.011 | 0.011 | 0 | 0.4 | 0.45 | 0.44 | 0.43 | 0.38 | 0.22 |
| 2 | 236040005 | rs6711272 | 0.139 | 0.417 | -0.782 | 1.405 | -0.007 | -0.002 | -0.002 | 0 | 0.26 | 0.29 | 0.17 | 0.32 | 0.18 | 0.36 |
| 2 | 236040155 | rs4663185 | 0.068 | 0.411 | -1.35 | 1.317 | 0.011 | 0.042 | 0.042 | 0 | 0.26 | 0.34 | 0.37 | 0.39 | 0.3 | 0.19 |
| 2 | 236040935 | rs4536605 | -0.535 | -0.067 | -1.712 | 1.307 | 0.038 | -0.058 | -0.058 | 0.82 | 0.27 | 0.1 | 0.19 | 0.38 | 0.19 | 0.35 |
| 2 | 236044868 | rs10199636 | -0.68 | 0.356 | -1.084 | 1.456 | 0 | 0.021 | 0.021 | 0 | 0.19 | 0.31 | 0.22 | 0.31 | 0.2 | 0.33 |
| 2 | 236046265 | rs10205899 | -1.17 | 0.15 | -0.686 | 1.761 | 0.002 | -0.004 | -0.004 | 0.016 | 0.37 | 0.68 | 0.27 | 0.46 | 0.33 | 0.38 |
| 2 | 236047007 | rs12478507 | -0.963 | -0.339 | -0.077 | 1.525 | 0.008 | 0.004 | 0.004 | 0 | 0.4 | 0.4 | 0.26 | 0.36 | 0.34 | 0.42 |
| 2 | 236047489 | rs6431343 | 0.54 | 0.003 | -1.134 | 1.301 | 0.05 | 0.001 | 0.001 | 0 | 0.27 | 0.35 | 0.5 | 0.3 | 0.34 | 0.26 |
| 2 | 236052078 | rs11693123 | 0.896 | -0.072 | -1.183 | 1.07 | 0.001 | 0.001 | 0.001 | 0 | 0.17 | 0.17 | 0.09 | 0.13 | 0.15 | 0.1 |
| 2 | 236055911 | rs7605399 | -0.755 | -0.187 | -1.212 | 0.985 | 0.038 | -0.013 | -0.013 | 1.938 | 0.47 | 0.79 | 0.36 | 0.45 | 0.4 | 0.47 |
| 2 | 236058975 | rs6431344 | -1.019 | -0.527 | -1.399 | 0.754 | -0.014 | -0.024 | -0.024 | 0 | 0.71 | 0.97 | 0.8 | 0.78 | 0.72 | 0.68 |
| 2 | 236063063 | rs6746703 | -0.42 | -0.56 | -1.142 | 0.824 | -0.031 | -0.001 | -0.001 | 0 | 0.25 | 0.4 | 0.18 | 0.25 | 0.19 | 0.32 |
| 2 | 236068095 | rs3924658 | -0.318 | -0.267 | -2.058 | 0.562 | 0.061 | 0.021 | 0.021 | 0 | 0.63 | 0.88 | 0.75 | 0.74 | 0.69 | 0.66 |
| 2 | 236070038 | rs13013452 | 0.407 | 0.109 | -2.645 | 0.107 | -0.069 | 0.047 | 0.047 | 0 | 0.34 | 0.05 | 0.26 | 0.19 | 0.33 | 0.36 |
| 2 | 236075756 | rs7594585 | -0.262 | 0.064 | -2.353 | 0.398 | -0.004 | -0.017 | -0.017 | 0.531 | 0.2 | 0.2 | 0.22 | 0.27 | 0.22 | 0.32 |
| 2 | 236079419 | rs3935483 | 0.005 | 0.202 | -2.001 | 0.278 | 0.029 | 0.067 | 0.067 | 0 | 0.5 | 0.3 | 0.42 | 0.34 | 0.41 | 0.54 |
| 2 | 236079843 | rs12464740 | -0.3 | 0.19 | -1.952 | 0.321 | -0.001 | 0.011 | 0.011 | 0 | 0.56 | 0.33 | 0.52 | 0.38 | 0.52 | 0.63 |
| 2 | 236080409 | rs6741321 | 0.412 | 0.054 | -1.921 | 0.266 | 0.005 | -0.01 | -0.01 | 0 | 0.12 | 0.09 | 0.12 | 0.23 | 0.1 | 0.22 |
| 2 | 236080737 | rs13416085 | -0.059 | 0.237 | -0.915 | 0.039 | -0.049 | -0.139 | -0.139 | 0 | 0.25 | 0.47 | 0.18 | 0.02 | 0.16 | 0.16 |
| 2 | 236085432 | rs10221738 | -0.537 | 0.299 | -0.941 | 0.095 | -0.008 | -0.122 | -0.122 | 0.28 | 0.3 | 0.41 | 0.27 | 0.07 | 0.27 | 0.25 |
| 2 | 236090823 | rs4332874 | -0.672 | 0.509 | -0.974 | 0.592 | -0.033 | -0.061 | -0.061 | 1.326 | 0.81 | 0.89 | 0.74 | 0.49 | 0.68 | 0.8 |
| 2 | 236094177 | rs6719254 | -0.613 | 0.341 | -0.102 | 0.138 | 0.031 | 0.017 | 0.017 | 0.164 | 0.3 | 0.56 | 0.44 | 0.64 | 0.44 | 0.31 |
| 2 | 236097368 | rs6741387 | -0.971 | -0.386 | 0.762 | -0.361 | -0.028 | -0.058 | -0.058 | 2.619 | 0.14 | 0.3 | 0.14 | 0.05 | 0.08 | 0.09 |
| 2 | 236097739 | rs6744631 | -0.584 | -0.597 | 0.967 | -0.479 | -0.007 | -0.023 | -0.023 | 0 | 0.21 | 0.3 | 0.15 | 0.07 | 0.14 | 0.15 |
| 2 | 236098533 | rs6748417 | -1.485 | -0.598 | 0.815 | -0.417 | -0.037 | -0.071 | -0.071 | 0 | 0.32 | 0.55 | 0.27 | 0.12 | 0.21 | 0.24 |
| 2 | 236100615 | rs4663552 | -1.025 | -0.629 | 0.467 | -0.402 | 0.033 | 0.073 | 0.073 | 0.185 | 0.27 | 0.17 | 0.23 | 0.08 | 0.18 | 0.22 |
| 2 | 236100852 | rs4663553 | -1.376 | -0.783 | 0.817 | -0.473 | -0.034 | -0.064 | -0.064 | 0.037 | 0.28 | 0.67 | 0.19 | 0.06 | 0.11 | 0.14 |
| 2 | 236100957 | rs10929104 | 1.418 | -0.619 | 0.749 | -0.462 | 0.017 | 0.025 | 0.025 | 1.976 | 0.65 | 0.25 | 0.71 | 0.87 | 0.77 | 0.76 |
| 2 | 236108448 | rs6414036 | 0.111 | -0.631 | 0.847 | -0.391 | 0.003 | -0.027 | -0.027 | 0 | 0.79 | 0.81 | 0.86 | 1 | 0.9 | 0.87 |
| 2 | 236114611 | rs891717 | 0.248 | -0.669 | 1.167 | -0.41 | -0.057 | -0.111 | -0.111 | 0 | 0.78 | 0.63 | 0.85 | 1 | 0.9 | 0.87 |
| 2 | 236115607 | rs4663554 | 1.529 | -0.686 | 1.473 | -0.469 | 0.013 | 0.015 | 0.015 | 3.658 | 0.66 | 0.26 | 0.72 | 0.86 | 0.77 | 0.8 |
| 2 | 236120110 | rs2316033 | 0.059 | -0.693 | 1.382 | -0.541 | -0.052 | -0.086 | -0.086 | 0 | 0.8 | 0.73 | 0.86 | 1 | 0.92 | 0.89 |
| 2 | 236124555 | rs2116397 | 1.164 | -0.723 | 1.728 | -0.565 | -0.049 | -0.089 | -0.089 | 1.182 | 0.72 | 0.55 | 0.71 | 0.92 | 0.81 | 0.84 |
| 2 | 236128951 | rs1991705 | 0.07 | -0.75 | 1.528 | -0.583 | -0.047 | -0.081 | -0.081 | 0.013 | 0.8 | 0.73 | 0.82 | 1 | 0.92 | 0.89 |
| 2 | 236140707 | rs4663557 | 1.526 | -0.611 | 1.495 | -0.442 | 0.033 | 0.057 | 0.057 | 1.573 | 0.63 | 0.25 | 0.69 | 0.81 | 0.73 | 0.78 |
| 2 | 236146526 | rs6431355 | 0.408 | -0.722 | 1.85 | -0.507 | -0.035 | -0.095 | -0.095 | 0 | 0.76 | 0.69 | 0.8 | 1 | 0.87 | 0.86 |
| 2 | 236148186 | rs9678801 | 0.382 | -0.733 | 1.876 | -0.512 | -0.028 | -0.085 | -0.085 | 0.363 | 0.76 | 0.71 | 0.8 | 1 | 0.87 | 0.86 |
| 2 | 236149692 | rs2060126 | 1.194 | -0.671 | 2.233 | -0.526 | 0.032 | 0.039 | 0.039 | 1.302 | 0.69 | 0.26 | 0.74 | 0.88 | 0.82 | 0.81 |
| 2 | 236150508 | rs10173831 | 0.371 | -0.668 | 2.023 | -0.532 | -0.02 | -0.072 | -0.072 | 0 | 0.76 | 0.72 | 0.8 | 1 | 0.87 | 0.84 |
| 2 | 236152504 | rs11682403 | 0.27 | -0.668 | 1.966 | -0.532 | 0.005 | -0.03 | -0.03 | 0 | 0.77 | 0.8 | 0.81 | 1 | 0.87 | 0.86 |
| 2 | 236156105 | rs11687316 | 1.424 | -0.451 | 0.424 | -0.438 | -0.16 | -0.165 | -0.165 | 0.068 | 0.29 | 0.02 | 0.28 | 0.45 | 0.43 | 0.3 |
| 2 | 236166480 | rs7595069 | 0.894 | -0.599 | 0.948 | -0.488 | -0.012 | -0.031 | -0.031 | 0 | 0.66 | 0.39 | 0.73 | 0.88 | 0.76 | 0.81 |
| 2 | 236167636 | rs12052405 | 0.24 | -0.403 | 0.679 | -0.411 | 0.008 | -0.001 | -0.001 | 0 | 0.07 | 0.08 | 0.05 | 0.07 | 0.1 | 0.05 |
| 2 | 236167826 | rs12052330 | 0.76 | -0.383 | 0.258 | -0.399 | -0.032 | -0.018 | -0.018 | 0 | 0.06 | 0.01 | 0.04 | 0.07 | 0.1 | 0.03 |
| 2 | 236172291 | rs10929113 | 0.626 | -0.581 | 0.254 | -0.496 | 0.067 | 0.097 | 0.097 | 1.51 | 0.69 | 0.15 | 0.73 | 0.88 | 0.8 | 0.84 |
| 2 | 236173098 | rs1431793 | 0.42 | -0.636 | 0.851 | -0.556 | -0.016 | -0.06 | -0.06 | 0 | 0.73 | 0.68 | 0.74 | 0.93 | 0.77 | 0.8 |
| 2 | 236174591 | rs4663189 | 1.225 | -0.833 | -0.043 | -0.958 | 0.002 | 0.027 | 0.027 | 0 | 0.2 | 0.12 | 0.25 | 0.14 | 0.18 | 0.29 |
| 2 | 236174661 | rs4663190 | -0.599 | -1.199 | -0.026 | -0.897 | 0.105 | 0.03 | 0.03 | 0.619 | 0.47 | 0.76 | 0.59 | 0.34 | 0.28 | 0.42 |
| 2 | 236174758 | rs4663191 | 0.305 | -0.561 | 0.478 | -1.75 | -0.021 | -0.1 | -0.1 | 0.168 | 0.68 | 0.53 | 0.68 | 0.94 | 0.76 | 0.78 |
| 2 | 236175412 | rs1541888 | 0.587 | -0.788 | 0.784 | -1.609 | -0.034 | -0.023 | -0.023 | 1.956 | 0.25 | 0.48 | 0.29 | 0.2 | 0.2 | 0.34 |
| 2 | 236178354 | rs1816370 | 0.261 | -0.772 | 0.671 | -1.657 | 0.077 | 0.119 | 0.119 | 1.111 | 0.55 | 0.19 | 0.5 | 0.78 | 0.68 | 0.56 |
| 2 | 236179155 | rs1431794 | 1.403 | -0.752 | 0.742 | -1.646 | 0.002 | -0.001 | -0.001 | 0 | 0.2 | 0.17 | 0.26 | 0.21 | 0.19 | 0.29 |
| 2 | 236180188 | rs1431795 | -0.534 | -0.751 | 0.867 | -1.638 | -0.057 | -0.162 | -0.162 | 0.092 | 0.77 | 0.49 | 0.77 | 1 | 0.87 | 0.87 |
| 2 | 236186583 | rs6710167 | 0.233 | -0.505 | 0.599 | -1.555 | 0.024 | 0.067 | 0.067 | 1.136 | 0.5 | 0.4 | 0.44 | 0.72 | 0.56 | 0.51 |
| 2 | 236188646 | rs1897332 | 0.682 | -0.487 | 0.313 | -1.543 | -0.004 | 0.004 | 0.004 | 0 | 0.1 | 0.06 | 0.08 | 0.07 | 0.12 | 0.06 |
| 2 | 236188718 | rs2163565 | -0.523 | -0.479 | 0.258 | -1.546 | -0.003 | -0.031 | -0.031 | 0 | 0.62 | 0.61 | 0.55 | 0.9 | 0.72 | 0.6 |
| 2 | 236189461 | rs11689488 | 0.732 | -0.296 | -0.466 | -1.371 | -0.125 | 0.064 | 0.064 | 0.677 | 0.41 | 0.04 | 0.41 | 0.72 | 0.51 | 0.47 |
| 2 | 236190199 | rs4527164 | 0.416 | -0.178 | -0.311 | -1.391 | 0.005 | 0.075 | 0.075 | 2.355 | 0.46 | 0.27 | 0.43 | 0.76 | 0.54 | 0.51 |
| 2 | 236194197 | rs4663563 | 0.937 | 0.319 | -0.294 | -0.812 | -0.048 | -0.057 | -0.057 | 0 | 0.14 | 0.01 | 0.32 | 0.19 | 0.17 | 0.16 |
| 2 | 236194711 | rs1367882 | 1.239 | 0.138 | 0.56 | -0.71 | 0.015 | 0.136 | 0.136 | 1.386 | 0.6 | 0.17 | 0.76 | 0.86 | 0.65 | 0.76 |
| 2 | 236197888 | rs13422090 | -0.836 | 0.159 | 0.83 | -0.68 | 0.04 | -0.04 | -0.04 | 0 | 0.14 | 0.37 | 0.11 | 0.07 | 0.16 | 0.1 |
| 2 | 236200056 | rs10929123 | 1.301 | 0.149 | 0.797 | -0.705 | 0.006 | 0.017 | 0.017 | 0.982 | 0.61 | 0.33 | 0.78 | 0.86 | 0.65 | 0.75 |
| 2 | 236200453 | rs6718069 | -1.075 | 0.793 | 0.413 | -0.363 | 0.022 | -0.034 | -0.034 | 0 | 0.14 | 0.28 | 0.1 | 0.07 | 0.16 | 0.1 |
| 2 | 236203142 | rs7566072 | 0.649 | 0.591 | 0.199 | -0.316 | -0.006 | -0.085 | -0.085 | 0.303 | 0.74 | 0.42 | 0.85 | 0.93 | 0.76 | 0.82 |
| 2 | 236203747 | rs11901273 | -0.443 | 0.834 | -0.328 | 0.034 | -0.002 | 0.012 | 0.012 | 0 | 0.28 | 0.67 | 0.33 | 0.59 | 0.26 | 0.26 |
| 2 | 236205639 | rs4561612 | 0.624 | 0.807 | -0.283 | -0.2 | 0.002 | -0.006 | -0.006 | 0 | 0.6 | 0.46 | 0.63 | 0.35 | 0.6 | 0.54 |
| 2 | 236206703 | rs7340415 | -0.959 | 0.847 | -0.682 | -0.359 | -0.014 | 0.034 | 0.034 | 0 | 0.39 | 0.73 | 0.42 | 0.7 | 0.4 | 0.38 |
| 2 | 236208067 | rs11691751 | -0.59 | 0.832 | -0.445 | -0.681 | 0.009 | -0.058 | -0.058 | 0 | 0.37 | 0.47 | 0.4 | 0.18 | 0.42 | 0.29 |

| | | | | | | | | | | | | | | | | |
|---|---|---|---|---|---|---|---|---|---|---|---|---|---|---|---|---|
| 2 | 236209918 | rs13019316 | -0.523 | 0.744 | -0.621 | -0.843 | -0.01 | 0.054 | 0.054 | 0 | 0.41 | 0.29 | 0.4 | 0.17 | 0.43 | 0.41 |
| 2 | 236212004 | rs1367876 | 0.413 | 0.844 | -0.889 | -0.111 | -0.011 | -0.02 | -0.02 | 0.173 | 0.17 | 0.11 | 0.19 | 0.55 | 0.22 | 0.21 |
| 2 | 236212448 | rs2316019 | -1.656 | 0.391 | -1.607 | -0.691 | -0.052 | 0.172 | 0.172 | 2.778 | 0.66 | 0.99 | 0.75 | 0.91 | 0.7 | 0.68 |
| 2 | 236214007 | rs13020554 | 0.03 | 0.687 | -0.989 | -0.44 | 0 | -0.001 | -0.001 | 0 | 0.24 | 0.22 | 0.37 | 0.17 | 0.28 | 0.19 |
| 2 | 236214712 | rs16772 | -1.305 | 0.607 | -0.515 | -0.424 | 0.014 | 0.048 | 0.048 | 4.626 | 0.51 | 0.6 | 0.63 | 0.78 | 0.61 | 0.57 |
| 2 | 236218851 | rs2116399 | -0.754 | 0.78 | -0.754 | -0.177 | -0.022 | 0.037 | 0.037 | 1.67 | 0.4 | 0.3 | 0.55 | 0.72 | 0.53 | 0.43 |
| 2 | 236224692 | rs12620850 | 0.416 | 0.814 | -0.875 | 0.012 | -0.04 | -0.085 | -0.085 | 0.011 | 0.18 | 0.06 | 0.19 | 0.55 | 0.23 | 0.27 |
| 2 | 236228143 | rs2163566 | -1.016 | 0.713 | -0.624 | -0.228 | 0.003 | -0.036 | -0.036 | 0 | 0.44 | 0.52 | 0.59 | 0.75 | 0.54 | 0.46 |
| 2 | 236228570 | rs1864713 | -0.219 | 0.646 | -1.017 | 0.097 | -0.017 | 0.007 | 0.007 | 0.268 | 0.21 | 0.13 | 0.36 | 0.17 | 0.27 | 0.16 |
| 2 | 236234442 | rs10929124 | -0.194 | 0.764 | -1.17 | -0.042 | -0.105 | 0.017 | 0.017 | 0 | 0.23 | 0.02 | 0.45 | 0.15 | 0.33 | 0.24 |
| 2 | 236236199 | rs10803637 | -1.126 | 0.306 | -0.834 | -0.187 | 0.009 | 0.017 | 0.017 | 0 | 0.63 | 0.74 | 0.81 | 0.75 | 0.69 | 0.69 |
| 2 | 236241723 | rs10929125 | -0.205 | 0.27 | -0.848 | -0.102 | -0.042 | 0.044 | 0.044 | 0 | 0.27 | 0.18 | 0.45 | 0.18 | 0.35 | 0.27 |
| 2 | 236242266 | rs4381737 | -1.227 | 0.013 | -0.245 | -0.031 | 0.003 | -0.017 | -0.017 | 0 | 0.38 | 0.49 | 0.54 | 0.29 | 0.4 | 0.46 |
| 2 | 236242437 | rs6730368 | 0.35 | 0.058 | -0.518 | -0.065 | -0.055 | 0.051 | 0.051 | 0 | 0.24 | 0.12 | 0.42 | 0.15 | 0.33 | 0.25 |
| 2 | 236244276 | rs2217538 | 0.165 | 0.101 | -0.513 | -0.048 | -0.068 | 0.04 | 0.04 | 0 | 0.26 | 0.12 | 0.45 | 0.18 | 0.36 | 0.27 |
| 2 | 236248116 | rs4444469 | -0.505 | 0.05 | -0.427 | -0.066 | -0.029 | 0.051 | 0.051 | 0 | 0.32 | 0.21 | 0.5 | 0.2 | 0.38 | 0.37 |
| 2 | 236248940 | rs6728599 | -1.448 | -0.054 | -0.64 | -0.066 | -0.1 | -0.178 | -0.178 | 0 | 0.75 | 0.98 | 0.74 | 0.48 | 0.7 | 0.76 |
| 2 | 236250584 | rs4663195 | -1.324 | -0.024 | -0.26 | -0.056 | -0.01 | -0.033 | -0.033 | 0 | 0.74 | 0.82 | 0.72 | 0.48 | 0.7 | 0.75 |
| 2 | 236258449 | rs12692166 | 0.3 | -0.445 | -0.183 | -0.136 | 0.014 | 0.017 | 0.017 | 0 | 0.45 | 0.52 | 0.27 | 0.26 | 0.34 | 0.42 |
| 2 | 236261524 | rs11680844 | 0.883 | -0.243 | -0.625 | -0.232 | 0.015 | 0.023 | 0.023 | 0 | 0.36 | 0.26 | 0.18 | 0.2 | 0.29 | 0.3 |
| 2 | 236265496 | rs13386664 | 0.859 | -0.06 | -1.307 | -0.097 | 0.003 | 0.007 | 0.007 | 0 | 0.28 | 0.31 | 0.3 | 0.54 | 0.35 | 0.27 |
| 2 | 236267189 | rs7597280 | -0.821 | 0.2 | -1.726 | -0.467 | 0.01 | 0.038 | 0.038 | 0 | 0.67 | 0.93 | 0.84 | 0.8 | 0.72 | 0.71 |
| 2 | 236270060 | rs4663576 | -0.424 | 0.263 | -1.037 | 0.113 | -0.074 | 0.132 | 0.132 | 0 | 0.53 | 0.91 | 0.51 | 0.77 | 0.49 | 0.59 |
| 2 | 236276440 | rs1878393 | -0.238 | 0.674 | -0.757 | 0.344 | -0.002 | 0.019 | 0.019 | 0 | 0.45 | 0.6 | 0.42 | 0.75 | 0.46 | 0.48 |
| 2 | 236282912 | rs10803639 | -0.782 | 0.816 | -0.495 | 0.774 | 0.002 | 0.007 | 0.007 | 0 | 0.21 | 0.33 | 0.28 | 0.3 | 0.24 | 0.24 |
| 2 | 236287266 | rs10190128 | -1.162 | 0.497 | 0.486 | 0.52 | 0.014 | 0.022 | 0.022 | 0 | 0.23 | 0.62 | 0.31 | 0.3 | 0.26 | 0.31 |
| 2 | 236301375 | rs10201495 | -1.212 | 0.025 | -0.851 | 0.205 | -0.127 | -0.09 | -0.09 | 0.011 | 0.58 | 0.95 | 0.56 | 0.47 | 0.5 | 0.63 |
| 2 | 236301786 | rs10196107 | -1.158 | 0.015 | 0.187 | 0.262 | -0.002 | -0.003 | -0.003 | 0 | 0.29 | 0.71 | 0.32 | 0.29 | 0.27 | 0.35 |
| 2 | 236312290 | rs7578935 | -0.913 | 0.667 | 0.505 | 0.297 | 0.008 | 0.041 | 0.041 | 0 | 0.19 | 0.34 | 0.11 | 0.31 | 0.17 | 0.24 |
| 2 | 236320610 | rs6431373 | -0.435 | 0.974 | 0.562 | 0.215 | 0.019 | 0.019 | 0.019 | 0 | 0.14 | 0.17 | 0.09 | 0.27 | 0.16 | 0.24 |
| 2 | 236322756 | rs11681481 | -0.724 | 0.815 | 0.739 | -0.046 | 0.044 | -0.078 | -0.078 | 0 | 0.17 | 0.43 | 0.17 | 0.03 | 0.19 | 0.15 |
| 2 | 236327921 | rs4663581 | 0.57 | 0.153 | 0.511 | -0.054 | 0.016 | 0.02 | 0.02 | 0 | 0.3 | 0.22 | 0.28 | 0.18 | 0.21 | 0.29 |
| 2 | 236336402 | rs13428454 | -1.34 | 0.292 | 0.748 | 0.042 | 0.048 | 0.006 | 0.006 | 0.675 | 0.3 | 0.57 | 0.27 | 0.34 | 0.37 | 0.39 |
| 2 | 236340067 | rs11691754 | -0.73 | 0.407 | 0.295 | 0.064 | -0.087 | -0.034 | -0.034 | 0 | 0.25 | 0.14 | 0.24 | 0.32 | 0.38 | 0.35 |
| 2 | 236354344 | rs6726521 | -1.097 | 0.1 | 0.372 | -0.109 | 0.019 | -0.011 | -0.011 | 0 | 0.33 | 0.51 | 0.34 | 0.31 | 0.39 | 0.33 |
| 2 | 236355415 | rs7607788 | 0.413 | -0.19 | 0.014 | -0.283 | -0.005 | -0.002 | -0.002 | 0 | 0.25 | 0.23 | 0.21 | 0.21 | 0.19 | 0.25 |
| 2 | 236357372 | rs12469570 | 0.591 | -0.277 | 0.077 | -0.344 | 0.021 | -0.031 | -0.031 | 0 | 0.68 | 0.39 | 0.64 | 0.77 | 0.6 | 0.66 |
| 2 | 236358786 | rs2316426 | 0.038 | -0.134 | 0.1 | -0.418 | 0.037 | -0.045 | -0.045 | 0.475 | 0.21 | 0.45 | 0.12 | 0.18 | 0.22 | 0.18 |
| 2 | 236361738 | rs7355273 | 0.845 | -0.047 | -0.601 | -0.404 | -0.002 | -0.001 | -0.001 | 0 | 0.66 | 0.3 | 0.64 | 0.77 | 0.6 | 0.66 |
| 2 | 236363806 | rs4663587 | 0.062 | -0.287 | -0.448 | -0.217 | -0.049 | 0.035 | 0.035 | 0 | 0.91 | 0.99 | 0.78 | 0.99 | 0.82 | 0.89 |
| 2 | 236367367 | rs12474898 | -0.239 | -0.765 | -0.134 | -0.03 | 0.008 | 0.003 | 0.003 | 1.825 | 0.23 | 0.19 | 0.21 | 0.21 | 0.17 | 0.23 |
| 2 | 236369719 | rs10929133 | -0.255 | -0.77 | -0.21 | -0.033 | 0.002 | 0.001 | 0.001 | 1.927 | 0.23 | 0.24 | 0.21 | 0.21 | 0.17 | 0.23 |
| 2 | 236377559 | rs4663588 | 0.285 | -0.296 | -0.348 | -0.556 | -0.004 | -0.095 | -0.095 | 1.973 | 0.64 | 0.62 | 0.7 | 0.93 | 0.67 | 0.71 |
| 2 | 236387136 | rs4663591 | 0.356 | -0.474 | -0.463 | -0.211 | -0.029 | 0.043 | 0.043 | 0 | 0.23 | 0.44 | 0.23 | 0.24 | 0.16 | 0.18 |
| 2 | 236394141 | rs1464158 | 0.801 | -0.622 | -0.713 | -0.389 | 0.003 | 0.029 | 0.029 | 0 | 0.56 | 0.31 | 0.5 | 0.69 | 0.56 | 0.6 |
| 2 | 236400824 | rs1071983 | -0.321 | -0.197 | -1.12 | -0.179 | -0.05 | 0.074 | 0.074 | 0.128 | 0.79 | 0.93 | 0.76 | 0.93 | 0.73 | 0.77 |
| 2 | 236401204 | rs4663596 | -0.608 | 0.131 | -1.05 | -0.02 | -0.041 | 0.019 | 0.019 | 0 | 0.93 | 0.99 | 0.9 | 0.99 | 0.84 | 0.96 |
| 2 | 236401474 | rs34568733 | 0.738 | -0.148 | -0.861 | 0.027 | -0.027 | 0.022 | 0.022 | 0 | 0.43 | 0.3 | 0.48 | 0.69 | 0.51 | 0.53 |
| 2 | 236410779 | rs10204812 | 0.039 | -0.149 | -0.529 | 0.453 | 0 | -0.001 | -0.001 | 0 | 0.69 | 0.73 | 0.75 | 0.65 | 0.68 | 0.7 |
| 2 | 236416245 | rs885539 | 0.257 | -0.354 | 0.018 | 0.512 | 0.001 | -0.004 | -0.004 | 0 | 0.65 | 0.43 | 0.7 | 0.68 | 0.65 | 0.72 |
| 2 | 236422975 | rs7559293 | 1.212 | -0.393 | 0.263 | 0.528 | -0.035 | -0.012 | -0.012 | 0 | 0.4 | 0.25 | 0.56 | 0.42 | 0.49 | 0.51 |
| 2 | 236424382 | rs3795897 | 0.679 | -0.445 | 0.434 | 0.425 | -0.016 | 0.024 | 0.024 | 0.038 | 0.17 | 0.25 | 0.15 | 0.23 | 0.13 | 0.14 |
| 2 | 236429510 | rs4663604 | -0.865 | -0.204 | 0.281 | 0.362 | -0.032 | 0.013 | 0.013 | 0 | 0.95 | 0.99 | 0.9 | 0.99 | 0.84 | 0.97 |
| 2 | 236430846 | rs6731720 | 0.142 | -0.219 | 0.404 | 0.367 | 0.077 | 0.029 | 0.029 | 0 | 0.92 | 0.75 | 0.88 | 0.91 | 0.84 | 0.9 |
| 2 | 236430910 | rs10460277 | 0.337 | -0.262 | 0.452 | 0.362 | 0.037 | -0.054 | -0.054 | 0 | 0.81 | 0.58 | 0.85 | 0.91 | 0.79 | 0.85 |
| 2 | 236435574 | rs2316436 | 0.529 | -0.241 | 0.084 | 0.465 | 0.003 | 0.016 | 0.016 | 0 | 0.58 | 0.64 | 0.72 | 0.65 | 0.62 | 0.65 |
| 2 | 236447354 | rs7569111 | 0.546 | 0.284 | 0.057 | 0.719 | 0.017 | 0 | 0 | 0 | 0.88 | 0.85 | 0.79 | 0.86 | 0.74 | 0.77 |
| 2 | 236452412 | rs6717219 | -0.904 | 0.091 | 0.664 | 0.822 | 0.046 | 0.102 | 0.102 | 0.241 | 0.06 | 0.24 | 0.1 | 0.18 | 0.07 | 0.19 |
| 2 | 236457888 | rs12475206 | -0.305 | -0.379 | 0.767 | 0.7 | -0.008 | 0.006 | 0.006 | 0 | 0.3 | 0.34 | 0.22 | 0.46 | 0.2 | 0.45 |
| 2 | 236464232 | rs13431390 | 0.225 | -0.449 | 0.697 | 0.659 | -0.04 | 0.012 | 0.012 | 0 | 0.26 | 0.35 | 0.15 | 0.32 | 0.13 | 0.26 |
| 2 | 236468798 | rs4663609 | 0.885 | 0.117 | 0.39 | 0.749 | 0.056 | 0.027 | 0.027 | 2.202 | 0.83 | 0.41 | 0.76 | 0.74 | 0.74 | 0.71 |
| 2 | 236471151 | rs6718006 | 0.42 | -0.188 | -0.211 | 0.614 | -0.043 | 0.006 | 0.006 | 0 | 0.4 | 0.25 | 0.54 | 0.33 | 0.45 | 0.32 |
| 2 | 236476847 | rs10208267 | -0.801 | -0.447 | 0.461 | 0.975 | -0.017 | 0.271 | 0.271 | 1.798 | 0.06 | 0.24 | 0.02 | 0.34 | 0 | 0.11 |
| 2 | 236478656 | rs12151399 | -0.035 | -0.488 | 0.106 | 0.35 | -0.007 | -0.032 | -0.032 | 0 | 0.47 | 0.47 | 0.26 | 0.16 | 0.35 | 0.45 |
| 2 | 236478926 | rs4571013 | -1.016 | 0.095 | -0.263 | 0.324 | -0.166 | -0.141 | -0.141 | 0 | 0.63 | 0.96 | 0.46 | 0.51 | 0.52 | 0.58 |
| 2 | 236484484 | rs13385267 | -0.296 | 0.04 | -0.075 | 0.348 | 0.001 | 0.31 | 0.31 | 1.859 | 0.07 | 0.31 | 0.03 | 0.33 | 0 | 0.11 |
| 2 | 236492988 | rs6431389 | 0.061 | 0.092 | 0.134 | 0.12 | 0.006 | 0 | 0 | 0 | 0.5 | 0.45 | 0.36 | 0.17 | 0.38 | 0.45 |
| 2 | 236494162 | rs7562244 | -0.781 | 0.512 | 0.467 | 0.616 | 0.11 | -0.009 | -0.009 | 0 | 0.07 | 0.37 | 0.08 | 0 | 0.14 | 0.02 |
| 2 | 236504782 | rs7595184 | 0.639 | 0.363 | -0.378 | 0.584 | -0.009 | 0.007 | 0.007 | 0 | 0.16 | 0.18 | 0.09 | 0.21 | 0.1 | 0.31 |

| | | | | | | | | | | | | | | | | |
|---|---|---|---|---|---|---|---|---|---|---|---|---|---|---|---|---|
| 2 | 236508275 | rs11675035 | -1.242 | 0.298 | -0.823 | 0.663 | -0.001 | -0.108 | -0.108 | 0 | 0.33 | 0.85 | 0.22 | 0.56 | 0.27 | 0.44 |
| 2 | 236508340 | rs6731727 | -1.318 | 0.679 | -0.457 | 0.873 | 0.029 | 0.123 | 0.123 | 0.568 | 0.17 | 0.57 | 0.13 | 0.35 | 0.17 | 0.13 |
| 2 | 236514769 | rs11676226 | 0.213 | 1.043 | -1.921 | 0.249 | -0.037 | 0.027 | 0.027 | 0 | 0.6 | 0.82 | 0.46 | 0.34 | 0.57 | 0.56 |
| 2 | 236521330 | rs6431390 | 0.626 | 1.164 | -1.835 | 0.125 | -0.041 | -0.022 | -0.022 | 0.084 | 0.9 | 0.99 | 0.71 | 0.88 | 0.88 | 0.89 |
| 2 | 236522161 | rs7607661 | -0.426 | 1.092 | -1.644 | -0.453 | -0.003 | -0.01 | -0.01 | 0 | 0.22 | 0.18 | 0.19 | 0 | 0.27 | 0.11 |
| 2 | 236525826 | rs7571980 | -0.146 | 1.522 | -1.971 | -0.033 | -0.069 | -0.029 | -0.029 | 0 | 0.8 | 0.96 | 0.74 | 0.81 | 0.73 | 0.73 |
| 2 | 236532308 | rs13411636 | -0.036 | 1.163 | -1.703 | -0.287 | -0.001 | -0.024 | -0.024 | 0 | 0.17 | 0.14 | 0.13 | 0 | 0.19 | 0.12 |
| 2 | 236543711 | rs10177178 | 0.462 | 0.501 | -1.391 | -0.148 | -0.003 | -0.007 | -0.007 | 0.046 | 0.34 | 0.33 | 0.31 | 0.4 | 0.34 | 0.52 |
| 2 | 236544481 | rs11680070 | 0.592 | 0.612 | -1.496 | -0.239 | 0.001 | 0.01 | 0.01 | 0 | 0.28 | 0.33 | 0.28 | 0.4 | 0.27 | 0.51 |
| 2 | 236546636 | rs12328363 | -0.748 | 0.369 | -1.05 | -0.316 | -0.014 | 0 | 0 | 0 | 0.77 | 0.85 | 0.7 | 0.81 | 0.69 | 0.74 |
| 2 | 236547496 | rs2316908 | -0.108 | -0.071 | -1.147 | -0.278 | -0.012 | 0 | 0 | 0.198 | 0.69 | 0.23 | 0.64 | 0.68 | 0.67 | 0.62 |
| 2 | 236550704 | rs2103279 | -0.797 | 0.043 | -0.823 | -0.249 | 0.015 | -0.002 | -0.002 | 1.499 | 0.77 | 0.74 | 0.68 | 0.8 | 0.69 | 0.74 |
| 2 | 236551136 | rs2103280 | -0.602 | 0.052 | -0.545 | -0.267 | 0.048 | 0.01 | 0.01 | 0 | 0.75 | 0.57 | 0.67 | 0.75 | 0.69 | 0.68 |
| 2 | 236553171 | rs12991774 | -0.006 | 0.047 | -0.755 | -0.276 | 0.048 | 0.049 | 0.049 | 0.004 | 0.48 | 0.32 | 0.4 | 0.4 | 0.41 | 0.23 |
| 2 | 236560230 | rs12474016 | -0.411 | 0.364 | -0.912 | -0.251 | -0.027 | 0.008 | 0.008 | 1.688 | 0.76 | 0.86 | 0.77 | 0.81 | 0.73 | 0.7 |
| 2 | 236566346 | rs2873308 | -0.022 | -0.213 | -0.468 | -0.511 | -0.029 | -0.114 | -0.114 | 1.012 | 0.27 | 0.41 | 0.21 | 0.03 | 0.27 | 0.07 |
| 2 | 236587761 | rs12472001 | 0.459 | -0.262 | -1.083 | -0.304 | 0.021 | -0.022 | -0.022 | 15.939 | 0.27 | 0.14 | 0.38 | 0.32 | 0.2 | 0.08 |
| 2 | 236588800 | rs10929146 | 0.506 | -0.228 | -0.716 | -0.195 | -0.005 | 0.004 | 0.004 | 0 | 0.26 | 0.27 | 0.38 | 0.39 | 0.19 | 0.18 |
| 2 | 236590828 | rs4233623 | -0.782 | 0.425 | -1.212 | -0.111 | -0.136 | -0.039 | -0.039 | 0 | 0.76 | 0.97 | 0.76 | 0.81 | 0.67 | 0.64 |
| 2 | 236607924 | rs1881191 | -0.57 | 0.734 | -0.904 | -0.089 | -0.043 | 0.009 | 0.009 | 1.078 | 0.73 | 0.85 | 0.72 | 0.77 | 0.63 | 0.54 |
| 2 | 236611263 | rs10166654 | 1.335 | 0.351 | -0.601 | 0.31 | 0.002 | -0.023 | -0.023 | 0 | 0.2 | 0.14 | 0.13 | 0.42 | 0.16 | 0.37 |
| 2 | 236617595 | rs3754670 | 0.894 | 0.308 | -0.476 | 0.314 | -0.016 | 0.115 | 0.115 | 0 | 0.23 | 0.47 | 0.15 | 0.42 | 0.17 | 0.37 |
| 2 | 236617736 | rs3754669 | 0.813 | 0.27 | -1.55 | 0.292 | -0.002 | -0.095 | -0.095 | 0 | 0.21 | 0.03 | 0.32 | 0.28 | 0.14 | 0.07 |
| 2 | 236618226 | rs10190094 | 0.824 | 0.733 | -1.221 | 0.395 | -0.021 | -0.044 | -0.044 | 0 | 0.22 | 0.13 | 0.18 | 0.42 | 0.26 | 0.38 |
| 2 | 236618425 | rs3820779 | -0.06 | 0.717 | -1.585 | 0.448 | -0.007 | -0.015 | -0.015 | 0 | 0.56 | 0.7 | 0.57 | 0.54 | 0.44 | 0.54 |
| 2 | 236618642 | rs1568069 | 0.115 | 0.339 | -1.271 | -0.002 | -0.011 | -0.021 | -0.021 | 0.045 | 0.37 | 0.57 | 0.52 | 0.76 | 0.32 | 0.53 |
| 2 | 236619385 | rs3768969 | -0.13 | 0.798 | -1.606 | 0.07 | 0.032 | 0.033 | 0.033 | 0 | 0.5 | 0.8 | 0.35 | 0.46 | 0.39 | 0.64 |
| 2 | 236627544 | rs1728289 | -0.182 | 0.918 | -0.748 | -0.568 | -0.006 | -0.128 | -0.128 | 0 | 0.76 | 0.62 | 0.83 | 0.99 | 0.78 | 0.8 |
| 2 | 236627892 | rs3768968 | 1.327 | 0.921 | -1.319 | -0.442 | 0.015 | -0.087 | -0.087 | 0.466 | 0.29 | 0.08 | 0.23 | 0.46 | 0.24 | 0.33 |
| 2 | 236631040 | rs2675124 | 1.137 | 1.094 | -1.705 | -0.486 | -0.018 | -0.115 | -0.115 | 3.175 | 0.17 | 0.02 | 0.39 | 0.37 | 0.18 | 0.09 |
| 2 | 236641479 | rs2675144 | -0.305 | 1.024 | -1.633 | -0.536 | -0.113 | -0.133 | -0.133 | 0.439 | 0.41 | 0.07 | 0.56 | 0.52 | 0.53 | 0.42 |
| 2 | 236655152 | rs2696398 | -0.618 | 1.261 | -0.647 | -0.836 | 0.002 | -0.15 | -0.15 | 0.515 | 0.75 | 0.54 | 0.82 | 0.99 | 0.77 | 0.8 |
| 2 | 236655526 | rs3738996 | 0.121 | 1.582 | -0.978 | -0.329 | -0.118 | 0.041 | 0.041 | 0.222 | 0.23 | 0.04 | 0.17 | 0.15 | 0.34 | 0.31 |
| 2 | 236670936 | rs6715112 | 0.248 | 1.396 | -0.398 | -0.06 | 0.005 | 0.014 | 0.014 | 0 | 0.21 | 0.27 | 0.41 | 0.34 | 0.25 | 0.1 |
| 2 | 236679586 | rs1710822 | 0.208 | 1.359 | -0.213 | -0.141 | 0.007 | 0.022 | 0.022 | 0 | 0.21 | 0.3 | 0.42 | 0.32 | 0.25 | 0.1 |
| 2 | 236691982 | rs1878154 | 0.376 | 1.338 | -0.038 | -0.151 | -0.025 | -0.043 | -0.043 | 0 | 0.19 | 0.12 | 0.4 | 0.32 | 0.25 | 0.1 |
| 2 | 236696691 | rs12472150 | 0.268 | 1.316 | 0.017 | -0.162 | -0.002 | -0.002 | -0.002 | 0.528 | 0.2 | 0.19 | 0.41 | 0.32 | 0.25 | 0.1 |
| 2 | 236700821 | rs12471567 | 0.248 | 0.795 | -0.052 | -0.683 | -0.01 | 0.024 | 0.024 | 0 | 0.57 | 0.36 | 0.69 | 0.83 | 0.5 | 0.54 |
| 2 | 236701122 | rs11685098 | -0.82 | 0.74 | 0.079 | -0.706 | -0.013 | -0.106 | -0.106 | 0 | 0.8 | 0.64 | 0.85 | 0.96 | 0.8 | 0.82 |
| 2 | 236708166 | rs2292708 | 0.77 | 1.069 | 0.011 | -0.432 | -0.1 | -0.08 | -0.08 | 0 | 0.19 | 0.03 | 0.41 | 0.27 | 0.26 | 0.1 |
| 2 | 236709123 | rs6760477 | -1.081 | 0.99 | 0.047 | -0.492 | -0.007 | -0.003 | -0.003 | 0 | 0.44 | 0.48 | 0.6 | 0.45 | 0.56 | 0.39 |
| 2 | 236717001 | rs7565206 | 0.301 | 1.053 | -0.094 | -0.549 | -0.052 | 0.05 | 0.05 | 0.074 | 0.21 | 0.12 | 0.16 | 0.12 | 0.3 | 0.28 |
| 2 | 236731624 | rs952608 | 0.562 | 0.972 | -0.219 | -0.484 | 0.008 | 0.007 | 0.007 | 0 | 0.21 | 0.25 | 0.43 | 0.31 | 0.27 | 0.11 |
| 2 | 236734213 | rs883521 | -0.719 | 0.777 | -0.425 | -0.444 | -0.007 | -0.006 | -0.006 | 0 | 0.45 | 0.5 | 0.61 | 0.43 | 0.57 | 0.39 |
| 2 | 236737133 | rs12478435 | 0.082 | 0.692 | -0.863 | -0.589 | -0.02 | 0.072 | 0.072 | 0.068 | 0.74 | 0.22 | 0.79 | 0.94 | 0.72 | 0.78 |
| 2 | 236738151 | rs12463531 | 0.89 | 0.643 | -1.139 | -0.504 | -0.074 | 0.149 | 0.149 | 0 | 0.58 | 0.14 | 0.65 | 0.78 | 0.52 | 0.52 |
| 2 | 236762851 | rs4625852 | -0.084 | 0.712 | -0.471 | 0.215 | -0.024 | -0.035 | -0.035 | 0 | 0.14 | 0.03 | 0.13 | 0.17 | 0.2 | 0.23 |
| 2 | 236793864 | rs13017704 | 0.64 | 0.596 | 0.811 | 0.147 | 0.036 | 0.007 | 0.007 | 0.031 | 0.47 | 0.27 | 0.37 | 0.54 | 0.36 | 0.67 |
| 2 | 236796548 | rs4663627 | 0.68 | 0.58 | 0.774 | 0.034 | 0.052 | 0.017 | 0.017 | 0.076 | 0.47 | 0.14 | 0.36 | 0.58 | 0.36 | 0.67 |
| 2 | 236801872 | rs4663629 | -0.203 | 1.044 | 0.996 | -0.17 | 0.005 | -0.057 | -0.057 | 0 | 0.79 | 0.73 | 0.85 | 0.99 | 0.75 | 0.91 |
| 2 | 236806685 | rs2873334 | 0.095 | 1.035 | 0.895 | -0.17 | 0.02 | -0.112 | -0.112 | 0 | 0.77 | 0.61 | 0.83 | 0.95 | 0.74 | 0.8 |
| 2 | 236827369 | rs11690665 | -0.422 | 0.996 | 0.837 | 0.53 | 0.004 | 0.005 | 0.005 | 0 | 0.28 | 0.42 | 0.48 | 0.4 | 0.37 | 0.13 |
| 2 | 236832074 | rs12471554 | -0.106 | 1.012 | 0.078 | 0.516 | -0.09 | -0.097 | -0.097 | 0 | 0.25 | 0.08 | 0.45 | 0.38 | 0.36 | 0.13 |
| 2 | 236832573 | rs7571708 | -0.618 | 1.012 | 0.248 | 0.528 | -0.005 | -0.005 | -0.005 | 0 | 0.32 | 0.75 | 0.5 | 0.4 | 0.38 | 0.24 |
| 2 | 236834160 | rs1962299 | -0.47 | 1.182 | -0.19 | 0.611 | 0.032 | 0.035 | 0.035 | 0 | 0.29 | 0.47 | 0.48 | 0.4 | 0.38 | 0.24 |
| 2 | 236837395 | rs12162297 | 0.085 | 1.04 | -1.324 | 0.746 | -0.092 | -0.097 | -0.097 | 0.05 | 0.34 | 0.1 | 0.36 | 0.43 | 0.37 | 0.41 |
| 2 | 236838338 | rs10211191 | -0.155 | 0.414 | -0.769 | 0.337 | -0.031 | -0.025 | -0.025 | 0 | 0.26 | 0.39 | 0.15 | 0.15 | 0.19 | 0.42 |
| 2 | 236844942 | rs4663634 | -0.355 | 0.772 | -0.997 | 0.567 | -0.121 | -0.117 | -0.117 | 3.131 | 0.66 | 0.93 | 0.55 | 0.49 | 0.54 | 0.82 |
| 2 | 236850240 | rs1962443 | -0.454 | 0.754 | -0.847 | 0.553 | -0.088 | -0.085 | -0.085 | 0 | 0.65 | 0.88 | 0.54 | 0.48 | 0.54 | 0.78 |
| 2 | 236859195 | rs13418964 | -0.34 | 0.74 | -0.757 | 0.534 | -0.079 | -0.071 | -0.071 | 0 | 0.66 | 0.88 | 0.54 | 0.49 | 0.53 | 0.79 |
| 2 | 236872445 | rs6717245 | -0.832 | 0.418 | 0.409 | 0.478 | 0.106 | 0.071 | 0.071 | 0.259 | 0.05 | 0.43 | 0.04 | 0 | 0.02 | 0.11 |
| 2 | 236877473 | rs13428207 | -1.097 | 0.39 | 0.56 | 0.471 | 0.089 | 0.061 | 0.061 | 0.54 | 0.05 | 0.47 | 0.04 | 0 | 0.02 | 0.11 |
| 2 | 236896521 | rs1453117 | -0.277 | 0.666 | 0.393 | 0.459 | 0.003 | 0.004 | 0.004 | 0.345 | 0.64 | 0.59 | 0.53 | 0.5 | 0.56 | 0.71 |
| 2 | 236898903 | rs10182721 | -0.27 | 0.659 | 0.39 | 0.456 | 0.003 | 0.004 | 0.004 | 0.261 | 0.64 | 0.59 | 0.53 | 0.5 | 0.56 | 0.7 |
| 2 | 236905308 | rs1530940 | 0.201 | 0.636 | 0.08 | 0.433 | 0.003 | 0 | 0 | 0 | 0.36 | 0.4 | 0.22 | 0.32 | 0.31 | 0.43 |
| 2 | 236915245 | rs1377377 | 0.457 | 0.199 | 0.09 | 0.304 | -0.016 | -0.006 | -0.006 | 0 | 0.24 | 0.29 | 0.17 | 0.19 | 0.19 | 0.3 |
| 2 | 236920673 | rs7598567 | -0.916 | 0.396 | -0.806 | 0.321 | -0.04 | -0.039 | -0.039 | 0 | 0.71 | 0.84 | 0.45 | 0.52 | 0.6 | 0.83 |
| 2 | 236922737 | rs7420415 | 0.125 | 0.343 | -0.743 | 0.296 | -0.004 | 0.015 | 0.015 | 0 | 0.46 | 0.29 | 0.26 | 0.34 | 0.42 | 0.53 |
| 2 | 236926347 | rs7565059 | -0.829 | 0.481 | -1.101 | 0.415 | -0.056 | -0.053 | -0.053 | 0 | 0.71 | 0.87 | 0.45 | 0.52 | 0.61 | 0.83 |

| | | | | | | | | | | | | | | | | |
|---|---|---|---|---|---|---|---|---|---|---|---|---|---|---|---|---|
| 2 | 236946552 | rs13024834 | 0.368 | 0.294 | -1.588 | 0.562 | -0.071 | -0.098 | -0.098 | 0 | 0.35 | 0.07 | 0.18 | 0.36 | 0.3 | 0.49 |
| 2 | 236949119 | rs6431421 | -0.239 | -0.036 | -0.595 | 0.452 | -0.065 | -0.033 | -0.033 | 0.901 | 0.25 | 0.48 | 0.18 | 0.16 | 0.19 | 0.28 |
| 2 | 236951982 | rs6739594 | 0.009 | 0.972 | -1.468 | 0.593 | -0.057 | 0.015 | 0.015 | 1.058 | 0.73 | 0.91 | 0.73 | 0.74 | 0.66 | 0.6 |
| 2 | 236954470 | rs3768951 | -0.732 | 0.334 | -0.909 | 1.201 | -0.013 | 0.04 | 0.04 | 0 | 0.09 | 0.16 | 0.13 | 0.19 | 0.06 | 0.1 |
| 2 | 236956749 | rs3768948 | -0.657 | 0.387 | -1.031 | 1.249 | 0.007 | -0.025 | -0.025 | 0 | 0.08 | 0.05 | 0.11 | 0.18 | 0.07 | 0.1 |
| 2 | 236958097 | rs13398226 | -0.011 | -0.011 | -0.614 | 1.267 | 0.004 | -0.005 | -0.005 | 0 | 0.37 | 0.34 | 0.52 | 0.49 | 0.35 | 0.38 |
| 2 | 236958688 | rs7587505 | -0.665 | -0.154 | -1.498 | 0.686 | -0.035 | 0.045 | 0.045 | 0 | 0.55 | 0.93 | 0.6 | 0.64 | 0.46 | 0.41 |
| 2 | 236967850 | rs2084934 | -0.2 | -0.964 | -1.33 | 0.596 | 0.028 | -0.051 | -0.051 | 0 | 0.31 | 0.74 | 0.39 | 0.46 | 0.21 | 0.39 |
| 2 | 236970210 | rs6749870 | 0.058 | -0.968 | -1.163 | 0.366 | 0.002 | -0.018 | -0.018 | 0 | 0.69 | 0.72 | 0.76 | 0.92 | 0.73 | 0.88 |
| 2 | 236972635 | rs3768945 | 0.32 | -1.124 | -0.795 | 0.63 | -0.009 | 0.007 | 0.007 | 0 | 0.27 | 0.3 | 0.29 | 0.29 | 0.21 | 0.34 |
| 2 | 236976106 | rs7581447 | 1.137 | -0.836 | -1.123 | 0.759 | -0.11 | -0.101 | -0.101 | 0 | 0.39 | 0.03 | 0.36 | 0.45 | 0.51 | 0.5 |
| 2 | 236988413 | rs1530933 | -0.996 | -0.757 | 0.076 | 0.984 | 0.074 | 0.243 | 0.243 | 0 | 0.05 | 0.31 | 0.13 | 0.16 | 0.05 | 0.06 |
| 2 | 236990057 | rs3768939 | -0.054 | -0.964 | 0.098 | 0.897 | -0.03 | 0.018 | 0.018 | 0 | 0.3 | 0.33 | 0.27 | 0.35 | 0.18 | 0.33 |
| 2 | 237007049 | rs9287596 | -1.029 | -0.184 | 0.25 | 0.668 | 0.028 | 0.114 | 0.114 | 0.009 | 0.1 | 0.32 | 0.16 | 0.17 | 0.08 | 0.07 |
| 2 | 237008172 | rs6431424 | -1.576 | -0.159 | 0.039 | 0.686 | 0.101 | 0.268 | 0.268 | 0.075 | 0.06 | 0.41 | 0.13 | 0.16 | 0.05 | 0.1 |
| 2 | 237010612 | rs4663638 | -0.618 | -0.299 | -1.539 | 0.108 | 0.006 | 0.017 | 0.017 | 0 | 0.54 | 0.78 | 0.5 | 0.41 | 0.45 | 0.5 |
| 2 | 237018510 | rs3768926 | -0.904 | -0.262 | -1.464 | -0.373 | -0.02 | 0.033 | 0.033 | 0 | 0.58 | 0.81 | 0.41 | 0.33 | 0.46 | 0.45 |
| 2 | 237027593 | rs3768921 | -0.989 | 0.077 | -0.619 | -0.143 | 0.02 | 0.038 | 0.038 | 0.28 | 0.48 | 0.22 | 0.31 | 0.32 | 0.42 | 0.44 |
| 2 | 237029104 | rs2278884 | -1.024 | 0.273 | -0.065 | 0.354 | -0.005 | -0.019 | -0.019 | 0 | 0.32 | 0.15 | 0.23 | 0.32 | 0.3 | 0.38 |
| 2 | 237040797 | rs6742722 | -1.518 | 0.338 | 0.557 | 0.374 | 0.002 | -0.013 | -0.013 | 1.38 | 0.22 | 0.77 | 0.11 | 0.04 | 0.14 | 0.12 |
| 2 | 237040812 | rs12620543 | -1.177 | 0.304 | 0.235 | 0.582 | -0.022 | -0.039 | -0.039 | 2.257 | 0.09 | 0.02 | 0.05 | 0.24 | 0.11 | 0.26 |
| 2 | 237048179 | rs6431431 | -2.242 | 0.275 | 0.195 | 0.529 | 0.009 | -0.029 | -0.029 | 0 | 0.32 | 0.9 | 0.17 | 0.28 | 0.25 | 0.36 |
| 2 | 237052475 | rs6753712 | -1.468 | 0.417 | 1.007 | 0.681 | 0.029 | 0.066 | 0.066 | 0 | 0.13 | 0.28 | 0.09 | 0.23 | 0.17 | 0.25 |
| 2 | 237061954 | rs4377285 | -1.21 | 0.449 | 0.751 | 0.701 | 0.003 | 0.013 | 0.013 | 0 | 0.12 | 0.14 | 0.09 | 0.23 | 0.16 | 0.25 |
| 2 | 237065805 | rs1316457 | -1.14 | 0.452 | 0.734 | 0.708 | 0.001 | 0.011 | 0.011 | 0 | 0.12 | 0.14 | 0.09 | 0.24 | 0.16 | 0.25 |
| 2 | 237073776 | rs6752442 | -1.993 | 0.365 | 0.6 | 0.587 | -0.011 | -0.03 | -0.03 | 0 | 0.32 | 0.84 | 0.2 | 0.29 | 0.3 | 0.39 |
| 2 | 237083077 | rs1921660 | -2.14 | 0.332 | 0.794 | 0.477 | -0.006 | 0.026 | 0.026 | 3.636 | 0.34 | 0.87 | 0.23 | 0.24 | 0.32 | 0.38 |
| 2 | 237091317 | rs10177160 | -0.811 | 0.236 | 1.722 | 0.304 | -0.007 | -0.09 | -0.09 | 0.026 | 0.15 | 0.25 | 0.08 | 0 | 0.13 | 0.12 |
| 2 | 237099410 | rs13386897 | -0.334 | 0.307 | 1.271 | 0.342 | -0.005 | 0.018 | 0.018 | 0 | 0.11 | 0.06 | 0.06 | 0 | 0.13 | 0.13 |
| 2 | 237100952 | rs7589642 | -0.398 | 0.294 | 1.271 | 0.322 | -0.004 | 0.019 | 0.019 | 0 | 0.12 | 0.06 | 0.06 | 0 | 0.14 | 0.13 |
| 2 | 237103548 | rs10177825 | -1.201 | 0.176 | 2.283 | 0.273 | -0.008 | -0.134 | -0.134 | 8.711 | 0.17 | 0.44 | 0.1 | 0 | 0.15 | 0.13 |
| 2 | 237104664 | rs12329227 | -0.862 | 0.279 | 2.283 | 0.301 | -0.002 | -0.077 | -0.077 | 0 | 0.14 | 0.23 | 0.08 | 0 | 0.14 | 0.12 |
| 2 | 237104704 | rs12328175 | -0.862 | 0.279 | 2.283 | 0.301 | -0.002 | -0.077 | -0.077 | 5.05 | 0.14 | 0.23 | 0.08 | 0 | 0.14 | 0.12 |
| 2 | 237105981 | rs7564581 | -1.497 | 0.234 | 2.535 | 0.209 | -0.019 | -0.105 | -0.105 | 0 | 0.22 | 0.65 | 0.13 | 0 | 0.16 | 0.12 |
| 2 | 237110321 | rs10929164 | -0.625 | 0.38 | 2.219 | 0.485 | 0.017 | 0.042 | 0.042 | 0 | 0.13 | 0.16 | 0.13 | 0.24 | 0.14 | 0.26 |
| 2 | 237113030 | rs12472312 | -0.625 | 0.38 | 2.219 | 0.485 | 0.017 | 0.042 | 0.042 | 0 | 0.13 | 0.17 | 0.13 | 0.24 | 0.14 | 0.26 |
| 2 | 237119400 | rs7584752 | -2.378 | 0.281 | 1.844 | 0.424 | -0.034 | 0.022 | 0.022 | 3.286 | 0.37 | 0.92 | 0.27 | 0.24 | 0.31 | 0.45 |
| 2 | 237124209 | rs6431439 | -2.397 | 0.376 | 1.809 | 0.673 | -0.072 | -0.03 | -0.03 | 0 | 0.37 | 0.95 | 0.36 | 0.26 | 0.32 | 0.43 |
| 2 | 237125171 | rs6737755 | -1.074 | 0.386 | 2.484 | 0.661 | 0.059 | 0.138 | 0.138 | 0 | 0.16 | 0.52 | 0.24 | 0.24 | 0.16 | 0.24 |
| 2 | 237127095 | rs7591668 | -2.537 | 0.38 | 2.164 | 0.653 | -0.073 | -0.051 | -0.051 | 1.067 | 0.37 | 0.95 | 0.36 | 0.28 | 0.32 | 0.43 |
| 2 | 237129535 | rs10929177 | -0.57 | 0.508 | 2.508 | 0.573 | 0 | -0.024 | -0.024 | 0 | 0.16 | 0.17 | 0.09 | 0.04 | 0.15 | 0.15 |
| 2 | 237148600 | rs2219063 | 0.627 | 0.46 | 2.226 | 0.732 | -0.023 | 0.023 | 0.023 | 0 | 0.81 | 0.63 | 0.78 | 0.79 | 0.81 | 0.8 |
| 2 | 237149941 | rs6756597 | 2.026 | 0.556 | 1.804 | 0.768 | -0.011 | -0.011 | -0.011 | 0 | 0.52 | 0.3 | 0.48 | 0.57 | 0.51 | 0.41 |
| 2 | 237150311 | rs6431443 | -0.557 | 0.548 | 1.326 | 0.763 | -0.013 | -0.014 | -0.014 | 3.736 | 0.14 | 0.11 | 0.24 | 0.23 | 0.21 | 0.27 |
| 2 | 237156549 | rs13390159 | -1.336 | -0.029 | 1.936 | 0.312 | -0.014 | -0.055 | -0.055 | 0.001 | 0.23 | 0.6 | 0.14 | 0.05 | 0.16 | 0.19 |
| 2 | 237183463 | rs1377445 | 1.472 | -0.265 | 2.41 | 1.034 | -0.009 | 0 | 0 | 0.377 | 0.77 | 0.29 | 0.76 | 0.78 | 0.83 | 0.77 |
| 2 | 237187934 | rs10195546 | -1.808 | -0.178 | 2.189 | 1.068 | -0.007 | 0.075 | 0.075 | 0.325 | 0.07 | 0.33 | 0.04 | 0.06 | 0.01 | 0.09 |
| 2 | 237189869 | rs10184445 | -1.808 | -0.178 | 2.189 | 1.068 | -0.007 | 0.075 | 0.075 | 0.325 | 0.07 | 0.33 | 0.04 | 0.06 | 0.01 | 0.09 |
| 2 | 237201756 | rs10206697 | -0.756 | -0.172 | 1.436 | 1.306 | -0.023 | 0.058 | 0.058 | 0.03 | 0.21 | 0.38 | 0.13 | 0.31 | 0.21 | 0.15 |
| 2 | 237212899 | rs7560409 | -1.942 | -0.146 | 2.556 | 1.646 | 0.019 | 0.262 | 0.262 | 0.652 | 0.07 | 0.41 | 0.05 | 0.23 | 0.02 | 0.03 |
| 2 | 237214545 | rs935925 | -2.699 | -0.272 | 2.553 | 1.855 | -0.044 | 0.104 | 0.104 | 6.751 | 0.11 | 0.77 | 0.15 | 0.33 | 0.02 | 0.04 |
| 2 | 237233178 | rs6431452 | -0.864 | 0.304 | 2.044 | 0.953 | 0.011 | -0.053 | -0.053 | 0 | 0.14 | 0.23 | 0.08 | 0.01 | 0.15 | 0.13 |
| 2 | 237233920 | rs7608734 | -1.567 | 0.233 | 2.919 | 1.19 | 0.004 | 0.01 | 0.01 | 0.562 | 0.18 | 0.64 | 0.13 | 0.17 | 0.17 | 0.15 |
| 2 | 237238496 | rs2046284 | -0.874 | 0.625 | 2.654 | 0.738 | 0.024 | -0.053 | -0.053 | 0 | 0.14 | 0.23 | 0.11 | 0.01 | 0.19 | 0.14 |
| 2 | 237241640 | rs6714545 | -0.487 | 0.115 | 2.124 | 0.419 | -0.022 | -0.005 | -0.005 | 0.051 | 0.19 | 0.27 | 0.25 | 0.15 | 0.15 | 0.12 |
| 2 | 237241852 | rs6704601 | -0.487 | 0.115 | 2.124 | 0.419 | -0.022 | -0.005 | -0.005 | 0.005 | 0.19 | 0.27 | 0.25 | 0.15 | 0.15 | 0.12 |
| 2 | 237242097 | rs6431457 | -1.521 | 0.087 | 2.042 | 0.419 | -0.002 | -0.053 | -0.053 | 0 | 0.33 | 0.5 | 0.35 | 0.16 | 0.33 | 0.25 |
| 2 | 237258309 | rs6748498 | -0.563 | 0.137 | 1.902 | 0.469 | -0.04 | -0.006 | -0.006 | 0 | 0.2 | 0.42 | 0.26 | 0.15 | 0.16 | 0.11 |
| 2 | 237272056 | rs6724356 | -0.429 | 0.408 | 1.272 | 0.461 | -0.017 | 0.015 | 0.015 | 0 | 0.14 | 0.11 | 0.1 | 0.01 | 0.2 | 0.14 |
| 2 | 237274657 | rs7585637 | -0.673 | 0.146 | 1.083 | 0.302 | 0.016 | 0.031 | 0.031 | 0 | 0.19 | 0.09 | 0.24 | 0.06 | 0.17 | 0.11 |
| 2 | 237279707 | rs1108890 | -1.786 | 0.429 | 1.242 | 0.178 | 0.007 | -0.029 | -0.029 | 0 | 0.37 | 0.5 | 0.38 | 0.19 | 0.42 | 0.26 |
| 2 | 237300677 | rs10204742 | -0.814 | 0.585 | 1.461 | 0.356 | 0.01 | -0.036 | -0.036 | 0 | 0.07 | 0.18 | 0.05 | 0 | 0.06 | 0.01 |
| 2 | 237305200 | rs11885437 | -0.556 | 0.588 | 1.396 | 0.357 | 0.003 | -0.031 | -0.031 | 0 | 0.06 | 0.13 | 0.05 | 0 | 0.06 | 0.01 |
| 2 | 237314747 | rs6431461 | -1.654 | 0.744 | 1.295 | 0.306 | 0.012 | -0.034 | -0.034 | 0 | 0.37 | 0.52 | 0.39 | 0.18 | 0.43 | 0.28 |
| 2 | 237344926 | rs1384579 | 0.073 | 0.757 | 0.917 | 0.301 | -0.004 | -0.015 | -0.015 | 0.101 | 0.12 | 0.12 | 0.09 | 0 | 0.19 | 0.12 |
| 2 | 237353017 | rs11897843 | -0.823 | 0.7 | 0.751 | 0.327 | -0.007 | -0.005 | -0.005 | 0 | 0.25 | 0.37 | 0.3 | 0.18 | 0.23 | 0.15 |
| 2 | 237371776 | rs900439 | -0.72 | 0.777 | 0.608 | 0.347 | -0.001 | -0.001 | -0.001 | 0 | 0.24 | 0.26 | 0.28 | 0.18 | 0.23 | 0.15 |
| 2 | 237382889 | rs2317639 | 0.073 | 0.856 | 0.425 | 0.328 | 0.003 | -0.026 | -0.026 | 0.629 | 0.12 | 0.14 | 0.08 | 0 | 0.19 | 0.12 |
| 2 | 237396935 | rs6751722 | -0.995 | 0.707 | 0.095 | 0.494 | 0.013 | -0.03 | -0.03 | 0 | 0.36 | 0.49 | 0.37 | 0.18 | 0.42 | 0.28 |

| | | | | | | | | | | | | | | | | |
|---|---|---|---|---|---|---|---|---|---|---|---|---|---|---|---|---|
| 2 | 237403115 | rs2317644 | -0.816 | 0.306 | -1.193 | 0.566 | -0.053 | -0.019 | -0.019 | 3.38 | 0.54 | 0.96 | 0.54 | 0.35 | 0.39 | 0.32 |
| 2 | 237405689 | rs6709076 | -1.222 | -0.106 | 0.826 | 0.932 | 0.065 | 0.482 | 0.482 | 0.214 | 0.07 | 0.47 | 0.26 | 0.33 | 0.03 | 0.06 |
| 2 | 237415521 | rs4663650 | -2.291 | -0.475 | 1.91 | 1.062 | -0.011 | 0.019 | 0.019 | 0.874 | 0.17 | 0.76 | 0.32 | 0.38 | 0.08 | 0.09 |
| 2 | 237433532 | rs1986988 | -2.527 | -0.478 | 2.465 | 0.959 | -0.014 | 0.021 | 0.021 | 0.552 | 0.18 | 0.73 | 0.25 | 0.35 | 0.1 | 0.14 |
| 2 | 237436404 | rs13392175 | -1.573 | -0.482 | 3.053 | 0.954 | -0.016 | -0.034 | -0.034 | 0.237 | 0.09 | 0.27 | 0.05 | 0.05 | 0.05 | 0.02 |
| 2 | 237439358 | rs10169073 | -2.991 | -0.483 | 2.843 | 0.963 | -0.015 | 0.022 | 0.022 | 0.379 | 0.18 | 0.73 | 0.24 | 0.32 | 0.09 | 0.14 |
| 2 | 237440052 | rs6431467 | -2.597 | -0.47 | 2.946 | 0.976 | 0.005 | 0.244 | 0.244 | 0.148 | 0.1 | 0.54 | 0.19 | 0.3 | 0.04 | 0.11 |
| 2 | 237442621 | rs4663654 | -1.594 | -0.455 | 2.554 | 0.983 | -0.002 | 0.013 | 0.013 | 0 | 0.06 | 0.04 | 0.16 | 0.3 | 0.04 | 0.11 |
| 2 | 237446265 | rs6431470 | -2.565 | -0.503 | 2.775 | 0.635 | -0.042 | 0.012 | 0.012 | 0.505 | 0.16 | 0.62 | 0.12 | 0.18 | 0.08 | 0.11 |
| 2 | 237453049 | rs4265959 | -2.36 | -0.353 | 2.458 | 0.591 | 0.016 | 0.036 | 0.036 | 0.535 | 0.13 | 0.52 | 0.17 | 0.16 | 0.13 | 0.16 |
| 2 | 237460365 | rs7582574 | -2.634 | -0.378 | 2.071 | 0.645 | -0.011 | 0.017 | 0.017 | 1.158 | 0.15 | 0.65 | 0.18 | 0.17 | 0.13 | 0.17 |
| 2 | 237464991 | rs2317643 | -2.359 | -0.266 | 2.123 | 1.024 | 0.008 | 0.185 | 0.185 | 0.956 | 0.11 | 0.56 | 0.17 | 0.27 | 0.06 | 0.16 |
| 2 | 237468542 | rs4663655 | -2.554 | -0.217 | 1.551 | 1.045 | -0.002 | 0.033 | 0.033 | 1.337 | 0.16 | 0.73 | 0.23 | 0.29 | 0.15 | 0.19 |
| 2 | 237481969 | rs10183022 | -1.077 | -0.672 | -0.313 | 0.098 | -0.082 | -0.003 | -0.003 | 0 | 0.56 | 0.97 | 0.37 | 0.37 | 0.39 | 0.48 |
| 2 | 237486160 | rs9808217 | -0.169 | -1.137 | 0.704 | -0.312 | -0.016 | -0.083 | -0.083 | 0 | 0.25 | 0.35 | 0.08 | 0 | 0.16 | 0.11 |
| 2 | 237486787 | rs1106166 | -1.158 | -1.193 | 1.415 | -0.29 | 0.088 | 0.013 | 0.013 | 0.022 | 0.07 | 0.31 | 0.09 | 0.01 | 0.06 | 0.14 |
| 2 | 237487588 | rs10179774 | -0.183 | -1.242 | 1.153 | -0.415 | 0.013 | 0.032 | 0.032 | 0 | 0.25 | 0.15 | 0.11 | 0.05 | 0.19 | 0.18 |
| 2 | 237488450 | rs7559855 | -1.445 | -1.204 | 1.184 | -0.392 | 0.086 | -0.013 | -0.013 | 0 | 0.08 | 0.43 | 0.1 | 0 | 0.06 | 0.14 |
| 2 | 237489904 | rs1045879 | -1.192 | -1.182 | 0.412 | -0.38 | 0.023 | 0.061 | 0.061 | 0 | 0.36 | 0.74 | 0.22 | 0.05 | 0.25 | 0.32 |
| 2 | 237497237 | rs11901793 | -1.32 | -1.594 | -0.081 | -0.074 | 0.084 | 0.036 | 0.036 | 0.121 | 0.37 | 0.93 | 0.2 | 0.27 | 0.21 | 0.29 |
| 2 | 237509207 | rs896543 | -1.104 | -1.544 | 0.323 | -0.449 | 0.066 | 0.074 | 0.074 | 0 | 0.34 | 0.76 | 0.16 | 0.2 | 0.2 | 0.25 |
| 2 | 237517937 | rs6431476 | -1.319 | -1.873 | 0.293 | -0.741 | 0.068 | 0.086 | 0.086 | 0 | 0.4 | 0.71 | 0.17 | 0.21 | 0.21 | 0.36 |
| 2 | 237523328 | rs11691569 | -0.764 | -1.88 | 0.673 | -0.766 | 0.03 | 0.044 | 0.044 | 0 | 0.31 | 0.19 | 0.1 | 0.15 | 0.2 | 0.23 |
| 2 | 237525893 | rs6736272 | -1.298 | -1.96 | 0.904 | -0.785 | 0.025 | 0.02 | 0.02 | 0.09 | 0.36 | 0.65 | 0.11 | 0.09 | 0.18 | 0.19 |
| 2 | 237531744 | rs7588292 | -1.82 | -1.862 | 1.216 | -0.7 | -0.036 | 0.024 | 0.024 | 0.543 | 0.11 | 0.45 | 0.05 | 0.09 | 0.03 | 0.06 |
| 2 | 237537935 | rs7594454 | -1.465 | -1.839 | 0.784 | -0.751 | 0.089 | 0.135 | 0.135 | 0 | 0.41 | 0.75 | 0.16 | 0.16 | 0.22 | 0.26 |
| 2 | 237545101 | rs1796443 | 0.681 | -1.964 | 0.889 | -1.035 | -0.026 | -0.019 | -0.019 | 0 | 0.85 | 0.77 | 0.93 | 0.89 | 0.87 | 0.94 |
| 2 | 237546708 | rs1796438 | 0.921 | -2.43 | 0.705 | -1.287 | 0.075 | 0.092 | 0.092 | 0 | 0.58 | 0.33 | 0.86 | 0.88 | 0.75 | 0.8 |
| 2 | 237550197 | rs2952665 | 1.256 | -2.151 | 0.641 | -1.119 | 0.069 | 0.098 | 0.098 | 0 | 0.55 | 0.35 | 0.74 | 0.91 | 0.71 | 0.72 |
| 2 | 237553041 | rs1669776 | 0.86 | -1.896 | 1.003 | -1.009 | -0.029 | -0.018 | -0.018 | 0 | 0.78 | 0.66 | 0.87 | 0.84 | 0.8 | 0.87 |
| 2 | 237561751 | rs2600737 | 1.227 | -1.839 | 1.071 | -0.852 | 0.004 | 0.003 | 0.003 | 0 | 0.56 | 0.58 | 0.8 | 0.77 | 0.64 | 0.72 |
| 2 | 237566300 | rs2600733 | 0.573 | -1.835 | 1.058 | -0.856 | -0.029 | -0.014 | -0.014 | 0 | 0.79 | 0.63 | 0.87 | 0.84 | 0.8 | 0.87 |
| 2 | 237566315 | rs2600732 | 1.051 | -1.782 | 0.659 | -0.846 | 0.003 | -0.001 | -0.001 | 0.861 | 0.52 | 0.53 | 0.78 | 0.8 | 0.63 | 0.73 |
| 2 | 237568742 | rs2600726 | 0.933 | -1.499 | 0.537 | -0.817 | 0 | -0.006 | -0.006 | 0 | 0.51 | 0.49 | 0.77 | 0.8 | 0.6 | 0.71 |
| 2 | 237573900 | rs1869093 | 0.94 | -1.494 | 0.186 | -0.881 | 0.038 | 0.056 | 0.056 | 0.208 | 0.52 | 0.39 | 0.77 | 0.82 | 0.6 | 0.71 |
| 2 | 237575172 | rs3109647 | -1 | -1.328 | 0.248 | -1.074 | -0.033 | -0.054 | -0.054 | 0 | 0.29 | 0.47 | 0.18 | 0.18 | 0.24 | 0.13 |
| 2 | 237578513 | rs2100335 | -0.769 | -1.357 | 0.66 | -1.025 | -0.032 | -0.026 | -0.026 | 0 | 0.24 | 0.45 | 0.17 | 0.2 | 0.18 | 0.11 |
| 2 | 237581541 | rs10181573 | 0.102 | -1.484 | 0.623 | -1.043 | -0.035 | 0.004 | 0.004 | 0 | 0.16 | 0.29 | 0.09 | 0.18 | 0.08 | 0.08 |
| 2 | 237583477 | rs11695375 | -0.818 | -1.516 | -0.648 | -1.493 | -0.008 | -0.05 | -0.05 | 0 | 0.36 | 0.39 | 0.24 | 0.11 | 0.34 | 0.15 |
| 2 | 237585368 | rs2720107 | -1.053 | -1.641 | -0.86 | -1.669 | -0.011 | -0.024 | -0.024 | 0 | 0.31 | 0.62 | 0.22 | 0.18 | 0.25 | 0.16 |
| 2 | 237586032 | rs12613625 | -0.465 | -1.636 | -0.861 | -1.669 | -0.018 | -0.051 | -0.051 | 0 | 0.27 | 0.33 | 0.16 | 0.11 | 0.24 | 0.09 |
| 2 | 237590339 | rs2720100 | -2.004 | -1.267 | -1.592 | -1.443 | -0.093 | 0.054 | 0.054 | 0 | 0.59 | 0.96 | 0.39 | 0.3 | 0.49 | 0.27 |
| 2 | 237592752 | rs11889670 | -0.302 | -0.785 | -0.709 | -1.182 | 0.001 | -0.025 | -0.025 | 0 | 0.09 | 0.11 | 0.08 | 0 | 0.1 | 0.07 |
| 2 | 237594511 | rs1105711 | -0.895 | -1.297 | -0.753 | -1.182 | -0.001 | -0.001 | -0.001 | 0.255 | 0.32 | 0.64 | 0.23 | 0.24 | 0.28 | 0.14 |
| 2 | 237595384 | rs2720099 | -1.652 | -1.009 | -1.41 | -0.948 | 0.03 | 0.018 | 0.018 | 0.269 | 0.47 | 0.88 | 0.37 | 0.4 | 0.43 | 0.21 |
| 2 | 237595867 | rs1869091 | -1.451 | -1.047 | -1.278 | -1.178 | 0.024 | 0.106 | 0.106 | 0.35 | 0.44 | 0.82 | 0.35 | 0.24 | 0.4 | 0.21 |
| 2 | 237597846 | rs2720098 | -1.867 | -0.941 | -0.862 | -1.019 | -0.007 | 0.025 | 0.025 | 1.105 | 0.56 | 0.74 | 0.43 | 0.36 | 0.54 | 0.26 |
| 2 | 237598117 | rs2720096 | -1.673 | -1.15 | -0.041 | -1.386 | 0.001 | 0 | 0 | 0 | 0.3 | 0.57 | 0.22 | 0.3 | 0.38 | 0.14 |
| 2 | 237598883 | rs2600776 | -2.215 | -1.223 | 0.343 | -1.661 | 0 | -0.023 | -0.023 | 0 | 0.35 | 0.44 | 0.26 | 0.2 | 0.45 | 0.19 |
| 2 | 237604353 | rs2600791 | -1.312 | -0.973 | 0.105 | -1.53 | 0.005 | 0.056 | 0.056 | 1.542 | 0.13 | 0.05 | 0.06 | 0.05 | 0.13 | 0.12 |
| 2 | 237609667 | rs2167580 | -1.949 | -1.066 | 0.266 | -1.582 | 0.049 | 0.12 | 0.12 | 2.092 | 0.36 | 0.19 | 0.22 | 0.11 | 0.3 | 0.2 |
| 2 | 237611437 | rs1869092 | -2.026 | -1.072 | 0.332 | -1.585 | 0.029 | 0.046 | 0.046 | 1.875 | 0.38 | 0.29 | 0.23 | 0.11 | 0.3 | 0.2 |
| 2 | 237611604 | rs7604535 | -1.118 | -0.798 | 0.817 | -1.483 | 0.014 | 0.048 | 0.048 | 0 | 0.17 | 0.36 | 0.15 | 0.18 | 0.22 | 0.09 |
| 2 | 237613894 | rs2720156 | -2.096 | -1.021 | 0.665 | -1.462 | -0.01 | -0.03 | -0.03 | 0 | 0.37 | 0.49 | 0.24 | 0.23 | 0.41 | 0.2 |
| 2 | 237614291 | rs2600783 | -1.179 | -0.629 | 1.064 | -1.306 | -0.008 | -0.063 | -0.063 | 0.02 | 0.15 | 0.41 | 0.15 | 0.06 | 0.16 | 0.11 |
| 2 | 237626734 | rs7585432 | -1.654 | -0.934 | 0.999 | -1.484 | -0.053 | -0.135 | -0.135 | 0.173 | 0.25 | 0.54 | 0.13 | 0 | 0.14 | 0.07 |
| 2 | 237629101 | rs10929183 | -1.314 | -0.981 | 1.137 | -1.509 | -0.056 | -0.118 | -0.118 | 2.17 | 0.21 | 0.36 | 0.1 | 0 | 0.1 | 0.07 |
| 2 | 237640659 | rs896542 | -2.471 | -0.959 | 0.293 | -1.487 | 0.051 | 0.149 | 0.149 | 0 | 0.51 | 0.87 | 0.44 | 0.25 | 0.47 | 0.3 |
| 2 | 237642373 | rs2067620 | -1.097 | -1.025 | 0.447 | -1.525 | -0.031 | -0.138 | -0.138 | 0 | 0.18 | 0.49 | 0.12 | 0 | 0.13 | 0.04 |
| 2 | 237644199 | rs7556982 | -2.535 | -0.989 | 0.279 | -1.491 | 0.052 | 0.145 | 0.145 | 0 | 0.51 | 0.87 | 0.44 | 0.25 | 0.48 | 0.3 |
| 2 | 237647424 | rs11679123 | -1.088 | -0.973 | 0.395 | -1.824 | -0.02 | -0.129 | -0.129 | 0 | 0.17 | 0.42 | 0.12 | 0 | 0.13 | 0.04 |
| 2 | 237656626 | rs6738577 | -0.94 | -0.732 | -0.367 | -1.473 | -0.018 | -0.041 | -0.041 | 0.834 | 0.15 | 0.24 | 0.1 | 0.01 | 0.09 | 0.09 |
| 2 | 237664197 | rs6743039 | -1.995 | -0.703 | -1.35 | -1.438 | 0.001 | 0.165 | 0.165 | 1.119 | 0.47 | 0.95 | 0.37 | 0.24 | 0.4 | 0.3 |
| 2 | 237673053 | rs7573720 | -1.206 | -0.331 | -0.655 | -1.266 | -0.003 | 0.019 | 0.019 | 0 | 0.28 | 0.21 | 0.24 | 0.22 | 0.31 | 0.21 |
| 2 | 237680816 | rs6733916 | -0.993 | -0.435 | -0.752 | -1.308 | 0.058 | 0.059 | 0.059 | 0.071 | 0.3 | 0.17 | 0.17 | 0.15 | 0.2 | 0.23 |
| 2 | 237683005 | rs11694626 | -1.364 | -0.093 | -0.286 | -1.374 | -0.006 | -0.044 | -0.044 | 1.525 | 0.38 | 0.47 | 0.27 | 0.17 | 0.38 | 0.33 |
| 2 | 237690418 | rs999031 | 1.02 | 0.304 | -0.313 | -1.377 | 0.006 | -0.087 | -0.087 | 0.092 | 0.73 | 0.35 | 0.77 | 0.98 | 0.7 | 0.8 |
| 2 | 237692770 | rs10929189 | 0.383 | 0.24 | -0.602 | -0.99 | 0.001 | -0.018 | -0.018 | 0 | 0.66 | 0.7 | 0.76 | 0.85 | 0.66 | 0.8 |
| 2 | 237697699 | rs1991761 | 1.09 | 0.062 | -0.189 | -0.932 | -0.031 | -0.094 | -0.094 | 1.076 | 0.83 | 0.35 | 0.91 | 1 | 0.93 | 0.95 |

| | | | | | | | | | | | | | | | | |
|---|---|---|---|---|---|---|---|---|---|---|---|---|---|---|---|---|
| 2 | 237698676 | rs1897464 | 1.694 | -0.35 | -0.822 | -0.621 | 0.073 | 0.02 | 0.02 | 5.557 | 0.68 | 0.11 | 0.72 | 0.76 | 0.77 | 0.79 |
| 2 | 237706939 | rs934397 | 1.08 | 0.301 | -0.758 | -0.616 | 0.005 | -0.027 | -0.027 | 1.667 | 0.81 | 0.27 | 0.82 | 0.95 | 0.79 | 0.87 |
| 2 | 237707551 | rs10209000 | -0.778 | 0.27 | -0.756 | -0.448 | 0.006 | 0.002 | 0.002 | 0 | 0.14 | 0.15 | 0.16 | 0.1 | 0.1 | 0.09 |
| 2 | 237708511 | rs13396122 | -0.792 | 0.263 | -0.76 | -0.452 | 0.001 | -0.003 | -0.003 | 1.01 | 0.14 | 0.16 | 0.16 | 0.2 | 0.11 | 0.1 |
| 2 | 237714186 | rs6431484 | 1.48 | 0.334 | -0.924 | -0.563 | -0.041 | 0.068 | 0.068 | 4.675 | 0.71 | 0.14 | 0.66 | 0.85 | 0.69 | 0.78 |
| 2 | 237715095 | rs12992344 | 1.652 | 0.073 | -1.079 | -0.572 | -0.028 | 0.053 | 0.053 | 3.343 | 0.66 | 0.11 | 0.66 | 0.76 | 0.68 | 0.73 |
| 2 | 237720634 | rs10929194 | 1.503 | -0.242 | -0.987 | -0.897 | 0.001 | 0.094 | 0.094 | 3.077 | 0.65 | 0.16 | 0.7 | 0.83 | 0.71 | 0.71 |
| 2 | 237723694 | rs13000320 | -2.244 | -0.164 | 0.256 | -0.919 | -0.03 | -0.043 | -0.043 | 0.047 | 0.19 | 0.57 | 0.2 | 0.23 | 0.16 | 0.23 |
| 2 | 237729467 | rs744296 | -1.78 | -0.216 | 0.575 | -0.784 | -0.024 | 0.007 | 0.007 | 0 | 0.08 | 0.19 | 0.12 | 0.18 | 0.04 | 0.07 |
| 2 | 237731854 | rs12329181 | -2.537 | -0.175 | 1.355 | -0.847 | -0.011 | 0.006 | 0.006 | 0.575 | 0.1 | 0.47 | 0.06 | 0.07 | 0.04 | 0.06 |
| 2 | 237736216 | rs7584906 | -1.373 | 0.189 | 1.417 | -0.921 | 0.008 | 0 | 0 | 0 | 0.09 | 0.08 | 0.11 | 0.04 | 0.19 | 0.11 |
| 2 | 237736561 | rs10170633 | -2.452 | 0.262 | 1.553 | -0.961 | -0.002 | -0.001 | -0.001 | 0 | 0.1 | 0.4 | 0.06 | 0.07 | 0.05 | 0.06 |
| 2 | 237740508 | rs10166334 | -2.316 | 0.263 | 1.59 | -0.964 | 0.14 | -0.03 | -0.03 | 0.154 | 0.12 | 0.38 | 0.13 | 0.06 | 0.2 | 0.11 |
| 2 | 237742697 | rs7563419 | -1.597 | 0.207 | 1.509 | -0.963 | -0.015 | -0.011 | -0.011 | 0 | 0.07 | 0.15 | 0.03 | 0.05 | 0.03 | 0.06 |
| 2 | 237751111 | rs10193277 | -0.343 | 0.116 | 1.188 | -1.231 | 0.003 | 0 | 0 | 0.016 | 0.08 | 0.07 | 0.09 | 0 | 0.05 | 0.05 |
| 2 | 237752054 | rs1000007 | -2.961 | 0.176 | 1.285 | -1.207 | 0.09 | -0.063 | -0.063 | 0 | 0.19 | 0.53 | 0.17 | 0.1 | 0.24 | 0.16 |
| 2 | 237752442 | rs11688999 | -1.413 | 0.296 | 1.085 | -1.158 | -0.018 | 0.016 | 0.016 | 0 | 0.1 | 0.06 | 0.11 | 0.02 | 0.2 | 0.1 |
| 2 | 237755950 | rs10186949 | -2.34 | 0.319 | 0.829 | -1.506 | 0.017 | -0.065 | -0.065 | 0.398 | 0.34 | 0.6 | 0.29 | 0.11 | 0.36 | 0.23 |
| 2 | 237757204 | rs11691692 | -2.03 | 0.411 | 1.057 | -1.439 | 0.052 | -0.07 | -0.07 | 0 | 0.17 | 0.28 | 0.2 | 0.03 | 0.24 | 0.16 |
| 2 | 237758257 | rs4503948 | 0.647 | 0.272 | 0.216 | -1.596 | -0.016 | 0.064 | 0.064 | 0.654 | 0.57 | 0.66 | 0.54 | 0.76 | 0.47 | 0.7 |
| 2 | 237758706 | rs13029887 | -0.377 | -0.157 | 0.079 | -1.532 | -0.003 | -0.022 | -0.022 | 0 | 0.1 | 0.11 | 0.11 | 0.02 | 0.08 | 0.1 |
| 2 | 237760783 | rs6431485 | -0.126 | -0.161 | -0.032 | -1.556 | -0.003 | 0.038 | 0.038 | 0 | 0.69 | 0.84 | 0.66 | 0.79 | 0.66 | 0.81 |
| 2 | 237765359 | rs7581272 | -1.364 | -0.294 | 0.549 | -1.753 | -0.002 | -0.116 | -0.116 | 0 | 0.27 | 0.43 | 0.22 | 0.04 | 0.23 | 0.2 |
| 2 | 237770442 | rs7578264 | -0.257 | -0.418 | 0.622 | -1.713 | 0.001 | -0.008 | -0.008 | 0.02 | 0.07 | 0.38 | 0.04 | 0 | 0.01 | 0 |
| 2 | 237771012 | rs11690379 | -0.679 | -0.367 | 0.823 | -1.809 | -0.007 | 0.002 | 0.002 | 2.782 | 0.21 | 0.16 | 0.19 | 0.04 | 0.23 | 0.19 |
| 2 | 237771589 | rs2318127 | -1.581 | -0.298 | 0.62 | -1.772 | -0.007 | -0.128 | -0.128 | 7.663 | 0.29 | 0.56 | 0.23 | 0.04 | 0.23 | 0.19 |
| 2 | 237772825 | rs6431487 | -1.716 | -0.449 | 0.099 | -1.618 | 0.014 | 0.046 | 0.046 | 4.218 | 0.32 | 0.77 | 0.33 | 0.17 | 0.24 | 0.34 |
| 2 | 237776239 | rs11680023 | 1.809 | -0.482 | -0.326 | -1.58 | -0.116 | -0.025 | -0.025 | 0 | 0.36 | 0.03 | 0.35 | 0.61 | 0.43 | 0.41 |
| 2 | 237780124 | rs3888913 | 1.216 | -0.493 | 0.018 | -1.479 | 0.008 | 0.109 | 0.109 | 1.756 | 0.52 | 0.18 | 0.49 | 0.65 | 0.55 | 0.55 |
| 2 | 237780727 | rs4074882 | -0.884 | -0.487 | -0.112 | -1.467 | -0.001 | 0.012 | 0.012 | 0.089 | 0.78 | 0.85 | 0.72 | 0.82 | 0.75 | 0.83 |
| 2 | 237794438 | rs6707132 | 1.073 | -0.47 | 0.06 | -1.441 | 0.007 | 0.034 | 0.034 | 1.304 | 0.53 | 0.32 | 0.49 | 0.66 | 0.55 | 0.54 |
| 2 | 237794545 | rs30105 | -0.902 | -0.254 | -0.359 | -1.338 | -0.01 | 0.01 | 0.01 | 0 | 0.79 | 0.89 | 0.73 | 0.84 | 0.75 | 0.83 |
| 2 | 237799960 | rs6737351 | 0.174 | -0.742 | -0.512 | -1.614 | -0.042 | 0.101 | 0.101 | 0.12 | 0.62 | 0.81 | 0.63 | 0.82 | 0.52 | 0.73 |
| 2 | 237803300 | rs10192099 | 0.199 | -0.761 | -0.453 | -1.623 | -0.037 | 0.096 | 0.096 | 0.142 | 0.62 | 0.79 | 0.63 | 0.82 | 0.52 | 0.73 |
| 2 | 237805870 | rs10169105 | 0.387 | -1.069 | -0.214 | -1.86 | -0.013 | -0.044 | -0.044 | 0 | 0.13 | 0.17 | 0.06 | 0.04 | 0.08 | 0.1 |
| 2 | 237806193 | rs2195869 | 0.059 | -1.034 | -0.293 | -1.829 | -0.037 | 0.097 | 0.097 | 0.141 | 0.62 | 0.79 | 0.63 | 0.82 | 0.52 | 0.73 |
| 2 | 237807135 | rs4663234 | 0.068 | -0.993 | -0.291 | -1.824 | -0.037 | 0.115 | 0.115 | 0.207 | 0.6 | 0.79 | 0.63 | 0.83 | 0.51 | 0.71 |
| 2 | 237810403 | rs6431488 | 0.129 | -0.967 | -0.293 | -1.824 | -0.041 | 0.131 | 0.131 | 0.244 | 0.6 | 0.81 | 0.63 | 0.85 | 0.51 | 0.72 |
| 2 | 237810566 | rs12463648 | 1.758 | -1.133 | 0.459 | -1.75 | -0.012 | 0.021 | 0.021 | 0.305 | 0.37 | 0.3 | 0.53 | 0.76 | 0.41 | 0.61 |
| 2 | 237811515 | rs4663236 | 0.899 | -1.046 | 0.36 | -1.725 | 0.003 | 0.053 | 0.053 | 0 | 0.47 | 0.63 | 0.57 | 0.79 | 0.43 | 0.62 |
| 2 | 237817522 | rs10803649 | 0.372 | -0.782 | 0.602 | -1.598 | -0.004 | 0.018 | 0.018 | 0 | 0.58 | 0.61 | 0.53 | 0.71 | 0.53 | 0.59 |
| 2 | 237821589 | rs7603272 | 1.036 | -0.857 | 0.532 | -1.732 | 0.075 | 0.143 | 0.143 | 1.228 | 0.61 | 0.2 | 0.79 | 0.91 | 0.77 | 0.76 |
| 2 | 237829432 | rs1107194 | 0.272 | -0.836 | 0.868 | -1.703 | -0.041 | -0.058 | -0.058 | 0 | 0.2 | 0.29 | 0.08 | 0.05 | 0.09 | 0.12 |
| 2 | 237831359 | rs7565151 | 0.135 | -0.797 | 1.042 | -1.694 | -0.002 | -0.081 | -0.081 | 0.031 | 0.15 | 0.31 | 0.1 | 0.03 | 0.14 | 0.17 |
| 2 | 237840220 | rs10164931 | -0.191 | -0.811 | 0.63 | -1.641 | -0.082 | -0.097 | -0.097 | 0.932 | 0.23 | 0.45 | 0.1 | 0.08 | 0.09 | 0.12 |
| 2 | 237840593 | rs10167711 | -0.191 | -0.811 | 0.63 | -1.641 | -0.082 | -0.097 | -0.097 | 0 | 0.23 | 0.46 | 0.1 | 0.08 | 0.09 | 0.12 |
| 2 | 237841705 | rs10168628 | -0.553 | -0.333 | 0.17 | -1.391 | -0.051 | -0.004 | -0.004 | 1.05 | 0.74 | 0.84 | 0.63 | 0.67 | 0.64 | 0.7 |
| 2 | 237841816 | rs10203831 | -0.921 | -0.506 | 0.337 | -1.468 | -0.012 | -0.049 | -0.049 | 0 | 0.39 | 0.58 | 0.27 | 0.08 | 0.23 | 0.31 |
| 2 | 237843486 | rs4541209 | 1.454 | -0.403 | 0.21 | -1.43 | 0.003 | -0.004 | -0.004 | 0 | 0.39 | 0.4 | 0.38 | 0.59 | 0.42 | 0.4 |
| 2 | 237843723 | rs1865671 | -1.107 | -0.463 | -0.054 | -1.452 | -0.083 | 0.008 | 0.008 | 0 | 0.81 | 0.95 | 0.69 | 0.87 | 0.7 | 0.84 |
| 2 | 237848049 | rs1865672 | -0.633 | -0.471 | 0.224 | -1.495 | -0.014 | -0.067 | -0.067 | 0 | 0.36 | 0.35 | 0.25 | 0.05 | 0.23 | 0.3 |
| 2 | 237848512 | rs1435846 | 1.088 | 0 | 0.35 | -1.222 | -0.002 | 0.001 | 0.001 | 0 | 0.38 | 0.35 | 0.38 | 0.59 | 0.44 | 0.39 |
| 2 | 237848643 | rs1836711 | 0.348 | -0.097 | 0.115 | -1.294 | -0.002 | -0.007 | -0.007 | 0 | 0.51 | 0.49 | 0.46 | 0.63 | 0.56 | 0.55 |
| 2 | 237851338 | rs729454 | 1.124 | -0.086 | -0.522 | -1.033 | -0.052 | -0.01 | -0.01 | 0 | 0.36 | 0.12 | 0.36 | 0.58 | 0.43 | 0.4 |
| 2 | 237851380 | rs893417 | 1.084 | -0.074 | -0.49 | -1.094 | -0.032 | 0.013 | 0.013 | 0 | 0.37 | 0.18 | 0.36 | 0.62 | 0.43 | 0.4 |
| 2 | 237857351 | rs6733928 | 0.833 | -0.14 | -0.51 | -0.735 | -0.008 | 0.004 | 0.004 | 0.008 | 0.43 | 0.1 | 0.41 | 0.6 | 0.4 | 0.5 |
| 2 | 237863149 | rs1435847 | -0.469 | -0.288 | -0.215 | -0.766 | -0.038 | 0 | 0 | 0.934 | 0.75 | 0.76 | 0.59 | 0.83 | 0.59 | 0.81 |
| 2 | 237864981 | rs6431501 | -0.34 | -0.49 | 0.715 | -0.733 | 0.035 | -0.033 | -0.033 | 3.627 | 0.1 | 0.26 | 0.06 | 0.02 | 0.1 | 0.1 |
| 2 | 237865757 | rs6748006 | 0.56 | -0.743 | 1.013 | -0.878 | -0.014 | -0.013 | -0.013 | 0 | 0.22 | 0.23 | 0.1 | 0.18 | 0.1 | 0.19 |
| 2 | 237865903 | rs6748162 | 0.669 | -0.782 | 0.95 | -0.897 | 0.019 | 0.018 | 0.018 | 0.294 | 0.2 | 0.16 | 0.09 | 0.18 | 0.1 | 0.19 |
| 2 | 237866575 | rs2060518 | -1.004 | -0.812 | 0.841 | -0.911 | 0.031 | 0.041 | 0.041 | 3.457 | 0.35 | 0.67 | 0.18 | 0.24 | 0.2 | 0.3 |
| 2 | 237868232 | rs11679109 | -0.791 | -0.808 | 0.98 | -0.909 | -0.018 | -0.032 | -0.032 | 0 | 0.33 | 0.56 | 0.17 | 0.24 | 0.2 | 0.3 |
| 2 | 237873958 | rs7600637 | -0.381 | -0.866 | 0.715 | -0.886 | -0.017 | -0.013 | -0.013 | 0 | 0.28 | 0.62 | 0.13 | 0.21 | 0.1 | 0.2 |
| 2 | 237877769 | rs7562412 | -0.834 | -0.791 | 0.508 | -0.997 | 0.056 | 0.129 | 0.129 | 0.732 | 0.39 | 0.75 | 0.28 | 0.19 | 0.24 | 0.36 |
| 2 | 237885874 | rs2701323 | -0.453 | -0.747 | 0.827 | -0.928 | -0.055 | -0.016 | -0.016 | 0 | 0.76 | 0.8 | 0.6 | 0.81 | 0.6 | 0.82 |
| 2 | 237895243 | rs9646714 | -0.276 | -0.782 | 1.505 | -0.86 | -0.022 | -0.053 | -0.053 | 0 | 0.35 | 0.46 | 0.25 | 0.23 | 0.24 | 0.33 |
| 2 | 237895703 | rs9287613 | -0.587 | -0.798 | 1.421 | -0.868 | 0.031 | 0.055 | 0.055 | 1.555 | 0.38 | 0.68 | 0.27 | 0.23 | 0.24 | 0.37 |
| 2 | 237897671 | rs13024150 | -0.058 | -0.788 | 1.387 | -0.866 | 0.006 | 0.01 | 0.01 | 0.128 | 0.33 | 0.27 | 0.24 | 0.2 | 0.23 | 0.33 |
| 2 | 237901771 | rs11899301 | -0.202 | -0.778 | 0.876 | -0.87 | -0.017 | -0.053 | -0.053 | 0.091 | 0.34 | 0.4 | 0.24 | 0.2 | 0.24 | 0.33 |

| | | | | | | | | | | | | | | | | |
|---|---|---|---|---|---|---|---|---|---|---|---|---|---|---|---|---|
| 2 | 237902511 | rs7584451 | 1.005 | -0.801 | -0.092 | -0.89 | 0.03 | 0.038 | 0.038 | 0.067 | 0.23 | 0.13 | 0.17 | 0.16 | 0.13 | 0.25 |
| 2 | 237906903 | rs2701336 | 1.541 | -0.622 | -0.578 | -0.812 | -0.046 | -0.114 | -0.114 | 0.121 | 0.38 | 0.03 | 0.32 | 0.59 | 0.36 | 0.47 |
| 2 | 237909007 | rs2701337 | -0.235 | -0.193 | -0.186 | -0.716 | 0.007 | 0 | 0 | 0.266 | 0.77 | 0.72 | 0.59 | 0.81 | 0.61 | 0.84 |
| 2 | 237911082 | rs755512 | 0.651 | 0.017 | -0.671 | -0.781 | -0.033 | 0.018 | 0.018 | 0 | 0.07 | 0.01 | 0.06 | 0.03 | 0.11 | 0.08 |
| 2 | 237916980 | rs2701339 | 0.736 | -0.41 | -0.733 | -0.61 | 0.002 | -0.079 | -0.079 | 0 | 0.36 | 0.05 | 0.37 | 0.6 | 0.32 | 0.41 |
| 2 | 237917604 | rs2573718 | -0.475 | 0.407 | -0.243 | -0.744 | 0.05 | -0.039 | -0.039 | 0.063 | 0.88 | 0.82 | 0.82 | 1 | 0.71 | 0.92 |
| 2 | 237933966 | rs2318131 | 0.271 | -0.112 | -0.165 | -0.624 | 0.002 | -0.008 | -0.008 | 2.837 | 0.38 | 0.31 | 0.38 | 0.38 | 0.36 | 0.34 |
| 2 | 237943585 | rs7591958 | 0.08 | -0.075 | -0.215 | -0.642 | 0 | -0.001 | -0.001 | 0 | 0.4 | 0.37 | 0.39 | 0.38 | 0.37 | 0.35 |
| 2 | 237944903 | rs13383408 | -0.763 | -0.686 | -0.208 | -0.512 | -0.029 | -0.03 | -0.03 | 0.137 | 0.09 | 0.14 | 0.04 | 0 | 0.02 | 0.08 |
| 2 | 237947288 | rs7576705 | -0.08 | -0.375 | -0.591 | -0.32 | 0.076 | -0.018 | -0.018 | 0.1 | 0.53 | 0.25 | 0.69 | 0.42 | 0.67 | 0.46 |
| 2 | 237957754 | rs7597414 | -1.366 | -0.1 | 0.756 | -0.884 | 0.003 | -0.114 | -0.114 | 0 | 0.22 | 0.44 | 0.12 | 0.06 | 0.19 | 0.09 |
| 2 | 237958171 | rs7597880 | -0.859 | -0.049 | 0.98 | -0.915 | -0.02 | -0.056 | -0.056 | 0 | 0.16 | 0.24 | 0.07 | 0.01 | 0.09 | 0.06 |
| 2 | 237958255 | rs6721793 | -2.026 | -0.322 | 1.573 | -1.02 | 0.064 | -0.057 | -0.057 | 0.394 | 0.13 | 0.55 | 0.09 | 0.05 | 0.13 | 0.05 |
| 2 | 237978320 | rs1986830 | -1.905 | -0.315 | 1.311 | -1.038 | -0.002 | -0.038 | -0.038 | 0 | 0.27 | 0.75 | 0.15 | 0.06 | 0.21 | 0.1 |
| 2 | 237980920 | rs7593758 | -1.988 | -0.318 | 1.556 | -1.039 | -0.007 | -0.062 | -0.062 | 0 | 0.28 | 0.72 | 0.15 | 0.06 | 0.21 | 0.1 |
| 2 | 237991777 | rs13404290 | -1.695 | -0.446 | 0.517 | -1.27 | -0.137 | 0.105 | 0.105 | 2.026 | 0.55 | 0.97 | 0.36 | 0.29 | 0.51 | 0.49 |
| 2 | 237992613 | rs7577630 | -1.671 | -0.446 | 0.772 | -1.27 | -0.059 | 0.136 | 0.136 | 1.912 | 0.55 | 0.9 | 0.36 | 0.29 | 0.51 | 0.49 |
| 2 | 237998872 | rs3748990 | 0.272 | 0.011 | 1.468 | -1.077 | -0.043 | -0.005 | -0.005 | 0 | 0.19 | 0.03 | 0.15 | 0.18 | 0.26 | 0.21 |
| 2 | 238032491 | rs10209493 | -1.394 | -0.156 | 2.246 | -0.944 | -0.053 | -0.109 | -0.109 | 0 | 0.17 | 0.6 | 0.09 | 0.02 | 0.07 | 0.06 |
| 2 | 238052911 | rs4549025 | 0.651 | -0.063 | 1.957 | -0.926 | 0.018 | -0.021 | -0.021 | 2.371 | 0.94 | 0.94 | 0.97 | 1 | 0.9 | 0.97 |
| 2 | 238065463 | rs10172321 | -1.765 | -0.182 | 2.252 | -0.991 | -0.065 | -0.114 | -0.114 | 0.037 | 0.21 | 0.64 | 0.1 | 0.02 | 0.08 | 0.08 |
| 2 | 238070572 | rs4575672 | 2.157 | -0.178 | 2.182 | -0.994 | -0.005 | -0.063 | -0.063 | 0 | 0.73 | 0.28 | 0.86 | 0.98 | 0.81 | 0.89 |
| 2 | 238086404 | rs10165956 | -1.664 | -0.23 | 2.163 | -0.958 | -0.011 | -0.019 | -0.019 | 0.091 | 0.06 | 0.24 | 0.04 | 0 | 0.02 | 0.02 |
| 2 | 238089055 | rs13420519 | -1.058 | -0.06 | 1.713 | -0.872 | -0.032 | -0.092 | -0.092 | 0.572 | 0.14 | 0.38 | 0.06 | 0 | 0.06 | 0.06 |
| 2 | 238100486 | rs4527163 | 1.675 | 0.145 | 1.412 | -0.788 | 0.055 | -0.063 | -0.063 | 0.307 | 0.89 | 0.67 | 0.93 | 1 | 0.88 | 0.96 |
| 2 | 238102905 | rs4663242 | 1.766 | 0.053 | 0.669 | -0.962 | -0.006 | 0.063 | 0.063 | 0.958 | 0.72 | 0.17 | 0.8 | 0.98 | 0.76 | 0.92 |
| 2 | 238114043 | rs6711456 | 1.82 | 0.182 | 0.897 | -0.621 | 0.001 | -0.014 | -0.014 | 0 | 0.8 | 0.26 | 0.79 | 0.94 | 0.81 | 0.91 |
| 2 | 238118447 | rs10929217 | 0.574 | 0.423 | 0.365 | -0.255 | 0.036 | -0.024 | -0.024 | 0 | 0.43 | 0.19 | 0.55 | 0.47 | 0.4 | 0.32 |
| 2 | 238120185 | rs6714480 | 0.973 | 0.827 | 0.907 | -0.187 | 0.091 | -0.056 | -0.056 | 0 | 0.9 | 0.72 | 0.84 | 0.99 | 0.84 | 0.94 |
| 2 | 238137652 | rs10180362 | -0.995 | 0.328 | 0.825 | 0.075 | 0.03 | 0.035 | 0.035 | 0 | 0.06 | 0.22 | 0.1 | 0.06 | 0.06 | 0.06 |
| 2 | 238140743 | rs10195752 | -0.918 | 0.341 | 0.854 | 0.065 | 0.032 | -0.027 | -0.027 | 0.202 | 0.06 | 0.22 | 0.1 | 0.01 | 0.07 | 0.06 |
| 2 | 238143803 | rs4490127 | -0.789 | 0.181 | 0.569 | 0.002 | -0.131 | -0.144 | -0.144 | 0 | 0.55 | 0.95 | 0.52 | 0.52 | 0.51 | 0.65 |
| 2 | 238153766 | rs4530312 | 0.203 | 0.066 | 0.875 | 0.035 | -0.047 | -0.05 | -0.05 | 0.109 | 0.4 | 0.14 | 0.36 | 0.46 | 0.44 | 0.59 |
| 2 | 238172570 | rs6431534 | -0.527 | 0.333 | -0.284 | 0.185 | -0.034 | 0.006 | 0.006 | 0 | 0.51 | 0.83 | 0.51 | 0.47 | 0.51 | 0.65 |
| 2 | 238173270 | rs12328617 | -0.648 | 0.51 | -0.25 | 0.062 | 0.033 | -0.025 | -0.025 | 0 | 0.06 | 0.22 | 0.1 | 0.01 | 0.07 | 0.04 |
| 2 | 238178706 | rs6728906 | -0.518 | 0.451 | -1.157 | 0.229 | -0.066 | -0.053 | -0.053 | 0 | 0.54 | 0.89 | 0.58 | 0.53 | 0.51 | 0.66 |
| 2 | 238192587 | rs11687617 | -0.343 | 0.146 | -0.151 | 0.732 | -0.013 | 0.06 | 0.06 | 0 | 0.2 | 0.25 | 0.12 | 0.32 | 0.19 | 0.28 |
| 2 | 238192670 | rs4663708 | -0.794 | 0.198 | -0.687 | 0.676 | 0.001 | 0.005 | 0.005 | 0 | 0.47 | 0.49 | 0.62 | 0.61 | 0.46 | 0.67 |
| 2 | 238195820 | rs4274551 | 0.592 | 0.288 | -1.07 | 0.682 | -0.085 | -0.081 | -0.081 | 0 | 0.23 | 0.02 | 0.41 | 0.28 | 0.25 | 0.37 |
| 2 | 238197191 | rs6756356 | -1.136 | 0.413 | -1.413 | 0.492 | -0.129 | -0.046 | -0.046 | 0 | 0.64 | 0.98 | 0.79 | 0.65 | 0.59 | 0.74 |
| 2 | 238197739 | rs4496283 | -0.952 | 0.384 | -1.012 | 0.351 | -0.05 | -0.008 | -0.008 | 0 | 0.61 | 0.86 | 0.74 | 0.6 | 0.58 | 0.7 |
| 2 | 238205060 | rs11676186 | -0.013 | 0.538 | -0.434 | 0.278 | 0.008 | -0.056 | -0.056 | 2.378 | 0.11 | 0.22 | 0.11 | 0 | 0.11 | 0.03 |
| 2 | 238208652 | rs1908970 | -0.229 | 0.453 | -0.94 | 0.574 | -0.005 | -0.007 | -0.007 | 0 | 0.3 | 0.19 | 0.46 | 0.32 | 0.32 | 0.44 |
| 2 | 238209382 | rs4663248 | -0.259 | 0.047 | -0.688 | 0.639 | -0.026 | -0.001 | -0.001 | 0 | 0.36 | 0.36 | 0.31 | 0.37 | 0.24 | 0.39 |
| 2 | 238214039 | rs10803657 | -0.251 | 0.851 | -0.588 | 0.68 | -0.004 | -0.004 | -0.004 | 0 | 0.68 | 0.77 | 0.59 | 0.62 | 0.68 | 0.59 |
| 2 | 238214198 | rs13426046 | -0.439 | 0.912 | -0.645 | 0.537 | 0.005 | -0.019 | -0.019 | 0.051 | 0.34 | 0.51 | 0.35 | 0.31 | 0.38 | 0.37 |
| 2 | 238214279 | rs12477269 | -0.439 | 0.912 | -0.645 | 0.537 | 0.005 | -0.019 | -0.019 | 0 | 0.34 | 0.51 | 0.35 | 0.31 | 0.38 | 0.37 |
| 2 | 238214571 | rs13404258 | 0.611 | 1.05 | -0.367 | 0.325 | 0 | -0.059 | -0.059 | 0.417 | 0.13 | 0.23 | 0.12 | 0 | 0.14 | 0.06 |
| 2 | 238215341 | rs6732495 | 0.26 | 0.953 | -0.443 | 0.259 | 0.015 | 0.024 | 0.024 | 0.186 | 0.17 | 0.27 | 0.21 | 0.31 | 0.22 | 0.25 |
| 2 | 238215479 | rs6729500 | 0.487 | 0.923 | -0.288 | 0.25 | 0.004 | -0.078 | -0.078 | 0 | 0.13 | 0.31 | 0.12 | 0 | 0.15 | 0.06 |
| 2 | 238217353 | rs4663714 | -0.384 | 0.921 | -0.145 | 0.254 | -0.012 | -0.018 | -0.018 | 0 | 0.21 | 0.16 | 0.22 | 0.31 | 0.23 | 0.31 |
| 2 | 238222143 | rs4392201 | -0.954 | 0.675 | -0.082 | 0.344 | -0.023 | -0.021 | -0.021 | 0 | 0.3 | 0.59 | 0.28 | 0.25 | 0.24 | 0.29 |
| 2 | 238227776 | rs10929225 | -0.245 | 0.492 | -0.288 | 0.589 | -0.034 | 0.034 | 0.034 | 0.285 | 0.18 | 0.37 | 0.17 | 0.25 | 0.1 | 0.21 |
| 2 | 238227919 | rs6759927 | 1.17 | 0.523 | -0.749 | 0.543 | 0.005 | -0.027 | -0.027 | 0 | 0.32 | 0.14 | 0.24 | 0.31 | 0.33 | 0.22 |
| 2 | 238231803 | rs12465759 | -0.571 | 0.405 | -0.054 | 0.625 | -0.019 | 0.015 | 0.015 | 0.322 | 0.18 | 0.28 | 0.16 | 0.26 | 0.09 | 0.21 |
| 2 | 238236921 | rs6720773 | 0.841 | 0.977 | -0.029 | 0.673 | 0.014 | -0.002 | -0.002 | 0 | 0.49 | 0.42 | 0.35 | 0.49 | 0.44 | 0.3 |
| 2 | 238244963 | rs6728818 | -0.208 | 1.473 | 0.079 | 0.448 | -0.013 | -0.024 | -0.024 | 0 | 0.73 | 0.81 | 0.55 | 0.54 | 0.69 | 0.51 |
| 2 | 238253065 | rs2291795 | 0.406 | 1.443 | -0.08 | 0.476 | -0.012 | 0.007 | 0.007 | 0 | 0.15 | 0.17 | 0.12 | 0.19 | 0.09 | 0.18 |
| 2 | 238253930 | rs2256485 | 0.445 | 1.433 | -0.22 | 0.234 | -0.003 | 0.026 | 0.026 | 0 | 0.12 | 0.08 | 0.17 | 0.05 | 0.13 | 0.12 |
| 2 | 238255686 | rs7586225 | 0.406 | 1.551 | 0.009 | 0.212 | -0.006 | -0.005 | -0.005 | 0 | 0.79 | 0.77 | 0.66 | 0.73 | 0.69 | 0.65 |
| 2 | 238256171 | rs2645767 | -0.945 | 1.273 | 0.142 | 0.13 | -0.012 | -0.052 | -0.052 | 0.956 | 0.18 | 0.31 | 0.21 | 0.06 | 0.15 | 0.14 |
| 2 | 238257013 | rs2646265 | 0.417 | 1.283 | 0.072 | 0.139 | -0.003 | 0.026 | 0.026 | 0 | 0.12 | 0.08 | 0.17 | 0.05 | 0.13 | 0.12 |
| 2 | 238257896 | rs2645766 | -0.878 | 1.345 | 0.323 | 0.161 | -0.009 | -0.104 | -0.104 | 0 | 0.21 | 0.44 | 0.23 | 0.07 | 0.19 | 0.18 |
| 2 | 238260610 | rs3790998 | -0.114 | 1.692 | 0.042 | 0.356 | -0.018 | -0.016 | -0.016 | 4.902 | 0.84 | 0.85 | 0.68 | 0.73 | 0.7 | 0.66 |
| 2 | 238261112 | rs4663726 | 1.131 | 1.489 | 0.184 | 0.19 | 0.001 | -0.002 | -0.002 | 0 | 0.61 | 0.52 | 0.55 | 0.66 | 0.59 | 0.45 |
| 2 | 238265531 | rs11677456 | 1.789 | 0.904 | 0.26 | 0.437 | -0.002 | 0 | 0 | 0 | 0.43 | 0.39 | 0.34 | 0.39 | 0.42 | 0.29 |
| 2 | 238267058 | rs4663731 | 1.111 | 1.089 | 0.771 | 0.515 | 0.053 | 0.057 | 0.057 | 0.31 | 0.7 | 0.56 | 0.5 | 0.45 | 0.61 | 0.47 |
| 2 | 238267717 | rs2646254 | -1.13 | 0.727 | 1.175 | 0.681 | -0.036 | 0.035 | 0.035 | 0.015 | 0.23 | 0.54 | 0.25 | 0.35 | 0.15 | 0.27 |
| 2 | 238269120 | rs12622722 | 1.463 | 1.079 | 0.808 | 0.501 | -0.004 | -0.007 | -0.007 | 0 | 0.58 | 0.33 | 0.36 | 0.4 | 0.49 | 0.37 |

| | | | | | | | | | | | | | | | | |
|---|---|---|---|---|---|---|---|---|---|---|---|---|---|---|---|---|
| 2 | 238270726 | rs7597795 | -1.316 | 0.638 | 1.263 | 0.154 | -0.018 | -0.12 | -0.12 | 0 | 0.23 | 0.45 | 0.17 | 0.09 | 0.18 | 0.21 |
| 2 | 238271284 | rs10167850 | -0.813 | 0.581 | 1.289 | 0.137 | -0.025 | -0.119 | -0.119 | 0.014 | 0.2 | 0.37 | 0.14 | 0.05 | 0.14 | 0.18 |
| 2 | 238276335 | rs2645771 | -1.078 | 0.778 | 1.193 | 0.429 | 0.004 | 0.003 | 0.003 | 0 | 0.22 | 0.33 | 0.31 | 0.3 | 0.22 | 0.3 |
| 2 | 238277795 | rs2646260 | -1.711 | 0.666 | 1.282 | 0.326 | -0.001 | -0.008 | -0.008 | 0 | 0.27 | 0.64 | 0.35 | 0.3 | 0.25 | 0.32 |
| 2 | 238283605 | rs12622093 | -0.609 | 0.532 | 1.269 | 0.271 | -0.032 | 0.007 | 0.007 | 0 | 0.23 | 0.11 | 0.25 | 0.27 | 0.25 | 0.25 |
| 2 | 238294434 | rs10929230 | -1.225 | 0.575 | 1.337 | 0.298 | -0.006 | -0.014 | -0.014 | 0.008 | 0.3 | 0.63 | 0.36 | 0.31 | 0.27 | 0.31 |
| 2 | 238301184 | rs1874573 | 0.343 | 0.236 | 0.751 | 0.127 | -0.01 | 0.012 | 0.012 | 0.63 | 0.69 | 0.77 | 0.71 | 0.82 | 0.69 | 0.62 |
| 2 | 238311612 | rs2645779 | -1.406 | -0.298 | 1.538 | 0.128 | -0.006 | -0.008 | -0.008 | 0 | 0.3 | 0.66 | 0.37 | 0.33 | 0.27 | 0.31 |
| 2 | 238317822 | rs2645782 | 0.825 | 0.029 | 1.539 | -0.012 | 0 | 0.003 | 0.003 | 0 | 0.55 | 0.44 | 0.52 | 0.66 | 0.55 | 0.41 |
| 2 | 238318315 | rs4663747 | -0.243 | 0.019 | 0.788 | -0.042 | 0.048 | 0.008 | 0.008 | 0.093 | 0.32 | 0.2 | 0.35 | 0.29 | 0.23 | 0.37 |
| 2 | 238320371 | rs7579936 | 0.986 | -0.066 | 0.634 | 0.276 | -0.023 | 0.041 | 0.041 | 0 | 0.46 | 0.14 | 0.49 | 0.62 | 0.52 | 0.33 |
| 2 | 238321317 | rs12476069 | -0.238 | -0.015 | 0.433 | 0.341 | -0.045 | 0.034 | 0.034 | 0 | 0.93 | 0.99 | 0.95 | 0.98 | 0.85 | 0.81 |
| 2 | 238326433 | rs3863910 | -0.746 | -0.898 | 0.896 | 0.605 | 0.029 | 0.019 | 0.019 | 0.073 | 0.45 | 0.7 | 0.45 | 0.35 | 0.34 | 0.47 |
| 2 | 238334279 | rs6717825 | -1.696 | -0.734 | 1.962 | 0.908 | -0.002 | 0 | 0 | 0 | 0.1 | 0.14 | 0.17 | 0.13 | 0.08 | 0.11 |
| 2 | 238336802 | rs6719451 | -2.827 | -0.994 | 2.027 | 0.905 | 0.044 | 0.02 | 0.02 | 0.015 | 0.26 | 0.82 | 0.26 | 0.23 | 0.12 | 0.31 |
| 2 | 238349940 | rs7588511 | -2.192 | -0.763 | 2.878 | 1.05 | -0.062 | -0.046 | -0.046 | 0 | 0.14 | 0.48 | 0.06 | 0.06 | 0.04 | 0.1 |
| 2 | 238351337 | rs12471077 | -1.992 | -0.387 | 3.002 | 1.19 | 0.02 | 0.006 | 0.006 | 0 | 0.07 | 0.13 | 0.19 | 0.13 | 0.12 | 0.16 |
| 2 | 238356795 | rs7602387 | -3.111 | -0.655 | 3.153 | 1.157 | 0.027 | 0.025 | 0.025 | 0.441 | 0.29 | 0.83 | 0.32 | 0.23 | 0.21 | 0.36 |
| 2 | 238365528 | rs12469894 | -3.098 | -0.538 | 3.574 | 1.155 | 0.005 | 0.008 | 0.008 | 0.117 | 0.29 | 0.74 | 0.34 | 0.23 | 0.22 | 0.35 |
| 2 | 238368751 | rs10445707 | -2.996 | -0.544 | 3.949 | 1.154 | -0.009 | -0.017 | -0.017 | 0.382 | 0.28 | 0.63 | 0.32 | 0.23 | 0.22 | 0.35 |
| 2 | 238378476 | rs7582828 | -2.926 | -0.559 | 3.993 | 1.163 | 0.009 | -0.021 | -0.021 | 3.516 | 0.24 | 0.59 | 0.31 | 0.21 | 0.22 | 0.32 |
| 2 | 238380272 | rs10929235 | -2.035 | -0.583 | 3.822 | 1.193 | -0.009 | 0 | 0 | 0.05 | 0.15 | 0.15 | 0.25 | 0.15 | 0.18 | 0.21 |
| 2 | 238387228 | rs7584330 | -3.264 | -0.651 | 3.792 | 1.258 | 0.014 | 0.007 | 0.007 | 0.129 | 0.31 | 0.74 | 0.34 | 0.25 | 0.21 | 0.36 |
| 2 | 238395871 | rs1574529 | -2.055 | -0.695 | 3.792 | 1.319 | -0.025 | 0.012 | 0.012 | 0 | 0.13 | 0.31 | 0.22 | 0.15 | 0.08 | 0.15 |
| 2 | 238413338 | rs7569427 | -2.493 | -0.718 | 2.87 | 1.445 | -0.04 | 0.019 | 0.019 | 0.235 | 0.19 | 0.72 | 0.27 | 0.27 | 0.08 | 0.26 |
| 2 | 238417728 | rs6431544 | -2.692 | -0.623 | 2.769 | 1.451 | 0.001 | -0.003 | -0.003 | 0 | 0.25 | 0.7 | 0.31 | 0.28 | 0.2 | 0.27 |
| 2 | 238427194 | rs3751109 | -2.762 | -0.387 | 3.269 | 1.451 | 0.069 | 0.027 | 0.027 | 0.079 | 0.11 | 0.37 | 0.1 | 0.08 | 0.14 | 0.11 |
| 2 | 238427251 | rs3751107 | -2.762 | -0.393 | 3.267 | 1.496 | 0.069 | 0.027 | 0.027 | 0.079 | 0.11 | 0.37 | 0.1 | 0.08 | 0.14 | 0.11 |
| 2 | 238433955 | rs6705903 | -3.635 | -0.376 | 3.851 | 1.772 | 0.071 | 0.182 | 0.182 | 0.568 | 0.05 | 0.46 | 0.07 | 0.14 | 0.02 | 0.13 |
| 2 | 238434249 | rs2292881 | -1.438 | -0.633 | 3.842 | 1.717 | 0.021 | -0.011 | -0.011 | 1.446 | 0.16 | 0.07 | 0.2 | 0.19 | 0.07 | 0.24 |
| 2 | 238435398 | rs7598559 | -2.68 | -0.615 | 3.95 | 1.7 | 0.048 | 0.011 | 0.011 | 0.001 | 0.11 | 0.23 | 0.08 | 0.07 | 0.14 | 0.09 |
| 2 | 238443226 | rs2292884 | -2.962 | -0.624 | 3.33 | 1.735 | -0.01 | 0.002 | 0.002 | 0 | 0.3 | 0.62 | 0.31 | 0.27 | 0.21 | 0.34 |
| 2 | 238443542 | rs9646713 | -2.436 | -0.593 | 2.86 | 1.663 | 0.063 | 0.019 | 0.019 | 0 | 0.11 | 0.35 | 0.09 | 0.08 | 0.14 | 0.13 |
| 2 | 238445593 | rs13418586 | -2.367 | -0.513 | 1.612 | 1.654 | 0.023 | -0.031 | -0.031 | 0 | 0.33 | 0.78 | 0.34 | 0.41 | 0.23 | 0.46 |
| 2 | 238447168 | rs13425580 | -2.577 | -0.504 | 1.185 | 1.64 | 0.028 | -0.079 | -0.079 | 0.058 | 0.35 | 0.92 | 0.36 | 0.41 | 0.23 | 0.46 |
| 2 | 238449007 | rs3817362 | -2.044 | -0.698 | 1.281 | 1.592 | 0.052 | -0.027 | -0.027 | 0 | 0.28 | 0.81 | 0.3 | 0.39 | 0.12 | 0.38 |
| 2 | 238454154 | rs867306 | -2.36 | -0.078 | 1.965 | 1.482 | 0.048 | 0.064 | 0.064 | 0 | 0.1 | 0.28 | 0.08 | 0.1 | 0.1 | 0.06 |
| 2 | 238457684 | rs10173589 | -2.416 | 0.047 | 2.354 | 1.33 | 0.037 | -0.022 | -0.022 | 0.458 | 0.15 | 0.35 | 0.11 | 0.13 | 0.17 | 0.12 |
| 2 | 238458180 | rs2292877 | -3.193 | -0.111 | 3.152 | 1.266 | -0.004 | -0.051 | -0.051 | 0 | 0.12 | 0.59 | 0.07 | 0.07 | 0.07 | 0.05 |
| 2 | 238462725 | rs2325814 | -2.607 | -0.366 | 2.896 | 2.001 | -0.005 | 0.013 | 0.013 | 0.646 | 0.21 | 0.69 | 0.27 | 0.39 | 0.12 | 0.29 |
| 2 | 238467914 | rs1320203 | -2.658 | -0.391 | 3.278 | 2.001 | -0.036 | 0.163 | 0.163 | 0.11 | 0.19 | 0.47 | 0.25 | 0.39 | 0.11 | 0.27 |
| 2 | 238476173 | rs11896561 | -2.498 | -0.457 | 3.02 | 1.997 | -0.029 | 0.099 | 0.099 | 0.126 | 0.17 | 0.35 | 0.23 | 0.38 | 0.09 | 0.2 |
| 2 | 238480686 | rs10183301 | -3.068 | 0.109 | 2.225 | 1.962 | 0.013 | -0.003 | -0.003 | 0 | 0.24 | 0.88 | 0.2 | 0.27 | 0.17 | 0.37 |
| 2 | 238482636 | rs10190103 | -2.286 | 0.139 | 2.366 | 1.924 | 0.026 | -0.017 | -0.017 | 0.795 | 0.05 | 0.16 | 0.05 | 0.03 | 0.08 | 0.17 |
| 2 | 238483729 | rs2280289 | -2.701 | 0.276 | 2.443 | 2.127 | -0.005 | 0.014 | 0.014 | 1.122 | 0.21 | 0.71 | 0.28 | 0.41 | 0.14 | 0.21 |
| 2 | 238491003 | rs2325810 | -1.444 | -0.089 | 2.04 | 1.726 | 0.007 | 0.009 | 0.009 | 12.847 | 0.14 | 0.07 | 0.07 | 0.09 | 0.09 | 0.11 |
| 2 | 238493118 | rs6431547 | -2.778 | -0.086 | 2.357 | 1.843 | -0.001 | -0.002 | -0.002 | 0 | 0.3 | 0.66 | 0.26 | 0.26 | 0.23 | 0.38 |
| 2 | 238494742 | rs3751112 | -2.333 | -0.011 | 2.454 | 1.842 | 0.029 | 0.027 | 0.027 | 0 | 0.18 | 0.55 | 0.21 | 0.17 | 0.16 | 0.29 |
| 2 | 238495843 | rs6733415 | -2.276 | 0.002 | 2.461 | 1.846 | 0.033 | 0.034 | 0.034 | 0 | 0.18 | 0.55 | 0.2 | 0.17 | 0.15 | 0.29 |
| 2 | 238498545 | rs9287621 | -1.85 | -0.013 | 2.481 | 1.853 | -0.019 | 0.083 | 0.083 | 0 | 0.14 | 0.54 | 0.11 | 0.16 | 0.07 | 0.19 |
| 2 | 238498996 | rs6760287 | -1.388 | 0.067 | 1.772 | 1.868 | -0.003 | -0.004 | -0.004 | 0.804 | 0.07 | 0.07 | 0.08 | 0.13 | 0.11 | 0.1 |
| 2 | 238499444 | rs2292873 | -1.333 | 0.043 | 1.772 | 1.872 | -0.003 | -0.004 | -0.004 | 0 | 0.07 | 0.07 | 0.06 | 0.12 | 0.11 | 0.1 |
| 2 | 238499690 | rs2292871 | -3.479 | 0.017 | 1.814 | 1.905 | -0.005 | -0.021 | -0.021 | 0 | 0.39 | 0.82 | 0.35 | 0.42 | 0.34 | 0.51 |
| 2 | 238499727 | rs6755617 | -1.423 | 0.025 | 1.784 | 1.902 | -0.003 | -0.005 | -0.005 | 0 | 0.07 | 0.07 | 0.07 | 0.12 | 0.11 | 0.1 |
| 2 | 238500046 | rs6742960 | -2.08 | 0.04 | 1.889 | 1.818 | -0.025 | 0.054 | 0.054 | 0.04 | 0.23 | 0.57 | 0.19 | 0.29 | 0.18 | 0.28 |
| 2 | 238500211 | rs6431548 | -2.471 | 0.056 | 0.873 | 1.755 | -0.001 | -0.017 | -0.017 | 0 | 0.39 | 0.79 | 0.35 | 0.42 | 0.34 | 0.51 |
| 2 | 238502591 | rs10929236 | -1.653 | 0.393 | 0.496 | 2.169 | -0.015 | -0.072 | -0.072 | 0 | 0.25 | 0.11 | 0.23 | 0.52 | 0.24 | 0.3 |
| 2 | 238502987 | rs13392999 | -2.023 | 0.384 | 1.109 | 2.67 | -0.001 | 0.128 | 0.128 | 0 | 0.12 | 0.18 | 0.18 | 0.54 | 0.12 | 0.24 |
| 2 | 238503066 | rs6710977 | -2.885 | 0.649 | 1.851 | 2.463 | 0.038 | 0.145 | 0.145 | 0.007 | 0.24 | 0.44 | 0.32 | 0.49 | 0.31 | 0.36 |
| 2 | 238504191 | rs12467097 | -2.131 | 1.22 | 1.951 | 2.068 | -0.007 | 0.03 | 0.03 | 0.544 | 0.26 | 0.35 | 0.18 | 0.43 | 0.19 | 0.3 |
| 2 | 238505096 | rs12464142 | -1.144 | 1.205 | 1.829 | 1.995 | 0.006 | -0.011 | -0.011 | 0 | 0.11 | 0.05 | 0.09 | 0.36 | 0.07 | 0.22 |
| 2 | 238505625 | rs6709393 | -2.154 | 1.321 | 2.567 | 1.877 | 0.042 | -0.153 | -0.153 | 0 | 0.25 | 0.52 | 0.24 | 0.04 | 0.3 | 0.2 |
| 2 | 238508099 | rs1316476 | -2.31 | 1.26 | 2.758 | 2.121 | 0.017 | -0.007 | -0.007 | 0 | 0.22 | 0.38 | 0.27 | 0.21 | 0.25 | 0.35 |
| 2 | 238509273 | rs7585075 | -0.355 | 1.47 | 1.983 | 1.863 | -0.016 | 0.013 | 0.013 | 0 | 0.09 | 0.09 | 0.07 | 0 | 0.12 | 0.05 |
| 2 | 238509820 | rs10929238 | -0.118 | 1.434 | 1.9 | 1.911 | -0.037 | 0.017 | 0.017 | 0.024 | 0.06 | 0.01 | 0.1 | 0.04 | 0.16 | 0.12 |
| 2 | 238513839 | rs4663262 | -1.344 | 1.438 | 2.064 | 1.976 | -0.001 | -0.001 | -0.001 | 0 | 0.13 | 0.12 | 0.13 | 0.17 | 0.06 | 0.22 |
| 2 | 238521322 | rs10199579 | -1.869 | 1.71 | 2.41 | 1.771 | 0.046 | -0.066 | -0.066 | 1.121 | 0.18 | 0.3 | 0.19 | 0.04 | 0.31 | 0.16 |
| 2 | 238522744 | rs4278886 | -0.908 | 0.982 | 1.293 | 1.205 | -0.006 | 0.017 | 0.017 | 0.856 | 0.47 | 0.29 | 0.48 | 0.47 | 0.5 | 0.55 |
| 2 | 238524267 | rs13414926 | -2.535 | 1.298 | 0.925 | 1.185 | -0.073 | 0.035 | 0.035 | 0.671 | 0.67 | 0.97 | 0.69 | 0.82 | 0.61 | 0.79 |

| | | | | | | | | | | | | | | | | |
|---|---|---|---|---|---|---|---|---|---|---|---|---|---|---|---|---|
| 2 | 238524416 | rs13430910 | -0.968 | 1.278 | 1.338 | 0.94 | 0.017 | -0.073 | -0.073 | 0 | 0.14 | 0.23 | 0.11 | 0 | 0.16 | 0.05 |
| 2 | 238526411 | rs12464145 | -1.104 | 1.148 | 1.771 | 1.092 | -0.047 | 0.028 | 0.028 | 0.117 | 0.17 | 0.48 | 0.16 | 0.17 | 0.07 | 0.22 |
| 2 | 238532983 | rs6740174 | 0.38 | 1.414 | 1.672 | 1.197 | 0.026 | -0.049 | -0.049 | 0 | 0.3 | 0.12 | 0.32 | 0.43 | 0.22 | 0.38 |
| 2 | 238538597 | rs4663263 | -0.582 | 1.745 | 1.739 | 1.208 | -0.007 | -0.003 | -0.003 | 0 | 0.38 | 0.33 | 0.42 | 0.47 | 0.4 | 0.49 |
| 2 | 238538613 | rs4663264 | -0.753 | 1.749 | 1.724 | 1.21 | -0.002 | -0.001 | -0.001 | 0 | 0.4 | 0.38 | 0.43 | 0.47 | 0.4 | 0.49 |
| 2 | 238549369 | rs13416014 | -1.476 | 1.858 | 1.642 | 1.01 | 0.05 | -0.065 | -0.065 | 0 | 0.19 | 0.31 | 0.18 | 0.05 | 0.29 | 0.21 |
| 2 | 238552006 | rs11894086 | -0.82 | 1.57 | 1.935 | 1.371 | 0 | -0.001 | -0.001 | 0 | 0.44 | 0.43 | 0.45 | 0.49 | 0.37 | 0.5 |
| 2 | 238552973 | rs11895665 | -1.746 | 1.682 | 2.271 | 1.542 | 0.122 | 0.307 | 0.307 | 0 | 0.05 | 0.29 | 0.13 | 0.2 | 0.09 | 0.1 |
| 2 | 238555352 | rs6757425 | -1.408 | 1.349 | 1.724 | 1.188 | 0.003 | 0.027 | 0.027 | 0.8 | 0.55 | 0.8 | 0.59 | 0.64 | 0.54 | 0.63 |
| 2 | 238555898 | rs4663774 | -1.416 | 1.348 | 1.717 | 1.195 | 0.002 | 0.027 | 0.027 | 0.742 | 0.55 | 0.81 | 0.59 | 0.64 | 0.54 | 0.63 |
| 2 | 238561083 | rs2029771 | -1.51 | 1.275 | 1.714 | 1.143 | 0.004 | -0.001 | -0.001 | 0 | 0.54 | 0.72 | 0.61 | 0.54 | 0.54 | 0.64 |
| 2 | 238561249 | rs2029770 | 0.047 | 1.236 | 1.743 | 1.057 | -0.051 | -0.01 | -0.01 | 0.166 | 0.34 | 0.52 | 0.34 | 0.31 | 0.18 | 0.36 |
| 2 | 238564791 | rs13408642 | -0.308 | 1.262 | 1.798 | 1.047 | -0.003 | -0.003 | -0.003 | 0 | 0.38 | 0.36 | 0.39 | 0.34 | 0.33 | 0.48 |
| 2 | 238569354 | rs12995190 | 0.575 | 1.658 | 2.035 | 0.827 | -0.031 | 0.018 | 0.018 | 0.061 | 0.06 | 0.01 | 0.08 | 0.04 | 0.16 | 0.12 |
| 2 | 238570232 | rs6742060 | -1.049 | 1.171 | 2.289 | 0.765 | 0.024 | -0.099 | -0.099 | 0.098 | 0.14 | 0.41 | 0.12 | 0 | 0.13 | 0.05 |
| 2 | 238571031 | rs7597491 | 0.803 | 1.295 | 1.869 | 0.776 | 0.004 | -0.032 | -0.032 | 0 | 0.82 | 0.75 | 0.82 | 0.98 | 0.76 | 0.84 |
| 2 | 238571074 | rs7597504 | 1.856 | 1.305 | 1.919 | 0.844 | 0.01 | -0.003 | -0.003 | 0 | 0.74 | 0.28 | 0.68 | 0.8 | 0.68 | 0.73 |
| 2 | 238571499 | rs903375 | 0.868 | 1.263 | 1.285 | 0.677 | 0.007 | -0.034 | -0.034 | 0.186 | 0.82 | 0.73 | 0.81 | 0.97 | 0.76 | 0.83 |
| 2 | 238574426 | rs6758509 | 0.413 | 0.93 | 0.681 | 0.718 | -0.004 | 0.008 | 0.008 | 0 | 0.88 | 0.9 | 0.82 | 0.92 | 0.89 | 0.82 |
| 2 | 238576444 | rs4663779 | 1.872 | 0.912 | 1.101 | 0.804 | -0.005 | 0 | 0 | 0 | 0.68 | 0.29 | 0.56 | 0.71 | 0.68 | 0.63 |
| 2 | 238576689 | rs10171943 | -2.049 | 0.585 | 1.555 | 1.131 | 0.022 | 0.111 | 0.111 | 0 | 0.06 | 0.16 | 0.11 | 0.18 | 0.08 | 0.1 |
| 2 | 238578536 | rs6733552 | 0.688 | 0.548 | 1.148 | 0.964 | 0.001 | -0.015 | -0.015 | 0.922 | 0.87 | 0.84 | 0.9 | 0.94 | 0.89 | 0.83 |
| 2 | 238581009 | rs7562075 | 1.731 | 0.508 | 1.473 | 1.223 | 0.019 | 0.041 | 0.041 | 0 | 0.76 | 0.66 | 0.68 | 0.66 | 0.7 | 0.64 |
| 2 | 238589737 | rs12995157 | 0.93 | 0.5 | 1.27 | 1.022 | -0.01 | 0.009 | 0.009 | 0 | 0.82 | 0.86 | 0.8 | 0.85 | 0.79 | 0.73 |
| 2 | 238591672 | rs10929245 | 1.513 | 0.417 | 2.122 | 1.053 | 0.015 | -0.016 | -0.016 | 0 | 0.78 | 0.5 | 0.76 | 0.8 | 0.78 | 0.7 |
| 2 | 238592234 | rs6743986 | 0.843 | 0.478 | 1.779 | 1.054 | -0.004 | 0.002 | 0.002 | 0 | 0.82 | 0.84 | 0.8 | 0.86 | 0.79 | 0.73 |
| 2 | 238603174 | rs10171166 | -0.616 | -0.764 | 1.903 | 1.16 | -0.015 | 0 | 0 | 0 | 0.29 | 0.67 | 0.18 | 0.29 | 0.22 | 0.33 |
| 2 | 238604650 | rs11680933 | 0.255 | -0.731 | 1.651 | 1.177 | 0.012 | -0.011 | -0.011 | 0.229 | 0.23 | 0.18 | 0.23 | 0.29 | 0.21 | 0.31 |
| 2 | 238621790 | rs3769090 | 0.279 | -0.74 | 1.538 | 1.142 | 0 | 0 | 0 | 0 | 0.23 | 0.17 | 0.23 | 0.26 | 0.21 | 0.32 |
| 2 | 238623413 | rs2047169 | -0.296 | -0.731 | 1.201 | 1.155 | -0.041 | 0.01 | 0.01 | 1.054 | 0.27 | 0.53 | 0.26 | 0.32 | 0.21 | 0.34 |
| 2 | 238628509 | rs6740706 | -0.12 | -0.584 | 0.738 | 1.346 | -0.052 | -0.062 | -0.062 | 0 | 0.9 | 0.99 | 0.76 | 0.77 | 0.84 | 0.82 |
| 2 | 238630236 | rs1565853 | 1.057 | -0.789 | 0.89 | 1.197 | 0.007 | 0.002 | 0.002 | 0.464 | 0.58 | 0.36 | 0.48 | 0.4 | 0.59 | 0.42 |
| 2 | 238632377 | rs3769085 | -0.852 | -1.54 | 0.516 | 0.986 | -0.002 | -0.001 | -0.001 | 0 | 0.28 | 0.31 | 0.32 | 0.26 | 0.22 | 0.32 |
| 2 | 238635137 | rs10177762 | -1.006 | -2.206 | 0.738 | 1.002 | 0.011 | -0.007 | -0.007 | 0.032 | 0.33 | 0.34 | 0.23 | 0.42 | 0.21 | 0.4 |
| 2 | 238640263 | rs10187629 | -1.666 | -2.109 | 0.622 | 1.044 | 0.044 | 0.11 | 0.11 | 2.226 | 0.08 | 0.44 | 0.07 | 0.07 | 0.05 | 0.02 |
| 2 | 238641521 | rs3769080 | 0.642 | -2.136 | 0.143 | 1.043 | 0.014 | -0.016 | -0.016 | 0 | 0.17 | 0.1 | 0.17 | 0.24 | 0.13 | 0.24 |
| 2 | 238645035 | rs3820809 | -1.855 | -1.815 | 0.246 | 0.764 | -0.042 | 0.002 | 0.002 | 1.587 | 0.2 | 0.61 | 0.13 | 0.18 | 0.13 | 0.17 |
| 2 | 238648521 | rs3769071 | -0.009 | -1.507 | -0.638 | 0.786 | 0.012 | -0.008 | -0.008 | 0 | 0.08 | 0 | 0.03 | 0.06 | 0.05 | 0.04 |
| 2 | 238650986 | rs11693524 | -0.7 | -1.509 | -0.478 | 0.773 | 0.008 | 0.004 | 0.004 | 2.007 | 0.1 | 0.06 | 0.03 | 0.07 | 0.05 | 0.04 |
| 2 | 238656212 | rs3754713 | -1.484 | -1.942 | -0.239 | 1.052 | -0.034 | 0.015 | 0.015 | 1.527 | 0.36 | 0.5 | 0.31 | 0.53 | 0.25 | 0.38 |
| 2 | 238660169 | rs12613388 | -0.979 | -1.826 | -0.388 | 1.156 | 0.01 | -0.012 | -0.012 | 0 | 0.11 | 0.08 | 0.1 | 0.2 | 0.08 | 0.11 |
| 2 | 238661832 | rs733579 | -1.243 | -1.902 | -0.308 | 1.149 | 0.009 | -0.013 | -0.013 | 0.02 | 0.31 | 0.27 | 0.29 | 0.5 | 0.24 | 0.38 |
| 2 | 238663503 | rs3754706 | -0.125 | -1.921 | -0.313 | 1.143 | 0.007 | -0.082 | -0.082 | 0 | 0.21 | 0.1 | 0.25 | 0.44 | 0.2 | 0.35 |
| 2 | 238664010 | rs3769058 | -1.364 | -1.922 | -0.267 | 1.141 | -0.002 | 0 | 0 | 0 | 0.32 | 0.34 | 0.3 | 0.5 | 0.25 | 0.39 |
| 2 | 238668783 | rs3213869 | 0.466 | -1.805 | -0.226 | 1.095 | 0.009 | -0.011 | -0.011 | 0 | 0.07 | 0 | 0.03 | 0.06 | 0.05 | 0.03 |
| 2 | 238671940 | rs15232 | -2.784 | -1.823 | -0.2 | 1.118 | -0.01 | 0.145 | 0.145 | 0 | 0.08 | 0.32 | 0.09 | 0.14 | 0.03 | 0.08 |
| 2 | 238672290 | rs3739040 | -2.451 | -1.813 | -0.187 | 1.08 | 0.023 | 0.145 | 0.145 | 0.511 | 0.05 | 0.32 | 0.02 | 0.08 | 0.01 | 0.01 |
| 2 | 238677614 | rs12617354 | -0.241 | -1.803 | -0.321 | 1.078 | -0.001 | -0.001 | -0.001 | 0.012 | 0.09 | 0.09 | 0.04 | 0.14 | 0.06 | 0.06 |
| 2 | 238682770 | rs2279642 | -1.258 | -1.9 | -0.345 | 1.048 | 0.029 | -0.036 | -0.036 | 0.004 | 0.32 | 0.23 | 0.3 | 0.51 | 0.25 | 0.4 |
| 2 | 238688305 | rs2117364 | -0.377 | -1.984 | -0.319 | 1.046 | -0.011 | 0.04 | 0.04 | 2.48 | 0.23 | 0.29 | 0.21 | 0.41 | 0.18 | 0.26 |
| 2 | 238694777 | rs11694275 | -1.638 | -1.898 | -0.557 | 0.766 | 0.015 | 0.037 | 0.037 | 0.895 | 0.15 | 0.08 | 0.07 | 0.06 | 0.11 | 0.06 |
| 2 | 238697636 | rs2325994 | -2.204 | -1.893 | -0.521 | 0.815 | -0.023 | -0.006 | -0.006 | 0 | 0.12 | 0.4 | 0.2 | 0.07 | 0.07 | 0.11 |
| 2 | 238712175 | rs11676396 | -2.072 | -1.833 | -0.664 | 0.846 | -0.05 | 0.016 | 0.016 | 0.295 | 0.22 | 0.42 | 0.21 | 0.2 | 0.14 | 0.18 |
| 2 | 238714830 | rs10202875 | -1.716 | -1.676 | -0.641 | 0.677 | -0.026 | -0.056 | -0.056 | 0.092 | 0.18 | 0.34 | 0.09 | 0.06 | 0.13 | 0.08 |
| 2 | 238718850 | rs1030744 | -0.019 | -1.605 | -1.186 | 0.365 | 0.013 | 0.081 | 0.081 | 0 | 0.26 | 0.11 | 0.27 | 0.07 | 0.22 | 0.22 |
| 2 | 238721240 | rs12474573 | -0.033 | -1.628 | -1.186 | 0.344 | 0.015 | 0.082 | 0.082 | 0 | 0.26 | 0.11 | 0.26 | 0.06 | 0.22 | 0.21 |
| 2 | 238721315 | rs4663791 | -0.325 | -1.764 | -0.997 | 0.789 | 0.045 | -0.041 | -0.041 | 0 | 0.28 | 0.11 | 0.19 | 0.38 | 0.17 | 0.33 |
| 2 | 238731931 | rs4663794 | -0.328 | -1.774 | -1.025 | 0.503 | -0.016 | 0.012 | 0.012 | 6.909 | 0.3 | 0.32 | 0.2 | 0.44 | 0.17 | 0.31 |
| 2 | 238733016 | rs4663795 | 0.295 | -1.422 | -0.823 | 0.659 | -0.026 | 0.047 | 0.047 | 10.116 | 0.23 | 0.3 | 0.18 | 0.39 | 0.14 | 0.27 |
| 2 | 238746880 | rs1198819 | -0.491 | -0.579 | -0.949 | 0.252 | -0.001 | -0.013 | -0.013 | 0 | 0.57 | 0.5 | 0.62 | 0.69 | 0.48 | 0.66 |
| 2 | 238760118 | rs1198823 | -0.451 | -0.838 | -0.99 | -0.267 | 0.005 | -0.005 | -0.005 | 0 | 0.43 | 0.4 | 0.45 | 0.14 | 0.35 | 0.44 |
| 2 | 238769680 | rs3769050 | -0.624 | -1.705 | -1.842 | -0.199 | 0.157 | 0.111 | 0.111 | 2.736 | 0.49 | 0.89 | 0.38 | 0.27 | 0.31 | 0.39 |
| 2 | 238779534 | rs3769036 | -0.075 | -2.077 | -0.405 | -0.017 | -0.02 | -0.004 | -0.004 | 2.021 | 0.19 | 0.24 | 0.13 | 0.13 | 0.14 | 0.16 |
| 2 | 238780317 | rs1469772 | -1.056 | -2.855 | -0.795 | -0.4 | 0.139 | 0.048 | 0.048 | 1.44 | 0.52 | 0.8 | 0.37 | 0.31 | 0.3 | 0.4 |
| 2 | 238787247 | rs13405506 | -0.75 | -2.96 | -0.165 | -0.244 | -0.048 | -0.008 | -0.008 | 0 | 0.29 | 0.42 | 0.18 | 0.27 | 0.13 | 0.2 |
| 2 | 238789044 | rs302679 | -0.721 | -1.893 | -1.131 | -0.564 | -0.063 | 0.063 | 0.063 | 0.242 | 0.84 | 0.98 | 0.71 | 0.95 | 0.78 | 0.71 |
| 2 | 238792133 | rs6728476 | -0.977 | -2.432 | -0.919 | -0.484 | 0.116 | 0.036 | 0.036 | 1.338 | 0.5 | 0.77 | 0.34 | 0.46 | 0.29 | 0.47 |
| 2 | 238793068 | rs1198826 | 1.187 | -2.05 | -1.06 | -0.287 | -0.031 | -0.038 | -0.038 | 0 | 0.35 | 0.26 | 0.4 | 0.46 | 0.5 | 0.29 |
| 2 | 238795602 | rs302673 | 0.67 | -1.967 | -0.786 | -0.162 | 0.026 | -0.007 | -0.007 | 0.595 | 0.55 | 0.35 | 0.53 | 0.49 | 0.65 | 0.47 |

| | | | | | | | | | | | | | | | | |
|---|---|---|---|---|---|---|---|---|---|---|---|---|---|---|---|---|
| 2 | 238800874 | rs13382236 | -0.568 | -2.085 | -0.734 | 0.067 | -0.065 | -0.001 | -0.001 | 0 | 0.23 | 0.31 | 0.13 | 0.23 | 0.07 | 0.27 |
| 2 | 238801786 | rs6746771 | -0.787 | -1.981 | -0.772 | 0.026 | -0.028 | -0.004 | -0.004 | 0.296 | 0.26 | 0.28 | 0.14 | 0.22 | 0.1 | 0.28 |
| 2 | 238803478 | rs11675972 | -0.128 | -0.799 | -0.837 | -0.123 | -0.005 | -0.008 | -0.008 | 0 | 0.85 | 0.87 | 0.92 | 0.93 | 0.9 | 0.72 |
| 2 | 238804069 | rs7603344 | -0.533 | -0.416 | -1.017 | -0.027 | -0.014 | -0.003 | -0.003 | 1.257 | 0.26 | 0.15 | 0.3 | 0.3 | 0.29 | 0.33 |
| 2 | 238804355 | rs6431563 | -0.179 | 0.219 | -0.954 | 0.014 | 0.001 | -0.007 | -0.007 | 0.051 | 0.8 | 0.82 | 0.87 | 0.89 | 0.8 | 0.69 |
| 2 | 238806370 | rs4663269 | 1.292 | -0.032 | -1.041 | -0.067 | -0.001 | 0.003 | 0.003 | 0 | 0.52 | 0.5 | 0.57 | 0.58 | 0.51 | 0.39 |
| 2 | 238809725 | rs1867934 | 0.226 | -0.033 | -1.201 | -0.256 | -0.007 | -0.039 | -0.039 | 0 | 0.79 | 0.76 | 0.88 | 0.95 | 0.84 | 0.71 |
| 2 | 238810763 | rs6717794 | 0.421 | 0.448 | -1.507 | -0.363 | 0.011 | 0.057 | 0.057 | 0 | 0.59 | 0.73 | 0.82 | 0.88 | 0.68 | 0.51 |
| 2 | 238811312 | rs11680171 | -0.292 | 0.669 | -1.603 | 0.178 | 0.005 | -0.018 | -0.018 | 0.35 | 0.45 | 0.29 | 0.37 | 0.47 | 0.35 | 0.53 |
| 2 | 238812164 | rs4663805 | -0.01 | 0.662 | -1.693 | 0.059 | 0.004 | 0.006 | 0.006 | 0 | 0.85 | 0.94 | 0.86 | 0.92 | 0.86 | 0.86 |
| 2 | 238815878 | rs1584243 | 0.006 | 0.685 | -1.707 | 0.032 | 0.005 | 0.01 | 0.01 | 0 | 0.84 | 0.94 | 0.9 | 0.93 | 0.87 | 0.86 |
| 2 | 238816201 | rs1080519 | -0.191 | 0.419 | -1.495 | -0.149 | 0.002 | 0.003 | 0.003 | 0 | 0.28 | 0.21 | 0.21 | 0.16 | 0.24 | 0.37 |
| 2 | 238822331 | rs10185142 | 0.327 | 0.266 | -1.328 | -0.138 | -0.005 | -0.012 | -0.012 | 0 | 0.35 | 0.42 | 0.21 | 0.21 | 0.32 | 0.31 |
| 2 | 238825809 | rs6729271 | -0.313 | 0.385 | -1.354 | -0.296 | -0.033 | 0.011 | 0.011 | 0.26 | 0.68 | 0.77 | 0.39 | 0.25 | 0.54 | 0.48 |
| 2 | 238829613 | rs6738488 | 0.035 | 0.127 | -1.424 | -0.483 | 0.002 | 0.001 | 0.001 | 0 | 0.15 | 0.13 | 0.08 | 0.14 | 0.12 | 0.15 |
| 2 | 238833264 | rs6707038 | -0.028 | 0.429 | -1.63 | 0.116 | -0.005 | -0.004 | -0.004 | 0 | 0.64 | 0.66 | 0.63 | 0.65 | 0.58 | 0.57 |
| 2 | 238836195 | rs6719401 | -0.283 | 0.022 | -1.909 | -0.11 | 0.012 | 0.04 | 0.04 | 0 | 0.17 | 0.13 | 0.12 | 0.02 | 0.16 | 0.15 |
| 2 | 238840859 | rs7559272 | -0.868 | 0.118 | -2.218 | 0.394 | -0.006 | -0.135 | -0.135 | 0.442 | 0.36 | 0.14 | 0.3 | 0.53 | 0.3 | 0.42 |
| 2 | 238851442 | rs4302179 | 0.361 | -0.385 | -0.869 | 0.379 | 0.014 | -0.045 | -0.045 | 0.263 | 0.4 | 0.44 | 0.5 | 0.22 | 0.57 | 0.32 |
| 2 | 238851935 | rs6713087 | 0.069 | -0.605 | -1.015 | 0.789 | -0.008 | 0.044 | 0.044 | 0 | 0.53 | 0.76 | 0.59 | 0.43 | 0.62 | 0.43 |
| 2 | 238863455 | rs6707140 | -0.475 | -0.274 | -0.553 | 1.18 | 0.014 | 0.011 | 0.011 | 0 | 0.52 | 0.36 | 0.38 | 0.55 | 0.4 | 0.42 |
| 2 | 238867059 | rs1446927 | -0.075 | -0.65 | -0.535 | 1.265 | 0.016 | 0.023 | 0.023 | 0.383 | 0.7 | 0.84 | 0.78 | 0.76 | 0.77 | 0.59 |
| 2 | 238867240 | rs1446928 | 0.083 | -0.524 | -0.056 | 1.402 | -0.001 | 0 | 0 | 0 | 0.4 | 0.45 | 0.35 | 0.54 | 0.35 | 0.38 |
| 2 | 238869884 | rs6723077 | 0.77 | -0.608 | -0.021 | 1.263 | -0.047 | 0.014 | 0.014 | 0.762 | 0.4 | 0.17 | 0.45 | 0.4 | 0.48 | 0.31 |
| 2 | 238870583 | rs1446929 | 0.066 | -0.578 | 0.225 | 1.306 | 0.001 | -0.001 | -0.001 | 0 | 0.39 | 0.38 | 0.34 | 0.37 | 0.35 | 0.33 |
| 2 | 238875491 | rs35277801 | 0.104 | -0.573 | 0.249 | 1.303 | -0.002 | -0.01 | -0.01 | 0 | 0.39 | 0.24 | 0.34 | 0.37 | 0.37 | 0.33 |
| 2 | 238879165 | rs6717405 | -1.287 | -0.54 | 0.318 | 1.238 | -0.008 | 0.036 | 0.036 | 0.102 | 0.83 | 0.93 | 0.82 | 0.97 | 0.84 | 0.68 |
| 2 | 238885728 | rs6708774 | -0.093 | -0.58 | 0.315 | 1.25 | 0 | -0.001 | -0.001 | 0 | 0.43 | 0.46 | 0.35 | 0.7 | 0.37 | 0.39 |
| 2 | 238888974 | rs895574 | 0.077 | -0.553 | 0.162 | 1.151 | 0.006 | -0.014 | -0.014 | 4.693 | 0.41 | 0.28 | 0.34 | 0.5 | 0.38 | 0.36 |
| 2 | 238906511 | rs10166061 | -0.9 | -0.672 | 0.143 | 0.686 | -0.003 | 0.016 | 0.016 | 0.117 | 0.83 | 0.9 | 0.82 | 0.97 | 0.84 | 0.69 |
| 2 | 238908120 | rs4663826 | -0.881 | -0.672 | 0.187 | 0.686 | -0.001 | 0.001 | 0.001 | 0.115 | 0.83 | 0.89 | 0.82 | 0.97 | 0.84 | 0.69 |
| 2 | 238923645 | rs2121473 | -0.881 | -0.67 | 0.071 | 0.687 | -0.001 | -0.002 | -0.002 | 0 | 0.83 | 0.89 | 0.82 | 0.97 | 0.84 | 0.69 |
| 2 | 238943966 | rs4663281 | -0.778 | -0.641 | -0.022 | 0.702 | 0 | -0.016 | -0.016 | 0.074 | 0.82 | 0.82 | 0.81 | 0.97 | 0.84 | 0.69 |
| 2 | 238964927 | rs821487 | -0.808 | -0.619 | -0.073 | 0.712 | -0.001 | 0.001 | 0.001 | 0.183 | 0.82 | 0.85 | 0.82 | 0.97 | 0.84 | 0.69 |
| 2 | 238975740 | rs2249435 | 1.126 | -0.765 | -0.515 | 0.688 | -0.003 | 0.004 | 0.004 | 0.018 | 0.22 | 0.01 | 0.1 | 0.16 | 0.14 | 0.14 |
| 2 | 238990388 | rs3210400 | 1.743 | -0.783 | -0.493 | 0.629 | -0.147 | 0.019 | 0.019 | 1.261 | 0.28 | 0.04 | 0.44 | 0.23 | 0.46 | 0.23 |
| 2 | 238994676 | rs2264132 | -0.546 | -0.881 | -0.18 | 0.717 | -0.004 | 0.043 | 0.043 | 0 | 0.78 | 0.62 | 0.76 | 0.76 | 0.8 | 0.65 |
| 2 | 238997956 | rs1374386 | -0.333 | -0.817 | -0.717 | 0.582 | 0.017 | -0.072 | -0.072 | 0.933 | 0.63 | 0.81 | 0.75 | 0.51 | 0.71 | 0.65 |
| 2 | 238999883 | rs1128552 | 0.272 | -0.693 | -0.554 | 0.523 | 0 | 0.01 | 0.01 | 0 | 0.25 | 0.18 | 0.23 | 0.07 | 0.22 | 0.39 |
| 2 | 239007806 | rs1054640 | 0.034 | -0.695 | -0.689 | 0.797 | 0.019 | -0.036 | -0.036 | 0.022 | 0.61 | 0.74 | 0.73 | 0.52 | 0.69 | 0.62 |
| 2 | 239008203 | rs1562337 | 0.886 | -0.294 | -0.738 | 0.458 | 0.018 | 0.067 | 0.067 | 0 | 0.47 | 0.31 | 0.48 | 0.16 | 0.62 | 0.51 |
| 2 | 239011252 | rs10210979 | 0.145 | -0.329 | -0.774 | 0.86 | -0.031 | 0.31 | 0.31 | 0 | 0.13 | 0.38 | 0.08 | 0.42 | 0.06 | 0.1 |
| 2 | 239013662 | rs7565244 | -0.243 | 0.121 | -1.891 | 0.57 | -0.03 | -0.035 | -0.035 | 0 | 0.73 | 0.92 | 0.77 | 0.73 | 0.76 | 0.71 |
| 2 | 239020497 | rs4663290 | -1.12 | 0.379 | -1.692 | 0.534 | 0.002 | 0 | 0 | 0.014 | 0.82 | 0.87 | 0.79 | 0.82 | 0.86 | 0.84 |
| 2 | 239021376 | rs12692210 | -0.424 | 0.498 | -1.728 | 0.45 | -0.002 | 0.012 | 0.012 | 3.756 | 0.69 | 0.79 | 0.62 | 0.79 | 0.71 | 0.51 |
| 2 | 239021690 | rs11898344 | -0.004 | 0.565 | -0.678 | 1.055 | 0.034 | 0.059 | 0.059 | 0.047 | 0.12 | 0.29 | 0.14 | 0.23 | 0.13 | 0.14 |
| 2 | 239023528 | rs10186514 | 0.12 | 0.477 | -0.781 | 1.043 | -0.046 | 0.143 | 0.143 | 0 | 0.22 | 0.47 | 0.18 | 0.42 | 0.12 | 0.15 |
| 2 | 239025630 | rs11904159 | -0.339 | 0.84 | -0.899 | 0.984 | 0.03 | 0.053 | 0.053 | 0.104 | 0.12 | 0.29 | 0.14 | 0.23 | 0.13 | 0.14 |
| 2 | 239025645 | rs11897736 | -0.401 | 0.839 | -0.899 | 0.984 | 0.028 | 0.051 | 0.051 | 0.024 | 0.12 | 0.29 | 0.14 | 0.23 | 0.13 | 0.15 |
| 2 | 239035853 | rs12479385 | -0.402 | 0.698 | -1.895 | 0.483 | -0.009 | 0.084 | 0.084 | 0.014 | 0.46 | 0.84 | 0.51 | 0.68 | 0.4 | 0.62 |
| 2 | 239040708 | rs11695472 | 0.132 | 1.017 | -2.255 | 0.197 | -0.09 | 0.013 | 0.013 | 1.218 | 0.79 | 0.98 | 0.78 | 0.93 | 0.71 | 0.82 |
| 2 | 239041826 | rs935412 | 0.008 | 0.581 | -2.007 | 0.35 | 0.008 | -0.009 | -0.009 | 0.002 | 0.43 | 0.65 | 0.42 | 0.53 | 0.35 | 0.47 |
| 2 | 239047020 | rs11688382 | 0.467 | 0.208 | -1.531 | 0.021 | 0.007 | 0.008 | 0.008 | 0 | 0.21 | 0.22 | 0.09 | 0.05 | 0.12 | 0.12 |
| 2 | 239047910 | rs13421970 | 0.355 | 0.168 | -1.26 | 0.001 | -0.012 | -0.028 | -0.028 | 0 | 0.22 | 0.29 | 0.09 | 0.05 | 0.12 | 0.12 |
| 2 | 239051550 | rs935411 | -0.214 | 0.583 | -1.466 | 0.27 | -0.032 | -0.029 | -0.029 | 0 | 0.91 | 0.98 | 0.89 | 0.93 | 0.86 | 0.96 |
| 2 | 239054724 | rs10192297 | 0.538 | 0.257 | -1.114 | 0.157 | 0.004 | 0.004 | 0.004 | 3.273 | 0.7 | 0.7 | 0.69 | 0.8 | 0.7 | 0.82 |
| 2 | 239055828 | rs7602358 | -0.003 | 0.45 | -1.505 | 0.087 | -0.044 | 0.029 | 0.029 | 4.294 | 0.77 | 0.97 | 0.81 | 0.91 | 0.75 | 0.85 |
| 2 | 239059484 | rs7605194 | 0.133 | 0.594 | -1.569 | 0.011 | -0.063 | 0.069 | 0.069 | 2.993 | 0.75 | 0.97 | 0.78 | 0.93 | 0.7 | 0.85 |
| 2 | 239059902 | rs7588480 | 0.129 | 0.475 | -1.447 | -0.007 | -0.004 | 0.06 | 0.06 | 5.949 | 0.76 | 0.9 | 0.8 | 0.92 | 0.75 | 0.86 |
| 2 | 239060766 | rs7601275 | -0.318 | 0.412 | -0.891 | 0.308 | 0.009 | 0.005 | 0.005 | 1.7 | 0.45 | 0.69 | 0.53 | 0.6 | 0.38 | 0.51 |
| 2 | 239066379 | rs1982338 | -0.126 | -0.183 | -0.541 | 0.275 | -0.001 | 0 | 0 | 10.002 | 0.32 | 0.36 | 0.36 | 0.39 | 0.25 | 0.2 |
| 2 | 239071623 | rs7564476 | 0.097 | 0.079 | -0.113 | 0.118 | 0.065 | 0.018 | 0.018 | 0 | 0.84 | 0.65 | 0.78 | 0.8 | 0.78 | 0.8 |
| 2 | 239075812 | rs3795905 | 1.718 | -0.021 | -0.298 | 0.04 | -0.02 | 0.027 | 0.027 | 3.071 | 0.28 | 0.19 | 0.26 | 0.2 | 0.34 | 0.35 |
| 2 | 239075935 | rs3820817 | 0.272 | -0.179 | -0.141 | 0.078 | 0.022 | 0.001 | 0.001 | 1.844 | 0.7 | 0.63 | 0.63 | 0.67 | 0.6 | 0.6 |
| 2 | 239079542 | rs2278737 | 1.325 | -0.102 | -0.312 | 0.123 | -0.047 | -0.006 | -0.006 | 0 | 0.15 | 0.02 | 0.15 | 0.1 | 0.2 | 0.12 |
| 2 | 239082976 | rs2880132 | -0.754 | -0.169 | -0.155 | 0.09 | 0 | 0.001 | 0.001 | 1.293 | 0.56 | 0.46 | 0.52 | 0.57 | 0.45 | 0.37 |
| 2 | 239083077 | rs6722284 | 1.145 | -0.503 | -0.219 | -0.133 | -0.03 | -0.069 | -0.069 | 0 | 0.13 | 0.26 | 0.05 | 0 | 0.05 | 0.02 |
| 2 | 239087912 | rs12731 | -1.095 | -0.477 | -0.485 | -0.116 | -0.002 | -0.006 | -0.006 | 2.31 | 0.57 | 0.53 | 0.52 | 0.57 | 0.45 | 0.37 |
| 2 | 239089035 | rs10187898 | 1.142 | -0.473 | -0.494 | -0.17 | -0.031 | -0.066 | -0.066 | 0 | 0.13 | 0.25 | 0.04 | 0 | 0.04 | 0.02 |

| | | | | | | | | | | | | | | | | |
|---|---|---|---|---|---|---|---|---|---|---|---|---|---|---|---|---|
| 2 | 239089151 | rs2880131 | 0.354 | -0.275 | -1.189 | -0.039 | 0.022 | -0.045 | -0.045 | 0 | 0.28 | 0.17 | 0.31 | 0.47 | 0.21 | 0.24 |
| 2 | 239093788 | rs6431588 | -1.426 | -0.222 | -1.343 | -0.062 | 0.019 | -0.032 | -0.032 | 4.328 | 0.9 | 0.91 | 0.89 | 0.99 | 0.83 | 0.89 |
| 2 | 239095422 | rs12472274 | 0.3 | -0.335 | -1.379 | -0.019 | 0.021 | -0.042 | -0.042 | 0 | 0.28 | 0.16 | 0.31 | 0.47 | 0.21 | 0.24 |
| 2 | 239097140 | rs2305171 | 1.104 | 0.051 | -2.035 | 0.069 | -0.006 | -0.116 | -0.116 | 0 | 0.18 | 0.02 | 0.27 | 0.36 | 0.16 | 0.1 |
| 2 | 239102856 | rs3795903 | 1.199 | 0.809 | -1.388 | -0.338 | -0.016 | 0.027 | 0.027 | 2.756 | 0.29 | 0.19 | 0.26 | 0.19 | 0.34 | 0.35 |
| 2 | 239113794 | rs11695186 | -0.076 | 0.671 | -0.951 | 0.052 | 0.026 | -0.029 | -0.029 | 0 | 0.87 | 0.85 | 0.84 | 0.97 | 0.79 | 0.75 |
| 2 | 239120652 | rs2341697 | -0.475 | -0.448 | -0.916 | 0.318 | 0.036 | 0.024 | 0.024 | 5.935 | 0.46 | 0.65 | 0.44 | 0.66 | 0.27 | 0.28 |
| 2 | 239133324 | rs2305174 | -0.62 | -0.603 | -0.727 | 0.512 | -0.011 | -0.002 | -0.002 | 0.471 | 0.28 | 0.35 | 0.19 | 0.25 | 0.19 | 0.31 |
| 2 | 239134056 | rs7572285 | -0.049 | -0.51 | -1.157 | -0.027 | -0.002 | -0.011 | -0.011 | 0 | 0.39 | 0.39 | 0.35 | 0.25 | 0.4 | 0.29 |
| 2 | 239138259 | rs11694250 | 1.03 | -0.571 | -1.404 | -0.069 | -0.087 | 0.067 | 0.067 | 0 | 0.26 | 0.02 | 0.24 | 0.12 | 0.34 | 0.19 |
| 2 | 239144542 | rs6744132 | -1.205 | -1.094 | -0.734 | -0.322 | -0.025 | 0.001 | 0.001 | 0.872 | 0.63 | 0.71 | 0.44 | 0.35 | 0.54 | 0.53 |
| 2 | 239145960 | rs4663865 | 0.97 | -1.071 | -0.713 | -0.251 | 0.001 | 0.003 | 0.003 | 0 | 0.13 | 0.11 | 0.05 | 0.06 | 0.1 | 0.13 |
| 2 | 239150815 | rs4663866 | 1.01 | -1.078 | -0.731 | 0.294 | 0.001 | 0.007 | 0.007 | 0 | 0.13 | 0.1 | 0.05 | 0.06 | 0.1 | 0.13 |
| 2 | 239155053 | rs934945 | 1.488 | -1.181 | -1.267 | 0.571 | 0.023 | -0.056 | -0.056 | 0 | 0.26 | 0.01 | 0.32 | 0.26 | 0.18 | 0.2 |
| 2 | 239155732 | rs880140 | 2.03 | -0.747 | -0.606 | 0.603 | -0.005 | 0.003 | 0.003 | 12.888 | 0.08 | 0.07 | 0.05 | 0.1 | 0.05 | 0.12 |
| 2 | 239163744 | rs744837 | -1.047 | -0.772 | -0.266 | 0.591 | 0.018 | 0.134 | 0.134 | 0 | 0.05 | 0.26 | 0.04 | 0.11 | 0.03 | 0.12 |
| 2 | 239165128 | rs6431590 | -1.727 | -0.711 | -0.121 | 0.605 | 0.012 | 0.011 | 0.011 | 0.411 | 0.38 | 0.72 | 0.31 | 0.33 | 0.34 | 0.38 |
| 2 | 239165636 | rs2304670 | 1.039 | -0.653 | -0.51 | 0.622 | 0 | 0.003 | 0.003 | 0 | 0.13 | 0.11 | 0.06 | 0.06 | 0.1 | 0.12 |
| 2 | 239165663 | rs2304669 | 0.998 | -0.661 | -0.522 | 0.664 | -0.018 | -0.016 | -0.016 | 0 | 0.05 | 0.02 | 0.15 | 0.14 | 0.16 | 0.05 |
| 2 | 239172863 | rs7570188 | 1.225 | -0.714 | -0.486 | 0.568 | 0.001 | -0.002 | -0.002 | 0 | 0.12 | 0.09 | 0.05 | 0.13 | 0.06 | 0.2 |
| 2 | 239176386 | rs3739064 | 0.101 | -0.604 | -0.729 | 0.614 | -0.014 | 0 | 0 | 0 | 0.18 | 0.12 | 0.19 | 0.18 | 0.25 | 0.17 |
| 2 | 239177073 | rs11894535 | -0.56 | -0.621 | -0.52 | 0.62 | 0.001 | -0.002 | -0.002 | 0 | 0.26 | 0.7 | 0.25 | 0.26 | 0.23 | 0.26 |
| 2 | 239184581 | rs10462023 | 0.915 | 0.121 | -1.379 | 0.821 | -0.141 | -0.045 | -0.045 | 0 | 0.22 | 0.04 | 0.25 | 0.23 | 0.36 | 0.31 |
| 2 | 239188479 | rs11892306 | -0.484 | -0.047 | -0.936 | 0.561 | -0.002 | -0.001 | -0.001 | 0.552 | 0.35 | 0.58 | 0.27 | 0.32 | 0.32 | 0.38 |
| 2 | 239202131 | rs13002160 | -0.757 | -0.026 | -1.446 | 0.578 | 0.028 | 0.015 | 0.015 | 1.087 | 0.38 | 0.8 | 0.3 | 0.32 | 0.32 | 0.38 |
| 2 | 239211380 | rs12692218 | -0.393 | 0.371 | -1.222 | 0.567 | 0.005 | 0.005 | 0.005 | 0 | 0.36 | 0.39 | 0.32 | 0.56 | 0.4 | 0.54 |
| 2 | 239215687 | rs7582286 | -0.964 | 0.553 | -1.779 | 0.466 | 0.008 | 0.025 | 0.025 | 0.072 | 0.61 | 0.96 | 0.57 | 0.71 | 0.69 | 0.7 |
| 2 | 239229854 | rs6706911 | -0.229 | 1.252 | -0.967 | 0.752 | 0.038 | 0.077 | 0.077 | 0 | 0.81 | 0.72 | 0.73 | 0.49 | 0.74 | 0.64 |
| 2 | 239230657 | rs7598777 | 1.5 | 1.217 | -1.222 | 0.178 | 0.004 | 0.071 | 0.071 | 6.685 | 0.24 | 0.11 | 0.2 | 0.1 | 0.25 | 0.26 |
| 2 | 239233732 | rs4663304 | -0.548 | 1.533 | -1.027 | 0.371 | 0.028 | 0.073 | 0.073 | 0 | 0.82 | 0.73 | 0.74 | 0.49 | 0.75 | 0.64 |
| 2 | 239239612 | rs1079435 | -0.369 | 1.54 | -1.046 | 0.381 | 0.039 | 0.081 | 0.081 | 0 | 0.81 | 0.71 | 0.72 | 0.48 | 0.74 | 0.64 |
| 2 | 239240370 | rs13424837 | -0.062 | 1.247 | -0.488 | 0.285 | -0.018 | 0.046 | 0.046 | 0.732 | 0.09 | 0.33 | 0.08 | 0.1 | 0.06 | 0.12 |
| 2 | 239253836 | rs4663305 | 1.24 | 1.056 | -0.807 | 0.006 | -0.001 | -0.024 | -0.024 | 0.641 | 0.32 | 0.31 | 0.26 | 0.1 | 0.29 | 0.26 |
| 2 | 239260601 | rs2572858 | 0.549 | 1.456 | -1.439 | 0.16 | -0.028 | 0.015 | 0.015 | 0.356 | 0.38 | 0.03 | 0.36 | 0.24 | 0.32 | 0.22 |
| 2 | 239279384 | rs821808 | 0.628 | 1.365 | -1.42 | 0.173 | -0.008 | 0.017 | 0.017 | 0.374 | 0.37 | 0.03 | 0.35 | 0.24 | 0.3 | 0.22 |
| 2 | 239295700 | rs509063 | -0.519 | 1.717 | -0.484 | 0.744 | 0.027 | 0.064 | 0.064 | 0 | 0.82 | 0.75 | 0.74 | 0.53 | 0.76 | 0.67 |
| 2 | 239314634 | rs541864 | -0.136 | 1.211 | -0.708 | 0.627 | 0.001 | 0.021 | 0.021 | 0 | 0.8 | 0.82 | 0.85 | 0.51 | 0.81 | 0.65 |
| 2 | 239318016 | rs13392396 | -0.94 | 0.81 | -0.415 | 0.359 | -0.023 | 0.004 | 0.004 | 0.244 | 0.14 | 0.66 | 0.14 | 0.14 | 0.12 | 0.16 |
| 2 | 239318063 | rs502932 | 0.464 | 0.649 | -1.014 | 0.061 | -0.002 | -0.012 | -0.012 | 0.01 | 0.36 | 0.02 | 0.44 | 0.26 | 0.29 | 0.21 |
| 2 | 239318280 | rs7561954 | -0.711 | 1.221 | -0.778 | 0.237 | 0.041 | 0.061 | 0.061 | 0.476 | 0.13 | 0.65 | 0.16 | 0.18 | 0.18 | 0.17 |
| 2 | 239322485 | rs10084192 | -0.31 | 1.164 | -0.668 | 0.207 | -0.027 | 0.047 | 0.047 | 0.414 | 0.11 | 0.63 | 0.12 | 0.14 | 0.06 | 0.12 |
| 2 | 239324624 | rs494819 | -0.054 | 1.195 | -0.884 | 0.017 | 0 | 0.007 | 0.007 | 0 | 0.74 | 0.76 | 0.8 | 0.5 | 0.74 | 0.59 |
| 2 | 239327783 | rs510902 | 0.557 | 1.32 | -0.927 | 0.085 | 0.001 | 0.008 | 0.008 | 0 | 0.69 | 0.33 | 0.69 | 0.45 | 0.69 | 0.58 |
| 2 | 239328519 | rs484968 | -0.345 | 1.327 | -0.391 | 0.106 | 0.003 | 0.011 | 0.011 | 0 | 0.77 | 0.8 | 0.8 | 0.55 | 0.75 | 0.7 |
| 2 | 239343044 | rs579345 | -0.645 | 1.167 | -0.258 | 0.12 | -0.014 | -0.033 | -0.033 | 0 | 0.8 | 0.88 | 0.81 | 0.58 | 0.78 | 0.7 |
| 2 | 239344070 | rs509903 | 0.02 | 1.143 | -0.121 | 0.105 | 0.002 | 0.015 | 0.015 | 0 | 0.75 | 0.73 | 0.8 | 0.51 | 0.74 | 0.61 |
| 2 | 239348995 | rs579861 | -0.457 | 1.153 | -0.089 | 0.239 | -0.002 | -0.001 | -0.001 | 0 | 0.79 | 0.82 | 0.81 | 0.54 | 0.78 | 0.7 |
| 2 | 239355561 | rs2278768 | -0.317 | 1.04 | 0.212 | 0.209 | -0.013 | 0.092 | 0.092 | 0.215 | 0.1 | 0.49 | 0.1 | 0.14 | 0.06 | 0.12 |
| 2 | 239361687 | rs7601182 | 0.118 | 1.061 | 0.162 | 0.198 | -0.006 | 0.072 | 0.072 | 0 | 0.09 | 0.42 | 0.1 | 0.1 | 0.06 | 0.12 |
| 2 | 239362938 | rs7605254 | 1.086 | 0.995 | -0.006 | 0.027 | -0.007 | -0.017 | -0.017 | 0 | 0.36 | 0.44 | 0.3 | 0.15 | 0.31 | 0.31 |
| 2 | 239369556 | rs4663309 | 2.143 | 1.013 | -0.592 | -0.048 | -0.004 | 0.103 | 0.103 | 4.034 | 0.26 | 0.05 | 0.21 | 0.06 | 0.24 | 0.2 |
| 2 | 239369790 | rs479276 | -0.416 | 1.451 | -0.084 | 0.347 | 0.021 | 0.038 | 0.038 | 0.029 | 0.42 | 0.25 | 0.47 | 0.33 | 0.35 | 0.31 |
| 2 | 239398809 | rs523437 | -0.089 | 1.422 | -0.411 | 0.403 | -0.041 | 0.027 | 0.027 | 0.203 | 0.35 | 0.07 | 0.44 | 0.25 | 0.34 | 0.24 |
| 2 | 239400430 | rs580209 | -0.143 | 1.449 | 0.204 | 0.422 | -0.004 | 0.037 | 0.037 | 0.075 | 0.36 | 0.18 | 0.45 | 0.25 | 0.34 | 0.24 |
| 2 | 239406446 | rs892839 | -1.44 | 1.619 | 0.522 | 0.562 | 0.075 | 0.075 | 0.075 | 0 | 0.05 | 0.14 | 0.06 | 0.13 | 0.14 | 0.1 |
| 2 | 239413172 | rs12692222 | -2.167 | 1.381 | 0.319 | 0.533 | 0.008 | 0.001 | 0.001 | 0.816 | 0.17 | 0.81 | 0.14 | 0.22 | 0.09 | 0.21 |
| 2 | 239419603 | rs6739251 | -2.401 | 1.576 | 0.725 | 0.604 | 0.027 | 0.098 | 0.098 | 0 | 0.15 | 0.6 | 0.15 | 0.28 | 0.18 | 0.23 |
| 2 | 239434892 | rs1039887 | -1.817 | 1.515 | 0.594 | 0.598 | 0.048 | 0.144 | 0.144 | 1.145 | 0.12 | 0.57 | 0.11 | 0.24 | 0.16 | 0.19 |
| 2 | 239437342 | rs9287630 | -0.927 | 1.462 | 0.3 | 0.614 | -0.018 | 0.083 | 0.083 | 0.422 | 0.09 | 0.45 | 0.05 | 0.14 | 0.04 | 0.14 |
| 2 | 239440110 | rs4663901 | 0.541 | 1.69 | -0.861 | 0.672 | -0.046 | -0.087 | -0.087 | 0 | 0.19 | 0.01 | 0.36 | 0.24 | 0.21 | 0.12 |
| 2 | 239443158 | rs9287631 | -0.702 | 1.851 | -0.416 | 0.762 | -0.087 | -0.042 | -0.042 | 0 | 0.52 | 0.93 | 0.63 | 0.56 | 0.53 | 0.52 |
| 2 | 239455842 | rs2166855 | -0.455 | 2.288 | -0.105 | 1.041 | -0.077 | -0.083 | -0.083 | 0 | 0.88 | 0.99 | 0.86 | 0.63 | 0.8 | 0.72 |
| 2 | 239466567 | rs10200433 | -0.224 | 1.969 | 0.393 | 0.763 | -0.002 | -0.001 | -0.001 | 0 | 0.43 | 0.46 | 0.56 | 0.39 | 0.42 | 0.33 |
| 2 | 239473435 | rs4663907 | 0.657 | 1.814 | 0.73 | 0.69 | 0.017 | 0.021 | 0.021 | 0 | 0.83 | 0.8 | 0.82 | 0.69 | 0.79 | 0.61 |
| 2 | 239478248 | rs11692233 | 0.697 | 1.424 | 0.793 | 0.686 | 0.03 | 0.11 | 0.11 | 0 | 0.83 | 0.48 | 0.83 | 0.73 | 0.85 | 0.68 |
| 2 | 239482459 | rs6728391 | 0.081 | 1.05 | 0.455 | 0 | -0.006 | -0.024 | -0.024 | 0 | 0.23 | 0.23 | 0.14 | 0.08 | 0.2 | 0.23 |
| 2 | 239486710 | rs4528715 | -1.225 | 1.217 | 0.584 | -0.076 | 0.011 | 0.026 | 0.026 | 1.089 | 0.51 | 0.56 | 0.38 | 0.37 | 0.47 | 0.58 |
| 2 | 239489888 | rs6738503 | -1.346 | 1.198 | 0.513 | -0.102 | 0.003 | 0.009 | 0.009 | 1.021 | 0.53 | 0.56 | 0.38 | 0.37 | 0.46 | 0.58 |
| 2 | 239491250 | rs2334029 | -0.699 | 1.201 | 0.578 | -0.107 | -0.004 | 0.002 | 0.002 | 0 | 0.3 | 0.43 | 0.25 | 0.29 | 0.26 | 0.35 |

| | | | | | | | | | | | | | | | | |
|---|---|---|---|---|---|---|---|---|---|---|---|---|---|---|---|---|
| 2 | 239492898 | rs2136772 | 0.004 | 1.093 | 0.515 | -0.126 | -0.025 | 0.003 | 0.003 | 0 | 0.22 | 0.3 | 0.19 | 0.23 | 0.14 | 0.23 |
| 2 | 239494246 | rs6710937 | -0.382 | 1.195 | 0.409 | -0.119 | 0.003 | 0.033 | 0.033 | 1.179 | 0.31 | 0.24 | 0.19 | 0.13 | 0.32 | 0.35 |
| 2 | 239495554 | rs10188868 | -0.465 | 1.539 | 0.134 | -0.294 | 0.021 | -0.013 | -0.013 | 9.204 | 0.36 | 0.49 | 0.54 | 0.71 | 0.45 | 0.5 |
| 2 | 239503407 | rs6716187 | -1.506 | 1.608 | 0.106 | -0.302 | -0.008 | 0.101 | 0.101 | 1.191 | 0.58 | 0.7 | 0.73 | 0.94 | 0.57 | 0.74 |
| 2 | 239504607 | rs11687086 | 0.558 | 1.689 | -0.29 | -0.301 | 0.007 | 0.012 | 0.012 | 0.084 | 0.24 | 0.17 | 0.16 | 0.08 | 0.23 | 0.23 |
| 2 | 239506809 | rs6731687 | -1.958 | 1.706 | -0.301 | -0.261 | 0.021 | 0.103 | 0.103 | 3.59 | 0.65 | 0.83 | 0.79 | 1 | 0.7 | 0.85 |
| 2 | 239508012 | rs986846 | -0.494 | 1.722 | -0.336 | -0.253 | 0.003 | -0.003 | -0.003 | 0 | 0.37 | 0.38 | 0.57 | 0.71 | 0.45 | 0.5 |
| 2 | 239509030 | rs7586445 | 0.485 | 1.562 | -1.237 | -0.819 | -0.036 | -0.018 | -0.018 | 0 | 0.06 | 0.01 | 0.05 | 0.05 | 0.13 | 0.11 |
| 2 | 239509174 | rs6414045 | -1.577 | 1.327 | -1.047 | -0.665 | 0.02 | 0.103 | 0.103 | 3.69 | 0.65 | 0.83 | 0.79 | 1 | 0.7 | 0.85 |
| 2 | 239514022 | rs2055863 | -0.439 | 1.096 | -1.005 | 0.081 | -0.008 | 0.065 | 0.065 | 0.913 | 0.32 | 0.17 | 0.24 | 0.13 | 0.37 | 0.35 |
| 2 | 239524082 | rs1027928 | -0.514 | 0.641 | -0.911 | 1.054 | 0.008 | -0.042 | -0.042 | 0.42 | 0.21 | 0.16 | 0.35 | 0.54 | 0.17 | 0.45 |
| 2 | 239524556 | rs7425902 | -1.292 | 0.806 | -0.914 | 0.547 | -0.028 | 0.05 | 0.05 | 0.488 | 0.54 | 0.74 | 0.66 | 0.7 | 0.49 | 0.68 |
| 2 | 239525212 | rs11888417 | -1.15 | 0.527 | -0.192 | 0.765 | 0.027 | 0.03 | 0.03 | 0.281 | 0.48 | 0.53 | 0.32 | 0.26 | 0.32 | 0.52 |
| 2 | 239533669 | rs1039888 | -0.734 | 0.351 | 0.637 | 1.051 | -0.004 | 0.006 | 0.006 | 0 | 0.26 | 0.27 | 0.19 | 0.47 | 0.19 | 0.36 |
| 2 | 239536797 | rs11689108 | -0.152 | 0.56 | 0.35 | 0.845 | 0.003 | 0.063 | 0.063 | 0.018 | 0.22 | 0.12 | 0.13 | 0.04 | 0.21 | 0.19 |
| 2 | 239547028 | rs13383709 | -0.487 | 0.431 | 0.324 | 1.018 | -0.008 | -0.011 | -0.011 | 0 | 0.27 | 0.13 | 0.4 | 0.29 | 0.28 | 0.3 |
| 2 | 239549312 | rs13004019 | -0.505 | -0.009 | 0.793 | 0.538 | -0.048 | 0.101 | 0.101 | 0 | 0.22 | 0.49 | 0.18 | 0.46 | 0.14 | 0.28 |
| 2 | 239553803 | rs12613966 | -0.745 | -0.012 | 0.452 | 0.548 | -0.039 | 0.086 | 0.086 | 0 | 0.24 | 0.54 | 0.21 | 0.48 | 0.17 | 0.35 |
| 2 | 239554578 | rs1995888 | -0.244 | -0.016 | 0.38 | 0.547 | -0.006 | 0.068 | 0.068 | 0 | 0.27 | 0.12 | 0.14 | 0.09 | 0.27 | 0.27 |
| 2 | 239565575 | rs10181391 | -1.068 | -0.129 | 0.508 | 0.555 | 0.005 | 0.014 | 0.014 | 0 | 0.34 | 0.28 | 0.21 | 0.15 | 0.33 | 0.35 |
| 2 | 239567125 | rs4994756 | -2.099 | -0.156 | 0.497 | 0.545 | 0 | -0.055 | -0.055 | 0 | 0.54 | 0.82 | 0.38 | 0.56 | 0.45 | 0.62 |
| 2 | 239574950 | rs7582271 | -0.296 | -0.199 | 0.576 | 0.581 | -0.04 | 0.131 | 0.131 | 0 | 0.2 | 0.46 | 0.16 | 0.41 | 0.12 | 0.27 |
| 2 | 239600087 | rs11694455 | -0.38 | -0.005 | 0.01 | 0.458 | -0.008 | 0.079 | 0.079 | 0 | 0.28 | 0.12 | 0.17 | 0.13 | 0.29 | 0.29 |
| 2 | 239600686 | rs7594129 | -1.875 | 0.005 | -0.036 | 0.471 | 0.025 | -0.024 | -0.024 | 0 | 0.53 | 0.77 | 0.38 | 0.56 | 0.44 | 0.62 |
| 2 | 239604970 | rs6431620 | -0.863 | -0.07 | -0.016 | 0.371 | 0.002 | -0.005 | -0.005 | 0 | 0.29 | 0.33 | 0.17 | 0.11 | 0.25 | 0.25 |
| 2 | 239610287 | rs4663946 | -1.919 | 0.206 | -0.315 | 0.417 | 0.026 | -0.035 | -0.035 | 0 | 0.52 | 0.83 | 0.35 | 0.56 | 0.42 | 0.55 |
| 2 | 239625150 | rs13392601 | -0.928 | 0.349 | -0.578 | 0.312 | 0.01 | 0.038 | 0.038 | 0 | 0.29 | 0.24 | 0.17 | 0.13 | 0.27 | 0.31 |
| 2 | 239632318 | rs7571108 | -0.248 | 0.394 | -0.948 | 0.3 | 0.025 | 0.065 | 0.065 | 0.567 | 0.19 | 0.08 | 0.07 | 0.05 | 0.11 | 0.09 |
| 2 | 239634596 | rs10191860 | 0.124 | 0.575 | -1.18 | 0.611 | -0.06 | -0.123 | -0.123 | 0 | 0.81 | 0.98 | 0.86 | 0.58 | 0.79 | 0.74 |
| 2 | 239636856 | rs7580245 | 1.485 | 0.805 | -1.169 | 0.443 | 0.016 | -0.004 | -0.004 | 0 | 0.5 | 0.18 | 0.63 | 0.46 | 0.56 | 0.49 |
| 2 | 239636949 | rs6740301 | -1.474 | 0.746 | -0.398 | 0.375 | 0.025 | -0.008 | -0.008 | 0 | 0.12 | 0.32 | 0.1 | 0.1 | 0.14 | 0.17 |
| 2 | 239637228 | rs11690021 | -0.705 | 0.765 | -0.698 | 0.407 | -0.001 | 0 | 0 | 1.89 | 0.08 | 0.03 | 0.06 | 0.08 | 0.09 | 0.09 |
| 2 | 239637675 | rs10929290 | 1.662 | 0.982 | -0.793 | 0.446 | 0.005 | -0.008 | -0.008 | 0 | 0.5 | 0.18 | 0.63 | 0.45 | 0.55 | 0.49 |
| 2 | 239638153 | rs6716943 | -1.499 | 1.449 | -0.578 | -0.025 | 0.005 | -0.052 | -0.052 | 0 | 0.27 | 0.38 | 0.18 | 0.11 | 0.26 | 0.27 |
| 2 | 239638300 | rs4545955 | 0.217 | 1.023 | -1.121 | 0.165 | -0.024 | 0.007 | 0.007 | 0 | 0.81 | 0.99 | 0.84 | 0.86 | 0.82 | 0.7 |
| 2 | 239638681 | rs6720396 | -1.942 | 0.804 | -0.847 | -0.235 | -0.007 | -0.049 | -0.049 | 0.704 | 0.38 | 0.57 | 0.32 | 0.07 | 0.34 | 0.28 |
| 2 | 239638781 | rs11674589 | -1.269 | 0.431 | -0.997 | 0.12 | -0.017 | 0.052 | 0.052 | 3.067 | 0.23 | 0.12 | 0.17 | 0.09 | 0.25 | 0.31 |
| 2 | 239646391 | rs12692242 | -0.384 | 0.188 | -0.17 | 1.378 | 0.015 | -0.005 | -0.005 | 0 | 0.32 | 0.18 | 0.2 | 0.38 | 0.28 | 0.42 |
| 2 | 239646465 | rs4663955 | -1.519 | 0.32 | 0.12 | 2.023 | 0.001 | 0.06 | 0.06 | 1.227 | 0.48 | 0.37 | 0.42 | 0.7 | 0.48 | 0.69 |
| 2 | 239646682 | rs11673727 | -1.201 | 0.137 | 0.572 | 1.951 | 0.014 | -0.021 | -0.021 | 1.902 | 0.08 | 0.16 | 0.09 | 0.02 | 0.13 | 0.08 |
| 2 | 239646741 | rs6745975 | -0.871 | 0.108 | 0.51 | 2.175 | -0.002 | 0.001 | 0.001 | 2.451 | 0.41 | 0.45 | 0.3 | 0.5 | 0.41 | 0.52 |
| 2 | 239665710 | rs4470287 | 1.531 | 0.102 | 0.721 | 1.72 | -0.002 | -0.017 | -0.017 | 0 | 0.46 | 0.42 | 0.5 | 0.27 | 0.43 | 0.26 |
| 2 | 239670186 | rs4663957 | 1.209 | -0.037 | 0.793 | 1.729 | 0.001 | 0.009 | 0.009 | 0.535 | 0.49 | 0.5 | 0.51 | 0.38 | 0.46 | 0.28 |
| 2 | 239671606 | rs4643485 | 1.269 | 0.025 | 0.753 | 1.759 | 0.007 | 0.035 | 0.035 | 0.467 | 0.5 | 0.55 | 0.52 | 0.38 | 0.46 | 0.28 |
| 2 | 239673336 | rs11676424 | -0.271 | 0.09 | 0.257 | 1.587 | 0.005 | -0.007 | -0.007 | 0 | 0.06 | 0.06 | 0.07 | 0.02 | 0.11 | 0.07 |
| 2 | 239674329 | rs12466434 | 0.983 | -0.12 | 0.427 | 2.129 | 0.011 | 0.004 | 0.004 | 2.707 | 0.5 | 0.39 | 0.59 | 0.51 | 0.54 | 0.49 |
| 2 | 239674876 | rs6431623 | 1.643 | -0.022 | -0.033 | 1.602 | -0.024 | 0.044 | 0.044 | 0.054 | 0.38 | 0.16 | 0.44 | 0.3 | 0.41 | 0.23 |
| 2 | 239680992 | rs9287636 | -0.098 | -0.473 | 0.456 | 1.858 | 0 | -0.001 | -0.001 | 0 | 0.72 | 0.7 | 0.79 | 0.79 | 0.71 | 0.67 |
| 2 | 239684289 | rs9711993 | 0.711 | -0.326 | 0.612 | 2.281 | -0.008 | 0.052 | 0.052 | 0 | 0.8 | 0.71 | 0.79 | 0.57 | 0.85 | 0.7 |
| 2 | 239686723 | rs12614110 | 1.03 | -0.267 | 0.02 | 2.272 | 0.002 | -0.009 | -0.009 | 0 | 0.78 | 0.78 | 0.74 | 0.49 | 0.81 | 0.67 |
| 2 | 239693565 | rs4073407 | 0.184 | -0.057 | -0.423 | 2.324 | -0.041 | -0.1 | -0.1 | 0.031 | 0.85 | 1 | 0.81 | 0.72 | 0.85 | 0.77 |
| 2 | 239697444 | rs9711321 | 0.948 | 0.062 | -0.216 | 1.909 | -0.007 | 0.224 | 0.224 | 1.047 | 0.49 | 0.19 | 0.34 | 0.19 | 0.51 | 0.29 |
| 2 | 239710123 | rs10929296 | -0.021 | -0.137 | 0.116 | 2.766 | -0.018 | -0.047 | -0.047 | 0 | 0.88 | 0.94 | 0.8 | 0.73 | 0.88 | 0.8 |
| 2 | 239719697 | rs12990932 | -0.686 | -0.676 | 0.878 | 1.86 | 0.003 | 0.006 | 0.006 | 0.008 | 0.11 | 0.28 | 0.06 | 0.14 | 0.11 | 0.17 |
| 2 | 239722254 | rs9710693 | 1.038 | -0.713 | 1.232 | 1.893 | -0.014 | -0.014 | -0.014 | 0 | 0.72 | 0.59 | 0.53 | 0.26 | 0.72 | 0.61 |
| 2 | 239722506 | rs9287645 | 1.183 | -0.738 | 0.525 | 1.263 | 0.017 | 0.1 | 0.1 | 0.731 | 0.49 | 0.69 | 0.39 | 0.02 | 0.51 | 0.26 |
| 2 | 239722948 | rs3934973 | 1.014 | -0.849 | 1.133 | 2.309 | 0.014 | -0.006 | -0.006 | 0 | 0.59 | 0.37 | 0.57 | 0.5 | 0.68 | 0.41 |
| 2 | 239723593 | rs10929297 | -0.008 | -0.721 | 0.403 | 2.638 | 0.017 | -0.059 | -0.059 | 2.187 | 0.89 | 0.98 | 0.82 | 0.6 | 0.9 | 0.74 |
| 2 | 239723729 | rs10929298 | -0.283 | -0.706 | 0.378 | 2.535 | 0.005 | -0.066 | -0.066 | 0 | 0.9 | 1 | 0.82 | 0.72 | 0.91 | 0.78 |
| 2 | 239731651 | rs12471757 | 0.837 | -0.631 | 0.585 | 1.966 | 0.025 | -0.007 | -0.007 | 0 | 0.63 | 0.74 | 0.6 | 0.38 | 0.66 | 0.36 |
| 2 | 239734678 | rs9752491 | -0.472 | -0.041 | 0.443 | 1.513 | 0.008 | 0.007 | 0.007 | 1.358 | 0.23 | 0.1 | 0.16 | 0.21 | 0.22 | 0.34 |
| 2 | 239736364 | rs9750952 | -0.486 | -0.041 | 0.426 | 1.513 | 0.009 | 0.008 | 0.008 | 1.401 | 0.23 | 0.1 | 0.16 | 0.21 | 0.22 | 0.34 |
| 2 | 239749411 | rs12613833 | -0.542 | 0.026 | 0.133 | 1.279 | 0.01 | -0.038 | -0.038 | 0 | 0.24 | 0.13 | 0.34 | 0.41 | 0.23 | 0.35 |
| 2 | 239799547 | rs3923767 | -0.271 | 0.555 | -0.85 | 0.37 | 0.012 | 0.063 | 0.063 | 0 | 0.42 | 0.23 | 0.44 | 0.18 | 0.41 | 0.35 |
| 2 | 239832812 | rs3935423 | 0.258 | 1.255 | -0.692 | 0.65 | 0.015 | 0.004 | 0.004 | 1.064 | 0.68 | 0.61 | 0.52 | 0.69 | 0.54 | 0.71 |
| 2 | 239833716 | rs3934910 | -0.388 | 0.893 | -0.167 | 1.463 | 0.004 | 0.187 | 0.187 | 0 | 0.06 | 0.15 | 0.09 | 0.33 | 0.06 | 0.08 |
| 2 | 239834334 | rs6736036 | -1.054 | 0.576 | -0.117 | 1.314 | -0.01 | 0.017 | 0.017 | 0 | 0.17 | 0.28 | 0.11 | 0.25 | 0.11 | 0.21 |
| 2 | 239835211 | rs4078406 | -0.599 | 0.128 | -0.48 | 0.604 | -0.002 | 0.032 | 0.032 | 1.675 | 0.5 | 0.64 | 0.65 | 0.63 | 0.54 | 0.43 |
| 2 | 239841003 | rs11124234 | -0.364 | 0.039 | -1.109 | 0.389 | -0.03 | -0.065 | -0.065 | 0 | 0.39 | 0.21 | 0.52 | 0.5 | 0.42 | 0.47 |

| | | | | | | | | | | | | | | | | |
|---|---|---|---|---|---|---|---|---|---|---|---|---|---|---|---|---|
| 2 | 239841135 | rs11674577 | -0.174 | -0.387 | -0.942 | 0.643 | -0.026 | -0.046 | -0.046 | 0.922 | 0.14 | 0.01 | 0.09 | 0.18 | 0.14 | 0.22 |
| 2 | 239843067 | rs7587437 | -0.523 | -0.747 | -0.017 | 0.589 | -0.059 | 0.234 | 0.234 | 0.97 | 0.13 | 0.38 | 0.1 | 0.35 | 0.05 | 0.12 |
| 2 | 239844842 | rs11901161 | -1.205 | -0.715 | -0.146 | 0.471 | -0.061 | 0.107 | 0.107 | 0 | 0.28 | 0.46 | 0.19 | 0.54 | 0.19 | 0.35 |
| 2 | 239846337 | rs7569771 | -1.178 | -0.725 | -0.176 | 0.587 | -0.026 | 0.052 | 0.052 | 0 | 0.27 | 0.57 | 0.21 | 0.5 | 0.22 | 0.34 |
| 2 | 239847178 | rs13383477 | 0.094 | -0.932 | -1.047 | -0.124 | 0.015 | 0.082 | 0.082 | 0 | 0.36 | 0.75 | 0.21 | 0.08 | 0.31 | 0.15 |
| 2 | 239847530 | rs7558224 | -1.527 | -1.054 | -0.788 | 0.304 | 0.03 | -0.049 | -0.049 | 0 | 0.3 | 0.75 | 0.24 | 0.52 | 0.22 | 0.36 |
| 2 | 239849075 | rs7571467 | 0.001 | -1.225 | -1.137 | 0.196 | -0.031 | 0.016 | 0.016 | 0 | 0.67 | 0.88 | 0.75 | 0.72 | 0.62 | 0.56 |
| 2 | 239850578 | rs13400762 | -1.519 | -1.111 | -1.073 | 0.511 | -0.077 | 0.022 | 0.022 | 0.997 | 0.83 | 0.99 | 0.85 | 0.9 | 0.75 | 0.78 |
| 2 | 239854389 | rs4852014 | 1.258 | -1.255 | -0.953 | 0.637 | 0.016 | 0.045 | 0.045 | 0.118 | 0.52 | 0.18 | 0.64 | 0.37 | 0.54 | 0.45 |
| 2 | 239857995 | rs9636382 | 0.334 | -0.948 | -0.695 | 0.428 | -0.01 | 0.025 | 0.025 | 0 | 0.65 | 0.74 | 0.74 | 0.73 | 0.6 | 0.56 |
| 2 | 239858277 | rs11124185 | -0.848 | -0.724 | -0.265 | 0.418 | -0.008 | 0.015 | 0.015 | 0.019 | 0.79 | 0.8 | 0.83 | 0.9 | 0.73 | 0.77 |
| 2 | 239861787 | rs4377355 | -1.413 | -0.405 | -0.215 | 0.396 | -0.007 | 0.008 | 0.008 | 0 | 0.94 | 0.94 | 0.9 | 0.97 | 0.83 | 0.84 |
| 2 | 239864826 | rs4521068 | 0.468 | -0.699 | -0.46 | 0.653 | -0.002 | -0.106 | -0.106 | 0.668 | 0.22 | 0.03 | 0.15 | 0.36 | 0.21 | 0.16 |
| 2 | 239865136 | rs4353673 | 0.366 | -0.901 | 0.249 | 0.428 | -0.019 | -0.029 | -0.029 | 2.968 | 0.37 | 0.48 | 0.19 | 0.22 | 0.28 | 0.22 |
| 2 | 239871074 | rs4599143 | 1.673 | -0.926 | 0.791 | 0.372 | -0.012 | -0.012 | -0.012 | 0 | 0.31 | 0.19 | 0.53 | 0.38 | 0.3 | 0.45 |
| 2 | 239871193 | rs7599854 | 1.396 | -1.068 | 0.998 | 0.3 | -0.001 | 0.025 | 0.025 | 0 | 0.27 | 0.2 | 0.14 | 0.04 | 0.26 | 0.12 |
| 2 | 239877356 | rs4368353 | -0.037 | -1.017 | 0.938 | 0.375 | -0.032 | 0.072 | 0.072 | 0 | 0.09 | 0.22 | 0.04 | 0.17 | 0.03 | 0.09 |
| 2 | 239877738 | rs4591362 | 0.944 | -0.893 | 0.506 | 0.504 | -0.022 | -0.088 | -0.088 | 0 | 0.2 | 0.01 | 0.15 | 0.29 | 0.2 | 0.11 |
| 2 | 239878079 | rs4073751 | -1.306 | -0.953 | 0.83 | 0.352 | 0.058 | -0.015 | -0.015 | 0 | 0.93 | 0.86 | 0.89 | 0.95 | 0.81 | 0.83 |
| 2 | 239879183 | rs4302232 | 0.714 | -1.015 | 0.364 | 0.348 | -0.012 | 0.062 | 0.062 | 0 | 0.51 | 0.19 | 0.68 | 0.67 | 0.5 | 0.56 |
| 2 | 239887435 | rs13006827 | 0.339 | -0.503 | 0.556 | -0.076 | 0 | -0.002 | -0.002 | 0 | 0.69 | 0.67 | 0.66 | 0.61 | 0.62 | 0.7 |
| 2 | 239890643 | rs4851976 | 0.458 | -0.518 | 0.656 | -0.373 | 0.003 | 0 | 0 | 0 | 0.69 | 0.58 | 0.71 | 0.66 | 0.61 | 0.7 |
| 2 | 239890696 | rs4852066 | -1.257 | -0.444 | 0.512 | -0.458 | 0.019 | -0.021 | -0.021 | 0 | 0.93 | 0.92 | 0.9 | 0.98 | 0.82 | 0.85 |
| 2 | 239892918 | rs6543544 | -0.79 | -0.618 | 0.711 | -0.568 | -0.002 | -0.001 | -0.001 | 0.001 | 0.52 | 0.45 | 0.3 | 0.43 | 0.46 | 0.28 |
| 2 | 239893009 | rs11679161 | 0.54 | -0.604 | 0.204 | -0.542 | -0.019 | -0.101 | -0.101 | 0.195 | 0.2 | 0.01 | 0.16 | 0.32 | 0.21 | 0.15 |
| 2 | 239896861 | rs10199914 | 0.757 | -0.478 | 0.73 | -0.58 | 0 | -0.015 | -0.015 | 0 | 0.4 | 0.32 | 0.59 | 0.55 | 0.36 | 0.57 |
| 2 | 239897650 | rs11124207 | -0.389 | -0.4 | 0.572 | -0.544 | -0.002 | -0.001 | -0.001 | 1.111 | 0.53 | 0.49 | 0.31 | 0.43 | 0.46 | 0.29 |
| 2 | 239899327 | rs4852079 | 0.69 | -0.218 | 0.922 | -0.623 | 0.071 | 0 | 0 | 0 | 0.86 | 0.58 | 0.86 | 0.9 | 0.8 | 0.84 |
| 2 | 239908773 | rs11678706 | -1.136 | -1.049 | 0.478 | -0.927 | -0.042 | -0.073 | -0.073 | 0.344 | 0.3 | 0.57 | 0.22 | 0.09 | 0.17 | 0.2 |
| 2 | 239913858 | rs13429449 | -0.044 | -1.117 | 0.618 | -0.955 | -0.017 | -0.026 | -0.026 | 0 | 0.05 | 0.18 | 0.02 | 0.01 | 0.01 | 0.06 |
| 2 | 239916135 | rs12328525 | -1.188 | -1.124 | 0.404 | -0.962 | -0.017 | -0.035 | -0.035 | 0 | 0.31 | 0.64 | 0.23 | 0.1 | 0.17 | 0.2 |
| 2 | 239920634 | rs6744347 | -1.039 | -0.449 | 0.338 | -0.696 | -0.002 | -0.047 | -0.047 | 0 | 0.3 | 0.45 | 0.38 | 0.16 | 0.31 | 0.23 |
| 2 | 239923651 | rs13418083 | -0.678 | -0.833 | 0.443 | -0.443 | 0.006 | 0.064 | 0.064 | 0 | 0.06 | 0.28 | 0.03 | 0.05 | 0.02 | 0.06 |
| 2 | 239935543 | rs3922831 | -1.055 | -1.294 | -0.045 | -1.199 | -0.021 | -0.101 | -0.101 | 0 | 0.34 | 0.32 | 0.17 | 0.01 | 0.22 | 0.18 |
| 2 | 239938387 | rs7581424 | -0.361 | -1.011 | -1.48 | -0.97 | 0.021 | 0.011 | 0.011 | 0 | 0.32 | 0.14 | 0.37 | 0.3 | 0.31 | 0.34 |
| 2 | 239944933 | rs13405988 | -0.969 | -0.635 | -1.217 | -0.853 | 0.049 | 0.113 | 0.113 | 0 | 0.51 | 0.74 | 0.52 | 0.3 | 0.45 | 0.41 |
| 2 | 239949681 | rs4075018 | -0.769 | -0.623 | -0.675 | -0.888 | -0.02 | -0.101 | -0.101 | 0 | 0.27 | 0.64 | 0.18 | 0.01 | 0.2 | 0.16 |
| 2 | 239952664 | rs11689632 | -0.452 | -0.341 | -1.305 | -0.189 | 0.035 | -0.038 | -0.038 | 0 | 0.4 | 0.25 | 0.35 | 0.45 | 0.3 | 0.35 |
| 2 | 239954199 | rs4851966 | 0.278 | 0.661 | -0.752 | -0.306 | 0.021 | -0.018 | -0.018 | 0.016 | 0.63 | 0.52 | 0.49 | 0.7 | 0.57 | 0.65 |
| 2 | 239957392 | rs1868318 | 0.996 | 0.706 | -1.228 | -0.325 | 0.058 | -0.015 | -0.015 | 3.096 | 0.43 | 0.28 | 0.3 | 0.56 | 0.34 | 0.48 |
| 2 | 239961615 | rs1470603 | -0.087 | 1.625 | -1.111 | -0.603 | -0.01 | -0.014 | -0.014 | 0 | 0.73 | 0.88 | 0.59 | 0.74 | 0.74 | 0.81 |
| 2 | 239964373 | rs11124176 | -0.244 | 1.998 | -0.448 | -0.378 | 0.072 | -0.1 | -0.1 | 0 | 0.16 | 0.56 | 0.21 | 0.04 | 0.25 | 0.16 |
| 2 | 239964748 | rs11124179 | -1.102 | 1.933 | -0.524 | -0.434 | 0.077 | -0.132 | -0.132 | 0 | 0.24 | 0.58 | 0.31 | 0.04 | 0.4 | 0.25 |
| 2 | 239970360 | rs895808 | -1.147 | 1.928 | -0.61 | 0.109 | 0.011 | 0.003 | 0.003 | 0 | 0.29 | 0.29 | 0.48 | 0.3 | 0.37 | 0.28 |
| 2 | 239972561 | rs13393217 | -0.376 | 1.915 | -0.81 | 0.251 | 0.02 | -0.028 | -0.028 | 0 | 0.07 | 0.08 | 0.11 | 0 | 0.15 | 0.07 |
| 2 | 239983350 | rs4852013 | -0.922 | 1.239 | -1.26 | 0.374 | -0.01 | -0.077 | -0.077 | 0.367 | 0.19 | 0.04 | 0.38 | 0.31 | 0.16 | 0.2 |
| 2 | 240001331 | rs3791370 | -1.343 | 1.686 | -0.566 | 0.403 | -0.005 | -0.003 | -0.003 | 0 | 0.35 | 0.34 | 0.6 | 0.39 | 0.43 | 0.34 |
| 2 | 240016177 | rs2291188 | -0.951 | 1.976 | -0.693 | 0.714 | -0.001 | -0.021 | -0.021 | 0 | 0.21 | 0.17 | 0.29 | 0.3 | 0.2 | 0.21 |
| 2 | 240020622 | rs4852019 | -1.381 | 1.573 | -0.637 | -0.156 | -0.002 | 0.025 | 0.025 | 0 | 0.51 | 0.72 | 0.61 | 0.4 | 0.52 | 0.49 |
| 2 | 240024654 | rs6543510 | 0.413 | 1.46 | 0.015 | -0.204 | -0.004 | -0.047 | -0.047 | 0 | 0.22 | 0.22 | 0.12 | 0.03 | 0.2 | 0.22 |
| 2 | 240025761 | rs3791399 | -1.408 | 1.29 | 0.246 | -0.276 | 0 | 0.001 | 0.001 | 0 | 0.39 | 0.42 | 0.55 | 0.38 | 0.44 | 0.4 |
| 2 | 240030484 | rs3791406 | 0.872 | 0.976 | 0.022 | 0.2 | 0 | -0.002 | -0.002 | 0 | 0.68 | 0.3 | 0.67 | 0.54 | 0.67 | 0.78 |
| 2 | 240032214 | rs3791414 | -0.646 | 1.016 | -0.018 | 0.176 | 0.012 | 0.011 | 0.011 | 0 | 0.85 | 0.78 | 0.9 | 0.82 | 0.82 | 0.92 |
| 2 | 240038687 | rs3791423 | 0.378 | 0.964 | -0.277 | 0.178 | -0.013 | -0.013 | -0.013 | 5.66 | 0.06 | 0.01 | 0.25 | 0.06 | 0.1 | 0.04 |
| 2 | 240038996 | rs3791424 | 1.635 | 0.61 | -0.509 | 0.048 | 0.004 | 0.003 | 0.003 | 0.342 | 0.53 | 0.17 | 0.38 | 0.5 | 0.51 | 0.7 |
| 2 | 240039119 | rs3791425 | -0.864 | 0.774 | -0.272 | 0.164 | 0.003 | -0.003 | -0.003 | 0 | 0.86 | 0.83 | 0.9 | 0.88 | 0.82 | 0.92 |
| 2 | 240040446 | rs3791428 | -0.643 | 0.646 | -0.174 | 0.252 | -0.002 | -0.05 | -0.05 | 0 | 0.21 | 0.11 | 0.23 | 0.32 | 0.21 | 0.17 |
| 2 | 240045350 | rs3791446 | -1.633 | 0.423 | 0.416 | 0.482 | -0.025 | 0.172 | 0.172 | 0 | 0.14 | 0.61 | 0.25 | 0.36 | 0.07 | 0.17 |
| 2 | 240048581 | rs3791449 | -0.644 | 0.475 | 0.679 | 0.227 | -0.007 | 0.057 | 0.057 | 0 | 0.12 | 0.36 | 0.2 | 0.11 | 0.07 | 0.17 |
| 2 | 240048911 | rs2898716 | 0.891 | 0.702 | 0.365 | 0.068 | -0.021 | -0.016 | -0.016 | 0 | 0.73 | 0.54 | 0.7 | 0.8 | 0.76 | 0.76 |
| 2 | 240053966 | rs12469994 | 1.094 | 0.815 | -0.563 | -0.02 | -0.035 | -0.015 | -0.015 | 0 | 0.32 | 0.11 | 0.37 | 0.72 | 0.32 | 0.49 |
| 2 | 240059625 | rs3791471 | -0.2 | 0.379 | 0.031 | 0.335 | 0.045 | -0.057 | -0.057 | 0.276 | 0.14 | 0.41 | 0.24 | 0.04 | 0.18 | 0.05 |
| 2 | 240060488 | rs3791474 | -0.553 | 0.317 | -0.524 | 0.61 | 0.014 | 0.022 | 0.022 | 0 | 0.35 | 0.33 | 0.27 | 0.16 | 0.23 | 0.34 |
| 2 | 240061735 | rs3791476 | -0.36 | 0.485 | -0.157 | 1.502 | 0.011 | -0.01 | -0.01 | 6.761 | 0.12 | 0.04 | 0.08 | 0.13 | 0.07 | 0.14 |
| 2 | 240065733 | rs3791480 | 0.941 | 1.076 | 0.709 | 1.68 | 0.06 | 0.17 | 0.17 | 0 | 0.82 | 0.54 | 0.59 | 0.66 | 0.76 | 0.92 |
| 2 | 240066417 | rs2290087 | 0.261 | 1.04 | -0.283 | 1.559 | -0.035 | -0.056 | -0.056 | 0 | 0.87 | 0.98 | 0.71 | 0.85 | 0.86 | 0.95 |
| 2 | 240075035 | rs3791498 | 0.905 | 0.9 | 0.374 | 1.311 | 0.004 | 0.026 | 0.026 | 1.043 | 0.69 | 0.51 | 0.41 | 0.46 | 0.71 | 0.79 |
| 2 | 240075545 | rs3791500 | 0.902 | 1.188 | 1.068 | 1.394 | -0.001 | -0.001 | -0.001 | 0 | 0.69 | 0.69 | 0.35 | 0.27 | 0.6 | 0.76 |
| 2 | 240077371 | rs3791504 | 1.124 | 1.189 | 0.878 | 1.403 | 0.024 | -0.042 | -0.042 | 3.554 | 0.67 | 0.46 | 0.34 | 0.27 | 0.6 | 0.76 |

| | | | | | | | | | | | | | | | | |
|---|---|---|---|---|---|---|---|---|---|---|---|---|---|---|---|---|
| 2 | 240085795 | rs2290089 | 0.929 | 1.107 | 1.185 | 1.399 | 0.003 | 0.023 | 0.023 | 1.842 | 0.68 | 0.54 | 0.41 | 0.46 | 0.71 | 0.72 |
| 2 | 240086416 | rs3828219 | 1.14 | 1.131 | 1.118 | 1.453 | 0.016 | -0.017 | -0.017 | 0 | 0.66 | 0.43 | 0.34 | 0.33 | 0.6 | 0.7 |
| 2 | 240095180 | rs552954 | 1.146 | 1.147 | 1.148 | 1.471 | 0.015 | -0.029 | -0.029 | 0 | 0.66 | 0.42 | 0.34 | 0.28 | 0.6 | 0.71 |
| 2 | 240096690 | rs6543514 | -0.224 | 1.143 | 0.85 | 1.905 | -0.008 | -0.015 | -0.015 | 0 | 0.15 | 0.1 | 0.28 | 0.28 | 0.15 | 0.22 |
| 2 | 240101713 | rs3791536 | -0.712 | 1.048 | 0.759 | 1.968 | -0.004 | -0.045 | -0.045 | 0 | 0.21 | 0.1 | 0.37 | 0.31 | 0.17 | 0.22 |
| 2 | 240101940 | rs592695 | 0.432 | 0.464 | 1.045 | 1.881 | -0.001 | 0.001 | 0.001 | 0.325 | 0.85 | 0.79 | 0.79 | 0.82 | 0.87 | 0.94 |
| 2 | 240104346 | rs506324 | 0.824 | 0.631 | 1.475 | 1.991 | 0.031 | 0.088 | 0.088 | 0.563 | 0.83 | 0.69 | 0.71 | 0.66 | 0.76 | 0.92 |
| 2 | 240105617 | rs843469 | 0.054 | 0.548 | 1.822 | 1.87 | 0.007 | 0.029 | 0.029 | 0.011 | 0.88 | 0.78 | 0.8 | 0.86 | 0.88 | 0.97 |
| 2 | 240108117 | rs1962113 | -0.703 | 0.479 | 1.423 | 1.861 | -0.017 | -0.035 | -0.035 | 2.045 | 0.23 | 0.17 | 0.44 | 0.48 | 0.28 | 0.23 |
| 2 | 240109568 | rs3828222 | -0.416 | 0.488 | 1.411 | 1.919 | -0.007 | -0.056 | -0.056 | 1.104 | 0.21 | 0.12 | 0.36 | 0.32 | 0.16 | 0.21 |
| 2 | 240110346 | rs4852032 | -0.678 | 0.495 | 1.427 | 1.915 | -0.017 | -0.033 | -0.033 | 0 | 0.23 | 0.17 | 0.44 | 0.48 | 0.28 | 0.23 |
| 2 | 240112972 | rs3791556 | 0.105 | 0.458 | 1.445 | 1.894 | -0.008 | -0.024 | -0.024 | 0 | 0.17 | 0.13 | 0.27 | 0.28 | 0.14 | 0.21 |
| 2 | 240114137 | rs753302 | 0.83 | -0.21 | 1.165 | 1.155 | -0.029 | -0.032 | -0.032 | 0.116 | 0.23 | 0.25 | 0.17 | 0.13 | 0.17 | 0.22 |
| 2 | 240116159 | rs6739116 | 0.718 | -0.168 | 1.17 | 1.147 | 0.002 | -0.035 | -0.035 | 0 | 0.23 | 0.27 | 0.22 | 0.13 | 0.28 | 0.22 |
| 2 | 240116168 | rs625839 | 1.126 | -0.163 | 0.905 | 1.172 | -0.048 | 0.128 | 0.128 | 1.573 | 0.45 | 0.14 | 0.28 | 0.3 | 0.48 | 0.52 |
| 2 | 240116841 | rs631292 | -0.551 | 0.3 | 1.644 | 1.695 | -0.001 | 0.002 | 0.002 | 0 | 0.91 | 0.9 | 0.83 | 0.84 | 0.9 | 0.96 |
| 2 | 240123729 | rs3828225 | -0.398 | 0.368 | 1.741 | 1.391 | 0.059 | 0.016 | 0.016 | 0.134 | 0.07 | 0.21 | 0.17 | 0.08 | 0.13 | 0.03 |
| 2 | 240127024 | rs678228 | 1.668 | 0.417 | 1.636 | 1.177 | -0.001 | -0.001 | -0.001 | 0 | 0.65 | 0.3 | 0.5 | 0.4 | 0.6 | 0.73 |
| 2 | 240132668 | rs3791599 | -0.236 | 0.311 | 1.548 | 0.96 | -0.003 | -0.007 | -0.007 | 0.67 | 0.15 | 0.12 | 0.11 | 0.22 | 0.16 | 0.2 |
| 2 | 240141383 | rs519165 | 1.524 | 0.067 | 1.112 | 0.38 | -0.009 | 0.12 | 0.12 | 2.03 | 0.4 | 0.16 | 0.21 | 0.04 | 0.44 | 0.38 |
| 2 | 240142118 | rs606531 | -0.342 | 0.215 | 1.705 | 1.389 | 0 | -0.01 | -0.01 | 0 | 0.92 | 0.93 | 0.83 | 0.88 | 0.9 | 0.96 |
| 2 | 240144903 | rs291330 | 2.013 | 0.267 | 1.841 | 1.182 | -0.01 | -0.046 | -0.046 | 0 | 0.69 | 0.24 | 0.52 | 0.39 | 0.63 | 0.8 |
| 2 | 240145376 | rs291333 | 1.27 | 0.293 | 1.809 | 1.23 | 0.024 | 0.004 | 0.004 | 0 | 0.83 | 0.77 | 0.8 | 0.76 | 0.73 | 0.94 |
| 2 | 240147847 | rs291335 | 2.076 | 0.312 | 1.804 | 1.239 | -0.005 | -0.024 | -0.024 | 0 | 0.69 | 0.27 | 0.52 | 0.39 | 0.64 | 0.79 |
| 2 | 240151759 | rs11691856 | -0.255 | 0.442 | 1.803 | 1.245 | 0.025 | 0.005 | 0.005 | 0 | 0.08 | 0.15 | 0.1 | 0.13 | 0.15 | 0.02 |
| 2 | 240164690 | rs6712996 | -2.046 | -0.2 | 1.563 | 1.413 | -0.007 | 0.092 | 0.092 | 0.083 | 0.23 | 0.5 | 0.36 | 0.4 | 0.24 | 0.24 |
| 2 | 240169077 | rs7573680 | -2.176 | -0.167 | 1.181 | 1.431 | -0.011 | 0.063 | 0.063 | 0.603 | 0.24 | 0.59 | 0.36 | 0.49 | 0.24 | 0.23 |
| 2 | 240176033 | rs1403607 | -0.771 | -0.306 | 1.071 | 1.449 | -0.009 | 0.091 | 0.091 | 0 | 0.11 | 0.15 | 0.24 | 0.37 | 0.09 | 0.16 |
| 2 | 240180336 | rs7595357 | -1.592 | -0.281 | 0.695 | 1.276 | -0.02 | 0.174 | 0.174 | 1.118 | 0.19 | 0.49 | 0.32 | 0.47 | 0.15 | 0.23 |
| 2 | 240182666 | rs12615371 | -1.148 | -0.54 | 0.185 | 1.173 | -0.051 | 0.118 | 0.118 | 2.711 | 0.18 | 0.25 | 0.25 | 0.49 | 0.1 | 0.29 |
| 2 | 240188918 | rs10208713 | -0.247 | -1.035 | -0.133 | 0.075 | -0.001 | -0.039 | -0.039 | 0.602 | 0.32 | 0.24 | 0.13 | 0.06 | 0.21 | 0.24 |
| 2 | 240193889 | rs6705378 | -1.013 | -0.802 | -0.013 | -0.003 | 0.038 | 0.001 | 0.001 | 0.064 | 0.53 | 0.66 | 0.45 | 0.56 | 0.4 | 0.57 |
| 2 | 240199015 | rs11683046 | 0.446 | -0.877 | 0.311 | -0.33 | 0.003 | -0.017 | -0.017 | 0 | 0.28 | 0.19 | 0.12 | 0.03 | 0.22 | 0.22 |
| 2 | 240201641 | rs13411439 | 0.194 | -0.927 | 0.719 | -0.352 | -0.021 | -0.158 | -0.158 | 0 | 0.3 | 0.41 | 0.14 | 0.03 | 0.22 | 0.22 |
| 2 | 240201797 | rs13414769 | -0.954 | -1.026 | 1.186 | -0.332 | 0.062 | -0.01 | -0.01 | 0.47 | 0.47 | 0.67 | 0.42 | 0.53 | 0.35 | 0.53 |
| 2 | 240207019 | rs6739632 | -0.247 | -1.052 | 1.03 | -0.246 | -0.024 | -0.098 | -0.098 | 0.011 | 0.35 | 0.42 | 0.2 | 0.11 | 0.27 | 0.23 |
| 2 | 240208154 | rs2048765 | 0.49 | -1.046 | 0.995 | -0.133 | 0.011 | 0 | 0 | 0 | 0.11 | 0.08 | 0.21 | 0.41 | 0.08 | 0.31 |
| 2 | 240216021 | rs6737742 | 0.451 | -1.037 | 1.11 | -0.215 | 0.003 | -0.027 | -0.027 | 0 | 0.28 | 0.21 | 0.12 | 0.02 | 0.23 | 0.22 |
| 2 | 240217116 | rs11883623 | 0.116 | -0.402 | 0.598 | -0.49 | -0.003 | 0.001 | 0.001 | 0 | 0.42 | 0.31 | 0.42 | 0.34 | 0.38 | 0.39 |
| 2 | 240221868 | rs13389628 | -0.279 | -0.497 | 0.739 | -0.462 | -0.035 | 0.119 | 0.119 | 0.097 | 0.16 | 0.28 | 0.25 | 0.5 | 0.1 | 0.31 |
| 2 | 240231126 | rs13405743 | 0.889 | -0.477 | 0.581 | -0.428 | 0.016 | -0.026 | -0.026 | 0.697 | 0.11 | 0.06 | 0.21 | 0.4 | 0.08 | 0.31 |
| 2 | 240241255 | rs925738 | 0.744 | -0.051 | 0.191 | -0.92 | -0.025 | 0.073 | 0.073 | 0 | 0.38 | 0.2 | 0.24 | 0.07 | 0.43 | 0.27 |
| 2 | 240244132 | rs11675637 | 1.198 | -0.065 | 0.223 | -0.884 | -0.003 | 0.016 | 0.016 | 0 | 0.13 | 0.12 | 0.3 | 0.32 | 0.14 | 0.17 |
| 2 | 240245917 | rs10182074 | -0.3 | -0.126 | 0.283 | -0.962 | -0.006 | -0.005 | -0.005 | 0 | 0.56 | 0.65 | 0.49 | 0.57 | 0.53 | 0.57 |
| 2 | 240248919 | rs4851972 | 1.09 | -0.056 | -0.137 | -0.903 | -0.003 | 0.013 | 0.013 | 0 | 0.14 | 0.12 | 0.3 | 0.32 | 0.15 | 0.18 |
| 2 | 240260505 | rs10174432 | -0.748 | -0.116 | -0.224 | -0.938 | -0.008 | -0.033 | -0.033 | 0 | 0.65 | 0.59 | 0.76 | 0.8 | 0.65 | 0.72 |
| 2 | 240262287 | rs908265 | -0.587 | -0.2 | -0.445 | -0.944 | 0.003 | -0.011 | -0.011 | 0 | 0.64 | 0.66 | 0.57 | 0.6 | 0.66 | 0.66 |
| 2 | 240262426 | rs2017669 | 0.32 | -0.772 | -0.209 | -0.682 | -0.028 | 0.233 | 0.233 | 0 | 0.13 | 0.39 | 0.23 | 0.41 | 0.08 | 0.31 |
| 2 | 240263584 | rs10211084 | -1.154 | -0.671 | -0.623 | -1.485 | -0.013 | -0.036 | -0.036 | 8.796 | 0.77 | 0.68 | 0.86 | 0.92 | 0.8 | 0.83 |
| 2 | 240282208 | rs12471054 | 0.7 | -0.027 | -0.791 | -0.731 | 0.02 | -0.043 | -0.043 | 0.02 | 0.11 | 0.04 | 0.38 | 0.67 | 0.08 | 0.37 |
| 2 | 240288261 | rs10197739 | -0.128 | 0.1 | -0.596 | -0.704 | -0.002 | 0.062 | 0.062 | 2.243 | 0.06 | 0.24 | 0.05 | 0.1 | 0.02 | 0.01 |
| 2 | 240288363 | rs10164440 | -0.165 | 0.099 | -0.597 | -0.705 | -0.003 | 0.061 | 0.061 | 2.306 | 0.06 | 0.24 | 0.05 | 0.1 | 0.02 | 0.01 |
| 2 | 240294969 | rs2411843 | -0.69 | 0.373 | -1.477 | -0.947 | 0.028 | 0.07 | 0.07 | 0 | 0.51 | 0.67 | 0.75 | 0.89 | 0.56 | 0.63 |
| 2 | 240295613 | rs2176046 | 0.244 | 0.292 | -1.962 | -0.028 | 0.017 | -0.059 | -0.059 | 0.7 | 0.11 | 0.03 | 0.37 | 0.63 | 0.07 | 0.4 |
| 2 | 240299620 | rs11680658 | 0.264 | 0.528 | -2.029 | -0.424 | -0.005 | 0.008 | 0.008 | 0.019 | 0.36 | 0.14 | 0.53 | 0.73 | 0.35 | 0.69 |
| 2 | 240307801 | rs908259 | -0.177 | 0.761 | -1.606 | -0.329 | -0.003 | 0.084 | 0.084 | 0 | 0.78 | 0.9 | 0.88 | 0.93 | 0.78 | 0.86 |
| 2 | 240318755 | rs843462 | -0.619 | 1.247 | -1.328 | 0.512 | -0.049 | -0.02 | -0.02 | 0.983 | 0.11 | 0.01 | 0.1 | 0.1 | 0.19 | 0.03 |
| 2 | 240327863 | rs843458 | -1.019 | 1.178 | -1.052 | 0.509 | -0.004 | 0.001 | 0.001 | 0 | 0.13 | 0.12 | 0.12 | 0.12 | 0.21 | 0.04 |
| 2 | 240330994 | rs843451 | 0.317 | 1.406 | -0.637 | 0.605 | 0.043 | 0.032 | 0.032 | 0 | 0.72 | 0.49 | 0.73 | 0.69 | 0.7 | 0.64 |
| 2 | 240331004 | rs3947630 | 1.03 | 0.808 | -0.794 | 0.229 | 0.014 | -0.001 | -0.001 | 0 | 0.47 | 0.25 | 0.52 | 0.57 | 0.46 | 0.58 |
| 2 | 240331971 | rs709274 | -0.392 | 0.832 | -0.314 | 0.325 | 0.039 | 0.026 | 0.026 | 0 | 0.85 | 0.75 | 0.85 | 0.82 | 0.79 | 0.71 |
| 2 | 240332234 | rs6738626 | 0.464 | 0.51 | -0.57 | 0.188 | 0.032 | 0.028 | 0.028 | 0 | 0.11 | 0.06 | 0.05 | 0 | 0.05 | 0.02 |
| 2 | 240333979 | rs843455 | -0.6 | 0.654 | -0.467 | 0.28 | 0.001 | -0.001 | -0.001 | 0 | 0.87 | 0.87 | 0.85 | 0.87 | 0.8 | 0.95 |
| 2 | 240336735 | rs13010856 | 0.484 | 0.482 | -0.407 | 0.257 | -0.033 | 0.03 | 0.03 | 0 | 0.09 | 0.22 | 0.07 | 0.12 | 0.02 | 0.05 |
| 2 | 240338904 | rs6543534 | 0.25 | 0.812 | -0.925 | -0.023 | -0.052 | -0.026 | -0.026 | 1.559 | 0.75 | 0.87 | 0.76 | 0.76 | 0.69 | 0.76 |
| 2 | 240342708 | rs7568826 | 0.263 | 0.828 | -0.938 | -0.031 | -0.056 | -0.024 | -0.024 | 0 | 0.75 | 0.88 | 0.76 | 0.76 | 0.69 | 0.76 |
| 2 | 240343513 | rs4851975 | 0.994 | 0.71 | -0.297 | -0.036 | -0.003 | 0.019 | 0.019 | 0 | 0.49 | 0.38 | 0.56 | 0.63 | 0.49 | 0.61 |
| 2 | 240350457 | rs7582778 | -0.053 | 0.71 | -0.646 | -0.045 | -0.005 | 0.002 | 0.002 | 0 | 0.11 | 0.09 | 0.12 | 0.1 | 0.14 | 0.1 |
| 2 | 240351323 | rs6757498 | 1.078 | 0.418 | -0.591 | 0.073 | -0.039 | -0.031 | -0.031 | 0.257 | 0.42 | 0.19 | 0.42 | 0.53 | 0.46 | 0.63 |

| | | | | | | | | | | | | | | | | |
|---|---|---|---|---|---|---|---|---|---|---|---|---|---|---|---|---|
| 2 | 240355738 | rs1564976 | -0.405 | 0.281 | -0.005 | 0.049 | 0.001 | 0.013 | 0.013 | 0 | 0.77 | 0.71 | 0.74 | 0.72 | 0.73 | 0.89 |
| 2 | 240359131 | rs6747515 | 0.69 | 0.447 | 0.05 | 0.189 | -0.02 | -0.016 | -0.016 | 0 | 0.43 | 0.28 | 0.43 | 0.52 | 0.46 | 0.62 |
| 2 | 240383056 | rs6743573 | -1.42 | 0.097 | 0.583 | 0.191 | -0.023 | -0.003 | -0.003 | 0.824 | 0.18 | 0.59 | 0.12 | 0.13 | 0.12 | 0.11 |
| 2 | 240385971 | rs1872618 | -1.301 | 0.431 | 0.071 | 0.131 | 0 | 0.008 | 0.008 | 0.257 | 0.72 | 0.73 | 0.62 | 0.51 | 0.73 | 0.48 |
| 2 | 240391590 | rs6756261 | 0.051 | 0.455 | -0.076 | -0.014 | 0.009 | 0.015 | 0.015 | 0.405 | 0.54 | 0.31 | 0.53 | 0.4 | 0.61 | 0.38 |
| 2 | 240394854 | rs6710481 | 0.304 | 0.002 | -0.243 | -0.029 | -0.02 | 0.005 | 0.005 | 0.549 | 0.16 | 0.2 | 0.14 | 0.15 | 0.09 | 0.11 |
| 2 | 240397496 | rs13430446 | -0.041 | 0.543 | -0.541 | -0.13 | -0.001 | -0.003 | -0.003 | 0 | 0.22 | 0.19 | 0.25 | 0.21 | 0.23 | 0.21 |
| 2 | 240397644 | rs13430569 | -0.054 | 0.548 | -0.54 | -0.128 | -0.001 | -0.002 | -0.002 | 0 | 0.22 | 0.19 | 0.25 | 0.21 | 0.23 | 0.21 |
| 2 | 240399614 | rs6726998 | -1.133 | 0.397 | -0.832 | 0.172 | 0.009 | -0.065 | -0.065 | 0 | 0.7 | 0.87 | 0.63 | 0.48 | 0.72 | 0.52 |
| 2 | 240401778 | rs11894982 | -0.602 | 0.278 | -0.629 | -0.48 | 0.002 | 0.002 | 0.002 | 0.363 | 0.6 | 0.64 | 0.73 | 0.69 | 0.64 | 0.78 |
| 2 | 240406374 | rs6716571 | -0.041 | 0.413 | -0.418 | -0.219 | -0.001 | 0 | 0 | 1.532 | 0.22 | 0.26 | 0.22 | 0.19 | 0.21 | 0.22 |
| 2 | 240409324 | rs6543540 | -0.875 | 0.169 | -0.344 | -0.366 | -0.007 | 0 | 0 | 0 | 0.7 | 0.65 | 0.78 | 0.71 | 0.73 | 0.79 |
| 2 | 240415571 | rs7597221 | -0.25 | -0.225 | -0.425 | -0.286 | -0.001 | -0.002 | -0.002 | 0 | 0.47 | 0.47 | 0.55 | 0.7 | 0.36 | 0.56 |
| 2 | 240417788 | rs1486323 | -0.016 | -0.385 | -0.612 | -0.359 | 0.02 | 0.035 | 0.035 | 0 | 0.39 | 0.28 | 0.54 | 0.71 | 0.34 | 0.54 |
| 2 | 240418660 | rs6743931 | -0.105 | 0.529 | 0.339 | -0.356 | 0.02 | -0.089 | -0.089 | 0 | 0.16 | 0.23 | 0.13 | 0.01 | 0.16 | 0.16 |
| 2 | 240420675 | rs1796449 | 0.307 | 0.718 | 0.323 | 0.145 | -0.015 | -0.017 | -0.017 | 0 | 0.44 | 0.15 | 0.55 | 0.63 | 0.46 | 0.54 |
| 2 | 240423291 | rs13415709 | -0.389 | 0.888 | 0.501 | 0.104 | 0.048 | -0.075 | -0.075 | 0 | 0.13 | 0.24 | 0.11 | 0.02 | 0.14 | 0.16 |
| 2 | 240424704 | rs13429807 | 0.244 | 1.08 | 0.565 | 0.223 | 0.081 | -0.051 | -0.051 | 0 | 0.09 | 0.23 | 0.1 | 0.01 | 0.12 | 0.14 |
| 2 | 240425643 | rs7562005 | -1.26 | 0.554 | 0.611 | 0.556 | 0.013 | 0.328 | 0.328 | 0 | 0.06 | 0.2 | 0.18 | 0.31 | 0.05 | 0.07 |
| 2 | 240426414 | rs12622672 | 0.708 | 0.777 | -0.023 | 0.182 | 0.014 | -0.009 | -0.009 | 0 | 0.38 | 0.31 | 0.4 | 0.62 | 0.34 | 0.41 |
| 2 | 240426607 | rs2630754 | 0.218 | 1.422 | -0.737 | -0.476 | -0.036 | 0.031 | 0.031 | 0.206 | 0.16 | 0.1 | 0.19 | 0.06 | 0.26 | 0.29 |
| 2 | 240426756 | rs1684274 | -0.344 | 0.647 | -0.426 | -0.366 | -0.007 | 0.043 | 0.043 | 1.84 | 0.66 | 0.81 | 0.8 | 0.97 | 0.65 | 0.77 |
| 2 | 240436304 | rs1564974 | -1.326 | 1.061 | 0.387 | 0.761 | 0.018 | 0.015 | 0.015 | 0.02 | 0.37 | 0.52 | 0.38 | 0.38 | 0.45 | 0.43 |
| 2 | 240437810 | rs6718108 | -0.795 | 1.293 | 0.698 | 0.891 | 0.006 | 0.001 | 0.001 | 0.278 | 0.23 | 0.23 | 0.32 | 0.24 | 0.34 | 0.37 |
| 2 | 240441158 | rs10174165 | -0.834 | 1.055 | 1.156 | 0.84 | -0.005 | 0.026 | 0.026 | 0.093 | 0.14 | 0.31 | 0.06 | 0.13 | 0.09 | 0.07 |
| 2 | 240442314 | rs11675968 | -0.245 | 1.058 | 0.998 | 0.753 | -0.001 | 0.007 | 0.007 | 0 | 0.05 | 0.03 | 0.02 | 0.02 | 0.06 | 0.03 |
| 2 | 240443992 | rs6727608 | -1.485 | 1.234 | 1.41 | 0.784 | 0.019 | 0.074 | 0.074 | 0 | 0.16 | 0.34 | 0.23 | 0.25 | 0.15 | 0.1 |
| 2 | 240445491 | rs11682976 | -0.313 | 1.231 | 1.076 | 0.765 | -0.014 | 0.024 | 0.024 | 0 | 0.06 | 0.01 | 0.02 | 0.02 | 0.06 | 0.03 |
| 2 | 240448123 | rs7573739 | -0.958 | 1.177 | 1.579 | 0.746 | 0.032 | -0.013 | -0.013 | 0 | 0.07 | 0.3 | 0.04 | 0.02 | 0.06 | 0.03 |
| 2 | 240452008 | rs874014 | -0.423 | 1.141 | 1.278 | 0.791 | 0.012 | 0.097 | 0.097 | 0 | 0.11 | 0.2 | 0.19 | 0.23 | 0.1 | 0.07 |
| 2 | 240452167 | rs7568886 | -0.899 | 0.957 | 1.157 | 0.593 | -0.008 | 0.027 | 0.027 | 0 | 0.15 | 0.33 | 0.07 | 0.14 | 0.1 | 0.07 |
| 2 | 240453552 | rs7592100 | -1.433 | 1.052 | 0.262 | 0.333 | -0.019 | -0.001 | -0.001 | 0 | 0.4 | 0.66 | 0.4 | 0.51 | 0.47 | 0.44 |
| 2 | 240457989 | rs731154 | -0.925 | 0.93 | -0.036 | -0.059 | -0.005 | 0.003 | 0.003 | 0.393 | 0.26 | 0.24 | 0.23 | 0.18 | 0.34 | 0.34 |
| 2 | 240458353 | rs731153 | 0.7 | 0.92 | 0.098 | -0.076 | 0.014 | -0.015 | -0.015 | 0 | 0.76 | 0.65 | 0.78 | 0.83 | 0.69 | 0.69 |
| 2 | 240463531 | rs11694791 | 0.287 | 0.672 | -0.193 | -0.76 | -0.003 | -0.125 | -0.125 | 0 | 0.23 | 0.4 | 0.13 | 0.01 | 0.19 | 0.13 |
| 2 | 240467554 | rs7571643 | -0.954 | 0.504 | -0.839 | -1.156 | -0.004 | 0.005 | 0.005 | 0 | 0.63 | 0.66 | 0.49 | 0.21 | 0.57 | 0.55 |
| 2 | 240471568 | rs6543547 | -0.218 | 0.433 | -0.581 | -0.905 | -0.014 | -0.074 | -0.074 | 0.856 | 0.19 | 0.36 | 0.17 | 0.08 | 0.16 | 0.15 |
| 2 | 240471992 | rs10206786 | 1.142 | 0.149 | -1.737 | -0.588 | 0.015 | 0.027 | 0.027 | 0 | 0.49 | 0.42 | 0.43 | 0.56 | 0.52 | 0.34 |
| 2 | 240473961 | rs10177901 | -0.3 | 0.452 | -1.489 | -0.889 | -0.004 | 0.006 | 0.006 | 1.185 | 0.19 | 0.16 | 0.18 | 0.12 | 0.26 | 0.26 |
| 2 | 240474582 | rs1872621 | 0.565 | 0.615 | -1.928 | -0.64 | 0 | -0.016 | -0.016 | 0.109 | 0.57 | 0.47 | 0.68 | 0.81 | 0.58 | 0.54 |
| 2 | 240476642 | rs1796453 | -0.867 | 0.936 | -2.6 | -0.048 | -0.057 | 0.023 | 0.023 | 0 | 0.16 | 0.01 | 0.16 | 0.09 | 0.22 | 0.24 |
| 2 | 240479364 | rs7596550 | -0.358 | 0.832 | -2.839 | 0.41 | -0.045 | -0.089 | -0.089 | 0.36 | 0.46 | 0.92 | 0.48 | 0.46 | 0.43 | 0.53 |
| 2 | 240482579 | rs11683432 | -0.129 | 0.713 | -2.719 | 0.265 | -0.018 | 0.041 | 0.041 | 0 | 0.2 | 0.05 | 0.19 | 0.13 | 0.23 | 0.07 |
| 2 | 240482934 | rs9636383 | 0.242 | 0.998 | -2.946 | 0.576 | -0.034 | -0.016 | -0.016 | 0 | 0.25 | 0.18 | 0.35 | 0.25 | 0.31 | 0.3 |
| 2 | 240484543 | rs9636384 | 0.245 | 0.935 | -2.972 | 0.593 | -0.037 | -0.007 | -0.007 | 0 | 0.26 | 0.18 | 0.33 | 0.25 | 0.32 | 0.3 |
| 2 | 240484995 | rs10202203 | 0.265 | 0.694 | -2.119 | 0.327 | -0.002 | -0.007 | -0.007 | 0 | 0.21 | 0.19 | 0.2 | 0.13 | 0.23 | 0.07 |
| 2 | 240489258 | rs4328655 | -0.312 | 0.283 | -1.915 | 0.817 | 0.015 | 0.015 | 0.015 | 1.859 | 0.52 | 0.45 | 0.63 | 0.61 | 0.62 | 0.47 |
| 2 | 240490213 | rs11124208 | -0.426 | 0.325 | -2.003 | 0.919 | -0.009 | -0.031 | -0.031 | 0 | 0.7 | 0.87 | 0.76 | 0.63 | 0.7 | 0.55 |
| 2 | 240491666 | rs1882676 | -0.443 | -0.204 | -1.661 | 1.095 | 0.007 | -0.007 | -0.007 | 0.7 | 0.31 | 0.3 | 0.27 | 0.41 | 0.2 | 0.34 |
| 2 | 240499524 | rs6543553 | -0.704 | -0.03 | -1.667 | 1.606 | 0.007 | -0.055 | -0.055 | 0.253 | 0.84 | 0.93 | 0.82 | 0.63 | 0.87 | 0.78 |
| 2 | 240500294 | rs11683242 | 0.131 | -0.002 | -1.662 | 1.249 | -0.003 | 0.02 | 0.02 | 5.651 | 0.16 | 0.11 | 0.21 | 0.07 | 0.16 | 0.11 |
| 2 | 240500437 | rs10180205 | 0.933 | 0.26 | -1.248 | 1.047 | -0.007 | 0.106 | 0.106 | 0 | 0.49 | 0.32 | 0.35 | 0.17 | 0.5 | 0.37 |
| 2 | 240502117 | rs7570018 | 0.242 | 0.197 | -0.474 | 1.292 | 0.02 | 0.079 | 0.079 | 0.207 | 0.13 | 0.24 | 0.23 | 0.32 | 0.17 | 0.24 |
| 2 | 240503373 | rs13390776 | 1.049 | 0.015 | -0.444 | 1.077 | -0.014 | 0.117 | 0.117 | 0 | 0.48 | 0.3 | 0.35 | 0.18 | 0.5 | 0.36 |
| 2 | 240504931 | rs13021357 | 0.443 | 0.056 | -0.26 | 1.014 | 0.001 | -0.001 | -0.001 | 0 | 0.17 | 0.15 | 0.21 | 0.07 | 0.15 | 0.11 |
| 2 | 240506093 | rs10189961 | -0.151 | 0.111 | 0.133 | 1.285 | 0 | 0.007 | 0.007 | 0.184 | 0.72 | 0.69 | 0.73 | 0.58 | 0.7 | 0.57 |
| 2 | 240507331 | rs2008714 | -0.022 | 0.086 | 0.123 | 0.914 | 0.002 | 0.02 | 0.02 | 0.095 | 0.71 | 0.67 | 0.72 | 0.52 | 0.68 | 0.57 |
| 2 | 240516749 | rs11685345 | 1.025 | -0.484 | -0.203 | 0.45 | 0.009 | 0.02 | 0.02 | 0 | 0.48 | 0.46 | 0.37 | 0.18 | 0.56 | 0.39 |
| 2 | 240517194 | rs7606519 | -0.185 | -0.503 | -0.218 | 0.787 | 0.05 | -0.016 | -0.016 | 0.786 | 0.71 | 0.86 | 0.76 | 0.63 | 0.78 | 0.59 |
| 2 | 240518771 | rs2028192 | 0.016 | -0.437 | -0.124 | 0.82 | 0.009 | -0.015 | -0.015 | 0 | 0.69 | 0.77 | 0.72 | 0.55 | 0.71 | 0.56 |
| 2 | 240520878 | rs7557121 | -0.66 | -0.44 | -0.251 | 0.81 | -0.002 | -0.028 | -0.028 | 0 | 0.3 | 0.23 | 0.39 | 0.48 | 0.33 | 0.36 |
| 2 | 240524052 | rs7593722 | -0.756 | -0.494 | -0.395 | 0.837 | 0.014 | -0.067 | -0.067 | 0 | 0.78 | 0.89 | 0.76 | 0.59 | 0.82 | 0.72 |
| 2 | 240529494 | rs12712292 | -1.061 | -0.606 | -0.346 | 0.809 | -0.005 | 0.001 | 0.001 | 0.684 | 0.8 | 0.81 | 0.78 | 0.67 | 0.89 | 0.76 |
| 2 | 240537244 | rs7604209 | -1.138 | -0.559 | -0.711 | 0.802 | 0.034 | -0.064 | -0.064 | 0 | 0.81 | 0.93 | 0.79 | 0.67 | 0.88 | 0.76 |
| 2 | 240539765 | rs1030891 | 0.273 | -0.537 | -0.616 | 0.733 | 0.001 | 0.002 | 0.002 | 5.687 | 0.19 | 0.18 | 0.36 | 0.38 | 0.16 | 0.17 |
| 2 | 240547172 | rs7568812 | 0.13 | -0.483 | -0.729 | 0.533 | 0.022 | -0.047 | -0.047 | 0 | 0.68 | 0.82 | 0.73 | 0.56 | 0.69 | 0.56 |
| 2 | 240554036 | rs754700 | 0.608 | -0.908 | -0.851 | -0.979 | 0.012 | -0.029 | -0.029 | 0 | 0.37 | 0.44 | 0.32 | 0.16 | 0.48 | 0.36 |
| 2 | 240562386 | rs3916020 | -0.472 | -1.186 | -0.764 | -0.826 | 0.027 | 0.016 | 0.016 | 0.522 | 0.66 | 0.8 | 0.47 | 0.26 | 0.75 | 0.58 |
| 2 | 240568714 | rs6760787 | -0.809 | -1.008 | -0.862 | -1.233 | 0.011 | 0.036 | 0.036 | 0.058 | 0.5 | 0.57 | 0.58 | 0.83 | 0.44 | 0.62 |

| | | | | | | | | | | | | | | | | |
|---|---|---|---|---|---|---|---|---|---|---|---|---|---|---|---|---|
| 2 | 240574731 | rs10165876 | -0.209 | -1.149 | -0.851 | -1.196 | 0.015 | 0.021 | 0.021 | 0 | 0.4 | 0.32 | 0.53 | 0.76 | 0.36 | 0.57 |
| 2 | 240581797 | rs13419075 | -0.26 | -0.956 | -0.068 | -0.908 | -0.03 | -0.043 | -0.043 | 0.418 | 0.15 | 0.26 | 0.09 | 0.04 | 0.09 | 0.15 |
| 2 | 240581960 | rs13024940 | -0.635 | -0.339 | -0.016 | -0.527 | 0.018 | 0.016 | 0.016 | 0 | 0.08 | 0.23 | 0.05 | 0.07 | 0.07 | 0.05 |
| 2 | 240586407 | rs9646967 | 0.129 | 0.101 | -0.333 | -0.712 | 0.002 | 0.006 | 0.006 | 0 | 0.33 | 0.26 | 0.55 | 0.76 | 0.31 | 0.44 |
| 2 | 240589914 | rs12612406 | -1.158 | 0.125 | -0.407 | -0.746 | 0.006 | 0.058 | 0.058 | 1.054 | 0.49 | 0.37 | 0.62 | 0.84 | 0.46 | 0.63 |
| 2 | 240601166 | rs6543570 | -0.22 | 0.067 | -0.783 | -0.544 | -0.019 | -0.032 | -0.032 | 0 | 0.38 | 0.48 | 0.58 | 0.75 | 0.32 | 0.47 |
| 2 | 240605763 | rs1465820 | -0.679 | 0.278 | -0.574 | -0.354 | -0.002 | -0.01 | -0.01 | 0.9 | 0.48 | 0.5 | 0.61 | 0.77 | 0.41 | 0.63 |
| 2 | 240618118 | rs11124220 | -1.018 | 0.567 | -0.349 | 0.23 | 0 | -0.011 | -0.011 | 1.805 | 0.38 | 0.34 | 0.59 | 0.84 | 0.41 | 0.63 |
| 2 | 240618610 | rs920247 | -0.853 | 0.579 | -0.362 | 0.537 | -0.003 | -0.003 | -0.003 | 0 | 0.35 | 0.35 | 0.55 | 0.73 | 0.34 | 0.6 |
| 2 | 240620658 | rs12712294 | -1.215 | 0.424 | -0.308 | 0.554 | 0.01 | 0.027 | 0.027 | 0.042 | 0.42 | 0.37 | 0.57 | 0.8 | 0.36 | 0.68 |
| 2 | 240620843 | rs6749693 | -1.295 | 0.55 | -0.429 | 0.867 | 0.001 | 0 | 0 | 0 | 0.36 | 0.29 | 0.21 | 0.36 | 0.35 | 0.42 |
| 2 | 240622298 | rs13026302 | -1.624 | 0.96 | -0.672 | 0.977 | 0.01 | 0.014 | 0.014 | 0 | 0.13 | 0.17 | 0.09 | 0.18 | 0.17 | 0.19 |
| 2 | 240625255 | rs1020080 | -1.17 | 0.779 | -0.873 | 0.81 | 0.002 | 0.02 | 0.02 | 0 | 0.14 | 0.12 | 0.12 | 0.1 | 0.14 | 0.16 |
| 2 | 240626443 | rs1437739 | -0.751 | 0.803 | -0.938 | 0.822 | -0.031 | 0.015 | 0.015 | 0 | 0.12 | 0 | 0.11 | 0.1 | 0.14 | 0.16 |
| 2 | 240626750 | rs7566783 | -1.133 | 0.638 | -0.844 | 0.892 | -0.002 | -0.002 | -0.002 | 0 | 0.28 | 0.26 | 0.23 | 0.29 | 0.27 | 0.39 |
| 2 | 240630573 | rs6711207 | -1.311 | 0.506 | -0.119 | 0.982 | 0.015 | 0.051 | 0.051 | 0 | 0.11 | 0.31 | 0.06 | 0.19 | 0.08 | 0.15 |
| 2 | 240632788 | rs1437730 | 0.006 | 0.534 | -1.622 | -0.275 | -0.107 | 0.26 | 0.26 | 0.173 | 0.39 | 0.06 | 0.25 | 0.03 | 0.49 | 0.22 |
| 2 | 240634587 | rs4852110 | -0.407 | -0.109 | -1.81 | 0.304 | 0.049 | 0.016 | 0.016 | 0 | 0.61 | 0.17 | 0.72 | 0.59 | 0.71 | 0.55 |
| 2 | 240634833 | rs4852111 | -0.377 | -0.12 | -1.818 | 0.554 | 0.053 | -0.017 | -0.017 | 0.518 | 0.61 | 0.17 | 0.67 | 0.53 | 0.7 | 0.51 |
| 2 | 240640112 | rs12470565 | -0.121 | 0.022 | -1.453 | 0.481 | 0.017 | 0.039 | 0.039 | 0 | 0.46 | 0.39 | 0.4 | 0.24 | 0.38 | 0.43 |
| 2 | 240642799 | rs4852113 | -0.892 | 0.013 | -0.887 | 0.519 | 0.003 | 0.042 | 0.042 | 0.04 | 0.47 | 0.73 | 0.65 | 0.59 | 0.58 | 0.39 |
| 2 | 240644043 | rs6724893 | 0.518 | -0.521 | -1.311 | 0.479 | -0.038 | -0.008 | -0.008 | 0 | 0.83 | 0.65 | 0.89 | 0.84 | 0.92 | 0.6 |
| 2 | 240644907 | rs11891511 | 1.404 | -0.527 | -1.809 | 0.523 | -0.006 | 0.003 | 0.003 | 0 | 0.23 | 0.26 | 0.15 | 0.31 | 0.22 | 0.15 |
| 2 | 240647402 | rs6734016 | 0.242 | -0.604 | -2.072 | 0.476 | 0.051 | 0.107 | 0.107 | 0.031 | 0.53 | 0.84 | 0.54 | 0.73 | 0.6 | 0.33 |
| 2 | 240649925 | rs10865083 | 0.906 | -0.553 | -1.28 | 1.147 | 0.053 | 0.032 | 0.032 | 0 | 0.67 | 0.86 | 0.62 | 0.77 | 0.74 | 0.51 |
| 2 | 240654444 | rs11124222 | -0.474 | -0.097 | -1.133 | 1.455 | 0.016 | 0.281 | 0.281 | 0 | 0.05 | 0.17 | 0.14 | 0.28 | 0.03 | 0.03 |
| 2 | 240655344 | rs11682702 | 0.068 | 0.03 | -1.81 | 0.56 | -0.001 | 0.005 | 0.005 | 0 | 0.11 | 0.02 | 0.05 | 0.1 | 0.09 | 0.2 |
| 2 | 240660209 | rs4852118 | -0.048 | -0.036 | -1.686 | 0.623 | 0.02 | -0.003 | -0.003 | 0.324 | 0.14 | 0.03 | 0.26 | 0.08 | 0.13 | 0.09 |
| 2 | 240661843 | rs12990084 | -0.033 | 0.077 | -1.758 | 0.661 | 0.009 | -0.01 | -0.01 | 0 | 0.13 | 0.01 | 0.09 | 0.09 | 0.13 | 0.14 |
| 2 | 240670045 | rs1530009 | 0.737 | 0.098 | -2 | 0.652 | 0.003 | -0.015 | -0.015 | 0.004 | 0.78 | 0.92 | 0.81 | 0.8 | 0.78 | 0.67 |
| 2 | 240671768 | rs731161 | 0.9 | 0.213 | -2.618 | -0.06 | -0.02 | 0.131 | 0.131 | 0 | 0.37 | 0.17 | 0.19 | 0.17 | 0.36 | 0.36 |
| 2 | 240672353 | rs748946 | 0.079 | 0.037 | -2.815 | 0.771 | 0.039 | -0.067 | -0.067 | 2.905 | 0.81 | 0.92 | 0.78 | 0.48 | 0.86 | 0.71 |
| 2 | 240675093 | rs11681707 | 0.106 | 0.075 | -2.769 | 0.745 | 0 | -0.023 | -0.023 | 0.245 | 0.78 | 0.94 | 0.81 | 0.8 | 0.79 | 0.67 |
| 2 | 240678769 | rs11897501 | -0.418 | -0.079 | -1.85 | 0.598 | 0.017 | 0.001 | 0.001 | 0 | 0.53 | 0.61 | 0.64 | 0.45 | 0.61 | 0.44 |
| 2 | 240678849 | rs4852123 | 0.918 | 0.454 | -1.67 | 0.165 | 0.019 | -0.001 | -0.001 | 0 | 0.12 | 0.12 | 0.3 | 0.16 | 0.2 | 0.14 |
| 2 | 240680022 | rs4852124 | 0.633 | -0.104 | -1.38 | 0.106 | 0.06 | 0.118 | 0.118 | 0 | 0.5 | 0.35 | 0.51 | 0.34 | 0.57 | 0.51 |
| 2 | 240682076 | rs4851992 | -0.309 | 0.411 | -1.75 | -0.022 | -0.085 | 0.026 | 0.026 | 0.078 | 0.78 | 0.93 | 0.64 | 0.82 | 0.66 | 0.81 |
| 2 | 240683327 | rs11124223 | -0.405 | -0.218 | -1.067 | -0.192 | -0.017 | -0.002 | -0.002 | 0.004 | 0.31 | 0.45 | 0.37 | 0.69 | 0.26 | 0.38 |
| 2 | 240683576 | rs2028191 | -0.18 | -0.276 | -1.021 | 0.163 | -0.011 | 0.032 | 0.032 | 0 | 0.28 | 0.42 | 0.36 | 0.55 | 0.25 | 0.21 |
| 2 | 240683658 | rs2028190 | 2.059 | -0.342 | -1.073 | 0.301 | 0.012 | -0.005 | -0.005 | 0.013 | 0.11 | 0.03 | 0.12 | 0.1 | 0.12 | 0.09 |
| 2 | 240684263 | rs12468632 | 2.109 | -0.343 | -1.073 | 0.302 | 0.014 | 0.022 | 0.022 | 0 | 0.11 | 0.03 | 0.07 | 0.06 | 0.12 | 0.09 |
| 2 | 240684619 | rs6543576 | -0.813 | -0.393 | -0.953 | 0.322 | -0.012 | 0.304 | 0.304 | 0 | 0.09 | 0.36 | 0.17 | 0.59 | 0.05 | 0.23 |
| 2 | 240685573 | rs6748389 | 1.035 | -0.496 | -1.143 | 0.031 | -0.003 | 0.035 | 0.035 | 0 | 0.54 | 0.57 | 0.39 | 0.14 | 0.46 | 0.36 |
| 2 | 240686528 | rs6748893 | -0.547 | -0.423 | -1.17 | 0.023 | -0.077 | 0.029 | 0.029 | 0.384 | 0.74 | 0.87 | 0.63 | 0.78 | 0.63 | 0.68 |
| 2 | 240687289 | rs10207060 | -0.877 | -0.641 | -0.648 | -0.229 | 0.022 | 0.067 | 0.067 | 0.18 | 0.52 | 0.63 | 0.54 | 0.78 | 0.46 | 0.51 |
| 2 | 240688338 | rs11904223 | 0.717 | -0.534 | -0.33 | -0.098 | 0.003 | 0.002 | 0.002 | 0.022 | 0.58 | 0.57 | 0.45 | 0.46 | 0.51 | 0.58 |
| 2 | 240689097 | rs7602963 | 1.434 | -0.491 | -0.386 | -0.066 | 0.029 | 0.055 | 0.055 | 0 | 0.43 | 0.29 | 0.27 | 0.23 | 0.36 | 0.46 |
| 2 | 240699174 | rs4852126 | -0.256 | -0.337 | 0.062 | 0.287 | 0.068 | 0.587 | 0.587 | 0 | 0.07 | 0.41 | 0.17 | 0.45 | 0.04 | 0.06 |
| 2 | 240699992 | rs6730007 | 0.28 | -0.376 | -0.156 | 0.098 | 0.015 | 0.008 | 0.008 | 0 | 0.68 | 0.63 | 0.56 | 0.52 | 0.58 | 0.48 |
| 2 | 240703201 | rs13014729 | 1.973 | -0.6 | -0.394 | 0.039 | 0.011 | -0.025 | -0.025 | 0 | 0.27 | 0.22 | 0.25 | 0.36 | 0.18 | 0.2 |
| 2 | 240705043 | rs12614692 | 1.249 | -0.408 | -0.494 | -0.054 | -0.007 | -0.014 | -0.014 | 0.622 | 0.1 | 0.08 | 0.09 | 0.14 | 0.14 | 0.19 |
| 2 | 240705460 | rs2412030 | -0.713 | -0.421 | -0.379 | -0.117 | 0.002 | 0.034 | 0.034 | 0 | 0.53 | 0.62 | 0.42 | 0.29 | 0.53 | 0.46 |
| 2 | 240709286 | rs10195015 | -0.8 | -0.389 | -0.488 | -0.118 | -0.046 | -0.021 | -0.021 | 0 | 0.81 | 0.89 | 0.67 | 0.65 | 0.72 | 0.67 |
| 2 | 240709496 | rs10181238 | 0.99 | -0.487 | -0.31 | -0.189 | -0.005 | 0.06 | 0.06 | 0 | 0.27 | 0.16 | 0.23 | 0.09 | 0.28 | 0.19 |
| 2 | 240711141 | rs2412033 | -0.153 | -0.43 | -0.285 | -0.138 | -0.032 | 0.004 | 0.004 | 0 | 0.68 | 0.81 | 0.6 | 0.59 | 0.59 | 0.58 |
| 2 | 240720131 | rs1316924 | 0.826 | -0.366 | -0.183 | -0.079 | -0.011 | 0.002 | 0.002 | 0 | 0.44 | 0.45 | 0.33 | 0.45 | 0.33 | 0.42 |
| 2 | 240721728 | rs1574030 | -0.871 | -0.164 | -0.216 | 0.013 | 0.046 | 0.013 | 0.013 | 0 | 0.83 | 0.69 | 0.66 | 0.68 | 0.75 | 0.8 |
| 2 | 240722820 | rs2412042 | 0.582 | 0.1 | -0.659 | 0.083 | 0.037 | 0.055 | 0.055 | 0 | 0.48 | 0.29 | 0.39 | 0.57 | 0.42 | 0.47 |
| 2 | 240727833 | rs10206264 | 2.169 | -0.077 | -0.695 | 0.075 | -0.003 | -0.027 | -0.027 | 0.432 | 0.23 | 0.25 | 0.15 | 0.04 | 0.16 | 0.17 |
| 2 | 240728483 | rs11679150 | 0.085 | -0.023 | -1.22 | -0.074 | -0.026 | -0.01 | -0.01 | 0 | 0.29 | 0.05 | 0.17 | 0.22 | 0.31 | 0.46 |
| 2 | 240728640 | rs11695034 | 2.678 | 0.168 | -1.199 | 0.106 | -0.005 | 0.09 | 0.09 | 0 | 0.2 | 0.06 | 0.12 | 0.04 | 0.15 | 0.17 |
| 2 | 240729948 | rs6543586 | -0.948 | 0.45 | -0.757 | 0.377 | -0.005 | -0.006 | -0.006 | 0.11 | 0.86 | 0.9 | 0.67 | 0.67 | 0.73 | 0.79 |
| 2 | 240730845 | rs7572106 | -0.58 | -0.063 | -0.458 | 0.108 | -0.002 | -0.023 | -0.023 | 0 | 0.38 | 0.39 | 0.29 | 0.12 | 0.33 | 0.32 |
| 2 | 240732594 | rs11124232 | -0.588 | 0.157 | -0.808 | 0.571 | 0.006 | -0.024 | -0.024 | 0 | 0.17 | 0.25 | 0.12 | 0.1 | 0.19 | 0.2 |
| 2 | 240732640 | rs12712298 | -1.57 | 0.624 | -1.16 | 0.846 | -0.014 | -0.012 | -0.012 | 0 | 0.95 | 0.98 | 0.95 | 0.92 | 0.91 | 0.94 |
| 2 | 240733032 | rs11124233 | -0.596 | 0.472 | -1.2 | 1.164 | -0.018 | 0.021 | 0.021 | 0 | 0.29 | 0.41 | 0.23 | 0.37 | 0.19 | 0.36 |
| 2 | 240733607 | rs7606359 | -0.752 | 0.906 | -1.494 | 1.059 | -0.008 | -0.005 | -0.005 | 0 | 0.89 | 0.93 | 0.92 | 0.65 | 0.83 | 0.79 |
| 2 | 240736174 | rs13000485 | 0.342 | 0.815 | -1.406 | 0.649 | -0.016 | 0.095 | 0.095 | 0 | 0.47 | 0.28 | 0.39 | 0.2 | 0.45 | 0.49 |
| 2 | 240736419 | rs12612507 | -0.936 | 0.795 | -1.76 | 0.458 | -0.023 | 0.066 | 0.066 | 0.466 | 0.47 | 0.86 | 0.55 | 0.67 | 0.46 | 0.59 |

| | | | | | | | | | | | | | | | | |
|---|---|---|---|---|---|---|---|---|---|---|---|---|---|---|---|---|
| 2 | 240736739 | rs6750193 | 0.442 | 0.666 | -1.148 | 0.496 | -0.001 | -0.032 | -0.032 | 0 | 0.12 | 0.29 | 0.12 | 0.14 | 0.1 | 0.08 |
| 2 | 240737309 | rs10167014 | 0.59 | 0.692 | -1.273 | 0.509 | -0.004 | -0.025 | -0.025 | 0.197 | 0.12 | 0.19 | 0.11 | 0.14 | 0.1 | 0.08 |
| 2 | 240739225 | rs4852139 | -0.429 | 0.764 | -1.814 | 0.777 | -0.015 | -0.019 | -0.019 | 0 | 0.71 | 0.92 | 0.72 | 0.69 | 0.71 | 0.76 |
| 2 | 240740612 | rs4241236 | -0.121 | 0.717 | -1.253 | 0.783 | -0.001 | -0.005 | -0.005 | 0 | 0.47 | 0.57 | 0.5 | 0.53 | 0.51 | 0.45 |
| 2 | 240745718 | rs4852140 | 0.121 | 0.554 | -1.284 | 0.916 | 0.003 | -0.023 | -0.023 | 0.078 | 0.74 | 0.78 | 0.67 | 0.48 | 0.79 | 0.63 |
| 2 | 240751267 | rs2412077 | 0.357 | 0.48 | -0.911 | 1.06 | -0.014 | -0.113 | -0.113 | 0 | 0.15 | 0.04 | 0.27 | 0.47 | 0.12 | 0.27 |
| 2 | 240752779 | rs11124235 | 0.68 | 0.671 | -1.356 | 0.273 | -0.029 | 0.019 | 0.019 | 0.007 | 0.35 | 0.2 | 0.36 | 0.34 | 0.37 | 0.18 |
| 2 | 240754930 | rs11680915 | 0.441 | 0.704 | -1.741 | 0.117 | -0.073 | -0.041 | -0.041 | 0.274 | 0.15 | 0.02 | 0.18 | 0.16 | 0.23 | 0.23 |
| 2 | 240755338 | rs10196581 | -0.467 | 0.007 | -1.564 | -0.134 | 0.028 | -0.005 | -0.005 | 0 | 0.39 | 0.65 | 0.38 | 0.57 | 0.26 | 0.46 |
| 2 | 240765562 | rs12987919 | 0.563 | 0.517 | -1.386 | 0.012 | 0.034 | -0.002 | -0.002 | 0 | 0.53 | 0.27 | 0.6 | 0.56 | 0.63 | 0.49 |
| 2 | 240773871 | rs13014066 | 0.242 | 0.43 | -1.234 | -0.329 | 0.027 | 0.13 | 0.13 | 0.195 | 0.59 | 0.29 | 0.75 | 0.89 | 0.67 | 0.63 |
| 2 | 240777836 | rs11894116 | -0.409 | 0.393 | -1.366 | -0.303 | -0.004 | -0.146 | -0.146 | 0.86 | 0.77 | 0.51 | 0.82 | 0.98 | 0.8 | 0.78 |
| 2 | 240780714 | rs2412024 | 0.091 | 0.511 | -1.755 | 0.282 | 0.002 | 0.004 | 0.004 | 0.581 | 0.42 | 0.31 | 0.45 | 0.38 | 0.42 | 0.27 |
| 2 | 240827284 | rs9288729 | -0.022 | 0.737 | -2.369 | 0.218 | -0.16 | -0.072 | -0.072 | 0 | 0.37 | 0.04 | 0.35 | 0.58 | 0.46 | 0.37 |
| 2 | 240830028 | rs9677897 | -0.219 | 0.444 | -1.337 | -0.013 | 0.006 | 0.015 | 0.015 | 0 | 0.49 | 0.58 | 0.48 | 0.64 | 0.55 | 0.4 |
| 2 | 240831679 | rs10933613 | -0.084 | 0.95 | -1.137 | -0.064 | 0.011 | 0.01 | 0.01 | 0 | 0.09 | 0.1 | 0.24 | 0.16 | 0.15 | 0.19 |
| 2 | 240837872 | rs11676967 | -0.143 | 0.942 | -1.166 | -0.15 | 0.006 | 0.001 | 0.001 | 0 | 0.09 | 0.1 | 0.25 | 0.12 | 0.16 | 0.18 |
| 2 | 240839969 | rs7599067 | -0.812 | 0.327 | -1.172 | -0.509 | -0.001 | -0.045 | -0.045 | 0 | 0.85 | 0.82 | 0.88 | 0.99 | 0.86 | 0.82 |
| 2 | 240842503 | rs11684477 | 0.407 | 0.68 | -1.104 | -0.136 | -0.001 | -0.004 | -0.004 | 0 | 0.37 | 0.33 | 0.61 | 0.52 | 0.38 | 0.43 |
| 2 | 240844350 | rs13402622 | 0.069 | 0.213 | -1.044 | -0.644 | -0.009 | -0.09 | -0.09 | 3.575 | 0.65 | 0.61 | 0.81 | 0.94 | 0.74 | 0.66 |
| 2 | 240851475 | rs10933571 | -0.175 | 0.683 | -1.318 | 0.138 | -0.011 | -0.005 | -0.005 | 0 | 0.74 | 0.8 | 0.76 | 0.74 | 0.67 | 0.81 |
| 2 | 240851523 | rs11686460 | 0.041 | 0.629 | -0.979 | 0.114 | 0.003 | 0.047 | 0.047 | 0 | 0.33 | 0.38 | 0.4 | 0.48 | 0.34 | 0.38 |
| 2 | 240857988 | rs10181407 | 0.28 | 0.506 | -0.539 | 0.037 | -0.02 | -0.044 | -0.044 | 0 | 0.16 | 0.5 | 0.13 | 0.08 | 0.14 | 0.16 |
| 2 | 240858334 | rs10933578 | 0.28 | 0.439 | -0.124 | 0.004 | -0.02 | -0.044 | -0.044 | 0.018 | 0.16 | 0.5 | 0.13 | 0.08 | 0.14 | 0.16 |
| 2 | 240858490 | rs12474521 | -1.201 | 0.217 | -0.939 | -0.313 | -0.043 | 0.005 | 0.005 | 0 | 0.71 | 0.94 | 0.79 | 0.76 | 0.72 | 0.62 |
| 2 | 240862048 | rs11900622 | 0.12 | 0.127 | -0.933 | 0.016 | 0.006 | 0.004 | 0.004 | 0.052 | 0.43 | 0.66 | 0.33 | 0.4 | 0.42 | 0.32 |
| 2 | 240864044 | rs4411726 | 0.263 | 0.131 | -0.903 | 0.017 | 0.001 | 0 | 0 | 0.029 | 0.42 | 0.62 | 0.32 | 0.4 | 0.42 | 0.32 |
| 2 | 240864911 | rs10178738 | 0.112 | 0.231 | -0.82 | 0.283 | 0 | 0.011 | 0.011 | 0 | 0.22 | 0.18 | 0.31 | 0.35 | 0.22 | 0.27 |
| 2 | 240870970 | rs4538238 | 1.134 | 0.263 | -1.128 | 0.385 | -0.025 | -0.053 | -0.053 | 0.203 | 0.1 | 0.02 | 0.19 | 0.22 | 0.08 | 0.12 |
| 2 | 240871018 | rs4505564 | -1.451 | 0.299 | -1.482 | 0.086 | -0.034 | -0.006 | -0.006 | 0 | 0.75 | 0.95 | 0.82 | 0.78 | 0.77 | 0.61 |
| 2 | 240871893 | rs4853981 | 0.211 | 0.35 | -0.885 | 0.004 | 0.009 | -0.005 | -0.005 | 0.115 | 0.45 | 0.34 | 0.33 | 0.48 | 0.46 | 0.33 |
| 2 | 240874059 | rs4996007 | -0.302 | 0.264 | -0.089 | -0.131 | -0.003 | -0.058 | -0.058 | 0 | 0.34 | 0.19 | 0.37 | 0.45 | 0.31 | 0.38 |
| 2 | 240875852 | rs13019067 | 0.773 | 0.387 | -0.002 | -0.395 | -0.008 | -0.01 | -0.01 | 0 | 0.28 | 0.25 | 0.27 | 0.34 | 0.35 | 0.23 |
| 2 | 240877650 | rs13405880 | 0.132 | 0.58 | 0.507 | -0.341 | 0.047 | -0.034 | -0.034 | 0 | 0.06 | 0.13 | 0.15 | 0 | 0.09 | 0.03 |
| 2 | 240878374 | rs6437330 | -0.063 | 0.368 | 0.617 | -0.433 | -0.006 | 0.03 | 0.03 | 0 | 0.35 | 0.44 | 0.44 | 0.47 | 0.32 | 0.34 |
| 2 | 240879532 | rs4312524 | -1.38 | 0.318 | 0.57 | -0.459 | 0.006 | 0.058 | 0.058 | 0.017 | 0.74 | 0.88 | 0.85 | 0.81 | 0.77 | 0.65 |
| 2 | 240879623 | rs6437332 | -0.488 | 0.306 | 0.589 | -0.439 | -0.004 | -0.007 | -0.007 | 0.23 | 0.41 | 0.57 | 0.58 | 0.48 | 0.41 | 0.37 |
| 2 | 240880447 | rs4356684 | -0.378 | 0.547 | 0.959 | 0.069 | 0.004 | -0.006 | -0.006 | 0 | 0.39 | 0.44 | 0.42 | 0.34 | 0.45 | 0.3 |
| 2 | 240886298 | rs3923002 | -0.306 | -0.146 | 1.194 | -0.043 | -0.004 | -0.032 | -0.032 | 0.183 | 0.09 | 0.13 | 0.16 | 0 | 0.06 | 0.03 |
| 2 | 240888514 | rs12695027 | -0.621 | -0.172 | 1.51 | -0.026 | 0.002 | 0.017 | 0.017 | 1.822 | 0.51 | 0.47 | 0.62 | 0.68 | 0.53 | 0.41 |
| 2 | 240890343 | rs4853978 | -1.699 | -0.284 | 1.097 | 0.004 | 0.009 | 0.042 | 0.042 | 0.596 | 0.7 | 0.87 | 0.77 | 0.78 | 0.73 | 0.59 |
| 2 | 240898578 | rs4854069 | -0.64 | 0.325 | 1.459 | 0.375 | 0 | 0 | 0 | 0 | 0.43 | 0.56 | 0.46 | 0.54 | 0.55 | 0.4 |
| 2 | 240900097 | rs13396556 | 0.845 | 0.305 | 1.458 | 0.413 | 0.01 | -0.017 | -0.017 | 0.306 | 0.15 | 0.1 | 0.18 | 0.19 | 0.13 | 0.14 |
| 2 | 240900121 | rs13424612 | 1.345 | 0.389 | 1.325 | 0.468 | -0.037 | -0.038 | -0.038 | 0 | 0.21 | 0.1 | 0.28 | 0.4 | 0.35 | 0.16 |
| 2 | 240906028 | rs4853976 | -0.234 | 0.377 | 1.318 | 0.462 | 0.031 | 0.036 | 0.036 | 0.847 | 0.33 | 0.54 | 0.47 | 0.51 | 0.41 | 0.27 |
| 2 | 240912712 | rs4149558 | -1.755 | 0.369 | 1.204 | 0.453 | 0.044 | 0.083 | 0.083 | 0 | 0.64 | 0.87 | 0.78 | 0.78 | 0.73 | 0.59 |
| 2 | 240915307 | rs4149556 | 1.218 | 0.397 | 1.216 | 0.462 | -0.027 | -0.028 | -0.028 | 0.482 | 0.22 | 0.13 | 0.29 | 0.41 | 0.35 | 0.16 |
| 2 | 240920291 | rs967476 | 1.014 | 0.365 | 0.742 | 0.48 | -0.001 | -0.001 | -0.001 | 0 | 0.13 | 0.13 | 0.11 | 0.1 | 0.16 | 0.17 |
| 2 | 240921463 | rs4149551 | 1.176 | 0.368 | 0.444 | 0.482 | -0.018 | -0.02 | -0.02 | 0.264 | 0.22 | 0.16 | 0.29 | 0.44 | 0.35 | 0.22 |
| 2 | 240922822 | rs958960 | -1.536 | 0.36 | 0.468 | 0.477 | 0.018 | 0.072 | 0.072 | 0 | 0.62 | 0.78 | 0.75 | 0.78 | 0.7 | 0.58 |
| 2 | 240923146 | rs6716293 | 0.122 | 0.364 | 0.462 | 0.479 | -0.008 | -0.007 | -0.007 | 0 | 0.31 | 0.3 | 0.44 | 0.44 | 0.41 | 0.25 |
| 2 | 240931266 | rs4149549 | 0.499 | 0.388 | 0.466 | 0.457 | -0.002 | 0.017 | 0.017 | 0 | 0.19 | 0.32 | 0.21 | 0.19 | 0.16 | 0.17 |
| 2 | 240933308 | rs4149547 | 1.334 | 0.416 | 0.432 | 0.481 | -0.045 | -0.047 | -0.047 | 0.002 | 0.21 | 0.09 | 0.28 | 0.41 | 0.35 | 0.16 |
| 2 | 240940636 | rs10193614 | 0.967 | 0.413 | 0.282 | 0.368 | -0.013 | 0.015 | 0.015 | 0 | 0.16 | 0.22 | 0.18 | 0.19 | 0.13 | 0.16 |
| 2 | 240947066 | rs3792089 | 1.117 | 0.378 | -0.111 | 0.324 | -0.01 | -0.005 | -0.005 | 0 | 0.12 | 0.02 | 0.1 | 0.09 | 0.16 | 0.15 |
| 2 | 240954110 | rs2289407 | 1.229 | 0.48 | -0.232 | 0.364 | -0.017 | -0.019 | -0.019 | 0.515 | 0.21 | 0.16 | 0.29 | 0.46 | 0.35 | 0.17 |
| 2 | 240954972 | rs6735806 | -0.743 | 0.016 | -0.577 | -0.191 | -0.013 | -0.046 | -0.046 | 0 | 0.24 | 0.4 | 0.27 | 0.1 | 0.22 | 0.2 |
| 2 | 240957801 | rs10804402 | 0.07 | 0.434 | -0.386 | 0.244 | -0.004 | -0.006 | -0.006 | 0 | 0.15 | 0.27 | 0.12 | 0.1 | 0.16 | 0.17 |
| 2 | 240961728 | rs2083411 | -0.563 | 0.464 | -1.149 | 0.227 | 0.049 | 0.126 | 0.126 | 0.207 | 0.56 | 0.8 | 0.63 | 0.79 | 0.68 | 0.56 |
| 2 | 240967561 | rs6709170 | -0.35 | 0.912 | -0.708 | 0.62 | 0.001 | -0.005 | -0.005 | 0.005 | 0.41 | 0.56 | 0.51 | 0.69 | 0.51 | 0.41 |
| 2 | 240968700 | rs13409568 | -0.449 | 0.669 | -0.261 | 1.024 | -0.01 | 0.008 | 0.008 | 0 | 0.18 | 0.22 | 0.18 | 0.25 | 0.13 | 0.16 |
| 2 | 240971450 | rs874060 | 0.068 | 1.062 | -0.101 | 0.984 | -0.014 | 0.004 | 0.004 | 0.112 | 0.58 | 0.46 | 0.63 | 0.54 | 0.63 | 0.5 |
| 2 | 240971519 | rs10933552 | -0.088 | 1.942 | -0.908 | 0.613 | -0.123 | 0.073 | 0.073 | 1.129 | 0.33 | 0.04 | 0.31 | 0.17 | 0.39 | 0.42 |
| 2 | 240972120 | rs10933554 | 0.385 | 2.025 | -0.516 | 0.87 | 0.006 | 0 | 0 | 1.778 | 0.49 | 0.54 | 0.34 | 0.5 | 0.45 | 0.2 |
| 2 | 240972417 | rs11683773 | -0.105 | 2.26 | -0.833 | 0.732 | -0.028 | 0.053 | 0.053 | 2.332 | 0.55 | 0.88 | 0.45 | 0.71 | 0.61 | 0.43 |
| 2 | 240976581 | rs13415895 | -0.516 | 2.303 | -0.418 | 0.388 | 0.021 | 0.016 | 0.016 | 0 | 0.1 | 0.13 | 0.1 | 0.21 | 0.17 | 0.23 |
| 2 | 240980394 | rs1004897 | -1.16 | 1.99 | -0.143 | -0.095 | 0.016 | -0.059 | -0.059 | 0 | 0.27 | 0.35 | 0.31 | 0.08 | 0.29 | 0.33 |
| 2 | 240981179 | rs6731916 | -0.976 | 2.149 | -0.494 | 0.002 | -0.082 | 0.064 | 0.064 | 2.767 | 0.62 | 0.93 | 0.59 | 0.78 | 0.63 | 0.59 |
| 2 | 240981262 | rs6728493 | 0.391 | 1.899 | -0.281 | 0.034 | -0.015 | 0.001 | 0.001 | 1.511 | 0.4 | 0.45 | 0.38 | 0.58 | 0.33 | 0.32 |

| | | | | | | | | | | | | | | | | |
|---|---|---|---|---|---|---|---|---|---|---|---|---|---|---|---|---|
| 2 | 240982791 | rs10171741 | -0.539 | 1.708 | -0.583 | -0.15 | -0.025 | 0.08 | 0.08 | 2.255 | 0.59 | 0.77 | 0.55 | 0.78 | 0.59 | 0.57 |
| 2 | 240990568 | rs7587455 | -1.375 | 1.142 | -0.125 | -0.227 | -0.027 | -0.115 | -0.115 | 0.04 | 0.17 | 0.63 | 0.2 | 0.01 | 0.08 | 0.15 |
| 2 | 240993114 | rs10933566 | 0.547 | 1.548 | -0.895 | -0.048 | -0.051 | 0.011 | 0.011 | 2.06 | 0.37 | 0.09 | 0.29 | 0.64 | 0.47 | 0.36 |
| 2 | 240993699 | rs6739025 | -0.296 | 1.241 | 0.236 | 0.015 | 0.034 | -0.024 | -0.024 | 0 | 0.06 | 0.24 | 0.04 | 0 | 0.05 | 0.01 |
| 2 | 240995216 | rs12471071 | -0.658 | 1.386 | -0.947 | -0.234 | -0.052 | 0.142 | 0.142 | 0.473 | 0.52 | 0.92 | 0.61 | 0.75 | 0.49 | 0.57 |
| 2 | 240996042 | rs4854045 | -0.069 | 1.28 | -1.026 | -0.375 | -0.02 | 0.056 | 0.056 | 0 | 0.1 | 0 | 0.13 | 0 | 0.07 | 0.22 |
| 2 | 240997420 | rs7569052 | -1.088 | 1.29 | -0.783 | -0.435 | -0.048 | 0.108 | 0.108 | 0.057 | 0.56 | 0.84 | 0.63 | 0.75 | 0.53 | 0.59 |
| 2 | 241000862 | rs6437268 | -0.561 | 1.252 | -0.182 | -0.401 | 0.056 | 0.058 | 0.058 | 0 | 0.47 | 0.79 | 0.51 | 0.63 | 0.36 | 0.56 |
| 2 | 241003331 | rs7568323 | -0.136 | 1.28 | -0.139 | -0.402 | 0.024 | -0.057 | -0.057 | 0 | 0.12 | 0.17 | 0.14 | 0 | 0.17 | 0.02 |
| 2 | 241006933 | rs2165216 | -0.699 | 1.105 | 0.175 | -0.43 | 0.003 | 0.055 | 0.055 | 0 | 0.51 | 0.35 | 0.59 | 0.75 | 0.52 | 0.58 |
| 2 | 241009262 | rs6724603 | -0.699 | 1.105 | 0.175 | -0.43 | 0.003 | 0.055 | 0.055 | 0 | 0.51 | 0.35 | 0.59 | 0.75 | 0.52 | 0.58 |
| 2 | 241019477 | rs2099728 | -1.435 | 1.108 | 0.074 | -0.421 | -0.088 | 0.095 | 0.095 | 0.265 | 0.58 | 0.9 | 0.65 | 0.75 | 0.53 | 0.59 |
| 2 | 241021460 | rs2368502 | -0.951 | 1.156 | 0.059 | -0.393 | -0.001 | 0.003 | 0.003 | 0 | 0.53 | 0.42 | 0.61 | 0.75 | 0.52 | 0.58 |
| 2 | 241023450 | rs4854039 | -0.997 | 1.151 | 0.189 | -0.396 | 0.001 | -0.014 | -0.014 | 0.032 | 0.53 | 0.48 | 0.62 | 0.75 | 0.52 | 0.58 |
| 2 | 241026828 | rs12479164 | 1.132 | 1.094 | -0.59 | -0.28 | -0.005 | -0.072 | -0.072 | 0.365 | 0.28 | 0.05 | 0.31 | 0.63 | 0.28 | 0.34 |
| 2 | 241046988 | rs1992307 | -1.208 | 1.516 | -0.843 | -0.222 | -0.112 | -0.019 | -0.019 | 0 | 0.59 | 0.97 | 0.71 | 0.64 | 0.58 | 0.58 |
| 2 | 241051887 | rs2369089 | -0.036 | 1.226 | -0.384 | 0.067 | -0.003 | 0.014 | 0.014 | 0 | 0.06 | 0.1 | 0.14 | 0.09 | 0.05 | 0.08 |
| 2 | 241052745 | rs9784121 | -1.172 | 1.156 | -0.761 | -0.059 | -0.125 | 0.064 | 0.064 | 1.435 | 0.6 | 0.94 | 0.67 | 0.72 | 0.56 | 0.51 |
| 2 | 241057425 | rs1866645 | 0.078 | 1.146 | -0.545 | -0.225 | -0.008 | 0.109 | 0.109 | 2.423 | 0.45 | 0.18 | 0.52 | 0.7 | 0.45 | 0.47 |
| 2 | 241061741 | rs10166127 | -0.188 | 1.143 | 0.313 | -0.298 | 0.032 | -0.06 | -0.06 | 0 | 0.1 | 0.36 | 0.1 | 0 | 0.11 | 0.05 |
| 2 | 241063482 | rs4854026 | -0.398 | 1.114 | 0.687 | -0.185 | 0.002 | -0.046 | -0.046 | 0 | 0.09 | 0.38 | 0.17 | 0 | 0.04 | 0.12 |
| 2 | 241066280 | rs4292120 | -1.155 | 1.129 | 0.495 | -0.176 | -0.153 | 0.055 | 0.055 | 1.504 | 0.61 | 0.97 | 0.67 | 0.73 | 0.57 | 0.56 |
| 2 | 241069831 | rs7567372 | 0.821 | 1.016 | 0.451 | 0.256 | -0.009 | -0.03 | -0.03 | 0 | 0.34 | 0.41 | 0.3 | 0.18 | 0.32 | 0.31 |
| 2 | 241073170 | rs747741 | 0.753 | 1.02 | 0.504 | 0.257 | -0.018 | -0.051 | -0.051 | 0.003 | 0.35 | 0.51 | 0.31 | 0.18 | 0.32 | 0.31 |
| 2 | 241073450 | rs3749078 | 0.818 | 1.026 | 0.372 | 0.265 | -0.009 | -0.03 | -0.03 | 0 | 0.34 | 0.41 | 0.3 | 0.18 | 0.32 | 0.31 |
| 2 | 241076471 | rs4853964 | -0.942 | 1.193 | 0.397 | 0.582 | -0.005 | 0.051 | 0.051 | 0.455 | 0.28 | 0.43 | 0.34 | 0.38 | 0.22 | 0.24 |
| 2 | 241077054 | rs12473823 | -0.992 | 1.148 | 0.077 | 0.156 | -0.078 | 0.085 | 0.085 | 0.824 | 0.6 | 0.89 | 0.65 | 0.72 | 0.54 | 0.57 |
| 2 | 241078945 | rs903759 | 1.056 | 0.952 | 0.489 | 0.354 | -0.003 | -0.011 | -0.011 | 0.017 | 0.31 | 0.31 | 0.31 | 0.16 | 0.34 | 0.29 |
| 2 | 241079382 | rs2291769 | 0.59 | 1.421 | 0.538 | 0.741 | 0.002 | 0.002 | 0.002 | 0.193 | 0.1 | 0.09 | 0.12 | 0.17 | 0.18 | 0.13 |
| 2 | 241081598 | rs4853963 | -1.887 | 1.481 | 0.417 | 0.559 | -0.007 | 0.033 | 0.033 | 1.258 | 0.42 | 0.68 | 0.5 | 0.71 | 0.42 | 0.39 |
| 2 | 241083287 | rs6437294 | -0.888 | 1.47 | 0.456 | 0.491 | 0.023 | 0.018 | 0.018 | 0.119 | 0.22 | 0.46 | 0.3 | 0.23 | 0.25 | 0.21 |
| 2 | 241085044 | rs13028057 | -0.386 | 1.413 | -0.288 | 0.695 | 0.002 | -0.032 | -0.032 | 0.003 | 0.26 | 0.26 | 0.24 | 0.43 | 0.22 | 0.19 |
| 2 | 241085226 | rs12695007 | -0.684 | 1.213 | -0.684 | 0.01 | -0.004 | 0.017 | 0.017 | 1.249 | 0.42 | 0.64 | 0.41 | 0.65 | 0.42 | 0.38 |
| 2 | 241092808 | rs903756 | -0.265 | 1.189 | -0.581 | -0.004 | 0.005 | 0.002 | 0.002 | 0.108 | 0.31 | 0.35 | 0.31 | 0.37 | 0.31 | 0.3 |
| 2 | 241094602 | rs903757 | -0.596 | 1.111 | -1.209 | -0.342 | -0.04 | -0.063 | -0.063 | 0 | 0.75 | 0.89 | 0.6 | 0.46 | 0.69 | 0.53 |
| 2 | 241100910 | rs903751 | -0.535 | 0.446 | -0.846 | -0.43 | 0.057 | 0.1 | 0.1 | 0.005 | 0.72 | 0.85 | 0.8 | 0.96 | 0.81 | 0.81 |
| 2 | 241103210 | rs13017098 | -0.49 | 0.652 | -0.529 | -0.323 | 0.003 | 0.031 | 0.031 | 0 | 0.55 | 0.73 | 0.51 | 0.43 | 0.47 | 0.47 |
| 2 | 241108454 | rs6719685 | -0.455 | 0.838 | -0.795 | -0.182 | -0.012 | 0.061 | 0.061 | 0 | 0.54 | 0.73 | 0.53 | 0.44 | 0.5 | 0.48 |
| 2 | 241108545 | rs6736600 | -0.412 | 1.181 | -0.372 | 0.438 | 0.02 | 0.012 | 0.012 | 0 | 0.28 | 0.52 | 0.32 | 0.43 | 0.31 | 0.31 |
| 2 | 241112958 | rs7576471 | -0.81 | 0.362 | -0.756 | 1.21 | -0.003 | 0.008 | 0.008 | 0.031 | 0.18 | 0.26 | 0.21 | 0.43 | 0.1 | 0.24 |
| 2 | 241113483 | rs13424628 | -1.469 | 0.821 | -0.472 | 0.708 | 0.001 | -0.006 | -0.006 | 0 | 0.12 | 0.18 | 0.11 | 0.07 | 0.14 | 0.12 |
| 2 | 241113670 | rs13424851 | -1.105 | 0.674 | -0.091 | 0.759 | 0.008 | 0 | 0 | 0 | 0.35 | 0.54 | 0.3 | 0.35 | 0.4 | 0.23 |
| 2 | 241114873 | rs7592376 | -0.988 | 0.362 | 0.218 | 0.881 | 0.052 | 0.124 | 0.124 | 0 | 0.12 | 0.28 | 0.17 | 0.35 | 0.24 | 0.15 |
| 2 | 241120829 | rs11695283 | -1.068 | 0.316 | 0.164 | 0.847 | 0.01 | 0.039 | 0.039 | 0 | 0.13 | 0.18 | 0.16 | 0.35 | 0.24 | 0.14 |
| 2 | 241121347 | rs11677535 | -1.518 | 0.22 | 0.513 | 0.8 | 0.036 | 0.112 | 0.112 | 0 | 0.16 | 0.34 | 0.18 | 0.36 | 0.24 | 0.14 |
| 2 | 241123603 | rs7597331 | -0.311 | 0.178 | 0.36 | 0.946 | -0.055 | -0.055 | -0.055 | 0 | 0.09 | 0.02 | 0.15 | 0.28 | 0.23 | 0.1 |
| 2 | 241123808 | rs7587174 | -1.312 | 0.2 | 0.42 | 0.923 | 0.019 | 0.062 | 0.062 | 0 | 0.16 | 0.28 | 0.18 | 0.36 | 0.24 | 0.15 |
| 2 | 241133660 | rs4853987 | -1.065 | -0.458 | 0.03 | 0.352 | 0.032 | 0.029 | 0.029 | 0.115 | 0.44 | 0.63 | 0.27 | 0.34 | 0.34 | 0.21 |
| 2 | 241140571 | rs4854097 | -1.094 | -0.466 | 0.153 | 0.364 | 0.02 | 0.007 | 0.007 | 0.369 | 0.45 | 0.62 | 0.28 | 0.38 | 0.34 | 0.21 |
| 2 | 241141196 | rs6437301 | 0.619 | -0.558 | -0.312 | 0.288 | 0.009 | -0.009 | -0.009 | 0.022 | 0.15 | 0.09 | 0.04 | 0 | 0.08 | 0.04 |
| 2 | 241144009 | rs869953 | -1.064 | -0.318 | -0.483 | 0.45 | 0.078 | 0.031 | 0.031 | 0.741 | 0.47 | 0.78 | 0.29 | 0.38 | 0.34 | 0.21 |
| 2 | 241152102 | rs6437302 | 0.033 | -0.133 | -0.504 | 1.149 | -0.001 | 0.055 | 0.055 | 0 | 0.86 | 0.84 | 0.68 | 0.58 | 0.88 | 0.89 |
| 2 | 241155550 | rs868379 | 0.372 | -0.575 | -0.71 | 0.28 | 0.021 | 0.025 | 0.025 | 0.72 | 0.26 | 0.27 | 0.09 | 0.05 | 0.15 | 0.08 |
| 2 | 241162084 | rs7592097 | 0.456 | 0.322 | -1.406 | 0.472 | -0.04 | -0.033 | -0.033 | 0 | 0.24 | 0.13 | 0.36 | 0.36 | 0.36 | 0.41 |
| 2 | 241167913 | rs7420256 | -0.034 | 0.272 | -1.092 | 0.884 | -0.03 | 0.042 | 0.042 | 0 | 0.31 | 0.48 | 0.38 | 0.43 | 0.23 | 0.23 |
| 2 | 241168633 | rs4854090 | -0.026 | 0.941 | -1.283 | 0.891 | -0.018 | -0.017 | -0.017 | 0 | 0.79 | 0.91 | 0.82 | 0.77 | 0.72 | 0.71 |
| 2 | 241173445 | rs10204130 | -0.799 | 0.88 | -0.934 | 0.898 | -0.004 | -0.008 | -0.008 | 0 | 0.17 | 0.19 | 0.15 | 0.26 | 0.24 | 0.09 |
| 2 | 241173514 | rs903760 | -0.416 | 0.571 | -0.686 | 0.531 | 0.003 | -0.023 | -0.023 | 0 | 0.12 | 0.2 | 0.09 | 0.02 | 0.1 | 0.09 |
| 2 | 241176022 | rs4854083 | -1.146 | 0.238 | 0.022 | 0.949 | -0.038 | 0.132 | 0.132 | 0.789 | 0.19 | 0.5 | 0.4 | 0.39 | 0.17 | 0.26 |
| 2 | 241188859 | rs10933584 | 1.27 | 0.031 | -0.549 | -0.255 | 0.012 | 0 | 0 | 0 | 0.28 | 0.18 | 0.15 | 0.01 | 0.19 | 0.09 |
| 2 | 241191425 | rs2279687 | -0.544 | 0.096 | 0.225 | 0.497 | -0.015 | 0.157 | 0.157 | 0 | 0.12 | 0.31 | 0.12 | 0.41 | 0.1 | 0.29 |
| 2 | 241193279 | rs12695010 | 0.16 | 0.155 | 0.211 | 0.199 | 0.023 | 0.021 | 0.021 | 0 | 0.49 | 0.37 | 0.32 | 0.42 | 0.42 | 0.44 |
| 2 | 241193615 | rs1565985 | -0.5 | 0.774 | -0.636 | -0.265 | -0.072 | 0.121 | 0.121 | 0 | 0.78 | 0.98 | 0.86 | 0.99 | 0.75 | 0.85 |
| 2 | 241194081 | rs1009283 | 0.303 | 0.846 | -0.024 | 0.428 | 0.027 | 0.04 | 0.04 | 0 | 0.23 | 0.29 | 0.29 | 0.43 | 0.33 | 0.34 |
| 2 | 241200086 | rs12105315 | -1.025 | 0.356 | 0.338 | 0.279 | 0.018 | -0.003 | -0.003 | 0 | 0.05 | 0.24 | 0.04 | 0 | 0.03 | 0.03 |
| 2 | 241206028 | rs6437317 | 0.667 | 0.174 | 0.278 | 0.207 | -0.012 | -0.023 | -0.023 | 0 | 0.77 | 0.66 | 0.63 | 0.85 | 0.8 | 0.81 |
| 2 | 241209928 | rs1495921 | -0.387 | 0.189 | 0.238 | 0.198 | -0.003 | 0.017 | 0.017 | 0.289 | 0.84 | 0.9 | 0.9 | 1 | 0.83 | 0.89 |
| 2 | 241213217 | rs10170009 | 0.559 | 0.197 | 0.279 | 0.245 | -0.01 | -0.018 | -0.018 | 0.052 | 0.77 | 0.66 | 0.63 | 0.84 | 0.8 | 0.81 |
| 2 | 241216359 | rs1495922 | 0.796 | 0.553 | -0.005 | 0.493 | 0.028 | 0.02 | 0.02 | 0 | 0.48 | 0.61 | 0.33 | 0.41 | 0.4 | 0.39 |

| | | | | | | | | | | | | | | | | |
|---|---|---|---|---|---|---|---|---|---|---|---|---|---|---|---|---|
| 2 | 241216673 | rs10933587 | -0.112 | 0.8 | -0.021 | 0.58 | -0.019 | -0.014 | -0.014 | 0 | 0.33 | 0.28 | 0.31 | 0.42 | 0.41 | 0.43 |
| 2 | 241224940 | rs7589895 | 0.291 | 0.429 | 0.287 | 0.348 | -0.001 | -0.005 | -0.005 | 3.109 | 0.54 | 0.47 | 0.36 | 0.26 | 0.41 | 0.53 |
| 2 | 241225364 | rs4417745 | 0.197 | 0.363 | 0.504 | 0.662 | 0.001 | 0.069 | 0.069 | 0 | 0.08 | 0.18 | 0.05 | 0.22 | 0.08 | 0.02 |
| 2 | 241227452 | rs13418001 | 0.542 | 0.299 | 0.418 | 0.383 | 0.015 | 0.031 | 0.031 | 0.693 | 0.49 | 0.42 | 0.34 | 0.26 | 0.38 | 0.5 |
| 2 | 241228985 | rs7562354 | 0.884 | 0.269 | 0.153 | 0.325 | 0.023 | 0.122 | 0.122 | 1.022 | 0.44 | 0.17 | 0.29 | 0.22 | 0.36 | 0.43 |
| 2 | 241231989 | rs1106154 | 1.093 | 0.081 | 0.145 | -0.025 | 0.016 | 0.149 | 0.149 | 5.919 | 0.37 | 0.17 | 0.24 | 0.03 | 0.38 | 0.17 |
| 2 | 241233933 | rs4854033 | 0.114 | 0.171 | 0.692 | 0.455 | -0.017 | 0.066 | 0.066 | 0.091 | 0.77 | 0.69 | 0.84 | 0.49 | 0.84 | 0.62 |
| 2 | 241239377 | rs7607116 | -0.171 | 0.027 | 1.166 | 0.655 | -0.026 | 0.047 | 0.047 | 0.416 | 0.81 | 0.75 | 0.88 | 0.71 | 0.92 | 0.67 |
| 2 | 241239911 | rs11685834 | 0.177 | 0.034 | 0.944 | 0.608 | -0.032 | -0.02 | -0.02 | 0 | 0.36 | 0.28 | 0.59 | 0.45 | 0.45 | 0.43 |
| 2 | 241247280 | rs7420288 | 0.051 | -0.133 | 1.056 | 0.671 | -0.047 | 0.047 | 0.047 | 0.728 | 0.78 | 0.64 | 0.86 | 0.7 | 0.92 | 0.64 |
| 2 | 241256535 | rs4676363 | -0.443 | 0.152 | 0.544 | 0.624 | -0.024 | -0.016 | -0.016 | 0 | 0.64 | 0.72 | 0.42 | 0.54 | 0.57 | 0.55 |
| 2 | 241261657 | rs2352821 | 0.6 | 0.333 | 0.67 | 0.259 | -0.005 | 0.057 | 0.057 | 2.142 | 0.41 | 0.24 | 0.27 | 0.23 | 0.47 | 0.21 |
| 2 | 241264925 | rs13026586 | -2.687 | 0.258 | 1.573 | 1.067 | 0.018 | 0.167 | 0.167 | 0.168 | 0.1 | 0.56 | 0.07 | 0.2 | 0.08 | 0.05 |
| 2 | 241271462 | rs12997738 | -2.224 | 0.313 | 1.461 | 1.097 | 0.061 | 0.231 | 0.231 | 0.021 | 0.08 | 0.41 | 0.06 | 0.2 | 0.09 | 0.03 |
| 2 | 241281421 | rs2352838 | 0.752 | 0.31 | 1.565 | 1.318 | -0.002 | 0.044 | 0.044 | 0.316 | 0.85 | 0.81 | 0.72 | 0.28 | 0.85 | 0.65 |
| 2 | 241282787 | rs7579238 | 1.442 | 0.181 | 1.782 | 1.253 | -0.024 | 0.061 | 0.061 | 0.684 | 0.81 | 0.65 | 0.71 | 0.28 | 0.85 | 0.62 |
| 2 | 241291165 | rs6437325 | 0.201 | 0.361 | 1.084 | 2.086 | -0.002 | 0.043 | 0.043 | 0.046 | 0.87 | 0.89 | 0.86 | 0.65 | 0.89 | 0.64 |
| 2 | 241291745 | rs10178014 | -0.671 | 0.819 | 0.775 | 1.072 | 0.026 | 0.064 | 0.064 | 1.372 | 0.26 | 0.58 | 0.34 | 0.54 | 0.32 | 0.32 |
| 2 | 241291822 | rs10933595 | 1.05 | 0.761 | 0.569 | 0.87 | -0.012 | -0.002 | -0.002 | 0 | 0.62 | 0.32 | 0.54 | 0.39 | 0.59 | 0.33 |
| 2 | 241297036 | rs7609518 | -0.382 | -0.137 | -0.155 | 0.149 | 0.002 | -0.087 | -0.087 | 0 | 0.75 | 0.92 | 0.7 | 0.38 | 0.76 | 0.6 |
| 2 | 241300692 | rs11679243 | 0.105 | 0.117 | 0.119 | -0.304 | 0.03 | 0.046 | 0.046 | 0 | 0.25 | 0.41 | 0.3 | 0.34 | 0.26 | 0.28 |
| 2 | 241303596 | rs4676439 | -1.04 | -0.102 | 0.27 | -0.276 | 0.001 | -0.014 | -0.014 | 3.095 | 0.8 | 0.83 | 0.87 | 0.96 | 0.8 | 0.88 |
| 2 | 241307312 | rs4571052 | -0.69 | -0.092 | 0.276 | -0.161 | 0.029 | -0.002 | -0.002 | 0 | 0.72 | 0.8 | 0.69 | 0.27 | 0.78 | 0.62 |
| 2 | 241309040 | rs7586891 | 0.035 | -0.105 | 0.108 | -0.565 | 0.004 | 0.018 | 0.018 | 0 | 0.33 | 0.37 | 0.31 | 0.39 | 0.3 | 0.35 |
| 2 | 241313129 | rs10165842 | -1.032 | -0.176 | -0.602 | -0.465 | 0.031 | -0.129 | -0.129 | 0 | 0.72 | 0.92 | 0.7 | 0.44 | 0.78 | 0.61 |
| 2 | 241315601 | rs6735344 | 1.002 | -0.015 | -0.524 | -0.933 | -0.1 | 0.133 | 0.133 | 0 | 0.25 | 0.06 | 0.31 | 0.04 | 0.4 | 0.22 |
| 2 | 241322107 | rs1316929 | 0.185 | -0.142 | 0.704 | -0.276 | -0.012 | -0.029 | -0.029 | 0 | 0.17 | 0.35 | 0.1 | 0.09 | 0.11 | 0.22 |
| 2 | 241334340 | rs6748992 | -0.083 | -0.123 | 0.345 | -0.262 | -0.008 | -0.038 | -0.038 | 0 | 0.07 | 0.22 | 0.04 | 0 | 0.05 | 0.02 |
| 2 | 241335924 | rs12614493 | -0.668 | -0.204 | 0.292 | -0.241 | -0.008 | 0.015 | 0.015 | 0.747 | 0.28 | 0.51 | 0.36 | 0.29 | 0.25 | 0.16 |
| 2 | 241338785 | rs2352852 | 0.336 | -0.125 | -0.085 | -0.186 | -0.005 | -0.025 | -0.025 | 0 | 0.06 | 0.1 | 0.03 | 0 | 0.05 | 0.02 |
| 2 | 241345108 | rs7570885 | -1.335 | -0.305 | -0.314 | -0.308 | 0.048 | 0.035 | 0.035 | 3.34 | 0.49 | 0.83 | 0.49 | 0.38 | 0.38 | 0.44 |
| 2 | 241349531 | rs1317979 | 0.791 | -0.349 | -0.143 | -0.377 | -0.007 | -0.02 | -0.02 | 0 | 0.18 | 0.12 | 0.29 | 0.29 | 0.18 | 0.13 |
| 2 | 241359366 | rs2078402 | -0.833 | -0.422 | 0.111 | -0.405 | 0.006 | 0 | 0 | 1.246 | 0.41 | 0.58 | 0.44 | 0.38 | 0.33 | 0.4 |
| 2 | 241359706 | rs1316479 | 0.07 | -0.409 | 0.181 | -0.396 | -0.043 | -0.076 | -0.076 | 0 | 0.22 | 0.45 | 0.14 | 0.08 | 0.14 | 0.27 |
| 2 | 241362669 | rs6750380 | -1.096 | -0.261 | 0.129 | -0.418 | 0.02 | 0.013 | 0.013 | 1.785 | 0.45 | 0.77 | 0.5 | 0.38 | 0.39 | 0.41 |
| 2 | 241367384 | rs6437342 | 0.545 | -0.347 | 0.08 | -0.267 | -0.019 | -0.032 | -0.032 | 0 | 0.17 | 0.35 | 0.1 | 0.08 | 0.11 | 0.23 |
| 2 | 241371065 | rs6707262 | -1.223 | -0.176 | -0.939 | -0.141 | 0.038 | 0.024 | 0.024 | 3.502 | 0.5 | 0.86 | 0.49 | 0.38 | 0.39 | 0.44 |
| 2 | 241372002 | rs7604524 | -1.214 | -0.177 | -0.935 | -0.135 | 0.041 | 0.031 | 0.031 | 3.293 | 0.5 | 0.85 | 0.49 | 0.38 | 0.39 | 0.44 |
| 2 | 241373531 | rs6716265 | -1.225 | -0.167 | -1.042 | -0.111 | 0.039 | 0.009 | 0.009 | 3.512 | 0.5 | 0.87 | 0.5 | 0.39 | 0.39 | 0.44 |
| 2 | 241373780 | rs9973786 | -0.565 | -0.234 | -0.209 | 0.03 | -0.031 | -0.049 | -0.049 | 0.004 | 0.25 | 0.6 | 0.16 | 0.08 | 0.15 | 0.3 |
| 2 | 241382083 | rs7577243 | 0.202 | 0.278 | -0.577 | 0.119 | -0.005 | -0.01 | -0.01 | 0 | 0.56 | 0.48 | 0.57 | 0.37 | 0.61 | 0.31 |
| 2 | 241385787 | rs13424854 | -0.361 | 0.259 | -0.664 | -0.134 | 0.001 | 0.004 | 0.004 | 0 | 0.2 | 0.21 | 0.13 | 0.07 | 0.17 | 0.26 |
| 2 | 241385823 | rs7589322 | -0.192 | 0.192 | -1.081 | 0.391 | 0.009 | -0.065 | -0.065 | 0.705 | 0.84 | 0.96 | 0.7 | 0.39 | 0.83 | 0.72 |
| 2 | 241390002 | rs3828334 | 0.581 | 0.276 | -1.131 | -0.665 | -0.002 | 0.038 | 0.038 | 0 | 0.33 | 0.24 | 0.22 | 0.06 | 0.32 | 0.13 |
| 2 | 241390516 | rs3828336 | 0.293 | 0.484 | -1.409 | -0.989 | -0.011 | 0.039 | 0.039 | 0 | 0.28 | 0.18 | 0.17 | 0.08 | 0.31 | 0.32 |
| 2 | 241395503 | rs2292832 | 0.088 | 0.236 | -0.774 | -0.493 | 0.006 | 0.001 | 0.001 | 0 | 0.66 | 0.73 | 0.69 | 0.36 | 0.72 | 0.55 |
| 2 | 241403014 | rs881029 | 0.777 | 0.075 | -0.755 | -1.319 | -0.003 | -0.083 | -0.083 | 0.271 | 0.23 | 0.36 | 0.16 | 0.02 | 0.2 | 0.16 |
| 2 | 241403957 | rs12695020 | -0.336 | 0.649 | -0.953 | -1.064 | 0.005 | 0.015 | 0.015 | 0 | 0.58 | 0.5 | 0.46 | 0.31 | 0.58 | 0.34 |
| 2 | 241404499 | rs2228327 | 0.319 | 0.641 | -0.583 | -1.29 | -0.003 | -0.036 | -0.036 | 0 | 0.35 | 0.31 | 0.24 | 0.09 | 0.32 | 0.26 |
| 2 | 241405528 | rs2228331 | -1.184 | 0.603 | -0.852 | -1.261 | 0.001 | -0.067 | -0.067 | 0.524 | 0.69 | 0.92 | 0.65 | 0.34 | 0.7 | 0.62 |
| 2 | 241408125 | rs2280261 | -0.704 | 0.592 | -0.972 | -1.122 | -0.011 | 0.003 | 0.003 | 0 | 0.63 | 0.78 | 0.55 | 0.31 | 0.6 | 0.54 |
| 2 | 241418137 | rs13005412 | -0.826 | 0.845 | -0.911 | -1.53 | 0.006 | -0.058 | -0.058 | 0 | 0.69 | 0.49 | 0.62 | 0.81 | 0.68 | 0.77 |
| 2 | 241419842 | rs6739196 | -0.072 | 0.837 | -1.148 | -1.585 | 0.001 | -0.019 | -0.019 | 0.9 | 0.26 | 0.29 | 0.26 | 0.06 | 0.31 | 0.24 |
| 2 | 241421011 | rs11693855 | 1.468 | 0.789 | -1.188 | -1.52 | 0.029 | -0.04 | -0.04 | 0 | 0.32 | 0.11 | 0.3 | 0.68 | 0.28 | 0.38 |
| 2 | 241421980 | rs7340350 | -0.058 | 0.846 | -1.176 | -1.509 | 0.001 | -0.019 | -0.019 | 0 | 0.26 | 0.29 | 0.26 | 0.06 | 0.31 | 0.24 |
| 2 | 241430049 | rs5004380 | 2.707 | 0.726 | -1.212 | -1.483 | -0.009 | -0.02 | -0.02 | 0 | 0.1 | 0.02 | 0.06 | 0.07 | 0.09 | 0.14 |
| 2 | 241435839 | rs4234126 | 1.451 | 0.737 | -1.24 | -1.515 | 0.017 | -0.038 | -0.038 | 0.156 | 0.32 | 0.12 | 0.3 | 0.68 | 0.3 | 0.38 |
| 2 | 241437578 | rs13005508 | 1.115 | 0.899 | -0.786 | -1.389 | 0.016 | 0.084 | 0.084 | 0 | 0.43 | 0.15 | 0.36 | 0.71 | 0.39 | 0.52 |
| 2 | 241438491 | rs4305277 | 1.732 | 0.925 | -0.581 | -1.885 | -0.008 | 0.016 | 0.016 | 0 | 0.14 | 0.32 | 0.24 | 0.17 | 0.1 | 0.06 |
| 2 | 241440020 | rs12986801 | 0.172 | 0.972 | -0.639 | -1.916 | -0.011 | 0.069 | 0.069 | 0 | 0.56 | 0.32 | 0.51 | 0.78 | 0.53 | 0.69 |
| 2 | 241442591 | rs10180897 | 0.188 | 0.917 | -0.091 | -1.615 | 0 | 0.006 | 0.006 | 0 | 0.63 | 0.4 | 0.6 | 0.78 | 0.64 | 0.68 |
| 2 | 241447428 | rs4676436 | 1.015 | 1.227 | -0.137 | -1.248 | 0.019 | -0.024 | -0.024 | 0 | 0.08 | 0.16 | 0.14 | 0.03 | 0.11 | 0.08 |
| 2 | 241452068 | rs12695024 | 0.136 | 0.831 | -0.329 | -0.989 | 0.013 | -0.049 | -0.049 | 0.699 | 0.7 | 0.55 | 0.62 | 0.79 | 0.68 | 0.77 |
| 2 | 241453126 | rs6730110 | -0.945 | 1.2 | -1.113 | -0.836 | -0.004 | -0.004 | -0.004 | 0 | 0.84 | 0.84 | 0.86 | 0.97 | 0.78 | 0.83 |
| 2 | 241456297 | rs4676432 | -0.466 | 0.875 | -1.475 | -0.866 | 0.007 | 0.046 | 0.046 | 0 | 0.73 | 0.85 | 0.65 | 0.88 | 0.74 | 0.83 |
| 2 | 241456401 | rs4676431 | 0.55 | 0.658 | -1.5 | -1.118 | -0.002 | -0.004 | -0.004 | 0 | 0.21 | 0.22 | 0.14 | 0.12 | 0.14 | 0.17 |
| 2 | 241458609 | rs10165759 | -0.481 | 1.524 | -1.357 | -0.97 | 0.017 | -0.042 | -0.042 | 0 | 0.69 | 0.6 | 0.52 | 0.86 | 0.63 | 0.75 |
| 2 | 241462456 | rs6728549 | 1.034 | 1.628 | -1.414 | -0.779 | -0.009 | -0.083 | -0.083 | 0 | 0.24 | 0.09 | 0.25 | 0.71 | 0.24 | 0.35 |
| 2 | 241463453 | rs3821348 | 0.682 | 2.065 | -0.905 | -0.27 | -0.009 | -0.004 | -0.004 | 0.177 | 0.15 | 0.11 | 0.14 | 0.1 | 0.18 | 0.3 |

| | | | | | | | | | | | | | | | | |
|---|---|---|---|---|---|---|---|---|---|---|---|---|---|---|---|---|
| 2 | 241466032 | rs10933612 | -0.485 | 2.036 | -0.943 | -0.318 | 0.093 | 0.004 | 0.004 | 0.265 | 0.82 | 0.66 | 0.6 | 0.85 | 0.7 | 0.84 |
| 2 | 241466210 | rs7603521 | 0.675 | 2.144 | -2.07 | -0.323 | -0.009 | -0.004 | -0.004 | 0.497 | 0.15 | 0.11 | 0.14 | 0.1 | 0.17 | 0.3 |
| 2 | 241477073 | rs4676430 | -0.226 | 1.967 | -1.713 | -0.096 | -0.035 | -0.015 | -0.015 | 0 | 0.84 | 0.85 | 0.62 | 0.83 | 0.7 | 0.84 |
| 2 | 241506669 | rs11683693 | 0.045 | 0.396 | -1.789 | -0.671 | 0.028 | 0.084 | 0.084 | 0.18 | 0.23 | 0.06 | 0.11 | 0.01 | 0.16 | 0.07 |
| 2 | 241511566 | rs4676423 | -0.233 | 0.742 | -1.158 | -0.553 | 0.016 | -0.045 | -0.045 | 0 | 0.87 | 0.79 | 0.88 | 0.99 | 0.85 | 0.9 |
| 2 | 241512271 | rs2001821 | -0.09 | 0.716 | -2.211 | -0.154 | -0.049 | -0.157 | -0.157 | 4.838 | 0.33 | 0.04 | 0.25 | 0.66 | 0.33 | 0.37 |
| 2 | 241512470 | rs2001823 | -0.187 | -0.221 | -1.344 | -1.182 | -0.023 | -0.127 | -0.127 | 0.103 | 0.64 | 0.45 | 0.77 | 0.98 | 0.71 | 0.87 |
| 2 | 241512983 | rs2001544 | -0.081 | 0.015 | -1.53 | -0.269 | -0.048 | -0.067 | -0.067 | 0.24 | 0.21 | 0.08 | 0.18 | 0.37 | 0.27 | 0.37 |
| 2 | 241515252 | rs2953146 | -0.393 | -0.738 | -1.129 | -1.172 | 0.027 | 0.037 | 0.037 | 0.751 | 0.68 | 0.77 | 0.86 | 0.96 | 0.78 | 0.72 |
| 2 | 241518989 | rs2975748 | -0.463 | -0.713 | -1.19 | -1.147 | 0.04 | 0.085 | 0.085 | 1.019 | 0.69 | 0.84 | 0.87 | 0.96 | 0.79 | 0.73 |
| 2 | 241525316 | rs2953172 | -0.308 | -0.629 | -1.423 | -1.24 | 0.02 | 0.095 | 0.095 | 0.004 | 0.66 | 0.82 | 0.81 | 0.91 | 0.73 | 0.69 |
| 2 | 241525691 | rs3749164 | 0.945 | -0.551 | -1.552 | -1.184 | -0.026 | -0.003 | -0.003 | 0 | 0.08 | 0.02 | 0.09 | 0.07 | 0.11 | 0.12 |
| 2 | 241530750 | rs2975758 | -0.974 | -0.721 | -1.286 | -0.96 | -0.005 | -0.043 | -0.043 | 0 | 0.74 | 0.76 | 0.89 | 0.98 | 0.84 | 0.81 |
| 2 | 241531174 | rs3792267 | 1.668 | -0.524 | -2.651 | -1.051 | -0.034 | 0.024 | 0.024 | 0 | 0.21 | 0.16 | 0.28 | 0.08 | 0.27 | 0.18 |
| 2 | 241531479 | rs3792269 | -0.062 | -0.465 | -2.415 | -0.814 | -0.027 | 0.003 | 0.003 | 0 | 0.13 | 0.03 | 0.13 | 0.1 | 0.16 | 0.21 |
| 2 | 241531737 | rs2975762 | 0.506 | -0.588 | -2.516 | -0.902 | 0.053 | 0.08 | 0.08 | 0 | 0.48 | 0.29 | 0.57 | 0.65 | 0.61 | 0.54 |
| 2 | 241536819 | rs3828345 | -0.216 | -0.021 | -3.103 | -0.665 | -0.024 | 0.007 | 0.007 | 0 | 0.13 | 0.03 | 0.13 | 0.09 | 0.16 | 0.21 |
| 2 | 241541640 | rs4234123 | -0.354 | 0.082 | -2.83 | -0.332 | 0.02 | 0.156 | 0.156 | 0.195 | 0.63 | 0.88 | 0.81 | 0.89 | 0.69 | 0.6 |
| 2 | 241545401 | rs2953161 | -0.378 | 0.668 | -1.983 | 0.539 | -0.04 | -0.029 | -0.029 | 0.636 | 0.3 | 0.18 | 0.36 | 0.64 | 0.47 | 0.61 |
| 2 | 241551402 | rs2953158 | -0.267 | 0.378 | -1.684 | 0.983 | 0.009 | -0.012 | -0.012 | 0.554 | 0.15 | 0.1 | 0.2 | 0.3 | 0.11 | 0.16 |
| 2 | 241558936 | rs10933621 | -0.191 | 1.062 | -1.897 | 0.415 | -0.038 | 0.041 | 0.041 | 0 | 0.26 | 0.15 | 0.3 | 0.08 | 0.29 | 0.26 |
| 2 | 241561088 | rs4676411 | -0.137 | 0.796 | -1.782 | 0.839 | -0.001 | -0.002 | -0.002 | 0.138 | 0.32 | 0.3 | 0.35 | 0.48 | 0.36 | 0.39 |
| 2 | 241561774 | rs11676358 | -0.613 | 0.041 | -2.686 | 1.009 | 0.012 | -0.084 | -0.084 | 0.546 | 0.22 | 0.01 | 0.16 | 0.34 | 0.16 | 0.28 |
| 2 | 241562515 | rs2975778 | -0.002 | 0.727 | -2.955 | 0.999 | -0.049 | -0.033 | -0.033 | 0.122 | 0.83 | 0.97 | 0.87 | 0.84 | 0.78 | 0.83 |
| 2 | 241562542 | rs11887524 | 0.249 | 0.565 | -2.767 | 0.597 | 0.034 | 0.062 | 0.062 | 0 | 0.5 | 0.65 | 0.43 | 0.28 | 0.57 | 0.62 |
| 2 | 241563169 | rs2975779 | 0.092 | 0.451 | -3.137 | 0.504 | 0.035 | -0.088 | -0.088 | 0.05 | 0.36 | 0.02 | 0.26 | 0.41 | 0.25 | 0.38 |
| 2 | 241563739 | rs4676410 | -0.115 | 1.332 | -1.537 | 0.767 | 0.072 | 0.155 | 0.155 | 0 | 0.14 | 0.34 | 0.4 | 0.29 | 0.2 | 0.14 |
| 2 | 241566012 | rs2953153 | -0.251 | 1.551 | -1.041 | 0.249 | 0.003 | 0.029 | 0.029 | 0.109 | 0.55 | 0.28 | 0.39 | 0.71 | 0.56 | 0.8 |
| 2 | 241566429 | rs2975781 | 0.205 | 1.978 | -1.327 | 0.168 | -0.022 | 0.021 | 0.021 | 0 | 0.48 | 0.08 | 0.33 | 0.7 | 0.45 | 0.66 |
| 2 | 241566871 | rs12477906 | -1.4 | 1.757 | -1.13 | 0.117 | -0.008 | -0.089 | -0.089 | 1.946 | 0.68 | 0.61 | 0.79 | 1 | 0.74 | 0.92 |
| 2 | 241569329 | rs2975785 | -0.298 | 1.74 | -1.008 | 0.189 | -0.013 | 0.056 | 0.056 | 0.049 | 0.53 | 0.18 | 0.37 | 0.71 | 0.53 | 0.79 |
| 2 | 241570249 | rs3749172 | -0.392 | 1.727 | -0.981 | 0.207 | -0.003 | 0.042 | 0.042 | 0.119 | 0.55 | 0.19 | 0.39 | 0.71 | 0.56 | 0.8 |
| 2 | 241571784 | rs33986981 | 0.289 | 1.454 | -1.082 | 0.19 | -0.072 | -0.083 | -0.083 | 0 | 0.14 | 0.01 | 0.13 | 0.31 | 0.22 | 0.29 |
| 2 | 241576626 | rs4335944 | -1.298 | 0.752 | -0.709 | -0.386 | -0.015 | -0.131 | -0.131 | 0 | 0.7 | 0.55 | 0.77 | 1 | 0.74 | 0.93 |
| 2 | 241583676 | rs4676403 | -1.199 | 1.363 | 0.112 | 0.271 | 0.04 | 0.075 | 0.075 | 0 | 0.17 | 0.38 | 0.41 | 0.29 | 0.2 | 0.28 |
| 2 | 241584823 | rs10933626 | -0.451 | 1.21 | 0.269 | -0.029 | -0.001 | -0.041 | -0.041 | 0 | 0.57 | 0.52 | 0.63 | 0.9 | 0.6 | 0.78 |
| 2 | 241585281 | rs4676402 | 0.127 | 1.269 | 0.232 | 0.018 | 0.053 | 0.008 | 0.008 | 0 | 0.08 | 0.17 | 0.15 | 0.09 | 0.13 | 0.11 |
| 2 | 241586810 | rs6724516 | -1.106 | 0.942 | 0.078 | -0.088 | 0.022 | 0.014 | 0.014 | 0.797 | 0.66 | 0.68 | 0.78 | 0.99 | 0.72 | 0.9 |
| 2 | 241591644 | rs6437356 | -1.253 | 0.958 | -0.252 | -0.089 | 0.032 | 0.106 | 0.106 | 1.648 | 0.67 | 0.84 | 0.78 | 0.99 | 0.72 | 0.9 |
| 2 | 241593944 | rs2060188 | -0.302 | 1.185 | -0.039 | 0.202 | 0.002 | -0.018 | -0.018 | 0.016 | 0.55 | 0.54 | 0.4 | 0.76 | 0.54 | 0.76 |
| 2 | 241596432 | rs4676345 | 0.107 | 1.234 | -0.971 | 0.436 | -0.014 | -0.053 | -0.053 | 0.265 | 0.84 | 0.89 | 0.86 | 0.63 | 0.78 | 0.71 |
| 2 | 241604696 | rs10203408 | 0.149 | 1.228 | -0.969 | 0.296 | -0.01 | -0.016 | -0.016 | 1.318 | 0.54 | 0.77 | 0.73 | 0.58 | 0.53 | 0.37 |
| 2 | 241609579 | rs7587179 | -0.069 | 1.964 | -0.386 | 0.452 | 0.038 | -0.001 | -0.001 | 0 | 0.07 | 0.11 | 0.14 | 0.08 | 0.13 | 0.12 |
| 2 | 241623894 | rs4676385 | 0.324 | 1.321 | -0.442 | 0.622 | 0.002 | 0.029 | 0.029 | 1.613 | 0.86 | 0.84 | 0.86 | 0.66 | 0.8 | 0.67 |
| 2 | 241626057 | rs10804412 | 0.037 | 0.335 | -0.343 | 0.336 | -0.005 | -0.001 | -0.001 | 4.011 | 0.61 | 0.62 | 0.73 | 0.64 | 0.56 | 0.51 |
| 2 | 241627928 | rs1560619 | -0.169 | 0.427 | -0.596 | 0.434 | -0.023 | -0.011 | -0.011 | 0.053 | 0.66 | 0.73 | 0.77 | 0.65 | 0.58 | 0.54 |
| 2 | 241642478 | rs12695029 | -0.542 | 0.364 | -0.752 | 0.451 | -0.021 | 0.023 | 0.023 | 0.182 | 0.61 | 0.83 | 0.78 | 0.68 | 0.63 | 0.43 |
| 2 | 241643000 | rs4676381 | -0.712 | 0.326 | -0.784 | 0.447 | -0.025 | -0.002 | -0.002 | 0 | 0.63 | 0.85 | 0.79 | 0.66 | 0.65 | 0.45 |
| 2 | 241643273 | rs4676380 | -0.137 | 0.203 | 0.075 | 0.369 | 0.001 | 0.002 | 0.002 | 0.641 | 0.56 | 0.39 | 0.74 | 0.64 | 0.64 | 0.43 |
| 2 | 241644175 | rs1864869 | -0.304 | 0.226 | -0.175 | 0.056 | 0.018 | 0.046 | 0.046 | 1.51 | 0.26 | 0.42 | 0.51 | 0.65 | 0.31 | 0.34 |
| 2 | 241644934 | rs1583259 | -0.782 | 0.521 | -0.702 | 0.361 | -0.012 | 0 | 0 | 0.517 | 0.63 | 0.83 | 0.79 | 0.66 | 0.66 | 0.46 |
| 2 | 241647342 | rs1001495 | -0.972 | 0.88 | 0.048 | 0.678 | 0.109 | 0.219 | 0.219 | 1.251 | 0.1 | 0.47 | 0.31 | 0.23 | 0.09 | 0.14 |
| 2 | 241652395 | rs755302 | 0.625 | 0.422 | -0.907 | 0.151 | -0.035 | 0.081 | 0.081 | 0 | 0.23 | 0.06 | 0.2 | 0 | 0.29 | 0.07 |
| 2 | 241652703 | rs755300 | -0.516 | 0.363 | -0.846 | 0.441 | -0.002 | 0.039 | 0.039 | 0 | 0.37 | 0.7 | 0.57 | 0.24 | 0.39 | 0.19 |
| 2 | 241653287 | rs4613 | -0.672 | 0.266 | -1.02 | 0.561 | 0.013 | 0.052 | 0.052 | 0 | 0.4 | 0.74 | 0.57 | 0.28 | 0.39 | 0.19 |
| 2 | 241655087 | rs3732339 | -1.297 | 0.221 | -0.448 | 1.26 | 0.005 | 0.17 | 0.17 | 2.299 | 0.07 | 0.28 | 0.33 | 0.24 | 0.04 | 0.05 |
| 2 | 241659876 | rs6735606 | -0.274 | 0.284 | -1.394 | 1.118 | 0.01 | -0.007 | -0.007 | 0 | 0.12 | 0.06 | 0.15 | 0.13 | 0.08 | 0.12 |
| 2 | 241666628 | rs3772047 | -0.492 | 1.033 | -1.747 | 0.86 | -0.045 | 0.023 | 0.023 | 0 | 0.21 | 0.1 | 0.1 | 0.13 | 0.25 | 0.15 |
| 2 | 241672740 | rs3772050 | -1.121 | 0.73 | -2.341 | 0.763 | 0.014 | -0.075 | -0.075 | 0 | 0.67 | 0.92 | 0.64 | 0.69 | 0.74 | 0.5 |
| 2 | 241675969 | rs11886658 | 0.928 | 1.297 | -1.666 | 0.363 | -0.013 | 0.016 | 0.016 | 0 | 0.71 | 0.75 | 0.84 | 0.86 | 0.66 | 0.74 |
| 2 | 241677264 | rs7578279 | -0.948 | 1.273 | -0.842 | 0.345 | 0.001 | 0.003 | 0.003 | 0.048 | 0.42 | 0.59 | 0.45 | 0.44 | 0.54 | 0.29 |
| 2 | 241678528 | rs3772054 | -0.133 | 1.626 | 0.047 | 0.438 | 0.053 | 0.018 | 0.018 | 0.017 | 0.26 | 0.45 | 0.33 | 0.31 | 0.3 | 0.19 |
| 2 | 241680633 | rs4414678 | 1.008 | 1.297 | -0.067 | 0.15 | -0.001 | -0.001 | -0.001 | 0 | 0.53 | 0.48 | 0.47 | 0.56 | 0.36 | 0.58 |
| 2 | 241681764 | rs3755534 | -1.326 | 1.048 | 0.582 | 0.435 | -0.011 | -0.03 | -0.03 | 0.024 | 0.19 | 0.41 | 0.1 | 0.12 | 0.15 | 0.1 |
| 2 | 241693240 | rs12624059 | -0.882 | 1.257 | 0.315 | 0.613 | -0.02 | -0.05 | -0.05 | 0 | 0.23 | 0.16 | 0.25 | 0.57 | 0.28 | 0.29 |
| 2 | 241695096 | rs4676447 | -2.13 | 1.331 | 0.324 | 0.382 | 0.032 | 0.045 | 0.045 | 0 | 0.52 | 0.74 | 0.51 | 0.7 | 0.58 | 0.42 |
| 2 | 241698298 | rs7598218 | -2.015 | 1.397 | 0.278 | 0.378 | -0.001 | -0.025 | -0.025 | 0 | 0.38 | 0.72 | 0.36 | 0.63 | 0.38 | 0.39 |
| 2 | 241699883 | rs3772059 | 0.6 | 1.727 | 0.139 | 0.751 | -0.044 | 0.019 | 0.019 | 0.062 | 0.16 | 0.06 | 0.16 | 0.14 | 0.22 | 0.07 |
| 2 | 241708675 | rs2288753 | -1.178 | 1.382 | 1.015 | 1.035 | 0.014 | 0.047 | 0.047 | 0 | 0.23 | 0.26 | 0.27 | 0.57 | 0.28 | 0.29 |

| | | | | | | | | | | | | | | | |
|---|---|---|---|---|---|---|---|---|---|---|---|---|---|---|---|
| 2 | 241722445 | rs2288750 | -0.801 | 1.471 | 1.171 | 1.205 | 0.037 | 0.109 | 0.109 | 0 | 0.17 | 0.49 | 0.25 | 0.34 | 0.19 | 0.1 |
| 2 | 241734754 | rs734586 | -1.436 | 1.11 | 0.574 | 1.097 | -0.057 | -0.023 | -0.023 | 0 | 0.41 | 0.81 | 0.42 | 0.5 | 0.48 | 0.29 |
| 2 | 241740907 | rs4676442 | -1.539 | 0.967 | 0.898 | 1.231 | -0.002 | -0.029 | -0.029 | 0 | 0.36 | 0.7 | 0.35 | 0.56 | 0.37 | 0.34 |
| 2 | 241741977 | rs7599943 | -1.75 | 0.956 | 0.855 | 1.226 | -0.002 | -0.068 | -0.068 | 0 | 0.37 | 0.79 | 0.37 | 0.58 | 0.37 | 0.37 |
| 2 | 241744372 | rs4455151 | -0.85 | 1.142 | 1.341 | 1.249 | 0.02 | 0.029 | 0.029 | 0 | 0.27 | 0.57 | 0.29 | 0.4 | 0.31 | 0.2 |
| 2 | 241753815 | rs4129943 | 0.341 | 1.149 | 1.787 | 0.952 | 0.02 | -0.009 | -0.009 | 0 | 0.76 | 0.65 | 0.83 | 0.76 | 0.7 | 0.79 |
| 2 | 241762579 | rs6745084 | -0.882 | 1.269 | 1.876 | 0.832 | 0.015 | -0.042 | -0.042 | 0 | 0.27 | 0.43 | 0.25 | 0.17 | 0.31 | 0.22 |
| 2 | 241763168 | rs4499423 | 0.243 | 1.283 | 1.804 | 0.833 | 0.008 | -0.005 | -0.005 | 0 | 0.76 | 0.72 | 0.83 | 0.76 | 0.7 | 0.79 |
| 2 | 241767075 | rs12465902 | 0.4 | 1.304 | 1.639 | 0.84 | 0.025 | -0.022 | -0.022 | 0.946 | 0.74 | 0.49 | 0.81 | 0.76 | 0.7 | 0.79 |
| 2 | 241769281 | rs4234121 | 1.505 | 0.557 | 1.284 | 1.057 | 0.038 | -0.002 | -0.002 | 0 | 0.47 | 0.2 | 0.57 | 0.5 | 0.39 | 0.53 |
| 2 | 241791028 | rs6719991 | -0.367 | 0.873 | 1.172 | 1.004 | 0.03 | -0.011 | -0.011 | 0.166 | 0.26 | 0.3 | 0.21 | 0.24 | 0.33 | 0.25 |
| 2 | 241792959 | rs11680616 | -0.135 | 0.515 | 1.019 | 0.983 | -0.002 | 0.001 | 0.001 | 0 | 0.18 | 0.19 | 0.15 | 0.09 | 0.21 | 0.11 |
| 2 | 241794655 | rs10933638 | -1.936 | 0.113 | 0.658 | 0.895 | 0.077 | -0.053 | -0.053 | 0.412 | 0.6 | 0.86 | 0.47 | 0.51 | 0.71 | 0.53 |
| 2 | 241800707 | rs7600847 | 0.012 | 0.195 | 0.735 | 0.858 | -0.088 | 0.006 | 0.006 | 0.069 | 0.27 | 0.11 | 0.23 | 0.24 | 0.39 | 0.27 |
| 2 | 241804078 | rs4129624 | 0.302 | 0.037 | 1.332 | 0.907 | -0.012 | 0.01 | 0.01 | 0 | 0.13 | 0.21 | 0.07 | 0.19 | 0.1 | 0.15 |
| 2 | 241805851 | rs7340215 | -1.202 | 0.019 | 1.606 | 0.896 | -0.003 | -0.004 | -0.004 | 0.049 | 0.46 | 0.51 | 0.39 | 0.33 | 0.61 | 0.38 |
| 2 | 241806305 | rs11886288 | 0.225 | -0.177 | 1.43 | 0.963 | -0.012 | 0.011 | 0.011 | 0 | 0.13 | 0.21 | 0.07 | 0.19 | 0.1 | 0.13 |
| 2 | 241816545 | rs6732985 | -0.034 | 0.025 | 1.084 | 1.247 | -0.075 | 0.005 | 0.005 | 0.005 | 0.26 | 0.13 | 0.24 | 0.26 | 0.42 | 0.26 |
| 2 | 241817516 | rs4426527 | 0.617 | -0.023 | 1.062 | 1.307 | -0.05 | 0.028 | 0.028 | 0 | 0.16 | 0.04 | 0.16 | 0.09 | 0.2 | 0.11 |
| 2 | 241818279 | rs4273214 | 0.078 | -0.034 | 1.061 | 1.235 | -0.084 | 0.009 | 0.009 | 0.022 | 0.27 | 0.12 | 0.24 | 0.26 | 0.42 | 0.27 |
| 2 | 241818527 | rs4344931 | -2.114 | -0.139 | 0.982 | 1.257 | 0.092 | 0.005 | 0.005 | 0 | 0.61 | 0.89 | 0.6 | 0.77 | 0.72 | 0.53 |
| 2 | 241819526 | rs4675792 | -1.059 | -0.147 | 1.306 | 1.26 | 0.033 | -0.013 | -0.013 | 0 | 0.44 | 0.33 | 0.49 | 0.59 | 0.63 | 0.39 |
| 2 | 241820501 | rs11886874 | 0.175 | -0.132 | 1.11 | 1.216 | -0.017 | 0.011 | 0.011 | 0 | 0.14 | 0.29 | 0.08 | 0.18 | 0.1 | 0.14 |
| 2 | 241821504 | rs9288742 | -0.276 | -0.095 | 1.137 | 1.194 | 0.037 | -0.017 | -0.017 | 0 | 0.31 | 0.51 | 0.27 | 0.26 | 0.42 | 0.27 |
| 2 | 241824566 | rs12994627 | 0.722 | -0.094 | 1.115 | 1.174 | -0.031 | 0.045 | 0.045 | 0 | 0.15 | 0.06 | 0.14 | 0 | 0.2 | 0.06 |
| 2 | 241824647 | rs10177846 | -0.407 | -0.103 | 1.137 | 1.219 | 0.031 | 0.034 | 0.034 | 0 | 0.32 | 0.54 | 0.38 | 0.57 | 0.42 | 0.32 |
| 2 | 241825253 | rs13025495 | 0.723 | -0.098 | 1.114 | 1.209 | -0.031 | 0.045 | 0.045 | 0 | 0.15 | 0.06 | 0.14 | 0 | 0.2 | 0.06 |
| 2 | 241826050 | rs13397515 | -0.637 | 0.171 | 1.464 | 1.575 | 0.029 | 0.033 | 0.033 | 0 | 0.32 | 0.54 | 0.38 | 0.57 | 0.42 | 0.32 |
| 2 | 241826139 | rs13422330 | -0.637 | 0.171 | 1.464 | 1.575 | 0.029 | 0.033 | 0.033 | 0 | 0.32 | 0.54 | 0.38 | 0.57 | 0.42 | 0.32 |
| 2 | 241835543 | rs10206101 | -1.396 | 0.231 | 0.026 | 1.383 | 0 | 0.016 | 0.016 | 0.29 | 0.45 | 0.78 | 0.35 | 0.44 | 0.53 | 0.42 |
| 2 | 241846140 | rs4076342 | -0.473 | 0.798 | 0.119 | 1.384 | -0.02 | -0.016 | -0.016 | 0 | 0.27 | 0.24 | 0.46 | 0.38 | 0.48 | 0.32 |
| 2 | 241859255 | rs4675790 | 0.054 | 0.613 | 0.372 | 1.072 | -0.05 | -0.044 | -0.044 | 0 | 0.73 | 0.81 | 0.56 | 0.5 | 0.54 | 0.62 |
| 2 | 241876580 | rs4675841 | 1.07 | 0.253 | 0.072 | 0.739 | 0.129 | 0.141 | 0.141 | 0.022 | 0.49 | 0.16 | 0.37 | 0.34 | 0.35 | 0.43 |
| 2 | 241879702 | rs11684777 | 0.2 | 0.184 | 0.647 | 0.68 | 0.011 | -0.06 | -0.06 | 0.112 | 0.15 | 0.21 | 0.14 | 0 | 0.18 | 0.04 |
| 2 | 241891799 | rs4464265 | -0.605 | -0.234 | 0.524 | 0.96 | -0.001 | 0 | 0 | 0 | 0.46 | 0.45 | 0.59 | 0.5 | 0.64 | 0.44 |
| 2 | 241894333 | rs4430951 | -0.923 | -0.193 | -0.042 | 0.978 | 0.128 | 0.14 | 0.14 | 0.012 | 0.51 | 0.84 | 0.63 | 0.66 | 0.65 | 0.57 |
| 2 | 241898520 | rs11886565 | -0.706 | 0.032 | -0.226 | 0.732 | 0.009 | -0.001 | -0.001 | 0 | 0.46 | 0.6 | 0.6 | 0.5 | 0.64 | 0.44 |
| 2 | 241911027 | rs4234091 | -1.031 | 0.186 | -0.159 | 0.905 | -0.007 | 0.016 | 0.016 | 0.021 | 0.25 | 0.7 | 0.35 | 0.33 | 0.19 | 0.17 |
| 2 | 241913553 | rs4675839 | -0.237 | 0.034 | -0.416 | -0.134 | 0.011 | 0.024 | 0.024 | 1.379 | 0.49 | 0.57 | 0.68 | 0.92 | 0.54 | 0.68 |
| 2 | 241914341 | rs10804401 | -0.927 | -0.236 | -1.083 | -0.441 | -0.012 | 0.178 | 0.178 | 1.952 | 0.63 | 0.9 | 0.78 | 0.93 | 0.65 | 0.75 |
| 2 | 241922182 | rs4234090 | 0.628 | 0.131 | -0.947 | 0.244 | -0.01 | -0.01 | -0.01 | 0 | 0.33 | 0.3 | 0.36 | 0.54 | 0.41 | 0.28 |
| 2 | 241927005 | rs7425888 | 0.204 | 0.348 | -0.387 | 0.761 | -0.001 | 0.001 | 0.001 | 8.398 | 0.12 | 0.09 | 0.14 | 0.09 | 0.12 | 0.18 |
| 2 | 241933700 | rs4602221 | -0.369 | 0.237 | -0.513 | 0.671 | -0.001 | 0.035 | 0.035 | 0 | 0.1 | 0.05 | 0.08 | 0.01 | 0.12 | 0.07 |
| 2 | 241943575 | rs4553820 | -1.19 | 0.489 | -0.157 | 0.954 | 0.006 | 0.055 | 0.055 | 2.817 | 0.26 | 0.53 | 0.39 | 0.4 | 0.27 | 0.47 |
| 2 | 241947122 | rs4675833 | 0.467 | 0.735 | -0.919 | 0.394 | -0.028 | 0.048 | 0.048 | 2.77 | 0.81 | 0.92 | 0.8 | 0.92 | 0.75 | 0.75 |
| 2 | 241957677 | rs4497870 | 0.17 | 0.298 | -0.955 | -0.148 | 0.004 | -0.004 | -0.004 | 0 | 0.3 | 0.21 | 0.22 | 0.06 | 0.31 | 0.22 |
| 2 | 241961346 | rs7573476 | -0.168 | 0.497 | -0.837 | 0.136 | -0.001 | -0.001 | -0.001 | 0 | 0.14 | 0.18 | 0.09 | 0.16 | 0.16 | 0.14 |
| 2 | 241968979 | rs4234102 | -1.069 | -0.173 | -1.062 | 0.856 | -0.076 | -0.007 | -0.007 | 0 | 0.65 | 0.85 | 0.66 | 0.69 | 0.56 | 0.67 |
| 2 | 241969033 | rs7562180 | 0.772 | -0.407 | -1.033 | 0.719 | 0.007 | 0.014 | 0.014 | 0 | 0.32 | 0.2 | 0.24 | 0.1 | 0.33 | 0.25 |
| 2 | 241971828 | rs2058061 | -0.892 | 0.039 | -1.426 | 1.095 | -0.086 | 0.033 | 0.033 | 0 | 0.65 | 0.87 | 0.66 | 0.75 | 0.56 | 0.68 |
| 2 | 241974289 | rs2074822 | -0.791 | -0.127 | -1.401 | 1.118 | -0.055 | 0.014 | 0.014 | 0 | 0.63 | 0.85 | 0.65 | 0.67 | 0.5 | 0.67 |
| 2 | 241979550 | rs7571117 | -0.249 | -0.131 | -0.78 | 1.115 | 0.003 | 0.012 | 0.012 | 0 | 0.56 | 0.61 | 0.35 | 0.32 | 0.46 | 0.42 |
| 2 | 241981906 | rs1123650 | 0.522 | -0.163 | -0.846 | 1.084 | -0.001 | -0.014 | -0.014 | 0 | 0.11 | 0.05 | 0.08 | 0.19 | 0.12 | 0.13 |
| 2 | 241986030 | rs6723732 | 0.934 | -0.301 | -0.633 | 0.74 | 0.015 | 0.081 | 0.081 | 0 | 0.31 | 0.14 | 0.21 | 0.06 | 0.3 | 0.21 |
| 2 | 241989723 | rs13426107 | -0.998 | -0.305 | 0.051 | 1.258 | -0.02 | 0.223 | 0.223 | 0.056 | 0.09 | 0.26 | 0.23 | 0.32 | 0.05 | 0.22 |
| 2 | 241990727 | rs7609450 | 0.95 | -0.324 | -0.61 | 0.609 | 0.027 | 0.084 | 0.084 | 0 | 0.3 | 0.13 | 0.19 | 0.04 | 0.27 | 0.21 |
| 2 | 241993027 | rs6715941 | -1.124 | 0.264 | -0.96 | 0.823 | -0.039 | -0.042 | -0.042 | 0.021 | 0.92 | 0.99 | 0.92 | 0.87 | 0.9 | 0.83 |
| 2 | 241995014 | rs10183172 | -0.918 | -0.043 | -0.816 | 0.834 | -0.088 | 0.067 | 0.067 | 0 | 0.65 | 0.85 | 0.69 | 0.8 | 0.52 | 0.67 |
| 2 | 242006736 | rs7559967 | -0.868 | -0.398 | 0.136 | 1.287 | -0.022 | 0.08 | 0.08 | 0 | 0.22 | 0.56 | 0.3 | 0.38 | 0.14 | 0.24 |
| 2 | 242011084 | rs17440466 | -0.392 | -0.437 | -0.351 | 1.255 | -0.042 | -0.031 | -0.031 | 0 | 0.15 | 0.02 | 0.08 | 0.19 | 0.17 | 0.1 |
| 2 | 242013387 | rs13030338 | 1.035 | -1.091 | -0.146 | 0.644 | 0.043 | 0.098 | 0.098 | 0 | 0.29 | 0.12 | 0.18 | 0.05 | 0.25 | 0.2 |
| 2 | 242014364 | rs12694993 | -1.069 | -1.267 | -0.272 | 0.883 | -0.027 | -0.097 | -0.097 | 0.135 | 0.61 | 0.85 | 0.53 | 0.55 | 0.43 | 0.51 |
| 2 | 242015650 | rs13411510 | -1.113 | -1.236 | 0.498 | 1.04 | -0.017 | 0.039 | 0.039 | 0.882 | 0.29 | 0.57 | 0.35 | 0.5 | 0.19 | 0.22 |
| 2 | 242021742 | rs2108485 | -1.256 | -1.471 | 0.136 | 1.078 | 0.032 | -0.038 | -0.038 | 4.218 | 0.3 | 0.75 | 0.35 | 0.55 | 0.17 | 0.17 |
| 2 | 242025882 | rs10188678 | -0.473 | -1.61 | 0.954 | 1.299 | -0.03 | 0.027 | 0.027 | 0.003 | 0.19 | 0.37 | 0.1 | 0.21 | 0.1 | 0.08 |
| 2 | 242026649 | rs13411556 | -0.473 | -1.61 | 0.954 | 1.299 | -0.03 | 0.027 | 0.027 | 0.003 | 0.19 | 0.37 | 0.1 | 0.21 | 0.1 | 0.08 |
| 2 | 242028282 | rs10933529 | -1.714 | -1.617 | 0.905 | 1.302 | 0.031 | -0.039 | -0.039 | 0 | 0.3 | 0.75 | 0.35 | 0.55 | 0.17 | 0.17 |
| 2 | 242028741 | rs1062748 | 1.32 | -1.842 | 0.6 | 1 | 0.055 | 0.099 | 0.099 | 0 | 0.26 | 0.1 | 0.14 | 0.05 | 0.18 | 0.26 |
| 2 | 242031090 | rs16843241 | 1.202 | -1.754 | 0.622 | 1.056 | -0.025 | -0.057 | -0.057 | 0.398 | 0.1 | 0 | 0.07 | 0.21 | 0.1 | 0.08 |

| | | | | | | | | | | | | | | | | |
|---|---|---|---|---|---|---|---|---|---|---|---|---|---|---|---|---|
| 2 | 242031281 | rs16843242 | 0.482 | -1.863 | 0.927 | 1.006 | -0.001 | 0.002 | 0.002 | 0 | 0.13 | 0.1 | 0.09 | 0.21 | 0.1 | 0.08 |
| 2 | 242032540 | rs3815291 | -1.407 | -1.842 | 1.284 | 1.024 | 0.026 | -0.021 | -0.021 | 0 | 0.57 | 0.87 | 0.49 | 0.6 | 0.35 | 0.43 |
| 2 | 242032709 | rs16843245 | -0.174 | -1.636 | 1.404 | 1.323 | -0.004 | -0.003 | -0.003 | 0 | 0.1 | 0.14 | 0.1 | 0.12 | 0.06 | 0.04 |
| 2 | 242032762 | rs3815292 | -0.174 | -1.636 | 1.404 | 1.328 | -0.004 | -0.002 | -0.002 | 0 | 0.1 | 0.14 | 0.1 | 0.13 | 0.06 | 0.04 |
| 2 | 242036797 | rs2286323 | 0.816 | -1.653 | 1.393 | 1.313 | -0.001 | -0.002 | -0.002 | 0 | 0.11 | 0.09 | 0.08 | 0.21 | 0.1 | 0.08 |
| 2 | 242039198 | rs3796093 | -0.174 | -1.64 | 1.317 | 1.26 | -0.004 | -0.003 | -0.003 | 0.214 | 0.1 | 0.14 | 0.1 | 0.12 | 0.06 | 0.04 |
| 2 | 242046785 | rs1131293 | -0.623 | -1.564 | 1.345 | 1.295 | -0.019 | 0.156 | 0.156 | 0 | 0.18 | 0.58 | 0.26 | 0.43 | 0.11 | 0.13 |
| 2 | 242051777 | rs2074824 | -0.593 | -1.552 | 1.351 | 1.301 | -0.019 | 0.159 | 0.159 | 0 | 0.18 | 0.58 | 0.26 | 0.43 | 0.11 | 0.13 |
| 2 | 242053823 | rs10187363 | -1.041 | -1.936 | 1.089 | 1.025 | -0.042 | 0.075 | 0.075 | 0.059 | 0.26 | 0.44 | 0.32 | 0.55 | 0.14 | 0.17 |
| 2 | 242062162 | rs2302050 | -0.211 | -1.689 | 1.147 | 1.195 | 0.002 | 0.31 | 0.31 | 0 | 0.15 | 0.48 | 0.25 | 0.44 | 0.1 | 0.13 |
| 2 | 242065498 | rs740393 | -1.842 | -1.704 | 1.296 | 0.786 | -0.023 | 0.106 | 0.106 | 1.805 | 0.15 | 0.63 | 0.27 | 0.36 | 0.07 | 0.09 |
| 2 | 242065624 | rs2240541 | -1.419 | -1.662 | 0.99 | 0.784 | 0.005 | -0.005 | -0.005 | 0.002 | 0.13 | 0.42 | 0.12 | 0.12 | 0.07 | 0.04 |
| 2 | 242066314 | rs2240542 | -0.795 | -1.759 | 0.036 | 0.659 | 0.027 | 0.01 | 0.01 | 0 | 0.44 | 0.63 | 0.42 | 0.41 | 0.26 | 0.36 |
| 2 | 242066830 | rs2240544 | -0.719 | -0.981 | -0.186 | 1 | -0.004 | -0.003 | -0.003 | 0 | 0.09 | 0.1 | 0.1 | 0.12 | 0.06 | 0.03 |
| 2 | 242068968 | rs2058062 | 0.094 | -0.758 | -0.684 | 1.009 | 0.006 | -0.015 | -0.015 | 0 | 0.13 | 0.13 | 0.32 | 0.67 | 0.12 | 0.25 |
| 2 | 242074402 | rs4258787 | -0.366 | -0.724 | -0.37 | 0.396 | -0.002 | -0.006 | -0.006 | 0 | 0.58 | 0.45 | 0.62 | 0.91 | 0.46 | 0.6 |
| 2 | 242075192 | rs2108484 | -0.476 | -0.786 | 0.036 | 0.364 | 0.008 | -0.087 | -0.087 | 0 | 0.6 | 0.53 | 0.62 | 0.91 | 0.46 | 0.6 |
| 2 | 242077496 | rs1470414 | 0.537 | -1.198 | -0.202 | 0.196 | 0.013 | 0.181 | 0.181 | 0.168 | 0.42 | 0.12 | 0.3 | 0.17 | 0.37 | 0.21 |
| 2 | 242079553 | rs10933532 | 0.569 | -1.214 | -0.156 | 0.379 | -0.02 | 0.049 | 0.049 | 2.047 | 0.14 | 0.29 | 0.2 | 0.26 | 0.09 | 0.26 |
| 2 | 242082262 | rs6709462 | -0.612 | -1.226 | -0.128 | 0.396 | -0.002 | -0.011 | -0.011 | 0 | 0.58 | 0.45 | 0.62 | 0.91 | 0.46 | 0.6 |
| 2 | 242082480 | rs758068 | -0.228 | -1.224 | -0.202 | 0.408 | 0.004 | 0.06 | 0.06 | 0 | 0.54 | 0.33 | 0.46 | 0.33 | 0.45 | 0.45 |
| 2 | 242088921 | rs3806599 | -0.868 | -0.447 | -0.347 | -0.119 | 0.008 | -0.043 | -0.043 | 0 | 0.81 | 0.7 | 0.85 | 0.93 | 0.8 | 0.74 |
| 2 | 242090639 | rs7568781 | -0.862 | -0.447 | -0.352 | 0.011 | 0.008 | -0.046 | -0.046 | 5.071 | 0.81 | 0.7 | 0.85 | 0.93 | 0.8 | 0.74 |
| 2 | 242092483 | rs4675993 | -0.026 | -0.485 | -0.437 | -0.051 | 0.001 | -0.012 | -0.012 | 0.214 | 0.59 | 0.54 | 0.48 | 0.71 | 0.46 | 0.52 |
| 2 | 242113054 | rs6710177 | -0.451 | -0.473 | -0.334 | 0.066 | 0.007 | 0.101 | 0.101 | 0 | 0.78 | 0.59 | 0.67 | 0.63 | 0.78 | 0.65 |
| 2 | 242113819 | rs7562408 | -1.054 | -0.529 | -0.135 | 0.009 | 0.016 | 0.536 | 0.536 | 0.916 | 0.06 | 0.37 | 0.14 | 0.5 | 0 | 0.13 |
| 2 | 242114037 | rs7565364 | -0.441 | -0.581 | -0.384 | -0.193 | 0.005 | -0.008 | -0.008 | 0 | 0.39 | 0.56 | 0.45 | 0.66 | 0.43 | 0.48 |
| 2 | 242116247 | rs12987697 | -0.525 | -0.581 | -0.387 | -0.215 | 0.004 | -0.057 | -0.057 | 0 | 0.4 | 0.56 | 0.59 | 0.86 | 0.43 | 0.55 |
| 2 | 242119419 | rs11685235 | 0.917 | -0.362 | -0.416 | 0.018 | 0.001 | -0.004 | -0.004 | 0 | 0.23 | 0.23 | 0.27 | 0.13 | 0.33 | 0.14 |
| 2 | 242121720 | rs2268904 | -0.802 | -0.391 | -0.123 | 0.044 | 0.048 | 0.675 | 0.675 | 1.606 | 0.06 | 0.41 | 0.14 | 0.48 | 0 | 0.12 |
| 2 | 242121781 | rs2268905 | 0.511 | -0.491 | -0.789 | -0.136 | -0.031 | -0.013 | -0.013 | 0 | 0.52 | 0.09 | 0.43 | 0.6 | 0.45 | 0.49 |
| 2 | 242122391 | rs3868 | -0.516 | -0.305 | -0.471 | -0.063 | 0 | -0.019 | -0.019 | 0 | 0.79 | 0.79 | 0.87 | 0.93 | 0.79 | 0.75 |
| 2 | 242128714 | rs1444997 | -2.028 | -0.284 | -0.723 | 0.011 | -0.047 | -0.01 | -0.01 | 0 | 0.9 | 0.99 | 0.92 | 0.93 | 0.86 | 0.79 |
| 2 | 242137702 | rs16843438 | 0.736 | -0.108 | -0.071 | 0.005 | -0.007 | -0.035 | -0.035 | 0 | 0.15 | 0.25 | 0.08 | 0.06 | 0.12 | 0.21 |
| 2 | 242138065 | rs16843440 | 1.326 | 0.551 | -0.612 | -0.597 | -0.036 | 0.027 | 0.027 | 0 | 0.14 | 0.09 | 0.18 | 0.01 | 0.28 | 0.13 |
| 2 | 242138334 | rs4675824 | -0.83 | 0.039 | -0.597 | -0.42 | 0.01 | 0.024 | 0.024 | 0 | 0.79 | 0.83 | 0.88 | 0.92 | 0.82 | 0.77 |
| 2 | 242141719 | rs2074840 | 0.258 | 0.085 | -0.903 | -1.674 | -0.003 | -0.056 | -0.056 | 0 | 0.6 | 0.48 | 0.51 | 0.08 | 0.68 | 0.33 |
| 2 | 242143741 | rs11683405 | 0.078 | 0.521 | -1.756 | -1.3 | -0.01 | 0.062 | 0.062 | 1.235 | 0.44 | 0.71 | 0.54 | 0.78 | 0.47 | 0.61 |
| 2 | 242144828 | rs2012160 | 0.19 | 0.573 | -1.76 | -0.881 | 0.007 | 0.061 | 0.061 | 0.194 | 0.7 | 0.82 | 0.84 | 0.92 | 0.74 | 0.75 |
| 2 | 242151585 | rs7593101 | -0.135 | 0.869 | -1.342 | -0.713 | -0.002 | -0.032 | -0.032 | 0 | 0.24 | 0.27 | 0.2 | 0 | 0.24 | 0.05 |
| 2 | 242163359 | rs7590653 | 0.044 | 0.902 | -1.245 | -0.715 | -0.004 | -0.018 | -0.018 | 0.622 | 0.22 | 0.22 | 0.19 | 0 | 0.23 | 0.05 |
| 2 | 242164562 | rs7568928 | 0.475 | 0.589 | -2.114 | -0.46 | 0.01 | 0.055 | 0.055 | 0 | 0.28 | 0.1 | 0.2 | 0.18 | 0.27 | 0.22 |
| 2 | 242166550 | rs1677 | -1.072 | 0.109 | -1.379 | -0.894 | -0.004 | -0.105 | -0.105 | 0 | 0.79 | 0.5 | 0.85 | 0.96 | 0.81 | 0.7 |
| 2 | 242167337 | rs1064767 | 1.061 | 0.182 | -1.532 | -0.813 | 0 | 0.007 | 0.007 | 0.22 | 0.17 | 0.16 | 0.12 | 0.6 | 0.2 | 0.28 |
| 2 | 242168976 | rs15129 | 0.901 | 0.125 | -1.198 | -0.314 | -0.001 | 0.006 | 0.006 | 0.17 | 0.17 | 0.2 | 0.38 | 0.75 | 0.2 | 0.37 |
| 2 | 242173882 | rs3755326 | 2.02 | -0.334 | -1.792 | -0.728 | 0.036 | 0.058 | 0.058 | 0 | 0.16 | 0.01 | 0.03 | 0.07 | 0.09 | 0.1 |
| 2 | 242173916 | rs3755325 | 0.243 | -0.396 | -1.433 | -0.759 | 0.013 | 0.119 | 0.119 | 0 | 0.46 | 0.27 | 0.44 | 0.16 | 0.52 | 0.23 |
| 2 | 242175331 | rs6437249 | -0.88 | -0.484 | -0.797 | -0.951 | 0.002 | -0.043 | -0.043 | 0 | 0.73 | 0.71 | 0.8 | 0.94 | 0.72 | 0.61 |
| 2 | 242177565 | rs2289795 | 0.447 | -0.563 | -0.915 | -0.807 | 0.04 | 0.007 | 0.007 | 0 | 0.35 | 0.17 | 0.17 | 0.66 | 0.29 | 0.36 |
| 2 | 242192848 | rs7578199 | 1.986 | -0.27 | -0.947 | -0.917 | -0.006 | -0.02 | -0.02 | 0 | 0.23 | 0.18 | 0.19 | 0 | 0.27 | 0.05 |
| 2 | 242197095 | rs4675973 | 0.508 | -0.61 | -0.412 | -0.607 | -0.002 | -0.002 | -0.002 | 0 | 0.32 | 0.37 | 0.44 | 0.72 | 0.28 | 0.47 |
| 2 | 242197173 | rs4675971 | -0.289 | -0.992 | -0.36 | -0.574 | -0.001 | 0.004 | 0.004 | 0.006 | 0.22 | 0.23 | 0.43 | 0.39 | 0.15 | 0.27 |
| 2 | 242204286 | rs3771343 | 0.202 | -0.767 | -0.397 | -0.519 | 0.005 | -0.054 | -0.054 | 0.82 | 0.43 | 0.15 | 0.24 | 0.51 | 0.38 | 0.43 |
| 2 | 242212672 | rs6705421 | -0.732 | -0.767 | -0.24 | -0.586 | -0.009 | 0.039 | 0.039 | 0 | 0.56 | 0.37 | 0.67 | 0.76 | 0.53 | 0.62 |
| 2 | 242239918 | rs7559564 | 1.03 | -0.822 | -0.457 | -0.621 | 0.025 | 0.108 | 0.108 | 0 | 0.28 | 0.09 | 0.12 | 0.1 | 0.19 | 0.19 |
| 2 | 242245570 | rs6757876 | 0.833 | -0.681 | -0.318 | -0.466 | -0.008 | -0.025 | -0.025 | 0 | 0.11 | 0.08 | 0.42 | 0.25 | 0.15 | 0.19 |
| 2 | 242253995 | rs10185319 | -2.107 | -0.739 | -0.09 | -0.636 | 0.001 | -0.024 | -0.024 | 3.339 | 0.83 | 0.79 | 0.87 | 0.94 | 0.81 | 0.69 |
| 2 | 242259614 | rs6745662 | -1.63 | -0.718 | -0.083 | -0.596 | 0.004 | 0.011 | 0.011 | 3.296 | 0.82 | 0.75 | 0.86 | 0.76 | 0.78 | 0.68 |
| 2 | 242262986 | rs12694997 | 1.956 | -0.791 | -0.181 | -0.717 | -0.024 | 0.037 | 0.037 | 0 | 0.23 | 0.12 | 0.17 | 0 | 0.27 | 0.05 |
| 2 | 242286747 | rs10933553 | 0.972 | -0.889 | -0.075 | -0.731 | 0.018 | 0.025 | 0.025 | 0 | 0.3 | 0.19 | 0.14 | 0.09 | 0.19 | 0.19 |
| 2 | 242287487 | rs12619647 | 0.146 | -0.87 | -0.149 | -0.7 | -0.021 | -0.024 | -0.024 | 0 | 0.25 | 0.16 | 0.53 | 0.67 | 0.32 | 0.43 |
| 2 | 242291743 | rs7568 | 1.317 | -0.907 | -0.191 | -0.549 | -0.024 | -0.088 | -0.088 | 0 | 0.16 | 0.07 | 0.12 | 0.42 | 0.19 | 0.25 |
| 2 | 242315636 | rs3771551 | 2.023 | -0.957 | -0.202 | -0.66 | 0.037 | 0.042 | 0.042 | 0 | 0.18 | 0.08 | 0.05 | 0.07 | 0.09 | 0.1 |
| 2 | 242322667 | rs6723363 | -3.295 | -0.831 | -0.274 | -0.648 | -0.043 | 0.039 | 0.039 | 1.008 | 0.93 | 0.99 | 0.95 | 1 | 0.86 | 0.88 |
| 2 | 242344333 | rs11681497 | 2.166 | -0.906 | -0.223 | -0.652 | -0.002 | -0.009 | -0.009 | 1.951 | 0.11 | 0.11 | 0.08 | 0.02 | 0.1 | 0.08 |
| 2 | 242350937 | rs2055566 | -3.489 | -0.742 | -0.244 | -0.615 | -0.031 | 0.035 | 0.035 | 0.417 | 0.93 | 0.99 | 0.96 | 1 | 0.91 | 0.89 |
| 2 | 242351379 | rs16843642 | 1.301 | -1.009 | 0.02 | -0.491 | -0.009 | -0.026 | -0.026 | 0 | 0.15 | 0.09 | 0.12 | 0.41 | 0.19 | 0.25 |
| 2 | 242353704 | rs3771561 | 0.78 | -1.026 | 0.241 | -0.496 | 0 | 0.033 | 0.033 | 0 | 0.17 | 0.18 | 0.13 | 0.41 | 0.19 | 0.25 |
| 2 | 242360245 | rs1006309 | -1.462 | -0.83 | 0.16 | -0.451 | 0.002 | 0.017 | 0.017 | 3.08 | 0.81 | 0.71 | 0.86 | 0.76 | 0.79 | 0.67 |

| | | | | | | | | | | | | | | | | |
|---|---|---|---|---|---|---|---|---|---|---|---|---|---|---|---|---|
| 2 | 242371101 | rs757978 | 1.71 | -0.711 | 0.005 | -0.363 | -0.002 | -0.009 | -0.009 | 0.35 | 0.11 | 0.11 | 0.08 | 0.02 | 0.1 | 0.08 |
| 2 | 242394141 | rs764081 | 0.024 | -0.968 | -0.132 | -0.336 | -0.013 | -0.005 | -0.005 | 0 | 0.39 | 0.43 | 0.47 | 0.62 | 0.3 | 0.43 |
| 2 | 242399445 | rs3771583 | 2.201 | -0.965 | -0.401 | -0.922 | 0.047 | 0.053 | 0.053 | 0 | 0.15 | 0.04 | 0.04 | 0.08 | 0.07 | 0.09 |
| 2 | 242400597 | rs13388333 | 0.184 | -0.468 | 0.165 | -0.769 | -0.012 | 0.015 | 0.015 | 0 | 0.29 | 0.26 | 0.3 | 0.11 | 0.38 | 0.14 |
| 2 | 242409933 | rs11682333 | 0.153 | -0.462 | 0.119 | -0.69 | 0.008 | -0.008 | -0.008 | 0 | 0.3 | 0.36 | 0.58 | 0.25 | 0.39 | 0.23 |
| 2 | 242424110 | rs4675947 | 0.153 | -0.461 | 0.12 | -0.671 | 0.008 | -0.008 | -0.008 | 0 | 0.3 | 0.36 | 0.58 | 0.25 | 0.39 | 0.23 |
| 2 | 242426399 | rs11689496 | 1.254 | -0.504 | 0 | -0.715 | 0 | -0.038 | -0.038 | 0 | 0.15 | 0.18 | 0.1 | 0.03 | 0.12 | 0.11 |
| 2 | 242432142 | rs4075818 | 0.788 | -0.2 | -0.541 | -0.553 | -0.038 | -0.026 | -0.026 | 0.292 | 0.09 | 0.01 | 0.14 | 0.11 | 0.15 | 0.09 |
| 2 | 242433348 | rs2018761 | -1.541 | -0.407 | -0.307 | -0.489 | 0.001 | -0.023 | -0.023 | 0.844 | 0.83 | 0.79 | 0.87 | 0.94 | 0.81 | 0.68 |
| 2 | 242444173 | rs7596240 | 0.152 | 0.29 | 0.284 | 0.01 | 0.02 | -0.041 | -0.041 | 0 | 0.25 | 0.37 | 0.2 | 0.08 | 0.3 | 0.1 |
| 2 | 242445336 | rs1995496 | -0.391 | -0.227 | -0.312 | 0.163 | 0 | 0.004 | 0.004 | 0.213 | 0.46 | 0.58 | 0.68 | 0.29 | 0.51 | 0.34 |
| 2 | 242448642 | rs4675809 | -0.424 | -0.376 | -0.143 | 0.956 | -0.01 | 0.039 | 0.039 | 0 | 0.1 | 0.16 | 0.38 | 0.26 | 0.08 | 0.19 |
| 2 | 242457864 | rs4381775 | -0.609 | -0.522 | -0.614 | 0.769 | 0.027 | 0.023 | 0.023 | 0 | 0.15 | 0.1 | 0.15 | 0.12 | 0.12 | 0.1 |
| 2 | 242459618 | rs4675807 | -1.064 | -0.646 | -0.63 | 0.733 | 0.036 | -0.014 | -0.014 | 0 | 0.3 | 0.21 | 0.47 | 0.36 | 0.22 | 0.29 |
| 2 | 242471244 | rs10933562 | -0.765 | -0.354 | 0.177 | 1.014 | -0.043 | 0.121 | 0.121 | 0.216 | 0.17 | 0.6 | 0.43 | 0.38 | 0.12 | 0.18 |
| 2 | 242476987 | rs13001905 | -1.33 | 0.025 | -0.442 | 0.63 | 0.015 | -0.021 | -0.021 | 1.934 | 0.39 | 0.82 | 0.58 | 0.41 | 0.38 | 0.24 |
| 2 | 242477457 | rs2174276 | -0.119 | 0.092 | -0.07 | 0.83 | -0.03 | 0.133 | 0.133 | 0 | 0.16 | 0.3 | 0.41 | 0.4 | 0.14 | 0.2 |
| 2 | 242480111 | rs2055567 | -0.243 | 0.72 | -0.378 | 0.344 | -0.005 | -0.022 | -0.022 | 0 | 0.2 | 0.21 | 0.17 | 0.01 | 0.27 | 0.04 |
| 2 | 242486521 | rs10175075 | -0.535 | 0.579 | -0.535 | 0.74 | 0 | 0 | 0 | 0.427 | 0.44 | 0.56 | 0.34 | 0.48 | 0.48 | 0.32 |
| 2 | 242490658 | rs13408032 | -0.558 | -0.126 | -0.036 | 1.145 | -0.058 | 0.125 | 0.125 | 0 | 0.19 | 0.46 | 0.28 | 0.33 | 0.14 | 0.3 |
| 2 | 242492540 | rs2379195 | -0.617 | 0.427 | -1.047 | 0.737 | -0.105 | 0.045 | 0.045 | 0.542 | 0.71 | 0.94 | 0.77 | 0.87 | 0.64 | 0.73 |
| 2 | 242493511 | rs4675801 | 0.006 | -0.001 | -0.565 | 0.728 | 0 | 0.002 | 0.002 | 0.52 | 0.48 | 0.55 | 0.63 | 0.49 | 0.5 | 0.58 |
| 2 | 242496325 | rs7572476 | 0.006 | -0.001 | -0.565 | 0.725 | 0 | 0.002 | 0.002 | 0.52 | 0.48 | 0.55 | 0.62 | 0.49 | 0.5 | 0.58 |
| 2 | 242502642 | rs4675800 | -0.298 | 0.093 | -0.592 | 0.155 | -0.001 | -0.021 | -0.021 | 0 | 0.61 | 0.54 | 0.7 | 0.87 | 0.6 | 0.7 |
| 2 | 242502956 | rs12479254 | -0.269 | 0.055 | -0.81 | 0.586 | -0.005 | 0.005 | 0.005 | 0 | 0.65 | 0.72 | 0.75 | 0.71 | 0.64 | 0.69 |
| 2 | 242503543 | rs12469165 | -0.987 | 0.271 | -1.094 | 0.467 | -0.05 | 0.046 | 0.046 | 0.073 | 0.73 | 0.89 | 0.79 | 0.87 | 0.68 | 0.74 |
| 2 | 242508407 | rs7558279 | -0.61 | 0.661 | -0.595 | 0.989 | 0 | -0.001 | -0.001 | 1.036 | 0.41 | 0.61 | 0.23 | 0.44 | 0.35 | 0.22 |
| 2 | 242509012 | rs7558942 | -0.676 | 0.369 | -0.455 | 0.905 | -0.003 | -0.009 | -0.009 | 0.142 | 0.54 | 0.49 | 0.61 | 0.73 | 0.59 | 0.63 |
| 2 | 242511083 | rs13004470 | -0.191 | -0.318 | -0.298 | 1.186 | 0.004 | 0.006 | 0.006 | 0.322 | 0.25 | 0.22 | 0.13 | 0.15 | 0.17 | 0.07 |
| 2 | 242511217 | rs13416312 | 0.121 | -0.32 | -0.335 | 1.151 | -0.004 | -0.002 | -0.002 | 0 | 0.22 | 0.22 | 0.09 | 0.15 | 0.12 | 0.06 |
| 2 | 242515096 | rs6716290 | -1.158 | -0.434 | -0.37 | 1.5 | 0.021 | -0.001 | -0.001 | 2.058 | 0.41 | 0.3 | 0.33 | 0.43 | 0.31 | 0.49 |
| 2 | 242515197 | rs7420703 | -0.038 | -0.435 | -0.371 | 1.5 | 0.01 | -0.03 | -0.03 | 0 | 0.16 | 0.09 | 0.2 | 0.35 | 0.14 | 0.42 |
| 2 | 242516105 | rs6717283 | -0.004 | -0.434 | -0.37 | 1.492 | 0.004 | 0.001 | 0.001 | 0.322 | 0.25 | 0.22 | 0.13 | 0.09 | 0.17 | 0.07 |
| 2 | 242517966 | rs10201535 | -1.348 | -0.444 | -0.391 | 1.544 | -0.013 | -0.002 | -0.002 | 0 | 0.43 | 0.43 | 0.41 | 0.63 | 0.33 | 0.54 |
| 2 | 242521405 | rs10191556 | -0.038 | -0.427 | -0.458 | 1.557 | 0.01 | -0.03 | -0.03 | 0 | 0.16 | 0.09 | 0.2 | 0.35 | 0.14 | 0.42 |
| 2 | 242522140 | rs4675925 | -0.758 | -0.039 | -0.326 | 1.301 | -0.014 | -0.002 | -0.002 | 0 | 0.43 | 0.44 | 0.41 | 0.63 | 0.33 | 0.54 |
| 2 | 242665008 | rs12048 | -0.834 | -0.001 | 0.119 | 1.731 | 0.014 | 0.382 | 0.382 | 1.079 | 0.08 | 0.38 | 0.18 | 0.31 | 0.04 | 0.16 |
| 2 | 242667894 | rs7588212 | -0.388 | 0.379 | -1.255 | 0.591 | -0.003 | 0.138 | 0.138 | 1.791 | 0.51 | 0.76 | 0.49 | 0.81 | 0.52 | 0.82 |
| 2 | 242710662 | rs10755057 | -0.035 | 0.66 | -2.002 | 0.767 | -0.084 | -0.063 | -0.063 | 0 | 0.6 | 0.9 | 0.52 | 0.59 | 0.52 | 0.43 |
| 2 | 242710935 | rs10933568 | 0.041 | 0.706 | -1.992 | 0.884 | -0.161 | -0.16 | -0.16 | 0.02 | 0.6 | 0.96 | 0.53 | 0.54 | 0.52 | 0.43 |
| 2 | 242712870 | rs7563672 | 0.449 | 0.309 | -1.252 | 0.718 | 0.007 | 0.005 | 0.005 | 0 | 0.44 | 0.55 | 0.28 | 0.4 | 0.39 | 0.29 |
| 2 | 242725752 | rs9973663 | -0.31 | 0.486 | -1.164 | 0.211 | 0.004 | -0.064 | -0.064 | 0 | 0.16 | 0.27 | 0.13 | 0.01 | 0.19 | 0.08 |
| 2 | 242743943 | rs741776 | 0.867 | 1.113 | -1.335 | 0.386 | -0.008 | -0.019 | -0.019 | 3.216 | 0.4 | 0.43 | 0.32 | 0.25 | 0.33 | 0.23 |
| 2 | 242750743 | rs28528975 | 0.024 | -0.8 | -1.224 | 0.291 | -0.031 | 0.014 | 0.014 | 0 | 0.38 | 0.41 | 0.44 | 0.52 | 0.29 | 0.37 |
| 3 | 247473 | rs5024851 | 1.44 | -0.624 | -0.315 | -1.395 | -0.003 | 0.076 | 0.076 | 0 | 0.1 | 0.13 | 0.18 | 0.39 | 0.1 | 0.31 |
| 3 | 253799 | rs6768035 | 0.877 | -0.989 | -0.161 | -1.455 | 0.002 | -0.003 | -0.003 | 0 | 0.65 | 0.38 | 0.65 | 0.59 | 0.73 | 0.51 |
| 3 | 261442 | rs459362 | 1.23 | -1.512 | 0.149 | -0.833 | -0.002 | 0.1 | 0.1 | 0 | 0.1 | 0.12 | 0.18 | 0.46 | 0.1 | 0.33 |
| 3 | 264293 | rs331855 | -0.57 | -2.799 | 0.461 | -1.312 | 0.015 | 0 | 0 | 0 | 0.5 | 0.46 | 0.46 | 0.45 | 0.38 | 0.64 |
| 3 | 269208 | rs634896 | 1.33 | -2.779 | 0.02 | -1.028 | 0.006 | 0 | 0 | 6.653 | 0.1 | 0.07 | 0.14 | 0.37 | 0.09 | 0.31 |
| 3 | 270521 | rs3872649 | 0.089 | -2.885 | 0.238 | -1.088 | -0.042 | 0.19 | 0.19 | 0 | 0.82 | 0.5 | 0.82 | 0.63 | 0.88 | 0.65 |
| 3 | 285316 | rs455635 | -1.745 | -2.796 | 0.195 | -0.804 | -0.042 | -0.02 | -0.02 | 0 | 0.64 | 0.83 | 0.44 | 0.63 | 0.48 | 0.75 |
| 3 | 287419 | rs455357 | -0.718 | -2.82 | 0.221 | -0.871 | 0.037 | -0.013 | -0.013 | 0 | 0.54 | 0.66 | 0.35 | 0.45 | 0.4 | 0.69 |
| 3 | 289311 | rs3864046 | -0.584 | -2.683 | 0.088 | -0.764 | -0.01 | 0.015 | 0.015 | 0 | 0.21 | 0.27 | 0.2 | 0.38 | 0.16 | 0.35 |
| 3 | 292992 | rs162738 | -0.65 | -2.531 | -0.386 | -0.662 | 0.024 | 0.222 | 0.222 | 1.292 | 0.52 | 0.86 | 0.38 | 0.18 | 0.4 | 0.36 |
| 3 | 294397 | rs331885 | -0.374 | -2.366 | -0.069 | -0.627 | 0.022 | -0.069 | -0.069 | 0.187 | 0.18 | 0.11 | 0.2 | 0.38 | 0.15 | 0.37 |
| 3 | 297321 | rs445658 | -0.435 | -2.368 | 0.029 | -0.628 | 0.016 | -0.037 | -0.037 | 2.885 | 0.18 | 0.15 | 0.2 | 0.38 | 0.15 | 0.37 |
| 3 | 298171 | rs432685 | -0.611 | -2.386 | 0.314 | -0.683 | 0.039 | -0.002 | -0.002 | 0 | 0.51 | 0.37 | 0.36 | 0.45 | 0.4 | 0.65 |
| 3 | 310616 | rs3956142 | -1.137 | -2.549 | 0.85 | -0.794 | -0.015 | 0.01 | 0.01 | 0 | 0.35 | 0.42 | 0.27 | 0.39 | 0.25 | 0.39 |
| 3 | 310747 | rs17016917 | 0.339 | -2.459 | 0.369 | -1.378 | 0.009 | 0.001 | 0.001 | 0 | 0.34 | 0.27 | 0.18 | 0.08 | 0.29 | 0.34 |
| 3 | 312349 | rs6764363 | -0.096 | -2.372 | 0.473 | -1.319 | 0.004 | -0.015 | -0.015 | 0 | 0.41 | 0.37 | 0.26 | 0.09 | 0.37 | 0.35 |
| 3 | 315016 | rs332453 | -1.718 | -2.137 | -0.007 | -1.226 | -0.079 | 0.196 | 0.196 | 0.006 | 0.66 | 0.91 | 0.68 | 0.99 | 0.49 | 0.8 |
| 3 | 316417 | rs11707796 | 0.178 | -1.815 | 0.621 | -0.838 | 0.024 | 0.041 | 0.041 | 0.074 | 0.37 | 0.28 | 0.23 | 0.11 | 0.29 | 0.35 |
| 3 | 320201 | rs11929364 | -0.061 | -1.195 | 1.403 | 0.174 | -0.005 | 0.33 | 0.33 | 0 | 0.11 | 0.37 | 0.16 | 0.4 | 0.08 | 0.29 |
| 3 | 323615 | rs1909383 | 0.316 | -1.202 | 1.181 | -0.251 | 0.017 | 0.022 | 0.022 | 0 | 0.33 | 0.26 | 0.2 | 0.07 | 0.27 | 0.33 |
| 3 | 325410 | rs425366 | -1.534 | -1.128 | 0.874 | -0.192 | -0.033 | -0.069 | -0.069 | 3.904 | 0.6 | 0.9 | 0.43 | 0.47 | 0.44 | 0.68 |
| 3 | 330037 | rs3933034 | -0.457 | -1.104 | 1.1 | -0.162 | 0.001 | 0 | 0 | 0.04 | 0.46 | 0.45 | 0.37 | 0.46 | 0.38 | 0.62 |
| 3 | 336028 | rs11717061 | 0.288 | -1.018 | 0.964 | -0.19 | 0.012 | 0.018 | 0.018 | 0 | 0.34 | 0.29 | 0.23 | 0.1 | 0.28 | 0.33 |
| 3 | 337727 | rs331892 | -1.14 | -0.679 | 1.072 | 0.236 | -0.043 | -0.072 | -0.072 | 0.209 | 0.62 | 0.9 | 0.45 | 0.47 | 0.46 | 0.75 |
| 3 | 340440 | rs332479 | -1.134 | -0.723 | 1.288 | 0.218 | -0.098 | -0.114 | -0.114 | 0 | 0.62 | 0.93 | 0.45 | 0.47 | 0.46 | 0.75 |

| | | | | | | | | | | | | | | | | |
|---|---|---|---|---|---|---|---|---|---|---|---|---|---|---|---|---|
| 3 | 340919 | rs193081 | -0.855 | -0.698 | 1.702 | 0.271 | -0.039 | 0.225 | 0.225 | 0 | 0.17 | 0.47 | 0.2 | 0.36 | 0.11 | 0.34 |
| 3 | 342281 | rs12637355 | -0.489 | -0.614 | 1.212 | 0.384 | -0.014 | 0.061 | 0.061 | 0 | 0.16 | 0.24 | 0.17 | 0.34 | 0.12 | 0.34 |
| 3 | 343022 | rs13093895 | 0.01 | -0.448 | 1.042 | 0.374 | 0.025 | 0.056 | 0.056 | 0 | 0.44 | 0.31 | 0.25 | 0.17 | 0.35 | 0.39 |
| 3 | 348599 | rs17023537 | -0.59 | 0.272 | 1.342 | 1.225 | 0.009 | -0.027 | -0.027 | 0 | 0.12 | 0.07 | 0.13 | 0.33 | 0.1 | 0.31 |
| 3 | 353314 | rs2648466 | -1.288 | 0.291 | 1.699 | 1.207 | -0.015 | 0.055 | 0.055 | 0 | 0.16 | 0.3 | 0.19 | 0.3 | 0.15 | 0.4 |
| 3 | 354495 | rs381357 | -2.036 | 0.96 | 1.965 | 1.075 | 0.005 | 0.03 | 0.03 | 0 | 0.26 | 0.62 | 0.27 | 0.31 | 0.26 | 0.46 |
| 3 | 359454 | rs4685578 | -0.743 | 0.986 | 2.015 | 0.988 | 0.115 | -0.035 | -0.035 | 0 | 0.1 | 0.41 | 0.09 | 0.01 | 0.11 | 0.05 |
| 3 | 361508 | rs2272522 | -1.601 | 0.927 | 1.961 | 1.067 | 0.011 | 0.064 | 0.064 | 0 | 0.22 | 0.56 | 0.24 | 0.31 | 0.2 | 0.34 |
| 3 | 366039 | rs162416 | -2.235 | 0.819 | 1.297 | 1.068 | -0.015 | -0.098 | -0.098 | 0.555 | 0.32 | 0.91 | 0.34 | 0.35 | 0.28 | 0.49 |
| 3 | 372448 | rs330893 | 0.711 | 0.687 | 1.861 | 1.313 | -0.012 | 0.087 | 0.087 | 0 | 0.87 | 0.77 | 0.85 | 0.7 | 0.89 | 0.73 |
| 3 | 373693 | rs4685598 | 1.823 | 0.734 | 2.101 | 1.254 | -0.017 | 0.043 | 0.043 | 0.614 | 0.8 | 0.34 | 0.75 | 0.69 | 0.8 | 0.57 |
| 3 | 374168 | rs7630509 | 1.976 | 0.659 | 2.084 | 1.161 | -0.02 | 0.072 | 0.072 | 0.23 | 0.79 | 0.43 | 0.75 | 0.69 | 0.79 | 0.59 |
| 3 | 374525 | rs330895 | -1.587 | 0.772 | 1.354 | 1.193 | -0.01 | 0.01 | 0.01 | 0 | 0.06 | 0.04 | 0.07 | 0.01 | 0.11 | 0.15 |
| 3 | 374788 | rs1975192 | -0.389 | 0.765 | 1.426 | 1.21 | 0.001 | 0.004 | 0.004 | 0 | 0.11 | 0.09 | 0.14 | 0.3 | 0.1 | 0.27 |
| 3 | 374869 | rs6800364 | 1.578 | 0.766 | 1.494 | 1.208 | -0.001 | 0.088 | 0.088 | 0 | 0.82 | 0.64 | 0.77 | 0.7 | 0.8 | 0.6 |
| 3 | 375590 | rs11129453 | 1.675 | 1.164 | 0.854 | 0.673 | 0.008 | -0.032 | -0.032 | 0.031 | 0.89 | 0.82 | 0.9 | 0.99 | 0.84 | 0.82 |
| 3 | 378069 | rs7649544 | 1.4 | 1.095 | 0.831 | 0.767 | 0.018 | -0.017 | -0.017 | 0 | 0.93 | 0.83 | 0.92 | 0.99 | 0.89 | 0.84 |
| 3 | 381302 | rs1353811 | 1.167 | 0.983 | 0.741 | 1.081 | -0.008 | 0.077 | 0.077 | 0 | 0.72 | 0.48 | 0.67 | 0.61 | 0.71 | 0.53 |
| 3 | 387597 | rs9637430 | 1.136 | 1.285 | 0.831 | 0.925 | 0.01 | 0.058 | 0.058 | 0.351 | 0.74 | 0.38 | 0.75 | 0.62 | 0.7 | 0.58 |
| 3 | 392578 | rs3773395 | -0.999 | 1.583 | 0.842 | 0.884 | -0.02 | -0.029 | -0.029 | 0.001 | 0.19 | 0.14 | 0.21 | 0.38 | 0.32 | 0.42 |
| 3 | 397958 | rs1874947 | 1.223 | 1.326 | 0.328 | 0.644 | -0.073 | -0.063 | -0.063 | 0.507 | 0.61 | 0.11 | 0.68 | 0.62 | 0.59 | 0.46 |
| 3 | 398423 | rs7628476 | 1.031 | 1.359 | 0.914 | 0.656 | -0.012 | -0.011 | -0.011 | 0 | 0.65 | 0.28 | 0.7 | 0.63 | 0.59 | 0.46 |
| 3 | 398934 | rs9841789 | 0.899 | 1.405 | 0.85 | 0.712 | 0.021 | 0.021 | 0.021 | 0 | 0.66 | 0.36 | 0.71 | 0.61 | 0.6 | 0.47 |
| 3 | 414050 | rs9876565 | -0.562 | 0.734 | 0.061 | 0.262 | -0.002 | 0.021 | 0.021 | 0 | 0.73 | 0.67 | 0.56 | 0.49 | 0.74 | 0.83 |
| 3 | 416441 | rs3773384 | -0.575 | 0.367 | -0.373 | -0.075 | -0.003 | -0.003 | -0.003 | 1.145 | 0.39 | 0.63 | 0.33 | 0.38 | 0.37 | 0.55 |
| 3 | 416943 | rs3773383 | 0.661 | 0.387 | -0.034 | -0.023 | 0.043 | 0.057 | 0.057 | 0.062 | 0.21 | 0.31 | 0.21 | 0.35 | 0.3 | 0.48 |
| 3 | 417207 | rs3821619 | -0.073 | 0.339 | -0.064 | -0.218 | 0.014 | 0.013 | 0.013 | 0.295 | 0.27 | 0.27 | 0.25 | 0.38 | 0.37 | 0.51 |
| 3 | 417994 | rs930237 | -0.179 | 0.069 | 0.329 | -0.218 | -0.049 | -0.106 | -0.106 | 0.025 | 0.83 | 0.58 | 0.87 | 1 | 0.94 | 0.96 |
| 3 | 418089 | rs3773380 | -0.315 | 0.345 | -0.291 | -0.023 | 0.013 | 0.004 | 0.004 | 0 | 0.35 | 0.54 | 0.3 | 0.35 | 0.36 | 0.52 |
| 3 | 419022 | rs2387181 | -0.606 | 0.491 | -0.374 | 0.025 | 0.008 | 0.006 | 0.006 | 0 | 0.51 | 0.56 | 0.44 | 0.5 | 0.52 | 0.66 |
| 3 | 419143 | rs3773377 | 0.277 | 0.81 | -0.496 | 0.329 | -0.005 | -0.004 | -0.004 | 0 | 0.61 | 0.62 | 0.64 | 0.5 | 0.57 | 0.46 |
| 3 | 422639 | rs2196039 | -0.107 | 0.689 | -1.214 | 0.46 | -0.056 | -0.105 | -0.105 | 1.294 | 0.29 | 0.03 | 0.34 | 0.58 | 0.28 | 0.4 |
| 3 | 423018 | rs11706690 | 0.089 | 0.844 | -1.15 | 0.048 | -0.054 | -0.002 | -0.002 | 0 | 0.61 | 0.12 | 0.51 | 0.63 | 0.66 | 0.74 |
| 3 | 430682 | rs4685668 | 0.953 | 0.26 | 0.556 | -0.038 | -0.01 | -0.061 | -0.061 | 1.008 | 0.89 | 0.46 | 0.93 | 1 | 0.96 | 0.97 |
| 3 | 433955 | rs11129762 | 0.914 | 0.328 | 1.051 | 0.028 | 0.031 | 0.009 | 0.009 | 2.345 | 0.93 | 0.53 | 0.95 | 1 | 0.98 | 0.99 |
| 3 | 436885 | rs1865164 | 1.224 | 0.239 | 1.08 | 0.022 | -0.018 | -0.041 | -0.041 | 0 | 0.91 | 0.71 | 0.96 | 1 | 0.98 | 0.97 |
| 3 | 438253 | rs3773376 | -0.189 | 0.7 | 0.317 | 0.629 | -0.064 | -0.118 | -0.118 | 0 | 0.16 | 0.01 | 0.22 | 0.52 | 0.2 | 0.31 |
| 3 | 448527 | rs2117046 | 1.594 | 0.609 | 1.482 | -0.117 | 0.042 | 0.007 | 0.007 | 5.095 | 0.93 | 0.59 | 0.95 | 1 | 0.98 | 0.98 |
| 3 | 449246 | rs9820546 | -1.59 | 0.643 | 1.521 | -0.102 | 0.047 | 0.014 | 0.014 | 1.678 | 0.05 | 0.35 | 0.04 | 0 | 0.02 | 0.02 |
| 3 | 461969 | rs9878298 | -0.284 | 0.78 | 0.773 | -0.767 | -0.002 | -0.017 | -0.017 | 0 | 0.53 | 0.56 | 0.59 | 0.75 | 0.43 | 0.65 |
| 3 | 464068 | rs7627995 | 0.198 | 0.946 | 0.783 | -1.033 | 0.034 | 0.052 | 0.052 | 0.17 | 0.5 | 0.33 | 0.43 | 0.25 | 0.63 | 0.37 |
| 3 | 467157 | rs4494911 | 0.724 | 1.609 | 0.221 | -0.983 | -0.039 | 0.012 | 0.012 | 0 | 0.32 | 0.06 | 0.47 | 0.71 | 0.32 | 0.48 |
| 3 | 469463 | rs1155327 | 0.21 | 0.348 | 0.302 | -1.29 | 0.001 | -0.001 | -0.001 | 0 | 0.53 | 0.42 | 0.42 | 0.26 | 0.61 | 0.38 |
| 3 | 471460 | rs13079540 | 0.518 | 0.179 | 0.187 | -1.397 | 0.002 | -0.014 | -0.014 | 0.002 | 0.43 | 0.41 | 0.39 | 0.25 | 0.52 | 0.33 |
| 3 | 489181 | rs1988234 | 0.646 | -0.296 | 0.84 | -0.983 | -0.046 | -0.083 | -0.083 | 0 | 0.84 | 0.6 | 0.92 | 0.96 | 0.93 | 0.85 |
| 3 | 504756 | rs9311359 | -1.59 | 0.127 | 1.041 | -0.554 | -0.003 | 0.043 | 0.043 | 0 | 0.08 | 0.28 | 0.08 | 0.12 | 0.03 | 0.17 |
| 3 | 529666 | rs11130113 | -0.005 | 0.26 | 0.78 | -1.03 | 0.017 | 0.049 | 0.049 | 3.666 | 0.5 | 0.58 | 0.54 | 0.7 | 0.47 | 0.53 |
| 3 | 543373 | rs7426480 | -0.465 | 0.114 | 1.058 | -0.998 | -0.001 | 0.006 | 0.006 | 0.303 | 0.64 | 0.67 | 0.69 | 0.71 | 0.68 | 0.62 |
| 3 | 546738 | rs17267716 | -0.066 | 0.131 | 0.325 | -0.973 | 0.022 | 0.045 | 0.045 | 0.783 | 0.42 | 0.74 | 0.34 | 0.29 | 0.31 | 0.4 |
| 3 | 552113 | rs4072082 | 0.091 | 0.105 | 0.436 | -0.949 | -0.01 | -0.019 | -0.019 | 0 | 0.4 | 0.52 | 0.31 | 0.3 | 0.29 | 0.39 |
| 3 | 565961 | rs1531913 | -0.268 | -0.102 | -0.438 | -0.524 | 0.001 | 0.053 | 0.053 | 0 | 0.52 | 0.82 | 0.49 | 0.38 | 0.51 | 0.47 |
| 3 | 580021 | rs7616596 | -0.229 | -0.475 | 0.023 | -0.52 | 0.03 | 0.03 | 0.03 | 0.469 | 0.58 | 0.78 | 0.54 | 0.32 | 0.62 | 0.48 |
| 3 | 581027 | rs9835014 | -1.083 | -0.584 | 0.528 | -0.536 | -0.01 | -0.03 | -0.03 | 1.024 | 0.07 | 0.2 | 0.05 | 0.01 | 0.05 | 0.02 |
| 3 | 586487 | rs17208795 | 0.602 | -0.837 | 0.983 | -0.673 | 0.032 | 0.064 | 0.064 | 0 | 0.46 | 0.21 | 0.44 | 0.31 | 0.56 | 0.44 |
| 3 | 595870 | rs9882746 | -3.127 | -0.743 | 2.401 | -0.603 | -0.042 | -0.068 | -0.068 | 0 | 0.15 | 0.66 | 0.11 | 0.01 | 0.06 | 0.05 |
| 3 | 597064 | rs9816725 | -1.868 | -0.597 | 2.134 | -0.536 | -0.013 | -0.031 | -0.031 | 0.025 | 0.07 | 0.18 | 0.06 | 0.01 | 0.05 | 0.03 |
| 3 | 598777 | rs9837987 | -1.962 | -0.6 | 2.173 | -0.537 | -0.006 | -0.028 | -0.028 | 0 | 0.07 | 0.29 | 0.06 | 0.01 | 0.05 | 0.03 |
| 3 | 602689 | rs2729048 | -2.464 | -0.608 | 2.301 | -0.539 | -0.024 | -0.039 | -0.039 | 0 | 0.07 | 0.26 | 0.05 | 0 | 0.03 | 0.03 |
| 3 | 609566 | rs2729257 | -2.288 | -0.492 | 2.189 | -0.472 | 0.027 | 0.01 | 0.01 | 0.686 | 0.06 | 0.29 | 0.06 | 0.01 | 0.04 | 0.03 |
| 3 | 668832 | rs9844190 | -2.492 | -0.34 | 2.383 | -0.512 | 0.058 | -0.026 | -0.026 | 1.832 | 0.1 | 0.43 | 0.09 | 0.01 | 0.07 | 0.04 |
| 3 | 669478 | rs9826534 | -3.181 | -0.487 | 2.132 | -0.581 | -0.01 | -0.041 | -0.041 | 0 | 0.15 | 0.77 | 0.14 | 0.01 | 0.09 | 0.05 |
| 3 | 671993 | rs9809898 | -1.766 | -0.355 | 2.158 | -0.517 | 0.044 | 0.012 | 0.012 | 2.616 | 0.07 | 0.4 | 0.07 | 0 | 0.03 | 0.03 |
| 3 | 673704 | rs9311812 | -2.539 | -0.289 | 1.758 | -0.526 | -0.002 | -0.077 | -0.077 | 0 | 0.13 | 0.58 | 0.12 | 0.01 | 0.1 | 0.05 |
| 3 | 674242 | rs9824721 | -1.955 | -0.153 | 1.563 | -0.455 | 0.054 | -0.042 | -0.042 | 0 | 0.1 | 0.49 | 0.1 | 0.01 | 0.08 | 0.05 |
| 3 | 677112 | rs7647780 | -0.712 | -0.569 | 0.263 | -0.771 | 0.022 | 0.041 | 0.041 | 0 | 0.24 | 0.12 | 0.11 | 0.07 | 0.18 | 0.26 |
| 3 | 678347 | rs1351865 | 0.324 | -0.124 | 0.164 | -0.53 | 0.006 | 0.053 | 0.053 | 0 | 0.51 | 0.35 | 0.61 | 0.67 | 0.51 | 0.59 |
| 3 | 684744 | rs9631481 | -0.23 | -0.127 | 0.677 | -0.456 | -0.001 | -0.001 | -0.001 | 0.117 | 0.46 | 0.49 | 0.39 | 0.35 | 0.51 | 0.41 |
| 3 | 698777 | rs9311883 | -1.968 | 0.521 | 1.043 | -0.019 | 0.036 | 0.042 | 0.042 | 0.79 | 0.14 | 0.35 | 0.13 | 0.21 | 0.14 | 0.19 |
| 3 | 700642 | rs2729262 | -1.991 | 0.603 | 1.171 | 0.039 | 0.022 | 0.031 | 0.031 | 1.904 | 0.14 | 0.27 | 0.19 | 0.22 | 0.15 | 0.2 |

| | | | | | | | | | | | | | | | | |
|---|---|---|---|---|---|---|---|---|---|---|---|---|---|---|---|---|
| 3 | 701825 | rs2627980 | -2.328 | 0.569 | 1.175 | 0.026 | 0.024 | 0.031 | 0.031 | 1.542 | 0.15 | 0.31 | 0.2 | 0.21 | 0.15 | 0.22 |
| 3 | 703725 | rs1905605 | -1.815 | 0.529 | 1.217 | 0.083 | -0.002 | 0.043 | 0.043 | 0.579 | 0.09 | 0.25 | 0.06 | 0.15 | 0.05 | 0.17 |
| 3 | 707654 | rs10049372 | 1.821 | 0.219 | 1.653 | -0.118 | 0.028 | -0.022 | -0.022 | 0.738 | 0.83 | 0.62 | 0.79 | 0.88 | 0.82 | 0.82 |
| 3 | 710880 | rs7615785 | 0.51 | 0.133 | 0.896 | -0.188 | -0.08 | 0 | 0 | 0 | 0.23 | 0.02 | 0.32 | 0.16 | 0.27 | 0.13 |
| 3 | 712551 | rs2729152 | 2.081 | -0.168 | 2.36 | 0.442 | 0.033 | 0.203 | 0.203 | 0 | 0.94 | 0.71 | 0.96 | 0.77 | 0.97 | 0.85 |
| 3 | 715829 | rs13085985 | -2.075 | -0.253 | 2.309 | 0.425 | -0.032 | 0.056 | 0.056 | 0.441 | 0.11 | 0.22 | 0.05 | 0.28 | 0.07 | 0.27 |
| 3 | 722279 | rs2627965 | 2.352 | -0.272 | 2.18 | 0.686 | -0.07 | 0.166 | 0.166 | 2.164 | 0.85 | 0.47 | 0.91 | 0.66 | 0.93 | 0.73 |
| 3 | 722831 | rs2627964 | 2.293 | -0.26 | 2.289 | 0.695 | 0.015 | 0.32 | 0.32 | 0.254 | 0.92 | 0.51 | 0.93 | 0.71 | 0.97 | 0.83 |
| 3 | 729510 | rs13066393 | -2.538 | -0.266 | 2.501 | 0.717 | 0.014 | 0.306 | 0.306 | 0.432 | 0.09 | 0.48 | 0.08 | 0.29 | 0.03 | 0.17 |
| 3 | 730719 | rs1905603 | -3.62 | -0.218 | 2.703 | 0.674 | 0 | 0 | 0 | 3.34 | 0.24 | 0.7 | 0.26 | 0.42 | 0.21 | 0.33 |
| 3 | 732114 | rs1872300 | 3.114 | -0.386 | 2.545 | 0.659 | -0.06 | 0.176 | 0.176 | 3.124 | 0.86 | 0.49 | 0.9 | 0.67 | 0.93 | 0.8 |
| 3 | 732750 | rs2729164 | -0.94 | -0.173 | 2.188 | 0.529 | 0.001 | -0.005 | -0.005 | 2.61 | 0.12 | 0.14 | 0.15 | 0.03 | 0.13 | 0.07 |
| 3 | 746092 | rs1484515 | 2.704 | -0.424 | 1.632 | 0.647 | -0.059 | 0.177 | 0.177 | 3.335 | 0.85 | 0.5 | 0.82 | 0.67 | 0.92 | 0.71 |
| 3 | 761907 | rs6789520 | -2.31 | -0.183 | 1.624 | 0.739 | -0.006 | 0.279 | 0.279 | 3.067 | 0.1 | 0.53 | 0.07 | 0.29 | 0.02 | 0.15 |
| 3 | 778709 | rs13320566 | -1.374 | -0.147 | 0.832 | 0.228 | 0.001 | 0.001 | 0.001 | 0 | 0.24 | 0.28 | 0.29 | 0.11 | 0.23 | 0.14 |
| 3 | 779167 | rs9849397 | -1.343 | -0.17 | 0.936 | 0.218 | -0.005 | -0.031 | -0.031 | 0 | 0.25 | 0.36 | 0.3 | 0.11 | 0.23 | 0.14 |
| 3 | 789427 | rs7621645 | -1.762 | 0.026 | 1.063 | 0.37 | -0.006 | 0.004 | 0.004 | 0.293 | 0.34 | 0.61 | 0.38 | 0.38 | 0.36 | 0.41 |
| 3 | 792207 | rs4254643 | -0.86 | -0.018 | 0.939 | 0.278 | -0.002 | 0.021 | 0.021 | 0 | 0.17 | 0.16 | 0.21 | 0.06 | 0.18 | 0.09 |
| 3 | 800548 | rs2689860 | 1.71 | -0.211 | 1.299 | 0.244 | 0.014 | -0.001 | -0.001 | 0 | 0.64 | 0.29 | 0.56 | 0.61 | 0.6 | 0.58 |
| 3 | 809720 | rs11714239 | -1.864 | -0.185 | 1.541 | 0.256 | -0.005 | 0.001 | 0.001 | 0.051 | 0.34 | 0.59 | 0.42 | 0.39 | 0.39 | 0.43 |
| 3 | 810051 | rs7432355 | -1.06 | -0.18 | 1.441 | 0.263 | 0.006 | -0.008 | -0.008 | 0 | 0.07 | 0.24 | 0.12 | 0.04 | 0.06 | 0.04 |
| 3 | 813254 | rs4283555 | -1.169 | -0.126 | 0.955 | 0.312 | -0.003 | -0.031 | -0.031 | 0.067 | 0.25 | 0.13 | 0.28 | 0.34 | 0.33 | 0.38 |
| 3 | 818135 | rs17047823 | -0.019 | -0.03 | 0.897 | 0.436 | -0.016 | -0.047 | -0.047 | 0 | 0.1 | 0.03 | 0.06 | 0.29 | 0.14 | 0.28 |
| 3 | 834393 | rs13099887 | 0.446 | -0.919 | 1.095 | -0.128 | -0.008 | -0.025 | -0.025 | 0 | 0.15 | 0.1 | 0.05 | 0 | 0.08 | 0.17 |
| 3 | 837481 | rs1357614 | 0.718 | -0.947 | 0.789 | -0.047 | -0.028 | -0.052 | -0.052 | 0 | 0.09 | 0.03 | 0.05 | 0.24 | 0.13 | 0.26 |
| 3 | 840276 | rs17486433 | 1.756 | -0.132 | 0.247 | 0.231 | -0.032 | -0.021 | -0.021 | 0.108 | 0.35 | 0.13 | 0.18 | 0.37 | 0.3 | 0.3 |
| 3 | 843748 | rs1523903 | -0.2 | -0.371 | 0.265 | 0.158 | 0.013 | 0.116 | 0.116 | 0 | 0.32 | 0.09 | 0.36 | 0.1 | 0.33 | 0.32 |
| 3 | 845999 | rs6769721 | -2.094 | 0.049 | 0.317 | 0.183 | -0.043 | -0.005 | -0.005 | 0 | 0.65 | 0.8 | 0.76 | 0.62 | 0.64 | 0.67 |
| 3 | 850922 | rs6786909 | -0.402 | 0.194 | 0.204 | 0.195 | 0.025 | 0.116 | 0.116 | 0 | 0.33 | 0.13 | 0.38 | 0.1 | 0.33 | 0.32 |
| 3 | 857325 | rs7629783 | -0.306 | 0.197 | 0.164 | 0.185 | 0.015 | 0.128 | 0.128 | 0 | 0.32 | 0.1 | 0.37 | 0.1 | 0.33 | 0.32 |
| 3 | 857458 | rs1403919 | 0.241 | 0.199 | 0.316 | 0.273 | -0.043 | -0.094 | -0.094 | 0.141 | 0.25 | 0.11 | 0.36 | 0.52 | 0.31 | 0.33 |
| 3 | 864513 | rs11914853 | -0.362 | 0.14 | 0.372 | 0.216 | 0.024 | 0.085 | 0.085 | 0 | 0.33 | 0.17 | 0.38 | 0.09 | 0.33 | 0.32 |
| 3 | 865779 | rs13076532 | 0.558 | 0.073 | 0.339 | 0.208 | -0.012 | 0.037 | 0.037 | 0 | 0.19 | 0.1 | 0.32 | 0.09 | 0.25 | 0.15 |
| 3 | 874972 | rs13062025 | 1.062 | 0.156 | 0.261 | 0.248 | 0.017 | 0.013 | 0.013 | 0 | 0.14 | 0.09 | 0.06 | 0.01 | 0.08 | 0.18 |
| 3 | 877099 | rs1464700 | -1.866 | 0.183 | 0.61 | 0.368 | -0.018 | -0.001 | -0.001 | 0.827 | 0.63 | 0.74 | 0.75 | 0.61 | 0.63 | 0.7 |
| 3 | 877925 | rs13066227 | 0.101 | 0.196 | 0.68 | 0.429 | 0.012 | -0.007 | -0.007 | 0.186 | 0.3 | 0.2 | 0.34 | 0.3 | 0.25 | 0.25 |
| 3 | 880532 | rs17486713 | 1.743 | 0.455 | 0.808 | 0.717 | -0.073 | 0.002 | 0.002 | 0 | 0.35 | 0.15 | 0.24 | 0.36 | 0.37 | 0.37 |
| 3 | 880594 | rs10510199 | 0.858 | 0.552 | 1.095 | 1.162 | -0.008 | -0.029 | -0.029 | 6.891 | 0.16 | 0.11 | 0.29 | 0.32 | 0.17 | 0.09 |
| 3 | 883851 | rs4684585 | 0.913 | 0.586 | 1.427 | 1.137 | 0.012 | 0.004 | 0.004 | 0 | 0.56 | 0.41 | 0.57 | 0.45 | 0.62 | 0.52 |
| 3 | 885794 | rs4686226 | 0.616 | 0.712 | 0.976 | 1.142 | 0.008 | 0 | 0 | 0 | 0.59 | 0.65 | 0.59 | 0.45 | 0.61 | 0.52 |
| 3 | 892152 | rs12639429 | -0.75 | 1.599 | 1.022 | 1.917 | 0.01 | 0 | 0 | 0.01 | 0.4 | 0.26 | 0.4 | 0.56 | 0.39 | 0.48 |
| 3 | 898661 | rs35964523 | -0.854 | 0.502 | 0.463 | 1.148 | 0.023 | 0.003 | 0.003 | 0 | 0.43 | 0.29 | 0.41 | 0.55 | 0.38 | 0.48 |
| 3 | 915389 | rs1464701 | 0.246 | 0.307 | -0.033 | 0.793 | -0.032 | -0.008 | -0.008 | 0 | 0.74 | 0.66 | 0.87 | 0.77 | 0.81 | 0.6 |
| 3 | 927360 | rs6788253 | 0.427 | 0.595 | -1.027 | 0.537 | 0.017 | 0.068 | 0.068 | 0 | 0.4 | 0.2 | 0.28 | 0.26 | 0.33 | 0.31 |
| 3 | 932212 | rs13090889 | -0.038 | 0.218 | -0.992 | 0.124 | 0.013 | 0.127 | 0.127 | 0 | 0.45 | 0.19 | 0.64 | 0.78 | 0.61 | 0.65 |
| 3 | 940698 | rs2313182 | 0.888 | 1.044 | -1.282 | 0.904 | -0.045 | -0.073 | -0.073 | 1.089 | 0.25 | 0.12 | 0.4 | 0.33 | 0.32 | 0.27 |
| 3 | 941639 | rs9830614 | -1.438 | 0.364 | -1.08 | 0.597 | 0.013 | 0.087 | 0.087 | 0 | 0.58 | 0.79 | 0.7 | 0.8 | 0.63 | 0.74 |
| 3 | 942672 | rs17049673 | -1.064 | 0.304 | -0.282 | 1.035 | 0.002 | -0.003 | -0.003 | 0.624 | 0.18 | 0.14 | 0.11 | 0.27 | 0.09 | 0.34 |
| 3 | 949345 | rs9812345 | -0.528 | 0.342 | 0.101 | 0.611 | -0.04 | -0.065 | -0.065 | 0.755 | 0.12 | 0.18 | 0.03 | 0.01 | 0.03 | 0.1 |
| 3 | 958096 | rs9838891 | -1.194 | 1.36 | -0.131 | 0.942 | -0.005 | 0.013 | 0.013 | 0 | 0.26 | 0.19 | 0.19 | 0.21 | 0.26 | 0.2 |
| 3 | 963381 | rs1357616 | 0.805 | 1.184 | 0.306 | 0.944 | 0.004 | -0.007 | -0.007 | 1.03 | 0.75 | 0.76 | 0.82 | 0.8 | 0.71 | 0.79 |
| 3 | 967323 | rs1464705 | 1.205 | 1.217 | -0.323 | 1.211 | -0.086 | -0.052 | -0.052 | 0 | 0.23 | 0.02 | 0.41 | 0.22 | 0.29 | 0.2 |
| 3 | 970300 | rs4684618 | 0.416 | 1.111 | -0.09 | 1.1 | -0.001 | -0.02 | -0.02 | 0 | 0.87 | 0.89 | 0.95 | 0.99 | 0.93 | 0.87 |
| 3 | 975836 | rs11920873 | 1.21 | 1.54 | 0.215 | 1.6 | -0.007 | 0.01 | 0.01 | 0 | 0.72 | 0.75 | 0.81 | 0.78 | 0.69 | 0.75 |
| 3 | 977968 | rs17435539 | 1.228 | 1.382 | 0.551 | 1.603 | 0.003 | -0.004 | -0.004 | 0 | 0.43 | 0.3 | 0.32 | 0.39 | 0.39 | 0.34 |
| 3 | 979343 | rs4280626 | -0.208 | 1.326 | 0.466 | 1.559 | -0.003 | -0.024 | -0.024 | 0 | 0.9 | 0.91 | 0.97 | 0.99 | 0.95 | 0.91 |
| 3 | 983297 | rs4686323 | 0.794 | 1.408 | 0.802 | 1.709 | 0.002 | -0.003 | -0.003 | 0 | 0.73 | 0.7 | 0.79 | 0.76 | 0.69 | 0.76 |
| 3 | 984435 | rs11713167 | 1.462 | 1.402 | 0.678 | 1.769 | 0.022 | 0.072 | 0.072 | 0.01 | 0.39 | 0.21 | 0.24 | 0.09 | 0.3 | 0.13 |
| 3 | 985070 | rs13092875 | 1.362 | 1.373 | 0.638 | 1.874 | -0.077 | -0.039 | -0.039 | 0 | 0.25 | 0.02 | 0.46 | 0.22 | 0.31 | 0.23 |
| 3 | 986199 | rs4452318 | 1.408 | 1.216 | 0.618 | 1.63 | -0.002 | -0.013 | -0.013 | 2.688 | 0.39 | 0.22 | 0.28 | 0.38 | 0.37 | 0.31 |
| 3 | 986917 | rs4686326 | 0.635 | 1.428 | 0.98 | 1.681 | -0.005 | -0.002 | -0.002 | 0 | 0.31 | 0.28 | 0.52 | 0.68 | 0.39 | 0.64 |
| 3 | 991967 | rs10510202 | 0.283 | 0.937 | 0.893 | 1.354 | 0.001 | -0.002 | -0.002 | 8.452 | 0.39 | 0.4 | 0.48 | 0.46 | 0.35 | 0.39 |
| 3 | 992289 | rs11914673 | 0.204 | 0.895 | 1.073 | 0.552 | 0.011 | 0.018 | 0.018 | 0 | 0.49 | 0.39 | 0.5 | 0.69 | 0.39 | 0.71 |
| 3 | 994203 | rs6771833 | 0.677 | 0.605 | 0.528 | 0.635 | 0.009 | 0.153 | 0.153 | 0 | 0.36 | 0.12 | 0.23 | 0.08 | 0.31 | 0.16 |
| 3 | 994754 | rs4303857 | 0.191 | 0.748 | 0.792 | 0.647 | 0.083 | 0.087 | 0.087 | 0 | 0.85 | 0.64 | 0.79 | 0.74 | 0.76 | 0.9 |
| 3 | 995334 | rs12629825 | 0.265 | 0.883 | 0.26 | 0.725 | -0.022 | -0.023 | -0.023 | 0.089 | 0.12 | 0.09 | 0.18 | 0.25 | 0.23 | 0.1 |
| 3 | 995511 | rs12630927 | -0.948 | 0.807 | 0.214 | 0.617 | -0.021 | 0.003 | 0.003 | 0 | 0.21 | 0.16 | 0.2 | 0.24 | 0.29 | 0.2 |
| 3 | 996315 | rs7629228 | -0.661 | 0.704 | 0.333 | 0.651 | -0.007 | -0.015 | -0.015 | 0 | 0.1 | 0.15 | 0.05 | 0.03 | 0.06 | 0.08 |
| 3 | 998513 | rs11131168 | 0.625 | 0.722 | 0.09 | 0.752 | -0.005 | 0.007 | 0.007 | 0 | 0.39 | 0.31 | 0.5 | 0.65 | 0.41 | 0.66 |

| | | | | | | | | | | | | | | | | |
|---|---|---|---|---|---|---|---|---|---|---|---|---|---|---|---|---|
| 3 | 1005874 | rs12634464 | -1.027 | 0.815 | 0.243 | 1.29 | -0.008 | 0.021 | 0.021 | 0 | 0.24 | 0.22 | 0.17 | 0.17 | 0.27 | 0.22 |
| 3 | 1014666 | rs4594610 | -0.878 | 0.461 | 0.854 | 2.335 | -0.006 | 0.042 | 0.042 | 0 | 0.1 | 0.17 | 0.35 | 0.25 | 0.08 | 0.29 |
| 3 | 1016458 | rs9855434 | -1.158 | 0.438 | 1.137 | 2.354 | -0.01 | 0.122 | 0.122 | 0 | 0.11 | 0.27 | 0.36 | 0.26 | 0.08 | 0.29 |
| 3 | 1020877 | rs6781767 | 2.084 | 0.53 | 1.779 | 2.989 | 0.001 | 0.29 | 0.29 | 2.158 | 0.88 | 0.57 | 0.84 | 0.57 | 0.92 | 0.77 |
| 3 | 1030428 | rs7641637 | 1.914 | 0.82 | 2.068 | 3.29 | 0.052 | 0.372 | 0.372 | 1.852 | 0.91 | 0.64 | 0.84 | 0.56 | 0.92 | 0.81 |
| 3 | 1033790 | rs4401351 | 2.287 | 0.804 | 2.137 | 3.282 | 0.056 | 0.408 | 0.408 | 3.179 | 0.91 | 0.56 | 0.84 | 0.56 | 0.92 | 0.81 |
| 3 | 1034188 | rs9312040 | 2.226 | 0.82 | 2.26 | 3.465 | 0.054 | 0.392 | 0.392 | 2.891 | 0.91 | 0.58 | 0.84 | 0.57 | 0.92 | 0.82 |
| 3 | 1035503 | rs4686368 | 2.286 | 0.487 | 1.86 | 2.686 | 0.046 | 0.408 | 0.408 | 3.179 | 0.91 | 0.56 | 0.84 | 0.56 | 0.92 | 0.81 |
| 3 | 1036877 | rs7644439 | 1.919 | 0.557 | 1.576 | 2.714 | 0.04 | 0.343 | 0.343 | 1.852 | 0.91 | 0.65 | 0.84 | 0.56 | 0.92 | 0.81 |
| 3 | 1051760 | rs4234583 | 1.786 | 0.642 | 1.325 | 2.603 | 0.011 | 0.183 | 0.183 | 0.111 | 0.91 | 0.83 | 0.83 | 0.59 | 0.91 | 0.78 |
| 3 | 1059654 | rs4266176 | 1.45 | 0.675 | 1.314 | 2.49 | 0.023 | 0.215 | 0.215 | 0 | 0.93 | 0.85 | 0.84 | 0.57 | 0.9 | 0.83 |
| 3 | 1061761 | rs4686384 | -1.249 | 0.684 | 1.256 | 1.724 | 0.001 | 0.004 | 0.004 | 0 | 0.5 | 0.5 | 0.55 | 0.74 | 0.59 | 0.74 |
| 3 | 1063826 | rs6793028 | -1.435 | 0.711 | 1.327 | 1.677 | 0.016 | 0.087 | 0.087 | 0 | 0.52 | 0.66 | 0.57 | 0.77 | 0.59 | 0.75 |
| 3 | 1070039 | rs17032482 | -0.351 | 0.796 | 1.515 | 1.464 | 0.009 | -0.071 | -0.071 | 0 | 0.34 | 0.37 | 0.26 | 0.09 | 0.42 | 0.27 |
| 3 | 1079024 | rs7631088 | -0.937 | 0.796 | 1.181 | 1.789 | 0.006 | 0.007 | 0.007 | 0 | 0.41 | 0.53 | 0.43 | 0.5 | 0.51 | 0.42 |
| 3 | 1082282 | rs12634679 | -1.363 | 0.54 | 0.815 | 1.267 | 0.047 | 0.118 | 0.118 | 0 | 0.51 | 0.69 | 0.56 | 0.77 | 0.59 | 0.75 |
| 3 | 1090602 | rs1504058 | -0.123 | 1.424 | 0.436 | 1.081 | -0.003 | 0.001 | 0.001 | 0 | 0.26 | 0.26 | 0.22 | 0.13 | 0.4 | 0.19 |
| 3 | 1092494 | rs12496517 | -1.761 | 0.828 | 0.87 | 1.871 | 0.036 | 0.244 | 0.244 | 1.697 | 0.07 | 0.2 | 0.16 | 0.41 | 0.08 | 0.15 |
| 3 | 1096770 | rs6442164 | -1.307 | 0.51 | 1.09 | 1.627 | 0.017 | 0.069 | 0.069 | 2.533 | 0.11 | 0.49 | 0.1 | 0.11 | 0.07 | 0.08 |
| 3 | 1106163 | rs6442167 | -0.705 | 0.653 | -0.118 | 1.412 | 0.006 | -0.035 | -0.035 | 0 | 0.23 | 0.13 | 0.4 | 0.31 | 0.22 | 0.42 |
| 3 | 1117233 | rs2729117 | 0.678 | -0.009 | 0.54 | 1.035 | -0.017 | 0.017 | 0.017 | 0 | 0.65 | 0.57 | 0.58 | 0.48 | 0.72 | 0.46 |
| 3 | 1118365 | rs2047887 | -0.472 | -0.643 | 0.717 | 0.493 | -0.028 | -0.03 | -0.03 | 0.003 | 0.18 | 0.26 | 0.11 | 0.07 | 0.09 | 0.13 |
| 3 | 1122000 | rs11719195 | 0.448 | 0.609 | 0.792 | 0.44 | 0.004 | -0.014 | -0.014 | 0.063 | 0.09 | 0.09 | 0.1 | 0.01 | 0.11 | 0.05 |
| 3 | 1122130 | rs2729124 | 0.551 | 0.114 | 1.476 | 0.944 | -0.01 | 0.02 | 0.02 | 0 | 0.72 | 0.63 | 0.63 | 0.57 | 0.74 | 0.51 |
| 3 | 1122456 | rs2729125 | 0.571 | 0.34 | 1.612 | 1.009 | 0.061 | 0.02 | 0.02 | 0 | 0.93 | 0.76 | 0.92 | 0.9 | 0.86 | 0.96 |
| 3 | 1125605 | rs2729130 | 1.014 | 0.46 | 1.741 | 1.126 | 0.086 | 0.332 | 0.332 | 0.402 | 0.9 | 0.63 | 0.75 | 0.6 | 0.89 | 0.8 |
| 3 | 1125688 | rs2729131 | 1.701 | 0.402 | 1.602 | 1.065 | 0.049 | 0.254 | 0.254 | 0.629 | 0.87 | 0.58 | 0.73 | 0.6 | 0.87 | 0.79 |
| 3 | 1127033 | rs2655198 | 0.492 | -0.301 | 1.073 | 0.335 | -0.004 | 0.006 | 0.006 | 0 | 0.87 | 0.86 | 0.89 | 0.9 | 0.82 | 0.92 |
| 3 | 1128074 | rs2729139 | 1.267 | -0.428 | 0.687 | 0.358 | 0.032 | -0.008 | -0.008 | 0 | 0.57 | 0.31 | 0.53 | 0.51 | 0.61 | 0.44 |
| 3 | 1128245 | rs17033401 | -0.481 | -0.82 | 0.726 | 0.869 | 0.035 | -0.042 | -0.042 | 0 | 0.19 | 0.12 | 0.27 | 0.38 | 0.12 | 0.42 |
| 3 | 1134445 | rs6807118 | -2.081 | 0.026 | 2.402 | 0.273 | -0.003 | -0.043 | -0.043 | 0 | 0.1 | 0.58 | 0.07 | 0 | 0.06 | 0.07 |
| 3 | 1139641 | rs9881241 | -2.244 | -0.1 | 2.294 | 0.21 | 0.005 | -0.041 | -0.041 | 0.037 | 0.09 | 0.53 | 0.06 | 0 | 0.06 | 0.07 |
| 3 | 1143424 | rs9878022 | -2.434 | 0.253 | 1.85 | 0.212 | -0.008 | -0.048 | -0.048 | 0.01 | 0.11 | 0.77 | 0.09 | 0 | 0.09 | 0.08 |
| 3 | 1149544 | rs6442206 | -2.631 | 0.511 | 1.788 | 0.179 | -0.003 | -0.028 | -0.028 | 0 | 0.12 | 0.78 | 0.1 | 0 | 0.09 | 0.07 |
| 3 | 1154559 | rs1504076 | 1.711 | 0.578 | 1.919 | 0.203 | 0.034 | -0.033 | -0.033 | 0.333 | 0.93 | 0.8 | 0.94 | 1 | 0.91 | 0.93 |
| 3 | 1159787 | rs13069565 | 1.209 | 0.604 | 1.06 | 0.518 | -0.001 | -0.002 | -0.002 | 1.887 | 0.66 | 0.34 | 0.54 | 0.62 | 0.64 | 0.59 |
| 3 | 1162041 | rs2872338 | 1.289 | 0.788 | 1.468 | 0.406 | 0.079 | -0.009 | -0.009 | 0.467 | 0.93 | 0.71 | 0.93 | 1 | 0.91 | 0.93 |
| 3 | 1164309 | rs11128538 | 0.57 | 0.172 | 0.855 | 0.424 | 0 | 0 | 0 | 0.774 | 0.65 | 0.59 | 0.77 | 0.72 | 0.69 | 0.64 |
| 3 | 1164339 | rs6782048 | 0.904 | 1.136 | 0.486 | 0.701 | 0.011 | 0.049 | 0.049 | 0 | 0.54 | 0.6 | 0.54 | 0.3 | 0.44 | 0.37 |
| 3 | 1165633 | rs996424 | 0.385 | 1.523 | -0.307 | 0.927 | -0.09 | -0.138 | -0.138 | 0 | 0.66 | 0.95 | 0.56 | 0.48 | 0.63 | 0.48 |
| 3 | 1169496 | rs4324457 | 1.037 | 0.992 | 0.049 | 0.038 | 0.002 | 0.012 | 0.012 | 0 | 0.4 | 0.51 | 0.4 | 0.48 | 0.48 | 0.35 |
| 3 | 1170794 | rs7646994 | 1.202 | 0.944 | 0.543 | 0.088 | 0.002 | 0.02 | 0.02 | 0.001 | 0.38 | 0.48 | 0.38 | 0.46 | 0.45 | 0.31 |
| 3 | 1173131 | rs2130276 | 0.623 | 0.478 | 0.493 | -0.246 | -0.001 | -0.028 | -0.028 | 0.912 | 0.14 | 0.12 | 0.06 | 0 | 0.08 | 0.11 |
| 3 | 1175539 | rs4481144 | -0.441 | 0.486 | 0.409 | -0.296 | -0.04 | -0.082 | -0.082 | 0 | 0.67 | 0.9 | 0.55 | 0.48 | 0.65 | 0.51 |
| 3 | 1175575 | rs4481145 | 0.98 | 0.427 | 0.657 | 0.043 | 0 | 0.003 | 0.003 | 0 | 0.5 | 0.48 | 0.42 | 0.39 | 0.53 | 0.41 |
| 3 | 1175950 | rs7620637 | -0.142 | 0.387 | 0.638 | 0.055 | -0.006 | -0.02 | -0.02 | 0.881 | 0.65 | 0.74 | 0.54 | 0.48 | 0.65 | 0.5 |
| 3 | 1183679 | rs1032669 | -1.618 | 0.873 | 0.231 | 0.239 | -0.029 | -0.043 | -0.043 | 0 | 0.62 | 0.79 | 0.63 | 0.52 | 0.52 | 0.68 |
| 3 | 1186687 | rs902118 | -1.498 | 0.913 | 0.542 | 0.253 | -0.005 | -0.007 | -0.007 | 0 | 0.61 | 0.7 | 0.61 | 0.51 | 0.5 | 0.68 |
| 3 | 1187510 | rs1504073 | -0.62 | 1.086 | 0.433 | 0.262 | 0.04 | 0.001 | 0.001 | 0 | 0.43 | 0.41 | 0.47 | 0.47 | 0.32 | 0.59 |
| 3 | 1191305 | rs3772374 | 0.206 | 1.195 | 0.586 | 0.844 | 0.014 | 0.183 | 0.183 | 0.193 | 0.05 | 0.15 | 0.13 | 0.28 | 0.08 | 0.1 |
| 3 | 1191638 | rs17034364 | -0.773 | 1.349 | 0.87 | 0.545 | 0.058 | -0.046 | -0.046 | 0.689 | 0.13 | 0.36 | 0.13 | 0.03 | 0.14 | 0.04 |
| 3 | 1192619 | rs2036236 | 1.739 | 0.975 | 0.247 | 0.188 | -0.015 | -0.011 | -0.011 | 0 | 0.57 | 0.32 | 0.38 | 0.46 | 0.5 | 0.49 |
| 3 | 1192829 | rs1846466 | -1.318 | 0.596 | 0.759 | 0.453 | 0.092 | -0.034 | -0.034 | 0 | 0.15 | 0.57 | 0.15 | 0.06 | 0.16 | 0.05 |
| 3 | 1197281 | rs17034421 | -0.654 | 0.509 | 0.714 | 0.751 | 0.08 | 0.387 | 0.387 | 0.95 | 0.07 | 0.36 | 0.12 | 0.3 | 0.1 | 0.13 |
| 3 | 1202295 | rs11921911 | -0.325 | 0.34 | 0.616 | 0.699 | 0.064 | 0.275 | 0.275 | 0.055 | 0.08 | 0.3 | 0.15 | 0.3 | 0.14 | 0.14 |
| 3 | 1204591 | rs3772371 | -0.372 | 0.328 | 0.604 | 0.687 | 0.049 | 0.201 | 0.201 | 0.152 | 0.08 | 0.29 | 0.14 | 0.25 | 0.14 | 0.14 |
| 3 | 1206696 | rs3772369 | -1.926 | -0.222 | -0.428 | 0.608 | -0.058 | 0.055 | 0.055 | 0.118 | 0.57 | 0.8 | 0.68 | 0.72 | 0.46 | 0.67 |
| 3 | 1207591 | rs1391920 | -0.336 | -0.335 | -0.403 | 0.576 | 0.055 | -0.032 | -0.032 | 2.503 | 0.35 | 0.15 | 0.43 | 0.43 | 0.19 | 0.47 |
| 3 | 1215013 | rs7632166 | -2.163 | -0.17 | -0.564 | 0.4 | -0.006 | 0.008 | 0.008 | 0 | 0.42 | 0.63 | 0.63 | 0.69 | 0.47 | 0.61 |
| 3 | 1218279 | rs9823491 | -2.081 | -0.201 | -1.39 | 0.412 | -0.084 | 0.045 | 0.045 | 0.008 | 0.6 | 0.88 | 0.69 | 0.75 | 0.51 | 0.72 |
| 3 | 1218873 | rs1499100 | -1.147 | -0.351 | -1.279 | 0.464 | 0.009 | 0.064 | 0.064 | 0.41 | 0.47 | 0.87 | 0.62 | 0.69 | 0.37 | 0.63 |
| 3 | 1220546 | rs12106966 | 0.552 | -0.343 | -1.178 | 0.73 | 0.004 | -0.02 | -0.02 | 0.746 | 0.1 | 0.06 | 0.32 | 0.44 | 0.07 | 0.33 |
| 3 | 1222180 | rs6806307 | -1.163 | -0.308 | -0.949 | 0.637 | 0.01 | 0.004 | 0.004 | 0 | 0.45 | 0.37 | 0.51 | 0.45 | 0.34 | 0.52 |
| 3 | 1226953 | rs6772304 | 0.92 | 0.039 | -1.316 | 0.922 | 0.017 | 0.008 | 0.008 | 0.399 | 0.08 | 0.04 | 0.32 | 0.47 | 0.04 | 0.32 |
| 3 | 1227382 | rs1499097 | -0.817 | 0.081 | -1.584 | 0.849 | -0.055 | -0.087 | -0.087 | 0 | 0.17 | 0.02 | 0.17 | 0.24 | 0.25 | 0.21 |
| 3 | 1227622 | rs1846463 | -1.747 | -0.069 | -1.986 | 0.382 | -0.028 | -0.059 | -0.059 | 0 | 0.55 | 0.91 | 0.37 | 0.41 | 0.46 | 0.46 |
| 3 | 1228607 | rs3772360 | -0.807 | 0.07 | -1.735 | 0.61 | -0.088 | -0.135 | -0.135 | 0.372 | 0.21 | 0.07 | 0.45 | 0.5 | 0.26 | 0.38 |
| 3 | 1230424 | rs6442271 | 0.411 | -0.733 | -0.678 | 0.105 | -0.111 | -0.157 | -0.157 | 0 | 0.24 | 0.5 | 0.07 | 0 | 0.04 | 0.12 |
| 3 | 1236435 | rs2886164 | -1.255 | -0.432 | -0.842 | 0.184 | 0.1 | 0.189 | 0.189 | 0.022 | 0.49 | 0.78 | 0.53 | 0.85 | 0.36 | 0.75 |

| | | | | | | | | | | | | | | | | |
|---|---|---|---|---|---|---|---|---|---|---|---|---|---|---|---|---|
| 3 | 1238287 | rs1353821 | -0.246 | -0.412 | -0.431 | 0.671 | -0.012 | 0 | 0 | 0 | 0.31 | 0.64 | 0.21 | 0.3 | 0.18 | 0.3 |
| 3 | 1238954 | rs17440919 | 0.296 | 0.29 | -0.341 | 0.63 | 0.067 | 0 | 0 | 0 | 0.06 | 0.12 | 0.08 | 0.03 | 0.13 | 0.03 |
| 3 | 1247015 | rs11128578 | -1.06 | -0.01 | -0.269 | 0.591 | 0 | 0.007 | 0.007 | 0 | 0.43 | 0.57 | 0.31 | 0.32 | 0.39 | 0.35 |
| 3 | 1247669 | rs11919828 | -0.967 | -0.03 | 0.19 | 0.504 | -0.001 | -0.004 | -0.004 | 0.594 | 0.42 | 0.42 | 0.29 | 0.3 | 0.39 | 0.35 |
| 3 | 1250310 | rs10510165 | -0.568 | -0.158 | 0.2 | 0.417 | -0.004 | -0.01 | -0.01 | 1.324 | 0.36 | 0.55 | 0.26 | 0.28 | 0.31 | 0.34 |
| 3 | 1250356 | rs17035484 | -0.564 | -0.361 | 0.2 | 0.361 | -0.005 | 0.107 | 0.107 | 0 | 0.12 | 0.33 | 0.11 | 0.25 | 0.12 | 0.19 |
| 3 | 1250634 | rs17035505 | 1.042 | -0.313 | 0.104 | 0.324 | 0.006 | -0.016 | -0.016 | 1.044 | 0.19 | 0.14 | 0.06 | 0 | 0.07 | 0.13 |
| 3 | 1253821 | rs17035611 | 1.105 | -0.178 | -0.107 | 0.621 | 0.01 | 0.01 | 0.01 | 0 | 0.08 | 0.04 | 0.31 | 0.5 | 0.06 | 0.33 |
| 3 | 1256852 | rs1499102 | -0.603 | -0.114 | 0.06 | 0.156 | 0 | -0.027 | -0.027 | 2.075 | 0.35 | 0.62 | 0.22 | 0.05 | 0.29 | 0.18 |
| 3 | 1257668 | rs7644569 | -0.519 | -0.363 | -0.047 | 0.103 | 0.046 | 0.036 | 0.036 | 0.016 | 0.33 | 0.75 | 0.17 | 0.01 | 0.17 | 0.16 |
| 3 | 1260932 | rs6777489 | -0.119 | -0.184 | 0.132 | 0.214 | 0.009 | -0.049 | -0.049 | 0 | 0.11 | 0.28 | 0.08 | 0.01 | 0.1 | 0.03 |
| 3 | 1262474 | rs2013828 | -1.016 | -0.245 | 0.162 | 0.563 | -0.008 | 0.002 | 0.002 | 0 | 0.43 | 0.47 | 0.48 | 0.51 | 0.34 | 0.56 |
| 3 | 1270698 | rs3796176 | -0.826 | -0.272 | -0.167 | 0.124 | 0.014 | 0.056 | 0.056 | 1.332 | 0.4 | 0.64 | 0.24 | 0.04 | 0.3 | 0.23 |
| 3 | 1272373 | rs9844484 | -0.986 | -0.075 | -0.061 | 0.461 | -0.006 | 0 | 0 | 0 | 0.43 | 0.52 | 0.39 | 0.51 | 0.25 | 0.57 |
| 3 | 1273621 | rs749993 | -1.019 | 0.326 | 0.178 | 0.843 | 0.032 | -0.017 | -0.017 | 0 | 0.14 | 0.19 | 0.13 | 0.04 | 0.21 | 0.1 |
| 3 | 1274590 | rs3772341 | 0.798 | 0.256 | -0.011 | 0.965 | 0.011 | 0.015 | 0.015 | 0 | 0.08 | 0.04 | 0.09 | 0.47 | 0.06 | 0.36 |
| 3 | 1276092 | rs11925058 | -1.365 | 0.504 | 0.069 | 0.951 | 0.049 | 0.071 | 0.071 | 0 | 0.3 | 0.48 | 0.34 | 0.45 | 0.36 | 0.51 |
| 3 | 1286804 | rs13315256 | -1.966 | 0.193 | -0.159 | 0.443 | -0.004 | -0.087 | -0.087 | 0 | 0.44 | 0.51 | 0.62 | 0.91 | 0.42 | 0.68 |
| 3 | 1289346 | rs3772340 | -1.642 | 0.137 | 0.011 | 1.069 | -0.007 | 0.006 | 0.006 | 0.006 | 0.33 | 0.44 | 0.39 | 0.59 | 0.25 | 0.5 |
| 3 | 1289452 | rs17036322 | 0.387 | 0.199 | -0.024 | 1.063 | 0.002 | -0.005 | -0.005 | 0 | 0.06 | 0.04 | 0.18 | 0.16 | 0.05 | 0.03 |
| 3 | 1289570 | rs9844920 | -1.199 | 0.167 | 0.009 | 1.072 | -0.001 | -0.003 | -0.003 | 0 | 0.17 | 0.25 | 0.14 | 0.12 | 0.16 | 0.11 |
| 3 | 1291744 | rs6794274 | -0.907 | 0.135 | -0.043 | 1.131 | -0.001 | 0.009 | 0.009 | 0 | 0.1 | 0.11 | 0.23 | 0.26 | 0.11 | 0.05 |
| 3 | 1293342 | rs9837484 | -1.425 | 0.111 | 0.062 | 1.08 | -0.02 | 0.05 | 0.05 | 0 | 0.25 | 0.39 | 0.17 | 0.43 | 0.17 | 0.44 |
| 3 | 1293401 | rs3913305 | -0.537 | 0.092 | -0.243 | 0.858 | 0 | -0.002 | -0.002 | 0 | 0.4 | 0.37 | 0.58 | 0.3 | 0.45 | 0.41 |
| 3 | 1293647 | rs3772337 | -0.967 | 0.089 | -0.458 | 0.836 | -0.002 | 0.002 | 0.002 | 0 | 0.14 | 0.11 | 0.09 | 0.12 | 0.16 | 0.11 |
| 3 | 1294736 | rs3772336 | 0.478 | 0.059 | -0.443 | 1.043 | 0.005 | 0.006 | 0.006 | 0 | 0.1 | 0.05 | 0.07 | 0.31 | 0.05 | 0.36 |
| 3 | 1296579 | rs17036648 | 0.516 | 0.045 | -0.431 | 0.958 | 0.008 | -0.008 | -0.008 | 0 | 0.1 | 0.05 | 0.07 | 0.3 | 0.05 | 0.34 |
| 3 | 1297263 | rs4082640 | -1.651 | -0.05 | -0.255 | 0.804 | -0.004 | 0.021 | 0.021 | 0 | 0.29 | 0.33 | 0.22 | 0.43 | 0.24 | 0.45 |
| 3 | 1298121 | rs3864028 | -1.079 | -0.045 | -0.197 | 0.819 | -0.012 | 0.092 | 0.092 | 0 | 0.2 | 0.35 | 0.18 | 0.41 | 0.16 | 0.39 |
| 3 | 1299077 | rs17036749 | -1.126 | 0.077 | -0.034 | 0.188 | 0.014 | 0.021 | 0.021 | 0.608 | 0.11 | 0.36 | 0.11 | 0.11 | 0.11 | 0.06 |
| 3 | 1299571 | rs3772334 | 0.087 | 0.099 | -0.363 | -0.227 | 0.001 | 0.042 | 0.042 | 0.165 | 0.41 | 0.24 | 0.58 | 0.7 | 0.39 | 0.64 |
| 3 | 1300820 | rs10510172 | 0.363 | -0.096 | -0.453 | 0.065 | -0.02 | 0 | 0 | 0 | 0.1 | 0 | 0.05 | 0.04 | 0.08 | 0.07 |
| 3 | 1301453 | rs12489745 | -0.952 | -0.123 | -0.246 | 0.044 | 0.002 | 0.007 | 0.007 | 0.575 | 0.11 | 0.25 | 0.11 | 0.1 | 0.11 | 0.06 |
| 3 | 1302694 | rs3872628 | -1.36 | -0.105 | -0.264 | 0.046 | -0.001 | -0.003 | -0.003 | 0 | 0.21 | 0.27 | 0.17 | 0.14 | 0.19 | 0.13 |
| 3 | 1310741 | rs9872414 | -0.51 | -0.166 | -0.44 | 0.019 | 0.025 | -0.012 | -0.012 | 0.066 | 0.43 | 0.71 | 0.4 | 0.51 | 0.33 | 0.4 |
| 3 | 1310929 | rs4085434 | -1.087 | -0.083 | -0.577 | -0.092 | -0.007 | 0.128 | 0.128 | 0 | 0.55 | 0.78 | 0.68 | 0.83 | 0.46 | 0.73 |
| 3 | 1313477 | rs3772324 | 0.463 | -0.175 | -0.283 | -0.089 | -0.043 | 0.075 | 0.075 | 0 | 0.31 | 0.52 | 0.29 | 0.51 | 0.18 | 0.27 |
| 3 | 1314793 | rs3772320 | -0.708 | -0.152 | -0.93 | -0.341 | 0.077 | 0.242 | 0.242 | 0 | 0.5 | 0.88 | 0.64 | 0.83 | 0.37 | 0.64 |
| 3 | 1315199 | rs10510169 | 0.412 | 0.036 | -0.971 | -0.384 | -0.025 | 0.019 | 0.019 | 0 | 0.11 | 0.02 | 0.09 | 0.04 | 0.12 | 0.09 |
| 3 | 1315900 | rs3772318 | -2.029 | 0.118 | -0.994 | -0.445 | -0.068 | 0.117 | 0.117 | 0 | 0.66 | 0.89 | 0.73 | 0.87 | 0.54 | 0.8 |
| 3 | 1316860 | rs7647469 | 0.062 | -0.462 | -0.538 | -0.368 | -0.005 | 0 | 0 | 0 | 0.58 | 0.48 | 0.65 | 0.47 | 0.71 | 0.68 |
| 3 | 1316884 | rs10510189 | -0.145 | -0.389 | -1.214 | -0.524 | 0.004 | 0.042 | 0.042 | 0 | 0.09 | 0.02 | 0.06 | 0 | 0.07 | 0.1 |
| 3 | 1317289 | rs12374025 | 0.724 | -0.39 | -1.109 | -0.562 | 0.046 | 0.145 | 0.145 | 0 | 0.47 | 0.23 | 0.39 | 0.17 | 0.63 | 0.3 |
| 3 | 1317710 | rs4286412 | 0.355 | -0.39 | -1.084 | -0.53 | 0.092 | 0.203 | 0.203 | 0 | 0.51 | 0.18 | 0.4 | 0.17 | 0.68 | 0.37 |
| 3 | 1317869 | rs3821435 | 1.106 | -0.299 | -1.073 | -0.503 | -0.041 | 0.115 | 0.115 | 0 | 0.4 | 0.22 | 0.32 | 0.14 | 0.56 | 0.27 |
| 3 | 1319496 | rs7646412 | -0.41 | -0.842 | -1.439 | -0.208 | 0.068 | -0.103 | -0.103 | 0.276 | 0.7 | 0.93 | 0.73 | 0.51 | 0.84 | 0.75 |
| 3 | 1319698 | rs6442316 | -0.924 | -0.899 | -1.456 | -0.239 | 0.064 | -0.128 | -0.128 | 0 | 0.74 | 0.95 | 0.78 | 0.52 | 0.86 | 0.77 |
| 3 | 1319738 | rs6442317 | -0.164 | -0.7 | -1.517 | -0.274 | 0.066 | -0.103 | -0.103 | 0 | 0.66 | 0.92 | 0.71 | 0.51 | 0.78 | 0.67 |
| 3 | 1320271 | rs3732517 | 0.033 | -0.508 | -1.212 | -0.676 | 0.044 | 0.064 | 0.064 | 0 | 0.56 | 0.7 | 0.45 | 0.17 | 0.72 | 0.36 |
| 3 | 1321825 | rs17037117 | -0.433 | 0.112 | -0.659 | -0.408 | -0.006 | -0.063 | -0.063 | 0 | 0.16 | 0.61 | 0.1 | 0.03 | 0.14 | 0.14 |
| 3 | 1322978 | rs9815996 | -0.2 | -0.167 | -1.27 | -0.17 | 0.007 | -0.01 | -0.01 | 2.758 | 0.21 | 0.18 | 0.14 | 0.28 | 0.16 | 0.38 |
| 3 | 1323078 | rs6806144 | 0.581 | 1.109 | -1.376 | -1.201 | -0.013 | 0.014 | 0.014 | 0 | 0.3 | 0.3 | 0.28 | 0.12 | 0.42 | 0.16 |
| 3 | 1323929 | rs6798638 | -0.11 | 0.96 | -1.678 | -1.239 | 0.021 | 0.11 | 0.11 | 0 | 0.61 | 0.86 | 0.79 | 0.84 | 0.67 | 0.71 |
| 3 | 1324048 | rs6786142 | 0.265 | 1.17 | -1.577 | -1.024 | -0.004 | 0.049 | 0.049 | 0 | 0.54 | 0.74 | 0.63 | 0.67 | 0.55 | 0.59 |
| 3 | 1325955 | rs4684862 | -0.328 | 0.685 | -1.492 | -1.069 | 0.069 | 0.09 | 0.09 | 0 | 0.66 | 0.87 | 0.81 | 0.82 | 0.8 | 0.72 |
| 3 | 1327522 | rs6763212 | -0.975 | 0.671 | -1.42 | -1.075 | 0.007 | -0.002 | -0.002 | 0 | 0.76 | 0.78 | 0.94 | 0.97 | 0.88 | 0.86 |
| 3 | 1329953 | rs17037184 | 0.316 | 0.953 | -1.107 | -0.785 | 0 | 0.015 | 0.015 | 0 | 0.05 | 0.07 | 0.14 | 0.17 | 0.08 | 0.12 |
| 3 | 1331135 | rs3772313 | 0.439 | 1.213 | -0.974 | -1.098 | 0 | 0.004 | 0.004 | 0 | 0.47 | 0.48 | 0.57 | 0.67 | 0.48 | 0.6 |
| 3 | 1333797 | rs174489 | 0.007 | 0.826 | -0.521 | -1.357 | 0.045 | 0.085 | 0.085 | 0 | 0.6 | 0.72 | 0.8 | 0.89 | 0.73 | 0.75 |
| 3 | 1334530 | rs10510207 | 0.337 | 1.245 | -0.344 | -1.422 | 0.024 | 0.025 | 0.025 | 0.127 | 0.06 | 0.16 | 0.06 | 0.11 | 0.07 | 0.03 |
| 3 | 1334767 | rs262806 | -0.133 | 1.315 | -0.445 | -1.565 | 0.03 | 0.106 | 0.106 | 0 | 0.61 | 0.75 | 0.81 | 0.93 | 0.72 | 0.79 |
| 3 | 1335476 | rs17271469 | -0.707 | 1.74 | -0.332 | -1.529 | -0.005 | -0.001 | -0.001 | 0 | 0.2 | 0.2 | 0.17 | 0.2 | 0.21 | 0.1 |
| 3 | 1335751 | rs183795 | -1.01 | 1.281 | 0.023 | -1.708 | 0.017 | -0.022 | -0.022 | 0 | 0.29 | 0.54 | 0.22 | 0.2 | 0.26 | 0.12 |
| 3 | 1336137 | rs961102 | 0.985 | 1.244 | -0.543 | -1.86 | -0.096 | 0.136 | 0.136 | 1.397 | 0.43 | 0.07 | 0.66 | 0.8 | 0.55 | 0.73 |
| 3 | 1336397 | rs17271519 | -0.639 | 1.065 | -0.431 | -1.8 | -0.017 | 0.011 | 0.011 | 0 | 0.26 | 0.19 | 0.18 | 0.2 | 0.25 | 0.11 |
| 3 | 1337162 | rs2063243 | 1.088 | 0.872 | -0.492 | -1.821 | -0.097 | 0.104 | 0.104 | 1.439 | 0.4 | 0.07 | 0.66 | 0.8 | 0.53 | 0.67 |
| 3 | 1337203 | rs565499 | -0.722 | 0.863 | -0.449 | -1.848 | 0.046 | 0.095 | 0.095 | 0 | 0.66 | 0.81 | 0.89 | 1 | 0.79 | 0.83 |
| 3 | 1338259 | rs1353825 | -0.595 | 1.212 | -0.195 | -1.294 | -0.009 | -0.002 | -0.002 | 1.284 | 0.19 | 0.19 | 0.17 | 0.19 | 0.22 | 0.1 |
| 3 | 1342013 | rs17037357 | -1.389 | 0.757 | 0.153 | -1.122 | -0.01 | -0.045 | -0.045 | 1.89 | 0.15 | 0.64 | 0.1 | 0.08 | 0.11 | 0.09 |

| | | | | | | | | | | | | | | | | |
|---|---|---|---|---|---|---|---|---|---|---|---|---|---|---|---|---|
| 3 | 1343839 | rs7627835 | 0.01 | 0.839 | -0.47 | -0.926 | 0 | 0.007 | 0.007 | 0 | 0.06 | 0.1 | 0.04 | 0.13 | 0.06 | 0.03 |
| 3 | 1344922 | rs3898715 | -0.28 | 0.633 | -0.773 | -1.582 | 0.015 | 0.093 | 0.093 | 0.949 | 0.67 | 0.21 | 0.81 | 1 | 0.73 | 0.89 |
| 3 | 1345563 | rs411331 | -0.363 | 0.634 | -0.207 | -1.577 | -0.006 | -0.098 | -0.098 | 0.287 | 0.68 | 0.43 | 0.83 | 1 | 0.73 | 0.89 |
| 3 | 1345601 | rs9823418 | -0.326 | 0.64 | -0.631 | -1.575 | 0.011 | 0.016 | 0.016 | 0.922 | 0.68 | 0.31 | 0.82 | 1 | 0.73 | 0.89 |
| 3 | 1347567 | rs1858066 | 0.317 | 1.03 | -1.62 | -1.464 | -0.081 | -0.024 | -0.024 | 0 | 0.25 | 0.02 | 0.21 | 0.19 | 0.27 | 0.3 |
| 3 | 1349915 | rs4684882 | -0.432 | 0.554 | -1.811 | -1.707 | -0.025 | 0.146 | 0.146 | 0.436 | 0.53 | 0.94 | 0.66 | 0.8 | 0.47 | 0.62 |
| 3 | 1350134 | rs1846485 | -0.268 | 0.213 | -1.342 | -1.67 | -0.022 | -0.013 | -0.013 | 0 | 0.22 | 0.68 | 0.16 | 0.16 | 0.14 | 0.12 |
| 3 | 1350458 | rs11128627 | 1.095 | 0.217 | -1.586 | -1.665 | -0.058 | -0.112 | -0.112 | 0.017 | 0.29 | 0.03 | 0.48 | 0.63 | 0.32 | 0.5 |
| 3 | 1355266 | rs155409 | -0.7 | -0.152 | -1.505 | -1.679 | 0.059 | 0.04 | 0.04 | 0.927 | 0.24 | 0.91 | 0.17 | 0.15 | 0.15 | 0.12 |
| 3 | 1356306 | rs155227 | -0.766 | 0.293 | -1.691 | -1.885 | -0.028 | 0.103 | 0.103 | 0.984 | 0.57 | 0.94 | 0.62 | 0.77 | 0.41 | 0.65 |
| 3 | 1358927 | rs262809 | 0.142 | 0.029 | -1.101 | -1.858 | 0.045 | 0.014 | 0.014 | 1.392 | 0.37 | 0.79 | 0.51 | 0.69 | 0.27 | 0.48 |
| 3 | 1362072 | rs17443601 | 1.114 | 0.562 | -1.46 | -2.003 | -0.069 | -0.049 | -0.049 | 0 | 0.14 | 0.02 | 0.21 | 0.22 | 0.26 | 0.25 |
| 3 | 1363099 | rs11927748 | 0.419 | -0.018 | -0.483 | -1.915 | -0.011 | -0.003 | -0.003 | 0.453 | 0.33 | 0.36 | 0.48 | 0.69 | 0.27 | 0.48 |
| 3 | 1366340 | rs265785 | 0.311 | -0.013 | -0.244 | -1.871 | -0.003 | -0.019 | -0.019 | 0 | 0.19 | 0.32 | 0.12 | 0.08 | 0.14 | 0.16 |
| 3 | 1366558 | rs17444084 | 1.565 | -0.004 | -0.436 | -1.87 | -0.033 | 0.008 | 0.008 | 0 | 0.12 | 0.01 | 0.08 | 0.08 | 0.1 | 0.1 |
| 3 | 1367042 | rs17037701 | 1.72 | 0.002 | -0.39 | -1.877 | -0.007 | -0.024 | -0.024 | 0 | 0.05 | 0.09 | 0.02 | 0 | 0.04 | 0.06 |
| 3 | 1367192 | rs265782 | -0.935 | -0.012 | -0.352 | -1.934 | 0.017 | 0.097 | 0.097 | 0.204 | 0.55 | 0.79 | 0.6 | 0.77 | 0.41 | 0.65 |
| 3 | 1367204 | rs265781 | -0.527 | 0.006 | -0.415 | -1.917 | 0.009 | 0.011 | 0.011 | 0.084 | 0.5 | 0.42 | 0.56 | 0.77 | 0.39 | 0.64 |
| 3 | 1368840 | rs265777 | 1.214 | 0.305 | -0.686 | -1.564 | -0.004 | 0.022 | 0.022 | 0 | 0.12 | 0.06 | 0.07 | 0.08 | 0.09 | 0.1 |
| 3 | 1369428 | rs265774 | 1.099 | 0.238 | -0.694 | -1.564 | -0.013 | 0.025 | 0.025 | 0 | 0.13 | 0.06 | 0.08 | 0.08 | 0.1 | 0.1 |
| 3 | 1374539 | rs3772295 | 0.073 | 0.174 | -0.368 | -1.476 | -0.002 | 0.005 | 0.005 | 0 | 0.21 | 0.4 | 0.2 | 0.21 | 0.17 | 0.12 |
| 3 | 1375734 | rs7628540 | 1.315 | -0.084 | -0.65 | -1.433 | 0 | 0.011 | 0.011 | 0 | 0.16 | 0.14 | 0.42 | 0.56 | 0.16 | 0.46 |
| 3 | 1376235 | rs265766 | -0.7 | -0.167 | -0.97 | -1.537 | 0.013 | 0.034 | 0.034 | 0.079 | 0.42 | 0.68 | 0.64 | 0.77 | 0.37 | 0.64 |
| 3 | 1376610 | rs987963 | 1.178 | 0.098 | -0.905 | -1.248 | 0.004 | -0.044 | -0.044 | 0 | 0.16 | 0.1 | 0.41 | 0.56 | 0.16 | 0.46 |
| 3 | 1380095 | rs1353823 | 0.144 | 0.52 | -0.816 | -1.234 | 0.054 | 0.084 | 0.084 | 0.237 | 0.1 | 0.4 | 0.13 | 0.14 | 0.1 | 0.03 |
| 3 | 1381174 | rs1479533 | 0.788 | 1.074 | -2.13 | -1.518 | -0.068 | -0.054 | -0.054 | 0 | 0.18 | 0.02 | 0.16 | 0.22 | 0.26 | 0.22 |
| 3 | 1382151 | rs2055738 | 0.023 | 0.738 | -2.3 | -1.583 | 0.01 | 0.156 | 0.156 | 0.062 | 0.48 | 0.15 | 0.32 | 0.21 | 0.58 | 0.28 |
| 3 | 1386467 | rs155399 | 0.281 | 0.892 | -1.835 | -1.156 | -0.023 | -0.015 | -0.015 | 0 | 0.2 | 0.16 | 0.19 | 0.21 | 0.28 | 0.2 |
| 3 | 1387438 | rs6794738 | 0.075 | 0.364 | -2.031 | -1.866 | -0.01 | 0.077 | 0.077 | 0 | 0.44 | 0.7 | 0.34 | 0.12 | 0.5 | 0.17 |
| 3 | 1388589 | rs11928188 | 0.943 | -0.207 | -1.873 | -1.708 | 0.001 | 0.002 | 0.002 | 0 | 0.19 | 0.13 | 0.1 | 0.07 | 0.14 | 0.08 |
| 3 | 1389117 | rs17038032 | 0.978 | -0.206 | -1.873 | -1.708 | 0.001 | 0.002 | 0.002 | 0 | 0.19 | 0.16 | 0.11 | 0.07 | 0.14 | 0.16 |
| 3 | 1389720 | rs17785924 | 1.476 | -0.251 | -1.927 | -1.685 | -0.001 | 0.025 | 0.025 | 0 | 0.14 | 0.07 | 0.09 | 0.07 | 0.09 | 0.1 |
| 3 | 1390985 | rs1499214 | 0.865 | -0.173 | -2.063 | -1.872 | 0 | -0.007 | -0.007 | 0 | 0.33 | 0.66 | 0.25 | 0.11 | 0.34 | 0.13 |
| 3 | 1391317 | rs1846484 | -1.926 | -0.106 | -2.105 | -1.862 | 0.004 | -0.017 | -0.017 | 0 | 0.83 | 0.88 | 0.58 | 0.41 | 0.85 | 0.56 |
| 3 | 1392751 | rs3796174 | -0.05 | -0.105 | -2.132 | -1.907 | -0.026 | 0.16 | 0.16 | 0 | 0.55 | 0.81 | 0.38 | 0.16 | 0.52 | 0.31 |
| 3 | 1395906 | rs7648067 | 0.719 | 0.004 | -1.755 | -1.86 | 0.002 | -0.009 | -0.009 | 0.148 | 0.36 | 0.64 | 0.26 | 0.08 | 0.37 | 0.14 |
| 3 | 1395966 | rs2133452 | 0.911 | 0.006 | -2.08 | -1.812 | -0.076 | -0.076 | -0.076 | 4.165 | 0.24 | 0.03 | 0.17 | 0.26 | 0.3 | 0.28 |
| 3 | 1398990 | rs269672 | -0.313 | -0.087 | -2.096 | -1.87 | -0.09 | 0.018 | 0.018 | 0 | 0.6 | 0.94 | 0.73 | 0.75 | 0.6 | 0.65 |
| 3 | 1400096 | rs1551995 | 0.123 | -0.097 | -2.094 | -1.876 | -0.081 | 0.018 | 0.018 | 0 | 0.39 | 0.07 | 0.27 | 0.25 | 0.4 | 0.33 |
| 3 | 1401122 | rs155895 | -1.135 | -0.027 | -2.046 | -1.822 | -0.081 | -0.061 | -0.061 | 1.688 | 0.76 | 0.98 | 0.83 | 0.77 | 0.7 | 0.73 |
| 3 | 1401808 | rs9837911 | 0.083 | 0.008 | -1.539 | -2.006 | 0.007 | 0.071 | 0.071 | 0 | 0.41 | 0.74 | 0.29 | 0.09 | 0.42 | 0.2 |
| 3 | 1404038 | rs9829288 | 0.397 | 0.09 | -1.439 | -1.897 | -0.002 | -0.069 | -0.069 | 0 | 0.36 | 0.58 | 0.26 | 0.08 | 0.37 | 0.14 |
| 3 | 1404179 | rs9848880 | 0.372 | 0.1 | -1.496 | -1.891 | -0.002 | -0.047 | -0.047 | 0 | 0.36 | 0.62 | 0.26 | 0.09 | 0.37 | 0.14 |
| 3 | 1405829 | rs10510209 | 0.89 | 0.351 | -1.649 | -1.428 | -0.037 | 0.037 | 0.037 | 0 | 0.16 | 0 | 0.12 | 0.07 | 0.11 | 0.11 |
| 3 | 1408344 | rs13063912 | 0.737 | 0.436 | -1.117 | -1.525 | -0.001 | -0.042 | -0.042 | 0.124 | 0.21 | 0.21 | 0.14 | 0 | 0.21 | 0.07 |
| 3 | 1415195 | rs3765098 | -0.63 | 0.427 | -1.08 | -1.586 | -0.013 | 0.108 | 0.108 | 0 | 0.54 | 0.77 | 0.39 | 0.17 | 0.48 | 0.29 |
| 3 | 1418753 | rs17038365 | 0.119 | 0.565 | -1.019 | -1.447 | 0 | -0.007 | -0.007 | 0 | 0.07 | 0.1 | 0.07 | 0.03 | 0.08 | 0.07 |
| 3 | 1419010 | rs155856 | -1.091 | 0.664 | -1.267 | -1.308 | -0.059 | -0.045 | -0.045 | 0 | 0.79 | 0.95 | 0.83 | 0.77 | 0.72 | 0.76 |
| 3 | 1419563 | rs155384 | -0.74 | 0.727 | -0.779 | -1.399 | 0.058 | -0.022 | -0.022 | 5.317 | 0.19 | 0.52 | 0.22 | 0.15 | 0.28 | 0.14 |
| 3 | 1420284 | rs3843386 | 0.818 | 0.17 | -0.842 | -1.404 | 0.02 | -0.017 | -0.017 | 0.172 | 0.38 | 0.26 | 0.47 | 0.58 | 0.32 | 0.43 |
| 3 | 1420519 | rs155387 | -0.258 | 0.291 | -0.868 | -1.449 | -0.002 | 0.021 | 0.021 | 0 | 0.36 | 0.71 | 0.32 | 0.19 | 0.34 | 0.25 |
| 3 | 1421673 | rs155390 | -0.675 | 0.499 | -1.169 | -1.294 | -0.022 | -0.003 | -0.003 | 0.982 | 0.73 | 0.81 | 0.74 | 0.75 | 0.65 | 0.68 |
| 3 | 1424745 | rs4684146 | 0.641 | 0.36 | -0.88 | -1.435 | 0 | -0.009 | -0.009 | 6.568 | 0.24 | 0.23 | 0.17 | 0.06 | 0.16 | 0.12 |
| 3 | 1424850 | rs2291100 | 0.981 | 0.357 | -1.105 | -1.469 | -0.101 | -0.096 | -0.096 | 0 | 0.27 | 0.04 | 0.28 | 0.35 | 0.39 | 0.35 |
| 3 | 1425506 | rs3845158 | 1.315 | 0.261 | -0.801 | -1.457 | 0.006 | -0.023 | -0.023 | 0 | 0.26 | 0.21 | 0.38 | 0.59 | 0.23 | 0.39 |
| 3 | 1425997 | rs3915436 | -0.176 | 0.358 | -0.9 | -1.429 | -0.002 | 0.004 | 0.004 | 0 | 0.54 | 0.44 | 0.58 | 0.65 | 0.44 | 0.57 |
| 3 | 1427769 | rs9821129 | 0.453 | 0.37 | -1.074 | -1.561 | 0.001 | 0.043 | 0.043 | 0 | 0.23 | 0.12 | 0.15 | 0.05 | 0.18 | 0.17 |
| 3 | 1440124 | rs3916111 | -0.541 | 0.356 | -0.664 | -1.396 | -0.01 | -0.14 | -0.14 | 0 | 0.81 | 0.49 | 0.86 | 1 | 0.83 | 0.93 |
| 3 | 1440777 | rs7622631 | 0.131 | 0.434 | -1.087 | -1.204 | -0.005 | -0.03 | -0.03 | 0 | 0.57 | 0.53 | 0.7 | 0.82 | 0.66 | 0.81 |
| 3 | 1441376 | rs9875177 | 0.217 | 1.033 | -1.042 | -0.562 | -0.013 | 0.024 | 0.024 | 0 | 0.55 | 0.71 | 0.59 | 0.68 | 0.45 | 0.6 |
| 3 | 1444243 | rs13326014 | -1.104 | 1.631 | -0.783 | -0.969 | 0.005 | -0.052 | -0.052 | 0 | 0.23 | 0.24 | 0.23 | 0.01 | 0.31 | 0.18 |
| 3 | 1446510 | rs6791266 | -0.871 | 1.556 | -0.864 | -1.272 | -0.001 | -0.025 | -0.025 | 0.469 | 0.19 | 0.19 | 0.2 | 0.01 | 0.25 | 0.16 |
| 3 | 1448615 | rs7644659 | -0.172 | 1.898 | -1.054 | -1.235 | -0.036 | 0.044 | 0.044 | 1.081 | 0.14 | 0.06 | 0.16 | 0.01 | 0.22 | 0.13 |
| 3 | 1449068 | rs12497579 | -0.47 | 1.814 | -1.432 | -1.782 | -0.002 | 0.013 | 0.013 | 0 | 0.15 | 0.09 | 0.1 | 0 | 0.16 | 0.11 |
| 3 | 1449151 | rs155863 | -1.385 | 1.417 | -0.822 | -1.379 | -0.018 | 0.045 | 0.045 | 0 | 0.43 | 0.67 | 0.54 | 0.21 | 0.5 | 0.37 |
| 3 | 1453696 | rs155871 | -0.939 | 1.468 | -0.748 | -1.615 | -0.012 | 0.04 | 0.04 | 0 | 0.42 | 0.3 | 0.53 | 0.16 | 0.48 | 0.33 |
| 3 | 1460921 | rs155879 | 0.917 | 1.001 | -0.355 | -1.427 | 0.048 | 0.069 | 0.069 | 0.178 | 0.87 | 0.59 | 0.71 | 0.85 | 0.86 | 0.9 |
| 3 | 1461498 | rs9810930 | 0.508 | 1.21 | -1.006 | -1.525 | 0.002 | -0.005 | -0.005 | 1.088 | 0.88 | 0.87 | 0.87 | 0.96 | 0.84 | 0.9 |
| 3 | 1463373 | rs9817190 | 1.385 | 1.142 | -0.792 | -1.492 | 0.004 | 0.002 | 0.002 | 0 | 0.81 | 0.7 | 0.65 | 0.82 | 0.77 | 0.84 |

| | | | | | | | | | | | | | | | | |
|---|---|---|---|---|---|---|---|---|---|---|---|---|---|---|---|---|
| 3 | 1466413 | rs1382862 | 1.475 | 1.357 | -1.091 | -1.817 | -0.006 | 0.032 | 0.032 | 0 | 0.55 | 0.61 | 0.46 | 0.82 | 0.45 | 0.61 |
| 3 | 1466592 | rs6777408 | 0.611 | 1.362 | -1.177 | -1.857 | -0.07 | 0.108 | 0.108 | 2.36 | 0.2 | 0.04 | 0.16 | 0 | 0.28 | 0.17 |
| 3 | 1467145 | rs17038909 | -0.059 | 1.322 | -1.129 | -1.808 | 0.003 | -0.016 | -0.016 | 0 | 0.06 | 0.05 | 0.22 | 0.15 | 0.08 | 0.06 |
| 3 | 1467675 | rs7626838 | 1.005 | 1.322 | -1.111 | -1.812 | 0.001 | -0.018 | -0.018 | 0 | 0.82 | 0.8 | 0.86 | 0.96 | 0.81 | 0.86 |
| 3 | 1470616 | rs17038965 | -0.803 | 1.358 | -1.098 | -1.789 | 0 | 0.003 | 0.003 | 0 | 0.1 | 0.12 | 0.23 | 0.13 | 0.13 | 0.14 |
| 3 | 1470892 | rs17038979 | 0.831 | 0.954 | -1.589 | -2.07 | 0.004 | 0.005 | 0.005 | 0.02 | 0.14 | 0.12 | 0.06 | 0.04 | 0.1 | 0.04 |
| 3 | 1471838 | rs155829 | -0.328 | 0.99 | -1.596 | -2.038 | 0 | 0.003 | 0.003 | 0 | 0.08 | 0.04 | 0.05 | 0.03 | 0.07 | 0.05 |
| 3 | 1474128 | rs155834 | -1.036 | 1.03 | -1.583 | -2.02 | -0.002 | 0.001 | 0.001 | 0 | 0.94 | 0.93 | 0.93 | 0.99 | 0.92 | 0.98 |
| 3 | 1478453 | rs1037386 | 1.689 | 0.982 | -1.5 | -2.076 | 0 | -0.003 | -0.003 | 0 | 0.43 | 0.44 | 0.39 | 0.78 | 0.36 | 0.6 |
| 3 | 1479813 | rs155844 | -1.639 | 1.186 | -1.706 | -1.946 | 0.002 | -0.001 | -0.001 | 0 | 0.23 | 0.23 | 0.37 | 0.19 | 0.27 | 0.19 |
| 3 | 1480396 | rs149216 | -0.781 | 1.18 | -1.731 | -1.942 | -0.001 | 0 | 0 | 0 | 0.13 | 0.12 | 0.25 | 0.13 | 0.13 | 0.12 |
| 3 | 1480622 | rs1530894 | 0.908 | 1.173 | -1.729 | -1.947 | 0 | -0.001 | -0.001 | 0 | 0.82 | 0.81 | 0.69 | 0.86 | 0.8 | 0.86 |
| 3 | 1480753 | rs155847 | -0.967 | 1.16 | -1.771 | -1.978 | -0.036 | 0.084 | 0.084 | 0 | 0.38 | 0.22 | 0.48 | 0.16 | 0.48 | 0.31 |
| 3 | 1483973 | rs17039234 | -0.389 | 1.269 | -1.703 | -2.134 | -0.002 | -0.013 | -0.013 | 0 | 0.29 | 0.23 | 0.24 | 0.04 | 0.35 | 0.29 |
| 3 | 1486518 | rs4553960 | -0.834 | 0.787 | -2.128 | -1.921 | 0.038 | 0.035 | 0.035 | 0.909 | 0.53 | 0.21 | 0.56 | 0.35 | 0.6 | 0.42 |
| 3 | 1486580 | rs11929623 | -0.192 | 0.765 | -2.12 | -1.922 | -0.001 | -0.015 | -0.015 | 0 | 0.09 | 0.06 | 0.04 | 0 | 0.08 | 0.09 |
| 3 | 1486965 | rs12486002 | -1.301 | 0.774 | -2.103 | -1.915 | 0.02 | -0.014 | -0.014 | 1.469 | 0.6 | 0.29 | 0.62 | 0.38 | 0.67 | 0.47 |
| 3 | 1492235 | rs12488675 | 0.062 | 0.399 | -1.776 | -1.805 | 0.019 | -0.027 | -0.027 | 2.332 | 0.08 | 0.13 | 0.09 | 0.02 | 0.12 | 0.1 |
| 3 | 1494264 | rs17039471 | 0.202 | 0.354 | -1.791 | -1.804 | 0.002 | -0.003 | -0.003 | 0.233 | 0.07 | 0.09 | 0.06 | 0.03 | 0.07 | 0.05 |
| 3 | 1494483 | rs718558 | 1.114 | 0.33 | -2.015 | -1.911 | 0.006 | 0.001 | 0.001 | 0.064 | 0.43 | 0.7 | 0.43 | 0.64 | 0.39 | 0.58 |
| 3 | 1495223 | rs2122993 | 0.417 | 0.287 | -1.904 | -1.944 | 0.001 | -0.036 | -0.036 | 0 | 0.56 | 0.59 | 0.68 | 0.91 | 0.52 | 0.66 |
| 3 | 1495993 | rs12489351 | -0.513 | 0.394 | -1.839 | -1.794 | -0.021 | 0.013 | 0.013 | 0 | 0.12 | 0.05 | 0.12 | 0.05 | 0.15 | 0.07 |
| 3 | 1496821 | rs17786098 | -1.147 | 0.373 | -1.834 | -1.809 | -0.003 | 0.028 | 0.028 | 0 | 0.18 | 0.12 | 0.15 | 0.05 | 0.19 | 0.1 |
| 3 | 1498979 | rs9825372 | 1.606 | 0.546 | -1.777 | -1.696 | -0.005 | 0.029 | 0.029 | 0.049 | 0.32 | 0.58 | 0.38 | 0.6 | 0.3 | 0.55 |
| 3 | 1500042 | rs13082351 | -0.596 | 0.664 | -1.757 | -1.758 | 0.001 | 0.005 | 0.005 | 0 | 0.3 | 0.2 | 0.19 | 0.06 | 0.28 | 0.17 |
| 3 | 1500657 | rs17193526 | -0.036 | 0.549 | -1.851 | -1.479 | -0.039 | -0.005 | -0.005 | 0 | 0.1 | 0.01 | 0.08 | 0.05 | 0.13 | 0.05 |
| 3 | 1503156 | rs17193533 | 0.317 | 0.471 | -1.869 | -1.532 | 0.002 | -0.036 | -0.036 | 0 | 0.25 | 0.18 | 0.16 | 0.02 | 0.23 | 0.14 |
| 3 | 1503929 | rs6442449 | -1.216 | 0.51 | -1.961 | -1.48 | 0 | 0.001 | 0.001 | 0.044 | 0.64 | 0.34 | 0.55 | 0.39 | 0.62 | 0.43 |
| 3 | 1504132 | rs6442451 | -1.216 | 0.51 | -1.961 | -1.48 | 0 | 0.001 | 0.001 | 0.044 | 0.64 | 0.34 | 0.55 | 0.39 | 0.62 | 0.43 |
| 3 | 1506235 | rs2167470 | 0.482 | 0.402 | -2.174 | -1.665 | 0.046 | -0.079 | -0.079 | 0 | 0.2 | 0.07 | 0.24 | 0.3 | 0.13 | 0.1 |
| 3 | 1507454 | rs13097651 | 0.73 | 0.83 | -1.571 | -1.426 | 0.002 | 0.004 | 0.004 | 0.011 | 0.36 | 0.58 | 0.46 | 0.67 | 0.37 | 0.58 |
| 3 | 1510262 | rs10865730 | -0.245 | 0.824 | -1.558 | -1.456 | 0.01 | 0.055 | 0.055 | 0 | 0.74 | 0.88 | 0.85 | 0.98 | 0.75 | 0.86 |
| 3 | 1510310 | rs11919047 | -0.303 | 1.179 | -1.3 | -0.509 | -0.002 | 0 | 0 | 0 | 0.38 | 0.31 | 0.38 | 0.32 | 0.38 | 0.28 |
| 3 | 1511108 | rs9856251 | 0.037 | 0.935 | -1.171 | -0.17 | 0.005 | -0.014 | -0.014 | 0 | 0.05 | 0.08 | 0.03 | 0.02 | 0.05 | 0.03 |
| 3 | 1525557 | rs2648451 | -0.138 | 0.761 | -1.198 | 0.052 | 0.003 | 0.035 | 0.035 | 0 | 0.48 | 0.32 | 0.49 | 0.33 | 0.52 | 0.3 |
| 3 | 1528588 | rs172171 | -0.476 | 0.486 | -0.958 | 0.113 | -0.004 | -0.011 | -0.011 | 0.097 | 0.35 | 0.41 | 0.23 | 0.22 | 0.34 | 0.27 |
| 3 | 1530388 | rs6771147 | 0.074 | 0.869 | -1.346 | 0.437 | -0.006 | -0.006 | -0.006 | 0 | 0.18 | 0.12 | 0.39 | 0.19 | 0.22 | 0.27 |
| 3 | 1532531 | rs17040275 | -0.083 | 0.137 | -1.122 | 0.067 | -0.034 | -0.036 | -0.036 | 0 | 0.28 | 0.48 | 0.27 | 0.3 | 0.26 | 0.22 |
| 3 | 1535741 | rs6781581 | -0.747 | -0.221 | -1.454 | -0.205 | 0.009 | 0.012 | 0.012 | 0 | 0.48 | 0.62 | 0.36 | 0.42 | 0.39 | 0.34 |
| 3 | 1536945 | rs7622999 | 0.44 | -0.229 | -1.434 | -0.183 | -0.006 | 0.023 | 0.023 | 0 | 0.14 | 0.1 | 0.09 | 0.01 | 0.14 | 0.11 |
| 3 | 1547688 | rs17040559 | 0.284 | -0.724 | -1.262 | -0.135 | -0.039 | -0.024 | -0.024 | 0 | 0.27 | 0.48 | 0.2 | 0.34 | 0.24 | 0.11 |
| 3 | 1548001 | rs1482272 | 1.342 | -0.29 | -1.585 | -0.548 | -0.064 | 0.059 | 0.059 | 0 | 0.12 | 0.03 | 0.12 | 0.01 | 0.27 | 0.14 |
| 3 | 1553690 | rs6796218 | -0.59 | -1.203 | -1.287 | -0.247 | -0.014 | -0.045 | -0.045 | 0.099 | 0.42 | 0.53 | 0.27 | 0.27 | 0.37 | 0.22 |
| 3 | 1556680 | rs9813890 | -0.906 | -1.262 | -1.31 | -0.306 | -0.007 | -0.02 | -0.02 | 0.625 | 0.44 | 0.56 | 0.34 | 0.33 | 0.38 | 0.34 |
| 3 | 1557413 | rs11128726 | 0.587 | -1.572 | -1.315 | -0.514 | -0.029 | 0.041 | 0.041 | 1.554 | 0.13 | 0.39 | 0.11 | 0.25 | 0.1 | 0.1 |
| 3 | 1557494 | rs11128727 | -1.203 | -1.542 | -1.278 | -0.461 | 0 | -0.004 | -0.004 | 0 | 0.44 | 0.59 | 0.34 | 0.33 | 0.38 | 0.34 |
| 3 | 1563444 | rs13066435 | -0.523 | -1.358 | -1.471 | -0.485 | 0.01 | 0.065 | 0.065 | 0 | 0.21 | 0.1 | 0.16 | 0.09 | 0.16 | 0.22 |
| 3 | 1575889 | rs1969290 | -0.557 | -1.255 | -1.268 | -0.288 | -0.029 | 0.004 | 0.004 | 0 | 0.14 | 0.02 | 0.12 | 0.12 | 0.11 | 0.18 |
| 3 | 1576946 | rs1000130 | -1.479 | -1.736 | -0.709 | 0.292 | -0.01 | -0.003 | -0.003 | 0 | 0.43 | 0.59 | 0.3 | 0.47 | 0.33 | 0.35 |
| 3 | 1577232 | rs1499258 | -1.454 | -1.711 | -0.689 | 0.307 | -0.012 | -0.004 | -0.004 | 1.545 | 0.42 | 0.59 | 0.3 | 0.46 | 0.33 | 0.35 |
| 3 | 1579270 | rs9814262 | -0.475 | -1.63 | -0.888 | 0.093 | 0.002 | 0.007 | 0.007 | 0 | 0.13 | 0.12 | 0.06 | 0.09 | 0.09 | 0.05 |
| 3 | 1587151 | rs2728007 | 0.103 | -1.705 | -0.766 | 0.436 | -0.011 | 0.088 | 0.088 | 0 | 0.9 | 0.89 | 0.83 | 0.44 | 0.96 | 0.75 |
| 3 | 1587860 | rs400048 | -1.434 | -1.83 | -0.719 | 0.316 | 0.036 | 0.06 | 0.06 | 0.024 | 0.46 | 0.69 | 0.39 | 0.85 | 0.34 | 0.48 |
| 3 | 1588919 | rs443052 | 0.664 | -1.752 | -0.833 | 0.389 | 0.005 | -0.022 | -0.022 | 0 | 0.82 | 0.88 | 0.79 | 0.45 | 0.88 | 0.68 |
| 3 | 1590441 | rs9878958 | -0.007 | -1.593 | -0.9 | -0.298 | -0.035 | -0.018 | -0.018 | 0 | 0.25 | 0.42 | 0.16 | 0.22 | 0.21 | 0.16 |
| 3 | 1590775 | rs9817135 | -0.456 | -0.85 | -1.106 | 0.024 | -0.004 | 0.006 | 0.006 | 0 | 0.08 | 0.1 | 0.1 | 0.16 | 0.04 | 0.14 |
| 3 | 1593450 | rs1587386 | -1.116 | -0.842 | -1.226 | 0.023 | -0.001 | -0.007 | -0.007 | 0.115 | 0.16 | 0.07 | 0.13 | 0.2 | 0.12 | 0.21 |
| 3 | 1594653 | rs2314764 | -0.269 | -0.132 | -0.959 | 0.191 | 0 | -0.001 | -0.001 | 0.717 | 0.07 | 0.08 | 0.11 | 0.11 | 0.04 | 0.15 |
| 3 | 1597057 | rs11712183 | -0.851 | -0.42 | -0.865 | -0.112 | -0.004 | 0.003 | 0.003 | 0 | 0.32 | 0.64 | 0.32 | 0.39 | 0.27 | 0.3 |
| 3 | 1597746 | rs471449 | 0.792 | -0.325 | -1.758 | -0.939 | -0.097 | 0.166 | 0.166 | 0.28 | 0.33 | 0.08 | 0.22 | 0.01 | 0.44 | 0.37 |
| 3 | 1597795 | rs471342 | 0.483 | -0.634 | -1.442 | -0.937 | -0.024 | 0.202 | 0.202 | 0 | 0.17 | 0.55 | 0.3 | 0.53 | 0.11 | 0.16 |
| 3 | 1600348 | rs440961 | -0.169 | -0.724 | -1.615 | -1.267 | -0.008 | -0.02 | -0.02 | 0 | 0.44 | 0.52 | 0.64 | 0.75 | 0.37 | 0.48 |
| 3 | 1600422 | rs403961 | -0.239 | -0.762 | -1.484 | -1.283 | 0.015 | 0.039 | 0.039 | 1.303 | 0.47 | 0.6 | 0.65 | 0.82 | 0.38 | 0.5 |
| 3 | 1603110 | rs13072051 | 1.321 | -0.751 | -1.709 | -1.333 | -0.129 | 0.151 | 0.151 | 0.001 | 0.29 | 0.07 | 0.21 | 0.07 | 0.43 | 0.35 |
| 3 | 1605071 | rs409430 | -0.371 | -1.013 | -1.36 | -1.189 | -0.029 | -0.025 | -0.025 | 0 | 0.71 | 0.62 | 0.85 | 0.84 | 0.79 | 0.84 |
| 3 | 1610284 | rs428179 | -0.57 | -1.022 | -1.727 | -1.107 | 0.012 | 0.1 | 0.1 | 0 | 0.63 | 0.93 | 0.63 | 0.76 | 0.71 | 0.64 |
| 3 | 1618188 | rs9820473 | 0.104 | -0.651 | -1.131 | -1.317 | 0.004 | 0.011 | 0.011 | 0 | 0.47 | 0.45 | 0.34 | 0.16 | 0.59 | 0.45 |
| 3 | 1619821 | rs2128252 | -0.155 | -0.595 | -1.328 | -1.277 | 0.003 | 0.002 | 0.002 | 0 | 0.5 | 0.53 | 0.35 | 0.22 | 0.6 | 0.47 |
| 3 | 1622574 | rs576453 | 0.389 | -0.865 | -0.651 | -0.642 | -0.049 | -0.029 | -0.029 | 0.12 | 0.66 | 0.5 | 0.83 | 0.78 | 0.78 | 0.82 |

| | | | | | | | | | | | | | | | | |
|---|---|---|---|---|---|---|---|---|---|---|---|---|---|---|---|---|
| 3 | 1627108 | rs12490663 | -0.315 | -0.276 | 0.055 | -1.149 | -0.008 | -0.076 | -0.076 | 0.136 | 0.31 | 0.46 | 0.39 | 0.15 | 0.31 | 0.19 |
| 3 | 1631903 | rs3968917 | 0.811 | 0 | -0.346 | -1.8 | 0.032 | 0.109 | 0.109 | 0 | 0.41 | 0.25 | 0.42 | 0.83 | 0.34 | 0.63 |
| 3 | 1632353 | rs4684256 | 1.89 | 0.239 | -0.465 | -1.779 | -0.07 | 0.071 | 0.071 | 0 | 0.15 | 0.05 | 0.12 | 0.01 | 0.27 | 0.12 |
| 3 | 1632567 | rs9835271 | 0.062 | -0.091 | -0.435 | -1.981 | 0.04 | -0.067 | -0.067 | 0 | 0.3 | 0.53 | 0.39 | 0.16 | 0.41 | 0.27 |
| 3 | 1633254 | rs9840070 | 0.533 | -0.625 | 0.367 | -1.525 | -0.011 | -0.084 | -0.084 | 0 | 0.2 | 0.26 | 0.14 | 0 | 0.19 | 0.07 |
| 3 | 1633542 | rs9857772 | -0.691 | -0.776 | 0.091 | -1.458 | -0.011 | -0.07 | -0.07 | 0 | 0.33 | 0.54 | 0.4 | 0.12 | 0.32 | 0.2 |
| 3 | 1636410 | rs9836895 | 0.175 | -0.953 | -0.069 | -1.147 | 0.015 | -0.025 | -0.025 | 0.005 | 0.13 | 0.36 | 0.26 | 0.11 | 0.12 | 0.13 |
| 3 | 1636865 | rs4685256 | 0.494 | -0.978 | -1.239 | -0.339 | 0.022 | -0.026 | -0.026 | 0 | 0.72 | 0.28 | 0.64 | 0.55 | 0.82 | 0.82 |
| 3 | 1638054 | rs12495819 | 0.973 | -0.815 | -1.259 | -0.425 | -0.006 | 0.084 | 0.084 | 0 | 0.18 | 0.05 | 0.15 | 0 | 0.2 | 0.05 |
| 3 | 1638613 | rs6779648 | 0.35 | -0.273 | -1.001 | -0.033 | 0.076 | 0.017 | 0.017 | 0.118 | 0.52 | 0.24 | 0.49 | 0.58 | 0.62 | 0.71 |
| 3 | 1647955 | rs280791 | 0.224 | 0.029 | -0.167 | 1.022 | -0.002 | 0.078 | 0.078 | 0 | 0.86 | 0.76 | 0.79 | 0.41 | 0.86 | 0.8 |
| 3 | 1648276 | rs2046239 | 0.537 | -0.18 | 0.388 | 1.104 | -0.016 | 0.041 | 0.041 | 0 | 0.79 | 0.71 | 0.76 | 0.73 | 0.82 | 0.8 |
| 3 | 1651487 | rs13073236 | 1.291 | -0.263 | 0.67 | 1.035 | -0.02 | 0.01 | 0.01 | 0.047 | 0.72 | 0.44 | 0.7 | 0.36 | 0.79 | 0.76 |
| 3 | 1654454 | rs401306 | 0.442 | -0.317 | 0.517 | 1.123 | -0.001 | -0.081 | -0.081 | 0.005 | 0.87 | 0.97 | 0.91 | 0.63 | 0.91 | 0.93 |
| 3 | 1655984 | rs417975 | -0.528 | -0.307 | 0.516 | 1.11 | -0.002 | 0.004 | 0.004 | 0.128 | 0.9 | 0.99 | 0.94 | 0.95 | 0.94 | 0.95 |
| 3 | 1656488 | rs2648447 | -0.232 | -0.327 | 1.037 | 1.121 | 0.007 | 0.16 | 0.16 | 0 | 0.87 | 0.62 | 0.81 | 0.78 | 0.88 | 0.85 |
| 3 | 1659713 | rs280123 | 0.777 | -0.292 | 1.038 | 1.143 | 0.011 | 0.268 | 0.268 | 0.452 | 0.95 | 0.79 | 0.95 | 0.69 | 0.97 | 0.97 |
| 3 | 1661120 | rs449622 | -1.537 | -0.298 | 1.064 | 1.152 | 0.02 | 0.318 | 0.318 | 1.76 | 0.07 | 0.34 | 0.06 | 0.35 | 0.03 | 0.04 |
| 3 | 1668488 | rs10510212 | 0.774 | -0.593 | 0.434 | 0.954 | -0.007 | -0.05 | -0.05 | 0.141 | 0.12 | 0 | 0.26 | 0.17 | 0.1 | 0.19 |
| 3 | 1669025 | rs364432 | -0.118 | 0.322 | 0.478 | 0.697 | -0.044 | 0.012 | 0.012 | 0 | 0.09 | 0.01 | 0.11 | 0.04 | 0.14 | 0.06 |
| 3 | 1669178 | rs1564585 | 0.663 | 0.231 | 0.568 | 0.711 | 0.005 | 0.115 | 0.115 | 0 | 0.12 | 0.28 | 0.17 | 0.2 | 0.11 | 0.14 |
| 3 | 1670301 | rs4143127 | 0.408 | 0.33 | -0.077 | 0.284 | 0.016 | -0.003 | -0.003 | 0.033 | 0.32 | 0.22 | 0.37 | 0.35 | 0.25 | 0.51 |
| 3 | 1670852 | rs1353813 | 0.81 | 0.321 | 0.093 | 0.439 | 0.007 | 0.042 | 0.042 | 0.025 | 0.75 | 0.47 | 0.68 | 0.4 | 0.73 | 0.78 |
| 3 | 1673173 | rs9838484 | 0.562 | 0.416 | 0.041 | 0.225 | 0.024 | 0.047 | 0.047 | 0 | 0.77 | 0.58 | 0.7 | 0.76 | 0.74 | 0.8 |
| 3 | 1674210 | rs459439 | 0.285 | 0.348 | -0.245 | 0.049 | -0.02 | 0.011 | 0.011 | 0 | 0.61 | 0.76 | 0.49 | 0.63 | 0.58 | 0.44 |
| 3 | 1684810 | rs1878627 | 1.101 | 0.208 | 0.358 | -0.241 | -0.022 | -0.022 | -0.022 | 0.486 | 0.87 | 0.28 | 0.85 | 0.94 | 0.93 | 0.96 |
| 3 | 1691064 | rs1878178 | 0.594 | 0.464 | -0.339 | -0.693 | -0.061 | 0.182 | 0.182 | 0.645 | 0.29 | 0.05 | 0.15 | 0.01 | 0.32 | 0.15 |
| 3 | 1691090 | rs1914006 | 0.345 | 0.457 | 0.303 | -0.635 | 0.001 | -0.004 | -0.004 | 0.08 | 0.57 | 0.36 | 0.7 | 0.92 | 0.59 | 0.81 |
| 3 | 1695712 | rs4685326 | 0.197 | 0.328 | 0.329 | -0.444 | -0.006 | 0.008 | 0.008 | 0.052 | 0.91 | 0.74 | 0.89 | 0.94 | 0.95 | 0.97 |
| 3 | 1704667 | rs7645123 | 0.052 | 0.725 | 0.079 | -0.516 | -0.012 | 0.024 | 0.024 | 0.07 | 0.36 | 0.26 | 0.2 | 0.01 | 0.37 | 0.16 |
| 3 | 1707030 | rs13078044 | -0.159 | 0.693 | -0.6 | 0.394 | -0.009 | 0.006 | 0.006 | 0 | 0.4 | 0.26 | 0.2 | 0.36 | 0.39 | 0.27 |
| 3 | 1707513 | rs9864375 | -0.124 | 0.828 | -0.585 | 0.369 | 0.001 | -0.002 | -0.002 | 0 | 0.55 | 0.48 | 0.67 | 0.63 | 0.53 | 0.73 |
| 3 | 1710306 | rs1320119 | -1.178 | 1.136 | -0.099 | 0.191 | 0.07 | 0.036 | 0.036 | 0.84 | 0.13 | 0.46 | 0.36 | 0.13 | 0.22 | 0.23 |
| 3 | 1710731 | rs1317443 | 0.876 | 0.596 | -1.326 | -0.243 | 0.013 | -0.067 | -0.067 | 0.136 | 0.3 | 0.04 | 0.2 | 0.66 | 0.2 | 0.22 |
| 3 | 1711415 | rs1579005 | 0.079 | 0.719 | -0.635 | 0.103 | -0.006 | -0.007 | -0.007 | 0 | 0.07 | 0.06 | 0.28 | 0.13 | 0.19 | 0.18 |
| 3 | 1712108 | rs4292218 | -0.57 | 0.802 | -0.711 | -0.271 | 0.001 | -0.004 | -0.004 | 0 | 0.31 | 0.4 | 0.45 | 0.29 | 0.35 | 0.58 |
| 3 | 1712410 | rs4298033 | -0.398 | 1.042 | -0.647 | -0.075 | 0.037 | 0.022 | 0.022 | 0.045 | 0.1 | 0.25 | 0.31 | 0.13 | 0.2 | 0.18 |
| 3 | 1717845 | rs7636546 | 0.196 | 0.606 | -0.98 | -0.391 | 0.008 | 0.048 | 0.048 | 0.017 | 0.22 | 0.12 | 0.17 | 0.14 | 0.17 | 0.42 |
| 3 | 1720667 | rs9815216 | -1.108 | 1.102 | -0.172 | -0.714 | 0.043 | -0.109 | -0.109 | 0 | 0.85 | 0.67 | 0.91 | 1 | 0.86 | 0.91 |
| 3 | 1722181 | rs6801074 | 0.023 | 1.302 | -0.06 | -0.219 | 0.069 | 0.099 | 0.099 | 0.055 | 0.09 | 0.32 | 0.36 | 0.19 | 0.19 | 0.18 |
| 3 | 1722281 | rs12498043 | 0.405 | 1.299 | -0.06 | -0.23 | 0.011 | -0.005 | -0.005 | 0 | 0.75 | 0.35 | 0.55 | 0.8 | 0.66 | 0.73 |
| 3 | 1731173 | rs9831634 | -1.062 | 1.137 | -0.013 | -0.308 | 0.043 | -0.115 | -0.115 | 5.275 | 0.85 | 0.62 | 0.91 | 1 | 0.86 | 0.91 |
| 3 | 1738391 | rs9845938 | -0.066 | 1.339 | -0.179 | -0.604 | 0.042 | -0.163 | -0.163 | 0 | 0.76 | 0.62 | 0.83 | 1 | 0.72 | 0.88 |
| 3 | 1739724 | rs1996439 | 0.913 | 1.104 | 0.167 | -0.588 | 0.007 | 0.032 | 0.032 | 0 | 0.51 | 0.3 | 0.29 | 0.55 | 0.42 | 0.32 |
| 3 | 1740069 | rs4410431 | -0.264 | 1.009 | 0.132 | -0.629 | 0.006 | -0.034 | -0.034 | 0.285 | 0.83 | 0.8 | 0.82 | 0.91 | 0.79 | 0.96 |
| 3 | 1740624 | rs4684296 | 0.806 | 0.914 | 0.118 | -0.648 | 0.007 | 0.006 | 0.006 | 0 | 0.56 | 0.32 | 0.35 | 0.56 | 0.54 | 0.35 |
| 3 | 1744782 | rs2872742 | 0.136 | 0.993 | 0.024 | -0.519 | 0.052 | 0.116 | 0.116 | 0.32 | 0.11 | 0.33 | 0.36 | 0.23 | 0.2 | 0.21 |
| 3 | 1745013 | rs2314772 | -0.455 | 0.907 | 0.214 | -0.819 | 0.024 | -0.046 | -0.046 | 0 | 0.83 | 0.67 | 0.82 | 0.91 | 0.8 | 0.97 |
| 3 | 1752478 | rs11709143 | 1.807 | 0.534 | -0.121 | -0.771 | 0.01 | -0.051 | -0.051 | 0 | 0.33 | 0.18 | 0.2 | 0.48 | 0.24 | 0.23 |
| 3 | 1754872 | rs7630359 | -0.821 | 0.921 | -0.002 | -0.642 | -0.016 | 0.023 | 0.023 | 0.193 | 0.45 | 0.67 | 0.68 | 0.64 | 0.54 | 0.6 |
| 3 | 1756682 | rs12489545 | 0.476 | 0.951 | 0.056 | -0.623 | -0.015 | -0.048 | -0.048 | 0 | 0.27 | 0.23 | 0.48 | 0.54 | 0.34 | 0.52 |
| 3 | 1757919 | rs6795466 | 0.322 | 0.947 | 0.097 | -0.624 | 0.002 | 0.014 | 0.014 | 0.022 | 0.28 | 0.37 | 0.5 | 0.55 | 0.34 | 0.54 |
| 3 | 1764453 | rs4143121 | -1.188 | 0.842 | 0.015 | -0.608 | 0.034 | 0.031 | 0.031 | 0.017 | 0.56 | 0.73 | 0.75 | 0.66 | 0.68 | 0.7 |
| 3 | 1770321 | rs17043113 | 1.214 | 0.8 | 0.012 | -0.64 | 0.039 | -0.063 | -0.063 | 0.934 | 0.14 | 0.27 | 0.09 | 0.03 | 0.15 | 0.11 |
| 3 | 1771749 | rs6442664 | -2.066 | 0.827 | -0.156 | -0.702 | 0 | 0 | 0 | 0 | 0.65 | 0.7 | 0.79 | 0.68 | 0.73 | 0.76 |
| 3 | 1772270 | rs7648926 | 0.648 | 1.004 | -0.5 | -0.641 | 0.002 | -0.024 | -0.024 | 2.51 | 0.25 | 0.25 | 0.45 | 0.55 | 0.26 | 0.51 |
| 3 | 1772716 | rs13090057 | 0.487 | 1.171 | -0.709 | -0.609 | -0.085 | 0.033 | 0.033 | 6.346 | 0.16 | 0.01 | 0.2 | 0.06 | 0.28 | 0.09 |
| 3 | 1779650 | rs6768566 | -0.045 | 1.113 | -0.673 | -0.602 | -0.025 | 0.042 | 0.042 | 0 | 0.18 | 0.07 | 0.19 | 0.08 | 0.22 | 0.14 |
| 3 | 1781739 | rs9856348 | 1.574 | 1.022 | -0.166 | -0.633 | 0.051 | -0.055 | -0.055 | 0 | 0.12 | 0.23 | 0.08 | 0.02 | 0.15 | 0.09 |
| 3 | 1782524 | rs6808990 | -1.906 | 0.7 | -0.337 | -0.751 | 0.033 | -0.002 | -0.002 | 0.003 | 0.65 | 0.86 | 0.81 | 0.68 | 0.77 | 0.7 |
| 3 | 1783334 | rs13096082 | 0.944 | 0.749 | -0.391 | -0.743 | -0.053 | 0.004 | 0.004 | 0 | 0.14 | 0.01 | 0.16 | 0.09 | 0.2 | 0.07 |
| 3 | 1783833 | rs6776792 | -0.209 | 0.805 | -0.183 | -0.748 | -0.002 | 0.013 | 0.013 | 0 | 0.36 | 0.51 | 0.55 | 0.58 | 0.39 | 0.53 |
| 3 | 1783885 | rs9880247 | 1.613 | 0.741 | 0.118 | -0.491 | 0.048 | -0.044 | -0.044 | 0 | 0.12 | 0.2 | 0.08 | 0.02 | 0.15 | 0.09 |
| 3 | 1787581 | rs9883944 | -0.868 | 0.701 | 0.172 | -0.538 | 0.005 | -0.042 | -0.042 | 0 | 0.34 | 0.36 | 0.28 | 0.12 | 0.39 | 0.22 |
| 3 | 1789514 | rs4482650 | 1.311 | 1.018 | 0.335 | -0.401 | 0.034 | -0.047 | -0.047 | 0 | 0.13 | 0.21 | 0.09 | 0.02 | 0.15 | 0.09 |
| 3 | 1790254 | rs12638360 | 0.416 | 0.909 | -0.002 | -0.448 | -0.056 | 0.022 | 0.022 | 4.325 | 0.17 | 0.01 | 0.17 | 0.08 | 0.21 | 0.09 |
| 3 | 1790316 | rs4597690 | -1.827 | 0.898 | -0.07 | -0.389 | 0.013 | 0.01 | 0.01 | 0 | 0.59 | 0.84 | 0.72 | 0.67 | 0.7 | 0.68 |
| 3 | 1791236 | rs4432626 | -0.443 | 0.995 | 0.349 | -0.134 | 0.019 | 0.038 | 0.038 | 0 | 0.32 | 0.47 | 0.45 | 0.56 | 0.42 | 0.51 |
| 3 | 1794194 | rs13316141 | 1.211 | 1.068 | 0.307 | -0.337 | 0.035 | -0.047 | -0.047 | 0.186 | 0.12 | 0.21 | 0.08 | 0.02 | 0.14 | 0.08 |

| | | | | | | | | | | | | | | | | |
|---|---|---|---|---|---|---|---|---|---|---|---|---|---|---|---|---|
| 3 | 1807252 | rs17043656 | 0.326 | 0.79 | -0.049 | 0.046 | 0.006 | 0.061 | 0.061 | 0 | 0.17 | 0.25 | 0.37 | 0.58 | 0.26 | 0.41 |
| 3 | 1807913 | rs17043661 | -1.223 | 0.652 | -0.381 | -0.169 | 0.016 | 0.067 | 0.067 | 0.17 | 0.54 | 0.81 | 0.71 | 0.7 | 0.66 | 0.68 |
| 3 | 1810799 | rs7632433 | -0.823 | 0.517 | 0.298 | 0.516 | -0.005 | 0.042 | 0.042 | 0.031 | 0.23 | 0.34 | 0.44 | 0.6 | 0.29 | 0.43 |
| 3 | 1811952 | rs6772544 | -0.611 | 0.594 | -0.027 | 0.518 | -0.001 | -0.005 | -0.005 | 0 | 0.22 | 0.2 | 0.44 | 0.6 | 0.29 | 0.43 |
| 3 | 1814351 | rs9848095 | -0.413 | 0.706 | 0.129 | 0.033 | -0.004 | -0.059 | -0.059 | 0 | 0.13 | 0.19 | 0.11 | 0 | 0.13 | 0.06 |
| 3 | 1816205 | rs11720928 | -0.294 | 0.743 | 0.088 | 0.047 | 0.001 | -0.035 | -0.035 | 0 | 0.11 | 0.14 | 0.11 | 0 | 0.14 | 0.07 |
| 3 | 1820437 | rs11927938 | -1.54 | 0.614 | 0.235 | 0.573 | -0.001 | -0.001 | -0.001 | 0 | 0.25 | 0.69 | 0.44 | 0.24 | 0.32 | 0.27 |
| 3 | 1821213 | rs13069133 | 0.668 | 0.75 | -0.231 | 0.286 | -0.039 | 0.033 | 0.033 | 8.29 | 0.12 | 0.01 | 0.13 | 0.04 | 0.18 | 0.04 |
| 3 | 1823910 | rs6780487 | -1.643 | 0.706 | -0.132 | 0.393 | -0.025 | 0.007 | 0.007 | 1.924 | 0.38 | 0.75 | 0.54 | 0.29 | 0.49 | 0.31 |
| 3 | 1826067 | rs4603939 | -0.809 | 0.673 | 0.073 | 0.418 | 0.023 | 0.09 | 0.09 | 1.062 | 0.15 | 0.58 | 0.32 | 0.24 | 0.18 | 0.21 |
| 3 | 1827927 | rs17786361 | 0.62 | 0.627 | -0.922 | -0.24 | -0.04 | 0.07 | 0.07 | 0 | 0.21 | 0.02 | 0.19 | 0.07 | 0.17 | 0.18 |
| 3 | 1829830 | rs9861760 | -1.655 | 1.09 | -0.172 | 0.466 | 0 | 0.056 | 0.056 | 1.851 | 0.42 | 0.76 | 0.63 | 0.66 | 0.48 | 0.56 |
| 3 | 1833607 | rs6771229 | -0.447 | 1.03 | 0.674 | -0.015 | 0.004 | -0.065 | -0.065 | 0 | 0.18 | 0.43 | 0.15 | 0.05 | 0.19 | 0.04 |
| 3 | 1833898 | rs6771572 | -1.143 | 0.968 | 0.85 | -0.156 | 0.004 | -0.047 | -0.047 | 0 | 0.36 | 0.41 | 0.35 | 0.06 | 0.41 | 0.13 |
| 3 | 1834534 | rs10510216 | 0.171 | 1.043 | 0.485 | -0.084 | -0.08 | 0.065 | 0.065 | 0.75 | 0.21 | 0.02 | 0.21 | 0.06 | 0.27 | 0.06 |
| 3 | 1836718 | rs13085953 | -1.393 | 0.95 | 0.913 | 0.27 | -0.002 | -0.005 | -0.005 | 0 | 0.46 | 0.49 | 0.63 | 0.28 | 0.57 | 0.32 |
| 3 | 1840129 | rs6442690 | -1.467 | 0.932 | 0.887 | 0.254 | 0.011 | 0.036 | 0.036 | 0.368 | 0.47 | 0.67 | 0.68 | 0.29 | 0.58 | 0.33 |
| 3 | 1841292 | rs17044045 | -0.13 | 1.039 | 0.975 | 0.429 | 0.003 | 0.019 | 0.019 | 0 | 0.1 | 0.1 | 0.28 | 0.23 | 0.17 | 0.21 |
| 3 | 1852696 | rs6789391 | -0.988 | 1.119 | 0.796 | 0.31 | 0.027 | -0.024 | -0.024 | 0 | 0.33 | 0.42 | 0.53 | 0.23 | 0.44 | 0.3 |
| 3 | 1854498 | rs6796167 | -0.997 | 1.116 | 0.795 | 0.309 | 0.026 | -0.024 | -0.024 | 0.003 | 0.33 | 0.42 | 0.53 | 0.23 | 0.44 | 0.3 |
| 3 | 1855755 | rs8179883 | -0.619 | 1.146 | 0.752 | 0.426 | 0.021 | 0.049 | 0.049 | 0 | 0.12 | 0.22 | 0.3 | 0.22 | 0.18 | 0.24 |
| 3 | 1856689 | rs6442696 | 0.11 | 0.365 | 0.147 | 0.903 | -0.013 | -0.076 | -0.076 | 0 | 0.83 | 0.96 | 0.79 | 0.73 | 0.84 | 0.71 |
| 3 | 1857515 | rs9833248 | 0.118 | 0.42 | 0.595 | 0.876 | -0.005 | 0.006 | 0.006 | 0 | 0.23 | 0.18 | 0.21 | 0.11 | 0.26 | 0.07 |
| 3 | 1863397 | rs17044201 | -0.266 | 0.503 | 0.871 | 1.033 | 0.078 | 0.229 | 0.229 | 0.408 | 0.07 | 0.33 | 0.27 | 0.21 | 0.11 | 0.22 |
| 3 | 1865505 | rs9838423 | -0.511 | 0.408 | 1.197 | 1.027 | 0.027 | 0.017 | 0.017 | 0 | 0.3 | 0.48 | 0.47 | 0.35 | 0.36 | 0.32 |
| 3 | 1867614 | rs4684321 | -0.102 | 0.231 | 1.299 | 1.128 | -0.005 | 0.017 | 0.017 | 0 | 0.9 | 0.9 | 0.87 | 0.72 | 0.95 | 0.75 |
| 3 | 1871275 | rs7647217 | -1.32 | 0.369 | 1.347 | 0.986 | 0.009 | 0.013 | 0.013 | 0.109 | 0.21 | 0.44 | 0.38 | 0.18 | 0.24 | 0.39 |
| 3 | 1872403 | rs12488941 | 0.44 | 0.444 | 1.116 | 0.933 | -0.082 | 0.067 | 0.067 | 0 | 0.24 | 0.02 | 0.2 | 0.1 | 0.27 | 0.08 |
| 3 | 1874179 | rs4365613 | -1.168 | 0.257 | 0.897 | 0.618 | -0.003 | 0 | 0 | 0 | 0.23 | 0.2 | 0.36 | 0.21 | 0.28 | 0.32 |
| 3 | 1882191 | rs17044355 | -0.008 | 0.525 | 1.138 | 0.875 | 0.018 | 0.09 | 0.09 | 0.171 | 0.07 | 0.14 | 0.25 | 0.21 | 0.11 | 0.23 |
| 3 | 1882270 | rs6807710 | -1.241 | 0.483 | 1.247 | 0.994 | 0 | -0.002 | -0.002 | 0.19 | 0.46 | 0.43 | 0.59 | 0.37 | 0.52 | 0.41 |
| 3 | 1884441 | rs6781373 | -0.037 | 0.524 | 1.161 | 1.019 | 0.021 | 0.1 | 0.1 | 0.202 | 0.07 | 0.18 | 0.25 | 0.21 | 0.11 | 0.24 |
| 3 | 1884889 | rs7625240 | -0.762 | 0.511 | 0.787 | 0.923 | -0.013 | 0.086 | 0.086 | 0 | 0.39 | 0.2 | 0.33 | 0.16 | 0.4 | 0.18 |
| 3 | 1884957 | rs7635437 | -0.652 | 0.531 | 0.657 | 0.933 | -0.038 | 0.123 | 0.123 | 0 | 0.38 | 0.14 | 0.32 | 0.16 | 0.4 | 0.18 |
| 3 | 1889450 | rs6442698 | 0.221 | 0.612 | 0.634 | 0.972 | -0.035 | 0.052 | 0.052 | 0 | 0.27 | 0.09 | 0.22 | 0.13 | 0.28 | 0.08 |
| 3 | 1889552 | rs1948181 | 0.322 | 0.606 | 0.533 | 0.969 | -0.084 | 0.036 | 0.036 | 0 | 0.26 | 0.02 | 0.21 | 0.13 | 0.28 | 0.08 |
| 3 | 1894541 | rs2727965 | 1.097 | 0.538 | 0.859 | 1.133 | 0.001 | 0.013 | 0.013 | 0 | 0.56 | 0.33 | 0.42 | 0.39 | 0.57 | 0.48 |
| 3 | 1895810 | rs2029359 | -0.453 | 0.531 | 0.908 | 1.166 | 0 | -0.003 | -0.003 | 0 | 0.35 | 0.34 | 0.34 | 0.41 | 0.32 | 0.3 |
| 3 | 1899796 | rs2731513 | 1.292 | 0.19 | 0.273 | 0.742 | -0.014 | 0.038 | 0.038 | 0 | 0.69 | 0.43 | 0.49 | 0.52 | 0.71 | 0.45 |
| 3 | 1906972 | rs2035667 | 0.312 | 0.462 | 0.31 | 1.12 | -0.045 | 0.039 | 0.039 | 0 | 0.24 | 0.05 | 0.21 | 0.14 | 0.27 | 0.08 |
| 3 | 1908074 | rs9878659 | -0.993 | 0.386 | 0.388 | 1.184 | 0.002 | 0.048 | 0.048 | 0.013 | 0.42 | 0.67 | 0.61 | 0.8 | 0.42 | 0.52 |
| 3 | 1909126 | rs4685452 | -0.939 | 0.48 | -0.125 | 1.177 | 0.002 | 0.007 | 0.007 | 0 | 0.41 | 0.64 | 0.61 | 0.64 | 0.41 | 0.57 |
| 3 | 1910527 | rs7631181 | -0.383 | 0.051 | -0.488 | 1.218 | -0.007 | 0.003 | 0.003 | 0 | 0.14 | 0.13 | 0.08 | 0.2 | 0.06 | 0.3 |
| 3 | 1913371 | rs17193911 | 0.212 | 0.562 | -1.051 | 0.698 | -0.056 | 0.136 | 0.136 | 0 | 0.23 | 0.03 | 0.18 | 0.01 | 0.24 | 0.07 |
| 3 | 1913585 | rs7631764 | -0.703 | 1.142 | -1.255 | 0.979 | -0.014 | -0.046 | -0.046 | 0 | 0.24 | 0.09 | 0.4 | 0.32 | 0.25 | 0.54 |
| 3 | 1913776 | rs7610626 | -1.54 | 0.957 | -1.389 | 1.265 | -0.101 | -0.014 | -0.014 | 0.023 | 0.57 | 0.93 | 0.66 | 0.63 | 0.5 | 0.65 |
| 3 | 1917909 | rs6442700 | -1.548 | 0.787 | -1.307 | 1.078 | -0.097 | 0.085 | 0.085 | 0 | 0.57 | 0.92 | 0.68 | 0.72 | 0.5 | 0.66 |
| 3 | 1918209 | rs6803270 | 0.408 | 1.189 | -0.617 | 0.786 | -0.017 | 0.033 | 0.033 | 0 | 0.21 | 0.08 | 0.2 | 0.11 | 0.22 | 0.06 |
| 3 | 1925851 | rs6442701 | -0.66 | 0.803 | 0.362 | 0.591 | 0 | -0.006 | -0.006 | 0.116 | 0.35 | 0.37 | 0.51 | 0.29 | 0.34 | 0.37 |
| 3 | 1926058 | rs6799163 | -0.72 | 0.775 | 0.786 | 0.587 | 0.002 | -0.03 | -0.03 | 0.256 | 0.36 | 0.48 | 0.52 | 0.28 | 0.34 | 0.37 |
| 3 | 1929666 | rs11713534 | 0.397 | 0.766 | 0.652 | 0.501 | -0.005 | 0.046 | 0.046 | 0.58 | 0.23 | 0.11 | 0.18 | 0.09 | 0.22 | 0.07 |
| 3 | 1932203 | rs4685462 | -1.453 | 1.179 | 0.613 | 0.502 | -0.015 | 0.058 | 0.058 | 2.096 | 0.47 | 0.82 | 0.64 | 0.4 | 0.52 | 0.4 |
| 3 | 1933927 | rs4317081 | -0.65 | 1.282 | 0.926 | 0.616 | 0 | 0.002 | 0.002 | 0.01 | 0.36 | 0.57 | 0.53 | 0.44 | 0.35 | 0.36 |
| 3 | 1936873 | rs12637547 | -1.35 | 1.24 | 0.731 | 0.568 | -0.008 | 0.056 | 0.056 | 0.455 | 0.45 | 0.68 | 0.62 | 0.29 | 0.51 | 0.39 |
| 3 | 1946687 | rs4387949 | -0.412 | 1.115 | 1.107 | 0.65 | 0.041 | 0.176 | 0.176 | 0 | 0.1 | 0.3 | 0.34 | 0.33 | 0.1 | 0.23 |
| 3 | 1949117 | rs1825946 | -1.356 | 0.682 | 0.898 | 0.678 | -0.025 | 0.07 | 0.07 | 0 | 0.23 | 0.36 | 0.39 | 0.49 | 0.15 | 0.49 |
| 3 | 1949227 | rs11710889 | 0.971 | 0.66 | 0.102 | 0.726 | 0.032 | -0.022 | -0.022 | 0 | 0.09 | 0.03 | 0.02 | 0.14 | 0.03 | 0.18 |
| 3 | 1949294 | rs13087010 | 0.81 | 1.16 | -0.106 | 0.687 | -0.068 | 0.059 | 0.059 | 1.045 | 0.21 | 0.02 | 0.17 | 0.08 | 0.22 | 0.07 |
| 3 | 1949594 | rs9821217 | -1.612 | 1.204 | 0 | 0.714 | -0.001 | -0.011 | -0.011 | 0 | 0.54 | 0.83 | 0.7 | 0.6 | 0.55 | 0.6 |
| 3 | 1962536 | rs4685465 | -0.342 | 1.058 | -0.304 | 0.78 | 0.054 | 0.2 | 0.2 | 0.026 | 0.1 | 0.34 | 0.3 | 0.32 | 0.1 | 0.23 |
| 3 | 1968641 | rs1488749 | -1.549 | 1.077 | -1.606 | 0.374 | 0.014 | -0.082 | -0.082 | 2.332 | 0.6 | 0.85 | 0.74 | 0.55 | 0.64 | 0.63 |
| 3 | 1971616 | rs10510222 | 0.043 | 1.213 | -1.219 | 0.156 | -0.026 | 0.05 | 0.05 | 0.41 | 0.23 | 0.1 | 0.19 | 0.1 | 0.26 | 0.08 |
| 3 | 1981567 | rs1488757 | -0.262 | 0.701 | -0.975 | 0.844 | -0.003 | 0.006 | 0.006 | 0 | 0.08 | 0.04 | 0.26 | 0.28 | 0.09 | 0.23 |
| 3 | 1982410 | rs17005753 | -0.716 | 0.295 | -1.332 | 0.624 | -0.014 | -0.081 | -0.081 | 1.719 | 0.14 | 0.01 | 0.35 | 0.3 | 0.11 | 0.35 |
| 3 | 1985238 | rs6783551 | -1.092 | -0.15 | -2.102 | 0.306 | 0.051 | -0.066 | -0.066 | 0 | 0.48 | 0.94 | 0.59 | 0.52 | 0.36 | 0.55 |
| 3 | 1985637 | rs17005777 | -0.343 | -0.025 | -1.359 | 0.345 | 0.007 | 0.035 | 0.035 | 0 | 0.1 | 0.14 | 0.34 | 0.23 | 0.11 | 0.3 |
| 3 | 1989416 | rs2600029 | 0.696 | 0.203 | -1.88 | -0.061 | -0.027 | -0.043 | -0.043 | 0 | 0.25 | 0.11 | 0.19 | 0.51 | 0.3 | 0.27 |
| 3 | 1996786 | rs6763575 | 0.571 | -0.426 | -1.115 | -0.295 | 0.015 | 0.011 | 0.011 | 2.566 | 0.19 | 0.14 | 0.11 | 0.11 | 0.1 | 0.06 |
| 3 | 1997837 | rs9847882 | -0.323 | -0.05 | -0.562 | -0.62 | 0.001 | -0.046 | -0.046 | 0 | 0.21 | 0.34 | 0.13 | 0.09 | 0.23 | 0.15 |

| | | | | | | | | | | | | | | | | |
|---|---|---|---|---|---|---|---|---|---|---|---|---|---|---|---|---|
| 3 | 2000736 | rs7646172 | 0.203 | -0.598 | -1.122 | -0.906 | 0.055 | 0.081 | 0.081 | 4.713 | 0.22 | 0.06 | 0.1 | 0.04 | 0.12 | 0.07 |
| 3 | 2001802 | rs1488745 | -1.27 | -0.482 | -1.327 | -0.651 | 0.023 | 0.104 | 0.104 | 3.581 | 0.5 | 0.9 | 0.59 | 0.31 | 0.44 | 0.51 |
| 3 | 2001904 | rs1488744 | 0.088 | -0.61 | -0.118 | -0.723 | -0.038 | -0.038 | -0.038 | 0.234 | 0.16 | 0.44 | 0.08 | 0.1 | 0.09 | 0.17 |
| 3 | 2004251 | rs17005939 | -0.079 | -0.485 | -0.462 | -0.671 | -0.063 | -0.064 | -0.064 | 0 | 0.15 | 0.03 | 0.37 | 0.25 | 0.25 | 0.3 |
| 3 | 2004374 | rs6763978 | 0.448 | -0.431 | -0.431 | -0.912 | 0.023 | 0.071 | 0.071 | 1.589 | 0.2 | 0.07 | 0.14 | 0.05 | 0.15 | 0.09 |
| 3 | 2004442 | rs2129995 | 0.769 | -0.484 | -0.4 | -0.879 | 0.047 | 0.063 | 0.063 | 2.582 | 0.18 | 0.06 | 0.11 | 0.05 | 0.09 | 0.05 |
| 3 | 2006604 | rs6771006 | 1.401 | -0.444 | -0.567 | -1.01 | 0.005 | 0.049 | 0.049 | 0 | 0.1 | 0 | 0.08 | 0.04 | 0.06 | 0.06 |
| 3 | 2008523 | rs2600033 | -1.643 | -0.37 | -0.335 | -1.082 | 0.049 | 0.116 | 0.116 | 2.731 | 0.48 | 0.78 | 0.56 | 0.3 | 0.41 | 0.52 |
| 3 | 2008771 | rs11128918 | 1.603 | -0.31 | -0.358 | -1.106 | -0.04 | 0.003 | 0.003 | 0 | 0.07 | 0.01 | 0.08 | 0.03 | 0.15 | 0.03 |
| 3 | 2011341 | rs1387092 | -1.702 | -0.485 | 0.672 | -1.235 | 0.004 | -0.009 | -0.009 | 0 | 0.43 | 0.63 | 0.29 | 0.11 | 0.4 | 0.32 |
| 3 | 2017591 | rs1852447 | -1.496 | -0.647 | 0.851 | -1.025 | 0.001 | 0.002 | 0.002 | 0.381 | 0.37 | 0.62 | 0.48 | 0.27 | 0.36 | 0.46 |
| 3 | 2023020 | rs2729298 | -1.06 | -0.619 | 0.875 | -0.962 | -0.016 | 0 | 0 | 0 | 0.25 | 0.63 | 0.39 | 0.24 | 0.19 | 0.43 |
| 3 | 2025196 | rs2729306 | -1.474 | -0.627 | 0.731 | -0.975 | 0.077 | 0.083 | 0.083 | 2.637 | 0.43 | 0.78 | 0.49 | 0.28 | 0.29 | 0.49 |
| 3 | 2025910 | rs2729308 | -0.751 | -0.579 | 0.915 | -0.929 | -0.007 | -0.005 | -0.005 | 0 | 0.29 | 0.64 | 0.44 | 0.22 | 0.19 | 0.3 |
| 3 | 2028829 | rs2634113 | -1.383 | -0.622 | 1.118 | -1.016 | 0.008 | 0.027 | 0.027 | 0.405 | 0.48 | 0.58 | 0.55 | 0.3 | 0.44 | 0.51 |
| 3 | 2029483 | rs9857735 | -0.815 | -0.307 | 1.295 | -1.019 | 0.002 | -0.032 | -0.032 | 0 | 0.36 | 0.53 | 0.51 | 0.24 | 0.34 | 0.33 |
| 3 | 2029929 | rs2600046 | -0.435 | -0.431 | 0.995 | -1.092 | -0.007 | 0.036 | 0.036 | 0 | 0.17 | 0.3 | 0.33 | 0.21 | 0.14 | 0.25 |
| 3 | 2030468 | rs6802020 | -0.605 | -0.497 | 0.65 | -1.113 | -0.007 | -0.1 | -0.1 | 0.083 | 0.33 | 0.42 | 0.22 | 0.1 | 0.32 | 0.28 |
| 3 | 2031953 | rs2600048 | 1.674 | -0.505 | 0.567 | -1.325 | -0.003 | 0.034 | 0.034 | 0.195 | 0.46 | 0.36 | 0.43 | 0.69 | 0.52 | 0.37 |
| 3 | 2032413 | rs7433476 | -0.505 | -0.496 | 0.563 | -1.324 | -0.011 | -0.121 | -0.121 | 1.174 | 0.34 | 0.45 | 0.22 | 0.1 | 0.31 | 0.29 |
| 3 | 2032902 | rs6791289 | 1.14 | -0.52 | 0.394 | -1.314 | -0.014 | -0.026 | -0.026 | 0 | 0.12 | 0.15 | 0.04 | 0.06 | 0.08 | 0.2 |
| 3 | 2033356 | rs2729312 | 0.763 | -0.306 | -0.008 | -1.181 | 0 | 0.015 | 0.015 | 0.145 | 0.55 | 0.64 | 0.49 | 0.72 | 0.56 | 0.37 |
| 3 | 2035573 | rs2634135 | -0.323 | -0.353 | -0.162 | -1.103 | -0.005 | 0 | 0 | 0 | 0.8 | 0.71 | 0.65 | 0.83 | 0.84 | 0.66 |
| 3 | 2035726 | rs2727950 | 0.611 | -0.03 | -0.747 | -0.562 | 0.002 | 0.035 | 0.035 | 0.506 | 0.46 | 0.29 | 0.43 | 0.65 | 0.53 | 0.36 |
| 3 | 2038356 | rs2727952 | -0.072 | 0.641 | -1.161 | 0.129 | -0.005 | -0.027 | -0.027 | 0 | 0.26 | 0.2 | 0.2 | 0.54 | 0.26 | 0.4 |
| 3 | 2047418 | rs2600062 | -0.202 | 0.681 | -1.204 | -0.131 | -0.001 | 0.005 | 0.005 | 0.298 | 0.59 | 0.66 | 0.54 | 0.62 | 0.57 | 0.42 |
| 3 | 2050777 | rs2727956 | 0.279 | 0.657 | -1.201 | 0.183 | 0.001 | 0.007 | 0.007 | 0 | 0.23 | 0.22 | 0.28 | 0.52 | 0.27 | 0.27 |
| 3 | 2057749 | rs17786584 | 1.38 | -0.083 | -1.533 | -0.03 | 0.023 | 0.032 | 0.032 | 1.383 | 0.1 | 0.01 | 0.06 | 0.05 | 0.03 | 0.05 |
| 3 | 2058537 | rs9835207 | 0.101 | -0.401 | -1.458 | -0.171 | -0.015 | -0.014 | -0.014 | 0 | 0.58 | 0.46 | 0.7 | 0.7 | 0.66 | 0.62 |
| 3 | 2059893 | rs4593008 | 1.35 | -0.072 | -1.944 | -0.148 | -0.047 | -0.018 | -0.018 | 0 | 0.26 | 0.07 | 0.39 | 0.26 | 0.32 | 0.23 |
| 3 | 2064647 | rs2729319 | -1.682 | 0.021 | -2.03 | -0.094 | -0.101 | -0.017 | -0.017 | 1.148 | 0.77 | 0.98 | 0.79 | 0.78 | 0.71 | 0.8 |
| 3 | 2066897 | rs4522753 | 0.702 | 0.082 | -2.077 | -0.242 | -0.103 | -0.026 | -0.026 | 0 | 0.27 | 0.05 | 0.27 | 0.26 | 0.36 | 0.24 |
| 3 | 2070846 | rs7616568 | -0.08 | -0.065 | -1.657 | -0.255 | 0.002 | -0.003 | -0.003 | 0 | 0.46 | 0.4 | 0.56 | 0.51 | 0.42 | 0.46 |
| 3 | 2071931 | rs2729324 | 1.355 | -0.14 | -1.347 | -0.152 | -0.012 | 0.015 | 0.015 | 0.155 | 0.27 | 0.45 | 0.2 | 0.24 | 0.26 | 0.31 |
| 3 | 2074515 | rs9878112 | 1.036 | -0.224 | -1.288 | 0.256 | -0.006 | 0.009 | 0.009 | 0 | 0.19 | 0.28 | 0.11 | 0.24 | 0.15 | 0.17 |
| 3 | 2076026 | rs4685479 | -0.512 | -0.234 | -1.296 | 0.226 | 0.003 | 0.008 | 0.008 | 0 | 0.61 | 0.57 | 0.66 | 0.53 | 0.58 | 0.51 |
| 3 | 2078499 | rs4434131 | -0.023 | -0.469 | -1.127 | 0.257 | -0.017 | -0.014 | -0.014 | 0 | 0.39 | 0.25 | 0.34 | 0.44 | 0.43 | 0.38 |
| 3 | 2079706 | rs2729021 | -0.063 | -1.674 | -0.211 | 0.325 | -0.086 | 0.153 | 0.153 | 0.367 | 0.15 | 0.54 | 0.08 | 0.23 | 0.06 | 0.22 |
| 3 | 2079789 | rs4575885 | 0.715 | -1.649 | -0.296 | 0.334 | -0.047 | -0.077 | -0.077 | 0 | 0.22 | 0.02 | 0.22 | 0.28 | 0.23 | 0.23 |
| 3 | 2080561 | rs6767146 | 1.531 | -1.81 | 0.592 | 0.413 | -0.078 | 0.047 | 0.047 | 0.05 | 0.73 | 0.4 | 0.69 | 0.67 | 0.86 | 0.65 |
| 3 | 2082992 | rs6793934 | 0.838 | -1.651 | 0.71 | 0.533 | 0.006 | 0.126 | 0.126 | 2.841 | 0.92 | 0.63 | 0.94 | 0.88 | 0.97 | 0.98 |
| 3 | 2087260 | rs2729330 | -1.697 | -1.676 | 0.568 | 0.494 | -0.082 | 0.062 | 0.062 | 0 | 0.26 | 0.57 | 0.29 | 0.33 | 0.14 | 0.35 |
| 3 | 2089027 | rs2729265 | -1.488 | -1.325 | 0.544 | 0.154 | -0.045 | 0.05 | 0.05 | 0.001 | 0.17 | 0.34 | 0.26 | 0.22 | 0.12 | 0.14 |
| 3 | 2095542 | rs9858266 | -0.656 | -1.003 | 0 | -0.603 | -0.004 | -0.013 | -0.013 | 0.166 | 0.47 | 0.45 | 0.42 | 0.17 | 0.36 | 0.47 |
| 3 | 2101328 | rs17786608 | 0.715 | -1.136 | -0.125 | -0.737 | -0.035 | 0.157 | 0.157 | 0 | 0.27 | 0.06 | 0.2 | 0 | 0.27 | 0.15 |
| 3 | 2103131 | rs6777569 | -1.262 | -1.05 | 0.013 | -0.781 | 0.022 | 0.056 | 0.056 | 0 | 0.52 | 0.37 | 0.45 | 0.17 | 0.42 | 0.49 |
| 3 | 2104048 | rs17006968 | -1.174 | -1.001 | 0.535 | -0.219 | -0.034 | 0.004 | 0.004 | 0 | 0.19 | 0.22 | 0.21 | 0.17 | 0.09 | 0.32 |
| 3 | 2109880 | rs9825844 | 0.907 | -1.007 | 0.466 | -0.208 | -0.036 | 0.005 | 0.005 | 0 | 0.08 | 0.11 | 0.02 | 0.06 | 0.02 | 0.21 |
| 3 | 2114929 | rs9880266 | -1.774 | -1.015 | 0.462 | -0.219 | -0.056 | -0.013 | -0.013 | 2.474 | 0.25 | 0.39 | 0.25 | 0.17 | 0.15 | 0.34 |
| 3 | 2119361 | rs17007173 | -1.874 | -0.467 | 0.198 | -0.499 | -0.013 | -0.019 | -0.019 | 0 | 0.17 | 0.27 | 0.24 | 0.1 | 0.13 | 0.13 |
| 3 | 2122311 | rs9854866 | -1.78 | -0.29 | 0.302 | -0.476 | 0.022 | -0.069 | -0.069 | 0 | 0.12 | 0.41 | 0.23 | 0 | 0.1 | 0.07 |
| 3 | 2127845 | rs13079877 | -0.798 | 0.087 | -0.522 | -0.326 | -0.016 | -0.002 | -0.002 | 0 | 0.56 | 0.69 | 0.48 | 0.47 | 0.49 | 0.59 |
| 3 | 2127911 | rs3935773 | 0.744 | 0.324 | -0.557 | -0.367 | -0.022 | 0.109 | 0.109 | 0 | 0.23 | 0.09 | 0.14 | 0 | 0.21 | 0.15 |
| 3 | 2130963 | rs6791544 | 0.334 | 0.76 | -0.3 | -0.017 | -0.002 | 0.01 | 0.01 | 0.249 | 0.25 | 0.22 | 0.26 | 0.46 | 0.23 | 0.43 |
| 3 | 2133377 | rs2729042 | -0.242 | 0.614 | -0.796 | 0.033 | -0.001 | -0.013 | -0.013 | 0.857 | 0.63 | 0.59 | 0.62 | 0.86 | 0.68 | 0.72 |
| 3 | 2134765 | rs2729283 | -0.217 | 0.669 | -0.773 | 0.058 | 0 | -0.011 | -0.011 | 0 | 0.63 | 0.59 | 0.62 | 0.86 | 0.68 | 0.72 |
| 3 | 2135638 | rs2729284 | 0.748 | 0.376 | -0.729 | -0.044 | -0.003 | 0.001 | 0.001 | 0.159 | 0.38 | 0.34 | 0.37 | 0.4 | 0.48 | 0.3 |
| 3 | 2137538 | rs2729287 | 0.36 | 0.531 | -0.776 | -0.311 | -0.002 | -0.021 | -0.021 | 0 | 0.66 | 0.61 | 0.65 | 0.88 | 0.74 | 0.74 |
| 3 | 2145996 | rs2727926 | -0.512 | 1.057 | -0.912 | 0.24 | 0 | 0.006 | 0.006 | 0.044 | 0.43 | 0.4 | 0.52 | 0.48 | 0.52 | 0.45 |
| 3 | 2146283 | rs10470643 | -0.628 | 1.794 | -0.358 | -0.06 | 0.002 | -0.003 | -0.003 | 2.44 | 0.42 | 0.42 | 0.48 | 0.47 | 0.45 | 0.43 |
| 3 | 2146692 | rs4459884 | -0.154 | 2.716 | -0.512 | -0.133 | -0.011 | -0.015 | -0.015 | 0 | 0.18 | 0.11 | 0.27 | 0.45 | 0.21 | 0.24 |
| 3 | 2146913 | rs4461405 | -0.224 | 2.752 | -0.36 | -0.141 | 0.027 | 0.055 | 0.055 | 0 | 0.18 | 0.23 | 0.28 | 0.45 | 0.21 | 0.24 |
| 3 | 2150400 | rs2729005 | -1.317 | 3.043 | -0.853 | -0.176 | 0.007 | 0 | 0 | 0 | 0.49 | 0.46 | 0.7 | 0.55 | 0.61 | 0.42 |
| 3 | 2152370 | rs17007773 | -0.057 | 3.065 | -0.835 | 0.388 | 0.022 | 0.047 | 0.047 | 0 | 0.19 | 0.22 | 0.28 | 0.45 | 0.21 | 0.25 |
| 3 | 2157357 | rs2729007 | -0.158 | 2.723 | -1.184 | 0.221 | -0.063 | -0.018 | -0.018 | 0 | 0.15 | 0.03 | 0.29 | 0.1 | 0.23 | 0.06 |
| 3 | 2160106 | rs9842506 | -1.066 | 2.188 | -1.573 | 0.353 | -0.045 | 0 | 0 | 0 | 0.25 | 0.1 | 0.37 | 0.15 | 0.31 | 0.13 |
| 3 | 2161513 | rs2727905 | -1.391 | 1.363 | -1.335 | 0.649 | 0 | -0.007 | -0.007 | 0 | 0.52 | 0.51 | 0.64 | 0.71 | 0.48 | 0.6 |
| 3 | 2165384 | rs4552334 | -0.727 | 1.216 | -1.373 | 1.084 | -0.002 | 0.012 | 0.012 | 0 | 0.07 | 0.08 | 0.09 | 0.2 | 0.06 | 0.11 |
| 3 | 2169803 | rs4129976 | -0.183 | 0.938 | -1.407 | 1.214 | 0.023 | 0.003 | 0.003 | 0 | 0.76 | 0.9 | 0.83 | 0.74 | 0.85 | 0.62 |

| | | | | | | | | | | | | | | | | |
|---|---|---|---|---|---|---|---|---|---|---|---|---|---|---|---|---|
| 3 | 2170519 | rs3924068 | -0.397 | 0.965 | -1.424 | 1.243 | 0 | -0.012 | -0.012 | 0 | 0.06 | 0.03 | 0.08 | 0.21 | 0.06 | 0.12 |
| 3 | 2175896 | rs6786748 | -0.322 | 0.941 | -1.437 | 1.189 | 0.019 | -0.014 | -0.014 | 0.389 | 0.77 | 0.93 | 0.83 | 0.74 | 0.85 | 0.61 |
| 3 | 2183750 | rs2727918 | -1.43 | 0.681 | -1.257 | 0.839 | 0 | -0.001 | -0.001 | 0 | 0.56 | 0.55 | 0.65 | 0.56 | 0.5 | 0.56 |
| 3 | 2184749 | rs2727922 | -1.165 | 0.642 | -1.272 | 0.895 | 0 | 0.003 | 0.003 | 0 | 0.52 | 0.44 | 0.63 | 0.56 | 0.5 | 0.56 |
| 3 | 2186551 | rs4685485 | 0.478 | 0.441 | -1.182 | 0.776 | -0.026 | -0.008 | -0.008 | 0 | 0.6 | 0.57 | 0.71 | 0.6 | 0.79 | 0.52 |
| 3 | 2194511 | rs2729018 | -1.484 | 0.26 | -1.118 | 0.733 | -0.009 | 0.025 | 0.025 | 0 | 0.57 | 0.66 | 0.65 | 0.69 | 0.49 | 0.61 |
| 3 | 2195312 | rs7649507 | 0.445 | -0.15 | -2.255 | 0.755 | -0.039 | -0.141 | -0.141 | 0 | 0.27 | 0.05 | 0.17 | 0.52 | 0.3 | 0.36 |
| 3 | 2195364 | rs2727925 | -0.073 | -0.994 | -1.89 | 0.646 | 0.058 | -0.017 | -0.017 | 0 | 0.63 | 0.75 | 0.72 | 0.56 | 0.81 | 0.5 |
| 3 | 2197796 | rs13327252 | -0.082 | -0.34 | -1.847 | 0.549 | -0.035 | 0.032 | 0.032 | 0 | 0.42 | 0.27 | 0.63 | 0.58 | 0.48 | 0.46 |
| 3 | 2197958 | rs17008491 | 2.951 | -0.396 | -1.561 | 0.644 | -0.014 | 0.027 | 0.027 | 0 | 0.06 | 0.08 | 0.02 | 0.16 | 0.02 | 0.1 |
| 3 | 2198673 | rs17194140 | 0.898 | 0.052 | -1.987 | 0.649 | -0.11 | -0.058 | -0.058 | 0 | 0.18 | 0.01 | 0.16 | 0.13 | 0.31 | 0.16 |
| 3 | 2200851 | rs9310645 | 0.04 | 0.05 | -1.747 | 0.21 | -0.007 | -0.008 | -0.008 | 0 | 0.34 | 0.4 | 0.49 | 0.74 | 0.24 | 0.47 |
| 3 | 2201147 | rs17786674 | 0.99 | 0.334 | -1.814 | 0.317 | -0.11 | 0.041 | 0.041 | 0 | 0.18 | 0.01 | 0.16 | 0.06 | 0.31 | 0.16 |
| 3 | 2206373 | rs2727927 | -0.341 | -0.01 | -2.005 | 0.445 | 0.01 | -0.007 | -0.007 | 0.378 | 0.62 | 0.64 | 0.75 | 0.58 | 0.71 | 0.59 |
| 3 | 2206702 | rs2320375 | 0.428 | 0.104 | -1.971 | 0.472 | -0.087 | -0.08 | -0.08 | 0 | 0.25 | 0.12 | 0.47 | 0.38 | 0.38 | 0.32 |
| 3 | 2208203 | rs4685491 | -0.271 | -0.298 | -1.549 | 0.043 | -0.003 | -0.009 | -0.009 | 1.031 | 0.43 | 0.5 | 0.55 | 0.69 | 0.35 | 0.56 |
| 3 | 2217011 | rs4684337 | 1.004 | -0.468 | -1.926 | -0.525 | -0.168 | 0.174 | 0.174 | 0 | 0.28 | 0.03 | 0.24 | 0.02 | 0.42 | 0.2 |
| 3 | 2223722 | rs7613165 | 0.553 | -0.539 | -1.639 | -0.144 | 0.012 | -0.036 | -0.036 | 0 | 0.3 | 0.19 | 0.3 | 0.49 | 0.24 | 0.39 |
| 3 | 2227373 | rs17009194 | 0.42 | -0.522 | -1.32 | -0.161 | -0.017 | 0.031 | 0.031 | 0.56 | 0.32 | 0.42 | 0.32 | 0.49 | 0.24 | 0.39 |
| 3 | 2231148 | rs2171475 | 0.994 | -0.482 | -1.635 | -0.162 | -0.006 | -0.035 | -0.035 | 0 | 0.13 | 0.09 | 0.27 | 0.25 | 0.13 | 0.22 |
| 3 | 2232424 | rs11720157 | 0.172 | -0.986 | -1.514 | -0.183 | 0.04 | 0.105 | 0.105 | 0.349 | 0.49 | 0.28 | 0.53 | 0.28 | 0.55 | 0.42 |
| 3 | 2233338 | rs2320378 | -0.93 | -0.221 | -0.912 | -0.621 | -0.009 | -0.02 | -0.02 | 0 | 0.79 | 0.58 | 0.82 | 0.77 | 0.77 | 0.81 |
| 3 | 2247857 | rs7641285 | -0.127 | -0.325 | -1.329 | -0.595 | 0.005 | 0.02 | 0.02 | 0 | 0.69 | 0.71 | 0.72 | 0.83 | 0.73 | 0.68 |
| 3 | 2254736 | rs1153535 | -0.411 | -0.557 | -1.509 | -0.812 | 0.049 | 0.065 | 0.065 | 0.039 | 0.68 | 0.78 | 0.58 | 0.87 | 0.81 | 0.76 |
| 3 | 2255344 | rs1153533 | 0.091 | -0.22 | -1.41 | -0.51 | 0.013 | 0.003 | 0.003 | 0.18 | 0.59 | 0.66 | 0.51 | 0.63 | 0.72 | 0.58 |
| 3 | 2256286 | rs1178517 | -0.165 | -0.527 | -1.766 | -0.59 | 0.046 | 0.005 | 0.005 | 0.022 | 0.55 | 0.67 | 0.47 | 0.45 | 0.7 | 0.61 |
| 3 | 2256426 | rs17010381 | 1.764 | -0.18 | -1.699 | -0.325 | -0.031 | -0.042 | -0.042 | 0 | 0.08 | 0.01 | 0.06 | 0.24 | 0.09 | 0.18 |
| 3 | 2260651 | rs1183149 | -0.381 | 0.941 | -1.159 | -0.72 | -0.003 | -0.006 | -0.006 | 0 | 0.36 | 0.39 | 0.32 | 0.4 | 0.43 | 0.31 |
| 3 | 2264609 | rs4685500 | 1.69 | 1.16 | -1.124 | -1.056 | -0.073 | 0.05 | 0.05 | 0 | 0.14 | 0.02 | 0.13 | 0.03 | 0.22 | 0.09 |
| 3 | 2266520 | rs1178484 | -0.244 | 0.934 | -0.874 | -0.985 | 0.034 | -0.048 | -0.048 | 0 | 0.09 | 0.35 | 0.08 | 0 | 0.12 | 0.08 |
| 3 | 2267782 | rs1153524 | 0.092 | 1.069 | -1.201 | -0.953 | -0.014 | 0.011 | 0.011 | 0.22 | 0.06 | 0.03 | 0.04 | 0 | 0.11 | 0.05 |
| 3 | 2270812 | rs7613221 | 0.71 | 0.546 | -1.208 | -1.26 | -0.002 | -0.042 | -0.042 | 0.007 | 0.33 | 0.55 | 0.23 | 0.18 | 0.32 | 0.33 |
| 3 | 2273611 | rs6442720 | 0.332 | -0.022 | -1.168 | -1.182 | 0.007 | 0.037 | 0.037 | 0 | 0.48 | 0.57 | 0.37 | 0.24 | 0.55 | 0.43 |
| 3 | 2277951 | rs17010936 | 1.141 | -0.236 | -1.297 | -1.15 | 0.065 | -0.039 | -0.039 | 0.485 | 0.31 | 0.08 | 0.4 | 0.35 | 0.19 | 0.37 |
| 3 | 2278069 | rs6795508 | 0.208 | 0.491 | -0.178 | -0.566 | -0.019 | -0.05 | -0.05 | 2.891 | 0.79 | 0.82 | 0.78 | 0.59 | 0.73 | 0.8 |
| 3 | 2281628 | rs1153520 | -0.689 | 0.528 | 0.036 | -0.567 | -0.023 | -0.057 | -0.057 | 0 | 0.84 | 0.89 | 0.82 | 0.56 | 0.8 | 0.85 |
| 3 | 2292019 | rs1523320 | 0.924 | 0.569 | -0.003 | -0.751 | -0.005 | -0.007 | -0.007 | 0.056 | 0.6 | 0.55 | 0.64 | 0.53 | 0.54 | 0.7 |
| 3 | 2302790 | rs1476184 | 1.414 | 0.409 | 0.129 | -0.705 | 0 | 0 | 0 | 0 | 0.52 | 0.46 | 0.6 | 0.53 | 0.51 | 0.66 |
| 3 | 2308394 | rs4685504 | 0.657 | 0.36 | -0.198 | -0.708 | 0 | 0.001 | 0.001 | 2.651 | 0.6 | 0.37 | 0.62 | 0.53 | 0.57 | 0.69 |
| 3 | 2309379 | rs1868218 | 0.913 | 0.483 | -0.129 | -0.647 | -0.008 | 0.027 | 0.027 | 3.155 | 0.66 | 0.38 | 0.71 | 0.55 | 0.71 | 0.75 |
| 3 | 2314035 | rs11917323 | 0.776 | 0.515 | -0.181 | -0.654 | -0.003 | -0.004 | -0.004 | 0 | 0.58 | 0.35 | 0.61 | 0.53 | 0.56 | 0.69 |
| 3 | 2315400 | rs2044602 | 1.204 | 0.645 | -0.145 | -0.579 | -0.002 | -0.002 | -0.002 | 0 | 0.53 | 0.3 | 0.61 | 0.53 | 0.55 | 0.67 |
| 3 | 2316806 | rs1880042 | 0.186 | 0.602 | -0.059 | -0.526 | 0.012 | 0.079 | 0.079 | 3.785 | 0.72 | 0.43 | 0.72 | 0.56 | 0.72 | 0.77 |
| 3 | 2325103 | rs11923606 | 0.104 | 0.671 | -0.518 | -0.454 | 0.014 | 0.079 | 0.079 | 6.371 | 0.23 | 0.33 | 0.25 | 0.45 | 0.24 | 0.2 |
| 3 | 2327135 | rs9841294 | 0.401 | 0.781 | -0.694 | -0.053 | -0.002 | -0.001 | -0.001 | 0 | 0.52 | 0.4 | 0.61 | 0.54 | 0.53 | 0.67 |
| 3 | 2331459 | rs1851612 | 0.478 | 0.885 | -0.704 | 0.011 | -0.001 | 0 | 0 | 1.558 | 0.49 | 0.4 | 0.61 | 0.54 | 0.52 | 0.65 |
| 3 | 2332104 | rs9310707 | 0.729 | 1.147 | -0.711 | 0.168 | -0.001 | -0.003 | -0.003 | 0 | 0.31 | 0.33 | 0.49 | 0.53 | 0.37 | 0.57 |
| 3 | 2332139 | rs1720199 | -1.08 | 0.622 | -0.803 | 0.018 | -0.014 | 0.035 | 0.035 | 0 | 0.88 | 0.95 | 0.92 | 0.96 | 0.85 | 0.95 |
| 3 | 2335062 | rs11915941 | 0.882 | 0.512 | -0.736 | 0.114 | -0.007 | 0.002 | 0.002 | 0 | 0.13 | 0.08 | 0.07 | 0.07 | 0.14 | 0.07 |
| 3 | 2338231 | rs1720201 | -0.548 | 0.427 | -0.108 | 0.329 | 0.043 | 0.114 | 0.114 | 0 | 0.24 | 0.42 | 0.3 | 0.59 | 0.28 | 0.2 |
| 3 | 2340743 | rs1178487 | -1.199 | 0.352 | 0.107 | 0.162 | -0.025 | 0.041 | 0.041 | 0 | 0.43 | 0.71 | 0.46 | 0.65 | 0.5 | 0.31 |
| 3 | 2342825 | rs1178491 | -0.741 | -0.381 | 0.584 | 0.001 | 0 | 0.005 | 0.005 | 0 | 0.41 | 0.59 | 0.38 | 0.61 | 0.43 | 0.25 |
| 3 | 2343448 | rs1178495 | -0.435 | -0.223 | 1.018 | 0.22 | -0.021 | 0.084 | 0.084 | 0 | 0.23 | 0.33 | 0.27 | 0.49 | 0.24 | 0.18 |
| 3 | 2345773 | rs1566383 | -1.575 | -0.279 | 1 | 0.141 | -0.011 | 0.105 | 0.105 | 0 | 0.51 | 0.72 | 0.48 | 0.66 | 0.52 | 0.34 |
| 3 | 2347409 | rs12632736 | 0.753 | -0.084 | 1.193 | 0.114 | 0.016 | -0.011 | -0.011 | 0.184 | 0.13 | 0.06 | 0.17 | 0.49 | 0.1 | 0.11 |
| 3 | 2347522 | rs1626056 | -0.461 | -0.04 | 1.194 | 0.14 | -0.014 | 0.064 | 0.064 | 0.025 | 0.23 | 0.29 | 0.27 | 0.49 | 0.24 | 0.18 |
| 3 | 2348159 | rs1666349 | -1.126 | -0.021 | 1.163 | 0.107 | 0.013 | 0.006 | 0.006 | 0 | 0.45 | 0.67 | 0.46 | 0.55 | 0.48 | 0.29 |
| 3 | 2350984 | rs1153483 | -1.073 | 0.016 | 1.252 | 0.196 | 0.002 | 0.113 | 0.113 | 0 | 0.25 | 0.48 | 0.29 | 0.53 | 0.27 | 0.19 |
| 3 | 2353515 | rs1967597 | -0.261 | -0.168 | 0.527 | 0.213 | -0.028 | 0.09 | 0.09 | 0.695 | 0.19 | 0.25 | 0.24 | 0.48 | 0.15 | 0.17 |
| 3 | 2355167 | rs11129096 | 0.311 | 0.005 | 0.286 | 0.208 | 0 | -0.003 | -0.003 | 0.306 | 0.64 | 0.67 | 0.69 | 0.48 | 0.61 | 0.75 |
| 3 | 2357019 | rs984085 | 0.276 | -0.068 | -0.081 | -0.444 | 0.027 | 0.003 | 0.003 | 0 | 0.35 | 0.19 | 0.3 | 0.71 | 0.3 | 0.45 |
| 3 | 2357058 | rs12494110 | 1.223 | 0.076 | 0.187 | -0.819 | -0.031 | 0.075 | 0.075 | 0 | 0.3 | 0.2 | 0.47 | 0.13 | 0.35 | 0.34 |
| 3 | 2357190 | rs12490811 | 0.673 | -0.034 | 0.173 | -0.828 | -0.033 | 0.173 | 0.173 | 0 | 0.46 | 0.2 | 0.56 | 0.2 | 0.53 | 0.41 |
| 3 | 2364025 | rs12633765 | 0.592 | -0.355 | 0.061 | -0.988 | 0.022 | 0.052 | 0.052 | 0.191 | 0.31 | 0.1 | 0.26 | 0.81 | 0.25 | 0.44 |
| 3 | 2364600 | rs1145044 | -0.347 | 0.084 | 0.56 | -0.884 | 0.07 | -0.062 | -0.062 | 0 | 0.89 | 0.67 | 0.85 | 1 | 0.86 | 0.92 |
| 3 | 2368791 | rs12491715 | -0.018 | 0.023 | 0.531 | -0.903 | 0.008 | 0.07 | 0.07 | 0 | 0.39 | 0.19 | 0.32 | 0.82 | 0.35 | 0.5 |
| 3 | 2375542 | rs1178536 | 0.067 | 0.03 | 0.711 | -0.892 | 0.069 | -0.079 | -0.079 | 0 | 0.88 | 0.56 | 0.84 | 1 | 0.85 | 0.92 |
| 3 | 2376679 | rs1178542 | -0.443 | -0.046 | 0.629 | -0.866 | -0.01 | -0.058 | -0.058 | 0 | 0.73 | 0.62 | 0.83 | 0.95 | 0.77 | 0.86 |
| 3 | 2384160 | rs1720180 | -1.358 | -0.099 | 0.647 | -0.846 | 0.022 | -0.026 | -0.026 | 0.319 | 0.93 | 0.77 | 0.93 | 1 | 0.95 | 0.95 |

| | | | | | | | | | | | | | | | | |
|---|---|---|---|---|---|---|---|---|---|---|---|---|---|---|---|---|
| 3 | 2390612 | rs1685495 | 0.482 | -0.127 | 0.623 | -0.889 | 0.001 | 0.051 | 0.051 | 0 | 0.51 | 0.48 | 0.54 | 0.17 | 0.52 | 0.43 |
| 3 | 2391940 | rs1685496 | 1.361 | -0.109 | 0.626 | -0.88 | -0.003 | -0.008 | -0.008 | 0 | 0.31 | 0.35 | 0.45 | 0.13 | 0.34 | 0.35 |
| 3 | 2399760 | rs1502574 | 0.476 | -0.485 | 0.762 | -0.738 | -0.012 | -0.06 | -0.06 | 0.146 | 0.26 | 0.36 | 0.17 | 0.05 | 0.23 | 0.13 |
| 3 | 2400967 | rs6785626 | -0.866 | -0.551 | 0.652 | -0.755 | 0.003 | -0.081 | -0.081 | 0 | 0.63 | 0.42 | 0.48 | 0.88 | 0.61 | 0.65 |
| 3 | 2403275 | rs6442726 | -1.52 | -0.449 | 0.499 | -0.698 | -0.054 | 0.115 | 0.115 | 0 | 0.71 | 0.87 | 0.57 | 0.87 | 0.66 | 0.65 |
| 3 | 2408010 | rs4685518 | 0.099 | -0.349 | 0.439 | -0.392 | -0.002 | -0.001 | -0.001 | 0 | 0.37 | 0.37 | 0.29 | 0.32 | 0.36 | 0.43 |
| 3 | 2409519 | rs9876065 | -1.662 | -0.316 | -0.043 | -0.455 | -0.046 | 0.073 | 0.073 | 0 | 0.73 | 0.85 | 0.53 | 0.88 | 0.62 | 0.63 |
| 3 | 2409532 | rs9876071 | -0.254 | -0.237 | 0.04 | -0.293 | -0.001 | -0.001 | -0.001 | 0.23 | 0.45 | 0.52 | 0.35 | 0.49 | 0.42 | 0.53 |
| 3 | 2412714 | rs12631357 | 0.903 | 0.161 | -0.773 | -0.333 | -0.014 | 0.035 | 0.035 | 0 | 0.19 | 0.02 | 0.08 | 0.05 | 0.17 | 0.08 |
| 3 | 2412847 | rs6792956 | -0.42 | 0.295 | -0.698 | -0.272 | 0.003 | 0.009 | 0.009 | 0.141 | 0.4 | 0.54 | 0.33 | 0.39 | 0.4 | 0.44 |
| 3 | 2415367 | rs9879546 | -1.336 | -0.087 | -1.107 | -0.655 | 0.011 | -0.077 | -0.077 | 0.008 | 0.63 | 0.57 | 0.44 | 0.88 | 0.55 | 0.58 |
| 3 | 2418270 | rs17013470 | -0.098 | 0.482 | -1.99 | 0.325 | -0.004 | 0 | 0 | 0 | 0.12 | 0.1 | 0.09 | 0 | 0.14 | 0.07 |
| 3 | 2420109 | rs1827452 | 0.487 | 0.751 | -1.799 | 0.725 | -0.03 | 0.049 | 0.049 | 0 | 0.06 | 0 | 0.11 | 0.5 | 0.08 | 0.09 |
| 3 | 2421201 | rs17013501 | -0.25 | 0.123 | -1.46 | 0.464 | 0.003 | -0.002 | -0.002 | 0 | 0.33 | 0.28 | 0.23 | 0.31 | 0.29 | 0.41 |
| 3 | 2426255 | rs11715813 | -0.629 | 0.523 | -1.84 | 0.239 | -0.046 | -0.01 | -0.01 | 0.142 | 0.18 | 0.01 | 0.08 | 0.1 | 0.18 | 0.1 |
| 3 | 2426632 | rs9854419 | 0.013 | 0.107 | -1.692 | -0.893 | 0.012 | 0.007 | 0.007 | 1.1 | 0.55 | 0.61 | 0.28 | 0.09 | 0.38 | 0.42 |
| 3 | 2427030 | rs1827451 | -0.564 | -0.497 | -1.255 | -0.202 | -0.02 | 0.014 | 0.014 | 0 | 0.28 | 0.37 | 0.23 | 0.61 | 0.17 | 0.44 |
| 3 | 2427838 | rs12636837 | -0.32 | -0.229 | -1.642 | -0.86 | -0.005 | 0.049 | 0.049 | 0 | 0.22 | 0.14 | 0.16 | 0.07 | 0.23 | 0.12 |
| 3 | 2428151 | rs1032784 | 0.3 | 0.058 | -1.627 | -0.938 | 0 | -0.002 | -0.002 | 0 | 0.24 | 0.18 | 0.39 | 0.25 | 0.22 | 0.24 |
| 3 | 2429860 | rs1393483 | -0.781 | 0.245 | -1.624 | -1.3 | -0.001 | -0.004 | -0.004 | 1.177 | 0.89 | 0.92 | 0.93 | 0.98 | 0.91 | 0.96 |
| 3 | 2431117 | rs7373598 | 0.352 | 0.321 | -0.987 | -1.251 | -0.002 | 0.007 | 0.007 | 0.088 | 0.39 | 0.35 | 0.36 | 0.65 | 0.43 | 0.57 |
| 3 | 2431546 | rs1604553 | 0.467 | 1.104 | -1.22 | -0.377 | -0.039 | -0.034 | -0.034 | 0 | 0.08 | 0 | 0.1 | 0.56 | 0.13 | 0.34 |
| 3 | 2435268 | rs13067318 | -0.335 | 0.718 | -0.594 | -1.009 | 0 | -0.023 | -0.023 | 0 | 0.55 | 0.55 | 0.54 | 0.17 | 0.46 | 0.37 |
| 3 | 2436134 | rs6787193 | 0.306 | 1.117 | -1.226 | -1.168 | -0.07 | 0.077 | 0.077 | 0.025 | 0.2 | 0.01 | 0.16 | 0.07 | 0.28 | 0.1 |
| 3 | 2459320 | rs3937932 | 0.962 | 0.753 | -0.72 | -1.278 | 0.002 | -0.02 | -0.02 | 0.098 | 0.29 | 0.32 | 0.41 | 0.2 | 0.28 | 0.26 |
| 3 | 2462852 | rs13084976 | 0.404 | 0.496 | -0.888 | -1.309 | -0.084 | 0.075 | 0.075 | 1.248 | 0.2 | 0.01 | 0.17 | 0.08 | 0.32 | 0.11 |
| 3 | 2464724 | rs17194371 | 0.455 | 0.575 | -0.922 | -1.233 | -0.036 | 0.062 | 0.062 | 0.88 | 0.19 | 0.01 | 0.14 | 0.08 | 0.23 | 0.11 |
| 3 | 2472932 | rs2134357 | -0.841 | 0.421 | -0.616 | -1.439 | 0.012 | 0.029 | 0.029 | 0 | 0.53 | 0.37 | 0.6 | 0.23 | 0.62 | 0.37 |
| 3 | 2473281 | rs2616575 | 0.704 | 0.444 | -0.602 | -1.438 | 0 | -0.009 | -0.009 | 0.052 | 0.33 | 0.3 | 0.43 | 0.21 | 0.31 | 0.28 |
| 3 | 2476543 | rs2648690 | 0.278 | 0.448 | -0.495 | -1.542 | 0.005 | -0.009 | -0.009 | 0 | 0.93 | 0.88 | 0.93 | 0.96 | 0.88 | 0.85 |
| 3 | 2476848 | rs9878431 | 0.747 | 0.432 | -0.463 | -1.576 | -0.005 | -0.012 | -0.012 | 0 | 0.44 | 0.55 | 0.35 | 0.77 | 0.31 | 0.61 |
| 3 | 2483305 | rs2648694 | 1.653 | 0.404 | -0.432 | -1.582 | -0.03 | 0.027 | 0.027 | 0 | 0.26 | 0.13 | 0.4 | 0.2 | 0.3 | 0.18 |
| 3 | 2484953 | rs2174017 | 1.446 | 0.087 | -0.422 | -1.344 | -0.014 | 0.007 | 0.007 | 8.626 | 0.33 | 0.44 | 0.29 | 0.61 | 0.27 | 0.53 |
| 3 | 2485134 | rs1948796 | 1.307 | -0.065 | -0.397 | -1.344 | -0.009 | 0.005 | 0.005 | 8.495 | 0.34 | 0.44 | 0.29 | 0.62 | 0.28 | 0.53 |
| 3 | 2486040 | rs2653495 | -0.18 | -0.077 | -0.624 | -1.343 | 0.035 | 0.086 | 0.086 | 1.295 | 0.51 | 0.29 | 0.59 | 0.23 | 0.62 | 0.38 |
| 3 | 2487038 | rs12054152 | 1.484 | -0.074 | -0.627 | -1.288 | -0.006 | -0.002 | -0.002 | 0 | 0.09 | 0.08 | 0.05 | 0.14 | 0.02 | 0.05 |
| 3 | 2487697 | rs2174018 | -1.445 | 0.008 | -0.62 | -1.349 | -0.018 | 0.02 | 0.02 | 0 | 0.7 | 0.77 | 0.55 | 0.8 | 0.65 | 0.79 |
| 3 | 2487786 | rs2134356 | 0.409 | -0.003 | -0.644 | -1.359 | 0.03 | 0.045 | 0.045 | 0.385 | 0.45 | 0.66 | 0.36 | 0.77 | 0.31 | 0.61 |
| 3 | 2488631 | rs17194406 | 0.381 | 0.11 | -0.79 | -1.27 | -0.087 | 0.14 | 0.14 | 0.669 | 0.23 | 0.02 | 0.18 | 0.02 | 0.34 | 0.11 |
| 3 | 2494395 | rs9841813 | -1.277 | 0.079 | -0.475 | -1.093 | -0.047 | 0.051 | 0.051 | 0 | 0.69 | 0.84 | 0.56 | 0.78 | 0.65 | 0.79 |
| 3 | 2494746 | rs990204 | 1.18 | 0.048 | -0.308 | -1.091 | -0.004 | -0.001 | -0.001 | 0.179 | 0.36 | 0.47 | 0.33 | 0.62 | 0.35 | 0.58 |
| 3 | 2495330 | rs6442731 | 1.208 | 0.126 | -0.314 | -1.045 | -0.003 | 0.004 | 0.004 | 7.483 | 0.36 | 0.55 | 0.33 | 0.61 | 0.36 | 0.55 |
| 3 | 2499708 | rs9310741 | -1.417 | 0.145 | -0.802 | -0.863 | -0.024 | 0.034 | 0.034 | 0 | 0.71 | 0.88 | 0.59 | 0.79 | 0.72 | 0.7 |
| 3 | 2502638 | rs9860735 | 0.084 | 0.191 | -0.733 | -0.841 | 0.011 | 0.049 | 0.049 | 0.334 | 0.49 | 0.64 | 0.42 | 0.76 | 0.45 | 0.6 |
| 3 | 2510409 | rs6802636 | -0.921 | 0.147 | -1.267 | -0.772 | 0.002 | 0.004 | 0.004 | 2.731 | 0.3 | 0.25 | 0.22 | 0.16 | 0.29 | 0.12 |
| 3 | 2519703 | rs17014863 | 0.346 | 0.484 | -1.784 | -0.853 | -0.007 | 0.052 | 0.052 | 0.736 | 0.17 | 0.08 | 0.12 | 0.02 | 0.19 | 0.08 |
| 3 | 2519956 | rs11129155 | 1.273 | 0.649 | -1.975 | -0.713 | -0.059 | 0.107 | 0.107 | 4.94 | 0.26 | 0.03 | 0.16 | 0.08 | 0.27 | 0.24 |
| 3 | 2529000 | rs11713873 | 0.508 | 0.552 | -1.895 | -0.363 | -0.004 | -0.064 | -0.064 | 0.258 | 0.38 | 0.08 | 0.44 | 0.39 | 0.32 | 0.29 |
| 3 | 2532330 | rs7651861 | 0.899 | 0.35 | -1.924 | -0.437 | -0.071 | -0.101 | -0.101 | 0.4 | 0.35 | 0.05 | 0.46 | 0.4 | 0.37 | 0.29 |
| 3 | 2533117 | rs17015059 | 0.323 | 0.912 | -0.571 | -0.46 | 0.11 | -0.007 | -0.007 | 0 | 0.07 | 0.29 | 0.1 | 0.03 | 0.16 | 0.03 |
| 3 | 2534560 | rs1125102 | 0.727 | 0.731 | -0.233 | -0.567 | 0.004 | -0.014 | -0.014 | 0 | 0.42 | 0.19 | 0.49 | 0.4 | 0.37 | 0.29 |
| 3 | 2537916 | rs17194476 | 0.23 | 0.656 | -0.572 | -0.871 | -0.029 | 0.086 | 0.086 | 0 | 0.14 | 0.01 | 0.11 | 0 | 0.15 | 0.09 |
| 3 | 2538769 | rs4571230 | -0.934 | 0.519 | 0.223 | -0.858 | 0.001 | 0.02 | 0.02 | 2.926 | 0.69 | 0.64 | 0.72 | 0.43 | 0.68 | 0.5 |
| 3 | 2543302 | rs9828150 | -0.142 | 0.454 | 0.129 | -0.744 | -0.004 | -0.006 | -0.006 | 0 | 0.6 | 0.39 | 0.63 | 0.42 | 0.53 | 0.48 |
| 3 | 2552565 | rs749613 | 0.911 | 0.25 | -0.412 | -0.703 | -0.06 | -0.094 | -0.094 | 4.409 | 0.43 | 0.08 | 0.5 | 0.43 | 0.39 | 0.33 |
| 3 | 2555483 | rs1382875 | -1.238 | 0.248 | 0.642 | -1.199 | -0.001 | -0.085 | -0.085 | 0.068 | 0.15 | 0.53 | 0.14 | 0.03 | 0.15 | 0.13 |
| 3 | 2559451 | rs7626087 | -0.94 | 0.498 | 0.246 | -1.132 | -0.001 | -0.06 | -0.06 | 0 | 0.19 | 0.27 | 0.14 | 0.01 | 0.2 | 0.2 |
| 3 | 2561556 | rs17620999 | -0.826 | 0.504 | 0.07 | -1.13 | -0.001 | -0.018 | -0.018 | 2.722 | 0.18 | 0.2 | 0.13 | 0.01 | 0.2 | 0.2 |
| 3 | 2563341 | rs1842220 | -0.776 | 0.317 | 0 | -0.963 | 0 | -0.007 | -0.007 | 0.788 | 0.69 | 0.7 | 0.75 | 0.86 | 0.72 | 0.76 |
| 3 | 2566466 | rs1382876 | -0.027 | 0.579 | -0.523 | -0.882 | 0 | 0.038 | 0.038 | 8.369 | 0.54 | 0.4 | 0.61 | 0.75 | 0.49 | 0.68 |
| 3 | 2572993 | rs13089709 | -0.709 | 0.72 | -0.684 | -0.497 | -0.039 | 0.069 | 0.069 | 0.015 | 0.15 | 0.03 | 0.11 | 0.03 | 0.17 | 0.1 |
| 3 | 2573148 | rs13093102 | -0.568 | 0.717 | -0.685 | -0.481 | -0.039 | 0.065 | 0.065 | 0.015 | 0.15 | 0.03 | 0.1 | 0.03 | 0.17 | 0.1 |
| 3 | 2573461 | rs17015816 | -1.303 | 0.704 | -0.449 | -0.166 | -0.001 | 0.003 | 0.003 | 0 | 0.21 | 0.21 | 0.18 | 0.56 | 0.21 | 0.44 |
| 3 | 2573531 | rs12636309 | 0.169 | 0.657 | -0.245 | 0.023 | 0.012 | 0.302 | 0.302 | 0 | 0.06 | 0.18 | 0.08 | 0.53 | 0.04 | 0.35 |
| 3 | 2578402 | rs724707 | 0.528 | 0.404 | -0.769 | -0.863 | 0.027 | 0.108 | 0.108 | 0.261 | 0.5 | 0.27 | 0.6 | 0.23 | 0.56 | 0.45 |
| 3 | 2579127 | rs6793017 | -0.799 | 0.619 | -1.647 | -0.719 | 0.008 | 0.013 | 0.013 | 0 | 0.73 | 0.77 | 0.75 | 0.8 | 0.77 | 0.83 |
| 3 | 2582549 | rs13058742 | 0.083 | 0.747 | -2.082 | -0.7 | -0.056 | 0.071 | 0.071 | 0 | 0.12 | 0 | 0.09 | 0.01 | 0.16 | 0.11 |
| 3 | 2583496 | rs11705951 | 1.295 | 0.619 | -2.14 | -0.716 | -0.046 | -0.016 | -0.016 | 0 | 0.27 | 0.1 | 0.43 | 0.22 | 0.32 | 0.15 |
| 3 | 2589322 | rs6763008 | -0.559 | 0.169 | -2.164 | -1.166 | 0.018 | 0.05 | 0.05 | 0 | 0.7 | 0.85 | 0.73 | 0.8 | 0.74 | 0.79 |

| | | | | | | | | | | | | | | | |
|---|---|---|---|---|---|---|---|---|---|---|---|---|---|---|---|
| 3 | 2593544 | rs2728054 | 0.665 | 1.285 | -1.921 | -0.463 | -0.069 | -0.054 | -0.054 | 0 | 0.25 | 0.06 | 0.4 | 0.21 | 0.3 | 0.16 |
| 3 | 2596726 | rs4685527 | -0.279 | 1.197 | -0.989 | 0.003 | -0.016 | 0.054 | 0.054 | 0 | 0.1 | 0.02 | 0.06 | 0.01 | 0.11 | 0.09 |
| 3 | 2600804 | rs2046037 | -1.037 | 1.386 | -0.362 | 0.595 | 0.018 | 0.072 | 0.072 | 0 | 0.16 | 0.17 | 0.18 | 0.59 | 0.24 | 0.34 |
| 3 | 2603228 | rs9859658 | -0.92 | 1.085 | -0.305 | -0.519 | -0.009 | -0.059 | -0.059 | 0.017 | 0.12 | 0.21 | 0.05 | 0 | 0.06 | 0.04 |
| 3 | 2603517 | rs2728011 | 0.744 | 1.253 | -0.244 | -0.383 | 0.024 | -0.031 | -0.031 | 0 | 0.84 | 0.73 | 0.83 | 0.92 | 0.77 | 0.95 |
| 3 | 2608506 | rs13081043 | -0.971 | 1.292 | -0.94 | -0.751 | -0.029 | 0.063 | 0.063 | 0.081 | 0.25 | 0.1 | 0.17 | 0.08 | 0.28 | 0.18 |
| 3 | 2609657 | rs11718965 | 0.087 | 1.544 | -0.725 | -1.352 | 0.002 | -0.045 | -0.045 | 0 | 0.21 | 0.25 | 0.09 | 0.02 | 0.2 | 0.13 |
| 3 | 2613001 | rs2616585 | 0.854 | 1.502 | -0.607 | -1.566 | 0.002 | -0.045 | -0.045 | 0 | 0.59 | 0.5 | 0.73 | 0.87 | 0.51 | 0.69 |
| 3 | 2614626 | rs11707897 | 0.253 | 1.197 | -1.026 | -1.307 | -0.061 | 0.099 | 0.099 | 0 | 0.18 | 0.03 | 0.08 | 0.03 | 0.19 | 0.14 |
| 3 | 2616290 | rs2728055 | -0.33 | 0.787 | -0.767 | -0.775 | -0.003 | -0.001 | -0.001 | 0 | 0.58 | 0.57 | 0.63 | 0.5 | 0.67 | 0.45 |
| 3 | 2617404 | rs13314366 | 0.402 | 0.293 | -1.151 | -1.172 | 0.02 | 0.085 | 0.085 | 0 | 0.48 | 0.26 | 0.66 | 0.64 | 0.57 | 0.4 |
| 3 | 2617954 | rs2728056 | 0.584 | 0.06 | -1.514 | -1.364 | -0.024 | 0.111 | 0.111 | 0.003 | 0.49 | 0.07 | 0.67 | 0.69 | 0.57 | 0.46 |
| 3 | 2620454 | rs12715075 | 0.207 | -0.092 | -0.551 | -1.398 | -0.013 | -0.057 | -0.057 | 0.015 | 0.29 | 0.35 | 0.21 | 0.09 | 0.22 | 0.26 |
| 3 | 2621687 | rs1466459 | 0.718 | -0.223 | -0.334 | -1.3 | 0.018 | 0.052 | 0.052 | 0 | 0.55 | 0.3 | 0.71 | 0.7 | 0.61 | 0.51 |
| 3 | 2624673 | rs7609846 | -0.372 | -0.05 | -0.233 | -1.408 | -0.007 | 0.001 | 0.001 | 0 | 0.14 | 0.11 | 0.07 | 0.01 | 0.18 | 0.07 |
| 3 | 2627817 | rs2616570 | 1.351 | -0.254 | -0.127 | -0.728 | 0.003 | 0.018 | 0.018 | 0 | 0.47 | 0.27 | 0.47 | 0.53 | 0.53 | 0.38 |
| 3 | 2629909 | rs2728060 | 0.471 | -0.637 | 0.643 | -0.407 | 0.057 | 0.213 | 0.213 | 4.35 | 0.94 | 0.63 | 0.77 | 0.84 | 0.98 | 0.93 |
| 3 | 2635044 | rs17564391 | 0.243 | -0.761 | 0.248 | -0.603 | -0.003 | -0.002 | -0.002 | 0 | 0.26 | 0.24 | 0.11 | 0.21 | 0.17 | 0.31 |
| 3 | 2636747 | rs4553961 | -0.514 | -0.739 | 0.312 | -0.56 | -0.006 | -0.006 | -0.006 | 0 | 0.37 | 0.36 | 0.23 | 0.28 | 0.26 | 0.5 |
| 3 | 2639146 | rs9819935 | -1.285 | -0.067 | 0.593 | -0.113 | -0.012 | 0.107 | 0.107 | 0.002 | 0.11 | 0.39 | 0.24 | 0.16 | 0.04 | 0.08 |
| 3 | 2639829 | rs2728063 | -0.396 | 0.645 | -0.764 | -0.402 | -0.017 | 0.039 | 0.039 | 0 | 0.23 | 0.09 | 0.18 | 0.17 | 0.24 | 0.15 |
| 3 | 2649873 | rs1382866 | 1.264 | 1.387 | 0.14 | -0.763 | 0.023 | -0.041 | -0.041 | 0 | 0.72 | 0.44 | 0.52 | 0.86 | 0.66 | 0.72 |
| 3 | 2651236 | rs12487827 | 1.556 | 1.059 | -0.735 | -1.127 | -0.049 | 0.246 | 0.246 | 3.16 | 0.4 | 0.05 | 0.24 | 0.1 | 0.36 | 0.35 |
| 3 | 2653577 | rs1020997 | 0.164 | 1.105 | -0.042 | -1.075 | 0.009 | -0.034 | -0.034 | 0 | 0.62 | 0.54 | 0.61 | 0.2 | 0.54 | 0.55 |
| 3 | 2655046 | rs12494352 | 1.049 | 1.132 | 0.376 | -1.134 | 0.01 | 0.096 | 0.096 | 2.02 | 0.49 | 0.37 | 0.34 | 0.1 | 0.5 | 0.37 |
| 3 | 2655555 | rs2600305 | -0.517 | 1.188 | 0.577 | -1.081 | -0.001 | -0.033 | -0.033 | 2.48 | 0.19 | 0.31 | 0.36 | 0.11 | 0.19 | 0.22 |
| 3 | 2657255 | rs13068877 | 1.029 | 1.005 | 0.251 | -0.784 | 0.004 | 0.16 | 0.16 | 1.367 | 0.47 | 0.24 | 0.34 | 0.17 | 0.5 | 0.37 |
| 3 | 2665743 | rs2170004 | -0.588 | 0.699 | 0.585 | -0.379 | -0.005 | 0.018 | 0.018 | 0 | 0.42 | 0.71 | 0.38 | 0.71 | 0.44 | 0.46 |
| 3 | 2667189 | rs1479546 | -0.125 | 0.659 | 0.706 | -0.382 | -0.006 | 0.001 | 0.001 | 0 | 0.37 | 0.36 | 0.34 | 0.7 | 0.44 | 0.46 |
| 3 | 2667880 | rs1904396 | -0.397 | 0.728 | 0.654 | -0.395 | -0.014 | 0.022 | 0.022 | 0 | 0.4 | 0.39 | 0.43 | 0.77 | 0.49 | 0.47 |
| 3 | 2668594 | rs6803088 | -1.417 | 0.811 | 1.174 | -0.166 | 0.009 | 0.029 | 0.029 | 0 | 0.17 | 0.53 | 0.39 | 0.16 | 0.17 | 0.17 |
| 3 | 2670624 | rs2600318 | -1.819 | 0.897 | 1.222 | -0.335 | 0.012 | -0.03 | -0.03 | 0 | 0.07 | 0.21 | 0.17 | 0 | 0.05 | 0.02 |
| 3 | 2672936 | rs6770875 | -0.245 | 1.113 | 1.163 | -0.304 | 0.014 | 0.002 | 0.002 | 0.233 | 0.37 | 0.46 | 0.48 | 0.69 | 0.47 | 0.4 |
| 3 | 2675155 | rs2619579 | 1.261 | 0.9 | 1.193 | -0.571 | -0.003 | -0.057 | -0.057 | 0 | 0.8 | 0.58 | 0.79 | 0.9 | 0.79 | 0.75 |
| 3 | 2675189 | rs9849237 | -1.702 | 1.165 | 1.363 | -0.404 | 0.022 | -0.049 | -0.049 | 0 | 0.17 | 0.46 | 0.3 | 0.1 | 0.18 | 0.23 |
| 3 | 2677960 | rs17017710 | -0.48 | 1.047 | 1.028 | -0.288 | 0.002 | -0.004 | -0.004 | 0 | 0.1 | 0.16 | 0.14 | 0.09 | 0.13 | 0.2 |
| 3 | 2682000 | rs1479545 | -1.046 | 0.66 | 0.474 | -0.513 | 0.049 | 0.083 | 0.083 | 4.127 | 0.55 | 0.82 | 0.7 | 0.8 | 0.65 | 0.62 |
| 3 | 2684357 | rs2728076 | -0.496 | 0.794 | 0.579 | -0.493 | 0.001 | 0.012 | 0.012 | 0.119 | 0.48 | 0.62 | 0.48 | 0.71 | 0.54 | 0.37 |
| 3 | 2687527 | rs1479544 | -0.337 | 0.883 | 0.332 | -0.474 | 0.029 | -0.012 | -0.012 | 0 | 0.11 | 0.25 | 0.25 | 0.09 | 0.13 | 0.22 |
| 3 | 2688399 | rs6442742 | -0.252 | 0.868 | 0.327 | -0.339 | -0.033 | 0.003 | 0.003 | 3.572 | 0.42 | 0.3 | 0.41 | 0.66 | 0.48 | 0.35 |
| 3 | 2690405 | rs1600359 | -0.321 | 1.07 | -0.139 | -0.267 | 0.037 | 0.08 | 0.08 | 3.026 | 0.45 | 0.25 | 0.64 | 0.74 | 0.6 | 0.41 |
| 3 | 2695813 | rs11129229 | 0.029 | 1.064 | -0.741 | -0.179 | -0.16 | -0.069 | -0.069 | 0 | 0.4 | 0.03 | 0.44 | 0.66 | 0.49 | 0.34 |
| 3 | 2697187 | rs9942085 | -2.371 | 1.107 | 1.475 | 0.029 | 0.044 | -0.036 | -0.036 | 0 | 0.18 | 0.37 | 0.28 | 0.11 | 0.22 | 0.15 |
| 3 | 2698276 | rs4583635 | -1.486 | 0.756 | 1.243 | -0.017 | 0.01 | -0.013 | -0.013 | 0.01 | 0.11 | 0.19 | 0.19 | 0.06 | 0.13 | 0.06 |
| 3 | 2700315 | rs7637377 | -2.275 | 0.684 | 1.938 | -0.042 | 0.042 | -0.038 | -0.038 | 1.553 | 0.13 | 0.54 | 0.16 | 0.05 | 0.11 | 0.06 |
| 3 | 2701863 | rs6442743 | -2.046 | 0.497 | 1.683 | -0.085 | -0.001 | 0.004 | 0.004 | 0 | 0.4 | 0.6 | 0.51 | 0.36 | 0.35 | 0.61 |
| 3 | 2701890 | rs17018135 | -0.563 | 0.512 | 1.479 | -0.064 | -0.001 | -0.001 | -0.001 | 0 | 0.06 | 0.06 | 0.15 | 0.06 | 0.07 | 0.05 |
| 3 | 2705062 | rs4234549 | -1.109 | 0.414 | 1.448 | -0.096 | 0.017 | -0.013 | -0.013 | 0 | 0.19 | 0.09 | 0.37 | 0.23 | 0.15 | 0.36 |
| 3 | 2706233 | rs7618043 | -0.589 | 0.474 | 1.939 | -0.131 | 0.012 | -0.021 | -0.021 | 0.015 | 0.12 | 0.18 | 0.08 | 0.06 | 0.1 | 0.19 |
| 3 | 2706571 | rs17018270 | -1.737 | 0.464 | 2.103 | -0.152 | 0.004 | 0.001 | 0.001 | 0 | 0.23 | 0.21 | 0.4 | 0.24 | 0.18 | 0.38 |
| 3 | 2706671 | rs6442744 | -2.267 | 0.462 | 2.013 | -0.144 | -0.003 | -0.023 | -0.023 | 0 | 0.35 | 0.48 | 0.48 | 0.3 | 0.28 | 0.58 |
| 3 | 2715025 | rs6804966 | -2.824 | 0.532 | 1.74 | -0.422 | -0.007 | 0.054 | 0.054 | 0 | 0.44 | 0.65 | 0.52 | 0.27 | 0.38 | 0.62 |
| 3 | 2716743 | rs2600274 | -1.902 | 0.603 | 1.911 | -0.468 | 0.064 | -0.137 | -0.137 | 0.118 | 0.2 | 0.49 | 0.11 | 0.04 | 0.19 | 0.23 |
| 3 | 2717559 | rs1914011 | -0.436 | 0.549 | 1.758 | -0.334 | 0 | -0.008 | -0.008 | 0 | 0.11 | 0.14 | 0.21 | 0.06 | 0.11 | 0.09 |
| 3 | 2720391 | rs2600280 | -2.31 | 0.483 | 1.526 | -0.318 | 0.004 | 0.033 | 0.033 | 0 | 0.42 | 0.65 | 0.49 | 0.3 | 0.33 | 0.58 |
| 3 | 2720586 | rs1516393 | -1.555 | 0.546 | 1.088 | -0.423 | 0.013 | -0.051 | -0.051 | 0 | 0.19 | 0.29 | 0.2 | 0.09 | 0.2 | 0.14 |
| 3 | 2724599 | rs2675308 | -1.784 | 0.519 | 0.291 | -0.102 | -0.006 | 0.026 | 0.026 | 0.006 | 0.41 | 0.8 | 0.5 | 0.38 | 0.33 | 0.59 |
| 3 | 2725785 | rs1878181 | -1.535 | 0.208 | 0.688 | 0.194 | -0.002 | 0.042 | 0.042 | 0 | 0.26 | 0.39 | 0.41 | 0.4 | 0.19 | 0.5 |
| 3 | 2727523 | rs1914009 | 1.368 | 0.365 | -0.367 | -0.095 | 0.002 | -0.017 | -0.017 | 0 | 0.52 | 0.12 | 0.56 | 0.44 | 0.63 | 0.21 |
| 3 | 2728391 | rs1516390 | 1.505 | 0.422 | -0.425 | -0.174 | -0.021 | -0.017 | -0.017 | 0.203 | 0.49 | 0.13 | 0.54 | 0.44 | 0.57 | 0.19 |
| 3 | 2728533 | rs2728020 | -1.089 | 0.134 | 0.511 | -0.038 | 0.012 | 0.029 | 0.029 | 1.371 | 0.1 | 0.52 | 0.14 | 0.04 | 0.05 | 0.07 |
| 3 | 2730587 | rs729752 | -1.438 | 0.527 | -0.152 | -0.108 | -0.052 | -0.072 | -0.072 | 0 | 0.22 | 0.04 | 0.25 | 0.33 | 0.24 | 0.49 |
| 3 | 2731319 | rs7646667 | -1.841 | 0.4 | 0.065 | -0.093 | -0.02 | -0.03 | -0.03 | 0 | 0.2 | 0.15 | 0.29 | 0.32 | 0.21 | 0.47 |
| 3 | 2732575 | rs932157 | 0.342 | 0.393 | 0.732 | -0.161 | -0.003 | -0.037 | -0.037 | 0 | 0.74 | 0.42 | 0.74 | 0.81 | 0.76 | 0.75 |
| 3 | 2733172 | rs2675304 | -0.176 | 0.391 | 0.873 | -0.157 | 0 | -0.003 | -0.003 | 2.788 | 0.78 | 0.75 | 0.86 | 0.81 | 0.78 | 0.76 |
| 3 | 2734207 | rs1024147 | 2.092 | 0.16 | 0.767 | -0.254 | -0.001 | -0.003 | -0.003 | 0 | 0.46 | 0.4 | 0.47 | 0.48 | 0.51 | 0.22 |
| 3 | 2734701 | rs965396 | 1.421 | 0.217 | 0.624 | -0.171 | -0.009 | -0.009 | -0.009 | 0 | 0.56 | 0.64 | 0.59 | 0.46 | 0.54 | 0.27 |
| 3 | 2738457 | rs1949343 | -1.292 | 0.197 | 0.851 | 0.267 | -0.007 | -0.022 | -0.022 | 0.225 | 0.18 | 0.16 | 0.2 | 0.36 | 0.16 | 0.46 |
| 3 | 2738701 | rs1516398 | -0.719 | 0.31 | 0.787 | 0.2 | -0.033 | 0.006 | 0.006 | 0.021 | 0.81 | 0.91 | 0.86 | 0.81 | 0.75 | 0.79 |

| | | | | | | | | | | | | | | | | |
|---|---|---|---|---|---|---|---|---|---|---|---|---|---|---|---|---|
| 3 | 2738721 | rs2728027 | -1.436 | 0.307 | 0.864 | 0.202 | 0.003 | 0.016 | 0.016 | 0.008 | 0.2 | 0.26 | 0.3 | 0.36 | 0.18 | 0.48 |
| 3 | 2740275 | rs2728030 | -0.594 | 0.32 | 0.531 | 0.229 | -0.036 | 0.008 | 0.008 | 0.068 | 0.81 | 0.92 | 0.86 | 0.82 | 0.75 | 0.8 |
| 3 | 2743441 | rs9811943 | -0.449 | 0.212 | 0.55 | 0.282 | -0.02 | -0.044 | -0.044 | 0 | 0.08 | 0.25 | 0.13 | 0 | 0.03 | 0.04 |
| 3 | 2743688 | rs7645216 | 1.521 | 0.159 | 0.612 | 0.265 | 0.004 | -0.001 | -0.001 | 0 | 0.51 | 0.39 | 0.46 | 0.46 | 0.53 | 0.28 |
| 3 | 2744803 | rs2600289 | -0.403 | 0.407 | 0.29 | 0.296 | -0.015 | -0.037 | -0.037 | 0 | 0.18 | 0.11 | 0.24 | 0.33 | 0.16 | 0.45 |
| 3 | 2746456 | rs13080275 | 1.083 | 0.511 | 0.739 | -0.027 | -0.004 | 0.012 | 0.012 | 0 | 0.64 | 0.32 | 0.68 | 0.79 | 0.66 | 0.62 |
| 3 | 2748076 | rs2728034 | -1.403 | -0.073 | 0.75 | 0.453 | 0.008 | -0.052 | -0.052 | 0 | 0.12 | 0.32 | 0.14 | 0.03 | 0.07 | 0.18 |
| 3 | 2754005 | rs2675289 | -1.476 | -0.587 | 0.616 | 1.232 | -0.06 | 0.184 | 0.184 | 0 | 0.2 | 0.45 | 0.3 | 0.48 | 0.13 | 0.34 |
| 3 | 2754700 | rs2728041 | -1.872 | -0.708 | 1.106 | 1.208 | -0.042 | 0.117 | 0.117 | 0.029 | 0.2 | 0.48 | 0.25 | 0.42 | 0.19 | 0.27 |
| 3 | 2755983 | rs6442749 | 1.667 | -0.808 | 1.01 | 1.283 | 0 | -0.001 | -0.001 | 0 | 0.7 | 0.41 | 0.71 | 0.78 | 0.75 | 0.67 |
| 3 | 2756407 | rs2728047 | 1.995 | -0.966 | 0.966 | 1.307 | 0.001 | -0.001 | -0.001 | 0 | 0.66 | 0.33 | 0.55 | 0.35 | 0.7 | 0.45 |
| 3 | 2757448 | rs1400197 | -0.631 | -0.96 | 1.134 | 1.373 | -0.006 | -0.009 | -0.009 | 0.02 | 0.06 | 0.23 | 0.02 | 0 | 0.01 | 0.02 |
| 3 | 2758388 | rs2063895 | -2.313 | -0.918 | 1.128 | 1.416 | -0.039 | 0.118 | 0.118 | 0.316 | 0.18 | 0.62 | 0.25 | 0.42 | 0.14 | 0.26 |
| 3 | 2759283 | rs12494838 | -1.738 | -0.518 | 1.033 | 1.612 | -0.002 | 0.08 | 0.08 | 1.647 | 0.27 | 0.46 | 0.34 | 0.59 | 0.28 | 0.47 |
| 3 | 2760177 | rs9310832 | -0.386 | -0.469 | 1.13 | 1.682 | 0.018 | 0.061 | 0.061 | 1.243 | 0.13 | 0.23 | 0.13 | 0.17 | 0.14 | 0.21 |
| 3 | 2765358 | rs7628663 | -1.299 | -0.253 | 0.524 | 1.621 | 0.006 | 0.18 | 0.18 | 0 | 0.09 | 0.23 | 0.2 | 0.44 | 0.1 | 0.23 |
| 3 | 2768820 | rs2320830 | -0.754 | -0.095 | -0.014 | 1.629 | -0.001 | 0.069 | 0.069 | 0 | 0.08 | 0.14 | 0.18 | 0.47 | 0.09 | 0.25 |
| 3 | 2770509 | rs17019466 | -1.413 | -0.358 | 0.101 | 0.833 | -0.011 | -0.006 | -0.006 | 0 | 0.1 | 0.14 | 0.11 | 0.05 | 0.07 | 0.05 |
| 3 | 2770914 | rs1033237 | 1.274 | -0.336 | 0.853 | 0.602 | 0.015 | -0.042 | -0.042 | 0.004 | 0.91 | 0.68 | 0.94 | 0.99 | 0.93 | 0.96 |
| 3 | 2772862 | rs11129265 | -0.135 | -0.285 | -0.02 | 0.359 | -0.029 | 0.042 | 0.042 | 0 | 0.39 | 0.22 | 0.44 | 0.29 | 0.55 | 0.28 |
| 3 | 2773049 | rs7626866 | -0.418 | -1.681 | -0.012 | 0.353 | 0.001 | -0.006 | -0.006 | 0.627 | 0.66 | 0.71 | 0.75 | 0.48 | 0.73 | 0.59 |
| 3 | 2773343 | rs12638165 | 0.552 | -2.208 | 0.048 | 0.194 | 0.037 | -0.009 | -0.009 | 0 | 0.22 | 0.05 | 0.27 | 0.19 | 0.14 | 0.28 |
| 3 | 2774996 | rs7611703 | 0.806 | -2.507 | -0.382 | 0.278 | 0.173 | 0.086 | 0.086 | 0.427 | 0.5 | 0.12 | 0.55 | 0.66 | 0.72 | 0.49 |
| 3 | 2776203 | rs12638512 | 0.724 | -2.561 | -0.39 | 0.277 | 0.18 | 0.079 | 0.079 | 0.377 | 0.51 | 0.12 | 0.63 | 0.66 | 0.73 | 0.54 |
| 3 | 2777262 | rs1973683 | -0.2 | -2.628 | 0.037 | 0.206 | -0.087 | -0.049 | -0.049 | 0.55 | 0.77 | 0.4 | 0.92 | 0.86 | 0.92 | 0.84 |
| 3 | 2779119 | rs7637988 | 0.481 | -2.632 | -0.227 | 0.404 | 0.155 | -0.007 | -0.007 | 0.367 | 0.54 | 0.22 | 0.57 | 0.57 | 0.78 | 0.62 |
| 3 | 2780113 | rs7648792 | 0.744 | -2.604 | -0.27 | 0.421 | 0.168 | -0.017 | -0.017 | 0 | 0.51 | 0.13 | 0.55 | 0.55 | 0.72 | 0.57 |
| 3 | 2780449 | rs2320958 | -0.011 | -1.899 | 0.013 | 0.666 | 0 | -0.001 | -0.001 | 1.292 | 0.27 | 0.22 | 0.37 | 0.3 | 0.2 | 0.31 |
| 3 | 2783549 | rs12631108 | 0.2 | -1.86 | -0.252 | 0.523 | 0.022 | 0.013 | 0.013 | 0 | 0.25 | 0.12 | 0.35 | 0.18 | 0.19 | 0.27 |
| 3 | 2783884 | rs17019645 | -0.084 | -1.665 | -0.001 | 0.222 | -0.01 | -0.025 | -0.025 | 0 | 0.83 | 0.8 | 0.93 | 0.89 | 0.85 | 0.84 |
| 3 | 2786502 | rs9822614 | -0.176 | -0.784 | 0.263 | 0.69 | -0.002 | 0.023 | 0.023 | 0 | 0.21 | 0.3 | 0.23 | 0.39 | 0.19 | 0.3 |
| 3 | 2790335 | rs7614036 | -0.69 | -0.675 | 0.716 | 0.651 | -0.001 | 0.035 | 0.035 | 0.474 | 0.25 | 0.62 | 0.21 | 0.36 | 0.22 | 0.35 |
| 3 | 2796414 | rs3897737 | 0.763 | -0.798 | 0.751 | 0.705 | -0.005 | 0.012 | 0.012 | 0 | 0.66 | 0.44 | 0.75 | 0.54 | 0.7 | 0.57 |
| 3 | 2798951 | rs6810122 | 1.226 | -1.029 | 0.543 | 0.698 | 0.031 | -0.032 | -0.032 | 0 | 0.67 | 0.25 | 0.78 | 0.54 | 0.78 | 0.59 |
| 3 | 2801981 | rs6442753 | -0.585 | -1.437 | 0.415 | 0.539 | -0.016 | 0.007 | 0.007 | 0 | 0.39 | 0.5 | 0.42 | 0.51 | 0.27 | 0.38 |
| 3 | 2803270 | rs6772481 | 0.698 | -1.045 | -0.283 | 0.314 | 0.03 | -0.06 | -0.06 | 0 | 0.26 | 0.11 | 0.37 | 0.36 | 0.2 | 0.27 |
| 3 | 2803304 | rs1400202 | 1.349 | -0.93 | -0.322 | 0.27 | -0.018 | 0.141 | 0.141 | 0 | 0.43 | 0.22 | 0.4 | 0.19 | 0.56 | 0.32 |
| 3 | 2803671 | rs2222104 | 0.638 | -0.821 | 0.122 | 0.423 | 0.004 | -0.008 | -0.008 | 0.351 | 0.7 | 0.35 | 0.77 | 0.55 | 0.77 | 0.59 |
| 3 | 2805947 | rs11129289 | 0.601 | -0.613 | 0.025 | 0.362 | 0.005 | -0.008 | -0.008 | 0 | 0.7 | 0.35 | 0.77 | 0.56 | 0.77 | 0.62 |
| 3 | 2806287 | rs11718833 | 1.248 | -0.442 | -0.685 | 0.514 | -0.057 | 0.062 | 0.062 | 1.007 | 0.41 | 0.15 | 0.4 | 0.3 | 0.58 | 0.34 |
| 3 | 2813968 | rs12330913 | 0.063 | -0.626 | 0.847 | 1.08 | -0.041 | 0.085 | 0.085 | 0 | 0.82 | 0.64 | 0.86 | 0.68 | 0.92 | 0.74 |
| 3 | 2815150 | rs3935671 | 1.34 | -0.816 | 0.977 | 1.057 | 0.002 | -0.004 | -0.004 | 0 | 0.68 | 0.34 | 0.78 | 0.55 | 0.77 | 0.58 |
| 3 | 2815263 | rs3935673 | 0.773 | -1.09 | 1.27 | 1.116 | -0.043 | 0.079 | 0.079 | 0 | 0.74 | 0.44 | 0.82 | 0.56 | 0.85 | 0.63 |
| 3 | 2821345 | rs955340 | -1.078 | -1.021 | 1.139 | 1.165 | -0.034 | 0.067 | 0.067 | 0 | 0.27 | 0.43 | 0.18 | 0.42 | 0.2 | 0.41 |
| 3 | 2821443 | rs955341 | 0.758 | -0.535 | 0.788 | 0.845 | -0.003 | 0.005 | 0.005 | 0 | 0.08 | 0.14 | 0.03 | 0.13 | 0.08 | 0.09 |
| 3 | 2824558 | rs1474021 | 0.604 | -0.443 | 0.958 | 0.627 | 0.018 | -0.045 | -0.045 | 0 | 0.83 | 0.5 | 0.9 | 0.9 | 0.86 | 0.88 |
| 3 | 2826373 | rs7646671 | 0.193 | -0.584 | 1.116 | 0.933 | -0.015 | 0.066 | 0.066 | 0 | 0.13 | 0.28 | 0.14 | 0.23 | 0.1 | 0.16 |
| 3 | 2826568 | rs956764 | 0.609 | -0.357 | 1.059 | 1.085 | 0.024 | 0.009 | 0.009 | 0.038 | 0.94 | 0.83 | 0.93 | 0.97 | 0.95 | 0.99 |
| 3 | 2831511 | rs2728090 | 0.649 | -0.564 | 0.49 | 0.858 | 0 | 0.001 | 0.001 | 0 | 0.38 | 0.38 | 0.47 | 0.36 | 0.44 | 0.46 |
| 3 | 2832318 | rs7615939 | -0.271 | -0.567 | 0.586 | 0.799 | 0 | 0.002 | 0.002 | 0 | 0.55 | 0.45 | 0.46 | 0.64 | 0.51 | 0.53 |
| 3 | 2835642 | rs7635694 | -0.065 | -0.219 | 0.73 | 0.15 | 0.013 | -0.019 | -0.019 | 0 | 0.35 | 0.49 | 0.35 | 0.26 | 0.37 | 0.25 |
| 3 | 2836270 | rs7628425 | 0.532 | -0.29 | 0.643 | 0.129 | -0.004 | 0.011 | 0.011 | 0 | 0.3 | 0.25 | 0.28 | 0.23 | 0.32 | 0.23 |
| 3 | 2836943 | rs2728095 | 0.352 | -0.327 | 0.578 | 0.144 | -0.003 | 0.024 | 0.024 | 0 | 0.6 | 0.33 | 0.62 | 0.74 | 0.62 | 0.75 |
| 3 | 2839444 | rs7647953 | 0.319 | -0.24 | 0.103 | -0.154 | -0.056 | 0.09 | 0.09 | 0 | 0.32 | 0.18 | 0.25 | 0.11 | 0.38 | 0.23 |
| 3 | 2844404 | rs2675317 | 0.114 | -0.112 | -0.74 | -0.163 | 0.004 | -0.031 | -0.031 | 0 | 0.32 | 0.36 | 0.28 | 0.08 | 0.38 | 0.23 |
| 3 | 2844653 | rs17020557 | 0.487 | -0.329 | -1.091 | 0.073 | 0.02 | -0.057 | -0.057 | 0 | 0.14 | 0 | 0.05 | 0.18 | 0.08 | 0.12 |
| 3 | 2844699 | rs2675316 | 0.587 | -0.026 | -0.814 | -0.119 | 0.008 | -0.022 | -0.022 | 0 | 0.26 | 0.28 | 0.24 | 0.07 | 0.33 | 0.23 |
| 3 | 2845843 | rs2728089 | -0.343 | -0.335 | -1.49 | -0.157 | -0.008 | -0.01 | -0.01 | 0 | 0.65 | 0.4 | 0.74 | 0.76 | 0.76 | 0.74 |
| 3 | 2846316 | rs975334 | -0.056 | 0.318 | -2.212 | 0.018 | 0.064 | 0.133 | 0.133 | 0 | 0.48 | 0.76 | 0.63 | 0.79 | 0.39 | 0.58 |
| 3 | 2851955 | rs4685564 | 0.327 | 0.741 | -1.851 | 0.91 | -0.093 | -0.011 | -0.011 | 0.129 | 0.17 | 0.01 | 0.16 | 0.1 | 0.26 | 0.18 |
| 3 | 2856723 | rs11129308 | -0.572 | -0.194 | -1.017 | 1.32 | -0.003 | 0.013 | 0.013 | 0 | 0.14 | 0.17 | 0.13 | 0.4 | 0.09 | 0.21 |
| 3 | 2862414 | rs11129309 | 0.219 | 0.389 | -1.047 | 0.613 | -0.001 | -0.003 | -0.003 | 0.982 | 0.75 | 0.74 | 0.62 | 0.79 | 0.75 | 0.72 |
| 3 | 2863279 | rs6772294 | -1.222 | 0.589 | -1.227 | 0.357 | -0.006 | 0.005 | 0.005 | 0 | 0.08 | 0.07 | 0.12 | 0.01 | 0.13 | 0.06 |
| 3 | 2867637 | rs1400207 | -0.291 | 0.422 | -1.854 | 0.059 | -0.046 | 0.075 | 0.075 | 0.19 | 0.62 | 0.81 | 0.65 | 0.75 | 0.59 | 0.6 |
| 3 | 2868988 | rs925820 | -0.416 | 0.591 | -1.889 | 0.03 | -0.027 | 0.07 | 0.07 | 0.026 | 0.63 | 0.81 | 0.68 | 0.76 | 0.64 | 0.62 |
| 3 | 2869449 | rs6793237 | 0.285 | 0.48 | -1.501 | 0.548 | -0.015 | -0.004 | -0.004 | 1.239 | 0.37 | 0.53 | 0.36 | 0.3 | 0.31 | 0.28 |
| 3 | 2872068 | rs6776496 | -0.172 | 0.114 | -1.358 | 0.755 | -0.016 | 0.179 | 0.179 | 0 | 0.05 | 0.11 | 0.19 | 0.43 | 0.05 | 0.13 |
| 3 | 2872154 | rs925819 | -0.831 | -0.105 | -1.51 | 0.639 | -0.008 | 0.036 | 0.036 | 0 | 0.11 | 0.11 | 0.2 | 0.42 | 0.08 | 0.18 |
| 3 | 2874048 | rs9859742 | -1.213 | -0.196 | -1.824 | 0.347 | -0.023 | 0.036 | 0.036 | 0 | 0.67 | 0.77 | 0.68 | 0.84 | 0.62 | 0.64 |

| | | | | | | | | | | | | | | | | |
|---|---|---|---|---|---|---|---|---|---|---|---|---|---|---|---|---|
| 3 | 2874906 | rs12636999 | -0.399 | -0.284 | -1.877 | 0.321 | 0.082 | 0.127 | 0.127 | 0 | 0.52 | 0.75 | 0.57 | 0.79 | 0.36 | 0.48 |
| 3 | 2877547 | rs9310863 | 0.038 | -0.295 | -1.791 | 0.341 | -0.01 | 0 | 0 | 0 | 0.44 | 0.53 | 0.35 | 0.38 | 0.29 | 0.33 |
| 3 | 2877859 | rs13093332 | 0.712 | 0.065 | -1.929 | 0.201 | -0.09 | 0.008 | 0.008 | 0.101 | 0.15 | 0.01 | 0.12 | 0.05 | 0.28 | 0.16 |
| 3 | 2881134 | rs9810902 | -0.226 | -1.291 | -1.118 | 0.547 | -0.012 | 0.003 | 0.003 | 0.031 | 0.29 | 0.32 | 0.26 | 0.31 | 0.2 | 0.23 |
| 3 | 2881575 | rs6781149 | -0.543 | -0.61 | -0.742 | 0.942 | -0.052 | 0.061 | 0.061 | 1.148 | 0.27 | 0.39 | 0.24 | 0.48 | 0.14 | 0.3 |
| 3 | 2886527 | rs17584516 | -0.429 | 0.457 | -0.992 | 0.481 | -0.006 | 0.016 | 0.016 | 0 | 0.27 | 0.23 | 0.31 | 0.21 | 0.31 | 0.23 |
| 3 | 2893155 | rs7636234 | -0.398 | 0.409 | -0.932 | 0.451 | -0.008 | 0.018 | 0.018 | 0 | 0.27 | 0.22 | 0.3 | 0.17 | 0.29 | 0.23 |
| 3 | 2895684 | rs11713158 | -0.202 | 0.441 | -0.684 | 0.93 | -0.001 | 0.013 | 0.013 | 0 | 0.8 | 0.78 | 0.84 | 0.64 | 0.77 | 0.83 |
| 3 | 2895719 | rs17584783 | 0.259 | 0.405 | -0.663 | 0.669 | 0 | 0 | 0 | 0 | 0.53 | 0.46 | 0.54 | 0.39 | 0.43 | 0.44 |
| 3 | 2895810 | rs11712375 | -0.541 | 0.398 | -0.711 | 1.086 | -0.002 | -0.01 | -0.01 | 0 | 0.82 | 0.88 | 0.71 | 0.53 | 0.77 | 0.83 |
| 3 | 2896952 | rs9827210 | -0.018 | 0.557 | -0.559 | 0.022 | 0.013 | -0.026 | -0.026 | 0 | 0.38 | 0.59 | 0.31 | 0.22 | 0.42 | 0.46 |
| 3 | 2897056 | rs9867619 | -0.185 | 0.424 | -0.662 | -0.061 | 0 | 0.003 | 0.003 | 0.195 | 0.45 | 0.51 | 0.31 | 0.32 | 0.43 | 0.49 |
| 3 | 2898037 | rs11715403 | 0.328 | 0.015 | -0.574 | -1.062 | 0.022 | 0.029 | 0.029 | 0 | 0.23 | 0.16 | 0.14 | 0.07 | 0.17 | 0.15 |
| 3 | 2899575 | rs4684365 | -0.181 | 0.665 | -0.411 | -1.35 | 0.002 | 0.003 | 0.003 | 0 | 0.36 | 0.3 | 0.34 | 0.13 | 0.34 | 0.3 |
| 3 | 2899654 | rs4234553 | -0.854 | 0.61 | -0.471 | -1.51 | 0.004 | -0.011 | -0.011 | 0.018 | 0.88 | 0.84 | 0.91 | 0.96 | 0.84 | 0.85 |
| 3 | 2901647 | rs10510238 | 1.44 | 0.826 | -0.514 | -1.486 | -0.018 | 0.015 | 0.015 | 0 | 0.11 | 0.06 | 0.18 | 0.05 | 0.13 | 0.1 |
| 3 | 2902492 | rs17584923 | 1.057 | 0.92 | -0.508 | -2.077 | -0.001 | -0.029 | -0.029 | 0.445 | 0.2 | 0.23 | 0.13 | 0.04 | 0.24 | 0.35 |
| 3 | 2902594 | rs6768322 | 1.412 | 0.722 | -0.453 | -2.117 | -0.024 | -0.037 | -0.037 | 0 | 0.23 | 0.26 | 0.16 | 0.08 | 0.18 | 0.16 |
| 3 | 2903786 | rs11924729 | 1.63 | 0.574 | -0.626 | -2.128 | -0.006 | -0.005 | -0.005 | 0 | 0.21 | 0.2 | 0.12 | 0.08 | 0.11 | 0.16 |
| 3 | 2903983 | rs10510239 | 1.091 | 0.449 | -0.821 | -2.01 | 0.012 | -0.014 | -0.014 | 0 | 0.26 | 0.16 | 0.29 | 0.64 | 0.21 | 0.22 |
| 3 | 2906446 | rs9812746 | -0.746 | 0.603 | -1.027 | -2.126 | -0.032 | 0.009 | 0.009 | 0.41 | 0.77 | 0.83 | 0.75 | 0.8 | 0.7 | 0.79 |
| 3 | 2907097 | rs10510241 | 0.944 | 0.532 | -0.998 | -2.227 | -0.002 | -0.002 | -0.002 | 0 | 0.32 | 0.29 | 0.45 | 0.68 | 0.29 | 0.28 |
| 3 | 2907746 | rs4343610 | -0.496 | 0.613 | -0.862 | -2.119 | -0.014 | 0.006 | 0.006 | 1.604 | 0.76 | 0.76 | 0.74 | 0.8 | 0.7 | 0.79 |
| 3 | 2908694 | rs6789163 | 0.056 | 0.252 | -1.056 | -1.735 | -0.003 | -0.005 | -0.005 | 0 | 0.4 | 0.38 | 0.51 | 0.7 | 0.35 | 0.34 |
| 3 | 2910122 | rs7637662 | 0.382 | 0.476 | -0.838 | -1.554 | -0.004 | -0.054 | -0.054 | 1.242 | 0.57 | 0.49 | 0.55 | 0.9 | 0.43 | 0.39 |
| 3 | 2912699 | rs4234555 | -0.244 | 0.44 | -1.007 | -0.949 | -0.003 | 0.033 | 0.033 | 0 | 0.9 | 0.96 | 0.96 | 0.97 | 0.89 | 0.86 |
| 3 | 2912864 | rs4608676 | 0.517 | 0.017 | -1.049 | -0.638 | 0.002 | 0.022 | 0.022 | 0.157 | 0.11 | 0.1 | 0.22 | 0.64 | 0.07 | 0.15 |
| 3 | 2913017 | rs4684366 | 1.127 | -0.184 | -1.116 | -1.312 | 0.018 | 0.057 | 0.057 | 0 | 0.33 | 0.19 | 0.3 | 0.12 | 0.29 | 0.31 |
| 3 | 2913680 | rs9310866 | 0.403 | -0.123 | -1.115 | -1.275 | 0.011 | -0.019 | -0.019 | 0.044 | 0.77 | 0.85 | 0.73 | 0.29 | 0.8 | 0.66 |
| 3 | 2918045 | rs9847797 | -0.234 | -0.432 | -0.477 | -1.092 | 0 | -0.008 | -0.008 | 0 | 0.49 | 0.48 | 0.53 | 0.78 | 0.38 | 0.5 |
| 3 | 2919516 | rs4637279 | 0.468 | -0.223 | -0.739 | -1.251 | 0.026 | 0.066 | 0.066 | 0 | 0.35 | 0.21 | 0.31 | 0.14 | 0.29 | 0.31 |
| 3 | 2921515 | rs6442763 | 0.109 | 0.223 | -0.702 | -1.32 | 0.005 | 0.01 | 0.01 | 0.767 | 0.52 | 0.55 | 0.45 | 0.13 | 0.46 | 0.37 |
| 3 | 2924686 | rs4685575 | -0.417 | 0.479 | -0.877 | -0.903 | -0.018 | 0.053 | 0.053 | 0.525 | 0.55 | 0.67 | 0.57 | 0.77 | 0.47 | 0.54 |
| 3 | 2925030 | rs17646346 | 0.718 | -0.176 | -1.292 | -1.035 | 0.016 | 0.024 | 0.024 | 0 | 0.19 | 0.16 | 0.08 | 0.08 | 0.1 | 0.21 |
| 3 | 2926235 | rs17646423 | -0.16 | 0.236 | -1.008 | -0.869 | 0.004 | 0.025 | 0.025 | 0.001 | 0.37 | 0.17 | 0.46 | 0.73 | 0.35 | 0.33 |
| 3 | 2930995 | rs13315983 | -0.06 | 0.334 | -0.592 | -0.838 | -0.018 | -0.058 | -0.058 | 0.103 | 0.14 | 0.31 | 0.14 | 0.06 | 0.11 | 0.1 |
| 3 | 2932365 | rs13061928 | 0.895 | 1.017 | -1.1 | -0.759 | -0.053 | -0.054 | -0.054 | 0 | 0.1 | 0.01 | 0.17 | 0.13 | 0.18 | 0.14 |
| 3 | 2932614 | rs17586083 | -0.535 | 0.388 | -0.968 | -0.584 | -0.007 | -0.016 | -0.016 | 0 | 0.41 | 0.25 | 0.55 | 0.58 | 0.41 | 0.44 |
| 3 | 2933942 | rs9859594 | 0.435 | 0.206 | -1.039 | -0.675 | -0.019 | -0.068 | -0.068 | 0.736 | 0.12 | 0.25 | 0.08 | 0 | 0.08 | 0.09 |
| 3 | 2936576 | rs17021982 | 0.624 | 0.542 | -1.748 | -0.35 | 0.005 | 0.028 | 0.028 | 0 | 0.17 | 0.07 | 0.05 | 0.1 | 0.11 | 0.17 |
| 3 | 2938734 | rs17022006 | 0.175 | 0.556 | -1.529 | -0.348 | -0.026 | -0.052 | -0.052 | 0.583 | 0.34 | 0.1 | 0.41 | 0.59 | 0.31 | 0.42 |
| 3 | 2939763 | rs7639614 | -0.387 | 0.502 | -0.704 | -0.481 | 0.004 | 0.007 | 0.007 | 0 | 0.12 | 0.1 | 0.07 | 0.05 | 0.11 | 0.1 |
| 3 | 2943790 | rs7635644 | -0.679 | 0.678 | -0.314 | -0.436 | 0.002 | -0.035 | -0.035 | 0.485 | 0.16 | 0.26 | 0.11 | 0.11 | 0.16 | 0.14 |
| 3 | 2946457 | rs17022164 | 0.045 | 0.698 | -0.356 | -0.877 | 0.001 | -0.007 | -0.007 | 0 | 0.4 | 0.43 | 0.46 | 0.7 | 0.41 | 0.46 |
| 3 | 2946688 | rs1566365 | -0.864 | 0.328 | -0.358 | -1.454 | -0.01 | 0.041 | 0.041 | 0.538 | 0.65 | 0.84 | 0.65 | 0.73 | 0.65 | 0.67 |
| 3 | 2947575 | rs1499135 | 0.287 | 0.241 | -0.062 | -1.44 | -0.01 | -0.068 | -0.068 | 0.858 | 0.11 | 0.34 | 0.11 | 0 | 0.09 | 0.07 |
| 3 | 2950536 | rs11129336 | -0.149 | 0.21 | 0.111 | -1.454 | 0.002 | 0.051 | 0.051 | 0 | 0.49 | 0.63 | 0.56 | 0.68 | 0.49 | 0.52 |
| 3 | 2952214 | rs1499133 | 0.337 | 0.274 | -0.013 | -1.401 | 0.002 | -0.012 | -0.012 | 0.148 | 0.39 | 0.44 | 0.47 | 0.67 | 0.39 | 0.45 |
| 3 | 2962596 | rs6779224 | -0.11 | 0.026 | -0.394 | -1.395 | 0 | 0 | 0 | 0 | 0.54 | 0.54 | 0.55 | 0.68 | 0.54 | 0.58 |
| 3 | 2963983 | rs6771937 | -0.612 | 0.152 | -0.381 | -1.401 | -0.007 | -0.008 | -0.008 | 0 | 0.63 | 0.61 | 0.67 | 0.68 | 0.69 | 0.65 |
| 3 | 2966604 | rs6781913 | 0.94 | 0.61 | -0.61 | -1.76 | -0.014 | 0.12 | 0.12 | 0 | 0.5 | 0.27 | 0.51 | 0.72 | 0.44 | 0.49 |
| 3 | 2966836 | rs1499139 | 0.048 | 0.625 | -0.525 | -1.68 | -0.001 | 0.016 | 0.016 | 1.706 | 0.77 | 0.62 | 0.71 | 0.73 | 0.74 | 0.75 |
| 3 | 2974333 | rs9310883 | 0.96 | 0.794 | -0.869 | -1.178 | -0.002 | -0.059 | -0.059 | 0.803 | 0.43 | 0.13 | 0.49 | 0.53 | 0.38 | 0.44 |
| 3 | 2975082 | rs11129351 | 1.404 | 0.708 | -0.25 | -0.783 | -0.053 | -0.099 | -0.099 | 0 | 0.39 | 0.11 | 0.53 | 0.48 | 0.45 | 0.51 |
| 3 | 2975573 | rs6442766 | 1.063 | 1.501 | 0.953 | 0.175 | 0 | 0.002 | 0.002 | 0 | 0.73 | 0.66 | 0.71 | 0.56 | 0.68 | 0.66 |
| 3 | 2976094 | rs6789338 | 1.606 | 1.565 | 1.116 | 0.207 | 0.014 | 0.022 | 0.022 | 0 | 0.64 | 0.4 | 0.61 | 0.56 | 0.56 | 0.59 |
| 3 | 2981182 | rs6779663 | 0.738 | 2.133 | 0.721 | -0.105 | 0.07 | 0.043 | 0.043 | 0.737 | 0.84 | 0.57 | 0.74 | 0.76 | 0.74 | 0.77 |
| 3 | 2982630 | rs6786174 | -0.432 | 1.82 | -0.342 | 0.155 | -0.008 | 0.004 | 0.004 | 0 | 0.07 | 0.06 | 0.09 | 0 | 0.12 | 0.08 |
| 3 | 2983948 | rs6787320 | 0.177 | 1.488 | -0.366 | 0.177 | -0.002 | 0.001 | 0.001 | 0.209 | 0.9 | 0.84 | 0.8 | 0.76 | 0.81 | 0.79 |
| 3 | 2988333 | rs34063866 | -1.276 | 0.85 | -0.226 | -0.207 | -0.005 | -0.058 | -0.058 | 0 | 0.1 | 0.26 | 0.08 | 0 | 0.09 | 0.08 |
| 3 | 2991620 | rs10510242 | -0.91 | 1.05 | -0.861 | -0.141 | -0.003 | 0.028 | 0.028 | 0 | 0.11 | 0.07 | 0.09 | 0 | 0.1 | 0.1 |
| 3 | 2993366 | rs344405 | 0.427 | 1.236 | -0.679 | 0.145 | -0.003 | 0.011 | 0.011 | 0 | 0.56 | 0.57 | 0.63 | 0.65 | 0.51 | 0.75 |
| 3 | 2996963 | rs344410 | 0.773 | 1.639 | -0.591 | -0.092 | -0.001 | 0.038 | 0.038 | 0.009 | 0.56 | 0.59 | 0.63 | 0.71 | 0.48 | 0.76 |
| 3 | 2999717 | rs9864546 | -0.299 | 1.606 | -0.465 | 0.014 | -0.007 | 0.008 | 0.008 | 0 | 0.34 | 0.32 | 0.26 | 0.32 | 0.34 | 0.2 |
| 3 | 3000908 | rs751056 | -0.299 | 1.115 | -0.07 | 0.769 | -0.005 | 0.207 | 0.207 | 0.037 | 0.13 | 0.29 | 0.26 | 0.39 | 0.15 | 0.24 |
| 3 | 3003178 | rs17023680 | -0.094 | 1.274 | -0.769 | 0.614 | -0.036 | -0.013 | -0.013 | 0 | 0.27 | 0.09 | 0.16 | 0.28 | 0.25 | 0.12 |
| 3 | 3005616 | rs7633715 | -1.028 | 1.483 | -0.488 | 0.625 | 0.003 | 0.059 | 0.059 | 1.256 | 0.44 | 0.24 | 0.43 | 0.7 | 0.55 | 0.33 |
| 3 | 3006381 | rs12715157 | -1.052 | 1.338 | 0.128 | 0.611 | -0.001 | -0.001 | -0.001 | 0 | 0.47 | 0.47 | 0.35 | 0.44 | 0.47 | 0.26 |
| 3 | 3006444 | rs10510243 | -0.772 | 1.017 | 0.315 | 0.318 | -0.017 | -0.02 | -0.02 | 1.289 | 0.16 | 0.22 | 0.14 | 0.12 | 0.12 | 0.09 |

| | | | | | | | | | | | | | | | | |
|---|---|---|---|---|---|---|---|---|---|---|---|---|---|---|---|---|
| 3 | 3014060 | rs340804 | -0.484 | 1.155 | 0.2 | 0.071 | 0.004 | -0.01 | -0.01 | 0.273 | 0.31 | 0.32 | 0.25 | 0.27 | 0.32 | 0.33 |
| 3 | 3016114 | rs17023926 | -0.067 | 0.914 | 0.406 | -0.171 | -0.001 | -0.023 | -0.023 | 0.076 | 0.12 | 0.16 | 0.1 | 0.01 | 0.13 | 0.04 |
| 3 | 3017397 | rs168165 | -1.359 | 0.693 | 0.287 | -0.339 | -0.003 | 0.03 | 0.03 | 2.473 | 0.36 | 0.78 | 0.4 | 0.3 | 0.33 | 0.31 |
| 3 | 3019437 | rs9834836 | -1.38 | 0.678 | 0.343 | -0.301 | -0.003 | 0.03 | 0.03 | 1.043 | 0.36 | 0.82 | 0.42 | 0.31 | 0.33 | 0.34 |
| 3 | 3020113 | rs7356030 | -0.273 | 0.749 | 0.786 | -0.281 | 0.012 | 0.044 | 0.044 | 1.573 | 0.21 | 0.45 | 0.22 | 0.28 | 0.2 | 0.26 |
| 3 | 3020199 | rs7619485 | 0.784 | 0.744 | 0.245 | -0.326 | -0.039 | 0.047 | 0.047 | 0 | 0.08 | 0.01 | 0.09 | 0.01 | 0.12 | 0.04 |
| 3 | 3026162 | rs4685589 | 0.035 | 0.676 | 0.258 | -0.499 | -0.047 | 0.049 | 0.049 | 0.348 | 0.22 | 0.05 | 0.13 | 0.14 | 0.23 | 0.21 |
| 3 | 3028102 | rs340817 | -1.687 | 0.766 | 0.136 | -0.542 | -0.055 | 0.155 | 0.155 | 0.266 | 0.5 | 0.88 | 0.47 | 0.3 | 0.46 | 0.4 |
| 3 | 3029483 | rs2055318 | -0.61 | 0.823 | 0.433 | -0.546 | 0.009 | -0.064 | -0.064 | 0 | 0.19 | 0.36 | 0.28 | 0.05 | 0.19 | 0.11 |
| 3 | 3031308 | rs7613278 | -0.726 | 0.815 | 0.43 | -0.549 | 0.007 | -0.064 | -0.064 | 0 | 0.19 | 0.36 | 0.28 | 0.05 | 0.2 | 0.11 |
| 3 | 3033626 | rs9853379 | -1.268 | 0.835 | 0.287 | -0.443 | -0.008 | 0.053 | 0.053 | 0.017 | 0.47 | 0.67 | 0.46 | 0.31 | 0.46 | 0.41 |
| 3 | 3039150 | rs17650371 | 0.193 | 1.124 | -0.395 | -0.274 | -0.032 | 0.038 | 0.038 | 0 | 0.12 | 0.05 | 0.11 | 0 | 0.18 | 0.07 |
| 3 | 3041148 | rs13060035 | 1.614 | 0.922 | -1.301 | 0.03 | -0.043 | -0.075 | -0.075 | 0 | 0.19 | 0.01 | 0.32 | 0.21 | 0.24 | 0.22 |
| 3 | 3041819 | rs149467 | 0.36 | 0.969 | -0.481 | -0.151 | 0.005 | 0.009 | 0.009 | 0.207 | 0.14 | 0.12 | 0.09 | 0.24 | 0.14 | 0.22 |
| 3 | 3046754 | rs6799919 | -0.968 | 0.373 | -0.096 | -0.563 | -0.006 | -0.072 | -0.072 | 0 | 0.25 | 0.69 | 0.31 | 0.03 | 0.21 | 0.14 |
| 3 | 3047566 | rs163554 | 0.626 | 0.744 | -0.016 | 0.18 | 0.002 | 0.042 | 0.042 | 0 | 0.88 | 0.8 | 0.89 | 0.73 | 0.85 | 0.88 |
| 3 | 3048519 | rs10510248 | -0.241 | 0.451 | 0.29 | 0.048 | 0.003 | 0.061 | 0.061 | 0 | 0.18 | 0.29 | 0.17 | 0.31 | 0.23 | 0.11 |
| 3 | 3049520 | rs13088151 | 0.597 | 0.557 | -0.722 | -0.414 | -0.049 | 0.105 | 0.105 | 3.474 | 0.3 | 0.05 | 0.22 | 0.14 | 0.25 | 0.14 |
| 3 | 3052679 | rs9310923 | 0.023 | -0.113 | -0.366 | -0.427 | -0.007 | -0.053 | -0.053 | 0 | 0.13 | 0.2 | 0.1 | 0.01 | 0.11 | 0.15 |
| 3 | 3057903 | rs3907591 | -0.277 | 0.08 | -0.095 | -0.324 | 0.033 | 0.031 | 0.031 | 2.065 | 0.5 | 0.32 | 0.39 | 0.41 | 0.57 | 0.28 |
| 3 | 3062242 | rs2018016 | 0.683 | 0.206 | -0.055 | -0.349 | 0 | -0.036 | -0.036 | 0 | 0.09 | 0.16 | 0.08 | 0.01 | 0.09 | 0.08 |
| 3 | 3066139 | rs3911919 | -0.416 | 0.205 | 0.352 | -0.424 | 0.017 | -0.097 | -0.097 | 0 | 0.15 | 0.33 | 0.19 | 0.01 | 0.18 | 0.11 |
| 3 | 3067850 | rs7629924 | 0.938 | 0.031 | 0.257 | -0.557 | -0.032 | -0.03 | -0.03 | 1.481 | 0.23 | 0.3 | 0.15 | 0.1 | 0.1 | 0.07 |
| 3 | 3068148 | rs13084872 | 0.728 | 0.121 | -0.004 | -0.453 | -0.038 | -0.022 | -0.022 | 0 | 0.23 | 0.15 | 0.21 | 0.32 | 0.37 | 0.18 |
| 3 | 3069063 | rs9874811 | 1.117 | 0.05 | 0.014 | -0.435 | 0.005 | -0.025 | -0.025 | 0 | 0.06 | 0.07 | 0.11 | 0 | 0.09 | 0.02 |
| 3 | 3069773 | rs9837045 | -0.077 | 0.057 | 0.007 | -0.432 | -0.028 | -0.001 | -0.001 | 0 | 0.32 | 0.27 | 0.28 | 0.32 | 0.47 | 0.26 |
| 3 | 3070471 | rs7649063 | -0.792 | 0.205 | 0.18 | -0.301 | -0.005 | -0.001 | -0.001 | 0 | 0.56 | 0.53 | 0.43 | 0.42 | 0.58 | 0.33 |
| 3 | 3073509 | rs1039255 | -0.773 | 0.227 | -0.263 | -0.258 | -0.003 | -0.001 | -0.001 | 0.187 | 0.55 | 0.56 | 0.42 | 0.43 | 0.57 | 0.36 |
| 3 | 3075096 | rs4685592 | -1.159 | 0.331 | -0.733 | -0.133 | 0.006 | -0.012 | -0.012 | 0 | 0.62 | 0.71 | 0.54 | 0.46 | 0.66 | 0.39 |
| 3 | 3075224 | rs7646097 | -1.159 | 0.331 | -0.733 | -0.133 | 0.006 | -0.012 | -0.012 | 0.301 | 0.62 | 0.71 | 0.54 | 0.46 | 0.66 | 0.39 |
| 3 | 3076218 | rs9845519 | -0.158 | 0.317 | -0.728 | -0.032 | -0.022 | -0.024 | -0.024 | 0 | 0.45 | 0.23 | 0.39 | 0.52 | 0.47 | 0.25 |
| 3 | 3076443 | rs6803232 | -0.707 | -0.033 | -1.149 | 0.19 | 0.006 | -0.011 | -0.011 | 0.036 | 0.62 | 0.71 | 0.54 | 0.45 | 0.66 | 0.39 |
| 3 | 3079127 | rs3749363 | 0.347 | -0.25 | -1.418 | 0.055 | 0.05 | 0.032 | 0.032 | 4.294 | 0.2 | 0.05 | 0.12 | 0.11 | 0.11 | 0.07 |
| 3 | 3080200 | rs7617493 | 0.068 | 0.354 | -0.962 | 0.179 | 0 | 0.002 | 0.002 | 1.184 | 0.6 | 0.6 | 0.39 | 0.51 | 0.47 | 0.34 |
| 3 | 3080452 | rs7625369 | 0.146 | 1.24 | -0.904 | -0.362 | -0.007 | 0.003 | 0.003 | 0 | 0.13 | 0.12 | 0.12 | 0.01 | 0.16 | 0.07 |
| 3 | 3080509 | rs7648232 | -0.392 | 0.237 | -0.328 | -0.456 | -0.025 | -0.009 | -0.009 | 0 | 0.25 | 0.37 | 0.32 | 0.19 | 0.17 | 0.19 |
| 3 | 3081516 | rs339287 | -0.553 | 0.064 | 0.043 | -0.572 | -0.035 | 0.034 | 0.034 | 0 | 0.2 | 0.51 | 0.14 | 0.19 | 0.14 | 0.1 |
| 3 | 3081544 | rs339286 | 0.119 | 0.026 | 0.292 | -0.657 | -0.007 | -0.081 | -0.081 | 0 | 0.12 | 0.37 | 0.11 | 0 | 0.1 | 0.05 |
| 3 | 3081825 | rs339285 | -1.016 | -0.17 | -0.159 | -0.703 | 0 | -0.011 | -0.011 | 0 | 0.91 | 0.97 | 0.88 | 0.88 | 0.91 | 0.8 |
| 3 | 3081959 | rs339284 | 0.611 | -0.151 | 0.089 | -0.689 | -0.011 | -0.068 | -0.068 | 0 | 0.11 | 0.29 | 0.1 | 0 | 0.1 | 0.05 |
| 3 | 3085780 | rs4685597 | 1.051 | -0.188 | -0.342 | -0.973 | 0.007 | 0.001 | 0.001 | 1.157 | 0.33 | 0.27 | 0.21 | 0.11 | 0.27 | 0.17 |
| 3 | 3086195 | rs163563 | -1.275 | 0.04 | -0.495 | -0.761 | -0.014 | -0.02 | -0.02 | 0 | 0.91 | 0.97 | 0.88 | 0.87 | 0.9 | 0.8 |
| 3 | 3091331 | rs7649378 | 0.686 | 0.083 | -0.352 | -0.838 | -0.003 | 0.001 | 0.001 | 0 | 0.64 | 0.47 | 0.63 | 0.67 | 0.65 | 0.65 |
| 3 | 3092546 | rs11920374 | 0.969 | 0.403 | -0.51 | -0.876 | -0.038 | -0.062 | -0.062 | 1.77 | 0.31 | 0.15 | 0.34 | 0.56 | 0.3 | 0.47 |
| 3 | 3093366 | rs6776572 | 0.136 | -0.225 | -0.321 | -0.856 | 0.006 | 0.095 | 0.095 | 0.197 | 0.46 | 0.32 | 0.55 | 0.78 | 0.5 | 0.49 |
| 3 | 3094216 | rs149150 | 0.193 | -0.293 | -0.05 | -0.867 | -0.008 | -0.05 | -0.05 | 0 | 0.44 | 0.54 | 0.32 | 0.12 | 0.41 | 0.23 |
| 3 | 3094450 | rs163567 | 0.712 | -0.023 | -0.043 | -0.839 | 0.016 | -0.074 | -0.074 | 0.663 | 0.12 | 0.32 | 0.12 | 0 | 0.15 | 0.06 |
| 3 | 3094469 | rs163568 | -0.131 | 0.037 | -0.553 | -0.723 | 0.025 | 0.043 | 0.043 | 0.109 | 0.52 | 0.66 | 0.36 | 0.28 | 0.45 | 0.27 |
| 3 | 3095404 | rs163573 | 0.135 | 0.382 | 0.112 | -0.312 | 0.031 | -0.069 | -0.069 | 0.648 | 0.11 | 0.36 | 0.1 | 0 | 0.13 | 0.06 |
| 3 | 3096927 | rs163574 | -0.044 | 0.322 | -0.554 | 0.127 | 0.01 | 0.017 | 0.017 | 0 | 0.9 | 0.8 | 0.87 | 0.87 | 0.91 | 0.72 |
| 3 | 3099460 | rs11708737 | -0.344 | 0.493 | -1.68 | 0.201 | 0.002 | -0.013 | -0.013 | 1.712 | 0.28 | 0.1 | 0.13 | 0.2 | 0.19 | 0.14 |
| 3 | 3100615 | rs4054727 | 0.083 | 0.66 | -1.819 | 0.345 | 0.002 | 0.003 | 0.003 | 0 | 0.55 | 0.21 | 0.5 | 0.56 | 0.58 | 0.36 |
| 3 | 3100787 | rs163577 | -0.241 | 0.368 | -1.333 | -0.037 | -0.001 | -0.005 | -0.005 | 0 | 0.16 | 0.13 | 0.11 | 0.07 | 0.11 | 0.2 |
| 3 | 3101825 | rs9310935 | -0.227 | 0.68 | -1.233 | -0.198 | -0.001 | 0.028 | 0.028 | 6.551 | 0.49 | 0.49 | 0.52 | 0.76 | 0.49 | 0.5 |
| 3 | 3102867 | rs3913308 | 0.226 | 0.716 | -0.727 | 0.262 | 0 | -0.01 | -0.01 | 0 | 0.36 | 0.43 | 0.22 | 0.3 | 0.33 | 0.3 |
| 3 | 3103110 | rs163582 | 0.304 | 1.147 | -0.888 | 0.067 | 0.016 | 0.023 | 0.023 | 0.127 | 0.7 | 0.64 | 0.52 | 0.48 | 0.57 | 0.65 |
| 3 | 3104303 | rs7629198 | -0.66 | 1.144 | -0.926 | -0.323 | 0.005 | -0.03 | -0.03 | 0.051 | 0.12 | 0.27 | 0.1 | 0.06 | 0.1 | 0.1 |
| 3 | 3104758 | rs340826 | 0.419 | 1.16 | -1.014 | -0.262 | 0.016 | 0.03 | 0.03 | 0 | 0.68 | 0.57 | 0.5 | 0.5 | 0.59 | 0.67 |
| 3 | 3104951 | rs17606924 | 0.184 | 1.369 | -0.979 | -0.34 | -0.011 | 0.01 | 0.01 | 0 | 0.08 | 0.04 | 0.06 | 0.05 | 0.13 | 0.09 |
| 3 | 3107691 | rs6794523 | -0.015 | 0.811 | -0.783 | -0.557 | -0.001 | -0.002 | -0.002 | 1.179 | 0.45 | 0.44 | 0.25 | 0.3 | 0.41 | 0.43 |
| 3 | 3108083 | rs340828 | 0.172 | 0.816 | -1.108 | -0.744 | 0 | -0.001 | -0.001 | 1.161 | 0.35 | 0.34 | 0.36 | 0.25 | 0.42 | 0.38 |
| 3 | 3108711 | rs17659192 | 1.05 | 0.673 | -1.297 | -0.609 | -0.003 | 0.002 | 0.002 | 0 | 0.08 | 0.05 | 0.05 | 0.06 | 0.1 | 0.06 |
| 3 | 3109444 | rs340831 | -0.753 | 0.441 | -1.344 | -0.632 | -0.051 | -0.092 | -0.092 | 0 | 0.74 | 0.89 | 0.6 | 0.44 | 0.69 | 0.74 |
| 3 | 3111436 | rs17659353 | 0.977 | 0.393 | -1.306 | -0.985 | -0.058 | 0.047 | 0.047 | 0.02 | 0.15 | 0.02 | 0.1 | 0.06 | 0.24 | 0.1 |
| 3 | 3111458 | rs340833 | -0.333 | 0.085 | -1.074 | -1.112 | 0 | -0.004 | -0.004 | 0 | 0.47 | 0.48 | 0.44 | 0.25 | 0.51 | 0.44 |
| 3 | 3115514 | rs340807 | 0.985 | 0.483 | -1.019 | -0.95 | -0.014 | -0.02 | -0.02 | 1.18 | 0.25 | 0.36 | 0.31 | 0.17 | 0.23 | 0.33 |
| 3 | 3116438 | rs2290611 | 1.135 | 0.76 | -1.185 | -0.796 | -0.057 | 0.042 | 0.042 | 0 | 0.13 | 0.02 | 0.09 | 0.05 | 0.24 | 0.1 |
| 3 | 3117700 | rs3792424 | 0.366 | 0.63 | -0.699 | -0.573 | 0 | -0.056 | -0.056 | 0.551 | 0.11 | 0.24 | 0.06 | 0.03 | 0.09 | 0.08 |
| 3 | 3120571 | rs4322988 | 0.582 | 0.639 | -1.531 | -0.064 | 0.003 | -0.009 | -0.009 | 0 | 0.15 | 0.14 | 0.12 | 0.27 | 0.09 | 0.26 |

| | | | | | | | | | | | | | | | | |
|---|---|---|---|---|---|---|---|---|---|---|---|---|---|---|---|---|
| 3 | 3121848 | rs340805 | 0.647 | 0.879 | -1.521 | -0.215 | 0.004 | -0.081 | -0.081 | 0.88 | 0.29 | 0.49 | 0.17 | 0.09 | 0.3 | 0.17 |
| 3 | 3122827 | rs6809408 | 0.331 | 1.491 | -2.194 | 0.284 | -0.003 | 0.05 | 0.05 | 0 | 0.33 | 0.22 | 0.38 | 0.17 | 0.37 | 0.34 |
| 3 | 3124182 | rs3804803 | 0.277 | 2.01 | -1.98 | 1.38 | -0.002 | -0.021 | -0.021 | 0 | 0.12 | 0.07 | 0.11 | 0.28 | 0.11 | 0.27 |
| 3 | 3126050 | rs163551 | 0.352 | 2.302 | -1.801 | 1.324 | -0.014 | -0.085 | -0.085 | 1.573 | 0.71 | 0.88 | 0.65 | 0.55 | 0.72 | 0.78 |
| 3 | 3127519 | rs3804800 | -1.339 | 3.193 | -1.169 | 1.256 | 0.056 | 0.016 | 0.016 | 0.13 | 0.09 | 0.14 | 0.3 | 0.12 | 0.21 | 0.09 |
| 3 | 3127730 | rs163549 | 0.701 | 2.846 | -1.329 | 1.447 | -0.005 | -0.006 | -0.006 | 0 | 0.79 | 0.81 | 0.55 | 0.71 | 0.63 | 0.65 |
| 3 | 3129349 | rs334807 | -0.208 | 1.41 | -1.313 | 0.74 | -0.011 | -0.043 | -0.043 | 0 | 0.55 | 0.5 | 0.5 | 0.86 | 0.64 | 0.72 |
| 3 | 3129632 | rs3804797 | -1.484 | 1.461 | -1.512 | 1.768 | 0.005 | -0.032 | -0.032 | 0 | 0.14 | 0.12 | 0.12 | 0.36 | 0.14 | 0.37 |
| 3 | 3130645 | rs3804795 | -1.076 | 1.428 | -1.476 | 1.564 | -0.013 | -0.056 | -0.056 | 0 | 0.09 | 0 | 0.07 | 0.24 | 0.08 | 0.31 |
| 3 | 3133791 | rs7647903 | -0.896 | 2.24 | -1.431 | 0.749 | -0.007 | 0.005 | 0.005 | 0 | 0.14 | 0.15 | 0.18 | 0.07 | 0.21 | 0.12 |
| 3 | 3133896 | rs7640267 | -1.214 | 1.834 | -1.579 | 0.626 | -0.008 | 0.031 | 0.031 | 0 | 0.06 | 0 | 0.03 | 0 | 0.05 | 0.02 |
| 3 | 3139957 | rs2290610 | -0.175 | 1.339 | -1.464 | 0.568 | 0.009 | 0.003 | 0.003 | 0 | 0.44 | 0.4 | 0.4 | 0.41 | 0.35 | 0.58 |
| 3 | 3142423 | rs6793085 | -0.195 | 0.897 | -1.054 | 0.102 | 0.001 | -0.001 | -0.001 | 0.132 | 0.52 | 0.46 | 0.47 | 0.53 | 0.49 | 0.76 |
| 3 | 3142943 | rs6808378 | -1.021 | 0.809 | -1.046 | -0.032 | -0.009 | -0.01 | -0.01 | 0 | 0.81 | 0.85 | 0.84 | 0.76 | 0.76 | 0.91 |
| 3 | 3147196 | rs17884458 | 0.208 | 0.224 | -0.811 | -0.539 | -0.009 | -0.074 | -0.074 | 0 | 0.26 | 0.33 | 0.24 | 0.04 | 0.21 | 0.13 |
| 3 | 3147676 | rs3856848 | 0.204 | 0.576 | -0.773 | -0.628 | 0.007 | -0.054 | -0.054 | 0 | 0.2 | 0.28 | 0.24 | 0.03 | 0.21 | 0.16 |
| 3 | 3148458 | rs334782 | -0.383 | 0.5 | -0.677 | -0.598 | 0.007 | -0.008 | -0.008 | 0 | 0.31 | 0.36 | 0.5 | 0.24 | 0.39 | 0.29 |
| 3 | 3149568 | rs3804791 | 0.583 | 0.079 | -0.671 | -0.839 | -0.021 | 0.005 | 0.005 | 0 | 0.37 | 0.22 | 0.24 | 0.37 | 0.4 | 0.26 |
| 3 | 3150267 | rs11713419 | 0.219 | 0.231 | -0.225 | -0.764 | 0.056 | 0.077 | 0.077 | 0.053 | 0.1 | 0.31 | 0.13 | 0.15 | 0.13 | 0.1 |
| 3 | 3151668 | rs4054760 | 0.175 | 0.173 | -0.272 | -0.993 | -0.007 | -0.005 | -0.005 | 1.648 | 0.22 | 0.27 | 0.3 | 0.02 | 0.28 | 0.18 |
| 3 | 3151759 | rs2290608 | 1.006 | 0.697 | -0.314 | -0.441 | -0.048 | 0.031 | 0.031 | 0 | 0.25 | 0.09 | 0.17 | 0.2 | 0.3 | 0.18 |
| 3 | 3153069 | rs334788 | 0.303 | 0.168 | 0.701 | -0.131 | 0.157 | 0.238 | 0.238 | 0.368 | 0.94 | 0.65 | 0.9 | 0.81 | 0.93 | 0.89 |
| 3 | 3153143 | rs334789 | 0.955 | 0.105 | 0.142 | -0.235 | -0.039 | 0.016 | 0.016 | 0 | 0.31 | 0.21 | 0.24 | 0.3 | 0.45 | 0.25 |
| 3 | 3154931 | rs334793 | -0.401 | 0.036 | 0.383 | -0.277 | 0.016 | 0.074 | 0.074 | 0.068 | 0.95 | 0.86 | 0.93 | 0.84 | 0.95 | 0.9 |
| 3 | 3155229 | rs9814648 | 0.184 | -0.37 | 0.407 | -0.415 | -0.003 | -0.06 | -0.06 | 0 | 0.32 | 0.38 | 0.32 | 0.04 | 0.26 | 0.2 |
| 3 | 3156066 | rs7618222 | 1.323 | -0.362 | -0.295 | -0.245 | 0.033 | -0.051 | -0.051 | 0 | 0.22 | 0.11 | 0.33 | 0.34 | 0.15 | 0.37 |
| 3 | 3156811 | rs2540078 | 0.209 | -0.049 | -0.407 | -0.799 | -0.001 | 0.002 | 0.002 | 0 | 0.43 | 0.44 | 0.37 | 0.45 | 0.56 | 0.33 |
| 3 | 3157051 | rs1153462 | 0.733 | 0.157 | -0.915 | -0.528 | -0.043 | -0.008 | -0.008 | 0.193 | 0.33 | 0.21 | 0.24 | 0.43 | 0.48 | 0.28 |
| 3 | 3157618 | rs9808896 | -0.546 | -0.108 | -0.426 | -0.487 | -0.005 | -0.055 | -0.055 | 0 | 0.25 | 0.26 | 0.2 | 0.02 | 0.18 | 0.16 |
| 3 | 3165907 | rs1278150 | -0.057 | 0.175 | -0.324 | 0.165 | 0.003 | 0.005 | 0.005 | 0.006 | 0.5 | 0.61 | 0.62 | 0.52 | 0.54 | 0.64 |
| 3 | 3184442 | rs3804783 | -1.013 | -0.859 | 1.691 | -0.26 | 0 | -0.001 | -0.001 | 0 | 0.14 | 0.13 | 0.2 | 0.05 | 0.12 | 0.15 |
| 3 | 3189279 | rs1705805 | 1.47 | -0.945 | 2.335 | -0.381 | -0.051 | -0.079 | -0.079 | 0.5 | 0.78 | 0.43 | 0.76 | 0.95 | 0.84 | 0.83 |
| 3 | 3191426 | rs1045433 | -1.011 | -0.987 | 2.222 | -0.395 | 0.01 | 0.01 | 0.01 | 0 | 0.15 | 0.12 | 0.2 | 0.05 | 0.13 | 0.18 |
| 3 | 3195574 | rs3846135 | -0.909 | -0.795 | 2.182 | -0.376 | 0.006 | 0.006 | 0.006 | 0 | 0.14 | 0.12 | 0.2 | 0.05 | 0.12 | 0.15 |
| 3 | 3199635 | rs1669338 | 1.586 | -1.003 | 2.383 | -0.399 | -0.039 | -0.056 | -0.056 | 0 | 0.78 | 0.36 | 0.75 | 0.93 | 0.85 | 0.84 |
| 3 | 3207754 | rs711619 | 1.872 | -0.972 | 2.29 | -0.387 | 0 | -0.031 | -0.031 | 0.021 | 0.9 | 0.46 | 0.94 | 0.98 | 0.95 | 0.97 |
| 3 | 3208731 | rs4685611 | -1.194 | -1.228 | 2.018 | -0.388 | 0.013 | 0.012 | 0.012 | 0.685 | 0.2 | 0.16 | 0.22 | 0.12 | 0.16 | 0.28 |
| 3 | 3220571 | rs711612 | 0.827 | -0.813 | 2.272 | -0.208 | 0.042 | 0.033 | 0.033 | 2.149 | 0.94 | 0.6 | 0.97 | 1 | 0.99 | 1 |
| 3 | 3232895 | rs9861982 | -1.652 | -0.892 | 2.228 | -0.245 | 0.028 | 0.014 | 0.014 | 0.122 | 0.06 | 0.34 | 0.02 | 0 | 0.02 | 0.03 |
| 3 | 3233271 | rs7624992 | -1.854 | -0.294 | 2.182 | -0.471 | 0.052 | -0.083 | -0.083 | 0 | 0.17 | 0.5 | 0.12 | 0.04 | 0.17 | 0.09 |
| 3 | 3238654 | rs11129466 | -0.575 | -0.33 | 1.833 | -0.419 | 0.05 | -0.02 | -0.02 | 0 | 0.09 | 0.19 | 0.09 | 0.04 | 0.11 | 0.04 |
| 3 | 3238726 | rs7640918 | -1.059 | -0.437 | 0.932 | -0.316 | 0.031 | 0 | 0 | 0 | 0.23 | 0.11 | 0.45 | 0.75 | 0.18 | 0.54 |
| 3 | 3239120 | rs1672735 | 1.049 | -0.424 | 1.042 | -0.294 | 0.013 | -0.065 | -0.065 | 0 | 0.83 | 0.36 | 0.88 | 0.96 | 0.86 | 0.93 |
| 3 | 3240356 | rs4109078 | -1.327 | -0.64 | 0.491 | -0.01 | 0.036 | 0.002 | 0.002 | 0 | 0.24 | 0.12 | 0.46 | 0.76 | 0.18 | 0.55 |
| 3 | 3242027 | rs1629634 | 1.129 | -0.429 | 0.824 | 0.206 | -0.021 | -0.014 | -0.014 | 0 | 0.78 | 0.69 | 0.7 | 0.86 | 0.82 | 0.77 |
| 3 | 3245509 | rs6768756 | 0.616 | 0.125 | 0.649 | 0.943 | 0.003 | 0.008 | 0.008 | 0.04 | 0.78 | 0.78 | 0.75 | 0.71 | 0.78 | 0.73 |
| 3 | 3245960 | rs11717189 | 1.416 | -0.211 | 0.811 | 1.376 | -0.004 | -0.013 | -0.013 | 8.417 | 0.65 | 0.57 | 0.46 | 0.31 | 0.67 | 0.45 |
| 3 | 3249106 | rs4685616 | -1.137 | -0.314 | 1.089 | 1.623 | -0.018 | 0.221 | 0.221 | 0 | 0.06 | 0.15 | 0.24 | 0.44 | 0.06 | 0.22 |
| 3 | 3252500 | rs6771981 | -1.023 | 0.038 | 0.892 | 0.6 | 0.05 | 0.074 | 0.074 | 0.86 | 0.05 | 0.28 | 0.11 | 0.07 | 0.03 | 0.11 |
| 3 | 3258800 | rs11915101 | -1.028 | 0.135 | 0.322 | 0.342 | -0.003 | -0.028 | -0.028 | 0.027 | 0.1 | 0.14 | 0.06 | 0 | 0.06 | 0.09 |
| 3 | 3260833 | rs3856858 | -0.123 | 0.069 | 0.172 | 0.218 | -0.001 | -0.001 | -0.001 | 0.021 | 0.64 | 0.62 | 0.55 | 0.62 | 0.64 | 0.69 |
| 3 | 3261014 | rs12488686 | 0.197 | -0.228 | 0.147 | 0.541 | 0.023 | 0.018 | 0.018 | 0 | 0.54 | 0.23 | 0.47 | 0.43 | 0.59 | 0.71 |
| 3 | 3261438 | rs13080820 | 0.362 | 0.1 | -0.217 | -0.22 | -0.005 | 0.074 | 0.074 | 0 | 0.36 | 0.16 | 0.24 | 0.17 | 0.36 | 0.44 |
| 3 | 3267027 | rs13074852 | 0.85 | 0.451 | -0.47 | -0.483 | -0.018 | 0.009 | 0.009 | 0 | 0.23 | 0.12 | 0.2 | 0.19 | 0.26 | 0.34 |
| 3 | 3276014 | rs12715200 | -0.3 | 0.397 | -0.029 | -0.428 | 0.003 | 0.019 | 0.019 | 0 | 0.39 | 0.39 | 0.29 | 0.27 | 0.37 | 0.47 |
| 3 | 3277275 | rs3846142 | -0.006 | 0.383 | -0.098 | -0.466 | -0.005 | 0.069 | 0.069 | 0 | 0.36 | 0.15 | 0.27 | 0.22 | 0.37 | 0.43 |
| 3 | 3278662 | rs6762199 | -1.25 | 0.108 | -0.189 | -0.771 | -0.003 | -0.011 | -0.011 | 0 | 0.6 | 0.68 | 0.48 | 0.46 | 0.58 | 0.57 |
| 3 | 3280819 | rs3843388 | -1.867 | 0.089 | -0.108 | -1.243 | -0.003 | -0.031 | -0.031 | 0 | 0.89 | 0.87 | 0.95 | 0.99 | 0.9 | 0.93 |
| 3 | 3283582 | rs12489660 | 0.556 | 0.115 | -0.272 | -1.165 | -0.008 | 0.007 | 0.007 | 0 | 0.31 | 0.3 | 0.27 | 0.25 | 0.35 | 0.36 |
| 3 | 3287422 | rs11129510 | -0.378 | 0.135 | -0.161 | -1.181 | 0 | -0.011 | -0.011 | 0 | 0.46 | 0.53 | 0.36 | 0.25 | 0.49 | 0.48 |
| 3 | 3288938 | rs3914771 | 1.005 | -0.214 | -0.955 | -0.955 | -0.015 | 0.009 | 0.009 | 0 | 0.25 | 0.16 | 0.2 | 0.24 | 0.27 | 0.38 |
| 3 | 3298121 | rs7622674 | -0.42 | -0.43 | -0.343 | -0.817 | -0.035 | 0.006 | 0.006 | 0 | 0.25 | 0.4 | 0.16 | 0.22 | 0.2 | 0.18 |
| 3 | 3302387 | rs711591 | -1.703 | -0.61 | -0.173 | -0.972 | -0.015 | -0.047 | -0.047 | 0 | 0.91 | 0.8 | 0.96 | 1 | 0.95 | 0.96 |
| 3 | 3304982 | rs3914769 | 1.521 | -0.483 | -0.529 | -1.114 | -0.003 | -0.002 | -0.002 | 0.248 | 0.27 | 0.25 | 0.46 | 0.53 | 0.38 | 0.32 |
| 3 | 3305737 | rs711588 | 1.102 | -0.195 | -0.411 | -0.993 | -0.033 | 0.017 | 0.017 | 0 | 0.28 | 0.08 | 0.28 | 0.24 | 0.29 | 0.42 |
| 3 | 3306941 | rs4685622 | 0.228 | -0.154 | -0.044 | -0.548 | -0.019 | -0.011 | -0.011 | 0.621 | 0.42 | 0.32 | 0.56 | 0.57 | 0.48 | 0.43 |
| 3 | 3307984 | rs3846145 | -0.214 | -0.195 | -0.611 | -0.604 | 0.006 | 0.018 | 0.018 | 0.031 | 0.49 | 0.65 | 0.59 | 0.61 | 0.46 | 0.44 |
| 3 | 3311720 | rs1153452 | 0.266 | -0.458 | -0.313 | -0.457 | -0.02 | -0.007 | -0.007 | 0 | 0.78 | 0.67 | 0.88 | 0.84 | 0.85 | 0.92 |
| 3 | 3312795 | rs1153451 | 0.163 | -0.434 | -0.375 | -0.456 | -0.016 | -0.007 | -0.007 | 0.005 | 0.78 | 0.72 | 0.89 | 0.85 | 0.85 | 0.92 |

| | | | | | | | | | | | | | | | | |
|---|---|---|---|---|---|---|---|---|---|---|---|---|---|---|---|---|
| 3 | 3315364 | rs13077321 | -0.423 | -0.741 | -0.483 | -0.472 | -0.025 | 0.019 | 0.019 | 0 | 0.17 | 0.28 | 0.09 | 0.21 | 0.11 | 0.2 |
| 3 | 3315392 | rs7612426 | -0.865 | -0.259 | -0.898 | -0.837 | -0.033 | -0.026 | -0.026 | 1.064 | 0.71 | 0.84 | 0.56 | 0.42 | 0.61 | 0.71 |
| 3 | 3319387 | rs9865956 | -0.401 | -1.213 | -0.681 | -1.069 | -0.032 | -0.003 | -0.003 | 0 | 0.23 | 0.34 | 0.11 | 0.2 | 0.13 | 0.23 |
| 3 | 3321124 | rs9828666 | -0.685 | -1.694 | -1.176 | -1.707 | 0.098 | 0.108 | 0.108 | 1.952 | 0.42 | 0.79 | 0.2 | 0.2 | 0.26 | 0.35 |
| 3 | 3323892 | rs2322041 | -0.361 | -1.144 | -0.69 | -1.709 | -0.008 | -0.023 | -0.023 | 0 | 0.27 | 0.39 | 0.2 | 0.16 | 0.25 | 0.29 |
| 3 | 3326866 | rs9817602 | -0.541 | -0.957 | -0.614 | -1.877 | 0.021 | 0.036 | 0.036 | 0 | 0.43 | 0.63 | 0.25 | 0.15 | 0.35 | 0.4 |
| 3 | 3330438 | rs2322044 | -0.964 | -1.025 | -0.708 | -1.92 | 0.061 | 0.181 | 0.181 | 0 | 0.5 | 0.83 | 0.3 | 0.18 | 0.41 | 0.46 |
| 3 | 3332441 | rs13073417 | -0.707 | -0.788 | -0.259 | -1.712 | -0.007 | -0.005 | -0.005 | 0.051 | 0.16 | 0.28 | 0.1 | 0.16 | 0.13 | 0.22 |
| 3 | 3340244 | rs1356615 | -0.864 | -0.763 | -0.005 | -1.757 | 0.005 | -0.01 | -0.01 | 0 | 0.16 | 0.35 | 0.11 | 0.15 | 0.15 | 0.23 |
| 3 | 3341324 | rs12497287 | -0.879 | -0.979 | 0.49 | -1.986 | -0.009 | -0.105 | -0.105 | 1.092 | 0.3 | 0.53 | 0.23 | 0.11 | 0.28 | 0.29 |
| 3 | 3343318 | rs13091627 | 0.313 | -0.927 | 0.263 | -1.932 | -0.001 | -0.021 | -0.021 | 0.615 | 0.1 | 0.14 | 0.06 | 0.01 | 0.1 | 0.08 |
| 3 | 3343531 | rs13065382 | 0.313 | -0.927 | 0.263 | -1.932 | -0.001 | -0.021 | -0.021 | 0 | 0.1 | 0.14 | 0.06 | 0.01 | 0.1 | 0.07 |
| 3 | 3344733 | rs6442791 | -1.854 | -0.183 | -0.104 | -2.311 | -0.007 | 0.02 | 0.02 | 0 | 0.7 | 0.82 | 0.6 | 0.31 | 0.72 | 0.76 |
| 3 | 3345004 | rs17029941 | -0.052 | 0.027 | -0.197 | -2.506 | 0.004 | -0.039 | -0.039 | 0.332 | 0.42 | 0.59 | 0.36 | 0.03 | 0.41 | 0.3 |
| 3 | 3345098 | rs1159832 | 0.074 | 0.095 | -0.451 | -2.458 | -0.033 | 0.018 | 0.018 | 0.969 | 0.27 | 0.15 | 0.23 | 0.27 | 0.31 | 0.46 |
| 3 | 3345465 | rs17671139 | -0.431 | -0.164 | -0.457 | -2.455 | 0.001 | 0.03 | 0.03 | 0 | 0.29 | 0.15 | 0.25 | 0.27 | 0.27 | 0.4 |
| 3 | 3346138 | rs10510257 | 0.075 | -0.224 | -0.344 | -2.497 | 0.007 | 0.034 | 0.034 | 0 | 0.26 | 0.17 | 0.24 | 0.18 | 0.22 | 0.22 |
| 3 | 3347090 | rs4684384 | -1.469 | -0.21 | -0.04 | -2.347 | 0.005 | -0.015 | -0.015 | 0 | 0.61 | 0.59 | 0.53 | 0.32 | 0.59 | 0.67 |
| 3 | 3347282 | rs4684385 | -0.509 | -0.248 | -0.257 | -2.208 | -0.006 | 0.048 | 0.048 | 0 | 0.33 | 0.18 | 0.28 | 0.28 | 0.32 | 0.41 |
| 3 | 3349500 | rs9866082 | -0.363 | -0.282 | 0.27 | -1.548 | -0.005 | -0.101 | -0.101 | 0 | 0.38 | 0.45 | 0.33 | 0.03 | 0.39 | 0.25 |
| 3 | 3355740 | rs17030006 | 0.413 | -0.653 | 0.57 | -0.939 | -0.001 | -0.038 | -0.038 | 0.144 | 0.22 | 0.25 | 0.2 | 0.01 | 0.19 | 0.15 |
| 3 | 3363379 | rs1995196 | 0.006 | -0.347 | 0.429 | 0.012 | -0.011 | 0.019 | 0.019 | 0.039 | 0.19 | 0.21 | 0.31 | 0.68 | 0.12 | 0.17 |
| 3 | 3364182 | rs7648378 | -0.725 | -0.606 | 0.383 | 0.54 | -0.024 | 0.012 | 0.012 | 0.211 | 0.3 | 0.41 | 0.19 | 0.58 | 0.19 | 0.21 |
| 3 | 3365596 | rs9849296 | -1.008 | -0.166 | 0.26 | 1.212 | 0.015 | -0.003 | -0.003 | 0 | 0.38 | 0.7 | 0.4 | 0.56 | 0.33 | 0.31 |
| 3 | 3370924 | rs6768984 | 0.151 | 0.159 | 1.117 | 2.061 | 0.023 | 0.039 | 0.039 | 0.004 | 0.79 | 0.6 | 0.8 | 0.69 | 0.73 | 0.85 |
| 3 | 3376689 | rs1499088 | 0.523 | 0.193 | 0.766 | 1.519 | -0.002 | 0.002 | 0.002 | 0 | 0.55 | 0.54 | 0.45 | 0.39 | 0.47 | 0.55 |
| 3 | 3376821 | rs1499087 | 1.098 | 0.105 | 0.931 | 1.583 | -0.003 | 0.122 | 0.122 | 0 | 0.49 | 0.23 | 0.41 | 0.29 | 0.45 | 0.47 |
| 3 | 3378900 | rs1499085 | -0.217 | 0.175 | 0.435 | 1.689 | 0.016 | 0.036 | 0.036 | 0 | 0.8 | 0.74 | 0.78 | 0.67 | 0.71 | 0.81 |
| 3 | 3383955 | rs10510260 | 0.53 | 0.25 | 0.163 | 1.598 | 0.002 | -0.015 | -0.015 | 0 | 0.26 | 0.2 | 0.37 | 0.39 | 0.28 | 0.34 |
| 3 | 3385806 | rs9861093 | -0.446 | 0.246 | 0.037 | 1.541 | 0.006 | 0.094 | 0.094 | 1.238 | 0.05 | 0.26 | 0.05 | 0.09 | 0.01 | 0.07 |
| 3 | 3388486 | rs1472473 | -0.215 | 0.657 | -0.869 | 1.513 | 0.021 | 0.011 | 0.011 | 0.023 | 0.81 | 0.71 | 0.82 | 0.8 | 0.74 | 0.87 |
| 3 | 3396403 | rs4142860 | 0.367 | 0.704 | -0.882 | 1.164 | -0.018 | 0.014 | 0.014 | 0.241 | 0.58 | 0.72 | 0.49 | 0.41 | 0.49 | 0.56 |
| 3 | 3402228 | rs1508741 | 0.273 | 0.588 | -1.479 | 1.198 | -0.056 | -0.011 | -0.011 | 0 | 0.57 | 0.83 | 0.48 | 0.43 | 0.47 | 0.54 |
| 3 | 3409961 | rs1508733 | -0.332 | 1.069 | -1.307 | 0.973 | -0.089 | -0.032 | -0.032 | 1.787 | 0.85 | 0.98 | 0.83 | 0.87 | 0.75 | 0.88 |
| 3 | 3414282 | rs2279975 | 0.229 | 0.403 | -1.073 | 1.196 | 0.016 | -0.029 | -0.029 | 0 | 0.33 | 0.27 | 0.39 | 0.54 | 0.29 | 0.4 |
| 3 | 3414586 | rs2279977 | 0.391 | 0.372 | -0.417 | 1.057 | -0.003 | 0.033 | 0.033 | 0 | 0.44 | 0.31 | 0.39 | 0.34 | 0.44 | 0.47 |
| 3 | 3416536 | rs1905764 | 0.161 | 0.187 | -0.202 | 1.613 | 0.008 | -0.057 | -0.057 | 0 | 0.12 | 0 | 0.18 | 0.21 | 0.09 | 0.1 |
| 3 | 3419135 | rs4685631 | 1.11 | 0.366 | 0.382 | 1.688 | -0.009 | 0.002 | 0.002 | 0.368 | 0.67 | 0.53 | 0.66 | 0.57 | 0.69 | 0.71 |
| 3 | 3419888 | rs9882221 | 1.973 | 0.438 | 0.436 | 1.241 | 0.01 | 0.085 | 0.085 | 0.305 | 0.37 | 0.2 | 0.18 | 0.03 | 0.32 | 0.38 |
| 3 | 3421278 | rs979152 | 0.612 | 0.491 | 0.554 | 1.293 | -0.006 | -0.053 | -0.053 | 0.378 | 0.72 | 0.58 | 0.7 | 0.9 | 0.75 | 0.81 |
| 3 | 3423954 | rs9838824 | 1.875 | 0.435 | 0.692 | 1.743 | -0.012 | -0.029 | -0.029 | 0 | 0.4 | 0.43 | 0.2 | 0.18 | 0.32 | 0.42 |
| 3 | 3424418 | rs1355070 | -0.2 | 0.723 | 1.366 | 2.296 | -0.015 | 0.019 | 0.019 | 0.227 | 0.28 | 0.24 | 0.38 | 0.12 | 0.37 | 0.21 |
| 3 | 3425382 | rs978762 | 0.083 | 0.649 | 1.077 | 2.287 | -0.082 | 0.072 | 0.072 | 0.979 | 0.25 | 0.04 | 0.35 | 0.12 | 0.31 | 0.21 |
| 3 | 3426420 | rs17030873 | -0.468 | 0.646 | 1.12 | 2.385 | -0.038 | -0.077 | -0.077 | 0 | 0.29 | 0.13 | 0.35 | 0.61 | 0.35 | 0.35 |
| 3 | 3428056 | rs13100659 | -0.783 | 0.485 | 1.137 | 2.554 | -0.004 | 0.151 | 0.151 | 0 | 0.06 | 0.09 | 0.05 | 0.48 | 0.06 | 0.14 |
| 3 | 3428614 | rs17628085 | 0.227 | 0.558 | 1.047 | 2.489 | -0.097 | 0.061 | 0.061 | 0.934 | 0.24 | 0.02 | 0.3 | 0.12 | 0.3 | 0.2 |
| 3 | 3429642 | rs9870347 | -2.168 | 0.175 | 2.259 | 3.425 | 0.002 | -0.046 | -0.046 | 3.105 | 0.28 | 0.79 | 0.23 | 0.55 | 0.26 | 0.29 |
| 3 | 3431858 | rs9876606 | -1.213 | 0.162 | 2.358 | 3.373 | 0.007 | -0.083 | -0.083 | 0 | 0.21 | 0.48 | 0.16 | 0.08 | 0.2 | 0.15 |
| 3 | 3434542 | rs11129561 | -1.955 | 0.179 | 2.466 | 3.326 | 0 | -0.012 | -0.012 | 0 | 0.27 | 0.73 | 0.23 | 0.55 | 0.26 | 0.28 |
| 3 | 3434942 | rs11926725 | -1.129 | 0.197 | 2.276 | 3.298 | 0.007 | 0.005 | 0.005 | 0 | 0.21 | 0.42 | 0.17 | 0.22 | 0.21 | 0.18 |
| 3 | 3448534 | rs1355071 | -1.74 | 0.195 | 2.311 | 3.309 | -0.002 | -0.016 | -0.016 | 0 | 0.28 | 0.46 | 0.31 | 0.79 | 0.27 | 0.36 |
| 3 | 3463165 | rs17031295 | -2.714 | 0.14 | 2.256 | 3.074 | 0.085 | 0.585 | 0.585 | 0.192 | 0.05 | 0.36 | 0.13 | 0.29 | 0.03 | 0.07 |
| 3 | 3475578 | rs9820648 | -1.59 | 0.192 | 2.002 | 2.764 | 0.001 | 0.063 | 0.063 | 0 | 0.25 | 0.45 | 0.19 | 0.45 | 0.24 | 0.3 |
| 3 | 3475859 | rs13088831 | -1.591 | 0.194 | 2.125 | 2.765 | 0.005 | 0.095 | 0.095 | 0 | 0.25 | 0.52 | 0.19 | 0.45 | 0.24 | 0.3 |
| 3 | 3496754 | rs1396415 | -1.178 | 0.103 | 1.663 | 2.751 | -0.001 | 0.069 | 0.069 | 0 | 0.21 | 0.3 | 0.25 | 0.59 | 0.21 | 0.26 |
| 3 | 3502091 | rs12486658 | -1.833 | 0.15 | 0.916 | 2.551 | -0.001 | -0.07 | -0.07 | 0 | 0.31 | 0.84 | 0.33 | 0.6 | 0.29 | 0.3 |
| 3 | 3527685 | rs7633914 | -1.65 | 0.481 | 1.613 | 2.331 | 0.036 | 0.103 | 0.103 | 0.763 | 0.19 | 0.49 | 0.19 | 0.29 | 0.19 | 0.17 |
| 3 | 3530023 | rs1967251 | 1.192 | -0.007 | 1.549 | 2.094 | -0.015 | -0.014 | -0.014 | 0 | 0.77 | 0.68 | 0.74 | 0.85 | 0.8 | 0.82 |
| 3 | 3530439 | rs9311069 | 1.158 | 0.029 | 1.551 | 2.11 | -0.013 | -0.012 | -0.012 | 0 | 0.77 | 0.68 | 0.74 | 0.85 | 0.8 | 0.83 |
| 3 | 3535441 | rs7649856 | 1.471 | 0.04 | 1.05 | 1.846 | -0.007 | -0.016 | -0.016 | 2.107 | 0.73 | 0.7 | 0.64 | 0.16 | 0.76 | 0.53 |
| 3 | 3540936 | rs9871391 | 1.46 | -0.064 | 0.406 | 2.152 | 0.011 | -0.071 | -0.071 | 0 | 0.8 | 0.89 | 0.69 | 0.64 | 0.85 | 0.78 |
| 3 | 3541722 | rs10510265 | -1.813 | 0.1 | 1.092 | 2.266 | -0.013 | 0.145 | 0.145 | 0 | 0.22 | 0.42 | 0.26 | 0.52 | 0.18 | 0.29 |
| 3 | 3547685 | rs1508729 | 1.442 | 0.214 | 0.361 | 2.178 | -0.037 | -0.032 | -0.032 | 0 | 0.86 | 0.99 | 0.78 | 0.86 | 0.84 | 0.9 |
| 3 | 3548182 | rs2874731 | -1.166 | 0.132 | 0.435 | 2.077 | 0 | -0.021 | -0.021 | 0 | 0.06 | 0.09 | 0.04 | 0 | 0.03 | 0.05 |
| 3 | 3548524 | rs17032533 | -0.75 | -0.033 | 0.803 | 2.077 | -0.009 | -0.042 | -0.042 | 1.08 | 0.09 | 0.2 | 0.04 | 0.02 | 0.07 | 0.01 |
| 3 | 3551353 | rs9311083 | 1.129 | 0.158 | 0.23 | 2.28 | -0.039 | -0.011 | -0.011 | 0 | 0.88 | 0.99 | 0.79 | 0.92 | 0.85 | 0.94 |
| 3 | 3552756 | rs10510266 | 1.585 | 0.186 | 0.271 | 2.276 | -0.01 | 0.003 | 0.003 | 0.022 | 0.87 | 0.9 | 0.76 | 0.9 | 0.84 | 0.9 |
| 3 | 3555312 | rs7634908 | 0.956 | 0.22 | 0.045 | 2.211 | -0.037 | 0.007 | 0.007 | 0 | 0.88 | 0.99 | 0.79 | 0.93 | 0.85 | 0.91 |
| 3 | 3556881 | rs9876329 | 1.47 | 0.372 | 0.514 | 1.887 | 0.009 | 0.119 | 0.119 | 0.33 | 0.75 | 0.5 | 0.56 | 0.47 | 0.76 | 0.64 |

| | | | | | | | | | | | | | | | | |
|---|---|---|---|---|---|---|---|---|---|---|---|---|---|---|---|---|
| 3 | 3557902 | rs6789036 | 1.329 | 0.08 | 0.058 | 1.757 | -0.018 | 0.019 | 0.019 | 0.042 | 0.66 | 0.47 | 0.58 | 0.53 | 0.72 | 0.67 |
| 3 | 3558481 | rs7644488 | -1.227 | -0.315 | -0.721 | 1.765 | 0.027 | -0.04 | -0.04 | 0 | 0.23 | 0.82 | 0.35 | 0.39 | 0.16 | 0.2 |
| 3 | 3558889 | rs2322212 | -0.438 | 0.442 | -1.388 | -0.154 | -0.049 | 0.127 | 0.127 | 0 | 0.39 | 0.19 | 0.26 | 0.1 | 0.43 | 0.31 |
| 3 | 3560143 | rs988071 | -0.755 | 0.176 | -1.552 | -0.423 | 0.009 | 0.127 | 0.127 | 0.019 | 0.48 | 0.82 | 0.52 | 0.71 | 0.44 | 0.46 |
| 3 | 3564116 | rs1508735 | -0.501 | 0.05 | -0.931 | -0.684 | -0.03 | -0.012 | -0.012 | 0.035 | 0.33 | 0.68 | 0.42 | 0.68 | 0.25 | 0.32 |
| 3 | 3564622 | rs17632978 | -0.097 | 0.316 | -1.054 | -0.594 | 0.022 | -0.034 | -0.034 | 0 | 0.13 | 0.12 | 0.1 | 0.03 | 0.2 | 0.15 |
| 3 | 3569967 | rs4685648 | -0.714 | 0.138 | -1.356 | -0.134 | -0.067 | -0.108 | -0.108 | 0 | 0.18 | 0.03 | 0.27 | 0.53 | 0.24 | 0.3 |
| 3 | 3572242 | rs330243 | 0.313 | -0.472 | -0.718 | -0.339 | -0.024 | -0.016 | -0.016 | 0 | 0.74 | 0.56 | 0.78 | 0.81 | 0.79 | 0.79 |
| 3 | 3572511 | rs330246 | 0.478 | -0.054 | -1.301 | -0.609 | -0.075 | -0.087 | -0.087 | 0.033 | 0.75 | 0.92 | 0.7 | 0.41 | 0.65 | 0.58 |
| 3 | 3576925 | rs4684395 | -0.906 | -0.273 | -1.219 | -0.726 | 0.002 | 0.008 | 0.008 | 0 | 0.85 | 0.88 | 0.93 | 0.94 | 0.86 | 0.84 |
| 3 | 3580449 | rs11129648 | 0.675 | -0.193 | -1.317 | -0.714 | 0.002 | 0.012 | 0.012 | 0 | 0.62 | 0.48 | 0.64 | 0.4 | 0.62 | 0.5 |
| 3 | 3581077 | rs2060642 | -0.038 | -0.348 | -1.302 | -0.698 | -0.048 | 0.092 | 0.092 | 0 | 0.73 | 0.48 | 0.72 | 0.43 | 0.8 | 0.65 |
| 3 | 3582105 | rs9879339 | 0.798 | -0.497 | -0.95 | -0.623 | -0.006 | 0.058 | 0.058 | 0.179 | 0.47 | 0.34 | 0.47 | 0.21 | 0.48 | 0.35 |
| 3 | 3583063 | rs723168 | 0.979 | -0.035 | -1.286 | -0.412 | -0.032 | 0.085 | 0.085 | 0.318 | 0.41 | 0.18 | 0.44 | 0.21 | 0.43 | 0.33 |
| 3 | 3585219 | rs17633588 | -0.655 | 0.01 | -0.827 | 0.169 | 0.001 | -0.002 | -0.002 | 0 | 0.16 | 0.14 | 0.18 | 0.19 | 0.16 | 0.16 |
| 3 | 3587732 | rs2054873 | 0.797 | -0.213 | -0.935 | -0.475 | 0.024 | 0.012 | 0.012 | 1.998 | 0.77 | 0.21 | 0.86 | 1 | 0.87 | 0.98 |
| 3 | 3588042 | rs17684824 | -0.506 | 0.723 | -0.955 | 0.276 | -0.037 | -0.042 | -0.042 | 0 | 0.08 | 0.01 | 0.1 | 0.17 | 0.13 | 0.12 |
| 3 | 3599937 | rs9811783 | -1.204 | 1.389 | 0.574 | 0.192 | 0.111 | 0.083 | 0.083 | 0.384 | 0.1 | 0.35 | 0.11 | 0.13 | 0.14 | 0.1 |
| 3 | 3612598 | rs9879901 | -0.8 | 0.735 | 0.755 | 0.237 | -0.013 | -0.002 | -0.002 | 0 | 0.21 | 0.24 | 0.16 | 0.18 | 0.14 | 0.09 |
| 3 | 3614887 | rs9815663 | -1.167 | 1.09 | 0.632 | 0.178 | -0.001 | -0.001 | -0.001 | 0.573 | 0.25 | 0.24 | 0.18 | 0.18 | 0.2 | 0.13 |
| 3 | 3617037 | rs1584572 | -1.054 | 1.357 | 0.795 | 0.177 | -0.015 | -0.007 | -0.007 | 0.006 | 0.24 | 0.29 | 0.17 | 0.18 | 0.17 | 0.11 |
| 3 | 3619829 | rs6778553 | -1.807 | 2.173 | 1.143 | 0.071 | 0.084 | -0.002 | -0.002 | 2.544 | 0.18 | 0.44 | 0.2 | 0.14 | 0.19 | 0.14 |
| 3 | 3623171 | rs13318643 | -1.866 | 2.066 | 2.107 | 0.043 | 0.067 | -0.002 | -0.002 | 0 | 0.18 | 0.57 | 0.2 | 0.13 | 0.2 | 0.11 |
| 3 | 3623315 | rs7620027 | -2.088 | 1.973 | 2.189 | 0.033 | 0.004 | -0.006 | -0.006 | 0 | 0.19 | 0.64 | 0.18 | 0.13 | 0.17 | 0.11 |
| 3 | 3627205 | rs17634664 | 0.145 | 1.996 | 2.032 | 0.065 | -0.033 | -0.038 | -0.038 | 0 | 0.08 | 0.01 | 0.1 | 0.13 | 0.11 | 0.1 |
| 3 | 3628512 | rs715882 | -1.903 | 1.952 | 2.517 | 0.031 | 0.019 | -0.055 | -0.055 | 0 | 0.1 | 0.55 | 0.08 | 0 | 0.06 | 0.01 |
| 3 | 3628667 | rs1375915 | -1.722 | 1.871 | 2.823 | -0.005 | -0.052 | -0.076 | -0.076 | 0.049 | 0.12 | 0.4 | 0.06 | 0.01 | 0.03 | 0.04 |
| 3 | 3628981 | rs10865864 | 0.034 | 2.466 | 1.775 | 0.341 | -0.061 | -0.028 | -0.028 | 0.201 | 0.15 | 0.03 | 0.13 | 0.22 | 0.24 | 0.17 |
| 3 | 3629346 | rs9814693 | -0.501 | 2.024 | 2.354 | 0.387 | 0.09 | 0.072 | 0.072 | 0 | 0.1 | 0.24 | 0.15 | 0.13 | 0.15 | 0.1 |
| 3 | 3635340 | rs6804997 | 0.496 | 1.687 | 1.712 | 0.669 | 0.008 | 0.036 | 0.036 | 0 | 0.85 | 0.7 | 0.79 | 0.79 | 0.88 | 0.85 |
| 3 | 3635641 | rs6794507 | 0.634 | 1.624 | 1.684 | 0.642 | 0.002 | 0.028 | 0.028 | 0 | 0.84 | 0.69 | 0.78 | 0.79 | 0.88 | 0.85 |
| 3 | 3639025 | rs9647362 | -1.523 | 1.619 | 1.637 | 0.532 | 0.054 | -0.019 | -0.019 | 0.333 | 0.2 | 0.45 | 0.2 | 0.11 | 0.2 | 0.15 |
| 3 | 3643141 | rs1349312 | 1.752 | 1.577 | 1.642 | 0.517 | -0.003 | -0.007 | -0.007 | 0 | 0.73 | 0.38 | 0.73 | 0.76 | 0.83 | 0.82 |
| 3 | 3644552 | rs1349309 | 2.162 | 1.448 | 1.376 | 0.472 | -0.059 | -0.053 | -0.053 | 0.309 | 0.62 | 0.07 | 0.6 | 0.64 | 0.72 | 0.71 |
| 3 | 3648076 | rs1450088 | 0.128 | 0.765 | 2.064 | 0.42 | -0.007 | 0.087 | 0.087 | 0 | 0.89 | 0.73 | 0.84 | 0.77 | 0.94 | 0.88 |
| 3 | 3648342 | rs1450087 | 1.255 | 0.745 | 2.344 | 0.409 | -0.033 | 0.079 | 0.079 | 1.106 | 0.85 | 0.44 | 0.82 | 0.76 | 0.94 | 0.87 |
| 3 | 3649300 | rs2129896 | 1.875 | 0.77 | 2.01 | 0.402 | 0.013 | 0.003 | 0.003 | 1.365 | 0.76 | 0.22 | 0.75 | 0.76 | 0.88 | 0.84 |
| 3 | 3650239 | rs9837561 | -1.462 | 0.849 | 2.064 | 0.328 | 0.013 | -0.073 | -0.073 | 0 | 0.12 | 0.36 | 0.07 | 0 | 0.06 | 0.03 |
| 3 | 3655251 | rs9881476 | 0.045 | 0.84 | 1.626 | 0.378 | -0.003 | -0.011 | -0.011 | 0 | 0.06 | 0.11 | 0.03 | 0.01 | 0.02 | 0.02 |
| 3 | 3657425 | rs4684400 | -0.939 | 0.244 | 1.115 | 0.158 | 0.002 | -0.002 | -0.002 | 0 | 0.54 | 0.48 | 0.48 | 0.51 | 0.53 | 0.5 |
| 3 | 3658840 | rs1601875 | 0.653 | 0.26 | 1.379 | 0.287 | -0.003 | 0.043 | 0.043 | 0.2 | 0.49 | 0.64 | 0.48 | 0.28 | 0.51 | 0.42 |
| 3 | 3659605 | rs1386948 | 0.271 | 0.238 | 1.226 | 0.332 | -0.004 | 0.028 | 0.028 | 0 | 0.76 | 0.7 | 0.72 | 0.58 | 0.8 | 0.77 |
| 3 | 3659759 | rs2035663 | 0.902 | 0.253 | 1.413 | 0.313 | 0.023 | 0.038 | 0.038 | 0 | 0.66 | 0.56 | 0.57 | 0.47 | 0.67 | 0.62 |
| 3 | 3659802 | rs2129895 | 0.803 | 0.288 | 1.253 | 0.312 | -0.001 | -0.002 | -0.002 | 0 | 0.67 | 0.68 | 0.61 | 0.47 | 0.68 | 0.62 |
| 3 | 3662149 | rs1002912 | 1.2 | 0.286 | 1.145 | 0.345 | -0.031 | -0.029 | -0.029 | 0 | 0.07 | 0.01 | 0.1 | 0.09 | 0.1 | 0.11 |
| 3 | 3663278 | rs13084784 | 0.429 | 0.179 | 1.132 | 0.313 | -0.043 | -0.052 | -0.052 | 0 | 0.33 | 0.09 | 0.28 | 0.39 | 0.35 | 0.3 |
| 3 | 3667994 | rs4685669 | 0.167 | 0.174 | 1.336 | 0.308 | -0.007 | -0.001 | -0.001 | 0.005 | 0.35 | 0.31 | 0.32 | 0.39 | 0.38 | 0.29 |
| 3 | 3669247 | rs9813139 | -0.418 | 0.214 | 1.332 | 0.27 | 0.025 | 0.013 | 0.013 | 0 | 0.13 | 0.16 | 0.15 | 0.12 | 0.13 | 0.17 |
| 3 | 3669386 | rs7616995 | 1.584 | 0.16 | 1.174 | 0.191 | -0.014 | 0.027 | 0.027 | 0.205 | 0.37 | 0.21 | 0.38 | 0.26 | 0.43 | 0.38 |
| 3 | 3671084 | rs11713638 | 0.922 | 0.331 | 0.895 | 0.413 | -0.001 | -0.004 | -0.004 | 0 | 0.48 | 0.52 | 0.44 | 0.27 | 0.48 | 0.43 |
| 3 | 3671849 | rs901859 | 0.472 | 0.545 | 0.34 | 0.748 | 0.015 | 0.019 | 0.019 | 0 | 0.88 | 0.8 | 0.86 | 0.85 | 0.86 | 0.83 |
| 3 | 3673608 | rs2200706 | 0.833 | 0.525 | 0.167 | 0.713 | -0.02 | -0.037 | -0.037 | 0 | 0.07 | 0 | 0.1 | 0.15 | 0.1 | 0.11 |
| 3 | 3673983 | rs6774407 | 1.412 | 0.406 | 0.428 | 0.617 | 0.007 | 0.053 | 0.053 | 0.102 | 0.87 | 0.7 | 0.83 | 0.8 | 0.93 | 0.84 |
| 3 | 3681179 | rs4685672 | -0.336 | 0.468 | -0.09 | 0.369 | -0.005 | 0.005 | 0.005 | 0 | 0.42 | 0.37 | 0.37 | 0.42 | 0.41 | 0.31 |
| 3 | 3685016 | rs12493103 | 0.572 | 0.332 | -0.414 | 0.633 | -0.018 | -0.043 | -0.043 | 0.114 | 0.08 | 0.01 | 0.09 | 0.24 | 0.1 | 0.13 |
| 3 | 3685593 | rs12491030 | -0.045 | 0.17 | -0.262 | 0.442 | -0.007 | 0.027 | 0.027 | 0 | 0.39 | 0.26 | 0.31 | 0.32 | 0.4 | 0.3 |
| 3 | 3686414 | rs13075893 | 1.392 | -0.407 | -0.415 | 0.164 | -0.044 | -0.048 | -0.048 | 2.328 | 0.35 | 0.18 | 0.5 | 0.39 | 0.44 | 0.43 |
| 3 | 3687827 | rs1873026 | -0.403 | -0.298 | -0.438 | 0.078 | -0.004 | 0.013 | 0.013 | 0 | 0.62 | 0.71 | 0.69 | 0.68 | 0.61 | 0.71 |
| 3 | 3689371 | rs7609583 | 1.21 | -0.105 | -0.696 | 0.262 | -0.021 | -0.021 | -0.021 | 0.121 | 0.08 | 0 | 0.09 | 0.11 | 0.1 | 0.1 |
| 3 | 3690193 | rs17034929 | -0.312 | -0.314 | -0.361 | 0.066 | -0.013 | -0.044 | -0.044 | 0.35 | 0.12 | 0.2 | 0.06 | 0.05 | 0.05 | 0.12 |
| 3 | 3690221 | rs17293711 | 0.549 | -0.356 | -0.77 | 0.191 | -0.021 | -0.059 | -0.059 | 0 | 0.1 | 0 | 0.11 | 0.25 | 0.12 | 0.14 |
| 3 | 3690348 | rs2874754 | 0.964 | -0.794 | -1.062 | -0.194 | -0.013 | -0.023 | -0.023 | 0.383 | 0.47 | 0.25 | 0.58 | 0.43 | 0.54 | 0.46 |
| 3 | 3692598 | rs1873023 | -0.511 | -0.266 | -0.826 | -0.207 | -0.011 | -0.051 | -0.051 | 0 | 0.15 | 0.21 | 0.08 | 0.04 | 0.1 | 0.12 |
| 3 | 3696902 | rs4685676 | 0.106 | 0.282 | -1.113 | 0.115 | -0.008 | 0.03 | 0.03 | 0 | 0.4 | 0.28 | 0.34 | 0.29 | 0.42 | 0.38 |
| 3 | 3699454 | rs6801861 | 0.672 | 0.74 | -1.383 | 0.587 | -0.055 | -0.093 | -0.093 | 0 | 0.31 | 0.16 | 0.49 | 0.48 | 0.4 | 0.43 |
| 3 | 3700675 | rs6781946 | -0.679 | 1.002 | -0.959 | 0.777 | -0.009 | -0.007 | -0.007 | 5.616 | 0.43 | 0.63 | 0.34 | 0.6 | 0.38 | 0.49 |
| 3 | 3702536 | rs17035163 | -0.261 | 0.213 | -0.618 | 1.079 | 0.017 | -0.003 | -0.003 | 0.18 | 0.16 | 0.1 | 0.07 | 0.16 | 0.09 | 0.19 |
| 3 | 3702601 | rs9872637 | 0.266 | 0.71 | -0.025 | 1.201 | -0.006 | -0.01 | -0.01 | 0 | 0.24 | 0.16 | 0.25 | 0.35 | 0.27 | 0.25 |
| 3 | 3706957 | rs7643899 | -0.758 | 0.547 | -0.212 | 1.576 | -0.021 | -0.054 | -0.054 | 0 | 0.23 | 0.1 | 0.31 | 0.39 | 0.24 | 0.33 |

| | | | | | | | | | | | | | | | | |
|---|---|---|---|---|---|---|---|---|---|---|---|---|---|---|---|---|
| 3 | 3708002 | rs983154 | -1.405 | 0.39 | 0.321 | 1.426 | 0 | 0.037 | 0.037 | 0 | 0.5 | 0.59 | 0.57 | 0.72 | 0.46 | 0.56 |
| 3 | 3709134 | rs6791862 | 0.515 | 0.479 | -0.004 | 1.502 | -0.001 | -0.002 | -0.002 | 0.018 | 0.23 | 0.19 | 0.23 | 0.35 | 0.22 | 0.25 |
| 3 | 3709508 | rs6442820 | -0.071 | 0.616 | -0.676 | 1.522 | 0.004 | -0.004 | -0.004 | 0 | 0.3 | 0.73 | 0.29 | 0.36 | 0.22 | 0.25 |
| 3 | 3712668 | rs6799467 | -0.812 | 0.608 | -0.933 | 0.735 | -0.004 | 0.087 | 0.087 | 0.081 | 0.5 | 0.85 | 0.59 | 0.69 | 0.43 | 0.52 |
| 3 | 3723840 | rs711570 | -0.858 | 0.259 | -0.046 | 1.17 | -0.023 | 0.03 | 0.03 | 0 | 0.21 | 0.35 | 0.32 | 0.22 | 0.15 | 0.2 |
| 3 | 3726156 | rs9809849 | 0.67 | 0.478 | -0.955 | 0.715 | -0.05 | 0.083 | 0.083 | 0 | 0.39 | 0.12 | 0.38 | 0.27 | 0.41 | 0.24 |
| 3 | 3726784 | rs1485273 | 0.652 | 0.493 | -0.859 | 0.695 | -0.033 | 0.077 | 0.077 | 0 | 0.39 | 0.15 | 0.38 | 0.27 | 0.41 | 0.24 |
| 3 | 3727589 | rs1485272 | 0.137 | 0.264 | -0.287 | 1.444 | -0.005 | -0.001 | -0.001 | 2.329 | 0.35 | 0.27 | 0.29 | 0.38 | 0.38 | 0.36 |
| 3 | 3728697 | rs17035515 | 0.405 | 0.628 | -0.374 | 1.716 | -0.001 | -0.003 | -0.003 | 0 | 0.29 | 0.26 | 0.15 | 0.21 | 0.28 | 0.32 |
| 3 | 3729229 | rs6801538 | -0.408 | 0.658 | -0.135 | 1.783 | 0.009 | 0.044 | 0.044 | 0.306 | 0.06 | 0.12 | 0.15 | 0.17 | 0.1 | 0.22 |
| 3 | 3731439 | rs711565 | 0.233 | 0.58 | -0.392 | 1.716 | -0.001 | -0.06 | -0.06 | 0 | 0.12 | 0.03 | 0.14 | 0.17 | 0.12 | 0.18 |
| 3 | 3735454 | rs1921019 | 0.468 | 0.804 | 0.089 | 1.265 | -0.005 | -0.008 | -0.008 | 0 | 0.27 | 0.27 | 0.15 | 0.21 | 0.26 | 0.31 |
| 3 | 3736295 | rs1312821 | 0.819 | 0.59 | -0.109 | 1.219 | -0.04 | -0.004 | -0.004 | 2.501 | 0.54 | 0.22 | 0.51 | 0.44 | 0.51 | 0.42 |
| 3 | 3740417 | rs10510174 | 0.07 | 0.362 | 0.052 | 0.664 | -0.004 | 0.02 | 0.02 | 0 | 0.29 | 0.15 | 0.15 | 0.22 | 0.28 | 0.32 |
| 3 | 3740692 | rs13081238 | 0.176 | 0.489 | 0.143 | 0.733 | 0 | -0.009 | -0.009 | 1.678 | 0.28 | 0.25 | 0.14 | 0.21 | 0.26 | 0.31 |
| 3 | 3742077 | rs6796414 | -1.429 | 0.454 | 0.143 | 0.9 | -0.002 | 0.017 | 0.017 | 0.735 | 0.08 | 0.17 | 0.17 | 0.12 | 0.07 | 0.23 |
| 3 | 3743714 | rs769773 | -0.394 | 0.75 | -0.846 | 0.821 | -0.039 | -0.038 | -0.038 | 0 | 0.1 | 0.01 | 0.08 | 0.12 | 0.13 | 0.21 |
| 3 | 3745964 | rs7611251 | 0.097 | 0.623 | -0.803 | 0.511 | -0.001 | -0.003 | -0.003 | 0.031 | 0.44 | 0.44 | 0.43 | 0.23 | 0.46 | 0.4 |
| 3 | 3749521 | rs1596153 | 0.784 | 0.552 | -0.985 | 0.821 | 0.007 | 0.061 | 0.061 | 0 | 0.76 | 0.45 | 0.76 | 0.6 | 0.76 | 0.58 |
| 3 | 3755608 | rs769787 | 1.094 | -0.221 | -1.412 | 0.639 | 0.014 | -0.021 | -0.021 | 0 | 0.65 | 0.31 | 0.71 | 0.51 | 0.73 | 0.57 |
| 3 | 3755909 | rs17035891 | 1.188 | -0.102 | -1.424 | 0.778 | -0.037 | -0.02 | -0.02 | 0 | 0.06 | 0.01 | 0.12 | 0.27 | 0.13 | 0.05 |
| 3 | 3757926 | rs769782 | 1.156 | -0.32 | -1.555 | 0.59 | 0.003 | 0.028 | 0.028 | 0 | 0.33 | 0.12 | 0.31 | 0.26 | 0.33 | 0.25 |
| 3 | 3759119 | rs769781 | 0.986 | -0.332 | -1.531 | 0.614 | 0.012 | 0.027 | 0.027 | 0 | 0.35 | 0.19 | 0.32 | 0.27 | 0.33 | 0.25 |
| 3 | 3761042 | rs4638948 | 0.778 | -1.129 | -1.615 | 0.507 | -0.099 | -0.028 | -0.028 | 0.093 | 0.38 | 0.11 | 0.45 | 0.35 | 0.45 | 0.6 |
| 3 | 3767186 | rs7643736 | -1.348 | -1.164 | -1.306 | 0.6 | -0.004 | 0.008 | 0.008 | 0 | 0.35 | 0.61 | 0.29 | 0.49 | 0.31 | 0.43 |
| 3 | 3769493 | rs733765 | 0.668 | -1.304 | -1.225 | 0.498 | -0.028 | -0.019 | -0.019 | 0.403 | 0.77 | 0.37 | 0.87 | 0.86 | 0.85 | 0.62 |
| 3 | 3769568 | rs12631298 | -0.282 | -1.298 | -1.137 | 0.531 | -0.001 | 0.002 | 0.002 | 0 | 0.1 | 0.11 | 0.05 | 0.14 | 0.09 | 0.3 |
| 3 | 3771470 | rs17361688 | -0.829 | -1.337 | -1.154 | 0.538 | 0.001 | -0.002 | -0.002 | 0 | 0.16 | 0.17 | 0.09 | 0.21 | 0.11 | 0.3 |
| 3 | 3772538 | rs769801 | 1.383 | -1.346 | -1.281 | 0.527 | 0.018 | 0.015 | 0.015 | 0 | 0.3 | 0.2 | 0.28 | 0.25 | 0.26 | 0.23 |
| 3 | 3773725 | rs769799 | -1.547 | -1.347 | -1.319 | 0.532 | -0.051 | 0.026 | 0.026 | 0 | 0.66 | 0.8 | 0.57 | 0.76 | 0.58 | 0.66 |
| 3 | 3775107 | rs769798 | -0.873 | -1.358 | -1.309 | 0.461 | 0.005 | -0.01 | -0.01 | 0.056 | 0.59 | 0.68 | 0.44 | 0.45 | 0.45 | 0.61 |
| 3 | 3775161 | rs6442828 | -1.214 | -1.406 | -1.275 | 0.602 | -0.044 | -0.023 | -0.023 | 0 | 0.27 | 0.59 | 0.15 | 0.19 | 0.16 | 0.37 |
| 3 | 3776465 | rs6783172 | 0.807 | -1.377 | -1.234 | 0.562 | -0.024 | -0.09 | -0.09 | 0.089 | 0.13 | 0.42 | 0.07 | 0 | 0.07 | 0.08 |
| 3 | 3776669 | rs769795 | 1.75 | -1.383 | -1.262 | 0.588 | 0.011 | -0.018 | -0.018 | 3.691 | 0.66 | 0.3 | 0.72 | 0.52 | 0.72 | 0.6 |
| 3 | 3777132 | rs769792 | -0.534 | -1.373 | -1.258 | 0.578 | 0.006 | 0.043 | 0.043 | 0 | 0.52 | 0.39 | 0.5 | 0.73 | 0.5 | 0.56 |
| 3 | 3780218 | rs17036173 | -0.935 | -1.229 | 0.03 | 1.125 | -0.003 | -0.001 | -0.001 | 0 | 0.11 | 0.13 | 0.08 | 0.14 | 0.06 | 0.24 |
| 3 | 3780926 | rs769621 | 0.84 | -1.094 | 0.152 | 1.26 | -0.004 | 0.026 | 0.026 | 0 | 0.79 | 0.36 | 0.79 | 0.71 | 0.81 | 0.88 |
| 3 | 3781719 | rs6805805 | -0.11 | -1.113 | 0.146 | 1.194 | -0.041 | -0.1 | -0.1 | 0 | 0.14 | 0.51 | 0.08 | 0 | 0.06 | 0.08 |
| 3 | 3781877 | rs711562 | 1.248 | -1.053 | -0.271 | 1.257 | -0.002 | 0.009 | 0.009 | 0 | 0.75 | 0.3 | 0.76 | 0.61 | 0.77 | 0.64 |
| 3 | 3786517 | rs17362851 | 0.37 | -0.951 | -0.25 | 1.276 | -0.001 | 0 | 0 | 0 | 0.09 | 0.07 | 0.05 | 0.09 | 0.05 | 0.04 |
| 3 | 3788916 | rs769618 | 1.051 | -0.932 | -0.442 | 1.148 | 0.003 | -0.011 | -0.011 | 2.133 | 0.64 | 0.26 | 0.71 | 0.59 | 0.67 | 0.58 |
| 3 | 3792945 | rs1073657 | -1.211 | -0.939 | -0.038 | 1.57 | -0.011 | -0.028 | -0.028 | 0 | 0.14 | 0.07 | 0.16 | 0.38 | 0.18 | 0.09 |
| 3 | 3796045 | rs711544 | 0.939 | -0.759 | 0.326 | 1.763 | 0.005 | 0.054 | 0.054 | 0 | 0.81 | 0.43 | 0.8 | 0.7 | 0.82 | 0.88 |
| 3 | 3802465 | rs711540 | 1.045 | -1.754 | 0.061 | 1.701 | 0.025 | 0.088 | 0.088 | 0 | 0.51 | 0.23 | 0.5 | 0.27 | 0.58 | 0.4 |
| 3 | 3802980 | rs2119809 | 0.568 | -1.852 | -0.031 | 2.337 | -0.002 | 0.003 | 0.003 | 0 | 0.05 | 0 | 0.14 | 0.32 | 0.02 | 0.02 |
| 3 | 3804410 | rs809954 | 0.434 | -2.019 | 0.701 | 2.307 | -0.001 | 0.001 | 0.001 | 0 | 0.64 | 0.43 | 0.69 | 0.59 | 0.63 | 0.48 |
| 3 | 3804555 | rs811272 | 1.307 | -2.207 | 1.202 | 2.336 | -0.063 | 0.174 | 0.174 | 0 | 0.82 | 0.57 | 0.75 | 0.57 | 0.92 | 0.73 |
| 3 | 3805331 | rs3931232 | 0.725 | -2.192 | 1.403 | 2.398 | -0.04 | 0.107 | 0.107 | 0 | 0.87 | 0.64 | 0.83 | 0.78 | 0.94 | 0.74 |
| 3 | 3807031 | rs1065344 | -0.421 | -2.08 | 1.155 | 2.27 | -0.006 | -0.014 | -0.014 | 0 | 0.29 | 0.28 | 0.29 | 0.52 | 0.34 | 0.27 |
| 3 | 3812310 | rs11708888 | -0.092 | -1.84 | 1.36 | 2.127 | -0.002 | 0 | 0 | 0 | 0.3 | 0.29 | 0.26 | 0.3 | 0.35 | 0.31 |
| 3 | 3820517 | rs6442831 | -0.389 | -1.563 | 1.146 | 1.586 | -0.002 | -0.014 | -0.014 | 0 | 0.46 | 0.52 | 0.48 | 0.76 | 0.42 | 0.61 |
| 3 | 3822559 | rs769630 | 0.808 | -1.396 | -0.057 | 1.662 | 0.005 | 0.108 | 0.108 | 0 | 0.47 | 0.22 | 0.43 | 0.24 | 0.57 | 0.38 |
| 3 | 3830641 | rs12636288 | -1.312 | -1.253 | 1.081 | 2.476 | -0.042 | 0.188 | 0.188 | 1.216 | 0.09 | 0.34 | 0.17 | 0.29 | 0.02 | 0.05 |
| 3 | 3840471 | rs711582 | 1.318 | -0.924 | 1.197 | 2.465 | 0.015 | 0.26 | 0.26 | 0 | 0.89 | 0.57 | 0.76 | 0.67 | 0.93 | 0.68 |
| 3 | 3843436 | rs17037086 | -1.53 | -0.908 | 0.846 | 2.55 | -0.002 | 0.096 | 0.096 | 0 | 0.09 | 0.16 | 0.21 | 0.36 | 0.08 | 0.33 |
| 3 | 3845835 | rs998655 | 0.844 | -1.854 | 0.688 | 1.854 | 0.038 | 0.043 | 0.043 | 0 | 0.51 | 0.34 | 0.59 | 0.23 | 0.73 | 0.36 |
| 3 | 3845970 | rs10510276 | -0.857 | -2.348 | 1.069 | 2.432 | -0.043 | 0.133 | 0.133 | 0.021 | 0.19 | 0.47 | 0.12 | 0.39 | 0.1 | 0.49 |
| 3 | 3846204 | rs10510277 | -0.068 | -2.626 | 0.943 | 2.444 | 0.019 | 0.003 | 0.003 | 0 | 0.22 | 0.08 | 0.17 | 0.22 | 0.12 | 0.15 |
| 3 | 3846805 | rs1488454 | 0.376 | -3.182 | 0.545 | 2.026 | 0.033 | 0.036 | 0.036 | 0.091 | 0.72 | 0.86 | 0.79 | 0.78 | 0.83 | 0.85 |
| 3 | 3847031 | rs10780025 | -1.299 | -3.243 | 0.732 | 2.024 | -0.04 | 0.046 | 0.046 | 0.074 | 0.31 | 0.6 | 0.26 | 0.55 | 0.17 | 0.51 |
| 3 | 3850233 | rs1561801 | 1.44 | -3.176 | 0.661 | 2.017 | 0.016 | 0.014 | 0.014 | 0 | 0.57 | 0.34 | 0.63 | 0.24 | 0.78 | 0.37 |
| 3 | 3850309 | rs1976077 | -0.261 | -3.167 | 0.199 | 1.937 | 0.026 | -0.041 | -0.041 | 0.231 | 0.22 | 0.13 | 0.24 | 0.37 | 0.11 | 0.18 |
| 3 | 3852890 | rs711580 | 0.773 | -3.182 | 0.922 | 1.822 | -0.046 | 0.075 | 0.075 | 0.08 | 0.77 | 0.46 | 0.88 | 0.55 | 0.89 | 0.62 |
| 3 | 3858932 | rs711578 | 1.919 | -3.232 | 0.361 | 1.389 | -0.029 | 0.066 | 0.066 | 0 | 0.39 | 0.13 | 0.45 | 0.2 | 0.64 | 0.33 |
| 3 | 3860491 | rs711576 | 0.27 | -3.045 | 1.384 | 1.274 | -0.016 | -0.098 | -0.098 | 0.074 | 0.8 | 0.65 | 0.85 | 0.95 | 0.87 | 0.9 |
| 3 | 3860968 | rs9834164 | 0.289 | -3.056 | 1.566 | 1.282 | 0.024 | -0.053 | -0.053 | 0 | 0.87 | 0.6 | 0.88 | 0.95 | 0.92 | 0.91 |
| 3 | 3863319 | rs696487 | 0.156 | -3.039 | 1.696 | 1.284 | 0.012 | -0.049 | -0.049 | 0.022 | 0.88 | 0.7 | 0.89 | 0.95 | 0.93 | 0.91 |
| 3 | 3864892 | rs17037382 | 0.28 | -3.202 | 1.966 | 1.364 | 0.01 | -0.003 | -0.003 | 0 | 0.12 | 0.27 | 0.15 | 0.14 | 0.1 | 0.19 |
| 3 | 3866316 | rs1825912 | 1.453 | -3.277 | 1.834 | 1.573 | -0.006 | -0.003 | -0.003 | 0 | 0.65 | 0.39 | 0.6 | 0.63 | 0.77 | 0.69 |

| | | | | | | | | | | | | | | | | |
|---|---|---|---|---|---|---|---|---|---|---|---|---|---|---|---|---|
| 3 | 3878883 | rs13083087 | 1.493 | -3.298 | 1.836 | 1.648 | -0.005 | -0.004 | -0.004 | 0 | 0.64 | 0.39 | 0.6 | 0.65 | 0.76 | 0.7 |
| 3 | 3880784 | rs7610133 | -1.361 | -3.743 | 1.8 | 1.692 | -0.005 | 0.001 | 0.001 | 0.659 | 0.36 | 0.63 | 0.41 | 0.62 | 0.25 | 0.51 |
| 3 | 3882283 | rs6442836 | 0.925 | -3.633 | 1.856 | 1.679 | 0.007 | 0.142 | 0.142 | 0 | 0.95 | 0.78 | 0.98 | 0.87 | 1 | 0.95 |
| 3 | 3885609 | rs17037518 | -0.818 | -3.802 | 1.857 | 1.443 | -0.033 | 0.004 | 0.004 | 0.019 | 0.15 | 0.26 | 0.2 | 0.19 | 0.08 | 0.12 |
| 3 | 3886919 | rs3749348 | 0.225 | -3.772 | 2.376 | 1.553 | 0.003 | 0.066 | 0.066 | 0.017 | 0.13 | 0.29 | 0.15 | 0.26 | 0.1 | 0.21 |
| 3 | 3887876 | rs2120609 | 1.47 | -3.789 | 2.43 | 1.558 | -0.006 | -0.005 | -0.005 | 0 | 0.57 | 0.43 | 0.57 | 0.25 | 0.73 | 0.45 |
| 3 | 3888894 | rs1872418 | -0.241 | -3.611 | 2.218 | 1.559 | -0.041 | 0.021 | 0.021 | 0 | 0.09 | 0.15 | 0.16 | 0.19 | 0.01 | 0.06 |
| 3 | 3891187 | rs4685690 | -0.957 | -3.55 | 2.205 | 1.567 | -0.04 | 0.092 | 0.092 | 0 | 0.21 | 0.52 | 0.31 | 0.45 | 0.11 | 0.26 |
| 3 | 3892716 | rs1994274 | 0.211 | -3.319 | 1.89 | 1.509 | -0.002 | 0.016 | 0.016 | 0 | 0.87 | 0.9 | 0.9 | 0.95 | 0.9 | 0.9 |
| 3 | 3892727 | rs4685692 | 0.281 | -3.189 | 1.862 | 1.562 | 0.005 | 0.049 | 0.049 | 1.268 | 0.13 | 0.29 | 0.14 | 0.26 | 0.1 | 0.21 |
| 3 | 3899791 | rs6776332 | 1.656 | -3.232 | 1.53 | 1.289 | 0.118 | 0.142 | 0.142 | 0.605 | 0.48 | 0.16 | 0.5 | 0.19 | 0.71 | 0.4 |
| 3 | 3900135 | rs13075986 | -1.409 | -3.116 | 1.881 | 1.391 | -0.001 | -0.001 | -0.001 | 0 | 0.5 | 0.55 | 0.4 | 0.66 | 0.36 | 0.64 |
| 3 | 3915789 | rs2596890 | 1.733 | -2.925 | 2.443 | 1.369 | -0.051 | 0.064 | 0.064 | 0.942 | 0.89 | 0.38 | 0.85 | 0.83 | 0.99 | 0.94 |
| 3 | 3915943 | rs2633820 | 0.833 | -2.912 | 2.553 | 1.375 | -0.034 | 0.068 | 0.068 | 0 | 0.91 | 0.75 | 0.89 | 0.83 | 0.99 | 0.94 |
| 3 | 3916426 | rs2633822 | 1.228 | -2.912 | 2.664 | 1.412 | -0.026 | 0.134 | 0.134 | 0.315 | 0.9 | 0.6 | 0.88 | 0.82 | 0.99 | 0.94 |
| 3 | 3920844 | rs1488445 | 0.731 | -3.021 | 2.33 | 1.158 | -0.038 | 0.01 | 0.01 | 0 | 0.89 | 0.59 | 0.96 | 0.96 | 0.97 | 0.93 |
| 3 | 3941431 | rs923092 | 0.479 | -3.069 | 1.752 | 1.451 | -0.002 | -0.007 | -0.007 | 0 | 0.13 | 0.13 | 0.16 | 0.25 | 0.11 | 0.2 |
| 3 | 3945544 | rs4685707 | -0.677 | -2.594 | 1.481 | 1.381 | 0.013 | -0.075 | -0.075 | 1.02 | 0.38 | 0.16 | 0.36 | 0.54 | 0.33 | 0.41 |
| 3 | 3948722 | rs9850979 | 0.227 | -2.308 | 1.568 | 1.49 | 0.001 | 0.019 | 0.019 | 0.204 | 0.14 | 0.2 | 0.16 | 0.28 | 0.11 | 0.22 |
| 3 | 3958764 | rs6785370 | 0.406 | -2.299 | 1.388 | 1.479 | 0 | 0.005 | 0.005 | 0.441 | 0.13 | 0.16 | 0.16 | 0.26 | 0.11 | 0.2 |
| 3 | 3960052 | rs4349487 | 0.764 | -2.393 | 1.527 | 1.488 | -0.03 | 0.079 | 0.079 | 0.625 | 0.76 | 0.48 | 0.72 | 0.67 | 0.87 | 0.72 |
| 3 | 3965271 | rs11716741 | 0.002 | -2.533 | 1.023 | 1.557 | 0.001 | -0.002 | -0.002 | 0 | 0.21 | 0.18 | 0.15 | 0.47 | 0.2 | 0.29 |
| 3 | 3967467 | rs1385464 | 1.71 | -2.818 | 1.209 | 1.212 | 0.013 | 0.127 | 0.127 | 0 | 0.51 | 0.27 | 0.45 | 0.19 | 0.58 | 0.36 |
| 3 | 3968208 | rs10865884 | 2.079 | -2.997 | 1.171 | 1.211 | 0.026 | 0.134 | 0.134 | 0 | 0.45 | 0.22 | 0.44 | 0.16 | 0.62 | 0.33 |
| 3 | 3968439 | rs10780027 | 1.326 | -2.985 | 1.201 | 1.225 | 0.028 | 0.068 | 0.068 | 0.002 | 0.58 | 0.27 | 0.47 | 0.19 | 0.66 | 0.42 |
| 3 | 3971502 | rs1825503 | 1.318 | -3.007 | 1.157 | 1.18 | 0.03 | 0.082 | 0.082 | 0 | 0.58 | 0.27 | 0.47 | 0.16 | 0.66 | 0.42 |
| 3 | 3972118 | rs9866915 | 0.116 | -2.971 | 1.099 | 1.417 | 0.002 | -0.008 | -0.008 | 0 | 0.21 | 0.16 | 0.15 | 0.42 | 0.2 | 0.3 |
| 3 | 4004427 | rs17749633 | 1.817 | -2.673 | 0.932 | 1.281 | 0.026 | 0.184 | 0.184 | 0 | 0.45 | 0.2 | 0.45 | 0.13 | 0.62 | 0.33 |
| 3 | 4011307 | rs4685723 | 1.655 | -2.754 | 0.868 | 1.267 | 0.069 | 0.181 | 0.181 | 0 | 0.48 | 0.22 | 0.45 | 0.15 | 0.65 | 0.37 |
| 3 | 4014660 | rs9845164 | -0.973 | -2.79 | 1.295 | 1.397 | 0.011 | -0.014 | -0.014 | 0 | 0.4 | 0.37 | 0.45 | 0.45 | 0.29 | 0.37 |
| 3 | 4015109 | rs317572 | 2.04 | -2.922 | 1.329 | 1.378 | 0.032 | 0.149 | 0.149 | 0 | 0.44 | 0.22 | 0.46 | 0.14 | 0.63 | 0.32 |
| 3 | 4015439 | rs9850111 | -0.008 | -2.382 | 1.494 | 1.795 | -0.018 | 0.08 | 0.08 | 0.01 | 0.19 | 0.36 | 0.34 | 0.39 | 0.12 | 0.26 |
| 3 | 4015564 | rs317570 | -0.091 | -2.354 | 1.368 | 1.77 | -0.026 | 0.115 | 0.115 | 0.02 | 0.8 | 0.48 | 0.65 | 0.61 | 0.87 | 0.74 |
| 3 | 4020725 | rs2279751 | -0.624 | -2.306 | 0.701 | 1.856 | 0.011 | -0.038 | -0.038 | 0 | 0.13 | 0.07 | 0.08 | 0.42 | 0.11 | 0.31 |
| 3 | 4023232 | rs17039519 | -0.121 | -2.294 | 0.675 | 2.044 | 0.011 | -0.044 | -0.044 | 0.308 | 0.11 | 0.03 | 0.07 | 0.37 | 0.09 | 0.29 |
| 3 | 4025152 | rs17039568 | -0.361 | -2.43 | 0.638 | 1.689 | -0.018 | 0.048 | 0.048 | 0.03 | 0.2 | 0.32 | 0.35 | 0.41 | 0.13 | 0.28 |
| 3 | 4025464 | rs17039576 | 0.148 | -0.234 | -0.174 | 1.08 | -0.026 | -0.032 | -0.032 | 0.002 | 0.2 | 0.12 | 0.27 | 0.4 | 0.29 | 0.34 |
| 3 | 4026680 | rs1074650 | 0.115 | -0.218 | 0.192 | 0.856 | -0.005 | 0.041 | 0.041 | 0.002 | 0.17 | 0.23 | 0.45 | 0.39 | 0.15 | 0.21 |
| 3 | 4031736 | rs1110798 | -0.84 | -0.301 | 0.15 | 0.704 | -0.005 | 0.034 | 0.034 | 0 | 0.25 | 0.34 | 0.42 | 0.41 | 0.2 | 0.29 |
| 3 | 4035443 | rs4685724 | 0.057 | -0.29 | 0.038 | 0.761 | -0.004 | 0.056 | 0.056 | 0 | 0.17 | 0.25 | 0.42 | 0.41 | 0.13 | 0.25 |
| 3 | 4041690 | rs317544 | 0.745 | -0.435 | -0.105 | 0.117 | 0.017 | -0.027 | -0.027 | 0 | 0.67 | 0.26 | 0.46 | 0.51 | 0.75 | 0.71 |
| 3 | 4045707 | rs7615618 | -0.581 | -0.52 | 0.022 | 0.126 | -0.016 | -0.047 | -0.047 | 0 | 0.85 | 0.62 | 0.89 | 0.98 | 0.89 | 0.94 |
| 3 | 4046691 | rs317588 | -1.052 | -0.496 | -0.255 | 0.097 | 0.004 | -0.037 | -0.037 | 0 | 0.34 | 0.71 | 0.53 | 0.48 | 0.29 | 0.34 |
| 3 | 4050668 | rs317592 | -0.366 | -0.494 | -0.157 | -0.041 | -0.005 | -0.015 | -0.015 | 0 | 0.86 | 0.72 | 0.9 | 0.96 | 0.89 | 0.93 |
| 3 | 4052274 | rs317593 | 1.288 | -0.665 | -0.181 | -0.193 | -0.009 | -0.034 | -0.034 | 1.393 | 0.7 | 0.43 | 0.82 | 0.93 | 0.75 | 0.79 |
| 3 | 4061316 | rs317614 | -1.215 | -0.622 | -0.097 | -0.218 | 0.018 | -0.05 | -0.05 | 2.855 | 0.91 | 0.77 | 0.92 | 1 | 0.91 | 0.99 |
| 3 | 4061814 | rs317615 | 1.298 | -0.662 | -0.106 | -0.225 | -0.01 | -0.047 | -0.047 | 0 | 0.71 | 0.43 | 0.83 | 0.95 | 0.76 | 0.85 |
| 3 | 4062565 | rs11928296 | -0.775 | -0.67 | -0.148 | -0.229 | -0.007 | -0.059 | -0.059 | 0 | 0.16 | 0.29 | 0.07 | 0.03 | 0.12 | 0.13 |
| 3 | 4062753 | rs317616 | 2.115 | -0.341 | -0.944 | -0.516 | -0.127 | 0.073 | 0.073 | 0 | 0.31 | 0.05 | 0.24 | 0.13 | 0.41 | 0.25 |
| 3 | 4066869 | rs317623 | 0.372 | -0.953 | -0.29 | -0.629 | -0.031 | -0.053 | -0.053 | 0.028 | 0.79 | 0.51 | 0.87 | 0.94 | 0.84 | 0.87 |
| 3 | 4070251 | rs6806504 | -0.115 | -0.99 | -0.495 | -0.594 | 0.021 | -0.033 | -0.033 | 0 | 0.29 | 0.09 | 0.16 | 0.39 | 0.24 | 0.43 |
| 3 | 4070649 | rs6806903 | -0.915 | -0.873 | -0.386 | -0.618 | 0.059 | -0.009 | -0.009 | 0 | 0.42 | 0.2 | 0.2 | 0.44 | 0.31 | 0.54 |
| 3 | 4071151 | rs1092733 | -0.93 | -0.863 | -0.376 | -0.617 | 0.012 | 0 | 0 | 1.007 | 0.43 | 0.55 | 0.26 | 0.43 | 0.31 | 0.53 |
| 3 | 4074033 | rs317605 | 0.714 | -0.594 | -0.296 | -0.683 | -0.005 | -0.006 | -0.006 | 1.378 | 0.58 | 0.53 | 0.74 | 0.62 | 0.65 | 0.51 |
| 3 | 4074425 | rs10865894 | -0.506 | -0.94 | -0.847 | -0.581 | 0.032 | -0.001 | -0.001 | 0 | 0.35 | 0.15 | 0.17 | 0.4 | 0.27 | 0.48 |
| 3 | 4076977 | rs7624815 | 1.051 | -0.168 | -0.66 | -0.717 | -0.028 | -0.039 | -0.039 | 0.988 | 0.36 | 0.27 | 0.49 | 0.61 | 0.37 | 0.32 |
| 3 | 4078654 | rs1987888 | 0.64 | 0.077 | -0.274 | -1.098 | 0.042 | 0.075 | 0.075 | 0 | 0.52 | 0.35 | 0.7 | 0.73 | 0.64 | 0.5 |
| 3 | 4088576 | rs1087817 | -0.031 | 0.557 | 0 | -0.775 | 0.005 | -0.097 | -0.097 | 0 | 0.14 | 0.56 | 0.12 | 0 | 0.11 | 0.08 |
| 3 | 4094293 | rs317530 | 0.889 | 0.646 | 0.098 | -0.834 | 0.012 | -0.119 | -0.119 | 0.115 | 0.75 | 0.42 | 0.79 | 1 | 0.7 | 0.87 |
| 3 | 4096255 | rs317528 | -0.114 | 0.509 | -0.208 | -0.467 | 0.027 | -0.097 | -0.097 | 0.052 | 0.87 | 0.57 | 0.89 | 1 | 0.88 | 0.92 |
| 3 | 4103926 | rs795727 | 0.12 | 0.48 | -0.769 | -0.057 | -0.001 | 0.085 | 0.085 | 0.756 | 0.54 | 0.76 | 0.68 | 0.76 | 0.5 | 0.58 |
| 3 | 4107619 | rs795729 | -0.892 | 0.463 | -0.409 | 0.554 | 0 | -0.001 | -0.001 | 0 | 0.2 | 0.19 | 0.23 | 0.2 | 0.2 | 0.11 |
| 3 | 4108261 | rs666412 | -0.229 | 0.294 | -1.531 | 0.346 | -0.046 | -0.012 | -0.012 | 0 | 0.32 | 0.03 | 0.25 | 0.26 | 0.27 | 0.21 |
| 3 | 4109700 | rs795734 | -0.488 | -0.228 | -0.435 | 1.025 | -0.004 | 0.003 | 0.003 | 0 | 0.36 | 0.4 | 0.45 | 0.43 | 0.31 | 0.33 |
| 3 | 4110799 | rs711676 | 0.776 | -0.348 | -0.293 | 0.92 | -0.015 | 0.121 | 0.121 | 0 | 0.88 | 0.59 | 0.77 | 0.82 | 0.92 | 0.85 |
| 3 | 4122269 | rs2648481 | -0.228 | -0.204 | -0.594 | 0.98 | -0.063 | -0.009 | -0.009 | 9.373 | 0.3 | 0.03 | 0.25 | 0.26 | 0.27 | 0.22 |
| 3 | 4132271 | rs11918722 | -0.737 | -0.237 | 0.314 | 0.866 | 0.056 | 0.228 | 0.228 | 1.06 | 0.06 | 0.45 | 0.2 | 0.15 | 0.04 | 0.13 |
| 3 | 4135452 | rs663377 | 0.418 | 0.168 | 0.458 | 1.086 | 0.001 | 0.004 | 0.004 | 2.791 | 0.13 | 0.13 | 0.12 | 0.22 | 0.18 | 0.28 |
| 3 | 4145581 | rs17039906 | -0.228 | 0.037 | 0.434 | 0.889 | 0.033 | 0.169 | 0.169 | 0.046 | 0.06 | 0.31 | 0.19 | 0.14 | 0.04 | 0.14 |

| | | | | | | | | | | | | | | | | |
|---|---|---|---|---|---|---|---|---|---|---|---|---|---|---|---|---|
| 3 | 4148058 | rs2600151 | -0.089 | -1.311 | 0.421 | 0.499 | -0.001 | 0 | 0 | 0.176 | 0.75 | 0.76 | 0.84 | 0.68 | 0.77 | 0.59 |
| 3 | 4157503 | rs13084851 | 0.597 | -0.889 | 0.429 | 0.574 | -0.004 | 0.008 | 0.008 | 0.714 | 0.42 | 0.63 | 0.57 | 0.26 | 0.49 | 0.36 |
| 3 | 4158345 | rs17702174 | 1.369 | -0.947 | 0.36 | 0.554 | -0.04 | 0.085 | 0.085 | 0.332 | 0.34 | 0.18 | 0.36 | 0.12 | 0.41 | 0.22 |
| 3 | 4164486 | rs795306 | -0.214 | -1.063 | 0.579 | 0.871 | -0.017 | 0.026 | 0.026 | 0.331 | 0.79 | 0.67 | 0.86 | 0.67 | 0.83 | 0.59 |
| 3 | 4168641 | rs795301 | 0.674 | -1.099 | 0.363 | 0.759 | -0.048 | -0.087 | -0.087 | 0.158 | 0.33 | 0.06 | 0.27 | 0.4 | 0.3 | 0.23 |
| 3 | 4170874 | rs592767 | -0.034 | -0.989 | 0.499 | 0.795 | -0.014 | 0.024 | 0.024 | 0.046 | 0.78 | 0.68 | 0.84 | 0.66 | 0.79 | 0.6 |
| 3 | 4180069 | rs2587924 | -0.006 | -1.053 | 0.662 | 0.813 | -0.005 | 0.007 | 0.007 | 0 | 0.4 | 0.44 | 0.46 | 0.55 | 0.34 | 0.38 |
| 3 | 4185026 | rs7632212 | 1.113 | 0.118 | 0.74 | 0.56 | -0.008 | 0.049 | 0.049 | 0.018 | 0.37 | 0.28 | 0.4 | 0.11 | 0.47 | 0.21 |
| 3 | 4190452 | rs2686702 | 0.578 | 0.217 | 0.737 | 0.637 | -0.006 | -0.015 | -0.015 | 0 | 0.35 | 0.23 | 0.29 | 0.39 | 0.32 | 0.23 |
| 3 | 4193500 | rs2629276 | 0.176 | 0.248 | 0.75 | 0.683 | 0 | 0.01 | 0.01 | 0 | 0.75 | 0.67 | 0.83 | 0.68 | 0.75 | 0.59 |
| 3 | 4205623 | rs1351950 | 1.171 | 0.178 | 0.264 | 0.225 | 0.016 | 0.06 | 0.06 | 0 | 0.35 | 0.19 | 0.32 | 0.18 | 0.32 | 0.23 |
| 3 | 4212007 | rs1017903 | 2.045 | 0.184 | 0.243 | 0.186 | -0.036 | -0.084 | -0.084 | 3.202 | 0.23 | 0.05 | 0.23 | 0.39 | 0.24 | 0.21 |
| 3 | 4229796 | rs1160604 | 0.793 | 0.174 | 0.288 | 0.397 | -0.017 | 0.052 | 0.052 | 0 | 0.38 | 0.25 | 0.38 | 0.22 | 0.44 | 0.24 |
| 3 | 4230372 | rs2629243 | -0.748 | 0.034 | 0.425 | 0.559 | -0.009 | 0.04 | 0.04 | 0 | 0.84 | 0.69 | 0.89 | 0.77 | 0.87 | 0.66 |
| 3 | 4230956 | rs2248840 | 0.708 | 0.062 | 0.379 | 0.562 | -0.001 | 0.001 | 0.001 | 0 | 0.36 | 0.38 | 0.31 | 0.4 | 0.32 | 0.23 |
| 3 | 4245833 | rs7624988 | -0.934 | 0.021 | 0.247 | 0.55 | 0 | 0.001 | 0.001 | 0 | 0.65 | 0.65 | 0.7 | 0.6 | 0.68 | 0.76 |
| 3 | 4247185 | rs7630606 | 0.476 | -0.039 | 0.279 | 0.539 | -0.02 | 0.1 | 0.1 | 0 | 0.45 | 0.26 | 0.44 | 0.22 | 0.55 | 0.26 |
| 3 | 4250309 | rs1116129 | -0.614 | -0.122 | 0.306 | 0.54 | 0 | 0 | 0 | 0 | 0.61 | 0.63 | 0.66 | 0.61 | 0.59 | 0.74 |
| 3 | 4253219 | rs2587940 | 0.39 | -0.11 | 0.334 | 0.573 | -0.01 | 0.025 | 0.025 | 0.005 | 0.16 | 0.3 | 0.11 | 0.22 | 0.12 | 0.37 |
| 3 | 4260946 | rs313681 | -0.432 | -0.091 | 0.486 | 0.417 | -0.007 | 0 | 0 | 0 | 0.59 | 0.68 | 0.66 | 0.6 | 0.57 | 0.73 |
| 3 | 4261274 | rs313682 | -0.459 | -0.054 | 0.413 | 0.426 | -0.003 | -0.001 | -0.001 | 0 | 0.59 | 0.64 | 0.65 | 0.6 | 0.56 | 0.73 |
| 3 | 4266844 | rs313691 | -0.059 | -0.947 | 0.07 | -0.034 | -0.077 | 0.035 | 0.035 | 0 | 0.25 | 0.49 | 0.13 | 0.28 | 0.11 | 0.42 |
| 3 | 4272253 | rs1242799 | 0.122 | -0.954 | 0.215 | -0.046 | -0.034 | 0.017 | 0.017 | 0 | 0.23 | 0.34 | 0.12 | 0.28 | 0.11 | 0.43 |
| 3 | 4272845 | rs1242807 | -0.67 | -0.584 | 0.026 | -0.134 | -0.01 | -0.008 | -0.008 | 0.095 | 0.59 | 0.74 | 0.52 | 0.42 | 0.57 | 0.66 |
| 3 | 4273489 | rs1242810 | -0.987 | -0.629 | -0.06 | -0.08 | -0.126 | -0.106 | -0.106 | 0 | 0.63 | 0.97 | 0.7 | 0.57 | 0.61 | 0.75 |
| 3 | 4274646 | rs13065523 | 0.846 | -0.557 | 0.054 | -0.772 | -0.03 | 0.047 | 0.047 | 0 | 0.33 | 0.21 | 0.39 | 0.14 | 0.46 | 0.23 |
| 3 | 4275803 | rs1153572 | -0.814 | -0.562 | 0.12 | -0.77 | -0.025 | -0.021 | -0.021 | 0 | 0.59 | 0.81 | 0.52 | 0.42 | 0.57 | 0.66 |
| 3 | 4276680 | rs4279083 | 0.723 | -0.335 | 0.032 | -0.781 | -0.03 | 0.047 | 0.047 | 0 | 0.33 | 0.21 | 0.38 | 0.14 | 0.46 | 0.23 |
| 3 | 4282950 | rs1876613 | 0.598 | -0.331 | 0.017 | -0.773 | -0.005 | 0.001 | 0.001 | 0 | 0.34 | 0.31 | 0.39 | 0.14 | 0.46 | 0.23 |
| 3 | 4286700 | rs13063552 | 0.489 | -0.392 | -0.079 | -0.855 | -0.003 | -0.002 | -0.002 | 0.302 | 0.36 | 0.35 | 0.39 | 0.14 | 0.46 | 0.26 |
| 3 | 4304897 | rs3912462 | 0.645 | -0.282 | -0.124 | -0.779 | -0.014 | 0.019 | 0.019 | 0 | 0.34 | 0.27 | 0.39 | 0.14 | 0.46 | 0.23 |
| 3 | 4313485 | rs310694 | 0.235 | -0.621 | -0.002 | -0.743 | -0.005 | 0 | 0 | 0.17 | 0.24 | 0.24 | 0.11 | 0.28 | 0.11 | 0.43 |
| 3 | 4317445 | rs314416 | -0.113 | -0.403 | -0.099 | -1.08 | -0.004 | -0.039 | -0.039 | 0 | 0.53 | 0.48 | 0.56 | 0.87 | 0.46 | 0.71 |
| 3 | 4328131 | rs310731 | -1.617 | -0.312 | -0.537 | -1.113 | -0.031 | 0.045 | 0.045 | 0 | 0.9 | 0.97 | 0.95 | 1 | 0.89 | 0.96 |
| 3 | 4328411 | rs997828 | 1.762 | -0.435 | -0.654 | -1.194 | -0.074 | -0.133 | -0.133 | 0 | 0.27 | 0.04 | 0.42 | 0.58 | 0.33 | 0.31 |
| 3 | 4331654 | rs13068964 | 0.657 | -0.913 | -0.055 | -1.039 | 0.025 | -0.066 | -0.066 | 0 | 0.34 | 0.44 | 0.39 | 0.12 | 0.43 | 0.24 |
| 3 | 4333294 | rs13101121 | 0.713 | -0.925 | 0.049 | -1.062 | -0.001 | -0.013 | -0.013 | 0 | 0.34 | 0.34 | 0.38 | 0.12 | 0.43 | 0.24 |
| 3 | 4335805 | rs2322679 | 0.79 | -0.947 | -0.063 | -1.068 | -0.051 | 0.082 | 0.082 | 0 | 0.33 | 0.15 | 0.36 | 0.12 | 0.43 | 0.24 |
| 3 | 4336589 | rs310727 | -0.731 | -0.976 | -0.321 | -1.028 | -0.017 | 0 | 0 | 0 | 0.58 | 0.73 | 0.49 | 0.42 | 0.51 | 0.65 |
| 3 | 4337312 | rs182568 | 0.412 | -1.072 | -1.067 | -0.959 | -0.038 | 0.009 | 0.009 | 0.354 | 0.23 | 0.31 | 0.1 | 0.29 | 0.08 | 0.4 |
| 3 | 4340847 | rs310709 | -1.061 | -0.915 | -1.189 | -0.967 | -0.08 | -0.039 | -0.039 | 0.156 | 0.61 | 0.86 | 0.51 | 0.42 | 0.55 | 0.65 |
| 3 | 4342274 | rs310711 | -0.083 | -1.138 | -0.795 | -0.446 | -0.014 | 0.001 | 0.001 | 0.124 | 0.23 | 0.25 | 0.11 | 0.3 | 0.12 | 0.41 |
| 3 | 4345766 | rs13099918 | 0.033 | -0.258 | -1.131 | -0.646 | -0.08 | 0.097 | 0.097 | 0.823 | 0.24 | 0.07 | 0.3 | 0.08 | 0.39 | 0.13 |
| 3 | 4351751 | rs310724 | 0.329 | -1.209 | -0.589 | -0.65 | 0.01 | 0.005 | 0.005 | 0 | 0.34 | 0.67 | 0.21 | 0.28 | 0.2 | 0.49 |
| 3 | 4353466 | rs7632080 | 0.205 | -1.196 | -0.782 | -0.945 | 0.003 | 0.035 | 0.035 | 0 | 0.19 | 0.12 | 0.1 | 0.01 | 0.18 | 0.1 |
| 3 | 4354697 | rs6801634 | 1.798 | -1.491 | -0.687 | -0.979 | 0.033 | 0.036 | 0.036 | 0 | 0.21 | 0.1 | 0.12 | 0.1 | 0.16 | 0.14 |
| 3 | 4356985 | rs9836925 | 0.141 | -1.436 | -0.341 | -0.962 | 0.04 | 0.04 | 0.04 | 0.094 | 0.38 | 0.71 | 0.21 | 0.1 | 0.22 | 0.2 |
| 3 | 4361483 | rs9866572 | -0.087 | -1.203 | -0.477 | -0.801 | 0.082 | 0.139 | 0.139 | 0.067 | 0.46 | 0.77 | 0.29 | 0.11 | 0.36 | 0.24 |
| 3 | 4365560 | rs711673 | -0.918 | -1.097 | -0.945 | -0.696 | 0.024 | 0.034 | 0.034 | 0.09 | 0.56 | 0.96 | 0.33 | 0.3 | 0.39 | 0.59 |
| 3 | 4372513 | rs794205 | -1.401 | -0.667 | -0.496 | -0.632 | -0.04 | -0.021 | -0.021 | 4.365 | 0.73 | 0.76 | 0.51 | 0.38 | 0.64 | 0.71 |
| 3 | 4376030 | rs308739 | -0.349 | -1.249 | -0.203 | -0.372 | -0.016 | 0.017 | 0.017 | 0 | 0.11 | 0.17 | 0.06 | 0.19 | 0.05 | 0.34 |
| 3 | 4378142 | rs894013 | 0.702 | -1.168 | -0.128 | 0.03 | -0.001 | 0.001 | 0.001 | 0 | 0.45 | 0.58 | 0.58 | 0.47 | 0.5 | 0.37 |
| 3 | 4380416 | rs308726 | 0.552 | -1.064 | -0.052 | 0.126 | 0 | 0.009 | 0.009 | 0 | 0.07 | 0.08 | 0.05 | 0.2 | 0.05 | 0.3 |
| 3 | 4383915 | rs17030493 | 0.627 | -0.92 | -0.131 | -0.25 | -0.044 | -0.064 | -0.064 | 0 | 0.28 | 0.32 | 0.15 | 0.1 | 0.19 | 0.15 |
| 3 | 4384362 | rs809437 | -0.693 | -0.355 | -0.421 | -0.045 | -0.06 | -0.03 | -0.03 | 13.511 | 0.8 | 0.94 | 0.78 | 0.76 | 0.73 | 0.85 |
| 3 | 4392786 | rs11928114 | 0.653 | -1.021 | 0.208 | -0.295 | -0.03 | -0.074 | -0.074 | 0 | 0.24 | 0.38 | 0.15 | 0.08 | 0.19 | 0.15 |
| 3 | 4393415 | rs2582349 | -0.506 | -1.38 | 0.471 | -0.235 | -0.013 | -0.012 | -0.012 | 0 | 0.41 | 0.56 | 0.2 | 0.3 | 0.28 | 0.49 |
| 3 | 4393684 | rs1687858 | -1.3 | -1.46 | -0.23 | 0.101 | -0.047 | -0.133 | -0.133 | 0 | 0.58 | 0.95 | 0.35 | 0.54 | 0.42 | 0.64 |
| 3 | 4398276 | rs1688411 | 0.251 | -1.817 | 0.426 | 0.266 | -0.051 | -0.024 | -0.024 | 2.56 | 0.31 | 0.42 | 0.2 | 0.24 | 0.17 | 0.42 |
| 3 | 4398825 | rs17040478 | -0.562 | -1.861 | 0.713 | 0.438 | -0.003 | 0.069 | 0.069 | 0 | 0.1 | 0.13 | 0.1 | 0.31 | 0.07 | 0.12 |
| 3 | 4398881 | rs1688413 | -0.98 | -1.773 | 0.86 | 0.443 | 0.084 | -0.019 | -0.019 | 0 | 0.45 | 0.78 | 0.35 | 0.56 | 0.31 | 0.56 |
| 3 | 4400735 | rs2819563 | -0.53 | -1.779 | 1.016 | -0.052 | -0.082 | -0.001 | -0.001 | 0 | 0.15 | 0.36 | 0.11 | 0.17 | 0.02 | 0.29 |
| 3 | 4402649 | rs1688416 | 0.302 | -1.748 | 1.013 | -0.054 | -0.036 | -0.008 | -0.008 | 0 | 0.84 | 0.64 | 0.87 | 0.83 | 0.92 | 0.69 |
| 3 | 4403537 | rs4685744 | 0.434 | -1.314 | 0.653 | -0.108 | -0.016 | 0.04 | 0.04 | 0 | 0.48 | 0.18 | 0.52 | 0.59 | 0.51 | 0.46 |
| 3 | 4403614 | rs2819562 | 0.922 | -1.277 | 0.626 | -0.096 | -0.011 | 0.013 | 0.013 | 0 | 0.35 | 0.28 | 0.33 | 0.24 | 0.41 | 0.24 |
| 3 | 4403767 | rs2819561 | -1.042 | -1.267 | 0.63 | -0.092 | -0.012 | 0.013 | 0.013 | 0 | 0.65 | 0.72 | 0.67 | 0.76 | 0.59 | 0.76 |
| 3 | 4403837 | rs2633852 | -1.331 | -1.286 | 0.459 | -0.1 | -0.098 | 0.019 | 0.019 | 0 | 0.67 | 0.92 | 0.69 | 0.76 | 0.59 | 0.76 |
| 3 | 4410534 | rs11915920 | 0.357 | -1.063 | 0.491 | -0.007 | -0.023 | 0.032 | 0.032 | 0 | 0.47 | 0.19 | 0.51 | 0.58 | 0.51 | 0.42 |
| 3 | 4411691 | rs1687894 | 0.677 | -1 | 0.924 | -0.024 | 0.065 | 0.067 | 0.067 | 0 | 0.58 | 0.23 | 0.69 | 0.66 | 0.7 | 0.48 |

| | | | | | | | | | | | | | | | | |
|---|---|---|---|---|---|---|---|---|---|---|---|---|---|---|---|---|
| 3 | 4414011 | rs1968930 | 0.905 | -1.014 | 1.267 | -0.267 | -0.022 | -0.054 | -0.054 | 0 | 0.23 | 0.29 | 0.12 | 0.07 | 0.17 | 0.16 |
| 3 | 4415276 | rs793938 | 0.381 | -1.059 | 1.164 | -0.267 | 0.058 | 0.077 | 0.077 | 0 | 0.63 | 0.2 | 0.7 | 0.74 | 0.74 | 0.51 |
| 3 | 4416407 | rs10494173 | 0.362 | -0.928 | 1.185 | -0.278 | -0.019 | -0.126 | -0.126 | 0 | 0.27 | 0.46 | 0.16 | 0.07 | 0.23 | 0.17 |
| 3 | 4420255 | rs9864595 | 0.626 | -0.736 | 0.778 | -0.195 | 0.01 | 0.017 | 0.017 | 0 | 0.27 | 0.19 | 0.14 | 0.07 | 0.24 | 0.18 |
| 3 | 4420713 | rs807785 | 0.407 | -0.759 | 0.823 | -0.148 | -0.019 | -0.035 | -0.035 | 0 | 0.59 | 0.5 | 0.7 | 0.72 | 0.68 | 0.46 |
| 3 | 4429632 | rs9990134 | 0.277 | 0.004 | 0.275 | -0.22 | -0.012 | -0.05 | -0.05 | 7.525 | 0.27 | 0.26 | 0.13 | 0.07 | 0.21 | 0.18 |
| 3 | 4430848 | rs7641926 | -0.062 | 0.361 | -0.449 | 0.604 | 0.004 | -0.021 | -0.021 | 0.301 | 0.58 | 0.65 | 0.56 | 0.49 | 0.6 | 0.4 |
| 3 | 4431225 | rs12489356 | -0.336 | 0.518 | 0.039 | 0.873 | 0.003 | 0.03 | 0.03 | 0 | 0.18 | 0.32 | 0.31 | 0.25 | 0.13 | 0.28 |
| 3 | 4431356 | rs9990361 | -0.606 | 1.211 | -0.238 | 0.762 | -0.01 | 0.003 | 0.003 | 0 | 0.22 | 0.18 | 0.36 | 0.22 | 0.35 | 0.18 |
| 3 | 4432033 | rs964403 | 0.794 | 0.284 | -0.298 | 0.73 | 0.051 | 0.016 | 0.016 | 0 | 0.41 | 0.21 | 0.33 | 0.4 | 0.35 | 0.29 |
| 3 | 4432116 | rs3864051 | 0.847 | 0.276 | -0.314 | 0.683 | 0.056 | 0.02 | 0.02 | 0 | 0.41 | 0.17 | 0.32 | 0.39 | 0.35 | 0.29 |
| 3 | 4436172 | rs794187 | 0.274 | 0.834 | 0.391 | 0.359 | 0.119 | -0.114 | -0.114 | 0 | 0.19 | 0.35 | 0.3 | 0.01 | 0.34 | 0.13 |
| 3 | 4437358 | rs794185 | -0.915 | 0.504 | -0.33 | 0.473 | 0.083 | -0.006 | -0.006 | 0.004 | 0.3 | 0.5 | 0.47 | 0.32 | 0.43 | 0.26 |
| 3 | 4437484 | rs11714594 | 1.283 | 0.257 | -0.682 | 0.336 | 0.053 | 0.142 | 0.142 | 0.039 | 0.38 | 0.09 | 0.19 | 0.14 | 0.32 | 0.26 |
| 3 | 4451420 | rs17040638 | 1.082 | -0.045 | -0.877 | 0.973 | 0.037 | -0.061 | -0.061 | 0 | 0.13 | 0.05 | 0.2 | 0.32 | 0.08 | 0.3 |
| 3 | 4452737 | rs813811 | -0.622 | 0.514 | -0.473 | 0.269 | 0.005 | -0.003 | -0.003 | 0 | 0.63 | 0.66 | 0.63 | 0.44 | 0.69 | 0.46 |
| 3 | 4461324 | rs6782372 | 0.211 | 0.649 | -0.629 | 0.493 | -0.001 | 0.004 | 0.004 | 0 | 0.08 | 0.07 | 0.05 | 0.22 | 0.07 | 0.08 |
| 3 | 4462774 | rs711663 | 0.178 | 0.568 | -0.615 | 0.427 | 0.045 | 0.053 | 0.053 | 0.659 | 0.93 | 0.57 | 0.94 | 0.99 | 0.99 | 0.93 |
| 3 | 4468080 | rs17685501 | 0.862 | 0.725 | -1.234 | 0.267 | -0.004 | 0.013 | 0.013 | 0 | 0.4 | 0.12 | 0.33 | 0.39 | 0.41 | 0.31 |
| 3 | 4468598 | rs2322683 | 0.446 | 0.913 | -1.009 | -0.159 | 0.124 | -0.128 | -0.128 | 2.819 | 0.78 | 0.5 | 0.67 | 0.98 | 0.65 | 0.82 |
| 3 | 4469192 | rs793391 | 0.298 | 1.325 | -2.248 | 0.01 | -0.116 | 0.096 | 0.096 | 0 | 0.15 | 0.01 | 0.26 | 0 | 0.33 | 0.11 |
| 3 | 4477911 | rs11713682 | 0.816 | 0.866 | -1.935 | 0.34 | -0.002 | 0.083 | 0.083 | 0 | 0.33 | 0.09 | 0.27 | 0.17 | 0.35 | 0.23 |
| 3 | 4486303 | rs12634249 | -0.411 | 1.745 | -1.286 | 1.094 | -0.036 | -0.033 | -0.033 | 0.344 | 0.14 | 0.04 | 0.33 | 0.19 | 0.26 | 0.26 |
| 3 | 4490513 | rs711635 | -1.34 | 1.43 | -0.681 | 0.551 | 0.06 | -0.018 | -0.018 | 0 | 0.27 | 0.39 | 0.37 | 0.21 | 0.39 | 0.2 |
| 3 | 4491126 | rs17040706 | -0.443 | 1.178 | -1.053 | 0.687 | -0.002 | -0.009 | -0.009 | 0 | 0.09 | 0.05 | 0.15 | 0.21 | 0.08 | 0.1 |
| 3 | 4504624 | rs9834719 | -0.206 | 1.196 | -1.029 | 0.721 | -0.001 | 0.009 | 0.009 | 0 | 0.08 | 0.07 | 0.14 | 0.23 | 0.08 | 0.09 |
| 3 | 4513912 | rs1825916 | -1.368 | 1.27 | -0.658 | 0.579 | 0.1 | -0.004 | -0.004 | 0.004 | 0.27 | 0.5 | 0.45 | 0.25 | 0.38 | 0.2 |
| 3 | 4515191 | rs17040736 | -0.816 | 0.684 | -1.271 | 0.355 | 0.001 | -0.018 | -0.018 | 0 | 0.14 | 0.07 | 0.23 | 0.25 | 0.13 | 0.12 |
| 3 | 4524176 | rs13327148 | -0.202 | 0.969 | -1.19 | 0.519 | 0.009 | 0.057 | 0.057 | 0 | 0.07 | 0.16 | 0.22 | 0.23 | 0.08 | 0.09 |
| 3 | 4527619 | rs304082 | -0.037 | 1.073 | -1.331 | 0.384 | -0.107 | 0.091 | 0.091 | 0 | 0.15 | 0.01 | 0.19 | 0 | 0.29 | 0.1 |
| 3 | 4531076 | rs957384 | 0.108 | 1.062 | -1.303 | 0.426 | 0.002 | 0.035 | 0.035 | 0 | 0.06 | 0.1 | 0.2 | 0.22 | 0.08 | 0.08 |
| 3 | 4532469 | rs304079 | -1.247 | 0.709 | -1.139 | 0.388 | 0.044 | -0.004 | -0.004 | 0 | 0.27 | 0.42 | 0.45 | 0.23 | 0.38 | 0.2 |
| 3 | 4534629 | rs304077 | -1.177 | 0.646 | -1.134 | 0.418 | 0.028 | -0.007 | -0.007 | 0.026 | 0.26 | 0.36 | 0.44 | 0.23 | 0.38 | 0.2 |
| 3 | 4536386 | rs304073 | -1.146 | 0.729 | -1.598 | 0.444 | -0.032 | 0.007 | 0.007 | 0 | 0.25 | 0.2 | 0.43 | 0.22 | 0.38 | 0.2 |
| 3 | 4536403 | rs2291597 | 1.058 | 0.116 | -2.232 | 0.914 | 0.01 | -0.075 | -0.075 | 2.472 | 0.11 | 0.01 | 0.1 | 0.3 | 0.07 | 0.35 |
| 3 | 4538738 | rs11714599 | 1.143 | 0.286 | -2.492 | 0.288 | 0.034 | -0.036 | -0.036 | 0.045 | 0.18 | 0.02 | 0.07 | 0.2 | 0.1 | 0.27 |
| 3 | 4540226 | rs2131458 | -0.557 | 1.071 | -1.65 | 0.476 | -0.06 | -0.023 | -0.023 | 0 | 0.17 | 0.05 | 0.34 | 0.2 | 0.27 | 0.15 |
| 3 | 4560263 | rs304010 | 0.834 | 0.83 | -1.608 | 0.391 | -0.043 | 0.004 | 0.004 | 0 | 0.77 | 0.84 | 0.59 | 0.8 | 0.63 | 0.81 |
| 3 | 4562667 | rs1038639 | 1.37 | 0.669 | -1.903 | 0.335 | -0.161 | -0.027 | -0.027 | 0 | 0.64 | 0.06 | 0.5 | 0.7 | 0.57 | 0.76 |
| 3 | 4565456 | rs4684418 | -0.496 | 0.912 | -1.321 | 0.456 | -0.047 | -0.034 | -0.034 | 0 | 0.13 | 0.05 | 0.28 | 0.19 | 0.28 | 0.09 |
| 3 | 4565559 | rs7639802 | 0.394 | 0.681 | -1.495 | 0.501 | 0.019 | -0.017 | -0.017 | 0 | 0.15 | 0.12 | 0.06 | 0.11 | 0.12 | 0.03 |
| 3 | 4566033 | rs304023 | 0.835 | 0.687 | -1.109 | 0.41 | -0.017 | -0.005 | -0.005 | 1.776 | 0.64 | 0.68 | 0.61 | 0.7 | 0.53 | 0.84 |
| 3 | 4571671 | rs6796205 | -1.218 | 0.798 | -0.516 | 0.504 | 0.013 | 0.031 | 0.031 | 0 | 0.24 | 0.6 | 0.3 | 0.26 | 0.27 | 0.17 |
| 3 | 4572505 | rs6784152 | -1.023 | 0.781 | -0.259 | 0.381 | 0.018 | 0.029 | 0.029 | 0 | 0.22 | 0.58 | 0.23 | 0.22 | 0.24 | 0.16 |
| 3 | 4577500 | rs304053 | 0.62 | 0.101 | -0.867 | -0.035 | 0.079 | 0.09 | 0.09 | 0 | 0.55 | 0.24 | 0.78 | 0.75 | 0.68 | 0.7 |
| 3 | 4578306 | rs304051 | -0.856 | 0.223 | -0.861 | -0.085 | 0.059 | 0.111 | 0.111 | 0 | 0.48 | 0.76 | 0.25 | 0.25 | 0.38 | 0.36 |
| 3 | 4579561 | rs6783234 | -0.411 | 0.104 | 0.006 | -0.101 | -0.029 | -0.051 | -0.051 | 0.619 | 0.21 | 0.44 | 0.13 | 0.14 | 0.16 | 0.24 |
| 3 | 4582227 | rs304048 | 0.541 | -0.176 | 0.363 | 0.169 | -0.021 | 0.122 | 0.122 | 0 | 0.84 | 0.55 | 0.9 | 0.69 | 0.87 | 0.71 |
| 3 | 4583026 | rs184231 | 1.352 | -0.535 | 0.422 | -0.03 | -0.091 | 0.058 | 0.058 | 0 | 0.76 | 0.47 | 0.89 | 0.69 | 0.88 | 0.66 |
| 3 | 4584044 | rs304003 | 0.655 | -0.313 | 0.415 | 0.121 | -0.028 | 0.144 | 0.144 | 0 | 0.83 | 0.46 | 0.88 | 0.69 | 0.87 | 0.72 |
| 3 | 4585549 | rs304006 | -0.281 | 0.426 | -0.316 | 0.118 | 0.008 | 0.015 | 0.015 | 0 | 0.5 | 0.35 | 0.44 | 0.33 | 0.52 | 0.31 |
| 3 | 4588013 | rs7616234 | 0.038 | 0.662 | -0.736 | 0.042 | -0.033 | -0.081 | -0.081 | 3.134 | 0.12 | 0.02 | 0.07 | 0.32 | 0.12 | 0.2 |
| 3 | 4588214 | rs4684419 | 0.876 | 0.801 | -0.537 | -0.309 | -0.012 | -0.004 | -0.004 | 0 | 0.44 | 0.33 | 0.64 | 0.55 | 0.52 | 0.48 |
| 3 | 4589707 | rs12639183 | 0.854 | 0.515 | -0.781 | -0.308 | 0.002 | 0.008 | 0.008 | 0.14 | 0.17 | 0.1 | 0.09 | 0.13 | 0.16 | 0.11 |
| 3 | 4591247 | rs6764083 | -0.102 | 0.393 | -0.723 | -0.495 | 0.001 | 0.004 | 0.004 | 0 | 0.77 | 0.76 | 0.81 | 0.71 | 0.78 | 0.7 |
| 3 | 4593188 | rs902982 | 1.282 | 0.417 | -0.874 | -0.629 | -0.039 | 0.002 | 0.002 | 0 | 0.44 | 0.22 | 0.65 | 0.6 | 0.55 | 0.51 |
| 3 | 4594882 | rs11130059 | 0.729 | 0.963 | -1.438 | -0.315 | -0.049 | 0.063 | 0.063 | 0 | 0.2 | 0.02 | 0.1 | 0.08 | 0.21 | 0.13 |
| 3 | 4602847 | rs9822773 | 1.241 | 1.256 | -1.417 | -0.36 | 0.012 | 0.025 | 0.025 | 0 | 0.46 | 0.66 | 0.67 | 0.63 | 0.48 | 0.52 |
| 3 | 4604297 | rs9832410 | 0.167 | 1.187 | -0.745 | 0.195 | 0.007 | 0.026 | 0.026 | 0 | 0.79 | 0.69 | 0.82 | 0.71 | 0.82 | 0.72 |
| 3 | 4607085 | rs4684420 | 0.596 | 1.08 | -0.754 | 0.255 | 0.001 | 0 | 0 | 0 | 0.67 | 0.64 | 0.76 | 0.66 | 0.69 | 0.63 |
| 3 | 4609648 | rs11923960 | 0.141 | 0.815 | -0.941 | 0.356 | 0.001 | 0.011 | 0.011 | 0 | 0.56 | 0.4 | 0.66 | 0.65 | 0.65 | 0.56 |
| 3 | 4613805 | rs11130069 | 0.076 | 0.769 | -0.705 | 0.47 | -0.003 | -0.003 | -0.003 | 0 | 0.91 | 0.92 | 0.93 | 0.86 | 0.9 | 0.93 |
| 3 | 4613873 | rs12488824 | 0.846 | 0.412 | -1.771 | 0.011 | -0.099 | 0.09 | 0.09 | 0 | 0.32 | 0.03 | 0.33 | 0.16 | 0.34 | 0.15 |
| 3 | 4615243 | rs6774037 | 0.255 | 0.082 | -0.181 | 0.435 | 0.04 | 0.039 | 0.039 | 0 | 0.89 | 0.73 | 0.91 | 0.86 | 0.89 | 0.93 |
| 3 | 4616061 | rs6442887 | 1.128 | -0.61 | 0.248 | 0.621 | 0 | -0.001 | -0.001 | 0 | 0.69 | 0.29 | 0.77 | 0.38 | 0.8 | 0.47 |
| 3 | 4621315 | rs6442888 | -0.716 | -0.279 | 0.383 | 0.477 | 0.025 | 0.025 | 0.025 | 0 | 0.13 | 0.34 | 0.1 | 0.14 | 0.12 | 0.07 |
| 3 | 4622135 | rs6764946 | 0.897 | -0.3 | 0.364 | 0.464 | 0.041 | 0.032 | 0.032 | 0 | 0.86 | 0.57 | 0.88 | 0.84 | 0.86 | 0.91 |
| 3 | 4626197 | rs6804671 | 1.319 | -0.577 | -0.195 | 0.488 | 0.041 | 0.002 | 0.002 | 0 | 0.6 | 0.28 | 0.53 | 0.38 | 0.71 | 0.41 |
| 3 | 4628541 | rs4685771 | -0.244 | -0.521 | 0.18 | 0.697 | -0.056 | 0.193 | 0.193 | 0 | 0.18 | 0.38 | 0.18 | 0.41 | 0.11 | 0.46 |

| | | | | | | | | | | | | | | | | |
|---|---|---|---|---|---|---|---|---|---|---|---|---|---|---|---|---|
| 3 | 4628859 | rs12633680 | 0.774 | -0.368 | 0.272 | 0.763 | -0.015 | 0.035 | 0.035 | 0 | 0.72 | 0.44 | 0.73 | 0.43 | 0.75 | 0.43 |
| 3 | 4636260 | rs9856060 | 0.142 | -0.287 | -0.393 | 0.905 | -0.024 | 0.091 | 0.091 | 0 | 0.18 | 0.27 | 0.25 | 0.41 | 0.14 | 0.43 |
| 3 | 4637939 | rs17041030 | 0.673 | -0.176 | -1.312 | 0.221 | -0.001 | -0.022 | -0.022 | 0.192 | 0.39 | 0.49 | 0.41 | 0.22 | 0.38 | 0.22 |
| 3 | 4639163 | rs9871881 | 0.125 | -0.441 | -1.015 | 0.552 | -0.024 | 0.13 | 0.13 | 0 | 0.77 | 0.53 | 0.7 | 0.41 | 0.82 | 0.48 |
| 3 | 4643555 | rs2322735 | 0.387 | -0.33 | -1.63 | 0.577 | -0.013 | 0.043 | 0.043 | 0.027 | 0.73 | 0.64 | 0.75 | 0.47 | 0.78 | 0.46 |
| 3 | 4644803 | rs6782233 | 0.44 | -0.362 | -1.62 | 0.627 | -0.013 | 0.05 | 0.05 | 0 | 0.72 | 0.64 | 0.75 | 0.49 | 0.77 | 0.47 |
| 3 | 4645916 | rs4684427 | -0.453 | 0.314 | -2.498 | 0.621 | -0.086 | -0.028 | -0.028 | 0 | 0.62 | 0.85 | 0.65 | 0.61 | 0.51 | 0.67 |
| 3 | 4647026 | rs2101694 | -0.112 | 0.671 | -1.537 | 0.436 | -0.039 | 0.022 | 0.022 | 0 | 0.71 | 0.86 | 0.72 | 0.76 | 0.66 | 0.85 |
| 3 | 4664754 | rs931503 | -0.476 | -0.159 | -1.151 | 1.01 | 0.016 | -0.028 | -0.028 | 0 | 0.2 | 0.14 | 0.14 | 0.48 | 0.12 | 0.49 |
| 3 | 4665228 | rs7638474 | -0.715 | 0.335 | -1.077 | 0.517 | -0.001 | -0.008 | -0.008 | 0 | 0.08 | 0.04 | 0.16 | 0.08 | 0.08 | 0.07 |
| 3 | 4665565 | rs2029665 | -0.48 | 0.508 | -1.529 | 0.01 | -0.085 | 0.128 | 0.128 | 0.34 | 0.6 | 0.89 | 0.58 | 0.84 | 0.56 | 0.76 |
| 3 | 4669692 | rs2307067 | -1.217 | 0.377 | -0.062 | 0.492 | -0.016 | 0.043 | 0.043 | 0 | 0.08 | 0.16 | 0.12 | 0.12 | 0.04 | 0.08 |
| 3 | 4671633 | rs902984 | -1.711 | 0.265 | 0.841 | 0.454 | -0.012 | 0.119 | 0.119 | 0.535 | 0.1 | 0.43 | 0.14 | 0.14 | 0.04 | 0.08 |
| 3 | 4676285 | rs9818111 | -1.912 | 0.205 | 1.167 | 0.457 | -0.006 | 0.15 | 0.15 | 0.674 | 0.1 | 0.52 | 0.15 | 0.15 | 0.04 | 0.06 |
| 3 | 4682229 | rs4073665 | 1.518 | 0.096 | 0.999 | 0.201 | -0.012 | -0.002 | -0.002 | 0 | 0.8 | 0.74 | 0.82 | 0.85 | 0.88 | 0.89 |
| 3 | 4685084 | rs4685785 | 1.397 | 0.97 | 1.804 | -0.138 | 0.057 | -0.167 | -0.167 | 1.984 | 0.78 | 0.59 | 0.83 | 1 | 0.76 | 0.96 |
| 3 | 4686814 | rs4594601 | -1.59 | 1.074 | 1.861 | -0.142 | 0.006 | -0.125 | -0.125 | 0 | 0.2 | 0.33 | 0.12 | 0 | 0.16 | 0.03 |
| 3 | 4688615 | rs2686605 | -1.447 | 1.055 | 1.638 | -0.146 | -0.014 | -0.008 | -0.008 | 0 | 0.18 | 0.12 | 0.13 | 0 | 0.24 | 0.04 |
| 3 | 4693087 | rs4685793 | 1.009 | 0.724 | 1.836 | -0.118 | 0.024 | -0.122 | -0.122 | 0 | 0.83 | 0.64 | 0.88 | 1 | 0.85 | 0.97 |
| 3 | 4693553 | rs3796336 | 1.637 | 0.664 | 1.44 | 0.011 | -0.002 | -0.013 | -0.013 | 0.333 | 0.72 | 0.41 | 0.72 | 0.85 | 0.75 | 0.88 |
| 3 | 4695995 | rs17041113 | -1.266 | 0.413 | 1.429 | 0.409 | -0.021 | 0.134 | 0.134 | 3.066 | 0.07 | 0.36 | 0.13 | 0.14 | 0.01 | 0.03 |
| 3 | 4696646 | rs17709863 | -1.097 | 0.891 | 0.712 | 0.694 | -0.007 | -0.009 | -0.009 | 0 | 0.4 | 0.63 | 0.45 | 0.5 | 0.42 | 0.38 |
| 3 | 4712234 | rs1994500 | -0.166 | 0.172 | 0.964 | -0.182 | -0.009 | -0.025 | -0.025 | 9.114 | 0.43 | 0.53 | 0.46 | 0.23 | 0.57 | 0.26 |
| 3 | 4712413 | rs2306875 | -0.431 | 0.058 | 0.865 | -0.119 | 0.016 | 0.019 | 0.019 | 0 | 0.61 | 0.71 | 0.65 | 0.37 | 0.68 | 0.64 |
| 3 | 4713162 | rs3792490 | -0.549 | 0.212 | 0.746 | -0.134 | 0.025 | 0.03 | 0.03 | 0 | 0.62 | 0.76 | 0.67 | 0.37 | 0.69 | 0.65 |
| 3 | 4716214 | rs3804984 | -0.832 | -0.633 | 0.641 | -0.212 | -0.007 | -0.037 | -0.037 | 0 | 0.36 | 0.38 | 0.42 | 0.83 | 0.38 | 0.63 |
| 3 | 4716811 | rs2306877 | -0.129 | -0.705 | 0.618 | -0.253 | -0.041 | 0 | 0 | 0 | 0.23 | 0.39 | 0.32 | 0.29 | 0.2 | 0.45 |
| 3 | 4718896 | rs4685797 | 1.327 | -1.147 | 0.34 | -0.572 | -0.027 | 0.042 | 0.042 | 0 | 0.36 | 0.31 | 0.34 | 0.08 | 0.48 | 0.21 |
| 3 | 4719236 | rs4684433 | -0.54 | -1.38 | 0.342 | -0.546 | -0.002 | -0.004 | -0.004 | 0 | 0.35 | 0.3 | 0.33 | 0.72 | 0.35 | 0.61 |
| 3 | 4719426 | rs4684436 | -1.005 | -1.41 | 0.343 | -0.599 | 0.033 | 0.045 | 0.045 | 0 | 0.74 | 0.84 | 0.77 | 0.9 | 0.86 | 0.84 |
| 3 | 4720206 | rs17041165 | 0.641 | -1.404 | 0.461 | -0.492 | -0.036 | 0.003 | 0.003 | 0 | 0.2 | 0.27 | 0.21 | 0.24 | 0.15 | 0.41 |
| 3 | 4720224 | rs13325766 | 0.949 | -1.639 | -0.089 | -0.665 | 0.047 | 0.088 | 0.088 | 0 | 0.26 | 0.05 | 0.22 | 0.08 | 0.15 | 0.17 |
| 3 | 4720566 | rs34917285 | 0.698 | -1.697 | -0.167 | -0.676 | 0.048 | 0.08 | 0.08 | 0 | 0.28 | 0.11 | 0.23 | 0.09 | 0.15 | 0.18 |
| 3 | 4721377 | rs6786687 | -0.596 | -1.609 | -0.153 | -0.549 | 0.001 | 0.039 | 0.039 | 0 | 0.84 | 0.78 | 0.87 | 0.47 | 0.81 | 0.79 |
| 3 | 4722532 | rs9866640 | 0.849 | -1.711 | -0.514 | -0.838 | 0.036 | 0.134 | 0.134 | 0 | 0.31 | 0.02 | 0.24 | 0.09 | 0.18 | 0.18 |
| 3 | 4722956 | rs4684438 | 0.666 | -1.447 | 0.073 | -0.998 | -0.035 | -0.007 | -0.007 | 0.503 | 0.2 | 0.34 | 0.23 | 0.21 | 0.16 | 0.4 |
| 3 | 4724744 | rs6773688 | 0.194 | -0.955 | -0.172 | -1.536 | -0.001 | 0.036 | 0.036 | 0.02 | 0.43 | 0.37 | 0.41 | 0.17 | 0.53 | 0.22 |
| 3 | 4726947 | rs733018 | -0.396 | -0.656 | -0.09 | -2.331 | -0.023 | -0.124 | -0.124 | 0 | 0.24 | 0.39 | 0.22 | 0.02 | 0.21 | 0.11 |
| 3 | 4728804 | rs11709648 | 0.209 | 0.162 | -0.632 | -2.222 | -0.07 | 0.005 | 0.005 | 0 | 0.27 | 0.07 | 0.41 | 0.82 | 0.31 | 0.51 |
| 3 | 4731593 | rs6803046 | -0.133 | 0.67 | 0.023 | -2.374 | -0.023 | 0.034 | 0.034 | 0 | 0.32 | 0.25 | 0.39 | 0.08 | 0.5 | 0.21 |
| 3 | 4732040 | rs6803471 | -0.133 | 0.67 | 0.023 | -2.367 | -0.023 | 0.034 | 0.034 | 0 | 0.32 | 0.25 | 0.39 | 0.08 | 0.51 | 0.21 |
| 3 | 4734046 | rs12715419 | -0.118 | 0.673 | 0.007 | -2.37 | -0.052 | 0.08 | 0.08 | 0 | 0.32 | 0.18 | 0.39 | 0.08 | 0.5 | 0.21 |
| 3 | 4738774 | rs3749382 | -0.001 | 0.665 | 0.175 | -2.413 | -0.033 | 0.036 | 0.036 | 0 | 0.31 | 0.25 | 0.4 | 0.08 | 0.49 | 0.21 |
| 3 | 4741821 | rs9311395 | -0.244 | 0.501 | 0.25 | -2.438 | -0.025 | 0.042 | 0.042 | 0 | 0.33 | 0.29 | 0.41 | 0.08 | 0.51 | 0.21 |
| 3 | 4742030 | rs7652028 | 0.285 | 0.719 | 0.308 | -2.518 | -0.031 | -0.135 | -0.135 | 0.657 | 0.22 | 0.43 | 0.2 | 0.02 | 0.17 | 0.13 |
| 3 | 4742779 | rs6774180 | -0.326 | 0.826 | 0.422 | -2.462 | 0.198 | -0.054 | -0.054 | 0.462 | 0.18 | 0.48 | 0.25 | 0.07 | 0.35 | 0.08 |
| 3 | 4744047 | rs9831844 | 0.404 | 0.887 | -0.112 | -2.435 | -0.028 | 0.014 | 0.014 | 0 | 0.14 | 0.11 | 0.22 | 0.07 | 0.34 | 0.08 |
| 3 | 4747247 | rs9867580 | 0.467 | 0.893 | -0.132 | -2.412 | -0.042 | 0.011 | 0.011 | 0 | 0.14 | 0.09 | 0.22 | 0.08 | 0.34 | 0.09 |
| 3 | 4748675 | rs17711545 | 0.993 | 0.743 | -0.254 | -2.474 | -0.024 | 0.104 | 0.104 | 0 | 0.19 | 0.01 | 0.17 | 0.01 | 0.16 | 0.13 |
| 3 | 4750489 | rs3804992 | -0.302 | 0.925 | -0.11 | -2.348 | -0.116 | 0.097 | 0.097 | 0 | 0.3 | 0.07 | 0.38 | 0.09 | 0.49 | 0.18 |
| 3 | 4750674 | rs3804993 | 0.59 | 0.585 | -0.242 | -2.544 | -0.004 | 0.007 | 0.007 | 0 | 0.27 | 0.22 | 0.39 | 0.8 | 0.27 | 0.51 |
| 3 | 4751252 | rs7631664 | -1.352 | 0.559 | -0.119 | -2.555 | 0.029 | 0.024 | 0.024 | 0 | 0.58 | 0.4 | 0.78 | 0.89 | 0.77 | 0.69 |
| 3 | 4751802 | rs3816835 | -0.209 | 0.843 | -0.201 | -2.411 | -0.088 | 0.096 | 0.096 | 0 | 0.31 | 0.11 | 0.39 | 0.09 | 0.5 | 0.18 |
| 3 | 4753178 | rs17041262 | 1.072 | 0.402 | -0.021 | -2.316 | 0.003 | 0.014 | 0.014 | 0 | 0.17 | 0.13 | 0.17 | 0.01 | 0.15 | 0.09 |
| 3 | 4754125 | rs9880562 | -0.719 | 0.256 | -0.02 | -2.421 | 0.019 | 0.053 | 0.053 | 0 | 0.41 | 0.33 | 0.46 | 0.12 | 0.59 | 0.22 |
| 3 | 4755979 | rs722368 | -0.919 | 0.11 | -0.013 | -2.465 | -0.014 | -0.047 | -0.047 | 0 | 0.44 | 0.54 | 0.5 | 0.15 | 0.59 | 0.3 |
| 3 | 4761477 | rs9859826 | -1.468 | -0.166 | -0.135 | -2.494 | -0.058 | -0.096 | -0.096 | 0 | 0.63 | 0.56 | 0.81 | 0.92 | 0.8 | 0.77 |
| 3 | 4765872 | rs7651990 | 0.664 | 0.468 | -0.357 | -2.188 | -0.002 | 0.004 | 0.004 | 0 | 0.2 | 0.24 | 0.35 | 0.76 | 0.2 | 0.46 |
| 3 | 4773465 | rs9824272 | -0.819 | 1.622 | 0.436 | -1.982 | -0.022 | 0.015 | 0.015 | 0.053 | 0.27 | 0.21 | 0.32 | 0.16 | 0.46 | 0.25 |
| 3 | 4777544 | rs931389 | 0.811 | 1.862 | 0.114 | -1.804 | -0.016 | 0.034 | 0.034 | 0.011 | 0.66 | 0.26 | 0.59 | 0.88 | 0.59 | 0.76 |
| 3 | 4780074 | rs3804999 | -1.205 | 2.31 | 0.602 | -1.33 | 0.058 | -0.059 | -0.059 | 0.013 | 0.19 | 0.34 | 0.31 | 0.07 | 0.29 | 0.19 |
| 3 | 4781349 | rs13092274 | 0.31 | 2.252 | 0.429 | -1.565 | 0.011 | -0.116 | -0.116 | 0.399 | 0.76 | 0.62 | 0.79 | 1 | 0.74 | 0.88 |
| 3 | 4783407 | rs11714054 | 0.14 | 2.546 | 0.326 | -1.546 | 0.049 | -0.026 | -0.026 | 0 | 0.11 | 0.17 | 0.14 | 0.03 | 0.25 | 0.09 |
| 3 | 4785230 | rs12638018 | 1.254 | 1.35 | -0.057 | -1.522 | 0.002 | 0.046 | 0.046 | 0 | 0.57 | 0.3 | 0.54 | 0.92 | 0.43 | 0.59 |
| 3 | 4786040 | rs1866999 | 1.118 | 1.52 | -0.002 | -1.459 | -0.002 | 0.023 | 0.023 | 0 | 0.59 | 0.3 | 0.54 | 0.92 | 0.43 | 0.6 |
| 3 | 4789935 | rs6786487 | 0.328 | 0.707 | -0.124 | -0.925 | -0.028 | -0.08 | -0.08 | 0.186 | 0.82 | 0.73 | 0.87 | 1 | 0.88 | 0.89 |
| 3 | 4794099 | rs1866997 | 1.419 | 1.623 | 0.108 | 0.489 | 0.073 | 0.222 | 0.222 | 0.02 | 0.92 | 0.77 | 0.88 | 0.63 | 0.86 | 0.75 |
| 3 | 4794380 | rs901849 | 1.257 | 2.472 | 0.294 | 0.363 | 0.124 | 0.105 | 0.105 | 0 | 0.77 | 0.58 | 0.66 | 0.56 | 0.56 | 0.6 |
| 3 | 4794677 | rs6785305 | -0.931 | 1.794 | -0.303 | 0.566 | -0.029 | -0.037 | -0.037 | 0.419 | 0.37 | 0.16 | 0.4 | 0.4 | 0.53 | 0.43 |

| | | | | | | | | | | | | | | | | |
|---|---|---|---|---|---|---|---|---|---|---|---|---|---|---|---|---|
| 3 | 4794911 | rs746039 | -0.36 | 1.108 | -0.985 | -0.106 | 0.016 | 0.082 | 0.082 | 0 | 0.73 | 0.89 | 0.84 | 0.99 | 0.77 | 0.85 |
| 3 | 4794980 | rs17041369 | 0.259 | 0.904 | -0.908 | 0.268 | 0.078 | 0.113 | 0.113 | 0.326 | 0.08 | 0.48 | 0.15 | 0.1 | 0.07 | 0.14 |
| 3 | 4796424 | rs3805005 | -0.051 | 0.804 | -0.826 | 0.646 | 0.07 | 0.456 | 0.456 | 0 | 0.09 | 0.41 | 0.26 | 0.41 | 0.09 | 0.22 |
| 3 | 4796807 | rs11706308 | -0.427 | 1.026 | -1.108 | 0.352 | -0.043 | 0.192 | 0.192 | 0 | 0.39 | 0.13 | 0.37 | 0.07 | 0.57 | 0.33 |
| 3 | 4797041 | rs11710644 | 0.785 | 1.278 | -1.159 | 0.377 | -0.088 | 0.011 | 0.011 | 0 | 0.12 | 0.03 | 0.18 | 0.06 | 0.34 | 0.14 |
| 3 | 4798840 | rs2015714 | 0.845 | 0.985 | -1.147 | 0.426 | 0.002 | 0.002 | 0.002 | 0 | 0.52 | 0.42 | 0.56 | 0.57 | 0.39 | 0.42 |
| 3 | 4802573 | rs9831960 | 0.619 | 0.871 | -1.078 | 0.559 | 0.026 | -0.013 | -0.013 | 0.72 | 0.58 | 0.3 | 0.52 | 0.48 | 0.66 | 0.45 |
| 3 | 4804000 | rs10510297 | 0.184 | 0.972 | -1.147 | 0.516 | 0.022 | -0.007 | -0.007 | 0.052 | 0.16 | 0.15 | 0.24 | 0.06 | 0.36 | 0.2 |
| 3 | 4804081 | rs3792499 | 1.466 | 0.905 | -1.212 | 0.52 | 0.019 | -0.075 | -0.075 | 0 | 0.43 | 0.04 | 0.31 | 0.44 | 0.31 | 0.31 |
| 3 | 4805489 | rs3805009 | -0.363 | 0.843 | -1.087 | 0.393 | 0.022 | 0.034 | 0.034 | 0 | 0.71 | 0.77 | 0.83 | 0.9 | 0.78 | 0.67 |
| 3 | 4807021 | rs3805011 | 0.773 | 1.018 | -1.101 | 0.359 | 0.026 | 0.059 | 0.059 | 0 | 0.53 | 0.63 | 0.58 | 0.85 | 0.38 | 0.48 |
| 3 | 4807357 | rs11713227 | 0.526 | 1.083 | -1.111 | 0.382 | 0.129 | -0.022 | -0.022 | 0.207 | 0.14 | 0.27 | 0.21 | 0.06 | 0.35 | 0.14 |
| 3 | 4807550 | rs3805012 | -0.036 | 0.732 | -1.244 | 0.547 | 0.02 | -0.008 | -0.008 | 0 | 0.67 | 0.75 | 0.74 | 0.59 | 0.74 | 0.55 |
| 3 | 4808703 | rs2291863 | 1.313 | 0.782 | -1.723 | 0.112 | 0.093 | -0.051 | -0.051 | 0 | 0.42 | 0.09 | 0.3 | 0.44 | 0.28 | 0.3 |
| 3 | 4809969 | rs17041401 | -0.418 | 1.656 | -1.474 | -0.129 | 0.077 | -0.016 | -0.016 | 0.566 | 0.18 | 0.24 | 0.22 | 0.09 | 0.43 | 0.18 |
| 3 | 4812298 | rs3792501 | 1.249 | 0.679 | -2.071 | -0.268 | 0.055 | -0.136 | -0.136 | 0 | 0.42 | 0.04 | 0.31 | 0.52 | 0.31 | 0.4 |
| 3 | 4818127 | rs17729477 | 0.062 | 0.866 | -1.588 | -0.958 | -0.02 | 0.111 | 0.111 | 0 | 0.19 | 0.06 | 0.14 | 0 | 0.2 | 0.1 |
| 3 | 4818792 | rs4684443 | 0.273 | 0.796 | -1.318 | -0.916 | -0.005 | -0.106 | -0.106 | 0 | 0.75 | 0.62 | 0.77 | 0.98 | 0.76 | 0.8 |
| 3 | 4827533 | rs1018109 | 0.537 | 0.933 | -1.202 | -0.809 | 0.001 | -0.057 | -0.057 | 0 | 0.64 | 0.54 | 0.75 | 0.91 | 0.64 | 0.67 |
| 3 | 4829575 | rs17786144 | 0.613 | 1.122 | -1.357 | -0.736 | -0.032 | 0.031 | 0.031 | 0 | 0.06 | 0.01 | 0.07 | 0 | 0.12 | 0.02 |
| 3 | 4831141 | rs7652212 | 0.176 | -0.064 | -1.134 | -0.607 | -0.021 | -0.049 | -0.049 | 1.005 | 0.9 | 0.8 | 0.96 | 1 | 0.96 | 0.98 |
| 3 | 4837269 | rs11713160 | 0.525 | 0.2 | -1.052 | -0.662 | 0.002 | -0.044 | -0.044 | 0.015 | 0.6 | 0.54 | 0.67 | 0.93 | 0.5 | 0.66 |
| 3 | 4838298 | rs931388 | -0.167 | 0.052 | -1.119 | 0.009 | -0.035 | 0.046 | 0.046 | 1.108 | 0.25 | 0.48 | 0.29 | 0.34 | 0.18 | 0.44 |
| 3 | 4838819 | rs3805018 | -0.745 | 0.183 | -1.311 | 0.147 | 0.006 | 0.002 | 0.002 | 0 | 0.52 | 0.55 | 0.5 | 0.45 | 0.57 | 0.68 |
| 3 | 4841657 | rs4142942 | 0.181 | -0.593 | -0.983 | 0.487 | -0.068 | 0.072 | 0.072 | 0.236 | 0.74 | 0.53 | 0.7 | 0.56 | 0.85 | 0.53 |
| 3 | 4842231 | rs2291862 | -0.526 | -0.392 | -1.742 | 0.215 | -0.003 | -0.032 | -0.032 | 0.014 | 0.33 | 0.12 | 0.36 | 0.38 | 0.31 | 0.24 |
| 3 | 4842488 | rs2291861 | 0.378 | -0.272 | -1.075 | 0.088 | 0.008 | 0.008 | 0.008 | 0 | 0.46 | 0.46 | 0.53 | 0.35 | 0.63 | 0.39 |
| 3 | 4847009 | rs2119802 | 0.479 | -0.548 | -0.459 | -0.316 | -0.037 | -0.083 | -0.083 | 0.015 | 0.69 | 0.44 | 0.8 | 0.94 | 0.79 | 0.73 |
| 3 | 4852238 | rs874131 | 0.626 | -0.643 | 0.25 | -0.361 | -0.042 | -0.086 | -0.086 | 0 | 0.72 | 0.44 | 0.79 | 0.93 | 0.83 | 0.73 |
| 3 | 4852446 | rs7612703 | -0.057 | -0.898 | 0.61 | -0.503 | -0.004 | -0.028 | -0.028 | 0 | 0.12 | 0.12 | 0.12 | 0.04 | 0.11 | 0.24 |
| 3 | 4852734 | rs878135 | 0.686 | -0.694 | -0.425 | -0.325 | -0.087 | -0.076 | -0.076 | 0.845 | 0.41 | 0.14 | 0.44 | 0.54 | 0.45 | 0.43 |
| 3 | 4855348 | rs731915 | 0.917 | -0.776 | -0.131 | -0.186 | -0.008 | -0.021 | -0.021 | 2.119 | 0.31 | 0.22 | 0.42 | 0.48 | 0.4 | 0.29 |
| 3 | 4856180 | rs711631 | 0.054 | -1.219 | 0.262 | -0.586 | 0.003 | -0.026 | -0.026 | 1.663 | 0.72 | 0.68 | 0.82 | 0.96 | 0.84 | 0.7 |
| 3 | 4856234 | rs901854 | 0.517 | -0.678 | -0.132 | -0.493 | 0 | 0.019 | 0.019 | 0 | 0.57 | 0.53 | 0.53 | 0.72 | 0.59 | 0.57 |
| 3 | 4856430 | rs3816252 | 1.478 | -0.152 | -0.279 | -0.198 | -0.004 | -0.006 | -0.006 | 0 | 0.29 | 0.2 | 0.36 | 0.5 | 0.4 | 0.33 |
| 3 | 4861697 | rs9838930 | 0.484 | -0.411 | -0.152 | -0.274 | 0.007 | 0.019 | 0.019 | 0.205 | 0.42 | 0.32 | 0.64 | 0.72 | 0.62 | 0.46 |
| 3 | 4861853 | rs9858750 | -0.36 | -0.827 | -0.182 | -0.473 | -0.058 | -0.07 | -0.07 | 0 | 0.81 | 0.3 | 0.89 | 0.97 | 0.94 | 0.74 |
| 3 | 4863387 | rs1873020 | -0.075 | -0.236 | -0.2 | 0.078 | -0.042 | 0.006 | 0.006 | 0 | 0.37 | 0.22 | 0.42 | 0.32 | 0.42 | 0.39 |
| 3 | 4864491 | rs1546228 | -1.15 | -0.205 | -0.044 | 0.173 | -0.013 | -0.018 | -0.018 | 0 | 0.71 | 0.8 | 0.57 | 0.38 | 0.62 | 0.72 |
| 3 | 4865354 | rs9883291 | -1.434 | -0.158 | -0.023 | 0.103 | -0.011 | -0.017 | -0.017 | 0.643 | 0.73 | 0.81 | 0.59 | 0.51 | 0.63 | 0.73 |
| 3 | 4867097 | rs2291860 | 0.555 | -0.061 | -0.202 | 0.13 | -0.027 | -0.024 | -0.024 | 0.01 | 0.15 | 0.12 | 0.29 | 0.22 | 0.23 | 0.15 |
| 3 | 4868669 | rs17041493 | 0.874 | -0.328 | -0.125 | -0.053 | 0.007 | 0.005 | 0.005 | 0 | 0.21 | 0.13 | 0.11 | 0.04 | 0.14 | 0.25 |
| 3 | 4868927 | rs17786757 | 1.179 | -0.252 | -0.487 | -0.019 | 0.002 | 0.071 | 0.071 | 0 | 0.19 | 0.01 | 0.11 | 0.04 | 0.14 | 0.22 |
| 3 | 4869590 | rs3805025 | 0.456 | 0.202 | -0.057 | 0.196 | 0.017 | 0.017 | 0.017 | 0.085 | 0.15 | 0.22 | 0.24 | 0.24 | 0.22 | 0.16 |
| 3 | 4871598 | rs7432768 | 0.214 | -0.068 | -0.147 | 0.03 | -0.006 | 0.099 | 0.099 | 0.294 | 0.49 | 0.26 | 0.61 | 0.77 | 0.52 | 0.6 |
| 3 | 4872198 | rs4685829 | -0.118 | 0.107 | -0.194 | 0.085 | -0.003 | 0.029 | 0.029 | 0.048 | 0.48 | 0.47 | 0.36 | 0.19 | 0.46 | 0.37 |
| 3 | 4873374 | rs3828439 | 0.465 | 0.832 | -0.396 | 0.68 | -0.004 | -0.008 | -0.008 | 0.485 | 0.26 | 0.18 | 0.36 | 0.53 | 0.33 | 0.35 |
| 3 | 4874498 | rs13096481 | 0.548 | 0.906 | -0.692 | 0.504 | -0.015 | -0.034 | -0.034 | 0 | 0.25 | 0.14 | 0.34 | 0.49 | 0.32 | 0.34 |
| 3 | 4874791 | rs11708908 | -0.564 | 1.361 | -0.636 | 0.268 | 0.005 | -0.022 | -0.022 | 0.013 | 0.09 | 0.14 | 0.09 | 0.02 | 0.1 | 0.09 |
| 3 | 4876661 | rs11711879 | -0.195 | 1.36 | -1.093 | 0.624 | 0.02 | 0.022 | 0.022 | 0 | 0.38 | 0.49 | 0.51 | 0.57 | 0.49 | 0.42 |
| 3 | 4877199 | rs12493413 | -0.352 | 2.024 | -1.798 | 0.183 | -0.126 | 0.011 | 0.011 | 0.02 | 0.19 | 0.02 | 0.17 | 0.13 | 0.38 | 0.09 |
| 3 | 4877348 | rs3805032 | 0.325 | 0.759 | -1.172 | 0.238 | -0.003 | -0.003 | -0.003 | 0 | 0.65 | 0.64 | 0.75 | 0.72 | 0.73 | 0.69 |
| 3 | 4880266 | rs3805034 | 0.303 | 0.557 | -1.141 | 0.15 | 0.005 | -0.004 | -0.004 | 0 | 0.86 | 0.88 | 0.82 | 0.85 | 0.91 | 0.95 |
| 3 | 4880300 | rs3805035 | -0.092 | 0.669 | -1.093 | 0.264 | -0.001 | 0.003 | 0.003 | 0.06 | 0.9 | 0.88 | 0.82 | 0.83 | 0.92 | 0.97 |
| 3 | 4882700 | rs3805036 | 0.098 | 0.471 | -1.189 | -0.329 | 0.007 | 0.044 | 0.044 | 0 | 0.51 | 0.39 | 0.45 | 0.56 | 0.56 | 0.51 |
| 3 | 4891407 | rs1827101 | 0.317 | 0.071 | -0.656 | -0.245 | -0.006 | 0.048 | 0.048 | 0.261 | 0.87 | 0.79 | 0.78 | 0.76 | 0.9 | 0.97 |
| 3 | 4893623 | rs994541 | 0.777 | -0.218 | -0.336 | -0.336 | -0.04 | 0.055 | 0.055 | 0.925 | 0.81 | 0.53 | 0.75 | 0.72 | 0.89 | 0.74 |
| 3 | 4896495 | rs4684448 | 0.727 | 0.079 | -0.632 | -0.643 | 0.004 | 0.023 | 0.023 | 0 | 0.51 | 0.35 | 0.5 | 0.69 | 0.56 | 0.59 |
| 3 | 4899009 | rs9880505 | -0.069 | 0.126 | -0.675 | -0.572 | -0.006 | 0.069 | 0.069 | 0.869 | 0.14 | 0.31 | 0.22 | 0.25 | 0.1 | 0.05 |
| 3 | 4905201 | rs2859808 | -0.206 | -0.037 | -0.721 | -0.641 | 0 | -0.005 | -0.005 | 0 | 0.65 | 0.68 | 0.73 | 0.94 | 0.66 | 0.64 |
| 3 | 4906628 | rs6794744 | -0.28 | -0.243 | -1.004 | -0.197 | -0.013 | 0.071 | 0.071 | 1.411 | 0.14 | 0.31 | 0.22 | 0.25 | 0.09 | 0.02 |
| 3 | 4922720 | rs4685840 | -0.145 | 0.261 | -1.623 | -0.475 | 0.02 | 0.075 | 0.075 | 0 | 0.73 | 0.84 | 0.83 | 0.94 | 0.73 | 0.86 |
| 3 | 4927281 | rs4685841 | 0.129 | 1.17 | -1.787 | 0.275 | -0.062 | -0.102 | -0.102 | 0 | 0.75 | 0.92 | 0.72 | 0.52 | 0.68 | 0.61 |
| 3 | 4928098 | rs876349 | -0.092 | -0.492 | -1.274 | -0.826 | -0.03 | -0.07 | -0.07 | 0 | 0.67 | 0.46 | 0.85 | 0.95 | 0.81 | 0.85 |
| 3 | 4928159 | rs3856883 | -0.058 | -0.557 | -1.211 | -0.853 | -0.02 | -0.053 | -0.053 | 0.091 | 0.66 | 0.45 | 0.84 | 0.95 | 0.81 | 0.81 |
| 3 | 4928944 | rs3762781 | 0.369 | -0.197 | -1.21 | -1.261 | 0.011 | 0.061 | 0.061 | 0.001 | 0.47 | 0.39 | 0.7 | 0.85 | 0.57 | 0.46 |
| 3 | 4936864 | rs6807140 | -0.124 | 0.137 | -1.779 | -1.594 | -0.011 | 0.04 | 0.04 | 0 | 0.44 | 0.39 | 0.39 | 0.1 | 0.54 | 0.24 |
| 3 | 4936913 | rs6794962 | -0.252 | 0.279 | -1.908 | -1.774 | 0.03 | -0.056 | -0.056 | 1.356 | 0.24 | 0.36 | 0.34 | 0.06 | 0.4 | 0.16 |
| 3 | 4940736 | rs6808127 | 0.151 | 0.171 | -1.038 | -1.085 | -0.004 | 0.029 | 0.029 | 0 | 0.33 | 0.6 | 0.26 | 0.47 | 0.26 | 0.51 |

| | | | | | | | | | | | | | | | | |
|---|---|---|---|---|---|---|---|---|---|---|---|---|---|---|---|---|
| 3 | 4941141 | rs9836040 | 1.354 | 0.505 | -1.131 | -1.415 | -0.008 | 0.096 | 0.096 | 0 | 0.25 | 0.07 | 0.14 | 0.06 | 0.23 | 0.12 |
| 3 | 4941214 | rs11922612 | 1.189 | 0.333 | -0.742 | -1.08 | -0.003 | -0.027 | -0.027 | 0 | 0.08 | 0.11 | 0.05 | 0.01 | 0.05 | 0.02 |
| 3 | 4946028 | rs1514453 | 0.62 | 0.428 | -0.768 | -1.087 | -0.028 | 0.077 | 0.077 | 0.033 | 0.37 | 0.23 | 0.35 | 0.06 | 0.49 | 0.17 |
| 3 | 4947829 | rs17041710 | -0.278 | 0.327 | 0.271 | 0.113 | -0.001 | 0.004 | 0.004 | 0 | 0.29 | 0.32 | 0.33 | 0.65 | 0.23 | 0.51 |
| 3 | 4952779 | rs6777136 | -0.077 | 0.637 | 0.113 | 0.394 | -0.007 | -0.012 | -0.012 | 0 | 0.26 | 0.27 | 0.34 | 0.62 | 0.29 | 0.46 |
| 3 | 4969387 | rs10049082 | -0.774 | 0.737 | 0.122 | 0.306 | -0.018 | 0.01 | 0.01 | 0 | 0.33 | 0.3 | 0.4 | 0.68 | 0.4 | 0.6 |
| 3 | 4973586 | rs4496464 | 0.722 | 0.513 | 0.795 | 0.872 | 0.015 | 0.153 | 0.153 | 0.018 | 0.88 | 0.61 | 0.9 | 0.83 | 0.89 | 0.81 |
| 3 | 4975451 | rs17041758 | -0.277 | 0.889 | 0.819 | 0.737 | 0.064 | -0.026 | -0.026 | 0 | 0.1 | 0.27 | 0.09 | 0.08 | 0.09 | 0.1 |
| 3 | 4978097 | rs6797654 | 0.819 | 0.503 | 0.905 | 0.876 | 0.03 | 0.146 | 0.146 | 0 | 0.88 | 0.58 | 0.9 | 0.84 | 0.89 | 0.81 |
| 3 | 4978355 | rs6442919 | -0.212 | 0.665 | 0.523 | 0.977 | 0.021 | 0.082 | 0.082 | 0 | 0.92 | 0.82 | 0.93 | 0.85 | 0.88 | 0.87 |
| 3 | 4979672 | rs9311438 | 0.446 | -0.08 | -0.317 | 0.437 | 0.029 | 0.045 | 0.045 | 0 | 0.42 | 0.3 | 0.36 | 0.22 | 0.33 | 0.32 |
| 3 | 4982356 | rs1955179 | 0.05 | -0.284 | -0.136 | 0.741 | -0.001 | 0.001 | 0.001 | 0 | 0.54 | 0.53 | 0.58 | 0.41 | 0.44 | 0.27 |
| 3 | 4983263 | rs4386489 | 0.278 | -0.225 | -0.016 | 0.826 | -0.001 | -0.004 | -0.004 | 0.537 | 0.49 | 0.48 | 0.44 | 0.38 | 0.43 | 0.27 |
| 3 | 4991139 | rs7635516 | 0.23 | 0.031 | 0.038 | 0.433 | -0.012 | 0.032 | 0.032 | 0 | 0.42 | 0.25 | 0.46 | 0.67 | 0.46 | 0.58 |
| 3 | 4991538 | rs9836876 | -0.775 | -0.362 | 0.809 | 0.683 | -0.005 | 0.029 | 0.029 | 0 | 0.12 | 0.26 | 0.19 | 0.15 | 0.12 | 0.19 |
| 3 | 4994341 | rs7638383 | -0.083 | -0.516 | 0.208 | 0.044 | 0.009 | 0.038 | 0.038 | 0 | 0.44 | 0.35 | 0.35 | 0.11 | 0.38 | 0.36 |
| 3 | 4994471 | rs7628494 | 0.674 | -0.697 | 0.704 | 0.692 | -0.017 | 0.067 | 0.067 | 0 | 0.8 | 0.49 | 0.76 | 0.76 | 0.81 | 0.81 |
| 3 | 5003570 | rs7640124 | 0.162 | -0.554 | 0.363 | 0.855 | -0.02 | 0.074 | 0.074 | 0 | 0.92 | 0.78 | 0.85 | 0.82 | 0.95 | 0.86 |
| 3 | 5013876 | rs6442925 | -0.641 | -0.713 | 0.095 | 0.069 | -0.005 | -0.027 | -0.027 | 0 | 0.75 | 0.71 | 0.82 | 0.92 | 0.85 | 0.72 |
| 3 | 5016106 | rs1588794 | -0.363 | -0.232 | 0.546 | 0.753 | 0.003 | 0.06 | 0.06 | 0 | 0.09 | 0.13 | 0.18 | 0.24 | 0.11 | 0.05 |
| 3 | 5019936 | rs1110261 | -1.218 | -0.305 | 0.517 | 0.367 | -0.019 | 0.01 | 0.01 | 0 | 0.2 | 0.32 | 0.28 | 0.17 | 0.22 | 0.19 |
| 3 | 5024771 | rs908078 | -0.682 | -0.151 | 0.111 | 0.383 | 0.015 | -0.02 | -0.02 | 0 | 0.17 | 0.09 | 0.24 | 0.19 | 0.15 | 0.17 |
| 3 | 5026008 | rs11130215 | -1.327 | -0.155 | 0.805 | 0.288 | -0.009 | 0.019 | 0.019 | 0 | 0.19 | 0.38 | 0.27 | 0.16 | 0.2 | 0.17 |
| 3 | 5029354 | rs11920786 | -1.051 | -0.737 | 0.988 | 0.335 | -0.041 | 0.028 | 0.028 | 0.015 | 0.17 | 0.31 | 0.24 | 0.16 | 0.14 | 0.16 |
| 3 | 5029776 | rs11918262 | -0.301 | -1.172 | 0.592 | -0.078 | -0.014 | -0.046 | -0.046 | 0 | 0.71 | 0.61 | 0.8 | 0.91 | 0.82 | 0.71 |
| 3 | 5030961 | rs2137947 | 0.09 | -1.25 | 0.48 | 0.191 | 0.013 | 0.007 | 0.007 | 0.226 | 0.67 | 0.26 | 0.68 | 0.66 | 0.78 | 0.67 |
| 3 | 5035285 | rs9311445 | -0.157 | -0.866 | 0.089 | 0.063 | -0.001 | -0.001 | -0.001 | 0 | 0.68 | 0.61 | 0.68 | 0.64 | 0.69 | 0.69 |
| 3 | 5037061 | rs6442928 | 0.869 | -0.893 | -0.205 | 0.05 | -0.02 | -0.018 | -0.018 | 0 | 0.44 | 0.24 | 0.39 | 0.5 | 0.5 | 0.48 |
| 3 | 5041215 | rs908072 | 0.871 | -1.372 | 0.189 | 0.77 | -0.046 | 0.117 | 0.117 | 0 | 0.81 | 0.63 | 0.69 | 0.5 | 0.85 | 0.68 |
| 3 | 5044464 | rs962837 | 1.506 | -1.431 | -0.778 | 0.642 | 0.048 | -0.097 | -0.097 | 0 | 0.63 | 0.11 | 0.53 | 0.45 | 0.69 | 0.61 |
| 3 | 5053815 | rs6782157 | -1.749 | -1.268 | -0.309 | 0.916 | 0.005 | -0.007 | -0.007 | 0.34 | 0.25 | 0.2 | 0.27 | 0.37 | 0.22 | 0.19 |
| 3 | 5054180 | rs11130232 | -1.252 | -1.175 | 0.148 | 1.399 | -0.024 | 0.067 | 0.067 | 0 | 0.13 | 0.19 | 0.18 | 0.37 | 0.12 | 0.15 |
| 3 | 5059576 | rs2671750 | 0.94 | -0.822 | 0.337 | 1.331 | -0.053 | 0.147 | 0.147 | 0 | 0.79 | 0.43 | 0.67 | 0.52 | 0.84 | 0.64 |
| 3 | 5062609 | rs9821267 | 0.087 | 0.285 | -0.905 | 0.793 | -0.009 | 0.018 | 0.018 | 0 | 0.42 | 0.35 | 0.38 | 0.28 | 0.47 | 0.31 |
| 3 | 5064834 | rs2606776 | -0.624 | 0.426 | -1.241 | 1.347 | -0.013 | -0.048 | -0.048 | 0 | 0.12 | 0.05 | 0.17 | 0.28 | 0.14 | 0.34 |
| 3 | 5069843 | rs11716050 | 0.293 | 0.161 | -1.15 | 0.719 | 0.023 | 0.028 | 0.028 | 0 | 0.39 | 0.28 | 0.38 | 0.3 | 0.33 | 0.35 |
| 3 | 5077330 | rs11927386 | -0.458 | 0.379 | -1.151 | 0.528 | -0.003 | 0.05 | 0.05 | 0 | 0.23 | 0.07 | 0.16 | 0.15 | 0.23 | 0.17 |
| 3 | 5078255 | rs9858342 | 0.801 | 0.319 | -0.097 | 0.801 | 0.019 | 0.133 | 0.133 | 0.155 | 0.87 | 0.53 | 0.9 | 0.78 | 0.9 | 0.79 |
| 3 | 5082256 | rs17790604 | -0.29 | 0.108 | -0.424 | 0.497 | -0.015 | 0.009 | 0.009 | 0.101 | 0.17 | 0.06 | 0.12 | 0.1 | 0.19 | 0.15 |
| 3 | 5082955 | rs7616918 | -0.264 | -0.475 | -0.615 | 0.372 | 0.038 | -0.002 | -0.002 | 0 | 0.49 | 0.22 | 0.5 | 0.41 | 0.44 | 0.51 |
| 3 | 5085165 | rs7639315 | 1.104 | -0.589 | -0.733 | 0.259 | 0.015 | -0.075 | -0.075 | 1.498 | 0.29 | 0.03 | 0.35 | 0.3 | 0.23 | 0.35 |
| 3 | 5085835 | rs11918600 | -0.15 | -0.327 | -0.607 | 0.133 | -0.014 | 0.013 | 0.013 | 0.021 | 0.19 | 0.08 | 0.13 | 0.11 | 0.23 | 0.16 |
| 3 | 5091740 | rs17041935 | -0.035 | -0.396 | 0.43 | -0.148 | 0.024 | -0.004 | -0.004 | 0 | 0.07 | 0.31 | 0.03 | 0 | 0.02 | 0.13 |
| 3 | 5092475 | rs6809156 | 0.895 | -0.596 | -0.042 | -0.068 | 0.035 | -0.008 | -0.008 | 0.794 | 0.32 | 0.2 | 0.33 | 0.3 | 0.23 | 0.35 |
| 3 | 5093704 | rs2570014 | -0.78 | -0.274 | 0.18 | -0.61 | 0.015 | -0.125 | -0.125 | 0.281 | 0.25 | 0.54 | 0.17 | 0.04 | 0.26 | 0.27 |
| 3 | 5093821 | rs2606768 | 0.123 | -0.194 | -0.107 | -0.573 | -0.055 | 0.091 | 0.091 | 0 | 0.18 | 0.05 | 0.13 | 0.03 | 0.24 | 0.16 |
| 3 | 5093858 | rs17791096 | 0.043 | -0.136 | 0.013 | -0.43 | -0.015 | 0.089 | 0.089 | 0.663 | 0.48 | 0.16 | 0.48 | 0.34 | 0.49 | 0.51 |
| 3 | 5097993 | rs6442929 | 0.01 | -0.26 | -0.179 | -0.629 | 0.022 | 0.025 | 0.025 | 0 | 0.43 | 0.61 | 0.33 | 0.23 | 0.31 | 0.47 |
| 3 | 5098705 | rs6773089 | -0.051 | -0.145 | -0.502 | -0.694 | 0.054 | 0.067 | 0.067 | 0 | 0.45 | 0.64 | 0.34 | 0.23 | 0.32 | 0.47 |
| 3 | 5099122 | rs6806899 | -0.214 | -0.229 | -0.49 | -0.682 | 0.054 | 0.063 | 0.063 | 0 | 0.45 | 0.64 | 0.34 | 0.23 | 0.32 | 0.47 |
| 3 | 5100291 | rs17734825 | 0.377 | 0.212 | -1.227 | -0.322 | 0.067 | 0.066 | 0.066 | 0 | 0.35 | 0.11 | 0.29 | 0.22 | 0.29 | 0.37 |
| 3 | 5102235 | rs980730 | 0.356 | 0.319 | -1.133 | -0.26 | 0.027 | 0.055 | 0.055 | 0 | 0.35 | 0.18 | 0.31 | 0.22 | 0.32 | 0.37 |
| 3 | 5103712 | rs6780573 | 0.345 | 0.405 | -1.391 | -0.222 | 0.027 | 0.054 | 0.054 | 0 | 0.36 | 0.19 | 0.31 | 0.22 | 0.32 | 0.37 |
| 3 | 5107167 | rs2570008 | -0.128 | 0.716 | -1.538 | 0.148 | -0.044 | 0.093 | 0.093 | 1.168 | 0.54 | 0.18 | 0.43 | 0.29 | 0.55 | 0.59 |
| 3 | 5115253 | rs9828404 | -1.038 | 0.769 | -1.078 | -0.033 | 0 | 0.002 | 0.002 | 0 | 0.13 | 0.07 | 0.15 | 0 | 0.11 | 0.11 |
| 3 | 5115389 | rs9867418 | -1.431 | 1.738 | -0.512 | 0.764 | 0.051 | 0.066 | 0.066 | 0 | 0.12 | 0.26 | 0.23 | 0.18 | 0.18 | 0.18 |
| 3 | 5126391 | rs925665 | 0.38 | 1.586 | -1.073 | 0.8 | -0.015 | -0.054 | -0.054 | 0 | 0.3 | 0.12 | 0.27 | 0.52 | 0.31 | 0.28 |
| 3 | 5129769 | rs11927981 | -0.66 | 1.513 | -0.12 | 0.904 | 0.099 | 0.136 | 0.136 | 1.352 | 0.11 | 0.49 | 0.33 | 0.18 | 0.16 | 0.17 |
| 3 | 5131363 | rs3864053 | 0.712 | 1.243 | -0.64 | 0.523 | -0.028 | 0.036 | 0.036 | 0.161 | 0.32 | 0.07 | 0.27 | 0.18 | 0.31 | 0.39 |
| 3 | 5131948 | rs4685865 | -1.017 | 1 | -0.376 | 0.399 | 0.003 | 0.005 | 0.005 | 0 | 0.77 | 0.74 | 0.87 | 0.88 | 0.8 | 0.88 |
| 3 | 5134494 | rs4685867 | 0.14 | 1.002 | 0.066 | 0.496 | 0 | 0.009 | 0.009 | 0.006 | 0.65 | 0.2 | 0.55 | 0.72 | 0.64 | 0.72 |
| 3 | 5135111 | rs2671762 | 0.095 | 1.14 | -0.027 | 0.589 | 0.044 | 0.029 | 0.029 | 0 | 0.91 | 0.8 | 0.82 | 0.91 | 0.84 | 0.84 |
| 3 | 5135571 | rs3864055 | 0.662 | 1.191 | -1.005 | 0.763 | -0.042 | -0.001 | -0.001 | 0 | 0.29 | 0.11 | 0.21 | 0.26 | 0.3 | 0.34 |
| 3 | 5136564 | rs17042007 | 0.424 | 1.589 | -0.515 | 1.27 | -0.095 | -0.146 | -0.146 | 0 | 0.28 | 0.01 | 0.34 | 0.44 | 0.32 | 0.35 |
| 3 | 5140364 | rs2671758 | 0.21 | 1.036 | 0.087 | 1.027 | -0.003 | 0.002 | 0.002 | 2.128 | 0.58 | 0.39 | 0.36 | 0.34 | 0.51 | 0.47 |
| 3 | 5147044 | rs2600100 | 1.352 | 0.885 | 0.54 | 1.114 | 0.059 | 0.085 | 0.085 | 2.012 | 0.92 | 0.59 | 0.87 | 0.91 | 0.92 | 0.82 |
| 3 | 5164769 | rs6772815 | -1.545 | 0.578 | 0.361 | 1.243 | -0.008 | 0.076 | 0.076 | 0 | 0.06 | 0.19 | 0.13 | 0.14 | 0.03 | 0.08 |
| 3 | 5167618 | rs4499578 | -0.858 | 0.625 | 0.288 | 1.003 | -0.003 | 0.002 | 0.002 | 0 | 0.26 | 0.58 | 0.3 | 0.31 | 0.29 | 0.28 |
| 3 | 5174332 | rs13072633 | -1.199 | 0.667 | 0.191 | 0.875 | -0.002 | -0.002 | -0.002 | 0 | 0.28 | 0.61 | 0.32 | 0.31 | 0.32 | 0.31 |

| | | | | | | | | | | | | | | | | |
|---|---|---|---|---|---|---|---|---|---|---|---|---|---|---|---|---|
| 3 | 5191756 | rs11707685 | -1.161 | 0.768 | 0.655 | 0.909 | 0 | 0 | 0 | 0 | 0.28 | 0.67 | 0.33 | 0.32 | 0.33 | 0.31 |
| 3 | 5193135 | rs4054851 | -0.589 | 0.702 | 1.584 | 0.861 | -0.002 | -0.022 | -0.022 | 0.012 | 0.17 | 0.27 | 0.22 | 0.15 | 0.17 | 0.18 |
| 3 | 5195655 | rs10428151 | -1.537 | 0.71 | 1.319 | 0.866 | 0 | -0.001 | -0.001 | 0.701 | 0.26 | 0.65 | 0.31 | 0.32 | 0.29 | 0.28 |
| 3 | 5197539 | rs11925441 | -1.533 | 0.734 | 1.156 | 0.501 | -0.003 | -0.011 | -0.011 | 0 | 0.28 | 0.6 | 0.32 | 0.2 | 0.3 | 0.19 |
| 3 | 5203776 | rs11130291 | -1.557 | 0.722 | 1.035 | 0.567 | -0.001 | -0.001 | -0.001 | 0 | 0.27 | 0.65 | 0.33 | 0.32 | 0.33 | 0.31 |
| 3 | 5232074 | rs343419 | 2.046 | 0.661 | 1.134 | 0.474 | -0.004 | -0.006 | -0.006 | 0 | 0.86 | 0.32 | 0.9 | 0.87 | 0.86 | 0.75 |
| 3 | 5245146 | rs341981 | 0.15 | 0.802 | -0.233 | 0.567 | -0.087 | -0.145 | -0.145 | 0 | 0.35 | 0.05 | 0.31 | 0.46 | 0.36 | 0.33 |
| 3 | 5256169 | rs341987 | 1.392 | 1.074 | 0.092 | 0.571 | -0.002 | 0.017 | 0.017 | 0 | 0.68 | 0.33 | 0.76 | 0.79 | 0.65 | 0.64 |
| 3 | 5258431 | rs370476 | 0.003 | 1.175 | 0.263 | 0.534 | 0.007 | -0.028 | -0.028 | 0 | 0.83 | 0.76 | 0.86 | 0.94 | 0.81 | 0.8 |
| 3 | 5260130 | rs461077 | 1.335 | 0.863 | 0.646 | 0.685 | 0.058 | 0.112 | 0.112 | 0.102 | 0.95 | 0.63 | 0.96 | 0.94 | 0.96 | 0.85 |
| 3 | 5260299 | rs464392 | 0.594 | 0.959 | 0.146 | 0.674 | 0.002 | -0.002 | -0.002 | 0 | 0.59 | 0.48 | 0.56 | 0.6 | 0.52 | 0.67 |
| 3 | 5267251 | rs463156 | 0.103 | 0.711 | -0.651 | 0.78 | -0.016 | -0.046 | -0.046 | 0 | 0.63 | 0.75 | 0.63 | 0.47 | 0.58 | 0.54 |
| 3 | 5268162 | rs341973 | 0.447 | 0.649 | -0.366 | 0.69 | -0.006 | 0.021 | 0.021 | 0.045 | 0.53 | 0.68 | 0.59 | 0.39 | 0.48 | 0.42 |
| 3 | 5273247 | rs341967 | 0.715 | 0.38 | -0.62 | 1.101 | 0.001 | -0.027 | -0.027 | 0.227 | 0.75 | 0.8 | 0.73 | 0.52 | 0.77 | 0.63 |
| 3 | 5273528 | rs3846158 | 1.576 | -0.064 | -1.136 | 0.707 | 0.051 | 0.068 | 0.068 | 0.005 | 0.54 | 0.18 | 0.61 | 0.37 | 0.63 | 0.28 |
| 3 | 5277162 | rs458381 | -1.348 | 0.147 | -0.253 | 0.707 | -0.022 | 0.027 | 0.027 | 1.824 | 0.15 | 0.57 | 0.09 | 0.17 | 0.1 | 0.29 |
| 3 | 5280161 | rs6793708 | 1.033 | -0.068 | -0.789 | 0.567 | 0.02 | 0.008 | 0.008 | 0 | 0.57 | 0.67 | 0.55 | 0.42 | 0.65 | 0.35 |
| 3 | 5280271 | rs6778534 | 1.254 | -0.101 | -0.562 | 0.394 | 0.014 | 0.034 | 0.034 | 0.094 | 0.5 | 0.58 | 0.49 | 0.39 | 0.55 | 0.26 |
| 3 | 5285286 | rs7373728 | 1.309 | -0.913 | 0.602 | 0.595 | 0.002 | -0.001 | -0.001 | 0.02 | 0.55 | 0.39 | 0.51 | 0.44 | 0.65 | 0.43 |
| 3 | 5287207 | rs9822748 | 1.323 | -0.928 | 0.681 | 0.592 | 0.022 | -0.002 | -0.002 | 0 | 0.55 | 0.32 | 0.51 | 0.44 | 0.66 | 0.43 |
| 3 | 5289558 | rs9833854 | 1.32 | -0.924 | 0.776 | 0.588 | 0.017 | -0.001 | -0.001 | 0 | 0.55 | 0.35 | 0.52 | 0.43 | 0.66 | 0.44 |
| 3 | 5290450 | rs466718 | 0.562 | -1.83 | 0.911 | -0.051 | -0.081 | -0.092 | -0.092 | 0 | 0.8 | 0.6 | 0.9 | 0.95 | 0.92 | 0.74 |
| 3 | 5302911 | rs6775103 | 0.978 | -1.807 | 0.821 | -0.084 | -0.015 | -0.052 | -0.052 | 0.214 | 0.76 | 0.67 | 0.85 | 0.93 | 0.83 | 0.67 |
| 3 | 5305473 | rs6776517 | 2.046 | -1.853 | 0.657 | 0.098 | 0.059 | 0.088 | 0.088 | 0.012 | 0.52 | 0.24 | 0.39 | 0.31 | 0.59 | 0.46 |
| 3 | 5307598 | rs342003 | -1.479 | -1.876 | 1.144 | 0.568 | -0.042 | -0.019 | -0.019 | 0 | 0.26 | 0.53 | 0.15 | 0.19 | 0.14 | 0.19 |
| 3 | 5312162 | rs12485270 | -0.759 | -1.935 | 0.944 | 0.609 | 0 | -0.048 | -0.048 | 0 | 0.19 | 0.13 | 0.37 | 0.44 | 0.19 | 0.24 |
| 3 | 5312819 | rs342007 | 2.002 | -1.96 | 1.045 | 0.557 | 0.048 | 0.03 | 0.03 | 0 | 0.56 | 0.35 | 0.48 | 0.34 | 0.68 | 0.58 |
| 3 | 5314243 | rs12497785 | -0.669 | -1.616 | 0.819 | 0.551 | -0.005 | -0.021 | -0.021 | 0 | 0.18 | 0.15 | 0.39 | 0.48 | 0.21 | 0.25 |
| 3 | 5316226 | rs7638016 | 0.377 | -1.819 | 1.26 | 0.46 | -0.029 | 0.054 | 0.054 | 0 | 0.07 | 0.17 | 0.01 | 0.16 | 0.01 | 0.09 |
| 3 | 5317068 | rs11130309 | 1.463 | -1.815 | 1.282 | 0.25 | -0.014 | -0.045 | -0.045 | 0.017 | 0.65 | 0.54 | 0.7 | 0.82 | 0.68 | 0.59 |
| 3 | 5318004 | rs11708761 | 1.323 | -1.798 | 0.987 | 0.242 | -0.004 | -0.014 | -0.014 | 1.576 | 0.65 | 0.42 | 0.69 | 0.82 | 0.67 | 0.59 |
| 3 | 5319430 | rs1874972 | 1.915 | -1.946 | 0.893 | 0.252 | 0.061 | 0.039 | 0.039 | 0 | 0.55 | 0.24 | 0.67 | 0.66 | 0.65 | 0.5 |
| 3 | 5327270 | rs924358 | 1.564 | -1.791 | 0.932 | 0.225 | 0.003 | 0.006 | 0.006 | 0 | 0.63 | 0.37 | 0.68 | 0.82 | 0.65 | 0.59 |
| 3 | 5332265 | rs9311480 | 1.677 | -1.854 | 0.647 | 0.418 | 0.053 | 0.041 | 0.041 | 0 | 0.55 | 0.16 | 0.61 | 0.68 | 0.64 | 0.49 |
| 3 | 5340428 | rs343401 | 1.218 | -2.057 | 1.328 | 0.321 | -0.04 | -0.042 | -0.042 | 0 | 0.69 | 0.55 | 0.75 | 0.83 | 0.79 | 0.69 |
| 3 | 5343801 | rs1995205 | 2.633 | -2.254 | 0.937 | 0.149 | -0.083 | 0.275 | 0.275 | 0 | 0.41 | 0.07 | 0.28 | 0.04 | 0.5 | 0.28 |
| 3 | 5345166 | rs11130330 | 2.011 | -2.484 | 1.098 | 0.296 | 0.023 | 0.048 | 0.048 | 0 | 0.57 | 0.34 | 0.37 | 0.24 | 0.59 | 0.39 |
| 3 | 5345191 | rs595615 | -0.61 | -2.199 | 1.01 | 0.422 | -0.026 | -0.027 | -0.027 | 2.351 | 0.06 | 0 | 0.05 | 0.1 | 0.06 | 0.11 |
| 3 | 5348688 | rs17042217 | -0.105 | -2.187 | 1.175 | 0.395 | 0.014 | 0.014 | 0.014 | 0 | 0.14 | 0.08 | 0.1 | 0.07 | 0.08 | 0.1 |
| 3 | 5348717 | rs17042221 | -0.497 | -2.213 | 1.265 | 0.374 | 0.016 | -0.006 | -0.006 | 0.042 | 0.13 | 0.08 | 0.13 | 0.13 | 0.11 | 0.16 |
| 3 | 5349208 | rs6803503 | -0.548 | -2.091 | 1.716 | 0.442 | -0.009 | 0.01 | 0.01 | 0 | 0.12 | 0.25 | 0.13 | 0.14 | 0.11 | 0.18 |
| 3 | 5349336 | rs6803605 | 1.033 | -2.073 | 1.707 | 0.583 | 0.003 | -0.005 | -0.005 | 0.122 | 0.7 | 0.71 | 0.56 | 0.34 | 0.75 | 0.63 |
| 3 | 5352849 | rs13068773 | 2.199 | -2.178 | 1.024 | 0.466 | 0.01 | 0.134 | 0.134 | 0 | 0.45 | 0.26 | 0.28 | 0.18 | 0.51 | 0.29 |
| 3 | 5355556 | rs6768764 | 1.754 | -2.182 | 0.907 | 0.271 | 0.071 | 0.021 | 0.021 | 0 | 0.54 | 0.27 | 0.62 | 0.62 | 0.65 | 0.55 |
| 3 | 5357532 | rs398708 | 0.316 | -1.826 | 1.289 | 0.431 | -0.015 | 0.014 | 0.014 | 0 | 0.89 | 0.76 | 0.92 | 0.85 | 0.9 | 0.82 |
| 3 | 5362294 | rs343431 | 1.342 | -1.867 | 1.118 | 0.396 | -0.029 | 0.022 | 0.022 | 0 | 0.72 | 0.59 | 0.75 | 0.66 | 0.78 | 0.63 |
| 3 | 5365063 | rs4684476 | 1.335 | -1.688 | 0.594 | 0.489 | 0.015 | -0.01 | -0.01 | 0 | 0.71 | 0.3 | 0.73 | 0.66 | 0.78 | 0.64 |
| 3 | 5367819 | rs392405 | -0.853 | -1.375 | 0.672 | 0.76 | -0.038 | 0.057 | 0.057 | 0 | 0.19 | 0.41 | 0.19 | 0.21 | 0.12 | 0.19 |
| 3 | 5368801 | rs2322977 | -0.909 | -1.049 | -0.039 | 0.848 | -0.013 | 0.016 | 0.016 | 0 | 0.33 | 0.58 | 0.41 | 0.57 | 0.29 | 0.35 |
| 3 | 5373391 | rs1114430 | 0.888 | -1.46 | 0.227 | 0.575 | -0.049 | -0.04 | -0.04 | 0 | 0.75 | 0.52 | 0.83 | 0.82 | 0.84 | 0.88 |
| 3 | 5375206 | rs7629289 | 1.094 | -2.92 | -0.145 | 0.98 | 0.097 | -0.013 | -0.013 | 0 | 0.55 | 0.26 | 0.58 | 0.46 | 0.71 | 0.45 |
| 3 | 5377364 | rs9871505 | 0.656 | -3.082 | 0.484 | 1.24 | -0.031 | 0.094 | 0.094 | 0 | 0.78 | 0.54 | 0.73 | 0.61 | 0.82 | 0.64 |
| 3 | 5380959 | rs9836671 | -0.097 | -2.849 | 0.742 | 1.227 | 0.01 | 0.034 | 0.034 | 0 | 0.07 | 0.23 | 0.07 | 0.19 | 0.09 | 0.2 |
| 3 | 5391554 | rs6794390 | 1.179 | -2.707 | 0.209 | 1.178 | -0.001 | -0.002 | -0.002 | 0 | 0.46 | 0.47 | 0.42 | 0.34 | 0.57 | 0.26 |
| 3 | 5400168 | rs4234565 | -0.61 | -2.871 | -0.378 | 1.656 | -0.002 | 0.009 | 0.009 | 0 | 0.14 | 0.2 | 0.34 | 0.21 | 0.11 | 0.2 |
| 3 | 5403935 | rs9851025 | 1.559 | -2.852 | -0.871 | 1.53 | -0.043 | 0.014 | 0.014 | 2.378 | 0.42 | 0.04 | 0.39 | 0.33 | 0.58 | 0.25 |
| 3 | 5410364 | rs4143244 | 0.321 | -2.727 | -0.575 | 1.418 | -0.003 | 0.001 | 0.001 | 5.945 | 0.62 | 0.6 | 0.72 | 0.46 | 0.7 | 0.45 |
| 3 | 5411891 | rs13075306 | 1.119 | -2.398 | -1.265 | 1.3 | -0.026 | 0.039 | 0.039 | 2.7 | 0.44 | 0.03 | 0.4 | 0.28 | 0.59 | 0.27 |
| 3 | 5420067 | rs17042380 | -0.843 | -1.751 | -0.398 | 1.746 | -0.014 | 0.008 | 0.008 | 0.054 | 0.1 | 0.21 | 0.05 | 0.13 | 0.07 | 0.12 |
| 3 | 5420175 | rs7649517 | -2.005 | -1.452 | -0.314 | 2.31 | -0.003 | 0.031 | 0.031 | 0 | 0.31 | 0.41 | 0.35 | 0.47 | 0.25 | 0.35 |
| 3 | 5426928 | rs17042385 | -0.235 | -1.416 | -0.444 | 2.239 | -0.001 | 0.001 | 0.001 | 0 | 0.08 | 0.06 | 0.04 | 0.06 | 0.06 | 0.07 |
| 3 | 5427679 | rs9842582 | -0.427 | -1.51 | -0.474 | 2.347 | 0 | 0 | 0 | 0.522 | 0.09 | 0.13 | 0.06 | 0.07 | 0.08 | 0.07 |
| 3 | 5428371 | rs4685917 | -1.372 | -1.43 | -0.127 | 2.572 | -0.026 | 0.046 | 0.046 | 0 | 0.23 | 0.46 | 0.27 | 0.34 | 0.2 | 0.44 |
| 3 | 5430579 | rs9830212 | -2.329 | -1.474 | -0.206 | 2.563 | -0.005 | 0.003 | 0.003 | 0 | 0.36 | 0.47 | 0.52 | 0.58 | 0.33 | 0.59 |
| 3 | 5432114 | rs9311513 | -1.64 | -1.541 | -0.356 | 2.327 | 0.002 | -0.013 | -0.013 | 0 | 0.23 | 0.22 | 0.27 | 0.29 | 0.2 | 0.26 |
| 3 | 5432511 | rs17042413 | -1.629 | -1.539 | -0.355 | 2.321 | 0.002 | -0.013 | -0.013 | 0.668 | 0.23 | 0.22 | 0.27 | 0.29 | 0.2 | 0.26 |
| 3 | 5434207 | rs9855025 | -1.151 | -1.374 | -0.203 | 2.275 | -0.003 | 0.013 | 0.013 | 0 | 0.17 | 0.13 | 0.1 | 0.09 | 0.14 | 0.12 |
| 3 | 5434345 | rs4685920 | -1.68 | -1.508 | -0.083 | 2.341 | -0.014 | 0.037 | 0.037 | 0 | 0.29 | 0.43 | 0.46 | 0.45 | 0.24 | 0.47 |
| 3 | 5434964 | rs13325114 | -1.474 | -1.501 | -0.082 | 2.173 | -0.015 | 0.035 | 0.035 | 0.948 | 0.14 | 0.24 | 0.12 | 0.26 | 0.07 | 0.1 |

| | | | | | | | | | | | | | | | | |
|---|---|---|---|---|---|---|---|---|---|---|---|---|---|---|---|---|
| 3 | 5436077 | rs13319992 | -1.031 | -0.833 | -0.058 | 1.952 | -0.022 | 0.048 | 0.048 | 0 | 0.24 | 0.3 | 0.11 | 0.34 | 0.17 | 0.32 |
| 3 | 5436345 | rs4425237 | 0.09 | -0.65 | -0.547 | 1.631 | -0.033 | 0.023 | 0.023 | 0 | 0.14 | 0.01 | 0.14 | 0.1 | 0.13 | 0.11 |
| 3 | 5438374 | rs4452317 | -0.061 | -0.643 | -0.492 | 1.535 | -0.022 | 0.002 | 0.002 | 0 | 0.14 | 0.02 | 0.07 | 0.12 | 0.13 | 0.12 |
| 3 | 5440984 | rs7624105 | 0.176 | -0.64 | -0.496 | 1.537 | -0.03 | -0.004 | -0.004 | 0.146 | 0.14 | 0.01 | 0.07 | 0.12 | 0.14 | 0.12 |
| 3 | 5442857 | rs7635144 | -1.644 | -0.494 | -0.304 | 1.43 | -0.007 | 0.03 | 0.03 | 0 | 0.56 | 0.56 | 0.56 | 0.66 | 0.5 | 0.77 |
| 3 | 5447838 | rs1403067 | 0.946 | -0.565 | -0.236 | 1.638 | -0.006 | -0.003 | -0.003 | 0 | 0.6 | 0.42 | 0.8 | 0.66 | 0.69 | 0.59 |
| 3 | 5449157 | rs4685929 | -0.063 | -0.631 | -0.25 | 1.582 | 0.001 | 0.053 | 0.053 | 0 | 0.23 | 0.3 | 0.41 | 0.35 | 0.24 | 0.38 |
| 3 | 5450470 | rs9880190 | 0.927 | -0.514 | -0.129 | 1.417 | -0.004 | -0.005 | -0.005 | 0 | 0.61 | 0.51 | 0.81 | 0.69 | 0.69 | 0.6 |
| 3 | 5461226 | rs7653222 | 1.345 | -0.808 | -0.376 | 1.32 | -0.025 | 0.031 | 0.031 | 0 | 0.41 | 0.27 | 0.35 | 0.35 | 0.49 | 0.22 |
| 3 | 5461734 | rs6810210 | -0.02 | -0.921 | -0.195 | 1.387 | 0.036 | 0.242 | 0.242 | 0.654 | 0.05 | 0.32 | 0.06 | 0.2 | 0.03 | 0.24 |
| 3 | 5462347 | rs10510305 | 0.294 | -0.963 | -1.037 | 1.084 | 0.001 | -0.005 | -0.005 | 2.609 | 0.18 | 0.02 | 0.05 | 0.12 | 0.14 | 0.23 |
| 3 | 5462541 | rs4684484 | 1.366 | -0.775 | -0.917 | 0.871 | -0.072 | 0.024 | 0.024 | 0 | 0.38 | 0.13 | 0.37 | 0.34 | 0.48 | 0.21 |
| 3 | 5468243 | rs2875002 | -1.466 | -0.249 | -1.009 | 0.411 | -0.001 | 0 | 0 | 0.073 | 0.61 | 0.65 | 0.59 | 0.52 | 0.55 | 0.65 |
| 3 | 5469862 | rs6798021 | -1.166 | -0.126 | -0.845 | 0.368 | -0.006 | -0.006 | -0.006 | 0 | 0.56 | 0.65 | 0.56 | 0.49 | 0.49 | 0.61 |
| 3 | 5470493 | rs10428066 | 0.306 | -0.1 | -1.204 | 0.413 | -0.028 | -0.07 | -0.07 | 0 | 0.2 | 0.01 | 0.12 | 0.21 | 0.19 | 0.28 |
| 3 | 5475904 | rs4600807 | 0.546 | -0.734 | -1.131 | 0.509 | 0.011 | -0.075 | -0.075 | 1.105 | 0.18 | 0 | 0.1 | 0.18 | 0.11 | 0.27 |
| 3 | 5476231 | rs6786147 | 0.723 | -0.741 | -1.029 | 0.379 | -0.003 | 0.002 | 0.002 | 0 | 0.17 | 0.11 | 0.23 | 0.16 | 0.17 | 0.2 |
| 3 | 5478213 | rs4132225 | -1.506 | -0.781 | -0.916 | 0.34 | -0.007 | -0.01 | -0.01 | 0 | 0.55 | 0.42 | 0.45 | 0.47 | 0.46 | 0.62 |
| 3 | 5479366 | rs17803128 | -0.679 | -0.823 | -1.084 | 0.406 | 0.056 | -0.037 | -0.037 | 3.25 | 0.34 | 0.14 | 0.18 | 0.31 | 0.24 | 0.39 |
| 3 | 5490708 | rs11130418 | -1.441 | -0.846 | -1.088 | 0.143 | 0.001 | -0.006 | -0.006 | 0 | 0.55 | 0.64 | 0.53 | 0.48 | 0.43 | 0.62 |
| 3 | 5491322 | rs17803320 | 0.498 | -0.177 | -1.739 | 0.476 | -0.007 | 0.01 | 0.01 | 0 | 0.18 | 0.11 | 0.23 | 0.12 | 0.19 | 0.22 |
| 3 | 5492484 | rs17746899 | 0.377 | -0.226 | -1.847 | 0.572 | 0.015 | -0.066 | -0.066 | 0 | 0.18 | 0.01 | 0.1 | 0.22 | 0.12 | 0.28 |
| 3 | 5493089 | rs17042584 | -0.508 | 0.532 | -0.964 | 0.335 | 0.004 | 0.001 | 0.001 | 0 | 0.25 | 0.31 | 0.37 | 0.27 | 0.26 | 0.27 |
| 3 | 5493389 | rs17042585 | -0.594 | 0.351 | -1.066 | 0.684 | 0.004 | -0.001 | -0.001 | 0 | 0.12 | 0.1 | 0.08 | 0.12 | 0.11 | 0.12 |
| 3 | 5497476 | rs6773522 | 0.12 | 0.671 | -1.009 | 0.506 | 0 | -0.002 | -0.002 | 0.01 | 0.47 | 0.56 | 0.49 | 0.68 | 0.49 | 0.48 |
| 3 | 5497517 | rs9858005 | -0.091 | 0.68 | -1.076 | -0.014 | -0.006 | 0.072 | 0.072 | 0 | 0.35 | 0.21 | 0.39 | 0.15 | 0.33 | 0.34 |
| 3 | 5500322 | rs7621563 | -0.699 | 0.135 | -0.731 | -0.04 | 0.001 | 0.001 | 0.001 | 0 | 0.43 | 0.61 | 0.36 | 0.72 | 0.34 | 0.61 |
| 3 | 5501375 | rs9850819 | -0.066 | -0.129 | -0.615 | 0.756 | 0.013 | -0.039 | -0.039 | 1.19 | 0.27 | 0.16 | 0.25 | 0.38 | 0.23 | 0.36 |
| 3 | 5506706 | rs6775400 | -1.175 | -0.242 | -0.226 | 1.069 | -0.024 | 0.094 | 0.094 | 0.109 | 0.2 | 0.6 | 0.28 | 0.44 | 0.16 | 0.32 |
| 3 | 5508001 | rs7627560 | 0.308 | -0.258 | -0.079 | 1.143 | -0.001 | -0.009 | -0.009 | 0 | 0.11 | 0.14 | 0.12 | 0.09 | 0.09 | 0.14 |
| 3 | 5511975 | rs2565614 | 1.636 | -0.718 | 0.217 | 0.875 | -0.002 | 0.092 | 0.092 | 0.913 | 0.85 | 0.52 | 0.9 | 0.75 | 0.89 | 0.83 |
| 3 | 5512154 | rs9880886 | 1.313 | -0.864 | -0.052 | 1.082 | 0.008 | -0.027 | -0.027 | 0 | 0.16 | 0.06 | 0.08 | 0.17 | 0.12 | 0.25 |
| 3 | 5512860 | rs4685941 | -0.839 | -0.442 | -0.002 | 0.873 | 0.004 | 0.042 | 0.042 | 0 | 0.06 | 0.18 | 0.22 | 0.11 | 0.07 | 0.06 |
| 3 | 5515770 | rs13097571 | 1.18 | 0.103 | -0.899 | 0.102 | -0.134 | 0.037 | 0.037 | 0 | 0.22 | 0.02 | 0.25 | 0.1 | 0.32 | 0.1 |
| 3 | 5515986 | rs9809817 | 1.068 | 0.409 | -0.24 | 0.498 | 0.014 | -0.064 | -0.064 | 0 | 0.17 | 0.11 | 0.15 | 0.3 | 0.13 | 0.26 |
| 3 | 5518671 | rs9311538 | -0.722 | 0.446 | 0.096 | 0.452 | -0.001 | 0.001 | 0.001 | 0 | 0.35 | 0.44 | 0.3 | 0.4 | 0.28 | 0.48 |
| 3 | 5522945 | rs2565612 | -0.641 | 0.898 | -0.036 | 0.205 | 0 | -0.064 | -0.064 | 0.167 | 0.81 | 0.79 | 0.88 | 1 | 0.8 | 0.92 |
| 3 | 5525968 | rs2686534 | 0.721 | 1.06 | -0.281 | 0.443 | -0.003 | -0.019 | -0.019 | 0 | 0.43 | 0.47 | 0.59 | 0.72 | 0.42 | 0.57 |
| 3 | 5528884 | rs6442950 | -1.725 | 0.783 | 0.667 | 1.518 | -0.011 | 0.006 | 0.006 | 0.897 | 0.53 | 0.66 | 0.43 | 0.58 | 0.49 | 0.62 |
| 3 | 5532378 | rs17042689 | -0.524 | -0.433 | 0.91 | 2.128 | -0.003 | 0.017 | 0.017 | 0.409 | 0.24 | 0.24 | 0.17 | 0.49 | 0.2 | 0.41 |
| 3 | 5533023 | rs941061 | -1.674 | 0.022 | 0.699 | 1.799 | -0.001 | 0.027 | 0.027 | 0 | 0.22 | 0.6 | 0.22 | 0.34 | 0.2 | 0.28 |
| 3 | 5540508 | rs1429997 | -1.631 | 0.23 | 0.21 | 1.074 | 0.009 | -0.059 | -0.059 | 0 | 0.23 | 0.42 | 0.18 | 0.1 | 0.23 | 0.24 |
| 3 | 5541735 | rs9311542 | 0.704 | -0.115 | 0.183 | 1.615 | -0.006 | 0.029 | 0.029 | 0 | 0.43 | 0.25 | 0.49 | 0.31 | 0.47 | 0.26 |
| 3 | 5542079 | rs4143287 | -0.891 | -0.114 | 0.364 | 1.646 | -0.001 | 0.012 | 0.012 | 0 | 0.65 | 0.64 | 0.65 | 0.52 | 0.68 | 0.52 |
| 3 | 5542369 | rs4143286 | -0.222 | -0.106 | 0.403 | 1.533 | -0.006 | 0.018 | 0.018 | 0 | 0.08 | 0.12 | 0.02 | 0.15 | 0.05 | 0.11 |
| 3 | 5542461 | rs13098133 | -1.401 | -0.062 | 0.272 | 1.55 | 0.002 | -0.019 | -0.019 | 0 | 0.2 | 0.27 | 0.12 | 0.12 | 0.19 | 0.17 |
| 3 | 5543950 | rs9874195 | 0.574 | -0.081 | 0.202 | 1.574 | -0.031 | 0.02 | 0.02 | 0 | 0.47 | 0.22 | 0.53 | 0.39 | 0.52 | 0.32 |
| 3 | 5544395 | rs882507 | -0.305 | -0.044 | 0.25 | 1.609 | -0.001 | -0.001 | -0.001 | 0.532 | 0.58 | 0.44 | 0.6 | 0.43 | 0.62 | 0.48 |
| 3 | 5548037 | rs12715476 | 0.188 | -0.281 | 0.048 | 1.138 | 0.013 | 0.023 | 0.023 | 0 | 0.5 | 0.42 | 0.54 | 0.39 | 0.56 | 0.33 |
| 3 | 5549546 | rs7645616 | 0.852 | 0.316 | 0.181 | 1.325 | -0.047 | 0.012 | 0.012 | 0 | 0.41 | 0.15 | 0.49 | 0.32 | 0.49 | 0.26 |
| 3 | 5549658 | rs7611671 | -0.519 | 0.315 | 0.182 | 1.328 | -0.001 | 0 | 0 | 0 | 0.58 | 0.47 | 0.6 | 0.36 | 0.62 | 0.41 |
| 3 | 5551114 | rs1367141 | -0.187 | 0.323 | 0.033 | 1.372 | 0.001 | 0.006 | 0.006 | 0 | 0.53 | 0.46 | 0.57 | 0.37 | 0.6 | 0.39 |
| 3 | 5561429 | rs17042804 | -0.383 | 0.394 | -0.114 | 1.305 | 0.002 | -0.024 | -0.024 | 0.37 | 0.06 | 0.1 | 0.04 | 0 | 0.05 | 0.04 |
| 3 | 5563371 | rs17749328 | -0.483 | 0.754 | -0.556 | 1.035 | 0.005 | -0.025 | -0.025 | 0 | 0.1 | 0.19 | 0.07 | 0.04 | 0.1 | 0.11 |
| 3 | 5564422 | rs9823228 | 0.15 | 0.771 | -1.172 | 1.237 | -0.031 | 0.05 | 0.05 | 0 | 0.46 | 0.24 | 0.52 | 0.32 | 0.53 | 0.32 |
| 3 | 5589870 | rs7633722 | -1.478 | 0.702 | 0.635 | 2.554 | 0.034 | 0.413 | 0.413 | 1.314 | 0.13 | 0.48 | 0.2 | 0.47 | 0.15 | 0.19 |
| 3 | 5592749 | rs1820920 | -1.244 | 0.776 | 0.44 | 2.355 | -0.01 | -0.01 | -0.01 | 0.374 | 0.06 | 0.06 | 0.07 | 0.12 | 0.13 | 0.14 |
| 3 | 5598666 | rs1430005 | 0.834 | 0.586 | 1.301 | 2.035 | -0.005 | 0.085 | 0.085 | 1.137 | 0.78 | 0.5 | 0.66 | 0.65 | 0.78 | 0.75 |
| 3 | 5599714 | rs1568996 | 0.682 | 0.595 | 1.628 | 1.956 | 0.011 | -0.069 | -0.069 | 0.226 | 0.87 | 0.64 | 0.88 | 0.99 | 0.84 | 0.97 |
| 3 | 5605274 | rs961245 | 2.248 | 1.177 | 2.506 | 1.844 | 0.023 | 0.051 | 0.051 | 0 | 0.77 | 0.46 | 0.63 | 0.65 | 0.73 | 0.72 |
| 3 | 5605715 | rs2033299 | 0.875 | 1.164 | 2.603 | 1.832 | 0.007 | -0.013 | -0.013 | 0 | 0.87 | 0.82 | 0.84 | 0.9 | 0.83 | 0.94 |
| 3 | 5607482 | rs1430012 | 0.454 | 1.258 | 2.565 | 1.825 | 0.005 | -0.026 | -0.026 | 0 | 0.89 | 0.86 | 0.89 | 0.99 | 0.82 | 0.97 |
| 3 | 5607784 | rs1430010 | 1.901 | 1.307 | 2.593 | 1.909 | 0.046 | 0.117 | 0.117 | 0.014 | 0.81 | 0.51 | 0.77 | 0.67 | 0.76 | 0.89 |
| 3 | 5610345 | rs6775123 | 0.668 | 1.627 | 2.255 | 1.75 | -0.029 | 0.042 | 0.042 | 0 | 0.89 | 0.96 | 0.88 | 0.99 | 0.81 | 0.95 |
| 3 | 5610543 | rs6787951 | 1.45 | 1.455 | 1.905 | 1.34 | -0.003 | -0.026 | -0.026 | 0 | 0.75 | 0.85 | 0.73 | 0.67 | 0.73 | 0.88 |
| 3 | 5615030 | rs6771379 | 2.011 | 0.727 | 2.264 | 0.809 | -0.011 | -0.093 | -0.093 | 1.43 | 0.63 | 0.53 | 0.77 | 0.88 | 0.67 | 0.85 |
| 3 | 5621655 | rs6776939 | 1.844 | 0.308 | 2.29 | 0.67 | -0.007 | -0.103 | -0.103 | 0.069 | 0.67 | 0.53 | 0.78 | 0.92 | 0.69 | 0.85 |
| 3 | 5625761 | rs6765805 | 1.946 | 0.146 | 1.961 | 1.12 | 0.004 | -0.009 | -0.009 | 0.708 | 0.55 | 0.36 | 0.54 | 0.51 | 0.5 | 0.47 |
| 3 | 5626851 | rs1864427 | 1.554 | 0.324 | 1.621 | 0.423 | -0.001 | 0.011 | 0.011 | 0 | 0.65 | 0.75 | 0.79 | 0.99 | 0.64 | 0.83 |

| | | | | | | | | | | | | | | | | |
|---|---|---|---|---|---|---|---|---|---|---|---|---|---|---|---|---|
| 3 | 5630590 | rs7628449 | 1.165 | 1.237 | 1.479 | 0.349 | -0.002 | -0.018 | -0.018 | 0 | 0.68 | 0.72 | 0.75 | 0.9 | 0.62 | 0.7 |
| 3 | 5631147 | rs4685951 | 1.275 | 1.356 | 0.56 | -0.148 | -0.054 | 0.033 | 0.033 | 0 | 0.19 | 0.05 | 0.23 | 0.15 | 0.22 | 0.18 |
| 3 | 5631528 | rs7615007 | 1.364 | 1.303 | 1.32 | 0.247 | 0.002 | -0.047 | -0.047 | 0 | 0.64 | 0.62 | 0.73 | 0.86 | 0.62 | 0.69 |
| 3 | 5631999 | rs1864429 | 1.056 | 1.433 | 1.344 | 0.32 | -0.018 | 0.046 | 0.046 | 0 | 0.68 | 0.81 | 0.74 | 0.86 | 0.62 | 0.7 |
| 3 | 5639540 | rs17042994 | 0.105 | 1.196 | 1.292 | 0.671 | 0.003 | -0.021 | -0.021 | 0 | 0.09 | 0 | 0.05 | 0.07 | 0.08 | 0.1 |
| 3 | 5641955 | rs4637282 | 1.546 | 1.474 | 1.603 | 0.539 | 0.015 | -0.037 | -0.037 | 0 | 0.71 | 0.55 | 0.75 | 0.78 | 0.69 | 0.7 |
| 3 | 5654382 | rs6786916 | 1.858 | 1.548 | 1.699 | 0.528 | 0.003 | -0.039 | -0.039 | 0 | 0.62 | 0.5 | 0.72 | 0.78 | 0.63 | 0.67 |
| 3 | 5661801 | rs2014115 | -2.569 | 2.072 | 1.937 | 1.062 | 0.064 | -0.016 | -0.016 | 0.942 | 0.25 | 0.49 | 0.27 | 0.24 | 0.33 | 0.35 |
| 3 | 5668014 | rs6442965 | 1.337 | 2.059 | 1.989 | 1.012 | 0.063 | -0.039 | -0.039 | 0 | 0.79 | 0.65 | 0.77 | 0.85 | 0.69 | 0.71 |
| 3 | 5669804 | rs7630031 | 0.97 | 2.058 | 1.922 | 1.019 | 0.016 | -0.011 | -0.011 | 0.004 | 0.81 | 0.83 | 0.79 | 0.85 | 0.7 | 0.72 |
| 3 | 5670133 | rs7630333 | 2.249 | 2.058 | 1.922 | 1.039 | 0.059 | -0.021 | -0.021 | 0 | 0.76 | 0.56 | 0.75 | 0.78 | 0.68 | 0.7 |
| 3 | 5674506 | rs1452709 | 1.207 | 2.096 | 1.88 | 0.823 | 0.015 | -0.012 | -0.012 | 0.229 | 0.8 | 0.8 | 0.79 | 0.85 | 0.7 | 0.67 |
| 3 | 5676938 | rs6776116 | 1.773 | 2.234 | 1.714 | 0.871 | 0.056 | -0.054 | -0.054 | 0 | 0.78 | 0.63 | 0.8 | 0.86 | 0.68 | 0.7 |
| 3 | 5679633 | rs996046 | 1.801 | 2.2 | 1.698 | 0.918 | 0.036 | -0.013 | -0.013 | 0.04 | 0.71 | 0.68 | 0.72 | 0.72 | 0.61 | 0.7 |
| 3 | 5686466 | rs11914737 | 1.769 | 2.293 | 1.69 | 0.886 | 0.033 | -0.056 | -0.056 | 0 | 0.72 | 0.6 | 0.77 | 0.84 | 0.65 | 0.71 |
| 3 | 5687674 | rs9284892 | -0.225 | 2.238 | 1.72 | 0.978 | -0.015 | 0.01 | 0.01 | 0.196 | 0.1 | 0.18 | 0.09 | 0.14 | 0.06 | 0.03 |
| 3 | 5688639 | rs7643083 | 0.945 | 2.282 | 1.736 | 0.966 | 0.053 | -0.019 | -0.019 | 0.258 | 0.82 | 0.75 | 0.81 | 0.85 | 0.67 | 0.73 |
| 3 | 5689472 | rs11130514 | 2.143 | 2.286 | 1.685 | 0.746 | 0.055 | -0.027 | -0.027 | 2.48 | 0.7 | 0.52 | 0.7 | 0.72 | 0.61 | 0.71 |
| 3 | 5692747 | rs6794753 | -1.193 | 1.922 | 1.591 | 0.766 | 0.117 | -0.012 | -0.012 | 0 | 0.17 | 0.31 | 0.2 | 0.17 | 0.31 | 0.25 |
| 3 | 5697751 | rs9854936 | -1.813 | 1.943 | 1.116 | 0.773 | 0.025 | -0.049 | -0.049 | 0 | 0.29 | 0.55 | 0.23 | 0.16 | 0.34 | 0.27 |
| 3 | 5700132 | rs9841751 | -1.968 | 1.952 | 0.991 | 0.779 | 0.011 | -0.026 | -0.026 | 0 | 0.3 | 0.58 | 0.24 | 0.17 | 0.35 | 0.27 |
| 3 | 5713757 | rs13100888 | -1.436 | 1.717 | 0.469 | 0.92 | 0.011 | -0.005 | -0.005 | 0 | 0.27 | 0.32 | 0.24 | 0.3 | 0.3 | 0.27 |
| 3 | 5714159 | rs4283565 | -1.618 | 1.768 | 0.515 | 1.081 | 0.008 | 0.004 | 0.004 | 0 | 0.28 | 0.33 | 0.27 | 0.37 | 0.32 | 0.27 |
| 3 | 5715540 | rs9876922 | -0.807 | 1.876 | 0.559 | 1.351 | 0.008 | -0.004 | -0.004 | 0 | 0.17 | 0.17 | 0.16 | 0.17 | 0.24 | 0.24 |
| 3 | 5716365 | rs6791910 | -1.717 | 1.563 | 0.913 | 1.166 | 0.003 | 0.009 | 0.009 | 0 | 0.33 | 0.52 | 0.31 | 0.42 | 0.32 | 0.29 |
| 3 | 5719861 | rs1545506 | -1.398 | 1.63 | 0.674 | 1.414 | 0.013 | -0.004 | -0.004 | 0 | 0.38 | 0.4 | 0.46 | 0.62 | 0.48 | 0.61 |
| 3 | 5727171 | rs6796220 | -1.725 | 1.417 | 1.165 | 2.322 | 0.009 | 0.082 | 0.082 | 0 | 0.18 | 0.35 | 0.37 | 0.63 | 0.2 | 0.22 |
| 3 | 5729325 | rs11717279 | -0.693 | 1.522 | 0.829 | 2.095 | 0.015 | 0.101 | 0.101 | 0 | 0.13 | 0.18 | 0.24 | 0.39 | 0.18 | 0.2 |
| 3 | 5730884 | rs7428299 | -1.635 | 1.785 | 0.663 | 1.938 | 0.005 | 0.019 | 0.019 | 0 | 0.19 | 0.18 | 0.25 | 0.36 | 0.26 | 0.26 |
| 3 | 5733933 | rs7612973 | -0.699 | 1.623 | 0.499 | 1.859 | -0.003 | 0.01 | 0.01 | 0 | 0.13 | 0.11 | 0.22 | 0.32 | 0.18 | 0.19 |
| 3 | 5752152 | rs6776169 | 0.232 | 0.987 | 0.763 | 0.901 | -0.01 | -0.047 | -0.047 | 0.035 | 0.1 | 0.17 | 0.03 | 0 | 0.06 | 0.01 |
| 3 | 5760464 | rs4143283 | -0.662 | 1.235 | 0.438 | 1.087 | -0.04 | -0.065 | -0.065 | 0 | 0.2 | 0.11 | 0.39 | 0.46 | 0.32 | 0.42 |
| 3 | 5773145 | rs9311642 | -1.531 | 0.864 | 1.052 | 0.988 | 0 | -0.001 | -0.001 | 0 | 0.37 | 0.66 | 0.54 | 0.66 | 0.42 | 0.42 |
| 3 | 5776857 | rs2470644 | -1.437 | 0.207 | 1.654 | 0.985 | -0.017 | -0.001 | -0.001 | 1.506 | 0.18 | 0.7 | 0.16 | 0.19 | 0.07 | 0.02 |
| 3 | 5777222 | rs1452713 | -0.301 | 0.255 | 1.873 | 1.004 | -0.038 | 0.035 | 0.035 | 1.324 | 0.13 | 0.29 | 0.11 | 0.19 | 0.06 | 0.02 |
| 3 | 5786854 | rs12636531 | -0.929 | 0.56 | 1.354 | 1.007 | -0.008 | -0.087 | -0.087 | 3.618 | 0.34 | 0.12 | 0.3 | 0.5 | 0.34 | 0.4 |
| 3 | 5787176 | rs12629173 | -1.759 | 0.779 | 1.802 | 0.824 | 0 | -0.007 | -0.007 | 0.233 | 0.48 | 0.46 | 0.5 | 0.81 | 0.51 | 0.66 |
| 3 | 5788862 | rs17043429 | -1.179 | 0.726 | 1.8 | 0.865 | 0.019 | 0.017 | 0.017 | 0.009 | 0.28 | 0.32 | 0.34 | 0.57 | 0.4 | 0.59 |
| 3 | 5789030 | rs1978772 | -2.177 | 0.681 | 2.037 | 0.833 | -0.005 | 0.171 | 0.171 | 0 | 0.51 | 0.77 | 0.52 | 0.81 | 0.51 | 0.66 |
| 3 | 5789520 | rs1836579 | -1.314 | 0.78 | 2.044 | 0.882 | 0.007 | 0.016 | 0.016 | 0 | 0.39 | 0.31 | 0.41 | 0.7 | 0.37 | 0.41 |
| 3 | 5789768 | rs1435361 | -0.66 | 0.783 | 2.017 | 0.89 | 0.041 | 0.081 | 0.081 | 0 | 0.13 | 0.23 | 0.18 | 0.3 | 0.17 | 0.26 |
| 3 | 5790183 | rs997500 | -1.811 | 0.777 | 2.065 | 0.88 | -0.009 | 0.033 | 0.033 | 0 | 0.47 | 0.6 | 0.42 | 0.61 | 0.48 | 0.64 |
| 3 | 5790348 | rs997499 | -1.009 | 0.838 | 1.307 | 0.911 | 0.005 | -0.016 | -0.016 | 0.566 | 0.36 | 0.27 | 0.31 | 0.51 | 0.34 | 0.4 |
| 3 | 5791627 | rs2444229 | -0.39 | 0.909 | 1.311 | 1.005 | 0.007 | 0.055 | 0.055 | 0 | 0.19 | 0.22 | 0.25 | 0.44 | 0.26 | 0.34 |
| 3 | 5792135 | rs7632996 | 0.031 | 0.859 | 0.978 | 0.928 | 0.005 | 0 | 0 | 0 | 0.1 | 0.14 | 0.09 | 0.1 | 0.14 | 0.25 |
| 3 | 5796218 | rs2470485 | -1.857 | 0.96 | 0.675 | 0.865 | 0 | -0.009 | -0.009 | 0.187 | 0.49 | 0.47 | 0.5 | 0.8 | 0.53 | 0.67 |
| 3 | 5796253 | rs10461037 | -0.157 | 0.962 | 0.504 | 0.89 | -0.008 | -0.023 | -0.023 | 0.593 | 0.18 | 0.13 | 0.23 | 0.43 | 0.25 | 0.35 |
| 3 | 5796661 | rs2444218 | -0.908 | 1.193 | 0.165 | 0.297 | 0 | -0.005 | -0.005 | 0.189 | 0.28 | 0.27 | 0.17 | 0.18 | 0.24 | 0.3 |
| 3 | 5797319 | rs1823765 | -0.159 | 0.613 | 0.252 | -0.101 | -0.021 | -0.031 | -0.031 | 0 | 0.2 | 0.23 | 0.1 | 0.08 | 0.11 | 0.08 |
| 3 | 5798232 | rs7609894 | -0.681 | 0.963 | -0.106 | 0.012 | 0.001 | 0.015 | 0.015 | 0 | 0.49 | 0.48 | 0.43 | 0.77 | 0.5 | 0.68 |
| 3 | 5798892 | rs1452708 | -0.24 | 0.96 | -0.605 | 0.235 | 0 | -0.007 | -0.007 | 0 | 0.46 | 0.39 | 0.61 | 0.78 | 0.54 | 0.58 |
| 3 | 5801886 | rs1836577 | 0.049 | 0.516 | -0.945 | 0.567 | 0.033 | -0.005 | -0.005 | 0 | 0.48 | 0.38 | 0.55 | 0.48 | 0.55 | 0.6 |
| 3 | 5802109 | rs17043484 | -0.704 | 0.26 | -0.707 | 0.6 | 0 | -0.001 | -0.001 | 0.701 | 0.2 | 0.2 | 0.21 | 0.24 | 0.14 | 0.13 |
| 3 | 5806750 | rs13088667 | -0.335 | 0.738 | 0.012 | 0.738 | -0.051 | -0.036 | -0.036 | 1.235 | 0.26 | 0.18 | 0.3 | 0.34 | 0.53 | 0.39 |
| 3 | 5807895 | rs4685983 | -1.128 | -0.355 | -0.021 | 0.573 | 0.098 | 0.004 | 0.004 | 0.704 | 0.48 | 0.71 | 0.64 | 0.5 | 0.73 | 0.62 |
| 3 | 5808682 | rs6779060 | -0.526 | -0.31 | 0.394 | 0.23 | -0.016 | -0.075 | -0.075 | 0.266 | 0.17 | 0.6 | 0.23 | 0.04 | 0.12 | 0.03 |
| 3 | 5813102 | rs9863945 | -0.618 | -0.348 | 0.594 | 0.23 | 0.022 | -0.005 | -0.005 | 0 | 0.08 | 0.52 | 0.06 | 0 | 0.01 | 0 |
| 3 | 5822491 | rs9864734 | 0.329 | -0.281 | 0.316 | 0.33 | -0.038 | -0.032 | -0.032 | 0 | 0.22 | 0.17 | 0.28 | 0.34 | 0.52 | 0.41 |
| 3 | 5823074 | rs4574278 | -1.172 | -0.38 | 0.081 | 0.349 | 0.029 | -0.043 | -0.043 | 0 | 0.4 | 0.8 | 0.52 | 0.41 | 0.64 | 0.44 |
| 3 | 5825005 | rs17233863 | 0.767 | -0.882 | -0.144 | 0.073 | 0.019 | -0.015 | -0.015 | 0 | 0.58 | 0.24 | 0.59 | 0.63 | 0.36 | 0.45 |
| 3 | 5825657 | rs1897958 | 0.846 | -1.087 | -0.202 | 0.012 | 0.054 | 0.001 | 0.001 | 0.014 | 0.54 | 0.24 | 0.55 | 0.6 | 0.31 | 0.43 |
| 3 | 5826383 | rs6442990 | 0.913 | -0.674 | -0.069 | 0.023 | -0.006 | -0.054 | -0.054 | 0.699 | 0.82 | 0.33 | 0.79 | 0.92 | 0.8 | 0.89 |
| 3 | 5826871 | rs11711086 | -0.175 | -0.173 | -0.509 | 0.115 | -0.104 | -0.055 | -0.055 | 0 | 0.29 | 0.1 | 0.25 | 0.32 | 0.48 | 0.47 |
| 3 | 5829838 | rs2470626 | 0.852 | -0.742 | 0.125 | 0.041 | 0.007 | 0.003 | 0.003 | 0.229 | 0.55 | 0.58 | 0.53 | 0.65 | 0.37 | 0.44 |
| 3 | 5830444 | rs4685985 | -1.149 | -1.087 | 0.315 | 0.029 | 0.024 | 0.005 | 0.005 | 0.628 | 0.49 | 0.46 | 0.59 | 0.42 | 0.78 | 0.65 |
| 3 | 5831036 | rs2437219 | 0.557 | 0.804 | 0.167 | -0.122 | -0.023 | 0.002 | 0.002 | 0.52 | 0.63 | 0.71 | 0.63 | 0.69 | 0.44 | 0.44 |
| 3 | 5832156 | rs1836581 | -0.613 | 0.742 | 0.607 | -0.335 | 0.044 | -0.138 | -0.138 | 1.173 | 0.2 | 0.46 | 0.26 | 0.01 | 0.23 | 0.05 |
| 3 | 5835743 | rs17043597 | 1.183 | 0.85 | 0.708 | -0.395 | -0.002 | -0.002 | -0.002 | 0 | 0.6 | 0.44 | 0.53 | 0.73 | 0.37 | 0.47 |
| 3 | 5838500 | rs17043606 | 0.403 | 0.881 | 0.708 | -0.408 | 0.019 | 0.003 | 0.003 | 0 | 0.07 | 0.13 | 0.14 | 0.07 | 0.14 | 0.11 |

| | | | | | | | | | | | | | | | | |
|---|---|---|---|---|---|---|---|---|---|---|---|---|---|---|---|---|
| 3 | 5838677 | rs2117636 | 0.531 | 2.069 | 0.458 | -0.47 | -0.078 | -0.097 | -0.097 | 0 | 0.18 | 0.01 | 0.16 | 0.22 | 0.27 | 0.31 |
| 3 | 5839900 | rs2437220 | -1.391 | 1.862 | 0.979 | -0.394 | 0.134 | -0.06 | -0.06 | 0 | 0.18 | 0.41 | 0.35 | 0.09 | 0.35 | 0.14 |
| 3 | 5841018 | rs7641425 | -0.647 | 1.476 | 0.627 | -0.524 | -0.08 | -0.006 | -0.006 | 0.222 | 0.31 | 0.13 | 0.25 | 0.23 | 0.43 | 0.49 |
| 3 | 5847377 | rs2244301 | 1.315 | 1.011 | 0.718 | -0.213 | -0.003 | -0.011 | -0.011 | 0.434 | 0.57 | 0.34 | 0.56 | 0.58 | 0.43 | 0.43 |
| 3 | 5848216 | rs9871134 | -0.99 | 1.356 | 0.468 | -0.16 | -0.034 | 0.044 | 0.044 | 0.096 | 0.34 | 0.26 | 0.33 | 0.13 | 0.47 | 0.42 |
| 3 | 5849483 | rs17043611 | 1.765 | 0.134 | 0.462 | -0.272 | -0.009 | 0 | 0 | 0 | 0.44 | 0.53 | 0.38 | 0.69 | 0.2 | 0.36 |
| 3 | 5851892 | rs2004019 | 1.48 | 0.274 | 0.537 | -0.264 | 0.038 | 0.018 | 0.018 | 0 | 0.51 | 0.39 | 0.38 | 0.7 | 0.23 | 0.38 |
| 3 | 5853299 | rs6781156 | -0.213 | 0.849 | -0.217 | -0.297 | -0.097 | 0.002 | 0.002 | 1.698 | 0.3 | 0.08 | 0.24 | 0.21 | 0.42 | 0.44 |
| 3 | 5853752 | rs7648808 | -0.13 | 0.749 | -0.184 | -0.295 | -0.09 | -0.006 | -0.006 | 0 | 0.28 | 0.1 | 0.24 | 0.22 | 0.43 | 0.47 |
| 3 | 5853915 | rs1435368 | 2.174 | 0.147 | -0.756 | -0.183 | 0.048 | -0.147 | -0.147 | 0.257 | 0.36 | 0.04 | 0.33 | 0.52 | 0.23 | 0.32 |
| 3 | 5856373 | rs1529860 | -0.141 | 0.826 | 0.107 | -0.325 | 0.044 | -0.033 | -0.033 | 0 | 0.17 | 0.35 | 0.34 | 0.08 | 0.23 | 0.15 |
| 3 | 5858488 | rs7609993 | 0.069 | 1.31 | 0.193 | -0.357 | 0.075 | 0 | 0 | 0 | 0.09 | 0.17 | 0.16 | 0.06 | 0.16 | 0.14 |
| 3 | 5862244 | rs717186 | 0.657 | 1.199 | -0.204 | -0.341 | -0.035 | -0.014 | -0.014 | 0.017 | 0.21 | 0.07 | 0.17 | 0.19 | 0.28 | 0.34 |
| 3 | 5862369 | rs717187 | -0.383 | 1.092 | 0.053 | -0.301 | 0.009 | -0.012 | -0.012 | 0.105 | 0.2 | 0.23 | 0.32 | 0.1 | 0.29 | 0.15 |
| 3 | 5867895 | rs962062 | 2.461 | 0.613 | 0.185 | -0.52 | -0.002 | -0.002 | -0.002 | 0.129 | 0.34 | 0.32 | 0.32 | 0.64 | 0.18 | 0.32 |
| 3 | 5879366 | rs9811522 | 0.389 | 0.889 | 0.148 | -0.483 | -0.009 | 0.014 | 0.014 | 0 | 0.18 | 0.13 | 0.15 | 0.07 | 0.18 | 0.15 |
| 3 | 5880322 | rs2262396 | 1.031 | 0.878 | 0.162 | -0.516 | 0.009 | -0.031 | -0.031 | 0.066 | 0.67 | 0.45 | 0.57 | 0.84 | 0.65 | 0.71 |
| 3 | 5881534 | rs2572645 | -0.306 | 0.874 | 0.105 | -0.494 | 0.016 | -0.022 | -0.022 | 0 | 0.85 | 0.66 | 0.73 | 0.91 | 0.83 | 0.87 |
| 3 | 5881625 | rs4684510 | 0.47 | 0.922 | -0.054 | -0.473 | 0 | -0.001 | -0.001 | 0.053 | 0.13 | 0.15 | 0.26 | 0.09 | 0.17 | 0.13 |
| 3 | 5882683 | rs977905 | -1.657 | 0.743 | -0.311 | -0.431 | 0.016 | 0.003 | 0.003 | 0 | 0.55 | 0.62 | 0.6 | 0.36 | 0.64 | 0.63 |
| 3 | 5883135 | rs4685991 | 0.614 | 0.645 | -0.409 | -0.079 | 0.001 | -0.005 | -0.005 | 0 | 0.58 | 0.53 | 0.55 | 0.71 | 0.43 | 0.52 |
| 3 | 5888012 | rs7624621 | -0.277 | 0.328 | -0.536 | -0.161 | 0.004 | 0.008 | 0.008 | 2.536 | 0.28 | 0.23 | 0.27 | 0.14 | 0.22 | 0.15 |
| 3 | 5888521 | rs1159474 | -1.793 | 0.6 | -1.016 | -0.281 | 0.027 | 0.078 | 0.078 | 0.255 | 0.57 | 0.77 | 0.6 | 0.23 | 0.58 | 0.51 |
| 3 | 5888909 | rs748191 | -0.268 | 0.849 | -0.698 | -0.132 | -0.01 | 0.011 | 0.011 | 1.502 | 0.79 | 0.84 | 0.75 | 0.94 | 0.76 | 0.66 |
| 3 | 5890053 | rs10510319 | 0.487 | 0.073 | -0.865 | 0.063 | -0.021 | -0.012 | -0.012 | 0 | 0.2 | 0.02 | 0.27 | 0.12 | 0.18 | 0.12 |
| 3 | 5890820 | rs1836586 | -0.604 | -0.039 | -0.403 | 0.563 | -0.003 | 0.001 | 0.001 | 3.288 | 0.29 | 0.3 | 0.38 | 0.36 | 0.24 | 0.14 |
| 3 | 5891648 | rs2572690 | 0.32 | -0.447 | -0.022 | 1.082 | -0.008 | 0.128 | 0.128 | 0 | 0.07 | 0.17 | 0.13 | 0.28 | 0.05 | 0.02 |
| 3 | 5894166 | rs17215755 | -0.464 | -0.713 | -0.554 | 1.323 | 0.011 | -0.053 | -0.053 | 0 | 0.15 | 0.04 | 0.24 | 0.32 | 0.11 | 0.34 |
| 3 | 5894565 | rs7629530 | -0.71 | -0.864 | -0.116 | 1.252 | -0.01 | 0.025 | 0.025 | 0.032 | 0.17 | 0.18 | 0.25 | 0.32 | 0.11 | 0.34 |
| 3 | 5895027 | rs4685993 | -0.716 | -0.874 | -0.099 | 1.248 | -0.007 | 0.021 | 0.021 | 0.045 | 0.17 | 0.17 | 0.26 | 0.32 | 0.1 | 0.35 |
| 3 | 5896152 | rs1960988 | -1.685 | -0.938 | -0.796 | 0.798 | 0.006 | -0.04 | -0.04 | 0 | 0.59 | 0.8 | 0.64 | 0.45 | 0.56 | 0.6 |
| 3 | 5897751 | rs7427632 | 1.101 | -0.785 | -0.542 | 0.551 | 0.003 | 0 | 0 | 0 | 0.15 | 0.13 | 0.17 | 0.04 | 0.09 | 0.1 |
| 3 | 5898038 | rs1075699 | -1.211 | -0.782 | -0.364 | 0.74 | 0.004 | -0.025 | -0.025 | 1.253 | 0.41 | 0.68 | 0.48 | 0.51 | 0.36 | 0.26 |
| 3 | 5899967 | rs722615 | 1.053 | -1.046 | -0.231 | 0.698 | 0.014 | 0.009 | 0.009 | 0.406 | 0.64 | 0.26 | 0.64 | 0.69 | 0.71 | 0.61 |
| 3 | 5901230 | rs1369404 | 0.895 | -1.429 | -0.123 | 0.601 | 0.034 | 0.066 | 0.066 | 0.102 | 0.19 | 0.09 | 0.19 | 0.07 | 0.09 | 0.12 |
| 3 | 5902056 | rs11130652 | 0.817 | -1.584 | 0.199 | 0.537 | 0.032 | 0.027 | 0.027 | 0 | 0.55 | 0.3 | 0.5 | 0.63 | 0.7 | 0.6 |
| 3 | 5902298 | rs11710507 | 1.787 | -1.167 | -0.082 | 0.811 | -0.036 | -0.02 | -0.02 | 0 | 0.06 | 0.01 | 0.04 | 0.04 | 0.13 | 0.02 |
| 3 | 5904553 | rs2730313 | 0.067 | -1.415 | 0.151 | 0.86 | -0.017 | -0.004 | -0.004 | 0.063 | 0.24 | 0.33 | 0.26 | 0.22 | 0.1 | 0.12 |
| 3 | 5907475 | rs2572683 | -0.994 | -1.595 | 0.47 | 1.134 | -0.049 | 0.178 | 0.178 | 0.017 | 0.17 | 0.48 | 0.16 | 0.48 | 0.08 | 0.15 |
| 3 | 5907990 | rs1865533 | 0.89 | -1.498 | -0.344 | 0.45 | -0.015 | 0.015 | 0.015 | 0 | 0.06 | 0.06 | 0.02 | 0.14 | 0.01 | 0.08 |
| 3 | 5913291 | rs1469805 | 1.573 | -1.642 | -0.956 | 0.201 | -0.126 | -0.015 | -0.015 | 0 | 0.41 | 0.07 | 0.57 | 0.64 | 0.52 | 0.52 |
| 3 | 5925079 | rs2730320 | -1.789 | -1.511 | -0.59 | -0.1 | -0.001 | -0.022 | -0.022 | 0 | 0.95 | 0.94 | 0.97 | 1 | 0.95 | 0.99 |
| 3 | 5925737 | rs2730321 | -0.21 | -1.71 | -0.522 | -0.309 | 0.001 | 0.033 | 0.033 | 0 | 0.73 | 0.75 | 0.74 | 0.91 | 0.77 | 0.71 |
| 3 | 5926673 | rs6791121 | -0.491 | -1.763 | -0.049 | -0.311 | 0.011 | 0.003 | 0.003 | 0.245 | 0.19 | 0.44 | 0.1 | 0.16 | 0.18 | 0.1 |
| 3 | 5927739 | rs2323505 | 1.496 | -1.804 | 0.036 | -0.304 | 0.01 | 0.004 | 0.004 | 0 | 0.06 | 0.05 | 0.04 | 0.05 | 0.13 | 0.02 |
| 3 | 5928176 | rs2572667 | -0.188 | -1.941 | -0.089 | -0.388 | -0.039 | 0.019 | 0.019 | 0 | 0.8 | 0.82 | 0.75 | 0.92 | 0.67 | 0.72 |
| 3 | 5928190 | rs2572666 | 1.926 | -1.832 | -0.263 | -0.372 | -0.001 | -0.023 | -0.023 | 0.477 | 0.05 | 0.06 | 0.04 | 0 | 0.05 | 0.01 |
| 3 | 5930619 | rs13073377 | 2.262 | -1.889 | -0.417 | -0.135 | -0.16 | -0.156 | -0.156 | 0 | 0.38 | 0.04 | 0.54 | 0.49 | 0.49 | 0.48 |
| 3 | 5931884 | rs6790270 | -0.379 | -2.169 | -0.213 | -0.45 | 0.014 | -0.058 | -0.058 | 0 | 0.17 | 0.39 | 0.09 | 0.11 | 0.18 | 0.1 |
| 3 | 5933395 | rs2730328 | -0.977 | -2.21 | -0.191 | -0.455 | 0.005 | -0.003 | -0.003 | 0.884 | 0.37 | 0.64 | 0.17 | 0.44 | 0.14 | 0.25 |
| 3 | 5935361 | rs4685995 | 0.809 | -2.018 | -0.242 | -0.487 | -0.03 | -0.06 | -0.06 | 0.435 | 0.63 | 0.51 | 0.84 | 0.77 | 0.73 | 0.82 |
| 3 | 5936245 | rs11130673 | 2.158 | -1.859 | -0.564 | -0.526 | -0.142 | -0.15 | -0.15 | 2.306 | 0.36 | 0.06 | 0.51 | 0.48 | 0.44 | 0.45 |
| 3 | 5938105 | rs12233495 | 1.48 | -1.933 | -0.496 | -0.48 | 0.005 | 0.042 | 0.042 | 0 | 0.06 | 0.02 | 0.04 | 0.28 | 0.01 | 0.09 |
| 3 | 5939108 | rs13088715 | 1.643 | -1.876 | -0.503 | -0.676 | -0.079 | 0.162 | 0.162 | 0 | 0.42 | 0.09 | 0.55 | 0.76 | 0.44 | 0.54 |
| 3 | 5939435 | rs9869388 | 0.399 | -1.877 | -0.324 | -0.697 | -0.006 | -0.086 | -0.086 | 0.532 | 0.61 | 0.51 | 0.65 | 0.89 | 0.63 | 0.65 |
| 3 | 5940480 | rs9846727 | -0.652 | -1.906 | -0.124 | -0.592 | 0.005 | -0.062 | -0.062 | 0 | 0.19 | 0.41 | 0.1 | 0.13 | 0.2 | 0.11 |
| 3 | 5943854 | rs7623234 | -1.897 | -1.491 | -0.3 | -0.552 | 0 | -0.011 | -0.011 | 0 | 0.94 | 0.96 | 0.95 | 1 | 0.93 | 0.98 |
| 3 | 5944905 | rs9754628 | 0.085 | -1.691 | -0.183 | -0.628 | 0.018 | -0.091 | -0.091 | 0 | 0.19 | 0.35 | 0.27 | 0.02 | 0.24 | 0.24 |
| 3 | 5956207 | rs9845152 | -1.54 | -1.648 | -0.643 | -0.404 | -0.019 | -0.05 | -0.05 | 0 | 0.58 | 0.81 | 0.44 | 0.52 | 0.49 | 0.54 |
| 3 | 5962755 | rs9810413 | -1.625 | -1.54 | -0.517 | -0.448 | -0.016 | -0.042 | -0.042 | 0 | 0.56 | 0.88 | 0.39 | 0.48 | 0.45 | 0.54 |
| 3 | 5965125 | rs925856 | -0.498 | -1.095 | -0.051 | -0.405 | 0.043 | -0.037 | -0.037 | 0 | 0.92 | 0.85 | 0.92 | 0.97 | 0.83 | 0.96 |
| 3 | 5967249 | rs1568082 | 0.181 | -1.37 | 0.434 | -0.44 | 0.007 | -0.025 | -0.025 | 0 | 0.88 | 0.68 | 0.93 | 0.93 | 0.91 | 0.98 |
| 3 | 5968314 | rs1969304 | -0.272 | -1.482 | 0.229 | -0.358 | 0 | 0.005 | 0.005 | 0 | 0.88 | 0.9 | 0.94 | 0.74 | 0.92 | 0.87 |
| 3 | 5973939 | rs1517383 | -0.825 | -1.795 | 0.409 | -0.694 | 0 | -0.022 | -0.022 | 0 | 0.31 | 0.7 | 0.32 | 0.13 | 0.37 | 0.31 |
| 3 | 5975204 | rs6768719 | 1.182 | -2.025 | 0.567 | -0.791 | -0.005 | -0.001 | -0.001 | 0 | 0.57 | 0.45 | 0.83 | 0.55 | 0.73 | 0.69 |
| 3 | 5990857 | rs17043965 | -0.112 | -1.829 | 0.525 | -0.753 | -0.002 | -0.008 | -0.008 | 0 | 0.24 | 0.23 | 0.26 | 0.28 | 0.26 | 0.36 |
| 3 | 5991376 | rs9875406 | -0.675 | -1.834 | 0.656 | -0.741 | 0.083 | 0.027 | 0.027 | 0 | 0.08 | 0.31 | 0.06 | 0.1 | 0.12 | 0.03 |
| 3 | 5992687 | rs1400671 | -1.929 | -1.906 | 0.272 | -0.772 | -0.024 | -0.045 | -0.045 | 0 | 0.58 | 0.87 | 0.39 | 0.48 | 0.45 | 0.55 |
| 3 | 6002693 | rs6767324 | -1.004 | -1.765 | 0.489 | -0.721 | -0.005 | -0.017 | -0.017 | 0.753 | 0.34 | 0.44 | 0.32 | 0.26 | 0.35 | 0.39 |

| | | | | | | | | | | | | | | | | |
|---|---|---|---|---|---|---|---|---|---|---|---|---|---|---|---|---|
| 3 | 6002768 | rs6807385 | -0.59 | -1.694 | 0.377 | -0.65 | -0.007 | 0.001 | 0.001 | 2.301 | 0.28 | 0.34 | 0.28 | 0.26 | 0.28 | 0.37 |
| 3 | 6005401 | rs6768819 | -0.425 | -1.621 | 0.379 | -0.646 | -0.02 | -0.007 | -0.007 | 0 | 0.27 | 0.22 | 0.28 | 0.29 | 0.36 | 0.39 |
| 3 | 6008503 | rs11130721 | 0.596 | -1.442 | 0.435 | -0.516 | 0.094 | 0.046 | 0.046 | 0.114 | 0.93 | 0.67 | 0.94 | 0.9 | 0.88 | 0.98 |
| 3 | 6009647 | rs4684515 | 2.049 | -1.596 | -0.355 | -0.516 | -0.163 | -0.134 | -0.134 | 0 | 0.35 | 0.06 | 0.55 | 0.54 | 0.49 | 0.35 |
| 3 | 6010772 | rs2048923 | -0.429 | -1.482 | 0.273 | -0.43 | -0.002 | 0 | 0 | 0.39 | 0.26 | 0.31 | 0.26 | 0.25 | 0.24 | 0.41 |
| 3 | 6014629 | rs11927011 | -0.472 | -1.288 | 0.162 | -0.723 | -0.003 | -0.038 | -0.038 | 0 | 0.08 | 0.19 | 0.07 | 0 | 0.08 | 0.06 |
| 3 | 6014760 | rs13325414 | 0.737 | -1.443 | -0.223 | -0.794 | 0.006 | 0.048 | 0.048 | 0 | 0.19 | 0.07 | 0.08 | 0.13 | 0.16 | 0.12 |
| 3 | 6016209 | rs17031549 | -0.325 | -1.288 | 0.332 | -0.542 | -0.005 | -0.003 | -0.003 | 0 | 0.22 | 0.24 | 0.21 | 0.18 | 0.18 | 0.38 |
| 3 | 6018739 | rs4686007 | 0.738 | -1.268 | 0.537 | -0.6 | -0.029 | -0.028 | -0.028 | 0 | 0.72 | 0.6 | 0.77 | 0.8 | 0.79 | 0.53 |
| 3 | 6018903 | rs4686008 | -0.088 | -1.065 | 0.262 | -0.489 | -0.007 | -0.003 | -0.003 | 0 | 0.79 | 0.74 | 0.79 | 0.82 | 0.82 | 0.62 |
| 3 | 6020600 | rs6799932 | -0.177 | -0.784 | 0.056 | -0.369 | 0.013 | -0.023 | -0.023 | 3.131 | 0.06 | 0.11 | 0.07 | 0 | 0.09 | 0.07 |
| 3 | 6027244 | rs9835716 | 1.055 | -1.582 | -0.025 | -0.517 | 0.057 | 0.042 | 0.042 | 0 | 0.54 | 0.35 | 0.77 | 0.68 | 0.75 | 0.69 |
| 3 | 6027486 | rs7615041 | -0.552 | -1.397 | 0.028 | -0.636 | 0.012 | 0.008 | 0.008 | 0 | 0.33 | 0.26 | 0.13 | 0.28 | 0.19 | 0.24 |
| 3 | 6031098 | rs9828567 | -0.864 | -1.491 | 0.059 | -0.592 | 0.003 | 0.006 | 0.006 | 0 | 0.38 | 0.33 | 0.22 | 0.27 | 0.32 | 0.28 |
| 3 | 6032360 | rs9834359 | 0.731 | -1.503 | -0.104 | -0.593 | 0.004 | 0.008 | 0.008 | 0 | 0.62 | 0.68 | 0.78 | 0.73 | 0.69 | 0.73 |
| 3 | 6042658 | rs1517378 | -1.692 | -1.729 | -0.791 | -0.705 | 0.072 | 0.138 | 0.138 | 0.023 | 0.52 | 0.87 | 0.29 | 0.29 | 0.32 | 0.51 |
| 3 | 6061053 | rs6770246 | -0.446 | -1.534 | -0.307 | -0.788 | 0.02 | -0.115 | -0.115 | 0.506 | 0.17 | 0.43 | 0.14 | 0.01 | 0.15 | 0.23 |
| 3 | 6068860 | rs9853582 | 0.473 | -1.922 | -0.762 | -1.071 | 0.08 | 0.1 | 0.1 | 0 | 0.26 | 0.07 | 0.19 | 0.1 | 0.14 | 0.22 |
| 3 | 6070049 | rs11923634 | -0.639 | -1.21 | -0.56 | -0.71 | 0.089 | -0.1 | -0.1 | 0.027 | 0.19 | 0.47 | 0.17 | 0.06 | 0.24 | 0.26 |
| 3 | 6081827 | rs1033251 | -1.252 | -0.618 | -1.424 | -0.114 | -0.071 | -0.007 | -0.007 | 0 | 0.6 | 0.9 | 0.42 | 0.38 | 0.43 | 0.63 |
| 3 | 6082552 | rs1914413 | 0.605 | -0.236 | -0.765 | 0.432 | 0.002 | 0.001 | 0.001 | 0.561 | 0.59 | 0.41 | 0.37 | 0.37 | 0.63 | 0.46 |
| 3 | 6088622 | rs2102811 | -1.448 | -0.034 | -0.081 | 0.783 | 0.004 | 0.094 | 0.094 | 0.932 | 0.25 | 0.46 | 0.39 | 0.53 | 0.21 | 0.38 |
| 3 | 6089091 | rs3941735 | -1.026 | -0.08 | 0.083 | 0.741 | -0.007 | -0.038 | -0.038 | 0 | 0.19 | 0.15 | 0.35 | 0.54 | 0.17 | 0.36 |
| 3 | 6089961 | rs9870341 | -0.593 | -0.134 | 0.183 | 0.07 | -0.001 | -0.151 | -0.151 | 0.925 | 0.27 | 0.47 | 0.26 | 0.02 | 0.24 | 0.19 |
| 3 | 6092724 | rs9844813 | 1.855 | -0.314 | -0.111 | 0.12 | -0.019 | 0.033 | 0.033 | 0 | 0.54 | 0.21 | 0.49 | 0.4 | 0.6 | 0.52 |
| 3 | 6094373 | rs1607193 | 0.844 | -0.257 | 0.112 | -0.021 | 0.022 | -0.101 | -0.101 | 0 | 0.75 | 0.57 | 0.82 | 0.95 | 0.7 | 0.88 |
| 3 | 6094918 | rs9832245 | 2.584 | -0.29 | -0.167 | 0.059 | -0.027 | 0.039 | 0.039 | 0 | 0.42 | 0.19 | 0.3 | 0.34 | 0.47 | 0.37 |
| 3 | 6095736 | rs1517375 | 1.43 | -0.031 | 0.176 | 0.176 | -0.012 | -0.067 | -0.067 | 0 | 0.71 | 0.46 | 0.83 | 0.88 | 0.73 | 0.9 |
| 3 | 6097432 | rs17287953 | -0.098 | 0.284 | -0.525 | 0.434 | -0.025 | -0.054 | -0.054 | 0 | 0.1 | 0.02 | 0.27 | 0.26 | 0.09 | 0.14 |
| 3 | 6099472 | rs6773693 | -0.199 | 0.36 | -0.184 | -0.085 | 0.002 | -0.052 | -0.052 | 0 | 0.17 | 0.2 | 0.09 | 0.01 | 0.15 | 0.05 |
| 3 | 6102823 | rs9865909 | -1.856 | 0.113 | -0.365 | 0.087 | -0.006 | 0.022 | 0.022 | 0 | 0.54 | 0.69 | 0.67 | 0.63 | 0.51 | 0.59 |
| 3 | 6104625 | rs908603 | 0.665 | 0.043 | 0.019 | 0.079 | -0.037 | -0.031 | -0.031 | 0 | 0.82 | 0.5 | 0.87 | 0.88 | 0.9 | 0.82 |
| 3 | 6105186 | rs973062 | -0.159 | -0.092 | -0.485 | -0.141 | -0.022 | -0.083 | -0.083 | 0 | 0.32 | 0.15 | 0.29 | 0.43 | 0.26 | 0.34 |
| 3 | 6106251 | rs6792001 | 0.432 | 0.313 | -0.249 | 0.029 | 0.033 | 0.022 | 0.022 | 0.042 | 0.83 | 0.47 | 0.83 | 0.79 | 0.82 | 0.82 |
| 3 | 6109111 | rs7643770 | 0.987 | 0.207 | 0.086 | -0.049 | -0.002 | -0.003 | -0.003 | 0 | 0.57 | 0.48 | 0.6 | 0.41 | 0.64 | 0.52 |
| 3 | 6110764 | rs13058925 | -0.081 | 0.727 | -0.424 | 0.161 | -0.05 | -0.084 | -0.084 | 0 | 0.74 | 0.85 | 0.68 | 0.56 | 0.72 | 0.69 |
| 3 | 6113062 | rs1517379 | 1.65 | 0.575 | -0.347 | -0.029 | 0.004 | 0.008 | 0.008 | 0 | 0.42 | 0.35 | 0.25 | 0.33 | 0.44 | 0.41 |
| 3 | 6114930 | rs17234241 | -0.15 | -0.061 | -0.464 | 0.151 | 0.024 | 0.091 | 0.091 | 0.106 | 0.16 | 0.01 | 0.27 | 0.03 | 0.12 | 0.11 |
| 3 | 6119146 | rs925852 | 0.212 | 0.527 | -0.975 | 0.639 | -0.043 | -0.1 | -0.1 | 0.034 | 0.73 | 0.9 | 0.72 | 0.61 | 0.73 | 0.73 |
| 3 | 6119416 | rs925850 | 0.829 | 0.495 | -0.697 | 0.681 | -0.02 | -0.018 | -0.018 | 0 | 0.66 | 0.72 | 0.62 | 0.44 | 0.62 | 0.64 |
| 3 | 6120442 | rs868243 | 0.882 | 0.557 | -0.52 | 0.593 | -0.005 | 0.002 | 0.002 | 0 | 0.53 | 0.69 | 0.36 | 0.54 | 0.49 | 0.57 |
| 3 | 6121501 | rs13088182 | 0.81 | 0.587 | -0.497 | 0.622 | -0.023 | -0.014 | -0.014 | 0.378 | 0.55 | 0.76 | 0.38 | 0.54 | 0.5 | 0.59 |
| 3 | 6131198 | rs7609589 | 1.025 | 0.361 | -0.138 | 0.499 | 0 | 0 | 0 | 0.271 | 0.49 | 0.47 | 0.41 | 0.39 | 0.65 | 0.56 |
| 3 | 6131889 | rs990156 | 0.681 | 1.538 | 0.093 | 1.357 | -0.001 | -0.022 | -0.022 | 0 | 0.75 | 0.75 | 0.71 | 0.44 | 0.78 | 0.67 |
| 3 | 6132069 | rs990155 | 0.609 | 1.549 | 0.028 | 1.346 | -0.002 | -0.023 | -0.023 | 0 | 0.76 | 0.76 | 0.71 | 0.44 | 0.78 | 0.67 |
| 3 | 6132140 | rs7624190 | 0.842 | 1.536 | 0.035 | 1.344 | 0 | -0.001 | -0.001 | 3.206 | 0.74 | 0.72 | 0.7 | 0.44 | 0.78 | 0.67 |
| 3 | 6134046 | rs9311784 | 0.581 | 1.634 | -0.009 | 1.06 | -0.012 | -0.015 | -0.015 | 0 | 0.76 | 0.76 | 0.69 | 0.58 | 0.7 | 0.71 |
| 3 | 6134126 | rs9311785 | 0.602 | 1.715 | -0.209 | 1.118 | 0.01 | 0.017 | 0.017 | 0 | 0.75 | 0.72 | 0.69 | 0.59 | 0.7 | 0.71 |
| 3 | 6137379 | rs1504036 | -0.255 | 1.123 | -0.586 | 0.953 | 0.008 | -0.028 | -0.028 | 1.779 | 0.64 | 0.74 | 0.51 | 0.53 | 0.64 | 0.67 |
| 3 | 6138104 | rs17215874 | -0.46 | 0.957 | 0.126 | 1.157 | 0.018 | 0.19 | 0.19 | 0.03 | 0.11 | 0.27 | 0.1 | 0.38 | 0.09 | 0.2 |
| 3 | 6139380 | rs17288023 | -0.928 | 0.521 | -0.013 | 1.721 | 0.017 | -0.025 | -0.025 | 0 | 0.11 | 0.05 | 0.08 | 0.32 | 0.06 | 0.16 |
| 3 | 6139410 | rs2134776 | -0.547 | 0.941 | -0.069 | 1.389 | -0.015 | -0.017 | -0.017 | 0.452 | 0.61 | 0.79 | 0.52 | 0.65 | 0.57 | 0.63 |
| 3 | 6140244 | rs6805381 | 0.118 | 1.156 | -0.042 | 0.738 | 0.038 | -0.026 | -0.026 | 0 | 0.23 | 0.4 | 0.24 | 0.2 | 0.28 | 0.16 |
| 3 | 6145045 | rs6801950 | 0.163 | 1.333 | 0.011 | 0.598 | -0.006 | -0.019 | -0.019 | 0 | 0.22 | 0.25 | 0.2 | 0.02 | 0.31 | 0.15 |
| 3 | 6145135 | rs6443023 | -0.877 | 1.161 | -0.033 | 0.77 | -0.003 | -0.017 | -0.017 | 0 | 0.57 | 0.75 | 0.47 | 0.51 | 0.55 | 0.64 |
| 3 | 6154425 | rs6443025 | -0.257 | 0.138 | 0.594 | 0.521 | -0.046 | -0.071 | -0.071 | 0 | 0.34 | 0.41 | 0.19 | 0.07 | 0.18 | 0.35 |
| 3 | 6156260 | rs1604809 | -0.99 | 0.502 | 0.087 | 0.563 | 0.067 | 0.041 | 0.041 | 0.808 | 0.49 | 0.76 | 0.32 | 0.41 | 0.37 | 0.52 |
| 3 | 6162218 | rs1504037 | 0.026 | 0.32 | 0.064 | 0.66 | 0.016 | -0.018 | -0.018 | 0 | 0.11 | 0.07 | 0.03 | 0.16 | 0.08 | 0.15 |
| 3 | 6165284 | rs1504041 | -0.889 | 0.275 | -0.006 | 0.545 | 0.098 | 0.037 | 0.037 | 0.015 | 0.49 | 0.79 | 0.31 | 0.4 | 0.34 | 0.53 |
| 3 | 6169451 | rs34615664 | -1.185 | 0.262 | -0.018 | 0.541 | 0.074 | 0.018 | 0.018 | 0.752 | 0.52 | 0.81 | 0.33 | 0.41 | 0.37 | 0.53 |
| 3 | 6178557 | rs17288044 | 0.081 | 0.035 | -0.011 | 0.583 | 0.017 | -0.029 | -0.029 | 0.679 | 0.11 | 0.05 | 0.04 | 0.17 | 0.08 | 0.18 |
| 3 | 6181122 | rs9811754 | -1.55 | 0.256 | -0.307 | 0.303 | -0.073 | -0.077 | -0.077 | 0 | 0.64 | 0.9 | 0.43 | 0.45 | 0.57 | 0.63 |
| 3 | 6182793 | rs7638535 | 0.28 | -0.151 | 0.121 | 0.007 | 0.001 | -0.042 | -0.042 | 0 | 0.23 | 0.49 | 0.18 | 0.18 | 0.2 | 0.12 |
| 3 | 6184718 | rs1504030 | 0.053 | -0.294 | 0.391 | -0.11 | 0.034 | -0.044 | -0.044 | 0.013 | 0.33 | 0.39 | 0.28 | 0.21 | 0.45 | 0.23 |
| 3 | 6185577 | rs17234332 | 0.168 | -0.22 | 0.341 | -0.082 | -0.007 | 0.031 | 0.031 | 0.516 | 0.25 | 0.1 | 0.14 | 0.15 | 0.23 | 0.23 |
| 3 | 6187051 | rs7651296 | -0.36 | -0.342 | 0.329 | -0.107 | -0.006 | 0.058 | 0.058 | 0 | 0.06 | 0.09 | 0.07 | 0.25 | 0.06 | 0.16 |
| 3 | 6187451 | rs9820630 | -0.866 | -0.334 | 0.362 | -0.134 | -0.085 | 0.033 | 0.033 | 0.022 | 0.2 | 0.47 | 0.15 | 0.25 | 0.11 | 0.26 |
| 3 | 6189342 | rs2134773 | 1.5 | -0.311 | 0.123 | -0.113 | -0.015 | 0.025 | 0.025 | 0 | 0.15 | 0.06 | 0.1 | 0 | 0.15 | 0.05 |
| 3 | 6189636 | rs2174262 | -1.848 | -0.272 | 0.043 | -0.084 | -0.07 | -0.036 | -0.036 | 0.015 | 0.62 | 0.88 | 0.42 | 0.42 | 0.56 | 0.62 |

| | | | | | | | | | | | | | | | | |
|---|---|---|---|---|---|---|---|---|---|---|---|---|---|---|---|---|
| 3 | 6197199 | rs7624103 | -2.295 | -0.652 | -0.04 | -0.275 | -0.119 | -0.178 | -0.178 | 2.665 | 0.7 | 0.97 | 0.49 | 0.54 | 0.63 | 0.71 |
| 3 | 6197932 | rs9311817 | -0.799 | -1.098 | 0.341 | -1.099 | 0.017 | 0.065 | 0.065 | 0.444 | 0.36 | 0.71 | 0.25 | 0.09 | 0.3 | 0.37 |
| 3 | 6199170 | rs13091404 | -2.013 | -1.01 | 0.325 | -0.985 | -0.087 | 0.001 | 0.001 | 0 | 0.64 | 0.9 | 0.4 | 0.28 | 0.56 | 0.62 |
| 3 | 6201267 | rs17215939 | 0.401 | -1.076 | 0.36 | -0.932 | 0.012 | -0.018 | -0.018 | 3.029 | 0.1 | 0.08 | 0.04 | 0.18 | 0.08 | 0.19 |
| 3 | 6205835 | rs9860865 | -0.202 | -0.907 | 0.433 | -0.918 | -0.002 | -0.006 | -0.006 | 0 | 0.07 | 0.35 | 0.03 | 0 | 0.01 | 0.03 |
| 3 | 6206636 | rs13080576 | 0.996 | -0.746 | -0.048 | -0.917 | 0.012 | -0.045 | -0.045 | 0.689 | 0.16 | 0.15 | 0.12 | 0.03 | 0.25 | 0.15 |
| 3 | 6206901 | rs17215946 | -0.293 | -1.024 | -0.062 | -0.743 | 0.017 | -0.005 | -0.005 | 0 | 0.06 | 0.04 | 0.04 | 0.21 | 0.01 | 0.27 |
| 3 | 6209095 | rs6798664 | -1.055 | -0.961 | -0.091 | -0.772 | 0.07 | 0.066 | 0.066 | 0.353 | 0.48 | 0.69 | 0.27 | 0.24 | 0.3 | 0.47 |
| 3 | 6213788 | rs1398788 | -0.371 | -1.08 | -0.12 | -0.645 | -0.044 | 0.067 | 0.067 | 0.495 | 0.09 | 0.24 | 0.13 | 0.22 | 0.02 | 0.25 |
| 3 | 6215292 | rs7636014 | 0.441 | -0.768 | -0.93 | -0.645 | -0.074 | 0.025 | 0.025 | 0.055 | 0.21 | 0.05 | 0.19 | 0.1 | 0.31 | 0.19 |
| 3 | 6215556 | rs7613556 | 0.481 | -0.241 | -0.677 | -0.623 | 0.009 | -0.012 | -0.012 | 0 | 0.2 | 0.11 | 0.12 | 0 | 0.13 | 0.06 |
| 3 | 6219430 | rs7625781 | -0.344 | 0.256 | -0.514 | 0.065 | -0.006 | 0.003 | 0.003 | 0 | 0.59 | 0.64 | 0.48 | 0.32 | 0.52 | 0.57 |
| 3 | 6224872 | rs1394156 | -0.39 | 0.341 | -1.062 | 0.055 | -0.04 | 0.012 | 0.012 | 0 | 0.6 | 0.78 | 0.49 | 0.3 | 0.51 | 0.57 |
| 3 | 6225798 | rs6782704 | -1.209 | -0.389 | -0.204 | -0.384 | -0.037 | -0.079 | -0.079 | 0.015 | 0.14 | 0.27 | 0.09 | 0.01 | 0.08 | 0.09 |
| 3 | 6226626 | rs13078519 | -1.042 | -0.423 | -0.871 | -0.716 | 0.011 | 0.083 | 0.083 | 0 | 0.21 | 0.08 | 0.31 | 0.01 | 0.26 | 0.23 |
| 3 | 6230553 | rs12374158 | -1.716 | -0.556 | 0.044 | -0.699 | 0.016 | -0.037 | -0.037 | 0.239 | 0.08 | 0.3 | 0.22 | 0.01 | 0.06 | 0.06 |
| 3 | 6231998 | rs9839456 | -1.581 | -0.657 | -0.091 | -0.749 | -0.003 | -0.022 | -0.022 | 0 | 0.1 | 0.16 | 0.22 | 0.05 | 0.08 | 0.09 |
| 3 | 6236038 | rs6790803 | -1.833 | -0.977 | 0.223 | -0.543 | 0.03 | 0.034 | 0.034 | 0 | 0.44 | 0.66 | 0.46 | 0.27 | 0.41 | 0.35 |
| 3 | 6236442 | rs13059452 | -1.338 | -1.359 | 0.196 | -0.884 | 0.052 | 0.072 | 0.072 | 0 | 0.33 | 0.2 | 0.35 | 0.06 | 0.28 | 0.3 |
| 3 | 6237572 | rs6797204 | -1.317 | -1.383 | 0.194 | -0.884 | 0.056 | 0.082 | 0.082 | 0 | 0.32 | 0.19 | 0.36 | 0.06 | 0.29 | 0.3 |
| 3 | 6240746 | rs7648545 | -1.278 | -1.316 | -0.088 | -0.844 | 0.074 | 0.138 | 0.138 | 0 | 0.32 | 0.14 | 0.34 | 0.06 | 0.28 | 0.28 |
| 3 | 6246651 | rs182286 | 1.285 | -0.522 | -0.466 | -0.303 | 0.012 | -0.046 | -0.046 | 0 | 0.42 | 0.23 | 0.28 | 0.5 | 0.32 | 0.35 |
| 3 | 6248988 | rs9850025 | -1.188 | -0.681 | 0.149 | -0.148 | 0 | 0.002 | 0.002 | 0 | 0.45 | 0.6 | 0.65 | 0.59 | 0.6 | 0.61 |
| 3 | 6249846 | rs6780164 | -1.179 | -0.618 | 0.306 | -0.145 | 0.002 | 0.003 | 0.003 | 0 | 0.45 | 0.6 | 0.65 | 0.6 | 0.6 | 0.62 |
| 3 | 6251762 | rs9837289 | -0.456 | -0.802 | 0.902 | -0.169 | 0.017 | 0.012 | 0.012 | 0 | 0.33 | 0.36 | 0.53 | 0.5 | 0.46 | 0.48 |
| 3 | 6255997 | rs6774280 | -0.49 | -0.848 | 0.813 | -0.301 | -0.002 | -0.019 | -0.019 | 2.197 | 0.13 | 0.18 | 0.29 | 0.07 | 0.18 | 0.17 |
| 3 | 6261246 | rs9848532 | -0.406 | -0.476 | 1.173 | -0.112 | 0.031 | 0.023 | 0.023 | 0 | 0.3 | 0.42 | 0.36 | 0.59 | 0.4 | 0.44 |
| 3 | 6262786 | rs11715135 | -0.69 | -0.452 | 1.219 | -0.13 | -0.004 | -0.025 | -0.025 | 0.883 | 0.14 | 0.22 | 0.29 | 0.12 | 0.18 | 0.19 |
| 3 | 6265920 | rs9870867 | -0.152 | -0.346 | 1.556 | 0.011 | 0.016 | 0.019 | 0.019 | 0 | 0.27 | 0.3 | 0.34 | 0.53 | 0.38 | 0.42 |
| 3 | 6267517 | rs286589 | -1.742 | -0.626 | 1.392 | -0.174 | -0.016 | 0.03 | 0.03 | 0 | 0.47 | 0.75 | 0.63 | 0.66 | 0.57 | 0.6 |
| 3 | 6272664 | rs4686025 | -1.358 | -0.262 | 1.474 | 0.065 | 0.016 | 0.006 | 0.006 | 0 | 0.39 | 0.55 | 0.59 | 0.57 | 0.54 | 0.56 |
| 3 | 6273813 | rs17044410 | -0.231 | -0.228 | 1.204 | 0.087 | 0.006 | 0.009 | 0.009 | 0 | 0.27 | 0.27 | 0.33 | 0.48 | 0.36 | 0.4 |
| 3 | 6279652 | rs1603866 | -1.086 | -0.263 | 1.089 | 0.06 | -0.001 | -0.008 | -0.008 | 2.204 | 0.1 | 0.13 | 0.25 | 0.1 | 0.15 | 0.14 |
| 3 | 6281402 | rs6776326 | -0.429 | -0.281 | 1.081 | 0.062 | -0.005 | -0.006 | -0.006 | 0 | 0.07 | 0.1 | 0.06 | 0.06 | 0.09 | 0.1 |
| 3 | 6287176 | rs6796959 | -1.017 | -0.101 | 0.911 | 0.103 | -0.02 | -0.012 | -0.012 | 0.061 | 0.34 | 0.27 | 0.52 | 0.51 | 0.47 | 0.47 |
| 3 | 6289556 | rs4686031 | -0.687 | 0.03 | 0.665 | 0.064 | -0.005 | 0.012 | 0.012 | 0.05 | 0.07 | 0.04 | 0.21 | 0.04 | 0.1 | 0.07 |
| 3 | 6291637 | rs6767746 | -1.947 | -0.116 | 1.128 | 0.004 | -0.009 | -0.054 | -0.054 | 0.04 | 0.12 | 0.23 | 0.25 | 0.04 | 0.16 | 0.16 |
| 3 | 6292920 | rs9818235 | -1.497 | -0.276 | 1.086 | -0.018 | -0.01 | -0.003 | -0.003 | 0.146 | 0.39 | 0.66 | 0.57 | 0.56 | 0.49 | 0.5 |
| 3 | 6293907 | rs2082938 | -1.816 | -0.255 | 0.728 | -0.004 | -0.028 | 0.003 | 0.003 | 0.059 | 0.44 | 0.76 | 0.62 | 0.56 | 0.56 | 0.59 |
| 3 | 6298349 | rs7636434 | -0.441 | -0.152 | 0.949 | 0.025 | 0.012 | 0.012 | 0.012 | 0 | 0.27 | 0.29 | 0.34 | 0.52 | 0.39 | 0.4 |
| 3 | 6302110 | rs7629918 | 0.302 | -0.054 | 0.808 | -0.045 | -0.012 | 0.005 | 0.005 | 0.023 | 0.16 | 0.12 | 0.12 | 0.02 | 0.24 | 0.16 |
| 3 | 6303579 | rs17272131 | -0.54 | -0.263 | 0.854 | 0.062 | -0.01 | -0.01 | -0.01 | 0.162 | 0.29 | 0.26 | 0.34 | 0.52 | 0.37 | 0.41 |
| 3 | 6307998 | rs1846439 | -1.555 | -0.189 | -0.017 | 0.122 | -0.028 | 0.012 | 0.012 | 0 | 0.46 | 0.79 | 0.54 | 0.57 | 0.53 | 0.6 |
| 3 | 6320151 | rs7625439 | -0.866 | -0.222 | 0.38 | 0.113 | -0.003 | -0.013 | -0.013 | 1.387 | 0.07 | 0.26 | 0.04 | 0.04 | 0.05 | 0.03 |
| 3 | 6322086 | rs3967024 | -1.283 | -0.267 | 0.263 | 0.488 | -0.034 | -0.019 | -0.019 | 0 | 0.4 | 0.78 | 0.41 | 0.41 | 0.48 | 0.52 |
| 3 | 6323162 | rs4686038 | -1.867 | -0.346 | 0.197 | 0.437 | -0.084 | -0.042 | -0.042 | 0.038 | 0.48 | 0.94 | 0.58 | 0.58 | 0.53 | 0.6 |
| 3 | 6328081 | rs1595039 | -1.198 | -0.302 | 0.252 | 0.47 | -0.038 | -0.034 | -0.034 | 0 | 0.4 | 0.78 | 0.46 | 0.55 | 0.48 | 0.55 |
| 3 | 6328641 | rs6780639 | -0.889 | -0.276 | 0.562 | 0.265 | -0.013 | -0.016 | -0.016 | 0 | 0.34 | 0.27 | 0.37 | 0.51 | 0.41 | 0.46 |
| 3 | 6342860 | rs157301 | -0.418 | -0.854 | -0.455 | 0.176 | -0.032 | -0.002 | -0.002 | 0 | 0.39 | 0.24 | 0.31 | 0.4 | 0.46 | 0.39 |
| 3 | 6343913 | rs1369621 | -1.87 | -0.987 | -0.726 | 0.046 | -0.095 | -0.094 | -0.094 | 0 | 0.57 | 0.95 | 0.62 | 0.57 | 0.56 | 0.67 |
| 3 | 6345015 | rs2117784 | -0.022 | -0.931 | -0.636 | 0.169 | -0.023 | -0.023 | -0.023 | 0 | 0.33 | 0.25 | 0.3 | 0.47 | 0.39 | 0.47 |
| 3 | 6354049 | rs11925550 | -0.271 | -0.88 | -0.287 | 0.15 | -0.036 | -0.059 | -0.059 | 0 | 0.1 | 0.19 | 0.08 | 0 | 0.09 | 0.11 |
| 3 | 6354598 | rs7613956 | -0.288 | -0.647 | -0.62 | 0.319 | 0.004 | 0.024 | 0.024 | 0 | 0.08 | 0.24 | 0.21 | 0.06 | 0.05 | 0.05 |
| 3 | 6370544 | rs6791211 | -1.277 | -0.579 | -1.587 | 0.325 | -0.103 | -0.048 | -0.048 | 1.243 | 0.55 | 0.95 | 0.62 | 0.6 | 0.56 | 0.66 |
| 3 | 6379792 | rs157297 | 0.833 | -0.17 | -1.787 | -0.111 | -0.116 | -0.111 | -0.111 | 0 | 0.5 | 0.02 | 0.41 | 0.52 | 0.54 | 0.39 |
| 3 | 6389075 | rs6800214 | 1.078 | 0.213 | -1.878 | -0.314 | -0.028 | -0.083 | -0.083 | 0.081 | 0.41 | 0.14 | 0.33 | 0.5 | 0.41 | 0.33 |
| 3 | 6389368 | rs1471968 | 0.645 | 0.532 | -1.276 | -0.402 | -0.009 | -0.005 | -0.005 | 0 | 0.55 | 0.5 | 0.51 | 0.64 | 0.65 | 0.49 |
| 3 | 6390337 | rs1435723 | -0.496 | 0.369 | -1.066 | -0.389 | 0.006 | 0.003 | 0.003 | 0 | 0.73 | 0.74 | 0.83 | 0.76 | 0.83 | 0.73 |
| 3 | 6391818 | rs1488419 | -0.232 | 0.571 | -0.844 | -0.28 | 0.033 | 0.05 | 0.05 | 0 | 0.65 | 0.79 | 0.78 | 0.82 | 0.73 | 0.61 |
| 3 | 6391939 | rs6806982 | 0.081 | 0.651 | -0.568 | -0.271 | 0.051 | 0.093 | 0.093 | 1.343 | 0.07 | 0.35 | 0.23 | 0.06 | 0.05 | 0.09 |
| 3 | 6394012 | rs9845211 | -0.407 | 0.978 | -0.908 | -0.515 | 0.024 | -0.012 | -0.012 | 0 | 0.21 | 0.2 | 0.24 | 0.15 | 0.38 | 0.18 |
| 3 | 6395238 | rs1488416 | 0.099 | 1.043 | -0.685 | -0.108 | 0.084 | 0.011 | 0.011 | 0 | 0.14 | 0.27 | 0.19 | 0.15 | 0.24 | 0.16 |
| 3 | 6395994 | rs9856021 | 0.829 | 0.854 | -0.805 | -0.125 | 0.001 | 0.04 | 0.04 | 0 | 0.51 | 0.45 | 0.59 | 0.67 | 0.49 | 0.46 |
| 3 | 6399073 | rs17044744 | 0.004 | 0.94 | -1.164 | -0.093 | 0.087 | 0.01 | 0.01 | 0 | 0.14 | 0.28 | 0.19 | 0.16 | 0.25 | 0.16 |
| 3 | 6399317 | rs6443047 | 0.809 | 0.087 | -0.949 | -0.112 | 0.015 | 0.015 | 0.015 | 0 | 0.56 | 0.38 | 0.54 | 0.75 | 0.68 | 0.51 |
| 3 | 6401420 | rs7651125 | 0.862 | -0.102 | -1.286 | -0.079 | 0.047 | 0.037 | 0.037 | 0 | 0.53 | 0.31 | 0.53 | 0.74 | 0.68 | 0.47 |
| 3 | 6402036 | rs17288121 | 0.236 | -0.086 | -1.5 | -0.083 | 0 | 0.001 | 0.001 | 0 | 0.06 | 0.08 | 0.13 | 0.07 | 0.05 | 0.06 |
| 3 | 6406809 | rs2217845 | 1.452 | 0.085 | -1.614 | -0.107 | -0.122 | -0.045 | -0.045 | 0 | 0.42 | 0.06 | 0.51 | 0.62 | 0.51 | 0.36 |
| 3 | 6409144 | rs6796561 | 0.328 | 0.144 | -1.518 | -0.351 | -0.012 | -0.019 | -0.019 | 0 | 0.59 | 0.4 | 0.75 | 0.83 | 0.72 | 0.52 |

| | | | | | | | | | | | | | | | | |
|---|---|---|---|---|---|---|---|---|---|---|---|---|---|---|---|---|
| 3 | 6410960 | rs9879475 | 1.542 | 0.085 | -1.545 | -0.221 | -0.12 | -0.113 | -0.113 | 2.091 | 0.41 | 0.06 | 0.39 | 0.5 | 0.49 | 0.35 |
| 3 | 6415990 | rs17288128 | 0.377 | 0.793 | -1.311 | 0.107 | 0 | 0.026 | 0.026 | 0 | 0.07 | 0.08 | 0.25 | 0.21 | 0.06 | 0.06 |
| 3 | 6416044 | rs1352400 | 0.326 | 0.843 | -1.292 | 0.082 | 0.088 | -0.008 | -0.008 | 0 | 0.13 | 0.27 | 0.13 | 0.13 | 0.18 | 0.13 |
| 3 | 6418430 | rs157463 | 0.536 | -0.486 | -2.186 | 0.21 | 0.023 | 0.009 | 0.009 | 0 | 0.15 | 0.02 | 0.09 | 0.12 | 0.09 | 0.2 |
| 3 | 6419564 | rs4686053 | -0.155 | -0.018 | -1.896 | 0.452 | -0.001 | 0.036 | 0.036 | 0 | 0.07 | 0.12 | 0.25 | 0.22 | 0.05 | 0.06 |
| 3 | 6420269 | rs1435716 | 0.207 | -0.011 | -1.967 | -0.486 | 0.032 | 0.043 | 0.043 | 0 | 0.65 | 0.78 | 0.81 | 0.83 | 0.73 | 0.6 |
| 3 | 6422814 | rs904460 | -0.061 | 0.63 | -2.363 | -0.811 | -0.014 | 0.221 | 0.221 | 0.298 | 0.49 | 0.2 | 0.69 | 0.91 | 0.53 | 0.7 |
| 3 | 6423320 | rs156168 | -0.349 | 0.806 | -2.823 | 0.032 | -0.052 | -0.102 | -0.102 | 1.19 | 0.82 | 0.99 | 0.74 | 0.62 | 0.8 | 0.78 |
| 3 | 6425168 | rs156166 | -0.068 | 0.643 | -3.254 | -1.125 | -0.062 | 0.168 | 0.168 | 0 | 0.38 | 0.17 | 0.28 | 0.04 | 0.43 | 0.17 |
| 3 | 6426631 | rs156164 | -0.309 | -0.008 | -3.594 | -0.684 | 0.088 | -0.046 | -0.046 | 0.209 | 0.5 | 0.91 | 0.34 | 0.45 | 0.42 | 0.59 |
| 3 | 6428207 | rs17600119 | 0.232 | -0.029 | -2.984 | -0.8 | 0.006 | 0.037 | 0.037 | 0.1 | 0.14 | 0.05 | 0.1 | 0.04 | 0.12 | 0.08 |
| 3 | 6435584 | rs458669 | 0.164 | -0.067 | -2.298 | -1.143 | 0 | -0.058 | -0.058 | 0 | 0.69 | 0.68 | 0.79 | 0.95 | 0.68 | 0.79 |
| 3 | 6436873 | rs12487852 | -0.344 | -0.908 | -2.191 | -1.214 | -0.004 | -0.024 | -0.024 | 0 | 0.28 | 0.26 | 0.19 | 0.12 | 0.24 | 0.2 |
| 3 | 6437832 | rs474423 | -0.947 | -0.869 | -1.568 | -0.715 | 0.007 | 0.106 | 0.106 | 0 | 0.54 | 0.77 | 0.34 | 0.26 | 0.44 | 0.55 |
| 3 | 6438854 | rs411041 | -0.972 | -0.871 | -1.675 | -0.699 | -0.003 | 0.111 | 0.111 | 0 | 0.54 | 0.82 | 0.35 | 0.27 | 0.45 | 0.55 |
| 3 | 6457290 | rs156175 | 0.756 | -0.097 | -1.959 | -0.093 | -0.084 | -0.075 | -0.075 | 0 | 0.18 | 0.02 | 0.29 | 0.3 | 0.26 | 0.17 |
| 3 | 6458324 | rs155514 | 0.394 | -0.717 | -1.941 | -0.182 | 0.032 | 0.051 | 0.051 | 0 | 0.6 | 0.12 | 0.68 | 0.69 | 0.69 | 0.64 |
| 3 | 6462066 | rs156672 | 0.257 | -1.033 | -1.213 | -0.584 | -0.014 | -0.069 | -0.069 | 0.102 | 0.26 | 0.3 | 0.17 | 0.04 | 0.18 | 0.15 |
| 3 | 6462422 | rs156495 | 0.497 | -0.728 | -2.187 | -0.312 | -0.078 | -0.141 | -0.141 | 0.185 | 0.26 | 0.05 | 0.4 | 0.51 | 0.31 | 0.43 |
| 3 | 6462629 | rs9814586 | 1.159 | -0.996 | -2.124 | -0.276 | -0.104 | -0.141 | -0.141 | 0.116 | 0.41 | 0.03 | 0.35 | 0.43 | 0.46 | 0.4 |
| 3 | 6464294 | rs155421 | 0.09 | -0.153 | -1.418 | 0.191 | -0.053 | -0.079 | -0.079 | 0 | 0.71 | 0.89 | 0.56 | 0.5 | 0.61 | 0.57 |
| 3 | 6474068 | rs6780751 | -0.282 | -0.49 | -0.645 | -0.127 | 0.005 | -0.003 | -0.003 | 0 | 0.36 | 0.43 | 0.48 | 0.42 | 0.47 | 0.44 |
| 3 | 6475006 | rs156488 | -1.357 | -1.268 | -0.871 | -0.104 | -0.024 | 0.027 | 0.027 | 0 | 0.68 | 0.79 | 0.69 | 0.84 | 0.62 | 0.73 |
| 3 | 6475608 | rs17216007 | 0.815 | -1.274 | -0.854 | -0.03 | 0.005 | -0.006 | -0.006 | 0 | 0.18 | 0.16 | 0.11 | 0.35 | 0.09 | 0.13 |
| 3 | 6478949 | rs2124811 | 0.024 | -0.616 | -0.517 | -0.038 | -0.026 | 0.009 | 0.009 | 0.065 | 0.35 | 0.3 | 0.48 | 0.42 | 0.47 | 0.44 |
| 3 | 6481625 | rs17216035 | 0.863 | -0.403 | -0.496 | 0.175 | 0.01 | 0.005 | 0.005 | 0 | 0.17 | 0.19 | 0.34 | 0.37 | 0.29 | 0.22 |
| 3 | 6483288 | rs267485 | 0.028 | -1.556 | -0.339 | 0.359 | -0.043 | 0.032 | 0.032 | 0 | 0.31 | 0.33 | 0.2 | 0.42 | 0.15 | 0.29 |
| 3 | 6485741 | rs266410 | 0.51 | -0.845 | -0.418 | 0.287 | -0.007 | 0 | 0 | 0 | 0.22 | 0.24 | 0.22 | 0.17 | 0.2 | 0.17 |
| 3 | 6486592 | rs17824615 | 0.076 | -0.612 | -0.184 | 0.544 | -0.008 | 0.019 | 0.019 | 0 | 0.12 | 0.09 | 0.15 | 0.05 | 0.16 | 0.09 |
| 3 | 6488102 | rs266413 | 1.022 | -1.541 | 0.355 | 0.291 | -0.036 | 0.004 | 0.004 | 0 | 0.73 | 0.56 | 0.74 | 0.75 | 0.78 | 0.59 |
| 3 | 6491868 | rs267501 | 0.886 | -1.56 | 0.344 | 0.315 | -0.012 | 0.044 | 0.044 | 0 | 0.07 | 0.09 | 0.12 | 0.2 | 0.04 | 0.15 |
| 3 | 6493262 | rs9855624 | -1.536 | -0.971 | 0.515 | 0.425 | -0.018 | 0.001 | 0.001 | 0 | 0.42 | 0.73 | 0.62 | 0.63 | 0.48 | 0.6 |
| 3 | 6496362 | rs267509 | -0.827 | -1.06 | 0.575 | 0.412 | 0.006 | 0.018 | 0.018 | 0 | 0.31 | 0.44 | 0.49 | 0.64 | 0.34 | 0.53 |
| 3 | 6497148 | rs7621178 | -1.453 | -1.599 | 0.845 | 0.661 | -0.046 | 0.023 | 0.023 | 0 | 0.29 | 0.53 | 0.33 | 0.3 | 0.22 | 0.43 |
| 3 | 6507290 | rs263705 | -0.762 | -1.592 | 0.752 | 0.782 | -0.018 | 0.065 | 0.065 | 0 | 0.92 | 0.87 | 0.88 | 0.8 | 0.95 | 0.85 |
| 3 | 6508807 | rs8180087 | 0.364 | -1.986 | 0.692 | 0.695 | -0.016 | 0.037 | 0.037 | 0.641 | 0.14 | 0.16 | 0.06 | 0.28 | 0.09 | 0.1 |
| 3 | 6516544 | rs154961 | -1.314 | -1.653 | 1.028 | 0.489 | -0.043 | 0.017 | 0.017 | 0 | 0.28 | 0.54 | 0.33 | 0.29 | 0.21 | 0.38 |
| 3 | 6518182 | rs155203 | 0.159 | -1.614 | 0.819 | 0.478 | -0.013 | 0.048 | 0.048 | 0 | 0.09 | 0.12 | 0.18 | 0.27 | 0.05 | 0.18 |
| 3 | 6522043 | rs9849820 | -1.033 | -1.062 | 1.092 | 0.527 | -0.012 | 0.031 | 0.031 | 0 | 0.21 | 0.48 | 0.3 | 0.26 | 0.18 | 0.36 |
| 3 | 6534200 | rs9810593 | 0.092 | -1.021 | 1.094 | 0.531 | -0.027 | -0.009 | -0.009 | 0 | 0.07 | 0.21 | 0.07 | 0.05 | 0.01 | 0.18 |
| 3 | 6538290 | rs1902540 | -1.098 | -1.03 | 1.216 | 0.523 | -0.008 | 0.046 | 0.046 | 0 | 0.2 | 0.55 | 0.31 | 0.25 | 0.18 | 0.37 |
| 3 | 6546891 | rs9834048 | -0.656 | -0.977 | 0.885 | 0.431 | 0 | -0.044 | -0.044 | 0 | 0.17 | 0.25 | 0.17 | 0.06 | 0.17 | 0.24 |
| 3 | 6563176 | rs2034870 | -1.133 | -0.957 | 1.175 | 0.55 | -0.001 | 0.045 | 0.045 | 0 | 0.21 | 0.57 | 0.32 | 0.27 | 0.2 | 0.36 |
| 3 | 6565877 | rs10510338 | 0.266 | -1.008 | 1.251 | 0.553 | -0.022 | -0.004 | -0.004 | 1.735 | 0.06 | 0.18 | 0.07 | 0.05 | 0.01 | 0.18 |
| 3 | 6598777 | rs9864202 | -1.178 | -0.978 | 1.224 | 0.542 | 0.003 | 0.055 | 0.055 | 0 | 0.2 | 0.49 | 0.32 | 0.28 | 0.2 | 0.36 |
| 3 | 6616306 | rs6765070 | 0.218 | -0.33 | 0.495 | 0.67 | -0.051 | -0.045 | -0.045 | 0.245 | 0.22 | 0.14 | 0.35 | 0.47 | 0.34 | 0.38 |
| 3 | 6627756 | rs352747 | -0.129 | -1.28 | 0.189 | 0.32 | -0.009 | -0.002 | -0.002 | 0 | 0.72 | 0.76 | 0.88 | 0.71 | 0.87 | 0.77 |
| 3 | 6634839 | rs6791299 | -0.737 | -0.156 | 0.264 | 0.736 | -0.041 | -0.016 | -0.016 | 0.383 | 0.33 | 0.25 | 0.46 | 0.47 | 0.5 | 0.44 |
| 3 | 6649648 | rs1499064 | -0.27 | 0.063 | 0.378 | 0.812 | -0.013 | -0.022 | -0.022 | 0 | 0.29 | 0.26 | 0.38 | 0.49 | 0.34 | 0.41 |
| 3 | 6659001 | rs164723 | -0.685 | 0.043 | 0.364 | 0.835 | -0.039 | -0.024 | -0.024 | 0.054 | 0.38 | 0.25 | 0.47 | 0.5 | 0.51 | 0.47 |
| 3 | 6659688 | rs352278 | -0.645 | 0.063 | 0.218 | 0.833 | -0.054 | -0.035 | -0.035 | 0 | 0.37 | 0.22 | 0.46 | 0.5 | 0.51 | 0.47 |
| 3 | 6665147 | rs352284 | -0.795 | 0.056 | 0.395 | 0.864 | -0.024 | -0.003 | -0.003 | 0 | 0.4 | 0.29 | 0.47 | 0.46 | 0.51 | 0.45 |
| 3 | 6665520 | rs13101172 | 0.704 | -0.239 | 0.338 | 1.045 | -0.003 | 0.001 | 0.001 | 0.376 | 0.09 | 0.07 | 0.03 | 0.19 | 0.05 | 0.08 |
| 3 | 6675631 | rs587284 | -0.775 | 0.759 | 0.78 | 1.245 | 0.042 | 0.043 | 0.043 | 0 | 0.14 | 0.22 | 0.29 | 0.26 | 0.24 | 0.18 |
| 3 | 6677459 | rs682889 | -1.484 | 0.698 | 0.791 | 1.105 | 0.012 | 0.019 | 0.019 | 0 | 0.44 | 0.37 | 0.65 | 0.72 | 0.59 | 0.6 |
| 3 | 6678484 | rs9811958 | -0.727 | 0.71 | 0.808 | 1.116 | 0 | 0 | 0 | 0 | 0.35 | 0.37 | 0.55 | 0.72 | 0.43 | 0.57 |
| 3 | 6680311 | rs9850138 | -1.1 | 0.704 | 0.627 | 1.285 | 0.12 | 0.376 | 0.376 | 0.142 | 0.06 | 0.38 | 0.21 | 0.3 | 0.08 | 0.15 |
| 3 | 6683606 | rs6802713 | -0.217 | 0.688 | 0.557 | 1.282 | -0.033 | -0.027 | -0.027 | 0 | 0.3 | 0.18 | 0.36 | 0.43 | 0.35 | 0.42 |
| 3 | 6685044 | rs352276 | -1.364 | 0.694 | 0.467 | 1.144 | 0.002 | 0.003 | 0.003 | 0 | 0.46 | 0.6 | 0.67 | 0.72 | 0.59 | 0.6 |
| 3 | 6689058 | rs6796951 | -0.81 | 0.854 | 0.523 | 1.383 | 0.026 | 0.005 | 0.005 | 0 | 0.34 | 0.47 | 0.56 | 0.7 | 0.43 | 0.43 |
| 3 | 6693609 | rs17045515 | -0.511 | 0.817 | 0.315 | 1.523 | 0.117 | 0.441 | 0.441 | 0.029 | 0.05 | 0.29 | 0.2 | 0.33 | 0.08 | 0.14 |
| 3 | 6698506 | rs10490893 | -0.62 | 0.816 | 0.323 | 1.522 | 0.134 | 0.483 | 0.483 | 0.029 | 0.05 | 0.35 | 0.21 | 0.33 | 0.08 | 0.14 |
| 3 | 6718995 | rs7610613 | -0.656 | 0.818 | 0.189 | 1.525 | 0.13 | 0.483 | 0.483 | 0.19 | 0.06 | 0.37 | 0.21 | 0.33 | 0.08 | 0.14 |
| 3 | 6721335 | rs9841570 | -0.499 | 0.829 | 0.074 | 1.53 | 0.111 | 0.436 | 0.436 | 0.214 | 0.05 | 0.31 | 0.2 | 0.33 | 0.08 | 0.14 |
| 3 | 6722500 | rs183004 | 0.026 | 0.973 | -1.037 | 0.808 | -0.052 | 0.054 | 0.054 | 0 | 0.09 | 0 | 0.09 | 0 | 0.16 | 0.03 |
| 3 | 6723880 | rs1403146 | -0.593 | 0.792 | -0.758 | 1.096 | -0.003 | -0.005 | -0.005 | 0 | 0.28 | 0.3 | 0.35 | 0.42 | 0.34 | 0.36 |
| 3 | 6725076 | rs12631413 | -0.178 | 1.155 | -1.123 | 1.268 | -0.055 | -0.076 | -0.076 | 0.908 | 0.15 | 0.01 | 0.09 | 0.33 | 0.18 | 0.2 |
| 3 | 6725175 | rs17045707 | -0.847 | 1.561 | -1.013 | 0.831 | -0.02 | -0.002 | -0.002 | 0.135 | 0.29 | 0.25 | 0.28 | 0.31 | 0.41 | 0.23 |
| 3 | 6728385 | rs6787150 | -0.358 | 0.971 | -1.074 | 0.533 | 0.005 | 0.052 | 0.052 | 0 | 0.53 | 0.75 | 0.61 | 0.38 | 0.51 | 0.37 |

| | | | | | | | | | | | | | | | | |
|---|---|---|---|---|---|---|---|---|---|---|---|---|---|---|---|---|
| 3 | 6734031 | rs11929315 | -0.486 | -0.001 | -1.003 | 0.597 | -0.002 | -0.001 | -0.001 | 0 | 0.84 | 0.84 | 0.89 | 0.86 | 0.89 | 0.82 |
| 3 | 6738287 | rs10490892 | 1.089 | 0.076 | -0.814 | 0.495 | -0.001 | -0.014 | -0.014 | 0 | 0.12 | 0.15 | 0.11 | 0 | 0.15 | 0.02 |
| 3 | 6738389 | rs9811534 | 0.717 | -0.022 | -0.595 | 0.412 | -0.001 | 0.007 | 0.007 | 0 | 0.46 | 0.5 | 0.47 | 0.72 | 0.52 | 0.66 |
| 3 | 6745703 | rs9838132 | 0.74 | 0.075 | -0.678 | 0.379 | 0 | 0.031 | 0.031 | 0 | 0.25 | 0.14 | 0.31 | 0.14 | 0.22 | 0.14 |
| 3 | 6748069 | rs9990237 | 0.889 | 0.042 | -0.61 | 0.287 | 0 | -0.052 | -0.052 | 0.511 | 0.13 | 0.23 | 0.11 | 0 | 0.15 | 0.02 |
| 3 | 6748249 | rs9311962 | -0.545 | -0.119 | 0.103 | 0.514 | -0.001 | -0.002 | -0.002 | 0 | 0.42 | 0.56 | 0.46 | 0.33 | 0.39 | 0.25 |
| 3 | 6751477 | rs9841590 | -0.151 | -0.044 | -0.301 | 0.255 | 0.001 | 0.026 | 0.026 | 0 | 0.38 | 0.34 | 0.42 | 0.14 | 0.37 | 0.16 |
| 3 | 6751582 | rs9841750 | 0.785 | -0.052 | -0.287 | 0.249 | 0.002 | 0.016 | 0.016 | 0 | 0.26 | 0.22 | 0.31 | 0.14 | 0.22 | 0.14 |
| 3 | 6752246 | rs9829430 | -0.15 | -0.037 | -0.316 | 0.265 | 0.001 | 0.026 | 0.026 | 0 | 0.38 | 0.34 | 0.42 | 0.14 | 0.37 | 0.16 |
| 3 | 6752294 | rs9829452 | -0.15 | -0.037 | -0.316 | 0.253 | 0.001 | 0.026 | 0.026 | 0 | 0.38 | 0.34 | 0.42 | 0.14 | 0.37 | 0.16 |
| 3 | 6753259 | rs9851886 | 0.871 | 0.029 | -0.616 | 0.407 | -0.003 | 0.052 | 0.052 | 0 | 0.25 | 0.11 | 0.31 | 0.14 | 0.22 | 0.14 |
| 3 | 6760499 | rs7638636 | 0.844 | 0.051 | -0.774 | 0.65 | -0.076 | -0.001 | -0.001 | 0 | 0.41 | 0.15 | 0.42 | 0.44 | 0.51 | 0.55 |
| 3 | 6764788 | rs460579 | 0.358 | -0.454 | -0.289 | 0.256 | -0.001 | 0.086 | 0.086 | 0.426 | 0.27 | 0.09 | 0.14 | 0.11 | 0.23 | 0.15 |
| 3 | 6766178 | rs978658 | 1.12 | -0.605 | 0.238 | 0.398 | 0.035 | 0.043 | 0.043 | 1.381 | 0.51 | 0.26 | 0.56 | 0.62 | 0.63 | 0.58 |
| 3 | 6766266 | rs978659 | 0.905 | -0.677 | 0.881 | 0.395 | -0.003 | -0.018 | -0.018 | 0.218 | 0.55 | 0.52 | 0.58 | 0.79 | 0.62 | 0.67 |
| 3 | 6766899 | rs345246 | -0.063 | -0.627 | 1.093 | 0.438 | -0.006 | -0.002 | -0.002 | 0.161 | 0.85 | 0.77 | 0.76 | 0.89 | 0.89 | 0.83 |
| 3 | 6770752 | rs345233 | -0.017 | -0.624 | 1.133 | 0.409 | -0.016 | -0.018 | -0.018 | 0.897 | 0.31 | 0.41 | 0.17 | 0.27 | 0.24 | 0.24 |
| 3 | 6770925 | rs345232 | 0.464 | -0.617 | 1.126 | 0.402 | 0.002 | 0.065 | 0.065 | 0.45 | 0.27 | 0.11 | 0.14 | 0.11 | 0.23 | 0.15 |
| 3 | 6771475 | rs164030 | -0.159 | -0.538 | 1.24 | 0.376 | 0 | 0.001 | 0.001 | 0 | 0.87 | 0.89 | 0.78 | 0.92 | 0.89 | 0.84 |
| 3 | 6781329 | rs12638623 | -1.495 | -0.49 | 0.937 | 0.533 | 0.025 | 0.186 | 0.186 | 0.892 | 0.11 | 0.51 | 0.21 | 0.25 | 0.07 | 0.22 |
| 3 | 6781909 | rs7629489 | 0.302 | -0.464 | 0.474 | 0.214 | 0.007 | 0.035 | 0.035 | 0.72 | 0.61 | 0.74 | 0.64 | 0.85 | 0.67 | 0.7 |
| 3 | 6785413 | rs12638350 | -0.743 | -0.384 | 0.331 | 0.676 | 0.001 | 0.006 | 0.006 | 0 | 0.05 | 0.05 | 0.12 | 0.16 | 0.04 | 0.04 |
| 3 | 6785430 | rs2633748 | 0.895 | -0.6 | 0.643 | 0.538 | -0.011 | 0.02 | 0.02 | 0 | 0.85 | 0.51 | 0.79 | 0.88 | 0.91 | 0.82 |
| 3 | 6788580 | rs4686091 | -1.418 | -0.497 | 0.526 | 0.587 | 0.01 | 0.156 | 0.156 | 0.137 | 0.07 | 0.26 | 0.22 | 0.22 | 0.04 | 0.15 |
| 3 | 6789265 | rs12491369 | 1.363 | -0.671 | -0.388 | 0.297 | -0.061 | -0.051 | -0.051 | 0 | 0.47 | 0.05 | 0.35 | 0.56 | 0.59 | 0.52 |
| 3 | 6792400 | rs163972 | 1.435 | -0.28 | 0.482 | 0.315 | 0.022 | 0.015 | 0.015 | 0 | 0.87 | 0.63 | 0.76 | 0.87 | 0.88 | 0.84 |
| 3 | 6794477 | rs163970 | 0.036 | -0.247 | 0.11 | 0.309 | -0.001 | 0 | 0 | 0 | 0.94 | 0.9 | 0.94 | 0.95 | 0.95 | 0.96 |
| 3 | 6795490 | rs163968 | 0.058 | -0.261 | -0.077 | 0 | -0.011 | 0.045 | 0.045 | 0 | 0.16 | 0.08 | 0.18 | 0.02 | 0.15 | 0.12 |
| 3 | 6795710 | rs171121 | -0.089 | -0.233 | -0.461 | -0.368 | -0.057 | 0.158 | 0.158 | 0.907 | 0.3 | 0.08 | 0.28 | 0.02 | 0.34 | 0.27 |
| 3 | 6796046 | rs7612734 | 1.034 | -0.251 | 0.222 | -0.188 | 0.001 | -0.013 | -0.013 | 1.828 | 0.56 | 0.47 | 0.52 | 0.8 | 0.53 | 0.49 |
| 3 | 6801391 | rs345200 | -0.073 | -0.141 | 0.088 | -0.238 | 0.006 | -0.047 | -0.047 | 0 | 0.3 | 0.34 | 0.3 | 0.08 | 0.34 | 0.37 |
| 3 | 6804839 | rs4686092 | 0.935 | -0.148 | -0.155 | 0.162 | -0.018 | 0.067 | 0.067 | 0.228 | 0.49 | 0.17 | 0.43 | 0.66 | 0.52 | 0.4 |
| 3 | 6805407 | rs378246 | -1.176 | -0.304 | 0.083 | 0.146 | -0.015 | -0.014 | -0.014 | 0 | 0.17 | 0.29 | 0.11 | 0.08 | 0.14 | 0.16 |
| 3 | 6805700 | rs168324 | -1.123 | -0.28 | 0.117 | 0.154 | -0.017 | 0.024 | 0.024 | 0 | 0.17 | 0.31 | 0.19 | 0.13 | 0.15 | 0.17 |
| 3 | 6813105 | rs11131052 | 0.917 | 0.009 | 0.095 | 0.622 | -0.013 | -0.006 | -0.006 | 0 | 0.5 | 0.18 | 0.43 | 0.59 | 0.51 | 0.43 |
| 3 | 6816887 | rs345206 | 0.175 | 0.047 | 0.138 | 0.64 | -0.001 | -0.001 | -0.001 | 0.121 | 0.61 | 0.35 | 0.52 | 0.65 | 0.64 | 0.53 |
| 3 | 6821304 | rs466544 | -0.28 | 0.163 | 0.554 | 0.964 | -0.005 | -0.007 | -0.007 | 0 | 0.58 | 0.5 | 0.62 | 0.66 | 0.65 | 0.65 |
| 3 | 6824734 | rs345215 | -0.766 | 0.2 | 0.187 | 0.785 | 0.004 | 0.057 | 0.057 | 0 | 0.63 | 0.74 | 0.73 | 0.88 | 0.68 | 0.7 |
| 3 | 6825162 | rs164026 | -0.903 | 0.404 | 0.252 | 1.113 | 0 | -0.004 | -0.004 | 0 | 0.12 | 0.1 | 0.14 | 0.06 | 0.14 | 0.14 |
| 3 | 6826255 | rs1400160 | -1.132 | 0.782 | 0.4 | 1.227 | 0.001 | 0.056 | 0.056 | 0 | 0.16 | 0.37 | 0.28 | 0.26 | 0.15 | 0.15 |
| 3 | 6827949 | rs345222 | -0.429 | 0.795 | -0.04 | 1.315 | 0.007 | 0.008 | 0.008 | 0 | 0.61 | 0.44 | 0.55 | 0.49 | 0.57 | 0.65 |
| 3 | 6832712 | rs12635360 | 0.028 | 0.935 | -0.117 | 0.801 | -0.007 | 0.079 | 0.079 | 0 | 0.54 | 0.3 | 0.41 | 0.23 | 0.48 | 0.58 |
| 3 | 6840697 | rs163541 | -1.199 | 1.201 | 0.238 | 1.095 | 0.041 | 0.092 | 0.092 | 0 | 0.08 | 0.19 | 0.13 | 0.2 | 0.11 | 0.08 |
| 3 | 6848338 | rs2046428 | -0.635 | 1.197 | -0.143 | 0.73 | -0.013 | -0.006 | -0.006 | 0 | 0.43 | 0.64 | 0.48 | 0.43 | 0.5 | 0.51 |
| 3 | 6852800 | rs12634144 | -0.337 | 1.473 | -0.381 | 0.942 | -0.017 | -0.024 | -0.024 | 0 | 0.36 | 0.26 | 0.42 | 0.43 | 0.46 | 0.47 |
| 3 | 6853593 | rs6808904 | -0.588 | 1.556 | -0.291 | 0.93 | -0.011 | -0.011 | -0.011 | 0.294 | 0.39 | 0.34 | 0.45 | 0.47 | 0.49 | 0.51 |
| 3 | 6861715 | rs1400286 | -0.467 | 1.544 | -0.593 | 1.071 | 0 | 0.001 | 0.001 | 0 | 0.06 | 0.11 | 0.17 | 0.06 | 0.05 | 0.04 |
| 3 | 6863068 | rs1516560 | -0.183 | 1.758 | -0.74 | 0.96 | -0.01 | -0.006 | -0.006 | 0 | 0.28 | 0.23 | 0.24 | 0.34 | 0.39 | 0.35 |
| 3 | 6867518 | rs11131055 | -0.239 | 1.575 | -0.738 | 0.88 | -0.008 | -0.008 | -0.008 | 0 | 0.29 | 0.25 | 0.24 | 0.37 | 0.38 | 0.38 |
| 3 | 6869166 | rs17046139 | -0.309 | 0.846 | -0.724 | 0.443 | 0 | -0.002 | -0.002 | 0 | 0.11 | 0.11 | 0.04 | 0.03 | 0.08 | 0.06 |
| 3 | 6874819 | rs437555 | -0.586 | 1.19 | -1.374 | 0.431 | -0.026 | -0.039 | -0.039 | 0.36 | 0.6 | 0.9 | 0.57 | 0.64 | 0.59 | 0.38 |
| 3 | 6879727 | rs184653 | -0.233 | 1.569 | -1.588 | 0.236 | 0.004 | 0.024 | 0.024 | 0.899 | 0.5 | 0.88 | 0.55 | 0.62 | 0.55 | 0.3 |
| 3 | 6882212 | rs342016 | 0.249 | 2.063 | -0.743 | 0.298 | 0.059 | 0.002 | 0.002 | 1.215 | 0.16 | 0.58 | 0.17 | 0.17 | 0.17 | 0.06 |
| 3 | 6889799 | rs342042 | -0.032 | 2.208 | -0.955 | 0.309 | 0.075 | -0.013 | -0.013 | 0.153 | 0.18 | 0.63 | 0.23 | 0.17 | 0.29 | 0.09 |
| 3 | 6893897 | rs7651971 | -0.264 | 1.363 | -1.345 | 0.43 | -0.005 | -0.039 | -0.039 | 0 | 0.43 | 0.19 | 0.4 | 0.49 | 0.43 | 0.33 |
| 3 | 6903297 | rs3749380 | 0.317 | 0.838 | -0.989 | 0.362 | 0.031 | 0.043 | 0.043 | 0.055 | 0.51 | 0.36 | 0.45 | 0.32 | 0.41 | 0.58 |
| 3 | 6908535 | rs342026 | 0.241 | 0.665 | -0.707 | 0.68 | -0.006 | 0.023 | 0.023 | 2.203 | 0.66 | 0.61 | 0.62 | 0.45 | 0.68 | 0.67 |
| 3 | 6910125 | rs17046224 | 0.185 | 0.715 | -0.888 | 0.77 | -0.005 | -0.032 | -0.032 | 0.049 | 0.08 | 0.05 | 0.13 | 0.17 | 0.08 | 0.1 |
| 3 | 6911400 | rs456835 | 0.68 | 0.658 | -0.795 | 0.037 | 0.011 | 0.083 | 0.083 | 0.591 | 0.48 | 0.27 | 0.24 | 0.31 | 0.44 | 0.55 |
| 3 | 6912613 | rs6773334 | 0.879 | 0.606 | -0.845 | -0.234 | -0.008 | 0.005 | 0.005 | 0 | 0.07 | 0.03 | 0.09 | 0 | 0.1 | 0.04 |
| 3 | 6916027 | rs17046239 | 0.873 | 0.275 | -0.923 | -0.226 | -0.001 | 0 | 0 | 0 | 0.06 | 0.06 | 0.03 | 0.1 | 0.06 | 0.07 |
| 3 | 6919079 | rs465152 | 0.542 | 0.37 | -0.561 | -0.322 | -0.001 | -0.009 | -0.009 | 0 | 0.48 | 0.49 | 0.31 | 0.2 | 0.45 | 0.51 |
| 3 | 6920419 | rs455860 | 1.046 | 0.704 | -0.44 | -0.195 | -0.002 | 0.078 | 0.078 | 0.366 | 0.42 | 0.24 | 0.21 | 0.2 | 0.37 | 0.46 |
| 3 | 6921906 | rs339805 | 0.056 | 0.816 | -0.533 | 0.104 | -0.001 | 0 | 0 | 0.92 | 0.61 | 0.61 | 0.47 | 0.29 | 0.58 | 0.59 |
| 3 | 6922000 | rs191443 | -0.27 | 0.691 | -0.452 | 0.186 | -0.001 | -0.001 | -0.001 | 0 | 0.65 | 0.61 | 0.52 | 0.29 | 0.63 | 0.64 |
| 3 | 6923647 | rs7651591 | -0.784 | 0.72 | 0.04 | 0.59 | -0.009 | 0.022 | 0.022 | 0.594 | 0.13 | 0.32 | 0.18 | 0.1 | 0.12 | 0.1 |
| 3 | 6925815 | rs12487836 | -0.684 | 0.923 | -0.331 | 0.303 | -0.001 | -0.004 | -0.004 | 0 | 0.09 | 0.06 | 0.08 | 0 | 0.1 | 0.08 |
| 3 | 6939834 | rs3846161 | -0.723 | 1.216 | -0.211 | -0.392 | 0.008 | -0.062 | -0.062 | 0.056 | 0.2 | 0.25 | 0.2 | 0.01 | 0.22 | 0.1 |
| 3 | 6942072 | rs9856655 | -0.215 | 1.31 | -0.219 | -0.41 | -0.002 | -0.011 | -0.011 | 0 | 0.08 | 0.05 | 0.06 | 0 | 0.13 | 0.05 |

| | | | | | | | | | | | | | | | | |
|---|---|---|---|---|---|---|---|---|---|---|---|---|---|---|---|---|
| 3 | 6950189 | rs7644436 | -1.135 | 0.915 | 0.321 | 0.199 | -0.018 | 0.04 | 0.04 | 0.34 | 0.25 | 0.59 | 0.17 | 0.32 | 0.19 | 0.12 |
| 3 | 6952375 | rs983534 | -1.382 | 0.842 | 0.129 | 0.694 | 0.025 | -0.026 | -0.026 | 0 | 0.31 | 0.81 | 0.28 | 0.57 | 0.22 | 0.26 |
| 3 | 6954873 | rs11921083 | -1.854 | 0.636 | 0.812 | 0.56 | 0.029 | 0.02 | 0.02 | 0.503 | 0.42 | 0.83 | 0.42 | 0.59 | 0.34 | 0.31 |
| 3 | 6955371 | rs9859343 | 0.798 | 0.592 | 1.194 | 0.529 | 0 | -0.028 | -0.028 | 0 | 0.07 | 0.09 | 0.03 | 0 | 0.06 | 0.01 |
| 3 | 6955734 | rs9311966 | -2.568 | 0.599 | 1.135 | 0.515 | -0.029 | 0.072 | 0.072 | 0.722 | 0.5 | 0.87 | 0.48 | 0.59 | 0.46 | 0.37 |
| 3 | 6959514 | rs10490858 | -0.003 | 0.554 | 1.716 | 0.455 | 0.004 | 0.341 | 0.341 | 0.043 | 0.07 | 0.31 | 0.07 | 0.32 | 0.02 | 0.05 |
| 3 | 6959982 | rs10490857 | 0.453 | 0.69 | 1.103 | 0.179 | -0.016 | 0.013 | 0.013 | 0 | 0.08 | 0.02 | 0.06 | 0 | 0.12 | 0.06 |
| 3 | 6963757 | rs182885 | -2.197 | 1.139 | 0.575 | -0.051 | 0.021 | 0.017 | 0.017 | 0 | 0.61 | 0.77 | 0.69 | 0.67 | 0.64 | 0.53 |
| 3 | 6966491 | rs339015 | -1.912 | 1.228 | 0.648 | -0.059 | -0.03 | 0.06 | 0.06 | 0 | 0.53 | 0.84 | 0.55 | 0.66 | 0.51 | 0.47 |
| 3 | 6968207 | rs2116711 | 0.296 | 1.224 | 0.633 | -0.089 | -0.002 | -0.035 | -0.035 | 0.375 | 0.11 | 0.12 | 0.07 | 0 | 0.1 | 0.07 |
| 3 | 6971081 | rs1516561 | -0.791 | 1.239 | 0.62 | -0.104 | 0 | -0.033 | -0.033 | 0 | 0.21 | 0.22 | 0.22 | 0.01 | 0.21 | 0.12 |
| 3 | 6971349 | rs17693618 | 0.802 | 1.375 | 0.05 | -0.123 | -0.017 | 0.026 | 0.026 | 0 | 0.09 | 0.04 | 0.08 | 0 | 0.11 | 0.05 |
| 3 | 6971989 | rs339022 | -0.472 | 1.326 | 0.097 | 0.074 | -0.019 | -0.023 | -0.023 | 0 | 0.27 | 0.14 | 0.37 | 0.32 | 0.31 | 0.36 |
| 3 | 6973116 | rs339020 | 0.341 | 1.263 | 0.279 | 0.248 | -0.005 | -0.023 | -0.023 | 0 | 0.18 | 0.13 | 0.31 | 0.32 | 0.2 | 0.29 |
| 3 | 6980293 | rs17046415 | -0.187 | 1.414 | 0.608 | 0.311 | 0.001 | -0.005 | -0.005 | 0.887 | 0.13 | 0.17 | 0.06 | 0.13 | 0.15 | 0.09 |
| 3 | 6980388 | rs17046418 | 0.582 | 1.489 | 0.611 | -0.006 | 0.004 | -0.037 | -0.037 | 0 | 0.1 | 0.15 | 0.09 | 0 | 0.11 | 0.05 |
| 3 | 6981134 | rs687338 | -0.269 | 1.286 | 0.551 | -0.039 | -0.001 | -0.018 | -0.018 | 0.209 | 0.27 | 0.17 | 0.34 | 0.34 | 0.26 | 0.3 |
| 3 | 6994953 | rs17694321 | -0.851 | 1.379 | 0.568 | -0.188 | 0.001 | -0.061 | -0.061 | 0 | 0.26 | 0.25 | 0.19 | 0 | 0.27 | 0.13 |
| 3 | 6995426 | rs2163735 | 0.29 | 1.316 | 0.398 | 0.302 | -0.003 | 0.044 | 0.044 | 0.263 | 0.12 | 0.11 | 0.09 | 0.34 | 0.1 | 0.07 |
| 3 | 6998093 | rs17046457 | 0.501 | 1.298 | 0.288 | -0.117 | 0 | -0.022 | -0.022 | 0 | 0.11 | 0.1 | 0.09 | 0 | 0.11 | 0.05 |
| 3 | 6999617 | rs163326 | -1.36 | 1.338 | 0.254 | 0.002 | -0.005 | 0.031 | 0.031 | 0 | 0.58 | 0.8 | 0.57 | 0.68 | 0.55 | 0.53 |
| 3 | 7003389 | rs693200 | -0.542 | 1.229 | 0.596 | -0.428 | -0.008 | -0.07 | -0.07 | 0 | 0.27 | 0.57 | 0.26 | 0.12 | 0.26 | 0.16 |
| 3 | 7003531 | rs1667850 | -0.393 | 1.349 | 0.124 | -0.376 | -0.003 | -0.048 | -0.048 | 0 | 0.26 | 0.44 | 0.26 | 0.12 | 0.26 | 0.16 |
| 3 | 7005021 | rs1516570 | -0.751 | 2.107 | -0.243 | -0.14 | 0.014 | -0.009 | -0.009 | 1.153 | 0.26 | 0.26 | 0.3 | 0.2 | 0.33 | 0.18 |
| 3 | 7005445 | rs478250 | -0.818 | 2.06 | -0.524 | -0.505 | 0.012 | -0.038 | -0.038 | 0 | 0.28 | 0.41 | 0.26 | 0.13 | 0.31 | 0.2 |
| 3 | 7006880 | rs4686101 | 0.141 | 1.592 | -0.921 | -1.024 | -0.001 | -0.02 | -0.02 | 0 | 0.49 | 0.51 | 0.43 | 0.19 | 0.44 | 0.27 |
| 3 | 7010081 | rs480409 | 0.792 | 1.431 | -1.476 | -0.43 | 0.015 | -0.075 | -0.075 | 1.881 | 0.2 | 0.08 | 0.34 | 0.63 | 0.23 | 0.44 |
| 3 | 7011534 | rs781393 | 0.169 | 2.297 | -1.496 | -0.892 | -0.005 | -0.026 | -0.026 | 0 | 0.81 | 0.78 | 0.73 | 0.99 | 0.78 | 0.83 |
| 3 | 7011962 | rs484283 | -0.423 | 2.328 | -1.726 | -0.5 | -0.027 | 0.029 | 0.029 | 0 | 0.94 | 0.95 | 0.92 | 1 | 0.89 | 0.96 |
| 3 | 7012842 | rs6796324 | 0.343 | 1.969 | -1.785 | -0.341 | -0.049 | -0.058 | -0.058 | 1.154 | 0.12 | 0.01 | 0.07 | 0.15 | 0.14 | 0.18 |
| 3 | 7013463 | rs1154352 | -0.242 | 1.546 | -1.244 | -0.657 | 0.011 | -0.043 | -0.043 | 0 | 0.66 | 0.57 | 0.71 | 0.85 | 0.64 | 0.92 |
| 3 | 7017919 | rs9883142 | 0.128 | 1.336 | -1.421 | 0.509 | 0.002 | 0.008 | 0.008 | 0 | 0.28 | 0.27 | 0.29 | 0.39 | 0.31 | 0.24 |
| 3 | 7018932 | rs781388 | 0.152 | 0.764 | -2.252 | 0.776 | -0.051 | -0.027 | -0.027 | 0.144 | 0.73 | 0.87 | 0.65 | 0.64 | 0.62 | 0.61 |
| 3 | 7020832 | rs17046516 | 0.728 | 0.243 | -2.309 | 0.839 | 0.065 | 0.016 | 0.016 | 2.203 | 0.26 | 0.06 | 0.22 | 0.21 | 0.15 | 0.25 |
| 3 | 7022968 | rs34165704 | 0.422 | 1.162 | -2.047 | 1.07 | -0.031 | -0.012 | -0.012 | 4.797 | 0.11 | 0.05 | 0.08 | 0.15 | 0.13 | 0.12 |
| 3 | 7024127 | rs712752 | 0.382 | 0.89 | -1.646 | 1.221 | -0.025 | -0.02 | -0.02 | 0 | 0.82 | 0.91 | 0.72 | 0.77 | 0.72 | 0.75 |
| 3 | 7026341 | rs670764 | 0.819 | 0.563 | -1.657 | 0.824 | 0.036 | 0.019 | 0.019 | 0 | 0.51 | 0.83 | 0.59 | 0.59 | 0.57 | 0.47 |
| 3 | 7026535 | rs13317247 | -0.171 | 0.25 | -1.435 | 0.852 | -0.001 | -0.001 | -0.001 | 0.535 | 0.15 | 0.11 | 0.14 | 0.15 | 0.19 | 0.17 |
| 3 | 7026696 | rs576913 | -0.103 | -0.124 | -1.361 | 0.771 | 0.062 | 0.002 | 0.002 | 0.533 | 0.27 | 0.05 | 0.22 | 0.22 | 0.17 | 0.3 |
| 3 | 7033701 | rs2324046 | -0.834 | 0.786 | -0.528 | 0.711 | 0.005 | 0.075 | 0.075 | 0 | 0.39 | 0.23 | 0.22 | 0.21 | 0.34 | 0.36 |
| 3 | 7035024 | rs1154357 | 1.181 | 0.489 | -0.602 | 0.653 | -0.036 | -0.072 | -0.072 | 0 | 0.48 | 0.1 | 0.52 | 0.46 | 0.57 | 0.38 |
| 3 | 7038214 | rs17752444 | 0.552 | 0.75 | -0.81 | 0.453 | -0.075 | -0.004 | -0.004 | 0 | 0.12 | 0.01 | 0.11 | 0.11 | 0.2 | 0.26 |
| 3 | 7039450 | rs2014195 | -0.703 | 1.054 | -0.817 | 0.398 | -0.076 | -0.081 | -0.081 | 0 | 0.21 | 0.01 | 0.25 | 0.29 | 0.24 | 0.25 |
| 3 | 7045455 | rs1499161 | 1.5 | 0.899 | -0.74 | 0.51 | -0.084 | -0.023 | -0.023 | 0.066 | 0.48 | 0.1 | 0.53 | 0.53 | 0.56 | 0.38 |
| 3 | 7052542 | rs1472476 | -0.02 | 0.809 | 0.268 | 0.39 | -0.009 | 0.02 | 0.02 | 0 | 0.27 | 0.23 | 0.19 | 0.18 | 0.26 | 0.4 |
| 3 | 7052752 | rs1532544 | -0.951 | 0.809 | 0.631 | 0.457 | -0.029 | 0.023 | 0.023 | 0.016 | 0.21 | 0.57 | 0.26 | 0.3 | 0.16 | 0.18 |
| 3 | 7057753 | rs7618803 | -1.134 | 0.912 | 0.311 | 0.427 | -0.007 | 0.013 | 0.013 | 0 | 0.47 | 0.34 | 0.43 | 0.49 | 0.45 | 0.61 |
| 3 | 7063064 | rs12639259 | -0.131 | 0.496 | -0.101 | 0.516 | 0.025 | -0.042 | -0.042 | 0 | 0.15 | 0.09 | 0.21 | 0.31 | 0.13 | 0.2 |
| 3 | 7066180 | rs7629851 | -0.097 | 0.917 | -0.217 | 0.42 | -0.032 | 0.057 | 0.057 | 0 | 0.33 | 0.22 | 0.22 | 0.18 | 0.34 | 0.44 |
| 3 | 7068147 | rs1021687 | -0.396 | 1.245 | 0.121 | 0.58 | -0.02 | 0.015 | 0.015 | 0 | 0.48 | 0.25 | 0.42 | 0.48 | 0.47 | 0.6 |
| 3 | 7072823 | rs1566366 | 0.521 | 1.246 | -0.896 | 0.734 | -0.063 | -0.083 | -0.083 | 0.006 | 0.44 | 0.07 | 0.49 | 0.48 | 0.48 | 0.36 |
| 3 | 7083387 | rs6764486 | -1.174 | 1.781 | 0.307 | 1.212 | 0.095 | 0.111 | 0.111 | 0 | 0.13 | 0.3 | 0.26 | 0.34 | 0.23 | 0.21 |
| 3 | 7083814 | rs1353832 | 0.859 | 0.749 | -0.266 | 1.181 | 0.019 | -0.026 | -0.026 | 0 | 0.72 | 0.8 | 0.68 | 0.7 | 0.75 | 0.62 |
| 3 | 7089629 | rs10510344 | -1.04 | -0.08 | 0.759 | 1.075 | -0.059 | 0.022 | 0.022 | 0 | 0.15 | 0.35 | 0.07 | 0.13 | 0.06 | 0.07 |
| 3 | 7090422 | rs7428271 | -0.978 | -0.206 | 0.181 | 0.734 | -0.043 | -0.031 | -0.031 | 1.265 | 0.59 | 0.81 | 0.49 | 0.48 | 0.48 | 0.55 |
| 3 | 7091501 | rs17234879 | 0.77 | -0.475 | -0.136 | 0.37 | -0.032 | 0.001 | 0.001 | 0 | 0.23 | 0.04 | 0.1 | 0.23 | 0.22 | 0.22 |
| 3 | 7093533 | rs10510347 | -0.008 | -0.574 | 0.12 | 0.221 | -0.026 | -0.014 | -0.014 | 0 | 0.15 | 0.17 | 0.03 | 0.05 | 0.05 | 0.05 |
| 3 | 7093553 | rs17046730 | -0.695 | -0.665 | 0.135 | 0.213 | -0.068 | -0.008 | -0.008 | 0 | 0.19 | 0.33 | 0.07 | 0.13 | 0.07 | 0.08 |
| 3 | 7104505 | rs4686115 | 0.185 | -0.92 | 0.412 | 0.066 | 0.009 | -0.027 | -0.027 | 0 | 0.2 | 0.38 | 0.3 | 0.13 | 0.2 | 0.22 |
| 3 | 7105375 | rs9311975 | -0.469 | -1.015 | 0.464 | 0.082 | 0 | -0.013 | -0.013 | 0 | 0.43 | 0.55 | 0.41 | 0.32 | 0.42 | 0.44 |
| 3 | 7105647 | rs1391946 | 0.121 | -1.028 | 0.424 | 0.092 | 0.012 | -0.033 | -0.033 | 1.366 | 0.2 | 0.45 | 0.31 | 0.13 | 0.2 | 0.22 |
| 3 | 7110451 | rs11917382 | 0.099 | -1.092 | 0.224 | 0.17 | -0.051 | 0.017 | 0.017 | 0 | 0.15 | 0.29 | 0.07 | 0.13 | 0.06 | 0.08 |
| 3 | 7111392 | rs11919292 | -0.106 | -1.121 | 0.22 | 0.156 | -0.057 | 0.004 | 0.004 | 0 | 0.18 | 0.31 | 0.08 | 0.13 | 0.07 | 0.08 |
| 3 | 7112861 | rs1874965 | 0.41 | -1.156 | 0.585 | 0.138 | -0.006 | 0.005 | 0.005 | 0.641 | 0.17 | 0.17 | 0.28 | 0.13 | 0.2 | 0.22 |
| 3 | 7115713 | rs17046762 | 0.057 | -1.139 | 0.487 | 0.185 | -0.059 | 0.012 | 0.012 | 0 | 0.17 | 0.31 | 0.07 | 0.13 | 0.07 | 0.07 |
| 3 | 7116696 | rs6772333 | -0.292 | -0.981 | 0.343 | 0.201 | -0.011 | -0.007 | -0.007 | 0 | 0.41 | 0.42 | 0.18 | 0.33 | 0.28 | 0.32 |
| 3 | 7120529 | rs9848973 | -1.492 | -0.749 | -0.354 | 0.233 | -0.129 | -0.109 | -0.109 | 0.105 | 0.65 | 0.92 | 0.5 | 0.45 | 0.5 | 0.55 |
| 3 | 7122638 | rs9311976 | -0.845 | -1.022 | 0.408 | 0.031 | -0.007 | -0.007 | -0.007 | 0 | 0.37 | 0.47 | 0.37 | 0.23 | 0.28 | 0.33 |
| 3 | 7125230 | rs9875041 | -0.638 | -1.073 | 0.528 | 0.135 | 0.001 | 0.026 | 0.026 | 0 | 0.48 | 0.76 | 0.47 | 0.44 | 0.44 | 0.49 |

| | | | | | | | | | | | | | | | | |
|---|---|---|---|---|---|---|---|---|---|---|---|---|---|---|---|---|
| 3 | 7125901 | rs6785425 | -0.463 | -1.184 | 0.758 | 0.12 | -0.073 | -0.001 | -0.001 | 0.091 | 0.2 | 0.58 | 0.25 | 0.13 | 0.06 | 0.09 |
| 3 | 7131501 | rs13072518 | -1.562 | -1.095 | 0.405 | 0.108 | -0.093 | -0.076 | -0.076 | 0.099 | 0.63 | 0.9 | 0.5 | 0.46 | 0.49 | 0.54 |
| 3 | 7142227 | rs9837834 | -0.552 | -0.929 | 0.726 | 0.106 | 0.003 | 0.005 | 0.005 | 0 | 0.44 | 0.71 | 0.45 | 0.43 | 0.41 | 0.46 |
| 3 | 7148907 | rs1499153 | -0.243 | -0.86 | 0.797 | 0.025 | -0.002 | 0 | 0 | 0 | 0.39 | 0.64 | 0.41 | 0.43 | 0.35 | 0.46 |
| 3 | 7151125 | rs1995200 | -0.432 | -1.293 | 0.733 | -0.182 | -0.05 | 0.015 | 0.015 | 0 | 0.23 | 0.51 | 0.31 | 0.21 | 0.16 | 0.25 |
| 3 | 7152640 | rs9876724 | -1.429 | -1.197 | 0.187 | -0.209 | -0.035 | -0.081 | -0.081 | 0 | 0.56 | 0.93 | 0.47 | 0.45 | 0.41 | 0.55 |
| 3 | 7153652 | rs6778030 | -1.568 | -0.845 | -0.435 | -0.194 | -0.086 | -0.094 | -0.094 | 1.192 | 0.63 | 0.9 | 0.5 | 0.49 | 0.53 | 0.58 |
| 3 | 7157733 | rs7646844 | -0.341 | -1.093 | 0.171 | -0.488 | -0.001 | -0.006 | -0.006 | 0 | 0.33 | 0.28 | 0.14 | 0.11 | 0.21 | 0.25 |
| 3 | 7161276 | rs11710715 | -1.584 | -1.057 | -0.017 | -0.396 | -0.107 | -0.046 | -0.046 | 0 | 0.61 | 0.92 | 0.38 | 0.42 | 0.49 | 0.52 |
| 3 | 7167442 | rs11915246 | 0.276 | -0.929 | 0.473 | -0.38 | 0.013 | 0 | 0 | 0 | 0.26 | 0.44 | 0.22 | 0.3 | 0.28 | 0.26 |
| 3 | 7167701 | rs11131063 | 0.474 | -0.957 | 0.569 | -0.382 | -0.011 | 0.007 | 0.007 | 0 | 0.23 | 0.23 | 0.14 | 0.22 | 0.27 | 0.22 |
| 3 | 7170011 | rs6766686 | -0.611 | -1.069 | 0.588 | -0.43 | 0.027 | 0.025 | 0.025 | 1.013 | 0.36 | 0.63 | 0.23 | 0.19 | 0.22 | 0.29 |
| 3 | 7172280 | rs6443091 | -1.481 | -1.015 | 0.571 | -0.419 | -0.01 | 0.015 | 0.015 | 1.535 | 0.59 | 0.65 | 0.29 | 0.33 | 0.48 | 0.48 |
| 3 | 7172721 | rs1874981 | 0.826 | -1.178 | 0.607 | -0.506 | 0.008 | 0.031 | 0.031 | 0 | 0.16 | 0.04 | 0.08 | 0.1 | 0.16 | 0.2 |
| 3 | 7173002 | rs1878164 | -1.33 | -0.881 | 0.638 | -0.345 | -0.01 | 0.016 | 0.016 | 1.497 | 0.59 | 0.65 | 0.35 | 0.33 | 0.48 | 0.48 |
| 3 | 7175878 | rs9880404 | 0.262 | -0.76 | 0.667 | -0.284 | 0.01 | 0 | 0 | 0 | 0.25 | 0.39 | 0.23 | 0.29 | 0.28 | 0.25 |
| 3 | 7176252 | rs17824908 | 0.454 | -0.758 | 0.618 | -0.284 | -0.025 | 0.012 | 0.012 | 0 | 0.23 | 0.16 | 0.13 | 0.21 | 0.27 | 0.22 |
| 3 | 7182917 | rs1568075 | -0.921 | -0.839 | 0.482 | -0.188 | -0.001 | 0.016 | 0.016 | 0 | 0.54 | 0.35 | 0.26 | 0.33 | 0.47 | 0.46 |
| 3 | 7188006 | rs3828472 | 0.648 | -0.761 | 0.366 | -0.269 | -0.036 | 0.013 | 0.013 | 0 | 0.22 | 0.14 | 0.17 | 0.21 | 0.32 | 0.23 |
| 3 | 7188116 | rs3749448 | -0.19 | -0.857 | 0.404 | 0.213 | -0.042 | -0.034 | -0.034 | 0 | 0.34 | 0.45 | 0.16 | 0.24 | 0.21 | 0.26 |
| 3 | 7208213 | rs9819783 | -1.623 | -0.506 | -0.373 | 0.206 | -0.076 | -0.093 | -0.093 | 0 | 0.63 | 0.95 | 0.42 | 0.44 | 0.56 | 0.54 |
| 3 | 7208953 | rs1516302 | -0.413 | 0.499 | -0.053 | -0.183 | 0.001 | -0.042 | -0.042 | 0.127 | 0.69 | 0.63 | 0.81 | 0.9 | 0.7 | 0.76 |
| 3 | 7211490 | rs17824974 | 0.399 | 0.583 | -0.236 | -0.111 | -0.049 | 0.034 | 0.034 | 0 | 0.26 | 0.14 | 0.17 | 0.19 | 0.35 | 0.25 |
| 3 | 7212607 | rs17288561 | 0.59 | 0.442 | -0.107 | 0.027 | -0.011 | -0.011 | -0.011 | 0.13 | 0.44 | 0.54 | 0.64 | 0.67 | 0.35 | 0.5 |
| 3 | 7214563 | rs965170 | 0.24 | 0.444 | -0.921 | 0.008 | -0.009 | 0.012 | 0.012 | 0 | 0.25 | 0.19 | 0.14 | 0.18 | 0.29 | 0.24 |
| 3 | 7214945 | rs11131064 | 0.903 | 0.491 | -1.371 | 0.188 | -0.031 | -0.051 | -0.051 | 0 | 0.37 | 0.05 | 0.56 | 0.61 | 0.37 | 0.43 |
| 3 | 7218175 | rs1400166 | -0.596 | 1.161 | -0.085 | 0.526 | 0.073 | 0.015 | 0.015 | 1.68 | 0.22 | 0.54 | 0.24 | 0.22 | 0.32 | 0.25 |
| 3 | 7222094 | rs11715681 | -0.058 | 1.344 | -0.068 | 0.504 | -0.049 | 0.005 | 0.005 | 0.71 | 0.17 | 0.09 | 0.17 | 0.15 | 0.36 | 0.26 |
| 3 | 7222236 | rs6784317 | -0.863 | 1.331 | 0.262 | 0.54 | 0.124 | 0.003 | 0.003 | 2.012 | 0.21 | 0.51 | 0.19 | 0.2 | 0.34 | 0.28 |
| 3 | 7225763 | rs6795397 | -0.021 | 1.114 | 0.182 | 0.542 | -0.052 | 0.005 | 0.005 | 0.64 | 0.16 | 0.09 | 0.17 | 0.15 | 0.36 | 0.25 |
| 3 | 7228869 | rs2063880 | -0.905 | 0.524 | 0.021 | 0.513 | 0.043 | -0.038 | -0.038 | 0 | 0.6 | 0.96 | 0.46 | 0.4 | 0.68 | 0.51 |
| 3 | 7229067 | rs17216426 | 1.415 | 0.344 | -0.17 | 1.147 | -0.041 | -0.066 | -0.066 | 0 | 0.11 | 0.01 | 0.16 | 0.22 | 0.13 | 0.18 |
| 3 | 7231390 | rs2063882 | 1.026 | 0.445 | 0.258 | 0.64 | 0.009 | 0.034 | 0.034 | 0 | 0.37 | 0.29 | 0.23 | 0.2 | 0.32 | 0.25 |
| 3 | 7232606 | rs7634808 | 0.669 | 0.429 | 0.692 | 0.485 | 0.008 | 0.018 | 0.018 | 0 | 0.41 | 0.64 | 0.25 | 0.19 | 0.33 | 0.27 |
| 3 | 7233935 | rs1963265 | 1.045 | 0.402 | 0.488 | 0.478 | 0.013 | 0.034 | 0.034 | 0 | 0.36 | 0.29 | 0.2 | 0.12 | 0.31 | 0.24 |
| 3 | 7247989 | rs7621313 | -0.42 | 0.206 | 0.078 | 0.747 | 0.074 | 0.036 | 0.036 | 0.029 | 0.59 | 0.81 | 0.43 | 0.34 | 0.69 | 0.51 |
| 3 | 7257630 | rs1543146 | -0.967 | 0.787 | 0.893 | 1.075 | 0.004 | -0.001 | -0.001 | 0 | 0.27 | 0.67 | 0.25 | 0.25 | 0.41 | 0.33 |
| 3 | 7259313 | rs13070476 | 0.124 | 1.375 | 0.544 | 1.063 | -0.053 | -0.007 | -0.007 | 0 | 0.14 | 0.02 | 0.12 | 0.15 | 0.23 | 0.16 |
| 3 | 7259684 | rs7652154 | -0.997 | 1.254 | 0.744 | 1.075 | 0.129 | 0.002 | 0.002 | 2.118 | 0.24 | 0.44 | 0.24 | 0.24 | 0.41 | 0.31 |
| 3 | 7261964 | rs9851849 | 0.174 | 0.306 | 0.742 | 1.396 | -0.011 | 0.015 | 0.015 | 0 | 0.61 | 0.5 | 0.59 | 0.5 | 0.66 | 0.5 |
| 3 | 7262014 | rs6797852 | -0.605 | 0.111 | 0.639 | 1.066 | -0.027 | -0.004 | -0.004 | 2.33 | 0.73 | 0.7 | 0.7 | 0.77 | 0.86 | 0.78 |
| 3 | 7284458 | rs870440 | 0.226 | -0.008 | 0.587 | 0.784 | 0.025 | -0.041 | -0.041 | 0.009 | 0.87 | 0.73 | 0.91 | 0.98 | 0.82 | 0.87 |
| 3 | 7292607 | rs9860580 | 0.457 | 0.021 | 0.184 | 0.954 | 0.008 | 0.03 | 0.03 | 0 | 0.5 | 0.38 | 0.32 | 0.44 | 0.42 | 0.43 |
| 3 | 7293469 | rs2055737 | 1.626 | -0.111 | -0.436 | 0.668 | -0.037 | 0.093 | 0.093 | 0 | 0.28 | 0.03 | 0.21 | 0.11 | 0.27 | 0.2 |
| 3 | 7299331 | rs9873905 | 0.826 | -0.005 | -0.04 | 0.673 | 0.001 | 0.005 | 0.005 | 0 | 0.42 | 0.26 | 0.29 | 0.44 | 0.39 | 0.34 |
| 3 | 7302907 | rs1396409 | 0.255 | 0.152 | -0.759 | 0.856 | -0.117 | -0.119 | -0.119 | 0.235 | 0.39 | 0.03 | 0.55 | 0.41 | 0.42 | 0.47 |
| 3 | 7309658 | rs4095095 | 1.316 | 0.314 | -0.545 | 0.75 | -0.083 | 0.024 | 0.024 | 0.133 | 0.23 | 0.03 | 0.37 | 0.14 | 0.29 | 0.18 |
| 3 | 7311647 | rs6763969 | -0.236 | 0.586 | 0.085 | 0.677 | -0.005 | 0.005 | 0.005 | 0 | 0.89 | 0.89 | 0.91 | 0.98 | 0.81 | 0.88 |
| 3 | 7320514 | rs6443099 | 0.635 | 0.561 | 0.343 | 0.811 | 0 | -0.012 | -0.012 | 0 | 0.38 | 0.32 | 0.56 | 0.49 | 0.38 | 0.45 |
| 3 | 7336452 | rs2133450 | 0.275 | 0.535 | 0.535 | 0.807 | -0.009 | -0.015 | -0.015 | 1.017 | 0.46 | 0.22 | 0.32 | 0.49 | 0.45 | 0.4 |
| 3 | 7340164 | rs2291867 | -1.002 | 0.656 | 0.469 | 0.617 | -0.027 | -0.108 | -0.108 | 0 | 0.62 | 0.92 | 0.46 | 0.53 | 0.63 | 0.53 |
| 3 | 7348129 | rs17047071 | 0.614 | 0.727 | 0.543 | 0.695 | 0 | -0.001 | -0.001 | 0.011 | 0.12 | 0.11 | 0.07 | 0.14 | 0.12 | 0.14 |
| 3 | 7349755 | rs17697287 | 0.404 | 0.498 | 0.801 | 0.675 | -0.019 | -0.006 | -0.006 | 0 | 0.46 | 0.16 | 0.31 | 0.47 | 0.45 | 0.39 |
| 3 | 7351373 | rs984123 | -1.29 | 0.505 | 1.328 | 0.652 | 0.028 | 0.035 | 0.035 | 3.731 | 0.08 | 0.45 | 0.05 | 0.05 | 0.02 | 0.03 |
| 3 | 7353942 | rs2136150 | -0.416 | 0.483 | 1.126 | 0.651 | -0.066 | -0.068 | -0.068 | 0 | 0.56 | 0.86 | 0.38 | 0.52 | 0.46 | 0.42 |
| 3 | 7357086 | rs2875237 | -0.341 | 0.571 | 1.434 | 0.82 | 0.068 | 0.018 | 0.018 | 0.065 | 0.07 | 0.2 | 0.08 | 0.07 | 0.15 | 0.11 |
| 3 | 7358711 | rs13082785 | 0.414 | 0.521 | 1.089 | 0.68 | -0.016 | -0.026 | -0.026 | 0 | 0.42 | 0.16 | 0.28 | 0.47 | 0.42 | 0.36 |
| 3 | 7359949 | rs1396410 | 0.619 | 0.452 | 0.901 | 0.716 | -0.03 | 0 | 0 | 0 | 0.39 | 0.14 | 0.27 | 0.39 | 0.41 | 0.33 |
| 3 | 7363887 | rs6443104 | -0.449 | 0.808 | 0.505 | 0.666 | -0.022 | -0.076 | -0.076 | 0.053 | 0.57 | 0.91 | 0.44 | 0.54 | 0.59 | 0.48 |
| 3 | 7364572 | rs1508708 | -0.199 | 0.279 | 0.765 | 0.685 | 0.011 | -0.005 | -0.005 | 0 | 0.51 | 0.71 | 0.35 | 0.52 | 0.44 | 0.4 |
| 3 | 7368673 | rs9867566 | 0.146 | 0.444 | 0.224 | 0.757 | -0.001 | -0.001 | -0.001 | 0 | 0.46 | 0.48 | 0.33 | 0.48 | 0.43 | 0.37 |
| 3 | 7371835 | rs1499210 | -0.105 | 0.454 | -0.014 | 0.781 | 0.004 | 0.004 | 0.004 | 0 | 0.5 | 0.56 | 0.36 | 0.48 | 0.46 | 0.39 |
| 3 | 7371919 | rs17652614 | -0.096 | 0.454 | -0.164 | 0.715 | -0.001 | -0.002 | -0.002 | 0.139 | 0.51 | 0.57 | 0.37 | 0.52 | 0.49 | 0.42 |
| 3 | 7381271 | rs10510364 | -0.105 | -0.45 | 0.063 | 0.964 | 0.002 | -0.003 | -0.003 | 0.325 | 0.2 | 0.19 | 0.17 | 0.29 | 0.12 | 0.11 |
| 3 | 7382827 | rs1508722 | -0.308 | -0.436 | -0.034 | 0.607 | 0 | -0.002 | -0.002 | 0 | 0.38 | 0.38 | 0.41 | 0.36 | 0.39 | 0.46 |
| 3 | 7386444 | rs1499199 | -0.377 | -0.672 | -0.077 | 0.501 | 0.002 | 0.014 | 0.014 | 0.12 | 0.4 | 0.33 | 0.41 | 0.35 | 0.39 | 0.46 |
| 3 | 7389984 | rs6791111 | -1.346 | -0.665 | -0.001 | 0.297 | -0.047 | 0.004 | 0.004 | 0 | 0.6 | 0.85 | 0.61 | 0.64 | 0.5 | 0.6 |
| 3 | 7390577 | rs7628504 | 0.662 | -0.44 | -0.079 | 0.18 | -0.021 | -0.076 | -0.076 | 1.683 | 0.2 | 0.04 | 0.27 | 0.35 | 0.22 | 0.33 |
| 3 | 7390780 | rs7617297 | -0.67 | -0.305 | 0.068 | 0.173 | -0.001 | -0.003 | -0.003 | 0 | 0.31 | 0.35 | 0.37 | 0.33 | 0.31 | 0.44 |

| | | | | | | | | | | | | | | | | |
|---|---|---|---|---|---|---|---|---|---|---|---|---|---|---|---|---|
| 3 | 7391647 | rs9830108 | 0.269 | -0.255 | 0.337 | 0.011 | 0.004 | -0.045 | -0.045 | 0.019 | 0.1 | 0.15 | 0.08 | 0 | 0.08 | 0.11 |
| 3 | 7394557 | rs2324148 | 0.111 | -0.258 | 0.299 | 0.01 | 0.017 | -0.063 | -0.063 | 0 | 0.11 | 0.27 | 0.09 | 0 | 0.08 | 0.12 |
| 3 | 7398367 | rs6779412 | 0.867 | -0.254 | 0.081 | 0.007 | 0.007 | -0.029 | -0.029 | 0 | 0.05 | 0.16 | 0.05 | 0 | 0.06 | 0.02 |
| 3 | 7398982 | rs6803602 | -0.467 | -0.379 | -0.089 | 0.061 | -0.002 | 0.012 | 0.012 | 0.75 | 0.27 | 0.21 | 0.35 | 0.24 | 0.31 | 0.35 |
| 3 | 7399749 | rs6783210 | 0.952 | -0.085 | 0.274 | 0.078 | 0.001 | -0.027 | -0.027 | 0.541 | 0.05 | 0.11 | 0.05 | 0 | 0.06 | 0.02 |
| 3 | 7405044 | rs13091584 | -1.102 | -0.224 | 0.311 | 0.156 | -0.005 | -0.001 | -0.001 | 0.975 | 0.56 | 0.62 | 0.51 | 0.55 | 0.49 | 0.57 |
| 3 | 7406960 | rs12491212 | 0.543 | -0.133 | -0.362 | 0.153 | -0.001 | -0.011 | -0.011 | 0 | 0.08 | 0.06 | 0.04 | 0 | 0.07 | 0.09 |
| 3 | 7411730 | rs7647001 | -0.23 | -0.157 | -0.497 | 0.228 | 0.014 | -0.006 | -0.006 | 0 | 0.46 | 0.24 | 0.43 | 0.55 | 0.42 | 0.44 |
| 3 | 7415452 | rs10510366 | -0.812 | -0.222 | 0.231 | 0.426 | 0.014 | -0.12 | -0.12 | 0 | 0.17 | 0.55 | 0.14 | 0 | 0.15 | 0.15 |
| 3 | 7419156 | rs9883258 | -0.57 | -0.63 | -0.049 | 1.192 | 0 | 0.007 | 0.007 | 0 | 0.23 | 0.2 | 0.2 | 0.49 | 0.24 | 0.3 |
| 3 | 7423494 | rs9863764 | -0.649 | -1.15 | 0.207 | 1.193 | -0.06 | 0.08 | 0.08 | 0 | 0.23 | 0.46 | 0.17 | 0.35 | 0.12 | 0.16 |
| 3 | 7425658 | rs13064153 | -1.003 | -1.121 | 0.525 | 0.881 | 0.017 | -0.014 | -0.014 | 0 | 0.08 | 0.24 | 0.06 | 0.02 | 0.06 | 0.07 |
| 3 | 7426507 | rs9856461 | -0.391 | -1.115 | 0.285 | 0.561 | 0 | 0 | 0 | 0 | 0.64 | 0.57 | 0.45 | 0.4 | 0.62 | 0.49 |
| 3 | 7428082 | rs1995194 | 0.158 | -0.978 | 0.233 | 0.503 | -0.011 | 0.011 | 0.011 | 0 | 0.24 | 0.25 | 0.16 | 0.33 | 0.15 | 0.18 |
| 3 | 7438708 | rs13066750 | 1.197 | -0.653 | -0.35 | 0.464 | -0.043 | 0.127 | 0.127 | 3.991 | 0.36 | 0.15 | 0.24 | 0.06 | 0.4 | 0.17 |
| 3 | 7443688 | rs712767 | -0.135 | -0.56 | -0.197 | 0.663 | -0.007 | -0.022 | -0.022 | 0.372 | 0.62 | 0.7 | 0.45 | 0.45 | 0.56 | 0.44 |
| 3 | 7451750 | rs17700470 | 1.417 | -0.466 | -0.456 | 0.521 | -0.033 | 0.066 | 0.066 | 6.121 | 0.3 | 0.2 | 0.26 | 0.11 | 0.37 | 0.14 |
| 3 | 7456558 | rs712773 | -0.33 | -0.441 | -0.173 | 1.022 | 0.002 | -0.004 | -0.004 | 1.507 | 0.38 | 0.36 | 0.27 | 0.44 | 0.29 | 0.49 |
| 3 | 7461505 | rs3804979 | 1.247 | -0.463 | -0.076 | 0.949 | 0.007 | -0.084 | -0.084 | 0 | 0.4 | 0.51 | 0.31 | 0.11 | 0.42 | 0.24 |
| 3 | 7463316 | rs712776 | -0.595 | -0.773 | -0.253 | 1.074 | -0.007 | -0.025 | -0.025 | 0 | 0.78 | 0.85 | 0.58 | 0.57 | 0.72 | 0.73 |
| 3 | 7468858 | rs712778 | -0.65 | -0.92 | -0.158 | 1.134 | 0.008 | -0.006 | -0.006 | 0 | 0.42 | 0.56 | 0.29 | 0.45 | 0.3 | 0.47 |
| 3 | 7476597 | rs712779 | -0.379 | -1.069 | -0.182 | 1.188 | -0.045 | 0.079 | 0.079 | 0.605 | 0.76 | 0.43 | 0.81 | 0.62 | 0.85 | 0.83 |
| 3 | 7479889 | rs11712265 | 0.551 | -1.071 | -0.365 | 1.136 | -0.001 | 0.039 | 0.039 | 0 | 0.19 | 0.03 | 0.11 | 0.1 | 0.16 | 0.33 |
| 3 | 7479947 | rs712781 | -1.038 | -1.473 | -0.264 | 1.182 | -0.022 | 0.052 | 0.052 | 0.031 | 0.81 | 0.67 | 0.85 | 0.69 | 0.85 | 0.84 |
| 3 | 7481801 | rs712782 | 0.564 | -1.47 | -0.25 | 0.977 | -0.024 | 0.061 | 0.061 | 0.058 | 0.21 | 0.33 | 0.16 | 0.37 | 0.15 | 0.18 |
| 3 | 7481854 | rs17717959 | 1.713 | -1.219 | -0.256 | 0.952 | -0.113 | 0.172 | 0.172 | 5.606 | 0.36 | 0.11 | 0.28 | 0.09 | 0.45 | 0.21 |
| 3 | 7484471 | rs779867 | 0.047 | -1.248 | -0.21 | 1.01 | -0.002 | 0 | 0 | 0 | 0.64 | 0.64 | 0.76 | 0.56 | 0.73 | 0.55 |
| 3 | 7486556 | rs3804972 | 1.639 | -1.038 | -0.272 | 0.492 | -0.101 | 0.176 | 0.176 | 4.987 | 0.36 | 0.12 | 0.27 | 0.07 | 0.44 | 0.19 |
| 3 | 7490129 | rs13096022 | 1.533 | -0.989 | -0.174 | 0.555 | -0.073 | 0.141 | 0.141 | 4.651 | 0.37 | 0.15 | 0.27 | 0.09 | 0.45 | 0.23 |
| 3 | 7493351 | rs712793 | 0.791 | -1.112 | -0.215 | 0.839 | -0.008 | -0.006 | -0.006 | 0 | 0.57 | 0.57 | 0.46 | 0.58 | 0.6 | 0.38 |
| 3 | 7496400 | rs6771374 | 1.279 | -0.969 | -0.766 | 0.636 | -0.071 | 0.218 | 0.218 | 3.496 | 0.4 | 0.13 | 0.31 | 0.09 | 0.51 | 0.3 |
| 3 | 7498258 | rs779724 | -0.12 | -0.935 | -0.597 | 0.793 | -0.009 | 0.011 | 0.011 | 0.325 | 0.73 | 0.57 | 0.56 | 0.66 | 0.73 | 0.66 |
| 3 | 7498342 | rs779723 | 0.907 | -0.825 | -0.775 | 0.885 | 0 | 0.011 | 0.011 | 0.002 | 0.2 | 0.26 | 0.38 | 0.29 | 0.21 | 0.22 |
| 3 | 7501010 | rs1375919 | -1.034 | -0.814 | -0.792 | 0.907 | -0.017 | 0.11 | 0.11 | 0 | 0.82 | 0.71 | 0.84 | 0.55 | 0.85 | 0.84 |
| 3 | 7504862 | rs3792476 | 0.396 | -0.637 | -1.598 | 0.647 | 0.023 | 0.066 | 0.066 | 1.726 | 0.58 | 0.28 | 0.43 | 0.21 | 0.63 | 0.57 |
| 3 | 7505468 | rs1868617 | 0.065 | -0.025 | -1.373 | 1.206 | 0.009 | 0.01 | 0.01 | 0 | 0.17 | 0.11 | 0.11 | 0.13 | 0.13 | 0.32 |
| 3 | 7507463 | rs779716 | 0.079 | 0.021 | -1.305 | 1.338 | -0.005 | 0.01 | 0.01 | 0.003 | 0.2 | 0.25 | 0.37 | 0.29 | 0.2 | 0.25 |
| 3 | 7508573 | rs17718789 | -0.754 | 0.256 | -2.069 | 1.024 | 0.003 | 0.012 | 0.012 | 0 | 0.22 | 0.15 | 0.16 | 0.13 | 0.2 | 0.39 |
| 3 | 7508738 | rs17656831 | -0.745 | 0.155 | -2.081 | 1.186 | 0.005 | -0.014 | -0.014 | 0 | 0.22 | 0.14 | 0.16 | 0.33 | 0.19 | 0.39 |
| 3 | 7509265 | rs17718867 | -0.61 | 0.277 | -2.54 | 0.228 | 0 | 0.056 | 0.056 | 0 | 0.21 | 0.05 | 0.22 | 0.12 | 0.19 | 0.35 |
| 3 | 7510856 | rs7623046 | 0.211 | 0.617 | -3 | 0.378 | -0.096 | -0.056 | -0.056 | 0.203 | 0.56 | 0.96 | 0.63 | 0.5 | 0.52 | 0.54 |
| 3 | 7510891 | rs7623055 | 0.108 | 0.657 | -3.378 | 0.18 | 0.06 | -0.056 | -0.056 | 1.47 | 0.52 | 0.94 | 0.55 | 0.5 | 0.38 | 0.56 |
| 3 | 7511375 | rs779710 | -0.295 | 0.745 | -2.381 | 0.307 | -0.025 | -0.022 | -0.022 | 0 | 0.37 | 0.71 | 0.36 | 0.44 | 0.39 | 0.41 |
| 3 | 7512728 | rs779708 | -0.759 | 0.373 | -2.537 | 0.514 | -0.045 | 0.036 | 0.036 | 0 | 0.62 | 0.97 | 0.72 | 0.71 | 0.68 | 0.59 |
| 3 | 7516145 | rs3804959 | -0.705 | 0.676 | -1.194 | 0.85 | -0.003 | -0.017 | -0.017 | 0 | 0.31 | 0.72 | 0.35 | 0.41 | 0.3 | 0.32 |
| 3 | 7519813 | rs752299 | -0.499 | 1.173 | -0.73 | 1.001 | 0.022 | 0.057 | 0.057 | 0 | 0.27 | 0.42 | 0.31 | 0.4 | 0.28 | 0.28 |
| 3 | 7523588 | rs1083800 | -0.393 | 1.259 | -0.835 | 1.017 | 0.033 | 0.07 | 0.07 | 0 | 0.26 | 0.4 | 0.32 | 0.42 | 0.29 | 0.26 |
| 3 | 7528503 | rs7374553 | 0.135 | 0.904 | -0.64 | 0.839 | -0.003 | 0.001 | 0.001 | 0 | 0.45 | 0.51 | 0.41 | 0.52 | 0.4 | 0.33 |
| 3 | 7529555 | rs9870680 | -0.114 | 1.502 | -0.623 | 0.564 | -0.013 | 0.015 | 0.015 | 0 | 0.36 | 0.35 | 0.49 | 0.71 | 0.48 | 0.36 |
| 3 | 7531715 | rs1450101 | 0.476 | 1.995 | -0.591 | 0.768 | -0.003 | -0.041 | -0.041 | 0 | 0.28 | 0.11 | 0.3 | 0.62 | 0.29 | 0.28 |
| 3 | 7533393 | rs12494654 | -0.482 | 1.836 | 0.006 | 0.304 | -0.002 | -0.006 | -0.006 | 0.38 | 0.42 | 0.44 | 0.49 | 0.72 | 0.45 | 0.41 |
| 3 | 7533704 | rs10222352 | -0.882 | 1.424 | 0.067 | 0.071 | 0 | -0.003 | -0.003 | 3.265 | 0.5 | 0.58 | 0.62 | 0.75 | 0.6 | 0.51 |
| 3 | 7537468 | rs6766032 | 0.027 | 1.391 | 0.261 | 0.074 | 0.007 | 0.016 | 0.016 | 0.749 | 0.06 | 0.22 | 0.08 | 0.07 | 0.05 | 0.02 |
| 3 | 7539974 | rs3804945 | -1.237 | 0.871 | 0.484 | -0.394 | -0.022 | 0.085 | 0.085 | 1.748 | 0.61 | 0.85 | 0.71 | 0.81 | 0.67 | 0.71 |
| 3 | 7544835 | rs7641476 | 0.642 | 0.805 | 0.478 | 0.066 | -0.011 | 0 | 0 | 0 | 0.36 | 0.45 | 0.36 | 0.66 | 0.34 | 0.31 |
| 3 | 7548067 | rs11131078 | 1.379 | 1.035 | 0.286 | 0.023 | -0.022 | -0.074 | -0.074 | 0.004 | 0.29 | 0.1 | 0.29 | 0.59 | 0.31 | 0.29 |
| 3 | 7551114 | rs924353 | 0.632 | 1.082 | 0.299 | -0.166 | -0.024 | 0.019 | 0.019 | 0 | 0.37 | 0.13 | 0.43 | 0.66 | 0.43 | 0.39 |
| 3 | 7553544 | rs779727 | -1.328 | 1.001 | 0.223 | -0.113 | -0.027 | -0.053 | -0.053 | 0 | 0.71 | 0.91 | 0.69 | 0.38 | 0.68 | 0.67 |
| 3 | 7557251 | rs17047503 | -0.872 | 0.875 | 0.435 | -0.208 | -0.016 | -0.044 | -0.044 | 0 | 0.17 | 0.3 | 0.16 | 0.04 | 0.17 | 0.15 |
| 3 | 7557427 | rs6774660 | -0.508 | 0.949 | 0.679 | -0.24 | 0.009 | -0.008 | -0.008 | 0 | 0.13 | 0.18 | 0.25 | 0.06 | 0.28 | 0.21 |
| 3 | 7557492 | rs1145136 | 1.083 | 0.489 | 0.7 | -0.088 | -0.011 | 0.034 | 0.034 | 1.92 | 0.38 | 0.2 | 0.34 | 0.31 | 0.33 | 0.25 |
| 3 | 7558907 | rs6778646 | 0.142 | 0.502 | 0.826 | -0.144 | -0.004 | -0.009 | -0.009 | 0.019 | 0.1 | 0.1 | 0.12 | 0.03 | 0.11 | 0.09 |
| 3 | 7562364 | rs1106486 | -1.104 | 0.519 | 0.824 | 0.146 | -0.004 | 0.01 | 0.01 | 1.952 | 0.61 | 0.75 | 0.61 | 0.64 | 0.67 | 0.72 |
| 3 | 7565092 | rs965595 | 0.318 | 0.51 | 0.674 | 0.145 | -0.002 | -0.008 | -0.008 | 0 | 0.45 | 0.51 | 0.42 | 0.36 | 0.37 | 0.32 |
| 3 | 7576514 | rs3804928 | 0.282 | 0.41 | 0.212 | -0.101 | -0.053 | 0.014 | 0.014 | 0 | 0.17 | 0.02 | 0.21 | 0.13 | 0.22 | 0.22 |
| 3 | 7578531 | rs779746 | -0.558 | 0.359 | 0.311 | 0.158 | -0.019 | -0.003 | -0.003 | 0.073 | 0.56 | 0.7 | 0.48 | 0.49 | 0.41 | 0.52 |
| 3 | 7580979 | rs3804925 | 1.356 | 0.417 | 0.182 | 0.151 | -0.045 | -0.006 | -0.006 | 0.642 | 0.1 | 0.01 | 0.1 | 0.09 | 0.11 | 0.12 |
| 3 | 7583801 | rs779740 | 0.347 | 0.305 | 0.442 | 0.24 | -0.005 | 0.012 | 0.012 | 0.815 | 0.45 | 0.4 | 0.46 | 0.61 | 0.53 | 0.55 |
| 3 | 7591405 | rs17047582 | -0.274 | 0.355 | 0.189 | 0.344 | -0.054 | -0.037 | -0.037 | 0.009 | 0.18 | 0.51 | 0.11 | 0.12 | 0.11 | 0.23 |

| | | | | | | | | | | | | | | | | |
|---|---|---|---|---|---|---|---|---|---|---|---|---|---|---|---|---|
| 3 | 7599918 | rs779718 | -0.373 | 0.844 | -0.221 | 0.264 | -0.015 | -0.005 | -0.005 | 0.078 | 0.62 | 0.8 | 0.61 | 0.58 | 0.68 | 0.72 |
| 3 | 7603708 | rs1485179 | -0.923 | 1.119 | 0.496 | -0.189 | -0.02 | 0.015 | 0.015 | 0 | 0.4 | 0.7 | 0.5 | 0.28 | 0.47 | 0.5 |
| 3 | 7607033 | rs1569285 | 0.566 | 0.687 | 0.784 | -0.278 | 0.015 | -0.035 | -0.035 | 0 | 0.65 | 0.45 | 0.62 | 0.81 | 0.61 | 0.62 |
| 3 | 7607994 | rs9832297 | -0.974 | 0.713 | 0.76 | -0.438 | 0.003 | -0.012 | -0.012 | 0.361 | 0.38 | 0.6 | 0.45 | 0.23 | 0.42 | 0.47 |
| 3 | 7619230 | rs3804908 | 0.095 | 0.332 | 0.536 | -0.577 | 0.023 | 0.026 | 0.026 | 0.007 | 0.47 | 0.53 | 0.26 | 0.29 | 0.29 | 0.31 |
| 3 | 7620382 | rs7614915 | -0.084 | 0.706 | 0.315 | -0.598 | -0.033 | -0.015 | -0.015 | 1.183 | 0.13 | 0.06 | 0.26 | 0.14 | 0.29 | 0.17 |
| 3 | 7620789 | rs1485175 | -0.722 | 0.649 | 0.476 | -0.598 | -0.002 | -0.003 | -0.003 | 0.367 | 0.59 | 0.57 | 0.51 | 0.44 | 0.58 | 0.51 |
| 3 | 7621158 | rs1485172 | -0.624 | 0.629 | 0.523 | -0.607 | 0.001 | 0.002 | 0.002 | 0 | 0.58 | 0.54 | 0.51 | 0.43 | 0.58 | 0.44 |
| 3 | 7621512 | rs9826424 | 0.347 | 0.632 | 0.527 | -0.604 | -0.014 | -0.015 | -0.015 | 0.684 | 0.46 | 0.48 | 0.26 | 0.29 | 0.29 | 0.27 |
| 3 | 7624469 | rs11717471 | -0.304 | 1.274 | 0.27 | -0.299 | -0.033 | -0.015 | -0.015 | 1.023 | 0.13 | 0.06 | 0.26 | 0.14 | 0.29 | 0.19 |
| 3 | 7639708 | rs3804883 | 0.198 | 0.338 | 0.638 | -0.412 | 0.007 | 0.007 | 0.007 | 0.743 | 0.46 | 0.48 | 0.26 | 0.3 | 0.3 | 0.34 |
| 3 | 7641515 | rs3804880 | 0.271 | 0.282 | 0.426 | -0.531 | -0.017 | 0.01 | 0.01 | 0 | 0.13 | 0.24 | 0.09 | 0.11 | 0.08 | 0.14 |
| 3 | 7641740 | rs17666242 | 1.298 | 0.342 | -0.489 | -0.529 | 0.012 | 0.154 | 0.154 | 1.171 | 0.3 | 0.08 | 0.15 | 0.09 | 0.22 | 0.16 |
| 3 | 7645217 | rs11713266 | -0.608 | 0.928 | 0.037 | -0.008 | 0.012 | 0.079 | 0.079 | 0 | 0.11 | 0.21 | 0.16 | 0.25 | 0.18 | 0.17 |
| 3 | 7651185 | rs1485171 | -0.419 | 0.851 | -0.128 | -0.344 | 0.003 | 0.004 | 0.004 | 0 | 0.12 | 0.16 | 0.17 | 0.13 | 0.18 | 0.14 |
| 3 | 7660133 | rs7640685 | -0.867 | 0.989 | -0.673 | -0.59 | -0.03 | -0.053 | -0.053 | 0 | 0.65 | 0.87 | 0.56 | 0.43 | 0.65 | 0.53 |
| 3 | 7663718 | rs12107224 | 0.535 | 1.137 | -0.448 | -0.931 | 0.001 | -0.027 | -0.027 | 0.195 | 0.1 | 0.19 | 0.08 | 0.04 | 0.07 | 0.08 |
| 3 | 7665463 | rs3804869 | 1.093 | 1.119 | -0.463 | -1.114 | 0.003 | -0.001 | -0.001 | 0 | 0.08 | 0.08 | 0.1 | 0.09 | 0.15 | 0.04 |
| 3 | 7666413 | rs6799023 | 1.329 | 0.939 | -0.386 | -1.218 | 0.026 | -0.021 | -0.021 | 0 | 0.67 | 0.5 | 0.65 | 0.74 | 0.58 | 0.67 |
| 3 | 7667909 | rs3804867 | 0.887 | 1.192 | -0.196 | -0.827 | -0.002 | -0.001 | -0.001 | 0 | 0.08 | 0.08 | 0.1 | 0.09 | 0.14 | 0.04 |
| 3 | 7672002 | rs3792451 | 1.344 | 1.053 | -0.375 | -0.919 | 0.029 | 0.138 | 0.138 | 0.838 | 0.36 | 0.13 | 0.18 | 0.11 | 0.27 | 0.25 |
| 3 | 7674009 | rs6443118 | 0.708 | 1.145 | -0.421 | -0.777 | -0.003 | -0.002 | -0.002 | 1.616 | 0.79 | 0.79 | 0.74 | 0.78 | 0.68 | 0.76 |
| 3 | 7677215 | rs3864067 | 0.714 | 1.144 | -0.447 | -0.778 | 0.009 | 0.002 | 0.002 | 0 | 0.79 | 0.72 | 0.73 | 0.78 | 0.68 | 0.83 |
| 3 | 7678134 | rs17047754 | 1.292 | 1.143 | -0.482 | -0.827 | 0.024 | 0.058 | 0.058 | 0.652 | 0.38 | 0.24 | 0.22 | 0.14 | 0.28 | 0.24 |
| 3 | 7679361 | rs9816982 | 0.252 | 1.595 | -0.311 | -0.559 | 0.005 | -0.001 | -0.001 | 0.337 | 0.38 | 0.43 | 0.5 | 0.61 | 0.39 | 0.52 |
| 3 | 7680452 | rs1872396 | -0.581 | 1.841 | -0.856 | -0.574 | -0.023 | -0.024 | -0.024 | 0 | 0.9 | 0.96 | 0.85 | 0.86 | 0.83 | 0.88 |
| 3 | 7681695 | rs9819314 | 0.239 | 1.86 | -0.85 | -0.663 | 0.005 | 0.001 | 0.001 | 0 | 0.84 | 0.79 | 0.74 | 0.78 | 0.68 | 0.83 |
| 3 | 7683434 | rs11918634 | 0.557 | 1.211 | -1.232 | -0.349 | -0.025 | -0.087 | -0.087 | 0 | 0.25 | 0.07 | 0.31 | 0.56 | 0.26 | 0.33 |
| 3 | 7685663 | rs17673467 | 0.139 | 0.996 | -0.854 | -0.997 | 0.004 | 0.082 | 0.082 | 0.753 | 0.47 | 0.31 | 0.33 | 0.14 | 0.44 | 0.24 |
| 3 | 7688564 | rs4686146 | -0.533 | 1.553 | -0.546 | -0.964 | 0.028 | -0.059 | -0.059 | 0.546 | 0.25 | 0.36 | 0.28 | 0.05 | 0.31 | 0.26 |
| 3 | 7688702 | rs3804850 | 0.401 | 1.078 | -0.908 | -0.987 | -0.001 | 0.003 | 0.003 | 3.333 | 0.23 | 0.14 | 0.2 | 0.17 | 0.22 | 0.18 |
| 3 | 7690534 | rs7635518 | 0.337 | 1.454 | -0.584 | -1.123 | -0.017 | -0.001 | -0.001 | 0 | 0.18 | 0.13 | 0.16 | 0 | 0.24 | 0.16 |
| 3 | 7691088 | rs13321020 | -0.583 | 1.124 | -0.219 | -0.997 | -0.024 | 0.027 | 0.027 | 0 | 0.09 | 0.31 | 0.15 | 0.1 | 0.06 | 0.14 |
| 3 | 7694402 | rs9877193 | -0.883 | 0.133 | -0.107 | -1.388 | 0.002 | 0.017 | 0.017 | 0.056 | 0.38 | 0.69 | 0.36 | 0.22 | 0.35 | 0.36 |
| 3 | 7695640 | rs3792439 | -0.783 | 0.324 | 0.008 | -1.335 | 0 | 0 | 0 | 0.206 | 0.55 | 0.59 | 0.52 | 0.28 | 0.53 | 0.5 |
| 3 | 7697810 | rs4414838 | -1.294 | -0.167 | -0.079 | -1.314 | -0.001 | -0.01 | -0.01 | 0 | 0.65 | 0.82 | 0.6 | 0.35 | 0.62 | 0.57 |
| 3 | 7699439 | rs3804849 | 1.203 | -0.504 | -0.203 | -1.375 | -0.01 | -0.036 | -0.036 | 0 | 0.31 | 0.15 | 0.38 | 0.61 | 0.36 | 0.4 |
| 3 | 7701440 | rs3864074 | 0.258 | -0.709 | 0.171 | -1.511 | 0.008 | 0.09 | 0.09 | 1.068 | 0.47 | 0.26 | 0.44 | 0.15 | 0.49 | 0.49 |
| 3 | 7708008 | rs3864076 | -0.323 | -0.312 | 0.169 | -1.585 | -0.006 | 0.055 | 0.055 | 1.729 | 0.52 | 0.71 | 0.44 | 0.29 | 0.47 | 0.49 |
| 3 | 7710922 | rs9870241 | -0.241 | -0.408 | 0.536 | -1.537 | -0.014 | 0.014 | 0.014 | 0 | 0.07 | 0.27 | 0.08 | 0.08 | 0.05 | 0.04 |
| 3 | 7716889 | rs162810 | 0.121 | -0.428 | 0.382 | -1.624 | -0.002 | 0.053 | 0.053 | 1.887 | 0.48 | 0.31 | 0.56 | 0.74 | 0.53 | 0.55 |
| 3 | 7720805 | rs162805 | 0.694 | -0.469 | 0.375 | -1.65 | -0.023 | -0.011 | -0.011 | 0.475 | 0.1 | 0.14 | 0.06 | 0.06 | 0.05 | 0.02 |
| 3 | 7721613 | rs162802 | 0.363 | -0.732 | 0.532 | -1.891 | -0.007 | 0.114 | 0.114 | 0.077 | 0.47 | 0.3 | 0.4 | 0.12 | 0.47 | 0.38 |
| 3 | 7721997 | rs2280739 | -0.942 | -0.68 | 0.717 | -1.758 | 0.011 | -0.01 | -0.01 | 0.262 | 0.08 | 0.22 | 0.11 | 0.03 | 0.07 | 0.1 |
| 3 | 7730014 | rs1504045 | 1.181 | -0.586 | 0.474 | -2.008 | -0.005 | 0.003 | 0.003 | 1.285 | 0.27 | 0.33 | 0.41 | 0.74 | 0.3 | 0.39 |
| 3 | 7732499 | rs162782 | 0.213 | -0.538 | 0.336 | -1.913 | -0.002 | 0.009 | 0.009 | 0 | 0.92 | 0.88 | 0.91 | 0.86 | 0.92 | 0.84 |
| 3 | 7739919 | rs162772 | -1.135 | -0.714 | 0.311 | -2.251 | -0.023 | 0.061 | 0.061 | 2.092 | 0.6 | 0.71 | 0.63 | 0.9 | 0.54 | 0.67 |
| 3 | 7741272 | rs162769 | -0.812 | -0.876 | 0.527 | -2.192 | -0.064 | 0.016 | 0.016 | 0 | 0.2 | 0.35 | 0.15 | 0.2 | 0.1 | 0.22 |
| 3 | 7741897 | rs162209 | -0.434 | -0.79 | 0.559 | -2.216 | 0.019 | 0.03 | 0.03 | 0.016 | 0.49 | 0.58 | 0.54 | 0.78 | 0.42 | 0.58 |
| 3 | 7745206 | rs10510371 | 1.514 | -0.784 | 0.553 | -2.126 | -0.034 | 0.007 | 0.007 | 1.014 | 0.08 | 0.26 | 0.05 | 0.06 | 0.02 | 0.01 |
| 3 | 7748843 | rs1504047 | 0.168 | -1.146 | 0.139 | -2.366 | 0.003 | 0.025 | 0.025 | 0 | 0.46 | 0.37 | 0.49 | 0.68 | 0.44 | 0.48 |
| 3 | 7753473 | rs332936 | 0.42 | -1.041 | -0.17 | -2.501 | 0.018 | 0.279 | 0.279 | 0.503 | 0.47 | 0.13 | 0.4 | 0.06 | 0.54 | 0.38 |
| 3 | 7757870 | rs3804820 | -0.222 | -0.71 | 0.225 | -2.121 | -0.008 | -0.005 | -0.005 | 2.475 | 0.08 | 0.12 | 0.08 | 0.05 | 0.05 | 0.14 |
| 3 | 7763386 | rs329037 | 0.186 | -0.327 | -0.345 | -2.358 | 0.023 | 0.083 | 0.083 | 0 | 0.4 | 0.21 | 0.3 | 0.16 | 0.35 | 0.35 |
| 3 | 7769282 | rs162724 | -0.23 | -0.076 | 0.455 | -1.921 | 0.006 | -0.069 | -0.069 | 0.156 | 0.18 | 0.54 | 0.19 | 0.06 | 0.16 | 0.14 |
| 3 | 7770147 | rs162722 | 0.732 | -0.135 | 0.505 | -1.937 | -0.001 | -0.001 | -0.001 | 0.003 | 0.74 | 0.3 | 0.77 | 0.85 | 0.77 | 0.8 |
| 3 | 7775834 | rs17047874 | 0.217 | 0.091 | 0.419 | -1.874 | 0.051 | -0.081 | -0.081 | 0 | 0.16 | 0.33 | 0.17 | 0.05 | 0.23 | 0.17 |
| 3 | 7782371 | rs9826579 | -0.929 | 0.027 | -0.048 | -1.876 | -0.001 | -0.001 | -0.001 | 0 | 0.24 | 0.69 | 0.24 | 0.16 | 0.23 | 0.19 |
| 3 | 7784626 | rs6806317 | -0.901 | 0.03 | -0.018 | -1.873 | 0.001 | -0.012 | -0.012 | 0 | 0.24 | 0.67 | 0.23 | 0.16 | 0.23 | 0.19 |
| 3 | 7785618 | rs323482 | -1.928 | 0.239 | -0.606 | -0.084 | -0.068 | -0.102 | -0.102 | 0 | 0.86 | 0.99 | 0.87 | 0.61 | 0.79 | 0.83 |
| 3 | 7797492 | rs161885 | -0.945 | -0.188 | -0.594 | -0.878 | 0.004 | 0.021 | 0.021 | 1.743 | 0.61 | 0.83 | 0.59 | 0.35 | 0.62 | 0.67 |
| 3 | 7805209 | rs1020102 | 0.47 | -0.206 | 0.681 | -0.523 | 0.026 | 0.005 | 0.005 | 1.192 | 0.14 | 0.26 | 0.13 | 0.14 | 0.17 | 0.17 |
| 3 | 7811984 | rs725382 | -0.801 | -0.437 | 0.357 | -0.626 | -0.001 | -0.032 | -0.032 | 0.728 | 0.25 | 0.65 | 0.19 | 0.14 | 0.2 | 0.2 |
| 3 | 7813017 | rs6443123 | -0.491 | -0.43 | 0.488 | -0.622 | 0.017 | -0.055 | -0.055 | 3.543 | 0.23 | 0.48 | 0.16 | 0.14 | 0.2 | 0.19 |
| 3 | 7817657 | rs9866595 | -0.851 | -0.196 | 0.051 | -0.522 | -0.002 | -0.019 | -0.019 | 0 | 0.25 | 0.69 | 0.2 | 0.14 | 0.2 | 0.19 |
| 3 | 7820444 | rs7634325 | -0.587 | -0.14 | 0.116 | -0.475 | -0.002 | -0.02 | -0.02 | 0 | 0.23 | 0.66 | 0.19 | 0.12 | 0.2 | 0.22 |
| 3 | 7820659 | rs9841316 | -0.071 | -0.235 | 0.162 | -0.826 | -0.002 | -0.009 | -0.009 | 0.053 | 0.68 | 0.66 | 0.72 | 0.75 | 0.68 | 0.63 |
| 3 | 7821284 | rs11708550 | 0.013 | -0.56 | 0.374 | -0.715 | -0.032 | 0.035 | 0.035 | 0.042 | 0.13 | 0.28 | 0.24 | 0.21 | 0.08 | 0.15 |
| 3 | 7821610 | rs161909 | 0.377 | -0.884 | 0.401 | -1.264 | -0.035 | -0.063 | -0.063 | 0 | 0.22 | 0.39 | 0.15 | 0.08 | 0.17 | 0.3 |

| | | | | | | | | | | | | | | | | |
|---|---|---|---|---|---|---|---|---|---|---|---|---|---|---|---|---|
| 3 | 7822471 | rs45840 | -0.444 | -0.901 | -0.142 | -1.464 | -0.047 | 0.083 | 0.083 | 0.84 | 0.76 | 0.9 | 0.74 | 0.96 | 0.69 | 0.76 |
| 3 | 7823055 | rs162211 | 0.911 | -1.258 | 0.401 | -1.558 | -0.007 | -0.064 | -0.064 | 0 | 0.61 | 0.47 | 0.65 | 0.86 | 0.63 | 0.62 |
| 3 | 7823658 | rs161445 | 0.329 | -1.258 | 0.508 | -1.57 | 0.009 | -0.115 | -0.115 | 0.617 | 0.69 | 0.52 | 0.7 | 0.95 | 0.68 | 0.73 |
| 3 | 7829298 | rs11131090 | 1.791 | -1.263 | 0.188 | -1.616 | -0.076 | 0.08 | 0.08 | 0 | 0.4 | 0.16 | 0.49 | 0.77 | 0.47 | 0.36 |
| 3 | 7829411 | rs161449 | 1.206 | -1.398 | 0.164 | -1.43 | 0.019 | 0.043 | 0.043 | 1.769 | 0.19 | 0.03 | 0.12 | 0.08 | 0.17 | 0.26 |
| 3 | 7829821 | rs9814244 | -0.645 | -1.063 | 0.779 | -1.116 | -0.071 | -0.027 | -0.027 | 0 | 0.16 | 0.51 | 0.1 | 0.1 | 0.06 | 0.14 |
| 3 | 7835289 | rs161926 | -1.028 | -0.759 | 0.165 | -1.12 | -0.002 | 0.025 | 0.025 | 0 | 0.88 | 0.94 | 0.87 | 0.96 | 0.88 | 0.89 |
| 3 | 7835759 | rs2174263 | 1.482 | -0.792 | 0.084 | -1.113 | 0.012 | 0.001 | 0.001 | 0 | 0.07 | 0 | 0.05 | 0.05 | 0.04 | 0.09 |
| 3 | 7838242 | rs161927 | -0.321 | -0.754 | 0.596 | -1.121 | -0.018 | -0.06 | -0.06 | 1.243 | 0.84 | 0.61 | 0.85 | 0.95 | 0.88 | 0.89 |
| 3 | 7839477 | rs162212 | 0.083 | -0.884 | 0.158 | -1.375 | 0.011 | 0.044 | 0.044 | 0 | 0.34 | 0.22 | 0.24 | 0.08 | 0.31 | 0.43 |
| 3 | 7839716 | rs9815643 | -0.113 | -0.235 | 0.348 | -0.969 | 0.05 | 0.055 | 0.055 | 0.551 | 0.06 | 0.22 | 0.05 | 0.09 | 0.06 | 0.07 |
| 3 | 7840303 | rs6772809 | 0.422 | -0.234 | 0.493 | -1.151 | 0.004 | 0.018 | 0.018 | 0 | 0.51 | 0.57 | 0.58 | 0.88 | 0.51 | 0.46 |
| 3 | 7840349 | rs1488472 | 0.067 | -0.574 | 0.357 | -1.1 | -0.013 | -0.002 | -0.002 | 0 | 0.15 | 0.27 | 0.12 | 0.14 | 0.1 | 0.15 |
| 3 | 7840740 | rs9829343 | -0.05 | -0.958 | 0.197 | -1.242 | 0.022 | 0.106 | 0.106 | 0 | 0.59 | 0.78 | 0.74 | 0.9 | 0.62 | 0.56 |
| 3 | 7841254 | rs17677071 | 0.058 | -0.701 | 0.482 | -0.3 | 0.029 | 0.092 | 0.092 | 0.166 | 0.2 | 0.48 | 0.27 | 0.26 | 0.23 | 0.27 |
| 3 | 7843597 | rs11921735 | 0.217 | -0.698 | 0.509 | -0.293 | 0.021 | 0.08 | 0.08 | 0.08 | 0.19 | 0.42 | 0.25 | 0.26 | 0.23 | 0.27 |
| 3 | 7843657 | rs9845228 | 0.031 | -0.809 | 0.083 | -0.246 | -0.018 | -0.001 | -0.001 | 0 | 0.23 | 0.25 | 0.17 | 0.23 | 0.11 | 0.21 |
| 3 | 7844612 | rs11706770 | 0.965 | -0.693 | 0.015 | -0.314 | 0.04 | 0.014 | 0.014 | 2.648 | 0.5 | 0.31 | 0.51 | 0.51 | 0.58 | 0.43 |
| 3 | 7844685 | rs11131091 | -0.367 | -0.61 | -0.074 | -0.268 | 0.002 | 0.006 | 0.006 | 0.471 | 0.78 | 0.72 | 0.7 | 0.75 | 0.72 | 0.67 |
| 3 | 7845802 | rs9843022 | 0.56 | -0.701 | -0.042 | -0.41 | 0.008 | 0.005 | 0.005 | 0 | 0.21 | 0.16 | 0.17 | 0.13 | 0.14 | 0.18 |
| 3 | 7846000 | rs13075346 | 1.4 | -0.247 | -0.306 | -0.406 | -0.041 | -0.034 | -0.034 | 0.046 | 0.37 | 0.2 | 0.4 | 0.51 | 0.4 | 0.32 |
| 3 | 7847183 | rs9865636 | 0.437 | -0.111 | -0.479 | -0.484 | 0 | -0.008 | -0.008 | 0.012 | 0.31 | 0.27 | 0.3 | 0.13 | 0.32 | 0.27 |
| 3 | 7849434 | rs2062850 | 0.277 | 0.284 | -0.401 | -0.322 | 0 | -0.031 | -0.031 | 0.013 | 0.81 | 0.75 | 0.83 | 0.99 | 0.78 | 0.88 |
| 3 | 7850284 | rs6780542 | -0.318 | 0.438 | -0.555 | 0.096 | 0.006 | 0.004 | 0.004 | 3.755 | 0.51 | 0.55 | 0.45 | 0.5 | 0.42 | 0.54 |
| 3 | 7852651 | rs17048002 | 1.154 | 0.528 | -0.576 | 0.242 | -0.004 | 0.026 | 0.026 | 0.983 | 0.24 | 0.3 | 0.34 | 0.46 | 0.19 | 0.12 |
| 3 | 7853659 | rs161894 | -0.043 | 0.29 | -0.97 | -0.151 | -0.051 | 0.052 | 0.052 | 1.534 | 0.38 | 0.21 | 0.44 | 0.27 | 0.52 | 0.48 |
| 3 | 7856288 | rs161898 | 1.127 | 0.731 | -1.166 | -0.309 | -0.033 | 0.044 | 0.044 | 0 | 0.08 | 0.02 | 0.09 | 0 | 0.13 | 0.07 |
| 3 | 7856781 | rs11706204 | 1.885 | 0.815 | -1.338 | -0.461 | -0.007 | -0.033 | -0.033 | 0 | 0.11 | 0.05 | 0.16 | 0.26 | 0.12 | 0.25 |
| 3 | 7858048 | rs162216 | -0.96 | 0.12 | -1.017 | -0.525 | 0.001 | 0.003 | 0.003 | 0 | 0.75 | 0.66 | 0.86 | 0.81 | 0.78 | 0.72 |
| 3 | 7858150 | rs4686162 | 1.597 | 0.198 | -0.988 | -0.813 | -0.032 | 0.045 | 0.045 | 1.559 | 0.14 | 0.04 | 0.13 | 0 | 0.21 | 0.17 |
| 3 | 7859835 | rs11918125 | 1.022 | -0.19 | -0.817 | -0.666 | -0.014 | -0.005 | -0.005 | 0 | 0.13 | 0.14 | 0.08 | 0.09 | 0.05 | 0.12 |
| 3 | 7860093 | rs161901 | -0.888 | -0.21 | -0.724 | -0.693 | 0.01 | 0.047 | 0.047 | 0.676 | 0.5 | 0.33 | 0.52 | 0.35 | 0.58 | 0.61 |
| 3 | 7860223 | rs11923266 | 0.149 | -0.395 | -0.418 | -0.583 | -0.055 | -0.029 | -0.029 | 0 | 0.16 | 0.37 | 0.1 | 0.08 | 0.07 | 0.1 |
| 3 | 7861685 | rs161906 | 0.247 | -0.115 | -1.072 | -0.617 | -0.09 | -0.034 | -0.034 | 0 | 0.3 | 0.08 | 0.35 | 0.33 | 0.44 | 0.41 |
| 3 | 7861851 | rs161455 | 0.011 | -0.114 | -0.695 | -0.632 | -0.007 | 0.001 | 0.001 | 0.635 | 0.32 | 0.26 | 0.37 | 0.32 | 0.44 | 0.41 |
| 3 | 7862620 | rs162705 | -1.203 | -0.322 | -0.934 | -0.871 | -0.001 | -0.002 | -0.002 | 0.045 | 0.55 | 0.51 | 0.5 | 0.41 | 0.62 | 0.63 |
| 3 | 7863283 | rs162708 | 0.422 | -0.267 | -1.18 | -0.837 | -0.081 | -0.031 | -0.031 | 0 | 0.3 | 0.09 | 0.36 | 0.33 | 0.44 | 0.41 |
| 3 | 7866171 | rs162713 | -0.98 | -0.228 | -0.81 | -0.661 | 0.003 | 0 | 0 | 0 | 0.57 | 0.61 | 0.52 | 0.45 | 0.64 | 0.69 |
| 3 | 7866522 | rs7651284 | 1.231 | -0.426 | -0.983 | -0.933 | -0.014 | -0.032 | -0.032 | 0 | 0.19 | 0.22 | 0.09 | 0.05 | 0.11 | 0.18 |
| 3 | 7870246 | rs2129922 | 1.899 | -0.104 | -1.295 | -0.638 | 0 | -0.01 | -0.01 | 0 | 0.24 | 0.25 | 0.36 | 0.44 | 0.19 | 0.14 |
| 3 | 7871402 | rs1542699 | 2.354 | 0.002 | -1.723 | -0.6 | -0.019 | -0.082 | -0.082 | 0.004 | 0.21 | 0.11 | 0.35 | 0.44 | 0.2 | 0.1 |
| 3 | 7873577 | rs17678083 | 1.727 | 0.64 | -1.486 | -0.672 | -0.02 | 0.026 | 0.026 | 0.047 | 0.1 | 0.03 | 0.09 | 0 | 0.14 | 0.1 |
| 3 | 7874159 | rs12494397 | 0.015 | 0.373 | -1.417 | -0.528 | 0.008 | 0.044 | 0.044 | 0.224 | 0.19 | 0.05 | 0.11 | 0.08 | 0.17 | 0.16 |
| 3 | 7881737 | rs4686168 | -0.238 | 0.369 | -1.403 | -0.474 | -0.017 | -0.035 | -0.035 | 0.106 | 0.71 | 0.81 | 0.7 | 0.61 | 0.64 | 0.63 |
| 3 | 7881985 | rs7628601 | 1.123 | 0.137 | -0.941 | -0.873 | 0 | 0.002 | 0.002 | 0 | 0.39 | 0.43 | 0.5 | 0.5 | 0.38 | 0.3 |
| 3 | 7883440 | rs1386981 | -0.494 | 0.127 | -1.088 | -0.758 | -0.003 | -0.037 | -0.037 | 0 | 0.71 | 0.81 | 0.7 | 0.61 | 0.67 | 0.61 |
| 3 | 7889657 | rs901877 | -0.099 | 0.075 | -0.757 | -0.85 | -0.001 | -0.002 | -0.002 | 0 | 0.66 | 0.67 | 0.66 | 0.61 | 0.64 | 0.63 |
| 3 | 7891245 | rs370017 | 1.263 | -0.006 | -0.715 | -1.152 | -0.001 | 0.022 | 0.022 | 0.809 | 0.33 | 0.53 | 0.44 | 0.43 | 0.32 | 0.17 |
| 3 | 7891262 | rs391664 | -0.673 | 0.008 | -0.813 | -1.309 | -0.001 | -0.02 | -0.02 | 0 | 0.44 | 0.42 | 0.31 | 0.15 | 0.42 | 0.48 |
| 3 | 7895790 | rs326419 | 1.682 | 0.048 | -0.762 | -1.302 | -0.021 | 0.048 | 0.048 | 0 | 0.09 | 0.02 | 0.08 | 0 | 0.11 | 0.08 |
| 3 | 7895867 | rs326418 | 0.126 | 0.085 | -0.807 | -1.351 | -0.001 | -0.066 | -0.066 | 0.01 | 0.28 | 0.38 | 0.23 | 0.08 | 0.28 | 0.28 |
| 3 | 7896679 | rs11718722 | 1.93 | 0.123 | -0.743 | -1.287 | -0.007 | 0.015 | 0.015 | 0.039 | 0.13 | 0.09 | 0.11 | 0.03 | 0.16 | 0.17 |
| 3 | 7897016 | rs326416 | -1.93 | 0.093 | -0.752 | -1.234 | -0.004 | -0.047 | -0.047 | 0 | 0.81 | 0.9 | 0.78 | 0.65 | 0.81 | 0.75 |
| 3 | 7897220 | rs11719723 | 1.862 | 0.151 | -0.592 | -1.265 | -0.007 | 0.015 | 0.015 | 0.039 | 0.13 | 0.09 | 0.11 | 0.03 | 0.16 | 0.17 |
| 3 | 7897821 | rs4684558 | 1.597 | 0.151 | -0.576 | -1.265 | 0.013 | -0.058 | -0.058 | 0 | 0.15 | 0.25 | 0.12 | 0.03 | 0.16 | 0.17 |
| 3 | 7898720 | rs13323237 | 1.331 | 0.185 | -0.654 | -1.134 | -0.001 | -0.008 | -0.008 | 0 | 0.17 | 0.19 | 0.14 | 0.08 | 0.17 | 0.2 |
| 3 | 7903022 | rs9311998 | 1.298 | 0.189 | -0.658 | -1.126 | -0.001 | -0.011 | -0.011 | 0.005 | 0.17 | 0.2 | 0.14 | 0.08 | 0.16 | 0.2 |
| 3 | 7903040 | rs326414 | 0.211 | 0.132 | -0.672 | -1.057 | 0.005 | 0.017 | 0.017 | 4.267 | 0.5 | 0.69 | 0.58 | 0.47 | 0.51 | 0.38 |
| 3 | 7905524 | rs9867817 | 2.665 | 0.157 | -0.695 | -0.957 | 0.004 | 0.052 | 0.052 | 1.097 | 0.22 | 0.34 | 0.35 | 0.44 | 0.2 | 0.09 |
| 3 | 7905601 | rs9829153 | 2.665 | 0.157 | -0.695 | -0.958 | 0.004 | 0.051 | 0.051 | 1.097 | 0.22 | 0.34 | 0.35 | 0.44 | 0.2 | 0.09 |
| 3 | 7906607 | rs227 | 0.027 | 0.098 | -0.643 | -1.031 | -0.011 | -0.039 | -0.039 | 0 | 0.33 | 0.4 | 0.21 | 0.18 | 0.28 | 0.37 |
| 3 | 7908883 | rs1488496 | -2.805 | 0.006 | -0.691 | -0.914 | -0.003 | -0.002 | -0.002 | 0 | 0.92 | 0.91 | 0.95 | 0.93 | 0.93 | 0.97 |
| 3 | 7909277 | rs411165 | -0.367 | -0.005 | -0.857 | -1.002 | 0.008 | -0.027 | -0.027 | 0.51 | 0.6 | 0.78 | 0.64 | 0.57 | 0.6 | 0.5 |
| 3 | 7910902 | rs450138 | -0.178 | -0.031 | -0.529 | -1.001 | -0.002 | 0.001 | 0.001 | 1.278 | 0.58 | 0.6 | 0.62 | 0.57 | 0.6 | 0.5 |
| 3 | 7912645 | rs9838153 | 0.187 | 0.033 | -0.441 | -1.162 | 0.007 | 0.043 | 0.043 | 0 | 0.31 | 0.22 | 0.19 | 0.12 | 0.3 | 0.36 |
| 3 | 7913209 | rs11131095 | 2.193 | 0.06 | -0.433 | -1.132 | 0.016 | 0.038 | 0.038 | 0.037 | 0.27 | 0.39 | 0.42 | 0.45 | 0.3 | 0.14 |
| 3 | 7915149 | rs397864 | -0.059 | 0.031 | -0.298 | -1.276 | -0.002 | -0.024 | -0.024 | 0 | 0.32 | 0.34 | 0.2 | 0.12 | 0.3 | 0.36 |
| 3 | 7916083 | rs13072357 | 0.732 | -0.016 | -0.497 | -1.328 | 0.001 | 0.005 | 0.005 | 1.429 | 0.13 | 0.11 | 0.09 | 0.03 | 0.13 | 0.16 |
| 3 | 7920512 | rs17690708 | 2.479 | 0.004 | -0.555 | -1.415 | -0.008 | 0.002 | 0.002 | 0.156 | 0.1 | 0.06 | 0.09 | 0 | 0.14 | 0.1 |

| | | | | | | | | | | | | | | | | |
|---|---|---|---|---|---|---|---|---|---|---|---|---|---|---|---|---|
| 3 | 7922313 | rs1074506 | 0.758 | -0.072 | -0.491 | -1.424 | 0.003 | 0.006 | 0.006 | 5.523 | 0.41 | 0.55 | 0.54 | 0.5 | 0.43 | 0.29 |
| 3 | 7925552 | rs453203 | -0.917 | -0.153 | -0.623 | -1.344 | 0.02 | -0.041 | -0.041 | 1.651 | 0.69 | 0.81 | 0.73 | 0.58 | 0.74 | 0.6 |
| 3 | 7949418 | rs2054884 | 0.465 | -0.13 | -0.5 | -1.35 | 0.011 | 0 | 0 | 0 | 0.49 | 0.45 | 0.6 | 0.5 | 0.57 | 0.39 |
| 3 | 7950172 | rs1386984 | 1.49 | -0.124 | -0.526 | -1.378 | 0.009 | 0.009 | 0.009 | 0 | 0.37 | 0.42 | 0.51 | 0.47 | 0.44 | 0.24 |
| 3 | 7951021 | rs1488487 | 0.965 | -0.139 | -0.558 | -1.35 | -0.007 | -0.07 | -0.07 | 0.608 | 0.19 | 0.34 | 0.13 | 0.09 | 0.17 | 0.21 |
| 3 | 7952801 | rs6802222 | -0.877 | -0.132 | -0.655 | -1.224 | 0.019 | -0.088 | -0.088 | 0.144 | 0.7 | 0.9 | 0.73 | 0.58 | 0.74 | 0.61 |
| 3 | 7955693 | rs6787368 | 0.552 | 0.078 | -0.737 | -0.992 | -0.007 | 0.055 | 0.055 | 1.668 | 0.13 | 0.03 | 0.09 | 0.03 | 0.13 | 0.16 |
| 3 | 7962060 | rs7625080 | 1.65 | 0.198 | -0.511 | -0.934 | 0.01 | 0.032 | 0.032 | 0.155 | 0.28 | 0.37 | 0.43 | 0.47 | 0.3 | 0.15 |
| 3 | 7962284 | rs1825924 | -0.281 | 0.206 | -0.422 | -1.164 | 0.003 | 0.013 | 0.013 | 0.584 | 0.73 | 0.68 | 0.71 | 0.62 | 0.68 | 0.65 |
| 3 | 7963677 | rs4686194 | 0.277 | 0.111 | -0.53 | -1.233 | 0.01 | -0.004 | -0.004 | 0 | 0.6 | 0.7 | 0.67 | 0.6 | 0.62 | 0.56 |
| 3 | 7963833 | rs3919962 | -0.193 | 0.122 | -0.555 | -1.23 | 0.001 | -0.011 | -0.011 | 0 | 0.64 | 0.78 | 0.7 | 0.63 | 0.64 | 0.66 |
| 3 | 7965419 | rs17048365 | 1.697 | 0.11 | -0.624 | -1.132 | 0.003 | -0.057 | -0.057 | 2.053 | 0.2 | 0.03 | 0.15 | 0.26 | 0.17 | 0.25 |
| 3 | 7966941 | rs10510378 | 1.331 | 0.516 | -0.335 | -1.027 | 0.016 | -0.034 | -0.034 | 0.77 | 0.08 | 0.11 | 0.09 | 0 | 0.12 | 0.05 |
| 3 | 7970956 | rs4686197 | 0.464 | 0.496 | 0.138 | -1.199 | -0.003 | 0.004 | 0.004 | 0 | 0.47 | 0.43 | 0.56 | 0.59 | 0.46 | 0.36 |
| 3 | 7971351 | rs17048378 | 1.692 | 0.438 | 0.152 | -1.386 | 0.003 | 0.008 | 0.008 | 0 | 0.16 | 0.13 | 0.09 | 0.1 | 0.14 | 0.16 |
| 3 | 7971904 | rs1488484 | -0.34 | 0.439 | 0.141 | -1.373 | 0 | 0 | 0 | 4.657 | 0.54 | 0.54 | 0.65 | 0.6 | 0.58 | 0.42 |
| 3 | 7972660 | rs9873903 | 0.33 | 0.447 | -0.16 | -1.322 | -0.003 | 0.005 | 0.005 | 0 | 0.47 | 0.43 | 0.56 | 0.6 | 0.46 | 0.36 |
| 3 | 7973163 | rs1488482 | -1.407 | 0.436 | -0.31 | -1.287 | 0.032 | -0.02 | -0.02 | 0 | 0.7 | 0.89 | 0.81 | 0.68 | 0.76 | 0.72 |
| 3 | 7982083 | rs10510380 | 1.328 | 0.481 | -0.233 | -1.126 | -0.013 | -0.041 | -0.041 | 0 | 0.3 | 0.22 | 0.46 | 0.45 | 0.33 | 0.2 |
| 3 | 7982827 | rs4591494 | 2.84 | 0.493 | -0.238 | -1.169 | -0.035 | -0.097 | -0.097 | 0 | 0.2 | 0.09 | 0.37 | 0.45 | 0.22 | 0.12 |
| 3 | 7983344 | rs6443144 | -0.923 | 0.487 | -0.095 | -0.909 | -0.01 | 0.04 | 0.04 | 2.192 | 0.4 | 0.68 | 0.35 | 0.23 | 0.44 | 0.51 |
| 3 | 7986779 | rs6443145 | -0.153 | 0.413 | 0.018 | -0.933 | 0.036 | -0.063 | -0.063 | 0 | 0.23 | 0.41 | 0.25 | 0.08 | 0.3 | 0.34 |
| 3 | 7987521 | rs9850126 | 0.235 | -0.017 | -0.112 | -0.726 | 0.002 | 0.004 | 0.004 | 0 | 0.56 | 0.6 | 0.69 | 0.67 | 0.6 | 0.52 |
| 3 | 7989144 | rs17048416 | 0.448 | 0.785 | 0.035 | -0.801 | 0.024 | -0.071 | -0.071 | 0 | 0.14 | 0.27 | 0.12 | 0.01 | 0.16 | 0.2 |
| 3 | 7990493 | rs1385437 | 0.379 | 0.784 | -0.643 | -0.767 | -0.009 | 0.019 | 0.019 | 0.8 | 0.58 | 0.38 | 0.62 | 0.83 | 0.49 | 0.49 |
| 3 | 7990913 | rs17048429 | 0.04 | 0.441 | -0.613 | -0.276 | 0.001 | -0.001 | -0.001 | 0.536 | 0.23 | 0.2 | 0.19 | 0.27 | 0.17 | 0.23 |
| 3 | 7990995 | rs1485155 | -1.113 | 1.087 | -0.22 | -0.253 | 0 | -0.005 | -0.005 | 0.046 | 0.11 | 0.15 | 0.09 | 0.06 | 0.08 | 0.14 |
| 3 | 7991133 | rs9855700 | 0.114 | 1.234 | -0.257 | -0.333 | -0.01 | 0.051 | 0.051 | 0.145 | 0.22 | 0.13 | 0.15 | 0.03 | 0.22 | 0.23 |
| 3 | 7991981 | rs6764556 | -0.119 | 0.932 | 0.496 | 0.372 | -0.012 | 0.057 | 0.057 | 0 | 0.77 | 0.65 | 0.65 | 0.43 | 0.81 | 0.61 |
| 3 | 7996336 | rs4686203 | -1.349 | 0.777 | 0.629 | 0.005 | -0.001 | -0.001 | -0.001 | 0 | 0.63 | 0.67 | 0.7 | 0.7 | 0.7 | 0.71 |
| 3 | 7996852 | rs9846963 | -0.265 | 1.198 | 0.756 | -0.047 | 0.027 | 0.014 | 0.014 | 0 | 0.28 | 0.39 | 0.46 | 0.67 | 0.34 | 0.44 |
| 3 | 7997953 | rs10510382 | 0.606 | 1.302 | 0.559 | 0.195 | -0.015 | 0.034 | 0.034 | 0 | 0.21 | 0.11 | 0.12 | 0.13 | 0.23 | 0.28 |
| 3 | 7998294 | rs6785352 | -0.058 | 1.297 | 0.446 | -0.201 | 0.077 | 0.053 | 0.053 | 0 | 0.06 | 0.21 | 0.12 | 0.1 | 0.1 | 0.06 |
| 3 | 7998577 | rs9818263 | -0.865 | 1.239 | 0.42 | -0.2 | -0.001 | -0.004 | -0.004 | 0 | 0.58 | 0.61 | 0.6 | 0.61 | 0.6 | 0.66 |
| 3 | 8004252 | rs13074331 | -0.23 | 0.934 | 0.105 | -0.046 | 0.003 | 0.058 | 0.058 | 0 | 0.73 | 0.54 | 0.6 | 0.42 | 0.74 | 0.55 |
| 3 | 8012506 | rs6443147 | -0.906 | 0.726 | -0.663 | 0.04 | 0.016 | 0.029 | 0.029 | 1.162 | 0.64 | 0.8 | 0.72 | 0.71 | 0.69 | 0.7 |
| 3 | 8016833 | rs6798912 | -0.528 | 1.011 | 0.021 | 0.099 | 0 | -0.023 | -0.023 | 0 | 0.39 | 0.46 | 0.33 | 0.18 | 0.43 | 0.27 |
| 3 | 8016955 | rs1825500 | 0.083 | 0.759 | 0.106 | -0.16 | 0.005 | 0.007 | 0.007 | 0.103 | 0.26 | 0.25 | 0.15 | 0.08 | 0.21 | 0.16 |
| 3 | 8017681 | rs17048457 | -0.755 | 1.003 | 0.487 | -0.016 | 0.002 | -0.023 | -0.023 | 0.451 | 0.07 | 0.08 | 0.05 | 0 | 0.08 | 0.05 |
| 3 | 8018306 | rs6805008 | 0.021 | 0.864 | 0.717 | -0.042 | -0.006 | -0.03 | -0.03 | 1.089 | 0.23 | 0.27 | 0.15 | 0.07 | 0.18 | 0.17 |
| 3 | 8019240 | rs17048460 | 0.825 | 1.013 | 0.593 | 0.354 | -0.006 | -0.008 | -0.008 | 0.063 | 0.14 | 0.11 | 0.18 | 0.23 | 0.19 | 0.14 |
| 3 | 8024019 | rs1385436 | -0.34 | 0.809 | 0.689 | 0.343 | 0 | 0 | 0 | 0 | 0.22 | 0.22 | 0.24 | 0.23 | 0.28 | 0.22 |
| 3 | 8027826 | rs6793461 | -1.524 | 0.619 | 0.768 | 0.57 | -0.026 | 0.069 | 0.069 | 0.954 | 0.56 | 0.84 | 0.57 | 0.73 | 0.54 | 0.62 |
| 3 | 8028886 | rs17048476 | 0.198 | 0.565 | 0.909 | 0.581 | -0.017 | -0.066 | -0.066 | 0 | 0.27 | 0.4 | 0.18 | 0.08 | 0.21 | 0.26 |
| 3 | 8030610 | rs9858956 | -0.803 | 1.063 | 0.987 | 1.22 | -0.001 | -0.038 | -0.038 | 0 | 0.07 | 0.15 | 0.04 | 0 | 0.07 | 0.06 |
| 3 | 8030809 | rs9859247 | -1.19 | 1.075 | 0.786 | 1.259 | 0.017 | 0.031 | 0.031 | 0 | 0.28 | 0.4 | 0.39 | 0.65 | 0.33 | 0.36 |
| 3 | 8041502 | rs3912448 | 0.107 | 1.002 | 0.244 | 1.152 | -0.011 | -0.009 | -0.009 | 0 | 0.15 | 0.13 | 0.18 | 0.21 | 0.21 | 0.13 |
| 3 | 8043370 | rs9880106 | -1.012 | 0.842 | 0.647 | 0.908 | 0.034 | 0.01 | 0.01 | 0 | 0.19 | 0.43 | 0.27 | 0.14 | 0.26 | 0.28 |
| 3 | 8068419 | rs512156 | -0.52 | 0.735 | 0.952 | 1.381 | 0.037 | 0.023 | 0.023 | 0 | 0.15 | 0.28 | 0.25 | 0.14 | 0.24 | 0.21 |
| 3 | 8070268 | rs471886 | -1.407 | 0.617 | 0.85 | 1.284 | 0.015 | 0.106 | 0.106 | 0 | 0.22 | 0.44 | 0.39 | 0.4 | 0.28 | 0.48 |
| 3 | 8078583 | rs516334 | 0.73 | 0.065 | 0.641 | 0.336 | -0.001 | -0.026 | -0.026 | 1.122 | 0.7 | 0.59 | 0.67 | 0.86 | 0.67 | 0.56 |
| 3 | 8078915 | rs492476 | -1.305 | 0.113 | -0.289 | 0.148 | 0.003 | 0.065 | 0.065 | 0.617 | 0.61 | 0.71 | 0.74 | 0.84 | 0.68 | 0.85 |
| 3 | 8090865 | rs1488342 | 0.289 | 0.045 | -0.144 | 0.136 | 0.01 | 0.014 | 0.014 | 0 | 0.32 | 0.45 | 0.39 | 0.69 | 0.35 | 0.42 |
| 3 | 8092805 | rs547906 | -0.451 | 0.002 | -0.228 | 0.121 | -0.003 | 0.011 | 0.011 | 0 | 0.28 | 0.21 | 0.37 | 0.22 | 0.33 | 0.41 |
| 3 | 8093466 | rs882955 | -0.231 | -0.055 | -0.266 | 0.096 | -0.013 | 0.008 | 0.008 | 0 | 0.26 | 0.15 | 0.36 | 0.22 | 0.33 | 0.37 |
| 3 | 8095274 | rs523678 | -1.34 | 0.175 | -0.306 | 0.182 | 0.001 | 0.062 | 0.062 | 0.506 | 0.62 | 0.72 | 0.79 | 0.92 | 0.69 | 0.85 |
| 3 | 8095826 | rs1111569 | -0.36 | 0.403 | -0.134 | 0.651 | 0 | 0.001 | 0.001 | 0 | 0.34 | 0.5 | 0.41 | 0.69 | 0.37 | 0.49 |
| 3 | 8098653 | rs507837 | -0.822 | 0.75 | -0.411 | 0.891 | -0.011 | -0.009 | -0.009 | 0 | 0.22 | 0.16 | 0.33 | 0.24 | 0.3 | 0.29 |
| 3 | 8111267 | rs9852161 | -0.249 | -0.276 | -0.551 | 0.182 | -0.004 | -0.014 | -0.014 | 0 | 0.18 | 0.22 | 0.11 | 0.05 | 0.09 | 0.18 |
| 3 | 8113242 | rs2633640 | 0.41 | 0.503 | -0.261 | 0.986 | -0.043 | -0.031 | -0.031 | 0 | 0.8 | 0.94 | 0.69 | 0.76 | 0.72 | 0.74 |
| 3 | 8113760 | rs1610182 | 0.932 | 0.456 | 0.321 | 0.911 | 0.03 | 0.024 | 0.024 | 0.775 | 0.76 | 0.65 | 0.64 | 0.7 | 0.68 | 0.69 |
| 3 | 8117179 | rs7632952 | -0.678 | 0.554 | -0.425 | 0.994 | -0.048 | -0.044 | -0.044 | 0.188 | 0.2 | 0.05 | 0.34 | 0.25 | 0.29 | 0.26 |
| 3 | 8131758 | rs472126 | -0.649 | 0.492 | -0.246 | 0.553 | 0.001 | -0.039 | -0.039 | 0.074 | 0.35 | 0.46 | 0.48 | 0.74 | 0.42 | 0.59 |
| 3 | 8143674 | rs6785623 | 1.229 | 0.661 | 0.043 | 0.825 | 0.003 | 0.063 | 0.063 | 0.325 | 0.54 | 0.22 | 0.33 | 0.33 | 0.47 | 0.38 |
| 3 | 8145637 | rs931298 | 0.077 | 0.682 | 0.308 | 1.019 | 0.018 | 0.061 | 0.061 | 0.105 | 0.74 | 0.66 | 0.66 | 0.57 | 0.63 | 0.53 |
| 3 | 8146961 | rs491726 | 0.338 | 0.827 | 0.38 | 0.647 | 0.071 | -0.056 | -0.056 | 0 | 0.9 | 0.7 | 0.9 | 0.99 | 0.91 | 0.9 |
| 3 | 8147718 | rs13097609 | 0.499 | 0.587 | 0.423 | 0.682 | 0.03 | -0.024 | -0.024 | 0 | 0.93 | 0.7 | 0.93 | 0.99 | 0.96 | 0.94 |
| 3 | 8148696 | rs6779176 | 1.347 | 0.863 | 0.35 | 0.626 | 0.001 | 0.058 | 0.058 | 0.085 | 0.87 | 0.62 | 0.83 | 0.79 | 0.92 | 0.89 |
| 3 | 8152315 | rs1850267 | 0.699 | 0.93 | 0.323 | 0.821 | 0.011 | 0.008 | 0.008 | 0 | 0.66 | 0.54 | 0.61 | 0.4 | 0.57 | 0.44 |

| | | | | | | | | | | | | | | | | |
|---|---|---|---|---|---|---|---|---|---|---|---|---|---|---|---|---|
| 3 | 8155409 | rs908481 | -0.564 | 1.019 | 0.514 | 0.754 | 0.004 | 0.028 | 0.028 | 0 | 0.29 | 0.41 | 0.35 | 0.6 | 0.35 | 0.49 |
| 3 | 8160017 | rs17048651 | -0.585 | 0.674 | 0.291 | 0.759 | -0.004 | -0.007 | -0.007 | 0 | 0.11 | 0.05 | 0.16 | 0.14 | 0.11 | 0.07 |
| 3 | 8160466 | rs7645904 | 0.688 | 0.434 | 0.564 | 0.513 | -0.006 | 0.02 | 0.02 | 0 | 0.72 | 0.58 | 0.6 | 0.34 | 0.71 | 0.46 |
| 3 | 8162513 | rs6790853 | -0.036 | 0.362 | 0.698 | 0.992 | -0.013 | 0.097 | 0.097 | 0 | 0.11 | 0.28 | 0.18 | 0.21 | 0.08 | 0.26 |
| 3 | 8164718 | rs1878175 | 0.437 | 0.368 | 0.711 | 0.99 | -0.003 | 0.084 | 0.084 | 0 | 0.87 | 0.63 | 0.8 | 0.78 | 0.9 | 0.7 |
| 3 | 8173161 | rs2729668 | 1.472 | 0.331 | 0.92 | 0.989 | -0.013 | 0.061 | 0.061 | 0.382 | 0.75 | 0.47 | 0.64 | 0.6 | 0.79 | 0.63 |
| 3 | 8177234 | rs4684581 | -0.964 | 0.362 | 0.922 | 0.932 | -0.008 | 0.019 | 0.019 | 0 | 0.25 | 0.36 | 0.38 | 0.68 | 0.26 | 0.5 |
| 3 | 8182914 | rs11715664 | 0.166 | 0.353 | 0.783 | 0.962 | -0.006 | -0.029 | -0.029 | 0 | 0.15 | 0.08 | 0.2 | 0.47 | 0.18 | 0.24 |
| 3 | 8184528 | rs465632 | 1.035 | 0.336 | 0.792 | 0.91 | -0.003 | 0.017 | 0.017 | 0 | 0.73 | 0.58 | 0.6 | 0.32 | 0.72 | 0.45 |
| 3 | 8187422 | rs6789772 | -0.384 | 0.201 | 0.664 | 0.932 | -0.002 | 0.012 | 0.012 | 0 | 0.12 | 0.17 | 0.16 | 0.2 | 0.11 | 0.07 |
| 3 | 8193426 | rs13061566 | -0.337 | 0.263 | 0.603 | 0.931 | 0 | -0.004 | -0.004 | 0 | 0.12 | 0.12 | 0.16 | 0.2 | 0.11 | 0.07 |
| 3 | 8194734 | rs9681162 | -1.074 | 0.307 | 0.623 | 0.848 | -0.001 | -0.002 | -0.002 | 0.345 | 0.27 | 0.25 | 0.39 | 0.68 | 0.29 | 0.3 |
| 3 | 8195676 | rs6796806 | -0.112 | 0.255 | 0.713 | 0.951 | -0.008 | 0.1 | 0.1 | 0 | 0.1 | 0.29 | 0.15 | 0.19 | 0.07 | 0.27 |
| 3 | 8200271 | rs6764798 | -0.391 | 0.224 | 0.731 | 0.946 | 0.003 | 0.138 | 0.138 | 0 | 0.11 | 0.41 | 0.16 | 0.19 | 0.07 | 0.26 |
| 3 | 8205397 | rs522949 | -1.734 | 0.288 | 0.362 | 0.643 | -0.008 | 0.019 | 0.019 | 0 | 0.36 | 0.53 | 0.44 | 0.52 | 0.3 | 0.46 |
| 3 | 8207706 | rs619481 | -0.546 | 0.287 | 0.445 | 0.682 | 0.007 | 0.002 | 0.002 | 0 | 0.12 | 0.18 | 0.14 | 0.12 | 0.12 | 0.11 |
| 3 | 8208054 | rs521085 | -2.453 | 0.26 | 0.246 | 0.5 | -0.002 | 0.003 | 0.003 | 0.27 | 0.42 | 0.53 | 0.47 | 0.52 | 0.37 | 0.5 |
| 3 | 8210196 | rs17048809 | -0.558 | 0.474 | 0.396 | 0.575 | -0.026 | 0.038 | 0.038 | 0 | 0.15 | 0.27 | 0.15 | 0.2 | 0.1 | 0.3 |
| 3 | 8210421 | rs662340 | -1.556 | 0.565 | 0.36 | 0.5 | 0.016 | 0.022 | 0.022 | 0.257 | 0.3 | 0.54 | 0.37 | 0.66 | 0.31 | 0.36 |
| 3 | 8212635 | rs9312005 | -1.412 | 0.52 | 0.402 | 0.496 | 0.014 | 0.035 | 0.035 | 0.084 | 0.28 | 0.53 | 0.36 | 0.66 | 0.28 | 0.32 |
| 3 | 8212656 | rs13318406 | -0.177 | 0.546 | 0.386 | 0.607 | 0.004 | 0.137 | 0.137 | 0 | 0.18 | 0.36 | 0.23 | 0.47 | 0.17 | 0.24 |
| 3 | 8217162 | rs17704732 | 0.184 | 0.584 | 0.204 | 0.682 | -0.007 | -0.007 | -0.007 | 0 | 0.09 | 0.06 | 0.11 | 0.12 | 0.11 | 0.07 |
| 3 | 8230904 | rs13088606 | -0.166 | 0.382 | 0.435 | 0.689 | -0.022 | 0.057 | 0.057 | 0 | 0.13 | 0.23 | 0.08 | 0.2 | 0.08 | 0.28 |
| 3 | 8233002 | rs390783 | 0.716 | 0.302 | 0.482 | 0.646 | -0.041 | 0.076 | 0.076 | 0 | 0.84 | 0.65 | 0.9 | 0.74 | 0.91 | 0.68 |
| 3 | 8235545 | rs9843014 | -0.42 | 0.314 | 0.476 | 0.656 | -0.027 | 0.096 | 0.096 | 0 | 0.13 | 0.29 | 0.08 | 0.26 | 0.08 | 0.28 |
| 3 | 8238326 | rs3912613 | -0.068 | 0.892 | -0.134 | 0.148 | -0.077 | -0.093 | -0.093 | 0 | 0.27 | 0.08 | 0.42 | 0.59 | 0.36 | 0.31 |
| 3 | 8238518 | rs4686232 | 0.386 | 0.6 | 0.093 | 0.202 | -0.011 | -0.038 | -0.038 | 0 | 0.22 | 0.14 | 0.33 | 0.47 | 0.27 | 0.24 |
| 3 | 8240382 | rs1488793 | -0.99 | 0.636 | 0.171 | 0.095 | -0.005 | -0.009 | -0.009 | 0 | 0.39 | 0.55 | 0.43 | 0.74 | 0.34 | 0.56 |
| 3 | 8244910 | rs6443171 | 0.025 | 0.435 | -0.599 | -0.406 | -0.007 | 0.022 | 0.022 | 0 | 0.56 | 0.64 | 0.52 | 0.74 | 0.53 | 0.65 |
| 3 | 8253480 | rs9882319 | -0.045 | 0.563 | -0.784 | -0.412 | 0 | -0.017 | -0.017 | 0 | 0.55 | 0.49 | 0.52 | 0.69 | 0.56 | 0.65 |
| 3 | 8260685 | rs6775971 | 1.166 | 0.213 | -0.736 | -0.529 | -0.034 | -0.011 | -0.011 | 0.48 | 0.28 | 0.41 | 0.16 | 0.21 | 0.18 | 0.29 |
| 3 | 8272008 | rs984475 | -1.186 | 0.632 | -1.567 | -0.826 | 0.003 | 0.053 | 0.053 | 0 | 0.77 | 0.91 | 0.87 | 0.99 | 0.78 | 0.79 |
| 3 | 8273312 | rs7636883 | 0.399 | 1.284 | -1.58 | -0.414 | -0.03 | -0.022 | -0.022 | 0 | 0.17 | 0.11 | 0.26 | 0.25 | 0.23 | 0.13 |
| 3 | 8275541 | rs17048879 | 0.835 | 0.553 | -1.572 | -0.284 | -0.039 | -0.112 | -0.112 | 0.97 | 0.31 | 0.09 | 0.43 | 0.52 | 0.38 | 0.37 |
| 3 | 8278215 | rs6443178 | 0.412 | 0.713 | -1.024 | -0.602 | -0.009 | 0.037 | 0.037 | 0 | 0.56 | 0.59 | 0.57 | 0.72 | 0.53 | 0.64 |
| 3 | 8281680 | rs7622900 | 0.407 | 1.32 | -0.489 | -0.058 | -0.001 | 0 | 0 | 0.153 | 0.8 | 0.82 | 0.74 | 0.82 | 0.71 | 0.82 |
| 3 | 8291309 | rs9869998 | 0.819 | 0.993 | -0.323 | -0.08 | 0.046 | -0.015 | -0.015 | 1.958 | 0.69 | 0.55 | 0.66 | 0.73 | 0.63 | 0.71 |
| 3 | 8296654 | rs6799128 | -0.808 | 0.071 | -0.766 | -0.067 | -0.038 | 0.052 | 0.052 | 0.41 | 0.46 | 0.83 | 0.5 | 0.37 | 0.46 | 0.47 |
| 3 | 8297505 | rs9882964 | -0.31 | 0.163 | -0.237 | -0.054 | 0.005 | -0.018 | -0.018 | 0.062 | 0.23 | 0.36 | 0.35 | 0.17 | 0.26 | 0.34 |
| 3 | 8302786 | rs1121119 | -0.728 | 0.259 | -1.084 | -0.014 | -0.021 | 0.021 | 0.021 | 0.126 | 0.4 | 0.72 | 0.48 | 0.35 | 0.44 | 0.42 |
| 3 | 8310990 | rs1979582 | -0.125 | 0.09 | -0.885 | 0.104 | -0.002 | -0.001 | -0.001 | 0.256 | 0.38 | 0.41 | 0.32 | 0.33 | 0.38 | 0.45 |
| 3 | 8314924 | rs923381 | -0.671 | -0.02 | -0.878 | 0.044 | -0.003 | 0.002 | 0.002 | 0 | 0.62 | 0.61 | 0.48 | 0.53 | 0.65 | 0.62 |
| 3 | 8316110 | rs17763297 | -0.027 | 0.077 | -1.254 | 0.095 | 0 | 0 | 0 | 0 | 0.38 | 0.39 | 0.31 | 0.34 | 0.39 | 0.45 |
| 3 | 8319151 | rs13081504 | 1.249 | 0.58 | -1.541 | 0.075 | -0.062 | -0.022 | -0.022 | 0 | 0.2 | 0.03 | 0.13 | 0.17 | 0.22 | 0.11 |
| 3 | 8322531 | rs1027107 | -1.134 | 0.494 | -1.424 | -0.062 | -0.015 | -0.029 | -0.029 | 0 | 0.74 | 0.83 | 0.72 | 0.58 | 0.69 | 0.69 |
| 3 | 8323456 | rs12494294 | 0.939 | 0.095 | -1.33 | 0.554 | -0.056 | -0.057 | -0.057 | 0 | 0.21 | 0.03 | 0.14 | 0.22 | 0.25 | 0.12 |
| 3 | 8323828 | rs4686251 | 0.033 | -0.363 | -0.723 | 0.488 | 0.006 | -0.006 | -0.006 | 0 | 0.26 | 0.24 | 0.26 | 0.3 | 0.16 | 0.3 |
| 3 | 8324589 | rs9312013 | -0.498 | 0.98 | -0.452 | 0.19 | 0.018 | -0.024 | -0.024 | 0 | 0.07 | 0.24 | 0.17 | 0.01 | 0.05 | 0.05 |
| 3 | 8327085 | rs6799462 | -0.079 | 1.226 | -1.226 | 0.268 | -0.008 | 0.01 | 0.01 | 0 | 0.31 | 0.21 | 0.21 | 0.2 | 0.35 | 0.23 |
| 3 | 8327822 | rs4686252 | -0.07 | 1.881 | -2.003 | 0.705 | -0.148 | -0.147 | -0.147 | 0.195 | 0.76 | 0.97 | 0.55 | 0.49 | 0.56 | 0.68 |
| 3 | 8327887 | rs11709567 | 0.243 | 0.949 | -2.212 | 0.117 | 0.012 | 0.085 | 0.085 | 1.057 | 0.33 | 0.1 | 0.17 | 0.16 | 0.33 | 0.25 |
| 3 | 8328967 | rs17763559 | 0.428 | 0.801 | -1.937 | 0.256 | 0 | 0 | 0 | 0 | 0.08 | 0.07 | 0.06 | 0.06 | 0.07 | 0.04 |
| 3 | 8330426 | rs17048945 | 0.284 | 0.684 | -1.761 | 0.056 | -0.006 | -0.02 | -0.02 | 1.026 | 0.09 | 0.15 | 0.06 | 0.02 | 0.06 | 0.04 |
| 3 | 8330534 | rs6763794 | 0.335 | 0.519 | -1.821 | -0.083 | 0.027 | 0.084 | 0.084 | 0.803 | 0.35 | 0.14 | 0.18 | 0.15 | 0.33 | 0.25 |
| 3 | 8336967 | rs4487223 | -0.814 | 0.34 | -0.976 | -0.08 | -0.022 | -0.017 | -0.017 | 0.217 | 0.66 | 0.58 | 0.69 | 0.71 | 0.79 | 0.63 |
| 3 | 8340430 | rs7635918 | 0.999 | 0.669 | -1.093 | -0.138 | 0.036 | 0.039 | 0.039 | 0 | 0.31 | 0.06 | 0.17 | 0.17 | 0.29 | 0.27 |
| 3 | 8342581 | rs17763954 | 0.317 | 1.578 | -1.081 | -0.151 | 0 | 0.001 | 0.001 | 0 | 0.12 | 0.11 | 0.1 | 0.05 | 0.18 | 0.06 |
| 3 | 8343977 | rs7619176 | -0.058 | 1.525 | -0.796 | -0.148 | 0 | 0.006 | 0.006 | 0 | 0.46 | 0.41 | 0.31 | 0.22 | 0.48 | 0.33 |
| 3 | 8344541 | rs754808 | 0.655 | 1.544 | -0.846 | -0.07 | -0.05 | -0.067 | -0.067 | 0.021 | 0.17 | 0.1 | 0.35 | 0.48 | 0.3 | 0.29 |
| 3 | 8348688 | rs901045 | 1.428 | 1.309 | -0.977 | -0.307 | 0.026 | 0.042 | 0.042 | 0 | 0.25 | 0.13 | 0.13 | 0.09 | 0.22 | 0.19 |
| 3 | 8358721 | rs6793282 | -0.275 | 1.259 | -0.983 | -0.388 | 0.008 | 0.025 | 0.025 | 0 | 0.49 | 0.43 | 0.56 | 0.65 | 0.6 | 0.54 |
| 3 | 8359650 | rs6771359 | -1.164 | 0.96 | -1.08 | -0.424 | -0.027 | -0.016 | -0.016 | 0 | 0.64 | 0.58 | 0.68 | 0.7 | 0.8 | 0.61 |
| 3 | 8361157 | rs12492127 | 1.445 | 1.015 | -1.067 | -0.379 | 0.024 | 0.041 | 0.041 | 0 | 0.25 | 0.13 | 0.14 | 0.08 | 0.22 | 0.17 |
| 3 | 8369647 | rs1485222 | -1.08 | 1.068 | -0.846 | -0.201 | -0.032 | -0.024 | -0.024 | 0.077 | 0.66 | 0.54 | 0.56 | 0.7 | 0.77 | 0.57 |
| 3 | 8379719 | rs4684591 | 0.914 | 1.08 | -0.722 | -0.128 | -0.011 | -0.004 | -0.004 | 0 | 0.34 | 0.39 | 0.44 | 0.32 | 0.24 | 0.43 |
| 3 | 8380711 | rs9855812 | -0.702 | 1.707 | -0.991 | 0.026 | -0.071 | -0.096 | -0.096 | 0 | 0.84 | 0.94 | 0.77 | 0.5 | 0.72 | 0.75 |
| 3 | 8382513 | rs1905772 | 0.027 | 1.181 | -0.644 | -0.405 | -0.002 | 0.025 | 0.025 | 0 | 0.36 | 0.27 | 0.22 | 0.16 | 0.39 | 0.27 |
| 3 | 8384132 | rs11924632 | 0.624 | 1.217 | -0.684 | -0.435 | -0.002 | 0.068 | 0.068 | 0 | 0.29 | 0.13 | 0.17 | 0.09 | 0.32 | 0.21 |
| 3 | 8384934 | rs9877426 | -0.139 | 1.014 | -0.22 | 0.282 | 0.016 | 0.022 | 0.022 | 0 | 0.08 | 0.18 | 0.06 | 0.07 | 0.07 | 0.06 |

| | | | | | | | | | | | | | | | | |
|---|---|---|---|---|---|---|---|---|---|---|---|---|---|---|---|---|
| 3 | 8390226 | rs1385469 | 0.408 | 1.316 | -0.504 | 0.215 | -0.018 | -0.005 | -0.005 | 0 | 0.89 | 0.95 | 0.79 | 0.92 | 0.83 | 0.88 |
| 3 | 8392254 | rs2062371 | -0.422 | 0.795 | -0.151 | -0.422 | 0.001 | 0.002 | 0.002 | 0.013 | 0.43 | 0.42 | 0.46 | 0.79 | 0.33 | 0.58 |
| 3 | 8403739 | rs359030 | -0.427 | 0.474 | 0.01 | -0.388 | 0.001 | -0.002 | -0.002 | 0 | 0.09 | 0.1 | 0.17 | 0.09 | 0.1 | 0.1 |
| 3 | 8404733 | rs359029 | -0.536 | 0.299 | 0.578 | -0.348 | -0.003 | 0.046 | 0.046 | 0 | 0.1 | 0.3 | 0.19 | 0.1 | 0.09 | 0.1 |
| 3 | 8406919 | rs358994 | -1.146 | 0.091 | 0.518 | -0.44 | -0.03 | 0.008 | 0.008 | 0 | 0.13 | 0.29 | 0.2 | 0.1 | 0.09 | 0.11 |
| 3 | 8410755 | rs358988 | 0.252 | 0.061 | 0.588 | -0.539 | -0.002 | -0.013 | -0.013 | 0 | 0.56 | 0.46 | 0.53 | 0.21 | 0.64 | 0.43 |
| 3 | 8412570 | rs358985 | 0.488 | 0.032 | 0.446 | -0.55 | 0.025 | 0.102 | 0.102 | 0.932 | 0.52 | 0.3 | 0.5 | 0.19 | 0.66 | 0.42 |
| 3 | 8421679 | rs1994267 | -0.781 | -0.055 | 0.102 | -0.468 | -0.029 | -0.002 | -0.002 | 0.015 | 0.9 | 1 | 0.91 | 0.91 | 0.92 | 0.96 |
| 3 | 8422531 | rs924366 | 0.185 | 0.03 | 0.778 | -0.497 | -0.002 | 0.055 | 0.055 | 0 | 0.58 | 0.71 | 0.55 | 0.21 | 0.66 | 0.41 |
| 3 | 8423035 | rs1385472 | 0.914 | -0.084 | 0.392 | 0.286 | -0.019 | -0.024 | -0.024 | 0 | 0.11 | 0.05 | 0.08 | 0.44 | 0.13 | 0.3 |
| 3 | 8423182 | rs2648762 | -0.11 | -0.093 | 0.386 | 0.124 | -0.002 | 0.001 | 0.001 | 0 | 0.09 | 0.12 | 0.18 | 0.08 | 0.1 | 0.1 |
| 3 | 8424159 | rs2648758 | -0.048 | -0.087 | 0.312 | 0.127 | 0.001 | -0.003 | -0.003 | 0 | 0.09 | 0.1 | 0.17 | 0.08 | 0.1 | 0.1 |
| 3 | 8426122 | rs359003 | 0.565 | -0.027 | 0.145 | 0.243 | -0.002 | -0.03 | -0.03 | 0 | 0.08 | 0.01 | 0.16 | 0.1 | 0.09 | 0.1 |
| 3 | 8428999 | rs1994268 | 1.209 | 0.067 | 0.326 | 0.166 | -0.016 | 0.099 | 0.099 | 0.008 | 0.44 | 0.24 | 0.34 | 0.11 | 0.56 | 0.32 |
| 3 | 8434822 | rs359045 | -0.84 | 0.439 | 0.6 | 0.502 | -0.012 | -0.001 | -0.001 | 0 | 0.43 | 0.5 | 0.43 | 0.44 | 0.31 | 0.4 |
| 3 | 8442494 | rs1161171 | 0.241 | -0.739 | 0.902 | 0.629 | -0.014 | 0.015 | 0.015 | 0.042 | 0.67 | 0.6 | 0.68 | 0.58 | 0.78 | 0.69 |
| 3 | 8445729 | rs359025 | -0.143 | -0.75 | 1.034 | 0.645 | 0.018 | -0.041 | -0.041 | 0 | 0.69 | 0.82 | 0.71 | 0.61 | 0.78 | 0.7 |
| 3 | 8451805 | rs359017 | 1.173 | -0.606 | 0.188 | 0.62 | 0.053 | 0.167 | 0.167 | 0.926 | 0.51 | 0.19 | 0.51 | 0.17 | 0.67 | 0.4 |
| 3 | 8460083 | rs1018114 | 0.436 | -0.853 | 0.299 | 0.662 | -0.024 | 0.016 | 0.016 | 0.161 | 0.65 | 0.47 | 0.67 | 0.6 | 0.79 | 0.71 |
| 3 | 8465869 | rs6798206 | -0.987 | -0.541 | 0.579 | 0.5 | -0.027 | -0.021 | -0.021 | 0 | 0.07 | 0.27 | 0.03 | 0.03 | 0.01 | 0.01 |
| 3 | 8467511 | rs359009 | 0.021 | -1.089 | -0.231 | 0.247 | 0.038 | -0.02 | -0.02 | 0 | 0.53 | 0.86 | 0.5 | 0.55 | 0.62 | 0.52 |
| 3 | 8469784 | rs359006 | 1.319 | -0.871 | 0.07 | -0.049 | -0.046 | 0.06 | 0.06 | 0 | 0.33 | 0.21 | 0.32 | 0.07 | 0.53 | 0.23 |
| 3 | 8472855 | rs181559 | 0.659 | -1.447 | -0.055 | -0.364 | -0.003 | -0.02 | -0.02 | 0 | 0.34 | 0.63 | 0.37 | 0.1 | 0.53 | 0.2 |
| 3 | 8473745 | rs13100943 | 0.74 | -1.733 | -0.216 | 0.173 | 0.009 | -0.015 | -0.015 | 0 | 0.16 | 0.13 | 0.12 | 0.39 | 0.11 | 0.31 |
| 3 | 8473812 | rs13084939 | 0.03 | -1.788 | -0.146 | 0.149 | -0.008 | 0.012 | 0.012 | 0.033 | 0.19 | 0.22 | 0.13 | 0.39 | 0.11 | 0.31 |
| 3 | 8475557 | rs12630404 | 0.144 | -1.663 | -0.087 | 0.27 | 0 | 0.006 | 0.006 | 0.035 | 0.18 | 0.19 | 0.14 | 0.44 | 0.13 | 0.32 |
| 3 | 8475598 | rs11715302 | 0.526 | -1.786 | -0.46 | -0.017 | 0.067 | -0.034 | -0.034 | 0.47 | 0.43 | 0.12 | 0.48 | 0.47 | 0.31 | 0.43 |
| 3 | 8487491 | rs1161149 | -0.488 | -1.832 | -0.44 | 0.047 | 0.056 | -0.027 | -0.027 | 0.671 | 0.53 | 0.87 | 0.5 | 0.52 | 0.66 | 0.51 |
| 3 | 8490576 | rs358271 | 0.907 | -1.533 | 0.184 | -0.499 | 0.05 | -0.13 | -0.13 | 0 | 0.32 | 0.45 | 0.33 | 0.07 | 0.54 | 0.21 |
| 3 | 8492028 | rs358268 | 0.851 | -1.557 | 0.17 | -0.494 | 0.025 | -0.076 | -0.076 | 0 | 0.33 | 0.54 | 0.34 | 0.07 | 0.53 | 0.2 |
| 3 | 8494223 | rs358265 | 0.688 | -1.358 | 0.116 | -0.616 | 0.019 | -0.087 | -0.087 | 1.189 | 0.34 | 0.55 | 0.37 | 0.1 | 0.53 | 0.21 |
| 3 | 8495335 | rs11718095 | 0.837 | -1.295 | 0.102 | -0.631 | 0.039 | -0.113 | -0.113 | 1.213 | 0.33 | 0.53 | 0.34 | 0.05 | 0.5 | 0.2 |
| 3 | 8495425 | rs7626393 | -0.531 | -1.693 | -0.308 | -0.646 | -0.004 | 0.015 | 0.015 | 0 | 0.77 | 0.81 | 0.83 | 0.88 | 0.77 | 0.81 |
| 3 | 8501886 | rs17049161 | 0.229 | -2.052 | -0.546 | -0.594 | 0.064 | -0.001 | -0.001 | 0 | 0.26 | 0.06 | 0.23 | 0.23 | 0.14 | 0.33 |
| 3 | 8509477 | rs17049172 | 0.292 | -2.214 | -0.498 | -0.639 | 0.029 | -0.009 | -0.009 | 5.825 | 0.23 | 0.18 | 0.24 | 0.25 | 0.12 | 0.34 |
| 3 | 8509497 | rs17049174 | -0.329 | -2.195 | -0.34 | -0.535 | 0.021 | -0.031 | -0.031 | 0 | 0.21 | 0.12 | 0.23 | 0.55 | 0.14 | 0.24 |
| 3 | 8512668 | rs9864415 | 1.065 | -1.118 | -0.126 | -1.285 | -0.046 | 0.05 | 0.05 | 0.38 | 0.32 | 0.24 | 0.31 | 0.05 | 0.49 | 0.21 |
| 3 | 8519329 | rs6777857 | -0.014 | -1.307 | 0.001 | -1.339 | 0.003 | -0.029 | -0.029 | 0 | 0.47 | 0.52 | 0.39 | 0.09 | 0.58 | 0.42 |
| 3 | 8521841 | rs17049179 | 0.198 | -1.232 | -0.116 | -1.206 | -0.006 | -0.005 | -0.005 | 1.332 | 0.12 | 0.16 | 0.06 | 0.08 | 0.06 | 0.2 |
| 3 | 8524321 | rs10510390 | -0.072 | -1.223 | -0.148 | -1.192 | 0.012 | 0.014 | 0.014 | 0 | 0.17 | 0.15 | 0.07 | 0.08 | 0.09 | 0.2 |
| 3 | 8524337 | rs9811322 | -0.313 | -1.191 | -0.121 | -1.254 | 0.007 | -0.003 | -0.003 | 0 | 0.51 | 0.51 | 0.38 | 0.13 | 0.58 | 0.42 |
| 3 | 8528431 | rs7647092 | 0.615 | -0.618 | -0.295 | -1.117 | -0.019 | 0.001 | 0.001 | 0.456 | 0.33 | 0.31 | 0.3 | 0.05 | 0.5 | 0.22 |
| 3 | 8542278 | rs3755789 | 0.632 | -0.614 | -0.375 | -1.058 | -0.023 | 0.01 | 0.01 | 0.892 | 0.33 | 0.3 | 0.29 | 0.05 | 0.51 | 0.21 |
| 3 | 8543757 | rs2324625 | 0.617 | -0.569 | -0.328 | -1.084 | -0.026 | 0.026 | 0.026 | 0 | 0.35 | 0.3 | 0.3 | 0.06 | 0.51 | 0.23 |
| 3 | 8546102 | rs9312023 | 0.079 | -0.818 | -0.358 | -1.155 | -0.001 | 0.058 | 0.058 | 0 | 0.43 | 0.32 | 0.32 | 0.08 | 0.56 | 0.38 |
| 3 | 8548827 | rs6798584 | 0.171 | -0.624 | -0.029 | -1.171 | -0.001 | -0.055 | -0.055 | 0.002 | 0.45 | 0.45 | 0.3 | 0.08 | 0.51 | 0.4 |
| 3 | 8549590 | rs12485277 | 1.045 | -0.484 | -0.585 | -1.023 | -0.034 | -0.035 | -0.035 | 0.082 | 0.09 | 0.01 | 0.08 | 0.1 | 0.09 | 0.08 |
| 3 | 8551112 | rs9866032 | -1.148 | -0.778 | -0.609 | -0.854 | -0.023 | 0.037 | 0.037 | 0.17 | 0.67 | 0.95 | 0.68 | 0.78 | 0.68 | 0.72 |
| 3 | 8551649 | rs17049223 | 0.101 | -0.413 | -0.105 | -1.072 | -0.009 | -0.023 | -0.023 | 0 | 0.07 | 0.14 | 0.01 | 0.02 | 0.03 | 0.17 |
| 3 | 8554052 | rs17049231 | 0.641 | -0.404 | -0.301 | -1.053 | 0.011 | 0.014 | 0.014 | 0 | 0.06 | 0.06 | 0.01 | 0.01 | 0.02 | 0.16 |
| 3 | 8557559 | rs3774220 | -0.132 | -0.473 | -0.096 | -1.186 | 0.001 | -0.055 | -0.055 | 0.293 | 0.45 | 0.49 | 0.3 | 0.07 | 0.51 | 0.4 |
| 3 | 8561732 | rs9312024 | 0.546 | -0.429 | -0.101 | -1.172 | -0.034 | 0.052 | 0.052 | 0 | 0.36 | 0.26 | 0.28 | 0.06 | 0.49 | 0.22 |
| 3 | 8563144 | rs9856112 | 0.537 | -0.426 | -0.111 | -1.088 | -0.016 | -0.024 | -0.024 | 0 | 0.07 | 0.17 | 0.02 | 0.01 | 0.02 | 0.16 |
| 3 | 8568468 | rs9827267 | 0.16 | -0.503 | -0.026 | -1.182 | -0.002 | -0.024 | -0.024 | 0.254 | 0.41 | 0.4 | 0.3 | 0.06 | 0.52 | 0.23 |
| 3 | 8572373 | rs931517 | -0.224 | -0.425 | -0.102 | -1.042 | -0.001 | -0.024 | -0.024 | 0 | 0.48 | 0.49 | 0.32 | 0.11 | 0.52 | 0.4 |
| 3 | 8573095 | rs6801316 | -1.208 | -0.685 | -0.321 | -1.288 | -0.034 | 0.025 | 0.025 | 0 | 0.8 | 0.89 | 0.83 | 0.89 | 0.72 | 0.81 |
| 3 | 8575411 | rs6795922 | -0.461 | 0.119 | -0.234 | -1.187 | -0.02 | -0.053 | -0.053 | 0 | 0.14 | 0.23 | 0.06 | 0.02 | 0.1 | 0.24 |
| 3 | 8576401 | rs10510392 | -0.233 | 0.134 | -0.284 | -1.18 | 0.01 | 0.018 | 0.018 | 0.045 | 0.13 | 0.11 | 0.06 | 0.02 | 0.11 | 0.24 |
| 3 | 8579850 | rs6784926 | 0.36 | 0.173 | -0.329 | -1.194 | -0.011 | 0.001 | 0.001 | 0.013 | 0.34 | 0.33 | 0.25 | 0.05 | 0.41 | 0.16 |
| 3 | 8579952 | rs6809788 | -0.419 | 0.238 | -0.369 | -1.009 | -0.001 | -0.034 | -0.034 | 1.784 | 0.47 | 0.5 | 0.32 | 0.08 | 0.52 | 0.39 |
| 3 | 8592351 | rs17049242 | 0.071 | 0.183 | -0.107 | -0.53 | -0.009 | -0.026 | -0.026 | 0 | 0.15 | 0.27 | 0.1 | 0.06 | 0.11 | 0.08 |
| 3 | 8592573 | rs1545416 | -1.313 | -0.005 | -0.029 | -0.791 | -0.033 | -0.085 | -0.085 | 0 | 0.31 | 0.51 | 0.17 | 0.08 | 0.23 | 0.32 |
| 3 | 8593038 | rs1971883 | -0.56 | 0.091 | -0.207 | -0.723 | -0.005 | -0.018 | -0.018 | 0 | 0.15 | 0.16 | 0.07 | 0.03 | 0.12 | 0.25 |
| 3 | 8597771 | rs7653602 | -0.89 | 0.127 | -0.054 | -0.708 | -0.016 | -0.061 | -0.061 | 0.151 | 0.26 | 0.4 | 0.15 | 0.08 | 0.23 | 0.32 |
| 3 | 8601334 | rs6443195 | -1.12 | 0.135 | -0.236 | 0.324 | -0.012 | 0.031 | 0.031 | 0 | 0.27 | 0.4 | 0.35 | 0.42 | 0.2 | 0.37 |
| 3 | 8602939 | rs9820959 | 0.273 | 0.066 | -0.199 | 0.281 | -0.01 | -0.045 | -0.045 | 0 | 0.14 | 0.21 | 0.06 | 0.02 | 0.09 | 0.1 |
| 3 | 8603243 | rs165177 | -1.152 | -0.026 | -1.148 | 0.168 | -0.077 | -0.067 | -0.067 | 0 | 0.75 | 0.99 | 0.79 | 0.72 | 0.74 | 0.73 |
| 3 | 8603483 | rs1876611 | -1.421 | -0.137 | -0.325 | 0.333 | -0.015 | 0.051 | 0.051 | 0 | 0.3 | 0.54 | 0.38 | 0.44 | 0.22 | 0.37 |
| 3 | 8603936 | rs17049256 | 0.293 | -0.035 | -0.463 | 0.422 | -0.007 | -0.06 | -0.06 | 0 | 0.15 | 0.35 | 0.08 | 0.05 | 0.1 | 0.1 |

| | | | | | | | | | | | | | | | |
|---|---|---|---|---|---|---|---|---|---|---|---|---|---|---|---|
| 3 | 8605570 | rs12489970 | -0.225 | 0.151 | -0.626 | 0.811 | 0 | -0.046 | -0.046 | 1.221 | 0.12 | 0.04 | 0.28 | 0.38 | 0.09 | 0.27 |
| 3 | 8606839 | rs1027327 | -0.735 | 0.013 | -0.45 | 0.946 | -0.002 | 0.03 | 0.03 | 0 | 0.11 | 0.16 | 0.25 | 0.33 | 0.09 | 0.23 |
| 3 | 8608920 | rs171408 | -0.334 | 0.027 | -0.131 | 0.54 | -0.014 | -0.007 | -0.007 | 0.376 | 0.71 | 0.6 | 0.77 | 0.73 | 0.79 | 0.75 |
| 3 | 8610581 | rs6764383 | -0.163 | 0.873 | -0.356 | 0.839 | -0.03 | -0.025 | -0.025 | 0 | 0.24 | 0.15 | 0.41 | 0.38 | 0.41 | 0.43 |
| 3 | 8615277 | rs164966 | -0.82 | 0.674 | -0.401 | 0.767 | -0.026 | -0.004 | -0.004 | 0 | 0.73 | 0.51 | 0.76 | 0.74 | 0.8 | 0.76 |
| 3 | 8621403 | rs956377 | -0.343 | 0.876 | -0.395 | 0.816 | -0.001 | -0.001 | -0.001 | 0 | 0.24 | 0.22 | 0.42 | 0.39 | 0.42 | 0.43 |
| 3 | 8631179 | rs355122 | 0.675 | 1.045 | -0.608 | 0.694 | 0.013 | 0.03 | 0.03 | 1.509 | 0.46 | 0.3 | 0.37 | 0.35 | 0.42 | 0.33 |
| 3 | 8631614 | rs4686276 | -0.112 | 1.304 | -0.477 | 0.829 | 0.033 | 0.028 | 0.028 | 0 | 0.23 | 0.26 | 0.39 | 0.35 | 0.35 | 0.38 |
| 3 | 8632542 | rs355123 | -0.226 | 0.799 | -0.552 | 0.771 | -0.004 | 0.005 | 0.005 | 0.419 | 0.62 | 0.56 | 0.54 | 0.49 | 0.71 | 0.62 |
| 3 | 8633172 | rs355124 | 0.22 | 0.648 | -0.391 | 0.648 | 0 | -0.002 | -0.002 | 0 | 0.46 | 0.55 | 0.39 | 0.31 | 0.38 | 0.31 |
| 3 | 8636293 | rs13316251 | 0.975 | 1.078 | -0.205 | 0.478 | 0 | 0.005 | 0.005 | 0 | 0.06 | 0.06 | 0.15 | 0.19 | 0.04 | 0.11 |
| 3 | 8637456 | rs355127 | -0.108 | 0.642 | -0.185 | -0.078 | 0 | -0.049 | -0.049 | 0 | 0.32 | 0.59 | 0.3 | 0.14 | 0.32 | 0.18 |
| 3 | 8637715 | rs9864298 | 1.165 | 0.769 | -0.232 | -0.39 | 0.009 | -0.076 | -0.076 | 0 | 0.16 | 0.31 | 0.13 | 0.01 | 0.16 | 0.04 |
| 3 | 8639150 | rs415926 | 0.341 | 0.836 | -0.747 | -0.349 | 0 | -0.074 | -0.074 | 0 | 0.16 | 0.23 | 0.1 | 0 | 0.16 | 0.12 |
| 3 | 8639568 | rs371734 | -0.123 | 0.725 | -0.663 | -0.344 | 0.003 | -0.086 | -0.086 | 0 | 0.29 | 0.35 | 0.21 | 0.01 | 0.32 | 0.15 |
| 3 | 8640309 | rs6763944 | -0.633 | 1.097 | -1.432 | -0.126 | -0.071 | -0.069 | -0.069 | 0.283 | 0.79 | 0.98 | 0.75 | 0.74 | 0.73 | 0.79 |
| 3 | 8645381 | rs7631882 | 0.404 | 1.085 | -1.163 | -0.213 | -0.008 | 0.029 | 0.029 | 0.28 | 0.6 | 0.64 | 0.64 | 0.78 | 0.59 | 0.73 |
| 3 | 8645666 | rs9878578 | 0.197 | 1.088 | -1.108 | -0.405 | -0.001 | -0.004 | -0.004 | 8.251 | 0.64 | 0.64 | 0.67 | 0.82 | 0.62 | 0.74 |
| 3 | 8646024 | rs430312 | -0.288 | 1.07 | -1.253 | -0.525 | 0 | 0.002 | 0.002 | 0.265 | 0.34 | 0.36 | 0.32 | 0.17 | 0.32 | 0.25 |
| 3 | 8646095 | rs4613438 | 0.576 | 0.741 | -0.983 | -0.144 | 0.001 | 0 | 0 | 0 | 0.46 | 0.51 | 0.44 | 0.54 | 0.38 | 0.56 |
| 3 | 8646781 | rs355090 | 0.3 | 0.945 | -1.14 | -0.224 | -0.025 | 0.054 | 0.054 | 0.248 | 0.41 | 0.26 | 0.36 | 0.28 | 0.49 | 0.25 |
| 3 | 8646870 | rs355091 | 0.275 | 0.802 | -1.045 | -0.217 | -0.01 | 0.049 | 0.049 | 0 | 0.45 | 0.34 | 0.39 | 0.28 | 0.53 | 0.29 |
| 3 | 8647709 | rs355097 | 0.369 | 0.64 | -0.223 | 0.114 | 0.014 | -0.016 | -0.016 | 0 | 0.83 | 0.57 | 0.76 | 0.88 | 0.87 | 0.82 |
| 3 | 8648021 | rs715709 | 0.473 | 0.626 | -0.948 | 0.022 | 0 | -0.021 | -0.021 | 0 | 0.36 | 0.17 | 0.44 | 0.42 | 0.35 | 0.37 |
| 3 | 8648067 | rs48221 | 0.168 | 0.631 | -0.67 | 0.074 | 0.015 | 0.047 | 0.047 | 0.334 | 0.52 | 0.31 | 0.56 | 0.68 | 0.58 | 0.62 |
| 3 | 8648159 | rs355099 | -0.502 | 0.129 | 0.187 | 0.247 | -0.039 | -0.016 | -0.016 | 0.059 | 0.84 | 0.49 | 0.81 | 0.88 | 0.93 | 0.88 |
| 3 | 8648296 | rs9857383 | -0.275 | 0.817 | -0.112 | 0.283 | -0.007 | 0.034 | 0.034 | 0 | 0.39 | 0.3 | 0.46 | 0.24 | 0.48 | 0.41 |
| 3 | 8648496 | rs10510393 | 0.268 | 0.569 | 0.617 | 0.543 | -0.015 | -0.04 | -0.04 | 1 | 0.13 | 0.2 | 0.09 | 0.02 | 0.09 | 0.11 |
| 3 | 8648561 | rs355101 | 0.304 | 0.544 | 0.653 | 0.897 | -0.031 | 0.042 | 0.042 | 0 | 0.79 | 0.5 | 0.8 | 0.77 | 0.85 | 0.83 |
| 3 | 8649260 | rs903115 | 0.813 | 1.764 | 0.719 | 1.299 | 0.063 | 0.07 | 0.07 | 3.223 | 0.75 | 0.46 | 0.58 | 0.6 | 0.6 | 0.74 |
| 3 | 8649462 | rs2324663 | 1.422 | 1.589 | 0.686 | 1.437 | -0.04 | -0.033 | -0.033 | 0 | 0.66 | 0.27 | 0.52 | 0.59 | 0.54 | 0.61 |
| 3 | 8649639 | rs873227 | -1.344 | 1.121 | 0.873 | 1.447 | 0.027 | 0.165 | 0.165 | 0 | 0.13 | 0.38 | 0.19 | 0.29 | 0.11 | 0.18 |
| 3 | 8651999 | rs2014165 | -0.948 | 0.818 | 0.313 | 1.178 | -0.004 | 0.017 | 0.017 | 0 | 0.11 | 0.13 | 0.24 | 0.23 | 0.09 | 0.15 |
| 3 | 8653169 | rs9312026 | -1.022 | 0.753 | 0.725 | 0.984 | 0.043 | 0.068 | 0.068 | 0 | 0.08 | 0.42 | 0.12 | 0.07 | 0.06 | 0.11 |
| 3 | 8659459 | rs2324666 | 0.489 | 1.337 | -0.287 | 0.978 | -0.062 | -0.061 | -0.061 | 0.204 | 0.76 | 0.88 | 0.61 | 0.67 | 0.66 | 0.65 |
| 3 | 8660008 | rs903116 | 1.156 | 0.965 | -0.638 | 0.389 | 0.032 | -0.019 | -0.019 | 0.041 | 0.52 | 0.17 | 0.35 | 0.49 | 0.42 | 0.47 |
| 3 | 8667735 | rs355057 | -0.286 | 0.866 | -0.391 | -0.019 | 0.002 | -0.008 | -0.008 | 0.309 | 0.35 | 0.35 | 0.36 | 0.39 | 0.3 | 0.36 |
| 3 | 8667896 | rs355058 | -0.076 | 1.071 | 0.092 | -0.294 | 0.05 | -0.115 | -0.115 | 0 | 0.83 | 0.63 | 0.82 | 1 | 0.81 | 0.85 |
| 3 | 8670161 | rs1965975 | 0.717 | 0.299 | -0.311 | 0.091 | -0.002 | -0.024 | -0.024 | 0 | 0.38 | 0.24 | 0.27 | 0.49 | 0.4 | 0.42 |
| 3 | 8672045 | rs9873950 | -1.32 | 0.068 | 0.541 | -0.163 | 0.052 | 0.048 | 0.048 | 4.915 | 0.06 | 0.49 | 0.06 | 0 | 0 | 0.01 |
| 3 | 8674360 | rs11710750 | 0.498 | 0.491 | -0.169 | -0.464 | -0.002 | -0.002 | -0.002 | 0 | 0.62 | 0.4 | 0.75 | 0.88 | 0.62 | 0.74 |
| 3 | 8674677 | rs4686281 | 0.835 | 0.023 | -0.322 | -0.24 | 0.006 | 0.011 | 0.011 | 0 | 0.65 | 0.29 | 0.62 | 0.76 | 0.74 | 0.65 |
| 3 | 8675539 | rs2276800 | -0.583 | -0.653 | -0.267 | -0.329 | 0.004 | 0.004 | 0.004 | 1.771 | 0.32 | 0.68 | 0.27 | 0.2 | 0.18 | 0.2 |
| 3 | 8675778 | rs11131140 | 0.999 | 0.619 | 0.078 | 0.358 | 0.02 | 0 | 0 | 0 | 0.85 | 0.74 | 0.72 | 0.79 | 0.81 | 0.83 |
| 3 | 8683848 | rs6768226 | -0.525 | -0.221 | -0.205 | -0.039 | -0.003 | -0.005 | -0.005 | 0.144 | 0.31 | 0.67 | 0.2 | 0.22 | 0.22 | 0.28 |
| 3 | 8684764 | rs164956 | -0.946 | -0.568 | -0.799 | -0.633 | 0.043 | 0.075 | 0.075 | 0.642 | 0.49 | 0.66 | 0.3 | 0.25 | 0.38 | 0.43 |
| 3 | 8687453 | rs4684602 | 1.063 | -0.641 | -1.219 | -0.692 | 0.026 | 0.047 | 0.047 | 0 | 0.51 | 0.22 | 0.51 | 0.59 | 0.53 | 0.54 |
| 3 | 8687561 | rs164961 | 0.444 | -0.606 | -1.374 | -0.872 | -0.013 | 0.061 | 0.061 | 0.972 | 0.19 | 0.01 | 0.11 | 0.07 | 0.17 | 0.17 |
| 3 | 8692449 | rs370508 | -0.093 | -0.671 | -0.276 | -0.673 | 0.012 | 0.056 | 0.056 | 0 | 0.87 | 0.67 | 0.79 | 0.86 | 0.87 | 0.88 |
| 3 | 8693750 | rs6795577 | 0.783 | 0.063 | -0.162 | -0.886 | -0.002 | -0.021 | -0.021 | 0 | 0.19 | 0.13 | 0.09 | 0.01 | 0.17 | 0.08 |
| 3 | 8694057 | rs3774209 | 1.851 | -0.037 | -0.064 | -0.861 | -0.017 | -0.009 | -0.009 | 0 | 0.36 | 0.24 | 0.52 | 0.59 | 0.38 | 0.45 |
| 3 | 8696316 | rs420867 | -0.071 | -0.756 | 0.17 | -0.567 | 0 | 0.006 | 0.006 | 0.718 | 0.24 | 0.15 | 0.15 | 0.25 | 0.24 | 0.26 |
| 3 | 8697372 | rs6785267 | 1.047 | -0.754 | 0.068 | -0.662 | -0.001 | 0 | 0 | 0 | 0.55 | 0.46 | 0.53 | 0.59 | 0.53 | 0.52 |
| 3 | 8702963 | rs412508 | 0.142 | -0.631 | 0.09 | -0.667 | 0.003 | -0.007 | -0.007 | 2.316 | 0.83 | 0.74 | 0.81 | 0.87 | 0.81 | 0.81 |
| 3 | 8709205 | rs2172557 | -0.446 | -0.44 | -0.434 | -0.598 | 0.011 | -0.042 | -0.042 | 0 | 0.15 | 0.35 | 0.13 | 0.06 | 0.16 | 0.17 |
| 3 | 8710291 | rs11131142 | -0.895 | -0.346 | -0.555 | -0.652 | 0.004 | -0.05 | -0.05 | 0 | 0.2 | 0.31 | 0.18 | 0.06 | 0.22 | 0.18 |
| 3 | 8711098 | rs12490191 | 0.628 | -0.362 | -1.101 | -0.435 | 0.009 | 0.024 | 0.024 | 0 | 0.52 | 0.35 | 0.57 | 0.62 | 0.52 | 0.5 |
| 3 | 8711575 | rs11916383 | 0.729 | -0.285 | -1.2 | -0.39 | 0.017 | 0.037 | 0.037 | 0 | 0.51 | 0.31 | 0.56 | 0.62 | 0.52 | 0.47 |
| 3 | 8712227 | rs164954 | -0.415 | 0.146 | -0.851 | 0.12 | 0.062 | 0.071 | 0.071 | 0 | 0.49 | 0.68 | 0.3 | 0.33 | 0.37 | 0.41 |
| 3 | 8717341 | rs11131143 | -0.9 | 0.764 | -0.437 | 0.923 | 0.024 | 0.07 | 0.07 | 0 | 0.1 | 0.21 | 0.07 | 0.26 | 0.13 | 0.18 |
| 3 | 8729551 | rs350811 | 0.29 | 0.014 | -0.526 | 0.699 | -0.026 | 0.015 | 0.015 | 0 | 0.72 | 0.6 | 0.76 | 0.68 | 0.74 | 0.8 |
| 3 | 8734471 | rs6766036 | 0.061 | 0.769 | -1.207 | 0.147 | -0.018 | -0.065 | -0.065 | 0.973 | 0.31 | 0.13 | 0.31 | 0.38 | 0.26 | 0.31 |
| 3 | 8745085 | rs4686294 | -1.141 | 0.735 | 0.042 | -0.129 | -0.006 | 0.011 | 0.011 | 0.77 | 0.1 | 0.18 | 0.19 | 0.09 | 0.1 | 0.06 |
| 3 | 8745493 | rs164459 | -1.133 | 1.457 | 0.448 | -1.039 | 0.043 | -0.116 | -0.116 | 0.59 | 0.27 | 0.55 | 0.26 | 0.03 | 0.35 | 0.19 |
| 3 | 8748648 | rs164462 | 0.342 | 1.44 | 0.205 | -1.089 | 0.015 | 0.02 | 0.02 | 0.471 | 0.55 | 0.69 | 0.65 | 0.62 | 0.44 | 0.39 |
| 3 | 8749012 | rs164464 | -0.057 | 2.068 | 0.478 | -1.093 | -0.059 | 0.071 | 0.071 | 0 | 0.88 | 0.99 | 0.92 | 1 | 0.82 | 0.86 |
| 3 | 8749619 | rs9822843 | -1.421 | 1.748 | 0.985 | -1.013 | 0.026 | -0.023 | -0.023 | 2.15 | 0.05 | 0.18 | 0.05 | 0 | 0.05 | 0.01 |
| 3 | 8751490 | rs12497498 | -0.517 | 2.019 | 0.538 | -1.079 | -0.021 | 0.009 | 0.009 | 0 | 0.12 | 0.06 | 0.13 | 0 | 0.21 | 0.07 |
| 3 | 8751984 | rs17049459 | -0.566 | 1.785 | 1.037 | -1.115 | 0.076 | -0.105 | -0.105 | 0 | 0.15 | 0.42 | 0.14 | 0 | 0.17 | 0.07 |

| | | | | | | | | | | | | | | | | |
|---|---|---|---|---|---|---|---|---|---|---|---|---|---|---|---|---|
| 3 | 8752296 | rs765450 | -0.984 | 0.845 | 0.676 | -0.997 | -0.013 | -0.105 | -0.105 | 0.66 | 0.21 | 0.38 | 0.13 | 0 | 0.19 | 0.13 |
| 3 | 8752675 | rs7615168 | -0.6 | 1.351 | 0.741 | -0.635 | 0.034 | -0.076 | -0.076 | 0 | 0.12 | 0.36 | 0.07 | 0 | 0.13 | 0.09 |
| 3 | 8754169 | rs7634632 | 0.358 | 1.416 | 0.116 | -0.487 | 0 | 0.013 | 0.013 | 0 | 0.47 | 0.42 | 0.39 | 0.39 | 0.5 | 0.48 |
| 3 | 8754314 | rs6443203 | -0.619 | 1.511 | -0.031 | -0.446 | 0.029 | -0.009 | -0.009 | 0 | 0.6 | 0.9 | 0.48 | 0.39 | 0.64 | 0.58 |
| 3 | 8754577 | rs6774842 | -0.587 | 1.353 | -0.207 | -0.418 | 0.027 | 0.004 | 0.004 | 0 | 0.59 | 0.79 | 0.47 | 0.38 | 0.65 | 0.58 |
| 3 | 8754648 | rs6798626 | -1.211 | 1.742 | 0.166 | -0.23 | 0.037 | -0.072 | -0.072 | 0 | 0.11 | 0.35 | 0.07 | 0 | 0.14 | 0.09 |
| 3 | 8767642 | rs17362990 | -1.166 | 2.26 | -0.5 | -0.172 | -0.003 | -0.011 | -0.011 | 0 | 0.1 | 0.12 | 0.2 | 0.01 | 0.18 | 0.08 |
| 3 | 8771402 | rs11914885 | 0.235 | 1.015 | -0.603 | -0.035 | 0.01 | 0.001 | 0.001 | 3.078 | 0.49 | 0.5 | 0.47 | 0.5 | 0.31 | 0.47 |
| 3 | 8771776 | rs10490801 | 0.375 | 0.288 | -0.437 | -0.344 | 0.041 | -0.016 | -0.016 | 0 | 0.4 | 0.21 | 0.43 | 0.49 | 0.28 | 0.46 |
| 3 | 8773131 | rs237860 | 0.55 | 0.675 | -1.081 | 0.517 | -0.053 | -0.075 | -0.075 | 0.149 | 0.74 | 0.82 | 0.57 | 0.44 | 0.65 | 0.58 |
| 3 | 8773285 | rs237862 | 0.687 | -0.509 | -1.59 | 0.116 | -0.049 | 0.05 | 0.05 | 0 | 0.32 | 0.12 | 0.23 | 0.19 | 0.42 | 0.25 |
| 3 | 8773552 | rs237864 | -0.774 | 0.351 | -1.681 | 0.378 | -0.017 | -0.015 | -0.015 | 0.127 | 0.11 | 0.07 | 0.08 | 0.2 | 0.15 | 0.15 |
| 3 | 8773950 | rs2268484 | 0.056 | -0.656 | -1.556 | 0.696 | -0.003 | 0.003 | 0.003 | 0 | 0.34 | 0.65 | 0.45 | 0.42 | 0.22 | 0.34 |
| 3 | 8773968 | rs237865 | -0.387 | 0.711 | -1.407 | 0.934 | -0.004 | 0.005 | 0.005 | 0 | 0.43 | 0.49 | 0.47 | 0.42 | 0.39 | 0.34 |
| 3 | 8774129 | rs237866 | -0.084 | 1.251 | -1.939 | 0.552 | -0.086 | 0.004 | 0.004 | 0 | 0.35 | 0.08 | 0.29 | 0.27 | 0.43 | 0.22 |
| 3 | 8774285 | rs2268485 | -0.013 | 1.046 | -1.949 | 0.454 | -0.076 | 0.028 | 0.028 | 0.014 | 0.32 | 0.06 | 0.27 | 0.22 | 0.37 | 0.19 |
| 3 | 8775077 | rs2270463 | 0.428 | 0.566 | -1.46 | 0.671 | 0.017 | -0.023 | -0.023 | 1.061 | 0.27 | 0.23 | 0.24 | 0.42 | 0.22 | 0.21 |
| 3 | 8775223 | rs6793441 | -0.288 | 0.729 | -0.842 | 0.074 | -0.007 | -0.017 | -0.017 | 0 | 0.35 | 0.42 | 0.36 | 0.21 | 0.35 | 0.22 |
| 3 | 8775379 | rs2072582 | 0.918 | 0.901 | -1.009 | 0.397 | -0.002 | -0.004 | -0.004 | 0 | 0.17 | 0.16 | 0.16 | 0.41 | 0.19 | 0.28 |
| 3 | 8775457 | rs2072583 | 0.475 | 0.55 | -0.478 | -0.032 | 0.007 | 0.012 | 0.012 | 0 | 0.3 | 0.22 | 0.22 | 0.23 | 0.27 | 0.19 |
| 3 | 8775589 | rs1974763 | 2.134 | 1.155 | -0.545 | -0.045 | -0.03 | 0.025 | 0.025 | 0 | 0.05 | 0.01 | 0.06 | 0.01 | 0.13 | 0.03 |
| 3 | 8775661 | rs1008642 | 0.13 | 0.615 | 0.008 | -0.099 | -0.018 | 0.004 | 0.004 | 0 | 0.33 | 0.62 | 0.31 | 0.37 | 0.27 | 0.18 |
| 3 | 8777037 | rs11713611 | 1.113 | 0.823 | 0.24 | -0.185 | 0.005 | -0.011 | -0.011 | 0 | 0.07 | 0.07 | 0.07 | 0.01 | 0.14 | 0.03 |
| 3 | 8778162 | rs237868 | 0.176 | 0.287 | 0.362 | -0.291 | -0.008 | -0.005 | -0.005 | 0.201 | 0.36 | 0.38 | 0.31 | 0.22 | 0.26 | 0.24 |
| 3 | 8779066 | rs237870 | -0.197 | 0.55 | -0.04 | -0.381 | 0.01 | 0.033 | 0.033 | 2.704 | 0.38 | 0.73 | 0.31 | 0.2 | 0.29 | 0.27 |
| 3 | 8781331 | rs11916022 | 0.718 | 0.775 | 0.177 | -0.679 | 0.015 | -0.112 | -0.112 | 0 | 0.22 | 0.32 | 0.14 | 0.01 | 0.24 | 0.07 |
| 3 | 8782406 | rs237875 | -0.513 | 0.708 | -0.276 | -0.468 | 0 | 0.001 | 0.001 | 2.033 | 0.51 | 0.55 | 0.55 | 0.41 | 0.5 | 0.48 |
| 3 | 8786487 | rs6777088 | -0.086 | 0.446 | -0.186 | -0.944 | 0 | -0.088 | -0.088 | 0 | 0.26 | 0.61 | 0.18 | 0.01 | 0.25 | 0.08 |
| 3 | 8787220 | rs13087941 | -0.233 | 0.746 | 0.106 | -0.79 | 0.119 | -0.051 | -0.051 | 0 | 0.1 | 0.28 | 0.13 | 0.01 | 0.23 | 0.05 |
| 3 | 8788096 | rs13093809 | 1.01 | 0.841 | -0.557 | -0.765 | -0.037 | 0.032 | 0.032 | 0 | 0.06 | 0.01 | 0.08 | 0.01 | 0.16 | 0.02 |
| 3 | 8788336 | rs7629329 | 0.009 | 0.533 | -0.88 | -0.778 | -0.011 | 0.007 | 0.007 | 0.008 | 0.32 | 0.19 | 0.41 | 0.3 | 0.35 | 0.16 |
| 3 | 8792728 | rs2324728 | 0.174 | 0.803 | -0.449 | -0.866 | 0.015 | -0.109 | -0.109 | 0.308 | 0.22 | 0.58 | 0.17 | 0.01 | 0.25 | 0.06 |
| 3 | 8793585 | rs237884 | -0.06 | 0.757 | -0.538 | -0.873 | 0.002 | -0.102 | -0.102 | 0 | 0.76 | 0.41 | 0.82 | 0.99 | 0.73 | 0.93 |
| 3 | 8794545 | rs1042778 | -0.544 | 1.106 | -0.734 | -0.685 | -0.005 | -0.004 | -0.004 | 0 | 0.33 | 0.68 | 0.46 | 0.08 | 0.38 | 0.39 |
| 3 | 8795543 | rs237885 | -0.553 | 0.701 | -0.399 | 0.175 | 0.001 | 0.009 | 0.009 | 0 | 0.5 | 0.53 | 0.57 | 0.28 | 0.51 | 0.56 |
| 3 | 8796547 | rs11706648 | 0.022 | 0.828 | -0.673 | -0.151 | -0.024 | 0.03 | 0.03 | 0.007 | 0.34 | 0.13 | 0.44 | 0.27 | 0.32 | 0.4 |
| 3 | 8797042 | rs237887 | -1.237 | 1.047 | -0.563 | -0.215 | 0.016 | -0.013 | -0.013 | 0 | 0.53 | 0.76 | 0.63 | 0.43 | 0.54 | 0.61 |
| 3 | 8797085 | rs2268490 | 0.791 | 0.411 | -0.801 | -0.435 | -0.015 | 0.044 | 0.044 | 0 | 0.2 | 0.26 | 0.23 | 0.5 | 0.14 | 0.15 |
| 3 | 8797095 | rs237888 | -0.404 | 0.851 | -0.392 | -0.277 | 0.143 | 0.041 | 0.041 | 0.004 | 0.06 | 0.39 | 0.06 | 0 | 0.06 | 0.01 |
| 3 | 8798181 | rs918316 | 0.317 | 0.574 | -0.679 | -0.037 | -0.015 | 0.116 | 0.116 | 0 | 0.1 | 0.24 | 0.07 | 0.23 | 0.05 | 0.14 |
| 3 | 8798586 | rs4686301 | 0.265 | 0.969 | -1.09 | -0.198 | -0.027 | 0.056 | 0.056 | 0 | 0.33 | 0.1 | 0.43 | 0.2 | 0.31 | 0.39 |
| 3 | 8800398 | rs2268491 | 0.62 | 0.33 | -0.539 | 0.452 | -0.029 | 0.071 | 0.071 | 0 | 0.15 | 0.27 | 0.22 | 0.34 | 0.11 | 0.1 |
| 3 | 8802483 | rs237889 | -1.148 | 0.768 | -0.846 | 0.342 | -0.021 | -0.032 | -0.032 | 0 | 0.6 | 0.91 | 0.8 | 0.59 | 0.6 | 0.57 |
| 3 | 8802851 | rs11131149 | -0.657 | 1.201 | -0.356 | -0.177 | 0.017 | -0.018 | -0.018 | 0.115 | 0.29 | 0.66 | 0.46 | 0.16 | 0.38 | 0.27 |
| 3 | 8807535 | rs2268495 | -0.336 | 1.072 | 0.07 | -0.156 | 0.039 | -0.008 | -0.008 | 0 | 0.15 | 0.32 | 0.33 | 0.18 | 0.21 | 0.16 |
| 3 | 8808285 | rs237897 | -1.762 | 1.124 | -0.747 | -0.502 | 0.02 | 0.104 | 0.104 | 0.863 | 0.56 | 0.83 | 0.66 | 0.32 | 0.59 | 0.53 |
| 3 | 8808515 | rs237899 | -0.892 | 1.127 | -0.497 | -0.431 | -0.003 | 0.001 | 0.001 | 0 | 0.37 | 0.34 | 0.31 | 0.14 | 0.37 | 0.35 |
| 3 | 8810896 | rs2301261 | -0.18 | 0.916 | -0.308 | -0.107 | -0.021 | 0.004 | 0.004 | 0 | 0.1 | 0.22 | 0.11 | 0.09 | 0.09 | 0.07 |
| 3 | 8813494 | rs6777726 | 0.526 | 0.989 | -0.531 | 0.04 | -0.008 | 0.031 | 0.031 | 0.014 | 0.07 | 0.11 | 0.07 | 0.12 | 0.07 | 0.06 |
| 3 | 8813927 | rs180789 | -0.405 | 1.3 | -1.061 | -0.256 | -0.078 | 0.107 | 0.107 | 0 | 0.24 | 0.03 | 0.2 | 0.06 | 0.26 | 0.25 |
| 3 | 8820732 | rs75775 | 0.322 | 1.256 | -0.852 | 0.563 | -0.018 | -0.025 | -0.025 | 0 | 0.9 | 0.94 | 0.75 | 0.64 | 0.84 | 0.88 |
| 3 | 8820740 | rs9860869 | -0.808 | 0.915 | -0.279 | -0.306 | -0.005 | -0.068 | -0.068 | 0 | 0.15 | 0.6 | 0.12 | 0.07 | 0.1 | 0.06 |
| 3 | 8821742 | rs237925 | -0.091 | 1.08 | -1.009 | -0.558 | -0.087 | 0.122 | 0.122 | 0 | 0.25 | 0.02 | 0.24 | 0.05 | 0.32 | 0.24 |
| 3 | 8826396 | rs6793234 | -1.414 | 0.617 | -0.353 | -0.317 | -0.002 | -0.001 | -0.001 | 0 | 0.2 | 0.74 | 0.13 | 0.16 | 0.11 | 0.18 |
| 3 | 8829872 | rs6791619 | -0.69 | 0.378 | -0.945 | -0.365 | 0.039 | -0.013 | -0.013 | 0 | 0.39 | 0.89 | 0.35 | 0.35 | 0.31 | 0.38 |
| 3 | 8833878 | rs237852 | 1.07 | 0.544 | -0.609 | -0.445 | -0.003 | 0.154 | 0.154 | 0 | 0.49 | 0.24 | 0.4 | 0.19 | 0.5 | 0.5 |
| 3 | 8834933 | rs3901926 | -0.688 | 0.623 | 0.092 | -0.218 | 0.039 | 0.01 | 0.01 | 0.05 | 0.06 | 0.31 | 0.03 | 0.01 | 0.02 | 0.1 |
| 3 | 8836063 | rs9817913 | -0.277 | 0.648 | -0.241 | -0.207 | -0.01 | -0.028 | -0.028 | 0 | 0.05 | 0.15 | 0.03 | 0.01 | 0.02 | 0.1 |
| 3 | 8841641 | rs237837 | 0.059 | 0.745 | -0.345 | -0.076 | 0.024 | 0.122 | 0.122 | 0 | 0.85 | 0.65 | 0.67 | 0.54 | 0.79 | 0.86 |
| 3 | 8848916 | rs1714299 | 1.5 | 1.061 | -0.67 | -0.271 | -0.103 | 0.112 | 0.112 | 0 | 0.43 | 0.02 | 0.28 | 0.19 | 0.42 | 0.48 |
| 3 | 8850723 | rs1643027 | 0.35 | 1.16 | 0.211 | 0.254 | 0.03 | 0.147 | 0.147 | 0 | 0.87 | 0.66 | 0.68 | 0.54 | 0.81 | 0.88 |
| 3 | 8854675 | rs1076508 | 0.824 | 0.629 | 0.259 | 0.649 | 0.045 | -0.044 | -0.044 | 1.64 | 0.16 | 0.02 | 0.14 | 0.19 | 0.07 | 0.16 |
| 3 | 8857963 | rs2241330 | -0.529 | 0.638 | 0.541 | 0.604 | -0.025 | -0.003 | -0.003 | 0 | 0.23 | 0.69 | 0.13 | 0.2 | 0.1 | 0.23 |
| 3 | 8862925 | rs345065 | 0.568 | 0.866 | 0.502 | 0.629 | -0.01 | -0.018 | -0.018 | 0 | 0.77 | 0.79 | 0.62 | 0.47 | 0.69 | 0.77 |
| 3 | 8864726 | rs3864087 | -2.152 | 0.789 | 0.557 | 0.651 | 0.016 | 0.156 | 0.156 | 1.46 | 0.08 | 0.67 | 0.1 | 0.2 | 0.04 | 0.17 |
| 3 | 8881649 | rs124844 | 1.676 | 0.803 | 0.376 | 0.581 | -0.02 | 0.134 | 0.134 | 0 | 0.53 | 0.07 | 0.5 | 0.27 | 0.59 | 0.55 |
| 3 | 8881713 | rs83622 | 0.628 | 0.959 | 0.651 | 0.77 | 0.001 | 0.009 | 0.009 | 0 | 0.77 | 0.75 | 0.58 | 0.47 | 0.68 | 0.78 |
| 3 | 8883603 | rs17049639 | -1.618 | 0.927 | 0.804 | 0.214 | 0.03 | -0.033 | -0.033 | 0 | 0.08 | 0.57 | 0.11 | 0 | 0.1 | 0.03 |
| 3 | 8883878 | rs9846430 | -1.255 | 0.902 | 0.63 | 0.332 | -0.004 | 0.024 | 0.024 | 0 | 0.38 | 0.32 | 0.48 | 0.73 | 0.38 | 0.4 |

| | | | | | | | | | | | | | | | | |
|---|---|---|---|---|---|---|---|---|---|---|---|---|---|---|---|---|
| 3 | 8884204 | rs9867782 | -1.084 | 1.001 | 0.565 | 0.452 | -0.012 | -0.002 | -0.002 | 0 | 0.24 | 0.23 | 0.46 | 0.73 | 0.35 | 0.37 |
| 3 | 8889809 | rs7634338 | 0.551 | 0.457 | 1.262 | 0.412 | -0.071 | -0.093 | -0.093 | 4.465 | 0.14 | 0.31 | 0.04 | 0 | 0.02 | 0.01 |
| 3 | 8895531 | rs6799969 | -2.051 | 1.81 | 1.832 | 1.159 | 0.041 | 0.036 | 0.036 | 0.371 | 0.28 | 0.48 | 0.38 | 0.67 | 0.42 | 0.24 |
| 3 | 8896114 | rs9827966 | -2.341 | 1.857 | 2.188 | 1.54 | 0.057 | 0.204 | 0.204 | 0 | 0.08 | 0.36 | 0.09 | 0.21 | 0.08 | 0.07 |
| 3 | 8896462 | rs1005988 | -1.929 | 1.885 | 1.953 | 1.504 | 0.029 | 0.052 | 0.052 | 0.338 | 0.26 | 0.41 | 0.35 | 0.46 | 0.39 | 0.22 |
| 3 | 8903387 | rs4686311 | -1.383 | 1.914 | 1.211 | 1.545 | -0.008 | -0.014 | -0.014 | 0 | 0.2 | 0.18 | 0.29 | 0.45 | 0.32 | 0.19 |
| 3 | 8905897 | rs9835766 | -2.162 | 1.599 | 1.75 | 1.365 | 0.018 | 0.048 | 0.048 | 0 | 0.17 | 0.54 | 0.13 | 0.21 | 0.16 | 0.18 |
| 3 | 8906295 | rs9816708 | -1.848 | 1.644 | 1.967 | 1.222 | 0.042 | -0.091 | -0.091 | 0 | 0.13 | 0.48 | 0.1 | 0.01 | 0.12 | 0.12 |
| 3 | 8909647 | rs10510394 | -2.107 | 1.982 | 1.939 | 1.596 | 0.047 | 0.134 | 0.134 | 0 | 0.2 | 0.44 | 0.28 | 0.45 | 0.25 | 0.13 |
| 3 | 8912662 | rs6798600 | -2.58 | 1.921 | 1.739 | 1.567 | -0.002 | -0.003 | -0.003 | 7.077 | 0.25 | 0.7 | 0.33 | 0.64 | 0.27 | 0.15 |
| 3 | 8923840 | rs4389469 | -1.572 | 2.124 | 1.948 | 1.324 | 0.029 | 0.021 | 0.021 | 0 | 0.31 | 0.6 | 0.32 | 0.44 | 0.37 | 0.22 |
| 3 | 8932189 | rs607531 | 2.458 | 1.346 | 2.782 | 1.641 | 0.005 | -0.028 | -0.028 | 1.823 | 0.72 | 0.21 | 0.67 | 0.36 | 0.76 | 0.76 |
| 3 | 8933946 | rs686195 | 2.458 | 1.346 | 2.788 | 1.641 | 0.005 | -0.026 | -0.026 | 0 | 0.72 | 0.23 | 0.67 | 0.36 | 0.76 | 0.76 |
| 3 | 8945081 | rs11711315 | -1.475 | 1.351 | 2.791 | 1.681 | -0.018 | 0.051 | 0.051 | 0 | 0.13 | 0.25 | 0.08 | 0.23 | 0.08 | 0.16 |
| 3 | 8947601 | rs547904 | 2.058 | 1.37 | 2.861 | 1.626 | 0.005 | 0.035 | 0.035 | 0.034 | 0.77 | 0.66 | 0.71 | 0.36 | 0.76 | 0.77 |
| 3 | 8955389 | rs373572 | 2.042 | 1.389 | 2.872 | 1.653 | 0.005 | 0.037 | 0.037 | 0.041 | 0.78 | 0.65 | 0.71 | 0.36 | 0.76 | 0.77 |
| 3 | 8956371 | rs575559 | 2.683 | 1.352 | 3.145 | 1.636 | -0.006 | 0.036 | 0.036 | 1.32 | 0.72 | 0.34 | 0.68 | 0.36 | 0.76 | 0.76 |
| 3 | 8956418 | rs574776 | 2.052 | 1.393 | 2.959 | 1.654 | 0.012 | 0.069 | 0.069 | 0.06 | 0.77 | 0.63 | 0.71 | 0.36 | 0.76 | 0.77 |
| 3 | 8975200 | rs408483 | 1.94 | 1.399 | 2.947 | 1.657 | 0.008 | 0.041 | 0.041 | 0.012 | 0.78 | 0.68 | 0.72 | 0.36 | 0.76 | 0.77 |
| 3 | 8975742 | rs447423 | 2.223 | 1.371 | 2.989 | 1.647 | 0.009 | 0.064 | 0.064 | 0.112 | 0.77 | 0.58 | 0.7 | 0.36 | 0.76 | 0.77 |
| 3 | 8978476 | rs11709327 | -1.162 | 1.364 | 2.876 | 1.708 | -0.01 | 0.038 | 0.038 | 0 | 0.11 | 0.19 | 0.08 | 0.23 | 0.08 | 0.15 |
| 3 | 8980898 | rs341769 | 1.87 | 1.337 | 2.943 | 1.714 | 0.039 | 0.146 | 0.146 | 0.001 | 0.86 | 0.67 | 0.77 | 0.59 | 0.84 | 0.93 |
| 3 | 8982228 | rs7641235 | -2.468 | 1.299 | 2.613 | 1.579 | -0.024 | 0.026 | 0.026 | 0.227 | 0.18 | 0.67 | 0.14 | 0.23 | 0.11 | 0.19 |
| 3 | 8989806 | rs11719775 | -1.268 | 1.34 | 2.532 | 1.564 | 0 | -0.037 | -0.037 | 0 | 0.12 | 0.19 | 0.08 | 0.02 | 0.1 | 0.16 |
| 3 | 8996747 | rs9827283 | -2.358 | 1.334 | 1.58 | 1.627 | -0.002 | 0 | 0 | 1.836 | 0.2 | 0.76 | 0.15 | 0.23 | 0.11 | 0.19 |
| 3 | 9012903 | rs369651 | 2.056 | 1.373 | 1.298 | 1.56 | -0.001 | -0.005 | -0.005 | 0.633 | 0.71 | 0.3 | 0.7 | 0.32 | 0.71 | 0.79 |
| 3 | 9017261 | rs813606 | 0.285 | 1.603 | 0.938 | 1.909 | -0.058 | -0.074 | -0.074 | 0 | 0.88 | 0.97 | 0.81 | 0.57 | 0.82 | 0.94 |
| 3 | 9024017 | rs341791 | 1.797 | 1.211 | 1.519 | 1.679 | -0.013 | 0.043 | 0.043 | 0 | 0.73 | 0.63 | 0.7 | 0.55 | 0.73 | 0.8 |
| 3 | 9024521 | rs1043951 | -1.017 | 1.282 | 1.081 | 1.196 | 0.017 | -0.039 | -0.039 | 0 | 0.11 | 0.33 | 0.1 | 0.02 | 0.09 | 0.13 |
| 3 | 9034145 | rs413789 | 0.371 | 1.158 | -0.264 | 1.116 | -0.008 | -0.007 | -0.007 | 0 | 0.62 | 0.76 | 0.66 | 0.36 | 0.55 | 0.65 |
| 3 | 9039469 | rs523684 | 0.203 | 1.116 | -0.435 | 1.208 | -0.05 | -0.07 | -0.07 | 1.246 | 0.64 | 0.86 | 0.65 | 0.39 | 0.54 | 0.65 |
| 3 | 9041299 | rs486012 | 0.336 | 1.044 | -0.207 | 1.168 | -0.019 | -0.029 | -0.029 | 0 | 0.63 | 0.83 | 0.65 | 0.39 | 0.54 | 0.67 |
| 3 | 9048152 | rs1161765 | -0.329 | 1.077 | -0.494 | 1.298 | -0.015 | -0.076 | -0.076 | 0.424 | 0.84 | 0.97 | 0.81 | 0.69 | 0.81 | 0.92 |
| 3 | 9051633 | rs1532999 | 0.574 | 1.173 | -0.371 | 1.238 | -0.035 | 0.061 | 0.061 | 0 | 0.19 | 0.07 | 0.14 | 0.02 | 0.27 | 0.22 |
| 3 | 9052494 | rs383563 | 0.839 | 1.106 | -0.223 | 1.434 | -0.006 | -0.001 | -0.001 | 0 | 0.62 | 0.72 | 0.66 | 0.62 | 0.53 | 0.68 |
| 3 | 9062544 | rs6443218 | -0.16 | 0.077 | -0.107 | 0.67 | -0.012 | -0.124 | -0.124 | 0 | 0.32 | 0.47 | 0.19 | 0.05 | 0.29 | 0.26 |
| 3 | 9066156 | rs2662089 | 0.511 | -0.064 | -0.416 | 1.118 | 0.001 | 0.008 | 0.008 | 0 | 0.51 | 0.54 | 0.56 | 0.38 | 0.41 | 0.52 |
| 3 | 9066853 | rs421041 | 0.017 | -0.006 | -0.689 | 1.102 | 0.001 | -0.005 | -0.005 | 0 | 0.79 | 0.8 | 0.79 | 0.7 | 0.76 | 0.88 |
| 3 | 9067684 | rs412680 | 0.395 | -0.021 | -1.143 | 0.99 | -0.005 | -0.017 | -0.017 | 0.194 | 0.61 | 0.7 | 0.4 | 0.4 | 0.59 | 0.64 |
| 3 | 9069133 | rs2252592 | 0.397 | 0.092 | -0.933 | 0.616 | 0.001 | 0.003 | 0.003 | 0 | 0.19 | 0.19 | 0.39 | 0.27 | 0.17 | 0.25 |
| 3 | 9072286 | rs1513315 | -0.364 | -0.335 | -1.199 | -0.247 | 0.017 | 0.002 | 0.002 | 0.108 | 0.25 | 0.22 | 0.14 | 0 | 0.16 | 0.08 |
| 3 | 9074843 | rs6777129 | -0.392 | 0.208 | -1.266 | 0.003 | -0.004 | 0.023 | 0.023 | 1.186 | 0.45 | 0.41 | 0.37 | 0.68 | 0.43 | 0.39 |
| 3 | 9074865 | rs6762129 | 0.203 | 0.333 | -1.434 | 0.293 | -0.009 | 0.01 | 0.01 | 2.61 | 0.32 | 0.25 | 0.3 | 0.3 | 0.34 | 0.23 |
| 3 | 9077896 | rs9875468 | -0.445 | 0.133 | -1.033 | 0.554 | -0.006 | 0.149 | 0.149 | 0 | 0.16 | 0.37 | 0.1 | 0.38 | 0.1 | 0.16 |
| 3 | 9078201 | rs2670009 | 0.36 | 0.18 | -1.726 | 0.139 | 0.001 | -0.027 | -0.027 | 0 | 0.21 | 0.09 | 0.34 | 0.26 | 0.25 | 0.37 |
| 3 | 9078581 | rs2662088 | 0.227 | 0.597 | -2.134 | -0.19 | -0.06 | 0.005 | 0.005 | 1.297 | 0.38 | 0.06 | 0.49 | 0.34 | 0.41 | 0.35 |
| 3 | 9079291 | rs17049971 | 0.565 | 0.458 | -1.728 | -0.067 | 0.007 | -0.013 | -0.013 | 0 | 0.06 | 0.05 | 0.03 | 0.09 | 0.05 | 0.11 |
| 3 | 9079714 | rs867528 | 1.531 | 0.3 | -1.193 | -0.428 | -0.005 | -0.101 | -0.101 | 0 | 0.15 | 0.29 | 0.09 | 0 | 0.1 | 0.02 |
| 3 | 9080869 | rs578911 | 0.183 | 0.188 | -1.757 | -0.493 | 0.001 | -0.012 | -0.012 | 0 | 0.35 | 0.34 | 0.36 | 0.49 | 0.32 | 0.47 |
| 3 | 9080893 | rs639057 | -0.604 | 0.445 | -2.002 | -0.813 | -0.039 | -0.06 | -0.06 | 0 | 0.78 | 0.88 | 0.63 | 0.62 | 0.68 | 0.81 |
| 3 | 9081339 | rs1162326 | 0.291 | 0.444 | -1.967 | -0.873 | 0 | 0 | 0 | 0 | 0.56 | 0.49 | 0.5 | 0.54 | 0.53 | 0.68 |
| 3 | 9082851 | rs2670008 | 2.094 | 0.475 | -1.975 | -0.792 | -0.006 | 0.036 | 0.036 | 0.218 | 0.21 | 0.13 | 0.14 | 0.04 | 0.21 | 0.21 |
| 3 | 9087655 | rs624528 | -0.327 | 0.416 | -1.795 | -0.367 | 0.028 | -0.025 | -0.025 | 0.033 | 0.43 | 0.72 | 0.44 | 0.51 | 0.35 | 0.48 |
| 3 | 9087967 | rs2670003 | 0.129 | 0.571 | -1.782 | -0.338 | 0.003 | 0.065 | 0.065 | 0 | 0.74 | 0.47 | 0.63 | 0.43 | 0.7 | 0.69 |
| 3 | 9089384 | rs1622607 | 0.159 | 0.556 | -1.81 | -0.141 | -0.003 | -0.005 | -0.005 | 0.449 | 0.36 | 0.34 | 0.42 | 0.37 | 0.37 | 0.43 |
| 3 | 9090673 | rs2670002 | -0.556 | 0.445 | -1.906 | 0.112 | 0.021 | 0.007 | 0.007 | 0.021 | 0.46 | 0.39 | 0.28 | 0.48 | 0.36 | 0.57 |
| 3 | 9090905 | rs611406 | -0.095 | 0.309 | -2.087 | -0.229 | -0.001 | -0.001 | -0.001 | 0 | 0.34 | 0.34 | 0.31 | 0.35 | 0.35 | 0.41 |
| 3 | 9091946 | rs2662090 | -0.532 | -0.233 | -2.015 | -0.68 | -0.008 | 0.043 | 0.043 | 0 | 0.51 | 0.32 | 0.56 | 0.66 | 0.5 | 0.66 |
| 3 | 9092661 | rs7609881 | 1.475 | 0.071 | -2.002 | -0.555 | 0.02 | 0.037 | 0.037 | 0 | 0.11 | 0.04 | 0.05 | 0.07 | 0.07 | 0.12 |
| 3 | 9094531 | rs13091604 | 1.046 | 0.51 | -1.664 | -0.454 | -0.019 | 0.078 | 0.078 | 0 | 0.26 | 0.13 | 0.16 | 0.03 | 0.26 | 0.19 |
| 3 | 9099653 | rs2664095 | -0.104 | 0.011 | -1.132 | 0.291 | -0.002 | -0.001 | -0.001 | 0.211 | 0.7 | 0.37 | 0.67 | 0.69 | 0.74 | 0.84 |
| 3 | 9101597 | rs7644129 | 1.158 | 0.175 | -0.614 | -0.131 | -0.004 | -0.071 | -0.071 | 0 | 0.14 | 0.28 | 0.06 | 0 | 0.1 | 0.02 |
| 3 | 9101668 | rs6443219 | -0.298 | 0.047 | -1.407 | 0.411 | -0.016 | -0.124 | -0.124 | 0.971 | 0.8 | 0.94 | 0.73 | 0.58 | 0.73 | 0.69 |
| 3 | 9104321 | rs4686320 | 0.436 | 0.423 | -1.377 | 0.052 | -0.022 | -0.037 | -0.037 | 0.121 | 0.34 | 0.17 | 0.46 | 0.49 | 0.35 | 0.42 |
| 3 | 9106893 | rs2669995 | 1.438 | 0.703 | -1.23 | -0.054 | -0.023 | -0.033 | -0.033 | 0 | 0.06 | 0.01 | 0.07 | 0.13 | 0.13 | 0.13 |
| 3 | 9107475 | rs7632256 | 1.291 | 0.508 | -1.145 | -0.732 | 0.001 | 0.098 | 0.098 | 0.564 | 0.36 | 0.14 | 0.23 | 0.09 | 0.38 | 0.34 |
| 3 | 9107979 | rs6787607 | 0.683 | 0.292 | -0.599 | -0.488 | -0.006 | -0.055 | -0.055 | 0 | 0.17 | 0.34 | 0.09 | 0.08 | 0.12 | 0.13 |
| 3 | 9108056 | rs6772477 | 1.553 | 0.344 | -1.033 | -0.202 | 0.001 | -0.019 | -0.019 | 0 | 0.14 | 0.15 | 0.06 | 0 | 0.11 | 0.02 |
| 3 | 9111221 | rs1162395 | 0.069 | 0.125 | -0.98 | 0.368 | -0.012 | 0.058 | 0.058 | 0 | 0.75 | 0.39 | 0.62 | 0.58 | 0.77 | 0.7 |

| | | | | | | | | | | | | | | | | |
|---|---|---|---|---|---|---|---|---|---|---|---|---|---|---|---|---|
| 3 | 9111847 | rs13088083 | 1.405 | 0.004 | -1.345 | -0.251 | 0.005 | 0.007 | 0.007 | 0 | 0.2 | 0.12 | 0.1 | 0.07 | 0.13 | 0.17 |
| 3 | 9112746 | rs1001603 | 0.659 | 0.394 | -1.114 | 0.33 | 0.013 | 0.046 | 0.046 | 0 | 0.55 | 0.26 | 0.43 | 0.35 | 0.57 | 0.55 |
| 3 | 9112876 | rs1543143 | 0.732 | 0.624 | -1.529 | 0.638 | -0.018 | 0.015 | 0.015 | 0 | 0.19 | 0.06 | 0.28 | 0.1 | 0.22 | 0.16 |
| 3 | 9113057 | rs2675174 | -0.934 | 0.776 | -1.303 | 1.024 | 0 | -0.053 | -0.053 | 1.764 | 0.84 | 0.89 | 0.81 | 0.67 | 0.86 | 0.83 |
| 3 | 9114759 | rs6793348 | -0.261 | 1.071 | -0.678 | 0.891 | -0.011 | -0.001 | -0.001 | 0 | 0.22 | 0.18 | 0.25 | 0.27 | 0.26 | 0.25 |
| 3 | 9118069 | rs9846342 | -0.55 | 0.751 | -0.229 | 0.813 | 0.012 | 0.05 | 0.05 | 0 | 0.07 | 0.25 | 0.07 | 0.11 | 0.07 | 0.12 |
| 3 | 9119182 | rs1996299 | 0.511 | 1.003 | -1.148 | 0.572 | -0.054 | -0.043 | -0.043 | 0 | 0.75 | 0.89 | 0.61 | 0.65 | 0.68 | 0.63 |
| 3 | 9120004 | rs2669989 | 1.027 | 0.257 | -0.476 | 0.485 | 0 | 0.005 | 0.005 | 0 | 0.89 | 0.89 | 0.89 | 0.78 | 0.88 | 0.85 |
| 3 | 9133882 | rs2664077 | 0.179 | -0.358 | -0.175 | -0.261 | -0.001 | 0.01 | 0.01 | 0 | 0.46 | 0.57 | 0.33 | 0.35 | 0.41 | 0.38 |
| 3 | 9136060 | rs531719 | 0.806 | -0.293 | -0.191 | -0.324 | -0.022 | 0.016 | 0.016 | 0 | 0.32 | 0.18 | 0.23 | 0.35 | 0.33 | 0.37 |
| 3 | 9142890 | rs386423 | 0.618 | -0.204 | -0.16 | -0.376 | -0.048 | 0.032 | 0.032 | 0 | 0.33 | 0.13 | 0.28 | 0.34 | 0.39 | 0.36 |
| 3 | 9147178 | rs1861941 | -0.109 | -0.443 | -0.18 | -0.307 | 0.006 | 0.001 | 0.001 | 0 | 0.54 | 0.61 | 0.5 | 0.49 | 0.57 | 0.54 |
| 3 | 9148183 | rs372576 | 0.123 | -0.335 | -0.811 | -0.209 | 0.013 | 0.003 | 0.003 | 0 | 0.24 | 0.14 | 0.25 | 0.15 | 0.23 | 0.2 |
| 3 | 9149099 | rs2664073 | -0.526 | -0.348 | -0.754 | -0.099 | -0.01 | -0.056 | -0.056 | 4.873 | 0.64 | 0.84 | 0.61 | 0.5 | 0.64 | 0.56 |
| 3 | 9156033 | rs391923 | -0.041 | -0.49 | -0.542 | -0.12 | 0.02 | 0 | 0 | 0 | 0.22 | 0.15 | 0.22 | 0.14 | 0.19 | 0.16 |
| 3 | 9157737 | rs17050039 | 0.799 | -0.453 | -0.563 | -0.194 | -0.001 | -0.016 | -0.016 | 0 | 0.07 | 0.09 | 0.07 | 0 | 0.07 | 0.01 |
| 3 | 9158138 | rs422552 | -0.439 | -0.399 | -0.14 | -0.107 | -0.013 | -0.015 | -0.015 | 0.486 | 0.3 | 0.39 | 0.3 | 0.15 | 0.26 | 0.17 |
| 3 | 9160465 | rs2543491 | 0.469 | -0.221 | -0.046 | 0.089 | 0.014 | -0.003 | -0.003 | 0 | 0.48 | 0.35 | 0.51 | 0.5 | 0.56 | 0.47 |
| 3 | 9161375 | rs2664122 | -0.044 | -0.33 | 0.232 | -0.048 | -0.01 | 0.002 | 0.002 | 0 | 0.58 | 0.54 | 0.63 | 0.58 | 0.63 | 0.6 |
| 3 | 9162915 | rs13078449 | -0.811 | -0.262 | 0.369 | -0.013 | 0 | 0 | 0 | 0 | 0.51 | 0.53 | 0.48 | 0.51 | 0.45 | 0.53 |
| 3 | 9163206 | rs6781236 | -0.814 | -0.262 | 0.37 | -0.013 | 0 | 0 | 0 | 1.394 | 0.51 | 0.53 | 0.48 | 0.51 | 0.44 | 0.53 |
| 3 | 9172578 | rs2287492 | -0.814 | -0.73 | -0.343 | -0.134 | -0.004 | -0.023 | -0.023 | 0 | 0.43 | 0.83 | 0.43 | 0.48 | 0.43 | 0.43 |
| 3 | 9181403 | rs2668361 | 1.109 | -1.006 | -0.579 | -0.15 | 0.001 | -0.05 | -0.05 | 0.657 | 0.46 | 0.09 | 0.46 | 0.51 | 0.57 | 0.44 |
| 3 | 9181903 | rs2543494 | -0.106 | -0.186 | 0.07 | 0.155 | -0.016 | 0.034 | 0.034 | 0 | 0.69 | 0.55 | 0.71 | 0.6 | 0.69 | 0.64 |
| 3 | 9187812 | rs9827505 | -0.216 | -0.079 | -0.938 | 0.163 | -0.001 | 0 | 0 | 0 | 0.13 | 0.13 | 0.22 | 0.13 | 0.13 | 0.17 |
| 3 | 9188615 | rs2257056 | 1.144 | -0.062 | -1.369 | 0.209 | -0.011 | 0.026 | 0.026 | 1.888 | 0.5 | 0.09 | 0.43 | 0.43 | 0.51 | 0.49 |
| 3 | 9193324 | rs2675186 | 0.074 | 0.084 | -1.36 | 0.368 | 0.008 | -0.054 | -0.054 | 0 | 0.71 | 0.81 | 0.74 | 0.57 | 0.72 | 0.69 |
| 3 | 9194156 | rs2543499 | -0.124 | 0.16 | -1.482 | 0.391 | -0.003 | -0.085 | -0.085 | 0.263 | 0.73 | 0.9 | 0.76 | 0.57 | 0.72 | 0.69 |
| 3 | 9195331 | rs6762693 | -0.939 | 0.311 | -0.697 | 0.064 | 0.019 | -0.016 | -0.016 | 0 | 0.16 | 0.33 | 0.25 | 0.12 | 0.17 | 0.15 |
| 3 | 9197231 | rs2675181 | 0.144 | 0.388 | -1.83 | 0.317 | -0.008 | -0.033 | -0.033 | 0 | 0.67 | 0.82 | 0.65 | 0.36 | 0.66 | 0.58 |
| 3 | 9200375 | rs17530358 | 0.559 | 0.113 | -2.473 | -0.159 | -0.045 | 0.055 | 0.055 | 0 | 0.25 | 0.02 | 0.36 | 0.09 | 0.26 | 0.24 |
| 3 | 9201696 | rs17194525 | 0.401 | 0.011 | -2.12 | -0.204 | 0.009 | 0.056 | 0.056 | 0 | 0.27 | 0.11 | 0.36 | 0.09 | 0.27 | 0.24 |
| 3 | 9207615 | rs2664107 | -0.188 | -0.065 | -2.992 | -0.42 | -0.008 | 0.02 | 0.02 | 0.166 | 0.56 | 0.08 | 0.56 | 0.33 | 0.64 | 0.61 |
| 3 | 9211356 | rs986119 | -0.039 | -0.165 | -2.702 | -0.409 | -0.053 | 0.025 | 0.025 | 0 | 0.36 | 0.15 | 0.25 | 0.29 | 0.41 | 0.38 |
| 3 | 9215296 | rs396323 | -0.33 | 0.246 | -3.051 | -0.096 | -0.115 | -0.115 | -0.115 | 0.291 | 0.78 | 0.96 | 0.78 | 0.62 | 0.65 | 0.74 |
| 3 | 9218727 | rs400052 | 0.172 | -0.496 | -2.554 | -0.481 | 0 | -0.005 | -0.005 | 0 | 0.42 | 0.64 | 0.42 | 0.56 | 0.36 | 0.43 |
| 3 | 9219440 | rs452714 | -0.146 | -0.474 | -2.618 | -0.551 | 0.003 | 0.006 | 0.006 | 0 | 0.31 | 0.34 | 0.27 | 0.38 | 0.36 | 0.3 |
| 3 | 9224175 | rs802792 | 0.844 | -0.697 | -2.625 | -0.501 | -0.003 | 0.035 | 0.035 | 0 | 0.3 | 0.55 | 0.34 | 0.4 | 0.28 | 0.28 |
| 3 | 9225376 | rs418207 | -0.009 | -0.339 | -2.038 | -0.303 | 0.051 | 0.015 | 0.015 | 0 | 0.27 | 0.5 | 0.29 | 0.33 | 0.38 | 0.27 |
| 3 | 9227644 | rs1145150 | 0.314 | -0.434 | -1.744 | 0.2 | 0.028 | 0.083 | 0.083 | 0 | 0.33 | 0.16 | 0.41 | 0.17 | 0.28 | 0.3 |
| 3 | 9229433 | rs1154386 | -0.742 | -0.567 | -1.712 | -0.01 | 0.008 | 0.027 | 0.027 | 0.045 | 0.51 | 0.6 | 0.51 | 0.23 | 0.45 | 0.45 |
| 3 | 9229760 | rs9839438 | 0.013 | 0.238 | -1.427 | 0.328 | 0.01 | -0.051 | -0.051 | 0 | 0.18 | 0.41 | 0.12 | 0.1 | 0.18 | 0.15 |
| 3 | 9230006 | rs10490820 | 0.768 | 0.428 | -1.641 | 0.366 | -0.017 | 0.009 | 0.009 | 0 | 0.06 | 0.03 | 0.11 | 0.38 | 0.1 | 0.1 |
| 3 | 9232109 | rs6765880 | -0.192 | 0.354 | -1.312 | -0.484 | 0.015 | -0.05 | -0.05 | 0 | 0.18 | 0.44 | 0.12 | 0.1 | 0.18 | 0.15 |
| 3 | 9234763 | rs1145153 | 0.253 | 0.557 | -1.868 | -0.107 | -0.035 | 0.097 | 0.097 | 0 | 0.22 | 0.03 | 0.3 | 0.05 | 0.23 | 0.22 |
| 3 | 9250653 | rs6797462 | -0.634 | 1.025 | -1.016 | 0.779 | 0.01 | 0.058 | 0.058 | 0 | 0.22 | 0.24 | 0.22 | 0.52 | 0.24 | 0.18 |
| 3 | 9250949 | rs441300 | -0.727 | 0.96 | -0.645 | 0.216 | -0.006 | -0.001 | -0.001 | 0.727 | 0.42 | 0.48 | 0.58 | 0.6 | 0.37 | 0.44 |
| 3 | 9252998 | rs12638703 | 0.34 | 0.819 | -0.145 | 0.585 | 0.004 | 0.08 | 0.08 | 0 | 0.15 | 0.19 | 0.16 | 0.49 | 0.14 | 0.14 |
| 3 | 9253191 | rs1154392 | 1.066 | 1.225 | -0.459 | -0.308 | 0.01 | 0.063 | 0.063 | 0 | 0.37 | 0.23 | 0.25 | 0.06 | 0.36 | 0.36 |
| 3 | 9256398 | rs4684627 | 0.702 | 1.166 | -0.785 | -0.494 | -0.084 | 0.118 | 0.118 | 0 | 0.27 | 0.02 | 0.41 | 0.09 | 0.29 | 0.26 |
| 3 | 9257096 | rs2675170 | -0.887 | 0.992 | -1.078 | -0.824 | -0.001 | -0.019 | -0.019 | 0.046 | 0.74 | 0.78 | 0.8 | 0.5 | 0.76 | 0.84 |
| 3 | 9257313 | rs1162435 | -0.106 | 1.019 | -0.725 | -0.835 | 0.004 | 0.036 | 0.036 | 0 | 0.16 | 0.24 | 0.14 | 0.36 | 0.18 | 0.26 |
| 3 | 9260103 | rs4108605 | 0.947 | 0.785 | -0.423 | -1 | -0.001 | -0.012 | -0.012 | 0.062 | 0.29 | 0.28 | 0.44 | 0.11 | 0.3 | 0.29 |
| 3 | 9262650 | rs1995994 | 0.265 | 0.7 | -0.295 | -0.974 | -0.001 | 0 | 0 | 0.029 | 0.33 | 0.34 | 0.28 | 0.7 | 0.33 | 0.22 |
| 3 | 9265620 | rs1028154 | 1.595 | 0.276 | -0.637 | -1.349 | 0.003 | -0.026 | -0.026 | 0.359 | 0.17 | 0.15 | 0.14 | 0.04 | 0.21 | 0.2 |
| 3 | 9266056 | rs2700331 | 0.18 | 0.17 | -0.602 | -1.356 | -0.007 | -0.031 | -0.031 | 0 | 0.63 | 0.58 | 0.73 | 0.8 | 0.63 | 0.5 |
| 3 | 9271859 | rs2700326 | -0.124 | 0.43 | -0.665 | -1.359 | 0.001 | -0.012 | -0.012 | 0 | 0.54 | 0.48 | 0.57 | 0.3 | 0.54 | 0.63 |
| 3 | 9272213 | rs1162707 | 0.485 | 0.391 | -0.562 | -1.385 | 0.005 | 0.024 | 0.024 | 0 | 0.12 | 0.22 | 0.09 | 0.15 | 0.12 | 0.23 |
| 3 | 9272907 | rs9845704 | 0.572 | 0.437 | -0.544 | -1.258 | 0.003 | -0.002 | -0.002 | 0 | 0.18 | 0.22 | 0.16 | 0.2 | 0.2 | 0.21 |
| 3 | 9274895 | rs1162704 | 1.318 | 0.416 | -0.561 | -1.586 | 0.01 | 0.043 | 0.043 | 0 | 0.08 | 0.19 | 0.08 | 0.15 | 0.09 | 0.21 |
| 3 | 9276083 | rs938116 | 0.479 | 0.4 | -0.695 | -1.628 | 0.008 | 0.101 | 0.101 | 0.007 | 0.45 | 0.25 | 0.49 | 0.12 | 0.45 | 0.41 |
| 3 | 9278830 | rs2137215 | -0.833 | 0.42 | -0.834 | -1.412 | -0.001 | -0.002 | -0.002 | 0 | 0.71 | 0.71 | 0.73 | 0.31 | 0.72 | 0.8 |
| 3 | 9287737 | rs9825800 | -0.271 | 0.469 | -0.711 | -1.531 | 0.001 | -0.068 | -0.068 | 0.183 | 0.6 | 0.48 | 0.61 | 0.15 | 0.59 | 0.57 |
| 3 | 9288545 | rs2137216 | -0.331 | 0.565 | -0.983 | -1.074 | 0.021 | 0.099 | 0.099 | 0 | 0.06 | 0.15 | 0.07 | 0.2 | 0.09 | 0.19 |
| 3 | 9291441 | rs17744749 | 1.301 | 0.255 | -1.235 | -1.379 | 0.028 | 0.048 | 0.048 | 0.03 | 0.11 | 0.03 | 0.15 | 0.01 | 0.06 | 0.12 |
| 3 | 9291861 | rs11131163 | 1.139 | 0.46 | -1.591 | -1.643 | -0.082 | 0.117 | 0.117 | 0 | 0.28 | 0.02 | 0.29 | 0.09 | 0.3 | 0.3 |
| 3 | 9292368 | rs2648521 | -0.271 | 0.264 | -1.115 | -1.361 | 0.025 | 0.121 | 0.121 | 0.019 | 0.07 | 0.18 | 0.07 | 0.22 | 0.09 | 0.15 |
| 3 | 9293722 | rs1512053 | 0.457 | -0.06 | -1.155 | -1.507 | 0.034 | 0.043 | 0.043 | 0 | 0.15 | 0.05 | 0.18 | 0.04 | 0.1 | 0.16 |
| 3 | 9294880 | rs2600175 | -0.645 | 0.435 | -0.717 | -1.337 | 0.009 | 0.018 | 0.018 | 0 | 0.65 | 0.52 | 0.68 | 0.39 | 0.65 | 0.77 |

| | | | | | | | | | | | | | | | | |
|---|---|---|---|---|---|---|---|---|---|---|---|---|---|---|---|---|
| 3 | 9299830 | rs9861610 | 0.106 | 0.502 | -0.583 | -1.606 | -0.004 | -0.055 | -0.055 | 0 | 0.15 | 0.22 | 0.12 | 0 | 0.13 | 0.16 |
| 3 | 9300870 | rs10222369 | -1.282 | 0.49 | -0.626 | -1.609 | -0.003 | -0.052 | -0.052 | 0.333 | 0.2 | 0.39 | 0.23 | 0.07 | 0.17 | 0.25 |
| 3 | 9302032 | rs2648512 | -0.619 | 0.63 | -1.086 | -1.849 | -0.013 | -0.03 | -0.03 | 0 | 0.59 | 0.47 | 0.67 | 0.23 | 0.68 | 0.64 |
| 3 | 9309199 | rs17744934 | 0.592 | 1.033 | -1.957 | -2.044 | -0.047 | 0.099 | 0.099 | 0 | 0.33 | 0.11 | 0.36 | 0.16 | 0.34 | 0.29 |
| 3 | 9312227 | rs2600162 | -1.401 | 0.796 | -1.664 | -2.142 | -0.02 | -0.03 | -0.03 | 0 | 0.64 | 0.48 | 0.72 | 0.23 | 0.72 | 0.66 |
| 3 | 9312398 | rs2600164 | 0.515 | 0.891 | -1.904 | -2.264 | -0.038 | 0.081 | 0.081 | 0 | 0.19 | 0.04 | 0.16 | 0.01 | 0.26 | 0.16 |
| 3 | 9312536 | rs13063276 | 0.308 | 0.787 | -1.861 | -2.316 | -0.066 | 0.118 | 0.118 | 0 | 0.4 | 0.16 | 0.51 | 0.19 | 0.43 | 0.39 |
| 3 | 9313668 | rs7618964 | 0.507 | 0.762 | -1.806 | -2.258 | -0.045 | 0.124 | 0.124 | 0 | 0.71 | 0.92 | 0.67 | 0.94 | 0.64 | 0.6 |
| 3 | 9313707 | rs2324763 | 1.058 | 0.823 | -1.692 | -2.138 | -0.001 | 0.003 | 0.003 | 0 | 0.07 | 0.05 | 0.16 | 0.02 | 0.08 | 0.12 |
| 3 | 9317758 | rs2600192 | -0.563 | 0.748 | -1.563 | -2.2 | 0 | -0.045 | -0.045 | 0 | 0.32 | 0.35 | 0.36 | 0.06 | 0.37 | 0.38 |
| 3 | 9318416 | rs17050210 | -0.744 | 0.737 | -1.549 | -2.12 | -0.006 | -0.037 | -0.037 | 0 | 0.59 | 0.49 | 0.62 | 0.21 | 0.58 | 0.53 |
| 3 | 9327287 | rs12631463 | 1.179 | 0.944 | -1.834 | -1.944 | -0.02 | 0.011 | 0.011 | 0 | 0.15 | 0.05 | 0.08 | 0.13 | 0.17 | 0.22 |
| 3 | 9334620 | rs2648568 | -1.313 | 0.869 | -1.829 | -2.043 | -0.015 | -0.031 | -0.031 | 0 | 0.63 | 0.47 | 0.6 | 0.22 | 0.7 | 0.63 |
| 3 | 9339644 | rs2648563 | -0.029 | 1.476 | -1.915 | -1.76 | -0.001 | -0.036 | -0.036 | 0 | 0.21 | 0.17 | 0.16 | 0.01 | 0.22 | 0.15 |
| 3 | 9340435 | rs4686339 | -0.067 | 1.249 | -2.047 | -1.158 | -0.103 | 0.049 | 0.049 | 1.342 | 0.2 | 0.01 | 0.31 | 0.06 | 0.29 | 0.16 |
| 3 | 9350167 | rs17050220 | -0.425 | -0.005 | -1.748 | -1.038 | 0.001 | 0 | 0 | 0.177 | 0.28 | 0.69 | 0.18 | 0.11 | 0.15 | 0.19 |
| 3 | 9350287 | rs17050221 | -0.046 | -0.142 | -0.927 | -1.13 | -0.052 | -0.109 | -0.109 | 2.798 | 0.22 | 0.41 | 0.08 | 0.03 | 0.13 | 0.12 |
| 3 | 9352319 | rs11707847 | -0.896 | 0.209 | -1.552 | -1.108 | 0.028 | 0.04 | 0.04 | 0.103 | 0.36 | 0.72 | 0.33 | 0.21 | 0.24 | 0.3 |
| 3 | 9357390 | rs7613074 | -0.688 | 1.351 | -2.313 | -1.003 | -0.072 | -0.008 | -0.008 | 0.857 | 0.89 | 0.99 | 0.85 | 0.9 | 0.75 | 0.92 |
| 3 | 9358616 | rs3852050 | 0.644 | 0.743 | -2.336 | 0.183 | -0.001 | 0.041 | 0.041 | 0 | 0.07 | 0.07 | 0.09 | 0.29 | 0.07 | 0.14 |
| 3 | 9363272 | rs4686342 | -0.131 | 0.393 | -3.094 | -0.246 | -0.017 | 0.148 | 0.148 | 0 | 0.57 | 0.82 | 0.67 | 0.84 | 0.4 | 0.62 |
| 3 | 9363925 | rs2324797 | 0.552 | 0.399 | -3.044 | -0.025 | -0.002 | -0.112 | -0.112 | 0 | 0.37 | 0.1 | 0.44 | 0.49 | 0.3 | 0.38 |
| 3 | 9366920 | rs4053780 | 0.67 | 0.418 | -2.952 | -0.162 | -0.014 | -0.044 | -0.044 | 0.021 | 0.34 | 0.2 | 0.33 | 0.53 | 0.32 | 0.44 |
| 3 | 9367619 | rs2648558 | -0.177 | 1.542 | -3.116 | -0.122 | -0.077 | 0.017 | 0.017 | 0 | 0.85 | 0.99 | 0.81 | 0.91 | 0.74 | 0.88 |
| 3 | 9374170 | rs10510398 | 0.039 | 1.238 | -3.176 | -0.318 | -0.046 | 0.129 | 0.129 | 0 | 0.24 | 0.03 | 0.28 | 0.01 | 0.28 | 0.21 |
| 3 | 9376490 | rs2600168 | -0.074 | 0.639 | -2.778 | -0.157 | -0.074 | -0.003 | -0.003 | 0 | 0.83 | 0.98 | 0.82 | 0.88 | 0.75 | 0.9 |
| 3 | 9381122 | rs2648549 | 0.98 | 0.613 | -1.99 | -0.131 | -0.033 | -0.045 | -0.045 | 0.446 | 0.62 | 0.22 | 0.48 | 0.53 | 0.57 | 0.73 |
| 3 | 9381250 | rs2728944 | 0.635 | 0.568 | -1.013 | -0.087 | 0.057 | -0.057 | -0.057 | 0 | 0.74 | 0.56 | 0.59 | 0.89 | 0.65 | 0.82 |
| 3 | 9389223 | rs2648545 | 2.022 | 0.035 | -1.583 | -0.377 | -0.096 | -0.098 | -0.098 | 3.357 | 0.44 | 0.03 | 0.39 | 0.45 | 0.43 | 0.51 |
| 3 | 9389577 | rs2728939 | -0.403 | 0.639 | -0.788 | 0.066 | -0.04 | 0.005 | 0.005 | 0 | 0.89 | 0.96 | 0.85 | 0.93 | 0.74 | 0.9 |
| 3 | 9391300 | rs2728937 | -0.545 | 0.226 | -0.817 | -0.067 | -0.062 | 0.053 | 0.053 | 0 | 0.91 | 0.99 | 0.87 | 1 | 0.76 | 0.91 |
| 3 | 9396030 | rs13068999 | 0.021 | 0.044 | -0.163 | 0.089 | -0.011 | -0.005 | -0.005 | 1.709 | 0.14 | 0.34 | 0.28 | 0.1 | 0.1 | 0.08 |
| 3 | 9396051 | rs13069009 | 0.026 | 0.044 | -0.188 | -0.155 | -0.011 | -0.042 | -0.042 | 0 | 0.14 | 0.29 | 0.25 | 0.04 | 0.1 | 0.08 |
| 3 | 9399887 | rs2728950 | 0.572 | 0.277 | -0.105 | -0.061 | 0.107 | 0.157 | 0.157 | 0 | 0.85 | 0.63 | 0.8 | 0.66 | 0.73 | 0.86 |
| 3 | 9404877 | rs2258768 | 1.317 | 0 | -0.082 | -0.163 | 0.043 | 0.057 | 0.057 | 0 | 0.7 | 0.54 | 0.6 | 0.58 | 0.6 | 0.79 |
| 3 | 9404887 | rs2258767 | -0.461 | 0.003 | 0.021 | -0.134 | 0.012 | 0.269 | 0.269 | 1.045 | 0.09 | 0.32 | 0.09 | 0.34 | 0.05 | 0.06 |
| 3 | 9406836 | rs1054975 | 1.426 | -0.004 | 0.009 | -0.141 | 0.044 | 0.058 | 0.058 | 0 | 0.7 | 0.55 | 0.59 | 0.58 | 0.6 | 0.79 |
| 3 | 9416832 | rs2255558 | 1.575 | 0.078 | 0.222 | -0.121 | 0.023 | 0.033 | 0.033 | 0.021 | 0.69 | 0.45 | 0.58 | 0.57 | 0.61 | 0.79 |
| 3 | 9418510 | rs2728945 | -0.517 | 0.226 | 0.178 | -0.108 | -0.03 | 0.025 | 0.025 | 0 | 0.86 | 0.95 | 0.77 | 0.94 | 0.81 | 0.92 |
| 3 | 9421024 | rs2596917 | -0.52 | 0.221 | 0.2 | -0.13 | 0.009 | 0.262 | 0.262 | 1.021 | 0.09 | 0.32 | 0.09 | 0.35 | 0.05 | 0.06 |
| 3 | 9426224 | rs1129174 | 0.377 | 0.24 | 0.132 | -0.19 | -0.035 | 0.028 | 0.028 | 0 | 0.13 | 0.04 | 0.22 | 0.06 | 0.19 | 0.07 |
| 3 | 9428270 | rs2251361 | 1.698 | -0.083 | 0.284 | -0.413 | -0.027 | -0.03 | -0.03 | 0 | 0.61 | 0.31 | 0.5 | 0.52 | 0.54 | 0.72 |
| 3 | 9435240 | rs2596914 | 0.632 | -0.259 | 0.548 | 0.083 | 0.016 | 0.325 | 0.325 | 0.381 | 0.9 | 0.63 | 0.91 | 0.65 | 0.94 | 0.94 |
| 3 | 9469655 | rs904467 | 0.936 | -0.262 | 0.239 | -0.064 | -0.075 | 0.023 | 0.023 | 0 | 0.18 | 0.01 | 0.33 | 0.07 | 0.28 | 0.28 |
| 3 | 9479142 | rs2442825 | 1.476 | -0.302 | 0.346 | -0.11 | -0.11 | -0.097 | -0.097 | 0 | 0.51 | 0.02 | 0.31 | 0.49 | 0.48 | 0.51 |
| 3 | 9480906 | rs2596937 | -2.125 | -0.359 | 1.717 | 0.628 | 0.098 | -0.001 | -0.001 | 0.07 | 0.13 | 0.47 | 0.24 | 0.08 | 0.18 | 0.13 |
| 3 | 9495385 | rs2648580 | 2.165 | -0.418 | 1.424 | 0.736 | 0.017 | 0.136 | 0.136 | 2.061 | 0.79 | 0.44 | 0.68 | 0.55 | 0.78 | 0.82 |
| 3 | 9502463 | rs9845420 | -2.211 | -0.287 | 1.672 | 0.052 | 0.052 | 0 | 0 | 1.135 | 0.13 | 0.55 | 0.24 | 0.08 | 0.13 | 0.13 |
| 3 | 9518511 | rs2279440 | -2.019 | -0.38 | 1.61 | 0.337 | 0.002 | -0.008 | -0.008 | 0 | 0.11 | 0.19 | 0.19 | 0.06 | 0.11 | 0.12 |
| 3 | 9533974 | rs17050395 | -2.052 | -0.367 | 1.308 | 0.329 | 0.061 | -0.044 | -0.044 | 0.525 | 0.15 | 0.48 | 0.24 | 0.07 | 0.17 | 0.13 |
| 3 | 9539655 | rs6774483 | -1.866 | -0.448 | 1.127 | 0.378 | -0.013 | -0.039 | -0.039 | 0.109 | 0.16 | 0.65 | 0.23 | 0.07 | 0.14 | 0.13 |
| 3 | 9539684 | rs2596915 | 1.694 | -0.512 | 0.902 | 0.827 | -0.011 | 0.052 | 0.052 | 2.927 | 0.75 | 0.41 | 0.71 | 0.51 | 0.79 | 0.82 |
| 3 | 9540323 | rs3732975 | -1.007 | 0.212 | 0.524 | 0.529 | 0.039 | -0.024 | -0.024 | 0 | 0.11 | 0.27 | 0.2 | 0.06 | 0.16 | 0.09 |
| 3 | 9544554 | rs2200233 | 1.139 | 0.23 | -0.274 | 0.566 | -0.005 | -0.03 | -0.03 | 0 | 0.67 | 0.28 | 0.6 | 0.54 | 0.66 | 0.66 |
| 3 | 9546173 | rs2596936 | 1.005 | -0.531 | 0.172 | 0.562 | -0.039 | 0.079 | 0.079 | 0.92 | 0.77 | 0.42 | 0.82 | 0.58 | 0.83 | 0.84 |
| 3 | 9547959 | rs2625889 | 0.699 | -0.239 | 0.222 | -0.261 | 0.005 | -0.046 | -0.046 | 0 | 0.7 | 0.65 | 0.72 | 0.91 | 0.64 | 0.66 |
| 3 | 9555554 | rs6807946 | -0.615 | -0.383 | 0.356 | 0.053 | -0.024 | 0.259 | 0.259 | 0.023 | 0.13 | 0.37 | 0.11 | 0.41 | 0.08 | 0.05 |
| 3 | 9556914 | rs9818611 | -1.816 | -0.613 | 0.482 | -0.1 | 0.002 | -0.006 | -0.006 | 0 | 0.45 | 0.59 | 0.38 | 0.49 | 0.47 | 0.42 |
| 3 | 9561205 | rs2633770 | -1.696 | -0.635 | 0.402 | -0.09 | 0.013 | -0.019 | -0.019 | 0.966 | 0.46 | 0.7 | 0.39 | 0.49 | 0.47 | 0.43 |
| 3 | 9568136 | rs9846242 | -0.775 | -0.683 | 0.263 | -0.255 | -0.004 | 0.042 | 0.042 | 0 | 0.31 | 0.23 | 0.25 | 0.08 | 0.35 | 0.33 |
| 3 | 9592982 | rs7649416 | -0.472 | -0.639 | 0.481 | -0.031 | 0.005 | 0.023 | 0.023 | 0 | 0.19 | 0.17 | 0.16 | 0.07 | 0.19 | 0.21 |
| 3 | 9595374 | rs2648593 | -1.483 | -0.674 | 0.772 | -0.006 | -0.007 | 0.014 | 0.014 | 0 | 0.32 | 0.61 | 0.28 | 0.47 | 0.28 | 0.26 |
| 3 | 9605874 | rs9836700 | -0.34 | -0.596 | 0.599 | -0.106 | 0 | -0.004 | -0.004 | 0 | 0.18 | 0.19 | 0.17 | 0.07 | 0.19 | 0.22 |
| 3 | 9612856 | rs2542353 | -0.794 | -0.391 | 1.051 | 0.363 | -0.002 | 0.302 | 0.302 | 0 | 0.13 | 0.35 | 0.09 | 0.41 | 0.09 | 0.05 |
| 3 | 9614001 | rs2633788 | -0.933 | -0.442 | 0.959 | 0.341 | -0.009 | 0.304 | 0.304 | 0.025 | 0.14 | 0.37 | 0.09 | 0.42 | 0.09 | 0.05 |
| 3 | 9616140 | rs2633789 | -1.108 | -0.453 | 0.825 | 0.336 | -0.021 | 0.247 | 0.247 | 0.031 | 0.14 | 0.49 | 0.1 | 0.42 | 0.09 | 0.06 |
| 3 | 9617634 | rs2542361 | -0.196 | -0.159 | -0.121 | -0.908 | -0.002 | -0.017 | -0.017 | 0 | 0.72 | 0.71 | 0.73 | 0.91 | 0.72 | 0.78 |
| 3 | 9635471 | rs2648573 | -1.024 | 0.512 | -0.113 | -1.093 | 0.027 | -0.131 | -0.131 | 3.615 | 0.27 | 0.41 | 0.17 | 0.01 | 0.33 | 0.23 |
| 3 | 9640458 | rs7634260 | -0.711 | 0.18 | 0.098 | -0.732 | 0.014 | -0.035 | -0.035 | 0.178 | 0.11 | 0.29 | 0.1 | 0.04 | 0.09 | 0.1 |

| | | | | | | | | | | | | | | | | |
|---|---|---|---|---|---|---|---|---|---|---|---|---|---|---|---|---|
| 3 | 9649470 | rs7645857 | -0.815 | 0.097 | -0.133 | -0.755 | -0.014 | -0.024 | -0.024 | 0.011 | 0.11 | 0.21 | 0.12 | 0.07 | 0.09 | 0.19 |
| 3 | 9652168 | rs2542391 | 0.331 | 0.276 | -0.911 | -1.396 | -0.038 | 0.088 | 0.088 | 0 | 0.8 | 0.95 | 0.88 | 0.99 | 0.75 | 0.8 |
| 3 | 9660064 | rs6443251 | -0.792 | 0.505 | -0.543 | -1.828 | 0.003 | -0.034 | -0.034 | 0 | 0.21 | 0.29 | 0.2 | 0.03 | 0.25 | 0.14 |
| 3 | 9673018 | rs7618981 | -0.818 | 0.959 | -0.192 | -2.334 | 0.008 | -0.1 | -0.1 | 2.849 | 0.33 | 0.55 | 0.24 | 0.05 | 0.3 | 0.2 |
| 3 | 9687811 | rs787839 | -0.706 | 0.857 | -0.225 | -2.31 | -0.064 | 0.06 | 0.06 | 0 | 0.89 | 0.99 | 0.9 | 0.99 | 0.79 | 0.92 |
| 3 | 9702755 | rs6809387 | -0.395 | 0.588 | -0.139 | -2.285 | -0.007 | -0.012 | -0.012 | 3.72 | 0.13 | 0.53 | 0.15 | 0.07 | 0.1 | 0.08 |
| 3 | 9709043 | rs9816328 | -0.677 | 0.564 | 0.457 | -2.376 | 0.001 | -0.033 | -0.033 | 0 | 0.14 | 0.49 | 0.15 | 0.05 | 0.11 | 0.09 |
| 3 | 9727675 | rs787837 | -0.821 | 0.244 | -0.091 | -2.123 | -0.012 | -0.015 | -0.015 | 0 | 0.42 | 0.49 | 0.35 | 0.3 | 0.32 | 0.41 |
| 3 | 9730522 | rs3868042 | 1.062 | 0.25 | -0.109 | -2.036 | 0.008 | -0.017 | -0.017 | 0 | 0.06 | 0.21 | 0.16 | 0 | 0.03 | 0.04 |
| 3 | 9734855 | rs4684652 | 0.175 | 0.234 | -0.142 | -2.052 | 0.001 | -0.016 | -0.016 | 0 | 0.11 | 0.27 | 0.13 | 0.04 | 0.1 | 0.06 |
| 3 | 9735085 | rs7615134 | -1.027 | 0.278 | -0.472 | -2.042 | -0.019 | -0.065 | -0.065 | 0.003 | 0.18 | 0.51 | 0.32 | 0.06 | 0.15 | 0.1 |
| 3 | 9735683 | rs11131183 | 0.524 | 0.575 | -1.37 | -1.848 | 0 | -0.003 | -0.003 | 0 | 0.09 | 0.12 | 0.11 | 0.04 | 0.1 | 0.06 |
| 3 | 9735908 | rs159147 | 0.076 | 0.938 | -1.868 | -1.281 | -0.051 | 0.006 | 0.006 | 0 | 0.37 | 0.12 | 0.28 | 0.71 | 0.38 | 0.48 |
| 3 | 9745043 | rs7618350 | 0.228 | 1.118 | -1.17 | -0.909 | -0.064 | 0.015 | 0.015 | 0 | 0.89 | 0.99 | 0.89 | 0.96 | 0.76 | 0.92 |
| 3 | 9748636 | rs468056 | -0.74 | 0.861 | -0.977 | -0.965 | -0.004 | 0.006 | 0.006 | 0.001 | 0.26 | 0.21 | 0.33 | 0.05 | 0.33 | 0.13 |
| 3 | 9752333 | rs293777 | -0.487 | 0.326 | -0.539 | -0.541 | -0.006 | -0.03 | -0.03 | 0 | 0.17 | 0.21 | 0.12 | 0.04 | 0.11 | 0.14 |
| 3 | 9765677 | rs293779 | -0.848 | 0.42 | -0.434 | -0.681 | -0.005 | -0.09 | -0.09 | 0.915 | 0.33 | 0.58 | 0.38 | 0.07 | 0.28 | 0.3 |
| 3 | 9772070 | rs125700 | 0.176 | 0.179 | -0.769 | 0.732 | 0.006 | -0.046 | -0.046 | 0 | 0.89 | 0.99 | 0.88 | 0.36 | 0.88 | 0.71 |
| 3 | 9775160 | rs159146 | -0.935 | 0.639 | -0.091 | 0.441 | 0.075 | 0.003 | 0.003 | 0 | 0.13 | 0.37 | 0.16 | 0.09 | 0.23 | 0.13 |
| 3 | 9778315 | rs159157 | -0.172 | 0.428 | -0.428 | 1.351 | -0.006 | -0.032 | -0.032 | 0 | 0.91 | 0.99 | 0.93 | 0.68 | 0.89 | 0.82 |
| 3 | 9788168 | rs2269112 | 0.58 | -0.139 | -1.077 | 0.784 | 0.017 | -0.024 | -0.024 | 0.064 | 0.24 | 0.02 | 0.16 | 0.17 | 0.15 | 0.28 |
| 3 | 9797723 | rs2075747 | 0.191 | -0.038 | -0.379 | 1.042 | 0.023 | -0.01 | -0.01 | 0.138 | 0.31 | 0.23 | 0.22 | 0.4 | 0.17 | 0.33 |
| 3 | 9799113 | rs293795 | -0.584 | 0.107 | 0.036 | 0.71 | 0.109 | 0.008 | 0.008 | 0 | 0.08 | 0.33 | 0.14 | 0.04 | 0.15 | 0.13 |
| 3 | 9803627 | rs293794 | -0.158 | 0.236 | 0.092 | 0.78 | 0.157 | 0.05 | 0.05 | 0 | 0.05 | 0.25 | 0.13 | 0.05 | 0.14 | 0.12 |
| 3 | 9809082 | rs293796 | -0.242 | -0.102 | -0.215 | 0.974 | -0.008 | -0.053 | -0.053 | 0 | 0.92 | 1 | 0.93 | 0.75 | 0.92 | 0.82 |
| 3 | 9823302 | rs6443264 | 0.367 | -0.251 | 0.442 | 0.941 | 0.008 | -0.013 | -0.013 | 0 | 0.3 | 0.28 | 0.3 | 0.55 | 0.2 | 0.34 |
| 3 | 9824651 | rs6443265 | 1.02 | -0.222 | 0.964 | 0.831 | 0.043 | 0.09 | 0.09 | 0.06 | 0.81 | 0.52 | 0.61 | 0.7 | 0.77 | 0.68 |
| 3 | 9831970 | rs11711241 | 1.868 | -0.343 | 0.513 | 0.424 | -0.135 | 0.205 | 0.205 | 0 | 0.34 | 0.06 | 0.24 | 0.03 | 0.43 | 0.19 |
| 3 | 9835140 | rs159143 | 0.811 | -0.642 | 0.876 | 0.71 | -0.032 | 0.049 | 0.049 | 0.025 | 0.83 | 0.64 | 0.71 | 0.75 | 0.87 | 0.81 |
| 3 | 9836386 | rs11131187 | 1.531 | -0.639 | 0.763 | 0.725 | -0.003 | 0.01 | 0.01 | 0.173 | 0.76 | 0.34 | 0.59 | 0.7 | 0.75 | 0.69 |
| 3 | 9839182 | rs4686371 | 0.604 | -0.51 | 0.629 | 0.87 | 0.027 | -0.07 | -0.07 | 0 | 0.29 | 0.17 | 0.29 | 0.53 | 0.2 | 0.34 |
| 3 | 9840983 | rs6774930 | 1.118 | -0.617 | 0.968 | 0.927 | 0.028 | 0.094 | 0.094 | 0.737 | 0.8 | 0.48 | 0.62 | 0.68 | 0.77 | 0.67 |
| 3 | 9860880 | rs3806667 | 0.056 | -0.66 | 0.76 | 1.099 | -0.002 | 0.014 | 0.014 | 0 | 0.89 | 0.86 | 0.74 | 0.75 | 0.9 | 0.78 |
| 3 | 9863233 | rs6773564 | 0.962 | -0.767 | 0.765 | 0.924 | 0.003 | -0.002 | -0.002 | 0 | 0.49 | 0.48 | 0.43 | 0.2 | 0.69 | 0.32 |
| 3 | 9867625 | rs3732527 | 0.811 | -0.496 | 0.736 | 1.194 | 0.005 | 0.003 | 0.003 | 0 | 0.55 | 0.56 | 0.44 | 0.21 | 0.72 | 0.4 |
| 3 | 9870414 | rs17050593 | 0.674 | -0.376 | 0.79 | 1.549 | 0.014 | -0.14 | -0.14 | 0 | 0.26 | 0.03 | 0.19 | 0.44 | 0.16 | 0.36 |
| 3 | 9871030 | rs2290305 | 0.6 | -0.288 | 0.906 | 1.213 | -0.003 | -0.008 | -0.008 | 0.462 | 0.58 | 0.56 | 0.46 | 0.22 | 0.73 | 0.42 |
| 3 | 9876919 | rs1057278 | -0.34 | 0.031 | 0.809 | 1.26 | 0.072 | 0.017 | 0.017 | 0 | 0.07 | 0.26 | 0.11 | 0.06 | 0.12 | 0.08 |
| 3 | 9877452 | rs4686377 | 0.484 | -0.113 | 0.227 | 1.33 | 0.024 | -0.126 | -0.126 | 1.977 | 0.27 | 0.08 | 0.28 | 0.51 | 0.18 | 0.34 |
| 3 | 9877481 | rs9878419 | 0.528 | -0.008 | 0.334 | 1.111 | 0.008 | 0.027 | 0.027 | 0 | 0.53 | 0.37 | 0.35 | 0.16 | 0.61 | 0.37 |
| 3 | 9883021 | rs2426 | 1.661 | 0.805 | -0.191 | 2.322 | -0.001 | 0.043 | 0.043 | 0 | 0.87 | 0.88 | 0.69 | 0.39 | 0.87 | 0.73 |
| 3 | 9888026 | rs9819655 | 0.504 | 0.845 | -0.502 | 1.752 | -0.019 | -0.008 | -0.008 | 0 | 0.95 | 1 | 0.93 | 0.69 | 0.92 | 0.79 |
| 3 | 9898280 | rs279565 | -1.294 | 1.419 | 0.059 | 1.145 | 0.099 | 0.019 | 0.019 | 0.105 | 0.06 | 0.23 | 0.15 | 0.07 | 0.16 | 0.08 |
| 3 | 9898582 | rs192495 | 0.904 | 0.238 | -0.274 | 0.436 | -0.005 | -0.002 | -0.002 | 0 | 0.66 | 0.39 | 0.54 | 0.27 | 0.76 | 0.62 |
| 3 | 9901479 | rs1695432 | 0.447 | 0.503 | -0.52 | -0.884 | 0.001 | -0.012 | -0.012 | 0 | 0.6 | 0.6 | 0.4 | 0.21 | 0.54 | 0.5 |
| 3 | 9907397 | rs375024 | 0.778 | -0.104 | -0.532 | -1.635 | -0.001 | -0.015 | -0.015 | 0 | 0.48 | 0.48 | 0.41 | 0.1 | 0.51 | 0.35 |
| 3 | 9908525 | rs2479 | -0.739 | 0.208 | -0.213 | -1.203 | -0.001 | 0.003 | 0.003 | 0.302 | 0.81 | 0.81 | 0.68 | 0.74 | 0.81 | 0.81 |
| 3 | 9917654 | rs462378 | -0.621 | 0.225 | 0.02 | -1.145 | 0.005 | 0.037 | 0.037 | 0.233 | 0.8 | 0.63 | 0.67 | 0.75 | 0.81 | 0.81 |
| 3 | 9926348 | rs11918795 | 0.504 | 0.552 | -0.498 | -1.314 | 0.006 | -0.061 | -0.061 | 0 | 0.12 | 0.2 | 0.05 | 0.01 | 0.1 | 0.11 |
| 3 | 9926423 | rs11922283 | 0.504 | 0.552 | -0.545 | -1.314 | 0.001 | -0.044 | -0.044 | 0.323 | 0.12 | 0.17 | 0.05 | 0.01 | 0.1 | 0.11 |
| 3 | 9934231 | rs279554 | -0.263 | 0.888 | -0.364 | -1.159 | 0 | -0.003 | -0.003 | 0.806 | 0.68 | 0.68 | 0.63 | 0.72 | 0.71 | 0.68 |
| 3 | 9934605 | rs279553 | -0.263 | 0.888 | -0.364 | -1.159 | 0 | -0.003 | -0.003 | 0 | 0.68 | 0.68 | 0.63 | 0.72 | 0.71 | 0.68 |
| 3 | 9935263 | rs279543 | 0.982 | 0.91 | -0.467 | -1.193 | 0.006 | 0.063 | 0.063 | 0 | 0.49 | 0.42 | 0.46 | 0.1 | 0.55 | 0.36 |
| 3 | 9939620 | rs11706730 | 1.247 | 0.981 | -0.657 | -1.107 | -0.021 | -0.083 | -0.083 | 0 | 0.14 | 0.02 | 0.16 | 0.62 | 0.15 | 0.26 |
| 3 | 9939828 | rs12053867 | 1.225 | 0.984 | -0.656 | -1.106 | -0.021 | -0.084 | -0.084 | 0 | 0.14 | 0.02 | 0.16 | 0.62 | 0.15 | 0.26 |
| 3 | 9942234 | rs395042 | -0.082 | 1.079 | -0.115 | -1.669 | 0 | -0.015 | -0.015 | 0 | 0.54 | 0.61 | 0.48 | 0.1 | 0.55 | 0.41 |
| 3 | 9943355 | rs4686381 | -0.09 | 1.405 | -0.339 | -1.365 | -0.04 | 0.029 | 0.029 | 0 | 0.08 | 0.02 | 0.12 | 0.02 | 0.19 | 0.12 |
| 3 | 9952706 | rs279572 | 0.238 | 0.607 | -0.094 | -1.487 | 0.007 | 0.038 | 0.038 | 0.67 | 0.51 | 0.64 | 0.47 | 0.09 | 0.55 | 0.33 |
| 3 | 9956279 | rs455863 | 0.585 | 0.73 | 0.171 | -1.433 | 0.006 | 0.051 | 0.051 | 0 | 0.48 | 0.41 | 0.46 | 0.09 | 0.54 | 0.32 |
| 3 | 9957898 | rs8883 | 0.293 | 0.857 | -0.293 | -1.317 | 0.006 | 0.025 | 0.025 | 0.039 | 0.52 | 0.64 | 0.48 | 0.08 | 0.54 | 0.34 |
| 3 | 9962816 | rs376511 | -0.384 | 0.884 | -0.439 | -1.239 | 0 | -0.048 | -0.048 | 1.08 | 0.48 | 0.45 | 0.54 | 0.9 | 0.44 | 0.64 |
| 3 | 9969328 | rs7627880 | -0.347 | 0.899 | -0.353 | -1.301 | 0 | -0.046 | -0.046 | 0.962 | 0.38 | 0.42 | 0.55 | 0.87 | 0.44 | 0.52 |
| 3 | 9977244 | rs3894571 | -0.501 | 0.404 | -0.662 | -0.992 | -0.046 | -0.011 | -0.011 | 0.122 | 0.57 | 0.82 | 0.57 | 0.45 | 0.49 | 0.6 |
| 3 | 9979660 | rs2302786 | -1.341 | -0.04 | -0.345 | -1.368 | 0.028 | 0.079 | 0.079 | 0 | 0.33 | 0.89 | 0.34 | 0.16 | 0.22 | 0.24 |
| 3 | 9985457 | rs9878047 | -1.112 | -0.06 | -0.178 | -1.346 | 0.021 | 0.039 | 0.039 | 0 | 0.3 | 0.81 | 0.33 | 0.16 | 0.21 | 0.24 |
| 3 | 9985656 | rs3774207 | -1.119 | -0.075 | -0.184 | -1.353 | 0.015 | 0.031 | 0.031 | 0 | 0.29 | 0.81 | 0.33 | 0.16 | 0.21 | 0.24 |
| 3 | 9996426 | rs3846167 | -1.122 | -0.081 | 0.18 | -1.518 | 0.007 | 0.013 | 0.013 | 0 | 0.3 | 0.75 | 0.32 | 0.13 | 0.21 | 0.25 |
| 3 | 10001772 | rs17050660 | -0.496 | -0.138 | 0.803 | -1.566 | -0.011 | -0.028 | -0.028 | 0 | 0.2 | 0.33 | 0.22 | 0.11 | 0.15 | 0.17 |
| 3 | 10009919 | rs13063098 | -0.72 | -0.18 | 0.846 | -1.584 | -0.019 | -0.038 | -0.038 | 0 | 0.22 | 0.36 | 0.23 | 0.11 | 0.15 | 0.17 |

| | | | | | | | | | | | | | | | | |
|---|---|---|---|---|---|---|---|---|---|---|---|---|---|---|---|---|
| 3 | 10010308 | rs13063929 | -0.918 | -0.177 | 0.865 | -1.583 | -0.027 | -0.046 | -0.046 | 0 | 0.23 | 0.38 | 0.23 | 0.12 | 0.15 | 0.17 |
| 3 | 10013273 | rs3732968 | -0.932 | -0.168 | 0.866 | -1.582 | -0.03 | -0.046 | -0.046 | 0.375 | 0.23 | 0.39 | 0.23 | 0.11 | 0.15 | 0.17 |
| 3 | 10027724 | rs17535749 | 0.297 | -0.177 | 0.834 | -1.622 | -0.01 | 0.024 | 0.024 | 0 | 0.07 | 0.03 | 0.07 | 0 | 0.08 | 0.04 |
| 3 | 10029289 | rs3755783 | -0.488 | -0.194 | 1.119 | -1.635 | -0.002 | -0.003 | -0.003 | 0 | 0.2 | 0.17 | 0.22 | 0.13 | 0.15 | 0.23 |
| 3 | 10045703 | rs6786636 | -1.526 | -0.206 | 0.989 | -1.652 | 0.026 | 0.059 | 0.059 | 0 | 0.31 | 0.85 | 0.33 | 0.14 | 0.21 | 0.25 |
| 3 | 10045734 | rs17050704 | -1.477 | -0.188 | 1.014 | -1.644 | 0.005 | 0.011 | 0.011 | 0 | 0.29 | 0.74 | 0.32 | 0.14 | 0.21 | 0.25 |
| 3 | 10046703 | rs2272120 | -1.477 | -0.188 | 1.014 | -1.644 | 0.005 | 0.011 | 0.011 | 0 | 0.29 | 0.74 | 0.32 | 0.14 | 0.21 | 0.24 |
| 3 | 10048601 | rs7638846 | -1.132 | -0.193 | 1.031 | -1.647 | -0.001 | -0.003 | -0.003 | 1.222 | 0.26 | 0.7 | 0.3 | 0.12 | 0.21 | 0.18 |
| 3 | 10049287 | rs2272118 | -1.476 | -0.186 | 1.029 | -1.644 | 0.005 | 0.011 | 0.011 | 0 | 0.29 | 0.74 | 0.32 | 0.14 | 0.21 | 0.25 |
| 3 | 10063960 | rs7611218 | -0.467 | -0.299 | 1.044 | -1.692 | -0.012 | -0.058 | -0.058 | 0 | 0.2 | 0.46 | 0.17 | 0.05 | 0.14 | 0.14 |
| 3 | 10085035 | rs803335 | -0.967 | -0.186 | 0.828 | -1.943 | -0.022 | 0.053 | 0.053 | 0.083 | 0.9 | 0.99 | 0.92 | 1 | 0.91 | 0.92 |
| 3 | 10122575 | rs6442150 | -0.465 | -0.316 | 0.983 | -1.857 | -0.015 | -0.058 | -0.058 | 0.094 | 0.21 | 0.35 | 0.16 | 0.05 | 0.14 | 0.14 |
| 3 | 10124120 | rs6789156 | -0.88 | -0.333 | 1.01 | -1.851 | -0.027 | -0.057 | -0.057 | 0 | 0.23 | 0.5 | 0.19 | 0.07 | 0.14 | 0.21 |
| 3 | 10126312 | rs13319597 | -0.47 | -0.329 | 1.012 | -1.849 | -0.016 | -0.059 | -0.059 | 0 | 0.21 | 0.47 | 0.17 | 0.05 | 0.14 | 0.14 |
| 3 | 10133949 | rs9811771 | -0.869 | -0.236 | 1.142 | -1.849 | -0.02 | -0.042 | -0.042 | 0 | 0.23 | 0.54 | 0.2 | 0.07 | 0.14 | 0.21 |
| 3 | 10135577 | rs1552244 | -0.961 | -0.244 | 1.142 | -1.853 | -0.018 | -0.038 | -0.038 | 0.089 | 0.24 | 0.55 | 0.2 | 0.07 | 0.14 | 0.21 |
| 3 | 10138189 | rs2272124 | -0.486 | -0.224 | 0.652 | -1.844 | -0.014 | -0.04 | -0.04 | 3.132 | 0.21 | 0.52 | 0.2 | 0.07 | 0.14 | 0.19 |
| 3 | 10140115 | rs2272123 | -0.423 | -0.224 | 0.632 | -1.844 | -0.014 | -0.071 | -0.071 | 0.054 | 0.21 | 0.4 | 0.17 | 0.05 | 0.14 | 0.13 |
| 3 | 10140696 | rs7647987 | -0.469 | -0.219 | 0.609 | -1.845 | -0.014 | -0.041 | -0.041 | 0 | 0.21 | 0.52 | 0.2 | 0.07 | 0.14 | 0.18 |
| 3 | 10142452 | rs6792811 | -0.042 | -0.02 | -0.161 | -1.988 | -0.124 | 0.03 | 0.03 | 4.831 | 0.67 | 0.96 | 0.48 | 0.22 | 0.61 | 0.46 |
| 3 | 10142949 | rs3172417 | 1.797 | 0.163 | 0.001 | -1.811 | -0.138 | 0.168 | 0.168 | 6.97 | 0.37 | 0.04 | 0.25 | 0.17 | 0.45 | 0.24 |
| 3 | 10143770 | rs12152512 | -1.1 | -0.258 | 1.209 | -1.271 | -0.015 | -0.042 | -0.042 | 0 | 0.21 | 0.52 | 0.2 | 0.07 | 0.14 | 0.19 |
| 3 | 10167112 | rs17032426 | -0.684 | -0.192 | 0.617 | -1.051 | 0.005 | -0.075 | -0.075 | 0.085 | 0.13 | 0.34 | 0.15 | 0.02 | 0.1 | 0.16 |
| 3 | 10168647 | rs6785976 | -0.684 | -0.192 | 0.603 | -1.051 | 0.005 | -0.075 | -0.075 | 0.006 | 0.13 | 0.34 | 0.15 | 0.02 | 0.1 | 0.16 |
| 3 | 10168785 | rs6442154 | -0.684 | -0.192 | 0.603 | -1.051 | 0.005 | -0.075 | -0.075 | 0 | 0.13 | 0.34 | 0.15 | 0.02 | 0.1 | 0.16 |
| 3 | 10169819 | rs17032440 | -0.684 | -0.192 | 0.603 | -1.051 | 0.005 | -0.075 | -0.075 | 0.631 | 0.13 | 0.34 | 0.15 | 0.02 | 0.1 | 0.16 |
| 3 | 10182562 | rs779803 | 1.106 | -0.043 | -0.442 | -0.98 | 0.022 | 0.059 | 0.059 | 0.007 | 0.6 | 0.27 | 0.61 | 0.82 | 0.7 | 0.58 |
| 3 | 10183337 | rs779805 | 1.113 | -0.044 | -1.06 | -0.981 | 0.028 | 0.125 | 0.125 | 0.029 | 0.6 | 0.16 | 0.6 | 0.82 | 0.7 | 0.58 |
| 3 | 10191943 | rs1642742 | 1.135 | -0.002 | -1.011 | -0.953 | 0.031 | 0.091 | 0.091 | 0 | 0.59 | 0.21 | 0.61 | 0.82 | 0.7 | 0.58 |
| 3 | 10195485 | rs1642739 | 0.035 | 0.004 | -1.075 | -0.96 | 0.007 | 0.044 | 0.044 | 0.11 | 0.83 | 0.95 | 0.83 | 0.89 | 0.87 | 0.83 |
| 3 | 10202884 | rs457414 | 0.633 | 0.277 | -0.917 | -0.737 | 0.031 | 0.083 | 0.083 | 0.007 | 0.59 | 0.22 | 0.61 | 0.81 | 0.69 | 0.59 |
| 3 | 10211918 | rs3844278 | -1.117 | 0.817 | -0.159 | -0.627 | 0.042 | -0.051 | -0.051 | 0 | 0.09 | 0.28 | 0.33 | 0.01 | 0.14 | 0.1 |
| 3 | 10212258 | rs3844280 | -1.275 | 0.766 | 0.022 | -0.65 | 0.047 | -0.054 | -0.054 | 0.148 | 0.09 | 0.31 | 0.34 | 0.01 | 0.14 | 0.1 |
| 3 | 10212494 | rs708030 | -0.353 | 0.23 | -0.478 | -1.668 | 0.018 | 0.14 | 0.14 | 1.536 | 0.52 | 0.71 | 0.56 | 0.11 | 0.48 | 0.4 |
| 3 | 10215950 | rs2619508 | 0.335 | 0.085 | -0.784 | -2.099 | 0.009 | 0.083 | 0.083 | 2.216 | 0.49 | 0.31 | 0.29 | 0.11 | 0.56 | 0.27 |
| 3 | 10219508 | rs2306696 | -0.252 | 0.368 | -1.205 | -2.119 | -0.039 | 0.046 | 0.046 | 0 | 0.17 | 0.08 | 0.35 | 0.1 | 0.21 | 0.22 |
| 3 | 10220830 | rs17610830 | -0.504 | 0.101 | -0.808 | -2.107 | -0.015 | 0.083 | 0.083 | 1.372 | 0.41 | 0.23 | 0.45 | 0.2 | 0.41 | 0.43 |
| 3 | 10226144 | rs1177614 | -0.73 | 0.103 | -0.459 | -1.774 | -0.001 | 0 | 0 | 0 | 0.73 | 0.69 | 0.7 | 0.24 | 0.77 | 0.6 |
| 3 | 10234065 | rs1144911 | 1.433 | -0.605 | -0.533 | -2.034 | -0.008 | -0.004 | -0.004 | 0 | 0.26 | 0.15 | 0.22 | 0 | 0.37 | 0.09 |
| 3 | 10237297 | rs263408 | 0.451 | -0.664 | -0.828 | -1.88 | -0.014 | 0.04 | 0.04 | 0 | 0.88 | 0.99 | 0.94 | 0.93 | 0.92 | 0.86 |
| 3 | 10238124 | rs696356 | 0.886 | -0.627 | -0.379 | -1.667 | -0.022 | 0.024 | 0.024 | 0.07 | 0.21 | 0.33 | 0.22 | 0.73 | 0.14 | 0.32 |
| 3 | 10241987 | rs263410 | -1.042 | -0.634 | -0.311 | -1.099 | -0.022 | 0.061 | 0.061 | 0.05 | 0.79 | 0.67 | 0.78 | 0.35 | 0.85 | 0.63 |
| 3 | 10242243 | rs263411 | -0.845 | -0.701 | -0.532 | -1.483 | 0 | 0.008 | 0.008 | 0 | 0.6 | 0.62 | 0.59 | 0.27 | 0.54 | 0.5 |
| 3 | 10242671 | rs263412 | -0.704 | -0.585 | -0.999 | -1.302 | -0.015 | -0.053 | -0.053 | 0 | 0.11 | 0.01 | 0.09 | 0.17 | 0.1 | 0.14 |
| 3 | 10242711 | rs263413 | -0.443 | -0.933 | -0.746 | -1.27 | -0.007 | 0.014 | 0.014 | 0 | 0.43 | 0.35 | 0.54 | 0.29 | 0.45 | 0.43 |
| 3 | 10245029 | rs380545 | -0.052 | -0.876 | -1.059 | -0.825 | -0.005 | -0.061 | -0.061 | 2.338 | 0.11 | 0.01 | 0.05 | 0.2 | 0.09 | 0.15 |
| 3 | 10249720 | rs708033 | -1.157 | -1.265 | -0.723 | -0.952 | -0.01 | 0.033 | 0.033 | 0.239 | 0.8 | 0.75 | 0.76 | 0.44 | 0.85 | 0.62 |
| 3 | 10249984 | rs7373858 | 0.433 | -1.386 | -0.373 | -1.349 | -0.014 | 0.02 | 0.02 | 0.041 | 0.34 | 0.24 | 0.45 | 0.13 | 0.39 | 0.28 |
| 3 | 10250783 | rs11465886 | -0.228 | -1.516 | -0.252 | -1.376 | -0.027 | -0.001 | -0.001 | 0 | 0.08 | 0.25 | 0.05 | 0.08 | 0.02 | 0.07 |
| 3 | 10254797 | rs11465897 | 1.525 | -1.472 | -0.326 | -1.568 | -0.009 | 0.006 | 0.006 | 0 | 0.28 | 0.23 | 0.2 | 0.01 | 0.35 | 0.12 |
| 3 | 10258493 | rs2302862 | 0.25 | -1.678 | -0.385 | -1.332 | 0.003 | 0.002 | 0.002 | 0 | 0.64 | 0.65 | 0.67 | 0.14 | 0.74 | 0.4 |
| 3 | 10264299 | rs813591 | 0.262 | -1.727 | -0.357 | -1.219 | -0.03 | 0.035 | 0.035 | 0.086 | 0.24 | 0.33 | 0.27 | 0.64 | 0.15 | 0.46 |
| 3 | 10272775 | rs4684672 | 0.229 | -1.606 | -0.472 | -1.488 | -0.048 | 0.11 | 0.11 | 0 | 0.39 | 0.22 | 0.49 | 0.06 | 0.45 | 0.3 |
| 3 | 10275475 | rs776514 | -1.024 | -1.856 | -0.265 | -1.49 | -0.002 | -0.02 | -0.02 | 0 | 0.6 | 0.55 | 0.74 | 0.76 | 0.56 | 0.74 |
| 3 | 10277866 | rs11706450 | -0.411 | -1.731 | -0.544 | -1.436 | 0.042 | 0.136 | 0.136 | 0 | 0.51 | 0.23 | 0.54 | 0.28 | 0.54 | 0.43 |
| 3 | 10278980 | rs11711184 | -0.458 | -1.708 | -0.528 | -1.425 | 0.037 | 0.118 | 0.118 | 0 | 0.51 | 0.25 | 0.55 | 0.28 | 0.54 | 0.43 |
| 3 | 10285888 | rs242724 | -2.186 | -1.414 | -0.53 | -1.431 | -0.019 | 0.041 | 0.041 | 0 | 0.75 | 0.88 | 0.82 | 0.99 | 0.65 | 0.92 |
| 3 | 10292140 | rs2270454 | 0.242 | -1.565 | 0.155 | -0.92 | -0.033 | 0.023 | 0.023 | 0.038 | 0.21 | 0.29 | 0.24 | 0.66 | 0.1 | 0.53 |
| 3 | 10295883 | rs2241308 | -0.324 | -1.288 | -0.034 | -1.099 | -0.008 | 0.008 | 0.008 | 0 | 0.26 | 0.34 | 0.5 | 0.66 | 0.19 | 0.58 |
| 3 | 10302056 | rs2241314 | 0.278 | -1.385 | -0.187 | -1.089 | -0.029 | 0.014 | 0.014 | 0.05 | 0.21 | 0.28 | 0.23 | 0.65 | 0.1 | 0.53 |
| 3 | 10302172 | rs394558 | -1.209 | -1.225 | -0.608 | -1.532 | -0.004 | -0.029 | -0.029 | 0 | 0.57 | 0.37 | 0.42 | 0.96 | 0.43 | 0.81 |
| 3 | 10308326 | rs1642974 | -0.557 | -1.414 | -0.8 | -1.592 | 0.005 | 0.018 | 0.018 | 0.004 | 0.43 | 0.35 | 0.36 | 0.66 | 0.34 | 0.65 |
| 3 | 10311939 | rs2075352 | 0.101 | -1.56 | -0.35 | -1.593 | -0.007 | 0.062 | 0.062 | 0 | 0.13 | 0.26 | 0.08 | 0.28 | 0.1 | 0.12 |
| 3 | 10312507 | rs2075353 | 0.676 | -1.552 | -0.555 | -1.565 | -0.014 | -0.061 | -0.061 | 0.272 | 0.11 | 0.01 | 0.06 | 0.29 | 0.1 | 0.12 |
| 3 | 10315103 | rs164938 | -0.556 | -1.787 | -0.407 | -1.821 | 0.005 | 0.041 | 0.041 | 0 | 0.45 | 0.62 | 0.6 | 0.7 | 0.41 | 0.71 |
| 3 | 10320865 | rs2075355 | 0.633 | -1.917 | -0.561 | -1.975 | -0.001 | 0.002 | 0.002 | 0 | 0.61 | 0.63 | 0.78 | 0.7 | 0.63 | 0.69 |
| 3 | 10321734 | rs514658 | 0.322 | -1.919 | -0.564 | -1.978 | 0.006 | -0.097 | -0.097 | 6.054 | 0.28 | 0.36 | 0.16 | 0.03 | 0.27 | 0.19 |
| 3 | 10322521 | rs2072578 | 0.856 | -1.757 | -0.761 | -1.85 | -0.014 | -0.056 | -0.056 | 0.205 | 0.11 | 0.01 | 0.06 | 0.27 | 0.1 | 0.11 |
| 3 | 10325950 | rs11718213 | 0.856 | -1.757 | -0.761 | -1.85 | -0.014 | -0.056 | -0.056 | 0.019 | 0.11 | 0.01 | 0.06 | 0.27 | 0.1 | 0.11 |

| | | | | | | | | | | | | | | | |
|---|---|---|---|---|---|---|---|---|---|---|---|---|---|---|---|
| 3 | 10326169 | rs171407 | -2.35 | -1.773 | -0.777 | -2.147 | -0.027 | 0.061 | 0.061 | 0.05 | 0.66 | 0.81 | 0.74 | 0.94 | 0.55 | 0.83 |
| 3 | 10328250 | rs35683 | -1.659 | -1.238 | -0.749 | -1.979 | -0.005 | -0.012 | -0.012 | 3.002 | 0.59 | 0.63 | 0.67 | 0.97 | 0.55 | 0.84 |
| 3 | 10328782 | rs35682 | -1.608 | -1.365 | -0.744 | -1.947 | -0.004 | -0.025 | -0.025 | 0.323 | 0.58 | 0.62 | 0.67 | 0.97 | 0.54 | 0.84 |
| 3 | 10329377 | rs35681 | -1.504 | -1.296 | -0.79 | -1.815 | 0 | -0.061 | -0.061 | 0.095 | 0.57 | 0.53 | 0.66 | 0.95 | 0.54 | 0.78 |
| 3 | 10330377 | rs42451 | 0.09 | -0.793 | -0.888 | -1.055 | -0.006 | 0.015 | 0.015 | 0 | 0.28 | 0.21 | 0.17 | 0.14 | 0.29 | 0.19 |
| 3 | 10331457 | rs696217 | 0.503 | -0.575 | -0.987 | -0.614 | -0.016 | -0.033 | -0.033 | 0 | 0.07 | 0.01 | 0.05 | 0.19 | 0.09 | 0.09 |
| 3 | 10332365 | rs26802 | -1.045 | -0.548 | -0.879 | -1.188 | 0.013 | 0.051 | 0.051 | 0 | 0.43 | 0.32 | 0.3 | 0.09 | 0.34 | 0.38 |
| 3 | 10335145 | rs10490815 | 0.142 | -0.042 | -0.583 | -0.579 | 0 | 0.001 | 0.001 | 0 | 0.29 | 0.28 | 0.49 | 0.57 | 0.3 | 0.49 |
| 3 | 10335786 | rs2619507 | -0.348 | -0.439 | -0.963 | -0.669 | -0.004 | 0.02 | 0.02 | 0 | 0.22 | 0.22 | 0.16 | 0.44 | 0.17 | 0.16 |
| 3 | 10341082 | rs12630739 | 1.105 | -0.24 | -0.69 | -0.794 | -0.001 | 0 | 0 | 0.156 | 0.22 | 0.19 | 0.44 | 0.53 | 0.25 | 0.47 |
| 3 | 10342262 | rs149162 | -2.631 | -0.384 | -0.98 | -1.203 | -0.004 | 0.027 | 0.027 | 0 | 0.71 | 0.8 | 0.81 | 0.98 | 0.64 | 0.85 |
| 3 | 10342876 | rs696221 | -1.8 | -0.309 | -1.084 | -1.295 | -0.006 | 0.117 | 0.117 | 0.308 | 0.6 | 0.79 | 0.7 | 0.98 | 0.55 | 0.81 |
| 3 | 10344111 | rs696220 | -0.558 | -0.154 | -0.686 | -1.11 | -0.003 | -0.007 | -0.007 | 0 | 0.39 | 0.37 | 0.56 | 0.93 | 0.39 | 0.6 |
| 3 | 10345921 | rs40515 | 0.009 | -0.21 | -0.699 | -1.122 | -0.002 | -0.072 | -0.072 | 0 | 0.28 | 0.28 | 0.22 | 0.01 | 0.23 | 0.17 |
| 3 | 10346421 | rs697231 | 1.012 | -0.149 | -0.877 | -0.914 | -0.012 | -0.068 | -0.068 | 0.624 | 0.13 | 0.04 | 0.09 | 0.39 | 0.12 | 0.12 |
| 3 | 10351830 | rs35668 | -1.736 | -0.205 | -0.906 | -0.978 | -0.061 | -0.055 | -0.055 | 0 | 0.61 | 0.89 | 0.74 | 0.58 | 0.52 | 0.73 |
| 3 | 10352703 | rs3952701 | 1.23 | -0.129 | -0.826 | -0.837 | -0.006 | -0.053 | -0.053 | 0.019 | 0.22 | 0.14 | 0.43 | 0.53 | 0.25 | 0.47 |
| 3 | 10356314 | rs154236 | -1.836 | -0.035 | -0.362 | -0.566 | -0.007 | 0.058 | 0.058 | 0.055 | 0.7 | 0.79 | 0.8 | 0.95 | 0.64 | 0.84 |
| 3 | 10368102 | rs41147 | -0.875 | 0.059 | -0.832 | -0.33 | -0.021 | 0.006 | 0.006 | 0 | 0.57 | 0.71 | 0.7 | 0.6 | 0.49 | 0.71 |
| 3 | 10369180 | rs1719571 | -0.689 | 0.278 | -0.33 | 0.397 | 0.011 | 0.072 | 0.072 | 0 | 0.18 | 0.29 | 0.13 | 0.35 | 0.15 | 0.14 |
| 3 | 10372840 | rs26310 | -0.988 | -0.008 | -0.656 | 0.638 | -0.014 | 0.023 | 0.023 | 0 | 0.35 | 0.49 | 0.5 | 0.56 | 0.33 | 0.4 |
| 3 | 10382638 | rs709641 | -0.316 | 0.193 | -1.623 | 0.557 | -0.053 | -0.048 | -0.048 | 0.405 | 0.52 | 0.15 | 0.61 | 0.5 | 0.46 | 0.6 |
| 3 | 10383517 | rs171406 | -0.303 | 0.115 | -1.393 | 0.4 | -0.003 | -0.009 | -0.009 | 1.301 | 0.49 | 0.24 | 0.59 | 0.45 | 0.44 | 0.58 |
| 3 | 10383996 | rs241508 | -0.407 | 0.113 | -1.286 | 0.458 | 0.009 | 0.02 | 0.02 | 0 | 0.51 | 0.38 | 0.58 | 0.52 | 0.42 | 0.67 |
| 3 | 10390603 | rs703913 | -0.137 | 0.421 | -1.573 | 0.718 | -0.046 | -0.089 | -0.089 | 0 | 0.12 | 0.01 | 0.09 | 0.37 | 0.16 | 0.21 |
| 3 | 10400643 | rs28113 | -1.129 | 0.234 | -1.338 | 0.13 | -0.004 | -0.019 | -0.019 | 0 | 0.62 | 0.41 | 0.69 | 0.95 | 0.58 | 0.85 |
| 3 | 10400959 | rs187675 | -0.37 | 0.209 | -1.23 | 0.452 | 0.002 | 0.054 | 0.054 | 0 | 0.15 | 0.23 | 0.12 | 0.41 | 0.16 | 0.22 |
| 3 | 10402103 | rs2278556 | 0.109 | 0.089 | -1.585 | 0.453 | -0.004 | -0.032 | -0.032 | 0.151 | 0.42 | 0.16 | 0.55 | 0.52 | 0.38 | 0.54 |
| 3 | 10404739 | rs775022 | -0.413 | 0.049 | -1.847 | 0.62 | 0.01 | -0.027 | -0.027 | 0.131 | 0.46 | 0.17 | 0.51 | 0.57 | 0.42 | 0.58 |
| 3 | 10413601 | rs2289273 | 0.334 | 0.085 | -1.786 | 0.654 | -0.039 | -0.081 | -0.081 | 0.005 | 0.12 | 0.01 | 0.04 | 0.32 | 0.14 | 0.2 |
| 3 | 10413715 | rs2289274 | -0.227 | -0.158 | -1.878 | 0.369 | -0.004 | -0.01 | -0.01 | 0.041 | 0.29 | 0.13 | 0.33 | 0.33 | 0.3 | 0.36 |
| 3 | 10416225 | rs3774170 | 0.771 | -0.05 | -1.873 | 0.558 | -0.034 | -0.072 | -0.072 | 0 | 0.19 | 0.02 | 0.21 | 0.25 | 0.17 | 0.24 |
| 3 | 10416287 | rs3774169 | 0.362 | -0.008 | -1.713 | 0.576 | -0.024 | -0.011 | -0.011 | 0 | 0.07 | 0.01 | 0.11 | 0.31 | 0.07 | 0.17 |
| 3 | 10417929 | rs741495 | -0.983 | -0.262 | -1.694 | 0.119 | 0.001 | 0.045 | 0.045 | 0 | 0.44 | 0.2 | 0.31 | 0.7 | 0.4 | 0.61 |
| 3 | 10419173 | rs3774165 | 0.606 | -0.24 | -1.494 | 0.245 | 0.003 | -0.054 | -0.054 | 0.566 | 0.27 | 0.09 | 0.22 | 0.39 | 0.22 | 0.35 |
| 3 | 10420087 | rs751122 | -0.982 | -0.358 | -1.177 | 0.176 | 0.004 | 0.009 | 0.009 | 0 | 0.46 | 0.44 | 0.32 | 0.7 | 0.37 | 0.6 |
| 3 | 10420875 | rs4684686 | -0.068 | -0.507 | -0.3 | 0.11 | -0.008 | -0.002 | -0.002 | 2.161 | 0.33 | 0.34 | 0.33 | 0.65 | 0.27 | 0.46 |
| 3 | 10427190 | rs725027 | -0.988 | -0.417 | -0.584 | -0.05 | 0.021 | 0.165 | 0.165 | 0 | 0.47 | 0.71 | 0.45 | 0.97 | 0.4 | 0.68 |
| 3 | 10427434 | rs2075315 | -0.244 | -0.383 | -0.556 | 0.153 | 0.087 | 0.475 | 0.475 | 0 | 0.07 | 0.38 | 0.15 | 0.28 | 0.05 | 0.15 |
| 3 | 10430862 | rs6442169 | -2.551 | -0.204 | -1.389 | -0.279 | -0.106 | 0.16 | 0.16 | 0 | 0.77 | 0.98 | 0.82 | 0.99 | 0.62 | 0.92 |
| 3 | 10434224 | rs3774161 | 0.814 | -0.166 | -1.141 | 0.058 | -0.034 | -0.058 | -0.058 | 0 | 0.09 | 0 | 0.06 | 0.31 | 0.12 | 0.2 |
| 3 | 10434941 | rs1317122 | -0.118 | -0.291 | -1.121 | -0.084 | -0.001 | 0.012 | 0.012 | 0 | 0.36 | 0.23 | 0.29 | 0.72 | 0.34 | 0.51 |
| 3 | 10437280 | rs1438168 | -0.229 | -0.263 | -1.262 | -0.034 | 0.012 | -0.038 | -0.038 | 0.091 | 0.35 | 0.19 | 0.24 | 0.43 | 0.28 | 0.37 |
| 3 | 10442439 | rs9647359 | 0.827 | -0.225 | -1.237 | -0.104 | 0 | -0.015 | -0.015 | 0.193 | 0.06 | 0.06 | 0.03 | 0 | 0.05 | 0.05 |
| 3 | 10443090 | rs11710165 | 0.77 | -0.314 | -1.513 | -0.091 | -0.029 | -0.117 | -0.117 | 0 | 0.25 | 0.04 | 0.21 | 0.4 | 0.23 | 0.3 |
| 3 | 10446826 | rs2160871 | 0.927 | -0.449 | -1.696 | -0.459 | -0.01 | 0.092 | 0.092 | 0 | 0.22 | 0.1 | 0.31 | 0.01 | 0.17 | 0.17 |
| 3 | 10446960 | rs929708 | -0.557 | -0.467 | -1.483 | -0.389 | 0.002 | 0.025 | 0.025 | 0 | 0.37 | 0.26 | 0.41 | 0.3 | 0.33 | 0.38 |
| 3 | 10447146 | rs3774155 | 1.18 | -0.43 | -1.454 | -0.292 | -0.015 | -0.07 | -0.07 | 0.054 | 0.23 | 0.08 | 0.2 | 0.38 | 0.22 | 0.3 |
| 3 | 10447277 | rs1370712 | -0.696 | -0.588 | -1.234 | -0.487 | -0.019 | -0.044 | -0.044 | 0 | 0.12 | 0.57 | 0.1 | 0.02 | 0.06 | 0.05 |
| 3 | 10448426 | rs4684689 | -1.868 | -0.445 | -1.367 | -0.551 | -0.077 | -0.005 | -0.005 | 0 | 0.68 | 0.88 | 0.73 | 0.69 | 0.53 | 0.78 |
| 3 | 10448772 | rs6786239 | 1.919 | -0.459 | -1.433 | -0.655 | 0 | 0.073 | 0.073 | 0 | 0.15 | 0.05 | 0.25 | 0.01 | 0.12 | 0.15 |
| 3 | 10449459 | rs34903 | -0.659 | -0.342 | -1.167 | -0.582 | -0.023 | 0.069 | 0.069 | 0 | 0.53 | 0.25 | 0.63 | 0.67 | 0.43 | 0.66 |
| 3 | 10452231 | rs34907 | -0.782 | -0.299 | -1.042 | -0.668 | -0.002 | 0.008 | 0.008 | 0 | 0.43 | 0.37 | 0.42 | 0.96 | 0.4 | 0.66 |
| 3 | 10453218 | rs34909 | 0.565 | -0.261 | -1.075 | -0.385 | -0.006 | -0.019 | -0.019 | 0 | 0.29 | 0.16 | 0.32 | 0.63 | 0.28 | 0.44 |
| 3 | 10457572 | rs11719939 | -0.1 | -0.393 | -0.996 | -0.717 | 0.003 | -0.013 | -0.013 | 0.789 | 0.31 | 0.31 | 0.37 | 0.01 | 0.26 | 0.26 |
| 3 | 10459934 | rs7649656 | -0.498 | -0.481 | -0.638 | -0.747 | -0.008 | -0.077 | -0.077 | 0 | 0.36 | 0.56 | 0.42 | 0.03 | 0.23 | 0.23 |
| 3 | 10460174 | rs34918 | 0.981 | -0.312 | -0.811 | -0.453 | -0.046 | -0.088 | -0.088 | 0.271 | 0.25 | 0.03 | 0.17 | 0.39 | 0.24 | 0.31 |
| 3 | 10461163 | rs3774146 | -0.308 | -0.236 | -0.512 | -0.411 | 0.014 | -0.041 | -0.041 | 0 | 0.11 | 0.53 | 0.1 | 0.01 | 0.06 | 0.05 |
| 3 | 10463569 | rs34921 | -1.036 | -0.154 | -0.887 | -0.283 | -0.049 | 0.026 | 0.026 | 0 | 0.62 | 0.81 | 0.72 | 0.69 | 0.5 | 0.72 |
| 3 | 10468280 | rs2470450 | -0.719 | -0.209 | -0.385 | -0.222 | 0.002 | -0.002 | -0.002 | 0 | 0.57 | 0.52 | 0.53 | 0.4 | 0.45 | 0.59 |
| 3 | 10468723 | rs495435 | -1.663 | -0.174 | -0.692 | -0.203 | -0.005 | 0.002 | 0.002 | 0 | 0.68 | 0.71 | 0.72 | 0.71 | 0.55 | 0.81 |
| 3 | 10470828 | rs523180 | 0.697 | -0.219 | -1.413 | -0.914 | -0.08 | -0.085 | -0.085 | 0.004 | 0.28 | 0.04 | 0.24 | 0.68 | 0.32 | 0.46 |
| 3 | 10472916 | rs4684693 | -0.217 | -0.309 | -0.829 | -1.135 | 0.004 | 0.099 | 0.099 | 0 | 0.29 | 0.14 | 0.34 | 0 | 0.23 | 0.25 |
| 3 | 10474137 | rs719318 | -1.107 | -0.416 | -0.224 | -0.829 | -0.052 | -0.117 | -0.117 | 0.417 | 0.19 | 0.51 | 0.14 | 0.02 | 0.07 | 0.1 |
| 3 | 10475438 | rs34856 | -0.295 | 0.49 | -0.682 | -0.493 | 0.005 | -0.023 | -0.023 | 0 | 0.93 | 0.94 | 0.96 | 1 | 0.93 | 0.96 |
| 3 | 10479259 | rs34859 | 0.264 | 0.62 | -1.319 | -0.303 | -0.084 | -0.148 | -0.148 | 0 | 0.26 | 0.02 | 0.2 | 0.55 | 0.27 | 0.41 |
| 3 | 10479545 | rs42307 | 0.783 | 0.433 | -0.315 | -0.173 | 0.009 | 0.092 | 0.092 | 0 | 0.83 | 0.72 | 0.79 | 0.66 | 0.86 | 0.71 |
| 3 | 10483958 | rs39667 | 0.826 | 0.34 | -0.227 | -0.25 | 0 | 0.002 | 0.002 | 0 | 0.81 | 0.79 | 0.75 | 0.73 | 0.84 | 0.77 |
| 3 | 10484726 | rs17603886 | 0.888 | 0.329 | -0.276 | -0.193 | -0.003 | 0.003 | 0.003 | 0 | 0.06 | 0.02 | 0.04 | 0.02 | 0.04 | 0.01 |

| | | | | | | | | | | | | | | | | |
|---|---|---|---|---|---|---|---|---|---|---|---|---|---|---|---|---|
| 3 | 10488362 | rs34866 | 2.024 | 0.014 | 0.798 | -0.399 | 0.004 | -0.008 | -0.008 | 0.074 | 0.73 | 0.29 | 0.68 | 0.7 | 0.81 | 0.67 |
| 3 | 10488638 | rs682575 | 0.918 | -0.012 | 1.438 | -0.398 | 0.015 | 0.086 | 0.086 | 0.398 | 0.85 | 0.63 | 0.76 | 0.74 | 0.88 | 0.8 |
| 3 | 10492652 | rs40509 | 1.863 | 0.089 | 0.821 | -0.454 | 0.002 | -0.005 | -0.005 | 0.411 | 0.73 | 0.2 | 0.67 | 0.71 | 0.8 | 0.72 |
| 3 | 10493434 | rs714737 | 0.846 | 0.03 | 0.14 | -0.843 | 0.003 | 0.013 | 0.013 | 0 | 0.17 | 0.13 | 0.28 | 0.11 | 0.17 | 0.17 |
| 3 | 10498165 | rs520006 | 0.298 | 0.061 | 0.605 | -0.777 | 0.055 | 0.104 | 0.104 | 0 | 0.87 | 0.55 | 0.81 | 0.77 | 0.88 | 0.87 |
| 3 | 10501498 | rs34870 | 1.801 | 0.212 | -0.26 | -1.018 | -0.105 | -0.004 | -0.004 | 0 | 0.49 | 0.08 | 0.34 | 0.57 | 0.53 | 0.45 |
| 3 | 10505649 | rs1009826 | 0.29 | 0.424 | 0.312 | -1.315 | -0.004 | -0.059 | -0.059 | 0 | 0.23 | 0.4 | 0.32 | 0.1 | 0.23 | 0.19 |
| 3 | 10508130 | rs1009825 | 2.456 | 0.569 | -0.212 | -1.56 | -0.071 | 0.128 | 0.128 | 0 | 0.23 | 0.06 | 0.18 | 0.02 | 0.36 | 0.08 |
| 3 | 10508371 | rs34871 | 1.48 | 0.499 | -0.064 | -1.545 | 0.003 | -0.008 | -0.008 | 0.254 | 0.69 | 0.46 | 0.65 | 0.71 | 0.75 | 0.69 |
| 3 | 10508495 | rs34872 | -0.747 | 0.629 | -0.01 | -1.529 | -0.021 | 0.06 | 0.06 | 0 | 0.59 | 0.75 | 0.59 | 0.74 | 0.52 | 0.72 |
| 3 | 10510287 | rs17032939 | 2.03 | 0.616 | 0.182 | -1.669 | 0.006 | -0.086 | -0.086 | 0 | 0.25 | 0.33 | 0.2 | 0.02 | 0.36 | 0.08 |
| 3 | 10510618 | rs7644022 | 0.64 | 0.573 | 0.086 | -1.702 | -0.003 | 0.065 | 0.065 | 0 | 0.37 | 0.73 | 0.32 | 0.08 | 0.43 | 0.24 |
| 3 | 10511602 | rs11928209 | -0.236 | 0.79 | -0.13 | -1.751 | -0.009 | 0.026 | 0.026 | 0 | 0.32 | 0.24 | 0.41 | 0.14 | 0.35 | 0.29 |
| 3 | 10512428 | rs17032943 | 2.423 | 0.828 | -0.415 | -1.87 | 0.001 | -0.043 | -0.043 | 0 | 0.2 | 0.25 | 0.16 | 0.01 | 0.28 | 0.06 |
| 3 | 10512455 | rs602709 | 0.393 | 0.386 | -0.116 | -1.621 | -0.001 | -0.011 | -0.011 | 0.053 | 0.64 | 0.41 | 0.54 | 0.83 | 0.65 | 0.68 |
| 3 | 10514660 | rs34878 | 0.41 | 0.17 | -0.158 | -1.465 | 0.044 | 0.231 | 0.231 | 0 | 0.51 | 0.78 | 0.51 | 0.05 | 0.61 | 0.31 |
| 3 | 10517836 | rs13064158 | 0.729 | 0.032 | -0.402 | -1.01 | 0.029 | -0.066 | -0.066 | 0 | 0.3 | 0.1 | 0.19 | 0.48 | 0.2 | 0.43 |
| 3 | 10519680 | rs2111751 | -0.796 | 0.219 | -0.503 | -1.107 | 0.01 | -0.052 | -0.052 | 0.762 | 0.5 | 0.44 | 0.53 | 0.99 | 0.38 | 0.74 |
| 3 | 10521905 | rs7623572 | -0.258 | 0.169 | -0.59 | -0.895 | 0 | -0.068 | -0.068 | 0 | 0.14 | 0.03 | 0.25 | 0.4 | 0.13 | 0.17 |
| 3 | 10522678 | rs34881 | 0.888 | 0.235 | -0.562 | -1.056 | -0.006 | -0.027 | -0.027 | 0 | 0.44 | 0.36 | 0.43 | 0.01 | 0.55 | 0.21 |
| 3 | 10524037 | rs34888 | 0.117 | 0.424 | -0.36 | -0.664 | 0.001 | -0.004 | -0.004 | 0.917 | 0.6 | 0.38 | 0.67 | 0.47 | 0.68 | 0.43 |
| 3 | 10525057 | rs34890 | -0.132 | 0.324 | -0.477 | -0.91 | 0.006 | -0.001 | -0.001 | 0 | 0.62 | 0.66 | 0.74 | 0.41 | 0.67 | 0.47 |
| 3 | 10525096 | rs735931 | 0.297 | 0.49 | -1.379 | -1.202 | -0.065 | 0.024 | 0.024 | 0.075 | 0.47 | 0.87 | 0.52 | 0.41 | 0.5 | 0.36 |
| 3 | 10525187 | rs34891 | -0.763 | 0.348 | -0.886 | -0.813 | 0.001 | -0.004 | -0.004 | 0 | 0.33 | 0.24 | 0.49 | 0.36 | 0.31 | 0.3 |
| 3 | 10525228 | rs39670 | -0.143 | 0.523 | -0.649 | -0.818 | -0.001 | -0.002 | -0.002 | 0.362 | 0.23 | 0.21 | 0.34 | 0.16 | 0.22 | 0.21 |
| 3 | 10525646 | rs34892 | 0.344 | 0.503 | -0.59 | -0.507 | -0.001 | -0.045 | -0.045 | 0.615 | 0.92 | 0.77 | 0.95 | 1 | 0.93 | 0.95 |
| 3 | 10526050 | rs1009965 | -0.29 | 0.749 | -1.156 | -0.277 | -0.019 | -0.009 | -0.009 | 0.392 | 0.05 | 0.01 | 0.07 | 0.05 | 0.07 | 0.07 |
| 3 | 10526438 | rs28280 | 0.12 | 0.524 | -1.034 | -0.442 | -0.01 | -0.019 | -0.019 | 0 | 0.59 | 0.59 | 0.6 | 0.28 | 0.68 | 0.5 |
| 3 | 10527775 | rs7619994 | 0.167 | 1.063 | -1.835 | -1.203 | -0.091 | 0.158 | 0.158 | 0 | 0.35 | 0.14 | 0.25 | 0.02 | 0.5 | 0.16 |
| 3 | 10527891 | rs741496 | 0.218 | 0.19 | -1.601 | -1.167 | -0.005 | -0.013 | -0.013 | 0 | 0.6 | 0.39 | 0.56 | 0.23 | 0.68 | 0.46 |
| 3 | 10528910 | rs808933 | -1.507 | 0.285 | -2.114 | -1.009 | -0.136 | 0.182 | 0.182 | 0.079 | 0.58 | 0.98 | 0.65 | 0.84 | 0.53 | 0.76 |
| 3 | 10529067 | rs17032981 | -0.192 | -0.152 | -1.567 | -1.235 | 0.002 | -0.021 | -0.021 | 2.511 | 0.17 | 0.17 | 0.07 | 0 | 0.1 | 0.11 |
| 3 | 10529069 | rs17032984 | -0.194 | 0.517 | -0.554 | -0.51 | 0.014 | 0.003 | 0.003 | 0 | 0.11 | 0.21 | 0.06 | 0.11 | 0.1 | 0.14 |
| 3 | 10532556 | rs2430890 | -0.089 | 0.568 | -0.54 | 0.552 | 0.003 | 0.029 | 0.029 | 0 | 0.17 | 0.25 | 0.32 | 0.27 | 0.18 | 0.39 |
| 3 | 10535771 | rs6807064 | -1.11 | 0.375 | -0.19 | 0.28 | -0.023 | -0.039 | -0.039 | 2.428 | 0.34 | 0.52 | 0.4 | 0.23 | 0.29 | 0.34 |
| 3 | 10536005 | rs11719906 | -0.875 | 0.491 | -0.108 | 0.287 | -0.013 | 0.011 | 0.011 | 0 | 0.32 | 0.28 | 0.39 | 0.24 | 0.41 | 0.34 |
| 3 | 10539482 | rs9873870 | -1.3 | 0.727 | 0.144 | 1.25 | 0.004 | 0.001 | 0.001 | 0 | 0.32 | 0.62 | 0.41 | 0.42 | 0.36 | 0.35 |
| 3 | 10553635 | rs7629412 | -1.173 | 0.591 | 0.545 | 2.193 | 0.084 | 0.44 | 0.44 | 0 | 0.06 | 0.4 | 0.13 | 0.28 | 0.03 | 0.03 |
| 3 | 10553949 | rs7629702 | -0.594 | 0.544 | 0.641 | 2.055 | -0.026 | -0.027 | -0.027 | 0 | 0.05 | 0.1 | 0.03 | 0 | 0.02 | 0.01 |
| 3 | 10554558 | rs4684704 | -1.286 | 1.031 | 0.432 | 2.346 | -0.011 | -0.009 | -0.009 | 0 | 0.15 | 0.14 | 0.24 | 0.33 | 0.28 | 0.41 |
| 3 | 10556104 | rs882527 | -1.362 | 0.997 | 0.396 | 2.423 | -0.023 | -0.02 | -0.02 | 0 | 0.15 | 0.13 | 0.24 | 0.35 | 0.27 | 0.41 |
| 3 | 10560942 | rs9875875 | -2.038 | 0.108 | 0.423 | 2.4 | -0.016 | 0.031 | 0.031 | 0 | 0.13 | 0.16 | 0.14 | 0.29 | 0.08 | 0.11 |
| 3 | 10566940 | rs4468954 | -1.65 | 0.243 | 0.413 | 2.45 | -0.025 | 0.07 | 0.07 | 0.086 | 0.08 | 0.2 | 0.04 | 0.14 | 0.03 | 0.11 |
| 3 | 10573967 | rs6442177 | -1.766 | 0.352 | 0.009 | 2.05 | -0.011 | 0.03 | 0.03 | 0 | 0.36 | 0.28 | 0.36 | 0.22 | 0.39 | 0.48 |
| 3 | 10574672 | rs4501094 | -1.909 | 0.551 | -0.005 | 2.301 | -0.007 | -0.002 | -0.002 | 0.189 | 0.17 | 0.17 | 0.33 | 0.49 | 0.25 | 0.44 |
| 3 | 10576987 | rs12714854 | -2.528 | 0.251 | 0.211 | 2.209 | -0.046 | 0.041 | 0.041 | 0 | 0.21 | 0.39 | 0.24 | 0.58 | 0.14 | 0.24 |
| 3 | 10579834 | rs9873122 | -2.656 | 0.242 | 0.207 | 2.127 | -0.051 | 0.113 | 0.113 | 0 | 0.22 | 0.4 | 0.23 | 0.42 | 0.13 | 0.21 |
| 3 | 10588369 | rs754354 | -2.272 | 0.455 | 0.175 | 1.989 | -0.003 | 0.003 | 0.003 | 0.66 | 0.42 | 0.4 | 0.55 | 0.61 | 0.5 | 0.6 |
| 3 | 10589024 | rs13316275 | -1.339 | 0.405 | 0.217 | 1.919 | -0.023 | -0.011 | -0.011 | 0.007 | 0.09 | 0.18 | 0.09 | 0.05 | 0.05 | 0.08 |
| 3 | 10590736 | rs4684044 | -2.476 | 0.427 | 0.198 | 2.055 | 0.005 | 0.005 | 0.005 | 0 | 0.44 | 0.53 | 0.54 | 0.52 | 0.47 | 0.6 |
| 3 | 10594207 | rs6810046 | -1.741 | 0.397 | 0.215 | 2.038 | -0.003 | -0.003 | -0.003 | 0.011 | 0.36 | 0.35 | 0.46 | 0.51 | 0.42 | 0.53 |
| 3 | 10594938 | rs754016 | -0.605 | 0.397 | 0.266 | 2.023 | -0.029 | -0.018 | -0.018 | 0 | 0.07 | 0.15 | 0.04 | 0.03 | 0.03 | 0.06 |
| 3 | 10595393 | rs750423 | -1.8 | 0.322 | 0.278 | 2.065 | -0.03 | -0.039 | -0.039 | 0 | 0.34 | 0.26 | 0.44 | 0.53 | 0.41 | 0.54 |
| 3 | 10599028 | rs4383496 | -2.918 | 0.229 | 0.284 | 1.957 | -0.003 | 0.029 | 0.029 | 0.064 | 0.52 | 0.6 | 0.59 | 0.76 | 0.51 | 0.68 |
| 3 | 10600232 | rs6806666 | 0.008 | 0.479 | 0.135 | 2.057 | -0.006 | -0.004 | -0.004 | 0 | 0.09 | 0.05 | 0.1 | 0.16 | 0.13 | 0.15 |
| 3 | 10600451 | rs11917518 | -0.947 | 0.031 | 0.102 | 1.89 | 0.002 | -0.003 | -0.003 | 0 | 0.19 | 0.17 | 0.18 | 0.27 | 0.14 | 0.1 |
| 3 | 10605733 | rs17509502 | -0.533 | 0.138 | 0.107 | 1.855 | -0.005 | 0.001 | 0.001 | 0.248 | 0.11 | 0.09 | 0.14 | 0.17 | 0.13 | 0.28 |
| 3 | 10608755 | rs4234502 | -2.285 | 0.049 | 0.123 | 1.82 | -0.006 | -0.024 | -0.024 | 0 | 0.38 | 0.43 | 0.38 | 0.22 | 0.3 | 0.47 |
| 3 | 10611407 | rs7629381 | -2.525 | 0.075 | 0.129 | 1.837 | -0.005 | 0.004 | 0.004 | 0 | 0.4 | 0.46 | 0.46 | 0.49 | 0.33 | 0.5 |
| 3 | 10611507 | rs7629483 | -0.989 | 0.477 | 0.186 | 2.057 | -0.008 | 0.003 | 0.003 | 0.579 | 0.11 | 0.09 | 0.21 | 0.17 | 0.17 | 0.31 |
| 3 | 10617048 | rs11914694 | -1.676 | 0.349 | 0.209 | 2.123 | 0.005 | 0.203 | 0.203 | 0.572 | 0.06 | 0.18 | 0.11 | 0.29 | 0.04 | 0.06 |
| 3 | 10621793 | rs11709049 | -2.446 | 0.247 | 0.144 | 1.141 | -0.001 | 0.049 | 0.049 | 0.116 | 0.52 | 0.65 | 0.6 | 0.88 | 0.49 | 0.7 |
| 3 | 10622168 | rs13059702 | -1.553 | 0.204 | 0.188 | 1.196 | 0.002 | 0.037 | 0.037 | 0 | 0.35 | 0.19 | 0.42 | 0.7 | 0.36 | 0.41 |
| 3 | 10622448 | rs9829797 | -2.662 | 0.184 | 0.195 | 1.18 | -0.004 | 0.064 | 0.064 | 0.145 | 0.52 | 0.68 | 0.61 | 0.91 | 0.5 | 0.74 |
| 3 | 10622941 | rs6803992 | -2.286 | 0.144 | 0.196 | 1.193 | 0.006 | 0.035 | 0.035 | 0.21 | 0.49 | 0.62 | 0.56 | 0.86 | 0.46 | 0.64 |
| 3 | 10623391 | rs4684717 | -2.087 | 0.175 | -0.071 | 1.896 | -0.04 | 0.076 | 0.076 | 0 | 0.12 | 0.29 | 0.1 | 0.27 | 0.06 | 0.14 |
| 3 | 10624908 | rs4622889 | -1.823 | 0.287 | -0.273 | 1.947 | 0.019 | 0.022 | 0.022 | 0 | 0.33 | 0.2 | 0.36 | 0.69 | 0.3 | 0.4 |
| 3 | 10625996 | rs7613069 | -1.7 | 0.228 | 0.41 | 2.316 | -0.053 | 0.096 | 0.096 | 0 | 0.11 | 0.35 | 0.06 | 0.22 | 0.03 | 0.08 |
| 3 | 10630258 | rs17033143 | -1.029 | 0.26 | -0.163 | 2.311 | 0.014 | -0.059 | -0.059 | 1.153 | 0.15 | 0.04 | 0.09 | 0.24 | 0.09 | 0.16 |

| | | | | | | | | | | | | | | | | |
|---|---|---|---|---|---|---|---|---|---|---|---|---|---|---|---|---|
| 3 | 10632832 | rs4269071 | 0.874 | -0.01 | 0.64 | 2.187 | -0.027 | 0.09 | 0.09 | 0 | 0.87 | 0.64 | 0.88 | 0.75 | 0.91 | 0.83 |
| 3 | 10633169 | rs9839935 | 0.904 | 0.013 | 0.629 | 2.184 | -0.026 | 0.089 | 0.089 | 0 | 0.86 | 0.64 | 0.87 | 0.75 | 0.9 | 0.83 |
| 3 | 10633335 | rs6799104 | -1.367 | 0.062 | 0.435 | 1.795 | 0.011 | 0.019 | 0.019 | 0 | 0.41 | 0.33 | 0.4 | 0.72 | 0.31 | 0.49 |
| 3 | 10642523 | rs9822272 | -1.121 | 0.677 | 0.807 | 2.418 | 0.003 | -0.004 | -0.004 | 0 | 0.24 | 0.19 | 0.1 | 0.27 | 0.13 | 0.24 |
| 3 | 10647143 | rs1962654 | 0.96 | 0.68 | 0.828 | 2.295 | 0 | 0.002 | 0.002 | 0 | 0.93 | 0.92 | 0.88 | 0.84 | 0.92 | 0.92 |
| 3 | 10649468 | rs12487529 | 2.676 | 0.678 | 0.975 | 1.896 | 0.008 | 0.117 | 0.117 | 0.003 | 0.46 | 0.23 | 0.33 | 0.03 | 0.57 | 0.24 |
| 3 | 10653320 | rs9834762 | -1.951 | 0.709 | 1.127 | 2.002 | 0.029 | 0.053 | 0.053 | 0.046 | 0.45 | 0.67 | 0.49 | 0.77 | 0.3 | 0.57 |
| 3 | 10654539 | rs9858042 | 1.367 | 0.719 | 1.085 | 2.238 | 0.011 | 0.057 | 0.057 | 0 | 0.91 | 0.77 | 0.81 | 0.8 | 0.87 | 0.82 |
| 3 | 10654982 | rs7632035 | 1.217 | 0.628 | 1.052 | 2.368 | -0.007 | 0.024 | 0.024 | 0 | 0.76 | 0.63 | 0.73 | 0.67 | 0.78 | 0.62 |
| 3 | 10656823 | rs749477 | 1.982 | 0.671 | 1.243 | 1.794 | 0.002 | -0.011 | -0.011 | 0 | 0.56 | 0.44 | 0.55 | 0.12 | 0.62 | 0.34 |
| 3 | 10657877 | rs6809504 | 1.619 | 0.72 | 1.378 | 1.937 | -0.001 | -0.008 | -0.008 | 0 | 0.62 | 0.65 | 0.6 | 0.19 | 0.65 | 0.51 |
| 3 | 10658014 | rs6772499 | 1.579 | 0.238 | 1.397 | 2.074 | -0.021 | -0.057 | -0.057 | 1.091 | 0.67 | 0.54 | 0.64 | 0.19 | 0.72 | 0.54 |
| 3 | 10659768 | rs930273 | 1.169 | 0.014 | 0.861 | 2.571 | -0.016 | 0.269 | 0.269 | 0 | 0.84 | 0.51 | 0.79 | 0.41 | 0.86 | 0.74 |
| 3 | 10666072 | rs1865395 | -1.373 | 0.718 | 0.171 | 2.698 | 0.029 | 0.136 | 0.136 | 0 | 0.13 | 0.2 | 0.17 | 0.54 | 0.2 | 0.19 |
| 3 | 10666144 | rs17033257 | -0.465 | -0.021 | 0.12 | 2.17 | 0.015 | -0.023 | -0.023 | 0 | 0.16 | 0.08 | 0.1 | 0.21 | 0.1 | 0.19 |
| 3 | 10666664 | rs4684731 | -1.647 | 0.429 | 0.171 | 1.931 | -0.002 | 0.014 | 0.014 | 0.102 | 0.07 | 0.1 | 0.12 | 0.19 | 0.07 | 0.24 |
| 3 | 10666716 | rs1865396 | -1.084 | 0.548 | 0.076 | 1.862 | 0.002 | 0.101 | 0.101 | 0 | 0.11 | 0.24 | 0.13 | 0.29 | 0.08 | 0.17 |
| 3 | 10669760 | rs11128520 | 0.466 | 0.725 | -0.486 | 1.286 | -0.001 | 0.003 | 0.003 | 0 | 0.64 | 0.43 | 0.53 | 0.61 | 0.65 | 0.49 |
| 3 | 10671218 | rs1000772 | -0.986 | 0.469 | -0.374 | 2.07 | -0.018 | 0.04 | 0.04 | 0.198 | 0.23 | 0.45 | 0.39 | 0.67 | 0.19 | 0.41 |
| 3 | 10675684 | rs892608 | -0.144 | 1.137 | -1.916 | 1.202 | -0.152 | -0.008 | -0.008 | 0 | 0.32 | 0.04 | 0.49 | 0.3 | 0.5 | 0.31 |
| 3 | 10676407 | rs879595 | -0.69 | -0.258 | -2.814 | 1.095 | -0.001 | 0.074 | 0.074 | 3.097 | 0.6 | 0.93 | 0.76 | 0.76 | 0.67 | 0.62 |
| 3 | 10676736 | rs11707497 | 0.037 | -0.777 | -2.953 | 0.915 | 0.002 | 0.002 | 0.002 | 0 | 0.07 | 0.06 | 0.05 | 0.05 | 0.04 | 0.17 |
| 3 | 10677196 | rs879594 | 0.296 | -1.144 | -1.707 | 1.113 | -0.07 | 0.129 | 0.129 | 0.057 | 0.2 | 0.54 | 0.17 | 0.37 | 0.08 | 0.16 |
| 3 | 10680590 | rs7631749 | 0.391 | -1.313 | -1.574 | 1.039 | -0.056 | 0.161 | 0.161 | 0.102 | 0.17 | 0.58 | 0.15 | 0.37 | 0.06 | 0.15 |
| 3 | 10680667 | rs7620122 | -0.465 | -1.336 | -1.586 | 1.029 | -0.052 | 0.076 | 0.076 | 0.835 | 0.23 | 0.62 | 0.19 | 0.41 | 0.1 | 0.34 |
| 3 | 10685297 | rs13090409 | -0.029 | -1.141 | -1.393 | 0.997 | -0.009 | 0.03 | 0.03 | 0.623 | 0.2 | 0.77 | 0.18 | 0.39 | 0.09 | 0.2 |
| 3 | 10686376 | rs9841925 | -0.593 | -1.166 | -1.415 | 0.984 | 0.013 | -0.02 | -0.02 | 2.007 | 0.24 | 0.8 | 0.19 | 0.41 | 0.12 | 0.29 |
| 3 | 10688397 | rs13065552 | -1.406 | -1.367 | -1.497 | 0.932 | -0.008 | 0.03 | 0.03 | 5.048 | 0.68 | 0.98 | 0.82 | 0.81 | 0.74 | 0.8 |
| 3 | 10695666 | rs9942062 | 0.074 | -1.335 | -1.456 | 0.929 | -0.064 | 0.112 | 0.112 | 0.189 | 0.21 | 0.58 | 0.17 | 0.36 | 0.09 | 0.16 |
| 3 | 10696036 | rs11128522 | 0.638 | -1.491 | -1.605 | 0.793 | -0.028 | 0.022 | 0.022 | 0 | 0.38 | 0.12 | 0.56 | 0.36 | 0.59 | 0.41 |
| 3 | 10706597 | rs11719375 | -1.36 | -1.529 | -1.65 | 0.864 | 0.091 | -0.117 | -0.117 | 0 | 0.35 | 0.93 | 0.33 | 0.51 | 0.18 | 0.48 |
| 3 | 10707421 | rs745643 | 1.087 | -1.528 | -1.661 | 0.844 | -0.104 | 0.016 | 0.016 | 0 | 0.33 | 0.06 | 0.49 | 0.3 | 0.56 | 0.32 |
| 3 | 10707861 | rs892605 | -0.861 | -1.53 | -1.627 | 0.861 | 0.025 | 0.04 | 0.04 | 3.585 | 0.63 | 0.91 | 0.73 | 0.75 | 0.7 | 0.71 |
| 3 | 10709750 | rs6770332 | -0.015 | -1.457 | -1.447 | 1.043 | -0.069 | 0.091 | 0.091 | 0.087 | 0.21 | 0.58 | 0.17 | 0.34 | 0.08 | 0.15 |
| 3 | 10719530 | rs11920812 | -0.496 | -1.74 | -0.597 | 0.965 | -0.024 | 0.08 | 0.08 | 0.08 | 0.19 | 0.29 | 0.21 | 0.43 | 0.12 | 0.21 |
| 3 | 10720167 | rs4234504 | -1.263 | -1.74 | -1.452 | 0.521 | -0.033 | -0.068 | -0.068 | 0 | 0.84 | 0.98 | 0.88 | 0.78 | 0.83 | 0.89 |
| 3 | 10723886 | rs6442199 | -0.151 | -2.526 | -1.086 | 0.225 | 0.01 | 0 | 0 | 0.056 | 0.4 | 0.62 | 0.26 | 0.38 | 0.24 | 0.25 |
| 3 | 10738839 | rs2571211 | 1.199 | -2.833 | -1.551 | 0.03 | -0.129 | -0.009 | -0.009 | 0 | 0.29 | 0.06 | 0.5 | 0.26 | 0.58 | 0.32 |
| 3 | 10746274 | rs973188 | -1.168 | -3.006 | -1.471 | 0.292 | 0.014 | -0.072 | -0.072 | 0.168 | 0.82 | 0.98 | 0.87 | 0.72 | 0.84 | 0.86 |
| 3 | 10751976 | rs12497240 | 0.259 | -3.447 | -1.696 | 0.079 | 0.002 | -0.061 | -0.061 | 0 | 0.23 | 0.02 | 0.14 | 0.27 | 0.2 | 0.12 |
| 3 | 10753490 | rs2596852 | -0.24 | -3.512 | -1.006 | 0.297 | -0.007 | 0.047 | 0.047 | 0 | 0.77 | 0.63 | 0.79 | 0.67 | 0.8 | 0.72 |
| 3 | 10763460 | rs6798029 | 0.083 | -3.672 | -0.779 | 0.301 | 0.017 | 0.009 | 0.009 | 0 | 0.45 | 0.56 | 0.28 | 0.38 | 0.23 | 0.49 |
| 3 | 10764022 | rs962835 | 0.281 | -3.741 | -0.774 | 0.311 | -0.05 | -0.023 | -0.023 | 0.325 | 0.42 | 0.46 | 0.23 | 0.32 | 0.2 | 0.31 |
| 3 | 10771533 | rs9826914 | 0.125 | -3.654 | -0.889 | 0.398 | -0.025 | -0.007 | -0.007 | 0 | 0.43 | 0.52 | 0.23 | 0.35 | 0.21 | 0.29 |
| 3 | 10774759 | rs1682825 | -1.061 | -2.958 | -1.136 | 0.64 | -0.001 | -0.038 | -0.038 | 0 | 0.82 | 0.89 | 0.81 | 0.65 | 0.81 | 0.77 |
| 3 | 10777632 | rs17028925 | 0.345 | -2.887 | -0.854 | 0.545 | 0.001 | 0.036 | 0.036 | 0 | 0.21 | 0.26 | 0.25 | 0.38 | 0.21 | 0.39 |
| 3 | 10780817 | rs171239 | 0.55 | -2.935 | -1.581 | 0.403 | -0.076 | -0.003 | -0.003 | 0 | 0.37 | 0.05 | 0.49 | 0.31 | 0.58 | 0.31 |
| 3 | 10783572 | rs1560922 | -1.085 | -2.926 | -0.197 | 0.607 | 0.052 | 0.159 | 0.159 | 4.137 | 0.07 | 0.58 | 0.1 | 0.09 | 0.02 | 0.12 |
| 3 | 10784877 | rs9830378 | -1.123 | -2.933 | -0.2 | 0.604 | 0.05 | 0.156 | 0.156 | 4.508 | 0.07 | 0.58 | 0.1 | 0.09 | 0.02 | 0.12 |
| 3 | 10785012 | rs350665 | 0.508 | -2.743 | -1.016 | 0.518 | -0.035 | 0.092 | 0.092 | 0 | 0.42 | 0.05 | 0.51 | 0.25 | 0.64 | 0.35 |
| 3 | 10786186 | rs9877719 | -0.222 | -2.956 | -0.339 | 0.41 | -0.01 | -0.004 | -0.004 | 0.631 | 0.3 | 0.66 | 0.26 | 0.16 | 0.16 | 0.29 |
| 3 | 10789999 | rs1682821 | -0.307 | -2.859 | -0.081 | 0.451 | -0.052 | -0.016 | -0.016 | 0.181 | 0.81 | 0.53 | 0.84 | 0.91 | 0.9 | 0.82 |
| 3 | 10791676 | rs2629126 | 0.157 | -3.273 | -0.424 | 0.269 | -0.001 | -0.001 | -0.001 | 0.023 | 0.35 | 0.33 | 0.25 | 0.61 | 0.22 | 0.44 |
| 3 | 10792193 | rs3856784 | 0.019 | -3.362 | -0.776 | 0.509 | 0.086 | -0.025 | -0.025 | 0 | 0.57 | 0.82 | 0.73 | 0.54 | 0.7 | 0.61 |
| 3 | 10796183 | rs350658 | 0.676 | -3.559 | -0.151 | 0.446 | -0.05 | -0.027 | -0.027 | 0 | 0.68 | 0.43 | 0.82 | 0.86 | 0.84 | 0.73 |
| 3 | 10797131 | rs350660 | 1.063 | -3.417 | -0.323 | 1.075 | 0.077 | -0.049 | -0.049 | 1.273 | 0.66 | 0.21 | 0.76 | 0.66 | 0.81 | 0.67 |
| 3 | 10801121 | rs3900078 | -0.156 | -3.439 | 0.073 | 0.969 | -0.019 | 0.005 | 0.005 | 0 | 0.08 | 0.14 | 0.07 | 0.07 | 0.05 | 0.07 |
| 3 | 10801251 | rs3900076 | -0.942 | -3.765 | 0.496 | 0.884 | -0.029 | 0.012 | 0.012 | 0.031 | 0.27 | 0.61 | 0.25 | 0.28 | 0.13 | 0.2 |
| 3 | 10801693 | rs13078886 | -0.645 | -3.772 | 0.859 | 0.912 | -0.007 | 0.075 | 0.075 | 0 | 0.1 | 0.21 | 0.1 | 0.18 | 0.07 | 0.13 |
| 3 | 10802061 | rs7635599 | 0.595 | -3.746 | 0.766 | 0.874 | -0.072 | -0.038 | -0.038 | 0.247 | 0.17 | 0.37 | 0.14 | 0.07 | 0.05 | 0.07 |
| 3 | 10802648 | rs350663 | -1.669 | -3.966 | -0.179 | 0.655 | -0.118 | 0.119 | 0.119 | 0.17 | 0.7 | 0.97 | 0.46 | 0.87 | 0.37 | 0.79 |
| 3 | 10807667 | rs9820724 | -0.852 | -3.928 | 0.058 | 0.754 | 0.028 | -0.014 | -0.014 | 0.004 | 0.3 | 0.73 | 0.29 | 0.29 | 0.16 | 0.2 |
| 3 | 10810225 | rs350667 | 1.598 | -4.187 | -0.352 | 1.16 | 0.075 | -0.05 | -0.05 | 0 | 0.27 | 0.12 | 0.17 | 0.39 | 0.13 | 0.34 |
| 3 | 10815485 | rs2581204 | 0.021 | -3.563 | -0.116 | 0.952 | 0.043 | 0.005 | 0.005 | 0.791 | 0.48 | 0.63 | 0.31 | 0.45 | 0.22 | 0.42 |
| 3 | 10817070 | rs2629123 | -1.319 | -3.548 | -0.441 | 0.926 | -0.033 | 0.051 | 0.051 | 0 | 0.66 | 0.8 | 0.41 | 0.75 | 0.37 | 0.71 |
| 3 | 10818641 | rs13064472 | -0.916 | -3.116 | -0.167 | 0.929 | 0.026 | 0.095 | 0.095 | 0 | 0.19 | 0.37 | 0.12 | 0.3 | 0.16 | 0.29 |
| 3 | 10819226 | rs11707734 | 0.506 | -2.929 | -1.12 | 0.819 | 0.006 | 0.014 | 0.014 | 0 | 0.07 | 0.01 | 0.03 | 0.04 | 0.04 | 0.12 |
| 3 | 10820067 | rs2596834 | -1.623 | -2.943 | -1.162 | 0.792 | -0.153 | 0.048 | 0.048 | 0 | 0.73 | 0.97 | 0.48 | 0.81 | 0.42 | 0.77 |
| 3 | 10820971 | rs9871974 | -0.021 | -2.201 | -0.382 | 1.051 | 0.001 | -0.003 | -0.003 | 0 | 0.13 | 0.1 | 0.06 | 0.14 | 0.08 | 0.19 |

| | | | | | | | | | | | | | | | | |
|---|---|---|---|---|---|---|---|---|---|---|---|---|---|---|---|---|
| 3 | 10821474 | rs2342375 | -0.872 | -2.223 | -0.492 | 0.979 | 0.041 | -0.018 | -0.018 | 0.056 | 0.39 | 0.76 | 0.29 | 0.35 | 0.25 | 0.37 |
| 3 | 10822795 | rs7635344 | -0.874 | -0.607 | 0.343 | 0.976 | 0.055 | 0.069 | 0.069 | 0 | 0.09 | 0.24 | 0.16 | 0.11 | 0.16 | 0.06 |
| 3 | 10823781 | rs6442205 | -1.417 | -0.966 | 0.723 | 0.664 | -0.017 | -0.03 | -0.03 | 0.717 | 0.13 | 0.4 | 0.15 | 0.06 | 0.12 | 0.12 |
| 3 | 10825837 | rs2017903 | 0.279 | -0.59 | 0.412 | 0.244 | -0.004 | 0.015 | 0.015 | 0.604 | 0.6 | 0.39 | 0.38 | 0.76 | 0.41 | 0.66 |
| 3 | 10830031 | rs1601365 | 0.189 | -0.607 | 0.512 | 0.236 | -0.002 | 0.002 | 0.002 | 0.657 | 0.6 | 0.4 | 0.38 | 0.76 | 0.41 | 0.66 |
| 3 | 10831167 | rs2596842 | 0.232 | -0.656 | 0.69 | 0.214 | 0.008 | -0.043 | -0.043 | 0 | 0.61 | 0.53 | 0.39 | 0.76 | 0.41 | 0.66 |
| 3 | 10831291 | rs1485134 | 0.884 | -1.348 | 0 | 0.534 | 0.097 | -0.014 | -0.014 | 0.363 | 0.45 | 0.18 | 0.28 | 0.51 | 0.31 | 0.51 |
| 3 | 10833964 | rs1905752 | 0.052 | -1.117 | -0.188 | 0.514 | 0.006 | 0.033 | 0.033 | 0 | 0.15 | 0.25 | 0.11 | 0.25 | 0.1 | 0.15 |
| 3 | 10835441 | rs1905755 | 0.144 | -1.118 | -0.472 | 0.514 | 0.001 | 0.011 | 0.011 | 0 | 0.15 | 0.18 | 0.1 | 0.25 | 0.1 | 0.16 |
| 3 | 10840684 | rs2600078 | -1.715 | -1.049 | -0.702 | 0.456 | -0.156 | 0.121 | 0.121 | 0 | 0.76 | 0.96 | 0.55 | 0.93 | 0.54 | 0.87 |
| 3 | 10848410 | rs2171003 | 1.249 | -1.468 | -1.042 | 0.371 | 0.083 | -0.046 | -0.046 | 3.439 | 0.39 | 0.17 | 0.24 | 0.48 | 0.26 | 0.4 |
| 3 | 10852828 | rs874589 | 1.548 | -1.741 | -1.265 | 0.766 | 0.053 | -0.086 | -0.086 | 0 | 0.35 | 0.05 | 0.15 | 0.39 | 0.25 | 0.39 |
| 3 | 10856773 | rs2581219 | -1.723 | -1.531 | -1.228 | 0.699 | -0.18 | 0.129 | 0.129 | 0.097 | 0.75 | 0.98 | 0.55 | 0.93 | 0.55 | 0.85 |
| 3 | 10857067 | rs2600072 | 0.646 | -1.613 | -0.96 | 0.815 | 0.09 | -0.01 | -0.01 | 0 | 0.45 | 0.19 | 0.28 | 0.51 | 0.31 | 0.51 |
| 3 | 10860297 | rs2254931 | 0.952 | -2.061 | -0.97 | 0.612 | 0.09 | -0.03 | -0.03 | 0 | 0.39 | 0.16 | 0.24 | 0.51 | 0.22 | 0.48 |
| 3 | 10864445 | rs9835618 | -1.16 | -1.166 | -0.881 | 0.483 | -0.006 | 0.005 | 0.005 | 0.51 | 0.25 | 0.64 | 0.33 | 0.25 | 0.23 | 0.26 |
| 3 | 10872262 | rs971930 | -0.668 | -0.741 | -1.224 | 0.356 | -0.062 | 0.067 | 0.067 | 0 | 0.66 | 0.89 | 0.62 | 0.84 | 0.41 | 0.83 |
| 3 | 10877611 | rs1809529 | -0.007 | -1.105 | -0.707 | 0.69 | 0.059 | 0.014 | 0.014 | 0.271 | 0.53 | 0.78 | 0.31 | 0.44 | 0.36 | 0.66 |
| 3 | 10878049 | rs6442209 | 0.855 | -0.887 | -0.427 | 0.614 | -0.004 | -0.039 | -0.039 | 0.023 | 0.08 | 0.16 | 0.04 | 0.01 | 0.04 | 0.11 |
| 3 | 10885920 | rs2304725 | 0.918 | -1.059 | -0.434 | 0.98 | 0.039 | 0.009 | 0.009 | 0 | 0.46 | 0.3 | 0.22 | 0.43 | 0.33 | 0.58 |
| 3 | 10887967 | rs2880586 | 1.439 | -1.287 | -0.653 | 0.811 | 0.03 | -0.008 | -0.008 | 0.237 | 0.37 | 0.28 | 0.16 | 0.39 | 0.23 | 0.42 |
| 3 | 10888402 | rs3774121 | 0.411 | -1.278 | -0.665 | 0.894 | 0.004 | 0.006 | 0.006 | 0 | 0.11 | 0.12 | 0.08 | 0.16 | 0.11 | 0.22 |
| 3 | 10891237 | rs3774120 | 1.74 | -1.308 | -0.76 | 0.91 | 0.047 | -0.011 | -0.011 | 0 | 0.35 | 0.02 | 0.13 | 0.27 | 0.23 | 0.37 |
| 3 | 10892030 | rs7631448 | -1.529 | -1.24 | -0.909 | 1.07 | -0.156 | -0.024 | -0.024 | 0 | 0.74 | 0.96 | 0.65 | 0.78 | 0.53 | 0.85 |
| 3 | 10893705 | rs3774119 | 0.785 | -1.327 | -0.607 | 1.147 | -0.007 | 0.026 | 0.026 | 0 | 0.09 | 0.11 | 0.27 | 0.23 | 0.05 | 0.08 |
| 3 | 10897423 | rs11710402 | 1.923 | -1.356 | -0.686 | 1.112 | 0.048 | -0.004 | -0.004 | 0 | 0.35 | 0.02 | 0.13 | 0.27 | 0.23 | 0.37 |
| 3 | 10901959 | rs1609480 | -0.537 | -1.284 | 0.45 | 1.184 | -0.013 | -0.006 | -0.006 | 0 | 0.66 | 0.7 | 0.55 | 0.66 | 0.42 | 0.79 |
| 3 | 10902353 | rs999448 | 0.881 | -1.244 | 0.456 | 1.265 | -0.011 | -0.007 | -0.007 | 0.045 | 0.09 | 0.18 | 0.05 | 0.11 | 0.05 | 0.12 |
| 3 | 10902862 | rs4684739 | 0.994 | -1.227 | 0.34 | 1.266 | 0.043 | 0.017 | 0.017 | 0 | 0.46 | 0.3 | 0.21 | 0.41 | 0.32 | 0.6 |
| 3 | 10907556 | rs6809562 | -0.073 | -0.965 | 0.353 | 1.335 | 0.004 | 0.063 | 0.063 | 0.508 | 0.08 | 0.25 | 0.11 | 0.13 | 0.11 | 0.06 |
| 3 | 10911151 | rs6442211 | -1.11 | -0.667 | -0.576 | 1.338 | -0.101 | -0.002 | -0.002 | 0.158 | 0.72 | 0.9 | 0.66 | 0.79 | 0.54 | 0.87 |
| 3 | 10913765 | rs4684742 | 0.289 | -1.11 | -0.171 | 1.417 | -0.002 | -0.005 | -0.005 | 0 | 0.54 | 0.54 | 0.51 | 0.63 | 0.37 | 0.63 |
| 3 | 10916332 | rs966030 | 0.119 | -1.107 | -0.197 | 1.411 | 0.018 | 0.068 | 0.068 | 0 | 0.11 | 0.31 | 0.09 | 0.21 | 0.09 | 0.24 |
| 3 | 10917457 | rs1610180 | -0.833 | -1.156 | 0.002 | 1.344 | -0.058 | 0.039 | 0.039 | 0.073 | 0.67 | 0.82 | 0.58 | 0.74 | 0.43 | 0.79 |
| 3 | 10923193 | rs3774108 | -0.196 | -1.773 | -0.92 | 1.614 | 0 | -0.004 | -0.004 | 0.35 | 0.74 | 0.7 | 0.86 | 0.59 | 0.8 | 0.6 |
| 3 | 10923941 | rs3821767 | 0.024 | -0.896 | -0.731 | 1.996 | 0.001 | 0.052 | 0.052 | 0 | 0.12 | 0.18 | 0.41 | 0.24 | 0.11 | 0.16 |
| 3 | 10933887 | rs2655278 | 1.904 | -1.094 | -1.235 | 1.726 | 0.016 | -0.004 | -0.004 | 0 | 0.34 | 0.03 | 0.14 | 0.26 | 0.23 | 0.25 |
| 3 | 10937056 | rs1807101 | -0.032 | -1.057 | -0.999 | 1.646 | 0.012 | -0.036 | -0.036 | 0.015 | 0.59 | 0.56 | 0.65 | 0.72 | 0.44 | 0.64 |
| 3 | 10938584 | rs1994260 | 1.668 | -0.963 | -1.146 | 1.796 | 0.026 | 0.014 | 0.014 | 1.007 | 0.35 | 0.03 | 0.13 | 0.26 | 0.23 | 0.25 |
| 3 | 10941102 | rs958469 | 0.484 | -0.341 | -0.471 | 1.977 | 0.029 | 0.147 | 0.147 | 0 | 0.07 | 0.33 | 0.32 | 0.15 | 0.04 | 0.14 |
| 3 | 10942903 | rs1960131 | -0.678 | 0.015 | -0.523 | 1.733 | 0.021 | -0.022 | -0.022 | 0 | 0.23 | 0.43 | 0.21 | 0.21 | 0.23 | 0.26 |
| 3 | 10945374 | rs901029 | 0.685 | 0.34 | -0.44 | 1.782 | 0 | 0 | 0 | 0.103 | 0.05 | 0.08 | 0.11 | 0.07 | 0.07 | 0.02 |
| 3 | 10948354 | rs3821763 | 0.775 | 0.347 | -0.597 | 1.471 | -0.002 | -0.005 | -0.005 | 0 | 0.05 | 0.06 | 0.1 | 0.07 | 0.07 | 0.02 |
| 3 | 10948486 | rs3821761 | 0.303 | 0.337 | -0.326 | 1.512 | 0.029 | 0.147 | 0.147 | 0 | 0.07 | 0.31 | 0.32 | 0.15 | 0.04 | 0.14 |
| 3 | 10948776 | rs2581210 | -0.856 | 0.419 | -1.212 | 1.072 | -0.087 | 0.062 | 0.062 | 0 | 0.74 | 0.93 | 0.79 | 0.86 | 0.6 | 0.85 |
| 3 | 10950051 | rs9868537 | -0.522 | 0.615 | -0.513 | 1.152 | 0.075 | 0.028 | 0.028 | 0 | 0.11 | 0.37 | 0.13 | 0.09 | 0.09 | 0.17 |
| 3 | 10953549 | rs2581206 | 0.47 | 0.111 | -1.502 | 1.253 | 0.039 | -0.025 | -0.025 | 0 | 0.45 | 0.24 | 0.53 | 0.5 | 0.37 | 0.43 |
| 3 | 10955024 | rs6442214 | 0.652 | 0.324 | -1.97 | 0.997 | -0.016 | -0.017 | -0.017 | 0.116 | 0.07 | 0.03 | 0.33 | 0.14 | 0.1 | 0.12 |
| 3 | 10957703 | rs1843278 | 1.608 | -0.024 | -2.013 | 0.535 | 0.056 | 0.152 | 0.152 | 0.531 | 0.31 | 0.03 | 0.12 | 0.14 | 0.2 | 0.24 |
| 3 | 10958276 | rs3774084 | 0.535 | 1.041 | -1.337 | 1.079 | -0.125 | 0.027 | 0.027 | 0.076 | 0.22 | 0.01 | 0.19 | 0.13 | 0.37 | 0.15 |
| 3 | 10960562 | rs6779181 | -0.598 | 0.672 | -0.634 | 1.444 | -0.009 | 0.03 | 0.03 | 0.173 | 0.06 | 0.22 | 0.09 | 0.08 | 0.06 | 0.02 |
| 3 | 10966711 | rs2581215 | -0.537 | 0.782 | -0.287 | 1.505 | 0.018 | -0.028 | -0.028 | 2.265 | 0.37 | 0.48 | 0.37 | 0.23 | 0.54 | 0.24 |
| 3 | 10967342 | rs17033655 | 0.875 | 0.883 | -0.402 | 1.59 | 0.004 | 0.002 | 0.002 | 0 | 0.1 | 0.16 | 0.07 | 0.12 | 0.1 | 0.17 |
| 3 | 10967739 | rs2272395 | -0.058 | 1.002 | -0.481 | 1.551 | 0.03 | -0.066 | -0.066 | 0.405 | 0.33 | 0.46 | 0.31 | 0.17 | 0.46 | 0.21 |
| 3 | 10969663 | rs9878482 | -0.576 | 0.885 | -0.461 | 1.94 | 0.023 | 0.194 | 0.194 | 0 | 0.19 | 0.51 | 0.19 | 0.51 | 0.18 | 0.32 |
| 3 | 10970053 | rs2581208 | 0.547 | 1.025 | -0.611 | 1.745 | 0.013 | 0.029 | 0.029 | 0 | 0.78 | 0.7 | 0.54 | 0.69 | 0.74 | 0.65 |
| 3 | 10970207 | rs2581207 | -0.383 | 0.999 | -0.606 | 1.659 | 0.011 | 0.021 | 0.021 | 0 | 0.84 | 0.71 | 0.86 | 0.81 | 0.84 | 0.81 |
| 3 | 10971358 | rs1881364 | 1.388 | 0.822 | -0.793 | 1.545 | 0.074 | -0.005 | -0.005 | 0 | 0.43 | 0.24 | 0.22 | 0.45 | 0.27 | 0.44 |
| 3 | 10971499 | rs1881365 | -1.462 | 0.913 | -0.656 | 1.443 | 0.077 | -0.019 | -0.019 | 0 | 0.55 | 0.74 | 0.76 | 0.47 | 0.73 | 0.56 |
| 3 | 10972387 | rs6791465 | 0.422 | 1.117 | -0.32 | 0.984 | 0.017 | -0.042 | -0.042 | 0 | 0.27 | 0.3 | 0.23 | 0.09 | 0.39 | 0.19 |
| 3 | 10974051 | rs1881367 | -0.079 | 1.608 | -0.45 | 1.616 | 0.008 | 0.023 | 0.023 | 0 | 0.86 | 0.78 | 0.9 | 0.82 | 0.88 | 0.81 |
| 3 | 10974427 | rs2697159 | 0.979 | 1.554 | -0.82 | 1.322 | -0.047 | 0.069 | 0.069 | 0.219 | 0.56 | 0.25 | 0.66 | 0.72 | 0.53 | 0.58 |
| 3 | 10975745 | rs2272399 | 1.657 | 1.381 | -0.816 | 1.212 | 0.091 | -0.063 | -0.063 | 0.071 | 0.45 | 0.21 | 0.25 | 0.58 | 0.32 | 0.41 |
| 3 | 10976876 | rs2245532 | -1.106 | 1.174 | -0.367 | 0.123 | 0.007 | -0.048 | -0.048 | 1.632 | 0.35 | 0.54 | 0.3 | 0.1 | 0.43 | 0.24 |
| 3 | 10977277 | rs11128532 | -0.712 | 1.494 | -0.22 | 0.281 | 0.021 | -0.071 | -0.071 | 0 | 0.29 | 0.51 | 0.23 | 0.09 | 0.35 | 0.22 |
| 3 | 10986153 | rs11128534 | -0.683 | 1.581 | 0.26 | 0.329 | 0.027 | -0.073 | -0.073 | 0.927 | 0.26 | 0.51 | 0.22 | 0.08 | 0.31 | 0.21 |
| 3 | 10995671 | rs2930144 | -0.468 | 1.265 | 0.303 | 0.432 | -0.003 | -0.011 | -0.011 | 0 | 0.18 | 0.29 | 0.11 | 0.17 | 0.14 | 0.19 |
| 3 | 10996556 | rs1919078 | -0.839 | 1.728 | 0.506 | 0.03 | -0.001 | -0.023 | -0.023 | 0.522 | 0.28 | 0.25 | 0.22 | 0.05 | 0.35 | 0.28 |
| 3 | 10997255 | rs2697152 | -0.61 | 1.513 | 0.929 | 0.379 | 0.038 | -0.053 | -0.053 | 0 | 0.34 | 0.52 | 0.26 | 0.23 | 0.41 | 0.32 |

| | | | | | | | | | | | | | | | | |
|---|---|---|---|---|---|---|---|---|---|---|---|---|---|---|---|---|
| 3 | 10998663 | rs6796285 | -2.197 | 1.223 | 1.716 | 1.618 | 0.052 | 0.225 | 0.225 | 0 | 0.12 | 0.48 | 0.13 | 0.32 | 0.09 | 0.22 |
| 3 | 10999494 | rs4610207 | -1.439 | 1.42 | 1.446 | 1.251 | 0.054 | -0.016 | -0.016 | 0 | 0.08 | 0.21 | 0.07 | 0.04 | 0.08 | 0.07 |
| 3 | 11010671 | rs1881363 | -2.465 | 1.439 | 0.8 | 1.549 | 0.059 | 0.233 | 0.233 | 1.512 | 0.19 | 0.5 | 0.42 | 0.42 | 0.21 | 0.35 |
| 3 | 11013380 | rs1004999 | -1.493 | 1.024 | 0.94 | 1.746 | 0.047 | 0.264 | 0.264 | 8.411 | 0.07 | 0.45 | 0.05 | 0.17 | 0.02 | 0.13 |
| 3 | 11014727 | rs2950799 | -1.296 | 1.023 | 0.673 | 0.973 | -0.023 | -0.02 | -0.02 | 0 | 0.09 | 0.05 | 0.33 | 0.24 | 0.16 | 0.17 |
| 3 | 11017136 | rs1710896 | -1.183 | 0.81 | 0.777 | 1.07 | 0.051 | 0.205 | 0.205 | 2.818 | 0.12 | 0.45 | 0.3 | 0.24 | 0.13 | 0.13 |
| 3 | 11019209 | rs1710884 | -0.496 | 0.65 | -1.222 | 0.262 | -0.065 | 0.009 | 0.009 | 0 | 0.21 | 0.01 | 0.23 | 0.15 | 0.27 | 0.12 |
| 3 | 11020033 | rs2928084 | -0.631 | -0.896 | -0.877 | -0.32 | -0.032 | -0.044 | -0.044 | 0.623 | 0.27 | 0.31 | 0.2 | 0.05 | 0.15 | 0.19 |
| 3 | 11021002 | rs17529534 | -0.409 | -0.2 | -1.181 | -0.147 | -0.03 | 0.001 | 0.001 | 0 | 0.08 | 0.01 | 0.04 | 0.05 | 0.06 | 0.18 |
| 3 | 11021097 | rs2930149 | 0.068 | 0.454 | -1.23 | -0.124 | -0.013 | 0.003 | 0.003 | 0.014 | 0.69 | 0.69 | 0.74 | 0.69 | 0.51 | 0.65 |
| 3 | 11022665 | rs1710895 | 0.168 | -0.675 | -1.647 | 0.143 | -0.018 | 0.016 | 0.016 | 0.387 | 0.32 | 0.33 | 0.44 | 0.45 | 0.23 | 0.28 |
| 3 | 11022826 | rs956053 | 1.339 | -0.979 | -1.882 | 0.035 | -0.008 | 0.016 | 0.016 | 0 | 0.33 | 0.04 | 0.26 | 0.24 | 0.28 | 0.37 |
| 3 | 11023672 | rs2697153 | -1.362 | -0.433 | -1.619 | 0.364 | -0.15 | -0.033 | -0.033 | 0.021 | 0.8 | 0.99 | 0.82 | 0.81 | 0.61 | 0.88 |
| 3 | 11036150 | rs2697150 | 0.063 | -0.592 | -1.441 | 0.643 | -0.024 | 0.019 | 0.019 | 0.254 | 0.31 | 0.35 | 0.43 | 0.43 | 0.23 | 0.27 |
| 3 | 11036480 | rs2697149 | 0.026 | -0.627 | -1.457 | 0.627 | -0.024 | 0.018 | 0.018 | 0.277 | 0.32 | 0.35 | 0.43 | 0.43 | 0.23 | 0.28 |
| 3 | 11036624 | rs2601126 | 0.438 | -0.451 | -1.358 | 0.637 | 0.009 | 0.008 | 0.008 | 0 | 0.48 | 0.59 | 0.39 | 0.38 | 0.38 | 0.61 |
| 3 | 11038807 | rs1710885 | -0.247 | -0.466 | -1.122 | 0.931 | 0 | 0.005 | 0.005 | 0.389 | 0.23 | 0.22 | 0.34 | 0.51 | 0.17 | 0.27 |
| 3 | 11039960 | rs1710887 | 0.143 | 0.279 | -1.315 | 0.835 | -0.003 | -0.03 | -0.03 | 0 | 0.62 | 0.79 | 0.76 | 0.46 | 0.65 | 0.68 |
| 3 | 11040439 | rs9990174 | 0.727 | 0.331 | -1.054 | 0.658 | 0.004 | 0.038 | 0.038 | 0 | 0.34 | 0.2 | 0.45 | 0.24 | 0.33 | 0.41 |
| 3 | 11041606 | rs1568072 | 1.025 | 0.616 | -1.001 | 0.858 | -0.062 | -0.064 | -0.064 | 0 | 0.1 | 0.01 | 0.1 | 0.26 | 0.22 | 0.07 |
| 3 | 11041870 | rs1728811 | -1.573 | 0.284 | -0.908 | 0.815 | -0.009 | -0.02 | -0.02 | 2.004 | 0.43 | 0.75 | 0.34 | 0.52 | 0.42 | 0.42 |
| 3 | 11045827 | rs1710892 | -1.446 | 0.238 | -0.252 | 0.731 | -0.008 | -0.018 | -0.018 | 0 | 0.41 | 0.73 | 0.33 | 0.5 | 0.41 | 0.42 |
| 3 | 11049517 | rs11707097 | -0.677 | 0.016 | -0.317 | 0.747 | 0.005 | 0.009 | 0.009 | 0 | 0.22 | 0.13 | 0.14 | 0.18 | 0.15 | 0.27 |
| 3 | 11053242 | rs2944366 | 1.23 | -0.057 | -0.36 | 0.778 | 0.001 | -0.049 | -0.049 | 0 | 0.29 | 0.14 | 0.22 | 0.36 | 0.3 | 0.31 |
| 3 | 11053389 | rs2930154 | -1.469 | 0.064 | -0.387 | 0.753 | -0.061 | -0.059 | -0.059 | 0.063 | 0.9 | 0.99 | 0.85 | 0.77 | 0.79 | 0.94 |
| 3 | 11053517 | rs6778281 | -0.245 | -0.024 | -0.162 | 0.714 | 0.007 | 0.024 | 0.024 | 0 | 0.09 | 0.27 | 0.07 | 0.08 | 0.07 | 0.05 |
| 3 | 11055624 | rs2933308 | 0.323 | -0.027 | -0.521 | 0.755 | -0.003 | -0.045 | -0.045 | 0 | 0.4 | 0.16 | 0.28 | 0.47 | 0.4 | 0.4 |
| 3 | 11058359 | rs2697142 | 1.721 | 0.07 | -0.466 | 0.652 | -0.001 | 0.016 | 0.016 | 0 | 0.31 | 0.25 | 0.46 | 0.15 | 0.32 | 0.31 |
| 3 | 11059847 | rs2928078 | 0.303 | -1.053 | -0.501 | 0.802 | 0.007 | 0.011 | 0.011 | 0 | 0.49 | 0.71 | 0.68 | 0.46 | 0.55 | 0.42 |
| 3 | 11062062 | rs3774070 | -1.559 | -1.197 | 0.238 | 1.293 | -0.033 | 0.019 | 0.019 | 0 | 0.17 | 0.46 | 0.09 | 0.13 | 0.07 | 0.19 |
| 3 | 11062934 | rs2950798 | 0.694 | -1.027 | -0.26 | 1.317 | -0.025 | -0.053 | -0.053 | 0 | 0.09 | 0.01 | 0.09 | 0.25 | 0.09 | 0.04 |
| 3 | 11066055 | rs11708202 | -1.711 | -1.165 | 0.24 | 0.837 | 0.001 | 0.007 | 0.007 | 0 | 0.22 | 0.17 | 0.09 | 0.16 | 0.16 | 0.29 |
| 3 | 11066388 | rs9879137 | -1.644 | -0.802 | 0.34 | 1.023 | -0.009 | 0.073 | 0.073 | 0 | 0.13 | 0.45 | 0.08 | 0.14 | 0.05 | 0.16 |
| 3 | 11066670 | rs10510403 | -1.227 | -0.299 | -0.778 | 1.216 | 0 | -0.004 | -0.004 | 0 | 0.2 | 0.16 | 0.1 | 0.24 | 0.15 | 0.3 |
| 3 | 11073249 | rs2272403 | -0.767 | -0.913 | -0.691 | 1.263 | -0.04 | 0.024 | 0.024 | 0.977 | 0.29 | 0.67 | 0.2 | 0.4 | 0.12 | 0.3 |
| 3 | 11077898 | rs2697135 | -0.046 | -0.052 | -1.212 | 1.366 | -0.038 | -0.05 | -0.05 | 1.366 | 0.09 | 0 | 0.1 | 0.15 | 0.14 | 0.15 |
| 3 | 11079070 | rs2675165 | -0.112 | -0.123 | -1.088 | 1.339 | -0.015 | -0.026 | -0.026 | 0 | 0.1 | 0.04 | 0.1 | 0.15 | 0.14 | 0.14 |
| 3 | 11079256 | rs2675166 | -0.006 | -0.119 | -1.119 | 1.337 | -0.038 | -0.05 | -0.05 | 0.211 | 0.09 | 0 | 0.1 | 0.15 | 0.14 | 0.15 |
| 3 | 11079520 | rs2944367 | -0.349 | 0.32 | -1.276 | 1.452 | -0.005 | -0.005 | -0.005 | 3.563 | 0.17 | 0.14 | 0.15 | 0.23 | 0.22 | 0.16 |
| 3 | 11080169 | rs1062246 | -0.365 | -0.596 | -1.708 | 0.982 | 0.055 | 0 | 0 | 0.213 | 0.46 | 0.68 | 0.42 | 0.54 | 0.37 | 0.47 |
| 3 | 11086001 | rs12186082 | 0.112 | 0.327 | -1.596 | 0.923 | -0.036 | -0.028 | -0.028 | 1.136 | 0.15 | 0.09 | 0.18 | 0.21 | 0.26 | 0.24 |
| 3 | 11086548 | rs1400155 | 0.564 | 0.62 | -1.834 | 0.874 | -0.062 | 0.017 | 0.017 | 0.032 | 0.14 | 0.01 | 0.1 | 0.07 | 0.23 | 0.12 |
| 3 | 11087044 | rs11719708 | -0.771 | -0.288 | -1.009 | 0.993 | 0.005 | 0 | 0 | 0 | 0.39 | 0.65 | 0.3 | 0.38 | 0.38 | 0.32 |
| 3 | 11092264 | rs11711623 | -0.324 | -0.445 | -0.69 | 1.002 | 0.006 | 0.018 | 0.018 | 0 | 0.2 | 0.36 | 0.19 | 0.24 | 0.2 | 0.28 |
| 3 | 11092828 | rs11128537 | -0.67 | -0.479 | -1.27 | 0.87 | 0.027 | -0.016 | -0.016 | 0 | 0.41 | 0.71 | 0.36 | 0.44 | 0.33 | 0.49 |
| 3 | 11096975 | rs6763039 | -0.365 | -0.039 | -1.44 | 0.274 | 0.01 | 0.031 | 0.031 | 0 | 0.39 | 0.3 | 0.31 | 0.25 | 0.34 | 0.36 |
| 3 | 11107761 | rs2930142 | -0.214 | 0.063 | -2.215 | 0.224 | 0.031 | -0.083 | -0.083 | 0 | 0.42 | 0.89 | 0.58 | 0.53 | 0.36 | 0.21 |
| 3 | 11115921 | rs12495227 | 1.049 | 0.727 | -2.429 | -0.183 | -0.025 | 0.031 | 0.031 | 0 | 0.12 | 0.06 | 0.07 | 0 | 0.15 | 0.1 |
| 3 | 11118696 | rs17033921 | 0.429 | 0.153 | -0.985 | 0.3 | 0.005 | 0.205 | 0.205 | 0 | 0.1 | 0.25 | 0.12 | 0.27 | 0.06 | 0.03 |
| 3 | 11118817 | rs2675152 | 0.842 | -0.177 | -0.853 | 0.094 | -0.007 | -0.003 | -0.003 | 0 | 0.21 | 0.15 | 0.42 | 0.26 | 0.26 | 0.2 |
| 3 | 11125942 | rs7618934 | -0.447 | 0.05 | -0.801 | -0.22 | 0.015 | -0.137 | -0.137 | 0 | 0.19 | 0.54 | 0.17 | 0.01 | 0.17 | 0.06 |
| 3 | 11129858 | rs349714 | 1.042 | 0.052 | -0.929 | -0.201 | -0.038 | -0.037 | -0.037 | 0 | 0.51 | 0.08 | 0.31 | 0.46 | 0.52 | 0.72 |
| 3 | 11130020 | rs349715 | -1.825 | -0.046 | -1.285 | -0.585 | -0.128 | 0.16 | 0.16 | 0 | 0.75 | 0.99 | 0.86 | 1 | 0.71 | 0.84 |
| 3 | 11130342 | rs17033937 | -0.858 | 0.292 | 0.282 | 0.16 | 0 | 0.014 | 0.014 | 0 | 0.21 | 0.24 | 0.2 | 0.29 | 0.19 | 0.08 |
| 3 | 11134319 | rs9873230 | -0.025 | 0.341 | 0.286 | 0.139 | 0.011 | -0.001 | -0.001 | 0 | 0.06 | 0.39 | 0.05 | 0 | 0.02 | 0.01 |
| 3 | 11153613 | rs9990176 | -1.767 | 0.16 | 0.424 | 0.604 | -0.012 | 0.012 | 0.012 | 0 | 0.29 | 0.6 | 0.25 | 0.39 | 0.24 | 0.31 |
| 3 | 11154466 | rs349707 | 0.476 | 0.154 | 0.297 | 0.554 | -0.001 | -0.006 | -0.006 | 0 | 0.28 | 0.2 | 0.49 | 0.29 | 0.31 | 0.15 |
| 3 | 11157856 | rs7625056 | -1.068 | 0.123 | 0.272 | 0.575 | -0.035 | 0 | 0 | 3.571 | 0.14 | 0.48 | 0.11 | 0.12 | 0.07 | 0.09 |
| 3 | 11161938 | rs6810358 | -0.021 | 0.222 | -0.027 | 0.655 | -0.004 | 0.013 | 0.013 | 0.85 | 0.35 | 0.4 | 0.55 | 0.39 | 0.34 | 0.24 |
| 3 | 11163408 | rs17034004 | -0.706 | 0.16 | -0.067 | 0.633 | 0.003 | 0.011 | 0.011 | 0.044 | 0.16 | 0.19 | 0.16 | 0.27 | 0.18 | 0.23 |
| 3 | 11165909 | rs4487224 | -0.071 | 0.219 | -0.109 | 0.63 | -0.004 | -0.006 | -0.006 | 0 | 0.35 | 0.39 | 0.55 | 0.28 | 0.34 | 0.15 |
| 3 | 11168333 | rs3915855 | -1.335 | 0.097 | -0.718 | 0.588 | -0.114 | -0.011 | -0.011 | 0 | 0.58 | 0.93 | 0.76 | 0.67 | 0.54 | 0.47 |
| 3 | 11174324 | rs729327 | -1.092 | -0.128 | 0.27 | 0.721 | -0.023 | 0.033 | 0.033 | 0.104 | 0.11 | 0.18 | 0.08 | 0.23 | 0.07 | 0.18 |
| 3 | 11176867 | rs6778270 | -1.062 | 0.171 | 0.113 | 0.721 | -0.001 | 0.003 | 0.003 | 0 | 0.11 | 0.1 | 0.1 | 0.21 | 0.1 | 0.19 |
| 3 | 11200110 | rs6796787 | -1.791 | 0.423 | 0.6 | 0.733 | 0.021 | 0.078 | 0.078 | 0 | 0.19 | 0.38 | 0.18 | 0.29 | 0.16 | 0.26 |
| 3 | 11205350 | rs2171544 | 0.117 | 0.463 | -0.147 | 0.418 | -0.02 | 0.075 | 0.075 | 1.206 | 0.24 | 0.09 | 0.41 | 0.09 | 0.26 | 0.13 |
| 3 | 11206322 | rs17034063 | -1.704 | 0.295 | 0.775 | 1.032 | -0.009 | 0.031 | 0.031 | 0 | 0.16 | 0.21 | 0.15 | 0.29 | 0.12 | 0.25 |
| 3 | 11206966 | rs11929549 | -2.288 | 0.15 | 0.83 | 1.073 | -0.004 | 0.016 | 0.016 | 0.625 | 0.23 | 0.66 | 0.24 | 0.37 | 0.18 | 0.27 |
| 3 | 11211338 | rs17034071 | -1.331 | 0.121 | 0.67 | 0.912 | -0.003 | -0.021 | -0.021 | 0.244 | 0.11 | 0.06 | 0.13 | 0.23 | 0.09 | 0.24 |

| | | | | | | | | | | | | | | | |
|---|---|---|---|---|---|---|---|---|---|---|---|---|---|---|---|
| 3 | 11211697 | rs4684761 | -1.53 | -0.251 | 0.794 | 0.834 | -0.013 | 0.034 | 0.034 | 0 | 0.11 | 0.14 | 0.13 | 0.23 | 0.07 | 0.23 |
| 3 | 11213268 | rs6442233 | -1.779 | 0.1 | 1.029 | 0.268 | -0.018 | -0.065 | -0.065 | 0 | 0.16 | 0.5 | 0.11 | 0.07 | 0.09 | 0.02 |
| 3 | 11213477 | rs6442234 | -1.861 | 0.099 | 1.014 | 0.268 | -0.022 | -0.061 | -0.061 | 0 | 0.17 | 0.56 | 0.11 | 0.07 | 0.09 | 0.02 |
| 3 | 11219002 | rs17034093 | -1.18 | 0.351 | 0.589 | 0.189 | 0.02 | -0.085 | -0.085 | 0.01 | 0.13 | 0.42 | 0.08 | 0 | 0.07 | 0.01 |
| 3 | 11220278 | rs13077064 | -1.096 | 0.386 | 0.242 | 0.205 | 0.01 | -0.081 | -0.081 | 0.712 | 0.12 | 0.35 | 0.08 | 0 | 0.08 | 0.01 |
| 3 | 11227460 | rs168333 | -0.041 | 0.225 | -0.997 | 0.31 | 0.076 | 0.073 | 0.073 | 0 | 0.5 | 0.24 | 0.69 | 0.34 | 0.65 | 0.53 |
| 3 | 11230464 | rs1552498 | -0.226 | 1.426 | -0.96 | 0.37 | 0.017 | 0.009 | 0.009 | 0.021 | 0.1 | 0.14 | 0.38 | 0.18 | 0.16 | 0.24 |
| 3 | 11234129 | rs238615 | 0.55 | 0.284 | -0.75 | 0.385 | -0.011 | 0.002 | 0.002 | 0 | 0.44 | 0.43 | 0.26 | 0.44 | 0.31 | 0.5 |
| 3 | 11234185 | rs7639145 | -0.876 | 0.954 | -0.996 | 0.134 | 0.015 | 0.015 | 0.015 | 0 | 0.39 | 0.55 | 0.63 | 0.56 | 0.5 | 0.44 |
| 3 | 11239076 | rs7630582 | -0.02 | 1.188 | -1.01 | 0.403 | -0.012 | -0.018 | -0.018 | 0 | 0.16 | 0.16 | 0.41 | 0.34 | 0.22 | 0.29 |
| 3 | 11239366 | rs9822871 | 0.582 | 1.082 | -0.899 | 0.326 | -0.022 | -0.008 | -0.008 | 4.466 | 0.15 | 0.1 | 0.2 | 0.17 | 0.29 | 0.09 |
| 3 | 11249695 | rs13061242 | 0.746 | -0.157 | -0.687 | 0.265 | -0.003 | -0.001 | -0.001 | 0 | 0.31 | 0.27 | 0.22 | 0.3 | 0.21 | 0.39 |
| 3 | 11249717 | rs7625756 | 0.715 | -0.17 | -0.487 | 0.259 | -0.022 | -0.005 | -0.005 | 0 | 0.32 | 0.33 | 0.22 | 0.3 | 0.21 | 0.39 |
| 3 | 11253721 | rs443137 | 0.737 | 0.177 | -0.898 | 0.46 | -0.083 | -0.071 | -0.071 | 0 | 0.21 | 0.06 | 0.21 | 0.37 | 0.36 | 0.13 |
| 3 | 11254071 | rs430353 | 0.023 | 0.032 | -0.541 | 0.346 | -0.002 | -0.002 | -0.002 | 0 | 0.55 | 0.46 | 0.46 | 0.68 | 0.61 | 0.54 |
| 3 | 11254494 | rs617156 | -1.078 | -0.364 | -0.412 | 0.074 | -0.026 | -0.069 | -0.069 | 0 | 0.89 | 0.68 | 0.93 | 1 | 0.94 | 0.95 |
| 3 | 11260221 | rs13353463 | 0.075 | -0.267 | -0.48 | 0.222 | 0.007 | 0.018 | 0.018 | 0 | 0.58 | 0.6 | 0.47 | 0.69 | 0.61 | 0.54 |
| 3 | 11265322 | rs6773737 | 1.205 | -0.471 | -0.094 | 0.355 | -0.001 | -0.001 | -0.001 | 0 | 0.34 | 0.27 | 0.21 | 0.3 | 0.23 | 0.41 |
| 3 | 11274354 | rs346087 | -0.423 | -0.579 | 0.143 | 0.299 | 0.003 | -0.002 | -0.002 | 0 | 0.66 | 0.69 | 0.53 | 0.68 | 0.68 | 0.6 |
| 3 | 11275244 | rs12496673 | 1.733 | -0.59 | 0.175 | 0.299 | -0.011 | -0.029 | -0.029 | 0 | 0.08 | 0.11 | 0.03 | 0 | 0.04 | 0.06 |
| 3 | 11275481 | rs6773307 | 0.994 | -0.568 | 0.229 | 0.165 | -0.001 | 0.045 | 0.045 | 0.423 | 0.11 | 0.21 | 0.34 | 0.31 | 0.12 | 0.34 |
| 3 | 11284950 | rs1666988 | -0.237 | -0.58 | 0.213 | 0.219 | -0.001 | -0.001 | -0.001 | 0.005 | 0.65 | 0.58 | 0.51 | 0.72 | 0.68 | 0.58 |
| 3 | 11290122 | rs347591 | 0.314 | -0.614 | 0.296 | 0.206 | -0.011 | -0.018 | -0.018 | 0 | 0.57 | 0.44 | 0.48 | 0.72 | 0.64 | 0.52 |
| 3 | 11292682 | rs346076 | 0.543 | -0.707 | 0.277 | 0.327 | 0.015 | 0.012 | 0.012 | 0 | 0.55 | 0.3 | 0.47 | 0.64 | 0.63 | 0.51 |
| 3 | 11300707 | rs901865 | -1.009 | -0.889 | 0.17 | 0 | 0 | -0.05 | -0.05 | 0 | 0.79 | 0.67 | 0.87 | 0.91 | 0.81 | 0.93 |
| 3 | 11302884 | rs1809049 | 0.728 | -0.931 | 0.128 | 0.263 | -0.014 | 0.049 | 0.049 | 0 | 0.17 | 0.34 | 0.39 | 0.28 | 0.14 | 0.37 |
| 3 | 11303874 | rs346070 | -1.457 | -0.88 | -0.028 | 0.257 | -0.007 | 0.018 | 0.018 | 0 | 0.82 | 0.89 | 0.88 | 1 | 0.81 | 0.93 |
| 3 | 11309274 | rs367745 | 0.766 | -0.84 | -0.042 | 0.317 | 0.01 | 0.007 | 0.007 | 0 | 0.55 | 0.31 | 0.47 | 0.64 | 0.64 | 0.54 |
| 3 | 11350553 | rs2305686 | 0.863 | -0.816 | -0.083 | 0.384 | 0.004 | -0.018 | -0.018 | 0 | 0.18 | 0.19 | 0.39 | 0.29 | 0.18 | 0.36 |
| 3 | 11352585 | rs11927249 | 1.192 | -0.886 | 0.016 | 0.299 | -0.031 | -0.047 | -0.047 | 0 | 0.09 | 0.15 | 0.02 | 0 | 0.03 | 0.04 |
| 3 | 11360997 | rs2594992 | 1.567 | -0.605 | -0.402 | 0.58 | -0.109 | -0.101 | -0.101 | 0 | 0.17 | 0.01 | 0.23 | 0.33 | 0.35 | 0.11 |
| 3 | 11381973 | rs3816380 | 0.59 | -1.112 | 0.036 | 0.453 | 0.002 | -0.009 | -0.009 | 0 | 0.18 | 0.2 | 0.39 | 0.3 | 0.18 | 0.35 |
| 3 | 11393268 | rs11128547 | 0.554 | -1.075 | -0.107 | 0.194 | 0.002 | -0.01 | -0.01 | 0 | 0.18 | 0.2 | 0.39 | 0.3 | 0.18 | 0.35 |
| 3 | 11397417 | rs2594972 | 0.081 | -1.224 | -0.233 | -0.003 | 0.001 | 0 | 0 | 0 | 0.65 | 0.6 | 0.49 | 0.69 | 0.63 | 0.58 |
| 3 | 11403441 | rs2248971 | 0.734 | -1.257 | -0.25 | -0.028 | 0.002 | 0.003 | 0.003 | 0 | 0.53 | 0.36 | 0.45 | 0.69 | 0.59 | 0.53 |
| 3 | 11418924 | rs12636305 | 0.226 | -1.184 | -0.26 | 0.106 | 0 | 0.002 | 0.002 | 0 | 0.21 | 0.28 | 0.41 | 0.29 | 0.18 | 0.35 |
| 3 | 11461863 | rs9826803 | 0.184 | -1.247 | -0.192 | -0.032 | -0.001 | -0.002 | -0.002 | 0 | 0.64 | 0.53 | 0.47 | 0.71 | 0.63 | 0.58 |
| 3 | 11474238 | rs6442253 | -1.451 | -1.199 | -0.062 | -0.134 | 0.008 | -0.056 | -0.056 | 0 | 0.84 | 0.79 | 0.89 | 1 | 0.81 | 0.94 |
| 3 | 11475297 | rs9816564 | 1.081 | -1.552 | -0.302 | -0.171 | 0.027 | 0.032 | 0.032 | 0 | 0.5 | 0.3 | 0.43 | 0.69 | 0.57 | 0.52 |
| 3 | 11483769 | rs2442781 | 0.093 | -1.541 | -0.316 | -0.242 | -0.006 | -0.083 | -0.083 | 0 | 0.7 | 0.65 | 0.82 | 0.98 | 0.73 | 0.87 |
| 3 | 11483793 | rs2442782 | 0.696 | -1.578 | -0.374 | -0.223 | 0 | 0.003 | 0.003 | 0 | 0.19 | 0.26 | 0.4 | 0.29 | 0.17 | 0.35 |
| 3 | 11487506 | rs2454512 | -1.603 | -1.497 | -0.453 | -0.239 | -0.021 | 0.032 | 0.032 | 0 | 0.84 | 0.89 | 0.88 | 1 | 0.8 | 0.94 |
| 3 | 11493926 | rs2442772 | -0.634 | -1.527 | -0.382 | -0.181 | -0.052 | 0.019 | 0.019 | 0 | 0.33 | 0.53 | 0.45 | 0.33 | 0.23 | 0.42 |
| 3 | 11497325 | rs2454505 | -0.653 | -1.539 | -0.399 | -0.188 | -0.054 | 0.017 | 0.017 | 0 | 0.34 | 0.54 | 0.46 | 0.33 | 0.24 | 0.42 |
| 3 | 11509668 | rs9818393 | 2.035 | -1.659 | -1.002 | -0.371 | -0.124 | 0.003 | 0.003 | 0 | 0.19 | 0.01 | 0.23 | 0.12 | 0.35 | 0.13 |
| 3 | 11510568 | rs2442780 | -1.825 | -1.569 | -0.721 | -0.251 | -0.015 | 0.015 | 0.015 | 0 | 0.84 | 0.88 | 0.88 | 1 | 0.8 | 0.94 |
| 3 | 11530121 | rs34621655 | 0.661 | -1.493 | -0.741 | -0.197 | 0.011 | 0.01 | 0.01 | 0 | 0.53 | 0.34 | 0.44 | 0.67 | 0.59 | 0.53 |
| 3 | 11539955 | rs11128552 | 0.78 | -1.511 | -1.066 | -0.078 | 0.007 | -0.014 | -0.014 | 1.452 | 0.19 | 0.2 | 0.39 | 0.31 | 0.15 | 0.34 |
| 3 | 11547936 | rs13086126 | 0.827 | -1.55 | -1.207 | -0.135 | 0.02 | -0.071 | -0.071 | 0 | 0.2 | 0.09 | 0.36 | 0.31 | 0.17 | 0.38 |
| 3 | 11551200 | rs2442795 | -1.312 | -1.447 | -1.219 | -0.246 | -0.012 | 0.059 | 0.059 | 0 | 0.81 | 0.93 | 0.88 | 0.97 | 0.79 | 0.92 |
| 3 | 11552027 | rs2447607 | 0.956 | -1.616 | -1.062 | -0.278 | 0.011 | 0.011 | 0.011 | 0 | 0.5 | 0.4 | 0.46 | 0.66 | 0.58 | 0.5 |
| 3 | 11562627 | rs2344826 | -1.23 | -1.587 | -1.332 | -0.434 | -0.018 | 0.009 | 0.009 | 7.464 | 0.93 | 0.99 | 0.95 | 0.95 | 0.91 | 0.81 |
| 3 | 11564940 | rs9840982 | 1.167 | -1.565 | -1.14 | -0.337 | -0.006 | 0.019 | 0.019 | 0 | 0.19 | 0.11 | 0.12 | 0.09 | 0.21 | 0.08 |
| 3 | 11568144 | rs9861286 | -0.136 | -1.613 | -1.263 | -0.355 | -0.024 | -0.113 | -0.113 | 0 | 0.3 | 0.5 | 0.18 | 0.09 | 0.25 | 0.12 |
| 3 | 11578151 | rs6442259 | -0.696 | -1.194 | -1.208 | 0.006 | 0.008 | 0.003 | 0.003 | 0 | 0.45 | 0.66 | 0.55 | 0.38 | 0.41 | 0.46 |
| 3 | 11579865 | rs4684787 | 1.619 | -0.55 | -0.995 | 0.086 | 0.015 | 0.026 | 0.026 | 0 | 0.23 | 0.23 | 0.23 | 0.44 | 0.3 | 0.17 |
| 3 | 11591200 | rs9985349 | 1.202 | -0.611 | -1.094 | 0.038 | -0.02 | 0.024 | 0.024 | 0 | 0.16 | 0.11 | 0.13 | 0 | 0.22 | 0.07 |
| 3 | 11593013 | rs2437685 | 0.156 | -0.677 | -1.007 | 0.059 | 0.001 | -0.056 | -0.056 | 0.598 | 0.22 | 0.33 | 0.16 | 0.01 | 0.23 | 0.07 |
| 3 | 11597277 | rs9855986 | 0.4 | -1.099 | -1.113 | 0.124 | -0.04 | -0.053 | -0.053 | 3.78 | 0.1 | 0.2 | 0.05 | 0 | 0.03 | 0.03 |
| 3 | 11600563 | rs7617620 | -0.491 | -0.603 | -1.914 | 0.358 | -0.036 | 0.016 | 0.016 | 0 | 0.73 | 0.9 | 0.72 | 0.81 | 0.67 | 0.57 |
| 3 | 11602131 | rs17034661 | 0.468 | -1.266 | -1.55 | 0.802 | 0.005 | -0.046 | -0.046 | 5.221 | 0.19 | 0.07 | 0.31 | 0.29 | 0.16 | 0.32 |
| 3 | 11605373 | rs9832407 | -0.085 | -0.994 | -1.296 | 0.503 | 0.003 | 0.036 | 0.036 | 0 | 0.19 | 0.13 | 0.12 | 0.09 | 0.19 | 0.1 |
| 3 | 11609198 | rs2443728 | 0.584 | -0.912 | -1.309 | 0.368 | -0.025 | 0.061 | 0.061 | 0 | 0.15 | 0.05 | 0.11 | 0 | 0.18 | 0.09 |
| 3 | 11610729 | rs730178 | 0.3 | -1.228 | -0.884 | 0.447 | -0.036 | 0.031 | 0.031 | 0 | 0.2 | 0.39 | 0.31 | 0.21 | 0.15 | 0.33 |
| 3 | 11611768 | rs2596814 | 0.201 | -1.325 | -1.382 | 0.65 | 0.001 | -0.004 | -0.004 | 0 | 0.31 | 0.28 | 0.28 | 0.4 | 0.23 | 0.35 |
| 3 | 11614969 | rs7640358 | 0.953 | -1.321 | -1.191 | 0.606 | -0.019 | 0.037 | 0.037 | 0 | 0.18 | 0.32 | 0.3 | 0.21 | 0.15 | 0.33 |
| 3 | 11615207 | rs2574706 | -0.143 | -0.821 | -1.815 | 0.703 | 0.001 | -0.003 | -0.003 | 0 | 0.32 | 0.29 | 0.29 | 0.37 | 0.24 | 0.35 |
| 3 | 11617308 | rs2004837 | -0.245 | 0.135 | -2.182 | 0.455 | -0.034 | -0.012 | -0.012 | 0 | 0.93 | 1 | 0.86 | 0.95 | 0.89 | 0.84 |
| 3 | 11619958 | rs11712666 | 0.813 | -0.033 | -1.959 | 0.545 | -0.13 | -0.004 | -0.004 | 0 | 0.3 | 0.04 | 0.28 | 0.3 | 0.45 | 0.23 |

| | | | | | | | | | | | | | | | | |
|---|---|---|---|---|---|---|---|---|---|---|---|---|---|---|---|---|
| 3 | 11625202 | rs6766130 | 0.423 | -1.062 | -1.057 | 0.806 | 0.011 | -0.001 | -0.001 | 0 | 0.61 | 0.59 | 0.53 | 0.47 | 0.73 | 0.41 |
| 3 | 11627016 | rs12493225 | 0.531 | -1.235 | -0.993 | 1.26 | 0.001 | 0.002 | 0.002 | 0 | 0.92 | 0.95 | 0.81 | 0.82 | 0.95 | 0.77 |
| 3 | 11627554 | rs2447617 | 1.292 | -2.083 | -1.405 | 0.993 | 0.096 | 0.027 | 0.027 | 0.39 | 0.51 | 0.21 | 0.44 | 0.42 | 0.67 | 0.31 |
| 3 | 11627978 | rs2574726 | -0.139 | -2.03 | -1.328 | 0.982 | 0 | 0.008 | 0.008 | 0 | 0.08 | 0.05 | 0.19 | 0.17 | 0.05 | 0.23 |
| 3 | 11628813 | rs6442262 | -0.468 | -2.487 | -0.975 | 1.061 | 0 | -0.001 | -0.001 | 0.007 | 0.16 | 0.18 | 0.07 | 0.24 | 0.08 | 0.2 |
| 3 | 11630533 | rs3856796 | -0.088 | -2.489 | -0.186 | 1.063 | 0 | 0.012 | 0.012 | 0 | 0.92 | 0.93 | 0.81 | 0.82 | 0.95 | 0.78 |
| 3 | 11630569 | rs3893266 | -1.026 | -2.643 | 0.306 | 0.951 | -0.033 | 0.013 | 0.013 | 0 | 0.2 | 0.34 | 0.11 | 0.24 | 0.08 | 0.2 |
| 3 | 11633951 | rs17776482 | 0.356 | -2.699 | 0.127 | 1.046 | -0.013 | -0.034 | -0.034 | 0.573 | 0.15 | 0.02 | 0.22 | 0.19 | 0.16 | 0.25 |
| 3 | 11633979 | rs17034742 | 0.338 | -2.76 | 0.141 | 1.015 | 0.002 | 0.002 | 0.002 | 0 | 0.07 | 0.05 | 0.02 | 0.19 | 0.04 | 0.14 |
| 3 | 11635631 | rs9870506 | -0.851 | -2.764 | 0.226 | 0.97 | -0.036 | -0.037 | -0.037 | 1.022 | 0.24 | 0.47 | 0.3 | 0.18 | 0.19 | 0.26 |
| 3 | 11640601 | rs2442776 | 1.42 | -2.625 | 0.243 | 1.186 | -0.003 | 0.005 | 0.005 | 0.038 | 0.76 | 0.75 | 0.75 | 0.65 | 0.87 | 0.63 |
| 3 | 11643465 | rs2276749 | -0.717 | -2.613 | 0.17 | 1.258 | -0.001 | -0.01 | -0.01 | 0 | 0.94 | 1 | 0.89 | 0.94 | 0.95 | 0.83 |
| 3 | 11647173 | rs892935 | 0.008 | -2.6 | 0.202 | 1.285 | -0.003 | 0.01 | 0.01 | 0 | 0.93 | 0.93 | 0.83 | 0.89 | 0.95 | 0.82 |
| 3 | 11647301 | rs892936 | -0.205 | -2.596 | 0.189 | 1.286 | 0 | 0.002 | 0.002 | 0 | 0.94 | 0.96 | 0.83 | 0.89 | 0.95 | 0.83 |
| 3 | 11656182 | rs6769370 | -1.42 | -2.975 | 0.222 | 1.088 | -0.042 | 0.023 | 0.023 | 0 | 0.23 | 0.58 | 0.17 | 0.3 | 0.09 | 0.2 |
| 3 | 11657278 | rs17034817 | -1.107 | -2.981 | 0.379 | 1.07 | -0.032 | 0.027 | 0.027 | 0 | 0.21 | 0.32 | 0.14 | 0.3 | 0.09 | 0.19 |
| 3 | 11662122 | rs2574716 | -2.478 | -3.055 | 0.034 | 1.014 | 0.067 | 0.006 | 0.006 | 0.003 | 0.55 | 0.86 | 0.59 | 0.6 | 0.34 | 0.72 |
| 3 | 11662320 | rs7651349 | 2.46 | -2.692 | -0.307 | 1.076 | -0.151 | 0.01 | 0.01 | 0 | 0.37 | 0.03 | 0.33 | 0.26 | 0.51 | 0.17 |
| 3 | 11662730 | rs9865799 | -0.633 | -2.59 | -0.123 | 0.672 | -0.03 | -0.018 | -0.018 | 0.595 | 0.15 | 0.22 | 0.09 | 0.07 | 0.05 | 0.09 |
| 3 | 11663976 | rs9871522 | -0.633 | -2.59 | -0.123 | 0.672 | -0.03 | -0.018 | -0.018 | 0.013 | 0.15 | 0.22 | 0.09 | 0.07 | 0.05 | 0.09 |
| 3 | 11664859 | rs6766610 | 1.54 | -2.578 | 0.02 | 0.722 | 0.008 | 0.006 | 0.006 | 0 | 0.49 | 0.54 | 0.48 | 0.52 | 0.53 | 0.29 |
| 3 | 11667123 | rs2616551 | -0.983 | -2.66 | -0.129 | 0.662 | 0.021 | 0.008 | 0.008 | 0.232 | 0.28 | 0.24 | 0.38 | 0.29 | 0.19 | 0.47 |
| 3 | 11667766 | rs2574711 | -0.29 | -2.666 | -0.155 | 0.615 | -0.003 | -0.002 | -0.002 | 0.527 | 0.12 | 0.16 | 0.17 | 0.11 | 0.06 | 0.21 |
| 3 | 11669322 | rs2616548 | -1.727 | -2.831 | -0.149 | 0.595 | -0.056 | -0.061 | -0.061 | 0.033 | 0.29 | 0.53 | 0.25 | 0.13 | 0.11 | 0.28 |
| 3 | 11670573 | rs2447613 | 0.189 | -2.461 | -0.318 | 0.727 | -0.042 | -0.019 | -0.019 | 0.274 | 0.18 | 0.08 | 0.13 | 0.17 | 0.27 | 0.21 |
| 3 | 11671484 | rs6767949 | 0.868 | -2.557 | -0.131 | 0.545 | -0.024 | -0.044 | -0.044 | 3.375 | 0.11 | 0.1 | 0.03 | 0 | 0.04 | 0.06 |
| 3 | 11672805 | rs6783054 | 1.141 | -1.717 | -0.368 | 0.809 | -0.004 | -0.006 | -0.006 | 0.205 | 0.48 | 0.42 | 0.42 | 0.49 | 0.5 | 0.3 |
| 3 | 11674229 | rs9863261 | 1.444 | -1.604 | -0.868 | 0.804 | -0.058 | -0.018 | -0.018 | 0 | 0.44 | 0.19 | 0.38 | 0.42 | 0.5 | 0.22 |
| 3 | 11675789 | rs6782029 | -1.329 | -1.393 | -0.783 | 0.916 | 0.009 | -0.043 | -0.043 | 0 | 0.28 | 0.19 | 0.4 | 0.39 | 0.23 | 0.5 |
| 3 | 11677077 | rs11128560 | 0.37 | -1.251 | -0.774 | 0.882 | -0.039 | -0.043 | -0.043 | 0 | 0.09 | 0 | 0.19 | 0.18 | 0.11 | 0.24 |
| 3 | 11679509 | rs11128564 | -1.402 | -1.497 | -0.294 | 0.899 | -0.014 | 0.017 | 0.017 | 0.57 | 0.21 | 0.31 | 0.31 | 0.32 | 0.14 | 0.37 |
| 3 | 11681487 | rs2616543 | 0.841 | -1.387 | -0.187 | 0.581 | -0.009 | -0.004 | -0.004 | 1.032 | 0.57 | 0.41 | 0.68 | 0.61 | 0.63 | 0.62 |
| 3 | 11682129 | rs2443704 | 0.029 | -1.283 | -0.252 | 0.552 | -0.013 | -0.008 | -0.008 | 0.198 | 0.9 | 0.89 | 0.87 | 0.95 | 0.94 | 0.88 |
| 3 | 11690450 | rs7613596 | 1.302 | -1.418 | -0.357 | 0.519 | -0.001 | -0.001 | -0.001 | 0 | 0.42 | 0.41 | 0.47 | 0.59 | 0.51 | 0.4 |
| 3 | 11690639 | rs7646184 | -0.111 | -1.428 | -0.451 | 0.611 | -0.01 | -0.01 | -0.01 | 0 | 0.8 | 0.81 | 0.81 | 0.8 | 0.74 | 0.8 |
| 3 | 11692869 | rs6804121 | -0.099 | -1.725 | -0.242 | 0.577 | -0.014 | 0.001 | 0.001 | 0.169 | 0.12 | 0.14 | 0.13 | 0.1 | 0.07 | 0.14 |
| 3 | 11693869 | rs17034986 | 0.343 | -1.907 | -0.38 | 0.36 | 0.006 | 0.009 | 0.009 | 0.298 | 0.13 | 0.11 | 0.06 | 0.03 | 0.1 | 0.11 |
| 3 | 11694467 | rs4684792 | 0.159 | -1.849 | -0.449 | 0.823 | -0.009 | -0.036 | -0.036 | 0 | 0.77 | 0.79 | 0.8 | 0.62 | 0.75 | 0.62 |
| 3 | 11698214 | rs3846114 | -0.177 | -1.493 | 0.007 | 0.707 | -0.007 | 0.006 | 0.006 | 0.21 | 0.13 | 0.21 | 0.16 | 0.1 | 0.12 | 0.17 |
| 3 | 11700713 | rs2346021 | 1.348 | -1.623 | -0.144 | 0.628 | 0.032 | -0.012 | -0.012 | 0.108 | 0.57 | 0.24 | 0.59 | 0.5 | 0.6 | 0.49 |
| 3 | 11702456 | rs7650685 | 0.86 | -1.55 | -0.057 | 0.685 | -0.006 | 0.01 | 0.01 | 4.553 | 0.65 | 0.54 | 0.65 | 0.51 | 0.66 | 0.51 |
| 3 | 11703635 | rs11715990 | -0.134 | -1.668 | 0.121 | 0.845 | 0.001 | 0.02 | 0.02 | 0 | 0.83 | 0.78 | 0.77 | 0.75 | 0.8 | 0.81 |
| 3 | 11710471 | rs17035071 | -0.804 | -1.418 | -0.27 | 0.652 | -0.01 | -0.011 | -0.011 | 0 | 0.25 | 0.35 | 0.3 | 0.27 | 0.18 | 0.35 |
| 3 | 11711190 | rs3856804 | -1.039 | -1.39 | -0.34 | 0.663 | -0.003 | -0.009 | -0.009 | 0 | 0.3 | 0.35 | 0.33 | 0.28 | 0.26 | 0.36 |
| 3 | 11714521 | rs7627256 | 0.896 | -1.428 | -0.98 | 0.884 | -0.026 | -0.036 | -0.036 | 0.046 | 0.41 | 0.19 | 0.46 | 0.46 | 0.53 | 0.44 |
| 3 | 11714890 | rs6442274 | 0.392 | -1.64 | -0.619 | 0.924 | 0.003 | 0.001 | 0.001 | 0.118 | 0.57 | 0.42 | 0.58 | 0.58 | 0.61 | 0.53 |
| 3 | 11730767 | rs13097188 | 0.168 | -1.465 | -0.559 | 0.864 | 0.001 | 0.001 | 0.001 | 0 | 0.14 | 0.14 | 0.27 | 0.12 | 0.09 | 0.1 |
| 3 | 11731029 | rs1039201 | 0.923 | -2.078 | -0.69 | 1.069 | 0.027 | 0.056 | 0.056 | 0 | 0.53 | 0.31 | 0.49 | 0.31 | 0.61 | 0.43 |
| 3 | 11731454 | rs4143285 | -0.984 | -2.232 | -0.442 | 1.311 | 0 | 0.008 | 0.008 | 0 | 0.81 | 0.78 | 0.88 | 0.69 | 0.79 | 0.79 |
| 3 | 11732276 | rs6782324 | 0.632 | -2.376 | -0.079 | 0.989 | -0.007 | 0.008 | 0.008 | 0 | 0.61 | 0.44 | 0.53 | 0.53 | 0.67 | 0.67 |
| 3 | 11736267 | rs9835698 | -0.305 | -2.327 | -0.387 | 1.069 | 0.001 | -0.04 | -0.04 | 2.966 | 0.72 | 0.77 | 0.72 | 0.53 | 0.69 | 0.6 |
| 3 | 11740887 | rs6786278 | 0.241 | -2.48 | -0.479 | 0.983 | 0.002 | -0.003 | -0.003 | 1.715 | 0.13 | 0.11 | 0.07 | 0.21 | 0.08 | 0.1 |
| 3 | 11744584 | rs6781438 | 1.421 | -2.252 | -0.417 | 1.108 | -0.013 | 0.019 | 0.019 | 0 | 0.4 | 0.28 | 0.4 | 0.32 | 0.53 | 0.39 |
| 3 | 11746737 | rs6788714 | 0.199 | -2.119 | -0.63 | 1.027 | 0.021 | -0.03 | -0.03 | 0 | 0.6 | 0.71 | 0.74 | 0.53 | 0.61 | 0.58 |
| 3 | 11758513 | rs4499579 | 0.916 | -1.65 | -0.615 | 0.997 | -0.005 | 0.003 | 0.003 | 0 | 0.39 | 0.35 | 0.42 | 0.36 | 0.53 | 0.34 |
| 3 | 11763001 | rs11128570 | 0.652 | -1.807 | -1.049 | 1.277 | 0.059 | 0.015 | 0.015 | 0.037 | 0.51 | 0.21 | 0.54 | 0.44 | 0.58 | 0.54 |
| 3 | 11767553 | rs7621951 | 0.272 | -1.664 | -0.956 | 1.232 | 0.004 | 0.001 | 0.001 | 0 | 0.31 | 0.23 | 0.22 | 0.27 | 0.35 | 0.35 |
| 3 | 11772784 | rs4684078 | -0.798 | -1.694 | -0.911 | 1.205 | 0.022 | 0.043 | 0.043 | 0 | 0.49 | 0.39 | 0.44 | 0.67 | 0.44 | 0.52 |
| 3 | 11772815 | rs4684079 | -0.826 | -1.364 | -0.504 | 1.589 | 0.004 | -0.004 | -0.004 | 0.06 | 0.18 | 0.16 | 0.22 | 0.4 | 0.1 | 0.16 |
| 3 | 11774148 | rs6810096 | -0.718 | -0.91 | -0.233 | 1.396 | -0.01 | 0.05 | 0.05 | 0 | 0.18 | 0.25 | 0.26 | 0.27 | 0.13 | 0.19 |
| 3 | 11778256 | rs4684800 | -0.23 | 0.258 | -0.507 | 1.048 | -0.011 | 0.007 | 0.007 | 0 | 0.71 | 0.76 | 0.63 | 0.82 | 0.66 | 0.7 |
| 3 | 11779370 | rs6442280 | -0.222 | -0.11 | 0.107 | 0.892 | 0.047 | 0.067 | 0.067 | 0.798 | 0.06 | 0.42 | 0.03 | 0.02 | 0.01 | 0 |
| 3 | 11781002 | rs9814925 | 0.221 | 0.03 | -0.054 | 1.139 | 0.018 | 0.004 | 0.004 | 0 | 0.64 | 0.56 | 0.41 | 0.65 | 0.6 | 0.58 |
| 3 | 11781682 | rs9815906 | -0.52 | 0.293 | -0.897 | 0.594 | -0.002 | -0.032 | -0.032 | 1.538 | 0.87 | 0.85 | 0.78 | 1 | 0.92 | 0.87 |
| 3 | 11781717 | rs17035113 | 0.582 | 0.579 | -1.077 | 1.026 | -0.028 | -0.03 | -0.03 | 1.883 | 0.05 | 0.01 | 0.19 | 0.13 | 0.07 | 0.13 |
| 3 | 11785147 | rs370234 | -0.912 | 0.369 | -1.568 | 0.827 | -0.009 | 0.059 | 0.059 | 0.022 | 0.57 | 0.93 | 0.58 | 0.66 | 0.6 | 0.68 |
| 3 | 11790188 | rs6800849 | 0.404 | 0.693 | -1.287 | 0.854 | -0.012 | 0.008 | 0.008 | 0 | 0.17 | 0.1 | 0.14 | 0.15 | 0.21 | 0.16 |
| 3 | 11790710 | rs301554 | -0.255 | 0.352 | -0.739 | 1.102 | -0.002 | -0.001 | -0.001 | 0 | 0.36 | 0.59 | 0.39 | 0.33 | 0.37 | 0.45 |
| 3 | 11791680 | rs301556 | -0.518 | 0.383 | -0.373 | 1.021 | 0.003 | -0.004 | -0.004 | 0 | 0.33 | 0.43 | 0.31 | 0.37 | 0.34 | 0.35 |

| | | | | | | | | | | | | | | | | |
|---|---|---|---|---|---|---|---|---|---|---|---|---|---|---|---|---|
| 3 | 11792304 | rs9310393 | -0.069 | 0.342 | -0.582 | 1.127 | -0.006 | 0.011 | 0.011 | 0.097 | 0.07 | 0.11 | 0.09 | 0.15 | 0.06 | 0.19 |
| 3 | 11792931 | rs301557 | 0.071 | 0.386 | -0.574 | 0.936 | -0.003 | 0.003 | 0.003 | 0 | 0.25 | 0.25 | 0.2 | 0.22 | 0.26 | 0.17 |
| 3 | 11793217 | rs2133065 | -0.011 | 0.344 | -0.576 | 1.028 | -0.002 | 0.005 | 0.005 | 0 | 0.49 | 0.6 | 0.42 | 0.44 | 0.51 | 0.51 |
| 3 | 11794478 | rs9836764 | 0.3 | 0.071 | -0.593 | 0.671 | -0.02 | -0.046 | -0.046 | 0.398 | 0.33 | 0.47 | 0.29 | 0.17 | 0.26 | 0.22 |
| 3 | 11796487 | rs7647310 | -0.355 | 0.348 | -0.897 | 0.8 | -0.002 | -0.003 | -0.003 | 0 | 0.56 | 0.66 | 0.44 | 0.48 | 0.57 | 0.59 |
| 3 | 11797832 | rs17661673 | 0.77 | 0.694 | -1.277 | 0.576 | -0.088 | 0.01 | 0.01 | 0 | 0.18 | 0.01 | 0.13 | 0.13 | 0.26 | 0.32 |
| 3 | 11797881 | rs11923454 | 0.833 | 0.106 | -0.952 | 0.25 | 0.001 | 0.014 | 0.014 | 0 | 0.21 | 0.11 | 0.1 | 0.01 | 0.19 | 0.16 |
| 3 | 11798039 | rs301551 | -1.15 | -0.169 | -1.265 | 0.527 | 0.039 | -0.044 | -0.044 | 0 | 0.75 | 0.87 | 0.65 | 0.67 | 0.82 | 0.74 |
| 3 | 11798681 | rs301552 | 0.472 | 0.44 | -0.311 | 0.738 | 0.016 | 0.014 | 0.014 | 0.138 | 0.18 | 0.25 | 0.19 | 0.26 | 0.23 | 0.08 |
| 3 | 11801890 | rs17610125 | 0.474 | 0.503 | -0.72 | 0.803 | -0.031 | 0.009 | 0.009 | 0 | 0.33 | 0.15 | 0.31 | 0.3 | 0.39 | 0.47 |
| 3 | 11803927 | rs301564 | -0.347 | 0.778 | -0.085 | 0.82 | 0.051 | 0.04 | 0.04 | 0.298 | 0.19 | 0.42 | 0.2 | 0.26 | 0.24 | 0.07 |
| 3 | 11811671 | rs301567 | -0.003 | -0.135 | -0.727 | 0.614 | 0 | -0.001 | -0.001 | 0 | 0.63 | 0.34 | 0.79 | 0.57 | 0.72 | 0.74 |
| 3 | 11812866 | rs904267 | -0.156 | -0.194 | -0.783 | -0.568 | 0.026 | -0.028 | -0.028 | 0.456 | 0.37 | 0.51 | 0.51 | 0.32 | 0.44 | 0.29 |
| 3 | 11818619 | rs301571 | -0.248 | -1.08 | -1.504 | -0.655 | -0.064 | 0.136 | 0.136 | 0.602 | 0.46 | 0.04 | 0.52 | 0.15 | 0.44 | 0.33 |
| 3 | 11819683 | rs4684807 | -1.43 | -0.224 | -0.166 | -0.33 | 0.005 | 0.023 | 0.023 | 0 | 0.48 | 0.65 | 0.69 | 0.75 | 0.48 | 0.64 |
| 3 | 11830536 | rs7627219 | -1.584 | -0.164 | -0.124 | -0.299 | 0.017 | 0.079 | 0.079 | 0 | 0.5 | 0.69 | 0.7 | 0.75 | 0.49 | 0.64 |
| 3 | 11833473 | rs1498091 | -0.482 | -0.197 | 0.172 | -0.408 | 0.004 | -0.004 | -0.004 | 0.001 | 0.3 | 0.43 | 0.58 | 0.63 | 0.34 | 0.35 |
| 3 | 11836386 | rs7629273 | -1.898 | -0.172 | 0.211 | -0.355 | 0 | 0.179 | 0.179 | 0 | 0.52 | 0.84 | 0.71 | 0.75 | 0.48 | 0.64 |
| 3 | 11840903 | rs1846206 | -0.964 | -0.174 | 0.836 | 0.014 | 0 | -0.003 | -0.003 | 0 | 0.4 | 0.39 | 0.54 | 0.3 | 0.39 | 0.44 |
| 3 | 11847081 | rs6442288 | -0.879 | -0.322 | 1.094 | 0.083 | -0.028 | 0.448 | 0.448 | 0 | 0.09 | 0.44 | 0.14 | 0.45 | 0.02 | 0.18 |
| 3 | 11860657 | rs4143293 | -0.352 | -0.326 | 1.166 | 0.087 | 0.003 | 0.522 | 0.522 | 0 | 0.07 | 0.4 | 0.14 | 0.45 | 0.02 | 0.18 |
| 3 | 11861966 | rs9820117 | 0.242 | -0.353 | 0.613 | 0.037 | -0.016 | 0.032 | 0.032 | 0 | 0.05 | 0.16 | 0.05 | 0.12 | 0.01 | 0.15 |
| 3 | 11866819 | rs7617855 | -0.035 | -0.16 | 0.541 | 0.013 | -0.001 | 0.485 | 0.485 | 0 | 0.07 | 0.36 | 0.14 | 0.44 | 0.02 | 0.17 |
| 3 | 11871898 | rs1857961 | -0.78 | -0.215 | -0.225 | -0.263 | -0.004 | 0.018 | 0.018 | 0 | 0.4 | 0.28 | 0.52 | 0.29 | 0.4 | 0.46 |
| 3 | 11874762 | rs12494203 | 0.103 | 0.041 | -1.125 | -0.604 | -0.056 | 0.064 | 0.064 | 0 | 0.17 | 0.06 | 0.32 | 0.01 | 0.26 | 0.14 |
| 3 | 11878732 | rs2173331 | -0.827 | 0.21 | -0.879 | -0.121 | 0.017 | 0.199 | 0.199 | 0 | 0.52 | 0.85 | 0.68 | 0.77 | 0.43 | 0.63 |
| 3 | 11880816 | rs303862 | 0.359 | 0.639 | -0.543 | 0.061 | 0.001 | -0.002 | -0.002 | 0 | 0.56 | 0.58 | 0.5 | 0.61 | 0.55 | 0.47 |
| 3 | 11886451 | rs380999 | 0.908 | -0.048 | -0.852 | -0.383 | 0.035 | 0.097 | 0.097 | 0.052 | 0.3 | 0.16 | 0.19 | 0.02 | 0.27 | 0.19 |
| 3 | 11888556 | rs440746 | 0.019 | 0.167 | -0.099 | -0.12 | 0.007 | -0.039 | -0.039 | 0.02 | 0.6 | 0.54 | 0.42 | 0.24 | 0.46 | 0.46 |
| 3 | 11889365 | rs430373 | 0.181 | 0.093 | -0.368 | -0.148 | -0.004 | 0.05 | 0.05 | 0 | 0.57 | 0.34 | 0.4 | 0.26 | 0.46 | 0.45 |
| 3 | 11889787 | rs4684823 | 0.443 | -0.154 | -0.727 | 0.58 | -0.033 | -0.025 | -0.025 | 0 | 0.25 | 0.13 | 0.3 | 0.32 | 0.33 | 0.26 |
| 3 | 11891529 | rs9819059 | -0.247 | 0.17 | -0.582 | 0.537 | 0 | -0.002 | -0.002 | 0.586 | 0.46 | 0.58 | 0.5 | 0.53 | 0.41 | 0.32 |
| 3 | 11895450 | rs1605404 | -0.026 | 0.418 | -0.508 | 0.753 | 0.004 | -0.002 | -0.002 | 0 | 0.46 | 0.43 | 0.45 | 0.48 | 0.39 | 0.31 |
| 3 | 11898859 | rs2133064 | -0.796 | 0.528 | -0.513 | 1.057 | 0.004 | 0.028 | 0.028 | 0.14 | 0.05 | 0.12 | 0.07 | 0.14 | 0.04 | 0.02 |
| 3 | 11912635 | rs6772434 | -0.704 | -0.017 | -0.68 | 0.994 | -0.01 | 0.009 | 0.009 | 0 | 0.23 | 0.21 | 0.16 | 0.34 | 0.14 | 0.28 |
| 3 | 11913188 | rs51578 | -0.285 | 0.205 | -1.2 | 0.95 | -0.01 | -0.042 | -0.042 | 0 | 0.77 | 0.87 | 0.64 | 0.63 | 0.75 | 0.66 |
| 3 | 11917239 | rs12631353 | 0.481 | 0.461 | -0.903 | 0.075 | 0.002 | 0.001 | 0.001 | 0 | 0.23 | 0.33 | 0.17 | 0.23 | 0.25 | 0.11 |
| 3 | 11920880 | rs425456 | 1.049 | 0.852 | -1.151 | -0.398 | -0.049 | 0.126 | 0.126 | 0 | 0.28 | 0.13 | 0.25 | 0.01 | 0.32 | 0.23 |
| 3 | 11931002 | rs7626703 | -0.572 | 0.416 | -1.136 | -0.276 | -0.02 | 0.067 | 0.067 | 0.233 | 0.54 | 0.75 | 0.45 | 0.7 | 0.5 | 0.42 |
| 3 | 11932455 | rs9869051 | 0.011 | -0.07 | -0.994 | -0.355 | -0.009 | 0.018 | 0.018 | 0.027 | 0.28 | 0.35 | 0.26 | 0.41 | 0.21 | 0.29 |
| 3 | 11940847 | rs366436 | -0.379 | -0.106 | -0.942 | -0.343 | -0.003 | 0.004 | 0.004 | 0 | 0.76 | 0.76 | 0.67 | 0.71 | 0.78 | 0.62 |
| 3 | 11942561 | rs377818 | -0.451 | 0.141 | -1.077 | -0.373 | -0.004 | -0.011 | -0.011 | 0.592 | 0.46 | 0.63 | 0.33 | 0.49 | 0.48 | 0.23 |
| 3 | 11943587 | rs1973443 | 0.562 | -0.081 | -0.544 | -0.22 | -0.017 | -0.011 | -0.011 | 0 | 0.19 | 0.31 | 0.1 | 0.16 | 0.12 | 0.05 |
| 3 | 11945285 | rs448244 | -0.482 | 0.071 | -0.93 | -0.214 | -0.019 | -0.022 | -0.022 | 0 | 0.59 | 0.79 | 0.51 | 0.46 | 0.57 | 0.39 |
| 3 | 11945930 | rs431470 | 0.266 | -0.144 | 0.184 | 0.211 | -0.018 | 0.036 | 0.036 | 0 | 0.09 | 0.2 | 0.05 | 0.15 | 0.05 | 0.03 |
| 3 | 11948258 | rs13083227 | -0.037 | 0.383 | -0.017 | 0.469 | -0.004 | 0.004 | 0.004 | 0 | 0.26 | 0.22 | 0.2 | 0.24 | 0.32 | 0.25 |
| 3 | 11948434 | rs17035391 | -0.1 | -0.126 | -0.121 | 1.242 | -0.002 | -0.06 | -0.06 | 0 | 0.13 | 0 | 0.13 | 0.2 | 0.11 | 0.1 |
| 3 | 11951282 | rs9855914 | -0.414 | -0.187 | 0.213 | 1.185 | -0.012 | 0.007 | 0.007 | 0.206 | 0.18 | 0.71 | 0.18 | 0.24 | 0.17 | 0.15 |
| 3 | 11953848 | rs9830204 | -0.483 | 0.064 | 0.231 | 1.395 | -0.015 | 0.015 | 0.015 | 0 | 0.3 | 0.54 | 0.29 | 0.37 | 0.31 | 0.26 |
| 3 | 11954129 | rs9834366 | -0.606 | 0.124 | -0.214 | 1.371 | -0.01 | 0.013 | 0.013 | 0 | 0.32 | 0.65 | 0.34 | 0.44 | 0.3 | 0.25 |
| 3 | 11957005 | rs9849971 | -0.805 | 0.149 | -0.847 | 1.419 | 0.007 | -0.008 | -0.008 | 0.129 | 0.12 | 0.13 | 0.17 | 0.25 | 0.08 | 0.08 |
| 3 | 11960565 | rs9812594 | -1.261 | 0.117 | -0.962 | 0.999 | 0.006 | 0.003 | 0.003 | 0.008 | 0.36 | 0.37 | 0.42 | 0.34 | 0.34 | 0.23 |
| 3 | 11969777 | rs2881489 | 0.365 | 0.029 | -1.677 | 1.273 | 0.031 | -0.078 | -0.078 | 0 | 0.72 | 0.84 | 0.7 | 0.53 | 0.78 | 0.47 |
| 3 | 11970658 | rs13091458 | -0.14 | 0.304 | -1.355 | 0.936 | -0.008 | 0.003 | 0.003 | 0 | 0.17 | 0.14 | 0.14 | 0.15 | 0.23 | 0.1 |
| 3 | 11978271 | rs2600251 | 1.412 | 0.701 | -0.937 | 0.86 | 0.003 | 0.003 | 0.003 | 0 | 0.6 | 0.43 | 0.42 | 0.58 | 0.55 | 0.71 |
| 3 | 11979988 | rs1018183 | 0.849 | 0.504 | -0.942 | 1.03 | -0.003 | -0.032 | -0.032 | 0 | 0.78 | 0.85 | 0.59 | 0.75 | 0.78 | 0.81 |
| 3 | 11983591 | rs9823022 | 0.236 | 0.48 | -0.795 | 0.949 | -0.011 | 0.012 | 0.012 | 0 | 0.07 | 0.14 | 0.12 | 0.11 | 0.05 | 0.08 |
| 3 | 11992489 | rs378603 | 0.801 | 0.308 | -0.507 | 1.031 | 0.011 | 0.062 | 0.062 | 0 | 0.8 | 0.69 | 0.65 | 0.72 | 0.78 | 0.83 |
| 3 | 11993206 | rs11918200 | 0.77 | 0.361 | -0.468 | 1.046 | -0.004 | 0.006 | 0.006 | 0 | 0.06 | 0.09 | 0.12 | 0.11 | 0.05 | 0.08 |
| 3 | 11997619 | rs305496 | 1.517 | 0.323 | -0.44 | 1.066 | -0.007 | 0.095 | 0.095 | 0 | 0.73 | 0.5 | 0.52 | 0.6 | 0.73 | 0.76 |
| 3 | 11998856 | rs305494 | 0.765 | 0.353 | -0.417 | 0.991 | 0.028 | 0.095 | 0.095 | 0 | 0.81 | 0.61 | 0.65 | 0.74 | 0.78 | 0.85 |
| 3 | 11999083 | rs4684091 | -0.792 | 0.165 | -1.239 | 0.245 | 0.001 | -0.001 | -0.001 | 0 | 0.44 | 0.56 | 0.56 | 0.38 | 0.38 | 0.26 |
| 3 | 12007733 | rs10780003 | -0.305 | 0.592 | -1.29 | 0.362 | 0.014 | 0.001 | 0.001 | 0.148 | 0.38 | 0.45 | 0.34 | 0.35 | 0.44 | 0.19 |
| 3 | 12007910 | rs438129 | -0.261 | 0.566 | -1.365 | 0.177 | -0.016 | 0.027 | 0.027 | 0 | 0.83 | 0.94 | 0.67 | 0.93 | 0.78 | 0.8 |
| 3 | 12009829 | rs11706089 | -0.227 | 0.65 | -1.301 | 0.245 | 0.012 | 0.001 | 0.001 | 0.12 | 0.38 | 0.43 | 0.34 | 0.35 | 0.44 | 0.19 |
| 3 | 12010027 | rs11708773 | 0.435 | 0.728 | -1.36 | 0.186 | -0.008 | -0.013 | -0.013 | 0.286 | 0.15 | 0.11 | 0.17 | 0.2 | 0.19 | 0.13 |
| 3 | 12011284 | rs6767822 | 0.72 | 0.83 | -1.302 | 0.237 | 0.013 | -0.02 | -0.02 | 0 | 0.23 | 0.33 | 0.17 | 0.15 | 0.26 | 0.07 |
| 3 | 12020115 | rs305516 | -0.234 | 0.223 | -1.13 | 0.331 | -0.022 | 0.049 | 0.049 | 0 | 0.27 | 0.46 | 0.24 | 0.34 | 0.24 | 0.16 |
| 3 | 12021458 | rs305512 | 0.001 | 0.214 | -1.139 | 0.324 | 0.006 | -0.057 | -0.057 | 0.073 | 0.25 | 0.1 | 0.22 | 0.34 | 0.25 | 0.17 |

| | | | | | | | | | | | | | | | | |
|---|---|---|---|---|---|---|---|---|---|---|---|---|---|---|---|---|
| 3 | 12022084 | rs9841769 | 1.096 | 0.379 | -1.295 | 0.32 | 0.001 | -0.004 | -0.004 | 0 | 0.56 | 0.41 | 0.41 | 0.59 | 0.55 | 0.64 |
| 3 | 12024543 | rs305508 | -0.514 | 0.072 | -0.991 | 0.633 | -0.024 | 0.051 | 0.051 | 0 | 0.28 | 0.5 | 0.25 | 0.36 | 0.24 | 0.16 |
| 3 | 12025330 | rs4684093 | 0.005 | 0.608 | -1.618 | 0.886 | -0.061 | -0.13 | -0.13 | 0.088 | 0.24 | 0.04 | 0.24 | 0.63 | 0.29 | 0.56 |
| 3 | 12025633 | rs3905903 | -0.87 | 0.549 | -0.809 | 0.938 | -0.003 | 0.022 | 0.022 | 0.027 | 0.15 | 0.19 | 0.19 | 0.28 | 0.14 | 0.12 |
| 3 | 12025779 | rs3905902 | -0.811 | 0.395 | -0.252 | 0.661 | -0.016 | 0.004 | 0.004 | 0.091 | 0.14 | 0.57 | 0.16 | 0.14 | 0.08 | 0.09 |
| 3 | 12028346 | rs4234509 | 0.854 | 1.22 | -0.885 | 0.436 | -0.004 | -0.01 | -0.01 | 0 | 0.72 | 0.31 | 0.75 | 0.66 | 0.71 | 0.75 |
| 3 | 12032812 | rs2600268 | 0.038 | 1.364 | 0.176 | 0.393 | 0.161 | 0.037 | 0.037 | 0 | 0.94 | 0.69 | 0.89 | 0.98 | 0.85 | 0.9 |
| 3 | 12036495 | rs2600272 | 1.014 | 1.235 | 0.59 | 0.426 | -0.015 | 0.075 | 0.075 | 0.777 | 0.1 | 0.2 | 0.06 | 0.18 | 0.07 | 0.04 |
| 3 | 12042835 | rs17669026 | 0.902 | 0.736 | -0.271 | -0.252 | 0.004 | 0.066 | 0.066 | 0.02 | 0.19 | 0.03 | 0.14 | 0.04 | 0.18 | 0.08 |
| 3 | 12049218 | rs2623882 | -0.392 | 0.729 | 0.037 | -0.207 | -0.03 | 0.029 | 0.029 | 0.822 | 0.23 | 0.58 | 0.17 | 0.32 | 0.13 | 0.14 |
| 3 | 12076475 | rs373827 | -0.982 | 1.154 | 0.134 | -0.141 | 0.02 | -0.001 | -0.001 | 0.764 | 0.95 | 0.95 | 0.92 | 0.98 | 0.85 | 0.89 |
| 3 | 12085149 | rs307582 | 0.431 | 1.136 | 0.153 | -0.111 | 0.134 | 0.198 | 0.198 | 0 | 0.92 | 0.72 | 0.87 | 0.79 | 0.82 | 0.86 |
| 3 | 12117220 | rs1994077 | 0.36 | 1.175 | -0.255 | -0.342 | -0.038 | -0.043 | -0.043 | 0.549 | 0.12 | 0.02 | 0.1 | 0.17 | 0.12 | 0.14 |
| 3 | 12145239 | rs160113 | 0.172 | 1.18 | -0.157 | -0.464 | 0.006 | -0.003 | -0.003 | 1.151 | 0.85 | 0.83 | 0.84 | 0.88 | 0.79 | 0.8 |
| 3 | 12150014 | rs308961 | 1.439 | 0.96 | -0.573 | -0.586 | -0.02 | 0.136 | 0.136 | 3.606 | 0.39 | 0.17 | 0.43 | 0.08 | 0.4 | 0.21 |
| 3 | 12156316 | rs11720130 | 0.573 | 1.039 | -0.183 | -0.477 | 0.007 | -0.002 | -0.002 | 0 | 0.12 | 0.16 | 0.11 | 0.12 | 0.12 | 0.14 |
| 3 | 12159294 | rs160208 | 0.161 | 1.038 | -0.033 | -0.417 | 0.141 | 0.257 | 0.257 | 0.089 | 0.94 | 0.66 | 0.9 | 0.82 | 0.87 | 0.9 |
| 3 | 12176975 | rs17035883 | 0.999 | 1.037 | -0.071 | -0.423 | -0.006 | -0.009 | -0.009 | 0 | 0.1 | 0.15 | 0.08 | 0.1 | 0.06 | 0.09 |
| 3 | 12177641 | rs92090 | 1.037 | 0.976 | -0.128 | -0.454 | -0.001 | -0.001 | -0.001 | 0 | 0.78 | 0.29 | 0.76 | 0.69 | 0.75 | 0.75 |
| 3 | 12188243 | rs1867012 | 0.324 | 0.945 | -0.065 | -0.454 | 0.049 | 0.024 | 0.024 | 1.058 | 0.06 | 0.23 | 0.06 | 0.02 | 0.06 | 0.05 |
| 3 | 12194085 | rs99365 | -0.006 | 1.007 | 0.03 | -0.396 | 0.122 | 0.236 | 0.236 | 0 | 0.94 | 0.72 | 0.9 | 0.82 | 0.87 | 0.9 |
| 3 | 12194584 | rs17035943 | 1.121 | 0.899 | -0.13 | -0.421 | -0.008 | -0.013 | -0.013 | 0 | 0.1 | 0.15 | 0.08 | 0.09 | 0.06 | 0.09 |
| 3 | 12200906 | rs3755724 | 0.474 | 1.115 | -0.447 | -0.424 | -0.05 | -0.062 | -0.062 | 0 | 0.36 | 0.12 | 0.31 | 0.66 | 0.36 | 0.56 |
| 3 | 12224151 | rs310762 | 1.053 | 1.466 | -0.49 | -0.413 | 0.007 | 0.029 | 0.029 | 0.196 | 0.48 | 0.2 | 0.54 | 0.66 | 0.4 | 0.61 |
| 3 | 12230704 | rs310763 | -0.28 | 1.666 | -0.351 | -0.286 | -0.047 | 0.009 | 0.009 | 0 | 0.85 | 0.94 | 0.86 | 0.88 | 0.8 | 0.8 |
| 3 | 12233482 | rs310766 | 0.399 | 1.664 | -0.284 | -0.313 | 0.064 | 0.113 | 0.113 | 0.086 | 0.81 | 0.52 | 0.79 | 0.7 | 0.76 | 0.77 |
| 3 | 12239931 | rs13098031 | 1.001 | 1.36 | -0.778 | -0.556 | -0.042 | 0.082 | 0.082 | 0 | 0.21 | 0.01 | 0.17 | 0.03 | 0.27 | 0.11 |
| 3 | 12272199 | rs6775191 | 0.299 | 1.347 | -0.319 | -0.569 | 0 | -0.006 | -0.006 | 0 | 0.28 | 0.22 | 0.21 | 0.05 | 0.32 | 0.13 |
| 3 | 12272683 | rs310752 | 0.867 | 1.297 | -0.448 | -0.48 | -0.026 | 0.048 | 0.048 | 0.021 | 0.52 | 0.22 | 0.57 | 0.67 | 0.45 | 0.66 |
| 3 | 12281063 | rs310757 | 0.24 | 1.324 | -0.103 | -0.22 | -0.019 | 0.006 | 0.006 | 0 | 0.62 | 0.36 | 0.58 | 0.74 | 0.46 | 0.75 |
| 3 | 12281103 | rs167466 | -0.256 | 1.355 | 0.129 | -0.498 | 0.022 | -0.063 | -0.063 | 0.287 | 0.68 | 0.63 | 0.64 | 0.93 | 0.54 | 0.83 |
| 3 | 12286720 | rs9850825 | -0.013 | 1.34 | 0.236 | -0.511 | 0.014 | -0.059 | -0.059 | 0 | 0.33 | 0.53 | 0.37 | 0.07 | 0.44 | 0.17 |
| 3 | 12293739 | rs7610451 | 0.841 | 1.224 | -0.005 | 0.011 | 0.017 | -0.009 | -0.009 | 0.113 | 0.52 | 0.29 | 0.55 | 0.55 | 0.4 | 0.66 |
| 3 | 12296469 | rs9875338 | 0.048 | 1.381 | 0.02 | 0.039 | 0.011 | -0.054 | -0.054 | 0.063 | 0.32 | 0.39 | 0.27 | 0.07 | 0.4 | 0.21 |
| 3 | 12296540 | rs7641345 | -1.411 | 0.993 | -0.217 | 0.181 | 0.016 | 0.014 | 0.014 | 0 | 0.8 | 0.9 | 0.75 | 0.85 | 0.87 | 0.88 |
| 3 | 12302462 | rs9878908 | 0.281 | 1.65 | -0.527 | 0.726 | 0.012 | 0.09 | 0.09 | 0 | 0.92 | 0.91 | 0.85 | 0.6 | 0.81 | 0.84 |
| 3 | 12314846 | rs4279078 | 0.139 | 1.437 | -0.561 | 0.741 | 0.001 | 0.106 | 0.106 | 0 | 0.05 | 0.04 | 0.13 | 0.38 | 0.11 | 0.07 |
| 3 | 12319058 | rs2920499 | -0.537 | 1.153 | -1.124 | 0.245 | -0.009 | 0.06 | 0.06 | 0.323 | 0.44 | 0.34 | 0.47 | 0.79 | 0.47 | 0.75 |
| 3 | 12322628 | rs2960421 | 1.465 | 1.159 | -1.425 | 0.115 | -0.037 | -0.118 | -0.118 | 0.2 | 0.21 | 0.02 | 0.17 | 0.4 | 0.18 | 0.51 |
| 3 | 12323413 | rs2920500 | -0.471 | 0.817 | -1.261 | -0.271 | -0.005 | 0.121 | 0.121 | 0.699 | 0.47 | 0.27 | 0.66 | 0.91 | 0.52 | 0.81 |
| 3 | 12327431 | rs13076933 | 1.051 | 1.043 | -1.752 | -0.27 | -0.061 | 0.076 | 0.076 | 0 | 0.2 | 0.01 | 0.16 | 0.03 | 0.26 | 0.11 |
| 3 | 12329783 | rs17036160 | 1.411 | 0.737 | -1.323 | 0.105 | -0.029 | -0.007 | -0.007 | 1.098 | 0.06 | 0 | 0.11 | 0.02 | 0.12 | 0.11 |
| 3 | 12334416 | rs2972164 | -0.419 | 0.521 | -1.549 | -0.243 | -0.05 | 0.251 | 0.251 | 1.205 | 0.46 | 0.17 | 0.65 | 0.92 | 0.52 | 0.81 |
| 3 | 12338115 | rs6768587 | -0.993 | 0.821 | -0.808 | 0.401 | 0 | 0.017 | 0.017 | 0 | 0.24 | 0.3 | 0.4 | 0.49 | 0.26 | 0.27 |
| 3 | 12340267 | rs9817428 | -1.069 | 0.796 | -0.597 | 0.39 | 0 | 0.056 | 0.056 | 0 | 0.24 | 0.37 | 0.41 | 0.49 | 0.26 | 0.28 |
| 3 | 12341406 | rs12631028 | -1.024 | 0.783 | -0.678 | 0.419 | 0 | 0.011 | 0.011 | 0 | 0.24 | 0.28 | 0.4 | 0.48 | 0.26 | 0.28 |
| 3 | 12341830 | rs12636461 | -0.947 | 0.818 | -0.662 | 0.435 | 0 | 0.014 | 0.014 | 0 | 0.23 | 0.28 | 0.4 | 0.48 | 0.26 | 0.28 |
| 3 | 12341996 | rs13076055 | -0.921 | 0.816 | -0.663 | 0.434 | 0 | 0.016 | 0.016 | 0 | 0.23 | 0.28 | 0.4 | 0.48 | 0.26 | 0.28 |
| 3 | 12343117 | rs11128596 | -1.005 | 0.809 | -0.667 | 0.418 | 0 | 0.017 | 0.017 | 0 | 0.24 | 0.29 | 0.4 | 0.49 | 0.26 | 0.28 |
| 3 | 12344441 | rs7620165 | 0.762 | 0.711 | -0.733 | 0.156 | -0.048 | 0.091 | 0.091 | 0 | 0.24 | 0.07 | 0.19 | 0.04 | 0.33 | 0.12 |
| 3 | 12344636 | rs11128597 | -0.972 | 0.585 | -0.564 | 0.365 | 0 | 0.012 | 0.012 | 0 | 0.24 | 0.28 | 0.4 | 0.49 | 0.26 | 0.28 |
| 3 | 12345971 | rs17793693 | 0.748 | 0.627 | -0.662 | -0.062 | -0.012 | 0.051 | 0.051 | 0 | 0.11 | 0.03 | 0.2 | 0 | 0.1 | 0.07 |
| 3 | 12346738 | rs10510410 | -1.246 | 0.602 | -0.394 | 0.284 | 0.002 | 0.018 | 0.018 | 0 | 0.23 | 0.32 | 0.37 | 0.31 | 0.26 | 0.23 |
| 3 | 12349490 | rs17036242 | -1.311 | 0.442 | -0.001 | 0.231 | 0 | 0.018 | 0.018 | 0.075 | 0.23 | 0.33 | 0.37 | 0.31 | 0.26 | 0.23 |
| 3 | 12349924 | rs13061415 | -1.284 | 0.426 | -0.008 | 0.224 | 0.001 | 0.021 | 0.021 | 0.021 | 0.23 | 0.33 | 0.37 | 0.31 | 0.25 | 0.23 |
| 3 | 12351574 | rs12629240 | -1.283 | 0.429 | -0.389 | 0.261 | 0 | -0.014 | -0.014 | 0 | 0.23 | 0.22 | 0.36 | 0.31 | 0.25 | 0.23 |
| 3 | 12352468 | rs9310401 | -1.341 | 0.426 | -0.38 | 0.26 | 0 | 0.006 | 0.006 | 0.384 | 0.23 | 0.3 | 0.37 | 0.31 | 0.26 | 0.23 |
| 3 | 12352971 | rs11715073 | -1.254 | 0.427 | -0.387 | 0.261 | -0.001 | -0.014 | -0.014 | 0 | 0.23 | 0.21 | 0.36 | 0.31 | 0.26 | 0.23 |
| 3 | 12353326 | rs11128598 | -1.254 | 0.413 | -0.39 | 0.257 | -0.001 | -0.014 | -0.014 | 0 | 0.23 | 0.21 | 0.36 | 0.31 | 0.26 | 0.23 |
| 3 | 12354411 | rs17036281 | 0.754 | 0.42 | -0.393 | 0.252 | -0.01 | 0.046 | 0.046 | 0 | 0.11 | 0.03 | 0.2 | 0 | 0.1 | 0.07 |
| 3 | 12356158 | rs12496505 | -1.262 | 0.454 | -0.369 | 0.272 | -0.002 | -0.015 | -0.015 | 0 | 0.23 | 0.21 | 0.36 | 0.31 | 0.26 | 0.23 |
| 3 | 12356196 | rs12487012 | -1.255 | 0.453 | -0.37 | 0.272 | -0.002 | -0.015 | -0.015 | 0 | 0.23 | 0.21 | 0.36 | 0.31 | 0.26 | 0.23 |
| 3 | 12356605 | rs13070963 | -1.263 | 0.454 | -0.37 | 0.272 | -0.002 | -0.015 | -0.015 | 0 | 0.23 | 0.21 | 0.36 | 0.31 | 0.26 | 0.23 |
| 3 | 12359049 | rs2067819 | -0.779 | 0.453 | -0.361 | 0.269 | -0.001 | 0.001 | 0.001 | 0 | 0.19 | 0.22 | 0.34 | 0.03 | 0.22 | 0.18 |
| 3 | 12360214 | rs12636454 | -1.279 | 0.476 | -0.377 | 0.289 | -0.001 | 0.005 | 0.005 | 0.336 | 0.23 | 0.3 | 0.37 | 0.31 | 0.25 | 0.23 |
| 3 | 12360769 | rs11128599 | -1.179 | 0.481 | -0.417 | 0.291 | -0.001 | -0.018 | -0.018 | 0.617 | 0.23 | 0.2 | 0.36 | 0.31 | 0.25 | 0.23 |
| 3 | 12363637 | rs12493718 | -1.18 | 0.489 | -0.397 | 0.295 | -0.001 | -0.015 | -0.015 | 0 | 0.23 | 0.21 | 0.36 | 0.31 | 0.25 | 0.23 |
| 3 | 12365308 | rs13083375 | 1.424 | 0.524 | -0.432 | 0.273 | -0.029 | -0.007 | -0.007 | 1.108 | 0.06 | 0.01 | 0.11 | 0.03 | 0.12 | 0.11 |
| 3 | 12369401 | rs13064760 | 1.425 | 0.525 | -0.432 | 0.273 | -0.029 | -0.007 | -0.007 | 0 | 0.06 | 0 | 0.11 | 0.03 | 0.12 | 0.11 |

| | | | | | | | | | | | | | | | | |
|---|---|---|---|---|---|---|---|---|---|---|---|---|---|---|---|---|
| 3 | 12370737 | rs17793951 | 0.791 | 0.57 | -0.642 | 0.082 | -0.062 | 0.106 | 0.106 | 0 | 0.24 | 0.05 | 0.2 | 0.04 | 0.33 | 0.12 |
| 3 | 12372074 | rs4145574 | -1.082 | 0.529 | -0.51 | 0.15 | -0.002 | -0.014 | -0.014 | 0 | 0.23 | 0.22 | 0.36 | 0.31 | 0.26 | 0.23 |
| 3 | 12372386 | rs4145573 | -1.082 | 0.529 | -0.51 | 0.15 | -0.002 | -0.014 | -0.014 | 1.201 | 0.23 | 0.22 | 0.36 | 0.31 | 0.26 | 0.23 |
| 3 | 12372882 | rs2028760 | -1.068 | 0.528 | -0.51 | 0.15 | -0.002 | -0.014 | -0.014 | 0 | 0.23 | 0.22 | 0.36 | 0.31 | 0.26 | 0.23 |
| 3 | 12375391 | rs1122648 | -1.158 | 0.525 | -0.467 | 0.149 | 0 | 0.01 | 0.01 | 0.239 | 0.23 | 0.31 | 0.37 | 0.31 | 0.26 | 0.23 |
| 3 | 12381165 | rs11128601 | -1.088 | 0.532 | -0.55 | 0.151 | -0.002 | -0.017 | -0.017 | 0 | 0.23 | 0.21 | 0.36 | 0.31 | 0.26 | 0.23 |
| 3 | 12383714 | rs7638903 | 0.024 | 0.524 | -0.403 | 0.133 | 0.005 | -0.014 | -0.014 | 0 | 0.09 | 0.18 | 0.14 | 0.03 | 0.12 | 0.13 |
| 3 | 12386337 | rs4684847 | 1.55 | 0.545 | -0.427 | 0.143 | -0.029 | -0.008 | -0.008 | 1.017 | 0.06 | 0.01 | 0.11 | 0.03 | 0.12 | 0.12 |
| 3 | 12388563 | rs10510418 | 0.739 | 0.516 | -0.566 | 0.29 | -0.065 | 0.006 | 0.006 | 0 | 0.25 | 0.05 | 0.22 | 0.18 | 0.32 | 0.16 |
| 3 | 12390135 | rs12497191 | 0.018 | 0.353 | -0.182 | 0.417 | -0.014 | -0.062 | -0.062 | 0.05 | 0.14 | 0.04 | 0.23 | 0.28 | 0.14 | 0.11 |
| 3 | 12390484 | rs17036328 | -0.321 | 0.341 | -0.151 | 0.4 | 0.002 | -0.016 | -0.016 | 0 | 0.1 | 0.18 | 0.13 | 0.03 | 0.12 | 0.13 |
| 3 | 12391207 | rs6802898 | -0.788 | 0.365 | -0.114 | 0.408 | -0.002 | -0.03 | -0.03 | 0 | 0.12 | 0.67 | 0.18 | 0.03 | 0.12 | 0.13 |
| 3 | 12391583 | rs2197423 | -0.05 | 0.348 | -0.122 | 0.413 | 0.005 | -0.014 | -0.014 | 0 | 0.09 | 0.18 | 0.13 | 0.03 | 0.12 | 0.13 |
| 3 | 12391813 | rs7647481 | -0.05 | 0.35 | -0.123 | 0.415 | 0.005 | -0.014 | -0.014 | 0 | 0.09 | 0.18 | 0.13 | 0.03 | 0.12 | 0.13 |
| 3 | 12394840 | rs1899951 | -0.655 | 0.357 | -0.177 | 0.418 | -0.003 | -0.03 | -0.03 | 0 | 0.12 | 0.67 | 0.18 | 0.03 | 0.12 | 0.13 |
| 3 | 12395645 | rs4684848 | -0.655 | 0.357 | -0.181 | 0.418 | -0.003 | -0.03 | -0.03 | 0 | 0.12 | 0.67 | 0.18 | 0.03 | 0.12 | 0.13 |
| 3 | 12395958 | rs7615916 | -1.5 | 0.335 | -0.198 | 0.435 | 0.003 | -0.008 | -0.008 | 0 | 0.26 | 0.71 | 0.41 | 0.3 | 0.26 | 0.24 |
| 3 | 12399270 | rs7627605 | -1.331 | 0.026 | -0.01 | 0.463 | -0.018 | -0.02 | -0.02 | 0 | 0.06 | 0.66 | 0.07 | 0 | 0 | 0.01 |
| 3 | 12399407 | rs12629751 | 0.097 | 0.112 | -0.135 | 0.561 | 0.012 | -0.046 | -0.046 | 0 | 0.12 | 0.04 | 0.06 | 0.28 | 0.1 | 0.09 |
| 3 | 12402474 | rs1373641 | -0.488 | 0.518 | -0.319 | 0.141 | -0.022 | 0.035 | 0.035 | 0 | 0.74 | 0.85 | 0.78 | 0.86 | 0.66 | 0.83 |
| 3 | 12402601 | rs1373640 | -0.887 | 0.47 | -0.298 | 0.12 | -0.063 | 0.038 | 0.038 | 0 | 0.77 | 0.95 | 0.79 | 0.86 | 0.66 | 0.84 |
| 3 | 12408278 | rs2938397 | 0.039 | 0.334 | -0.343 | 0.498 | -0.005 | -0.015 | -0.015 | 0.826 | 0.69 | 0.74 | 0.74 | 0.58 | 0.62 | 0.79 |
| 3 | 12410349 | rs17817469 | 0.465 | 0.09 | -0.72 | -0.038 | 0.001 | -0.002 | -0.002 | 0.715 | 0.15 | 0.11 | 0.25 | 0 | 0.15 | 0.1 |
| 3 | 12412115 | rs2921188 | -1.18 | 0.074 | -0.319 | -0.018 | -0.063 | 0.027 | 0.027 | 0 | 0.77 | 0.95 | 0.78 | 0.86 | 0.66 | 0.84 |
| 3 | 12413339 | rs2120825 | 1.662 | 0.136 | -0.439 | -0.088 | -0.027 | 0.005 | 0.005 | 0 | 0.06 | 0.01 | 0.11 | 0.02 | 0.11 | 0.14 |
| 3 | 12414420 | rs6809832 | 0.308 | 0.075 | -0.323 | -0.04 | 0.002 | 0.018 | 0.018 | 0 | 0.16 | 0.09 | 0.25 | 0 | 0.15 | 0.1 |
| 3 | 12418612 | rs2028759 | 1.169 | 0.037 | -0.364 | -0.046 | -0.037 | -0.021 | -0.021 | 0 | 0.51 | 0.18 | 0.44 | 0.57 | 0.47 | 0.68 |
| 3 | 12419475 | rs2921190 | 0.729 | 0.04 | -0.321 | -0.051 | -0.022 | -0.01 | -0.01 | 0 | 0.57 | 0.74 | 0.51 | 0.58 | 0.47 | 0.69 |
| 3 | 12423266 | rs4135263 | 0.162 | -0.111 | -0.385 | 0.013 | 0.003 | 0.005 | 0.005 | 0 | 0.16 | 0.11 | 0.25 | 0 | 0.15 | 0.1 |
| 3 | 12424793 | rs2972162 | 0.72 | -0.113 | -0.386 | 0.013 | -0.023 | -0.012 | -0.012 | 0 | 0.57 | 0.74 | 0.51 | 0.58 | 0.47 | 0.69 |
| 3 | 12426936 | rs10510419 | 0.156 | -0.089 | -0.374 | 0.023 | 0.004 | 0.009 | 0.009 | 0 | 0.16 | 0.11 | 0.25 | 0 | 0.15 | 0.1 |
| 3 | 12429468 | rs2938395 | -0.401 | -0.073 | -0.369 | 0.03 | -0.025 | -0.056 | -0.056 | 2.381 | 0.72 | 0.85 | 0.75 | 0.58 | 0.62 | 0.79 |
| 3 | 12430075 | rs2959268 | -0.543 | -0.073 | -0.37 | 0.028 | -0.047 | -0.085 | -0.085 | 0.228 | 0.73 | 0.9 | 0.76 | 0.58 | 0.62 | 0.79 |
| 3 | 12431568 | rs2938394 | -0.478 | -0.074 | -0.353 | 0.028 | -0.047 | -0.089 | -0.089 | 0.342 | 0.72 | 0.9 | 0.76 | 0.58 | 0.62 | 0.79 |
| 3 | 12434608 | rs2938392 | 1.288 | -0.085 | -0.363 | 0.026 | -0.029 | -0.005 | -0.005 | 0.009 | 0.5 | 0.18 | 0.44 | 0.58 | 0.47 | 0.68 |
| 3 | 12439387 | rs2938387 | -0.457 | -0.087 | -0.349 | 0.126 | -0.058 | -0.114 | -0.114 | 0.022 | 0.72 | 0.92 | 0.76 | 0.57 | 0.62 | 0.78 |
| 3 | 12441507 | rs2921193 | 0.836 | -0.076 | -0.332 | 0.029 | -0.021 | -0.008 | -0.008 | 0 | 0.56 | 0.73 | 0.51 | 0.58 | 0.47 | 0.68 |
| 3 | 12442833 | rs2959272 | 0.845 | -0.078 | -0.518 | 0.024 | -0.009 | -0.002 | -0.002 | 0 | 0.56 | 0.64 | 0.5 | 0.57 | 0.47 | 0.68 |
| 3 | 12443657 | rs1875796 | 1.169 | -0.126 | -0.563 | 0.004 | -0.044 | -0.015 | -0.015 | 0 | 0.51 | 0.16 | 0.44 | 0.58 | 0.47 | 0.69 |
| 3 | 12443844 | rs4135275 | 1.807 | -0.221 | -1.018 | -0.176 | -0.065 | -0.123 | -0.123 | 2.464 | 0.2 | 0.04 | 0.23 | 0.49 | 0.19 | 0.49 |
| 3 | 12445807 | rs1151996 | -0.559 | -0.149 | -0.692 | -0.111 | -0.061 | -0.103 | -0.103 | 3.037 | 0.73 | 0.91 | 0.76 | 0.58 | 0.62 | 0.79 |
| 3 | 12446004 | rs1151997 | 1.049 | -0.138 | -0.678 | -0.103 | -0.046 | -0.019 | -0.019 | 0 | 0.52 | 0.17 | 0.45 | 0.58 | 0.47 | 0.69 |
| 3 | 12446492 | rs1151998 | 1.058 | -0.138 | -0.679 | -0.103 | -0.046 | -0.018 | -0.018 | 0 | 0.52 | 0.17 | 0.45 | 0.58 | 0.47 | 0.68 |
| 3 | 12447153 | rs1151999 | -1.141 | -0.13 | -0.676 | -0.091 | -0.047 | -0.015 | -0.015 | 0.84 | 0.48 | 0.83 | 0.55 | 0.42 | 0.53 | 0.32 |
| 3 | 12449528 | rs796313 | -0.792 | -0.013 | -0.602 | -0.046 | -0.037 | -0.018 | -0.018 | 0 | 0.43 | 0.22 | 0.49 | 0.42 | 0.53 | 0.31 |
| 3 | 12449866 | rs4135284 | 0.518 | 0.013 | -0.673 | -0.056 | 0.002 | 0.002 | 0.002 | 0 | 0.15 | 0.11 | 0.25 | 0 | 0.15 | 0.1 |
| 3 | 12449963 | rs1822825 | -1.125 | 0.045 | -0.855 | -0.041 | -0.046 | -0.019 | -0.019 | 0.277 | 0.48 | 0.83 | 0.55 | 0.42 | 0.53 | 0.31 |
| 3 | 12450354 | rs709149 | 0.379 | 0.183 | -0.807 | 0.104 | -0.049 | -0.087 | -0.087 | 0.197 | 0.27 | 0.11 | 0.24 | 0.42 | 0.38 | 0.22 |
| 3 | 12454999 | rs709151 | 0.476 | 0.273 | -1.127 | 0.179 | -0.062 | -0.107 | -0.107 | 0.057 | 0.25 | 0.09 | 0.23 | 0.42 | 0.35 | 0.16 |
| 3 | 12461615 | rs709156 | 0.672 | 0.253 | -1.136 | 0.125 | -0.054 | 0.031 | 0.031 | 0 | 0.23 | 0.08 | 0.2 | 0.13 | 0.31 | 0.12 |
| 3 | 12462024 | rs709157 | 1.23 | 0.252 | -1.144 | -0.099 | -0.067 | 0.086 | 0.086 | 0 | 0.2 | 0.03 | 0.17 | 0.02 | 0.3 | 0.07 |
| 3 | 12463176 | rs709158 | 0.622 | 0.267 | -1.062 | -0.069 | -0.054 | -0.097 | -0.097 | 0.001 | 0.25 | 0.1 | 0.23 | 0.41 | 0.35 | 0.16 |
| 3 | 12465488 | rs1175541 | 0.64 | 0.362 | -1.046 | -0.021 | -0.062 | -0.108 | -0.108 | 0 | 0.25 | 0.09 | 0.23 | 0.41 | 0.35 | 0.16 |
| 3 | 12466214 | rs1175542 | -1.077 | 0.296 | -1.062 | -0.209 | -0.041 | -0.013 | -0.013 | 2.199 | 0.48 | 0.83 | 0.56 | 0.41 | 0.53 | 0.31 |
| 3 | 12466490 | rs1177809 | 0.429 | 0.392 | -0.987 | 0.003 | -0.055 | -0.104 | -0.104 | 0 | 0.27 | 0.1 | 0.23 | 0.39 | 0.35 | 0.17 |
| 3 | 12471362 | rs6794024 | -0.386 | -0.016 | -0.065 | -0.352 | -0.016 | -0.136 | -0.136 | 0.663 | 0.21 | 0.36 | 0.28 | 0 | 0.17 | 0.14 |
| 3 | 12471862 | rs1152001 | -1.219 | -0.11 | -0.95 | -0.447 | 0.02 | 0.046 | 0.046 | 1.746 | 0.7 | 0.83 | 0.84 | 0.9 | 0.77 | 0.77 |
| 3 | 12471871 | rs1152002 | -0.176 | 0.834 | -0.625 | 0.152 | 0.025 | 0.025 | 0.025 | 0 | 0.34 | 0.42 | 0.4 | 0.43 | 0.49 | 0.2 |
| 3 | 12475088 | rs7626560 | -0.316 | 0.708 | -0.67 | -0.266 | -0.009 | -0.074 | -0.074 | 0 | 0.17 | 0.16 | 0.26 | 0 | 0.15 | 0.13 |
| 3 | 12475557 | rs3856806 | -0.003 | 1.18 | -0.983 | 0.385 | -0.014 | -0.023 | -0.023 | 0 | 0.08 | 0.05 | 0.11 | 0.21 | 0.12 | 0.16 |
| 3 | 12475994 | rs4135300 | 0.458 | 1.431 | -0.992 | 0.231 | -0.035 | -0.038 | -0.038 | 1.665 | 0.06 | 0.01 | 0.11 | 0.19 | 0.09 | 0.04 |
| 3 | 12478817 | rs9833097 | -0.5 | 0.769 | 0.09 | -0.497 | -0.002 | -0.029 | -0.029 | 0.031 | 0.1 | 0.1 | 0.2 | 0 | 0.1 | 0.08 |
| 3 | 12481375 | rs4498025 | -0.284 | 1.383 | -0.591 | -0.882 | -0.083 | 0.017 | 0.017 | 0 | 0.23 | 0.01 | 0.2 | 0.14 | 0.24 | 0.1 |
| 3 | 12483104 | rs1152004 | 0.503 | 1.811 | -0.369 | -1.269 | 0.036 | -0.009 | -0.009 | 0 | 0.11 | 0.19 | 0.19 | 0.08 | 0.24 | 0.16 |
| 3 | 12483451 | rs1152005 | -2.076 | 0.975 | -0.993 | -1.871 | -0.061 | -0.154 | -0.154 | 4.238 | 0.78 | 0.47 | 0.91 | 1 | 0.9 | 0.95 |
| 3 | 12484849 | rs4684104 | 0.008 | 0.85 | -2.501 | -1.784 | -0.074 | 0.287 | 0.287 | 0.797 | 0.56 | 0.06 | 0.5 | 0.92 | 0.56 | 0.71 |
| 3 | 12486964 | rs10440120 | -0.082 | 1.018 | -1.859 | -1.498 | 0.015 | 0.011 | 0.011 | 0.872 | 0.23 | 0.71 | 0.17 | 0.03 | 0.15 | 0.11 |
| 3 | 12487047 | rs1152006 | -1.178 | 1.539 | -2.087 | -0.94 | 0.076 | -0.051 | -0.051 | 0 | 0.54 | 0.85 | 0.73 | 0.53 | 0.65 | 0.46 |
| 3 | 12487511 | rs1186464 | 0.816 | 1.624 | -2.065 | -1.129 | -0.023 | 0.023 | 0.023 | 0 | 0.07 | 0.03 | 0.12 | 0.06 | 0.14 | 0.1 |

| | | | | | | | | | | | | | | | | |
|---|---|---|---|---|---|---|---|---|---|---|---|---|---|---|---|---|
| 3 | 12487612 | rs1152008 | -0.486 | 1.708 | -1.929 | -1.371 | 0.005 | 0.012 | 0.012 | 0 | 0.39 | 0.61 | 0.46 | 0.09 | 0.4 | 0.27 |
| 3 | 12487650 | rs6806708 | 0.255 | 1.636 | -1.864 | -1.311 | -0.004 | -0.061 | -0.061 | 0 | 0.31 | 0.56 | 0.34 | 0.03 | 0.26 | 0.18 |
| 3 | 12487827 | rs13088214 | 0.283 | 1.637 | -1.874 | -1.311 | -0.01 | -0.083 | -0.083 | 0.227 | 0.31 | 0.53 | 0.34 | 0.03 | 0.26 | 0.18 |
| 3 | 12488027 | rs7618046 | -0.669 | 1.991 | -1.927 | -1.12 | -0.005 | 0.012 | 0.012 | 0 | 0.46 | 0.56 | 0.59 | 0.43 | 0.51 | 0.38 |
| 3 | 12493347 | rs9855622 | 0.837 | 2.173 | -2.031 | -1.973 | 0.014 | -0.014 | -0.014 | 0 | 0.09 | 0.13 | 0.12 | 0.09 | 0.11 | 0.04 |
| 3 | 12494794 | rs2249407 | -0.04 | 1.841 | -2.348 | -2.286 | -0.004 | 0.234 | 0.234 | 0.066 | 0.46 | 0.2 | 0.53 | 0.95 | 0.54 | 0.73 |
| 3 | 12496461 | rs9872031 | 0.341 | 1.588 | -2.251 | -2.205 | -0.038 | 0.168 | 0.168 | 0 | 0.4 | 0.17 | 0.42 | 0.91 | 0.4 | 0.64 |
| 3 | 12500651 | rs9809905 | -0.737 | 1.554 | -1.882 | -1.774 | -0.002 | 0.236 | 0.236 | 0.053 | 0.42 | 0.83 | 0.49 | 0.11 | 0.37 | 0.22 |
| 3 | 12507401 | rs17036788 | -1.201 | 1.935 | -0.564 | -1.492 | 0.045 | -0.041 | -0.041 | 0.556 | 0.13 | 0.39 | 0.2 | 0.07 | 0.18 | 0.12 |
| 3 | 12507564 | rs1699348 | -1.612 | 2.148 | -0.349 | -1.902 | -0.001 | -0.001 | -0.001 | 0.078 | 0.4 | 0.64 | 0.63 | 0.15 | 0.56 | 0.27 |
| 3 | 12511695 | rs9820178 | -0.392 | 2.525 | -0.01 | -1.971 | 0.028 | -0.066 | -0.066 | 0.059 | 0.17 | 0.38 | 0.13 | 0.04 | 0.19 | 0.1 |
| 3 | 12521180 | rs709162 | -1.781 | 2.358 | -0.076 | -2.02 | -0.006 | 0.06 | 0.06 | 1.229 | 0.39 | 0.73 | 0.56 | 0.13 | 0.55 | 0.18 |
| 3 | 12521698 | rs1797886 | -1.332 | 2.44 | 0.031 | -2.051 | 0.019 | -0.053 | -0.053 | 2.272 | 0.33 | 0.6 | 0.37 | 0.11 | 0.49 | 0.15 |
| 3 | 12525656 | rs3806663 | -0.707 | 2.225 | 0.074 | -1.966 | 0.007 | -0.027 | -0.027 | 0 | 0.06 | 0.13 | 0.19 | 0.02 | 0.06 | 0.03 |
| 3 | 12526006 | rs3806662 | -0.648 | 2.233 | 0.082 | -1.963 | 0.007 | -0.024 | -0.024 | 0 | 0.05 | 0.12 | 0.19 | 0.02 | 0.06 | 0.03 |
| 3 | 12538926 | rs299648 | -1.406 | 2.388 | 0.132 | -2.043 | 0.003 | -0.024 | -0.024 | 0 | 0.33 | 0.64 | 0.38 | 0.11 | 0.49 | 0.16 |
| 3 | 12573518 | rs299642 | -0.999 | 2.349 | 0.156 | -1.996 | 0.045 | -0.077 | -0.077 | 0 | 0.3 | 0.53 | 0.31 | 0.13 | 0.4 | 0.16 |
| 3 | 12574512 | rs299643 | -1.709 | 2.302 | -0.03 | -2.101 | -0.052 | 0.081 | 0.081 | 0.107 | 0.36 | 0.77 | 0.44 | 0.13 | 0.48 | 0.18 |
| 3 | 12576846 | rs299629 | -0.991 | 2.298 | -0.006 | -2.104 | 0.008 | -0.029 | -0.029 | 0 | 0.29 | 0.64 | 0.31 | 0.11 | 0.39 | 0.14 |
| 3 | 12578104 | rs116002 | -2.26 | 1.845 | -0.31 | -2.354 | -0.103 | 0.269 | 0.269 | 1.619 | 0.46 | 0.96 | 0.64 | 0.15 | 0.54 | 0.22 |
| 3 | 12581592 | rs13069599 | -1.532 | 1.767 | -0.171 | -2.303 | -0.002 | -0.071 | -0.071 | 0 | 0.14 | 0.27 | 0.26 | 0.02 | 0.13 | 0.06 |
| 3 | 12587133 | rs11706408 | 0.109 | 1.838 | -0.262 | -2.255 | 0.005 | -0.017 | -0.017 | 0.034 | 0.05 | 0.08 | 0.07 | 0 | 0.08 | 0.02 |
| 3 | 12598963 | rs2255648 | -2.053 | 1.899 | -0.384 | -2.321 | -0.076 | 0.22 | 0.22 | 0.594 | 0.41 | 0.85 | 0.56 | 0.1 | 0.49 | 0.18 |
| 3 | 12607929 | rs2454427 | 1.386 | 1.96 | -0.486 | -2.327 | -0.071 | 0.023 | 0.023 | 0 | 0.11 | 0.01 | 0.16 | 0.06 | 0.22 | 0.05 |
| 3 | 12613866 | rs2648307 | -2.559 | 1.987 | -0.705 | -2.53 | -0.002 | 0.347 | 0.347 | 1.819 | 0.49 | 0.96 | 0.66 | 0.13 | 0.62 | 0.28 |
| 3 | 12615984 | rs713178 | -1.547 | 2.009 | -0.489 | -2.449 | -0.009 | 0.002 | 0.002 | 0 | 0.18 | 0.12 | 0.35 | 0.04 | 0.25 | 0.16 |
| 3 | 12616533 | rs3773341 | 0.251 | 1.497 | -0.458 | -2.041 | 0.036 | 0.001 | 0.001 | 0 | 0.43 | 0.64 | 0.3 | 0.56 | 0.27 | 0.61 |
| 3 | 12618262 | rs9855183 | -0.698 | 1.883 | 0.476 | -2.285 | 0.068 | -0.035 | -0.035 | 0 | 0.12 | 0.44 | 0.07 | 0.03 | 0.11 | 0.06 |
| 3 | 12619083 | rs2344825 | -0.994 | 1.932 | 0.444 | -2.241 | 0.023 | -0.066 | -0.066 | 2.969 | 0.28 | 0.37 | 0.47 | 0.09 | 0.37 | 0.18 |
| 3 | 12624070 | rs15997 | -2.512 | 1.872 | 0.548 | -2.311 | 0.023 | 0.097 | 0.097 | 0.427 | 0.46 | 0.7 | 0.63 | 0.12 | 0.59 | 0.26 |
| 3 | 12624321 | rs5746255 | -1.703 | 2.051 | 0.696 | -2.311 | 0.037 | -0.09 | -0.09 | 0 | 0.19 | 0.42 | 0.21 | 0.03 | 0.22 | 0.09 |
| 3 | 12624763 | rs7956 | -1.59 | 2.071 | 0.669 | -2.302 | 0.044 | -0.088 | -0.088 | 0 | 0.18 | 0.4 | 0.21 | 0.03 | 0.22 | 0.09 |
| 3 | 12626516 | rs3729931 | -2.441 | 2.048 | 0.61 | -2.33 | -0.004 | -0.02 | -0.02 | 2.823 | 0.32 | 0.65 | 0.47 | 0.06 | 0.38 | 0.21 |
| 3 | 12635706 | rs1532533 | -1.444 | 1.44 | 0.581 | -2.293 | 0.055 | -0.086 | -0.086 | 0 | 0.32 | 0.55 | 0.38 | 0.11 | 0.45 | 0.16 |
| 3 | 12649576 | rs2442812 | -1.387 | 1.446 | 0.594 | -2.29 | 0.056 | -0.09 | -0.09 | 0 | 0.31 | 0.48 | 0.38 | 0.11 | 0.45 | 0.16 |
| 3 | 12653233 | rs3730269 | -1.477 | 1.553 | 0.58 | -2.301 | 0.012 | -0.085 | -0.085 | 2.396 | 0.18 | 0.4 | 0.2 | 0.03 | 0.19 | 0.09 |
| 3 | 12654653 | rs3773353 | -1.273 | 1.556 | 0.56 | -2.3 | -0.08 | 0.02 | 0.02 | 0 | 0.88 | 0.99 | 0.89 | 0.94 | 0.76 | 0.95 |
| 3 | 12655689 | rs9809501 | 0.104 | 1.557 | 0.652 | -2.306 | 0.003 | -0.028 | -0.028 | 0.181 | 0.11 | 0.16 | 0.12 | 0.03 | 0.11 | 0.06 |
| 3 | 12662370 | rs11705805 | 2.756 | 1.438 | 0.279 | -2.338 | -0.051 | 0.267 | 0.267 | 1.256 | 0.53 | 0.03 | 0.35 | 0.86 | 0.41 | 0.72 |
| 3 | 12663233 | rs7643321 | -0.989 | 1.48 | 0.423 | -2.24 | 0.001 | -0.07 | -0.07 | 0 | 0.17 | 0.33 | 0.19 | 0.03 | 0.18 | 0.08 |
| 3 | 12673275 | rs11128607 | -1.425 | 1.476 | 0.418 | -2.239 | 0.013 | -0.085 | -0.085 | 0 | 0.18 | 0.4 | 0.2 | 0.03 | 0.19 | 0.09 |
| 3 | 12674172 | rs13099808 | -0.016 | 1.464 | 0.399 | -2.245 | -0.012 | 0.024 | 0.024 | 0 | 0.11 | 0.05 | 0.22 | 0.01 | 0.14 | 0.1 |
| 3 | 12697498 | rs4684871 | 1.188 | 1.484 | 0.405 | -2.27 | 0.063 | -0.098 | -0.098 | 0.279 | 0.69 | 0.48 | 0.68 | 0.91 | 0.57 | 0.86 |
| 3 | 12702562 | rs6795441 | 1.844 | 1.529 | 0.296 | -2.233 | -0.002 | -0.056 | -0.056 | 0.792 | 0.58 | 0.43 | 0.45 | 0.9 | 0.43 | 0.76 |
| 3 | 12709167 | rs7617041 | -2.202 | 1.369 | 0.274 | -2.2 | 0.005 | -0.103 | -0.103 | 0.682 | 0.3 | 0.42 | 0.42 | 0.04 | 0.33 | 0.19 |
| 3 | 12717321 | rs13061104 | -1.262 | 1.393 | 0.26 | -2.173 | 0.002 | -0.082 | -0.082 | 0 | 0.23 | 0.3 | 0.35 | 0.04 | 0.25 | 0.17 |
| 3 | 12722870 | rs6442323 | 1.367 | 1.139 | 0.022 | -2.142 | -0.008 | 0.012 | 0.012 | 0.86 | 0.6 | 0.37 | 0.44 | 0.89 | 0.49 | 0.75 |
| 3 | 12727962 | rs9874315 | 1.519 | 0.968 | -0.107 | -2.098 | 0.022 | 0.117 | 0.117 | 0.794 | 0.55 | 0.26 | 0.37 | 0.89 | 0.41 | 0.73 |
| 3 | 12742433 | rs13072983 | -0.488 | -0.121 | 0.175 | -1.59 | -0.009 | 0.02 | 0.02 | 0.015 | 0.11 | 0.08 | 0.23 | 0.01 | 0.14 | 0.1 |
| 3 | 12763841 | rs6442325 | -1.534 | -0.583 | -0.616 | -1.499 | 0.072 | 0.037 | 0.037 | 0.144 | 0.57 | 0.96 | 0.78 | 0.32 | 0.72 | 0.38 |
| 3 | 12776428 | rs3732687 | 0.073 | -0.13 | 0.111 | -1.448 | 0.054 | -0.033 | -0.033 | 0 | 0.27 | 0.42 | 0.34 | 0.2 | 0.39 | 0.2 |
| 3 | 12778794 | rs6442327 | -0.89 | -0.743 | 0.226 | -1.34 | -0.011 | 0.162 | 0.162 | 0.126 | 0.47 | 0.84 | 0.55 | 0.24 | 0.55 | 0.27 |
| 3 | 12788938 | rs1467025 | -0.328 | -0.317 | 0.342 | -1.179 | 0.023 | -0.036 | -0.036 | 0 | 0.36 | 0.4 | 0.42 | 0.2 | 0.48 | 0.23 |
| 3 | 12797747 | rs6783990 | -0.212 | -1.222 | 0.833 | -0.728 | 0.032 | 0.169 | 0.169 | 0 | 0.08 | 0.31 | 0.05 | 0.17 | 0.04 | 0.1 |
| 3 | 12803792 | rs9876483 | -0.4 | -1.435 | 0.706 | -0.945 | -0.013 | 0.111 | 0.111 | 1.585 | 0.38 | 0.78 | 0.43 | 0.16 | 0.58 | 0.2 |
| 3 | 12805910 | rs9812144 | -1.282 | -1.548 | 0.535 | -1.011 | 0.119 | 0.097 | 0.097 | 0 | 0.5 | 0.9 | 0.51 | 0.33 | 0.68 | 0.34 |
| 3 | 12806080 | rs7626609 | -1.109 | -1.516 | 0.7 | -1.003 | 0.058 | 0.064 | 0.064 | 0 | 0.45 | 0.82 | 0.46 | 0.33 | 0.63 | 0.32 |
| 3 | 12806123 | rs7626624 | -0.667 | -1.139 | 0.669 | -0.848 | -0.007 | -0.005 | -0.005 | 0 | 0.12 | 0.14 | 0.08 | 0.08 | 0.1 | 0.08 |
| 3 | 12818103 | rs17037220 | -0.148 | -1.016 | 0.803 | -0.652 | -0.005 | -0.059 | -0.059 | 0 | 0.37 | 0.51 | 0.42 | 0.18 | 0.6 | 0.21 |
| 3 | 12824363 | rs3901666 | -1.057 | -1.288 | 0.003 | -0.592 | 0.035 | -0.033 | -0.033 | 0 | 0.52 | 0.98 | 0.5 | 0.44 | 0.64 | 0.36 |
| 3 | 12836792 | rs4034942 | 0.051 | -1.258 | 0.048 | -0.593 | 0.005 | -0.024 | -0.024 | 0 | 0.37 | 0.52 | 0.42 | 0.33 | 0.58 | 0.25 |
| 3 | 12839915 | rs6806919 | -1.141 | -1.335 | 0.017 | -0.569 | -0.029 | 0.006 | 0.006 | 0 | 0.2 | 0.7 | 0.13 | 0.27 | 0.07 | 0.17 |
| 3 | 12842223 | rs4642101 | -0.827 | -1.178 | -0.633 | -1.135 | 0.109 | 0.102 | 0.102 | 0 | 0.49 | 0.73 | 0.48 | 0.29 | 0.69 | 0.36 |
| 3 | 12845158 | rs9835677 | -0.126 | -1.015 | -1.506 | -1.065 | 0.007 | 0.017 | 0.017 | 0 | 0.38 | 0.33 | 0.42 | 0.18 | 0.65 | 0.26 |
| 3 | 12848822 | rs11718898 | -1.091 | -1.016 | -1.527 | -1.059 | 0.085 | 0.062 | 0.062 | 0 | 0.55 | 0.97 | 0.51 | 0.29 | 0.7 | 0.37 |
| 3 | 12856856 | rs2305398 | -1.038 | -0.683 | -1.647 | -0.814 | 0.049 | 0.103 | 0.103 | 0.101 | 0.51 | 0.97 | 0.5 | 0.29 | 0.66 | 0.31 |
| 3 | 12857493 | rs2305397 | -0.382 | 0.77 | -0.81 | -0.606 | 0.001 | -0.008 | -0.008 | 0 | 0.32 | 0.32 | 0.37 | 0.18 | 0.58 | 0.19 |
| 3 | 12860988 | rs906671 | -0.095 | 0.737 | -1.083 | -0.793 | 0.01 | 0.069 | 0.069 | 0 | 0.54 | 0.34 | 0.47 | 0.82 | 0.54 | 0.64 |
| 3 | 12861586 | rs4684883 | -0.024 | 0.917 | -1.592 | -0.902 | -0.033 | 0.059 | 0.059 | 0 | 0.19 | 0.05 | 0.3 | 0.12 | 0.18 | 0.18 |

| | | | | | | | | | | | | | | | | |
|---|---|---|---|---|---|---|---|---|---|---|---|---|---|---|---|---|
| 3 | 12861608 | rs3817121 | 0.222 | 0.905 | -1.602 | -0.855 | -0.046 | 0.034 | 0.034 | 0 | 0.17 | 0.03 | 0.29 | 0.12 | 0.17 | 0.17 |
| 3 | 12862257 | rs7616406 | -0.621 | 0.904 | -1.176 | -0.77 | 0.016 | 0.029 | 0.029 | 5.538 | 0.58 | 0.71 | 0.57 | 0.31 | 0.62 | 0.51 |
| 3 | 12864386 | rs6442333 | -0.724 | 0.951 | -1.152 | 0.102 | 0.007 | -0.132 | -0.132 | 0 | 0.62 | 0.93 | 0.62 | 0.5 | 0.65 | 0.54 |
| 3 | 12864518 | rs6800812 | 0.207 | 1.349 | -0.013 | -0.086 | -0.006 | 0.028 | 0.028 | 1.9 | 0.45 | 0.37 | 0.46 | 0.22 | 0.48 | 0.3 |
| 3 | 12866725 | rs7629133 | -0.81 | 1.434 | 0.01 | -0.185 | 0.021 | -0.016 | -0.016 | 0.238 | 0.59 | 0.69 | 0.58 | 0.49 | 0.66 | 0.55 |
| 3 | 12872442 | rs9813534 | -0.305 | 1.163 | 0.588 | 0.03 | 0.024 | -0.006 | -0.006 | 0.608 | 0.12 | 0.6 | 0.11 | 0.09 | 0.1 | 0.23 |
| 3 | 12873265 | rs3732680 | -1.173 | 0.914 | 0.066 | 0.041 | 0.019 | -0.103 | -0.103 | 0 | 0.73 | 0.89 | 0.65 | 0.54 | 0.76 | 0.69 |
| 3 | 12878465 | rs3773304 | -0.027 | 1.109 | 0.39 | 0.028 | -0.001 | 0 | 0 | 0 | 0.54 | 0.44 | 0.52 | 0.44 | 0.56 | 0.4 |
| 3 | 12898941 | rs906673 | -0.013 | 0.95 | -0.142 | -0.2 | 0.023 | 0.056 | 0.056 | 0.072 | 0.54 | 0.27 | 0.51 | 0.35 | 0.61 | 0.43 |
| 3 | 12899890 | rs7645264 | -0.692 | 1.02 | 0.481 | -0.135 | 0.046 | -0.016 | -0.016 | 0.366 | 0.15 | 0.35 | 0.11 | 0.1 | 0.14 | 0.26 |
| 3 | 12902199 | rs1848466 | -0.333 | 1.02 | 0.356 | -0.141 | 0.004 | -0.009 | -0.009 | 0 | 0.12 | 0.16 | 0.07 | 0.09 | 0.12 | 0.24 |
| 3 | 12908510 | rs2600228 | -0.16 | 1.318 | 0.154 | -0.333 | 0.005 | 0.001 | 0.001 | 0 | 0.58 | 0.37 | 0.5 | 0.44 | 0.59 | 0.64 |
| 3 | 12914900 | rs361218 | -0.769 | 1.374 | 0.31 | -0.303 | 0.004 | -0.007 | -0.007 | 0 | 0.68 | 0.72 | 0.6 | 0.45 | 0.7 | 0.67 |
| 3 | 12916106 | rs6808492 | -1.645 | 1.144 | 0.281 | -0.444 | -0.008 | -0.039 | -0.039 | 0.496 | 0.11 | 0.27 | 0.05 | 0.01 | 0.05 | 0.03 |
| 3 | 12917207 | rs434460 | 0.019 | 1.848 | -0.071 | -0.659 | -0.001 | 0.079 | 0.079 | 0.22 | 0.57 | 0.28 | 0.46 | 0.25 | 0.56 | 0.51 |
| 3 | 12920171 | rs361225 | -1.36 | 1.722 | 0.031 | -0.462 | 0.041 | -0.043 | -0.043 | 0.014 | 0.73 | 0.85 | 0.64 | 0.42 | 0.78 | 0.73 |
| 3 | 12923197 | rs2569991 | -0.831 | 1.711 | 0.339 | -0.272 | 0.016 | -0.013 | -0.013 | 0 | 0.67 | 0.71 | 0.62 | 0.44 | 0.75 | 0.71 |
| 3 | 12926041 | rs2600227 | -0.791 | 1.244 | -0.269 | -0.696 | -0.112 | 0.136 | 0.136 | 0 | 0.58 | 0.96 | 0.6 | 0.76 | 0.56 | 0.59 |
| 3 | 12928762 | rs2600226 | -0.68 | 1.398 | 0.414 | -0.559 | -0.009 | 0.001 | 0.001 | 0 | 0.64 | 0.49 | 0.57 | 0.36 | 0.66 | 0.48 |
| 3 | 12940346 | rs14191 | -1.716 | 1.293 | 0.088 | 0.004 | 0.043 | -0.155 | -0.155 | 0 | 0.73 | 0.98 | 0.66 | 0.44 | 0.79 | 0.71 |
| 3 | 12942328 | rs3773309 | 0.111 | 1.538 | 0.209 | -0.202 | 0.011 | -0.002 | -0.002 | 0 | 0.08 | 0.1 | 0.07 | 0.08 | 0.13 | 0.24 |
| 3 | 12943608 | rs4547681 | -0.121 | 1.49 | 0.299 | -0.221 | 0.083 | 0.01 | 0.01 | 0.143 | 0.08 | 0.31 | 0.09 | 0.08 | 0.12 | 0.21 |
| 3 | 12946142 | rs3773312 | 0.188 | 1.542 | 0.533 | 0.481 | 0.044 | 0.144 | 0.144 | 0 | 0.05 | 0.14 | 0.09 | 0.15 | 0.12 | 0.03 |
| 3 | 12946433 | rs2600323 | 1.434 | 1.349 | 0.014 | 0.722 | 0.001 | -0.138 | -0.138 | 0.028 | 0.29 | 0.02 | 0.33 | 0.46 | 0.26 | 0.3 |
| 3 | 12946690 | rs734390 | 1.517 | 1.316 | -0.534 | -0.078 | -0.114 | 0.085 | 0.085 | 0 | 0.21 | 0.02 | 0.32 | 0.06 | 0.29 | 0.2 |
| 3 | 12949028 | rs734525 | -1.595 | 0.729 | -0.349 | 0.051 | 0.052 | -0.147 | -0.147 | 0 | 0.71 | 0.96 | 0.66 | 0.49 | 0.78 | 0.7 |
| 3 | 12955156 | rs2117011 | 1.008 | 0.889 | -0.246 | 0.059 | -0.017 | 0.003 | 0.003 | 0 | 0.08 | 0.06 | 0.07 | 0.07 | 0.12 | 0.22 |
| 3 | 12956905 | rs17037462 | -0.312 | 0.848 | 0.028 | 0.063 | 0.044 | 0.154 | 0.154 | 0 | 0.09 | 0.22 | 0.11 | 0.16 | 0.14 | 0.03 |
| 3 | 12959342 | rs2596901 | -1.453 | 0.599 | -0.336 | 0.092 | 0.06 | -0.094 | -0.094 | 2.444 | 0.7 | 0.9 | 0.66 | 0.49 | 0.78 | 0.71 |
| 3 | 12963216 | rs2600221 | -1.454 | 0.625 | -0.401 | 0.05 | 0.054 | -0.099 | -0.099 | 0 | 0.7 | 0.92 | 0.66 | 0.49 | 0.78 | 0.71 |
| 3 | 12970494 | rs2600208 | -0.231 | 0.569 | -0.148 | -0.457 | 0.017 | 0.025 | 0.025 | 0 | 0.55 | 0.35 | 0.53 | 0.35 | 0.64 | 0.45 |
| 3 | 12974143 | rs2596899 | -1.536 | 0.291 | -0.565 | -0.302 | 0.013 | -0.155 | -0.155 | 0 | 0.72 | 0.99 | 0.66 | 0.43 | 0.76 | 0.71 |
| 3 | 12979173 | rs939899 | 0.774 | 0.459 | 0.009 | -0.679 | -0.003 | 0.011 | 0.011 | 0.948 | 0.08 | 0.12 | 0.06 | 0.15 | 0.09 | 0.12 |
| 3 | 12981873 | rs2012023 | 0.743 | 0.1 | 0.121 | -1.284 | -0.022 | -0.127 | -0.127 | 0 | 0.26 | 0.4 | 0.21 | 0.03 | 0.25 | 0.22 |
| 3 | 12991383 | rs2569997 | 0.063 | 0.729 | 0.394 | -0.998 | 0 | -0.016 | -0.016 | 0 | 0.52 | 0.5 | 0.51 | 0.85 | 0.47 | 0.68 |
| 3 | 12999160 | rs2600327 | -1.657 | 0.817 | -0.346 | -1.307 | -0.084 | 0.08 | 0.08 | 0 | 0.79 | 0.98 | 0.69 | 0.93 | 0.72 | 0.85 |
| 3 | 13001642 | rs1529756 | 0.998 | 1.597 | 0.041 | -0.816 | -0.047 | -0.027 | -0.027 | 0.14 | 0.07 | 0.01 | 0.08 | 0.09 | 0.11 | 0.2 |
| 3 | 13001931 | rs765248 | 0.605 | 1.603 | 0.268 | -0.944 | -0.027 | 0.101 | 0.101 | 0 | 0.38 | 0.21 | 0.44 | 0.11 | 0.45 | 0.17 |
| 3 | 13005609 | rs747831 | -1.347 | 1.567 | 0.447 | -0.931 | 0.011 | -0.054 | -0.054 | 1.093 | 0.82 | 0.69 | 0.89 | 0.96 | 0.8 | 0.83 |
| 3 | 13006661 | rs1010412 | 1.532 | 1.617 | -0.296 | -0.842 | 0.03 | -0.039 | -0.039 | 0.508 | 0.26 | 0.05 | 0.33 | 0.76 | 0.21 | 0.33 |
| 3 | 13012816 | rs17774704 | 0.775 | 1.959 | -0.048 | -0.646 | 0.015 | 0.001 | 0.001 | 0 | 0.08 | 0.12 | 0.08 | 0.09 | 0.11 | 0.2 |
| 3 | 13015655 | rs4684892 | 0.309 | 1.611 | -0.053 | -0.875 | -0.012 | 0.078 | 0.078 | 0 | 0.4 | 0.26 | 0.44 | 0.11 | 0.46 | 0.17 |
| 3 | 13018488 | rs11922549 | 0.345 | 1.175 | 0.247 | -0.56 | 0.056 | -0.002 | -0.002 | 0 | 0.06 | 0.15 | 0.1 | 0.04 | 0.14 | 0.01 |
| 3 | 13025899 | rs360745 | 0.971 | 1.31 | -0.305 | -0.571 | -0.025 | 0.137 | 0.137 | 0 | 0.33 | 0.11 | 0.36 | 0.12 | 0.34 | 0.19 |
| 3 | 13032126 | rs6442344 | -0.512 | 1.105 | 0.474 | -0.353 | 0.069 | -0.013 | -0.013 | 1.36 | 0.13 | 0.41 | 0.14 | 0.05 | 0.17 | 0.13 |
| 3 | 13036932 | rs360836 | 0.52 | 1.048 | 0.258 | -0.43 | -0.023 | -0.114 | -0.114 | 0 | 0.21 | 0.38 | 0.11 | 0.02 | 0.17 | 0.19 |
| 3 | 13042982 | rs428862 | 0.033 | 1.065 | -0.463 | -0.13 | -0.002 | -0.019 | -0.019 | 0 | 0.34 | 0.63 | 0.23 | 0.09 | 0.34 | 0.34 |
| 3 | 13045495 | rs361021 | 0.28 | 1.606 | -0.501 | 0.664 | -0.005 | 0.007 | 0.007 | 0 | 0.56 | 0.56 | 0.55 | 0.6 | 0.59 | 0.66 |
| 3 | 13045939 | rs892272 | -0.513 | 2.09 | -0.477 | 1.128 | -0.012 | -0.012 | -0.012 | 0 | 0.06 | 0.05 | 0.07 | 0.16 | 0.09 | 0.15 |
| 3 | 13047602 | rs360833 | 0.528 | 1.441 | 0.018 | 1.304 | 0.019 | 0.121 | 0.121 | 0.022 | 0.78 | 0.62 | 0.59 | 0.5 | 0.76 | 0.59 |
| 3 | 13052644 | rs11712537 | -0.048 | 0.964 | 0.253 | 0.629 | 0 | -0.009 | -0.009 | 0 | 0.06 | 0.04 | 0.02 | 0 | 0.07 | 0.05 |
| 3 | 13053835 | rs17037535 | 0.541 | 0.136 | -0.989 | 0.44 | 0.005 | -0.147 | -0.147 | 0 | 0.28 | 0.01 | 0.19 | 0.42 | 0.18 | 0.24 |
| 3 | 13053937 | rs360830 | 0.895 | 0.262 | -1.437 | -0.443 | 0.015 | 0.1 | 0.1 | 0.028 | 0.27 | 0.1 | 0.18 | 0.07 | 0.28 | 0.19 |
| 3 | 13055576 | rs458459 | -0.911 | 0.581 | -1.162 | 0.284 | -0.005 | -0.073 | -0.073 | 0 | 0.67 | 0.91 | 0.7 | 0.47 | 0.74 | 0.74 |
| 3 | 13057011 | rs4684895 | -0.086 | 0.798 | -0.116 | 0.511 | -0.001 | 0.044 | 0.044 | 0.031 | 0.15 | 0.28 | 0.22 | 0.29 | 0.15 | 0.24 |
| 3 | 13057791 | rs9830953 | -0.302 | 1.185 | -0.161 | 0.305 | 0.018 | -0.015 | -0.015 | 0 | 0.09 | 0.24 | 0.1 | 0.04 | 0.1 | 0.16 |
| 3 | 13067049 | rs522491 | 0.505 | 0.55 | -0.516 | 0.393 | -0.012 | 0.017 | 0.017 | 2.339 | 0.65 | 0.45 | 0.54 | 0.49 | 0.7 | 0.48 |
| 3 | 13069494 | rs2881979 | 0.169 | 0.82 | -0.288 | -0.062 | 0.003 | -0.03 | -0.03 | 0 | 0.07 | 0.1 | 0.05 | 0 | 0.08 | 0.02 |
| 3 | 13070113 | rs380499 | -0.374 | 0.862 | -0.528 | -0.278 | 0.004 | -0.071 | -0.071 | 0 | 0.83 | 0.72 | 0.87 | 0.99 | 0.81 | 0.92 |
| 3 | 13070799 | rs361052 | 0.518 | 0.74 | -0.236 | 0.048 | 0.024 | 0.019 | 0.019 | 0.548 | 0.81 | 0.49 | 0.8 | 0.81 | 0.81 | 0.81 |
| 3 | 13072590 | rs6790533 | 0.23 | 0.434 | -1.307 | -0.41 | -0.06 | 0.11 | 0.11 | 0.064 | 0.63 | 0.82 | 0.58 | 0.8 | 0.57 | 0.85 |
| 3 | 13075760 | rs892276 | 1.319 | -0.3 | -1.49 | -0.697 | -0.024 | -0.019 | -0.019 | 0 | 0.32 | 0.17 | 0.52 | 0.57 | 0.35 | 0.39 |
| 3 | 13076226 | rs11128630 | 0.977 | -0.809 | -1.57 | -0.914 | 0.017 | 0.009 | 0.009 | 0.186 | 0.23 | 0.09 | 0.16 | 0.21 | 0.23 | 0.25 |
| 3 | 13078387 | rs6442346 | -0.725 | -0.89 | -1.471 | -0.963 | -0.065 | 0.089 | 0.089 | 0.396 | 0.68 | 0.85 | 0.76 | 0.81 | 0.63 | 0.87 |
| 3 | 13080976 | rs9841967 | 0.621 | -0.922 | -0.619 | -0.984 | 0 | 0 | 0 | 0.022 | 0.27 | 0.28 | 0.17 | 0.22 | 0.23 | 0.37 |
| 3 | 13081885 | rs4684901 | -0.718 | -0.924 | -0.831 | -0.99 | -0.07 | 0.085 | 0.085 | 0.245 | 0.68 | 0.86 | 0.76 | 0.81 | 0.63 | 0.87 |
| 3 | 13084130 | rs6810137 | -0.578 | -1.081 | -0.493 | -1.106 | 0.018 | 0.017 | 0.017 | 0 | 0.41 | 0.6 | 0.22 | 0.23 | 0.28 | 0.47 |
| 3 | 13088331 | rs360901 | 1.348 | -1.062 | -0.644 | -1.144 | -0.022 | 0.009 | 0.009 | 0.827 | 0.37 | 0.19 | 0.57 | 0.62 | 0.39 | 0.39 |
| 3 | 13093188 | rs360894 | 0.335 | -0.973 | -0.427 | -1.203 | 0.001 | -0.03 | -0.03 | 0 | 0.64 | 0.38 | 0.73 | 0.84 | 0.63 | 0.76 |

| | | | | | | | | | | | | | | | | |
|---|---|---|---|---|---|---|---|---|---|---|---|---|---|---|---|---|
| 3 | 13095532 | rs6784345 | 0.501 | -1.11 | -0.323 | -1.128 | -0.048 | -0.08 | -0.08 | 0 | 0.13 | 0.35 | 0.06 | 0.01 | 0.04 | 0.1 |
| 3 | 13100249 | rs6765254 | 0.304 | -1.258 | -0.497 | -1.28 | -0.037 | -0.041 | -0.041 | 0 | 0.35 | 0.37 | 0.21 | 0.21 | 0.26 | 0.36 |
| 3 | 13101791 | rs360762 | 1.048 | -1.168 | -0.415 | -1.368 | -0.012 | 0.031 | 0.031 | 0 | 0.41 | 0.27 | 0.56 | 0.64 | 0.4 | 0.51 |
| 3 | 13104266 | rs360757 | -0.843 | -1.018 | -0.388 | -1.322 | 0.029 | -0.044 | -0.044 | 0 | 0.75 | 0.59 | 0.77 | 0.85 | 0.66 | 0.85 |
| 3 | 13106302 | rs360875 | 0.385 | -1.05 | -0.499 | -1.268 | 0.001 | 0.057 | 0.057 | 0 | 0.5 | 0.56 | 0.63 | 0.64 | 0.45 | 0.49 |
| 3 | 13106888 | rs360874 | 0.648 | -0.999 | -0.849 | -1.24 | -0.017 | 0.073 | 0.073 | 0.001 | 0.46 | 0.25 | 0.6 | 0.64 | 0.44 | 0.49 |
| 3 | 13106959 | rs13061267 | 1.855 | -0.342 | -1.347 | -0.474 | -0.038 | -0.042 | -0.042 | 0 | 0.13 | 0.07 | 0.4 | 0.39 | 0.19 | 0.22 |
| 3 | 13107882 | rs360871 | 2.286 | -0.425 | -1.388 | -0.623 | -0.013 | 0.038 | 0.038 | 0 | 0.07 | 0 | 0.03 | 0 | 0.07 | 0.01 |
| 3 | 13110476 | rs1157649 | 1.864 | -0.385 | -1.358 | -0.374 | -0.039 | -0.046 | -0.046 | 1.599 | 0.13 | 0.07 | 0.39 | 0.37 | 0.19 | 0.23 |
| 3 | 13110536 | rs417123 | 0.312 | -0.368 | -1.367 | -0.336 | 0.019 | 0.039 | 0.039 | 0 | 0.34 | 0.54 | 0.48 | 0.51 | 0.35 | 0.37 |
| 3 | 13112931 | rs413245 | 0.175 | -0.293 | -1.142 | -0.545 | 0 | -0.003 | -0.003 | 1.117 | 0.51 | 0.56 | 0.61 | 0.63 | 0.49 | 0.53 |
| 3 | 13116061 | rs11718794 | 1.08 | -0.323 | -1.416 | -0.325 | 0.026 | -0.015 | -0.015 | 0 | 0.23 | 0.04 | 0.14 | 0.21 | 0.21 | 0.23 |
| 3 | 13117870 | rs460970 | 0.14 | -0.097 | -0.979 | -0.645 | 0.041 | -0.043 | -0.043 | 0 | 0.74 | 0.6 | 0.73 | 0.78 | 0.67 | 0.68 |
| 3 | 13117999 | rs444944 | -0.435 | -0.267 | -1.233 | -0.759 | -0.015 | 0.032 | 0.032 | 0 | 0.57 | 0.24 | 0.34 | 0.32 | 0.47 | 0.52 |
| 3 | 13118489 | rs4684905 | 0.183 | -0.241 | -1.119 | -0.662 | 0.006 | 0.006 | 0.006 | 0 | 0.44 | 0.41 | 0.54 | 0.6 | 0.4 | 0.57 |
| 3 | 13123355 | rs2648699 | -0.505 | -0.308 | -1.787 | -0.682 | 0.002 | 0.021 | 0.021 | 0 | 0.64 | 0.8 | 0.72 | 0.74 | 0.67 | 0.81 |
| 3 | 13124572 | rs607829 | 0.224 | -0.12 | -1.583 | -1.104 | 0.001 | -0.038 | -0.038 | 0 | 0.18 | 0.22 | 0.11 | 0.01 | 0.19 | 0.04 |
| 3 | 13126434 | rs636377 | -0.686 | -0.08 | -1.683 | -1.088 | -0.009 | 0.015 | 0.015 | 0 | 0.83 | 0.88 | 0.91 | 0.99 | 0.82 | 0.96 |
| 3 | 13134777 | rs521573 | -0.019 | -0.491 | -1.54 | -1.062 | 0.012 | 0.036 | 0.036 | 0 | 0.49 | 0.7 | 0.6 | 0.6 | 0.49 | 0.58 |
| 3 | 13135575 | rs662634 | 0.129 | -0.72 | -1.726 | -1.094 | 0.006 | 0.022 | 0.022 | 0 | 0.19 | 0.19 | 0.12 | 0.07 | 0.18 | 0.15 |
| 3 | 13137229 | rs3856816 | 1.11 | -0.476 | -1.907 | -0.892 | 0.004 | 0.08 | 0.08 | 0 | 0.29 | 0.44 | 0.46 | 0.51 | 0.31 | 0.42 |
| 3 | 13140472 | rs165569 | 1.115 | -0.199 | -1.866 | -0.92 | 0 | -0.037 | -0.037 | 0 | 0.09 | 0.18 | 0.06 | 0 | 0.1 | 0.05 |
| 3 | 13140732 | rs165567 | -1.421 | -0.077 | -2.079 | -0.837 | 0.033 | -0.012 | -0.012 | 1.505 | 0.87 | 0.8 | 0.86 | 0.89 | 0.75 | 0.87 |
| 3 | 13140835 | rs360749 | 0.047 | -0.693 | -1.971 | -0.679 | 0.018 | 0.019 | 0.019 | 0 | 0.47 | 0.71 | 0.58 | 0.58 | 0.43 | 0.61 |
| 3 | 13144375 | rs3856817 | 0.077 | -1.222 | -1.922 | -0.455 | -0.003 | 0 | 0 | 0 | 0.24 | 0.19 | 0.19 | 0.36 | 0.11 | 0.38 |
| 3 | 13146033 | rs2686749 | 0.877 | -1.292 | -2.506 | -0.702 | 0.042 | 0.05 | 0.05 | 0 | 0.6 | 0.67 | 0.83 | 0.74 | 0.75 | 0.65 |
| 3 | 13146594 | rs2686748 | -0.456 | -0.758 | -2.379 | -0.473 | 0.039 | 0.079 | 0.079 | 0 | 0.28 | 0.15 | 0.1 | 0.13 | 0.18 | 0.17 |
| 3 | 13148872 | rs17833015 | 0.087 | -0.098 | -2.148 | 0.004 | -0.01 | -0.002 | -0.002 | 0 | 0.15 | 0 | 0.05 | 0.13 | 0.09 | 0.15 |
| 3 | 13149343 | rs4684909 | 0.692 | 0.83 | -1.99 | 0.243 | -0.026 | -0.043 | -0.043 | 1.196 | 0.25 | 0.18 | 0.48 | 0.38 | 0.35 | 0.19 |
| 3 | 13149782 | rs636758 | -0.578 | 0.071 | -1.792 | -0.235 | 0.032 | 0.051 | 0.051 | 1.284 | 0.51 | 0.37 | 0.3 | 0.34 | 0.35 | 0.52 |
| 3 | 13150039 | rs17833070 | -0.641 | 0.654 | -1.447 | 0.329 | -0.001 | 0.001 | 0.001 | 0.042 | 0.11 | 0.09 | 0.07 | 0.13 | 0.08 | 0.09 |
| 3 | 13154366 | rs360741 | -0.334 | 0.774 | -2.117 | -0.102 | -0.022 | 0.119 | 0.119 | 0 | 0.43 | 0.83 | 0.37 | 0.26 | 0.43 | 0.47 |
| 3 | 13157050 | rs6800859 | -0.372 | 0.381 | -1.751 | -0.309 | 0.011 | 0.008 | 0.008 | 0.139 | 0.39 | 0.65 | 0.3 | 0.36 | 0.3 | 0.51 |
| 3 | 13164783 | rs1615157 | -0.841 | 0.208 | -1.481 | -0.778 | -0.015 | 0.019 | 0.019 | 0.916 | 0.55 | 0.74 | 0.36 | 0.43 | 0.44 | 0.64 |
| 3 | 13165857 | rs7622345 | 0.505 | -0.086 | -0.804 | -0.961 | 0.004 | -0.003 | -0.003 | 0.029 | 0.29 | 0.22 | 0.24 | 0.35 | 0.25 | 0.47 |
| 3 | 13166918 | rs360738 | 0.407 | -0.252 | -0.808 | -0.998 | -0.045 | 0.039 | 0.039 | 0 | 0.18 | 0.02 | 0.07 | 0.09 | 0.16 | 0.18 |
| 3 | 13168301 | rs11128635 | -0.37 | 0.364 | -0.815 | -0.979 | 0.009 | 0.004 | 0.004 | 0.225 | 0.4 | 0.67 | 0.3 | 0.39 | 0.32 | 0.49 |
| 3 | 13178145 | rs358386 | 0.358 | 0.376 | -0.765 | -0.896 | -0.005 | 0.001 | 0.001 | 1.336 | 0.55 | 0.62 | 0.67 | 0.55 | 0.55 | 0.36 |
| 3 | 13178247 | rs2004402 | 0.421 | 0.363 | -0.926 | -0.878 | 0 | 0.001 | 0.001 | 6.614 | 0.28 | 0.24 | 0.25 | 0.37 | 0.29 | 0.47 |
| 3 | 13180372 | rs358380 | -0.097 | 0.43 | -1.513 | -0.971 | -0.075 | 0.064 | 0.064 | 0 | 0.78 | 0.99 | 0.91 | 0.92 | 0.79 | 0.81 |
| 3 | 13182935 | rs165435 | -0.886 | 0.338 | -1.369 | -0.911 | 0.005 | -0.005 | -0.005 | 0.828 | 0.55 | 0.73 | 0.38 | 0.46 | 0.5 | 0.68 |
| 3 | 13186109 | rs358374 | 0.214 | 0.57 | -1.303 | -0.963 | -0.028 | 0.085 | 0.085 | 0 | 0.79 | 0.94 | 0.9 | 0.93 | 0.81 | 0.81 |
| 3 | 13192364 | rs11719364 | 0.124 | 0.214 | -1.299 | -0.656 | 0.002 | 0.001 | 0.001 | 1.394 | 0.54 | 0.41 | 0.47 | 0.42 | 0.59 | 0.49 |
| 3 | 13192688 | rs358345 | 0.345 | 0.153 | -1.137 | -0.484 | 0 | -0.001 | -0.001 | 0 | 0.29 | 0.31 | 0.44 | 0.48 | 0.25 | 0.29 |
| 3 | 13207536 | rs357161 | 0.304 | 0.282 | -1.048 | -0.595 | 0.002 | -0.001 | -0.001 | 0.067 | 0.47 | 0.48 | 0.53 | 0.47 | 0.4 | 0.42 |
| 3 | 13208328 | rs11715528 | 0.198 | 0.398 | -1.605 | -0.333 | -0.038 | -0.039 | -0.039 | 0 | 0.33 | 0.19 | 0.34 | 0.41 | 0.41 | 0.34 |
| 3 | 13213525 | rs9875054 | -0.775 | 0.03 | -1.426 | 0.152 | 0.039 | -0.009 | -0.009 | 0 | 0.44 | 0.82 | 0.29 | 0.43 | 0.33 | 0.5 |
| 3 | 13214507 | rs9822701 | -0.403 | 0.165 | -0.528 | -0.034 | -0.032 | -0.123 | -0.123 | 2.142 | 0.22 | 0.48 | 0.14 | 0.05 | 0.16 | 0.09 |
| 3 | 13215984 | rs165265 | -0.353 | 0.872 | -0.727 | 0.7 | 0.018 | 0.032 | 0.032 | 0 | 0.34 | 0.51 | 0.45 | 0.51 | 0.37 | 0.38 |
| 3 | 13218281 | rs357135 | 0.241 | 0.221 | -0.864 | 0.683 | 0.012 | -0.004 | -0.004 | 0 | 0.54 | 0.37 | 0.57 | 0.49 | 0.63 | 0.67 |
| 3 | 13218414 | rs357136 | -0.413 | -0.251 | -0.941 | 0.49 | 0.006 | 0.007 | 0.007 | 0.018 | 0.8 | 0.89 | 0.78 | 0.79 | 0.82 | 0.84 |
| 3 | 13219337 | rs357139 | 0.299 | -0.318 | -0.728 | 0.55 | -0.014 | 0.026 | 0.026 | 0.783 | 0.67 | 0.87 | 0.73 | 0.68 | 0.71 | 0.77 |
| 3 | 13226080 | rs358597 | 0.035 | -0.505 | -0.359 | 0.509 | -0.001 | -0.001 | -0.001 | 1.659 | 0.7 | 0.67 | 0.74 | 0.75 | 0.78 | 0.77 |
| 3 | 13226100 | rs358598 | -0.205 | -0.473 | -0.441 | 0.523 | 0.003 | 0 | 0 | 2.463 | 0.73 | 0.9 | 0.76 | 0.75 | 0.78 | 0.81 |
| 3 | 13226945 | rs17037831 | 0.216 | -0.244 | -0.308 | 0.787 | -0.001 | 0.024 | 0.024 | 0 | 0.11 | 0.13 | 0.1 | 0.28 | 0.1 | 0.12 |
| 3 | 13231519 | rs582111 | -0.203 | 0.033 | -0.884 | 0.354 | 0.005 | 0.007 | 0.007 | 0.026 | 0.42 | 0.48 | 0.5 | 0.47 | 0.53 | 0.45 |
| 3 | 13232319 | rs1124781 | -0.501 | 0.198 | -0.608 | 0.651 | -0.002 | -0.016 | -0.016 | 0.414 | 0.16 | 0.12 | 0.14 | 0.36 | 0.16 | 0.16 |
| 3 | 13232556 | rs242519 | 0.867 | 0.024 | -0.616 | 0.56 | 0 | 0 | 0 | 0.464 | 0.54 | 0.52 | 0.59 | 0.36 | 0.61 | 0.62 |
| 3 | 13233385 | rs11717784 | -0.922 | 0.2 | -0.996 | 0.586 | 0.004 | 0.056 | 0.056 | 0 | 0.21 | 0.27 | 0.16 | 0.35 | 0.22 | 0.21 |
| 3 | 13233451 | rs357119 | 0.17 | 0.156 | -1.621 | 0.502 | -0.014 | -0.045 | -0.045 | 0.289 | 0.63 | 0.88 | 0.7 | 0.43 | 0.62 | 0.69 |
| 3 | 13244384 | rs4684917 | -1.14 | 0.486 | -1.672 | 0.118 | -0.076 | 0.15 | 0.15 | 0.129 | 0.64 | 0.95 | 0.65 | 0.87 | 0.59 | 0.72 |
| 3 | 13250682 | rs17037891 | 0.347 | 0.319 | -1.348 | 0.63 | 0.007 | -0.011 | -0.011 | 0 | 0.12 | 0.09 | 0.12 | 0.29 | 0.12 | 0.1 |
| 3 | 13255140 | rs9814792 | -0.724 | 0.232 | -0.668 | 0.515 | -0.004 | -0.007 | -0.007 | 0 | 0.35 | 0.68 | 0.44 | 0.51 | 0.37 | 0.46 |
| 3 | 13257064 | rs3852028 | -0.882 | 0.215 | -0.648 | 0.483 | -0.001 | -0.001 | -0.001 | 0.472 | 0.37 | 0.61 | 0.43 | 0.51 | 0.38 | 0.46 |
| 3 | 13257109 | rs6762472 | 0.42 | 0.048 | -0.075 | 0.788 | -0.014 | 0.14 | 0.14 | 0 | 0.05 | 0.24 | 0.02 | 0.21 | 0 | 0.23 |
| 3 | 13257231 | rs2349558 | 0.451 | 0.133 | 0.164 | 0.803 | -0.019 | -0.011 | -0.011 | 0 | 0.19 | 0.08 | 0.25 | 0.18 | 0.27 | 0.15 |
| 3 | 13257512 | rs6442359 | -0.39 | -0.066 | 0.459 | 0.879 | -0.028 | 0.043 | 0.043 | 0 | 0.13 | 0.34 | 0.09 | 0.21 | 0.07 | 0.25 |
| 3 | 13258548 | rs12632098 | -1.068 | -0.079 | 0.5 | 0.884 | -0.033 | 0.117 | 0.117 | 0 | 0.18 | 0.43 | 0.18 | 0.33 | 0.12 | 0.31 |
| 3 | 13262586 | rs9873778 | -0.698 | -0.12 | 0.46 | 0.552 | -0.033 | 0.118 | 0.118 | 0 | 0.16 | 0.41 | 0.17 | 0.33 | 0.11 | 0.31 |

| | | | | | | | | | | | | | | | |
|---|---|---|---|---|---|---|---|---|---|---|---|---|---|---|---|
| 3 | 13263342 | rs17037914 | -0.841 | -0.125 | 0.253 | 0.562 | -0.032 | 0.077 | 0.077 | 0 | 0.18 | 0.37 | 0.17 | 0.33 | 0.12 | 0.31 |
| 3 | 13265680 | rs17080939 | -0.878 | -0.12 | 0.155 | 0.564 | -0.039 | 0.1 | 0.1 | 0 | 0.18 | 0.47 | 0.18 | 0.33 | 0.12 | 0.31 |
| 3 | 13267771 | rs6770415 | 0.292 | -0.11 | -0.435 | -0.305 | -0.005 | 0.007 | 0.007 | 0 | 0.25 | 0.25 | 0.21 | 0.35 | 0.17 | 0.22 |
| 3 | 13270827 | rs895752 | -0.903 | 0.277 | -0.925 | -1.023 | -0.01 | 0.105 | 0.105 | 0 | 0.64 | 0.79 | 0.66 | 0.85 | 0.59 | 0.7 |
| 3 | 13273299 | rs4684126 | -0.444 | 0.402 | -0.947 | -0.845 | 0.005 | 0.023 | 0.023 | 0 | 0.54 | 0.53 | 0.58 | 0.85 | 0.51 | 0.67 |
| 3 | 13275607 | rs9310419 | -0.451 | 0.552 | -1.358 | -0.312 | 0.017 | 0.192 | 0.192 | 0 | 0.51 | 0.79 | 0.51 | 0.82 | 0.39 | 0.58 |
| 3 | 13277637 | rs530922 | 0.284 | 0.173 | -1.144 | 0.375 | -0.023 | -0.026 | -0.026 | 1.173 | 0.55 | 0.22 | 0.53 | 0.57 | 0.53 | 0.59 |
| 3 | 13282611 | rs12495466 | -0.724 | -0.364 | -0.418 | 0.217 | -0.011 | -0.031 | -0.031 | 0 | 0.07 | 0.22 | 0.05 | 0 | 0.02 | 0.04 |
| 3 | 13284218 | rs17037953 | -0.024 | -0.486 | -0.056 | 0.037 | -0.008 | -0.038 | -0.038 | 0.813 | 0.09 | 0.13 | 0.04 | 0 | 0.08 | 0.01 |
| 3 | 13284746 | rs7631991 | -1.866 | -0.663 | 0.564 | 0.169 | -0.043 | -0.051 | -0.051 | 0 | 0.19 | 0.63 | 0.13 | 0.06 | 0.11 | 0.07 |
| 3 | 13287362 | rs6442363 | -1.624 | -0.7 | 0.809 | 0.659 | -0.04 | 0.025 | 0.025 | 1.594 | 0.19 | 0.53 | 0.14 | 0.26 | 0.13 | 0.19 |
| 3 | 13287930 | rs9875796 | -1.072 | -0.702 | 1.057 | 0.471 | -0.045 | -0.091 | -0.091 | 0 | 0.16 | 0.46 | 0.08 | 0.05 | 0.1 | 0.14 |
| 3 | 13288579 | rs13316861 | -1.266 | -0.704 | 0.814 | 0.47 | -0.032 | -0.058 | -0.058 | 0 | 0.17 | 0.59 | 0.1 | 0.05 | 0.1 | 0.14 |
| 3 | 13289938 | rs7647863 | -1.416 | -0.759 | 0.833 | 0.459 | -0.035 | -0.063 | -0.063 | 0.138 | 0.17 | 0.6 | 0.11 | 0.05 | 0.09 | 0.13 |
| 3 | 13290146 | rs11128638 | 0.528 | -1.009 | 0.389 | 1.185 | 0.015 | -0.003 | -0.003 | 0 | 0.07 | 0.01 | 0.15 | 0.36 | 0.03 | 0.11 |
| 3 | 13296300 | rs6785220 | -0.389 | -0.617 | 0.261 | 0.521 | 0.002 | 0.003 | 0.003 | 0 | 0.1 | 0.12 | 0.15 | 0.16 | 0.16 | 0.15 |
| 3 | 13299104 | rs6805636 | -1.967 | -1.286 | -0.362 | 0.127 | 0.037 | 0.061 | 0.061 | 0.076 | 0.37 | 0.79 | 0.29 | 0.27 | 0.29 | 0.4 |
| 3 | 13301696 | rs6784815 | -1.944 | -1.73 | -0.234 | 0.175 | 0.031 | 0.034 | 0.034 | 0 | 0.32 | 0.77 | 0.26 | 0.27 | 0.25 | 0.39 |
| 3 | 13305786 | rs581086 | -0.019 | -1.591 | 0.316 | -0.646 | 0.002 | -0.093 | -0.093 | 0 | 0.63 | 0.47 | 0.76 | 0.96 | 0.66 | 0.79 |
| 3 | 13305930 | rs648445 | -0.675 | -1.582 | 0.178 | -0.684 | -0.016 | 0.033 | 0.033 | 0 | 0.7 | 0.78 | 0.8 | 0.93 | 0.7 | 0.79 |
| 3 | 13315549 | rs6765843 | 0.166 | -1.232 | 1.06 | -0.355 | 0.019 | -0.031 | -0.031 | 0.002 | 0.08 | 0.31 | 0.07 | 0 | 0.06 | 0.01 |
| 3 | 13321852 | rs357116 | 0.566 | -1.679 | 0.77 | -0.131 | 0.002 | -0.006 | -0.006 | 2.058 | 0.47 | 0.35 | 0.59 | 0.55 | 0.57 | 0.54 |
| 3 | 13325922 | rs357111 | -0.441 | -1.891 | 0.777 | -0.492 | -0.023 | 0.037 | 0.037 | 0 | 0.63 | 0.77 | 0.75 | 0.97 | 0.59 | 0.77 |
| 3 | 13327947 | rs357113 | 1.488 | -1.982 | 0.614 | -0.498 | -0.027 | 0.017 | 0.017 | 0 | 0.29 | 0.16 | 0.39 | 0.28 | 0.35 | 0.33 |
| 3 | 13337476 | rs357125 | -0.732 | -1.976 | 0.641 | -0.562 | -0.026 | 0.036 | 0.036 | 0 | 0.63 | 0.77 | 0.74 | 0.98 | 0.58 | 0.78 |
| 3 | 13337553 | rs7648762 | -0.155 | -2.132 | 0.924 | -0.374 | 0.018 | -0.03 | -0.03 | 0 | 0.27 | 0.22 | 0.26 | 0.42 | 0.21 | 0.3 |
| 3 | 13340010 | rs357121 | 0.289 | -1.96 | 1.012 | -0.305 | 0.005 | -0.005 | -0.005 | 2.16 | 0.41 | 0.68 | 0.5 | 0.58 | 0.4 | 0.52 |
| 3 | 13343360 | rs184793 | 1.249 | -1.926 | 0.486 | -0.457 | -0.026 | 0.03 | 0.03 | 0 | 0.32 | 0.11 | 0.39 | 0.27 | 0.34 | 0.37 |
| 3 | 13345450 | rs580438 | 1.076 | -1.919 | 0.562 | -0.454 | -0.001 | 0.027 | 0.027 | 0 | 0.33 | 0.2 | 0.4 | 0.29 | 0.35 | 0.38 |
| 3 | 13357986 | rs354476 | -0.269 | -1.922 | 0.517 | -0.354 | 0 | -0.006 | -0.006 | 0 | 0.54 | 0.53 | 0.62 | 0.7 | 0.51 | 0.67 |
| 3 | 13361287 | rs354478 | -0.225 | -1.924 | 0.516 | -0.354 | 0 | -0.003 | -0.003 | 0 | 0.56 | 0.57 | 0.62 | 0.7 | 0.51 | 0.67 |
| 3 | 13361391 | rs354479 | -2.214 | -1.962 | 0.403 | -0.442 | -0.098 | 0.127 | 0.127 | 0 | 0.8 | 0.97 | 0.85 | 0.99 | 0.73 | 0.88 |
| 3 | 13366545 | rs6801423 | 0.682 | -1.98 | 0.476 | -0.434 | -0.003 | 0.033 | 0.033 | 0 | 0.18 | 0.22 | 0.17 | 0.29 | 0.16 | 0.2 |
| 3 | 13367406 | rs665064 | 0.54 | -1.989 | 0.514 | -0.441 | 0.013 | 0.04 | 0.04 | 0.167 | 0.47 | 0.18 | 0.58 | 0.67 | 0.48 | 0.62 |
| 3 | 13368892 | rs2271509 | 0.066 | -1.908 | 0.335 | -0.405 | -0.001 | -0.005 | -0.005 | 0.026 | 0.45 | 0.46 | 0.38 | 0.3 | 0.49 | 0.33 |
| 3 | 13379452 | rs3773301 | 0.517 | -1.955 | 0.14 | -0.368 | -0.002 | 0.019 | 0.019 | 0 | 0.18 | 0.2 | 0.17 | 0.29 | 0.16 | 0.19 |
| 3 | 13380613 | rs17038034 | 1.369 | -1.979 | 0.227 | -0.386 | 0.012 | 0.005 | 0.005 | 0 | 0.06 | 0.19 | 0.03 | 0.03 | 0.03 | 0.04 |
| 3 | 13395475 | rs2280085 | 0.514 | -1.944 | -0.331 | -0.355 | -0.004 | 0.027 | 0.027 | 0 | 0.18 | 0.21 | 0.17 | 0.29 | 0.17 | 0.19 |
| 3 | 13395579 | rs2280084 | -0.896 | -1.987 | -0.542 | -0.355 | -0.039 | -0.066 | -0.066 | 0 | 0.57 | 0.9 | 0.45 | 0.4 | 0.55 | 0.44 |
| 3 | 13400275 | rs17038082 | 0.3 | -2.001 | -0.522 | -0.367 | -0.01 | 0 | 0 | 0 | 0.05 | 0.07 | 0.02 | 0.07 | 0.02 | 0.09 |
| 3 | 13407556 | rs3732671 | -0.951 | -2.003 | -0.527 | -0.369 | -0.041 | -0.072 | -0.072 | 0 | 0.57 | 0.9 | 0.45 | 0.41 | 0.54 | 0.44 |
| 3 | 13423110 | rs6771144 | -0.96 | -1.956 | -0.526 | -0.368 | -0.05 | -0.072 | -0.072 | 1.242 | 0.58 | 0.9 | 0.44 | 0.4 | 0.54 | 0.44 |
| 3 | 13425416 | rs13062319 | 2.037 | -2.051 | -0.524 | -0.45 | -0.095 | 0.123 | 0.123 | 0 | 0.2 | 0.04 | 0.15 | 0.01 | 0.27 | 0.12 |
| 3 | 13426610 | rs9837443 | -0.288 | -2.03 | -0.339 | -0.203 | 0.007 | 0.019 | 0.019 | 0.18 | 0.53 | 0.82 | 0.43 | 0.34 | 0.53 | 0.36 |
| 3 | 13433158 | rs9845966 | -0.189 | -1.903 | -0.279 | -0.17 | 0.014 | 0.03 | 0.03 | 0.198 | 0.52 | 0.83 | 0.4 | 0.32 | 0.52 | 0.35 |
| 3 | 13436336 | rs7622028 | -0.155 | -1.899 | -0.273 | -0.167 | 0.021 | 0.036 | 0.036 | 0.093 | 0.51 | 0.8 | 0.4 | 0.32 | 0.52 | 0.35 |
| 3 | 13447791 | rs9817861 | -0.735 | -1.737 | -0.181 | -0.471 | -0.01 | -0.024 | -0.024 | 0 | 0.27 | 0.66 | 0.15 | 0.09 | 0.14 | 0.2 |
| 3 | 13449517 | rs9828122 | -0.052 | -1.731 | -0.138 | -0.473 | -0.035 | -0.103 | -0.103 | 0 | 0.21 | 0.56 | 0.13 | 0.03 | 0.12 | 0.12 |
| 3 | 13456803 | rs9817742 | -0.052 | -1.731 | -0.138 | -0.473 | -0.035 | -0.103 | -0.103 | 4.248 | 0.21 | 0.56 | 0.13 | 0.03 | 0.12 | 0.12 |
| 3 | 13476507 | rs6799613 | -0.531 | -1.585 | -0.396 | -0.405 | -0.023 | -0.044 | -0.044 | 0 | 0.28 | 0.63 | 0.15 | 0.09 | 0.14 | 0.19 |
| 3 | 13478434 | rs365691 | -0.967 | -1.382 | -0.526 | -0.275 | -0.036 | -0.11 | -0.11 | 0 | 0.65 | 0.9 | 0.45 | 0.44 | 0.59 | 0.51 |
| 3 | 13480595 | rs2675239 | 0.568 | -1.025 | -0.352 | -0.228 | -0.043 | -0.022 | -0.022 | 0 | 0.38 | 0.24 | 0.3 | 0.35 | 0.45 | 0.31 |
| 3 | 13499551 | rs468113 | 0.578 | -0.894 | -0.608 | -0.188 | 0.016 | 0.01 | 0.01 | 0 | 0.38 | 0.47 | 0.31 | 0.35 | 0.44 | 0.32 |
| 3 | 13503058 | rs468141 | 1.08 | -0.869 | -0.677 | 0.29 | 0.009 | 0.03 | 0.03 | 0 | 0.27 | 0.35 | 0.23 | 0.36 | 0.29 | 0.28 |
| 3 | 13504556 | rs240743 | 1.064 | -0.885 | -0.973 | 0.3 | 0.016 | 0.05 | 0.05 | 0 | 0.27 | 0.39 | 0.23 | 0.36 | 0.29 | 0.29 |
| 3 | 13504610 | rs352385 | -0.485 | -0.33 | -1.981 | 0.8 | -0.093 | 0.041 | 0.041 | 0 | 0.62 | 0.89 | 0.52 | 0.67 | 0.48 | 0.67 |
| 3 | 13505180 | rs3846119 | 0.073 | -0.609 | -1.831 | 0.95 | 0.006 | -0.011 | -0.011 | 0 | 0.13 | 0.09 | 0.17 | 0.28 | 0.06 | 0.27 |
| 3 | 13515826 | rs12636440 | 0.248 | -0.498 | -1.81 | 0.977 | 0.018 | -0.021 | -0.021 | 0 | 0.12 | 0.07 | 0.17 | 0.3 | 0.06 | 0.28 |
| 3 | 13517751 | rs2675234 | 0.76 | -0.649 | -1.753 | 0.612 | -0.011 | -0.002 | -0.002 | 3.018 | 0.46 | 0.72 | 0.43 | 0.36 | 0.43 | 0.45 |
| 3 | 13517761 | rs2655215 | 0.743 | -0.651 | -1.754 | 0.611 | -0.01 | -0.001 | -0.001 | 0.339 | 0.46 | 0.72 | 0.43 | 0.37 | 0.43 | 0.45 |
| 3 | 13519938 | rs2655217 | -0.388 | -0.486 | -2.22 | 0.925 | -0.108 | 0.004 | 0.004 | 0 | 0.65 | 0.94 | 0.65 | 0.67 | 0.53 | 0.73 |
| 3 | 13523010 | rs2655218 | -0.371 | -0.437 | -2.195 | 0.947 | -0.093 | 0.014 | 0.014 | 0 | 0.65 | 0.93 | 0.66 | 0.67 | 0.54 | 0.73 |
| 3 | 13523057 | rs3732670 | -0.483 | -0.509 | -1.493 | 1.204 | -0.01 | 0.005 | 0.005 | 0 | 0.13 | 0.11 | 0.17 | 0.3 | 0.06 | 0.27 |
| 3 | 13527134 | rs2597515 | 0.175 | 0.004 | -1.964 | 0.796 | 0 | -0.001 | -0.001 | 1.045 | 0.51 | 0.83 | 0.48 | 0.37 | 0.46 | 0.43 |
| 3 | 13528498 | rs468513 | 0.363 | -0.51 | -1.721 | 0.377 | 0.005 | 0.016 | 0.016 | 0 | 0.63 | 0.66 | 0.8 | 0.81 | 0.68 | 0.67 |
| 3 | 13533582 | rs7634112 | -0.566 | -1.387 | -1.586 | 0.512 | 0.002 | -0.002 | -0.002 | 0 | 0.25 | 0.2 | 0.2 | 0.33 | 0.13 | 0.33 |
| 3 | 13535631 | rs6796154 | 0.223 | -1.291 | -1.128 | 0.324 | 0.081 | -0.019 | -0.019 | 0 | 0.06 | 0.25 | 0.06 | 0.01 | 0.11 | 0.06 |
| 3 | 13537240 | rs2655222 | -0.151 | -1.275 | -1.071 | 0.769 | 0.011 | -0.034 | -0.034 | 0 | 0.71 | 0.78 | 0.77 | 0.59 | 0.78 | 0.79 |
| 3 | 13537336 | rs1116048 | -0.321 | -1.276 | -1.095 | 0.769 | 0.017 | -0.081 | -0.081 | 0 | 0.73 | 0.85 | 0.78 | 0.59 | 0.8 | 0.8 |

| | | | | | | | | | | | | | | | | |
|---|---|---|---|---|---|---|---|---|---|---|---|---|---|---|---|---|
| 3 | 13538012 | rs2290193 | -1.197 | -1.516 | -0.759 | 0.737 | -0.002 | 0.001 | 0.001 | 0 | 0.34 | 0.64 | 0.29 | 0.61 | 0.26 | 0.41 |
| 3 | 13538054 | rs2290194 | -0.361 | -1.389 | -0.864 | 0.8 | 0.033 | 0.039 | 0.039 | 0.076 | 0.34 | 0.16 | 0.25 | 0.22 | 0.22 | 0.25 |
| 3 | 13540334 | rs2675231 | -0.413 | -1.625 | -0.897 | 0.678 | -0.005 | 0.036 | 0.036 | 0 | 0.82 | 0.98 | 0.9 | 0.91 | 0.87 | 0.89 |
| 3 | 13540641 | rs17038236 | 0.519 | -1.502 | -0.72 | 0.716 | 0.088 | -0.015 | -0.015 | 1.031 | 0.06 | 0.28 | 0.07 | 0.01 | 0.12 | 0.06 |
| 3 | 13541294 | rs9838980 | -1.09 | -1.522 | -0.792 | 0.786 | 0.017 | 0.164 | 0.164 | 0 | 0.17 | 0.53 | 0.13 | 0.36 | 0.17 | 0.23 |
| 3 | 13548520 | rs1356234 | 0.869 | -1.529 | -0.714 | 0.315 | -0.009 | 0.072 | 0.072 | 1.681 | 0.73 | 0.4 | 0.72 | 0.55 | 0.74 | 0.77 |
| 3 | 13551420 | rs2655225 | -0.059 | -1.558 | -0.628 | 0.541 | -0.024 | 0.123 | 0.123 | 3.945 | 0.81 | 0.61 | 0.79 | 0.54 | 0.86 | 0.85 |
| 3 | 13551810 | rs2675229 | -0.822 | -1.541 | -0.663 | 0.32 | 0.01 | 0.026 | 0.026 | 1.18 | 0.86 | 0.89 | 0.92 | 1 | 0.9 | 0.99 |
| 3 | 13555665 | rs2655226 | 0.665 | -1.751 | -0.743 | 0.272 | -0.018 | 0.03 | 0.03 | 4.753 | 0.73 | 0.6 | 0.76 | 0.55 | 0.82 | 0.79 |
| 3 | 13559567 | rs2630441 | 0.358 | -1.188 | -1.067 | 0.15 | 0.01 | -0.045 | -0.045 | 0 | 0.61 | 0.76 | 0.71 | 0.48 | 0.7 | 0.64 |
| 3 | 13561326 | rs2244564 | 0.418 | -1.079 | -0.81 | 0.082 | -0.014 | -0.018 | -0.018 | 0 | 0.67 | 0.63 | 0.74 | 0.78 | 0.79 | 0.79 |
| 3 | 13562685 | rs2597520 | 0.511 | -1.061 | -1.287 | -0.083 | -0.002 | 0 | 0 | 0.061 | 0.52 | 0.55 | 0.62 | 0.51 | 0.55 | 0.66 |
| 3 | 13564048 | rs2731343 | 0.743 | -0.931 | -0.64 | 0.498 | -0.024 | -0.028 | -0.028 | 0 | 0.67 | 0.61 | 0.74 | 0.74 | 0.76 | 0.77 |
| 3 | 13571228 | rs2630452 | -0.435 | -0.584 | -1.469 | 0.084 | 0.001 | 0.012 | 0.012 | 2.321 | 0.86 | 0.87 | 0.93 | 1 | 0.89 | 0.98 |
| 3 | 13571802 | rs2630453 | 0.312 | -0.386 | -1.609 | 0.379 | 0.027 | 0.052 | 0.052 | 1.403 | 0.54 | 0.25 | 0.51 | 0.3 | 0.61 | 0.6 |
| 3 | 13578054 | rs9849584 | -0.324 | 0.073 | -1.544 | 0.676 | 0.025 | -0.036 | -0.036 | 1.477 | 0.37 | 0.16 | 0.38 | 0.44 | 0.35 | 0.37 |
| 3 | 13579099 | rs9854903 | -0.512 | 0.224 | -0.774 | 0.407 | 0.018 | -0.025 | -0.025 | 0.167 | 0.05 | 0.19 | 0.05 | 0 | 0.06 | 0.01 |
| 3 | 13579299 | rs6808244 | -0.4 | 0.015 | -0.413 | 0.808 | -0.006 | 0.048 | 0.048 | 0 | 0.08 | 0.17 | 0.14 | 0.15 | 0.08 | 0.11 |
| 3 | 13582768 | rs9881100 | 0.535 | -0.066 | -0.522 | 0.168 | -0.002 | -0.058 | -0.058 | 0 | 0.81 | 0.76 | 0.85 | 1 | 0.8 | 0.97 |
| 3 | 13583528 | rs896056 | 0.535 | -0.066 | -0.522 | 0.168 | -0.002 | -0.058 | -0.058 | 0 | 0.81 | 0.82 | 0.86 | 1 | 0.8 | 0.98 |
| 3 | 13584980 | rs17178474 | 0.601 | -0.695 | -0.824 | -0.095 | 0.015 | 0.012 | 0.012 | 0 | 0.15 | 0.08 | 0.08 | 0.09 | 0.1 | 0.26 |
| 3 | 13585313 | rs6775085 | 1.146 | -0.313 | -0.479 | 0.038 | 0.002 | -0.001 | -0.001 | 0 | 0.38 | 0.39 | 0.39 | 0.47 | 0.39 | 0.43 |
| 3 | 13587987 | rs2016058 | 0.463 | -0.675 | 0.412 | 0.122 | -0.017 | -0.146 | -0.146 | 0.817 | 0.77 | 0.45 | 0.82 | 1 | 0.79 | 0.93 |
| 3 | 13592321 | rs2122499 | -0.031 | -0.589 | -0.309 | 0.163 | 0 | -0.028 | -0.028 | 0 | 0.81 | 0.8 | 0.85 | 1 | 0.8 | 0.97 |
| 3 | 13600056 | rs11712980 | 0.507 | -0.576 | -0.515 | 0.357 | 0.003 | -0.013 | -0.013 | 0 | 0.08 | 0.05 | 0.12 | 0.15 | 0.08 | 0.12 |
| 3 | 13600524 | rs10865714 | 0.246 | -0.521 | -0.164 | 0.139 | -0.004 | -0.066 | -0.066 | 0 | 0.8 | 0.72 | 0.84 | 1 | 0.79 | 0.93 |
| 3 | 13612936 | rs3732666 | -1.412 | -1.319 | -0.225 | 0.508 | 0.04 | -0.013 | -0.013 | 0 | 0.34 | 0.76 | 0.28 | 0.55 | 0.19 | 0.41 |
| 3 | 13614411 | rs3773300 | -0.793 | -1.382 | 0.387 | 0.558 | -0.014 | 0.098 | 0.098 | 0 | 0.14 | 0.44 | 0.19 | 0.19 | 0.09 | 0.12 |
| 3 | 13617049 | rs1028157 | 0.206 | -1.328 | 0.199 | 0.567 | 0.016 | 0.001 | 0.001 | 0.256 | 0.13 | 0.05 | 0.05 | 0.1 | 0.05 | 0.16 |
| 3 | 13621320 | rs9862523 | -0.555 | -1.373 | 0.464 | 0.551 | -0.043 | -0.014 | -0.014 | 0 | 0.16 | 0.24 | 0.06 | 0.1 | 0.05 | 0.21 |
| 3 | 13622987 | rs7615835 | -0.827 | -0.775 | -0.499 | -0.187 | -0.087 | 0.136 | 0.136 | 0 | 0.69 | 0.9 | 0.65 | 0.99 | 0.56 | 0.82 |
| 3 | 13626592 | rs3773283 | -1.491 | -0.508 | 0.269 | 1 | -0.011 | 0.036 | 0.036 | 0 | 0.3 | 0.47 | 0.28 | 0.51 | 0.24 | 0.42 |
| 3 | 13630307 | rs3773282 | -1.715 | -0.474 | 0.354 | 1.086 | -0.026 | 0.029 | 0.029 | 0 | 0.32 | 0.57 | 0.28 | 0.51 | 0.23 | 0.44 |
| 3 | 13637116 | rs2350310 | -1.51 | 0.131 | 0.527 | 1.31 | -0.016 | 0.201 | 0.201 | 0 | 0.18 | 0.48 | 0.21 | 0.4 | 0.13 | 0.19 |
| 3 | 13640933 | rs11128646 | -1.178 | 0.411 | 0.29 | 1.154 | 0.011 | 0.026 | 0.026 | 0.185 | 0.49 | 0.62 | 0.48 | 0.62 | 0.51 | 0.53 |
| 3 | 13642946 | rs1878173 | -1.167 | 0.449 | 0.222 | 1.254 | 0.009 | 0.038 | 0.038 | 0.261 | 0.48 | 0.66 | 0.5 | 0.62 | 0.5 | 0.53 |
| 3 | 13644176 | rs6771311 | -0.307 | 0.406 | 0.551 | 1.329 | 0.008 | -0.01 | -0.01 | 0 | 0.36 | 0.4 | 0.39 | 0.31 | 0.4 | 0.4 |
| 3 | 13646247 | rs6765409 | 0.217 | 0.504 | 0.492 | 1.352 | -0.019 | 0.001 | 0.001 | 0.45 | 0.28 | 0.2 | 0.35 | 0.3 | 0.33 | 0.33 |
| 3 | 13647142 | rs17038334 | -1.021 | 0.161 | 1.411 | 1.666 | 0.014 | 0.155 | 0.155 | 0 | 0.11 | 0.26 | 0.11 | 0.31 | 0.1 | 0.1 |
| 3 | 13649339 | rs6442403 | -2.14 | 0.191 | 1.443 | 0.608 | 0 | -0.024 | -0.024 | 0.009 | 0.25 | 0.52 | 0.16 | 0.18 | 0.21 | 0.22 |
| 3 | 13649762 | rs2276747 | -1.894 | 0.197 | 0.995 | 0.609 | -0.002 | -0.008 | -0.008 | 0 | 0.22 | 0.66 | 0.18 | 0.16 | 0.18 | 0.16 |
| 3 | 13650434 | rs6803572 | -0.792 | 0.115 | 0.939 | 0.442 | 0.033 | -0.047 | -0.047 | 2.759 | 0.11 | 0.29 | 0.05 | 0.03 | 0.1 | 0.11 |
| 3 | 13653400 | rs9835273 | -1.243 | 0.205 | 0.005 | 0.501 | -0.018 | -0.097 | -0.097 | 0.078 | 0.22 | 0.53 | 0.12 | 0.05 | 0.15 | 0.2 |
| 3 | 13659897 | rs4596126 | -1.508 | 0.169 | -1.155 | 0.714 | -0.015 | 0.012 | 0.012 | 3.473 | 0.29 | 0.92 | 0.21 | 0.2 | 0.22 | 0.25 |
| 3 | 13660809 | rs2280305 | -1.354 | -0.062 | -0.742 | 1.382 | 0.006 | -0.008 | -0.008 | 0.058 | 0.19 | 0.18 | 0.17 | 0.36 | 0.11 | 0.25 |
| 3 | 13664936 | rs3773260 | -0.321 | 0.547 | -1.023 | 0.859 | -0.012 | 0.01 | 0.01 | 0 | 0.2 | 0.16 | 0.18 | 0.12 | 0.27 | 0.23 |
| 3 | 13665410 | rs2176621 | 1.159 | 0.761 | -1.16 | 0.202 | 0.037 | 0.114 | 0.114 | 0 | 0.48 | 0.23 | 0.6 | 0.75 | 0.42 | 0.39 |
| 3 | 13666263 | rs6773406 | 0.203 | 0.741 | -0.148 | 0.585 | 0.01 | -0.013 | -0.013 | 0 | 0.7 | 0.61 | 0.79 | 0.76 | 0.69 | 0.66 |
| 3 | 13670536 | rs9843344 | 0.17 | 0.731 | -0.089 | 0.382 | -0.013 | -0.002 | -0.002 | 0 | 0.73 | 0.76 | 0.83 | 0.99 | 0.69 | 0.77 |
| 3 | 13670776 | rs4684968 | 0.143 | 0.366 | -0.326 | 0.147 | -0.014 | 0.021 | 0.021 | 0 | 0.69 | 0.76 | 0.83 | 0.99 | 0.68 | 0.77 |
| 3 | 13675694 | rs3773254 | 1.057 | 0.13 | -0.334 | 0.17 | 0.029 | 0.034 | 0.034 | 0 | 0.48 | 0.31 | 0.64 | 0.65 | 0.44 | 0.41 |
| 3 | 13676250 | rs11711559 | 0.547 | 0.162 | -0.146 | 0.334 | 0.018 | 0.069 | 0.069 | 0 | 0.18 | 0.27 | 0.16 | 0.34 | 0.23 | 0.33 |
| 3 | 13676940 | rs9838270 | 0.518 | 0.341 | -0.66 | -0.075 | 0.003 | 0.056 | 0.056 | 0 | 0.84 | 0.92 | 0.89 | 0.99 | 0.86 | 0.88 |
| 3 | 13677751 | rs3816411 | 0.46 | -0.338 | -0.153 | 0.516 | -0.004 | -0.002 | -0.002 | 0 | 0.62 | 0.54 | 0.67 | 0.6 | 0.63 | 0.5 |
| 3 | 13679044 | rs4684148 | 0.029 | -1.033 | -1.02 | 0.189 | 0.069 | 0.024 | 0.024 | 0 | 0.45 | 0.69 | 0.37 | 0.64 | 0.32 | 0.34 |
| 3 | 13679203 | rs1061375 | 0.037 | -1.047 | -0.963 | 0.184 | 0.1 | 0.028 | 0.028 | 0 | 0.45 | 0.76 | 0.39 | 0.64 | 0.32 | 0.34 |
| 3 | 13679335 | rs1061376 | 0.035 | -1.045 | -0.965 | 0.184 | 0.069 | 0.023 | 0.023 | 0 | 0.45 | 0.69 | 0.38 | 0.64 | 0.32 | 0.34 |
| 3 | 13680861 | rs7610284 | -0.412 | -0.263 | -1.042 | -0.352 | 0.028 | 0.044 | 0.044 | 0 | 0.44 | 0.63 | 0.52 | 0.79 | 0.33 | 0.59 |
| 3 | 13686178 | rs9882140 | -0.839 | 0.054 | -0.918 | -0.102 | -0.011 | 0.043 | 0.043 | 0 | 0.76 | 0.82 | 0.79 | 0.92 | 0.72 | 0.78 |
| 3 | 13689452 | rs745940 | 0.58 | -0.065 | -0.815 | 0.092 | 0.024 | -0.001 | -0.001 | 0.976 | 0.15 | 0.21 | 0.17 | 0.12 | 0.2 | 0.15 |
| 3 | 13695304 | rs2974386 | 0.262 | -0.341 | -1.135 | 0.198 | -0.002 | 0.082 | 0.082 | 0 | 0.52 | 0.73 | 0.66 | 0.72 | 0.49 | 0.57 |
| 3 | 13701622 | rs9848698 | -0.881 | -0.244 | -0.902 | 0.149 | -0.009 | 0.003 | 0.003 | 0 | 0.61 | 0.63 | 0.47 | 0.65 | 0.5 | 0.58 |
| 3 | 13705345 | rs6793679 | 0.738 | -0.763 | -0.766 | 0.152 | -0.009 | 0.009 | 0.009 | 0 | 0.12 | 0.15 | 0.09 | 0.13 | 0.07 | 0.13 |
| 3 | 13705569 | rs7650215 | -0.707 | -1.051 | -0.735 | -0.014 | -0.035 | -0.005 | -0.005 | 0 | 0.3 | 0.56 | 0.3 | 0.71 | 0.14 | 0.32 |
| 3 | 13707300 | rs2956507 | -2.575 | -0.925 | -1.131 | -0.245 | -0.128 | -0.046 | -0.046 | 0 | 0.75 | 0.98 | 0.62 | 0.77 | 0.62 | 0.67 |
| 3 | 13709181 | rs7618109 | -0.299 | -1.022 | -0.623 | 0.091 | -0.056 | 0.058 | 0.058 | 0.096 | 0.25 | 0.5 | 0.22 | 0.58 | 0.13 | 0.3 |
| 3 | 13710562 | rs3934833 | 2.361 | -0.57 | -0.817 | -0.043 | -0.057 | 0.043 | 0.043 | 0 | 0.1 | 0.03 | 0.08 | 0.01 | 0.19 | 0.11 |
| 3 | 13711684 | rs9310429 | 0.748 | -0.953 | -0.68 | -0.002 | -0.059 | 0.126 | 0.126 | 0 | 0.19 | 0.36 | 0.16 | 0.39 | 0.11 | 0.16 |
| 3 | 13717252 | rs12495328 | 1.56 | -0.96 | -0.479 | -0.264 | -0.008 | -0.058 | -0.058 | 5.485 | 0.2 | 0.25 | 0.16 | 0.05 | 0.17 | 0.12 |

| | | | | | | | | | | | | | | | | |
|---|---|---|---|---|---|---|---|---|---|---|---|---|---|---|---|---|
| 3 | 13720097 | rs6775581 | -2.062 | -1.042 | -0.706 | -0.21 | -0.037 | 0.04 | 0.04 | 0 | 0.65 | 0.81 | 0.61 | 0.76 | 0.57 | 0.6 |
| 3 | 13722366 | rs7636218 | 0.725 | -0.763 | -0.359 | 0.004 | -0.018 | 0.042 | 0.042 | 0 | 0.12 | 0.18 | 0.09 | 0.2 | 0.07 | 0.14 |
| 3 | 13726306 | rs13062655 | 1.514 | -0.724 | -0.851 | -0.598 | 0.011 | 0.01 | 0.01 | 0.282 | 0.23 | 0.17 | 0.16 | 0.05 | 0.18 | 0.07 |
| 3 | 13726888 | rs6794344 | -1.032 | -0.842 | -0.794 | -0.655 | -0.014 | 0.028 | 0.028 | 0 | 0.56 | 0.65 | 0.53 | 0.64 | 0.51 | 0.56 |
| 3 | 13728715 | rs9823655 | -1.189 | -0.711 | -0.771 | -0.672 | 0.006 | 0.037 | 0.037 | 0 | 0.55 | 0.7 | 0.5 | 0.64 | 0.46 | 0.49 |
| 3 | 13728876 | rs9823857 | 0.416 | -0.918 | -0.633 | -0.538 | -0.082 | 0.094 | 0.094 | 0.044 | 0.25 | 0.43 | 0.28 | 0.58 | 0.12 | 0.4 |
| 3 | 13730682 | rs6795216 | -1.087 | -0.731 | -1.008 | -0.441 | -0.035 | 0.025 | 0.025 | 0 | 0.58 | 0.76 | 0.54 | 0.64 | 0.51 | 0.57 |
| 3 | 13731856 | rs9848869 | -0.614 | -0.573 | -0.641 | -0.112 | 0 | 0.001 | 0.001 | 0.215 | 0.35 | 0.59 | 0.39 | 0.59 | 0.32 | 0.53 |
| 3 | 13731948 | rs9811564 | -0.438 | -0.68 | -0.532 | -0.129 | -0.053 | 0.035 | 0.035 | 0 | 0.3 | 0.55 | 0.31 | 0.58 | 0.17 | 0.41 |
| 3 | 13732454 | rs4684981 | 0.291 | -0.308 | -0.296 | -0.177 | 0.066 | -0.01 | -0.01 | 0 | 0.17 | 0.41 | 0.19 | 0.13 | 0.22 | 0.12 |
| 3 | 13736633 | rs4234514 | 0.631 | -1.322 | -0.943 | -0.24 | 0.029 | -0.001 | -0.001 | 0 | 0.11 | 0.03 | 0.09 | 0.11 | 0.04 | 0.22 |
| 3 | 13737000 | rs4450798 | 0.686 | -1.452 | -0.896 | 0.145 | 0.002 | 0.011 | 0.011 | 0 | 0.19 | 0.22 | 0.2 | 0.25 | 0.21 | 0.24 |
| 3 | 13741016 | rs4234515 | 0.535 | -1.771 | -1.718 | -0.56 | -0.055 | 0.084 | 0.084 | 0.011 | 0.3 | 0.04 | 0.27 | 0.16 | 0.29 | 0.3 |
| 3 | 13744059 | rs11714418 | 2.062 | -1.744 | -1.731 | -0.587 | -0.084 | 0.079 | 0.079 | 0 | 0.18 | 0.01 | 0.18 | 0.05 | 0.26 | 0.08 |
| 3 | 13747337 | rs13082757 | 0.866 | -2.125 | -1.131 | -0.458 | 0 | 0.009 | 0.009 | 0 | 0.49 | 0.38 | 0.49 | 0.48 | 0.52 | 0.47 |
| 3 | 13750083 | rs4684988 | 0.652 | -2.15 | -1.241 | -0.449 | -0.069 | 0.072 | 0.072 | 0.009 | 0.3 | 0.04 | 0.28 | 0.16 | 0.3 | 0.3 |
| 3 | 13756808 | rs4234517 | -0.782 | -2.088 | -0.924 | -0.349 | -0.011 | 0.023 | 0.023 | 0 | 0.32 | 0.62 | 0.33 | 0.45 | 0.22 | 0.45 |
| 3 | 13763808 | rs4611801 | 1.481 | -1.858 | -0.116 | -0.205 | 0.032 | 0.162 | 0.162 | 0 | 0.15 | 0.31 | 0.19 | 0.36 | 0.16 | 0.21 |
| 3 | 13765391 | rs2974384 | -1.013 | -2.365 | -0.388 | -0.542 | -0.004 | 0.007 | 0.007 | 0.752 | 0.52 | 0.65 | 0.5 | 0.5 | 0.48 | 0.56 |
| 3 | 13795677 | rs7634577 | -0.1 | -2.506 | -0.451 | -0.559 | 0 | -0.001 | -0.001 | 0 | 0.55 | 0.5 | 0.53 | 0.54 | 0.54 | 0.7 |
| 3 | 13799270 | rs7612896 | 0.911 | -3.055 | -0.902 | -0.835 | -0.021 | 0.027 | 0.027 | 0 | 0.21 | 0.22 | 0.16 | 0.41 | 0.12 | 0.16 |
| 3 | 13803506 | rs7631252 | 2.157 | -2.745 | -1.159 | -0.702 | 0.009 | 0.092 | 0.092 | 0 | 0.13 | 0.2 | 0.17 | 0.36 | 0.15 | 0.19 |
| 3 | 13804145 | rs6786420 | -1.544 | -2.8 | -1.432 | -0.805 | -0.031 | 0.096 | 0.096 | 0 | 0.75 | 0.92 | 0.8 | 0.94 | 0.69 | 0.91 |
| 3 | 13805593 | rs7637646 | -0.672 | -2.798 | -1.375 | -0.787 | 0.003 | 0.006 | 0.006 | 0 | 0.6 | 0.56 | 0.61 | 0.58 | 0.54 | 0.72 |
| 3 | 13807781 | rs4684157 | 1.135 | -2.311 | -1.605 | -0.91 | -0.001 | 0.039 | 0.039 | 0 | 0.29 | 0.38 | 0.35 | 0.39 | 0.28 | 0.24 |
| 3 | 13807961 | rs7611685 | -0.932 | -2.016 | -2.275 | -1.225 | -0.029 | 0.143 | 0.143 | 0.436 | 0.59 | 0.8 | 0.56 | 0.81 | 0.47 | 0.39 |
| 3 | 13810670 | rs4525847 | -1.836 | -1.128 | -2.082 | -0.711 | -0.069 | 0.109 | 0.109 | 0 | 0.77 | 0.96 | 0.81 | 0.95 | 0.7 | 0.91 |
| 3 | 13811824 | rs9872360 | -0.746 | -1.311 | -2.099 | -0.733 | -0.03 | 0.122 | 0.122 | 0 | 0.67 | 0.9 | 0.71 | 0.9 | 0.66 | 0.88 |
| 3 | 13813545 | rs4685020 | 1.15 | -0.926 | -1.911 | -0.539 | 0.01 | -0.002 | -0.002 | 0.016 | 0.1 | 0.01 | 0.1 | 0.04 | 0.04 | 0.03 |
| 3 | 13813806 | rs6777806 | -1.506 | -1.019 | -1.957 | -0.201 | -0.074 | -0.139 | -0.139 | 0 | 0.69 | 0.99 | 0.71 | 0.56 | 0.67 | 0.81 |
| 3 | 13816624 | rs11716601 | 0.731 | -0.524 | -1.034 | -0.636 | 0.028 | -0.062 | -0.062 | 0 | 0.21 | 0.3 | 0.24 | 0.07 | 0.27 | 0.28 |
| 3 | 13816690 | rs11708722 | 0.674 | -0.532 | -0.966 | -0.639 | 0.036 | -0.076 | -0.076 | 0.852 | 0.22 | 0.34 | 0.25 | 0.07 | 0.27 | 0.28 |
| 3 | 13817382 | rs6791421 | 0.43 | -0.921 | -1.569 | -0.294 | -0.002 | 0.004 | 0.004 | 0 | 0.22 | 0.21 | 0.22 | 0.14 | 0.21 | 0.16 |
| 3 | 13818385 | rs4591488 | -0.964 | -1.084 | -2.042 | -0.018 | -0.052 | -0.186 | -0.186 | 0 | 0.66 | 0.99 | 0.61 | 0.5 | 0.66 | 0.79 |
| 3 | 13823342 | rs6793656 | 1.363 | -0.678 | -1.59 | -0.704 | -0.019 | 0.027 | 0.027 | 0 | 0.15 | 0.09 | 0.07 | 0.08 | 0.16 | 0.12 |
| 3 | 13824469 | rs9310432 | -0.257 | -0.915 | -1.259 | -0.984 | -0.017 | 0.051 | 0.051 | 0.056 | 0.41 | 0.65 | 0.36 | 0.22 | 0.45 | 0.65 |
| 3 | 13825666 | rs13097641 | 0.019 | -1.016 | -0.792 | -0.812 | 0.011 | -0.056 | -0.056 | 0 | 0.3 | 0.54 | 0.32 | 0.13 | 0.32 | 0.55 |
| 3 | 13826854 | rs729639 | -0.523 | -0.943 | -1.575 | -0.605 | -0.046 | 0.057 | 0.057 | 0 | 0.61 | 0.98 | 0.56 | 0.25 | 0.62 | 0.71 |
| 3 | 13829565 | rs13079210 | -0.128 | -0.087 | -0.644 | 0.079 | 0.012 | -0.011 | -0.011 | 0 | 0.24 | 0.29 | 0.25 | 0.21 | 0.29 | 0.57 |
| 3 | 13835087 | rs6442409 | -0.453 | -0.47 | -1.375 | 0.249 | -0.103 | -0.128 | -0.128 | 0 | 0.71 | 0.99 | 0.57 | 0.43 | 0.66 | 0.8 |
| 3 | 13835823 | rs4685026 | 0.397 | -0.375 | -0.703 | -0.472 | -0.005 | -0.036 | -0.036 | 0.309 | 0.44 | 0.54 | 0.24 | 0.22 | 0.38 | 0.42 |
| 3 | 13835944 | rs12495493 | 0.061 | 0.384 | -0.501 | -0.067 | 0.009 | -0.01 | -0.01 | 0 | 0.18 | 0.19 | 0.29 | 0.07 | 0.25 | 0.31 |
| 3 | 13843635 | rs4685033 | 0.579 | 0 | -0.963 | -0.211 | 0.005 | -0.007 | -0.007 | 0 | 0.38 | 0.34 | 0.2 | 0.13 | 0.31 | 0.42 |
| 3 | 13848486 | rs7617314 | 0.251 | 0.08 | -1.064 | 0.263 | -0.007 | -0.004 | -0.004 | 0 | 0.45 | 0.49 | 0.33 | 0.48 | 0.39 | 0.49 |
| 3 | 13850666 | rs2053101 | 0.367 | 0.278 | -1.566 | 0.344 | -0.077 | 0.034 | 0.034 | 0 | 0.17 | 0.02 | 0.33 | 0.07 | 0.25 | 0.32 |
| 3 | 13850702 | rs17038609 | -0.269 | -0.317 | -1.224 | 0.708 | 0.026 | 0.051 | 0.051 | 0 | 0.05 | 0 | 0.05 | 0.38 | 0.01 | 0.07 |
| 3 | 13855350 | rs11720627 | 0.019 | 0.064 | -0.961 | -0.62 | 0.003 | -0.016 | -0.016 | 4.444 | 0.38 | 0.35 | 0.21 | 0.12 | 0.3 | 0.43 |
| 3 | 13857577 | rs3762725 | -0.601 | 0.533 | -0.841 | -0.305 | 0.002 | -0.026 | -0.026 | 0 | 0.22 | 0.24 | 0.35 | 0.05 | 0.26 | 0.24 |
| 3 | 13857892 | rs2163910 | -0.753 | 0.245 | -0.816 | 0.619 | -0.003 | 0.043 | 0.043 | 0 | 0.11 | 0.12 | 0.2 | 0.65 | 0.1 | 0.37 |
| 3 | 13859173 | rs3796316 | -0.28 | 0.269 | -1.792 | -0.044 | -0.035 | 0.079 | 0.079 | 0 | 0.57 | 0.84 | 0.44 | 0.29 | 0.43 | 0.59 |
| 3 | 13859332 | rs3796314 | 0.73 | -0.184 | -2.062 | -0.583 | 0.003 | 0.071 | 0.071 | 0 | 0.19 | 0.04 | 0.09 | 0.06 | 0.16 | 0.14 |
| 3 | 13864338 | rs873853 | 1.022 | 0.235 | -1.368 | -0.38 | 0.012 | 0.109 | 0.109 | 0.092 | 0.15 | 0.21 | 0.18 | 0.61 | 0.18 | 0.15 |
| 3 | 13865075 | rs9840696 | -1.113 | 0.209 | -1.601 | -0.583 | -0.091 | 0.093 | 0.093 | 0 | 0.71 | 0.91 | 0.64 | 0.91 | 0.62 | 0.76 |
| 3 | 13866499 | rs4685039 | 1.069 | 0.219 | -1.27 | -0.39 | 0.003 | 0.053 | 0.053 | 0.06 | 0.15 | 0.14 | 0.17 | 0.61 | 0.18 | 0.15 |
| 3 | 13867255 | rs12492620 | 0.551 | -0.491 | -1.366 | -0.841 | 0.009 | 0.041 | 0.041 | 0 | 0.25 | 0.14 | 0.24 | 0.08 | 0.22 | 0.14 |
| 3 | 13870052 | rs934453 | -0.202 | -0.679 | -0.577 | -0.916 | -0.007 | -0.04 | -0.04 | 0 | 0.17 | 0.28 | 0.12 | 0.09 | 0.11 | 0.36 |
| 3 | 13872270 | rs12634816 | -0.193 | -0.838 | -0.354 | -0.626 | 0.008 | -0.004 | -0.004 | 0 | 0.34 | 0.28 | 0.35 | 0.67 | 0.31 | 0.21 |
| 3 | 13873451 | rs734176 | 1.672 | -0.812 | -0.497 | -0.582 | -0.019 | -0.054 | -0.054 | 0 | 0.13 | 0.05 | 0.2 | 0.6 | 0.17 | 0.14 |
| 3 | 13878889 | rs12634112 | 0.255 | -1.033 | -0.116 | -0.625 | -0.029 | -0.026 | -0.026 | 0.047 | 0.22 | 0.34 | 0.16 | 0.09 | 0.13 | 0.06 |
| 3 | 13879616 | rs11922919 | 1.1 | -1.1 | -0.044 | -0.786 | 0 | 0.002 | 0.002 | 0.026 | 0.2 | 0.16 | 0.14 | 0.18 | 0.19 | 0.42 |
| 3 | 13883360 | rs9863149 | -0.207 | -0.866 | 0.157 | -0.676 | 0 | 0.001 | 0.001 | 0 | 0.42 | 0.49 | 0.29 | 0.27 | 0.32 | 0.48 |
| 3 | 13890485 | rs12492784 | -0.283 | -0.456 | -0.155 | -0.406 | 0.016 | 0.037 | 0.037 | 0 | 0.48 | 0.54 | 0.29 | 0.17 | 0.37 | 0.26 |
| 3 | 13892543 | rs17038710 | 1.28 | -0.317 | -0.911 | 0.106 | -0.033 | -0.099 | -0.099 | 0 | 0.14 | 0.03 | 0.18 | 0.56 | 0.17 | 0.19 |
| 3 | 13896284 | rs12639607 | 1.237 | -0.331 | -0.917 | 0.052 | -0.032 | -0.1 | -0.1 | 0.467 | 0.14 | 0.03 | 0.19 | 0.58 | 0.17 | 0.19 |
| 3 | 13897245 | rs1433354 | -0.558 | -0.735 | -0.703 | -0.248 | 0.014 | 0.017 | 0.017 | 1.092 | 0.38 | 0.66 | 0.52 | 0.83 | 0.28 | 0.47 |
| 3 | 13903303 | rs13069140 | 0.189 | -0.437 | -0.58 | -0.272 | 0.006 | 0.002 | 0.002 | 0 | 0.06 | 0.1 | 0.18 | 0.08 | 0.11 | 0.08 |
| 3 | 13903474 | rs7641735 | 0.015 | -0.893 | -0.275 | -0.215 | -0.078 | 0.002 | 0.002 | 0.836 | 0.21 | 0.45 | 0.18 | 0.17 | 0.1 | 0.25 |
| 3 | 13905000 | rs9828013 | -0.576 | -0.124 | -1.121 | -0.348 | -0.076 | 0.106 | 0.106 | 0 | 0.62 | 0.85 | 0.51 | 0.84 | 0.48 | 0.62 |
| 3 | 13906838 | rs6442414 | 0.286 | 0.37 | -0.67 | -0.916 | -0.006 | 0.029 | 0.029 | 0.8 | 0.6 | 0.72 | 0.72 | 0.92 | 0.56 | 0.88 |

| | | | | | | | | | | | | | | | | |
|---|---|---|---|---|---|---|---|---|---|---|---|---|---|---|---|---|
| 3 | 13907100 | rs6442416 | 0.551 | 0.184 | -0.034 | -1.091 | 0.002 | -0.058 | -0.058 | 0 | 0.65 | 0.39 | 0.71 | 0.91 | 0.61 | 0.87 |
| 3 | 13908071 | rs9864031 | 0.782 | 0.236 | 0.277 | -1.03 | -0.028 | -0.058 | -0.058 | 0 | 0.25 | 0.29 | 0.15 | 0.09 | 0.18 | 0.04 |
| 3 | 13912488 | rs12635960 | 0.077 | 0.381 | 0.471 | -0.846 | -0.004 | 0.009 | 0.009 | 0.878 | 0.58 | 0.66 | 0.73 | 0.92 | 0.56 | 0.87 |
| 3 | 13912705 | rs11128662 | 0.479 | 0.357 | 0.314 | -0.401 | -0.005 | 0.053 | 0.053 | 1.159 | 0.52 | 0.34 | 0.6 | 0.76 | 0.5 | 0.64 |
| 3 | 13915465 | rs4684166 | 0.485 | 0.323 | 0.215 | -0.42 | -0.008 | 0.102 | 0.102 | 0.922 | 0.52 | 0.27 | 0.6 | 0.77 | 0.52 | 0.63 |
| 3 | 13922735 | rs4685048 | 0.46 | 0.373 | 0.411 | -0.33 | 0.008 | 0.048 | 0.048 | 0.028 | 0.51 | 0.38 | 0.62 | 0.77 | 0.47 | 0.58 |
| 3 | 13929756 | rs13077668 | 0.893 | 0.659 | 0.171 | 0.056 | 0.056 | 0.003 | 0.003 | 0 | 0.42 | 0.13 | 0.27 | 0.39 | 0.36 | 0.3 |
| 3 | 13930133 | rs1077524 | 0.509 | 0.277 | 0.944 | 0.424 | -0.002 | -0.006 | -0.006 | 0 | 0.34 | 0.3 | 0.38 | 0.41 | 0.33 | 0.52 |
| 3 | 13976830 | rs11128668 | -0.203 | 0.91 | 1.633 | 0.368 | -0.03 | 0.033 | 0.033 | 0 | 0.18 | 0.08 | 0.14 | 0 | 0.25 | 0.14 |
| 3 | 13977365 | rs12490233 | -0.69 | 0.839 | 1.916 | 0.425 | 0.014 | 0.013 | 0.013 | 0 | 0.49 | 0.43 | 0.53 | 0.43 | 0.57 | 0.61 |
| 3 | 13978803 | rs6776709 | 0.787 | 0.809 | 1.931 | 0.429 | -0.005 | -0.011 | -0.011 | 0 | 0.3 | 0.29 | 0.37 | 0.43 | 0.32 | 0.48 |
| 3 | 14018793 | rs11718142 | -0.721 | 0.812 | 2.183 | 0.447 | 0.001 | 0.001 | 0.001 | 0 | 0.49 | 0.52 | 0.53 | 0.43 | 0.57 | 0.59 |
| 3 | 14042534 | rs4685060 | 1.252 | 0.788 | 2.048 | 0.43 | -0.026 | -0.046 | -0.046 | 0.12 | 0.26 | 0.15 | 0.35 | 0.42 | 0.3 | 0.33 |
| 3 | 14048394 | rs4684175 | 1.082 | 0.819 | 1.932 | 0.445 | -0.009 | -0.019 | -0.019 | 0 | 0.27 | 0.23 | 0.36 | 0.43 | 0.3 | 0.33 |
| 3 | 14084157 | rs2168718 | -0.427 | 0.415 | 1.555 | 0.659 | 0.011 | 0.01 | 0.01 | 0 | 0.48 | 0.43 | 0.53 | 0.43 | 0.57 | 0.59 |
| 3 | 14136506 | rs2733566 | -0.924 | 0.482 | 0.869 | 0.369 | 0.035 | 0.054 | 0.054 | 0 | 0.53 | 0.81 | 0.69 | 0.64 | 0.57 | 0.62 |
| 3 | 14155453 | rs9856296 | 0.386 | 1.145 | 1.027 | 0.902 | -0.003 | 0 | 0 | 0 | 0.2 | 0.22 | 0.34 | 0.44 | 0.25 | 0.4 |
| 3 | 14158220 | rs2731313 | -1.161 | 1.34 | 1.241 | 0.544 | 0.006 | 0.001 | 0.001 | 0 | 0.38 | 0.44 | 0.51 | 0.44 | 0.5 | 0.54 |
| 3 | 14166872 | rs2733580 | -1.296 | 1.604 | 1.469 | 0.85 | 0.058 | 0.136 | 0.136 | 0 | 0.13 | 0.45 | 0.26 | 0.22 | 0.13 | 0.11 |
| 3 | 14168288 | rs2733584 | -1.918 | 1.632 | 1.331 | 0.981 | 0.02 | 0.192 | 0.192 | 0 | 0.18 | 0.45 | 0.37 | 0.59 | 0.16 | 0.22 |
| 3 | 14172827 | rs17038942 | 0.624 | 1.65 | 1.351 | 0.934 | 0.016 | -0.036 | -0.036 | 3.394 | 0.18 | 0.3 | 0.25 | 0.06 | 0.23 | 0.29 |
| 3 | 14174146 | rs4685075 | -1.179 | 1.659 | 1.093 | 0.968 | 0.002 | -0.005 | -0.005 | 0.044 | 0.36 | 0.62 | 0.59 | 0.68 | 0.4 | 0.48 |
| 3 | 14177954 | rs2733587 | -0.141 | 1.827 | 0.703 | 1.045 | -0.035 | -0.02 | -0.02 | 0 | 0.07 | 0.01 | 0.17 | 0.39 | 0.12 | 0.1 |
| 3 | 14183758 | rs1043953 | -0.041 | 1.669 | 0.683 | 0.866 | -0.062 | 0.059 | 0.059 | 0 | 0.12 | 0.05 | 0.13 | 0 | 0.22 | 0.09 |
| 3 | 14184669 | rs8516 | -0.196 | 1.706 | 0.866 | 0.841 | -0.025 | 0.019 | 0.019 | 0 | 0.12 | 0.09 | 0.14 | 0 | 0.22 | 0.09 |
| 3 | 14184719 | rs10468 | -0.891 | 1.616 | 1.14 | 0.789 | -0.005 | -0.013 | -0.013 | 0 | 0.16 | 0.14 | 0.15 | 0 | 0.25 | 0.11 |
| 3 | 14186823 | rs2470458 | -0.215 | 1.648 | 1.057 | 0.799 | -0.022 | 0.016 | 0.016 | 0 | 0.12 | 0.1 | 0.14 | 0 | 0.22 | 0.09 |
| 3 | 14188700 | rs2733533 | -1.26 | 1.735 | 1.515 | 0.665 | 0.012 | -0.044 | -0.044 | 0 | 0.32 | 0.38 | 0.39 | 0.04 | 0.46 | 0.37 |
| 3 | 14194094 | rs3731149 | 0.78 | 1.756 | 1.528 | 0.637 | 0.008 | -0.029 | -0.029 | 0.088 | 0.16 | 0.23 | 0.25 | 0.03 | 0.22 | 0.26 |
| 3 | 14199887 | rs2228000 | -0.718 | 1.775 | 1.136 | 0.834 | -0.062 | -0.067 | -0.067 | 0.996 | 0.16 | 0.09 | 0.25 | 0.41 | 0.26 | 0.2 |
| 3 | 14199908 | rs2227999 | 1.435 | 1.768 | 1.131 | 0.771 | -0.015 | 0.03 | 0.03 | 0 | 0.06 | 0.01 | 0.02 | 0 | 0.06 | 0.03 |
| 3 | 14201352 | rs3731125 | 0.324 | 1.792 | 1.158 | 0.675 | 0.006 | -0.011 | -0.011 | 0 | 0.06 | 0.06 | 0.05 | 0.01 | 0.09 | 0.13 |
| 3 | 14211101 | rs2733537 | -1.266 | 1.837 | 1.334 | 0.879 | -0.042 | -0.033 | -0.033 | 0 | 0.22 | 0.16 | 0.3 | 0.42 | 0.35 | 0.34 |
| 3 | 14217576 | rs1350344 | -0.792 | 1.808 | 1.465 | 0.685 | 0 | -0.03 | -0.03 | 0 | 0.32 | 0.32 | 0.39 | 0.03 | 0.46 | 0.37 |
| 3 | 14228726 | rs2733541 | -1.116 | 1.844 | 0.962 | 0.681 | 0.041 | -0.013 | -0.013 | 0 | 0.35 | 0.51 | 0.54 | 0.24 | 0.47 | 0.43 |
| 3 | 14240535 | rs2733553 | -0.993 | 1.25 | 0.177 | 0.342 | -0.001 | -0.018 | -0.018 | 0.021 | 0.87 | 0.88 | 0.94 | 1 | 0.88 | 0.93 |
| 3 | 14243183 | rs4685078 | 0.657 | 1.468 | -0.047 | 0.56 | -0.029 | 0.018 | 0.018 | 0 | 0.13 | 0.04 | 0.23 | 0.06 | 0.22 | 0.27 |
| 3 | 14249532 | rs4458353 | 0.849 | 0.796 | 0.003 | 0.664 | -0.011 | 0.033 | 0.033 | 0 | 0.53 | 0.28 | 0.32 | 0.35 | 0.48 | 0.35 |
| 3 | 14253587 | rs4428160 | 0.719 | 0.768 | 0.321 | 0.445 | 0 | -0.005 | -0.005 | 0 | 0.56 | 0.48 | 0.5 | 0.55 | 0.53 | 0.46 |
| 3 | 14253947 | rs4685080 | -0.112 | 0.883 | 0.44 | 0.193 | 0.005 | -0.045 | -0.045 | 2.049 | 0.8 | 0.79 | 0.88 | 0.99 | 0.79 | 0.8 |
| 3 | 14255758 | rs9855053 | -0.172 | 0.655 | 0.865 | 0.564 | 0.013 | 0.032 | 0.032 | 0.219 | 0.24 | 0.51 | 0.4 | 0.24 | 0.23 | 0.24 |
| 3 | 14257187 | rs12491506 | 0.127 | 0.83 | 0.865 | 0.551 | 0.008 | -0.022 | -0.022 | 2.104 | 0.74 | 0.62 | 0.64 | 0.9 | 0.69 | 0.8 |
| 3 | 14258107 | rs9869943 | 0.778 | 1.018 | 0.428 | 0.722 | -0.03 | -0.031 | -0.031 | 0 | 0.14 | 0.07 | 0.14 | 0.21 | 0.17 | 0.13 |
| 3 | 14259640 | rs9876159 | -0.402 | 0.581 | 0.915 | 0.37 | -0.008 | 0.038 | 0.038 | 0 | 0.83 | 0.63 | 0.69 | 0.82 | 0.86 | 0.8 |
| 3 | 14263206 | rs12630964 | 1.427 | -0.089 | 0.145 | 0.148 | -0.011 | 0.051 | 0.051 | 0 | 0.29 | 0.12 | 0.2 | 0.16 | 0.32 | 0.14 |
| 3 | 14263701 | rs7626307 | 0.024 | -0.007 | 0.184 | 0.086 | -0.011 | -0.002 | -0.002 | 0.056 | 0.57 | 0.68 | 0.72 | 0.38 | 0.57 | 0.43 |
| 3 | 14265600 | rs9833599 | 0.943 | 0.073 | -0.117 | -0.176 | -0.001 | 0.022 | 0.022 | 0 | 0.2 | 0.17 | 0.23 | 0.05 | 0.21 | 0.21 |
| 3 | 14272914 | rs4684185 | 0.445 | 0.028 | -0.087 | -0.14 | -0.013 | 0.031 | 0.031 | 0 | 0.31 | 0.23 | 0.4 | 0.83 | 0.3 | 0.51 |
| 3 | 14275457 | rs6762079 | 0.632 | -0.181 | 0.034 | -0.395 | 0.002 | 0.011 | 0.011 | 0 | 0.25 | 0.21 | 0.26 | 0.04 | 0.23 | 0.23 |
| 3 | 14276982 | rs17226476 | 2.137 | -0.227 | 0.089 | -0.637 | 0.002 | 0.032 | 0.032 | 0 | 0.22 | 0.15 | 0.14 | 0.11 | 0.24 | 0.08 |
| 3 | 14278009 | rs6782386 | -1.937 | -0.201 | -0.091 | -0.656 | -0.011 | 0.048 | 0.048 | 0 | 0.89 | 0.99 | 0.89 | 0.98 | 0.88 | 0.87 |
| 3 | 14280171 | rs12493406 | 1.27 | -0.215 | 0.087 | -0.665 | -0.006 | 0.005 | 0.005 | 0 | 0.09 | 0.05 | 0.07 | 0 | 0.11 | 0.05 |
| 3 | 14282367 | rs4684187 | -0.469 | -0.226 | 0.325 | -0.671 | -0.002 | 0.002 | 0.002 | 0 | 0.8 | 0.87 | 0.82 | 0.98 | 0.77 | 0.82 |
| 3 | 14283861 | rs7618619 | 0.944 | -0.262 | 0.173 | -0.269 | -0.061 | -0.087 | -0.087 | 1.087 | 0.29 | 0.11 | 0.17 | 0.67 | 0.3 | 0.43 |
| 3 | 14285488 | rs17305345 | 1.902 | -0.338 | 0.117 | -0.304 | -0.05 | 0.052 | 0.052 | 0.014 | 0.17 | 0.02 | 0.22 | 0.04 | 0.2 | 0.22 |
| 3 | 14287703 | rs10510427 | 1.081 | 0.056 | 0.298 | -0.185 | -0.007 | 0.005 | 0.005 | 0 | 0.09 | 0.05 | 0.07 | 0 | 0.12 | 0.05 |
| 3 | 14290893 | rs1871856 | -0.277 | -0.092 | 0.3 | -0.107 | -0.002 | -0.061 | -0.061 | 0 | 0.56 | 0.53 | 0.67 | 0.87 | 0.53 | 0.74 |
| 3 | 14293622 | rs12631029 | 0.406 | 0.033 | -0.033 | 0.075 | -0.011 | 0.032 | 0.032 | 0 | 0.31 | 0.23 | 0.4 | 0.83 | 0.3 | 0.51 |
| 3 | 14294172 | rs2128163 | 0.051 | 0.099 | -0.634 | 0.504 | -0.006 | -0.015 | -0.015 | 0 | 0.58 | 0.66 | 0.7 | 0.45 | 0.58 | 0.44 |
| 3 | 14298264 | rs2128162 | -0.554 | 0.414 | -1.29 | 0.18 | -0.001 | 0.123 | 0.123 | 0 | 0.65 | 0.81 | 0.64 | 0.94 | 0.65 | 0.64 |
| 3 | 14302245 | rs7644094 | 0.514 | 0.05 | -0.894 | 0.376 | -0.001 | -0.001 | -0.001 | 0.526 | 0.33 | 0.35 | 0.42 | 0.28 | 0.33 | 0.2 |
| 3 | 14304145 | rs17227017 | 1.766 | 0.137 | -1.389 | -0.268 | 0.006 | 0.039 | 0.039 | 0 | 0.21 | 0.1 | 0.12 | 0.11 | 0.21 | 0.09 |
| 3 | 14305341 | rs13072044 | 0.668 | 0.144 | -1.363 | -0.253 | -0.001 | -0.013 | -0.013 | 0.552 | 0.1 | 0.13 | 0.08 | 0 | 0.12 | 0.05 |
| 3 | 14306081 | rs754020 | -0.254 | 0.125 | -1.443 | -0.256 | -0.002 | 0.024 | 0.024 | 0 | 0.57 | 0.4 | 0.47 | 0.24 | 0.58 | 0.42 |
| 3 | 14306114 | rs900186 | 0.777 | -0.171 | -1.599 | 0.005 | -0.004 | 0.037 | 0.037 | 0 | 0.27 | 0.18 | 0.27 | 0.13 | 0.26 | 0.28 |
| 3 | 14306278 | rs1038299 | 0.282 | -0.122 | -1.479 | -0.042 | 0.007 | 0.131 | 0.131 | 0 | 0.48 | 0.26 | 0.39 | 0.15 | 0.47 | 0.37 |
| 3 | 14308098 | rs6769584 | 0.481 | -0.506 | -1.343 | 0.86 | 0.022 | -0.026 | -0.026 | 0.311 | 0.15 | 0.1 | 0.06 | 0.32 | 0.1 | 0.2 |
| 3 | 14308924 | rs17039221 | 0.573 | 0.261 | -1.115 | 0.486 | 0.004 | 0.028 | 0.028 | 0 | 0.2 | 0.13 | 0.12 | 0.1 | 0.21 | 0.09 |
| 3 | 14311273 | rs9881693 | -0.004 | 0.562 | -0.823 | 1.279 | 0 | 0.016 | 0.016 | 0.099 | 0.82 | 0.83 | 0.82 | 0.62 | 0.8 | 0.78 |

| | | | | | | | | | | | | | | | | |
|---|---|---|---|---|---|---|---|---|---|---|---|---|---|---|---|---|
| 3 | 14318853 | rs1806947 | 0.329 | 0.331 | -0.78 | 0.811 | 0.002 | 0.003 | 0.003 | 0 | 0.34 | 0.35 | 0.21 | 0.19 | 0.31 | 0.25 |
| 3 | 14321177 | rs1482038 | 0.015 | 0.145 | -0.921 | 0.868 | 0.025 | 0.042 | 0.042 | 0 | 0.38 | 0.73 | 0.38 | 0.21 | 0.31 | 0.31 |
| 3 | 14322924 | rs11711684 | 0.758 | 0.369 | -1.278 | 1.224 | -0.021 | -0.067 | -0.067 | 0 | 0.14 | 0.03 | 0.13 | 0.25 | 0.13 | 0.15 |
| 3 | 14330296 | rs7610197 | -0.296 | 0.199 | -0.596 | 0.849 | -0.005 | -0.046 | -0.046 | 0 | 0.65 | 0.54 | 0.76 | 0.84 | 0.64 | 0.76 |
| 3 | 14331328 | rs6796741 | -0.043 | 0.267 | -0.802 | 0.889 | -0.007 | -0.016 | -0.016 | 0 | 0.61 | 0.45 | 0.62 | 0.68 | 0.63 | 0.67 |
| 3 | 14345707 | rs9883639 | -0.294 | 0.233 | -1.771 | 0.721 | -0.024 | 0.009 | 0.009 | 0.56 | 0.39 | 0.12 | 0.47 | 0.67 | 0.36 | 0.46 |
| 3 | 14346445 | rs6794448 | -0.163 | 0.007 | -1.254 | 0.955 | -0.01 | 0.034 | 0.034 | 0 | 0.92 | 0.88 | 0.93 | 0.84 | 0.95 | 0.9 |
| 3 | 14347085 | rs9837707 | 0.041 | -0.022 | -1.189 | 0.922 | -0.015 | 0.073 | 0.073 | 0 | 0.91 | 0.85 | 0.86 | 0.81 | 0.95 | 0.86 |
| 3 | 14350659 | rs4684204 | -0.732 | 0.166 | -1.537 | 0.932 | 0.002 | -0.036 | -0.036 | 0 | 0.82 | 0.91 | 0.71 | 0.72 | 0.82 | 0.76 |
| 3 | 14355700 | rs4684205 | -0.187 | 0.508 | -1.8 | 0.69 | 0.005 | -0.008 | -0.008 | 3.98 | 0.52 | 0.8 | 0.58 | 0.45 | 0.51 | 0.61 |
| 3 | 14357107 | rs12491174 | 1.355 | 0.442 | -1.84 | 0.148 | -0.003 | 0.02 | 0.02 | 0.461 | 0.18 | 0.1 | 0.12 | 0 | 0.2 | 0.06 |
| 3 | 14359685 | rs6442441 | -0.353 | 0.42 | -1.733 | 0.117 | 0.002 | 0.025 | 0.025 | 0 | 0.49 | 0.75 | 0.39 | 0.38 | 0.44 | 0.36 |
| 3 | 14360339 | rs9842431 | 1.23 | 0.406 | -1.35 | 0.205 | -0.013 | -0.022 | -0.022 | 0.372 | 0.14 | 0.1 | 0.12 | 0.21 | 0.14 | 0.1 |
| 3 | 14362548 | rs9815883 | 1.927 | 0.318 | -0.963 | 0.231 | -0.01 | -0.044 | -0.044 | 1.357 | 0.08 | 0.02 | 0.09 | 0.21 | 0.07 | 0.08 |
| 3 | 14365266 | rs9851340 | -0.15 | 0.303 | -0.814 | 0.155 | -0.02 | 0.049 | 0.049 | 0.46 | 0.21 | 0.53 | 0.21 | 0.23 | 0.13 | 0.17 |
| 3 | 14369344 | rs4685122 | 0.346 | 0.552 | -1.137 | 0.238 | -0.014 | 0.016 | 0.016 | 0.649 | 0.37 | 0.17 | 0.31 | 0.34 | 0.34 | 0.45 |
| 3 | 14374615 | rs4684207 | -0.638 | 0.672 | -1.272 | 0.434 | -0.011 | -0.034 | -0.034 | 0.551 | 0.71 | 0.83 | 0.65 | 0.6 | 0.66 | 0.61 |
| 3 | 14377469 | rs9310445 | 0.814 | 0.686 | -1.332 | 0.289 | -0.016 | -0.018 | -0.018 | 0 | 0.31 | 0.17 | 0.35 | 0.3 | 0.37 | 0.17 |
| 3 | 14377609 | rs874969 | 1.501 | 0.627 | -1.345 | 0.262 | 0.007 | -0.014 | -0.014 | 0.556 | 0.11 | 0.1 | 0.17 | 0.27 | 0.08 | 0.11 |
| 3 | 14377632 | rs874967 | -0.195 | 0.654 | -1.378 | 0.203 | -0.003 | 0.03 | 0.03 | 0 | 0.61 | 0.74 | 0.48 | 0.33 | 0.59 | 0.49 |
| 3 | 14377925 | rs900176 | -0.091 | 0.579 | -0.891 | 0.143 | 0.006 | 0.008 | 0.008 | 0.289 | 0.45 | 0.39 | 0.44 | 0.6 | 0.33 | 0.52 |
| 3 | 14379479 | rs900179 | -1.725 | 0.622 | -1.153 | 0.271 | -0.05 | -0.046 | -0.046 | 0 | 0.79 | 0.96 | 0.7 | 0.76 | 0.73 | 0.76 |
| 3 | 14380774 | rs13091718 | -0.263 | 0.39 | -1.077 | 0.177 | 0.003 | 0.004 | 0.004 | 0 | 0.56 | 0.58 | 0.45 | 0.55 | 0.57 | 0.43 |
| 3 | 14380985 | rs12638446 | 0.892 | 0.412 | -0.847 | 0.079 | 0.005 | -0.033 | -0.033 | 0 | 0.33 | 0.43 | 0.26 | 0.17 | 0.38 | 0.25 |
| 3 | 14381907 | rs9861586 | 1.1 | 0.302 | -0.859 | 0.147 | 0.003 | 0.006 | 0.006 | 0 | 0.2 | 0.18 | 0.16 | 0.13 | 0.13 | 0.3 |
| 3 | 14382283 | rs9810264 | 1.828 | 0.429 | -0.947 | 0.396 | 0.003 | -0.04 | -0.04 | 0 | 0.1 | 0.04 | 0.1 | 0.21 | 0.07 | 0.09 |
| 3 | 14382866 | rs9828655 | 1.956 | 0.415 | -0.886 | 0.333 | 0 | 0.02 | 0.02 | 0.164 | 0.08 | 0.09 | 0.17 | 0.27 | 0.08 | 0.12 |
| 3 | 14383816 | rs7639399 | 0.256 | 0.37 | -0.926 | 0.008 | 0.001 | -0.003 | -0.003 | 0 | 0.45 | 0.6 | 0.39 | 0.46 | 0.43 | 0.32 |
| 3 | 14385116 | rs12163644 | 1.988 | 0.356 | -1.064 | -0.008 | -0.049 | -0.061 | -0.061 | 0 | 0.11 | 0.01 | 0.08 | 0.2 | 0.12 | 0.1 |
| 3 | 14385513 | rs9844200 | -0.142 | 0.312 | -0.989 | -0.172 | -0.002 | 0.006 | 0.006 | 0 | 0.49 | 0.62 | 0.45 | 0.46 | 0.51 | 0.33 |
| 3 | 14385901 | rs12497942 | 1.198 | 0.273 | -1.038 | -0.247 | -0.001 | 0.003 | 0.003 | 0 | 0.1 | 0.07 | 0.1 | 0 | 0.14 | 0.02 |
| 3 | 14386387 | rs13061796 | 1.619 | 0.212 | -0.758 | -0.164 | -0.007 | -0.142 | -0.142 | 0 | 0.23 | 0.41 | 0.13 | 0 | 0.23 | 0.12 |
| 3 | 14387154 | rs749855 | -1.168 | 0.26 | -1.061 | -0.086 | -0.007 | -0.021 | -0.021 | 0 | 0.72 | 0.79 | 0.64 | 0.57 | 0.67 | 0.59 |
| 3 | 14387203 | rs11711461 | 0.223 | 0.24 | -1.057 | -0.094 | -0.011 | -0.053 | -0.053 | 0 | 0.32 | 0.19 | 0.36 | 0.46 | 0.33 | 0.22 |
| 3 | 14388278 | rs4685137 | -0.893 | 0.34 | -0.963 | 0.175 | -0.006 | -0.019 | -0.019 | 0 | 0.72 | 0.78 | 0.64 | 0.57 | 0.67 | 0.59 |
| 3 | 14390507 | rs12639065 | -0.6 | 0.127 | -0.701 | 0.084 | -0.001 | 0.005 | 0.005 | 0 | 0.58 | 0.51 | 0.4 | 0.5 | 0.58 | 0.57 |
| 3 | 14393625 | rs999969 | 0.712 | 0.003 | -0.817 | -0.125 | 0.002 | -0.009 | -0.009 | 0 | 0.35 | 0.36 | 0.23 | 0.32 | 0.37 | 0.45 |
| 3 | 14393742 | rs893081 | 0.618 | 0.069 | -1.022 | 0.058 | 0 | -0.009 | -0.009 | 0 | 0.23 | 0.13 | 0.24 | 0.21 | 0.21 | 0.14 |
| 3 | 14395876 | rs1384106 | 1.95 | 0.315 | -1.002 | 0.112 | -0.021 | -0.018 | -0.018 | 0 | 0.08 | 0.01 | 0.05 | 0.12 | 0.08 | 0.19 |
| 3 | 14399756 | rs9853336 | 0.836 | 0.127 | -0.764 | 0.106 | -0.001 | 0 | 0 | 0 | 0.22 | 0.26 | 0.28 | 0.21 | 0.24 | 0.18 |
| 3 | 14400077 | rs13081924 | 2.005 | 0.013 | -1.071 | 0.144 | -0.039 | -0.099 | -0.099 | 2.586 | 0.16 | 0 | 0.19 | 0.25 | 0.14 | 0.19 |
| 3 | 14401563 | rs6796254 | 0.765 | 0.129 | -0.555 | 0.063 | -0.008 | -0.041 | -0.041 | 0.176 | 0.31 | 0.51 | 0.19 | 0.21 | 0.29 | 0.24 |
| 3 | 14402491 | rs9878149 | -1.665 | 0.203 | -0.636 | 0.088 | -0.007 | 0.006 | 0.006 | 0 | 0.78 | 0.84 | 0.71 | 0.77 | 0.74 | 0.79 |
| 3 | 14402962 | rs13095528 | 0.176 | -0.092 | -0.453 | -0.039 | -0.001 | -0.001 | -0.001 | 0 | 0.47 | 0.46 | 0.44 | 0.39 | 0.43 | 0.47 |
| 3 | 14405954 | rs1550161 | -1.514 | 0.206 | -0.449 | 0.111 | 0 | 0 | 0 | 0 | 0.77 | 0.76 | 0.69 | 0.68 | 0.69 | 0.78 |
| 3 | 14406331 | rs1550160 | 1.27 | -0.011 | -0.396 | -0.015 | -0.011 | -0.128 | -0.128 | 0 | 0.25 | 0.38 | 0.13 | 0 | 0.23 | 0.11 |
| 3 | 14406623 | rs1550158 | -0.127 | 0.111 | -0.602 | 0.182 | 0.012 | 0.012 | 0.012 | 0 | 0.65 | 0.56 | 0.55 | 0.55 | 0.6 | 0.55 |
| 3 | 14407960 | rs1871852 | 0.029 | 0.106 | -0.639 | 0.242 | 0.003 | -0.001 | -0.001 | 0 | 0.45 | 0.56 | 0.44 | 0.4 | 0.39 | 0.41 |
| 3 | 14408618 | rs2118105 | 0.013 | 0.021 | -0.618 | 0.459 | -0.001 | 0.003 | 0.003 | 0.131 | 0.27 | 0.24 | 0.28 | 0.39 | 0.22 | 0.36 |
| 3 | 14409443 | rs17307846 | 0.131 | 0.363 | -1.03 | -0.05 | 0 | -0.008 | -0.008 | 0 | 0.62 | 0.76 | 0.65 | 0.63 | 0.66 | 0.58 |
| 3 | 14409794 | rs7643329 | 1.098 | 0.817 | -0.614 | -0.516 | -0.021 | 0.05 | 0.05 | 0 | 0.44 | 0.21 | 0.45 | 0.64 | 0.44 | 0.45 |
| 3 | 14411655 | rs1865884 | 0.172 | 0.651 | -0.483 | -0.822 | 0.011 | 0.029 | 0.029 | 0.133 | 0.67 | 0.76 | 0.65 | 0.75 | 0.67 | 0.75 |
| 3 | 14421239 | rs10446439 | 0.787 | 0.847 | 0.304 | -0.114 | -0.002 | -0.023 | -0.023 | 0 | 0.92 | 0.81 | 0.97 | 0.95 | 0.94 | 0.88 |
| 3 | 14427076 | rs1038300 | -1.485 | 1.202 | 0.297 | -0.015 | 0.022 | 0.049 | 0.049 | 0 | 0.27 | 0.45 | 0.39 | 0.36 | 0.31 | 0.38 |
| 3 | 14427543 | rs1038301 | -0.32 | 1.484 | -0.183 | -0.015 | -0.008 | 0.005 | 0.005 | 0 | 0.08 | 0.03 | 0.06 | 0 | 0.11 | 0.04 |
| 3 | 14428788 | rs4685148 | 0.214 | 1.492 | -0.272 | 0.099 | -0.011 | -0.009 | -0.009 | 4.334 | 0.93 | 0.92 | 0.95 | 0.88 | 0.91 | 0.92 |
| 3 | 14441382 | rs2033688 | 1.182 | 1.238 | -0.211 | -0.072 | -0.017 | -0.006 | -0.006 | 0 | 0.62 | 0.22 | 0.65 | 0.65 | 0.62 | 0.65 |
| 3 | 14441957 | rs4685153 | 1.013 | 1.647 | -0.634 | 0.157 | 0.004 | 0.003 | 0.003 | 0 | 0.67 | 0.59 | 0.69 | 0.65 | 0.63 | 0.65 |
| 3 | 14446545 | rs1156568 | 0.846 | 1.403 | -0.903 | 0.299 | -0.013 | -0.01 | -0.01 | 0 | 0.93 | 0.92 | 0.95 | 0.85 | 0.91 | 0.93 |
| 3 | 14451837 | rs9810939 | -0.246 | 0.918 | -1.118 | -1.023 | -0.001 | 0.133 | 0.133 | 0.266 | 0.51 | 0.71 | 0.48 | 0.05 | 0.51 | 0.25 |
| 3 | 14452800 | rs9835759 | -0.292 | 0.487 | -0.146 | -1.222 | -0.023 | -0.085 | -0.085 | 1.933 | 0.14 | 0.37 | 0.06 | 0 | 0.07 | 0.08 |
| 3 | 14454286 | rs6765103 | -1.137 | 0.556 | -0.484 | -1.591 | -0.005 | -0.087 | -0.087 | 1.262 | 0.27 | 0.39 | 0.15 | 0.02 | 0.24 | 0.16 |
| 3 | 14459473 | rs4331657 | -0.76 | 0.362 | -0.48 | -1.597 | 0.003 | 0.017 | 0.017 | 3.141 | 0.21 | 0.12 | 0.11 | 0.05 | 0.21 | 0.11 |
| 3 | 14459623 | rs1157564 | 1.028 | 0.234 | -0.112 | -2.027 | 0 | -0.051 | -0.051 | 0 | 0.6 | 0.48 | 0.62 | 0.91 | 0.55 | 0.74 |
| 3 | 14460915 | rs12638941 | 1.15 | 0.041 | -0.606 | -2.108 | 0.008 | 0.214 | 0.214 | 0.055 | 0.55 | 0.16 | 0.57 | 0.91 | 0.55 | 0.67 |
| 3 | 14463458 | rs4685158 | 0.596 | -0.037 | -0.359 | -2.043 | 0.013 | 0.009 | 0.009 | 0.382 | 0.68 | 0.28 | 0.69 | 0.96 | 0.72 | 0.76 |
| 3 | 14468767 | rs13058781 | -0.733 | 0.624 | -0.075 | -1.492 | 0.026 | -0.023 | -0.023 | 0 | 0.26 | 0.38 | 0.21 | 0.22 | 0.31 | 0.34 |
| 3 | 14469704 | rs6774615 | -0.083 | 0.495 | -0.424 | -1.641 | -0.031 | -0.048 | -0.048 | 0 | 0.15 | 0.01 | 0.09 | 0.21 | 0.17 | 0.27 |
| 3 | 14470510 | rs13081322 | -1.18 | 0.426 | -0.109 | -1.428 | 0.009 | -0.012 | -0.012 | 0 | 0.28 | 0.37 | 0.22 | 0.28 | 0.32 | 0.34 |

| | | | | | | | | | | | | | | | | |
|---|---|---|---|---|---|---|---|---|---|---|---|---|---|---|---|---|
| 3 | 14470911 | rs748098 | -1.247 | 0.303 | -0.381 | -1.154 | -0.001 | 0.011 | 0.011 | 0 | 0.49 | 0.75 | 0.44 | 0.42 | 0.49 | 0.55 |
| 3 | 14471838 | rs11919568 | 1.125 | 0.389 | -1.039 | -0.982 | 0.003 | -0.003 | -0.003 | 0 | 0.21 | 0.19 | 0.33 | 0.13 | 0.27 | 0.22 |
| 3 | 14472004 | rs4684217 | -0.543 | 1.04 | -0.622 | -0.937 | 0.001 | -0.004 | -0.004 | 0 | 0.44 | 0.45 | 0.5 | 0.41 | 0.54 | 0.57 |
| 3 | 14473278 | rs6801669 | 0.385 | 1.028 | -0.37 | -0.927 | 0.006 | 0.002 | 0.002 | 0 | 0.22 | 0.24 | 0.18 | 0.27 | 0.27 | 0.35 |
| 3 | 14476457 | rs2164576 | 0.668 | 0.8 | -0.128 | -0.987 | 0 | 0.001 | 0.001 | 0 | 0.24 | 0.24 | 0.22 | 0.12 | 0.21 | 0.22 |
| 3 | 14476497 | rs4685160 | -2.132 | 0.76 | -0.321 | -0.926 | 0.015 | -0.088 | -0.088 | 0 | 0.7 | 0.92 | 0.75 | 0.54 | 0.75 | 0.79 |
| 3 | 14476628 | rs4684218 | 0.989 | 0.825 | -0.189 | -0.9 | 0.011 | -0.015 | -0.015 | 0 | 0.22 | 0.25 | 0.33 | 0.15 | 0.28 | 0.22 |
| 3 | 14480205 | rs9683047 | -0.031 | 0.617 | -0.144 | -0.812 | 0.001 | -0.006 | -0.006 | 0 | 0.46 | 0.5 | 0.55 | 0.27 | 0.49 | 0.37 |
| 3 | 14480778 | rs6790433 | -0.176 | 0.842 | -0.017 | -0.652 | 0.016 | 0.003 | 0.003 | 0 | 0.23 | 0.37 | 0.19 | 0.27 | 0.27 | 0.35 |
| 3 | 14482180 | rs4684220 | -0.05 | 0.532 | -0.324 | 0.214 | -0.001 | -0.001 | -0.001 | 0 | 0.41 | 0.37 | 0.64 | 0.29 | 0.49 | 0.4 |
| 3 | 14482659 | rs6766142 | -0.393 | 0.199 | -0.149 | 0.336 | -0.004 | -0.001 | -0.001 | 0 | 0.42 | 0.4 | 0.59 | 0.45 | 0.55 | 0.47 |
| 3 | 14483826 | rs7638413 | -1.211 | -0.079 | -0.642 | -0.077 | -0.008 | -0.001 | -0.001 | 0 | 0.79 | 0.88 | 0.83 | 0.88 | 0.78 | 0.84 |
| 3 | 14484113 | rs7628386 | -0.182 | -0.431 | -0.212 | 0.233 | 0.013 | -0.037 | -0.037 | 0 | 0.32 | 0.72 | 0.24 | 0.53 | 0.22 | 0.32 |
| 3 | 14489370 | rs2304511 | -0.7 | -0.684 | 0.625 | -0.145 | -0.007 | -0.022 | -0.022 | 0.208 | 0.25 | 0.34 | 0.21 | 0.06 | 0.24 | 0.22 |
| 3 | 14489694 | rs6765627 | -1.172 | -0.764 | 0.975 | -0.082 | -0.002 | 0.03 | 0.03 | 4.639 | 0.28 | 0.37 | 0.2 | 0.45 | 0.26 | 0.46 |
| 3 | 14494855 | rs3773182 | -0.732 | -0.511 | 1.091 | 0.039 | 0.002 | -0.013 | -0.013 | 0 | 0.37 | 0.46 | 0.28 | 0.39 | 0.43 | 0.51 |
| 3 | 14496074 | rs11919644 | -0.916 | -0.549 | 1.257 | 0.021 | -0.001 | -0.012 | -0.012 | 0 | 0.39 | 0.57 | 0.29 | 0.4 | 0.43 | 0.52 |
| 3 | 14496162 | rs11128699 | -0.878 | -0.549 | 1.275 | 0.021 | 0 | -0.024 | -0.024 | 0 | 0.39 | 0.52 | 0.29 | 0.4 | 0.43 | 0.52 |
| 3 | 14496974 | rs3773180 | -0.688 | -0.548 | 1.33 | 0.022 | 0 | -0.001 | -0.001 | 0 | 0.37 | 0.38 | 0.28 | 0.39 | 0.42 | 0.48 |
| 3 | 14497019 | rs3773179 | 2.242 | -0.698 | 1.063 | -0.058 | 0.003 | 0.018 | 0.018 | 2.1 | 0.13 | 0.01 | 0.13 | 0.09 | 0.07 | 0.13 |
| 3 | 14498844 | rs3773177 | -1.087 | 0.065 | 2.085 | 0.453 | -0.001 | 0.001 | 0.001 | 4.892 | 0.37 | 0.38 | 0.28 | 0.4 | 0.43 | 0.49 |
| 3 | 14500744 | rs3773176 | -2.083 | -0.011 | 1.707 | 0.479 | 0.009 | 0.027 | 0.027 | 6.658 | 0.5 | 0.88 | 0.35 | 0.45 | 0.46 | 0.61 |
| 3 | 14501153 | rs9833294 | -1.289 | 0.077 | 1.659 | 0.515 | 0 | -0.023 | -0.023 | 0 | 0.39 | 0.52 | 0.29 | 0.4 | 0.42 | 0.52 |
| 3 | 14501174 | rs12489456 | 0.188 | 0.179 | 0.781 | 0.552 | -0.039 | -0.059 | -0.059 | 0 | 0.13 | 0.04 | 0.11 | 0.38 | 0.18 | 0.34 |
| 3 | 14506279 | rs6763180 | -1.714 | 0.168 | 0.836 | 0.567 | 0.011 | 0.024 | 0.024 | 4.658 | 0.48 | 0.76 | 0.34 | 0.45 | 0.46 | 0.57 |
| 3 | 14506503 | rs7610741 | -1.714 | 0.168 | 0.836 | 0.567 | 0.011 | 0.024 | 0.024 | 4.658 | 0.48 | 0.76 | 0.34 | 0.45 | 0.46 | 0.57 |
| 3 | 14506634 | rs7611000 | -1.714 | 0.168 | 0.836 | 0.567 | 0.011 | 0.024 | 0.024 | 4.658 | 0.48 | 0.76 | 0.34 | 0.45 | 0.46 | 0.57 |
| 3 | 14508311 | rs2289129 | -1.041 | -0.299 | 0.502 | 0.334 | 0.014 | -0.028 | -0.028 | 1.363 | 0.09 | 0.03 | 0.03 | 0.05 | 0.05 | 0.11 |
| 3 | 14509865 | rs7642354 | -0.875 | -0.084 | 0.976 | 0.264 | -0.016 | -0.089 | -0.089 | 0 | 0.27 | 0.64 | 0.19 | 0.01 | 0.23 | 0.2 |
| 3 | 14510861 | rs3773174 | -1.039 | 0.217 | 0.633 | 0.936 | -0.018 | 0.081 | 0.081 | 0 | 0.09 | 0.2 | 0.06 | 0.18 | 0.05 | 0.16 |
| 3 | 14511699 | rs3773172 | -1.128 | 0.15 | 0.611 | 0.934 | -0.022 | 0.081 | 0.081 | 0 | 0.09 | 0.18 | 0.05 | 0.16 | 0.05 | 0.16 |
| 3 | 14512093 | rs3773171 | -0.761 | 0.162 | 0.533 | 0.94 | -0.011 | 0.038 | 0.038 | 0 | 0.08 | 0.09 | 0.04 | 0.16 | 0.05 | 0.16 |
| 3 | 14512731 | rs9826660 | -1.381 | 0.15 | 0.66 | 0.944 | -0.021 | 0.131 | 0.131 | 0 | 0.1 | 0.28 | 0.06 | 0.16 | 0.05 | 0.16 |
| 3 | 14513695 | rs2289128 | -0.761 | 0.179 | 0.538 | 0.951 | -0.01 | 0.036 | 0.036 | 0 | 0.08 | 0.09 | 0.04 | 0.16 | 0.05 | 0.16 |
| 3 | 14517115 | rs9834678 | -1.102 | 0.153 | 0.427 | 0.89 | -0.014 | 0.123 | 0.123 | 0 | 0.1 | 0.3 | 0.06 | 0.16 | 0.06 | 0.15 |
| 3 | 14519102 | rs9845444 | -0.941 | 0.162 | 0.31 | 0.908 | -0.015 | 0.138 | 0.138 | 0 | 0.09 | 0.3 | 0.06 | 0.15 | 0.05 | 0.16 |
| 3 | 14522194 | rs11710744 | -0.576 | 0.645 | -0.228 | 0.299 | -0.007 | -0.07 | -0.07 | 0 | 0.25 | 0.29 | 0.26 | 0.01 | 0.25 | 0.12 |
| 3 | 14524270 | rs3773165 | -0.861 | 1.345 | -0.348 | 1.45 | 0.051 | 0.146 | 0.146 | 3.538 | 0.12 | 0.32 | 0.21 | 0.37 | 0.15 | 0.24 |
| 3 | 14526031 | rs3773162 | -0.713 | 1.657 | -1.424 | 1.462 | -0.08 | -0.111 | -0.111 | 0 | 0.19 | 0.03 | 0.2 | 0.4 | 0.33 | 0.22 |
| 3 | 14526538 | rs1062540 | -0.306 | 1.252 | -2.06 | 0.46 | -0.106 | 0.14 | 0.14 | 2.445 | 0.37 | 0.05 | 0.39 | 0.13 | 0.56 | 0.19 |
| 3 | 14527363 | rs2327896 | -0.249 | 0.369 | -0.97 | 0.063 | -0.032 | -0.072 | -0.072 | 0.246 | 0.88 | 0.68 | 0.93 | 0.99 | 0.95 | 0.84 |
| 3 | 14528004 | rs2341984 | -0.221 | -0.288 | -1.437 | -0.273 | -0.031 | -0.075 | -0.075 | 0.128 | 0.59 | 0.51 | 0.8 | 0.85 | 0.75 | 0.63 |
| 3 | 14528028 | rs2341985 | 0.433 | -0.09 | -1.106 | 0.344 | 0.019 | 0.068 | 0.068 | 0.062 | 0.12 | 0.23 | 0.22 | 0.23 | 0.16 | 0.2 |
| 3 | 14528256 | rs4685165 | -0.129 | -0.429 | -1.108 | -0.127 | -0.023 | -0.06 | -0.06 | 0.058 | 0.58 | 0.51 | 0.79 | 0.85 | 0.75 | 0.63 |
| 3 | 14530721 | rs9036 | 0.544 | -0.387 | -1.478 | -0.097 | 0.029 | 0.084 | 0.084 | 0.422 | 0.25 | 0.09 | 0.14 | 0.12 | 0.18 | 0.18 |
| 3 | 14531382 | rs6794801 | -0.617 | -0.141 | -1.143 | 0.471 | 0.008 | 0.042 | 0.042 | 0.861 | 0.29 | 0.42 | 0.35 | 0.48 | 0.27 | 0.4 |
| 3 | 14531786 | rs11718660 | -0.169 | 0.447 | -2.152 | 0.199 | -0.115 | -0.124 | -0.124 | 0.025 | 0.45 | 0.04 | 0.45 | 0.46 | 0.5 | 0.46 |
| 3 | 14545776 | rs6800512 | -0.028 | 1.449 | -1.49 | -0.084 | -0.072 | -0.029 | -0.029 | 0 | 0.73 | 0.87 | 0.69 | 0.66 | 0.55 | 0.81 |
| 3 | 14547802 | rs4234525 | 0.195 | 1.178 | -0.94 | -0.208 | -0.008 | -0.002 | -0.002 | 0 | 0.7 | 0.67 | 0.65 | 0.66 | 0.53 | 0.81 |
| 3 | 14550500 | rs9878341 | 0.602 | 0.485 | -0.613 | -0.629 | -0.018 | -0.075 | -0.075 | 0 | 0.14 | 0.24 | 0.09 | 0.01 | 0.1 | 0.14 |
| 3 | 14555216 | rs4685171 | 0.194 | 0.574 | -0.038 | -0.577 | 0.024 | -0.054 | -0.054 | 0.023 | 0.65 | 0.59 | 0.67 | 0.77 | 0.55 | 0.75 |
| 3 | 14555811 | rs4613440 | 0.194 | 0.595 | -0.038 | -0.577 | 0.025 | -0.054 | -0.054 | 0 | 0.65 | 0.59 | 0.67 | 0.77 | 0.55 | 0.75 |
| 3 | 14562127 | rs1106333 | 0.361 | 0.248 | 0.01 | -0.431 | -0.068 | -0.03 | -0.03 | 0 | 0.14 | 0.37 | 0.06 | 0.06 | 0.04 | 0.16 |
| 3 | 14567666 | rs4684230 | 0.281 | 0.944 | -0.493 | -0.48 | 0.04 | 0.032 | 0.032 | 1.157 | 0.71 | 0.48 | 0.64 | 0.61 | 0.6 | 0.8 |
| 3 | 14568071 | rs11921520 | 0.21 | 0.761 | -0.361 | -0.601 | -0.003 | -0.075 | -0.075 | 0.061 | 0.2 | 0.29 | 0.12 | 0.02 | 0.18 | 0.16 |
| 3 | 14569560 | rs2090700 | 1.143 | 1.003 | -0.696 | -0.52 | 0.004 | 0.037 | 0.037 | 1.337 | 0.51 | 0.19 | 0.52 | 0.57 | 0.43 | 0.65 |
| 3 | 14569733 | rs2102916 | 2.05 | 0.862 | -1.405 | -0.685 | -0.043 | -0.158 | -0.158 | 0.391 | 0.21 | 0.02 | 0.32 | 0.43 | 0.21 | 0.19 |
| 3 | 14571310 | rs4684231 | 0.832 | 0.819 | -1.084 | -0.803 | -0.017 | 0.029 | 0.029 | 1.904 | 0.53 | 0.19 | 0.55 | 0.57 | 0.47 | 0.65 |
| 3 | 14575244 | rs17317288 | 0.339 | 1.093 | -0.861 | -0.729 | -0.001 | -0.035 | -0.035 | 1.659 | 0.2 | 0.19 | 0.1 | 0.02 | 0.18 | 0.2 |
| 3 | 14575333 | rs17317323 | 0.339 | 1.088 | -0.874 | -0.763 | -0.001 | -0.032 | -0.032 | 0.007 | 0.2 | 0.19 | 0.1 | 0.02 | 0.18 | 0.2 |
| 3 | 14578162 | rs4685194 | 0.753 | 0.84 | -1.043 | -0.522 | 0 | -0.027 | -0.027 | 0 | 0.31 | 0.23 | 0.42 | 0.56 | 0.34 | 0.34 |
| 3 | 14580152 | rs1949539 | 0.604 | 0.772 | -0.854 | -0.881 | -0.001 | -0.027 | -0.027 | 1.255 | 0.12 | 0.11 | 0.03 | 0.01 | 0.07 | 0.1 |
| 3 | 14581594 | rs6442461 | -0.789 | 0.951 | -0.779 | -1.022 | 0.001 | -0.011 | -0.011 | 0.133 | 0.4 | 0.54 | 0.23 | 0.2 | 0.39 | 0.33 |
| 3 | 14582669 | rs11128705 | -1.046 | 1.119 | -0.712 | -0.759 | 0.004 | -0.001 | -0.001 | 0 | 0.81 | 0.77 | 0.73 | 0.77 | 0.71 | 0.89 |
| 3 | 14582923 | rs11709154 | -0.37 | 0.806 | -0.676 | -1.04 | -0.006 | -0.046 | -0.046 | 0 | 0.54 | 0.43 | 0.31 | 0.11 | 0.38 | 0.53 |
| 3 | 14582986 | rs9883238 | 0.003 | -0.229 | -0.797 | -1.02 | 0.024 | 0.01 | 0.01 | 0.575 | 0.46 | 0.52 | 0.25 | 0.07 | 0.22 | 0.48 |
| 3 | 14583194 | rs13062253 | 0.489 | 0.792 | -0.584 | -0.103 | 0.002 | -0.003 | -0.003 | 0 | 0.58 | 0.35 | 0.62 | 0.6 | 0.54 | 0.67 |
| 3 | 14591346 | rs9310449 | -0.486 | 0.789 | -0.342 | 0.245 | 0.013 | 0.019 | 0.019 | 2.198 | 0.84 | 0.77 | 0.72 | 0.74 | 0.83 | 0.93 |
| 3 | 14596864 | rs9834063 | 0.168 | 0.372 | -1.211 | -0.129 | 0.001 | 0.063 | 0.063 | 0.29 | 0.26 | 0.1 | 0.22 | 0.09 | 0.26 | 0.35 |

| | | | | | | | | | | | | | | | | |
|---|---|---|---|---|---|---|---|---|---|---|---|---|---|---|---|---|
| 3 | 14598843 | rs11128708 | 1.433 | 0.365 | -1.614 | 0.221 | -0.071 | -0.099 | -0.099 | 0 | 0.34 | 0.05 | 0.29 | 0.38 | 0.4 | 0.41 |
| 3 | 14598965 | rs4449299 | 0.252 | 0.551 | -1.122 | 0.075 | 0.003 | 0.058 | 0.058 | 1.01 | 0.26 | 0.13 | 0.22 | 0.09 | 0.24 | 0.33 |
| 3 | 14604672 | rs7647806 | 0.513 | 0.57 | -0.24 | 0.154 | -0.003 | 0.003 | 0.003 | 0 | 0.57 | 0.63 | 0.45 | 0.65 | 0.53 | 0.62 |
| 3 | 14605987 | rs7629002 | 0.395 | 0.452 | 0.101 | 0.001 | 0.031 | 0.017 | 0.017 | 0.33 | 0.72 | 0.59 | 0.56 | 0.69 | 0.7 | 0.73 |
| 3 | 14606700 | rs11128709 | 0.682 | 0.62 | -0.005 | -0.005 | 0.002 | -0.002 | -0.002 | 0 | 0.11 | 0.09 | 0.18 | 0.08 | 0.15 | 0.22 |
| 3 | 14607976 | rs7613527 | 0.677 | 0.522 | -0.018 | 0.172 | 0.007 | 0.034 | 0.034 | 0 | 0.2 | 0.11 | 0.16 | 0.15 | 0.19 | 0.19 |
| 3 | 14609339 | rs13090193 | -1.25 | 0.559 | -0.022 | 0.182 | 0.005 | 0.012 | 0.012 | 2.705 | 0.86 | 0.84 | 0.74 | 0.73 | 0.84 | 0.94 |
| 3 | 14610761 | rs11714664 | -0.165 | 0.669 | -0.326 | 0.158 | -0.009 | 0.025 | 0.025 | 0.506 | 0.28 | 0.17 | 0.29 | 0.08 | 0.32 | 0.33 |
| 3 | 14613814 | rs9868402 | 0.189 | 0.842 | -0.16 | 0.53 | 0.039 | -0.005 | -0.005 | 0 | 0.12 | 0.23 | 0.18 | 0.08 | 0.15 | 0.21 |
| 3 | 14621295 | rs4684234 | 0.875 | 0.365 | -1.281 | 0.745 | -0.06 | -0.044 | -0.044 | 0 | 0.35 | 0.05 | 0.28 | 0.32 | 0.37 | 0.35 |
| 3 | 14623302 | rs4685197 | 0.432 | 0.233 | 0.038 | 0.963 | -0.002 | 0.066 | 0.066 | 3.831 | 0.79 | 0.46 | 0.67 | 0.58 | 0.83 | 0.87 |
| 3 | 14628404 | rs6801503 | -0.219 | 0.227 | -0.388 | 0.491 | 0 | 0.01 | 0.01 | 0 | 0.53 | 0.44 | 0.47 | 0.65 | 0.49 | 0.44 |
| 3 | 14630164 | rs17039877 | 0.174 | -0.017 | -0.226 | 0.253 | -0.009 | -0.041 | -0.041 | 0.877 | 0.1 | 0.26 | 0.06 | 0.04 | 0.06 | 0.06 |
| 3 | 14632542 | rs4685199 | -0.294 | -0.174 | -0.402 | 0.472 | -0.009 | -0.004 | -0.004 | 0 | 0.55 | 0.36 | 0.48 | 0.62 | 0.5 | 0.42 |
| 3 | 14632711 | rs4462948 | -0.382 | -0.502 | -0.766 | 0.601 | -0.004 | 0.001 | 0.001 | 0 | 0.24 | 0.3 | 0.25 | 0.34 | 0.21 | 0.13 |
| 3 | 14635048 | rs17318350 | 0.947 | -0.385 | -1.108 | 0.361 | -0.009 | -0.011 | -0.011 | 0 | 0.3 | 0.05 | 0.22 | 0.25 | 0.28 | 0.29 |
| 3 | 14637075 | rs4685201 | -0.834 | -0.407 | -0.429 | 0.214 | -0.048 | 0.014 | 0.014 | 0.857 | 0.62 | 0.81 | 0.53 | 0.7 | 0.51 | 0.5 |
| 3 | 14638566 | rs7615913 | -0.517 | -0.612 | 0.455 | 0.17 | -0.034 | -0.048 | -0.048 | 0 | 0.16 | 0.53 | 0.1 | 0.05 | 0.07 | 0.09 |
| 3 | 14641414 | rs4298030 | 0.683 | -0.607 | 0.152 | -0.209 | 0.002 | -0.019 | -0.019 | 0.003 | 0.29 | 0.28 | 0.12 | 0.05 | 0.15 | 0.23 |
| 3 | 14641812 | rs4511884 | -0.465 | -0.245 | 0.053 | -0.083 | -0.002 | -0.001 | -0.001 | 0 | 0.49 | 0.5 | 0.38 | 0.47 | 0.38 | 0.42 |
| 3 | 14642278 | rs12497747 | -0.097 | -0.031 | 0.108 | 0.029 | -0.012 | -0.038 | -0.038 | 0 | 0.18 | 0.11 | 0.25 | 0.42 | 0.23 | 0.18 |
| 3 | 14642891 | rs12631299 | -0.596 | 0.222 | 0.047 | -0.017 | 0.003 | -0.001 | -0.001 | 0 | 0.59 | 0.49 | 0.47 | 0.63 | 0.48 | 0.55 |
| 3 | 14648833 | rs6798424 | 0.045 | 0.523 | 0.102 | -0.176 | 0.043 | -0.015 | -0.015 | 0 | 0.06 | 0.19 | 0.07 | 0.01 | 0.09 | 0.1 |
| 3 | 14655725 | rs12495033 | -0.226 | 0.416 | -0.05 | -0.468 | 0.004 | 0.005 | 0.005 | 0.028 | 0.44 | 0.6 | 0.29 | 0.13 | 0.37 | 0.41 |
| 3 | 14657598 | rs12493810 | -0.304 | 0.538 | -0.003 | -0.093 | 0.003 | 0.009 | 0.009 | 0 | 0.27 | 0.31 | 0.33 | 0.51 | 0.32 | 0.26 |
| 3 | 14659515 | rs12495599 | -0.561 | 0.486 | -0.603 | -0.061 | -0.036 | -0.047 | -0.047 | 0.39 | 0.64 | 0.78 | 0.49 | 0.57 | 0.54 | 0.53 |
| 3 | 14661876 | rs4583633 | -0.591 | 1.353 | -0.398 | 0.303 | 0.012 | 0.015 | 0.015 | 2.043 | 0.31 | 0.33 | 0.49 | 0.54 | 0.45 | 0.29 |
| 3 | 14665333 | rs11717103 | -0.304 | 1.763 | -0.447 | 0.527 | 0.043 | 0.056 | 0.056 | 5.335 | 0.26 | 0.44 | 0.47 | 0.52 | 0.41 | 0.29 |
| 3 | 14665953 | rs4685204 | -0.968 | 0.762 | -0.788 | -0.464 | 0.042 | 0.072 | 0.072 | 0.44 | 0.58 | 0.78 | 0.79 | 0.94 | 0.72 | 0.68 |
| 3 | 14666117 | rs1401892 | -0.21 | 1.042 | -0.72 | 0.341 | -0.002 | -0.007 | -0.007 | 0 | 0.21 | 0.27 | 0.2 | 0.18 | 0.18 | 0.22 |
| 3 | 14670155 | rs10865731 | -0.537 | 1.362 | -1.596 | 0.388 | -0.088 | -0.12 | -0.12 | 0 | 0.33 | 0.09 | 0.29 | 0.44 | 0.4 | 0.51 |
| 3 | 14670563 | rs7644101 | 0.327 | 1.266 | -0.663 | 0.401 | 0.023 | 0.132 | 0.132 | 1.887 | 0.86 | 0.68 | 0.63 | 0.56 | 0.83 | 0.83 |
| 3 | 14672311 | rs7640236 | -0.003 | 1.207 | -0.51 | 0.406 | 0.005 | 0.051 | 0.051 | 0 | 0.89 | 0.87 | 0.66 | 0.55 | 0.84 | 0.83 |
| 3 | 14676294 | rs17252904 | 1.403 | 0.147 | -0.94 | -0.44 | 0.037 | 0.057 | 0.057 | 0 | 0.26 | 0.13 | 0.11 | 0.04 | 0.17 | 0.09 |
| 3 | 14678299 | rs3846121 | 0.534 | 0.088 | 0.01 | -0.25 | -0.015 | -0.019 | -0.019 | 0.389 | 0.34 | 0.61 | 0.34 | 0.19 | 0.23 | 0.19 |
| 3 | 14679273 | rs13088257 | 0.879 | 0.51 | -0.499 | -0.169 | -0.1 | -0.121 | -0.121 | 0.003 | 0.22 | 0.04 | 0.23 | 0.39 | 0.33 | 0.37 |
| 3 | 14688071 | rs9310457 | -0.032 | 0.367 | 0.104 | -0.135 | -0.006 | -0.016 | -0.016 | 0.426 | 0.6 | 0.52 | 0.6 | 0.63 | 0.61 | 0.65 |
| 3 | 14689393 | rs11708596 | 0.918 | 0.182 | 0.129 | -0.393 | 0.021 | 0.057 | 0.057 | 0 | 0.22 | 0.06 | 0.15 | 0.05 | 0.17 | 0.19 |
| 3 | 14691111 | rs11928962 | 0.752 | 0.262 | 0.008 | -0.341 | 0.003 | -0.001 | -0.001 | 0 | 0.55 | 0.4 | 0.41 | 0.48 | 0.55 | 0.56 |
| 3 | 14692314 | rs7637233 | 0.807 | 0.257 | -0.213 | -0.342 | 0.018 | 0.005 | 0.005 | 0.104 | 0.52 | 0.24 | 0.38 | 0.48 | 0.54 | 0.55 |
| 3 | 14697590 | rs13091225 | 1.066 | 0.529 | 0.04 | -0.204 | -0.046 | -0.102 | -0.102 | 0 | 0.6 | 0.12 | 0.37 | 0.46 | 0.55 | 0.67 |
| 3 | 14700046 | rs4684236 | -0.482 | 0.529 | 0.669 | -0.193 | -0.001 | 0 | 0 | 0 | 0.28 | 0.27 | 0.43 | 0.23 | 0.34 | 0.26 |
| 3 | 14702351 | rs17040040 | 0.896 | 0.542 | 0.721 | -0.233 | 0.005 | -0.037 | -0.037 | 0 | 0.12 | 0.16 | 0.1 | 0.01 | 0.12 | 0.05 |
| 3 | 14702982 | rs2276754 | -0.376 | 0.537 | 0.799 | -0.214 | 0.097 | 0.241 | 0.241 | 0.214 | 0.92 | 0.62 | 0.83 | 0.69 | 0.91 | 0.89 |
| 3 | 14706848 | rs3773487 | -0.579 | 0.526 | 0.768 | -0.235 | -0.001 | -0.001 | -0.001 | 0 | 0.28 | 0.27 | 0.42 | 0.23 | 0.34 | 0.23 |
| 3 | 14708918 | rs2280383 | -0.247 | 0.531 | 0.785 | -0.243 | 0.104 | 0.257 | 0.257 | 0.312 | 0.92 | 0.57 | 0.82 | 0.68 | 0.91 | 0.89 |
| 3 | 14709935 | rs3773490 | 1.451 | 0.574 | 0.79 | -0.25 | -0.029 | -0.071 | -0.071 | 0 | 0.61 | 0.16 | 0.38 | 0.46 | 0.56 | 0.66 |
| 3 | 14710588 | rs4684237 | 0.69 | 0.563 | 0.767 | -0.26 | 0 | -0.013 | -0.013 | 0.091 | 0.16 | 0.11 | 0.33 | 0.21 | 0.22 | 0.17 |
| 3 | 14712473 | rs1901 | -0.767 | 0.512 | 0.719 | -0.219 | -0.006 | -0.025 | -0.025 | 0 | 0.32 | 0.46 | 0.46 | 0.23 | 0.35 | 0.23 |
| 3 | 14713058 | rs11717438 | -0.778 | 0.512 | 0.703 | -0.206 | -0.006 | -0.023 | -0.023 | 0 | 0.32 | 0.47 | 0.46 | 0.23 | 0.35 | 0.23 |
| 3 | 14713306 | rs11709673 | -0.303 | 0.517 | 0.667 | -0.206 | -0.001 | -0.001 | -0.001 | 1.223 | 0.28 | 0.27 | 0.42 | 0.23 | 0.34 | 0.22 |
| 3 | 14716371 | rs2975 | 1.29 | 0.518 | 0.544 | -0.202 | -0.055 | -0.112 | -0.112 | 0 | 0.6 | 0.1 | 0.37 | 0.46 | 0.56 | 0.66 |
| 3 | 14721074 | rs1344783 | -0.139 | 0.432 | 0.652 | -0.125 | 0.112 | 0.271 | 0.271 | 0.363 | 0.92 | 0.52 | 0.82 | 0.68 | 0.91 | 0.89 |
| 3 | 14723750 | rs6442468 | -1.108 | 0.603 | 0.752 | -0.11 | 0.067 | 0.179 | 0.179 | 0 | 0.94 | 0.78 | 0.84 | 0.69 | 0.91 | 0.89 |
| 3 | 14724413 | rs9821143 | -0.392 | 0.622 | 0.549 | -0.269 | 0.002 | -0.003 | -0.003 | 0 | 0.29 | 0.28 | 0.44 | 0.23 | 0.37 | 0.22 |
| 3 | 14728880 | rs753412 | -0.11 | 0.276 | 0.164 | -0.215 | 0.004 | 0.027 | 0.027 | 0 | 0.87 | 0.84 | 0.83 | 0.71 | 0.87 | 0.73 |
| 3 | 14730457 | rs871971 | -0.129 | 0.275 | 0.127 | -0.216 | -0.001 | 0.001 | 0.001 | 0 | 0.87 | 0.89 | 0.83 | 0.71 | 0.87 | 0.73 |
| 3 | 14730645 | rs871972 | -0.054 | 0.668 | -0.13 | -0.193 | 0.041 | 0.013 | 0.013 | 0 | 0.24 | 0.39 | 0.38 | 0.25 | 0.3 | 0.19 |
| 3 | 14731808 | rs1432601 | 1.567 | 0.424 | -0.438 | -0.237 | 0.055 | 0.004 | 0.004 | 0 | 0.46 | 0.14 | 0.25 | 0.41 | 0.36 | 0.46 |
| 3 | 14742513 | rs9860601 | 0.606 | 0.304 | 0.148 | 0.246 | -0.004 | 0.05 | 0.05 | 0 | 0.13 | 0.23 | 0.26 | 0.26 | 0.12 | 0.14 |
| 3 | 14745383 | rs4685211 | 1.471 | 0.434 | -0.093 | 0.449 | 0.001 | 0.008 | 0.008 | 0 | 0.47 | 0.42 | 0.46 | 0.32 | 0.42 | 0.28 |
| 3 | 14745571 | rs11128716 | 1.029 | 0.471 | -0.076 | 0.437 | -0.022 | 0.015 | 0.015 | 0 | 0.13 | 0.07 | 0.16 | 0.04 | 0.2 | 0.07 |
| 3 | 14749248 | rs9822010 | 0.002 | 0.345 | -0.032 | 0.43 | -0.002 | 0.03 | 0.03 | 0 | 0.85 | 0.76 | 0.83 | 0.72 | 0.88 | 0.74 |
| 3 | 14750914 | rs7614045 | 1.05 | 0.338 | 0.005 | 0.426 | 0.004 | 0.002 | 0.002 | 0 | 0.68 | 0.6 | 0.65 | 0.68 | 0.66 | 0.66 |
| 3 | 14755572 | rs6765537 | 1.512 | 0.46 | -0.012 | 0.525 | 0.001 | 0.008 | 0.008 | 0 | 0.47 | 0.42 | 0.45 | 0.32 | 0.42 | 0.28 |
| 3 | 14758055 | rs13075489 | -0.136 | 0.634 | -0.552 | 0.527 | 0.006 | 0.076 | 0.076 | 0 | 0.88 | 0.78 | 0.81 | 0.65 | 0.89 | 0.76 |
| 3 | 14765122 | rs13316767 | 0.509 | 0.437 | -1.108 | 0.221 | -0.024 | 0.015 | 0.015 | 0 | 0.13 | 0.07 | 0.16 | 0.04 | 0.2 | 0.07 |
| 3 | 14768534 | rs7636830 | -0.872 | 0.397 | -1.072 | 0.22 | -0.001 | -0.006 | -0.006 | 0.007 | 0.39 | 0.35 | 0.39 | 0.4 | 0.47 | 0.46 |
| 3 | 14768855 | rs4312639 | -0.203 | 0.005 | -0.988 | 0.464 | 0.008 | -0.017 | -0.017 | 0.66 | 0.25 | 0.18 | 0.23 | 0.36 | 0.27 | 0.39 |

| | | | | | | | | | | | | | | | |
|---|---|---|---|---|---|---|---|---|---|---|---|---|---|---|---|
| 3 | 14773267 | rs12496846 | -0.234 | 0.032 | -0.977 | 0.456 | 0.009 | -0.015 | -0.015 | 0 | 0.26 | 0.18 | 0.23 | 0.36 | 0.27 | 0.39 |
| 3 | 14782794 | rs7610462 | -0.869 | -0.037 | -0.599 | 0.143 | -0.029 | -0.03 | -0.03 | 2.283 | 0.11 | 0.25 | 0.03 | 0.03 | 0.02 | 0.15 |
| 3 | 14786651 | rs2568848 | 0.462 | 0.445 | -1.118 | 0.327 | -0.025 | -0.004 | -0.004 | 0.513 | 0.69 | 0.85 | 0.64 | 0.72 | 0.63 | 0.74 |
| 3 | 14796144 | rs17040258 | 0.204 | 0.147 | -1.263 | 0.599 | -0.021 | -0.061 | -0.061 | 0 | 0.11 | 0 | 0.14 | 0.19 | 0.07 | 0.11 |
| 3 | 14798380 | rs11128718 | 0.074 | 0.092 | -1.376 | 0.75 | -0.001 | -0.015 | -0.015 | 0 | 0.17 | 0.15 | 0.22 | 0.34 | 0.17 | 0.45 |
| 3 | 14807375 | rs11128719 | 0.107 | -0.502 | -1.502 | 0.163 | 0.008 | 0.008 | 0.008 | 0 | 0.64 | 0.68 | 0.71 | 0.76 | 0.72 | 0.7 |
| 3 | 14810290 | rs903056 | 0.978 | -0.397 | -2.272 | 0.194 | -0.108 | -0.026 | -0.026 | 0 | 0.4 | 0.06 | 0.43 | 0.33 | 0.52 | 0.2 |
| 3 | 14810854 | rs1687282 | -0.152 | -0.713 | -2.123 | 0.185 | -0.027 | -0.061 | -0.061 | 0.003 | 0.73 | 0.81 | 0.6 | 0.41 | 0.66 | 0.56 |
| 3 | 14811160 | rs2568856 | -0.154 | -0.713 | -2.002 | 0.139 | -0.01 | -0.021 | -0.021 | 0.002 | 0.73 | 0.74 | 0.59 | 0.4 | 0.66 | 0.55 |
| 3 | 14812118 | rs7651825 | 1.335 | -0.353 | -2.53 | 0.005 | -0.119 | -0.006 | -0.006 | 0 | 0.31 | 0.03 | 0.37 | 0.22 | 0.42 | 0.13 |
| 3 | 14812460 | rs6442476 | 0.308 | -0.244 | -2.273 | 0.285 | -0.1 | 0.025 | 0.025 | 0.289 | 0.59 | 0.9 | 0.52 | 0.29 | 0.53 | 0.43 |
| 3 | 14814156 | rs3796306 | 0.928 | 0.423 | -1.742 | 1.032 | -0.045 | -0.041 | -0.041 | 0 | 0.08 | 0 | 0.18 | 0.33 | 0.14 | 0.19 |
| 3 | 14815753 | rs829182 | 0.381 | -0.4 | -1.354 | 0.785 | 0.004 | 0.002 | 0.002 | 0.663 | 0.56 | 0.53 | 0.54 | 0.47 | 0.63 | 0.49 |
| 3 | 14816252 | rs829183 | 0.247 | -0.064 | -1.048 | 0.796 | -0.011 | 0.009 | 0.009 | 0 | 0.73 | 0.64 | 0.76 | 0.68 | 0.78 | 0.56 |
| 3 | 14817362 | rs864361 | -0.268 | 0.471 | -1.032 | 0.748 | 0 | -0.001 | -0.001 | 0.105 | 0.46 | 0.58 | 0.37 | 0.4 | 0.39 | 0.64 |
| 3 | 14817654 | rs853334 | 0.214 | 0.128 | -1.275 | 0.762 | 0.014 | -0.009 | -0.009 | 0 | 0.56 | 0.31 | 0.51 | 0.53 | 0.63 | 0.51 |
| 3 | 14817727 | rs3899614 | 0.142 | 0.419 | -1.761 | 0.438 | 0.037 | 0.066 | 0.066 | 0 | 0.51 | 0.77 | 0.64 | 0.6 | 0.46 | 0.57 |
| 3 | 14818231 | rs829185 | 0.185 | 0.796 | -1.6 | -0.358 | -0.011 | 0.011 | 0.011 | 0 | 0.61 | 0.64 | 0.67 | 0.88 | 0.44 | 0.71 |
| 3 | 14818512 | rs13066658 | 0.923 | 0.938 | -2.423 | -0.577 | -0.069 | 0.059 | 0.059 | 0.73 | 0.24 | 0.1 | 0.2 | 0.18 | 0.35 | 0.16 |
| 3 | 14818716 | rs829186 | -1.241 | 0.711 | -2.001 | 0.221 | 0.01 | 0.147 | 0.147 | 0 | 0.91 | 0.85 | 0.72 | 0.56 | 0.88 | 0.87 |
| 3 | 14818821 | rs9816383 | 0.697 | -0.464 | -1.758 | -0.375 | -0.065 | -0.097 | -0.097 | 0.15 | 0.28 | 0.46 | 0.17 | 0.05 | 0.13 | 0.15 |
| 3 | 14818830 | rs829188 | -0.971 | -0.161 | -1.876 | -0.027 | -0.027 | -0.023 | -0.023 | 0 | 0.67 | 0.74 | 0.52 | 0.38 | 0.55 | 0.72 |
| 3 | 14819652 | rs13072398 | 1.083 | -0.247 | -2.126 | -0.593 | -0.069 | 0.095 | 0.095 | 0 | 0.3 | 0.11 | 0.22 | 0.19 | 0.4 | 0.25 |
| 3 | 14822488 | rs9828560 | 0.874 | -0.268 | -1.976 | -0.595 | -0.002 | -0.005 | -0.005 | 0 | 0.32 | 0.32 | 0.26 | 0.19 | 0.4 | 0.18 |
| 3 | 14824030 | rs11920440 | 1.612 | -0.025 | -2.485 | -0.459 | -0.069 | 0.063 | 0.063 | 0.179 | 0.24 | 0.11 | 0.21 | 0.18 | 0.37 | 0.17 |
| 3 | 14828174 | rs829193 | 1.4 | 0.019 | -2.669 | -0.358 | -0.058 | -0.032 | -0.032 | 0 | 0.16 | 0.01 | 0.22 | 0.13 | 0.23 | 0.31 |
| 3 | 14829349 | rs13100512 | 1.368 | 0.024 | -2.256 | -0.116 | -0.061 | 0.048 | 0.048 | 0.478 | 0.25 | 0.11 | 0.21 | 0.2 | 0.37 | 0.17 |
| 3 | 14830469 | rs13074606 | 1.243 | 0.015 | -2.255 | -0.117 | -0.059 | 0.055 | 0.055 | 0.19 | 0.27 | 0.11 | 0.22 | 0.21 | 0.37 | 0.19 |
| 3 | 14837473 | rs1689549 | 0.936 | -0.392 | -1.941 | 0.7 | -0.032 | -0.053 | -0.053 | 0 | 0.09 | 0.01 | 0.11 | 0.37 | 0.12 | 0.17 |
| 3 | 14841602 | rs192676 | 0.562 | -0.411 | -1.952 | 0.666 | -0.028 | -0.079 | -0.079 | 0.021 | 0.11 | 0.01 | 0.11 | 0.37 | 0.12 | 0.19 |
| 3 | 14845673 | rs9875241 | 1.658 | -0.378 | -1.923 | 0.477 | 0.001 | -0.011 | -0.011 | 0.569 | 0.06 | 0.08 | 0.14 | 0.01 | 0.07 | 0.02 |
| 3 | 14846758 | rs1687310 | -0.907 | -0.939 | -2.014 | 0.253 | 0.042 | 0.004 | 0.004 | 0 | 0.46 | 0.74 | 0.51 | 0.55 | 0.34 | 0.4 |
| 3 | 14849002 | rs1159749 | 0.694 | -0.395 | -1.731 | 0.519 | -0.022 | -0.07 | -0.07 | 0 | 0.11 | 0.01 | 0.07 | 0.31 | 0.11 | 0.08 |
| 3 | 14855787 | rs752078 | 0.395 | -0.534 | -1.385 | 0.129 | 0.002 | -0.025 | -0.025 | 0 | 0.25 | 0.48 | 0.3 | 0.14 | 0.26 | 0.3 |
| 3 | 14856838 | rs731580 | 1.132 | -0.364 | -1.995 | 0.143 | -0.129 | -0.099 | -0.099 | 0 | 0.34 | 0.02 | 0.37 | 0.35 | 0.43 | 0.3 |
| 3 | 14857773 | rs6442478 | -0.699 | -0.368 | -1.574 | 0.262 | -0.002 | -0.069 | -0.069 | 0 | 0.47 | 0.92 | 0.5 | 0.53 | 0.42 | 0.48 |
| 3 | 14865367 | rs10510432 | 0.163 | -0.236 | -0.992 | 0.924 | -0.013 | -0.037 | -0.037 | 1.531 | 0.11 | 0.04 | 0.15 | 0.39 | 0.14 | 0.14 |
| 3 | 14871766 | rs744892 | 0.353 | -0.224 | -1.058 | 0.962 | -0.021 | -0.048 | -0.048 | 0 | 0.1 | 0.02 | 0.14 | 0.39 | 0.13 | 0.14 |
| 3 | 14873794 | rs7653784 | -0.118 | -0.751 | -1.482 | 0.711 | 0.02 | -0.036 | -0.036 | 0.484 | 0.69 | 0.89 | 0.68 | 0.66 | 0.74 | 0.72 |
| 3 | 14877535 | rs1077535 | 0.983 | -1.005 | -1.044 | 0.333 | 0.014 | 0.029 | 0.029 | 0 | 0.5 | 0.53 | 0.48 | 0.26 | 0.55 | 0.54 |
| 3 | 14880436 | rs9846767 | -0.109 | -0.597 | -1.356 | 0.31 | 0.004 | 0.003 | 0.003 | 0 | 0.08 | 0.05 | 0.03 | 0 | 0.08 | 0.04 |
| 3 | 14882553 | rs4684241 | -0.643 | -0.75 | -1.507 | 0.209 | -0.016 | 0.063 | 0.063 | 0.554 | 0.8 | 0.92 | 0.87 | 0.97 | 0.8 | 0.85 |
| 3 | 14886796 | rs1389300 | 0.683 | -0.943 | -1.397 | 0.303 | 0.001 | 0.028 | 0.028 | 0.26 | 0.6 | 0.77 | 0.7 | 0.67 | 0.61 | 0.72 |
| 3 | 14890939 | rs6789614 | 1.579 | -0.961 | -1.208 | 0.373 | 0.006 | -0.016 | -0.016 | 0 | 0.44 | 0.57 | 0.47 | 0.29 | 0.49 | 0.56 |
| 3 | 14892502 | rs17040381 | 0.544 | -0.937 | -1.088 | 0.518 | 0.003 | -0.073 | -0.073 | 0.027 | 0.11 | 0 | 0.15 | 0.22 | 0.09 | 0.06 |
| 3 | 14894140 | rs4685218 | -0.194 | -0.898 | -0.972 | 0.342 | -0.008 | -0.045 | -0.045 | 0 | 0.13 | 0.12 | 0.06 | 0.01 | 0.1 | 0.07 |
| 3 | 14898136 | rs2467473 | -0.647 | -0.9 | -1.065 | 0.637 | 0.005 | -0.008 | -0.008 | 0.089 | 0.21 | 0.14 | 0.25 | 0.37 | 0.17 | 0.2 |
| 3 | 14911406 | rs1656442 | -0.394 | -0.823 | -1.25 | 0.305 | -0.016 | 0.09 | 0.09 | 0.996 | 0.77 | 0.91 | 0.87 | 0.98 | 0.77 | 0.81 |
| 3 | 14914385 | rs1625709 | -0.549 | -1.057 | -0.917 | 0.466 | -0.029 | 0.058 | 0.058 | 0 | 0.19 | 0.28 | 0.24 | 0.38 | 0.13 | 0.19 |
| 3 | 14917526 | rs17040476 | 0.574 | -0.985 | -1.086 | 0.518 | 0.004 | -0.074 | -0.074 | 0 | 0.12 | 0 | 0.15 | 0.22 | 0.09 | 0.06 |
| 3 | 14920105 | rs293924 | -0.039 | -0.966 | -1.042 | 0.451 | 0.001 | 0.005 | 0.005 | 0 | 0.16 | 0.12 | 0.23 | 0.38 | 0.13 | 0.17 |
| 3 | 14923230 | rs13062241 | 1.534 | -1.098 | -0.981 | 0.321 | 0.007 | -0.015 | -0.015 | 0.01 | 0.42 | 0.56 | 0.47 | 0.29 | 0.49 | 0.56 |
| 3 | 14923396 | rs6804922 | 1.015 | -1.009 | -1.48 | 0.321 | 0.016 | 0.046 | 0.046 | 0.749 | 0.51 | 0.66 | 0.52 | 0.3 | 0.55 | 0.58 |
| 3 | 14925256 | rs903055 | -0.199 | -0.768 | -1.553 | 0.248 | -0.006 | 0.076 | 0.076 | 1.118 | 0.75 | 0.89 | 0.86 | 0.97 | 0.76 | 0.81 |
| 3 | 14925821 | rs17040499 | 0.666 | -0.998 | -1.605 | 0.539 | -0.024 | -0.057 | -0.057 | 0 | 0.09 | 0.01 | 0.11 | 0.31 | 0.1 | 0.11 |
| 3 | 14926229 | rs13080281 | 0.596 | -1.121 | -1.476 | 0.599 | 0.025 | -0.031 | -0.031 | 0 | 0.66 | 0.8 | 0.69 | 0.6 | 0.73 | 0.65 |
| 3 | 14928335 | rs2087919 | 0.683 | -0.775 | -1.477 | 0.591 | 0.004 | -0.007 | -0.007 | 0.492 | 0.65 | 0.7 | 0.68 | 0.6 | 0.7 | 0.66 |
| 3 | 14932156 | rs293927 | -0.614 | -0.717 | -1.665 | 0.563 | -0.001 | 0.065 | 0.065 | 0.186 | 0.78 | 0.91 | 0.87 | 0.97 | 0.79 | 0.8 |
| 3 | 14935189 | rs293926 | -0.495 | -0.74 | -1.668 | 0.535 | -0.001 | 0.075 | 0.075 | 0.408 | 0.78 | 0.91 | 0.86 | 0.97 | 0.79 | 0.78 |
| 3 | 14939479 | rs2164356 | 0.616 | -0.668 | -1.622 | 0.655 | 0.007 | -0.019 | -0.019 | 0.685 | 0.67 | 0.76 | 0.69 | 0.59 | 0.71 | 0.63 |
| 3 | 14941152 | rs293909 | 0.452 | -0.477 | -1.622 | 0.933 | 0.008 | -0.007 | -0.007 | 0.298 | 0.06 | 0.04 | 0.07 | 0.19 | 0.04 | 0.1 |
| 3 | 14941447 | rs293908 | -0.404 | -0.462 | -1.626 | 0.755 | -0.006 | 0.068 | 0.068 | 0 | 0.12 | 0.14 | 0.18 | 0.38 | 0.08 | 0.16 |
| 3 | 14942024 | rs2307088 | 0.656 | -0.449 | -1.532 | 0.356 | 0 | 0.052 | 0.052 | 0.491 | 0.46 | 0.33 | 0.48 | 0.3 | 0.53 | 0.53 |
| 3 | 14960062 | rs293915 | 0.748 | -0.312 | -1.722 | 0.365 | -0.051 | -0.01 | -0.01 | 0 | 0.38 | 0.2 | 0.56 | 0.4 | 0.5 | 0.58 |
| 3 | 14960170 | rs293916 | -0.189 | -0.565 | -2.113 | 0.19 | 0.004 | 0.037 | 0.037 | 0 | 0.83 | 0.99 | 0.94 | 0.91 | 0.88 | 0.91 |
| 3 | 14960204 | rs2117776 | 1.869 | -0.616 | -2.149 | 0.354 | -0.033 | -0.048 | -0.048 | 0 | 0.14 | 0.05 | 0.14 | 0.28 | 0.19 | 0.12 |
| 3 | 14964296 | rs7644443 | 0.146 | -0.527 | -1.713 | 0.194 | -0.023 | -0.009 | -0.009 | 0.032 | 0.28 | 0.62 | 0.21 | 0.22 | 0.17 | 0.18 |
| 3 | 14965466 | rs4685234 | 0.438 | 0.052 | -1.763 | 0.643 | 0 | -0.019 | -0.019 | 0 | 0.66 | 0.83 | 0.76 | 0.62 | 0.68 | 0.78 |
| 3 | 14966318 | rs2013496 | 0.301 | 0.18 | -1.242 | 0.27 | -0.007 | 0.042 | 0.042 | 0 | 0.92 | 0.99 | 0.96 | 1 | 0.93 | 0.95 |

| | | | | | | | | | | | | | | | | |
|---|---|---|---|---|---|---|---|---|---|---|---|---|---|---|---|---|
| 3 | 14969197 | rs12491077 | -0.224 | 0.265 | -0.756 | 0.67 | -0.003 | 0.055 | 0.055 | 0 | 0.07 | 0.19 | 0.12 | 0.16 | 0.08 | 0.1 |
| 3 | 14981262 | rs6442490 | -0.657 | 0.093 | -1.153 | 0.174 | 0.019 | 0.025 | 0.025 | 2.367 | 0.34 | 0.77 | 0.27 | 0.26 | 0.25 | 0.27 |
| 3 | 14985201 | rs9663 | 0.563 | 0.026 | -0.833 | -0.008 | 0 | -0.02 | -0.02 | 0.295 | 0.1 | 0.09 | 0.05 | 0 | 0.09 | 0.04 |
| 3 | 14999621 | rs13097792 | -0.126 | -0.143 | -0.071 | -0.029 | 0.008 | -0.017 | -0.017 | 0.133 | 0.13 | 0.03 | 0.12 | 0.14 | 0.15 | 0.12 |
| 3 | 15042036 | rs6805692 | -1.527 | 0.096 | 0.225 | -0.284 | -0.002 | -0.057 | -0.057 | 0 | 0.28 | 0.43 | 0.22 | 0.12 | 0.29 | 0.17 |
| 3 | 15051128 | rs2164358 | -1.037 | -0.365 | -0.212 | -0.675 | 0.011 | 0.021 | 0.021 | 0.037 | 0.93 | 0.99 | 0.95 | 0.98 | 0.93 | 0.96 |
| 3 | 15055905 | rs11716145 | 0.149 | -0.369 | -0.204 | -0.643 | -0.039 | -0.024 | -0.024 | 0 | 0.12 | 0 | 0.12 | 0.12 | 0.18 | 0.12 |
| 3 | 15058956 | rs3773480 | 0.149 | -0.369 | -0.204 | -0.643 | -0.039 | -0.024 | -0.024 | 0 | 0.12 | 0 | 0.12 | 0.12 | 0.18 | 0.12 |
| 3 | 15061708 | rs12489827 | 1.46 | 0.481 | -0.839 | -0.82 | -0.06 | -0.018 | -0.018 | 0 | 0.11 | 0 | 0.21 | 0.11 | 0.15 | 0.1 |
| 3 | 15074769 | rs17477815 | 1.68 | -0.156 | -0.973 | -0.858 | -0.002 | 0 | 0 | 0 | 0.1 | 0.01 | 0.11 | 0.08 | 0.12 | 0.18 |
| 3 | 15075857 | rs6790230 | -1.388 | 0.105 | -0.603 | -0.608 | 0 | -0.037 | -0.037 | 0.069 | 0.33 | 0.52 | 0.29 | 0.23 | 0.32 | 0.28 |
| 3 | 15081352 | rs6442494 | -0.126 | 0.089 | -0.581 | -0.632 | 0.006 | -0.025 | -0.025 | 0.126 | 0.17 | 0.31 | 0.13 | 0.11 | 0.12 | 0.15 |
| 3 | 15091248 | rs10510437 | -0.018 | 0.118 | -0.801 | -0.592 | -0.014 | 0.007 | 0.007 | 0.431 | 0.15 | 0.03 | 0.14 | 0.12 | 0.2 | 0.13 |
| 3 | 15100801 | rs13068576 | -0.025 | 0.129 | -0.923 | -0.587 | -0.014 | 0.007 | 0.007 | 0.495 | 0.16 | 0.03 | 0.14 | 0.12 | 0.2 | 0.13 |
| 3 | 15102111 | rs11925733 | 1.494 | 0.145 | -0.877 | -0.623 | 0.002 | -0.026 | -0.026 | 0.016 | 0.09 | 0.09 | 0.05 | 0 | 0.09 | 0.04 |
| 3 | 15102538 | rs11717371 | -0.013 | 0.133 | -0.897 | -0.612 | -0.014 | 0.007 | 0.007 | 0.495 | 0.16 | 0.03 | 0.14 | 0.12 | 0.2 | 0.13 |
| 3 | 15106915 | rs11706367 | -0.473 | 0.005 | -0.745 | -0.563 | -0.007 | 0.028 | 0.028 | 0 | 0.23 | 0.13 | 0.18 | 0.12 | 0.26 | 0.17 |
| 3 | 15113123 | rs1464390 | -1.407 | -0.183 | -0.587 | -0.629 | -0.002 | -0.011 | -0.011 | 3.277 | 0.34 | 0.63 | 0.29 | 0.23 | 0.3 | 0.28 |
| 3 | 15113519 | rs9820194 | -1.07 | -0.161 | -0.538 | -0.619 | -0.001 | 0.01 | 0.01 | 0 | 0.31 | 0.32 | 0.26 | 0.23 | 0.3 | 0.28 |
| 3 | 15115270 | rs3831 | -0.408 | -0.161 | -0.549 | -0.619 | -0.012 | 0.038 | 0.038 | 0 | 0.23 | 0.1 | 0.18 | 0.12 | 0.26 | 0.17 |
| 3 | 15115478 | rs9830744 | -0.005 | -0.145 | -0.516 | -0.591 | -0.004 | 0.006 | 0.006 | 0.413 | 0.07 | 0.16 | 0.08 | 0.11 | 0.03 | 0.11 |
| 3 | 15124802 | rs480644 | -1.951 | 0.188 | -0.866 | -0.689 | 0.002 | -0.061 | -0.061 | 0 | 0.69 | 0.88 | 0.72 | 0.46 | 0.72 | 0.66 |
| 3 | 15147989 | rs11712564 | -0.962 | 0.364 | -0.726 | -0.573 | 0 | -0.025 | -0.025 | 0 | 0.31 | 0.53 | 0.28 | 0.25 | 0.3 | 0.28 |
| 3 | 15152062 | rs6808450 | -0.235 | 0.265 | -1.166 | -0.549 | -0.002 | -0.002 | -0.002 | 0 | 0.46 | 0.47 | 0.36 | 0.54 | 0.44 | 0.33 |
| 3 | 15152762 | rs6763162 | 1.525 | 0.369 | -0.888 | -0.564 | -0.006 | -0.009 | -0.009 | 0.101 | 0.12 | 0.07 | 0.07 | 0 | 0.13 | 0.05 |
| 3 | 15153519 | rs9845522 | -0.934 | 0.4 | -1.203 | -0.54 | -0.001 | -0.029 | -0.029 | 0 | 0.32 | 0.55 | 0.28 | 0.23 | 0.3 | 0.28 |
| 3 | 15155121 | rs6442499 | -1.355 | 0.381 | -1.237 | -0.548 | 0.005 | 0.022 | 0.022 | 1.246 | 0.34 | 0.75 | 0.31 | 0.23 | 0.3 | 0.28 |
| 3 | 15158151 | rs7639103 | -0.691 | 0.906 | -1.311 | -0.451 | 0.017 | -0.075 | -0.075 | 0 | 0.28 | 0.44 | 0.21 | 0.12 | 0.3 | 0.17 |
| 3 | 15160002 | rs6798961 | 0.252 | 0.875 | -1.444 | -0.338 | -0.004 | 0.001 | 0.001 | 0 | 0.13 | 0.06 | 0.12 | 0.12 | 0.17 | 0.13 |
| 3 | 15160885 | rs7624183 | 0.376 | 1.131 | -1.354 | -0.211 | 0.053 | -0.051 | -0.051 | 0 | 0.19 | 0.46 | 0.17 | 0.12 | 0.17 | 0.16 |
| 3 | 15162577 | rs7625117 | -1.371 | 0.918 | -1.654 | -0.005 | -0.031 | -0.028 | -0.028 | 0 | 0.91 | 0.98 | 0.88 | 0.88 | 0.89 | 0.82 |
| 3 | 15173478 | rs17081778 | 0.447 | 0.787 | -1.365 | 0.103 | -0.027 | -0.031 | -0.031 | 1.361 | 0.09 | 0 | 0.1 | 0.12 | 0.14 | 0.12 |
| 3 | 15185634 | rs3864041 | 0.064 | 0.658 | -1.163 | 0.153 | -0.011 | -0.009 | -0.009 | 0.448 | 0.44 | 0.66 | 0.38 | 0.53 | 0.39 | 0.32 |
| 3 | 15192306 | rs353088 | 0.067 | 0.258 | -0.527 | -0.334 | 0.01 | 0.052 | 0.052 | 0 | 0.39 | 0.26 | 0.31 | 0.15 | 0.36 | 0.26 |
| 3 | 15192862 | rs9858968 | -0.737 | 0.686 | -0.457 | 0.536 | 0.024 | 0.24 | 0.24 | 0 | 0.17 | 0.46 | 0.27 | 0.47 | 0.14 | 0.32 |
| 3 | 15194136 | rs1549983 | 0.761 | 0.773 | -1.046 | 0.196 | -0.006 | -0.009 | -0.009 | 0.281 | 0.59 | 0.27 | 0.43 | 0.53 | 0.59 | 0.49 |
| 3 | 15194949 | rs353091 | -1.173 | -0.382 | -0.677 | 0.255 | 0 | 0 | 0 | 0.381 | 0.33 | 0.66 | 0.48 | 0.25 | 0.31 | 0.3 |
| 3 | 15203172 | rs4685240 | 0.705 | 0.073 | 0.24 | 0.071 | 0.007 | -0.038 | -0.038 | 0 | 0.07 | 0.21 | 0.1 | 0 | 0.08 | 0.07 |
| 3 | 15211394 | rs9827530 | -1.633 | 0.146 | 0.451 | 0.214 | 0.034 | 0.169 | 0.169 | 0 | 0.22 | 0.53 | 0.36 | 0.48 | 0.24 | 0.39 |
| 3 | 15211967 | rs7636348 | -1.657 | 0.124 | 0.451 | 0.199 | 0.032 | 0.162 | 0.162 | 0 | 0.22 | 0.53 | 0.36 | 0.48 | 0.24 | 0.4 |
| 3 | 15215008 | rs4685241 | -1.657 | 0.124 | 0.451 | 0.199 | 0.032 | 0.162 | 0.162 | 0 | 0.22 | 0.53 | 0.36 | 0.48 | 0.24 | 0.4 |
| 3 | 15216429 | rs7639618 | -0.886 | 0.128 | 0.499 | 0.202 | 0.015 | 0.096 | 0.096 | 0 | 0.13 | 0.21 | 0.25 | 0.48 | 0.16 | 0.33 |
| 3 | 15216615 | rs7639807 | 0.817 | 0.027 | 0.459 | 0.155 | 0.008 | -0.037 | -0.037 | 0.182 | 0.07 | 0.2 | 0.1 | 0 | 0.08 | 0.06 |
| 3 | 15216791 | rs7651842 | 0.817 | 0.027 | 0.459 | 0.155 | 0.008 | -0.037 | -0.037 | 0 | 0.07 | 0.2 | 0.1 | 0 | 0.08 | 0.06 |
| 3 | 15224983 | rs353093 | 1.451 | -0.184 | 0.235 | 0.114 | 0 | 0.017 | 0.017 | 3.08 | 0.2 | 0.19 | 0.23 | 0.4 | 0.19 | 0.13 |
| 3 | 15259461 | rs17539401 | 1.111 | -0.336 | 0.327 | 0.125 | -0.008 | 0.02 | 0.02 | 0 | 0.07 | 0.02 | 0.1 | 0 | 0.1 | 0.08 |
| 3 | 15276541 | rs17539942 | 1.159 | -0.389 | 0.251 | 0.082 | -0.019 | 0.034 | 0.034 | 0 | 0.07 | 0.01 | 0.09 | 0 | 0.1 | 0.08 |
| 3 | 15295364 | rs1318937 | -1.245 | -0.292 | 0.402 | 0.555 | -0.011 | 0.214 | 0.214 | 0 | 0.17 | 0.55 | 0.27 | 0.43 | 0.13 | 0.25 |
| 3 | 15299227 | rs6771876 | -1.008 | -0.184 | 0.352 | 0.12 | 0.008 | 0.06 | 0.06 | 4.383 | 0.07 | 0.57 | 0.05 | 0.08 | 0.01 | 0.07 |
| 3 | 15301363 | rs2291853 | -0.75 | -0.03 | -1.075 | -0.018 | -0.087 | -0.056 | -0.056 | 0.469 | 0.67 | 0.89 | 0.49 | 0.62 | 0.57 | 0.77 |
| 3 | 15311325 | rs1287467 | -0.019 | -0.136 | -0.047 | -0.561 | 0 | -0.004 | -0.004 | 0 | 0.66 | 0.62 | 0.47 | 0.19 | 0.69 | 0.53 |
| 3 | 15315029 | rs4036836 | -1.526 | -0.066 | 0.035 | -0.272 | 0.001 | 0.011 | 0.011 | 0.709 | 0.29 | 0.84 | 0.26 | 0.25 | 0.32 | 0.38 |
| 3 | 15318013 | rs988682 | -1.345 | 0.086 | 0.225 | -0.216 | 0.001 | -0.003 | -0.003 | 0.134 | 0.28 | 0.66 | 0.25 | 0.28 | 0.31 | 0.4 |
| 3 | 15322501 | rs6772096 | -1.36 | -0.515 | 0.531 | 0.14 | -0.036 | 0.042 | 0.042 | 0 | 0.22 | 0.51 | 0.15 | 0.29 | 0.19 | 0.31 |
| 3 | 15326527 | rs2729689 | 0.614 | -0.629 | 0.103 | -0.199 | 0.047 | 0.125 | 0.125 | 1.639 | 0.54 | 0.24 | 0.51 | 0.08 | 0.68 | 0.45 |
| 3 | 15329619 | rs2633979 | 0.631 | -0.648 | -0.026 | -0.208 | 0.051 | 0.158 | 0.158 | 1.707 | 0.54 | 0.22 | 0.49 | 0.08 | 0.68 | 0.45 |
| 3 | 15332310 | rs1106130 | -0.243 | -0.695 | -0.049 | 0.078 | 0.011 | -0.022 | -0.022 | 0 | 0.7 | 0.81 | 0.61 | 0.36 | 0.78 | 0.71 |
| 3 | 15332733 | rs17040960 | -0.975 | -0.372 | 0.96 | 0.811 | -0.002 | 0.015 | 0.015 | 0 | 0.1 | 0.12 | 0.06 | 0.24 | 0.09 | 0.22 |
| 3 | 15333551 | rs7636739 | -1.687 | -0.36 | 0.533 | 0.816 | -0.021 | 0.018 | 0.018 | 2.028 | 0.16 | 0.75 | 0.13 | 0.24 | 0.1 | 0.24 |
| 3 | 15340663 | rs6442511 | -1.507 | -0.111 | 0.815 | 0.991 | -0.011 | 0.128 | 0.128 | 0 | 0.14 | 0.59 | 0.13 | 0.29 | 0.13 | 0.21 |
| 3 | 15340747 | rs6442512 | -1.409 | -0.118 | 1.221 | 1.037 | 0.024 | 0.256 | 0.256 | 0 | 0.1 | 0.4 | 0.1 | 0.29 | 0.1 | 0.2 |
| 3 | 15342850 | rs9831807 | -0.863 | -0.036 | 0.622 | 0.99 | 0.004 | -0.006 | -0.006 | 0 | 0.39 | 0.57 | 0.39 | 0.61 | 0.35 | 0.43 |
| 3 | 15345924 | rs9828229 | -0.678 | -0.115 | 0.929 | 1.04 | 0 | 0.007 | 0.007 | 0 | 0.35 | 0.36 | 0.36 | 0.6 | 0.34 | 0.41 |
| 3 | 15348823 | rs6801433 | -0.868 | -0.505 | 0.67 | 0.724 | -0.001 | 0 | 0 | 0 | 0.42 | 0.43 | 0.4 | 0.57 | 0.39 | 0.46 |
| 3 | 15349552 | rs2010265 | -0.961 | -0.744 | 0.827 | 0.444 | -0.018 | -0.048 | -0.048 | 0 | 0.08 | 0.25 | 0.06 | 0.01 | 0.04 | 0.11 |
| 3 | 15351965 | rs6793635 | -0.833 | -0.812 | 0.041 | 0.624 | 0.019 | 0.026 | 0.026 | 0 | 0.52 | 0.61 | 0.36 | 0.4 | 0.43 | 0.43 |
| 3 | 15354031 | rs2729683 | -0.375 | 0.286 | -0.302 | 0.629 | -0.037 | -0.043 | -0.043 | 0.006 | 0.88 | 0.93 | 0.74 | 0.67 | 0.81 | 0.64 |
| 3 | 15354258 | rs2688672 | -0.004 | 0.242 | -0.327 | 0.605 | -0.043 | -0.061 | -0.061 | 0 | 0.86 | 0.93 | 0.74 | 0.67 | 0.8 | 0.64 |
| 3 | 15354946 | rs2729682 | 1.73 | 0.168 | -0.208 | 0.517 | -0.04 | 0.012 | 0.012 | 0.162 | 0.33 | 0.17 | 0.36 | 0.27 | 0.37 | 0.25 |

| | | | | | | | | | | | | | | | | |
|---|---|---|---|---|---|---|---|---|---|---|---|---|---|---|---|---|
| 3 | 15359819 | rs12638270 | 1.614 | 0.165 | -0.209 | 0.512 | -0.016 | 0.013 | 0.013 | 0.119 | 0.34 | 0.21 | 0.38 | 0.3 | 0.37 | 0.25 |
| 3 | 15371003 | rs9854364 | 0.453 | -0.532 | -0.238 | 0.523 | 0.013 | 0.02 | 0.02 | 0 | 0.22 | 0.14 | 0.2 | 0.13 | 0.21 | 0.19 |
| 3 | 15371819 | rs3773464 | -1.554 | -0.315 | -0.273 | 0.516 | -0.01 | 0.004 | 0.004 | 0 | 0.63 | 0.63 | 0.59 | 0.67 | 0.53 | 0.74 |
| 3 | 15375822 | rs17041032 | -0.319 | -0.57 | -0.629 | 0.418 | -0.005 | -0.022 | -0.022 | 0 | 0.08 | 0.11 | 0.05 | 0 | 0.04 | 0.09 |
| 3 | 15380888 | rs17482502 | 0.694 | -0.443 | -0.688 | 0.748 | 0 | -0.018 | -0.018 | 0 | 0.16 | 0.04 | 0.08 | 0.24 | 0.15 | 0.1 |
| 3 | 15381404 | rs17041034 | 0.1 | -0.263 | -0.531 | 1.007 | -0.002 | 0.023 | 0.023 | 0 | 0.06 | 0.09 | 0.18 | 0.14 | 0.04 | 0.08 |
| 3 | 15391894 | rs11926398 | -0.751 | -0.403 | -0.937 | 0.822 | 0.097 | 0.015 | 0.015 | 0 | 0.5 | 0.8 | 0.46 | 0.64 | 0.35 | 0.37 |
| 3 | 15410365 | rs6766086 | -0.652 | -0.79 | -0.235 | 0.718 | -0.021 | -0.006 | -0.006 | 0 | 0.27 | 0.58 | 0.2 | 0.33 | 0.15 | 0.17 |
| 3 | 15427117 | rs2344675 | -0.606 | -0.445 | -0.958 | -0.014 | 0.004 | -0.04 | -0.04 | 0 | 0.82 | 0.74 | 0.76 | 0.95 | 0.78 | 0.9 |
| 3 | 15429168 | rs2219251 | -0.632 | -0.429 | -1.532 | -0.189 | 0.013 | 0.001 | 0.001 | 0 | 0.52 | 0.69 | 0.48 | 0.57 | 0.47 | 0.45 |
| 3 | 15431443 | rs6807885 | 0.074 | -0.745 | -0.755 | -0.296 | 0.007 | 0.033 | 0.033 | 1.274 | 0.1 | 0.39 | 0.1 | 0.08 | 0.05 | 0.07 |
| 3 | 15448448 | rs6764006 | 0.784 | -0.768 | -0.185 | -0.291 | -0.007 | 0.003 | 0.003 | 0.086 | 0.07 | 0.13 | 0.08 | 0.07 | 0.04 | 0.07 |
| 3 | 15450400 | rs13075694 | -0.831 | -0.736 | -0.25 | -0.363 | 0.013 | 0.001 | 0.001 | 0 | 0.52 | 0.69 | 0.48 | 0.57 | 0.47 | 0.45 |
| 3 | 15455717 | rs2290537 | 0.205 | -0.771 | -0.228 | -0.401 | 0.007 | 0.001 | 0.001 | 0.061 | 0.4 | 0.26 | 0.37 | 0.49 | 0.42 | 0.38 |
| 3 | 15456836 | rs1869853 | 0.091 | -0.73 | 0.163 | -0.393 | -0.002 | 0.01 | 0.01 | 0.411 | 0.09 | 0.24 | 0.09 | 0.06 | 0.05 | 0.07 |
| 3 | 15459328 | rs11919472 | -0.383 | -0.731 | 0.248 | -0.393 | 0.014 | 0.039 | 0.039 | 0.969 | 0.1 | 0.36 | 0.1 | 0.08 | 0.05 | 0.07 |
| 3 | 15469001 | rs2290536 | -1.802 | -0.7 | -0.23 | -0.43 | -0.054 | 0.065 | 0.065 | 0 | 0.87 | 0.99 | 0.79 | 0.98 | 0.82 | 0.92 |
| 3 | 15477228 | rs7647032 | 2.098 | -0.581 | -0.198 | -0.269 | -0.081 | -0.152 | -0.152 | 0 | 0.3 | 0.04 | 0.28 | 0.38 | 0.31 | 0.45 |
| 3 | 15480726 | rs17041129 | 0.364 | -0.71 | 0.481 | 0.144 | -0.007 | 0.002 | 0.002 | 0.086 | 0.07 | 0.13 | 0.08 | 0.06 | 0.05 | 0.07 |
| 3 | 15483830 | rs1048608 | 0.629 | -0.771 | 0.378 | 0.08 | -0.043 | -0.02 | -0.02 | 0 | 0.24 | 0.13 | 0.28 | 0.29 | 0.33 | 0.28 |
| 3 | 15491756 | rs3274 | -0.507 | -0.924 | 0.44 | 0.02 | 0.004 | -0.003 | -0.003 | 0.103 | 0.39 | 0.35 | 0.35 | 0.5 | 0.39 | 0.36 |
| 3 | 15492636 | rs10154896 | 0.522 | -0.825 | 0.583 | 0.033 | 0.006 | -0.021 | -0.021 | 0 | 0.05 | 0.17 | 0.04 | 0 | 0.03 | 0.02 |
| 3 | 15495926 | rs1079067 | 0.707 | -0.809 | 0.571 | 0.037 | -0.028 | -0.02 | -0.02 | 3.174 | 0.22 | 0.15 | 0.26 | 0.31 | 0.31 | 0.26 |
| 3 | 15500295 | rs12634346 | -0.918 | -0.942 | 0.589 | 0.147 | 0.001 | 0 | 0 | 0 | 0.47 | 0.47 | 0.52 | 0.6 | 0.44 | 0.42 |
| 3 | 15500933 | rs13060973 | -0.776 | -0.981 | 0.667 | 0.13 | 0.008 | -0.002 | -0.002 | 0 | 0.45 | 0.38 | 0.51 | 0.6 | 0.44 | 0.42 |
| 3 | 15501686 | rs17041167 | 1.448 | -1.001 | 0.568 | 0.122 | 0.01 | 0.002 | 0.002 | 0.013 | 0.16 | 0.11 | 0.08 | 0.19 | 0.09 | 0.1 |
| 3 | 15503055 | rs4395387 | -1.603 | -0.996 | 0.638 | 0.099 | 0.016 | -0.008 | -0.008 | 0 | 0.53 | 0.77 | 0.57 | 0.6 | 0.46 | 0.43 |
| 3 | 15512276 | rs2305618 | -0.184 | -0.874 | 0.53 | 0.046 | -0.023 | -0.04 | -0.04 | 0.011 | 0.09 | 0.21 | 0.05 | 0.01 | 0.03 | 0.02 |
| 3 | 15516910 | rs3816526 | -0.365 | -0.779 | 0.476 | 0.058 | -0.001 | -0.004 | -0.004 | 0 | 0.37 | 0.33 | 0.34 | 0.5 | 0.4 | 0.36 |
| 3 | 15518889 | rs13074638 | -0.732 | -0.793 | 0.563 | 0.05 | 0.002 | -0.002 | -0.002 | 0 | 0.41 | 0.58 | 0.36 | 0.52 | 0.4 | 0.37 |
| 3 | 15519262 | rs3773448 | -0.677 | -0.699 | -0.048 | 0.083 | 0 | 0 | 0 | 0 | 0.4 | 0.4 | 0.35 | 0.5 | 0.4 | 0.36 |
| 3 | 15520639 | rs2290538 | 0.698 | -0.489 | -0.244 | 0.178 | -0.028 | -0.021 | -0.021 | 0 | 0.22 | 0.15 | 0.25 | 0.3 | 0.3 | 0.26 |
| 3 | 15520838 | rs750387 | -0.055 | -0.749 | 0.027 | 0.389 | -0.003 | -0.016 | -0.016 | 0 | 0.07 | 0.08 | 0.03 | 0 | 0.03 | 0.02 |
| 3 | 15530595 | rs6773439 | -0.749 | -0.935 | 0.131 | 0.497 | -0.06 | 0.07 | 0.07 | 0 | 0.28 | 0.48 | 0.26 | 0.52 | 0.17 | 0.25 |
| 3 | 15537992 | rs9847828 | -1.425 | -0.573 | 0.146 | 0.445 | 0 | 0.037 | 0.037 | 0 | 0.26 | 0.6 | 0.41 | 0.42 | 0.21 | 0.27 |
| 3 | 15542290 | rs6777812 | -0.779 | -0.615 | 0.308 | 0.393 | -0.039 | -0.06 | -0.06 | 0 | 0.29 | 0.44 | 0.15 | 0.2 | 0.22 | 0.17 |
| 3 | 15545105 | rs9823378 | -0.621 | -0.626 | 0.391 | 0.386 | -0.03 | -0.053 | -0.053 | 0.444 | 0.27 | 0.4 | 0.15 | 0.19 | 0.22 | 0.17 |
| 3 | 15547970 | rs983289 | -1.972 | -0.676 | 0.433 | 0.368 | 0.007 | -0.009 | -0.009 | 0 | 0.39 | 0.63 | 0.47 | 0.62 | 0.33 | 0.37 |
| 3 | 15555270 | rs2124493 | 0.524 | -0.688 | 0.429 | 0.371 | 0.022 | 0.008 | 0.008 | 0 | 0.17 | 0.08 | 0.07 | 0.19 | 0.15 | 0.11 |
| 3 | 15559169 | rs12629790 | -0.998 | -0.562 | 0.389 | 0.447 | 0.016 | 0.127 | 0.127 | 0 | 0.1 | 0.2 | 0.32 | 0.43 | 0.11 | 0.21 |
| 3 | 15564453 | rs989682 | -0.846 | -0.647 | -0.04 | 0.12 | -0.001 | 0.001 | 0.001 | 0 | 0.33 | 0.62 | 0.3 | 0.52 | 0.28 | 0.32 |
| 3 | 15589591 | rs12633887 | -1.511 | -1.065 | -0.316 | -0.086 | 0.034 | -0.021 | -0.021 | 0 | 0.42 | 0.7 | 0.47 | 0.61 | 0.32 | 0.39 |
| 3 | 15599405 | rs11706669 | 1.285 | -1.176 | -0.149 | -0.044 | 0.006 | 0.001 | 0.001 | 0 | 0.14 | 0.08 | 0.07 | 0.17 | 0.12 | 0.12 |
| 3 | 15607002 | rs10510440 | 0.727 | -1.197 | -0.108 | 0.295 | 0 | 0.048 | 0.048 | 0 | 0.08 | 0.11 | 0.15 | 0.36 | 0.08 | 0.18 |
| 3 | 15610070 | rs905650 | 1.364 | -1.012 | -0.003 | 0.322 | 0.01 | 0.001 | 0.001 | 0 | 0.13 | 0.07 | 0.06 | 0.17 | 0.12 | 0.13 |
| 3 | 15648004 | rs7651039 | -1.522 | -0.76 | -0.312 | 0.073 | -0.061 | -0.001 | -0.001 | 0 | 0.57 | 0.9 | 0.63 | 0.68 | 0.52 | 0.51 |
| 3 | 15663060 | rs2455826 | -0.025 | -0.806 | 0.191 | 0.314 | 0.004 | 0.006 | 0.006 | 0 | 0.3 | 0.26 | 0.17 | 0.23 | 0.25 | 0.19 |
| 3 | 15666856 | rs2455823 | -0.421 | -0.717 | 0.239 | 0.376 | -0.002 | 0.007 | 0.007 | 0 | 0.33 | 0.37 | 0.3 | 0.56 | 0.34 | 0.36 |
| 3 | 15671026 | rs9310479 | -0.776 | -0.61 | 0.746 | 0.423 | 0.13 | 0.347 | 0.347 | 0 | 0.05 | 0.27 | 0.16 | 0.33 | 0.1 | 0.17 |
| 3 | 15685137 | rs7640807 | -0.901 | -0.876 | 0.337 | 0.319 | 0.027 | -0.009 | -0.009 | 0.46 | 0.4 | 0.83 | 0.51 | 0.67 | 0.35 | 0.43 |
| 3 | 15686776 | rs3817641 | -0.274 | -0.802 | 0.352 | 0.503 | 0.001 | 0.067 | 0.067 | 0.32 | 0.05 | 0.07 | 0.13 | 0.38 | 0.09 | 0.17 |
| 3 | 15689118 | rs2470531 | 1.03 | -0.964 | 0.173 | 0.167 | 0.039 | -0.045 | -0.045 | 0 | 0.55 | 0.13 | 0.64 | 0.37 | 0.63 | 0.59 |
| 3 | 15690025 | rs2455853 | -0.201 | -0.979 | 0.47 | 0.159 | 0.003 | 0.003 | 0.003 | 0 | 0.68 | 0.71 | 0.83 | 0.77 | 0.74 | 0.8 |
| 3 | 15694088 | rs12490588 | 0.642 | -0.899 | 0.463 | 0.215 | 0.015 | 0.12 | 0.12 | 0.019 | 0.07 | 0.13 | 0.14 | 0.4 | 0.1 | 0.19 |
| 3 | 15696909 | rs2470532 | 1.026 | -0.387 | 0.092 | -0.571 | 0.004 | -0.001 | -0.001 | 0 | 0.5 | 0.11 | 0.6 | 0.35 | 0.57 | 0.51 |
| 3 | 15701589 | rs2455800 | 0.53 | -0.496 | 0.36 | -0.572 | 0.001 | -0.001 | -0.001 | 0 | 0.54 | 0.44 | 0.63 | 0.37 | 0.58 | 0.52 |
| 3 | 15701857 | rs2455801 | 0.555 | -0.352 | 0.274 | -0.568 | 0.005 | -0.001 | -0.001 | 0 | 0.54 | 0.36 | 0.63 | 0.37 | 0.58 | 0.52 |
| 3 | 15706124 | rs924753 | 0.407 | -0.36 | 0.401 | -0.616 | -0.004 | -0.002 | -0.002 | 0 | 0.55 | 0.46 | 0.55 | 0.35 | 0.61 | 0.5 |
| 3 | 15707445 | rs2455807 | -0.035 | -0.42 | 0.694 | -0.707 | 0.011 | 0.016 | 0.016 | 0 | 0.63 | 0.71 | 0.81 | 0.94 | 0.7 | 0.74 |
| 3 | 15716898 | rs2470537 | 0.656 | -0.234 | 0.488 | -0.772 | 0.012 | 0.002 | 0.002 | 0.024 | 0.53 | 0.34 | 0.63 | 0.35 | 0.6 | 0.48 |
| 3 | 15719279 | rs17041442 | 3.204 | 0.012 | -0.331 | -1.104 | -0.041 | -0.095 | -0.095 | 0 | 0.15 | 0 | 0.09 | 0.21 | 0.15 | 0.2 |
| 3 | 15737610 | rs2470547 | 0.57 | -0.028 | -0.05 | -1.124 | 0.028 | -0.005 | -0.005 | 0.326 | 0.52 | 0.16 | 0.61 | 0.35 | 0.6 | 0.48 |
| 3 | 15737698 | rs2470549 | 0.57 | -0.028 | -0.05 | -1.124 | 0.028 | -0.005 | -0.005 | 0 | 0.52 | 0.16 | 0.61 | 0.35 | 0.6 | 0.48 |
| 3 | 15752830 | rs2455833 | 0.821 | 0 | 0.221 | -1.103 | 0.02 | -0.019 | -0.019 | 0.731 | 0.51 | 0.1 | 0.6 | 0.35 | 0.6 | 0.48 |
| 3 | 15759491 | rs2470518 | 0.163 | 0.002 | 0.626 | -1.155 | -0.01 | -0.079 | -0.079 | 2.728 | 0.61 | 0.54 | 0.79 | 0.91 | 0.69 | 0.72 |
| 3 | 15797454 | rs4128307 | 0.734 | 0.071 | 0.77 | -1 | 0.011 | 0.192 | 0.192 | 0 | 0.09 | 0.17 | 0.16 | 0.58 | 0.09 | 0.22 |
| 3 | 15800740 | rs7617039 | 0.84 | 0.015 | 0.72 | -1.112 | 0.022 | 0.013 | 0.013 | 0.019 | 0.51 | 0.19 | 0.61 | 0.33 | 0.6 | 0.5 |
| 3 | 15805224 | rs6442549 | -1.2 | -0.249 | 0.758 | -1.41 | -0.007 | 0.031 | 0.031 | 0 | 0.81 | 0.89 | 0.87 | 0.94 | 0.77 | 0.84 |
| 3 | 15810605 | rs6442551 | 0.088 | -0.194 | 0.836 | -1.341 | 0.012 | 0.037 | 0.037 | 0 | 0.64 | 0.75 | 0.81 | 0.94 | 0.69 | 0.74 |

| | | | | | | | | | | | | | | | | |
|---|---|---|---|---|---|---|---|---|---|---|---|---|---|---|---|---|
| 3 | 15872730 | rs11923217 | 0.399 | -0.141 | 0.545 | -1.294 | -0.01 | -0.078 | -0.078 | 0 | 0.61 | 0.53 | 0.79 | 0.91 | 0.69 | 0.72 |
| 3 | 15873052 | rs11128759 | 1.315 | 0.022 | 0.265 | -1.053 | 0.002 | 0.125 | 0.125 | 0.226 | 0.08 | 0.09 | 0.15 | 0.56 | 0.09 | 0.22 |
| 3 | 15906997 | rs13093693 | 2.053 | 0.333 | -0.151 | -1.311 | -0.044 | 0.045 | 0.045 | 0 | 0.12 | 0.04 | 0.23 | 0 | 0.23 | 0.09 |
| 3 | 15908301 | rs17041602 | 0.342 | 0.097 | 0.096 | -1.307 | -0.006 | 0.043 | 0.043 | 0.933 | 0.23 | 0.3 | 0.21 | 0.59 | 0.18 | 0.3 |
| 3 | 15927193 | rs6442553 | -0.856 | 0.097 | -0.29 | -1.403 | -0.004 | 0.032 | 0.032 | 0 | 0.78 | 0.88 | 0.87 | 0.93 | 0.78 | 0.84 |
| 3 | 15935841 | rs753196 | 0.288 | 0.065 | 0.004 | -1.414 | -0.017 | 0.123 | 0.123 | 0.575 | 0.24 | 0.43 | 0.22 | 0.59 | 0.19 | 0.3 |
| 3 | 15939142 | rs4684263 | 0.963 | 0.557 | 0.194 | -1.843 | -0.071 | 0.096 | 0.096 | 0 | 0.19 | 0.04 | 0.24 | 0 | 0.27 | 0.12 |
| 3 | 15939311 | rs1466072 | -0.241 | 0.089 | 0.792 | -1.334 | -0.003 | 0.029 | 0.029 | 0 | 0.78 | 0.87 | 0.87 | 0.93 | 0.78 | 0.84 |
| 3 | 15963163 | rs4685277 | -0.119 | 0.14 | 0.291 | -1.303 | -0.001 | -0.021 | -0.021 | 0 | 0.78 | 0.78 | 0.87 | 0.93 | 0.8 | 0.84 |
| 3 | 15963751 | rs6442560 | 0.144 | 0.158 | 0.118 | -1.292 | -0.008 | -0.081 | -0.081 | 0 | 0.75 | 0.62 | 0.85 | 0.92 | 0.8 | 0.82 |
| 3 | 15978955 | rs9310490 | -0.528 | 0.105 | -0.162 | -0.865 | 0.001 | 0.091 | 0.091 | 0.017 | 0.19 | 0.54 | 0.22 | 0.66 | 0.15 | 0.24 |
| 3 | 15981306 | rs6769112 | 1.1 | 0.363 | -0.612 | -1.248 | -0.065 | 0.081 | 0.081 | 0 | 0.17 | 0.05 | 0.27 | 0.02 | 0.29 | 0.11 |
| 3 | 15981946 | rs6442562 | -0.643 | -0.008 | 0.043 | -0.629 | 0.006 | 0.116 | 0.116 | 0 | 0.19 | 0.53 | 0.22 | 0.66 | 0.15 | 0.24 |
| 3 | 15990231 | rs6775519 | -1.172 | -0.206 | -0.314 | -0.864 | 0.011 | -0.002 | -0.002 | 0.361 | 0.34 | 0.72 | 0.29 | 0.71 | 0.25 | 0.37 |
| 3 | 16000477 | rs35067774 | 0.255 | -0.375 | 0.258 | -0.478 | 0.005 | 0.211 | 0.211 | 0 | 0.13 | 0.37 | 0.18 | 0.63 | 0.11 | 0.27 |
| 3 | 16004488 | rs1869627 | -0.085 | -0.507 | -0.091 | -0.865 | 0 | -0.025 | -0.025 | 0.015 | 0.62 | 0.55 | 0.63 | 0.26 | 0.62 | 0.45 |
| 3 | 16009044 | rs1454776 | 0.948 | -0.56 | -0.545 | -0.893 | -0.015 | 0.107 | 0.107 | 0 | 0.45 | 0.26 | 0.54 | 0.24 | 0.53 | 0.33 |
| 3 | 16018470 | rs6442565 | -0.197 | -0.506 | -0.624 | -0.938 | 0 | 0.094 | 0.094 | 0 | 0.63 | 0.86 | 0.77 | 0.86 | 0.65 | 0.62 |
| 3 | 16019023 | rs6806574 | 1.016 | -0.504 | -0.729 | -0.955 | 0.011 | 0.012 | 0.012 | 0 | 0.11 | 0.06 | 0.04 | 0 | 0.07 | 0.09 |
| 3 | 16036830 | rs6770617 | -0.871 | -0.386 | -0.21 | -0.803 | -0.001 | 0.001 | 0.001 | 0 | 0.74 | 0.79 | 0.8 | 0.85 | 0.73 | 0.71 |
| 3 | 16038735 | rs6801560 | -0.569 | -0.369 | -0.076 | -0.787 | 0.002 | -0.02 | -0.02 | 0 | 0.72 | 0.67 | 0.77 | 0.84 | 0.71 | 0.67 |
| 3 | 16042244 | rs17041715 | 0.45 | -0.385 | -0.07 | -0.736 | -0.007 | 0.085 | 0.085 | 0.031 | 0.14 | 0.24 | 0.18 | 0.6 | 0.11 | 0.27 |
| 3 | 16052206 | rs2346283 | -0.443 | -0.117 | -0.109 | -0.585 | 0.007 | -0.013 | -0.013 | 0 | 0.72 | 0.6 | 0.71 | 0.81 | 0.71 | 0.67 |
| 3 | 16068354 | rs9815997 | 0.879 | -0.279 | -1.091 | -0.746 | -0.033 | 0.08 | 0.08 | 0 | 0.39 | 0.24 | 0.47 | 0.21 | 0.49 | 0.3 |
| 3 | 16073266 | rs2730367 | 0.002 | -0.261 | -0.992 | -0.655 | -0.042 | 0.029 | 0.029 | 0 | 0.3 | 0.53 | 0.23 | 0.62 | 0.2 | 0.36 |
| 3 | 16077847 | rs2730335 | 0.082 | -0.19 | -1.396 | -0.617 | -0.022 | 0.01 | 0.01 | 0 | 0.31 | 0.59 | 0.24 | 0.62 | 0.2 | 0.36 |
| 3 | 16082493 | rs1598255 | -0.14 | 0.683 | -1.586 | -0.106 | -0.011 | 0.01 | 0.01 | 0 | 0.74 | 0.86 | 0.73 | 0.82 | 0.7 | 0.7 |
| 3 | 16090103 | rs17041810 | -0.238 | 1.032 | -2.008 | 0.097 | -0.055 | -0.024 | -0.024 | 0.029 | 0.25 | 0.01 | 0.29 | 0.15 | 0.25 | 0.38 |
| 3 | 16093524 | rs2603764 | -0.297 | 1.006 | -1.652 | -0.051 | -0.002 | 0.003 | 0.003 | 0 | 0.12 | 0.11 | 0.26 | 0.1 | 0.12 | 0.1 |
| 3 | 16093862 | rs6766571 | 0.148 | 1.3 | -1.495 | 0.213 | 0.004 | -0.028 | -0.028 | 0 | 0.24 | 0.12 | 0.4 | 0.31 | 0.22 | 0.36 |
| 3 | 16096066 | rs897204 | -0.664 | 0.84 | -1.14 | 0.318 | 0.011 | 0.03 | 0.03 | 0 | 0.43 | 0.66 | 0.62 | 0.66 | 0.39 | 0.58 |
| 3 | 16098470 | rs7628888 | -0.931 | 0.79 | -1.059 | 0.448 | 0.012 | 0.035 | 0.035 | 0 | 0.46 | 0.66 | 0.62 | 0.68 | 0.4 | 0.56 |
| 3 | 16100768 | rs744243 | 1.162 | 0.633 | -1.413 | -0.316 | 0.005 | 0.012 | 0.012 | 0 | 0.22 | 0.14 | 0.11 | 0 | 0.2 | 0.09 |
| 3 | 16101104 | rs1454769 | 0.654 | 0.505 | -0.977 | 0.543 | -0.055 | -0.036 | -0.036 | 0.277 | 0.24 | 0.02 | 0.24 | 0.17 | 0.21 | 0.34 |
| 3 | 16101186 | rs1454768 | -0.337 | 0.542 | -0.866 | 0.233 | -0.009 | -0.01 | -0.01 | 1.603 | 0.35 | 0.58 | 0.53 | 0.66 | 0.28 | 0.49 |
| 3 | 16105565 | rs4434130 | -0.676 | 0.394 | -0.024 | -0.071 | -0.005 | -0.006 | -0.006 | 0 | 0.42 | 0.55 | 0.53 | 0.38 | 0.35 | 0.43 |
| 3 | 16109760 | rs6442578 | -0.776 | 0.388 | -0.024 | -0.074 | 0.008 | 0.01 | 0.01 | 0 | 0.42 | 0.65 | 0.54 | 0.38 | 0.35 | 0.43 |
| 3 | 16110666 | rs1454765 | -0.135 | 0.418 | 0.081 | -0.089 | -0.006 | -0.001 | -0.001 | 0 | 0.34 | 0.37 | 0.47 | 0.34 | 0.27 | 0.42 |
| 3 | 16122017 | rs11916777 | -0.332 | 0.707 | -0.432 | 0.167 | -0.005 | -0.008 | -0.008 | 0 | 0.4 | 0.54 | 0.53 | 0.38 | 0.35 | 0.43 |
| 3 | 16133383 | rs10865738 | -0.375 | 0.803 | -0.812 | 0.202 | -0.001 | -0.002 | -0.002 | 0 | 0.4 | 0.46 | 0.53 | 0.38 | 0.35 | 0.43 |
| 3 | 16139112 | rs4234534 | -0.002 | 0.94 | -1.048 | 0.167 | 0 | 0 | 0 | 0.278 | 0.56 | 0.48 | 0.38 | 0.32 | 0.59 | 0.48 |
| 3 | 16140475 | rs12494204 | 0.337 | 1.255 | -1.405 | 0.22 | -0.025 | 0.011 | 0.011 | 0 | 0.07 | 0.01 | 0.07 | 0.05 | 0.11 | 0.06 |
| 3 | 16140831 | rs9861567 | -0.536 | 0.99 | -1.154 | 0.531 | -0.001 | 0.011 | 0.011 | 1.342 | 0.73 | 0.71 | 0.58 | 0.7 | 0.75 | 0.68 |
| 3 | 16141464 | rs4504158 | 0.338 | 0.95 | -1.818 | 0.162 | -0.016 | -0.026 | -0.026 | 0 | 0.28 | 0.1 | 0.17 | 0.26 | 0.29 | 0.3 |
| 3 | 16146243 | rs12714923 | 0.253 | 1.275 | -1.123 | -0.096 | 0.004 | -0.046 | -0.046 | 0.406 | 0.14 | 0.14 | 0.09 | 0 | 0.17 | 0.12 |
| 3 | 16146964 | rs4419364 | -0.687 | 1.452 | -1.232 | 0.347 | -0.006 | -0.007 | -0.007 | 0.107 | 0.43 | 0.64 | 0.51 | 0.44 | 0.55 | 0.48 |
| 3 | 16147937 | rs4643692 | -1.103 | 1.143 | -1.649 | 0.353 | -0.024 | -0.064 | -0.064 | 0 | 0.83 | 0.95 | 0.69 | 0.74 | 0.81 | 0.8 |
| 3 | 16148056 | rs6776952 | -0.494 | 1.6 | -1.58 | 0.274 | -0.034 | -0.035 | -0.035 | 0.374 | 0.78 | 0.9 | 0.63 | 0.71 | 0.72 | 0.76 |
| 3 | 16150852 | rs4423748 | -0.139 | 0.929 | -1.69 | -0.43 | 0.009 | 0.069 | 0.069 | 0.388 | 0.5 | 0.63 | 0.51 | 0.68 | 0.56 | 0.54 |
| 3 | 16151388 | rs6783201 | 0.495 | 1.154 | -1.084 | -0.121 | 0.004 | 0.015 | 0.015 | 0.105 | 0.25 | 0.26 | 0.31 | 0.37 | 0.31 | 0.22 |
| 3 | 16160879 | rs13060631 | -0.523 | 1.103 | -1.077 | -0.114 | 0.006 | 0.055 | 0.055 | 0.225 | 0.49 | 0.63 | 0.51 | 0.63 | 0.56 | 0.61 |
| 3 | 16162533 | rs4234535 | 1.69 | 0.953 | -1.181 | -0.415 | 0.024 | 0.055 | 0.055 | 1.736 | 0.25 | 0.17 | 0.11 | 0.03 | 0.15 | 0.15 |
| 3 | 16162613 | rs4423749 | 0.691 | 1.137 | -1.09 | -0.304 | -0.004 | 0.019 | 0.019 | 0 | 0.25 | 0.39 | 0.2 | 0.31 | 0.26 | 0.3 |
| 3 | 16163432 | rs4078148 | 1.423 | 1.105 | -1.04 | -0.686 | -0.005 | -0.003 | -0.003 | 0 | 0.13 | 0.1 | 0.06 | 0.07 | 0.08 | 0.21 |
| 3 | 16163689 | rs6782512 | 1.411 | 1.462 | -1.18 | -0.859 | -0.006 | 0.004 | 0.004 | 0 | 0.07 | 0.06 | 0.05 | 0 | 0.1 | 0.05 |
| 3 | 16164695 | rs9868739 | 1.573 | 1.122 | -0.863 | -0.764 | 0.004 | -0.058 | -0.058 | 0 | 0.15 | 0.21 | 0.11 | 0 | 0.18 | 0.1 |
| 3 | 16165418 | rs4685307 | -0.355 | 0.918 | -1.161 | -0.313 | -0.005 | 0.084 | 0.084 | 0.947 | 0.53 | 0.63 | 0.51 | 0.78 | 0.57 | 0.61 |
| 3 | 16165607 | rs2346489 | 1.094 | 1.132 | -1.13 | 0.557 | -0.038 | -0.023 | -0.023 | 0 | 0.15 | 0.05 | 0.31 | 0.17 | 0.19 | 0.22 |
| 3 | 16168677 | rs17041904 | -0.013 | 0.839 | -0.615 | 0.973 | -0.012 | 0.104 | 0.104 | 0 | 0.14 | 0.22 | 0.17 | 0.48 | 0.1 | 0.28 |
| 3 | 16169280 | rs6442581 | -0.058 | 1.176 | -0.698 | 0.386 | 0.009 | 0.007 | 0.007 | 0 | 0.68 | 0.63 | 0.49 | 0.58 | 0.57 | 0.56 |
| 3 | 16169491 | rs7624406 | -0.806 | 1.325 | -0.531 | 0.329 | -0.007 | 0.011 | 0.011 | 0 | 0.78 | 0.84 | 0.71 | 0.8 | 0.74 | 0.64 |
| 3 | 16169628 | rs17041907 | 0.637 | 1.262 | -0.55 | 0.69 | 0 | 0.046 | 0.046 | 0 | 0.05 | 0.08 | 0.03 | 0.22 | 0.05 | 0.08 |
| 3 | 16169676 | rs2730332 | -0.509 | 1.644 | -1.057 | -0.088 | 0 | 0.038 | 0.038 | 0 | 0.43 | 0.63 | 0.53 | 0.76 | 0.59 | 0.5 |
| 3 | 16169710 | rs17272271 | 1.317 | 1.433 | -0.994 | 0.079 | 0.009 | 0.016 | 0.016 | 0 | 0.2 | 0.07 | 0.23 | 0.18 | 0.14 | 0.32 |
| 3 | 16172032 | rs924935 | -0.46 | 1.443 | -0.657 | 0.018 | 0.001 | 0.008 | 0.008 | 0 | 0.47 | 0.53 | 0.52 | 0.74 | 0.58 | 0.49 |
| 3 | 16173235 | rs1505611 | 0.22 | 1.345 | -0.462 | 0.196 | -0.001 | -0.005 | -0.005 | 0.275 | 0.43 | 0.67 | 0.48 | 0.51 | 0.42 | 0.29 |
| 3 | 16177592 | rs4685308 | 1.774 | 1.323 | -0.873 | 0.126 | 0.016 | 0.149 | 0.149 | 0.356 | 0.29 | 0.03 | 0.16 | 0.05 | 0.17 | 0.15 |
| 3 | 16179863 | rs2730338 | -0.646 | 1.374 | -0.818 | 0.283 | -0.002 | -0.024 | -0.024 | 0 | 0.75 | 0.79 | 0.57 | 0.52 | 0.77 | 0.61 |
| 3 | 16181586 | rs13079803 | 0.183 | 1.261 | -0.797 | 0.245 | -0.005 | 0.005 | 0.005 | 0 | 0.4 | 0.33 | 0.31 | 0.3 | 0.48 | 0.4 |

| | | | | | | | | | | | | | | | | |
|---|---|---|---|---|---|---|---|---|---|---|---|---|---|---|---|---|
| 3 | 16185929 | rs12485734 | 0.077 | 1.262 | -0.757 | 0.246 | 0.003 | -0.008 | -0.008 | 0.002 | 0.41 | 0.52 | 0.33 | 0.3 | 0.48 | 0.4 |
| 3 | 16186303 | rs13089442 | 0.227 | 1.319 | -0.813 | 0.271 | 0.006 | -0.01 | -0.01 | 2.303 | 0.4 | 0.45 | 0.32 | 0.3 | 0.48 | 0.4 |
| 3 | 16189021 | rs10510446 | 1.346 | 0.991 | -0.868 | 0.518 | 0.018 | 0.003 | 0.003 | 0 | 0.17 | 0.04 | 0.21 | 0.18 | 0.1 | 0.31 |
| 3 | 16192143 | rs6771632 | 0.037 | 1.863 | -0.504 | 0.704 | 0.054 | 0.048 | 0.048 | 0 | 0.69 | 0.44 | 0.57 | 0.54 | 0.51 | 0.73 |
| 3 | 16196224 | rs6775152 | 0.096 | 1.876 | -0.523 | 0.833 | 0.014 | 0.053 | 0.053 | 0 | 0.24 | 0.33 | 0.37 | 0.53 | 0.33 | 0.33 |
| 3 | 16196660 | rs13095794 | 0.363 | 1.993 | -0.631 | 0.691 | -0.043 | -0.035 | -0.035 | 0 | 0.26 | 0.16 | 0.34 | 0.3 | 0.37 | 0.22 |
| 3 | 16197119 | rs4685313 | 0.384 | 2.002 | -0.749 | 0.695 | -0.074 | -0.062 | -0.062 | 0.43 | 0.26 | 0.1 | 0.33 | 0.29 | 0.37 | 0.22 |
| 3 | 16197294 | rs7624605 | 0.127 | 1.964 | -0.196 | 0.897 | 0.002 | 0.047 | 0.047 | 0 | 0.21 | 0.32 | 0.19 | 0.3 | 0.24 | 0.27 |
| 3 | 16199034 | rs2881410 | -0.041 | 1.863 | -0.201 | 0.894 | 0.035 | 0.038 | 0.038 | 0 | 0.78 | 0.66 | 0.76 | 0.69 | 0.64 | 0.68 |
| 3 | 16213362 | rs2657595 | -0.607 | 1.337 | -0.962 | -0.071 | -0.004 | 0.16 | 0.16 | 0 | 0.58 | 0.81 | 0.79 | 0.91 | 0.62 | 0.72 |
| 3 | 16213680 | rs2730350 | -0.256 | 2.18 | -0.222 | 0.344 | 0.066 | 0.056 | 0.056 | 0 | 0.22 | 0.36 | 0.22 | 0.31 | 0.34 | 0.29 |
| 3 | 16215261 | rs6804033 | -0.287 | 1.711 | -0.299 | 0.344 | -0.009 | 0.024 | 0.024 | 0 | 0.36 | 0.53 | 0.41 | 0.47 | 0.29 | 0.55 |
| 3 | 16220352 | rs4685320 | 0.769 | 2.026 | -0.73 | 0.196 | -0.032 | -0.037 | -0.037 | 1.468 | 0.55 | 0.22 | 0.45 | 0.52 | 0.47 | 0.4 |
| 3 | 16222989 | rs2047975 | 0.742 | 1.579 | -0.848 | -0.104 | -0.024 | -0.02 | -0.02 | 0 | 0.21 | 0.11 | 0.18 | 0.24 | 0.23 | 0.15 |
| 3 | 16224346 | rs17041971 | 0.926 | 1.797 | -0.403 | -0.282 | -0.01 | -0.022 | -0.022 | 0 | 0.53 | 0.24 | 0.53 | 0.53 | 0.5 | 0.39 |
| 3 | 16228798 | rs2102303 | 0.643 | 1.471 | -0.22 | -0.725 | -0.007 | 0.008 | 0.008 | 0 | 0.69 | 0.31 | 0.69 | 0.76 | 0.58 | 0.75 |
| 3 | 16234295 | rs9310495 | 0.225 | 1.497 | 0.289 | -0.695 | -0.008 | -0.081 | -0.081 | 0 | 0.13 | 0.36 | 0.08 | 0 | 0.07 | 0.05 |
| 3 | 16235042 | rs924931 | 0.831 | 1.559 | 0.351 | -0.76 | -0.009 | -0.008 | -0.008 | 0 | 0.61 | 0.71 | 0.54 | 0.58 | 0.55 | 0.51 |
| 3 | 16235398 | rs13074942 | -0.428 | 1.471 | 0.424 | -0.681 | 0.002 | -0.006 | -0.006 | 0 | 0.32 | 0.25 | 0.36 | 0.42 | 0.27 | 0.41 |
| 3 | 16239063 | rs6799256 | -0.6 | 1.48 | 0.459 | -0.759 | -0.011 | -0.113 | -0.113 | 0 | 0.18 | 0.41 | 0.12 | 0 | 0.13 | 0.08 |
| 3 | 16242101 | rs3796301 | 0.475 | 1.361 | 0.424 | -0.662 | -0.001 | 0 | 0 | 0 | 0.16 | 0.16 | 0.23 | 0.19 | 0.11 | 0.28 |
| 3 | 16243596 | rs11711765 | 1.569 | 1.554 | 0.301 | -0.724 | -0.017 | 0.016 | 0.016 | 0 | 0.48 | 0.26 | 0.48 | 0.58 | 0.5 | 0.47 |
| 3 | 16245397 | rs6804047 | -0.219 | 1.537 | 0.401 | -0.762 | 0.004 | 0 | 0 | 0 | 0.24 | 0.23 | 0.19 | 0.18 | 0.14 | 0.28 |
| 3 | 16251995 | rs6807124 | 1.146 | 1.456 | 0.156 | -0.822 | -0.002 | -0.002 | -0.002 | 0.872 | 0.54 | 0.51 | 0.6 | 0.58 | 0.55 | 0.51 |
| 3 | 16253421 | rs6442586 | -0.141 | 1.354 | 0.049 | -0.788 | -0.042 | -0.003 | -0.003 | 0 | 0.23 | 0.35 | 0.18 | 0.2 | 0.11 | 0.28 |
| 3 | 16256677 | rs1039636 | 1.752 | 1.674 | -0.386 | -0.703 | 0.011 | -0.002 | -0.002 | 0 | 0.45 | 0.23 | 0.44 | 0.57 | 0.43 | 0.44 |
| 3 | 16258093 | rs2089249 | 0.491 | 2.295 | -0.578 | -0.568 | -0.028 | 0.021 | 0.021 | 0 | 0.91 | 0.94 | 0.89 | 1 | 0.78 | 0.92 |
| 3 | 16258650 | rs905945 | 0.776 | 1.555 | -0.462 | -0.462 | -0.051 | 0.048 | 0.048 | 0.034 | 0.85 | 0.93 | 0.85 | 1 | 0.72 | 0.88 |
| 3 | 16258891 | rs905946 | 1.006 | 1.449 | -0.417 | -0.44 | -0.022 | 0.011 | 0.011 | 0.161 | 0.74 | 0.81 | 0.78 | 0.81 | 0.67 | 0.65 |
| 3 | 16261139 | rs13063387 | 0.087 | 0.615 | -0.023 | -0.086 | -0.001 | 0 | 0 | 0 | 0.17 | 0.14 | 0.19 | 0.2 | 0.15 | 0.11 |
| 3 | 16261420 | rs2271077 | 0.333 | 0.751 | -0.234 | -0.228 | -0.01 | 0.01 | 0.01 | 0 | 0.12 | 0.15 | 0.07 | 0.19 | 0.06 | 0.23 |
| 3 | 16269066 | rs14210 | -0.008 | 1.265 | -0.188 | -0.433 | -0.001 | -0.029 | -0.029 | 0 | 0.15 | 0.14 | 0.11 | 0.04 | 0.15 | 0.05 |
| 3 | 16269555 | rs842280 | 0.035 | 1.595 | -0.157 | -0.214 | 0 | -0.016 | -0.016 | 0 | 0.05 | 0.07 | 0.04 | 0 | 0.07 | 0.06 |
| 3 | 16272184 | rs6776381 | -0.917 | 1.935 | 0.217 | -0.146 | 0.03 | -0.072 | -0.072 | 0.135 | 0.16 | 0.27 | 0.21 | 0.04 | 0.22 | 0.11 |
| 3 | 16273558 | rs9848911 | -1.657 | 2.011 | 0.096 | -0.14 | -0.008 | 0.022 | 0.022 | 0.14 | 0.34 | 0.73 | 0.38 | 0.21 | 0.37 | 0.23 |
| 3 | 16275500 | rs6442589 | -0.19 | 1.763 | 0.181 | -0.089 | 0.046 | -0.054 | -0.054 | 0 | 0.09 | 0.23 | 0.1 | 0 | 0.12 | 0.04 |
| 3 | 16277043 | rs2672564 | 0.214 | 1.604 | -0.447 | 0.368 | -0.017 | -0.025 | -0.025 | 0 | 0.78 | 0.85 | 0.74 | 0.65 | 0.74 | 0.72 |
| 3 | 16278197 | rs9837613 | -0.455 | 0.989 | -0.168 | 0.156 | 0.005 | -0.062 | -0.062 | 0 | 0.11 | 0.3 | 0.14 | 0 | 0.09 | 0.07 |
| 3 | 16285941 | rs842261 | -0.122 | 1.144 | -0.928 | 0.748 | -0.018 | -0.049 | -0.049 | 0 | 0.29 | 0.17 | 0.34 | 0.49 | 0.34 | 0.34 |
| 3 | 16288232 | rs842249 | 0.095 | 1.215 | -0.963 | 1.358 | -0.019 | -0.009 | -0.009 | 0 | 0.71 | 0.82 | 0.67 | 0.61 | 0.65 | 0.74 |
| 3 | 16288762 | rs842248 | 0.127 | 1.163 | -0.936 | 1.326 | -0.019 | -0.055 | -0.055 | 0 | 0.7 | 0.81 | 0.66 | 0.52 | 0.66 | 0.66 |
| 3 | 16290167 | rs2063777 | 0.173 | 1.302 | -0.342 | 0.857 | 0.014 | -0.026 | -0.026 | 0 | 0.14 | 0.21 | 0.08 | 0.05 | 0.16 | 0.05 |
| 3 | 16293151 | rs6763776 | 0.696 | 0.65 | -0.4 | 0.867 | -0.009 | -0.004 | -0.004 | 0 | 0.25 | 0.25 | 0.4 | 0.26 | 0.32 | 0.35 |
| 3 | 16294425 | rs6442592 | 0.452 | 0.641 | 0.091 | 0.926 | 0.011 | 0.059 | 0.059 | 0 | 0.15 | 0.25 | 0.11 | 0.19 | 0.16 | 0.13 |
| 3 | 16295635 | rs712874 | 0.498 | 0.225 | -0.093 | 0.664 | 0.01 | 0.06 | 0.06 | 0 | 0.42 | 0.19 | 0.4 | 0.36 | 0.41 | 0.33 |
| 3 | 16299968 | rs842252 | 1.183 | 0.051 | -0.18 | 0.571 | -0.034 | 0.011 | 0.011 | 0.127 | 0.32 | 0.13 | 0.36 | 0.33 | 0.38 | 0.3 |
| 3 | 16301148 | rs10087 | 0.479 | -0.061 | -0.116 | 0.523 | -0.028 | -0.03 | -0.03 | 0 | 0.41 | 0.26 | 0.47 | 0.48 | 0.43 | 0.53 |
| 3 | 16301959 | rs552 | 0.286 | -0.08 | 0.13 | 0.532 | 0.006 | 0.048 | 0.048 | 0 | 0.18 | 0.52 | 0.14 | 0.17 | 0.16 | 0.13 |
| 3 | 16310379 | rs6804208 | 0.32 | -0.069 | 0.3 | 0.536 | 0.007 | 0.046 | 0.046 | 0 | 0.17 | 0.55 | 0.14 | 0.17 | 0.16 | 0.13 |
| 3 | 16312564 | rs842259 | 0.482 | -0.099 | 0.268 | 0.52 | -0.026 | -0.031 | -0.031 | 0 | 0.41 | 0.26 | 0.47 | 0.48 | 0.43 | 0.54 |
| 3 | 16327855 | rs842274 | 0.403 | -0.098 | 0.266 | 0.521 | -0.017 | -0.02 | -0.02 | 0 | 0.42 | 0.31 | 0.47 | 0.48 | 0.43 | 0.53 |
| 3 | 16327909 | rs6777976 | 0.988 | -0.102 | 0.321 | 0.52 | 0.029 | 0.07 | 0.07 | 0 | 0.14 | 0.27 | 0.11 | 0.17 | 0.16 | 0.13 |
| 3 | 16328356 | rs842273 | -0.548 | -0.104 | 0.183 | 0.495 | -0.018 | -0.02 | -0.02 | 1.005 | 0.58 | 0.7 | 0.52 | 0.52 | 0.56 | 0.47 |
| 3 | 16332988 | rs842270 | -0.565 | 0.019 | 0.164 | 0.48 | -0.017 | -0.021 | -0.021 | 0 | 0.59 | 0.69 | 0.53 | 0.52 | 0.57 | 0.47 |
| 3 | 16357708 | rs1050604 | 0.819 | -0.019 | 0.233 | 0.442 | -0.024 | -0.025 | -0.025 | 0 | 0.11 | 0.14 | 0.04 | 0.02 | 0.03 | 0.03 |
| 3 | 16358584 | rs690241 | -0.536 | 0.189 | -0.177 | 0.586 | -0.029 | -0.038 | -0.038 | 0 | 0.59 | 0.74 | 0.53 | 0.51 | 0.56 | 0.47 |
| 3 | 16364919 | rs690216 | -0.543 | 0.121 | -0.235 | 0.614 | -0.026 | -0.036 | -0.036 | 0 | 0.59 | 0.74 | 0.54 | 0.51 | 0.56 | 0.47 |
| 3 | 16365223 | rs483682 | 1.046 | 0.158 | -0.231 | 0.67 | -0.025 | -0.071 | -0.071 | 0 | 0.32 | 0.16 | 0.38 | 0.47 | 0.37 | 0.39 |
| 3 | 16370495 | rs690606 | -0.609 | 0.155 | -0.266 | 0.788 | 0.003 | -0.006 | -0.006 | 0 | 0.66 | 0.68 | 0.64 | 0.44 | 0.67 | 0.42 |
| 3 | 16370675 | rs581152 | -0.715 | 0.181 | -0.281 | 0.757 | 0.002 | -0.011 | -0.011 | 0.091 | 0.67 | 0.69 | 0.67 | 0.56 | 0.68 | 0.51 |
| 3 | 16376279 | rs538525 | 1.038 | 0.487 | -0.507 | 0.555 | -0.061 | 0.021 | 0.021 | 0.284 | 0.33 | 0.14 | 0.46 | 0.3 | 0.42 | 0.31 |
| 3 | 16377067 | rs690522 | 1.091 | 0.652 | -0.603 | 0.548 | -0.079 | 0.002 | 0.002 | 0.994 | 0.31 | 0.1 | 0.45 | 0.3 | 0.41 | 0.31 |
| 3 | 16381111 | rs490059 | -0.658 | 0.479 | -0.493 | 0.296 | 0.008 | 0.024 | 0.024 | 0 | 0.65 | 0.73 | 0.78 | 0.7 | 0.69 | 0.5 |
| 3 | 16385395 | rs1060699 | 0.227 | 0.322 | -0.375 | 0.293 | 0 | 0.001 | 0.001 | 0 | 0.11 | 0.09 | 0.15 | 0.11 | 0.12 | 0.05 |
| 3 | 16386354 | rs690023 | 1.086 | 0.217 | 0.083 | 0.458 | -0.006 | -0.007 | -0.007 | 0.667 | 0.31 | 0.3 | 0.48 | 0.53 | 0.4 | 0.39 |
| 3 | 16388390 | rs7622858 | 0.109 | 0.325 | 0.117 | 0.074 | -0.007 | 0.021 | 0.021 | 0 | 0.27 | 0.18 | 0.26 | 0.11 | 0.3 | 0.12 |
| 3 | 16391908 | rs542591 | -0.663 | 0.393 | -0.41 | 0.266 | 0.013 | 0.006 | 0.006 | 0 | 0.66 | 0.74 | 0.69 | 0.65 | 0.73 | 0.54 |
| 3 | 16395513 | rs574083 | 0.096 | 0.512 | -0.84 | 0.649 | -0.02 | 0.002 | 0.002 | 0 | 0.39 | 0.24 | 0.42 | 0.57 | 0.44 | 0.44 |
| 3 | 16400787 | rs6787251 | -0.1 | 0.094 | -1.056 | 0.256 | 0.005 | 0.007 | 0.007 | 0 | 0.22 | 0.17 | 0.25 | 0.11 | 0.2 | 0.26 |

| | | | | | | | | | | | | | | | | |
|---|---|---|---|---|---|---|---|---|---|---|---|---|---|---|---|---|
| 3 | 16401146 | rs689691 | 0.334 | 0.15 | -1.088 | 0.185 | -0.017 | 0.01 | 0.01 | 0.703 | 0.42 | 0.27 | 0.33 | 0.49 | 0.5 | 0.45 |
| 3 | 16403766 | rs690343 | -0.255 | -0.057 | -0.613 | 0.083 | -0.005 | -0.004 | -0.004 | 0 | 0.53 | 0.48 | 0.4 | 0.5 | 0.57 | 0.51 |
| 3 | 16405740 | rs574141 | 1.193 | 0.697 | -1.304 | -0.066 | -0.111 | -0.015 | -0.015 | 0 | 0.18 | 0.02 | 0.16 | 0.19 | 0.31 | 0.16 |
| 3 | 16405868 | rs11916806 | 0.821 | 0.085 | -0.563 | 0.036 | -0.012 | -0.021 | -0.021 | 0.009 | 0.11 | 0.25 | 0.09 | 0.08 | 0.07 | 0.06 |
| 3 | 16407973 | rs526100 | 0.25 | -0.253 | -0.828 | -0.112 | -0.001 | 0 | 0 | 0.012 | 0.61 | 0.38 | 0.59 | 0.65 | 0.62 | 0.65 |
| 3 | 16408457 | rs9835911 | -0.598 | -1.274 | -1.291 | -0.657 | 0.102 | 0.121 | 0.121 | 0.799 | 0.48 | 0.81 | 0.48 | 0.24 | 0.31 | 0.38 |
| 3 | 16408489 | rs12635698 | -0.384 | -1.719 | -1.211 | -0.868 | -0.05 | -0.1 | -0.1 | 0.068 | 0.28 | 0.58 | 0.16 | 0.03 | 0.13 | 0.15 |
| 3 | 16408534 | rs552666 | -1.592 | -1.781 | -1.576 | -0.913 | -0.026 | -0.148 | -0.148 | 0.787 | 0.77 | 0.94 | 0.73 | 0.54 | 0.77 | 0.57 |
| 3 | 16408665 | rs553681 | 1.65 | -1.715 | -1.576 | -1 | -0.039 | 0.002 | 0.002 | 0 | 0.27 | 0.13 | 0.26 | 0.3 | 0.45 | 0.17 |
| 3 | 16408778 | rs689953 | 0.16 | -1.89 | -1.652 | -1.042 | 0 | -0.001 | -0.001 | 0 | 0.44 | 0.53 | 0.38 | 0.32 | 0.55 | 0.28 |
| 3 | 16408949 | rs556322 | -0.2 | -1.742 | -1.554 | -0.885 | 0.015 | -0.04 | -0.04 | 0 | 0.62 | 0.83 | 0.64 | 0.51 | 0.65 | 0.44 |
| 3 | 16411531 | rs17042188 | 0.548 | -1.458 | -1.147 | -0.678 | -0.013 | -0.041 | -0.041 | 0 | 0.08 | 0.21 | 0.04 | 0 | 0.03 | 0.03 |
| 3 | 16417485 | rs4685330 | 1.352 | -1.573 | -2.033 | -0.719 | 0.018 | -0.001 | -0.001 | 0 | 0.2 | 0.1 | 0.29 | 0.19 | 0.15 | 0.28 |
| 3 | 16420179 | rs483050 | -0.343 | -1.579 | -2.341 | -0.745 | 0.034 | 0.033 | 0.033 | 0 | 0.59 | 0.83 | 0.47 | 0.36 | 0.64 | 0.39 |
| 3 | 16420794 | rs842416 | 0.028 | -1.303 | -2.037 | -0.758 | 0.009 | -0.033 | -0.033 | 0 | 0.65 | 0.76 | 0.69 | 0.54 | 0.71 | 0.43 |
| 3 | 16422989 | rs9310496 | 1.662 | -0.963 | -2.193 | -0.834 | 0.016 | 0.01 | 0.01 | 0.353 | 0.24 | 0.1 | 0.33 | 0.19 | 0.2 | 0.25 |
| 3 | 16424008 | rs1464171 | 0.764 | -0.97 | -2.122 | -0.964 | 0.011 | 0.024 | 0.024 | 1.346 | 0.36 | 0.27 | 0.41 | 0.21 | 0.29 | 0.3 |
| 3 | 16425121 | rs548740 | 0.097 | -0.897 | -2.643 | -0.882 | 0.014 | -0.061 | -0.061 | 0 | 0.33 | 0.19 | 0.3 | 0.47 | 0.29 | 0.58 |
| 3 | 16425349 | rs690219 | 0.295 | -0.857 | -2.61 | -0.865 | -0.006 | 0.001 | 0.001 | 0 | 0.34 | 0.69 | 0.31 | 0.33 | 0.42 | 0.11 |
| 3 | 16426427 | rs471336 | 0.542 | -1.011 | -2.611 | -0.897 | 0.043 | 0.067 | 0.067 | 0 | 0.41 | 0.21 | 0.41 | 0.2 | 0.3 | 0.47 |
| 3 | 16426891 | rs17042200 | 1.469 | -1.182 | -2.086 | -0.807 | -0.024 | -0.088 | -0.088 | 0 | 0.17 | 0.07 | 0.22 | 0.46 | 0.19 | 0.36 |
| 3 | 16428593 | rs712877 | -0.046 | -1.03 | -2.027 | -0.914 | 0.007 | -0.012 | -0.012 | 0.1 | 0.17 | 0.14 | 0.08 | 0.01 | 0.1 | 0.23 |
| 3 | 16431421 | rs523909 | 0.192 | -0.664 | -1.921 | -0.278 | -0.017 | -0.056 | -0.056 | 0 | 0.24 | 0.08 | 0.23 | 0.27 | 0.23 | 0.26 |
| 3 | 16433301 | rs7624893 | 1.128 | -0.203 | -1.694 | -0.138 | -0.006 | -0.056 | -0.056 | 0.974 | 0.18 | 0.63 | 0.12 | 0.08 | 0.13 | 0.03 |
| 3 | 16433519 | rs1864259 | 1.249 | 0.221 | -1.929 | 0.182 | -0.018 | -0.061 | -0.061 | 4.268 | 0.16 | 0.07 | 0.2 | 0.28 | 0.18 | 0.23 |
| 3 | 16436725 | rs12488145 | 0.13 | 0.242 | -1.981 | 0.17 | -0.047 | -0.124 | -0.124 | 0.527 | 0.26 | 0.11 | 0.3 | 0.49 | 0.3 | 0.4 |
| 3 | 16438270 | rs11715320 | 0.646 | 0.311 | -1.764 | 0.159 | -0.015 | -0.031 | -0.031 | 0 | 0.1 | 0.04 | 0.09 | 0.22 | 0.12 | 0.17 |
| 3 | 16440244 | rs842496 | 0.16 | 0.07 | -1.824 | -0.293 | 0.022 | 0.013 | 0.013 | 0.837 | 0.54 | 0.28 | 0.62 | 0.42 | 0.62 | 0.47 |
| 3 | 16441428 | rs6442600 | -1.184 | 0.315 | -1.753 | -0.155 | -0.028 | -0.02 | -0.02 | 0.311 | 0.64 | 0.8 | 0.5 | 0.6 | 0.56 | 0.64 |
| 3 | 16450872 | rs1546650 | -0.783 | -0.002 | -1.75 | -0.176 | -0.001 | 0.01 | 0.01 | 0.75 | 0.57 | 0.8 | 0.56 | 0.59 | 0.55 | 0.57 |
| 3 | 16455710 | rs12486361 | -0.501 | 0.675 | -0.808 | 0.323 | 0.021 | 0.005 | 0.005 | 0.123 | 0.39 | 0.53 | 0.45 | 0.58 | 0.44 | 0.45 |
| 3 | 16457444 | rs4685336 | -0.89 | 1.176 | -0.519 | 0.313 | 0 | 0 | 0 | 0 | 0.27 | 0.31 | 0.34 | 0.24 | 0.26 | 0.33 |
| 3 | 16458929 | rs7626299 | -0.928 | 1.086 | -0.511 | 0.321 | 0 | 0 | 0 | 0 | 0.27 | 0.31 | 0.35 | 0.24 | 0.28 | 0.33 |
| 3 | 16461063 | rs7632675 | -0.96 | 0.974 | -0.322 | 0.028 | -0.009 | -0.05 | -0.05 | 0 | 0.22 | 0.31 | 0.14 | 0.05 | 0.17 | 0.2 |
| 3 | 16463271 | rs17042258 | 0.145 | 1.175 | -0.596 | 0.142 | 0.008 | 0.008 | 0.008 | 0 | 0.13 | 0.14 | 0.25 | 0.16 | 0.17 | 0.17 |
| 3 | 16470125 | rs17042260 | -0.631 | 0.507 | -0.029 | -0.096 | -0.044 | -0.096 | -0.096 | 0.273 | 0.14 | 0.37 | 0.09 | 0.01 | 0.05 | 0.17 |
| 3 | 16470197 | rs17042262 | -1.035 | 0.631 | -0.241 | -0.097 | -0.008 | -0.091 | -0.091 | 1.649 | 0.2 | 0.33 | 0.14 | 0.02 | 0.16 | 0.23 |
| 3 | 16472764 | rs11720129 | -0.324 | 1.272 | -0.783 | 0.353 | 0.006 | -0.019 | -0.019 | 0 | 0.09 | 0.1 | 0.07 | 0.02 | 0.11 | 0.06 |
| 3 | 16474332 | rs4685337 | 0.083 | 1.529 | -1.44 | 0.755 | -0.071 | -0.054 | -0.054 | 0 | 0.7 | 0.83 | 0.57 | 0.52 | 0.53 | 0.61 |
| 3 | 16475215 | rs1517516 | -0.55 | 1.228 | -1.272 | 0.844 | 0.002 | -0.003 | -0.003 | 0 | 0.27 | 0.22 | 0.23 | 0.31 | 0.26 | 0.26 |
| 3 | 16480827 | rs9817739 | -1.023 | 2.273 | -0.769 | 1.165 | 0.031 | 0.038 | 0.038 | 0 | 0.17 | 0.29 | 0.26 | 0.25 | 0.21 | 0.08 |
| 3 | 16485219 | rs7644810 | 1.334 | 2.233 | -0.907 | 1.242 | 0.011 | 0.04 | 0.04 | 0 | 0.7 | 0.59 | 0.64 | 0.6 | 0.67 | 0.78 |
| 3 | 16491898 | rs9867980 | -0.614 | 2.492 | -0.974 | 1.205 | -0.014 | 0 | 0 | 0 | 0.87 | 0.91 | 0.85 | 0.88 | 0.78 | 0.86 |
| 3 | 16495468 | rs4684281 | -1.399 | 1.911 | -0.807 | 1.236 | -0.005 | 0.073 | 0.073 | 0 | 0.17 | 0.33 | 0.22 | 0.28 | 0.13 | 0.1 |
| 3 | 16499003 | rs11710881 | -0.52 | 2.38 | -1.051 | 1.169 | -0.054 | -0.003 | -0.003 | 0 | 0.09 | 0.01 | 0.09 | 0.05 | 0.15 | 0.03 |
| 3 | 16504557 | rs17042294 | -1.606 | 2.17 | -0.782 | 1.353 | 0.001 | 0.026 | 0.026 | 0 | 0.14 | 0.23 | 0.11 | 0.27 | 0.13 | 0.11 |
| 3 | 16505411 | rs17042300 | 0.007 | 2.131 | -0.789 | 1.358 | -0.001 | 0.033 | 0.033 | 0 | 0.06 | 0.06 | 0.04 | 0.22 | 0.02 | 0.08 |
| 3 | 16506665 | rs12487874 | 0.903 | 1.715 | -1.536 | 1.181 | -0.09 | -0.013 | -0.013 | 0 | 0.19 | 0.02 | 0.16 | 0.1 | 0.24 | 0.17 |
| 3 | 16508057 | rs6767890 | -1.268 | 1.541 | -1.235 | 1.185 | 0.002 | 0.045 | 0.045 | 0 | 0.15 | 0.26 | 0.12 | 0.27 | 0.14 | 0.11 |
| 3 | 16516102 | rs9848484 | 1.026 | 0.691 | -1.14 | 0.355 | -0.008 | -0.032 | -0.032 | 0.033 | 0.22 | 0.22 | 0.21 | 0.04 | 0.16 | 0.23 |
| 3 | 16518270 | rs7621664 | -0.491 | 0.648 | -0.962 | 0.391 | 0 | -0.002 | -0.002 | 0.142 | 0.1 | 0.15 | 0.08 | 0.05 | 0.12 | 0.03 |
| 3 | 16520435 | rs11128791 | 0.097 | 0.507 | -0.647 | 0.743 | 0.007 | 0.006 | 0.006 | 0 | 0.66 | 0.57 | 0.59 | 0.5 | 0.51 | 0.66 |
| 3 | 16532132 | rs10865744 | 0.855 | -0.149 | -0.427 | 0.769 | -0.031 | -0.004 | -0.004 | 0 | 0.06 | 0.01 | 0.05 | 0.04 | 0.09 | 0.02 |
| 3 | 16535097 | rs10454232 | 1.233 | -0.239 | -0.404 | 0.567 | -0.002 | -0.034 | -0.034 | 0 | 0.21 | 0.22 | 0.22 | 0.04 | 0.18 | 0.24 |
| 3 | 16535229 | rs3815962 | 0.132 | -0.411 | -0.355 | 0.865 | 0.012 | 0.011 | 0.011 | 0 | 0.46 | 0.36 | 0.46 | 0.39 | 0.35 | 0.48 |
| 3 | 16543322 | rs17272530 | -0.869 | -0.462 | -0.237 | 1.012 | -0.001 | -0.003 | -0.003 | 0 | 0.56 | 0.59 | 0.6 | 0.55 | 0.45 | 0.55 |
| 3 | 16543983 | rs2881512 | -0.729 | -0.42 | -0.286 | 1.021 | 0 | 0.005 | 0.005 | 0 | 0.55 | 0.51 | 0.59 | 0.55 | 0.45 | 0.56 |
| 3 | 16547665 | rs9854409 | -1.973 | -0.307 | 0.192 | 1.039 | 0.03 | 0.207 | 0.207 | 2.179 | 0.13 | 0.52 | 0.18 | 0.3 | 0.11 | 0.07 |
| 3 | 16548760 | rs9859948 | -0.84 | -0.538 | 0.064 | 1.005 | 0.002 | 0.002 | 0.002 | 0 | 0.51 | 0.55 | 0.58 | 0.61 | 0.42 | 0.54 |
| 3 | 16551795 | rs11128793 | -0.155 | -0.487 | -0.711 | 0.972 | 0.002 | 0.001 | 0.001 | 0 | 0.43 | 0.56 | 0.45 | 0.31 | 0.32 | 0.48 |
| 3 | 16553680 | rs2881514 | -0.947 | -0.403 | -1.278 | 1.001 | 0.005 | -0.002 | -0.002 | 0 | 0.53 | 0.79 | 0.6 | 0.6 | 0.43 | 0.54 |
| 3 | 16563873 | rs9310499 | -0.165 | -0.751 | -0.97 | 1.245 | -0.001 | 0.003 | 0.003 | 0 | 0.3 | 0.34 | 0.37 | 0.34 | 0.28 | 0.3 |
| 3 | 16565559 | rs17042365 | -0.049 | -0.707 | -0.977 | 1.322 | -0.005 | -0.006 | -0.006 | 0 | 0.29 | 0.34 | 0.35 | 0.23 | 0.23 | 0.28 |
| 3 | 16567340 | rs13093768 | 1.066 | -0.693 | -1.245 | 1.293 | -0.019 | 0.07 | 0.07 | 0 | 0.19 | 0.05 | 0.2 | 0.04 | 0.17 | 0.24 |
| 3 | 16569746 | rs6774181 | -0.923 | -0.953 | -1.388 | 1.246 | 0.032 | 0.005 | 0.005 | 0.04 | 0.47 | 0.8 | 0.57 | 0.61 | 0.37 | 0.5 |
| 3 | 16571573 | rs17042372 | 0.35 | -0.029 | -1.494 | 0.895 | -0.032 | -0.03 | -0.03 | 0 | 0.05 | 0.01 | 0.06 | 0.16 | 0.1 | 0.03 |
| 3 | 16572975 | rs13093250 | 0.156 | -0.07 | -1.536 | 0.684 | -0.047 | 0.063 | 0.063 | 0 | 0.23 | 0.04 | 0.23 | 0.09 | 0.22 | 0.25 |
| 3 | 16573140 | rs13097853 | 0.673 | -0.089 | -1.51 | 0.65 | -0.019 | 0.071 | 0.071 | 0 | 0.19 | 0.05 | 0.2 | 0.04 | 0.17 | 0.24 |
| 3 | 16578457 | rs3856833 | -0.149 | -0.203 | -1.326 | 0.662 | -0.002 | -0.003 | -0.003 | 2.964 | 0.31 | 0.3 | 0.37 | 0.34 | 0.3 | 0.3 |

| | | | | | | | | | | | | | | | | |
|---|---|---|---|---|---|---|---|---|---|---|---|---|---|---|---|---|
| 3 | 16585132 | rs2042826 | -0.767 | -0.002 | -1.389 | 0.7 | -0.042 | -0.007 | -0.007 | 0 | 0.86 | 0.95 | 0.83 | 0.87 | 0.75 | 0.84 |
| 3 | 16585193 | rs6799971 | 0.732 | -0.522 | -1.505 | 0.52 | -0.005 | -0.001 | -0.001 | 0 | 0.32 | 0.29 | 0.24 | 0.29 | 0.31 | 0.28 |
| 3 | 16586222 | rs10510450 | 0.635 | -1.313 | -1.654 | 0.636 | -0.015 | 0 | 0 | 0 | 0.37 | 0.58 | 0.26 | 0.3 | 0.17 | 0.39 |
| 3 | 16588273 | rs6442608 | 0.346 | -1.386 | -1.804 | 0.713 | 0.098 | -0.023 | -0.023 | 0 | 0.4 | 0.67 | 0.29 | 0.5 | 0.18 | 0.39 |
| 3 | 16590910 | rs6793330 | -0.327 | -0.91 | -1.763 | 0.852 | 0.097 | -0.06 | -0.06 | 0 | 0.54 | 0.88 | 0.42 | 0.55 | 0.37 | 0.47 |
| 3 | 16592134 | rs17200369 | 0.03 | -0.862 | -1.717 | 0.077 | 0.154 | 0.108 | 0.108 | 0.133 | 0.49 | 0.77 | 0.52 | 0.73 | 0.28 | 0.56 |
| 3 | 16594218 | rs6763943 | -0.107 | -0.387 | -1.504 | 0.184 | -0.046 | 0.019 | 0.019 | 0 | 0.76 | 0.83 | 0.71 | 0.83 | 0.58 | 0.8 |
| 3 | 16596267 | rs2347309 | -0.151 | -0.893 | -1.162 | 0.006 | 0.006 | -0.001 | -0.001 | 0 | 0.61 | 0.69 | 0.64 | 0.57 | 0.67 | 0.54 |
| 3 | 16602491 | rs11916740 | -1.489 | -0.587 | -1.04 | 0.059 | -0.042 | -0.011 | -0.011 | 0.003 | 0.86 | 0.95 | 0.83 | 0.87 | 0.76 | 0.85 |
| 3 | 16603877 | rs12497831 | 0.818 | -0.748 | -0.938 | -0.173 | -0.05 | -0.001 | -0.001 | 0 | 0.24 | 0.27 | 0.19 | 0.28 | 0.06 | 0.3 |
| 3 | 16617862 | rs12635244 | 1.075 | -0.22 | -0.752 | 0.04 | 0.013 | 0.164 | 0.164 | 0 | 0.07 | 0.09 | 0.24 | 0.48 | 0.1 | 0.18 |
| 3 | 16619147 | rs9882426 | -0.59 | -0.219 | -0.79 | 0.007 | 0.009 | 0.003 | 0.003 | 0.199 | 0.62 | 0.71 | 0.64 | 0.62 | 0.69 | 0.56 |
| 3 | 16625048 | rs10510452 | 1.359 | -0.334 | -0.9 | -0.11 | -0.094 | 0.066 | 0.066 | 0 | 0.26 | 0.06 | 0.19 | 0.09 | 0.3 | 0.23 |
| 3 | 16627187 | rs17200390 | 1.13 | -0.284 | -0.947 | -0.129 | -0.044 | 0.007 | 0.007 | 0 | 0.08 | 0.01 | 0.11 | 0.05 | 0.16 | 0.05 |
| 3 | 16627806 | rs1808518 | 0.844 | -0.303 | -0.895 | -0.125 | 0.026 | 0.235 | 0.235 | 0.049 | 0.08 | 0.18 | 0.25 | 0.48 | 0.1 | 0.19 |
| 3 | 16628023 | rs10510453 | 1.265 | -0.318 | -0.919 | -0.135 | -0.053 | -0.118 | -0.118 | 0.102 | 0.19 | 0.37 | 0.09 | 0.01 | 0.14 | 0.09 |
| 3 | 16638775 | rs2217204 | -0.794 | -0.324 | -0.826 | -0.054 | 0.028 | 0.049 | 0.049 | 0.256 | 0.63 | 0.85 | 0.77 | 0.77 | 0.69 | 0.61 |
| 3 | 16639754 | rs3773827 | -0.336 | -0.31 | -0.947 | -0.322 | 0.005 | 0.016 | 0.016 | 0.312 | 0.54 | 0.6 | 0.52 | 0.3 | 0.59 | 0.43 |
| 3 | 16643940 | rs3773832 | 0.881 | -0.139 | -1.535 | -0.209 | -0.014 | 0.008 | 0.008 | 0 | 0.27 | 0.2 | 0.33 | 0.24 | 0.3 | 0.29 |
| 3 | 16646194 | rs2162616 | 0.873 | -0.059 | -1.518 | -0.184 | -0.011 | 0.007 | 0.007 | 0 | 0.27 | 0.22 | 0.33 | 0.25 | 0.3 | 0.29 |
| 3 | 16656331 | rs4685366 | 0.647 | 0.71 | -1.405 | 0.565 | 0.019 | 0.094 | 0.094 | 0 | 0.5 | 0.66 | 0.42 | 0.25 | 0.44 | 0.38 |
| 3 | 16662488 | rs6797529 | 0.225 | 0.611 | -1.64 | 0.763 | 0.014 | 0.002 | 0.002 | 0.045 | 0.65 | 0.83 | 0.65 | 0.62 | 0.69 | 0.57 |
| 3 | 16663683 | rs7623201 | -0.471 | 0.288 | -1.665 | 0.916 | 0.035 | -0.04 | -0.04 | 0 | 0.28 | 0.19 | 0.22 | 0.4 | 0.16 | 0.4 |
| 3 | 16664444 | rs7625875 | 0.473 | 1.559 | -1.946 | 0.257 | -0.053 | 0.05 | 0.05 | 0 | 0.75 | 0.9 | 0.77 | 0.9 | 0.69 | 0.67 |
| 3 | 16664547 | rs2059742 | -1.024 | 2.564 | -1.427 | 0.352 | -0.01 | 0.007 | 0.007 | 0.013 | 0.16 | 0.14 | 0.27 | 0.07 | 0.3 | 0.19 |
| 3 | 16667968 | rs7635873 | 0.092 | 2.253 | -1.25 | 0.783 | -0.016 | -0.065 | -0.065 | 3.63 | 0.65 | 0.89 | 0.59 | 0.64 | 0.66 | 0.65 |
| 3 | 16670272 | rs17272614 | 0.029 | 2.236 | -1.19 | 0.743 | -0.004 | 0.001 | 0.001 | 0.007 | 0.16 | 0.16 | 0.17 | 0.16 | 0.2 | 0.3 |
| 3 | 16671425 | rs1991419 | 0.939 | 1.375 | -1.147 | 0.339 | 0.026 | 0.004 | 0.004 | 0 | 0.11 | 0.28 | 0.14 | 0.09 | 0.13 | 0.08 |
| 3 | 16674026 | rs9809465 | 1.325 | 1.159 | -1.028 | 0.315 | -0.014 | 0.003 | 0.003 | 0 | 0.26 | 0.25 | 0.17 | 0.29 | 0.18 | 0.22 |
| 3 | 16675127 | rs7647846 | 0.038 | 1.24 | -1.073 | 0.283 | -0.003 | 0.002 | 0.002 | 0 | 0.42 | 0.41 | 0.34 | 0.55 | 0.37 | 0.53 |
| 3 | 16676779 | rs17200418 | 0.952 | 1.657 | -1.056 | -0.051 | -0.017 | 0.013 | 0.013 | 0 | 0.09 | 0.04 | 0.06 | 0 | 0.12 | 0.2 |
| 3 | 16677519 | rs7612566 | -0.548 | 1.61 | -0.796 | 0.024 | 0.015 | -0.007 | -0.007 | 0 | 0.27 | 0.32 | 0.25 | 0.25 | 0.32 | 0.38 |
| 3 | 16680339 | rs1035847 | 0.977 | 0.815 | -0.752 | -0.184 | -0.015 | -0.03 | -0.03 | 0 | 0.28 | 0.33 | 0.18 | 0.13 | 0.21 | 0.17 |
| 3 | 16680667 | rs11128801 | 0.43 | 0.98 | -0.961 | -0.198 | -0.019 | -0.056 | -0.056 | 0.13 | 0.29 | 0.42 | 0.3 | 0.2 | 0.24 | 0.26 |
| 3 | 16681999 | rs4685368 | -1.321 | 1.301 | -1.011 | 0.324 | -0.055 | -0.1 | -0.1 | 0 | 0.87 | 0.99 | 0.82 | 0.59 | 0.82 | 0.82 |
| 3 | 16684658 | rs10510457 | 1.457 | 0.418 | -1.443 | -0.666 | 0.01 | 0.158 | 0.158 | 0.003 | 0.35 | 0.11 | 0.18 | 0.01 | 0.3 | 0.24 |
| 3 | 16686195 | rs6802075 | 1.94 | 0.198 | -1.06 | -0.533 | -0.054 | 0.003 | 0.003 | 0 | 0.15 | 0.27 | 0.08 | 0.12 | 0.05 | 0.07 |
| 3 | 16687037 | rs10510458 | 1.517 | 0.426 | -1.121 | -0.627 | -0.004 | -0.004 | -0.004 | 0.042 | 0.11 | 0.1 | 0.09 | 0.01 | 0.13 | 0.19 |
| 3 | 16687807 | rs6806077 | -2.472 | 0.333 | -1.138 | -0.265 | -0.013 | 0.017 | 0.017 | 0 | 0.92 | 0.96 | 0.83 | 0.65 | 0.87 | 0.83 |
| 3 | 16687844 | rs17042532 | 2.217 | 0.234 | -1.119 | -0.182 | -0.003 | -0.014 | -0.014 | 0 | 0.05 | 0 | 0.09 | 0.26 | 0.04 | 0.08 |
| 3 | 16690419 | rs6767531 | 2.044 | 0.151 | -1.033 | -0.285 | -0.048 | 0.004 | 0.004 | 0 | 0.14 | 0.22 | 0.07 | 0.13 | 0.06 | 0.07 |
| 3 | 16692542 | rs10510459 | 0.042 | 0.558 | -0.911 | -0.941 | 0.009 | -0.045 | -0.045 | 0.273 | 0.29 | 0.38 | 0.28 | 0.08 | 0.35 | 0.33 |
| 3 | 16696226 | rs6782488 | 0.465 | 0.466 | -0.891 | -0.83 | -0.017 | 0.15 | 0.15 | 0 | 0.12 | 0.26 | 0.27 | 0.44 | 0.1 | 0.18 |
| 3 | 16699558 | rs9811661 | 0.835 | 0.482 | -1.189 | -1.812 | -0.001 | -0.072 | -0.072 | 1.109 | 0.51 | 0.49 | 0.31 | 0.06 | 0.37 | 0.32 |
| 3 | 16699931 | rs977676 | -1.255 | 0.144 | -1.031 | -1.513 | -0.021 | -0.051 | -0.051 | 0 | 0.86 | 0.96 | 0.74 | 0.28 | 0.83 | 0.76 |
| 3 | 16700334 | rs12494691 | 0.712 | 0.254 | -0.753 | -1.348 | 0.021 | -0.011 | -0.011 | 0 | 0.22 | 0.35 | 0.32 | 0.21 | 0.27 | 0.23 |
| 3 | 16701670 | rs7615979 | 1.221 | -0.297 | -0.62 | -1.672 | -0.057 | -0.091 | -0.091 | 2.736 | 0.16 | 0.27 | 0.07 | 0.03 | 0.06 | 0.11 |
| 3 | 16702160 | rs767375 | 0.21 | 0.458 | -1.251 | -1.47 | -0.021 | 0.112 | 0.112 | 0 | 0.38 | 0.25 | 0.27 | 0.04 | 0.37 | 0.21 |
| 3 | 16710679 | rs9813959 | -0.269 | -0.243 | -0.389 | -1.196 | -0.025 | -0.11 | -0.11 | 0.526 | 0.25 | 0.54 | 0.19 | 0.06 | 0.21 | 0.3 |
| 3 | 16711443 | rs9818882 | -1.189 | 0.012 | -0.737 | -0.553 | -0.011 | -0.008 | -0.008 | 0 | 0.88 | 0.89 | 0.84 | 0.52 | 0.82 | 0.74 |
| 3 | 16727170 | rs2114775 | 0.785 | -0.146 | -0.513 | -0.661 | 0.004 | -0.008 | -0.008 | 0.297 | 0.61 | 0.35 | 0.64 | 0.47 | 0.61 | 0.45 |
| 3 | 16734396 | rs2060598 | 0.652 | -0.201 | -0.719 | -1.194 | 0.003 | -0.011 | -0.011 | 0 | 0.24 | 0.22 | 0.34 | 0.44 | 0.19 | 0.21 |
| 3 | 16737497 | rs4260417 | 1.222 | -0.273 | -0.299 | -1.288 | 0.011 | -0.039 | -0.039 | 0.366 | 0.08 | 0.18 | 0.07 | 0.01 | 0.1 | 0.18 |
| 3 | 16739637 | rs1449536 | -1.032 | -0.394 | -0.342 | -1.346 | 0.001 | -0.037 | -0.037 | 0 | 0.79 | 0.87 | 0.81 | 0.52 | 0.78 | 0.7 |
| 3 | 16741105 | rs10510461 | 1.322 | -0.377 | -0.313 | -1.985 | -0.017 | -0.124 | -0.124 | 0.016 | 0.31 | 0.4 | 0.21 | 0.02 | 0.26 | 0.15 |
| 3 | 16743520 | rs7636863 | 0.036 | -0.375 | -0.386 | -1.968 | 0.006 | 0.062 | 0.062 | 0 | 0.54 | 0.63 | 0.41 | 0.05 | 0.55 | 0.46 |
| 3 | 16744246 | rs7617386 | 0.704 | -0.364 | -0.362 | -1.991 | -0.006 | -0.122 | -0.122 | 1.093 | 0.39 | 0.52 | 0.27 | 0.03 | 0.36 | 0.33 |
| 3 | 16744285 | rs11711008 | 1.43 | -0.219 | -0.573 | -1.674 | 0 | -0.015 | -0.015 | 0 | 0.15 | 0.11 | 0.13 | 0.02 | 0.18 | 0.13 |
| 3 | 16747387 | rs12634883 | 0.368 | -0.663 | -0.422 | -1.329 | -0.003 | 0.003 | 0.003 | 0 | 0.29 | 0.28 | 0.34 | 0.4 | 0.2 | 0.26 |
| 3 | 16748531 | rs9818121 | 1.942 | -0.256 | -0.708 | -1.283 | 0.008 | 0.007 | 0.007 | 0 | 0.11 | 0.11 | 0.05 | 0 | 0.07 | 0.04 |
| 3 | 16748611 | rs9876033 | 0.751 | -0.183 | -0.63 | -1.35 | 0 | -0.028 | -0.028 | 0 | 0.19 | 0.2 | 0.1 | 0.01 | 0.17 | 0.22 |
| 3 | 16749024 | rs9860475 | 0.8 | -0.174 | -0.822 | -1.346 | 0 | 0.016 | 0.016 | 0 | 0.18 | 0.1 | 0.09 | 0.02 | 0.17 | 0.22 |
| 3 | 16758156 | rs2122258 | 0.062 | -0.12 | -0.644 | -0.979 | -0.003 | -0.006 | -0.006 | 0 | 0.63 | 0.34 | 0.55 | 0.44 | 0.57 | 0.54 |
| 3 | 16759275 | rs2347646 | 0.596 | -0.222 | -1.108 | -1.061 | 0.019 | -0.085 | -0.085 | 0 | 0.32 | 0.14 | 0.3 | 0.41 | 0.24 | 0.26 |
| 3 | 16759632 | rs895945 | -0.468 | -0.346 | -0.802 | -1.605 | 0 | 0 | 0 | 0 | 0.67 | 0.62 | 0.61 | 0.37 | 0.62 | 0.59 |
| 3 | 16760709 | rs1375637 | 0.649 | -0.358 | -0.67 | -1.582 | -0.001 | -0.072 | -0.072 | 0.097 | 0.37 | 0.55 | 0.26 | 0.02 | 0.34 | 0.32 |
| 3 | 16761693 | rs4684295 | 0.406 | -0.237 | -0.823 | -1.636 | -0.031 | 0.038 | 0.038 | 0 | 0.45 | 0.2 | 0.45 | 0.33 | 0.42 | 0.39 |
| 3 | 16762185 | rs7618461 | 0.655 | -0.175 | -0.392 | -1.723 | 0.002 | -0.123 | -0.123 | 0 | 0.38 | 0.41 | 0.25 | 0.02 | 0.41 | 0.36 |
| 3 | 16764156 | rs6778810 | 0.586 | -0.329 | -0.832 | -1.658 | -0.044 | -0.027 | -0.027 | 0.11 | 0.35 | 0.15 | 0.41 | 0.33 | 0.35 | 0.34 |

| | | | | | | | | | | | | | | | | |
|---|---|---|---|---|---|---|---|---|---|---|---|---|---|---|---|---|
| 3 | 16764234 | rs6778843 | -1.792 | -0.245 | -0.664 | -1.47 | -0.004 | 0.008 | 0.008 | 0 | 0.87 | 0.88 | 0.81 | 0.38 | 0.84 | 0.75 |
| 3 | 16771147 | rs2044739 | -0.017 | -0.398 | -0.387 | -1.577 | -0.012 | 0.041 | 0.041 | 0.14 | 0.74 | 0.4 | 0.73 | 0.37 | 0.79 | 0.7 |
| 3 | 16778701 | rs9829793 | 1.808 | -0.337 | -0.465 | -1.702 | -0.01 | 0.009 | 0.009 | 0.179 | 0.16 | 0.17 | 0.16 | 0.01 | 0.19 | 0.14 |
| 3 | 16778771 | rs9849824 | 1.113 | -0.602 | -0.511 | -1.753 | -0.039 | 0.093 | 0.093 | 0.589 | 0.4 | 0.28 | 0.37 | 0.02 | 0.5 | 0.32 |
| 3 | 16778910 | rs17200488 | 1.714 | -0.703 | -0.519 | -1.748 | 0.013 | -0.041 | -0.041 | 0.123 | 0.19 | 0.08 | 0.24 | 0.32 | 0.15 | 0.2 |
| 3 | 16783061 | rs978664 | 0.938 | -0.62 | -0.586 | -1.733 | -0.018 | 0.117 | 0.117 | 0.78 | 0.42 | 0.29 | 0.45 | 0.05 | 0.56 | 0.35 |
| 3 | 16784389 | rs9833158 | 0.545 | -0.586 | -0.502 | -1.636 | 0.04 | 0.126 | 0.126 | 0.862 | 0.52 | 0.32 | 0.49 | 0.05 | 0.63 | 0.4 |
| 3 | 16785673 | rs2167062 | -0.74 | -0.216 | 0.366 | -0.889 | 0.007 | 0.136 | 0.136 | 0 | 0.83 | 0.65 | 0.78 | 0.39 | 0.82 | 0.74 |
| 3 | 16791642 | rs9842776 | 1.672 | -0.301 | -0.211 | -1.065 | 0.001 | 0.109 | 0.109 | 0 | 0.34 | 0.12 | 0.38 | 0.07 | 0.33 | 0.21 |
| 3 | 16793773 | rs6442631 | 0.261 | -0.516 | 0.107 | -1.193 | 0.013 | -0.052 | -0.052 | 0 | 0.62 | 0.54 | 0.55 | 0.11 | 0.51 | 0.44 |
| 3 | 16801747 | rs9829920 | -1.53 | -0.498 | 0.216 | -0.916 | 0.002 | 0.053 | 0.053 | 0 | 0.87 | 0.85 | 0.81 | 0.39 | 0.84 | 0.76 |
| 3 | 16809683 | rs6808150 | -1.013 | -0.559 | 0.27 | -0.922 | 0.007 | 0.067 | 0.067 | 0 | 0.85 | 0.76 | 0.72 | 0.37 | 0.78 | 0.73 |
| 3 | 16811826 | rs2242003 | 1.814 | -0.571 | 0.106 | -1.012 | -0.013 | 0.124 | 0.124 | 0 | 0.34 | 0.11 | 0.38 | 0.08 | 0.34 | 0.2 |
| 3 | 16812671 | rs1375629 | -1.421 | -0.297 | 0.166 | -0.696 | -0.004 | -0.014 | -0.014 | 1.549 | 0.87 | 0.89 | 0.81 | 0.4 | 0.84 | 0.76 |
| 3 | 16812771 | rs2218736 | -0.797 | -0.349 | 0.212 | -0.718 | -0.006 | 0.111 | 0.111 | 0.664 | 0.84 | 0.73 | 0.79 | 0.4 | 0.84 | 0.76 |
| 3 | 16817585 | rs12634691 | -0.63 | -1.11 | 0.663 | -0.045 | 0.043 | 0.091 | 0.091 | 1.143 | 0.28 | 0.52 | 0.36 | 0.51 | 0.39 | 0.38 |
| 3 | 16832828 | rs6442641 | -1.11 | -1.047 | 0.701 | 0.156 | 0.013 | 0.187 | 0.187 | 0.389 | 0.06 | 0.26 | 0.12 | 0.25 | 0.07 | 0.05 |
| 3 | 16838450 | rs9866541 | -1.624 | -1.3 | 0.482 | -0.109 | -0.04 | -0.054 | -0.054 | 0.441 | 0.55 | 0.88 | 0.47 | 0.54 | 0.53 | 0.58 |
| 3 | 16843044 | rs10510465 | -1.64 | -1.317 | 1.125 | -0.22 | 0.031 | 0.215 | 0.215 | 1.025 | 0.07 | 0.31 | 0.12 | 0.24 | 0.07 | 0.05 |
| 3 | 16850944 | rs7625399 | 0.281 | -1.392 | 1.034 | -0.42 | -0.022 | -0.086 | -0.086 | 1.619 | 0.15 | 0.27 | 0.07 | 0 | 0.08 | 0.07 |
| 3 | 16854379 | rs1823368 | -0.869 | -1.277 | 1.129 | -0.428 | 0.017 | -0.009 | -0.009 | 0 | 0.24 | 0.47 | 0.23 | 0.23 | 0.23 | 0.28 |
| 3 | 16860151 | rs6782026 | 0.118 | -1.28 | 0.944 | -0.562 | -0.021 | 0.038 | 0.038 | 0.975 | 0.18 | 0.07 | 0.18 | 0.01 | 0.21 | 0.26 |
| 3 | 16860824 | rs9836131 | -0.192 | -1.26 | 1.044 | -0.538 | 0.014 | -0.09 | -0.09 | 0 | 0.21 | 0.3 | 0.2 | 0.01 | 0.23 | 0.29 |
| 3 | 16862873 | rs9846711 | -1.285 | -1.484 | 0.972 | -0.498 | -0.001 | -0.003 | -0.003 | 0 | 0.39 | 0.56 | 0.35 | 0.3 | 0.36 | 0.4 |
| 3 | 16866253 | rs9867227 | -0.833 | -1.206 | 1.084 | -0.356 | 0.029 | -0.105 | -0.105 | 0 | 0.24 | 0.48 | 0.3 | 0.04 | 0.28 | 0.25 |
| 3 | 16874088 | rs13327361 | -0.787 | -1.199 | 0.799 | -0.183 | -0.011 | -0.023 | -0.023 | 0 | 0.36 | 0.51 | 0.27 | 0.3 | 0.3 | 0.32 |
| 3 | 16878403 | rs1866860 | 0.491 | -1.226 | 0.478 | -0.051 | -0.032 | 0.061 | 0.061 | 0 | 0.14 | 0.24 | 0.06 | 0.27 | 0.06 | 0.07 |
| 3 | 16879208 | rs1031300 | -1.018 | -0.808 | -0.576 | 0.014 | 0.011 | 0.029 | 0.029 | 0.031 | 0.44 | 0.77 | 0.49 | 0.39 | 0.43 | 0.41 |
| 3 | 16885235 | rs6796746 | -0.175 | -1.032 | -0.359 | -0.239 | 0.014 | 0.013 | 0.013 | 0 | 0.19 | 0.13 | 0.17 | 0.07 | 0.08 | 0.12 |
| 3 | 16891389 | rs9877897 | -1.201 | -0.88 | 0.221 | -0.271 | -0.006 | -0.054 | -0.054 | 0 | 0.31 | 0.43 | 0.37 | 0.14 | 0.31 | 0.34 |
| 3 | 16892058 | rs17042807 | 1.191 | -1.358 | -0.201 | -0.093 | 0.011 | -0.064 | -0.064 | 0 | 0.1 | 0 | 0.12 | 0.38 | 0.05 | 0.11 |
| 3 | 16894193 | rs2347655 | -1.056 | -1.091 | 0.022 | -0.602 | -0.005 | -0.05 | -0.05 | 0 | 0.31 | 0.43 | 0.37 | 0.15 | 0.31 | 0.34 |
| 3 | 16899031 | rs11929434 | -0.188 | -1.067 | 0.049 | -0.635 | 0.005 | 0.004 | 0.004 | 0 | 0.22 | 0.17 | 0.28 | 0.09 | 0.15 | 0.2 |
| 3 | 16900224 | rs9822053 | -0.654 | -1.049 | 0.162 | -0.631 | 0.062 | -0.012 | -0.012 | 0.084 | 0.09 | 0.24 | 0.08 | 0.05 | 0.16 | 0.15 |
| 3 | 16902837 | rs10510466 | -1.082 | -1.084 | 0.175 | -0.646 | -0.017 | -0.076 | -0.076 | 0.23 | 0.29 | 0.42 | 0.26 | 0.11 | 0.25 | 0.29 |
| 3 | 16911330 | rs9852648 | -1.3 | -1.026 | -0.279 | -0.613 | 0.092 | 0.007 | 0.007 | 0 | 0.11 | 0.33 | 0.2 | 0.08 | 0.23 | 0.21 |
| 3 | 16914975 | rs9852010 | -0.736 | -1.053 | -0.25 | -0.624 | 0.036 | -0.016 | -0.016 | 0 | 0.09 | 0.2 | 0.08 | 0.05 | 0.16 | 0.15 |
| 3 | 16917553 | rs12485389 | 0.161 | -0.988 | -0.316 | -0.68 | 0.047 | 0.071 | 0.071 | 0 | 0.2 | 0.03 | 0.16 | 0.06 | 0.09 | 0.14 |
| 3 | 16919573 | rs9878006 | -0.631 | -0.932 | -0.276 | -0.671 | 0.033 | -0.013 | -0.013 | 0.451 | 0.09 | 0.17 | 0.08 | 0.05 | 0.16 | 0.14 |
| 3 | 16923765 | rs1992380 | -0.711 | -1.126 | -0.127 | -0.846 | -0.034 | -0.059 | -0.059 | 0 | 0.27 | 0.41 | 0.3 | 0.09 | 0.17 | 0.23 |
| 3 | 16929109 | rs6788010 | -0.478 | -0.859 | -0.115 | -0.628 | -0.011 | -0.003 | -0.003 | 0 | 0.11 | 0.15 | 0.26 | 0.09 | 0.07 | 0.17 |
| 3 | 16949514 | rs1348109 | -0.998 | -0.729 | -0.522 | -0.676 | 0.004 | 0.014 | 0.014 | 1.579 | 0.48 | 0.36 | 0.52 | 0.6 | 0.46 | 0.51 |
| 3 | 16951768 | rs6799397 | 0.093 | -0.735 | -0.654 | -0.675 | 0.001 | -0.039 | -0.039 | 0 | 0.33 | 0.18 | 0.23 | 0.51 | 0.32 | 0.32 |
| 3 | 16959319 | rs1441039 | -0.747 | -0.729 | -0.658 | -0.687 | 0.012 | -0.008 | -0.008 | 0.104 | 0.42 | 0.28 | 0.47 | 0.59 | 0.4 | 0.48 |
| 3 | 16965021 | rs1025818 | -0.197 | -0.674 | -0.65 | -0.651 | -0.01 | -0.047 | -0.047 | 0.749 | 0.36 | 0.19 | 0.26 | 0.53 | 0.37 | 0.33 |
| 3 | 16970781 | rs9879150 | -0.194 | -0.622 | -0.609 | -0.617 | -0.005 | -0.026 | -0.026 | 0 | 0.34 | 0.25 | 0.33 | 0.58 | 0.38 | 0.41 |
| 3 | 16974610 | rs9880307 | -0.962 | -0.829 | -0.629 | -0.734 | 0.008 | 0.006 | 0.006 | 0 | 0.47 | 0.4 | 0.5 | 0.6 | 0.44 | 0.5 |
| 3 | 16974628 | rs9825009 | 0.176 | -0.934 | -0.638 | -0.511 | 0.04 | -0.07 | -0.07 | 1.14 | 0.28 | 0.18 | 0.18 | 0.53 | 0.19 | 0.3 |
| 3 | 16980586 | rs1372071 | 0.078 | -0.999 | -0.662 | -0.497 | 0.031 | -0.037 | -0.037 | 0 | 0.29 | 0.21 | 0.18 | 0.52 | 0.17 | 0.34 |
| 3 | 16985272 | rs7639882 | -0.182 | -1.168 | -0.647 | -0.973 | 0.029 | -0.002 | -0.002 | 0 | 0.46 | 0.56 | 0.27 | 0.53 | 0.29 | 0.47 |
| 3 | 17002619 | rs6769249 | -0.772 | -1.047 | -0.711 | -1.314 | -0.005 | 0.006 | 0.006 | 0.064 | 0.34 | 0.66 | 0.41 | 0.09 | 0.39 | 0.36 |
| 3 | 17015935 | rs34322012 | 0.826 | -1.199 | -0.798 | -1.291 | 0.05 | -0.127 | -0.127 | 1.439 | 0.29 | 0.09 | 0.19 | 0.55 | 0.2 | 0.3 |
| 3 | 17016794 | rs1806555 | -1.164 | -1.187 | -0.743 | -1.294 | -0.02 | -0.017 | -0.017 | 0 | 0.61 | 0.68 | 0.49 | 0.61 | 0.54 | 0.65 |
| 3 | 17020573 | rs9870977 | -0.352 | -1.148 | 0.051 | -1.393 | 0.003 | -0.044 | -0.044 | 0 | 0.3 | 0.35 | 0.37 | 0.09 | 0.33 | 0.32 |
| 3 | 17049965 | rs9876768 | 0.317 | -1.148 | -0.031 | -1.325 | -0.026 | 0.01 | 0.01 | 0 | 0.13 | 0.07 | 0.28 | 0.08 | 0.21 | 0.13 |
| 3 | 17051328 | rs7653834 | -0.257 | -1.312 | 0.084 | -1.381 | 0.002 | -0.001 | -0.001 | 0 | 0.48 | 0.51 | 0.29 | 0.55 | 0.32 | 0.53 |
| 3 | 17067085 | rs6442662 | 0.753 | -1.337 | 0.082 | -1.405 | -0.022 | -0.094 | -0.094 | 0 | 0.18 | 0.27 | 0.09 | 0.01 | 0.12 | 0.24 |
| 3 | 17074871 | rs12108065 | 0.014 | -1.194 | 0.088 | -1.351 | -0.006 | 0 | 0 | 0 | 0.46 | 0.51 | 0.29 | 0.55 | 0.3 | 0.49 |
| 3 | 17081577 | rs10510469 | 1.598 | -0.343 | -0.524 | -1.839 | -0.065 | 0.089 | 0.089 | 0.686 | 0.18 | 0.03 | 0.17 | 0 | 0.24 | 0.13 |
| 3 | 17083709 | rs6762816 | -0.863 | -1.03 | -0.162 | -1.474 | -0.02 | -0.006 | -0.006 | 0 | 0.61 | 0.65 | 0.59 | 0.63 | 0.53 | 0.68 |
| 3 | 17099309 | rs7610811 | -0.957 | -1.029 | -0.241 | -1.455 | -0.043 | -0.014 | -0.014 | 0 | 0.62 | 0.75 | 0.59 | 0.64 | 0.53 | 0.67 |
| 3 | 17106027 | rs4334629 | -0.749 | -0.646 | -0.041 | -1.324 | -0.012 | -0.004 | -0.004 | 0 | 0.53 | 0.66 | 0.46 | 0.58 | 0.51 | 0.58 |
| 3 | 17120206 | rs13062572 | 0.068 | 0.347 | 0.551 | -0.849 | 0.111 | -0.027 | -0.027 | 0 | 0.06 | 0.17 | 0.08 | 0.01 | 0.19 | 0.06 |
| 3 | 17121692 | rs6772001 | 0.509 | 0.157 | 0.522 | -0.883 | -0.028 | -0.107 | -0.107 | 0 | 0.16 | 0.43 | 0.11 | 0.03 | 0.1 | 0.15 |
| 3 | 17131501 | rs11549706 | 0.621 | 0.295 | -0.077 | -0.874 | -0.052 | 0.04 | 0.04 | 0.735 | 0.08 | 0.01 | 0.07 | 0.01 | 0.2 | 0.06 |
| 3 | 17132005 | rs11549705 | 0.76 | 0.166 | 0.065 | -0.908 | -0.021 | -0.066 | -0.066 | 0 | 0.15 | 0.22 | 0.09 | 0.03 | 0.1 | 0.15 |
| 3 | 17135184 | rs9881375 | 0.145 | 0.394 | -0.274 | -0.684 | 0.023 | -0.088 | -0.088 | 0.033 | 0.27 | 0.47 | 0.29 | 0.1 | 0.32 | 0.26 |
| 3 | 17135597 | rs6765957 | -0.818 | -0.071 | -0.687 | -0.749 | -0.012 | 0.03 | 0.03 | 0 | 0.51 | 0.72 | 0.49 | 0.61 | 0.51 | 0.56 |
| 3 | 17147963 | rs4684309 | -0.477 | 0.288 | -0.474 | 0.157 | 0.003 | -0.024 | -0.024 | 0.083 | 0.26 | 0.22 | 0.22 | 0.52 | 0.22 | 0.31 |

| | | | | | | | | | | | | | | | | |
|---|---|---|---|---|---|---|---|---|---|---|---|---|---|---|---|---|
| 3 | 17168021 | rs11710739 | -0.57 | 0.589 | 0.085 | -0.311 | 0.007 | -0.047 | -0.047 | 0.529 | 0.16 | 0.33 | 0.2 | 0.09 | 0.1 | 0.08 |
| 3 | 17173415 | rs4077616 | -0.486 | 0.862 | 0.712 | -0.312 | 0.032 | 0.01 | 0.01 | 0 | 0.1 | 0.25 | 0.06 | 0.07 | 0.1 | 0.13 |
| 3 | 17175592 | rs11716921 | -0.45 | 0.661 | 0.397 | -0.267 | 0.003 | -0.027 | -0.027 | 0.33 | 0.14 | 0.23 | 0.19 | 0.09 | 0.1 | 0.08 |
| 3 | 17178779 | rs4072191 | -0.952 | 1.109 | 0.24 | -0.245 | 0.01 | -0.013 | -0.013 | 0 | 0.37 | 0.54 | 0.38 | 0.24 | 0.43 | 0.24 |
| 3 | 17181317 | rs4685433 | -0.796 | 0.509 | -0.25 | -0.711 | 0.021 | 0.073 | 0.073 | 0 | 0.59 | 0.69 | 0.58 | 0.85 | 0.64 | 0.63 |
| 3 | 17182697 | rs4234541 | -0.138 | 0.046 | -0.002 | 0.174 | -0.002 | 0 | 0 | 0 | 0.42 | 0.42 | 0.4 | 0.57 | 0.44 | 0.47 |
| 3 | 17185728 | rs9813123 | -0.612 | -0.249 | 0.291 | 0.156 | -0.014 | 0.022 | 0.022 | 1.992 | 0.05 | 0.21 | 0.13 | 0.07 | 0.01 | 0.05 |
| 3 | 17187647 | rs17043133 | -0.713 | -0.263 | 0.475 | 0.372 | -0.005 | 0.054 | 0.054 | 0 | 0.14 | 0.34 | 0.19 | 0.2 | 0.1 | 0.16 |
| 3 | 17188392 | rs6778609 | -0.324 | -0.129 | -0.142 | 0.135 | 0 | 0.096 | 0.096 | 0 | 0.51 | 0.36 | 0.47 | 0.76 | 0.53 | 0.62 |
| 3 | 17190589 | rs17043135 | -0.126 | -0.309 | 0.197 | 0.858 | -0.005 | 0.052 | 0.052 | 0 | 0.16 | 0.2 | 0.16 | 0.3 | 0.12 | 0.2 |
| 3 | 17191688 | rs12632495 | -1.031 | -0.768 | 0.261 | 0.424 | -0.012 | 0.013 | 0.013 | 1.247 | 0.35 | 0.58 | 0.4 | 0.52 | 0.22 | 0.47 |
| 3 | 17191874 | rs11927361 | -0.069 | -0.657 | 0.606 | 0.131 | 0.008 | -0.016 | -0.016 | 0 | 0.08 | 0.29 | 0.06 | 0.02 | 0.03 | 0.12 |
| 3 | 17193348 | rs11720982 | 0.523 | -0.267 | -0.048 | -0.314 | 0.001 | 0.011 | 0.011 | 0 | 0.49 | 0.54 | 0.45 | 0.38 | 0.51 | 0.49 |
| 3 | 17194071 | rs13062346 | -1.174 | -0.322 | -0.603 | -0.261 | -0.017 | 0.022 | 0.022 | 0 | 0.79 | 0.88 | 0.83 | 0.93 | 0.68 | 0.87 |
| 3 | 17196846 | rs17043152 | 0.54 | -0.356 | -0.745 | -0.247 | 0.03 | -0.052 | -0.052 | 0 | 0.23 | 0.15 | 0.35 | 0.55 | 0.14 | 0.33 |
| 3 | 17198623 | rs4685439 | -1.105 | -0.018 | -0.736 | -0.383 | -0.039 | 0.041 | 0.041 | 0 | 0.81 | 0.91 | 0.83 | 0.93 | 0.68 | 0.88 |
| 3 | 17200091 | rs7610560 | -1.47 | 0.09 | -0.751 | -0.396 | -0.077 | 0.041 | 0.041 | 0 | 0.83 | 0.98 | 0.85 | 0.94 | 0.69 | 0.88 |
| 3 | 17202334 | rs13078644 | -1.172 | 0.032 | -0.733 | -0.364 | -0.033 | 0.038 | 0.038 | 0 | 0.81 | 0.92 | 0.84 | 0.94 | 0.69 | 0.88 |
| 3 | 17208267 | rs1138454 | 0.851 | -0.272 | -0.839 | -0.548 | -0.023 | 0.096 | 0.096 | 0.722 | 0.37 | 0.21 | 0.31 | 0.08 | 0.39 | 0.38 |
| 3 | 17213357 | rs4685442 | 0.136 | -0.347 | -0.884 | -0.484 | 0.042 | -0.033 | -0.033 | 0.266 | 0.32 | 0.21 | 0.37 | 0.58 | 0.2 | 0.4 |
| 3 | 17214070 | rs2470584 | 1.084 | -0.241 | -0.792 | -0.656 | -0.009 | 0.052 | 0.052 | 0.02 | 0.31 | 0.17 | 0.29 | 0.07 | 0.32 | 0.31 |
| 3 | 17221252 | rs1463222 | 0.346 | -0.089 | -0.401 | -0.549 | 0.006 | 0.087 | 0.087 | 0.483 | 0.47 | 0.69 | 0.39 | 0.25 | 0.46 | 0.35 |
| 3 | 17230884 | rs4685443 | 0.843 | 0.135 | -0.627 | -0.102 | -0.03 | -0.169 | -0.169 | 0 | 0.22 | 0.01 | 0.28 | 0.46 | 0.17 | 0.27 |
| 3 | 17266023 | rs1375817 | 0.352 | 0.117 | 0.358 | -0.252 | -0.001 | -0.043 | -0.043 | 0 | 0.44 | 0.42 | 0.33 | 0.04 | 0.45 | 0.29 |
| 3 | 17276455 | rs2733498 | 0.353 | 0.112 | 0.351 | -0.254 | -0.001 | -0.031 | -0.031 | 0 | 0.44 | 0.39 | 0.33 | 0.04 | 0.45 | 0.29 |
| 3 | 17295308 | rs2348004 | 0.251 | 0.102 | 0.428 | -0.245 | 0.001 | 0.02 | 0.02 | 0.585 | 0.46 | 0.62 | 0.4 | 0.22 | 0.47 | 0.35 |
| 3 | 17302558 | rs2596648 | 0.196 | 0.164 | 0.423 | -0.804 | 0.003 | 0.064 | 0.064 | 0.556 | 0.47 | 0.67 | 0.41 | 0.23 | 0.47 | 0.35 |
| 3 | 17315758 | rs2084572 | 0.336 | 0.205 | 0.349 | -1.059 | -0.001 | -0.043 | -0.043 | 0 | 0.44 | 0.42 | 0.33 | 0.02 | 0.44 | 0.29 |
| 3 | 17322397 | rs2733524 | 0.253 | 0.204 | 0.353 | -1.059 | 0 | -0.086 | -0.086 | 0.079 | 0.45 | 0.47 | 0.33 | 0.04 | 0.44 | 0.3 |
| 3 | 17324733 | rs767685 | 0.344 | 0.233 | 0.25 | -1.052 | -0.002 | -0.015 | -0.015 | 0 | 0.44 | 0.36 | 0.33 | 0.02 | 0.44 | 0.29 |
| 3 | 17346355 | rs283943 | -0.453 | 0.289 | 0.084 | -1.031 | -0.001 | -0.118 | -0.118 | 2.368 | 0.56 | 0.5 | 0.66 | 0.96 | 0.56 | 0.69 |
| 3 | 17364413 | rs283932 | -0.621 | 0.432 | 0.035 | -0.893 | -0.016 | -0.002 | -0.002 | 0 | 0.76 | 0.82 | 0.79 | 0.72 | 0.66 | 0.76 |
| 3 | 17394615 | rs1374197 | 0.271 | 0.228 | 0.082 | -1.143 | 0.007 | 0.201 | 0.201 | 0.773 | 0.49 | 0.75 | 0.37 | 0.08 | 0.47 | 0.38 |
| 3 | 17398386 | rs9850127 | 0.361 | 0.224 | 0.123 | -1.166 | 0.008 | 0.143 | 0.143 | 0.637 | 0.48 | 0.72 | 0.37 | 0.06 | 0.47 | 0.37 |
| 3 | 17399281 | rs13318886 | 0.325 | 0.236 | 0.13 | -1.161 | 0.003 | 0.234 | 0.234 | 0.754 | 0.49 | 0.78 | 0.37 | 0.07 | 0.47 | 0.38 |
| 3 | 17406273 | rs6801690 | 0.386 | 0.089 | -0.013 | -1.217 | -0.007 | 0.094 | 0.094 | 0.337 | 0.43 | 0.71 | 0.36 | 0.07 | 0.46 | 0.38 |
| 3 | 17452925 | rs1446300 | -0.693 | 0.136 | -0.086 | -1.056 | -0.107 | -0.014 | -0.014 | 0 | 0.73 | 0.98 | 0.8 | 0.77 | 0.65 | 0.83 |
| 3 | 17453982 | rs9824191 | 1.747 | 0.133 | 0.021 | -1.517 | -0.01 | 0.001 | 0.001 | 0 | 0.17 | 0.13 | 0.14 | 0.02 | 0.22 | 0.19 |
| 3 | 17464862 | rs13317478 | 1.018 | 0.095 | 0.222 | -1.513 | -0.002 | -0.018 | -0.018 | 0 | 0.31 | 0.24 | 0.24 | 0.02 | 0.33 | 0.27 |
| 3 | 17486643 | rs17043638 | 0.65 | 0.194 | 0.241 | -1.399 | 0.013 | 0.01 | 0.01 | 1.16 | 0.3 | 0.21 | 0.44 | 0.7 | 0.22 | 0.45 |
| 3 | 17532320 | rs13095571 | 0.041 | 0.334 | 0.466 | -1.557 | -0.001 | -0.067 | -0.067 | 0 | 0.42 | 0.41 | 0.33 | 0.02 | 0.44 | 0.3 |
| 3 | 17627877 | rs9681332 | 1.374 | 0.495 | -0.046 | -1.188 | 0.013 | -0.035 | -0.035 | 0 | 0.07 | 0.16 | 0.07 | 0 | 0.11 | 0.02 |
| 3 | 17661762 | rs9310527 | -0.12 | 1.005 | -0.389 | -1.532 | -0.041 | 0.222 | 0.222 | 0 | 0.46 | 0.85 | 0.45 | 0.08 | 0.49 | 0.39 |
| 3 | 17671037 | rs4470554 | -0.248 | 0.599 | 0.187 | -1.559 | 0 | -0.104 | -0.104 | 0 | 0.58 | 0.53 | 0.67 | 0.98 | 0.56 | 0.7 |
| 3 | 17679106 | rs9855515 | -0.817 | 0.577 | 0.242 | -1.606 | 0.001 | -0.014 | -0.014 | 0.02 | 0.66 | 0.74 | 0.75 | 0.98 | 0.67 | 0.73 |
| 3 | 17697368 | rs11128838 | -0.208 | 0.631 | 0.225 | -1.404 | -0.007 | 0.193 | 0.193 | 0 | 0.54 | 0.21 | 0.62 | 0.93 | 0.53 | 0.62 |
| 3 | 17721297 | rs10510477 | -0.416 | 0.64 | 0.309 | -1.409 | 0 | -0.097 | -0.097 | 0.081 | 0.57 | 0.44 | 0.66 | 0.98 | 0.56 | 0.7 |
| 3 | 17736035 | rs4908959 | -0.477 | 0.64 | 0.198 | -1.409 | 0.001 | -0.138 | -0.138 | 0 | 0.58 | 0.48 | 0.67 | 0.98 | 0.56 | 0.7 |
| 3 | 17756467 | rs4909012 | -0.477 | 0.654 | 0.179 | -1.409 | 0.002 | -0.134 | -0.134 | 0 | 0.58 | 0.48 | 0.67 | 0.98 | 0.56 | 0.7 |
| 3 | 17765959 | rs10433559 | 1.103 | 0.478 | -0.425 | -0.683 | -0.028 | -0.139 | -0.139 | 0 | 0.21 | 0.01 | 0.26 | 0.4 | 0.15 | 0.26 |
| 3 | 17788430 | rs4312680 | -0.454 | 0.431 | -0.209 | -0.906 | -0.003 | 0.15 | 0.15 | 0 | 0.54 | 0.24 | 0.63 | 0.93 | 0.53 | 0.62 |
| 3 | 17791027 | rs11708664 | 0.846 | 0.804 | -0.364 | -0.44 | -0.097 | -0.056 | -0.056 | 0 | 0.16 | 0.01 | 0.14 | 0.22 | 0.31 | 0.14 |
| 3 | 17812688 | rs11921152 | -0.407 | 0.795 | 0.195 | -0.557 | -0.001 | -0.07 | -0.07 | 0 | 0.44 | 0.5 | 0.62 | 0.98 | 0.48 | 0.62 |
| 3 | 17814553 | rs11128857 | -0.115 | 0.795 | 0.184 | -0.546 | -0.009 | 0.026 | 0.026 | 0 | 0.42 | 0.3 | 0.59 | 0.93 | 0.46 | 0.54 |
| 3 | 17856532 | rs12492361 | 0.768 | 0.794 | -0.463 | -0.695 | 0.009 | -0.03 | -0.03 | 0 | 0.36 | 0.37 | 0.23 | 0.01 | 0.28 | 0.18 |
| 3 | 17868000 | rs10510480 | 1.178 | 0.977 | -0.294 | 0.273 | -0.028 | -0.069 | -0.069 | 0 | 0.11 | 0 | 0.24 | 0.36 | 0.09 | 0.2 |
| 3 | 17877828 | rs17200979 | -0.624 | 0.869 | 0.084 | -0.253 | -0.007 | 0.027 | 0.027 | 0 | 0.44 | 0.56 | 0.6 | 0.78 | 0.48 | 0.57 |
| 3 | 17886959 | rs9862290 | -0.464 | 1.01 | -0.201 | -1.099 | 0.006 | -0.09 | -0.09 | 0 | 0.43 | 0.44 | 0.61 | 0.91 | 0.46 | 0.55 |
| 3 | 17892481 | rs6577599 | 0.292 | 1.029 | -0.604 | -0.415 | -0.019 | 0.051 | 0.051 | 0 | 0.68 | 0.88 | 0.59 | 0.26 | 0.68 | 0.54 |
| 3 | 17915113 | rs12493277 | -0.702 | 1.466 | -0.542 | -0.296 | -0.034 | -0.024 | -0.024 | 0 | 0.07 | 0.01 | 0.04 | 0.07 | 0.08 | 0.06 |
| 3 | 17941042 | rs11128872 | -1.53 | 1.26 | 0.325 | -0.384 | 0.008 | 0.005 | 0.005 | 0.084 | 0.09 | 0.17 | 0.21 | 0.1 | 0.07 | 0.07 |
| 3 | 17950027 | rs12492931 | -1.17 | 1.285 | 0.318 | -0.373 | 0.003 | 0.001 | 0.001 | 0 | 0.07 | 0.11 | 0.21 | 0.1 | 0.07 | 0.07 |
| 3 | 17957737 | rs1430893 | -0.74 | 1.287 | 0.255 | -0.378 | -0.014 | 0.012 | 0.012 | 0 | 0.06 | 0.02 | 0.2 | 0.05 | 0.07 | 0.07 |
| 3 | 17963984 | rs1346887 | -1.271 | 1.281 | 0.412 | -0.354 | 0.011 | 0.012 | 0.012 | 0 | 0.07 | 0.16 | 0.23 | 0.09 | 0.07 | 0.06 |
| 3 | 17967203 | rs9814647 | -0.978 | 1.346 | 0.64 | -0.624 | 0.011 | -0.08 | -0.08 | 0.078 | 0.35 | 0.45 | 0.59 | 0.86 | 0.38 | 0.51 |
| 3 | 17977785 | rs11928474 | -0.265 | 1.404 | 0.548 | -0.619 | 0.001 | -0.005 | -0.005 | 0 | 0.28 | 0.25 | 0.36 | 0.77 | 0.31 | 0.46 |
| 3 | 17984441 | rs17044152 | -1.643 | 1.382 | 0.524 | -0.702 | 0.002 | 0.002 | 0.002 | 0 | 0.11 | 0.11 | 0.26 | 0.12 | 0.12 | 0.07 |
| 3 | 17987400 | rs6577604 | -0.891 | 1.354 | 0.502 | -0.771 | 0.003 | -0.036 | -0.036 | 0 | 0.36 | 0.32 | 0.58 | 0.86 | 0.38 | 0.53 |
| 3 | 17989553 | rs4128378 | -1.393 | 1.334 | 0.458 | -0.816 | 0.006 | -0.104 | -0.104 | 0 | 0.4 | 0.37 | 0.63 | 0.93 | 0.43 | 0.55 |

| | | | | | | | | | | | | | | | | |
|---|---|---|---|---|---|---|---|---|---|---|---|---|---|---|---|---|
| 3 | 17995710 | rs4908948 | -1.559 | 1.043 | 0.858 | -0.681 | 0.019 | -0.019 | -0.019 | 0 | 0.1 | 0.33 | 0.24 | 0.05 | 0.07 | 0.05 |
| 3 | 18010703 | rs6577606 | -0.065 | 1.471 | 0.368 | -0.617 | -0.04 | 0.047 | 0.047 | 0 | 0.26 | 0.13 | 0.54 | 0.84 | 0.32 | 0.45 |
| 3 | 18031227 | rs7429951 | -0.512 | 1.248 | 0.64 | -0.723 | -0.002 | -0.001 | -0.001 | 0 | 0.32 | 0.26 | 0.58 | 0.83 | 0.36 | 0.49 |
| 3 | 18083383 | rs4130063 | -1.65 | 1.786 | 0.952 | -0.576 | 0.06 | 0.056 | 0.056 | 1.146 | 0.1 | 0.27 | 0.09 | 0.12 | 0.11 | 0.06 |
| 3 | 18089105 | rs7620376 | 0.896 | 1.686 | 0.731 | 0.455 | -0.014 | 0.12 | 0.12 | 1.012 | 0.1 | 0.19 | 0.12 | 0.39 | 0.06 | 0.05 |
| 3 | 18095128 | rs7432188 | 0.409 | 1.938 | 0.402 | 0.483 | -0.088 | -0.128 | -0.128 | 0 | 0.16 | 0.02 | 0.44 | 0.47 | 0.28 | 0.42 |
| 3 | 18095177 | rs9838434 | -0.978 | 1.636 | 0.793 | -0.159 | 0.02 | 0.034 | 0.034 | 0.128 | 0.05 | 0.15 | 0.06 | 0.08 | 0.05 | 0.02 |
| 3 | 18117085 | rs9861146 | -2.085 | 1.835 | 1.394 | -0.246 | 0.109 | 0.036 | 0.036 | 3.016 | 0.14 | 0.47 | 0.14 | 0.12 | 0.16 | 0.06 |
| 3 | 18117117 | rs9824023 | -1.424 | 1.81 | 1.216 | -0.247 | 0.066 | 0.003 | 0.003 | 3.452 | 0.1 | 0.39 | 0.07 | 0.04 | 0.07 | 0.04 |
| 3 | 18133992 | rs9854107 | -0.255 | 2.057 | 0.847 | -0.575 | -0.013 | 0.038 | 0.038 | 0 | 0.29 | 0.19 | 0.56 | 0.81 | 0.34 | 0.5 |
| 3 | 18134253 | rs1355725 | -0.322 | 2.022 | 0.835 | -0.615 | -0.007 | 0.052 | 0.052 | 0 | 0.3 | 0.22 | 0.56 | 0.83 | 0.34 | 0.5 |
| 3 | 18138740 | rs955673 | -0.65 | 2.126 | 1.18 | -0.252 | 0.088 | 0.05 | 0.05 | 0 | 0.05 | 0.16 | 0.09 | 0.08 | 0.11 | 0.02 |
| 3 | 18148270 | rs6577615 | -0.869 | 1.82 | -0.368 | -0.864 | -0.042 | 0.115 | 0.115 | 0 | 0.81 | 0.99 | 0.88 | 1 | 0.82 | 0.92 |
| 3 | 18168803 | rs7627406 | -0.422 | 1.998 | 0.398 | -0.413 | 0.047 | 0.023 | 0.023 | 2.152 | 0.06 | 0.23 | 0.04 | 0.04 | 0.05 | 0.04 |
| 3 | 18190493 | rs13060927 | -0.262 | 1.85 | -0.107 | -0.709 | -0.009 | 0.001 | 0.001 | 0 | 0.11 | 0.04 | 0.09 | 0.08 | 0.13 | 0.16 |
| 3 | 18193653 | rs11713547 | 0.813 | 1.791 | -0.177 | -0.726 | 0.009 | 0.023 | 0.023 | 0 | 0.07 | 0.01 | 0.01 | 0 | 0.04 | 0.13 |
| 3 | 18201859 | rs938182 | -0.674 | 1.517 | -0.337 | -1.549 | 0.006 | -0.048 | -0.048 | 0 | 0.5 | 0.49 | 0.68 | 0.93 | 0.59 | 0.67 |
| 3 | 18206165 | rs2102623 | 0.649 | 1.857 | -0.737 | -1.282 | -0.027 | -0.007 | -0.007 | 0 | 0.05 | 0.01 | 0.05 | 0.09 | 0.09 | 0.07 |
| 3 | 18227421 | rs9809856 | -0.002 | 2.418 | -0.681 | -1.214 | -0.018 | 0.025 | 0.025 | 0.014 | 0.38 | 0.28 | 0.61 | 0.84 | 0.46 | 0.5 |
| 3 | 18230131 | rs2001792 | 1.056 | 1.938 | -1.11 | -1.58 | 0.051 | 0.153 | 0.153 | 0 | 0.32 | 0.08 | 0.11 | 0 | 0.26 | 0.2 |
| 3 | 18235724 | rs13076743 | 1.584 | 1.983 | -1.119 | -1.522 | -0.05 | -0.017 | -0.017 | 0 | 0.11 | 0.01 | 0.11 | 0.08 | 0.15 | 0.05 |
| 3 | 18250509 | rs7618405 | 1.744 | 1.781 | -1.19 | -1.476 | 0.042 | 0.101 | 0.101 | 0 | 0.25 | 0.08 | 0.11 | 0.07 | 0.21 | 0.21 |
| 3 | 18261521 | rs6796378 | 0.479 | 2.462 | -0.805 | -1.004 | 0.014 | -0.01 | -0.01 | 0.009 | 0.27 | 0.3 | 0.53 | 0.76 | 0.33 | 0.44 |
| 3 | 18271059 | rs13092526 | 0.114 | 2.559 | -0.719 | -0.905 | -0.011 | 0.003 | 0.003 | 0 | 0.17 | 0.11 | 0.15 | 0.17 | 0.22 | 0.11 |
| 3 | 18302624 | rs9845097 | -0.681 | 2.393 | -0.7 | -0.169 | 0.013 | 0.009 | 0.009 | 0 | 0.34 | 0.37 | 0.58 | 0.63 | 0.51 | 0.53 |
| 3 | 18308311 | rs9876469 | 0.272 | 2.571 | -0.722 | -0.165 | -0.022 | 0.006 | 0.006 | 0 | 0.17 | 0.1 | 0.16 | 0.17 | 0.26 | 0.11 |
| 3 | 18308436 | rs7432753 | 1.094 | 2.574 | -0.654 | -0.139 | 0.035 | 0.084 | 0.084 | 0.716 | 0.16 | 0.2 | 0.41 | 0.46 | 0.25 | 0.41 |
| 3 | 18313624 | rs9837205 | -0.69 | 2.519 | -0.538 | -0.167 | 0.018 | 0.01 | 0.01 | 0 | 0.36 | 0.44 | 0.58 | 0.65 | 0.51 | 0.53 |
| 3 | 18314866 | rs7653460 | 0.867 | 2.369 | -0.747 | -0.375 | 0.004 | 0.001 | 0.001 | 0 | 0.05 | 0.03 | 0.05 | 0.1 | 0.09 | 0.06 |
| 3 | 18318129 | rs7427439 | -1.246 | 2.237 | -0.593 | -0.632 | 0.017 | -0.002 | -0.002 | 0 | 0.65 | 0.71 | 0.86 | 0.93 | 0.75 | 0.64 |
| 3 | 18318327 | rs4128294 | 1.005 | 2.857 | -0.481 | -0.017 | -0.052 | 0.006 | 0.006 | 0.68 | 0.11 | 0.01 | 0.1 | 0.06 | 0.17 | 0.05 |
| 3 | 18319785 | rs7428577 | -0.03 | 2.28 | -0.443 | 0.003 | -0.031 | 0.007 | 0.007 | 0 | 0.17 | 0.07 | 0.16 | 0.17 | 0.26 | 0.11 |
| 3 | 18335741 | rs7374717 | 0.57 | 1.65 | -0.241 | 0.212 | -0.001 | 0.027 | 0.027 | 0 | 0.12 | 0.09 | 0.38 | 0.47 | 0.12 | 0.31 |
| 3 | 18336700 | rs6807560 | 0.276 | 1.855 | -0.297 | 0.177 | -0.048 | -0.02 | -0.02 | 0 | 0.3 | 0.15 | 0.27 | 0.3 | 0.49 | 0.16 |
| 3 | 18353002 | rs11921478 | 0.269 | 1.725 | -0.296 | 0.142 | -0.041 | -0.018 | -0.018 | 0.478 | 0.3 | 0.16 | 0.27 | 0.3 | 0.47 | 0.15 |
| 3 | 18380444 | rs7433691 | 0.743 | 1.215 | -0.226 | 0.187 | -0.002 | 0.004 | 0.004 | 0 | 0.13 | 0.09 | 0.42 | 0.57 | 0.14 | 0.33 |
| 3 | 18383022 | rs9872636 | -1.034 | 1.135 | -0.183 | -0.029 | 0.031 | 0.03 | 0.03 | 0.146 | 0.49 | 0.6 | 0.76 | 0.92 | 0.65 | 0.57 |
| 3 | 18385732 | rs7434191 | 0.27 | 1.165 | -0.821 | 0.2 | -0.017 | -0.006 | -0.006 | 0 | 0.31 | 0.23 | 0.29 | 0.3 | 0.52 | 0.16 |
| 3 | 18389210 | rs3935618 | 0.816 | 1.058 | -1.219 | 0.47 | -0.021 | -0.083 | -0.083 | 0 | 0.12 | 0.01 | 0.4 | 0.48 | 0.13 | 0.3 |
| 3 | 18391478 | rs12330375 | -0.325 | 0.713 | -0.996 | -0.388 | 0.024 | 0.081 | 0.081 | 0 | 0.43 | 0.31 | 0.69 | 0.75 | 0.63 | 0.44 |
| 3 | 18400203 | rs7635386 | 0.235 | 1.145 | -0.959 | 0.106 | -0.031 | -0.072 | -0.072 | 0.016 | 0.28 | 0.11 | 0.2 | 0.37 | 0.39 | 0.21 |
| 3 | 18400621 | rs4129096 | 1.025 | 0.545 | -1.527 | -0.423 | -0.004 | -0.055 | -0.055 | 0 | 0.3 | 0.11 | 0.51 | 0.39 | 0.27 | 0.31 |
| 3 | 18401567 | rs11927883 | 0.512 | -0.335 | -0.712 | -0.498 | -0.009 | -0.025 | -0.025 | 0.097 | 0.08 | 0.16 | 0.03 | 0.04 | 0.05 | 0.18 |
| 3 | 18409077 | rs9841621 | -0.949 | -0.422 | 0.117 | -0.304 | 0.022 | 0.075 | 0.075 | 0 | 0.07 | 0.33 | 0.06 | 0.16 | 0.02 | 0.25 |
| 3 | 18409579 | rs11720566 | 0.726 | -0.285 | -0.515 | -0.3 | -0.054 | -0.099 | -0.099 | 10.491 | 0.26 | 0.08 | 0.2 | 0.36 | 0.39 | 0.21 |
| 3 | 18412358 | rs9714119 | 0.218 | -0.659 | -0.298 | -0.487 | -0.001 | -0.001 | -0.001 | 0 | 0.44 | 0.53 | 0.62 | 0.54 | 0.39 | 0.61 |
| 3 | 18415632 | rs9310553 | 0.146 | -0.456 | -0.452 | -0.592 | 0.002 | 0.003 | 0.003 | 0 | 0.58 | 0.61 | 0.73 | 0.92 | 0.57 | 0.69 |
| 3 | 18415702 | rs9809323 | -1.224 | -0.543 | -0.416 | -0.66 | 0.025 | 0.064 | 0.064 | 0 | 0.77 | 0.86 | 0.86 | 0.93 | 0.83 | 0.85 |
| 3 | 18419859 | rs10460976 | 0.288 | -0.434 | -0.414 | -0.615 | 0.001 | -0.015 | -0.015 | 0.029 | 0.56 | 0.58 | 0.73 | 0.92 | 0.57 | 0.69 |
| 3 | 18421990 | rs11128887 | 0.966 | -0.064 | -0.925 | -0.454 | 0.01 | -0.047 | -0.047 | 0 | 0.18 | 0.19 | 0.12 | 0.01 | 0.25 | 0.16 |
| 3 | 18427477 | rs9756797 | 0.425 | 0.284 | -1.31 | 0.187 | -0.033 | -0.021 | -0.021 | 0.055 | 0.35 | 0.2 | 0.54 | 0.4 | 0.35 | 0.42 |
| 3 | 18439331 | rs11712525 | 1.03 | 0.318 | -1.572 | 0.224 | -0.001 | -0.016 | -0.016 | 0 | 0.16 | 0.11 | 0.14 | 0.39 | 0.21 | 0.14 |
| 3 | 18453843 | rs9713269 | -0.143 | 0.431 | -1.448 | 0.456 | -0.052 | -0.031 | -0.031 | 0 | 0.35 | 0.16 | 0.54 | 0.4 | 0.35 | 0.41 |
| 3 | 18465443 | rs6762753 | 0.527 | -0.439 | -0.915 | 0.914 | 0.006 | -0.007 | -0.007 | 0.015 | 0.76 | 0.78 | 0.9 | 0.64 | 0.85 | 0.82 |
| 3 | 18468523 | rs6550461 | -0.361 | -0.789 | -0.611 | 1.026 | -0.006 | 0.009 | 0.009 | 0 | 0.2 | 0.28 | 0.13 | 0.22 | 0.18 | 0.15 |
| 3 | 18472769 | rs7642423 | -0.949 | -1.303 | -0.313 | 1.044 | 0.005 | -0.002 | -0.002 | 0.722 | 0.36 | 0.64 | 0.33 | 0.6 | 0.27 | 0.39 |
| 3 | 18482558 | rs1328542 | 0.899 | -0.823 | -0.544 | 0.728 | -0.029 | -0.033 | -0.033 | 0 | 0.26 | 0.1 | 0.43 | 0.32 | 0.25 | 0.33 |
| 3 | 18495301 | rs714141 | -1.264 | -0.764 | -0.382 | 0.653 | 0.035 | 0.01 | 0.01 | 7.174 | 0.48 | 0.66 | 0.59 | 0.49 | 0.36 | 0.39 |
| 3 | 18507382 | rs6550850 | 1.122 | -0.506 | -0.355 | 0.618 | -0.024 | 0.013 | 0.013 | 0.358 | 0.19 | 0.05 | 0.14 | 0.23 | 0.17 | 0.31 |
| 3 | 18512239 | rs1513763 | -0.983 | -0.887 | 0.558 | 0.928 | -0.039 | -0.01 | -0.01 | 0 | 0.24 | 0.42 | 0.24 | 0.15 | 0.17 | 0.18 |
| 3 | 18521394 | rs2076802 | -0.781 | -0.203 | 0.707 | 1.221 | 0.003 | -0.005 | -0.005 | 0 | 0.17 | 0.15 | 0.3 | 0.24 | 0.12 | 0.14 |
| 3 | 18536553 | rs1822120 | -1.604 | -0.359 | 1.01 | 1.312 | -0.007 | 0.009 | 0.009 | 0 | 0.32 | 0.34 | 0.41 | 0.35 | 0.29 | 0.37 |
| 3 | 18538587 | rs17029286 | 0.039 | -0.391 | 0.947 | 1.319 | -0.002 | 0.014 | 0.014 | 0 | 0.13 | 0.06 | 0.08 | 0.05 | 0.15 | 0.14 |
| 3 | 18549716 | rs4973727 | -0.246 | -0.394 | 0.898 | 1.338 | -0.007 | 0.014 | 0.014 | 0.016 | 0.13 | 0.14 | 0.27 | 0.25 | 0.08 | 0.13 |
| 3 | 18557288 | rs6807937 | -1.829 | -0.394 | 0.988 | 1.27 | -0.001 | 0 | 0 | 0.933 | 0.37 | 0.61 | 0.55 | 0.46 | 0.31 | 0.4 |
| 3 | 18581848 | rs7633075 | -1.346 | -0.755 | 1.358 | 1.356 | -0.053 | 0.101 | 0.101 | 1.992 | 0.21 | 0.46 | 0.43 | 0.35 | 0.12 | 0.24 |
| 3 | 18585365 | rs1877648 | -1.521 | -0.252 | 0.808 | 1.46 | -0.008 | 0.228 | 0.228 | 0 | 0.11 | 0.39 | 0.31 | 0.27 | 0.07 | 0.17 |
| 3 | 18589151 | rs17044604 | -0.241 | 0.128 | -0.276 | 1.658 | -0.006 | 0.112 | 0.112 | 0 | 0.07 | 0.12 | 0.26 | 0.27 | 0.05 | 0.1 |
| 3 | 18602534 | rs717793 | -0.63 | 0.173 | -0.352 | 1.181 | -0.004 | 0.009 | 0.009 | 1 | 0.36 | 0.64 | 0.57 | 0.43 | 0.33 | 0.4 |

| | | | | | | | | | | | | | | | | |
|---|---|---|---|---|---|---|---|---|---|---|---|---|---|---|---|---|
| 3 | 18603113 | rs6785093 | -0.681 | 0.006 | -0.055 | 1.13 | -0.016 | 0.049 | 0.049 | 0.009 | 0.25 | 0.5 | 0.37 | 0.39 | 0.21 | 0.23 |
| 3 | 18604810 | rs189221 | -0.433 | -0.308 | -0.66 | 0.082 | 0.009 | 0.064 | 0.064 | 0 | 0.57 | 0.67 | 0.58 | 0.88 | 0.59 | 0.72 |
| 3 | 18606228 | rs13074328 | 0.186 | 0.265 | -0.803 | 0.432 | -0.026 | 0.025 | 0.025 | 0.003 | 0.16 | 0.29 | 0.14 | 0.19 | 0.12 | 0.15 |
| 3 | 18607171 | rs336629 | 0.239 | 0.918 | -1.534 | 0.483 | -0.121 | 0.042 | 0.042 | 0 | 0.26 | 0.06 | 0.26 | 0.17 | 0.4 | 0.29 |
| 3 | 18607538 | rs336630 | 0.336 | 0.987 | -1.643 | 0.512 | -0.153 | 0.026 | 0.026 | 0 | 0.25 | 0.03 | 0.25 | 0.17 | 0.4 | 0.29 |
| 3 | 18609345 | rs9838549 | -0.525 | 0.841 | -0.982 | 0.577 | -0.001 | -0.015 | -0.015 | 0 | 0.38 | 0.5 | 0.31 | 0.26 | 0.37 | 0.32 |
| 3 | 18609478 | rs9876904 | -0.836 | 0.877 | -1.431 | 0.879 | 0.02 | 0.024 | 0.024 | 1.725 | 0.4 | 0.85 | 0.57 | 0.66 | 0.38 | 0.38 |
| 3 | 18609712 | rs17044638 | -0.364 | 1.477 | -1.185 | 1.791 | -0.028 | -0.011 | -0.011 | 0 | 0.07 | 0.01 | 0.1 | 0.41 | 0.13 | 0.13 |
| 3 | 18610179 | rs455932 | -0.259 | 1.783 | -1.591 | 1.188 | -0.08 | -0.016 | -0.016 | 2.098 | 0.31 | 0.08 | 0.28 | 0.27 | 0.4 | 0.3 |
| 3 | 18610677 | rs334958 | -0.385 | 1.752 | -1.636 | 1.174 | -0.095 | -0.024 | -0.024 | 0 | 0.32 | 0.07 | 0.28 | 0.28 | 0.4 | 0.31 |
| 3 | 18616112 | rs336639 | 0.353 | 1.83 | -1.617 | 1.118 | -0.082 | 0.146 | 0.146 | 0 | 0.26 | 0.06 | 0.19 | 0.03 | 0.31 | 0.19 |
| 3 | 18617152 | rs4364185 | -1.25 | 1.557 | -0.786 | 1.065 | -0.004 | 0.014 | 0.014 | 0.636 | 0.32 | 0.63 | 0.55 | 0.61 | 0.35 | 0.31 |
| 3 | 18617627 | rs996000 | -1.415 | 1.511 | -0.708 | 1.045 | 0 | -0.002 | -0.002 | 0.088 | 0.33 | 0.67 | 0.55 | 0.61 | 0.36 | 0.31 |
| 3 | 18621139 | rs462296 | -1.728 | 1.659 | -0.611 | 1.148 | 0.011 | 0.004 | 0.004 | 0.383 | 0.59 | 0.73 | 0.75 | 0.63 | 0.67 | 0.5 |
| 3 | 18623462 | rs2361571 | -0.412 | 1.601 | 0.03 | 0.42 | 0.005 | 0.033 | 0.033 | 0.084 | 0.21 | 0.28 | 0.43 | 0.34 | 0.25 | 0.17 |
| 3 | 18624900 | rs1849590 | -0.414 | 1.547 | 0.126 | 0.37 | 0.009 | -0.048 | -0.048 | 0 | 0.13 | 0.25 | 0.29 | 0.06 | 0.16 | 0.05 |
| 3 | 18630689 | rs336616 | -0.616 | 1.628 | 0.138 | 0.539 | -0.048 | -0.026 | -0.026 | 0 | 0.41 | 0.19 | 0.42 | 0.44 | 0.5 | 0.44 |
| 3 | 18645611 | rs6762250 | -1.27 | 1.602 | 0.551 | 0.497 | 0.04 | 0.022 | 0.022 | 0.408 | 0.12 | 0.37 | 0.08 | 0.12 | 0.13 | 0.14 |
| 3 | 18650015 | rs336610 | -1.194 | 1.58 | 0.698 | 0.457 | -0.001 | -0.001 | -0.001 | 0 | 0.51 | 0.54 | 0.43 | 0.43 | 0.56 | 0.47 |
| 3 | 18662477 | rs1440968 | 0.213 | 1.572 | 0.533 | 0.409 | -0.002 | -0.016 | -0.016 | 0 | 0.06 | 0.09 | 0.02 | 0 | 0.02 | 0.01 |
| 3 | 18673993 | rs6792314 | 0.383 | 1.683 | 0.46 | 0.329 | -0.025 | 0.055 | 0.055 | 0 | 0.3 | 0.16 | 0.22 | 0.04 | 0.34 | 0.22 |
| 3 | 18674920 | rs4280655 | -1.664 | 1.607 | 0.373 | 0.338 | 0.007 | 0.03 | 0.03 | 1.604 | 0.26 | 0.61 | 0.49 | 0.45 | 0.34 | 0.31 |
| 3 | 18676151 | rs6802224 | -0.853 | 1.587 | 0.393 | 0.375 | -0.022 | 0.057 | 0.057 | 0 | 0.45 | 0.26 | 0.36 | 0.31 | 0.46 | 0.34 |
| 3 | 18677075 | rs7641940 | -1.403 | 1.564 | 0.42 | 0.378 | 0.026 | -0.009 | -0.009 | 0 | 0.17 | 0.61 | 0.36 | 0.17 | 0.24 | 0.18 |
| 3 | 18695490 | rs9848614 | -1.862 | 1.553 | 0.256 | 0.387 | 0.044 | -0.067 | -0.067 | 2.041 | 0.61 | 0.85 | 0.72 | 0.49 | 0.7 | 0.53 |
| 3 | 18704719 | rs11128900 | -1.083 | 1.586 | 0.519 | 0.472 | 0.023 | 0.014 | 0.014 | 0 | 0.5 | 0.43 | 0.43 | 0.44 | 0.56 | 0.47 |
| 3 | 18709522 | rs12106877 | -1.564 | 2.094 | 0.321 | 0.316 | 0.005 | 0.006 | 0.006 | 0.301 | 0.29 | 0.34 | 0.32 | 0.41 | 0.36 | 0.35 |
| 3 | 18711400 | rs13070928 | -1.154 | 2.144 | 0.341 | 0.354 | -0.001 | -0.001 | -0.001 | 0.017 | 0.25 | 0.27 | 0.23 | 0.28 | 0.25 | 0.22 |
| 3 | 18712885 | rs9883391 | -0.14 | 2.122 | 0.333 | 0.317 | -0.001 | -0.051 | -0.051 | 0 | 0.32 | 0.57 | 0.42 | 0.08 | 0.34 | 0.17 |
| 3 | 18737796 | rs9826966 | -2.121 | 2.088 | 0.198 | 0.361 | -0.012 | -0.172 | -0.172 | 4.053 | 0.62 | 0.98 | 0.75 | 0.5 | 0.7 | 0.53 |
| 3 | 18754697 | rs4390955 | -2.052 | 2.056 | 0.236 | 0.381 | 0.021 | -0.126 | -0.126 | 2.521 | 0.61 | 0.91 | 0.74 | 0.5 | 0.7 | 0.53 |
| 3 | 18758419 | rs13098781 | -0.665 | 2.108 | 0.439 | 0.541 | 0.011 | -0.003 | -0.003 | 0.079 | 0.17 | 0.28 | 0.15 | 0.17 | 0.17 | 0.1 |
| 3 | 18762932 | rs4260464 | -0.665 | 2.02 | 0.486 | 0.466 | 0.011 | -0.005 | -0.005 | 1.513 | 0.17 | 0.28 | 0.15 | 0.16 | 0.17 | 0.1 |
| 3 | 18772770 | rs4626115 | 0.621 | 2.095 | 0.23 | 0.507 | -0.011 | -0.007 | -0.007 | 0 | 0.09 | 0.05 | 0.1 | 0.15 | 0.13 | 0.09 |
| 3 | 18779294 | rs13068695 | -0.089 | 2.207 | 0.64 | 0.473 | 0.012 | 0.02 | 0.02 | 0 | 0.13 | 0.16 | 0.15 | 0.15 | 0.17 | 0.13 |
| 3 | 18781973 | rs9876614 | -1.543 | 2.263 | 0.257 | 0.572 | 0.004 | 0.007 | 0.007 | 0.611 | 0.26 | 0.24 | 0.34 | 0.45 | 0.41 | 0.28 |
| 3 | 18784423 | rs6809854 | 0.865 | 2.179 | -0.225 | 0.463 | 0.002 | 0.009 | 0.009 | 0 | 0.22 | 0.21 | 0.31 | 0.07 | 0.2 | 0.15 |
| 3 | 18800046 | rs9824781 | -0.533 | 2.249 | -0.246 | 0.616 | 0.019 | 0.024 | 0.024 | 0 | 0.14 | 0.17 | 0.2 | 0.29 | 0.23 | 0.22 |
| 3 | 18802882 | rs6550631 | -1.284 | 2.174 | -0.317 | 0.605 | 0.012 | 0.047 | 0.047 | 0.188 | 0.5 | 0.8 | 0.63 | 0.37 | 0.56 | 0.43 |
| 3 | 18808580 | rs4370047 | -0.864 | 1.957 | -0.341 | 0.442 | -0.003 | 0.067 | 0.067 | 0 | 0.46 | 0.65 | 0.53 | 0.24 | 0.46 | 0.3 |
| 3 | 18812072 | rs7644430 | 0.559 | 1.716 | -0.993 | 0.52 | -0.001 | 0.004 | 0.004 | 0 | 0.26 | 0.24 | 0.34 | 0.07 | 0.26 | 0.16 |
| 3 | 18820405 | rs13091481 | 0.348 | 1.22 | -1.458 | 0.885 | -0.021 | -0.049 | -0.049 | 1.723 | 0.08 | 0.01 | 0.08 | 0.16 | 0.09 | 0.05 |
| 3 | 18824298 | rs7428918 | -0.881 | 1.112 | -1.217 | 0.719 | 0.048 | -0.069 | -0.069 | 0 | 0.6 | 0.87 | 0.73 | 0.54 | 0.69 | 0.52 |
| 3 | 18835650 | rs6797445 | -0.948 | 1.49 | -0.887 | 0.301 | -0.051 | -0.114 | -0.114 | 0 | 0.64 | 0.93 | 0.65 | 0.48 | 0.63 | 0.65 |
| 3 | 18838880 | rs11926588 | -0.461 | 1.771 | -0.379 | 0.184 | 0.002 | 0.038 | 0.038 | 0 | 0.53 | 0.79 | 0.52 | 0.4 | 0.46 | 0.47 |
| 3 | 18840933 | rs7640928 | 0.184 | 1.78 | 0.266 | 0.066 | -0.035 | -0.026 | -0.026 | 1.09 | 0.41 | 0.48 | 0.36 | 0.24 | 0.24 | 0.31 |
| 3 | 18847711 | rs9832559 | -0.417 | 1.994 | 0.091 | -0.011 | 0.04 | -0.029 | -0.029 | 0 | 0.07 | 0.17 | 0.07 | 0.01 | 0.09 | 0.03 |
| 3 | 18848311 | rs9817260 | -0.597 | 1.979 | 0.085 | -0.018 | 0.034 | -0.032 | -0.032 | 0.403 | 0.07 | 0.16 | 0.07 | 0.01 | 0.09 | 0.04 |
| 3 | 18857812 | rs7428376 | 0.528 | 0.298 | -0.081 | -0.334 | 0.063 | 0.142 | 0.142 | 0.057 | 0.34 | 0.08 | 0.3 | 0.08 | 0.25 | 0.29 |
| 3 | 18861144 | rs7427170 | 0.551 | 0.307 | -0.077 | -0.33 | 0.061 | 0.14 | 0.14 | 0.057 | 0.34 | 0.08 | 0.3 | 0.08 | 0.25 | 0.28 |
| 3 | 18865304 | rs6779916 | -0.439 | 0.373 | 0.316 | -0.345 | 0.059 | -0.022 | -0.022 | 0.021 | 0.07 | 0.19 | 0.06 | 0.01 | 0.08 | 0.03 |
| 3 | 18866539 | rs9830798 | -1.808 | 0.36 | -0.092 | -0.287 | -0.002 | -0.006 | -0.006 | 0 | 0.28 | 0.76 | 0.29 | 0.32 | 0.25 | 0.34 |
| 3 | 18875039 | rs7430675 | 0.647 | 0.353 | -0.254 | -0.308 | 0.06 | 0.139 | 0.139 | 0.055 | 0.34 | 0.08 | 0.3 | 0.08 | 0.25 | 0.28 |
| 3 | 18877426 | rs9881034 | -1.156 | 0.81 | -0.326 | -0.045 | -0.123 | -0.064 | -0.064 | 0 | 0.62 | 0.93 | 0.6 | 0.41 | 0.5 | 0.62 |
| 3 | 18882582 | rs9859674 | -1.174 | 1.279 | 0.257 | 0.084 | 0.045 | 0.112 | 0.112 | 0.016 | 0.09 | 0.19 | 0.06 | 0.26 | 0.13 | 0.18 |
| 3 | 18883999 | rs6808949 | -0.835 | 1.205 | -0.137 | 0.049 | 0.015 | -0.038 | -0.038 | 1.559 | 0.08 | 0.18 | 0.1 | 0.01 | 0.07 | 0.07 |
| 3 | 18885718 | rs9820501 | -0.302 | 0.756 | -0.516 | 0.068 | 0.021 | 0.035 | 0.035 | 0 | 0.48 | 0.59 | 0.42 | 0.33 | 0.41 | 0.47 |
| 3 | 18887173 | rs7617199 | 0.789 | 0.616 | -0.507 | -0.024 | -0.001 | -0.006 | -0.006 | 0.489 | 0.86 | 0.84 | 0.9 | 0.95 | 0.84 | 0.9 |
| 3 | 18887390 | rs7617388 | 0.588 | 1.348 | -0.609 | -0.224 | 0.012 | -0.018 | -0.018 | 0 | 0.94 | 0.89 | 0.82 | 1 | 0.89 | 0.95 |
| 3 | 18889373 | rs7623645 | -0.21 | 0.627 | -0.635 | -0.875 | -0.006 | -0.039 | -0.039 | 0.879 | 0.4 | 0.55 | 0.29 | 0.08 | 0.26 | 0.33 |
| 3 | 18897425 | rs6789997 | 0.725 | 0.131 | -0.745 | -0.498 | 0.034 | 0.037 | 0.037 | 0 | 0.19 | 0.1 | 0.19 | 0.07 | 0.1 | 0.17 |
| 3 | 18902798 | rs4521250 | -1.931 | 0.1 | -0.886 | -0.353 | -0.102 | -0.103 | -0.103 | 0 | 0.6 | 0.93 | 0.56 | 0.41 | 0.49 | 0.48 |
| 3 | 18909119 | rs6792900 | -1.353 | 0.048 | -0.429 | -0.291 | 0.016 | 0.021 | 0.021 | 0.017 | 0.44 | 0.69 | 0.43 | 0.29 | 0.31 | 0.42 |
| 3 | 18914471 | rs6794710 | -0.062 | 0.427 | -0.173 | -0.101 | 0.008 | -0.148 | -0.148 | 0 | 0.19 | 0.45 | 0.09 | 0 | 0.14 | 0.09 |
| 3 | 18921772 | rs4973706 | -0.57 | 0.424 | -0.497 | -0.04 | 0.01 | 0.008 | 0.008 | 0.037 | 0.25 | 0.62 | 0.26 | 0.24 | 0.2 | 0.22 |
| 3 | 18928109 | rs9860501 | -0.577 | 0.026 | -0.367 | -0.4 | -0.024 | -0.06 | -0.06 | 0 | 0.39 | 0.48 | 0.3 | 0.16 | 0.26 | 0.31 |
| 3 | 18929861 | rs9847294 | -0.107 | 0.314 | -0.505 | -0.646 | 0 | -0.007 | -0.007 | 0 | 0.5 | 0.47 | 0.6 | 0.8 | 0.59 | 0.64 |
| 3 | 18929887 | rs6789426 | 0.618 | 1.646 | -0.173 | -0.935 | -0.015 | 0.013 | 0.013 | 1.851 | 0.72 | 0.24 | 0.64 | 0.81 | 0.64 | 0.71 |
| 3 | 18929996 | rs9860355 | -1.554 | 1.647 | 0.606 | -1.237 | 0.085 | -0.098 | -0.098 | 0 | 0.14 | 0.53 | 0.23 | 0 | 0.14 | 0.11 |

| | | | | | | | | | | | | | | | | |
|---|---|---|---|---|---|---|---|---|---|---|---|---|---|---|---|---|
| 3 | 18930125 | rs4401379 | -0.781 | 0.743 | 0.584 | -0.913 | -0.019 | -0.008 | -0.008 | 0 | 0.17 | 0.28 | 0.09 | 0.13 | 0.11 | 0.13 |
| 3 | 18930274 | rs4637309 | -1.311 | 0.03 | 0.675 | -0.547 | -0.051 | 0.011 | 0.011 | 0 | 0.3 | 0.44 | 0.32 | 0.29 | 0.17 | 0.34 |
| 3 | 18931796 | rs11717739 | 0.71 | 0.939 | 0.437 | -0.698 | -0.016 | 0.067 | 0.067 | 0 | 0.48 | 0.32 | 0.52 | 0.74 | 0.52 | 0.63 |
| 3 | 18939227 | rs11710762 | -0.868 | 1.554 | 0.883 | 0.144 | -0.008 | 0.006 | 0.006 | 0 | 0.1 | 0.11 | 0.07 | 0.06 | 0.19 | 0.21 |
| 3 | 18946037 | rs7614379 | -1.417 | 1.049 | 1.078 | 0.196 | -0.021 | -0.046 | -0.046 | 0.998 | 0.14 | 0.26 | 0.13 | 0.03 | 0.07 | 0.06 |
| 3 | 18957376 | rs13089058 | -0.486 | 0.842 | 1.091 | 0.325 | -0.007 | -0.013 | -0.013 | 2.186 | 0.1 | 0.11 | 0.04 | 0.03 | 0.06 | 0.04 |
| 3 | 18959956 | rs7428879 | -1.763 | 0.976 | 1.166 | 0.397 | 0.005 | 0.002 | 0.002 | 0.061 | 0.31 | 0.52 | 0.26 | 0.27 | 0.27 | 0.35 |
| 3 | 18960276 | rs6805368 | -0.869 | 0.985 | 0.915 | 0.429 | -0.055 | 0.028 | 0.028 | 0 | 0.2 | 0.31 | 0.19 | 0.2 | 0.09 | 0.16 |
| 3 | 18990356 | rs9872071 | -0.464 | 1.456 | 0.039 | 0.096 | -0.04 | 0.003 | 0.003 | 0 | 0.08 | 0.04 | 0.05 | 0.07 | 0.18 | 0.17 |
| 3 | 19017070 | rs4410460 | -0.816 | 0.913 | 0.057 | 0.223 | 0.006 | -0.014 | -0.014 | 0 | 0.32 | 0.44 | 0.25 | 0.27 | 0.31 | 0.45 |
| 3 | 19040447 | rs4408886 | 0.223 | 0.524 | 0.478 | 0.071 | -0.005 | -0.1 | -0.1 | 0.423 | 0.84 | 0.54 | 0.87 | 0.99 | 0.88 | 0.86 |
| 3 | 19041089 | rs4973721 | -0.641 | 0.568 | 0.329 | 0.167 | 0 | -0.004 | -0.004 | 0 | 0.11 | 0.08 | 0.13 | 0.14 | 0.11 | 0.14 |
| 3 | 19043554 | rs6807205 | -0.128 | 0.654 | 0.747 | 0.089 | -0.002 | -0.057 | -0.057 | 0.284 | 0.88 | 0.84 | 0.91 | 1 | 0.88 | 0.87 |
| 3 | 19047145 | rs7427678 | 0.369 | 0.649 | 0.914 | 0.086 | 0 | -0.098 | -0.098 | 0 | 0.86 | 0.69 | 0.9 | 1 | 0.88 | 0.87 |
| 3 | 19055818 | rs9861405 | 1.128 | 0.539 | 0.743 | 0.276 | -0.02 | -0.045 | -0.045 | 1.382 | 0.8 | 0.54 | 0.85 | 0.9 | 0.84 | 0.82 |
| 3 | 19057380 | rs7426697 | 0.062 | 0.579 | 0.688 | 0.284 | -0.003 | -0.063 | -0.063 | 0 | 0.87 | 0.7 | 0.89 | 0.99 | 0.88 | 0.85 |
| 3 | 19112134 | rs4101508 | -0.148 | 1.646 | 0.533 | 0.425 | -0.028 | -0.03 | -0.03 | 0 | 0.06 | 0.01 | 0.17 | 0.16 | 0.15 | 0.05 |
| 3 | 19115356 | rs9832959 | -0.628 | 0.44 | 0.756 | -0.052 | -0.011 | -0.101 | -0.101 | 0 | 0.16 | 0.38 | 0.12 | 0.01 | 0.13 | 0.15 |
| 3 | 19121644 | rs9837132 | -0.331 | 0.487 | 0.292 | -0.063 | -0.007 | -0.069 | -0.069 | 0.067 | 0.14 | 0.23 | 0.11 | 0.01 | 0.13 | 0.13 |
| 3 | 19123609 | rs6773955 | -0.461 | -0.469 | 0.297 | -0.297 | -0.06 | -0.117 | -0.117 | 0.53 | 0.2 | 0.32 | 0.08 | 0 | 0.08 | 0.11 |
| 3 | 19224498 | rs997384 | -0.351 | -0.209 | -0.216 | 0.578 | -0.011 | -0.093 | -0.093 | 0 | 0.8 | 0.96 | 0.83 | 0.61 | 0.8 | 0.77 |
| 3 | 19237783 | rs4129625 | 0.332 | -0.739 | -0.268 | 0.715 | 0.013 | -0.011 | -0.011 | 0 | 0.12 | 0.06 | 0.09 | 0.18 | 0.07 | 0.13 |
| 3 | 19282990 | rs9814599 | 0.274 | -1.212 | 0.52 | 0.737 | -0.066 | 0.115 | 0.115 | 0.645 | 0.15 | 0.57 | 0.19 | 0.21 | 0.06 | 0.17 |
| 3 | 19339747 | rs4264750 | 0.738 | 0.585 | -0.424 | -0.12 | 0.014 | 0.088 | 0.088 | 0 | 0.48 | 0.31 | 0.42 | 0.16 | 0.52 | 0.29 |
| 3 | 19341221 | rs11709636 | 1.478 | 0.232 | -0.19 | -0.611 | -0.035 | -0.051 | -0.051 | 0.262 | 0.3 | 0.44 | 0.25 | 0.13 | 0.23 | 0.16 |
| 3 | 19348681 | rs9845765 | 2.203 | -0.005 | -0.191 | -0.865 | -0.01 | -0.087 | -0.087 | 0.189 | 0.25 | 0.25 | 0.18 | 0 | 0.24 | 0.12 |
| 3 | 19352451 | rs6782326 | 0.673 | -0.139 | -0.845 | -0.366 | 0.05 | 0.043 | 0.043 | 0 | 0.43 | 0.71 | 0.33 | 0.31 | 0.36 | 0.35 |
| 3 | 19358167 | rs7618209 | 0.429 | 0.187 | -1.052 | -0.512 | -0.071 | 0.068 | 0.068 | 0 | 0.23 | 0.1 | 0.32 | 0.06 | 0.34 | 0.18 |
| 3 | 19371580 | rs1372555 | 0.208 | 0.558 | -1.047 | -0.494 | -0.036 | 0.011 | 0.011 | 0 | 0.37 | 0.18 | 0.47 | 0.68 | 0.39 | 0.5 |
| 3 | 19371858 | rs2034007 | 1.165 | 0.404 | -1.286 | 0.129 | -0.001 | -0.016 | -0.016 | 0 | 0.27 | 0.23 | 0.34 | 0.56 | 0.28 | 0.42 |
| 3 | 19375595 | rs2053505 | 0.847 | 0.29 | -1.297 | 0.102 | -0.04 | -0.096 | -0.096 | 0.027 | 0.3 | 0.1 | 0.39 | 0.55 | 0.27 | 0.45 |
| 3 | 19375791 | rs2053506 | -0.101 | 0.483 | -1.171 | 0.292 | -0.081 | 0.062 | 0.062 | 0 | 0.75 | 0.92 | 0.75 | 0.86 | 0.64 | 0.8 |
| 3 | 19382787 | rs2034008 | 1.035 | -0.79 | -0.598 | 0.582 | -0.017 | 0.02 | 0.02 | 0 | 0.07 | 0.11 | 0.06 | 0.13 | 0.01 | 0.08 |
| 3 | 19385317 | rs9865301 | 0.351 | -0.795 | -0.535 | 0.623 | -0.008 | 0.022 | 0.022 | 0 | 0.1 | 0.16 | 0.05 | 0.19 | 0.09 | 0.2 |
| 3 | 19385436 | rs12488588 | 0.579 | -0.888 | -0.759 | 0.482 | 0.012 | 0.019 | 0.019 | 0.25 | 0.14 | 0.08 | 0.05 | 0.1 | 0.1 | 0.06 |
| 3 | 19400348 | rs9851592 | -1.539 | -0.796 | -0.604 | 0.556 | -0.03 | 0.022 | 0.022 | 0.625 | 0.31 | 0.46 | 0.17 | 0.41 | 0.2 | 0.31 |
| 3 | 19401472 | rs17005867 | 0.657 | -0.657 | -0.756 | 0.572 | -0.011 | 0.033 | 0.033 | 0 | 0.09 | 0.16 | 0.04 | 0.19 | 0.08 | 0.18 |
| 3 | 19407103 | rs12485979 | 0.22 | -0.663 | -0.85 | 0.486 | -0.004 | -0.003 | -0.003 | 0.003 | 0.16 | 0.18 | 0.11 | 0.12 | 0.1 | 0.11 |
| 3 | 19408279 | rs1372553 | -0.822 | -0.691 | -0.84 | 0.609 | -0.001 | 0.011 | 0.011 | 0 | 0.14 | 0.18 | 0.12 | 0.31 | 0.12 | 0.25 |
| 3 | 19408549 | rs7640623 | 0.887 | -0.946 | -0.947 | 0.102 | 0.009 | -0.002 | -0.002 | 0.121 | 0.43 | 0.54 | 0.29 | 0.12 | 0.33 | 0.24 |
| 3 | 19409153 | rs7645658 | -0.829 | -0.629 | -1.149 | 0.38 | -0.034 | -0.039 | -0.039 | 0 | 0.66 | 0.76 | 0.47 | 0.54 | 0.55 | 0.55 |
| 3 | 19409996 | rs9875843 | -0.408 | -0.725 | -1.058 | 0.317 | -0.012 | 0.002 | 0.002 | 2.602 | 0.22 | 0.29 | 0.12 | 0.24 | 0.12 | 0.16 |
| 3 | 19410169 | rs9310595 | 0.111 | -0.703 | -1.19 | 0.134 | 0.042 | 0.062 | 0.062 | 0 | 0.53 | 0.66 | 0.32 | 0.31 | 0.41 | 0.41 |
| 3 | 19419822 | rs11914956 | 0.171 | -0.336 | -1.508 | 0.386 | -0.001 | 0.034 | 0.034 | 0 | 0.16 | 0.05 | 0.1 | 0.1 | 0.17 | 0.07 |
| 3 | 19420687 | rs9846656 | -0.644 | -0.367 | -1.425 | 0.392 | 0.008 | 0.006 | 0.006 | 0 | 0.23 | 0.18 | 0.17 | 0.23 | 0.19 | 0.14 |
| 3 | 19423561 | rs2197126 | -1.168 | 0.583 | -1.862 | 0.987 | -0.123 | -0.139 | -0.139 | 0 | 0.77 | 0.97 | 0.64 | 0.53 | 0.67 | 0.59 |
| 3 | 19428402 | rs7630132 | 0.244 | -0.561 | -1.549 | 1.042 | -0.028 | -0.011 | -0.011 | 0.098 | 0.57 | 0.71 | 0.46 | 0.4 | 0.5 | 0.51 |
| 3 | 19433647 | rs7616342 | -0.286 | -0.516 | -1.584 | 0.973 | -0.045 | -0.06 | -0.06 | 2.114 | 0.64 | 0.83 | 0.53 | 0.4 | 0.59 | 0.54 |
| 3 | 19434568 | rs7635327 | 2.536 | -0.897 | -1.333 | 0.363 | 0.005 | -0.004 | -0.004 | 1.301 | 0.29 | 0.19 | 0.17 | 0.01 | 0.26 | 0.13 |
| 3 | 19436030 | rs17570934 | 2.525 | -0.923 | -1.08 | 0.344 | 0 | -0.051 | -0.051 | 0.655 | 0.3 | 0.25 | 0.18 | 0.01 | 0.26 | 0.13 |
| 3 | 19436270 | rs10470605 | -0.735 | -0.931 | -1.074 | 0.62 | -0.001 | 0.085 | 0.085 | 0 | 0.28 | 0.47 | 0.37 | 0.44 | 0.27 | 0.39 |
| 3 | 19436587 | rs10470642 | -1.801 | -0.731 | -1.473 | 1.03 | -0.11 | -0.089 | -0.089 | 0 | 0.79 | 0.98 | 0.73 | 0.63 | 0.7 | 0.62 |
| 3 | 19437414 | rs6551250 | -0.131 | -0.663 | -1.432 | 1.074 | -0.025 | -0.009 | -0.009 | 0.084 | 0.62 | 0.71 | 0.5 | 0.35 | 0.56 | 0.54 |
| 3 | 19438345 | rs7647575 | -0.189 | -0.658 | -1.42 | 1.072 | -0.013 | -0.027 | -0.027 | 0 | 0.62 | 0.71 | 0.59 | 0.44 | 0.6 | 0.52 |
| 3 | 19441126 | rs1348232 | 0.359 | -0.694 | -1.452 | 1.082 | 0.023 | 0.048 | 0.048 | 0.431 | 0.52 | 0.69 | 0.43 | 0.32 | 0.43 | 0.44 |
| 3 | 19449092 | rs7611730 | -0.397 | -0.434 | -1.04 | 2.063 | -0.01 | 0 | 0 | 0 | 0.7 | 0.7 | 0.54 | 0.28 | 0.64 | 0.53 |
| 3 | 19450214 | rs6799035 | -0.34 | -0.435 | -1.041 | 2.105 | -0.005 | -0.008 | -0.008 | 0 | 0.71 | 0.7 | 0.62 | 0.37 | 0.68 | 0.54 |
| 3 | 19457107 | rs1867001 | -1.276 | -0.409 | -1.217 | 2.068 | -0.114 | -0.127 | -0.127 | 0.042 | 0.78 | 0.97 | 0.64 | 0.59 | 0.68 | 0.58 |
| 3 | 19459577 | rs4142701 | -1.5 | -0.407 | -1.217 | 2.066 | -0.111 | -0.056 | -0.056 | 0 | 0.79 | 0.98 | 0.72 | 0.7 | 0.7 | 0.61 |
| 3 | 19462931 | rs9310596 | -1.183 | -0.408 | -1.216 | 2.085 | -0.132 | -0.105 | -0.105 | 0.415 | 0.77 | 0.97 | 0.62 | 0.63 | 0.65 | 0.59 |
| 3 | 19473190 | rs6549867 | -0.379 | -0.417 | -1.14 | 2.156 | -0.023 | -0.021 | -0.021 | 0 | 0.71 | 0.74 | 0.61 | 0.36 | 0.66 | 0.55 |
| 3 | 19477536 | rs10510487 | -0.003 | -0.517 | -1.106 | 2.273 | 0.005 | -0.002 | -0.002 | 0.243 | 0.05 | 0.06 | 0.05 | 0.08 | 0.04 | 0.07 |
| 3 | 19482975 | rs1348229 | -1.268 | -0.293 | -1.18 | 2.245 | -0.123 | -0.134 | -0.134 | 0 | 0.78 | 0.97 | 0.63 | 0.58 | 0.66 | 0.58 |
| 3 | 19497353 | rs17005983 | 0.002 | -0.405 | -0.802 | 2.113 | -0.025 | 0.114 | 0.114 | 0.021 | 0.06 | 0.19 | 0.07 | 0.25 | 0.01 | 0.06 |
| 3 | 19555271 | rs6549944 | -1.501 | -0.252 | -0.986 | 1.932 | -0.088 | -0.096 | -0.096 | 0 | 0.79 | 0.98 | 0.74 | 0.69 | 0.71 | 0.7 |
| 3 | 19557968 | rs7372627 | 0.393 | -0.879 | -0.649 | 1.674 | -0.002 | -0.005 | -0.005 | 0.457 | 0.61 | 0.61 | 0.6 | 0.41 | 0.61 | 0.44 |
| 3 | 19562271 | rs9853477 | 1.414 | -1.139 | -0.689 | 1.365 | 0.01 | 0.057 | 0.057 | 1.72 | 0.35 | 0.21 | 0.27 | 0.1 | 0.37 | 0.19 |
| 3 | 19562980 | rs6770402 | -0.231 | -0.831 | -1.059 | 1.575 | 0 | -0.001 | -0.001 | 0 | 0.07 | 0.08 | 0.06 | 0.09 | 0.05 | 0.09 |
| 3 | 19568009 | rs6549948 | -1.471 | -0.768 | -1.109 | 1.593 | -0.099 | -0.027 | -0.027 | 0 | 0.79 | 0.97 | 0.74 | 0.75 | 0.7 | 0.6 |

| | | | | | | | | | | | | | | | | |
|---|---|---|---|---|---|---|---|---|---|---|---|---|---|---|---|---|
| 3 | 19578530 | rs17006052 | -0.017 | -0.979 | -1.039 | 1.608 | 0.002 | -0.01 | -0.01 | 0.071 | 0.06 | 0.05 | 0.06 | 0.09 | 0.05 | 0.09 |
| 3 | 19580707 | rs908216 | -1.31 | -0.915 | -0.973 | 1.503 | -0.026 | 0.013 | 0.013 | 0 | 0.78 | 0.87 | 0.73 | 0.78 | 0.69 | 0.6 |
| 3 | 19592688 | rs7626907 | -0.767 | -1.388 | -0.768 | 1.699 | -0.013 | 0.026 | 0.026 | 0 | 0.1 | 0.16 | 0.07 | 0.25 | 0.03 | 0.05 |
| 3 | 19595902 | rs7628760 | -1.482 | -1.3 | -0.812 | 1.607 | 0.013 | -0.025 | -0.025 | 0 | 0.27 | 0.22 | 0.33 | 0.61 | 0.23 | 0.28 |
| 3 | 19606024 | rs17006069 | -0.698 | -1.031 | -0.629 | 1.402 | -0.005 | 0.037 | 0.037 | 0.374 | 0.11 | 0.13 | 0.14 | 0.37 | 0.08 | 0.15 |
| 3 | 19614280 | rs17543807 | 1.895 | -1.223 | -0.994 | 0.894 | -0.026 | 0.153 | 0.153 | 0 | 0.23 | 0.01 | 0.12 | 0 | 0.23 | 0.1 |
| 3 | 19618837 | rs13075095 | 1.974 | -1.143 | -0.958 | 0.93 | -0.029 | 0.147 | 0.147 | 0.369 | 0.22 | 0.01 | 0.12 | 0 | 0.23 | 0.1 |
| 3 | 19640745 | rs9873476 | 1.129 | -0.594 | -0.559 | 1.24 | -0.007 | 0.135 | 0.135 | 0.967 | 0.21 | 0.04 | 0.1 | 0 | 0.2 | 0.06 |
| 3 | 19653387 | rs6804525 | 0.934 | -0.523 | 0.086 | 1.61 | 0.016 | 0.014 | 0.014 | 1.98 | 0.38 | 0.27 | 0.28 | 0.25 | 0.35 | 0.21 |
| 3 | 19683373 | rs6804427 | 0.986 | 0.82 | 0.834 | 2.208 | 0.015 | 0.031 | 0.031 | 0 | 0.65 | 0.54 | 0.54 | 0.41 | 0.62 | 0.42 |
| 3 | 19684181 | rs17006125 | -0.917 | 0.941 | 0.778 | 2.292 | -0.003 | 0 | 0 | 0 | 0.08 | 0.1 | 0.11 | 0.08 | 0.07 | 0.1 |
| 3 | 19711814 | rs11532602 | -1.564 | 0.953 | 0.66 | 2.309 | -0.011 | 0.207 | 0.207 | 2.204 | 0.07 | 0.24 | 0.05 | 0.28 | 0.01 | 0.05 |
| 3 | 19751987 | rs6784806 | -0.623 | 1.014 | 0.327 | 1.969 | -0.111 | -0.022 | -0.022 | 0.44 | 0.8 | 0.97 | 0.71 | 0.77 | 0.7 | 0.54 |
| 3 | 19754793 | rs2062978 | -2.358 | 0.097 | 1.484 | 2.44 | -0.03 | 0.257 | 0.257 | 4.327 | 0.14 | 0.42 | 0.16 | 0.38 | 0.08 | 0.12 |
| 3 | 19782050 | rs17006216 | -1.103 | 0.455 | 0.456 | 1.238 | -0.001 | 0 | 0 | 0.018 | 0.09 | 0.08 | 0.07 | 0.1 | 0.08 | 0.08 |
| 3 | 19783277 | rs17006217 | -1.665 | 0.479 | 0.5 | 1.274 | -0.005 | 0.014 | 0.014 | 0.03 | 0.16 | 0.23 | 0.16 | 0.17 | 0.12 | 0.24 |
| 3 | 19788213 | rs10510491 | -1.448 | 0.124 | 0.517 | 1.042 | -0.003 | 0.024 | 0.024 | 0.644 | 0.12 | 0.36 | 0.09 | 0.11 | 0.08 | 0.14 |
| 3 | 19788618 | rs4858568 | -1.233 | -0.087 | 0.283 | 1.212 | -0.04 | 0.09 | 0.09 | 0.328 | 0.16 | 0.55 | 0.21 | 0.18 | 0.12 | 0.13 |
| 3 | 19799545 | rs11706482 | -0.049 | 0.093 | -1.143 | 1.184 | -0.035 | -0.133 | -0.133 | 2.291 | 0.27 | 0.06 | 0.35 | 0.43 | 0.3 | 0.19 |
| 3 | 19804889 | rs3931311 | -0.265 | -0.334 | -0.419 | 1.002 | 0.002 | -0.021 | -0.021 | 0 | 0.28 | 0.23 | 0.37 | 0.43 | 0.29 | 0.19 |
| 3 | 19817805 | rs1506095 | 0.366 | 0.963 | 0.598 | 0.42 | -0.022 | 0.019 | 0.019 | 0 | 0.11 | 0.09 | 0.21 | 0.03 | 0.25 | 0.14 |
| 3 | 19837587 | rs1506097 | -0.829 | 0.53 | 0.558 | 0.429 | 0.002 | 0.002 | 0.002 | 0 | 0.46 | 0.73 | 0.62 | 0.42 | 0.55 | 0.34 |
| 3 | 19849542 | rs269392 | -1.291 | 0.015 | -0.615 | -0.384 | -0.082 | 0.027 | 0.027 | 0 | 0.77 | 0.98 | 0.85 | 0.85 | 0.76 | 0.86 |
| 3 | 19875252 | rs11926759 | -0.003 | 0.422 | -0.012 | -0.399 | 0.019 | -0.033 | -0.033 | 0 | 0.16 | 0.2 | 0.22 | 0.02 | 0.25 | 0.13 |
| 3 | 19907859 | rs405571 | 1.029 | -0.088 | 0.204 | -0.62 | -0.002 | -0.007 | -0.007 | 0 | 0.19 | 0.2 | 0.09 | 0.08 | 0.1 | 0.2 |
| 3 | 19910769 | rs17006314 | 0.394 | -0.061 | 0.297 | -0.619 | 0.007 | -0.02 | -0.02 | 0 | 0.15 | 0.15 | 0.21 | 0.02 | 0.25 | 0.13 |
| 3 | 19920147 | rs12632924 | 1.426 | -0.14 | -0.111 | -0.901 | -0.046 | 0.01 | 0.01 | 0 | 0.22 | 0.01 | 0.13 | 0.14 | 0.21 | 0.14 |
| 3 | 19923822 | rs2306774 | 0.842 | -0.227 | 0.224 | -0.584 | -0.003 | -0.005 | -0.005 | 0 | 0.3 | 0.07 | 0.2 | 0.26 | 0.3 | 0.3 |
| 3 | 19930743 | rs11925477 | 0.9 | -0.104 | 0.508 | -0.622 | -0.01 | -0.023 | -0.023 | 0 | 0.19 | 0.24 | 0.09 | 0.08 | 0.1 | 0.2 |
| 3 | 19939707 | rs2931383 | -0.506 | -0.055 | 0.658 | -0.595 | -0.033 | 0.059 | 0.059 | 0 | 0.37 | 0.81 | 0.38 | 0.17 | 0.38 | 0.43 |
| 3 | 19939928 | rs2931384 | -0.454 | -0.045 | 0.629 | -0.59 | -0.022 | 0.039 | 0.039 | 0 | 0.37 | 0.78 | 0.37 | 0.17 | 0.38 | 0.43 |
| 3 | 19946057 | rs3916092 | -0.534 | 0.706 | 0.838 | -0.539 | 0.199 | -0.003 | -0.003 | 0 | 0.09 | 0.44 | 0.18 | 0.01 | 0.18 | 0.08 |
| 3 | 19946992 | rs2931394 | -0.623 | 0.002 | 0.13 | -0.713 | -0.005 | -0.023 | -0.023 | 0.274 | 0.84 | 0.88 | 0.61 | 0.3 | 0.78 | 0.66 |
| 3 | 19947296 | rs2929377 | -0.193 | -0.231 | 0.075 | -1.075 | 0.021 | -0.05 | -0.05 | 0.411 | 0.33 | 0.46 | 0.33 | 0.13 | 0.4 | 0.35 |
| 3 | 19948053 | rs2929374 | -0.201 | -0.229 | 0.077 | -1.074 | 0.02 | -0.049 | -0.049 | 0.248 | 0.33 | 0.46 | 0.33 | 0.13 | 0.4 | 0.35 |
| 3 | 19962044 | rs13078867 | 0.778 | -0.236 | -0.447 | -1.282 | 0.007 | 0.022 | 0.022 | 0 | 0.29 | 0.07 | 0.18 | 0.21 | 0.25 | 0.28 |
| 3 | 19968899 | rs2929358 | 0.756 | -0.264 | -0.259 | -1.463 | -0.001 | -0.002 | -0.002 | 0 | 0.18 | 0.23 | 0.12 | 0.11 | 0.14 | 0.22 |
| 3 | 19975139 | rs17180771 | 1.696 | -0.1 | -0.412 | -1.519 | -0.069 | 0.038 | 0.038 | 0 | 0.1 | 0.01 | 0.2 | 0.01 | 0.24 | 0.12 |
| 3 | 19975215 | rs17795400 | 0.938 | -0.28 | -0.52 | -1.444 | 0.012 | 0.014 | 0.014 | 0 | 0.28 | 0.08 | 0.17 | 0.22 | 0.24 | 0.27 |
| 3 | 19975582 | rs2929349 | -0.312 | -0.258 | 0.329 | -1.125 | -0.005 | -0.003 | -0.003 | 0 | 0.57 | 0.49 | 0.5 | 0.35 | 0.63 | 0.62 |
| 3 | 19976253 | rs6550227 | 0.444 | -0.21 | 0.1 | -1.148 | 0.005 | 0.001 | 0.001 | 0 | 0.38 | 0.36 | 0.45 | 0.65 | 0.34 | 0.38 |
| 3 | 19978971 | rs2929347 | -0.502 | -0.125 | -0.16 | -1.072 | 0.002 | 0.001 | 0.001 | 0 | 0.59 | 0.6 | 0.52 | 0.35 | 0.63 | 0.61 |
| 3 | 19982135 | rs2170342 | 0.589 | -0.108 | -0.016 | -1.117 | -0.015 | -0.021 | -0.021 | 1.197 | 0.19 | 0.34 | 0.14 | 0.12 | 0.16 | 0.27 |
| 3 | 19990813 | rs11128928 | 0.691 | 0.483 | -0.078 | -1.195 | -0.012 | -0.003 | -0.003 | 0.062 | 0.12 | 0.06 | 0.14 | 0.15 | 0.16 | 0.22 |
| 3 | 19994475 | rs4858660 | 0.458 | 0.141 | -0.355 | -1.804 | 0.005 | 0.026 | 0.026 | 0 | 0.52 | 0.57 | 0.46 | 0.17 | 0.57 | 0.42 |
| 3 | 19996373 | rs11128929 | 1.356 | 0.157 | -0.368 | -1.785 | -0.013 | -0.004 | -0.004 | 0 | 0.12 | 0.06 | 0.14 | 0.15 | 0.16 | 0.22 |
| 3 | 19996508 | rs6778866 | 1.542 | 0.197 | -0.33 | -1.816 | -0.014 | -0.005 | -0.005 | 0 | 0.18 | 0.38 | 0.15 | 0.16 | 0.16 | 0.26 |
| 3 | 19999504 | rs17181547 | 2.916 | 0.286 | -0.522 | -1.957 | -0.027 | 0.113 | 0.113 | 0 | 0.2 | 0.04 | 0.1 | 0 | 0.19 | 0.05 |
| 3 | 20002394 | rs12486167 | 2.451 | 0.243 | -0.518 | -1.948 | -0.067 | 0.056 | 0.056 | 0 | 0.11 | 0.01 | 0.2 | 0.01 | 0.22 | 0.11 |
| 3 | 20003611 | rs4241539 | 2.245 | 0.292 | -0.471 | -1.928 | 0.008 | -0.021 | -0.021 | 0 | 0.12 | 0.1 | 0.21 | 0.01 | 0.22 | 0.11 |
| 3 | 20009231 | rs7634934 | 1.173 | 0.203 | -0.293 | -1.916 | 0.01 | -0.001 | -0.001 | 0.86 | 0.13 | 0.17 | 0.15 | 0.15 | 0.16 | 0.22 |
| 3 | 20009298 | rs2127956 | -0.862 | 0.187 | -0.399 | -1.935 | 0.025 | 0.013 | 0.013 | 0 | 0.65 | 0.74 | 0.61 | 0.31 | 0.73 | 0.64 |
| 3 | 20010886 | rs4398451 | 1.592 | 0.185 | -0.406 | -1.939 | -0.029 | 0.041 | 0.041 | 0 | 0.33 | 0.18 | 0.31 | 0.01 | 0.41 | 0.16 |
| 3 | 20020154 | rs1143451 | 1.145 | 0.208 | -0.504 | -1.848 | -0.001 | -0.001 | -0.001 | 0.009 | 0.13 | 0.14 | 0.15 | 0.15 | 0.16 | 0.22 |
| 3 | 20020443 | rs9835991 | 1.718 | 0.006 | -0.644 | -1.916 | 0.03 | 0.051 | 0.051 | 0 | 0.23 | 0.13 | 0.09 | 0 | 0.15 | 0.06 |
| 3 | 20023990 | rs13085694 | -0.094 | 0.278 | -0.414 | -1.493 | 0.005 | -0.009 | -0.009 | 0.296 | 0.32 | 0.56 | 0.3 | 0.31 | 0.32 | 0.49 |
| 3 | 20033962 | rs2929416 | 2.64 | 0.432 | -1.283 | -1.31 | -0.014 | 0.027 | 0.027 | 0 | 0.08 | 0 | 0.05 | 0.01 | 0.09 | 0.05 |
| 3 | 20042834 | rs4103004 | 0.946 | 0.549 | -1.053 | -1.052 | -0.015 | 0.045 | 0.045 | 0 | 0.35 | 0.32 | 0.28 | 0.22 | 0.36 | 0.27 |
| 3 | 20062060 | rs2929387 | -0.758 | 0.331 | -0.844 | -0.249 | 0 | -0.001 | -0.001 | 0 | 0.87 | 0.87 | 0.72 | 0.41 | 0.88 | 0.77 |
| 3 | 20065662 | rs2948091 | -1.127 | 0.225 | -0.682 | -0.457 | 0.001 | 0.031 | 0.031 | 0 | 0.82 | 0.62 | 0.83 | 0.79 | 0.8 | 0.67 |
| 3 | 20079878 | rs12639078 | 0.145 | 0.198 | -1.869 | -0.382 | 0.012 | -0.069 | -0.069 | 0 | 0.33 | 0.11 | 0.33 | 0.36 | 0.29 | 0.27 |
| 3 | 20080132 | rs17626938 | 0.958 | -0.015 | -1.792 | -1.176 | 0.046 | 0.051 | 0.051 | 0 | 0.21 | 0.02 | 0.08 | 0.11 | 0.12 | 0.19 |
| 3 | 20083881 | rs1915919 | 0.175 | 1.066 | -0.85 | -0.785 | 0.016 | -0.119 | -0.119 | 0 | 0.36 | 0.39 | 0.33 | 0.04 | 0.4 | 0.2 |
| 3 | 20090908 | rs6808352 | -0.325 | 0.821 | -0.933 | 0.103 | 0.002 | 0.003 | 0.003 | 0.371 | 0.43 | 0.6 | 0.46 | 0.39 | 0.33 | 0.37 |
| 3 | 20096735 | rs2929401 | -0.345 | 1.023 | -0.821 | 0.049 | 0.001 | -0.003 | -0.003 | 0 | 0.39 | 0.41 | 0.37 | 0.56 | 0.42 | 0.6 |
| 3 | 20098119 | rs2948085 | -1.309 | 0.732 | -1.12 | -0.581 | 0.071 | 0.115 | 0.115 | 0 | 0.7 | 0.91 | 0.81 | 0.96 | 0.85 | 0.91 |
| 3 | 20107401 | rs7627847 | -0.155 | 1.691 | -0.969 | -0.075 | 0 | 0.002 | 0.002 | 0 | 0.41 | 0.38 | 0.27 | 0.42 | 0.44 | 0.31 |
| 3 | 20108546 | rs1124376 | 1.511 | 1.853 | -0.808 | 0.054 | -0.01 | 0.091 | 0.091 | 0 | 0.17 | 0.02 | 0.11 | 0.01 | 0.21 | 0.1 |
| 3 | 20116934 | rs4858754 | -1.075 | 2.127 | 0.201 | 0.572 | 0.062 | 0.044 | 0.044 | 0 | 0.29 | 0.52 | 0.45 | 0.41 | 0.4 | 0.53 |

| | | | | | | | | | | | | | | | | |
|---|---|---|---|---|---|---|---|---|---|---|---|---|---|---|---|---|
| 3 | 20121828 | rs11714225 | -1.034 | 1.358 | -0.148 | 0.693 | -0.001 | 0 | 0 | 0 | 0.26 | 0.26 | 0.3 | 0.42 | 0.28 | 0.27 |
| 3 | 20122315 | rs11128934 | -1.034 | 1.358 | -0.145 | 0.693 | 0 | 0.002 | 0.002 | 0 | 0.26 | 0.27 | 0.3 | 0.42 | 0.28 | 0.27 |
| 3 | 20125393 | rs11128935 | -1.508 | 1.349 | -0.057 | 0.619 | -0.008 | -0.011 | -0.011 | 0.346 | 0.38 | 0.76 | 0.45 | 0.59 | 0.37 | 0.65 |
| 3 | 20125788 | rs7623123 | -0.994 | 1.35 | -0.097 | 0.754 | 0.02 | 0.053 | 0.053 | 0 | 0.09 | 0.19 | 0.12 | 0.23 | 0.11 | 0.08 |
| 3 | 20136959 | rs12636856 | -0.51 | 1.147 | -0.096 | 0.929 | -0.007 | 0.128 | 0.128 | 0 | 0.1 | 0.17 | 0.06 | 0.38 | 0.08 | 0.15 |
| 3 | 20137895 | rs7641684 | -0.452 | 1.137 | 0.508 | 0.999 | 0.017 | 0.047 | 0.047 | 0 | 0.12 | 0.26 | 0.19 | 0.21 | 0.14 | 0.39 |
| 3 | 20140129 | rs11715447 | 1.697 | 0.996 | 0.154 | 0.46 | 0.002 | 0.038 | 0.038 | 0 | 0.13 | 0.03 | 0.08 | 0 | 0.14 | 0.05 |
| 3 | 20141522 | rs13323791 | -0.702 | 1.023 | 0.474 | 0.6 | 0.031 | 0.208 | 0.208 | 0.017 | 0.14 | 0.41 | 0.22 | 0.41 | 0.15 | 0.4 |
| 3 | 20152563 | rs17006610 | 0.814 | 0.768 | 0.702 | 0.339 | -0.019 | -0.115 | -0.115 | 0.084 | 0.2 | 0.31 | 0.14 | 0.01 | 0.21 | 0.1 |
| 3 | 20152772 | rs11923928 | -0.272 | 1.163 | 0.276 | 0.075 | -0.013 | -0.004 | -0.004 | 0 | 0.22 | 0.12 | 0.2 | 0.19 | 0.21 | 0.24 |
| 3 | 20156302 | rs2365794 | -1.609 | 1.289 | 0.182 | 0.095 | -0.013 | 0.014 | 0.014 | 0 | 0.57 | 0.81 | 0.58 | 0.61 | 0.61 | 0.74 |
| 3 | 20156873 | rs6771737 | -0.673 | 1.238 | 0.306 | 0.103 | -0.003 | -0.031 | -0.031 | 0.093 | 0.43 | 0.44 | 0.34 | 0.19 | 0.43 | 0.34 |
| 3 | 20162216 | rs7617840 | -0.832 | 1.262 | 0.216 | 0.068 | 0.002 | 0.002 | 0.002 | 0 | 0.44 | 0.52 | 0.37 | 0.4 | 0.44 | 0.35 |
| 3 | 20167830 | rs13097409 | 1.101 | 1.497 | 0.231 | 0.247 | -0.052 | 0.047 | 0.047 | 0 | 0.11 | 0.01 | 0.17 | 0.01 | 0.15 | 0.08 |
| 3 | 20183231 | rs17006645 | 0.686 | 1.299 | 0.61 | 0.435 | 0.012 | -0.023 | -0.023 | 0 | 0.07 | 0.03 | 0.09 | 0.2 | 0.04 | 0.35 |
| 3 | 20187080 | rs2661377 | -0.271 | 1.43 | 1.076 | 0.22 | 0.024 | -0.019 | -0.019 | 0 | 0.06 | 0.1 | 0.09 | 0.02 | 0.1 | 0.03 |
| 3 | 20188301 | rs10510498 | 1.118 | 1.327 | 0.933 | 0.37 | 0.004 | -0.013 | -0.013 | 0.077 | 0.05 | 0.03 | 0.07 | 0.13 | 0.03 | 0.15 |
| 3 | 20194423 | rs4858770 | -0.141 | 2.143 | 1.215 | 0.461 | 0.074 | 0.22 | 0.22 | 0 | 0.07 | 0.18 | 0.29 | 0.42 | 0.17 | 0.33 |
| 3 | 20196571 | rs7620481 | -0.141 | 2.218 | 1.188 | 0.442 | 0.068 | 0.208 | 0.208 | 0 | 0.07 | 0.17 | 0.29 | 0.42 | 0.16 | 0.33 |
| 3 | 20200232 | rs10865760 | -0.635 | 2.126 | 1.283 | 0.497 | 0.005 | -0.002 | -0.002 | 0.562 | 0.25 | 0.22 | 0.3 | 0.2 | 0.27 | 0.2 |
| 3 | 20200636 | rs1828079 | -1.059 | 2.069 | 1.405 | 0.448 | 0.006 | 0.059 | 0.059 | 0 | 0.31 | 0.48 | 0.4 | 0.53 | 0.29 | 0.36 |
| 3 | 20226659 | rs17006708 | -0.607 | 2.057 | 1.411 | 0.496 | 0.013 | -0.001 | -0.001 | 0 | 0.25 | 0.26 | 0.29 | 0.2 | 0.27 | 0.2 |
| 3 | 20231519 | rs6550413 | 0.124 | 2.119 | 1.351 | 0.563 | 0.084 | 0.231 | 0.231 | 0 | 0.07 | 0.18 | 0.29 | 0.41 | 0.17 | 0.33 |
| 3 | 20237433 | rs1395985 | -0.908 | 2.011 | 1.325 | 0.416 | 0.014 | 0.024 | 0.024 | 0 | 0.3 | 0.4 | 0.38 | 0.33 | 0.31 | 0.34 |
| 3 | 20264950 | rs952466 | 2.223 | 0.768 | 0.404 | -0.254 | 0.044 | 0.019 | 0.019 | 0 | 0.22 | 0.11 | 0.14 | 0.2 | 0.12 | 0.07 |
| 3 | 20266516 | rs2365799 | -0.631 | 0.86 | 0.524 | -0.002 | 0.058 | 0.02 | 0.02 | 0 | 0.32 | 0.47 | 0.54 | 0.58 | 0.44 | 0.54 |
| 3 | 20287463 | rs12638108 | 2.125 | 0.705 | 0.205 | -0.024 | 0.024 | 0.012 | 0.012 | 0 | 0.22 | 0.15 | 0.15 | 0.2 | 0.11 | 0.07 |
| 3 | 20292220 | rs1605519 | -0.747 | 0.838 | 0.058 | -0.122 | -0.012 | 0.212 | 0.212 | 0 | 0.6 | 0.9 | 0.78 | 0.91 | 0.59 | 0.74 |
| 3 | 20292978 | rs4858166 | -0.459 | 0.91 | 0.179 | -0.034 | 0.016 | 0.134 | 0.134 | 0 | 0.55 | 0.83 | 0.71 | 0.78 | 0.56 | 0.6 |
| 3 | 20301354 | rs12489472 | 1.865 | 1.219 | 0.173 | -0.144 | 0.009 | 0.058 | 0.058 | 0 | 0.16 | 0.05 | 0.1 | 0 | 0.18 | 0.11 |
| 3 | 20301653 | rs1466988 | -1.502 | 1.141 | 0.311 | 0.088 | 0.079 | 0.001 | 0.001 | 0 | 0.67 | 0.84 | 0.76 | 0.67 | 0.81 | 0.76 |
| 3 | 20306520 | rs9860301 | 0.375 | 1.36 | 0.945 | 0.106 | -0.002 | -0.014 | -0.014 | 0 | 0.38 | 0.39 | 0.29 | 0.11 | 0.45 | 0.27 |
| 3 | 20307061 | rs9827605 | -1.31 | 1.197 | 0.659 | 0.225 | 0.077 | 0.005 | 0.005 | 0 | 0.67 | 0.81 | 0.76 | 0.66 | 0.81 | 0.76 |
| 3 | 20339157 | rs9838186 | 0.013 | 1.731 | 0.829 | 0.443 | 0.079 | 0.204 | 0.204 | 0 | 0.08 | 0.22 | 0.25 | 0.39 | 0.17 | 0.32 |
| 3 | 20345376 | rs6801666 | -0.232 | 1.171 | 0.717 | 0.188 | 0.002 | 0.008 | 0.008 | 0 | 0.47 | 0.46 | 0.49 | 0.63 | 0.56 | 0.61 |
| 3 | 20348723 | rs9820730 | 0.672 | 1.582 | 0.312 | 0.363 | -0.088 | 0.051 | 0.051 | 0 | 0.34 | 0.07 | 0.23 | 0.21 | 0.42 | 0.25 |
| 3 | 20352766 | rs9871864 | 0.17 | 1.587 | 0.568 | 0.403 | -0.008 | 0.014 | 0.014 | 0 | 0.39 | 0.31 | 0.28 | 0.25 | 0.45 | 0.29 |
| 3 | 20355533 | rs4241504 | -0.437 | 1.599 | 0.416 | 0.283 | 0.001 | 0.001 | 0.001 | 0 | 0.53 | 0.45 | 0.48 | 0.26 | 0.62 | 0.39 |
| 3 | 20358655 | rs4858183 | -1.475 | 2.401 | 1.296 | 0.707 | 0.064 | 0.059 | 0.059 | 0 | 0.28 | 0.48 | 0.5 | 0.42 | 0.36 | 0.45 |
| 3 | 20362439 | rs6809146 | -0.81 | 2.356 | 0.706 | 0.708 | 0.121 | 0.081 | 0.081 | 0.156 | 0.08 | 0.41 | 0.12 | 0.06 | 0.1 | 0.07 |
| 3 | 20365258 | rs11709686 | -0.013 | 2.336 | -0.197 | 0.661 | -0.029 | 0.056 | 0.056 | 0 | 0.16 | 0.07 | 0.19 | 0.01 | 0.18 | 0.11 |
| 3 | 20367901 | rs7624305 | -1.364 | 1.7 | 0.23 | 1.09 | 0.017 | 0.122 | 0.122 | 0 | 0.12 | 0.26 | 0.23 | 0.39 | 0.13 | 0.33 |
| 3 | 20371453 | rs7619947 | -1.305 | 1.719 | 0.186 | 1.099 | 0.017 | 0.124 | 0.124 | 0 | 0.12 | 0.25 | 0.23 | 0.39 | 0.13 | 0.33 |
| 3 | 20374929 | rs6784144 | -1.228 | 1.674 | -0.141 | 1.33 | 0.001 | 0.039 | 0.039 | 0 | 0.07 | 0.07 | 0.16 | 0.3 | 0.08 | 0.24 |
| 3 | 20375219 | rs11915718 | -0.545 | 2.109 | -0.062 | 1.041 | 0.014 | -0.006 | -0.006 | 3.485 | 0.05 | 0.09 | 0.08 | 0.02 | 0.12 | 0.03 |
| 3 | 20375546 | rs11128951 | -1.563 | 2.331 | -0.312 | 1.342 | -0.011 | -0.018 | -0.018 | 0 | 0.14 | 0.13 | 0.2 | 0.35 | 0.19 | 0.25 |
| 3 | 20378153 | rs1356792 | 0.958 | 1.674 | -0.77 | 0.899 | 0.087 | 0.1 | 0.1 | 0 | 0.5 | 0.24 | 0.3 | 0.35 | 0.4 | 0.27 |
| 3 | 20391205 | rs1464294 | 0.859 | 1.41 | -0.169 | 0.974 | -0.004 | 0.052 | 0.052 | 0.016 | 0.72 | 0.39 | 0.69 | 0.5 | 0.76 | 0.41 |
| 3 | 20401474 | rs1520055 | -1.193 | 1.179 | 0.117 | 0.694 | -0.015 | -0.051 | -0.051 | 0.581 | 0.89 | 0.87 | 0.95 | 1 | 0.92 | 0.88 |
| 3 | 20403193 | rs9842115 | 0.871 | 1.755 | 0.294 | 0.775 | 0 | -0.001 | -0.001 | 0 | 0.63 | 0.43 | 0.39 | 0.44 | 0.54 | 0.37 |
| 3 | 20414230 | rs9854943 | 1.122 | 0.996 | -0.65 | 0.476 | -0.057 | -0.038 | -0.038 | 0.659 | 0.43 | 0.13 | 0.5 | 0.46 | 0.46 | 0.3 |
| 3 | 20417593 | rs17805021 | 1.754 | 1.132 | -0.818 | 0.496 | -0.017 | -0.013 | -0.013 | 0 | 0.07 | 0.03 | 0.04 | 0.08 | 0.08 | 0.03 |
| 3 | 20418070 | rs1879231 | -0.318 | 1.097 | -0.6 | 0.45 | 0.014 | 0.097 | 0.097 | 0 | 0.07 | 0.13 | 0.17 | 0.33 | 0.11 | 0.13 |
| 3 | 20419583 | rs6768597 | 1.804 | 1.191 | -0.854 | 0.225 | -0.096 | -0.044 | -0.044 | 0.024 | 0.14 | 0.01 | 0.31 | 0.1 | 0.26 | 0.16 |
| 3 | 20421544 | rs17006915 | 1.733 | 0.661 | -0.476 | -0.087 | -0.012 | -0.07 | -0.07 | 0.962 | 0.17 | 0.19 | 0.09 | 0 | 0.14 | 0.07 |
| 3 | 20431544 | rs6770717 | -0.927 | 0.541 | -0.712 | 0.052 | -0.008 | 0.024 | 0.024 | 0 | 0.81 | 0.87 | 0.84 | 0.96 | 0.73 | 0.81 |
| 3 | 20433212 | rs11128953 | 0.044 | 0.534 | -0.778 | 0.106 | -0.023 | 0.051 | 0.051 | 0 | 0.74 | 0.86 | 0.81 | 0.88 | 0.66 | 0.78 |
| 3 | 20436149 | rs9857925 | 2.119 | 0.359 | -0.67 | 0.261 | 0.055 | -0.031 | -0.031 | 0 | 0.3 | 0.08 | 0.18 | 0.4 | 0.2 | 0.14 |
| 3 | 20436493 | rs1401921 | -0.866 | 0.428 | -0.59 | 0.179 | -0.002 | -0.004 | -0.004 | 0 | 0.81 | 0.81 | 0.84 | 0.96 | 0.73 | 0.81 |
| 3 | 20440199 | rs17006942 | 1.294 | 0.368 | -0.661 | 0.361 | 0.019 | -0.015 | -0.015 | 0 | 0.36 | 0.28 | 0.26 | 0.52 | 0.24 | 0.35 |
| 3 | 20446843 | rs1357502 | 0.108 | 0.915 | -0.585 | 0.193 | -0.062 | 0.103 | 0.103 | 0 | 0.38 | 0.2 | 0.44 | 0.1 | 0.47 | 0.36 |
| 3 | 20450399 | rs6769110 | -1.385 | -0.307 | -0.536 | 0.289 | 0.06 | 0.095 | 0.095 | 0 | 0.54 | 0.72 | 0.7 | 0.73 | 0.59 | 0.71 |
| 3 | 20456375 | rs6765440 | -1.199 | -0.291 | -0.519 | 0.769 | 0.055 | 0 | 0 | 0 | 0.52 | 0.7 | 0.61 | 0.5 | 0.56 | 0.58 |
| 3 | 20459302 | rs9811678 | -1.196 | -0.352 | -0.376 | 0.742 | 0.043 | 0.001 | 0.001 | 0 | 0.52 | 0.68 | 0.61 | 0.5 | 0.56 | 0.58 |
| 3 | 20462528 | rs4858199 | 0.231 | -0.092 | 0.467 | 0.679 | 0.042 | 0.026 | 0.026 | 0 | 0.31 | 0.4 | 0.53 | 0.34 | 0.44 | 0.43 |
| 3 | 20465084 | rs6801593 | 2.412 | 0.137 | 0.155 | 0.543 | -0.113 | -0.051 | -0.051 | 0 | 0.15 | 0.01 | 0.37 | 0.1 | 0.32 | 0.14 |
| 3 | 20476362 | rs13070421 | 0.373 | -0.178 | 0.323 | 0.558 | -0.007 | 0.014 | 0.014 | 0 | 0.22 | 0.28 | 0.17 | 0.39 | 0.15 | 0.28 |
| 3 | 20477223 | rs939299 | 0.261 | -0.07 | 0.319 | 0.506 | 0.051 | -0.027 | -0.027 | 0.067 | 0.29 | 0.38 | 0.44 | 0.11 | 0.41 | 0.29 |
| 3 | 20480253 | rs12492284 | 1.812 | -0.155 | 0.355 | 0.664 | -0.001 | 0 | 0 | 0 | 0.13 | 0.1 | 0.07 | 0.22 | 0.11 | 0.07 |

| | | | | | | | | | | | | | | | | |
|---|---|---|---|---|---|---|---|---|---|---|---|---|---|---|---|---|
| 3 | 20496022 | rs6793867 | -0.781 | -0.243 | 0.547 | 0.656 | -0.037 | 0.029 | 0.029 | 0 | 0.28 | 0.49 | 0.19 | 0.4 | 0.16 | 0.41 |
| 3 | 20498236 | rs7633394 | -2.061 | -0.531 | 0.543 | 0.561 | 0.038 | 0.104 | 0.104 | 0 | 0.61 | 0.88 | 0.73 | 0.77 | 0.61 | 0.76 |
| 3 | 20498673 | rs9880609 | 1.58 | -0.533 | 0.613 | 0.574 | -0.012 | 0.049 | 0.049 | 0 | 0.08 | 0.14 | 0.11 | 0.23 | 0.06 | 0.15 |
| 3 | 20501403 | rs1474180 | 1.128 | -0.518 | 0.618 | 0.527 | -0.006 | 0.028 | 0.028 | 0 | 0.09 | 0.2 | 0.1 | 0.18 | 0.04 | 0.22 |
| 3 | 20501468 | rs17806455 | 2.377 | -0.443 | 0.41 | 0.503 | -0.004 | 0 | 0 | 0 | 0.08 | 0.1 | 0.04 | 0 | 0.08 | 0.03 |
| 3 | 20503129 | rs6762123 | 0.819 | -0.527 | 0.345 | 0.558 | -0.027 | 0.005 | 0.005 | 0 | 0.18 | 0.29 | 0.09 | 0.22 | 0.11 | 0.18 |
| 3 | 20507621 | rs6550532 | -0.032 | -0.529 | 0.28 | 0.576 | 0 | 0 | 0 | 0 | 0.34 | 0.38 | 0.51 | 0.27 | 0.45 | 0.4 |
| 3 | 20511825 | rs4858211 | -0.401 | -0.509 | 0.035 | 0.601 | 0.007 | -0.005 | -0.005 | 0 | 0.38 | 0.57 | 0.53 | 0.29 | 0.46 | 0.5 |
| 3 | 20512027 | rs6550534 | -0.45 | -0.5 | 0.039 | 0.605 | 0.006 | -0.005 | -0.005 | 0 | 0.39 | 0.57 | 0.54 | 0.29 | 0.46 | 0.51 |
| 3 | 20512768 | rs1523347 | -0.007 | -0.413 | 0.06 | 0.632 | -0.013 | 0.036 | 0.036 | 0 | 0.23 | 0.29 | 0.23 | 0.46 | 0.2 | 0.23 |
| 3 | 20513525 | rs1520045 | 1.149 | -0.111 | -0.109 | 0.578 | -0.001 | 0.002 | 0.002 | 0 | 0.08 | 0.12 | 0.09 | 0.17 | 0.04 | 0.22 |
| 3 | 20515558 | rs610310 | 1.373 | -0.119 | -0.358 | 0.571 | 0.002 | -0.011 | -0.011 | 0 | 0.15 | 0.09 | 0.09 | 0.22 | 0.14 | 0.08 |
| 3 | 20517023 | rs615861 | -1.014 | -0.121 | -0.24 | 0.631 | 0.016 | -0.001 | -0.001 | 0 | 0.46 | 0.63 | 0.6 | 0.47 | 0.52 | 0.56 |
| 3 | 20517372 | rs617621 | 1.347 | -0.197 | -0.306 | 0.692 | 0 | -0.002 | -0.002 | 0 | 0.15 | 0.11 | 0.09 | 0.22 | 0.14 | 0.08 |
| 3 | 20524946 | rs9968234 | 0.376 | -0.312 | -0.013 | 0.833 | -0.002 | 0.063 | 0.063 | 0 | 0.11 | 0.37 | 0.17 | 0.17 | 0.04 | 0.22 |
| 3 | 20527329 | rs689307 | -1.994 | -0.767 | -0.527 | 0.407 | -0.005 | 0.053 | 0.053 | 0 | 0.85 | 0.95 | 0.92 | 1 | 0.81 | 0.85 |
| 3 | 20527630 | rs582714 | -1.853 | -0.951 | -0.579 | 0.567 | 0.032 | 0.038 | 0.038 | 0 | 0.63 | 0.9 | 0.75 | 0.77 | 0.64 | 0.77 |
| 3 | 20540687 | rs11919213 | 0.244 | -0.697 | -0.273 | 0.547 | -0.003 | 0.066 | 0.066 | 0 | 0.12 | 0.4 | 0.17 | 0.18 | 0.04 | 0.23 |
| 3 | 20552038 | rs1587978 | -1.363 | -0.693 | -0.555 | 0.183 | 0.011 | 0.004 | 0.004 | 0 | 0.81 | 0.85 | 0.92 | 0.8 | 0.87 | 0.92 |
| 3 | 20552462 | rs7620685 | 2.378 | -0.762 | -0.799 | 0.085 | -0.123 | 0.011 | 0.011 | 0.008 | 0.16 | 0.01 | 0.23 | 0.05 | 0.34 | 0.14 |
| 3 | 20559052 | rs661669 | 0.783 | -0.836 | -0.669 | 0.059 | -0.001 | -0.001 | -0.001 | 0 | 0.54 | 0.54 | 0.38 | 0.57 | 0.5 | 0.57 |
| 3 | 20567783 | rs594583 | 0.484 | -1.091 | -0.462 | 0.085 | -0.049 | -0.002 | -0.002 | 0 | 0.7 | 0.56 | 0.75 | 0.63 | 0.83 | 0.68 |
| 3 | 20570373 | rs1403651 | 0.126 | -0.689 | -0.241 | 0.261 | -0.007 | 0.061 | 0.061 | 0 | 0.11 | 0.34 | 0.17 | 0.18 | 0.04 | 0.26 |
| 3 | 20575986 | rs1506048 | -0.271 | -1.615 | -0.844 | -0.253 | 0.018 | 0.015 | 0.015 | 0 | 0.62 | 0.66 | 0.71 | 0.67 | 0.67 | 0.58 |
| 3 | 20577473 | rs1523351 | 0.821 | -0.985 | -0.733 | -0.378 | 0.05 | -0.045 | -0.045 | 2.656 | 0.27 | 0.33 | 0.42 | 0.1 | 0.37 | 0.3 |
| 3 | 20578998 | rs10440098 | 0.821 | -0.985 | -0.754 | -0.378 | 0.044 | -0.04 | -0.04 | 0 | 0.27 | 0.32 | 0.43 | 0.1 | 0.37 | 0.3 |
| 3 | 20580104 | rs1027777 | 1.028 | -0.405 | -0.844 | -0.144 | 0 | -0.003 | -0.003 | 0 | 0.22 | 0.19 | 0.41 | 0.05 | 0.36 | 0.18 |
| 3 | 20583201 | rs1506049 | 1.609 | -0.951 | -1.14 | 0.404 | 0.016 | -0.047 | -0.047 | 0 | 0.12 | 0.04 | 0.13 | 0.25 | 0.08 | 0.11 |
| 3 | 20583674 | rs996482 | 0.024 | -1.273 | -0.841 | 0.207 | -0.001 | 0.009 | 0.009 | 0 | 0.3 | 0.57 | 0.23 | 0.34 | 0.3 | 0.33 |
| 3 | 20601131 | rs1113746 | -2.264 | -1.419 | -1.202 | -0.114 | 0.018 | 0.012 | 0.012 | 0 | 0.78 | 0.88 | 0.92 | 0.8 | 0.83 | 0.81 |
| 3 | 20604370 | rs1506057 | 1.998 | -1.285 | -0.883 | -0.067 | 0 | -0.008 | -0.008 | 0 | 0.16 | 0.17 | 0.14 | 0.28 | 0.15 | 0.19 |
| 3 | 20605188 | rs13062445 | 2.477 | -1.259 | -1.08 | -0.819 | -0.017 | 0.079 | 0.079 | 0.283 | 0.13 | 0.02 | 0.06 | 0 | 0.11 | 0.06 |
| 3 | 20605302 | rs4292253 | 2.819 | -0.703 | -1.069 | -0.756 | -0.124 | 0.034 | 0.034 | 0 | 0.17 | 0.01 | 0.36 | 0.04 | 0.31 | 0.15 |
| 3 | 20611348 | rs9815914 | 0.801 | -1.295 | 0.264 | -0.46 | -0.001 | 0.01 | 0.01 | 0 | 0.08 | 0.26 | 0.04 | 0.04 | 0.03 | 0.08 |
| 3 | 20619752 | rs11720317 | 0.661 | -0.946 | 0.221 | -0.438 | 0.051 | -0.045 | -0.045 | 0 | 0.1 | 0.25 | 0.08 | 0.01 | 0.15 | 0.08 |
| 3 | 20621602 | rs1605193 | -0.125 | -1.153 | -0.055 | -0.476 | -0.008 | -0.031 | -0.031 | 0.36 | 0.25 | 0.34 | 0.25 | 0.17 | 0.26 | 0.3 |
| 3 | 20624780 | rs2055652 | -1.058 | -1.544 | -0.759 | -0.322 | 0.017 | 0.023 | 0.023 | 0 | 0.72 | 0.82 | 0.89 | 0.77 | 0.78 | 0.73 |
| 3 | 20627558 | rs17007204 | 1.138 | -1.14 | -0.379 | 0.087 | 0.032 | 0.01 | 0.01 | 0 | 0.13 | 0.1 | 0.16 | 0.14 | 0.08 | 0.15 |
| 3 | 20638228 | rs6769686 | -0.742 | -0.613 | 0.268 | 0.063 | -0.018 | -0.105 | -0.105 | 0.108 | 0.15 | 0.43 | 0.09 | 0.01 | 0.09 | 0.13 |
| 3 | 20642753 | rs9859271 | -1.135 | -0.738 | 0.145 | 0.713 | -0.031 | 0.112 | 0.112 | 0 | 0.23 | 0.41 | 0.25 | 0.34 | 0.16 | 0.22 |
| 3 | 20643480 | rs6766464 | -1.05 | -0.443 | -0.387 | 0.459 | -0.001 | 0.009 | 0.009 | 0.116 | 0.29 | 0.66 | 0.41 | 0.53 | 0.29 | 0.37 |
| 3 | 20645063 | rs1021937 | -1.37 | -0.91 | -0.968 | -0.017 | 0.027 | 0.031 | 0.031 | 0 | 0.76 | 0.88 | 0.91 | 0.8 | 0.82 | 0.83 |
| 3 | 20645600 | rs6784556 | 0.517 | -1.192 | -0.997 | -0.454 | 0.004 | 0.01 | 0.01 | 0 | 0.25 | 0.23 | 0.21 | 0.23 | 0.24 | 0.31 |
| 3 | 20651459 | rs1121305 | -0.534 | -1.222 | -1.111 | -0.554 | 0.007 | 0.04 | 0.04 | 0 | 0.35 | 0.26 | 0.27 | 0.24 | 0.35 | 0.42 |
| 3 | 20667279 | rs1500412 | 3.115 | -0.809 | -1.506 | -0.668 | -0.077 | 0.036 | 0.036 | 0 | 0.12 | 0.01 | 0.24 | 0.03 | 0.22 | 0.12 |
| 3 | 20667710 | rs12496130 | 2.232 | -1.702 | -1.576 | -0.603 | -0.031 | 0.046 | 0.046 | 0 | 0.08 | 0.02 | 0.07 | 0 | 0.13 | 0.06 |
| 3 | 20669485 | rs9826976 | 0.549 | -1.889 | -0.785 | -0.014 | 0.051 | 0.164 | 0.164 | 0 | 0.16 | 0.35 | 0.27 | 0.38 | 0.22 | 0.3 |
| 3 | 20674832 | rs4337653 | -1.431 | -2.048 | -1.137 | 0.031 | 0.031 | -0.142 | -0.142 | 0 | 0.66 | 0.97 | 0.8 | 0.55 | 0.74 | 0.75 |
| 3 | 20682440 | rs9839718 | 1.717 | -1.919 | -0.776 | 0.15 | -0.008 | 0.002 | 0.002 | 0 | 0.12 | 0.13 | 0.13 | 0.16 | 0.09 | 0.14 |
| 3 | 20688879 | rs11714736 | 0.635 | -1.85 | -0.46 | 0.129 | 0.027 | 0.04 | 0.04 | 0 | 0.16 | 0.27 | 0.26 | 0.26 | 0.21 | 0.27 |
| 3 | 20690491 | rs17007299 | 0.994 | -1.846 | -0.274 | 0.173 | -0.006 | 0.003 | 0.003 | 0 | 0.13 | 0.09 | 0.23 | 0.36 | 0.21 | 0.26 |
| 3 | 20692368 | rs4462985 | 0.458 | -1.858 | -0.588 | 0.05 | 0.018 | 0.013 | 0.013 | 0 | 0.34 | 0.38 | 0.54 | 0.4 | 0.49 | 0.42 |
| 3 | 20693062 | rs6781212 | 1.989 | -1.96 | -0.679 | 0.075 | -0.013 | 0.002 | 0.002 | 0.013 | 0.11 | 0.12 | 0.1 | 0.15 | 0.06 | 0.13 |
| 3 | 20695652 | rs1392573 | -1 | -2.029 | -0.51 | -0.03 | 0.044 | -0.009 | -0.009 | 0.026 | 0.62 | 0.79 | 0.75 | 0.63 | 0.73 | 0.74 |
| 3 | 20696379 | rs904827 | -1.069 | -2.064 | -0.795 | -0.045 | 0.061 | -0.067 | -0.067 | 0 | 0.63 | 0.97 | 0.78 | 0.64 | 0.74 | 0.73 |
| 3 | 20703952 | rs7647157 | -1.464 | -2.032 | -0.779 | -0.031 | 0.03 | -0.126 | -0.126 | 0.021 | 0.66 | 0.97 | 0.79 | 0.53 | 0.74 | 0.74 |
| 3 | 20705014 | rs4858248 | -0.023 | -1.932 | -0.469 | -0.115 | -0.003 | 0.027 | 0.027 | 0 | 0.45 | 0.39 | 0.46 | 0.26 | 0.49 | 0.46 |
| 3 | 20707404 | rs9841080 | -0.748 | -1.45 | 0.034 | 0.021 | 0.013 | 0.014 | 0.014 | 0 | 0.35 | 0.54 | 0.39 | 0.45 | 0.44 | 0.44 |
| 3 | 20708397 | rs7623569 | 1.046 | -1.135 | -0.097 | 0.208 | 0.023 | 0.158 | 0.158 | 0 | 0.12 | 0.26 | 0.27 | 0.37 | 0.17 | 0.17 |
| 3 | 20710994 | rs1566479 | 1.991 | -0.979 | -0.179 | -0.711 | -0.1 | 0.064 | 0.064 | 0 | 0.16 | 0.01 | 0.26 | 0.03 | 0.24 | 0.17 |
| 3 | 20711653 | rs6807165 | 0.477 | -1.372 | 0.069 | -0.694 | 0.018 | -0.043 | -0.043 | 0 | 0.31 | 0.36 | 0.36 | 0.11 | 0.42 | 0.33 |
| 3 | 20713455 | rs2886695 | 0.126 | -0.985 | 0.024 | -0.151 | 0.015 | 0.048 | 0.048 | 0.576 | 0.2 | 0.31 | 0.29 | 0.3 | 0.26 | 0.25 |
| 3 | 20718124 | rs7630274 | -0.124 | -1.151 | -0.664 | -0.334 | -0.01 | 0.019 | 0.019 | 1.988 | 0.28 | 0.65 | 0.38 | 0.5 | 0.24 | 0.35 |
| 3 | 20720138 | rs1500422 | 1.172 | -0.763 | -1.692 | -0.947 | -0.13 | 0.144 | 0.144 | 0.011 | 0.25 | 0.04 | 0.3 | 0.03 | 0.35 | 0.19 |
| 3 | 20725113 | rs17203361 | 0.639 | -0.818 | -1.493 | -1.022 | -0.024 | 0.018 | 0.018 | 0 | 0.31 | 0.26 | 0.35 | 0.07 | 0.41 | 0.25 |
| 3 | 20725743 | rs11716554 | 0.33 | -1.032 | -1.598 | -1.079 | 0.02 | 0.007 | 0.007 | 0 | 0.23 | 0.21 | 0.18 | 0.18 | 0.18 | 0.19 |
| 3 | 20726723 | rs9866656 | -0.669 | -1.042 | -1.486 | -1.083 | 0 | -0.002 | -0.002 | 0 | 0.55 | 0.62 | 0.55 | 0.24 | 0.58 | 0.44 |
| 3 | 20727885 | rs17007409 | 1.877 | -1.043 | -1.485 | -1.055 | -0.002 | -0.001 | -0.001 | 0 | 0.11 | 0.1 | 0.1 | 0.18 | 0.05 | 0.14 |
| 3 | 20729091 | rs4858253 | 1.145 | -0.75 | -1.486 | -1.005 | -0.01 | -0.009 | -0.009 | 0 | 0.27 | 0.26 | 0.34 | 0.03 | 0.35 | 0.21 |

| | | | | | | | | | | | | | | | | |
|---|---|---|---|---|---|---|---|---|---|---|---|---|---|---|---|---|
| 3 | 20731156 | rs6790689 | 0.084 | -0.301 | -1.392 | -0.556 | 0.001 | 0 | 0 | 1.686 | 0.32 | 0.39 | 0.33 | 0.36 | 0.34 | 0.27 |
| 3 | 20736153 | rs931717 | -1.502 | -0.017 | -1.531 | -0.351 | 0.014 | 0.017 | 0.017 | 0 | 0.85 | 1 | 0.95 | 0.87 | 0.92 | 0.89 |
| 3 | 20736328 | rs12489804 | 0.891 | 0.494 | -1.128 | -0.405 | 0.028 | -0.04 | -0.04 | 0.022 | 0.11 | 0.22 | 0.08 | 0.03 | 0.13 | 0.14 |
| 3 | 20744029 | rs12491632 | 1.112 | -0.251 | -1.626 | 0.124 | -0.015 | 0.039 | 0.039 | 0 | 0.07 | 0.01 | 0.05 | 0 | 0.1 | 0.05 |
| 3 | 20751876 | rs9839849 | 0.536 | -0.412 | -1.208 | 0.56 | -0.009 | -0.025 | -0.025 | 1.163 | 0.6 | 0.64 | 0.59 | 0.21 | 0.66 | 0.53 |
| 3 | 20752472 | rs4858256 | 0.414 | -0.431 | -1.2 | 0.651 | -0.001 | 0.018 | 0.018 | 1.432 | 0.14 | 0.15 | 0.21 | 0.36 | 0.15 | 0.15 |
| 3 | 20752492 | rs4858257 | 0.147 | -0.542 | -1.249 | 0.829 | -0.011 | 0.017 | 0.017 | 0.142 | 0.73 | 0.8 | 0.8 | 0.57 | 0.81 | 0.63 |
| 3 | 20754342 | rs1983020 | -0.465 | -0.091 | -1.533 | 0.349 | 0.022 | 0.032 | 0.032 | 0.199 | 0.22 | 0.1 | 0.15 | 0.13 | 0.14 | 0.16 |
| 3 | 20761438 | rs870016 | -0.485 | -0.875 | -1.364 | 0.799 | -0.02 | 0.014 | 0.014 | 0 | 0.26 | 0.25 | 0.27 | 0.52 | 0.14 | 0.37 |
| 3 | 20762123 | rs873977 | -0.196 | -1.26 | -1.429 | 1.268 | -0.051 | 0.071 | 0.071 | 0 | 0.16 | 0.24 | 0.24 | 0.3 | 0.07 | 0.21 |
| 3 | 20764228 | rs985115 | -0.32 | -1.371 | -1.494 | 1.296 | 0.026 | -0.039 | -0.039 | 0 | 0.09 | 0.02 | 0.03 | 0.2 | 0.05 | 0.16 |
| 3 | 20764696 | rs6789500 | -1.474 | -1.407 | -1.572 | 0.934 | 0.005 | -0.005 | -0.005 | 0 | 0.34 | 0.29 | 0.32 | 0.58 | 0.25 | 0.51 |
| 3 | 20766050 | rs2366312 | 1.048 | -1.822 | -2.158 | 0.548 | 0.012 | -0.069 | -0.069 | 2.467 | 0.12 | 0.02 | 0.18 | 0.38 | 0.09 | 0.1 |
| 3 | 20766560 | rs17810229 | 0.572 | -0.677 | -1.835 | -0.679 | 0.004 | -0.047 | -0.047 | 0.03 | 0.21 | 0.31 | 0.18 | 0 | 0.31 | 0.14 |
| 3 | 20777360 | rs9839419 | -0.349 | -0.63 | -1.55 | -0.416 | 0.033 | -0.072 | -0.072 | 0 | 0.31 | 0.18 | 0.24 | 0.51 | 0.23 | 0.37 |
| 3 | 20777750 | rs2068188 | 0.161 | -0.699 | -1.584 | -0.4 | 0.036 | -0.068 | -0.068 | 0 | 0.22 | 0.11 | 0.18 | 0.42 | 0.14 | 0.28 |
| 3 | 20781002 | rs11918349 | -0.67 | -0.378 | -1.385 | -0.466 | -0.023 | -0.023 | -0.023 | 0 | 0.36 | 0.28 | 0.43 | 0.56 | 0.42 | 0.49 |
| 3 | 20788581 | rs964911 | -1.531 | -0.76 | -1.297 | -1.06 | 0.006 | -0.028 | -0.028 | 0.765 | 0.63 | 0.82 | 0.62 | 0.63 | 0.67 | 0.66 |
| 3 | 20788875 | rs964910 | 2.412 | -0.853 | -1.367 | -1.024 | 0.001 | -0.063 | -0.063 | 0 | 0.11 | 0.02 | 0.15 | 0.37 | 0.07 | 0.14 |
| 3 | 20790181 | rs4857969 | -1.042 | -1.025 | -0.696 | -1.25 | 0.012 | 0.009 | 0.009 | 0 | 0.51 | 0.57 | 0.55 | 0.63 | 0.58 | 0.64 |
| 3 | 20798214 | rs7630411 | 0.694 | -0.701 | -0.714 | -1.116 | 0.015 | 0.012 | 0.012 | 0 | 0.11 | 0.06 | 0.04 | 0.04 | 0.07 | 0.14 |
| 3 | 20803801 | rs9990378 | 1.484 | -0.46 | -0.585 | -1.187 | 0.105 | -0.008 | -0.008 | 0 | 0.06 | 0.11 | 0.19 | 0.04 | 0.18 | 0.15 |
| 3 | 20806820 | rs9990230 | -1.004 | -0.471 | -0.767 | -1.402 | 0.032 | 0.059 | 0.059 | 0 | 0.54 | 0.31 | 0.7 | 0.98 | 0.64 | 0.84 |
| 3 | 20807454 | rs13070736 | 0.48 | -0.502 | -1.183 | -1.231 | 0.061 | -0.104 | -0.104 | 4.273 | 0.23 | 0.07 | 0.25 | 0.39 | 0.16 | 0.34 |
| 3 | 20815289 | rs1392574 | -1.733 | 0.102 | -0.789 | -1.542 | -0.002 | 0.025 | 0.025 | 0 | 0.73 | 0.84 | 0.84 | 0.98 | 0.76 | 0.9 |
| 3 | 20817921 | rs1500418 | -0.484 | 0.216 | -0.799 | -1.435 | -0.01 | -0.13 | -0.13 | 0 | 0.55 | 0.48 | 0.72 | 0.98 | 0.63 | 0.84 |
| 3 | 20819623 | rs1500415 | -0.591 | 0.325 | -0.905 | -1.354 | 0.007 | -0.047 | -0.047 | 0 | 0.54 | 0.4 | 0.7 | 0.98 | 0.62 | 0.81 |
| 3 | 20827150 | rs1827226 | -0.09 | 0.278 | -1.239 | -1.281 | 0.024 | 0.159 | 0.159 | 0 | 0.47 | 0.22 | 0.5 | 0.93 | 0.44 | 0.66 |
| 3 | 20834135 | rs1392580 | -0.876 | 0.843 | -1.229 | -0.265 | 0.018 | 0.034 | 0.034 | 0.6 | 0.5 | 0.84 | 0.63 | 0.52 | 0.58 | 0.65 |
| 3 | 20835200 | rs17007652 | -0.01 | 1.864 | -0.691 | -0.613 | 0.088 | -0.012 | -0.012 | 0.845 | 0.08 | 0.15 | 0.22 | 0.04 | 0.18 | 0.19 |
| 3 | 20839740 | rs1500427 | -0.503 | 1.339 | -0.716 | -0.599 | 0.093 | -0.023 | -0.023 | 0 | 0.14 | 0.25 | 0.28 | 0.08 | 0.29 | 0.28 |
| 3 | 20856539 | rs4858270 | -0.344 | 0.572 | -1.575 | -0.834 | -0.023 | 0.101 | 0.101 | 0.311 | 0.72 | 0.94 | 0.86 | 0.9 | 0.71 | 0.8 |
| 3 | 20863963 | rs6769651 | -0.36 | 0.591 | -1.654 | -0.915 | -0.022 | 0.148 | 0.148 | 0.027 | 0.72 | 0.95 | 0.88 | 0.96 | 0.74 | 0.83 |
| 3 | 20882268 | rs17007701 | 0.481 | -0.051 | -1.907 | -1.139 | 0.001 | 0 | 0 | 0 | 0.11 | 0.11 | 0.07 | 0.1 | 0.13 | 0.12 |
| 3 | 20883163 | rs1995191 | 1.078 | -0.472 | -1.686 | -0.562 | -0.007 | 0.088 | 0.088 | 0 | 0.09 | 0.08 | 0.22 | 0.39 | 0.07 | 0.14 |
| 3 | 20886763 | rs1032665 | 0.021 | -0.45 | -1.667 | -0.579 | -0.004 | 0.032 | 0.032 | 0 | 0.2 | 0.19 | 0.29 | 0.49 | 0.19 | 0.26 |
| 3 | 20893060 | rs4858276 | 1.037 | -0.626 | -1.289 | -0.735 | -0.031 | -0.029 | -0.029 | 0.488 | 0.35 | 0.22 | 0.39 | 0.45 | 0.39 | 0.43 |
| 3 | 20898944 | rs1391857 | 0.893 | -0.638 | -1.308 | -0.917 | -0.049 | -0.026 | -0.026 | 0.279 | 0.36 | 0.2 | 0.34 | 0.41 | 0.39 | 0.43 |
| 3 | 20907259 | rs12486758 | 0.986 | -0.547 | -0.872 | -1.058 | -0.011 | -0.05 | -0.05 | 0 | 0.11 | 0.2 | 0.07 | 0 | 0.08 | 0.02 |
| 3 | 20912122 | rs6807580 | -0.905 | -0.533 | -1.312 | -1.147 | -0.002 | 0.043 | 0.043 | 0 | 0.75 | 0.87 | 0.89 | 0.95 | 0.76 | 0.76 |
| 3 | 20919920 | rs4549291 | -0.869 | -0.532 | -1.285 | -1.148 | 0 | 0.03 | 0.03 | 0 | 0.74 | 0.83 | 0.88 | 0.95 | 0.76 | 0.76 |
| 3 | 20930467 | rs17811881 | 0.222 | -0.896 | -1.198 | -0.731 | 0.001 | 0.01 | 0.01 | 0 | 0.13 | 0.09 | 0.07 | 0.1 | 0.15 | 0.13 |
| 3 | 20934441 | rs13093247 | 0.494 | -0.83 | -1.293 | -0.747 | -0.032 | -0.01 | -0.01 | 0.352 | 0.12 | 0 | 0.07 | 0.08 | 0.17 | 0.09 |
| 3 | 20934496 | rs13093281 | 0.306 | -0.818 | -1.037 | -0.742 | -0.001 | -0.006 | -0.006 | 0 | 0.13 | 0.15 | 0.09 | 0.08 | 0.17 | 0.1 |
| 3 | 20939291 | rs1580800 | 0.61 | -0.656 | -1.583 | -0.988 | 0.002 | 0.025 | 0.025 | 1.671 | 0.15 | 0.09 | 0.08 | 0 | 0.12 | 0.03 |
| 3 | 20944820 | rs1498994 | 0.066 | -0.43 | -1.591 | -0.476 | 0.014 | 0.002 | 0.002 | 0 | 0.45 | 0.17 | 0.33 | 0.42 | 0.53 | 0.5 |
| 3 | 20946983 | rs1498997 | -0.343 | 0.232 | -1.468 | -0.107 | -0.003 | -0.019 | -0.019 | 1.669 | 0.93 | 0.9 | 0.93 | 0.99 | 0.94 | 0.93 |
| 3 | 20949819 | rs4858280 | -0.053 | 0.618 | -1.821 | 0.128 | -0.105 | -0.072 | -0.072 | 0 | 0.32 | 0.08 | 0.23 | 0.42 | 0.42 | 0.43 |
| 3 | 20970042 | rs10510509 | -0.332 | 0.445 | -1.373 | -0.087 | -0.009 | 0.015 | 0.015 | 5.881 | 0.08 | 0.05 | 0.05 | 0 | 0.1 | 0.03 |
| 3 | 20972475 | rs9864991 | 0.443 | -0.122 | -1.459 | 0.158 | 0.012 | 0.034 | 0.034 | 0 | 0.43 | 0.32 | 0.48 | 0.35 | 0.62 | 0.3 |
| 3 | 20973020 | rs1498999 | -0.469 | -0.319 | -0.849 | 0.378 | 0.021 | 0.008 | 0.008 | 0 | 0.52 | 0.54 | 0.47 | 0.6 | 0.33 | 0.64 |
| 3 | 20979796 | rs4482681 | 0.455 | -0.54 | -0.643 | 0.198 | 0.023 | 0.023 | 0.023 | 0.071 | 0.47 | 0.45 | 0.51 | 0.38 | 0.67 | 0.34 |
| 3 | 20980214 | rs1587379 | 0.696 | -0.717 | -0.869 | 0.191 | -0.005 | -0.008 | -0.008 | 0.038 | 0.92 | 0.96 | 0.95 | 0.93 | 0.93 | 0.84 |
| 3 | 20984404 | rs9862856 | -0.223 | -1.312 | -0.842 | 0.391 | 0.011 | 0.003 | 0.003 | 0.877 | 0.45 | 0.51 | 0.35 | 0.15 | 0.3 | 0.31 |
| 3 | 20984501 | rs9863026 | -0.275 | -1.3 | -0.873 | 0.412 | 0.018 | 0.017 | 0.017 | 1.076 | 0.46 | 0.54 | 0.42 | 0.16 | 0.3 | 0.31 |
| 3 | 20984552 | rs12636704 | -0.193 | -0.832 | -0.896 | 0.887 | 0.014 | 0.026 | 0.026 | 0 | 0.09 | 0.07 | 0.13 | 0.43 | 0.04 | 0.34 |
| 3 | 20994957 | rs9851291 | -0.705 | -0.716 | -0.994 | 0.306 | 0.021 | 0.049 | 0.049 | 0 | 0.54 | 0.67 | 0.4 | 0.68 | 0.38 | 0.65 |
| 3 | 20995849 | rs2201910 | 0.227 | -1.063 | -0.662 | 0.005 | 0 | 0.001 | 0.001 | 0.012 | 0.52 | 0.56 | 0.62 | 0.33 | 0.61 | 0.43 |
| 3 | 20997613 | rs2366681 | 0.265 | -1.269 | -0.838 | 0.214 | 0.013 | -0.029 | -0.029 | 0.616 | 0.69 | 0.72 | 0.79 | 0.56 | 0.79 | 0.49 |
| 3 | 20999820 | rs882238 | -0.299 | -1.233 | -0.773 | 0.305 | 0.006 | -0.014 | -0.014 | 0.767 | 0.79 | 0.77 | 0.84 | 0.57 | 0.86 | 0.54 |
| 3 | 21000069 | rs882239 | -0.312 | -1.239 | -0.772 | 0.306 | 0.006 | -0.014 | -0.014 | 0 | 0.79 | 0.77 | 0.84 | 0.57 | 0.86 | 0.54 |
| 3 | 21001290 | rs13073034 | 0.365 | -1.145 | -0.776 | 0.002 | -0.002 | 0.045 | 0.045 | 0 | 0.18 | 0.06 | 0.08 | 0.07 | 0.18 | 0.13 |
| 3 | 21006294 | rs13061016 | 0.131 | -1.189 | -0.245 | -0.029 | -0.004 | -0.04 | -0.04 | 0.257 | 0.2 | 0.22 | 0.09 | 0.07 | 0.18 | 0.13 |
| 3 | 21008710 | rs4462987 | 1.734 | -0.86 | -0.388 | 0.145 | 0.032 | 0 | 0 | 0 | 0.23 | 0.29 | 0.28 | 0.23 | 0.41 | 0.1 |
| 3 | 21020875 | rs9829660 | -0.667 | -0.77 | -0.342 | 0.743 | 0.016 | 0.023 | 0.023 | 0 | 0.64 | 0.49 | 0.63 | 0.55 | 0.55 | 0.76 |
| 3 | 21027150 | rs1497524 | 0.124 | -1.096 | -0.732 | 0.697 | -0.024 | -0.022 | -0.022 | 0 | 0.44 | 0.18 | 0.23 | 0.41 | 0.4 | 0.61 |
| 3 | 21027910 | rs7625611 | -1.757 | -1.141 | -0.549 | 0.731 | -0.013 | 0.01 | 0.01 | 0 | 0.72 | 0.73 | 0.73 | 0.79 | 0.6 | 0.88 |
| 3 | 21030097 | rs1846077 | -0.825 | -1.289 | -0.723 | 0.69 | 0.005 | 0.007 | 0.007 | 0.009 | 0.53 | 0.37 | 0.55 | 0.59 | 0.45 | 0.78 |
| 3 | 21031454 | rs13096029 | -0.207 | -1.392 | -0.636 | 0.813 | -0.002 | 0.003 | 0.003 | 0.129 | 0.1 | 0.05 | 0.05 | 0.05 | 0.11 | 0.11 |

| | | | | | | | | | | | | | | | | |
|---|---|---|---|---|---|---|---|---|---|---|---|---|---|---|---|---|
| 3 | 21035183 | rs12491034 | 0.675 | -1.364 | -0.71 | 0.84 | -0.048 | -0.015 | -0.015 | 0 | 0.4 | 0.16 | 0.23 | 0.38 | 0.39 | 0.54 |
| 3 | 21038610 | rs13073288 | 0.906 | -1.25 | -0.677 | 0.802 | -0.01 | 0.025 | 0.025 | 0 | 0.05 | 0.01 | 0.03 | 0.01 | 0.07 | 0.04 |
| 3 | 21042010 | rs13059917 | 0.61 | -1.354 | -0.794 | 0.787 | -0.049 | -0.009 | -0.009 | 0.318 | 0.41 | 0.15 | 0.24 | 0.38 | 0.42 | 0.54 |
| 3 | 21043491 | rs1497504 | -0.335 | -1.733 | -0.394 | 0.829 | -0.031 | 0.041 | 0.041 | 0 | 0.19 | 0.38 | 0.11 | 0.19 | 0.14 | 0.19 |
| 3 | 21049245 | rs6793362 | 0.226 | -1.531 | -0.313 | 0.512 | -0.007 | 0.011 | 0.011 | 1.154 | 0.46 | 0.33 | 0.26 | 0.39 | 0.44 | 0.6 |
| 3 | 21052737 | rs12633333 | 0.386 | -1.515 | -0.505 | 0.538 | -0.032 | -0.029 | -0.029 | 0 | 0.45 | 0.19 | 0.26 | 0.43 | 0.44 | 0.61 |
| 3 | 21062584 | rs4306882 | -1.395 | -1.7 | 0.129 | 0.402 | 0.016 | -0.029 | -0.029 | 0 | 0.71 | 0.63 | 0.74 | 0.81 | 0.62 | 0.89 |
| 3 | 21063578 | rs7625772 | -1.493 | -1.718 | 0.183 | 0.396 | 0.005 | -0.011 | -0.011 | 0 | 0.72 | 0.73 | 0.77 | 0.82 | 0.64 | 0.89 |
| 3 | 21065052 | rs13065045 | 0.239 | -1.801 | -0.312 | 0.616 | -0.034 | 0.052 | 0.052 | 1.679 | 0.45 | 0.22 | 0.26 | 0.31 | 0.45 | 0.59 |
| 3 | 21065354 | rs1497516 | -1.864 | -1.775 | -0.378 | 0.582 | -0.13 | 0.014 | 0.014 | 1.34 | 0.75 | 0.98 | 0.8 | 0.81 | 0.62 | 0.89 |
| 3 | 21066829 | rs17008011 | 0.758 | -1.557 | -0.171 | 0.613 | 0 | -0.014 | -0.014 | 0 | 0.06 | 0.07 | 0.03 | 0.01 | 0.06 | 0.04 |
| 3 | 21072043 | rs958635 | -0.325 | -1.574 | -0.107 | 0.656 | 0.005 | 0.013 | 0.013 | 0.816 | 0.52 | 0.39 | 0.29 | 0.36 | 0.5 | 0.64 |
| 3 | 21083986 | rs1532461 | 0.231 | -1.763 | -0.413 | 0.524 | -0.033 | 0.042 | 0.042 | 0 | 0.41 | 0.21 | 0.25 | 0.26 | 0.42 | 0.56 |
| 3 | 21084555 | rs1391142 | 1.295 | -1.77 | -0.461 | 0.518 | -0.027 | 0.027 | 0.027 | 0.081 | 0.3 | 0.16 | 0.2 | 0.21 | 0.31 | 0.48 |
| 3 | 21086158 | rs924224 | 0.147 | -1.939 | -0.29 | 0.572 | 0.018 | -0.025 | -0.025 | 0 | 0.28 | 0.2 | 0.47 | 0.55 | 0.18 | 0.32 |
| 3 | 21087219 | rs10510510 | 0.41 | -1.497 | -0.357 | 0.525 | 0.003 | 0.017 | 0.017 | 0 | 0.07 | 0.22 | 0.04 | 0.05 | 0.05 | 0.05 |
| 3 | 21097113 | rs9825481 | -0.064 | -1.572 | -0.463 | 0.595 | 0.011 | 0.013 | 0.013 | 1.07 | 0.51 | 0.33 | 0.27 | 0.39 | 0.48 | 0.63 |
| 3 | 21099489 | rs7652147 | -1.671 | -1.617 | -0.34 | 0.561 | -0.016 | 0.018 | 0.018 | 1.2 | 0.73 | 0.76 | 0.76 | 0.81 | 0.63 | 0.89 |
| 3 | 21103323 | rs12637202 | 0.684 | -1.649 | -0.314 | 0.684 | -0.041 | -0.005 | -0.005 | 0.208 | 0.39 | 0.14 | 0.21 | 0.34 | 0.37 | 0.55 |
| 3 | 21106656 | rs4858289 | -0.419 | -1.293 | 0.116 | 1.063 | -0.082 | 0.116 | 0.116 | 0 | 0.22 | 0.44 | 0.48 | 0.42 | 0.14 | 0.26 |
| 3 | 21112698 | rs1908988 | 0.515 | -0.618 | -0.265 | 0.557 | -0.01 | 0.005 | 0.005 | 0.718 | 0.3 | 0.28 | 0.24 | 0.23 | 0.43 | 0.17 |
| 3 | 21125033 | rs9810583 | 1.375 | -0.481 | -0.221 | 0.558 | -0.006 | 0.004 | 0.004 | 2.419 | 0.22 | 0.24 | 0.21 | 0.18 | 0.33 | 0.09 |
| 3 | 21127149 | rs11128984 | -0.446 | -0.792 | 0.066 | 0.968 | -0.012 | -0.011 | -0.011 | 0.009 | 0.42 | 0.72 | 0.67 | 0.49 | 0.47 | 0.39 |
| 3 | 21128190 | rs1846072 | 1.158 | -0.72 | 0.103 | 1.043 | -0.007 | 0.022 | 0.022 | 0.354 | 0.09 | 0.09 | 0.37 | 0.25 | 0.03 | 0.22 |
| 3 | 21134566 | rs1497517 | 1.366 | -1 | -0.098 | 0.988 | -0.01 | -0.06 | -0.06 | 0 | 0.1 | 0.01 | 0.08 | 0.31 | 0.1 | 0.13 |
| 3 | 21135162 | rs17008182 | -0.327 | -0.759 | 0.287 | 0.527 | -0.001 | -0.001 | -0.001 | 0 | 0.64 | 0.32 | 0.38 | 0.56 | 0.61 | 0.68 |
| 3 | 21136757 | rs9883374 | -0.313 | -0.731 | 0.127 | 0.466 | -0.001 | -0.001 | -0.001 | 0 | 0.43 | 0.6 | 0.66 | 0.5 | 0.5 | 0.4 |
| 3 | 21165530 | rs1497502 | 0.235 | -0.933 | -0.15 | 0.583 | -0.039 | -0.033 | -0.033 | 0 | 0.35 | 0.22 | 0.58 | 0.48 | 0.45 | 0.35 |
| 3 | 21182719 | rs13060855 | -0.054 | -0.994 | -0.344 | 0.529 | -0.008 | -0.011 | -0.011 | 1.235 | 0.38 | 0.31 | 0.58 | 0.51 | 0.44 | 0.36 |
| 3 | 21193366 | rs7622113 | -0.269 | -1.258 | -0.906 | 0.27 | -0.031 | -0.009 | -0.009 | 0.022 | 0.43 | 0.14 | 0.59 | 0.46 | 0.47 | 0.44 |
| 3 | 21205670 | rs1073891 | 1.464 | -1.178 | -1.029 | 0.194 | -0.081 | 0.008 | 0.008 | 0.148 | 0.19 | 0.05 | 0.18 | 0.21 | 0.33 | 0.12 |
| 3 | 21214596 | rs13094843 | -0.595 | -1.003 | -0.901 | -0.035 | 0.005 | -0.008 | -0.008 | 0 | 0.55 | 0.23 | 0.64 | 0.49 | 0.58 | 0.49 |
| 3 | 21218130 | rs9879165 | 0.47 | -0.628 | -0.45 | -0.236 | -0.02 | 0.058 | 0.058 | 0 | 0.12 | 0.04 | 0.05 | 0 | 0.12 | 0.04 |
| 3 | 21241148 | rs4858297 | -0.468 | -0.938 | -0.784 | -0.345 | 0.048 | 0.097 | 0.097 | 0 | 0.54 | 0.72 | 0.63 | 0.81 | 0.39 | 0.69 |
| 3 | 21248642 | rs11720452 | 1.525 | -0.454 | -1.167 | -0.853 | -0.127 | 0.034 | 0.034 | 0.17 | 0.23 | 0.02 | 0.17 | 0.12 | 0.32 | 0.11 |
| 3 | 21251319 | rs17008237 | 1.279 | -1.333 | -0.873 | -0.56 | 0 | 0.14 | 0.14 | 0 | 0.24 | 0.41 | 0.12 | 0.53 | 0.23 | 0.21 |
| 3 | 21257309 | rs1436831 | -0.71 | -1.322 | -1.016 | -0.597 | 0.037 | 0.13 | 0.13 | 0.1 | 0.55 | 0.75 | 0.64 | 0.86 | 0.42 | 0.72 |
| 3 | 21259194 | rs10510512 | 1.118 | -1.692 | -1.104 | -0.689 | 0.009 | 0.008 | 0.008 | 0 | 0.14 | 0.1 | 0.06 | 0 | 0.11 | 0.05 |
| 3 | 21259435 | rs7627715 | 0.074 | -1.743 | -1.163 | -0.609 | 0.033 | 0.04 | 0.04 | 0 | 0.23 | 0.16 | 0.09 | 0.06 | 0.15 | 0.27 |
| 3 | 21260370 | rs10510511 | 1.707 | -1.21 | -1.116 | -0.308 | 0.013 | -0.011 | -0.011 | 0 | 0.05 | 0 | 0.41 | 0.26 | 0.04 | 0.22 |
| 3 | 21265540 | rs9831928 | 0.765 | -1.152 | -1.016 | -0.555 | -0.033 | 0.07 | 0.07 | 0.064 | 0.31 | 0.16 | 0.21 | 0.13 | 0.35 | 0.16 |
| 3 | 21265871 | rs13063191 | 0.053 | -1.27 | -1.084 | -0.531 | -0.021 | 0.085 | 0.085 | 0 | 0.39 | 0.16 | 0.24 | 0.2 | 0.4 | 0.38 |
| 3 | 21270162 | rs6550577 | -1.436 | -1.163 | -1.075 | -0.656 | -0.001 | -0.016 | -0.016 | 0.218 | 0.89 | 0.89 | 0.87 | 1 | 0.82 | 0.92 |
| 3 | 21270688 | rs13061121 | 1.021 | -1.178 | -1.156 | -0.43 | -0.039 | 0.037 | 0.037 | 0.085 | 0.31 | 0.13 | 0.21 | 0.19 | 0.35 | 0.16 |
| 3 | 21274335 | rs1865912 | 0.294 | -1.357 | -0.867 | -0.442 | -0.024 | 0.014 | 0.014 | 0 | 0.25 | 0.4 | 0.53 | 0.27 | 0.24 | 0.35 |
| 3 | 21275538 | rs13326143 | -0.315 | -1.409 | -0.68 | -0.491 | -0.004 | -0.033 | -0.033 | 0.996 | 0.56 | 0.47 | 0.65 | 0.81 | 0.48 | 0.74 |
| 3 | 21278077 | rs6770301 | 1.054 | -1.405 | -0.784 | -0.462 | 0.03 | -0.068 | -0.068 | 0.032 | 0.32 | 0.13 | 0.13 | 0.53 | 0.26 | 0.39 |
| 3 | 21280306 | rs7615883 | 1.391 | -1.365 | -0.637 | -0.489 | -0.003 | -0.043 | -0.043 | 1.669 | 0.18 | 0.19 | 0.08 | 0 | 0.16 | 0.16 |
| 3 | 21280562 | rs1992102 | -0.367 | -1.325 | -0.64 | -0.487 | -0.007 | 0.009 | 0.009 | 0 | 0.81 | 0.83 | 0.84 | 0.94 | 0.77 | 0.71 |
| 3 | 21281019 | rs12638167 | 1.654 | -1.01 | -0.494 | -0.335 | 0.007 | -0.069 | -0.069 | 0 | 0.24 | 0.07 | 0.1 | 0.47 | 0.21 | 0.2 |
| 3 | 21281355 | rs13074521 | 1.042 | -0.768 | 0.122 | -0.238 | 0.001 | 0.033 | 0.033 | 0 | 0.55 | 0.13 | 0.3 | 0.67 | 0.57 | 0.5 |
| 3 | 21282042 | rs17008294 | 0.41 | -0.132 | 0.799 | 0.61 | -0.02 | 0.03 | 0.03 | 1.416 | 0.09 | 0.29 | 0.05 | 0.05 | 0.02 | 0.05 |
| 3 | 21289232 | rs4858303 | -0.505 | 0.516 | 0.081 | 0.684 | 0.001 | -0.018 | -0.018 | 0 | 0.89 | 0.89 | 0.87 | 1 | 0.82 | 0.93 |
| 3 | 21290055 | rs6785004 | -0.642 | 0.522 | 0.073 | 0.691 | -0.007 | 0.159 | 0.159 | 0 | 0.07 | 0.26 | 0.45 | 0.27 | 0.06 | 0.21 |
| 3 | 21290238 | rs1436842 | -0.36 | 0.806 | -0.585 | 0.422 | 0 | -0.016 | -0.016 | 0 | 0.89 | 0.9 | 0.87 | 1 | 0.82 | 0.93 |
| 3 | 21299605 | rs4858305 | -0.189 | 0.126 | -0.922 | 0.771 | -0.036 | -0.026 | -0.026 | 0 | 0.65 | 0.81 | 0.76 | 0.52 | 0.59 | 0.7 |
| 3 | 21308534 | rs9310639 | 0.688 | -0.19 | -1.001 | 0.578 | 0.022 | -0.053 | -0.053 | 0 | 0.3 | 0.16 | 0.12 | 0.49 | 0.22 | 0.22 |
| 3 | 21310668 | rs2118127 | -0.368 | -0.218 | -0.978 | 0.561 | -0.018 | 0.006 | 0.006 | 0 | 0.88 | 0.92 | 0.88 | 1 | 0.81 | 0.89 |
| 3 | 21310889 | rs4045832 | 0.336 | -0.467 | 0.037 | 0.552 | 0.027 | 0.023 | 0.023 | 0 | 0.34 | 0.21 | 0.53 | 0.75 | 0.26 | 0.39 |
| 3 | 21311787 | rs6763061 | -1.256 | -0.441 | 0.006 | 0.563 | 0.002 | 0.202 | 0.202 | 0 | 0.08 | 0.27 | 0.46 | 0.28 | 0.08 | 0.22 |
| 3 | 21313148 | rs6550581 | -0.061 | -0.081 | 0.428 | 0.288 | 0.015 | 0 | 0 | 0 | 0.44 | 0.64 | 0.23 | 0.6 | 0.35 | 0.45 |
| 3 | 21315193 | rs4045949 | -0.245 | -0.069 | 0.407 | 0.304 | -0.003 | 0.091 | 0.091 | 0 | 0.06 | 0.08 | 0.43 | 0.28 | 0.06 | 0.21 |
| 3 | 21317215 | rs7621020 | -0.356 | -0.123 | 0.573 | 0.22 | 0.026 | 0.082 | 0.082 | 0 | 0.48 | 0.68 | 0.64 | 0.86 | 0.39 | 0.62 |
| 3 | 21320118 | rs9859926 | 0.554 | -0.035 | 0.433 | 0.27 | -0.005 | 0.025 | 0.025 | 0 | 0.36 | 0.44 | 0.17 | 0.53 | 0.3 | 0.29 |
| 3 | 21329182 | rs4453865 | 0.778 | 0.022 | 0.185 | 0.174 | 0.021 | 0.047 | 0.047 | 0 | 0.34 | 0.15 | 0.53 | 0.78 | 0.27 | 0.42 |
| 3 | 21329581 | rs6795699 | -0.893 | -0.073 | 0.646 | 0.122 | 0.003 | -0.015 | -0.015 | 0.067 | 0.14 | 0.28 | 0.08 | 0.08 | 0.13 | 0.22 |
| 3 | 21331233 | rs1836950 | 0.732 | -0.056 | 0.642 | 0.008 | -0.002 | -0.009 | -0.009 | 3.125 | 0.42 | 0.37 | 0.28 | 0.12 | 0.46 | 0.26 |
| 3 | 21333846 | rs2335191 | 0.964 | -0.057 | 0.608 | 0.005 | 0.018 | 0.044 | 0.044 | 0 | 0.34 | 0.14 | 0.53 | 0.79 | 0.27 | 0.43 |
| 3 | 21337754 | rs1436821 | 0.811 | -0.016 | 0.739 | 0.029 | 0.015 | -0.06 | -0.06 | 0 | 0.76 | 0.57 | 0.81 | 0.91 | 0.73 | 0.69 |

| | | | | | | | | | | | | | | | | |
|---|---|---|---|---|---|---|---|---|---|---|---|---|---|---|---|---|
| 3 | 21337956 | rs1436822 | -0.714 | 0.072 | 0.703 | -0.003 | 0.003 | -0.021 | -0.021 | 0 | 0.89 | 0.9 | 0.89 | 1 | 0.82 | 0.93 |
| 3 | 21345639 | rs680612 | -0.824 | 0.063 | 0.665 | 0.199 | -0.023 | 0.015 | 0.015 | 0 | 0.9 | 0.89 | 0.89 | 1 | 0.82 | 0.81 |
| 3 | 21351637 | rs13080495 | -0.501 | -0.068 | 0.76 | 0.239 | -0.031 | -0.011 | -0.011 | 0 | 0.12 | 0.33 | 0.07 | 0.06 | 0.06 | 0.19 |
| 3 | 21356406 | rs1396003 | 0 | 0.05 | 0.603 | 0.24 | 0.003 | -0.005 | -0.005 | 0 | 0.47 | 0.53 | 0.62 | 0.86 | 0.36 | 0.63 |
| 3 | 21358252 | rs2661406 | -1.124 | 0.01 | 0.74 | 0.194 | -0.016 | 0.063 | 0.063 | 0 | 0.6 | 0.74 | 0.73 | 0.88 | 0.54 | 0.75 |
| 3 | 21359356 | rs1395998 | -0.855 | -0.002 | 0.78 | 0.265 | -0.019 | 0.035 | 0.035 | 0 | 0.13 | 0.39 | 0.1 | 0.12 | 0.08 | 0.24 |
| 3 | 21361784 | rs13059346 | 1.183 | -0.036 | 0.428 | 0.203 | 0.021 | -0.012 | -0.012 | 0 | 0.06 | 0 | 0.02 | 0.06 | 0.04 | 0.16 |
| 3 | 21365934 | rs10154837 | 1.326 | -0.001 | 0.435 | 0.177 | 0.011 | -0.067 | -0.067 | 0.18 | 0.3 | 0.1 | 0.12 | 0.46 | 0.24 | 0.23 |
| 3 | 21369586 | rs1395993 | -0.288 | 0.217 | 0.58 | 0.078 | 0.045 | 0.119 | 0.119 | 0 | 0.49 | 0.67 | 0.62 | 0.87 | 0.37 | 0.68 |
| 3 | 21373177 | rs2661388 | 0.294 | 0.26 | 0.573 | 0.078 | -0.005 | -0.009 | -0.009 | 2.085 | 0.1 | 0.09 | 0.09 | 0 | 0.14 | 0.07 |
| 3 | 21383828 | rs2731945 | -0.837 | 0.443 | 0.419 | 0.011 | -0.022 | 0.077 | 0.077 | 0 | 0.61 | 0.79 | 0.74 | 0.87 | 0.56 | 0.74 |
| 3 | 21385878 | rs2731946 | -0.949 | 0.422 | 0.778 | 0.048 | 0.082 | -0.092 | -0.092 | 0 | 0.13 | 0.48 | 0.15 | 0 | 0.15 | 0.11 |
| 3 | 21388267 | rs9820497 | 1.188 | 0.275 | 0.688 | 0.184 | 0.009 | -0.055 | -0.055 | 1.741 | 0.33 | 0.15 | 0.14 | 0.48 | 0.27 | 0.3 |
| 3 | 21389545 | rs2731948 | -0.789 | 0.331 | 0.696 | 0.162 | 0.079 | -0.085 | -0.085 | 0 | 0.13 | 0.48 | 0.13 | 0 | 0.15 | 0.05 |
| 3 | 21392307 | rs6770065 | 1.407 | 0.272 | 0.397 | 0.198 | 0.003 | -0.071 | -0.071 | 0 | 0.31 | 0.1 | 0.13 | 0.46 | 0.25 | 0.24 |
| 3 | 21393248 | rs13070110 | -0.133 | 0.436 | 0.412 | 0.289 | -0.014 | -0.006 | -0.006 | 0.105 | 0.1 | 0.06 | 0.15 | 0.09 | 0.13 | 0.18 |
| 3 | 21396096 | rs2731938 | -0.579 | 0.549 | 0.922 | 0.448 | 0.106 | 0.071 | 0.071 | 0 | 0.15 | 0.35 | 0.39 | 0.23 | 0.24 | 0.19 |
| 3 | 21397755 | rs13095327 | -0.013 | 0.555 | 0.93 | 0.411 | 0.046 | -0.061 | -0.061 | 0.226 | 0.12 | 0.2 | 0.14 | 0 | 0.22 | 0.06 |
| 3 | 21406128 | rs776650 | -0.952 | 0.442 | 0.943 | 0.426 | -0.022 | 0.12 | 0.12 | 0 | 0.6 | 0.85 | 0.7 | 0.87 | 0.57 | 0.73 |
| 3 | 21411341 | rs1395991 | -1.915 | 0.416 | 1.12 | 0.502 | 0.071 | 0.308 | 0.308 | 0.275 | 0.08 | 0.45 | 0.3 | 0.3 | 0.05 | 0.16 |
| 3 | 21424685 | rs341832 | -0.344 | 0.266 | 0.946 | 0.242 | 0.02 | -0.064 | -0.064 | 0.505 | 0.16 | 0.21 | 0.17 | 0 | 0.23 | 0.12 |
| 3 | 21427018 | rs9835915 | 0.809 | 0.133 | 0.526 | 0.35 | 0.034 | -0.05 | -0.05 | 0.395 | 0.39 | 0.19 | 0.25 | 0.53 | 0.3 | 0.4 |
| 3 | 21433365 | rs341859 | -1.224 | 0.152 | 0.845 | 0.38 | 0.022 | -0.076 | -0.076 | 0 | 0.23 | 0.46 | 0.3 | 0.09 | 0.27 | 0.22 |
| 3 | 21433422 | rs9844163 | 1.56 | 0.111 | 0.784 | 0.499 | 0.011 | -0.049 | -0.049 | 0 | 0.31 | 0.1 | 0.14 | 0.38 | 0.26 | 0.24 |
| 3 | 21433969 | rs768737 | -1.465 | 0.067 | 0.832 | 0.537 | 0.026 | 0.304 | 0.304 | 0.861 | 0.06 | 0.26 | 0.28 | 0.36 | 0.03 | 0.23 |
| 3 | 21435383 | rs9872314 | 1.457 | 0.092 | 0.234 | 0.583 | 0.011 | -0.069 | -0.069 | 0 | 0.32 | 0.11 | 0.13 | 0.42 | 0.27 | 0.28 |
| 3 | 21435989 | rs9873271 | 0.463 | 0.096 | 0.322 | 0.484 | -0.026 | 0.027 | 0.027 | 0 | 0.08 | 0.24 | 0.14 | 0.09 | 0.05 | 0.16 |
| 3 | 21437406 | rs442931 | -0.901 | 0.216 | -0.091 | 0.476 | -0.073 | 0.139 | 0.139 | 0 | 0.63 | 0.89 | 0.72 | 0.87 | 0.58 | 0.74 |
| 3 | 21443008 | rs411616 | -0.955 | 0.216 | -0.103 | 0.476 | -0.084 | 0.137 | 0.137 | 0 | 0.63 | 0.92 | 0.72 | 0.87 | 0.58 | 0.74 |
| 3 | 21447492 | rs4857993 | -0.86 | 0.234 | 0.052 | 0.609 | -0.018 | 0.126 | 0.126 | 0 | 0.07 | 0.16 | 0.27 | 0.34 | 0.03 | 0.17 |
| 3 | 21449109 | rs341864 | -0.799 | 0.245 | -0.17 | 0.371 | -0.054 | 0.14 | 0.14 | 0 | 0.62 | 0.87 | 0.72 | 0.87 | 0.57 | 0.71 |
| 3 | 21450656 | rs1865716 | 1.382 | 0.243 | -0.28 | 0.653 | 0.006 | -0.039 | -0.039 | 0 | 0.32 | 0.1 | 0.14 | 0.36 | 0.27 | 0.24 |
| 3 | 21456170 | rs376703 | -0.933 | 0.243 | 0.366 | 0.652 | 0.02 | 0.114 | 0.114 | 0 | 0.23 | 0.45 | 0.5 | 0.4 | 0.24 | 0.33 |
| 3 | 21458362 | rs450092 | -0.51 | 0.855 | -0.979 | 0.55 | -0.015 | -0.007 | -0.007 | 0 | 0.78 | 0.95 | 0.8 | 0.77 | 0.76 | 0.64 |
| 3 | 21460201 | rs9700 | 0.541 | 0.712 | -0.62 | 0.41 | 0.007 | 0.02 | 0.02 | 0 | 0.5 | 0.3 | 0.36 | 0.53 | 0.51 | 0.44 |
| 3 | 21467663 | rs341844 | 0.7 | 1.414 | -0.294 | 0.098 | 0.055 | -0.058 | -0.058 | 0 | 0.72 | 0.5 | 0.53 | 0.84 | 0.61 | 0.76 |
| 3 | 21474671 | rs7614523 | 0.091 | 0.12 | -0.327 | 0.374 | -0.051 | 0.025 | 0.025 | 0 | 0.2 | 0.31 | 0.15 | 0.33 | 0.13 | 0.29 |
| 3 | 21475640 | rs6806788 | 0.297 | 0.083 | -0.336 | 0.39 | -0.073 | 0.034 | 0.034 | 0 | 0.18 | 0.3 | 0.14 | 0.33 | 0.09 | 0.24 |
| 3 | 21482318 | rs9879872 | -0.125 | 0.175 | -0.426 | 0.444 | -0.071 | 0.051 | 0.051 | 0 | 0.22 | 0.52 | 0.16 | 0.38 | 0.15 | 0.4 |
| 3 | 21483086 | rs3821733 | -0.524 | 0.427 | -0.498 | 0.248 | -0.03 | -0.001 | -0.001 | 0 | 0.28 | 0.59 | 0.34 | 0.36 | 0.2 | 0.35 |
| 3 | 21484762 | rs411674 | 0.078 | 0.471 | -0.566 | 0.109 | -0.001 | -0.016 | -0.016 | 0.356 | 0.94 | 0.94 | 0.96 | 1 | 0.95 | 0.97 |
| 3 | 21487927 | rs12495770 | 0.267 | 0.744 | -0.969 | 0.216 | -0.067 | -0.009 | -0.009 | 0 | 0.27 | 0.1 | 0.34 | 0.21 | 0.38 | 0.26 |
| 3 | 21490145 | rs424695 | -1.159 | 0.815 | -1.215 | 0.174 | 0.03 | -0.008 | -0.008 | 0.363 | 0.57 | 0.75 | 0.69 | 0.61 | 0.62 | 0.72 |
| 3 | 21491327 | rs451582 | -0.96 | 0.81 | -1.215 | 0.179 | 0.027 | -0.01 | -0.01 | 0 | 0.54 | 0.68 | 0.55 | 0.54 | 0.58 | 0.55 |
| 3 | 21491425 | rs6776560 | 1.507 | 0.894 | -1.112 | 0.19 | 0.008 | -0.02 | -0.02 | 0.228 | 0.06 | 0.08 | 0.05 | 0 | 0.07 | 0.01 |
| 3 | 21491939 | rs424842 | 0.793 | 0.719 | -1.102 | 0.241 | -0.006 | -0.02 | -0.02 | 0 | 0.91 | 0.76 | 0.8 | 0.93 | 0.93 | 0.8 |
| 3 | 21498649 | rs17273488 | 0.904 | 0.828 | -1.445 | 0.29 | 0.011 | -0.017 | -0.017 | 0.164 | 0.42 | 0.2 | 0.28 | 0.4 | 0.39 | 0.26 |
| 3 | 21499213 | rs800621 | -0.663 | 1.263 | -1.538 | 0.159 | 0.012 | 0.026 | 0.026 | 0 | 0.45 | 0.69 | 0.54 | 0.45 | 0.41 | 0.61 |
| 3 | 21500324 | rs399821 | 0.713 | 1.236 | -1.574 | 0.195 | -0.013 | -0.008 | -0.008 | 0 | 0.55 | 0.3 | 0.57 | 0.45 | 0.52 | 0.43 |
| 3 | 21500623 | rs388647 | 0.034 | 2.068 | -1.292 | 0.104 | -0.018 | 0.035 | 0.035 | 0 | 0.19 | 0.11 | 0.2 | 0.05 | 0.2 | 0.14 |
| 3 | 21501593 | rs711735 | 0.408 | 2.274 | -1.221 | 0.729 | -0.041 | -0.018 | -0.018 | 0.75 | 0.81 | 0.89 | 0.78 | 0.84 | 0.73 | 0.85 |
| 3 | 21505299 | rs17008871 | 0.173 | 1.603 | -1.237 | 0.859 | -0.009 | -0.024 | -0.024 | 0 | 0.18 | 0.14 | 0.28 | 0.3 | 0.2 | 0.23 |
| 3 | 21505480 | rs163469 | -0.965 | 1.981 | -1.303 | 0.388 | -0.01 | 0.005 | 0.005 | 0 | 0.41 | 0.33 | 0.45 | 0.35 | 0.53 | 0.41 |
| 3 | 21506161 | rs1026687 | 0.668 | 1.705 | -1.352 | 0.29 | 0.002 | 0.022 | 0.022 | 0 | 0.59 | 0.71 | 0.52 | 0.67 | 0.42 | 0.49 |
| 3 | 21506347 | rs1026690 | 0.644 | 1.737 | -1.47 | 0.291 | 0.001 | 0.024 | 0.024 | 0 | 0.59 | 0.75 | 0.53 | 0.67 | 0.42 | 0.49 |
| 3 | 21507615 | rs163470 | -0.312 | 1.888 | -1.542 | 0.212 | -0.05 | 0.055 | 0.055 | 0 | 0.8 | 0.9 | 0.85 | 0.99 | 0.72 | 0.8 |
| 3 | 21508309 | rs17008884 | 0.938 | 1.765 | -1.549 | 0.234 | -0.008 | -0.02 | -0.02 | 0 | 0.17 | 0.13 | 0.27 | 0.28 | 0.21 | 0.22 |
| 3 | 21508433 | rs11914832 | 2.198 | 0.94 | -1.329 | -0.074 | 0.011 | 0 | 0 | 0.763 | 0.32 | 0.28 | 0.23 | 0.11 | 0.19 | 0.16 |
| 3 | 21509189 | rs7618153 | 0.567 | 1.185 | -1.553 | 0.063 | 0.008 | 0.027 | 0.027 | 0 | 0.59 | 0.73 | 0.53 | 0.67 | 0.41 | 0.48 |
| 3 | 21510700 | rs1863090 | 0.115 | 0.911 | -1.47 | -0.145 | -0.1 | 0.098 | 0.098 | 0 | 0.76 | 0.92 | 0.81 | 0.94 | 0.62 | 0.7 |
| 3 | 21512235 | rs1457587 | -0.297 | 0.881 | -1.191 | -0.161 | 0.012 | -0.065 | -0.065 | 0.546 | 0.37 | 0.5 | 0.56 | 0.83 | 0.39 | 0.49 |
| 3 | 21515705 | rs3860575 | 1.445 | 0.43 | -1.121 | -0.242 | -0.017 | -0.039 | -0.039 | 0.149 | 0.38 | 0.37 | 0.25 | 0.11 | 0.22 | 0.2 |
| 3 | 21517575 | rs2060530 | 0.739 | 0.871 | -0.56 | 0.818 | -0.04 | 0.194 | 0.194 | 0.598 | 0.1 | 0.26 | 0.16 | 0.37 | 0.03 | 0.15 |
| 3 | 21519145 | rs6800260 | -0.424 | 0.917 | -0.813 | 0.804 | -0.065 | -0.047 | -0.047 | 0.058 | 0.31 | 0.19 | 0.44 | 0.52 | 0.46 | 0.43 |
| 3 | 21522197 | rs9811744 | 0.101 | 0.166 | -0.237 | 1.424 | -0.044 | 0.239 | 0.239 | 0.606 | 0.1 | 0.33 | 0.17 | 0.37 | 0.03 | 0.15 |
| 3 | 21523342 | rs2194607 | -0.624 | 0.582 | -0.713 | 0.752 | -0.011 | -0.008 | -0.008 | 0 | 0.39 | 0.55 | 0.49 | 0.7 | 0.33 | 0.48 |
| 3 | 21523966 | rs167936 | -1.16 | 1.041 | -0.771 | 1.121 | -0.003 | -0.007 | -0.007 | 0.152 | 0.19 | 0.18 | 0.19 | 0.31 | 0.22 | 0.21 |
| 3 | 21525138 | rs9883288 | -1.021 | 0.691 | -0.689 | 0.817 | -0.005 | -0.014 | -0.014 | 0 | 0.09 | 0.14 | 0.08 | 0.03 | 0.07 | 0.08 |
| 3 | 21526766 | rs162410 | -0.725 | 1.517 | -0.346 | 1.049 | 0.025 | 0.025 | 0.025 | 0 | 0.09 | 0.14 | 0.28 | 0.23 | 0.18 | 0.08 |

| | | | | | | | | | | | | | | | | |
|---|---|---|---|---|---|---|---|---|---|---|---|---|---|---|---|---|
| 3 | 21528920 | rs161199 | -1.216 | 1.064 | -1.177 | -0.017 | 0.004 | 0.099 | 0.099 | 0.086 | 0.78 | 0.96 | 0.87 | 1 | 0.78 | 0.93 |
| 3 | 21529297 | rs7617195 | 0.926 | 0.849 | -0.38 | 0.041 | -0.003 | 0.044 | 0.044 | 0 | 0.5 | 0.37 | 0.47 | 0.63 | 0.5 | 0.61 |
| 3 | 21535708 | rs163480 | -0.529 | 0.76 | -0.348 | 0.141 | 0.01 | 0.019 | 0.019 | 0 | 0.7 | 0.79 | 0.84 | 0.87 | 0.75 | 0.77 |
| 3 | 21544296 | rs1012728 | 1.324 | 0.914 | -0.962 | 0.034 | -0.002 | -0.14 | -0.14 | 0 | 0.25 | 0.1 | 0.29 | 0.51 | 0.21 | 0.45 |
| 3 | 21546663 | rs233165 | -1.176 | 1.005 | -0.859 | 0.153 | 0.016 | 0.04 | 0.04 | 0 | 0.71 | 0.91 | 0.85 | 0.86 | 0.75 | 0.77 |
| 3 | 21547349 | rs920118 | 0.371 | 1.062 | -0.414 | 0.177 | -0.001 | 0.004 | 0.004 | 0 | 0.52 | 0.56 | 0.48 | 0.64 | 0.49 | 0.63 |
| 3 | 21547703 | rs233162 | -0.419 | 1.073 | -0.533 | 0.158 | 0.018 | 0.049 | 0.049 | 0 | 0.62 | 0.78 | 0.77 | 0.84 | 0.67 | 0.7 |
| 3 | 21550556 | rs9310648 | 0.07 | 0.752 | -0.5 | 0.248 | -0.004 | -0.007 | -0.007 | 0 | 0.56 | 0.57 | 0.56 | 0.69 | 0.58 | 0.69 |
| 3 | 21558771 | rs13323807 | 0.717 | 0.707 | -0.45 | 0.297 | 0.021 | -0.022 | -0.022 | 0.29 | 0.29 | 0.41 | 0.23 | 0.17 | 0.36 | 0.22 |
| 3 | 21562716 | rs7646837 | -0.292 | 0.616 | -0.448 | 0.306 | 0.01 | 0.015 | 0.015 | 0 | 0.58 | 0.75 | 0.56 | 0.68 | 0.58 | 0.69 |
| 3 | 21564349 | rs3860579 | 0.744 | 0.799 | -0.464 | 0.168 | 0.021 | -0.022 | -0.022 | 0 | 0.29 | 0.4 | 0.23 | 0.17 | 0.34 | 0.23 |
| 3 | 21567690 | rs233158 | -0.852 | 0.685 | -0.505 | 0.109 | -0.011 | -0.026 | -0.026 | 1.533 | 0.7 | 0.67 | 0.84 | 0.89 | 0.77 | 0.78 |
| 3 | 21569097 | rs233155 | -1.134 | 0.693 | -0.555 | 0.126 | 0.012 | 0.028 | 0.028 | 0 | 0.72 | 0.83 | 0.86 | 0.89 | 0.77 | 0.79 |
| 3 | 21571130 | rs233146 | 0.177 | 0.721 | -0.539 | 0.237 | 0.014 | 0.026 | 0.026 | 0 | 0.11 | 0.18 | 0.29 | 0.18 | 0.16 | 0.1 |
| 3 | 21572733 | rs4858328 | -0.049 | 0.738 | -0.7 | 0.238 | -0.003 | -0.005 | -0.005 | 0 | 0.59 | 0.48 | 0.55 | 0.71 | 0.6 | 0.68 |
| 3 | 21573453 | rs192058 | -0.449 | 0.646 | -0.825 | 0.705 | 0.022 | 0.04 | 0.04 | 0 | 0.13 | 0.29 | 0.3 | 0.18 | 0.16 | 0.1 |
| 3 | 21582201 | rs17008992 | 0.439 | 0.863 | -1.021 | 0.691 | -0.046 | -0.176 | -0.176 | 0 | 0.28 | 0.03 | 0.3 | 0.52 | 0.24 | 0.44 |
| 3 | 21582347 | rs162412 | -1.233 | 1.117 | -0.109 | 1.013 | 0.017 | 0.044 | 0.044 | 0 | 0.18 | 0.28 | 0.31 | 0.27 | 0.21 | 0.2 |
| 3 | 21590882 | rs13092094 | -0.602 | 0.939 | -0.329 | 1.146 | 0.013 | 0.024 | 0.024 | 0 | 0.11 | 0.17 | 0.13 | 0.18 | 0.15 | 0.13 |
| 3 | 21591377 | rs11708571 | 1.075 | 1.286 | -0.886 | 0.424 | -0.057 | 0.078 | 0.078 | 0 | 0.21 | 0.05 | 0.14 | 0 | 0.29 | 0.11 |
| 3 | 21602714 | rs11919873 | -0.174 | 1.108 | -1.205 | 0.873 | -0.062 | 0.044 | 0.044 | 0.067 | 0.66 | 0.87 | 0.65 | 0.74 | 0.61 | 0.69 |
| 3 | 21603613 | rs10460951 | -0.215 | 1.111 | -1.256 | 0.875 | -0.095 | 0.042 | 0.042 | 0.154 | 0.66 | 0.93 | 0.66 | 0.74 | 0.61 | 0.69 |
| 3 | 21613585 | rs1457586 | -0.399 | 1.081 | -1.243 | 0.943 | -0.03 | -0.061 | -0.061 | 0.164 | 0.87 | 0.98 | 0.8 | 0.74 | 0.9 | 0.79 |
| 3 | 21617726 | rs17009067 | 1.039 | 0.598 | -0.589 | 1.181 | 0.009 | 0.013 | 0.013 | 0 | 0.44 | 0.4 | 0.39 | 0.32 | 0.4 | 0.3 |
| 3 | 21622803 | rs12637139 | 1.925 | 0.138 | -1.41 | 1.139 | -0.001 | 0.018 | 0.018 | 0 | 0.28 | 0.03 | 0.16 | 0.2 | 0.24 | 0.19 |
| 3 | 21623631 | rs1841914 | 0.758 | 0.649 | -0.573 | 1.428 | 0.014 | 0.015 | 0.015 | 0 | 0.5 | 0.61 | 0.5 | 0.46 | 0.48 | 0.48 |
| 3 | 21625101 | rs4102411 | 0.801 | 0.836 | -0.659 | 1.694 | -0.019 | -0.046 | -0.046 | 0.132 | 0.77 | 0.87 | 0.66 | 0.6 | 0.76 | 0.67 |
| 3 | 21628130 | rs9822059 | -0.834 | 0.897 | -0.456 | 1.658 | 0.018 | 0.094 | 0.094 | 0 | 0.14 | 0.22 | 0.16 | 0.3 | 0.16 | 0.24 |
| 3 | 21628232 | rs9841949 | -1.055 | 0.854 | -0.4 | 1.639 | 0.016 | 0.102 | 0.102 | 0 | 0.16 | 0.29 | 0.17 | 0.3 | 0.16 | 0.24 |
| 3 | 21628851 | rs11708720 | -0.073 | 0.939 | -0.663 | 1.602 | -0.011 | -0.015 | -0.015 | 0 | 0.1 | 0.06 | 0.11 | 0.18 | 0.14 | 0.1 |
| 3 | 21629318 | rs1457588 | -0.602 | 0.787 | -0.603 | 1.427 | -0.02 | -0.051 | -0.051 | 0 | 0.89 | 0.98 | 0.8 | 0.77 | 0.92 | 0.77 |
| 3 | 21630297 | rs9879945 | 0.345 | 0.676 | -0.462 | 1.277 | -0.015 | 0.021 | 0.021 | 0 | 0.57 | 0.36 | 0.52 | 0.6 | 0.57 | 0.51 |
| 3 | 21631330 | rs1457589 | 0.581 | 0.684 | -0.376 | 1.238 | 0.016 | -0.05 | -0.05 | 0 | 0.31 | 0.53 | 0.26 | 0.17 | 0.35 | 0.26 |
| 3 | 21631666 | rs1382165 | 0.306 | 0.68 | -0.374 | 1.263 | -0.006 | 0.006 | 0.006 | 0 | 0.57 | 0.43 | 0.53 | 0.6 | 0.57 | 0.51 |
| 3 | 21634204 | rs17009100 | -0.887 | 0.711 | -0.404 | 1.336 | 0.005 | 0.127 | 0.127 | 0.599 | 0.15 | 0.31 | 0.14 | 0.31 | 0.13 | 0.25 |
| 3 | 21634367 | rs6776795 | -0.195 | 0.682 | -0.488 | 1.38 | 0 | 0 | 0 | 0.099 | 0.4 | 0.53 | 0.38 | 0.45 | 0.47 | 0.4 |
| 3 | 21635561 | rs10510514 | 1.046 | 0.713 | -0.498 | 1.4 | 0.022 | 0.09 | 0.09 | 0.016 | 0.45 | 0.19 | 0.41 | 0.31 | 0.44 | 0.29 |
| 3 | 21636543 | rs1382167 | 0.322 | 0.864 | -0.205 | 1.739 | -0.007 | 0.019 | 0.019 | 0.018 | 0.83 | 0.72 | 0.75 | 0.74 | 0.86 | 0.68 |
| 3 | 21637396 | rs6776046 | -0.768 | 0.964 | -0.169 | 1.598 | 0.01 | 0.055 | 0.055 | 0 | 0.11 | 0.13 | 0.13 | 0.28 | 0.14 | 0.16 |
| 3 | 21645400 | rs4858339 | -0.22 | 0.706 | -0.983 | 0.409 | -0.017 | 0.05 | 0.05 | 0 | 0.7 | 0.85 | 0.83 | 0.88 | 0.65 | 0.77 |
| 3 | 21645676 | rs6773155 | 0.082 | 0.57 | -0.915 | 0.7 | -0.013 | 0.029 | 0.029 | 0 | 0.61 | 0.74 | 0.69 | 0.71 | 0.54 | 0.71 |
| 3 | 21647537 | rs11918612 | 0.435 | -0.088 | -1.145 | 0.902 | -0.042 | -0.06 | -0.06 | 0 | 0.37 | 0.15 | 0.52 | 0.45 | 0.41 | 0.52 |
| 3 | 21647818 | rs6780177 | 1.064 | -0.601 | -0.333 | 0.818 | 0.013 | 0.035 | 0.035 | 0.705 | 0.15 | 0.36 | 0.14 | 0.18 | 0.12 | 0.07 |
| 3 | 21650537 | rs721623 | -0.181 | -0.374 | -0.314 | 0.737 | -0.003 | 0.025 | 0.025 | 0 | 0.51 | 0.54 | 0.65 | 0.67 | 0.47 | 0.59 |
| 3 | 21650648 | rs1824212 | -0.085 | -0.395 | -0.222 | 0.751 | -0.002 | 0.048 | 0.048 | 0 | 0.5 | 0.6 | 0.65 | 0.66 | 0.47 | 0.59 |
| 3 | 21651240 | rs6550614 | -0.287 | -0.33 | -0.281 | 0.743 | -0.01 | 0.066 | 0.066 | 0 | 0.52 | 0.67 | 0.66 | 0.67 | 0.48 | 0.6 |
| 3 | 21654918 | rs7629800 | 0.046 | -0.383 | -0.291 | 0.864 | 0.013 | 0.002 | 0.002 | 0.238 | 0.46 | 0.37 | 0.52 | 0.53 | 0.38 | 0.65 |
| 3 | 21655297 | rs11129012 | 0.935 | -0.318 | -0.508 | 1.251 | -0.037 | -0.01 | -0.01 | 0.192 | 0.23 | 0.06 | 0.28 | 0.22 | 0.25 | 0.29 |
| 3 | 21658582 | rs17029417 | 1.209 | -0.31 | -0.366 | 1.221 | 0.013 | 0.069 | 0.069 | 0.219 | 0.16 | 0.4 | 0.17 | 0.21 | 0.14 | 0.07 |
| 3 | 21661148 | rs17009163 | -0.172 | -0.137 | -1.16 | 1.068 | -0.007 | -0.065 | -0.065 | 3.088 | 0.35 | 0.15 | 0.48 | 0.43 | 0.32 | 0.49 |
| 3 | 21662267 | rs3849543 | 0.801 | 0.007 | -1.233 | 0.87 | -0.044 | 0.016 | 0.016 | 0 | 0.3 | 0.08 | 0.33 | 0.22 | 0.33 | 0.33 |
| 3 | 21663649 | rs1902754 | 0.276 | -0.062 | -0.631 | 1.306 | 0.002 | 0.011 | 0.011 | 3.03 | 0.11 | 0.11 | 0.2 | 0.3 | 0.1 | 0.22 |
| 3 | 21666926 | rs10780010 | -0.317 | -0.208 | -0.344 | 1.185 | -0.018 | 0.056 | 0.056 | 0 | 0.27 | 0.43 | 0.37 | 0.55 | 0.23 | 0.33 |
| 3 | 21668990 | rs12054239 | 0.391 | 0.103 | -0.86 | 1.28 | -0.005 | -0.053 | -0.053 | 0 | 0.1 | 0.01 | 0.21 | 0.31 | 0.09 | 0.22 |
| 3 | 21670384 | rs3911498 | 0.452 | 0.096 | -1.02 | 1.014 | -0.003 | -0.02 | -0.02 | 0 | 0.15 | 0.04 | 0.19 | 0.17 | 0.13 | 0.17 |
| 3 | 21670520 | rs3860583 | 0.299 | 0.292 | -0.616 | 0.531 | -0.003 | 0.014 | 0.014 | 0 | 0.48 | 0.52 | 0.48 | 0.29 | 0.52 | 0.42 |
| 3 | 21673600 | rs7619318 | -1.058 | 0.105 | -0.346 | 0.675 | 0.002 | -0.006 | -0.006 | 0.012 | 0.71 | 0.74 | 0.81 | 0.56 | 0.75 | 0.69 |
| 3 | 21684130 | rs1993507 | 1.116 | 0.451 | -0.819 | 0.52 | -0.083 | -0.066 | -0.066 | 0 | 0.27 | 0.01 | 0.38 | 0.25 | 0.33 | 0.4 |
| 3 | 21686466 | rs1387822 | 0.008 | -0.034 | 0.142 | 1.007 | -0.003 | 0 | 0 | 0 | 0.58 | 0.38 | 0.66 | 0.39 | 0.69 | 0.54 |
| 3 | 21689964 | rs1352646 | 0.291 | -0.217 | 0.119 | 0.966 | 0 | 0 | 0 | 0.471 | 0.55 | 0.37 | 0.66 | 0.37 | 0.67 | 0.51 |
| 3 | 21692359 | rs9864367 | -1.027 | -0.186 | 0.313 | 0.968 | -0.002 | 0.012 | 0.012 | 0 | 0.22 | 0.28 | 0.28 | 0.29 | 0.2 | 0.24 |
| 3 | 21698394 | rs2054942 | 0.833 | -0.127 | 0.496 | 1.078 | 0.008 | 0.063 | 0.063 | 0.125 | 0.48 | 0.24 | 0.49 | 0.23 | 0.56 | 0.45 |
| 3 | 21701712 | rs1320292 | -0.967 | -0.03 | 0.404 | 1.182 | 0.006 | -0.01 | -0.01 | 0 | 0.71 | 0.68 | 0.81 | 0.52 | 0.78 | 0.69 |
| 3 | 21706369 | rs3821396 | 0.765 | 0.231 | 0.235 | 1.176 | -0.029 | -0.028 | -0.028 | 0 | 0.05 | 0.01 | 0.15 | 0.15 | 0.12 | 0.07 |
| 3 | 21712555 | rs9839129 | -0.83 | -0.027 | 0.499 | 1.151 | 0.002 | -0.001 | -0.001 | 0 | 0.68 | 0.64 | 0.79 | 0.6 | 0.76 | 0.68 |
| 3 | 21714103 | rs986503 | -0.938 | -0.037 | 0.382 | 1.158 | 0.012 | -0.006 | -0.006 | 0 | 0.71 | 0.72 | 0.81 | 0.61 | 0.79 | 0.69 |
| 3 | 21714495 | rs9850039 | -0.612 | 0.355 | 0.155 | 1.37 | 0.008 | 0.038 | 0.038 | 0 | 0.13 | 0.18 | 0.2 | 0.31 | 0.19 | 0.14 |
| 3 | 21716235 | rs9860048 | -0.906 | 0.323 | 0.227 | 1.36 | 0.02 | 0.058 | 0.058 | 0 | 0.13 | 0.26 | 0.26 | 0.27 | 0.17 | 0.18 |
| 3 | 21718112 | rs9870622 | 1.145 | 0.324 | 0.042 | 1.309 | 0 | 0.009 | 0.009 | 0 | 0.06 | 0.07 | 0.04 | 0.11 | 0.07 | 0.07 |

| | | | | | | | | | | | | | | | | |
|---|---|---|---|---|---|---|---|---|---|---|---|---|---|---|---|---|
| 3 | 21720374 | rs10510515 | 1.424 | 0.339 | -0.082 | 1.315 | -0.016 | -0.033 | -0.033 | 0.083 | 0.06 | 0 | 0.03 | 0.11 | 0.07 | 0.07 |
| 3 | 21732114 | rs11710743 | -0.587 | 0.275 | 0.156 | 1.219 | -0.008 | -0.001 | -0.001 | 0 | 0.65 | 0.6 | 0.65 | 0.59 | 0.75 | 0.67 |
| 3 | 21734567 | rs9862240 | 0.142 | 0.512 | 0.108 | 1.347 | 0.042 | 0.072 | 0.072 | 0 | 0.07 | 0.18 | 0.19 | 0.2 | 0.14 | 0.07 |
| 3 | 21738751 | rs6790087 | 0.816 | 0.221 | -0.1 | 0.948 | -0.026 | 0.091 | 0.091 | 0.066 | 0.46 | 0.16 | 0.32 | 0.22 | 0.51 | 0.4 |
| 3 | 21742025 | rs12632528 | -0.08 | 0.174 | 0.019 | 1.041 | -0.005 | 0.025 | 0.025 | 0 | 0.12 | 0.21 | 0.14 | 0.18 | 0.1 | 0.17 |
| 3 | 21745040 | rs2336038 | 0.478 | 0.265 | -0.261 | 0.707 | -0.008 | 0.051 | 0.051 | 0.013 | 0.47 | 0.27 | 0.34 | 0.22 | 0.51 | 0.42 |
| 3 | 21745418 | rs4858344 | 0.523 | 0.3 | -0.513 | 0.718 | -0.017 | 0.077 | 0.077 | 0.066 | 0.46 | 0.2 | 0.34 | 0.23 | 0.51 | 0.43 |
| 3 | 21751433 | rs9858811 | 0.532 | -0.714 | -0.352 | 1.235 | -0.03 | 0.036 | 0.036 | 0 | 0.79 | 0.62 | 0.82 | 0.65 | 0.9 | 0.7 |
| 3 | 21751690 | rs1080020 | -0.126 | -0.851 | -0.39 | 1.013 | -0.014 | -0.028 | -0.028 | 0 | 0.38 | 0.62 | 0.28 | 0.2 | 0.22 | 0.39 |
| 3 | 21752050 | rs957589 | -0.473 | 0.037 | -0.685 | 0.969 | 0 | 0.03 | 0.03 | 0.255 | 0.47 | 0.67 | 0.39 | 0.37 | 0.4 | 0.63 |
| 3 | 21753272 | rs4858007 | 0.812 | 0.352 | -0.87 | 1.142 | -0.038 | -0.059 | -0.059 | 0 | 0.12 | 0.04 | 0.35 | 0.26 | 0.19 | 0.05 |
| 3 | 21755681 | rs4858348 | -1.288 | 0.205 | -0.823 | 1.113 | 0.002 | 0.004 | 0.004 | 0 | 0.64 | 0.76 | 0.77 | 0.73 | 0.64 | 0.73 |
| 3 | 21756281 | rs11129016 | -0.509 | 0.21 | -0.836 | 1.121 | -0.005 | 0.009 | 0.009 | 0.945 | 0.52 | 0.71 | 0.42 | 0.51 | 0.45 | 0.68 |
| 3 | 21756553 | rs12106696 | -0.515 | 0.189 | -0.757 | 0.932 | 0.002 | -0.003 | -0.003 | 0.447 | 0.53 | 0.67 | 0.66 | 0.56 | 0.47 | 0.48 |
| 3 | 21764453 | rs968140 | -1.157 | 0.035 | -0.774 | 0.812 | 0.001 | 0.008 | 0.008 | 0 | 0.64 | 0.76 | 0.77 | 0.78 | 0.63 | 0.73 |
| 3 | 21771422 | rs9882615 | 1.398 | -0.032 | -0.78 | 0.563 | 0.016 | 0.032 | 0.032 | 0.458 | 0.25 | 0.15 | 0.15 | 0.11 | 0.19 | 0.21 |
| 3 | 21771710 | rs4440141 | 1.231 | -0.115 | -0.851 | 0.278 | 0.007 | 0.022 | 0.022 | 1.105 | 0.14 | 0.07 | 0.14 | 0.08 | 0.11 | 0.15 |
| 3 | 21771769 | rs17009383 | 0.316 | 0.667 | -0.317 | 0.841 | -0.002 | -0.003 | -0.003 | 0.42 | 0.12 | 0.11 | 0.11 | 0.2 | 0.18 | 0.24 |
| 3 | 21772057 | rs12489188 | 0.011 | 0.56 | -0.023 | 0.946 | 0.002 | 0.002 | 0.002 | 0 | 0.63 | 0.31 | 0.72 | 0.69 | 0.66 | 0.66 |
| 3 | 21776124 | rs2878599 | -0.153 | 0.845 | 0.094 | 0.905 | -0.002 | -0.03 | -0.03 | 0 | 0.78 | 0.68 | 0.9 | 0.86 | 0.81 | 0.66 |
| 3 | 21777499 | rs6810155 | 1.195 | 0.821 | -0.452 | 0.838 | -0.015 | -0.027 | -0.027 | 0.131 | 0.05 | 0.11 | 0.02 | 0 | 0.01 | 0 |
| 3 | 21778143 | rs7625853 | 0.65 | 1.067 | -0.628 | 1.139 | 0.001 | -0.001 | -0.001 | 0 | 0.64 | 0.41 | 0.52 | 0.61 | 0.62 | 0.59 |
| 3 | 21779621 | rs13062599 | 0.882 | 1.189 | -1.246 | 1.111 | -0.005 | -0.02 | -0.02 | 0.188 | 0.11 | 0.16 | 0.07 | 0.06 | 0.08 | 0.17 |
| 3 | 21781295 | rs4858353 | 0.022 | 1.383 | -2.06 | 1.542 | -0.049 | -0.062 | -0.062 | 0 | 0.29 | 0.09 | 0.49 | 0.38 | 0.36 | 0.22 |
| 3 | 21783008 | rs7428718 | 0.605 | 1.44 | -1.43 | 1.518 | 0.024 | 0.035 | 0.035 | 6.204 | 0.3 | 0.15 | 0.25 | 0.19 | 0.23 | 0.17 |
| 3 | 21784011 | rs4858012 | -0.657 | 2.21 | -0.779 | 2.171 | -0.038 | -0.047 | -0.047 | 0 | 0.09 | 0.04 | 0.34 | 0.24 | 0.17 | 0.05 |
| 3 | 21784815 | rs7626282 | -0.6 | 1.944 | -0.159 | 2.216 | 0.103 | 0.227 | 0.227 | 0.005 | 0.1 | 0.39 | 0.12 | 0.31 | 0.13 | 0.21 |
| 3 | 21785822 | rs12630043 | -0.75 | 1.628 | -0.959 | 2.115 | 0.001 | 0.008 | 0.008 | 0 | 0.07 | 0.05 | 0.26 | 0.26 | 0.06 | 0.05 |
| 3 | 21786893 | rs6798573 | -0.101 | 1.452 | -1.061 | 0.896 | 0.004 | 0.014 | 0.014 | 0 | 0.58 | 0.64 | 0.77 | 0.76 | 0.66 | 0.58 |
| 3 | 21788882 | rs17009449 | -0.471 | 1.775 | -1.109 | 1.258 | -0.028 | -0.026 | -0.026 | 0 | 0.06 | 0 | 0.04 | 0.07 | 0.07 | 0.13 |
| 3 | 21789545 | rs1879166 | -0.881 | 2.018 | -1.104 | 2.068 | -0.017 | 0.057 | 0.057 | 0 | 0.06 | 0.05 | 0.18 | 0.38 | 0.12 | 0.19 |
| 3 | 21792882 | rs7634827 | 0.251 | 2.148 | -0.903 | 0.422 | -0.031 | 0.069 | 0.069 | 0 | 0.73 | 0.86 | 0.74 | 0.95 | 0.66 | 0.77 |
| 3 | 21794174 | rs13340131 | 0.862 | 1.432 | -0.775 | 0.624 | 0.001 | 0.036 | 0.036 | 2 | 0.08 | 0.06 | 0.14 | 0.32 | 0.06 | 0.08 |
| 3 | 21794874 | rs4261893 | 0.383 | 1.29 | -0.453 | -0.214 | 0.003 | 0.023 | 0.023 | 1.073 | 0.67 | 0.61 | 0.61 | 0.48 | 0.64 | 0.42 |
| 3 | 21795780 | rs9879065 | 0.423 | 1.261 | -0.421 | -0.226 | 0.002 | 0.019 | 0.019 | 0.966 | 0.67 | 0.61 | 0.61 | 0.48 | 0.64 | 0.42 |
| 3 | 21796865 | rs6805992 | -0.529 | 1.408 | -0.388 | -0.246 | -0.03 | -0.111 | -0.111 | 0 | 0.79 | 0.68 | 0.86 | 0.98 | 0.83 | 0.83 |
| 3 | 21797663 | rs7652766 | 0.55 | 1.367 | -0.161 | -0.258 | 0.037 | 0.016 | 0.016 | 0 | 0.06 | 0.2 | 0.11 | 0.09 | 0.07 | 0.19 |
| 3 | 21799428 | rs4334661 | 0.199 | 1.357 | -0.143 | -0.256 | -0.015 | -0.1 | -0.1 | 0.361 | 0.73 | 0.47 | 0.75 | 0.89 | 0.76 | 0.65 |
| 3 | 21799575 | rs9825051 | 0.199 | 1.357 | -0.142 | -0.256 | -0.014 | -0.096 | -0.096 | 0.004 | 0.73 | 0.46 | 0.75 | 0.89 | 0.76 | 0.65 |
| 3 | 21799771 | rs4588386 | 1.589 | 1.686 | -0.289 | -0.585 | 0.031 | 0.109 | 0.109 | 0 | 0.41 | 0.21 | 0.22 | 0.14 | 0.34 | 0.23 |
| 3 | 21802296 | rs1320288 | 1.13 | 1.848 | -0.311 | -0.558 | 0.031 | 0.014 | 0.014 | 1.758 | 0.5 | 0.35 | 0.36 | 0.45 | 0.4 | 0.32 |
| 3 | 21804309 | rs11708627 | 0.848 | 1.826 | -0.386 | -0.507 | -0.053 | 0.059 | 0.059 | 0 | 0.16 | 0.06 | 0.22 | 0.03 | 0.24 | 0.1 |
| 3 | 21805799 | rs13081154 | -0.969 | 1.625 | -0.174 | -0.626 | 0 | -0.011 | -0.011 | 0 | 0.94 | 0.97 | 0.97 | 1 | 0.94 | 0.99 |
| 3 | 21806432 | rs7619783 | 0.139 | 1.656 | -0.159 | -0.59 | 0.003 | 0.006 | 0.006 | 1.663 | 0.15 | 0.19 | 0.31 | 0.73 | 0.18 | 0.31 |
| 3 | 21807679 | rs12633231 | -0.104 | 1.634 | -0.264 | -0.544 | -0.06 | -0.086 | -0.086 | 0 | 0.23 | 0.12 | 0.38 | 0.43 | 0.35 | 0.33 |
| 3 | 21808848 | rs9864800 | 1.107 | 1.506 | -0.15 | -0.664 | 0.024 | 0.007 | 0.007 | 2.725 | 0.51 | 0.35 | 0.36 | 0.46 | 0.41 | 0.34 |
| 3 | 21809062 | rs4858017 | -0.905 | 1.472 | -0.08 | -0.77 | -0.001 | -0.014 | -0.014 | 0 | 0.46 | 0.55 | 0.63 | 0.76 | 0.52 | 0.47 |
| 3 | 21809702 | rs12715020 | -1.011 | 1.269 | -0.109 | -0.902 | -0.003 | 0.069 | 0.069 | 0 | 0.48 | 0.67 | 0.63 | 0.77 | 0.52 | 0.47 |
| 3 | 21809863 | rs4353825 | 0.665 | 1.448 | -0.236 | -0.888 | -0.058 | 0.07 | 0.07 | 0 | 0.17 | 0.06 | 0.22 | 0.02 | 0.25 | 0.12 |
| 3 | 21814858 | rs7628655 | -0.507 | 1.551 | -0.033 | -0.863 | 0.013 | 0.019 | 0.019 | 0 | 0.3 | 0.42 | 0.48 | 0.44 | 0.34 | 0.38 |
| 3 | 21816016 | rs6797528 | -0.274 | 1.547 | -0.038 | -0.887 | 0.035 | -0.055 | -0.055 | 0.056 | 0.23 | 0.38 | 0.34 | 0.11 | 0.29 | 0.27 |
| 3 | 21816589 | rs6798138 | 1.392 | 1.424 | 0.083 | -0.83 | -0.002 | 0.04 | 0.04 | 2.884 | 0.09 | 0.11 | 0.14 | 0.32 | 0.05 | 0.08 |
| 3 | 21817744 | rs2336046 | 0.34 | 1.424 | 0.093 | -0.884 | -0.01 | 0.113 | 0.113 | 0.141 | 0.2 | 0.31 | 0.24 | 0.41 | 0.2 | 0.27 |
| 3 | 21819469 | rs4858356 | -1.025 | 1.449 | -0.129 | -0.985 | 0 | -0.012 | -0.012 | 0 | 0.95 | 0.97 | 0.97 | 0.98 | 0.93 | 0.87 |
| 3 | 21819857 | rs4858357 | 1.276 | 1.497 | -0.199 | -0.928 | -0.011 | 0.14 | 0.14 | 0 | 0.49 | 0.23 | 0.33 | 0.25 | 0.45 | 0.4 |
| 3 | 21820331 | rs17621507 | 1.098 | 1.641 | -0.237 | -0.922 | -0.038 | 0.047 | 0.047 | 0 | 0.13 | 0.05 | 0.21 | 0.02 | 0.19 | 0.09 |
| 3 | 21821931 | rs17029431 | 0.991 | 1.592 | -0.054 | -0.763 | -0.007 | -0.017 | -0.017 | 0 | 0.63 | 0.31 | 0.59 | 0.51 | 0.6 | 0.4 |
| 3 | 21821970 | rs9877894 | -1.201 | 1.653 | 0.09 | -0.873 | 0.078 | -0.014 | -0.014 | 0 | 0.06 | 0.32 | 0.05 | 0 | 0.08 | 0.02 |
| 3 | 21823542 | rs4525889 | -0.891 | 1.16 | 0.225 | -1.084 | -0.011 | -0.064 | -0.064 | 0 | 0.38 | 0.51 | 0.48 | 0.8 | 0.34 | 0.59 |
| 3 | 21825247 | rs7645913 | -0.068 | 1.128 | 0.255 | -0.92 | -0.016 | 0.066 | 0.066 | 0.206 | 0.23 | 0.36 | 0.27 | 0.41 | 0.23 | 0.3 |
| 3 | 21825971 | rs12497298 | 1.87 | 1.008 | 0.071 | -1.245 | -0.013 | 0.114 | 0.114 | 0.244 | 0.4 | 0.18 | 0.23 | 0.16 | 0.4 | 0.23 |
| 3 | 21827159 | rs3111745 | 1.603 | 0.963 | 0.094 | -1.316 | -0.004 | 0.086 | 0.086 | 0 | 0.49 | 0.29 | 0.43 | 0.18 | 0.47 | 0.3 |
| 3 | 21828964 | rs3860586 | -0.27 | 0.786 | 0.159 | -1.147 | -0.008 | -0.009 | -0.009 | 0.014 | 0.3 | 0.34 | 0.37 | 0.7 | 0.27 | 0.41 |
| 3 | 21831337 | rs3933112 | 0.261 | 0.832 | 0.209 | -1.005 | -0.006 | 0.032 | 0.032 | 0.254 | 0.22 | 0.31 | 0.27 | 0.36 | 0.21 | 0.27 |
| 3 | 21833043 | rs17009597 | 1.718 | 0.784 | 0.157 | -1.018 | 0.01 | 0.016 | 0.016 | 4.555 | 0.09 | 0.1 | 0.11 | 0.29 | 0.04 | 0.13 |
| 3 | 21833212 | rs2946632 | 0.074 | 0.716 | 0.19 | -1.148 | -0.008 | 0.005 | 0.005 | 0 | 0.73 | 0.64 | 0.79 | 0.69 | 0.78 | 0.71 |
| 3 | 21835397 | rs4547720 | 0.847 | 0.874 | 0.071 | -1.193 | 0.002 | 0.01 | 0.01 | 0.007 | 0.65 | 0.57 | 0.57 | 0.54 | 0.58 | 0.56 |
| 3 | 21835655 | rs17009600 | 2.351 | 0.809 | 0.045 | -1.113 | 0.014 | 0.018 | 0.018 | 3.103 | 0.07 | 0.03 | 0.1 | 0.3 | 0.04 | 0.13 |
| 3 | 21836075 | rs7374877 | 1.258 | 0.669 | -0.184 | -1.448 | 0.003 | 0 | 0 | 0.002 | 0.46 | 0.45 | 0.41 | 0.52 | 0.37 | 0.45 |

| | | | | | | | | | | | | | | | | |
|---|---|---|---|---|---|---|---|---|---|---|---|---|---|---|---|---|
| 3 | 21837299 | rs1606850 | 1.378 | 0.594 | -0.099 | -1.446 | -0.003 | -0.037 | -0.037 | 0 | 0.4 | 0.49 | 0.31 | 0.22 | 0.34 | 0.32 |
| 3 | 21837635 | rs13093354 | 0.757 | 0.617 | -0.119 | -1.47 | -0.005 | 0.058 | 0.058 | 0.045 | 0.15 | 0.25 | 0.21 | 0.36 | 0.11 | 0.23 |
| 3 | 21837676 | rs1606852 | 2.073 | 0.674 | -0.228 | -1.471 | 0.003 | 0.007 | 0.007 | 0.041 | 0.32 | 0.25 | 0.2 | 0.15 | 0.27 | 0.21 |
| 3 | 21841043 | rs1356102 | 1.283 | 0.58 | -0.244 | -1.48 | -0.012 | 0.053 | 0.053 | 0 | 0.42 | 0.29 | 0.27 | 0.16 | 0.42 | 0.27 |
| 3 | 21841370 | rs9869184 | 0.388 | 0.542 | -0.387 | -1.477 | 0.007 | -0.058 | -0.058 | 0.101 | 0.25 | 0.15 | 0.24 | 0.37 | 0.22 | 0.32 |
| 3 | 21841904 | rs11129023 | 1.052 | 0.435 | -0.459 | -1.382 | 0 | -0.009 | -0.009 | 0.037 | 0.19 | 0.14 | 0.14 | 0.07 | 0.2 | 0.09 |
| 3 | 21844883 | rs13083531 | 1.63 | 0.503 | -0.203 | -1.162 | -0.018 | 0.072 | 0.072 | 0 | 0.32 | 0.17 | 0.22 | 0.16 | 0.31 | 0.24 |
| 3 | 21851803 | rs13071008 | 0.457 | 0.547 | 0.263 | -0.768 | -0.004 | -0.009 | -0.009 | 0 | 0.68 | 0.65 | 0.52 | 0.26 | 0.69 | 0.49 |
| 3 | 21861727 | rs2176081 | -0.104 | 0.332 | 0.131 | -0.341 | 0.005 | -0.039 | -0.039 | 0 | 0.69 | 0.73 | 0.78 | 0.82 | 0.68 | 0.71 |
| 3 | 21864270 | rs9833498 | -0.271 | 0.088 | -0.215 | 0.33 | -0.036 | 0.081 | 0.081 | 1.574 | 0.13 | 0.27 | 0.12 | 0.3 | 0.05 | 0.13 |
| 3 | 21865234 | rs2670259 | 0.024 | 1.09 | -0.246 | -0.166 | 0.011 | -0.044 | -0.044 | 0.278 | 0.27 | 0.34 | 0.19 | 0.16 | 0.26 | 0.2 |
| 3 | 21867328 | rs7614916 | -0.463 | 0.771 | -0.123 | 0.512 | 0.008 | 0.067 | 0.067 | 0 | 0.24 | 0.51 | 0.42 | 0.37 | 0.3 | 0.33 |
| 3 | 21867628 | rs2630811 | -0.05 | 0.169 | -0.267 | 0.204 | -0.003 | -0.007 | -0.007 | 0 | 0.73 | 0.71 | 0.8 | 0.91 | 0.79 | 0.85 |
| 3 | 21867786 | rs2670258 | 0.584 | 0.602 | 0.179 | 0.604 | 0.007 | 0.015 | 0.015 | 0 | 0.84 | 0.65 | 0.81 | 0.76 | 0.84 | 0.82 |
| 3 | 21868051 | rs2670256 | -0.377 | 1.054 | -0.415 | 0.783 | 0.009 | 0.016 | 0.016 | 5.623 | 0.36 | 0.43 | 0.49 | 0.56 | 0.4 | 0.33 |
| 3 | 21874628 | rs9821836 | -0.247 | 1.186 | -0.799 | 0.353 | -0.015 | 0.034 | 0.034 | 0 | 0.28 | 0.21 | 0.31 | 0.17 | 0.28 | 0.22 |
| 3 | 21879594 | rs2630770 | 0.649 | 1.12 | -0.905 | 0.752 | 0.05 | 0.026 | 0.026 | 0.21 | 0.48 | 0.61 | 0.32 | 0.4 | 0.39 | 0.4 |
| 3 | 21881162 | rs2670277 | -0.018 | 1.538 | -0.947 | 1.126 | -0.039 | 0.018 | 0.018 | 0.188 | 0.64 | 0.78 | 0.51 | 0.72 | 0.6 | 0.71 |
| 3 | 21882659 | rs3912513 | 0.639 | 1.66 | -0.947 | 1.087 | -0.012 | -0.017 | -0.017 | 2.328 | 0.16 | 0.17 | 0.18 | 0.32 | 0.21 | 0.29 |
| 3 | 21885965 | rs2630799 | 0.49 | 0.611 | -1.265 | 0.693 | -0.033 | 0.02 | 0.02 | 1.018 | 0.42 | 0.18 | 0.36 | 0.38 | 0.48 | 0.42 |
| 3 | 21890328 | rs2630803 | 1.572 | 0.562 | -0.344 | 0.675 | -0.01 | 0.118 | 0.118 | 2.401 | 0.28 | 0.05 | 0.19 | 0.07 | 0.28 | 0.13 |
| 3 | 21892549 | rs2630807 | 0.876 | 0.397 | 0.364 | 0.894 | -0.002 | 0.004 | 0.004 | 0.96 | 0.45 | 0.38 | 0.4 | 0.46 | 0.48 | 0.53 |
| 3 | 21893698 | rs17009807 | -0.229 | 0.327 | -0.357 | 1.068 | -0.008 | 0.001 | 0.001 | 0 | 0.07 | 0.1 | 0.05 | 0.07 | 0.04 | 0.05 |
| 3 | 21897140 | rs9310661 | 0.052 | 0.734 | -0.75 | 1.014 | 0.004 | 0.002 | 0.002 | 0 | 0.35 | 0.34 | 0.31 | 0.34 | 0.28 | 0.32 |
| 3 | 21897186 | rs2670283 | -0.552 | 1.187 | -0.926 | 1.004 | -0.018 | -0.019 | -0.019 | 0 | 0.93 | 0.98 | 0.82 | 0.88 | 0.88 | 0.91 |
| 3 | 21899165 | rs9822325 | -0.054 | 0.725 | -0.86 | 0.582 | -0.018 | 0.053 | 0.053 | 0 | 0.3 | 0.16 | 0.3 | 0.17 | 0.28 | 0.22 |
| 3 | 21901392 | rs1679236 | 0.213 | 0.532 | -0.052 | 1.246 | 0.008 | 0.12 | 0.12 | 0 | 0.91 | 0.84 | 0.66 | 0.81 | 0.92 | 0.93 |
| 3 | 21903434 | rs1684497 | -0.009 | 0.687 | 0.324 | 1.311 | 0 | 0 | 0 | 0.391 | 0.32 | 0.42 | 0.29 | 0.31 | 0.42 | 0.35 |
| 3 | 21903493 | rs12486502 | 1.26 | 0.444 | -0.016 | 1.42 | -0.015 | -0.044 | -0.044 | 0 | 0.08 | 0.01 | 0.04 | 0.17 | 0.08 | 0.17 |
| 3 | 21903726 | rs1352139 | -0.435 | 0.479 | 0.103 | 1.428 | 0.027 | 0.192 | 0.192 | 0 | 0.9 | 0.81 | 0.63 | 0.75 | 0.91 | 0.91 |
| 3 | 21904054 | rs12493709 | 1.829 | 0.042 | -0.285 | 1.161 | -0.02 | -0.014 | -0.014 | 0.065 | 0.12 | 0.04 | 0.08 | 0.15 | 0.11 | 0.1 |
| 3 | 21904132 | rs1352137 | -0.535 | 0.032 | -0.276 | 1.162 | -0.001 | 0.03 | 0.03 | 0.526 | 0.79 | 0.8 | 0.58 | 0.67 | 0.81 | 0.81 |
| 3 | 21905595 | rs2620535 | 1.32 | 0.066 | -0.175 | 1.094 | 0.034 | 0.088 | 0.088 | 0 | 0.38 | 0.22 | 0.21 | 0.12 | 0.3 | 0.3 |
| 3 | 21908453 | rs1291305 | -0.874 | 0.201 | -0.074 | 1.204 | 0.023 | 0.156 | 0.156 | 0 | 0.91 | 0.82 | 0.66 | 0.81 | 0.92 | 0.91 |
| 3 | 21909164 | rs1621132 | 0.469 | 0.266 | -0.015 | 1.146 | -0.009 | 0.001 | 0.001 | 1.097 | 0.32 | 0.4 | 0.29 | 0.32 | 0.42 | 0.35 |
| 3 | 21909399 | rs1679221 | 0.326 | 0.182 | -0.124 | 1.099 | 0.043 | 0.023 | 0.023 | 0.019 | 0.32 | 0.55 | 0.33 | 0.4 | 0.42 | 0.35 |
| 3 | 21919039 | rs779036 | -0.611 | -0.359 | -0.102 | 0.907 | 0.016 | 0.018 | 0.018 | 0 | 0.57 | 0.72 | 0.47 | 0.68 | 0.61 | 0.59 |
| 3 | 21920809 | rs705007 | -1.565 | -0.209 | -0.238 | 0.999 | -0.001 | 0.019 | 0.019 | 0 | 0.93 | 0.96 | 0.69 | 0.82 | 0.92 | 0.92 |
| 3 | 21921197 | rs779033 | 0.24 | -0.583 | -0.447 | 0.906 | -0.004 | 0.002 | 0.002 | 0 | 0.44 | 0.46 | 0.37 | 0.47 | 0.52 | 0.43 |
| 3 | 21921897 | rs779032 | -1.312 | -1.194 | -1.004 | 0.445 | 0.034 | -0.012 | -0.012 | 0 | 0.78 | 0.93 | 0.7 | 0.81 | 0.85 | 0.9 |
| 3 | 21923732 | rs3912211 | 1.138 | -1.158 | -0.935 | 0.566 | -0.002 | -0.042 | -0.042 | 0.049 | 0.1 | 0 | 0.04 | 0.15 | 0.06 | 0.19 |
| 3 | 21923838 | rs2673536 | 0.346 | -1.563 | -0.521 | 0.501 | 0.004 | -0.003 | -0.003 | 0 | 0.35 | 0.3 | 0.4 | 0.32 | 0.33 | 0.37 |
| 3 | 21925055 | rs259556 | -1.017 | -1.302 | -0.317 | -0.015 | 0.008 | 0.005 | 0.005 | 0 | 0.64 | 0.55 | 0.58 | 0.52 | 0.55 | 0.7 |
| 3 | 21927108 | rs259535 | 1.678 | -1.383 | -0.348 | -0.092 | -0.002 | 0.035 | 0.035 | 0 | 0.17 | 0.08 | 0.09 | 0 | 0.15 | 0.14 |
| 3 | 21928015 | rs2670253 | -0.104 | -1.424 | -0.124 | -0.053 | -0.001 | -0.001 | -0.001 | 0 | 0.48 | 0.46 | 0.57 | 0.52 | 0.4 | 0.56 |
| 3 | 21930938 | rs259532 | -0.064 | -1.479 | -0.094 | 0.108 | 0 | -0.001 | -0.001 | 1.427 | 0.3 | 0.24 | 0.42 | 0.32 | 0.21 | 0.4 |
| 3 | 21931417 | rs3849559 | 2.292 | -1.258 | -0.026 | -0.165 | 0.012 | -0.023 | -0.023 | 0 | 0.07 | 0.15 | 0.08 | 0.02 | 0.09 | 0.11 |
| 3 | 21932373 | rs259527 | 0.123 | -1.14 | -0.084 | -0.046 | 0.006 | 0.014 | 0.014 | 1.507 | 0.28 | 0.19 | 0.13 | 0.16 | 0.22 | 0.34 |
| 3 | 21932746 | rs2244460 | 1.072 | -1.287 | -0.033 | -0.212 | -0.019 | -0.036 | -0.036 | 0 | 0.31 | 0.4 | 0.18 | 0.13 | 0.24 | 0.28 |
| 3 | 21936532 | rs259498 | 0.036 | -1.174 | -0.018 | -0.183 | 0 | -0.023 | -0.023 | 5.649 | 0.48 | 0.51 | 0.26 | 0.13 | 0.39 | 0.41 |
| 3 | 21938226 | rs259493 | 1.414 | -1.025 | -0.062 | -0.125 | 0 | -0.021 | -0.021 | 0.004 | 0.18 | 0.13 | 0.09 | 0 | 0.15 | 0.14 |
| 3 | 21941259 | rs259489 | -0.498 | -0.992 | -0.117 | 0.032 | -0.004 | 0.013 | 0.013 | 1.032 | 0.36 | 0.47 | 0.51 | 0.4 | 0.32 | 0.42 |
| 3 | 21943191 | rs17641808 | 2.226 | -0.953 | -0.074 | 0.035 | 0.016 | 0.006 | 0.006 | 0 | 0.06 | 0.09 | 0.08 | 0.07 | 0.09 | 0.03 |
| 3 | 21943932 | rs259499 | 1.245 | -1.01 | -0.004 | 0.257 | -0.003 | 0.004 | 0.004 | 0 | 0.19 | 0.21 | 0.39 | 0.18 | 0.17 | 0.2 |
| 3 | 21944121 | rs259500 | -0.48 | -1.039 | -0.039 | 0.205 | -0.001 | -0.041 | -0.041 | 1.579 | 0.35 | 0.44 | 0.22 | 0.22 | 0.31 | 0.38 |
| 3 | 21947463 | rs259505 | 0.107 | -1.034 | -0.05 | 0.187 | 0.01 | 0.008 | 0.008 | 0 | 0.26 | 0.36 | 0.47 | 0.23 | 0.26 | 0.22 |
| 3 | 21947772 | rs17010054 | 1.829 | -1.019 | -0.134 | 0.109 | -0.012 | 0.007 | 0.007 | 0 | 0.1 | 0.12 | 0.04 | 0.14 | 0.06 | 0.22 |
| 3 | 21948672 | rs259506 | 1.34 | -1.002 | -0.334 | -0.043 | -0.002 | -0.058 | -0.058 | 0.374 | 0.19 | 0.2 | 0.1 | 0 | 0.16 | 0.14 |
| 3 | 21951024 | rs627341 | 1.139 | -0.898 | -0.859 | 0.118 | 0.029 | 0.031 | 0.031 | 1.001 | 0.29 | 0.18 | 0.16 | 0.16 | 0.23 | 0.28 |
| 3 | 21953644 | rs259545 | -1.258 | -0.56 | -0.849 | 0.358 | -0.012 | -0.003 | -0.003 | 0 | 0.79 | 0.79 | 0.77 | 0.74 | 0.66 | 0.86 |
| 3 | 21953965 | rs259544 | 0.161 | -0.824 | -0.893 | 0.216 | -0.025 | 0.03 | 0.03 | 0.585 | 0.29 | 0.4 | 0.43 | 0.34 | 0.22 | 0.41 |
| 3 | 21954050 | rs597902 | 0.924 | -0.71 | -1.629 | -0.234 | 0.026 | 0.079 | 0.079 | 0.031 | 0.25 | 0.06 | 0.15 | 0.07 | 0.19 | 0.07 |
| 3 | 21955198 | rs11719664 | 0.864 | 0.172 | -1.331 | 0.215 | -0.038 | -0.056 | -0.056 | 0 | 0.1 | 0 | 0.05 | 0.18 | 0.11 | 0.23 |
| 3 | 21955384 | rs259540 | 0.724 | -0.17 | -1.623 | -0.23 | -0.001 | 0.073 | 0.073 | 0 | 0.13 | 0.01 | 0.05 | 0 | 0.09 | 0.11 |
| 3 | 21955784 | rs10510520 | 0.32 | 0.438 | -1.727 | -0.017 | -0.041 | -0.043 | -0.043 | 0 | 0.2 | 0.1 | 0.22 | 0.34 | 0.35 | 0.32 |
| 3 | 21960467 | rs1352135 | -1.446 | -0.476 | -1.413 | -0.401 | -0.008 | -0.031 | -0.031 | 0 | 0.83 | 0.83 | 0.94 | 0.99 | 0.9 | 0.96 |
| 3 | 21960877 | rs1479989 | -0.521 | -0.621 | -1.19 | -0.504 | 0.034 | 0.013 | 0.013 | 1.122 | 0.43 | 0.24 | 0.44 | 0.6 | 0.67 | 0.69 |
| 3 | 21961267 | rs1684524 | -1.743 | -0.479 | -1.228 | -0.472 | 0.033 | 0.068 | 0.068 | 0 | 0.83 | 0.96 | 0.95 | 0.99 | 0.9 | 0.96 |
| 3 | 21967150 | rs259521 | 2.678 | -0.768 | -1.226 | -0.562 | 0.01 | -0.078 | -0.078 | 4.924 | 0.23 | 0.06 | 0.42 | 0.39 | 0.18 | 0.19 |

| | | | | | | | | | | | | | | | | |
|---|---|---|---|---|---|---|---|---|---|---|---|---|---|---|---|---|
| 3 | 21970405 | rs259480 | 2.086 | -0.87 | -0.678 | -0.727 | -0.016 | -0.032 | -0.032 | 0 | 0.1 | 0.13 | 0.04 | 0 | 0.04 | 0.08 |
| 3 | 21972835 | rs1684481 | 0.452 | -0.472 | -0.632 | -0.912 | 0.003 | -0.035 | -0.035 | 0.079 | 0.28 | 0.29 | 0.24 | 0.07 | 0.27 | 0.21 |
| 3 | 21973963 | rs665732 | -0.759 | -0.701 | -0.575 | -0.938 | -0.027 | -0.067 | -0.067 | 0 | 0.72 | 0.68 | 0.9 | 0.98 | 0.86 | 0.86 |
| 3 | 21976413 | rs1553084 | 0.056 | -0.421 | -0.346 | -0.774 | -0.004 | -0.003 | -0.003 | 0.09 | 0.33 | 0.27 | 0.36 | 0.4 | 0.49 | 0.42 |
| 3 | 21977471 | rs7651606 | 0.245 | -0.417 | -0.35 | -0.772 | -0.022 | -0.013 | -0.013 | 0.019 | 0.32 | 0.23 | 0.36 | 0.4 | 0.49 | 0.42 |
| 3 | 21983332 | rs9859150 | -0.593 | -0.45 | -0.137 | -0.738 | 0 | 0 | 0 | 0.6 | 0.42 | 0.38 | 0.42 | 0.58 | 0.6 | 0.67 |
| 3 | 21983554 | rs611146 | 2.396 | -0.526 | -0.164 | -0.753 | -0.004 | 0.003 | 0.003 | 0.096 | 0.28 | 0.34 | 0.45 | 0.4 | 0.18 | 0.19 |
| 3 | 21994215 | rs1486391 | -1.445 | -0.31 | -0.721 | -0.779 | 0.001 | 0.039 | 0.039 | 0 | 0.8 | 0.9 | 0.91 | 0.98 | 0.82 | 0.93 |
| 3 | 21996586 | rs716237 | 2.025 | -0.201 | -0.635 | -0.677 | -0.001 | -0.001 | -0.001 | 0 | 0.13 | 0.09 | 0.15 | 0.06 | 0.15 | 0.1 |
| 3 | 22000946 | rs12490009 | 2.053 | -0.038 | -0.801 | -0.556 | -0.051 | -0.055 | -0.055 | 0 | 0.1 | 0 | 0.07 | 0.19 | 0.16 | 0.23 |
| 3 | 22004875 | rs17583756 | 1.099 | -0.014 | -0.709 | -0.557 | 0.006 | 0.046 | 0.046 | 0 | 0.2 | 0.1 | 0.09 | 0.02 | 0.18 | 0.09 |
| 3 | 22005074 | rs259569 | -1.158 | -0.128 | -0.77 | -0.553 | 0.03 | 0.016 | 0.016 | 0.019 | 0.56 | 0.78 | 0.49 | 0.58 | 0.64 | 0.74 |
| 3 | 22006821 | rs677508 | -1.141 | 0.048 | -0.808 | -0.36 | -0.037 | 0.054 | 0.054 | 0 | 0.6 | 0.84 | 0.39 | 0.27 | 0.54 | 0.49 |
| 3 | 22009022 | rs6765798 | 0.681 | 0.433 | -0.46 | -0.023 | -0.013 | -0.015 | -0.015 | 0 | 0.17 | 0.11 | 0.19 | 0.32 | 0.29 | 0.33 |
| 3 | 22019794 | rs2291818 | 0.427 | 0.435 | -0.34 | -0.029 | 0.017 | 0.036 | 0.036 | 0 | 0.19 | 0.24 | 0.2 | 0.33 | 0.29 | 0.34 |
| 3 | 22021227 | rs11129036 | 0.626 | 0.469 | -0.645 | -0.014 | -0.007 | -0.006 | -0.006 | 0 | 0.18 | 0.13 | 0.19 | 0.33 | 0.29 | 0.33 |
| 3 | 22021394 | rs13066500 | -0.291 | 0.404 | -0.75 | -0.135 | -0.06 | -0.066 | -0.066 | 0 | 0.29 | 0.12 | 0.27 | 0.52 | 0.45 | 0.6 |
| 3 | 22022923 | rs12494033 | -0.291 | 0.389 | -0.753 | -0.135 | -0.057 | -0.064 | -0.064 | 2.127 | 0.29 | 0.12 | 0.27 | 0.52 | 0.44 | 0.6 |
| 3 | 22032335 | rs6790810 | 0.761 | -0.076 | -0.203 | -0.58 | -0.041 | -0.085 | -0.085 | 0.082 | 0.12 | 0.56 | 0.07 | 0 | 0.04 | 0.06 |
| 3 | 22033071 | rs7652319 | -0.352 | 0.118 | -0.637 | -0.548 | -0.001 | -0.002 | -0.002 | 0.386 | 0.31 | 0.73 | 0.2 | 0.2 | 0.27 | 0.36 |
| 3 | 22037999 | rs2055268 | -1.412 | 0.053 | -1.07 | -0.569 | -0.004 | 0.016 | 0.016 | 0.333 | 0.61 | 0.87 | 0.42 | 0.32 | 0.63 | 0.64 |
| 3 | 22047818 | rs1679254 | -1.164 | -0.337 | -0.76 | -0.659 | 0.008 | -0.011 | -0.011 | 0 | 0.61 | 0.32 | 0.39 | 0.42 | 0.71 | 0.72 |
| 3 | 22048024 | rs12494280 | 1.955 | -0.156 | -0.968 | -0.636 | -0.049 | -0.032 | -0.032 | 0.714 | 0.12 | 0 | 0.08 | 0.13 | 0.16 | 0.27 |
| 3 | 22048849 | rs1694020 | -0.577 | -0.193 | -0.973 | -0.723 | 0.023 | 0.068 | 0.068 | 0.03 | 0.45 | 0.75 | 0.28 | 0.21 | 0.38 | 0.43 |
| 3 | 22049529 | rs1679256 | 0.142 | 0.173 | -0.781 | -0.732 | 0.002 | 0.022 | 0.022 | 0 | 0.3 | 0.13 | 0.17 | 0.17 | 0.3 | 0.23 |
| 3 | 22055009 | rs779097 | -0.171 | -0.138 | -0.499 | -0.72 | -0.001 | 0.002 | 0.002 | 0 | 0.34 | 0.34 | 0.2 | 0.12 | 0.37 | 0.28 |
| 3 | 22055043 | rs2008355 | 1.509 | -0.211 | -0.592 | -0.58 | -0.016 | 0.028 | 0.028 | 1.703 | 0.13 | 0.05 | 0.12 | 0 | 0.15 | 0.07 |
| 3 | 22055968 | rs6770681 | -0.236 | -1.384 | -0.181 | -0.178 | -0.038 | 0.098 | 0.098 | 0 | 0.14 | 0.42 | 0.07 | 0.18 | 0.07 | 0.18 |
| 3 | 22057218 | rs259432 | 0.226 | -1.209 | -0.48 | -0.24 | -0.037 | 0.006 | 0.006 | 0 | 0.21 | 0.32 | 0.08 | 0.2 | 0.12 | 0.18 |
| 3 | 22063264 | rs259434 | -0.51 | -0.06 | -0.521 | -0.464 | -0.015 | 0.011 | 0.011 | 3.352 | 0.73 | 0.81 | 0.74 | 0.9 | 0.59 | 0.71 |
| 3 | 22063502 | rs259435 | 0.135 | -0.547 | -0.505 | -0.266 | 0.002 | 0.014 | 0.014 | 0.967 | 0.57 | 0.63 | 0.62 | 0.83 | 0.45 | 0.55 |
| 3 | 22076137 | rs11914361 | -0.435 | -1.027 | -0.464 | 0.262 | -0.019 | 0.004 | 0.004 | 0.583 | 0.3 | 0.33 | 0.19 | 0.28 | 0.2 | 0.41 |
| 3 | 22080819 | rs711688 | 1.355 | -0.625 | -1.11 | 0.575 | -0.044 | -0.042 | -0.042 | 0 | 0.26 | 0.06 | 0.37 | 0.44 | 0.29 | 0.22 |
| 3 | 22081161 | rs7623423 | 0.491 | -0.619 | -0.98 | 0.526 | -0.001 | -0.001 | -0.001 | 0 | 0.1 | 0.09 | 0.08 | 0.08 | 0.08 | 0.13 |
| 3 | 22081979 | rs166314 | -1.136 | -0.846 | -0.676 | 0.26 | -0.003 | -0.002 | -0.002 | 0 | 0.75 | 0.71 | 0.65 | 0.75 | 0.64 | 0.75 |
| 3 | 22083147 | rs711687 | -0.464 | -0.753 | -0.659 | 0.503 | -0.002 | 0.007 | 0.007 | 0.02 | 0.61 | 0.3 | 0.61 | 0.71 | 0.59 | 0.7 |
| 3 | 22083236 | rs6773103 | -0.495 | -0.909 | -0.72 | 0.758 | 0.003 | -0.01 | -0.01 | 0 | 0.33 | 0.27 | 0.26 | 0.39 | 0.3 | 0.48 |
| 3 | 22083587 | rs166315 | 1.044 | -0.873 | -0.711 | 0.732 | -0.001 | 0.021 | 0.021 | 0 | 0.33 | 0.17 | 0.37 | 0.38 | 0.3 | 0.23 |
| 3 | 22084139 | rs259456 | 1.236 | -0.853 | -0.736 | 0.739 | -0.019 | 0.016 | 0.016 | 0 | 0.31 | 0.09 | 0.35 | 0.38 | 0.3 | 0.23 |
| 3 | 22085607 | rs259458 | 1.5 | -0.85 | -0.683 | 0.736 | -0.013 | 0.03 | 0.03 | 0 | 0.29 | 0.16 | 0.35 | 0.33 | 0.29 | 0.22 |
| 3 | 22085809 | rs1117 | -0.921 | -0.808 | -0.464 | 0.797 | 0.011 | -0.02 | -0.02 | 0 | 0.65 | 0.49 | 0.63 | 0.75 | 0.59 | 0.71 |
| 3 | 22086438 | rs259460 | 0.593 | -0.81 | -0.982 | 0.819 | -0.024 | -0.018 | -0.018 | 0.098 | 0.45 | 0.17 | 0.45 | 0.52 | 0.43 | 0.51 |
| 3 | 22088181 | rs259463 | 1.542 | -0.741 | -1.048 | 0.809 | -0.031 | 0.044 | 0.044 | 0 | 0.29 | 0.06 | 0.34 | 0.33 | 0.29 | 0.21 |
| 3 | 22089184 | rs12715024 | -0.28 | -0.489 | -0.813 | 1.014 | 0.001 | -0.004 | -0.004 | 1.135 | 0.24 | 0.23 | 0.17 | 0.28 | 0.22 | 0.37 |
| 3 | 22100442 | rs779159 | 0.705 | -0.201 | -1.045 | 0.726 | 0.001 | -0.019 | -0.019 | 0.807 | 0.39 | 0.34 | 0.44 | 0.42 | 0.37 | 0.32 |
| 3 | 22101780 | rs779162 | 0.083 | 0.381 | -0.919 | 0.704 | 0.001 | 0 | 0 | 0 | 0.59 | 0.5 | 0.56 | 0.65 | 0.52 | 0.64 |
| 3 | 22105044 | rs779167 | -0.124 | 0.722 | -1.343 | 0.574 | -0.036 | -0.101 | -0.101 | 0.055 | 0.68 | 0.9 | 0.54 | 0.52 | 0.61 | 0.57 |
| 3 | 22106222 | rs11717259 | -0.041 | 0.816 | -1.264 | 0.538 | -0.039 | -0.087 | -0.087 | 0 | 0.67 | 0.87 | 0.53 | 0.51 | 0.6 | 0.56 |
| 3 | 22106715 | rs1463579 | 1.294 | 0.751 | -2.072 | -0.022 | -0.033 | 0.071 | 0.071 | 0 | 0.21 | 0.01 | 0.11 | 0.1 | 0.18 | 0.17 |
| 3 | 22108957 | rs4508805 | 1.049 | 0.533 | -1.527 | -0.168 | -0.011 | -0.003 | -0.003 | 0 | 0.11 | 0.21 | 0.05 | 0.09 | 0.07 | 0.07 |
| 3 | 22110442 | rs941349 | -0.531 | 0.474 | -1.27 | -0.244 | -0.008 | -0.004 | -0.004 | 0 | 0.43 | 0.82 | 0.42 | 0.4 | 0.41 | 0.38 |
| 3 | 22110667 | rs12497147 | -0.042 | 0.681 | -0.556 | -0.521 | 0.005 | -0.081 | -0.081 | 0 | 0.24 | 0.42 | 0.14 | 0.09 | 0.23 | 0.15 |
| 3 | 22111153 | rs11917648 | 0.013 | 0.201 | -0.316 | -0.218 | 0.012 | 0.038 | 0.038 | 0 | 0.09 | 0.26 | 0.07 | 0.13 | 0.08 | 0.12 |
| 3 | 22113690 | rs941348 | 1.102 | 0.175 | -0.458 | -0.19 | 0.005 | 0.025 | 0.025 | 0 | 0.26 | 0.18 | 0.14 | 0.18 | 0.21 | 0.27 |
| 3 | 22116971 | rs17010728 | -0.657 | -0.152 | 0.069 | -0.279 | -0.026 | 0.027 | 0.027 | 0.869 | 0.23 | 0.54 | 0.28 | 0.26 | 0.17 | 0.19 |
| 3 | 22121266 | rs6782385 | 0.396 | -0.014 | -0.073 | -0.429 | -0.005 | -0.001 | -0.001 | 0 | 0.15 | 0.19 | 0.11 | 0.13 | 0.13 | 0.12 |
| 3 | 22121708 | rs1479993 | -0.063 | 0.208 | -0.065 | -0.836 | -0.014 | 0.009 | 0.009 | 0.184 | 0.29 | 0.21 | 0.36 | 0.2 | 0.39 | 0.28 |
| 3 | 22122396 | rs2593321 | 0.736 | -0.069 | -0.393 | -1.099 | 0.032 | 0.062 | 0.062 | 0 | 0.36 | 0.22 | 0.21 | 0.17 | 0.29 | 0.34 |
| 3 | 22122803 | rs17010739 | 1.054 | -0.199 | -0.164 | -1.338 | 0.018 | -0.071 | -0.071 | 0 | 0.13 | 0.25 | 0.1 | 0 | 0.17 | 0.04 |
| 3 | 22129839 | rs2170091 | -0.315 | -0.211 | -0.128 | -1.342 | -0.001 | 0.004 | 0.004 | 4.226 | 0.45 | 0.54 | 0.5 | 0.33 | 0.51 | 0.39 |
| 3 | 22130392 | rs2695642 | 0.407 | -0.108 | -0.077 | -1.354 | 0.031 | 0.051 | 0.051 | 0 | 0.47 | 0.22 | 0.46 | 0.34 | 0.44 | 0.45 |
| 3 | 22131055 | rs2695641 | -0.518 | -0.12 | -0.35 | -1.274 | -0.002 | 0.017 | 0.017 | 1.481 | 0.67 | 0.56 | 0.58 | 0.38 | 0.67 | 0.61 |
| 3 | 22131305 | rs2695639 | -1.661 | -0.311 | -0.374 | -1.189 | 0.008 | -0.031 | -0.031 | 0.119 | 0.8 | 0.89 | 0.69 | 0.38 | 0.84 | 0.66 |
| 3 | 22132083 | rs2928351 | -0.261 | -0.241 | -0.261 | -1.208 | -0.014 | -0.018 | -0.018 | 0.356 | 0.66 | 0.58 | 0.44 | 0.23 | 0.7 | 0.54 |
| 3 | 22139413 | rs12491081 | -0.057 | 0.225 | -0.688 | -1 | 0.004 | -0.009 | -0.009 | 2.638 | 0.36 | 0.61 | 0.27 | 0.23 | 0.38 | 0.36 |
| 3 | 22139499 | rs1983489 | 1.096 | 0.303 | -0.698 | -1.018 | 0.012 | -0.021 | -0.021 | 0 | 0.19 | 0.19 | 0.12 | 0.05 | 0.25 | 0.17 |
| 3 | 22145834 | rs17010798 | 1.064 | 0.205 | -0.734 | -0.873 | -0.01 | -0.01 | -0.01 | 0.803 | 0.14 | 0.27 | 0.14 | 0.09 | 0.12 | 0.12 |
| 3 | 22147707 | rs779183 | 0.447 | -0.057 | -1.564 | -0.579 | 0.005 | 0.037 | 0.037 | 1.305 | 0.54 | 0.63 | 0.37 | 0.23 | 0.53 | 0.46 |
| 3 | 22147772 | rs779182 | -0.367 | -0.441 | -1.33 | -0.001 | -0.014 | 0.022 | 0.022 | 5.073 | 0.75 | 0.74 | 0.69 | 0.37 | 0.84 | 0.64 |

| | | | | | | | | | | | | | | | | |
|---|---|---|---|---|---|---|---|---|---|---|---|---|---|---|---|---|
| 3 | 22152886 | rs4858028 | 0.597 | -0.269 | -1.98 | -0.06 | -0.037 | -0.034 | -0.034 | 0 | 0.08 | 0.02 | 0.24 | 0.08 | 0.14 | 0.05 |
| 3 | 22153040 | rs9823158 | -0.176 | -0.259 | -1.765 | 0.057 | -0.008 | 0.009 | 0.009 | 5.302 | 0.75 | 0.74 | 0.66 | 0.33 | 0.84 | 0.65 |
| 3 | 22153897 | rs9310681 | -0.315 | 0.53 | -1.69 | -0.185 | 0.038 | 0.149 | 0.149 | 4.489 | 0.51 | 0.8 | 0.37 | 0.22 | 0.55 | 0.46 |
| 3 | 22159896 | rs6793553 | 0.104 | 0.76 | -1.595 | 0.212 | -0.01 | -0.013 | -0.013 | 3.398 | 0.63 | 0.78 | 0.62 | 0.44 | 0.6 | 0.54 |
| 3 | 22161079 | rs11710290 | 0.048 | 1.135 | -1.524 | -0.212 | -0.001 | 0.022 | 0.022 | 0 | 0.14 | 0.15 | 0.28 | 0.4 | 0.17 | 0.27 |
| 3 | 22161566 | rs6781787 | -0.187 | 0.882 | -2.305 | -0.369 | -0.012 | 0.01 | 0.01 | 0 | 0.78 | 0.9 | 0.83 | 0.8 | 0.73 | 0.81 |
| 3 | 22162351 | rs12493053 | 0.991 | 0.843 | -2.4 | -0.403 | -0.01 | -0.075 | -0.075 | 0.088 | 0.28 | 0.08 | 0.3 | 0.4 | 0.26 | 0.38 |
| 3 | 22163190 | rs6775657 | 0.492 | 1.095 | -2.03 | -0.326 | -0.005 | -0.04 | -0.04 | 0 | 0.69 | 0.78 | 0.59 | 0.45 | 0.64 | 0.57 |
| 3 | 22173759 | rs4858387 | -1.049 | 1.032 | -1.997 | -0.281 | -0.03 | -0.042 | -0.042 | 0 | 0.92 | 0.99 | 0.94 | 0.83 | 0.89 | 0.92 |
| 3 | 22175067 | rs1527637 | -0.263 | 0.655 | -1.501 | -0.759 | 0.002 | 0.032 | 0.032 | 0.681 | 0.51 | 0.39 | 0.61 | 0.81 | 0.49 | 0.69 |
| 3 | 22177068 | rs9812687 | -0.124 | 0.31 | -1.305 | -0.901 | -0.021 | -0.063 | -0.063 | 1.396 | 0.74 | 0.52 | 0.81 | 0.84 | 0.75 | 0.84 |
| 3 | 22182024 | rs11129047 | 0.682 | 0.331 | -1.657 | -0.779 | 0.007 | -0.079 | -0.079 | 0.382 | 0.36 | 0.09 | 0.33 | 0.41 | 0.3 | 0.41 |
| 3 | 22183438 | rs7618885 | 0.838 | 0.314 | -0.873 | -0.816 | -0.013 | -0.065 | -0.065 | 0 | 0.17 | 0.28 | 0.12 | 0 | 0.14 | 0.11 |
| 3 | 22186404 | rs7628166 | -0.642 | 0.406 | -0.825 | -0.715 | -0.005 | 0 | 0 | 0 | 0.41 | 0.48 | 0.4 | 0.43 | 0.35 | 0.4 |
| 3 | 22188382 | rs4858388 | 0.14 | 0.197 | -0.98 | -0.859 | -0.032 | 0.015 | 0.015 | 0.058 | 0.56 | 0.85 | 0.67 | 0.41 | 0.57 | 0.49 |
| 3 | 22188526 | rs11917423 | 1.639 | 0.471 | -0.612 | -0.76 | 0 | -0.081 | -0.081 | 0.056 | 0.24 | 0.31 | 0.25 | 0.03 | 0.25 | 0.11 |
| 3 | 22189491 | rs7639978 | 0.659 | 0.071 | -0.744 | -0.636 | -0.001 | 0.031 | 0.031 | 0 | 0.11 | 0.1 | 0.26 | 0.44 | 0.09 | 0.23 |
| 3 | 22190694 | rs17648344 | 0.648 | -0.193 | -0.905 | -0.718 | 0.021 | 0 | 0 | 0 | 0.25 | 0.25 | 0.1 | 0 | 0.18 | 0.14 |
| 3 | 22191086 | rs4858389 | 0.648 | -0.051 | -1.004 | -0.651 | 0.01 | 0.075 | 0.075 | 0 | 0.16 | 0.21 | 0.31 | 0.42 | 0.18 | 0.25 |
| 3 | 22192085 | rs9816170 | 0.273 | 0.074 | -1.348 | -0.429 | 0.003 | 0.003 | 0.003 | 0 | 0.5 | 0.46 | 0.53 | 0.42 | 0.54 | 0.52 |
| 3 | 22198015 | rs9864211 | -0.561 | 0.013 | -1.682 | 0.33 | -0.022 | -0.024 | -0.024 | 1.337 | 0.92 | 1 | 0.95 | 0.87 | 0.9 | 0.93 |
| 3 | 22199510 | rs6771467 | 0.012 | 0.203 | -1.432 | 0.461 | -0.024 | -0.059 | -0.059 | 0 | 0.23 | 0.1 | 0.23 | 0.37 | 0.24 | 0.38 |
| 3 | 22202153 | rs10513638 | 0.328 | 0.432 | -1.236 | 0.783 | 0.027 | 0.084 | 0.084 | 0.219 | 0.11 | 0.16 | 0.22 | 0.29 | 0.15 | 0.28 |
| 3 | 22203032 | rs6766674 | -0.057 | -0.271 | -1.636 | 0.342 | -0.008 | -0.003 | -0.003 | 0 | 0.69 | 0.63 | 0.85 | 0.76 | 0.77 | 0.83 |
| 3 | 22203294 | rs12637217 | -0.195 | 0.374 | -2.421 | 0.747 | -0.031 | -0.082 | -0.082 | 0 | 0.28 | 0.1 | 0.26 | 0.42 | 0.29 | 0.43 |
| 3 | 22203514 | rs17010924 | -0.831 | 0.607 | -2.216 | 1.1 | -0.045 | -0.048 | -0.048 | 0 | 0.08 | 0.01 | 0.12 | 0.27 | 0.14 | 0.08 |
| 3 | 22203636 | rs3932414 | -0.382 | -0.084 | -2.608 | 0.071 | -0.051 | 0.083 | 0.083 | 1.348 | 0.7 | 0.98 | 0.61 | 0.9 | 0.71 | 0.86 |
| 3 | 22204244 | rs9852181 | -0.508 | -0.99 | -1.436 | 0.074 | -0.075 | -0.051 | -0.051 | 0 | 0.23 | 0.54 | 0.14 | 0.1 | 0.11 | 0.13 |
| 3 | 22205505 | rs1965306 | -0.426 | -0.719 | -2.609 | 0.157 | -0.102 | -0.082 | -0.082 | 0.478 | 0.64 | 0.89 | 0.41 | 0.4 | 0.47 | 0.45 |
| 3 | 22207033 | rs9310685 | 0.328 | -1.591 | -2.321 | -0.504 | 0 | -0.002 | -0.002 | 0 | 0.22 | 0.2 | 0.16 | 0.07 | 0.21 | 0.2 |
| 3 | 22207701 | rs7626875 | 0.48 | -2.098 | -2.126 | -0.82 | -0.045 | -0.061 | -0.061 | 0 | 0.27 | 0.53 | 0.14 | 0.09 | 0.14 | 0.15 |
| 3 | 22214274 | rs6775362 | 0.64 | -2.111 | -2.162 | -0.791 | 0 | -0.008 | -0.008 | 0.084 | 0.22 | 0.24 | 0.17 | 0.07 | 0.21 | 0.2 |
| 3 | 22216840 | rs17029466 | -0.502 | -2.142 | -1.456 | -0.802 | 0.093 | 0.254 | 0.254 | 0 | 0.49 | 0.81 | 0.31 | 0.16 | 0.35 | 0.36 |
| 3 | 22229281 | rs9682433 | -0.95 | -2.099 | -1.339 | -0.93 | 0.013 | -0.006 | -0.006 | 0 | 0.61 | 0.56 | 0.39 | 0.32 | 0.5 | 0.45 |
| 3 | 22229497 | rs9310686 | -0.924 | -2.11 | -1.362 | -0.935 | 0.016 | -0.009 | -0.009 | 0 | 0.61 | 0.54 | 0.39 | 0.32 | 0.5 | 0.44 |
| 3 | 22231144 | rs10513640 | 0.98 | -2.146 | -1.518 | -0.871 | 0.004 | -0.024 | -0.024 | 0 | 0.15 | 0.11 | 0.11 | 0.23 | 0.13 | 0.1 |
| 3 | 22239509 | rs1948377 | -2.087 | -1.443 | -2.024 | -0.449 | -0.109 | -0.101 | -0.101 | 1.359 | 0.8 | 0.99 | 0.77 | 0.71 | 0.73 | 0.7 |
| 3 | 22244794 | rs7624502 | 1.029 | -1.558 | -1.868 | -0.649 | 0.009 | 0.008 | 0.008 | 0 | 0.17 | 0.12 | 0.11 | 0.12 | 0.13 | 0.11 |
| 3 | 22249364 | rs11709805 | 1.379 | -1.612 | -1.881 | -0.723 | 0.019 | 0.035 | 0.035 | 3.185 | 0.22 | 0.12 | 0.1 | 0.03 | 0.17 | 0.14 |
| 3 | 22253194 | rs2728981 | -0.461 | -1.579 | -1.618 | -0.709 | 0.012 | -0.025 | -0.025 | 0 | 0.61 | 0.56 | 0.39 | 0.21 | 0.5 | 0.44 |
| 3 | 22258978 | rs2638134 | 0.408 | -1.827 | -1.792 | -0.84 | 0.084 | 0.098 | 0.098 | 0.043 | 0.49 | 0.26 | 0.7 | 0.71 | 0.67 | 0.61 |
| 3 | 22261529 | rs2728983 | -0.935 | -1.838 | -1.62 | -0.852 | -0.047 | -0.087 | -0.087 | 0 | 0.73 | 0.53 | 0.85 | 0.9 | 0.82 | 0.84 |
| 3 | 22266124 | rs12635688 | 0.404 | -1.787 | -1.566 | -0.792 | 0.009 | 0.004 | 0.004 | 2.053 | 0.24 | 0.23 | 0.15 | 0.2 | 0.15 | 0.22 |
| 3 | 22267493 | rs7647595 | 1.582 | -1.589 | -1.835 | -0.802 | 0.011 | -0.085 | -0.085 | 2.611 | 0.17 | 0.27 | 0.16 | 0 | 0.19 | 0.08 |
| 3 | 22268086 | rs6791903 | 0.134 | -1.231 | -2.124 | -0.43 | 0.037 | 0.113 | 0.113 | 0.076 | 0.46 | 0.82 | 0.35 | 0.25 | 0.36 | 0.31 |
| 3 | 22276847 | rs2638123 | -1.112 | -1.101 | -1.851 | -0.292 | 0.03 | 0.062 | 0.062 | 3.555 | 0.78 | 0.93 | 0.9 | 0.97 | 0.83 | 0.86 |
| 3 | 22278497 | rs2035848 | 0.85 | -0.697 | -1.968 | 0.105 | -0.043 | -0.077 | -0.077 | 0.731 | 0.13 | 0.05 | 0.36 | 0.44 | 0.21 | 0.26 |
| 3 | 22280773 | rs2728951 | -0.92 | -1.159 | -1.931 | -0.476 | 0.009 | 0.123 | 0.123 | 2.772 | 0.78 | 0.99 | 0.9 | 0.98 | 0.83 | 0.83 |
| 3 | 22283291 | rs2728954 | -0.379 | -0.884 | -1.831 | 0.1 | -0.027 | 0.033 | 0.033 | 0.228 | 0.45 | 0.9 | 0.52 | 0.42 | 0.42 | 0.39 |
| 3 | 22284122 | rs2728955 | -0.129 | -0.912 | -0.671 | 0.023 | -0.004 | -0.017 | -0.017 | 0.69 | 0.36 | 0.46 | 0.43 | 0.33 | 0.36 | 0.38 |
| 3 | 22285562 | rs729286 | 0.22 | -0.944 | -0.509 | 0.014 | 0.002 | -0.003 | -0.003 | 0 | 0.32 | 0.29 | 0.33 | 0.31 | 0.36 | 0.26 |
| 3 | 22292516 | rs1388550 | -0.06 | -0.944 | -0.317 | 0.038 | -0.002 | 0.005 | 0.005 | 0 | 0.34 | 0.23 | 0.38 | 0.29 | 0.36 | 0.35 |
| 3 | 22300883 | rs2728988 | -0.026 | -0.908 | -0.062 | 0.183 | 0.001 | 0 | 0 | 0.465 | 0.35 | 0.4 | 0.37 | 0.38 | 0.36 | 0.41 |
| 3 | 22306489 | rs2638161 | -1.086 | -1.324 | -1.047 | 0.33 | -0.037 | -0.181 | -0.181 | 0 | 0.7 | 0.97 | 0.67 | 0.56 | 0.69 | 0.77 |
| 3 | 22318273 | rs2638145 | -0.529 | -0.244 | -0.926 | 0.217 | -0.026 | 0.06 | 0.06 | 0.583 | 0.45 | 0.86 | 0.48 | 0.39 | 0.44 | 0.41 |
| 3 | 22327742 | rs2638148 | -0.968 | -0.268 | -1.008 | 0.166 | -0.059 | 0.066 | 0.066 | 1.183 | 0.52 | 0.92 | 0.5 | 0.41 | 0.48 | 0.49 |
| 3 | 22330606 | rs1388549 | -0.154 | -0.396 | -0.313 | 0.105 | -0.003 | 0.008 | 0.008 | 2.194 | 0.39 | 0.3 | 0.28 | 0.29 | 0.42 | 0.4 |
| 3 | 22331129 | rs937105 | -0.771 | -0.363 | -0.347 | 0.155 | 0.021 | 0.017 | 0.017 | 0.344 | 0.1 | 0.51 | 0.09 | 0.1 | 0.05 | 0.07 |
| 3 | 22335953 | rs9856840 | -0.064 | -0.358 | -0.242 | 0.225 | 0.054 | 0.045 | 0.045 | 0.119 | 0.07 | 0.39 | 0.07 | 0.09 | 0.04 | 0.01 |
| 3 | 22337587 | rs7429312 | -0.497 | -0.367 | -0.273 | 0.222 | 0.018 | 0.013 | 0.013 | 0.319 | 0.1 | 0.5 | 0.09 | 0.1 | 0.05 | 0.07 |
| 3 | 22338009 | rs13082422 | -0.241 | -0.275 | -0.47 | 0.219 | 0.002 | -0.018 | -0.018 | 0.513 | 0.39 | 0.39 | 0.33 | 0.29 | 0.4 | 0.4 |
| 3 | 22338707 | rs4557183 | 0.525 | -0.151 | -0.594 | 0.207 | 0.005 | -0.023 | -0.023 | 0 | 0.31 | 0.62 | 0.32 | 0.27 | 0.28 | 0.34 |
| 3 | 22343571 | rs1874088 | -0.845 | -0.157 | -0.894 | 0.388 | -0.01 | -0.08 | -0.08 | 0 | 0.63 | 0.86 | 0.61 | 0.53 | 0.63 | 0.73 |
| 3 | 22348864 | rs11712617 | 1.584 | -0.144 | -1.195 | 0.117 | -0.071 | 0.036 | 0.036 | 0 | 0.15 | 0.03 | 0.23 | 0.1 | 0.28 | 0.09 |
| 3 | 22350042 | rs4858031 | -1.009 | -0.175 | -0.892 | 0.256 | 0.002 | -0.048 | -0.048 | 0 | 0.63 | 0.75 | 0.6 | 0.53 | 0.63 | 0.75 |
| 3 | 22351154 | rs9809134 | 1.037 | -0.374 | -0.732 | 0.094 | -0.004 | 0.036 | 0.036 | 12.202 | 0.05 | 0.22 | 0.03 | 0.1 | 0.01 | 0.01 |
| 3 | 22354943 | rs11917977 | 0.962 | -0.374 | -0.784 | 0.094 | -0.013 | 0.025 | 0.025 | 13.797 | 0.05 | 0.21 | 0.03 | 0.1 | 0.01 | 0.01 |
| 3 | 22356126 | rs9855468 | -1.199 | -0.278 | -1.272 | 0.521 | -0.01 | -0.076 | -0.076 | 2.507 | 0.66 | 0.92 | 0.7 | 0.65 | 0.67 | 0.76 |
| 3 | 22356930 | rs1389663 | -0.479 | 0.418 | -0.63 | -0.047 | -0.017 | 0.027 | 0.027 | 0 | 0.33 | 0.21 | 0.34 | 0.16 | 0.43 | 0.29 |

| | | | | | | | | | | | | | | | | |
|---|---|---|---|---|---|---|---|---|---|---|---|---|---|---|---|---|
| 3 | 22359166 | rs1494223 | -0.743 | 0.844 | -0.64 | -0.284 | 0.006 | 0.051 | 0.051 | 0 | 0.53 | 0.75 | 0.36 | 0.4 | 0.56 | 0.41 |
| 3 | 22363886 | rs6774197 | -0.059 | 0.969 | -0.366 | -0.205 | 0.02 | 0.002 | 0.002 | 1.103 | 0.23 | 0.45 | 0.2 | 0.25 | 0.29 | 0.19 |
| 3 | 22368095 | rs1157438 | -0.211 | 0.976 | -0.563 | -0.212 | 0.022 | 0.006 | 0.006 | 1.261 | 0.24 | 0.55 | 0.21 | 0.29 | 0.29 | 0.2 |
| 3 | 22376670 | rs9880225 | 1.307 | 0.754 | -0.476 | -0.338 | -0.013 | 0.075 | 0.075 | 0 | 0.22 | 0.11 | 0.12 | 0.01 | 0.23 | 0.14 |
| 3 | 22376894 | rs9880567 | 0.421 | 0.759 | -0.298 | -0.284 | 0.002 | 0.014 | 0.014 | 0 | 0.29 | 0.35 | 0.17 | 0.15 | 0.26 | 0.28 |
| 3 | 22381341 | rs9858479 | 1.255 | 0.802 | -0.443 | -0.344 | -0.003 | 0.046 | 0.046 | 0 | 0.22 | 0.15 | 0.13 | 0.01 | 0.23 | 0.15 |
| 3 | 22385375 | rs7623065 | -0.153 | 0.844 | -0.345 | -0.418 | 0.005 | 0.064 | 0.064 | 0 | 0.36 | 0.75 | 0.24 | 0.14 | 0.32 | 0.32 |
| 3 | 22393482 | rs1389661 | -0.379 | 0.833 | -0.557 | -0.263 | 0.018 | 0.068 | 0.068 | 0 | 0.37 | 0.83 | 0.22 | 0.24 | 0.29 | 0.32 |
| 3 | 22396657 | rs9854991 | 0.288 | 0.898 | -0.393 | -0.366 | -0.006 | -0.118 | -0.118 | 0 | 0.31 | 0.49 | 0.18 | 0.07 | 0.29 | 0.16 |
| 3 | 22401243 | rs9870287 | 1.136 | 0.915 | -1.12 | -0.11 | -0.01 | 0.052 | 0.052 | 0 | 0.22 | 0.13 | 0.26 | 0.1 | 0.26 | 0.13 |
| 3 | 22415767 | rs9842199 | 0.217 | 1.392 | -0.822 | 0.339 | -0.008 | 0.028 | 0.028 | 0.117 | 0.72 | 0.84 | 0.79 | 0.76 | 0.71 | 0.88 |
| 3 | 22416318 | rs6419844 | 0.468 | 1.439 | -0.824 | 0.366 | -0.012 | 0.031 | 0.031 | 0 | 0.69 | 0.8 | 0.63 | 0.7 | 0.67 | 0.79 |
| 3 | 22417757 | rs4858400 | -1.476 | 1.033 | -0.434 | 0.562 | 0.009 | 0.008 | 0.008 | 0.535 | 0.21 | 0.17 | 0.29 | 0.21 | 0.18 | 0.35 |
| 3 | 22422856 | rs3749374 | -1.998 | 1.365 | 0.466 | 1.121 | 0.011 | 0.082 | 0.082 | 1.452 | 0.24 | 0.55 | 0.22 | 0.43 | 0.25 | 0.34 |
| 3 | 22423131 | rs3749375 | 0.914 | 1.777 | 0.138 | 0.862 | 0.002 | -0.08 | -0.08 | 0.246 | 0.73 | 0.58 | 0.83 | 0.87 | 0.73 | 0.91 |
| 3 | 22426918 | rs17011280 | -1.116 | 1.822 | 0.172 | 1.185 | -0.009 | -0.004 | -0.004 | 0 | 0.08 | 0.08 | 0.06 | 0.24 | 0.12 | 0.16 |
| 3 | 22435661 | rs1494228 | 0.658 | 1.95 | 0.501 | 1.044 | 0.02 | 0.051 | 0.051 | 0 | 0.8 | 0.54 | 0.85 | 0.65 | 0.77 | 0.9 |
| 3 | 22441261 | rs1121249 | 2.465 | 1.891 | 0.436 | 1.011 | -0.022 | -0.052 | -0.052 | 0 | 0.58 | 0.21 | 0.64 | 0.43 | 0.54 | 0.58 |
| 3 | 22456933 | rs935736 | 2.444 | 1.923 | 0.463 | 1.021 | -0.042 | -0.078 | -0.078 | 0 | 0.59 | 0.17 | 0.63 | 0.43 | 0.54 | 0.59 |
| 3 | 22457663 | rs1548137 | -0.325 | 1.925 | 0.526 | 0.96 | -0.014 | -0.033 | -0.033 | 1.007 | 0.1 | 0.23 | 0.08 | 0.02 | 0.06 | 0.13 |
| 3 | 22460263 | rs6778816 | 1.846 | 1.974 | 0.577 | 0.983 | 0.01 | 0.023 | 0.023 | 0 | 0.7 | 0.49 | 0.72 | 0.46 | 0.6 | 0.8 |
| 3 | 22462205 | rs13061996 | 1.728 | 1.956 | 0.585 | 1.007 | -0.003 | 0.002 | 0.002 | 0 | 0.7 | 0.45 | 0.79 | 0.62 | 0.71 | 0.77 |
| 3 | 22465345 | rs7614013 | -2.121 | 1.972 | 0.56 | 1.229 | -0.001 | 0 | 0 | 0.539 | 0.2 | 0.25 | 0.2 | 0.21 | 0.23 | 0.23 |
| 3 | 22473092 | rs9861611 | -1.517 | 2.123 | 0.48 | 1.37 | -0.011 | -0.011 | -0.011 | 0 | 0.1 | 0.08 | 0.13 | 0.19 | 0.18 | 0.1 |
| 3 | 22473729 | rs17011371 | -0.973 | 1.635 | 0.841 | 1.24 | -0.015 | -0.011 | -0.011 | 0.014 | 0.08 | 0.14 | 0.08 | 0.07 | 0.06 | 0.02 |
| 3 | 22476773 | rs6789629 | -0.981 | 1.971 | 1.149 | 1.522 | 0.021 | 0.11 | 0.11 | 0 | 0.11 | 0.21 | 0.09 | 0.37 | 0.15 | 0.09 |
| 3 | 22482089 | rs10510529 | -1.467 | 2.265 | 0.819 | 1.233 | -0.015 | -0.013 | -0.013 | 0 | 0.1 | 0.09 | 0.13 | 0.18 | 0.18 | 0.08 |
| 3 | 22501079 | rs1349332 | -1.312 | 2.285 | 1.763 | 1.413 | 0.09 | 0.231 | 0.231 | 0 | 0.11 | 0.29 | 0.13 | 0.38 | 0.18 | 0.1 |
| 3 | 22504135 | rs1450348 | -2.165 | 2.233 | 1.444 | 1.329 | 0.021 | 0.077 | 0.077 | 0.121 | 0.24 | 0.54 | 0.2 | 0.39 | 0.25 | 0.3 |
| 3 | 22511091 | rs2018333 | -1.781 | 2.503 | 1.005 | 1.547 | 0.048 | 0.106 | 0.106 | 1.16 | 0.16 | 0.64 | 0.16 | 0.39 | 0.23 | 0.13 |
| 3 | 22513865 | rs1901104 | -1.212 | 2.437 | 0.672 | 1.078 | 0 | -0.003 | -0.003 | 0.072 | 0.22 | 0.22 | 0.2 | 0.13 | 0.25 | 0.21 |
| 3 | 22531603 | rs1349273 | -2.23 | 2.444 | 0.99 | 1.624 | -0.01 | -0.009 | -0.009 | 0 | 0.38 | 0.68 | 0.46 | 0.6 | 0.43 | 0.39 |
| 3 | 22531649 | rs7616263 | -0.448 | 2.534 | 1.24 | 1.596 | 0.014 | -0.03 | -0.03 | 0 | 0.11 | 0.28 | 0.12 | 0.09 | 0.12 | 0.03 |
| 3 | 22532660 | rs2122369 | -2.913 | 2.687 | 1.284 | 1.605 | -0.022 | 0.004 | 0.004 | 2.104 | 0.42 | 0.75 | 0.48 | 0.62 | 0.49 | 0.45 |
| 3 | 22535867 | rs17011474 | -0.727 | 2.708 | 1.487 | 1.627 | 0.005 | 0.053 | 0.053 | 0.43 | 0.12 | 0.18 | 0.24 | 0.47 | 0.19 | 0.13 |
| 3 | 22537634 | rs1449897 | -1.483 | 2.52 | 1.441 | 1.555 | 0.002 | -0.028 | -0.028 | 0 | 0.23 | 0.29 | 0.2 | 0.13 | 0.24 | 0.21 |
| 3 | 22544898 | rs1947238 | -2.832 | 2.741 | 0.853 | 1.52 | -0.068 | -0.03 | -0.03 | 0 | 0.46 | 0.96 | 0.53 | 0.65 | 0.53 | 0.48 |
| 3 | 22562237 | rs4392438 | -1.708 | 2.48 | 1.177 | 1.296 | -0.002 | -0.077 | -0.077 | 0 | 0.25 | 0.5 | 0.22 | 0.13 | 0.24 | 0.21 |
| 3 | 22562572 | rs2358691 | -1.749 | 2.482 | 1.177 | 1.297 | -0.004 | -0.078 | -0.078 | 0 | 0.25 | 0.5 | 0.23 | 0.13 | 0.24 | 0.21 |
| 3 | 22571896 | rs1466123 | 0.737 | 2.527 | 0.872 | 1.047 | 0.001 | -0.061 | -0.061 | 0 | 0.79 | 0.71 | 0.85 | 0.95 | 0.79 | 0.81 |
| 3 | 22574169 | rs2358693 | -1.171 | 2.732 | 0.983 | 1.8 | 0.016 | 0.01 | 0.01 | 0 | 0.17 | 0.24 | 0.15 | 0.27 | 0.24 | 0.11 |
| 3 | 22575592 | rs6769561 | -0.91 | 2.559 | 0.988 | 1.664 | 0.026 | -0.006 | -0.006 | 0.357 | 0.15 | 0.42 | 0.12 | 0.08 | 0.12 | 0.23 |
| 3 | 22576404 | rs11719717 | -1.449 | 2.868 | 0.86 | 1.887 | 0.034 | 0.057 | 0.057 | 0 | 0.28 | 0.44 | 0.37 | 0.56 | 0.38 | 0.18 |
| 3 | 22576545 | rs6785666 | -2.318 | 2.855 | 0.876 | 1.882 | -0.028 | 0.007 | 0.007 | 1.089 | 0.41 | 0.78 | 0.49 | 0.63 | 0.5 | 0.42 |
| 3 | 22581523 | rs12630192 | -1.74 | 2.643 | 0.733 | 2.092 | 0.035 | 0.059 | 0.059 | 0 | 0.28 | 0.43 | 0.37 | 0.56 | 0.38 | 0.18 |
| 3 | 22584574 | rs4479617 | -1.144 | 2.411 | 0.499 | 1.636 | 0.081 | 0.066 | 0.066 | 0.321 | 0.07 | 0.21 | 0.05 | 0.14 | 0.12 | 0.06 |
| 3 | 22587743 | rs17011536 | -0.899 | 1.987 | 0.346 | 1.5 | 0.018 | 0.049 | 0.049 | 0 | 0.08 | 0.18 | 0.07 | 0.15 | 0.09 | 0.21 |
| 3 | 22589584 | rs2122371 | 1.278 | 1.971 | 0.558 | 1.811 | 0.001 | 0.023 | 0.023 | 0.188 | 0.93 | 0.88 | 0.95 | 0.84 | 0.94 | 0.95 |
| 3 | 22593944 | rs13066782 | -0.316 | 1.98 | 0.556 | 1.793 | 0.011 | -0.023 | -0.023 | 0 | 0.06 | 0.18 | 0.04 | 0.02 | 0.06 | 0.1 |
| 3 | 22594055 | rs13071252 | -0.079 | 1.692 | 0.054 | 0.942 | 0 | -0.005 | -0.005 | 0 | 0.46 | 0.5 | 0.27 | 0.38 | 0.42 | 0.63 |
| 3 | 22594832 | rs17011585 | -1.171 | 1.659 | 0.188 | 0.944 | -0.002 | 0.014 | 0.014 | 0 | 0.11 | 0.22 | 0.05 | 0.17 | 0.09 | 0.14 |
| 3 | 22594900 | rs13095534 | -0.918 | 1.864 | 0.105 | 1.002 | 0.03 | 0.03 | 0.03 | 0 | 0.21 | 0.38 | 0.23 | 0.28 | 0.26 | 0.26 |
| 3 | 22597952 | rs6778123 | -0.594 | 1.783 | 0.081 | 0.992 | -0.006 | 0.001 | 0.001 | 0.615 | 0.57 | 0.54 | 0.42 | 0.52 | 0.6 | 0.66 |
| 3 | 22599912 | rs716736 | 0.335 | 1.725 | -0.069 | 1.035 | -0.001 | 0.001 | 0.001 | 0 | 0.09 | 0.06 | 0.07 | 0.05 | 0.11 | 0.09 |
| 3 | 22600086 | rs9824112 | 0.033 | 1.646 | 0.296 | 1.238 | -0.001 | 0.043 | 0.043 | 0 | 0.44 | 0.27 | 0.56 | 0.63 | 0.45 | 0.38 |
| 3 | 22600482 | rs9824519 | 0.918 | 1.636 | 0.309 | 1.26 | 0.015 | -0.021 | -0.021 | 0 | 0.33 | 0.26 | 0.42 | 0.51 | 0.27 | 0.34 |
| 3 | 22602018 | rs4303885 | -1.142 | 1.824 | 0.637 | 1.287 | 0.015 | 0.096 | 0.096 | 0.904 | 0.07 | 0.27 | 0.06 | 0.16 | 0.04 | 0.04 |
| 3 | 22608487 | rs9867308 | -1.055 | 1.954 | 0.507 | 1.281 | 0.035 | 0.054 | 0.054 | 0 | 0.21 | 0.35 | 0.29 | 0.34 | 0.27 | 0.26 |
| 3 | 22618845 | rs9310698 | 0.561 | 1.937 | 0.216 | 1.15 | -0.045 | -0.009 | -0.009 | 0.026 | 0.11 | 0.02 | 0.14 | 0.13 | 0.17 | 0.04 |
| 3 | 22623495 | rs6809842 | -1.341 | 1.865 | 0.446 | 1.284 | 0.037 | 0.111 | 0.111 | 0 | 0.13 | 0.39 | 0.15 | 0.25 | 0.14 | 0.16 |
| 3 | 22623604 | rs1531160 | -0.411 | 1.924 | 0.29 | 1.575 | -0.019 | -0.033 | -0.033 | 0.014 | 0.9 | 0.98 | 0.85 | 0.86 | 0.86 | 0.89 |
| 3 | 22626907 | rs1579984 | 0.425 | 1.84 | 0.353 | 1.597 | -0.001 | 0.018 | 0.018 | 0 | 0.86 | 0.85 | 0.83 | 0.72 | 0.85 | 0.88 |
| 3 | 22627609 | rs13070085 | -0.739 | 1.982 | 0.417 | 0.963 | 0.075 | 0.025 | 0.025 | 0 | 0.1 | 0.33 | 0.14 | 0.08 | 0.12 | 0.12 |
| 3 | 22628123 | rs6778150 | 0.064 | 1.728 | 0.593 | 1.124 | 0.019 | -0.001 | -0.001 | 0 | 0.12 | 0.24 | 0.16 | 0.14 | 0.14 | 0.04 |
| 3 | 22631593 | rs9829768 | 0.916 | 1.85 | 0.051 | 1.274 | 0.035 | 0.136 | 0.136 | 0 | 0.78 | 0.53 | 0.71 | 0.63 | 0.74 | 0.72 |
| 3 | 22632753 | rs6550703 | 0.474 | 1.568 | -0.6 | 1.167 | -0.001 | 0 | 0 | 0 | 0.73 | 0.71 | 0.65 | 0.55 | 0.67 | 0.71 |
| 3 | 22634352 | rs11921888 | 0.187 | 0.877 | -0.98 | 1.312 | -0.001 | 0 | 0 | 0.026 | 0.29 | 0.29 | 0.34 | 0.41 | 0.24 | 0.23 |
| 3 | 22644920 | rs1870480 | 0.137 | 1.067 | -1.11 | 0.676 | 0.032 | 0.064 | 0.064 | 0 | 0.5 | 0.65 | 0.42 | 0.31 | 0.57 | 0.55 |
| 3 | 22646735 | rs9882017 | -0.078 | 1.213 | -1.571 | 0.885 | -0.023 | 0.06 | 0.06 | 0 | 0.54 | 0.84 | 0.5 | 0.67 | 0.53 | 0.61 |

| | | | | | | | | | | | | | | | | |
|---|---|---|---|---|---|---|---|---|---|---|---|---|---|---|---|---|
| 3 | 22648091 | rs1349282 | 0.092 | 1.074 | -1.444 | 0.867 | 0.053 | 0.052 | 0.052 | 0.515 | 0.46 | 0.78 | 0.4 | 0.33 | 0.41 | 0.55 |
| 3 | 22648120 | rs1349283 | 0.205 | 0.922 | -1.318 | 0.905 | -0.001 | 0.003 | 0.003 | 0.751 | 0.59 | 0.52 | 0.63 | 0.52 | 0.49 | 0.53 |
| 3 | 22649910 | rs9310699 | -0.293 | 1.005 | -1.916 | 0.657 | -0.082 | -0.053 | -0.053 | 0 | 0.65 | 0.92 | 0.77 | 0.67 | 0.62 | 0.72 |
| 3 | 22675098 | rs1459522 | 0.241 | 0.63 | -1.844 | 0.541 | 0.012 | 0.13 | 0.13 | 0 | 0.45 | 0.78 | 0.41 | 0.24 | 0.47 | 0.52 |
| 3 | 22686512 | rs13096681 | 0.701 | 0.909 | -1.193 | 0.481 | 0.05 | 0.001 | 0.001 | 0.116 | 0.08 | 0.18 | 0.09 | 0.09 | 0.16 | 0.15 |
| 3 | 22694778 | rs6550706 | -0.684 | 0.773 | -1.372 | 0.626 | -0.079 | -0.057 | -0.057 | 1.188 | 0.66 | 0.93 | 0.77 | 0.67 | 0.62 | 0.73 |
| 3 | 22709578 | rs4858046 | -0.53 | 0.558 | -0.571 | 0.506 | -0.005 | -0.013 | -0.013 | 0 | 0.63 | 0.67 | 0.67 | 0.6 | 0.62 | 0.68 |
| 3 | 22717876 | rs4858429 | 1.443 | 0.411 | -0.699 | 0.445 | 0 | -0.024 | -0.024 | 0 | 0.13 | 0.06 | 0.33 | 0.42 | 0.09 | 0.17 |
| 3 | 22721090 | rs9877823 | -0.116 | 0.45 | -0.581 | 0.365 | -0.006 | -0.002 | -0.002 | 0 | 0.56 | 0.57 | 0.63 | 0.59 | 0.55 | 0.64 |
| 3 | 22722267 | rs2358930 | -0.098 | -0.189 | -0.744 | -0.15 | -0.005 | -0.009 | -0.009 | 0 | 0.41 | 0.34 | 0.53 | 0.56 | 0.42 | 0.61 |
| 3 | 22748006 | rs9847981 | -0.919 | -0.205 | -0.627 | -0.107 | -0.017 | 0.005 | 0.005 | 0.692 | 0.61 | 0.69 | 0.72 | 0.69 | 0.56 | 0.69 |
| 3 | 22756424 | rs1380366 | 0.261 | -0.192 | -0.719 | -0.119 | 0.006 | -0.01 | -0.01 | 0 | 0.34 | 0.3 | 0.45 | 0.49 | 0.26 | 0.46 |
| 3 | 22761145 | rs1471234 | 0.872 | -0.135 | -0.827 | -0.206 | -0.005 | 0.011 | 0.011 | 0 | 0.22 | 0.15 | 0.17 | 0.13 | 0.29 | 0.18 |
| 3 | 22766469 | rs1402366 | 1.884 | -0.378 | -0.718 | -0.23 | -0.01 | -0.032 | -0.032 | 0.023 | 0.14 | 0.14 | 0.1 | 0.03 | 0.13 | 0.03 |
| 3 | 22769681 | rs17012008 | 1.115 | -0.352 | -0.863 | -0.182 | 0.022 | 0.065 | 0.065 | 0.253 | 0.17 | 0.05 | 0.09 | 0.06 | 0.15 | 0.28 |
| 3 | 22804837 | rs909017 | -0.925 | 0.1 | -1.276 | -0.162 | -0.089 | -0.121 | -0.121 | 0 | 0.77 | 0.98 | 0.81 | 0.67 | 0.68 | 0.76 |
| 3 | 22815776 | rs2167175 | 2.286 | -0.19 | -0.799 | -0.373 | -0.008 | -0.04 | -0.04 | 3.598 | 0.08 | 0.11 | 0.06 | 0 | 0.08 | 0.02 |
| 3 | 22822569 | rs296165 | -0.192 | -0.295 | -0.668 | -0.037 | -0.011 | 0.049 | 0.049 | 0.104 | 0.84 | 0.66 | 0.8 | 0.76 | 0.86 | 0.86 |
| 3 | 22827129 | rs644642 | 0.751 | -0.294 | -0.771 | -0.24 | -0.001 | -0.002 | -0.002 | 0 | 0.53 | 0.41 | 0.56 | 0.47 | 0.48 | 0.53 |
| 3 | 22828137 | rs4858453 | 1.57 | -0.318 | -0.626 | -0.198 | -0.038 | 0.014 | 0.014 | 1.335 | 0.07 | 0.25 | 0.12 | 0.07 | 0 | 0.04 |
| 3 | 22833287 | rs296160 | -0.327 | -0.252 | -0.605 | -0.152 | -0.008 | 0.061 | 0.061 | 0.067 | 0.84 | 0.65 | 0.79 | 0.74 | 0.86 | 0.83 |
| 3 | 22833705 | rs296158 | -1.31 | -0.145 | -0.745 | -0.131 | 0.003 | 0.019 | 0.019 | 0 | 0.91 | 0.9 | 0.92 | 0.81 | 0.86 | 0.88 |
| 3 | 22840407 | rs33115 | 1.492 | 0.07 | -1.048 | -0.365 | -0.05 | -0.011 | -0.011 | 0.248 | 0.19 | 0.03 | 0.13 | 0.15 | 0.21 | 0.12 |
| 3 | 22841478 | rs978239 | -0.343 | -0.012 | -1.004 | -0.389 | -0.023 | -0.032 | -0.032 | 0 | 0.69 | 0.83 | 0.78 | 0.65 | 0.64 | 0.74 |
| 3 | 22841826 | rs33117 | 0.754 | -0.102 | -0.983 | -0.478 | -0.002 | -0.001 | -0.001 | 7.92 | 0.51 | 0.43 | 0.55 | 0.5 | 0.46 | 0.51 |
| 3 | 22858824 | rs4858052 | 1.571 | -0.861 | -1.083 | 0.174 | 0.009 | -0.009 | -0.009 | 0.001 | 0.11 | 0.1 | 0.3 | 0.29 | 0.04 | 0.12 |
| 3 | 22865347 | rs33119 | 1.514 | -0.773 | -1.34 | 0.003 | -0.028 | -0.017 | -0.017 | 0 | 0.19 | 0.04 | 0.12 | 0.16 | 0.19 | 0.13 |
| 3 | 22867428 | rs39637 | 1.757 | -1.029 | -1.263 | -0.172 | 0.003 | 0.068 | 0.068 | 0.175 | 0.34 | 0.14 | 0.2 | 0.18 | 0.35 | 0.4 |
| 3 | 22874584 | rs294790 | 1.038 | -0.757 | -1.055 | 0.05 | -0.012 | 0.079 | 0.079 | 0 | 0.52 | 0.22 | 0.32 | 0.33 | 0.54 | 0.49 |
| 3 | 22875581 | rs13100945 | 1.033 | -0.5 | -0.906 | 0.347 | 0.01 | 0.003 | 0.003 | 1.424 | 0.14 | 0.17 | 0.14 | 0.16 | 0.2 | 0.26 |
| 3 | 22884739 | rs3911122 | 0.948 | -0.449 | -0.861 | 0.433 | -0.007 | 0.033 | 0.033 | 0 | 0.18 | 0.25 | 0.37 | 0.3 | 0.14 | 0.14 |
| 3 | 22889452 | rs620918 | -0.52 | -0.373 | -1.299 | 0.115 | 0.001 | 0.01 | 0.01 | 0 | 0.48 | 0.54 | 0.64 | 0.68 | 0.42 | 0.53 |
| 3 | 22900401 | rs298419 | -1.271 | 0.336 | -1.674 | 0.473 | 0.02 | 0.025 | 0.025 | 0 | 0.93 | 0.91 | 0.94 | 0.85 | 0.87 | 0.94 |
| 3 | 22901845 | rs192722 | 1.141 | -0.093 | -2.16 | 0.41 | -0.02 | -0.01 | -0.01 | 0 | 0.17 | 0.05 | 0.1 | 0.14 | 0.17 | 0.11 |
| 3 | 22905895 | rs299070 | 0.302 | -0.241 | -2.017 | 0.139 | -0.007 | 0.107 | 0.107 | 0.357 | 0.4 | 0.71 | 0.31 | 0.18 | 0.43 | 0.39 |
| 3 | 22910437 | rs184174 | -1.222 | 0.723 | -1.81 | 0.752 | -0.066 | -0.066 | -0.066 | 0 | 0.89 | 0.99 | 0.9 | 0.76 | 0.79 | 0.9 |
| 3 | 22916401 | rs9876180 | 1.307 | -0.793 | -0.94 | 0.634 | 0.039 | 0.053 | 0.053 | 0.353 | 0.11 | 0.46 | 0.11 | 0.11 | 0.11 | 0.06 |
| 3 | 22920093 | rs11718139 | 0.942 | -0.856 | -1.373 | 0.899 | 0.012 | 0.013 | 0.013 | 0.056 | 0.1 | 0.16 | 0.15 | 0.15 | 0.15 | 0.22 |
| 3 | 22920246 | rs3911124 | 1.162 | -0.864 | -1.385 | 0.852 | 0.009 | 0.017 | 0.017 | 1.032 | 0.12 | 0.54 | 0.12 | 0.11 | 0.12 | 0.06 |
| 3 | 22929986 | rs637382 | 0.573 | -1.738 | -2.297 | 0.68 | 0.08 | -0.042 | -0.042 | 0 | 0.38 | 0.06 | 0.34 | 0.36 | 0.21 | 0.3 |
| 3 | 22931734 | rs580227 | -1.743 | -1.767 | -2.359 | 0.664 | -0.161 | -0.018 | -0.018 | 1.238 | 0.7 | 0.96 | 0.72 | 0.69 | 0.49 | 0.58 |
| 3 | 22939427 | rs6795012 | 0.59 | -1.904 | -2.185 | 0.434 | -0.013 | -0.018 | -0.018 | 0.279 | 0.16 | 0.67 | 0.13 | 0.11 | 0.14 | 0.08 |
| 3 | 22946172 | rs698224 | 0.697 | -1.999 | -2.335 | 0.691 | 0.075 | -0.025 | -0.025 | 6.413 | 0.36 | 0.06 | 0.33 | 0.31 | 0.19 | 0.29 |
| 3 | 22946545 | rs696500 | -0.609 | -1.785 | -2.183 | 0.722 | 0.057 | -0.02 | -0.02 | 0 | 0.54 | 0.69 | 0.54 | 0.49 | 0.32 | 0.41 |
| 3 | 22963690 | rs13058941 | 0.386 | -1.469 | -1.766 | 0.694 | 0.009 | 0.005 | 0.005 | 7.709 | 0.14 | 0.2 | 0.13 | 0.15 | 0.16 | 0.21 |
| 3 | 22964321 | rs6794105 | -0.547 | -1.78 | -1.955 | 0.662 | 0.05 | 0.002 | 0.002 | 0.19 | 0.5 | 0.33 | 0.46 | 0.47 | 0.36 | 0.5 |
| 3 | 22964868 | rs1708144 | -0.611 | -1.796 | -1.958 | 0.659 | 0.044 | -0.001 | -0.001 | 0.43 | 0.51 | 0.33 | 0.46 | 0.47 | 0.36 | 0.51 |
| 3 | 22966463 | rs1606478 | -0.948 | -1.865 | -1.993 | 0.574 | 0.019 | 0.002 | 0.002 | 0 | 0.55 | 0.34 | 0.55 | 0.53 | 0.36 | 0.56 |
| 3 | 22983152 | rs711704 | 0.04 | -2.701 | -1.829 | 0.972 | 0.096 | 0.095 | 0.095 | 0 | 0.37 | 0.09 | 0.33 | 0.24 | 0.2 | 0.3 |
| 3 | 22984643 | rs711705 | 1.366 | -3.02 | -2.046 | 0.82 | -0.016 | -0.003 | -0.003 | 0 | 0.13 | 0.02 | 0.08 | 0.08 | 0.12 | 0.12 |
| 3 | 22989819 | rs822780 | -0.685 | -2.924 | -1.779 | 0.941 | 0.13 | 0.051 | 0.051 | 1.955 | 0.47 | 0.23 | 0.4 | 0.39 | 0.26 | 0.49 |
| 3 | 22992191 | rs1824015 | -0.576 | -2.966 | -1.233 | 1.024 | -0.047 | 0.149 | 0.149 | 5.152 | 0.15 | 0.35 | 0.19 | 0.29 | 0.08 | 0.22 |
| 3 | 22994314 | rs822787 | -0.315 | -2.92 | -1.299 | 0.898 | 0.09 | -0.007 | -0.007 | 0 | 0.41 | 0.21 | 0.44 | 0.38 | 0.21 | 0.33 |
| 3 | 22997566 | rs711708 | -0.318 | -2.874 | -1.295 | 0.978 | 0.098 | -0.016 | -0.016 | 0 | 0.4 | 0.17 | 0.43 | 0.39 | 0.2 | 0.35 |
| 3 | 22998569 | rs711710 | -1.031 | -2.834 | -1.215 | 0.997 | 0.017 | 0.005 | 0.005 | 0.856 | 0.51 | 0.47 | 0.52 | 0.53 | 0.28 | 0.51 |
| 3 | 23002974 | rs822204 | 0.387 | -2.712 | -0.827 | 1.058 | -0.009 | 0.012 | 0.012 | 0 | 0.22 | 0.24 | 0.42 | 0.22 | 0.16 | 0.19 |
| 3 | 23003050 | rs822793 | -0.74 | -2.803 | -0.913 | 0.977 | -0.035 | 0.05 | 0.05 | 0 | 0.34 | 0.46 | 0.49 | 0.38 | 0.23 | 0.37 |
| 3 | 23005150 | rs711714 | 0.768 | -2.676 | -1.05 | 0.964 | 0.007 | 0.043 | 0.043 | 0 | 0.2 | 0.08 | 0.3 | 0.07 | 0.15 | 0.15 |
| 3 | 23010390 | rs711715 | -0.311 | -2.656 | -0.917 | 0.996 | 0.003 | -0.004 | -0.004 | 0 | 0.32 | 0.27 | 0.49 | 0.38 | 0.23 | 0.37 |
| 3 | 23010878 | rs6806064 | 0.618 | -2.785 | -0.253 | 0.729 | -0.027 | -0.046 | -0.046 | 0 | 0.18 | 0.39 | 0.1 | 0.13 | 0.14 | 0.08 |
| 3 | 23011352 | rs711718 | -0.959 | -2.285 | -0.231 | 0.772 | 0.029 | 0.013 | 0.013 | 0 | 0.52 | 0.67 | 0.61 | 0.51 | 0.41 | 0.46 |
| 3 | 23014385 | rs822202 | -0.125 | -2.288 | -0.432 | 0.741 | 0.033 | 0.015 | 0.015 | 0 | 0.35 | 0.24 | 0.38 | 0.26 | 0.22 | 0.28 |
| 3 | 23024216 | rs535689 | -1.568 | -1.136 | -0.012 | 0.9 | 0.013 | 0.016 | 0.016 | 0 | 0.75 | 0.76 | 0.75 | 0.58 | 0.64 | 0.75 |
| 3 | 23026888 | rs680930 | -0.9 | -1.241 | 0.191 | 1.097 | 0.021 | 0.043 | 0.043 | 0.324 | 0.69 | 0.6 | 0.63 | 0.46 | 0.59 | 0.66 |
| 3 | 23030914 | rs603056 | 0.71 | -1.595 | -0.083 | 0.609 | -0.033 | 0.093 | 0.093 | 0 | 0.33 | 0.18 | 0.43 | 0.04 | 0.34 | 0.38 |
| 3 | 23031200 | rs529219 | -1.747 | -1.271 | 0.073 | 0.979 | 0.017 | 0.019 | 0.019 | 0 | 0.75 | 0.73 | 0.75 | 0.6 | 0.65 | 0.75 |
| 3 | 23031894 | rs822774 | -0.133 | -1.625 | -0.005 | 0.65 | -0.004 | 0.048 | 0.048 | 4.324 | 0.57 | 0.33 | 0.54 | 0.28 | 0.42 | 0.6 |
| 3 | 23032225 | rs680110 | -0.323 | -1.792 | 0 | 0.587 | 0.001 | 0.004 | 0.004 | 0 | 0.49 | 0.61 | 0.56 | 0.25 | 0.5 | 0.48 |
| 3 | 23032988 | rs822770 | -1.004 | -1.539 | 0.074 | 0.846 | 0.015 | 0.029 | 0.029 | 0.676 | 0.69 | 0.52 | 0.62 | 0.47 | 0.59 | 0.71 |

| | | | | | | | | | | | | | | | | |
|---|---|---|---|---|---|---|---|---|---|---|---|---|---|---|---|---|
| 3 | 23041286 | rs6782300 | 0.069 | -1.558 | 0.145 | 0.669 | 0.019 | 0.027 | 0.027 | 0.729 | 0.18 | 0.44 | 0.13 | 0.2 | 0.17 | 0.12 |
| 3 | 23048040 | rs633464 | 1.697 | -1.467 | -0.048 | 0.707 | 0.016 | 0.017 | 0.017 | 0 | 0.24 | 0.27 | 0.26 | 0.41 | 0.37 | 0.21 |
| 3 | 23054508 | rs537660 | 1.921 | -1.489 | -0.252 | 0.704 | 0.004 | 0.003 | 0.003 | 0 | 0.23 | 0.21 | 0.23 | 0.37 | 0.36 | 0.2 |
| 3 | 23078311 | rs1817582 | 0.301 | -2.1 | -0.327 | 0.678 | -0.044 | -0.016 | -0.016 | 0 | 0.28 | 0.32 | 0.13 | 0.18 | 0.1 | 0.15 |
| 3 | 23081138 | rs4858469 | 1.401 | -2.05 | -0.79 | 0.748 | 0.073 | -0.008 | -0.008 | 0 | 0.22 | 0.08 | 0.09 | 0.22 | 0.08 | 0.15 |
| 3 | 23090051 | rs2359367 | -0.594 | -1.894 | -0.74 | 0.494 | 0.002 | -0.033 | -0.033 | 0 | 0.66 | 0.7 | 0.56 | 0.87 | 0.65 | 0.5 |
| 3 | 23093391 | rs10510535 | 2.021 | -1.821 | -1.682 | 0.331 | -0.022 | 0.079 | 0.079 | 0 | 0.13 | 0 | 0.26 | 0.01 | 0.1 | 0.13 |
| 3 | 23093574 | rs9310709 | 0.641 | -1.622 | -1.688 | 0.697 | -0.059 | -0.015 | -0.015 | 1.076 | 0.4 | 0.16 | 0.61 | 0.44 | 0.49 | 0.5 |
| 3 | 23097075 | rs861983 | 0.707 | -1.596 | -1.793 | 0.139 | -0.067 | 0.146 | 0.146 | 1.078 | 0.27 | 0.06 | 0.39 | 0.03 | 0.31 | 0.39 |
| 3 | 23107083 | rs13433892 | 1.548 | -1.033 | -1.72 | 0.456 | -0.019 | -0.001 | -0.001 | 0 | 0.31 | 0.25 | 0.27 | 0.35 | 0.44 | 0.21 |
| 3 | 23118908 | rs1485384 | 0.066 | -0.729 | -2.125 | 0.426 | 0.017 | -0.003 | -0.003 | 0 | 0.66 | 0.67 | 0.82 | 0.61 | 0.81 | 0.74 |
| 3 | 23120060 | rs9826441 | 0.082 | -0.539 | -2.84 | 0.061 | -0.083 | 0.015 | 0.015 | 0 | 0.7 | 0.97 | 0.78 | 0.81 | 0.67 | 0.81 |
| 3 | 23121967 | rs1385545 | -0.611 | -0.751 | -2.091 | 0.723 | 0.054 | 0.017 | 0.017 | 1.047 | 0.76 | 0.87 | 0.87 | 0.81 | 0.87 | 0.83 |
| 3 | 23125668 | rs9310710 | -0.1 | -0.309 | -2.491 | 0.93 | -0.011 | -0.102 | -0.102 | 0 | 0.6 | 0.96 | 0.7 | 0.52 | 0.67 | 0.65 |
| 3 | 23126649 | rs4467482 | -0.123 | -0.331 | -2.55 | 0.88 | 0.003 | -0.114 | -0.114 | 0 | 0.62 | 0.96 | 0.71 | 0.52 | 0.69 | 0.65 |
| 3 | 23139179 | rs6800955 | -0.23 | -0.35 | -2.525 | 1.034 | 0.025 | -0.135 | -0.135 | 0 | 0.58 | 0.95 | 0.71 | 0.56 | 0.67 | 0.71 |
| 3 | 23143047 | rs4591517 | 0.364 | 0.118 | -2.798 | 0.727 | -0.015 | 0.054 | 0.054 | 0 | 0.31 | 0.81 | 0.47 | 0.2 | 0.28 | 0.46 |
| 3 | 23143461 | rs7641032 | 1.246 | 0.298 | -2.801 | 1.125 | -0.119 | -0.165 | -0.165 | 0.117 | 0.24 | 0.03 | 0.33 | 0.49 | 0.38 | 0.24 |
| 3 | 23146772 | rs6807204 | 1.175 | 0.209 | -2.249 | 1.037 | -0.022 | -0.023 | -0.023 | 0 | 0.25 | 0.2 | 0.24 | 0.38 | 0.34 | 0.21 |
| 3 | 23147227 | rs12638485 | 1.282 | 0.51 | -2.113 | 0.834 | -0.002 | -0.001 | -0.001 | 0 | 0.13 | 0.1 | 0.08 | 0.15 | 0.16 | 0.13 |
| 3 | 23149728 | rs6550725 | 0.301 | 1.029 | -2.171 | 0.646 | -0.011 | -0.004 | -0.004 | 0 | 0.37 | 0.36 | 0.45 | 0.64 | 0.51 | 0.39 |
| 3 | 23153062 | rs7618603 | 0.484 | 0.991 | -1.642 | 0.866 | 0.001 | 0.045 | 0.045 | 0.415 | 0.18 | 0.5 | 0.35 | 0.18 | 0.13 | 0.45 |
| 3 | 23157067 | rs3922415 | 1.485 | 1.119 | -1.793 | 0.923 | -0.02 | -0.018 | -0.018 | 0 | 0.21 | 0.18 | 0.25 | 0.36 | 0.34 | 0.22 |
| 3 | 23158915 | rs7634983 | 1.836 | 0.841 | -1.91 | 0.986 | -0.062 | -0.075 | -0.075 | 0 | 0.19 | 0.05 | 0.2 | 0.36 | 0.3 | 0.22 |
| 3 | 23162240 | rs4390954 | 0.371 | 0.628 | -1.62 | 0.941 | -0.001 | -0.001 | -0.001 | 0.203 | 0.38 | 0.58 | 0.56 | 0.54 | 0.43 | 0.61 |
| 3 | 23175867 | rs4485741 | -0.591 | 0.427 | -2.099 | 0.815 | -0.02 | 0.074 | 0.074 | 8.371 | 0.55 | 0.89 | 0.67 | 0.68 | 0.58 | 0.74 |
| 3 | 23181557 | rs9825994 | 1.563 | 1.091 | -2.05 | 1.025 | -0.071 | -0.059 | -0.059 | 0 | 0.23 | 0.05 | 0.5 | 0.36 | 0.34 | 0.26 |
| 3 | 23186581 | rs7649830 | -0.556 | 0.924 | -1.228 | 1.315 | -0.023 | 0.13 | 0.13 | 0.767 | 0.12 | 0.62 | 0.17 | 0.23 | 0.09 | 0.26 |
| 3 | 23187407 | rs4422335 | -0.401 | 1.034 | -0.696 | 1.223 | 0.003 | 0.148 | 0.148 | 0.397 | 0.13 | 0.45 | 0.09 | 0.17 | 0.11 | 0.32 |
| 3 | 23187727 | rs4547724 | -0.613 | 0.681 | -1.433 | 0.875 | -0.007 | -0.09 | -0.09 | 0 | 0.39 | 0.85 | 0.3 | 0.45 | 0.34 | 0.48 |
| 3 | 23188043 | rs9879958 | 1.973 | 0.828 | -1.624 | 0.899 | -0.1 | -0.091 | -0.091 | 0.003 | 0.23 | 0.03 | 0.49 | 0.36 | 0.34 | 0.27 |
| 3 | 23193928 | rs4619807 | -0.777 | 0.674 | -1.566 | 0.731 | -0.051 | 0.089 | 0.089 | 1.494 | 0.62 | 0.98 | 0.8 | 0.81 | 0.68 | 0.75 |
| 3 | 23198484 | rs6780569 | -0.679 | 0.59 | -1.188 | 0.956 | -0.018 | 0.085 | 0.085 | 0.415 | 0.14 | 0.62 | 0.11 | 0.18 | 0.12 | 0.32 |
| 3 | 23203454 | rs7374732 | -0.74 | 0.68 | -1.417 | 0.924 | -0.037 | 0.001 | 0.001 | 1.369 | 0.61 | 0.94 | 0.72 | 0.67 | 0.65 | 0.72 |
| 3 | 23203734 | rs9867201 | 0.017 | 0.741 | -0.388 | 0.984 | 0.007 | 0.024 | 0.024 | 0 | 0.37 | 0.51 | 0.59 | 0.53 | 0.46 | 0.58 |
| 3 | 23206871 | rs9827762 | -0.818 | 0.621 | -0.249 | 1.024 | 0.016 | 0.266 | 0.266 | 0 | 0.14 | 0.49 | 0.18 | 0.32 | 0.14 | 0.35 |
| 3 | 23210370 | rs6550734 | 0.332 | 0.943 | -0.086 | 0.809 | 0 | 0.001 | 0.001 | 0 | 0.33 | 0.43 | 0.6 | 0.67 | 0.39 | 0.6 |
| 3 | 23214269 | rs6799137 | 0.566 | 0.988 | -0.459 | 0.757 | -0.012 | 0.002 | 0.002 | 0.063 | 0.26 | 0.25 | 0.24 | 0.26 | 0.31 | 0.17 |
| 3 | 23216324 | rs9831921 | -0.502 | 0.937 | -0.384 | 0.946 | -0.011 | 0.148 | 0.148 | 0.089 | 0.12 | 0.61 | 0.09 | 0.2 | 0.1 | 0.26 |
| 3 | 23217338 | rs3922891 | -2.353 | 0.974 | -0.52 | 0.886 | -0.05 | -0.067 | -0.067 | 0.839 | 0.78 | 0.98 | 0.55 | 0.67 | 0.74 | 0.72 |
| 3 | 23218126 | rs9842300 | 0.744 | 0.966 | -0.421 | 0.764 | -0.014 | 0.02 | 0.02 | 0.041 | 0.25 | 0.21 | 0.16 | 0.14 | 0.28 | 0.14 |
| 3 | 23221931 | rs11710420 | 0.588 | 0.397 | -0.766 | 0.504 | -0.019 | 0.059 | 0.059 | 2.142 | 0.33 | 0.09 | 0.2 | 0.2 | 0.29 | 0.26 |
| 3 | 23226298 | rs6550742 | 0.93 | 0.424 | -0.672 | 0.454 | 0.008 | 0.048 | 0.048 | 0 | 0.3 | 0.21 | 0.13 | 0.15 | 0.23 | 0.14 |
| 3 | 23234618 | rs3732395 | -0.069 | 0.277 | -0.484 | 0.634 | 0.003 | -0.037 | -0.037 | 0 | 0.42 | 0.2 | 0.24 | 0.39 | 0.38 | 0.54 |
| 3 | 23235424 | rs17012800 | -0.293 | 0.368 | -0.042 | 0.588 | -0.009 | 0.04 | 0.04 | 0 | 0.12 | 0.14 | 0.05 | 0.2 | 0.11 | 0.28 |
| 3 | 23249069 | rs4858483 | -1.617 | 0.786 | -0.393 | 0.383 | -0.018 | -0.019 | -0.019 | 0 | 0.69 | 0.88 | 0.48 | 0.65 | 0.66 | 0.68 |
| 3 | 23260467 | rs7617072 | -1.18 | 0.97 | 0.068 | 0.336 | -0.004 | 0.004 | 0.004 | 1.831 | 0.65 | 0.71 | 0.45 | 0.65 | 0.62 | 0.67 |
| 3 | 23261334 | rs17012829 | 0.322 | 0.93 | 0.068 | 0.41 | -0.005 | 0.067 | 0.067 | 0 | 0.09 | 0.13 | 0.05 | 0.19 | 0.09 | 0.24 |
| 3 | 23267063 | rs2132149 | 0.436 | 1.003 | 0.136 | 0.338 | -0.02 | -0.002 | -0.002 | 0 | 0.25 | 0.14 | 0.25 | 0.29 | 0.29 | 0.25 |
| 3 | 23280826 | rs2359637 | 0.038 | 0.989 | 0.252 | 0.336 | 0 | -0.006 | -0.006 | 0 | 0.4 | 0.36 | 0.2 | 0.35 | 0.36 | 0.42 |
| 3 | 23289892 | rs6792370 | -0.616 | 0.998 | 0.09 | 0.218 | -0.001 | 0.012 | 0.012 | 0 | 0.18 | 0.21 | 0.1 | 0.19 | 0.17 | 0.31 |
| 3 | 23297579 | rs6550748 | -1.894 | 1.171 | 0.421 | 0.413 | -0.017 | -0.015 | -0.015 | 0.002 | 0.69 | 0.88 | 0.48 | 0.65 | 0.66 | 0.67 |
| 3 | 23301213 | rs6771206 | 0.151 | 1.178 | 0.385 | 0.41 | 0.006 | -0.008 | -0.008 | 0.224 | 0.38 | 0.33 | 0.18 | 0.35 | 0.33 | 0.42 |
| 3 | 23317542 | rs7630344 | 0.803 | 1.227 | 0.663 | 0.452 | -0.002 | -0.002 | -0.002 | 0 | 0.38 | 0.58 | 0.68 | 0.48 | 0.46 | 0.36 |
| 3 | 23320855 | rs2132147 | -0.11 | 0.299 | 1.088 | 0.719 | 0.006 | -0.007 | -0.007 | 0 | 0.73 | 0.67 | 0.74 | 0.72 | 0.71 | 0.75 |
| 3 | 23336450 | rs7612463 | 0.39 | 0.307 | 1.119 | 0.765 | 0.007 | 0.06 | 0.06 | 0 | 0.12 | 0.2 | 0.08 | 0.19 | 0.13 | 0.26 |
| 3 | 23373052 | rs4858503 | 0.559 | 0.15 | 1.132 | 0.729 | -0.021 | 0.019 | 0.019 | 0 | 0.64 | 0.5 | 0.74 | 0.45 | 0.69 | 0.56 |
| 3 | 23382812 | rs1874503 | -0.866 | 0.706 | 0.963 | 0.892 | 0.008 | 0.023 | 0.023 | 3.49 | 0.79 | 0.9 | 0.87 | 0.83 | 0.83 | 0.79 |
| 3 | 23390301 | rs1353322 | 0.398 | 0.791 | 0.998 | 0.889 | 0.003 | 0.001 | 0.001 | 0 | 0.42 | 0.52 | 0.69 | 0.48 | 0.54 | 0.42 |
| 3 | 23403652 | rs9825778 | 1.175 | 0.845 | 0.781 | 0.895 | -0.141 | -0.059 | -0.059 | 0 | 0.35 | 0.03 | 0.53 | 0.35 | 0.43 | 0.39 |
| 3 | 23410933 | rs6550760 | -0.828 | 0.869 | 0.839 | 0.906 | 0.01 | 0.023 | 0.023 | 3.347 | 0.78 | 0.89 | 0.87 | 0.82 | 0.83 | 0.79 |
| 3 | 23417884 | rs6796840 | 0.904 | 0.911 | 0.962 | 0.923 | -0.004 | 0.003 | 0.003 | 0 | 0.37 | 0.31 | 0.56 | 0.35 | 0.43 | 0.38 |
| 3 | 23440644 | rs778506 | -0.789 | 1.016 | 0.916 | 0.928 | 0.007 | -0.011 | -0.011 | 0 | 0.62 | 0.59 | 0.44 | 0.65 | 0.57 | 0.62 |
| 3 | 23453798 | rs6807788 | 0.348 | 0.991 | 0.881 | 0.883 | 0.009 | -0.006 | -0.006 | 0 | 0.38 | 0.36 | 0.17 | 0.35 | 0.29 | 0.38 |
| 3 | 23454790 | rs1496653 | -0.767 | 0.972 | 0.929 | 0.934 | 0.054 | 0.096 | 0.096 | 0 | 0.16 | 0.39 | 0.14 | 0.2 | 0.22 | 0.27 |
| 3 | 23466101 | rs778493 | 0.306 | 1.007 | 0.629 | 0.901 | -0.063 | -0.028 | -0.028 | 0 | 0.23 | 0.07 | 0.24 | 0.29 | 0.28 | 0.25 |
| 3 | 23471072 | rs4340647 | -0.475 | 0.682 | 0.766 | 0.758 | 0.009 | 0.169 | 0.169 | 0 | 0.15 | 0.36 | 0.1 | 0.27 | 0.13 | 0.27 |
| 3 | 23474611 | rs778502 | 0.797 | 0.678 | 0.703 | 0.819 | 0.004 | -0.002 | -0.002 | 0 | 0.21 | 0.18 | 0.14 | 0.25 | 0.17 | 0.22 |
| 3 | 23483536 | rs13062101 | 0.149 | 0.675 | 0.734 | 0.791 | 0.014 | 0.019 | 0.019 | 0.389 | 0.48 | 0.38 | 0.36 | 0.48 | 0.42 | 0.36 |

| | | | | | | | | | | | | | | | | |
|---|---|---|---|---|---|---|---|---|---|---|---|---|---|---|---|---|
| 3 | 23484827 | rs704533 | -0.903 | 0.699 | 0.713 | 0.727 | 0.005 | -0.012 | -0.012 | 0 | 0.63 | 0.63 | 0.44 | 0.71 | 0.57 | 0.64 |
| 3 | 23494353 | rs7632099 | 1.687 | 0.711 | 0.656 | 0.727 | 0.031 | 0.061 | 0.061 | 0 | 0.24 | 0.14 | 0.1 | 0.08 | 0.16 | 0.11 |
| 3 | 23494496 | rs7610906 | -0.114 | 0.727 | 0.811 | 0.785 | -0.006 | 0.003 | 0.003 | 0.117 | 0.42 | 0.55 | 0.23 | 0.37 | 0.36 | 0.39 |
| 3 | 23495258 | rs2359768 | -0.044 | 0.792 | 0.891 | 0.85 | 0.006 | 0.154 | 0.154 | 0 | 0.13 | 0.31 | 0.08 | 0.27 | 0.11 | 0.26 |
| 3 | 23505899 | rs778472 | -1.104 | 0.827 | 0.777 | 0.835 | 0.003 | -0.008 | -0.008 | 0 | 0.39 | 0.59 | 0.36 | 0.63 | 0.41 | 0.55 |
| 3 | 23511338 | rs999764 | -0.207 | 0.682 | 0.76 | 0.838 | -0.005 | 0.023 | 0.023 | 0.717 | 0.31 | 0.36 | 0.2 | 0.44 | 0.27 | 0.46 |
| 3 | 23521204 | rs9862734 | 0.652 | 0.928 | 0.748 | 0.965 | 0 | -0.003 | -0.003 | 0 | 0.23 | 0.22 | 0.23 | 0.36 | 0.21 | 0.24 |
| 3 | 23522785 | rs1695183 | 0.535 | 0.898 | 0.688 | 0.913 | -0.023 | -0.01 | -0.01 | 0 | 0.23 | 0.43 | 0.17 | 0.25 | 0.19 | 0.22 |
| 3 | 23525625 | rs778521 | -0.508 | 0.981 | 0.314 | 0.925 | -0.036 | 0.01 | 0.01 | 0 | 0.61 | 0.8 | 0.33 | 0.6 | 0.46 | 0.6 |
| 3 | 23527471 | rs7641437 | 1.151 | 1.065 | 0.272 | 1.015 | -0.003 | 0.072 | 0.072 | 0 | 0.08 | 0.12 | 0.04 | 0.19 | 0.07 | 0.25 |
| 3 | 23545630 | rs7613091 | -0.429 | 1.321 | 0.075 | 0.476 | -0.008 | 0.014 | 0.014 | 0 | 0.6 | 0.67 | 0.4 | 0.72 | 0.48 | 0.62 |
| 3 | 23549075 | rs2133331 | -0.465 | 1.307 | 0.097 | 0.475 | -0.001 | -0.001 | -0.001 | 0 | 0.59 | 0.62 | 0.39 | 0.72 | 0.48 | 0.62 |
| 3 | 23564092 | rs17013412 | 0.419 | 1.161 | -0.175 | 0.475 | -0.006 | 0.003 | 0.003 | 0 | 0.37 | 0.42 | 0.17 | 0.37 | 0.27 | 0.38 |
| 3 | 23590068 | rs17013457 | 0.331 | 0.975 | -0.181 | -0.058 | -0.019 | 0.01 | 0.01 | 0.394 | 0.38 | 0.46 | 0.23 | 0.48 | 0.25 | 0.39 |
| 3 | 23609689 | rs7640308 | -0.826 | 1.127 | -0.793 | -0.035 | -0.067 | 0.041 | 0.041 | 0 | 0.65 | 0.88 | 0.46 | 0.72 | 0.58 | 0.64 |
| 3 | 23615762 | rs7609845 | 0.204 | 1.097 | -1.143 | -0.019 | -0.012 | 0.008 | 0.008 | 0.765 | 0.4 | 0.53 | 0.28 | 0.48 | 0.34 | 0.41 |
| 3 | 23635228 | rs11706875 | 0.842 | 1.123 | -1.299 | 0.004 | -0.014 | -0.001 | -0.001 | 1.963 | 0.22 | 0.1 | 0.3 | 0.24 | 0.24 | 0.23 |
| 3 | 23635636 | rs7649176 | -0.394 | 1.688 | -1.299 | -0.422 | -0.054 | -0.063 | -0.063 | 0 | 0.89 | 0.98 | 0.77 | 0.75 | 0.84 | 0.76 |
| 3 | 23638627 | rs7648443 | -0.337 | 1.175 | -0.953 | -0.085 | 0.003 | -0.009 | -0.009 | 1.016 | 0.6 | 0.58 | 0.53 | 0.72 | 0.49 | 0.63 |
| 3 | 23642466 | rs6787720 | 0.115 | 0.597 | -1.88 | 0.354 | -0.003 | 0.003 | 0.003 | 0.029 | 0.4 | 0.8 | 0.5 | 0.61 | 0.4 | 0.52 |
| 3 | 23648116 | rs11711194 | 0.056 | 1.279 | -1.569 | 0.817 | -0.089 | 0.005 | 0.005 | 1.171 | 0.66 | 0.92 | 0.46 | 0.69 | 0.57 | 0.63 |
| 3 | 23655487 | rs4518067 | 0.715 | 0.582 | -0.219 | 1.291 | -0.024 | 0.045 | 0.045 | 0.358 | 0.71 | 0.5 | 0.72 | 0.59 | 0.74 | 0.64 |
| 3 | 23657017 | rs7624577 | -1.387 | 1.344 | -0.015 | 1.858 | 0.027 | 0.065 | 0.065 | 0.043 | 0.3 | 0.43 | 0.37 | 0.45 | 0.31 | 0.37 |
| 3 | 23658030 | rs6773562 | -0.088 | 1.59 | 0.031 | 1.878 | 0.021 | -0.013 | -0.013 | 0.351 | 0.08 | 0.13 | 0.14 | 0.09 | 0.12 | 0.08 |
| 3 | 23658489 | rs4264676 | -0.083 | 1.532 | 0.069 | 2.037 | -0.027 | 0.1 | 0.1 | 0.296 | 0.13 | 0.29 | 0.13 | 0.29 | 0.06 | 0.21 |
| 3 | 23664029 | rs10084688 | -1.228 | 1.435 | -0.212 | 1.32 | 0 | 0.005 | 0.005 | 0 | 0.53 | 0.52 | 0.54 | 0.61 | 0.53 | 0.51 |
| 3 | 23666667 | rs12485752 | 0.277 | 1.201 | -0.682 | 1.574 | -0.05 | -0.053 | -0.053 | 0.407 | 0.08 | 0 | 0.1 | 0.17 | 0.13 | 0.07 |
| 3 | 23670620 | rs7639174 | -0.465 | 1.001 | -0.421 | 1.091 | -0.003 | -0.004 | -0.004 | 0 | 0.45 | 0.44 | 0.4 | 0.52 | 0.4 | 0.44 |
| 3 | 23680618 | rs7642418 | -1.425 | 1.296 | 0.214 | 1.424 | 0.043 | 0.075 | 0.075 | 0 | 0.11 | 0.3 | 0.17 | 0.15 | 0.14 | 0.18 |
| 3 | 23683479 | rs12108186 | -1.399 | 1.417 | 0.132 | 1.566 | -0.003 | 0.011 | 0.011 | 0 | 0.13 | 0.24 | 0.16 | 0.16 | 0.1 | 0.11 |
| 3 | 23683778 | rs9884096 | 2.133 | 2.025 | -0.064 | 1.454 | 0 | -0.001 | -0.001 | 0 | 0.81 | 0.76 | 0.74 | 0.77 | 0.78 | 0.73 |
| 3 | 23684312 | rs6793612 | 1.261 | 1.876 | -0.025 | 1.562 | 0.013 | 0.007 | 0.007 | 0.25 | 0.82 | 0.83 | 0.81 | 0.72 | 0.71 | 0.55 |
| 3 | 23687843 | rs9867427 | 0.987 | 1.478 | -1.048 | 0.985 | -0.047 | 0.074 | 0.074 | 0.948 | 0.32 | 0.16 | 0.26 | 0.22 | 0.38 | 0.13 |
| 3 | 23689201 | rs4072410 | -0.14 | 1.078 | -1.46 | 1.09 | 0.001 | -0.08 | -0.08 | 0.339 | 0.54 | 0.91 | 0.54 | 0.52 | 0.4 | 0.8 |
| 3 | 23695943 | rs4858524 | 1.401 | 0.86 | -0.219 | 0.866 | 0.001 | 0.003 | 0.003 | 0 | 0.67 | 0.63 | 0.61 | 0.55 | 0.63 | 0.56 |
| 3 | 23696842 | rs6768481 | -1.519 | 0.287 | 0.386 | 0.241 | -0.001 | -0.056 | -0.056 | 0.009 | 0.25 | 0.4 | 0.22 | 0.14 | 0.21 | 0.11 |
| 3 | 23698252 | rs11928764 | -1.23 | -0.216 | 0.388 | 0.161 | -0.025 | -0.044 | -0.044 | 0.187 | 0.25 | 0.4 | 0.28 | 0.09 | 0.19 | 0.18 |
| 3 | 23705238 | rs9830571 | -1.248 | -0.42 | 0.525 | 0.326 | -0.033 | 0.003 | 0.003 | 10.961 | 0.16 | 0.35 | 0.14 | 0.14 | 0.1 | 0.26 |
| 3 | 23705671 | rs7638256 | -0.61 | -0.6 | 0.554 | 0.114 | -0.035 | -0.011 | -0.011 | 0 | 0.16 | 0.49 | 0.24 | 0.09 | 0.1 | 0.11 |
| 3 | 23706368 | rs6807584 | -1.626 | -0.517 | -0.099 | 0.154 | 0.07 | 0.109 | 0.109 | 2.532 | 0.31 | 0.84 | 0.31 | 0.09 | 0.2 | 0.32 |
| 3 | 23713820 | rs7646855 | -1.775 | -0.319 | 0.023 | 0.099 | 0.042 | 0.031 | 0.031 | 1.389 | 0.34 | 0.78 | 0.38 | 0.2 | 0.24 | 0.41 |
| 3 | 23720288 | rs4994252 | -1.871 | -0.53 | 0.675 | -0.144 | -0.009 | -0.016 | -0.016 | 0 | 0.23 | 0.67 | 0.21 | 0.13 | 0.18 | 0.28 |
| 3 | 23725613 | rs4396825 | -0.312 | -0.296 | -0.611 | -1.164 | 0.009 | 0.151 | 0.151 | 0 | 0.54 | 0.77 | 0.69 | 0.9 | 0.43 | 0.64 |
| 3 | 23733192 | rs7633774 | -1.274 | -0.13 | -0.716 | -1.02 | -0.064 | 0.124 | 0.124 | 0 | 0.67 | 0.86 | 0.75 | 0.89 | 0.56 | 0.86 |
| 3 | 23734098 | rs4858527 | -1.565 | -0.116 | -0.823 | -0.988 | -0.083 | 0.135 | 0.135 | 0 | 0.69 | 0.92 | 0.77 | 0.9 | 0.6 | 0.88 |
| 3 | 23735174 | rs9812786 | -0.93 | -0.294 | -0.921 | -0.706 | -0.068 | 0.131 | 0.131 | 0 | 0.53 | 0.92 | 0.57 | 0.73 | 0.45 | 0.74 |
| 3 | 23742624 | rs7643401 | -0.729 | -0.463 | 0.302 | -0.508 | 0.008 | -0.044 | -0.044 | 0.125 | 0.09 | 0.3 | 0.06 | 0.01 | 0.05 | 0.03 |
| 3 | 23743503 | rs17013665 | -0.897 | -0.703 | 0.097 | -0.729 | -0.003 | -0.005 | -0.005 | 0 | 0.31 | 0.37 | 0.27 | 0.28 | 0.28 | 0.43 |
| 3 | 23744086 | rs6550783 | -0.673 | -0.518 | -0.554 | -1.018 | -0.042 | 0.055 | 0.055 | 2.946 | 0.59 | 0.82 | 0.47 | 0.3 | 0.5 | 0.49 |
| 3 | 23744103 | rs7637788 | 0.052 | -0.413 | -0.301 | -0.657 | -0.012 | 0.001 | 0.001 | 0 | 0.16 | 0.19 | 0.16 | 0.2 | 0.1 | 0.17 |
| 3 | 23748920 | rs7620545 | -0.512 | -0.039 | -1.079 | -0.579 | -0.091 | -0.002 | -0.002 | 0.127 | 0.62 | 0.9 | 0.62 | 0.33 | 0.45 | 0.48 |
| 3 | 23750521 | rs1037633 | -0.281 | -0.156 | -0.386 | -0.626 | 0.054 | -0.015 | -0.015 | 2.892 | 0.07 | 0.3 | 0.07 | 0 | 0.08 | 0.04 |
| 3 | 23757541 | rs4858533 | 0.859 | 0.752 | -1.385 | 0.226 | -0.075 | -0.137 | -0.137 | 0.373 | 0.31 | 0.05 | 0.25 | 0.46 | 0.35 | 0.3 |
| 3 | 23759239 | rs6784676 | -0.01 | 1.205 | -0.823 | 0.297 | -0.011 | 0.024 | 0.024 | 0.063 | 0.44 | 0.3 | 0.35 | 0.65 | 0.52 | 0.63 |
| 3 | 23759937 | rs6550788 | -0.733 | 0.914 | -1.162 | 0.066 | -0.072 | -0.065 | -0.065 | 0 | 0.8 | 0.98 | 0.63 | 0.76 | 0.79 | 0.79 |
| 3 | 23762371 | rs2061185 | 0.408 | 0.759 | -0.933 | 0.078 | -0.001 | -0.005 | -0.005 | 0.576 | 0.64 | 0.36 | 0.48 | 0.74 | 0.69 | 0.72 |
| 3 | 23766551 | rs9867568 | 0.501 | 0.938 | -0.882 | 0.38 | 0.003 | -0.017 | -0.017 | 3.622 | 0.39 | 0.21 | 0.28 | 0.43 | 0.39 | 0.45 |
| 3 | 23768866 | rs751481 | 0.553 | 0.63 | -0.035 | 0.125 | -0.001 | -0.004 | -0.004 | 0 | 0.46 | 0.38 | 0.34 | 0.54 | 0.49 | 0.55 |
| 3 | 23770133 | rs4510335 | -0.326 | 0.328 | -0.078 | 0.257 | -0.003 | -0.006 | -0.006 | 0.015 | 0.8 | 0.84 | 0.67 | 0.72 | 0.8 | 0.81 |
| 3 | 23784303 | rs6767354 | 0.514 | -0.159 | 0.471 | -0.313 | -0.044 | -0.038 | -0.038 | 0 | 0.26 | 0.52 | 0.25 | 0.16 | 0.19 | 0.11 |
| 3 | 23785060 | rs7651576 | 0.209 | -0.022 | 0.426 | -0.3 | -0.036 | -0.045 | -0.045 | 0 | 0.29 | 0.45 | 0.28 | 0.17 | 0.21 | 0.12 |
| 3 | 23785920 | rs17013704 | 0.422 | 0.175 | 0.159 | -0.186 | -0.025 | -0.032 | -0.032 | 0 | 0.27 | 0.42 | 0.28 | 0.18 | 0.22 | 0.14 |
| 3 | 23793276 | rs1993399 | -0.365 | 0.409 | -0.379 | -0.101 | 0.002 | -0.01 | -0.01 | 0 | 0.8 | 0.81 | 0.81 | 0.89 | 0.77 | 0.81 |
| 3 | 23793622 | rs6780385 | 0.128 | 0.035 | -0.77 | 0.273 | 0.002 | -0.005 | -0.005 | 0 | 0.38 | 0.35 | 0.28 | 0.55 | 0.31 | 0.33 |
| 3 | 23796713 | rs12485431 | -0.694 | 0.618 | -1.386 | 0.246 | -0.001 | 0 | 0 | 0 | 0.14 | 0.16 | 0.26 | 0.12 | 0.11 | 0.41 |
| 3 | 23801209 | rs17754530 | -0.487 | 0.721 | -1.396 | 0.208 | 0.001 | -0.001 | -0.001 | 0 | 0.12 | 0.14 | 0.26 | 0.1 | 0.11 | 0.4 |
| 3 | 23802647 | rs3845923 | 0.067 | 0.197 | -1.645 | 0.146 | -0.007 | -0.057 | -0.057 | 0 | 0.33 | 0.19 | 0.29 | 0.63 | 0.33 | 0.3 |
| 3 | 23804744 | rs6791919 | 0.604 | -0.006 | -1.226 | -0.266 | 0.011 | 0.021 | 0.021 | 0.011 | 0.81 | 0.89 | 0.76 | 0.92 | 0.89 | 0.63 |
| 3 | 23806758 | rs17754631 | -0.472 | 0.279 | -0.576 | -0.393 | 0.031 | -0.03 | -0.03 | 0.464 | 0.22 | 0.32 | 0.26 | 0.15 | 0.27 | 0.14 |

| | | | | | | | | | | | | | | | | |
|---|---|---|---|---|---|---|---|---|---|---|---|---|---|---|---|---|
| 3 | 23807550 | rs12497066 | -0.119 | 0.259 | -0.661 | -0.435 | 0.004 | 0 | 0 | 0.185 | 0.14 | 0.09 | 0.2 | 0.15 | 0.2 | 0.1 |
| 3 | 23822088 | rs6765263 | -1.161 | -0.284 | -1.562 | -0.818 | -0.002 | 0.112 | 0.112 | 1.638 | 0.53 | 0.81 | 0.55 | 0.26 | 0.49 | 0.57 |
| 3 | 23825088 | rs4535170 | -1.327 | -0.106 | -1.176 | -0.479 | 0.015 | 0.039 | 0.039 | 5.451 | 0.38 | 0.8 | 0.47 | 0.26 | 0.34 | 0.56 |
| 3 | 23825204 | rs9861357 | -0.582 | 0.043 | -0.732 | -0.329 | -0.018 | -0.007 | -0.007 | 0 | 0.08 | 0.15 | 0.17 | 0.05 | 0.05 | 0.32 |
| 3 | 23839884 | rs9851068 | -1.054 | 0.322 | -0.831 | -0.507 | -0.005 | 0.036 | 0.036 | 2.676 | 0.32 | 0.7 | 0.42 | 0.1 | 0.31 | 0.51 |
| 3 | 23853293 | rs4858547 | -1.153 | -0.042 | -0.015 | -0.57 | -0.011 | -0.064 | -0.064 | 0.368 | 0.32 | 0.46 | 0.32 | 0.17 | 0.27 | 0.3 |
| 3 | 23860415 | rs1471060 | -0.955 | 0.349 | -0.814 | -0.141 | -0.042 | -0.096 | -0.096 | 0 | 0.76 | 0.89 | 0.82 | 0.5 | 0.71 | 0.75 |
| 3 | 23862053 | rs6775367 | 0.967 | 0.331 | -0.714 | -0.158 | -0.003 | -0.004 | -0.004 | 0.153 | 0.46 | 0.5 | 0.51 | 0.33 | 0.44 | 0.48 |
| 3 | 23869248 | rs2168424 | 0.258 | 0.26 | -0.49 | -0.293 | -0.009 | 0.004 | 0.004 | 0 | 0.09 | 0.15 | 0.1 | 0.11 | 0.05 | 0.12 |
| 3 | 23871730 | rs6765292 | 0.436 | 0.233 | -0.369 | -0.38 | -0.007 | 0.016 | 0.016 | 10.547 | 0.19 | 0.08 | 0.19 | 0.05 | 0.23 | 0.18 |
| 3 | 23873602 | rs1473379 | 0.838 | 0.134 | -0.3 | -0.047 | 0.002 | 0.007 | 0.007 | 0.414 | 0.46 | 0.55 | 0.54 | 0.28 | 0.45 | 0.43 |
| 3 | 23875970 | rs12629302 | 1.522 | 0.255 | -0.143 | -0.086 | 0.001 | 0.05 | 0.05 | 0.017 | 0.37 | 0.17 | 0.33 | 0.26 | 0.39 | 0.18 |
| 3 | 23900401 | rs6779573 | 0.023 | 0.655 | 0.331 | -0.112 | 0.012 | 0 | 0 | 0 | 0.65 | 0.67 | 0.62 | 0.36 | 0.52 | 0.6 |
| 3 | 23904214 | rs9859567 | 0.013 | 0.586 | 0.82 | -0.227 | 0.036 | -0.041 | -0.041 | 0 | 0.08 | 0.26 | 0.17 | 0 | 0.07 | 0.2 |
| 3 | 23907775 | rs17787217 | 1.264 | 0.255 | 1 | -0.447 | 0.027 | 0.07 | 0.07 | 0.242 | 0.48 | 0.29 | 0.42 | 0.32 | 0.43 | 0.34 |
| 3 | 23909658 | rs7651185 | -0.885 | 0.752 | 0.773 | -0.418 | -0.024 | -0.053 | -0.053 | 0 | 0.78 | 0.86 | 0.83 | 0.46 | 0.73 | 0.82 |
| 3 | 23921062 | rs4858552 | 0.702 | 0.851 | 1.059 | -0.295 | 0.002 | -0.001 | -0.001 | 0 | 0.63 | 0.39 | 0.61 | 0.38 | 0.62 | 0.52 |
| 3 | 23921114 | rs4858553 | -1.134 | 0.866 | 1.041 | -0.275 | -0.031 | -0.052 | -0.052 | 0 | 0.8 | 0.88 | 0.82 | 0.42 | 0.72 | 0.84 |
| 3 | 23934115 | rs6789983 | 0.768 | 0.866 | 1.239 | -0.37 | 0.019 | -0.031 | -0.031 | 0 | 0.13 | 0.14 | 0.19 | 0.05 | 0.2 | 0.16 |
| 3 | 23934772 | rs3762775 | -0.054 | 0.878 | 1.31 | -0.378 | 0.036 | -0.04 | -0.04 | 0.103 | 0.08 | 0.28 | 0.18 | 0 | 0.07 | 0.21 |
| 3 | 23935839 | rs6778970 | 0.277 | 0.873 | 1.295 | -0.367 | 0.025 | -0.007 | -0.007 | 0 | 0.15 | 0.25 | 0.21 | 0.09 | 0.2 | 0.22 |
| 3 | 23936964 | rs11718625 | 0.971 | 0.984 | 1.269 | -0.415 | 0.011 | -0.014 | -0.014 | 0 | 0.09 | 0.08 | 0.16 | 0.05 | 0.15 | 0.16 |
| 3 | 23937789 | rs9876561 | -0.043 | 0.961 | 1.245 | -0.394 | 0.016 | -0.047 | -0.047 | 0 | 0.15 | 0.19 | 0.19 | 0.05 | 0.2 | 0.17 |
| 3 | 23957929 | rs2360610 | 1.459 | 0.93 | 0.997 | -0.015 | 0.044 | 0.019 | 0.019 | 0.556 | 0.48 | 0.21 | 0.44 | 0.43 | 0.41 | 0.36 |
| 3 | 23962646 | rs2001209 | -1.136 | 0.959 | 1.086 | -0.058 | 0 | -0.002 | -0.002 | 0 | 0.26 | 0.71 | 0.41 | 0.1 | 0.27 | 0.43 |
| 3 | 23976031 | rs4578971 | -0.042 | 0.985 | 1.156 | 0.006 | 0.022 | 0.033 | 0.033 | 0 | 0.71 | 0.62 | 0.65 | 0.45 | 0.66 | 0.58 |
| 3 | 23980440 | rs10865791 | -0.032 | 0.973 | 1.156 | 0.006 | 0.021 | 0.033 | 0.033 | 0 | 0.71 | 0.62 | 0.65 | 0.44 | 0.66 | 0.58 |
| 3 | 23982330 | rs13325754 | -0.178 | 0.746 | 1.277 | -0.116 | -0.002 | 0.012 | 0.012 | 0.313 | 0.17 | 0.11 | 0.19 | 0.05 | 0.2 | 0.18 |
| 3 | 23985809 | rs4131864 | -0.937 | 0.724 | 1.239 | -0.125 | -0.008 | -0.103 | -0.103 | 0 | 0.26 | 0.4 | 0.24 | 0.05 | 0.25 | 0.22 |
| 3 | 23985852 | rs4858564 | -0.089 | 0.784 | 1.23 | -0.043 | 0.032 | -0.04 | -0.04 | 0 | 0.08 | 0.28 | 0.18 | 0 | 0.07 | 0.2 |
| 3 | 23987946 | rs6770052 | -0.237 | 0.779 | 1.199 | -0.007 | 0.037 | 0.038 | 0.038 | 0.075 | 0.08 | 0.3 | 0.19 | 0.05 | 0.07 | 0.26 |
| 3 | 23990241 | rs11922577 | -0.918 | 0.744 | 1.099 | 0.022 | -0.007 | -0.075 | -0.075 | 0.171 | 0.26 | 0.4 | 0.24 | 0.07 | 0.25 | 0.23 |
| 3 | 23996371 | rs4858095 | -1.008 | 1.176 | 1.013 | 0.172 | 0.021 | 0.05 | 0.05 | 0 | 0.79 | 0.73 | 0.82 | 0.53 | 0.72 | 0.84 |
| 3 | 24009919 | rs4858098 | -1.011 | 1.217 | 0.947 | 0.195 | -0.068 | -0.118 | -0.118 | 0 | 0.78 | 0.94 | 0.84 | 0.49 | 0.72 | 0.84 |
| 3 | 24014314 | rs4858569 | -0.143 | 1.202 | 0.99 | 0.214 | -0.028 | -0.058 | -0.058 | 0.056 | 0.72 | 0.84 | 0.79 | 0.46 | 0.67 | 0.78 |
| 3 | 24019327 | rs4602333 | -0.26 | 1.328 | 1.232 | 0.321 | 0.039 | -0.039 | -0.039 | 0 | 0.08 | 0.29 | 0.18 | 0 | 0.07 | 0.2 |
| 3 | 24031090 | rs9310725 | -0.963 | 1.664 | 0.878 | 0.704 | -0.044 | -0.086 | -0.086 | 0 | 0.79 | 0.92 | 0.84 | 0.53 | 0.74 | 0.84 |
| 3 | 24034969 | rs6550825 | 0.385 | 1.867 | 1.079 | 0.669 | 0.003 | 0.013 | 0.013 | 0 | 0.67 | 0.42 | 0.65 | 0.48 | 0.67 | 0.61 |
| 3 | 24038687 | rs6550829 | -0.946 | 1.903 | 0.959 | 0.666 | -0.047 | -0.089 | -0.089 | 0 | 0.79 | 0.95 | 0.85 | 0.57 | 0.74 | 0.84 |
| 3 | 24044111 | rs12497944 | 0.238 | 1.7 | 0.771 | 0.311 | -0.024 | 0.059 | 0.059 | 0.369 | 0.17 | 0.05 | 0.22 | 0.05 | 0.23 | 0.21 |
| 3 | 24044400 | rs9883101 | 0.191 | 2.035 | 0.929 | 0.47 | 0.004 | 0.004 | 0.004 | 0 | 0.67 | 0.42 | 0.64 | 0.42 | 0.65 | 0.6 |
| 3 | 24050952 | rs4858571 | -0.071 | 1.842 | 1.168 | 0.505 | 0.061 | -0.002 | -0.002 | 0 | 0.07 | 0.24 | 0.2 | 0.05 | 0.07 | 0.16 |
| 3 | 24052025 | rs4356765 | -0.855 | 1.847 | 1.208 | 0.423 | 0.019 | -0.086 | -0.086 | 3.371 | 0.25 | 0.47 | 0.42 | 0.1 | 0.3 | 0.39 |
| 3 | 24053364 | rs9847444 | -1.361 | 1.889 | 0.944 | 0.526 | -0.055 | -0.112 | -0.112 | 0.189 | 0.8 | 0.97 | 0.86 | 0.55 | 0.76 | 0.86 |
| 3 | 24053792 | rs951015 | -1.412 | 1.774 | 1.215 | 0.443 | -0.003 | 0.018 | 0.018 | 0.201 | 0.29 | 0.73 | 0.46 | 0.18 | 0.32 | 0.45 |
| 3 | 24058855 | rs938078 | 0.323 | 1.776 | 1.038 | 0.198 | -0.016 | 0.047 | 0.047 | 0.139 | 0.17 | 0.08 | 0.21 | 0.05 | 0.23 | 0.23 |
| 3 | 24070055 | rs7431278 | -0.721 | 1.761 | 1.065 | 0.228 | 0.017 | -0.075 | -0.075 | 0 | 0.25 | 0.4 | 0.43 | 0.1 | 0.3 | 0.38 |
| 3 | 24081772 | rs11709614 | 1.551 | 1.573 | 0.311 | 0.064 | 0.035 | 0.185 | 0.185 | 0 | 0.4 | 0.17 | 0.22 | 0.07 | 0.36 | 0.27 |
| 3 | 24082776 | rs4858100 | 0.991 | 1.18 | 0.451 | 0.28 | -0.016 | 0.071 | 0.071 | 0 | 0.5 | 0.19 | 0.41 | 0.36 | 0.48 | 0.42 |
| 3 | 24085166 | rs1112195 | 1.15 | 1.277 | 0.376 | 0.324 | -0.039 | 0.166 | 0.166 | 0 | 0.47 | 0.18 | 0.37 | 0.22 | 0.46 | 0.41 |
| 3 | 24088171 | rs17014085 | 1.268 | 1.121 | 0.313 | 0.467 | -0.028 | -0.063 | -0.063 | 0 | 0.2 | 0.09 | 0.15 | 0.45 | 0.22 | 0.12 |
| 3 | 24089253 | rs11718240 | 0.833 | 0.95 | 0.468 | -0.021 | 0.003 | 0.012 | 0.012 | 0.784 | 0.13 | 0.13 | 0.18 | 0.06 | 0.15 | 0.21 |
| 3 | 24091463 | rs11927256 | -0.954 | 0.908 | 0.451 | -0.03 | -0.05 | 0.061 | 0.061 | 0 | 0.46 | 0.85 | 0.48 | 0.68 | 0.47 | 0.52 |
| 3 | 24092290 | rs11928242 | -0.928 | 0.907 | 0.484 | -0.03 | -0.045 | 0.061 | 0.061 | 0 | 0.46 | 0.83 | 0.47 | 0.68 | 0.47 | 0.53 |
| 3 | 24105626 | rs7634184 | -0.809 | 0.809 | 1.451 | -0.056 | -0.015 | -0.045 | -0.045 | 0 | 0.25 | 0.58 | 0.25 | 0.11 | 0.22 | 0.34 |
| 3 | 24106647 | rs2089848 | 0.713 | 0.804 | 1.382 | -0.059 | 0.006 | 0.015 | 0.015 | 0.555 | 0.14 | 0.13 | 0.18 | 0.05 | 0.15 | 0.21 |
| 3 | 24109112 | rs7612158 | -1.294 | 0.82 | 1.36 | -0.034 | 0.047 | 0.058 | 0.058 | 1.023 | 0.29 | 0.85 | 0.28 | 0.15 | 0.22 | 0.43 |
| 3 | 24109875 | rs7635612 | 0.129 | 0.871 | 1.425 | -0.018 | 0.031 | -0.016 | -0.016 | 0 | 0.1 | 0.31 | 0.06 | 0.05 | 0.07 | 0.13 |
| 3 | 24113697 | rs6801153 | 0.323 | 0.926 | 1.192 | 0.055 | 0.001 | -0.002 | -0.002 | 1.146 | 0.76 | 0.72 | 0.59 | 0.75 | 0.76 | 0.65 |
| 3 | 24145084 | rs6550848 | -0.895 | 0.975 | 1.23 | 0.007 | -0.006 | 0.017 | 0.017 | 0 | 0.47 | 0.73 | 0.46 | 0.63 | 0.46 | 0.5 |
| 3 | 24148622 | rs6769403 | 1.349 | 1.001 | 1.21 | 0.01 | -0.05 | -0.105 | -0.105 | 0 | 0.2 | 0.06 | 0.21 | 0.52 | 0.25 | 0.13 |
| 3 | 24153725 | rs11925334 | -0.849 | 0.897 | 1.372 | 0.019 | -0.022 | -0.046 | -0.046 | 0 | 0.27 | 0.62 | 0.26 | 0.12 | 0.22 | 0.36 |
| 3 | 24158397 | rs7428850 | 0.639 | 0.924 | 1.172 | 0.035 | 0.029 | 0.013 | 0.013 | 0 | 0.08 | 0.25 | 0.05 | 0.06 | 0.06 | 0.11 |
| 3 | 24158647 | rs17014136 | 0.567 | 0.91 | 1.01 | -0.067 | -0.019 | -0.04 | -0.04 | 0.007 | 0.16 | 0.29 | 0.2 | 0.05 | 0.15 | 0.25 |
| 3 | 24159387 | rs1349265 | 1.332 | 0.963 | 0.413 | -0.033 | 0.036 | -0.003 | -0.003 | 0.519 | 0.63 | 0.12 | 0.51 | 0.69 | 0.69 | 0.52 |
| 3 | 24163021 | rs844107 | -0.646 | 1.017 | 0.766 | 0.081 | 0 | 0 | 0 | 0.266 | 0.41 | 0.56 | 0.43 | 0.62 | 0.41 | 0.47 |
| 3 | 24163921 | rs826374 | 0.885 | 1.267 | 0.214 | 0.236 | -0.007 | -0.009 | -0.009 | 0 | 0.2 | 0.15 | 0.22 | 0.52 | 0.25 | 0.14 |
| 3 | 24168981 | rs13063628 | 0.802 | 0.82 | -0.271 | -0.215 | -0.016 | 0.008 | 0.008 | 5.63 | 0.1 | 0.06 | 0.06 | 0 | 0.14 | 0.06 |
| 3 | 24169361 | rs1667765 | 1.343 | 0.949 | -0.469 | -0.137 | -0.062 | -0.161 | -0.161 | 0.062 | 0.19 | 0.01 | 0.23 | 0.55 | 0.21 | 0.14 |

| | | | | | | | | | | | | | | | | |
|---|---|---|---|---|---|---|---|---|---|---|---|---|---|---|---|---|
| 3 | 24169493 | rs1868573 | -0.688 | 0.69 | 0.042 | -0.28 | -0.001 | -0.029 | -0.029 | 0 | 0.29 | 0.6 | 0.28 | 0.1 | 0.29 | 0.38 |
| 3 | 24169726 | rs826378 | -1.827 | 0.617 | -0.347 | -0.548 | -0.03 | -0.033 | -0.033 | 0 | 0.64 | 0.92 | 0.6 | 0.69 | 0.63 | 0.66 |
| 3 | 24173744 | rs1705734 | -1.221 | 0.338 | 0.186 | -0.163 | -0.002 | 0.049 | 0.049 | 0 | 0.52 | 0.68 | 0.52 | 0.66 | 0.52 | 0.51 |
| 3 | 24174159 | rs13066296 | 0.809 | 0.609 | -0.307 | -0.11 | -0.01 | -0.002 | -0.002 | 0 | 0.1 | 0.06 | 0.05 | 0 | 0.14 | 0.06 |
| 3 | 24186625 | rs10428188 | 0.158 | 0.397 | -0.304 | -0.103 | 0.006 | 0.008 | 0.008 | 0.393 | 0.18 | 0.18 | 0.19 | 0.06 | 0.15 | 0.25 |
| 3 | 24187084 | rs826385 | -0.049 | 0.435 | 0.037 | -0.076 | 0.026 | 0.031 | 0.031 | 0 | 0.33 | 0.46 | 0.33 | 0.61 | 0.36 | 0.26 |
| 3 | 24187500 | rs826386 | -0.351 | 0.141 | -0.304 | 0.243 | 0.004 | -0.002 | -0.002 | 0.061 | 0.4 | 0.35 | 0.47 | 0.61 | 0.37 | 0.38 |
| 3 | 24190184 | rs9844739 | -0.563 | -0.129 | -0.413 | 0.15 | 0.014 | 0.017 | 0.017 | 0.633 | 0.2 | 0.17 | 0.2 | 0.09 | 0.18 | 0.28 |
| 3 | 24191961 | rs1623905 | 0.326 | 0.357 | 0.299 | 0.604 | 0.032 | 0.052 | 0.052 | 0.301 | 0.82 | 0.66 | 0.65 | 0.69 | 0.8 | 0.75 |
| 3 | 24196196 | rs6774779 | -0.928 | 0.006 | 0.815 | 0.96 | -0.007 | 0.132 | 0.132 | 2.717 | 0.1 | 0.3 | 0.06 | 0.23 | 0.07 | 0.16 |
| 3 | 24197273 | rs1470661 | 0.102 | 0.039 | 0.81 | 0.917 | 0.001 | -0.001 | -0.001 | 0.372 | 0.13 | 0.13 | 0.32 | 0.17 | 0.16 | 0.17 |
| 3 | 24198726 | rs11919189 | -0.764 | -0.154 | 1.559 | 0.908 | -0.004 | 0.106 | 0.106 | 2.698 | 0.09 | 0.26 | 0.05 | 0.2 | 0.06 | 0.16 |
| 3 | 24200811 | rs9870693 | -0.069 | -0.265 | 0.867 | 0.261 | -0.002 | 0.022 | 0.022 | 0.22 | 0.32 | 0.26 | 0.3 | 0.12 | 0.35 | 0.37 |
| 3 | 24202111 | rs6777133 | -0.411 | -0.207 | 0.828 | 0.442 | 0.012 | 0.007 | 0.007 | 0 | 0.21 | 0.3 | 0.37 | 0.27 | 0.21 | 0.21 |
| 3 | 24202457 | rs17014251 | -0.297 | -0.13 | 0.773 | 0.462 | 0.006 | 0.013 | 0.013 | 3.215 | 0.11 | 0.17 | 0.07 | 0.13 | 0.1 | 0.12 |
| 3 | 24203529 | rs17014257 | -1.545 | -0.23 | 0.638 | 0.557 | 0.002 | -0.015 | -0.015 | 0 | 0.36 | 0.75 | 0.46 | 0.43 | 0.33 | 0.44 |
| 3 | 24208974 | rs2596625 | 0.634 | -0.115 | 0.644 | 0.554 | -0.005 | -0.006 | -0.006 | 0 | 0.13 | 0.11 | 0.32 | 0.24 | 0.15 | 0.16 |
| 3 | 24211739 | rs2683530 | -1.691 | -0.327 | 0.433 | 0.643 | 0.015 | 0.025 | 0.025 | 0 | 0.49 | 0.72 | 0.67 | 0.57 | 0.47 | 0.64 |
| 3 | 24212581 | rs2683529 | -0.395 | -0.185 | 0.551 | 0.667 | 0 | 0 | 0 | 0.658 | 0.23 | 0.23 | 0.39 | 0.32 | 0.27 | 0.23 |
| 3 | 24215973 | rs13320945 | -0.793 | -0.354 | 0.52 | 0.474 | -0.003 | -0.02 | -0.02 | 0 | 0.41 | 0.43 | 0.36 | 0.23 | 0.41 | 0.49 |
| 3 | 24216342 | rs7616636 | -0.135 | -0.292 | 0.509 | 0.413 | 0.006 | -0.073 | -0.073 | 1.17 | 0.25 | 0.44 | 0.15 | 0.1 | 0.23 | 0.27 |
| 3 | 24218879 | rs9862985 | -0.639 | -0.291 | 0.573 | 0.439 | 0.001 | -0.001 | -0.001 | 2.668 | 0.39 | 0.38 | 0.35 | 0.18 | 0.37 | 0.41 |
| 3 | 24219521 | rs4858582 | -1.697 | -0.183 | 0.654 | 0.577 | -0.002 | 0.016 | 0.016 | 0 | 0.52 | 0.73 | 0.68 | 0.44 | 0.54 | 0.72 |
| 3 | 24220643 | rs7652337 | -2.686 | -0.119 | 0.313 | 0.594 | -0.074 | -0.148 | -0.148 | 0 | 0.66 | 0.98 | 0.77 | 0.53 | 0.67 | 0.77 |
| 3 | 24220680 | rs7652347 | 0.272 | 0.105 | 0.852 | 0.492 | 0.006 | -0.083 | -0.083 | 0 | 0.2 | 0.33 | 0.12 | 0.02 | 0.19 | 0.11 |
| 3 | 24221506 | rs2596618 | 0.519 | 0.485 | 0.67 | 0.79 | -0.002 | 0.002 | 0.002 | 1.489 | 0.13 | 0.16 | 0.31 | 0.2 | 0.12 | 0.21 |
| 3 | 24222782 | rs2683541 | -0.6 | 0.517 | 0.798 | 0.879 | 0.03 | 0.061 | 0.061 | 0.574 | 0.19 | 0.38 | 0.36 | 0.31 | 0.2 | 0.35 |
| 3 | 24225870 | rs826216 | -1.373 | 0.566 | 0.801 | 0.954 | -0.008 | -0.011 | -0.011 | 0 | 0.33 | 0.71 | 0.48 | 0.39 | 0.35 | 0.54 |
| 3 | 24227175 | rs12639293 | -0.807 | 0.237 | 1.045 | 0.833 | -0.008 | -0.015 | -0.015 | 0 | 0.36 | 0.53 | 0.32 | 0.22 | 0.35 | 0.37 |
| 3 | 24228977 | rs844103 | -0.35 | 0.387 | 0.805 | 0.908 | -0.004 | 0.002 | 0.002 | 0 | 0.23 | 0.22 | 0.37 | 0.22 | 0.24 | 0.35 |
| 3 | 24232709 | rs7612541 | 0.257 | 0.203 | 1.02 | 0.977 | -0.016 | 0.021 | 0.021 | 0 | 0.16 | 0.23 | 0.23 | 0.22 | 0.14 | 0.2 |
| 3 | 24236613 | rs826229 | -0.401 | 0.327 | 0.75 | 0.952 | -0.016 | 0.008 | 0.008 | 0 | 0.24 | 0.21 | 0.39 | 0.21 | 0.28 | 0.29 |
| 3 | 24237950 | rs6797693 | -1.304 | 0.108 | 0.865 | 0.854 | 0.018 | -0.015 | -0.015 | 0.785 | 0.39 | 0.78 | 0.37 | 0.37 | 0.38 | 0.49 |
| 3 | 24240974 | rs9820526 | -0.138 | -0.067 | 1.367 | 1.096 | -0.017 | 0.195 | 0.195 | 0.054 | 0.14 | 0.47 | 0.24 | 0.26 | 0.12 | 0.18 |
| 3 | 24241978 | rs17014331 | 0.971 | -0.034 | 0.991 | 1.075 | 0.006 | -0.022 | -0.022 | 0 | 0.1 | 0.06 | 0.19 | 0.14 | 0.11 | 0.11 |
| 3 | 24253228 | rs826253 | -0.201 | -0.014 | 0.723 | 1.471 | -0.005 | -0.031 | -0.031 | 0.175 | 0.19 | 0.12 | 0.35 | 0.38 | 0.17 | 0.27 |
| 3 | 24258985 | rs851719 | -2.133 | 0.008 | 0.637 | 1.259 | -0.058 | -0.027 | -0.027 | 0 | 0.62 | 0.97 | 0.78 | 0.68 | 0.65 | 0.77 |
| 3 | 24259201 | rs4858106 | -0.107 | 0.218 | 0.958 | 1.118 | -0.037 | -0.02 | -0.02 | 0 | 0.23 | 0.51 | 0.14 | 0.14 | 0.2 | 0.21 |
| 3 | 24260059 | rs13097208 | -1.079 | 0.279 | 0.605 | 1.035 | 0.001 | -0.003 | -0.003 | 0 | 0.39 | 0.39 | 0.36 | 0.24 | 0.47 | 0.43 |
| 3 | 24262255 | rs826248 | -2.149 | 0.335 | 0.024 | 0.843 | -0.06 | -0.02 | -0.02 | 0 | 0.62 | 0.97 | 0.77 | 0.68 | 0.65 | 0.78 |
| 3 | 24263738 | rs6764136 | -0.698 | 0.23 | 0.483 | 0.844 | 0.001 | -0.002 | -0.002 | 0 | 0.34 | 0.29 | 0.27 | 0.43 | 0.32 | 0.39 |
| 3 | 24265142 | rs2683532 | -1.608 | 0.311 | 0.279 | 1.092 | 0.042 | 0.195 | 0.195 | 0 | 0.49 | 0.84 | 0.67 | 0.75 | 0.45 | 0.55 |
| 3 | 24266984 | rs6550853 | -1.201 | 0.573 | 0.717 | 1.225 | 0.017 | 0.044 | 0.044 | 0 | 0.28 | 0.58 | 0.32 | 0.43 | 0.31 | 0.42 |
| 3 | 24267140 | rs826223 | 0.312 | 0.587 | 0.639 | 1.408 | 0.008 | 0.121 | 0.121 | 0 | 0.12 | 0.23 | 0.35 | 0.36 | 0.11 | 0.16 |
| 3 | 24267842 | rs12491199 | -0.127 | 0.583 | 0.492 | 1.375 | 0.004 | 0.076 | 0.076 | 0 | 0.18 | 0.24 | 0.22 | 0.39 | 0.19 | 0.31 |
| 3 | 24268436 | rs826221 | 0.851 | 0.542 | -0.013 | 1.415 | -0.003 | 0.008 | 0.008 | 0 | 0.08 | 0.07 | 0.29 | 0.26 | 0.08 | 0.12 |
| 3 | 24269789 | rs826217 | -1.456 | 0.258 | 0.125 | 0.691 | 0.006 | 0.046 | 0.046 | 0 | 0.59 | 0.74 | 0.76 | 0.84 | 0.64 | 0.77 |
| 3 | 24270796 | rs1466122 | 0.42 | 0.358 | -0.09 | 0.653 | -0.002 | 0.013 | 0.013 | 0 | 0.57 | 0.67 | 0.4 | 0.68 | 0.47 | 0.65 |
| 3 | 24271599 | rs7619754 | 0.501 | 0.193 | -0.163 | 0.937 | -0.002 | 0.006 | 0.006 | 0 | 0.56 | 0.61 | 0.35 | 0.37 | 0.49 | 0.62 |
| 3 | 24271657 | rs9874037 | 1.032 | 0.21 | -0.383 | 0.617 | -0.004 | -0.035 | -0.035 | 2.168 | 0.11 | 0.11 | 0.07 | 0 | 0.12 | 0.03 |
| 3 | 24272388 | rs1466119 | -0.118 | 0.253 | -0.479 | 0.829 | -0.117 | -0.08 | -0.08 | 0 | 0.69 | 0.93 | 0.49 | 0.63 | 0.61 | 0.68 |
| 3 | 24273461 | rs7625900 | 0.053 | 0.45 | -0.247 | 0.728 | -0.012 | -0.003 | -0.003 | 0.261 | 0.64 | 0.7 | 0.44 | 0.62 | 0.56 | 0.67 |
| 3 | 24273528 | rs7625917 | 1 | 0.17 | -0.121 | 0.45 | -0.012 | -0.011 | -0.011 | 4.469 | 0.08 | 0.14 | 0.05 | 0.06 | 0.04 | 0.06 |
| 3 | 24273552 | rs17787283 | 2.026 | -0.301 | -0.406 | 0.225 | 0.004 | 0.106 | 0.106 | 0 | 0.3 | 0.1 | 0.17 | 0.13 | 0.28 | 0.29 |
| 3 | 24273978 | rs7617186 | -0.608 | -0.337 | 0.326 | 0.682 | -0.016 | 0.064 | 0.064 | 0 | 0.24 | 0.36 | 0.22 | 0.38 | 0.23 | 0.19 |
| 3 | 24274074 | rs4858110 | 0.991 | -0.349 | -0.47 | -0.045 | -0.016 | 0.015 | 0.015 | 0 | 0.58 | 0.31 | 0.32 | 0.39 | 0.49 | 0.61 |
| 3 | 24276575 | rs986575 | -0.45 | -0.5 | -0.49 | -0.1 | -0.046 | -0.07 | -0.07 | 0 | 0.29 | 0.44 | 0.16 | 0.13 | 0.21 | 0.27 |
| 3 | 24277179 | rs9829876 | 0.153 | 0.178 | -1.134 | 0.211 | 0.032 | 0.025 | 0.025 | 0 | 0.64 | 0.54 | 0.41 | 0.47 | 0.54 | 0.58 |
| 3 | 24277810 | rs11714355 | 0.31 | 0.071 | -2.126 | -0.692 | -0.036 | 0.187 | 0.187 | 0 | 0.39 | 0.08 | 0.23 | 0.13 | 0.37 | 0.33 |
| 3 | 24278313 | rs4858595 | -0.389 | -0.466 | -1.731 | -0.51 | 0.03 | 0.024 | 0.024 | 3.052 | 0.28 | 0.76 | 0.18 | 0.19 | 0.14 | 0.21 |
| 3 | 24278815 | rs1349264 | -0.382 | 1.288 | -1.936 | -0.387 | -0.053 | 0.008 | 0.008 | 0 | 0.92 | 0.99 | 0.67 | 0.93 | 0.8 | 0.84 |
| 3 | 24278912 | rs1449873 | 0.147 | 0.428 | -2.024 | -0.516 | 0.013 | 0.038 | 0.038 | 0.597 | 0.19 | 0.08 | 0.1 | 0.07 | 0.17 | 0.16 |
| 3 | 24281453 | rs10212213 | 0.143 | 0.877 | -2.311 | -0.608 | -0.08 | 0.038 | 0.038 | 4.777 | 0.72 | 0.94 | 0.54 | 0.83 | 0.62 | 0.71 |
| 3 | 24281488 | rs12386345 | 1.112 | 0.71 | -2.573 | -0.52 | -0.022 | 0.002 | 0.002 | 0 | 0.26 | 0.14 | 0.23 | 0.26 | 0.26 | 0.29 |
| 3 | 24281656 | rs11129141 | 0.919 | 0.187 | -1.604 | -0.413 | 0.002 | -0.007 | -0.007 | 0 | 0.34 | 0.33 | 0.22 | 0.53 | 0.29 | 0.36 |
| 3 | 24282982 | rs9849581 | 0.003 | 0.357 | -0.586 | -0.156 | -0.015 | -0.013 | -0.013 | 0 | 0.11 | 0.17 | 0.07 | 0.05 | 0.07 | 0.07 |
| 3 | 24284924 | rs9840296 | 0.527 | 0.625 | -0.389 | -0.202 | -0.001 | -0.019 | -0.019 | 0 | 0.61 | 0.48 | 0.46 | 0.78 | 0.57 | 0.66 |
| 3 | 24285768 | rs1562737 | -0.189 | 1.191 | -0.191 | 0.063 | -0.033 | 0.02 | 0.02 | 0 | 0.91 | 0.98 | 0.65 | 0.95 | 0.82 | 0.85 |
| 3 | 24286407 | rs9812189 | -0.933 | 1.171 | 0.511 | 0.757 | 0.051 | 0.164 | 0.164 | 3.922 | 0.14 | 0.56 | 0.18 | 0.33 | 0.12 | 0.15 |

| | | | | | | | | | | | | | | | | |
|---|---|---|---|---|---|---|---|---|---|---|---|---|---|---|---|---|
| 3 | 24292825 | rs2044756 | -0.338 | 1.21 | 0.034 | 0.737 | 0.012 | 0.003 | 0.003 | 0.28 | 0.1 | 0.15 | 0.14 | 0.16 | 0.1 | 0.11 |
| 3 | 24294854 | rs826227 | 1.246 | 1.375 | 0.303 | 0.784 | 0.07 | 0.133 | 0.133 | 0.036 | 0.76 | 0.52 | 0.48 | 0.54 | 0.71 | 0.65 |
| 3 | 24301536 | rs9873977 | 0.512 | 0.645 | 0.137 | 0.548 | 0.028 | 0.182 | 0.182 | 0 | 0.05 | 0.23 | 0.02 | 0.13 | 0.01 | 0.06 |
| 3 | 24306302 | rs826230 | -0.603 | 0.854 | -0.218 | 0.341 | -0.031 | 0.028 | 0.028 | 0 | 0.93 | 0.99 | 0.67 | 0.97 | 0.86 | 0.88 |
| 3 | 24307606 | rs826232 | 0.055 | 0.584 | -0.725 | -0.05 | 0.001 | 0.018 | 0.018 | 0.528 | 0.62 | 0.7 | 0.5 | 0.75 | 0.63 | 0.49 |
| 3 | 24307693 | rs862247 | -0.108 | 1.239 | -0.462 | 0.061 | -0.024 | 0.024 | 0.024 | 0 | 0.92 | 0.98 | 0.67 | 0.95 | 0.86 | 0.85 |
| 3 | 24313506 | rs1910854 | 0.146 | 0.974 | -0.659 | -0.189 | -0.029 | 0.089 | 0.089 | 6.195 | 0.17 | 0.01 | 0.1 | 0.04 | 0.19 | 0.1 |
| 3 | 24327131 | rs2360957 | 0.628 | 1.758 | -0.297 | 0.55 | -0.092 | -0.039 | -0.039 | 0 | 0.84 | 0.97 | 0.51 | 0.76 | 0.72 | 0.76 |
| 3 | 24336162 | rs1158265 | -0.688 | 1.456 | 0.547 | 0.671 | 0.036 | 0.035 | 0.035 | 0.246 | 0.26 | 0.39 | 0.27 | 0.38 | 0.3 | 0.36 |
| 3 | 24336257 | rs4404389 | 0.395 | 1.453 | 0.361 | 0.703 | 0.003 | -0.017 | -0.017 | 0 | 0.15 | 0.07 | 0.11 | 0.16 | 0.14 | 0.21 |
| 3 | 24337914 | rs9830674 | -1.191 | 1.277 | 0.91 | 0.828 | 0.057 | 0.038 | 0.038 | 0.491 | 0.18 | 0.62 | 0.24 | 0.28 | 0.26 | 0.19 |
| 3 | 24339734 | rs2167115 | -1.564 | 1.238 | 0.814 | 0.801 | -0.013 | -0.015 | -0.015 | 1.421 | 0.34 | 0.73 | 0.35 | 0.45 | 0.4 | 0.41 |
| 3 | 24344775 | rs17014491 | 0.49 | 1.202 | 0.752 | 0.789 | 0.002 | -0.02 | -0.02 | 0.526 | 0.15 | 0.07 | 0.11 | 0.19 | 0.14 | 0.2 |
| 3 | 24346109 | rs12485694 | -0.616 | 1.077 | 0.781 | 0.592 | 0.014 | 0.032 | 0.032 | 0 | 0.49 | 0.45 | 0.67 | 0.76 | 0.58 | 0.63 |
| 3 | 24346981 | rs9850879 | 0.139 | 1.14 | 1.005 | 0.653 | -0.002 | 0.04 | 0.04 | 0 | 0.33 | 0.48 | 0.49 | 0.45 | 0.37 | 0.33 |
| 3 | 24348350 | rs4575827 | 1.206 | 1.209 | 0.666 | 0.658 | -0.017 | -0.067 | -0.067 | 0.286 | 0.24 | 0.09 | 0.4 | 0.39 | 0.28 | 0.28 |
| 3 | 24349915 | rs4858604 | 1.022 | 1.357 | 0.525 | 0.684 | 0 | -0.002 | -0.002 | 0.466 | 0.27 | 0.28 | 0.43 | 0.4 | 0.29 | 0.3 |
| 3 | 24350811 | rs9310736 | -0.716 | 1.353 | 0.056 | 0.438 | 0.035 | 0.086 | 0.086 | 0 | 0.57 | 0.76 | 0.74 | 0.81 | 0.67 | 0.67 |
| 3 | 24351871 | rs7622481 | -0.386 | 1.276 | -0.019 | 0.458 | 0.021 | 0.021 | 0.021 | 0.195 | 0.21 | 0.26 | 0.23 | 0.33 | 0.26 | 0.34 |
| 3 | 24357082 | rs7610039 | 0.243 | 0.155 | -0.466 | -0.024 | -0.005 | 0.026 | 0.026 | 0 | 0.44 | 0.71 | 0.66 | 0.68 | 0.54 | 0.5 |
| 3 | 24357243 | rs7640580 | 0.103 | -0.096 | -0.477 | -0.266 | 0.016 | 0.102 | 0.102 | 0 | 0.6 | 0.83 | 0.75 | 0.8 | 0.63 | 0.69 |
| 3 | 24357286 | rs7610222 | -0.093 | -0.281 | -0.239 | -0.077 | 0.003 | 0.003 | 0.003 | 0 | 0.67 | 0.71 | 0.78 | 0.84 | 0.75 | 0.71 |
| 3 | 24358906 | rs6550857 | -1.321 | -0.234 | 0.457 | 0.418 | 0.03 | 0.095 | 0.095 | 2.149 | 0.06 | 0.37 | 0.05 | 0.03 | 0.02 | 0.03 |
| 3 | 24361096 | rs1857793 | -0.87 | -0.704 | -0.094 | 0.26 | 0.013 | 0.006 | 0.006 | 0 | 0.45 | 0.62 | 0.59 | 0.63 | 0.34 | 0.48 |
| 3 | 24364406 | rs1505296 | -0.149 | -1.104 | 0.045 | 0.499 | 0.007 | 0.057 | 0.057 | 0.178 | 0.21 | 0.46 | 0.42 | 0.33 | 0.17 | 0.36 |
| 3 | 24365661 | rs7649511 | -0.858 | -1.268 | -0.103 | 0.42 | -0.007 | -0.006 | -0.006 | 0 | 0.31 | 0.59 | 0.48 | 0.34 | 0.26 | 0.38 |
| 3 | 24370875 | rs9812034 | -1.44 | -1.388 | -0.24 | 0.291 | 0.053 | 0.03 | 0.03 | 0.006 | 0.47 | 0.72 | 0.6 | 0.63 | 0.35 | 0.48 |
| 3 | 24379641 | rs6787255 | -1.208 | -1.397 | -0.971 | 0.072 | 0.096 | 0.037 | 0.037 | 0.238 | 0.49 | 0.91 | 0.64 | 0.62 | 0.35 | 0.49 |
| 3 | 24379744 | rs7609823 | -0.412 | -1.472 | -0.627 | 0.068 | -0.006 | 0.008 | 0.008 | 0.38 | 0.23 | 0.66 | 0.46 | 0.34 | 0.18 | 0.33 |
| 3 | 24383347 | rs13318276 | -0.209 | -1.482 | -0.957 | 0.233 | 0.043 | -0.061 | -0.061 | 0 | 0.22 | 0.06 | 0.17 | 0.28 | 0.18 | 0.16 |
| 3 | 24384005 | rs9833191 | -1.128 | -1.035 | -1.743 | -0.56 | -0.123 | -0.082 | -0.082 | 0.069 | 0.7 | 0.92 | 0.74 | 0.61 | 0.58 | 0.69 |
| 3 | 24389619 | rs9809320 | 0.336 | -0.755 | -1.701 | -0.438 | 0.007 | -0.084 | -0.084 | 0 | 0.21 | 0.04 | 0.18 | 0.29 | 0.2 | 0.17 |
| 3 | 24390258 | rs7619474 | -1.258 | -0.593 | -1.597 | -0.306 | -0.092 | -0.077 | -0.077 | 0 | 0.75 | 0.93 | 0.77 | 0.64 | 0.66 | 0.69 |
| 3 | 24393296 | rs1394763 | -0.745 | -0.521 | -0.885 | -0.492 | -0.017 | -0.024 | -0.024 | 0 | 0.1 | 0.1 | 0.03 | 0 | 0.03 | 0.05 |
| 3 | 24401518 | rs1354601 | -0.437 | -0.234 | -0.473 | -0.359 | 0.031 | 0.016 | 0.016 | 0.053 | 0.07 | 0.36 | 0.05 | 0 | 0.02 | 0.01 |
| 3 | 24403210 | rs6781111 | 0.146 | 0.236 | -1.264 | -0.188 | 0.005 | 0.009 | 0.009 | 0.009 | 0.17 | 0.17 | 0.44 | 0.35 | 0.23 | 0.3 |
| 3 | 24409215 | rs6550858 | 0.589 | 0.14 | -1.848 | -0.688 | -0.038 | 0.121 | 0.121 | 0 | 0.25 | 0.07 | 0.11 | 0.01 | 0.25 | 0.21 |
| 3 | 24409823 | rs11924768 | -0.545 | -0.354 | -2.079 | -0.584 | -0.072 | 0.077 | 0.077 | 0.382 | 0.57 | 0.89 | 0.6 | 0.33 | 0.51 | 0.54 |
| 3 | 24410415 | rs6550859 | -0.447 | -0.524 | -2.393 | -0.616 | -0.124 | 0.001 | 0.001 | 0.985 | 0.58 | 0.95 | 0.61 | 0.32 | 0.52 | 0.55 |
| 3 | 24413479 | rs11919155 | -0.119 | -0.746 | -1.44 | -0.867 | 0.01 | -0.04 | -0.04 | 1.054 | 0.11 | 0.24 | 0.1 | 0.04 | 0.1 | 0.13 |
| 3 | 24419005 | rs12638970 | -0.125 | -1.206 | -1.658 | -0.948 | -0.001 | 0.014 | 0.014 | 0 | 0.2 | 0.15 | 0.12 | 0.06 | 0.18 | 0.13 |
| 3 | 24419559 | rs9809150 | -1.026 | -1.042 | -2.313 | -0.49 | 0.024 | -0.062 | -0.062 | 0 | 0.64 | 0.87 | 0.63 | 0.58 | 0.65 | 0.68 |
| 3 | 24422751 | rs9310740 | 0.761 | -1.223 | -1.639 | -0.506 | 0.049 | 0.04 | 0.04 | 1.437 | 0.32 | 0.48 | 0.45 | 0.49 | 0.43 | 0.49 |
| 3 | 24425496 | rs6771995 | 0.229 | -1.27 | -1.595 | -0.529 | 0.012 | 0.009 | 0.009 | 1.513 | 0.35 | 0.65 | 0.47 | 0.49 | 0.44 | 0.49 |
| 3 | 24427731 | rs11129151 | -0.175 | -1.224 | -1.497 | -0.463 | -0.045 | 0.004 | 0.004 | 3.226 | 0.87 | 0.63 | 0.91 | 0.91 | 0.96 | 0.94 |
| 3 | 24434524 | rs11129152 | 0.816 | -1.299 | -1.332 | -0.536 | -0.099 | -0.079 | -0.079 | 0 | 0.32 | 0.09 | 0.45 | 0.48 | 0.45 | 0.48 |
| 3 | 24441549 | rs4309672 | 0.637 | -1.407 | -1.165 | -0.55 | 0.033 | 0.02 | 0.02 | 1.217 | 0.35 | 0.45 | 0.48 | 0.48 | 0.47 | 0.48 |
| 3 | 24454727 | rs9882821 | 0.495 | -1.53 | -1.074 | -0.518 | -0.034 | -0.042 | -0.042 | 0.174 | 0.11 | 0.34 | 0.07 | 0.02 | 0.03 | 0.08 |
| 3 | 24456139 | rs4541352 | -1.234 | -1.669 | -1.191 | -0.705 | 0.012 | -0.08 | -0.08 | 0.993 | 0.66 | 0.88 | 0.66 | 0.5 | 0.69 | 0.69 |
| 3 | 24461325 | rs7626512 | 0.791 | -1.406 | -0.451 | -0.87 | 0.004 | -0.113 | -0.113 | 0 | 0.21 | 0.37 | 0.13 | 0.01 | 0.19 | 0.16 |
| 3 | 24469470 | rs9812620 | 0.761 | -1.172 | -0.261 | -0.87 | -0.012 | -0.118 | -0.118 | 0 | 0.28 | 0.38 | 0.14 | 0.01 | 0.19 | 0.14 |
| 3 | 24474053 | rs4630885 | -0.773 | -1.191 | 0.084 | -0.544 | -0.028 | 0.037 | 0.037 | 1.452 | 0.9 | 0.75 | 0.94 | 0.91 | 0.97 | 0.89 |
| 3 | 24475752 | rs9854304 | -0.327 | -1.234 | -0.153 | -0.092 | -0.005 | 0.007 | 0.007 | 0.827 | 0.63 | 0.62 | 0.61 | 0.55 | 0.67 | 0.55 |
| 3 | 24480540 | rs11714673 | 0.72 | -0.19 | -0.067 | 0.278 | -0.021 | 0.04 | 0.04 | 0 | 0.56 | 0.26 | 0.55 | 0.67 | 0.59 | 0.6 |
| 3 | 24480882 | rs11711197 | -0.707 | -0.228 | -0.104 | 0.169 | -0.017 | 0.095 | 0.095 | 0.734 | 0.9 | 0.59 | 0.93 | 0.87 | 0.96 | 0.85 |
| 3 | 24482318 | rs13077437 | 0.494 | 0.3 | 0.016 | 0.112 | -0.013 | -0.013 | -0.013 | 0 | 0.28 | 0.23 | 0.33 | 0.01 | 0.3 | 0.11 |
| 3 | 24483794 | rs9831450 | -0.022 | -0.129 | -0.136 | 0.833 | 0.01 | 0.016 | 0.016 | 0.292 | 0.56 | 0.58 | 0.59 | 0.29 | 0.61 | 0.42 |
| 3 | 24487630 | rs4858616 | 0.51 | -0.139 | -0.031 | 0.869 | -0.011 | 0.057 | 0.057 | 0 | 0.83 | 0.71 | 0.9 | 0.66 | 0.89 | 0.7 |
| 3 | 24490035 | rs17014722 | 0.803 | -0.173 | -0.187 | 0.887 | 0.016 | -0.013 | -0.013 | 0 | 0.07 | 0.01 | 0.03 | 0.06 | 0.03 | 0.11 |
| 3 | 24490129 | rs4563353 | 0.318 | 0.147 | 0.065 | 0.894 | 0.013 | 0.116 | 0.116 | 0 | 0.91 | 0.74 | 0.93 | 0.72 | 0.92 | 0.82 |
| 3 | 24503729 | rs2362185 | 0.858 | 0.309 | -0.354 | 0.275 | -0.001 | 0 | 0 | 0 | 0.34 | 0.24 | 0.32 | 0.34 | 0.33 | 0.33 |
| 3 | 24511270 | rs982952 | 0.993 | 0.345 | -0.326 | 0.281 | -0.001 | -0.002 | -0.002 | 0 | 0.32 | 0.25 | 0.31 | 0.34 | 0.32 | 0.33 |
| 3 | 24512956 | rs4353759 | 1.091 | 0.341 | -0.428 | 0.279 | -0.016 | -0.017 | -0.017 | 0 | 0.32 | 0.14 | 0.3 | 0.34 | 0.32 | 0.33 |
| 3 | 24516064 | rs9845983 | 0.773 | 0.457 | -0.12 | 0.168 | 0.014 | -0.024 | -0.024 | 0 | 0.06 | 0.17 | 0.03 | 0 | 0.05 | 0.06 |
| 3 | 24516898 | rs9310744 | 0.67 | 0.597 | -0.204 | 0.4 | 0.001 | 0.002 | 0.002 | 0 | 0.42 | 0.56 | 0.37 | 0.44 | 0.4 | 0.52 |
| 3 | 24520283 | rs6792725 | 0.14 | 0.394 | -0.813 | 0.751 | -0.019 | -0.119 | -0.119 | 0 | 0.74 | 0.91 | 0.56 | 0.49 | 0.67 | 0.75 |
| 3 | 24533278 | rs2196427 | -0.36 | 0.195 | -0.243 | 0.375 | -0.011 | 0.008 | 0.008 | 0.004 | 0.42 | 0.35 | 0.39 | 0.41 | 0.52 | 0.39 |
| 3 | 24538267 | rs3749376 | -0.602 | 0.041 | 0.252 | 0.058 | -0.019 | -0.027 | -0.027 | 0 | 0.17 | 0.45 | 0.28 | 0.1 | 0.12 | 0.17 |
| 3 | 24538838 | rs892940 | -1.099 | -0.13 | -0.31 | 0.212 | -0.013 | -0.019 | -0.019 | 0 | 0.59 | 0.82 | 0.5 | 0.49 | 0.58 | 0.54 |

| | | | | | | | | | | | | | | | | |
|---|---|---|---|---|---|---|---|---|---|---|---|---|---|---|---|---|
| 3 | 24554704 | rs10490840 | -0.268 | 0.325 | -0.397 | 0.017 | -0.027 | 0.001 | 0.001 | 0 | 0.35 | 0.26 | 0.37 | 0.39 | 0.47 | 0.21 |
| 3 | 24562444 | rs9868777 | -0.647 | 0.231 | -0.517 | 0.275 | 0.009 | 0.006 | 0.006 | 1.7 | 0.57 | 0.63 | 0.49 | 0.46 | 0.6 | 0.47 |
| 3 | 24570579 | rs2117876 | -0.62 | 0.764 | -0.286 | 0.4 | -0.021 | 0.026 | 0.026 | 14.959 | 0.94 | 0.99 | 0.95 | 1 | 0.9 | 0.99 |
| 3 | 24574154 | rs11709849 | 1.084 | 0.374 | -0.704 | 0.57 | 0.044 | 0.036 | 0.036 | 0 | 0.45 | 0.12 | 0.45 | 0.32 | 0.34 | 0.38 |
| 3 | 24578418 | rs9850443 | 0.268 | 0.125 | -0.322 | 0.697 | -0.026 | -0.056 | -0.056 | 0 | 0.63 | 0.74 | 0.62 | 0.47 | 0.44 | 0.67 |
| 3 | 24585522 | rs2196438 | -1.091 | 0.152 | 0.67 | 0.54 | -0.047 | -0.006 | -0.006 | 0 | 0.18 | 0.6 | 0.16 | 0.17 | 0.09 | 0.29 |
| 3 | 24590744 | rs9838720 | -1.142 | 0.127 | 1.064 | 0.529 | -0.035 | -0.005 | -0.005 | 0 | 0.19 | 0.63 | 0.16 | 0.17 | 0.09 | 0.29 |
| 3 | 24590841 | rs9838616 | -1.107 | 0.127 | 1.042 | 0.529 | -0.047 | -0.006 | -0.006 | 0 | 0.18 | 0.6 | 0.16 | 0.17 | 0.09 | 0.29 |
| 3 | 24593374 | rs9853927 | -0.625 | 0.121 | 1.36 | 0.533 | -0.002 | 0 | 0 | 0 | 0.14 | 0.11 | 0.12 | 0.16 | 0.09 | 0.25 |
| 3 | 24617220 | rs6781066 | -2.052 | -0.023 | 1.162 | 0.451 | -0.003 | 0.002 | 0.002 | 0.812 | 0.24 | 0.68 | 0.19 | 0.17 | 0.12 | 0.29 |
| 3 | 24617928 | rs6784489 | -0.023 | 0.556 | 0.861 | 0.579 | -0.081 | -0.038 | -0.038 | 0 | 0.86 | 0.98 | 0.81 | 0.73 | 0.73 | 0.88 |
| 3 | 24625873 | rs2043578 | -1.429 | 0.113 | 2.045 | 0.511 | -0.034 | -0.056 | -0.056 | 0 | 0.16 | 0.25 | 0.08 | 0.03 | 0.1 | 0.2 |
| 3 | 24633038 | rs9878606 | -1.921 | 0.123 | 1.46 | 0.521 | 0.009 | 0.005 | 0.005 | 1.227 | 0.21 | 0.7 | 0.18 | 0.17 | 0.11 | 0.25 |
| 3 | 24633757 | rs7632924 | -1.614 | 0.65 | 1.461 | 0.373 | -0.003 | -0.017 | -0.017 | 0 | 0.08 | 0.31 | 0.04 | 0.02 | 0.03 | 0.03 |
| 3 | 24639726 | rs9836150 | -1.552 | -0.246 | 0.779 | 0.487 | -0.029 | -0.055 | -0.055 | 0 | 0.1 | 0.25 | 0.05 | 0.02 | 0.05 | 0.04 |
| 3 | 24646076 | rs10440133 | -0.277 | -0.098 | 0.374 | 1.305 | -0.005 | -0.053 | -0.053 | 0 | 0.16 | 0.1 | 0.13 | 0.35 | 0.19 | 0.19 |
| 3 | 24646108 | rs10490835 | -0.858 | 0.637 | 0.412 | 1.275 | -0.005 | 0 | 0 | 0 | 0.08 | 0.06 | 0.14 | 0.12 | 0.19 | 0.31 |
| 3 | 24646154 | rs10490834 | -0.442 | 0.437 | 1.202 | 1.022 | 0.01 | -0.045 | -0.045 | 0 | 0.08 | 0.23 | 0.06 | 0 | 0.06 | 0.04 |
| 3 | 24652813 | rs17014951 | -1.351 | -0.359 | 0.778 | 1.089 | -0.001 | 0.057 | 0.057 | 0.635 | 0.25 | 0.63 | 0.27 | 0.5 | 0.27 | 0.44 |
| 3 | 24654338 | rs920117 | -0.413 | -0.391 | 0.949 | 1.142 | 0.024 | -0.046 | -0.046 | 0 | 0.14 | 0.06 | 0.09 | 0.37 | 0.12 | 0.26 |
| 3 | 24657675 | rs9819978 | -0.926 | -0.546 | 1.098 | 1.036 | 0.051 | 0.001 | 0.001 | 0 | 0.08 | 0.15 | 0.14 | 0.11 | 0.14 | 0.15 |
| 3 | 24658697 | rs7618297 | -2.208 | -0.265 | 0.942 | 1.15 | -0.013 | -0.091 | -0.091 | 0 | 0.34 | 0.87 | 0.32 | 0.48 | 0.33 | 0.49 |
| 3 | 24669437 | rs7613236 | -1.978 | -0.258 | 1.044 | 0.722 | -0.007 | 0.173 | 0.173 | 0.885 | 0.48 | 0.77 | 0.45 | 0.84 | 0.49 | 0.68 |
| 3 | 24669959 | rs2362758 | -2.149 | -0.265 | 0.976 | 0.723 | -0.013 | 0.229 | 0.229 | 1.738 | 0.5 | 0.81 | 0.46 | 0.85 | 0.51 | 0.7 |
| 3 | 24683601 | rs6550882 | -2.062 | -0.257 | 1.444 | 0.927 | 0.072 | 0.044 | 0.044 | 2.041 | 0.45 | 0.79 | 0.34 | 0.64 | 0.37 | 0.52 |
| 3 | 24684637 | rs7627134 | -0.498 | -0.3 | 1.636 | 1.266 | -0.011 | 0.024 | 0.024 | 0 | 0.14 | 0.19 | 0.09 | 0.32 | 0.1 | 0.26 |
| 3 | 24691022 | rs7610415 | -0.747 | -0.274 | 1.555 | 1.259 | -0.004 | 0.007 | 0.007 | 0 | 0.26 | 0.3 | 0.21 | 0.32 | 0.22 | 0.23 |
| 3 | 24694021 | rs7619728 | -0.246 | -0.116 | 1.58 | 0.866 | -0.005 | -0.047 | -0.047 | 0 | 0.1 | 0.14 | 0.06 | 0 | 0.07 | 0.06 |
| 3 | 24702734 | rs13433753 | -0.559 | -0.182 | 1.287 | 0.96 | -0.013 | 0.005 | 0.005 | 0.007 | 0.28 | 0.33 | 0.23 | 0.64 | 0.21 | 0.35 |
| 3 | 24705221 | rs9812637 | -1.158 | -0.119 | 0.504 | 0.904 | -0.02 | 0.093 | 0.093 | 1.333 | 0.47 | 0.81 | 0.46 | 0.72 | 0.5 | 0.67 |
| 3 | 24708172 | rs17015018 | -0.384 | -0.066 | 0.428 | 0.879 | 0.039 | -0.016 | -0.016 | 0 | 0.06 | 0.23 | 0.04 | 0.02 | 0.06 | 0.17 |
| 3 | 24708874 | rs6767967 | -0.89 | -0.107 | -0.563 | 0.99 | -0.006 | 0.094 | 0.094 | 0.962 | 0.49 | 0.73 | 0.46 | 0.73 | 0.53 | 0.7 |
| 3 | 24715105 | rs1984874 | -0.9 | -0.072 | -1.127 | 0.936 | -0.026 | 0.167 | 0.167 | 1.689 | 0.5 | 0.92 | 0.43 | 0.73 | 0.5 | 0.66 |
| 3 | 24716668 | rs2362755 | -0.519 | 0.496 | -0.446 | 1.065 | 0.02 | 0.025 | 0.025 | 0 | 0.36 | 0.51 | 0.34 | 0.45 | 0.47 | 0.51 |
| 3 | 24723928 | rs7628773 | 0.086 | 1.439 | -1.221 | 0.741 | -0.06 | -0.003 | -0.003 | 0.083 | 0.79 | 0.93 | 0.63 | 0.79 | 0.66 | 0.8 |
| 3 | 24724883 | rs4858630 | 0.478 | 0.784 | -1.674 | 0.549 | 0.02 | -0.04 | -0.04 | 0 | 0.4 | 0.25 | 0.32 | 0.48 | 0.33 | 0.53 |
| 3 | 24727377 | rs7629359 | -0.743 | 0.839 | -1.64 | 0.599 | -0.062 | 0.01 | 0.01 | 0.05 | 0.78 | 0.93 | 0.64 | 0.81 | 0.65 | 0.8 |
| 3 | 24727911 | rs11129159 | 0.738 | 0.428 | -2.151 | 0.406 | 0.003 | -0.027 | -0.027 | 0 | 0.28 | 0.22 | 0.25 | 0.42 | 0.23 | 0.39 |
| 3 | 24729694 | rs4365572 | 0.323 | -0.153 | -1.865 | 0.151 | 0.016 | -0.018 | -0.018 | 0 | 0.32 | 0.21 | 0.28 | 0.34 | 0.33 | 0.21 |
| 3 | 24729704 | rs4472000 | -0.131 | 0.099 | -1.613 | 0.11 | 0.014 | 0.002 | 0.002 | 0.119 | 0.37 | 0.26 | 0.3 | 0.35 | 0.37 | 0.34 |
| 3 | 24731037 | rs4600774 | -0.617 | 1.162 | -1.688 | -0.516 | -0.026 | 0.053 | 0.053 | 0.87 | 0.75 | 0.91 | 0.81 | 0.88 | 0.71 | 0.7 |
| 3 | 24732019 | rs4343572 | -0.17 | 1.07 | -1.007 | -0.674 | 0 | -0.017 | -0.017 | 0 | 0.7 | 0.7 | 0.72 | 0.89 | 0.65 | 0.63 |
| 3 | 24742308 | rs7643059 | -0.284 | 1.039 | -0.708 | -0.706 | 0.003 | -0.043 | -0.043 | 0 | 0.72 | 0.67 | 0.74 | 0.89 | 0.7 | 0.66 |
| 3 | 24746378 | rs9862855 | 0.377 | 1.099 | 0.023 | -0.615 | 0.008 | -0.035 | -0.035 | 0 | 0.59 | 0.5 | 0.53 | 0.71 | 0.53 | 0.59 |
| 3 | 24746454 | rs4558693 | 0.83 | 1.107 | 0.043 | -0.614 | 0 | -0.001 | -0.001 | 0 | 0.14 | 0.17 | 0.21 | 0.18 | 0.17 | 0.08 |
| 3 | 24746641 | rs3934760 | -1.312 | 1.115 | -0.055 | -0.617 | -0.008 | 0.039 | 0.039 | 0.272 | 0.78 | 0.9 | 0.84 | 0.92 | 0.76 | 0.73 |
| 3 | 24746852 | rs3934758 | 1.162 | 1.114 | 0.201 | -0.757 | 0.018 | 0.118 | 0.118 | 0 | 0.47 | 0.28 | 0.44 | 0.71 | 0.41 | 0.55 |
| 3 | 24746951 | rs3934759 | 1.424 | 1.025 | 0.143 | -0.835 | -0.004 | -0.023 | -0.023 | 0 | 0.28 | 0.19 | 0.31 | 0.56 | 0.25 | 0.39 |
| 3 | 24747030 | rs4073046 | -0.069 | 1.06 | 0.031 | -0.841 | -0.004 | 0.071 | 0.071 | 1.326 | 0.48 | 0.66 | 0.62 | 0.77 | 0.48 | 0.54 |
| 3 | 24748506 | rs4585139 | -0.26 | 0.94 | -0.085 | -0.791 | 0.007 | -0.092 | -0.092 | 0.116 | 0.63 | 0.59 | 0.73 | 0.91 | 0.61 | 0.68 |
| 3 | 24751005 | rs4077423 | 0.359 | 0.951 | -0.273 | -0.79 | -0.001 | -0.01 | -0.01 | 0.144 | 0.42 | 0.49 | 0.57 | 0.75 | 0.39 | 0.52 |
| 3 | 24751085 | rs4077422 | 1.164 | 0.919 | -0.308 | -0.731 | -0.005 | 0.021 | 0.021 | 0 | 0.3 | 0.48 | 0.38 | 0.56 | 0.25 | 0.45 |
| 3 | 24751734 | rs7651159 | 1.336 | 0.973 | -0.78 | -0.541 | -0.001 | 0.025 | 0.025 | 0 | 0.22 | 0.12 | 0.16 | 0.17 | 0.21 | 0.16 |
| 3 | 24752371 | rs13081423 | 0.696 | 1.007 | -0.56 | -0.17 | -0.052 | -0.042 | -0.042 | 0.053 | 0.13 | 0.01 | 0.2 | 0.19 | 0.17 | 0.08 |
| 3 | 24755790 | rs3935025 | 0.036 | 0.915 | -0.046 | -0.425 | 0.002 | 0.064 | 0.064 | 3.015 | 0.47 | 0.37 | 0.56 | 0.74 | 0.44 | 0.5 |
| 3 | 24756503 | rs3934713 | 0.003 | 0.947 | -0.165 | -0.388 | 0.001 | 0.02 | 0.02 | 2.804 | 0.48 | 0.45 | 0.57 | 0.74 | 0.44 | 0.49 |
| 3 | 24758368 | rs4858125 | -0.254 | 1.162 | -0.502 | -0.984 | 0 | 0.009 | 0.009 | 2.51 | 0.51 | 0.57 | 0.64 | 0.86 | 0.49 | 0.72 |
| 3 | 24761188 | rs4411831 | -0.209 | -0.312 | -1.319 | -0.76 | 0.048 | 0.049 | 0.049 | 0 | 0.48 | 0.69 | 0.33 | 0.43 | 0.42 | 0.52 |
| 3 | 24762070 | rs9864779 | -0.083 | -0.427 | -1.026 | -0.865 | -0.02 | -0.017 | -0.017 | 0.924 | 0.43 | 0.56 | 0.28 | 0.34 | 0.34 | 0.51 |
| 3 | 24764500 | rs9310754 | 0.916 | -0.127 | -1.385 | -0.699 | -0.029 | -0.026 | -0.026 | 0 | 0.31 | 0.23 | 0.54 | 0.47 | 0.37 | 0.29 |
| 3 | 24766256 | rs6794687 | 0.466 | -0.709 | -1.512 | -1.073 | 0 | -0.028 | -0.028 | 0 | 0.28 | 0.21 | 0.18 | 0.03 | 0.28 | 0.17 |
| 3 | 24767611 | rs4241531 | -1.216 | -0.421 | -1.717 | -0.433 | 0.002 | -0.042 | -0.042 | 2.76 | 0.73 | 0.85 | 0.77 | 0.57 | 0.76 | 0.69 |
| 3 | 24769500 | rs9882830 | 0.594 | -0.48 | -1.429 | -0.14 | -0.079 | -0.008 | -0.008 | 1.164 | 0.28 | 0.51 | 0.16 | 0.29 | 0.15 | 0.38 |
| 3 | 24771996 | rs4478032 | 0.302 | -0.897 | -1.256 | -0.369 | -0.078 | -0.007 | -0.007 | 0.247 | 0.27 | 0.48 | 0.18 | 0.27 | 0.16 | 0.28 |
| 3 | 24772115 | rs4487199 | -0.511 | -0.917 | -1.284 | -0.417 | -0.013 | -0.005 | -0.005 | 0 | 0.39 | 0.59 | 0.25 | 0.33 | 0.27 | 0.42 |
| 3 | 24773695 | rs9876486 | 1.334 | -0.178 | -1.223 | -0.207 | 0.003 | 0.014 | 0.014 | 0 | 0.12 | 0.07 | 0.07 | 0.06 | 0.11 | 0.17 |
| 3 | 24776203 | rs12486584 | 0.561 | -0.028 | -0.622 | 0.01 | 0.017 | 0.057 | 0.057 | 0.113 | 0.33 | 0.54 | 0.49 | 0.54 | 0.38 | 0.28 |
| 3 | 24778351 | rs7646431 | 0.82 | -0.254 | -0.713 | 0.401 | 0.015 | 0.014 | 0.014 | 0.07 | 0.3 | 0.33 | 0.21 | 0.2 | 0.22 | 0.2 |
| 3 | 24778690 | rs6766822 | 0.366 | 0.513 | -0.742 | 0.536 | -0.004 | -0.002 | -0.002 | 0 | 0.47 | 0.49 | 0.28 | 0.34 | 0.33 | 0.33 |

| | | | | | | | | | | | | | | | | |
|---|---|---|---|---|---|---|---|---|---|---|---|---|---|---|---|---|
| 3 | 24779648 | rs6772760 | -2.433 | 0.6 | -1.36 | 0.739 | -0.011 | -0.008 | -0.008 | 1.47 | 0.89 | 0.98 | 0.92 | 0.9 | 0.88 | 0.94 |
| 3 | 24780266 | rs4075988 | 1.485 | 0.816 | -1.255 | 0.813 | -0.006 | -0.01 | -0.01 | 0 | 0.15 | 0.11 | 0.2 | 0.37 | 0.15 | 0.27 |
| 3 | 24782701 | rs13433781 | 1.71 | 0.785 | -1.279 | 0.566 | 0.007 | -0.005 | -0.005 | 0 | 0.08 | 0.08 | 0.08 | 0.06 | 0.12 | 0.1 |
| 3 | 24784958 | rs11919098 | 1.905 | 0.527 | -1.317 | 0.715 | -0.019 | -0.054 | -0.054 | 0.002 | 0.13 | 0.04 | 0.15 | 0.27 | 0.11 | 0.23 |
| 3 | 24788904 | rs9853202 | -0.477 | 0.475 | -1.153 | 0.538 | -0.001 | -0.001 | -0.001 | 0.927 | 0.42 | 0.44 | 0.66 | 0.56 | 0.55 | 0.5 |
| 3 | 24790835 | rs9310755 | -0.439 | 0.168 | -1.096 | 0.625 | -0.001 | -0.002 | -0.002 | 0 | 0.38 | 0.33 | 0.33 | 0.37 | 0.36 | 0.44 |
| 3 | 24791375 | rs4858647 | 1.96 | -0.277 | -0.928 | 0.615 | -0.048 | 0.006 | 0.006 | 0 | 0.25 | 0.34 | 0.12 | 0.3 | 0.1 | 0.2 |
| 3 | 24791678 | rs11920796 | 1.828 | 0.254 | -0.899 | 0.37 | -0.018 | -0.057 | -0.057 | 0 | 0.14 | 0.04 | 0.16 | 0.27 | 0.11 | 0.23 |
| 3 | 24791970 | rs11920964 | 1.751 | 0.174 | -0.885 | 0.301 | -0.002 | -0.019 | -0.019 | 0 | 0.11 | 0.09 | 0.04 | 0 | 0.08 | 0.1 |
| 3 | 24791988 | rs9860870 | -0.076 | 0.301 | -0.956 | 0.396 | -0.001 | -0.001 | -0.001 | 0 | 0.28 | 0.31 | 0.25 | 0.34 | 0.25 | 0.4 |
| 3 | 24792424 | rs9310756 | 0.426 | 0.257 | -1.283 | 0.105 | -0.002 | 0.005 | 0.005 | 0 | 0.14 | 0.12 | 0.07 | 0.07 | 0.14 | 0.17 |
| 3 | 24793858 | rs7429711 | -0.31 | 0.867 | -1.106 | 0.62 | 0.016 | 0.005 | 0.005 | 0 | 0.33 | 0.36 | 0.57 | 0.6 | 0.46 | 0.52 |
| 3 | 24794908 | rs11707787 | 0.696 | 0.689 | -1.188 | 0.519 | -0.023 | -0.023 | -0.023 | 1.037 | 0.42 | 0.46 | 0.27 | 0.29 | 0.3 | 0.32 |
| 3 | 24795619 | rs6776123 | -1.678 | 1.74 | -1.411 | 0.991 | -0.023 | 0.024 | 0.024 | 0.225 | 0.8 | 0.95 | 0.86 | 0.88 | 0.8 | 0.88 |
| 3 | 24797615 | rs13088027 | -0.653 | 1.668 | -1.377 | 0.943 | -0.012 | 0.005 | 0.005 | 0.961 | 0.36 | 0.32 | 0.63 | 0.63 | 0.53 | 0.42 |
| 3 | 24798187 | rs4241536 | 0.683 | 2.319 | -1.028 | 1.249 | -0.111 | -0.114 | -0.114 | 0 | 0.17 | 0.04 | 0.47 | 0.53 | 0.31 | 0.28 |
| 3 | 24799541 | rs9845065 | 0.848 | 2.295 | -0.861 | 1.207 | 0.035 | -0.017 | -0.017 | 0 | 0.09 | 0.23 | 0.08 | 0.06 | 0.1 | 0.1 |
| 3 | 24800829 | rs7649758 | 0.142 | 2.577 | -0.872 | 1.488 | 0.13 | 0.144 | 0.144 | 0 | 0.12 | 0.25 | 0.39 | 0.35 | 0.29 | 0.15 |
| 3 | 24808506 | rs9827067 | 0.091 | 0.57 | -1.024 | 1.298 | -0.051 | -0.032 | -0.032 | 0 | 0.25 | 0.2 | 0.54 | 0.59 | 0.41 | 0.38 |
| 3 | 24812844 | rs4533581 | -0.531 | 0.472 | -0.926 | 1.18 | 0.005 | -0.017 | -0.017 | 0 | 0.75 | 0.68 | 0.8 | 0.81 | 0.72 | 0.82 |
| 3 | 24817052 | rs4276127 | -0.843 | 0.488 | -0.916 | 1.186 | -0.001 | 0.003 | 0.003 | 0.336 | 0.77 | 0.82 | 0.84 | 0.88 | 0.76 | 0.86 |
| 3 | 24818050 | rs11927156 | 0.901 | 0.394 | -0.961 | 1.199 | 0.025 | 0.025 | 0.025 | 0 | 0.5 | 0.53 | 0.27 | 0.21 | 0.31 | 0.44 |
| 3 | 24824302 | rs7640332 | 0.021 | 0.627 | -0.937 | 1.258 | -0.021 | -0.015 | -0.015 | 0.069 | 0.26 | 0.27 | 0.53 | 0.59 | 0.39 | 0.39 |
| 3 | 24825191 | rs4131189 | 1.053 | 0.68 | -1.189 | 1.227 | -0.111 | -0.071 | -0.071 | 0 | 0.17 | 0.04 | 0.47 | 0.25 | 0.31 | 0.29 |
| 3 | 24829088 | rs9881863 | 0.178 | 0.367 | -1.083 | 0.552 | 0.022 | -0.019 | -0.019 | 0.411 | 0.12 | 0.25 | 0.12 | 0.14 | 0.14 | 0.12 |
| 3 | 24829516 | rs9844787 | 0.456 | 0.223 | -1.382 | 0.664 | -0.006 | -0.007 | -0.007 | 0.026 | 0.51 | 0.52 | 0.3 | 0.3 | 0.35 | 0.47 |
| 3 | 24834378 | rs4361227 | -0.045 | 0.568 | -1.723 | 0.848 | -0.038 | -0.021 | -0.021 | 0.201 | 0.76 | 0.93 | 0.82 | 0.8 | 0.72 | 0.86 |
| 3 | 24836454 | rs4470447 | 0.441 | 1.063 | -1.56 | 1.083 | -0.079 | -0.069 | -0.069 | 0 | 0.13 | 0.03 | 0.44 | 0.3 | 0.32 | 0.34 |
| 3 | 24836522 | rs7617311 | 1.013 | -0.325 | -2.44 | 0.766 | 0.021 | -0.068 | -0.068 | 0.336 | 0.32 | 0.07 | 0.22 | 0.37 | 0.25 | 0.34 |
| 3 | 24838818 | rs6802797 | 0.245 | -0.988 | -2.011 | 0.514 | -0.036 | -0.03 | -0.03 | 0 | 0.66 | 0.36 | 0.46 | 0.54 | 0.54 | 0.55 |
| 3 | 24839916 | rs6550908 | -1.124 | -0.784 | -1.762 | 0.628 | -0.002 | 0.001 | 0.001 | 0.107 | 0.9 | 0.9 | 0.93 | 0.87 | 0.91 | 0.95 |
| 3 | 24841723 | rs6550911 | -0.326 | -1.092 | -1.885 | 0.391 | 0.016 | 0.012 | 0.012 | 0 | 0.8 | 0.89 | 0.9 | 0.86 | 0.84 | 0.93 |
| 3 | 24842231 | rs6797905 | 0.039 | -1.01 | -1.053 | 0.579 | 0.01 | 0.01 | 0.01 | 1.26 | 0.66 | 0.67 | 0.47 | 0.57 | 0.5 | 0.58 |
| 3 | 24847080 | rs6781264 | -0.25 | -1.043 | -0.657 | 0.737 | -0.003 | 0.002 | 0.002 | 1.45 | 0.87 | 0.82 | 0.9 | 0.87 | 0.87 | 0.93 |
| 3 | 24850546 | rs7623169 | 0.63 | -1.099 | -0.837 | 0.753 | -0.007 | -0.001 | -0.001 | 0.598 | 0.09 | 0.12 | 0.06 | 0.06 | 0.07 | 0.03 |
| 3 | 24852307 | rs11713190 | 0.564 | -1.008 | -1.101 | 0.838 | -0.03 | -0.067 | -0.067 | 0 | 0.33 | 0.08 | 0.23 | 0.38 | 0.3 | 0.36 |
| 3 | 24858630 | rs7429669 | 0.537 | -0.852 | 0.232 | 0.912 | 0.004 | 0.015 | 0.015 | 0.772 | 0.87 | 0.77 | 0.9 | 0.87 | 0.88 | 0.93 |
| 3 | 24883013 | rs6550921 | -0.989 | -0.772 | 0.196 | 1.163 | -0.028 | 0.11 | 0.11 | 0 | 0.19 | 0.34 | 0.18 | 0.34 | 0.14 | 0.13 |
| 3 | 24886249 | rs6774513 | -0.862 | -0.7 | 0.377 | 1.256 | -0.008 | 0.153 | 0.153 | 0.133 | 0.13 | 0.28 | 0.16 | 0.38 | 0.1 | 0.13 |
| 3 | 24888242 | rs1994909 | 0.191 | -1.317 | 0.266 | 0.516 | -0.02 | -0.019 | -0.019 | 2.452 | 0.85 | 0.72 | 0.89 | 0.92 | 0.9 | 0.96 |
| 3 | 24891014 | rs10865798 | -0.841 | -1.249 | -0.237 | -0.483 | 0.003 | 0.007 | 0.007 | 0 | 0.61 | 0.76 | 0.73 | 0.96 | 0.64 | 0.65 |
| 3 | 24891120 | rs9837889 | -0.519 | -1.649 | -0.304 | 0.156 | -0.063 | -0.007 | -0.007 | 0.808 | 0.32 | 0.56 | 0.16 | 0.68 | 0.17 | 0.27 |
| 3 | 24902942 | rs13064710 | 0.74 | -1.166 | -0.774 | 0.102 | -0.124 | 0.008 | 0.008 | 0.802 | 0.18 | 0.02 | 0.44 | 0.13 | 0.4 | 0.34 |
| 3 | 24903229 | rs7651101 | 0.123 | -1.964 | -0.05 | 0.58 | -0.06 | 0.042 | 0.042 | 0 | 0.14 | 0.26 | 0.12 | 0.26 | 0.05 | 0.05 |
| 3 | 24904904 | rs7610607 | -1.24 | -1.88 | -0.35 | 0.472 | 0.13 | 0.126 | 0.126 | 0.372 | 0.43 | 0.85 | 0.31 | 0.76 | 0.23 | 0.31 |
| 3 | 24908670 | rs9845715 | -0.094 | -1.704 | 0.184 | 0.931 | -0.017 | 0.148 | 0.148 | 0 | 0.06 | 0.21 | 0.04 | 0.18 | 0.02 | 0.04 |
| 3 | 24911917 | rs9310762 | -0.861 | -1.962 | 0.626 | 1.467 | -0.029 | 0.067 | 0.067 | 0.571 | 0.21 | 0.6 | 0.27 | 0.61 | 0.14 | 0.16 |
| 3 | 24913845 | rs2362771 | -1.774 | -1.915 | 0.189 | 1.14 | -0.053 | 0.151 | 0.151 | 0 | 0.63 | 0.93 | 0.76 | 0.87 | 0.65 | 0.67 |
| 3 | 24914550 | rs11129167 | -1.208 | -2.063 | 0.716 | 1.509 | 0.004 | 0.012 | 0.012 | 1.595 | 0.52 | 0.62 | 0.61 | 0.69 | 0.59 | 0.65 |
| 3 | 24921767 | rs11711907 | 0.564 | -2.069 | 0.441 | 1.646 | -0.015 | 0.025 | 0.025 | 0 | 0.06 | 0.14 | 0.02 | 0.15 | 0.03 | 0.1 |
| 3 | 24924495 | rs6550923 | 0.147 | -2.866 | -0.233 | 1.394 | 0.058 | 0.014 | 0.014 | 0.973 | 0.51 | 0.74 | 0.35 | 0.51 | 0.4 | 0.53 |
| 3 | 24927203 | rs10510551 | 0.662 | -2.528 | -0.042 | 1.504 | -0.166 | 0.067 | 0.067 | 1.147 | 0.28 | 0.03 | 0.43 | 0.14 | 0.43 | 0.42 |
| 3 | 24928351 | rs9819866 | -0.237 | -2.5 | 0.374 | 1.614 | -0.03 | 0.095 | 0.095 | 1.107 | 0.19 | 0.46 | 0.15 | 0.26 | 0.13 | 0.12 |
| 3 | 24931873 | rs9883880 | -1.623 | -2.498 | 0.198 | 1.583 | -0.042 | 0.169 | 0.169 | 0 | 0.62 | 0.92 | 0.76 | 0.88 | 0.64 | 0.67 |
| 3 | 24933325 | rs11711279 | 0.092 | -2.528 | 0.306 | 1.748 | -0.016 | 0.076 | 0.076 | 0 | 0.07 | 0.16 | 0.04 | 0.32 | 0.03 | 0.11 |
| 3 | 24942213 | rs2363518 | -1.396 | -2.216 | 0.038 | 1.09 | 0.015 | 0.132 | 0.132 | 0 | 0.61 | 0.85 | 0.77 | 0.88 | 0.68 | 0.65 |
| 3 | 24943995 | rs6769702 | -0.047 | -2.349 | 0.301 | 1.182 | 0.023 | 0.069 | 0.069 | 0.824 | 0.19 | 0.37 | 0.18 | 0.26 | 0.18 | 0.14 |
| 3 | 24945215 | rs6550926 | -1.706 | -2.647 | -0.771 | 0.775 | 0.015 | 0.119 | 0.119 | 0.931 | 0.65 | 0.88 | 0.79 | 0.87 | 0.71 | 0.7 |
| 3 | 24948515 | rs2363520 | 0.323 | -2.052 | -0.744 | 0.746 | -0.125 | 0.125 | 0.125 | 0 | 0.33 | 0.06 | 0.45 | 0.13 | 0.47 | 0.4 |
| 3 | 24951858 | rs4568101 | -0.785 | -2.134 | -0.401 | 0.893 | 0.02 | 0.009 | 0.009 | 0 | 0.53 | 0.57 | 0.64 | 0.38 | 0.65 | 0.55 |
| 3 | 24955790 | rs35342260 | -0.513 | -0.703 | -0.71 | 0.721 | -0.021 | -0.031 | -0.031 | 0 | 0.39 | 0.22 | 0.29 | 0.41 | 0.49 | 0.43 |
| 3 | 24961167 | rs6774983 | -1.293 | -1.152 | -1.031 | 0.41 | 0.016 | 0.168 | 0.168 | 0 | 0.52 | 0.85 | 0.42 | 0.76 | 0.55 | 0.64 |
| 3 | 24963629 | rs17015448 | 0.603 | -1.249 | -0.788 | 0.657 | 0.011 | -0.005 | -0.005 | 0.027 | 0.07 | 0.03 | 0.03 | 0.11 | 0.04 | 0.09 |
| 3 | 24964350 | rs6782972 | -1.077 | -1.012 | -0.149 | 0.72 | -0.009 | 0 | 0 | 0.047 | 0.3 | 0.55 | 0.22 | 0.31 | 0.22 | 0.23 |
| 3 | 24964676 | rs17015451 | -0.083 | -1.046 | 0.178 | 0.717 | 0.016 | 0.024 | 0.024 | 0 | 0.19 | 0.26 | 0.13 | 0.28 | 0.19 | 0.18 |
| 3 | 24966768 | rs10510553 | -0.292 | -1.147 | 0.283 | 0.79 | -0.054 | 0.044 | 0.044 | 0.011 | 0.14 | 0.36 | 0.11 | 0.21 | 0.06 | 0.11 |
| 3 | 24972416 | rs2363526 | -1.799 | -1.189 | 0.156 | 0.586 | -0.003 | 0.049 | 0.049 | 0.003 | 0.44 | 0.65 | 0.33 | 0.71 | 0.5 | 0.59 |
| 3 | 24972792 | rs2363527 | 0.803 | -1.017 | 0.195 | 0.619 | 0.013 | -0.015 | -0.015 | 0 | 0.15 | 0.17 | 0.11 | 0.1 | 0.16 | 0.11 |
| 3 | 24974882 | rs9878781 | -1.192 | -1.095 | 0.087 | 0.559 | 0.021 | 0.01 | 0.01 | 0 | 0.38 | 0.45 | 0.3 | 0.55 | 0.46 | 0.51 |

| | | | | | | | | | | | | | | | | |
|---|---|---|---|---|---|---|---|---|---|---|---|---|---|---|---|---|
| 3 | 24979142 | rs2363530 | -2.13 | -1.093 | -0.11 | 0.379 | -0.049 | 0.129 | 0.129 | 0 | 0.53 | 0.93 | 0.41 | 0.74 | 0.52 | 0.63 |
| 3 | 24981701 | rs7614855 | -1.404 | -0.675 | 0.135 | 0.646 | 0.002 | -0.008 | -0.008 | 0 | 0.41 | 0.52 | 0.32 | 0.71 | 0.48 | 0.58 |
| 3 | 24988244 | rs11129172 | 1.111 | -0.377 | -0.426 | 0.721 | -0.09 | -0.104 | -0.104 | 0.028 | 0.14 | 0.03 | 0.13 | 0.35 | 0.26 | 0.32 |
| 3 | 24991227 | rs17195094 | 0.994 | -0.49 | -0.254 | 0.432 | -0.022 | 0.016 | 0.016 | 0 | 0.12 | 0.07 | 0.09 | 0 | 0.16 | 0.07 |
| 3 | 24997512 | rs6780762 | -1.365 | -0.633 | -0.144 | 0.514 | 0.029 | 0.109 | 0.109 | 0 | 0.49 | 0.83 | 0.38 | 0.68 | 0.52 | 0.66 |
| 3 | 25004369 | rs12487571 | 0.642 | -0.537 | -0.366 | 0.221 | 0.057 | -0.002 | -0.002 | 0.013 | 0.48 | 0.32 | 0.58 | 0.51 | 0.41 | 0.51 |
| 3 | 25004487 | rs12490596 | 0.449 | -0.503 | -0.399 | 0.249 | 0.013 | 0.003 | 0.003 | 0.76 | 0.51 | 0.45 | 0.6 | 0.6 | 0.41 | 0.56 |
| 3 | 25006453 | rs4858670 | -0.387 | 0.012 | -0.857 | -0.21 | -0.033 | 0.088 | 0.088 | 0 | 0.68 | 0.82 | 0.81 | 0.86 | 0.69 | 0.85 |
| 3 | 25007199 | rs6550928 | -1.411 | -0.089 | -0.835 | -0.224 | 0.002 | 0 | 0 | 0.317 | 0.81 | 0.82 | 0.9 | 0.92 | 0.85 | 0.92 |
| 3 | 25009827 | rs9849459 | 0.56 | -0.14 | -1.128 | 0.248 | 0.029 | 0.008 | 0.008 | 0.196 | 0.5 | 0.58 | 0.62 | 0.54 | 0.42 | 0.51 |
| 3 | 25015541 | rs4502548 | -0.257 | 0.24 | -1.721 | 0.49 | -0.107 | -0.08 | -0.08 | 0 | 0.68 | 0.9 | 0.74 | 0.63 | 0.59 | 0.62 |
| 3 | 25021998 | rs12635887 | 0.555 | 0.658 | -1.443 | 0.846 | -0.074 | -0.083 | -0.083 | 0 | 0.15 | 0.04 | 0.16 | 0.3 | 0.28 | 0.32 |
| 3 | 25027071 | rs7639868 | 0.565 | 0.635 | -1.381 | 0.857 | -0.029 | -0.031 | -0.031 | 0.212 | 0.16 | 0.1 | 0.17 | 0.28 | 0.28 | 0.32 |
| 3 | 25033188 | rs12496073 | 0.549 | -0.48 | -1.692 | 0.56 | 0.006 | -0.09 | -0.09 | 0.441 | 0.32 | 0.15 | 0.5 | 0.52 | 0.32 | 0.46 |
| 3 | 25037457 | rs6771747 | 0.195 | -0.356 | -1.687 | 0.394 | 0.017 | 0.053 | 0.053 | 0.072 | 0.54 | 0.75 | 0.43 | 0.62 | 0.48 | 0.64 |
| 3 | 25046463 | rs12490228 | 1.603 | -0.149 | -1.899 | 0.389 | -0.09 | -0.053 | -0.053 | 0 | 0.19 | 0.02 | 0.36 | 0.2 | 0.26 | 0.22 |
| 3 | 25047254 | rs9842936 | -0.214 | -0.081 | -1.643 | 0.201 | -0.001 | -0.005 | -0.005 | 0 | 0.78 | 0.8 | 0.84 | 0.88 | 0.78 | 0.87 |
| 3 | 25047446 | rs4858678 | 1.691 | 0.035 | -1.84 | 0.511 | -0.09 | -0.085 | -0.085 | 0 | 0.19 | 0.03 | 0.37 | 0.25 | 0.26 | 0.22 |
| 3 | 25049310 | rs11129176 | 1.714 | 0.137 | -1.825 | 0.403 | -0.081 | -0.047 | -0.047 | 1.603 | 0.19 | 0.03 | 0.37 | 0.2 | 0.26 | 0.22 |
| 3 | 25050218 | rs9830542 | 0.688 | -0.588 | -1.822 | 0.51 | 0.06 | 0.015 | 0.015 | 0 | 0.41 | 0.66 | 0.27 | 0.35 | 0.24 | 0.27 |
| 3 | 25051076 | rs11711307 | 1.14 | 0.033 | -1.902 | 0.724 | -0.076 | -0.115 | -0.115 | 0 | 0.17 | 0.03 | 0.19 | 0.39 | 0.25 | 0.4 |
| 3 | 25052333 | rs6550934 | -0.034 | -0.603 | -1.839 | 0.441 | -0.036 | 0.023 | 0.023 | 0.514 | 0.56 | 0.82 | 0.47 | 0.59 | 0.52 | 0.56 |
| 3 | 25052754 | rs4241542 | 0.812 | 0.29 | -1.363 | 0.37 | -0.055 | -0.067 | -0.067 | 0.281 | 0.61 | 0.81 | 0.67 | 0.48 | 0.52 | 0.47 |
| 3 | 25053482 | rs4280597 | 0.89 | 0.28 | -0.84 | 0.298 | 0.005 | -0.017 | -0.017 | 0.333 | 0.69 | 0.69 | 0.75 | 0.79 | 0.69 | 0.78 |
| 3 | 25058757 | rs1021701 | 0.038 | 0.222 | -1.24 | 0.351 | -0.028 | 0.012 | 0.012 | 0 | 0.78 | 0.92 | 0.85 | 0.79 | 0.78 | 0.78 |
| 3 | 25059227 | rs1021702 | 1.376 | 0.106 | -0.863 | 0.219 | 0.065 | -0.002 | -0.002 | 0 | 0.48 | 0.21 | 0.53 | 0.4 | 0.43 | 0.46 |
| 3 | 25064354 | rs1604006 | 0.882 | 0.381 | -0.37 | -0.673 | 0.003 | -0.011 | -0.011 | 0.046 | 0.6 | 0.57 | 0.68 | 0.78 | 0.54 | 0.6 |
| 3 | 25065129 | rs873429 | -0.411 | -0.488 | -0.571 | -1.069 | -0.062 | 0.068 | 0.068 | 0 | 0.14 | 0.03 | 0.14 | 0 | 0.18 | 0.06 |
| 3 | 25068617 | rs1604003 | 1.665 | -0.753 | -0.321 | -1.21 | -0.002 | -0.034 | -0.034 | 1.138 | 0.66 | 0.64 | 0.57 | 0.88 | 0.69 | 0.69 |
| 3 | 25070898 | rs4482635 | -0.025 | -0.699 | -0.744 | -1.157 | 0.043 | 0.034 | 0.034 | 0.663 | 0.82 | 0.93 | 0.79 | 0.92 | 0.88 | 0.75 |
| 3 | 25074015 | rs1603996 | 0.534 | -0.538 | -0.556 | -1.028 | 0.015 | -0.116 | -0.116 | 0 | 0.73 | 0.55 | 0.69 | 0.94 | 0.74 | 0.87 |
| 3 | 25074806 | rs4858139 | 0.788 | -0.347 | -0.534 | -0.875 | -0.001 | -0.042 | -0.042 | 0 | 0.58 | 0.49 | 0.51 | 0.88 | 0.61 | 0.66 |
| 3 | 25077674 | rs7640521 | -0.397 | -0.914 | -0.414 | -0.565 | -0.012 | -0.003 | -0.003 | 0 | 0.26 | 0.27 | 0.16 | 0.24 | 0.18 | 0.22 |
| 3 | 25081857 | rs9847186 | -1.122 | -1.446 | -0.709 | -1.107 | 0.001 | 0.001 | 0.001 | 0 | 0.49 | 0.42 | 0.45 | 0.31 | 0.42 | 0.6 |
| 3 | 25084111 | rs6550938 | -1.559 | -1.945 | -0.878 | -1.115 | -0.083 | -0.074 | -0.074 | 1.58 | 0.64 | 0.91 | 0.52 | 0.42 | 0.48 | 0.76 |
| 3 | 25090904 | rs6786779 | 0.856 | -0.958 | 0.432 | -0.54 | -0.003 | -0.028 | -0.028 | 0 | 0.75 | 0.75 | 0.71 | 0.92 | 0.75 | 0.64 |
| 3 | 25091481 | rs6550939 | 0.836 | -0.96 | 0.482 | -0.54 | 0.002 | 0.003 | 0.003 | 0 | 0.75 | 0.83 | 0.71 | 0.92 | 0.75 | 0.64 |
| 3 | 25097736 | rs4858694 | 1.268 | -1.219 | 0.939 | -0.66 | -0.001 | -0.018 | -0.018 | 0.08 | 0.72 | 0.61 | 0.73 | 0.81 | 0.7 | 0.64 |
| 3 | 25099021 | rs11129178 | 1.758 | -1.497 | 0.982 | -0.713 | -0.029 | -0.009 | -0.009 | 0 | 0.66 | 0.56 | 0.62 | 0.69 | 0.71 | 0.46 |
| 3 | 25105266 | rs13327067 | -0.505 | -1.471 | 1.05 | -0.723 | 0.002 | -0.052 | -0.052 | 0.535 | 0.1 | 0.14 | 0.1 | 0 | 0.11 | 0.05 |
| 3 | 25106437 | rs6804842 | 2.479 | -1.508 | 0.885 | -0.725 | 0.013 | 0.02 | 0.02 | 0.317 | 0.54 | 0.36 | 0.54 | 0.65 | 0.56 | 0.39 |
| 3 | 25111128 | rs17015753 | -0.508 | -1.509 | 0.959 | -0.689 | -0.003 | 0.004 | 0.004 | 0 | 0.11 | 0.15 | 0.07 | 0.03 | 0.13 | 0.07 |
| 3 | 25111890 | rs6791983 | 0.908 | -1.542 | 1.013 | -0.644 | -0.004 | -0.008 | -0.008 | 0 | 0.76 | 0.72 | 0.67 | 0.83 | 0.75 | 0.67 |
| 3 | 25112171 | rs993804 | 0.908 | -1.542 | 1.012 | -0.644 | -0.004 | -0.008 | -0.008 | 0 | 0.76 | 0.72 | 0.67 | 0.83 | 0.75 | 0.67 |
| 3 | 25113247 | rs6772970 | -2.005 | -1.363 | 0.771 | -0.611 | -0.031 | -0.016 | -0.016 | 0 | 0.22 | 0.41 | 0.2 | 0.17 | 0.16 | 0.25 |
| 3 | 25113358 | rs6798255 | -2.092 | -1.436 | 0.764 | -0.611 | -0.048 | 0.009 | 0.009 | 0.878 | 0.22 | 0.43 | 0.2 | 0.24 | 0.15 | 0.25 |
| 3 | 25115413 | rs1574901 | -0.556 | -1.407 | 0.563 | -0.706 | -0.026 | -0.033 | -0.033 | 0 | 0.86 | 0.99 | 0.93 | 0.9 | 0.85 | 0.7 |
| 3 | 25115826 | rs7626290 | -1.899 | -1.401 | 0.677 | -0.724 | -0.037 | -0.076 | -0.076 | 0 | 0.2 | 0.51 | 0.16 | 0.11 | 0.13 | 0.12 |
| 3 | 25131827 | rs10212193 | -1.199 | -1.115 | 0.668 | -0.584 | -0.005 | 0.006 | 0.006 | 0 | 0.13 | 0.16 | 0.11 | 0.03 | 0.17 | 0.13 |
| 3 | 25133273 | rs4858703 | -0.415 | -1.202 | 0.614 | -0.556 | -0.028 | -0.035 | -0.035 | 0 | 0.86 | 0.99 | 0.91 | 0.89 | 0.84 | 0.67 |
| 3 | 25134883 | rs6801041 | -1.953 | -1.26 | 0.853 | -0.598 | -0.05 | -0.06 | -0.06 | 0 | 0.19 | 0.5 | 0.13 | 0.12 | 0.09 | 0.12 |
| 3 | 25140536 | rs1603987 | -0.277 | -1.208 | 0.634 | -0.488 | -0.01 | -0.01 | -0.01 | 0 | 0.86 | 0.95 | 0.91 | 0.9 | 0.84 | 0.67 |
| 3 | 25141824 | rs4353774 | 2.554 | -1.173 | 0.726 | -0.439 | 0.04 | 0.014 | 0.014 | 0.23 | 0.53 | 0.33 | 0.64 | 0.61 | 0.6 | 0.31 |
| 3 | 25143427 | rs4858142 | 1.556 | -0.962 | 0.753 | -0.414 | -0.025 | -0.032 | -0.032 | 0 | 0.71 | 0.59 | 0.79 | 0.79 | 0.75 | 0.58 |
| 3 | 25146601 | rs1580817 | -0.206 | -0.487 | 0.319 | -0.254 | -0.025 | -0.046 | -0.046 | 0 | 0.88 | 0.99 | 0.92 | 0.86 | 0.85 | 0.69 |
| 3 | 25148811 | rs1603986 | 1.779 | 0.195 | 0.535 | -0.604 | -0.004 | 0.002 | 0.002 | 0 | 0.6 | 0.62 | 0.72 | 0.88 | 0.54 | 0.53 |
| 3 | 25149547 | rs1603983 | 1.305 | 0.073 | 0.501 | -0.618 | -0.017 | -0.024 | -0.024 | 0 | 0.71 | 0.65 | 0.86 | 0.88 | 0.76 | 0.61 |
| 3 | 25152496 | rs1580816 | 0.503 | 0.079 | 0.604 | -0.597 | 0.005 | 0.063 | 0.063 | 0.052 | 0.93 | 0.76 | 0.96 | 0.92 | 0.95 | 0.94 |
| 3 | 25155831 | rs1909526 | 2.301 | 0.076 | 0.749 | -0.569 | -0.001 | 0.003 | 0.003 | 0.305 | 0.54 | 0.4 | 0.61 | 0.67 | 0.46 | 0.37 |
| 3 | 25156808 | rs11129180 | 1.675 | 0.157 | 0.768 | -0.682 | -0.007 | -0.013 | -0.013 | 0 | 0.67 | 0.54 | 0.75 | 0.75 | 0.66 | 0.46 |
| 3 | 25161230 | rs1580819 | 0.755 | 0.223 | 0.603 | -0.634 | 0.012 | 0.047 | 0.047 | 0 | 0.86 | 0.75 | 0.85 | 0.79 | 0.84 | 0.78 |
| 3 | 25171653 | rs17517019 | -0.335 | 0.006 | 0.543 | -0.47 | -0.019 | 0.063 | 0.063 | 0 | 0.17 | 0.02 | 0.08 | 0.05 | 0.18 | 0.32 |
| 3 | 25171988 | rs1587425 | 2.382 | -0.024 | 0.86 | -0.499 | 0 | 0.016 | 0.016 | 0 | 0.55 | 0.41 | 0.68 | 0.71 | 0.53 | 0.37 |
| 3 | 25183414 | rs7611139 | -1.58 | -0.031 | 0.885 | -0.557 | 0.001 | -0.034 | -0.034 | 0.141 | 0.27 | 0.23 | 0.16 | 0.05 | 0.29 | 0.38 |
| 3 | 25184389 | rs6764521 | 1.894 | -0.041 | 0.894 | -0.402 | 0 | 0 | 0 | 0 | 0.61 | 0.57 | 0.74 | 0.66 | 0.64 | 0.43 |
| 3 | 25193047 | rs904593 | 1.529 | -0.012 | 0.689 | -0.43 | -0.001 | -0.001 | -0.001 | 0.051 | 0.75 | 0.72 | 0.81 | 0.77 | 0.75 | 0.72 |
| 3 | 25208246 | rs13092408 | 2.741 | -0.138 | 0.865 | -0.51 | -0.003 | 0.06 | 0.06 | 0 | 0.48 | 0.3 | 0.64 | 0.69 | 0.49 | 0.41 |
| 3 | 25208341 | rs17577191 | -0.067 | -0.077 | 0.589 | -0.507 | -0.044 | 0.046 | 0.046 | 0.038 | 0.08 | 0.01 | 0.06 | 0 | 0.11 | 0.07 |
| 3 | 25212878 | rs321543 | -1.657 | 0.055 | 1.011 | -0.287 | -0.004 | -0.004 | -0.004 | 0 | 0.3 | 0.36 | 0.24 | 0.26 | 0.32 | 0.53 |

| | | | | | | | | | | | | | | | | |
|---|---|---|---|---|---|---|---|---|---|---|---|---|---|---|---|---|
| 3 | 25213659 | rs13068891 | -0.08 | 0.071 | 0.661 | -0.335 | -0.006 | 0.02 | 0.02 | 0 | 0.13 | 0.01 | 0.06 | 0.06 | 0.13 | 0.27 |
| 3 | 25228212 | rs6796899 | 1.473 | 0.116 | 0.736 | -0.307 | -0.001 | -0.001 | -0.001 | 0 | 0.77 | 0.74 | 0.81 | 0.77 | 0.76 | 0.73 |
| 3 | 25233025 | rs2649616 | -0.98 | 0.157 | 0.798 | -0.3 | 0.008 | 0.011 | 0.011 | 0 | 0.15 | 0.26 | 0.17 | 0.19 | 0.18 | 0.26 |
| 3 | 25240188 | rs321526 | -2.353 | 0.353 | 0.926 | -0.326 | 0 | 0.042 | 0.042 | 0 | 0.47 | 0.65 | 0.35 | 0.31 | 0.48 | 0.57 |
| 3 | 25252641 | rs1436244 | 0.872 | 0.422 | 0.906 | -0.002 | 0.016 | 0.015 | 0.015 | 0 | 0.86 | 0.74 | 0.86 | 0.83 | 0.83 | 0.74 |
| 3 | 25261724 | rs9830578 | 0.954 | 0.425 | 1.048 | -0.014 | 0.012 | 0.012 | 0.012 | 0 | 0.84 | 0.6 | 0.85 | 0.85 | 0.84 | 0.74 |
| 3 | 25267578 | rs9310773 | 0.741 | 0.416 | 0.917 | -0.017 | 0.007 | 0.005 | 0.005 | 0 | 0.85 | 0.72 | 0.85 | 0.84 | 0.84 | 0.73 |
| 3 | 25275474 | rs13092896 | -0.307 | 0.287 | 0.278 | -0.051 | 0.004 | 0.014 | 0.014 | 0 | 0.14 | 0.03 | 0.05 | 0.08 | 0.13 | 0.28 |
| 3 | 25290005 | rs1601161 | -0.31 | 0.29 | 0.276 | -0.047 | 0.001 | 0.021 | 0.021 | 0 | 0.14 | 0.03 | 0.06 | 0.07 | 0.14 | 0.28 |
| 3 | 25309054 | rs1483851 | 2.249 | 0.908 | 0.463 | -0.141 | 0.007 | 0.052 | 0.052 | 0.225 | 0.52 | 0.23 | 0.61 | 0.67 | 0.48 | 0.39 |
| 3 | 25315828 | rs1872144 | 0.489 | 0.976 | 0.615 | -0.24 | -0.003 | 0.001 | 0.001 | 0 | 0.83 | 0.84 | 0.85 | 0.84 | 0.82 | 0.71 |
| 3 | 25316314 | rs13084418 | -0.146 | 0.953 | 0.525 | -0.29 | 0.001 | 0.017 | 0.017 | 0 | 0.14 | 0.05 | 0.05 | 0.08 | 0.14 | 0.27 |
| 3 | 25317646 | rs13090773 | 2.662 | 1.294 | 0.589 | 0.684 | 0.001 | 0.041 | 0.041 | 0.643 | 0.49 | 0.1 | 0.58 | 0.39 | 0.48 | 0.37 |
| 3 | 25318014 | rs4681015 | 2.525 | 1.537 | 0.866 | 0.844 | 0.009 | 0.036 | 0.036 | 0.119 | 0.52 | 0.23 | 0.61 | 0.38 | 0.49 | 0.35 |
| 3 | 25319260 | rs322707 | 2.111 | 1.623 | 1.068 | 0.763 | -0.004 | 0.004 | 0.004 | 0 | 0.66 | 0.36 | 0.73 | 0.37 | 0.68 | 0.42 |
| 3 | 25320221 | rs4681044 | 1.383 | 1.867 | 0.749 | -0.402 | 0.008 | -0.003 | -0.003 | 0.053 | 0.78 | 0.77 | 0.77 | 0.84 | 0.69 | 0.68 |
| 3 | 25323954 | rs6550960 | -0.762 | 1.658 | -0.023 | -0.32 | -0.099 | 0.056 | 0.056 | 0 | 0.22 | 0.03 | 0.13 | 0.09 | 0.29 | 0.29 |
| 3 | 25326083 | rs13069258 | 1.059 | 1.306 | 0.959 | -0.265 | 0.011 | -0.005 | -0.005 | 0.047 | 0.82 | 0.65 | 0.84 | 0.85 | 0.82 | 0.71 |
| 3 | 25328838 | rs322683 | -1.474 | 0.766 | 0.959 | -0.777 | -0.004 | -0.007 | -0.007 | 0.914 | 0.42 | 0.56 | 0.3 | 0.24 | 0.35 | 0.57 |
| 3 | 25332423 | rs322673 | -1.253 | 0.703 | 0.99 | -1.255 | -0.027 | -0.067 | -0.067 | 0.622 | 0.28 | 0.48 | 0.21 | 0.13 | 0.25 | 0.49 |
| 3 | 25335519 | rs6810082 | -1.227 | 0.599 | 0.962 | -1.393 | -0.047 | -0.097 | -0.097 | 0.654 | 0.28 | 0.47 | 0.16 | 0.08 | 0.18 | 0.31 |
| 3 | 25337874 | rs977224 | 0.841 | 0.974 | 1.056 | -1.557 | 0.022 | -0.041 | -0.041 | 0 | 0.82 | 0.75 | 0.81 | 0.92 | 0.74 | 0.73 |
| 3 | 25340452 | rs2363596 | 1.36 | 0.989 | 1.088 | -1.563 | 0 | -0.012 | -0.012 | 0 | 0.67 | 0.7 | 0.71 | 0.88 | 0.57 | 0.47 |
| 3 | 25341796 | rs167732 | -1.765 | 0.888 | 1.186 | -1.588 | -0.012 | -0.05 | -0.05 | 1.477 | 0.42 | 0.51 | 0.27 | 0.12 | 0.35 | 0.56 |
| 3 | 25347555 | rs322688 | -0.811 | 0.928 | 0.98 | -1.535 | 0.003 | 0.018 | 0.018 | 0 | 0.22 | 0.13 | 0.17 | 0.12 | 0.24 | 0.49 |
| 3 | 25349414 | rs9809773 | 2.291 | 1.002 | 1.06 | -1.577 | -0.006 | 0.074 | 0.074 | 0.354 | 0.48 | 0.37 | 0.63 | 0.88 | 0.47 | 0.41 |
| 3 | 25349434 | rs322694 | -0.266 | 1.229 | 0.741 | -1.415 | 0.019 | -0.039 | -0.039 | 0 | 0.9 | 0.87 | 0.89 | 1 | 0.82 | 0.96 |
| 3 | 25349538 | rs322695 | -0.136 | 0.838 | 0.411 | -1.267 | -0.019 | 0.048 | 0.048 | 0 | 0.16 | 0.04 | 0.1 | 0.05 | 0.17 | 0.29 |
| 3 | 25354249 | rs322698 | -0.532 | 1.233 | 0.939 | -1.071 | 0.012 | 0.001 | 0.001 | 0 | 0.19 | 0.35 | 0.36 | 0.12 | 0.24 | 0.3 |
| 3 | 25354688 | rs322700 | -1.116 | 1.053 | 0.897 | -1.243 | -0.015 | -0.039 | -0.039 | 0.006 | 0.33 | 0.53 | 0.43 | 0.12 | 0.3 | 0.34 |
| 3 | 25359386 | rs9834664 | 1.24 | 1.254 | 0.743 | -1.341 | 0 | -0.007 | -0.007 | 0.007 | 0.62 | 0.4 | 0.51 | 0.88 | 0.57 | 0.64 |
| 3 | 25366205 | rs1561116 | -0.913 | 1.182 | 0.779 | -1.304 | -0.011 | -0.039 | -0.039 | 0 | 0.23 | 0.29 | 0.14 | 0.09 | 0.17 | 0.23 |
| 3 | 25367141 | rs17016204 | -0.918 | 1.166 | 0.963 | -1.313 | -0.018 | -0.059 | -0.059 | 1.512 | 0.23 | 0.32 | 0.14 | 0.09 | 0.17 | 0.23 |
| 3 | 25369060 | rs9832914 | -0.856 | 1.186 | 0.796 | -1.304 | -0.008 | -0.03 | -0.03 | 0 | 0.23 | 0.26 | 0.14 | 0.09 | 0.17 | 0.22 |
| 3 | 25377223 | rs322684 | -0.009 | 1.244 | 0.509 | -1.299 | -0.004 | -0.003 | -0.003 | 2.069 | 0.17 | 0.07 | 0.33 | 0.13 | 0.23 | 0.32 |
| 3 | 25378658 | rs1483836 | 2.44 | 1.012 | 0.139 | -1.322 | -0.069 | -0.043 | -0.043 | 0 | 0.38 | 0.07 | 0.34 | 0.69 | 0.38 | 0.36 |
| 3 | 25383258 | rs4681047 | -1.191 | 1.218 | 0.464 | -1.465 | -0.012 | 0.028 | 0.028 | 2.269 | 0.37 | 0.18 | 0.45 | 0.24 | 0.42 | 0.53 |
| 3 | 25386696 | rs6769438 | 1.068 | 1.079 | 0.739 | -1.441 | -0.013 | -0.049 | -0.049 | 0.063 | 0.11 | 0.16 | 0.06 | 0 | 0.07 | 0.01 |
| 3 | 25387962 | rs6775569 | -1.145 | 1.099 | 0.754 | -1.492 | -0.001 | 0.004 | 0.004 | 0.464 | 0.35 | 0.31 | 0.46 | 0.21 | 0.4 | 0.53 |
| 3 | 25388313 | rs12497123 | 0.364 | 1.139 | 0.721 | -1.473 | -0.034 | -0.03 | -0.03 | 0 | 0.16 | 0.03 | 0.33 | 0.12 | 0.23 | 0.3 |
| 3 | 25388569 | rs6787363 | 2.312 | 1.112 | 0.966 | -1.532 | -0.007 | 0.067 | 0.067 | 0.985 | 0.43 | 0.19 | 0.37 | 0.79 | 0.4 | 0.4 |
| 3 | 25389504 | rs11721271 | -0.278 | 1.055 | 1.121 | -1.441 | 0.006 | 0.001 | 0.001 | 0 | 0.08 | 0.12 | 0.08 | 0.09 | 0.11 | 0.21 |
| 3 | 25396760 | rs6550967 | 2.014 | 1.474 | 0.937 | -1.486 | 0.029 | 0.057 | 0.057 | 1.02 | 0.46 | 0.34 | 0.37 | 0.78 | 0.36 | 0.41 |
| 3 | 25405921 | rs1384899 | 1.032 | 0.642 | 1.023 | -1.503 | -0.014 | 0.034 | 0.034 | 0 | 0.66 | 0.73 | 0.53 | 0.85 | 0.58 | 0.65 |
| 3 | 25407268 | rs1531871 | 1.012 | 0.643 | 1.02 | -1.502 | -0.018 | 0.041 | 0.041 | 0 | 0.66 | 0.76 | 0.53 | 0.85 | 0.58 | 0.65 |
| 3 | 25407596 | rs7614710 | 2.129 | 0.633 | 1.321 | -1.493 | 0 | 0.004 | 0.004 | 0.784 | 0.45 | 0.36 | 0.39 | 0.76 | 0.4 | 0.39 |
| 3 | 25408748 | rs922943 | 1.874 | 0.63 | 1.33 | -1.476 | -0.007 | 0.05 | 0.05 | 0.168 | 0.52 | 0.3 | 0.41 | 0.76 | 0.46 | 0.41 |
| 3 | 25409842 | rs1483843 | -0.164 | 1.136 | 0.969 | -1.417 | 0.007 | -0.038 | -0.038 | 0 | 0.87 | 0.84 | 0.88 | 1 | 0.82 | 0.96 |
| 3 | 25413038 | rs873838 | 1.495 | 0.999 | 0.933 | -1.387 | -0.011 | 0.018 | 0.018 | 0.004 | 0.59 | 0.62 | 0.45 | 0.78 | 0.47 | 0.42 |
| 3 | 25418758 | rs9823425 | 2.018 | 0.773 | 0.975 | -1.364 | -0.002 | 0.022 | 0.022 | 0.219 | 0.47 | 0.37 | 0.4 | 0.76 | 0.45 | 0.44 |
| 3 | 25422696 | rs7628541 | 0.798 | 0.852 | 0.974 | -1.453 | -0.011 | -0.097 | -0.097 | 0 | 0.73 | 0.64 | 0.84 | 0.99 | 0.79 | 0.91 |
| 3 | 25423265 | rs9876345 | 2.121 | 0.794 | 0.868 | -1.388 | -0.002 | 0.013 | 0.013 | 0.243 | 0.47 | 0.43 | 0.41 | 0.73 | 0.44 | 0.42 |
| 3 | 25423431 | rs17523645 | 0.408 | 0.797 | 0.607 | -1.272 | -0.002 | -0.056 | -0.056 | 0 | 0.14 | 0.02 | 0.33 | 0.16 | 0.16 | 0.27 |
| 3 | 25426975 | rs11129198 | -1 | 0.919 | 0.556 | -1.196 | 0.018 | 0 | 0 | 0 | 0.12 | 0.22 | 0.1 | 0.1 | 0.18 | 0.24 |
| 3 | 25431389 | rs6784301 | 1.424 | 0.711 | 0.487 | -1.432 | 0.002 | -0.065 | -0.065 | 0 | 0.69 | 0.6 | 0.79 | 0.9 | 0.66 | 0.7 |
| 3 | 25432062 | rs1991786 | 2.22 | 0.679 | 0.319 | -1.452 | -0.022 | 0.024 | 0.024 | 0.262 | 0.43 | 0.31 | 0.4 | 0.72 | 0.42 | 0.39 |
| 3 | 25434491 | rs6794839 | -1.16 | 0.824 | 0.481 | -1.257 | -0.019 | -0.015 | -0.015 | 0 | 0.33 | 0.4 | 0.42 | 0.19 | 0.28 | 0.35 |
| 3 | 25438508 | rs13088742 | 0.024 | 1.228 | -0.127 | -0.836 | -0.004 | -0.064 | -0.064 | 0 | 0.15 | 0.02 | 0.33 | 0.18 | 0.17 | 0.29 |
| 3 | 25441977 | rs2116701 | -0.16 | 1.346 | 0.051 | -0.678 | -0.029 | 0.058 | 0.058 | 0 | 0.89 | 0.93 | 0.91 | 1 | 0.87 | 0.97 |
| 3 | 25443691 | rs6550971 | 1.103 | 1.368 | 0.181 | -0.707 | 0.014 | -0.009 | -0.009 | 0 | 0.73 | 0.64 | 0.55 | 0.82 | 0.67 | 0.76 |
| 3 | 25447603 | rs6783726 | -1.082 | 1.425 | -0.022 | -0.464 | 0.039 | 0.061 | 0.061 | 0 | 0.23 | 0.36 | 0.42 | 0.27 | 0.33 | 0.43 |
| 3 | 25449650 | rs4681060 | -0.331 | 1.717 | -0.825 | -0.526 | -0.003 | -0.002 | -0.002 | 0 | 0.09 | 0.05 | 0.16 | 0.12 | 0.14 | 0.21 |
| 3 | 25451077 | rs7620529 | 1.165 | 1.234 | -0.917 | -0.557 | -0.006 | 0.018 | 0.018 | 0 | 0.64 | 0.69 | 0.52 | 0.81 | 0.59 | 0.7 |
| 3 | 25452087 | rs13085878 | -0.163 | 1.443 | -1.577 | -0.453 | -0.006 | -0.017 | -0.017 | 0 | 0.16 | 0.08 | 0.33 | 0.19 | 0.22 | 0.25 |
| 3 | 25458100 | rs1897501 | 0.351 | 1.492 | -1.352 | -0.553 | -0.042 | 0.085 | 0.085 | 0 | 0.81 | 0.91 | 0.85 | 1 | 0.78 | 0.94 |
| 3 | 25460596 | rs6805482 | 0.15 | 1.155 | -0.969 | 0.118 | 0.009 | 0.014 | 0.014 | 0 | 0.62 | 0.44 | 0.61 | 0.45 | 0.65 | 0.7 |
| 3 | 25461807 | rs9865116 | 0.27 | 0.918 | -1.23 | -0.147 | 0 | 0.026 | 0.026 | 0.596 | 0.1 | 0.04 | 0.09 | 0.07 | 0.07 | 0.03 |
| 3 | 25461836 | rs6778608 | -0.045 | 1.351 | -1.438 | -0.444 | -0.009 | 0.008 | 0.008 | 0 | 0.81 | 0.87 | 0.74 | 0.87 | 0.75 | 0.75 |
| 3 | 25465008 | rs922939 | 0.957 | 0.802 | -0.595 | -0.73 | 0.004 | -0.056 | -0.056 | 0 | 0.65 | 0.54 | 0.69 | 0.82 | 0.65 | 0.71 |

| | | | | | | | | | | | | | | | | |
|---|---|---|---|---|---|---|---|---|---|---|---|---|---|---|---|---|
| 3 | 25472040 | rs755661 | -0.956 | 0.614 | -0.932 | -1.313 | 0.005 | 0.036 | 0.036 | 1.378 | 0.56 | 0.61 | 0.44 | 0.21 | 0.56 | 0.45 |
| 3 | 25477412 | rs7613553 | -0.364 | 0.646 | -0.887 | -1.563 | -0.034 | 0.076 | 0.076 | 0.176 | 0.38 | 0.24 | 0.26 | 0.11 | 0.46 | 0.33 |
| 3 | 25479778 | rs4416353 | -1.476 | 0.155 | -0.75 | -1.025 | 0.013 | 0.165 | 0.165 | 0 | 0.83 | 0.54 | 0.68 | 0.56 | 0.83 | 0.79 |
| 3 | 25479983 | rs9284856 | 0.398 | -0.37 | -1.27 | -1.017 | 0.019 | 0.018 | 0.018 | 0.083 | 0.19 | 0.14 | 0.15 | 0.09 | 0.12 | 0.16 |
| 3 | 25480108 | rs9284857 | 0.711 | -0.247 | -1.28 | -0.977 | -0.014 | -0.014 | -0.014 | 0 | 0.07 | 0.11 | 0.05 | 0.02 | 0.04 | 0.1 |
| 3 | 25480644 | rs6800566 | 0.435 | -0.017 | -1.622 | -1.079 | -0.01 | -0.02 | -0.02 | 0.15 | 0.3 | 0.15 | 0.2 | 0.32 | 0.34 | 0.26 |
| 3 | 25485756 | rs12631063 | 1.144 | 0.048 | -1.41 | -1.41 | 0.001 | -0.003 | -0.003 | 0.436 | 0.12 | 0.01 | 0.11 | 0.09 | 0.09 | 0.03 |
| 3 | 25489318 | rs4681063 | -0.29 | 0.189 | -1.054 | -1.599 | -0.021 | -0.064 | -0.064 | 0 | 0.37 | 0.55 | 0.38 | 0.17 | 0.31 | 0.28 |
| 3 | 25491968 | rs6793694 | 0.176 | 0.722 | -1.182 | -1.352 | -0.015 | 0.035 | 0.035 | 0 | 0.68 | 0.77 | 0.59 | 0.81 | 0.62 | 0.66 |
| 3 | 25493178 | rs13070407 | 0.334 | 0.886 | -1.364 | -1.409 | -0.02 | 0.042 | 0.042 | 0 | 0.23 | 0.07 | 0.28 | 0.18 | 0.26 | 0.3 |
| 3 | 25495637 | rs13314219 | -0.066 | 0.626 | -1.233 | -1.271 | -0.003 | 0.002 | 0.002 | 0 | 0.67 | 0.69 | 0.58 | 0.85 | 0.61 | 0.65 |
| 3 | 25496674 | rs957950 | -0.546 | 0.584 | -1.243 | -0.361 | -0.015 | -0.012 | -0.012 | 0 | 0.08 | 0.13 | 0.16 | 0.07 | 0.06 | 0.12 |
| 3 | 25500881 | rs7629902 | -0.048 | 0.99 | -1.188 | -0.291 | 0.002 | -0.056 | -0.056 | 0 | 0.13 | 0.22 | 0.1 | 0 | 0.13 | 0.11 |
| 3 | 25507030 | rs6778350 | -0.063 | 0.939 | -1.478 | 0.05 | -0.011 | -0.005 | -0.005 | 0 | 0.9 | 0.96 | 0.78 | 0.87 | 0.88 | 0.83 |
| 3 | 25508428 | rs7616467 | -0.376 | 0.99 | -1.202 | -0.014 | 0.008 | -0.032 | -0.032 | 0 | 0.32 | 0.49 | 0.36 | 0.16 | 0.38 | 0.36 |
| 3 | 25513209 | rs4393871 | 0.048 | 0.241 | -2.338 | 0.512 | 0.047 | -0.028 | -0.028 | 0.303 | 0.5 | 0.04 | 0.49 | 0.4 | 0.63 | 0.56 |
| 3 | 25513628 | rs7629478 | -0.545 | -0.456 | -1.305 | 1.072 | -0.032 | -0.004 | -0.004 | 0 | 0.34 | 0.41 | 0.4 | 0.34 | 0.24 | 0.33 |
| 3 | 25514276 | rs4607073 | 0.778 | 0.265 | -0.608 | 1.6 | -0.003 | -0.005 | -0.005 | 0 | 0.58 | 0.54 | 0.47 | 0.31 | 0.57 | 0.53 |
| 3 | 25515361 | rs6775425 | 0.889 | 0.77 | 0.2 | 2.255 | 0.032 | 0.155 | 0.155 | 0 | 0.84 | 0.69 | 0.59 | 0.58 | 0.76 | 0.76 |
| 3 | 25520582 | rs1529672 | -1.033 | 0.389 | 0.135 | 1.946 | -0.005 | 0.115 | 0.115 | 0.018 | 0.15 | 0.22 | 0.27 | 0.39 | 0.15 | 0.14 |
| 3 | 25521703 | rs2364120 | 0.596 | 0.719 | -0.176 | 1.428 | -0.037 | 0.028 | 0.028 | 0.097 | 0.94 | 1 | 0.92 | 1 | 0.9 | 0.99 |
| 3 | 25525545 | rs13314209 | -1.188 | 1.041 | 0.579 | 1.414 | 0.045 | 0.038 | 0.038 | 0.44 | 0.07 | 0.32 | 0.07 | 0.09 | 0.07 | 0.07 |
| 3 | 25528363 | rs1286661 | 1.543 | 1.031 | 1.251 | 1.387 | 0.083 | -0.017 | -0.017 | 0 | 0.9 | 0.55 | 0.88 | 0.96 | 0.88 | 0.99 |
| 3 | 25529280 | rs1286664 | -0.761 | 1.089 | 1.25 | 1.595 | 0.001 | 0.089 | 0.089 | 0.008 | 0.15 | 0.2 | 0.27 | 0.36 | 0.15 | 0.15 |
| 3 | 25529507 | rs7616278 | 2.259 | 0.696 | 0.885 | 1.293 | 0.002 | -0.003 | -0.003 | 0.419 | 0.57 | 0.31 | 0.45 | 0.44 | 0.58 | 0.63 |
| 3 | 25531265 | rs1286654 | -1.37 | 0.533 | 0.787 | 1.282 | 0 | -0.001 | -0.001 | 0 | 0.32 | 0.26 | 0.4 | 0.48 | 0.25 | 0.29 |
| 3 | 25532425 | rs11916491 | -0.609 | 0.512 | 0.755 | 1.245 | -0.022 | -0.017 | -0.017 | 0.59 | 0.21 | 0.34 | 0.2 | 0.19 | 0.17 | 0.21 |
| 3 | 25537503 | rs1286648 | -2.33 | 0.917 | 0.695 | 1.135 | -0.042 | 0.002 | 0.002 | 0.084 | 0.47 | 0.9 | 0.6 | 0.59 | 0.47 | 0.4 |
| 3 | 25538883 | rs2568875 | -1.895 | 0.952 | 0.837 | 1.152 | 0.008 | 0.092 | 0.092 | 0 | 0.26 | 0.58 | 0.37 | 0.41 | 0.26 | 0.17 |
| 3 | 25541371 | rs1406575 | -1.706 | 1.178 | 0.58 | 1.106 | 0.019 | -0.009 | -0.009 | 0 | 0.41 | 0.67 | 0.47 | 0.56 | 0.33 | 0.36 |
| 3 | 25548205 | rs1286644 | -2.122 | 1.294 | 0.158 | 1.096 | -0.119 | -0.077 | -0.077 | 0 | 0.46 | 0.94 | 0.59 | 0.58 | 0.44 | 0.39 |
| 3 | 25548555 | rs1560633 | 1.424 | 1.325 | 0.857 | 1.097 | 0.002 | 0.01 | 0.01 | 0 | 0.63 | 0.55 | 0.48 | 0.42 | 0.63 | 0.64 |
| 3 | 25554363 | rs1153589 | -1.924 | 1.005 | 0.651 | 1.246 | 0.001 | -0.047 | -0.047 | 0 | 0.25 | 0.81 | 0.46 | 0.48 | 0.21 | 0.16 |
| 3 | 25557684 | rs17591006 | -0.133 | 1.034 | 0.656 | 1.185 | -0.016 | 0.013 | 0.013 | 0 | 0.05 | 0.02 | 0.05 | 0 | 0.08 | 0.01 |
| 3 | 25559928 | rs1153594 | -0.866 | 0.941 | 1.603 | 1.323 | -0.021 | 0.096 | 0.096 | 0.116 | 0.16 | 0.33 | 0.27 | 0.35 | 0.11 | 0.13 |
| 3 | 25560231 | rs1435703 | -1.474 | 0.98 | 1.789 | 1.272 | 0.082 | -0.007 | -0.007 | 0 | 0.08 | 0.44 | 0.09 | 0 | 0.07 | 0.01 |
| 3 | 25561049 | rs1347458 | -2.481 | 0.962 | 1.818 | 1.291 | 0.046 | 0.061 | 0.061 | 0.18 | 0.1 | 0.56 | 0.19 | 0.12 | 0.08 | 0.02 |
| 3 | 25561488 | rs1153596 | -1.426 | 0.979 | 1.735 | 1.304 | -0.02 | 0.249 | 0.249 | 0.126 | 0.19 | 0.48 | 0.38 | 0.49 | 0.14 | 0.16 |
| 3 | 25564929 | rs6805350 | -0.586 | 1.037 | 1.665 | 1.207 | 0.032 | -0.025 | -0.025 | 0 | 0.06 | 0.21 | 0.06 | 0 | 0.06 | 0.01 |
| 3 | 25566195 | rs1286754 | -2.479 | 1.15 | 0.758 | 0.546 | -0.019 | -0.077 | -0.077 | 1.6 | 0.51 | 0.94 | 0.65 | 0.5 | 0.41 | 0.33 |
| 3 | 25567812 | rs1286757 | -2.53 | 1.179 | 0.685 | 0.402 | 0.011 | -0.018 | -0.018 | 0 | 0.52 | 0.8 | 0.64 | 0.5 | 0.42 | 0.37 |
| 3 | 25568871 | rs1435705 | 0.242 | 1.254 | 0.741 | 0.434 | 0.019 | 0.015 | 0.015 | 0 | 0.05 | 0.11 | 0.22 | 0.1 | 0.09 | 0.04 |
| 3 | 25569881 | rs17525900 | -0.494 | 1.257 | 0.785 | 0.427 | 0.033 | 0.011 | 0.011 | 0 | 0.07 | 0.14 | 0.23 | 0.1 | 0.11 | 0.08 |
| 3 | 25571725 | rs1153603 | -2.295 | 1.046 | 0.718 | 0.354 | 0.039 | -0.105 | -0.105 | 0 | 0.43 | 0.91 | 0.62 | 0.49 | 0.32 | 0.28 |
| 3 | 25573009 | rs1286760 | -1.049 | 0.884 | 0.763 | 0.119 | -0.029 | 0.109 | 0.109 | 1.606 | 0.23 | 0.48 | 0.21 | 0.37 | 0.15 | 0.17 |
| 3 | 25574186 | rs1153606 | -1.144 | 0.862 | 0.698 | 0.111 | -0.034 | 0.106 | 0.106 | 1.214 | 0.23 | 0.53 | 0.24 | 0.38 | 0.16 | 0.17 |
| 3 | 25575675 | rs1286665 | -1.915 | 0.93 | 0.453 | 0.091 | 0.03 | -0.002 | -0.002 | 0 | 0.42 | 0.71 | 0.33 | 0.39 | 0.27 | 0.29 |
| 3 | 25576865 | rs1992005 | 0.177 | 1.111 | 0.389 | 0.087 | 0.019 | 0.023 | 0.023 | 0 | 0.05 | 0.12 | 0.22 | 0.11 | 0.07 | 0.04 |
| 3 | 25578552 | rs1881703 | -2.385 | 2.057 | 0.185 | 0.079 | -0.057 | -0.035 | -0.035 | 0.013 | 0.49 | 0.91 | 0.73 | 0.54 | 0.47 | 0.4 |
| 3 | 25578892 | rs7616062 | -1.114 | 2.02 | 0.635 | 0.298 | 0.035 | 0.057 | 0.057 | 0.406 | 0.17 | 0.53 | 0.36 | 0.25 | 0.18 | 0.16 |
| 3 | 25578938 | rs7631979 | -2.07 | 1.924 | 0.359 | 0.053 | 0 | -0.002 | -0.002 | 0 | 0.4 | 0.39 | 0.38 | 0.3 | 0.38 | 0.26 |
| 3 | 25579915 | rs11707637 | -2.337 | 2.108 | 0.223 | 0.062 | 0.002 | -0.002 | -0.002 | 0 | 0.43 | 0.55 | 0.6 | 0.41 | 0.45 | 0.3 |
| 3 | 25581512 | rs1286771 | -0.338 | 1.903 | 0.711 | 0.332 | 0.012 | 0.034 | 0.034 | 0.427 | 0.13 | 0.35 | 0.13 | 0.14 | 0.11 | 0.12 |
| 3 | 25584358 | rs4681027 | 0.234 | 1.946 | 0.49 | 0.309 | -0.001 | -0.001 | -0.001 | 0 | 0.06 | 0.08 | 0.15 | 0.07 | 0.04 | 0.05 |
| 3 | 25585166 | rs1286769 | -2.571 | 2.016 | 0.221 | 0.465 | -0.03 | -0.026 | -0.026 | 0 | 0.48 | 0.93 | 0.71 | 0.56 | 0.43 | 0.36 |
| 3 | 25586106 | rs17016718 | -0.752 | 2.392 | 0.465 | 0.707 | 0.004 | -0.033 | -0.033 | 0 | 0.19 | 0.1 | 0.13 | 0.26 | 0.17 | 0.15 |
| 3 | 25593872 | rs1286765 | -2.524 | 2.693 | 0.499 | 0.763 | -0.121 | -0.056 | -0.056 | 5.329 | 0.46 | 0.95 | 0.74 | 0.59 | 0.48 | 0.36 |
| 3 | 25598909 | rs1286762 | -1.052 | 2.913 | 1.033 | 0.955 | 0.011 | 0.036 | 0.036 | 0.382 | 0.16 | 0.43 | 0.3 | 0.21 | 0.15 | 0.16 |
| 3 | 25600646 | rs17016738 | -1.69 | 2.806 | 0.874 | 0.594 | 0 | 0.004 | 0.004 | 0 | 0.19 | 0.19 | 0.36 | 0.36 | 0.25 | 0.21 |
| 3 | 25604599 | rs17016773 | -1.02 | 2.622 | 0.709 | 0.616 | -0.012 | -0.052 | -0.052 | 0 | 0.16 | 0.08 | 0.31 | 0.38 | 0.2 | 0.16 |
| 3 | 25605101 | rs17526839 | -0.094 | 2.723 | 0.89 | 0.529 | 0.019 | -0.022 | -0.022 | 0 | 0.05 | 0.07 | 0.05 | 0 | 0.08 | 0.01 |
| 3 | 25605886 | rs17016781 | -1.354 | 2.695 | 0.692 | 0.497 | 0.021 | -0.103 | -0.103 | 0 | 0.21 | 0.02 | 0.26 | 0.39 | 0.15 | 0.24 |
| 3 | 25608576 | rs1286729 | 0.037 | 2.929 | 0.928 | 0.627 | -0.034 | -0.016 | -0.016 | 5.352 | 0.06 | 0.01 | 0.13 | 0.09 | 0.1 | 0.08 |
| 3 | 25609011 | rs6795340 | -1.475 | 2.999 | 1.238 | 0.554 | -0.003 | 0.028 | 0.028 | 0.273 | 0.27 | 0.43 | 0.33 | 0.32 | 0.25 | 0.29 |
| 3 | 25611152 | rs4081949 | -1.422 | 2.793 | 1.396 | 0.468 | 0.054 | 0.066 | 0.066 | 0.065 | 0.05 | 0.24 | 0.16 | 0.07 | 0.04 | 0.15 |
| 3 | 25612565 | rs3773445 | -2.589 | 2.88 | 1.317 | 0.435 | 0.015 | 0.006 | 0.006 | 0 | 0.39 | 0.53 | 0.51 | 0.45 | 0.43 | 0.45 |
| 3 | 25613372 | rs1286733 | -1.824 | 2.902 | 1.426 | 0.422 | -0.017 | -0.004 | -0.004 | 0.331 | 0.31 | 0.3 | 0.34 | 0.35 | 0.36 | 0.27 |
| 3 | 25614532 | rs1286738 | -2.325 | 3.067 | 1.369 | 0.328 | 0.053 | -0.007 | -0.007 | 0 | 0.15 | 0.27 | 0.34 | 0.16 | 0.22 | 0.23 |
| 3 | 25617403 | rs17526942 | 0.045 | 3.076 | 1.174 | 0.323 | -0.002 | -0.002 | -0.002 | 0 | 0.06 | 0.04 | 0.14 | 0.1 | 0.1 | 0.08 |

| | | | | | | | | | | | | | | | | |
|---|---|---|---|---|---|---|---|---|---|---|---|---|---|---|---|---|
| 3 | 25617775 | rs3773439 | -2.394 | 3.004 | 1.273 | 0.332 | 0.012 | 0.01 | 0.01 | 0 | 0.35 | 0.54 | 0.48 | 0.43 | 0.35 | 0.4 |
| 3 | 25618159 | rs1656463 | -1.103 | 2.778 | 1.066 | 0.303 | -0.033 | 0.011 | 0.011 | 0 | 0.1 | 0.04 | 0.18 | 0.09 | 0.18 | 0.1 |
| 3 | 25620132 | rs3821629 | -1.069 | 2.056 | 1.213 | 0.321 | 0.021 | 0.057 | 0.057 | 0 | 0.22 | 0.36 | 0.31 | 0.33 | 0.22 | 0.3 |
| 3 | 25621302 | rs7634326 | -1.05 | 1.295 | 1.53 | -0.099 | 0.003 | -0.017 | -0.017 | 0 | 0.06 | 0.21 | 0.02 | 0 | 0.03 | 0.03 |
| 3 | 25623740 | rs868368 | -1.488 | 1.475 | 0.766 | 0.099 | -0.008 | 0.014 | 0.014 | 0 | 0.06 | 0.14 | 0.15 | 0.1 | 0.03 | 0.16 |
| 3 | 25624634 | rs11917304 | -2.142 | 1.557 | 0.755 | 0.069 | 0.028 | -0.027 | -0.027 | 0 | 0.13 | 0.29 | 0.24 | 0.1 | 0.13 | 0.2 |
| 3 | 25628749 | rs4681028 | -1.474 | 1.182 | 0.452 | 0.285 | 0.02 | 0.043 | 0.043 | 0 | 0.15 | 0.25 | 0.31 | 0.3 | 0.21 | 0.25 |
| 3 | 25630066 | rs6782425 | -2.204 | 0.804 | 0.73 | 0.204 | 0.008 | 0.049 | 0.049 | 0 | 0.21 | 0.42 | 0.37 | 0.32 | 0.21 | 0.27 |
| 3 | 25630857 | rs7621140 | -1.783 | 0.545 | 0.883 | 0.14 | -0.012 | -0.006 | -0.006 | 0 | 0.15 | 0.35 | 0.22 | 0.15 | 0.12 | 0.15 |
| 3 | 25632894 | rs2164360 | -1.309 | 0.723 | 0.691 | -0.107 | -0.025 | -0.023 | -0.023 | 0 | 0.16 | 0.1 | 0.21 | 0.17 | 0.2 | 0.17 |
| 3 | 25639394 | rs1058378 | -1.372 | 0.753 | 0.771 | -0.088 | 0.014 | 0.014 | 0.014 | 0.932 | 0.13 | 0.5 | 0.2 | 0.13 | 0.11 | 0.16 |
| 3 | 25645514 | rs1881708 | -2.293 | 0.623 | 0.574 | -0.139 | 0.007 | 0.004 | 0.004 | 0 | 0.31 | 0.51 | 0.4 | 0.31 | 0.31 | 0.33 |
| 3 | 25654677 | rs1001647 | -0.674 | 0.347 | 0.175 | -0.472 | 0.002 | -0.005 | -0.005 | 0 | 0.3 | 0.71 | 0.2 | 0.09 | 0.2 | 0.21 |
| 3 | 25681432 | rs2037860 | -1.742 | 0.5 | 1.018 | 0.027 | -0.002 | -0.01 | -0.01 | 0.007 | 0.13 | 0.21 | 0.2 | 0.13 | 0.13 | 0.15 |
| 3 | 25683930 | rs11705878 | -0.962 | 0.508 | 1.023 | 0.039 | 0.004 | 0.007 | 0.007 | 0 | 0.1 | 0.21 | 0.17 | 0.13 | 0.11 | 0.15 |
| 3 | 25706277 | rs10433573 | -2.171 | 0.612 | 0.655 | 0.063 | 0.013 | 0.014 | 0.014 | 0.049 | 0.28 | 0.5 | 0.4 | 0.31 | 0.3 | 0.33 |
| 3 | 25725070 | rs10446418 | -0.972 | 1.044 | -0.14 | 0.339 | -0.06 | -0.065 | -0.065 | 0 | 0.12 | 0.01 | 0.17 | 0.18 | 0.17 | 0.17 |
| 3 | 25730461 | rs1286841 | -1.739 | 0.215 | 1.039 | -0.061 | -0.001 | 0.031 | 0.031 | 0 | 0.14 | 0.25 | 0.25 | 0.25 | 0.17 | 0.28 |
| 3 | 25739241 | rs6785846 | 2.203 | -0.035 | 0.88 | -0.684 | 0.008 | 0.063 | 0.063 | 1.87 | 0.77 | 0.1 | 0.68 | 0.86 | 0.8 | 0.74 |
| 3 | 25755713 | rs2932015 | 2.341 | -0.238 | 0.771 | -0.776 | 0.025 | 0.071 | 0.071 | 0.946 | 0.74 | 0.08 | 0.66 | 0.84 | 0.8 | 0.69 |
| 3 | 25775356 | rs6550987 | 1.507 | -0.242 | 2.195 | -0.757 | -0.018 | -0.054 | -0.054 | 0 | 0.83 | 0.7 | 0.83 | 0.93 | 0.83 | 0.75 |
| 3 | 25795889 | rs6804099 | 2.199 | -0.255 | 2.883 | -0.763 | -0.016 | -0.061 | -0.061 | 0 | 0.78 | 0.32 | 0.8 | 0.93 | 0.81 | 0.73 |
| 3 | 25819628 | rs9877259 | 2.563 | -0.262 | 2.96 | -0.765 | 0.023 | 0.105 | 0.105 | 0.922 | 0.75 | 0.08 | 0.76 | 0.9 | 0.8 | 0.68 |
| 3 | 25826537 | rs9810848 | -2.015 | -0.207 | 3.241 | -0.823 | 0.09 | 0.042 | 0.042 | 0.372 | 0.05 | 0.35 | 0.03 | 0 | 0.02 | 0.02 |
| 3 | 25836023 | rs9831075 | 1.972 | -0.215 | 3.205 | -0.801 | -0.017 | -0.089 | -0.089 | 0 | 0.79 | 0.5 | 0.8 | 0.92 | 0.81 | 0.76 |
| 3 | 25836088 | rs9834955 | 1.901 | -0.215 | 3.206 | -0.801 | -0.014 | -0.081 | -0.081 | 0.117 | 0.8 | 0.52 | 0.81 | 0.92 | 0.81 | 0.76 |
| 3 | 25836733 | rs6550989 | 1.961 | -0.215 | 3.206 | -0.801 | -0.016 | -0.089 | -0.089 | 2.905 | 0.8 | 0.5 | 0.8 | 0.92 | 0.82 | 0.76 |
| 3 | 25838831 | rs6779291 | 1.925 | -0.216 | 3.147 | -0.805 | -0.01 | -0.039 | -0.039 | 0.071 | 0.8 | 0.54 | 0.72 | 0.86 | 0.81 | 0.75 |
| 3 | 25843732 | rs6789319 | -2.543 | -0.028 | 3.193 | -1.062 | 0.023 | -0.018 | -0.018 | 0 | 0.08 | 0.55 | 0.05 | 0.01 | 0.02 | 0.05 |
| 3 | 25846126 | rs6550990 | -2.763 | -0.012 | 2.502 | -1.294 | 0.014 | 0.049 | 0.049 | 0.833 | 0.23 | 0.83 | 0.16 | 0.03 | 0.16 | 0.28 |
| 3 | 25863660 | rs7626769 | 2.888 | -0.169 | 2.549 | -1.069 | 0.025 | 0.087 | 0.087 | 0 | 0.75 | 0.12 | 0.8 | 0.93 | 0.8 | 0.7 |
| 3 | 25872276 | rs4681079 | 2.389 | -0.168 | 2.991 | -1.043 | -0.013 | -0.089 | -0.089 | 0 | 0.8 | 0.38 | 0.83 | 0.94 | 0.81 | 0.74 |
| 3 | 25877906 | rs7340528 | -1.006 | -0.019 | 2.547 | -0.989 | -0.001 | -0.023 | -0.023 | 1.099 | 0.07 | 0.1 | 0.06 | 0.02 | 0.09 | 0.11 |
| 3 | 25885030 | rs11129210 | 2.265 | -0.234 | 2.974 | -1.065 | -0.029 | -0.102 | -0.102 | 0 | 0.81 | 0.52 | 0.86 | 0.95 | 0.83 | 0.74 |
| 3 | 25898495 | rs6810130 | 2.793 | -0.029 | 2.717 | -1.087 | 0.017 | 0.098 | 0.098 | 0.046 | 0.76 | 0.13 | 0.79 | 0.95 | 0.79 | 0.72 |
| 3 | 25913415 | rs4293672 | 1.809 | 0.157 | 3.073 | -1.041 | 0.013 | -0.075 | -0.075 | 1.463 | 0.84 | 0.62 | 0.84 | 0.95 | 0.8 | 0.73 |
| 3 | 25913883 | rs6551000 | 2.589 | 0.154 | 3.179 | -1.044 | 0.005 | 0.042 | 0.042 | 0 | 0.78 | 0.17 | 0.79 | 0.95 | 0.78 | 0.71 |
| 3 | 25913954 | rs6551001 | 1.754 | 0.317 | 2.947 | -1.004 | 0.019 | -0.08 | -0.08 | 0.023 | 0.84 | 0.64 | 0.84 | 0.97 | 0.8 | 0.78 |
| 3 | 25933984 | rs2968169 | 2.022 | 0.156 | 2.31 | -1.049 | -0.009 | -0.096 | -0.096 | 0 | 0.79 | 0.5 | 0.73 | 0.95 | 0.83 | 0.67 |
| 3 | 25944830 | rs4334607 | 2.068 | 0.404 | 2.384 | -1.1 | -0.01 | -0.094 | -0.094 | 0 | 0.79 | 0.5 | 0.72 | 0.93 | 0.83 | 0.67 |
| 3 | 25946030 | rs9874424 | 1.752 | 0.546 | 1.981 | -1.05 | 0.007 | -0.01 | -0.01 | 0 | 0.92 | 0.82 | 0.83 | 0.96 | 0.92 | 0.95 |
| 3 | 25949300 | rs9819197 | 2.232 | 0.496 | 1.967 | -1.063 | -0.002 | -0.011 | -0.011 | 0 | 0.88 | 0.81 | 0.81 | 0.96 | 0.9 | 0.94 |
| 3 | 25964812 | rs9872930 | -2.212 | 0.459 | 2.343 | -1.052 | 0.045 | -0.015 | -0.015 | 0 | 0.08 | 0.27 | 0.06 | 0.02 | 0.06 | 0.21 |
| 3 | 25965247 | rs6786329 | 1.72 | 0.382 | 1.952 | -1.289 | -0.045 | 0.056 | 0.056 | 0.011 | 0.88 | 0.98 | 0.78 | 0.97 | 0.84 | 0.94 |
| 3 | 25968912 | rs9856369 | 3.463 | 0.168 | 2.61 | -1.401 | 0 | -0.031 | -0.031 | 0 | 0.73 | 0.29 | 0.65 | 0.95 | 0.75 | 0.65 |
| 3 | 25985168 | rs6551009 | 3.369 | 0.081 | 2.664 | -1.349 | -0.002 | -0.038 | -0.038 | 0 | 0.75 | 0.3 | 0.68 | 0.94 | 0.79 | 0.65 |
| 3 | 25994433 | rs6551011 | 3.214 | -0.044 | 2.367 | -1.714 | 0.019 | 0.041 | 0.041 | 0 | 0.66 | 0.27 | 0.64 | 0.94 | 0.71 | 0.63 |
| 3 | 25994862 | rs9834799 | -2.194 | -0.046 | 2.583 | -1.722 | 0.025 | -0.056 | -0.056 | 0.012 | 0.1 | 0.29 | 0.07 | 0.03 | 0.08 | 0.21 |
| 3 | 25998294 | rs1426774 | 3.498 | -0.048 | 2.444 | -1.722 | 0.019 | 0.038 | 0.038 | 0 | 0.66 | 0.28 | 0.64 | 0.95 | 0.72 | 0.62 |
| 3 | 26002178 | rs12715076 | 2.435 | -0.04 | 2.551 | -1.682 | 0.015 | 0.073 | 0.073 | 1.429 | 0.74 | 0.85 | 0.72 | 0.96 | 0.76 | 0.83 |
| 3 | 26011981 | rs12715077 | 3.154 | -0.11 | 2.887 | -1.753 | 0.011 | -0.1 | -0.1 | 0.721 | 0.81 | 0.66 | 0.74 | 0.94 | 0.8 | 0.73 |
| 3 | 26014022 | rs12494445 | -2.07 | -0.171 | 2.9 | -1.73 | 0.016 | 0.007 | 0.007 | 0 | 0.18 | 0.14 | 0.1 | 0 | 0.13 | 0.11 |
| 3 | 26017607 | rs1426765 | 1.428 | -0.176 | 2.856 | -1.696 | -0.025 | 0.038 | 0.038 | 0.137 | 0.92 | 0.98 | 0.81 | 0.98 | 0.9 | 0.94 |
| 3 | 26018762 | rs6804277 | 1.772 | -0.176 | 2.911 | -1.696 | 0 | -0.012 | -0.012 | 0.04 | 0.91 | 0.9 | 0.8 | 0.98 | 0.9 | 0.94 |
| 3 | 26022980 | rs933944 | 2.152 | 0.092 | 2.713 | -1.89 | -0.024 | 0.105 | 0.105 | 2.611 | 0.76 | 0.91 | 0.71 | 0.98 | 0.74 | 0.78 |
| 3 | 26055831 | rs10470654 | 3.318 | -0.148 | 3.095 | -1.959 | 0.026 | 0.077 | 0.077 | 0 | 0.63 | 0.25 | 0.63 | 0.94 | 0.69 | 0.65 |
| 3 | 26069100 | rs1426763 | 1.883 | -0.109 | 3.067 | -1.943 | -0.064 | 0.147 | 0.147 | 5.752 | 0.77 | 0.98 | 0.74 | 0.97 | 0.75 | 0.79 |
| 3 | 26070745 | rs7613943 | 2.824 | -0.136 | 3.188 | -1.955 | -0.004 | -0.093 | -0.093 | 5.507 | 0.68 | 0.43 | 0.66 | 0.94 | 0.7 | 0.67 |
| 3 | 26076008 | rs17544105 | 3.257 | -0.154 | 3.374 | -1.963 | 0.011 | 0.121 | 0.121 | 0.627 | 0.67 | 0.16 | 0.63 | 0.95 | 0.7 | 0.69 |
| 3 | 26076713 | rs9867507 | 3.337 | -0.128 | 3.276 | -1.952 | 0 | -0.057 | -0.057 | 0 | 0.66 | 0.4 | 0.65 | 0.94 | 0.69 | 0.67 |
| 3 | 26085604 | rs9849293 | 3.316 | -0.127 | 3.253 | -1.951 | -0.001 | -0.065 | -0.065 | 0.7 | 0.66 | 0.41 | 0.65 | 0.94 | 0.69 | 0.67 |
| 3 | 26087857 | rs991623 | 1.976 | -0.121 | 3.192 | -1.949 | -0.022 | 0.1 | 0.1 | 4.696 | 0.77 | 0.92 | 0.73 | 0.98 | 0.75 | 0.8 |
| 3 | 26089949 | rs9833596 | -2.081 | -0.1 | 3.246 | -1.924 | -0.006 | -0.045 | -0.045 | 0 | 0.09 | 0.21 | 0.05 | 0.03 | 0.06 | 0.12 |
| 3 | 26106208 | rs12629334 | 2.695 | -0.1 | 3.238 | -1.94 | -0.011 | -0.131 | -0.131 | 0 | 0.7 | 0.47 | 0.68 | 0.98 | 0.73 | 0.79 |
| 3 | 26107171 | rs7372661 | -0.78 | -0.097 | 3.133 | -1.943 | -0.018 | 0.029 | 0.029 | 1.629 | 0.06 | 0.01 | 0.16 | 0 | 0.06 | 0.02 |
| 3 | 26111090 | rs6551031 | 2.964 | -0.12 | 3.229 | -1.937 | 0.002 | -0.03 | -0.03 | 0 | 0.68 | 0.32 | 0.64 | 0.94 | 0.7 | 0.68 |
| 3 | 26116590 | rs6772506 | -2.275 | -0.13 | 3.504 | -1.918 | -0.013 | -0.051 | -0.051 | 1.632 | 0.1 | 0.28 | 0.07 | 0.03 | 0.06 | 0.11 |
| 3 | 26117140 | rs9821503 | 1.875 | 0.184 | 3.535 | -1.733 | -0.026 | 0.087 | 0.087 | 4.41 | 0.78 | 0.92 | 0.73 | 0.98 | 0.75 | 0.8 |
| 3 | 26120360 | rs6551034 | 2.167 | 0.266 | 3.771 | -1.481 | -0.038 | 0.077 | 0.077 | 0 | 0.85 | 0.97 | 0.78 | 0.98 | 0.83 | 0.8 |

| | | | | | | | | | | | | | | | | |
|---|---|---|---|---|---|---|---|---|---|---|---|---|---|---|---|---|
| 3 | 26125050 | rs7641517 | 2.8 | 0.334 | 3.948 | -1.45 | 0.002 | -0.086 | -0.086 | 0 | 0.82 | 0.35 | 0.73 | 0.98 | 0.83 | 0.82 |
| 3 | 26131150 | rs6551040 | 2.387 | 0.342 | 4.122 | -1.401 | 0.035 | -0.094 | -0.094 | 0 | 0.84 | 0.59 | 0.73 | 0.96 | 0.83 | 0.82 |
| 3 | 26139590 | rs6419855 | 1.858 | 0.344 | 4.084 | -1.4 | 0.001 | -0.015 | -0.015 | 0 | 0.86 | 0.84 | 0.74 | 0.96 | 0.83 | 0.82 |
| 3 | 26140528 | rs9860291 | 1.901 | 0.343 | 4.09 | -1.4 | 0.003 | -0.021 | -0.021 | 0.111 | 0.86 | 0.83 | 0.74 | 0.96 | 0.83 | 0.81 |
| 3 | 26146045 | rs6784527 | 2.818 | 0.34 | 4.204 | -1.401 | 0.005 | -0.088 | -0.088 | 2.478 | 0.82 | 0.41 | 0.71 | 0.96 | 0.83 | 0.81 |
| 3 | 26146839 | rs6551046 | 1.861 | 0.358 | 4.182 | -1.393 | 0.006 | -0.03 | -0.03 | 0 | 0.86 | 0.81 | 0.74 | 0.96 | 0.83 | 0.82 |
| 3 | 26157145 | rs7430881 | 2.721 | 0.376 | 4.125 | -1.332 | 0.003 | -0.062 | -0.062 | 1.505 | 0.82 | 0.38 | 0.71 | 0.94 | 0.83 | 0.82 |
| 3 | 26160044 | rs8180001 | 2.966 | 0.374 | 4.118 | -1.336 | 0 | -0.035 | -0.035 | 0 | 0.8 | 0.27 | 0.68 | 0.92 | 0.8 | 0.7 |
| 3 | 26173223 | rs7638008 | 2.802 | 0.39 | 4.075 | -1.328 | 0.004 | -0.071 | -0.071 | 0 | 0.82 | 0.4 | 0.71 | 0.94 | 0.83 | 0.77 |
| 3 | 26182935 | rs9870231 | 1.721 | 0.6 | 3.557 | -1.253 | -0.019 | 0.025 | 0.025 | 0.035 | 0.87 | 0.93 | 0.75 | 0.93 | 0.83 | 0.75 |
| 3 | 26185773 | rs7629019 | 2.213 | 0.594 | 3.646 | -1.257 | 0.044 | -0.065 | -0.065 | 0 | 0.85 | 0.61 | 0.71 | 0.93 | 0.83 | 0.76 |
| 3 | 26193388 | rs2106590 | 2.874 | 0.466 | 3.264 | -1.24 | -0.001 | 0.007 | 0.007 | 0.28 | 0.8 | 0.16 | 0.66 | 0.91 | 0.82 | 0.69 |
| 3 | 26207425 | rs11129227 | -3.053 | 0.339 | 3.267 | -1.191 | 0 | -0.008 | -0.008 | 0 | 0.19 | 0.81 | 0.33 | 0.09 | 0.16 | 0.26 |
| 3 | 26224215 | rs6551069 | -3.024 | 0.354 | 3.273 | -1.184 | 0 | -0.009 | -0.009 | 0 | 0.19 | 0.81 | 0.33 | 0.09 | 0.16 | 0.25 |
| 3 | 26226918 | rs7633090 | 2.499 | 0.444 | 3.393 | -1.206 | 0.004 | -0.064 | -0.064 | 0 | 0.82 | 0.33 | 0.68 | 0.91 | 0.82 | 0.74 |
| 3 | 26247001 | rs17017671 | -1.406 | 0.496 | 3.524 | -1.228 | 0.029 | 0.01 | 0.01 | 0 | 0.07 | 0.34 | 0.04 | 0.01 | 0.03 | 0.03 |
| 3 | 26256357 | rs11928334 | -2.999 | 0.617 | 3.669 | -0.988 | 0 | -0.003 | -0.003 | 0.337 | 0.05 | 0.7 | 0.07 | 0 | 0 | 0 |
| 3 | 26256909 | rs2106585 | -2.737 | 0.649 | 3.566 | -1.003 | -0.002 | -0.029 | -0.029 | 0.669 | 0.18 | 0.78 | 0.33 | 0.1 | 0.16 | 0.26 |
| 3 | 26286986 | rs7633522 | -2.563 | 0.42 | 3.442 | -0.778 | 0.045 | 0.026 | 0.026 | 2.023 | 0.06 | 0.65 | 0.11 | 0 | 0.02 | 0.01 |
| 3 | 26297737 | rs9864307 | -2.369 | 0.479 | 3.117 | -1.135 | -0.003 | -0.033 | -0.033 | 0.036 | 0.23 | 0.81 | 0.36 | 0.08 | 0.17 | 0.26 |
| 3 | 26314903 | rs12490758 | -0.115 | 0.88 | 2.167 | -0.382 | -0.081 | -0.136 | -0.136 | 0.622 | 0.18 | 0.01 | 0.19 | 0.41 | 0.23 | 0.19 |
| 3 | 26342792 | rs7616136 | -2.388 | 0.941 | 2.534 | -0.25 | 0.003 | 0.014 | 0.014 | 0 | 0.24 | 0.84 | 0.36 | 0.1 | 0.17 | 0.24 |
| 3 | 26377577 | rs7372296 | -1.47 | 0.996 | 1.916 | -0.216 | 0.01 | -0.063 | -0.063 | 0 | 0.18 | 0.34 | 0.27 | 0.09 | 0.14 | 0.26 |
| 3 | 26396178 | rs1021795 | -1.692 | 0.723 | 1.578 | 0.176 | -0.011 | 0.018 | 0.018 | 0 | 0.13 | 0.4 | 0.08 | 0.14 | 0.09 | 0.16 |
| 3 | 26405240 | rs2174003 | -1.818 | 0.625 | 2.001 | -0.131 | 0.003 | -0.092 | -0.092 | 0.723 | 0.14 | 0.51 | 0.28 | 0 | 0.07 | 0.05 |
| 3 | 26406007 | rs4973792 | -1.146 | 0.634 | 1.958 | -0.13 | -0.006 | -0.05 | -0.05 | 2.219 | 0.11 | 0.17 | 0.25 | 0 | 0.06 | 0.05 |
| 3 | 26408174 | rs7651648 | 0.483 | 1.221 | 1.436 | 0.66 | -0.087 | -0.115 | -0.115 | 1.998 | 0.85 | 0.98 | 0.7 | 0.44 | 0.74 | 0.75 |
| 3 | 26410029 | rs10510573 | -1.545 | 0.51 | 0.714 | 0.346 | -0.003 | -0.009 | -0.009 | 0 | 0.39 | 0.63 | 0.44 | 0.2 | 0.29 | 0.34 |
| 3 | 26414083 | rs1160717 | 2.024 | 0.549 | 0.778 | 0.414 | -0.001 | 0.055 | 0.055 | 0 | 0.51 | 0.32 | 0.31 | 0.27 | 0.54 | 0.43 |
| 3 | 26418818 | rs1532719 | 0.047 | 0.623 | 0.595 | 0.479 | -0.038 | -0.047 | -0.047 | 0 | 0.88 | 0.95 | 0.73 | 0.46 | 0.8 | 0.76 |
| 3 | 26424850 | rs11713539 | -0.064 | 0.464 | 0.547 | 0.396 | -0.026 | -0.016 | -0.016 | 0 | 0.22 | 0.04 | 0.14 | 0.18 | 0.19 | 0.28 |
| 3 | 26425249 | rs1532718 | -0.347 | 0.494 | 0.788 | 0.345 | -0.001 | -0.018 | -0.018 | 0 | 0.1 | 0.11 | 0.25 | 0 | 0.06 | 0.05 |
| 3 | 26432346 | rs2134319 | 0.341 | 0.373 | 0.988 | 0.593 | -0.024 | -0.042 | -0.042 | 0.668 | 0.88 | 0.95 | 0.75 | 0.45 | 0.86 | 0.74 |
| 3 | 26439471 | rs7610718 | 1.314 | -0.142 | 0.989 | 0.583 | -0.004 | 0.012 | 0.012 | 0 | 0.75 | 0.33 | 0.67 | 0.44 | 0.8 | 0.66 |
| 3 | 26460080 | rs959685 | 2.005 | -0.247 | 0.539 | 0.55 | -0.071 | 0.055 | 0.055 | 0 | 0.44 | 0.12 | 0.48 | 0.31 | 0.49 | 0.4 |
| 3 | 26466967 | rs7631120 | -0.583 | -0.102 | 1.2 | 1.669 | -0.002 | 0.063 | 0.063 | 0 | 0.07 | 0.07 | 0.18 | 0.41 | 0.15 | 0.15 |
| 3 | 26469815 | rs9814029 | -1.414 | -0.393 | 1.54 | 1.553 | -0.037 | 0.002 | 0.002 | 0 | 0.27 | 0.57 | 0.22 | 0.25 | 0.2 | 0.21 |
| 3 | 26471909 | rs9310801 | -0.135 | -0.417 | 1.552 | 1.324 | -0.001 | -0.043 | -0.043 | 0 | 0.08 | 0.3 | 0.06 | 0 | 0.05 | 0.01 |
| 3 | 26480713 | rs9654031 | -1.016 | -0.401 | 1.216 | 1.331 | -0.032 | -0.079 | -0.079 | 0 | 0.12 | 0.44 | 0.09 | 0 | 0.07 | 0.01 |
| 3 | 26501241 | rs12495859 | -1.738 | -0.033 | 0.915 | 1.915 | 0 | -0.007 | -0.007 | 0 | 0.26 | 0.28 | 0.36 | 0.66 | 0.3 | 0.37 |
| 3 | 26503686 | rs13099283 | -0.561 | 0.028 | 0.883 | 2.116 | -0.033 | 0.013 | 0.013 | 0 | 0.06 | 0.02 | 0.19 | 0.41 | 0.15 | 0.16 |
| 3 | 26519005 | rs11712421 | -0.912 | -0.021 | 0.897 | 2.054 | 0.019 | -0.024 | -0.024 | 0 | 0.19 | 0.16 | 0.15 | 0.27 | 0.15 | 0.2 |
| 3 | 26522419 | rs12629714 | -2.203 | -0.089 | 0.953 | 2.017 | 0.018 | 0.006 | 0.006 | 0 | 0.37 | 0.31 | 0.39 | 0.68 | 0.35 | 0.49 |
| 3 | 26523363 | rs1158544 | -1.536 | -0.097 | 0.754 | 1.998 | 0.049 | -0.011 | -0.011 | 0 | 0.28 | 0.19 | 0.19 | 0.28 | 0.19 | 0.33 |
| 3 | 26531613 | rs2036430 | -1.788 | -0.079 | 0.811 | 2.007 | -0.031 | 0 | 0 | 0 | 0.3 | 0.38 | 0.21 | 0.3 | 0.19 | 0.32 |
| 3 | 26541692 | rs7642228 | -1.845 | 0.294 | 1.358 | 2.105 | -0.023 | 0.074 | 0.074 | 0 | 0.23 | 0.54 | 0.32 | 0.41 | 0.21 | 0.29 |
| 3 | 26542963 | rs7637842 | -1.683 | 0.388 | 1.097 | 2.033 | -0.019 | 0.147 | 0.147 | 0 | 0.22 | 0.53 | 0.32 | 0.52 | 0.22 | 0.3 |
| 3 | 26544115 | rs7651192 | -1.542 | 1.462 | 0.768 | 1.689 | 0.085 | 0.051 | 0.051 | 0 | 0.09 | 0.24 | 0.2 | 0.12 | 0.16 | 0.16 |
| 3 | 26547333 | rs1352718 | -1.062 | 1.263 | 0.506 | 1.964 | -0.002 | 0.112 | 0.112 | 0 | 0.18 | 0.25 | 0.27 | 0.51 | 0.2 | 0.28 |
| 3 | 26547966 | rs1847156 | 1.817 | 1.618 | 0.999 | 1.801 | 0.137 | 0.114 | 0.114 | 0 | 0.91 | 0.62 | 0.9 | 0.88 | 0.84 | 0.95 |
| 3 | 26548139 | rs1490687 | 1.152 | 1.65 | 0.61 | 1.516 | 0.006 | -0.006 | -0.006 | 0 | 0.87 | 0.86 | 0.85 | 0.93 | 0.8 | 0.93 |
| 3 | 26552078 | rs4973786 | -1.389 | 2.086 | 0.325 | 1.829 | -0.005 | -0.007 | -0.007 | 0.34 | 0.36 | 0.68 | 0.32 | 0.58 | 0.38 | 0.49 |
| 3 | 26557594 | rs7638812 | 1.235 | 1.974 | -0.009 | 1.371 | 0.005 | 0.06 | 0.06 | 0 | 0.54 | 0.29 | 0.55 | 0.28 | 0.42 | 0.4 |
| 3 | 26560272 | rs4973790 | 1.428 | 1.676 | -0.853 | 1.131 | -0.035 | 0.062 | 0.062 | 0 | 0.43 | 0.03 | 0.5 | 0.27 | 0.35 | 0.31 |
| 3 | 26560530 | rs1826219 | 0.868 | 2.144 | 0.61 | 2.036 | 0.002 | 0.091 | 0.091 | 0.24 | 0.77 | 0.45 | 0.75 | 0.55 | 0.75 | 0.64 |
| 3 | 26563909 | rs6763409 | -0.205 | 2.094 | 1.151 | 2.096 | -0.019 | 0.139 | 0.139 | 0.058 | 0.14 | 0.35 | 0.14 | 0.37 | 0.08 | 0.24 |
| 3 | 26581279 | rs13315748 | -0.131 | 2.189 | 1.372 | 2.17 | -0.02 | 0.08 | 0.08 | 0.017 | 0.13 | 0.26 | 0.13 | 0.3 | 0.08 | 0.24 |
| 3 | 26582570 | rs6789537 | -0.042 | 2.285 | 1.194 | 1.943 | 0.025 | -0.063 | -0.063 | 0 | 0.12 | 0.21 | 0.09 | 0.01 | 0.15 | 0.2 |
| 3 | 26594154 | rs7643655 | -0.623 | 2.144 | 1.219 | 2.114 | 0.023 | 0.106 | 0.106 | 0 | 0.05 | 0.16 | 0.18 | 0.16 | 0.12 | 0.02 |
| 3 | 26599551 | rs1602325 | -1.099 | 2.33 | 1.112 | 2.056 | 0.051 | 0.09 | 0.09 | 0.478 | 0.21 | 0.49 | 0.24 | 0.36 | 0.25 | 0.32 |
| 3 | 26613656 | rs9868148 | -0.267 | 2.143 | 0.874 | 2.006 | -0.02 | 0.208 | 0.208 | 0.177 | 0.14 | 0.45 | 0.15 | 0.35 | 0.08 | 0.22 |
| 3 | 26616411 | rs12491628 | -0.792 | 2.525 | 1.034 | 1.655 | 0.003 | 0.071 | 0.071 | 0 | 0.05 | 0.13 | 0.17 | 0.16 | 0.1 | 0.03 |
| 3 | 26622845 | rs13063381 | -0.59 | 2.609 | 0.975 | 1.602 | -0.008 | 0 | 0 | 0 | 0.07 | 0.03 | 0.1 | 0.06 | 0.19 | 0.07 |
| 3 | 26623735 | rs13069035 | -0.59 | 2.609 | 0.956 | 1.602 | -0.012 | 0 | 0 | 0 | 0.07 | 0.03 | 0.1 | 0.06 | 0.19 | 0.07 |
| 3 | 26629921 | rs17018362 | -0.831 | 2.605 | 0.982 | 1.6 | 0.016 | -0.001 | -0.001 | 0 | 0.07 | 0.09 | 0.1 | 0.06 | 0.19 | 0.07 |
| 3 | 26631359 | rs6551116 | -0.524 | 2.829 | 0.883 | 1.409 | 0.019 | -0.063 | -0.063 | 1.338 | 0.14 | 0.22 | 0.11 | 0.01 | 0.17 | 0.22 |
| 3 | 26633278 | rs957215 | -1.311 | 2.44 | 0.599 | 1.489 | 0.072 | 0.001 | 0.001 | 0 | 0.27 | 0.45 | 0.38 | 0.23 | 0.46 | 0.33 |
| 3 | 26639047 | rs990497 | 0.908 | 1.705 | 0.601 | 1.617 | 0.009 | 0.048 | 0.048 | 4.862 | 0.75 | 0.45 | 0.67 | 0.63 | 0.7 | 0.65 |
| 3 | 26639145 | rs6807559 | -0.503 | 0.975 | 0.347 | 1.166 | -0.019 | -0.007 | -0.007 | 3.026 | 0.26 | 0.39 | 0.22 | 0.3 | 0.21 | 0.33 |

| | | | | | | | | | | | | | | | | |
|---|---|---|---|---|---|---|---|---|---|---|---|---|---|---|---|---|
| 3 | 26651227 | rs1907158 | 1.664 | 0.488 | 0.198 | 1.08 | -0.051 | 0.052 | 0.052 | 1.586 | 0.49 | 0.13 | 0.46 | 0.32 | 0.5 | 0.4 |
| 3 | 26668383 | rs1586716 | 0.764 | 0.601 | 0.852 | 0.936 | 0 | -0.004 | -0.004 | 0.023 | 0.72 | 0.36 | 0.78 | 0.89 | 0.75 | 0.68 |
| 3 | 26682359 | rs6790894 | -0.748 | 0.523 | 0.967 | 1.143 | 0.023 | -0.022 | -0.022 | 0 | 0.2 | 0.35 | 0.2 | 0.22 | 0.23 | 0.26 |
| 3 | 26685101 | rs17018450 | -0.65 | 0.237 | 0.712 | 0.864 | -0.013 | -0.023 | -0.023 | 0 | 0.19 | 0.24 | 0.13 | 0.1 | 0.12 | 0.26 |
| 3 | 26685232 | rs7652737 | -1.103 | 0.021 | 0.807 | 1.219 | -0.028 | 0.054 | 0.054 | 0.181 | 0.24 | 0.39 | 0.26 | 0.45 | 0.14 | 0.3 |
| 3 | 26685397 | rs7652856 | 1.163 | 0.096 | 0.749 | 0.884 | 0 | -0.003 | -0.003 | 0 | 0.52 | 0.49 | 0.59 | 0.66 | 0.53 | 0.45 |
| 3 | 26685539 | rs12496948 | -0.985 | -0.004 | 0.769 | 0.911 | -0.01 | 0.002 | 0.002 | 0 | 0.26 | 0.38 | 0.26 | 0.33 | 0.21 | 0.43 |
| 3 | 26687117 | rs6767457 | -1.488 | -0.002 | 0.558 | 1.086 | -0.006 | 0.019 | 0.019 | 0.276 | 0.23 | 0.61 | 0.24 | 0.4 | 0.19 | 0.28 |
| 3 | 26687577 | rs17018468 | -0.279 | -0.014 | 0.387 | 1.101 | 0.025 | -0.02 | -0.02 | 0.002 | 0.19 | 0.14 | 0.12 | 0.26 | 0.09 | 0.26 |
| 3 | 26693662 | rs17018481 | 0.724 | 0.091 | 0.213 | 1.234 | -0.008 | -0.044 | -0.044 | 0 | 0.12 | 0.01 | 0.08 | 0.13 | 0.07 | 0.2 |
| 3 | 26696574 | rs6769536 | -0.74 | 0.074 | 0.494 | 1.118 | -0.004 | 0.074 | 0.074 | 0.646 | 0.18 | 0.58 | 0.29 | 0.39 | 0.16 | 0.21 |
| 3 | 26697920 | rs4327343 | 1.82 | 0.101 | 0.389 | 1.049 | -0.009 | 0.01 | 0.01 | 0.462 | 0.46 | 0.23 | 0.48 | 0.49 | 0.48 | 0.38 |
| 3 | 26698628 | rs6551125 | 0.482 | 0.117 | 0.318 | 1.037 | -0.02 | 0.126 | 0.126 | 2.969 | 0.64 | 0.83 | 0.78 | 0.9 | 0.66 | 0.6 |
| 3 | 26698857 | rs11720649 | 1.78 | 0.19 | -0.04 | 0.995 | -0.007 | 0.011 | 0.011 | 0.365 | 0.46 | 0.25 | 0.48 | 0.49 | 0.48 | 0.34 |
| 3 | 26704577 | rs4266153 | -0.769 | 0.446 | -0.458 | 0.6 | -0.008 | 0.02 | 0.02 | 0.008 | 0.79 | 0.86 | 0.86 | 1 | 0.77 | 0.85 |
| 3 | 26705637 | rs7616728 | -0.295 | 0.528 | -0.365 | 1.016 | 0.028 | 0.087 | 0.087 | 0.009 | 0.13 | 0.26 | 0.35 | 0.33 | 0.17 | 0.18 |
| 3 | 26714438 | rs11710809 | 1.384 | 0.364 | -1.2 | 1.02 | -0.073 | 0.042 | 0.042 | 0.881 | 0.39 | 0.1 | 0.34 | 0.26 | 0.43 | 0.31 |
| 3 | 26739650 | rs17018559 | -0.263 | 0.781 | -0.931 | 1.104 | -0.052 | -0.011 | -0.011 | 0 | 0.13 | 0 | 0.22 | 0.12 | 0.17 | 0.17 |
| 3 | 26749492 | rs17349384 | 1.393 | 0.586 | -1.043 | 0.813 | 0.007 | 0.15 | 0.15 | 0 | 0.43 | 0.18 | 0.29 | 0.12 | 0.35 | 0.31 |
| 3 | 26750891 | rs4473518 | -0.079 | 0.471 | -0.275 | 1.255 | -0.003 | 0.132 | 0.132 | 1.257 | 0.11 | 0.34 | 0.18 | 0.29 | 0.08 | 0.18 |
| 3 | 26763559 | rs1488239 | -0.357 | 0.635 | -0.555 | 1.358 | -0.031 | -0.041 | -0.041 | 0 | 0.2 | 0.04 | 0.27 | 0.24 | 0.22 | 0.26 |
| 3 | 26768818 | rs2129855 | 0.935 | 0.678 | -0.592 | 1.551 | -0.052 | -0.002 | -0.002 | 0 | 0.57 | 0.87 | 0.49 | 0.39 | 0.44 | 0.51 |
| 3 | 26777472 | rs1488233 | -1.584 | 0.66 | -0.597 | 1.508 | 0.006 | -0.046 | -0.046 | 0 | 0.34 | 0.75 | 0.47 | 0.52 | 0.31 | 0.47 |
| 3 | 26778662 | rs4088810 | -0.941 | 0.69 | -0.578 | 1.528 | -0.074 | -0.096 | -0.096 | 0.119 | 0.79 | 0.95 | 0.77 | 0.64 | 0.68 | 0.8 |
| 3 | 26802920 | rs9872064 | 1.867 | 0.597 | -0.93 | 1.302 | 0.023 | 0.018 | 0.018 | 0 | 0.45 | 0.22 | 0.34 | 0.32 | 0.37 | 0.36 |
| 3 | 26803095 | rs9858519 | 0.217 | 0.536 | -1.08 | 1.294 | -0.128 | -0.06 | -0.06 | 0 | 0.65 | 0.91 | 0.57 | 0.59 | 0.51 | 0.62 |
| 3 | 26803617 | rs17018637 | -1.401 | 0.694 | -0.755 | 1.195 | -0.014 | 0.002 | 0.002 | 1.949 | 0.19 | 0.7 | 0.23 | 0.27 | 0.14 | 0.26 |
| 3 | 26806383 | rs3912610 | 0.708 | 0.872 | -0.428 | 1.344 | 0.004 | 0.014 | 0.014 | 0 | 0.64 | 0.43 | 0.56 | 0.44 | 0.55 | 0.58 |
| 3 | 26815224 | rs6799773 | 0.27 | 1.081 | -0.591 | 1.28 | -0.082 | -0.07 | -0.07 | 0 | 0.7 | 0.89 | 0.57 | 0.4 | 0.61 | 0.67 |
| 3 | 26817659 | rs1018165 | 1.54 | 0.938 | -0.425 | 1.148 | -0.006 | 0.106 | 0.106 | 0 | 0.52 | 0.28 | 0.35 | 0.13 | 0.47 | 0.44 |
| 3 | 26835843 | rs6792668 | 0.836 | 0.989 | -0.369 | 1.188 | -0.026 | 0.026 | 0.026 | 0 | 0.57 | 0.74 | 0.51 | 0.39 | 0.52 | 0.6 |
| 3 | 26837389 | rs6790598 | 0.284 | 1.143 | -0.601 | 1.244 | -0.077 | -0.056 | -0.056 | 2.513 | 0.28 | 0.06 | 0.27 | 0.3 | 0.34 | 0.23 |
| 3 | 26843377 | rs1155107 | -0.023 | 1.106 | -0.561 | 1.254 | -0.016 | -0.042 | -0.042 | 0 | 0.88 | 0.94 | 0.8 | 0.7 | 0.88 | 0.85 |
| 3 | 26846174 | rs4267616 | 0.564 | 1.124 | -0.522 | 1.263 | -0.017 | -0.064 | -0.064 | 0 | 0.83 | 0.94 | 0.78 | 0.65 | 0.82 | 0.75 |
| 3 | 26846791 | rs1386874 | 1.783 | 1.05 | -0.506 | 1.224 | -0.036 | 0.181 | 0.181 | 0 | 0.46 | 0.15 | 0.33 | 0.15 | 0.44 | 0.43 |
| 3 | 26847447 | rs1488212 | 0.618 | 1.124 | -0.524 | 1.254 | -0.059 | 0.027 | 0.027 | 0 | 0.6 | 0.84 | 0.52 | 0.36 | 0.49 | 0.6 |
| 3 | 26847954 | rs9830761 | 0.004 | 0.908 | -0.39 | 1.306 | 0.022 | 0.131 | 0.131 | 0 | 0.86 | 0.71 | 0.79 | 0.67 | 0.87 | 0.84 |
| 3 | 26849303 | rs9820319 | 0.018 | 0.963 | -0.401 | 1.301 | -0.074 | -0.044 | -0.044 | 0 | 0.29 | 0.08 | 0.28 | 0.31 | 0.37 | 0.25 |
| 3 | 26850423 | rs17018761 | 0.253 | 0.385 | -0.181 | 1.32 | 0.044 | 0.131 | 0.131 | 0.795 | 0.08 | 0.37 | 0.18 | 0.2 | 0.07 | 0.12 |
| 3 | 26871939 | rs12631641 | 0.293 | 0.673 | -1.238 | 1.255 | -0.054 | -0.046 | -0.046 | 0 | 0.27 | 0.09 | 0.25 | 0.29 | 0.32 | 0.2 |
| 3 | 26872883 | rs17225714 | 1.437 | 0.722 | -1.271 | 1.199 | -0.027 | 0.161 | 0.161 | 0 | 0.45 | 0.14 | 0.32 | 0.15 | 0.42 | 0.43 |
| 3 | 26874626 | rs905936 | -0.313 | 0.8 | -0.709 | 1.183 | 0 | 0.001 | 0.001 | 0 | 0.29 | 0.32 | 0.27 | 0.32 | 0.32 | 0.21 |
| 3 | 26877202 | rs17018797 | -1.137 | 0.474 | -0.759 | 1.123 | -0.006 | 0.006 | 0.006 | 0 | 0.14 | 0.16 | 0.17 | 0.27 | 0.13 | 0.23 |
| 3 | 26877966 | rs7623573 | 0.484 | 0.643 | -0.725 | 1 | -0.007 | 0.065 | 0.065 | 0 | 0.68 | 0.49 | 0.62 | 0.47 | 0.7 | 0.65 |
| 3 | 26881106 | rs12635000 | -0.104 | 0.863 | -1.578 | 0.198 | 0.022 | 0.062 | 0.062 | 0 | 0.46 | 0.24 | 0.46 | 0.66 | 0.42 | 0.5 |
| 3 | 26885891 | rs11711355 | 0.194 | 0.862 | -1.766 | 0.216 | 0.014 | -0.034 | -0.034 | 1.172 | 0.38 | 0.15 | 0.32 | 0.57 | 0.34 | 0.33 |
| 3 | 26890400 | rs9871922 | 0.469 | 1.031 | -1.669 | -0.158 | 0.042 | 0.004 | 0.004 | 0 | 0.2 | 0.52 | 0.31 | 0.19 | 0.27 | 0.21 |
| 3 | 26892154 | rs9826565 | -0.744 | 0.928 | -1.776 | -0.253 | -0.027 | 0.117 | 0.117 | 0 | 0.63 | 0.87 | 0.78 | 0.84 | 0.65 | 0.57 |
| 3 | 26895022 | rs9310820 | 0.291 | 1.151 | -1.787 | -0.312 | -0.007 | -0.001 | -0.001 | 0 | 0.2 | 0.57 | 0.23 | 0.18 | 0.2 | 0.15 |
| 3 | 26897822 | rs9834454 | -0.412 | 1.131 | -1.707 | -0.177 | 0.006 | 0.069 | 0.069 | 0 | 0.62 | 0.78 | 0.78 | 0.78 | 0.66 | 0.56 |
| 3 | 26898377 | rs9858870 | 0.132 | 1.291 | -1.253 | 0.08 | 0.056 | 0.026 | 0.026 | 0 | 0.21 | 0.53 | 0.32 | 0.24 | 0.29 | 0.23 |
| 3 | 26901271 | rs1386884 | 0.25 | 0.865 | -1.459 | -0.087 | -0.004 | -0.026 | -0.026 | 2.383 | 0.06 | 0.1 | 0.04 | 0.04 | 0.06 | 0.03 |
| 3 | 26902598 | rs905933 | -0.42 | 1.083 | -1.918 | -0.039 | 0.003 | 0.103 | 0.103 | 0.871 | 0.7 | 0.88 | 0.82 | 0.87 | 0.74 | 0.72 |
| 3 | 26903228 | rs1488225 | 0.765 | 1.62 | -1.372 | 0.56 | -0.022 | -0.015 | -0.015 | 2.011 | 0.94 | 0.99 | 0.95 | 0.92 | 0.9 | 0.92 |
| 3 | 26903893 | rs11711747 | -0.535 | 1.418 | -0.796 | 1.168 | 0.011 | 0.104 | 0.104 | 0.154 | 0.19 | 0.37 | 0.2 | 0.48 | 0.21 | 0.21 |
| 3 | 26906882 | rs13067234 | 0.094 | 1.212 | -0.93 | 0.833 | -0.012 | -0.032 | -0.032 | 0.011 | 0.24 | 0.1 | 0.17 | 0.23 | 0.19 | 0.13 |
| 3 | 26907399 | rs9858938 | 0.602 | 0.841 | -1.098 | 0.948 | -0.049 | -0.042 | -0.042 | 0 | 0.44 | 0.12 | 0.63 | 0.46 | 0.58 | 0.52 |
| 3 | 26912568 | rs966763 | -0.549 | 0.715 | -0.342 | 0.927 | -0.018 | -0.006 | -0.006 | 0.927 | 0.71 | 0.47 | 0.83 | 0.71 | 0.77 | 0.68 |
| 3 | 26913820 | rs13077447 | 0.478 | 1.084 | -0.606 | 0.705 | -0.028 | -0.034 | -0.034 | 1.494 | 0.14 | 0.09 | 0.27 | 0.23 | 0.17 | 0.15 |
| 3 | 26947415 | rs955027 | 0.291 | 1.187 | -0.577 | 0.618 | -0.069 | -0.069 | -0.069 | 1.657 | 0.57 | 0.11 | 0.55 | 0.46 | 0.6 | 0.52 |
| 3 | 26947546 | rs17305779 | 1.103 | 1.363 | -0.621 | 0.692 | -0.047 | -0.049 | -0.049 | 0 | 0.08 | 0.01 | 0.21 | 0.24 | 0.17 | 0.08 |
| 3 | 26948644 | rs7615990 | -0.764 | 0.92 | 0.04 | 0.47 | -0.004 | -0.006 | -0.006 | 1.195 | 0.73 | 0.67 | 0.85 | 0.74 | 0.76 | 0.69 |
| 3 | 26948975 | rs7641122 | -0.52 | 0.924 | 0.08 | 0.469 | -0.006 | -0.006 | -0.006 | 0 | 0.71 | 0.41 | 0.82 | 0.72 | 0.75 | 0.66 |
| 3 | 26951226 | rs7629417 | -0.467 | 0.876 | -0.163 | 0.43 | -0.034 | -0.048 | -0.048 | 0 | 0.29 | 0.11 | 0.35 | 0.32 | 0.24 | 0.3 |
| 3 | 26951741 | rs1447639 | 1.257 | 0.906 | -0.143 | 0.419 | -0.047 | -0.063 | -0.063 | 0.265 | 0.4 | 0.13 | 0.46 | 0.45 | 0.52 | 0.37 |
| 3 | 26958417 | rs9310823 | 1.195 | 0.668 | -0.09 | 0.426 | -0.015 | -0.062 | -0.062 | 0 | 0.16 | 0.23 | 0.11 | 0 | 0.08 | 0.03 |
| 3 | 26963258 | rs1466957 | 0.644 | 0.78 | -0.146 | 0.572 | -0.075 | -0.024 | -0.024 | 0.03 | 0.55 | 0.11 | 0.53 | 0.42 | 0.57 | 0.51 |
| 3 | 26976847 | rs13065512 | -0.165 | 0.791 | -0.028 | 0.366 | 0.012 | 0.022 | 0.022 | 0.04 | 0.67 | 0.22 | 0.78 | 0.73 | 0.73 | 0.65 |
| 3 | 26979781 | rs2201978 | -0.183 | 0.798 | -0.019 | 0.369 | 0.011 | 0.023 | 0.023 | 0.062 | 0.67 | 0.24 | 0.78 | 0.73 | 0.73 | 0.64 |

| | | | | | | | | | | | | | | | | |
|---|---|---|---|---|---|---|---|---|---|---|---|---|---|---|---|---|
| 3 | 26982392 | rs1374797 | -0.291 | 0.828 | -0.089 | 0.382 | -0.004 | -0.007 | -0.007 | 0.207 | 0.68 | 0.4 | 0.78 | 0.73 | 0.73 | 0.64 |
| 3 | 26995291 | rs6762560 | -0.458 | 0.805 | -0.379 | 0.429 | -0.012 | -0.014 | -0.014 | 0 | 0.71 | 0.54 | 0.83 | 0.73 | 0.75 | 0.66 |
| 3 | 27009845 | rs13083708 | -0.733 | 0.845 | -1.068 | 0.439 | -0.001 | -0.002 | -0.002 | 1.005 | 0.73 | 0.74 | 0.84 | 0.75 | 0.76 | 0.68 |
| 3 | 27010197 | rs1868110 | -0.193 | 1.096 | -2.209 | 0.499 | -0.037 | -0.025 | -0.025 | 0 | 0.3 | 0.08 | 0.34 | 0.29 | 0.24 | 0.3 |
| 3 | 27010754 | rs13079997 | 0.333 | 0.771 | -1.657 | 0.697 | 0.05 | -0.042 | -0.042 | 0 | 0.49 | 0.1 | 0.67 | 0.45 | 0.65 | 0.44 |
| 3 | 27016393 | rs17019072 | -0.171 | 1.111 | -0.926 | 0.8 | 0 | 0.005 | 0.005 | 0.083 | 0.24 | 0.21 | 0.2 | 0.4 | 0.25 | 0.3 |
| 3 | 27017385 | rs1374799 | 0.139 | 1.027 | -0.446 | 0.146 | 0.003 | 0.009 | 0.009 | 0.806 | 0.69 | 0.65 | 0.6 | 0.39 | 0.67 | 0.42 |
| 3 | 27020726 | rs1374798 | -0.731 | 1.078 | -0.322 | -0.106 | -0.001 | -0.008 | -0.008 | 2.776 | 0.81 | 0.81 | 0.88 | 0.88 | 0.83 | 0.75 |
| 3 | 27024158 | rs7616789 | 0.492 | 0.784 | -0.653 | -0.707 | -0.003 | -0.044 | -0.044 | 0 | 0.45 | 0.51 | 0.27 | 0.09 | 0.34 | 0.26 |
| 3 | 27024947 | rs2044573 | 0.784 | 0.797 | -0.533 | -0.792 | 0.037 | -0.012 | -0.012 | 0 | 0.13 | 0.32 | 0.15 | 0.07 | 0.15 | 0.16 |
| 3 | 27025777 | rs9841676 | 0.827 | 0.809 | -0.775 | -0.994 | 0.024 | -0.039 | -0.039 | 0 | 0.13 | 0.24 | 0.12 | 0.04 | 0.15 | 0.16 |
| 3 | 27026003 | rs4973842 | 1.037 | 0.395 | -1.4 | -0.733 | -0.058 | -0.051 | -0.051 | 0.175 | 0.22 | 0.04 | 0.26 | 0.26 | 0.33 | 0.14 |
| 3 | 27026433 | rs11925608 | 0.337 | -0.301 | -0.707 | -1.044 | 0.045 | 0.078 | 0.078 | 2.962 | 0.32 | 0.19 | 0.12 | 0.02 | 0.19 | 0.1 |
| 3 | 27052635 | rs13320169 | -0.619 | 0.154 | 0.138 | -0.963 | -0.001 | 0.002 | 0.002 | 0 | 0.2 | 0.24 | 0.16 | 0.13 | 0.23 | 0.16 |
| 3 | 27053802 | rs1402991 | 0.253 | -0.071 | -0.329 | -1.418 | -0.06 | 0.016 | 0.016 | 0 | 0.3 | 0.2 | 0.57 | 0.76 | 0.46 | 0.51 |
| 3 | 27056797 | rs2177132 | -0.366 | -0.06 | 0.19 | -1.422 | -0.001 | -0.017 | -0.017 | 0 | 0.23 | 0.7 | 0.2 | 0.13 | 0.23 | 0.16 |
| 3 | 27056851 | rs1402989 | 0.17 | -0.282 | 0.139 | -1.498 | -0.095 | 0.051 | 0.051 | 0 | 0.32 | 0.14 | 0.58 | 0.79 | 0.48 | 0.52 |
| 3 | 27068426 | rs4973848 | 0.259 | -0.23 | 0.266 | -1.419 | 0.01 | -0.045 | -0.045 | 0 | 0.76 | 0.4 | 0.78 | 0.85 | 0.72 | 0.78 |
| 3 | 27072581 | rs6551161 | 0.136 | -0.241 | 0.269 | -1.416 | 0.005 | -0.049 | -0.049 | 0 | 0.77 | 0.41 | 0.8 | 0.87 | 0.75 | 0.83 |
| 3 | 27100979 | rs9855276 | 1.882 | -0.313 | 0.418 | -1.485 | -0.004 | -0.027 | -0.027 | 0.058 | 0.9 | 0.38 | 0.89 | 0.95 | 0.9 | 0.92 |
| 3 | 27114597 | rs12636719 | 0.645 | -0.275 | 0.137 | -1.58 | -0.165 | -0.05 | -0.05 | 0.42 | 0.26 | 0.02 | 0.4 | 0.74 | 0.39 | 0.45 |
| 3 | 27116147 | rs17019305 | 0.144 | -0.264 | 0.242 | -1.594 | -0.007 | 0.017 | 0.017 | 0 | 0.19 | 0.18 | 0.16 | 0.13 | 0.23 | 0.16 |
| 3 | 27123670 | rs1402980 | 0.301 | -0.316 | 0.226 | -1.651 | -0.195 | -0.016 | -0.016 | 0 | 0.31 | 0.02 | 0.45 | 0.73 | 0.46 | 0.47 |
| 3 | 27139429 | rs17019399 | -0.762 | -0.218 | 0.602 | -1.541 | 0.067 | 0.063 | 0.063 | 0.834 | 0.06 | 0.38 | 0.05 | 0.03 | 0.05 | 0.01 |
| 3 | 27147320 | rs13433970 | -0.275 | -0.241 | 0.327 | -1.597 | 0 | -0.004 | -0.004 | 2.575 | 0.17 | 0.27 | 0.15 | 0.16 | 0.21 | 0.19 |
| 3 | 27152266 | rs7628297 | -0.11 | -0.256 | 0.321 | -1.594 | -0.001 | -0.005 | -0.005 | 0 | 0.16 | 0.21 | 0.13 | 0.13 | 0.18 | 0.16 |
| 3 | 27188306 | rs1522166 | -0.356 | -0.388 | 0.277 | -1.631 | -0.001 | -0.012 | -0.012 | 0 | 0.17 | 0.26 | 0.14 | 0.15 | 0.18 | 0.15 |
| 3 | 27199604 | rs1522159 | 1.545 | -0.771 | 0.001 | -1.861 | 0.052 | 0.181 | 0.181 | 0 | 0.4 | 0.12 | 0.32 | 0.09 | 0.28 | 0.29 |
| 3 | 27207760 | rs7618127 | 0.997 | -0.769 | 0.115 | -1.88 | 0.042 | 0.095 | 0.095 | 0 | 0.47 | 0.66 | 0.41 | 0.09 | 0.35 | 0.4 |
| 3 | 27223710 | rs13098500 | 1.076 | -0.772 | 0.472 | -1.873 | 0.048 | 0.112 | 0.112 | 0 | 0.47 | 0.66 | 0.39 | 0.09 | 0.36 | 0.39 |
| 3 | 27253971 | rs1445111 | -0.366 | -0.516 | 0.332 | -1.823 | -0.182 | 0.01 | 0.01 | 0 | 0.67 | 0.97 | 0.58 | 0.26 | 0.56 | 0.55 |
| 3 | 27257026 | rs7652855 | -0.927 | -0.354 | 0.234 | -1.93 | -0.139 | -0.034 | -0.034 | 0 | 0.74 | 0.98 | 0.63 | 0.26 | 0.63 | 0.6 |
| 3 | 27257943 | rs11129270 | 0.027 | -0.438 | 0.271 | -1.919 | -0.105 | 0.074 | 0.074 | 0 | 0.64 | 0.9 | 0.57 | 0.24 | 0.55 | 0.54 |
| 3 | 27260283 | rs2370946 | 0.013 | -0.46 | 0.289 | -1.929 | -0.099 | 0.072 | 0.072 | 0 | 0.64 | 0.89 | 0.57 | 0.24 | 0.55 | 0.54 |
| 3 | 27273415 | rs2100006 | -0.231 | -0.499 | 0.219 | -1.923 | -0.189 | 0.037 | 0.037 | 0 | 0.65 | 0.97 | 0.61 | 0.24 | 0.54 | 0.49 |
| 3 | 27279235 | rs13075192 | -0.428 | -0.495 | 0.251 | -1.979 | -0.003 | 0.012 | 0.012 | 0 | 0.18 | 0.16 | 0.18 | 0.11 | 0.21 | 0.16 |
| 3 | 27279423 | rs4586769 | 1.635 | -0.603 | 0.376 | -1.968 | 0.052 | 0.138 | 0.138 | 0.054 | 0.39 | 0.14 | 0.35 | 0.11 | 0.28 | 0.29 |
| 3 | 27281769 | rs1158719 | 1.619 | -0.603 | 0.366 | -1.988 | 0.053 | 0.152 | 0.152 | 0.041 | 0.39 | 0.12 | 0.35 | 0.11 | 0.28 | 0.29 |
| 3 | 27296212 | rs2289803 | 0.744 | -0.609 | 0.344 | -1.935 | 0.082 | 0.24 | 0.24 | 0 | 0.51 | 0.92 | 0.46 | 0.21 | 0.35 | 0.37 |
| 3 | 27316919 | rs4973868 | 1.57 | -0.619 | 0.553 | -1.939 | 0.032 | 0.061 | 0.061 | 0.019 | 0.4 | 0.25 | 0.37 | 0.1 | 0.27 | 0.29 |
| 3 | 27326097 | rs11129280 | 1.666 | -0.619 | 0.536 | -1.939 | 0.051 | 0.148 | 0.148 | 0.043 | 0.39 | 0.12 | 0.35 | 0.1 | 0.27 | 0.29 |
| 3 | 27326451 | rs10510594 | 1.641 | -0.62 | 0.537 | -1.94 | 0.053 | 0.162 | 0.162 | 0.039 | 0.39 | 0.11 | 0.35 | 0.1 | 0.28 | 0.29 |
| 3 | 27332820 | rs10510592 | 1.666 | -0.636 | 0.529 | -1.947 | 0.051 | 0.148 | 0.148 | 0.052 | 0.39 | 0.12 | 0.35 | 0.1 | 0.27 | 0.28 |
| 3 | 27336213 | rs610118 | -1.367 | -0.777 | 0.858 | -1.924 | 0 | 0.015 | 0.015 | 2.214 | 0.1 | 0.63 | 0.1 | 0.11 | 0.08 | 0.08 |
| 3 | 27339326 | rs17317435 | 0.113 | -0.75 | 0.589 | -1.933 | -0.014 | 0.028 | 0.028 | 0 | 0.14 | 0.06 | 0.16 | 0.07 | 0.19 | 0.13 |
| 3 | 27349047 | rs4973760 | -1.522 | -0.743 | 0.262 | -1.926 | 0.005 | 0.039 | 0.039 | 0.935 | 0.25 | 0.85 | 0.26 | 0.18 | 0.26 | 0.21 |
| 3 | 27353716 | rs552647 | -0.075 | -0.597 | 0.196 | -1.899 | -0.191 | 0.034 | 0.034 | 0 | 0.64 | 0.97 | 0.61 | 0.27 | 0.53 | 0.49 |
| 3 | 27373066 | rs9864765 | 1.472 | -0.775 | 0.295 | -1.922 | 0.05 | 0.093 | 0.093 | 0 | 0.43 | 0.25 | 0.38 | 0.13 | 0.3 | 0.31 |
| 3 | 27389740 | rs12637322 | 0.679 | -0.73 | 0.18 | -1.972 | -0.159 | -0.03 | -0.03 | 0 | 0.28 | 0.02 | 0.34 | 0.73 | 0.41 | 0.44 |
| 3 | 27390490 | rs6551194 | 1.264 | -0.705 | 0.38 | -1.965 | -0.005 | 0.052 | 0.052 | 0 | 0.74 | 0.81 | 0.77 | 0.86 | 0.73 | 0.77 |
| 3 | 27395970 | rs7611368 | 1.158 | -0.667 | 0.551 | -1.901 | 0.014 | -0.06 | -0.06 | 0 | 0.85 | 0.47 | 0.89 | 0.92 | 0.89 | 0.88 |
| 3 | 27416013 | rs4973768 | 0.646 | -0.548 | 0.401 | -1.837 | -0.007 | 0.048 | 0.048 | 0.143 | 0.57 | 0.32 | 0.53 | 0.18 | 0.48 | 0.44 |
| 3 | 27421427 | rs11709954 | -0.144 | -0.025 | 0.494 | -1.799 | -0.037 | 0.046 | 0.046 | 0 | 0.15 | 0.03 | 0.15 | 0.06 | 0.21 | 0.19 |
| 3 | 27424718 | rs1472256 | -0.36 | -0.073 | 0.55 | -1.851 | -0.002 | -0.034 | -0.034 | 0 | 0.91 | 0.86 | 0.94 | 1 | 0.93 | 0.93 |
| 3 | 27427225 | rs1472254 | 0.353 | -0.2 | 0.436 | -1.946 | -0.07 | 0.1 | 0.1 | 0.144 | 0.62 | 0.79 | 0.58 | 0.19 | 0.52 | 0.51 |
| 3 | 27427521 | rs9823689 | 2.017 | -0.939 | 0.172 | -2.46 | -0.006 | -0.046 | -0.046 | 0.443 | 0.23 | 0.26 | 0.12 | 0.04 | 0.15 | 0.04 |
| 3 | 27431898 | rs13068734 | 1.293 | -0.886 | 0.092 | -2.497 | -0.024 | 0.024 | 0.024 | 0 | 0.06 | 0.01 | 0.09 | 0 | 0.12 | 0.06 |
| 3 | 27432779 | rs7644644 | 1.985 | -0.944 | 0.172 | -2.453 | 0.028 | 0.03 | 0.03 | 0 | 0.19 | 0.07 | 0.26 | 0.09 | 0.14 | 0.27 |
| 3 | 27432995 | rs2307032 | 1.856 | -1.028 | 0.001 | -1.847 | -0.119 | -0.144 | -0.144 | 0 | 0.19 | 0.01 | 0.32 | 0.57 | 0.34 | 0.18 |
| 3 | 27442336 | rs3736312 | 0.111 | -0.54 | 0.403 | -2.26 | -0.032 | 0.048 | 0.048 | 0 | 0.16 | 0.04 | 0.16 | 0.06 | 0.22 | 0.19 |
| 3 | 27458914 | rs923941 | 0.156 | -0.463 | 0.412 | -2.251 | -0.032 | 0.048 | 0.048 | 0 | 0.16 | 0.05 | 0.16 | 0.06 | 0.23 | 0.19 |
| 3 | 27472936 | rs3755652 | 0.156 | -0.463 | 0.412 | -2.251 | -0.032 | 0.048 | 0.048 | 0 | 0.16 | 0.05 | 0.16 | 0.06 | 0.23 | 0.19 |
| 3 | 27475351 | rs3278 | 1.946 | -0.481 | 0.457 | -2.25 | 0.016 | 0.038 | 0.038 | 0.005 | 0.23 | 0.15 | 0.11 | 0.04 | 0.15 | 0.04 |
| 3 | 27478899 | rs2029618 | 1.931 | -0.481 | 0.457 | -2.251 | 0.017 | 0.039 | 0.039 | 0.007 | 0.23 | 0.15 | 0.11 | 0.04 | 0.15 | 0.04 |
| 3 | 27484856 | rs9839331 | -0.554 | -0.44 | 0.451 | -2.221 | -0.002 | -0.033 | -0.033 | 0 | 0.91 | 0.86 | 0.94 | 1 | 0.93 | 0.93 |
| 3 | 27490194 | rs13096477 | 0.288 | -0.428 | 0.398 | -2.203 | -0.037 | 0.046 | 0.046 | 0 | 0.14 | 0.03 | 0.15 | 0.06 | 0.21 | 0.19 |
| 3 | 27529889 | rs9871261 | 0.853 | -0.561 | 0.478 | -2.221 | -0.016 | -0.044 | -0.044 | 0.167 | 0.44 | 0.46 | 0.41 | 0.14 | 0.3 | 0.32 |
| 3 | 27531796 | rs11129290 | 1.536 | -0.596 | 0.586 | -2.008 | 0.03 | 0.038 | 0.038 | 0 | 0.2 | 0.08 | 0.24 | 0.08 | 0.14 | 0.28 |
| 3 | 27547177 | rs6551205 | 2.157 | -0.674 | 0.559 | -1.799 | 0.014 | 0.008 | 0.008 | 0.373 | 0.2 | 0.15 | 0.09 | 0.12 | 0.12 | 0.05 |

| | | | | | | | | | | | | | | | | |
|---|---|---|---|---|---|---|---|---|---|---|---|---|---|---|---|---|
| 3 | 27548267 | rs3856675 | 1.606 | -0.744 | 0.492 | -1.811 | 0.03 | 0.042 | 0.042 | 0 | 0.2 | 0.08 | 0.24 | 0.07 | 0.14 | 0.27 |
| 3 | 27548813 | rs6774965 | 1.454 | -0.746 | 0.558 | -1.806 | -0.02 | -0.023 | -0.023 | 0 | 0.21 | 0.23 | 0.3 | 0.09 | 0.14 | 0.28 |
| 3 | 27549188 | rs820429 | 1.243 | -0.278 | 0.494 | -1.719 | -0.143 | -0.098 | -0.098 | 0 | 0.28 | 0.02 | 0.37 | 0.67 | 0.39 | 0.38 |
| 3 | 27552982 | rs6551207 | 0.297 | 0.139 | 1.663 | -1.213 | 0 | -0.023 | -0.023 | 0.038 | 0.9 | 0.89 | 0.93 | 1 | 0.91 | 0.92 |
| 3 | 27553839 | rs629462 | 1.498 | 0.133 | 1.676 | -1.205 | 0.002 | -0.061 | -0.061 | 0 | 0.85 | 0.57 | 0.89 | 0.92 | 0.87 | 0.89 |
| 3 | 27562613 | rs2643825 | -1.388 | 0.396 | 1.063 | -1.094 | -0.015 | 0.015 | 0.015 | 0 | 0.24 | 0.19 | 0.23 | 0.05 | 0.3 | 0.26 |
| 3 | 27572710 | rs1845700 | -1.552 | -0.135 | 1.079 | -0.929 | -0.001 | -0.036 | -0.036 | 0.127 | 0.26 | 0.66 | 0.24 | 0.13 | 0.26 | 0.25 |
| 3 | 27579916 | rs2724541 | -1.768 | -0.072 | 1.03 | -0.912 | -0.012 | 0.038 | 0.038 | 0.451 | 0.28 | 0.79 | 0.29 | 0.17 | 0.31 | 0.3 |
| 3 | 27588090 | rs13091835 | -0.452 | -0.211 | 1.107 | -0.62 | 0.01 | -0.035 | -0.035 | 0 | 0.16 | 0.22 | 0.2 | 0.06 | 0.24 | 0.19 |
| 3 | 27590422 | rs7429875 | -1.734 | -0.257 | 0.875 | -0.634 | -0.01 | 0.033 | 0.033 | 0.216 | 0.25 | 0.8 | 0.28 | 0.18 | 0.29 | 0.27 |
| 3 | 27590687 | rs3845994 | -1.216 | -0.3 | 0.917 | -0.595 | 0.002 | -0.042 | -0.042 | 0 | 0.21 | 0.64 | 0.23 | 0.13 | 0.25 | 0.2 |
| 3 | 27595912 | rs17019924 | -0.018 | -0.185 | 0.213 | -0.485 | -0.034 | -0.021 | -0.021 | 0 | 0.29 | 0.58 | 0.17 | 0.21 | 0.17 | 0.13 |
| 3 | 27602032 | rs9852113 | -1.507 | 0.784 | 0.771 | 0.029 | 0.037 | -0.031 | -0.031 | 0 | 0.19 | 0.58 | 0.23 | 0.16 | 0.27 | 0.22 |
| 3 | 27606268 | rs3892373 | -0.943 | 1.072 | -0.17 | 0.042 | 0.043 | -0.019 | -0.019 | 0 | 0.15 | 0.28 | 0.21 | 0.11 | 0.26 | 0.21 |
| 3 | 27612852 | rs12330312 | -0.112 | 0.449 | -0.621 | 0.315 | -0.018 | -0.012 | -0.012 | 0 | 0.43 | 0.52 | 0.43 | 0.72 | 0.32 | 0.46 |
| 3 | 27614316 | rs9854207 | -0.041 | -0.474 | -0.785 | 0.708 | 0.032 | -0.018 | -0.018 | 0 | 0.3 | 0.81 | 0.21 | 0.4 | 0.16 | 0.18 |
| 3 | 27615796 | rs2642925 | 0.277 | 0.467 | -1.324 | 0.254 | 0.011 | -0.004 | -0.004 | 0 | 0.22 | 0.2 | 0.22 | 0.11 | 0.38 | 0.25 |
| 3 | 27616054 | rs2643818 | -0.325 | 0.646 | -1.324 | 0.295 | -0.009 | 0.022 | 0.022 | 1.627 | 0.48 | 0.79 | 0.4 | 0.52 | 0.55 | 0.41 |
| 3 | 27620980 | rs13073001 | 1.038 | 0.481 | -1.293 | 0.307 | -0.029 | 0.039 | 0.039 | 4.824 | 0.26 | 0.67 | 0.19 | 0.43 | 0.17 | 0.13 |
| 3 | 27623969 | rs2642940 | 0.774 | 0.537 | -1.354 | 0.105 | -0.049 | 0.009 | 0.009 | 0 | 0.19 | 0.07 | 0.2 | 0.1 | 0.37 | 0.2 |
| 3 | 27635608 | rs2643842 | 0.598 | 0.523 | -1.045 | 0.287 | -0.049 | 0.021 | 0.021 | 0 | 0.28 | 0.57 | 0.19 | 0.35 | 0.18 | 0.15 |
| 3 | 27638500 | rs2642905 | -1.269 | 0.041 | -1.743 | -0.025 | -0.052 | 0.056 | 0.056 | 0 | 0.7 | 0.99 | 0.7 | 0.84 | 0.73 | 0.72 |
| 3 | 27646114 | rs2643823 | 0.46 | 0.264 | -1.357 | 0.036 | 0.041 | -0.025 | -0.025 | 0 | 0.31 | 0.81 | 0.22 | 0.37 | 0.21 | 0.18 |
| 3 | 27651544 | rs12715123 | -0.454 | 0.512 | -1.581 | -0.19 | 0.006 | 0 | 0 | 0 | 0.66 | 0.32 | 0.72 | 0.65 | 0.72 | 0.84 |
| 3 | 27651710 | rs13317121 | 0.65 | 0.532 | -1.702 | -0.174 | -0.045 | 0.019 | 0.019 | 0 | 0.22 | 0.1 | 0.2 | 0.11 | 0.35 | 0.23 |
| 3 | 27652096 | rs11710219 | -0.323 | 0.532 | -1.709 | -0.173 | 0.023 | -0.011 | -0.011 | 0 | 0.64 | 0.18 | 0.7 | 0.63 | 0.69 | 0.8 |
| 3 | 27652426 | rs6763146 | 0.535 | 0.435 | -1.574 | -0.04 | -0.026 | 0.014 | 0.014 | 0 | 0.22 | 0.16 | 0.21 | 0.11 | 0.35 | 0.23 |
| 3 | 27652901 | rs2642942 | -0.24 | 0.676 | -1.437 | 0.041 | -0.026 | -0.037 | -0.037 | 0 | 0.53 | 0.88 | 0.43 | 0.47 | 0.56 | 0.41 |
| 3 | 27663546 | rs3913110 | -0.04 | 0.461 | -0.624 | -0.216 | 0.001 | 0.001 | 0.001 | 0.139 | 0.59 | 0.62 | 0.56 | 0.6 | 0.68 | 0.57 |
| 3 | 27664144 | rs6762388 | 0.325 | 0.577 | -0.488 | -0.369 | 0 | 0.004 | 0.004 | 0 | 0.5 | 0.47 | 0.47 | 0.56 | 0.52 | 0.51 |
| 3 | 27666362 | rs6551219 | 0.211 | 0.527 | -0.37 | -0.373 | 0.002 | 0.014 | 0.014 | 0 | 0.52 | 0.59 | 0.48 | 0.56 | 0.56 | 0.54 |
| 3 | 27666794 | rs3846007 | 0.194 | 0.525 | -0.371 | -0.374 | 0.002 | 0.013 | 0.013 | 0 | 0.52 | 0.59 | 0.49 | 0.56 | 0.56 | 0.54 |
| 3 | 27677178 | rs2642911 | 0.007 | 0.29 | -0.567 | -0.257 | -0.006 | 0.009 | 0.009 | 0.413 | 0.4 | 0.24 | 0.44 | 0.4 | 0.35 | 0.46 |
| 3 | 27680063 | rs8179938 | -0.254 | -0.439 | 0.186 | -0.025 | -0.047 | 0.159 | 0.159 | 0.755 | 0.19 | 0.4 | 0.26 | 0.53 | 0.15 | 0.24 |
| 3 | 27690531 | rs2370990 | -0.858 | -0.377 | 0.449 | -0.19 | -0.023 | 0.045 | 0.045 | 0.152 | 0.27 | 0.38 | 0.33 | 0.56 | 0.27 | 0.27 |
| 3 | 27702442 | rs3856680 | -0.418 | -0.344 | 0.634 | -0.163 | -0.015 | 0.077 | 0.077 | 1.048 | 0.21 | 0.36 | 0.26 | 0.59 | 0.21 | 0.26 |
| 3 | 27706298 | rs9820465 | -0.494 | -0.356 | 0.641 | -0.171 | -0.02 | 0.111 | 0.111 | 1.492 | 0.22 | 0.5 | 0.27 | 0.6 | 0.2 | 0.25 |
| 3 | 27706375 | rs2643845 | 0.66 | -0.758 | -0.156 | -0.423 | -0.022 | -0.026 | -0.026 | 0 | 0.33 | 0.06 | 0.41 | 0.31 | 0.29 | 0.37 |
| 3 | 27709178 | rs6795883 | -0.743 | -0.735 | 0.771 | 0.326 | -0.023 | 0.035 | 0.035 | 1.398 | 0.22 | 0.64 | 0.27 | 0.61 | 0.16 | 0.25 |
| 3 | 27718345 | rs9836553 | -0.401 | -0.867 | 0.416 | 0.405 | -0.045 | 0.087 | 0.087 | 0.263 | 0.2 | 0.35 | 0.25 | 0.59 | 0.15 | 0.24 |
| 3 | 27721198 | rs6764447 | -1.121 | -0.893 | -0.271 | 0.296 | 0.037 | -0.021 | -0.021 | 0 | 0.29 | 0.77 | 0.36 | 0.63 | 0.22 | 0.28 |
| 3 | 27724852 | rs2724509 | -0.003 | -0.457 | -0.249 | -0.144 | 0.003 | 0.026 | 0.026 | 0.137 | 0.12 | 0.05 | 0.16 | 0.03 | 0.16 | 0.04 |
| 3 | 27725605 | rs9846205 | -1.615 | -0.497 | 0.579 | -0.28 | 0.093 | 0.042 | 0.042 | 1.532 | 0.06 | 0.41 | 0.05 | 0 | 0.04 | 0.02 |
| 3 | 27726951 | rs2642935 | -1.473 | -0.603 | 0.543 | -0.331 | -0.019 | -0.107 | -0.107 | 0 | 0.2 | 0.52 | 0.21 | 0.03 | 0.2 | 0.09 |
| 3 | 27756251 | rs2581199 | -1.123 | -0.585 | 0.455 | -0.257 | -0.01 | -0.03 | -0.03 | 0.472 | 0.18 | 0.3 | 0.18 | 0.06 | 0.17 | 0.09 |
| 3 | 27764623 | rs3806624 | -0.994 | -0.707 | -0.413 | -0.378 | -0.07 | 0.113 | 0.113 | 3.151 | 0.62 | 0.88 | 0.65 | 0.85 | 0.47 | 0.78 |
| 3 | 27782163 | rs12490648 | -0.042 | -1.029 | -0.74 | -0.456 | 0.016 | 0.003 | 0.003 | 0 | 0.13 | 0 | 0.04 | 0.09 | 0.08 | 0.17 |
| 3 | 27782360 | rs9878516 | -0.426 | -0.725 | -0.158 | -0.448 | 0.018 | -0.03 | -0.03 | 0 | 0.06 | 0.12 | 0.03 | 0 | 0.06 | 0.05 |
| 3 | 27783015 | rs13327021 | -0.379 | -1.004 | -0.073 | 0.188 | 0.048 | 0.022 | 0.022 | 1.14 | 0.49 | 0.26 | 0.53 | 0.46 | 0.38 | 0.69 |
| 3 | 27788780 | rs11129295 | -0.554 | -0.962 | 0.113 | 0.357 | 0.035 | 0.087 | 0.087 | 0.667 | 0.51 | 0.34 | 0.57 | 0.78 | 0.39 | 0.69 |
| 3 | 27797810 | rs12493245 | -0.258 | -0.937 | -0.058 | 0.32 | 0.025 | -0.002 | -0.002 | 0.258 | 0.46 | 0.33 | 0.54 | 0.45 | 0.35 | 0.68 |
| 3 | 27799034 | rs4680845 | -0.258 | -0.937 | -0.058 | 0.32 | 0.025 | -0.002 | -0.002 | 1.946 | 0.46 | 0.33 | 0.53 | 0.45 | 0.35 | 0.68 |
| 3 | 27801959 | rs17020288 | -0.061 | -0.725 | 0.084 | 0.435 | 0.011 | -0.012 | -0.012 | 0 | 0.41 | 0.3 | 0.51 | 0.46 | 0.33 | 0.66 |
| 3 | 27802978 | rs7633947 | 0.291 | -0.739 | 0.235 | 0.304 | -0.009 | -0.068 | -0.068 | 5.626 | 0.16 | 0.21 | 0.08 | 0 | 0.12 | 0.05 |
| 3 | 27804360 | rs6801242 | 0.102 | -0.73 | 0.234 | 0.308 | -0.006 | -0.061 | -0.061 | 0 | 0.16 | 0.21 | 0.08 | 0 | 0.13 | 0.07 |
| 3 | 27806147 | rs2371150 | -0.046 | -0.724 | 0.252 | 0.386 | 0.009 | -0.011 | -0.011 | 0 | 0.41 | 0.33 | 0.52 | 0.47 | 0.33 | 0.65 |
| 3 | 27812666 | rs9823274 | -0.396 | -0.736 | 0.089 | 0.412 | -0.001 | -0.006 | -0.006 | 4.806 | 0.45 | 0.53 | 0.56 | 0.79 | 0.35 | 0.65 |
| 3 | 27818453 | rs9872426 | -0.406 | -0.586 | -0.848 | 0.33 | 0.015 | -0.008 | -0.008 | 0 | 0.45 | 0.3 | 0.52 | 0.51 | 0.37 | 0.68 |
| 3 | 27819783 | rs7621606 | -1.262 | -0.531 | -0.945 | 0.281 | -0.096 | 0.147 | 0.147 | 1.407 | 0.62 | 0.84 | 0.66 | 0.84 | 0.5 | 0.75 |
| 3 | 27819955 | rs12633504 | 0.561 | -0.467 | -1.124 | 0.375 | 0.037 | 0.036 | 0.036 | 0 | 0.3 | 0.03 | 0.19 | 0.18 | 0.18 | 0.39 |
| 3 | 27822978 | rs9828781 | -0.487 | 0.174 | -0.769 | 0.097 | 0.047 | 0.134 | 0.134 | 0.567 | 0.14 | 0.3 | 0.32 | 0.32 | 0.18 | 0.26 |
| 3 | 27823887 | rs7627710 | 0.268 | 0.107 | -0.865 | -0.001 | -0.005 | 0 | 0 | 0.044 | 0.45 | 0.45 | 0.32 | 0.52 | 0.29 | 0.46 |
| 3 | 27827733 | rs9817112 | -0.651 | 0.393 | -1.476 | -0.937 | -0.063 | 0.117 | 0.117 | 1.159 | 0.62 | 0.8 | 0.66 | 0.85 | 0.5 | 0.75 |
| 3 | 27833251 | rs730510 | -0.443 | 0.426 | -1.272 | -0.593 | -0.001 | 0.02 | 0.02 | 0 | 0.38 | 0.33 | 0.45 | 0.19 | 0.34 | 0.36 |
| 3 | 27836045 | rs9873946 | -0.505 | 0.708 | -1.305 | -0.131 | 0.028 | 0.008 | 0.008 | 0 | 0.08 | 0.23 | 0.07 | 0.05 | 0.11 | 0.07 |
| 3 | 27839134 | rs2371117 | -0.066 | 0.64 | -2.481 | -0.341 | -0.074 | 0.087 | 0.087 | 2.97 | 0.63 | 0.98 | 0.69 | 0.81 | 0.61 | 0.69 |
| 3 | 27844548 | rs9851152 | 0.289 | -0.247 | -2.747 | -1.099 | 0.052 | 0.159 | 0.159 | 0 | 0.71 | 0.92 | 0.9 | 0.99 | 0.82 | 0.84 |
| 3 | 27846468 | rs4541368 | 0.518 | -0.011 | -2.489 | -0.766 | 0 | 0.017 | 0.017 | 0 | 0.28 | 0.26 | 0.34 | 0.51 | 0.26 | 0.28 |
| 3 | 27850725 | rs2371121 | -0.531 | -0.386 | -1.896 | -0.863 | 0.042 | 0.05 | 0.05 | 0 | 0.68 | 0.73 | 0.89 | 0.97 | 0.82 | 0.82 |

| | | | | | | | | | | | | | | | | |
|---|---|---|---|---|---|---|---|---|---|---|---|---|---|---|---|---|
| 3 | 27851720 | rs17020402 | 0.599 | -0.161 | -1.913 | -1.005 | 0 | 0.005 | 0.005 | 0 | 0.27 | 0.23 | 0.33 | 0.52 | 0.25 | 0.3 |
| 3 | 27866326 | rs7625647 | 0.772 | -0.853 | -1.523 | -1.635 | -0.035 | -0.021 | -0.021 | 0.477 | 0.3 | 0.28 | 0.45 | 0.45 | 0.48 | 0.47 |
| 3 | 27868849 | rs7621069 | 0.845 | -0.645 | -1.088 | -1.578 | -0.042 | -0.031 | -0.031 | 0.514 | 0.29 | 0.22 | 0.44 | 0.45 | 0.45 | 0.44 |
| 3 | 27875759 | rs9830335 | 0.538 | -0.077 | -0.762 | -1.659 | 0.056 | 0.079 | 0.079 | 0.386 | 0.25 | 0.48 | 0.38 | 0.46 | 0.34 | 0.43 |
| 3 | 27880417 | rs9810699 | 0.879 | -0.08 | -0.872 | -1.646 | -0.003 | -0.003 | -0.003 | 0 | 0.22 | 0.25 | 0.36 | 0.43 | 0.34 | 0.41 |
| 3 | 27887795 | rs17020449 | 1.164 | -0.224 | -0.685 | -1.693 | 0.006 | -0.006 | -0.006 | 2.113 | 0.1 | 0.06 | 0.06 | 0 | 0.08 | 0.08 |
| 3 | 27889035 | rs1542542 | 1.217 | 0.761 | -0.687 | -1.388 | -0.001 | -0.001 | -0.001 | 0 | 0.12 | 0.13 | 0.25 | 0.22 | 0.15 | 0.19 |
| 3 | 27903128 | rs11715548 | 0.94 | 0.838 | -0.71 | -1.383 | -0.006 | -0.007 | -0.007 | 0.56 | 0.11 | 0.11 | 0.12 | 0.2 | 0.19 | 0.22 |
| 3 | 27909537 | rs1080873 | 1.16 | 0.515 | -0.654 | -1.613 | -0.013 | -0.016 | -0.016 | 0 | 0.33 | 0.2 | 0.42 | 0.54 | 0.38 | 0.3 |
| 3 | 27916603 | rs11708641 | -0.47 | 1.037 | 0.042 | -1.059 | 0.03 | 0.043 | 0.043 | 0.059 | 0.24 | 0.37 | 0.37 | 0.45 | 0.34 | 0.43 |
| 3 | 27922474 | rs7623233 | 0.827 | 0.983 | 0.243 | -0.298 | 0.012 | 0.075 | 0.075 | 0 | 0.09 | 0.16 | 0.07 | 0.21 | 0.11 | 0.06 |
| 3 | 27924505 | rs2618101 | -0.356 | 0.845 | 0.276 | -0.396 | -0.017 | 0.011 | 0.011 | 0 | 0.19 | 0.52 | 0.15 | 0.22 | 0.19 | 0.12 |
| 3 | 27927647 | rs993710 | -0.095 | 0.938 | 0.077 | -0.506 | 0.012 | 0.005 | 0.005 | 0.531 | 0.21 | 0.31 | 0.29 | 0.27 | 0.2 | 0.21 |
| 3 | 27927730 | rs2581194 | -1.578 | 0.776 | -0.192 | -0.182 | 0.018 | 0.043 | 0.043 | 0.534 | 0.56 | 0.84 | 0.56 | 0.7 | 0.62 | 0.67 |
| 3 | 27929182 | rs2581188 | 1.168 | 0.477 | 0.119 | 0.068 | 0.021 | 0.084 | 0.084 | 0 | 0.08 | 0.14 | 0.07 | 0.2 | 0.1 | 0.05 |
| 3 | 27929871 | rs2581187 | -0.721 | 0.59 | 0.161 | -0.038 | -0.002 | 0.012 | 0.012 | 0 | 0.27 | 0.61 | 0.37 | 0.47 | 0.27 | 0.33 |
| 3 | 27931857 | rs12637187 | -0.394 | 0.622 | -0.241 | -0.02 | -0.016 | 0.014 | 0.014 | 0 | 0.24 | 0.16 | 0.16 | 0.18 | 0.3 | 0.38 |
| 3 | 27932447 | rs1382413 | 1.044 | 0.619 | -0.157 | 0.022 | 0.027 | 0.134 | 0.134 | 0.076 | 0.11 | 0.33 | 0.25 | 0.27 | 0.09 | 0.2 |
| 3 | 27934722 | rs2618108 | -1.746 | 0.66 | -0.871 | -0.015 | -0.048 | 0.043 | 0.043 | 0.882 | 0.53 | 0.88 | 0.55 | 0.65 | 0.54 | 0.65 |
| 3 | 27937774 | rs9878755 | 1.132 | 0.826 | -0.805 | 0.099 | 0.017 | 0.082 | 0.082 | 0 | 0.08 | 0.14 | 0.07 | 0.2 | 0.1 | 0.05 |
| 3 | 27938713 | rs17707359 | 0.493 | 0.814 | -1.48 | -0.25 | -0.075 | -0.056 | -0.056 | 0.042 | 0.14 | 0.02 | 0.11 | 0.18 | 0.23 | 0.3 |
| 3 | 27942217 | rs12496752 | 0.597 | 0.354 | -1.009 | -0.113 | -0.005 | -0.024 | -0.024 | 2.601 | 0.09 | 0.1 | 0.05 | 0 | 0.07 | 0.08 |
| 3 | 27945306 | rs9816729 | -0.924 | 0.427 | -1.305 | -0.037 | 0.028 | -0.043 | -0.043 | 0.036 | 0.38 | 0.81 | 0.44 | 0.47 | 0.33 | 0.36 |
| 3 | 27952483 | rs2581177 | -1.304 | 0.983 | -1.507 | -0.178 | -0.041 | 0.004 | 0.004 | 2.025 | 0.55 | 0.89 | 0.62 | 0.65 | 0.57 | 0.66 |
| 3 | 27953634 | rs1382417 | -0.643 | 1.047 | -0.812 | 0.111 | 0.013 | 0.081 | 0.081 | 0.113 | 0.27 | 0.44 | 0.42 | 0.61 | 0.31 | 0.31 |
| 3 | 27954037 | rs17020606 | -0.291 | 0.812 | -0.459 | 0.503 | 0.003 | 0.005 | 0.005 | 0.244 | 0.26 | 0.32 | 0.18 | 0.14 | 0.24 | 0.33 |
| 3 | 27954625 | rs1478556 | -0.676 | 1.158 | -0.137 | 1.284 | 0.045 | 0.315 | 0.315 | 0 | 0.15 | 0.38 | 0.34 | 0.53 | 0.19 | 0.22 |
| 3 | 27958681 | rs2253562 | -0.02 | 1.304 | -0.857 | 0.56 | -0.011 | 0.011 | 0.011 | 1.36 | 0.88 | 0.91 | 0.93 | 0.92 | 0.84 | 0.82 |
| 3 | 27959742 | rs12636813 | 0.478 | 0.577 | -1.772 | -0.185 | -0.007 | 0.207 | 0.207 | 0 | 0.52 | 0.16 | 0.62 | 0.85 | 0.55 | 0.58 |
| 3 | 27975263 | rs10510599 | 1.157 | 0.03 | -1.498 | 0.257 | -0.062 | -0.052 | -0.052 | 0 | 0.35 | 0.12 | 0.38 | 0.35 | 0.4 | 0.29 |
| 3 | 27975339 | rs1808099 | -0.357 | -0.325 | -1.052 | -0.384 | -0.006 | -0.011 | -0.011 | 0 | 0.8 | 0.88 | 0.84 | 0.99 | 0.78 | 0.81 |
| 3 | 27976461 | rs2127012 | 0.4 | -0.687 | -1.554 | -0.176 | -0.044 | -0.039 | -0.039 | 0 | 0.23 | 0.03 | 0.16 | 0.22 | 0.26 | 0.24 |
| 3 | 27976864 | rs7639905 | 0.824 | -0.778 | -1.414 | -0.093 | -0.032 | 0.179 | 0.179 | 0 | 0.5 | 0.12 | 0.6 | 0.76 | 0.49 | 0.51 |
| 3 | 27978145 | rs337426 | 0.637 | -1.213 | -0.457 | 0.491 | -0.008 | 0.004 | 0.004 | 0 | 0.61 | 0.47 | 0.51 | 0.57 | 0.64 | 0.53 |
| 3 | 27978488 | rs337427 | 0.485 | -1.048 | -0.644 | 0.429 | 0.024 | 0.021 | 0.021 | 0.156 | 0.27 | 0.31 | 0.34 | 0.37 | 0.38 | 0.26 |
| 3 | 27979646 | rs427532 | -0.761 | -1.259 | -0.289 | 0.356 | 0.086 | 0.111 | 0.111 | 0.576 | 0.51 | 0.69 | 0.58 | 0.7 | 0.69 | 0.56 |
| 3 | 27979804 | rs9832327 | 0.097 | -1.303 | -0.033 | 0.663 | 0.004 | 0.041 | 0.041 | 0 | 0.21 | 0.32 | 0.18 | 0.31 | 0.25 | 0.19 |
| 3 | 27981652 | rs1911153 | -1.64 | -1.552 | -0.299 | 0.177 | 0.048 | 0.062 | 0.062 | 0.122 | 0.72 | 0.78 | 0.66 | 0.91 | 0.83 | 0.84 |
| 3 | 27987104 | rs2199653 | -0.305 | -1.123 | -0.227 | 0.137 | 0.001 | 0.019 | 0.019 | 0 | 0.29 | 0.17 | 0.21 | 0.27 | 0.27 | 0.39 |
| 3 | 27991802 | rs404044 | 1.111 | -0.859 | -0.187 | 0.348 | -0.04 | -0.042 | -0.042 | 0 | 0.22 | 0.11 | 0.28 | 0.36 | 0.34 | 0.27 |
| 3 | 28006393 | rs2169823 | 0.476 | -0.39 | 0.148 | 0.756 | -0.017 | 0.002 | 0.002 | 0 | 0.08 | 0.02 | 0.1 | 0.08 | 0.11 | 0.13 |
| 3 | 28007315 | rs12630663 | -0.917 | -0.61 | 0.205 | 0.605 | -0.008 | -0.021 | -0.021 | 0 | 0.39 | 0.27 | 0.28 | 0.59 | 0.41 | 0.55 |
| 3 | 28007973 | rs4680886 | 0.869 | -0.54 | -0.006 | 0.774 | -0.052 | -0.082 | -0.082 | 0 | 0.13 | 0.02 | 0.12 | 0.31 | 0.22 | 0.22 |
| 3 | 28009822 | rs13318273 | -1.99 | -0.645 | 0.423 | 0.561 | -0.005 | 0.048 | 0.048 | 0 | 0.51 | 0.73 | 0.42 | 0.67 | 0.51 | 0.71 |
| 3 | 28011628 | rs6781471 | -1.98 | -0.659 | 0.424 | 0.561 | -0.005 | 0.048 | 0.048 | 0 | 0.51 | 0.73 | 0.41 | 0.67 | 0.51 | 0.71 |
| 3 | 28014302 | rs11710076 | -1.096 | -0.65 | 0.521 | 0.598 | -0.001 | -0.006 | -0.006 | 0 | 0.4 | 0.36 | 0.28 | 0.58 | 0.41 | 0.57 |
| 3 | 28014862 | rs11129306 | -0.477 | -0.659 | 0.478 | 0.603 | 0 | 0.051 | 0.051 | 0 | 0.22 | 0.34 | 0.18 | 0.39 | 0.25 | 0.36 |
| 3 | 28015470 | rs9815806 | 0.276 | -0.643 | 0.648 | 0.582 | 0.002 | 0.008 | 0.008 | 0 | 0.2 | 0.14 | 0.1 | 0.21 | 0.16 | 0.2 |
| 3 | 28017377 | rs11129307 | -0.015 | -0.536 | 0.644 | 0.687 | 0.002 | 0.021 | 0.021 | 0.489 | 0.18 | 0.2 | 0.17 | 0.38 | 0.24 | 0.33 |
| 3 | 28018581 | rs17020711 | -0.332 | -0.422 | -0.066 | 0.759 | -0.01 | -0.038 | -0.038 | 1.269 | 0.21 | 0.16 | 0.18 | 0.38 | 0.27 | 0.33 |
| 3 | 28020552 | rs12632457 | -0.845 | -0.432 | -0.026 | 0.647 | 0.016 | 0.067 | 0.067 | 0.201 | 0.3 | 0.44 | 0.31 | 0.46 | 0.38 | 0.48 |
| 3 | 28020954 | rs9637527 | -0.181 | -0.548 | -0.202 | 0.597 | -0.008 | -0.009 | -0.009 | 0 | 0.21 | 0.31 | 0.11 | 0.21 | 0.14 | 0.23 |
| 3 | 28026396 | rs12639500 | -0.412 | -0.392 | -0.86 | 0.772 | -0.018 | -0.123 | -0.123 | 0.869 | 0.26 | 0.07 | 0.17 | 0.42 | 0.26 | 0.24 |
| 3 | 28031787 | rs4680887 | -0.206 | -0.625 | -0.33 | 0.352 | 0.009 | 0.052 | 0.052 | 0 | 0.31 | 0.19 | 0.29 | 0.17 | 0.26 | 0.31 |
| 3 | 28032919 | rs2199654 | -1.332 | -0.848 | -0.122 | 0.442 | 0.006 | 0.013 | 0.013 | 0.856 | 0.52 | 0.6 | 0.45 | 0.62 | 0.44 | 0.55 |
| 3 | 28033233 | rs2219779 | -0.21 | -0.708 | -0.33 | 0.541 | -0.02 | 0.067 | 0.067 | 0 | 0.3 | 0.05 | 0.28 | 0.17 | 0.27 | 0.3 |
| 3 | 28035785 | rs1159028 | -0.128 | -0.686 | 0.038 | 0.582 | 0 | -0.002 | -0.002 | 0.071 | 0.3 | 0.25 | 0.32 | 0.24 | 0.27 | 0.4 |
| 3 | 28039368 | rs6804917 | -0.286 | -0.714 | 0.129 | 0.94 | -0.002 | 0.016 | 0.016 | 0 | 0.32 | 0.3 | 0.35 | 0.49 | 0.29 | 0.41 |
| 3 | 28043715 | rs9825129 | -0.183 | -0.742 | -0.311 | 0.547 | 0.035 | -0.013 | -0.013 | 0 | 0.4 | 0.29 | 0.49 | 0.48 | 0.26 | 0.38 |
| 3 | 28047508 | rs7645854 | 1.289 | 0.025 | -0.564 | 0.258 | -0.017 | 0.028 | 0.028 | 0.838 | 0.29 | 0.21 | 0.22 | 0.13 | 0.38 | 0.23 |
| 3 | 28049709 | rs17020780 | 1.1 | -0.002 | -0.542 | 0.372 | 0.009 | -0.011 | -0.011 | 0.246 | 0.1 | 0.08 | 0.06 | 0.25 | 0.07 | 0.32 |
| 3 | 28049737 | rs9857133 | -0.121 | 0.126 | -0.285 | 0.178 | 0.013 | 0.034 | 0.034 | 0.111 | 0.73 | 0.62 | 0.75 | 0.6 | 0.68 | 0.6 |
| 3 | 28049773 | rs13090784 | 0.952 | 0.29 | -0.119 | -0.241 | -0.012 | 0.016 | 0.016 | 0.616 | 0.29 | 0.22 | 0.22 | 0.14 | 0.37 | 0.23 |
| 3 | 28051829 | rs17020791 | 0.14 | -0.123 | 0.123 | -0.402 | -0.03 | -0.069 | -0.069 | 0 | 0.13 | 0.21 | 0.07 | 0 | 0.06 | 0.08 |
| 3 | 28057905 | rs1532244 | -0.517 | 0.307 | 0.276 | -0.061 | 0.043 | 0.047 | 0.047 | 0 | 0.11 | 0.34 | 0.22 | 0.09 | 0.11 | 0.16 |
| 3 | 28060456 | rs433317 | 0.581 | 0.152 | -0.263 | -0.215 | 0.008 | 0.003 | 0.003 | 0 | 0.51 | 0.45 | 0.56 | 0.53 | 0.45 | 0.43 |
| 3 | 28063788 | rs9310860 | 1.074 | 0.485 | -0.15 | -0.443 | 0.001 | -0.019 | -0.019 | 0 | 0.1 | 0.15 | 0.07 | 0.02 | 0.1 | 0.05 |
| 3 | 28064407 | rs365482 | -0.283 | 0.49 | -0.184 | -0.407 | -0.004 | 0.001 | 0.001 | 0 | 0.7 | 0.71 | 0.74 | 0.85 | 0.63 | 0.77 |
| 3 | 28071444 | rs669607 | 1.336 | 0.804 | -0.064 | -0.161 | -0.054 | 0.002 | 0.002 | 0 | 0.51 | 0.11 | 0.42 | 0.48 | 0.49 | 0.39 |

| | | | | | | | | | | | | | | | | |
|---|---|---|---|---|---|---|---|---|---|---|---|---|---|---|---|---|
| 3 | 28074885 | rs451375 | 0.059 | 0.831 | 0.283 | -0.149 | 0 | 0 | 0 | 0 | 0.72 | 0.67 | 0.72 | 0.59 | 0.72 | 0.6 |
| 3 | 28078786 | rs4680741 | 0.545 | 0.885 | 0.494 | -0.149 | 0.04 | 0.043 | 0.043 | 0.353 | 0.07 | 0.15 | 0.19 | 0.09 | 0.11 | 0.1 |
| 3 | 28078879 | rs11921176 | 1.085 | 0.845 | 0.336 | -0.179 | 0.006 | -0.014 | -0.014 | 0 | 0.1 | 0.14 | 0.07 | 0.03 | 0.12 | 0.06 |
| 3 | 28079085 | rs170934 | 1.339 | 0.749 | 0.231 | -0.178 | -0.022 | 0.025 | 0.025 | 0 | 0.51 | 0.16 | 0.43 | 0.47 | 0.49 | 0.38 |
| 3 | 28081442 | rs338610 | -0.13 | 0.271 | 0.588 | 0.067 | 0.006 | -0.01 | -0.01 | 0 | 0.73 | 0.72 | 0.74 | 0.52 | 0.78 | 0.59 |
| 3 | 28083823 | rs9840096 | -0.125 | 1.5 | 1.15 | -0.377 | -0.002 | -0.007 | -0.007 | 0 | 0.16 | 0.16 | 0.11 | 0.02 | 0.2 | 0.07 |
| 3 | 28096525 | rs6765138 | -1.863 | 1.36 | 1.226 | -0.403 | 0.001 | -0.008 | -0.008 | 0.024 | 0.28 | 0.65 | 0.36 | 0.11 | 0.32 | 0.19 |
| 3 | 28096687 | rs6777804 | -1.58 | 1.396 | 1.214 | -0.387 | 0.041 | -0.051 | -0.051 | 0 | 0.26 | 0.44 | 0.33 | 0.11 | 0.32 | 0.19 |
| 3 | 28110346 | rs9823676 | -0.362 | 1.281 | 0.025 | -0.557 | 0.01 | -0.035 | -0.035 | 0 | 0.16 | 0.2 | 0.12 | 0.02 | 0.2 | 0.07 |
| 3 | 28111191 | rs902979 | -1.321 | 1.258 | 0.045 | -0.571 | 0.017 | -0.096 | -0.096 | 0.989 | 0.21 | 0.4 | 0.16 | 0.03 | 0.22 | 0.07 |
| 3 | 28111840 | rs9849453 | -0.394 | 1.385 | -0.527 | -0.514 | -0.007 | 0.004 | 0.004 | 0 | 0.16 | 0.15 | 0.11 | 0.02 | 0.2 | 0.07 |
| 3 | 28122580 | rs2371258 | 0.052 | 1.259 | -1.41 | -0.098 | -0.036 | 0.008 | 0.008 | 2.006 | 0.79 | 0.97 | 0.79 | 0.84 | 0.75 | 0.9 |
| 3 | 28126595 | rs1156909 | -0.61 | 1.735 | -0.709 | -0.184 | 0.017 | -0.029 | -0.029 | 0 | 0.2 | 0.29 | 0.33 | 0.05 | 0.26 | 0.17 |
| 3 | 28132490 | rs9809389 | -0.292 | 1.547 | -1.056 | 0.129 | -0.02 | -0.025 | -0.025 | 0 | 0.31 | 0.22 | 0.45 | 0.4 | 0.37 | 0.52 |
| 3 | 28137183 | rs338616 | -0.727 | 1.355 | -0.711 | -0.135 | 0 | -0.007 | -0.007 | 0 | 0.72 | 0.71 | 0.74 | 0.86 | 0.69 | 0.91 |
| 3 | 28144414 | rs634274 | -1.118 | 1.506 | -0.701 | -0.192 | -0.007 | 0.022 | 0.022 | 1.35 | 0.75 | 0.93 | 0.81 | 0.86 | 0.74 | 0.92 |
| 3 | 28151568 | rs9816145 | -0.561 | 1.419 | -0.372 | -0.251 | 0.001 | -0.019 | -0.019 | 0.057 | 0.71 | 0.66 | 0.74 | 0.86 | 0.69 | 0.91 |
| 3 | 28163157 | rs7427073 | 0.215 | 1.383 | -0.314 | -0.267 | 0.001 | -0.027 | -0.027 | 0 | 0.3 | 0.37 | 0.26 | 0.14 | 0.31 | 0.09 |
| 3 | 28180249 | rs2371260 | 0.33 | 1.888 | -0.648 | -0.551 | 0.017 | -0.046 | -0.046 | 0 | 0.13 | 0.2 | 0.28 | 0.02 | 0.22 | 0.18 |
| 3 | 28193856 | rs925527 | 0.036 | 1.396 | -0.906 | -0.679 | -0.006 | -0.02 | -0.02 | 0.712 | 0.38 | 0.4 | 0.3 | 0.19 | 0.35 | 0.23 |
| 3 | 28193977 | rs2172437 | 0.287 | 1.395 | -0.855 | -0.68 | 0.003 | 0.009 | 0.009 | 0.002 | 0.36 | 0.27 | 0.27 | 0.19 | 0.35 | 0.23 |
| 3 | 28196462 | rs13095903 | 1.477 | 1.395 | -0.794 | -0.714 | 0.009 | 0.025 | 0.025 | 1.26 | 0.25 | 0.1 | 0.19 | 0.14 | 0.25 | 0.08 |
| 3 | 28197450 | rs11705740 | 0.205 | 1.375 | -1.043 | -0.738 | 0.004 | 0.013 | 0.013 | 0 | 0.37 | 0.26 | 0.27 | 0.18 | 0.35 | 0.25 |
| 3 | 28201121 | rs9876788 | -1.803 | 1.326 | -1.614 | -0.563 | -0.048 | -0.081 | -0.081 | 0 | 0.76 | 0.97 | 0.81 | 0.65 | 0.75 | 0.79 |
| 3 | 28203584 | rs11706637 | 0.718 | 0.92 | -1.226 | -0.349 | -0.056 | -0.137 | -0.137 | 0.768 | 0.2 | 0.03 | 0.19 | 0.39 | 0.17 | 0.34 |
| 3 | 28209580 | rs9882400 | -0.726 | 1.047 | -0.491 | -0.55 | 0.005 | -0.008 | -0.008 | 0 | 0.62 | 0.59 | 0.51 | 0.66 | 0.52 | 0.6 |
| 3 | 28211713 | rs17710733 | -0.373 | 0.611 | -0.326 | -0.52 | -0.006 | 0.02 | 0.02 | 0 | 0.11 | 0.06 | 0.07 | 0.05 | 0.1 | 0.17 |
| 3 | 28219791 | rs2063576 | -0.217 | 0.529 | 0.108 | -0.539 | -0.009 | -0.025 | -0.025 | 1.176 | 0.42 | 0.5 | 0.47 | 0.7 | 0.39 | 0.44 |
| 3 | 28226208 | rs11712251 | -0.251 | 0.513 | 0.283 | -0.546 | -0.006 | -0.017 | -0.017 | 0.023 | 0.43 | 0.54 | 0.47 | 0.7 | 0.39 | 0.43 |
| 3 | 28265642 | rs11718937 | 0.933 | 0.28 | 0.376 | -0.309 | -0.035 | -0.022 | -0.022 | 0 | 0.22 | 0.03 | 0.19 | 0.16 | 0.26 | 0.07 |
| 3 | 28270669 | rs6800659 | -0.692 | 0.529 | 0.799 | -0.604 | 0.016 | 0.023 | 0.023 | 0 | 0.32 | 0.57 | 0.56 | 0.56 | 0.35 | 0.48 |
| 3 | 28274233 | rs10510606 | 0.509 | 0.538 | 0.728 | -0.635 | 0.08 | -0.019 | -0.019 | 0.023 | 0.1 | 0.22 | 0.3 | 0.07 | 0.2 | 0.11 |
| 3 | 28280965 | rs1610238 | 1.135 | 0.618 | 0.619 | -0.596 | -0.031 | -0.018 | -0.018 | 1.234 | 0.22 | 0.03 | 0.2 | 0.16 | 0.26 | 0.07 |
| 3 | 28281551 | rs7645825 | 0.985 | 1.058 | 0.585 | -0.622 | -0.022 | -0.12 | -0.12 | 0 | 0.15 | 0.01 | 0.21 | 0.49 | 0.13 | 0.36 |
| 3 | 28296690 | rs17021298 | 0.588 | 1.043 | 0.615 | -0.589 | -0.04 | -0.104 | -0.104 | 4.315 | 0.17 | 0.04 | 0.24 | 0.48 | 0.18 | 0.38 |
| 3 | 28303111 | rs1609729 | -1.095 | 0.866 | 0.755 | -0.627 | 0.025 | 0.035 | 0.035 | 0 | 0.55 | 0.66 | 0.77 | 0.71 | 0.65 | 0.55 |
| 3 | 28308331 | rs11129315 | -1.12 | 0.891 | 0.808 | -0.576 | 0.026 | 0.039 | 0.039 | 0 | 0.56 | 0.68 | 0.77 | 0.7 | 0.65 | 0.56 |
| 3 | 28308752 | rs12107256 | 0.827 | 1.051 | 0.683 | -0.521 | 0.005 | 0.008 | 0.008 | 0.785 | 0.23 | 0.14 | 0.21 | 0.16 | 0.26 | 0.07 |
| 3 | 28332087 | rs11709034 | -0.355 | 0.868 | 0.946 | -0.554 | 0.001 | -0.011 | -0.011 | 0 | 0.43 | 0.57 | 0.56 | 0.22 | 0.52 | 0.3 |
| 3 | 28332601 | rs7624015 | 0.872 | 0.884 | 1.074 | -0.576 | -0.012 | 0.011 | 0.011 | 0 | 0.14 | 0.12 | 0.28 | 0.06 | 0.2 | 0.19 |
| 3 | 28334651 | rs12107556 | 0.872 | 0.884 | 1.074 | -0.576 | -0.012 | 0.011 | 0.011 | 0 | 0.14 | 0.12 | 0.29 | 0.06 | 0.21 | 0.19 |
| 3 | 28343334 | rs7641491 | -0.266 | 0.559 | 1.105 | -0.613 | 0.003 | -0.03 | -0.03 | 0.063 | 0.42 | 0.54 | 0.55 | 0.19 | 0.53 | 0.29 |
| 3 | 28358057 | rs11706332 | 1.008 | 0.59 | 0.985 | -0.555 | -0.027 | 0.017 | 0.017 | 0 | 0.24 | 0.07 | 0.22 | 0.16 | 0.29 | 0.11 |
| 3 | 28359788 | rs1599951 | -1.218 | 0.514 | 0.994 | -0.6 | -0.01 | -0.007 | -0.007 | 0 | 0.85 | 0.87 | 0.71 | 0.97 | 0.77 | 0.81 |
| 3 | 28363241 | rs7613767 | -1.942 | 0.254 | 0.827 | -0.629 | -0.03 | -0.024 | -0.024 | 0.799 | 0.7 | 0.97 | 0.83 | 0.74 | 0.73 | 0.77 |
| 3 | 28363559 | rs17357206 | 1.228 | 0.105 | 0.844 | -0.604 | -0.05 | 0.007 | 0.007 | 0 | 0.24 | 0.04 | 0.21 | 0.16 | 0.3 | 0.11 |
| 3 | 28384051 | rs12054402 | 1.099 | 0.065 | 0.831 | -0.583 | 0.034 | -0.117 | -0.117 | 0 | 0.14 | 0.01 | 0.19 | 0.48 | 0.08 | 0.35 |
| 3 | 28390041 | rs1870258 | 1.093 | 0.049 | 0.957 | -0.592 | -0.003 | -0.019 | -0.019 | 0 | 0.15 | 0.15 | 0.29 | 0.03 | 0.22 | 0.19 |
| 3 | 28390351 | rs1870259 | -0.298 | 0.037 | 1.106 | -0.595 | -0.001 | -0.008 | -0.008 | 0 | 0.45 | 0.58 | 0.56 | 0.18 | 0.52 | 0.3 |
| 3 | 28412966 | rs6551253 | -1.905 | 0.091 | 1.184 | -0.607 | -0.041 | -0.031 | -0.031 | 1.601 | 0.7 | 0.97 | 0.83 | 0.74 | 0.73 | 0.77 |
| 3 | 28426786 | rs17021402 | 0.681 | 0.236 | 1.412 | -0.564 | -0.001 | -0.022 | -0.022 | 0 | 0.29 | 0.36 | 0.27 | 0.15 | 0.31 | 0.11 |
| 3 | 28427029 | rs10510610 | 1.005 | 0.239 | 1.468 | -0.572 | 0.016 | -0.054 | -0.054 | 5.038 | 0.16 | 0.21 | 0.3 | 0.03 | 0.22 | 0.19 |
| 3 | 28429991 | rs13090439 | 0.756 | 0.243 | 1.495 | -0.571 | -0.001 | -0.012 | -0.012 | 0 | 0.28 | 0.29 | 0.26 | 0.15 | 0.3 | 0.11 |
| 3 | 28431394 | rs4680913 | -0.145 | 0.006 | 1.205 | -0.461 | -0.002 | 0 | 0 | 0 | 0.44 | 0.57 | 0.56 | 0.18 | 0.52 | 0.3 |
| 3 | 28505809 | rs9819034 | 0.594 | 0.219 | 1.178 | -0.311 | -0.01 | 0.035 | 0.035 | 0 | 0.06 | 0.21 | 0.03 | 0.06 | 0.03 | 0.09 |
| 3 | 28525740 | rs6805617 | -0.276 | 0.73 | 1.168 | -0.284 | 0 | 0.022 | 0.022 | 0 | 0.42 | 0.29 | 0.46 | 0.65 | 0.43 | 0.49 |
| 3 | 28533658 | rs13100556 | 0.532 | 0.69 | 1.158 | -0.231 | -0.005 | -0.045 | -0.045 | 0 | 0.16 | 0.09 | 0.23 | 0.47 | 0.16 | 0.38 |
| 3 | 28546773 | rs13061948 | -1.273 | 0.638 | 1.18 | -0.274 | 0.005 | -0.001 | -0.001 | 0 | 0.62 | 0.65 | 0.79 | 0.65 | 0.68 | 0.67 |
| 3 | 28549395 | rs11717248 | -0.346 | 0.62 | 1.053 | -0.224 | 0.001 | 0.008 | 0.008 | 0 | 0.42 | 0.3 | 0.46 | 0.62 | 0.43 | 0.48 |
| 3 | 28559193 | rs4234347 | 0.386 | 0.583 | 1.035 | -0.132 | -0.001 | -0.009 | -0.009 | 0.837 | 0.17 | 0.12 | 0.23 | 0.45 | 0.16 | 0.37 |
| 3 | 28567322 | rs963361 | -0.387 | 0.584 | 0.961 | -0.378 | 0.002 | 0.011 | 0.011 | 0 | 0.42 | 0.36 | 0.47 | 0.65 | 0.43 | 0.49 |
| 3 | 28587185 | rs6549859 | 1.257 | 0.294 | 0.927 | -0.428 | 0.001 | 0 | 0 | 0 | 0.36 | 0.35 | 0.2 | 0.32 | 0.29 | 0.32 |
| 3 | 28588561 | rs4640517 | 0.314 | 0.088 | 0.266 | -0.207 | 0.001 | -0.02 | -0.02 | 0 | 0.18 | 0.11 | 0.23 | 0.48 | 0.18 | 0.38 |
| 3 | 28590909 | rs7627575 | -1.06 | 0.119 | 0.113 | -0.334 | -0.049 | -0.016 | -0.016 | 0 | 0.76 | 0.98 | 0.8 | 0.84 | 0.73 | 0.9 |
| 3 | 28600137 | rs4680757 | 0.263 | 0.258 | 0.253 | -0.352 | 0.009 | 0.03 | 0.03 | 0 | 0.57 | 0.7 | 0.55 | 0.35 | 0.55 | 0.52 |
| 3 | 28613602 | rs4234341 | -0.646 | 0.169 | 0.243 | 0.022 | 0 | 0 | 0 | 0 | 0.76 | 0.77 | 0.77 | 0.84 | 0.73 | 0.89 |
| 3 | 28624327 | rs4133027 | -0.171 | 0.208 | 0.374 | -0.006 | -0.002 | -0.052 | -0.052 | 0.011 | 0.71 | 0.49 | 0.72 | 0.84 | 0.69 | 0.87 |
| 3 | 28629129 | rs6769096 | 0.707 | 0.177 | -0.591 | 0.035 | 0.025 | 0.006 | 0.006 | 0 | 0.35 | 0.22 | 0.2 | 0.34 | 0.28 | 0.31 |
| 3 | 28631155 | rs4356777 | 0.287 | 0.393 | -0.262 | 0.012 | 0.004 | -0.045 | -0.045 | 0 | 0.73 | 0.59 | 0.56 | 0.86 | 0.68 | 0.87 |

| | | | | | | | | | | | | | | | | |
|---|---|---|---|---|---|---|---|---|---|---|---|---|---|---|---|---|
| 3 | 28650238 | rs9839541 | -0.091 | -0.542 | -0.63 | 0.311 | 0.029 | 0.017 | 0.017 | 0 | 0.51 | 0.51 | 0.34 | 0.54 | 0.31 | 0.49 |
| 3 | 28653629 | rs4680766 | -0.168 | 0.699 | -0.717 | -0.038 | 0.016 | -0.013 | -0.013 | 0 | 0.86 | 0.79 | 0.82 | 0.88 | 0.77 | 0.79 |
| 3 | 28654162 | rs12487716 | 0.713 | -0.542 | -0.196 | 0.227 | 0.021 | 0.02 | 0.02 | 0 | 0.36 | 0.29 | 0.16 | 0.19 | 0.22 | 0.27 |
| 3 | 28654774 | rs7652119 | 0.757 | -0.62 | -0.243 | 0.192 | 0.033 | 0.031 | 0.031 | 0 | 0.35 | 0.24 | 0.15 | 0.19 | 0.21 | 0.27 |
| 3 | 28660626 | rs4306817 | -0.796 | -0.427 | -0.159 | 0.184 | 0 | -0.001 | -0.001 | 0 | 0.88 | 0.84 | 0.89 | 0.88 | 0.83 | 0.81 |
| 3 | 28669144 | rs6549866 | 0.843 | -0.518 | 0.321 | 0.313 | 0.005 | 0.008 | 0.008 | 0 | 0.14 | 0.08 | 0.17 | 0.36 | 0.09 | 0.25 |
| 3 | 28670884 | rs4494881 | 0.984 | -0.649 | 0.304 | 0.185 | -0.046 | 0.021 | 0.021 | 0 | 0.34 | 0.22 | 0.5 | 0.32 | 0.48 | 0.29 |
| 3 | 28671357 | rs4356778 | 0.499 | -0.612 | 0.642 | 0.24 | -0.017 | 0.089 | 0.089 | 0 | 0.72 | 0.56 | 0.68 | 0.53 | 0.73 | 0.54 |
| 3 | 28678285 | rs9830706 | 0.709 | -0.009 | 0.597 | 0.023 | 0.003 | 0.021 | 0.021 | 0 | 0.67 | 0.65 | 0.65 | 0.52 | 0.65 | 0.51 |
| 3 | 28681808 | rs6773699 | -0.785 | 0.019 | 0.68 | 0.053 | 0.008 | -0.001 | -0.001 | 0 | 0.89 | 0.9 | 0.88 | 0.9 | 0.79 | 0.79 |
| 3 | 28682296 | rs970624 | 0.234 | -0.154 | 0.745 | 0.064 | -0.025 | 0.046 | 0.046 | 0.201 | 0.19 | 0.2 | 0.22 | 0.36 | 0.1 | 0.27 |
| 3 | 28685997 | rs9841243 | 0.694 | -1.413 | 0.469 | -0.639 | -0.064 | -0.086 | -0.086 | 0 | 0.23 | 0.47 | 0.12 | 0.08 | 0.12 | 0.09 |
| 3 | 28689218 | rs1870816 | -0.586 | -1.479 | 0.065 | -0.687 | -0.086 | 0.004 | 0.004 | 0 | 0.74 | 0.95 | 0.81 | 0.77 | 0.66 | 0.59 |
| 3 | 28690589 | rs9310867 | 0.312 | -1.676 | 0.022 | 0.469 | -0.015 | -0.031 | -0.031 | 0 | 0.57 | 0.81 | 0.41 | 0.46 | 0.56 | 0.33 |
| 3 | 28699121 | rs1461803 | 0.596 | -1.613 | -0.135 | 0.292 | -0.009 | 0.055 | 0.055 | 0.013 | 0.48 | 0.27 | 0.62 | 0.66 | 0.5 | 0.49 |
| 3 | 28703364 | rs6549870 | 0.661 | -1.582 | -0.691 | 0.239 | -0.087 | 0.098 | 0.098 | 0.232 | 0.47 | 0.08 | 0.6 | 0.68 | 0.5 | 0.49 |
| 3 | 28704463 | rs4399848 | -1.434 | -1.316 | -0.541 | 0.312 | -0.023 | 0.027 | 0.027 | 0 | 0.93 | 0.99 | 0.94 | 0.98 | 0.91 | 0.83 |
| 3 | 28711912 | rs11715629 | -0.023 | -2.072 | -0.706 | 0.473 | 0.105 | -0.073 | -0.073 | 4.079 | 0.36 | 0.1 | 0.26 | 0.39 | 0.18 | 0.33 |
| 3 | 28712104 | rs7644788 | -0.491 | -1.059 | -0.905 | 0.293 | -0.105 | 0.036 | 0.036 | 0 | 0.69 | 0.92 | 0.77 | 0.75 | 0.56 | 0.54 |
| 3 | 28712239 | rs13075697 | 1.343 | -0.798 | -0.265 | 0.345 | 0.021 | 0.01 | 0.01 | 0 | 0.31 | 0.5 | 0.47 | 0.35 | 0.37 | 0.21 |
| 3 | 28712248 | rs10049305 | 0.336 | -1.159 | -0.698 | 0.449 | 0.033 | -0.07 | -0.07 | 0 | 0.19 | 0.08 | 0.22 | 0.35 | 0.12 | 0.29 |
| 3 | 28714733 | rs4680773 | 0.774 | -1.759 | -1.717 | 0.059 | -0.07 | -0.038 | -0.038 | 0.492 | 0.37 | 0.14 | 0.54 | 0.42 | 0.45 | 0.32 |
| 3 | 28720126 | rs17651822 | 1.39 | -1.889 | -1.771 | -0.077 | -0.038 | 0.012 | 0.012 | 0 | 0.15 | 0 | 0.07 | 0.1 | 0.15 | 0.2 |
| 3 | 28720221 | rs9875435 | -0.654 | -1.972 | -1.505 | 0.132 | -0.003 | 0.002 | 0.002 | 0 | 0.91 | 0.9 | 0.93 | 0.84 | 0.91 | 0.8 |
| 3 | 28721830 | rs4680704 | 0.226 | -2.278 | -1.008 | -0.037 | 0.008 | 0.014 | 0.014 | 0 | 0.59 | 0.71 | 0.77 | 0.71 | 0.66 | 0.47 |
| 3 | 28725054 | rs4680776 | 0.263 | -2.319 | -0.801 | 0.126 | -0.006 | 0.002 | 0.002 | 0 | 0.63 | 0.65 | 0.76 | 0.62 | 0.72 | 0.55 |
| 3 | 28733949 | rs9310869 | 1.57 | -2.086 | -0.589 | 0.249 | 0 | 0.002 | 0.002 | 0 | 0.12 | 0.13 | 0.1 | 0.18 | 0.13 | 0.19 |
| 3 | 28736622 | rs4398388 | 0.084 | -2.171 | -0.625 | 0.221 | -0.003 | 0.033 | 0.033 | 4.204 | 0.22 | 0.65 | 0.27 | 0.26 | 0.21 | 0.21 |
| 3 | 28737370 | rs4315650 | -1.537 | -2.127 | -0.595 | 0.189 | 0.008 | 0.066 | 0.066 | 0 | 0.95 | 0.87 | 0.94 | 0.84 | 0.95 | 0.83 |
| 3 | 28745091 | rs6549876 | 0.445 | -2.236 | -0.733 | 0.137 | 0.015 | -0.028 | -0.028 | 0 | 0.69 | 0.22 | 0.69 | 0.59 | 0.73 | 0.6 |
| 3 | 28745139 | rs6549877 | 0.098 | -2.23 | -0.338 | 0.156 | -0.023 | 0.074 | 0.074 | 3.789 | 0.86 | 0.48 | 0.79 | 0.76 | 0.87 | 0.79 |
| 3 | 28745247 | rs6549878 | -1.656 | -2.034 | -0.669 | 0.197 | -0.004 | 0.02 | 0.02 | 0 | 0.95 | 0.91 | 0.95 | 0.84 | 0.95 | 0.81 |
| 3 | 28747158 | rs1563981 | 1.116 | -2.113 | -1.032 | 0.013 | 0.026 | 0.028 | 0.028 | 0 | 0.16 | 0.08 | 0.07 | 0.05 | 0.09 | 0.09 |
| 3 | 28748823 | rs1461821 | -0.16 | -2.131 | -1.015 | 0.015 | -0.004 | -0.002 | -0.002 | 0.043 | 0.32 | 0.32 | 0.16 | 0.22 | 0.23 | 0.28 |
| 3 | 28749314 | rs1381392 | 1.005 | -1.619 | -0.963 | 0.218 | -0.061 | 0.03 | 0.03 | 0 | 0.18 | 0 | 0.09 | 0.1 | 0.19 | 0.26 |
| 3 | 28749348 | rs4298010 | 0.698 | -1.396 | -0.531 | 0.412 | 0.006 | 0.018 | 0.018 | 0 | 0.16 | 0.24 | 0.09 | 0.17 | 0.14 | 0.19 |
| 3 | 28750478 | rs17021778 | 0.918 | -0.429 | -1.012 | 0.473 | -0.054 | -0.054 | -0.054 | 0 | 0.15 | 0.02 | 0.09 | 0.17 | 0.16 | 0.21 |
| 3 | 28752588 | rs1461811 | 1.107 | -0.639 | -0.707 | 0.369 | 0.002 | 0.011 | 0.011 | 0 | 0.46 | 0.39 | 0.57 | 0.45 | 0.45 | 0.3 |
| 3 | 28755787 | rs4378932 | 0.045 | -0.772 | -0.309 | 0.384 | -0.003 | -0.019 | -0.019 | 0.32 | 0.66 | 0.63 | 0.71 | 0.78 | 0.65 | 0.66 |
| 3 | 28758394 | rs922287 | 0.449 | -0.795 | -0.34 | 0.409 | 0.007 | 0.005 | 0.005 | 0 | 0.59 | 0.4 | 0.65 | 0.63 | 0.6 | 0.51 |
| 3 | 28760544 | rs2888033 | 0.077 | -1.396 | -0.094 | 0.656 | -0.001 | 0.001 | 0.001 | 0 | 0.52 | 0.61 | 0.63 | 0.61 | 0.49 | 0.45 |
| 3 | 28763075 | rs17021808 | 1.008 | -1.288 | 0.15 | 0.77 | 0.008 | 0.079 | 0.079 | 0.014 | 0.06 | 0.22 | 0.06 | 0.15 | 0.05 | 0.16 |
| 3 | 28764272 | rs1381396 | 0.827 | -1.436 | -0.014 | 0.627 | -0.018 | 0.021 | 0.021 | 0.446 | 0.74 | 0.41 | 0.71 | 0.63 | 0.78 | 0.65 |
| 3 | 28765220 | rs6777958 | -0.754 | -1.47 | 0.128 | 0.647 | -0.008 | 0.008 | 0.008 | 0.019 | 0.88 | 0.89 | 0.92 | 0.85 | 0.9 | 0.83 |
| 3 | 28765289 | rs17021825 | 1.472 | -1.436 | 0.062 | 0.638 | -0.001 | 0.058 | 0.058 | 0 | 0.06 | 0.17 | 0.06 | 0.15 | 0.05 | 0.16 |
| 3 | 28765430 | rs6802374 | 0.33 | -1.348 | -0.12 | 0.552 | -0.033 | 0.045 | 0.045 | 0 | 0.8 | 0.63 | 0.77 | 0.78 | 0.83 | 0.81 |
| 3 | 28769680 | rs17653235 | 0.238 | -1.225 | -0.276 | 0.329 | -0.012 | 0.008 | 0.008 | 0 | 0.35 | 0.25 | 0.23 | 0.31 | 0.37 | 0.56 |
| 3 | 28770641 | rs12636605 | 1.188 | -1.132 | -0.385 | 0.291 | -0.13 | -0.014 | -0.014 | 0 | 0.25 | 0.01 | 0.14 | 0.17 | 0.32 | 0.36 |
| 3 | 28771502 | rs6549883 | -0.484 | -1.499 | 0.161 | 0.382 | -0.045 | 0.039 | 0.039 | 0 | 0.84 | 0.58 | 0.9 | 0.83 | 0.9 | 0.84 |
| 3 | 28774049 | rs13100124 | 0.786 | -1.005 | -0.298 | 0.567 | -0.005 | -0.007 | -0.007 | 0 | 0.31 | 0.34 | 0.3 | 0.28 | 0.26 | 0.08 |
| 3 | 28778641 | rs6804729 | 0.214 | -0.783 | 0.059 | 0.901 | 0.013 | 0.06 | 0.06 | 0.313 | 0.69 | 0.46 | 0.51 | 0.55 | 0.67 | 0.58 |
| 3 | 28780713 | rs7428667 | 1.181 | -1.122 | -0.249 | 0.593 | 0.008 | 0.007 | 0.007 | 0.133 | 0.24 | 0.19 | 0.12 | 0.09 | 0.13 | 0.2 |
| 3 | 28783357 | rs2199411 | 1.055 | -1.042 | -0.238 | 0.624 | -0.012 | 0.021 | 0.021 | 0 | 0.46 | 0.27 | 0.39 | 0.45 | 0.54 | 0.38 |
| 3 | 28786591 | rs11129330 | 0.267 | -0.935 | -0.472 | 0.765 | 0.001 | 0.001 | 0.001 | 0 | 0.3 | 0.25 | 0.17 | 0.23 | 0.17 | 0.36 |
| 3 | 28791586 | rs1461820 | 0.899 | -0.545 | -0.964 | 0.94 | -0.037 | -0.048 | -0.048 | 0 | 0.16 | 0.08 | 0.37 | 0.24 | 0.22 | 0.2 |
| 3 | 28796677 | rs1011416 | 0.643 | -0.762 | -1.108 | 0.902 | 0.044 | 0.01 | 0.01 | 0 | 0.29 | 0.02 | 0.13 | 0.26 | 0.17 | 0.39 |
| 3 | 28798512 | rs9841714 | 1.046 | -0.856 | -0.734 | 0.722 | -0.009 | -0.069 | -0.069 | 0.82 | 0.23 | 0.41 | 0.2 | 0.09 | 0.22 | 0.06 |
| 3 | 28798728 | rs2306082 | 1.698 | -0.876 | -0.784 | 0.966 | 0.002 | -0.001 | -0.001 | 0 | 0.06 | 0.03 | 0.06 | 0.22 | 0.06 | 0.17 |
| 3 | 28799676 | rs17021907 | 1.168 | -0.873 | -0.532 | 1.024 | 0.003 | -0.004 | -0.004 | 0.152 | 0.26 | 0.23 | 0.11 | 0.11 | 0.15 | 0.25 |
| 3 | 28800568 | rs1461818 | 0.82 | -0.831 | -0.519 | 1.109 | -0.028 | -0.043 | -0.043 | 0.002 | 0.18 | 0.12 | 0.41 | 0.32 | 0.27 | 0.22 |
| 3 | 28801369 | rs1007361 | 1.109 | -0.833 | -0.492 | 0.92 | -0.012 | -0.03 | -0.03 | 0.705 | 0.09 | 0.02 | 0.1 | 0.28 | 0.12 | 0.19 |
| 3 | 28801888 | rs7648722 | 1.123 | -0.872 | -0.401 | 0.905 | -0.006 | -0.007 | -0.007 | 0 | 0.23 | 0.34 | 0.26 | 0.27 | 0.22 | 0.07 |
| 3 | 28802104 | rs922285 | 0.112 | -0.745 | -0.634 | 0.979 | -0.028 | -0.044 | -0.044 | 0.3 | 0.24 | 0.14 | 0.46 | 0.53 | 0.33 | 0.38 |
| 3 | 28803294 | rs7610221 | -1.725 | -0.722 | -0.626 | 0.984 | -0.003 | 0.001 | 0.001 | 0 | 0.73 | 0.78 | 0.78 | 0.72 | 0.7 | 0.68 |
| 3 | 28807987 | rs1015856 | -1.735 | -0.717 | -0.623 | 0.85 | -0.002 | -0.002 | -0.002 | 0.076 | 0.73 | 0.78 | 0.84 | 0.91 | 0.7 | 0.69 |
| 3 | 28808844 | rs12634756 | -0.781 | -0.734 | -0.546 | 0.894 | 0.009 | 0.011 | 0.011 | 0 | 0.63 | 0.58 | 0.72 | 0.62 | 0.58 | 0.5 |
| 3 | 28810588 | rs7610366 | -1.727 | -0.739 | -0.572 | 0.871 | -0.002 | -0.001 | -0.001 | 2.979 | 0.73 | 0.78 | 0.84 | 0.9 | 0.69 | 0.69 |
| 3 | 28811329 | rs996646 | 0.101 | -0.755 | -0.604 | 0.879 | -0.033 | -0.051 | -0.051 | 0.18 | 0.24 | 0.13 | 0.47 | 0.53 | 0.33 | 0.38 |
| 3 | 28812010 | rs11920283 | -0.785 | -0.734 | -0.653 | 0.892 | 0.012 | 0.014 | 0.014 | 0 | 0.63 | 0.56 | 0.72 | 0.62 | 0.57 | 0.5 |

| | | | | | | | | | | | | | | | | |
|---|---|---|---|---|---|---|---|---|---|---|---|---|---|---|---|---|
| 3 | 28813677 | rs7647498 | -0.006 | -0.747 | -0.934 | 0.192 | 0.028 | 0.041 | 0.041 | 0 | 0.48 | 0.41 | 0.35 | 0.37 | 0.36 | 0.3 |
| 3 | 28816082 | rs17021931 | -0.099 | -0.392 | -0.62 | 0.165 | 0.004 | 0.041 | 0.041 | 0 | 0.49 | 0.38 | 0.58 | 0.63 | 0.47 | 0.62 |
| 3 | 28821970 | rs9862861 | -1.014 | -0.057 | -0.792 | 0.146 | -0.002 | -0.003 | -0.003 | 0 | 0.73 | 0.77 | 0.83 | 0.91 | 0.7 | 0.69 |
| 3 | 28823156 | rs7617605 | -0.403 | 0.421 | -1.034 | 0.269 | -0.014 | 0.023 | 0.023 | 0 | 0.25 | 0.17 | 0.2 | 0.14 | 0.32 | 0.27 |
| 3 | 28823318 | rs11915504 | -0.113 | 0.155 | -0.878 | 0.365 | -0.004 | -0.001 | -0.001 | 0 | 0.14 | 0.2 | 0.11 | 0.14 | 0.15 | 0.19 |
| 3 | 28823658 | rs11927465 | -0.175 | -0.12 | -0.937 | 0.48 | -0.005 | 0.002 | 0.002 | 0.435 | 0.31 | 0.24 | 0.34 | 0.37 | 0.38 | 0.28 |
| 3 | 28823801 | rs11923484 | -0.143 | -0.133 | -0.932 | 0.47 | -0.005 | 0.005 | 0.005 | 0.35 | 0.31 | 0.23 | 0.33 | 0.36 | 0.37 | 0.27 |
| 3 | 28826270 | rs9813026 | 0.509 | -0.748 | -0.74 | 0.498 | -0.014 | 0.035 | 0.035 | 0.01 | 0.07 | 0.27 | 0.05 | 0.04 | 0.02 | 0.01 |
| 3 | 28828525 | rs7633531 | 0.179 | -0.907 | -1.969 | 0.69 | 0.074 | -0.081 | -0.081 | 0 | 0.32 | 0.88 | 0.45 | 0.43 | 0.2 | 0.35 |
| 3 | 28830276 | rs13075403 | -0.97 | -0.13 | -1.701 | 0.416 | -0.059 | 0.076 | 0.076 | 1.897 | 0.68 | 0.93 | 0.86 | 0.85 | 0.69 | 0.68 |
| 3 | 28835662 | rs11922946 | -0.187 | -0.888 | -0.608 | 1.065 | -0.014 | 0.026 | 0.026 | 0 | 0.23 | 0.59 | 0.29 | 0.35 | 0.23 | 0.18 |
| 3 | 28837587 | rs7645110 | -1.182 | -1.278 | -1.312 | 0.803 | -0.113 | 0.048 | 0.048 | 0.056 | 0.65 | 0.92 | 0.71 | 0.82 | 0.52 | 0.64 |
| 3 | 28845890 | rs7631590 | 0.929 | -1.728 | -1.33 | 0.901 | 0.034 | -0.018 | -0.018 | 0.512 | 0.28 | 0.12 | 0.33 | 0.35 | 0.2 | 0.36 |
| 3 | 28849038 | rs1993761 | -0.59 | -1.786 | -1.325 | 0.878 | 0.049 | 0.042 | 0.042 | 0 | 0.52 | 0.72 | 0.62 | 0.7 | 0.42 | 0.54 |
| 3 | 28852080 | rs6762992 | -0.174 | -1.789 | -0.756 | 0.937 | -0.019 | -0.002 | -0.002 | 0.791 | 0.45 | 0.5 | 0.44 | 0.44 | 0.3 | 0.46 |
| 3 | 28859856 | rs9810180 | -1.48 | -1.479 | -1.056 | 0.749 | -0.177 | 0.002 | 0.002 | 0.072 | 0.68 | 0.97 | 0.72 | 0.83 | 0.53 | 0.65 |
| 3 | 28864888 | rs9857300 | -0.205 | -1.852 | -0.294 | 1.124 | -0.017 | -0.001 | -0.001 | 0 | 0.33 | 0.46 | 0.33 | 0.41 | 0.3 | 0.23 |
| 3 | 28866604 | rs9867716 | 0.715 | -1.947 | -0.309 | 1.028 | -0.03 | -0.001 | -0.001 | 1.443 | 0.12 | 0.22 | 0.05 | 0.13 | 0.07 | 0.07 |
| 3 | 28867765 | rs9873347 | -1.543 | -1.99 | -0.25 | 0.849 | -0.052 | 0.049 | 0.049 | 0 | 0.67 | 0.8 | 0.69 | 0.85 | 0.52 | 0.65 |
| 3 | 28868231 | rs1392301 | 1.268 | -2.13 | -0.51 | 0.964 | 0.025 | 0.058 | 0.058 | 0 | 0.25 | 0.08 | 0.25 | 0.1 | 0.15 | 0.27 |
| 3 | 28869948 | rs1353926 | -1.067 | -2.196 | -0.573 | 1.109 | 0.066 | 0.012 | 0.012 | 1.117 | 0.43 | 0.89 | 0.48 | 0.67 | 0.38 | 0.39 |
| 3 | 28871065 | rs12488415 | 0.848 | -2.194 | -0.69 | 1.067 | 0.011 | 0.062 | 0.062 | 0 | 0.28 | 0.03 | 0.27 | 0.19 | 0.19 | 0.32 |
| 3 | 28871192 | rs2221152 | 0.676 | -2.076 | -0.197 | 0.891 | -0.038 | -0.04 | -0.04 | 0 | 0.12 | 0.3 | 0.06 | 0.07 | 0.07 | 0.06 |
| 3 | 28877923 | rs885540 | -1.285 | -1.413 | -0.206 | 0.976 | -0.142 | 0.023 | 0.023 | 0 | 0.66 | 0.95 | 0.66 | 0.8 | 0.51 | 0.57 |
| 3 | 28878439 | rs6767753 | -1.173 | -1.347 | -0.156 | 0.49 | -0.036 | -0.039 | -0.039 | 0 | 0.65 | 0.8 | 0.64 | 0.64 | 0.51 | 0.54 |
| 3 | 28879197 | rs11129333 | -0.086 | -1.531 | -0.3 | 0.408 | -0.001 | 0 | 0 | 0 | 0.38 | 0.41 | 0.34 | 0.53 | 0.31 | 0.22 |
| 3 | 28882980 | rs7623722 | -0.292 | -0.641 | -0.509 | 0.736 | -0.002 | -0.003 | -0.003 | 0 | 0.61 | 0.62 | 0.61 | 0.63 | 0.47 | 0.5 |
| 3 | 28887146 | rs7614435 | -0.028 | -0.938 | -0.923 | -0.13 | 0 | -0.015 | -0.015 | 0.556 | 0.46 | 0.53 | 0.32 | 0.19 | 0.46 | 0.29 |
| 3 | 28890121 | rs1500007 | -0.547 | -0.468 | -1.343 | -0.002 | -0.076 | -0.093 | -0.093 | 0.659 | 0.71 | 0.88 | 0.66 | 0.48 | 0.61 | 0.69 |
| 3 | 28910107 | rs11713577 | -0.163 | -1.51 | -0.282 | -0.851 | -0.005 | -0.048 | -0.048 | 0.063 | 0.42 | 0.49 | 0.29 | 0.14 | 0.38 | 0.26 |
| 3 | 28912185 | rs9820080 | -0.589 | -1.476 | -0.254 | -0.992 | 0.006 | 0.063 | 0.063 | 0.459 | 0.5 | 0.39 | 0.35 | 0.14 | 0.48 | 0.32 |
| 3 | 28922565 | rs1392299 | -0.573 | -1.515 | 0.321 | -0.844 | 0.003 | 0.044 | 0.044 | 0.674 | 0.49 | 0.42 | 0.34 | 0.14 | 0.47 | 0.31 |
| 3 | 28926923 | rs6793728 | -0.1 | -1.461 | 0.293 | -0.865 | 0.004 | 0.02 | 0.02 | 1.76 | 0.41 | 0.35 | 0.27 | 0.06 | 0.38 | 0.2 |
| 3 | 28927195 | rs10510613 | -0.491 | -1.456 | -0.012 | -0.831 | -0.006 | 0.018 | 0.018 | 0 | 0.12 | 0.23 | 0.28 | 0.17 | 0.12 | 0.16 |
| 3 | 28928860 | rs2221154 | -1.223 | -1.471 | -0.411 | -0.564 | -0.058 | -0.049 | -0.049 | 0 | 0.58 | 0.88 | 0.59 | 0.45 | 0.51 | 0.48 |
| 3 | 28929230 | rs4680793 | 1.628 | -1.297 | -0.569 | -0.888 | -0.028 | -0.079 | -0.079 | 0 | 0.12 | 0.01 | 0.23 | 0.4 | 0.13 | 0.15 |
| 3 | 28936109 | rs9816337 | -1.373 | -1.344 | 0.194 | -1.709 | -0.015 | 0.124 | 0.124 | 0.075 | 0.54 | 0.74 | 0.57 | 0.23 | 0.48 | 0.36 |
| 3 | 28943520 | rs13078203 | 0.298 | -0.994 | -0.149 | -1.481 | -0.014 | 0.035 | 0.035 | 0 | 0.11 | 0.02 | 0.03 | 0.02 | 0.09 | 0.08 |
| 3 | 28960287 | rs11710772 | 0.893 | -0.687 | -0.255 | -1.439 | -0.058 | 0.062 | 0.062 | 0 | 0.13 | 0.04 | 0.12 | 0 | 0.21 | 0.06 |
| 3 | 28970909 | rs17022186 | -0.899 | -0.772 | -0.122 | -1.339 | -0.002 | -0.004 | -0.004 | 0 | 0.11 | 0.1 | 0.11 | 0.09 | 0.11 | 0.21 |
| 3 | 28971865 | rs2371470 | -1.075 | -0.786 | -0.363 | -1.401 | 0 | 0.058 | 0.058 | 0.022 | 0.46 | 0.57 | 0.45 | 0.24 | 0.43 | 0.47 |
| 3 | 28992861 | rs6765943 | -0.645 | -0.223 | -1.006 | -1.343 | 0 | -0.027 | -0.027 | 0 | 0.08 | 0.08 | 0.08 | 0 | 0.08 | 0.16 |
| 3 | 28993507 | rs6768946 | 0.966 | -0.484 | -1.141 | -1.408 | 0.035 | 0.228 | 0.228 | 0.01 | 0.52 | 0.13 | 0.69 | 0.85 | 0.62 | 0.54 |
| 3 | 28993693 | rs6793529 | 0.869 | -0.092 | -1.009 | -1.179 | 0.018 | 0.115 | 0.115 | 0.048 | 0.56 | 0.23 | 0.72 | 0.85 | 0.63 | 0.67 |
| 3 | 29016439 | rs2218746 | 0.23 | 0.326 | -0.63 | -0.493 | 0.001 | -0.005 | -0.005 | 0 | 0.54 | 0.43 | 0.69 | 0.85 | 0.59 | 0.67 |
| 3 | 29018088 | rs9870123 | -0.621 | 0.296 | -0.172 | 0.014 | -0.008 | -0.003 | -0.003 | 0 | 0.35 | 0.57 | 0.24 | 0.32 | 0.31 | 0.43 |
| 3 | 29020122 | rs11129339 | 0.335 | 0.058 | 0.487 | 0.797 | -0.023 | 0.067 | 0.067 | 2.881 | 0.16 | 0.23 | 0.13 | 0.36 | 0.12 | 0.17 |
| 3 | 29022934 | rs4680713 | 0.525 | 0.21 | 0.264 | 0.073 | -0.01 | 0.062 | 0.062 | 0 | 0.74 | 0.85 | 0.85 | 0.96 | 0.76 | 0.7 |
| 3 | 29026168 | rs1449688 | -0.604 | 0.38 | 0.75 | -0.024 | 0.003 | -0.044 | -0.044 | 0 | 0.16 | 0.22 | 0.11 | 0.04 | 0.14 | 0.25 |
| 3 | 29034403 | rs2167089 | 1.175 | 0.142 | 1.02 | -0.048 | 0 | -0.051 | -0.051 | 0 | 0.68 | 0.65 | 0.8 | 0.94 | 0.71 | 0.63 |
| 3 | 29035532 | rs13059806 | 1.763 | 0.101 | 0.953 | 0.301 | -0.029 | -0.021 | -0.021 | 0.036 | 0.48 | 0.13 | 0.6 | 0.5 | 0.54 | 0.41 |
| 3 | 29038253 | rs17022316 | 1.018 | 0.055 | 0.918 | 0.214 | 0.024 | -0.076 | -0.076 | 0 | 0.15 | 0.03 | 0.1 | 0.37 | 0.12 | 0.17 |
| 3 | 29041539 | rs4680800 | 1.341 | 0.152 | 1.035 | 0.24 | 0.009 | 0.003 | 0.003 | 0 | 0.52 | 0.42 | 0.67 | 0.57 | 0.58 | 0.46 |
| 3 | 29044226 | rs9828929 | -0.679 | -0.022 | 0.721 | 0.311 | 0.017 | -0.036 | -0.036 | 0 | 0.32 | 0.19 | 0.23 | 0.42 | 0.27 | 0.42 |
| 3 | 29047444 | rs6790709 | -0.822 | -0.254 | 1.316 | 0.353 | -0.018 | 0.015 | 0.015 | 4.072 | 0.38 | 0.39 | 0.43 | 0.46 | 0.28 | 0.44 |
| 3 | 29051075 | rs6793416 | -0.658 | -0.103 | 1.137 | 0.25 | -0.034 | 0.05 | 0.05 | 0.766 | 0.9 | 0.98 | 0.93 | 1 | 0.88 | 0.93 |
| 3 | 29051853 | rs1449692 | -1.42 | 0.174 | 1.315 | 0.205 | 0.041 | 0.021 | 0.021 | 0.092 | 0.49 | 0.68 | 0.53 | 0.53 | 0.39 | 0.57 |
| 3 | 29054809 | rs9854980 | -0.395 | 0.131 | 1.732 | 0.155 | -0.012 | -0.02 | -0.02 | 0 | 0.17 | 0.33 | 0.3 | 0.1 | 0.13 | 0.25 |
| 3 | 29059892 | rs9826412 | -0.466 | 0.12 | 1.954 | 0.153 | -0.011 | -0.024 | -0.024 | 0 | 0.18 | 0.38 | 0.32 | 0.1 | 0.13 | 0.26 |
| 3 | 29068750 | rs11927581 | -1.657 | 0.082 | 1.786 | 0.315 | 0.041 | 0.069 | 0.069 | 0 | 0.5 | 0.68 | 0.53 | 0.67 | 0.39 | 0.6 |
| 3 | 29086494 | rs17022500 | -0.029 | 0.036 | 1.826 | 0.36 | -0.001 | 0.002 | 0.002 | 0 | 0.16 | 0.16 | 0.28 | 0.23 | 0.12 | 0.28 |
| 3 | 29088999 | rs6796246 | 1.978 | 0.03 | 1.722 | 0.302 | -0.071 | 0.036 | 0.036 | 0 | 0.41 | 0.19 | 0.39 | 0.33 | 0.5 | 0.34 |
| 3 | 29090155 | rs2371578 | 0.968 | 0.261 | 1.733 | 0.202 | 0.005 | -0.035 | -0.035 | 0 | 0.19 | 0.1 | 0.14 | 0.31 | 0.17 | 0.19 |
| 3 | 29091210 | rs662120 | 0.068 | 0.457 | 1.742 | 0.004 | -0.002 | -0.002 | -0.002 | 0 | 0.15 | 0.37 | 0.31 | 0.1 | 0.12 | 0.24 |
| 3 | 29106733 | rs9850192 | 1.418 | 1.467 | 0.87 | -0.082 | -0.017 | 0.032 | 0.032 | 2.084 | 0.62 | 0.36 | 0.5 | 0.84 | 0.56 | 0.54 |
| 3 | 29114031 | rs527972 | 0.065 | 1.428 | 0.815 | 0.022 | 0.02 | 0.019 | 0.019 | 0 | 0.15 | 0.34 | 0.32 | 0.15 | 0.16 | 0.27 |
| 3 | 29123737 | rs569307 | -0.4 | 1.68 | 0.167 | -0.056 | 0.036 | 0.006 | 0.006 | 0 | 0.19 | 0.42 | 0.4 | 0.16 | 0.2 | 0.29 |
| 3 | 29130678 | rs9818278 | 0.491 | 1.554 | -0.038 | -0.03 | 0.014 | -0.036 | -0.036 | 0 | 0.24 | 0.15 | 0.15 | 0.43 | 0.19 | 0.21 |
| 3 | 29133039 | rs634868 | 0.401 | 1.628 | -0.086 | 0.008 | -0.027 | -0.014 | -0.014 | 0.141 | 0.61 | 0.73 | 0.73 | 0.56 | 0.58 | 0.67 |

| | | | | | | | | | | | | | | | | |
|---|---|---|---|---|---|---|---|---|---|---|---|---|---|---|---|---|
| 3 | 29133487 | rs512720 | -0.262 | 1.782 | -0.631 | 0.1 | -0.079 | 0.078 | 0.078 | 1.013 | 0.87 | 0.99 | 0.91 | 1 | 0.78 | 0.91 |
| 3 | 29139548 | rs474161 | -0.181 | 1.009 | -0.077 | 0.545 | 0.047 | 0.026 | 0.026 | 0 | 0.16 | 0.43 | 0.36 | 0.15 | 0.18 | 0.27 |
| 3 | 29140843 | rs1303912 | 0.633 | 0.444 | -1.001 | 0.656 | 0.018 | -0.056 | -0.056 | 0.162 | 0.36 | 0.1 | 0.32 | 0.39 | 0.34 | 0.31 |
| 3 | 29144250 | rs546270 | 1.106 | 0.523 | 0.132 | 0.644 | -0.006 | -0.001 | -0.001 | 0.522 | 0.81 | 0.43 | 0.81 | 0.79 | 0.82 | 0.85 |
| 3 | 29146469 | rs643454 | 0.518 | 0.258 | -0.77 | 0.695 | -0.091 | -0.052 | -0.052 | 0.006 | 0.34 | 0.12 | 0.27 | 0.44 | 0.46 | 0.38 |
| 3 | 29146855 | rs641725 | -0.718 | 0.032 | 0.352 | 0.618 | 0.009 | 0.001 | 0.001 | 0.039 | 0.51 | 0.63 | 0.4 | 0.59 | 0.6 | 0.46 |
| 3 | 29153605 | rs523660 | 0.994 | 0.057 | 0.479 | 0.655 | 0.009 | 0.083 | 0.083 | 0 | 0.89 | 0.69 | 0.89 | 0.79 | 0.93 | 0.91 |
| 3 | 29153755 | rs13059236 | -0.156 | 0.102 | 0.589 | 0.643 | 0.011 | -0.044 | -0.044 | 0.078 | 0.09 | 0.23 | 0.08 | 0.01 | 0.11 | 0.05 |
| 3 | 29158022 | rs604110 | 0.663 | 0.141 | 0.499 | 0.659 | -0.095 | -0.059 | -0.059 | 0 | 0.34 | 0.11 | 0.26 | 0.43 | 0.45 | 0.37 |
| 3 | 29160210 | rs11926062 | -0.378 | 0.365 | 0.975 | 0.632 | 0.06 | 0.072 | 0.072 | 0 | 0.05 | 0.27 | 0.09 | 0.07 | 0.04 | 0.06 |
| 3 | 29162760 | rs6788002 | -1.871 | 0.363 | 0.604 | 0.679 | 0.006 | 0.017 | 0.017 | 0 | 0.19 | 0.51 | 0.17 | 0.24 | 0.17 | 0.15 |
| 3 | 29178304 | rs9858901 | -1.585 | 0.354 | 0.55 | 0.66 | 0.003 | 0.03 | 0.03 | 1.413 | 0.16 | 0.65 | 0.18 | 0.16 | 0.16 | 0.12 |
| 3 | 29179292 | rs6779080 | -0.704 | 0.564 | -0.379 | 0.627 | -0.101 | -0.066 | -0.066 | 0 | 0.65 | 0.87 | 0.75 | 0.52 | 0.55 | 0.57 |
| 3 | 29183617 | rs17022765 | -0.883 | 0.496 | 0.156 | 0.472 | 0.057 | -0.074 | -0.074 | 0.375 | 0.13 | 0.25 | 0.11 | 0 | 0.19 | 0.08 |
| 3 | 29185516 | rs1878408 | 0.965 | 0.422 | -0.892 | 0.555 | 0.066 | 0.032 | 0.032 | 0.752 | 0.41 | 0.14 | 0.5 | 0.28 | 0.3 | 0.35 |
| 3 | 29192092 | rs4680807 | -1.07 | 0.614 | -0.725 | 0.726 | 0.008 | 0.001 | 0.001 | 0.423 | 0.22 | 0.69 | 0.22 | 0.21 | 0.25 | 0.14 |
| 3 | 29201460 | rs2102801 | -0.341 | 1.173 | -1.242 | 0.669 | -0.046 | -0.041 | -0.041 | 0.168 | 0.65 | 0.76 | 0.74 | 0.5 | 0.56 | 0.54 |
| 3 | 29212902 | rs12492761 | -1.186 | 1.336 | -0.832 | 0.481 | -0.009 | 0.009 | 0.009 | 0 | 0.11 | 0.07 | 0.06 | 0.02 | 0.15 | 0.12 |
| 3 | 29218259 | rs6780332 | -0.563 | 1.138 | -0.738 | 0.716 | 0 | 0.013 | 0.013 | 0 | 0.35 | 0.5 | 0.48 | 0.35 | 0.33 | 0.36 |
| 3 | 29223338 | rs13099375 | -0.066 | 0.895 | -1.324 | 0.399 | 0.008 | 0.04 | 0.04 | 0 | 0.26 | 0.12 | 0.36 | 0.11 | 0.22 | 0.18 |
| 3 | 29223853 | rs9883170 | -0.095 | 0.404 | -1.269 | 0.968 | -0.018 | 0.059 | 0.059 | 1.601 | 0.14 | 0.21 | 0.06 | 0.35 | 0.09 | 0.13 |
| 3 | 29224900 | rs1545106 | 0.225 | 1.426 | -1.287 | 1.288 | 0.015 | 0.061 | 0.061 | 0 | 0.91 | 0.86 | 0.93 | 0.67 | 0.84 | 0.77 |
| 3 | 29226447 | rs7645215 | 0.45 | 1.115 | -1.296 | 0.659 | 0.004 | -0.017 | -0.017 | 0 | 0.07 | 0.16 | 0.04 | 0.04 | 0.06 | 0.06 |
| 3 | 29230759 | rs4635667 | 0.074 | 1.195 | -1.626 | 0.518 | 0.005 | 0.068 | 0.068 | 0 | 0.3 | 0.09 | 0.39 | 0.11 | 0.26 | 0.2 |
| 3 | 29233149 | rs9831444 | 0.399 | 1.551 | -1.984 | 0.184 | -0.077 | 0.054 | 0.054 | 0 | 0.2 | 0.03 | 0.14 | 0.1 | 0.25 | 0.22 |
| 3 | 29236334 | rs1589059 | 0.053 | 0.812 | -1.794 | 0.106 | 0.015 | 0.053 | 0.053 | 0 | 0.44 | 0.29 | 0.47 | 0.19 | 0.38 | 0.34 |
| 3 | 29238081 | rs7643900 | -0.329 | 0.599 | -1.781 | 0.815 | 0.014 | -0.03 | -0.03 | 0 | 0.9 | 0.96 | 0.96 | 0.7 | 0.92 | 0.78 |
| 3 | 29239664 | rs1030842 | 0.238 | 0.706 | -1.422 | 0.447 | -0.001 | 0 | 0 | 0 | 0.58 | 0.57 | 0.55 | 0.41 | 0.49 | 0.42 |
| 3 | 29241894 | rs12634340 | -0.028 | 0.452 | -1.256 | 0.953 | 0.011 | -0.015 | -0.015 | 0 | 0.15 | 0.13 | 0.1 | 0.38 | 0.08 | 0.22 |
| 3 | 29244234 | rs2048896 | -0.426 | 0.767 | -1.064 | 0.963 | -0.016 | -0.006 | -0.006 | 0 | 0.7 | 0.76 | 0.63 | 0.67 | 0.58 | 0.6 |
| 3 | 29245064 | rs6805530 | 0.947 | 0.265 | -0.513 | 0.862 | 0.006 | 0.001 | 0.001 | 0 | 0.45 | 0.41 | 0.45 | 0.41 | 0.36 | 0.34 |
| 3 | 29248536 | rs1517150 | 0.263 | 0.387 | -0.298 | 0.762 | -0.001 | 0 | 0 | 0 | 0.57 | 0.54 | 0.54 | 0.58 | 0.49 | 0.44 |
| 3 | 29250696 | rs11129353 | 0.536 | 0.383 | 0.069 | 0.713 | 0.005 | -0.002 | -0.002 | 1.609 | 0.56 | 0.53 | 0.53 | 0.59 | 0.5 | 0.49 |
| 3 | 29251992 | rs7616684 | -0.593 | 0.38 | 0.017 | 0.732 | -0.027 | -0.01 | -0.01 | 0 | 0.69 | 0.77 | 0.62 | 0.67 | 0.58 | 0.59 |
| 3 | 29253107 | rs6773381 | 0.459 | 0.58 | -0.058 | 0.894 | 0.002 | -0.036 | -0.036 | 0 | 0.12 | 0.07 | 0.07 | 0.32 | 0.11 | 0.19 |
| 3 | 29254876 | rs11129354 | 0.616 | 0.578 | -0.027 | 0.788 | 0.005 | 0.002 | 0.002 | 0 | 0.5 | 0.45 | 0.47 | 0.42 | 0.41 | 0.37 |
| 3 | 29258177 | rs6790879 | 0.149 | 0.595 | -0.277 | 0.762 | -0.128 | 0.163 | 0.163 | 0 | 0.34 | 0.02 | 0.34 | 0.09 | 0.37 | 0.28 |
| 3 | 29258751 | rs2196554 | 0.191 | 0.642 | -0.133 | 0.946 | -0.003 | -0.002 | -0.002 | 0.412 | 0.62 | 0.6 | 0.54 | 0.58 | 0.52 | 0.54 |
| 3 | 29263650 | rs13098112 | 0.529 | 0.58 | -0.307 | 1.01 | -0.131 | 0.152 | 0.152 | 0 | 0.31 | 0.02 | 0.34 | 0.07 | 0.34 | 0.29 |
| 3 | 29265476 | rs6772594 | 1.348 | 0.491 | -0.093 | 1.142 | 0.007 | 0.002 | 0.002 | 0 | 0.41 | 0.36 | 0.27 | 0.37 | 0.3 | 0.29 |
| 3 | 29265952 | rs768303 | 0.5 | 0.637 | 0.093 | 1.477 | 0.005 | -0.005 | -0.005 | 0 | 0.57 | 0.52 | 0.36 | 0.39 | 0.48 | 0.45 |
| 3 | 29267732 | rs4496436 | 0.421 | 0.742 | 0.655 | 1.842 | 0.081 | 0.428 | 0.428 | 0 | 0.94 | 0.64 | 0.9 | 0.67 | 0.91 | 0.82 |
| 3 | 29293114 | rs12639059 | 0.224 | 1.285 | -0.939 | 1.43 | -0.082 | 0.008 | 0.008 | 0 | 0.28 | 0.02 | 0.44 | 0.14 | 0.29 | 0.25 |
| 3 | 29296192 | rs13072528 | 0.007 | 1.461 | -0.6 | 1.853 | -0.046 | -0.027 | -0.027 | 0 | 0.22 | 0.08 | 0.18 | 0.25 | 0.21 | 0.27 |
| 3 | 29296791 | rs7622721 | -1.058 | 1.037 | -0.269 | 1.546 | 0.011 | -0.003 | -0.003 | 0 | 0.34 | 0.59 | 0.3 | 0.76 | 0.36 | 0.53 |
| 3 | 29300050 | rs967220 | -0.648 | 0.555 | -0.591 | 1.713 | 0.004 | -0.006 | -0.006 | 0 | 0.29 | 0.26 | 0.19 | 0.6 | 0.21 | 0.31 |
| 3 | 29303962 | rs17706635 | 1.89 | 0.858 | -1.029 | 1.359 | -0.016 | 0.069 | 0.069 | 0 | 0.27 | 0.08 | 0.32 | 0.1 | 0.28 | 0.19 |
| 3 | 29304320 | rs17023089 | 0.924 | 0.852 | -0.974 | 1.392 | -0.014 | -0.029 | -0.029 | 0.312 | 0.05 | 0 | 0.05 | 0.16 | 0.06 | 0.04 |
| 3 | 29304503 | rs6798283 | 0.401 | 0.595 | -0.896 | 1.341 | 0.023 | 0.005 | 0.005 | 0 | 0.49 | 0.3 | 0.47 | 0.41 | 0.44 | 0.38 |
| 3 | 29305491 | rs987692 | -0.577 | 1.122 | -0.415 | 1.152 | 0.139 | 0.092 | 0.092 | 0 | 0.94 | 0.72 | 0.9 | 0.87 | 0.87 | 0.82 |
| 3 | 29305716 | rs987693 | -1.199 | 0.98 | -0.705 | 0.96 | 0.005 | 0.001 | 0.001 | 0 | 0.59 | 0.53 | 0.53 | 0.45 | 0.54 | 0.54 |
| 3 | 29307067 | rs9871008 | -0.378 | 1.215 | -0.946 | 1.052 | 0 | 0 | 0 | 0 | 0.19 | 0.21 | 0.12 | 0.14 | 0.14 | 0.13 |
| 3 | 29310744 | rs17023120 | -0.887 | 1.59 | -1.316 | 1.057 | -0.004 | -0.017 | -0.017 | 0 | 0.19 | 0.14 | 0.3 | 0.26 | 0.21 | 0.22 |
| 3 | 29310894 | rs13074088 | -1.073 | 1.98 | -1.62 | 0.796 | -0.032 | 0.02 | 0.02 | 1.955 | 0.19 | 0.1 | 0.15 | 0.13 | 0.25 | 0.19 |
| 3 | 29310929 | rs13094301 | -0.463 | 1.574 | -1.807 | 0.889 | -0.049 | -0.007 | -0.007 | 0.022 | 0.17 | 0.02 | 0.14 | 0.13 | 0.21 | 0.18 |
| 3 | 29312086 | rs9310891 | -0.208 | 1.559 | -1.396 | 0.824 | 0.013 | 0.048 | 0.048 | 1.059 | 0.77 | 0.56 | 0.81 | 0.66 | 0.77 | 0.59 |
| 3 | 29312287 | rs9812101 | 0.504 | 1.27 | -2.539 | 0.503 | -0.015 | 0.022 | 0.022 | 0 | 0.17 | 0.09 | 0.24 | 0 | 0.19 | 0.13 |
| 3 | 29312810 | rs10490855 | 0.003 | 0.734 | -2.378 | 1.387 | -0.015 | -0.122 | -0.122 | 0 | 0.37 | 0.05 | 0.35 | 0.43 | 0.35 | 0.26 |
| 3 | 29320833 | rs13079598 | -0.835 | 1.115 | -1.579 | 1.868 | -0.01 | -0.008 | -0.008 | 0 | 0.1 | 0.06 | 0.13 | 0.29 | 0.11 | 0.14 |
| 3 | 29322297 | rs2888099 | -0.799 | 1.025 | -1.129 | 1.635 | 0.012 | 0.032 | 0.032 | 0 | 0.17 | 0.32 | 0.27 | 0.27 | 0.16 | 0.31 |
| 3 | 29322363 | rs9874998 | -0.71 | 0.41 | -1.492 | 1.454 | 0.013 | -0.003 | -0.003 | 0.012 | 0.11 | 0.11 | 0.06 | 0.14 | 0.05 | 0.2 |
| 3 | 29325250 | rs4680816 | -0.414 | 0.818 | -1.927 | 0.888 | -0.046 | 0.034 | 0.034 | 4.567 | 0.2 | 0.06 | 0.3 | 0.12 | 0.24 | 0.19 |
| 3 | 29331859 | rs1025644 | 0.013 | 1.411 | -1.15 | 1.664 | 0.057 | 0.169 | 0.169 | 0 | 0.94 | 0.8 | 0.84 | 0.7 | 0.92 | 0.85 |
| 3 | 29334820 | rs17665167 | 1.241 | 0.867 | -1.441 | 0.492 | 0.004 | -0.012 | -0.012 | 0 | 0.29 | 0.21 | 0.14 | 0.06 | 0.23 | 0.14 |
| 3 | 29335046 | rs1020042 | 0.533 | 0.637 | -0.631 | 0.289 | -0.012 | -0.043 | -0.043 | 0 | 0.09 | 0.3 | 0.08 | 0.01 | 0.09 | 0.06 |
| 3 | 29335253 | rs1020043 | -0.157 | 1.03 | -0.579 | 0.376 | 0.003 | 0.019 | 0.019 | 0 | 0.67 | 0.6 | 0.5 | 0.52 | 0.65 | 0.65 |
| 3 | 29335426 | rs9878305 | 1.217 | 1.392 | -0.963 | 0.872 | -0.012 | -0.01 | -0.01 | 0.48 | 0.07 | 0.05 | 0.1 | 0.17 | 0.1 | 0.05 |
| 3 | 29337484 | rs17665280 | -0.11 | 1.095 | -0.64 | 0.644 | 0.009 | 0.026 | 0.026 | 0.022 | 0.35 | 0.26 | 0.4 | 0.19 | 0.35 | 0.27 |
| 3 | 29338610 | rs13093377 | 0.616 | 1.203 | -0.674 | 0.787 | 0.001 | 0.015 | 0.015 | 0 | 0.26 | 0.14 | 0.34 | 0.21 | 0.26 | 0.21 |

| | | | | | | | | | | | | | | | | |
|---|---|---|---|---|---|---|---|---|---|---|---|---|---|---|---|---|
| 3 | 29342717 | rs17023218 | 0.782 | 1.201 | -0.726 | 0.786 | -0.007 | 0.014 | 0.014 | 1.434 | 0.23 | 0.1 | 0.31 | 0.19 | 0.24 | 0.19 |
| 3 | 29343465 | rs9654034 | 0.209 | 1.159 | -0.767 | 0.877 | -0.003 | -0.013 | -0.013 | 0 | 0.2 | 0.21 | 0.18 | 0.3 | 0.22 | 0.38 |
| 3 | 29345181 | rs9865738 | 0.142 | 1.222 | -0.827 | 0.797 | -0.002 | 0 | 0 | 0 | 0.07 | 0.09 | 0.05 | 0.04 | 0.09 | 0.14 |
| 3 | 29347180 | rs2196550 | 1.15 | 1.151 | -0.996 | 0.726 | 0.019 | -0.008 | -0.008 | 0 | 0.2 | 0.08 | 0.28 | 0.22 | 0.14 | 0.18 |
| 3 | 29347933 | rs736357 | -0.112 | 0.617 | -0.774 | 0.452 | 0 | 0.005 | 0.005 | 0 | 0.54 | 0.55 | 0.37 | 0.22 | 0.55 | 0.31 |
| 3 | 29349070 | rs17023226 | 0.554 | 0.606 | -0.701 | 0.81 | -0.006 | 0.049 | 0.049 | 0.429 | 0.08 | 0.11 | 0.1 | 0.24 | 0.07 | 0.27 |
| 3 | 29349296 | rs9831559 | 2.085 | 0.64 | -0.743 | 0.404 | -0.012 | -0.028 | -0.028 | 0 | 0.25 | 0.3 | 0.16 | 0.15 | 0.22 | 0.12 |
| 3 | 29351830 | rs6791039 | 1.919 | 0.42 | -0.8 | -0.182 | -0.011 | -0.056 | -0.056 | 0.119 | 0.25 | 0.36 | 0.17 | 0.09 | 0.23 | 0.12 |
| 3 | 29352830 | rs9875819 | -0.357 | 0.708 | -1.327 | 0.004 | -0.067 | -0.105 | -0.105 | 0.011 | 0.23 | 0.03 | 0.21 | 0.35 | 0.27 | 0.33 |
| 3 | 29357063 | rs6549924 | 0.304 | 0.704 | -0.832 | -0.598 | 0.01 | 0.042 | 0.042 | 0 | 0.45 | 0.33 | 0.63 | 0.77 | 0.39 | 0.67 |
| 3 | 29357140 | rs7349464 | 2.345 | 0.815 | -0.948 | -0.826 | 0.001 | -0.004 | -0.004 | 1.166 | 0.14 | 0.12 | 0.07 | 0.02 | 0.12 | 0.06 |
| 3 | 29358171 | rs7624339 | -1.239 | 0.857 | -0.744 | -0.862 | -0.012 | -0.032 | -0.032 | 2.021 | 0.78 | 0.67 | 0.86 | 0.89 | 0.8 | 0.9 |
| 3 | 29358326 | rs9831361 | 1.597 | 0.858 | -0.747 | -0.87 | -0.019 | -0.024 | -0.024 | 0 | 0.19 | 0.33 | 0.11 | 0.11 | 0.15 | 0.1 |
| 3 | 29358602 | rs12495625 | 1.381 | 1.184 | -1.143 | -0.828 | -0.045 | 0.004 | 0.004 | 0 | 0.11 | 0.04 | 0.14 | 0.09 | 0.19 | 0.09 |
| 3 | 29360376 | rs10865825 | -0.152 | 1.155 | -1.12 | -0.927 | -0.044 | 0.072 | 0.072 | 1.799 | 0.69 | 0.8 | 0.76 | 0.88 | 0.58 | 0.77 |
| 3 | 29366164 | rs9839176 | -0.076 | 1.105 | -1.001 | -0.78 | -0.009 | 0.012 | 0.012 | 0 | 0.79 | 0.83 | 0.8 | 0.92 | 0.69 | 0.87 |
| 3 | 29367711 | rs12491584 | 0.665 | 0.784 | -0.757 | -0.749 | 0.01 | 0.042 | 0.042 | 0 | 0.45 | 0.34 | 0.63 | 0.8 | 0.39 | 0.69 |
| 3 | 29367784 | rs12494597 | 0.282 | 0.916 | -0.587 | -0.439 | -0.018 | 0.028 | 0.028 | 0.026 | 0.75 | 0.81 | 0.81 | 0.92 | 0.65 | 0.79 |
| 3 | 29369364 | rs6785458 | 0.705 | 0.66 | -0.558 | 0.307 | -0.007 | 0.166 | 0.166 | 0 | 0.06 | 0.16 | 0.1 | 0.25 | 0.04 | 0.27 |
| 3 | 29369685 | rs4380376 | -0.202 | 0.809 | -0.643 | -0.23 | 0.009 | -0.007 | -0.007 | 0 | 0.87 | 0.83 | 0.85 | 0.89 | 0.81 | 0.92 |
| 3 | 29370491 | rs9850492 | 0.678 | 0.139 | -0.65 | -0.235 | -0.001 | -0.006 | -0.006 | 0 | 0.66 | 0.63 | 0.82 | 0.71 | 0.67 | 0.62 |
| 3 | 29371024 | rs9814033 | 0.36 | 0.147 | -0.789 | -0.053 | 0.005 | -0.009 | -0.009 | 0 | 0.78 | 0.83 | 0.9 | 0.71 | 0.82 | 0.69 |
| 3 | 29372404 | rs6777437 | -0.232 | 0.253 | -0.46 | -0.259 | 0.01 | 0.043 | 0.043 | 0 | 0.48 | 0.69 | 0.69 | 0.68 | 0.58 | 0.6 |
| 3 | 29373274 | rs7374763 | 0.219 | 0.332 | -0.521 | -0.076 | 0.016 | -0.006 | -0.006 | 0 | 0.42 | 0.57 | 0.54 | 0.57 | 0.28 | 0.49 |
| 3 | 29373424 | rs6804900 | 2.036 | 0.539 | -0.786 | -0.06 | -0.04 | 0.067 | 0.067 | 0 | 0.12 | 0.01 | 0.07 | 0.02 | 0.16 | 0.06 |
| 3 | 29376271 | rs9882071 | 0.812 | 0.331 | -0.176 | 0.277 | 0.051 | 0.321 | 0.321 | 0 | 0.1 | 0.41 | 0.37 | 0.37 | 0.11 | 0.22 |
| 3 | 29376642 | rs9845524 | 0.322 | 0.086 | -0.533 | -0.271 | -0.004 | 0 | 0 | 0 | 0.39 | 0.36 | 0.3 | 0.41 | 0.25 | 0.48 |
| 3 | 29376770 | rs9845429 | 0.122 | 0.253 | -0.542 | -0.297 | -0.001 | 0.001 | 0.001 | 0 | 0.44 | 0.38 | 0.37 | 0.41 | 0.39 | 0.5 |
| 3 | 29377989 | rs6810211 | -0.397 | -0.29 | -0.343 | -0.103 | -0.027 | -0.01 | -0.01 | 0.078 | 0.72 | 0.59 | 0.71 | 0.71 | 0.81 | 0.62 |
| 3 | 29378090 | rs6773182 | 0.652 | -0.105 | -0.874 | -0.141 | -0.055 | -0.009 | -0.009 | 0 | 0.3 | 0.11 | 0.29 | 0.3 | 0.43 | 0.35 |
| 3 | 29381310 | rs13097390 | -1.695 | 0.152 | -0.714 | 0.093 | -0.014 | 0.016 | 0.016 | 0 | 0.87 | 0.98 | 0.94 | 0.9 | 0.91 | 0.82 |
| 3 | 29382541 | rs7431530 | 0.532 | 0.244 | -0.309 | 0.226 | -0.003 | 0.011 | 0.011 | 0.471 | 0.31 | 0.37 | 0.35 | 0.35 | 0.27 | 0.19 |
| 3 | 29385810 | rs6779814 | 1.187 | 0.281 | -0.722 | 0.278 | -0.004 | 0.003 | 0.003 | 0.425 | 0.17 | 0.18 | 0.13 | 0.23 | 0.14 | 0.13 |
| 3 | 29387083 | rs13090360 | -0.078 | 0.313 | -0.527 | 0.171 | -0.01 | 0.005 | 0.005 | 0 | 0.37 | 0.45 | 0.42 | 0.32 | 0.33 | 0.22 |
| 3 | 29388257 | rs4680822 | 1.569 | 0.533 | -0.637 | 0.325 | 0.001 | 0.027 | 0.027 | 0 | 0.15 | 0.24 | 0.25 | 0.14 | 0.14 | 0.1 |
| 3 | 29390775 | rs9863801 | 0.145 | 0.75 | -1.223 | 0.255 | 0.019 | 0.022 | 0.022 | 0 | 0.37 | 0.52 | 0.47 | 0.43 | 0.45 | 0.35 |
| 3 | 29391314 | rs9812752 | -0.452 | 0.708 | -1.274 | 0.074 | 0.006 | 0.014 | 0.014 | 2.056 | 0.52 | 0.59 | 0.66 | 0.62 | 0.58 | 0.48 |
| 3 | 29392163 | rs4129221 | 0.112 | 0.487 | -1.039 | 0.019 | -0.019 | 0.041 | 0.041 | 0 | 0.28 | 0.09 | 0.2 | 0.2 | 0.3 | 0.24 |
| 3 | 29394655 | rs7427457 | 0.02 | 0.476 | -0.966 | 0.014 | -0.005 | 0.038 | 0.038 | 0 | 0.28 | 0.15 | 0.21 | 0.2 | 0.3 | 0.24 |
| 3 | 29394725 | rs7427438 | 0.004 | 0.549 | -1.053 | 0.054 | 0.001 | -0.013 | -0.013 | 0 | 0.29 | 0.42 | 0.35 | 0.17 | 0.31 | 0.22 |
| 3 | 29395169 | rs9813870 | -0.207 | 0.359 | -0.629 | 0.58 | -0.006 | 0.014 | 0.014 | 0.473 | 0.31 | 0.4 | 0.37 | 0.31 | 0.27 | 0.21 |
| 3 | 29407073 | rs9849677 | 0.885 | 0.271 | -0.966 | 0.304 | -0.009 | 0.016 | 0.016 | 0 | 0.61 | 0.54 | 0.59 | 0.48 | 0.69 | 0.52 |
| 3 | 29407516 | rs6770055 | 0.172 | 0.435 | -1.403 | 0.417 | -0.052 | -0.125 | -0.125 | 0 | 0.19 | 0.02 | 0.29 | 0.4 | 0.19 | 0.27 |
| 3 | 29407918 | rs7651268 | 0.263 | 0.571 | -1.075 | 0.25 | -0.001 | 0 | 0 | 0 | 0.62 | 0.62 | 0.6 | 0.48 | 0.69 | 0.52 |
| 3 | 29412201 | rs4465907 | -0.1 | 0.973 | -1.546 | -0.855 | -0.026 | 0.046 | 0.046 | 0 | 0.37 | 0.24 | 0.28 | 0.04 | 0.46 | 0.22 |
| 3 | 29415650 | rs4645104 | -0.307 | 1.394 | -1.613 | -0.902 | -0.01 | 0.018 | 0.018 | 0 | 0.22 | 0.15 | 0.25 | 0.12 | 0.29 | 0.22 |
| 3 | 29420132 | rs4574243 | 0.441 | 0.996 | -2.515 | -1.2 | -0.108 | 0.247 | 0.247 | 0.043 | 0.45 | 0.05 | 0.24 | 0.15 | 0.49 | 0.41 |
| 3 | 29422762 | rs4287886 | 0.14 | 0.553 | -2.494 | -0.781 | 0.007 | -0.002 | -0.002 | 0.341 | 0.37 | 0.31 | 0.47 | 0.61 | 0.32 | 0.6 |
| 3 | 29424794 | rs13059377 | 0.815 | 0.537 | -2.559 | -0.948 | -0.02 | 0.028 | 0.028 | 0 | 0.2 | 0.04 | 0.19 | 0.1 | 0.22 | 0.19 |
| 3 | 29426601 | rs9819877 | -0.673 | 0.518 | -1.905 | -0.265 | 0.057 | 0.304 | 0.304 | 0.001 | 0.85 | 0.58 | 0.76 | 0.45 | 0.87 | 0.79 |
| 3 | 29426721 | rs9819772 | 0.809 | 0.471 | -1.974 | -0.391 | 0.016 | -0.021 | -0.021 | 0 | 0.22 | 0.44 | 0.16 | 0.22 | 0.22 | 0.3 |
| 3 | 29427676 | rs11713209 | 0.878 | 0.476 | -1.607 | -0.029 | -0.002 | 0.141 | 0.141 | 0 | 0.47 | 0.26 | 0.27 | 0.2 | 0.5 | 0.43 |
| 3 | 29428187 | rs4377455 | 0.183 | 0.469 | -1.569 | 0.037 | 0.002 | -0.007 | -0.007 | 0 | 0.64 | 0.52 | 0.57 | 0.34 | 0.64 | 0.6 |
| 3 | 29428510 | rs9830413 | -0.264 | 0.345 | -1.501 | 0.163 | 0.003 | 0.026 | 0.026 | 0 | 0.38 | 0.31 | 0.5 | 0.25 | 0.37 | 0.38 |
| 3 | 29431425 | rs4387928 | -0.676 | -0.223 | -0.531 | 0.433 | 0.043 | 0.012 | 0.012 | 5.087 | 0.42 | 0.74 | 0.44 | 0.64 | 0.35 | 0.53 |
| 3 | 29433430 | rs6549934 | -0.162 | -0.537 | 0.022 | 0.44 | -0.056 | 0.009 | 0.009 | 0.641 | 0.31 | 0.42 | 0.27 | 0.3 | 0.24 | 0.34 |
| 3 | 29436454 | rs9844602 | 0.653 | -0.058 | -0.57 | -0.117 | -0.024 | 0.003 | 0.003 | 1.119 | 0.37 | 0.15 | 0.42 | 0.31 | 0.38 | 0.35 |
| 3 | 29436928 | rs7636503 | 1.561 | -0.247 | -0.713 | 0.056 | 0.005 | 0.021 | 0.021 | 0 | 0.17 | 0.06 | 0.15 | 0.08 | 0.19 | 0.21 |
| 3 | 29437405 | rs4432591 | -2.26 | -0.095 | -0.688 | 0.093 | -0.063 | 0.088 | 0.088 | 0 | 0.79 | 0.93 | 0.86 | 0.97 | 0.72 | 0.89 |
| 3 | 29438500 | rs7642033 | -0.741 | -0.118 | -0.542 | 0.155 | 0.007 | -0.033 | -0.033 | 0 | 0.66 | 0.6 | 0.77 | 0.77 | 0.68 | 0.79 |
| 3 | 29441020 | rs7355803 | -0.568 | -0.362 | -0.838 | -0.029 | 0.011 | 0.009 | 0.009 | 0 | 0.45 | 0.64 | 0.55 | 0.25 | 0.38 | 0.37 |
| 3 | 29442294 | rs12491968 | 2.054 | -0.308 | -1.123 | 0.035 | -0.045 | 0.029 | 0.029 | 2.058 | 0.18 | 0.03 | 0.11 | 0.14 | 0.22 | 0.23 |
| 3 | 29443204 | rs7432004 | 2.083 | -0.315 | -1.136 | 0.032 | -0.069 | 0.014 | 0.014 | 2.284 | 0.18 | 0.01 | 0.11 | 0.14 | 0.22 | 0.23 |
| 3 | 29445931 | rs13079920 | 0.665 | -0.385 | -1.179 | 0.028 | -0.087 | -0.058 | -0.058 | 1.024 | 0.33 | 0.04 | 0.39 | 0.31 | 0.35 | 0.34 |
| 3 | 29446983 | rs4680830 | -2.392 | -0.37 | -1.173 | -0.048 | -0.065 | 0.083 | 0.083 | 0 | 0.79 | 0.94 | 0.85 | 0.96 | 0.72 | 0.89 |
| 3 | 29450307 | rs6772026 | -1.378 | -0.446 | -1.019 | 0.003 | -0.008 | 0.007 | 0.007 | 0.003 | 0.71 | 0.81 | 0.8 | 0.77 | 0.7 | 0.81 |
| 3 | 29450450 | rs4146795 | -2.391 | -0.348 | -1.118 | -0.041 | -0.062 | 0.082 | 0.082 | 0 | 0.79 | 0.93 | 0.85 | 0.97 | 0.72 | 0.9 |
| 3 | 29450982 | rs7622604 | -0.787 | -0.517 | -1.069 | 0.04 | 0.074 | 0.012 | 0.012 | 3.758 | 0.45 | 0.85 | 0.47 | 0.64 | 0.37 | 0.55 |
| 3 | 29455853 | rs4680724 | 1.245 | -0.355 | -1.116 | 0.151 | -0.026 | -0.072 | -0.072 | 0.183 | 0.14 | 0.01 | 0.28 | 0.17 | 0.13 | 0.11 |

| | | | | | | | | | | | | | | | | |
|---|---|---|---|---|---|---|---|---|---|---|---|---|---|---|---|---|
| 3 | 29456198 | rs1370796 | 1.098 | -0.28 | -1.024 | 0.266 | -0.047 | -0.056 | -0.056 | 1.108 | 0.25 | 0.13 | 0.25 | 0.43 | 0.33 | 0.44 |
| 3 | 29456714 | rs4680833 | 2.103 | -0.234 | -1.015 | 0.301 | -0.024 | 0.005 | 0.005 | 0 | 0.05 | 0.03 | 0.13 | 0.27 | 0.1 | 0.21 |
| 3 | 29458092 | rs1561374 | 0.376 | -0.519 | -0.569 | 0.163 | -0.011 | -0.022 | -0.022 | 0 | 0.41 | 0.52 | 0.28 | 0.29 | 0.39 | 0.45 |
| 3 | 29459035 | rs10222540 | -0.246 | -0.383 | -0.213 | 0.284 | 0.036 | 0.166 | 0.166 | 0.174 | 0.11 | 0.31 | 0.17 | 0.33 | 0.12 | 0.26 |
| 3 | 29459473 | rs9835541 | 1.287 | -0.446 | -0.333 | 0.135 | 0 | -0.007 | -0.007 | 0 | 0.23 | 0.25 | 0.14 | 0.22 | 0.24 | 0.28 |
| 3 | 29462116 | rs7640471 | 2.043 | -0.09 | -0.392 | 0.415 | -0.014 | 0.03 | 0.03 | 0.693 | 0.07 | 0.07 | 0.04 | 0.19 | 0.02 | 0.07 |
| 3 | 29465924 | rs2053288 | 0.274 | 0.08 | -0.405 | 0.388 | -0.007 | -0.023 | -0.023 | 0 | 0.44 | 0.53 | 0.33 | 0.23 | 0.41 | 0.43 |
| 3 | 29467931 | rs9861686 | -1.008 | 0.568 | -0.775 | 0.537 | -0.02 | 0.02 | 0.02 | 0.067 | 0.71 | 0.87 | 0.81 | 0.78 | 0.7 | 0.82 |
| 3 | 29472046 | rs7643656 | -0.612 | 0.496 | -0.749 | 0.55 | -0.045 | -0.053 | -0.053 | 0 | 0.66 | 0.82 | 0.68 | 0.5 | 0.6 | 0.61 |
| 3 | 29476428 | rs3773018 | 0.298 | 0.057 | -0.474 | 0.245 | -0.003 | -0.001 | -0.001 | 0 | 0.48 | 0.45 | 0.5 | 0.54 | 0.38 | 0.49 |
| 3 | 29476582 | rs3773019 | -0.258 | 0.194 | -0.359 | 0.317 | -0.002 | 0.028 | 0.028 | 1.083 | 0.54 | 0.54 | 0.64 | 0.84 | 0.5 | 0.66 |
| 3 | 29476873 | rs9863789 | 0.522 | 0.386 | -0.476 | 0.146 | 0.006 | 0.02 | 0.02 | 3.247 | 0.28 | 0.41 | 0.33 | 0.42 | 0.33 | 0.43 |
| 3 | 29479352 | rs9878948 | 0.506 | -0.384 | -0.78 | -0.239 | 0.025 | 0.057 | 0.057 | 0.634 | 0.36 | 0.22 | 0.21 | 0.11 | 0.31 | 0.29 |
| 3 | 29482046 | rs1438339 | -1.026 | -0.486 | -0.251 | 0.419 | -0.029 | 0.006 | 0.006 | 0.47 | 0.32 | 0.63 | 0.47 | 0.32 | 0.19 | 0.27 |
| 3 | 29482712 | rs17023487 | 0.575 | -0.205 | -0.076 | 0.683 | -0.002 | 0.011 | 0.011 | 0 | 0.11 | 0.19 | 0.2 | 0.19 | 0.07 | 0.15 |
| 3 | 29484510 | rs7632290 | -0.767 | -0.024 | 0.032 | 0.805 | 0 | 0.006 | 0.006 | 0 | 0.9 | 0.89 | 0.79 | 0.86 | 0.89 | 0.9 |
| 3 | 29485209 | rs9882397 | 1.082 | 0.167 | 0.09 | 0.786 | 0.005 | 0 | 0 | 0 | 0.36 | 0.38 | 0.41 | 0.39 | 0.42 | 0.45 |
| 3 | 29485303 | rs9882552 | -0.339 | 0.127 | 0.002 | 0.633 | 0 | 0 | 0 | 0.013 | 0.6 | 0.6 | 0.55 | 0.58 | 0.64 | 0.52 |
| 3 | 29486572 | rs17666790 | 0.578 | -0.004 | 0.081 | 0.597 | 0 | -0.004 | -0.004 | 1.415 | 0.46 | 0.57 | 0.39 | 0.54 | 0.44 | 0.6 |
| 3 | 29487637 | rs6784776 | 1.911 | 0.003 | 0.102 | 0.747 | 0.002 | -0.003 | -0.003 | 0 | 0.1 | 0.09 | 0.1 | 0.17 | 0.04 | 0.16 |
| 3 | 29489266 | rs4583597 | -0.887 | 0.04 | 0.129 | 0.75 | -0.001 | 0 | 0 | 0.359 | 0.69 | 0.62 | 0.64 | 0.74 | 0.68 | 0.67 |
| 3 | 29489607 | rs4334600 | -1.961 | 0.082 | -0.036 | 0.791 | 0.002 | -0.011 | -0.011 | 0 | 0.91 | 0.96 | 0.92 | 0.87 | 0.92 | 0.9 |
| 3 | 29490420 | rs5028435 | -0.961 | 0.079 | -0.007 | 0.772 | 0 | 0 | 0 | 0 | 0.7 | 0.69 | 0.65 | 0.75 | 0.68 | 0.68 |
| 3 | 29490600 | rs17023528 | 1.708 | 0.109 | 0.087 | 0.618 | -0.001 | -0.008 | -0.008 | 0 | 0.21 | 0.26 | 0.15 | 0.11 | 0.22 | 0.22 |
| 3 | 29491223 | rs7625682 | 1.837 | -0.13 | -0.092 | 0.56 | 0.008 | -0.012 | -0.012 | 3.266 | 0.09 | 0.07 | 0.09 | 0.17 | 0.04 | 0.16 |
| 3 | 29493019 | rs5028491 | -0.137 | -0.286 | 0.417 | 0.558 | -0.034 | 0.133 | 0.133 | 0 | 0.22 | 0.44 | 0.29 | 0.44 | 0.15 | 0.23 |
| 3 | 29493600 | rs17023546 | 1.879 | -0.168 | 0.529 | 0.334 | 0.007 | -0.01 | -0.01 | 0 | 0.11 | 0.22 | 0.21 | 0.09 | 0.13 | 0.06 |
| 3 | 29493709 | rs9310899 | -0.193 | -0.255 | 0.54 | 0.442 | 0.01 | 0.023 | 0.023 | 2.633 | 0.47 | 0.61 | 0.54 | 0.63 | 0.42 | 0.46 |
| 3 | 29494147 | rs9882137 | 1.6 | -0.225 | 0.614 | 0.46 | -0.009 | -0.03 | -0.03 | 0 | 0.08 | 0.27 | 0.06 | 0 | 0.03 | 0.01 |
| 3 | 29495442 | rs17667279 | 1.872 | -0.149 | 0.419 | 0.493 | -0.016 | 0.003 | 0.003 | 0 | 0.11 | 0.09 | 0.32 | 0.1 | 0.16 | 0.07 |
| 3 | 29497989 | rs9861677 | -1.583 | -0.266 | 0.389 | 0.432 | -0.033 | 0.076 | 0.076 | 0.603 | 0.57 | 0.82 | 0.73 | 0.72 | 0.55 | 0.52 |
| 3 | 29498007 | rs9824170 | -0.582 | -0.245 | 0.385 | 0.546 | -0.01 | 0.035 | 0.035 | 0 | 0.31 | 0.42 | 0.46 | 0.53 | 0.26 | 0.29 |
| 3 | 29498643 | rs7648007 | 0.544 | -0.085 | 0.42 | 0.714 | 0.011 | 0.172 | 0.172 | 0 | 0.09 | 0.27 | 0.17 | 0.27 | 0.06 | 0.07 |
| 3 | 29499253 | rs9310900 | 1.641 | -0.079 | 0.426 | 0.674 | 0.011 | 0.002 | 0.002 | 0.108 | 0.23 | 0.31 | 0.44 | 0.22 | 0.28 | 0.23 |
| 3 | 29499390 | rs9942107 | 1.804 | -0.093 | 0.431 | 0.675 | 0.007 | 0.001 | 0.001 | 0.32 | 0.22 | 0.31 | 0.34 | 0.2 | 0.25 | 0.22 |
| 3 | 29501137 | rs9853435 | 1.836 | -0.107 | 0.408 | 0.629 | -0.001 | 0.003 | 0.003 | 0 | 0.11 | 0.12 | 0.09 | 0.09 | 0.13 | 0.07 |
| 3 | 29502389 | rs9310901 | 0.522 | 0.203 | 0.482 | 0.631 | 0.008 | 0.035 | 0.035 | 1.04 | 0.33 | 0.4 | 0.53 | 0.46 | 0.37 | 0.32 |
| 3 | 29506471 | rs1438328 | 3.016 | 0.403 | 0.059 | 0.69 | -0.028 | -0.036 | -0.036 | 0 | 0.06 | 0.01 | 0.07 | 0.14 | 0.1 | 0.13 |
| 3 | 29510365 | rs9855960 | 1.111 | 0.368 | 0.29 | 0.672 | 0 | 0.017 | 0.017 | 2.703 | 0.3 | 0.3 | 0.42 | 0.45 | 0.32 | 0.28 |
| 3 | 29516639 | rs10510621 | 2.318 | 0.515 | 0.155 | 0.637 | 0.009 | 0.026 | 0.026 | 2.681 | 0.08 | 0.11 | 0.1 | 0.14 | 0.12 | 0.14 |
| 3 | 29517046 | rs7629495 | 2.051 | 0.22 | 0.24 | 0.647 | -0.009 | -0.002 | -0.002 | 0 | 0.21 | 0.19 | 0.44 | 0.22 | 0.3 | 0.23 |
| 3 | 29518724 | rs1866298 | -2.014 | 0.334 | -0.159 | 0.633 | -0.024 | 0.013 | 0.013 | 0.013 | 0.9 | 0.98 | 0.84 | 0.9 | 0.89 | 0.91 |
| 3 | 29520501 | rs1347772 | -0.921 | 0.465 | 0.17 | 0.71 | 0.007 | 0.042 | 0.042 | 0.741 | 0.55 | 0.71 | 0.72 | 0.7 | 0.59 | 0.51 |
| 3 | 29526264 | rs9837287 | -0.396 | -0.052 | 0.363 | 0.681 | -0.015 | 0.021 | 0.021 | 0 | 0.19 | 0.42 | 0.22 | 0.22 | 0.14 | 0.23 |
| 3 | 29529846 | rs3773020 | 0.581 | -0.255 | 0.359 | 0.709 | 0.02 | 0.026 | 0.026 | 0 | 0.05 | 0.19 | 0.12 | 0.05 | 0.03 | 0.05 |
| 3 | 29529907 | rs2371695 | -0.5 | -0.158 | 0.237 | 0.556 | 0.003 | -0.002 | -0.002 | 0.329 | 0.53 | 0.6 | 0.6 | 0.46 | 0.58 | 0.47 |
| 3 | 29530984 | rs3773025 | -0.13 | -2.769 | 0.724 | 0.614 | 0.016 | 0.006 | 0.006 | 0 | 0.05 | 0.19 | 0.12 | 0.03 | 0.03 | 0.04 |
| 3 | 29531222 | rs9815793 | -0.486 | -2.813 | 0.68 | 0.582 | -0.004 | -0.001 | -0.001 | 0 | 0.42 | 0.57 | 0.51 | 0.4 | 0.39 | 0.42 |
| 3 | 29531977 | rs3773026 | -0.151 | -2.702 | 0.58 | 0.475 | 0.016 | 0.006 | 0.006 | 0 | 0.05 | 0.2 | 0.12 | 0.03 | 0.03 | 0.04 |
| 3 | 29532038 | rs3773027 | -0.438 | -2.737 | 0.609 | 0.477 | -0.031 | 0.015 | 0.015 | 0 | 0.18 | 0.36 | 0.21 | 0.19 | 0.11 | 0.2 |
| 3 | 29534726 | rs12489361 | -0.156 | -2.723 | 0.328 | 0.553 | -0.008 | 0.03 | 0.03 | 0.312 | 0.36 | 0.5 | 0.56 | 0.5 | 0.37 | 0.37 |
| 3 | 29535987 | rs17023730 | 0.773 | -2.755 | 0.172 | 0.565 | 0.014 | -0.012 | -0.012 | 0 | 0.12 | 0.08 | 0.09 | 0.17 | 0.08 | 0.16 |
| 3 | 29538444 | rs35879 | -0.993 | -2.863 | 0.21 | 0.612 | 0.03 | 0.053 | 0.053 | 1.473 | 0.48 | 0.68 | 0.66 | 0.67 | 0.44 | 0.53 |
| 3 | 29540243 | rs9835577 | -0.536 | -2.573 | 0.963 | 0.863 | -0.039 | 0.004 | 0.004 | 0 | 0.16 | 0.29 | 0.11 | 0.17 | 0.08 | 0.19 |
| 3 | 29551939 | rs9867172 | 0.008 | -1.677 | 0.852 | 0.8 | 0.001 | 0.165 | 0.165 | 0 | 0.06 | 0.1 | 0.14 | 0.37 | 0.05 | 0.18 |
| 3 | 29554121 | rs3773050 | -0.823 | -1.773 | 1.085 | 0.741 | -0.003 | 0.247 | 0.247 | 0.003 | 0.13 | 0.27 | 0.21 | 0.47 | 0.1 | 0.28 |
| 3 | 29561962 | rs13081014 | -0.653 | -1.756 | 1.017 | 0.744 | -0.003 | 0.202 | 0.202 | 0 | 0.12 | 0.23 | 0.2 | 0.47 | 0.1 | 0.28 |
| 3 | 29573549 | rs35911 | -0.415 | -2.205 | 0.593 | 0.18 | -0.027 | 0.105 | 0.105 | 0.039 | 0.57 | 0.8 | 0.42 | 0.22 | 0.52 | 0.41 |
| 3 | 29576965 | rs9879838 | -1.116 | -2.344 | 1.477 | 0.367 | -0.037 | -0.094 | -0.094 | 0 | 0.14 | 0.5 | 0.13 | 0.01 | 0.11 | 0.14 |
| 3 | 29582808 | rs9883294 | -1.341 | -2.493 | 1.518 | 0.501 | -0.077 | 0.049 | 0.049 | 0.444 | 0.26 | 0.57 | 0.2 | 0.38 | 0.16 | 0.35 |
| 3 | 29594410 | rs35892 | 0.775 | -2.133 | 1.348 | 0.236 | -0.01 | 0.038 | 0.038 | 0 | 0.42 | 0.33 | 0.28 | 0.25 | 0.41 | 0.25 |
| 3 | 29597791 | rs35883 | -0.435 | -2.196 | 1.411 | 0.203 | -0.016 | 0.045 | 0.045 | 0 | 0.59 | 0.69 | 0.42 | 0.26 | 0.53 | 0.4 |
| 3 | 29598605 | rs7646981 | -0.717 | -2.113 | 0.939 | 0.258 | -0.051 | -0.096 | -0.096 | 0 | 0.17 | 0.34 | 0.14 | 0.01 | 0.12 | 0.14 |
| 3 | 29602141 | rs13070499 | -1.449 | -2.169 | 0.976 | 0.301 | -0.069 | -0.069 | -0.069 | 0.008 | 0.25 | 0.46 | 0.2 | 0.08 | 0.18 | 0.28 |
| 3 | 29605071 | rs7652067 | -1.443 | -2.006 | 0.804 | 0.5 | 0.016 | 0.293 | 0.293 | 0.985 | 0.1 | 0.42 | 0.18 | 0.36 | 0.08 | 0.19 |
| 3 | 29606334 | rs3773084 | -0.417 | -2.008 | 0.762 | 0.482 | -0.009 | -0.001 | -0.001 | 0 | 0.1 | 0.14 | 0.07 | 0.07 | 0.09 | 0.15 |
| 3 | 29607291 | rs3773086 | 0.368 | -2.001 | 0.707 | 0.454 | 0.002 | -0.009 | -0.009 | 0 | 0.11 | 0.1 | 0.11 | 0.01 | 0.09 | 0.13 |
| 3 | 29609040 | rs3773087 | -1.155 | -1.994 | 0.752 | 0.502 | -0.032 | 0.034 | 0.034 | 0 | 0.23 | 0.33 | 0.23 | 0.4 | 0.18 | 0.43 |
| 3 | 29620115 | rs3821575 | -1.575 | -1.976 | 0.746 | 0.466 | -0.029 | 0.028 | 0.028 | 0.278 | 0.29 | 0.59 | 0.35 | 0.45 | 0.23 | 0.45 |

| | | | | | | | | | | | | | | | | |
|---|---|---|---|---|---|---|---|---|---|---|---|---|---|---|---|---|
| 3 | 29620604 | rs13323338 | -1.121 | -1.716 | 0.758 | 0.344 | -0.138 | -0.129 | -0.129 | 0.638 | 0.73 | 0.98 | 0.64 | 0.68 | 0.64 | 0.7 |
| 3 | 29624609 | rs9881271 | -1.901 | -1.985 | 1.554 | 0.309 | -0.033 | 0.062 | 0.062 | 0 | 0.21 | 0.39 | 0.28 | 0.37 | 0.17 | 0.31 |
| 3 | 29629015 | rs9847589 | -1.102 | -1.974 | 1.342 | 0.336 | -0.012 | 0.009 | 0.009 | 0.054 | 0.17 | 0.19 | 0.17 | 0.33 | 0.13 | 0.29 |
| 3 | 29629048 | rs9851413 | -2.505 | -2.005 | 1.357 | 0.272 | -0.028 | 0.013 | 0.013 | 0 | 0.29 | 0.43 | 0.32 | 0.38 | 0.23 | 0.34 |
| 3 | 29633157 | rs17605724 | -1.232 | -1.998 | 1.342 | 0.267 | -0.004 | -0.033 | -0.033 | 0 | 0.09 | 0.13 | 0.03 | 0 | 0.07 | 0.04 |
| 3 | 29638641 | rs2034911 | -1.034 | -2.001 | 1.286 | 0.273 | -0.012 | 0.01 | 0.01 | 0.041 | 0.16 | 0.19 | 0.16 | 0.33 | 0.13 | 0.29 |
| 3 | 29639717 | rs9819610 | -1.035 | -2.003 | 1.285 | 0.272 | -0.012 | 0.01 | 0.01 | 0 | 0.16 | 0.19 | 0.16 | 0.33 | 0.12 | 0.29 |
| 3 | 29639871 | rs17546842 | -0.944 | -1.951 | 1.171 | 0.278 | 0 | -0.008 | -0.008 | 0 | 0.08 | 0.06 | 0.03 | 0 | 0.07 | 0.04 |
| 3 | 29650850 | rs12487575 | -2.543 | -1.969 | 1.242 | 0.239 | -0.028 | 0.007 | 0.007 | 0.006 | 0.3 | 0.54 | 0.31 | 0.38 | 0.23 | 0.34 |
| 3 | 29650928 | rs7638763 | -1.675 | -1.852 | 0.774 | 0.293 | -0.038 | 0.078 | 0.078 | 0 | 0.21 | 0.45 | 0.28 | 0.38 | 0.16 | 0.3 |
| 3 | 29656687 | rs7647230 | -2.253 | -1.689 | 0.121 | 0.338 | -0.032 | 0.008 | 0.008 | 0.136 | 0.29 | 0.41 | 0.25 | 0.37 | 0.23 | 0.32 |
| 3 | 29665461 | rs9829357 | -1.803 | -1.603 | -0.21 | 0.338 | -0.038 | -0.009 | -0.009 | 0 | 0.36 | 0.46 | 0.3 | 0.37 | 0.24 | 0.36 |
| 3 | 29676090 | rs1440493 | -1.291 | -1.215 | -0.546 | 0.433 | 0.009 | -0.028 | -0.028 | 0.014 | 0.27 | 0.27 | 0.19 | 0 | 0.2 | 0.17 |
| 3 | 29677273 | rs11721004 | -1.459 | -1.21 | -0.547 | 0.435 | 0.008 | -0.007 | -0.007 | 2.254 | 0.28 | 0.27 | 0.22 | 0.02 | 0.24 | 0.18 |
| 3 | 29683162 | rs9868264 | -0.869 | -1.254 | -0.521 | 0.506 | -0.016 | 0.183 | 0.183 | 0 | 0.06 | 0.12 | 0.11 | 0.37 | 0.03 | 0.17 |
| 3 | 29685871 | rs17613797 | -0.59 | -1.249 | -0.532 | 0.196 | 0.007 | -0.02 | -0.02 | 0.1 | 0.2 | 0.14 | 0.15 | 0 | 0.13 | 0.15 |
| 3 | 29692699 | rs9831085 | -1.714 | -1.264 | -0.569 | 0.234 | -0.016 | -0.006 | -0.006 | 0 | 0.35 | 0.4 | 0.33 | 0.37 | 0.26 | 0.36 |
| 3 | 29695588 | rs10510622 | -1.399 | -1.195 | -0.534 | 0.658 | -0.023 | 0.008 | 0.008 | 0 | 0.27 | 0.32 | 0.29 | 0.37 | 0.19 | 0.32 |
| 3 | 29700225 | rs9882829 | -0.936 | -1.144 | -0.647 | 0.235 | -0.009 | -0.005 | -0.005 | 0 | 0.23 | 0.18 | 0.18 | 0.21 | 0.16 | 0.29 |
| 3 | 29703236 | rs2888127 | -1.368 | -0.329 | -1.065 | 0.122 | -0.004 | 0.003 | 0.003 | 0 | 0.05 | 0.11 | 0.11 | 0.05 | 0.04 | 0.02 |
| 3 | 29707139 | rs7624637 | -0.89 | -0.911 | -1.438 | -0.28 | 0.052 | 0.071 | 0.071 | 0.379 | 0.24 | 0.09 | 0.23 | 0.03 | 0.15 | 0.13 |
| 3 | 29707569 | rs9845179 | -0.671 | -0.621 | -1.669 | 0.072 | -0.002 | -0.025 | -0.025 | 0 | 0.12 | 0.03 | 0.12 | 0.19 | 0.14 | 0.17 |
| 3 | 29707927 | rs9865546 | -0.803 | -0.517 | -1.646 | -0.59 | 0.012 | 0.022 | 0.022 | 0 | 0.22 | 0.14 | 0.17 | 0.01 | 0.16 | 0.14 |
| 3 | 29710221 | rs7622316 | -1.098 | -0.829 | -0.593 | -0.577 | -0.005 | 0.001 | 0.001 | 0.781 | 0.07 | 0.36 | 0.09 | 0.03 | 0.05 | 0.01 |
| 3 | 29710713 | rs7634507 | -0.444 | 0.256 | -0.736 | -0.876 | -0.008 | 0.004 | 0.004 | 0.27 | 0.81 | 0.87 | 0.68 | 0.81 | 0.65 | 0.72 |
| 3 | 29725062 | rs7647724 | -0.39 | -0.873 | -1.14 | -0.405 | -0.016 | -0.019 | -0.019 | 0 | 0.64 | 0.76 | 0.51 | 0.53 | 0.57 | 0.61 |
| 3 | 29728504 | rs1440501 | -0.421 | -0.891 | -0.671 | -0.402 | 0.002 | 0.001 | 0.001 | 0.391 | 0.64 | 0.69 | 0.5 | 0.54 | 0.56 | 0.63 |
| 3 | 29734821 | rs7626062 | 0.365 | -1.493 | -0.275 | -0.49 | -0.004 | 0.004 | 0.004 | 0 | 0.51 | 0.25 | 0.37 | 0.47 | 0.5 | 0.57 |
| 3 | 29735363 | rs9833942 | -0.501 | -1.45 | -0.15 | -0.48 | 0.011 | 0.011 | 0.011 | 0.286 | 0.63 | 0.64 | 0.5 | 0.53 | 0.57 | 0.59 |
| 3 | 29741400 | rs7620682 | -0.601 | -1.435 | -0.022 | -0.461 | -0.04 | -0.025 | -0.025 | 2.231 | 0.1 | 0.37 | 0.1 | 0.03 | 0.05 | 0.01 |
| 3 | 29745863 | rs9310908 | -0.558 | -1.465 | 0.129 | -0.484 | -0.038 | -0.026 | -0.026 | 0 | 0.1 | 0.39 | 0.11 | 0.03 | 0.05 | 0.01 |
| 3 | 29747197 | rs6782005 | -0.667 | -1.491 | 0.19 | -0.495 | -0.036 | -0.024 | -0.024 | 0 | 0.1 | 0.47 | 0.11 | 0.03 | 0.05 | 0.01 |
| 3 | 29748169 | rs12638297 | 1.234 | -0.907 | -0.101 | -0.583 | -0.009 | -0.014 | -0.014 | 0.395 | 0.11 | 0.03 | 0.19 | 0.16 | 0.15 | 0.16 |
| 3 | 29753431 | rs7427355 | -0.691 | -0.824 | 0.287 | -0.664 | -0.029 | -0.023 | -0.023 | 0 | 0.1 | 0.36 | 0.11 | 0.03 | 0.05 | 0.01 |
| 3 | 29761506 | rs1371844 | 0.693 | -1.048 | 0.219 | -1.003 | 0.023 | 0.047 | 0.047 | 0 | 0.28 | 0.38 | 0.21 | 0.42 | 0.29 | 0.41 |
| 3 | 29766410 | rs7650350 | 0.085 | -1.42 | 0.032 | -1.234 | 0.031 | 0.028 | 0.028 | 0.008 | 0.28 | 0.18 | 0.2 | 0.05 | 0.17 | 0.13 |
| 3 | 29772094 | rs9816394 | 0.536 | -1.383 | -0.149 | -1.218 | 0.047 | 0.06 | 0.06 | 0.96 | 0.23 | 0.11 | 0.14 | 0.01 | 0.12 | 0.12 |
| 3 | 29777050 | rs718634 | -0.759 | -1.086 | -0.103 | -1.036 | -0.025 | -0.022 | -0.022 | 0 | 0.63 | 0.74 | 0.45 | 0.46 | 0.52 | 0.57 |
| 3 | 29778243 | rs1866773 | 0.298 | -1.1 | -0.417 | -1.075 | -0.015 | -0.007 | -0.007 | 0 | 0.43 | 0.26 | 0.3 | 0.43 | 0.42 | 0.53 |
| 3 | 29784510 | rs1992352 | -0.811 | 0.04 | -0.283 | -1.307 | 0.054 | -0.015 | -0.015 | 0 | 0.94 | 0.91 | 0.89 | 0.98 | 0.84 | 0.93 |
| 3 | 29789353 | rs1371835 | -0.679 | -0.587 | -0.638 | -1.245 | 0 | 0.05 | 0.05 | 0.093 | 0.59 | 0.71 | 0.36 | 0.22 | 0.43 | 0.44 |
| 3 | 29790651 | rs6767352 | -0.611 | -0.491 | -0.526 | -1.171 | 0.027 | 0.079 | 0.079 | 0 | 0.53 | 0.7 | 0.39 | 0.24 | 0.41 | 0.43 |
| 3 | 29800913 | rs3773108 | -0.684 | -0.235 | -0.482 | -0.825 | 0.012 | 0.048 | 0.048 | 0 | 0.52 | 0.66 | 0.43 | 0.33 | 0.43 | 0.48 |
| 3 | 29808408 | rs7637304 | -0.063 | 0.342 | -0.729 | -0.325 | 0.001 | 0.002 | 0.002 | 0 | 0.27 | 0.24 | 0.22 | 0.21 | 0.23 | 0.27 |
| 3 | 29809958 | rs2083488 | -0.089 | 0.37 | -0.844 | -0.391 | 0.003 | 0.011 | 0.011 | 0 | 0.28 | 0.19 | 0.17 | 0.2 | 0.23 | 0.27 |
| 3 | 29817658 | rs2371816 | -0.291 | 0.607 | -0.465 | -0.662 | -0.001 | 0.021 | 0.021 | 0.608 | 0.44 | 0.42 | 0.56 | 0.8 | 0.55 | 0.55 |
| 3 | 29820062 | rs9817585 | -0.785 | 0.452 | -0.537 | -1.109 | 0.055 | 0.119 | 0.119 | 1.738 | 0.58 | 0.8 | 0.72 | 0.88 | 0.73 | 0.65 |
| 3 | 29831996 | rs7652078 | -0.831 | 1.206 | 0.38 | -0.287 | 0.016 | 0.073 | 0.073 | 0 | 0.08 | 0.17 | 0.11 | 0.18 | 0.08 | 0.09 |
| 3 | 29832462 | rs7640634 | -0.931 | 0.954 | 0.803 | -0.5 | 0.016 | -0.062 | -0.062 | 0 | 0.16 | 0.32 | 0.14 | 0.03 | 0.17 | 0.09 |
| 3 | 29835798 | rs7650407 | -1.466 | 0.94 | 0.44 | -0.472 | 0.011 | -0.009 | -0.009 | 0.068 | 0.23 | 0.46 | 0.24 | 0.21 | 0.25 | 0.19 |
| 3 | 29841445 | rs6784690 | -1.435 | 1.312 | 0.097 | -0.79 | -0.007 | 0.056 | 0.056 | 0 | 0.41 | 0.66 | 0.5 | 0.23 | 0.58 | 0.3 |
| 3 | 29843576 | rs10510627 | -1.231 | 1.551 | 0.594 | -0.886 | 0.059 | -0.028 | -0.028 | 0 | 0.3 | 0.46 | 0.38 | 0.22 | 0.45 | 0.27 |
| 3 | 29850829 | rs2371831 | -1.627 | 1.653 | 0.856 | -1.025 | 0.01 | -0.05 | -0.05 | 0 | 0.36 | 0.57 | 0.44 | 0.09 | 0.52 | 0.22 |
| 3 | 29853403 | rs10510628 | -1.154 | 1.893 | 0.921 | -1.025 | 0.025 | -0.032 | -0.032 | 4.083 | 0.24 | 0.24 | 0.33 | 0.06 | 0.41 | 0.17 |
| 3 | 29859930 | rs6790352 | -0.512 | 1.881 | 0.862 | -1.003 | 0.004 | -0.037 | -0.037 | 0 | 0.12 | 0.13 | 0.1 | 0 | 0.13 | 0.06 |
| 3 | 29860690 | rs13068149 | 0.277 | 1.953 | 0.631 | -0.976 | -0.059 | 0.049 | 0.049 | 0 | 0.14 | 0.04 | 0.24 | 0.03 | 0.32 | 0.08 |
| 3 | 29863163 | rs4680853 | -0.922 | 1.571 | 0.695 | -0.892 | 0.061 | -0.091 | -0.091 | 0 | 0.3 | 0.4 | 0.37 | 0.06 | 0.46 | 0.21 |
| 3 | 29867910 | rs9837143 | -1.646 | 1.523 | 0.739 | -0.911 | 0.011 | -0.033 | -0.033 | 0 | 0.38 | 0.52 | 0.47 | 0.25 | 0.53 | 0.28 |
| 3 | 29876233 | rs10446455 | -1.141 | 1.272 | 0.382 | -0.339 | 0 | 0.001 | 0.001 | 0 | 0.08 | 0.11 | 0.13 | 0.18 | 0.1 | 0.05 |
| 3 | 29882996 | rs7613154 | -0.869 | 1.435 | 0.212 | -0.551 | 0.001 | -0.008 | -0.008 | 0 | 0.39 | 0.58 | 0.43 | 0.28 | 0.59 | 0.44 |
| 3 | 29888528 | rs4680860 | -0.896 | 1.197 | 0.333 | -0.685 | -0.003 | -0.005 | -0.005 | 0 | 0.42 | 0.44 | 0.44 | 0.28 | 0.63 | 0.49 |
| 3 | 29891801 | rs7633961 | -0.757 | 2.043 | 0.809 | -0.319 | 0.102 | 0.047 | 0.047 | 0 | 0.21 | 0.58 | 0.33 | 0.26 | 0.39 | 0.29 |
| 3 | 29892306 | rs12494881 | -0.37 | 2.312 | 1.053 | -0.467 | 0.081 | -0.069 | -0.069 | 1.826 | 0.16 | 0.27 | 0.23 | 0.01 | 0.32 | 0.11 |
| 3 | 29901874 | rs17024502 | -0.069 | 1.31 | 1.179 | -0.054 | -0.01 | 0.023 | 0.023 | 0 | 0.11 | 0.16 | 0.12 | 0.15 | 0.11 | 0.13 |
| 3 | 29907310 | rs12490863 | -1.145 | 1.23 | 1.669 | 0.076 | -0.026 | 0.042 | 0.042 | 0 | 0.15 | 0.62 | 0.25 | 0.18 | 0.12 | 0.17 |
| 3 | 29907479 | rs4680733 | 0.477 | 1.513 | 0.752 | 0.668 | -0.032 | -0.007 | -0.007 | 4.936 | 0.06 | 0.01 | 0.17 | 0.36 | 0.11 | 0.11 |
| 3 | 29915860 | rs3773112 | -0.672 | 0.09 | 1.005 | -0.67 | -0.046 | -0.045 | -0.045 | 0.01 | 0.27 | 0.57 | 0.17 | 0.1 | 0.16 | 0.24 |
| 3 | 29918453 | rs7651150 | -0.774 | 0.096 | 0.968 | -0.656 | -0.009 | -0.006 | -0.006 | 0 | 0.27 | 0.67 | 0.26 | 0.16 | 0.18 | 0.28 |
| 3 | 29922333 | rs13324314 | 0.665 | 0.577 | 0.703 | -1.012 | 0.025 | -0.042 | -0.042 | 0.447 | 0.17 | 0.26 | 0.25 | 0.07 | 0.25 | 0.28 |

| | | | | | | | | | | | | | | | | |
|---|---|---|---|---|---|---|---|---|---|---|---|---|---|---|---|---|
| 3 | 29925518 | rs2218724 | -0.893 | 0.409 | 0.649 | -1.144 | 0.01 | -0.032 | -0.032 | 0.891 | 0.3 | 0.51 | 0.31 | 0.2 | 0.32 | 0.36 |
| 3 | 29928319 | rs9821576 | 0.686 | 0.487 | 0.526 | -1.113 | 0.022 | -0.043 | -0.043 | 0 | 0.18 | 0.26 | 0.26 | 0.07 | 0.25 | 0.29 |
| 3 | 29929324 | rs9826958 | 0.707 | 0.489 | 0.506 | -1.105 | 0.023 | -0.028 | -0.028 | 0 | 0.18 | 0.27 | 0.25 | 0.09 | 0.25 | 0.3 |
| 3 | 29931885 | rs1449292 | -0.962 | -0.717 | 0.099 | -1.741 | -0.007 | 0.043 | 0.043 | 0 | 0.58 | 0.66 | 0.4 | 0.18 | 0.46 | 0.51 |
| 3 | 29933717 | rs11916764 | 0.499 | -0.037 | 0.381 | -0.899 | 0.019 | -0.012 | -0.012 | 0 | 0.18 | 0.28 | 0.26 | 0.13 | 0.25 | 0.29 |
| 3 | 29937979 | rs17553785 | 1.51 | 0.352 | 0.319 | -1.124 | -0.022 | 0.028 | 0.028 | 0.024 | 0.05 | 0.01 | 0.03 | 0 | 0.07 | 0.07 |
| 3 | 29940074 | rs3773120 | 0.561 | -0.082 | 0.683 | -0.768 | 0.022 | -0.006 | -0.006 | 0 | 0.17 | 0.28 | 0.18 | 0.13 | 0.25 | 0.3 |
| 3 | 29954551 | rs9875937 | -0.349 | -0.877 | 0.4 | -0.376 | 0.006 | 0.008 | 0.008 | 0 | 0.09 | 0.1 | 0.03 | 0.04 | 0.05 | 0.04 |
| 3 | 29954690 | rs17024608 | 0.338 | -0.992 | 0.242 | -0.436 | 0.012 | 0.042 | 0.042 | 0.564 | 0.15 | 0.07 | 0.08 | 0 | 0.1 | 0.1 |
| 3 | 29956902 | rs6776043 | -0.876 | -0.426 | 0.929 | 0.43 | -0.02 | -0.038 | -0.038 | 0 | 0.1 | 0.28 | 0.04 | 0.04 | 0.04 | 0.04 |
| 3 | 29964567 | rs1868488 | 0.571 | -0.105 | 0.45 | 0.668 | -0.001 | -0.03 | -0.03 | 0 | 0.87 | 0.84 | 0.85 | 0.99 | 0.88 | 0.87 |
| 3 | 29968278 | rs6798671 | -0.124 | -0.032 | 0.014 | 0.993 | -0.009 | -0.008 | -0.008 | 0.231 | 0.16 | 0.08 | 0.2 | 0.22 | 0.2 | 0.2 |
| 3 | 29970919 | rs1449286 | 1.126 | -0.143 | 0.051 | 1.141 | 0.007 | -0.016 | -0.016 | 0 | 0.65 | 0.21 | 0.46 | 0.4 | 0.66 | 0.59 |
| 3 | 29982654 | rs9858736 | 0.757 | -0.019 | 0.547 | 0.77 | -0.005 | -0.065 | -0.065 | 0.681 | 0.84 | 0.63 | 0.85 | 0.95 | 0.9 | 0.85 |
| 3 | 29992733 | rs11717088 | -1.326 | -0.274 | 0.446 | 1.557 | -0.03 | 0.134 | 0.134 | 0.207 | 0.13 | 0.28 | 0.21 | 0.32 | 0.06 | 0.27 |
| 3 | 30011942 | rs1449281 | -0.738 | -0.427 | 0.556 | 1.506 | 0.003 | 0.026 | 0.026 | 0.346 | 0.22 | 0.28 | 0.24 | 0.26 | 0.23 | 0.11 |
| 3 | 30026811 | rs6792333 | -0.65 | -0.54 | 0.381 | 1.888 | -0.003 | 0.006 | 0.006 | 0.508 | 0.2 | 0.05 | 0.24 | 0.17 | 0.2 | 0.11 |
| 3 | 30029724 | rs1868491 | -1.111 | -0.739 | 0.591 | 1.919 | -0.003 | -0.023 | -0.023 | 0.04 | 0.21 | 0.15 | 0.33 | 0.35 | 0.24 | 0.19 |
| 3 | 30031420 | rs7647262 | -0.252 | -0.739 | 0.573 | 1.901 | -0.018 | -0.016 | -0.016 | 0 | 0.16 | 0.04 | 0.24 | 0.21 | 0.2 | 0.11 |
| 3 | 30032355 | rs3773124 | -1.699 | -0.858 | 0.781 | 1.845 | 0.005 | -0.015 | -0.015 | 0 | 0.26 | 0.23 | 0.34 | 0.38 | 0.25 | 0.21 |
| 3 | 30033043 | rs3773127 | -0.997 | -0.851 | 0.74 | 1.845 | -0.013 | -0.063 | -0.063 | 0 | 0.2 | 0.06 | 0.27 | 0.36 | 0.21 | 0.12 |
| 3 | 30044778 | rs6549963 | 0.886 | -0.728 | -0.419 | 1.366 | -0.105 | 0.027 | 0.027 | 0 | 0.33 | 0.06 | 0.21 | 0.21 | 0.41 | 0.26 |
| 3 | 30046191 | rs6549965 | -0.109 | -0.586 | -0.051 | 1.504 | 0.001 | 0.017 | 0.017 | 0 | 0.19 | 0.15 | 0.28 | 0.36 | 0.22 | 0.13 |
| 3 | 30049417 | rs3811723 | -0.635 | -0.668 | 0.18 | 1.428 | -0.008 | -0.009 | -0.009 | 0 | 0.23 | 0.25 | 0.27 | 0.21 | 0.21 | 0.11 |
| 3 | 30052399 | rs1857883 | -0.785 | -0.597 | 0.22 | 1.208 | -0.002 | -0.001 | -0.001 | 2.029 | 0.46 | 0.52 | 0.61 | 0.56 | 0.54 | 0.5 |
| 3 | 30059835 | rs6798324 | 0.923 | -0.598 | 0.174 | 1.164 | -0.016 | -0.015 | -0.015 | 0 | 0.15 | 0.05 | 0.23 | 0.2 | 0.19 | 0.08 |
| 3 | 30059851 | rs4680867 | 1.896 | -0.885 | -0.401 | 0.608 | -0.031 | -0.052 | -0.052 | 0 | 0.15 | 0.03 | 0.22 | 0.26 | 0.2 | 0.22 |
| 3 | 30059891 | rs1911070 | 0.971 | -0.807 | -0.032 | 0.744 | -0.013 | 0.01 | 0.01 | 0 | 0.23 | 0.17 | 0.26 | 0.19 | 0.31 | 0.3 |
| 3 | 30063238 | rs2062992 | -0.614 | -0.763 | 0.011 | 0.801 | 0.004 | -0.008 | -0.008 | 0.881 | 0.44 | 0.46 | 0.56 | 0.42 | 0.53 | 0.46 |
| 3 | 30064457 | rs1506283 | 1.075 | -0.809 | -0.025 | 0.818 | -0.016 | -0.017 | -0.017 | 0 | 0.15 | 0.05 | 0.23 | 0.2 | 0.19 | 0.08 |
| 3 | 30064795 | rs9847686 | -0.541 | -0.766 | 0.012 | 0.836 | -0.002 | -0.001 | -0.001 | 2.233 | 0.46 | 0.53 | 0.6 | 0.57 | 0.55 | 0.47 |
| 3 | 30066317 | rs2174583 | 0.257 | -0.758 | 0.012 | 0.802 | 0.015 | -0.04 | -0.04 | 2.302 | 0.29 | 0.42 | 0.34 | 0.21 | 0.34 | 0.38 |
| 3 | 30069870 | rs9854414 | -0.599 | -0.868 | -0.019 | 0.771 | -0.004 | 0.006 | 0.006 | 0 | 0.45 | 0.39 | 0.56 | 0.41 | 0.54 | 0.46 |
| 3 | 30074874 | rs4426642 | -0.4 | -0.847 | 0.008 | 0.619 | 0.01 | 0 | 0 | 0 | 0.4 | 0.47 | 0.48 | 0.4 | 0.56 | 0.33 |
| 3 | 30077482 | rs11706573 | 0.119 | -0.717 | -0.241 | 0.797 | 0.003 | 0.027 | 0.027 | 0 | 0.18 | 0.18 | 0.27 | 0.36 | 0.22 | 0.1 |
| 3 | 30086319 | rs6549972 | 0.334 | -0.888 | -0.562 | 0.446 | -0.033 | 0.048 | 0.048 | 5.574 | 0.19 | 0.06 | 0.15 | 0.08 | 0.27 | 0.09 |
| 3 | 30090772 | rs1506280 | 0.748 | -0.917 | -0.363 | 0.589 | -0.003 | 0 | 0 | 0 | 0.23 | 0.21 | 0.22 | 0.2 | 0.35 | 0.22 |
| 3 | 30109355 | rs950752 | 0.055 | -1.047 | 0.1 | 0.314 | 0.002 | 0.005 | 0.005 | 3.659 | 0.23 | 0.26 | 0.3 | 0.35 | 0.28 | 0.12 |
| 3 | 30111359 | rs7649490 | -0.464 | -1.305 | 0.559 | 0.405 | 0.04 | 0.003 | 0.003 | 0.273 | 0.48 | 0.7 | 0.54 | 0.45 | 0.63 | 0.31 |
| 3 | 30114498 | rs17024761 | 0.54 | -1.336 | 0.422 | -0.061 | 0 | -0.01 | -0.01 | 0.212 | 0.1 | 0.08 | 0.04 | 0.01 | 0.05 | 0.08 |
| 3 | 30118503 | rs1506292 | 1.502 | -1.223 | -0.051 | -0.029 | -0.09 | 0 | 0 | 3.555 | 0.18 | 0.04 | 0.25 | 0.14 | 0.34 | 0.17 |
| 3 | 30118718 | rs7649308 | -0.208 | -1.44 | 0.064 | -0.032 | 0 | 0 | 0 | 0 | 0.34 | 0.65 | 0.47 | 0.33 | 0.43 | 0.35 |
| 3 | 30118918 | rs7639172 | -0.685 | -1.652 | -0.219 | -0.071 | 0.047 | -0.021 | -0.021 | 0.677 | 0.57 | 0.78 | 0.61 | 0.42 | 0.67 | 0.46 |
| 3 | 30124940 | rs13079874 | -0.565 | -1.496 | -0.097 | -0.235 | 0 | 0.011 | 0.011 | 0 | 0.62 | 0.42 | 0.61 | 0.4 | 0.59 | 0.43 |
| 3 | 30126613 | rs6791084 | 0.232 | -1.354 | -0.175 | -0.03 | 0.021 | 0.015 | 0.015 | 0 | 0.38 | 0.59 | 0.51 | 0.58 | 0.54 | 0.46 |
| 3 | 30129784 | rs1347287 | -0.301 | -0.96 | 0.121 | 0.162 | 0.007 | 0.009 | 0.009 | 0 | 0.39 | 0.2 | 0.42 | 0.29 | 0.34 | 0.3 |
| 3 | 30134493 | rs7611558 | -0.372 | -1.151 | 0.075 | 0.075 | -0.001 | -0.009 | -0.009 | 0 | 0.55 | 0.74 | 0.56 | 0.58 | 0.6 | 0.57 |
| 3 | 30136362 | rs4426643 | -0.242 | -1.164 | 0.302 | -0.001 | 0.002 | 0 | 0 | 0.014 | 0.54 | 0.59 | 0.55 | 0.58 | 0.6 | 0.56 |
| 3 | 30137032 | rs9863443 | 1.61 | -1.152 | 0.311 | 0.019 | 0.016 | 0.048 | 0.048 | 0 | 0.26 | 0.4 | 0.33 | 0.44 | 0.34 | 0.34 |
| 3 | 30138470 | rs13088954 | -0.252 | -1.118 | 0.258 | 0.042 | 0.001 | 0.003 | 0.003 | 0.136 | 0.49 | 0.55 | 0.55 | 0.57 | 0.49 | 0.47 |
| 3 | 30139266 | rs9878212 | 0.233 | -1.088 | 0.294 | 0.051 | -0.008 | 0.001 | 0.001 | 0 | 0.38 | 0.55 | 0.36 | 0.41 | 0.4 | 0.4 |
| 3 | 30152143 | rs1032921 | -0.372 | -0.711 | 0.232 | 0.246 | 0 | 0.002 | 0.002 | 0 | 0.7 | 0.66 | 0.59 | 0.68 | 0.71 | 0.77 |
| 3 | 30153526 | rs11129385 | -2.093 | -0.647 | 0.14 | 0.147 | -0.024 | -0.003 | -0.003 | 0 | 0.85 | 0.95 | 0.8 | 0.85 | 0.8 | 0.85 |
| 3 | 30154862 | rs17024894 | -0.029 | -0.713 | 0.177 | 0.1 | -0.004 | -0.014 | -0.014 | 0 | 0.46 | 0.41 | 0.28 | 0.26 | 0.37 | 0.44 |
| 3 | 30156873 | rs1434009 | -1.975 | -0.448 | -0.201 | 0.169 | -0.014 | 0 | 0 | 0 | 0.85 | 0.94 | 0.82 | 0.86 | 0.8 | 0.84 |
| 3 | 30161722 | rs7619284 | -0.063 | -0.676 | -0.095 | 0.021 | -0.016 | -0.054 | -0.054 | 0 | 0.68 | 0.53 | 0.76 | 0.86 | 0.74 | 0.77 |
| 3 | 30163689 | rs11715853 | 0.509 | -0.724 | -0.154 | 0.109 | 0 | 0.006 | 0.006 | 0 | 0.34 | 0.28 | 0.26 | 0.35 | 0.31 | 0.38 |
| 3 | 30171659 | rs1032918 | -2.036 | -0.652 | -0.144 | -0.284 | -0.01 | 0.005 | 0.005 | 0 | 0.84 | 0.93 | 0.81 | 0.88 | 0.8 | 0.86 |
| 3 | 30172394 | rs4521180 | 0.815 | -0.841 | -0.057 | -0.215 | -0.033 | -0.014 | -0.014 | 0 | 0.15 | 0.21 | 0.05 | 0.05 | 0.06 | 0.09 |
| 3 | 30174833 | rs1434002 | 1.013 | -0.834 | -0.115 | -0.151 | -0.01 | -0.019 | -0.019 | 0 | 0.36 | 0.32 | 0.51 | 0.55 | 0.44 | 0.38 |
| 3 | 30175885 | rs6778417 | 0.797 | -0.806 | -0.157 | -0.151 | -0.002 | 0.003 | 0.003 | 0 | 0.32 | 0.25 | 0.24 | 0.26 | 0.31 | 0.37 |
| 3 | 30176020 | rs12493458 | 0.015 | -0.763 | -0.299 | -0.084 | 0.014 | 0.004 | 0.004 | 0 | 0.25 | 0.21 | 0.25 | 0.14 | 0.15 | 0.17 |
| 3 | 30181698 | rs11708467 | -0.532 | -0.613 | -0.472 | -0.036 | -0.001 | -0.002 | -0.002 | 0.077 | 0.72 | 0.73 | 0.76 | 0.83 | 0.75 | 0.77 |
| 3 | 30182375 | rs13079421 | 1.708 | -0.004 | -0.654 | 0.149 | -0.028 | -0.036 | -0.036 | 0 | 0.1 | 0.01 | 0.2 | 0.1 | 0.09 | 0.06 |
| 3 | 30188747 | rs1434019 | -1.78 | 0.097 | -0.736 | 0.098 | -0.021 | 0.004 | 0.004 | 0 | 0.85 | 0.95 | 0.81 | 0.88 | 0.8 | 0.88 |
| 3 | 30189304 | rs1991902 | 1.097 | 0.638 | -0.516 | 0.305 | -0.006 | -0.012 | -0.012 | 0 | 0.24 | 0.21 | 0.3 | 0.45 | 0.34 | 0.32 |
| 3 | 30192418 | rs4680877 | -0.008 | 0.677 | -0.64 | 0.34 | 0.004 | 0.01 | 0.01 | 0 | 0.37 | 0.47 | 0.52 | 0.56 | 0.43 | 0.37 |
| 3 | 30192906 | rs12629933 | 1.087 | 0.703 | -0.815 | 0.359 | -0.001 | 0 | 0 | 0 | 0.26 | 0.26 | 0.32 | 0.45 | 0.34 | 0.31 |
| 3 | 30193577 | rs4680879 | 0.504 | 0.802 | -1.273 | 0.156 | 0.006 | 0.015 | 0.015 | 0 | 0.39 | 0.53 | 0.52 | 0.56 | 0.44 | 0.38 |

| | | | | | | | | | | | | | | | | |
|---|---|---|---|---|---|---|---|---|---|---|---|---|---|---|---|---|
| 3 | 30197892 | rs7638300 | -0.153 | 0.983 | -1.469 | -0.086 | -0.034 | 0.033 | 0.033 | 0 | 0.17 | 0.07 | 0.16 | 0.1 | 0.24 | 0.16 |
| 3 | 30200658 | rs2371996 | 0.035 | 0.564 | -1.321 | -0.23 | -0.01 | 0.07 | 0.07 | 0 | 0.57 | 0.75 | 0.68 | 0.83 | 0.58 | 0.74 |
| 3 | 30207667 | rs1433986 | -0.575 | 0.052 | -0.585 | 0.551 | -0.01 | 0.026 | 0.026 | 0 | 0.26 | 0.63 | 0.41 | 0.33 | 0.25 | 0.32 |
| 3 | 30208036 | rs7611199 | 0.459 | 0.027 | -0.711 | 0.722 | -0.025 | -0.055 | -0.055 | 0.115 | 0.12 | 0.02 | 0.18 | 0.2 | 0.13 | 0.25 |
| 3 | 30211106 | rs2371997 | -1.095 | -0.432 | -0.7 | 0.391 | 0.01 | -0.005 | -0.005 | 0 | 0.47 | 0.66 | 0.61 | 0.45 | 0.44 | 0.57 |
| 3 | 30211349 | rs17025018 | -0.578 | -0.958 | -0.357 | 0.392 | -0.02 | 0.041 | 0.041 | 0 | 0.26 | 0.52 | 0.29 | 0.32 | 0.23 | 0.34 |
| 3 | 30225323 | rs2196131 | 0.454 | -0.977 | -0.601 | 0.468 | 0 | -0.066 | -0.066 | 0 | 0.17 | 0.07 | 0.2 | 0.32 | 0.16 | 0.33 |
| 3 | 30228083 | rs9846192 | -1.374 | -1.097 | -0.981 | -0.31 | -0.037 | 0.063 | 0.063 | 0 | 0.69 | 0.86 | 0.75 | 0.88 | 0.61 | 0.8 |
| 3 | 30228711 | rs9809165 | 0.778 | -1.071 | -0.76 | -0.073 | -0.015 | 0.022 | 0.022 | 0 | 0.24 | 0.15 | 0.15 | 0.18 | 0.26 | 0.24 |
| 3 | 30231416 | rs1897747 | 0.309 | -0.791 | 0.095 | -0.281 | -0.02 | -0.093 | -0.093 | 1.67 | 0.21 | 0.32 | 0.19 | 0 | 0.16 | 0.15 |
| 3 | 30234616 | rs1433989 | 0.794 | -0.918 | -0.285 | -0.267 | 0.008 | -0.011 | -0.011 | 1.507 | 0.18 | 0.14 | 0.17 | 0 | 0.16 | 0.15 |
| 3 | 30235221 | rs1036308 | -1.068 | -1.161 | -0.043 | -0.254 | -0.02 | 0.025 | 0.025 | 0.017 | 0.7 | 0.78 | 0.75 | 0.88 | 0.63 | 0.81 |
| 3 | 30243401 | rs9831993 | -0.051 | -1.155 | -0.132 | 0.005 | 0.002 | 0.006 | 0.006 | 0 | 0.5 | 0.5 | 0.65 | 0.81 | 0.46 | 0.65 |
| 3 | 30245137 | rs1560848 | -0.051 | -1.155 | -0.132 | 0.005 | 0.002 | 0.006 | 0.006 | 0 | 0.5 | 0.5 | 0.65 | 0.81 | 0.46 | 0.65 |
| 3 | 30246627 | rs755615 | -0.382 | -1.056 | -0.493 | -0.1 | -0.02 | 0.079 | 0.079 | 0 | 0.57 | 0.71 | 0.71 | 0.85 | 0.51 | 0.67 |
| 3 | 30252123 | rs17025096 | 0.936 | -1.042 | -0.56 | 0.232 | -0.006 | -0.029 | -0.029 | 0 | 0.18 | 0.16 | 0.35 | 0.33 | 0.19 | 0.32 |
| 3 | 30252515 | rs9851923 | -1.43 | -1.047 | -0.648 | 0.151 | -0.049 | 0.041 | 0.041 | 0.001 | 0.74 | 0.86 | 0.79 | 0.85 | 0.65 | 0.82 |
| 3 | 30258522 | rs6779483 | 0.422 | -1.473 | -0.161 | 0.504 | -0.017 | 0.026 | 0.026 | 0 | 0.31 | 0.43 | 0.23 | 0.42 | 0.25 | 0.27 |
| 3 | 30259240 | rs6549977 | -0.944 | -1.425 | -0.586 | 0.535 | 0.016 | -0.018 | -0.018 | 1.497 | 0.56 | 0.68 | 0.44 | 0.5 | 0.46 | 0.5 |
| 3 | 30261529 | rs13100297 | -0.834 | -1.158 | -0.423 | 0.417 | 0.038 | -0.006 | -0.006 | 0.984 | 0.52 | 0.62 | 0.41 | 0.5 | 0.42 | 0.49 |
| 3 | 30263217 | rs17025114 | -0.396 | -0.609 | -0.475 | 0.316 | -0.005 | -0.036 | -0.036 | 0 | 0.16 | 0.16 | 0.07 | 0 | 0.14 | 0.15 |
| 3 | 30267853 | rs1368857 | -1.03 | -0.254 | -0.243 | 1.297 | -0.025 | 0.1 | 0.1 | 0 | 0.18 | 0.3 | 0.18 | 0.43 | 0.13 | 0.27 |
| 3 | 30268246 | rs9864657 | -0.392 | 0.458 | -0.706 | 0.594 | -0.01 | 0.033 | 0.033 | 1.234 | 0.38 | 0.73 | 0.38 | 0.23 | 0.42 | 0.28 |
| 3 | 30268632 | rs892414 | -0.814 | 0.983 | -0.445 | 0.371 | 0.001 | -0.008 | -0.008 | 0 | 0.23 | 0.21 | 0.42 | 0.15 | 0.24 | 0.32 |
| 3 | 30268817 | rs1433997 | -1.034 | 0.663 | -0.193 | 0.74 | 0.003 | -0.001 | -0.001 | 0 | 0.3 | 0.35 | 0.47 | 0.26 | 0.29 | 0.36 |
| 3 | 30271646 | rs7644703 | -0.046 | 0.526 | -0.018 | 0.371 | -0.015 | -0.07 | -0.07 | 0.025 | 0.24 | 0.46 | 0.19 | 0.1 | 0.25 | 0.19 |
| 3 | 30271708 | rs7610877 | -0.901 | 0.23 | -0.606 | 0.706 | 0.026 | -0.031 | -0.031 | 0.58 | 0.44 | 0.77 | 0.48 | 0.47 | 0.39 | 0.5 |
| 3 | 30272520 | rs1433994 | -0.858 | 0.248 | -0.56 | 0.715 | 0.021 | -0.018 | -0.018 | 0 | 0.44 | 0.71 | 0.48 | 0.46 | 0.38 | 0.49 |
| 3 | 30275947 | rs1433993 | -0.821 | 0.284 | -0.799 | 0.577 | 0.006 | 0.014 | 0.014 | 0.282 | 0.55 | 0.75 | 0.68 | 0.55 | 0.53 | 0.53 |
| 3 | 30278290 | rs11926889 | -0.45 | -0.445 | -0.014 | 0.27 | -0.036 | -0.096 | -0.096 | 0 | 0.19 | 0.36 | 0.11 | 0.01 | 0.1 | 0.14 |
| 3 | 30285246 | rs6793027 | 0.269 | 0.15 | 0.578 | 1.136 | 0.032 | 0.202 | 0.202 | 0 | 0.2 | 0.51 | 0.19 | 0.49 | 0.24 | 0.22 |
| 3 | 30285495 | rs17564364 | 1.262 | 0.229 | -0.099 | 0.602 | -0.038 | 0.045 | 0.045 | 0 | 0.08 | 0.01 | 0.07 | 0 | 0.13 | 0.02 |
| 3 | 30303774 | rs2372079 | 0.815 | -0.554 | 0.028 | 0.57 | 0.021 | 0.014 | 0.014 | 0 | 0.15 | 0.06 | 0.07 | 0.11 | 0.09 | 0.22 |
| 3 | 30304226 | rs6797604 | -0.291 | -1.04 | 0.234 | 0.189 | -0.001 | 0.008 | 0.008 | 0 | 0.44 | 0.59 | 0.34 | 0.25 | 0.44 | 0.33 |
| 3 | 30307982 | rs9827620 | -0.884 | -1.039 | 0.27 | 0.222 | -0.011 | 0.022 | 0.022 | 0 | 0.57 | 0.65 | 0.41 | 0.36 | 0.53 | 0.56 |
| 3 | 30308287 | rs9869860 | -0.397 | -1.04 | 0.538 | 0.061 | 0.008 | 0.006 | 0.006 | 0 | 0.36 | 0.3 | 0.23 | 0.15 | 0.25 | 0.35 |
| 3 | 30308759 | rs9870410 | 1.751 | -1.085 | 0.52 | -0.032 | -0.005 | -0.035 | -0.035 | 0 | 0.09 | 0.14 | 0.04 | 0 | 0.06 | 0.06 |
| 3 | 30309850 | rs4955324 | -0.652 | -1.123 | 0.359 | 0.125 | 0.001 | 0.001 | 0.001 | 0 | 0.49 | 0.52 | 0.62 | 0.36 | 0.52 | 0.51 |
| 3 | 30309919 | rs4955322 | 0.683 | -1.133 | 0.351 | 0.11 | 0.007 | 0.03 | 0.03 | 0.046 | 0.2 | 0.14 | 0.15 | 0.05 | 0.16 | 0.13 |
| 3 | 30312764 | rs12631406 | 1.57 | -1.094 | 0.3 | 0.142 | 0.024 | -0.002 | -0.002 | 0 | 0.21 | 0.36 | 0.18 | 0.21 | 0.27 | 0.2 |
| 3 | 30313475 | rs4955166 | -0.233 | -1.126 | 0.281 | 0.128 | -0.029 | 0.056 | 0.056 | 0 | 0.28 | 0.13 | 0.44 | 0.15 | 0.25 | 0.29 |
| 3 | 30315877 | rs6549980 | -0.633 | -1.139 | 0.259 | 0.133 | 0.006 | 0.012 | 0.012 | 0 | 0.49 | 0.49 | 0.62 | 0.36 | 0.52 | 0.49 |
| 3 | 30316691 | rs7640965 | -1.773 | -1.125 | 0.174 | 0.441 | -0.006 | 0.01 | 0.01 | 0.069 | 0.67 | 0.84 | 0.73 | 0.72 | 0.62 | 0.75 |
| 3 | 30323253 | rs4955165 | 0.974 | -0.688 | 0.31 | 0.509 | -0.026 | -0.021 | -0.021 | 0 | 0.07 | 0.02 | 0.29 | 0.1 | 0.09 | 0.18 |
| 3 | 30328692 | rs7651821 | 0.051 | -0.767 | 0.387 | 0.535 | 0.004 | -0.008 | -0.008 | 0 | 0.22 | 0.24 | 0.18 | 0.32 | 0.16 | 0.13 |
| 3 | 30329559 | rs7643114 | 0.432 | -0.748 | 0.181 | 0.32 | 0.003 | 0.077 | 0.077 | 0.009 | 0.2 | 0.09 | 0.15 | 0.05 | 0.16 | 0.11 |
| 3 | 30332349 | rs2077164 | -0.681 | -0.677 | 0.394 | 0.537 | -0.026 | 0.009 | 0.009 | 0.992 | 0.43 | 0.79 | 0.55 | 0.58 | 0.43 | 0.53 |
| 3 | 30334755 | rs10510633 | 0.274 | -0.548 | 0.701 | 0.632 | 0.003 | -0.001 | -0.001 | 0.003 | 0.3 | 0.35 | 0.47 | 0.32 | 0.37 | 0.4 |
| 3 | 30341425 | rs9819616 | -0.08 | -0.642 | 0.355 | 0.617 | -0.001 | -0.009 | -0.009 | 0.382 | 0.35 | 0.7 | 0.25 | 0.43 | 0.34 | 0.35 |
| 3 | 30347455 | rs1495578 | 0.02 | -0.776 | 0.967 | 0.558 | 0.002 | 0 | 0 | 0 | 0.48 | 0.53 | 0.32 | 0.51 | 0.39 | 0.53 |
| 3 | 30348778 | rs6549982 | -0.921 | -0.323 | 0.661 | 0.462 | 0.014 | 0.13 | 0.13 | 0.082 | 0.64 | 0.91 | 0.77 | 0.82 | 0.66 | 0.83 |
| 3 | 30351574 | rs11929513 | 0.197 | -0.292 | 0.844 | 0.485 | 0.051 | 0.049 | 0.049 | 0.505 | 0.14 | 0.18 | 0.43 | 0.32 | 0.27 | 0.3 |
| 3 | 30354360 | rs11129397 | 0.476 | -0.467 | 1.205 | 0.405 | 0.013 | -0.001 | -0.001 | 0.226 | 0.45 | 0.42 | 0.31 | 0.5 | 0.39 | 0.53 |
| 3 | 30357427 | rs7632716 | -0.142 | -0.167 | 1.357 | 0.482 | 0.105 | -0.005 | -0.005 | 0 | 0.12 | 0.38 | 0.16 | 0.1 | 0.19 | 0.13 |
| 3 | 30361276 | rs2029828 | 0.959 | -0.048 | 1.327 | 0.537 | 0.017 | -0.002 | -0.002 | 0.496 | 0.08 | 0.09 | 0.13 | 0.1 | 0.19 | 0.13 |
| 3 | 30372053 | rs7634533 | -0.703 | -0.2 | 1.124 | 0.462 | 0.044 | 0.019 | 0.019 | 0.113 | 0.27 | 0.39 | 0.47 | 0.32 | 0.31 | 0.37 |
| 3 | 30381927 | rs6778682 | 0.436 | -0.171 | 1.175 | 0.368 | 0.098 | 0.002 | 0.002 | 0.175 | 0.1 | 0.28 | 0.14 | 0.11 | 0.19 | 0.19 |
| 3 | 30382276 | rs6781464 | 0.882 | -0.157 | 1.097 | 0.376 | 0.076 | -0.001 | -0.001 | 0 | 0.09 | 0.17 | 0.13 | 0.1 | 0.19 | 0.14 |
| 3 | 30384065 | rs12489521 | -1.851 | -0.376 | 0.569 | 0.181 | -0.005 | 0.064 | 0.064 | 0 | 0.77 | 0.96 | 0.9 | 0.92 | 0.79 | 0.89 |
| 3 | 30386860 | rs17623772 | 1.121 | -0.175 | 0.845 | 0.682 | -0.027 | -0.02 | -0.02 | 0 | 0.3 | 0.15 | 0.25 | 0.29 | 0.34 | 0.37 |
| 3 | 30387538 | rs17025329 | 1.225 | -0.221 | 1.071 | 0.631 | -0.016 | -0.049 | -0.049 | 0.227 | 0.12 | 0.17 | 0.04 | 0 | 0.05 | 0.08 |
| 3 | 30390867 | rs9838682 | -0.133 | -0.197 | 1.169 | 0.708 | 0.031 | 0.026 | 0.026 | 0.414 | 0.47 | 0.36 | 0.66 | 0.63 | 0.61 | 0.67 |
| 3 | 30393943 | rs7625884 | -1.257 | -0.04 | 1.112 | 0.688 | -0.013 | -0.028 | -0.028 | 0 | 0.34 | 0.74 | 0.52 | 0.54 | 0.32 | 0.45 |
| 3 | 30395951 | rs11129399 | 0.774 | -0.035 | 1.073 | 0.688 | -0.036 | 0.052 | 0.052 | 0.679 | 0.15 | 0.29 | 0.07 | 0.21 | 0.05 | 0.16 |
| 3 | 30399578 | rs11922492 | -1.169 | -0.017 | 0.972 | 0.66 | -0.01 | -0.019 | -0.019 | 0 | 0.33 | 0.71 | 0.52 | 0.54 | 0.32 | 0.45 |
| 3 | 30402863 | rs2132298 | 0.209 | -0.087 | 0.988 | 0.807 | -0.014 | 0.036 | 0.036 | 0.268 | 0.37 | 0.29 | 0.56 | 0.76 | 0.42 | 0.62 |
| 3 | 30404343 | rs9866764 | 0.433 | -0.254 | 1.376 | 1.245 | 0.06 | 0.152 | 0.152 | 0 | 0.06 | 0.19 | 0.3 | 0.24 | 0.08 | 0.16 |
| 3 | 30412176 | rs7649253 | -1.068 | -0.305 | 1.44 | 1.22 | -0.008 | 0.068 | 0.068 | 0.221 | 0.23 | 0.63 | 0.38 | 0.44 | 0.13 | 0.32 |
| 3 | 30414405 | rs1495591 | -1.002 | -0.208 | 1.085 | 1.264 | -0.009 | 0.103 | 0.103 | 0.241 | 0.22 | 0.61 | 0.38 | 0.43 | 0.13 | 0.32 |

| | | | | | | | | | | | | | | | | |
|---|---|---|---|---|---|---|---|---|---|---|---|---|---|---|---|---|
| 3 | 30415729 | rs13094467 | 0.525 | 0.05 | 0.239 | 0.503 | -0.018 | -0.051 | -0.051 | 1.822 | 0.13 | 0.17 | 0.04 | 0 | 0.05 | 0.08 |
| 3 | 30417471 | rs9812768 | 0.04 | 0.604 | -0.167 | 1.018 | 0.06 | 0.152 | 0.152 | 0 | 0.06 | 0.19 | 0.3 | 0.24 | 0.08 | 0.16 |
| 3 | 30425285 | rs6783860 | -0.223 | 0.118 | -0.582 | 0.559 | -0.004 | 0.01 | 0.01 | 0 | 0.59 | 0.75 | 0.46 | 0.59 | 0.58 | 0.66 |
| 3 | 30436528 | rs4518165 | 0.119 | -0.219 | -0.093 | 0.363 | -0.004 | -0.027 | -0.027 | 0 | 0.38 | 0.47 | 0.52 | 0.28 | 0.38 | 0.49 |
| 3 | 30443869 | rs7653262 | 0.107 | -0.793 | 0.075 | 0.266 | -0.003 | -0.002 | -0.002 | 0 | 0.45 | 0.48 | 0.73 | 0.64 | 0.64 | 0.56 |
| 3 | 30445021 | rs1495589 | 1.355 | -1.354 | -0.442 | -0.055 | 0.007 | 0.029 | 0.029 | 0 | 0.21 | 0.14 | 0.11 | 0.09 | 0.2 | 0.16 |
| 3 | 30447759 | rs9850570 | 1.355 | -1.354 | -0.442 | -0.055 | 0.007 | 0.029 | 0.029 | 0 | 0.21 | 0.14 | 0.11 | 0.09 | 0.2 | 0.15 |
| 3 | 30449446 | rs11129403 | 1.377 | -1.144 | -0.479 | 0.065 | -0.03 | 0.025 | 0.025 | 0.667 | 0.18 | 0.12 | 0.24 | 0.12 | 0.31 | 0.14 |
| 3 | 30460903 | rs4279142 | 1.328 | -1.143 | -0.321 | 0.24 | -0.017 | -0.001 | -0.001 | 0 | 0.23 | 0.14 | 0.45 | 0.27 | 0.3 | 0.33 |
| 3 | 30469927 | rs6549991 | -1.109 | -1.369 | -0.169 | 0.268 | -0.057 | 0.039 | 0.039 | 1.369 | 0.79 | 0.62 | 0.88 | 0.74 | 0.88 | 0.81 |
| 3 | 30478897 | rs4075749 | 0.465 | -0.522 | -0.477 | 0.356 | -0.071 | 0.084 | 0.084 | 0.71 | 0.23 | 0.1 | 0.25 | 0.11 | 0.32 | 0.17 |
| 3 | 30478985 | rs4075748 | 0.358 | -0.494 | -0.445 | 0.362 | 0.007 | 0.009 | 0.009 | 0.634 | 0.26 | 0.26 | 0.29 | 0.37 | 0.33 | 0.25 |
| 3 | 30482310 | rs13073411 | 0.447 | -0.231 | -0.832 | -0.282 | 0.003 | 0.037 | 0.037 | 0 | 0.2 | 0.11 | 0.1 | 0.11 | 0.2 | 0.14 |
| 3 | 30488348 | rs4325953 | -0.091 | -0.043 | -0.316 | -0.161 | 0.006 | -0.045 | -0.045 | 0 | 0.38 | 0.54 | 0.32 | 0.11 | 0.39 | 0.35 |
| 3 | 30510943 | rs4402966 | -0.518 | -0.079 | -0.934 | -0.122 | 0.018 | 0.14 | 0.14 | 0.132 | 0.51 | 0.76 | 0.42 | 0.8 | 0.51 | 0.63 |
| 3 | 30526856 | rs11718153 | -0.951 | -0.021 | -0.805 | 0.656 | -0.025 | 0.099 | 0.099 | 0 | 0.28 | 0.49 | 0.29 | 0.45 | 0.19 | 0.57 |
| 3 | 30528411 | rs6792212 | -0.042 | 0.406 | -0.487 | 1.272 | 0.022 | 0.287 | 0.287 | 0 | 0.11 | 0.33 | 0.1 | 0.36 | 0.07 | 0.24 |
| 3 | 30530017 | rs7642647 | -0.29 | 0.558 | -0.555 | 1.259 | 0.005 | 0.008 | 0.008 | 0.768 | 0.49 | 0.55 | 0.43 | 0.75 | 0.55 | 0.63 |
| 3 | 30530899 | rs11917602 | -0.194 | 0.809 | -0.704 | 1.023 | 0 | 0 | 0 | 0.141 | 0.45 | 0.49 | 0.39 | 0.4 | 0.52 | 0.52 |
| 3 | 30534343 | rs4533691 | -0.44 | 1.015 | -0.815 | 1.358 | -0.01 | -0.04 | -0.04 | 0.317 | 0.08 | 0.01 | 0.12 | 0.14 | 0.09 | 0.04 |
| 3 | 30534842 | rs7617478 | -0.935 | 1.182 | -0.656 | 1.241 | 0.001 | -0.003 | -0.003 | 0 | 0.07 | 0.1 | 0.05 | 0.05 | 0.09 | 0.14 |
| 3 | 30540436 | rs4331703 | -1.817 | 1.913 | -0.616 | 1.713 | -0.021 | -0.022 | -0.022 | 0 | 0.26 | 0.2 | 0.29 | 0.43 | 0.37 | 0.26 |
| 3 | 30543417 | rs4325955 | -1.505 | 2.119 | -0.544 | 1.817 | -0.002 | -0.003 | -0.003 | 0 | 0.2 | 0.18 | 0.22 | 0.24 | 0.26 | 0.21 |
| 3 | 30547439 | rs7641221 | -0.501 | 1.807 | -0.452 | 1.835 | 0.042 | 0.035 | 0.035 | 0 | 0.06 | 0.1 | 0.09 | 0.2 | 0.11 | 0.06 |
| 3 | 30554212 | rs9819666 | -1.822 | 1.74 | 0.12 | 1.901 | 0.061 | 0.103 | 0.103 | 0 | 0.25 | 0.48 | 0.32 | 0.52 | 0.31 | 0.3 |
| 3 | 30556353 | rs11922839 | -0.71 | 1.245 | 0.173 | 1.932 | -0.015 | -0.037 | -0.037 | 1.06 | 0.08 | 0.01 | 0.11 | 0.13 | 0.09 | 0.12 |
| 3 | 30557554 | rs9873583 | -1.607 | 1.147 | 0.292 | 1.879 | 0.063 | 0.126 | 0.126 | 0 | 0.2 | 0.48 | 0.28 | 0.44 | 0.22 | 0.26 |
| 3 | 30559496 | rs4305453 | -1.303 | 0.725 | 0.04 | 1.491 | 0 | -0.002 | -0.002 | 0 | 0.33 | 0.66 | 0.31 | 0.53 | 0.31 | 0.37 |
| 3 | 30567666 | rs4955159 | -0.548 | -0.08 | -0.728 | 0.307 | 0.018 | 0.085 | 0.085 | 0.865 | 0.73 | 0.87 | 0.69 | 1 | 0.78 | 0.88 |
| 3 | 30568804 | rs9878028 | -1.059 | -0.059 | 0.033 | 1.488 | -0.023 | 0.061 | 0.061 | 0 | 0.07 | 0.11 | 0.07 | 0.21 | 0.02 | 0.08 |
| 3 | 30570302 | rs9822117 | -1.415 | -0.272 | 0.563 | 1.462 | -0.05 | 0.117 | 0.117 | 0 | 0.13 | 0.31 | 0.14 | 0.31 | 0.06 | 0.26 |
| 3 | 30570719 | rs11716372 | 1.703 | -0.083 | 0.516 | 1.496 | -0.001 | 0.011 | 0.011 | 0.18 | 0.6 | 0.55 | 0.6 | 0.48 | 0.62 | 0.42 |
| 3 | 30571674 | rs2129843 | 0.793 | 0.226 | -0.02 | 1.507 | -0.042 | -0.026 | -0.026 | 0 | 0.73 | 0.87 | 0.74 | 0.73 | 0.68 | 0.67 |
| 3 | 30571768 | rs2171508 | 0.753 | 0.232 | -0.025 | 1.338 | -0.044 | 0.016 | 0.016 | 0 | 0.73 | 0.87 | 0.74 | 0.8 | 0.68 | 0.67 |
| 3 | 30586611 | rs12631194 | -0.205 | 0.188 | 0.247 | 1.336 | -0.006 | 0.032 | 0.032 | 0.023 | 0.1 | 0.18 | 0.09 | 0.2 | 0.09 | 0.07 |
| 3 | 30590213 | rs7616969 | 1.78 | -0.095 | 0.514 | 1.395 | -0.001 | 0.008 | 0.008 | 0.382 | 0.58 | 0.53 | 0.57 | 0.47 | 0.62 | 0.39 |
| 3 | 30591844 | rs7648418 | 0.523 | 0.003 | 0.319 | 1.154 | -0.024 | 0.003 | 0.003 | 0.206 | 0.81 | 0.86 | 0.78 | 0.86 | 0.76 | 0.71 |
| 3 | 30594168 | rs7613005 | 1.3 | -0.074 | 0.402 | 1.256 | -0.018 | -0.009 | -0.009 | 0 | 0.7 | 0.8 | 0.65 | 0.3 | 0.7 | 0.39 |
| 3 | 30594586 | rs973404 | 1.334 | -0.074 | 0.407 | 1.256 | -0.013 | -0.005 | -0.005 | 0 | 0.69 | 0.79 | 0.65 | 0.3 | 0.7 | 0.4 |
| 3 | 30606591 | rs2200685 | 1.844 | -0.126 | 1.21 | 0.614 | -0.002 | -0.011 | -0.011 | 0 | 0.7 | 0.33 | 0.7 | 0.72 | 0.75 | 0.53 |
| 3 | 30607113 | rs1431129 | 0.68 | -0.128 | 1.273 | 0.622 | -0.004 | -0.002 | -0.002 | 0.144 | 0.86 | 0.87 | 0.87 | 0.84 | 0.81 | 0.77 |
| 3 | 30610221 | rs7634675 | 1.805 | -0.147 | 1.625 | 0.611 | 0.003 | -0.002 | -0.002 | 0.978 | 0.75 | 0.7 | 0.75 | 0.72 | 0.76 | 0.53 |
| 3 | 30611095 | rs13321947 | -1.103 | -0.151 | 1.607 | 0.611 | -0.02 | -0.012 | -0.012 | 1.139 | 0.1 | 0.17 | 0.12 | 0.13 | 0.05 | 0.24 |
| 3 | 30615627 | rs9874154 | 0.783 | -0.152 | 1.507 | 0.604 | 0.008 | 0 | 0 | 0 | 0.84 | 0.79 | 0.86 | 0.85 | 0.81 | 0.77 |
| 3 | 30618613 | rs9814861 | 0.114 | -0.048 | 1.44 | 0.61 | -0.025 | 0.011 | 0.011 | 0 | 0.89 | 0.96 | 0.93 | 0.94 | 0.85 | 0.83 |
| 3 | 30625123 | rs7648529 | 2.246 | -0.042 | 1.591 | 0.616 | 0.002 | -0.004 | -0.004 | 0 | 0.74 | 0.68 | 0.74 | 0.72 | 0.76 | 0.54 |
| 3 | 30626595 | rs2129842 | 0.104 | 0 | 1.234 | 0.611 | -0.027 | 0.011 | 0.011 | 0 | 0.89 | 0.96 | 0.93 | 0.94 | 0.85 | 0.84 |
| 3 | 30629672 | rs4276229 | -0.528 | -0.065 | 1.206 | 0.606 | -0.014 | -0.003 | -0.003 | 0.354 | 0.07 | 0.1 | 0.11 | 0.12 | 0.04 | 0.16 |
| 3 | 30632802 | rs4955245 | 0.772 | 0.005 | 0.958 | 0.621 | -0.011 | -0.004 | -0.004 | 0 | 0.86 | 0.88 | 0.87 | 0.86 | 0.81 | 0.78 |
| 3 | 30647160 | rs3087465 | 2.08 | -0.187 | 1.1 | 0.361 | -0.002 | -0.005 | -0.005 | 0.309 | 0.71 | 0.27 | 0.71 | 0.76 | 0.77 | 0.55 |
| 3 | 30650064 | rs6550005 | 1.536 | 0.244 | 0.796 | 0.378 | 0.041 | -0.038 | -0.038 | 0.099 | 0.81 | 0.49 | 0.83 | 0.88 | 0.82 | 0.75 |
| 3 | 30650327 | rs6770038 | 0.939 | 0.384 | 0.885 | 0.441 | 0.078 | -0.009 | -0.009 | 0 | 0.84 | 0.5 | 0.83 | 0.9 | 0.83 | 0.83 |
| 3 | 30655481 | rs1835538 | 0.176 | 0.526 | -0.673 | 0.087 | -0.053 | 0.103 | 0.103 | 0 | 0.61 | 0.79 | 0.54 | 0.86 | 0.56 | 0.73 |
| 3 | 30663987 | rs9850060 | 0.016 | 0.365 | -0.271 | 0.631 | 0.002 | 0.012 | 0.012 | 2.443 | 0.27 | 0.24 | 0.35 | 0.18 | 0.24 | 0.2 |
| 3 | 30667425 | rs17025785 | 0.587 | 0.554 | -0.429 | -0.092 | -0.001 | 0 | 0 | 1.096 | 0.28 | 0.23 | 0.41 | 0.69 | 0.37 | 0.36 |
| 3 | 30668684 | rs4522809 | 0.108 | 0.293 | -0.317 | -0.523 | 0 | -0.005 | -0.005 | 0.152 | 0.55 | 0.5 | 0.51 | 0.29 | 0.48 | 0.47 |
| 3 | 30672362 | rs6809777 | 0.552 | 0.038 | -0.276 | -0.154 | 0 | 0.01 | 0.01 | 0 | 0.28 | 0.26 | 0.27 | 0.18 | 0.3 | 0.3 |
| 3 | 30674254 | rs12490899 | -0.571 | 0.077 | -0.305 | 0.61 | 0.003 | 0 | 0 | 0.722 | 0.07 | 0.05 | 0.1 | 0.28 | 0.04 | 0.16 |
| 3 | 30677269 | rs12487185 | 0.468 | 0.751 | -0.291 | 0.166 | -0.029 | -0.032 | -0.032 | 0 | 0.3 | 0.1 | 0.29 | 0.69 | 0.31 | 0.36 |
| 3 | 30677484 | rs11924422 | -0.348 | 0.627 | -0.133 | -0.038 | -0.003 | 0.007 | 0.007 | 0.022 | 0.6 | 0.63 | 0.6 | 0.72 | 0.6 | 0.61 |
| 3 | 30679612 | rs11129421 | -0.97 | 0.557 | -0.261 | -0.147 | -0.053 | 0.007 | 0.007 | 1.162 | 0.67 | 0.88 | 0.65 | 0.73 | 0.65 | 0.64 |
| 3 | 30680370 | rs3863057 | -1.15 | 0.474 | -0.46 | -0.149 | -0.081 | 0.015 | 0.015 | 3.083 | 0.72 | 0.94 | 0.66 | 0.8 | 0.68 | 0.81 |
| 3 | 30683506 | rs13075948 | 0.917 | 0.882 | -0.25 | -0.148 | 0.009 | 0.049 | 0.049 | 0 | 0.3 | 0.22 | 0.32 | 0.03 | 0.31 | 0.26 |
| 3 | 30686414 | rs1155705 | -0.266 | 1.164 | -0.025 | 0.56 | 0.002 | 0 | 0 | 0 | 0.32 | 0.34 | 0.3 | 0.71 | 0.34 | 0.37 |
| 3 | 30689755 | rs2082224 | -0.178 | 0.98 | -0.163 | 0.845 | 0 | -0.004 | -0.004 | 0 | 0.24 | 0.22 | 0.27 | 0.39 | 0.23 | 0.18 |
| 3 | 30690748 | rs2005061 | 0.261 | 0.957 | -0.089 | 1.035 | 0.003 | -0.02 | -0.02 | 0 | 0.14 | 0.12 | 0.22 | 0.37 | 0.12 | 0.2 |
| 3 | 30690911 | rs1078985 | 0.609 | 1.1 | -0.141 | 0.675 | -0.013 | 0.027 | 0.027 | 0 | 0.3 | 0.15 | 0.16 | 0.16 | 0.33 | 0.29 |
| 3 | 30693643 | rs1036097 | 0.805 | 1.016 | 0.157 | 0.677 | 0.011 | 0.001 | 0.001 | 0.009 | 0.44 | 0.31 | 0.5 | 0.46 | 0.42 | 0.44 |
| 3 | 30697661 | rs1019855 | 0.21 | 1.33 | 0.929 | 1.312 | -0.005 | 0.089 | 0.089 | 0.143 | 0.15 | 0.24 | 0.24 | 0.36 | 0.12 | 0.2 |

| | | | | | | | | | | | | | | | | |
|---|---|---|---|---|---|---|---|---|---|---|---|---|---|---|---|---|
| 3 | 30698977 | rs12490466 | 0.355 | 1.332 | 0.858 | 1.312 | -0.002 | 0.039 | 0.039 | 0.018 | 0.14 | 0.17 | 0.23 | 0.36 | 0.12 | 0.2 |
| 3 | 30700922 | rs995435 | 0.799 | 1.346 | 0.906 | 1.301 | 0.006 | 0.031 | 0.031 | 0.42 | 0.75 | 0.59 | 0.68 | 0.68 | 0.76 | 0.73 |
| 3 | 30702140 | rs991694 | 0.213 | 1.169 | 0.446 | 0.795 | -0.043 | 0.056 | 0.056 | 0 | 0.9 | 0.99 | 0.93 | 0.99 | 0.88 | 0.93 |
| 3 | 30704007 | rs6792117 | 0.732 | 1.34 | 0.567 | 0.701 | 0.012 | 0.069 | 0.069 | 4.868 | 0.51 | 0.35 | 0.44 | 0.13 | 0.53 | 0.35 |
| 3 | 30704619 | rs4583693 | 0.809 | 1.37 | 0.292 | 0.992 | -0.001 | -0.001 | -0.001 | 0.774 | 0.21 | 0.2 | 0.24 | 0.17 | 0.18 | 0.2 |
| 3 | 30708432 | rs749794 | 0.533 | 1.571 | 0.499 | 1.051 | 0.022 | 0.022 | 0.022 | 0.575 | 0.72 | 0.56 | 0.66 | 0.3 | 0.69 | 0.55 |
| 3 | 30712344 | rs3773644 | 0.919 | 1.008 | -0.191 | 1.216 | 0.012 | 0.049 | 0.049 | 6.141 | 0.5 | 0.29 | 0.28 | 0.42 | 0.42 | 0.48 |
| 3 | 30712827 | rs11466511 | 0.384 | 1.64 | -0.692 | 1.69 | 0.006 | 0.002 | 0.002 | 0.568 | 0.21 | 0.16 | 0.24 | 0.15 | 0.2 | 0.21 |
| 3 | 30716602 | rs3773649 | -0.273 | 2.337 | -0.776 | 1.629 | 0.001 | 0.001 | 0.001 | 0.068 | 0.17 | 0.18 | 0.41 | 0.33 | 0.31 | 0.22 |
| 3 | 30718942 | rs3773652 | -0.241 | 1.495 | -1.074 | 1.298 | 0.003 | -0.014 | -0.014 | 0 | 0.45 | 0.3 | 0.7 | 0.46 | 0.56 | 0.41 |
| 3 | 30720304 | rs2043136 | -0.463 | 0.454 | -0.912 | 1.059 | 0.018 | -0.017 | -0.017 | 0.069 | 0.66 | 0.7 | 0.85 | 0.55 | 0.76 | 0.54 |
| 3 | 30721201 | rs2372212 | -0.601 | 0.241 | -0.985 | 1.068 | 0.052 | -0.008 | -0.008 | 0 | 0.7 | 0.85 | 0.9 | 0.65 | 0.82 | 0.61 |
| 3 | 30723470 | rs1346907 | -0.701 | 0.05 | -1.217 | 0.82 | 0.006 | 0.046 | 0.046 | 0 | 0.46 | 0.79 | 0.61 | 0.66 | 0.54 | 0.58 |
| 3 | 30724436 | rs3773658 | 0.341 | 0.086 | -1.094 | 1.014 | 0.024 | -0.023 | -0.023 | 1.095 | 0.17 | 0.12 | 0.06 | 0.31 | 0.1 | 0.31 |
| 3 | 30725645 | rs876687 | 0.269 | 0.077 | -0.72 | 1.011 | 0.001 | -0.003 | -0.003 | 0 | 0.17 | 0.19 | 0.07 | 0.31 | 0.1 | 0.31 |
| 3 | 30725776 | rs876688 | 0.322 | 0.683 | -0.482 | 1.031 | 0.092 | 0.021 | 0.021 | 0 | 0.22 | 0.4 | 0.48 | 0.23 | 0.38 | 0.19 |
| 3 | 30729636 | rs9843942 | 0.013 | 0.144 | -0.813 | 0.501 | 0.029 | 0.022 | 0.022 | 0.631 | 0.49 | 0.58 | 0.51 | 0.37 | 0.36 | 0.47 |
| 3 | 30730239 | rs2276768 | -0.778 | 0.315 | -0.284 | 1.013 | -0.034 | 0.093 | 0.093 | 0.021 | 0.15 | 0.37 | 0.09 | 0.37 | 0.09 | 0.3 |
| 3 | 30730689 | rs3773662 | -0.015 | 0.258 | -0.079 | 0.877 | 0.002 | -0.022 | -0.022 | 0.053 | 0.07 | 0.17 | 0.16 | 0.02 | 0.05 | 0.11 |
| 3 | 30730872 | rs3773663 | -1.623 | 0.76 | -0.234 | 0.831 | 0.021 | 0.021 | 0.021 | 0.325 | 0.31 | 0.43 | 0.23 | 0.48 | 0.42 | 0.47 |
| 3 | 30732821 | rs2276767 | -0.232 | 1.224 | -0.488 | 0.682 | -0.051 | 0.008 | 0.008 | 0 | 0.15 | 0.07 | 0.14 | 0.11 | 0.33 | 0.16 |
| 3 | 30735937 | rs744751 | -0.113 | 1.383 | -0.49 | 0.679 | -0.053 | 0.009 | 0.009 | 0 | 0.15 | 0.07 | 0.14 | 0.11 | 0.33 | 0.17 |
| 3 | 30736551 | rs6550009 | -0.164 | 1.395 | -0.438 | 0.687 | -0.024 | 0.007 | 0.007 | 0 | 0.15 | 0.09 | 0.14 | 0.11 | 0.33 | 0.17 |
| 3 | 30740289 | rs795346 | -1.928 | 0.863 | -0.517 | 0.681 | -0.008 | -0.004 | -0.004 | 0 | 0.54 | 0.49 | 0.64 | 0.61 | 0.63 | 0.67 |
| 3 | 30741405 | rs795430 | -0.98 | 0.887 | -0.484 | 0.863 | 0.001 | -0.002 | -0.002 | 0 | 0.37 | 0.35 | 0.5 | 0.38 | 0.3 | 0.47 |
| 3 | 30742562 | rs304822 | -0.566 | 0.877 | -1.328 | 0.933 | -0.003 | -0.076 | -0.076 | 0.403 | 0.28 | 0.11 | 0.31 | 0.36 | 0.25 | 0.35 |
| 3 | 30745941 | rs795347 | -2.248 | 0.464 | -1.106 | 0.806 | 0.012 | -0.001 | -0.001 | 0 | 0.64 | 0.77 | 0.7 | 0.66 | 0.66 | 0.67 |
| 3 | 30746868 | rs6767233 | -0.346 | 0.874 | -1.085 | 0.748 | -0.025 | 0.004 | 0.004 | 0 | 0.19 | 0.15 | 0.21 | 0.19 | 0.36 | 0.22 |
| 3 | 30747697 | rs6795213 | -1.318 | 0.597 | -1.127 | 0.838 | -0.031 | -0.016 | -0.016 | 0 | 0.32 | 0.27 | 0.26 | 0.41 | 0.44 | 0.47 |
| 3 | 30749965 | rs6779998 | -2.027 | 0.367 | -1.074 | 0.791 | -0.024 | -0.019 | -0.019 | 0 | 0.41 | 0.76 | 0.31 | 0.51 | 0.45 | 0.56 |
| 3 | 30752261 | rs6776524 | 0.3 | -0.217 | -0.995 | 0.858 | 0.01 | -0.018 | -0.018 | 0 | 0.12 | 0.1 | 0.04 | 0.22 | 0.08 | 0.25 |
| 3 | 30753573 | rs6804368 | -1.692 | -0.074 | -0.651 | 0.819 | 0.005 | 0.002 | 0.002 | 0 | 0.36 | 0.57 | 0.26 | 0.43 | 0.41 | 0.51 |
| 3 | 30757269 | rs304824 | 0.119 | -0.07 | -0.703 | 0.804 | -0.024 | 0.012 | 0.012 | 0 | 0.22 | 0.05 | 0.31 | 0.15 | 0.22 | 0.14 |
| 3 | 30766438 | rs2372214 | -1.052 | -0.419 | -0.572 | 0.838 | 0.074 | -0.027 | -0.027 | 0 | 0.24 | 0.48 | 0.21 | 0.21 | 0.33 | 0.25 |
| 3 | 30769150 | rs17029870 | -0.296 | -1.798 | -0.59 | 0.879 | 0.022 | -0.025 | -0.025 | 0 | 0.15 | 0.1 | 0.04 | 0.22 | 0.08 | 0.24 |
| 3 | 30769931 | rs12054099 | -0.283 | -1.798 | -0.581 | 0.879 | 0.016 | -0.015 | -0.015 | 0.082 | 0.15 | 0.13 | 0.05 | 0.22 | 0.08 | 0.24 |
| 3 | 30774786 | rs7614821 | -1.267 | -1.565 | -0.499 | 0.874 | 0 | -0.001 | -0.001 | 0 | 0.27 | 0.66 | 0.19 | 0.3 | 0.27 | 0.29 |
| 3 | 30775466 | rs6783574 | -2.362 | -1.68 | -0.577 | 0.9 | 0.035 | -0.043 | -0.043 | 0 | 0.44 | 0.86 | 0.24 | 0.52 | 0.35 | 0.53 |
| 3 | 30775963 | rs812124 | 2.959 | -1.692 | -0.695 | 0.867 | -0.153 | -0.082 | -0.082 | 0 | 0.35 | 0.02 | 0.45 | 0.33 | 0.44 | 0.35 |
| 3 | 30780107 | rs6787491 | -0.47 | -1.936 | -0.343 | 0.81 | 0.014 | -0.015 | -0.015 | 0 | 0.15 | 0.13 | 0.05 | 0.22 | 0.08 | 0.23 |
| 3 | 30785840 | rs7610363 | -1.246 | -1.898 | -0.403 | 0.781 | 0.009 | 0 | 0 | 0 | 0.23 | 0.34 | 0.15 | 0.28 | 0.27 | 0.25 |
| 3 | 30788153 | rs6550013 | -2.242 | -1.946 | -0.441 | 0.733 | 0.009 | 0.002 | 0.002 | 0 | 0.41 | 0.72 | 0.24 | 0.42 | 0.37 | 0.48 |
| 3 | 30788621 | rs17026519 | -1.342 | -1.784 | -0.441 | 0.809 | -0.06 | 0.028 | 0.028 | 0 | 0.21 | 0.54 | 0.11 | 0.24 | 0.1 | 0.28 |
| 3 | 30788979 | rs13085035 | -0.509 | -1.512 | -0.702 | 0.833 | -0.04 | 0.003 | 0.003 | 0 | 0.18 | 0.07 | 0.13 | 0.19 | 0.27 | 0.19 |
| 3 | 30797125 | rs1875688 | -0.974 | -1.643 | -0.561 | 0.84 | -0.009 | 0.008 | 0.008 | 0.089 | 0.17 | 0.22 | 0.05 | 0.31 | 0.09 | 0.28 |
| 3 | 30800620 | rs795448 | -0.409 | -1.731 | -0.796 | 0.722 | -0.005 | 0.02 | 0.02 | 0 | 0.83 | 0.96 | 0.66 | 0.86 | 0.84 | 0.89 |
| 3 | 30802262 | rs7652095 | -1.8 | -1.834 | -0.558 | 0.662 | -0.004 | 0.006 | 0.006 | 0 | 0.38 | 0.47 | 0.21 | 0.5 | 0.36 | 0.5 |
| 3 | 30803040 | rs795443 | 2.291 | -1.661 | -0.596 | 0.692 | -0.002 | -0.001 | -0.001 | 0.796 | 0.43 | 0.37 | 0.45 | 0.36 | 0.47 | 0.38 |
| 3 | 30810912 | rs9844695 | -1.218 | -1.644 | -0.902 | 0.434 | -0.051 | -0.026 | -0.026 | 0 | 0.19 | 0.61 | 0.11 | 0.14 | 0.08 | 0.26 |
| 3 | 30811589 | rs2062775 | -1.218 | -1.644 | -0.902 | 0.434 | -0.051 | -0.026 | -0.026 | 0.002 | 0.19 | 0.61 | 0.11 | 0.14 | 0.08 | 0.26 |
| 3 | 30812487 | rs9854881 | -0.931 | -1.548 | -0.834 | 0.638 | -0.042 | 0.039 | 0.039 | 0 | 0.19 | 0.1 | 0.23 | 0.12 | 0.3 | 0.22 |
| 3 | 30814963 | rs9866328 | -0.146 | -1.443 | -0.791 | 0.67 | -0.013 | 0.013 | 0.013 | 0 | 0.11 | 0.06 | 0.1 | 0.01 | 0.21 | 0.12 |
| 3 | 30815257 | rs304819 | 0.009 | -1.597 | -0.631 | 0.947 | 0.008 | 0.014 | 0.014 | 0 | 0.81 | 0.91 | 0.73 | 0.83 | 0.84 | 0.9 |
| 3 | 30825421 | rs1159304 | -0.989 | -1.453 | -0.361 | 1.485 | 0.023 | 0.423 | 0.423 | 0.504 | 0.08 | 0.48 | 0.06 | 0.32 | 0.02 | 0.1 |
| 3 | 30828871 | rs1393748 | 1.934 | -1.599 | -0.526 | 1.325 | 0 | 0 | 0 | 0 | 0.52 | 0.52 | 0.49 | 0.57 | 0.53 | 0.45 |
| 3 | 30831150 | rs6805101 | -1.829 | -1.401 | -0.263 | 1.368 | -0.07 | 0.142 | 0.142 | 0.248 | 0.17 | 0.63 | 0.08 | 0.44 | 0.04 | 0.34 |
| 3 | 30833347 | rs13087758 | -0.086 | -1.117 | -0.925 | 1.321 | -0.026 | 0.01 | 0.01 | 0 | 0.14 | 0.06 | 0.19 | 0.14 | 0.24 | 0.15 |
| 3 | 30835061 | rs1503096 | 0.162 | -1.036 | -1.034 | 1.357 | 0.007 | 0.028 | 0.028 | 0 | 0.79 | 0.92 | 0.73 | 0.85 | 0.82 | 0.88 |
| 3 | 30835872 | rs1393753 | 0.15 | -1.029 | -1.03 | 1.296 | 0.007 | 0.035 | 0.035 | 0 | 0.79 | 0.92 | 0.73 | 0.87 | 0.82 | 0.88 |
| 3 | 30835984 | rs1393752 | 0.839 | -0.869 | -0.997 | 1.512 | 0.007 | 0.087 | 0.087 | 0 | 0.56 | 0.26 | 0.47 | 0.73 | 0.55 | 0.7 |
| 3 | 30836536 | rs4955328 | -1.408 | -0.778 | -0.008 | 2.195 | -0.029 | 0.191 | 0.191 | 0 | 0.09 | 0.23 | 0.09 | 0.46 | 0.03 | 0.29 |
| 3 | 30844147 | rs1158476 | -0.985 | -0.746 | 0.022 | 1.742 | -0.01 | 0.033 | 0.033 | 0 | 0.35 | 0.48 | 0.41 | 0.53 | 0.31 | 0.26 |
| 3 | 30845810 | rs9871174 | -1.52 | -1.09 | 0.429 | 1.985 | -0.016 | 0.137 | 0.137 | 1.407 | 0.1 | 0.53 | 0.12 | 0.19 | 0.02 | 0.24 |
| 3 | 30848365 | rs1393749 | -1.628 | -1.29 | 0.736 | 1.908 | 0.039 | 0.042 | 0.042 | 0.056 | 0.47 | 0.74 | 0.54 | 0.69 | 0.35 | 0.54 |
| 3 | 30852085 | rs9878903 | 0.115 | -1.376 | 0.832 | 1.864 | 0.003 | -0.008 | -0.008 | 0.939 | 0.12 | 0.13 | 0.11 | 0.05 | 0.17 | 0.09 |
| 3 | 30855276 | rs1158454 | -1.242 | -1.49 | 1.019 | 2.322 | 0.004 | 0.144 | 0.144 | 0.125 | 0.2 | 0.54 | 0.22 | 0.38 | 0.18 | 0.33 |
| 3 | 30858749 | rs9836482 | -0.426 | -1.491 | 0.937 | 2.25 | 0.01 | -0.032 | -0.032 | 0 | 0.15 | 0.22 | 0.13 | 0.06 | 0.17 | 0.09 |
| 3 | 30858939 | rs9856958 | -1.086 | -1.685 | 0.769 | 2.295 | 0.018 | -0.043 | -0.043 | 0 | 0.2 | 0.06 | 0.18 | 0.26 | 0.14 | 0.19 |
| 3 | 30862962 | rs9879113 | -1.034 | -1.679 | 0.769 | 2.295 | 0.017 | -0.043 | -0.043 | 0.031 | 0.19 | 0.06 | 0.18 | 0.26 | 0.15 | 0.19 |

| | | | | | | | | | | | | | | | | |
|---|---|---|---|---|---|---|---|---|---|---|---|---|---|---|---|---|
| 3 | 30870256 | rs11129425 | -1.859 | -1.766 | 0.858 | 2.243 | -0.056 | -0.004 | -0.004 | 0 | 0.29 | 0.46 | 0.33 | 0.29 | 0.16 | 0.23 |
| 3 | 30871224 | rs5004864 | -2.046 | -1.885 | 0.906 | 2.314 | -0.032 | 0.229 | 0.229 | 0 | 0.08 | 0.3 | 0.1 | 0.32 | 0.01 | 0.22 |
| 3 | 30871240 | rs1494730 | -2.282 | -1.845 | 0.746 | 2.184 | 0.06 | -0.024 | -0.024 | 1.828 | 0.35 | 0.76 | 0.43 | 0.61 | 0.17 | 0.45 |
| 3 | 30878291 | rs1389908 | -1.274 | -1.544 | 0.553 | 2.518 | 0.012 | -0.018 | -0.018 | 0 | 0.27 | 0.74 | 0.16 | 0.39 | 0.19 | 0.21 |
| 3 | 30884190 | rs13077553 | -0.063 | -1.469 | -0.087 | 2.083 | 0.008 | 0.01 | 0.01 | 0.486 | 0.25 | 0.19 | 0.23 | 0.21 | 0.22 | 0.38 |
| 3 | 30884289 | rs11709194 | -0.277 | -1.472 | -0.08 | 2.233 | 0.011 | -0.037 | -0.037 | 0.042 | 0.27 | 0.23 | 0.23 | 0.45 | 0.23 | 0.43 |
| 3 | 30885806 | rs1494738 | -1.242 | -1.355 | -0.011 | 1.894 | 0 | 0 | 0 | 0.4 | 0.43 | 0.55 | 0.52 | 0.62 | 0.33 | 0.56 |
| 3 | 30886154 | rs1494739 | -1.092 | -1.364 | -0.074 | 1.93 | -0.016 | -0.004 | -0.004 | 0 | 0.41 | 0.51 | 0.51 | 0.62 | 0.33 | 0.56 |
| 3 | 30889582 | rs903424 | -2.091 | -1.237 | -0.473 | 2.059 | -0.014 | 0.104 | 0.104 | 0 | 0.52 | 0.97 | 0.55 | 0.74 | 0.35 | 0.51 |
| 3 | 30893309 | rs6795046 | -1.861 | -1.221 | -0.347 | 2.471 | 0.044 | -0.011 | -0.011 | 0 | 0.49 | 0.93 | 0.53 | 0.61 | 0.33 | 0.45 |
| 3 | 30894470 | rs12486119 | -1.186 | -0.778 | 0.485 | 2.926 | 0.002 | -0.002 | -0.002 | 0.016 | 0.25 | 0.25 | 0.34 | 0.39 | 0.18 | 0.28 |
| 3 | 30903008 | rs6774917 | -1.03 | -0.269 | 0.631 | 3.239 | -0.002 | 0.034 | 0.034 | 0 | 0.17 | 0.21 | 0.31 | 0.49 | 0.15 | 0.18 |
| 3 | 30903823 | rs4955343 | -0.997 | -0.269 | 0.631 | 3.239 | -0.002 | 0.035 | 0.035 | 0 | 0.17 | 0.21 | 0.31 | 0.49 | 0.15 | 0.18 |
| 3 | 30904844 | rs4955348 | -1.717 | -0.189 | 0.847 | 3.259 | 0 | -0.002 | -0.002 | 0.01 | 0.31 | 0.37 | 0.45 | 0.67 | 0.3 | 0.37 |
| 3 | 30905384 | rs1389903 | -1.654 | -0.188 | 0.625 | 3.3 | -0.002 | -0.001 | -0.001 | 0.032 | 0.3 | 0.27 | 0.44 | 0.65 | 0.29 | 0.35 |
| 3 | 30905575 | rs1389904 | -0.587 | -0.132 | 0.659 | 3.113 | 0.003 | 0.02 | 0.02 | 0 | 0.15 | 0.22 | 0.15 | 0.17 | 0.14 | 0.17 |
| 3 | 30910474 | rs7609812 | -1.7 | -0.136 | 0.649 | 3.151 | -0.004 | -0.025 | -0.025 | 0 | 0.3 | 0.26 | 0.45 | 0.55 | 0.29 | 0.33 |
| 3 | 30912592 | rs17026715 | -0.934 | -0.193 | 0.696 | 3.201 | 0.001 | 0.002 | 0.002 | 0 | 0.17 | 0.2 | 0.31 | 0.37 | 0.13 | 0.15 |
| 3 | 30915905 | rs11718267 | -0.876 | 0.071 | 0.71 | 3.187 | 0.001 | -0.036 | -0.036 | 0 | 0.17 | 0.11 | 0.3 | 0.26 | 0.15 | 0.15 |
| 3 | 30919675 | rs4290836 | -0.967 | 0.065 | 0.8 | 3.231 | 0.001 | -0.056 | -0.056 | 0.452 | 0.17 | 0.12 | 0.36 | 0.43 | 0.16 | 0.19 |
| 3 | 30935761 | rs6794058 | -1.647 | 0.538 | 0.946 | 3.125 | 0.006 | 0.088 | 0.088 | 0 | 0.21 | 0.41 | 0.43 | 0.39 | 0.24 | 0.21 |
| 3 | 30940741 | rs4955170 | -0.925 | 0.131 | 0.845 | 3.161 | 0.001 | -0.014 | -0.014 | 0 | 0.17 | 0.17 | 0.36 | 0.39 | 0.16 | 0.19 |
| 3 | 30942285 | rs9682385 | -1.439 | 0.126 | 0.879 | 3.159 | -0.007 | 0.058 | 0.058 | 0 | 0.19 | 0.31 | 0.38 | 0.4 | 0.16 | 0.19 |
| 3 | 30943207 | rs11129429 | -0.783 | 0.19 | 0.573 | 3.218 | 0 | -0.007 | -0.007 | 0 | 0.16 | 0.16 | 0.29 | 0.27 | 0.16 | 0.14 |
| 3 | 30951066 | rs17432258 | -0.361 | 0.274 | 0.265 | 2.958 | -0.039 | -0.046 | -0.046 | 0 | 0.14 | 0.01 | 0.09 | 0.14 | 0.16 | 0.18 |
| 3 | 30951882 | rs9873423 | 0.163 | 0.271 | 0.443 | 3.061 | -0.016 | -0.033 | -0.033 | 0 | 0.85 | 0.94 | 0.89 | 0.75 | 0.81 | 0.77 |
| 3 | 30960588 | rs10510642 | -2.071 | -0.168 | 1.484 | 2.874 | -0.011 | 0.156 | 0.156 | 2.112 | 0.08 | 0.58 | 0.05 | 0.1 | 0.01 | 0.05 |
| 3 | 30970165 | rs11714737 | -0.512 | -0.193 | 1.29 | 3.001 | -0.013 | -0.003 | -0.003 | 0 | 0.13 | 0.06 | 0.22 | 0.11 | 0.16 | 0.09 |
| 3 | 30974374 | rs17026785 | -0.849 | -0.263 | 1.272 | 2.912 | -0.002 | -0.014 | -0.014 | 1.565 | 0.16 | 0.1 | 0.29 | 0.24 | 0.16 | 0.13 |
| 3 | 30975073 | rs6768230 | -2.151 | -0.267 | 1.571 | 2.695 | -0.014 | 0.149 | 0.149 | 2.162 | 0.08 | 0.59 | 0.05 | 0.1 | 0.01 | 0.05 |
| 3 | 30995986 | rs1494741 | -1.787 | -0.341 | 1.459 | 2.842 | -0.039 | 0.111 | 0.111 | 0.85 | 0.21 | 0.58 | 0.34 | 0.38 | 0.14 | 0.19 |
| 3 | 30996999 | rs9860804 | -2.02 | -0.255 | 1.288 | 2.687 | -0.029 | 0.073 | 0.073 | 0.123 | 0.25 | 0.58 | 0.38 | 0.38 | 0.17 | 0.21 |
| 3 | 31001640 | rs1494742 | -0.929 | -0.325 | 1.409 | 2.687 | 0.002 | -0.01 | -0.01 | 0.523 | 0.17 | 0.12 | 0.28 | 0.27 | 0.16 | 0.11 |
| 3 | 31008073 | rs6550032 | -1.983 | -0.361 | 1.726 | 2.695 | -0.028 | 0.084 | 0.084 | 0 | 0.24 | 0.46 | 0.37 | 0.38 | 0.18 | 0.21 |
| 3 | 31012087 | rs17433029 | -0.86 | -0.359 | 1.731 | 2.5 | 0.002 | -0.011 | -0.011 | 0 | 0.17 | 0.11 | 0.28 | 0.27 | 0.17 | 0.11 |
| 3 | 31015186 | rs1112792 | -1.921 | -0.376 | 1.833 | 2.605 | -0.029 | 0.065 | 0.065 | 0 | 0.23 | 0.45 | 0.37 | 0.35 | 0.17 | 0.22 |
| 3 | 31022013 | rs4955358 | -0.798 | -0.367 | 1.769 | 2.601 | 0.001 | -0.033 | -0.033 | 0 | 0.16 | 0.08 | 0.28 | 0.28 | 0.16 | 0.11 |
| 3 | 31026401 | rs6779686 | -2.131 | -0.389 | 1.905 | 2.656 | -0.032 | 0.059 | 0.059 | 0.014 | 0.25 | 0.54 | 0.37 | 0.36 | 0.17 | 0.22 |
| 3 | 31027815 | rs4572805 | -0.127 | 0.196 | 1.349 | 2.479 | -0.057 | -0.037 | -0.037 | 0 | 0.17 | 0.03 | 0.13 | 0.14 | 0.25 | 0.22 |
| 3 | 31028952 | rs1395163 | -1.861 | 0.183 | 1.483 | 2.139 | 0.009 | 0.089 | 0.089 | 1.44 | 0.07 | 0.36 | 0.08 | 0.06 | 0.01 | 0.11 |
| 3 | 31039428 | rs1876168 | -2.102 | -0.071 | 1.189 | 1.832 | 0.005 | 0.06 | 0.06 | 0 | 0.5 | 0.6 | 0.66 | 0.66 | 0.49 | 0.6 |
| 3 | 31040600 | rs13319756 | -1.163 | -0.087 | 1.204 | 1.848 | 0.014 | 0.05 | 0.05 | 0.319 | 0.06 | 0.24 | 0.03 | 0.04 | 0.01 | 0.09 |
| 3 | 31040707 | rs11129434 | -0.436 | -0.093 | 1.115 | 1.933 | -0.048 | -0.079 | -0.079 | 0 | 0.25 | 0.1 | 0.28 | 0.3 | 0.32 | 0.4 |
| 3 | 31042458 | rs1567025 | -2.253 | -0.104 | 1.125 | 1.911 | 0.004 | 0.097 | 0.097 | 0.002 | 0.51 | 0.68 | 0.66 | 0.66 | 0.49 | 0.61 |
| 3 | 31045654 | rs11918606 | -0.481 | -0.093 | 0.839 | 1.828 | -0.043 | -0.069 | -0.069 | 0 | 0.27 | 0.1 | 0.29 | 0.3 | 0.32 | 0.4 |
| 3 | 31050500 | rs1506200 | -0.616 | -0.12 | 0.835 | 1.828 | -0.055 | -0.071 | -0.071 | 0.572 | 0.29 | 0.1 | 0.29 | 0.3 | 0.34 | 0.4 |
| 3 | 31060089 | rs9844634 | -1.148 | -0.098 | 0.894 | 1.627 | -0.017 | 0.03 | 0.03 | 0 | 0.18 | 0.26 | 0.29 | 0.3 | 0.15 | 0.1 |
| 3 | 31064485 | rs17434315 | -1.389 | 0.404 | 0.197 | 1.602 | 0.001 | 0.041 | 0.041 | 0 | 0.25 | 0.36 | 0.4 | 0.34 | 0.24 | 0.21 |
| 3 | 31068091 | rs17026964 | 1.154 | 0.243 | -1.004 | 0.904 | -0.028 | -0.046 | -0.046 | 0 | 0.37 | 0.06 | 0.18 | 0.41 | 0.31 | 0.22 |
| 3 | 31079320 | rs294308 | 0.058 | -0.393 | -0.759 | 0.605 | 0.012 | 0.026 | 0.026 | 0.014 | 0.74 | 0.82 | 0.74 | 0.94 | 0.79 | 0.9 |
| 3 | 31079959 | rs294305 | 0.651 | -0.548 | -0.156 | 0.517 | 0.026 | 0.043 | 0.043 | 0.406 | 0.7 | 0.3 | 0.68 | 0.94 | 0.79 | 0.89 |
| 3 | 31083591 | rs294303 | 0.259 | -0.555 | -0.338 | 0.583 | -0.001 | 0.115 | 0.115 | 0 | 0.72 | 0.94 | 0.74 | 0.92 | 0.78 | 0.82 |
| 3 | 31089048 | rs11922205 | -0.759 | -0.19 | -0.172 | 0.708 | 0.001 | -0.019 | -0.019 | 0 | 0.24 | 0.24 | 0.39 | 0.53 | 0.25 | 0.58 |
| 3 | 31102035 | rs6550039 | -0.88 | -0.185 | 0.53 | 1.067 | -0.003 | 0.103 | 0.103 | 0.766 | 0.23 | 0.54 | 0.35 | 0.38 | 0.22 | 0.46 |
| 3 | 31104770 | rs1485924 | -0.132 | -0.174 | 0.508 | 1.079 | 0.025 | 0.186 | 0.186 | 0 | 0.18 | 0.43 | 0.33 | 0.36 | 0.21 | 0.39 |
| 3 | 31105718 | rs7614625 | 0.004 | -0.159 | 0.595 | 1.085 | 0.018 | 0.129 | 0.129 | 0 | 0.18 | 0.36 | 0.33 | 0.35 | 0.21 | 0.39 |
| 3 | 31110187 | rs13090678 | -0.985 | -0.204 | 0.717 | 1.063 | -0.01 | 0.063 | 0.063 | 1.075 | 0.25 | 0.61 | 0.36 | 0.38 | 0.22 | 0.46 |
| 3 | 31116647 | rs1385831 | -0.424 | -0.062 | 0.6 | 0.905 | 0.004 | 0.118 | 0.118 | 0.119 | 0.21 | 0.38 | 0.44 | 0.49 | 0.23 | 0.5 |
| 3 | 31126912 | rs294284 | -0.258 | -0.221 | -0.327 | 0.659 | -0.002 | 0.096 | 0.096 | 0.316 | 0.76 | 0.91 | 0.75 | 0.94 | 0.79 | 0.89 |
| 3 | 31128482 | rs294279 | -1.049 | -0.195 | -0.185 | 0.714 | 0.022 | 0.052 | 0.052 | 0 | 0.47 | 0.78 | 0.71 | 0.59 | 0.44 | 0.62 |
| 3 | 31135969 | rs294282 | -1.012 | -0.273 | -0.302 | 0.742 | 0.016 | 0.009 | 0.009 | 0 | 0.46 | 0.75 | 0.44 | 0.41 | 0.43 | 0.46 |
| 3 | 31144288 | rs294296 | -1.102 | -0.228 | -0.301 | 0.608 | 0.039 | 0.04 | 0.04 | 0.043 | 0.48 | 0.8 | 0.6 | 0.59 | 0.43 | 0.6 |
| 3 | 31149368 | rs1857876 | -0.345 | -0.195 | 0.09 | 0.373 | -0.01 | -0.043 | -0.043 | 0.562 | 0.29 | 0.37 | 0.24 | 0.13 | 0.25 | 0.18 |
| 3 | 31155829 | rs294288 | -2.109 | -0.011 | -0.821 | 0.157 | -0.088 | -0.045 | -0.045 | 0 | 0.68 | 0.98 | 0.77 | 0.73 | 0.71 | 0.82 |
| 3 | 31163822 | rs839581 | -0.78 | -0.927 | 0.297 | -0.028 | 0.007 | 0.006 | 0.006 | 0.608 | 0.35 | 0.63 | 0.24 | 0.12 | 0.22 | 0.23 |
| 3 | 31164966 | rs839584 | -1.462 | -0.824 | 0.461 | 0.035 | 0.024 | 0.038 | 0.038 | 0.015 | 0.51 | 0.73 | 0.58 | 0.57 | 0.44 | 0.66 |
| 3 | 31173171 | rs11925641 | 0.415 | -0.775 | 0.518 | 0.345 | -0.002 | -0.022 | -0.022 | 0 | 0.18 | 0.12 | 0.23 | 0.3 | 0.21 | 0.32 |
| 3 | 31175501 | rs7625163 | -1.374 | -0.804 | 0.399 | 0.093 | 0.046 | 0.012 | 0.012 | 0.091 | 0.48 | 0.9 | 0.56 | 0.55 | 0.38 | 0.55 |
| 3 | 31176875 | rs1625526 | -0.429 | -0.619 | 0.954 | 0.004 | -0.021 | -0.063 | -0.063 | 0 | 0.32 | 0.44 | 0.25 | 0.08 | 0.24 | 0.21 |

| | | | | | | | | | | | | | | | | |
|---|---|---|---|---|---|---|---|---|---|---|---|---|---|---|---|---|
| 3 | 31176949 | rs9826061 | 0.189 | -0.648 | 1.056 | 0.083 | 0.026 | 0.217 | 0.217 | 0 | 0.17 | 0.39 | 0.36 | 0.43 | 0.19 | 0.36 |
| 3 | 31177841 | rs1876167 | -0.13 | -0.628 | 0.985 | 0.095 | -0.002 | 0.002 | 0.002 | 0 | 0.19 | 0.19 | 0.35 | 0.46 | 0.21 | 0.4 |
| 3 | 31178850 | rs1395165 | -1.655 | -0.595 | 1.181 | 0.066 | 0.008 | 0.022 | 0.022 | 0 | 0.52 | 0.66 | 0.61 | 0.54 | 0.46 | 0.59 |
| 3 | 31179339 | rs1485916 | 0.278 | -0.487 | 1.156 | -0.007 | -0.002 | -0.038 | -0.038 | 0 | 0.22 | 0.29 | 0.15 | 0.02 | 0.19 | 0.09 |
| 3 | 31196694 | rs17027132 | 0.46 | -0.39 | 0.789 | 0.224 | -0.002 | 0.007 | 0.007 | 0.113 | 0.16 | 0.16 | 0.34 | 0.45 | 0.19 | 0.37 |
| 3 | 31202693 | rs1683756 | 0.034 | -0.372 | 0.985 | 0.004 | -0.017 | -0.098 | -0.098 | 0 | 0.25 | 0.6 | 0.15 | 0.01 | 0.18 | 0.08 |
| 3 | 31203987 | rs17643248 | -0.623 | -0.188 | 0.786 | 0.235 | 0 | -0.002 | -0.002 | 0 | 0.08 | 0.07 | 0.08 | 0.07 | 0.08 | 0.11 |
| 3 | 31219954 | rs12485312 | -1.089 | -0.52 | 1.044 | 0.194 | -0.011 | 0.045 | 0.045 | 0 | 0.27 | 0.42 | 0.43 | 0.56 | 0.25 | 0.51 |
| 3 | 31223049 | rs6791484 | -1.217 | -0.523 | 1.04 | 0.164 | -0.026 | 0.019 | 0.019 | 0 | 0.11 | 0.28 | 0.09 | 0.08 | 0.05 | 0.14 |
| 3 | 31229379 | rs2011774 | 0.502 | -0.443 | 0.664 | 0.293 | -0.01 | -0.027 | -0.027 | 0.108 | 0.15 | 0.08 | 0.34 | 0.46 | 0.19 | 0.36 |
| 3 | 31235468 | rs4515095 | 0.345 | -0.319 | 0.644 | -0.014 | -0.007 | 0.031 | 0.031 | 0 | 0.23 | 0.15 | 0.11 | 0.03 | 0.23 | 0.09 |
| 3 | 31238623 | rs9853319 | 0.431 | -0.048 | 0.557 | 0.378 | -0.006 | -0.015 | -0.015 | 0 | 0.15 | 0.12 | 0.34 | 0.42 | 0.19 | 0.37 |
| 3 | 31243972 | rs6778284 | -0.089 | -0.063 | 0.8 | 0.254 | 0.007 | 0.134 | 0.134 | 0.159 | 0.2 | 0.4 | 0.37 | 0.46 | 0.21 | 0.4 |
| 3 | 31252289 | rs11927074 | 0.299 | -0.029 | 0.269 | 0.269 | -0.003 | -0.009 | -0.009 | 0 | 0.18 | 0.15 | 0.35 | 0.46 | 0.21 | 0.4 |
| 3 | 31260498 | rs4955185 | -0.683 | -0.103 | 0.084 | 0.412 | 0.015 | 0.076 | 0.076 | 0 | 0.5 | 0.74 | 0.54 | 0.64 | 0.43 | 0.59 |
| 3 | 31262627 | rs4352403 | -0.275 | 0.348 | 0.422 | 0.93 | 0.002 | -0.004 | -0.004 | 0 | 0.42 | 0.64 | 0.29 | 0.48 | 0.43 | 0.36 |
| 3 | 31263782 | rs9838192 | -1.587 | 0.32 | 0.651 | 1.055 | -0.017 | 0.053 | 0.053 | 0 | 0.08 | 0.26 | 0.09 | 0.09 | 0.04 | 0.08 |
| 3 | 31273355 | rs4955188 | 0.876 | 0.937 | 0.528 | 1.039 | 0.027 | 0.014 | 0.014 | 0 | 0.83 | 0.72 | 0.82 | 0.84 | 0.82 | 0.83 |
| 3 | 31279217 | rs7646751 | -1.096 | 0.602 | 0.263 | 1.078 | -0.003 | 0.007 | 0.007 | 0 | 0.49 | 0.77 | 0.6 | 0.46 | 0.46 | 0.38 |
| 3 | 31279827 | rs4596184 | -0.221 | 0.695 | 0.625 | 1.322 | 0 | 0.03 | 0.03 | 0 | 0.15 | 0.23 | 0.16 | 0.14 | 0.13 | 0.13 |
| 3 | 31280168 | rs4317137 | -0.768 | 0.836 | 1.133 | 0.885 | -0.007 | 0.018 | 0.018 | 0 | 0.37 | 0.46 | 0.36 | 0.46 | 0.3 | 0.28 |
| 3 | 31285081 | rs13320098 | -1.358 | 0.714 | 1.753 | 0.882 | 0.102 | 0.029 | 0.029 | 0.116 | 0.13 | 0.42 | 0.25 | 0.12 | 0.17 | 0.15 |
| 3 | 31288543 | rs9876905 | -1.452 | 0.701 | 1.767 | 0.834 | 0.111 | 0.017 | 0.017 | 0.086 | 0.13 | 0.5 | 0.25 | 0.11 | 0.17 | 0.15 |
| 3 | 31292684 | rs9852655 | -1.806 | 0.298 | 1.5 | 0.808 | -0.002 | 0.006 | 0.006 | 0 | 0.28 | 0.7 | 0.47 | 0.19 | 0.27 | 0.26 |
| 3 | 31294492 | rs9843286 | -1.476 | 0.359 | 2.319 | 0.692 | 0.098 | -0.066 | -0.066 | 0.063 | 0.11 | 0.45 | 0.3 | 0 | 0.14 | 0.06 |
| 3 | 31304525 | rs6775336 | -1.687 | 0.268 | 2.595 | 0.665 | 0.065 | -0.068 | -0.068 | 0 | 0.11 | 0.36 | 0.3 | 0 | 0.15 | 0.06 |
| 3 | 31307907 | rs6810318 | -1.664 | -0.326 | 2.65 | 0.689 | -0.027 | -0.081 | -0.081 | 1.509 | 0.23 | 0.49 | 0.18 | 0.08 | 0.17 | 0.15 |
| 3 | 31312759 | rs6803159 | -0.73 | -0.63 | 2.218 | 0.563 | -0.01 | -0.005 | -0.005 | 0.038 | 0.05 | 0.08 | 0.02 | 0.04 | 0.01 | 0.09 |
| 3 | 31315991 | rs9845245 | -2.095 | -0.647 | 2.297 | 0.548 | -0.007 | -0.015 | -0.015 | 1.422 | 0.28 | 0.69 | 0.22 | 0.12 | 0.17 | 0.25 |
| 3 | 31317131 | rs7645512 | -2.074 | -0.648 | 2.278 | 0.548 | -0.003 | -0.013 | -0.013 | 0 | 0.28 | 0.69 | 0.22 | 0.09 | 0.17 | 0.23 |
| 3 | 31319355 | rs7652050 | -2.148 | -0.655 | 2.271 | 0.545 | -0.004 | -0.01 | -0.01 | 0 | 0.28 | 0.69 | 0.22 | 0.12 | 0.17 | 0.25 |
| 3 | 31325569 | rs6787074 | -0.326 | -0.637 | 2.215 | 0.499 | -0.006 | 0.045 | 0.045 | 0 | 0.15 | 0.04 | 0.14 | 0.06 | 0.12 | 0.14 |
| 3 | 31329185 | rs9811300 | -2.476 | -0.628 | 2.368 | 0.504 | -0.01 | -0.027 | -0.027 | 0 | 0.09 | 0.62 | 0.07 | 0.03 | 0.04 | 0.04 |
| 3 | 31332156 | rs6786337 | -2.179 | -0.664 | 2.205 | 0.62 | 0.019 | 0.019 | 0.019 | 0 | 0.29 | 0.73 | 0.25 | 0.18 | 0.18 | 0.27 |
| 3 | 31347242 | rs7623471 | -2.307 | -0.712 | 2.84 | 0.791 | 0.078 | 0.178 | 0.178 | 0 | 0.06 | 0.41 | 0.05 | 0.12 | 0.03 | 0.06 |
| 3 | 31348653 | rs7638704 | -1.892 | -0.798 | 2.911 | 0.829 | 0 | 0 | 0 | 0 | 0.23 | 0.71 | 0.2 | 0.18 | 0.15 | 0.21 |
| 3 | 31354395 | rs11921989 | -1.522 | -1.185 | 3.019 | 1.045 | -0.056 | -0.022 | -0.022 | 0 | 0.11 | 0.32 | 0.04 | 0.04 | 0.01 | 0.07 |
| 3 | 31364686 | rs9310966 | -3.308 | -1.126 | 3.408 | 1.058 | 0.033 | 0.123 | 0.123 | 0.357 | 0.09 | 0.5 | 0.13 | 0.18 | 0.08 | 0.14 |
| 3 | 31370819 | rs6767817 | -3.064 | -1.149 | 3.091 | 1.054 | -0.014 | 0 | 0 | 0 | 0.19 | 0.74 | 0.17 | 0.21 | 0.1 | 0.21 |
| 3 | 31383105 | rs1006965 | -0.785 | -1.183 | 2.969 | 1.029 | -0.009 | -0.004 | -0.004 | 0 | 0.09 | 0.09 | 0.03 | 0.03 | 0.04 | 0.08 |
| 3 | 31386232 | rs7619670 | -2.599 | -1.186 | 2.738 | 1.028 | -0.014 | 0 | 0 | 0.718 | 0.17 | 0.76 | 0.09 | 0.17 | 0.05 | 0.17 |
| 3 | 31392328 | rs9862255 | -2.812 | -1.197 | 2.724 | 1.596 | 0.002 | -0.006 | -0.006 | 0 | 0.19 | 0.77 | 0.23 | 0.29 | 0.1 | 0.21 |
| 3 | 31394943 | rs13434007 | 0.05 | -1.308 | 2.123 | 1.687 | -0.001 | -0.017 | -0.017 | 1.317 | 0.19 | 0.11 | 0.13 | 0.29 | 0.17 | 0.14 |
| 3 | 31404944 | rs17027453 | -0.407 | -1.37 | 3.33 | 1.883 | 0.002 | 0.088 | 0.088 | 0.282 | 0.22 | 0.36 | 0.3 | 0.5 | 0.19 | 0.21 |
| 3 | 31408977 | rs6788642 | -1.106 | -1.412 | 2.993 | 1.874 | 0.027 | -0.088 | -0.088 | 0.208 | 0.3 | 0.82 | 0.36 | 0.54 | 0.21 | 0.24 |
| 3 | 31419711 | rs10446456 | 0.022 | -0.998 | 3.183 | 1.784 | 0 | -0.003 | -0.003 | 0 | 0.2 | 0.13 | 0.15 | 0.31 | 0.17 | 0.12 |
| 3 | 31423893 | rs17027548 | -2.159 | -0.942 | 2.998 | 1.749 | 0.016 | 0.083 | 0.083 | 0.011 | 0.06 | 0.21 | 0.1 | 0.12 | 0.03 | 0.06 |
| 3 | 31431722 | rs12497875 | -0.661 | -1.217 | 2.884 | 1.712 | -0.015 | 0.036 | 0.036 | 0.642 | 0.26 | 0.61 | 0.26 | 0.43 | 0.19 | 0.17 |
| 3 | 31433893 | rs9845596 | -0.236 | -1.124 | 2.909 | 1.759 | -0.002 | 0.101 | 0.101 | 0.428 | 0.22 | 0.38 | 0.24 | 0.43 | 0.17 | 0.15 |
| 3 | 31445799 | rs9880831 | -1.738 | -1.059 | 2.646 | 1.786 | 0 | -0.012 | -0.012 | 0 | 0.38 | 0.69 | 0.56 | 0.5 | 0.34 | 0.28 |
| 3 | 31446097 | rs4254693 | -2.028 | -1.073 | 2.499 | 2.083 | -0.004 | 0.009 | 0.009 | 0.972 | 0.39 | 0.85 | 0.61 | 0.67 | 0.31 | 0.34 |
| 3 | 31447156 | rs9848966 | -1.7 | -1.043 | 2.774 | 2.088 | 0.035 | 0.026 | 0.026 | 0.069 | 0.14 | 0.49 | 0.3 | 0.07 | 0.11 | 0.13 |
| 3 | 31450478 | rs1495181 | -1.982 | -1.045 | 2.614 | 2.082 | -0.009 | -0.015 | -0.015 | 0.175 | 0.38 | 0.81 | 0.61 | 0.61 | 0.31 | 0.31 |
| 3 | 31451063 | rs9870013 | -1.941 | -1.03 | 2.744 | 2.117 | 0.025 | 0.127 | 0.127 | 0.129 | 0.15 | 0.57 | 0.38 | 0.23 | 0.12 | 0.17 |
| 3 | 31453696 | rs12490552 | -1.699 | -0.991 | 2.022 | 1.946 | 0 | -0.032 | -0.032 | 0 | 0.35 | 0.76 | 0.53 | 0.55 | 0.28 | 0.28 |
| 3 | 31456161 | rs9831111 | -2.063 | -0.779 | 2.027 | 1.647 | 0.013 | 0.035 | 0.035 | 0 | 0.15 | 0.28 | 0.35 | 0.13 | 0.14 | 0.13 |
| 3 | 31466659 | rs870387 | -0.309 | -1.229 | 1.361 | 1.495 | -0.005 | 0 | 0 | 0 | 0.35 | 0.37 | 0.34 | 0.64 | 0.28 | 0.34 |
| 3 | 31480114 | rs1495173 | -1.043 | -1.319 | 1.726 | 1.324 | 0.022 | 0.042 | 0.042 | 0.303 | 0.44 | 0.76 | 0.44 | 0.67 | 0.36 | 0.44 |
| 3 | 31481326 | rs9810311 | -0.717 | -1.419 | 2.049 | 1.392 | -0.006 | -0.009 | -0.009 | 0 | 0.4 | 0.52 | 0.42 | 0.64 | 0.35 | 0.34 |
| 3 | 31486740 | rs9829340 | -0.702 | -1.369 | 2.303 | 2.024 | 0.064 | -0.023 | -0.023 | 0.059 | 0.07 | 0.21 | 0.07 | 0.01 | 0.07 | 0.08 |
| 3 | 31488222 | rs6766414 | 0.19 | -1.482 | 1.799 | 2.054 | 0.013 | -0.035 | -0.035 | 0 | 0.3 | 0.15 | 0.25 | 0.35 | 0.28 | 0.17 |
| 3 | 31496145 | rs9849739 | -0.729 | -1.576 | 1.428 | 1.481 | -0.032 | -0.058 | -0.058 | 0 | 0.22 | 0.32 | 0.15 | 0.06 | 0.16 | 0.11 |
| 3 | 31503032 | rs6784586 | 0.609 | -1.623 | 0.499 | 1.46 | 0.022 | -0.042 | -0.042 | 0.116 | 0.3 | 0.16 | 0.26 | 0.38 | 0.26 | 0.18 |
| 3 | 31509373 | rs7646265 | -1.943 | -1.705 | 0.176 | 1.332 | -0.176 | 0.013 | 0.013 | 6.103 | 0.64 | 0.98 | 0.59 | 0.73 | 0.5 | 0.57 |
| 3 | 31510049 | rs12108038 | 0.783 | -1.724 | 0.2 | 1.316 | 0.024 | -0.035 | -0.035 | 0 | 0.29 | 0.07 | 0.25 | 0.32 | 0.25 | 0.18 |
| 3 | 31510258 | rs11129451 | 0.558 | -1.646 | 0.482 | 1.369 | 0.015 | -0.006 | -0.006 | 0 | 0.31 | 0.27 | 0.26 | 0.33 | 0.26 | 0.27 |
| 3 | 31514777 | rs9836051 | -1.908 | -1.669 | 0.966 | 0.874 | -0.013 | -0.01 | -0.01 | 0 | 0.31 | 0.52 | 0.23 | 0.3 | 0.24 | 0.27 |
| 3 | 31522212 | rs13319027 | -1.41 | -1.837 | 0.77 | 0.807 | -0.032 | -0.099 | -0.099 | 0 | 0.27 | 0.5 | 0.2 | 0.06 | 0.17 | 0.19 |
| 3 | 31527836 | rs9877296 | -1.045 | -1.908 | 0.872 | 0.855 | -0.051 | 0.02 | 0.02 | 2.195 | 0.23 | 0.4 | 0.15 | 0.3 | 0.14 | 0.2 |

| | | | | | | | | | | | | | | | | |
|---|---|---|---|---|---|---|---|---|---|---|---|---|---|---|---|---|
| 3 | 31527951 | rs9683182 | -0.44 | -1.968 | 0.086 | 0.833 | 0.005 | 0.01 | 0.01 | 0.121 | 0.42 | 0.38 | 0.42 | 0.32 | 0.36 | 0.28 |
| 3 | 31529089 | rs7639313 | -1.864 | -1.998 | -0.826 | 0.8 | -0.169 | -0.037 | -0.037 | 0 | 0.64 | 0.97 | 0.57 | 0.69 | 0.48 | 0.51 |
| 3 | 31534154 | rs9863982 | -2.455 | -2.196 | 0.161 | 1.121 | 0.049 | -0.038 | -0.038 | 0.328 | 0.33 | 0.92 | 0.22 | 0.36 | 0.21 | 0.3 |
| 3 | 31540876 | rs6793666 | -0.656 | -1.639 | 0.937 | 1.108 | 0.028 | 0.002 | 0.002 | 0.018 | 0.07 | 0.16 | 0.04 | 0.03 | 0.06 | 0.07 |
| 3 | 31541300 | rs12485487 | -0.015 | -1.972 | 0.626 | 1.044 | 0.026 | -0.036 | -0.036 | 0.114 | 0.27 | 0.14 | 0.26 | 0.31 | 0.21 | 0.19 |
| 3 | 31541869 | rs7641067 | -1.402 | -1.988 | 0.743 | 0.953 | 0.043 | 0.001 | 0.001 | 0.71 | 0.42 | 0.82 | 0.38 | 0.64 | 0.33 | 0.38 |
| 3 | 31549953 | rs12491561 | -0.275 | -2.004 | 1.046 | 0.657 | -0.004 | 0.004 | 0.004 | 0.043 | 0.3 | 0.33 | 0.3 | 0.37 | 0.24 | 0.25 |
| 3 | 31551780 | rs4390968 | -0.46 | -1.962 | 0.865 | 0.663 | -0.039 | 0.037 | 0.037 | 0.09 | 0.32 | 0.53 | 0.32 | 0.37 | 0.24 | 0.24 |
| 3 | 31554627 | rs6788100 | 0.234 | -1.774 | 0.534 | 0.634 | -0.018 | 0.002 | 0.002 | 0 | 0.31 | 0.37 | 0.35 | 0.31 | 0.19 | 0.18 |
| 3 | 31569962 | rs7641408 | 0.523 | -1.694 | 0.453 | 0.656 | -0.036 | 0.022 | 0.022 | 0 | 0.27 | 0.4 | 0.37 | 0.31 | 0.19 | 0.18 |
| 3 | 31573057 | rs9810072 | 0.009 | -1.568 | 0.34 | 0.318 | 0 | -0.001 | -0.001 | 0 | 0.1 | 0.09 | 0.05 | 0.05 | 0.1 | 0.11 |
| 3 | 31576435 | rs9877437 | -0.59 | -1.659 | 0.346 | 0.271 | -0.006 | -0.009 | -0.009 | 0 | 0.94 | 0.9 | 0.97 | 0.98 | 0.96 | 0.93 |
| 3 | 31593820 | rs9817014 | 0.416 | -1.761 | 0.084 | 0.426 | 0.079 | -0.028 | -0.028 | 2.393 | 0.58 | 0.22 | 0.45 | 0.38 | 0.7 | 0.71 |
| 3 | 31620122 | rs7649067 | -0.127 | -1.778 | 0.191 | 0.459 | -0.033 | 0.001 | 0.001 | 0 | 0.64 | 0.45 | 0.5 | 0.68 | 0.76 | 0.75 |
| 3 | 31644397 | rs4569693 | 0.374 | -1.753 | 0.05 | 0.371 | 0.072 | -0.028 | -0.028 | 0.011 | 0.59 | 0.22 | 0.63 | 0.39 | 0.71 | 0.72 |
| 3 | 31645684 | rs9839338 | -0.39 | -1.767 | 0.124 | 0.378 | -0.025 | 0.002 | 0.002 | 0 | 0.67 | 0.6 | 0.69 | 0.7 | 0.77 | 0.76 |
| 3 | 31652978 | rs13093994 | 1.135 | -1.683 | 0.013 | 0.406 | 0.061 | 0.03 | 0.03 | 0 | 0.48 | 0.1 | 0.57 | 0.34 | 0.61 | 0.61 |
| 3 | 31664171 | rs9681121 | 0.791 | -1.413 | 0.172 | 0.52 | -0.018 | 0.006 | 0.006 | 0 | 0.27 | 0.34 | 0.29 | 0.28 | 0.19 | 0.16 |
| 3 | 31672157 | rs7627284 | -0.156 | -1.42 | 0.169 | 0.477 | -0.048 | 0.002 | 0.002 | 0 | 0.66 | 0.51 | 0.68 | 0.7 | 0.77 | 0.76 |
| 3 | 31677395 | rs11129457 | -0.447 | -1.296 | -0.798 | 0.483 | -0.012 | -0.02 | -0.02 | 0 | 0.94 | 0.84 | 0.96 | 0.98 | 0.96 | 0.93 |
| 3 | 31678512 | rs9828641 | 0.073 | -1.285 | -0.793 | 0.488 | 0 | -0.002 | -0.002 | 0.247 | 0.1 | 0.09 | 0.05 | 0.05 | 0.09 | 0.11 |
| 3 | 31678613 | rs10026 | -0.082 | -1.415 | -0.81 | 0.513 | -0.017 | -0.026 | -0.026 | 0 | 0.93 | 0.82 | 0.96 | 0.98 | 0.96 | 0.92 |
| 3 | 31687232 | rs12491975 | 0.759 | -1.322 | -1.841 | 0.604 | 0.012 | -0.01 | -0.01 | 0 | 0.27 | 0.24 | 0.28 | 0.3 | 0.19 | 0.16 |
| 3 | 31688392 | rs9816273 | -0.087 | -0.67 | -2.462 | 0.886 | -0.003 | -0.002 | -0.002 | 0 | 0.1 | 0.1 | 0.04 | 0.04 | 0.07 | 0.1 |
| 3 | 31695245 | rs2124488 | -0.479 | -0.674 | -2.477 | 0.905 | 0.09 | 0.071 | 0.071 | 0.103 | 0.49 | 0.91 | 0.43 | 0.68 | 0.37 | 0.39 |
| 3 | 31698150 | rs9310975 | -0.101 | -0.512 | -2.384 | 0.926 | -0.002 | 0 | 0 | 0.817 | 0.1 | 0.1 | 0.05 | 0.05 | 0.08 | 0.1 |
| 3 | 31708743 | rs9866738 | -0.348 | 0.219 | -2.342 | 0.651 | 0.085 | 0.032 | 0.032 | 0.047 | 0.49 | 0.91 | 0.41 | 0.65 | 0.37 | 0.37 |
| 3 | 31712393 | rs3749405 | -0.354 | 0.245 | -2.325 | 0.632 | 0.095 | 0.082 | 0.082 | 0.062 | 0.49 | 0.85 | 0.41 | 0.68 | 0.37 | 0.37 |
| 3 | 31714174 | rs7643110 | -0.362 | 0.306 | -2.362 | 0.669 | 0.087 | 0.082 | 0.082 | 0.055 | 0.49 | 0.86 | 0.41 | 0.68 | 0.38 | 0.37 |
| 3 | 31717019 | rs1107643 | -0.402 | 0.418 | -2.234 | 0.705 | 0.03 | 0.049 | 0.049 | 0.223 | 0.51 | 0.83 | 0.43 | 0.65 | 0.44 | 0.41 |
| 3 | 31725316 | rs2278959 | -0.142 | 0.538 | -2.12 | 0.836 | -0.008 | -0.047 | -0.047 | 0 | 0.31 | 0.1 | 0.29 | 0.32 | 0.31 | 0.21 |
| 3 | 31725691 | rs2278960 | -0.391 | 0.765 | -2.136 | 0.266 | -0.112 | 0.044 | 0.044 | 0.046 | 0.61 | 0.94 | 0.48 | 0.7 | 0.51 | 0.5 |
| 3 | 31726314 | rs1902334 | 0.914 | 1.071 | -0.866 | 0.021 | 0.013 | -0.076 | -0.076 | 0 | 0.85 | 0.68 | 0.91 | 0.97 | 0.85 | 0.85 |
| 3 | 31726603 | rs6785222 | 1.047 | 1.3 | -0.585 | 0.327 | -0.007 | 0 | 0 | 0 | 0.6 | 0.39 | 0.57 | 0.33 | 0.51 | 0.46 |
| 3 | 31732743 | rs7649547 | 1.231 | 0.607 | 0.159 | 0.223 | -0.037 | -0.039 | -0.039 | 0 | 0.83 | 0.31 | 0.9 | 0.92 | 0.92 | 0.83 |
| 3 | 31734989 | rs11921928 | -0.838 | 0.831 | 0.218 | 0.646 | 0.001 | 0.031 | 0.031 | 0.048 | 0.16 | 0.22 | 0.44 | 0.58 | 0.21 | 0.24 |
| 3 | 31744233 | rs3805074 | 1.404 | 0.756 | 0.981 | 0.555 | -0.007 | 0.057 | 0.057 | 0.032 | 0.74 | 0.52 | 0.51 | 0.38 | 0.73 | 0.66 |
| 3 | 31752607 | rs11129458 | -0.472 | 0.806 | 0.674 | 0.606 | -0.023 | -0.066 | -0.066 | 0 | 0.14 | 0.06 | 0.41 | 0.61 | 0.2 | 0.21 |
| 3 | 31754127 | rs9310976 | 1.255 | 1.356 | 1.348 | 0.035 | 0.074 | -0.05 | -0.05 | 0 | 0.85 | 0.52 | 0.84 | 0.92 | 0.81 | 0.7 |
| 3 | 31755406 | rs9825159 | -1.623 | 1.294 | 1.369 | 0.046 | 0.076 | -0.051 | -0.051 | 0 | 0.14 | 0.48 | 0.15 | 0.05 | 0.17 | 0.22 |
| 3 | 31757599 | rs3792549 | -1.174 | 0.945 | 0.595 | 0.028 | -0.072 | -0.029 | -0.029 | 0 | 0.25 | 0.1 | 0.52 | 0.69 | 0.36 | 0.43 |
| 3 | 31760436 | rs3762794 | -0.55 | 1.066 | 0.638 | 0.15 | -0.068 | -0.073 | -0.073 | 0 | 0.2 | 0.1 | 0.49 | 0.61 | 0.34 | 0.38 |
| 3 | 31764508 | rs7613225 | -1.307 | 0.725 | 1.19 | 0.102 | 0.018 | 0.041 | 0.041 | 2.054 | 0.07 | 0.29 | 0.03 | 0.04 | 0.02 | 0.04 |
| 3 | 31770787 | rs7620402 | -1.079 | 1.332 | 0.964 | 0.07 | 0.044 | 0.038 | 0.038 | 0.266 | 0.24 | 0.37 | 0.51 | 0.59 | 0.38 | 0.37 |
| 3 | 31776980 | rs4374563 | -1.124 | 1.509 | 0.18 | 0.139 | 0.003 | 0.001 | 0.001 | 1.325 | 0.28 | 0.36 | 0.55 | 0.63 | 0.42 | 0.42 |
| 3 | 31783326 | rs9825170 | 0.962 | 1.009 | 0.145 | -0.405 | -0.004 | -0.009 | -0.009 | 0 | 0.56 | 0.5 | 0.31 | 0.3 | 0.4 | 0.4 |
| 3 | 31784179 | rs9826527 | -0.897 | 0.849 | 0.359 | -0.715 | -0.015 | 0.084 | 0.084 | 0 | 0.4 | 0.2 | 0.62 | 0.77 | 0.52 | 0.57 |
| 3 | 31785560 | rs7634193 | -0.347 | 0.845 | 0.946 | -0.84 | -0.008 | 0.004 | 0.004 | 0.18 | 0.3 | 0.26 | 0.59 | 0.75 | 0.44 | 0.51 |
| 3 | 31786371 | rs7614865 | -1.054 | -0.072 | 0.441 | -1.539 | -0.03 | -0.122 | -0.122 | 2.826 | 0.81 | 0.6 | 0.88 | 1 | 0.88 | 0.91 |
| 3 | 31789582 | rs2290532 | 0.336 | 0.287 | 0.188 | -1.354 | -0.163 | -0.003 | -0.003 | 0 | 0.29 | 0.05 | 0.56 | 0.74 | 0.44 | 0.52 |
| 3 | 31791851 | rs4955112 | 0.332 | 0.539 | 0.23 | -1.369 | -0.115 | 0.005 | 0.005 | 0.486 | 0.28 | 0.1 | 0.55 | 0.71 | 0.42 | 0.49 |
| 3 | 31792918 | rs12632849 | 0.525 | 0.904 | 0.195 | -1.139 | -0.125 | -0.104 | -0.104 | 0 | 0.26 | 0.05 | 0.45 | 0.6 | 0.35 | 0.37 |
| 3 | 31795570 | rs1902341 | -0.94 | 0.72 | 0.589 | -1.238 | -0.004 | 0.002 | 0.002 | 0 | 0.42 | 0.58 | 0.55 | 0.63 | 0.44 | 0.44 |
| 3 | 31804842 | rs2168422 | -0.375 | 0.804 | 0.463 | -1.231 | 0.002 | 0.006 | 0.006 | 0.007 | 0.29 | 0.7 | 0.27 | 0.18 | 0.23 | 0.23 |
| 3 | 31806188 | rs7621973 | -0.272 | 0.807 | 0.573 | -1.204 | -0.008 | -0.023 | -0.023 | 0 | 0.74 | 0.45 | 0.74 | 0.82 | 0.78 | 0.75 |
| 3 | 31807370 | rs11716090 | 0.595 | 0.756 | 0.373 | -1.232 | 0.013 | 0.061 | 0.061 | 0.755 | 0.66 | 0.12 | 0.69 | 0.81 | 0.73 | 0.69 |
| 3 | 31807576 | rs11716163 | 0.92 | 0.93 | 0.244 | -1.225 | -0.113 | -0.159 | -0.159 | 0.83 | 0.22 | 0.01 | 0.41 | 0.55 | 0.28 | 0.35 |
| 3 | 31807981 | rs1902338 | 1.026 | 1.018 | 1.066 | -1.263 | 0.016 | 0.061 | 0.061 | 0.795 | 0.66 | 0.12 | 0.7 | 0.81 | 0.74 | 0.69 |
| 3 | 31811625 | rs12497607 | 1.196 | 1.072 | 1.175 | -1.246 | 0.025 | 0.069 | 0.069 | 0.159 | 0.63 | 0.18 | 0.68 | 0.8 | 0.69 | 0.69 |
| 3 | 31817544 | rs2219250 | -0.001 | 1.132 | 1.191 | -1.233 | -0.004 | -0.008 | -0.008 | 0.682 | 0.75 | 0.62 | 0.76 | 0.82 | 0.78 | 0.76 |
| 3 | 31817884 | rs6796431 | -1.659 | 0.962 | 0.868 | -1.239 | -0.018 | -0.073 | -0.073 | 0.853 | 0.12 | 0.37 | 0.07 | 0.02 | 0.09 | 0.06 |
| 3 | 31827315 | rs4955200 | 0.798 | 1.056 | 0.452 | -1.306 | -0.002 | -0.032 | -0.032 | 3.29 | 0.67 | 0.37 | 0.7 | 0.8 | 0.69 | 0.68 |
| 3 | 31829150 | rs13094960 | 1.316 | 1.048 | 0.11 | -1.381 | 0.001 | 0.023 | 0.023 | 0.015 | 0.59 | 0.32 | 0.66 | 0.8 | 0.62 | 0.67 |
| 3 | 31831109 | rs6798458 | -1.349 | 1.045 | 0.111 | -1.38 | 0.005 | 0.026 | 0.026 | 0.015 | 0.39 | 0.69 | 0.33 | 0.2 | 0.36 | 0.3 |
| 3 | 31839135 | rs7653447 | 0.892 | 1.049 | -0.091 | -1.299 | 0 | 0.016 | 0.016 | 0.093 | 0.65 | 0.29 | 0.7 | 0.8 | 0.68 | 0.71 |
| 3 | 31839242 | rs7643025 | -0.443 | 1.105 | -0.182 | -1.169 | -0.014 | 0.009 | 0.009 | 0 | 0.77 | 0.88 | 0.77 | 0.82 | 0.75 | 0.76 |
| 3 | 31846251 | rs4955203 | 0.987 | 0.649 | -0.422 | -1.048 | 0.012 | 0.11 | 0.11 | 0.199 | 0.51 | 0.28 | 0.6 | 0.74 | 0.51 | 0.57 |
| 3 | 31847360 | rs4611855 | 0.473 | 1.289 | -0.783 | -0.594 | 0 | -0.007 | -0.007 | 0 | 0.64 | 0.67 | 0.71 | 0.77 | 0.61 | 0.64 |
| 3 | 31849818 | rs7646594 | -1.386 | 1.401 | -0.987 | -0.228 | -0.037 | 0.042 | 0.042 | 0 | 0.08 | 0.01 | 0.06 | 0.01 | 0.1 | 0.03 |

| | | | | | | | | | | | | | | | | |
|---|---|---|---|---|---|---|---|---|---|---|---|---|---|---|---|---|
| 3 | 31850663 | rs7649767 | -1.25 | 0.862 | -0.724 | -0.087 | -0.004 | -0.009 | -0.009 | 0.107 | 0.17 | 0.71 | 0.14 | 0.09 | 0.17 | 0.14 |
| 3 | 31851726 | rs6773772 | -0.683 | 0.684 | -0.53 | 0.164 | -0.007 | -0.003 | -0.003 | 0 | 0.32 | 0.35 | 0.28 | 0.29 | 0.3 | 0.25 |
| 3 | 31852083 | rs7653230 | -1.732 | 0.96 | 0.581 | -0.137 | 0.032 | -0.058 | -0.058 | 0 | 0.11 | 0.5 | 0.1 | 0 | 0.09 | 0.07 |
| 3 | 31856030 | rs13075708 | 0.942 | 2.26 | 1.031 | -0.455 | 0.083 | -0.073 | -0.073 | 0 | 0.83 | 0.71 | 0.83 | 0.96 | 0.73 | 0.85 |
| 3 | 31856491 | rs4591530 | 1.787 | 1.5 | 1.134 | -0.662 | 0.014 | -0.064 | -0.064 | 0 | 0.74 | 0.45 | 0.81 | 0.94 | 0.73 | 0.82 |
| 3 | 31857836 | rs13087711 | 1.692 | 1.205 | 0.758 | -0.678 | 0.007 | 0.076 | 0.076 | 0 | 0.49 | 0.36 | 0.58 | 0.7 | 0.51 | 0.62 |
| 3 | 31858879 | rs9877405 | 1.035 | 1.211 | 0.781 | -0.736 | 0.081 | -0.109 | -0.109 | 4.913 | 0.78 | 0.59 | 0.82 | 0.94 | 0.73 | 0.82 |
| 3 | 31860058 | rs4955208 | 1.068 | 1.212 | 0.807 | -0.736 | 0.079 | -0.111 | -0.111 | 0.112 | 0.78 | 0.57 | 0.82 | 0.94 | 0.73 | 0.82 |
| 3 | 31860451 | rs4955210 | 1.744 | 0.795 | -0.261 | -0.339 | -0.013 | 0.046 | 0.046 | 0 | 0.39 | 0.15 | 0.23 | 0.28 | 0.38 | 0.37 |
| 3 | 31862681 | rs4476542 | 0.086 | 0.559 | -0.314 | -0.167 | -0.024 | 0.023 | 0.023 | 0 | 0.27 | 0.08 | 0.24 | 0.19 | 0.28 | 0.21 |
| 3 | 31862952 | rs11720782 | -0.147 | 0.431 | -0.216 | -0.006 | 0.018 | -0.027 | -0.027 | 0.225 | 0.1 | 0.06 | 0.04 | 0.47 | 0.06 | 0.12 |
| 3 | 31873214 | rs9880436 | 1.347 | 0.564 | -0.021 | -0.09 | 0 | 0.042 | 0.042 | 0 | 0.49 | 0.35 | 0.56 | 0.32 | 0.52 | 0.51 |
| 3 | 31874014 | rs6807471 | 0.742 | 0.487 | -0.064 | -0.01 | -0.004 | 0.016 | 0.016 | 0 | 0.6 | 0.39 | 0.6 | 0.78 | 0.58 | 0.67 |
| 3 | 31876100 | rs4485758 | 1.084 | 0.663 | 0.053 | 0.102 | 0.022 | 0.072 | 0.072 | 0 | 0.75 | 0.64 | 0.79 | 0.47 | 0.72 | 0.75 |
| 3 | 31883771 | rs7611111 | -0.022 | 0.891 | 0.075 | 0.116 | -0.011 | -0.074 | -0.074 | 0 | 0.28 | 0.16 | 0.23 | 0.51 | 0.31 | 0.19 |
| 3 | 31885869 | rs7633447 | 0.192 | 1.185 | 0.076 | 0.156 | 0.106 | -0.006 | -0.006 | 0 | 0.86 | 0.71 | 0.83 | 0.9 | 0.78 | 0.9 |
| 3 | 31890725 | rs6550076 | 0.562 | 1.068 | 0.039 | 0.101 | 0.044 | -0.009 | -0.009 | 0 | 0.85 | 0.77 | 0.84 | 0.88 | 0.78 | 0.87 |
| 3 | 31891118 | rs12487988 | 0.837 | 0.502 | -0.339 | -0.053 | 0.01 | 0.013 | 0.013 | 0 | 0.51 | 0.42 | 0.55 | 0.34 | 0.41 | 0.57 |
| 3 | 31901521 | rs6776025 | -1.239 | 0.494 | 0.261 | 0.104 | -0.068 | -0.02 | -0.02 | 0 | 0.13 | 0.54 | 0.08 | 0.08 | 0.03 | 0.14 |
| 3 | 31901848 | rs6776254 | 0.423 | 1.625 | -0.386 | 0.097 | 0.029 | -0.008 | -0.008 | 0 | 0.86 | 0.81 | 0.57 | 0.91 | 0.78 | 0.92 |
| 3 | 31908377 | rs4452359 | -0.841 | 1.339 | -0.928 | 0.822 | -0.001 | 0 | 0 | 0 | 0.23 | 0.2 | 0.17 | 0.27 | 0.31 | 0.17 |
| 3 | 31919601 | rs13091847 | -0.735 | 0.797 | -1.396 | 1.036 | -0.005 | -0.037 | -0.037 | 0 | 0.17 | 0.02 | 0.08 | 0.13 | 0.14 | 0.12 |
| 3 | 31922955 | rs6550080 | -0.625 | 1.286 | -1.874 | 0.594 | -0.072 | 0.067 | 0.067 | 0 | 0.72 | 0.92 | 0.63 | 0.87 | 0.66 | 0.75 |
| 3 | 31926807 | rs4478116 | -0.584 | 1.267 | -0.77 | 1.072 | 0.074 | 0.171 | 0.171 | 0 | 0.05 | 0.25 | 0.23 | 0.17 | 0.08 | 0.05 |
| 3 | 31928640 | rs4282098 | 0.504 | 0.775 | -1.176 | 0.83 | -0.022 | -0.011 | -0.011 | 0 | 0.21 | 0.12 | 0.15 | 0.24 | 0.25 | 0.24 |
| 3 | 31933619 | rs4334675 | -0.654 | 0.609 | -0.736 | 0.537 | 0.004 | -0.005 | -0.005 | 0 | 0.15 | 0.25 | 0.11 | 0.13 | 0.13 | 0.14 |
| 3 | 31942810 | rs4478115 | 0.227 | 0.361 | -1.13 | 0.449 | -0.013 | -0.003 | -0.003 | 0.634 | 0.23 | 0.14 | 0.17 | 0.22 | 0.23 | 0.22 |
| 3 | 31944640 | rs9811517 | 0.385 | -0.199 | -0.836 | 0.191 | -0.023 | -0.021 | -0.021 | 0.695 | 0.1 | 0.21 | 0.06 | 0.06 | 0.07 | 0.11 |
| 3 | 31945442 | rs4334676 | 1.191 | -0.049 | -1.891 | -0.266 | -0.011 | 0.116 | 0.116 | 0 | 0.26 | 0.09 | 0.24 | 0.06 | 0.26 | 0.21 |
| 3 | 31946673 | rs11712058 | 0.498 | 0.121 | -1.638 | 0.049 | -0.014 | -0.007 | -0.007 | 0 | 0.23 | 0.1 | 0.16 | 0.22 | 0.23 | 0.22 |
| 3 | 31950590 | rs11129470 | -0.24 | -0.074 | -1.329 | -0.178 | 0.013 | 0.042 | 0.042 | 0 | 0.55 | 0.45 | 0.48 | 0.84 | 0.62 | 0.65 |
| 3 | 31951420 | rs13317583 | 0.668 | -0.032 | -0.921 | -0.028 | -0.007 | 0.009 | 0.009 | 0 | 0.15 | 0.41 | 0.29 | 0.12 | 0.12 | 0.18 |
| 3 | 31953943 | rs9849233 | -0.351 | -0.375 | -1.206 | 0.305 | -0.002 | 0.001 | 0.001 | 0 | 0.51 | 0.55 | 0.51 | 0.51 | 0.51 | 0.53 |
| 3 | 31954098 | rs6767086 | -0.04 | -0.411 | -1.19 | 0.326 | 0.002 | 0.005 | 0.005 | 0.098 | 0.28 | 0.53 | 0.32 | 0.24 | 0.25 | 0.27 |
| 3 | 31955806 | rs11924760 | 0.398 | -0.081 | -1.106 | 0.429 | 0.025 | 0.029 | 0.029 | 0 | 0.3 | 0.43 | 0.41 | 0.38 | 0.36 | 0.32 |
| 3 | 31956117 | rs9842494 | -0.352 | -0.648 | -1.254 | -0.121 | 0.05 | 0.104 | 0.104 | 0 | 0.54 | 0.69 | 0.69 | 0.92 | 0.65 | 0.71 |
| 3 | 31958529 | rs7641393 | -1.077 | -0.836 | -1.225 | 0.245 | -0.003 | 0.003 | 0.003 | 0 | 0.72 | 0.74 | 0.72 | 0.78 | 0.72 | 0.72 |
| 3 | 31959990 | rs11929134 | 1.162 | -0.712 | -0.975 | 0.207 | 0.022 | -0.007 | -0.007 | 0 | 0.12 | 0.3 | 0.23 | 0.09 | 0.12 | 0.2 |
| 3 | 31960866 | rs4444747 | 0.706 | -0.63 | -1.368 | 0.306 | -0.053 | -0.019 | -0.019 | 0 | 0.4 | 0.2 | 0.39 | 0.37 | 0.48 | 0.32 |
| 3 | 31962771 | rs4532171 | -1.405 | -0.625 | -1.261 | 0.036 | 0.004 | 0.009 | 0.009 | 0 | 0.73 | 0.74 | 0.77 | 0.96 | 0.78 | 0.85 |
| 3 | 31964898 | rs7644177 | -0.208 | -0.651 | -1.189 | 0.178 | 0.003 | -0.01 | -0.01 | 1.497 | 0.56 | 0.53 | 0.68 | 0.91 | 0.67 | 0.75 |
| 3 | 31965758 | rs4561865 | 0.161 | -0.698 | -1.205 | 0.258 | 0.004 | 0.015 | 0.015 | 0 | 0.51 | 0.38 | 0.62 | 0.75 | 0.6 | 0.64 |
| 3 | 31970818 | rs6796265 | -1.158 | -1.363 | -0.601 | 0.546 | 0 | -0.003 | -0.003 | 0 | 0.46 | 0.48 | 0.57 | 0.69 | 0.49 | 0.66 |
| 3 | 31978325 | rs11925027 | 0.66 | -1.37 | -0.476 | 0.554 | 0.014 | 0.002 | 0.002 | 0 | 0.13 | 0.29 | 0.25 | 0.11 | 0.13 | 0.28 |
| 3 | 31980606 | rs6800757 | -2.175 | -1.508 | -0.649 | 0.479 | -0.022 | 0.053 | 0.053 | 0 | 0.55 | 0.81 | 0.62 | 0.7 | 0.52 | 0.68 |
| 3 | 31981227 | rs12630254 | -0.279 | -1.288 | -0.612 | 0.425 | -0.075 | 0.075 | 0.075 | 0 | 0.27 | 0.05 | 0.17 | 0.16 | 0.27 | 0.18 |
| 3 | 31988094 | rs4073766 | -1.151 | -0.984 | 0.202 | 1.12 | -0.013 | 0.006 | 0.006 | 0 | 0.24 | 0.66 | 0.32 | 0.34 | 0.17 | 0.37 |
| 3 | 31989487 | rs9853939 | -2.045 | -0.911 | -0.293 | 1.041 | -0.064 | -0.062 | -0.062 | 0.86 | 0.51 | 0.92 | 0.5 | 0.54 | 0.46 | 0.59 |
| 3 | 31989920 | rs11916062 | 0.004 | -0.711 | 0.148 | 0.558 | 0.019 | -0.006 | -0.006 | 0 | 0.14 | 0.31 | 0.26 | 0.11 | 0.14 | 0.29 |
| 3 | 31999834 | rs11921166 | 0.038 | -0.703 | 0.077 | 0.55 | 0.017 | -0.009 | -0.009 | 0 | 0.14 | 0.29 | 0.25 | 0.11 | 0.14 | 0.3 |
| 3 | 32001042 | rs9847800 | -2.347 | -0.838 | -0.576 | 0.547 | -0.051 | 0.094 | 0.094 | 0 | 0.55 | 0.9 | 0.63 | 0.75 | 0.53 | 0.72 |
| 3 | 32001194 | rs9847964 | -1.015 | -0.79 | -0.332 | 0.592 | 0 | -0.004 | -0.004 | 0 | 0.32 | 0.36 | 0.21 | 0.18 | 0.31 | 0.22 |
| 3 | 32002469 | rs11916514 | -0.578 | -1.009 | -0.478 | 1.196 | -0.016 | -0.014 | -0.014 | 0 | 0.18 | 0.13 | 0.27 | 0.25 | 0.2 | 0.35 |
| 3 | 32003769 | rs12489744 | -1.914 | -1.455 | -0.297 | 1.251 | -0.022 | 0.065 | 0.065 | 0 | 0.2 | 0.66 | 0.2 | 0.53 | 0.11 | 0.25 |
| 3 | 32011584 | rs11719930 | 0.07 | -0.996 | -0.313 | 1.27 | -0.003 | 0.001 | 0.001 | 0 | 0.12 | 0.08 | 0.22 | 0.12 | 0.15 | 0.34 |
| 3 | 32016607 | rs7635119 | -1.668 | -1.224 | -0.136 | 1.279 | 0.012 | 0 | 0 | 0 | 0.3 | 0.7 | 0.35 | 0.31 | 0.21 | 0.39 |
| 3 | 32017806 | rs6769296 | -0.212 | -1.226 | 0.101 | 1.286 | 0.039 | 0.102 | 0.102 | 0.931 | 0.07 | 0.41 | 0.05 | 0.05 | 0.02 | 0.02 |
| 3 | 32018536 | rs12485651 | -0.459 | -1.249 | -0.208 | 1.422 | -0.022 | 0.046 | 0.046 | 0.095 | 0.05 | 0 | 0.12 | 0.47 | 0.09 | 0.18 |
| 3 | 32019685 | rs4383559 | -1.451 | -1.26 | -0.189 | 1.38 | -0.011 | -0.037 | -0.037 | 0.233 | 0.27 | 0.15 | 0.4 | 0.72 | 0.28 | 0.55 |
| 3 | 32033248 | rs11718700 | -1.866 | -1.171 | 0.026 | 1.33 | 0.031 | 0.003 | 0.003 | 3.798 | 0.31 | 0.8 | 0.36 | 0.3 | 0.19 | 0.36 |
| 3 | 32033850 | rs6806060 | -0.559 | -1.13 | 0.46 | 1.433 | 0.043 | 0.093 | 0.093 | 1.025 | 0.07 | 0.43 | 0.05 | 0.04 | 0.02 | 0.02 |
| 3 | 32048179 | rs13327469 | -0.824 | -1.108 | -0.439 | 1.45 | -0.004 | 0.037 | 0.037 | 0.014 | 0.18 | 0.06 | 0.16 | 0.13 | 0.16 | 0.1 |
| 3 | 32049091 | rs6772958 | -1.764 | -1.2 | 0.057 | 1.412 | -0.013 | 0.031 | 0.031 | 0 | 0.29 | 0.53 | 0.34 | 0.45 | 0.25 | 0.26 |
| 3 | 32062641 | rs4955124 | 1.099 | -1.607 | 0.039 | 1.548 | -0.043 | 0.098 | 0.098 | 1.1 | 0.79 | 0.45 | 0.78 | 0.43 | 0.9 | 0.79 |
| 3 | 32064607 | rs1597666 | 1.108 | -1.771 | -0.037 | 1.066 | 0.005 | -0.001 | -0.001 | 0 | 0.6 | 0.55 | 0.54 | 0.56 | 0.67 | 0.56 |
| 3 | 32068951 | rs12496020 | 1.107 | -1.904 | 0.242 | 0.775 | 0.016 | 0.018 | 0.018 | 0 | 0.69 | 0.33 | 0.78 | 0.78 | 0.81 | 0.87 |
| 3 | 32082789 | rs901089 | 0.394 | -1.721 | 0.674 | 0.693 | -0.021 | -0.153 | -0.153 | 0 | 0.77 | 0.58 | 0.83 | 1 | 0.81 | 0.91 |
| 3 | 32092872 | rs12488844 | 1.208 | -1.706 | 0.207 | 0.677 | 0.025 | 0.041 | 0.041 | 0 | 0.66 | 0.31 | 0.76 | 0.8 | 0.78 | 0.86 |
| 3 | 32101543 | rs1073208 | -0.161 | -1.637 | 0.816 | 0.679 | -0.044 | -0.078 | -0.078 | 0 | 0.81 | 0.58 | 0.92 | 0.9 | 0.91 | 0.97 |

| | | | | | | | | | | | | | | | | |
|---|---|---|---|---|---|---|---|---|---|---|---|---|---|---|---|---|
| 3 | 32107951 | rs11129479 | 1.358 | -1.564 | -0.124 | 1.223 | 0.028 | -0.068 | -0.068 | 0.016 | 0.59 | 0.13 | 0.62 | 0.42 | 0.69 | 0.61 |
| 3 | 32120106 | rs4955237 | 0.974 | -1.737 | 0.332 | 0.83 | 0.006 | 0.006 | 0.006 | 1.01 | 0.66 | 0.35 | 0.76 | 0.78 | 0.8 | 0.78 |
| 3 | 32121979 | rs1376605 | 0.927 | -1.777 | 0.285 | 1.001 | 0.013 | 0.012 | 0.012 | 0.909 | 0.66 | 0.33 | 0.72 | 0.82 | 0.84 | 0.79 |
| 3 | 32122731 | rs17028838 | -1.068 | -1.415 | 0.249 | 1.033 | 0 | 0 | 0 | 0.013 | 0.09 | 0.1 | 0.09 | 0.13 | 0.09 | 0.11 |
| 3 | 32125010 | rs7643045 | -0.581 | -1.668 | 0.413 | 0.974 | -0.006 | 0.018 | 0.018 | 0 | 0.25 | 0.38 | 0.32 | 0.74 | 0.25 | 0.56 |
| 3 | 32125402 | rs17028843 | -0.232 | -1.742 | 0.306 | 0.657 | -0.003 | -0.046 | -0.046 | 0 | 0.1 | 0.19 | 0.1 | 0 | 0.09 | 0.08 |
| 3 | 32125482 | rs4955238 | 1.844 | -2.102 | -0.317 | 0.455 | -0.144 | 0.152 | 0.152 | 3.514 | 0.39 | 0.05 | 0.39 | 0.12 | 0.53 | 0.27 |
| 3 | 32128803 | rs7632997 | 1.91 | -2.262 | -0.345 | 0.063 | -0.113 | 0.196 | 0.196 | 5.564 | 0.37 | 0.11 | 0.4 | 0.06 | 0.53 | 0.26 |
| 3 | 32129315 | rs4955239 | -0.598 | -2.302 | -0.036 | 0.69 | 0.014 | -0.025 | -0.025 | 0 | 0.81 | 0.91 | 0.81 | 0.72 | 0.88 | 0.91 |
| 3 | 32130942 | rs2341419 | -1.253 | -2.292 | 0.057 | 0.535 | 0.016 | 0.017 | 0.017 | 1.301 | 0.84 | 0.9 | 0.95 | 1 | 0.92 | 0.99 |
| 3 | 32141891 | rs6790564 | 0.091 | -2.282 | 0.053 | 0.956 | -0.009 | -0.034 | -0.034 | 4.633 | 0.17 | 0.14 | 0.14 | 0.49 | 0.2 | 0.48 |
| 3 | 32141967 | rs9874619 | 1.972 | -2.418 | -0.1 | 0.646 | -0.092 | 0.217 | 0.217 | 2.585 | 0.4 | 0.07 | 0.43 | 0.07 | 0.57 | 0.26 |
| 3 | 32146188 | rs1159444 | -0.563 | -1.606 | 1.168 | 1.226 | 0.016 | -0.071 | -0.071 | 0 | 0.12 | 0.26 | 0.09 | 0 | 0.11 | 0.07 |
| 3 | 32154324 | rs13061752 | -0.334 | -1.648 | 0.931 | 1.396 | -0.008 | -0.038 | -0.038 | 0 | 0.16 | 0.13 | 0.14 | 0.5 | 0.2 | 0.47 |
| 3 | 32154422 | rs13061945 | -1.882 | -1.689 | 1.206 | 1.318 | 0.003 | -0.007 | -0.007 | 0 | 0.33 | 0.65 | 0.27 | 0.64 | 0.33 | 0.59 |
| 3 | 32165335 | rs1839778 | -1.357 | -1.479 | 1.398 | 1.444 | -0.03 | 0.041 | 0.041 | 0 | 0.29 | 0.52 | 0.23 | 0.64 | 0.23 | 0.54 |
| 3 | 32181403 | rs12637969 | -1.21 | -1.28 | 1.597 | 1.375 | -0.026 | 0.041 | 0.041 | 0 | 0.27 | 0.41 | 0.3 | 0.63 | 0.23 | 0.45 |
| 3 | 32181761 | rs9835387 | -0.709 | -1.275 | 1.525 | 1.221 | 0.031 | -0.075 | -0.075 | 0 | 0.13 | 0.3 | 0.12 | 0 | 0.14 | 0.14 |
| 3 | 32187943 | rs9839669 | -1.603 | -1.254 | 1.296 | 1.309 | -0.004 | 0.008 | 0.008 | 0 | 0.37 | 0.57 | 0.39 | 0.49 | 0.34 | 0.54 |
| 3 | 32205940 | rs11129500 | -1.839 | -0.67 | 0.72 | 1.05 | -0.003 | 0.003 | 0.003 | 0 | 0.42 | 0.74 | 0.44 | 0.6 | 0.43 | 0.56 |
| 3 | 32208121 | rs11923184 | 0.124 | -0.668 | 0.859 | 1.014 | 0.002 | -0.018 | -0.018 | 0 | 0.08 | 0.08 | 0.07 | 0 | 0.12 | 0.11 |
| 3 | 32208431 | rs6802739 | -2.011 | -0.703 | 0.85 | 1.023 | -0.017 | 0 | 0 | 0 | 0.43 | 0.82 | 0.46 | 0.61 | 0.43 | 0.56 |
| 3 | 32208621 | rs2813 | -1.267 | -0.787 | 1.073 | 1.045 | -0.003 | 0.011 | 0.011 | 0 | 0.32 | 0.6 | 0.37 | 0.61 | 0.32 | 0.45 |
| 3 | 32209230 | rs3749278 | -1.829 | -0.793 | 1.044 | 1.005 | 0.005 | 0.001 | 0.001 | 0 | 0.41 | 0.7 | 0.43 | 0.61 | 0.4 | 0.55 |
| 3 | 32209927 | rs7363 | -1.815 | -0.988 | 0.861 | 0.846 | 0.025 | 0.024 | 0.024 | 0 | 0.47 | 0.71 | 0.45 | 0.6 | 0.43 | 0.59 |
| 3 | 32215390 | rs6550098 | -0.615 | -0.993 | 1.26 | 1.037 | -0.022 | 0.008 | 0.008 | 0 | 0.19 | 0.44 | 0.12 | 0.19 | 0.15 | 0.17 |
| 3 | 32216014 | rs6784980 | -0.383 | -0.975 | 1.18 | 1.045 | -0.014 | 0.002 | 0.002 | 0 | 0.17 | 0.31 | 0.1 | 0.19 | 0.15 | 0.18 |
| 3 | 32217789 | rs2060780 | -1.554 | -0.913 | 0.517 | 1.026 | -0.006 | -0.002 | -0.002 | 0 | 0.43 | 0.46 | 0.41 | 0.41 | 0.38 | 0.56 |
| 3 | 32222173 | rs17028998 | 0.015 | -0.996 | -0.223 | 1.168 | -0.003 | -0.044 | -0.044 | 0 | 0.1 | 0.01 | 0.04 | 0.33 | 0.08 | 0.31 |
| 3 | 32222434 | rs9827326 | -1.754 | -0.708 | -0.211 | 1.07 | 0.061 | 0.252 | 0.252 | 0 | 0.5 | 0.82 | 0.57 | 0.88 | 0.41 | 0.63 |
| 3 | 32223481 | rs9842565 | -0.409 | -0.752 | -0.088 | 0.931 | -0.017 | 0.082 | 0.082 | 0 | 0.23 | 0.49 | 0.25 | 0.39 | 0.19 | 0.38 |
| 3 | 32225203 | rs11922502 | 0.308 | -0.601 | -0.345 | 0.527 | -0.005 | -0.039 | -0.039 | 0 | 0.16 | 0.2 | 0.08 | 0 | 0.11 | 0.11 |
| 3 | 32225326 | rs7644491 | -0.889 | -0.02 | -0.827 | 0.921 | 0.053 | 0.04 | 0.04 | 0 | 0.49 | 0.73 | 0.44 | 0.61 | 0.41 | 0.6 |
| 3 | 32233643 | rs9867908 | -1.032 | 0.376 | -0.918 | 0.328 | 0 | 0.007 | 0.007 | 0 | 0.57 | 0.63 | 0.62 | 0.88 | 0.56 | 0.71 |
| 3 | 32234547 | rs11711134 | -0.271 | 0.48 | -1.416 | 0.767 | -0.117 | -0.128 | -0.128 | 0 | 0.3 | 0.08 | 0.3 | 0.41 | 0.38 | 0.46 |
| 3 | 32234614 | rs6550105 | -1.028 | 0.36 | -0.919 | 0.836 | 0.001 | 0.008 | 0.008 | 0 | 0.54 | 0.59 | 0.6 | 0.68 | 0.54 | 0.66 |
| 3 | 32235481 | rs11720070 | -0.766 | 0.056 | -0.92 | 0.967 | 0.027 | 0.037 | 0.037 | 0 | 0.42 | 0.74 | 0.52 | 0.31 | 0.36 | 0.41 |
| 3 | 32235747 | rs9878447 | 0.777 | -0.316 | 0.118 | 1.241 | -0.025 | 0.153 | 0.153 | 0.297 | 0.87 | 0.51 | 0.78 | 0.74 | 0.9 | 0.85 |
| 3 | 32236058 | rs4440157 | 0.088 | 0.31 | -0.133 | 1.213 | 0.033 | 0.101 | 0.101 | 0 | 0.9 | 0.73 | 0.85 | 0.82 | 0.89 | 0.94 |
| 3 | 32236999 | rs11713463 | -0.331 | -0.067 | -0.865 | 0.868 | 0.012 | 0.003 | 0.003 | 0 | 0.16 | 0.08 | 0.15 | 0.13 | 0.1 | 0.27 |
| 3 | 32239401 | rs4955252 | 0.63 | 0.117 | -1.686 | 0.288 | -0.094 | 0.19 | 0.19 | 0 | 0.43 | 0.13 | 0.39 | 0.19 | 0.53 | 0.33 |
| 3 | 32239659 | rs4245875 | -1.088 | 0.303 | -1.314 | 0.911 | 0.002 | -0.028 | -0.028 | 0.155 | 0.78 | 0.85 | 0.69 | 0.68 | 0.77 | 0.69 |
| 3 | 32241789 | rs4245876 | -1.273 | 0.149 | -1.158 | 0.551 | -0.006 | -0.018 | -0.018 | 0.191 | 0.92 | 0.89 | 0.94 | 1 | 0.96 | 0.98 |
| 3 | 32245255 | rs4078487 | 0.716 | 0.269 | -0.787 | 0.859 | 0.028 | 0.081 | 0.081 | 0 | 0.09 | 0.17 | 0.09 | 0.19 | 0.1 | 0.09 |
| 3 | 32245557 | rs11713777 | 0.1 | 0.252 | -0.753 | 0.349 | 0.001 | -0.043 | -0.043 | 0 | 0.13 | 0.25 | 0.08 | 0.02 | 0.1 | 0.09 |
| 3 | 32248962 | rs4955138 | -0.15 | 0.113 | -0.768 | 0.604 | 0 | 0.076 | 0.076 | 0.062 | 0.78 | 0.48 | 0.82 | 0.63 | 0.8 | 0.84 |
| 3 | 32250919 | rs9310986 | 0.742 | 0.011 | -1.487 | -0.303 | -0.048 | 0.166 | 0.166 | 0 | 0.37 | 0.11 | 0.24 | 0.17 | 0.41 | 0.29 |
| 3 | 32251471 | rs7616055 | 0.176 | 0.401 | -0.731 | 0.308 | -0.001 | -0.001 | -0.001 | 0.415 | 0.12 | 0.15 | 0.08 | 0.12 | 0.07 | 0.18 |
| 3 | 32258093 | rs4955261 | -0.469 | 0.417 | -0.63 | -0.095 | 0.009 | -0.053 | -0.053 | 0.768 | 0.7 | 0.61 | 0.78 | 0.83 | 0.68 | 0.77 |
| 3 | 32260154 | rs6800446 | -0.102 | 0.237 | -0.7 | 0.758 | 0.003 | 0.025 | 0.025 | 0 | 0.74 | 0.43 | 0.56 | 0.6 | 0.68 | 0.68 |
| 3 | 32260825 | rs9842135 | -0.654 | 0.013 | -1.328 | 0.408 | -0.043 | 0.025 | 0.025 | 0 | 0.69 | 0.92 | 0.84 | 0.74 | 0.66 | 0.7 |
| 3 | 32264265 | rs4245878 | 0.68 | -0.767 | -0.073 | 0.923 | -0.048 | 0.065 | 0.065 | 0.562 | 0.7 | 0.55 | 0.6 | 0.45 | 0.79 | 0.54 |
| 3 | 32264991 | rs4376071 | 1.192 | -0.945 | -0.009 | 0.339 | -0.027 | -0.014 | -0.014 | 0 | 0.67 | 0.49 | 0.61 | 0.67 | 0.72 | 0.59 |
| 3 | 32265784 | rs4381984 | 0.237 | -0.751 | 0.319 | 0.578 | -0.035 | 0.012 | 0.012 | 0.711 | 0.78 | 0.73 | 0.79 | 0.68 | 0.83 | 0.65 |
| 3 | 32267947 | rs4955265 | -0.108 | -0.776 | 0.309 | 0.516 | -0.036 | -0.078 | -0.078 | 0 | 0.82 | 0.73 | 0.85 | 0.92 | 0.86 | 0.68 |
| 3 | 32272534 | rs9833530 | 0.706 | -0.734 | 0.219 | 0.565 | -0.021 | 0.003 | 0.003 | 1.604 | 0.74 | 0.41 | 0.63 | 0.67 | 0.79 | 0.63 |
| 3 | 32275125 | rs6550108 | -0.075 | -0.546 | 0.322 | 0.086 | -0.032 | -0.062 | -0.062 | 1.304 | 0.86 | 0.66 | 0.92 | 0.91 | 0.9 | 0.73 |
| 3 | 32282686 | rs4075566 | -0.698 | -0.688 | 0.167 | -0.074 | -0.026 | -0.074 | -0.074 | 3.159 | 0.15 | 0.22 | 0.08 | 0 | 0.13 | 0.08 |
| 3 | 32285786 | rs4524313 | -1.062 | -0.232 | -0.133 | 0.439 | 0.001 | -0.015 | -0.015 | 0 | 0.23 | 0.16 | 0.48 | 0.61 | 0.22 | 0.25 |
| 3 | 32291203 | rs6550109 | 0.464 | 0.325 | -0.2 | 0.43 | 0.005 | 0.015 | 0.015 | 0 | 0.61 | 0.55 | 0.53 | 0.5 | 0.56 | 0.42 |
| 3 | 32299549 | rs12496256 | 0.296 | 0.398 | -0.229 | 0.398 | 0.003 | -0.018 | -0.018 | 0 | 0.67 | 0.75 | 0.64 | 0.55 | 0.67 | 0.51 |
| 3 | 32300636 | rs12497202 | -0.077 | 0.154 | -0.284 | -0.019 | -0.002 | 0.028 | 0.028 | 0.074 | 0.26 | 0.15 | 0.23 | 0.04 | 0.26 | 0.24 |
| 3 | 32303087 | rs13314943 | 0.199 | -0.062 | -0.336 | -0.402 | -0.001 | 0.107 | 0.107 | 1.408 | 0.79 | 0.36 | 0.69 | 0.49 | 0.77 | 0.81 |
| 3 | 32304103 | rs4245886 | 0.932 | 0.337 | -0.624 | -0.455 | -0.012 | 0.041 | 0.041 | 0 | 0.51 | 0.2 | 0.41 | 0.45 | 0.46 | 0.57 |
| 3 | 32312914 | rs7636660 | 0.645 | 0.349 | -0.365 | -0.44 | -0.003 | 0.012 | 0.012 | 0 | 0.55 | 0.35 | 0.46 | 0.45 | 0.51 | 0.57 |
| 3 | 32313676 | rs4955272 | 1.896 | 0.352 | -1.079 | -0.781 | -0.064 | 0.078 | 0.078 | 0 | 0.23 | 0.04 | 0.14 | 0.11 | 0.29 | 0.15 |
| 3 | 32316115 | rs13317353 | 0.131 | 0.633 | -0.187 | -0.279 | -0.002 | 0.034 | 0.034 | 1.001 | 0.22 | 0.23 | 0.18 | 0.46 | 0.2 | 0.21 |
| 3 | 32320408 | rs7644602 | 0.948 | 0.481 | -0.174 | -1.115 | 0 | 0.051 | 0.051 | 0 | 0.56 | 0.34 | 0.53 | 0.86 | 0.55 | 0.58 |
| 3 | 32324866 | rs6809090 | 0.297 | 0.923 | 0.088 | -0.92 | 0.005 | -0.052 | -0.052 | 0 | 0.69 | 0.58 | 0.59 | 0.87 | 0.65 | 0.67 |

| | | | | | | | | | | | | | | | | |
|---|---|---|---|---|---|---|---|---|---|---|---|---|---|---|---|---|
| 3 | 32325273 | rs4305450 | 0.079 | 1.024 | 0.084 | -0.845 | -0.007 | -0.054 | -0.054 | 0 | 0.25 | 0.3 | 0.14 | 0.02 | 0.21 | 0.28 |
| 3 | 32326122 | rs4325954 | -0.092 | 1.308 | 0.235 | -0.05 | 0.023 | -0.016 | -0.016 | 0 | 0.85 | 0.72 | 0.68 | 0.88 | 0.77 | 0.86 |
| 3 | 32329497 | rs4380451 | -0.241 | 1.309 | 0.156 | -0.13 | 0.029 | -0.035 | -0.035 | 1.721 | 0.17 | 0.31 | 0.32 | 0.1 | 0.26 | 0.14 |
| 3 | 32330809 | rs4484229 | -0.981 | 0.803 | 0.133 | -0.263 | 0 | -0.055 | -0.055 | 0 | 0.31 | 0.46 | 0.5 | 0.16 | 0.35 | 0.21 |
| 3 | 32334008 | rs11914777 | -1.718 | 1.028 | -0.049 | -0.149 | 0.007 | 0.081 | 0.081 | 0 | 0.62 | 0.76 | 0.75 | 0.8 | 0.63 | 0.56 |
| 3 | 32339493 | rs6793780 | -0.447 | 1.926 | 0.262 | 0.227 | 0.009 | -0.025 | -0.025 | 0 | 0.22 | 0.3 | 0.37 | 0.11 | 0.33 | 0.2 |
| 3 | 32343840 | rs9835916 | 0.353 | 1.425 | 0.452 | 0.815 | -0.001 | 0.118 | 0.118 | 0 | 0.09 | 0.17 | 0.11 | 0.4 | 0.11 | 0.05 |
| 3 | 32343855 | rs9853554 | -0.145 | 1.544 | 0.248 | 0.761 | 0.007 | -0.03 | -0.03 | 0.665 | 0.19 | 0.29 | 0.33 | 0.11 | 0.25 | 0.18 |
| 3 | 32344306 | rs6803740 | 0.73 | 1.572 | -0.389 | 0.779 | -0.011 | -0.019 | -0.019 | 0.087 | 0.2 | 0.11 | 0.13 | 0.26 | 0.18 | 0.27 |
| 3 | 32344588 | rs11129511 | -1.07 | 1.473 | -0.317 | 0.765 | -0.001 | 0.052 | 0.052 | 0 | 0.3 | 0.62 | 0.46 | 0.51 | 0.33 | 0.22 |
| 3 | 32345009 | rs12107664 | -1.362 | 1.403 | -0.599 | -0.202 | 0.023 | 0.125 | 0.125 | 0 | 0.51 | 0.74 | 0.6 | 0.76 | 0.54 | 0.5 |
| 3 | 32348195 | rs9875310 | -0.224 | 1.452 | -1.196 | 0.166 | 0.006 | -0.01 | -0.01 | 0.05 | 0.31 | 0.39 | 0.26 | 0.66 | 0.28 | 0.33 |
| 3 | 32348341 | rs12497324 | -0.495 | 1.279 | -1.568 | 0.289 | -0.011 | 0.035 | 0.035 | 0.043 | 0.13 | 0.07 | 0.18 | 0.02 | 0.16 | 0.12 |
| 3 | 32353131 | rs1370718 | -0.43 | 1.2 | -1.504 | 1.082 | -0.029 | -0.04 | -0.04 | 0 | 0.07 | 0.01 | 0.07 | 0.19 | 0.1 | 0.03 |
| 3 | 32354387 | rs1370717 | -0.279 | 0.866 | -2.041 | 0.402 | -0.005 | 0.136 | 0.136 | 2.563 | 0.47 | 0.17 | 0.54 | 0.71 | 0.45 | 0.49 |
| 3 | 32355865 | rs12629727 | -0.132 | 1.062 | -1.935 | 1.166 | -0.036 | -0.058 | -0.058 | 0 | 0.09 | 0.01 | 0.14 | 0.28 | 0.08 | 0.14 |
| 3 | 32358003 | rs17029201 | 0.39 | 0.559 | -2.158 | 0.239 | 0.011 | 0.018 | 0.018 | 1.303 | 0.14 | 0.06 | 0.09 | 0.12 | 0.11 | 0.19 |
| 3 | 32358666 | rs11129515 | -0.563 | 0.419 | -1.88 | -0.587 | 0.021 | -0.077 | -0.077 | 0 | 0.65 | 0.48 | 0.62 | 0.93 | 0.55 | 0.7 |
| 3 | 32359009 | rs2053280 | -0.411 | 0.495 | -2.321 | -0.514 | -0.011 | 0 | 0 | 0.053 | 0.56 | 0.79 | 0.52 | 0.41 | 0.44 | 0.48 |
| 3 | 32362998 | rs9833771 | 0.95 | 0.522 | -1.943 | -0.248 | -0.031 | 0.068 | 0.068 | 0.456 | 0.36 | 0.21 | 0.29 | 0.2 | 0.44 | 0.37 |
| 3 | 32364319 | rs1866259 | -0.385 | -0.054 | -1.004 | 0.176 | -0.001 | -0.005 | -0.005 | 0 | 0.35 | 0.3 | 0.49 | 0.74 | 0.31 | 0.43 |
| 3 | 32365449 | rs4955278 | 0.423 | -0.004 | -1.318 | 0.263 | -0.047 | -0.151 | -0.151 | 0 | 0.24 | 0.05 | 0.29 | 0.5 | 0.26 | 0.35 |
| 3 | 32365788 | rs1438182 | -0.044 | -0.172 | -0.35 | 0.576 | 0.015 | 0.05 | 0.05 | 0 | 0.13 | 0.04 | 0.03 | 0.04 | 0.1 | 0.1 |
| 3 | 32367230 | rs17029220 | -0.63 | 0.409 | -0.02 | 1.145 | -0.022 | 0.114 | 0.114 | 2.267 | 0.11 | 0.18 | 0.18 | 0.26 | 0.05 | 0.08 |
| 3 | 32373331 | rs7623462 | -1.096 | 1.202 | 0.04 | 0.483 | 0.023 | -0.055 | -0.055 | 0 | 0.12 | 0.22 | 0.19 | 0 | 0.14 | 0.18 |
| 3 | 32373530 | rs4293737 | -1.145 | 0.981 | 0.293 | 1.077 | -0.004 | 0.044 | 0.044 | 0.396 | 0.25 | 0.35 | 0.26 | 0.37 | 0.23 | 0.42 |
| 3 | 32376462 | rs9880262 | 1.1 | 1.513 | 0.385 | 1.188 | 0.023 | 0.009 | 0.009 | 0.059 | 0.64 | 0.49 | 0.31 | 0.61 | 0.49 | 0.52 |
| 3 | 32378093 | rs17029231 | 0.543 | 1.928 | 0.385 | 1.307 | -0.004 | -0.003 | -0.003 | 0 | 0.82 | 0.82 | 0.57 | 0.77 | 0.69 | 0.82 |
| 3 | 32382748 | rs7647165 | -1.016 | 1.182 | 0.198 | 1.133 | 0.006 | -0.01 | -0.01 | 0 | 0.39 | 0.59 | 0.38 | 0.46 | 0.36 | 0.41 |
| 3 | 32385571 | rs17029241 | 0.725 | 1.111 | 0.161 | 1.331 | 0.01 | -0.017 | -0.017 | 0.016 | 0.08 | 0.03 | 0.03 | 0.13 | 0.07 | 0.06 |
| 3 | 32389053 | rs17029245 | -0.581 | 1.096 | 0.575 | 1.33 | -0.003 | 0.001 | 0.001 | 0 | 0.18 | 0.17 | 0.09 | 0.19 | 0.14 | 0.14 |
| 3 | 32394968 | rs11129516 | 1.412 | 0.897 | 0.171 | 1.043 | 0.052 | -0.014 | -0.014 | 0.029 | 0.48 | 0.19 | 0.56 | 0.56 | 0.57 | 0.53 |
| 3 | 32396804 | rs6802418 | -0.451 | 1.643 | 0.89 | 1.335 | 0.237 | 0.229 | 0.229 | 0 | 0.09 | 0.31 | 0.25 | 0.34 | 0.24 | 0.11 |
| 3 | 32398024 | rs13075542 | -0.168 | 1.653 | 0.856 | 1.34 | 0.087 | 0.083 | 0.083 | 0 | 0.09 | 0.13 | 0.24 | 0.34 | 0.24 | 0.09 |
| 3 | 32398234 | rs13080060 | 0.637 | 1.652 | 0.86 | 1.314 | 0.108 | 0.068 | 0.068 | 0 | 0.06 | 0.11 | 0.22 | 0.22 | 0.21 | 0.07 |
| 3 | 32399181 | rs2289244 | 0.719 | 1.341 | 1.022 | 1.193 | -0.01 | -0.019 | -0.019 | 0.138 | 0.62 | 0.42 | 0.63 | 0.8 | 0.68 | 0.59 |
| 3 | 32399378 | rs17279970 | 1.283 | 1.346 | 0.878 | 1.246 | 0.028 | -0.007 | -0.007 | 0 | 0.53 | 0.27 | 0.39 | 0.47 | 0.43 | 0.49 |
| 3 | 32401965 | rs7617800 | -0.157 | 1.522 | 1.18 | 1.423 | 0.02 | 0.079 | 0.079 | 0 | 0.95 | 0.75 | 0.96 | 0.88 | 0.97 | 0.98 |
| 3 | 32401993 | rs11129517 | -0.449 | 1.669 | 0.969 | 1.298 | 0.183 | 0.082 | 0.082 | 0 | 0.11 | 0.28 | 0.28 | 0.22 | 0.27 | 0.1 |
| 3 | 32405158 | rs3844585 | 0.489 | 1.269 | 0.248 | 1.075 | 0.037 | 0.078 | 0.078 | 0 | 0.75 | 0.6 | 0.74 | 0.53 | 0.69 | 0.77 |
| 3 | 32405450 | rs3844586 | 1.486 | 0.998 | -0.857 | 0.854 | 0.026 | 0.056 | 0.056 | 0 | 0.45 | 0.23 | 0.38 | 0.34 | 0.4 | 0.41 |
| 3 | 32406584 | rs3853715 | -0.335 | 1.127 | -0.942 | 0.834 | -0.001 | 0.008 | 0.008 | 1.657 | 0.19 | 0.07 | 0.26 | 0.14 | 0.22 | 0.3 |
| 3 | 32407237 | rs7619214 | -0.292 | 1.705 | -0.693 | 1.067 | -0.016 | -0.012 | -0.012 | 0 | 0.08 | 0.04 | 0.17 | 0.24 | 0.15 | 0.08 |
| 3 | 32407482 | rs10510655 | 0.741 | 0.861 | -0.906 | 0.295 | -0.002 | 0.012 | 0.012 | 0 | 0.08 | 0.06 | 0.03 | 0.02 | 0.07 | 0.12 |
| 3 | 32411922 | rs11710558 | -0.043 | -0.336 | -1.565 | 0.117 | 0.024 | 0.013 | 0.013 | 0 | 0.3 | 0.14 | 0.31 | 0.26 | 0.23 | 0.22 |
| 3 | 32413126 | rs9837863 | 0.844 | -0.009 | -1.823 | -0.221 | -0.005 | 0.025 | 0.025 | 0 | 0.14 | 0.07 | 0.09 | 0.02 | 0.17 | 0.09 |
| 3 | 32413622 | rs17029340 | 1.01 | -0.005 | -1.922 | -0.079 | 0.002 | -0.006 | -0.006 | 0.357 | 0.09 | 0.03 | 0.07 | 0.12 | 0.09 | 0.1 |
| 3 | 32415114 | rs3867381 | 0.287 | 0.242 | -1.425 | 0.108 | -0.003 | -0.011 | -0.011 | 0.062 | 0.31 | 0.26 | 0.3 | 0.5 | 0.32 | 0.22 |
| 3 | 32417504 | rs3867382 | 1.24 | 0.015 | -1.552 | 0.201 | 0.001 | -0.007 | -0.007 | 0.274 | 0.08 | 0.03 | 0.21 | 0.13 | 0.08 | 0.08 |
| 3 | 32417644 | rs4364205 | -0.527 | -0.255 | -0.831 | 0.064 | 0.004 | 0 | 0 | 0 | 0.63 | 0.61 | 0.67 | 0.57 | 0.57 | 0.72 |
| 3 | 32419270 | rs17190433 | 0.891 | -0.085 | -0.537 | 0.092 | 0.022 | 0.011 | 0.011 | 0 | 0.19 | 0.29 | 0.26 | 0.17 | 0.23 | 0.37 |
| 3 | 32423251 | rs11924535 | -0.373 | -0.165 | -0.71 | 0.096 | 0.014 | 0.004 | 0.004 | 0 | 0.32 | 0.46 | 0.5 | 0.32 | 0.35 | 0.48 |
| 3 | 32426611 | rs6801981 | -0.291 | -1.124 | -0.522 | -0.031 | -0.056 | -0.044 | -0.044 | 0.946 | 0.18 | 0.47 | 0.28 | 0.13 | 0.09 | 0.16 |
| 3 | 32428470 | rs6808600 | 1.042 | -1.066 | -1.391 | 0.067 | 0.027 | -0.021 | -0.021 | 0.554 | 0.29 | 0.07 | 0.16 | 0.26 | 0.22 | 0.22 |
| 3 | 32439785 | rs347134 | -0.006 | -1.069 | -1.298 | 0.143 | 0.112 | 0.047 | 0.047 | 0.069 | 0.5 | 0.19 | 0.5 | 0.39 | 0.68 | 0.6 |
| 3 | 32442272 | rs6790078 | 1.218 | -0.005 | -1.543 | -0.013 | -0.053 | 0.057 | 0.057 | 0 | 0.16 | 0.07 | 0.1 | 0.02 | 0.24 | 0.12 |
| 3 | 32442673 | rs347133 | -1.063 | -0.423 | -0.958 | 0.214 | -0.008 | 0.004 | 0.004 | 0 | 0.63 | 0.63 | 0.67 | 0.53 | 0.75 | 0.74 |
| 3 | 32446152 | rs2343849 | -0.045 | -0.481 | -0.976 | 0.228 | 0.011 | -0.014 | -0.014 | 0.092 | 0.19 | 0.15 | 0.2 | 0.26 | 0.16 | 0.23 |
| 3 | 32446465 | rs4635747 | 0.177 | -0.524 | -0.877 | 0.204 | 0.005 | 0.004 | 0.004 | 0 | 0.17 | 0.16 | 0.09 | 0.15 | 0.1 | 0.17 |
| 3 | 32447305 | rs347128 | -0.066 | -0.53 | -0.801 | 0.177 | -0.006 | -0.002 | -0.002 | 0 | 0.48 | 0.5 | 0.58 | 0.38 | 0.65 | 0.58 |
| 3 | 32451345 | rs636006 | 0.258 | -0.564 | -0.665 | 0.444 | 0.013 | 0.013 | 0.013 | 0 | 0.34 | 0.4 | 0.48 | 0.36 | 0.44 | 0.49 |
| 3 | 32455081 | rs614004 | -0.491 | -0.58 | -0.689 | 0.4 | 0.002 | 0.002 | 0.002 | 0 | 0.44 | 0.44 | 0.55 | 0.47 | 0.53 | 0.6 |
| 3 | 32456946 | rs667181 | 0.093 | -0.603 | -0.634 | 0.354 | 0.008 | 0.006 | 0.006 | 0 | 0.36 | 0.41 | 0.49 | 0.35 | 0.45 | 0.5 |
| 3 | 32457198 | rs655754 | 0.613 | -0.563 | -1.125 | 0.018 | -0.014 | 0.007 | 0.007 | 0 | 0.29 | 0.23 | 0.35 | 0.23 | 0.38 | 0.43 |
| 3 | 32463161 | rs17029445 | 0.297 | -0.619 | -1.126 | 0.115 | 0.013 | 0.011 | 0.011 | 0 | 0.08 | 0.06 | 0.01 | 0.02 | 0.02 | 0.07 |
| 3 | 32463994 | rs17029455 | 1.276 | -0.149 | -0.99 | 0.371 | 0.004 | 0.003 | 0.003 | 0 | 0.07 | 0.03 | 0.04 | 0.05 | 0.05 | 0.03 |
| 3 | 32465430 | rs347158 | -0.151 | -0.032 | -0.845 | 0.351 | 0.003 | 0.015 | 0.015 | 0.038 | 0.5 | 0.48 | 0.46 | 0.38 | 0.51 | 0.64 |
| 3 | 32466107 | rs9811451 | -0.228 | 0.021 | -1.148 | 0.832 | -0.002 | -0.034 | -0.034 | 0 | 0.09 | 0.03 | 0.1 | 0.23 | 0.06 | 0.09 |
| 3 | 32466487 | rs635787 | 0.65 | 0.19 | -1.345 | 0.699 | -0.052 | -0.009 | -0.009 | 0 | 0.28 | 0.15 | 0.47 | 0.34 | 0.44 | 0.53 |

| | | | | | | | | | | | | | | | | |
|---|---|---|---|---|---|---|---|---|---|---|---|---|---|---|---|---|
| 3 | 32466797 | rs9853991 | 0.65 | 0.257 | -0.668 | 0.412 | 0 | -0.035 | -0.035 | 0.141 | 0.18 | 0.22 | 0.1 | 0.02 | 0.15 | 0.05 |
| 3 | 32468216 | rs347156 | -0.135 | -0.252 | -0.226 | 0.424 | -0.003 | -0.003 | -0.003 | 0 | 0.55 | 0.44 | 0.6 | 0.38 | 0.64 | 0.63 |
| 3 | 32472280 | rs6550133 | 0.737 | -0.628 | -0.141 | 0.68 | -0.01 | 0.011 | 0.011 | 0.341 | 0.18 | 0.24 | 0.14 | 0.16 | 0.17 | 0.14 |
| 3 | 32472324 | rs347153 | 0.3 | -0.55 | -0.048 | 0.688 | -0.006 | -0.006 | -0.006 | 0.047 | 0.56 | 0.51 | 0.61 | 0.39 | 0.65 | 0.63 |
| 3 | 32473441 | rs347151 | -0.368 | -0.566 | -0.237 | 0.779 | 0.003 | -0.022 | -0.022 | 0 | 0.71 | 0.75 | 0.71 | 0.49 | 0.76 | 0.73 |
| 3 | 32490894 | rs2304594 | 1.623 | -0.571 | -0.294 | 0.561 | 0.01 | -0.013 | -0.013 | 0 | 0.12 | 0.03 | 0.08 | 0.1 | 0.11 | 0.09 |
| 3 | 32491084 | rs2304593 | 2.106 | -0.539 | -0.289 | 0.537 | 0.006 | 0.013 | 0.013 | 1.002 | 0.09 | 0.03 | 0.05 | 0.05 | 0.08 | 0.04 |
| 3 | 32494837 | rs4955290 | 0.955 | -0.62 | -0.254 | 0.557 | 0.002 | 0.011 | 0.011 | 0 | 0.05 | 0.02 | 0.09 | 0.19 | 0.02 | 0.1 |
| 3 | 32497574 | rs3888882 | 1.08 | -0.391 | -0.056 | 0.333 | 0 | -0.016 | -0.016 | 0.283 | 0.18 | 0.2 | 0.1 | 0.02 | 0.17 | 0.07 |
| 3 | 32498088 | rs164215 | -0.446 | -0.459 | 0.169 | 0.374 | -0.001 | -0.001 | -0.001 | 0.464 | 0.49 | 0.45 | 0.38 | 0.49 | 0.4 | 0.32 |
| 3 | 32498154 | rs164214 | -0.431 | -0.576 | 0.19 | 0.142 | -0.001 | -0.002 | -0.002 | 0 | 0.49 | 0.46 | 0.38 | 0.47 | 0.41 | 0.31 |
| 3 | 32499727 | rs342738 | -0.977 | -0.593 | 0.209 | 0.144 | -0.008 | 0.002 | 0.002 | 0 | 0.55 | 0.6 | 0.43 | 0.46 | 0.46 | 0.37 |
| 3 | 32499736 | rs342737 | 0.989 | -0.932 | 0.235 | 0.296 | 0.007 | -0.007 | -0.007 | 0 | 0.21 | 0.28 | 0.19 | 0.26 | 0.19 | 0.13 |
| 3 | 32499817 | rs17029555 | 1.616 | -0.634 | 0.002 | 0.324 | 0.017 | -0.014 | -0.014 | 0.33 | 0.12 | 0.03 | 0.08 | 0.11 | 0.1 | 0.09 |
| 3 | 32502187 | rs342756 | -0.128 | -0.605 | 0.176 | 0.4 | -0.012 | 0.013 | 0.013 | 0 | 0.37 | 0.45 | 0.3 | 0.5 | 0.29 | 0.27 |
| 3 | 32511297 | rs369751 | -0.376 | 0.058 | 0.043 | 0.261 | 0.003 | 0.002 | 0.002 | 1.728 | 0.5 | 0.54 | 0.43 | 0.53 | 0.41 | 0.44 |
| 3 | 32523287 | rs17029597 | 0.458 | 0.169 | 0.484 | 0.665 | 0.003 | -0.004 | -0.004 | 0 | 0.06 | 0.05 | 0.05 | 0.06 | 0.08 | 0.02 |
| 3 | 32523777 | rs4796 | -0.925 | 0.04 | 0.705 | 0.519 | 0 | -0.002 | -0.002 | 0 | 0.41 | 0.64 | 0.4 | 0.53 | 0.36 | 0.33 |
| 3 | 32526153 | rs367841 | -0.696 | 0.057 | 0.803 | 0.665 | -0.002 | 0.014 | 0.014 | 0 | 0.37 | 0.54 | 0.36 | 0.48 | 0.33 | 0.28 |
| 3 | 32531442 | rs6793004 | 0.049 | 0.071 | 0.903 | 0.44 | 0.02 | -0.006 | -0.006 | 0 | 0.07 | 0.15 | 0.07 | 0.07 | 0.1 | 0.02 |
| 3 | 32533010 | rs7640978 | -0.877 | 0.061 | 0.94 | 0.434 | 0.034 | -0.008 | -0.008 | 0 | 0.1 | 0.32 | 0.09 | 0.07 | 0.11 | 0.03 |
| 3 | 32533964 | rs17279984 | 1.028 | 0.072 | 0.743 | 0.331 | 0.008 | 0.025 | 0.025 | 0.099 | 0.15 | 0.06 | 0.07 | 0.03 | 0.1 | 0.05 |
| 3 | 32535382 | rs9834932 | -0.582 | 0.119 | 0.741 | 0.368 | 0.015 | -0.016 | -0.016 | 0 | 0.09 | 0.23 | 0.08 | 0.07 | 0.11 | 0.03 |
| 3 | 32544941 | rs164208 | -1.671 | 0.208 | 0.585 | 0.391 | -0.009 | -0.012 | -0.012 | 0.958 | 0.61 | 0.7 | 0.53 | 0.54 | 0.56 | 0.45 |
| 3 | 32545691 | rs164207 | 0.482 | 0.2 | 0.491 | 0.298 | -0.006 | 0.012 | 0.012 | 0 | 0.26 | 0.22 | 0.19 | 0.22 | 0.24 | 0.21 |
| 3 | 32551946 | rs17201487 | -0.024 | 0.363 | 0.245 | 0.443 | -0.003 | 0.052 | 0.052 | 0.632 | 0.21 | 0.11 | 0.13 | 0.09 | 0.19 | 0.06 |
| 3 | 32561253 | rs7633407 | -0.689 | 0.348 | 0.345 | 0.458 | 0.001 | -0.006 | -0.006 | 0 | 0.38 | 0.66 | 0.4 | 0.44 | 0.35 | 0.39 |
| 3 | 32564484 | rs164219 | -1.862 | 0.297 | 0.307 | 0.235 | -0.009 | -0.026 | -0.026 | 0.787 | 0.59 | 0.76 | 0.53 | 0.5 | 0.55 | 0.44 |
| 3 | 32569096 | rs6782149 | -1.167 | 0.415 | -0.067 | 0.325 | -0.002 | 0 | 0 | 0 | 0.33 | 0.38 | 0.32 | 0.3 | 0.32 | 0.24 |
| 3 | 32572589 | rs11914299 | -0.054 | 0.3 | 0.079 | 0.437 | -0.02 | 0.093 | 0.093 | 0 | 0.12 | 0.3 | 0.2 | 0.22 | 0.12 | 0.17 |
| 3 | 32578505 | rs2303857 | 0.469 | 0.34 | -0.051 | 0.419 | -0.008 | 0.007 | 0.007 | 0 | 0.07 | 0.04 | 0.06 | 0.05 | 0.1 | 0.02 |
| 3 | 32599218 | rs9825747 | -0.228 | 0.254 | 0.287 | 0.765 | -0.019 | 0.095 | 0.095 | 0 | 0.12 | 0.29 | 0.2 | 0.22 | 0.12 | 0.17 |
| 3 | 32599534 | rs342731 | 0.083 | 0.263 | 0.333 | 0.732 | 0.006 | -0.013 | -0.013 | 0.687 | 0.24 | 0.26 | 0.19 | 0.2 | 0.23 | 0.19 |
| 3 | 32599845 | rs9868423 | -0.495 | -0.179 | 0.571 | 1.025 | -0.019 | 0.095 | 0.095 | 0 | 0.12 | 0.29 | 0.2 | 0.22 | 0.12 | 0.17 |
| 3 | 32606580 | rs12107280 | -0.495 | -0.179 | 0.571 | 1.025 | -0.019 | 0.095 | 0.095 | 0 | 0.12 | 0.29 | 0.2 | 0.22 | 0.12 | 0.17 |
| 3 | 32615669 | rs3853720 | -0.567 | 0.356 | 0.231 | 0.687 | -0.004 | -0.004 | -0.004 | 0.937 | 0.53 | 0.63 | 0.47 | 0.54 | 0.5 | 0.46 |
| 3 | 32637097 | rs10510656 | -0.937 | 0.492 | 0.466 | 0.679 | 0.02 | 0.014 | 0.014 | 0.015 | 0.21 | 0.55 | 0.14 | 0.21 | 0.18 | 0.17 |
| 3 | 32638175 | rs7641549 | -1.069 | 0.175 | 0.642 | 0.553 | 0.056 | 0.145 | 0.145 | 1.352 | 0.07 | 0.53 | 0.07 | 0.06 | 0.03 | 0.04 |
| 3 | 32648371 | rs9836624 | -1.896 | -0.084 | 0.816 | 0.602 | 0.003 | -0.004 | -0.004 | 0 | 0.29 | 0.87 | 0.19 | 0.31 | 0.22 | 0.29 |
| 3 | 32691140 | rs9847334 | -1.627 | -0.37 | 0.627 | 0.297 | 0.011 | 0.021 | 0.021 | 0 | 0.36 | 0.67 | 0.23 | 0.28 | 0.25 | 0.39 |
| 3 | 32699422 | rs7640848 | -1.699 | -0.356 | 0.538 | 0.993 | 0 | -0.003 | -0.003 | 1.631 | 0.39 | 0.6 | 0.39 | 0.54 | 0.34 | 0.49 |
| 3 | 32702348 | rs13098724 | -0.898 | -0.433 | 0.575 | 0.961 | -0.002 | 0.003 | 0.003 | 0 | 0.22 | 0.12 | 0.13 | 0.23 | 0.18 | 0.28 |
| 3 | 32704457 | rs9864339 | -1.489 | -0.404 | 0.699 | 1.005 | -0.006 | 0.067 | 0.067 | 0.188 | 0.2 | 0.58 | 0.26 | 0.36 | 0.18 | 0.24 |
| 3 | 32704743 | rs6550146 | -2.409 | -0.703 | -0.098 | 0.253 | -0.047 | -0.018 | -0.018 | 0 | 0.59 | 0.81 | 0.5 | 0.6 | 0.46 | 0.57 |
| 3 | 32714617 | rs7621006 | -1.611 | -0.575 | 0.098 | 0.374 | -0.007 | 0.052 | 0.052 | 0.577 | 0.23 | 0.6 | 0.31 | 0.42 | 0.2 | 0.28 |
| 3 | 32747644 | rs6809257 | -0.95 | -0.583 | 0.1 | 0.237 | -0.001 | -0.001 | -0.001 | 0 | 0.09 | 0.09 | 0.05 | 0.06 | 0.05 | 0.04 |
| 3 | 32763486 | rs12495733 | -0.079 | -0.581 | 0.561 | 0.287 | 0.052 | 0.239 | 0.239 | 0 | 0.09 | 0.35 | 0.2 | 0.26 | 0.11 | 0.21 |
| 3 | 32766905 | rs6550155 | -0.785 | -0.841 | 0.359 | 0.115 | -0.023 | 0.001 | 0.001 | 0 | 0.27 | 0.41 | 0.28 | 0.27 | 0.21 | 0.29 |
| 3 | 32809395 | rs9814104 | -0.915 | -0.747 | -0.005 | -0.148 | 0.017 | 0.052 | 0.052 | 0 | 0.22 | 0.13 | 0.13 | 0.06 | 0.13 | 0.1 |
| 3 | 32814699 | rs3763569 | -1.001 | -0.641 | -0.094 | -0.02 | -0.007 | 0.025 | 0.025 | 0 | 0.21 | 0.1 | 0.12 | 0.17 | 0.16 | 0.21 |
| 3 | 32821366 | rs9811760 | -0.274 | -0.573 | -0.139 | -0.077 | 0.008 | -0.003 | -0.003 | 0 | 0.16 | 0.19 | 0.11 | 0.18 | 0.15 | 0.22 |
| 3 | 32823463 | rs6804305 | 0.178 | -0.489 | -0.124 | -0.089 | 0.002 | 0.007 | 0.007 | 0 | 0.06 | 0.02 | 0.03 | 0 | 0.03 | 0 |
| 3 | 32832864 | rs871683 | -0.286 | -0.418 | -0.072 | 0.002 | 0.018 | -0.001 | -0.001 | 0 | 0.13 | 0.18 | 0.1 | 0.19 | 0.15 | 0.18 |
| 3 | 32835960 | rs11129533 | -1.106 | -0.381 | -0.506 | 0.192 | 0.002 | 0.007 | 0.007 | 0 | 0.31 | 0.62 | 0.33 | 0.51 | 0.32 | 0.49 |
| 3 | 32839607 | rs4909014 | 1.361 | -0.188 | -1.147 | 0.18 | -0.052 | -0.063 | -0.063 | 0 | 0.45 | 0.07 | 0.57 | 0.47 | 0.61 | 0.45 |
| 3 | 32844155 | rs9883625 | 0.761 | 0.041 | 0.095 | 0.356 | 0.022 | 0.097 | 0.097 | 0 | 0.88 | 0.69 | 0.79 | 0.75 | 0.84 | 0.75 |
| 3 | 32861628 | rs6796921 | -0.274 | 0.25 | -0.289 | -0.653 | -0.001 | 0.033 | 0.033 | 1.704 | 0.27 | 0.18 | 0.2 | 0.16 | 0.24 | 0.17 |
| 3 | 32864541 | rs4909038 | -0.756 | 0.563 | 0.678 | -0.596 | 0.048 | 0.007 | 0.007 | 0 | 0.14 | 0.39 | 0.21 | 0.11 | 0.18 | 0.26 |
| 3 | 32867104 | rs13060190 | 0.107 | 0.562 | 0.284 | -0.643 | -0.008 | 0.099 | 0.099 | 1.545 | 0.26 | 0.1 | 0.19 | 0.04 | 0.22 | 0.14 |
| 3 | 32869922 | rs13078069 | -0.251 | 0.345 | 0.317 | -0.727 | 0.002 | 0.067 | 0.067 | 3.643 | 0.29 | 0.18 | 0.19 | 0.16 | 0.23 | 0.17 |
| 3 | 32871036 | rs6809722 | 1.831 | 0.316 | 0.096 | -0.655 | -0.051 | 0.051 | 0.051 | 1.086 | 0.45 | 0.1 | 0.53 | 0.59 | 0.5 | 0.43 |
| 3 | 32876739 | rs7644738 | 0.577 | 0.359 | -0.614 | -0.911 | -0.033 | 0.081 | 0.081 | 0.02 | 0.22 | 0.09 | 0.19 | 0.04 | 0.24 | 0.12 |
| 3 | 32879663 | rs9832642 | -1.113 | -0.04 | -0.411 | -0.87 | 0.009 | -0.005 | -0.005 | 0 | 0.23 | 0.55 | 0.26 | 0.22 | 0.25 | 0.38 |
| 3 | 32880078 | rs12629685 | 1.526 | 0.026 | -0.73 | -0.92 | -0.086 | -0.032 | -0.032 | 0.075 | 0.39 | 0.04 | 0.47 | 0.59 | 0.41 | 0.45 |
| 3 | 32883596 | rs997521 | 0.711 | 0.458 | -0.288 | -0.944 | 0 | 0.002 | 0.002 | 0.836 | 0.52 | 0.46 | 0.67 | 0.62 | 0.57 | 0.67 |
| 3 | 32890314 | rs6550166 | 1.184 | 0.524 | 0.172 | -0.839 | 0.018 | -0.011 | -0.011 | 0.098 | 0.85 | 0.37 | 0.78 | 0.83 | 0.79 | 0.73 |
| 3 | 32898205 | rs4909018 | -0.093 | 0.531 | -0.542 | -1.035 | -0.001 | -0.055 | -0.055 | 0 | 0.46 | 0.53 | 0.62 | 0.84 | 0.5 | 0.7 |
| 3 | 32908632 | rs6550168 | -0.425 | 1.161 | -0.799 | 0.151 | 0.002 | 0 | 0 | 0.401 | 0.4 | 0.57 | 0.45 | 0.39 | 0.53 | 0.45 |
| 3 | 32913093 | rs6550169 | -0.006 | 1.201 | -1.432 | 0.327 | -0.037 | -0.056 | -0.056 | 0 | 0.57 | 0.84 | 0.44 | 0.47 | 0.52 | 0.43 |

| | | | | | | | | | | | | | | | | |
|---|---|---|---|---|---|---|---|---|---|---|---|---|---|---|---|---|
| 3 | 32917415 | rs12488341 | 0.879 | 0.886 | -1.563 | 0.199 | -0.047 | -0.153 | -0.153 | 0 | 0.26 | 0.02 | 0.43 | 0.55 | 0.3 | 0.51 |
| 3 | 32935126 | rs13433809 | 0.54 | 0.734 | -1.014 | -0.246 | -0.059 | 0.139 | 0.139 | 0 | 0.37 | 0.15 | 0.31 | 0.02 | 0.42 | 0.29 |
| 3 | 32937008 | rs11927331 | 0.539 | 0.769 | -0.391 | -0.15 | -0.003 | -0.013 | -0.013 | 0.003 | 0.36 | 0.45 | 0.27 | 0.24 | 0.33 | 0.28 |
| 3 | 32937596 | rs4909017 | -0.329 | 0.307 | -0.121 | 0.091 | 0.003 | -0.023 | -0.023 | 1.345 | 0.56 | 0.49 | 0.64 | 0.76 | 0.57 | 0.68 |
| 3 | 32949845 | rs6765751 | -0.532 | 0.65 | 0.445 | 0.527 | 0.004 | 0.073 | 0.073 | 0 | 0.12 | 0.19 | 0.1 | 0.3 | 0.09 | 0.15 |
| 3 | 32970737 | rs7651953 | 0.903 | 0.463 | 0.236 | 0.13 | -0.043 | 0.155 | 0.155 | 0 | 0.4 | 0.18 | 0.31 | 0.02 | 0.42 | 0.23 |
| 3 | 32979088 | rs6550177 | 0.376 | 0.751 | 0.515 | 0.388 | -0.001 | -0.004 | -0.004 | 0 | 0.49 | 0.6 | 0.36 | 0.16 | 0.44 | 0.34 |
| 3 | 32985486 | rs7428479 | 0.594 | 0.708 | 0.526 | 0.376 | -0.003 | 0.018 | 0.018 | 0 | 0.45 | 0.39 | 0.33 | 0.04 | 0.43 | 0.29 |
| 3 | 32990101 | rs7612287 | 0.741 | 0.707 | 0.507 | 0.382 | -0.003 | 0.042 | 0.042 | 0 | 0.45 | 0.39 | 0.34 | 0.17 | 0.44 | 0.29 |
| 3 | 32996509 | rs6770096 | -0.579 | 0.742 | 0.643 | 0.525 | 0.01 | 0.132 | 0.132 | 0 | 0.12 | 0.26 | 0.1 | 0.3 | 0.07 | 0.15 |
| 3 | 33005230 | rs6795737 | -0.342 | 0.798 | -0.613 | 0.599 | -0.022 | -0.054 | -0.054 | 0 | 0.61 | 0.83 | 0.46 | 0.47 | 0.52 | 0.49 |
| 3 | 33012309 | rs6550181 | -1.178 | 0.232 | 0.006 | 0.814 | -0.029 | 0.146 | 0.146 | 0.091 | 0.13 | 0.55 | 0.12 | 0.31 | 0.04 | 0.16 |
| 3 | 33013483 | rs4679081 | -0.157 | 0.801 | -0.683 | 0.494 | -0.001 | 0.044 | 0.044 | 0 | 0.47 | 0.31 | 0.57 | 0.66 | 0.53 | 0.58 |
| 3 | 33015469 | rs13314993 | -0.205 | 0.958 | -1.004 | 0.542 | -0.093 | -0.094 | -0.094 | 0 | 0.35 | 0.11 | 0.46 | 0.49 | 0.53 | 0.56 |
| 3 | 33018582 | rs11129541 | -0.188 | 1.029 | -0.743 | 0.54 | -0.064 | -0.065 | -0.065 | 0 | 0.34 | 0.17 | 0.44 | 0.49 | 0.51 | 0.56 |
| 3 | 33019374 | rs6764245 | 1.331 | 0.338 | -0.93 | -0.232 | 0.02 | 0.173 | 0.173 | 0 | 0.37 | 0.17 | 0.29 | 0.11 | 0.29 | 0.28 |
| 3 | 33019416 | rs4234236 | 0.365 | 0.207 | -0.435 | -0.062 | -0.013 | -0.105 | -0.105 | 0 | 0.77 | 0.43 | 0.83 | 1 | 0.83 | 0.93 |
| 3 | 33020035 | rs4678680 | 0.604 | 0.137 | -0.25 | -0.107 | -0.015 | -0.073 | -0.073 | 0 | 0.74 | 0.44 | 0.8 | 0.91 | 0.82 | 0.83 |
| 3 | 33024524 | rs4678710 | 0.984 | 0.042 | -0.038 | 0.027 | -0.036 | 0.086 | 0.086 | 0 | 0.75 | 0.39 | 0.78 | 0.49 | 0.87 | 0.75 |
| 3 | 33024773 | rs9854470 | 0.249 | 0.402 | -0.079 | 0.167 | 0.014 | -0.052 | -0.052 | 0 | 0.1 | 0.21 | 0.07 | 0 | 0.09 | 0.03 |
| 3 | 33025110 | rs7611260 | 0.408 | 0.938 | -0.918 | 0.145 | -0.129 | -0.181 | -0.181 | 0 | 0.29 | 0.03 | 0.36 | 0.47 | 0.38 | 0.45 |
| 3 | 33029986 | rs6793913 | -1.369 | 1.215 | -0.434 | 0.226 | 0.003 | 0.06 | 0.06 | 0 | 0.44 | 0.74 | 0.41 | 0.82 | 0.41 | 0.54 |
| 3 | 33038183 | rs12167 | 0.385 | 1.603 | -0.372 | 0.697 | -0.063 | -0.057 | -0.057 | 0 | 0.17 | 0.04 | 0.31 | 0.16 | 0.3 | 0.26 |
| 3 | 33043203 | rs9310999 | 0.152 | 1.8 | 0.066 | 0.815 | -0.007 | -0.007 | -0.007 | 3.181 | 0.14 | 0.13 | 0.33 | 0.16 | 0.27 | 0.22 |
| 3 | 33049152 | rs1133253 | 1.354 | 1.676 | 0.589 | 0.9 | -0.017 | -0.006 | -0.006 | 0.111 | 0.65 | 0.34 | 0.41 | 0.72 | 0.5 | 0.58 |
| 3 | 33062415 | rs6550191 | -1.486 | 1.395 | 1.355 | 1.066 | 0.001 | -0.028 | -0.028 | 0 | 0.25 | 0.34 | 0.24 | 0.04 | 0.35 | 0.2 |
| 3 | 33065339 | rs7613051 | -1.966 | 1.392 | 1.331 | 1.545 | 0.011 | 0.086 | 0.086 | 0 | 0.16 | 0.3 | 0.17 | 0.48 | 0.21 | 0.32 |
| 3 | 33070455 | rs9863198 | -1.9 | 0.856 | 1.595 | 1.472 | 0.003 | 0.016 | 0.016 | 0 | 0.27 | 0.38 | 0.26 | 0.38 | 0.25 | 0.25 |
| 3 | 33071627 | rs6762132 | -1.815 | 0.889 | 1.344 | 1.492 | 0.004 | -0.026 | -0.026 | 0 | 0.23 | 0.35 | 0.27 | 0.13 | 0.27 | 0.21 |
| 3 | 33082188 | rs6805519 | 0.738 | 0.861 | 1.211 | 1.567 | 0.036 | 0.234 | 0.234 | 0.006 | 0.89 | 0.6 | 0.85 | 0.67 | 0.91 | 0.85 |
| 3 | 33085549 | rs6773715 | -0.169 | 0.934 | 0.375 | 1.188 | -0.03 | 0.057 | 0.057 | 0 | 0.12 | 0.04 | 0.07 | 0 | 0.14 | 0.03 |
| 3 | 33087200 | rs6780220 | -2.366 | 0.823 | 0.727 | 1.691 | -0.01 | 0.147 | 0.147 | 0 | 0.13 | 0.19 | 0.14 | 0.57 | 0.13 | 0.29 |
| 3 | 33094524 | rs4257495 | -1.401 | 0.901 | 0.651 | 1.141 | -0.005 | 0.003 | 0.003 | 0 | 0.05 | 0.12 | 0.03 | 0.08 | 0.03 | 0.04 |
| 3 | 33099188 | rs4438612 | -1.68 | 0.993 | 0.721 | 1.009 | 0.004 | -0.014 | -0.014 | 0.075 | 0.27 | 0.32 | 0.39 | 0.17 | 0.32 | 0.27 |
| 3 | 33103028 | rs6550202 | 2.249 | 0.658 | 1.74 | 1.201 | 0.008 | 0.034 | 0.034 | 2.63 | 0.9 | 0.4 | 0.82 | 0.92 | 0.89 | 0.93 |
| 3 | 33123354 | rs9883177 | -1.33 | 0.603 | 1.232 | 1.554 | -0.021 | 0.133 | 0.133 | 0 | 0.16 | 0.52 | 0.1 | 0.37 | 0.1 | 0.21 |
| 3 | 33132144 | rs7612234 | 0.569 | 0.579 | 0.714 | 1.072 | -0.001 | -0.024 | -0.024 | 0.937 | 0.79 | 0.74 | 0.74 | 0.98 | 0.75 | 0.82 |
| 3 | 33138549 | rs7637099 | 1.812 | 0.586 | 0.201 | 1.215 | 0 | -0.103 | -0.103 | 0.448 | 0.6 | 0.07 | 0.57 | 0.49 | 0.65 | 0.54 |
| 3 | 33145481 | rs4678677 | 0.497 | 1.445 | 0.203 | 1.765 | -0.054 | -0.046 | -0.046 | 0 | 0.09 | 0.02 | 0.29 | 0.31 | 0.18 | 0.17 |
| 3 | 33146189 | rs4578976 | -0.661 | 0.711 | 0.944 | 1.781 | 0.036 | 0.261 | 0.261 | 0 | 0.1 | 0.29 | 0.13 | 0.39 | 0.09 | 0.17 |
| 3 | 33155264 | rs6809511 | -0.619 | 0.697 | 0.975 | 1.736 | 0.037 | 0.357 | 0.357 | 0 | 0.1 | 0.3 | 0.12 | 0.5 | 0.09 | 0.17 |
| 3 | 33158912 | rs4678686 | 0.168 | 0.638 | 0.795 | 1.349 | 0.001 | -0.05 | -0.05 | 0 | 0.84 | 0.76 | 0.8 | 0.96 | 0.86 | 0.87 |
| 3 | 33159266 | rs11129545 | 2.082 | 0.383 | 0.531 | 1.29 | 0.011 | -0.093 | -0.093 | 0 | 0.6 | 0.08 | 0.58 | 0.4 | 0.65 | 0.55 |
| 3 | 33161165 | rs4074418 | -1.11 | 0.503 | 1.256 | 1.43 | 0 | 0.209 | 0.209 | 0 | 0.13 | 0.33 | 0.14 | 0.58 | 0.1 | 0.2 |
| 3 | 33161898 | rs4076086 | -1.091 | 0.505 | 1.257 | 1.431 | 0 | 0.211 | 0.211 | 0 | 0.13 | 0.33 | 0.14 | 0.58 | 0.1 | 0.2 |
| 3 | 33167853 | rs6550209 | 0.596 | 0.579 | 0.913 | 1.035 | 0.001 | -0.045 | -0.045 | 0 | 0.81 | 0.77 | 0.77 | 0.96 | 0.82 | 0.86 |
| 3 | 33171659 | rs4392367 | 0.46 | 0.828 | 0.647 | 1.115 | -0.015 | 0.026 | 0.026 | 0 | 0.83 | 0.88 | 0.78 | 0.98 | 0.77 | 0.87 |
| 3 | 33182052 | rs7623768 | -0.641 | 0.928 | 0.687 | 1.481 | 0 | 0.01 | 0.01 | 0 | 0.12 | 0.11 | 0.18 | 0.23 | 0.18 | 0.2 |
| 3 | 33186356 | rs1127898 | -1.292 | 0.752 | 0.364 | 1.401 | 0.004 | 0.056 | 0.056 | 0 | 0.2 | 0.27 | 0.42 | 0.56 | 0.23 | 0.37 |
| 3 | 33187082 | rs12635415 | -0.51 | 0.702 | -0.044 | 1.294 | -0.006 | -0.041 | -0.041 | 0 | 0.13 | 0.05 | 0.17 | 0.22 | 0.16 | 0.25 |
| 3 | 33200507 | rs6781908 | -0.966 | 0.625 | 0.527 | 1.33 | -0.008 | 0.091 | 0.091 | 0.106 | 0.28 | 0.51 | 0.39 | 0.52 | 0.23 | 0.35 |
| 3 | 33201336 | rs12631761 | 0.052 | 0.568 | 0.1 | 1.252 | -0.011 | -0.052 | -0.052 | 0.097 | 0.14 | 0.03 | 0.17 | 0.19 | 0.15 | 0.23 |
| 3 | 33204539 | rs7635223 | -0.378 | 0.567 | 0.291 | 0.977 | 0.036 | -0.09 | -0.09 | 0.744 | 0.86 | 0.68 | 0.88 | 1 | 0.86 | 0.98 |
| 3 | 33206420 | rs11709566 | 1.135 | 0.51 | 0.587 | 1.001 | -0.009 | -0.026 | -0.026 | 0.097 | 0.6 | 0.27 | 0.51 | 0.47 | 0.64 | 0.63 |
| 3 | 33206687 | rs11709715 | -0.405 | 0.511 | 0.59 | 0.986 | 0.035 | -0.09 | -0.09 | 0.191 | 0.86 | 0.68 | 0.88 | 1 | 0.86 | 0.98 |
| 3 | 33207322 | rs9917752 | -0.467 | 0.524 | 0.557 | 1.014 | -0.001 | 0.067 | 0.067 | 0.097 | 0.26 | 0.41 | 0.37 | 0.53 | 0.22 | 0.35 |
| 3 | 33211735 | rs13068113 | 1.119 | 0.523 | 0.422 | 0.965 | -0.011 | -0.019 | -0.019 | 0.164 | 0.06 | 0.03 | 0.09 | 0.2 | 0.09 | 0.03 |
| 3 | 33213119 | rs4678497 | -1.153 | 0.483 | 0.519 | 0.904 | -0.001 | 0.007 | 0.007 | 0 | 0.39 | 0.38 | 0.54 | 0.72 | 0.38 | 0.59 |
| 3 | 33214676 | rs6793960 | 0.509 | 0.398 | 0.382 | 1.052 | 0.02 | 0.007 | 0.007 | 0.126 | 0.78 | 0.6 | 0.79 | 0.79 | 0.77 | 0.93 |
| 3 | 33224218 | rs7620019 | -0.475 | 0.266 | -0.095 | 1.014 | 0.011 | -0.052 | -0.052 | 0 | 0.19 | 0.07 | 0.18 | 0.22 | 0.17 | 0.28 |
| 3 | 33226976 | rs4621286 | -1.109 | 0.28 | -0.092 | 1.104 | 0.005 | -0.112 | -0.112 | 0.755 | 0.26 | 0.11 | 0.28 | 0.4 | 0.25 | 0.3 |
| 3 | 33229885 | rs7636399 | 1.098 | 0.313 | 0.262 | 1.181 | 0.002 | -0.008 | -0.008 | 1.676 | 0.59 | 0.64 | 0.61 | 0.6 | 0.61 | 0.68 |
| 3 | 33233524 | rs11710380 | 2.188 | 0.24 | 0.017 | 1.029 | -0.004 | 0.103 | 0.103 | 0 | 0.37 | 0.19 | 0.23 | 0.1 | 0.37 | 0.33 |
| 3 | 33238026 | rs4371466 | -0.589 | 0.418 | -0.279 | 1.233 | -0.007 | 0.023 | 0.023 | 1.091 | 0.13 | 0.19 | 0.12 | 0.23 | 0.11 | 0.05 |
| 3 | 33238293 | rs4234246 | 0.127 | 0.505 | -0.225 | 1.274 | -0.002 | 0.007 | 0.007 | 0.161 | 0.72 | 0.75 | 0.72 | 0.84 | 0.71 | 0.72 |
| 3 | 33240314 | rs4384906 | 0.793 | 0.581 | -0.219 | 1.303 | 0.009 | 0.076 | 0.076 | 0.049 | 0.07 | 0.17 | 0.11 | 0.18 | 0.09 | 0.02 |
| 3 | 33243386 | rs4389431 | 0.088 | 0.654 | -0.779 | 1.486 | -0.073 | -0.123 | -0.123 | 1.33 | 0.18 | 0.04 | 0.33 | 0.47 | 0.24 | 0.34 |
| 3 | 33249596 | rs12489019 | -0.075 | 0.287 | -0.068 | 1.049 | 0.016 | -0.004 | -0.004 | 0 | 0.05 | 0.03 | 0.01 | 0.05 | 0.02 | 0.03 |
| 3 | 33250531 | rs4678772 | 0.99 | 0.753 | 0.094 | 1.195 | -0.003 | -0.004 | -0.004 | 0 | 0.72 | 0.78 | 0.73 | 0.71 | 0.71 | 0.72 |

| | | | | | | | | | | | | | | | | |
|---|---|---|---|---|---|---|---|---|---|---|---|---|---|---|---|---|
| 3 | 33255592 | rs9637517 | 0.401 | 1.428 | -0.85 | 0.565 | -0.058 | 0.066 | 0.066 | 0 | 0.89 | 0.99 | 0.93 | 1 | 0.85 | 0.97 |
| 3 | 33257434 | rs4678778 | -1.391 | 1.314 | -0.886 | 1.569 | -0.003 | -0.002 | -0.002 | 3.75 | 0.08 | 0.07 | 0.33 | 0.34 | 0.13 | 0.24 |
| 3 | 33258380 | rs9832564 | -0.492 | 0.771 | -0.829 | 1.022 | -0.023 | 0.022 | 0.022 | 0.922 | 0.39 | 0.5 | 0.33 | 0.53 | 0.34 | 0.2 |
| 3 | 33264914 | rs10222597 | -0.61 | 1.095 | -0.866 | 1.12 | -0.007 | 0 | 0 | 0 | 0.07 | 0.06 | 0.33 | 0.32 | 0.12 | 0.23 |
| 3 | 33269181 | rs13085587 | -0.217 | 0.753 | -0.52 | 1.508 | 0.006 | -0.01 | -0.01 | 0.168 | 0.29 | 0.25 | 0.25 | 0.37 | 0.29 | 0.16 |
| 3 | 33271693 | rs4455284 | -1.649 | 0.659 | -0.439 | 1.374 | 0.013 | 0.035 | 0.035 | 0.204 | 0.49 | 0.59 | 0.66 | 0.86 | 0.47 | 0.43 |
| 3 | 33273478 | rs4678787 | -0.688 | 0.748 | -0.136 | 1.559 | 0.004 | 0.083 | 0.083 | 0 | 0.06 | 0.1 | 0.33 | 0.32 | 0.09 | 0.2 |
| 3 | 33277828 | rs6550215 | -1.559 | 0.439 | -0.138 | 1.719 | -0.018 | 0.085 | 0.085 | 0 | 0.29 | 0.52 | 0.53 | 0.58 | 0.29 | 0.35 |
| 3 | 33281293 | rs6800399 | -0.564 | 0.369 | -0.062 | 1.652 | -0.027 | 0.003 | 0.003 | 0 | 0.2 | 0.41 | 0.2 | 0.23 | 0.18 | 0.12 |
| 3 | 33289596 | rs7653239 | -1.138 | 0.246 | 0.168 | 1.721 | -0.015 | 0.082 | 0.082 | 0 | 0.14 | 0.25 | 0.1 | 0.26 | 0.09 | 0.05 |
| 3 | 33292542 | rs4678512 | -1.34 | 0.518 | 0.148 | 1.746 | 0.045 | 0.136 | 0.136 | 0 | 0.12 | 0.32 | 0.37 | 0.34 | 0.14 | 0.28 |
| 3 | 33295953 | rs9871600 | -2.271 | 0.439 | 0.173 | 1.296 | -0.005 | -0.003 | -0.003 | 0 | 0.37 | 0.57 | 0.59 | 0.74 | 0.34 | 0.41 |
| 3 | 33302952 | rs4572718 | 0.287 | 0.427 | -0.019 | 1.212 | -0.008 | -0.006 | -0.006 | 0 | 0.11 | 0.16 | 0.13 | 0.15 | 0.11 | 0.07 |
| 3 | 33315104 | rs9861219 | -2.028 | 0.765 | 0.349 | 1.066 | 0 | 0.006 | 0.006 | 0 | 0.34 | 0.57 | 0.57 | 0.72 | 0.33 | 0.41 |
| 3 | 33321616 | rs7626634 | -1.305 | 0.735 | 0.555 | 1.219 | -0.025 | -0.022 | -0.022 | 0 | 0.17 | 0.22 | 0.19 | 0.15 | 0.11 | 0.2 |
| 3 | 33321659 | rs4678848 | 0.439 | 0.73 | 0.593 | 1.23 | -0.006 | -0.006 | -0.006 | 0.784 | 0.09 | 0.11 | 0.12 | 0.12 | 0.07 | 0.12 |
| 3 | 33329435 | rs6777187 | -2.376 | 0.756 | 0.647 | 1.445 | 0 | 0.035 | 0.035 | 0 | 0.33 | 0.55 | 0.56 | 0.61 | 0.32 | 0.4 |
| 3 | 33341291 | rs10428231 | 0.357 | 0.734 | 0.588 | 1.362 | -0.004 | -0.007 | -0.007 | 0 | 0.09 | 0.12 | 0.12 | 0.12 | 0.08 | 0.08 |
| 3 | 33355978 | rs7640029 | 2.306 | 0.837 | -0.395 | 0.505 | -0.053 | 0.113 | 0.113 | 0.38 | 0.35 | 0.01 | 0.19 | 0.12 | 0.33 | 0.42 |
| 3 | 33373203 | rs17029976 | -1.305 | 0.706 | 0.37 | 0.794 | -0.025 | -0.021 | -0.021 | 0 | 0.18 | 0.23 | 0.2 | 0.15 | 0.11 | 0.16 |
| 3 | 33404914 | rs12486192 | -1.581 | 0.82 | 0.483 | 0.873 | 0.034 | 0.083 | 0.083 | 0 | 0.25 | 0.42 | 0.48 | 0.58 | 0.31 | 0.34 |
| 3 | 33413263 | rs4678929 | -0.391 | 0.728 | 0.541 | 0.886 | -0.02 | -0.016 | -0.016 | 0 | 0.13 | 0.1 | 0.34 | 0.46 | 0.21 | 0.24 |
| 3 | 33416140 | rs17805850 | 0.642 | 0.722 | 0.628 | 0.882 | 0.001 | 0.002 | 0.002 | 0 | 0.08 | 0.05 | 0.11 | 0.12 | 0.08 | 0.08 |
| 3 | 33419422 | rs2291897 | -0.868 | 0.73 | 0.631 | 0.887 | 0.055 | 0.14 | 0.14 | 0.018 | 0.16 | 0.37 | 0.37 | 0.46 | 0.21 | 0.24 |
| 3 | 33421178 | rs4045539 | -0.787 | 0.72 | 0.682 | 0.907 | 0.045 | 0.123 | 0.123 | 0 | 0.16 | 0.33 | 0.37 | 0.46 | 0.22 | 0.24 |
| 3 | 33429282 | rs13087731 | -1.585 | 0.83 | 0.4 | 0.713 | 0.013 | 0.008 | 0.008 | 0.266 | 0.33 | 0.39 | 0.46 | 0.47 | 0.4 | 0.34 |
| 3 | 33430093 | rs11260 | 0.878 | 0.854 | 0.434 | 0.749 | -0.009 | -0.001 | -0.001 | 0.221 | 0.05 | 0.04 | 0.07 | 0.03 | 0.01 | 0.05 |
| 3 | 33434831 | rs2293250 | -2.203 | 0.926 | 0.31 | 0.744 | -0.001 | -0.001 | -0.001 | 0 | 0.46 | 0.56 | 0.63 | 0.61 | 0.47 | 0.45 |
| 3 | 33440318 | rs17030583 | -1.33 | 0.858 | 0.43 | 0.826 | 0.035 | 0.076 | 0.076 | 0 | 0.21 | 0.37 | 0.4 | 0.46 | 0.24 | 0.28 |
| 3 | 33444286 | rs623244 | 1.734 | 0.778 | 0.628 | 0.845 | 0.012 | 0.067 | 0.067 | 0 | 0.69 | 0.5 | 0.44 | 0.39 | 0.69 | 0.62 |
| 3 | 33453720 | rs12493761 | -1.857 | 0.793 | 0.51 | 0.852 | 0.014 | 0.074 | 0.074 | 0 | 0.31 | 0.47 | 0.55 | 0.61 | 0.31 | 0.38 |
| 3 | 33457493 | rs11928797 | 0.513 | 0.912 | 0.394 | 0.846 | -0.034 | 0.046 | 0.046 | 0 | 0.08 | 0 | 0.08 | 0 | 0.1 | 0.26 |
| 3 | 33458266 | rs3736563 | -2.11 | 0.433 | 0.924 | 1.345 | 0.008 | 0.054 | 0.054 | 0 | 0.31 | 0.53 | 0.56 | 0.61 | 0.31 | 0.38 |
| 3 | 33468329 | rs9311014 | -2.441 | 0.433 | 0.672 | 1.347 | -0.001 | -0.006 | -0.006 | 0.006 | 0.33 | 0.66 | 0.57 | 0.61 | 0.32 | 0.39 |
| 3 | 33483744 | rs4678999 | -0.753 | 0.462 | 0.257 | 0.639 | 0.001 | 0.001 | 0.001 | 0 | 0.14 | 0.1 | 0.18 | 0.15 | 0.08 | 0.13 |
| 3 | 33509761 | rs7613545 | -0.77 | 0.462 | 0.149 | 0.632 | 0.01 | 0.004 | 0.004 | 0 | 0.14 | 0.08 | 0.18 | 0.15 | 0.08 | 0.13 |
| 3 | 33530636 | rs9838654 | -1.938 | 0.672 | 0.435 | 0.715 | 0.011 | 0.018 | 0.018 | 0 | 0.29 | 0.41 | 0.49 | 0.37 | 0.31 | 0.33 |
| 3 | 33531800 | rs6781106 | -2.093 | 0.755 | 0.369 | 0.538 | -0.006 | 0 | 0 | 0 | 0.36 | 0.33 | 0.53 | 0.37 | 0.42 | 0.35 |
| 3 | 33538041 | rs1052812 | -1.965 | 0.705 | 0.396 | 0.505 | -0.004 | -0.002 | -0.002 | 0 | 0.32 | 0.31 | 0.51 | 0.37 | 0.4 | 0.34 |
| 3 | 33543628 | rs1532590 | -2.105 | 0.71 | 0.446 | 0.496 | -0.028 | -0.009 | -0.009 | 0 | 0.33 | 0.22 | 0.51 | 0.37 | 0.38 | 0.32 |
| 3 | 33544110 | rs4679044 | -1.857 | 0.732 | 0.438 | 0.498 | -0.029 | -0.017 | -0.017 | 1.395 | 0.29 | 0.2 | 0.49 | 0.37 | 0.37 | 0.31 |
| 3 | 33545785 | rs6781484 | -1.429 | 0.761 | 0.438 | 0.516 | -0.044 | -0.031 | -0.031 | 0.061 | 0.22 | 0.12 | 0.38 | 0.27 | 0.3 | 0.24 |
| 3 | 33562114 | rs9862261 | -2.587 | 0.735 | 0.45 | 0.51 | -0.006 | 0 | 0 | 0 | 0.36 | 0.33 | 0.52 | 0.37 | 0.41 | 0.35 |
| 3 | 33570327 | rs9817055 | -2.485 | 0.75 | 0.53 | 0.561 | 0.01 | 0.017 | 0.017 | 0 | 0.28 | 0.39 | 0.49 | 0.36 | 0.3 | 0.33 |
| 3 | 33582575 | rs4678610 | 0.161 | 0.776 | 0.521 | 0.583 | 0 | -0.001 | -0.001 | 0 | 0.08 | 0.08 | 0.12 | 0.09 | 0.08 | 0.08 |
| 3 | 33587605 | rs12637850 | -1.614 | 1.118 | 0.518 | 0.604 | -0.051 | -0.031 | -0.031 | 0.916 | 0.21 | 0.12 | 0.39 | 0.27 | 0.33 | 0.23 |
| 3 | 33648715 | rs9856967 | -1.375 | 1.12 | 1.07 | 0.697 | 0.094 | 0.166 | 0.166 | 0 | 0.13 | 0.39 | 0.35 | 0.27 | 0.18 | 0.21 |
| 3 | 33653803 | rs6807732 | -2.666 | 1.13 | 1.122 | 0.625 | 0.026 | 0.001 | 0.001 | 0 | 0.36 | 0.44 | 0.55 | 0.37 | 0.45 | 0.34 |
| 3 | 33670344 | rs7374396 | -1.327 | 1.109 | 1.065 | 0.641 | -0.023 | -0.024 | -0.024 | 0 | 0.17 | 0.12 | 0.36 | 0.27 | 0.26 | 0.24 |
| 3 | 33683463 | rs9654036 | -1.015 | 1.021 | 1.042 | 0.617 | -0.022 | -0.048 | -0.048 | 0 | 0.15 | 0.29 | 0.15 | 0.09 | 0.11 | 0.12 |
| 3 | 33708114 | rs13100100 | -0.739 | 1.15 | 0.846 | 0.705 | -0.059 | 0.026 | 0.026 | 0.665 | 0.13 | 0.02 | 0.09 | 0.03 | 0.18 | 0.03 |
| 3 | 33736126 | rs13065450 | -0.939 | 1.025 | 0.881 | 0.629 | 0.023 | 0.03 | 0.03 | 0 | 0.11 | 0.14 | 0.34 | 0.27 | 0.23 | 0.21 |
| 3 | 33745827 | rs9839042 | -2.52 | 0.973 | 1.002 | 0.608 | 0.004 | 0.004 | 0.004 | 0 | 0.28 | 0.7 | 0.52 | 0.37 | 0.33 | 0.31 |
| 3 | 33767207 | rs4678614 | -1.822 | 1.378 | 0.661 | 0.948 | -0.019 | -0.017 | -0.017 | 0 | 0.2 | 0.16 | 0.39 | 0.29 | 0.34 | 0.22 |
| 3 | 33789382 | rs9858415 | -1.835 | 0.758 | 0.78 | 1.018 | 0.038 | 0.104 | 0.104 | 0 | 0.22 | 0.38 | 0.48 | 0.45 | 0.27 | 0.32 |
| 3 | 33794769 | rs4420815 | -0.819 | 0.762 | 0.356 | 1.065 | 0.006 | 0.044 | 0.044 | 0 | 0.11 | 0.12 | 0.33 | 0.34 | 0.18 | 0.23 |
| 3 | 33801024 | rs3934853 | -1.212 | 0.716 | 0.427 | 1.037 | 0.009 | 0.029 | 0.029 | 0 | 0.2 | 0.24 | 0.46 | 0.45 | 0.27 | 0.33 |
| 3 | 33811713 | rs7628348 | 0.665 | -0.304 | 0.263 | 1.02 | 0.005 | -0.022 | -0.022 | 0 | 0.82 | 0.92 | 0.87 | 0.84 | 0.86 | 0.88 |
| 3 | 33838124 | rs9843488 | -1.148 | -0.688 | 0.258 | 0.668 | -0.008 | 0.009 | 0.009 | 0 | 0.26 | 0.31 | 0.4 | 0.41 | 0.22 | 0.36 |
| 3 | 33838866 | rs13081248 | 0.098 | -0.84 | 0.03 | 0.309 | -0.011 | 0.041 | 0.041 | 0 | 0.1 | 0.02 | 0.11 | 0.02 | 0.09 | 0.04 |
| 3 | 33842470 | rs6772865 | -0.605 | -1.186 | 0.268 | 0.391 | -0.007 | 0.027 | 0.027 | 0 | 0.17 | 0.22 | 0.35 | 0.38 | 0.16 | 0.24 |
| 3 | 33844292 | rs6809978 | -1.747 | -1.079 | 0.237 | 0.358 | -0.008 | 0.013 | 0.013 | 0 | 0.25 | 0.3 | 0.4 | 0.41 | 0.23 | 0.34 |
| 3 | 33848798 | rs9853642 | -1.747 | -1.079 | 0.237 | 0.358 | -0.008 | 0.013 | 0.013 | 0 | 0.25 | 0.3 | 0.4 | 0.41 | 0.23 | 0.34 |
| 3 | 33857055 | rs11926545 | -1.964 | -1.242 | 1.077 | 0.286 | -0.02 | 0.018 | 0.018 | 0 | 0.28 | 0.65 | 0.44 | 0.41 | 0.23 | 0.34 |
| 3 | 33877626 | rs3792594 | -1.564 | -1.263 | 1.033 | 0.277 | -0.005 | 0.007 | 0.007 | 0 | 0.25 | 0.29 | 0.4 | 0.41 | 0.23 | 0.34 |
| 3 | 33882134 | rs7618512 | -1.97 | -1.278 | 1.33 | 0.27 | -0.032 | 0.031 | 0.031 | 0 | 0.28 | 0.58 | 0.44 | 0.41 | 0.23 | 0.35 |
| 3 | 33889784 | rs4678618 | -1.049 | -1.243 | 0.776 | 0.278 | 0.008 | -0.016 | -0.016 | 0 | 0.16 | 0.16 | 0.31 | 0.39 | 0.13 | 0.3 |
| 3 | 33893987 | rs9813017 | -0.404 | -1.243 | 0.779 | 0.278 | -0.008 | -0.037 | -0.037 | 0 | 0.09 | 0.14 | 0.09 | 0.02 | 0.1 | 0.05 |
| 3 | 33901914 | rs12493220 | 0.124 | -1.247 | 0.777 | 0.277 | -0.001 | 0.076 | 0.076 | 0 | 0.09 | 0.13 | 0.29 | 0.38 | 0.08 | 0.22 |

| | | | | | | | | | | | | | | | | |
|---|---|---|---|---|---|---|---|---|---|---|---|---|---|---|---|---|
| 3 | 33905566 | rs1127732 | -1.041 | -1.242 | 0.778 | 0.264 | 0.008 | -0.016 | -0.016 | 0 | 0.16 | 0.16 | 0.31 | 0.39 | 0.13 | 0.3 |
| 3 | 33907945 | rs3183987 | -1.814 | -1.21 | 0.754 | 0.295 | -0.005 | 0.007 | 0.007 | 0 | 0.25 | 0.29 | 0.4 | 0.41 | 0.23 | 0.34 |
| 3 | 33914508 | rs9311031 | -1.198 | -1.029 | 0.897 | 0.408 | -0.015 | 0.074 | 0.074 | 0 | 0.18 | 0.34 | 0.39 | 0.38 | 0.18 | 0.27 |
| 3 | 33917150 | rs6783500 | -1.213 | -1.052 | 0.737 | 0.377 | -0.026 | 0.067 | 0.067 | 0 | 0.19 | 0.38 | 0.39 | 0.36 | 0.17 | 0.25 |
| 3 | 33929604 | rs9826564 | -1.556 | -1.151 | 0.634 | -0.339 | -0.025 | -0.056 | -0.056 | 0 | 0.29 | 0.38 | 0.23 | 0.07 | 0.23 | 0.16 |
| 3 | 33936338 | rs9835248 | -1.916 | -1.521 | 0.442 | -0.509 | 0.054 | 0.065 | 0.065 | 0 | 0.48 | 0.67 | 0.42 | 0.27 | 0.4 | 0.25 |
| 3 | 33948370 | rs7642534 | -1.097 | -1.654 | 0.528 | -0.596 | -0.008 | -0.007 | -0.007 | 0 | 0.29 | 0.34 | 0.3 | 0.21 | 0.26 | 0.11 |
| 3 | 33956494 | rs13080359 | 0.17 | -1.739 | 0.054 | -0.509 | -0.017 | 0.012 | 0.012 | 0 | 0.07 | 0.17 | 0.04 | 0.04 | 0.04 | 0.06 |
| 3 | 33960238 | rs6780261 | -0.272 | -1.719 | 0.12 | -0.599 | 0.001 | -0.059 | -0.059 | 0.829 | 0.11 | 0.24 | 0.09 | 0.03 | 0.09 | 0.06 |
| 3 | 33963428 | rs9848590 | -0.589 | -1.881 | 0.187 | -0.635 | -0.069 | 0.021 | 0.021 | 0 | 0.19 | 0.47 | 0.25 | 0.2 | 0.1 | 0.08 |
| 3 | 33963571 | rs9848859 | -1.961 | -1.984 | -0.003 | -0.743 | 0.107 | 0.099 | 0.099 | 0 | 0.47 | 0.74 | 0.42 | 0.27 | 0.34 | 0.22 |
| 3 | 33965686 | rs10212462 | -1.01 | -1.68 | -0.163 | -0.461 | -0.062 | 0.009 | 0.009 | 0 | 0.25 | 0.41 | 0.26 | 0.24 | 0.15 | 0.15 |
| 3 | 33966294 | rs4626044 | -0.406 | -0.912 | -0.987 | -0.433 | 0.017 | 0.02 | 0.02 | 0 | 0.11 | 0.07 | 0.08 | 0 | 0.1 | 0.03 |
| 3 | 33970478 | rs7618730 | 0.598 | -0.066 | -1.274 | 0.14 | -0.03 | -0.037 | -0.037 | 0 | 0.75 | 0.85 | 0.6 | 0.67 | 0.66 | 0.52 |
| 3 | 33974172 | rs11916425 | 0.992 | 0.08 | -1.305 | -0.324 | -0.024 | 0.03 | 0.03 | 0 | 0.42 | 0.28 | 0.41 | 0.29 | 0.43 | 0.55 |
| 3 | 33974709 | rs9985486 | 0.21 | -0.304 | -1.111 | -0.024 | -0.01 | -0.03 | -0.03 | 0 | 0.64 | 0.74 | 0.62 | 0.45 | 0.65 | 0.67 |
| 3 | 33980567 | rs6776985 | -0.495 | -0.285 | -1.444 | -0.091 | -0.043 | 0.056 | 0.056 | 0 | 0.89 | 0.99 | 0.82 | 1 | 0.87 | 0.91 |
| 3 | 33983197 | rs11705838 | 1.253 | -0.346 | -1.393 | -0.044 | 0.015 | 0.045 | 0.045 | 0 | 0.48 | 0.31 | 0.44 | 0.33 | 0.55 | 0.64 |
| 3 | 33985368 | rs1112837 | 0.756 | 0.616 | -1.823 | 0.258 | -0.012 | -0.046 | -0.046 | 0 | 0.65 | 0.8 | 0.64 | 0.45 | 0.67 | 0.69 |
| 3 | 33986165 | rs7613581 | 0.605 | 0.647 | -1.334 | -0.015 | -0.003 | 0.005 | 0.005 | 0 | 0.89 | 0.92 | 0.82 | 1 | 0.87 | 0.91 |
| 3 | 33997849 | rs4678623 | -0.017 | 0.911 | -1.58 | 0.296 | -0.022 | 0.003 | 0.003 | 0.282 | 0.59 | 0.29 | 0.52 | 0.59 | 0.51 | 0.69 |
| 3 | 34001399 | rs12486443 | 0.248 | 0.677 | -2.366 | -0.595 | 0.041 | 0.154 | 0.154 | 0.233 | 0.66 | 0.16 | 0.67 | 0.96 | 0.7 | 0.9 |
| 3 | 34012971 | rs6550241 | -0.65 | 0.773 | -2.314 | -0.368 | -0.007 | -0.028 | -0.028 | 0 | 0.76 | 0.32 | 0.73 | 0.93 | 0.77 | 0.93 |
| 3 | 34013124 | rs6550242 | 1.149 | 0.62 | -2.403 | -0.33 | -0.057 | -0.06 | -0.06 | 0 | 0.48 | 0.09 | 0.45 | 0.51 | 0.41 | 0.63 |
| 3 | 34014928 | rs11717668 | 1.384 | 0.134 | -3.042 | -0.037 | -0.095 | -0.027 | -0.027 | 0 | 0.36 | 0.03 | 0.3 | 0.27 | 0.33 | 0.43 |
| 3 | 34015618 | rs6763221 | -1.385 | 0.113 | -1.978 | -0.21 | -0.034 | -0.08 | -0.08 | 0.02 | 0.79 | 0.51 | 0.85 | 0.93 | 0.82 | 0.93 |
| 3 | 34018833 | rs9850318 | 0.987 | 0.099 | -1.922 | -0.219 | -0.006 | -0.033 | -0.033 | 0 | 0.09 | 0.12 | 0.06 | 0.01 | 0.08 | 0.03 |
| 3 | 34021986 | rs9866710 | -0.875 | 0.103 | -1.914 | -0.217 | -0.004 | -0.02 | -0.02 | 0 | 0.76 | 0.31 | 0.75 | 0.93 | 0.78 | 0.93 |
| 3 | 34034343 | rs1109973 | -1.355 | 0.055 | -1.722 | -0.234 | -0.031 | -0.07 | -0.07 | 0 | 0.79 | 0.46 | 0.84 | 0.93 | 0.82 | 0.93 |
| 3 | 34039993 | rs11710273 | 1.245 | 0.027 | -1.884 | -0.181 | 0.003 | -0.054 | -0.054 | 0.532 | 0.11 | 0.02 | 0.14 | 0.24 | 0.08 | 0.2 |
| 3 | 34043325 | rs13074236 | 0.001 | -0.194 | -2.005 | -0.153 | -0.096 | -0.072 | -0.072 | 1.832 | 0.31 | 0.05 | 0.36 | 0.68 | 0.35 | 0.48 |
| 3 | 34044246 | rs4132079 | -0.14 | -0.187 | -1.941 | -0.125 | -0.004 | -0.006 | -0.006 | 0.026 | 0.29 | 0.26 | 0.29 | 0.45 | 0.35 | 0.31 |
| 3 | 34044916 | rs11709925 | 0.701 | -0.107 | -1.993 | -0.113 | -0.065 | -0.098 | -0.098 | 0 | 0.47 | 0.06 | 0.45 | 0.48 | 0.42 | 0.62 |
| 3 | 34051325 | rs12494973 | -0.726 | -0.126 | -1.842 | -0.124 | -0.006 | 0.004 | 0.004 | 0 | 0.4 | 0.34 | 0.44 | 0.68 | 0.44 | 0.48 |
| 3 | 34053980 | rs11714011 | 1.068 | -0.032 | -1.872 | -0.106 | 0.007 | -0.013 | -0.013 | 0 | 0.12 | 0.1 | 0.16 | 0.23 | 0.07 | 0.15 |
| 3 | 34059913 | rs4072314 | 0.719 | -0.105 | -1.908 | -0.211 | -0.004 | -0.033 | -0.033 | 0 | 0.1 | 0.13 | 0.07 | 0.01 | 0.1 | 0.03 |
| 3 | 34060079 | rs4072315 | 0.952 | -0.103 | -1.968 | -0.194 | -0.058 | -0.086 | -0.086 | 0.003 | 0.19 | 0.06 | 0.21 | 0.45 | 0.28 | 0.3 |
| 3 | 34061422 | rs4076661 | 0.376 | -0.192 | -2.035 | -0.27 | 0.054 | 0.066 | 0.066 | 0.056 | 0.54 | 0.11 | 0.52 | 0.69 | 0.6 | 0.75 |
| 3 | 34065749 | rs11129577 | -0.179 | 0.224 | -2.012 | -0.185 | -0.057 | -0.089 | -0.089 | 0.019 | 0.25 | 0.07 | 0.24 | 0.46 | 0.33 | 0.29 |
| 3 | 34077223 | rs4373023 | -0.64 | -0.088 | -1.903 | -0.405 | -0.014 | -0.077 | -0.077 | 3.021 | 0.72 | 0.37 | 0.8 | 0.98 | 0.76 | 0.92 |
| 3 | 34080270 | rs10865853 | 1.1 | -0.229 | -2.227 | -0.493 | 0.002 | -0.067 | -0.067 | 0 | 0.13 | 0.02 | 0.15 | 0.31 | 0.08 | 0.19 |
| 3 | 34080654 | rs6775954 | -0.172 | -0.289 | -2.242 | -0.533 | -0.007 | 0 | 0 | 0.684 | 0.56 | 0.34 | 0.6 | 0.53 | 0.52 | 0.66 |
| 3 | 34080929 | rs9815333 | 0.257 | -0.441 | -2.205 | -0.472 | -0.003 | 0.007 | 0.007 | 0.678 | 0.47 | 0.3 | 0.55 | 0.52 | 0.44 | 0.63 |
| 3 | 34084684 | rs6807347 | 0.178 | -0.346 | -1.817 | -0.367 | -0.003 | 0.084 | 0.084 | 0 | 0.15 | 0.23 | 0.26 | 0.31 | 0.12 | 0.2 |
| 3 | 34086193 | rs1992348 | 0.395 | -0.343 | -1.83 | -0.357 | 0.005 | 0.096 | 0.096 | 0 | 0.13 | 0.25 | 0.26 | 0.3 | 0.12 | 0.2 |
| 3 | 34100132 | rs2336036 | -0.118 | -0.554 | -1.966 | -0.81 | 0.034 | 0.163 | 0.163 | 0.902 | 0.66 | 0.12 | 0.67 | 0.98 | 0.71 | 0.91 |
| 3 | 34105739 | rs6794758 | 0.308 | -0.646 | -0.97 | -0.831 | -0.024 | -0.1 | -0.1 | 0.487 | 0.29 | 0.38 | 0.2 | 0.02 | 0.24 | 0.08 |
| 3 | 34106370 | rs2119266 | 0.847 | -0.491 | -1.733 | -0.848 | -0.076 | -0.133 | -0.133 | 0.008 | 0.23 | 0.03 | 0.25 | 0.41 | 0.25 | 0.3 |
| 3 | 34110517 | rs985463 | -0.334 | 0.163 | -1.399 | -0.807 | 0.002 | -0.014 | -0.014 | 0.455 | 0.66 | 0.27 | 0.65 | 0.51 | 0.67 | 0.7 |
| 3 | 34113178 | rs4678646 | -0.601 | 0.031 | -1.399 | -1.176 | -0.025 | 0.062 | 0.062 | 0 | 0.69 | 0.84 | 0.65 | 0.12 | 0.67 | 0.46 |
| 3 | 34114983 | rs7628777 | 0.801 | -0.853 | -0.519 | -1.623 | -0.016 | -0.095 | -0.095 | 0 | 0.27 | 0.3 | 0.18 | 0.01 | 0.23 | 0.08 |
| 3 | 34115997 | rs9813805 | 0.551 | -0.55 | -0.136 | -1.38 | 0.009 | -0.023 | -0.023 | 0 | 0.08 | 0.23 | 0.06 | 0.01 | 0.05 | 0.03 |
| 3 | 34119403 | rs9311044 | 0.287 | -0.45 | -0.335 | -0.977 | 0.002 | -0.02 | -0.02 | 0.533 | 0.35 | 0.23 | 0.31 | 0.44 | 0.31 | 0.31 |
| 3 | 34132701 | rs893923 | -0.186 | 0.361 | -0.057 | -0.807 | -0.008 | 0.055 | 0.055 | 0 | 0.71 | 0.6 | 0.72 | 0.55 | 0.71 | 0.64 |
| 3 | 34145597 | rs1440385 | 1.546 | 0.616 | -0.189 | -0.915 | -0.01 | -0.003 | -0.003 | 0 | 0.08 | 0.05 | 0.22 | 0.1 | 0.09 | 0.2 |
| 3 | 34147822 | rs6768921 | 1.571 | 0.439 | -0.209 | -0.971 | -0.018 | -0.061 | -0.061 | 0 | 0.23 | 0.1 | 0.26 | 0.43 | 0.25 | 0.28 |
| 3 | 34149458 | rs10049124 | 0.936 | 0.569 | 0.056 | -0.854 | -0.016 | -0.097 | -0.097 | 0 | 0.23 | 0.3 | 0.18 | 0.01 | 0.22 | 0.1 |
| 3 | 34155206 | rs1899082 | -0.453 | 0.504 | -0.13 | -0.764 | -0.007 | 0.034 | 0.034 | 0 | 0.72 | 0.63 | 0.73 | 0.55 | 0.71 | 0.63 |
| 3 | 34165785 | rs13315416 | 0.35 | 0.561 | -0.419 | -0.727 | -0.002 | 0.009 | 0.009 | 0.534 | 0.34 | 0.28 | 0.21 | 0.02 | 0.34 | 0.14 |
| 3 | 34184505 | rs6770257 | -0.161 | 0.782 | -0.228 | 0.366 | -0.002 | -0.048 | -0.048 | 0 | 0.13 | 0.16 | 0.09 | 0.01 | 0.12 | 0.07 |
| 3 | 34200942 | rs981120 | -0.704 | 0.975 | -0.251 | 0.512 | 0.003 | -0.006 | -0.006 | 0.01 | 0.09 | 0.14 | 0.21 | 0.1 | 0.09 | 0.19 |
| 3 | 34201975 | rs1348036 | 0.513 | 0.84 | -0.499 | 1.016 | -0.001 | 0.057 | 0.057 | 0 | 0.9 | 0.91 | 0.92 | 0.47 | 0.89 | 0.69 |
| 3 | 34204000 | rs2218182 | 1.445 | 0.95 | -0.258 | 1.028 | 0 | 0.004 | 0.004 | 0 | 0.8 | 0.83 | 0.58 | 0.34 | 0.78 | 0.44 |
| 3 | 34204828 | rs4678659 | -0.206 | 0.979 | -0.455 | 1.001 | -0.041 | -0.029 | -0.029 | 0 | 0.08 | 0 | 0.32 | 0.1 | 0.1 | 0.23 |
| 3 | 34212521 | rs9823543 | 0.718 | 0.912 | -0.396 | 0.771 | -0.012 | -0.024 | -0.024 | 0.017 | 0.82 | 0.89 | 0.6 | 0.35 | 0.79 | 0.49 |
| 3 | 34212616 | rs11708781 | 0.829 | 0.56 | -0.759 | -0.223 | -0.011 | 0.158 | 0.158 | 0 | 0.47 | 0.3 | 0.32 | 0.01 | 0.57 | 0.23 |
| 3 | 34216978 | rs991273 | 0.543 | 0.432 | -0.305 | -0.333 | -0.002 | 0 | 0 | 0.053 | 0.28 | 0.3 | 0.22 | 0.34 | 0.18 | 0.23 |
| 3 | 34217664 | rs7626572 | 0.53 | 0.607 | -0.613 | -0.933 | -0.013 | 0.153 | 0.153 | 0 | 0.47 | 0.29 | 0.32 | 0.01 | 0.57 | 0.23 |
| 3 | 34241686 | rs2053425 | 0.356 | 0.665 | -0.473 | -1.062 | 0 | -0.049 | -0.049 | 0 | 0.31 | 0.3 | 0.19 | 0.02 | 0.32 | 0.13 |

| | | | | | | | | | | | | | | | | |
|---|---|---|---|---|---|---|---|---|---|---|---|---|---|---|---|---|
| 3 | 34243929 | rs10510657 | 0.217 | 1.006 | 0.382 | 0.331 | 0.131 | 0.384 | 0.384 | 0 | 0.08 | 0.28 | 0.1 | 0.44 | 0.17 | 0.16 |
| 3 | 34245520 | rs7630319 | 0.967 | 1.041 | 0.038 | -0.098 | -0.01 | 0.003 | 0.003 | 0 | 0.12 | 0.08 | 0.11 | 0 | 0.14 | 0.07 |
| 3 | 34247545 | rs4678662 | 0.951 | 0.486 | -0.229 | -0.283 | -0.001 | 0.108 | 0.108 | 0 | 0.48 | 0.31 | 0.37 | 0.13 | 0.51 | 0.3 |
| 3 | 34247679 | rs6780603 | 0.293 | 0.271 | -0.401 | -0.191 | 0.028 | -0.042 | -0.042 | 0 | 0.27 | 0.13 | 0.41 | 0.4 | 0.21 | 0.47 |
| 3 | 34249893 | rs9985312 | 0.337 | 0.309 | -0.21 | -0.183 | 0.008 | 0.006 | 0.006 | 0 | 0.6 | 0.62 | 0.48 | 0.58 | 0.65 | 0.46 |
| 3 | 34252562 | rs6808824 | -0.623 | 0.343 | -0.007 | -0.223 | 0 | 0.012 | 0.012 | 0 | 0.43 | 0.34 | 0.65 | 0.41 | 0.41 | 0.6 |
| 3 | 34254285 | rs9810690 | 1.149 | 0.311 | 0.142 | -0.4 | 0.001 | 0.051 | 0.051 | 0 | 0.46 | 0.32 | 0.27 | 0.13 | 0.46 | 0.24 |
| 3 | 34264328 | rs7630757 | -0.496 | 0.137 | 0.097 | -0.359 | -0.002 | 0.006 | 0.006 | 0 | 0.39 | 0.41 | 0.48 | 0.22 | 0.36 | 0.39 |
| 3 | 34264385 | rs7630778 | -0.486 | 0.137 | 0.097 | -0.359 | -0.002 | 0.006 | 0.006 | 0 | 0.39 | 0.41 | 0.48 | 0.22 | 0.36 | 0.39 |
| 3 | 34270550 | rs4678665 | 0.226 | 0.215 | 0.213 | -0.363 | 0.011 | -0.017 | -0.017 | 0.124 | 0.65 | 0.45 | 0.55 | 0.25 | 0.63 | 0.52 |
| 3 | 34275465 | rs9859996 | 0.827 | 0.07 | 0.106 | -0.396 | 0.003 | -0.082 | -0.082 | 0 | 0.14 | 0.28 | 0.14 | 0 | 0.13 | 0.08 |
| 3 | 34276109 | rs12497198 | -0.905 | -0.073 | 0.067 | -0.309 | 0.041 | 0.285 | 0.285 | 0 | 0.89 | 0.66 | 0.91 | 0.48 | 0.86 | 0.84 |
| 3 | 34277749 | rs1401345 | 0.196 | -0.098 | 0.014 | -0.469 | 0.006 | -0.006 | -0.006 | 0 | 0.66 | 0.56 | 0.57 | 0.24 | 0.63 | 0.52 |
| 3 | 34290036 | rs7642509 | 0.89 | -0.079 | -0.622 | -0.479 | 0.005 | 0.078 | 0.078 | 0 | 0.5 | 0.27 | 0.43 | 0.24 | 0.51 | 0.41 |
| 3 | 34294343 | rs4577412 | 1.666 | -0.244 | -1.296 | -0.964 | -0.078 | 0.179 | 0.179 | 0.564 | 0.32 | 0.05 | 0.19 | 0 | 0.38 | 0.18 |
| 3 | 34299081 | rs1401340 | 1.128 | -0.303 | -1.3 | -0.863 | -0.033 | -0.013 | -0.013 | 0 | 0.21 | 0.08 | 0.34 | 0.23 | 0.22 | 0.33 |
| 3 | 34300719 | rs13096452 | 1.803 | -0.182 | -1.337 | -0.861 | -0.082 | 0.167 | 0.167 | 0.128 | 0.3 | 0.05 | 0.18 | 0 | 0.38 | 0.11 |
| 3 | 34307550 | rs7621054 | -0.5 | 0.211 | -0.624 | -0.22 | 0.052 | 0.343 | 0.343 | 0 | 0.87 | 0.57 | 0.89 | 0.47 | 0.85 | 0.8 |
| 3 | 34311764 | rs13067818 | 0.76 | 0.072 | -0.676 | -0.63 | -0.006 | -0.045 | -0.045 | 0 | 0.17 | 0.23 | 0.12 | 0 | 0.12 | 0.07 |
| 3 | 34314578 | rs12494048 | 0.954 | 0.087 | -0.812 | -0.502 | -0.077 | -0.012 | -0.012 | 0.743 | 0.18 | 0.02 | 0.26 | 0.16 | 0.21 | 0.32 |
| 3 | 34317979 | rs9838476 | -0.03 | 0.024 | -0.032 | 0.05 | 0.002 | 0.075 | 0.075 | 0 | 0.72 | 0.54 | 0.76 | 0.41 | 0.74 | 0.73 |
| 3 | 34319129 | rs2336569 | 0.165 | 0.094 | -0.626 | 0.081 | -0.004 | 0.017 | 0.017 | 0 | 0.7 | 0.39 | 0.74 | 0.41 | 0.74 | 0.73 |
| 3 | 34334484 | rs9834633 | -0.286 | -0.068 | -0.695 | -0.021 | -0.006 | 0.002 | 0.002 | 0 | 0.08 | 0.1 | 0.12 | 0.14 | 0.06 | 0.14 |
| 3 | 34352463 | rs9826990 | -1.102 | 0.044 | -0.8 | -1.038 | 0.007 | 0.053 | 0.053 | 1.349 | 0.59 | 0.73 | 0.7 | 0.97 | 0.59 | 0.73 |
| 3 | 34403194 | rs2069265 | 0.945 | 0.285 | -0.792 | -0.147 | -0.099 | -0.055 | -0.055 | 0.016 | 0.2 | 0.03 | 0.36 | 0.29 | 0.3 | 0.29 |
| 3 | 34408481 | rs7623112 | -0.196 | -0.021 | -0.2 | 0.025 | -0.007 | -0.011 | -0.011 | 0 | 0.36 | 0.31 | 0.53 | 0.52 | 0.41 | 0.55 |
| 3 | 34415876 | rs1191624 | -0.394 | -0.271 | -0.478 | -0.238 | -0.004 | -0.004 | -0.004 | 0.32 | 0.39 | 0.35 | 0.56 | 0.55 | 0.44 | 0.55 |
| 3 | 34416500 | rs13083647 | 0.646 | 0.234 | -0.992 | -0.269 | -0.046 | -0.083 | -0.083 | 0 | 0.22 | 0.07 | 0.15 | 0.42 | 0.29 | 0.16 |
| 3 | 34421745 | rs1919621 | -0.192 | 0.198 | -0.698 | -0.433 | 0.014 | 0.017 | 0.017 | 0 | 0.38 | 0.49 | 0.68 | 0.57 | 0.45 | 0.62 |
| 3 | 34446738 | rs17031657 | 1.577 | 0.016 | -0.948 | -0.301 | -0.061 | -0.124 | -0.124 | 0 | 0.15 | 0.01 | 0.11 | 0.48 | 0.18 | 0.12 |
| 3 | 34451940 | rs1534673 | 1.105 | -0.254 | -0.863 | -0.207 | 0.005 | -0.008 | -0.008 | 0.925 | 0.08 | 0.02 | 0.12 | 0.25 | 0.08 | 0.24 |
| 3 | 34465049 | rs9827159 | 1.18 | -0.353 | -0.715 | -0.637 | -0.005 | -0.076 | -0.076 | 0 | 0.37 | 0.42 | 0.27 | 0.02 | 0.27 | 0.27 |
| 3 | 34500346 | rs9875352 | 0.974 | -0.241 | -0.652 | -0.6 | 0.031 | 0.037 | 0.037 | 5.924 | 0.38 | 0.28 | 0.25 | 0.02 | 0.28 | 0.24 |
| 3 | 34502246 | rs4678469 | -0.008 | 0.011 | -0.891 | -0.534 | -0.001 | -0.003 | -0.003 | 0 | 0.35 | 0.34 | 0.52 | 0.49 | 0.36 | 0.57 |
| 3 | 34519341 | rs2336652 | -1.76 | 0.34 | -0.864 | -0.335 | -0.003 | -0.028 | -0.028 | 0 | 0.94 | 0.88 | 0.96 | 1 | 0.94 | 0.99 |
| 3 | 34533675 | rs1868528 | 0.362 | 0.185 | -1.021 | -0.203 | -0.036 | -0.072 | -0.072 | 0.806 | 0.25 | 0.11 | 0.17 | 0.48 | 0.31 | 0.19 |
| 3 | 34538558 | rs1919623 | 0.692 | 0.421 | -0.898 | -0.043 | 0.005 | 0.002 | 0.002 | 0.402 | 0.22 | 0.25 | 0.35 | 0.3 | 0.26 | 0.3 |
| 3 | 34542396 | rs1919617 | 0.349 | 0.234 | -0.615 | 0.565 | -0.034 | -0.059 | -0.059 | 4.168 | 0.21 | 0.1 | 0.17 | 0.46 | 0.3 | 0.17 |
| 3 | 34545151 | rs997334 | -0.021 | -0.074 | -0.458 | 0.046 | -0.025 | -0.04 | -0.04 | 0 | 0.33 | 0.43 | 0.37 | 0.27 | 0.25 | 0.36 |
| 3 | 34545267 | rs9881051 | 0.436 | -0.254 | -0.353 | 0.322 | -0.021 | 0.179 | 0.179 | 0.019 | 0.12 | 0.3 | 0.16 | 0.32 | 0.1 | 0.27 |
| 3 | 34546401 | rs17031805 | 1.009 | -0.066 | -0.529 | 0.185 | -0.038 | -0.047 | -0.047 | 0 | 0.2 | 0.1 | 0.16 | 0.41 | 0.3 | 0.16 |
| 3 | 34551563 | rs6777100 | 0.349 | -0.121 | -0.419 | 0.14 | -0.022 | 0.221 | 0.221 | 0.07 | 0.13 | 0.33 | 0.17 | 0.32 | 0.1 | 0.27 |
| 3 | 34556501 | rs1852808 | -0.323 | -0.123 | -0.448 | 0.07 | 0.014 | -0.026 | -0.026 | 0 | 0.33 | 0.39 | 0.32 | 0.72 | 0.39 | 0.43 |
| 3 | 34567536 | rs4678692 | -0.422 | -0.188 | -0.346 | 0.068 | 0.013 | 0.02 | 0.02 | 1.419 | 0.51 | 0.37 | 0.5 | 0.67 | 0.53 | 0.51 |
| 3 | 34568780 | rs17201571 | 1.232 | -0.334 | -0.443 | -0.023 | 0.042 | 0.036 | 0.036 | 0.604 | 0.2 | 0.07 | 0.07 | 0.15 | 0.1 | 0.13 |
| 3 | 34569954 | rs12632204 | 1.154 | -0.263 | -0.416 | 0.022 | -0.053 | -0.061 | -0.061 | 0 | 0.2 | 0.06 | 0.16 | 0.39 | 0.3 | 0.16 |
| 3 | 34580559 | rs4678471 | -0.599 | -0.424 | -0.417 | 0.021 | 0.05 | 0.025 | 0.025 | 0.275 | 0.46 | 0.72 | 0.52 | 0.6 | 0.35 | 0.63 |
| 3 | 34589121 | rs1402246 | -1.092 | 0.023 | -0.19 | 0.123 | -0.001 | 0.156 | 0.156 | 0.54 | 0.63 | 0.8 | 0.68 | 0.99 | 0.64 | 0.79 |
| 3 | 34592382 | rs6779484 | -1.183 | 0.14 | -0.202 | 0.158 | -0.037 | 0.18 | 0.18 | 0.659 | 0.64 | 0.85 | 0.68 | 0.99 | 0.61 | 0.78 |
| 3 | 34593849 | rs6783456 | 0.215 | 0.344 | 0.776 | 1.566 | 0.027 | 0.122 | 0.122 | 0.118 | 0.11 | 0.24 | 0.35 | 0.31 | 0.15 | 0.24 |
| 3 | 34610315 | rs9311059 | -0.386 | 0.135 | 0.624 | 1.065 | -0.01 | 0.038 | 0.038 | 0 | 0.13 | 0.27 | 0.1 | 0.19 | 0.1 | 0.28 |
| 3 | 34615754 | rs13074645 | -1.013 | 0.486 | 0.661 | 1.132 | 0.005 | 0.002 | 0.002 | 0 | 0.95 | 0.98 | 0.96 | 1 | 0.94 | 0.99 |
| 3 | 34622033 | rs9311060 | 0.536 | 0.437 | 0.775 | 1.193 | -0.013 | 0.03 | 0.03 | 0 | 0.85 | 0.71 | 0.87 | 0.82 | 0.84 | 0.7 |
| 3 | 34623437 | rs7628551 | -0.243 | 0.524 | 0.813 | 1.35 | -0.005 | 0.128 | 0.128 | 0 | 0.11 | 0.18 | 0.15 | 0.47 | 0.1 | 0.29 |
| 3 | 34654702 | rs7622138 | 0.016 | 0.653 | 1.01 | 1.265 | 0.042 | 0.048 | 0.048 | 0.015 | 0.12 | 0.32 | 0.3 | 0.16 | 0.14 | 0.23 |
| 3 | 34665118 | rs13075567 | 0.279 | 1.104 | 0.511 | 0.823 | -0.033 | -0.059 | -0.059 | 0 | 0.19 | 0.07 | 0.13 | 0.36 | 0.25 | 0.11 |
| 3 | 34676421 | rs9875943 | -0.094 | 1.268 | 1.004 | 1.225 | -0.004 | -0.032 | -0.032 | 0.036 | 0.9 | 0.9 | 0.93 | 1 | 0.89 | 0.98 |
| 3 | 34683998 | rs6790851 | -0.89 | 1.665 | 0.806 | 1.681 | 0.006 | 0.097 | 0.097 | 0 | 0.11 | 0.34 | 0.1 | 0.19 | 0.1 | 0.28 |
| 3 | 34698297 | rs13075731 | -1.182 | 1.696 | 0.362 | 1.696 | 0.03 | 0.047 | 0.047 | 0.028 | 0.13 | 0.3 | 0.12 | 0.18 | 0.18 | 0.29 |
| 3 | 34708263 | rs13061601 | -0.855 | 1.828 | -0.39 | 1.76 | -0.018 | -0.024 | -0.024 | 0 | 0.12 | 0.07 | 0.1 | 0.18 | 0.18 | 0.28 |
| 3 | 34765770 | rs951973 | -0.798 | 1.459 | -1.066 | 2.023 | -0.053 | -0.073 | -0.073 | 0 | 0.11 | 0.01 | 0.33 | 0.45 | 0.16 | 0.24 |
| 3 | 34814789 | rs11129615 | -0.911 | 1.264 | -0.553 | 1.809 | 0.005 | 0.082 | 0.082 | 0 | 0.21 | 0.36 | 0.38 | 0.5 | 0.19 | 0.34 |
| 3 | 34814806 | rs11129616 | -0.677 | 1.019 | -0.916 | 0.895 | 0.002 | 0 | 0 | 0 | 0.59 | 0.62 | 0.65 | 0.61 | 0.62 | 0.57 |
| 3 | 34816835 | rs653476 | 0.881 | 1.497 | -1.507 | 0.433 | -0.073 | 0.052 | 0.052 | 0.063 | 0.26 | 0.02 | 0.11 | 0.1 | 0.25 | 0.18 |
| 3 | 34817627 | rs667168 | -0.425 | 1.777 | -1.239 | 0.627 | -0.017 | 0.109 | 0.109 | 0.121 | 0.48 | 0.8 | 0.54 | 0.24 | 0.49 | 0.35 |
| 3 | 34820296 | rs591834 | -0.464 | 0.981 | -1.938 | 0.421 | -0.021 | -0.037 | -0.037 | 3.233 | 0.67 | 0.84 | 0.7 | 0.51 | 0.65 | 0.46 |
| 3 | 34822886 | rs7623820 | 0.548 | 0.916 | -1.391 | 0.369 | -0.01 | 0.023 | 0.023 | 0.207 | 0.4 | 0.62 | 0.34 | 0.14 | 0.44 | 0.22 |
| 3 | 34824952 | rs7635856 | -0.353 | 1.099 | -1.347 | 0.666 | 0 | -0.029 | -0.029 | 0 | 0.54 | 0.82 | 0.57 | 0.53 | 0.55 | 0.43 |
| 3 | 34834899 | rs1487993 | -0.143 | 1.19 | -1.355 | 0.663 | 0.013 | 0 | 0 | 0 | 0.37 | 0.26 | 0.5 | 0.66 | 0.38 | 0.34 |

| | | | | | | | | | | | | | | | | |
|---|---|---|---|---|---|---|---|---|---|---|---|---|---|---|---|---|
| 3 | 34834934 | rs511922 | 1.139 | 1.16 | -0.648 | 0.627 | -0.015 | 0.056 | 0.056 | 0.029 | 0.34 | 0.2 | 0.18 | 0.13 | 0.33 | 0.21 |
| 3 | 34838952 | rs522543 | -0.742 | 1.163 | -0.538 | 0.679 | -0.013 | 0.024 | 0.024 | 0.219 | 0.72 | 0.85 | 0.72 | 0.79 | 0.7 | 0.55 |
| 3 | 34841583 | rs6550303 | 0.371 | 1.147 | -0.562 | 0.727 | -0.004 | 0.022 | 0.022 | 0 | 0.13 | 0.19 | 0.24 | 0.37 | 0.11 | 0.21 |
| 3 | 34850303 | rs4643650 | 0.114 | 1.172 | -0.54 | 0.761 | -0.001 | 0.004 | 0.004 | 0 | 0.15 | 0.19 | 0.32 | 0.38 | 0.19 | 0.24 |
| 3 | 34850540 | rs564190 | 0.515 | 1.132 | -0.231 | 0.703 | 0.004 | 0.006 | 0.006 | 0.551 | 0.56 | 0.48 | 0.38 | 0.42 | 0.51 | 0.32 |
| 3 | 34852103 | rs6798946 | -0.441 | 1.083 | -0.096 | 0.597 | 0.039 | 0.016 | 0.016 | 0 | 0.06 | 0.28 | 0.06 | 0.01 | 0.05 | 0.01 |
| 3 | 34865842 | rs645648 | -0.585 | 1.044 | -0.206 | 0.585 | 0.006 | -0.02 | -0.02 | 0 | 0.72 | 0.64 | 0.7 | 0.79 | 0.7 | 0.58 |
| 3 | 34871929 | rs7612893 | -0.607 | 1.086 | -0.23 | 0.586 | 0.004 | -0.015 | -0.015 | 0 | 0.72 | 0.66 | 0.7 | 0.79 | 0.7 | 0.58 |
| 3 | 34876537 | rs613115 | 2.126 | 1.39 | -0.853 | 0.226 | -0.04 | 0.009 | 0.009 | 0.546 | 0.11 | 0.01 | 0.05 | 0.03 | 0.1 | 0.09 |
| 3 | 34877313 | rs627033 | 0.43 | 1.211 | -0.987 | 0.493 | -0.011 | -0.002 | -0.002 | 0.151 | 0.51 | 0.19 | 0.32 | 0.44 | 0.46 | 0.34 |
| 3 | 34882436 | rs11709953 | 1.267 | 1.062 | -0.875 | 0.65 | 0.027 | -0.059 | -0.059 | 0 | 0.2 | 0.08 | 0.16 | 0.28 | 0.15 | 0.11 |
| 3 | 34891189 | rs1916228 | -0.61 | 1.14 | -0.787 | 0.312 | 0.025 | 0.019 | 0.019 | 0 | 0.5 | 0.69 | 0.59 | 0.58 | 0.46 | 0.41 |
| 3 | 34898125 | rs6550306 | 0.682 | 1.152 | -0.44 | 0.295 | 0.018 | 0.06 | 0.06 | 0.564 | 0.1 | 0.23 | 0.17 | 0.27 | 0.15 | 0.12 |
| 3 | 34902705 | rs6792813 | -0.759 | 1.127 | -0.332 | 0.242 | 0.017 | 0.003 | 0.003 | 0 | 0.24 | 0.57 | 0.31 | 0.27 | 0.28 | 0.22 |
| 3 | 34905662 | rs7652586 | 1.177 | 1.087 | -0.419 | 0.229 | -0.026 | 0.053 | 0.053 | 0 | 0.1 | 0.01 | 0.09 | 0 | 0.11 | 0.1 |
| 3 | 34909205 | rs7620215 | 1.534 | 0.977 | -0.49 | 0.261 | 0.03 | -0.066 | -0.066 | 0.815 | 0.21 | 0.07 | 0.27 | 0.31 | 0.16 | 0.18 |
| 3 | 34910172 | rs4678714 | 1.376 | 1.048 | -0.401 | 0.282 | 0 | 0.05 | 0.05 | 0.043 | 0.06 | 0.12 | 0.07 | 0.27 | 0.08 | 0.1 |
| 3 | 34914885 | rs9311079 | 0.231 | 1.253 | -0.502 | 0.258 | -0.011 | -0.001 | -0.001 | 0.236 | 0.34 | 0.4 | 0.46 | 0.31 | 0.3 | 0.29 |
| 3 | 34915433 | rs2071489 | -0.456 | 1.198 | -0.517 | 0.203 | 0.01 | 0.005 | 0.005 | 0 | 0.46 | 0.67 | 0.64 | 0.58 | 0.45 | 0.41 |
| 3 | 34931283 | rs869226 | -0.618 | 1.155 | -0.506 | 0.196 | -0.004 | -0.001 | -0.001 | 0.302 | 0.22 | 0.28 | 0.2 | 0.27 | 0.2 | 0.2 |
| 3 | 34935472 | rs9647406 | 0.786 | 1.235 | -0.547 | 0.313 | -0.009 | -0.01 | -0.01 | 1.163 | 0.09 | 0.07 | 0.13 | 0.19 | 0.11 | 0.12 |
| 3 | 34937547 | rs9860323 | -1.339 | 1.246 | -0.585 | 0.223 | -0.012 | 0.058 | 0.058 | 0 | 0.58 | 0.78 | 0.78 | 0.78 | 0.56 | 0.54 |
| 3 | 34940000 | rs6774497 | -0.544 | 1.171 | -0.598 | 0.312 | 0.03 | 0.076 | 0.076 | 1.301 | 0.19 | 0.35 | 0.41 | 0.46 | 0.26 | 0.24 |
| 3 | 34972029 | rs4678492 | 0.42 | 1.208 | -0.555 | 0.142 | 0.008 | 0.024 | 0.024 | 0 | 0.51 | 0.52 | 0.41 | 0.22 | 0.56 | 0.49 |
| 3 | 34979273 | rs4678495 | 0.901 | 1.578 | -0.443 | 0.005 | 0.044 | -0.045 | -0.045 | 2.093 | 0.71 | 0.48 | 0.51 | 0.79 | 0.66 | 0.69 |
| 3 | 34991575 | rs10154831 | -0.146 | 1.634 | -0.325 | 0.034 | -0.037 | -0.028 | -0.028 | 0 | 0.86 | 0.94 | 0.84 | 0.78 | 0.78 | 0.85 |
| 3 | 34996151 | rs4678729 | -0.042 | 1.609 | -0.351 | 0.015 | 0.002 | 0.028 | 0.028 | 0 | 0.13 | 0.23 | 0.18 | 0.27 | 0.16 | 0.12 |
| 3 | 34997498 | rs11922997 | 0.214 | 1.542 | -0.631 | -0.29 | -0.002 | -0.141 | -0.141 | 2.723 | 0.25 | 0.43 | 0.36 | 0.02 | 0.23 | 0.18 |
| 3 | 35008453 | rs1463552 | 0.527 | 1.516 | -0.651 | -0.302 | 0.02 | -0.045 | -0.045 | 0.873 | 0.62 | 0.56 | 0.49 | 0.79 | 0.56 | 0.67 |
| 3 | 35008595 | rs9881697 | 1.282 | 1.407 | -1.03 | -0.176 | 0.039 | 0.016 | 0.016 | 0 | 0.47 | 0.25 | 0.31 | 0.52 | 0.41 | 0.53 |
| 3 | 35015022 | rs4678730 | 0.168 | 1.637 | -1.057 | 0.061 | -0.017 | -0.004 | -0.004 | 0 | 0.86 | 0.91 | 0.84 | 0.8 | 0.79 | 0.84 |
| 3 | 35038139 | rs11129629 | -0.19 | 1.636 | -1.076 | 0.027 | -0.008 | 0.004 | 0.004 | 0 | 0.91 | 0.93 | 0.88 | 0.93 | 0.87 | 0.93 |
| 3 | 35039861 | rs2359209 | 1.28 | 1.95 | -1.038 | 0.109 | -0.003 | -0.003 | -0.003 | 4.355 | 0.05 | 0.05 | 0.06 | 0.12 | 0.08 | 0.08 |
| 3 | 35040896 | rs2884658 | 1.11 | 1.33 | -1.2 | 0.281 | -0.036 | -0.045 | -0.045 | 0.001 | 0.59 | 0.18 | 0.6 | 0.45 | 0.56 | 0.55 |
| 3 | 35045331 | rs2035257 | 1.238 | 1.25 | -1.333 | 0.182 | -0.003 | -0.039 | -0.039 | 0.202 | 0.57 | 0.18 | 0.67 | 0.45 | 0.62 | 0.55 |
| 3 | 35046813 | rs6779123 | 1.033 | 1.277 | -1.009 | 0.308 | 0.032 | 0.103 | 0.103 | 0 | 0.07 | 0.15 | 0.08 | 0.28 | 0.08 | 0.08 |
| 3 | 35046864 | rs6792252 | 1.426 | 1.277 | -1.019 | 0.307 | -0.006 | 0.012 | 0.012 | 0 | 0.06 | 0.05 | 0.07 | 0.28 | 0.08 | 0.08 |
| 3 | 35054264 | rs9990367 | -0.772 | 1.145 | -0.826 | 0.225 | 0.034 | 0.11 | 0.11 | 1.217 | 0.12 | 0.46 | 0.16 | 0.19 | 0.12 | 0.23 |
| 3 | 35064339 | rs17032594 | 2.188 | 1.18 | -0.989 | 0.261 | 0.007 | 0.017 | 0.017 | 0 | 0.11 | 0.06 | 0.04 | 0.01 | 0.09 | 0.06 |
| 3 | 35064874 | rs2219794 | 0.083 | 1.289 | -0.987 | 0.359 | 0.025 | 0.029 | 0.029 | 0 | 0.84 | 0.77 | 0.8 | 0.65 | 0.77 | 0.84 |
| 3 | 35085414 | rs7620914 | -0.097 | 1.145 | -0.749 | 0.205 | -0.004 | 0.023 | 0.023 | 0 | 0.13 | 0.24 | 0.14 | 0.19 | 0.12 | 0.24 |
| 3 | 35089305 | rs2035260 | 0.853 | 0.957 | -1.692 | -0.201 | -0.052 | 0.032 | 0.032 | 0 | 0.23 | 0.02 | 0.22 | 0.12 | 0.24 | 0.24 |
| 3 | 35099044 | rs9854749 | -0.745 | 1.128 | -1.242 | 0.071 | 0.001 | -0.002 | -0.002 | 1.058 | 0.89 | 0.89 | 0.87 | 0.92 | 0.86 | 0.91 |
| 3 | 35100361 | rs1382552 | 0.099 | 0.823 | -1.177 | 0.071 | -0.017 | -0.018 | -0.018 | 0 | 0.25 | 0.43 | 0.18 | 0.2 | 0.21 | 0.3 |
| 3 | 35108589 | rs10433599 | 0.787 | 0.997 | -1.216 | 0.182 | -0.004 | 0.006 | 0.006 | 0 | 0.25 | 0.17 | 0.18 | 0.29 | 0.23 | 0.17 |
| 3 | 35108791 | rs10433542 | 1.151 | 0.957 | -1.236 | 0.235 | -0.006 | -0.004 | -0.004 | 0 | 0.22 | 0.13 | 0.17 | 0.29 | 0.2 | 0.16 |
| 3 | 35113640 | rs6785948 | 0.807 | 0.534 | -1.439 | 0.237 | 0.042 | -0.008 | -0.008 | 0.039 | 0.36 | 0.17 | 0.32 | 0.34 | 0.34 | 0.31 |
| 3 | 35128046 | rs1478838 | 2.089 | 0.361 | -1.533 | 0.235 | -0.027 | -0.07 | -0.07 | 0 | 0.13 | 0.02 | 0.17 | 0.22 | 0.16 | 0.11 |
| 3 | 35134831 | rs11129639 | -0.388 | 1.025 | -1.042 | 0.403 | -0.049 | 0.003 | 0.003 | 0 | 0.65 | 0.83 | 0.62 | 0.61 | 0.59 | 0.72 |
| 3 | 35142511 | rs11129640 | 1.453 | 0.886 | -0.723 | 0.389 | -0.006 | -0.007 | -0.007 | 0 | 0.23 | 0.22 | 0.24 | 0.27 | 0.23 | 0.4 |
| 3 | 35144053 | rs6786233 | 1.308 | 0.972 | -0.873 | 0.44 | 0.007 | 0.015 | 0.015 | 0.969 | 0.24 | 0.35 | 0.25 | 0.27 | 0.23 | 0.4 |
| 3 | 35167155 | rs4541358 | -0.238 | 1.17 | -1.916 | 0.55 | -0.036 | -0.008 | -0.008 | 1.035 | 0.64 | 0.78 | 0.62 | 0.58 | 0.59 | 0.71 |
| 3 | 35167376 | rs4133297 | 0.562 | 1.049 | -2.048 | 0.676 | 0.043 | 0.003 | 0.003 | 0 | 0.36 | 0.16 | 0.32 | 0.32 | 0.34 | 0.31 |
| 3 | 35170656 | rs4279040 | 0.531 | 1.729 | -1.42 | 0.919 | -0.041 | -0.011 | -0.011 | 0 | 0.86 | 0.96 | 0.79 | 0.87 | 0.79 | 0.88 |
| 3 | 35190794 | rs6803721 | 0.18 | 1.172 | -0.989 | 1.127 | -0.005 | -0.005 | -0.005 | 0 | 0.79 | 0.81 | 0.73 | 0.62 | 0.71 | 0.81 |
| 3 | 35231021 | rs13077466 | 0.259 | 1.259 | -0.875 | 1.213 | -0.003 | -0.003 | -0.003 | 0.023 | 0.16 | 0.14 | 0.25 | 0.31 | 0.25 | 0.13 |
| 3 | 35237877 | rs6788583 | -0.87 | 0.347 | -1.171 | 1.063 | -0.002 | 0.014 | 0.014 | 0 | 0.53 | 0.42 | 0.47 | 0.62 | 0.55 | 0.4 |
| 3 | 35240210 | rs6786768 | 0.469 | -0.087 | -1.734 | 1.016 | 0.054 | -0.062 | -0.062 | 0 | 0.17 | 0.05 | 0.04 | 0.25 | 0.09 | 0.18 |
| 3 | 35245127 | rs4346506 | -0.867 | 0.245 | -1.154 | 0.766 | -0.01 | 0.025 | 0.025 | 0 | 0.54 | 0.68 | 0.45 | 0.63 | 0.55 | 0.4 |
| 3 | 35245356 | rs11713784 | 1.143 | 0.678 | -1.467 | 0.319 | -0.051 | 0.06 | 0.06 | 0 | 0.1 | 0.01 | 0.07 | 0 | 0.15 | 0.08 |
| 3 | 35251044 | rs7647607 | 0.514 | -0.676 | -1.842 | 0.504 | 0.047 | 0.014 | 0.014 | 0 | 0.51 | 0.29 | 0.57 | 0.6 | 0.65 | 0.53 |
| 3 | 35261198 | rs28847946 | 1.294 | -0.849 | -1.754 | 0.639 | -0.024 | 0.007 | 0.007 | 0 | 0.38 | 0.25 | 0.4 | 0.37 | 0.46 | 0.37 |
| 3 | 35265216 | rs4078511 | 1.58 | -1.514 | -1.962 | 0.345 | 0.025 | -0.031 | -0.031 | 0 | 0.13 | 0.06 | 0.07 | 0.29 | 0.09 | 0.1 |
| 3 | 35265738 | rs6781932 | -0.538 | -1.543 | -2.107 | 0.1 | -0.005 | 0.033 | 0.033 | 0 | 0.79 | 0.91 | 0.74 | 0.91 | 0.8 | 0.76 |
| 3 | 35278091 | rs1376021 | 1.355 | -1.146 | -1.782 | 0.356 | -0.02 | -0.015 | -0.015 | 0.112 | 0.38 | 0.25 | 0.41 | 0.45 | 0.49 | 0.37 |
| 3 | 35278145 | rs17692875 | 0.413 | -0.776 | -1.84 | 0.163 | -0.006 | 0.004 | 0.004 | 0 | 0.16 | 0.06 | 0.16 | 0.15 | 0.13 | 0.2 |
| 3 | 35281232 | rs2222795 | 0.491 | -0.949 | -1.43 | 0.022 | 0.014 | 0.054 | 0.054 | 0.086 | 0.55 | 0.66 | 0.53 | 0.74 | 0.61 | 0.51 |
| 3 | 35281449 | rs1450322 | 1.079 | -1.707 | -1.234 | 0.524 | 0.021 | -0.022 | -0.022 | 0 | 0.12 | 0.06 | 0.07 | 0.2 | 0.08 | 0.09 |
| 3 | 35282759 | rs9811203 | -0.639 | -1.992 | -0.943 | 0.075 | -0.001 | 0 | 0 | 0 | 0.41 | 0.37 | 0.47 | 0.33 | 0.43 | 0.43 |

| | | | | | | | | | | | | | | | | |
|---|---|---|---|---|---|---|---|---|---|---|---|---|---|---|---|---|
| 3 | 35286114 | rs17744075 | 0.488 | -2.253 | -0.944 | -0.428 | -0.005 | -0.025 | -0.025 | 0 | 0.07 | 0 | 0.02 | 0.1 | 0.05 | 0.14 |
| 3 | 35286613 | rs17693202 | 0.112 | -2.585 | -0.606 | -0.69 | 0.006 | -0.058 | -0.058 | 0.221 | 0.16 | 0.25 | 0.28 | 0.07 | 0.18 | 0.22 |
| 3 | 35288963 | rs2060661 | 2.584 | -2.935 | -0.65 | -0.598 | -0.027 | -0.026 | -0.026 | 0 | 0.28 | 0.21 | 0.32 | 0.4 | 0.47 | 0.19 |
| 3 | 35290327 | rs1450310 | 1.331 | -3.254 | -0.629 | -0.701 | 0.073 | 0.083 | 0.083 | 0 | 0.5 | 0.35 | 0.52 | 0.67 | 0.66 | 0.54 |
| 3 | 35291791 | rs9680953 | 2.697 | -3.132 | -0.641 | -0.661 | -0.019 | -0.018 | -0.018 | 0.159 | 0.28 | 0.22 | 0.32 | 0.4 | 0.46 | 0.19 |
| 3 | 35298744 | rs4678757 | -1.063 | -3.224 | -0.607 | -0.723 | 0.041 | 0.019 | 0.019 | 1.201 | 0.79 | 0.9 | 0.87 | 0.84 | 0.91 | 0.9 |
| 3 | 35303387 | rs7434012 | 2.76 | -3.222 | -0.561 | -0.72 | -0.032 | -0.031 | -0.031 | 0.385 | 0.28 | 0.21 | 0.32 | 0.41 | 0.47 | 0.19 |
| 3 | 35305540 | rs1529049 | 0.917 | -3.176 | -0.393 | -0.714 | 0.005 | -0.012 | -0.012 | 0.329 | 0.22 | 0.1 | 0.21 | 0.24 | 0.2 | 0.32 |
| 3 | 35316680 | rs1450314 | -1.642 | -3.29 | -0.307 | -0.736 | 0.027 | 0.039 | 0.039 | 0.04 | 0.5 | 0.66 | 0.54 | 0.43 | 0.44 | 0.72 |
| 3 | 35317606 | rs6783653 | -0.403 | -3.235 | -0.147 | -0.715 | 0 | -0.036 | -0.036 | 0 | 0.22 | 0.29 | 0.27 | 0.16 | 0.22 | 0.34 |
| 3 | 35331357 | rs11129651 | -1.267 | -3.293 | -0.123 | -0.712 | -0.003 | -0.003 | -0.003 | 0 | 0.45 | 0.44 | 0.51 | 0.43 | 0.39 | 0.69 |
| 3 | 35334398 | rs17693703 | 0.409 | -3.318 | 0.011 | -0.757 | -0.002 | -0.024 | -0.024 | 0 | 0.16 | 0.16 | 0.22 | 0.05 | 0.15 | 0.23 |
| 3 | 35337113 | rs11927356 | -1.355 | -3.288 | -0.045 | -0.711 | -0.003 | -0.002 | -0.002 | 0 | 0.46 | 0.44 | 0.56 | 0.46 | 0.39 | 0.69 |
| 3 | 35342479 | rs7619621 | -0.954 | -3.33 | -0.105 | -0.675 | 0.031 | 0.023 | 0.023 | 1.737 | 0.79 | 0.87 | 0.92 | 0.87 | 0.91 | 0.91 |
| 3 | 35360207 | rs4678765 | 1.002 | -3.132 | -0.075 | -0.775 | 0.011 | -0.013 | -0.013 | 0.519 | 0.2 | 0.13 | 0.24 | 0.29 | 0.17 | 0.33 |
| 3 | 35364399 | rs7611068 | 0.232 | -3.054 | -0.028 | -0.943 | -0.006 | 0.05 | 0.05 | 0.147 | 0.17 | 0.07 | 0.21 | 0.03 | 0.18 | 0.24 |
| 3 | 35368482 | rs5007515 | -2.402 | -3.123 | 0.107 | -0.965 | 0.033 | 0.042 | 0.042 | 0 | 0.68 | 0.55 | 0.65 | 0.57 | 0.5 | 0.79 |
| 3 | 35371916 | rs6801705 | -0.774 | -3.012 | 0.063 | -0.969 | 0.001 | -0.025 | -0.025 | 0.259 | 0.24 | 0.27 | 0.26 | 0.13 | 0.24 | 0.35 |
| 3 | 35385225 | rs1455324 | 1.114 | -3.096 | -0.241 | -0.969 | 0.016 | -0.047 | -0.047 | 2.588 | 0.2 | 0.08 | 0.31 | 0.31 | 0.17 | 0.35 |
| 3 | 35386568 | rs7625092 | 0.662 | -3.212 | -0.006 | -1.029 | -0.018 | -0.015 | -0.015 | 0.651 | 0.24 | 0.25 | 0.08 | 0.11 | 0.09 | 0.09 |
| 3 | 35390220 | rs4086242 | -0.543 | -3.288 | 0.161 | -1.063 | -0.007 | -0.005 | -0.005 | 1.812 | 0.35 | 0.62 | 0.16 | 0.2 | 0.17 | 0.22 |
| 3 | 35390673 | rs7615737 | -0.083 | -3.295 | 0.18 | -1.07 | -0.063 | -0.069 | -0.069 | 0.485 | 0.3 | 0.58 | 0.13 | 0.11 | 0.12 | 0.14 |
| 3 | 35392137 | rs11705929 | -0.312 | -3.231 | 0.004 | -1.087 | 0.017 | -0.004 | -0.004 | 1.952 | 0.35 | 0.15 | 0.51 | 0.34 | 0.33 | 0.57 |
| 3 | 35393058 | rs12330405 | 0.519 | -3.323 | 0.123 | -1.046 | -0.026 | -0.027 | -0.027 | 0 | 0.25 | 0.29 | 0.1 | 0.11 | 0.1 | 0.11 |
| 3 | 35395020 | rs1003054 | -0.543 | -3.196 | -0.327 | -0.991 | 0.001 | 0.002 | 0.002 | 2.03 | 0.35 | 0.63 | 0.16 | 0.2 | 0.17 | 0.21 |
| 3 | 35397487 | rs2167176 | -2.441 | -2.78 | -0.233 | -0.793 | -0.061 | -0.068 | -0.068 | 0 | 0.73 | 0.86 | 0.68 | 0.55 | 0.51 | 0.81 |
| 3 | 35399185 | rs4678770 | 0.358 | -3.054 | -0.383 | -0.842 | 0.017 | -0.026 | -0.026 | 0.225 | 0.24 | 0.16 | 0.34 | 0.32 | 0.2 | 0.37 |
| 3 | 35405013 | rs9822492 | 0.316 | -3.056 | -0.106 | -0.927 | -0.024 | -0.019 | -0.019 | 8.104 | 0.23 | 0.24 | 0.17 | 0.11 | 0.09 | 0.14 |
| 3 | 35426559 | rs4678771 | -0.623 | -2.945 | -0.578 | -0.93 | 0.024 | 0.011 | 0.011 | 0 | 0.42 | 0.27 | 0.43 | 0.4 | 0.39 | 0.55 |
| 3 | 35435460 | rs814487 | 1.392 | -2.51 | -0.818 | -0.751 | 0.002 | 0.004 | 0.004 | 0 | 0.36 | 0.4 | 0.38 | 0.49 | 0.52 | 0.3 |
| 3 | 35439546 | rs814490 | 2.098 | -2.433 | -1.728 | -0.67 | -0.109 | -0.13 | -0.13 | 0.855 | 0.29 | 0.06 | 0.31 | 0.46 | 0.49 | 0.2 |
| 3 | 35442122 | rs1455326 | -1.082 | -2.593 | -1.569 | -0.692 | 0.061 | 0.036 | 0.036 | 0 | 0.53 | 0.25 | 0.54 | 0.42 | 0.39 | 0.61 |
| 3 | 35443653 | rs17033117 | 0.334 | -2.276 | -1.492 | -0.619 | 0.005 | -0.014 | -0.014 | 2.042 | 0.2 | 0.16 | 0.33 | 0.29 | 0.18 | 0.35 |
| 3 | 35447320 | rs9858821 | -0.33 | -1.918 | -1.512 | -0.852 | 0.021 | 0.068 | 0.068 | 0 | 0.18 | 0.07 | 0.17 | 0 | 0.15 | 0.19 |
| 3 | 35457227 | rs6784540 | -1.401 | -1.695 | -1.74 | -0.472 | -0.007 | -0.058 | -0.058 | 0 | 0.61 | 0.87 | 0.6 | 0.58 | 0.41 | 0.65 |
| 3 | 35457937 | rs4678773 | 1.375 | -1.594 | -1.921 | -0.532 | -0.038 | 0.011 | 0.011 | 0.018 | 0.11 | 0.01 | 0.13 | 0.07 | 0.14 | 0.16 |
| 3 | 35459355 | rs17033173 | 1.007 | -2.485 | -2.039 | -0.748 | 0.051 | 0.038 | 0.038 | 0.054 | 0.18 | 0.03 | 0.15 | 0.11 | 0.08 | 0.14 |
| 3 | 35459699 | rs12631043 | 0.936 | -2.609 | -1.587 | -0.706 | -0.01 | 0.049 | 0.049 | 0 | 0.12 | 0.16 | 0.26 | 0.38 | 0.09 | 0.22 |
| 3 | 35484172 | rs9869249 | 0.016 | -2.547 | -0.731 | -0.907 | -0.01 | -0.08 | -0.08 | 0.073 | 0.13 | 0.34 | 0.06 | 0 | 0.11 | 0.1 |
| 3 | 35486613 | rs13322664 | -1.771 | -2.59 | -0.85 | -0.868 | 0.001 | -0.039 | -0.039 | 0 | 0.61 | 0.83 | 0.55 | 0.58 | 0.42 | 0.66 |
| 3 | 35491431 | rs6794661 | 0.896 | -2.373 | -0.799 | -0.857 | -0.008 | 0.014 | 0.014 | 0.002 | 0.08 | 0.05 | 0.03 | 0 | 0.11 | 0.08 |
| 3 | 35494997 | rs17033255 | 0.379 | -2.484 | -0.631 | -0.663 | 0.005 | -0.006 | -0.006 | 0 | 0.2 | 0.19 | 0.2 | 0.23 | 0.17 | 0.18 |
| 3 | 35502060 | rs9870508 | -0.459 | -2.492 | -0.231 | -0.619 | -0.033 | -0.067 | -0.067 | 0.008 | 0.32 | 0.4 | 0.18 | 0.1 | 0.21 | 0.23 |
| 3 | 35512661 | rs6794749 | -0.633 | -2.37 | -0.174 | -1.635 | -0.006 | -0.027 | -0.027 | 0 | 0.34 | 0.37 | 0.2 | 0.1 | 0.23 | 0.27 |
| 3 | 35517986 | rs814494 | 0.574 | -1.275 | -0.344 | -1.712 | -0.001 | -0.001 | -0.001 | 0.526 | 0.58 | 0.61 | 0.55 | 0.65 | 0.57 | 0.58 |
| 3 | 35527409 | rs1523048 | 0.212 | -1.176 | -0.467 | -1.735 | -0.012 | 0.009 | 0.009 | 0 | 0.65 | 0.74 | 0.63 | 0.75 | 0.61 | 0.7 |
| 3 | 35528166 | rs13062354 | 1.004 | -0.745 | -0.712 | -1.55 | -0.028 | -0.005 | -0.005 | 0 | 0.39 | 0.26 | 0.5 | 0.63 | 0.5 | 0.52 |
| 3 | 35534318 | rs872133 | 1.339 | -0.63 | -0.666 | -1.486 | 0.002 | -0.003 | -0.003 | 0 | 0.08 | 0.09 | 0.14 | 0.07 | 0.13 | 0.15 |
| 3 | 35536237 | rs7650296 | 0.656 | -0.754 | -0.569 | -1.535 | 0.001 | 0.024 | 0.024 | 0 | 0.46 | 0.32 | 0.55 | 0.69 | 0.53 | 0.63 |
| 3 | 35537731 | rs13069451 | 0.654 | -0.751 | -0.562 | -1.534 | 0 | 0.005 | 0.005 | 0 | 0.46 | 0.38 | 0.56 | 0.69 | 0.53 | 0.63 |
| 3 | 35539447 | rs9866255 | 0.665 | -0.801 | -0.648 | -1.599 | -0.006 | 0.034 | 0.034 | 1.239 | 0.11 | 0.05 | 0.11 | 0.01 | 0.13 | 0.13 |
| 3 | 35542257 | rs4380362 | -0.167 | -0.814 | -0.655 | -1.599 | -0.014 | -0.014 | -0.014 | 0.015 | 0.58 | 0.47 | 0.67 | 0.69 | 0.66 | 0.76 |
| 3 | 35543442 | rs12629480 | 0.423 | -0.429 | -0.882 | -1.354 | -0.002 | 0.03 | 0.03 | 0 | 0.46 | 0.27 | 0.54 | 0.69 | 0.53 | 0.63 |
| 3 | 35552301 | rs2359669 | 1.779 | -0.363 | -1.217 | -1.344 | -0.064 | -0.088 | -0.088 | 0 | 0.28 | 0.11 | 0.35 | 0.58 | 0.37 | 0.38 |
| 3 | 35554832 | rs17033327 | -0.916 | -0.513 | -0.816 | -1.298 | 0.007 | 0.021 | 0.021 | 0 | 0.2 | 0.09 | 0.18 | 0.12 | 0.18 | 0.27 |
| 3 | 35559240 | rs10510663 | -0.022 | -0.524 | -0.812 | -1.294 | -0.013 | -0.006 | -0.006 | 0 | 0.08 | 0.05 | 0.05 | 0.06 | 0.04 | 0.1 |
| 3 | 35560971 | rs17033337 | -0.022 | -0.524 | -0.812 | -1.294 | -0.013 | -0.006 | -0.006 | 0.093 | 0.08 | 0.05 | 0.05 | 0.06 | 0.04 | 0.1 |
| 3 | 35565846 | rs17033355 | -0.737 | -0.069 | -0.51 | -1.109 | 0 | 0 | 0 | 0 | 0.16 | 0.12 | 0.17 | 0.12 | 0.16 | 0.26 |
| 3 | 35572379 | rs11917489 | -0.685 | -0.359 | -0.135 | -1.447 | -0.02 | -0.11 | -0.11 | 0.45 | 0.37 | 0.45 | 0.3 | 0.06 | 0.32 | 0.35 |
| 3 | 35572591 | rs12715275 | 0.156 | -0.193 | -0.154 | -1.434 | -0.001 | -0.013 | -0.013 | 0.152 | 0.68 | 0.71 | 0.68 | 0.6 | 0.69 | 0.72 |
| 3 | 35576342 | rs13077402 | 0.162 | -0.193 | 0.068 | -1.437 | 0 | -0.003 | -0.003 | 0 | 0.68 | 0.67 | 0.67 | 0.6 | 0.69 | 0.72 |
| 3 | 35577646 | rs6763494 | 2.111 | -0.22 | 0.194 | -1.462 | -0.035 | -0.064 | -0.064 | 0.329 | 0.32 | 0.15 | 0.37 | 0.54 | 0.37 | 0.38 |
| 3 | 35577952 | rs6550350 | -0.088 | -0.175 | 0.197 | -1.371 | -0.001 | -0.001 | -0.001 | 0.079 | 0.72 | 0.71 | 0.68 | 0.65 | 0.7 | 0.82 |
| 3 | 35580968 | rs6787053 | -0.122 | -0.149 | 0.142 | -1.375 | -0.011 | -0.018 | -0.018 | 0 | 0.72 | 0.77 | 0.69 | 0.65 | 0.7 | 0.82 |
| 3 | 35597986 | rs6796482 | 0.039 | -0.36 | 0.349 | -1.542 | -0.003 | -0.053 | -0.053 | 0 | 0.17 | 0.29 | 0.16 | 0.05 | 0.16 | 0.17 |
| 3 | 35600582 | rs6805985 | 0.344 | 0.099 | 0.584 | -1.28 | 0 | -0.004 | -0.004 | 0.096 | 0.67 | 0.66 | 0.67 | 0.6 | 0.69 | 0.72 |
| 3 | 35619559 | rs9870198 | 0.276 | 0.129 | 0.604 | -1.261 | -0.001 | -0.005 | -0.005 | 0 | 0.69 | 0.69 | 0.67 | 0.58 | 0.69 | 0.71 |
| 3 | 35630423 | rs9824956 | -0.613 | 0.164 | 0.645 | -1.348 | -0.01 | -0.092 | -0.092 | 0 | 0.31 | 0.39 | 0.29 | 0.07 | 0.3 | 0.33 |
| 3 | 35636709 | rs11129657 | 0.474 | 1.04 | 0.697 | -1.273 | 0.058 | -0.026 | -0.026 | 0.113 | 0.12 | 0.25 | 0.17 | 0.07 | 0.2 | 0.17 |

| | | | | | | | | | | | | | | | | |
|---|---|---|---|---|---|---|---|---|---|---|---|---|---|---|---|---|
| 3 | 35637909 | rs988174 | 0.285 | 0.954 | 0.603 | -1.278 | -0.001 | 0.005 | 0.005 | 0.105 | 0.69 | 0.64 | 0.68 | 0.57 | 0.69 | 0.7 |
| 3 | 35638443 | rs6779429 | 2.104 | 1.001 | 0.416 | -1.281 | 0.008 | -0.063 | -0.063 | 0 | 0.36 | 0.18 | 0.36 | 0.49 | 0.32 | 0.38 |
| 3 | 35649101 | rs9810198 | -1.315 | 1.613 | 0.261 | -1.044 | -0.02 | -0.037 | -0.037 | 0 | 0.9 | 0.98 | 0.91 | 0.88 | 0.86 | 0.93 |
| 3 | 35651949 | rs9881157 | 0.185 | 1.522 | 0.406 | -1.06 | -0.027 | -0.043 | -0.043 | 0 | 0.72 | 0.85 | 0.71 | 0.65 | 0.7 | 0.8 |
| 3 | 35661187 | rs13078110 | 0.161 | 1.655 | 0.819 | -0.907 | 0.066 | -0.011 | -0.011 | 0 | 0.12 | 0.29 | 0.17 | 0.11 | 0.21 | 0.17 |
| 3 | 35662850 | rs9856017 | 0.191 | 1.5 | 0.736 | -1.046 | 0.005 | -0.001 | -0.001 | 0 | 0.19 | 0.1 | 0.13 | 0 | 0.16 | 0.15 |
| 3 | 35667761 | rs10490871 | -1.018 | 1.484 | 0.882 | -1.032 | -0.004 | -0.068 | -0.068 | 0.341 | 0.37 | 0.49 | 0.33 | 0.16 | 0.36 | 0.41 |
| 3 | 35678494 | rs1542211 | 1.983 | 1.442 | 0.79 | -1.066 | 0.01 | -0.027 | -0.027 | 0.002 | 0.39 | 0.24 | 0.36 | 0.51 | 0.34 | 0.37 |
| 3 | 35683935 | rs1542212 | 1.749 | 1.454 | 0.832 | -1.094 | 0.01 | -0.005 | -0.005 | 0 | 0.43 | 0.32 | 0.45 | 0.5 | 0.38 | 0.43 |
| 3 | 35692163 | rs9311103 | -0.007 | 1.519 | 0.917 | -1.111 | 0.003 | 0.005 | 0.005 | 0 | 0.76 | 0.7 | 0.69 | 0.66 | 0.69 | 0.78 |
| 3 | 35695010 | rs17201627 | 0.842 | 1.616 | 0.69 | -1.013 | 0.036 | 0.048 | 0.048 | 0.351 | 0.15 | 0.01 | 0.04 | 0.09 | 0.08 | 0.15 |
| 3 | 35695512 | rs4678789 | 0.324 | 1.723 | 0.858 | -0.932 | 0.056 | -0.017 | -0.017 | 0 | 0.12 | 0.27 | 0.18 | 0.08 | 0.19 | 0.17 |
| 3 | 35701621 | rs6784162 | 0.853 | 1.771 | 0.881 | -0.942 | -0.023 | -0.009 | -0.009 | 0 | 0.6 | 0.76 | 0.65 | 0.57 | 0.57 | 0.6 |
| 3 | 35705729 | rs9859804 | 0.987 | 1.234 | 0.988 | -0.963 | 0 | 0 | 0 | 0 | 0.62 | 0.41 | 0.38 | 0.57 | 0.48 | 0.56 |
| 3 | 35711293 | rs17033657 | 0.341 | 1.371 | 1.159 | -0.837 | -0.003 | -0.013 | -0.013 | 0 | 0.05 | 0.15 | 0.06 | 0 | 0.03 | 0.07 |
| 3 | 35712071 | rs2280096 | 1.416 | 1.634 | 1.198 | -0.894 | 0.003 | 0.007 | 0.007 | 0 | 0.52 | 0.41 | 0.42 | 0.56 | 0.45 | 0.52 |
| 3 | 35713180 | rs2197732 | 1.296 | 1.698 | 1.187 | -0.899 | -0.004 | 0.001 | 0.001 | 0 | 0.56 | 0.41 | 0.43 | 0.56 | 0.5 | 0.56 |
| 3 | 35713837 | rs4678791 | 0.868 | 1.497 | 1.222 | -0.893 | 0 | -0.001 | -0.001 | 1.059 | 0.6 | 0.57 | 0.5 | 0.56 | 0.52 | 0.62 |
| 3 | 35717389 | rs17280201 | 1.901 | 1.5 | 0.976 | -0.896 | 0.017 | -0.03 | -0.03 | 0.571 | 0.45 | 0.22 | 0.32 | 0.48 | 0.4 | 0.43 |
| 3 | 35720567 | rs1513474 | -0.193 | 1.48 | 1.01 | -0.832 | -0.056 | -0.045 | -0.045 | 0.753 | 0.78 | 0.87 | 0.56 | 0.65 | 0.62 | 0.8 |
| 3 | 35724596 | rs9811585 | 1.521 | 1.481 | 1.035 | -0.83 | -0.001 | -0.001 | -0.001 | 0 | 0.48 | 0.48 | 0.39 | 0.48 | 0.37 | 0.47 |
| 3 | 35726748 | rs3821447 | 2.113 | 1.428 | 0.215 | -0.819 | 0.027 | -0.077 | -0.077 | 0 | 0.41 | 0.11 | 0.29 | 0.48 | 0.34 | 0.4 |
| 3 | 35729000 | rs4678796 | 0.59 | 0.88 | 0.375 | -1.581 | -0.001 | 0 | 0 | 0 | 0.09 | 0.08 | 0.1 | 0.08 | 0.1 | 0.12 |
| 3 | 35731725 | rs1513476 | -0.381 | 1.159 | 0.285 | -1.557 | -0.036 | 0.027 | 0.027 | 0 | 0.81 | 0.9 | 0.83 | 0.86 | 0.69 | 0.88 |
| 3 | 35731993 | rs13082065 | 1.693 | 1.23 | -0.042 | -1.455 | -0.006 | 0.105 | 0.105 | 0 | 0.44 | 0.16 | 0.57 | 0.69 | 0.41 | 0.48 |
| 3 | 35734231 | rs13094598 | 1.703 | 1.292 | -0.039 | -1.443 | 0.007 | 0.09 | 0.09 | 0 | 0.44 | 0.18 | 0.57 | 0.69 | 0.39 | 0.46 |
| 3 | 35743058 | rs7349512 | -0.735 | 1.297 | 0.072 | -1.445 | 0.003 | 0.019 | 0.019 | 0 | 0.3 | 0.24 | 0.17 | 0.18 | 0.26 | 0.34 |
| 3 | 35744679 | rs17033700 | 0.177 | 1.319 | -0.004 | -1.485 | 0.015 | 0.051 | 0.051 | 0 | 0.21 | 0.11 | 0.08 | 0.09 | 0.17 | 0.22 |
| 3 | 35751601 | rs7636980 | 1.231 | 1.34 | 0.03 | -1.469 | -0.002 | 0.078 | 0.078 | 0 | 0.52 | 0.31 | 0.61 | 0.77 | 0.49 | 0.58 |
| 3 | 35755869 | rs9872664 | 0.171 | 1.308 | -0.051 | -1.456 | 0.017 | 0.051 | 0.051 | 0 | 0.21 | 0.11 | 0.08 | 0.09 | 0.17 | 0.25 |
| 3 | 35762320 | rs1533273 | 0.46 | 1.444 | -0.08 | -1.209 | 0.012 | 0.004 | 0.004 | 0 | 0.08 | 0.12 | 0.1 | 0.08 | 0.1 | 0.13 |
| 3 | 35775249 | rs4678802 | 0.911 | 1.379 | -0.097 | -1.27 | -0.003 | 0.076 | 0.076 | 0 | 0.52 | 0.32 | 0.61 | 0.78 | 0.49 | 0.58 |
| 3 | 35778908 | rs11927763 | -0.609 | 1.888 | 0.195 | -1.054 | 0.049 | -0.051 | -0.051 | 0 | 0.1 | 0.19 | 0.11 | 0 | 0.18 | 0.05 |
| 3 | 35779750 | rs2012153 | -1.398 | 1.809 | 0.424 | -1.359 | 0.014 | -0.054 | -0.054 | 0.929 | 0.4 | 0.47 | 0.3 | 0.15 | 0.45 | 0.37 |
| 3 | 35784581 | rs13094224 | -1.009 | 1.504 | 0.689 | -1.366 | 0.001 | 0 | 0 | 0 | 0.33 | 0.25 | 0.28 | 0.15 | 0.3 | 0.35 |
| 3 | 35785608 | rs2305234 | 0.615 | 1.699 | 0.69 | -1.101 | 0.015 | 0.006 | 0.006 | 0 | 0.07 | 0.08 | 0.11 | 0.07 | 0.09 | 0.07 |
| 3 | 35799942 | rs6550369 | -1.565 | 1.894 | 0.81 | -1.216 | 0.052 | -0.024 | -0.024 | 2.842 | 0.2 | 0.35 | 0.21 | 0.15 | 0.27 | 0.24 |
| 3 | 35832877 | rs7621646 | -1.665 | 1.53 | 0.822 | -1.493 | -0.041 | 0.108 | 0.108 | 0 | 0.41 | 0.77 | 0.32 | 0.16 | 0.42 | 0.39 |
| 3 | 35840831 | rs4678808 | -1.266 | 1.515 | 0.889 | -1.519 | 0.002 | -0.025 | -0.025 | 1.482 | 0.34 | 0.6 | 0.29 | 0.07 | 0.4 | 0.25 |
| 3 | 35846872 | rs2670348 | 0.634 | 1.622 | 0.418 | -1.549 | -0.028 | -0.008 | -0.008 | 0 | 0.13 | 0.04 | 0.15 | 0.13 | 0.17 | 0.09 |
| 3 | 35859680 | rs9854002 | -2.086 | 1.111 | 0.8 | -1.506 | -0.051 | 0.153 | 0.153 | 0 | 0.43 | 0.82 | 0.39 | 0.16 | 0.43 | 0.41 |
| 3 | 35872376 | rs9857288 | -1.31 | 1.213 | 0.876 | -1.486 | 0.003 | -0.026 | -0.026 | 0.097 | 0.34 | 0.59 | 0.28 | 0.07 | 0.38 | 0.25 |
| 3 | 35877357 | rs9825404 | -1.311 | 1.076 | 0.874 | -1.487 | -0.005 | -0.008 | -0.008 | 0 | 0.34 | 0.62 | 0.28 | 0.07 | 0.38 | 0.25 |
| 3 | 35881597 | rs9682328 | -0.715 | 1.027 | 1.12 | -1.449 | 0.168 | -0.017 | -0.017 | 0 | 0.15 | 0.43 | 0.2 | 0.07 | 0.25 | 0.13 |
| 3 | 35891733 | rs6799609 | -2.109 | 0.884 | 0.8 | -1.516 | -0.081 | 0.188 | 0.188 | 0 | 0.42 | 0.89 | 0.33 | 0.17 | 0.4 | 0.42 |
| 3 | 35898774 | rs9847358 | -0.114 | 1.194 | 0.77 | -1.478 | -0.011 | -0.066 | -0.066 | 0 | 0.2 | 0.24 | 0.08 | 0 | 0.13 | 0.12 |
| 3 | 35902885 | rs1455495 | -0.596 | 1.434 | 0.296 | -1.255 | 0.071 | -0.025 | -0.025 | 0 | 0.14 | 0.2 | 0.18 | 0.06 | 0.24 | 0.13 |
| 3 | 35913342 | rs7644212 | -0.871 | 0.727 | -0.345 | -1.32 | -0.023 | 0.064 | 0.064 | 0 | 0.4 | 0.68 | 0.31 | 0.07 | 0.4 | 0.26 |
| 3 | 35913637 | rs7634955 | -0.805 | 0.42 | -0.629 | -1.183 | -0.011 | 0.087 | 0.087 | 0 | 0.42 | 0.71 | 0.33 | 0.15 | 0.36 | 0.28 |
| 3 | 35925872 | rs9869067 | -1.693 | -0.742 | -0.472 | -0.571 | 0.002 | 0.003 | 0.003 | 2.107 | 0.17 | 0.8 | 0.13 | 0.16 | 0.08 | 0.13 |
| 3 | 35936616 | rs11129672 | 0.889 | -1.467 | -1.484 | -0.339 | 0.109 | 0.026 | 0.026 | 0 | 0.52 | 0.14 | 0.55 | 0.45 | 0.61 | 0.6 |
| 3 | 35938795 | rs10049409 | -0.082 | -1.492 | -0.849 | -0.322 | 0.009 | -0.016 | -0.016 | 0 | 0.66 | 0.65 | 0.65 | 0.61 | 0.69 | 0.75 |
| 3 | 35950046 | rs7615307 | 0.823 | -1.387 | -1.493 | -0.3 | 0.091 | -0.003 | -0.003 | 1.014 | 0.52 | 0.09 | 0.55 | 0.45 | 0.62 | 0.6 |
| 3 | 35950281 | rs17033925 | -0.871 | -1.141 | -0.663 | -0.087 | 0.002 | -0.006 | -0.006 | 0.014 | 0.07 | 0.08 | 0.06 | 0.04 | 0.09 | 0.09 |
| 3 | 35954597 | rs1971222 | -0.546 | -1.425 | -0.242 | -0.008 | -0.038 | 0.004 | 0.004 | 0 | 0.16 | 0.27 | 0.06 | 0.14 | 0.06 | 0.12 |
| 3 | 35955145 | rs12636878 | -0.06 | -1.419 | -0.165 | 0.003 | -0.019 | 0.052 | 0.052 | 0 | 0.06 | 0.13 | 0.02 | 0.14 | 0.02 | 0.09 |
| 3 | 35967682 | rs6550376 | -0.105 | -1.797 | -0.766 | 0.185 | 0.021 | -0.063 | -0.063 | 0.605 | 0.84 | 0.95 | 0.78 | 0.61 | 0.87 | 0.82 |
| 3 | 35969079 | rs1016528 | 2.156 | -2.159 | -1.07 | -0.149 | 0.112 | 0.044 | 0.044 | 0.937 | 0.5 | 0.11 | 0.51 | 0.43 | 0.69 | 0.58 |
| 3 | 35973273 | rs11928185 | 0.559 | -1.952 | -0.463 | 0.181 | -0.053 | 0.049 | 0.049 | 0.595 | 0.88 | 0.35 | 0.92 | 0.86 | 0.98 | 0.91 |
| 3 | 35981900 | rs2123015 | 1.33 | -1.82 | -0.609 | -0.05 | 0.105 | -0.048 | -0.048 | 0 | 0.63 | 0.17 | 0.67 | 0.46 | 0.81 | 0.7 |
| 3 | 35982130 | rs6799705 | 0.309 | -0.898 | -0.098 | -0.088 | -0.013 | -0.049 | -0.049 | 0 | 0.09 | 0.15 | 0.04 | 0 | 0.04 | 0.03 |
| 3 | 35987444 | rs12486302 | -0.751 | -0.913 | -0.122 | 0.23 | 0.018 | -0.058 | -0.058 | 0 | 0.3 | 0.17 | 0.41 | 0.5 | 0.23 | 0.48 |
| 3 | 35988035 | rs10510669 | 0.149 | -0.752 | -0.421 | -0.008 | -0.018 | 0.011 | 0.011 | 0.163 | 0.11 | 0.03 | 0.07 | 0.09 | 0.11 | 0.09 |
| 3 | 35988262 | rs4234249 | -0.617 | -0.886 | -0.042 | -0.191 | 0.011 | -0.066 | -0.066 | 0.129 | 0.89 | 0.62 | 0.92 | 1 | 0.92 | 0.98 |
| 3 | 35992042 | rs1844064 | 0.098 | -0.975 | -0.323 | -0.016 | 0.006 | -0.017 | -0.017 | 0 | 0.22 | 0.15 | 0.42 | 0.37 | 0.18 | 0.37 |
| 3 | 35994851 | rs1488091 | 1.651 | -0.993 | -0.251 | -0.309 | 0.027 | 0.027 | 0.027 | 1.765 | 0.46 | 0.2 | 0.39 | 0.41 | 0.61 | 0.44 |
| 3 | 35996873 | rs1488092 | -0.98 | -0.931 | -0.363 | -0.263 | 0.01 | 0 | 0 | 0 | 0.4 | 0.32 | 0.5 | 0.6 | 0.3 | 0.54 |
| 3 | 36003489 | rs4538325 | 0.175 | -0.44 | -0.126 | -0.997 | 0.024 | -0.127 | -0.127 | 2.12 | 0.84 | 0.52 | 0.87 | 1 | 0.87 | 0.95 |
| 3 | 36003776 | rs2054856 | -0.039 | -0.156 | -0.438 | -0.583 | 0 | -0.032 | -0.032 | 0 | 0.25 | 0.12 | 0.3 | 0.66 | 0.25 | 0.44 |

| | | | | | | | | | | | | | | | | |
|---|---|---|---|---|---|---|---|---|---|---|---|---|---|---|---|---|
| 3 | 36007391 | rs6807845 | -1.977 | -0.722 | -0.394 | -0.864 | 0.007 | -0.017 | -0.017 | 0 | 0.58 | 0.47 | 0.54 | 0.75 | 0.49 | 0.6 |
| 3 | 36008561 | rs17827807 | 0.632 | -0.75 | -1.044 | -0.899 | -0.035 | 0.06 | 0.06 | 0 | 0.13 | 0.02 | 0.07 | 0.03 | 0.1 | 0.07 |
| 3 | 36008682 | rs10461001 | 0.566 | -0.726 | -0.997 | -0.862 | 0.02 | -0.072 | -0.072 | 2.302 | 0.21 | 0.07 | 0.25 | 0.67 | 0.2 | 0.43 |
| 3 | 36009058 | rs17033999 | 0.495 | -0.737 | -1.003 | -0.868 | 0.02 | -0.072 | -0.072 | 0 | 0.22 | 0.07 | 0.26 | 0.67 | 0.21 | 0.45 |
| 3 | 36012538 | rs6798316 | 0.781 | -0.729 | -1.034 | -0.889 | -0.02 | 0.039 | 0.039 | 0.434 | 0.13 | 0.01 | 0.13 | 0.04 | 0.12 | 0.05 |
| 3 | 36018767 | rs13071680 | 2.022 | -0.827 | -0.865 | -0.879 | -0.064 | 0.046 | 0.046 | 0.004 | 0.4 | 0.18 | 0.43 | 0.26 | 0.51 | 0.4 |
| 3 | 36021293 | rs6550381 | -1.471 | -0.406 | -0.728 | -0.252 | 0.006 | 0.009 | 0.009 | 0 | 0.4 | 0.33 | 0.45 | 0.71 | 0.38 | 0.52 |
| 3 | 36025977 | rs11129681 | -0.02 | -0.316 | -1.046 | -0.209 | 0.019 | -0.083 | -0.083 | 0 | 0.22 | 0.05 | 0.25 | 0.67 | 0.2 | 0.43 |
| 3 | 36026192 | rs4114674 | -1.433 | -0.26 | -1.054 | -0.199 | 0.01 | 0.019 | 0.019 | 0 | 0.39 | 0.28 | 0.43 | 0.71 | 0.38 | 0.51 |
| 3 | 36026320 | rs6788290 | -0.563 | 0.184 | -0.953 | -0.495 | -0.007 | 0.03 | 0.03 | 0.016 | 0.17 | 0.08 | 0.16 | 0.04 | 0.17 | 0.07 |
| 3 | 36033247 | rs9850040 | -1.022 | -0.05 | -0.449 | -0.662 | -0.011 | -0.093 | -0.093 | 0 | 0.23 | 0.27 | 0.13 | 0 | 0.17 | 0.08 |
| 3 | 36034590 | rs11714013 | 0.264 | 0.517 | -1.181 | -1.055 | -0.074 | 0.054 | 0.054 | 0 | 0.25 | 0.12 | 0.27 | 0.12 | 0.33 | 0.19 |
| 3 | 36044211 | rs968974 | -0.082 | -0.217 | -1.151 | -0.4 | 0.001 | -0.026 | -0.026 | 0.358 | 0.22 | 0.14 | 0.19 | 0.31 | 0.21 | 0.15 |
| 3 | 36047374 | rs4419340 | 0.134 | -0.38 | -0.406 | -0.475 | -0.008 | 0.013 | 0.013 | 0 | 0.76 | 0.63 | 0.78 | 0.69 | 0.79 | 0.85 |
| 3 | 36052065 | rs11927759 | 0.43 | -0.156 | -0.18 | -0.591 | -0.009 | 0.032 | 0.032 | 0 | 0.57 | 0.33 | 0.57 | 0.85 | 0.54 | 0.74 |
| 3 | 36068269 | rs11129685 | 2.07 | -0.062 | -0.408 | 0.003 | -0.014 | -0.12 | -0.12 | 0 | 0.31 | 0.01 | 0.29 | 0.35 | 0.28 | 0.43 |
| 3 | 36068686 | rs1839860 | 1.718 | -0.126 | -0.231 | -0.465 | -0.04 | -0.121 | -0.121 | 0 | 0.37 | 0.03 | 0.36 | 0.56 | 0.34 | 0.6 |
| 3 | 36069713 | rs9838039 | -0.22 | 0.005 | 0.133 | -0.568 | 0.019 | -0.06 | -0.06 | 0.549 | 0.33 | 0.5 | 0.35 | 0.14 | 0.35 | 0.24 |
| 3 | 36073516 | rs7611559 | 0.349 | 0.12 | 0.188 | -0.625 | 0.039 | -0.031 | -0.031 | 0.399 | 0.92 | 0.74 | 0.91 | 1 | 0.91 | 0.98 |
| 3 | 36079910 | rs12496323 | 0.952 | 0.12 | 0.201 | -0.802 | 0.02 | 0.104 | 0.104 | 0 | 0.5 | 0.29 | 0.53 | 0.88 | 0.47 | 0.7 |
| 3 | 36095941 | rs2359957 | 0.056 | 0.061 | -0.08 | -0.871 | -0.054 | 0.092 | 0.092 | 0 | 0.29 | 0.17 | 0.3 | 0.08 | 0.34 | 0.21 |
| 3 | 36097738 | rs4637238 | -0.626 | 0.264 | 0.064 | -0.626 | -0.001 | -0.032 | -0.032 | 0 | 0.46 | 0.49 | 0.43 | 0.11 | 0.48 | 0.28 |
| 3 | 36108994 | rs6795407 | -0.478 | 0.648 | 0.03 | -0.533 | -0.02 | 0.098 | 0.098 | 0 | 0.38 | 0.2 | 0.33 | 0.11 | 0.39 | 0.26 |
| 3 | 36109184 | rs1451794 | 0.779 | 0.749 | 0.063 | -0.394 | -0.001 | -0.001 | -0.001 | 0 | 0.07 | 0.06 | 0.12 | 0.03 | 0.09 | 0.08 |
| 3 | 36120024 | rs7613472 | 0.311 | 0.859 | -0.176 | -0.565 | -0.04 | 0.041 | 0.041 | 0 | 0.17 | 0.08 | 0.2 | 0.03 | 0.25 | 0.11 |
| 3 | 36123358 | rs4678516 | 1.184 | 0.495 | 0.077 | -0.749 | 0.018 | 0.083 | 0.083 | 0 | 0.45 | 0.28 | 0.52 | 0.89 | 0.44 | 0.7 |
| 3 | 36138227 | rs4678836 | 1.426 | 0.617 | 0.298 | -0.352 | 0.039 | -0.001 | -0.001 | 1.026 | 0.42 | 0.25 | 0.41 | 0.58 | 0.36 | 0.6 |
| 3 | 36138335 | rs4678837 | 0.879 | 0.621 | 0.303 | -0.362 | 0.027 | 0.073 | 0.073 | 0 | 0.49 | 0.39 | 0.54 | 0.89 | 0.45 | 0.71 |
| 3 | 36143350 | rs1375883 | 0.965 | 0.603 | 0.281 | -0.37 | 0.041 | 0.157 | 0.157 | 0 | 0.48 | 0.31 | 0.53 | 0.89 | 0.45 | 0.71 |
| 3 | 36148330 | rs1992991 | 0.448 | 0.602 | 0.269 | -0.353 | 0.003 | 0.108 | 0.108 | 0 | 0.06 | 0.14 | 0.13 | 0.31 | 0.09 | 0.12 |
| 3 | 36148866 | rs10510676 | 1.518 | 0.592 | 0.225 | -0.359 | 0.031 | -0.034 | -0.034 | 1.398 | 0.41 | 0.16 | 0.41 | 0.57 | 0.36 | 0.59 |
| 3 | 36152237 | rs10865872 | 1.463 | 0.616 | 0.215 | -0.348 | 0.041 | -0.012 | -0.012 | 0 | 0.42 | 0.2 | 0.41 | 0.58 | 0.36 | 0.59 |
| 3 | 36154313 | rs1449991 | -0.111 | 0.671 | 0.193 | -0.381 | -0.006 | -0.069 | -0.069 | 0 | 0.34 | 0.53 | 0.26 | 0.08 | 0.3 | 0.18 |
| 3 | 36160204 | rs9848822 | 1.222 | 0.75 | -0.274 | -0.334 | 0.026 | 0.007 | 0.007 | 0 | 0.45 | 0.65 | 0.46 | 0.58 | 0.36 | 0.6 |
| 3 | 36161094 | rs1449994 | 0.879 | 1.244 | 0.196 | 0.393 | -0.019 | -0.013 | -0.013 | 0 | 0.76 | 0.82 | 0.66 | 0.65 | 0.66 | 0.77 |
| 3 | 36170271 | rs7621073 | -0.613 | 0.138 | 0.552 | 0.997 | -0.001 | 0.075 | 0.075 | 0 | 0.06 | 0.1 | 0.13 | 0.31 | 0.09 | 0.12 |
| 3 | 36182324 | rs6773134 | 0.538 | 0.157 | 0.896 | 0.547 | 0.09 | 0.014 | 0.014 | 0.002 | 0.9 | 0.56 | 0.82 | 0.96 | 0.91 | 0.92 |
| 3 | 36189216 | rs4678850 | -0.825 | -0.448 | 0.594 | 0.78 | -0.002 | 0.005 | 0.005 | 0.435 | 0.24 | 0.27 | 0.27 | 0.38 | 0.25 | 0.22 |
| 3 | 36190651 | rs4678851 | -1.378 | -0.812 | 0.947 | 1.163 | 0.025 | -0.017 | -0.017 | 0 | 0.37 | 0.71 | 0.35 | 0.61 | 0.3 | 0.37 |
| 3 | 36213692 | rs1901030 | -1.222 | -0.822 | 0.811 | 1.109 | -0.022 | 0.001 | 0.001 | 0.36 | 0.34 | 0.46 | 0.31 | 0.38 | 0.29 | 0.25 |
| 3 | 36214197 | rs1450010 | -1.387 | -0.812 | 0.69 | 0.965 | -0.008 | 0.002 | 0.002 | 0 | 0.4 | 0.58 | 0.39 | 0.53 | 0.33 | 0.35 |
| 3 | 36216890 | rs4588310 | 1.437 | -0.413 | 0.457 | 0.932 | 0.001 | 0.001 | 0.001 | 0.049 | 0.56 | 0.53 | 0.53 | 0.57 | 0.57 | 0.62 |
| 3 | 36234539 | rs9881848 | -0.754 | -0.552 | 0.76 | 1.008 | 0.047 | -0.002 | -0.002 | 0 | 0.11 | 0.41 | 0.18 | 0.02 | 0.12 | 0.1 |
| 3 | 36235084 | rs6774457 | -1.598 | -0.742 | 0.87 | 0.997 | -0.015 | -0.06 | -0.06 | 0.876 | 0.15 | 0.54 | 0.21 | 0.03 | 0.13 | 0.11 |
| 3 | 36239406 | rs9870718 | -0.435 | -0.724 | 0.85 | 1.007 | 0.043 | 0.028 | 0.028 | 0 | 0.09 | 0.38 | 0.14 | 0.02 | 0.1 | 0.1 |
| 3 | 36240905 | rs10510678 | 0.177 | -0.764 | 0.748 | 1.245 | 0.005 | 0.036 | 0.036 | 0.162 | 0.08 | 0.08 | 0.13 | 0.35 | 0.15 | 0.12 |
| 3 | 36251715 | rs285317 | -1.625 | -0.71 | 0.507 | 1.161 | -0.012 | 0.005 | 0.005 | 0.15 | 0.28 | 0.67 | 0.27 | 0.28 | 0.22 | 0.26 |
| 3 | 36251807 | rs4678856 | 2.614 | -0.884 | 0.023 | 1.125 | -0.101 | 0.048 | 0.048 | 0.055 | 0.5 | 0.03 | 0.46 | 0.27 | 0.51 | 0.45 |
| 3 | 36254428 | rs285314 | -2.16 | -0.403 | 0.401 | 1.398 | 0.005 | -0.031 | -0.031 | 0.114 | 0.37 | 0.83 | 0.41 | 0.64 | 0.37 | 0.41 |
| 3 | 36254709 | rs285313 | -0.419 | -0.317 | 0.499 | 1.27 | -0.032 | -0.027 | -0.027 | 0 | 0.95 | 1 | 0.91 | 0.91 | 0.9 | 0.87 |
| 3 | 36267529 | rs285299 | -1.774 | -0.612 | 0.766 | 1.19 | -0.003 | 0.001 | 0.001 | 0.051 | 0.29 | 0.67 | 0.32 | 0.38 | 0.3 | 0.29 |
| 3 | 36270289 | rs1165955 | -0.433 | -0.573 | 0.655 | 1.027 | -0.011 | -0.054 | -0.054 | 0.944 | 0.15 | 0.4 | 0.15 | 0.02 | 0.14 | 0.15 |
| 3 | 36271512 | rs1165956 | -1.617 | -0.547 | -0.108 | 1.221 | 0.014 | -0.038 | -0.038 | 0.042 | 0.32 | 0.81 | 0.38 | 0.49 | 0.33 | 0.31 |
| 3 | 36285604 | rs17034665 | 0.001 | -0.406 | -0.219 | 1.355 | -0.031 | -0.041 | -0.041 | 0 | 0.08 | 0.02 | 0.13 | 0.35 | 0.16 | 0.15 |
| 3 | 36286389 | rs814010 | -2.028 | -0.599 | -0.159 | 1.468 | 0.017 | -0.036 | -0.036 | 0 | 0.36 | 0.84 | 0.38 | 0.66 | 0.36 | 0.4 |
| 3 | 36286503 | rs974622 | -1.668 | -0.63 | 0.439 | 1.479 | 0.013 | 0.091 | 0.091 | 0.173 | 0.17 | 0.27 | 0.2 | 0.63 | 0.22 | 0.27 |
| 3 | 36297407 | rs772622 | -2.082 | -0.635 | 0.427 | 1.457 | 0.021 | -0.038 | -0.038 | 0 | 0.33 | 0.78 | 0.37 | 0.66 | 0.33 | 0.38 |
| 3 | 36298453 | rs11129702 | -1.532 | -0.697 | 0.456 | 1.516 | -0.005 | 0.106 | 0.106 | 0 | 0.15 | 0.22 | 0.17 | 0.53 | 0.17 | 0.23 |
| 3 | 36301545 | rs772626 | -1.155 | -0.632 | 0.437 | 1.152 | -0.021 | 0 | 0 | 0.198 | 0.19 | 0.58 | 0.21 | 0.13 | 0.16 | 0.15 |
| 3 | 36307903 | rs285332 | -1.735 | -0.627 | 0.433 | 1.156 | -0.011 | 0.033 | 0.033 | 0 | 0.28 | 0.64 | 0.32 | 0.49 | 0.3 | 0.3 |
| 3 | 36311231 | rs6788863 | 1.522 | -0.433 | 0.679 | 1.086 | 0.005 | 0.026 | 0.026 | 0.182 | 0.71 | 0.38 | 0.64 | 0.46 | 0.62 | 0.67 |
| 3 | 36312234 | rs7647352 | 2.135 | -0.533 | 0.657 | 1.05 | -0.013 | -0.066 | -0.066 | 0 | 0.63 | 0.16 | 0.58 | 0.46 | 0.59 | 0.65 |
| 3 | 36314020 | rs285327 | 0.676 | -0.378 | 0.748 | 1.163 | 0.028 | 0.1 | 0.1 | 0.295 | 0.86 | 0.8 | 0.81 | 0.48 | 0.74 | 0.8 |
| 3 | 36317607 | rs285324 | -0.3 | -0.399 | 0.725 | 1.223 | -0.016 | -0.055 | -0.055 | 0 | 0.15 | 0.45 | 0.17 | 0.02 | 0.13 | 0.12 |
| 3 | 36329450 | rs12636822 | 0.061 | -0.16 | 0.174 | 1.52 | -0.033 | -0.035 | -0.035 | 0 | 0.08 | 0.01 | 0.13 | 0.41 | 0.16 | 0.16 |
| 3 | 36339375 | rs6782000 | -0.627 | -0.593 | 0.103 | 0.862 | 0 | -0.01 | -0.01 | 0.027 | 0.16 | 0.17 | 0.11 | 0.05 | 0.1 | 0.15 |
| 3 | 36344479 | rs1479151 | -1.176 | -0.609 | 0.097 | 1.082 | -0.001 | -0.032 | -0.032 | 0 | 0.24 | 0.18 | 0.4 | 0.49 | 0.24 | 0.31 |
| 3 | 36345965 | rs17034848 | -1.086 | -1.047 | 0.812 | 0.663 | 0.002 | -0.009 | -0.009 | 1.093 | 0.05 | 0.19 | 0.03 | 0 | 0.01 | 0.01 |
| 3 | 36348072 | rs9846752 | -0.074 | -1.565 | 0.533 | 0.354 | -0.026 | 0.005 | 0.005 | 0 | 0.23 | 0.42 | 0.21 | 0.16 | 0.16 | 0.19 |

| | | | | | | | | | | | | | | | | |
|---|---|---|---|---|---|---|---|---|---|---|---|---|---|---|---|---|
| 3 | 36352364 | rs2046000 | -0.029 | -2.066 | 0.391 | 0.611 | -0.001 | 0.005 | 0.005 | 0 | 0.84 | 0.84 | 0.74 | 0.77 | 0.85 | 0.79 |
| 3 | 36359873 | rs2045999 | 0.194 | -2.325 | -0.04 | 0.228 | 0.003 | 0.022 | 0.022 | 0 | 0.78 | 0.87 | 0.75 | 0.99 | 0.78 | 0.85 |
| 3 | 36360999 | rs1479146 | 0.995 | -2.392 | -0.078 | 0.02 | 0 | -0.04 | -0.04 | 0 | 0.7 | 0.77 | 0.7 | 0.99 | 0.71 | 0.8 |
| 3 | 36361555 | rs10510683 | 2.19 | -2.779 | -0.719 | -0.033 | -0.13 | -0.127 | -0.127 | 0 | 0.4 | 0.04 | 0.42 | 0.47 | 0.57 | 0.45 |
| 3 | 36378607 | rs12631282 | 0.53 | -2.225 | 0.32 | 0.085 | -0.013 | 0.007 | 0.007 | 0 | 0.16 | 0.19 | 0.13 | 0.12 | 0.08 | 0.18 |
| 3 | 36385822 | rs1351234 | -0.951 | -2.308 | 0.458 | 0.036 | -0.036 | 0.052 | 0.052 | 0 | 0.27 | 0.61 | 0.26 | 0.42 | 0.14 | 0.34 |
| 3 | 36387951 | rs7638034 | -1.093 | -2.292 | 0.458 | -0.064 | -0.028 | 0.036 | 0.036 | 0 | 0.28 | 0.61 | 0.29 | 0.42 | 0.16 | 0.35 |
| 3 | 36388644 | rs6807628 | -0.05 | -2.249 | 0.462 | -0.035 | -0.006 | 0.032 | 0.032 | 0 | 0.1 | 0.14 | 0.14 | 0.25 | 0.07 | 0.17 |
| 3 | 36391710 | rs1842149 | -0.815 | -2.269 | 0.434 | -0.077 | 0.001 | -0.052 | -0.052 | 0 | 0.88 | 0.85 | 0.85 | 0.99 | 0.87 | 0.89 |
| 3 | 36401634 | rs13066826 | -1.061 | -2.238 | 0.185 | -0.041 | -0.051 | 0.079 | 0.079 | 0 | 0.28 | 0.53 | 0.29 | 0.41 | 0.15 | 0.34 |
| 3 | 36403155 | rs12638953 | 0.348 | -1.998 | 0.325 | 0.062 | -0.013 | 0.006 | 0.006 | 0 | 0.15 | 0.19 | 0.12 | 0.13 | 0.08 | 0.16 |
| 3 | 36403951 | rs1993923 | -0.548 | -2.007 | 0.316 | 0.031 | 0.007 | -0.109 | -0.109 | 0 | 0.2 | 0.3 | 0.2 | 0.01 | 0.2 | 0.17 |
| 3 | 36403964 | rs9826783 | -0.026 | -1.845 | -0.232 | 0.165 | -0.035 | 0.046 | 0.046 | 0 | 0.17 | 0.39 | 0.16 | 0.16 | 0.09 | 0.16 |
| 3 | 36405074 | rs2127232 | -0.094 | -1.233 | -0.575 | 0.016 | -0.028 | 0.033 | 0.033 | 0 | 0.06 | 0 | 0.04 | 0 | 0.08 | 0.06 |
| 3 | 36409601 | rs2605393 | -1.004 | -1.195 | -0.458 | 0.096 | 0 | 0 | 0 | 0 | 0.34 | 0.65 | 0.38 | 0.3 | 0.28 | 0.35 |
| 3 | 36420628 | rs6550422 | -0.995 | -0.646 | -0.372 | 0.25 | -0.021 | 0.03 | 0.03 | 0 | 0.2 | 0.45 | 0.14 | 0.24 | 0.16 | 0.24 |
| 3 | 36425536 | rs1531134 | 0.633 | -0.413 | -0.961 | 0.316 | -0.046 | -0.102 | -0.102 | 0 | 0.17 | 0.01 | 0.25 | 0.2 | 0.22 | 0.19 |
| 3 | 36440897 | rs17035012 | 0.302 | -0.443 | -0.805 | 0.305 | 0 | 0.042 | 0.042 | 0 | 0.09 | 0.15 | 0.07 | 0.23 | 0.09 | 0.14 |
| 3 | 36450379 | rs728082 | -1.094 | -0.456 | -0.599 | 0.2 | -0.005 | 0.011 | 0.011 | 1.583 | 0.34 | 0.46 | 0.36 | 0.43 | 0.35 | 0.35 |
| 3 | 36459254 | rs4678875 | 0.128 | -0.387 | -0.625 | 0.187 | -0.008 | -0.031 | -0.031 | 0 | 0.21 | 0.17 | 0.26 | 0.2 | 0.26 | 0.21 |
| 3 | 36464275 | rs10514698 | 0.39 | -0.386 | -0.623 | 0.222 | -0.005 | -0.025 | -0.025 | 0 | 0.19 | 0.15 | 0.26 | 0.17 | 0.23 | 0.2 |
| 3 | 36466050 | rs6775506 | -1.682 | -0.717 | -0.802 | -0.256 | -0.067 | -0.036 | -0.036 | 0 | 0.58 | 0.88 | 0.56 | 0.45 | 0.55 | 0.58 |
| 3 | 36470307 | rs4678878 | -0.983 | -0.702 | -0.731 | -0.231 | -0.006 | 0.003 | 0.003 | 0.24 | 0.4 | 0.54 | 0.41 | 0.44 | 0.41 | 0.43 |
| 3 | 36473840 | rs11129714 | -0.54 | -0.494 | -1.28 | -0.002 | -0.001 | 0 | 0 | 0.354 | 0.31 | 0.38 | 0.34 | 0.21 | 0.32 | 0.29 |
| 3 | 36475542 | rs11921700 | 0.13 | -0.268 | -1.505 | -0.038 | 0 | -0.013 | -0.013 | 0.422 | 0.09 | 0.1 | 0.07 | 0.01 | 0.07 | 0.09 |
| 3 | 36477116 | rs1598529 | -0.365 | -0.069 | -1.291 | 0.304 | -0.001 | 0.009 | 0.009 | 3.058 | 0.47 | 0.66 | 0.49 | 0.45 | 0.48 | 0.5 |
| 3 | 36484375 | rs11717255 | 0.119 | 0.073 | -1.125 | 1.058 | 0.001 | 0 | 0 | 0 | 0.84 | 0.85 | 0.73 | 0.59 | 0.9 | 0.87 |
| 3 | 36484823 | rs3749279 | -0.303 | 0.665 | -1.005 | 0.828 | -0.005 | -0.013 | -0.013 | 0 | 0.16 | 0.1 | 0.26 | 0.18 | 0.2 | 0.17 |
| 3 | 36486859 | rs7610107 | -0.893 | 0.86 | -0.647 | 0.098 | 0.014 | -0.07 | -0.07 | 0 | 0.14 | 0.24 | 0.09 | 0 | 0.14 | 0.06 |
| 3 | 36488350 | rs870446 | -0.19 | 0.902 | -1.375 | -0.497 | -0.001 | 0.061 | 0.061 | 0.049 | 0.28 | 0.14 | 0.32 | 0.19 | 0.27 | 0.23 |
| 3 | 36488495 | rs870445 | -0.523 | 1.057 | -1.367 | -0.073 | 0.016 | -0.005 | -0.005 | 0 | 0.54 | 0.56 | 0.68 | 0.51 | 0.59 | 0.43 |
| 3 | 36490020 | rs4678879 | 0.402 | 0.81 | -1.689 | -0.303 | 0.002 | 0.101 | 0.101 | 0 | 0.34 | 0.14 | 0.35 | 0.19 | 0.3 | 0.24 |
| 3 | 36491641 | rs1456079 | -0.846 | 0.395 | -1.648 | -0.462 | -0.006 | -0.073 | -0.073 | 0 | 0.68 | 0.83 | 0.75 | 0.51 | 0.65 | 0.52 |
| 3 | 36493673 | rs897546 | 0.754 | 0.233 | -1.869 | -0.14 | -0.006 | -0.011 | -0.011 | 0.643 | 0.14 | 0.12 | 0.17 | 0.33 | 0.19 | 0.21 |
| 3 | 36500714 | rs3214011 | 1.331 | 0.981 | -1.429 | -0.624 | 0.012 | -0.022 | -0.022 | 0 | 0.07 | 0.1 | 0.1 | 0.02 | 0.11 | 0.05 |
| 3 | 36503067 | rs715660 | 1.089 | 0.882 | -1.55 | -0.899 | 0.054 | 0.125 | 0.125 | 0 | 0.41 | 0.16 | 0.33 | 0.21 | 0.3 | 0.25 |
| 3 | 36504281 | rs6767011 | -1.743 | 0.819 | -1.61 | -0.808 | -0.016 | -0.044 | -0.044 | 0 | 0.87 | 1 | 0.93 | 0.87 | 0.87 | 0.74 |
| 3 | 36512233 | rs7612223 | 1.899 | 1.208 | -1.083 | -1.009 | 0.084 | -0.05 | -0.05 | 3.75 | 0.09 | 0.2 | 0.11 | 0 | 0.15 | 0.07 |
| 3 | 36515716 | rs1456078 | 0.063 | 1.098 | -0.865 | -0.915 | -0.004 | 0.001 | 0.001 | 0 | 0.66 | 0.55 | 0.74 | 0.55 | 0.72 | 0.49 |
| 3 | 36517759 | rs11709139 | 0.002 | 1.101 | -0.906 | -0.855 | 0.003 | -0.002 | -0.002 | 0.713 | 0.67 | 0.6 | 0.75 | 0.56 | 0.72 | 0.49 |
| 3 | 36521264 | rs11708473 | 1.681 | 1.364 | -1.186 | -1.099 | 0.057 | 0.14 | 0.14 | 0.529 | 0.34 | 0.09 | 0.3 | 0.12 | 0.25 | 0.18 |
| 3 | 36523672 | rs6787386 | 0.111 | 1.379 | -0.964 | -0.873 | 0.03 | 0.014 | 0.014 | 0 | 0.65 | 0.57 | 0.53 | 0.53 | 0.5 | 0.52 |
| 3 | 36524206 | rs1456074 | -0.196 | 1.508 | -1.016 | -0.887 | -0.007 | -0.007 | -0.007 | 0 | 0.69 | 0.75 | 0.63 | 0.53 | 0.61 | 0.52 |
| 3 | 36524730 | rs4678880 | 1.81 | 0.994 | -1.194 | -0.758 | 0.052 | 0.038 | 0.038 | 0 | 0.19 | 0.02 | 0.18 | 0.11 | 0.1 | 0.13 |
| 3 | 36530418 | rs7645818 | -0.513 | 1.929 | -0.303 | -0.114 | 0.02 | 0.008 | 0.008 | 1.155 | 0.31 | 0.59 | 0.47 | 0.63 | 0.42 | 0.39 |
| 3 | 36534540 | rs2290528 | 1.069 | 1.184 | -0.552 | 0.32 | -0.001 | 0.008 | 0.008 | 0.382 | 0.11 | 0.13 | 0.15 | 0.31 | 0.12 | 0.19 |
| 3 | 36556725 | rs6797696 | -0.208 | 1.172 | -0.825 | -0.164 | -0.013 | 0.015 | 0.015 | 0 | 0.7 | 0.77 | 0.79 | 0.78 | 0.67 | 0.59 |
| 3 | 36563089 | rs6787556 | 0.703 | 1.149 | -0.512 | -0.482 | 0.017 | 0.002 | 0.002 | 0 | 0.44 | 0.36 | 0.51 | 0.52 | 0.4 | 0.24 |
| 3 | 36573091 | rs7647494 | 1.254 | 1.13 | -0.597 | -0.606 | -0.005 | -0.018 | -0.018 | 0 | 0.14 | 0.18 | 0.11 | 0.05 | 0.11 | 0.12 |
| 3 | 36588168 | rs17188371 | 2.071 | 1.331 | -0.339 | -0.821 | -0.01 | 0.008 | 0.008 | 0 | 0.12 | 0.08 | 0.09 | 0 | 0.14 | 0.07 |
| 3 | 36589040 | rs3772105 | 2.412 | 1.383 | 0.015 | -0.822 | 0.003 | 0.032 | 0.032 | 0.187 | 0.08 | 0.13 | 0.14 | 0.27 | 0.08 | 0.19 |
| 3 | 36590098 | rs4678888 | -2.575 | 1.653 | -0.194 | -0.873 | -0.033 | -0.051 | -0.051 | 0.047 | 0.88 | 0.99 | 0.92 | 0.84 | 0.87 | 0.74 |
| 3 | 36595444 | rs9817732 | 0.572 | 1.236 | -0.148 | -0.908 | -0.023 | 0.062 | 0.062 | 0 | 0.57 | 0.69 | 0.68 | 0.79 | 0.53 | 0.47 |
| 3 | 36596036 | rs1879545 | 1.16 | 1.24 | -0.327 | -0.896 | 0.019 | 0.017 | 0.017 | 0 | 0.46 | 0.25 | 0.51 | 0.52 | 0.44 | 0.29 |
| 3 | 36602353 | rs2361278 | -1.939 | 1.318 | -0.563 | -0.878 | -0.056 | -0.026 | -0.026 | 0 | 0.84 | 0.96 | 0.89 | 0.83 | 0.78 | 0.69 |
| 3 | 36608642 | rs11716101 | 1.676 | 1.314 | -0.235 | -0.78 | -0.004 | -0.008 | -0.008 | 0 | 0.13 | 0.11 | 0.1 | 0 | 0.14 | 0.07 |
| 3 | 36624641 | rs2176994 | -0.39 | 1.261 | -0.355 | -0.676 | -0.01 | 0.018 | 0.018 | 0 | 0.69 | 0.78 | 0.78 | 0.79 | 0.66 | 0.56 |
| 3 | 36643821 | rs7616895 | 1.277 | 0.741 | -0.896 | -0.593 | -0.017 | 0.143 | 0.143 | 0.085 | 0.32 | 0.04 | 0.17 | 0.1 | 0.29 | 0.19 |
| 3 | 36647490 | rs11928585 | 0.562 | 0.37 | 0.137 | 0.056 | 0 | 0.038 | 0.038 | 0 | 0.13 | 0.13 | 0.34 | 0.42 | 0.15 | 0.09 |
| 3 | 36670483 | rs17035343 | 0.181 | 0.299 | -0.056 | -0.73 | 0.014 | -0.077 | -0.077 | 0.047 | 0.17 | 0.31 | 0.18 | 0.01 | 0.19 | 0.12 |
| 3 | 36672645 | rs4678540 | 1.357 | 0.328 | -0.121 | -0.417 | 0.005 | 0.046 | 0.046 | 0.253 | 0.09 | 0.13 | 0.13 | 0.3 | 0.09 | 0.2 |
| 3 | 36674940 | rs6550429 | 1.359 | -0.058 | -0.638 | -0.781 | -0.012 | -0.032 | -0.032 | 0.183 | 0.34 | 0.2 | 0.46 | 0.49 | 0.34 | 0.26 |
| 3 | 36677431 | rs17035368 | 0.659 | 0.083 | -0.419 | -0.62 | 0.008 | -0.055 | -0.055 | 0.066 | 0.16 | 0.24 | 0.18 | 0.01 | 0.19 | 0.12 |
| 3 | 36680099 | rs950146 | 0.541 | -0.073 | -0.748 | -0.608 | -0.002 | 0.001 | 0.001 | 0 | 0.48 | 0.39 | 0.59 | 0.5 | 0.49 | 0.41 |
| 3 | 36680920 | rs1396028 | -0.45 | -0.509 | -0.56 | -0.721 | -0.019 | -0.015 | -0.015 | 0 | 0.32 | 0.58 | 0.27 | 0.3 | 0.21 | 0.33 |
| 3 | 36689486 | rs7617990 | 0.877 | -0.135 | -0.631 | -0.503 | -0.005 | 0.012 | 0.012 | 0.077 | 0.09 | 0.1 | 0.09 | 0.02 | 0.11 | 0.07 |
| 3 | 36690671 | rs1553657 | 0.406 | -0.8 | -0.146 | -0.631 | -0.033 | 0.049 | 0.049 | 0 | 0.17 | 0.26 | 0.16 | 0.3 | 0.09 | 0.22 |
| 3 | 36691744 | rs9843434 | 0.028 | -0.881 | -0.148 | -0.453 | 0.007 | 0.007 | 0.007 | 0.552 | 0.62 | 0.51 | 0.48 | 0.49 | 0.54 | 0.63 |
| 3 | 36694732 | rs2204039 | 0.002 | -0.834 | -0.453 | -0.64 | 0.009 | -0.01 | -0.01 | 0 | 0.27 | 0.16 | 0.22 | 0.38 | 0.22 | 0.32 |

| | | | | | | | | | | | | | | | | |
|---|---|---|---|---|---|---|---|---|---|---|---|---|---|---|---|---|
| 3 | 36701434 | rs13082991 | 2.171 | -0.869 | -0.812 | -0.736 | -0.079 | 0.087 | 0.087 | 0 | 0.27 | 0.05 | 0.22 | 0.11 | 0.28 | 0.25 |
| 3 | 36701507 | rs1396026 | -0.373 | -0.485 | -0.125 | -0.379 | 0 | 0.003 | 0.003 | 0 | 0.69 | 0.64 | 0.57 | 0.5 | 0.62 | 0.68 |
| 3 | 36703758 | rs1396029 | 1.517 | -0.389 | -0.455 | -0.253 | -0.012 | 0.062 | 0.062 | 0 | 0.11 | 0.16 | 0.13 | 0.3 | 0.07 | 0.2 |
| 3 | 36704321 | rs906475 | -1.72 | -0.375 | -1.047 | -0.197 | -0.064 | -0.036 | -0.036 | 0.019 | 0.81 | 0.99 | 0.89 | 0.8 | 0.75 | 0.78 |
| 3 | 36711297 | rs9869380 | 0.284 | -0.322 | -0.691 | -0.493 | 0 | 0.022 | 0.022 | 0.3 | 0.56 | 0.61 | 0.47 | 0.13 | 0.51 | 0.44 |
| 3 | 36715114 | rs9862308 | 0.616 | -0.248 | -0.599 | -0.076 | 0 | 0.015 | 0.015 | 0.019 | 0.11 | 0.09 | 0.11 | 0.02 | 0.12 | 0.08 |
| 3 | 36721944 | rs2361281 | 0.702 | -0.154 | -0.766 | -0.008 | -0.028 | 0.028 | 0.028 | 0.211 | 0.11 | 0.03 | 0.07 | 0.08 | 0.14 | 0.1 |
| 3 | 36730767 | rs12491128 | 0.863 | -0.277 | -0.246 | 0.433 | 0.002 | 0.001 | 0.001 | 0 | 0.1 | 0.1 | 0.13 | 0.32 | 0.07 | 0.2 |
| 3 | 36735419 | rs1507881 | -1.262 | 0.151 | 0.169 | -0.329 | -0.002 | -0.049 | -0.049 | 0 | 0.23 | 0.35 | 0.22 | 0.02 | 0.21 | 0.14 |
| 3 | 36741335 | rs4678545 | -1.068 | 0.174 | 0.303 | 0.053 | -0.029 | 0.114 | 0.114 | 0.607 | 0.17 | 0.4 | 0.33 | 0.29 | 0.14 | 0.13 |
| 3 | 36744766 | rs17035498 | -0.66 | 0.111 | 0.074 | -0.657 | -0.007 | -0.062 | -0.062 | 0.014 | 0.18 | 0.26 | 0.13 | 0 | 0.13 | 0.11 |
| 3 | 36748860 | rs7625360 | -0.783 | 0.904 | 0.144 | -0.764 | 0.019 | -0.044 | -0.044 | 0.022 | 0.18 | 0.27 | 0.16 | 0.09 | 0.25 | 0.13 |
| 3 | 36749177 | rs954493 | -1.07 | 0.922 | 0.521 | -0.922 | -0.002 | -0.043 | -0.043 | 0.588 | 0.27 | 0.38 | 0.16 | 0.14 | 0.21 | 0.16 |
| 3 | 36750445 | rs4678898 | 1.034 | 1.174 | 0.535 | -0.598 | 0.009 | 0 | 0 | 0 | 0.74 | 0.34 | 0.76 | 0.77 | 0.69 | 0.79 |
| 3 | 36751220 | rs12106858 | -1.322 | 1.204 | 0.764 | -0.625 | 0.107 | -0.012 | -0.012 | 0 | 0.08 | 0.51 | 0.08 | 0 | 0.09 | 0.01 |
| 3 | 36751494 | rs9842768 | -1.322 | 1.204 | 0.764 | -0.625 | 0.107 | -0.012 | -0.012 | 1.144 | 0.08 | 0.52 | 0.08 | 0 | 0.09 | 0.01 |
| 3 | 36752967 | rs12107083 | 0.126 | 0.55 | 0.298 | -0.499 | 0.003 | 0.011 | 0.011 | 0 | 0.82 | 0.77 | 0.83 | 0.77 | 0.78 | 0.8 |
| 3 | 36755703 | rs6803280 | 0.047 | 0.557 | 0.108 | -0.517 | -0.001 | 0 | 0 | 2.218 | 0.82 | 0.83 | 0.83 | 0.77 | 0.79 | 0.8 |
| 3 | 36757317 | rs13074517 | 0.345 | 0.147 | -0.992 | -0.551 | -0.044 | -0.087 | -0.087 | 0.19 | 0.4 | 0.09 | 0.41 | 0.56 | 0.41 | 0.49 |
| 3 | 36770832 | rs11717985 | 1.048 | 0.088 | -1.147 | -0.8 | -0.011 | -0.124 | -0.124 | 0 | 0.25 | 0.03 | 0.29 | 0.6 | 0.23 | 0.36 |
| 3 | 36775011 | rs6808243 | -0.84 | 0.87 | -0.423 | -0.418 | 0.017 | 0.091 | 0.091 | 0 | 0.88 | 0.67 | 0.88 | 0.82 | 0.85 | 0.84 |
| 3 | 36777041 | rs2063159 | -0.84 | 0.87 | -0.418 | -0.418 | 0.017 | 0.085 | 0.085 | 0 | 0.88 | 0.68 | 0.88 | 0.82 | 0.85 | 0.84 |
| 3 | 36779707 | rs3796187 | 1.14 | 0.799 | -0.948 | -0.629 | -0.113 | 0.077 | 0.077 | 0 | 0.31 | 0.07 | 0.38 | 0.17 | 0.41 | 0.3 |
| 3 | 36784164 | rs2135951 | -0.034 | 0.761 | -0.783 | -0.639 | 0.072 | 0.159 | 0.159 | 0 | 0.51 | 0.22 | 0.52 | 0.22 | 0.6 | 0.46 |
| 3 | 36789085 | rs11917941 | 0.748 | 1.175 | -0.801 | -0.307 | -0.061 | 0.067 | 0.067 | 0 | 0.33 | 0.14 | 0.4 | 0.17 | 0.41 | 0.31 |
| 3 | 36792606 | rs868488 | -0.178 | 1.122 | -0.661 | 0.024 | 0.002 | 0.041 | 0.041 | 0 | 0.84 | 0.79 | 0.84 | 0.7 | 0.8 | 0.82 |
| 3 | 36793391 | rs7615553 | 0.716 | 1.054 | -1.129 | -0.251 | -0.06 | 0.066 | 0.066 | 0 | 0.15 | 0.01 | 0.11 | 0.03 | 0.19 | 0.15 |
| 3 | 36794597 | rs7632683 | -0.027 | 1.237 | -0.222 | 0.485 | 0.022 | 0.191 | 0.191 | 0 | 0.86 | 0.66 | 0.83 | 0.62 | 0.83 | 0.83 |
| 3 | 36796647 | rs9820240 | 0.309 | 1.416 | -0.833 | -0.204 | -0.015 | 0.027 | 0.027 | 0 | 0.28 | 0.18 | 0.33 | 0.04 | 0.3 | 0.27 |
| 3 | 36798989 | rs2063162 | 0.441 | 0.815 | -0.182 | 0.521 | 0.007 | 0.148 | 0.148 | 0 | 0.83 | 0.54 | 0.78 | 0.74 | 0.83 | 0.85 |
| 3 | 36800257 | rs7620657 | 0.082 | 1.017 | -1.335 | 0.427 | -0.05 | -0.081 | -0.081 | 0 | 0.69 | 0.85 | 0.66 | 0.47 | 0.67 | 0.55 |
| 3 | 36800911 | rs7635239 | 0.952 | 0.728 | -1.647 | -0.308 | 0.022 | 0.1 | 0.1 | 0.851 | 0.34 | 0.19 | 0.32 | 0.07 | 0.3 | 0.26 |
| 3 | 36802741 | rs4678902 | -0.913 | 0.693 | -1.723 | 0.072 | -0.007 | -0.059 | -0.059 | 0 | 0.83 | 0.95 | 0.88 | 0.64 | 0.85 | 0.79 |
| 3 | 36803994 | rs4678550 | 0.553 | 0.762 | -1.417 | 0.108 | -0.002 | 0 | 0 | 0 | 0.14 | 0.17 | 0.21 | 0.17 | 0.16 | 0.24 |
| 3 | 36809119 | rs9311135 | -0.516 | 0.62 | -1.218 | 0.079 | 0.007 | 0.024 | 0.024 | 0.459 | 0.48 | 0.7 | 0.55 | 0.55 | 0.52 | 0.54 |
| 3 | 36811137 | rs9811881 | 0.87 | 0.768 | -0.492 | 0.364 | -0.002 | 0.136 | 0.136 | 0 | 0.14 | 0.26 | 0.19 | 0.32 | 0.16 | 0.15 |
| 3 | 36812570 | rs9876871 | -0.178 | 0.961 | -0.878 | 0.322 | 0.013 | 0.072 | 0.072 | 0 | 0.3 | 0.49 | 0.44 | 0.5 | 0.35 | 0.41 |
| 3 | 36812936 | rs9877525 | 0.434 | 0.669 | -1.049 | -0.004 | -0.006 | -0.005 | -0.005 | 3.399 | 0.6 | 0.43 | 0.64 | 0.31 | 0.67 | 0.59 |
| 3 | 36813568 | rs9882217 | 0.173 | 1.344 | -0.541 | 0.6 | -0.006 | 0.101 | 0.101 | 0 | 0.22 | 0.43 | 0.26 | 0.38 | 0.22 | 0.2 |
| 3 | 36815704 | rs9865184 | 0.027 | 1.134 | -0.365 | 0.27 | 0.002 | 0.043 | 0.043 | 0.547 | 0.34 | 0.49 | 0.45 | 0.52 | 0.36 | 0.42 |
| 3 | 36817025 | rs17035576 | 0.434 | 1.275 | -0.576 | 0.213 | 0.011 | 0.001 | 0.001 | 0.077 | 0.09 | 0.15 | 0.08 | 0.05 | 0.13 | 0.1 |
| 3 | 36825139 | rs1472862 | 1.165 | 0.868 | -1.023 | -0.085 | 0 | 0.098 | 0.098 | 0.053 | 0.34 | 0.15 | 0.29 | 0.08 | 0.34 | 0.23 |
| 3 | 36829980 | rs987190 | -0.462 | 0.836 | -0.959 | 0.152 | -0.007 | 0.012 | 0.012 | 0.361 | 0.76 | 0.71 | 0.81 | 0.63 | 0.82 | 0.75 |
| 3 | 36830901 | rs1507870 | 0.535 | 0.996 | -1.415 | -0.323 | -0.032 | -0.006 | -0.006 | 0 | 0.08 | 0.02 | 0.06 | 0.04 | 0.12 | 0.1 |
| 3 | 36831830 | rs11129732 | 0.251 | 0.894 | -1.2 | -0.186 | 0.004 | 0.008 | 0.008 | 0 | 0.31 | 0.32 | 0.42 | 0.62 | 0.34 | 0.4 |
| 3 | 36831937 | rs11713606 | 0.331 | 0.671 | -0.857 | 0.041 | -0.007 | -0.01 | -0.01 | 0 | 0.67 | 0.51 | 0.75 | 0.72 | 0.72 | 0.65 |
| 3 | 36842884 | rs9311137 | 0.529 | 0.748 | -1.457 | -0.381 | -0.012 | 0.083 | 0.083 | 0 | 0.74 | 0.88 | 0.64 | 0.11 | 0.75 | 0.61 |
| 3 | 36846786 | rs4678553 | 0.781 | 0.618 | -1.454 | -0.497 | -0.002 | 0.091 | 0.091 | 0 | 0.71 | 0.85 | 0.62 | 0.11 | 0.74 | 0.59 |
| 3 | 36847271 | rs9311138 | 0.561 | 0.421 | -0.672 | 0.4 | 0.015 | 0.153 | 0.153 | 0 | 0.9 | 0.74 | 0.81 | 0.71 | 0.91 | 0.82 |
| 3 | 36848743 | rs7646422 | 1.356 | 0.4 | -0.363 | 0.343 | -0.003 | -0.029 | -0.029 | 0 | 0.7 | 0.64 | 0.6 | 0.11 | 0.74 | 0.59 |
| 3 | 36854253 | rs925135 | 0.998 | 0.554 | -0.178 | -0.688 | 0.01 | 0.006 | 0.006 | 0 | 0.42 | 0.35 | 0.25 | 0.13 | 0.38 | 0.29 |
| 3 | 36855613 | rs4678554 | 0.588 | 0.492 | -0.13 | -0.736 | 0.003 | 0.077 | 0.077 | 0 | 0.71 | 0.88 | 0.62 | 0.18 | 0.74 | 0.59 |
| 3 | 36856030 | rs9834970 | -0.089 | 0.546 | -0.053 | -0.628 | 0.01 | 0.018 | 0.018 | 0 | 0.46 | 0.31 | 0.54 | 0.58 | 0.51 | 0.54 |
| 3 | 36856328 | rs906482 | 0.901 | 0.414 | 0.244 | -0.738 | 0.032 | 0.11 | 0.11 | 0 | 0.42 | 0.25 | 0.25 | 0.11 | 0.39 | 0.26 |
| 3 | 36859083 | rs1027840 | -0.445 | 0.417 | -0.058 | -0.714 | -0.041 | -0.066 | -0.066 | 0 | 0.88 | 0.97 | 0.81 | 0.73 | 0.84 | 0.8 |
| 3 | 36859705 | rs1553656 | 0.06 | 0.117 | 0.066 | -0.955 | 0.019 | -0.004 | -0.004 | 0 | 0.44 | 0.14 | 0.54 | 0.56 | 0.56 | 0.54 |
| 3 | 36860697 | rs1472864 | 0.799 | 0.073 | 0.432 | -0.984 | -0.004 | -0.024 | -0.024 | 0 | 0.4 | 0.41 | 0.22 | 0.15 | 0.27 | 0.26 |
| 3 | 36862288 | rs1507882 | 0.332 | 0.106 | 0.38 | -1.035 | -0.018 | 0.057 | 0.057 | 0 | 0.68 | 0.83 | 0.58 | 0.2 | 0.64 | 0.56 |
| 3 | 36862980 | rs4624519 | 0.893 | -0.012 | 0.596 | -1.093 | 0.062 | -0.132 | -0.132 | 0.012 | 0.3 | 0.5 | 0.36 | 0.06 | 0.37 | 0.3 |
| 3 | 36864160 | rs9917659 | 0.911 | -0.063 | 0.598 | -1.093 | 0.025 | 0.022 | 0.022 | 0 | 0.39 | 0.33 | 0.21 | 0.14 | 0.27 | 0.26 |
| 3 | 36869439 | rs4789 | -0.036 | 0.035 | 0.586 | -1.002 | 0.025 | -0.002 | -0.002 | 0 | 0.45 | 0.14 | 0.54 | 0.57 | 0.56 | 0.54 |
| 3 | 36878970 | rs4234258 | 0.713 | 0.164 | 0.646 | -1.101 | 0.039 | -0.106 | -0.106 | 0 | 0.31 | 0.57 | 0.38 | 0.06 | 0.37 | 0.3 |
| 3 | 36880684 | rs4431050 | 0.819 | 1.547 | 0.779 | -1.142 | 0.142 | 0.011 | 0.011 | 0 | 0.93 | 0.64 | 0.79 | 0.96 | 0.9 | 0.86 |
| 3 | 36883369 | rs9821223 | 0.755 | 1.023 | 0.696 | -1.132 | 0.046 | -0.106 | -0.106 | 0 | 0.29 | 0.44 | 0.36 | 0.06 | 0.37 | 0.29 |
| 3 | 36892717 | rs4441609 | -0.373 | 0.291 | 0.048 | -1.379 | 0.008 | 0 | 0 | 0 | 0.61 | 0.66 | 0.62 | 0.62 | 0.63 | 0.59 |
| 3 | 36893000 | rs4375995 | 1.255 | 0.268 | -0.272 | -1.251 | -0.049 | -0.07 | -0.07 | 0 | 0.11 | 0.01 | 0.2 | 0.54 | 0.16 | 0.22 |
| 3 | 36901580 | rs11129738 | 0.059 | 0.367 | -0.044 | -1.259 | 0.018 | -0.052 | -0.052 | 0 | 0.3 | 0.14 | 0.25 | 0.56 | 0.25 | 0.29 |
| 3 | 36921795 | rs4404400 | -0.335 | 0.353 | 0.03 | -1.243 | -0.022 | 0.022 | 0.022 | 0 | 0.34 | 0.53 | 0.29 | 0.56 | 0.25 | 0.29 |
| 3 | 36932232 | rs4602338 | 0.855 | 0.355 | -0.185 | -1.274 | -0.052 | -0.083 | -0.083 | 0.013 | 0.12 | 0.01 | 0.2 | 0.31 | 0.16 | 0.27 |

| | | | | | | | | | | | | | | | | |
|---|---|---|---|---|---|---|---|---|---|---|---|---|---|---|---|---|
| 3 | 36938180 | rs4328757 | -0.648 | 0.512 | 0.274 | -1.237 | 0.017 | -0.002 | -0.002 | 0 | 0.61 | 0.73 | 0.63 | 0.62 | 0.62 | 0.52 |
| 3 | 36943118 | rs11708164 | -0.462 | 0.112 | 0.456 | -0.926 | -0.008 | 0.007 | 0.007 | 0 | 0.33 | 0.36 | 0.27 | 0.56 | 0.24 | 0.29 |
| 3 | 36946818 | rs7611656 | -0.529 | 0.107 | 0.134 | -0.747 | -0.037 | 0.047 | 0.047 | 0 | 0.33 | 0.49 | 0.28 | 0.53 | 0.23 | 0.29 |
| 3 | 36958995 | rs6769400 | 0.244 | 0.535 | -0.422 | -1.084 | -0.032 | 0.061 | 0.061 | 0 | 0.37 | 0.27 | 0.38 | 0.08 | 0.44 | 0.25 |
| 3 | 36960660 | rs7622114 | -0.635 | 0.229 | -0.118 | -1.024 | -0.01 | -0.004 | -0.004 | 0 | 0.55 | 0.57 | 0.59 | 0.63 | 0.6 | 0.49 |
| 3 | 36962588 | rs11714248 | 0.978 | 0.668 | -0.502 | -1.137 | -0.109 | 0.106 | 0.106 | 0 | 0.27 | 0.09 | 0.35 | 0.08 | 0.41 | 0.24 |
| 3 | 36966067 | rs6806158 | -0.497 | 0.08 | 0.02 | -0.786 | 0.054 | 0.013 | 0.013 | 0 | 0.54 | 0.71 | 0.6 | 0.44 | 0.6 | 0.46 |
| 3 | 36968309 | rs9849555 | 0.341 | 0.001 | -0.201 | -0.953 | -0.006 | 0.055 | 0.055 | 0 | 0.11 | 0.17 | 0.15 | 0.35 | 0.08 | 0.21 |
| 3 | 36968347 | rs11926768 | 0.675 | -0.599 | -0.341 | -1.15 | -0.06 | 0.105 | 0.105 | 0 | 0.34 | 0.18 | 0.36 | 0.08 | 0.43 | 0.22 |
| 3 | 36981363 | rs7619689 | -0.666 | -0.482 | 0.087 | -1.045 | 0.044 | 0.016 | 0.016 | 0 | 0.53 | 0.69 | 0.6 | 0.43 | 0.59 | 0.46 |
| 3 | 36994234 | rs2058477 | 0.096 | -0.356 | 0.508 | -1.563 | -0.029 | -0.064 | -0.064 | 1.292 | 0.11 | 0.53 | 0.08 | 0 | 0.05 | 0.01 |
| 3 | 36998755 | rs6768108 | 1.2 | -0.281 | 0.158 | -1.58 | -0.127 | 0.129 | 0.129 | 0 | 0.28 | 0.06 | 0.35 | 0.08 | 0.43 | 0.21 |
| 3 | 37017024 | rs7611106 | 1.022 | -0.374 | 0.206 | -1.632 | -0.091 | 0.115 | 0.115 | 0 | 0.29 | 0.08 | 0.35 | 0.07 | 0.44 | 0.22 |
| 3 | 37025867 | rs6789043 | 0 | -0.531 | 0.543 | -1.723 | 0.002 | -0.081 | -0.081 | 0 | 0.41 | 0.51 | 0.41 | 0.07 | 0.47 | 0.22 |
| 3 | 37030175 | rs9311149 | -0.417 | -0.648 | 0.568 | -1.706 | 0.01 | 0.061 | 0.061 | 0 | 0.45 | 0.66 | 0.44 | 0.1 | 0.5 | 0.24 |
| 3 | 37032296 | rs1046512 | 1.116 | -0.582 | 0.45 | -1.64 | -0.123 | 0.143 | 0.143 | 0 | 0.29 | 0.06 | 0.35 | 0.07 | 0.44 | 0.21 |
| 3 | 37032370 | rs3172297 | 1.116 | -0.582 | 0.45 | -1.64 | -0.123 | 0.143 | 0.143 | 0 | 0.29 | 0.06 | 0.35 | 0.07 | 0.44 | 0.21 |
| 3 | 37034946 | rs1800734 | 0.703 | -1.56 | 0.164 | -1.828 | 0.069 | -0.041 | -0.041 | 0 | 0.36 | 0.15 | 0.3 | 0.55 | 0.27 | 0.39 |
| 3 | 37053120 | rs13320360 | 0.885 | -1.353 | 0.158 | -1.789 | -0.016 | -0.022 | -0.022 | 0 | 0.05 | 0.2 | 0.02 | 0 | 0 | 0.01 |
| 3 | 37053568 | rs1799977 | 2.231 | -0.93 | -0.052 | -1.903 | -0.098 | 0.133 | 0.133 | 0 | 0.23 | 0.04 | 0.17 | 0.03 | 0.33 | 0.13 |
| 3 | 37060459 | rs9857293 | 1.136 | -1.24 | 0.128 | -1.77 | -0.021 | -0.026 | -0.026 | 0 | 0.05 | 0.17 | 0.02 | 0 | 0 | 0.01 |
| 3 | 37074668 | rs3774332 | 1.482 | -1.398 | -0.064 | -1.859 | 0.006 | 0.02 | 0.02 | 0 | 0.07 | 0.02 | 0.04 | 0.04 | 0.07 | 0.06 |
| 3 | 37093064 | rs1558529 | 1.666 | -1.3 | 0.272 | -1.94 | -0.022 | 0.011 | 0.011 | 0 | 0.29 | 0.2 | 0.36 | 0.08 | 0.44 | 0.21 |
| 3 | 37095070 | rs10849 | -0.104 | -1.414 | 0.397 | -1.992 | 0.008 | 0.049 | 0.049 | 0 | 0.45 | 0.65 | 0.43 | 0.1 | 0.5 | 0.23 |
| 3 | 37107470 | rs2302503 | 0.734 | -1.291 | -0.065 | -1.954 | 0.016 | -0.084 | -0.084 | 0.314 | 0.35 | 0.39 | 0.38 | 0.08 | 0.44 | 0.22 |
| 3 | 37125699 | rs9846039 | 1.597 | -1.289 | -0.081 | -1.954 | -0.008 | -0.014 | -0.014 | 0 | 0.3 | 0.24 | 0.36 | 0.08 | 0.44 | 0.21 |
| 3 | 37136711 | rs9820888 | -1.693 | -1.36 | -0.299 | -1.871 | -0.05 | -0.06 | -0.06 | 2.285 | 0.9 | 1 | 0.8 | 0.71 | 0.84 | 0.69 |
| 3 | 37144535 | rs2302507 | 0.384 | -1.393 | -0.214 | -1.91 | 0.003 | -0.101 | -0.101 | 2.178 | 0.41 | 0.51 | 0.42 | 0.08 | 0.47 | 0.22 |
| 3 | 37147069 | rs17035950 | 2.394 | -1.392 | -0.223 | -1.898 | -0.014 | -0.028 | -0.028 | 0 | 0.06 | 0.12 | 0.04 | 0 | 0.02 | 0 |
| 3 | 37148958 | rs7643836 | 0.445 | -1.334 | -0.305 | -1.866 | 0.004 | -0.099 | -0.099 | 0 | 0.4 | 0.51 | 0.41 | 0.08 | 0.47 | 0.22 |
| 3 | 37153270 | rs3774322 | -1.658 | -0.969 | -0.451 | -1.63 | -0.051 | -0.063 | -0.063 | 0.03 | 0.9 | 1 | 0.8 | 0.71 | 0.84 | 0.69 |
| 3 | 37175960 | rs9832138 | 0.407 | -0.938 | -0.179 | -1.769 | 0.001 | -0.076 | -0.076 | 2.164 | 0.41 | 0.53 | 0.42 | 0.08 | 0.47 | 0.21 |
| 3 | 37181591 | rs6809297 | 1.274 | -0.872 | -0.13 | -1.745 | 0.022 | -0.068 | -0.068 | 2.811 | 0.3 | 0.32 | 0.38 | 0.08 | 0.44 | 0.2 |
| 3 | 37182198 | rs6550457 | 1.172 | -0.889 | -0.04 | -1.752 | 0.039 | -0.101 | -0.101 | 0 | 0.31 | 0.39 | 0.38 | 0.08 | 0.44 | 0.2 |
| 3 | 37184473 | rs336601 | 1.012 | -1.132 | -0.287 | -1.748 | -0.057 | -0.101 | -0.101 | 0.193 | 0.16 | 0.02 | 0.24 | 0.32 | 0.23 | 0.37 |
| 3 | 37185960 | rs10154916 | 1.219 | -0.787 | 0.237 | -1.884 | -0.023 | 0.01 | 0.01 | 0 | 0.3 | 0.2 | 0.36 | 0.07 | 0.44 | 0.2 |
| 3 | 37192199 | rs7630221 | 0.008 | -1.038 | 0.247 | -1.991 | 0.003 | -0.015 | -0.015 | 0 | 0.46 | 0.57 | 0.45 | 0.13 | 0.52 | 0.24 |
| 3 | 37196601 | rs9869432 | 1.086 | -0.677 | 0.451 | -1.86 | 0.02 | -0.073 | -0.073 | 4.139 | 0.31 | 0.32 | 0.38 | 0.07 | 0.44 | 0.21 |
| 3 | 37224099 | rs1053006 | 0.964 | -0.117 | 0.506 | -1.602 | -0.005 | 0.008 | 0.008 | 0 | 0.06 | 0.02 | 0.18 | 0.05 | 0.11 | 0.08 |
| 3 | 37256337 | rs7620618 | 0.064 | -1.013 | 0.394 | -1.534 | -0.043 | -0.083 | -0.083 | 0 | 0.12 | 0.41 | 0.07 | 0 | 0.04 | 0.02 |
| 3 | 37269039 | rs7639607 | 0.298 | -0.956 | 0.221 | -1.675 | 0.004 | -0.098 | -0.098 | 0 | 0.4 | 0.52 | 0.42 | 0.08 | 0.47 | 0.22 |
| 3 | 37276674 | rs2062587 | 0.133 | -0.537 | 0.331 | -1.602 | 0.016 | -0.111 | -0.111 | 1.562 | 0.4 | 0.52 | 0.37 | 0.08 | 0.41 | 0.22 |
| 3 | 37289234 | rs4678953 | 0.408 | -0.548 | 0.43 | -1.598 | 0.016 | -0.088 | -0.088 | 0 | 0.37 | 0.54 | 0.37 | 0.08 | 0.4 | 0.18 |
| 3 | 37289822 | rs336597 | -1.614 | -0.664 | 0.057 | -1.304 | -0.046 | -0.053 | -0.053 | 0 | 0.91 | 1 | 0.81 | 0.66 | 0.84 | 0.66 |
| 3 | 37297443 | rs11714766 | 1.637 | -0.718 | 0.25 | -1.466 | -0.089 | 0.087 | 0.087 | 0 | 0.24 | 0.11 | 0.3 | 0.08 | 0.36 | 0.16 |
| 3 | 37313088 | rs7651477 | -0.215 | -0.919 | 0.733 | -1.539 | -0.032 | 0.139 | 0.139 | 0.022 | 0.44 | 0.79 | 0.4 | 0.11 | 0.41 | 0.26 |
| 3 | 37332662 | rs9879975 | 1.92 | -0.93 | 0.822 | -1.508 | -0.005 | -0.029 | -0.029 | 0 | 0.06 | 0.13 | 0.04 | 0 | 0.04 | 0.01 |
| 3 | 37344180 | rs11129756 | 0.116 | -0.938 | 0.932 | -1.548 | 0.011 | -0.095 | -0.095 | 3.546 | 0.4 | 0.43 | 0.36 | 0.08 | 0.41 | 0.23 |
| 3 | 37346319 | rs4452278 | 1.112 | -0.954 | 0.9 | -1.558 | 0.017 | -0.062 | -0.062 | 0 | 0.31 | 0.32 | 0.34 | 0.08 | 0.39 | 0.16 |
| 3 | 37353242 | rs6550471 | -0.2 | -1.246 | 0.498 | -1.696 | -0.019 | 0.17 | 0.17 | 0.214 | 0.47 | 0.79 | 0.41 | 0.11 | 0.43 | 0.26 |
| 3 | 37366459 | rs11718848 | 1.862 | -1.239 | 0.279 | -1.693 | -0.06 | 0.058 | 0.058 | 0 | 0.25 | 0.14 | 0.3 | 0.08 | 0.36 | 0.16 |
| 3 | 37380334 | rs7612650 | -1.404 | -1.275 | -0.116 | -1.61 | -0.023 | -0.061 | -0.061 | 0 | 0.87 | 0.97 | 0.78 | 0.63 | 0.83 | 0.65 |
| 3 | 37403413 | rs4624520 | 0.488 | -1.16 | -0.095 | -1.586 | 0.014 | -0.114 | -0.114 | 4.931 | 0.41 | 0.5 | 0.37 | 0.08 | 0.41 | 0.23 |
| 3 | 37414439 | rs6773378 | -0.002 | -1.164 | -0.923 | -1.225 | 0.03 | -0.026 | -0.026 | 0 | 0.43 | 0.13 | 0.37 | 0.53 | 0.4 | 0.44 |
| 3 | 37453112 | rs13066950 | 0.448 | -0.786 | -0.69 | -0.97 | 0.035 | -0.045 | -0.045 | 0.54 | 0.39 | 0.12 | 0.35 | 0.52 | 0.36 | 0.37 |
| 3 | 37463903 | rs999516 | -0.006 | -0.357 | -0.158 | -1.141 | -0.002 | -0.029 | -0.029 | 0 | 0.11 | 0.13 | 0.09 | 0 | 0.09 | 0.02 |
| 3 | 37474256 | rs1029236 | -0.418 | -0.121 | 0.625 | -1.87 | 0.004 | -0.013 | -0.013 | 0 | 0.44 | 0.59 | 0.29 | 0.03 | 0.38 | 0.23 |
| 3 | 37474280 | rs17200475 | 1.693 | -0.243 | 0.211 | -1.949 | -0.067 | 0.113 | 0.113 | 0 | 0.21 | 0.04 | 0.15 | 0.03 | 0.25 | 0.15 |
| 3 | 37477958 | rs17200615 | 1.855 | -0.231 | 0.119 | -1.944 | -0.068 | 0.107 | 0.107 | 0 | 0.2 | 0.04 | 0.15 | 0.03 | 0.25 | 0.15 |
| 3 | 37483387 | rs13084370 | 0.563 | -0.228 | 0.259 | -1.955 | -0.03 | 0.074 | 0.074 | 0 | 0.31 | 0.17 | 0.24 | 0.03 | 0.34 | 0.16 |
| 3 | 37485641 | rs17812775 | 1.725 | -0.22 | 0.24 | -1.95 | -0.037 | 0.071 | 0.071 | 0 | 0.21 | 0.1 | 0.16 | 0.03 | 0.24 | 0.15 |
| 3 | 37487442 | rs17812793 | 0.63 | -0.387 | 0.288 | -1.789 | 0.002 | -0.002 | -0.002 | 0.223 | 0.11 | 0.07 | 0.08 | 0 | 0.09 | 0.02 |
| 3 | 37487580 | rs9844749 | 0.839 | -0.35 | 0.641 | -1.785 | 0.003 | -0.022 | -0.022 | 0.943 | 0.07 | 0.27 | 0.04 | 0 | 0.03 | 0.01 |
| 3 | 37489415 | rs7629063 | -0.783 | -0.383 | 0.584 | -1.811 | -0.035 | -0.114 | -0.114 | 0.002 | 0.21 | 0.32 | 0.12 | 0 | 0.13 | 0.07 |
| 3 | 37491401 | rs197723 | -0.247 | 0.295 | 0.291 | -0.652 | -0.001 | 0.021 | 0.021 | 0 | 0.84 | 0.82 | 0.72 | 0.52 | 0.83 | 0.66 |
| 3 | 37498210 | rs879904 | 0.924 | 0.455 | 0.155 | -0.726 | 0.028 | 0.045 | 0.045 | 0.091 | 0.43 | 0.24 | 0.34 | 0.38 | 0.39 | 0.37 |
| 3 | 37500658 | rs199279 | -0.386 | 0.56 | 0.329 | -0.068 | 0.001 | 0.056 | 0.056 | 0.07 | 0.88 | 0.86 | 0.75 | 0.51 | 0.86 | 0.67 |
| 3 | 37503831 | rs17036373 | 0.015 | 0.527 | 0.218 | -0.169 | 0.007 | -0.007 | -0.007 | 0 | 0.13 | 0.1 | 0.05 | 0 | 0.08 | 0.06 |
| 3 | 37508551 | rs149815 | 0.038 | 0.862 | 0.363 | 0.197 | 0.01 | 0.191 | 0.191 | 0.029 | 0.87 | 0.69 | 0.72 | 0.54 | 0.86 | 0.67 |

| | | | | | | | | | | | | | | | | |
|---|---|---|---|---|---|---|---|---|---|---|---|---|---|---|---|---|
| 3 | 37515341 | rs12019224 | 0.852 | 0.701 | -0.005 | -0.115 | 0.003 | -0.002 | -0.002 | 0 | 0.62 | 0.53 | 0.53 | 0.46 | 0.61 | 0.47 |
| 3 | 37515533 | rs169111 | 0.273 | 0.825 | 0.33 | -0.11 | 0.016 | -0.014 | -0.014 | 0 | 0.87 | 0.81 | 0.84 | 0.93 | 0.85 | 0.88 |
| 3 | 37515827 | rs197770 | 0.526 | 1.219 | -0.019 | -0.132 | -0.034 | -0.004 | -0.004 | 0 | 0.09 | 0.03 | 0.15 | 0.06 | 0.16 | 0.1 |
| 3 | 37516083 | rs267527 | -0.239 | 0.697 | -0.267 | -0.224 | -0.022 | 0.014 | 0.014 | 0 | 0.6 | 0.72 | 0.58 | 0.54 | 0.52 | 0.68 |
| 3 | 37516184 | rs2162683 | 0.899 | 1.084 | -0.251 | -0.543 | -0.025 | 0.053 | 0.053 | 0 | 0.26 | 0.16 | 0.23 | 0.09 | 0.31 | 0.16 |
| 3 | 37516206 | rs197766 | -0.933 | 0.415 | -0.112 | -0.436 | 0.032 | -0.017 | -0.017 | 0 | 0.59 | 0.68 | 0.62 | 0.61 | 0.7 | 0.47 |
| 3 | 37517462 | rs2212020 | -0.231 | 0.825 | -0.203 | -0.597 | 0.01 | -0.027 | -0.027 | 0 | 0.32 | 0.34 | 0.32 | 0.15 | 0.36 | 0.22 |
| 3 | 37520793 | rs399703 | -0.715 | 0.459 | -0.462 | -0.375 | 0.001 | -0.003 | -0.003 | 0 | 0.59 | 0.58 | 0.55 | 0.63 | 0.61 | 0.53 |
| 3 | 37521988 | rs17227650 | 1.304 | 0.559 | -0.753 | -0.336 | -0.007 | 0.027 | 0.027 | 0 | 0.12 | 0.05 | 0.1 | 0.09 | 0.13 | 0.08 |
| 3 | 37523079 | rs17227748 | 1.526 | 0.555 | -0.757 | -0.338 | -0.014 | 0.022 | 0.022 | 0 | 0.11 | 0.03 | 0.1 | 0.09 | 0.13 | 0.08 |
| 3 | 37523636 | rs391224 | -0.819 | 0.625 | -0.656 | -0.352 | 0.012 | -0.012 | -0.012 | 0.06 | 0.6 | 0.66 | 0.59 | 0.64 | 0.65 | 0.53 |
| 3 | 37525761 | rs10510691 | 1.338 | 0.744 | -0.613 | -0.304 | -0.004 | 0.026 | 0.026 | 0 | 0.12 | 0.06 | 0.1 | 0.08 | 0.13 | 0.08 |
| 3 | 37526013 | rs6550478 | -0.063 | 0.722 | -0.608 | -0.318 | 0.009 | 0 | 0 | 0 | 0.47 | 0.61 | 0.49 | 0.55 | 0.52 | 0.44 |
| 3 | 37526959 | rs17036423 | 0.117 | 1.368 | -0.254 | -0.272 | 0.001 | -0.014 | -0.014 | 0 | 0.15 | 0.19 | 0.14 | 0.08 | 0.17 | 0.08 |
| 3 | 37529636 | rs11716363 | 0.117 | 1.368 | -0.279 | -0.266 | 0.001 | -0.012 | -0.012 | 0.297 | 0.15 | 0.19 | 0.14 | 0.09 | 0.17 | 0.08 |
| 3 | 37535577 | rs174829 | -1.348 | 0.954 | -0.285 | -0.265 | 0.009 | 0 | 0 | 0 | 0.52 | 0.54 | 0.72 | 0.84 | 0.59 | 0.6 |
| 3 | 37537087 | rs267518 | -1.067 | 0.609 | -0.105 | -0.157 | -0.003 | 0.068 | 0.068 | 0 | 0.42 | 0.37 | 0.68 | 0.79 | 0.54 | 0.38 |
| 3 | 37548036 | rs155528 | 0.631 | 1.165 | -0.263 | 0.208 | -0.05 | -0.056 | -0.056 | 0 | 0.13 | 0.01 | 0.35 | 0.16 | 0.19 | 0.11 |
| 3 | 37550382 | rs928799 | -0.461 | 0.829 | 0.194 | -0.536 | -0.006 | -0.02 | -0.02 | 0 | 0.12 | 0.22 | 0.12 | 0.1 | 0.11 | 0.05 |
| 3 | 37560708 | rs3733138 | 0.635 | 0.85 | 0.127 | -0.517 | 0.01 | 0 | 0 | 0 | 0.08 | 0.03 | 0.07 | 0.09 | 0.07 | 0.05 |
| 3 | 37563713 | rs155525 | -2.059 | 0.868 | 0.275 | -0.61 | 0.009 | 0.035 | 0.035 | 0 | 0.49 | 0.65 | 0.71 | 0.86 | 0.55 | 0.58 |
| 3 | 37566946 | rs11915449 | -0.229 | 0.863 | 0.247 | -0.556 | -0.01 | -0.021 | -0.021 | 0 | 0.09 | 0.14 | 0.04 | 0.05 | 0.04 | 0.21 |
| 3 | 37573230 | rs267569 | -0.478 | 0.947 | 0.192 | -0.563 | -0.032 | 0.011 | 0.011 | 0.043 | 0.27 | 0.28 | 0.57 | 0.71 | 0.43 | 0.32 |
| 3 | 37576139 | rs267556 | -1.686 | 0.935 | 0.317 | -0.556 | -0.001 | -0.022 | -0.022 | 0 | 0.43 | 0.51 | 0.69 | 0.8 | 0.54 | 0.38 |
| 3 | 37576601 | rs267555 | -1.023 | 1.003 | 0.132 | -0.513 | -0.097 | 0.129 | 0.129 | 0.002 | 0.36 | 0.13 | 0.63 | 0.8 | 0.51 | 0.38 |
| 3 | 37582867 | rs2559894 | 1.027 | 1.41 | 0.137 | -0.197 | -0.041 | -0.019 | -0.019 | 0 | 0.08 | 0.02 | 0.08 | 0.08 | 0.18 | 0.14 |
| 3 | 37588447 | rs187892 | -0.297 | 0.406 | 0.347 | -0.536 | 0.012 | -0.008 | -0.008 | 0 | 0.31 | 0.38 | 0.59 | 0.72 | 0.46 | 0.34 |
| 3 | 37590136 | rs6785634 | -0.742 | 0.032 | 0.484 | -0.394 | -0.052 | -0.049 | -0.049 | 0.057 | 0.19 | 0.39 | 0.13 | 0.15 | 0.1 | 0.25 |
| 3 | 37597483 | rs1892803 | 0.445 | 0.104 | -0.244 | -0.34 | -0.06 | -0.053 | -0.053 | 0.066 | 0.8 | 0.44 | 0.85 | 0.85 | 0.9 | 0.74 |
| 3 | 37598382 | rs743395 | 0.823 | 0.552 | -0.528 | -0.509 | 0.026 | 0.091 | 0.091 | 0 | 0.5 | 0.29 | 0.26 | 0.1 | 0.42 | 0.42 |
| 3 | 37600470 | rs919113 | -0.639 | 1.318 | -0.57 | 0.441 | -0.008 | -0.008 | -0.008 | 0 | 0.3 | 0.26 | 0.51 | 0.52 | 0.41 | 0.23 |
| 3 | 37603526 | rs2555240 | -0.753 | 0.83 | -0.582 | -0.412 | 0.015 | 0.032 | 0.032 | 0 | 0.5 | 0.58 | 0.65 | 0.84 | 0.59 | 0.32 |
| 3 | 37604012 | rs9825420 | -0.286 | 1.416 | 0.226 | -0.385 | 0.074 | -0.033 | -0.033 | 2.156 | 0.1 | 0.35 | 0.12 | 0.05 | 0.12 | 0.08 |
| 3 | 37604095 | rs2555239 | -0.169 | 1.907 | -0.175 | 0.149 | -0.008 | -0.018 | -0.018 | 0 | 0.31 | 0.27 | 0.42 | 0.44 | 0.37 | 0.19 |
| 3 | 37606770 | rs17828523 | 0.115 | 1.87 | 0.329 | 0.1 | 0.037 | -0.026 | -0.026 | 0 | 0.09 | 0.21 | 0.09 | 0.05 | 0.12 | 0.08 |
| 3 | 37607725 | rs7642079 | -0.231 | 1.917 | 0.376 | 0.031 | 0.051 | -0.046 | -0.046 | 0 | 0.13 | 0.25 | 0.11 | 0.05 | 0.17 | 0.1 |
| 3 | 37611860 | rs2844363 | -0.919 | 1.337 | 0.22 | 0.124 | -0.001 | 0 | 0 | 0 | 0.46 | 0.54 | 0.55 | 0.49 | 0.56 | 0.31 |
| 3 | 37618011 | rs6419831 | -1.22 | 1.207 | 0.113 | -0.066 | 0.041 | 0.058 | 0.058 | 0 | 0.53 | 0.64 | 0.63 | 0.76 | 0.65 | 0.32 |
| 3 | 37633548 | rs2070481 | 0.307 | 1.395 | 0.051 | 0.406 | -0.02 | 0.03 | 0.03 | 0 | 0.14 | 0.3 | 0.35 | 0.2 | 0.14 | 0.06 |
| 3 | 37639213 | rs9823993 | -0.862 | 1.174 | -0.053 | 0.281 | -0.002 | -0.002 | -0.002 | 0.401 | 0.48 | 0.46 | 0.42 | 0.6 | 0.58 | 0.29 |
| 3 | 37644423 | rs7617277 | -2.051 | 0.977 | -0.599 | 0.115 | -0.016 | 0.101 | 0.101 | 0 | 0.7 | 0.97 | 0.82 | 0.91 | 0.77 | 0.51 |
| 3 | 37649328 | rs1573287 | -1.547 | 1.388 | -0.292 | 0.243 | 0.029 | 0.046 | 0.046 | 0 | 0.61 | 0.69 | 0.74 | 0.8 | 0.72 | 0.34 |
| 3 | 37650359 | rs4678977 | -0.842 | 1.357 | -0.262 | 0.264 | 0.013 | 0.011 | 0.011 | 0 | 0.45 | 0.33 | 0.42 | 0.61 | 0.57 | 0.28 |
| 3 | 37652245 | rs17036721 | 1.065 | 1.611 | -0.543 | 0.446 | -0.056 | -0.082 | -0.082 | 0 | 0.12 | 0 | 0.1 | 0.29 | 0.17 | 0.1 |
| 3 | 37652985 | rs6550491 | -1.732 | 0.623 | -0.443 | 0.02 | 0.032 | 0.044 | 0.044 | 0 | 0.62 | 0.7 | 0.74 | 0.8 | 0.73 | 0.35 |
| 3 | 37662867 | rs2027626 | -0.147 | 1.137 | -0.216 | 0.34 | -0.022 | -0.025 | -0.025 | 0 | 0.21 | 0.14 | 0.19 | 0.46 | 0.33 | 0.17 |
| 3 | 37663435 | rs2154762 | -0.678 | -0.182 | 0.014 | -0.106 | -0.015 | -0.024 | -0.024 | 0.042 | 0.24 | 0.28 | 0.2 | 0.11 | 0.2 | 0.09 |
| 3 | 37665385 | rs4678980 | -0.822 | -0.023 | -0.466 | -0.042 | 0.001 | -0.004 | -0.004 | 0 | 0.29 | 0.29 | 0.23 | 0.11 | 0.24 | 0.1 |
| 3 | 37671379 | rs2000494 | -0.67 | 0.285 | -0.429 | 0.321 | 0.002 | 0.001 | 0.001 | 0 | 0.22 | 0.16 | 0.17 | 0.22 | 0.22 | 0.21 |
| 3 | 37672777 | rs4678981 | -0.473 | 0.272 | -0.418 | 0.32 | 0.003 | 0 | 0 | 0 | 0.2 | 0.13 | 0.16 | 0.22 | 0.22 | 0.17 |
| 3 | 37682799 | rs11708693 | -0.465 | 0.261 | 0.01 | 0.494 | 0.009 | 0.016 | 0.016 | 0 | 0.23 | 0.31 | 0.38 | 0.34 | 0.3 | 0.23 |
| 3 | 37688152 | rs4678985 | -0.529 | 0.402 | 0.64 | 0.359 | -0.004 | -0.014 | -0.014 | 0 | 0.23 | 0.37 | 0.19 | 0.22 | 0.24 | 0.18 |
| 3 | 37692342 | rs6550492 | -1.816 | 0.093 | 0.994 | 0.254 | 0.002 | -0.005 | -0.005 | 0 | 0.49 | 0.69 | 0.59 | 0.56 | 0.54 | 0.41 |
| 3 | 37694036 | rs6798191 | -1.002 | 0.074 | 1.04 | 0.237 | 0.007 | -0.042 | -0.042 | 0 | 0.29 | 0.48 | 0.25 | 0.22 | 0.28 | 0.19 |
| 3 | 37705356 | rs17036765 | 0.639 | 0.071 | 0.887 | 0.271 | -0.02 | -0.025 | -0.025 | 0 | 0.08 | 0.02 | 0.11 | 0.15 | 0.12 | 0.13 |
| 3 | 37706267 | rs6550497 | -1.95 | 0.076 | 0.905 | 0.269 | 0.002 | -0.006 | -0.006 | 0.92 | 0.49 | 0.68 | 0.58 | 0.56 | 0.54 | 0.41 |
| 3 | 37716880 | rs2886141 | -2.028 | 0.065 | 0.803 | 0.275 | 0 | -0.005 | -0.005 | 0 | 0.5 | 0.76 | 0.59 | 0.55 | 0.54 | 0.42 |
| 3 | 37721795 | rs13097652 | -0.51 | 0.102 | 0.784 | 0.374 | -0.007 | 0.014 | 0.014 | 0 | 0.2 | 0.34 | 0.32 | 0.23 | 0.24 | 0.08 |
| 3 | 37726832 | rs6770747 | -0.35 | 0.368 | 0.703 | 0.552 | 0.006 | 0.015 | 0.015 | 0.277 | 0.09 | 0.13 | 0.08 | 0.21 | 0.12 | 0.25 |
| 3 | 37743968 | rs743393 | 0.707 | -0.33 | 0.463 | 0.332 | 0.016 | -0.03 | -0.03 | 0.166 | 0.69 | 0.9 | 0.65 | 0.68 | 0.64 | 0.85 |
| 3 | 37750624 | rs7373528 | 2.022 | -0.279 | 0.486 | 0.246 | 0.016 | -0.001 | -0.001 | 0 | 0.51 | 0.32 | 0.48 | 0.46 | 0.47 | 0.58 |
| 3 | 37757348 | rs6419835 | -0.847 | -0.312 | 0.171 | 0.234 | 0 | -0.007 | -0.007 | 0.053 | 0.86 | 0.99 | 0.95 | 0.91 | 0.86 | 0.94 |
| 3 | 37761233 | rs2844353 | -0.859 | -0.387 | 0.181 | 0.266 | -0.066 | -0.088 | -0.088 | 0 | 0.32 | 0.05 | 0.36 | 0.34 | 0.39 | 0.16 |
| 3 | 37765695 | rs762308 | -0.106 | -0.171 | 0.325 | 0.371 | -0.038 | -0.085 | -0.085 | 0 | 0.82 | 0.98 | 0.69 | 0.67 | 0.73 | 0.82 |
| 3 | 37775974 | rs17036845 | -0.976 | -0.059 | 1.101 | 0.542 | -0.008 | -0.025 | -0.025 | 0 | 0.29 | 0.35 | 0.17 | 0.18 | 0.26 | 0.24 |
| 3 | 37777368 | rs6792467 | -1.545 | 0.622 | 0.917 | 0.589 | -0.045 | -0.037 | -0.037 | 0.032 | 0.39 | 0.88 | 0.33 | 0.57 | 0.44 | 0.33 |
| 3 | 37780809 | rs13321209 | -0.365 | 0.479 | 1.12 | 0.5 | -0.004 | -0.043 | -0.043 | 0 | 0.24 | 0.36 | 0.16 | 0.1 | 0.24 | 0.1 |
| 3 | 37783832 | rs6808722 | -0.632 | 0.51 | 1.335 | 0.684 | -0.011 | 0.011 | 0.011 | 1.128 | 0.07 | 0.23 | 0.04 | 0.09 | 0.02 | 0.14 |
| 3 | 37792980 | rs2844377 | 1.348 | 0.542 | 0.842 | 0.684 | -0.077 | 0.05 | 0.05 | 0 | 0.52 | 0.08 | 0.6 | 0.33 | 0.45 | 0.64 |

| | | | | | | | | | | | | | | | | |
|---|---|---|---|---|---|---|---|---|---|---|---|---|---|---|---|---|
| 3 | 37796134 | rs9854824 | -1.061 | 0.572 | 0.962 | 0.576 | 0.013 | 0.027 | 0.027 | 0 | 0.39 | 0.66 | 0.27 | 0.19 | 0.35 | 0.26 |
| 3 | 37798504 | rs6782755 | 0.681 | 0.647 | 1.2 | 0.878 | 0.062 | 0.16 | 0.16 | 0 | 0.07 | 0.15 | 0.12 | 0.35 | 0.18 | 0.07 |
| 3 | 37804853 | rs9854372 | -1.071 | 0.606 | 0.98 | 0.829 | 0.013 | 0.029 | 0.029 | 0 | 0.39 | 0.67 | 0.27 | 0.19 | 0.35 | 0.26 |
| 3 | 37811992 | rs2685100 | -1.303 | 0.625 | 0.889 | 0.843 | -0.021 | -0.021 | -0.021 | 0 | 0.45 | 0.75 | 0.38 | 0.54 | 0.53 | 0.33 |
| 3 | 37812263 | rs2685099 | -0.564 | 0.618 | 1.157 | 0.869 | -0.002 | -0.037 | -0.037 | 0.362 | 0.32 | 0.37 | 0.22 | 0.1 | 0.34 | 0.12 |
| 3 | 37816249 | rs2364663 | -0.86 | 1.44 | 1.194 | 1.371 | 0.175 | 0.318 | 0.318 | 0.604 | 0.11 | 0.49 | 0.15 | 0.36 | 0.22 | 0.08 |
| 3 | 37825138 | rs2685113 | 0.899 | 1.61 | 1.024 | 1.015 | 0.195 | 0.277 | 0.277 | 0 | 0.87 | 0.46 | 0.8 | 0.58 | 0.73 | 0.9 |
| 3 | 37826174 | rs2685111 | 0.018 | -0.159 | 0.278 | -0.009 | 0.015 | 0.052 | 0.052 | 0 | 0.18 | 0.08 | 0.09 | 0 | 0.13 | 0.05 |
| 3 | 37826722 | rs2844397 | -0.404 | -0.13 | 0.485 | -0.102 | 0.003 | -0.03 | -0.03 | 0 | 0.21 | 0.21 | 0.1 | 0 | 0.15 | 0.18 |
| 3 | 37826845 | rs1984311 | 0.269 | -0.669 | -0.208 | -0.494 | 0.004 | -0.002 | -0.002 | 0.23 | 0.42 | 0.32 | 0.39 | 0.44 | 0.4 | 0.28 |
| 3 | 37827350 | rs2685109 | 1.005 | -0.689 | -0.418 | -0.509 | 0.008 | 0.047 | 0.047 | 0 | 0.16 | 0.07 | 0.07 | 0 | 0.12 | 0.05 |
| 3 | 37827376 | rs3845949 | 0.383 | -0.712 | -0.267 | -0.501 | 0.01 | -0.016 | -0.016 | 0.003 | 0.4 | 0.2 | 0.38 | 0.44 | 0.4 | 0.3 |
| 3 | 37832172 | rs2507948 | -0.955 | -0.635 | -0.106 | -0.397 | 0.005 | 0.003 | 0.003 | 0 | 0.63 | 0.58 | 0.5 | 0.45 | 0.55 | 0.48 |
| 3 | 37833387 | rs11129771 | -0.696 | -1.29 | 0.088 | -0.959 | -0.033 | -0.145 | -0.145 | 0 | 0.3 | 0.43 | 0.17 | 0 | 0.19 | 0.22 |
| 3 | 37837218 | rs7631600 | -0.374 | -1.685 | 0.34 | -0.693 | -0.056 | -0.028 | -0.028 | 0.844 | 0.13 | 0.33 | 0.09 | 0.05 | 0.06 | 0.03 |
| 3 | 37843817 | rs6550509 | -1.093 | -1.821 | -0.03 | -0.849 | -0.055 | -0.072 | -0.072 | 0 | 0.21 | 0.52 | 0.13 | 0.06 | 0.12 | 0.14 |
| 3 | 37845788 | rs17275376 | 0.751 | -1.576 | -0.636 | -0.856 | 0.015 | 0.076 | 0.076 | 0.015 | 0.14 | 0.02 | 0.04 | 0 | 0.09 | 0.04 |
| 3 | 37858771 | rs6782856 | -0.731 | -1.524 | -0.809 | -0.587 | 0.044 | 0.061 | 0.061 | 0.814 | 0.45 | 0.61 | 0.24 | 0.24 | 0.31 | 0.28 |
| 3 | 37862718 | rs6796546 | -1.119 | -1.294 | -1.537 | -0.137 | -0.071 | -0.083 | -0.083 | 0 | 0.62 | 0.88 | 0.49 | 0.52 | 0.42 | 0.49 |
| 3 | 37867504 | rs7617462 | -0.167 | -2.102 | -0.995 | -0.657 | 0.029 | 0.021 | 0.021 | 0.692 | 0.29 | 0.16 | 0.09 | 0.03 | 0.15 | 0.14 |
| 3 | 37870175 | rs13077801 | -0.338 | -1.944 | -0.803 | -0.334 | 0.012 | 0.003 | 0.003 | 1.155 | 0.31 | 0.19 | 0.13 | 0.28 | 0.21 | 0.23 |
| 3 | 37887360 | rs6794868 | -1.145 | -2.103 | -0.074 | -0.766 | -0.048 | -0.07 | -0.07 | 0 | 0.33 | 0.51 | 0.17 | 0.05 | 0.18 | 0.19 |
| 3 | 37891706 | rs6800173 | -1.752 | -1.973 | 0.147 | -0.71 | 0.024 | -0.047 | -0.047 | 0 | 0.1 | 0.51 | 0.09 | 0.01 | 0.05 | 0.09 |
| 3 | 37914922 | rs9877943 | -1.655 | -2.061 | 0.047 | -0.772 | -0.01 | -0.034 | -0.034 | 0 | 0.34 | 0.56 | 0.2 | 0.05 | 0.22 | 0.17 |
| 3 | 37940408 | rs11713329 | -0.342 | -1.576 | 0.439 | -0.556 | 0.015 | 0.017 | 0.017 | 0 | 0.18 | 0.11 | 0.08 | 0.06 | 0.12 | 0.09 |
| 3 | 37945154 | rs2012641 | -1.097 | -1.572 | 0.717 | -0.558 | -0.038 | -0.096 | -0.096 | 1.082 | 0.24 | 0.45 | 0.15 | 0.04 | 0.17 | 0.11 |
| 3 | 37949176 | rs6790217 | -1.861 | -1.592 | 0.933 | -0.635 | -0.041 | -0.099 | -0.099 | 0 | 0.33 | 0.45 | 0.17 | 0.07 | 0.22 | 0.22 |
| 3 | 37952594 | rs6792008 | -1.26 | -1.517 | 0.971 | -0.612 | -0.032 | -0.074 | -0.074 | 0.221 | 0.25 | 0.52 | 0.15 | 0.05 | 0.17 | 0.11 |
| 3 | 37957785 | rs11712794 | -0.847 | -1.517 | 1.141 | -0.611 | -0.015 | -0.038 | -0.038 | 3.302 | 0.22 | 0.23 | 0.13 | 0.05 | 0.16 | 0.11 |
| 3 | 37960591 | rs7633540 | -1.204 | -1.513 | 1.051 | -0.609 | -0.032 | -0.075 | -0.075 | 0 | 0.25 | 0.51 | 0.15 | 0.05 | 0.17 | 0.11 |
| 3 | 37969001 | rs9835886 | -1.985 | -1.523 | 0.898 | -0.621 | 0.001 | -0.017 | -0.017 | 0 | 0.34 | 0.59 | 0.19 | 0.05 | 0.23 | 0.22 |
| 3 | 37973595 | rs6779021 | -1.805 | -1.497 | 0.9 | -0.61 | -0.001 | -0.021 | -0.021 | 0 | 0.33 | 0.59 | 0.19 | 0.05 | 0.23 | 0.22 |
| 3 | 37976700 | rs2668069 | -0.698 | -0.431 | 0.343 | -0.004 | -0.062 | -0.064 | -0.064 | 0 | 0.87 | 0.97 | 0.61 | 0.79 | 0.73 | 0.86 |
| 3 | 37977714 | rs17037196 | -0.956 | -0.949 | 0.712 | -0.192 | 0.01 | -0.017 | -0.017 | 0 | 0.05 | 0.19 | 0.04 | 0 | 0.03 | 0.01 |
| 3 | 37982461 | rs9837352 | -0.919 | -1.061 | 0.343 | -0.184 | 0.002 | -0.027 | -0.027 | 0 | 0.06 | 0.17 | 0.04 | 0 | 0.04 | 0.01 |
| 3 | 37986249 | rs9844203 | -1.568 | -1.069 | -0.041 | -0.186 | -0.024 | -0.069 | -0.069 | 0.12 | 0.12 | 0.48 | 0.07 | 0 | 0.05 | 0.1 |
| 3 | 37996268 | rs11919078 | -1.199 | -1.073 | -0.079 | -0.179 | -0.062 | -0.08 | -0.08 | 5.162 | 0.25 | 0.48 | 0.13 | 0.05 | 0.13 | 0.18 |
| 3 | 38000885 | rs6787874 | -1.413 | -1.017 | -0.651 | 0.398 | 0.113 | -0.028 | -0.028 | 0 | 0.42 | 0.92 | 0.27 | 0.38 | 0.24 | 0.41 |
| 3 | 38004661 | rs6776745 | -1.368 | -0.268 | 0.052 | 0.225 | -0.041 | -0.082 | -0.082 | 0 | 0.25 | 0.37 | 0.12 | 0.05 | 0.14 | 0.24 |
| 3 | 38010708 | rs7372209 | -0.294 | 0.746 | -0.242 | 0.591 | -0.08 | -0.08 | -0.08 | 0.127 | 0.87 | 0.98 | 0.62 | 0.73 | 0.7 | 0.85 |
| 3 | 38013493 | rs6789468 | -1.555 | 0.569 | 0.204 | 0.22 | -0.022 | -0.065 | -0.065 | 0 | 0.26 | 0.55 | 0.16 | 0.05 | 0.17 | 0.25 |
| 3 | 38019428 | rs9826470 | -1.419 | 0.804 | 0.154 | 0.388 | -0.009 | -0.055 | -0.055 | 0 | 0.27 | 0.52 | 0.18 | 0.09 | 0.2 | 0.31 |
| 3 | 38022139 | rs11719578 | -0.674 | 0.591 | 0.131 | 0.413 | 0.003 | 0.015 | 0.015 | 0 | 0.2 | 0.13 | 0.11 | 0.08 | 0.14 | 0.27 |
| 3 | 38023553 | rs10095 | -0.715 | 0.589 | 0.286 | 0.412 | -0.006 | -0.026 | -0.026 | 0 | 0.2 | 0.23 | 0.12 | 0.08 | 0.14 | 0.27 |
| 3 | 38024408 | rs11129775 | -0.715 | 0.589 | 0.294 | 0.412 | -0.005 | -0.023 | -0.023 | 0 | 0.2 | 0.23 | 0.12 | 0.08 | 0.14 | 0.27 |
| 3 | 38026652 | rs11711286 | -0.878 | 0.582 | 0.325 | 0.409 | -0.008 | -0.029 | -0.029 | 0 | 0.21 | 0.26 | 0.12 | 0.09 | 0.14 | 0.3 |
| 3 | 38028675 | rs11129776 | -0.704 | 0.592 | 0.285 | 0.415 | -0.006 | -0.026 | -0.026 | 0.001 | 0.2 | 0.23 | 0.12 | 0.08 | 0.14 | 0.28 |
| 3 | 38034821 | rs6776766 | -0.059 | 0.789 | -0.095 | 0.699 | -0.081 | -0.091 | -0.091 | 0 | 0.85 | 0.98 | 0.63 | 0.72 | 0.7 | 0.83 |
| 3 | 38038982 | rs6809649 | -1.009 | 0.145 | 0.391 | 0.433 | -0.007 | -0.063 | -0.063 | 0.637 | 0.22 | 0.43 | 0.14 | 0.09 | 0.14 | 0.29 |
| 3 | 38043904 | rs9843296 | -0.539 | 0.127 | 0.412 | 0.423 | 0.015 | -0.033 | -0.033 | 0 | 0.18 | 0.35 | 0.13 | 0.09 | 0.13 | 0.21 |
| 3 | 38047680 | rs11707277 | 0.004 | 0.124 | 0.343 | 0.421 | -0.012 | 0.011 | 0.011 | 0 | 0.14 | 0.05 | 0.1 | 0.08 | 0.13 | 0.2 |
| 3 | 38047954 | rs9816693 | -0.996 | 0.157 | 0.247 | 0.437 | -0.008 | -0.063 | -0.063 | 0 | 0.22 | 0.42 | 0.14 | 0.09 | 0.14 | 0.29 |
| 3 | 38051941 | rs9857730 | -1.571 | 0.043 | 0.117 | 0.412 | -0.019 | -0.062 | -0.062 | 0.048 | 0.27 | 0.51 | 0.17 | 0.09 | 0.18 | 0.29 |
| 3 | 38057606 | rs9861030 | -1.898 | 0.247 | -0.104 | 0.608 | -0.013 | -0.037 | -0.037 | 0 | 0.1 | 0.15 | 0.04 | 0 | 0.04 | 0.09 |
| 3 | 38058838 | rs11922130 | -2.279 | 0.194 | -0.11 | 0.582 | -0.012 | -0.036 | -0.036 | 0.239 | 0.13 | 0.16 | 0.03 | 0 | 0.05 | 0.1 |
| 3 | 38062550 | rs3753169 | 1.469 | 0.961 | -0.209 | 0.87 | -0.01 | -0.01 | -0.01 | 0 | 0.75 | 0.78 | 0.59 | 0.66 | 0.65 | 0.78 |
| 3 | 38063372 | rs6599099 | 1.371 | 1.017 | -0.393 | 0.904 | -0.039 | -0.041 | -0.041 | 0 | 0.76 | 0.86 | 0.58 | 0.66 | 0.65 | 0.72 |
| 3 | 38073168 | rs6599013 | 1.048 | 1.308 | -0.207 | 1.028 | 0.045 | 0.04 | 0.04 | 0 | 0.74 | 0.64 | 0.55 | 0.66 | 0.63 | 0.78 |
| 3 | 38080101 | rs4389435 | -1.193 | 1.75 | -0.448 | 1.373 | 0.002 | 0.011 | 0.011 | 0 | 0.14 | 0.12 | 0.28 | 0.34 | 0.28 | 0.2 |
| 3 | 38086463 | rs11713291 | -0.063 | 1.135 | -0.929 | 1.016 | 0.001 | 0.015 | 0.015 | 0 | 0.13 | 0.03 | 0.05 | 0.06 | 0.08 | 0.2 |
| 3 | 38087123 | rs7630340 | -0.553 | 0.199 | -1.187 | 0.685 | 0.038 | 0.041 | 0.041 | 1.325 | 0.14 | 0.07 | 0.05 | 0.05 | 0.05 | 0.04 |
| 3 | 38106932 | rs9840792 | -1.823 | 0.754 | -0.646 | 0.979 | 0 | -0.009 | -0.009 | 0 | 0.39 | 0.52 | 0.44 | 0.62 | 0.36 | 0.28 |
| 3 | 38108513 | rs13083423 | 0.18 | 0.805 | -0.64 | 1.276 | -0.024 | -0.003 | -0.003 | 0 | 0.08 | 0.05 | 0.27 | 0.33 | 0.24 | 0.12 |
| 3 | 38109174 | rs6775473 | -2.023 | 0.803 | -0.579 | 1.251 | -0.001 | -0.01 | -0.01 | 0 | 0.39 | 0.55 | 0.44 | 0.62 | 0.36 | 0.28 |
| 3 | 38111270 | rs11928949 | -0.214 | 0.625 | -0.607 | 1.15 | 0.038 | 0.039 | 0.039 | 1.155 | 0.15 | 0.09 | 0.06 | 0.06 | 0.05 | 0.05 |
| 3 | 38134438 | rs11129778 | -0.477 | 0.589 | -0.788 | 1.125 | -0.03 | -0.013 | -0.013 | 0 | 0.95 | 0.99 | 0.95 | 0.93 | 0.93 | 0.95 |
| 3 | 38137624 | rs9825655 | -0.534 | 0.407 | -0.423 | 1.207 | -0.025 | 0.034 | 0.034 | 0 | 0.15 | 0.24 | 0.1 | 0.23 | 0.06 | 0.1 |
| 3 | 38140721 | rs1123569 | -1.18 | 0.366 | -0.325 | 1.207 | -0.081 | -0.009 | -0.009 | 0 | 0.3 | 0.46 | 0.17 | 0.27 | 0.12 | 0.14 |
| 3 | 38144162 | rs9311180 | -1.248 | 0.35 | -0.263 | 1.226 | -0.067 | -0.005 | -0.005 | 0.119 | 0.31 | 0.42 | 0.17 | 0.29 | 0.12 | 0.14 |

| | | | | | | | | | | | | | | | | |
|---|---|---|---|---|---|---|---|---|---|---|---|---|---|---|---|---|
| 3 | 38144875 | rs6599263 | -1.303 | 0.352 | -0.345 | 1.223 | -0.067 | -0.008 | -0.008 | 0 | 0.31 | 0.51 | 0.18 | 0.28 | 0.12 | 0.14 |
| 3 | 38151731 | rs12630114 | -1.886 | 0.547 | -0.382 | 0.959 | -0.001 | -0.003 | -0.003 | 0 | 0.4 | 0.57 | 0.45 | 0.61 | 0.36 | 0.25 |
| 3 | 38154167 | rs6790738 | -1.622 | 0.539 | -0.185 | 0.993 | 0 | -0.001 | -0.001 | 0 | 0.37 | 0.6 | 0.43 | 0.6 | 0.34 | 0.26 |
| 3 | 38161691 | rs11129780 | -1.044 | 0.162 | -0.101 | 0.522 | 0.045 | 0.051 | 0.051 | 0.001 | 0.26 | 0.14 | 0.08 | 0.02 | 0.13 | 0.06 |
| 3 | 38173733 | rs2239621 | -2.009 | 0.457 | 0.364 | 0.979 | -0.012 | -0.003 | -0.003 | 0.227 | 0.4 | 0.59 | 0.31 | 0.36 | 0.27 | 0.23 |
| 3 | 38178290 | rs7625290 | 0.055 | 0.276 | 0.325 | 0.923 | -0.001 | -0.014 | -0.014 | 0.227 | 0.11 | 0.09 | 0.03 | 0.01 | 0.05 | 0.01 |
| 3 | 38184021 | rs7744 | 0.688 | 0.551 | 0.242 | 1.296 | -0.018 | 0.047 | 0.047 | 0 | 0.06 | 0.03 | 0.15 | 0.33 | 0.14 | 0.1 |
| 3 | 38184370 | rs6853 | -1.214 | 0.385 | 0.279 | 1.18 | -0.031 | -0.067 | -0.067 | 1.506 | 0.27 | 0.29 | 0.1 | 0.02 | 0.13 | 0.06 |
| 3 | 38185174 | rs6767684 | -1.511 | 0.381 | 0.38 | 1.179 | -0.045 | -0.087 | -0.087 | 0.146 | 0.29 | 0.36 | 0.09 | 0.02 | 0.13 | 0.06 |
| 3 | 38206906 | rs9839010 | 1.13 | 0.588 | -0.246 | 0.627 | 0.01 | -0.026 | -0.026 | 0 | 0.64 | 0.56 | 0.44 | 0.24 | 0.47 | 0.52 |
| 3 | 38252406 | rs4955408 | -1.658 | 0.83 | -0.258 | 0.563 | -0.005 | -0.002 | -0.002 | 0 | 0.4 | 0.6 | 0.36 | 0.41 | 0.28 | 0.24 |
| 3 | 38261037 | rs1384006 | 1.137 | 0.766 | -0.179 | -0.311 | 0.017 | -0.038 | -0.038 | 0.064 | 0.64 | 0.51 | 0.47 | 0.29 | 0.48 | 0.55 |
| 3 | 38261807 | rs156260 | -0.479 | 0.806 | -0.083 | -0.318 | 0.042 | -0.04 | -0.04 | 0 | 0.85 | 0.69 | 0.86 | 0.94 | 0.83 | 0.85 |
| 3 | 38271881 | rs6599079 | -1.136 | 0.288 | -0.02 | -0.346 | -0.076 | -0.109 | -0.109 | 0.045 | 0.28 | 0.46 | 0.14 | 0.08 | 0.11 | 0.06 |
| 3 | 38283085 | rs9850171 | -0.674 | 0.263 | -0.025 | -0.346 | -0.039 | -0.052 | -0.052 | 0.048 | 0.24 | 0.29 | 0.12 | 0.08 | 0.1 | 0.05 |
| 3 | 38296817 | rs2011 | 2.689 | 0.332 | 0.01 | -0.364 | -0.051 | 0.089 | 0.089 | 0.393 | 0.35 | 0.01 | 0.32 | 0.21 | 0.35 | 0.47 |
| 3 | 38302137 | rs4955397 | 0.171 | 0.218 | 0.266 | -0.336 | 0 | -0.002 | -0.002 | 0.668 | 0.11 | 0.09 | 0.07 | 0.06 | 0.06 | 0.02 |
| 3 | 38304367 | rs6805200 | -0.953 | 0.188 | 0.281 | -0.352 | -0.054 | -0.07 | -0.07 | 0 | 0.25 | 0.36 | 0.13 | 0.08 | 0.1 | 0.14 |
| 3 | 38307510 | rs9842091 | 0.143 | 0.335 | 0.32 | -0.296 | -0.055 | -0.05 | -0.05 | 1.812 | 0.14 | 0.31 | 0.07 | 0.03 | 0.05 | 0.04 |
| 3 | 38307803 | rs4679027 | 0.398 | 0.321 | 0.263 | -0.156 | 0.008 | 0.012 | 0.012 | 0 | 0.1 | 0.06 | 0.05 | 0.07 | 0.05 | 0.01 |
| 3 | 38310091 | rs9311184 | 0.142 | 0.315 | 0.263 | -0.154 | -0.055 | -0.05 | -0.05 | 0 | 0.14 | 0.3 | 0.07 | 0.03 | 0.05 | 0.04 |
| 3 | 38311953 | rs1979845 | -1.229 | 0.291 | 0.309 | -0.173 | -0.072 | -0.078 | -0.078 | 0.172 | 0.27 | 0.53 | 0.15 | 0.09 | 0.11 | 0.05 |
| 3 | 38312372 | rs3805040 | 0.097 | 0.302 | 0.318 | -0.166 | -0.058 | -0.054 | -0.054 | 0 | 0.15 | 0.33 | 0.07 | 0.03 | 0.05 | 0.04 |
| 3 | 38313688 | rs9874338 | -1.202 | 0.304 | 0.297 | -0.166 | -0.078 | -0.085 | -0.085 | 0 | 0.27 | 0.49 | 0.14 | 0.09 | 0.12 | 0.05 |
| 3 | 38318025 | rs2236631 | 0.522 | 0.371 | 0.041 | -0.127 | -0.011 | -0.002 | -0.002 | 0 | 0.09 | 0 | 0.05 | 0.07 | 0.05 | 0.01 |
| 3 | 38322626 | rs2298419 | 2.369 | 0.341 | -0.13 | -0.203 | -0.014 | 0.117 | 0.117 | 0.229 | 0.37 | 0.05 | 0.31 | 0.22 | 0.37 | 0.46 |
| 3 | 38323289 | rs2298421 | 1.445 | 0.674 | 0.032 | -0.099 | 0.008 | 0.026 | 0.026 | 0.124 | 0.52 | 0.36 | 0.41 | 0.26 | 0.45 | 0.48 |
| 3 | 38327292 | rs4679028 | -1.644 | 0.833 | 0.271 | 0.144 | 0.006 | -0.031 | -0.031 | 0 | 0.28 | 0.56 | 0.23 | 0.25 | 0.24 | 0.17 |
| 3 | 38329091 | rs11708676 | 0.475 | 1.082 | -0.097 | 0.242 | -0.024 | 0.008 | 0.008 | 0 | 0.11 | 0.08 | 0.13 | 0.08 | 0.17 | 0.15 |
| 3 | 38342714 | rs4679029 | -0.066 | 0.832 | -0.034 | 0.316 | 0.006 | -0.003 | -0.003 | 0 | 0.1 | 0.07 | 0.06 | 0.16 | 0.06 | 0.02 |
| 3 | 38342864 | rs4679031 | -0.066 | 0.832 | -0.034 | 0.316 | 0.006 | -0.003 | -0.003 | 0 | 0.1 | 0.07 | 0.06 | 0.16 | 0.06 | 0.02 |
| 3 | 38347851 | rs2073714 | -0.066 | 1.039 | -0.169 | 0.377 | 0.006 | -0.003 | -0.003 | 1.934 | 0.1 | 0.07 | 0.06 | 0.16 | 0.06 | 0.02 |
| 3 | 38350134 | rs2070491 | 0.897 | 0.884 | -0.014 | 0.2 | -0.001 | -0.023 | -0.023 | 0 | 0.53 | 0.53 | 0.48 | 0.2 | 0.54 | 0.64 |
| 3 | 38350226 | rs818821 | -0.151 | 0.865 | 0.29 | 0.202 | -0.004 | 0.009 | 0.009 | 0.43 | 0.32 | 0.4 | 0.33 | 0.57 | 0.34 | 0.21 |
| 3 | 38350519 | rs818818 | 0.699 | 1.003 | 0.623 | 0.565 | -0.001 | 0.002 | 0.002 | 0 | 0.82 | 0.84 | 0.81 | 0.76 | 0.87 | 0.84 |
| 3 | 38350543 | rs818817 | 0.644 | 1.09 | 0.626 | 0.55 | -0.002 | -0.006 | -0.006 | 0.414 | 0.83 | 0.89 | 0.81 | 0.76 | 0.86 | 0.84 |
| 3 | 38350630 | rs704929 | 0.641 | 1.062 | 0.601 | 0.574 | -0.001 | 0.002 | 0.002 | 0 | 0.82 | 0.84 | 0.81 | 0.76 | 0.87 | 0.84 |
| 3 | 38351294 | rs818816 | 0.641 | 1.062 | 0.601 | 0.574 | -0.001 | 0.002 | 0.002 | 0 | 0.82 | 0.84 | 0.81 | 0.76 | 0.87 | 0.84 |
| 3 | 38352450 | rs818811 | 1.067 | 0.945 | 0.436 | 0.546 | 0.023 | -0.017 | -0.017 | 0 | 0.68 | 0.54 | 0.68 | 0.69 | 0.7 | 0.66 |
| 3 | 38355917 | rs194706 | -1.492 | 0.345 | 0.798 | 0.721 | -0.01 | 0.045 | 0.045 | 1.49 | 0.12 | 0.39 | 0.12 | 0.12 | 0.05 | 0.05 |
| 3 | 38357817 | rs2070492 | 0.174 | 0.144 | -0.049 | 0.432 | 0.007 | 0.001 | 0.001 | 0 | 0.12 | 0.07 | 0.15 | 0.16 | 0.11 | 0.06 |
| 3 | 38363393 | rs171248 | 0.759 | 1 | 0.466 | 1.107 | 0.067 | 0.038 | 0.038 | 0 | 0.73 | 0.57 | 0.59 | 0.56 | 0.58 | 0.76 |
| 3 | 38363837 | rs183574 | -0.653 | 0.994 | -0.112 | 1.105 | -0.077 | -0.052 | -0.052 | 0.56 | 0.25 | 0.09 | 0.37 | 0.44 | 0.41 | 0.24 |
| 3 | 38365841 | rs6792261 | 0.041 | -0.041 | -0.035 | 0.933 | -0.014 | 0.024 | 0.024 | 0.301 | 0.09 | 0.13 | 0.08 | 0.23 | 0.02 | 0.05 |
| 3 | 38370556 | rs196364 | -0.8 | 0.297 | -0.246 | 0.95 | -0.035 | 0 | 0 | 0 | 0.28 | 0.12 | 0.38 | 0.3 | 0.35 | 0.25 |
| 3 | 38375557 | rs696256 | 0.462 | -0.061 | 0.337 | 0.842 | -0.05 | 0.027 | 0.027 | 1.807 | 0.8 | 0.38 | 0.84 | 0.76 | 0.91 | 0.93 |
| 3 | 38386516 | rs580454 | 0.429 | 0.405 | 0.374 | 0.61 | -0.042 | -0.019 | -0.019 | 0.043 | 0.12 | 0.04 | 0.21 | 0.14 | 0.23 | 0.13 |
| 3 | 38388014 | rs2234600 | -0.103 | -0.635 | 0.876 | 0.601 | -0.016 | -0.024 | -0.024 | 0.33 | 0.93 | 0.73 | 0.95 | 1 | 0.98 | 0.97 |
| 3 | 38401677 | rs818851 | 0.44 | -0.851 | 0.98 | 0.447 | -0.057 | -0.118 | -0.118 | 0.054 | 0.82 | 0.63 | 0.92 | 1 | 0.92 | 0.96 |
| 3 | 38406478 | rs704941 | 1.419 | -0.83 | 1.103 | 0.794 | 0.046 | -0.05 | -0.05 | 0 | 0.63 | 0.23 | 0.57 | 0.46 | 0.76 | 0.76 |
| 3 | 38408361 | rs843890 | -0.342 | -0.73 | 1.304 | 0.83 | -0.026 | -0.079 | -0.079 | 1.109 | 0.87 | 0.71 | 0.94 | 1 | 0.94 | 0.98 |
| 3 | 38408742 | rs820292 | -0.013 | -0.753 | 1.606 | 1.045 | -0.043 | 0.101 | 0.101 | 4.416 | 0.85 | 0.44 | 0.87 | 0.75 | 0.94 | 0.93 |
| 3 | 38412303 | rs196381 | 0.223 | -0.665 | 0.705 | 0.787 | -0.008 | -0.006 | -0.006 | 0 | 0.13 | 0.09 | 0.24 | 0.12 | 0.14 | 0.14 |
| 3 | 38413880 | rs704952 | -0.979 | -0.695 | 0.764 | 0.618 | -0.009 | -0.039 | -0.039 | 0.049 | 0.89 | 0.85 | 0.96 | 1 | 0.94 | 0.98 |
| 3 | 38418005 | rs2070487 | -0.728 | -1.226 | 0.479 | 0.791 | -0.008 | 0.017 | 0.017 | 0 | 0.19 | 0.26 | 0.22 | 0.29 | 0.16 | 0.2 |
| 3 | 38419582 | rs169045 | 1.519 | -1.375 | -0.847 | 0.057 | -0.062 | 0.115 | 0.115 | 1.91 | 0.39 | 0.1 | 0.29 | 0.2 | 0.46 | 0.51 |
| 3 | 38421792 | rs196376 | 0.611 | -1.52 | -0.598 | 0.055 | 0 | 0.001 | 0.001 | 0.962 | 0.63 | 0.42 | 0.46 | 0.54 | 0.6 | 0.59 |
| 3 | 38424185 | rs151618 | 1.749 | -1.744 | -1.077 | -0.047 | -0.093 | -0.107 | -0.107 | 0.2 | 0.38 | 0.12 | 0.3 | 0.47 | 0.45 | 0.54 |
| 3 | 38424442 | rs564242 | -0.817 | -1.661 | -0.954 | -0.014 | -0.064 | -0.08 | -0.08 | 0.189 | 0.4 | 0.13 | 0.52 | 0.51 | 0.47 | 0.4 |
| 3 | 38425451 | rs17037638 | 0.204 | -1.516 | -0.093 | 0.021 | -0.022 | -0.058 | -0.058 | 1.526 | 0.14 | 0.21 | 0.05 | 0 | 0.06 | 0.03 |
| 3 | 38436175 | rs11710235 | -0.133 | -1.066 | -0.552 | 0.191 | -0.054 | 0.015 | 0.015 | 0 | 0.1 | 0.01 | 0.16 | 0.05 | 0.15 | 0.14 |
| 3 | 38437835 | rs2070988 | -0.993 | -1.455 | -0.593 | 0.41 | -0.014 | -0.071 | -0.071 | 2.349 | 0.32 | 0.15 | 0.45 | 0.55 | 0.35 | 0.37 |
| 3 | 38441543 | rs6809970 | 0.486 | -1.648 | -0.077 | 0.095 | -0.036 | -0.091 | -0.091 | 0.155 | 0.16 | 0.32 | 0.08 | 0 | 0.06 | 0.04 |
| 3 | 38445226 | rs1002675 | 1.81 | -1.892 | -0.022 | 0.318 | 0.015 | 0.103 | 0.103 | 0.071 | 0.46 | 0.25 | 0.33 | 0.26 | 0.57 | 0.54 |
| 3 | 38454931 | rs7373930 | 0.442 | -1.979 | -0.089 | 0.677 | -0.019 | -0.036 | -0.036 | 0 | 0.7 | 0.92 | 0.72 | 0.64 | 0.67 | 0.68 |
| 3 | 38455439 | rs7373805 | 1.88 | -1.98 | -0.079 | 0.673 | -0.015 | 0.127 | 0.127 | 0.27 | 0.44 | 0.13 | 0.32 | 0.26 | 0.57 | 0.54 |
| 3 | 38455704 | rs13074025 | 1.88 | -1.98 | -0.079 | 0.673 | -0.015 | 0.127 | 0.127 | 0.27 | 0.44 | 0.13 | 0.32 | 0.26 | 0.57 | 0.54 |
| 3 | 38456531 | rs4147025 | 1.88 | -1.991 | 0.069 | 0.716 | -0.015 | 0.127 | 0.127 | 0.27 | 0.44 | 0.13 | 0.32 | 0.26 | 0.57 | 0.54 |
| 3 | 38461279 | rs13085153 | 1.703 | -2.05 | 0.102 | 0.644 | 0.052 | 0.001 | 0.001 | 0.052 | 0.48 | 0.2 | 0.48 | 0.44 | 0.61 | 0.6 |

| | | | | | | | | | | | | | | | | |
|---|---|---|---|---|---|---|---|---|---|---|---|---|---|---|---|---|
| 3 | 38487802 | rs6599203 | 0.64 | -2.056 | 0.281 | 0.646 | 0.004 | -0.01 | -0.01 | 0 | 0.68 | 0.86 | 0.72 | 0.64 | 0.68 | 0.68 |
| 3 | 38501916 | rs2268755 | 1.395 | -1.708 | -0.002 | 0.642 | 0.055 | 0 | 0 | 0.046 | 0.48 | 0.2 | 0.48 | 0.44 | 0.61 | 0.6 |
| 3 | 38519424 | rs2070489 | 1.254 | -1.675 | 0.018 | 0.669 | 0.056 | 0.018 | 0.018 | 0.443 | 0.49 | 0.31 | 0.49 | 0.44 | 0.62 | 0.6 |
| 3 | 38524742 | rs1046048 | 1.651 | -1.781 | 0.002 | 0.836 | -0.009 | 0.132 | 0.132 | 1.254 | 0.45 | 0.13 | 0.32 | 0.26 | 0.58 | 0.55 |
| 3 | 38532511 | rs1058945 | 1.668 | -1.781 | 0.002 | 0.836 | -0.01 | 0.131 | 0.131 | 1.282 | 0.45 | 0.13 | 0.32 | 0.26 | 0.58 | 0.55 |
| 3 | 38541318 | rs2300669 | -1.43 | -1.876 | 0.66 | 0.835 | 0.02 | 0.008 | 0.008 | 0.059 | 0.49 | 0.57 | 0.5 | 0.56 | 0.39 | 0.39 |
| 3 | 38553965 | rs2186457 | -1.697 | -1.862 | 0.646 | 0.718 | 0.048 | 0.03 | 0.03 | 0.201 | 0.51 | 0.66 | 0.5 | 0.59 | 0.39 | 0.4 |
| 3 | 38556802 | rs11717146 | 0.564 | -1.891 | 0.154 | 0.631 | -0.01 | -0.018 | -0.018 | 0.105 | 0.08 | 0.02 | 0.06 | 0.11 | 0.08 | 0.09 |
| 3 | 38558210 | rs6599210 | 0.029 | -1.859 | 0.388 | 0.657 | 0.001 | -0.003 | -0.003 | 0 | 0.23 | 0.26 | 0.12 | 0.11 | 0.14 | 0.12 |
| 3 | 38559749 | rs2051211 | -0.822 | -1.672 | 0.008 | 0.773 | 0.003 | -0.008 | -0.008 | 0 | 0.23 | 0.14 | 0.22 | 0.3 | 0.25 | 0.24 |
| 3 | 38574491 | rs6792762 | -1.834 | -1.589 | -0.414 | 0.657 | 0.027 | 0.164 | 0.164 | 0.622 | 0.54 | 0.82 | 0.65 | 0.75 | 0.41 | 0.44 |
| 3 | 38574968 | rs6810361 | -1.12 | -1.916 | 0.214 | 1.076 | 0.009 | -0.01 | -0.01 | 0 | 0.34 | 0.71 | 0.47 | 0.49 | 0.21 | 0.23 |
| 3 | 38583025 | rs12634016 | -1.16 | -1.733 | 0.544 | 1.293 | -0.071 | 0.103 | 0.103 | 1.234 | 0.19 | 0.51 | 0.34 | 0.64 | 0.11 | 0.18 |
| 3 | 38589163 | rs11129795 | -0.459 | -1.594 | 0.307 | 0.646 | 0.001 | 0.031 | 0.031 | 0 | 0.27 | 0.18 | 0.2 | 0.11 | 0.26 | 0.14 |
| 3 | 38591689 | rs7429945 | -1.122 | -1.12 | -0.185 | 0.954 | 0.045 | 0.003 | 0.003 | 0.87 | 0.45 | 0.71 | 0.43 | 0.58 | 0.37 | 0.3 |
| 3 | 38592406 | rs1805126 | -1.119 | -1.118 | -0.193 | 0.955 | 0.044 | 0.003 | 0.003 | 0.682 | 0.45 | 0.7 | 0.43 | 0.58 | 0.36 | 0.3 |
| 3 | 38593393 | rs12053903 | -1.12 | -0.937 | -0.147 | 1.022 | 0.079 | 0.004 | 0.004 | 1.579 | 0.47 | 0.87 | 0.49 | 0.59 | 0.37 | 0.27 |
| 3 | 38599037 | rs6793245 | -1.017 | -0.905 | -0.278 | 0.339 | 0.039 | 0.017 | 0.017 | 0.391 | 0.46 | 0.64 | 0.33 | 0.38 | 0.36 | 0.23 |
| 3 | 38602855 | rs3935472 | -0.573 | -1.17 | -0.259 | 0.303 | -0.04 | -0.007 | -0.007 | 0.009 | 0.31 | 0.38 | 0.13 | 0.25 | 0.16 | 0.12 |
| 3 | 38606588 | rs7427447 | 0.793 | -0.193 | -0.566 | -0.176 | -0.002 | 0.062 | 0.062 | 0 | 0.58 | 0.31 | 0.56 | 0.87 | 0.59 | 0.6 |
| 3 | 38606714 | rs11129796 | -0.158 | -0.237 | -0.217 | -0.293 | -0.004 | -0.038 | -0.038 | 0 | 0.86 | 0.82 | 0.92 | 1 | 0.91 | 0.97 |
| 3 | 38609334 | rs9861242 | 0.866 | -0.196 | 0.069 | -0.491 | 0 | -0.119 | -0.119 | 0 | 0.74 | 0.57 | 0.72 | 0.96 | 0.77 | 0.76 |
| 3 | 38610471 | rs9833086 | 1.053 | 0.108 | 0.05 | -0.098 | 0.01 | -0.039 | -0.039 | 0 | 0.72 | 0.51 | 0.63 | 0.78 | 0.73 | 0.72 |
| 3 | 38619312 | rs7373819 | 1.486 | 0.509 | -0.642 | -0.074 | -0.068 | -0.023 | -0.023 | 0 | 0.39 | 0.09 | 0.4 | 0.38 | 0.46 | 0.42 |
| 3 | 38622467 | rs7430407 | -0.027 | 0.529 | -0.224 | -0.136 | 0.001 | -0.032 | -0.032 | 0 | 0.91 | 0.87 | 0.92 | 1 | 0.91 | 0.94 |
| 3 | 38624253 | rs3922844 | 0.797 | 0.412 | 0.099 | -0.167 | 0.001 | 0 | 0 | 0 | 0.69 | 0.34 | 0.65 | 0.88 | 0.71 | 0.71 |
| 3 | 38628642 | rs11711097 | 0.626 | 0.287 | 0.021 | -0.05 | 0.005 | -0.015 | -0.015 | 0 | 0.7 | 0.25 | 0.67 | 0.5 | 0.77 | 0.77 |
| 3 | 38629767 | rs7433889 | 0.175 | 0.449 | 0.01 | -0.108 | 0.003 | -0.032 | -0.032 | 0.972 | 0.9 | 0.85 | 0.91 | 1 | 0.89 | 0.94 |
| 3 | 38634142 | rs7374540 | 1.102 | 0.215 | -0.3 | -0.217 | 0.003 | -0.031 | -0.031 | 2.755 | 0.54 | 0.18 | 0.53 | 0.47 | 0.6 | 0.56 |
| 3 | 38635072 | rs9311194 | 0.669 | 0.178 | -0.385 | -0.316 | -0.002 | -0.007 | -0.007 | 0 | 0.18 | 0.14 | 0.19 | 0.1 | 0.19 | 0.19 |
| 3 | 38637160 | rs13062931 | 1.999 | 0.283 | -0.455 | -0.29 | -0.11 | -0.099 | -0.099 | 0 | 0.36 | 0.04 | 0.35 | 0.38 | 0.42 | 0.36 |
| 3 | 38641134 | rs9871385 | -1.041 | 0.457 | -0.303 | -0.365 | 0.01 | 0.04 | 0.04 | 0.18 | 0.39 | 0.27 | 0.34 | 0.12 | 0.36 | 0.4 |
| 3 | 38643279 | rs7427874 | -1.082 | 0.437 | -0.305 | -0.385 | 0.004 | 0.026 | 0.026 | 0.199 | 0.4 | 0.32 | 0.36 | 0.11 | 0.38 | 0.4 |
| 3 | 38645420 | rs1805124 | -0.368 | 0.137 | -0.053 | -0.321 | -0.014 | -0.029 | -0.029 | 0 | 0.25 | 0.31 | 0.23 | 0.1 | 0.22 | 0.27 |
| 3 | 38645999 | rs13084981 | 1.246 | 0.239 | -0.244 | -0.165 | -0.006 | -0.053 | -0.053 | 0.503 | 0.15 | 0.09 | 0.15 | 0.48 | 0.13 | 0.17 |
| 3 | 38647047 | rs11711602 | 0.532 | 0.361 | -0.134 | -0.35 | 0 | 0 | 0 | 0.101 | 0.17 | 0.12 | 0.19 | 0.1 | 0.18 | 0.19 |
| 3 | 38648062 | rs6599222 | 0.237 | 0.619 | 0.055 | -0.255 | 0.01 | -0.066 | -0.066 | 0 | 0.82 | 0.78 | 0.81 | 0.98 | 0.8 | 0.86 |
| 3 | 38650180 | rs9853984 | 1.206 | 0.5 | -0.569 | -0.218 | -0.03 | 0.02 | 0.02 | 0 | 0.39 | 0.17 | 0.38 | 0.34 | 0.41 | 0.37 |
| 3 | 38650460 | rs6599223 | -0.911 | 0.419 | -0.57 | 0.133 | -0.001 | 0.005 | 0.005 | 0 | 0.8 | 0.8 | 0.88 | 0.69 | 0.83 | 0.76 |
| 3 | 38653277 | rs6599225 | -0.743 | 0.407 | -0.294 | 0.126 | -0.003 | 0.042 | 0.042 | 0.018 | 0.78 | 0.64 | 0.87 | 0.68 | 0.83 | 0.76 |
| 3 | 38655643 | rs9856587 | -0.034 | 0.694 | -0.198 | -0.17 | 0.007 | -0.085 | -0.085 | 0.004 | 0.18 | 0.44 | 0.24 | 0.05 | 0.17 | 0.15 |
| 3 | 38656033 | rs6797133 | 0.219 | 0.876 | -0.536 | 0.114 | 0 | 0.011 | 0.011 | 0.08 | 0.57 | 0.68 | 0.6 | 0.37 | 0.59 | 0.51 |
| 3 | 38660936 | rs12498069 | -0.139 | 0.553 | -0.478 | 0.239 | 0 | -0.007 | -0.007 | 0 | 0.14 | 0.2 | 0.13 | 0.06 | 0.13 | 0.07 |
| 3 | 38661533 | rs9832895 | 0.186 | 0.449 | -0.797 | 0.326 | -0.012 | 0.021 | 0.021 | 0.019 | 0.44 | 0.3 | 0.42 | 0.39 | 0.47 | 0.54 |
| 3 | 38662961 | rs6599228 | -0.141 | 0.253 | -0.843 | 0.639 | 0.01 | 0.006 | 0.006 | 0.226 | 0.38 | 0.48 | 0.57 | 0.57 | 0.43 | 0.48 |
| 3 | 38664309 | rs3934936 | 0.206 | 0.197 | -0.869 | 0.321 | -0.001 | 0.002 | 0.002 | 0.233 | 0.36 | 0.35 | 0.3 | 0.22 | 0.33 | 0.33 |
| 3 | 38665202 | rs7624535 | -1.082 | 0.001 | -0.509 | 0.818 | 0.007 | -0.016 | -0.016 | 0 | 0.73 | 0.78 | 0.69 | 0.47 | 0.77 | 0.74 |
| 3 | 38672585 | rs6599229 | -0.651 | 0.619 | 0.011 | 0.517 | -0.005 | -0.002 | -0.002 | 0.03 | 0.45 | 0.53 | 0.53 | 0.56 | 0.57 | 0.52 |
| 3 | 38676686 | rs13060541 | -0.67 | 1.131 | -0.087 | 0.265 | 0.028 | -0.052 | -0.052 | 0 | 0.15 | 0.24 | 0.18 | 0.01 | 0.25 | 0.12 |
| 3 | 38680628 | rs7373102 | -1.201 | 0.511 | 0.196 | 0.467 | 0.028 | 0.039 | 0.039 | 0 | 0.51 | 0.76 | 0.5 | 0.58 | 0.52 | 0.54 |
| 3 | 38681394 | rs6763048 | -0.537 | 0.678 | -0.01 | 0.637 | 0.006 | -0.062 | -0.062 | 0.134 | 0.12 | 0.25 | 0.12 | 0.01 | 0.15 | 0.09 |
| 3 | 38686192 | rs7372712 | 1.074 | 0.796 | -0.391 | 0.805 | -0.024 | -0.073 | -0.073 | 0 | 0.16 | 0.06 | 0.2 | 0.49 | 0.17 | 0.34 |
| 3 | 38694261 | rs7373686 | 0.962 | 0.762 | -0.363 | 0.737 | -0.04 | -0.001 | -0.001 | 0 | 0.11 | 0.02 | 0.13 | 0.09 | 0.16 | 0.19 |
| 3 | 38695708 | rs7373934 | -0.593 | 0.683 | 0.177 | 0.762 | 0.015 | -0.036 | -0.036 | 0 | 0.21 | 0.41 | 0.25 | 0.15 | 0.25 | 0.31 |
| 3 | 38697339 | rs9861643 | -0.432 | 0.535 | -0.215 | 0.855 | 0.005 | 0.078 | 0.078 | 1.097 | 0.23 | 0.39 | 0.28 | 0.37 | 0.18 | 0.1 |
| 3 | 38703974 | rs7430660 | -0.488 | -0.003 | -0.736 | 1.002 | 0.001 | 0.003 | 0.003 | 0 | 0.18 | 0.38 | 0.18 | 0.17 | 0.18 | 0.19 |
| 3 | 38707352 | rs13061927 | 0.513 | 0.491 | -1.263 | 1.154 | -0.155 | 0.012 | 0.012 | 0.054 | 0.31 | 0.02 | 0.35 | 0.18 | 0.37 | 0.31 |
| 3 | 38711196 | rs6806438 | -1.365 | 0.62 | 0.121 | 1.718 | 0.021 | -0.054 | -0.054 | 2.201 | 0.17 | 0.35 | 0.16 | 0.08 | 0.21 | 0.21 |
| 3 | 38714476 | rs6773329 | -1.119 | 0.817 | -0.485 | 1.816 | 0.011 | 0.002 | 0.002 | 0 | 0.16 | 0.22 | 0.16 | 0.17 | 0.21 | 0.22 |
| 3 | 38714855 | rs9825287 | -0.019 | 0.952 | -0.621 | 1.458 | 0.003 | 0 | 0 | 0.658 | 0.06 | 0.09 | 0.07 | 0.06 | 0.05 | 0.12 |
| 3 | 38721160 | rs7372943 | -0.666 | 1.005 | -0.677 | 1.458 | 0.008 | 0.195 | 0.195 | 0 | 0.13 | 0.35 | 0.15 | 0.36 | 0.11 | 0.16 |
| 3 | 38721308 | rs7373154 | -0.827 | 0.97 | -0.763 | 1.355 | 0.013 | 0.077 | 0.077 | 0 | 0.25 | 0.4 | 0.29 | 0.44 | 0.29 | 0.35 |
| 3 | 38726685 | rs7374165 | -0.414 | 0.643 | -1.791 | 1.213 | -0.01 | -0.145 | -0.145 | 0 | 0.31 | 0.06 | 0.31 | 0.48 | 0.33 | 0.4 |
| 3 | 38730117 | rs6764249 | -0.489 | 1.751 | -1.072 | 1.527 | 0.035 | 0.144 | 0.144 | 0 | 0.12 | 0.22 | 0.22 | 0.35 | 0.17 | 0.11 |
| 3 | 38736230 | rs12639182 | 0.097 | 1.305 | -1.418 | 1.426 | 0.012 | -0.113 | -0.113 | 0.044 | 0.2 | 0.06 | 0.16 | 0.44 | 0.19 | 0.27 |
| 3 | 38737643 | rs7372391 | -1.482 | 1.201 | -0.833 | 1.148 | -0.001 | 0.105 | 0.105 | 0 | 0.18 | 0.34 | 0.21 | 0.31 | 0.16 | 0.12 |
| 3 | 38738717 | rs6599240 | 1.754 | 1.867 | -0.692 | 0.838 | -0.001 | -0.02 | -0.02 | 0 | 0.52 | 0.44 | 0.43 | 0.16 | 0.44 | 0.39 |
| 3 | 38742229 | rs11129799 | 0.856 | 1.611 | -0.656 | 0.813 | -0.024 | -0.031 | -0.031 | 0 | 0.86 | 0.91 | 0.78 | 0.74 | 0.79 | 0.76 |
| 3 | 38747735 | rs6599244 | -0.482 | 1.506 | -0.565 | 0.721 | 0.006 | 0.098 | 0.098 | 0 | 0.11 | 0.19 | 0.19 | 0.31 | 0.12 | 0.1 |

| | | | | | | | | | | | | | | | | |
|---|---|---|---|---|---|---|---|---|---|---|---|---|---|---|---|---|
| 3 | 38750375 | rs11129801 | 0.352 | 1.353 | -0.571 | 0.61 | 0 | 0.001 | 0.001 | 0.005 | 0.75 | 0.73 | 0.74 | 0.52 | 0.72 | 0.56 |
| 3 | 38754647 | rs6775197 | -1.051 | 0.929 | 0.424 | 0.553 | 0.03 | 0.047 | 0.047 | 4.231 | 0.07 | 0.29 | 0.09 | 0.04 | 0.02 | 0.09 |
| 3 | 38764998 | rs12632942 | 0.064 | 1.103 | 0.195 | 0.577 | -0.005 | -0.059 | -0.059 | 0.445 | 0.23 | 0.12 | 0.12 | 0.38 | 0.25 | 0.23 |
| 3 | 38766338 | rs9825762 | -0.56 | 0.859 | 0.241 | 0.026 | 0.004 | -0.09 | -0.09 | 0 | 0.19 | 0.27 | 0.2 | 0.01 | 0.2 | 0.18 |
| 3 | 38766675 | rs6795970 | -1.95 | 0.77 | 0.111 | 0.079 | -0.094 | 0.2 | 0.2 | 0.001 | 0.57 | 0.95 | 0.64 | 0.82 | 0.59 | 0.69 |
| 3 | 38766701 | rs6791171 | 0.241 | 0.837 | 0.371 | 0.09 | 0.087 | -0.033 | -0.033 | 0 | 0.07 | 0.2 | 0.14 | 0 | 0.13 | 0.09 |
| 3 | 38767315 | rs6801957 | -1.842 | 0.832 | 0.23 | 0.222 | -0.029 | 0.169 | 0.169 | 0 | 0.55 | 0.86 | 0.58 | 0.78 | 0.58 | 0.67 |
| 3 | 38768334 | rs7374804 | -0.017 | 0.83 | 0.298 | 0.205 | 0.004 | 0.09 | 0.09 | 0 | 0.05 | 0.11 | 0.1 | 0.26 | 0.06 | 0.15 |
| 3 | 38771464 | rs6780103 | -1.447 | 0.812 | 0.468 | 0.156 | -0.002 | -0.012 | -0.012 | 0 | 0.28 | 0.52 | 0.3 | 0.21 | 0.28 | 0.32 |
| 3 | 38777554 | rs10428132 | -1.876 | 0.862 | 0.401 | 0.16 | -0.043 | 0.202 | 0.202 | 0 | 0.55 | 0.88 | 0.57 | 0.79 | 0.58 | 0.67 |
| 3 | 38782026 | rs7615140 | -0.739 | 0.887 | 0.511 | 0.193 | -0.002 | -0.033 | -0.033 | 0 | 0.36 | 0.55 | 0.35 | 0.79 | 0.33 | 0.48 |
| 3 | 38793940 | rs7630989 | -0.545 | 0.89 | 0.392 | 0.218 | -0.003 | 0.133 | 0.133 | 2.629 | 0.08 | 0.21 | 0.09 | 0.2 | 0.03 | 0.14 |
| 3 | 38793989 | rs7617919 | 0.655 | 0.914 | 0.338 | 0.178 | -0.02 | -0.077 | -0.077 | 1.506 | 0.22 | 0.08 | 0.11 | 0.37 | 0.25 | 0.24 |
| 3 | 38794268 | rs7431144 | -1.352 | 0.923 | 0.377 | 0.188 | 0.016 | 0.284 | 0.284 | 4.086 | 0.12 | 0.36 | 0.24 | 0.41 | 0.08 | 0.25 |
| 3 | 38796985 | rs12630795 | 0.663 | 1.01 | 0.265 | 0.217 | -0.024 | -0.077 | -0.077 | 0.203 | 0.21 | 0.08 | 0.11 | 0.37 | 0.25 | 0.24 |
| 3 | 38798836 | rs6798015 | -2.021 | 0.734 | -0.145 | -0.332 | -0.056 | 0.107 | 0.107 | 0 | 0.64 | 0.89 | 0.6 | 0.79 | 0.63 | 0.71 |
| 3 | 38800763 | rs6809264 | -0.363 | 0.725 | 0.066 | -0.283 | 0.025 | -0.105 | -0.105 | 0 | 0.19 | 0.32 | 0.22 | 0.01 | 0.25 | 0.18 |
| 3 | 38800873 | rs9637536 | 0.549 | 0.712 | -0.063 | -0.262 | 0 | -0.001 | -0.001 | 0.28 | 0.17 | 0.17 | 0.15 | 0.1 | 0.15 | 0.15 |
| 3 | 38802251 | rs7641844 | -0.206 | 0.662 | -0.273 | -0.047 | -0.004 | -0.004 | -0.004 | 0 | 0.31 | 0.33 | 0.25 | 0.68 | 0.28 | 0.38 |
| 3 | 38804358 | rs7635221 | -0.424 | 0.739 | -0.421 | 0.023 | 0 | 0.095 | 0.095 | 0 | 0.08 | 0.23 | 0.04 | 0.16 | 0.04 | 0.05 |
| 3 | 38804588 | rs6599257 | -1.886 | 0.774 | -0.667 | -0.908 | -0.032 | 0.118 | 0.118 | 0 | 0.69 | 0.87 | 0.66 | 0.84 | 0.68 | 0.74 |
| 3 | 38812708 | rs4420804 | -0.88 | 0.506 | -0.261 | -0.554 | 0.005 | -0.009 | -0.009 | 0 | 0.37 | 0.46 | 0.5 | 0.31 | 0.38 | 0.44 |
| 3 | 38820774 | rs11927856 | -0.523 | 0.494 | -0.832 | -0.426 | 0.003 | 0.006 | 0.006 | 0 | 0.27 | 0.22 | 0.31 | 0.25 | 0.27 | 0.28 |
| 3 | 38830893 | rs4676595 | -0.86 | 0.556 | -0.532 | -0.246 | 0.004 | 0.024 | 0.024 | 0 | 0.79 | 0.81 | 0.82 | 0.54 | 0.75 | 0.84 |
| 3 | 38832997 | rs9815891 | 1.553 | 0.599 | -0.573 | -0.501 | 0.005 | 0.019 | 0.019 | 0 | 0.36 | 0.3 | 0.48 | 0.28 | 0.37 | 0.35 |
| 3 | 38835888 | rs11712354 | 0.092 | 0.398 | 0.394 | -0.517 | -0.004 | 0.038 | 0.038 | 0 | 0.14 | 0.04 | 0.12 | 0.07 | 0.13 | 0.16 |
| 3 | 38842912 | rs4676593 | -1.486 | 0.357 | 0.501 | -0.561 | -0.009 | -0.028 | -0.028 | 0 | 0.34 | 0.47 | 0.3 | 0.26 | 0.26 | 0.43 |
| 3 | 38843647 | rs11717455 | 0.128 | 0.394 | 0.391 | -0.578 | -0.005 | 0.038 | 0.038 | 0 | 0.14 | 0.04 | 0.12 | 0.07 | 0.13 | 0.15 |
| 3 | 38845645 | rs11922345 | 0.173 | 0.508 | 0.461 | -0.421 | 0.02 | -0.04 | -0.04 | 0.114 | 0.08 | 0.17 | 0.05 | 0 | 0.1 | 0.06 |
| 3 | 38847152 | rs7627881 | -0.881 | 0.471 | -0.014 | -0.211 | 0.008 | 0.077 | 0.077 | 0.053 | 0.43 | 0.76 | 0.37 | 0.26 | 0.36 | 0.48 |
| 3 | 38862148 | rs4130468 | -0.558 | 1.182 | -1.188 | 0.581 | -0.028 | 0.002 | 0.002 | 0 | 0.1 | 0.02 | 0.1 | 0.08 | 0.1 | 0.17 |
| 3 | 38863137 | rs4643645 | -0.219 | 1.213 | -1.245 | 0.154 | -0.003 | 0.059 | 0.059 | 0.003 | 0.48 | 0.35 | 0.47 | 0.72 | 0.5 | 0.56 |
| 3 | 38864387 | rs7635472 | -1.338 | 1.423 | -1.377 | 0.107 | -0.001 | -0.012 | -0.012 | 0 | 0.55 | 0.86 | 0.49 | 0.56 | 0.55 | 0.48 |
| 3 | 38864937 | rs4676592 | -0.755 | 2.116 | -0.821 | 0.266 | 0.025 | 0.007 | 0.007 | 0.04 | 0.09 | 0.14 | 0.11 | 0.16 | 0.13 | 0.14 |
| 3 | 38877793 | rs13062806 | -1.135 | 2.005 | -0.22 | 0.23 | 0.009 | -0.007 | -0.007 | 0.671 | 0.21 | 0.22 | 0.24 | 0.13 | 0.25 | 0.16 |
| 3 | 38885616 | rs13317208 | -0.7 | 1.789 | 0.566 | 0.09 | 0 | 0.009 | 0.009 | 0.112 | 0.23 | 0.32 | 0.14 | 0.23 | 0.24 | 0.2 |
| 3 | 38886413 | rs3923518 | -0.946 | 1.778 | 0.652 | 0.079 | -0.003 | 0.002 | 0.002 | 0 | 0.2 | 0.18 | 0.24 | 0.13 | 0.24 | 0.16 |
| 3 | 38888021 | rs4234133 | -2.207 | 1.757 | 0.849 | 0.085 | -0.008 | -0.008 | -0.008 | 0 | 0.47 | 0.8 | 0.44 | 0.51 | 0.49 | 0.4 |
| 3 | 38893532 | rs4561763 | -0.706 | 1.751 | 0.793 | 0.076 | 0.001 | 0.016 | 0.016 | 0.186 | 0.23 | 0.37 | 0.14 | 0.23 | 0.24 | 0.2 |
| 3 | 38895359 | rs4364115 | -2.105 | 1.731 | 0.71 | 0.054 | -0.009 | -0.01 | -0.01 | 0 | 0.47 | 0.8 | 0.44 | 0.51 | 0.49 | 0.4 |
| 3 | 38923700 | rs4315640 | -0.033 | 1.805 | 0.598 | -0.017 | -0.007 | 0 | 0 | 0.008 | 0.07 | 0.04 | 0.07 | 0 | 0.12 | 0.08 |
| 3 | 38928643 | rs9867831 | -0.211 | 1.68 | 0.719 | -0.078 | 0.016 | -0.03 | -0.03 | 2.474 | 0.08 | 0.3 | 0.06 | 0 | 0.05 | 0.03 |
| 3 | 38936829 | rs13099670 | -0.966 | 1.752 | 0.393 | -0.048 | -0.039 | 0.016 | 0.016 | 0 | 0.19 | 0.07 | 0.22 | 0.12 | 0.24 | 0.16 |
| 3 | 38945984 | rs11709492 | -1.251 | 1.406 | 0.914 | -0.065 | -0.027 | 0.005 | 0.005 | 0 | 0.25 | 0.39 | 0.14 | 0.24 | 0.22 | 0.25 |
| 3 | 38946416 | rs4553926 | -0.022 | 1.37 | 0.799 | -0.092 | -0.002 | -0.008 | -0.008 | 0 | 0.87 | 0.92 | 0.84 | 0.87 | 0.88 | 0.92 |
| 3 | 38948028 | rs9311199 | 0.126 | 1.357 | 0.948 | -0.098 | -0.002 | 0.001 | 0.001 | 0.072 | 0.86 | 0.85 | 0.84 | 0.87 | 0.88 | 0.92 |
| 3 | 38995532 | rs4076478 | -0.425 | 1.328 | 0.742 | -0.13 | 0.004 | -0.027 | -0.027 | 0 | 0.16 | 0.06 | 0.08 | 0.22 | 0.16 | 0.2 |
| 3 | 39005889 | rs4491835 | -3.574 | 1.694 | 0.542 | -0.235 | -0.055 | -0.116 | -0.116 | 0.839 | 0.56 | 0.97 | 0.51 | 0.53 | 0.6 | 0.49 |
| 3 | 39016788 | rs11710221 | -0.363 | 1.605 | 0.684 | -0.272 | 0.003 | -0.028 | -0.028 | 0 | 0.16 | 0.05 | 0.08 | 0.22 | 0.16 | 0.2 |
| 3 | 39021954 | rs12635888 | 0.344 | 1.588 | 0.657 | -0.289 | -0.012 | -0.04 | -0.04 | 0 | 0.1 | 0.03 | 0.05 | 0.19 | 0.11 | 0.12 |
| 3 | 39025135 | rs11929481 | -0.432 | 1.558 | 0.968 | -0.316 | 0 | -0.022 | -0.022 | 1.17 | 0.07 | 0.18 | 0.04 | 0.02 | 0.06 | 0.05 |
| 3 | 39028854 | rs4616582 | -2.509 | 1.547 | 0.834 | -0.336 | 0.029 | -0.112 | -0.112 | 0.406 | 0.4 | 0.92 | 0.35 | 0.49 | 0.31 | 0.28 |
| 3 | 39048112 | rs7615849 | -0.327 | 1.541 | 0.985 | -0.35 | 0.005 | -0.006 | -0.006 | 0 | 0.15 | 0.09 | 0.16 | 0.13 | 0.11 | 0.07 |
| 3 | 39051570 | rs4558686 | 0.403 | 1.539 | 0.901 | -0.35 | -0.028 | -0.061 | -0.061 | 0 | 0.1 | 0.01 | 0.05 | 0.19 | 0.12 | 0.12 |
| 3 | 39056526 | rs6796391 | -1.294 | 1.529 | 1.094 | -0.362 | 0.01 | -0.023 | -0.023 | 0 | 0.1 | 0.47 | 0.08 | 0.02 | 0.07 | 0.05 |
| 3 | 39059192 | rs4676660 | -4.048 | 1.696 | 0.845 | -0.414 | -0.058 | -0.14 | -0.14 | 0 | 0.57 | 0.97 | 0.51 | 0.51 | 0.61 | 0.47 |
| 3 | 39067925 | rs883349 | -2.839 | 1.235 | 0.957 | -0.309 | 0.038 | -0.099 | -0.099 | 0.297 | 0.44 | 0.93 | 0.37 | 0.51 | 0.37 | 0.35 |
| 3 | 39081171 | rs11925218 | -2.327 | 1.248 | 0.982 | -0.296 | 0.037 | -0.033 | -0.033 | 0 | 0.38 | 0.84 | 0.32 | 0.34 | 0.32 | 0.24 |
| 3 | 39091812 | rs6778963 | -2.527 | 1.249 | 0.978 | -0.295 | 0.027 | -0.109 | -0.109 | 0.122 | 0.4 | 0.92 | 0.36 | 0.49 | 0.34 | 0.28 |
| 3 | 39093326 | rs6599003 | -2.316 | 1.248 | 0.982 | -0.294 | 0.036 | -0.034 | -0.034 | 0 | 0.38 | 0.84 | 0.32 | 0.34 | 0.32 | 0.24 |
| 3 | 39100201 | rs6769051 | -2.795 | 1.376 | 0.94 | -0.328 | 0.018 | -0.091 | -0.091 | 0 | 0.42 | 0.92 | 0.37 | 0.5 | 0.37 | 0.29 |
| 3 | 39110041 | rs4676481 | -2.538 | 1.042 | 1.266 | -0.209 | 0.029 | -0.101 | -0.101 | 0 | 0.4 | 0.91 | 0.36 | 0.49 | 0.34 | 0.28 |
| 3 | 39111140 | rs2293312 | -2.452 | 1.042 | 1.258 | -0.214 | 0.037 | -0.033 | -0.033 | 0 | 0.38 | 0.84 | 0.32 | 0.34 | 0.32 | 0.24 |
| 3 | 39118104 | rs11707101 | -0.022 | 1.056 | 1.262 | -0.193 | -0.01 | -0.035 | -0.035 | 0 | 0.11 | 0.05 | 0.06 | 0.19 | 0.14 | 0.12 |
| 3 | 39124132 | rs12636980 | -1.862 | 1.053 | 1.463 | -0.2 | -0.01 | 0.021 | 0.021 | 0 | 0.29 | 0.39 | 0.25 | 0.32 | 0.26 | 0.19 |
| 3 | 39125293 | rs2056613 | -2.456 | 1.043 | 1.21 | -0.199 | 0.02 | -0.113 | -0.113 | 0 | 0.35 | 0.89 | 0.33 | 0.47 | 0.28 | 0.23 |
| 3 | 39137278 | rs1053489 | 0.221 | 0.979 | 1.236 | -0.131 | -0.012 | -0.04 | -0.04 | 0 | 0.1 | 0.03 | 0.06 | 0.19 | 0.12 | 0.12 |
| 3 | 39138840 | rs3732377 | 0.29 | 0.979 | 1.236 | -0.131 | -0.011 | -0.042 | -0.042 | 0.052 | 0.1 | 0.03 | 0.06 | 0.19 | 0.12 | 0.12 |
| 3 | 39142704 | rs2293307 | -1.182 | 1.027 | 1.454 | -0.502 | -0.021 | 0.011 | 0.011 | 1.622 | 0.18 | 0.48 | 0.2 | 0.13 | 0.12 | 0.07 |

| | | | | | | | | | | | | | | | |
|---|---|---|---|---|---|---|---|---|---|---|---|---|---|---|---|
| 3 | 39146429 | rs9809104 | -2.077 | 1.072 | 1.405 | -0.485 | 0.043 | -0.009 | -0.009 | 0 | 0.3 | 0.89 | 0.3 | 0.3 | 0.22 | 0.16 |
| 3 | 39150397 | rs3796127 | -0.484 | 1.061 | 1.483 | -0.399 | 0.01 | 0.025 | 0.025 | 0.349 | 0.13 | 0.36 | 0.1 | 0.11 | 0.12 | 0.11 |
| 3 | 39152345 | rs1112438 | -1.848 | 1.135 | 1.529 | -0.374 | -0.006 | 0.002 | 0.002 | 0.295 | 0.28 | 0.57 | 0.29 | 0.25 | 0.27 | 0.27 |
| 3 | 39161456 | rs1274972 | -2.612 | 1.152 | 1.246 | -0.372 | 0.041 | 0.043 | 0.043 | 0.725 | 0.4 | 0.85 | 0.34 | 0.27 | 0.33 | 0.32 |
| 3 | 39161464 | rs1274971 | -2.612 | 1.152 | 1.246 | -0.372 | 0.041 | 0.043 | 0.043 | 0.725 | 0.4 | 0.85 | 0.34 | 0.27 | 0.33 | 0.32 |
| 3 | 39163425 | rs2949565 | 2.473 | 1.152 | 1.246 | -0.372 | 0.041 | 0.043 | 0.043 | 0.725 | 0.6 | 0.15 | 0.66 | 0.73 | 0.67 | 0.68 |
| 3 | 39168283 | rs811971 | 1.62 | 1.144 | 1.295 | -0.358 | -0.001 | 0.003 | 0.003 | 1.461 | 0.73 | 0.29 | 0.68 | 0.62 | 0.75 | 0.79 |
| 3 | 39168757 | rs784505 | 2.791 | 1.188 | 0.681 | -0.348 | 0.026 | -0.049 | -0.049 | 0 | 0.57 | 0.07 | 0.62 | 0.57 | 0.65 | 0.65 |
| 3 | 39175004 | rs17791786 | -0.194 | 1.192 | 0.793 | -0.451 | 0.006 | 0.012 | 0.012 | 0 | 0.12 | 0.17 | 0.17 | 0.13 | 0.12 | 0.08 |
| 3 | 39180263 | rs704959 | 2.371 | 1.177 | 0.708 | -0.463 | 0.043 | 0.045 | 0.045 | 1.184 | 0.6 | 0.16 | 0.66 | 0.73 | 0.67 | 0.69 |
| 3 | 39183895 | rs784501 | 1.17 | 1.688 | 0.813 | -0.144 | 0.003 | 0.009 | 0.009 | 0 | 0.78 | 0.75 | 0.77 | 0.77 | 0.77 | 0.81 |
| 3 | 39184959 | rs1274958 | 1.089 | 1.708 | 0.69 | -0.135 | -0.001 | -0.002 | -0.002 | 2.799 | 0.79 | 0.8 | 0.77 | 0.77 | 0.77 | 0.81 |
| 3 | 39185248 | rs784519 | 0.156 | 1.697 | 0.469 | -0.171 | -0.002 | -0.005 | -0.005 | 1.568 | 0.9 | 0.93 | 0.94 | 0.89 | 0.89 | 0.89 |
| 3 | 39188182 | rs13084580 | -0.288 | 1.587 | 0.574 | -0.314 | 0.002 | 0.003 | 0.003 | 0 | 0.11 | 0.13 | 0.17 | 0.12 | 0.1 | 0.08 |
| 3 | 39188843 | rs885067 | -1.675 | 1.812 | 0.874 | -0.664 | 0.054 | -0.06 | -0.06 | 0 | 0.27 | 0.53 | 0.33 | 0.12 | 0.3 | 0.2 |
| 3 | 39191029 | rs1274963 | 0.582 | 1.75 | 0.843 | -0.673 | -0.011 | -0.002 | -0.002 | 0.794 | 0.84 | 0.86 | 0.87 | 1 | 0.8 | 0.88 |
| 3 | 39195964 | rs2280630 | -1.303 | 1.886 | 0.889 | -0.647 | 0.041 | 0.021 | 0.021 | 0 | 0.25 | 0.41 | 0.32 | 0.33 | 0.31 | 0.22 |
| 3 | 39196552 | rs811970 | 1.31 | 1.63 | 0.882 | -0.796 | 0 | -0.043 | -0.043 | 0.014 | 0.76 | 0.7 | 0.84 | 0.96 | 0.78 | 0.82 |
| 3 | 39198940 | rs784502 | 2.485 | 1.63 | 0.432 | -0.755 | 0.048 | 0.079 | 0.079 | 0.55 | 0.5 | 0.21 | 0.52 | 0.63 | 0.46 | 0.6 |
| 3 | 39202341 | rs11720125 | -0.126 | 1.894 | 0.375 | -0.685 | -0.047 | 0.01 | 0.01 | 0.017 | 0.1 | 0.01 | 0.09 | 0.06 | 0.21 | 0.12 |
| 3 | 39203895 | rs6801073 | 2.358 | 1.901 | 0.446 | -0.686 | 0.016 | 0.041 | 0.041 | 0.846 | 0.54 | 0.18 | 0.54 | 0.61 | 0.49 | 0.63 |
| 3 | 39208697 | rs9311208 | -1.447 | 1.943 | 0.814 | -0.616 | 0.089 | 0.034 | 0.034 | 0.194 | 0.17 | 0.47 | 0.16 | 0.24 | 0.23 | 0.15 |
| 3 | 39208934 | rs9869040 | -1.332 | 1.866 | 0.84 | -0.571 | 0.087 | 0.21 | 0.21 | 0.073 | 0.06 | 0.41 | 0.06 | 0.18 | 0.02 | 0.03 |
| 3 | 39214256 | rs4676609 | -0.473 | 2.246 | -0.02 | -0.629 | -0.055 | 0.056 | 0.056 | 0.973 | 0.15 | 0.05 | 0.11 | 0.03 | 0.18 | 0.13 |
| 3 | 39217854 | rs4075734 | 0.131 | 2.218 | 0.041 | -0.549 | -0.032 | 0 | 0 | 0.56 | 0.08 | 0.01 | 0.08 | 0.07 | 0.19 | 0.11 |
| 3 | 39226127 | rs3732383 | -0.378 | 2.147 | 0.003 | -0.575 | -0.058 | 0 | 0 | 0 | 0.13 | 0.01 | 0.14 | 0.14 | 0.22 | 0.18 |
| 3 | 39230997 | rs2271487 | -1.839 | 2.062 | 0.619 | -0.612 | 0.038 | -0.033 | -0.033 | 0.519 | 0.31 | 0.45 | 0.36 | 0.24 | 0.39 | 0.33 |
| 3 | 39231724 | rs11712101 | 2.581 | 1.797 | -0.312 | -0.715 | -0.035 | -0.05 | -0.05 | 0 | 0.43 | 0.03 | 0.48 | 0.6 | 0.41 | 0.47 |
| 3 | 39234170 | rs17729892 | 0.401 | 1.854 | 0.249 | -0.425 | -0.035 | 0.014 | 0.014 | 0.836 | 0.08 | 0.02 | 0.08 | 0.04 | 0.12 | 0.09 |
| 3 | 39240559 | rs7622284 | 0.626 | 1.499 | 0.578 | -0.361 | 0.005 | 0.009 | 0.009 | 0 | 0.87 | 0.79 | 0.91 | 0.86 | 0.88 | 0.93 |
| 3 | 39241437 | rs4676483 | 1.519 | 1.473 | 0.205 | -0.374 | 0.002 | 0.02 | 0.02 | 1.592 | 0.81 | 0.38 | 0.76 | 0.81 | 0.85 | 0.91 |
| 3 | 39248852 | rs12632771 | -0.488 | 1.623 | -0.55 | -0.452 | -0.005 | 0.002 | 0.002 | 0 | 0.1 | 0.08 | 0.07 | 0.06 | 0.11 | 0.04 |
| 3 | 39252669 | rs3938106 | 0.281 | 1.288 | -0.473 | -0.66 | -0.002 | -0.011 | -0.011 | 0.03 | 0.84 | 0.84 | 0.92 | 0.91 | 0.88 | 0.87 |
| 3 | 39257756 | rs17792215 | 0.146 | 1.829 | -0.821 | -0.619 | -0.036 | -0.025 | -0.025 | 0 | 0.08 | 0.02 | 0.1 | 0.16 | 0.11 | 0.12 |
| 3 | 39257892 | rs11707681 | -0.532 | 2.4 | -1.05 | -0.667 | -0.049 | 0.015 | 0.015 | 0 | 0.17 | 0.08 | 0.13 | 0.11 | 0.27 | 0.17 |
| 3 | 39265616 | rs4314106 | -0.456 | 1.988 | -0.791 | -0.848 | -0.006 | 0.004 | 0.004 | 0 | 0.11 | 0.09 | 0.08 | 0.06 | 0.11 | 0.06 |
| 3 | 39267352 | rs4676484 | -0.387 | 1.858 | -0.622 | -0.757 | -0.003 | -0.001 | -0.001 | 0.041 | 0.1 | 0.08 | 0.1 | 0.15 | 0.12 | 0.11 |
| 3 | 39280382 | rs4676617 | 1.61 | 1.482 | -1.21 | -1.085 | -0.054 | 0.155 | 0.155 | 0.669 | 0.48 | 0.13 | 0.62 | 0.76 | 0.54 | 0.62 |
| 3 | 39282923 | rs7619161 | 0.587 | 1.02 | 0.13 | -1.056 | 0.045 | -0.017 | -0.017 | 0 | 0.92 | 0.62 | 0.94 | 1 | 0.95 | 0.97 |
| 3 | 39286825 | rs7631529 | -0.083 | 0.782 | -0.183 | -1.148 | 0.034 | 0.033 | 0.033 | 0 | 0.15 | 0.06 | 0.03 | 0.01 | 0.06 | 0.08 |
| 3 | 39286934 | rs6599014 | -1.486 | 0.894 | 0.295 | -1.152 | -0.005 | -0.003 | -0.003 | 1.188 | 0.28 | 0.26 | 0.1 | 0.17 | 0.17 | 0.22 |
| 3 | 39291616 | rs9850385 | -1.629 | 0.918 | 0.388 | -1.14 | -0.026 | -0.011 | -0.011 | 0 | 0.3 | 0.4 | 0.12 | 0.23 | 0.19 | 0.24 |
| 3 | 39293804 | rs9861255 | -0.888 | 0.811 | 0.141 | -1.167 | -0.04 | -0.073 | -0.073 | 0 | 0.21 | 0.3 | 0.07 | 0.06 | 0.1 | 0.11 |
| 3 | 39300157 | rs17038640 | 2.374 | 0.882 | -1.439 | -1.441 | -0.038 | -0.091 | -0.091 | 0.488 | 0.36 | 0.03 | 0.47 | 0.62 | 0.34 | 0.47 |
| 3 | 39303147 | rs4676487 | 1.212 | 0.828 | -0.724 | -1.503 | -0.035 | 0.072 | 0.072 | 0 | 0.58 | 0.27 | 0.62 | 0.79 | 0.55 | 0.69 |
| 3 | 39304602 | rs11129819 | 0.015 | 0.899 | -0.664 | -1.614 | -0.015 | 0.011 | 0.011 | 0 | 0.19 | 0.11 | 0.24 | 0.03 | 0.29 | 0.13 |
| 3 | 39305304 | rs17038663 | 0.392 | 0.916 | -0.521 | -1.625 | 0.002 | -0.028 | -0.028 | 0 | 0.06 | 0.08 | 0.04 | 0 | 0.05 | 0.05 |
| 3 | 39306219 | rs11710546 | -0.064 | 0.933 | -0.721 | -1.438 | -0.023 | 0.031 | 0.031 | 0 | 0.19 | 0.09 | 0.24 | 0.03 | 0.29 | 0.13 |
| 3 | 39306605 | rs17038674 | 0.245 | 0.901 | -0.727 | -1.456 | 0.019 | 0.019 | 0.019 | 0 | 0.12 | 0.04 | 0.01 | 0.06 | 0.03 | 0.07 |
| 3 | 39306784 | rs1050592 | -0.3 | 1.548 | -0.631 | -1.191 | -0.023 | 0.033 | 0.033 | 0 | 0.19 | 0.09 | 0.24 | 0.03 | 0.29 | 0.13 |
| 3 | 39307162 | rs3732378 | 0.121 | 1.688 | -0.657 | -1.216 | -0.042 | 0.025 | 0.025 | 0 | 0.07 | 0.01 | 0.16 | 0.03 | 0.17 | 0.11 |
| 3 | 39307256 | rs3732379 | -0.242 | 1.593 | -0.616 | -1.184 | -0.023 | 0.031 | 0.031 | 0 | 0.19 | 0.09 | 0.24 | 0.03 | 0.29 | 0.13 |
| 3 | 39312941 | rs9868689 | -0.656 | 1.722 | -0.457 | -1.245 | 0.001 | -0.02 | -0.02 | 0 | 0.14 | 0.17 | 0.11 | 0.01 | 0.2 | 0.1 |
| 3 | 39314574 | rs4423707 | 0.511 | 1.588 | -0.248 | -1.35 | 0 | -0.006 | -0.006 | 0 | 0.1 | 0.14 | 0.09 | 0.04 | 0.08 | 0.12 |
| 3 | 39317913 | rs13062158 | 2.506 | 1.37 | -0.479 | -1.448 | 0.011 | -0.016 | -0.016 | 0 | 0.32 | 0.13 | 0.42 | 0.68 | 0.3 | 0.42 |
| 3 | 39318288 | rs2853712 | 1.423 | 1.282 | 0.203 | -1.417 | 0 | -0.007 | -0.007 | 0.325 | 0.56 | 0.41 | 0.66 | 0.83 | 0.56 | 0.7 |
| 3 | 39324283 | rs2853707 | 0.56 | 1.432 | 0.509 | -1.643 | 0.024 | -0.028 | -0.028 | 0 | 0.85 | 0.73 | 0.85 | 0.94 | 0.8 | 0.89 |
| 3 | 39331972 | rs2853705 | 0.258 | 1.479 | 0.541 | -1.758 | 0.018 | -0.044 | -0.044 | 0 | 0.87 | 0.81 | 0.86 | 0.99 | 0.8 | 0.93 |
| 3 | 39334720 | rs9825782 | -0.535 | 1.354 | 0.763 | -1.884 | 0 | -0.004 | -0.004 | 1.466 | 0.14 | 0.23 | 0.12 | 0.14 | 0.16 | 0.15 |
| 3 | 39335450 | rs10510699 | 0.491 | 1.525 | 0.495 | -1.942 | -0.008 | -0.001 | -0.001 | 3.51 | 0.12 | 0.06 | 0.11 | 0.08 | 0.12 | 0.13 |
| 3 | 39337570 | rs4473504 | 2.977 | 1.53 | 0.233 | -2.155 | -0.015 | 0.013 | 0.013 | 0.373 | 0.37 | 0.09 | 0.47 | 0.71 | 0.31 | 0.56 |
| 3 | 39337600 | rs4361224 | -0.436 | 1.646 | 0.515 | -2.182 | 0.016 | -0.121 | -0.121 | 0.174 | 0.25 | 0.36 | 0.21 | 0 | 0.34 | 0.12 |
| 3 | 39337941 | rs9842265 | -0.2 | 1.296 | 0.6 | -2.125 | -0.004 | -0.002 | -0.002 | 0 | 0.14 | 0.19 | 0.11 | 0.14 | 0.14 | 0.15 |
| 3 | 39340050 | rs7651347 | 0.003 | 1.22 | 0.494 | -2.157 | -0.04 | -0.14 | -0.14 | 1.345 | 0.21 | 0.39 | 0.13 | 0 | 0.17 | 0.04 |
| 3 | 39340292 | rs11719029 | 1.221 | 1.307 | -0.063 | -2.09 | -0.021 | -0.019 | -0.019 | 0 | 0.06 | 0 | 0.08 | 0.07 | 0.1 | 0.09 |
| 3 | 39357305 | rs2649747 | -1.37 | 1.22 | 0.065 | -2.117 | -0.01 | 0.004 | 0.004 | 4.434 | 0.31 | 0.29 | 0.31 | 0.23 | 0.44 | 0.35 |
| 3 | 39358757 | rs2649748 | -0.859 | 0.958 | -0.035 | -1.976 | -0.01 | -0.033 | -0.033 | 0 | 0.91 | 0.82 | 0.94 | 0.96 | 0.93 | 0.99 |
| 3 | 39360217 | rs1909560 | 2.395 | 0.992 | -0.696 | -1.828 | 0.001 | 0.004 | 0.004 | 0.096 | 0.37 | 0.09 | 0.46 | 0.7 | 0.3 | 0.56 |
| 3 | 39364602 | rs13077620 | 2.368 | 0.966 | -0.716 | -1.889 | -0.005 | 0.018 | 0.018 | 0.413 | 0.37 | 0.04 | 0.45 | 0.72 | 0.3 | 0.57 |

| | | | | | | | | | | | | | | | | |
|---|---|---|---|---|---|---|---|---|---|---|---|---|---|---|---|---|
| 3 | 39369682 | rs12107527 | 0.418 | 1.136 | -0.902 | -1.967 | -0.007 | -0.017 | -0.017 | 0 | 0.19 | 0.18 | 0.17 | 0 | 0.3 | 0.1 |
| 3 | 39369947 | rs7647254 | 0.744 | 1.177 | -1.298 | -1.946 | -0.044 | -0.02 | -0.02 | 0 | 0.11 | 0 | 0.11 | 0.06 | 0.11 | 0.13 |
| 3 | 39380545 | rs17038753 | -0.017 | 0.852 | -0.943 | -1.83 | -0.016 | -0.022 | -0.022 | 0.023 | 0.06 | 0.2 | 0.02 | 0 | 0.01 | 0.01 |
| 3 | 39381823 | rs2853717 | 0.365 | 1.237 | -1.642 | -1.698 | -0.028 | 0.033 | 0.033 | 0 | 0.18 | 0.1 | 0.16 | 0 | 0.3 | 0.1 |
| 3 | 39383876 | rs13088312 | 1.547 | 0.765 | -1.574 | -1.257 | -0.003 | 0.004 | 0.004 | 0.213 | 0.36 | 0.08 | 0.45 | 0.7 | 0.3 | 0.56 |
| 3 | 39422061 | rs3821374 | 0.691 | 0.663 | -0.682 | -1.053 | 0.018 | 0.06 | 0.06 | 0 | 0.75 | 0.84 | 0.79 | 0.92 | 0.78 | 0.77 |
| 3 | 39428241 | rs11129823 | -1.111 | 1.05 | -0.355 | -0.718 | -0.014 | 0 | 0 | 0.289 | 0.27 | 0.15 | 0.31 | 0.27 | 0.31 | 0.26 |
| 3 | 39437800 | rs9877539 | 0.726 | 0.622 | 0.635 | -0.628 | 0.003 | 0.094 | 0.094 | 1.725 | 0.94 | 0.73 | 0.98 | 0.93 | 1 | 0.83 |
| 3 | 39445780 | rs6806372 | -0.979 | 0.624 | 0.587 | -0.632 | -0.013 | -0.002 | -0.002 | 0 | 0.27 | 0.15 | 0.31 | 0.28 | 0.31 | 0.26 |
| 3 | 39448204 | rs3772146 | -1.088 | 0.765 | 0.562 | -0.637 | -0.018 | -0.002 | -0.002 | 0 | 0.27 | 0.16 | 0.32 | 0.28 | 0.33 | 0.26 |
| 3 | 39448581 | rs3772143 | -1.503 | 0.73 | 0.612 | -1.174 | -0.001 | 0.004 | 0.004 | 1.322 | 0.46 | 0.55 | 0.47 | 0.32 | 0.5 | 0.32 |
| 3 | 39448620 | rs11706951 | 1.15 | 0.731 | 0.538 | -1.188 | -0.027 | 0.079 | 0.079 | 0 | 0.13 | 0.01 | 0.09 | 0 | 0.16 | 0.03 |
| 3 | 39453057 | rs2269349 | 1.618 | 0.63 | 0.808 | -1.222 | 0.012 | 0.034 | 0.034 | 0.513 | 0.52 | 0.36 | 0.55 | 0.65 | 0.53 | 0.53 |
| 3 | 39463436 | rs6781117 | -0.747 | 0.607 | 0.85 | -1.193 | -0.007 | 0 | 0 | 1.006 | 0.27 | 0.2 | 0.32 | 0.28 | 0.31 | 0.26 |
| 3 | 39463986 | rs2039845 | -0.747 | 0.607 | 0.85 | -1.193 | -0.007 | 0 | 0 | 1.006 | 0.27 | 0.2 | 0.31 | 0.28 | 0.31 | 0.26 |
| 3 | 39468418 | rs521480 | 1.015 | 0.468 | 1.029 | -1.14 | -0.004 | 0.09 | 0.09 | 3.368 | 0.93 | 0.66 | 0.97 | 0.93 | 1 | 0.82 |
| 3 | 39475433 | rs6804150 | 1.04 | 0.464 | 1.029 | -1.189 | 0.001 | -0.006 | -0.006 | 0 | 0.14 | 0.08 | 0.1 | 0 | 0.15 | 0.03 |
| 3 | 39475596 | rs816513 | 1.003 | 0.437 | 1.355 | -1.106 | 0 | 0.103 | 0.103 | 3.144 | 0.93 | 0.64 | 0.97 | 0.93 | 1 | 0.81 |
| 3 | 39479881 | rs12495623 | -1.027 | 1.01 | 1.122 | -1.332 | -0.026 | 0.018 | 0.018 | 2.143 | 0.33 | 0.2 | 0.39 | 0.26 | 0.46 | 0.33 |
| 3 | 39480518 | rs1317871 | -0.212 | 1.063 | 1.097 | -1.423 | -0.019 | 0.024 | 0.024 | 1.331 | 0.17 | 0.09 | 0.22 | 0.04 | 0.25 | 0.19 |
| 3 | 39485162 | rs1278141 | 0.388 | 0.983 | 1.369 | -1.379 | 0.013 | 0.114 | 0.114 | 1.373 | 0.94 | 0.71 | 0.98 | 0.93 | 1 | 0.81 |
| 3 | 39489619 | rs630934 | 2.131 | 0.991 | 1.118 | -1.399 | -0.032 | 0.009 | 0.009 | 4.135 | 0.57 | 0.14 | 0.53 | 0.61 | 0.51 | 0.46 |
| 3 | 39489879 | rs6770798 | -1.362 | 1.123 | 0.942 | -1.382 | -0.025 | 0.018 | 0.018 | 2.055 | 0.33 | 0.21 | 0.39 | 0.28 | 0.46 | 0.33 |
| 3 | 39492594 | rs9852186 | -1.655 | 1.108 | 0.981 | -1.324 | 0.007 | -0.007 | -0.007 | 0 | 0.37 | 0.43 | 0.43 | 0.35 | 0.48 | 0.35 |
| 3 | 39493858 | rs9820623 | -1.628 | 1.109 | 0.968 | -1.342 | 0.007 | -0.007 | -0.007 | 0 | 0.37 | 0.43 | 0.43 | 0.35 | 0.48 | 0.35 |
| 3 | 39500200 | rs11709044 | -1.381 | 1.122 | 0.703 | -1.329 | -0.028 | 0.015 | 0.015 | 1.932 | 0.34 | 0.21 | 0.39 | 0.31 | 0.46 | 0.33 |
| 3 | 39502506 | rs1707981 | 2.149 | 1.302 | 0.23 | -1.311 | -0.092 | -0.079 | -0.079 | 2.239 | 0.4 | 0.03 | 0.42 | 0.59 | 0.35 | 0.42 |
| 3 | 39507945 | rs13092508 | -0.529 | 1.794 | 0.241 | -1.582 | -0.042 | 0.041 | 0.041 | 1.964 | 0.19 | 0.09 | 0.24 | 0.04 | 0.3 | 0.23 |
| 3 | 39509440 | rs1768190 | -2.145 | 1.226 | 0.27 | -1.487 | 0.007 | -0.006 | -0.006 | 0.178 | 0.61 | 0.93 | 0.6 | 0.38 | 0.69 | 0.6 |
| 3 | 39516485 | rs1768198 | -0.248 | 0.943 | 1.099 | -1.537 | -0.04 | -0.032 | -0.032 | 0 | 0.27 | 0.53 | 0.15 | 0.24 | 0.18 | 0.26 |
| 3 | 39521833 | rs1768206 | -0.031 | 0.931 | 1.322 | -1.548 | -0.015 | -0.023 | -0.023 | 0 | 0.24 | 0.33 | 0.13 | 0.19 | 0.18 | 0.09 |
| 3 | 39526874 | rs1464047 | -0.677 | 1.258 | 1.221 | -1.587 | -0.048 | 0.052 | 0.052 | 4.239 | 0.3 | 0.16 | 0.4 | 0.1 | 0.48 | 0.33 |
| 3 | 39530083 | rs545397 | -2.342 | 1.198 | 1.19 | -1.614 | -0.01 | -0.01 | -0.01 | 0 | 0.61 | 0.93 | 0.59 | 0.38 | 0.69 | 0.61 |
| 3 | 39534377 | rs500507 | -0.508 | 1.148 | 1.59 | -1.531 | -0.019 | -0.016 | -0.016 | 0 | 0.28 | 0.63 | 0.17 | 0.24 | 0.19 | 0.26 |
| 3 | 39535884 | rs535878 | -0.704 | 1.119 | 1.552 | -1.545 | 0.024 | 0.021 | 0.021 | 0.003 | 0.31 | 0.76 | 0.2 | 0.28 | 0.21 | 0.3 |
| 3 | 39536147 | rs2370969 | -0.572 | 1.17 | 1.478 | -1.501 | 0.026 | -0.039 | -0.039 | 0.397 | 0.27 | 0.33 | 0.43 | 0.13 | 0.42 | 0.27 |
| 3 | 39536524 | rs562545 | -1.988 | 0.635 | 1.764 | -1.073 | -0.078 | 0 | 0 | 0.169 | 0.58 | 0.96 | 0.52 | 0.38 | 0.58 | 0.59 |
| 3 | 39539526 | rs614359 | -0.603 | 0.225 | 2.462 | -0.894 | -0.03 | -0.017 | -0.017 | 0 | 0.26 | 0.49 | 0.22 | 0.26 | 0.19 | 0.3 |
| 3 | 39540727 | rs1417147 | -0.653 | 0.224 | 2.48 | -0.912 | -0.035 | -0.01 | -0.01 | 0.052 | 0.26 | 0.52 | 0.24 | 0.31 | 0.2 | 0.31 |
| 3 | 39544943 | rs17756489 | 1.632 | 0.299 | 2.14 | -0.8 | -0.048 | 0.023 | 0.023 | 0 | 0.45 | 0.08 | 0.48 | 0.6 | 0.44 | 0.43 |
| 3 | 39548420 | rs1768254 | -1.067 | -0.394 | 3.244 | -0.084 | 0.004 | 0.006 | 0.006 | 0 | 0.22 | 0.23 | 0.21 | 0.22 | 0.2 | 0.11 |
| 3 | 39554946 | rs552724 | -1.067 | -0.394 | 3.244 | -0.084 | 0.004 | 0.006 | 0.006 | 0 | 0.22 | 0.23 | 0.21 | 0.22 | 0.2 | 0.11 |
| 3 | 39555497 | rs816491 | -1.383 | -0.386 | 2.994 | -0.081 | -0.018 | -0.019 | -0.019 | 3.083 | 0.25 | 0.64 | 0.25 | 0.26 | 0.2 | 0.12 |
| 3 | 39556155 | rs816492 | -1.373 | -0.372 | 3 | -0.074 | -0.018 | -0.019 | -0.019 | 0.148 | 0.24 | 0.64 | 0.25 | 0.26 | 0.2 | 0.12 |
| 3 | 39556561 | rs1068953 | -1.115 | -0.368 | 2.918 | -0.073 | -0.018 | -0.026 | -0.026 | 0 | 0.23 | 0.43 | 0.23 | 0.23 | 0.2 | 0.11 |
| 3 | 39557120 | rs704962 | -1.428 | -0.371 | 2.924 | -0.075 | -0.017 | -0.018 | -0.018 | 0 | 0.26 | 0.67 | 0.26 | 0.26 | 0.21 | 0.14 |
| 3 | 39557238 | rs704963 | -1.218 | -0.108 | 2.169 | -0.026 | -0.018 | -0.026 | -0.026 | 0 | 0.23 | 0.43 | 0.23 | 0.23 | 0.2 | 0.11 |
| 3 | 39561389 | rs661422 | -0.655 | -0.112 | 2.132 | -0.033 | -0.003 | 0.011 | 0.011 | 0 | 0.07 | 0.09 | 0.1 | 0.21 | 0.03 | 0.06 |
| 3 | 39573800 | rs1615046 | -1.191 | -0.072 | 1.773 | -0.01 | -0.018 | -0.025 | -0.025 | 0 | 0.23 | 0.42 | 0.23 | 0.22 | 0.2 | 0.09 |
| 3 | 39590195 | rs1631776 | -1.302 | 0.075 | 0.806 | 0.017 | -0.033 | -0.039 | -0.039 | 0 | 0.28 | 0.52 | 0.25 | 0.23 | 0.21 | 0.1 |
| 3 | 39590291 | rs1768141 | -1.277 | 0.431 | 0.358 | 0.743 | -0.006 | 0.015 | 0.015 | 0 | 0.08 | 0.11 | 0.1 | 0.22 | 0.04 | 0.06 |
| 3 | 39609791 | rs845865 | 1.012 | 0.716 | 0.061 | 0.294 | -0.01 | 0.005 | 0.005 | 0 | 0.88 | 0.92 | 0.85 | 0.88 | 0.86 | 0.83 |
| 3 | 39613992 | rs816504 | 1.769 | 0.654 | 1.349 | 0.245 | 0.038 | 0.009 | 0.009 | 0 | 0.86 | 0.49 | 0.8 | 0.85 | 0.85 | 0.81 |
| 3 | 39630765 | rs6789275 | 1.949 | 0.587 | 2.353 | 0.415 | 0.009 | 0.011 | 0.011 | 1.368 | 0.86 | 0.33 | 0.78 | 0.81 | 0.86 | 0.83 |
| 3 | 39639754 | rs11129839 | -0.749 | 0.299 | 1.596 | 0.698 | 0.001 | -0.077 | -0.077 | 0 | 0.12 | 0.02 | 0.12 | 0.59 | 0.11 | 0.27 |
| 3 | 39640995 | rs2137562 | 1.583 | 0.123 | 2.198 | -0.308 | -0.01 | -0.079 | -0.079 | 0.3 | 0.71 | 0.47 | 0.79 | 0.86 | 0.79 | 0.76 |
| 3 | 39649777 | rs9872576 | 1.534 | 0.137 | 2.149 | -0.302 | -0.01 | -0.099 | -0.099 | 0 | 0.72 | 0.49 | 0.79 | 0.86 | 0.79 | 0.76 |
| 3 | 39654502 | rs4676503 | 1.139 | 0.128 | 2.003 | -0.302 | -0.001 | 0.054 | 0.054 | 0 | 0.76 | 0.87 | 0.83 | 0.92 | 0.81 | 0.84 |
| 3 | 39668202 | rs13091037 | 2.059 | -0.016 | 1.901 | -0.318 | 0.003 | -0.001 | -0.001 | 0.39 | 0.66 | 0.33 | 0.74 | 0.86 | 0.75 | 0.75 |
| 3 | 39670988 | rs9872714 | 0.961 | 0.108 | 1.556 | -0.268 | 0.01 | -0.041 | -0.041 | 0 | 0.93 | 0.83 | 0.94 | 1 | 0.95 | 0.98 |
| 3 | 39687488 | rs6776434 | 1.625 | -0.012 | 1.442 | -0.316 | -0.015 | -0.087 | -0.087 | 0.196 | 0.68 | 0.52 | 0.76 | 0.88 | 0.76 | 0.76 |
| 3 | 39694769 | rs1405796 | -0.711 | -0.029 | 0.872 | -0.314 | -0.034 | 0.028 | 0.028 | 0 | 0.1 | 0.01 | 0.13 | 0.06 | 0.12 | 0.13 |
| 3 | 39715764 | rs4314107 | 1.303 | 0.001 | 0.992 | -0.233 | -0.012 | -0.062 | -0.062 | 0 | 0.67 | 0.42 | 0.75 | 0.88 | 0.75 | 0.78 |
| 3 | 39749593 | rs9832461 | -1.296 | 0.739 | 1.075 | -0.077 | 0.028 | -0.052 | -0.052 | 1.353 | 0.21 | 0.38 | 0.18 | 0.13 | 0.24 | 0.21 |
| 3 | 39763962 | rs6778426 | -1.798 | 0.532 | 1.708 | -0.079 | 0 | 0.009 | 0.009 | 2.765 | 0.26 | 0.75 | 0.22 | 0.2 | 0.26 | 0.28 |
| 3 | 39776225 | rs17076570 | -0.811 | 0.497 | 2.136 | -0.12 | 0.004 | -0.013 | -0.013 | 1.402 | 0.13 | 0.15 | 0.13 | 0.05 | 0.2 | 0.15 |
| 3 | 39786363 | rs9849166 | -2.249 | 0.462 | 2.379 | -0.13 | -0.001 | -0.007 | -0.007 | 4.599 | 0.26 | 0.72 | 0.22 | 0.2 | 0.26 | 0.28 |
| 3 | 39791421 | rs9824054 | -2.251 | 0.432 | 2.253 | -0.112 | -0.002 | -0.022 | -0.022 | 0.019 | 0.26 | 0.63 | 0.21 | 0.2 | 0.25 | 0.25 |
| 3 | 39801185 | rs7632048 | -1.188 | 0.339 | 1.658 | -0.314 | -0.002 | -0.01 | -0.01 | 0 | 0.16 | 0.16 | 0.13 | 0.04 | 0.21 | 0.16 |

| | | | | | | | | | | | | | | | | |
|---|---|---|---|---|---|---|---|---|---|---|---|---|---|---|---|---|
| 3 | 39809582 | rs4676512 | -0.674 | -0.1 | 0.52 | -0.51 | -0.001 | 0.018 | 0.018 | 0 | 0.63 | 0.84 | 0.57 | 0.27 | 0.66 | 0.52 |
| 3 | 39809692 | rs12107380 | -0.765 | 0.377 | 0.807 | -0.578 | 0.003 | 0.001 | 0.001 | 0.019 | 0.16 | 0.19 | 0.09 | 0.18 | 0.18 | 0.22 |
| 3 | 39809870 | rs12108156 | -1.235 | 0.658 | 1.26 | -0.602 | 0.066 | 0.041 | 0.041 | 1.713 | 0.19 | 0.52 | 0.14 | 0.21 | 0.23 | 0.23 |
| 3 | 39812292 | rs4676517 | -0.963 | 0.549 | 1.126 | -0.374 | -0.003 | -0.008 | -0.008 | 0.074 | 0.64 | 0.84 | 0.57 | 0.31 | 0.67 | 0.52 |
| 3 | 39814559 | rs1918022 | -0.922 | 0.553 | 1.101 | -0.466 | 0.002 | 0.015 | 0.015 | 0 | 0.64 | 0.82 | 0.57 | 0.27 | 0.67 | 0.52 |
| 3 | 39817494 | rs1880787 | 0.234 | 0.74 | 0.868 | -0.879 | -0.011 | -0.015 | -0.015 | 0 | 0.14 | 0.2 | 0.1 | 0.05 | 0.08 | 0.08 |
| 3 | 39823517 | rs883035 | -0.144 | 0.749 | 0.635 | -0.88 | -0.026 | -0.02 | -0.02 | 0.173 | 0.12 | 0.08 | 0.08 | 0.18 | 0.18 | 0.22 |
| 3 | 39828262 | rs4676449 | 0.894 | 0.533 | 0.84 | -0.772 | -0.004 | 0.1 | 0.1 | 0 | 0.31 | 0.13 | 0.33 | 0.01 | 0.34 | 0.15 |
| 3 | 39837709 | rs13086503 | -0.638 | 0.867 | 0.43 | -0.559 | -0.002 | 0.014 | 0.014 | 0 | 0.54 | 0.58 | 0.5 | 0.24 | 0.52 | 0.47 |
| 3 | 39839388 | rs12493939 | 0.014 | 0.906 | -0.109 | -0.689 | 0.01 | 0.033 | 0.033 | 0.001 | 0.46 | 0.37 | 0.44 | 0.06 | 0.44 | 0.29 |
| 3 | 39844703 | rs9842160 | -1.152 | 1.285 | -0.516 | 0.13 | 0.023 | 0.022 | 0.022 | 0.059 | 0.07 | 0.13 | 0.1 | 0.17 | 0.14 | 0.29 |
| 3 | 39846240 | rs9848464 | 0.028 | 0.591 | -1.158 | 0.478 | -0.004 | 0.002 | 0.002 | 0 | 0.46 | 0.27 | 0.46 | 0.44 | 0.5 | 0.35 |
| 3 | 39847606 | rs9858242 | -0.051 | 0.79 | -1.606 | 0.631 | -0.05 | -0.023 | -0.023 | 0.041 | 0.84 | 0.99 | 0.9 | 0.88 | 0.8 | 0.85 |
| 3 | 39848341 | rs1918029 | 0.465 | 0.26 | -1.717 | 0.695 | -0.03 | 0.065 | 0.065 | 0 | 0.55 | 0.76 | 0.53 | 0.32 | 0.55 | 0.53 |
| 3 | 39849382 | rs9864768 | 0.923 | 0.195 | -1.536 | 0.401 | -0.001 | -0.007 | -0.007 | 0 | 0.18 | 0.22 | 0.16 | 0.08 | 0.18 | 0.32 |
| 3 | 39850446 | rs2049625 | 0.905 | 0.386 | -1.541 | 0.393 | -0.002 | 0.012 | 0.012 | 0 | 0.19 | 0.17 | 0.17 | 0.08 | 0.23 | 0.33 |
| 3 | 39852825 | rs871231 | -0.278 | 0.239 | -0.614 | 0.867 | -0.007 | 0.047 | 0.047 | 0 | 0.3 | 0.52 | 0.28 | 0.42 | 0.32 | 0.48 |
| 3 | 39862052 | rs9867310 | 1.006 | 0.24 | -0.643 | 0.899 | 0 | -0.002 | -0.002 | 0 | 0.19 | 0.23 | 0.19 | 0.14 | 0.23 | 0.33 |
| 3 | 39873484 | rs1918025 | 1.74 | 0.146 | -0.236 | 0.77 | 0.001 | 0.066 | 0.066 | 0 | 0.32 | 0.16 | 0.31 | 0.15 | 0.31 | 0.16 |
| 3 | 39873605 | rs17050883 | 1.568 | 0.142 | -0.168 | 0.785 | 0 | -0.012 | -0.012 | 0 | 0.07 | 0.14 | 0.05 | 0.03 | 0.07 | 0.08 |
| 3 | 39875648 | rs9821211 | 0.684 | 0.134 | 0.111 | 0.873 | -0.002 | 0.005 | 0.005 | 0 | 0.21 | 0.31 | 0.22 | 0.24 | 0.25 | 0.37 |
| 3 | 39885937 | rs2371091 | 0.248 | 0.223 | 0.041 | 0.899 | -0.001 | -0.008 | -0.008 | 1.59 | 0.27 | 0.37 | 0.24 | 0.18 | 0.3 | 0.42 |
| 3 | 39888305 | rs6786065 | -0.447 | 0.221 | 0.119 | 0.923 | -0.008 | -0.017 | -0.017 | 0 | 0.63 | 0.75 | 0.59 | 0.57 | 0.64 | 0.64 |
| 3 | 39889596 | rs930917 | -0.31 | 0.305 | 0.347 | 1.019 | -0.003 | -0.028 | -0.028 | 2.333 | 0.29 | 0.46 | 0.26 | 0.21 | 0.31 | 0.45 |
| 3 | 39898228 | rs11129852 | 0.415 | 0.06 | 0.396 | 0.683 | 0.028 | 0.008 | 0.008 | 0.029 | 0.53 | 0.75 | 0.46 | 0.44 | 0.42 | 0.46 |
| 3 | 39902642 | rs9832990 | 1.262 | -0.045 | 0.7 | 0.739 | 0.025 | 0.052 | 0.052 | 0.245 | 0.44 | 0.28 | 0.37 | 0.21 | 0.35 | 0.36 |
| 3 | 39907375 | rs4676456 | -0.762 | 0.058 | 0.74 | 0.875 | -0.046 | -0.036 | -0.036 | 0 | 0.65 | 0.8 | 0.55 | 0.58 | 0.59 | 0.64 |
| 3 | 39908845 | rs7617131 | -0.372 | 0.048 | 0.71 | 0.85 | -0.026 | -0.021 | -0.021 | 0 | 0.63 | 0.68 | 0.51 | 0.51 | 0.54 | 0.62 |
| 3 | 39909607 | rs7612386 | 1.195 | 0.013 | 0.641 | 0.826 | 0.015 | 0.026 | 0.026 | 0.101 | 0.46 | 0.37 | 0.39 | 0.21 | 0.35 | 0.37 |
| 3 | 39924901 | rs1598990 | 1.642 | 0.134 | 0.292 | 1.013 | -0.037 | -0.035 | -0.035 | 0 | 0.08 | 0 | 0.05 | 0.1 | 0.11 | 0.12 |
| 3 | 39931414 | rs13079551 | 1.914 | -0.17 | 0.511 | 0.669 | 0.021 | 0.051 | 0.051 | 0.209 | 0.31 | 0.06 | 0.29 | 0.19 | 0.26 | 0.18 |
| 3 | 39931515 | rs9827374 | -1.328 | -0.064 | 0.656 | 0.761 | 0.048 | 0.21 | 0.21 | 4.371 | 0.08 | 0.36 | 0.07 | 0.23 | 0.07 | 0.1 |
| 3 | 39938182 | rs12489300 | 1.642 | -0.347 | 0.308 | 0.6 | 0.021 | 0.032 | 0.032 | 0.048 | 0.33 | 0.17 | 0.31 | 0.2 | 0.26 | 0.18 |
| 3 | 39940997 | rs13060185 | 0.681 | -0.321 | 0.349 | 0.547 | 0.018 | 0.066 | 0.066 | 0.375 | 0.45 | 0.22 | 0.4 | 0.28 | 0.43 | 0.35 |
| 3 | 39943731 | rs11717059 | 0.582 | -0.333 | 0.358 | 0.58 | -0.003 | 0.004 | 0.004 | 0 | 0.13 | 0.12 | 0.09 | 0.09 | 0.17 | 0.17 |
| 3 | 39944942 | rs9860916 | -0.081 | -0.368 | 0.286 | 0.563 | 0.031 | 0.002 | 0.002 | 0.42 | 0.52 | 0.71 | 0.45 | 0.49 | 0.44 | 0.43 |
| 3 | 39947936 | rs7618607 | -0.039 | -0.37 | 0.268 | 0.562 | 0.034 | 0.004 | 0.004 | 0.72 | 0.52 | 0.71 | 0.45 | 0.49 | 0.44 | 0.41 |
| 3 | 39954073 | rs13086725 | 1.764 | -0.283 | 0.116 | 0.608 | 0.028 | 0.049 | 0.049 | 0.165 | 0.32 | 0.1 | 0.31 | 0.19 | 0.26 | 0.16 |
| 3 | 39954674 | rs1599902 | 0.462 | -0.212 | 0.154 | 0.743 | 0.021 | 0.009 | 0.009 | 0.218 | 0.5 | 0.53 | 0.44 | 0.49 | 0.44 | 0.37 |
| 3 | 39961966 | rs13060371 | 0.646 | -0.425 | -0.198 | 0.683 | -0.005 | -0.003 | -0.003 | 2.066 | 0.47 | 0.46 | 0.42 | 0.41 | 0.39 | 0.24 |
| 3 | 39963677 | rs6599061 | -1.038 | -0.229 | -0.694 | 0.615 | -0.126 | -0.157 | -0.157 | 0 | 0.75 | 0.98 | 0.58 | 0.59 | 0.67 | 0.68 |
| 3 | 39964074 | rs6776086 | 0.554 | -0.489 | -0.204 | 0.571 | -0.006 | -0.004 | -0.004 | 0 | 0.46 | 0.44 | 0.42 | 0.41 | 0.39 | 0.24 |
| 3 | 39966748 | rs11717034 | 1.173 | -0.622 | -0.12 | 0.599 | 0.015 | 0.1 | 0.1 | 1.268 | 0.38 | 0.12 | 0.35 | 0.18 | 0.36 | 0.15 |
| 3 | 39975119 | rs1585286 | 0.538 | -0.521 | -0.292 | 0.612 | -0.006 | -0.004 | -0.004 | 0.352 | 0.46 | 0.44 | 0.42 | 0.41 | 0.39 | 0.24 |
| 3 | 39975352 | rs9815380 | 0.542 | -0.518 | -0.291 | 0.62 | -0.007 | -0.004 | -0.004 | 0.503 | 0.46 | 0.44 | 0.42 | 0.41 | 0.39 | 0.24 |
| 3 | 39985393 | rs1598997 | -1.238 | 0.288 | 0.556 | 1.237 | -0.021 | 0.066 | 0.066 | 0.411 | 0.16 | 0.44 | 0.1 | 0.23 | 0.1 | 0.13 |
| 3 | 40003159 | rs17067271 | -0.722 | 0.881 | 0.581 | 1.369 | 0.001 | 0.002 | 0.002 | 0 | 0.19 | 0.2 | 0.13 | 0.26 | 0.22 | 0.33 |
| 3 | 40003898 | rs17183401 | 1.241 | 0.673 | 0.625 | 1.324 | -0.014 | 0.004 | 0.004 | 0.403 | 0.25 | 0.12 | 0.34 | 0.22 | 0.33 | 0.16 |
| 3 | 40004366 | rs1304138 | -0.61 | 0.299 | 0.687 | 1.407 | 0.041 | -0.016 | -0.016 | 1.647 | 0.45 | 0.81 | 0.44 | 0.46 | 0.38 | 0.41 |
| 3 | 40006045 | rs9820000 | 0.978 | -0.202 | 0.811 | 0.876 | -0.027 | -0.001 | -0.001 | 0 | 0.07 | 0.18 | 0.03 | 0.08 | 0.02 | 0.14 |
| 3 | 40008634 | rs9985399 | -1.198 | -0.219 | 0.702 | 0.874 | -0.09 | 0.042 | 0.042 | 0.343 | 0.19 | 0.55 | 0.08 | 0.26 | 0.06 | 0.25 |
| 3 | 40010608 | rs4549224 | -1.646 | 0.116 | 0.503 | 0.887 | -0.04 | -0.131 | -0.131 | 0 | 0.61 | 0.98 | 0.57 | 0.48 | 0.67 | 0.62 |
| 3 | 40014166 | rs9862625 | -1.17 | -0.047 | 1.092 | 0.747 | -0.066 | 0.063 | 0.063 | 0.167 | 0.16 | 0.57 | 0.08 | 0.25 | 0.06 | 0.25 |
| 3 | 40014390 | rs1585287 | -0.912 | 0.066 | 1.017 | 0.736 | 0 | 0.073 | 0.073 | 0.603 | 0.43 | 0.81 | 0.44 | 0.31 | 0.42 | 0.41 |
| 3 | 40023160 | rs7652142 | -0.609 | 0.057 | 1.047 | 0.604 | 0.001 | 0.048 | 0.048 | 0.089 | 0.41 | 0.66 | 0.42 | 0.3 | 0.42 | 0.41 |
| 3 | 40032958 | rs9851674 | 0.404 | -0.047 | 1.229 | 0.668 | -0.022 | 0.027 | 0.027 | 0.103 | 0.08 | 0.3 | 0.03 | 0.11 | 0.02 | 0.14 |
| 3 | 40033989 | rs11924220 | -0.998 | 0.606 | 0.886 | 0.724 | -0.014 | 0.066 | 0.066 | 0.224 | 0.09 | 0.3 | 0.06 | 0.13 | 0.04 | 0.06 |
| 3 | 40042376 | rs7611605 | -0.904 | 0.584 | 0.667 | 0.733 | -0.021 | 0.055 | 0.055 | 0.116 | 0.09 | 0.26 | 0.06 | 0.13 | 0.04 | 0.06 |
| 3 | 40050275 | rs6791790 | -0.904 | 0.584 | 0.667 | 0.733 | -0.021 | 0.055 | 0.055 | 0.116 | 0.09 | 0.26 | 0.06 | 0.14 | 0.04 | 0.06 |
| 3 | 40072811 | rs13092608 | 0.801 | 0.61 | 0.603 | 0.678 | -0.025 | 0.027 | 0.027 | 0.097 | 0.25 | 0.12 | 0.35 | 0.06 | 0.37 | 0.16 |
| 3 | 40080544 | rs7632391 | -0.347 | 0.608 | 0.39 | 0.717 | 0.004 | -0.015 | -0.015 | 0 | 0.34 | 0.34 | 0.4 | 0.2 | 0.41 | 0.22 |
| 3 | 40081119 | rs11707920 | 0.524 | 0.928 | 0.001 | 0.413 | -0.011 | 0.003 | 0.003 | 0 | 0.26 | 0.19 | 0.36 | 0.06 | 0.37 | 0.16 |
| 3 | 40086208 | rs9873761 | 0.95 | -0.717 | 0.134 | 1.002 | 0.049 | 0.016 | 0.016 | 0 | 0.42 | 0.21 | 0.28 | 0.37 | 0.26 | 0.3 |
| 3 | 40087181 | rs13326962 | 0.241 | -0.782 | 0.489 | 1.333 | -0.003 | -0.001 | -0.001 | 13.518 | 0.54 | 0.39 | 0.4 | 0.57 | 0.34 | 0.43 |
| 3 | 40088077 | rs2280319 | -0.266 | -0.791 | 0.514 | 1.307 | 0.005 | -0.011 | -0.011 | 0 | 0.1 | 0.07 | 0.11 | 0.19 | 0.08 | 0.09 |
| 3 | 40094795 | rs11920120 | 1.052 | -0.813 | 0.523 | 1.313 | 0.009 | 0.004 | 0.004 | 0 | 0.43 | 0.31 | 0.29 | 0.37 | 0.26 | 0.3 |
| 3 | 40096618 | rs6599077 | 1.06 | -0.811 | 0.524 | 1.314 | 0.008 | 0.004 | 0.004 | 0 | 0.43 | 0.31 | 0.29 | 0.37 | 0.26 | 0.3 |
| 3 | 40101795 | rs2887953 | -0.571 | -0.559 | 0.877 | 1.498 | 0.024 | -0.013 | -0.013 | 0.373 | 0.68 | 0.57 | 0.53 | 0.76 | 0.51 | 0.63 |
| 3 | 40105341 | rs4293655 | 0.45 | -0.585 | 1.043 | 1.493 | -0.009 | 0 | 0 | 0 | 0.54 | 0.44 | 0.4 | 0.56 | 0.34 | 0.44 |

| | | | | | | | | | | | | | | | | |
|---|---|---|---|---|---|---|---|---|---|---|---|---|---|---|---|---|
| 3 | 40107494 | rs17075172 | -0.154 | -0.578 | 1.03 | 1.496 | -0.004 | 0.007 | 0.007 | 0 | 0.1 | 0.12 | 0.12 | 0.19 | 0.08 | 0.09 |
| 3 | 40108336 | rs2371181 | -0.533 | -0.558 | 1.079 | 1.482 | 0.011 | -0.006 | -0.006 | 0 | 0.67 | 0.59 | 0.52 | 0.76 | 0.51 | 0.6 |
| 3 | 40109637 | rs17075188 | -1.455 | -0.619 | 0.96 | 1.53 | -0.001 | 0 | 0 | 0.507 | 0.24 | 0.28 | 0.24 | 0.4 | 0.25 | 0.3 |
| 3 | 40112540 | rs6786793 | 0.027 | -0.548 | 0.902 | 1.501 | -0.008 | -0.009 | -0.009 | 0.031 | 0.13 | 0.14 | 0.12 | 0.2 | 0.17 | 0.2 |
| 3 | 40113022 | rs10865893 | 1.498 | -0.583 | 1.517 | 1.506 | 0.004 | 0.045 | 0.045 | 0 | 0.73 | 0.45 | 0.73 | 0.6 | 0.75 | 0.7 |
| 3 | 40122623 | rs6803028 | 0.304 | -0.734 | 1.576 | 1.447 | 0.009 | -0.006 | -0.006 | 0 | 0.33 | 0.37 | 0.52 | 0.23 | 0.51 | 0.37 |
| 3 | 40129206 | rs9838815 | -0.672 | -0.816 | 1.647 | 1.472 | -0.002 | -0.001 | -0.001 | 0 | 0.44 | 0.49 | 0.6 | 0.43 | 0.58 | 0.49 |
| 3 | 40131246 | rs17075573 | -0.266 | -0.814 | 1.549 | 1.474 | 0 | -0.001 | -0.001 | 2.991 | 0.11 | 0.12 | 0.09 | 0.2 | 0.08 | 0.09 |
| 3 | 40135213 | rs4234131 | 1.482 | -0.832 | 1.595 | 1.459 | 0.093 | 0.011 | 0.011 | 0 | 0.42 | 0.14 | 0.26 | 0.37 | 0.25 | 0.3 |
| 3 | 40143794 | rs1799429 | 0.573 | -0.752 | 1.559 | 1.453 | -0.012 | 0.005 | 0.005 | 0 | 0.31 | 0.27 | 0.55 | 0.24 | 0.5 | 0.36 |
| 3 | 40149458 | rs1799414 | -0.179 | -0.767 | 1.436 | 1.435 | 0.011 | 0.012 | 0.012 | 0 | 0.87 | 0.75 | 0.87 | 0.8 | 0.83 | 0.82 |
| 3 | 40159756 | rs4530464 | -0.407 | -0.807 | 1.27 | 1.442 | 0.001 | -0.001 | -0.001 | 0.044 | 0.42 | 0.4 | 0.64 | 0.43 | 0.58 | 0.45 |
| 3 | 40165048 | rs939075 | 0.447 | -0.435 | 1.326 | 1.277 | 0.051 | 0.035 | 0.035 | 0 | 0.84 | 0.49 | 0.84 | 0.81 | 0.82 | 0.77 |
| 3 | 40174858 | rs9878321 | 0.855 | -0.8 | 1.171 | 1.631 | -0.004 | 0.001 | 0.001 | 0 | 0.51 | 0.46 | 0.34 | 0.46 | 0.27 | 0.58 |
| 3 | 40185635 | rs4132209 | 0.666 | -0.536 | 0.758 | 1.626 | -0.108 | 0.05 | 0.05 | 0.006 | 0.28 | 0.1 | 0.46 | 0.1 | 0.48 | 0.19 |
| 3 | 40188011 | rs1799428 | 1.617 | -0.75 | 1.058 | 1.634 | 0.003 | -0.028 | -0.028 | 0 | 0.7 | 0.27 | 0.73 | 0.43 | 0.74 | 0.56 |
| 3 | 40195206 | rs2679807 | -0.129 | -0.736 | 1.476 | 1.837 | 0.018 | -0.004 | -0.004 | 0 | 0.34 | 0.4 | 0.56 | 0.25 | 0.52 | 0.35 |
| 3 | 40202976 | rs1714412 | 0.394 | -0.821 | 1.385 | 2.011 | 0.018 | 0.045 | 0.045 | 0 | 0.83 | 0.67 | 0.83 | 0.72 | 0.8 | 0.8 |
| 3 | 40204178 | rs1714414 | -0.405 | -0.491 | 1.482 | 2.003 | 0.023 | -0.001 | -0.001 | 0.676 | 0.34 | 0.41 | 0.55 | 0.27 | 0.51 | 0.37 |
| 3 | 40214049 | rs2887961 | 0.056 | -0.251 | 0.724 | 1.721 | -0.005 | -0.004 | -0.004 | 0 | 0.68 | 0.34 | 0.45 | 0.68 | 0.49 | 0.73 |
| 3 | 40219446 | rs2255401 | 1.55 | -0.675 | 0.957 | 1.786 | -0.002 | 0.008 | 0.008 | 1.454 | 0.75 | 0.3 | 0.82 | 0.51 | 0.78 | 0.63 |
| 3 | 40226685 | rs2371176 | 1.734 | -0.8 | 1.989 | 1.54 | 0.035 | 0.12 | 0.12 | 0 | 0.9 | 0.61 | 0.9 | 0.8 | 0.94 | 0.85 |
| 3 | 40230406 | rs6769485 | 1.569 | -0.919 | 1.833 | 1.515 | -0.006 | 0.008 | 0.008 | 0 | 0.89 | 0.87 | 0.94 | 0.76 | 0.93 | 0.77 |
| 3 | 40231883 | rs1317317 | 1.989 | -0.7 | 2.2 | 1.494 | 0.018 | 0.051 | 0.051 | 0 | 0.89 | 0.75 | 0.92 | 0.8 | 0.9 | 0.84 |
| 3 | 40234083 | rs2261542 | 1.737 | -0.643 | 2.094 | 1.519 | 0.022 | 0.051 | 0.051 | 0 | 0.91 | 0.8 | 0.92 | 0.8 | 0.91 | 0.85 |
| 3 | 40235522 | rs2887962 | 0.762 | -0.644 | 2.086 | 1.52 | 0.003 | 0.035 | 0.035 | 0 | 0.93 | 0.89 | 0.94 | 0.8 | 0.94 | 0.86 |
| 3 | 40236723 | rs9311235 | 1.701 | -0.658 | 2.057 | 1.514 | 0.02 | 0.048 | 0.048 | 0 | 0.91 | 0.8 | 0.92 | 0.8 | 0.91 | 0.85 |
| 3 | 40237563 | rs2256927 | 2.156 | -0.661 | 2.053 | 1.512 | 0.031 | 0.075 | 0.075 | 0 | 0.88 | 0.51 | 0.89 | 0.8 | 0.9 | 0.85 |
| 3 | 40249244 | rs1344189 | -0.08 | -1.318 | 1.638 | 1.36 | -0.005 | -0.002 | -0.002 | 0.65 | 0.32 | 0.28 | 0.52 | 0.26 | 0.55 | 0.3 |
| 3 | 40257780 | rs1714408 | 0.034 | -1.325 | 1.627 | 1.352 | -0.119 | -0.077 | -0.077 | 1.239 | 0.3 | 0.06 | 0.4 | 0.26 | 0.55 | 0.3 |
| 3 | 40267321 | rs2679822 | 2.047 | -1.429 | 1.988 | 1.298 | -0.01 | -0.003 | -0.003 | 0 | 0.86 | 0.39 | 0.89 | 0.87 | 0.9 | 0.83 |
| 3 | 40268675 | rs9823575 | 0.109 | -1.596 | 2.103 | 1.208 | -0.025 | -0.019 | -0.019 | 0.078 | 0.06 | 0.12 | 0.02 | 0.04 | 0.01 | 0.07 |
| 3 | 40271697 | rs1317217 | 1.366 | -1.603 | 2.029 | 1.209 | 0.129 | -0.001 | -0.001 | 0.168 | 0.5 | 0.12 | 0.35 | 0.58 | 0.35 | 0.46 |
| 3 | 40275655 | rs2371136 | -1.272 | -1.57 | 2.085 | 1.203 | 0.016 | 0.043 | 0.043 | 0 | 0.05 | 0.2 | 0.05 | 0.08 | 0.04 | 0.1 |
| 3 | 40283255 | rs1880763 | 1.328 | -1.592 | 1.871 | 1.207 | 0.125 | 0 | 0 | 0.038 | 0.5 | 0.15 | 0.36 | 0.58 | 0.35 | 0.46 |
| 3 | 40285160 | rs7639223 | -0.036 | -1.77 | 1.899 | 1.26 | -0.006 | 0.004 | 0.004 | 0.189 | 0.65 | 0.75 | 0.47 | 0.76 | 0.43 | 0.69 |
| 3 | 40286017 | rs4618168 | -0.036 | -1.77 | 2.064 | 1.26 | -0.018 | 0.004 | 0.004 | 0.189 | 0.65 | 0.8 | 0.47 | 0.76 | 0.43 | 0.69 |
| 3 | 40293797 | rs7431934 | -0.174 | -1.776 | 1.976 | 1.28 | -0.002 | 0.002 | 0.002 | 0.232 | 0.35 | 0.31 | 0.54 | 0.24 | 0.58 | 0.31 |
| 3 | 40300405 | rs7625538 | -1.768 | -1.757 | 2.2 | 1.5 | -0.013 | 0.04 | 0.04 | 2.548 | 0.13 | 0.42 | 0.07 | 0.13 | 0.06 | 0.15 |
| 3 | 40301059 | rs9866106 | -0.569 | -1.758 | 2.257 | 1.483 | -0.029 | -0.003 | -0.003 | 0 | 0.07 | 0.18 | 0.02 | 0.06 | 0.01 | 0.08 |
| 3 | 40301178 | rs9883899 | -0.612 | -1.758 | 2.257 | 1.483 | -0.03 | -0.004 | -0.004 | 0 | 0.07 | 0.18 | 0.02 | 0.06 | 0.01 | 0.08 |
| 3 | 40309225 | rs9861194 | 1.192 | -2.062 | 1.994 | 1.334 | 0.11 | 0.001 | 0.001 | 0 | 0.49 | 0.21 | 0.35 | 0.6 | 0.34 | 0.44 |
| 3 | 40311325 | rs4974028 | 1.118 | -2.066 | 1.859 | 1.328 | -0.008 | 0.019 | 0.019 | 0 | 0.9 | 0.55 | 0.92 | 0.92 | 0.95 | 0.84 |
| 3 | 40326165 | rs11129874 | 0.104 | -2.073 | 1.746 | 1.324 | -0.086 | -0.052 | -0.052 | 0 | 0.34 | 0.07 | 0.52 | 0.25 | 0.56 | 0.33 |
| 3 | 40335292 | rs12633295 | -0.245 | -1.999 | 1.998 | 1.505 | 0.018 | 0.002 | 0.002 | 0.076 | 0.42 | 0.39 | 0.6 | 0.33 | 0.63 | 0.44 |
| 3 | 40342347 | rs9821440 | 0.405 | -1.972 | 1.847 | 1.487 | -0.014 | -0.005 | -0.005 | 0 | 0.55 | 0.47 | 0.38 | 0.67 | 0.36 | 0.54 |
| 3 | 40359977 | rs17078141 | 1.192 | -1.627 | 1.077 | 1.423 | 0.005 | -0.111 | -0.111 | 0.913 | 0.44 | 0.03 | 0.33 | 0.57 | 0.37 | 0.42 |
| 3 | 40362122 | rs11718621 | -0.107 | -2.144 | 1.408 | 1.637 | 0.01 | 0.01 | 0.01 | 0.288 | 0.4 | 0.13 | 0.57 | 0.24 | 0.61 | 0.35 |
| 3 | 40374551 | rs4974067 | 0.822 | -2.329 | 1.842 | 1.587 | 0.102 | 0 | 0 | 0.119 | 0.49 | 0.24 | 0.32 | 0.58 | 0.3 | 0.48 |
| 3 | 40378738 | rs2123999 | -0.267 | -2.318 | 1.809 | 1.593 | 0.092 | -0.059 | -0.059 | 0.987 | 0.43 | 0.15 | 0.62 | 0.39 | 0.68 | 0.46 |
| 3 | 40404770 | rs10510708 | 0.279 | -2.282 | 1.913 | 1.605 | 0.09 | -0.004 | -0.004 | 0.708 | 0.55 | 0.71 | 0.36 | 0.61 | 0.32 | 0.54 |
| 3 | 40424093 | rs9835219 | 1.032 | -2.272 | 2.027 | 1.657 | 0.004 | -0.001 | -0.001 | 0 | 0.45 | 0.42 | 0.32 | 0.52 | 0.3 | 0.38 |
| 3 | 40429183 | rs2305521 | 0.392 | -2.281 | 2.139 | 1.659 | 0.057 | -0.006 | -0.006 | 0 | 0.53 | 0.65 | 0.35 | 0.58 | 0.31 | 0.5 |
| 3 | 40458041 | rs9332450 | 0.495 | -2.503 | 2.223 | 1.659 | 0.06 | -0.004 | -0.004 | 0.089 | 0.52 | 0.63 | 0.35 | 0.56 | 0.31 | 0.48 |
| 3 | 40459532 | rs4973898 | -0.353 | -2.529 | 2.267 | 1.664 | 0.097 | -0.045 | -0.045 | 0 | 0.43 | 0.17 | 0.63 | 0.41 | 0.68 | 0.46 |
| 3 | 40459724 | rs13326389 | 1.249 | -2.542 | 2.266 | 1.664 | 0.096 | -0.048 | -0.048 | 0 | 0.42 | 0.07 | 0.29 | 0.49 | 0.29 | 0.37 |
| 3 | 40468206 | rs9873394 | -1.786 | -2.509 | 2.326 | 1.672 | -0.029 | 0.025 | 0.025 | 5.26 | 0.1 | 0.56 | 0.06 | 0.07 | 0.02 | 0.11 |
| 3 | 40468896 | rs1047855 | 1.384 | -2.519 | 2.249 | 1.668 | 0.071 | -0.079 | -0.079 | 0.158 | 0.41 | 0.04 | 0.29 | 0.49 | 0.29 | 0.37 |
| 3 | 40493218 | rs6801859 | -0.433 | -2.526 | 2.32 | 1.653 | -0.01 | 0 | 0 | 0 | 0.46 | 0.52 | 0.66 | 0.45 | 0.69 | 0.47 |
| 3 | 40494065 | rs11714912 | -0.334 | -2.513 | 2.016 | 1.656 | 0.092 | -0.057 | -0.057 | 0 | 0.43 | 0.14 | 0.62 | 0.41 | 0.68 | 0.46 |
| 3 | 40499273 | rs2276871 | -0.998 | -2.436 | 1.848 | 1.627 | 0.081 | 0 | 0 | 0 | 0.53 | 0.73 | 0.69 | 0.48 | 0.7 | 0.57 |
| 3 | 40503809 | rs11539046 | -1.153 | -2.421 | 1.846 | 1.539 | 0.074 | 0.012 | 0.012 | 0.01 | 0.55 | 0.75 | 0.69 | 0.51 | 0.7 | 0.63 |
| 3 | 40504326 | rs13073003 | -0.962 | -2.415 | 1.826 | 1.542 | -0.008 | -0.001 | -0.001 | 1.129 | 0.53 | 0.54 | 0.67 | 0.51 | 0.7 | 0.62 |
| 3 | 40519532 | rs4128740 | 1.204 | -2.298 | 0.72 | 1.6 | 0.09 | -0.033 | -0.033 | 0 | 0.41 | 0.08 | 0.29 | 0.48 | 0.29 | 0.37 |
| 3 | 40535809 | rs7645733 | 0.339 | -2.03 | 1.338 | 1.695 | 0.037 | -0.017 | -0.017 | 0 | 0.55 | 0.72 | 0.41 | 0.53 | 0.39 | 0.6 |
| 3 | 40536525 | rs9845644 | -0.194 | -2.172 | 1.349 | 1.834 | -0.003 | 0.025 | 0.025 | 0.548 | 0.63 | 0.88 | 0.48 | 0.75 | 0.4 | 0.74 |
| 3 | 40537829 | rs9851402 | 0.426 | -2.199 | 1.37 | 1.897 | 0.052 | -0.025 | -0.025 | 0.02 | 0.55 | 0.8 | 0.41 | 0.58 | 0.38 | 0.6 |
| 3 | 40540738 | rs6599106 | -2.543 | -2.13 | 2.116 | 1.805 | 0.019 | 0.147 | 0.147 | 0.19 | 0.06 | 0.35 | 0.08 | 0.12 | 0.02 | 0.03 |
| 3 | 40552916 | rs6790578 | 1.293 | -2.266 | 2.208 | 2.039 | 0.061 | 0.063 | 0.063 | 0.402 | 0.45 | 0.15 | 0.34 | 0.42 | 0.36 | 0.44 |

| | | | | | | | | | | | | | | | |
|---|---|---|---|---|---|---|---|---|---|---|---|---|---|---|---|
| 3 | 40556596 | rs9820580 | 0.984 | -2.186 | 2.474 | 1.983 | 0.06 | -0.005 | -0.005 | 0 | 0.49 | 0.27 | 0.36 | 0.56 | 0.37 | 0.61 |
| 3 | 40557328 | rs13095915 | -2.584 | -2.173 | 2.426 | 1.96 | 0.022 | 0.144 | 0.144 | 0.242 | 0.06 | 0.37 | 0.08 | 0.12 | 0.02 | 0.03 |
| 3 | 40565594 | rs6599108 | 0.516 | -2.18 | 2.47 | 1.943 | 0.046 | -0.037 | -0.037 | 0 | 0.56 | 0.8 | 0.42 | 0.56 | 0.39 | 0.62 |
| 3 | 40576061 | rs3924444 | -0.087 | -2.23 | 2.191 | 1.932 | 0.008 | 0.057 | 0.057 | 0.318 | 0.62 | 0.88 | 0.49 | 0.79 | 0.4 | 0.73 |
| 3 | 40579357 | rs4602359 | 1.059 | -2.223 | 2.175 | 1.937 | 0.068 | 0.01 | 0.01 | 0.011 | 0.48 | 0.26 | 0.36 | 0.58 | 0.37 | 0.59 |
| 3 | 40589558 | rs9810005 | -1.116 | -2.072 | 2.02 | 1.835 | 0.001 | 0.006 | 0.006 | 0 | 0.08 | 0.07 | 0.08 | 0.24 | 0.02 | 0.13 |
| 3 | 40595426 | rs11915318 | -0.004 | -2.025 | 2.064 | 1.753 | 0.019 | -0.005 | -0.005 | 0.047 | 0.6 | 0.8 | 0.43 | 0.5 | 0.4 | 0.53 |
| 3 | 40600392 | rs4974035 | -0.299 | -2.025 | 1.903 | 1.789 | -0.035 | -0.009 | -0.009 | 0 | 0.64 | 0.92 | 0.49 | 0.73 | 0.4 | 0.71 |
| 3 | 40617539 | rs6774841 | -0.195 | -1.804 | 1.706 | 1.226 | -0.021 | -0.063 | -0.063 | 0.177 | 0.62 | 0.91 | 0.5 | 0.67 | 0.4 | 0.55 |
| 3 | 40620339 | rs6808354 | -0.195 | -1.724 | 2.053 | 1.078 | -0.026 | -0.044 | -0.044 | 0 | 0.08 | 0.11 | 0.02 | 0 | 0.02 | 0.02 |
| 3 | 40626368 | rs12492728 | -0.134 | -1.672 | 1.669 | 1.167 | 0.009 | -0.012 | -0.012 | 0 | 0.61 | 0.78 | 0.51 | 0.64 | 0.4 | 0.58 |
| 3 | 40627829 | rs11708971 | 0.514 | -1.196 | 1.546 | 0.973 | 0.053 | -0.019 | -0.019 | 0 | 0.45 | 0.14 | 0.38 | 0.49 | 0.36 | 0.46 |
| 3 | 40631348 | rs6599111 | 0.521 | -1.153 | 1.632 | 0.992 | 0.039 | 0.009 | 0.009 | 0.468 | 0.47 | 0.39 | 0.4 | 0.49 | 0.37 | 0.5 |
| 3 | 40659555 | rs6772841 | -2.288 | -0.456 | 1.955 | 0.991 | 0.035 | 0.022 | 0.022 | 3.801 | 0.05 | 0.36 | 0.04 | 0.01 | 0.01 | 0.05 |
| 3 | 40668749 | rs4597685 | -2.014 | -0.509 | 1.935 | 0.958 | -0.023 | -0.031 | -0.031 | 0 | 0.11 | 0.44 | 0.06 | 0.06 | 0.04 | 0.09 |
| 3 | 40688939 | rs4077961 | -2.055 | -0.509 | 2.058 | 0.941 | -0.022 | -0.036 | -0.036 | 0 | 0.11 | 0.46 | 0.06 | 0.06 | 0.04 | 0.09 |
| 3 | 40697713 | rs4411862 | -2.286 | -0.429 | 2.132 | 1.151 | -0.008 | 0.058 | 0.058 | 0.438 | 0.12 | 0.48 | 0.11 | 0.13 | 0.07 | 0.12 |
| 3 | 40701424 | rs4513425 | -1.088 | -0.426 | 2.286 | 1.085 | -0.002 | -0.012 | -0.012 | 0 | 0.07 | 0.31 | 0.04 | 0.03 | 0.03 | 0.03 |
| 3 | 40708000 | rs9682256 | -2.466 | -0.392 | 2.145 | 1.116 | 0.008 | 0.06 | 0.06 | 1.818 | 0.05 | 0.17 | 0.07 | 0.06 | 0.04 | 0.09 |
| 3 | 40733620 | rs13067751 | -0.58 | -0.462 | 2.591 | 1.644 | -0.02 | -0.01 | -0.01 | 0 | 0.07 | 0 | 0.19 | 0.39 | 0.08 | 0.17 |
| 3 | 40744698 | rs7611057 | 2.52 | -0.82 | 3.365 | 0.449 | -0.045 | -0.019 | -0.019 | 0 | 0.82 | 0.31 | 0.86 | 0.91 | 0.9 | 0.89 |
| 3 | 40754437 | rs17080646 | -1.757 | -0.772 | 3.236 | 0.466 | -0.024 | -0.038 | -0.038 | 3.052 | 0.07 | 0.17 | 0.04 | 0 | 0.04 | 0.05 |
| 3 | 40762022 | rs4498012 | 2.07 | -0.774 | 3.224 | 0.47 | -0.042 | -0.07 | -0.07 | 0 | 0.86 | 0.49 | 0.92 | 0.96 | 0.93 | 0.91 |
| 3 | 40764236 | rs9875272 | 2.305 | -0.715 | 3.051 | 0.464 | -0.039 | -0.016 | -0.016 | 0 | 0.83 | 0.31 | 0.86 | 0.91 | 0.9 | 0.89 |
| 3 | 40809914 | rs655341 | 1.488 | -0.708 | 2.091 | 0.075 | -0.026 | -0.037 | -0.037 | 0.716 | 0.91 | 0.84 | 0.96 | 0.98 | 0.95 | 0.93 |
| 3 | 40816505 | rs6769201 | -1.184 | -0.705 | 2.018 | 0.076 | -0.011 | -0.025 | -0.025 | 0 | 0.08 | 0.09 | 0.03 | 0 | 0.05 | 0.07 |
| 3 | 40817096 | rs667189 | 1.618 | -0.758 | 2.022 | 0.01 | -0.062 | -0.116 | -0.116 | 0.044 | 0.84 | 0.46 | 0.92 | 0.98 | 0.92 | 0.91 |
| 3 | 40830766 | rs7613264 | -1.713 | -0.486 | 2.092 | 0.163 | -0.005 | -0.041 | -0.041 | 0 | 0.1 | 0.43 | 0.06 | 0 | 0.05 | 0.06 |
| 3 | 40885879 | rs627998 | -0.197 | 0.126 | 0.145 | 0.069 | -0.003 | 0.026 | 0.026 | 0 | 0.31 | 0.1 | 0.25 | 0.27 | 0.29 | 0.26 |
| 3 | 40899164 | rs7650433 | -0.844 | 1.589 | 0.315 | 0.012 | 0.004 | -0.02 | -0.02 | 0 | 0.1 | 0.14 | 0.1 | 0 | 0.18 | 0.04 |
| 3 | 40899921 | rs6807519 | -1.62 | 1.931 | -0.046 | 0.474 | -0.05 | -0.056 | -0.056 | 0 | 0.31 | 0.88 | 0.34 | 0.36 | 0.35 | 0.33 |
| 3 | 40900521 | rs9872703 | -1.278 | 1.456 | 0.263 | 0.155 | 0 | -0.025 | -0.025 | 3.627 | 0.11 | 0.1 | 0.07 | 0 | 0.1 | 0.03 |
| 3 | 40904201 | rs6768379 | -1.74 | 1.77 | 1.863 | 0.393 | 0.017 | 0.01 | 0.01 | 0 | 0.19 | 0.19 | 0.19 | 0.23 | 0.28 | 0.22 |
| 3 | 40911930 | rs9810950 | -2.608 | 1.502 | 1.952 | 0.397 | -0.083 | -0.103 | -0.103 | 0 | 0.32 | 0.96 | 0.36 | 0.39 | 0.33 | 0.36 |
| 3 | 40915216 | rs4973914 | -2.373 | 1.773 | 2.591 | 0.544 | 0.015 | 0.012 | 0.012 | 0 | 0.25 | 0.65 | 0.25 | 0.3 | 0.31 | 0.28 |
| 3 | 40917869 | rs1565981 | -0.974 | 1.787 | 2.514 | 0.534 | -0.029 | -0.054 | -0.054 | 0 | 0.12 | 0.03 | 0.08 | 0.25 | 0.14 | 0.2 |
| 3 | 40918851 | rs9837197 | -0.924 | 1.55 | 2.664 | 0.426 | 0.053 | -0.035 | -0.035 | 0 | 0.06 | 0.13 | 0.09 | 0 | 0.12 | 0.02 |
| 3 | 40939632 | rs17051772 | -2.257 | 1.274 | 2.1 | 0.402 | 0.001 | 0.021 | 0.021 | 3.487 | 0.17 | 0.7 | 0.2 | 0.22 | 0.17 | 0.19 |
| 3 | 40966907 | rs7629386 | -1.632 | 1.603 | 1.399 | 0.136 | 0.018 | -0.03 | -0.03 | 0 | 0.21 | 0.25 | 0.24 | 0.07 | 0.33 | 0.16 |
| 3 | 40982272 | rs6599129 | -2.274 | 1.354 | 0.363 | 0.21 | -0.016 | 0.026 | 0.026 | 0 | 0.39 | 0.71 | 0.38 | 0.34 | 0.4 | 0.37 |
| 3 | 40986300 | rs7625419 | -2.278 | 1.36 | 0.366 | 0.23 | -0.016 | 0.017 | 0.017 | 0 | 0.39 | 0.71 | 0.38 | 0.36 | 0.4 | 0.38 |
| 3 | 41010253 | rs13067838 | -0.855 | 0.933 | -0.316 | 0.474 | -0.072 | 0.036 | 0.036 | 0 | 0.16 | 0.01 | 0.1 | 0.08 | 0.2 | 0.14 |
| 3 | 41012473 | rs3828363 | -1.057 | 0.411 | 0.356 | 0.854 | -0.038 | 0.067 | 0.067 | 0 | 0.1 | 0.21 | 0.11 | 0.25 | 0.03 | 0.15 |
| 3 | 41019888 | rs4498013 | -0.125 | 0.611 | -0.334 | 0.711 | 0 | 0 | 0 | 0 | 0.53 | 0.53 | 0.44 | 0.6 | 0.54 | 0.39 |
| 3 | 41021263 | rs6789549 | -0.379 | 0.411 | -1.125 | 0.612 | 0.007 | 0.032 | 0.032 | 0 | 0.34 | 0.17 | 0.4 | 0.28 | 0.39 | 0.42 |
| 3 | 41025167 | rs6599130 | -0.97 | -0.154 | -0.77 | 1.078 | 0.043 | 0.018 | 0.018 | 0 | 0.49 | 0.76 | 0.52 | 0.48 | 0.47 | 0.61 |
| 3 | 41027338 | rs4276160 | -0.148 | -0.171 | -0.202 | 1.141 | 0.01 | 0.085 | 0.085 | 0 | 0.11 | 0.4 | 0.1 | 0.19 | 0.09 | 0.19 |
| 3 | 41029087 | rs11708803 | 0.141 | -0.418 | -0.471 | 1.27 | -0.011 | -0.058 | -0.058 | 0 | 0.33 | 0.09 | 0.4 | 0.36 | 0.37 | 0.39 |
| 3 | 41031039 | rs7653388 | -0.803 | -0.654 | 0.052 | 1.33 | -0.026 | 0.087 | 0.087 | 0.869 | 0.15 | 0.56 | 0.16 | 0.26 | 0.08 | 0.19 |
| 3 | 41032309 | rs6798084 | 0.236 | -0.871 | -0.504 | 0.922 | 0.011 | -0.011 | -0.011 | 0.148 | 0.23 | 0.16 | 0.12 | 0.01 | 0.2 | 0.22 |
| 3 | 41037617 | rs11716816 | 1.315 | -0.556 | -0.875 | 1.408 | 0.019 | 0.071 | 0.071 | 0 | 0.46 | 0.24 | 0.36 | 0.27 | 0.52 | 0.39 |
| 3 | 41039907 | rs6599132 | 0.282 | -0.336 | -1.207 | 1.137 | -0.013 | -0.003 | -0.003 | 0.1 | 0.43 | 0.65 | 0.42 | 0.46 | 0.43 | 0.54 |
| 3 | 41042846 | rs4393879 | -0.645 | -0.102 | -1.535 | 1.192 | -0.061 | -0.041 | -0.041 | 0 | 0.7 | 0.92 | 0.58 | 0.66 | 0.65 | 0.78 |
| 3 | 41055118 | rs9822113 | -0.349 | -0.214 | -1.257 | 0.821 | 0.004 | 0.007 | 0.007 | 0 | 0.5 | 0.34 | 0.47 | 0.52 | 0.49 | 0.55 |
| 3 | 41058826 | rs6802872 | -0.618 | -0.257 | -1.173 | 0.801 | -0.004 | -0.001 | -0.001 | 0.653 | 0.53 | 0.42 | 0.48 | 0.52 | 0.5 | 0.55 |
| 3 | 41059754 | rs17250766 | 0.616 | 0.238 | -1.57 | 0.777 | -0.006 | -0.014 | -0.014 | 0 | 0.26 | 0.19 | 0.26 | 0.31 | 0.24 | 0.3 |
| 3 | 41063002 | rs7651160 | 0.28 | 0.15 | -1.478 | 0.417 | 0.005 | 0.029 | 0.029 | 0 | 0.23 | 0.16 | 0.15 | 0.1 | 0.24 | 0.25 |
| 3 | 41063760 | rs440804 | 0.108 | 0.585 | -1.271 | 1.085 | -0.006 | -0.04 | -0.04 | 0.067 | 0.27 | 0.19 | 0.32 | 0.43 | 0.25 | 0.31 |
| 3 | 41065413 | rs409228 | -0.471 | -0.008 | -0.783 | 0.731 | -0.021 | -0.015 | -0.015 | 0 | 0.3 | 0.46 | 0.39 | 0.24 | 0.29 | 0.31 |
| 3 | 41066130 | rs764806 | 0.623 | -0.115 | -1.2 | 0.739 | -0.057 | -0.023 | -0.023 | 0 | 0.37 | 0.19 | 0.32 | 0.39 | 0.47 | 0.25 |
| 3 | 41066221 | rs4459874 | -0.258 | -0.831 | -0.742 | 0.648 | -0.032 | 0.099 | 0.099 | 0.023 | 0.72 | 0.55 | 0.6 | 0.5 | 0.77 | 0.64 |
| 3 | 41066528 | rs1544887 | 1.339 | -0.597 | -0.842 | 0.554 | -0.089 | -0.039 | -0.039 | 1.924 | 0.35 | 0.1 | 0.37 | 0.37 | 0.43 | 0.24 |
| 3 | 41067055 | rs9837626 | 0.672 | -0.686 | -0.612 | 0.655 | -0.034 | 0.113 | 0.113 | 0.332 | 0.22 | 0.34 | 0.28 | 0.41 | 0.16 | 0.33 |
| 3 | 41067696 | rs6809212 | -0.023 | -1.147 | -0.938 | 0.3 | -0.047 | -0.028 | -0.028 | 0 | 0.7 | 0.55 | 0.79 | 0.78 | 0.82 | 0.65 |
| 3 | 41068497 | rs4974056 | 1.113 | -0.439 | -0.801 | 0.375 | -0.023 | -0.027 | -0.027 | 0 | 0.39 | 0.34 | 0.45 | 0.47 | 0.46 | 0.32 |
| 3 | 41072131 | rs6782790 | 0.244 | -0.054 | -0.872 | 0.276 | -0.038 | 0.039 | 0.039 | 0 | 0.12 | 0.03 | 0.11 | 0 | 0.19 | 0.1 |
| 3 | 41085459 | rs106871 | 1.327 | -0.595 | -0.97 | 0.431 | -0.108 | -0.057 | -0.057 | 0 | 0.35 | 0.09 | 0.38 | 0.36 | 0.44 | 0.3 |
| 3 | 41086761 | rs453509 | 0.633 | -0.997 | -0.737 | 0.388 | 0.019 | 0.04 | 0.04 | 0.228 | 0.57 | 0.63 | 0.7 | 0.79 | 0.67 | 0.6 |
| 3 | 41093575 | rs408499 | 0.584 | -1.04 | -0.616 | 0.37 | -0.005 | -0.013 | -0.013 | 0 | 0.58 | 0.5 | 0.68 | 0.78 | 0.67 | 0.61 |

| | | | | | | | | | | | | | | | | |
|---|---|---|---|---|---|---|---|---|---|---|---|---|---|---|---|---|
| 3 | 41094080 | rs9284874 | 0.639 | -1.009 | -0.537 | 0.471 | -0.001 | 0.126 | 0.126 | 0.431 | 0.22 | 0.4 | 0.29 | 0.41 | 0.2 | 0.28 |
| 3 | 41097246 | rs367446 | -0.853 | -1.166 | -0.742 | 0.299 | 0.025 | 0.033 | 0.033 | 2.057 | 0.78 | 0.85 | 0.87 | 0.89 | 0.85 | 0.83 |
| 3 | 41100099 | rs9863665 | 0.569 | -1.177 | -0.699 | 0.38 | -0.005 | 0.119 | 0.119 | 0.773 | 0.22 | 0.51 | 0.32 | 0.43 | 0.22 | 0.29 |
| 3 | 41100200 | rs408277 | 1.679 | -1.043 | -0.857 | 0.369 | -0.118 | -0.041 | -0.041 | 0 | 0.35 | 0.09 | 0.38 | 0.36 | 0.46 | 0.31 |
| 3 | 41102711 | rs17052699 | -0.145 | -0.911 | -0.657 | 0.39 | 0 | -0.003 | -0.003 | 0.223 | 0.17 | 0.18 | 0.17 | 0.1 | 0.17 | 0.21 |
| 3 | 41103150 | rs6599141 | 0.234 | -1.441 | -0.438 | 0.535 | -0.006 | 0.088 | 0.088 | 1.248 | 0.23 | 0.54 | 0.32 | 0.43 | 0.22 | 0.29 |
| 3 | 41107173 | rs6788984 | -0.947 | -1.118 | -0.135 | 0.098 | -0.003 | -0.097 | -0.097 | 0 | 0.18 | 0.35 | 0.12 | 0.02 | 0.15 | 0.17 |
| 3 | 41109561 | rs370243 | 0.099 | -1.59 | -0.376 | 0.29 | -0.005 | -0.002 | -0.002 | 0 | 0.54 | 0.54 | 0.52 | 0.45 | 0.66 | 0.54 |
| 3 | 41114867 | rs384934 | 1.297 | -1.586 | -0.351 | 0.294 | -0.032 | 0.009 | 0.009 | 0.186 | 0.38 | 0.22 | 0.42 | 0.35 | 0.54 | 0.32 |
| 3 | 41115016 | rs384260 | -0.021 | -1.571 | -0.291 | 0.302 | -0.005 | -0.001 | -0.001 | 0 | 0.55 | 0.46 | 0.52 | 0.45 | 0.67 | 0.55 |
| 3 | 41118450 | rs442115 | 0.583 | -1.585 | -0.034 | 0.471 | 0.004 | -0.002 | -0.002 | 2.682 | 0.4 | 0.49 | 0.43 | 0.43 | 0.53 | 0.36 |
| 3 | 41119907 | rs431867 | 0.935 | -1.577 | -0.03 | 0.491 | -0.011 | 0.009 | 0.009 | 0 | 0.37 | 0.29 | 0.42 | 0.31 | 0.53 | 0.3 |
| 3 | 41122345 | rs383911 | 0.954 | -1.594 | -0.031 | 0.533 | -0.012 | 0.021 | 0.021 | 0 | 0.37 | 0.29 | 0.42 | 0.24 | 0.53 | 0.3 |
| 3 | 41123735 | rs10490823 | 1.208 | -1.656 | -0.087 | 0.503 | -0.081 | 0.035 | 0.035 | 0 | 0.34 | 0.11 | 0.4 | 0.24 | 0.53 | 0.29 |
| 3 | 41128564 | rs430727 | 0.433 | -1.803 | 0.492 | 0.45 | -0.001 | 0.013 | 0.013 | 0 | 0.42 | 0.64 | 0.45 | 0.36 | 0.54 | 0.36 |
| 3 | 41129297 | rs415997 | 1.061 | -1.774 | 0.431 | 0.438 | -0.024 | 0.033 | 0.033 | 0 | 0.36 | 0.23 | 0.41 | 0.24 | 0.53 | 0.29 |
| 3 | 41147757 | rs446717 | -0.471 | -1.744 | 0.568 | 0.496 | 0.001 | 0.046 | 0.046 | 0 | 0.56 | 0.19 | 0.53 | 0.63 | 0.47 | 0.65 |
| 3 | 41150549 | rs443824 | 0.774 | -2.117 | 0.191 | 0.109 | 0.03 | 0.005 | 0.005 | 1.486 | 0.21 | 0.01 | 0.11 | 0.15 | 0.14 | 0.19 |
| 3 | 41160573 | rs12485899 | 0.291 | -1.197 | 0.624 | 0.458 | -0.001 | 0.006 | 0.006 | 0 | 0.4 | 0.35 | 0.41 | 0.41 | 0.54 | 0.42 |
| 3 | 41161343 | rs4437116 | 0.635 | -1.229 | 0.548 | 0.249 | -0.023 | 0.053 | 0.053 | 0 | 0.36 | 0.25 | 0.4 | 0.18 | 0.52 | 0.34 |
| 3 | 41164908 | rs7642431 | 0.707 | -1.246 | 0.45 | 0.248 | -0.052 | 0.079 | 0.079 | 0 | 0.36 | 0.16 | 0.51 | 0.2 | 0.53 | 0.35 |
| 3 | 41168607 | rs7621892 | 0.236 | -1.509 | 0.7 | 0.143 | 0.011 | -0.012 | -0.012 | 0 | 0.38 | 0.53 | 0.42 | 0.37 | 0.52 | 0.39 |
| 3 | 41169335 | rs9883523 | -0.831 | -1.322 | 0.571 | 0.108 | 0.044 | 0.198 | 0.198 | 0.652 | 0.06 | 0.35 | 0.03 | 0.18 | 0 | 0.05 |
| 3 | 41177604 | rs375583 | -0.055 | -1.212 | -0.166 | 0.063 | -0.012 | 0.079 | 0.079 | 0 | 0.56 | 0.18 | 0.55 | 0.65 | 0.48 | 0.59 |
| 3 | 41190647 | rs1798796 | 1.726 | -0.965 | 0.842 | 0.315 | -0.015 | 0.003 | 0.003 | 3.867 | 0.89 | 0.42 | 0.67 | 0.92 | 0.96 | 0.91 |
| 3 | 41203895 | rs880026 | 0.11 | -0.849 | 0.097 | -0.069 | -0.043 | 0.021 | 0.021 | 0 | 0.3 | 0.15 | 0.33 | 0.24 | 0.48 | 0.4 |
| 3 | 41205419 | rs9864635 | -0.537 | -0.875 | 0.858 | -0.074 | 0.009 | 0.041 | 0.041 | 0 | 0.06 | 0.27 | 0.28 | 0.04 | 0.01 | 0.05 |
| 3 | 41206371 | rs17053666 | -1.115 | -0.987 | 0.965 | -0.119 | 0.013 | 0.015 | 0.015 | 0.11 | 0.05 | 0.3 | 0.04 | 0.03 | 0.01 | 0.05 |
| 3 | 41206998 | rs1016762 | 0.194 | -0.933 | 0.806 | -0.165 | -0.061 | 0.024 | 0.024 | 0 | 0.3 | 0.12 | 0.32 | 0.24 | 0.48 | 0.39 |
| 3 | 41210298 | rs1880479 | 1.413 | -1.001 | 0.949 | -0.139 | 0.025 | 0.026 | 0.026 | 1.068 | 0.93 | 0.56 | 0.95 | 0.97 | 0.99 | 0.95 |
| 3 | 41212753 | rs17252114 | 0.185 | -0.779 | 0.688 | -0.146 | -0.066 | 0.033 | 0.033 | 0 | 0.3 | 0.17 | 0.33 | 0.23 | 0.47 | 0.39 |
| 3 | 41220433 | rs6802052 | -1.53 | -1.079 | 0.9 | 0.246 | -0.037 | 0.144 | 0.144 | 0 | 0.1 | 0.33 | 0.29 | 0.19 | 0.02 | 0.14 |
| 3 | 41230654 | rs7630377 | 0.842 | -0.627 | 0.022 | 0.014 | -0.004 | 0.064 | 0.064 | 0 | 0.58 | 0.16 | 0.37 | 0.68 | 0.52 | 0.53 |
| 3 | 41240177 | rs3864004 | -0.277 | -0.31 | -0.329 | 0.122 | -0.017 | 0.016 | 0.016 | 0 | 0.33 | 0.3 | 0.34 | 0.26 | 0.43 | 0.43 |
| 3 | 41243742 | rs3915129 | -0.288 | -0.282 | -0.298 | 0.139 | -0.011 | 0.01 | 0.01 | 0 | 0.33 | 0.32 | 0.34 | 0.26 | 0.43 | 0.43 |
| 3 | 41279440 | rs4135385 | 0.877 | -0.868 | -0.807 | 0.005 | -0.052 | -0.155 | -0.155 | 0 | 0.2 | 0.01 | 0.28 | 0.53 | 0.24 | 0.19 |
| 3 | 41281388 | rs2953 | -0.449 | 0.185 | 0.36 | 0.389 | -0.001 | 0 | 0 | 0 | 0.33 | 0.34 | 0.34 | 0.28 | 0.43 | 0.43 |
| 3 | 41284536 | rs9883073 | -0.308 | 0.368 | -0.005 | 0.468 | -0.03 | 0.019 | 0.019 | 0 | 0.31 | 0.26 | 0.33 | 0.26 | 0.43 | 0.43 |
| 3 | 41287018 | rs11711946 | -0.311 | 0.453 | -0.497 | 0.501 | -0.08 | 0.008 | 0.008 | 0 | 0.3 | 0.15 | 0.33 | 0.28 | 0.43 | 0.43 |
| 3 | 41288466 | rs6769117 | 0.364 | 0.119 | -0.147 | 0.593 | 0.014 | 0.016 | 0.016 | 0 | 0.06 | 0.02 | 0.01 | 0 | 0.02 | 0 |
| 3 | 41292736 | rs9850653 | -1.217 | 0.338 | 0.671 | 1.046 | 0.003 | 0.166 | 0.166 | 0.254 | 0.11 | 0.48 | 0.23 | 0.23 | 0.06 | 0.16 |
| 3 | 41295767 | rs11129896 | -0.075 | 0.678 | 0.183 | 1.01 | -0.117 | -0.006 | -0.006 | 0 | 0.26 | 0.06 | 0.43 | 0.22 | 0.42 | 0.42 |
| 3 | 41297523 | rs9849022 | -1.353 | 0.555 | 0.297 | 0.921 | -0.007 | -0.013 | -0.013 | 0 | 0.39 | 0.71 | 0.56 | 0.45 | 0.48 | 0.6 |
| 3 | 41301649 | rs2293299 | -1.768 | 0.328 | 0.519 | 0.814 | -0.04 | 0.068 | 0.068 | 1.036 | 0.14 | 0.6 | 0.25 | 0.24 | 0.07 | 0.16 |
| 3 | 41303087 | rs9838277 | -1.417 | 0.369 | 0.535 | 0.832 | -0.028 | 0.12 | 0.12 | 0.196 | 0.13 | 0.52 | 0.24 | 0.24 | 0.07 | 0.16 |
| 3 | 41308449 | rs11720292 | -0.654 | 0.839 | -0.104 | 0.332 | -0.114 | 0.136 | 0.136 | 0 | 0.33 | 0.09 | 0.24 | 0.09 | 0.44 | 0.45 |
| 3 | 41312055 | rs9820765 | -0.375 | 1.115 | 1.306 | 0.776 | -0.007 | 0.002 | 0.002 | 0 | 0.28 | 0.22 | 0.22 | 0.09 | 0.36 | 0.38 |
| 3 | 41315914 | rs9311266 | -1.274 | 0.964 | 1.458 | 0.792 | -0.002 | -0.008 | -0.008 | 0 | 0.12 | 0.33 | 0.2 | 0.11 | 0.11 | 0.09 |
| 3 | 41318158 | rs1949836 | -1.583 | 0.953 | 1.297 | 0.796 | -0.069 | 0.004 | 0.004 | 0.571 | 0.45 | 0.88 | 0.48 | 0.37 | 0.47 | 0.58 |
| 3 | 41328182 | rs9849601 | -2.204 | 0.817 | 1.462 | 0.902 | -0.034 | 0.088 | 0.088 | 2.143 | 0.17 | 0.58 | 0.26 | 0.3 | 0.11 | 0.2 |
| 3 | 41339984 | rs17055247 | -1.274 | 0.883 | 1.462 | 0.806 | -0.006 | -0.015 | -0.015 | 0 | 0.11 | 0.26 | 0.2 | 0.09 | 0.11 | 0.09 |
| 3 | 41343854 | rs7637185 | -1.748 | 0.987 | 1.285 | 0.795 | -0.07 | 0.011 | 0.011 | 0.512 | 0.45 | 0.87 | 0.48 | 0.36 | 0.46 | 0.57 |
| 3 | 41345172 | rs4118041 | -0.27 | 0.938 | 1.522 | 0.785 | -0.007 | -0.001 | -0.001 | 1.252 | 0.28 | 0.21 | 0.22 | 0.07 | 0.35 | 0.39 |
| 3 | 41359534 | rs4973937 | -1.389 | 0.95 | 1.503 | 0.8 | -0.002 | -0.002 | -0.002 | 0 | 0.12 | 0.34 | 0.2 | 0.11 | 0.11 | 0.09 |
| 3 | 41364241 | rs6770252 | -1.773 | 0.914 | 1.367 | 0.751 | -0.088 | -0.006 | -0.006 | 0 | 0.43 | 0.88 | 0.48 | 0.37 | 0.45 | 0.55 |
| 3 | 41372090 | rs9820239 | -0.802 | 1.721 | 1.438 | 0.707 | -0.009 | 0.006 | 0.006 | 0 | 0.28 | 0.58 | 0.22 | 0.26 | 0.22 | 0.44 |
| 3 | 41373224 | rs9311268 | -1.878 | 1.729 | 1.432 | 0.685 | -0.067 | 0.014 | 0.014 | 0.26 | 0.42 | 0.81 | 0.48 | 0.35 | 0.44 | 0.55 |
| 3 | 41373677 | rs17055760 | -1.716 | 1.728 | 1.591 | 0.679 | 0.067 | -0.042 | -0.042 | 0 | 0.15 | 0.29 | 0.26 | 0.09 | 0.23 | 0.1 |
| 3 | 41374573 | rs6775739 | -1.036 | 1.689 | 1.651 | 0.694 | 0.01 | -0.004 | -0.004 | 0.355 | 0.1 | 0.28 | 0.19 | 0.09 | 0.11 | 0.09 |
| 3 | 41375033 | rs1123522 | -1.036 | 1.689 | 1.665 | 0.694 | 0.011 | -0.003 | -0.003 | 0 | 0.1 | 0.29 | 0.19 | 0.09 | 0.11 | 0.09 |
| 3 | 41375667 | rs6790569 | -1.014 | 1.707 | 1.604 | 0.687 | 0.041 | -0.005 | -0.005 | 0 | 0.31 | 0.49 | 0.28 | 0.26 | 0.34 | 0.46 |
| 3 | 41377373 | rs901688 | -0.659 | 1.616 | 1.662 | 0.653 | 0.03 | -0.06 | -0.06 | 0 | 0.27 | 0.34 | 0.21 | 0.07 | 0.33 | 0.36 |
| 3 | 41377828 | rs6773779 | -1.897 | 1.588 | 1.629 | 0.762 | -0.015 | 0.122 | 0.122 | 0.204 | 0.15 | 0.38 | 0.26 | 0.32 | 0.12 | 0.2 |
| 3 | 41377991 | rs6786558 | -1.906 | 1.521 | 1.598 | 0.676 | -0.019 | 0.088 | 0.088 | 0.043 | 0.16 | 0.38 | 0.28 | 0.3 | 0.13 | 0.21 |
| 3 | 41378621 | rs749932 | -2.057 | 1.464 | 1.48 | 0.639 | -0.073 | 0.02 | 0.02 | 0.306 | 0.46 | 0.89 | 0.51 | 0.38 | 0.47 | 0.59 |
| 3 | 41379517 | rs880603 | -0.41 | 1.552 | 1.612 | 0.661 | -0.001 | -0.011 | -0.011 | 0 | 0.25 | 0.24 | 0.2 | 0.07 | 0.33 | 0.36 |
| 3 | 41387911 | rs1386602 | -1.843 | 1.558 | 1.505 | 0.645 | 0.021 | -0.033 | -0.033 | 0 | 0.17 | 0.25 | 0.28 | 0.09 | 0.23 | 0.12 |
| 3 | 41397041 | rs9842536 | -0.642 | 1.672 | 1.384 | 0.743 | -0.001 | 0.009 | 0.009 | 0 | 0.26 | 0.36 | 0.19 | 0.28 | 0.23 | 0.45 |
| 3 | 41414528 | rs9843415 | -0.776 | 1.612 | 1.243 | 0.482 | -0.01 | 0.002 | 0.002 | 0 | 0.28 | 0.24 | 0.2 | 0.07 | 0.33 | 0.38 |

| | | | | | | | | | | | | | | | | |
|---|---|---|---|---|---|---|---|---|---|---|---|---|---|---|---|---|
| 3 | 41420792 | rs1386601 | -1.163 | 1.568 | 1.294 | 1.116 | 0.039 | 0.025 | 0.025 | 0 | 0.34 | 0.48 | 0.34 | 0.42 | 0.43 | 0.5 |
| 3 | 41421563 | rs13327260 | -1.338 | 1.506 | 1.204 | 0.833 | -0.001 | -0.018 | -0.018 | 1.867 | 0.11 | 0.13 | 0.2 | 0.05 | 0.11 | 0.05 |
| 3 | 41433086 | rs7618346 | -1.713 | 1.453 | 1.096 | 1.115 | -0.002 | -0.035 | -0.035 | 0.481 | 0.46 | 0.84 | 0.49 | 0.48 | 0.48 | 0.59 |
| 3 | 41438288 | rs9311274 | -1.792 | 1.439 | 1.101 | 1.124 | 0 | -0.034 | -0.034 | 0.007 | 0.46 | 0.86 | 0.49 | 0.48 | 0.48 | 0.57 |
| 3 | 41443267 | rs9824480 | -0.804 | 1.643 | 1.449 | 1.076 | 0.009 | 0.047 | 0.047 | 0 | 0.31 | 0.52 | 0.24 | 0.45 | 0.29 | 0.52 |
| 3 | 41448372 | rs9990047 | -2.021 | 1.657 | 1.458 | 1.061 | 0.047 | -0.065 | -0.065 | 0.626 | 0.16 | 0.36 | 0.25 | 0.06 | 0.2 | 0.05 |
| 3 | 41455363 | rs9832048 | -1.94 | 1.627 | 1.313 | 1.052 | -0.008 | -0.046 | -0.046 | 0.436 | 0.47 | 0.88 | 0.48 | 0.48 | 0.48 | 0.56 |
| 3 | 41457429 | rs17269115 | -0.313 | 1.648 | 1.583 | 1.073 | 0.015 | -0.053 | -0.053 | 0 | 0.26 | 0.29 | 0.17 | 0.08 | 0.26 | 0.42 |
| 3 | 41457654 | rs9866871 | -1.861 | 1.515 | 1.319 | 1.124 | -0.009 | -0.047 | -0.047 | 0 | 0.46 | 0.88 | 0.48 | 0.48 | 0.48 | 0.58 |
| 3 | 41458709 | rs9852385 | -1.691 | 1.515 | 1.369 | 0.472 | -0.013 | 0.083 | 0.083 | 0 | 0.42 | 0.7 | 0.43 | 0.13 | 0.46 | 0.46 |
| 3 | 41463200 | rs9818955 | -0.994 | 1.579 | 1.4 | 0.508 | 0.047 | -0.108 | -0.108 | 0.171 | 0.33 | 0.47 | 0.23 | 0.08 | 0.36 | 0.43 |
| 3 | 41471642 | rs13068880 | -1.965 | 1.567 | 1.242 | 0.589 | -0.005 | -0.024 | -0.024 | 2.778 | 0.1 | 0.17 | 0.21 | 0.03 | 0.09 | 0.06 |
| 3 | 41475001 | rs7623860 | -0.356 | 1.667 | 1.672 | 0.47 | 0.001 | -0.044 | -0.044 | 0 | 0.25 | 0.3 | 0.15 | 0.12 | 0.23 | 0.33 |
| 3 | 41476562 | rs11129907 | -2.344 | 1.729 | 1.81 | 1.079 | 0.067 | 0.104 | 0.104 | 0.06 | 0.15 | 0.29 | 0.31 | 0.56 | 0.25 | 0.24 |
| 3 | 41480047 | rs7627972 | -1.93 | 1.66 | 1.822 | 1.054 | -0.015 | 0.085 | 0.085 | 0 | 0.45 | 0.81 | 0.48 | 0.67 | 0.48 | 0.57 |
| 3 | 41483427 | rs7621118 | -1.746 | 1.8 | 1.748 | 1.116 | -0.01 | 0.029 | 0.029 | 0 | 0.41 | 0.68 | 0.46 | 0.67 | 0.48 | 0.57 |
| 3 | 41489604 | rs6599155 | -0.766 | 1.855 | 1.762 | 1.106 | 0.037 | -0.074 | -0.074 | 0 | 0.29 | 0.39 | 0.21 | 0.12 | 0.33 | 0.34 |
| 3 | 41489806 | rs1565958 | -2.238 | 1.885 | 1.591 | 1.18 | 0.027 | 0.167 | 0.167 | 0 | 0.13 | 0.3 | 0.25 | 0.56 | 0.15 | 0.23 |
| 3 | 41492630 | rs13059459 | -2.53 | 1.914 | 1.417 | 1.178 | 0.052 | 0.083 | 0.083 | 0 | 0.16 | 0.26 | 0.31 | 0.56 | 0.25 | 0.24 |
| 3 | 41494605 | rs6809441 | -0.752 | 1.692 | 1.281 | 1.112 | 0.026 | -0.042 | -0.042 | 0.052 | 0.29 | 0.35 | 0.2 | 0.13 | 0.33 | 0.36 |
| 3 | 41494618 | rs33916626 | -0.815 | 1.697 | 1.345 | 1.188 | 0.001 | -0.004 | -0.004 | 0 | 0.06 | 0.09 | 0.18 | 0.03 | 0.09 | 0.04 |
| 3 | 41496752 | rs2270605 | -0.709 | 1.686 | 1.323 | 1.146 | 0.027 | -0.043 | -0.043 | 0.771 | 0.29 | 0.36 | 0.2 | 0.13 | 0.33 | 0.36 |
| 3 | 41504134 | rs2291757 | -1.33 | 1.444 | 0.815 | 1.088 | 0.006 | 0.014 | 0.014 | 0 | 0.46 | 0.59 | 0.47 | 0.66 | 0.51 | 0.62 |
| 3 | 41531540 | rs1795317 | 1.766 | 1.339 | 1.277 | 1.573 | 0.04 | 0.341 | 0.341 | 0.074 | 0.93 | 0.71 | 0.91 | 0.56 | 0.95 | 0.81 |
| 3 | 41532550 | rs1495698 | -0.454 | 1.774 | 1.303 | 1.285 | 0.023 | -0.039 | -0.039 | 0.149 | 0.3 | 0.34 | 0.23 | 0.13 | 0.36 | 0.33 |
| 3 | 41537412 | rs4973881 | -1.074 | 1.703 | 1.372 | 1.273 | 0.016 | -0.043 | -0.043 | 0 | 0.37 | 0.48 | 0.4 | 0.17 | 0.44 | 0.4 |
| 3 | 41541196 | rs1495703 | 2.016 | 1.539 | 1.451 | 1.343 | 0.023 | 0.368 | 0.368 | 0.201 | 0.9 | 0.6 | 0.89 | 0.56 | 0.95 | 0.8 |
| 3 | 41542470 | rs9821623 | -0.177 | 1.899 | 1.499 | 1.254 | -0.004 | -0.039 | -0.039 | 0 | 0.27 | 0.34 | 0.18 | 0.13 | 0.26 | 0.31 |
| 3 | 41559534 | rs1630957 | 2.042 | 1.838 | 1.374 | 1.33 | 0.02 | 0.33 | 0.33 | 0.082 | 0.9 | 0.63 | 0.89 | 0.56 | 0.95 | 0.81 |
| 3 | 41567962 | rs953932 | -1.419 | 1.969 | 1.024 | 1.29 | 0.018 | -0.013 | -0.013 | 0.033 | 0.09 | 0.12 | 0.22 | 0.05 | 0.17 | 0.09 |
| 3 | 41570772 | rs7624356 | -0.257 | 1.932 | 1.055 | 1.172 | 0.002 | -0.067 | -0.067 | 0.005 | 0.27 | 0.43 | 0.18 | 0.12 | 0.27 | 0.31 |
| 3 | 41573511 | rs2700436 | 1.809 | 2.238 | 0.927 | 1.232 | -0.021 | -0.005 | -0.005 | 0 | 0.88 | 0.92 | 0.72 | 0.51 | 0.78 | 0.74 |
| 3 | 41584779 | rs1691961 | 1.67 | 2.095 | 0.936 | 1.184 | -0.01 | 0.02 | 0.02 | 0.121 | 0.57 | 0.26 | 0.51 | 0.38 | 0.51 | 0.42 |
| 3 | 41598769 | rs11926714 | 1.126 | 2.099 | 1.015 | 1.195 | -0.005 | -0.002 | -0.002 | 0.098 | 0.61 | 0.42 | 0.53 | 0.37 | 0.51 | 0.43 |
| 3 | 41599526 | rs1795355 | 1.654 | 2.078 | 1.019 | 1.188 | -0.028 | 0.011 | 0.011 | 0.092 | 0.57 | 0.19 | 0.51 | 0.37 | 0.5 | 0.42 |
| 3 | 41599834 | rs9311282 | -0.522 | 2.164 | 1.163 | 1.186 | 0.049 | -0.071 | -0.071 | 0.172 | 0.29 | 0.38 | 0.23 | 0.11 | 0.36 | 0.32 |
| 3 | 41606692 | rs3897976 | 0.635 | 2.195 | 0.67 | 0.996 | -0.018 | 0.002 | 0.002 | 0.029 | 0.1 | 0.02 | 0.12 | 0.08 | 0.11 | 0.09 |
| 3 | 41611228 | rs7627367 | 0.928 | 2.372 | 0.822 | 1.3 | -0.009 | -0.043 | -0.043 | 0 | 0.64 | 0.19 | 0.71 | 0.41 | 0.68 | 0.52 |
| 3 | 41613317 | rs1256388 | 2.18 | 2.431 | 0.881 | 1.289 | 0.08 | 0.191 | 0.191 | 0 | 0.83 | 0.6 | 0.69 | 0.49 | 0.77 | 0.74 |
| 3 | 41615845 | rs17058895 | -0.42 | 2.358 | 0.838 | 1.171 | -0.002 | -0.065 | -0.065 | 0.179 | 0.28 | 0.44 | 0.2 | 0.13 | 0.28 | 0.28 |
| 3 | 41623222 | rs9846536 | -1.702 | 2.094 | 0.865 | 0.509 | 0.026 | 0.044 | 0.044 | 0.806 | 0.06 | 0.52 | 0.04 | 0.01 | 0 | 0 |
| 3 | 41626139 | rs13086798 | -0.625 | 2.432 | 0.218 | 0.554 | -0.043 | 0 | 0 | 0 | 0.07 | 0.01 | 0.2 | 0.04 | 0.18 | 0.09 |
| 3 | 41634468 | rs4973975 | -0.975 | 2.185 | 0.212 | 0.404 | -0.056 | 0.081 | 0.081 | 0 | 0.36 | 0.23 | 0.38 | 0.15 | 0.46 | 0.41 |
| 3 | 41637388 | rs1108949 | -1.569 | 2.153 | 1.043 | 0.476 | -0.016 | -0.027 | -0.027 | 0.154 | 0.19 | 0.54 | 0.27 | 0.11 | 0.18 | 0.26 |
| 3 | 41638274 | rs9941986 | -2.524 | 2.118 | 0.171 | 0.079 | -0.033 | 0.077 | 0.077 | 0.059 | 0.35 | 0.83 | 0.38 | 0.16 | 0.33 | 0.34 |
| 3 | 41638984 | rs4973976 | -1.688 | 2.05 | 0.241 | 0.057 | 0.004 | 0.003 | 0.003 | 0 | 0.27 | 0.7 | 0.32 | 0.11 | 0.21 | 0.27 |
| 3 | 41645498 | rs4973889 | -2.259 | 2.081 | 0.022 | 0.054 | -0.041 | 0.069 | 0.069 | 0.054 | 0.35 | 0.83 | 0.4 | 0.17 | 0.34 | 0.34 |
| 3 | 41646670 | rs7629556 | -1.08 | 2.141 | 0.427 | 0.01 | 0.002 | -0.006 | -0.006 | 0 | 0.18 | 0.2 | 0.28 | 0.07 | 0.26 | 0.28 |
| 3 | 41651081 | rs13068312 | -1.174 | 2.126 | 0.428 | 0.032 | 0.001 | -0.004 | -0.004 | 0 | 0.18 | 0.2 | 0.29 | 0.07 | 0.28 | 0.28 |
| 3 | 41667545 | rs4973978 | -1.178 | 2.168 | 0.428 | 0.021 | 0.001 | -0.002 | -0.002 | 0 | 0.18 | 0.2 | 0.29 | 0.08 | 0.29 | 0.28 |
| 3 | 41676164 | rs974595 | 0.257 | 1.94 | 0.469 | 0.02 | -0.004 | -0.015 | -0.015 | 0 | 0.11 | 0.19 | 0.08 | 0.05 | 0.1 | 0.19 |
| 3 | 41681233 | rs9828875 | -2.016 | 1.922 | 0.173 | 0.004 | -0.04 | 0.105 | 0.105 | 0.056 | 0.35 | 0.83 | 0.4 | 0.12 | 0.33 | 0.34 |
| 3 | 41714831 | rs9876835 | -0.998 | 1.231 | 0.794 | 0.402 | -0.014 | 0.062 | 0.062 | 0 | 0.09 | 0.31 | 0.06 | 0.11 | 0.02 | 0.06 |
| 3 | 41716317 | rs994439 | -1.338 | 0.923 | -0.316 | 0.161 | -0.101 | 0.018 | 0.018 | 0 | 0.58 | 0.9 | 0.66 | 0.61 | 0.51 | 0.66 |
| 3 | 41717979 | rs1436850 | -1.361 | 0.255 | 0.594 | -0.023 | -0.045 | 0.04 | 0.04 | 0.695 | 0.13 | 0.42 | 0.07 | 0.12 | 0.03 | 0.06 |
| 3 | 41726062 | rs893118 | -0.19 | 0.266 | 0.512 | -0.051 | -0.021 | 0.063 | 0.063 | 3.905 | 0.06 | 0.24 | 0.02 | 0.08 | 0.01 | 0.01 |
| 3 | 41741896 | rs6781095 | -1.532 | 0.203 | 0.703 | -0.091 | -0.051 | -0.002 | -0.002 | 0 | 0.14 | 0.66 | 0.08 | 0.1 | 0.02 | 0.05 |
| 3 | 41760895 | rs1125203 | -0.262 | -0.08 | -0.126 | -0.473 | 0.042 | 0.054 | 0.054 | 0 | 0.37 | 0.73 | 0.18 | 0.15 | 0.19 | 0.18 |
| 3 | 41763648 | rs17060958 | -0.319 | -0.176 | 0.16 | -0.509 | -0.041 | -0.045 | -0.045 | 0 | 0.1 | 0.5 | 0.04 | 0 | 0.01 | 0 |
| 3 | 41764300 | rs9874975 | -0.289 | -0.147 | 0.021 | -0.502 | 0.046 | 0.059 | 0.059 | 0 | 0.37 | 0.73 | 0.18 | 0.15 | 0.19 | 0.18 |
| 3 | 41781118 | rs6787883 | 0.109 | -0.105 | 0.41 | -0.434 | -0.026 | -0.03 | -0.03 | 0 | 0.08 | 0.35 | 0.02 | 0 | 0 | 0 |
| 3 | 41786009 | rs6599175 | -0.236 | -0.109 | 0.101 | -0.485 | 0.042 | 0.055 | 0.055 | 0 | 0.37 | 0.71 | 0.17 | 0.15 | 0.19 | 0.18 |
| 3 | 41794937 | rs7650227 | -0.243 | -0.123 | 0.104 | -0.491 | 0.029 | 0.037 | 0.037 | 0 | 0.37 | 0.7 | 0.17 | 0.15 | 0.19 | 0.18 |
| 3 | 41810220 | rs13093299 | 1.836 | -0.538 | -0.379 | -0.321 | 0.012 | -0.036 | -0.036 | 0 | 0.15 | 0.04 | 0.26 | 0.17 | 0.16 | 0.16 |
| 3 | 41822054 | rs9857175 | -0.008 | -0.15 | -0.026 | -0.15 | 0.013 | 0.015 | 0.015 | 0 | 0.36 | 0.65 | 0.16 | 0.15 | 0.19 | 0.18 |
| 3 | 41824556 | rs6803652 | -0.116 | -0.148 | -0.077 | -0.16 | 0.03 | 0.038 | 0.038 | 0 | 0.37 | 0.71 | 0.17 | 0.15 | 0.19 | 0.18 |
| 3 | 41827063 | rs9311289 | 0.322 | 0.444 | -0.406 | 0.01 | -0.01 | -0.02 | -0.02 | 0 | 0.19 | 0.14 | 0.2 | 0.44 | 0.2 | 0.24 |
| 3 | 41829397 | rs4973892 | 0.741 | 0.625 | -0.75 | -0.131 | -0.059 | 0.039 | 0.039 | 0 | 0.38 | 0.07 | 0.55 | 0.29 | 0.45 | 0.46 |
| 3 | 41831203 | rs17215589 | 2.139 | 0.508 | -0.739 | -0.244 | -0.019 | 0.089 | 0.089 | 0 | 0.19 | 0.06 | 0.09 | 0 | 0.17 | 0.11 |

| | | | | | | | | | | | | | | | | |
|---|---|---|---|---|---|---|---|---|---|---|---|---|---|---|---|---|
| 3 | 41841716 | rs4973986 | -2.69 | 0.554 | -0.758 | -0.147 | -0.046 | -0.066 | -0.066 | 0 | 0.88 | 0.99 | 0.92 | 0.75 | 0.82 | 0.84 |
| 3 | 41851718 | rs6796210 | 0.157 | 0.146 | -0.582 | -0.21 | 0.027 | 0.033 | 0.033 | 0 | 0.36 | 0.71 | 0.18 | 0.15 | 0.19 | 0.18 |
| 3 | 41853161 | rs7632387 | 0.854 | 0.104 | -0.309 | -0.229 | -0.016 | -0.028 | -0.028 | 0.03 | 0.29 | 0.42 | 0.17 | 0.15 | 0.19 | 0.18 |
| 3 | 41855284 | rs6599179 | 0.151 | 0.113 | -0.341 | -0.211 | 0.026 | 0.032 | 0.032 | 0 | 0.36 | 0.71 | 0.18 | 0.15 | 0.19 | 0.18 |
| 3 | 41859973 | rs900569 | -0.514 | 0.288 | -0.581 | -0.335 | -0.183 | -0.153 | -0.153 | 2.766 | 0.66 | 0.97 | 0.68 | 0.4 | 0.51 | 0.62 |
| 3 | 41862873 | rs2128835 | 0.291 | 0.004 | -0.215 | -0.243 | -0.03 | -0.041 | -0.041 | 0 | 0.34 | 0.49 | 0.15 | 0.15 | 0.19 | 0.18 |
| 3 | 41877414 | rs3774372 | 1.101 | 0.044 | -0.253 | -0.225 | 0.002 | 0.005 | 0.005 | 0 | 0.27 | 0.2 | 0.14 | 0.15 | 0.18 | 0.18 |
| 3 | 41897482 | rs1994157 | 1.113 | 0.044 | -0.253 | -0.225 | 0.002 | 0.005 | 0.005 | 0 | 0.26 | 0.2 | 0.14 | 0.15 | 0.18 | 0.18 |
| 3 | 41910571 | rs1716670 | -2.022 | 0.257 | -0.277 | -0.177 | -0.054 | -0.061 | -0.061 | 2.52 | 0.87 | 0.98 | 0.9 | 0.74 | 0.78 | 0.83 |
| 3 | 41920335 | rs2030431 | -0.621 | 0.198 | -0.135 | -0.279 | -0.028 | -0.031 | -0.031 | 0 | 0.92 | 0.71 | 0.98 | 1 | 1 | 1 |
| 3 | 41925398 | rs1052501 | -0.236 | 0.201 | -0.194 | -0.282 | 0.011 | 0.013 | 0.013 | 0 | 0.64 | 0.3 | 0.82 | 0.85 | 0.81 | 0.82 |
| 3 | 41930219 | rs1717000 | -0.208 | 0.202 | -0.239 | -0.281 | 0.027 | 0.033 | 0.033 | 0 | 0.64 | 0.28 | 0.81 | 0.85 | 0.81 | 0.82 |
| 3 | 41941473 | rs1716697 | 0.754 | 0.234 | -0.18 | -0.185 | -0.067 | 0.08 | 0.08 | 1.986 | 0.85 | 0.52 | 0.92 | 0.86 | 0.98 | 0.93 |
| 3 | 41948665 | rs9838537 | 1.989 | 0.48 | -0.399 | -0.269 | -0.026 | 0.033 | 0.033 | 0 | 0.14 | 0.07 | 0.22 | 0.07 | 0.2 | 0.14 |
| 3 | 41952838 | rs9877875 | 0.736 | 0.302 | 0.063 | 0.218 | -0.011 | -0.007 | -0.007 | 1.328 | 0.07 | 0.12 | 0.04 | 0.03 | 0.02 | 0.06 |
| 3 | 41960006 | rs1716975 | -0.046 | 0.295 | -0.393 | 0.158 | 0.024 | 0.03 | 0.03 | 0 | 0.64 | 0.28 | 0.8 | 0.85 | 0.8 | 0.83 |
| 3 | 41961168 | rs1716979 | 0.417 | 0.429 | -0.466 | 0.208 | -0.123 | -0.156 | -0.156 | 0.927 | 0.25 | 0.03 | 0.26 | 0.6 | 0.37 | 0.36 |
| 3 | 41965234 | rs1615243 | 0.221 | -0.03 | 0.111 | -0.036 | 0.02 | 0.024 | 0.024 | 0 | 0.64 | 0.29 | 0.81 | 0.85 | 0.81 | 0.83 |
| 3 | 41965532 | rs1716984 | 0.286 | -0.028 | 0.014 | -0.035 | 0.047 | 0.061 | 0.061 | 0 | 0.63 | 0.21 | 0.8 | 0.85 | 0.81 | 0.83 |
| 3 | 41970925 | rs1716988 | 1.251 | 0.121 | -0.206 | 0.104 | -0.07 | -0.077 | -0.077 | 0 | 0.13 | 0.01 | 0.1 | 0.28 | 0.23 | 0.19 |
| 3 | 41996136 | rs2272007 | 0.606 | -1.001 | 0.609 | -0.43 | 0.027 | 0.032 | 0.032 | 0 | 0.63 | 0.28 | 0.81 | 0.85 | 0.81 | 0.83 |
| 3 | 41996275 | rs7651623 | 0.208 | -1.156 | 1.002 | -0.34 | -0.032 | -0.034 | -0.034 | 0 | 0.92 | 0.7 | 0.98 | 1 | 1 | 1 |
| 3 | 41998209 | rs4973899 | 1.405 | -1.169 | 0.593 | -0.351 | -0.068 | 0.036 | 0.036 | 1.777 | 0.84 | 0.39 | 0.91 | 0.85 | 0.98 | 0.93 |
| 3 | 42019565 | rs7634985 | 0.525 | -1.027 | 0.286 | -0.31 | -0.013 | -0.015 | -0.015 | 0 | 0.66 | 0.38 | 0.83 | 0.85 | 0.82 | 0.83 |
| 3 | 42032483 | rs794894 | 0.391 | -0.996 | 0.342 | -0.293 | -0.032 | -0.037 | -0.037 | 0 | 0.67 | 0.45 | 0.84 | 0.85 | 0.83 | 0.83 |
| 3 | 42043509 | rs7629767 | 0.169 | -1.017 | 0.219 | -0.311 | 0.009 | 0.011 | 0.011 | 0 | 0.25 | 0.17 | 0.13 | 0.15 | 0.17 | 0.17 |
| 3 | 42057101 | rs939561 | 0.28 | -0.869 | 0.023 | -0.302 | -0.013 | -0.017 | -0.017 | 1.567 | 0.67 | 0.38 | 0.83 | 0.86 | 0.82 | 0.83 |
| 3 | 42067377 | rs9877580 | -0.037 | -0.952 | 0.082 | -0.18 | 0 | -0.002 | -0.002 | 0 | 0.05 | 0.24 | 0.02 | 0 | 0.01 | 0 |
| 3 | 42067669 | rs17219347 | 1.003 | -1.351 | -0.35 | -0.509 | 0.032 | 0.061 | 0.061 | 0 | 0.16 | 0.05 | 0.04 | 0 | 0.08 | 0.1 |
| 3 | 42073957 | rs17066958 | -0.338 | -1.727 | 0.014 | -0.397 | -0.068 | -0.027 | -0.027 | 0 | 0.27 | 0.53 | 0.12 | 0.18 | 0.1 | 0.16 |
| 3 | 42076622 | rs904269 | -1.079 | -1.439 | -0.736 | -0.402 | -0.166 | -0.164 | -0.164 | 0 | 0.74 | 0.96 | 0.64 | 0.45 | 0.55 | 0.68 |
| 3 | 42076970 | rs12106811 | 0.777 | -1.714 | -0.336 | -0.8 | 0.005 | 0.029 | 0.029 | 0 | 0.18 | 0.44 | 0.27 | 0.17 | 0.17 | 0.11 |
| 3 | 42079111 | rs9847688 | -1.095 | -1.718 | -1.056 | -0.794 | 0.112 | 0.155 | 0.155 | 0 | 0.47 | 0.92 | 0.37 | 0.23 | 0.28 | 0.26 |
| 3 | 42090688 | rs975580 | 0.051 | -1.48 | -0.943 | -0.741 | -0.036 | 0.114 | 0.114 | 0.08 | 0.25 | 0.06 | 0.28 | 0.09 | 0.3 | 0.19 |
| 3 | 42091744 | rs7636449 | -1.062 | -1.262 | -0.084 | -0.028 | 0.085 | 0.074 | 0.074 | 0 | 0.37 | 0.78 | 0.17 | 0.18 | 0.19 | 0.21 |
| 3 | 42105675 | rs4974016 | 0.144 | -0.012 | 0.324 | 0.423 | 0.016 | 0.008 | 0.008 | 0 | 0.74 | 0.47 | 0.7 | 0.69 | 0.7 | 0.85 |
| 3 | 42110843 | rs5029848 | 1.039 | -0.151 | 0.286 | 0.132 | -0.014 | -0.011 | -0.011 | 0 | 0.62 | 0.29 | 0.6 | 0.43 | 0.57 | 0.7 |
| 3 | 42116757 | rs9861303 | 1.662 | -0.235 | 0.197 | 0.049 | -0.013 | 0.019 | 0.019 | 0.016 | 0.47 | 0.24 | 0.56 | 0.43 | 0.49 | 0.57 |
| 3 | 42119479 | rs1909116 | 0.012 | 0.178 | 0.559 | 0.296 | 0.039 | 0.091 | 0.091 | 0 | 0.75 | 0.56 | 0.69 | 0.58 | 0.68 | 0.81 |
| 3 | 42131415 | rs1995137 | 0.803 | 0.189 | 0.395 | 0.269 | 0.02 | 0.014 | 0.014 | 0 | 0.65 | 0.52 | 0.63 | 0.43 | 0.56 | 0.69 |
| 3 | 42131734 | rs2173336 | 0.014 | 0.324 | 0.109 | 0.212 | 0.006 | 0.011 | 0.011 | 0 | 0.75 | 0.78 | 0.7 | 0.59 | 0.65 | 0.82 |
| 3 | 42166169 | rs9311295 | 0.499 | 0.717 | 0.047 | 0.501 | 0.019 | 0.02 | 0.02 | 0.021 | 0.71 | 0.69 | 0.64 | 0.54 | 0.58 | 0.78 |
| 3 | 42177407 | rs1979789 | 1.16 | 0.662 | -0.478 | 0.446 | -0.004 | 0.003 | 0.003 | 1.691 | 0.54 | 0.43 | 0.58 | 0.43 | 0.51 | 0.62 |
| 3 | 42179679 | rs11129941 | 0.488 | 0.761 | -0.647 | 0.491 | 0.016 | 0.001 | 0.001 | 0.299 | 0.14 | 0.07 | 0.05 | 0.11 | 0.08 | 0.16 |
| 3 | 42180651 | rs2055285 | 0.655 | 0.85 | -0.65 | 0.255 | -0.002 | 0 | 0 | 0 | 0.55 | 0.49 | 0.6 | 0.54 | 0.51 | 0.68 |
| 3 | 42184173 | rs9872588 | -0.815 | 0.699 | -0.795 | 0.108 | -0.014 | 0.024 | 0.024 | 0 | 0.09 | 0.02 | 0.05 | 0.05 | 0.11 | 0.07 |
| 3 | 42192326 | rs4973900 | 0.11 | 0.207 | -0.034 | -0.829 | -0.021 | -0.144 | -0.144 | 0.711 | 0.75 | 0.49 | 0.83 | 0.98 | 0.78 | 0.72 |
| 3 | 42195217 | rs9682473 | 0.539 | -0.048 | -1.094 | -0.661 | 0.023 | -0.099 | -0.099 | 0 | 0.32 | 0.11 | 0.4 | 0.48 | 0.26 | 0.34 |
| 3 | 42208663 | rs9862639 | 0.205 | 0.474 | -0.497 | -1.209 | -0.001 | -0.057 | -0.057 | 0 | 0.57 | 0.4 | 0.73 | 0.95 | 0.59 | 0.54 |
| 3 | 42212734 | rs7633017 | 0.542 | 0.376 | -0.628 | -1.193 | -0.052 | 0.091 | 0.091 | 0 | 0.19 | 0.07 | 0.11 | 0.03 | 0.21 | 0.17 |
| 3 | 42221547 | rs6775303 | 1.126 | 0.519 | -0.732 | -1.291 | -0.048 | 0.054 | 0.054 | 0 | 0.12 | 0.02 | 0.07 | 0.03 | 0.14 | 0.09 |
| 3 | 42225951 | rs4682953 | -0.128 | 0.552 | -0.654 | -1.391 | -0.003 | -0.029 | -0.029 | 0 | 0.65 | 0.63 | 0.77 | 0.95 | 0.64 | 0.62 |
| 3 | 42226151 | rs4234445 | -0.363 | 0.412 | -0.58 | -1.336 | -0.003 | -0.073 | -0.073 | 0 | 0.68 | 0.65 | 0.82 | 0.98 | 0.7 | 0.62 |
| 3 | 42232783 | rs2276828 | 1.34 | 0.568 | -0.522 | -1.273 | -0.036 | -0.068 | -0.068 | 0 | 0.22 | 0.08 | 0.19 | 0.46 | 0.29 | 0.16 |
| 3 | 42233759 | rs6783599 | -1.105 | 0.487 | -0.612 | -1.273 | -0.004 | -0.067 | -0.067 | 0 | 0.77 | 0.72 | 0.86 | 0.98 | 0.77 | 0.71 |
| 3 | 42236415 | rs9851455 | -1.062 | 0.487 | -0.621 | -1.273 | -0.007 | -0.085 | -0.085 | 0 | 0.77 | 0.69 | 0.86 | 0.98 | 0.77 | 0.7 |
| 3 | 42251263 | rs2290134 | -0.206 | 0.707 | -0.467 | -1.276 | 0.001 | 0.005 | 0.005 | 0 | 0.53 | 0.43 | 0.65 | 0.5 | 0.5 | 0.56 |
| 3 | 42252141 | rs3196165 | -1.25 | 0.657 | -0.342 | -1.327 | 0.003 | 0.011 | 0.011 | 0 | 0.78 | 0.85 | 0.87 | 0.98 | 0.77 | 0.71 |
| 3 | 42252274 | rs9311311 | 0.835 | 0.711 | -0.331 | -1.272 | 0.009 | 0.078 | 0.078 | 0 | 0.25 | 0.42 | 0.22 | 0.48 | 0.27 | 0.15 |
| 3 | 42253664 | rs1046910 | 0.464 | 0.489 | -0.204 | -1.058 | 0.008 | -0.023 | -0.023 | 0 | 0.35 | 0.26 | 0.54 | 0.45 | 0.32 | 0.39 |
| 3 | 42254483 | rs10865916 | 0.35 | 0.578 | 0.095 | -0.991 | -0.007 | 0 | 0 | 0 | 0.41 | 0.37 | 0.3 | 0.47 | 0.46 | 0.32 |
| 3 | 42256517 | rs3774386 | 0.526 | 0.6 | -0.064 | -0.996 | 0.006 | -0.013 | -0.013 | 0 | 0.34 | 0.23 | 0.56 | 0.52 | 0.32 | 0.42 |
| 3 | 42257075 | rs3774388 | 0.934 | 0.498 | -0.745 | -0.801 | -0.068 | -0.016 | -0.016 | 0 | 0.32 | 0.12 | 0.21 | 0.36 | 0.37 | 0.21 |
| 3 | 42259299 | rs11928109 | 0.011 | 0.452 | -0.646 | -0.84 | 0.024 | 0.019 | 0.019 | 0.16 | 0.11 | 0.52 | 0.12 | 0.09 | 0.09 | 0.09 |
| 3 | 42264422 | rs3733054 | -0.145 | 0.451 | -0.667 | -0.807 | -0.024 | 0.014 | 0.014 | 0 | 0.44 | 0.71 | 0.33 | 0.45 | 0.45 | 0.31 |
| 3 | 42265166 | rs12487911 | -0.022 | 0.45 | -0.661 | -0.808 | 0.017 | 0.017 | 0.017 | 0.368 | 0.12 | 0.51 | 0.11 | 0.1 | 0.08 | 0.09 |
| 3 | 42265369 | rs3774391 | 0.832 | 0.648 | -1.074 | -0.718 | -0.091 | -0.028 | -0.028 | 0 | 0.32 | 0.08 | 0.21 | 0.35 | 0.37 | 0.22 |
| 3 | 42265634 | rs1043610 | 0.755 | 0.672 | -1.096 | -0.707 | -0.028 | 0.001 | 0.001 | 0.289 | 0.33 | 0.19 | 0.21 | 0.35 | 0.37 | 0.22 |
| 3 | 42266360 | rs3755809 | 0.403 | 0.116 | -1.393 | -0.422 | -0.017 | -0.001 | -0.001 | 0 | 0.21 | 0.07 | 0.19 | 0.15 | 0.21 | 0.27 |

| | | | | | | | | | | | | | | | | |
|---|---|---|---|---|---|---|---|---|---|---|---|---|---|---|---|---|
| 3 | 42270874 | rs11919824 | 0.07 | 0.07 | -1.343 | -1.09 | 0.003 | 0.035 | 0.035 | 0 | 0.35 | 0.29 | 0.3 | 0.11 | 0.3 | 0.45 |
| 3 | 42272118 | rs9879040 | -0.492 | -0.35 | -0.995 | -0.747 | -0.032 | -0.013 | -0.013 | 0.014 | 0.14 | 0.6 | 0.13 | 0.11 | 0.08 | 0.08 |
| 3 | 42274243 | rs12152415 | 0.365 | 0.187 | -1.521 | -0.268 | -0.029 | -0.085 | -0.085 | 0.173 | 0.13 | 0.01 | 0.26 | 0.61 | 0.15 | 0.2 |
| 3 | 42291664 | rs9682785 | 0.637 | -0.097 | -1.287 | -0.778 | 0.009 | -0.008 | -0.008 | 0 | 0.2 | 0.15 | 0.1 | 0.01 | 0.12 | 0.21 |
| 3 | 42293658 | rs17071704 | 0.16 | -0.189 | -0.704 | -0.505 | -0.003 | 0.028 | 0.028 | 0 | 0.14 | 0.16 | 0.16 | 0.59 | 0.13 | 0.22 |
| 3 | 42295194 | rs7628795 | -0.732 | -0.212 | -0.102 | -0.854 | 0.004 | 0.014 | 0.014 | 0 | 0.48 | 0.58 | 0.49 | 0.72 | 0.39 | 0.64 |
| 3 | 42295806 | rs9311317 | -0.875 | -0.473 | 0.181 | -1.143 | -0.025 | -0.035 | -0.035 | 0 | 0.28 | 0.38 | 0.28 | 0.79 | 0.24 | 0.33 |
| 3 | 42298550 | rs11129948 | -0.503 | -0.178 | 0.267 | -0.847 | 0.013 | 0.065 | 0.065 | 0 | 0.49 | 0.66 | 0.5 | 0.71 | 0.4 | 0.64 |
| 3 | 42299737 | rs3774396 | -1.351 | -0.195 | 1.041 | -0.352 | 0.038 | 0.199 | 0.199 | 5.994 | 0.05 | 0.34 | 0.08 | 0.13 | 0 | 0.03 |
| 3 | 42299870 | rs8192472 | 0.001 | -0.003 | 0.349 | -0.287 | 0.007 | -0.003 | -0.003 | 0 | 0.39 | 0.28 | 0.35 | 0.53 | 0.34 | 0.51 |
| 3 | 42303311 | rs10865918 | 1.06 | 0.229 | -0.359 | -0.394 | -0.133 | 0.091 | 0.091 | 0.022 | 0.31 | 0.06 | 0.23 | 0.16 | 0.41 | 0.21 |
| 3 | 42306294 | rs1799923 | -0.034 | 0.124 | -0.034 | -0.419 | -0.014 | 0.084 | 0.084 | 1.428 | 0.86 | 0.64 | 0.78 | 0.72 | 0.89 | 0.85 |
| 3 | 42306446 | rs11571842 | -0.107 | 0.127 | -0.05 | -0.42 | -0.001 | 0.002 | 0.002 | 0 | 0.55 | 0.53 | 0.55 | 0.55 | 0.49 | 0.64 |
| 3 | 42314587 | rs11129950 | 0.549 | -0.233 | -0.185 | -0.411 | 0.002 | 0.01 | 0.01 | 0 | 0.15 | 0.12 | 0.18 | 0.03 | 0.14 | 0.07 |
| 3 | 42320079 | rs4683161 | -0.036 | 0.005 | -0.401 | -0.172 | -0.003 | -0.003 | -0.003 | 0 | 0.52 | 0.44 | 0.52 | 0.55 | 0.45 | 0.59 |
| 3 | 42329986 | rs9311318 | -0.953 | 0.092 | -0.615 | -0.151 | -0.066 | 0.03 | 0.03 | 0.402 | 0.65 | 0.82 | 0.7 | 0.7 | 0.55 | 0.75 |
| 3 | 42331546 | rs9813958 | -0.408 | 0.229 | -0.181 | 0.213 | -0.004 | 0.022 | 0.022 | 0.187 | 0.12 | 0.28 | 0.17 | 0.15 | 0.1 | 0.15 |
| 3 | 42342619 | rs2888182 | -0.518 | 0.622 | -0.542 | 0.531 | 0.017 | 0.056 | 0.056 | 0.213 | 0.06 | 0.18 | 0.15 | 0.14 | 0.09 | 0.02 |
| 3 | 42343616 | rs9682293 | -0.413 | 0.599 | -1.184 | 0.912 | 0.022 | -0.001 | -0.001 | 0 | 0.56 | 0.36 | 0.62 | 0.5 | 0.67 | 0.46 |
| 3 | 42347132 | rs11129960 | 0.34 | 0.456 | -1.411 | 0.978 | -0.03 | 0.017 | 0.017 | 4.118 | 0.47 | 0.11 | 0.47 | 0.45 | 0.58 | 0.45 |
| 3 | 42347678 | rs11712777 | 0.993 | 0.795 | -1.767 | 0.879 | -0.125 | -0.023 | -0.023 | 0 | 0.27 | 0.04 | 0.18 | 0.26 | 0.39 | 0.25 |
| 3 | 42355010 | rs1442265 | 0.701 | 0.215 | -1.364 | 1.04 | -0.007 | -0.025 | -0.025 | 0.358 | 0.17 | 0.1 | 0.17 | 0.28 | 0.2 | 0.31 |
| 3 | 42356836 | rs659113 | 0.219 | 0.586 | -1.401 | 1.172 | -0.085 | -0.01 | -0.01 | 5.057 | 0.44 | 0.09 | 0.46 | 0.44 | 0.54 | 0.44 |
| 3 | 42360871 | rs474470 | -0.328 | 0.344 | -1.006 | 1.256 | 0.014 | 0.007 | 0.007 | 0 | 0.54 | 0.42 | 0.69 | 0.64 | 0.63 | 0.47 |
| 3 | 42362817 | rs678574 | -0.11 | 0.453 | -1.351 | 1.318 | 0.049 | 0.015 | 0.015 | 0 | 0.51 | 0.24 | 0.6 | 0.5 | 0.63 | 0.46 |
| 3 | 42371742 | rs683887 | -0.109 | 0.54 | -1.526 | 0.889 | 0.049 | 0.012 | 0.012 | 0 | 0.51 | 0.24 | 0.6 | 0.51 | 0.63 | 0.46 |
| 3 | 42378073 | rs6781092 | -0.49 | 0.747 | -1.154 | 1.117 | -0.004 | -0.013 | -0.013 | 0.111 | 0.28 | 0.29 | 0.29 | 0.57 | 0.34 | 0.31 |
| 3 | 42378179 | rs1019188 | -0.392 | 0.707 | -0.647 | 1.235 | 0.002 | 0.038 | 0.038 | 0 | 0.18 | 0.22 | 0.18 | 0.37 | 0.19 | 0.21 |
| 3 | 42380328 | rs1348247 | -0.071 | 0.509 | -0.783 | 0.975 | -0.001 | 0.006 | 0.006 | 0.424 | 0.26 | 0.31 | 0.23 | 0.37 | 0.28 | 0.3 |
| 3 | 42391480 | rs33510 | 1 | 0.088 | -1.101 | 0.507 | -0.055 | -0.111 | -0.111 | 0 | 0.24 | 0.03 | 0.41 | 0.38 | 0.22 | 0.26 |
| 3 | 42392724 | rs1442268 | 0.147 | 0.207 | -0.58 | 0.142 | -0.002 | -0.021 | -0.021 | 0 | 0.26 | 0.27 | 0.22 | 0.37 | 0.28 | 0.3 |
| 3 | 42394776 | rs33512 | -0.689 | 0.348 | -0.858 | -0.745 | 0.031 | -0.17 | -0.17 | 0.03 | 0.76 | 0.59 | 0.8 | 1 | 0.68 | 0.77 |
| 3 | 42399905 | rs13093997 | 0.152 | 0.315 | -1.287 | -0.222 | -0.009 | 0.021 | 0.021 | 0.152 | 0.26 | 0.21 | 0.21 | 0.18 | 0.28 | 0.29 |
| 3 | 42402393 | rs1442272 | 0.738 | 0.434 | -1.167 | -0.566 | -0.01 | -0.013 | -0.013 | 0.184 | 0.21 | 0.17 | 0.2 | 0 | 0.33 | 0.19 |
| 3 | 42402514 | rs33454 | -0.153 | -0.08 | -0.7 | -0.684 | -0.06 | -0.013 | -0.013 | 0.545 | 0.12 | 0.43 | 0.13 | 0.1 | 0.02 | 0.02 |
| 3 | 42404785 | rs26350 | 1.288 | -0.162 | -1.757 | -0.608 | -0.068 | -0.148 | -0.148 | 0.048 | 0.32 | 0.06 | 0.37 | 0.49 | 0.29 | 0.38 |
| 3 | 42404928 | rs26351 | 0.467 | 0.1 | -1.202 | -0.054 | 0.005 | 0.004 | 0.004 | 0 | 0.66 | 0.34 | 0.68 | 0.69 | 0.7 | 0.67 |
| 3 | 42405350 | rs26352 | 0.948 | -0.019 | -1.363 | -0.021 | -0.018 | -0.043 | -0.043 | 0 | 0.44 | 0.14 | 0.4 | 0.49 | 0.36 | 0.47 |
| 3 | 42408243 | rs33470 | -0.775 | -0.024 | -0.538 | -0.186 | -0.059 | -0.006 | -0.006 | 0 | 0.12 | 0.52 | 0.13 | 0.1 | 0.02 | 0.02 |
| 3 | 42408422 | rs33527 | -0.432 | 0.255 | -1.505 | -0.088 | -0.054 | -0.038 | -0.038 | 0 | 0.81 | 0.98 | 0.84 | 0.81 | 0.75 | 0.71 |
| 3 | 42413845 | rs33480 | 0.074 | -0.424 | -1.031 | -0.088 | -0.014 | -0.001 | -0.001 | 0 | 0.06 | 0.1 | 0.1 | 0.05 | 0.02 | 0.02 |
| 3 | 42415470 | rs33533 | -0.414 | -0.48 | -1.457 | -0.171 | -0.029 | -0.025 | -0.025 | 0 | 0.23 | 0.68 | 0.19 | 0.11 | 0.07 | 0.08 |
| 3 | 42415966 | rs33534 | 0.311 | 1.372 | -1.899 | 0.321 | -0.089 | -0.014 | -0.014 | 0 | 0.75 | 0.96 | 0.75 | 0.77 | 0.59 | 0.64 |
| 3 | 42417576 | rs33483 | -0.077 | 0.994 | -1.819 | 0.067 | -0.007 | 0.076 | 0.076 | 0 | 0.47 | 0.67 | 0.39 | 0.31 | 0.45 | 0.29 |
| 3 | 42417887 | rs33484 | 0.026 | 0.809 | -1.061 | -0.085 | -0.002 | 0.008 | 0.008 | 0 | 0.47 | 0.52 | 0.38 | 0.34 | 0.47 | 0.29 |
| 3 | 42417982 | rs33485 | -0.39 | 0.844 | -1.586 | -0.138 | -0.035 | 0.033 | 0.033 | 0 | 0.8 | 0.95 | 0.86 | 0.87 | 0.74 | 0.83 |
| 3 | 42418446 | rs28350 | -0.729 | 0.845 | -1.601 | -0.244 | -0.01 | 0.053 | 0.053 | 0 | 0.84 | 0.96 | 0.91 | 0.94 | 0.81 | 0.82 |
| 3 | 42419738 | rs33486 | 0.79 | 0.751 | -1.121 | -0.731 | 0 | -0.008 | -0.008 | 3.222 | 0.63 | 0.39 | 0.64 | 0.83 | 0.6 | 0.77 |
| 3 | 42423590 | rs33491 | 0.639 | 0.662 | -1.003 | -0.707 | 0.015 | 0.046 | 0.046 | 1.616 | 0.64 | 0.28 | 0.66 | 0.83 | 0.64 | 0.78 |
| 3 | 42427191 | rs33500 | -0.979 | 0.673 | -1.202 | -0.617 | 0.001 | 0.036 | 0.036 | 0 | 0.84 | 0.93 | 0.91 | 0.92 | 0.81 | 0.83 |
| 3 | 42428487 | rs11718659 | 0.679 | 0.426 | -1.489 | -0.124 | -0.077 | -0.139 | -0.139 | 0.868 | 0.29 | 0.07 | 0.42 | 0.48 | 0.33 | 0.55 |
| 3 | 42429817 | rs6792514 | -0.827 | 0.658 | -1.47 | -0.494 | 0.002 | 0.069 | 0.069 | 0 | 0.6 | 0.72 | 0.76 | 0.92 | 0.58 | 0.75 |
| 3 | 42433305 | rs17074351 | -0.506 | 0.544 | -1.015 | -0.372 | 0.025 | 0.047 | 0.047 | 0 | 0.24 | 0.31 | 0.28 | 0.38 | 0.27 | 0.17 |
| 3 | 42435119 | rs247421 | 1.126 | 0.373 | -1.369 | -0.869 | -0.005 | 0.066 | 0.066 | 0.609 | 0.28 | 0.15 | 0.14 | 0 | 0.25 | 0.08 |
| 3 | 42438076 | rs40415 | -1.168 | 0.329 | -1.248 | -0.33 | -0.039 | -0.07 | -0.07 | 1.533 | 0.65 | 0.87 | 0.52 | 0.43 | 0.58 | 0.42 |
| 3 | 42438754 | rs13054 | -1.168 | 0.329 | -1.248 | -0.33 | -0.039 | -0.07 | -0.07 | 0 | 0.65 | 0.87 | 0.53 | 0.44 | 0.58 | 0.43 |
| 3 | 42439589 | rs2041376 | -0.962 | 0.331 | -1.209 | -0.352 | 0.007 | -0.023 | -0.023 | 0 | 0.37 | 0.73 | 0.39 | 0.44 | 0.33 | 0.35 |
| 3 | 42448296 | rs33518 | -1.044 | 0.398 | -1.454 | -0.325 | -0.057 | -0.088 | -0.088 | 1.045 | 0.65 | 0.9 | 0.52 | 0.44 | 0.58 | 0.42 |
| 3 | 42448471 | rs2286720 | 1.554 | 0.528 | -1.466 | -0.238 | -0.038 | -0.042 | -0.042 | 0.121 | 0.08 | 0.01 | 0.05 | 0.28 | 0.11 | 0.13 |
| 3 | 42467919 | rs168208 | -0.998 | 0.355 | -1.416 | -0.2 | -0.036 | -0.078 | -0.078 | 0.105 | 0.65 | 0.89 | 0.53 | 0.45 | 0.6 | 0.36 |
| 3 | 42470077 | rs13317334 | -0.182 | 0.003 | -0.445 | 0.067 | 0.001 | 0.005 | 0.005 | 0 | 0.1 | 0.24 | 0.14 | 0.1 | 0.06 | 0.02 |
| 3 | 42470438 | rs6441802 | -0.453 | 0.005 | 0.047 | 0.076 | 0.013 | 0.02 | 0.02 | 0 | 0.21 | 0.3 | 0.18 | 0.28 | 0.24 | 0.16 |
| 3 | 42484782 | rs4682737 | 0.039 | 0.233 | -0.638 | 0.155 | -0.085 | -0.09 | -0.09 | 0.758 | 0.17 | 0.01 | 0.15 | 0.27 | 0.24 | 0.16 |
| 3 | 42484827 | rs10154877 | -0.501 | -0.216 | 0.006 | 0.024 | 0.002 | 0.006 | 0.006 | 0 | 0.1 | 0.24 | 0.14 | 0.1 | 0.06 | 0.03 |
| 3 | 42488962 | rs594254 | -0.281 | -0.532 | -0.377 | -0.088 | -0.005 | -0.009 | -0.009 | 1.063 | 0.38 | 0.55 | 0.27 | 0.24 | 0.27 | 0.4 |
| 3 | 42492513 | rs11717603 | 1.063 | -0.372 | -0.386 | 0.614 | -0.001 | 0.008 | 0.008 | 0 | 0.17 | 0.19 | 0.33 | 0.38 | 0.18 | 0.28 |
| 3 | 42493287 | rs397247 | 0.336 | -0.486 | -0.088 | 0.271 | -0.003 | -0.002 | -0.002 | 0 | 0.42 | 0.4 | 0.3 | 0.41 | 0.36 | 0.41 |
| 3 | 42494635 | rs6768356 | 1.227 | -0.665 | 0.068 | 0.172 | -0.001 | 0.004 | 0.004 | 0 | 0.33 | 0.28 | 0.18 | 0.26 | 0.3 | 0.26 |
| 3 | 42497914 | rs13324412 | 0.587 | -0.792 | -0.069 | 0.051 | -0.008 | 0.043 | 0.043 | 0 | 0.13 | 0.35 | 0.13 | 0.15 | 0.07 | 0.1 |

| | | | | | | | | | | | | | | | | |
|---|---|---|---|---|---|---|---|---|---|---|---|---|---|---|---|---|
| 3 | 42500125 | rs588801 | -1.438 | -0.81 | -0.071 | -0.005 | -0.044 | 0.104 | 0.104 | 0 | 0.72 | 0.9 | 0.76 | 0.93 | 0.62 | 0.79 |
| 3 | 42501338 | rs396426 | 0.609 | -0.789 | -0.096 | 0.037 | -0.001 | -0.009 | -0.009 | 0.433 | 0.23 | 0.22 | 0.43 | 0.5 | 0.25 | 0.44 |
| 3 | 42503482 | rs1508207 | 1.411 | -0.854 | -0.075 | 0.012 | -0.008 | 0.021 | 0.021 | 0 | 0.32 | 0.14 | 0.17 | 0.24 | 0.3 | 0.25 |
| 3 | 42515965 | rs17074496 | 0.259 | -0.662 | 0.241 | 0.097 | -0.033 | 0.061 | 0.061 | 0 | 0.14 | 0.44 | 0.12 | 0.15 | 0.05 | 0.1 |
| 3 | 42517802 | rs545499 | 0.232 | -0.466 | -0.345 | 0.186 | 0.006 | 0.026 | 0.026 | 0 | 0.07 | 0.11 | 0.11 | 0.11 | 0.07 | 0.16 |
| 3 | 42518379 | rs11129965 | 0.161 | -0.535 | -0.754 | 0.046 | 0.008 | 0.078 | 0.078 | 0 | 0.57 | 0.27 | 0.62 | 0.78 | 0.57 | 0.69 |
| 3 | 42521749 | rs4383487 | 0.169 | -0.459 | -0.836 | 0.233 | -0.006 | -0.046 | -0.046 | 1.195 | 0.25 | 0.18 | 0.36 | 0.48 | 0.27 | 0.41 |
| 3 | 42524866 | rs11714082 | 0.065 | -0.494 | -0.852 | 0.238 | -0.004 | -0.048 | -0.048 | 0.41 | 0.26 | 0.19 | 0.36 | 0.48 | 0.27 | 0.42 |
| 3 | 42528031 | rs601204 | 1.242 | -0.423 | -0.993 | 0.107 | 0.001 | 0.028 | 0.028 | 0 | 0.33 | 0.19 | 0.27 | 0.19 | 0.29 | 0.25 |
| 3 | 42529082 | rs11718502 | 1.14 | 0.002 | -1.027 | 0.389 | -0.025 | -0.059 | -0.059 | 0.172 | 0.16 | 0.11 | 0.24 | 0.43 | 0.2 | 0.23 |
| 3 | 42529750 | rs435698 | -0.601 | -0.049 | -0.478 | -0.083 | -0.045 | 0.101 | 0.101 | 0 | 0.73 | 0.91 | 0.75 | 0.89 | 0.65 | 0.81 |
| 3 | 42534042 | rs342520 | 0.057 | 0.144 | -0.29 | 0.078 | 0.002 | -0.025 | -0.025 | 0 | 0.6 | 0.51 | 0.64 | 0.73 | 0.59 | 0.69 |
| 3 | 42541887 | rs342543 | -0.108 | 0.204 | -0.206 | 0.185 | 0 | 0 | 0 | 0.252 | 0.49 | 0.51 | 0.66 | 0.5 | 0.58 | 0.44 |
| 3 | 42547127 | rs1876506 | -0.771 | -0.063 | 0.353 | 0.155 | 0.081 | 0.119 | 0.119 | 0.625 | 0.05 | 0.36 | 0.05 | 0.07 | 0.01 | 0.04 |
| 3 | 42549343 | rs399582 | -1.042 | -0.234 | -0.104 | -0.424 | 0.011 | -0.006 | -0.006 | 3.095 | 0.73 | 0.82 | 0.88 | 0.98 | 0.8 | 0.86 |
| 3 | 42550651 | rs12493291 | -0.007 | -0.377 | -0.539 | -0.285 | 0.074 | -0.018 | -0.018 | 0 | 0.55 | 0.22 | 0.64 | 0.57 | 0.67 | 0.64 |
| 3 | 42551373 | rs7627240 | 1.307 | -0.356 | -0.742 | -0.492 | -0.102 | 0.092 | 0.092 | 0.008 | 0.29 | 0.04 | 0.39 | 0.14 | 0.42 | 0.24 |
| 3 | 42553095 | rs1078008 | -0.806 | -0.232 | -0.502 | -0.474 | 0 | -0.021 | -0.021 | 0 | 0.67 | 0.38 | 0.83 | 0.91 | 0.79 | 0.82 |
| 3 | 42555524 | rs2278215 | 0.629 | -0.194 | -0.599 | 0.241 | -0.093 | -0.027 | -0.027 | 0 | 0.2 | 0.02 | 0.2 | 0.19 | 0.22 | 0.31 |
| 3 | 42557199 | rs2267535 | -0.25 | -0.327 | 0.137 | 0.445 | -0.033 | 0.148 | 0.148 | 0.626 | 0.16 | 0.4 | 0.22 | 0.37 | 0.12 | 0.18 |
| 3 | 42566142 | rs407931 | -0.853 | -0.035 | -0.566 | 0.302 | -0.006 | 0.02 | 0.02 | 0.449 | 0.26 | 0.69 | 0.27 | 0.44 | 0.21 | 0.28 |
| 3 | 42568440 | rs437876 | -0.22 | 0.191 | -1.09 | -0.171 | 0.015 | 0.158 | 0.158 | 0.09 | 0.58 | 0.89 | 0.47 | 0.84 | 0.37 | 0.66 |
| 3 | 42571729 | rs16810 | 0.203 | 0.622 | -0.435 | 0.236 | 0.002 | 0 | 0 | 0 | 0.55 | 0.53 | 0.53 | 0.57 | 0.45 | 0.6 |
| 3 | 42578509 | rs342511 | -0.144 | 0.782 | -0.767 | 0.057 | -0.073 | -0.006 | -0.006 | 0 | 0.67 | 0.87 | 0.59 | 0.68 | 0.52 | 0.66 |
| 3 | 42578768 | rs8913 | -0.248 | 0.949 | -0.838 | -0.062 | -0.103 | 0.02 | 0.02 | 1.692 | 0.72 | 0.92 | 0.61 | 0.81 | 0.54 | 0.7 |
| 3 | 42580706 | rs342508 | -0.707 | 0.19 | 0.083 | 0.312 | -0.025 | 0.178 | 0.178 | 1.538 | 0.2 | 0.5 | 0.22 | 0.54 | 0.16 | 0.33 |
| 3 | 42588822 | rs342521 | -1.331 | 0.602 | 0.033 | 0.031 | 0.001 | -0.024 | -0.024 | 0 | 0.36 | 0.8 | 0.3 | 0.58 | 0.34 | 0.51 |
| 3 | 42594019 | rs663258 | -0.393 | 0.667 | 0.329 | 0.064 | 0.014 | 0.215 | 0.215 | 0.526 | 0.19 | 0.48 | 0.23 | 0.47 | 0.2 | 0.26 |
| 3 | 42599091 | rs2271184 | -0.38 | 0.666 | 0.371 | 0.064 | 0.013 | 0.207 | 0.207 | 0.507 | 0.19 | 0.45 | 0.22 | 0.47 | 0.2 | 0.26 |
| 3 | 42599588 | rs6797545 | 0.81 | 0.654 | 0.428 | 0.007 | 0.007 | -0.075 | -0.075 | 0 | 0.13 | 0.26 | 0.05 | 0 | 0.11 | 0.15 |
| 3 | 42617654 | rs6793102 | 1.363 | 0.775 | -0.401 | -0.415 | -0.092 | -0.051 | -0.051 | 3.336 | 0.24 | 0.01 | 0.2 | 0.25 | 0.27 | 0.23 |
| 3 | 42630441 | rs2940407 | -0.079 | 0.697 | 0.287 | -0.054 | 0.016 | 0.198 | 0.198 | 0.003 | 0.17 | 0.4 | 0.21 | 0.47 | 0.18 | 0.26 |
| 3 | 42630979 | rs34801786 | -0.032 | 0.688 | 0.257 | -0.056 | -0.002 | -0.026 | -0.026 | 0 | 0.17 | 0.32 | 0.07 | 0.1 | 0.14 | 0.25 |
| 3 | 42636141 | rs4549 | -1.48 | 0.683 | 0.236 | -0.058 | 0.004 | -0.024 | -0.024 | 0 | 0.36 | 0.82 | 0.3 | 0.59 | 0.34 | 0.51 |
| 3 | 42637245 | rs637852 | 0.145 | 0.701 | 0.417 | -0.024 | 0.011 | 0.137 | 0.137 | 0.011 | 0.16 | 0.31 | 0.2 | 0.48 | 0.18 | 0.26 |
| 3 | 42638261 | rs13064311 | 0.445 | 0.691 | 0.045 | -0.029 | -0.033 | -0.06 | -0.06 | 0 | 0.13 | 0.06 | 0.18 | 0.48 | 0.18 | 0.26 |
| 3 | 42652674 | rs3774402 | 0.044 | 0.722 | 0.018 | -0.024 | 0.002 | -0.022 | -0.022 | 0 | 0.16 | 0.31 | 0.07 | 0.1 | 0.14 | 0.25 |
| 3 | 42668441 | rs168138 | -0.243 | 0.734 | -0.006 | -0.013 | 0.007 | 0.191 | 0.191 | 1.39 | 0.2 | 0.48 | 0.23 | 0.47 | 0.21 | 0.27 |
| 3 | 42672486 | rs1062051 | 0.032 | 0.835 | -0.09 | -0.026 | 0.015 | -0.024 | -0.024 | 0 | 0.16 | 0.32 | 0.07 | 0.1 | 0.15 | 0.25 |
| 3 | 42689777 | rs8351 | -1.003 | 0.589 | -0.69 | -0.032 | 0.003 | -0.025 | -0.025 | 0 | 0.37 | 0.82 | 0.31 | 0.58 | 0.35 | 0.52 |
| 3 | 42708632 | rs2290976 | 0.634 | 0.319 | -0.219 | -0.123 | -0.004 | -0.028 | -0.028 | 0 | 0.13 | 0.17 | 0.12 | 0.05 | 0.1 | 0.13 |
| 3 | 42711655 | rs339716 | -1 | 0.209 | -0.601 | -0.798 | -0.005 | -0.004 | -0.004 | 1.761 | 0.7 | 0.79 | 0.69 | 0.99 | 0.71 | 0.8 |
| 3 | 42712156 | rs339714 | 0.299 | 0.077 | -0.405 | -0.698 | -0.001 | -0.032 | -0.032 | 0 | 0.52 | 0.53 | 0.56 | 0.82 | 0.58 | 0.59 |
| 3 | 42713014 | rs9647353 | 1.327 | 0.198 | -0.967 | -0.631 | -0.057 | -0.003 | -0.003 | 0 | 0.23 | 0.05 | 0.22 | 0.22 | 0.27 | 0.25 |
| 3 | 42716120 | rs339712 | -0.565 | 0.222 | -0.086 | -0.817 | -0.006 | 0.014 | 0.014 | 0 | 0.7 | 0.8 | 0.69 | 0.99 | 0.71 | 0.8 |
| 3 | 42723605 | rs339708 | 0.567 | 0.023 | 0.129 | -0.024 | -0.005 | -0.011 | -0.011 | 0.252 | 0.13 | 0.15 | 0.1 | 0.07 | 0.09 | 0.13 |
| 3 | 42733468 | rs123509 | -0.496 | 0.636 | -0.492 | -0.26 | -0.042 | 0.057 | 0.057 | 0 | 0.85 | 0.94 | 0.87 | 1 | 0.77 | 0.89 |
| 3 | 42733667 | rs3846062 | 0.656 | -0.023 | 0.027 | -0.22 | 0 | 0.003 | 0.003 | 0.114 | 0.56 | 0.44 | 0.65 | 0.7 | 0.59 | 0.59 |
| 3 | 42735274 | rs339697 | 0.325 | -0.1 | -0.156 | -0.235 | -0.049 | 0.036 | 0.036 | 0.239 | 0.6 | 0.85 | 0.69 | 0.71 | 0.6 | 0.6 |
| 3 | 42737879 | rs339694 | 0.408 | -0.141 | 0.601 | -0.26 | -0.001 | 0.005 | 0.005 | 0.004 | 0.58 | 0.62 | 0.66 | 0.69 | 0.59 | 0.59 |
| 3 | 42738297 | rs2290973 | 1.028 | -0.65 | 0.028 | -0.47 | -0.02 | 0.03 | 0.03 | 0 | 0.43 | 0.2 | 0.6 | 0.66 | 0.54 | 0.51 |
| 3 | 42747548 | rs9836162 | 0.822 | -0.286 | -0.091 | -0.373 | 0.002 | 0.001 | 0.001 | 0 | 0.3 | 0.26 | 0.2 | 0.29 | 0.27 | 0.35 |
| 3 | 42750446 | rs2240860 | 0.873 | -0.42 | -0.136 | -0.603 | 0.005 | -0.004 | -0.004 | 0.004 | 0.27 | 0.22 | 0.21 | 0.3 | 0.19 | 0.3 |
| 3 | 42752061 | rs13072026 | 0.827 | -0.419 | -0.135 | -0.583 | 0.006 | -0.005 | -0.005 | 0 | 0.27 | 0.23 | 0.21 | 0.3 | 0.19 | 0.3 |
| 3 | 42753904 | rs339661 | -1.208 | -0.36 | -0.413 | -0.801 | -0.049 | 0.08 | 0.08 | 0 | 0.82 | 0.94 | 0.87 | 1 | 0.76 | 0.89 |
| 3 | 42754912 | rs3895740 | 0.054 | -0.387 | 0.105 | -0.757 | -0.01 | -0.009 | -0.009 | 0 | 0.58 | 0.62 | 0.41 | 0.47 | 0.46 | 0.61 |
| 3 | 42771791 | rs13085272 | 0.954 | -0.343 | -0.26 | -0.862 | 0.014 | 0.029 | 0.029 | 0 | 0.27 | 0.17 | 0.17 | 0.11 | 0.19 | 0.28 |
| 3 | 42781316 | rs11709500 | 1.407 | -0.341 | -0.331 | -0.852 | -0.004 | 0.047 | 0.047 | 0 | 0.28 | 0.14 | 0.17 | 0.16 | 0.27 | 0.28 |
| 3 | 42789316 | rs13095730 | 0.917 | -0.325 | -0.656 | -0.818 | 0.019 | 0.044 | 0.044 | 0 | 0.27 | 0.14 | 0.17 | 0.11 | 0.19 | 0.25 |
| 3 | 42799765 | rs17238798 | 1.569 | -0.126 | -0.717 | -0.712 | -0.081 | 0.021 | 0.021 | 0 | 0.25 | 0.01 | 0.16 | 0.17 | 0.27 | 0.28 |
| 3 | 42800291 | rs11915803 | 1.415 | -0.118 | -0.619 | -0.709 | -0.046 | 0.047 | 0.047 | 0 | 0.26 | 0.06 | 0.16 | 0.17 | 0.27 | 0.28 |
| 3 | 42808493 | rs339664 | -0.372 | -0.098 | -0.143 | -0.553 | -0.001 | -0.017 | -0.017 | 0 | 0.5 | 0.48 | 0.66 | 0.8 | 0.48 | 0.57 |
| 3 | 42819857 | rs2372110 | 1.218 | -0.16 | -0.312 | -0.466 | -0.087 | -0.151 | -0.151 | 0 | 0.18 | 0.02 | 0.43 | 0.51 | 0.28 | 0.3 |
| 3 | 42827437 | rs4683332 | 0.949 | -0.17 | -0.166 | -0.645 | -0.023 | -0.03 | -0.03 | 0 | 0.19 | 0.11 | 0.47 | 0.69 | 0.28 | 0.31 |
| 3 | 42828546 | rs11717346 | 0.389 | -0.183 | -0.137 | -0.668 | 0 | -0.01 | -0.01 | 0 | 0.49 | 0.53 | 0.34 | 0.2 | 0.52 | 0.43 |
| 3 | 42832979 | rs13061576 | 0.563 | -0.197 | -0.183 | -0.636 | 0.031 | 0.086 | 0.086 | 0 | 0.3 | 0.12 | 0.18 | 0.11 | 0.2 | 0.26 |
| 3 | 42855765 | rs4683335 | 0.585 | 0.414 | 0.001 | 0.027 | -0.022 | -0.027 | -0.027 | 0 | 0.18 | 0.11 | 0.47 | 0.7 | 0.28 | 0.31 |
| 3 | 42863739 | rs4682859 | 0.707 | 0.513 | -0.92 | -0.25 | -0.087 | -0.119 | -0.119 | 0.001 | 0.19 | 0.03 | 0.5 | 0.58 | 0.3 | 0.31 |
| 3 | 42865044 | rs7611888 | 0.786 | 0.523 | -0.93 | -0.245 | -0.097 | -0.128 | -0.128 | 0.111 | 0.19 | 0.01 | 0.5 | 0.58 | 0.3 | 0.31 |

| | | | | | | | | | | | | | | | | |
|---|---|---|---|---|---|---|---|---|---|---|---|---|---|---|---|---|
| 3 | 42869216 | rs7653015 | 0.428 | 0.847 | -0.466 | -0.621 | 0.033 | 0.016 | 0.016 | 0 | 0.13 | 0.27 | 0.16 | 0.15 | 0.18 | 0.1 |
| 3 | 42870395 | rs4396867 | 0.55 | 0.44 | -1.389 | -1.318 | -0.047 | 0.16 | 0.16 | 0 | 0.29 | 0.06 | 0.13 | 0.01 | 0.31 | 0.29 |
| 3 | 42872590 | rs7433284 | -0.115 | 0.061 | -1.478 | -1.126 | -0.076 | -0.06 | -0.06 | 0.058 | 0.35 | 0.08 | 0.55 | 0.62 | 0.4 | 0.49 |
| 3 | 42877413 | rs4682862 | -0.752 | -0.306 | -0.669 | -1.342 | -0.001 | -0.003 | -0.003 | 0 | 0.43 | 0.46 | 0.61 | 0.64 | 0.42 | 0.54 |
| 3 | 42880490 | rs9861236 | 1.332 | -0.427 | -0.639 | -1.405 | 0.006 | 0.003 | 0.003 | 0 | 0.12 | 0.1 | 0.03 | 0.03 | 0.07 | 0.12 |
| 3 | 42890731 | rs11129979 | -0.923 | -0.624 | -0.092 | -1.506 | 0.002 | 0.025 | 0.025 | 0 | 0.43 | 0.4 | 0.63 | 0.7 | 0.42 | 0.54 |
| 3 | 42895982 | rs11129980 | 0.958 | -0.622 | -0.371 | -1.496 | -0.097 | -0.125 | -0.125 | 0 | 0.23 | 0.03 | 0.52 | 0.59 | 0.32 | 0.3 |
| 3 | 42896639 | rs9815043 | -0.276 | -0.633 | -0.288 | -1.548 | -0.012 | 0.013 | 0.013 | 0 | 0.36 | 0.25 | 0.58 | 0.68 | 0.4 | 0.43 |
| 3 | 42898649 | rs17239701 | 1.371 | -0.631 | -0.329 | -1.523 | 0.015 | 0.05 | 0.05 | 0 | 0.12 | 0.03 | 0.03 | 0.03 | 0.07 | 0.12 |
| 3 | 42908775 | rs1366046 | 0 | -0.885 | -0.404 | -1.671 | -0.015 | 0.014 | 0.014 | 0 | 0.36 | 0.23 | 0.58 | 0.68 | 0.4 | 0.43 |
| 3 | 42909160 | rs3919627 | 0.018 | -0.885 | -0.416 | -1.671 | -0.017 | 0.014 | 0.014 | 0 | 0.36 | 0.22 | 0.58 | 0.68 | 0.4 | 0.43 |
| 3 | 42909687 | rs11715464 | 1.476 | -1.169 | -0.916 | -1.994 | -0.106 | 0.113 | 0.113 | 0 | 0.37 | 0.02 | 0.17 | 0.17 | 0.38 | 0.34 |
| 3 | 42910621 | rs7612912 | -0.091 | -0.692 | -0.377 | -1.716 | -0.005 | 0.016 | 0.016 | 0 | 0.38 | 0.31 | 0.59 | 0.7 | 0.4 | 0.52 |
| 3 | 42913845 | rs12494055 | -1.167 | -0.328 | -0.053 | -1.545 | 0.012 | 0.107 | 0.107 | 0 | 0.5 | 0.7 | 0.74 | 0.82 | 0.52 | 0.64 |
| 3 | 42914012 | rs3732860 | -1.162 | -0.321 | -0.05 | -1.542 | 0.012 | 0.106 | 0.106 | 0 | 0.5 | 0.7 | 0.74 | 0.82 | 0.52 | 0.64 |
| 3 | 42915385 | rs6782601 | 1.314 | -0.193 | -0.024 | -1.543 | 0.038 | -0.049 | -0.049 | 0 | 0.11 | 0.22 | 0.13 | 0.01 | 0.17 | 0.1 |
| 3 | 42915878 | rs735320 | 1.461 | -0.156 | 0.06 | -1.535 | 0.036 | -0.049 | -0.049 | 0.531 | 0.1 | 0.17 | 0.12 | 0 | 0.17 | 0.09 |
| 3 | 42917906 | rs734969 | -0.532 | -0.408 | 0.094 | -1.277 | 0 | 0.003 | 0.003 | 0 | 0.37 | 0.35 | 0.58 | 0.67 | 0.36 | 0.44 |
| 3 | 42918238 | rs760194 | -0.016 | -0.23 | 0.074 | -1.219 | -0.001 | 0.002 | 0.002 | 0 | 0.3 | 0.35 | 0.53 | 0.66 | 0.28 | 0.42 |
| 3 | 42922177 | rs11923822 | 1.234 | -0.339 | -0.106 | -1.278 | -0.089 | -0.128 | -0.128 | 0 | 0.19 | 0.01 | 0.48 | 0.58 | 0.26 | 0.3 |
| 3 | 42922697 | rs6771233 | -0.454 | -0.364 | -0.15 | -1.335 | 0.002 | 0.012 | 0.012 | 0 | 0.38 | 0.29 | 0.57 | 0.67 | 0.36 | 0.48 |
| 3 | 42924625 | rs11129981 | -0.27 | -0.428 | -0.132 | -1.229 | -0.004 | 0.006 | 0.006 | 0 | 0.36 | 0.27 | 0.56 | 0.67 | 0.35 | 0.46 |
| 3 | 42928515 | rs1427800 | 1.244 | -0.581 | 0.403 | -1.278 | 0.021 | 0.054 | 0.054 | 2.218 | 0.92 | 0.61 | 0.92 | 0.94 | 0.97 | 0.97 |
| 3 | 42946047 | rs7625959 | -0.059 | -0.262 | 0.705 | -1.579 | 0.057 | -0.076 | -0.076 | 0 | 0.15 | 0.39 | 0.15 | 0.04 | 0.19 | 0.13 |
| 3 | 42947793 | rs9832875 | 1.217 | -0.317 | 0.613 | -1.592 | 0.024 | -0.039 | -0.039 | 0 | 0.1 | 0.15 | 0.13 | 0 | 0.18 | 0.1 |
| 3 | 42954912 | rs11129987 | -0.14 | -0.342 | 0.567 | -1.609 | 0.054 | -0.108 | -0.108 | 0 | 0.83 | 0.47 | 0.82 | 0.96 | 0.79 | 0.87 |
| 3 | 42967597 | rs704901 | 1.36 | -0.133 | -0.33 | -1.282 | -0.089 | -0.129 | -0.129 | 0 | 0.18 | 0.01 | 0.41 | 0.57 | 0.25 | 0.32 |
| 3 | 42976797 | rs6768791 | 0.9 | 0.135 | -0.156 | -1.257 | 0 | 0.005 | 0.005 | 0 | 0.15 | 0.17 | 0.18 | 0.23 | 0.14 | 0.24 |
| 3 | 42994980 | rs11129989 | 1.197 | 0.222 | -0.665 | -1.232 | -0.001 | 0 | 0 | 0 | 0.14 | 0.13 | 0.16 | 0.17 | 0.14 | 0.24 |
| 3 | 43005979 | rs11919400 | 1.475 | -0.301 | -1.386 | -1.583 | -0.069 | 0.034 | 0.034 | 1.278 | 0.21 | 0.01 | 0.14 | 0.09 | 0.25 | 0.08 |
| 3 | 43032810 | rs13080201 | 0.032 | -0.871 | -0.89 | -1.629 | 0.016 | 0.029 | 0.029 | 0 | 0.33 | 0.26 | 0.17 | 0.17 | 0.25 | 0.2 |
| 3 | 43048159 | rs565521 | 1.352 | -0.487 | -0.865 | -1.529 | -0.002 | -0.008 | -0.008 | 2.267 | 0.36 | 0.31 | 0.51 | 0.57 | 0.4 | 0.35 |
| 3 | 43049011 | rs9874514 | -0.476 | -0.612 | -0.608 | -1.819 | -0.023 | 0.076 | 0.076 | 0 | 0.4 | 0.29 | 0.28 | 0.21 | 0.44 | 0.33 |
| 3 | 43050746 | rs17455760 | 1.08 | -0.531 | -0.773 | -1.737 | -0.06 | -0.006 | -0.006 | 0 | 0.22 | 0.01 | 0.12 | 0.17 | 0.23 | 0.19 |
| 3 | 43059578 | rs9828785 | 0.827 | -0.38 | -0.461 | -1.74 | 0.021 | -0.043 | -0.043 | 0 | 0.11 | 0.17 | 0.13 | 0 | 0.19 | 0.12 |
| 3 | 43068165 | rs9882262 | -0.99 | -0.47 | -0.197 | -1.656 | -0.004 | -0.013 | -0.013 | 0 | 0.61 | 0.52 | 0.45 | 0.34 | 0.59 | 0.68 |
| 3 | 43070847 | rs704903 | -0.958 | -0.514 | -0.304 | -1.665 | -0.003 | -0.002 | -0.002 | 0 | 0.77 | 0.88 | 0.84 | 0.76 | 0.78 | 0.8 |
| 3 | 43073453 | rs17075084 | 0.661 | -0.715 | -0.272 | -1.61 | -0.013 | -0.013 | -0.013 | 0 | 0.25 | 0.29 | 0.18 | 0.15 | 0.15 | 0.37 |
| 3 | 43073761 | rs658958 | 0.889 | -0.438 | -0.532 | -1.42 | 0 | 0.002 | 0.002 | 0.815 | 0.37 | 0.34 | 0.53 | 0.65 | 0.41 | 0.32 |
| 3 | 43074434 | rs2936817 | 0.88 | -0.437 | -0.532 | -1.419 | 0 | 0.002 | 0.002 | 0.846 | 0.38 | 0.34 | 0.53 | 0.65 | 0.41 | 0.32 |
| 3 | 43081964 | rs601055 | -0.292 | -0.73 | -0.347 | -1.618 | -0.031 | -0.03 | -0.03 | 0 | 0.69 | 0.41 | 0.79 | 0.83 | 0.83 | 0.59 |
| 3 | 43084131 | rs704912 | 1.437 | -0.757 | -0.315 | -1.687 | -0.007 | -0.017 | -0.017 | 0 | 0.33 | 0.25 | 0.5 | 0.58 | 0.39 | 0.32 |
| 3 | 43089086 | rs13087788 | -0.03 | -0.734 | -0.458 | -1.684 | -0.076 | 0.05 | 0.05 | 0.749 | 0.35 | 0.11 | 0.29 | 0.25 | 0.44 | 0.32 |
| 3 | 43095101 | rs536119 | 0.466 | -0.575 | -0.391 | -1.767 | -0.001 | -0.004 | -0.004 | 0 | 0.46 | 0.46 | 0.67 | 0.67 | 0.6 | 0.37 |
| 3 | 43097710 | rs664628 | 1.291 | -0.806 | -0.493 | -1.662 | -0.004 | -0.011 | -0.011 | 0 | 0.34 | 0.27 | 0.51 | 0.58 | 0.39 | 0.27 |
| 3 | 43097765 | rs3732858 | 0.707 | -1.283 | -0.325 | -1.3 | -0.007 | 0.007 | 0.007 | 7.395 | 0.22 | 0.03 | 0.16 | 0.15 | 0.17 | 0.38 |
| 3 | 43101132 | rs530842 | -0.221 | -0.745 | 0.432 | -1.14 | 0.008 | 0.01 | 0.01 | 1.083 | 0.93 | 0.57 | 0.96 | 1 | 0.99 | 0.98 |
| 3 | 43102416 | rs1075408 | 0.963 | -0.61 | -0.177 | -1.197 | -0.035 | -0.023 | -0.023 | 0 | 0.16 | 0.01 | 0.09 | 0.16 | 0.16 | 0.22 |
| 3 | 43102509 | rs9842444 | 1.287 | -0.273 | 0.403 | -1.099 | -0.018 | -0.02 | -0.02 | 0.005 | 0.05 | 0.19 | 0.02 | 0 | 0 | 0 |
| 3 | 43104383 | rs513875 | 0.375 | -0.094 | 0.059 | -1.219 | 0.011 | 0.062 | 0.062 | 0.615 | 0.66 | 0.24 | 0.72 | 0.87 | 0.69 | 0.77 |
| 3 | 43116170 | rs7619872 | 0.843 | -0.194 | 0.423 | -1.167 | -0.001 | -0.006 | -0.006 | 0 | 0.22 | 0.2 | 0.12 | 0.11 | 0.16 | 0.19 |
| 3 | 43121559 | rs604033 | -1.112 | -0.244 | 0.059 | -0.919 | 0.003 | -0.009 | -0.009 | 0 | 0.75 | 0.83 | 0.81 | 0.74 | 0.79 | 0.74 |
| 3 | 43121691 | rs9811883 | 1.364 | -0.277 | 0.088 | -0.933 | -0.027 | -0.029 | -0.029 | 0 | 0.06 | 0.18 | 0.01 | 0 | 0 | 0 |
| 3 | 43125481 | rs12497427 | 1.074 | -0.149 | -0.182 | -0.878 | -0.051 | -0.088 | -0.088 | 0 | 0.24 | 0.02 | 0.17 | 0.26 | 0.21 | 0.26 |
| 3 | 43132444 | rs488069 | -1.056 | -0.155 | -0.133 | -0.879 | 0.002 | -0.007 | -0.007 | 0 | 0.75 | 0.78 | 0.81 | 0.74 | 0.79 | 0.74 |
| 3 | 43156139 | rs7625703 | 0.945 | -0.559 | -0.239 | -0.818 | -0.021 | -0.049 | -0.049 | 0 | 0.36 | 0.21 | 0.47 | 0.57 | 0.39 | 0.47 |
| 3 | 43167735 | rs9840881 | 0.536 | -0.74 | -0.017 | -0.87 | -0.012 | -0.063 | -0.063 | 0.088 | 0.25 | 0.26 | 0.11 | 0.05 | 0.16 | 0.19 |
| 3 | 43170450 | rs12107602 | -0.927 | -0.99 | -0.565 | -0.623 | 0.006 | -0.018 | -0.018 | 0.015 | 0.75 | 0.89 | 0.82 | 0.75 | 0.8 | 0.74 |
| 3 | 43174229 | rs7648947 | 0.406 | -0.8 | -0.485 | -0.486 | -0.017 | -0.002 | -0.002 | 0 | 0.41 | 0.33 | 0.56 | 0.62 | 0.48 | 0.5 |
| 3 | 43176153 | rs1125065 | 1.003 | -0.577 | -0.901 | -0.46 | -0.007 | -0.063 | -0.063 | 0 | 0.23 | 0.05 | 0.15 | 0.25 | 0.18 | 0.24 |
| 3 | 43176831 | rs12635437 | 1.012 | -0.568 | -0.897 | -0.47 | -0.007 | -0.066 | -0.066 | 0 | 0.23 | 0.05 | 0.13 | 0.25 | 0.18 | 0.24 |
| 3 | 43179134 | rs2372335 | 0.885 | -0.092 | 0.187 | 0.237 | 0.001 | -0.002 | -0.002 | 0.136 | 0.56 | 0.5 | 0.4 | 0.42 | 0.51 | 0.59 |
| 3 | 43179422 | rs7622021 | 0.267 | 0.174 | 0.27 | 0.243 | -0.019 | -0.027 | -0.027 | 0 | 0.65 | 0.72 | 0.47 | 0.48 | 0.54 | 0.6 |
| 3 | 43184573 | rs9848231 | -0.15 | 0.292 | -0.374 | 0.282 | -0.034 | -0.036 | -0.036 | 0 | 0.07 | 0.2 | 0.02 | 0 | 0 | 0 |
| 3 | 43186244 | rs2372337 | -0.482 | 1.092 | -1.083 | 0.643 | -0.058 | -0.082 | -0.082 | 0 | 0.33 | 0.16 | 0.52 | 0.52 | 0.45 | 0.39 |
| 3 | 43196598 | rs1878722 | -0.311 | 1.142 | -0.656 | 0.789 | -0.01 | -0.018 | -0.018 | 0 | 0.27 | 0.23 | 0.47 | 0.56 | 0.34 | 0.39 |
| 3 | 43197817 | rs12489611 | 1.002 | 0.464 | 0.309 | 0.46 | 0.015 | -0.013 | -0.013 | 0.35 | 0.91 | 0.51 | 0.93 | 0.99 | 0.96 | 0.95 |
| 3 | 43199470 | rs2007700 | -1.204 | 1.226 | -0.278 | 0.771 | 0.035 | 0.029 | 0.029 | 0 | 0.09 | 0.18 | 0.15 | 0.18 | 0.19 | 0.2 |
| 3 | 43199941 | rs1106077 | -1.33 | 0.646 | -0.264 | 0.757 | -0.011 | 0.034 | 0.034 | 0 | 0.07 | 0.16 | 0.05 | 0.16 | 0.03 | 0.15 |

| | | | | | | | | | | | | | | | | |
|---|---|---|---|---|---|---|---|---|---|---|---|---|---|---|---|---|
| 3 | 43201561 | rs9855146 | 0.673 | -0.742 | -0.602 | 0.684 | 0.043 | -0.017 | -0.017 | 0 | 0.68 | 0.29 | 0.67 | 0.65 | 0.85 | 0.72 |
| 3 | 43203402 | rs11712506 | 0.732 | -0.033 | -1.648 | 0.215 | -0.124 | -0.014 | -0.014 | 0 | 0.44 | 0.08 | 0.58 | 0.61 | 0.48 | 0.58 |
| 3 | 43204084 | rs6766134 | 0.858 | 0.107 | -0.648 | -0.003 | -0.002 | -0.012 | -0.012 | 0.455 | 0.79 | 0.71 | 0.84 | 0.84 | 0.78 | 0.81 |
| 3 | 43205962 | rs2372342 | 1.043 | -0.069 | -0.639 | -0.022 | -0.028 | 0.001 | 0.001 | 0 | 0.85 | 0.75 | 0.9 | 0.84 | 0.93 | 0.84 |
| 3 | 43214092 | rs1814906 | -0.332 | -0.268 | -0.709 | -0.019 | -0.023 | 0.016 | 0.016 | 0 | 0.93 | 0.85 | 0.98 | 0.9 | 0.98 | 0.88 |
| 3 | 43217289 | rs13074033 | 1.652 | -0.097 | -0.577 | -0.094 | -0.003 | -0.008 | -0.008 | 0 | 0.71 | 0.63 | 0.81 | 0.75 | 0.73 | 0.69 |
| 3 | 43220849 | rs3843370 | 1.583 | 1.042 | -0.655 | -0.201 | 0.004 | -0.003 | -0.003 | 3.376 | 0.7 | 0.64 | 0.76 | 0.74 | 0.63 | 0.68 |
| 3 | 43222913 | rs13066654 | 1.332 | 0.715 | 0.263 | -0.098 | 0.045 | 0.001 | 0.001 | 0.004 | 0.81 | 0.49 | 0.75 | 0.83 | 0.77 | 0.83 |
| 3 | 43226220 | rs17075283 | -0.612 | 0.738 | 0.227 | -0.101 | 0.005 | -0.003 | -0.003 | 0 | 0.13 | 0.16 | 0.18 | 0.11 | 0.18 | 0.15 |
| 3 | 43233899 | rs7641667 | 1.558 | 0.656 | 0.397 | -0.045 | 0.035 | 0.025 | 0.025 | 0.022 | 0.74 | 0.44 | 0.66 | 0.67 | 0.67 | 0.68 |
| 3 | 43238338 | rs6767821 | -0.264 | 0.062 | 0.466 | 0.01 | -0.031 | 0.044 | 0.044 | 0 | 0.07 | 0.21 | 0.09 | 0.14 | 0.02 | 0.11 |
| 3 | 43238536 | rs17253286 | -0.406 | 0.132 | 0.097 | -0.318 | 0.007 | 0.031 | 0.031 | 0 | 0.19 | 0.11 | 0.1 | 0 | 0.16 | 0.11 |
| 3 | 43246004 | rs12635384 | -0.722 | 0.437 | 0.362 | 0.096 | 0.004 | 0.061 | 0.061 | 0 | 0.11 | 0.19 | 0.18 | 0.24 | 0.1 | 0.13 |
| 3 | 43247753 | rs9864154 | 1.473 | 0.137 | 0.04 | -0.069 | 0.039 | 0.036 | 0.036 | 1.854 | 0.64 | 0.14 | 0.68 | 0.73 | 0.74 | 0.72 |
| 3 | 43248393 | rs13080126 | -0.737 | 0.145 | 0.569 | -0.432 | -0.037 | -0.135 | -0.135 | 0 | 0.23 | 0.42 | 0.12 | 0.03 | 0.16 | 0.15 |
| 3 | 43250297 | rs9875582 | 1.384 | 0.68 | 0.327 | -0.347 | 0.003 | 0.016 | 0.016 | 1.508 | 0.77 | 0.17 | 0.57 | 0.85 | 0.79 | 0.81 |
| 3 | 43254107 | rs4683364 | -1.787 | 0.798 | 1.414 | -0.37 | 0.003 | -0.01 | -0.01 | 0 | 0.2 | 0.67 | 0.4 | 0.14 | 0.2 | 0.16 |
| 3 | 43256744 | rs6791526 | -1.303 | 0.805 | 2.261 | -0.38 | 0.053 | 0.008 | 0.008 | 0.437 | 0.12 | 0.4 | 0.29 | 0.08 | 0.16 | 0.05 |
| 3 | 43259636 | rs17075342 | -0.33 | 0.792 | 2.333 | -0.387 | 0.008 | 0.017 | 0.017 | 0 | 0.07 | 0.18 | 0.12 | 0.06 | 0.05 | 0.11 |
| 3 | 43266311 | rs7639584 | 2.15 | 0.83 | 2.353 | -0.39 | -0.001 | 0.003 | 0.003 | 1.402 | 0.79 | 0.22 | 0.58 | 0.85 | 0.8 | 0.84 |
| 3 | 43278884 | rs2372351 | -1.005 | 0.86 | 2.523 | -0.419 | 0.052 | -0.066 | -0.066 | 0.036 | 0.11 | 0.33 | 0.13 | 0 | 0.15 | 0.04 |
| 3 | 43294971 | rs6808396 | -0.682 | 1.073 | 2.304 | -0.325 | -0.005 | -0.008 | -0.008 | 0 | 0.09 | 0.1 | 0.11 | 0 | 0.15 | 0.04 |
| 3 | 43297045 | rs4682879 | -1.195 | 1.045 | 2.474 | -0.299 | 0.015 | 0.118 | 0.118 | 0 | 0.09 | 0.32 | 0.28 | 0.14 | 0.05 | 0.12 |
| 3 | 43300871 | rs7646966 | 1.978 | 1.067 | 2.245 | -0.332 | 0.024 | -0.023 | -0.023 | 0 | 0.86 | 0.34 | 0.7 | 0.92 | 0.84 | 0.95 |
| 3 | 43303296 | rs1320163 | 1.66 | 1.09 | 2.198 | -0.322 | 0.049 | -0.01 | -0.01 | 0 | 0.87 | 0.4 | 0.7 | 0.92 | 0.85 | 0.95 |
| 3 | 43334020 | rs13320571 | 2.085 | 0.838 | 2.322 | -0.402 | -0.008 | -0.007 | -0.007 | 0 | 0.85 | 0.23 | 0.82 | 0.93 | 0.94 | 0.87 |
| 3 | 43362597 | rs1018175 | 1.029 | 1.378 | 1.53 | -0.414 | -0.071 | -0.126 | -0.126 | 0 | 0.16 | 0.02 | 0.26 | 0.57 | 0.24 | 0.32 |
| 3 | 43383641 | rs7621298 | 0.305 | 1.046 | 0.662 | -1.183 | -0.074 | 0.157 | 0.157 | 0 | 0.45 | 0.15 | 0.44 | 0.79 | 0.51 | 0.65 |
| 3 | 43387934 | rs11130000 | 1.414 | 1.051 | 0.749 | -1.146 | -0.006 | 0.062 | 0.062 | 0 | 0.33 | 0.07 | 0.29 | 0.2 | 0.29 | 0.17 |
| 3 | 43396221 | rs17075597 | 1.165 | 1.031 | 1.188 | -1.047 | 0 | 0.004 | 0.004 | 0 | 0.07 | 0.09 | 0.12 | 0.07 | 0.05 | 0.12 |
| 3 | 43401268 | rs17075600 | -0.242 | 1.019 | 1.409 | -1.054 | 0.016 | 0.021 | 0.021 | 0 | 0.1 | 0.38 | 0.14 | 0.07 | 0.06 | 0.13 |
| 3 | 43417636 | rs17075631 | 1.604 | 1.236 | 0.908 | -1.004 | -0.074 | -0.108 | -0.108 | 0 | 0.14 | 0.02 | 0.28 | 0.56 | 0.25 | 0.3 |
| 3 | 43417792 | rs17643558 | 1.465 | 0.503 | 0.605 | -1.556 | -0.001 | 0.038 | 0.038 | 1.572 | 0.21 | 0.06 | 0.1 | 0.11 | 0.18 | 0.16 |
| 3 | 43433362 | rs11715896 | -0.228 | 0.492 | 0.864 | -1.694 | 0.002 | 0.022 | 0.022 | 0 | 0.46 | 0.38 | 0.41 | 0.19 | 0.42 | 0.22 |
| 3 | 43433557 | rs977911 | -1.11 | 0.45 | 1.034 | -1.667 | 0.002 | 0.037 | 0.037 | 0 | 0.93 | 0.76 | 0.88 | 0.94 | 0.97 | 0.89 |
| 3 | 43467895 | rs7650267 | -0.001 | 0.446 | 0.968 | -1.695 | 0.025 | -0.068 | -0.068 | 0.372 | 0.12 | 0.22 | 0.11 | 0 | 0.16 | 0.06 |
| 3 | 43477118 | rs4547674 | 0.713 | 0.31 | 0.833 | -1.74 | -0.005 | 0.076 | 0.076 | 0 | 0.47 | 0.38 | 0.62 | 0.83 | 0.54 | 0.67 |
| 3 | 43498248 | rs2372433 | 0.728 | 0.278 | 0.682 | -1.773 | -0.014 | 0.126 | 0.126 | 0 | 0.34 | 0.14 | 0.17 | 0.11 | 0.3 | 0.17 |
| 3 | 43499327 | rs17473616 | 1.752 | 0.36 | 0.76 | -1.796 | 0 | -0.016 | -0.016 | 0.019 | 0.05 | 0.04 | 0.04 | 0 | 0.05 | 0.05 |
| 3 | 43500735 | rs11709131 | 2.327 | 0.329 | 0.7 | -1.851 | -0.032 | 0.077 | 0.077 | 3.796 | 0.13 | 0 | 0.05 | 0 | 0.12 | 0.01 |
| 3 | 43500901 | rs17075727 | 1.816 | 0.498 | 0.911 | -1.434 | -0.065 | -0.132 | -0.132 | 1.342 | 0.16 | 0.02 | 0.2 | 0.55 | 0.22 | 0.31 |
| 3 | 43505561 | rs6775337 | 0.863 | 0.143 | 1.446 | -1.415 | -0.015 | 0.125 | 0.125 | 0 | 0.34 | 0.13 | 0.17 | 0.11 | 0.3 | 0.17 |
| 3 | 43532019 | rs7628317 | -1.109 | 0.124 | 1.735 | -1.416 | 0.007 | 0.05 | 0.05 | 0 | 0.93 | 0.73 | 0.87 | 0.94 | 0.97 | 0.91 |
| 3 | 43537085 | rs6802567 | -0.4 | 0.114 | 1.966 | -1.427 | 0.039 | -0.059 | -0.059 | 0.985 | 0.11 | 0.28 | 0.09 | 0.01 | 0.12 | 0.07 |
| 3 | 43564157 | rs4462942 | -1.455 | 0.112 | 2.018 | -1.428 | -0.002 | 0.036 | 0.036 | 0 | 0.94 | 0.8 | 0.88 | 0.94 | 0.97 | 0.91 |
| 3 | 43579340 | rs13092838 | 0.866 | 0.105 | 1.963 | -1.441 | -0.015 | 0.125 | 0.125 | 0 | 0.34 | 0.14 | 0.17 | 0.11 | 0.3 | 0.17 |
| 3 | 43591142 | rs12638282 | 0.748 | 0.099 | 1.516 | -1.363 | -0.041 | 0.184 | 0.184 | 0 | 0.46 | 0.19 | 0.61 | 0.83 | 0.55 | 0.68 |
| 3 | 43591327 | rs17409162 | 1.141 | 0.118 | 1.517 | -1.29 | 0 | -0.015 | -0.015 | 0 | 0.06 | 0.04 | 0.04 | 0 | 0.05 | 0.05 |
| 3 | 43593399 | rs4682689 | 0.874 | 0.095 | 1.498 | -1.272 | -0.003 | 0.037 | 0.037 | 0 | 0.07 | 0.24 | 0.12 | 0.06 | 0.03 | 0.09 |
| 3 | 43599924 | rs3772160 | 0.833 | 0.081 | 1.296 | -1.34 | -0.122 | 0.082 | 0.082 | 0 | 0.43 | 0.07 | 0.4 | 0.76 | 0.54 | 0.66 |
| 3 | 43600596 | rs2054554 | 0.091 | 0.079 | 1.484 | -1.345 | 0.001 | -0.011 | -0.011 | 0 | 0.53 | 0.48 | 0.74 | 0.88 | 0.58 | 0.76 |
| 3 | 43602803 | rs3772165 | 0.107 | 0.083 | 1.367 | -1.343 | 0.003 | 0.015 | 0.015 | 0 | 0.53 | 0.45 | 0.73 | 0.88 | 0.58 | 0.76 |
| 3 | 43607136 | rs7616708 | 0.134 | 0.084 | 1.158 | -1.343 | 0.007 | 0.098 | 0.098 | 0 | 0.52 | 0.35 | 0.72 | 0.88 | 0.58 | 0.77 |
| 3 | 43622656 | rs4682690 | 0.144 | 0.084 | 1.158 | -1.343 | 0.008 | 0.098 | 0.098 | 0 | 0.52 | 0.35 | 0.72 | 0.88 | 0.58 | 0.77 |
| 3 | 43627341 | rs3755603 | 1.576 | 0.073 | 1.192 | -1.343 | -0.012 | 0.008 | 0.008 | 0 | 0.06 | 0.1 | 0.11 | 0.06 | 0.03 | 0.09 |
| 3 | 43643284 | rs965928 | 1.04 | 0.075 | 1.071 | -1.359 | -0.011 | 0.081 | 0.081 | 0 | 0.07 | 0.18 | 0.31 | 0.12 | 0.03 | 0.1 |
| 3 | 43648035 | rs1482651 | -0.484 | 0.08 | 0.68 | -1.348 | -0.001 | -0.058 | -0.058 | 0 | 0.59 | 0.58 | 0.78 | 0.88 | 0.63 | 0.82 |
| 3 | 43668176 | rs9852783 | 0.702 | 0.279 | 0.141 | -1.254 | -0.108 | 0.088 | 0.088 | 0 | 0.44 | 0.09 | 0.39 | 0.76 | 0.52 | 0.65 |
| 3 | 43682052 | rs7614885 | -1.379 | 0.117 | 0.668 | -1.365 | -0.057 | 0.013 | 0.013 | 0.108 | 0.15 | 0.43 | 0.2 | 0.12 | 0.05 | 0.25 |
| 3 | 43683643 | rs7620653 | -1.379 | 0.117 | 0.668 | -1.365 | -0.057 | 0.013 | 0.013 | 0.108 | 0.15 | 0.43 | 0.2 | 0.12 | 0.05 | 0.25 |
| 3 | 43687273 | rs6793877 | -0.165 | 0.211 | 0.627 | -1.499 | -0.013 | 0.044 | 0.044 | 0 | 0.62 | 0.77 | 0.76 | 0.88 | 0.59 | 0.79 |
| 3 | 43688012 | rs9856765 | 0.474 | -0.118 | 1.406 | -0.633 | -0.001 | -0.023 | -0.023 | 0 | 0.06 | 0.05 | 0.04 | 0 | 0.05 | 0.05 |
| 3 | 43695469 | rs6782070 | -1.738 | -0.14 | 1.481 | -0.576 | -0.008 | 0.052 | 0.052 | 0.058 | 0.09 | 0.39 | 0.11 | 0.1 | 0.03 | 0.16 |
| 3 | 43717530 | rs4682900 | -0.221 | 0.062 | 0.887 | -0.67 | 0.006 | 0 | 0 | 0 | 0.88 | 0.85 | 0.91 | 0.88 | 0.85 | 0.86 |
| 3 | 43718504 | rs12054254 | -0.425 | 0.041 | 1.14 | -0.67 | -0.007 | 0.037 | 0.037 | 0 | 0.05 | 0.14 | 0.07 | 0.1 | 0.03 | 0.15 |
| 3 | 43747352 | rs3774792 | -0.256 | -0.002 | 0.986 | -0.679 | 0.007 | 0 | 0 | 0 | 0.88 | 0.85 | 0.91 | 0.88 | 0.85 | 0.86 |
| 3 | 43748755 | rs3774791 | -1.052 | -0.181 | 0.465 | -0.715 | -0.03 | 0.002 | 0.002 | 0 | 0.13 | 0.16 | 0.1 | 0.12 | 0.05 | 0.25 |
| 3 | 43755473 | rs3774789 | -0.44 | -0.144 | 0.482 | -0.699 | -0.007 | 0.048 | 0.048 | 0 | 0.06 | 0.14 | 0.07 | 0.1 | 0.03 | 0.15 |
| 3 | 43760062 | rs887472 | -0.44 | -0.144 | 0.482 | -0.699 | -0.007 | 0.048 | 0.048 | 0 | 0.06 | 0.15 | 0.07 | 0.1 | 0.03 | 0.15 |

| | | | | | | | | | | | | | | | | |
|---|---|---|---|---|---|---|---|---|---|---|---|---|---|---|---|---|
| 3 | 43761441 | rs3733156 | 1.724 | -0.225 | 0.473 | -0.822 | 0.041 | 0.047 | 0.047 | 1.879 | 0.63 | 0.17 | 0.7 | 0.76 | 0.73 | 0.55 |
| 3 | 43765737 | rs17075928 | -0.819 | -0.002 | 0.475 | -0.549 | -0.03 | 0.002 | 0.002 | 0 | 0.13 | 0.16 | 0.1 | 0.12 | 0.05 | 0.25 |
| 3 | 43772772 | rs6799877 | 1.099 | 0.124 | 0.516 | -0.554 | -0.029 | -0.001 | -0.001 | 0 | 0.85 | 0.75 | 0.88 | 0.87 | 0.92 | 0.72 |
| 3 | 43774426 | rs2372546 | 0.456 | 0.375 | 0.319 | -0.486 | 0.001 | 0.004 | 0.004 | 0 | 0.87 | 0.86 | 0.85 | 0.88 | 0.86 | 0.73 |
| 3 | 43782361 | rs6441791 | -0.884 | 0.688 | 0.691 | -0.649 | 0.031 | -0.049 | -0.049 | 0 | 0.18 | 0.48 | 0.15 | 0.1 | 0.18 | 0.16 |
| 3 | 43782668 | rs6441792 | -0.024 | 0.02 | 0.181 | -0.984 | -0.01 | -0.01 | -0.01 | 0.108 | 0.61 | 0.55 | 0.76 | 0.79 | 0.75 | 0.57 |
| 3 | 43784417 | rs17075971 | -0.309 | 0.633 | 0.585 | -0.603 | -0.007 | 0 | 0 | 0.04 | 0.12 | 0.1 | 0.11 | 0.11 | 0.21 | 0.14 |
| 3 | 43794560 | rs740836 | 1.392 | 0.096 | 0.739 | -0.41 | 0.018 | 0.078 | 0.078 | 0.013 | 0.86 | 0.63 | 0.84 | 0.79 | 0.85 | 0.78 |
| 3 | 43795005 | rs1078622 | 1.047 | -0.037 | 0.943 | -0.664 | 0.001 | 0.017 | 0.017 | 0 | 0.88 | 0.8 | 0.86 | 0.88 | 0.91 | 0.76 |
| 3 | 43795310 | rs1076413 | 1.047 | -0.037 | 0.943 | -0.664 | 0.001 | 0.017 | 0.017 | 0 | 0.88 | 0.8 | 0.86 | 0.88 | 0.91 | 0.76 |
| 3 | 43797458 | rs17255655 | 0.531 | -0.179 | 0.359 | -1.306 | 0.014 | 0.038 | 0.038 | 0.529 | 0.3 | 0.24 | 0.16 | 0.09 | 0.24 | 0.17 |
| 3 | 43798393 | rs17075984 | 0.597 | -0.004 | 0.265 | -1.105 | 0.008 | -0.032 | -0.032 | 3.219 | 0.06 | 0.1 | 0.05 | 0 | 0.05 | 0.11 |
| 3 | 43801941 | rs13100389 | 0.97 | -0.075 | -0.023 | -1.102 | -0.05 | 0.041 | 0.041 | 0 | 0.42 | 0.17 | 0.6 | 0.66 | 0.52 | 0.38 |
| 3 | 43803575 | rs9835853 | 1.437 | 0.176 | 0.332 | -1.758 | 0.002 | -0.035 | -0.035 | 1.276 | 0.74 | 0.34 | 0.8 | 0.91 | 0.74 | 0.8 |
| 3 | 43805178 | rs9637421 | -1.291 | 0.168 | 0.743 | -1.76 | 0.012 | -0.057 | -0.057 | 1.083 | 0.19 | 0.48 | 0.13 | 0.09 | 0.16 | 0.14 |
| 3 | 43805245 | rs9637422 | -0.957 | 0.192 | 0.795 | -1.738 | 0.033 | -0.014 | -0.014 | 0 | 0.13 | 0.34 | 0.1 | 0.08 | 0.13 | 0.06 |
| 3 | 43805528 | rs13069363 | -0.548 | 0.215 | 0.553 | -1.73 | 0 | -0.001 | -0.001 | 0 | 0.1 | 0.12 | 0.08 | 0.06 | 0.12 | 0.05 |
| 3 | 43807124 | rs7622411 | -0.266 | -0.446 | 0.161 | -2.081 | -0.003 | -0.029 | -0.029 | 0.176 | 0.27 | 0.35 | 0.21 | 0.13 | 0.22 | 0.19 |
| 3 | 43810452 | rs1558323 | 1.45 | -0.333 | -0.591 | -2.099 | -0.133 | 0.127 | 0.127 | 0.348 | 0.45 | 0.04 | 0.55 | 0.73 | 0.54 | 0.52 |
| 3 | 43822281 | rs13092458 | 1.358 | 0.299 | -1.149 | -2.142 | -0.106 | 0.015 | 0.015 | 0 | 0.2 | 0.02 | 0.17 | 0.12 | 0.32 | 0.23 |
| 3 | 43826250 | rs10490784 | 1.43 | -0.068 | -0.544 | -2.208 | -0.05 | 0.145 | 0.145 | 0 | 0.45 | 0.12 | 0.57 | 0.72 | 0.53 | 0.5 |
| 3 | 43826822 | rs4682696 | -0.477 | -0.186 | 0.118 | -2.323 | -0.003 | -0.025 | -0.025 | 0 | 0.41 | 0.39 | 0.35 | 0.22 | 0.37 | 0.41 |
| 3 | 43833177 | rs11130013 | 0.029 | -0.345 | -0.054 | -2.395 | 0.003 | 0.026 | 0.026 | 0 | 0.2 | 0.13 | 0.13 | 0.12 | 0.15 | 0.19 |
| 3 | 43836717 | rs4073774 | 1.119 | -0.315 | 0.309 | -2.418 | 0 | 0.04 | 0.04 | 0 | 0.6 | 0.3 | 0.65 | 0.79 | 0.6 | 0.66 |
| 3 | 43853667 | rs11706078 | -1.822 | -0.256 | 0.333 | -2.375 | -0.003 | 0.073 | 0.073 | 0 | 0.52 | 0.7 | 0.43 | 0.28 | 0.47 | 0.49 |
| 3 | 43854669 | rs4682910 | 0.248 | -0.217 | 0.161 | -2.374 | -0.015 | 0.054 | 0.054 | 0 | 0.22 | 0.06 | 0.22 | 0.12 | 0.27 | 0.16 |
| 3 | 43856009 | rs7625372 | -0.568 | -0.219 | 0.375 | -2.399 | -0.008 | 0.023 | 0.023 | 0 | 0.29 | 0.23 | 0.27 | 0.18 | 0.35 | 0.2 |
| 3 | 43857084 | rs4234436 | 1.15 | -0.468 | 1.21 | -2.082 | -0.02 | -0.066 | -0.066 | 0.115 | 0.7 | 0.51 | 0.8 | 0.84 | 0.8 | 0.67 |
| 3 | 43862251 | rs4392389 | 1.327 | -0.359 | 1.315 | -2.037 | 0.007 | 0.016 | 0.016 | 0 | 0.69 | 0.28 | 0.78 | 0.84 | 0.79 | 0.67 |
| 3 | 43866923 | rs4362698 | -1.385 | -0.234 | 0.41 | -1.983 | -0.002 | -0.03 | -0.03 | 0 | 0.94 | 0.88 | 0.96 | 1 | 0.96 | 0.89 |
| 3 | 43867027 | rs11708881 | -0.396 | -0.08 | -0.046 | -1.799 | 0.006 | -0.003 | -0.003 | 0 | 0.08 | 0.16 | 0.05 | 0.06 | 0.08 | 0.04 |
| 3 | 43870742 | rs4629304 | 0.992 | 0.016 | 0.005 | -1.76 | 0.008 | 0.019 | 0.019 | 0.061 | 0.12 | 0.07 | 0.09 | 0.07 | 0.07 | 0.15 |
| 3 | 43873105 | rs12493309 | 0.561 | 0.029 | -0.007 | -1.753 | 0 | -0.001 | -0.001 | 0 | 0.14 | 0.1 | 0.1 | 0.1 | 0.08 | 0.18 |
| 3 | 43874463 | rs4547675 | -0.011 | 0.212 | -0.389 | -1.613 | 0.014 | 0.002 | 0.002 | 0 | 0.07 | 0.16 | 0.05 | 0.06 | 0.08 | 0.04 |
| 3 | 43877835 | rs4502556 | 0.68 | 0.212 | -0.86 | -1.533 | -0.001 | -0.005 | -0.005 | 0 | 0.72 | 0.27 | 0.79 | 0.83 | 0.79 | 0.73 |
| 3 | 43880640 | rs4535201 | 0.642 | 0.26 | -1.637 | -1.668 | -0.09 | 0.042 | 0.042 | 1.669 | 0.29 | 0.03 | 0.17 | 0.17 | 0.34 | 0.32 |
| 3 | 43881519 | rs4419358 | 0.428 | 0.053 | -1.685 | -2.128 | 0.043 | 0.155 | 0.155 | 0.014 | 0.55 | 0.22 | 0.53 | 0.84 | 0.6 | 0.54 |
| 3 | 43892962 | rs7630503 | 1.334 | -0.152 | -0.92 | -2.067 | -0.012 | -0.061 | -0.061 | 0 | 0.1 | 0.22 | 0.06 | 0 | 0.06 | 0.07 |
| 3 | 43893418 | rs11713435 | 2.107 | 0.122 | -1.042 | -1.997 | -0.048 | -0.065 | -0.065 | 0 | 0.21 | 0.05 | 0.25 | 0.62 | 0.23 | 0.15 |
| 3 | 43896176 | rs11130015 | -0.294 | 0.02 | -0.708 | -2.124 | -0.003 | -0.027 | -0.027 | 0.808 | 0.3 | 0.28 | 0.38 | 0.12 | 0.3 | 0.31 |
| 3 | 43899743 | rs9989976 | 0.001 | -0.043 | -0.583 | -1.901 | 0 | -0.005 | -0.005 | 1.607 | 0.24 | 0.21 | 0.43 | 0.18 | 0.29 | 0.29 |
| 3 | 43902170 | rs9824340 | -0.715 | -0.287 | -0.423 | -2.019 | -0.006 | -0.028 | -0.028 | 0.235 | 0.63 | 0.48 | 0.58 | 0.32 | 0.67 | 0.64 |
| 3 | 43906996 | rs6766529 | -0.726 | -0.263 | -0.336 | -2.059 | 0.007 | -0.046 | -0.046 | 0 | 0.78 | 0.7 | 0.64 | 0.87 | 0.74 | 0.74 |
| 3 | 43910872 | rs9863510 | 0.219 | -0.315 | -0.136 | -2.042 | -0.002 | -0.035 | -0.035 | 0.077 | 0.46 | 0.53 | 0.27 | 0.2 | 0.41 | 0.41 |
| 3 | 43932045 | rs6789465 | 1.605 | -0.146 | -0.426 | -1.767 | -0.081 | -0.077 | -0.077 | 2.674 | 0.26 | 0.02 | 0.27 | 0.62 | 0.29 | 0.22 |
| 3 | 43940061 | rs9311344 | 0.388 | -0.112 | -0.152 | -1.959 | -0.008 | -0.055 | -0.055 | 0.247 | 0.23 | 0.3 | 0.11 | 0.06 | 0.18 | 0.15 |
| 3 | 43946897 | rs7612822 | -0.749 | -0.08 | -0.157 | -1.864 | 0.016 | -0.042 | -0.042 | 0.714 | 0.79 | 0.7 | 0.65 | 0.87 | 0.72 | 0.75 |
| 3 | 43953346 | rs4682921 | 1.578 | -0.057 | -0.643 | -1.636 | -0.078 | -0.014 | -0.014 | 0.576 | 0.19 | 0.01 | 0.14 | 0.18 | 0.23 | 0.28 |
| 3 | 43954893 | rs9861853 | -0.004 | -0.268 | -0.316 | -1.813 | -0.03 | -0.098 | -0.098 | 0 | 0.25 | 0.58 | 0.13 | 0.03 | 0.18 | 0.13 |
| 3 | 43959284 | rs7620256 | 0.867 | -0.154 | -0.653 | -1.787 | -0.022 | 0.04 | 0.04 | 0 | 0.21 | 0.12 | 0.33 | 0.14 | 0.28 | 0.28 |
| 3 | 43968796 | rs6441805 | -0.088 | -0.4 | -0.396 | -1.535 | 0.015 | 0.043 | 0.043 | 2.284 | 0.5 | 0.36 | 0.39 | 0.6 | 0.43 | 0.33 |
| 3 | 43989595 | rs9861276 | 1.261 | -0.308 | -0.456 | -1.434 | -0.022 | -0.044 | -0.044 | 1.767 | 0.28 | 0.09 | 0.28 | 0.57 | 0.27 | 0.21 |
| 3 | 43994607 | rs10212466 | -0.171 | -0.335 | -0.401 | -1.302 | 0.017 | 0.049 | 0.049 | 1.968 | 0.51 | 0.36 | 0.38 | 0.58 | 0.42 | 0.32 |
| 3 | 43994802 | rs6775579 | 0.315 | -0.578 | -0.423 | -1.423 | -0.012 | -0.04 | -0.04 | 0 | 0.16 | 0.18 | 0.08 | 0.02 | 0.11 | 0.1 |
| 3 | 43997666 | rs3843372 | 1.255 | -0.423 | -0.663 | -1.356 | -0.022 | -0.063 | -0.063 | 0 | 0.28 | 0.1 | 0.27 | 0.55 | 0.27 | 0.2 |
| 3 | 44003720 | rs1852037 | 1.222 | -0.434 | -0.676 | -1.361 | -0.018 | -0.058 | -0.058 | 0.017 | 0.28 | 0.12 | 0.27 | 0.55 | 0.27 | 0.2 |
| 3 | 44006666 | rs1404481 | -0.769 | -0.484 | -0.288 | -1.51 | -0.005 | 0.015 | 0.015 | 0.917 | 0.57 | 0.65 | 0.54 | 0.7 | 0.47 | 0.39 |
| 3 | 44015285 | rs6806526 | -0.138 | -0.493 | -0.342 | -1.489 | -0.008 | -0.037 | -0.037 | 0.062 | 0.23 | 0.26 | 0.12 | 0.04 | 0.16 | 0.14 |
| 3 | 44015668 | rs6794506 | 1.75 | -0.348 | -0.418 | -1.352 | -0.006 | -0.024 | -0.024 | 0 | 0.08 | 0.14 | 0.05 | 0.03 | 0.05 | 0.04 |
| 3 | 44045981 | rs3852012 | -1.656 | 0.064 | -1.226 | -1.421 | -0.076 | 0.002 | 0.002 | 0.159 | 0.79 | 0.97 | 0.82 | 0.83 | 0.71 | 0.67 |
| 3 | 44056898 | rs11718455 | 1.17 | -0.18 | -0.892 | -1.002 | -0.052 | -0.106 | -0.106 | 0.551 | 0.27 | 0.07 | 0.27 | 0.51 | 0.27 | 0.22 |
| 3 | 44065912 | rs7624276 | -0.725 | -0.596 | -0.381 | -1.215 | -0.012 | -0.03 | -0.03 | 0 | 0.29 | 0.65 | 0.28 | 0.14 | 0.2 | 0.21 |
| 3 | 44068577 | rs1608730 | -0.572 | -0.668 | -0.022 | -1.261 | 0.002 | -0.007 | -0.007 | 0.017 | 0.1 | 0.27 | 0.18 | 0.11 | 0.09 | 0.11 |
| 3 | 44074449 | rs7635417 | 0.284 | -0.659 | -0.005 | -1.255 | -0.04 | -0.096 | -0.096 | 0 | 0.19 | 0.38 | 0.1 | 0.04 | 0.11 | 0.11 |
| 3 | 44074503 | rs808719 | 0.815 | -0.488 | -0.303 | -1.052 | -0.017 | 0.019 | 0.019 | 0.283 | 0.43 | 0.28 | 0.44 | 0.36 | 0.51 | 0.56 |
| 3 | 44076205 | rs7639253 | 0.277 | -0.186 | 0.052 | -0.87 | -0.026 | -0.067 | -0.067 | 1.015 | 0.18 | 0.29 | 0.09 | 0.04 | 0.11 | 0.1 |
| 3 | 44079629 | rs17076342 | -0.977 | 0.349 | 0.144 | -0.506 | 0.004 | -0.007 | -0.007 | 0.11 | 0.09 | 0.21 | 0.17 | 0.1 | 0.09 | 0.09 |
| 3 | 44083188 | rs808725 | 0.009 | 0.427 | 0.04 | -0.179 | 0.002 | 0.004 | 0.004 | 1.323 | 0.7 | 0.61 | 0.61 | 0.47 | 0.66 | 0.65 |
| 3 | 44083462 | rs808726 | 0.106 | 0.07 | -0.276 | -0.295 | -0.085 | -0.096 | -0.096 | 0 | 0.68 | 0.95 | 0.58 | 0.42 | 0.65 | 0.66 |

| | | | | | | | | | | | | | | | | |
|---|---|---|---|---|---|---|---|---|---|---|---|---|---|---|---|---|
| 3 | 44084343 | rs6804085 | -0.952 | 0.124 | -0.068 | -0.518 | -0.023 | -0.091 | -0.091 | 0 | 0.28 | 0.56 | 0.29 | 0.15 | 0.22 | 0.2 |
| 3 | 44085852 | rs808727 | 1.113 | -0.122 | 0.025 | -0.359 | 0.01 | 0.039 | 0.039 | 0.897 | 0.51 | 0.3 | 0.48 | 0.37 | 0.6 | 0.55 |
| 3 | 44087312 | rs808729 | 1.113 | -0.118 | -0.011 | -0.368 | 0.014 | 0.062 | 0.062 | 0.963 | 0.5 | 0.21 | 0.47 | 0.37 | 0.6 | 0.55 |
| 3 | 44088348 | rs1034967 | -0.92 | -0.137 | 0.036 | -0.39 | 0.003 | -0.002 | -0.002 | 0 | 0.27 | 0.72 | 0.27 | 0.15 | 0.17 | 0.2 |
| 3 | 44091193 | rs4682931 | -0.763 | 0.021 | -0.057 | -0.328 | -0.022 | -0.046 | -0.046 | 0 | 0.26 | 0.63 | 0.26 | 0.15 | 0.17 | 0.19 |
| 3 | 44091635 | rs4682933 | -0.832 | 0.077 | -0.16 | -0.304 | -0.028 | -0.061 | -0.061 | 0 | 0.25 | 0.61 | 0.26 | 0.15 | 0.17 | 0.19 |
| 3 | 44093707 | rs808718 | -0.578 | 0.728 | -0.626 | 0.167 | -0.056 | -0.115 | -0.115 | 0.54 | 0.78 | 0.96 | 0.75 | 0.52 | 0.76 | 0.76 |
| 3 | 44097461 | rs12496134 | -1.04 | 0.554 | 0.093 | 0.045 | 0.024 | 0.01 | 0.01 | 0 | 0.09 | 0.28 | 0.18 | 0.12 | 0.09 | 0.1 |
| 3 | 44104875 | rs17076392 | -0.716 | 0.609 | -0.124 | 0.069 | 0.008 | 0 | 0 | 0 | 0.08 | 0.15 | 0.17 | 0.12 | 0.09 | 0.07 |
| 3 | 44105159 | rs1008369 | 0.792 | 0.632 | -1.347 | -0.427 | -0.164 | 0.13 | 0.13 | 4.465 | 0.44 | 0.02 | 0.42 | 0.21 | 0.49 | 0.31 |
| 3 | 44105198 | rs6766121 | -0.631 | 0.91 | -0.334 | -0.221 | 0.067 | 0.024 | 0.024 | 0 | 0.07 | 0.26 | 0.1 | 0.12 | 0.08 | 0.07 |
| 3 | 44106849 | rs17076399 | -0.011 | 0.288 | -0.044 | -0.466 | -0.008 | -0.063 | -0.063 | 0 | 0.12 | 0.26 | 0.09 | 0.02 | 0.11 | 0.06 |
| 3 | 44108553 | rs6804151 | -0.547 | 0.198 | -0.319 | -0.544 | -0.012 | -0.061 | -0.061 | 0 | 0.29 | 0.64 | 0.17 | 0.05 | 0.2 | 0.15 |
| 3 | 44111902 | rs9881814 | -0.223 | 0.198 | -0.067 | -0.531 | -0.032 | -0.134 | -0.134 | 0 | 0.26 | 0.47 | 0.14 | 0.06 | 0.19 | 0.15 |
| 3 | 44111945 | rs9844198 | -0.671 | 0.352 | -0.828 | -0.056 | 0.028 | 0.043 | 0.043 | 0.534 | 0.35 | 0.78 | 0.25 | 0.22 | 0.3 | 0.37 |
| 3 | 44119447 | rs4682935 | -0.1 | 0.92 | -0.679 | 0.6 | 0.016 | 0.062 | 0.062 | 1.637 | 0.79 | 0.72 | 0.67 | 0.54 | 0.77 | 0.77 |
| 3 | 44125372 | rs9839994 | -0.572 | 0.449 | -1.296 | 0.34 | 0.006 | 0.047 | 0.047 | 0.991 | 0.38 | 0.79 | 0.35 | 0.28 | 0.37 | 0.16 |
| 3 | 44134579 | rs6797157 | -0.639 | 0.291 | -1.432 | 0.403 | -0.08 | -0.064 | -0.064 | 0 | 0.55 | 0.9 | 0.45 | 0.48 | 0.54 | 0.53 |
| 3 | 44135356 | rs9882688 | -0.765 | -0.06 | 0.057 | 0.234 | -0.022 | -0.054 | -0.054 | 0 | 0.13 | 0.52 | 0.1 | 0.04 | 0.06 | 0.02 |
| 3 | 44140584 | rs2109409 | -0.534 | 0.195 | -0.35 | 0.631 | -0.003 | 0.01 | 0.01 | 0 | 0.34 | 0.4 | 0.45 | 0.41 | 0.33 | 0.55 |
| 3 | 44141157 | rs6800118 | 0.38 | 0.39 | -0.106 | 0.153 | 0.01 | -0.012 | -0.012 | 0 | 0.58 | 0.42 | 0.61 | 0.69 | 0.45 | 0.7 |
| 3 | 44146964 | rs6809200 | -1.162 | 0.593 | 0.776 | -0.419 | 0.045 | -0.07 | -0.07 | 0 | 0.11 | 0.54 | 0.09 | 0 | 0.1 | 0.07 |
| 3 | 44155415 | rs7621857 | -0.717 | 0.582 | 1.065 | -0.416 | 0.029 | 0.012 | 0.012 | 0.679 | 0.05 | 0.29 | 0.03 | 0 | 0.01 | 0.04 |
| 3 | 44160183 | rs17645573 | -0.317 | 0.623 | 0.826 | -0.403 | 0.026 | -0.034 | -0.034 | 0.604 | 0.06 | 0.18 | 0.06 | 0 | 0.08 | 0.02 |
| 3 | 44175887 | rs4234441 | 0.237 | 0.472 | -0.11 | -0.675 | -0.016 | -0.011 | -0.011 | 0 | 0.77 | 0.94 | 0.78 | 0.75 | 0.74 | 0.79 |
| 3 | 44177438 | rs11708137 | -0.877 | 0.423 | 0.278 | -0.727 | 0.031 | -0.067 | -0.067 | 0 | 0.11 | 0.29 | 0.08 | 0 | 0.1 | 0.07 |
| 3 | 44196660 | rs4682716 | 0.708 | 0.017 | -0.425 | -1.051 | 0.014 | 0.026 | 0.026 | 0.394 | 0.39 | 0.17 | 0.51 | 0.62 | 0.38 | 0.62 |
| 3 | 44198777 | rs4682938 | 0.325 | 0.169 | -0.069 | -1.019 | 0.006 | 0 | 0 | 0 | 0.86 | 0.88 | 0.86 | 0.85 | 0.77 | 0.84 |
| 3 | 44199758 | rs7618961 | -0.575 | 0.164 | 0.137 | -1.045 | 0.039 | -0.048 | -0.048 | 0.309 | 0.09 | 0.32 | 0.07 | 0 | 0.09 | 0.03 |
| 3 | 44206461 | rs6762587 | 0.127 | 0.063 | -0.778 | -0.851 | -0.003 | -0.012 | -0.012 | 0 | 0.79 | 0.91 | 0.8 | 0.77 | 0.79 | 0.81 |
| 3 | 44221458 | rs730004 | 0.218 | -0.324 | -0.81 | -1.708 | -0.027 | -0.103 | -0.103 | 0.005 | 0.37 | 0.46 | 0.25 | 0.05 | 0.26 | 0.19 |
| 3 | 44222317 | rs1110330 | 0.427 | 0.869 | -0.796 | -2.661 | -0.009 | 0.014 | 0.014 | 1.912 | 0.26 | 0.18 | 0.15 | 0.02 | 0.27 | 0.1 |
| 3 | 44222351 | rs1105023 | 1.096 | 0.715 | -0.842 | -2.85 | -0.001 | 0.166 | 0.166 | 0.12 | 0.5 | 0.21 | 0.61 | 0.82 | 0.52 | 0.76 |
| 3 | 44233507 | rs1565208 | 0.032 | 0.601 | -0.397 | -2.757 | 0.01 | 0.005 | 0.005 | 0 | 0.85 | 0.76 | 0.82 | 0.83 | 0.85 | 0.87 |
| 3 | 44251270 | rs9825080 | 1.814 | 0.242 | -0.923 | -3.111 | 0.029 | 0.089 | 0.089 | 0 | 0.38 | 0.17 | 0.24 | 0.04 | 0.31 | 0.26 |
| 3 | 44254630 | rs10510740 | 0.477 | 0.189 | -0.232 | -2.77 | -0.008 | -0.051 | -0.051 | 0.274 | 0.09 | 0.25 | 0.07 | 0 | 0.09 | 0.01 |
| 3 | 44282934 | rs9809107 | 1.006 | 0.549 | -0.226 | -2.924 | -0.001 | 0.075 | 0.075 | 0 | 0.51 | 0.33 | 0.59 | 0.81 | 0.52 | 0.75 |
| 3 | 44283808 | rs7645375 | -0.74 | 0.568 | -0.116 | -2.926 | 0.012 | 0.012 | 0.012 | 0.041 | 0.9 | 0.88 | 0.82 | 0.81 | 0.86 | 0.87 |
| 3 | 44284584 | rs9284879 | 1.395 | 0.516 | -0.09 | -3.038 | 0.002 | 0.123 | 0.123 | 0 | 0.5 | 0.26 | 0.59 | 0.81 | 0.52 | 0.75 |
| 3 | 44289015 | rs6785357 | 0.708 | 0.449 | 0.097 | -3.082 | -0.003 | 0.037 | 0.037 | 2.591 | 0.25 | 0.12 | 0.14 | 0.02 | 0.23 | 0.11 |
| 3 | 44309620 | rs6768749 | 1.582 | 0.499 | 0.023 | -3.093 | 0.001 | 0.122 | 0.122 | 0 | 0.5 | 0.26 | 0.59 | 0.81 | 0.52 | 0.75 |
| 3 | 44377959 | rs11707582 | -0.647 | 0.053 | 0.096 | -3.03 | -0.014 | -0.157 | -0.157 | 0.593 | 0.36 | 0.52 | 0.23 | 0.02 | 0.34 | 0.12 |
| 3 | 44379864 | rs2279454 | -0.462 | 0.136 | 0.088 | -3.036 | -0.001 | 0.007 | 0.007 | 0 | 0.89 | 0.84 | 0.84 | 0.83 | 0.9 | 0.88 |
| 3 | 44385502 | rs11706775 | -0.649 | 0.147 | 0.043 | -3.047 | -0.016 | -0.157 | -0.157 | 0 | 0.36 | 0.52 | 0.23 | 0.02 | 0.34 | 0.12 |
| 3 | 44450564 | rs6441841 | -0.396 | 0.15 | -0.103 | -2.945 | -0.017 | -0.06 | -0.06 | 0.007 | 0.9 | 0.74 | 0.93 | 1 | 0.9 | 0.99 |
| 3 | 44451894 | rs12488257 | 1.247 | 0.33 | -0.42 | -2.829 | 0.022 | 0.092 | 0.092 | 0.093 | 0.5 | 0.32 | 0.61 | 0.81 | 0.56 | 0.76 |
| 3 | 44461080 | rs3935210 | 0.356 | 0.485 | 0.104 | -3.081 | 0 | -0.113 | -0.113 | 0 | 0.74 | 0.6 | 0.82 | 1 | 0.66 | 0.93 |
| 3 | 44461904 | rs12492600 | 0.252 | 0.544 | 0.047 | -3.215 | 0.002 | -0.007 | -0.007 | 0 | 0.25 | 0.24 | 0.26 | 0.17 | 0.24 | 0.16 |
| 3 | 44468065 | rs9855433 | 1.524 | 0.474 | -0.025 | -3.222 | -0.004 | 0.117 | 0.117 | 0.458 | 0.45 | 0.23 | 0.55 | 0.83 | 0.42 | 0.8 |
| 3 | 44481153 | rs12637714 | 1.227 | 0.643 | -0.09 | -3.1 | -0.033 | -0.034 | -0.034 | 0 | 0.07 | 0.01 | 0.14 | 0.12 | 0.09 | 0.16 |
| 3 | 44485914 | rs4610267 | -0.103 | 0.561 | 0.087 | -3.202 | 0.009 | -0.101 | -0.101 | 0.073 | 0.78 | 0.68 | 0.84 | 1 | 0.68 | 0.95 |
| 3 | 44489880 | rs11710965 | 1.322 | 0.636 | 0.002 | -3.182 | -0.035 | 0.046 | 0.046 | 0.593 | 0.1 | 0.01 | 0.06 | 0 | 0.17 | 0.01 |
| 3 | 44520267 | rs13087917 | 0.883 | 0.489 | 0.057 | -3.2 | 0.027 | -0.058 | -0.058 | 0.009 | 0.12 | 0.18 | 0.1 | 0 | 0.23 | 0.02 |
| 3 | 44524674 | rs4234447 | -0.867 | 0.435 | 0.174 | -3.158 | 0 | 0 | 0 | 0 | 0.92 | 0.93 | 0.85 | 0.88 | 0.91 | 0.84 |
| 3 | 44528949 | rs12632439 | -0.683 | 0.427 | 0.212 | -3.161 | 0.003 | 0.004 | 0.004 | 0.046 | 0.92 | 0.89 | 0.84 | 0.88 | 0.91 | 0.84 |
| 3 | 44568820 | rs13072696 | 0.007 | 0.382 | 0.182 | -3.159 | 0.004 | -0.061 | -0.061 | 0.351 | 0.2 | 0.24 | 0.15 | 0 | 0.32 | 0.05 |
| 3 | 44577236 | rs2141574 | -0.808 | 0.268 | 0.326 | -3.212 | -0.009 | -0.145 | -0.145 | 0.67 | 0.28 | 0.43 | 0.17 | 0 | 0.34 | 0.06 |
| 3 | 44593523 | rs1010031 | -0.853 | 0.259 | 0.395 | -3.217 | -0.014 | -0.145 | -0.145 | 0 | 0.28 | 0.44 | 0.18 | 0 | 0.34 | 0.06 |
| 3 | 44593620 | rs6763832 | 1.182 | 0.215 | 0.429 | -3.392 | -0.004 | 0.002 | 0.002 | 2.557 | 0.17 | 0.11 | 0.1 | 0.04 | 0.15 | 0.04 |
| 3 | 44596149 | rs6441856 | -0.718 | 0.228 | 0.416 | -3.374 | -0.014 | -0.145 | -0.145 | 0.234 | 0.28 | 0.44 | 0.18 | 0 | 0.34 | 0.06 |
| 3 | 44596896 | rs6441857 | -0.718 | 0.228 | 0.416 | -3.374 | -0.014 | -0.145 | -0.145 | 0 | 0.28 | 0.44 | 0.18 | 0 | 0.34 | 0.06 |
| 3 | 44612050 | rs9873604 | 0.151 | 0.202 | 0.477 | -3.433 | 0.004 | -0.022 | -0.022 | 0 | 0.72 | 0.67 | 0.73 | 0.83 | 0.75 | 0.79 |
| 3 | 44613014 | rs2034475 | -0.285 | 0.206 | 0.513 | -3.428 | -0.001 | -0.094 | -0.094 | 0 | 0.24 | 0.3 | 0.16 | 0 | 0.32 | 0.06 |
| 3 | 44625624 | rs6441858 | -0.311 | 0.113 | 0.524 | -3.467 | -0.001 | -0.054 | -0.054 | 0 | 0.24 | 0.25 | 0.15 | 0 | 0.32 | 0.06 |
| 3 | 44632212 | rs939649 | -0.759 | 0.22 | 0.753 | -3.45 | 0.005 | 0.005 | 0.005 | 0 | 0.91 | 0.87 | 0.83 | 0.88 | 0.9 | 0.83 |
| 3 | 44634629 | rs10212298 | -0.343 | 0.241 | 0.735 | -3.444 | -0.003 | -0.111 | -0.111 | 0 | 0.24 | 0.35 | 0.16 | 0 | 0.31 | 0.06 |
| 3 | 44641826 | rs11711191 | -0.284 | 0.242 | 0.732 | -3.444 | -0.002 | -0.094 | -0.094 | 0.299 | 0.24 | 0.31 | 0.16 | 0 | 0.31 | 0.06 |
| 3 | 44643857 | rs13100919 | -0.284 | 0.242 | 0.732 | -3.444 | -0.002 | -0.094 | -0.094 | 0 | 0.24 | 0.31 | 0.16 | 0 | 0.31 | 0.06 |
| 3 | 44665285 | rs1551063 | -0.342 | 0.241 | 0.732 | -3.444 | -0.003 | -0.111 | -0.111 | 0 | 0.24 | 0.34 | 0.16 | 0 | 0.31 | 0.06 |

| | | | | | | | | | | | | | | | | |
|---|---|---|---|---|---|---|---|---|---|---|---|---|---|---|---|---|
| 3 | 44674664 | rs11710959 | -0.112 | 0.273 | 0.748 | -3.428 | 0 | -0.061 | -0.061 | 0 | 0.22 | 0.25 | 0.15 | 0 | 0.31 | 0.06 |
| 3 | 44678498 | rs12635624 | 1.925 | 0.249 | 0.743 | -3.457 | -0.008 | 0.129 | 0.129 | 0.211 | 0.45 | 0.22 | 0.52 | 0.84 | 0.42 | 0.73 |
| 3 | 44681393 | rs11717797 | -0.253 | 0.259 | 0.741 | -3.432 | 0 | -0.11 | -0.11 | 0 | 0.23 | 0.34 | 0.16 | 0 | 0.31 | 0.06 |
| 3 | 44687300 | rs1402752 | -0.668 | 0.263 | 0.683 | -3.417 | 0.006 | 0.006 | 0.006 | 0.02 | 0.91 | 0.85 | 0.8 | 0.88 | 0.89 | 0.83 |
| 3 | 44701738 | rs1402753 | -0.125 | 0.265 | 0.592 | -3.408 | 0 | -0.058 | -0.058 | 0.064 | 0.22 | 0.25 | 0.15 | 0 | 0.31 | 0.06 |
| 3 | 44704083 | rs11130032 | 1.682 | 0.376 | 0.415 | -3.373 | -0.005 | 0.128 | 0.128 | 0.019 | 0.46 | 0.24 | 0.53 | 0.83 | 0.43 | 0.73 |
| 3 | 44712097 | rs6809116 | 1.554 | 0.394 | 0.477 | -3.311 | -0.014 | 0.152 | 0.152 | 0 | 0.49 | 0.23 | 0.53 | 0.83 | 0.45 | 0.74 |
| 3 | 44743993 | rs6778410 | 1.622 | 0.473 | 0.315 | -3.303 | -0.022 | 0.165 | 0.165 | 0 | 0.48 | 0.21 | 0.53 | 0.84 | 0.45 | 0.74 |
| 3 | 44761729 | rs17076938 | 0.688 | 0.281 | 0.465 | -3.456 | 0.011 | -0.008 | -0.008 | 1.477 | 0.06 | 0.08 | 0.1 | 0.04 | 0.05 | 0.04 |
| 3 | 44763037 | rs7640654 | 1.699 | -0.046 | 0.381 | -3.656 | 0.017 | 0.164 | 0.164 | 0 | 0.52 | 0.21 | 0.53 | 0.84 | 0.52 | 0.7 |
| 3 | 44770943 | rs2257995 | 0.59 | -0.101 | 0.507 | -3.637 | 0 | -0.02 | -0.02 | 0 | 0.67 | 0.68 | 0.67 | 0.88 | 0.6 | 0.74 |
| 3 | 44771128 | rs2279908 | 0.729 | -0.119 | 0.449 | -3.644 | -0.001 | -0.033 | -0.033 | 0 | 0.66 | 0.61 | 0.66 | 0.88 | 0.6 | 0.74 |
| 3 | 44775962 | rs4682752 | 0.729 | -0.119 | 0.449 | -3.644 | -0.001 | -0.033 | -0.033 | 0 | 0.66 | 0.61 | 0.66 | 0.88 | 0.6 | 0.74 |
| 3 | 44777795 | rs11922470 | 0.599 | 0.004 | 0.466 | -3.611 | 0 | -0.023 | -0.023 | 0 | 0.67 | 0.66 | 0.66 | 0.86 | 0.59 | 0.74 |
| 3 | 44784591 | rs17554407 | 2.319 | 0.448 | 0.083 | -3.825 | -0.083 | -0.058 | -0.058 | 0 | 0.13 | 0.01 | 0.3 | 0.79 | 0.22 | 0.44 |
| 3 | 44795238 | rs4682993 | 0.772 | 0.477 | 0.483 | -3.611 | 0.012 | -0.008 | -0.008 | 0 | 0.06 | 0.08 | 0.09 | 0.04 | 0.05 | 0.04 |
| 3 | 44803130 | rs4682994 | 0.62 | 0.316 | 0.534 | -3.621 | 0.026 | -0.005 | -0.005 | 0 | 0.06 | 0.14 | 0.1 | 0.04 | 0.05 | 0.04 |
| 3 | 44810107 | rs6441867 | 0.723 | 0.309 | 0.466 | -3.63 | 0 | -0.024 | -0.024 | 0 | 0.67 | 0.68 | 0.67 | 0.88 | 0.59 | 0.74 |
| 3 | 44814073 | rs2372827 | 0.066 | 0.277 | 0.505 | -3.669 | 0.002 | -0.012 | -0.012 | 0 | 0.76 | 0.77 | 0.72 | 0.88 | 0.69 | 0.8 |
| 3 | 44818836 | rs6770793 | 0.872 | 0.298 | 0.483 | -3.732 | -0.028 | -0.034 | -0.034 | 0.4 | 0.07 | 0.24 | 0.03 | 0 | 0.01 | 0 |
| 3 | 44835573 | rs1463950 | 0.903 | 0.235 | 0.324 | -3.737 | 0.001 | -0.054 | -0.054 | 0 | 0.67 | 0.62 | 0.67 | 0.89 | 0.59 | 0.75 |
| 3 | 44854563 | rs3762636 | 1.76 | 0.335 | -0.163 | -3.703 | 0.007 | 0.172 | 0.172 | 0 | 0.53 | 0.2 | 0.53 | 0.85 | 0.52 | 0.71 |
| 3 | 44857004 | rs6792918 | 0.486 | 0.351 | -0.28 | -3.68 | 0.011 | -0.011 | -0.011 | 2.54 | 0.07 | 0.08 | 0.1 | 0.04 | 0.06 | 0.03 |
| 3 | 44868381 | rs9849390 | 0.964 | 0.371 | -0.749 | -3.621 | -0.001 | -0.019 | -0.019 | 0 | 0.09 | 0.09 | 0.05 | 0 | 0.09 | 0.05 |
| 3 | 44871505 | rs11710339 | 1.405 | 0.558 | -1.13 | -3.528 | -0.047 | 0.059 | 0.059 | 4.461 | 0.12 | 0.02 | 0.08 | 0 | 0.22 | 0.02 |
| 3 | 44883971 | rs10514710 | 1.672 | 0.186 | -1.465 | -3.436 | -0.049 | 0.129 | 0.129 | 0.034 | 0.24 | 0.01 | 0.16 | 0.04 | 0.22 | 0.13 |
| 3 | 44884647 | rs3804583 | 1.286 | 0.117 | -0.494 | -3.079 | 0.001 | 0.229 | 0.229 | 0 | 0.52 | 0.16 | 0.53 | 0.85 | 0.52 | 0.71 |
| 3 | 44892033 | rs3804580 | 1.118 | 0.332 | -0.288 | -2.992 | 0 | 0.125 | 0.125 | 0 | 0.55 | 0.23 | 0.55 | 0.85 | 0.52 | 0.71 |
| 3 | 44894445 | rs7645364 | 1.026 | 0.061 | 0.089 | -2.919 | -0.015 | -0.025 | -0.025 | 1.348 | 0.05 | 0.16 | 0.02 | 0 | 0.01 | 0 |
| 3 | 44907228 | rs7622843 | 0.278 | 0.436 | 0.101 | -2.838 | -0.002 | -0.092 | -0.092 | 0.263 | 0.75 | 0.71 | 0.85 | 1 | 0.69 | 0.92 |
| 3 | 44911097 | rs2888422 | 0.931 | 0.515 | -0.198 | -2.797 | 0.001 | -0.077 | -0.077 | 0 | 0.66 | 0.54 | 0.66 | 0.89 | 0.59 | 0.74 |
| 3 | 44912443 | rs1506659 | 0.193 | 0.533 | -0.131 | -2.718 | 0.002 | -0.099 | -0.099 | 4.256 | 0.78 | 0.7 | 0.86 | 1 | 0.7 | 0.93 |
| 3 | 44915992 | rs7621143 | -0.657 | 0.279 | -0.22 | -2.703 | -0.004 | -0.029 | -0.029 | 0 | 0.1 | 0.16 | 0.06 | 0 | 0.09 | 0.05 |
| 3 | 44917735 | rs12632450 | 0.374 | 0.255 | -0.801 | -3.457 | -0.007 | 0.066 | 0.066 | 0.471 | 0.33 | 0.17 | 0.42 | 0.83 | 0.29 | 0.54 |
| 3 | 44917850 | rs17076998 | 0.333 | 0.323 | -0.729 | -3.634 | -0.001 | 0 | 0 | 0 | 0.09 | 0.1 | 0.17 | 0.08 | 0.11 | 0.19 |
| 3 | 44922141 | rs1532898 | 0.473 | 0.377 | -0.778 | -3.588 | -0.008 | 0.08 | 0.08 | 0.471 | 0.33 | 0.16 | 0.42 | 0.83 | 0.29 | 0.54 |
| 3 | 44932270 | rs937837 | 1.445 | 0.351 | -1.216 | -3.521 | -0.084 | -0.009 | -0.009 | 0 | 0.22 | 0.01 | 0.3 | 0.79 | 0.22 | 0.48 |
| 3 | 44940413 | rs1109412 | -0.446 | 0.035 | -0.745 | -3.719 | 0.012 | 0.129 | 0.129 | 0 | 0.48 | 0.31 | 0.48 | 0.88 | 0.41 | 0.73 |
| 3 | 44942121 | rs4682756 | -0.397 | 0.103 | -0.325 | -3.825 | 0.006 | 0.007 | 0.007 | 0.199 | 0.5 | 0.57 | 0.51 | 0.82 | 0.4 | 0.61 |
| 3 | 44943389 | rs1995641 | -0.284 | 0.093 | -0.244 | -3.83 | 0.003 | -0.002 | -0.002 | 0.477 | 0.49 | 0.48 | 0.51 | 0.8 | 0.4 | 0.6 |
| 3 | 44945740 | rs937833 | -0.951 | -0.027 | -0.274 | -3.811 | 0.005 | -0.079 | -0.079 | 0 | 0.58 | 0.55 | 0.6 | 0.89 | 0.49 | 0.79 |
| 3 | 44948479 | rs3749195 | -0.769 | -0.035 | 0.105 | -3.717 | -0.009 | 0.042 | 0.042 | 0.194 | 0.56 | 0.67 | 0.58 | 0.89 | 0.45 | 0.76 |
| 3 | 44948674 | rs1395388 | 0.567 | -0.145 | 0.038 | -3.762 | 0.006 | 0 | 0 | 0 | 0.07 | 0.09 | 0.05 | 0 | 0.06 | 0.05 |
| 3 | 44954481 | rs6784318 | -0.721 | 0.221 | 0.2 | -3.65 | -0.004 | 0.067 | 0.067 | 0.285 | 0.57 | 0.69 | 0.58 | 0.89 | 0.43 | 0.76 |
| 3 | 44955530 | rs1007 | -0.985 | -0.008 | 0.16 | -3.656 | -0.029 | 0.068 | 0.068 | 0.718 | 0.61 | 0.76 | 0.6 | 0.91 | 0.46 | 0.76 |
| 3 | 44958125 | rs3749193 | 0.612 | -0.155 | 0.007 | -3.781 | -0.03 | 0.011 | 0.011 | 0 | 0.16 | 0.02 | 0.18 | 0.11 | 0.15 | 0.23 |
| 3 | 44986836 | rs3749189 | -0.293 | -0.236 | 0.469 | -3.817 | -0.02 | -0.044 | -0.044 | 0.001 | 0.21 | 0.31 | 0.22 | 0.11 | 0.16 | 0.23 |
| 3 | 44990471 | rs9810846 | -0.687 | -0.054 | 0.386 | -3.707 | -0.003 | -0.012 | -0.012 | 0 | 0.55 | 0.59 | 0.58 | 0.89 | 0.45 | 0.76 |
| 3 | 44995128 | rs17368353 | 1.911 | 0.373 | 0.144 | -3.931 | -0.053 | 0.107 | 0.107 | 0 | 0.25 | 0.05 | 0.28 | 0.09 | 0.28 | 0.18 |
| 3 | 45012782 | rs1533677 | -0.821 | 0.277 | 0.2 | -3.887 | -0.029 | 0.064 | 0.064 | 0.906 | 0.61 | 0.76 | 0.61 | 0.91 | 0.46 | 0.76 |
| 3 | 45016919 | rs17632716 | -0.281 | 0.429 | 0.292 | -3.939 | 0 | -0.052 | -0.052 | 0 | 0.56 | 0.53 | 0.6 | 0.88 | 0.5 | 0.76 |
| 3 | 45017905 | rs6441874 | 0.174 | 0.435 | 0.277 | -4.005 | 0.002 | -0.074 | -0.074 | 0 | 0.44 | 0.48 | 0.52 | 0.86 | 0.4 | 0.7 |
| 3 | 45022096 | rs11707321 | 2.158 | 0.738 | 0.049 | -4.005 | -0.031 | 0.037 | 0.037 | 0 | 0.08 | 0.01 | 0.05 | 0 | 0.15 | 0.01 |
| 3 | 45030749 | rs7631259 | 1.48 | 0.4 | 0.133 | -3.919 | -0.004 | -0.016 | -0.016 | 1.917 | 0.13 | 0.14 | 0.08 | 0.02 | 0.09 | 0.06 |
| 3 | 45034411 | rs4683017 | -0.859 | 0.478 | 0.041 | -3.903 | -0.016 | 0.033 | 0.033 | 1.94 | 0.63 | 0.73 | 0.63 | 0.9 | 0.51 | 0.76 |
| 3 | 45061073 | rs11924033 | 0.977 | 0.562 | 0.028 | -3.705 | -0.033 | -0.027 | -0.027 | 0 | 0.06 | 0.01 | 0.13 | 0.1 | 0.09 | 0.17 |
| 3 | 45062095 | rs17570449 | -0.119 | 0.406 | 0.231 | -3.893 | 0.003 | 0.013 | 0.013 | 1.549 | 0.49 | 0.58 | 0.54 | 0.86 | 0.4 | 0.7 |
| 3 | 45075932 | rs939310 | -0.113 | 0.398 | 0.145 | -3.894 | 0 | -0.036 | -0.036 | 0 | 0.48 | 0.58 | 0.54 | 0.87 | 0.39 | 0.71 |
| 3 | 45077123 | rs13963 | 0.263 | 0.387 | 0.102 | -3.899 | -0.002 | -0.025 | -0.025 | 0 | 0.42 | 0.4 | 0.51 | 0.86 | 0.38 | 0.7 |
| 3 | 45083741 | rs11924100 | 1.204 | 0.498 | 0.146 | -3.709 | -0.025 | -0.029 | -0.029 | 0 | 0.06 | 0.13 | 0.03 | 0.02 | 0.01 | 0 |
| 3 | 45083928 | rs10514712 | 1.71 | 0.658 | -0.114 | -3.862 | -0.017 | 0.1 | 0.1 | 0 | 0.23 | 0.05 | 0.27 | 0.09 | 0.25 | 0.2 |
| 3 | 45092806 | rs4323037 | -0.53 | 0.334 | 0.254 | -3.589 | 0.005 | 0.046 | 0.046 | 0.315 | 0.52 | 0.65 | 0.53 | 0.87 | 0.39 | 0.72 |
| 3 | 45096013 | rs1520077 | -1.234 | 0.336 | 0.223 | -3.522 | -0.024 | 0.028 | 0.028 | 0.839 | 0.65 | 0.78 | 0.61 | 0.89 | 0.48 | 0.78 |
| 3 | 45097509 | rs7618668 | -0.383 | 0.317 | 0.466 | -3.255 | 0.032 | 0.018 | 0.018 | 0 | 0.08 | 0.21 | 0.14 | 0.08 | 0.08 | 0.12 |
| 3 | 45101804 | rs4683033 | -0.555 | 0.612 | 0.148 | -3.42 | 0.015 | 0.003 | 0.003 | 0 | 0.49 | 0.55 | 0.5 | 0.87 | 0.35 | 0.71 |
| 3 | 45108163 | rs939313 | -1.264 | 1.107 | -0.163 | -3.31 | -0.006 | 0.014 | 0.014 | 0 | 0.64 | 0.71 | 0.64 | 0.88 | 0.6 | 0.78 |
| 3 | 45111239 | rs11130043 | -0.417 | 1.032 | -0.65 | -3.216 | 0.02 | 0.229 | 0.229 | 0.18 | 0.51 | 0.22 | 0.43 | 0.77 | 0.41 | 0.64 |
| 3 | 45114302 | rs12635851 | 0.496 | 0.967 | -0.455 | -2.674 | -0.057 | 0.019 | 0.019 | 0.111 | 0.27 | 0.02 | 0.3 | 0.75 | 0.23 | 0.51 |
| 3 | 45120140 | rs6801821 | 0.799 | 1.072 | -0.01 | -2.83 | 0.014 | 0.088 | 0.088 | 0 | 0.54 | 0.29 | 0.44 | 0.15 | 0.57 | 0.31 |

| | | | | | | | | | | | | | | | | |
|---|---|---|---|---|---|---|---|---|---|---|---|---|---|---|---|---|
| 3 | 45124169 | rs11063 | 1.136 | 1.144 | 0.254 | -2.614 | 0.029 | -0.024 | -0.024 | 0.355 | 0.87 | 0.31 | 0.82 | 0.89 | 0.83 | 0.88 |
| 3 | 45125566 | rs4682765 | 0.607 | 1.251 | 0.023 | -2.587 | -0.073 | 0.007 | 0.007 | 0.05 | 0.26 | 0.01 | 0.36 | 0.74 | 0.25 | 0.52 |
| 3 | 45125720 | rs4683038 | 1.957 | 1.058 | 1.172 | -1.806 | -0.002 | 0.032 | 0.032 | 1.116 | 0.76 | 0.18 | 0.75 | 0.87 | 0.74 | 0.8 |
| 3 | 45128097 | rs6441886 | 1.308 | 0.525 | 0.923 | -1.439 | 0.037 | 0.027 | 0.027 | 0.905 | 0.93 | 0.54 | 0.95 | 0.98 | 0.99 | 1 |
| 3 | 45130158 | rs6784502 | 1.053 | 0.774 | 0.692 | -1.451 | 0.037 | -0.056 | -0.056 | 0 | 0.89 | 0.54 | 0.9 | 0.97 | 0.9 | 0.93 |
| 3 | 45130725 | rs7618923 | -1.301 | 1.434 | 0.343 | -1.538 | 0.035 | -0.059 | -0.059 | 0 | 0.14 | 0.3 | 0.13 | 0.03 | 0.18 | 0.08 |
| 3 | 45131916 | rs11916758 | 0.499 | 1.18 | -0.828 | -1.346 | 0.043 | 0.113 | 0.113 | 0 | 0.48 | 0.28 | 0.32 | 0.23 | 0.41 | 0.41 |
| 3 | 45137087 | rs2037358 | 0.486 | 1.295 | -1.488 | -1.322 | -0.056 | 0.198 | 0.198 | 0 | 0.45 | 0.13 | 0.65 | 0.8 | 0.49 | 0.65 |
| 3 | 45138515 | rs17077267 | 0.338 | 1.467 | -0.866 | -1.287 | 0.042 | -0.053 | -0.053 | 0.673 | 0.1 | 0.22 | 0.09 | 0 | 0.15 | 0.06 |
| 3 | 45139101 | rs6775113 | -0.203 | 1.168 | -1.044 | -1.136 | 0.057 | -0.04 | -0.04 | 0 | 0.88 | 0.61 | 0.88 | 0.97 | 0.84 | 0.95 |
| 3 | 45140266 | rs1079472 | 1.083 | 0.715 | -1.955 | -0.849 | 0.002 | -0.016 | -0.016 | 0.267 | 0.18 | 0.03 | 0.17 | 0.12 | 0.18 | 0.12 |
| 3 | 45145019 | rs4579052 | -0.172 | 1.11 | -1.613 | 0.136 | -0.019 | -0.092 | -0.092 | 1.734 | 0.3 | 0.11 | 0.48 | 0.55 | 0.31 | 0.52 |
| 3 | 45148674 | rs6779858 | -0.3 | 0.852 | -1.216 | -0.349 | 0 | 0.015 | 0.015 | 0.029 | 0.58 | 0.63 | 0.69 | 0.83 | 0.56 | 0.79 |
| 3 | 45149708 | rs5007171 | -0.395 | 0.504 | -1.184 | 0.511 | 0.025 | -0.046 | -0.046 | 4.277 | 0.25 | 0.18 | 0.37 | 0.61 | 0.22 | 0.62 |
| 3 | 45153422 | rs6778556 | 0.214 | 1.567 | -0.259 | 0.879 | 0.112 | 0.217 | 0.217 | 1.18 | 0.89 | 0.7 | 0.82 | 0.65 | 0.83 | 0.87 |
| 3 | 45154357 | rs12494480 | 0.166 | 1.063 | -0.946 | 0.221 | -0.036 | -0.011 | -0.011 | 0 | 0.46 | 0.22 | 0.5 | 0.52 | 0.49 | 0.6 |
| 3 | 45154868 | rs4683045 | 0.549 | 1.3 | -1.41 | 0.091 | 0.003 | -0.116 | -0.116 | 0.265 | 0.25 | 0.09 | 0.31 | 0.44 | 0.24 | 0.46 |
| 3 | 45155294 | rs4683047 | 0.532 | 0.916 | -0.646 | -0.152 | 0.012 | 0.011 | 0.011 | 0.925 | 0.77 | 0.32 | 0.65 | 0.65 | 0.64 | 0.81 |
| 3 | 45157568 | rs7625872 | -0.571 | 0.114 | -0.02 | 0.37 | 0.088 | 0.254 | 0.254 | 0 | 0.09 | 0.32 | 0.18 | 0.32 | 0.15 | 0.11 |
| 3 | 45161070 | rs13325398 | 0.293 | 0.315 | -0.845 | 0.361 | 0.046 | 0.061 | 0.061 | 0 | 0.31 | 0.17 | 0.16 | 0.08 | 0.16 | 0.19 |
| 3 | 45162316 | rs9876540 | -0.758 | 1.398 | -0.757 | 1.328 | 0.053 | 0.164 | 0.164 | 0 | 0.06 | 0.14 | 0.14 | 0.31 | 0.15 | 0.1 |
| 3 | 45170095 | rs4682769 | 0.017 | 0.875 | -1.828 | 0.795 | -0.104 | -0.055 | -0.055 | 0 | 0.44 | 0.11 | 0.49 | 0.46 | 0.54 | 0.38 |
| 3 | 45170505 | rs11916555 | 0.007 | 0.763 | -1.652 | 0.799 | 0 | 0 | 0 | 1.082 | 0.13 | 0.1 | 0.08 | 0.13 | 0.11 | 0.11 |
| 3 | 45171167 | rs4683055 | -0.275 | 0.353 | -0.724 | 0.452 | 0.001 | 0.003 | 0.003 | 0 | 0.48 | 0.54 | 0.47 | 0.54 | 0.43 | 0.61 |
| 3 | 45177006 | rs10490812 | -0.52 | 0.28 | -0.26 | 0.011 | -0.009 | -0.05 | -0.05 | 0 | 0.18 | 0.26 | 0.15 | 0.07 | 0.15 | 0.15 |
| 3 | 45185519 | rs7633169 | 0.044 | 0.219 | 0.053 | 0.015 | -0.011 | -0.051 | -0.051 | 0 | 0.84 | 0.76 | 0.88 | 0.95 | 0.86 | 0.87 |
| 3 | 45189453 | rs7629057 | 0.633 | 0.276 | 0.022 | -0.145 | -0.003 | -0.031 | -0.031 | 0 | 0.65 | 0.57 | 0.64 | 0.75 | 0.63 | 0.76 |
| 3 | 45189580 | rs7636990 | 0.918 | 0.305 | -0.189 | 0.036 | -0.005 | -0.007 | -0.007 | 0 | 0.3 | 0.28 | 0.39 | 0.39 | 0.32 | 0.28 |
| 3 | 45192973 | rs4683068 | -0.193 | 0.095 | 0.201 | 0.032 | 0.018 | 0.267 | 0.267 | 0.249 | 0.91 | 0.65 | 0.87 | 0.71 | 0.92 | 0.73 |
| 3 | 45196743 | rs7651325 | 0.521 | 0.227 | -0.149 | -0.358 | -0.008 | 0.008 | 0.008 | 0 | 0.18 | 0.11 | 0.17 | 0.16 | 0.23 | 0.17 |
| 3 | 45199620 | rs4683070 | 0.886 | 0.274 | 0.037 | -0.151 | 0.005 | -0.006 | -0.006 | 0 | 0.21 | 0.18 | 0.27 | 0.26 | 0.21 | 0.3 |
| 3 | 45199631 | rs4683071 | -0.46 | 0.022 | -0.121 | -0.273 | 0.017 | 0.025 | 0.025 | 0 | 0.47 | 0.29 | 0.46 | 0.43 | 0.47 | 0.48 |
| 3 | 45203896 | rs4683072 | 0.343 | -0.119 | 0.05 | -0.407 | -0.01 | -0.018 | -0.018 | 0 | 0.55 | 0.53 | 0.59 | 0.72 | 0.62 | 0.55 |
| 3 | 45206115 | rs7618987 | -0.296 | 0.057 | 0.151 | -0.34 | 0.011 | 0.016 | 0.016 | 2.638 | 0.71 | 0.77 | 0.87 | 0.93 | 0.79 | 0.79 |
| 3 | 45211087 | rs9819089 | -0.271 | 0.076 | 0.203 | -0.271 | -0.014 | -0.017 | -0.017 | 0 | 0.35 | 0.43 | 0.32 | 0.24 | 0.27 | 0.43 |
| 3 | 45211399 | rs9857201 | -0.641 | 0.156 | 0.183 | -0.166 | -0.006 | -0.001 | -0.001 | 0 | 0.4 | 0.47 | 0.37 | 0.41 | 0.36 | 0.48 |
| 3 | 45212443 | rs4524316 | -0.76 | 0.576 | -0.178 | -0.303 | 0.015 | 0.012 | 0.012 | 2.188 | 0.4 | 0.45 | 0.4 | 0.47 | 0.45 | 0.49 |
| 3 | 45215474 | rs9816577 | 0.879 | 0.594 | -0.674 | -0.322 | 0 | 0.011 | 0.011 | 0 | 0.25 | 0.16 | 0.18 | 0.21 | 0.23 | 0.4 |
| 3 | 45215803 | rs9817295 | -0.057 | 0.734 | -0.487 | -0.264 | 0.002 | -0.006 | -0.006 | 0 | 0.71 | 0.35 | 0.71 | 0.78 | 0.7 | 0.8 |
| 3 | 45216928 | rs4683082 | 1.278 | 0.235 | -0.599 | -0.145 | 0.008 | -0.07 | -0.07 | 0.152 | 0.33 | 0.14 | 0.45 | 0.42 | 0.31 | 0.33 |
| 3 | 45217643 | rs10490814 | 1.381 | 0.472 | -0.513 | -0.182 | -0.002 | 0.015 | 0.015 | 0 | 0.13 | 0.07 | 0.11 | 0.04 | 0.15 | 0.12 |
| 3 | 45220085 | rs6767173 | -0.369 | 0.676 | -0.405 | -0.134 | -0.01 | 0.02 | 0.02 | 0.453 | 0.59 | 0.67 | 0.68 | 0.76 | 0.55 | 0.49 |
| 3 | 45221692 | rs7627767 | 0.463 | 0.811 | -0.854 | -0.006 | -0.051 | -0.041 | -0.041 | 0.016 | 0.39 | 0.13 | 0.53 | 0.56 | 0.48 | 0.42 |
| 3 | 45221916 | rs6441898 | -0.495 | 0.957 | -0.77 | 0.056 | 0.025 | 0.032 | 0.032 | 0 | 0.93 | 0.87 | 0.93 | 0.84 | 0.87 | 0.94 |
| 3 | 45223641 | rs12497339 | 0.835 | 0.407 | -1.057 | 0.045 | -0.04 | -0.109 | -0.109 | 0.558 | 0.34 | 0.07 | 0.47 | 0.43 | 0.36 | 0.36 |
| 3 | 45226973 | rs6799689 | 0.299 | -0.051 | -0.421 | -0.318 | -0.011 | 0.034 | 0.034 | 0.026 | 0.42 | 0.32 | 0.27 | 0.18 | 0.5 | 0.25 |
| 3 | 45227101 | rs17077435 | 1.515 | -0.112 | -0.568 | -0.391 | -0.004 | 0.021 | 0.021 | 0 | 0.13 | 0.07 | 0.1 | 0.04 | 0.13 | 0.1 |
| 3 | 45229193 | rs1017299 | 1.527 | -0.158 | -0.58 | -0.205 | -0.015 | -0.125 | -0.125 | 0.135 | 0.26 | 0.08 | 0.41 | 0.47 | 0.28 | 0.38 |
| 3 | 45230567 | rs9813119 | 1.02 | -0.248 | -0.222 | -0.4 | -0.005 | -0.033 | -0.033 | 0.369 | 0.07 | 0.23 | 0.03 | 0 | 0.03 | 0.01 |
| 3 | 45230989 | rs9831259 | 1.432 | -0.241 | -0.258 | -0.397 | -0.002 | -0.014 | -0.014 | 0 | 0.06 | 0.06 | 0.03 | 0 | 0.03 | 0.01 |
| 3 | 45237733 | rs7639787 | 0.383 | -0.398 | -0.354 | -0.365 | -0.018 | 0.023 | 0.023 | 0 | 0.41 | 0.29 | 0.33 | 0.36 | 0.51 | 0.33 |
| 3 | 45239835 | rs245215 | 1.248 | -0.299 | -0.908 | -0.537 | -0.123 | 0.181 | 0.181 | 0 | 0.31 | 0.06 | 0.21 | 0.04 | 0.43 | 0.18 |
| 3 | 45245744 | rs245207 | -0.65 | -0.793 | -0.36 | -0.413 | -0.03 | -0.094 | -0.094 | 1.037 | 0.79 | 0.41 | 0.86 | 0.97 | 0.87 | 0.9 |
| 3 | 45252540 | rs1559995 | 1.501 | -0.718 | -0.796 | -0.261 | -0.012 | -0.15 | -0.15 | 0.141 | 0.27 | 0.06 | 0.43 | 0.49 | 0.28 | 0.45 |
| 3 | 45253299 | rs33734 | 0.588 | -0.807 | -0.82 | -0.334 | -0.059 | 0.02 | 0.02 | 0 | 0.4 | 0.11 | 0.33 | 0.36 | 0.52 | 0.33 |
| 3 | 45257469 | rs28169 | 0.489 | -0.892 | -0.512 | -0.199 | 0.043 | 0.025 | 0.025 | 3.388 | 0.71 | 0.2 | 0.79 | 0.84 | 0.83 | 0.78 |
| 3 | 45262393 | rs33748 | -0.013 | -0.949 | 0.086 | -0.217 | -0.032 | 0.042 | 0.042 | 0 | 0.87 | 0.75 | 0.9 | 0.87 | 0.93 | 0.88 |
| 3 | 45263711 | rs4543049 | 0.653 | -0.88 | -0.242 | -0.171 | -0.012 | 0.061 | 0.061 | 0 | 0.07 | 0.12 | 0.06 | 0.13 | 0.04 | 0.11 |
| 3 | 45266555 | rs33749 | 0.48 | -0.9 | -0.413 | -0.195 | 0.039 | 0.022 | 0.022 | 2.836 | 0.71 | 0.24 | 0.8 | 0.84 | 0.83 | 0.78 |
| 3 | 45273915 | rs12493086 | 0.929 | -0.713 | -0.437 | 0.008 | 0.001 | 0 | 0 | 0.169 | 0.05 | 0.04 | 0.09 | 0.17 | 0.04 | 0.07 |
| 3 | 45281928 | rs12487673 | -0.663 | -0.687 | -0.105 | -0.073 | -0.018 | 0.055 | 0.055 | 0 | 0.18 | 0.24 | 0.18 | 0.3 | 0.12 | 0.19 |
| 3 | 45282691 | rs33770 | 0.93 | -0.67 | -0.336 | -0.136 | 0.054 | -0.03 | -0.03 | 0 | 0.62 | 0.15 | 0.68 | 0.68 | 0.75 | 0.71 |
| 3 | 45284315 | rs33773 | 1.045 | -0.704 | -0.857 | -0.33 | -0.111 | 0.1 | 0.1 | 0 | 0.33 | 0.1 | 0.23 | 0.18 | 0.47 | 0.26 |
| 3 | 45285743 | rs972850 | 1.15 | -0.767 | -0.444 | -0.28 | 0.004 | -0.012 | -0.012 | 0 | 0.31 | 0.28 | 0.46 | 0.51 | 0.29 | 0.45 |
| 3 | 45288534 | rs33779 | -0.627 | -0.898 | -0.086 | -0.3 | -0.023 | -0.033 | -0.033 | 0 | 0.77 | 0.51 | 0.78 | 0.82 | 0.83 | 0.8 |
| 3 | 45289715 | rs33780 | 1.066 | -0.845 | -0.127 | -0.35 | -0.079 | 0.085 | 0.085 | 0 | 0.34 | 0.13 | 0.24 | 0.18 | 0.47 | 0.25 |
| 3 | 45292244 | rs2888432 | -0.223 | -0.887 | 0.115 | 0.008 | -0.035 | 0.058 | 0.058 | 0 | 0.14 | 0.28 | 0.09 | 0.14 | 0.07 | 0.12 |
| 3 | 45293715 | rs4683095 | 1.733 | -0.686 | -0.455 | -0.121 | -0.02 | 0.045 | 0.045 | 0 | 0.15 | 0.01 | 0.22 | 0.06 | 0.19 | 0.07 |
| 3 | 45302410 | rs33797 | -0.332 | -0.57 | -0.058 | -0.075 | 0.027 | 0.027 | 0.027 | 0 | 0.6 | 0.28 | 0.45 | 0.37 | 0.67 | 0.48 |

| | | | | | | | | | | | | | | | | |
|---|---|---|---|---|---|---|---|---|---|---|---|---|---|---|---|---|
| 3 | 45304890 | rs9822723 | -0.119 | -0.009 | 0.19 | 0.461 | 0 | 0.027 | 0.027 | 0.266 | 0.09 | 0.15 | 0.11 | 0.2 | 0.08 | 0.1 |
| 3 | 45309500 | rs17077548 | -0.271 | -0.26 | 0.695 | 0.865 | 0.006 | 0.117 | 0.117 | 1.707 | 0.15 | 0.31 | 0.17 | 0.34 | 0.12 | 0.19 |
| 3 | 45309717 | rs11706314 | -0.581 | -0.207 | 0.733 | 0.777 | -0.003 | 0.009 | 0.009 | 0 | 0.06 | 0.08 | 0.03 | 0.14 | 0.04 | 0.11 |
| 3 | 45313505 | rs11923281 | -0.936 | -0.598 | 0.656 | 0.52 | 0.001 | -0.004 | -0.004 | 0 | 0.36 | 0.63 | 0.33 | 0.52 | 0.29 | 0.33 |
| 3 | 45316089 | rs33802 | 1.224 | -0.558 | 0.068 | 0.047 | -0.096 | 0.146 | 0.146 | 0 | 0.28 | 0.06 | 0.2 | 0.03 | 0.41 | 0.2 |
| 3 | 45324475 | rs6765891 | -0.752 | -0.648 | 0.591 | 0.294 | -0.005 | 0.055 | 0.055 | 2.963 | 0.33 | 0.48 | 0.32 | 0.52 | 0.29 | 0.33 |
| 3 | 45325521 | rs7650391 | -0.695 | -0.672 | 0.544 | 0.28 | -0.004 | 0.02 | 0.02 | 2.458 | 0.33 | 0.39 | 0.31 | 0.48 | 0.29 | 0.32 |
| 3 | 45333985 | rs7617429 | 0.138 | -0.536 | 0.627 | 0.502 | 0.007 | 0.081 | 0.081 | 1.59 | 0.15 | 0.33 | 0.26 | 0.3 | 0.12 | 0.19 |
| 3 | 45340998 | rs7620365 | -0.952 | -0.581 | 0.616 | 0.49 | -0.029 | 0.073 | 0.073 | 0.721 | 0.21 | 0.54 | 0.28 | 0.33 | 0.12 | 0.28 |
| 3 | 45360959 | rs6793847 | -1.329 | -0.094 | -0.028 | 0.143 | 0.01 | -0.002 | -0.002 | 0 | 0.53 | 0.76 | 0.5 | 0.57 | 0.42 | 0.47 |
| 3 | 45361222 | rs9867680 | 0.601 | -0.216 | 0.245 | 0.117 | -0.028 | -0.013 | -0.013 | 0 | 0.06 | 0.13 | 0.01 | 0.03 | 0 | 0.1 |
| 3 | 45362383 | rs11927517 | 0.558 | -0.326 | 0.311 | 0.099 | 0.001 | 0.01 | 0.01 | 0 | 0.3 | 0.22 | 0.21 | 0.24 | 0.3 | 0.18 |
| 3 | 45362601 | rs9821844 | -0.723 | -0.324 | 0.385 | 0.129 | -0.04 | -0.022 | -0.022 | 0.535 | 0.08 | 0.23 | 0.02 | 0.03 | 0 | 0.1 |
| 3 | 45367619 | rs11719661 | 0.234 | -0.271 | 0.341 | 0.156 | 0.003 | 0.04 | 0.04 | 1.269 | 0.05 | 0.12 | 0.03 | 0.12 | 0.04 | 0.03 |
| 3 | 45369114 | rs2372948 | -0.781 | -0.258 | 0.21 | 0.105 | 0.013 | -0.005 | -0.005 | 0 | 0.47 | 0.63 | 0.49 | 0.49 | 0.41 | 0.37 |
| 3 | 45370416 | rs1468696 | -1.389 | -0.176 | 0.093 | 0.119 | 0.017 | -0.007 | -0.007 | 0.012 | 0.52 | 0.7 | 0.49 | 0.52 | 0.42 | 0.47 |
| 3 | 45371605 | rs929578 | 0.925 | -0.198 | -0.449 | -0.06 | -0.017 | 0.09 | 0.09 | 0 | 0.25 | 0.05 | 0.15 | 0.08 | 0.25 | 0.16 |
| 3 | 45377695 | rs6785141 | -1.142 | 0.047 | -0.087 | -0.191 | 0.06 | -0.089 | -0.089 | 0.469 | 0.82 | 0.67 | 0.84 | 0.97 | 0.7 | 0.86 |
| 3 | 45381039 | rs4683098 | 0.724 | 0.211 | 0.058 | 0.067 | 0.018 | 0.001 | 0.001 | 2.546 | 0.67 | 0.42 | 0.65 | 0.66 | 0.66 | 0.6 |
| 3 | 45383854 | rs2110084 | 0.74 | 0.31 | -0.121 | 0.034 | 0.002 | 0.008 | 0.008 | 0 | 0.28 | 0.19 | 0.19 | 0.2 | 0.27 | 0.18 |
| 3 | 45385866 | rs6809473 | 1.612 | 0.529 | -0.486 | 0.079 | -0.041 | -0.109 | -0.109 | 0 | 0.37 | 0.06 | 0.46 | 0.41 | 0.38 | 0.42 |
| 3 | 45390944 | rs6768533 | 0.807 | 0.876 | 0.095 | -0.024 | 0.048 | -0.015 | -0.015 | 0 | 0.72 | 0.51 | 0.68 | 0.74 | 0.66 | 0.69 |
| 3 | 45392610 | rs1860928 | 0.802 | 0.859 | 0.065 | -0.144 | 0.05 | -0.037 | -0.037 | 0 | 0.72 | 0.52 | 0.79 | 0.78 | 0.64 | 0.68 |
| 3 | 45395685 | rs854221 | -0.949 | 0.23 | -0.03 | 0.052 | -0.028 | -0.01 | -0.01 | 0 | 0.21 | 0.56 | 0.13 | 0.19 | 0.15 | 0.22 |
| 3 | 45397986 | rs854222 | -1.068 | -0.517 | -0.961 | -0.094 | 0.056 | -0.046 | -0.046 | 0 | 0.65 | 0.82 | 0.65 | 0.42 | 0.78 | 0.63 |
| 3 | 45403666 | rs11709274 | 0.668 | -0.255 | -1.234 | -0.473 | -0.015 | -0.016 | -0.016 | 0 | 0.27 | 0.14 | 0.34 | 0.29 | 0.3 | 0.28 |
| 3 | 45403826 | rs6795131 | -1.157 | -0.275 | -0.807 | -0.536 | -0.013 | 0.004 | 0.004 | 1.157 | 0.61 | 0.43 | 0.6 | 0.42 | 0.78 | 0.63 |
| 3 | 45405467 | rs6793155 | -1.38 | -0.234 | -0.803 | -0.54 | -0.048 | 0.025 | 0.025 | 0.596 | 0.64 | 0.52 | 0.63 | 0.42 | 0.81 | 0.64 |
| 3 | 45406592 | rs13314784 | 0.064 | -0.063 | -1.033 | -0.494 | -0.013 | 0.012 | 0.012 | 0.314 | 0.34 | 0.29 | 0.26 | 0.13 | 0.48 | 0.36 |
| 3 | 45408398 | rs854226 | -1.021 | 0.33 | -0.97 | -0.455 | -0.023 | 0.057 | 0.057 | 0 | 0.54 | 0.78 | 0.54 | 0.31 | 0.53 | 0.56 |
| 3 | 45429891 | rs854187 | -0.504 | 0.425 | -0.502 | -0.355 | -0.003 | 0.052 | 0.052 | 0.103 | 0.1 | 0.41 | 0.08 | 0.12 | 0.05 | 0.06 |
| 3 | 45436473 | rs17637580 | 0.574 | 0.903 | -1.248 | -0.483 | -0.035 | 0.006 | 0.006 | 0 | 0.17 | 0.1 | 0.14 | 0.11 | 0.29 | 0.19 |
| 3 | 45436941 | rs17077759 | 0.759 | 0.81 | -1.193 | -0.641 | 0.005 | -0.038 | -0.038 | 1.069 | 0.13 | 0.18 | 0.14 | 0.01 | 0.18 | 0.14 |
| 3 | 45440944 | rs17575828 | 0.478 | 0.452 | -1.809 | -0.659 | -0.021 | 0.072 | 0.072 | 0 | 0.24 | 0.04 | 0.26 | 0.07 | 0.26 | 0.19 |
| 3 | 45448958 | rs6802884 | -0.612 | 0.049 | -1.447 | -0.693 | 0 | -0.003 | -0.003 | 0 | 0.23 | 0.27 | 0.22 | 0.13 | 0.17 | 0.28 |
| 3 | 45449373 | rs9830462 | -0.037 | 0.013 | -1.395 | -0.668 | -0.001 | 0 | 0 | 0 | 0.07 | 0.07 | 0.04 | 0.1 | 0.03 | 0.11 |
| 3 | 45452578 | rs854208 | -1.598 | -0.355 | -1.73 | -1.029 | 0.024 | 0.002 | 0.002 | 0.233 | 0.62 | 0.92 | 0.53 | 0.3 | 0.74 | 0.56 |
| 3 | 45458733 | rs17576289 | 2.48 | -1.015 | -2.22 | -0.514 | -0.019 | -0.069 | -0.069 | 1.387 | 0.16 | 0.01 | 0.21 | 0.25 | 0.15 | 0.08 |
| 3 | 45460460 | rs854200 | -0.878 | -0.556 | -1.651 | -0.79 | 0.061 | 0.116 | 0.116 | 0.002 | 0.54 | 0.75 | 0.48 | 0.19 | 0.7 | 0.44 |
| 3 | 45461483 | rs17638301 | 1.089 | -0.443 | -1.86 | -0.841 | -0.024 | 0.063 | 0.063 | 0 | 0.14 | 0.05 | 0.1 | 0.01 | 0.16 | 0.14 |
| 3 | 45466897 | rs12632096 | 0.092 | -0.407 | -1.779 | -0.801 | -0.037 | 0.046 | 0.046 | 0 | 0.37 | 0.2 | 0.32 | 0.19 | 0.54 | 0.3 |
| 3 | 45466958 | rs17638423 | 0.989 | -0.566 | -1.739 | -0.883 | -0.003 | 0.018 | 0.018 | 0 | 0.2 | 0.1 | 0.18 | 0.07 | 0.25 | 0.12 |
| 3 | 45475333 | rs4683113 | 1.422 | -0.533 | -2.019 | -0.665 | -0.04 | 0.069 | 0.069 | 0.167 | 0.46 | 0.09 | 0.53 | 0.69 | 0.55 | 0.57 |
| 3 | 45476040 | rs9825179 | 0.065 | -0.454 | -1.816 | -0.726 | 0.004 | 0.027 | 0.027 | 0 | 0.05 | 0.12 | 0.04 | 0.11 | 0.03 | 0.09 |
| 3 | 45476486 | rs2014378 | 1.219 | -0.723 | -1.94 | -0.74 | -0.028 | 0.059 | 0.059 | 0 | 0.14 | 0.05 | 0.11 | 0.01 | 0.16 | 0.16 |
| 3 | 45476694 | rs952621 | 0.43 | -0.756 | -1.843 | -0.677 | 0.016 | 0.014 | 0.014 | 0.152 | 0.23 | 0.12 | 0.17 | 0.17 | 0.19 | 0.12 |
| 3 | 45500950 | rs1949304 | 0.967 | -0.661 | -1.867 | -0.528 | 0.09 | 0.116 | 0.116 | 0.146 | 0.5 | 0.11 | 0.63 | 0.67 | 0.62 | 0.62 |
| 3 | 45506446 | rs3774694 | 0.956 | -0.159 | -1.572 | -0.231 | -0.054 | -0.02 | -0.02 | 0.343 | 0.16 | 0.04 | 0.16 | 0.2 | 0.31 | 0.2 |
| 3 | 45507779 | rs17638880 | -0.139 | -0.319 | -1.607 | -0.38 | -0.066 | -0.056 | -0.056 | 0.534 | 0.23 | 0.06 | 0.22 | 0.33 | 0.37 | 0.26 |
| 3 | 45517098 | rs1473443 | 0.491 | -0.635 | -1.178 | -0.562 | -0.045 | -0.048 | -0.048 | 0 | 0.88 | 0.36 | 0.91 | 0.98 | 0.95 | 0.93 |
| 3 | 45536988 | rs17639812 | 0.899 | -0.435 | -1.302 | -0.466 | -0.064 | -0.065 | -0.065 | 0 | 0.17 | 0.04 | 0.19 | 0.34 | 0.32 | 0.14 |
| 3 | 45545451 | rs12629394 | -0.44 | -1.234 | -0.765 | -1.194 | 0.01 | 0.063 | 0.063 | 0.1 | 0.37 | 0.2 | 0.26 | 0.19 | 0.33 | 0.27 |
| 3 | 45549028 | rs2128358 | 0.768 | -1.333 | -0.847 | -1.254 | 0.013 | 0.055 | 0.055 | 0 | 0.19 | 0.03 | 0.14 | 0.07 | 0.17 | 0.12 |
| 3 | 45557707 | rs11549809 | -0.231 | -1.283 | -0.681 | -1.116 | 0.013 | 0.064 | 0.064 | 0.086 | 0.37 | 0.2 | 0.25 | 0.19 | 0.33 | 0.27 |
| 3 | 45572257 | rs2118749 | 0.876 | -1.146 | -1.053 | -1.106 | 0.095 | 0.15 | 0.15 | 0.796 | 0.55 | 0.1 | 0.69 | 0.78 | 0.68 | 0.61 |
| 3 | 45594122 | rs267272 | 0.922 | -0.935 | -0.986 | -0.841 | 0.016 | 0.18 | 0.18 | 0.578 | 0.48 | 0.06 | 0.62 | 0.79 | 0.59 | 0.57 |
| 3 | 45598655 | rs10212409 | -0.619 | -0.714 | -0.452 | -0.771 | -0.003 | -0.005 | -0.005 | 0.307 | 0.06 | 0.09 | 0.05 | 0.03 | 0.06 | 0.03 |
| 3 | 45598744 | rs267280 | 0.639 | -0.602 | 0.115 | -0.956 | 0.005 | 0.065 | 0.065 | 0 | 0.52 | 0.35 | 0.63 | 0.88 | 0.54 | 0.62 |
| 3 | 45600019 | rs9819404 | -0.46 | -0.564 | 0.269 | -0.921 | -0.006 | -0.007 | -0.007 | 0 | 0.06 | 0.14 | 0.05 | 0.03 | 0.06 | 0.03 |
| 3 | 45601252 | rs267260 | 0.164 | -0.511 | 0.157 | -0.934 | 0 | -0.058 | -0.058 | 3.429 | 0.58 | 0.46 | 0.68 | 0.92 | 0.6 | 0.66 |
| 3 | 45602225 | rs13083568 | -0.113 | -0.43 | -0.16 | -0.937 | -0.008 | 0.096 | 0.096 | 0 | 0.37 | 0.22 | 0.27 | 0.08 | 0.37 | 0.34 |
| 3 | 45604131 | rs13075111 | 0.057 | -0.302 | 0.054 | -0.811 | -0.003 | 0.107 | 0.107 | 0.148 | 0.42 | 0.21 | 0.3 | 0.06 | 0.41 | 0.34 |
| 3 | 45604159 | rs13075121 | 0.057 | -0.302 | 0.054 | -0.811 | -0.003 | 0.107 | 0.107 | 0.148 | 0.42 | 0.21 | 0.3 | 0.06 | 0.41 | 0.33 |
| 3 | 45613379 | rs9823534 | -1.004 | -0.066 | 0.398 | 0.078 | -0.008 | 0.015 | 0.015 | 0 | 0.08 | 0.11 | 0.06 | 0.16 | 0.06 | 0.06 |
| 3 | 45616915 | rs11130070 | 1.408 | -0.12 | 0.858 | 0.273 | -0.004 | 0.189 | 0.189 | 1.538 | 0.89 | 0.48 | 0.85 | 0.74 | 0.93 | 0.92 |
| 3 | 45617040 | rs13066873 | 0.419 | -0.333 | 0.417 | -0.261 | -0.012 | 0.139 | 0.139 | 0 | 0.42 | 0.19 | 0.3 | 0.07 | 0.41 | 0.26 |
| 3 | 45617100 | rs9853624 | 0.561 | -0.168 | 0.29 | -0.163 | -0.024 | 0.051 | 0.051 | 0 | 0.43 | 0.24 | 0.54 | 0.68 | 0.51 | 0.65 |
| 3 | 45623809 | rs6799789 | -0.617 | -0.274 | 0.352 | 0.134 | -0.025 | 0.081 | 0.081 | 0 | 0.07 | 0.17 | 0.03 | 0.19 | 0.05 | 0.05 |
| 3 | 45624142 | rs267245 | 0.348 | -0.268 | 0.152 | -0.059 | -0.001 | -0.007 | -0.007 | 0 | 0.46 | 0.45 | 0.63 | 0.74 | 0.5 | 0.67 |

| | | | | | | | | | | | | | | | | |
|---|---|---|---|---|---|---|---|---|---|---|---|---|---|---|---|---|
| 3 | 45633834 | rs267218 | 0.476 | -0.122 | -0.464 | -0.015 | -0.037 | 0.043 | 0.043 | 0 | 0.42 | 0.17 | 0.55 | 0.65 | 0.5 | 0.64 |
| 3 | 45635274 | rs2294160 | 0.003 | -0.317 | -0.563 | 0.094 | -0.001 | 0.009 | 0.009 | 0 | 0.06 | 0.06 | 0.02 | 0.19 | 0.05 | 0.05 |
| 3 | 45638255 | rs2005227 | 1.035 | 0.486 | -0.825 | 0.59 | -0.003 | -0.08 | -0.08 | 1.071 | 0.31 | 0.08 | 0.34 | 0.4 | 0.26 | 0.48 |
| 3 | 45644183 | rs267232 | 0.258 | 1.114 | -0.563 | 0.536 | -0.005 | -0.006 | -0.006 | 0 | 0.1 | 0.08 | 0.14 | 0.22 | 0.22 | 0.11 |
| 3 | 45649417 | rs2578693 | -0.926 | 0.74 | -0.045 | 0.243 | -0.009 | -0.003 | -0.003 | 0 | 0.6 | 0.54 | 0.55 | 0.44 | 0.68 | 0.39 |
| 3 | 45649864 | rs2578692 | -0.869 | 0.694 | 0.104 | 0.213 | 0.026 | 0.042 | 0.042 | 0.901 | 0.19 | 0.31 | 0.2 | 0.38 | 0.29 | 0.16 |
| 3 | 45652275 | rs267210 | -0.401 | 0.684 | -0.174 | 0.224 | -0.015 | -0.022 | -0.022 | 0.464 | 0.17 | 0.15 | 0.16 | 0.38 | 0.27 | 0.16 |
| 3 | 45667387 | rs3774675 | -0.53 | 0.634 | -0.087 | 0.36 | 0.01 | 0.052 | 0.052 | 0.604 | 0.18 | 0.24 | 0.22 | 0.48 | 0.27 | 0.19 |
| 3 | 45667989 | rs4683131 | 1.003 | 0.487 | -0.194 | 0.181 | 0.021 | -0.039 | -0.039 | 0 | 0.4 | 0.17 | 0.36 | 0.46 | 0.34 | 0.59 |
| 3 | 45672871 | rs2012762 | -0.753 | 0.446 | -0.559 | -0.217 | -0.03 | 0.176 | 0.176 | 0.054 | 0.63 | 0.9 | 0.66 | 0.94 | 0.63 | 0.79 |
| 3 | 45674387 | rs3796362 | -0.199 | 0.523 | -0.626 | -0.123 | 0.01 | -0.003 | -0.003 | 0 | 0.13 | 0.09 | 0.07 | 0.17 | 0.1 | 0.08 |
| 3 | 45677398 | rs7648681 | -1.036 | 0.932 | -0.596 | -0.119 | -0.032 | -0.025 | -0.025 | 0 | 0.25 | 0.2 | 0.24 | 0.4 | 0.36 | 0.2 |
| 3 | 45706167 | rs12493946 | 1.256 | 0.648 | -0.32 | -0.454 | 0.017 | -0.011 | -0.011 | 0 | 0.36 | 0.27 | 0.38 | 0.52 | 0.25 | 0.57 |
| 3 | 45710349 | rs17078175 | 0.247 | 0.992 | -0.825 | -0.737 | -0.008 | -0.004 | -0.004 | 0 | 0.13 | 0.18 | 0.09 | 0.16 | 0.1 | 0.08 |
| 3 | 45711735 | rs1318992 | -1.139 | 1.098 | -1.071 | -0.767 | -0.017 | -0.016 | -0.016 | 0.347 | 0.9 | 0.95 | 0.86 | 0.78 | 0.76 | 0.89 |
| 3 | 45715767 | rs2742409 | 0.397 | 0.801 | -1.168 | -0.939 | -0.069 | 0.228 | 0.228 | 1.289 | 0.35 | 0.05 | 0.32 | 0.03 | 0.4 | 0.2 |
| 3 | 45719209 | rs2247771 | 0.406 | 0.807 | -1.145 | -0.937 | -0.059 | 0.213 | 0.213 | 1.147 | 0.34 | 0.05 | 0.32 | 0.02 | 0.38 | 0.2 |
| 3 | 45719561 | rs3796361 | 0.468 | 0.671 | -0.957 | -0.738 | 0.005 | -0.007 | -0.007 | 0 | 0.12 | 0.1 | 0.08 | 0.18 | 0.1 | 0.08 |
| 3 | 45723434 | rs17262464 | 1.419 | 0.515 | -1.192 | -1.079 | 0 | 0.118 | 0.118 | 0.097 | 0.21 | 0.01 | 0.12 | 0.02 | 0.16 | 0.1 |
| 3 | 45724610 | rs2742412 | -0.263 | 1.137 | -0.42 | -0.573 | 0.054 | 0.067 | 0.067 | 0.063 | 0.89 | 0.76 | 0.87 | 0.79 | 0.83 | 0.89 |
| 3 | 45725031 | rs2742413 | -0.141 | 1.193 | -0.549 | -0.507 | 0.003 | 0.005 | 0.005 | 0.299 | 0.94 | 0.9 | 0.95 | 0.91 | 0.92 | 0.98 |
| 3 | 45730117 | rs2742416 | -0.317 | 1.236 | -0.561 | -0.488 | 0.004 | 0.008 | 0.008 | 0.068 | 0.95 | 0.91 | 0.95 | 0.91 | 0.93 | 0.98 |
| 3 | 45732409 | rs2673051 | 0.042 | 1.203 | -0.342 | -0.692 | 0.002 | 0.049 | 0.049 | 0.011 | 0.49 | 0.42 | 0.45 | 0.26 | 0.56 | 0.32 |
| 3 | 45739807 | rs2673050 | 0.357 | 1.546 | -0.438 | -0.64 | -0.016 | 0.116 | 0.116 | 0 | 0.43 | 0.16 | 0.39 | 0.29 | 0.48 | 0.29 |
| 3 | 45754685 | rs2271619 | 0.372 | 0.649 | -0.18 | -0.601 | 0.033 | 0.025 | 0.025 | 0.253 | 0.1 | 0.21 | 0.11 | 0.12 | 0.12 | 0.09 |
| 3 | 45759654 | rs2673028 | 0.431 | 0.66 | -0.259 | -0.608 | -0.038 | 0.199 | 0.199 | 0.33 | 0.47 | 0.18 | 0.43 | 0.21 | 0.56 | 0.28 |
| 3 | 45762725 | rs13086080 | 0.602 | 0.692 | -0.357 | -0.623 | -0.046 | -0.043 | -0.043 | 0 | 0.1 | 0.01 | 0.1 | 0.2 | 0.18 | 0.08 |
| 3 | 45763635 | rs2673062 | -0.525 | 0.564 | -0.194 | -0.539 | 0.006 | 0.009 | 0.009 | 0 | 0.6 | 0.65 | 0.57 | 0.38 | 0.68 | 0.41 |
| 3 | 45792209 | rs12490575 | 1.454 | 0.489 | -0.934 | -0.012 | -0.038 | -0.072 | -0.072 | 0.872 | 0.12 | 0.01 | 0.16 | 0.25 | 0.14 | 0.08 |
| 3 | 45795498 | rs12495593 | 0.049 | 0.302 | -0.609 | -1.039 | 0.013 | -0.004 | -0.004 | 0 | 0.44 | 0.33 | 0.44 | 0.51 | 0.3 | 0.6 |
| 3 | 45796951 | rs2251109 | 0.764 | 0.524 | -1.035 | -1.466 | -0.122 | 0.133 | 0.133 | 0.755 | 0.37 | 0.09 | 0.29 | 0.2 | 0.49 | 0.23 |
| 3 | 45798226 | rs17213127 | 2.004 | 0.83 | -1.015 | -0.856 | -0.017 | 0.033 | 0.033 | 0 | 0.06 | 0 | 0.02 | 0 | 0.05 | 0.01 |
| 3 | 45803585 | rs2191028 | 0.584 | 0.878 | -0.632 | -0.846 | -0.002 | 0.022 | 0.022 | 0 | 0.35 | 0.29 | 0.27 | 0.19 | 0.37 | 0.19 |
| 3 | 45804069 | rs2286489 | 1.066 | 0.948 | -0.812 | -1.022 | 0.018 | 0.02 | 0.02 | 0 | 0.71 | 0.36 | 0.64 | 0.56 | 0.64 | 0.65 |
| 3 | 45804391 | rs2191027 | 1.569 | 0.919 | -0.88 | -1.154 | -0.075 | 0.159 | 0.159 | 0.375 | 0.27 | 0.02 | 0.19 | 0.02 | 0.3 | 0.15 |
| 3 | 45804734 | rs6770261 | -0.489 | 0.833 | -0.477 | -1.074 | 0.059 | -0.023 | -0.023 | 0 | 0.19 | 0.47 | 0.27 | 0.16 | 0.24 | 0.27 |
| 3 | 45808701 | rs10461016 | 1.099 | 0.881 | -0.703 | -1.06 | -0.029 | -0.032 | -0.032 | 0 | 0.07 | 0.02 | 0.08 | 0.17 | 0.1 | 0.15 |
| 3 | 45809273 | rs4299518 | 0.212 | 0.744 | -1.283 | -1.288 | -0.025 | 0.155 | 0.155 | 0 | 0.44 | 0.21 | 0.39 | 0.03 | 0.48 | 0.33 |
| 3 | 45809329 | rs4327428 | 0.855 | 0.892 | -1.322 | -0.977 | -0.012 | 0.016 | 0.016 | 0 | 0.07 | 0.07 | 0.14 | 0.02 | 0.11 | 0.13 |
| 3 | 45809843 | rs2108917 | 0.963 | 0.879 | -1.388 | -0.968 | 0 | -0.001 | -0.001 | 0 | 0.75 | 0.25 | 0.68 | 0.75 | 0.74 | 0.7 |
| 3 | 45810094 | rs720625 | -0.462 | 1.408 | -1.93 | -0.625 | 0.002 | 0.004 | 0.004 | 0 | 0.53 | 0.57 | 0.53 | 0.39 | 0.61 | 0.47 |
| 3 | 45813025 | rs758387 | -0.525 | 2.096 | -3.07 | -0.112 | 0.051 | 0.062 | 0.062 | 0.008 | 0.54 | 0.17 | 0.5 | 0.39 | 0.65 | 0.53 |
| 3 | 45814748 | rs17279465 | -0.056 | 1.643 | -2.958 | -0.401 | -0.087 | 0.266 | 0.266 | 0 | 0.41 | 0.09 | 0.36 | 0.03 | 0.44 | 0.34 |
| 3 | 45819837 | rs2531748 | -0.227 | 0.858 | -1.82 | 0.562 | 0.008 | -0.002 | -0.002 | 0 | 0.35 | 0.19 | 0.37 | 0.62 | 0.35 | 0.36 |
| 3 | 45821244 | rs13067466 | -0.379 | 1.308 | -2.089 | 0.386 | -0.069 | -0.019 | -0.019 | 3.099 | 0.17 | 0.02 | 0.18 | 0.23 | 0.19 | 0.14 |
| 3 | 45822437 | rs17078339 | -0.44 | 0.243 | -1.784 | 0.687 | 0.015 | -0.017 | -0.017 | 0 | 0.35 | 0.29 | 0.37 | 0.49 | 0.28 | 0.34 |
| 3 | 45828586 | rs2531747 | -0.983 | 0.958 | -1.927 | 0.679 | 0.012 | 0.033 | 0.033 | 0.045 | 0.53 | 0.77 | 0.75 | 0.68 | 0.57 | 0.71 |
| 3 | 45828841 | rs9848415 | 0.27 | 0.613 | -1.893 | 1.137 | 0.005 | -0.005 | -0.005 | 0.737 | 0.25 | 0.18 | 0.35 | 0.31 | 0.25 | 0.25 |
| 3 | 45829995 | rs2159272 | -0.816 | 0.85 | -2.223 | 0.585 | 0.005 | 0.013 | 0.013 | 0 | 0.44 | 0.74 | 0.67 | 0.65 | 0.45 | 0.61 |
| 3 | 45830886 | rs7644870 | 0.169 | 0.916 | -2.327 | 0.813 | 0.002 | -0.032 | -0.032 | 0 | 0.26 | 0.2 | 0.37 | 0.5 | 0.26 | 0.27 |
| 3 | 45831442 | rs12488144 | -0.111 | 1.194 | -2.358 | 1.083 | -0.054 | -0.023 | -0.023 | 0 | 0.14 | 0.02 | 0.12 | 0.12 | 0.17 | 0.11 |
| 3 | 45834811 | rs13064991 | 0.199 | 0.536 | -2.121 | 1.305 | 0.004 | -0.044 | -0.044 | 0 | 0.17 | 0.06 | 0.27 | 0.42 | 0.18 | 0.18 |
| 3 | 45837886 | rs2271615 | 0.121 | -0.035 | -2.67 | 0.283 | 0.001 | 0.077 | 0.077 | 0 | 0.22 | 0.01 | 0.11 | 0.08 | 0.16 | 0.11 |
| 3 | 45841068 | rs2531742 | 0.024 | 0.159 | -2.641 | 0.131 | -0.029 | 0 | 0 | 0 | 0.33 | 0.18 | 0.42 | 0.38 | 0.39 | 0.31 |
| 3 | 45845748 | rs17078346 | 0.966 | -0.658 | -2.245 | -0.816 | 0.007 | -0.017 | -0.017 | 0.739 | 0.2 | 0.2 | 0.07 | 0.02 | 0.1 | 0.32 |
| 3 | 45850520 | rs2531757 | 1.013 | -0.382 | -2.557 | -0.484 | -0.059 | -0.08 | -0.08 | 1.833 | 0.24 | 0.06 | 0.46 | 0.47 | 0.33 | 0.29 |
| 3 | 45853072 | rs11130076 | 0.546 | -0.907 | -2.561 | -0.926 | -0.085 | -0.04 | -0.04 | 0 | 0.41 | 0.1 | 0.57 | 0.62 | 0.49 | 0.41 |
| 3 | 45855172 | rs4683143 | -0.322 | -1.14 | -1.833 | -1.451 | -0.002 | -0.068 | -0.068 | 0 | 0.64 | 0.56 | 0.76 | 0.96 | 0.64 | 0.59 |
| 3 | 45856364 | rs2531756 | 0.657 | -1.23 | -1.877 | -1.543 | 0.011 | 0.028 | 0.028 | 0 | 0.1 | 0.05 | 0.05 | 0.04 | 0.09 | 0.09 |
| 3 | 45864481 | rs4683145 | -1.034 | -1.323 | -1.683 | -1.598 | 0.016 | -0.034 | -0.034 | 1.112 | 0.9 | 0.84 | 0.85 | 0.99 | 0.79 | 0.96 |
| 3 | 45865006 | rs1058961 | 0.24 | -1.357 | -1.66 | -1.585 | -0.003 | -0.009 | -0.009 | 0 | 0.67 | 0.29 | 0.69 | 0.69 | 0.67 | 0.81 |
| 3 | 45869972 | rs1129183 | 1.189 | -1.276 | -1.747 | -1.565 | 0.008 | 0.013 | 0.013 | 0 | 0.08 | 0.04 | 0.04 | 0.04 | 0.07 | 0.08 |
| 3 | 45871203 | rs2064061 | 1.359 | -1.174 | -1.337 | -1.471 | -0.033 | -0.004 | -0.004 | 0.62 | 0.41 | 0.27 | 0.59 | 0.65 | 0.49 | 0.42 |
| 3 | 45877712 | rs11130077 | 0.345 | -0.606 | -1.64 | -1.199 | -0.004 | -0.07 | -0.07 | 0 | 0.63 | 0.53 | 0.74 | 0.95 | 0.63 | 0.58 |
| 3 | 45900621 | rs7614952 | 0.247 | -0.438 | -1.456 | -1.014 | 0.057 | 0.078 | 0.078 | 0 | 0.83 | 0.68 | 0.77 | 0.74 | 0.72 | 0.93 |
| 3 | 45906356 | rs9856760 | -0.583 | -0.355 | -1.409 | -1.003 | 0.069 | -0.053 | -0.053 | 1.832 | 0.9 | 0.72 | 0.86 | 0.99 | 0.8 | 0.97 |
| 3 | 45908148 | rs7625839 | 0.413 | -0.537 | -1.615 | -0.984 | 0 | -0.001 | -0.001 | 0 | 0.57 | 0.53 | 0.67 | 0.7 | 0.56 | 0.55 |
| 3 | 45910870 | rs2191031 | 0.359 | -0.933 | -1.897 | -0.851 | 0.037 | -0.035 | -0.035 | 0 | 0.29 | 0.08 | 0.12 | 0.24 | 0.21 | 0.41 |
| 3 | 45923085 | rs1860264 | -1.051 | -0.813 | -2.116 | -1.082 | -0.027 | -0.065 | -0.065 | 0 | 0.58 | 0.91 | 0.34 | 0.37 | 0.38 | 0.6 |

| | | | | | | | | | | | | | | | | |
|---|---|---|---|---|---|---|---|---|---|---|---|---|---|---|---|---|
| 3 | 45924211 | rs9868455 | -1.011 | -1.132 | -1.71 | -1.241 | -0.018 | 0.01 | 0.01 | 0.397 | 0.35 | 0.47 | 0.23 | 0.36 | 0.24 | 0.21 |
| 3 | 45924791 | rs6441929 | -0.552 | -1.262 | -1.525 | -1.187 | -0.011 | 0.01 | 0.01 | 0 | 0.24 | 0.31 | 0.17 | 0.27 | 0.16 | 0.13 |
| 3 | 45927759 | rs7653372 | 1.013 | -0.996 | -1.245 | -1.013 | -0.051 | -0.171 | -0.171 | 0.016 | 0.24 | 0.44 | 0.11 | 0 | 0.14 | 0.39 |
| 3 | 45938939 | rs2236938 | 0.014 | -0.633 | -1.253 | -0.532 | -0.003 | 0.157 | 0.157 | 0 | 0.13 | 0.53 | 0.14 | 0.23 | 0.1 | 0.07 |
| 3 | 45948264 | rs4683147 | 0.942 | 0.31 | -2.327 | -0.687 | -0.072 | -0.106 | -0.106 | 0.437 | 0.33 | 0.08 | 0.54 | 0.63 | 0.39 | 0.32 |
| 3 | 45951678 | rs12493471 | -0.741 | 0.284 | -2.664 | -1.355 | -0.099 | 0.207 | 0.207 | 1.012 | 0.7 | 0.96 | 0.83 | 1 | 0.67 | 0.57 |
| 3 | 45956060 | rs4683148 | 1.208 | 0.088 | -2.796 | -1.161 | -0.076 | -0.085 | -0.085 | 0 | 0.33 | 0.11 | 0.54 | 0.65 | 0.41 | 0.36 |
| 3 | 45957689 | rs1488373 | -0.978 | 0.304 | -2.791 | -1.247 | -0.022 | 0.02 | 0.02 | 0 | 0.92 | 0.96 | 0.89 | 1 | 0.79 | 0.93 |
| 3 | 45960079 | rs1047444 | -1.014 | 0.33 | -2.806 | -1.247 | -0.035 | 0.033 | 0.033 | 0 | 0.92 | 0.97 | 0.9 | 1 | 0.79 | 0.93 |
| 3 | 45960851 | rs3796373 | -0.262 | -0.223 | -2.759 | -1.497 | 0.038 | -0.026 | -0.026 | 0 | 0.36 | 0.74 | 0.28 | 0.36 | 0.27 | 0.2 |
| 3 | 45961842 | rs1488374 | -1.096 | -0.165 | -2.724 | -1.507 | -0.036 | 0.034 | 0.034 | 0 | 0.92 | 0.96 | 0.89 | 1 | 0.8 | 0.93 |
| 3 | 45962752 | rs7652331 | -0.967 | -0.201 | -2.744 | -1.527 | -0.092 | 0.182 | 0.182 | 0.782 | 0.72 | 0.96 | 0.84 | 1 | 0.68 | 0.56 |
| 3 | 45962802 | rs6441934 | -1.611 | -0.012 | -2.681 | -1.439 | -0.04 | 0.035 | 0.035 | 0 | 0.93 | 0.99 | 0.91 | 1 | 0.82 | 0.97 |
| 3 | 45975983 | rs1072755 | 1.09 | -0.281 | -2.569 | -1.471 | -0.017 | -0.021 | -0.021 | 0 | 0.36 | 0.24 | 0.55 | 0.65 | 0.41 | 0.36 |
| 3 | 45980673 | rs17078449 | 0.627 | -0.457 | -2.257 | -1.585 | 0.001 | -0.028 | -0.028 | 1.644 | 0.21 | 0.16 | 0.17 | 0.3 | 0.21 | 0.15 |
| 3 | 45986623 | rs936939 | 0.627 | -0.473 | -1.972 | -1.581 | 0.001 | -0.027 | -0.027 | 0 | 0.21 | 0.16 | 0.17 | 0.3 | 0.21 | 0.15 |
| 3 | 45989044 | rs2234358 | 0.694 | -0.423 | -1.966 | -1.568 | -0.032 | -0.002 | -0.002 | 0 | 0.6 | 0.31 | 0.63 | 0.65 | 0.53 | 0.73 |
| 3 | 45993606 | rs4682799 | 0.865 | -0.409 | -2.053 | -1.577 | -0.038 | 0.001 | 0.001 | 0 | 0.58 | 0.26 | 0.61 | 0.65 | 0.53 | 0.73 |
| 3 | 46005478 | rs1545985 | -0.669 | -0.291 | -2.187 | -1.65 | -0.036 | 0.094 | 0.094 | 0.022 | 0.67 | 0.84 | 0.8 | 1 | 0.65 | 0.58 |
| 3 | 46008790 | rs3796375 | 1.477 | -0.288 | -2.236 | -1.404 | -0.103 | -0.089 | -0.089 | 1.976 | 0.32 | 0.09 | 0.56 | 0.66 | 0.43 | 0.37 |
| 3 | 46016944 | rs751553 | 1.477 | -0.279 | -2.252 | -1.414 | -0.111 | -0.096 | -0.096 | 4.639 | 0.32 | 0.06 | 0.56 | 0.66 | 0.43 | 0.37 |
| 3 | 46021218 | rs4682801 | -0.29 | -0.246 | -1.859 | -1.534 | 0.103 | -0.076 | -0.076 | 0 | 0.88 | 0.38 | 0.87 | 1 | 0.8 | 0.92 |
| 3 | 46025013 | rs4683159 | -0.5 | -0.482 | -1.972 | -1.652 | -0.005 | 0.018 | 0.018 | 0.194 | 0.65 | 0.28 | 0.79 | 1 | 0.67 | 0.56 |
| 3 | 46027410 | rs737452 | 1.632 | -0.437 | -2.026 | -1.503 | -0.115 | -0.112 | -0.112 | 0.593 | 0.32 | 0.05 | 0.56 | 0.65 | 0.43 | 0.37 |
| 3 | 46032847 | rs1873001 | 0.373 | -0.531 | -1.915 | -1.596 | 0.027 | 0.156 | 0.156 | 0 | 0.56 | 0.23 | 0.76 | 0.96 | 0.66 | 0.52 |
| 3 | 46034086 | rs3947589 | -0.043 | -0.476 | -1.665 | -1.491 | -0.064 | -0.053 | -0.053 | 2.122 | 0.12 | 0.55 | 0.07 | 0.04 | 0.02 | 0.04 |
| 3 | 46034791 | rs1392288 | 0.921 | -0.421 | -1.792 | -1.522 | 0 | -0.033 | -0.033 | 0.181 | 0.76 | 0.26 | 0.82 | 0.96 | 0.78 | 0.88 |
| 3 | 46047032 | rs2171529 | 0.476 | -0.73 | -2.042 | -1.676 | 0.039 | 0.17 | 0.17 | 0.021 | 0.53 | 0.27 | 0.75 | 0.95 | 0.63 | 0.51 |
| 3 | 46059139 | rs6764042 | -0.818 | -0.92 | -2.111 | -1.615 | 0.037 | -0.013 | -0.013 | 0.018 | 0.59 | 0.77 | 0.31 | 0.35 | 0.34 | 0.55 |
| 3 | 46061840 | rs13074382 | 1.744 | -1.001 | -2.064 | -1.537 | -0.053 | -0.072 | -0.072 | 0.053 | 0.23 | 0.44 | 0.09 | 0.05 | 0.12 | 0.35 |
| 3 | 46063329 | rs2230322 | 1.744 | -1.001 | -2.064 | -1.537 | -0.053 | -0.072 | -0.072 | 0 | 0.23 | 0.44 | 0.09 | 0.05 | 0.12 | 0.35 |
| 3 | 46072006 | rs13060901 | -0.809 | -0.919 | -2.1 | -1.503 | 0.035 | -0.012 | -0.012 | 5.731 | 0.59 | 0.76 | 0.31 | 0.35 | 0.34 | 0.55 |
| 3 | 46081179 | rs7638309 | -0.629 | -0.735 | -2.048 | -1.501 | -0.036 | 0.032 | 0.032 | 0 | 0.89 | 0.94 | 0.89 | 0.99 | 0.77 | 0.91 |
| 3 | 46090557 | rs7642229 | 0.749 | -0.931 | -2.067 | -1.554 | -0.056 | -0.063 | -0.063 | 0.176 | 0.32 | 0.47 | 0.13 | 0.09 | 0.14 | 0.39 |
| 3 | 46096385 | rs2036293 | -0.522 | -0.619 | -2.169 | -1.415 | -0.001 | 0 | 0 | 0 | 0.89 | 0.9 | 0.89 | 0.99 | 0.76 | 0.91 |
| 3 | 46100496 | rs2173640 | -0.835 | -0.748 | -2.227 | -1.474 | -0.03 | 0.07 | 0.07 | 1.927 | 0.68 | 0.84 | 0.82 | 0.94 | 0.64 | 0.56 |
| 3 | 46100691 | rs2036295 | -0.212 | -0.752 | -2.182 | -1.475 | 0.022 | 0.005 | 0.005 | 0.807 | 0.35 | 0.66 | 0.3 | 0.29 | 0.2 | 0.19 |
| 3 | 46101257 | rs4490403 | -0.918 | -0.737 | -2.203 | -1.471 | -0.034 | 0.077 | 0.077 | 1.959 | 0.68 | 0.85 | 0.82 | 0.94 | 0.64 | 0.56 |
| 3 | 46101371 | rs4490404 | -0.655 | -0.723 | -2.195 | -1.469 | -0.041 | 0.034 | 0.034 | 0 | 0.89 | 0.95 | 0.89 | 0.99 | 0.76 | 0.92 |
| 3 | 46101684 | rs1604068 | -0.728 | -0.648 | -2.246 | -1.514 | 0.085 | 0.033 | 0.033 | 3.236 | 0.39 | 0.8 | 0.32 | 0.29 | 0.24 | 0.2 |
| 3 | 46102620 | rs1500005 | 1.615 | -0.39 | -2.143 | -1.383 | -0.126 | -0.141 | -0.141 | 0 | 0.29 | 0.04 | 0.5 | 0.65 | 0.4 | 0.36 |
| 3 | 46103974 | rs9311384 | 0.469 | -0.198 | -1.911 | -1.349 | 0.044 | -0.053 | -0.053 | 0 | 0.84 | 0.71 | 0.86 | 0.99 | 0.73 | 0.91 |
| 3 | 46118749 | rs1388608 | -1.055 | -0.134 | -1.986 | -1.297 | -0.042 | 0.028 | 0.028 | 0 | 0.92 | 0.99 | 0.91 | 0.99 | 0.79 | 0.93 |
| 3 | 46119791 | rs13434336 | -0.418 | -0.554 | -2.169 | -1.346 | 0.069 | -0.002 | -0.002 | 0 | 0.56 | 0.84 | 0.38 | 0.32 | 0.33 | 0.55 |
| 3 | 46123333 | rs9838450 | 1.133 | -0.931 | -2.062 | -1.53 | -0.047 | -0.08 | -0.08 | 0 | 0.28 | 0.48 | 0.12 | 0.05 | 0.16 | 0.36 |
| 3 | 46125575 | rs9990060 | -0.632 | -0.845 | -2.023 | -1.479 | -0.015 | -0.006 | -0.006 | 0 | 0.37 | 0.48 | 0.31 | 0.28 | 0.24 | 0.2 |
| 3 | 46126467 | rs4683177 | -1.017 | -0.72 | -2.057 | -1.462 | -0.02 | 0.019 | 0.019 | 0 | 0.91 | 0.96 | 0.9 | 0.99 | 0.79 | 0.93 |
| 3 | 46127987 | rs902760 | -0.987 | -0.722 | -2.058 | -1.463 | -0.021 | 0.019 | 0.019 | 0 | 0.91 | 0.96 | 0.9 | 0.99 | 0.79 | 0.93 |
| 3 | 46128080 | rs751625 | -0.145 | -0.939 | -2.01 | -1.456 | 0.025 | 0 | 0 | 0.464 | 0.32 | 0.28 | 0.26 | 0.28 | 0.2 | 0.19 |
| 3 | 46130509 | rs10510747 | 0.12 | -0.937 | -2.06 | -1.456 | 0.045 | -0.025 | -0.025 | 0 | 0.29 | 0.14 | 0.25 | 0.28 | 0.2 | 0.19 |
| 3 | 46139499 | rs13086717 | 1.904 | 0.013 | -2.396 | -0.746 | 0 | -0.049 | -0.049 | 0 | 0.23 | 0.02 | 0.21 | 0.22 | 0.18 | 0.27 |
| 3 | 46140002 | rs9872121 | -1.022 | -0.634 | -2.332 | -1.112 | 0 | -0.01 | -0.01 | 0 | 0.21 | 0.77 | 0.13 | 0.04 | 0.06 | 0.04 |
| 3 | 46141844 | rs1491951 | 1.998 | -0.502 | -2.253 | -1.068 | -0.052 | -0.074 | -0.074 | 0.062 | 0.23 | 0.42 | 0.1 | 0.05 | 0.13 | 0.35 |
| 3 | 46143059 | rs6772051 | -0.004 | -0.163 | -2.423 | -1.042 | 0.005 | 0.024 | 0.024 | 0.198 | 0.48 | 0.68 | 0.62 | 0.7 | 0.48 | 0.41 |
| 3 | 46145295 | rs2888512 | -0.524 | -0.381 | -2.29 | -0.887 | 0.044 | 0.013 | 0.013 | 0.356 | 0.37 | 0.73 | 0.32 | 0.28 | 0.24 | 0.2 |
| 3 | 46150400 | rs1546077 | -0.658 | -0.028 | -2.331 | -0.757 | -0.029 | -0.079 | -0.079 | 0 | 0.61 | 0.94 | 0.41 | 0.33 | 0.37 | 0.56 |
| 3 | 46171700 | rs13320534 | 0.284 | -0.123 | -2.054 | -0.673 | -0.027 | 0.022 | 0.022 | 0.256 | 0.21 | 0.35 | 0.22 | 0.22 | 0.14 | 0.16 |
| 3 | 46173446 | rs6774742 | -0.561 | 0.281 | -1.788 | -1.151 | 0.007 | -0.065 | -0.065 | 2.778 | 0.67 | 0.48 | 0.8 | 0.95 | 0.67 | 0.58 |
| 3 | 46174694 | rs6789719 | -0.506 | 0.181 | -1.829 | -1.191 | 0.005 | -0.106 | -0.106 | 0 | 0.66 | 0.57 | 0.81 | 0.95 | 0.67 | 0.58 |
| 3 | 46180416 | rs10510749 | 1.427 | -0.026 | -1.735 | -1.244 | 0.027 | 0.033 | 0.033 | 0.045 | 0.2 | 0.11 | 0.06 | 0.04 | 0.09 | 0.29 |
| 3 | 46205686 | rs7616215 | 0.595 | 0.101 | -1.878 | -1.377 | 0 | -0.018 | -0.018 | 0 | 0.54 | 0.52 | 0.74 | 0.87 | 0.64 | 0.55 |
| 3 | 46214685 | rs2172246 | -0.235 | 0.143 | -1.896 | -1.35 | -0.05 | 0.043 | 0.043 | 0.164 | 0.82 | 0.93 | 0.85 | 0.92 | 0.74 | 0.84 |
| 3 | 46226646 | rs13061548 | 0.486 | 0.154 | -1.708 | -1.155 | -0.028 | -0.037 | -0.037 | 0 | 0.27 | 0.32 | 0.11 | 0.05 | 0.1 | 0.29 |
| 3 | 46242616 | rs3136673 | 1.413 | 0.097 | -1.79 | -1.115 | 0.041 | 0.052 | 0.052 | 0 | 0.21 | 0.08 | 0.09 | 0.04 | 0.1 | 0.29 |
| 3 | 46243175 | rs3136671 | 0.79 | 0.064 | -2.09 | -0.915 | -0.035 | -0.073 | -0.073 | 0.061 | 0.15 | 0.02 | 0.12 | 0.22 | 0.13 | 0.21 |
| 3 | 46263717 | rs11926063 | 0.612 | -0.198 | -1.642 | -1.127 | -0.057 | -0.062 | -0.062 | 0 | 0.25 | 0.38 | 0.12 | 0.04 | 0.09 | 0.28 |
| 3 | 46274109 | rs1979672 | 0.64 | 0.41 | -2.179 | -0.942 | 0.004 | -0.003 | -0.003 | 0 | 0.24 | 0.11 | 0.26 | 0.3 | 0.21 | 0.16 |
| 3 | 46274215 | rs1979671 | -0.5 | 0.559 | -2.32 | -0.929 | -0.079 | 0.023 | 0.023 | 0 | 0.82 | 0.95 | 0.8 | 0.89 | 0.7 | 0.79 |
| 3 | 46281744 | rs13096142 | 0.588 | -0.041 | -2.197 | -0.921 | 0.016 | 0.036 | 0.036 | 0 | 0.37 | 0.15 | 0.3 | 0.32 | 0.3 | 0.35 |

| | | | | | | | | | | | | | | | | |
|---|---|---|---|---|---|---|---|---|---|---|---|---|---|---|---|---|
| 3 | 46286663 | rs13073976 | 2.109 | 0.222 | -2.024 | -0.808 | -0.002 | 0.031 | 0.031 | 0.089 | 0.14 | 0 | 0.04 | 0.03 | 0.08 | 0.21 |
| 3 | 46295777 | rs6441948 | -0.21 | 0.895 | -1.642 | -0.85 | 0.08 | -0.043 | -0.043 | 0 | 0.8 | 0.66 | 0.66 | 0.9 | 0.57 | 0.73 |
| 3 | 46299765 | rs7652290 | 0.164 | 0.471 | -2.304 | -0.949 | 0.012 | 0.019 | 0.019 | 0 | 0.59 | 0.4 | 0.4 | 0.62 | 0.35 | 0.58 |
| 3 | 46304606 | rs1491962 | -0.309 | 0.566 | -2.248 | -0.924 | 0.07 | -0.042 | -0.042 | 4.253 | 0.79 | 0.41 | 0.65 | 0.9 | 0.57 | 0.73 |
| 3 | 46306700 | rs4987053 | 1.764 | 0.22 | -2.445 | -1.075 | 0.021 | 0.027 | 0.027 | 0 | 0.18 | 0.16 | 0.07 | 0.03 | 0.08 | 0.21 |
| 3 | 46310921 | rs4413345 | 0.419 | -0.106 | -2.689 | -1.32 | 0.102 | 0.085 | 0.085 | 0 | 0.5 | 0.37 | 0.35 | 0.63 | 0.28 | 0.56 |
| 3 | 46312539 | rs1027241 | -0.247 | 0.303 | -2.346 | -1.099 | 0.052 | -0.041 | -0.041 | 0.59 | 0.77 | 0.4 | 0.64 | 0.9 | 0.56 | 0.73 |
| 3 | 46313334 | rs9810242 | 1.558 | 0.146 | -2.263 | -0.932 | 0.017 | 0.021 | 0.021 | 0 | 0.18 | 0.15 | 0.07 | 0.03 | 0.08 | 0.21 |
| 3 | 46323859 | rs9839623 | -1.727 | 0.215 | -2.432 | -0.807 | -0.046 | -0.048 | -0.048 | 0 | 0.8 | 0.98 | 0.76 | 0.72 | 0.78 | 0.85 |
| 3 | 46336781 | rs7645716 | 0.338 | -0.012 | -2.42 | -0.843 | 0.064 | -0.001 | -0.001 | 0.321 | 0.39 | 0.23 | 0.33 | 0.6 | 0.27 | 0.38 |
| 3 | 46339812 | rs9990343 | 0.257 | 0.234 | -2.069 | -0.853 | 0.014 | -0.035 | -0.035 | 0 | 0.36 | 0.52 | 0.38 | 0.12 | 0.51 | 0.47 |
| 3 | 46345413 | rs4683211 | -1.277 | 0.087 | -2.238 | -0.668 | -0.004 | -0.005 | -0.005 | 0 | 0.76 | 0.87 | 0.72 | 0.72 | 0.77 | 0.85 |
| 3 | 46346025 | rs10510751 | 0.566 | -0.46 | -2.524 | -0.096 | -0.008 | 0.009 | 0.009 | 0 | 0.15 | 0.19 | 0.22 | 0.2 | 0.09 | 0.08 |
| 3 | 46346874 | rs11712150 | 1.044 | -0.101 | -3.169 | -0.263 | -0.069 | 0.035 | 0.035 | 0 | 0.17 | 0.04 | 0.19 | 0.1 | 0.3 | 0.25 |
| 3 | 46352355 | rs4513489 | 0.92 | -0.964 | -3.028 | -0.152 | 0.03 | -0.076 | -0.076 | 0.104 | 0.25 | 0.07 | 0.12 | 0.39 | 0.18 | 0.31 |
| 3 | 46352384 | rs6441961 | -0.501 | -0.653 | -2.695 | -0.13 | -0.006 | 0.016 | 0.016 | 0 | 0.63 | 0.89 | 0.68 | 0.69 | 0.7 | 0.65 |
| 3 | 46397039 | rs3092962 | 0.932 | -0.46 | -2.636 | 0.029 | -0.016 | -0.028 | -0.028 | 0 | 0.24 | 0.08 | 0.28 | 0.29 | 0.23 | 0.16 |
| 3 | 46413334 | rs2856762 | 1.684 | -0.683 | -2.145 | 0.212 | -0.015 | 0.028 | 0.028 | 0.394 | 0.13 | 0 | 0.05 | 0.03 | 0.09 | 0.2 |
| 3 | 46483717 | rs3828477 | -0.215 | -1.167 | -1.99 | 0.737 | 0.015 | -0.034 | -0.034 | 0 | 0.3 | 0.23 | 0.22 | 0.56 | 0.28 | 0.41 |
| 3 | 46498865 | rs11707471 | -1.033 | -1.954 | -2.509 | 0.56 | 0.085 | 0.038 | 0.038 | 0.222 | 0.48 | 0.96 | 0.49 | 0.65 | 0.35 | 0.53 |
| 3 | 46498975 | rs17078864 | 0.443 | -2.283 | -2.563 | 1.015 | 0.012 | -0.026 | -0.026 | 6.593 | 0.19 | 0.13 | 0.16 | 0.39 | 0.18 | 0.24 |
| 3 | 46500978 | rs17078876 | 0.595 | -2.345 | -1.638 | 1.31 | 0.014 | -0.043 | -0.043 | 0.025 | 0.18 | 0.09 | 0.15 | 0.28 | 0.18 | 0.24 |
| 3 | 46501213 | rs1126478 | -1.409 | -2.299 | -1.79 | 1.354 | 0.09 | 0.057 | 0.057 | 0.12 | 0.48 | 0.96 | 0.49 | 0.65 | 0.35 | 0.53 |
| 3 | 46501268 | rs1126477 | -0.77 | -2.156 | -1.475 | 1.171 | 0.064 | -0.033 | -0.033 | 4.206 | 0.35 | 0.77 | 0.42 | 0.42 | 0.26 | 0.37 |
| 3 | 46501931 | rs4683235 | 0.213 | -1.418 | -1.104 | 0.884 | -0.008 | 0.005 | 0.005 | 0.021 | 0.1 | 0.18 | 0.23 | 0.14 | 0.08 | 0.12 |
| 3 | 46503214 | rs13070740 | 1.725 | -1.121 | -1.272 | 0.799 | -0.013 | -0.03 | -0.03 | 1.063 | 0.09 | 0.01 | 0.04 | 0.06 | 0.08 | 0.14 |
| 3 | 46508524 | rs4683236 | 0.039 | -1.208 | -1.307 | 0.363 | -0.013 | -0.019 | -0.019 | 1.916 | 0.29 | 0.35 | 0.19 | 0.19 | 0.2 | 0.23 |
| 3 | 46513681 | rs1520484 | -0.451 | -1.571 | -1.186 | 0.488 | -0.01 | -0.002 | -0.002 | 0 | 0.38 | 0.58 | 0.41 | 0.41 | 0.25 | 0.34 |
| 3 | 46521392 | rs2139634 | 1.062 | -0.942 | -1.672 | -0.102 | -0.002 | -0.007 | -0.007 | 0 | 0.33 | 0.28 | 0.4 | 0.51 | 0.41 | 0.4 |
| 3 | 46527930 | rs1402151 | 1.111 | -1.108 | -1.95 | -0.073 | -0.095 | -0.171 | -0.171 | 0.114 | 0.32 | 0.03 | 0.38 | 0.49 | 0.42 | 0.39 |
| 3 | 46530169 | rs7644995 | -1.556 | -1.055 | -2.117 | -0.371 | -0.077 | 0.091 | 0.091 | 0 | 0.83 | 0.96 | 0.88 | 0.99 | 0.77 | 0.91 |
| 3 | 46538515 | rs883740 | 1.559 | -1.1 | -1.787 | 0.008 | -0.009 | 0.158 | 0.158 | 0.054 | 0.09 | 0.23 | 0.18 | 0.38 | 0.06 | 0.12 |
| 3 | 46541147 | rs7430431 | -0.625 | -1.698 | -1.927 | -0.366 | -0.028 | 0.118 | 0.118 | 0 | 0.57 | 0.78 | 0.69 | 0.95 | 0.55 | 0.73 |
| 3 | 46543617 | rs10514713 | 0.775 | -1.627 | -2.029 | -0.276 | -0.093 | -0.168 | -0.168 | 0.075 | 0.33 | 0.03 | 0.43 | 0.49 | 0.42 | 0.44 |
| 3 | 46549304 | rs2068099 | 1.995 | -1.393 | -2.123 | -0.151 | -0.03 | -0.028 | -0.028 | 0 | 0.17 | 0.02 | 0.15 | 0.19 | 0.2 | 0.24 |
| 3 | 46556835 | rs6808142 | -1.221 | -1.332 | -1.765 | -0.196 | -0.006 | -0.023 | -0.023 | 0 | 0.69 | 0.74 | 0.74 | 0.96 | 0.64 | 0.76 |
| 3 | 46556948 | rs17030627 | 0.472 | -1.298 | -1.974 | -0.033 | -0.099 | -0.13 | -0.13 | 0.779 | 0.42 | 0.04 | 0.47 | 0.52 | 0.5 | 0.47 |
| 3 | 46565341 | rs13093282 | 2.481 | -1.285 | -1.927 | 0.002 | -0.006 | -0.042 | -0.042 | 0.739 | 0.1 | 0.01 | 0.05 | 0.09 | 0.09 | 0.17 |
| 3 | 46569721 | rs6791703 | -0.218 | -1.283 | -1.768 | -0.016 | -0.008 | 0.028 | 0.028 | 0.441 | 0.54 | 0.34 | 0.56 | 0.61 | 0.58 | 0.64 |
| 3 | 46573318 | rs11713015 | 0.141 | -1.007 | -1.631 | -0.315 | 0.012 | 0.011 | 0.011 | 0 | 0.42 | 0.34 | 0.28 | 0.28 | 0.37 | 0.45 |
| 3 | 46574256 | rs9849400 | 0.141 | -1.007 | -1.631 | -0.315 | 0.012 | 0.011 | 0.011 | 0 | 0.42 | 0.34 | 0.28 | 0.28 | 0.37 | 0.45 |
| 3 | 46576081 | rs939421 | -0.638 | -0.936 | -2.05 | -0.272 | -0.007 | 0.031 | 0.031 | 0.642 | 0.74 | 0.57 | 0.52 | 0.32 | 0.73 | 0.69 |
| 3 | 46579125 | rs17220622 | 0.771 | -0.922 | -2.265 | -0.177 | -0.001 | 0.051 | 0.051 | 0 | 0.3 | 0.04 | 0.21 | 0.19 | 0.29 | 0.28 |
| 3 | 46586179 | rs7648827 | 1.255 | -0.418 | -1.505 | 0.032 | 0.016 | -0.015 | -0.015 | 0 | 0.12 | 0.07 | 0.05 | 0.09 | 0.09 | 0.17 |
| 3 | 46586622 | rs17286758 | 1.255 | -0.418 | -1.505 | 0.032 | 0.016 | -0.015 | -0.015 | 0 | 0.12 | 0.07 | 0.05 | 0.09 | 0.09 | 0.17 |
| 3 | 46588700 | rs9883208 | -0.016 | -0.232 | -1.32 | 0.099 | 0 | 0.047 | 0.047 | 0 | 0.49 | 0.56 | 0.32 | 0.13 | 0.44 | 0.41 |
| 3 | 46592530 | rs6774189 | 0.525 | -0.293 | -1.002 | 0.929 | -0.005 | 0.108 | 0.108 | 0 | 0.08 | 0.18 | 0.16 | 0.38 | 0.07 | 0.12 |
| 3 | 46593147 | rs1879325 | 0.117 | -0.012 | -0.978 | 0.505 | -0.001 | 0.002 | 0.002 | 1.916 | 0.65 | 0.35 | 0.52 | 0.43 | 0.63 | 0.67 |
| 3 | 46593906 | rs11919651 | 1.238 | 0.145 | -1.317 | 0.071 | -0.039 | 0.045 | 0.045 | 0 | 0.09 | 0.02 | 0.07 | 0 | 0.13 | 0.06 |
| 3 | 46598377 | rs13353497 | -0.174 | -0.294 | -1.098 | -0.086 | -0.005 | -0.019 | -0.019 | 0 | 0.44 | 0.46 | 0.51 | 0.78 | 0.37 | 0.48 |
| 3 | 46599937 | rs2280410 | 1.887 | -0.414 | -1.079 | 0.081 | 0.006 | -0.007 | -0.007 | 0.301 | 0.09 | 0.07 | 0.07 | 0.16 | 0.02 | 0.14 |
| 3 | 46599984 | rs2280411 | 0.352 | -0.356 | -1.126 | 0.106 | -0.004 | 0.002 | 0.002 | 1.06 | 0.35 | 0.4 | 0.44 | 0.62 | 0.35 | 0.36 |
| 3 | 46600520 | rs6783573 | 1.02 | -0.227 | -1.451 | -0.153 | -0.075 | 0.062 | 0.062 | 0.409 | 0.28 | 0.05 | 0.25 | 0.14 | 0.34 | 0.32 |
| 3 | 46605447 | rs17287098 | 1.518 | -0.263 | -1.182 | -0.16 | 0 | -0.025 | -0.025 | 0 | 0.16 | 0.17 | 0.14 | 0.07 | 0.16 | 0.13 |
| 3 | 46617502 | rs3806702 | 1.317 | -0.208 | -1.415 | -0.185 | -0.051 | 0.038 | 0.038 | 0.853 | 0.25 | 0.03 | 0.2 | 0.14 | 0.27 | 0.29 |
| 3 | 46620614 | rs11130097 | -0.34 | -0.464 | -1.258 | -0.028 | 0.003 | -0.002 | -0.002 | 0 | 0.57 | 0.49 | 0.49 | 0.42 | 0.55 | 0.59 |
| 3 | 46621215 | rs2280414 | 2.193 | -0.352 | -1.226 | 0.091 | 0.028 | -0.048 | -0.048 | 0.317 | 0.08 | 0.01 | 0.07 | 0.17 | 0.02 | 0.14 |
| 3 | 46626589 | rs9860011 | 0.788 | -0.281 | -0.966 | 0.161 | -0.004 | 0.144 | 0.144 | 0 | 0.1 | 0.25 | 0.07 | 0.37 | 0.06 | 0.17 |
| 3 | 46628443 | rs12637699 | 1.648 | -0.163 | -1.411 | -0.321 | -0.059 | 0.052 | 0.052 | 0.107 | 0.2 | 0.03 | 0.19 | 0.1 | 0.27 | 0.24 |
| 3 | 46634789 | rs17287344 | 1.159 | -0.175 | -1.388 | -0.259 | -0.009 | 0.02 | 0.02 | 0.256 | 0.07 | 0.02 | 0.07 | 0.03 | 0.09 | 0.02 |
| 3 | 46636120 | rs9880442 | -0.791 | -0.158 | -1.232 | -0.471 | -0.007 | -0.003 | -0.003 | 2.363 | 0.68 | 0.71 | 0.76 | 0.91 | 0.6 | 0.69 |
| 3 | 46636886 | rs11713702 | 1.525 | 0.168 | -1.616 | -0.348 | -0.04 | 0.047 | 0.047 | 0 | 0.1 | 0.02 | 0.07 | 0 | 0.13 | 0.06 |
| 3 | 46638971 | rs1520492 | 1.15 | -0.309 | -1.474 | 0.033 | 0.017 | -0.032 | -0.032 | 0 | 0.15 | 0.04 | 0.14 | 0.16 | 0.11 | 0.11 |
| 3 | 46644094 | rs1520485 | 1.251 | -0.381 | -1.486 | 0.128 | -0.004 | -0.024 | -0.024 | 0 | 0.22 | 0.12 | 0.2 | 0.26 | 0.27 | 0.19 |
| 3 | 46644862 | rs13076522 | 0.958 | -0.635 | -0.947 | 0.443 | 0.001 | -0.012 | -0.012 | 0 | 0.14 | 0.17 | 0.18 | 0.11 | 0.14 | 0.12 |
| 3 | 46644909 | rs6799581 | -0.462 | -0.607 | -0.934 | 0.47 | 0.01 | 0.013 | 0.013 | 0 | 0.29 | 0.2 | 0.32 | 0.26 | 0.25 | 0.24 |
| 3 | 46649700 | rs7640056 | -0.428 | -0.301 | -0.528 | 0.478 | -0.021 | 0.05 | 0.05 | 0 | 0.3 | 0.49 | 0.42 | 0.49 | 0.21 | 0.39 |
| 3 | 46652429 | rs6442009 | 0.738 | -0.223 | -0.516 | 0.405 | -0.002 | -0.01 | -0.01 | 0 | 0.57 | 0.39 | 0.5 | 0.51 | 0.61 | 0.7 |
| 3 | 46654132 | rs2139637 | 1.017 | -0.533 | -0.052 | 0.198 | -0.039 | -0.052 | -0.052 | 0 | 0.77 | 0.47 | 0.79 | 0.85 | 0.84 | 0.74 |

| | | | | | | | | | | | | | | | |
|---|---|---|---|---|---|---|---|---|---|---|---|---|---|---|---|
| 3 | 46655961 | rs10452028 | 0.215 | -0.157 | -0.859 | 0.148 | 0.007 | 0.046 | 0.046 | 0 | 0.12 | 0.05 | 0.11 | 0.03 | 0.08 | 0.08 |
| 3 | 46656351 | rs873382 | 0.215 | -0.157 | -0.859 | 0.148 | 0.007 | 0.046 | 0.046 | 0 | 0.12 | 0.05 | 0.11 | 0.03 | 0.08 | 0.08 |
| 3 | 46659860 | rs869701 | 0.248 | -0.093 | -0.774 | 0.308 | -0.001 | 0.011 | 0.011 | 0 | 0.49 | 0.4 | 0.51 | 0.32 | 0.54 | 0.45 |
| 3 | 46663372 | rs9817414 | -0.04 | -0.56 | -0.856 | 0.485 | -0.002 | -0.035 | -0.035 | 0 | 0.21 | 0.11 | 0.18 | 0.32 | 0.23 | 0.19 |
| 3 | 46666521 | rs9829227 | -0.714 | -0.713 | -0.575 | 0.507 | -0.01 | 0.076 | 0.076 | 0 | 0.09 | 0.13 | 0.09 | 0.32 | 0.07 | 0.13 |
| 3 | 46667110 | rs3796369 | -1.039 | -0.814 | -0.573 | 0.463 | -0.017 | 0.066 | 0.066 | 0 | 0.11 | 0.15 | 0.09 | 0.32 | 0.07 | 0.13 |
| 3 | 46668689 | rs1356875 | -0.487 | -0.794 | -0.444 | 0.28 | -0.001 | 0.001 | 0.001 | 0 | 0.65 | 0.65 | 0.58 | 0.61 | 0.69 | 0.59 |
| 3 | 46676589 | rs6805749 | 0.442 | -0.978 | -0.427 | 0.089 | 0.004 | 0.01 | 0.01 | 0 | 0.54 | 0.37 | 0.47 | 0.29 | 0.62 | 0.46 |
| 3 | 46677519 | rs1520489 | 0.571 | -1.312 | -0.058 | 0.276 | 0.001 | -0.012 | -0.012 | 0 | 0.45 | 0.43 | 0.53 | 0.61 | 0.5 | 0.53 |
| 3 | 46679675 | rs4234464 | 0.463 | -1.312 | 0.008 | 0.292 | -0.004 | 0.015 | 0.015 | 0 | 0.44 | 0.34 | 0.52 | 0.61 | 0.5 | 0.53 |
| 3 | 46680063 | rs11709020 | 1.104 | -1.288 | -0.043 | 0.286 | -0.013 | 0.007 | 0.007 | 0.181 | 0.37 | 0.24 | 0.37 | 0.27 | 0.46 | 0.41 |
| 3 | 46688498 | rs12491294 | -0.228 | -1.357 | -0.126 | 0.211 | -0.001 | 0.079 | 0.079 | 0 | 0.5 | 0.27 | 0.43 | 0.4 | 0.47 | 0.46 |
| 3 | 46688562 | rs7633770 | 1.408 | -1.339 | -0.257 | 0.34 | -0.043 | 0.004 | 0.004 | 0.639 | 0.31 | 0.15 | 0.29 | 0.23 | 0.43 | 0.28 |
| 3 | 46692379 | rs9861101 | 0.125 | -1.023 | -0.026 | 0.519 | -0.002 | 0.107 | 0.107 | 0 | 0.05 | 0.08 | 0.16 | 0.33 | 0.05 | 0.12 |
| 3 | 46700713 | rs6442013 | -0.418 | -1.301 | -0.077 | -0.179 | -0.069 | -0.122 | -0.122 | 0 | 0.84 | 0.56 | 0.92 | 0.99 | 0.94 | 0.9 |
| 3 | 46705369 | rs4682826 | 0.309 | -0.728 | -1.006 | 0.284 | -0.059 | -0.064 | -0.064 | 0 | 0.23 | 0.08 | 0.31 | 0.36 | 0.31 | 0.37 |
| 3 | 46711135 | rs3747519 | 0.319 | -1.03 | -0.846 | 0.102 | 0.001 | 0.051 | 0.051 | 0 | 0.44 | 0.28 | 0.34 | 0.33 | 0.46 | 0.35 |
| 3 | 46711507 | rs3827495 | 0.472 | -1.007 | -0.793 | -0.047 | -0.024 | -0.027 | -0.027 | 0 | 0.29 | 0.16 | 0.33 | 0.36 | 0.33 | 0.43 |
| 3 | 46711563 | rs11711074 | -0.46 | -1.514 | -0.242 | -0.069 | -0.015 | 0.012 | 0.012 | 0 | 0.73 | 0.61 | 0.68 | 0.68 | 0.78 | 0.75 |
| 3 | 46711850 | rs3210183 | -0.172 | -1.487 | -0.341 | -0.047 | -0.028 | 0.009 | 0.009 | 0 | 0.71 | 0.43 | 0.66 | 0.69 | 0.78 | 0.75 |
| 3 | 46718298 | rs6806400 | -1.639 | -0.998 | -0.993 | -0.133 | -0.02 | -0.063 | -0.063 | 0.06 | 0.77 | 0.96 | 0.81 | 0.7 | 0.76 | 0.84 |
| 3 | 46721211 | rs9817508 | 0.614 | -0.939 | -0.528 | -0.202 | -0.066 | -0.034 | -0.034 | 0 | 0.3 | 0.1 | 0.39 | 0.34 | 0.42 | 0.41 |
| 3 | 46721318 | rs6442019 | -0.757 | -1.039 | -0.47 | -0.238 | 0.015 | 0.009 | 0.009 | 0 | 0.63 | 0.83 | 0.76 | 0.65 | 0.67 | 0.67 |
| 3 | 46721519 | rs10451990 | -0.465 | -1.053 | -0.501 | -0.243 | -0.01 | -0.011 | -0.011 | 0.073 | 0.59 | 0.52 | 0.73 | 0.64 | 0.66 | 0.67 |
| 3 | 46721662 | rs6803988 | -1.511 | -0.976 | -0.893 | -0.189 | -0.02 | -0.06 | -0.06 | 0 | 0.74 | 0.92 | 0.8 | 0.66 | 0.74 | 0.8 |
| 3 | 46724569 | rs9847407 | 0.673 | -0.913 | -0.811 | -0.143 | -0.068 | -0.035 | -0.035 | 0 | 0.3 | 0.1 | 0.39 | 0.33 | 0.42 | 0.41 |
| 3 | 46728208 | rs7373358 | 0.006 | -1.025 | -0.667 | -0.155 | 0.018 | 0.007 | 0.007 | 0 | 0.33 | 0.73 | 0.37 | 0.29 | 0.25 | 0.25 |
| 3 | 46728659 | rs7633016 | -0.509 | -1.057 | -0.735 | -0.19 | 0.057 | 0.056 | 0.056 | 0 | 0.43 | 0.77 | 0.4 | 0.3 | 0.32 | 0.38 |
| 3 | 46730063 | rs12496585 | 0.414 | -1.039 | -0.456 | -0.153 | -0.027 | 0.001 | 0.001 | 0 | 0.28 | 0.42 | 0.34 | 0.29 | 0.24 | 0.25 |
| 3 | 46734441 | rs11718902 | 0.527 | -0.937 | -0.578 | -0.098 | -0.015 | -0.008 | -0.008 | 0 | 0.32 | 0.22 | 0.41 | 0.35 | 0.42 | 0.41 |
| 3 | 46735645 | rs6442020 | -0.304 | -0.833 | -0.718 | -0.391 | 0.039 | 0.06 | 0.06 | 0 | 0.48 | 0.68 | 0.42 | 0.31 | 0.39 | 0.39 |
| 3 | 46741299 | rs9867730 | -1.179 | -0.975 | -0.087 | 0.11 | 0.002 | 0.004 | 0.004 | 0 | 0.73 | 0.85 | 0.8 | 0.74 | 0.74 | 0.8 |
| 3 | 46742019 | rs6442021 | -0.068 | -1.211 | -0.066 | 0.374 | 0.009 | 0.002 | 0.002 | 0 | 0.54 | 0.6 | 0.61 | 0.5 | 0.6 | 0.55 |
| 3 | 46743895 | rs4683316 | -1.056 | -1.15 | 0.241 | 0.366 | -0.014 | 0.031 | 0.031 | 0 | 0.69 | 0.5 | 0.75 | 0.66 | 0.71 | 0.78 |
| 3 | 46744966 | rs7650366 | -0.479 | -1.37 | 0.099 | 0.275 | -0.016 | 0.005 | 0.005 | 0 | 0.25 | 0.34 | 0.26 | 0.3 | 0.16 | 0.35 |
| 3 | 46746975 | rs7426919 | 0.449 | -1.252 | 0.091 | 0.282 | -0.001 | -0.004 | -0.004 | 0.075 | 0.44 | 0.44 | 0.51 | 0.36 | 0.56 | 0.43 |
| 3 | 46748453 | rs12491225 | 0.505 | -1.217 | -0.42 | 0.297 | -0.011 | 0.051 | 0.051 | 0 | 0.44 | 0.16 | 0.49 | 0.36 | 0.56 | 0.43 |
| 3 | 46750202 | rs7628174 | 0.388 | -0.82 | 0.135 | 0.511 | -0.001 | 0 | 0 | 1.526 | 0.16 | 0.19 | 0.17 | 0.16 | 0.1 | 0.23 |
| 3 | 46753577 | rs9814780 | -1.916 | -0.806 | -0.294 | 0.501 | -0.021 | -0.05 | -0.05 | 0.089 | 0.89 | 0.99 | 0.89 | 0.83 | 0.89 | 0.84 |
| 3 | 46753976 | rs9815362 | -0.805 | -0.379 | 0.195 | 0.378 | -0.003 | -0.03 | -0.03 | 0 | 0.44 | 0.77 | 0.54 | 0.49 | 0.42 | 0.48 |
| 3 | 46763839 | rs7632176 | -0.939 | -0.542 | 1.255 | 0.398 | -0.008 | -0.004 | -0.004 | 0.954 | 0.19 | 0.27 | 0.19 | 0.15 | 0.14 | 0.24 |
| 3 | 46768026 | rs7623501 | -0.942 | -0.485 | 1.155 | 0.423 | -0.012 | 0 | 0 | 0 | 0.2 | 0.47 | 0.2 | 0.15 | 0.14 | 0.24 |
| 3 | 46778008 | rs6793661 | -0.892 | -0.466 | 1.159 | 0.242 | -0.011 | -0.026 | -0.026 | 0.093 | 0.19 | 0.45 | 0.21 | 0.12 | 0.15 | 0.25 |
| 3 | 46778988 | rs6763438 | -1.482 | -0.57 | 0.715 | 0.264 | -0.019 | -0.011 | -0.011 | 0 | 0.26 | 0.67 | 0.31 | 0.23 | 0.19 | 0.36 |
| 3 | 46781836 | rs6801459 | -1.068 | -0.128 | 0.743 | 0.3 | 0.024 | -0.03 | -0.03 | 0.369 | 0.26 | 0.53 | 0.25 | 0.25 | 0.29 | 0.34 |
| 3 | 46783432 | rs6810260 | -0.947 | -1.081 | 1.154 | 0.19 | -0.026 | -0.062 | -0.062 | 0 | 0.19 | 0.53 | 0.18 | 0.12 | 0.14 | 0.23 |
| 3 | 46790397 | rs7614762 | -1.74 | -1.187 | 0.474 | 0.157 | 0.037 | 0.027 | 0.027 | 0 | 0.31 | 0.82 | 0.34 | 0.24 | 0.19 | 0.38 |
| 3 | 46852679 | rs7647507 | -1.213 | -1.192 | 0.532 | 0.073 | -0.028 | -0.104 | -0.104 | 0.016 | 0.23 | 0.65 | 0.15 | 0.01 | 0.14 | 0.13 |
| 3 | 46866866 | rs6796502 | -1.137 | -1.295 | 0.407 | 0.363 | -0.014 | 0.005 | 0.005 | 0 | 0.13 | 0.18 | 0.21 | 0.14 | 0.09 | 0.24 |
| 3 | 46873957 | rs7636089 | -1.641 | -1.343 | 0.064 | 0.344 | 0.056 | 0.032 | 0.032 | 0 | 0.32 | 0.83 | 0.33 | 0.25 | 0.16 | 0.37 |
| 3 | 46879831 | rs12107311 | -1.641 | -1.319 | -0.001 | 0.363 | 0.064 | 0.031 | 0.031 | 0 | 0.32 | 0.83 | 0.33 | 0.27 | 0.16 | 0.37 |
| 3 | 46884552 | rs4683304 | -1.639 | -1.318 | -0.001 | 0.364 | 0.064 | 0.031 | 0.031 | 0.012 | 0.32 | 0.84 | 0.33 | 0.27 | 0.16 | 0.36 |
| 3 | 46889187 | rs6787229 | -1.441 | -1.091 | 0.566 | 0.446 | -0.021 | 0.005 | 0.005 | 0 | 0.15 | 0.26 | 0.12 | 0.15 | 0.1 | 0.26 |
| 3 | 46895333 | rs6781266 | -1.319 | -1.155 | 0.596 | 0.441 | -0.035 | 0.014 | 0.014 | 0 | 0.13 | 0.29 | 0.11 | 0.14 | 0.08 | 0.26 |
| 3 | 46899516 | rs1042973 | -1.396 | -1.408 | 0.687 | 0.342 | -0.053 | 0.001 | 0.001 | 0 | 0.23 | 0.32 | 0.14 | 0.24 | 0.11 | 0.4 |
| 3 | 46901654 | rs1531136 | -2.181 | -1.53 | 0.109 | 0.239 | 0.1 | 0.057 | 0.057 | 0.165 | 0.35 | 0.89 | 0.22 | 0.26 | 0.16 | 0.42 |
| 3 | 46902129 | rs2227294 | -2 | -1.511 | 0.155 | 0.981 | 0.102 | -0.072 | -0.072 | 0 | 0.33 | 0.89 | 0.3 | 0.39 | 0.14 | 0.42 |
| 3 | 46903043 | rs3792557 | -1.342 | -1.547 | 0.413 | 0.523 | -0.039 | -0.001 | -0.001 | 0 | 0.17 | 0.58 | 0.13 | 0.15 | 0.09 | 0.33 |
| 3 | 46903094 | rs3792558 | -1.586 | -1.341 | -0.061 | 0.591 | 0.078 | 0.036 | 0.036 | 0 | 0.32 | 0.82 | 0.2 | 0.26 | 0.13 | 0.42 |
| 3 | 46929546 | rs6442037 | 0.49 | -0.145 | -0.316 | 0.247 | -0.058 | 0.053 | 0.053 | 1.406 | 0.65 | 0.87 | 0.76 | 0.73 | 0.66 | 0.55 |
| 3 | 46944274 | rs1138518 | 0.914 | -0.025 | -0.331 | 0.74 | -0.001 | -0.04 | -0.04 | 0 | 0.61 | 0.82 | 0.65 | 0.44 | 0.64 | 0.6 |
| 3 | 46951935 | rs11719795 | 0.605 | -0.162 | -0.345 | 0.768 | 0.02 | -0.048 | -0.048 | 0.067 | 0.64 | 0.83 | 0.67 | 0.44 | 0.7 | 0.63 |
| 3 | 46952777 | rs12715420 | -0.625 | -0.538 | 0.365 | 0.535 | -0.003 | -0.019 | -0.019 | 0.178 | 0.06 | 0.27 | 0.05 | 0 | 0.02 | 0.02 |
| 3 | 46967040 | rs2290545 | 0.388 | -0.377 | -0.02 | 0.517 | -0.007 | -0.103 | -0.103 | 0 | 0.7 | 0.89 | 0.72 | 0.46 | 0.74 | 0.64 |
| 3 | 46973673 | rs4683297 | -1.135 | -0.344 | 0.096 | 0.528 | 0.015 | -0.027 | -0.027 | 0 | 0.39 | 0.2 | 0.35 | 0.56 | 0.33 | 0.38 |
| 3 | 46978923 | rs4683295 | -1.238 | 0.009 | -0.572 | 0.328 | 0.031 | -0.02 | -0.02 | 0 | 0.42 | 0.24 | 0.42 | 0.51 | 0.35 | 0.39 |
| 3 | 46982737 | rs4682844 | -0.947 | 0.176 | -0.97 | -0.158 | -0.008 | -0.02 | -0.02 | 0 | 0.48 | 0.2 | 0.37 | 0.48 | 0.41 | 0.43 |
| 3 | 46988561 | rs13092573 | 1.649 | 0.41 | -0.849 | -0.281 | -0.016 | -0.004 | -0.004 | 0 | 0.24 | 0.15 | 0.3 | 0.26 | 0.33 | 0.36 |
| 3 | 47024495 | rs11130112 | 0.469 | 0.337 | -0.577 | -0.179 | 0 | 0.002 | 0.002 | 0.252 | 0.52 | 0.56 | 0.6 | 0.46 | 0.59 | 0.57 |

| | | | | | | | | | | | | | | | | |
|---|---|---|---|---|---|---|---|---|---|---|---|---|---|---|---|---|
| 3 | 47052937 | rs1982510 | 0.74 | 0.322 | -0.661 | -0.467 | -0.012 | 0.011 | 0.011 | 0 | 0.19 | 0.14 | 0.11 | 0 | 0.19 | 0.08 |
| 3 | 47125385 | rs4082155 | 0.997 | 0.411 | -0.593 | -0.273 | -0.007 | -0.02 | -0.02 | 0 | 0.47 | 0.24 | 0.52 | 0.53 | 0.58 | 0.57 |
| 3 | 47126543 | rs4078466 | 0.739 | 0.397 | -0.498 | -0.279 | 0.001 | 0 | 0 | 0 | 0.5 | 0.54 | 0.55 | 0.53 | 0.58 | 0.57 |
| 3 | 47129903 | rs12631730 | -1.024 | 0.29 | -0.63 | 0.02 | -0.005 | 0.01 | 0.01 | 0.094 | 0.79 | 0.9 | 0.83 | 0.67 | 0.83 | 0.73 |
| 3 | 47162661 | rs6767907 | 0.444 | 0.304 | -0.599 | 0.049 | -0.011 | 0.05 | 0.05 | 0.384 | 0.56 | 0.78 | 0.68 | 0.66 | 0.61 | 0.63 |
| 3 | 47198826 | rs11708451 | 1.098 | 0.414 | -0.535 | 0.111 | -0.004 | -0.015 | -0.015 | 0.046 | 0.48 | 0.24 | 0.52 | 0.53 | 0.58 | 0.58 |
| 3 | 47214122 | rs7620997 | -0.234 | 0.458 | -0.604 | 0.11 | -0.014 | 0.03 | 0.03 | 0.02 | 0.41 | 0.21 | 0.3 | 0.34 | 0.37 | 0.36 |
| 3 | 47268306 | rs7632501 | -0.492 | 0.334 | -0.596 | 0.142 | -0.007 | 0.044 | 0.044 | 0.171 | 0.43 | 0.22 | 0.32 | 0.34 | 0.38 | 0.35 |
| 3 | 47285912 | rs11130123 | 1.014 | 0.312 | -0.522 | 0.151 | 0.017 | -0.01 | -0.01 | 0.009 | 0.49 | 0.24 | 0.52 | 0.54 | 0.6 | 0.59 |
| 3 | 47292183 | rs11130124 | 0.733 | 0.338 | -0.585 | 0.165 | 0 | -0.001 | -0.001 | 0 | 0.52 | 0.54 | 0.56 | 0.54 | 0.6 | 0.59 |
| 3 | 47306894 | rs11130126 | -0.482 | 0.361 | -0.765 | 0.136 | -0.009 | 0.044 | 0.044 | 0.119 | 0.43 | 0.22 | 0.31 | 0.34 | 0.38 | 0.34 |
| 3 | 47322781 | rs1076394 | -0.575 | 0.346 | -1.291 | 0.196 | -0.008 | -0.025 | -0.025 | 0 | 0.46 | 0.21 | 0.42 | 0.45 | 0.4 | 0.4 |
| 3 | 47328744 | rs10490798 | 1.226 | 0.353 | -1.361 | 0.159 | -0.023 | -0.098 | -0.098 | 0 | 0.19 | 0.01 | 0.16 | 0.31 | 0.16 | 0.25 |
| 3 | 47331136 | rs11710322 | -0.377 | 0.436 | -1.131 | 0.349 | -0.008 | -0.025 | -0.025 | 0 | 0.46 | 0.21 | 0.42 | 0.45 | 0.4 | 0.4 |
| 3 | 47351501 | rs11710121 | -0.051 | 0.446 | -1.147 | 0.347 | -0.004 | 0.033 | 0.033 | 0 | 0.41 | 0.2 | 0.3 | 0.33 | 0.35 | 0.36 |
| 3 | 47362080 | rs295433 | 1.741 | 0.529 | -1.392 | 0.266 | -0.008 | 0.001 | 0.001 | 0 | 0.27 | 0.2 | 0.27 | 0.28 | 0.35 | 0.36 |
| 3 | 47385585 | rs295458 | 0.942 | 0.205 | -0.805 | 0.337 | 0.014 | -0.003 | -0.003 | 0.18 | 0.49 | 0.23 | 0.52 | 0.55 | 0.6 | 0.58 |
| 3 | 47405456 | rs4858890 | -0.641 | 0.269 | -0.804 | 0.34 | -0.008 | -0.016 | -0.016 | 0.26 | 0.46 | 0.24 | 0.43 | 0.45 | 0.41 | 0.43 |
| 3 | 47415100 | rs1872164 | -0.644 | 0.185 | -0.624 | 0.603 | 0.003 | 0.004 | 0.004 | 0 | 0.46 | 0.27 | 0.43 | 0.53 | 0.4 | 0.44 |
| 3 | 47420627 | rs13075233 | 1.002 | 0.069 | -0.379 | 0.552 | 0.02 | 0.022 | 0.022 | 0 | 0.49 | 0.24 | 0.52 | 0.46 | 0.6 | 0.56 |
| 3 | 47425473 | rs11719877 | 1.002 | 0.089 | -0.379 | 0.552 | 0.023 | 0.022 | 0.022 | 0 | 0.49 | 0.24 | 0.52 | 0.46 | 0.6 | 0.56 |
| 3 | 47443262 | rs11705957 | -0.61 | 0.082 | -0.379 | 0.566 | 0.003 | 0.007 | 0.007 | 0 | 0.46 | 0.27 | 0.43 | 0.54 | 0.39 | 0.44 |
| 3 | 47452118 | rs6780013 | -0.585 | 0.069 | -0.429 | 0.726 | -0.001 | 0.009 | 0.009 | 0.015 | 0.43 | 0.25 | 0.32 | 0.4 | 0.38 | 0.39 |
| 3 | 47452786 | rs1531875 | 1.055 | 0.086 | -0.434 | 0.601 | 0.02 | 0.024 | 0.024 | 0 | 0.49 | 0.24 | 0.52 | 0.46 | 0.6 | 0.56 |
| 3 | 47455086 | rs4858868 | 1.055 | 0.086 | -0.434 | 0.601 | 0.02 | 0.024 | 0.024 | 0 | 0.49 | 0.24 | 0.52 | 0.46 | 0.6 | 0.56 |
| 3 | 47464122 | rs1038394 | 1.055 | 0.064 | -0.456 | 0.591 | 0.02 | 0.024 | 0.024 | 0 | 0.49 | 0.24 | 0.52 | 0.46 | 0.6 | 0.56 |
| 3 | 47468261 | rs11711824 | 1.05 | 0.058 | -0.423 | 0.588 | 0.021 | 0.025 | 0.025 | 0 | 0.49 | 0.24 | 0.52 | 0.46 | 0.6 | 0.56 |
| 3 | 47494157 | rs12490383 | 0.872 | -0.049 | -0.653 | 0.424 | -0.018 | -0.162 | -0.162 | 0 | 0.2 | 0.01 | 0.26 | 0.49 | 0.16 | 0.3 |
| 3 | 47511698 | rs17784714 | 1.29 | -0.01 | -0.641 | 0.371 | -0.078 | 0.098 | 0.098 | 0 | 0.16 | 0.01 | 0.09 | 0 | 0.19 | 0.08 |
| 3 | 47518266 | rs11716763 | -0.703 | 0.04 | -0.455 | 0.537 | 0.003 | 0.007 | 0.007 | 0 | 0.46 | 0.27 | 0.43 | 0.54 | 0.4 | 0.43 |
| 3 | 47528193 | rs11914913 | 1.111 | 0.053 | -0.418 | 0.543 | 0.02 | 0.023 | 0.023 | 0 | 0.49 | 0.26 | 0.52 | 0.46 | 0.6 | 0.56 |
| 3 | 47558448 | rs11711265 | 1.104 | 0.047 | -0.388 | 0.17 | 0.021 | 0.072 | 0.072 | 0 | 0.49 | 0.26 | 0.58 | 0.7 | 0.6 | 0.56 |
| 3 | 47575568 | rs4858833 | 0.391 | 0.056 | -0.018 | 0.387 | 0.071 | 0.067 | 0.067 | 3.629 | 0.95 | 0.5 | 0.95 | 1 | 1 | 1 |
| 3 | 47582724 | rs1564760 | 0.774 | -0.064 | -0.704 | 0.225 | 0.045 | 0.049 | 0.049 | 0 | 0.54 | 0.26 | 0.65 | 0.7 | 0.69 | 0.58 |
| 3 | 47586514 | rs9854395 | 0.344 | 0.302 | 0.227 | -0.088 | 0.054 | -0.07 | -0.07 | 0 | 0.14 | 0.44 | 0.09 | 0.02 | 0.13 | 0.05 |
| 3 | 47595879 | rs6785031 | 0.764 | -0.171 | -0.059 | -0.311 | 0 | -0.001 | -0.001 | 0.051 | 0.6 | 0.39 | 0.59 | 0.72 | 0.66 | 0.54 |
| 3 | 47598663 | rs11130138 | 0.764 | -0.171 | -0.059 | -0.311 | 0 | -0.001 | -0.001 | 0 | 0.6 | 0.39 | 0.59 | 0.72 | 0.66 | 0.54 |
| 3 | 47606215 | rs11716779 | 0.57 | -0.54 | 0.237 | -0.516 | -0.101 | 0.071 | 0.071 | 1.198 | 0.25 | 0.02 | 0.15 | 0.09 | 0.27 | 0.25 |
| 3 | 47613302 | rs869508 | -0.649 | -0.607 | 0.503 | -0.436 | -0.029 | 0.043 | 0.043 | 1.753 | 0.34 | 0.12 | 0.36 | 0.27 | 0.33 | 0.42 |
| 3 | 47618953 | rs3732530 | 1.428 | -0.64 | 0.691 | -0.076 | 0.006 | -0.002 | -0.002 | 0 | 0.59 | 0.36 | 0.48 | 0.6 | 0.66 | 0.54 |
| 3 | 47622210 | rs3755637 | 1.346 | -0.647 | 0.734 | -0.1 | -0.007 | -0.029 | -0.029 | 0.14 | 0.6 | 0.45 | 0.59 | 0.71 | 0.67 | 0.55 |
| 3 | 47652639 | rs1014228 | 1.444 | -0.671 | 0.733 | -0.101 | 0.006 | -0.001 | -0.001 | 0 | 0.59 | 0.36 | 0.48 | 0.6 | 0.67 | 0.55 |
| 3 | 47652795 | rs1014229 | 2.141 | -0.285 | 0.268 | -0.893 | -0.015 | -0.075 | -0.075 | 0.046 | 0.3 | 0.1 | 0.27 | 0.45 | 0.33 | 0.4 |
| 3 | 47672915 | rs17785248 | 1.319 | -0.228 | 0 | -0.833 | -0.095 | 0.058 | 0.058 | 1.133 | 0.26 | 0.02 | 0.15 | 0.11 | 0.26 | 0.24 |
| 3 | 47717626 | rs2291903 | 2.014 | -0.197 | 0.012 | -0.823 | 0 | -0.031 | -0.031 | 0 | 0.06 | 0 | 0.19 | 0.16 | 0.06 | 0.17 |
| 3 | 47777774 | rs4858825 | 2.152 | -0.292 | -0.226 | -0.952 | -0.114 | -0.094 | -0.094 | 0 | 0.23 | 0.01 | 0.28 | 0.25 | 0.34 | 0.15 |
| 3 | 47857575 | rs3888 | 0.096 | 0.035 | -0.03 | -0.809 | -0.007 | -0.029 | -0.029 | 3.035 | 0.61 | 0.47 | 0.6 | 0.71 | 0.68 | 0.56 |
| 3 | 47866161 | rs13072132 | 1.642 | 0.034 | -0.116 | -0.809 | -0.095 | 0.059 | 0.059 | 1.11 | 0.26 | 0.02 | 0.15 | 0.11 | 0.26 | 0.24 |
| 3 | 47868622 | rs3915654 | -0.043 | 0.164 | -0.047 | -0.755 | 0.004 | 0.013 | 0.013 | 0.199 | 0.6 | 0.37 | 0.59 | 0.71 | 0.68 | 0.56 |
| 3 | 47899878 | rs13066266 | 1.703 | -0.068 | -0.169 | -0.652 | -0.116 | -0.094 | -0.094 | 1.016 | 0.25 | 0.02 | 0.27 | 0.26 | 0.32 | 0.23 |
| 3 | 47936671 | rs777016 | -1.35 | -0.398 | 0.219 | -0.475 | 0.017 | -0.022 | -0.022 | 0.167 | 0.68 | 0.77 | 0.74 | 0.55 | 0.72 | 0.59 |
| 3 | 47943861 | rs319696 | -0.249 | -0.257 | 0.539 | -1.624 | -0.009 | 0.024 | 0.024 | 0.871 | 0.33 | 0.18 | 0.35 | 0.28 | 0.3 | 0.35 |
| 3 | 48016172 | rs9862913 | -0.642 | -0.268 | 0.592 | -1.657 | -0.005 | -0.03 | -0.03 | 0 | 0.37 | 0.54 | 0.4 | 0.28 | 0.32 | 0.35 |
| 3 | 48029118 | rs1013431 | 0.483 | -0.272 | 0.585 | -1.658 | -0.005 | -0.03 | -0.03 | 0 | 0.63 | 0.46 | 0.6 | 0.72 | 0.69 | 0.65 |
| 3 | 48074547 | rs5012971 | 0.682 | -0.258 | 0.183 | -1.651 | 0.002 | 0.006 | 0.006 | 0.362 | 0.62 | 0.37 | 0.59 | 0.72 | 0.68 | 0.65 |
| 3 | 48130893 | rs6442101 | 0.033 | -0.041 | -0.659 | -1.56 | -0.029 | 0.033 | 0.033 | 1.724 | 0.69 | 0.88 | 0.65 | 0.74 | 0.69 | 0.69 |
| 3 | 48138082 | rs9825114 | 0.051 | -0.24 | -0.631 | -1.528 | -0.039 | 0.037 | 0.037 | 1.591 | 0.68 | 0.88 | 0.64 | 0.74 | 0.67 | 0.67 |
| 3 | 48186267 | rs12496784 | 0.402 | -1.058 | -0.253 | -1.443 | -0.011 | -0.007 | -0.007 | 0 | 0.63 | 0.53 | 0.59 | 0.69 | 0.68 | 0.64 |
| 3 | 48193515 | rs7617773 | 0.225 | -1.021 | -0.356 | -1.426 | 0.012 | 0.004 | 0.004 | 1.967 | 0.65 | 0.77 | 0.62 | 0.69 | 0.68 | 0.64 |
| 3 | 48227462 | rs1380053 | 0.125 | -0.934 | -0.355 | -1.318 | -0.038 | 0.015 | 0.015 | 0 | 0.08 | 0.12 | 0.13 | 0.14 | 0.03 | 0.04 |
| 3 | 48268509 | rs2889854 | -1.813 | -0.85 | -0.655 | -1.611 | -0.09 | 0.096 | 0.096 | 0 | 0.85 | 0.99 | 0.88 | 1 | 0.76 | 0.96 |
| 3 | 48272536 | rs745116 | -0.466 | -0.99 | -0.021 | -1.049 | -0.003 | 0.026 | 0.026 | 0.212 | 0.3 | 0.28 | 0.22 | 0.12 | 0.29 | 0.27 |
| 3 | 48287879 | rs13059037 | 0.024 | -0.989 | -0.037 | -1.049 | -0.067 | 0.046 | 0.046 | 0.245 | 0.26 | 0.06 | 0.18 | 0.12 | 0.29 | 0.27 |
| 3 | 48297560 | rs11710861 | -0.389 | -0.989 | -0.025 | -1.049 | -0.004 | 0.024 | 0.024 | 0.217 | 0.29 | 0.26 | 0.22 | 0.12 | 0.29 | 0.27 |
| 3 | 48301614 | rs13068265 | 0.058 | -0.982 | -0.036 | -1.046 | -0.068 | 0.044 | 0.044 | 0.394 | 0.26 | 0.06 | 0.18 | 0.12 | 0.29 | 0.27 |
| 3 | 48303669 | rs11709691 | 0.058 | -0.982 | -0.036 | -1.046 | -0.068 | 0.044 | 0.044 | 0.394 | 0.26 | 0.06 | 0.18 | 0.12 | 0.29 | 0.27 |
| 3 | 48312229 | rs1045482 | -1.223 | -1.053 | 0.094 | -1.057 | -0.011 | -0.012 | -0.012 | 0 | 0.41 | 0.56 | 0.36 | 0.29 | 0.33 | 0.38 |
| 3 | 48312402 | rs3937 | -0.452 | -0.73 | -0.097 | -1.061 | -0.004 | 0.025 | 0.025 | 0.294 | 0.29 | 0.26 | 0.21 | 0.12 | 0.29 | 0.27 |
| 3 | 48335857 | rs3197223 | -0.041 | -0.561 | -0.57 | -0.978 | -0.064 | 0.042 | 0.042 | 0.212 | 0.26 | 0.07 | 0.18 | 0.12 | 0.28 | 0.28 |

| | | | | | | | | | | | | | | | | |
|---|---|---|---|---|---|---|---|---|---|---|---|---|---|---|---|---|
| 3 | 48338173 | rs4479622 | -0.042 | -0.554 | -0.568 | -1.013 | -0.064 | 0.046 | 0.046 | 0.196 | 0.26 | 0.07 | 0.18 | 0.12 | 0.28 | 0.28 |
| 3 | 48340045 | rs13085251 | -0.042 | -0.554 | -0.568 | -1.013 | -0.064 | 0.046 | 0.046 | 0.196 | 0.26 | 0.07 | 0.18 | 0.12 | 0.28 | 0.28 |
| 3 | 48360433 | rs11706277 | -0.033 | -0.548 | -0.565 | -1.01 | -0.064 | 0.046 | 0.046 | 0.162 | 0.26 | 0.07 | 0.18 | 0.12 | 0.28 | 0.28 |
| 3 | 48360992 | rs11718350 | 0.04 | -0.543 | -0.568 | -1.008 | -0.073 | 0.039 | 0.039 | 0.478 | 0.26 | 0.03 | 0.18 | 0.12 | 0.28 | 0.28 |
| 3 | 48362750 | rs11716371 | 0.034 | -0.544 | -0.563 | -1.008 | -0.066 | 0.043 | 0.043 | 0.434 | 0.26 | 0.06 | 0.18 | 0.12 | 0.28 | 0.27 |
| 3 | 48373214 | rs11710257 | -0.047 | -0.554 | -0.568 | -1.013 | -0.064 | 0.047 | 0.047 | 0.168 | 0.26 | 0.07 | 0.18 | 0.12 | 0.29 | 0.28 |
| 3 | 48381826 | rs13095891 | -0.724 | -0.962 | -0.311 | -1.194 | -0.046 | -0.004 | -0.004 | 4.371 | 0.13 | 0.2 | 0.06 | 0.15 | 0.05 | 0.13 |
| 3 | 48384388 | rs6779517 | -0.329 | -0.166 | -0.173 | -0.945 | -0.014 | 0.01 | 0.01 | 0.353 | 0.26 | 0.22 | 0.31 | 0.14 | 0.36 | 0.25 |
| 3 | 48396309 | rs11708617 | 0.297 | -0.884 | -0.847 | -0.721 | -0.137 | 0.011 | 0.011 | 0 | 0.28 | 0.04 | 0.36 | 0.18 | 0.45 | 0.28 |
| 3 | 48398586 | rs7630053 | -0.371 | -1.519 | -0.717 | -0.89 | -0.063 | 0.059 | 0.059 | 0 | 0.38 | 0.16 | 0.34 | 0.22 | 0.45 | 0.27 |
| 3 | 48407462 | rs4558783 | -0.24 | -1.045 | -0.479 | -0.746 | -0.042 | -0.012 | -0.012 | 0.13 | 0.66 | 0.76 | 0.64 | 0.64 | 0.52 | 0.71 |
| 3 | 48413537 | rs1459242 | -0.369 | -1.446 | 0.172 | -0.888 | 0.01 | -0.004 | -0.004 | 0 | 0.64 | 0.6 | 0.62 | 0.66 | 0.53 | 0.69 |
| 3 | 48418214 | rs2362452 | 0.484 | -1.994 | 0.489 | -1.04 | 0.011 | 0.076 | 0.076 | 0.19 | 0.5 | 0.25 | 0.58 | 0.66 | 0.52 | 0.69 |
| 3 | 48419897 | rs6442117 | -0.586 | -2.017 | 0.478 | -1.047 | 0.012 | -0.004 | -0.004 | 0 | 0.64 | 0.6 | 0.62 | 0.66 | 0.53 | 0.69 |
| 3 | 48436794 | rs1563736 | 0.449 | -1.931 | 0.516 | -1.015 | -0.012 | 0.003 | 0.003 | 0 | 0.36 | 0.3 | 0.37 | 0.34 | 0.46 | 0.31 |
| 3 | 48440963 | rs7642022 | 0.364 | -1.953 | 0.616 | -1.025 | -0.049 | -0.051 | -0.051 | 0.093 | 0.1 | 0.17 | 0.02 | 0 | 0 | 0 |
| 3 | 48449956 | rs10470686 | 0.428 | -1.872 | 0.284 | -0.989 | -0.012 | 0.003 | 0.003 | 0 | 0.36 | 0.3 | 0.37 | 0.34 | 0.46 | 0.31 |
| 3 | 48454468 | rs3214041 | 0.432 | -1.867 | 0.287 | -0.987 | -0.012 | 0.003 | 0.003 | 0 | 0.36 | 0.3 | 0.37 | 0.34 | 0.46 | 0.31 |
| 3 | 48455626 | rs6794875 | -0.019 | -1.913 | 0.271 | -1.015 | 0.004 | -0.014 | -0.014 | 0 | 0.41 | 0.51 | 0.32 | 0.32 | 0.47 | 0.31 |
| 3 | 48473204 | rs2290822 | 0.005 | -1.808 | 0.117 | -0.969 | 0.002 | -0.007 | -0.007 | 0 | 0.4 | 0.46 | 0.31 | 0.31 | 0.46 | 0.31 |
| 3 | 48474249 | rs2279077 | 0.388 | -1.805 | 0.109 | -0.968 | -0.023 | 0.018 | 0.018 | 0 | 0.37 | 0.24 | 0.29 | 0.31 | 0.46 | 0.31 |
| 3 | 48481647 | rs3774808 | 0.418 | -1.805 | 0.109 | -0.968 | -0.023 | 0.017 | 0.017 | 0 | 0.36 | 0.24 | 0.29 | 0.31 | 0.46 | 0.31 |
| 3 | 48483432 | rs9849509 | 0.014 | -1.791 | 0.123 | -0.961 | 0.002 | -0.007 | -0.007 | 0.341 | 0.4 | 0.46 | 0.31 | 0.31 | 0.46 | 0.31 |
| 3 | 48487048 | rs730566 | 1.329 | -2.087 | -0.108 | -1.237 | -0.033 | -0.07 | -0.07 | 0 | 0.24 | 0.07 | 0.4 | 0.61 | 0.25 | 0.36 |
| 3 | 48487353 | rs9838618 | 0.154 | -1.636 | 0.293 | -0.939 | -0.023 | 0.018 | 0.018 | 0 | 0.37 | 0.24 | 0.29 | 0.31 | 0.46 | 0.31 |
| 3 | 48505964 | rs2242150 | -0.098 | -1.611 | 0.268 | -0.928 | 0.002 | -0.007 | -0.007 | 0 | 0.4 | 0.47 | 0.31 | 0.31 | 0.46 | 0.31 |
| 3 | 48515079 | rs4858795 | 0.602 | -1.297 | -0.068 | -0.709 | 0.012 | 0.074 | 0.074 | 0.152 | 0.49 | 0.23 | 0.64 | 0.66 | 0.54 | 0.7 |
| 3 | 48520289 | rs9311428 | 0.095 | -0.602 | -0.431 | -0.38 | -0.005 | -0.001 | -0.001 | 0.032 | 0.32 | 0.31 | 0.3 | 0.34 | 0.44 | 0.25 |
| 3 | 48541016 | rs7434077 | -0.861 | -1.507 | -0.067 | -0.506 | -0.101 | -0.045 | -0.045 | 0 | 0.23 | 0.55 | 0.08 | 0.15 | 0.05 | 0.09 |
| 3 | 48556339 | rs2267846 | 0.782 | -1.464 | -0.782 | -0.563 | -0.038 | 0.062 | 0.062 | 0 | 0.28 | 0.06 | 0.17 | 0.15 | 0.26 | 0.34 |
| 3 | 48564507 | rs6806059 | -0.765 | -1.417 | 0.32 | -0.819 | -0.098 | 0.011 | 0.011 | 0.884 | 0.15 | 0.54 | 0.08 | 0.15 | 0.03 | 0.04 |
| 3 | 48583650 | rs9810991 | -0.893 | -1.533 | 0.115 | -0.869 | -0.116 | -0.004 | -0.004 | 0 | 0.17 | 0.47 | 0.07 | 0.14 | 0.03 | 0.04 |
| 3 | 48591481 | rs2267844 | -1.017 | -1.512 | -0.443 | -0.963 | -0.168 | 0.025 | 0.025 | 0 | 0.7 | 0.96 | 0.73 | 0.81 | 0.54 | 0.78 |
| 3 | 48599898 | rs9839959 | -0.487 | -1.249 | 0.708 | -0.462 | -0.055 | -0.058 | -0.058 | 4.42 | 0.1 | 0.22 | 0.03 | 0 | 0.01 | 0 |
| 3 | 48606701 | rs13060933 | -0.64 | -1.062 | 0.564 | -0.386 | -0.015 | 0.078 | 0.078 | 0 | 0.05 | 0.21 | 0.04 | 0.14 | 0.03 | 0.04 |
| 3 | 48618666 | rs1003649 | -1.152 | -1.256 | 0.351 | -0.475 | -0.112 | -0.001 | -0.001 | 0 | 0.16 | 0.47 | 0.08 | 0.13 | 0.03 | 0.04 |
| 3 | 48624611 | rs2255532 | -1.355 | -0.626 | -0.22 | -0.768 | 0.014 | 0.064 | 0.064 | 0.308 | 0.34 | 0.85 | 0.25 | 0.19 | 0.31 | 0.36 |
| 3 | 48628014 | rs2228561 | -0.13 | -0.415 | -0.214 | -0.54 | -0.032 | 0.001 | 0.001 | 0 | 0.06 | 0.03 | 0.07 | 0.04 | 0.12 | 0.25 |
| 3 | 48644202 | rs34591495 | -0.023 | -0.398 | -0.006 | -0.569 | -0.031 | -0.001 | -0.001 | 0 | 0.06 | 0.01 | 0.07 | 0.03 | 0.12 | 0.25 |
| 3 | 48651891 | rs1387005 | -0.424 | -0.66 | 0.617 | -0.595 | -0.023 | -0.026 | -0.026 | 0 | 0.07 | 0.22 | 0.03 | 0 | 0.01 | 0 |
| 3 | 48658467 | rs3895736 | -1.222 | -0.155 | 0.781 | -0.735 | 0.019 | 0.002 | 0.002 | 0 | 0.11 | 0.19 | 0.13 | 0.06 | 0.17 | 0.2 |
| 3 | 48659009 | rs9834639 | -1.346 | -0.25 | 1.06 | -0.772 | 0.034 | 0.011 | 0.011 | 0 | 0.08 | 0.27 | 0.09 | 0.02 | 0.08 | 0.02 |
| 3 | 48661985 | rs13063312 | -0.921 | -0.231 | 0.97 | -0.739 | 0.013 | 0.031 | 0.031 | 0 | 0.07 | 0.16 | 0.08 | 0.13 | 0.11 | 0.04 |
| 3 | 48663046 | rs7631574 | -2.132 | -0.345 | 1.201 | -0.786 | -0.041 | -0.013 | -0.013 | 0 | 0.16 | 0.54 | 0.13 | 0.13 | 0.12 | 0.04 |
| 3 | 48666923 | rs2276852 | -2.322 | -0.374 | 1.22 | -0.798 | -0.047 | -0.02 | -0.02 | 0.093 | 0.17 | 0.53 | 0.14 | 0.13 | 0.12 | 0.04 |
| 3 | 48668394 | rs3821876 | -0.891 | -0.267 | 1.093 | -0.754 | 0.013 | 0.03 | 0.03 | 0 | 0.07 | 0.16 | 0.08 | 0.13 | 0.11 | 0.04 |
| 3 | 48669447 | rs13324142 | -0.907 | -0.269 | 1.093 | -0.754 | 0.015 | 0.032 | 0.032 | 0 | 0.07 | 0.16 | 0.08 | 0.13 | 0.11 | 0.04 |
| 3 | 48669648 | rs2276850 | -1.487 | -0.397 | 1.055 | -0.811 | 0.002 | 0.016 | 0.016 | 0 | 0.09 | 0.16 | 0.08 | 0.13 | 0.11 | 0.04 |
| 3 | 48689192 | rs2286652 | -0.982 | -0.281 | 1.029 | -0.77 | 0.011 | 0.019 | 0.019 | 0 | 0.07 | 0.13 | 0.08 | 0.12 | 0.12 | 0.04 |
| 3 | 48689787 | rs12107418 | -2.132 | -0.373 | 1.191 | -0.809 | -0.032 | -0.011 | -0.011 | 0 | 0.16 | 0.48 | 0.13 | 0.12 | 0.13 | 0.04 |
| 3 | 48691316 | rs12107252 | -2.315 | -0.389 | 1.227 | -0.816 | -0.038 | -0.019 | -0.019 | 0.21 | 0.16 | 0.51 | 0.13 | 0.12 | 0.13 | 0.04 |
| 3 | 48705934 | rs13070798 | -1.082 | -0.783 | 1.112 | -1.07 | 0.042 | 0.064 | 0.064 | 0 | 0.08 | 0.26 | 0.09 | 0.12 | 0.12 | 0.04 |
| 3 | 48712791 | rs9836462 | -2.174 | -0.813 | 1 | -1.086 | -0.045 | -0.026 | -0.026 | 0.135 | 0.17 | 0.51 | 0.13 | 0.12 | 0.13 | 0.04 |
| 3 | 48730116 | rs12493578 | 0.846 | -0.533 | 0.347 | -1.851 | 0.046 | -0.116 | -0.116 | 0 | 0.73 | 0.56 | 0.75 | 0.95 | 0.65 | 0.93 |
| 3 | 48731487 | rs3172494 | 0.041 | -0.839 | 0.431 | -2.286 | 0.001 | -0.032 | -0.032 | 0 | 0.18 | 0.18 | 0.05 | 0 | 0.12 | 0.14 |
| 3 | 48735706 | rs11719291 | -0.42 | -0.729 | 0.603 | -2.169 | 0.064 | 0.02 | 0.02 | 0 | 0.08 | 0.3 | 0.09 | 0.05 | 0.13 | 0.04 |
| 3 | 48738678 | rs11706983 | -0.069 | -0.731 | 0.647 | -2.17 | 0.061 | 0.021 | 0.021 | 0 | 0.07 | 0.22 | 0.09 | 0.05 | 0.12 | 0.04 |
| 3 | 48749358 | rs9882549 | -0.416 | -0.73 | 0.623 | -2.17 | 0.063 | 0.019 | 0.019 | 0 | 0.08 | 0.27 | 0.09 | 0.05 | 0.12 | 0.04 |
| 3 | 48759562 | rs10865953 | -0.255 | -0.72 | 0.526 | -2.185 | -0.002 | -0.032 | -0.032 | 0.056 | 0.81 | 0.86 | 0.85 | 1 | 0.78 | 0.96 |
| 3 | 48787219 | rs6791542 | 0.956 | -0.994 | 0.594 | -2.331 | 0.017 | -0.093 | -0.093 | 0.867 | 0.69 | 0.45 | 0.74 | 0.95 | 0.66 | 0.92 |
| 3 | 48816725 | rs13074318 | 0.34 | -0.955 | 0.584 | -2.34 | -0.032 | -0.008 | -0.008 | 1.505 | 0.06 | 0 | 0.06 | 0.03 | 0.11 | 0.04 |
| 3 | 48819376 | rs12496747 | 1.491 | -1.4 | -0.155 | -2.661 | 0.014 | -0.076 | -0.076 | 0 | 0.16 | 0.01 | 0.38 | 0.74 | 0.07 | 0.35 |
| 3 | 48833615 | rs11713694 | 0.433 | -0.911 | 0.298 | -2.577 | -0.033 | -0.013 | -0.013 | 0 | 0.06 | 0.01 | 0.06 | 0.05 | 0.11 | 0.04 |
| 3 | 48844077 | rs11130176 | 0.567 | -0.935 | 0.402 | -2.582 | 0.031 | -0.086 | -0.086 | 0 | 0.72 | 0.55 | 0.76 | 0.95 | 0.67 | 0.92 |
| 3 | 48891275 | rs10780033 | 0.57 | -0.868 | 0.394 | -2.668 | 0.031 | -0.086 | -0.086 | 0 | 0.72 | 0.55 | 0.76 | 0.95 | 0.67 | 0.92 |
| 3 | 48893780 | rs9840684 | 0.546 | -0.868 | 0.404 | -2.669 | 0.027 | -0.08 | -0.08 | 0 | 0.73 | 0.61 | 0.77 | 0.95 | 0.67 | 0.92 |
| 3 | 48906469 | rs4974079 | 0.584 | -0.881 | 0.394 | -2.692 | 0.027 | -0.08 | -0.08 | 0 | 0.73 | 0.61 | 0.77 | 0.95 | 0.67 | 0.92 |
| 3 | 48937790 | rs11924597 | 0.607 | -0.874 | 0.381 | -2.706 | 0.031 | -0.086 | -0.086 | 0 | 0.72 | 0.55 | 0.76 | 0.95 | 0.67 | 0.92 |
| 3 | 48961726 | rs13087930 | 0.572 | -0.725 | 0.401 | -2.755 | -0.033 | -0.015 | -0.015 | 0.606 | 0.06 | 0.01 | 0.06 | 0.05 | 0.11 | 0.04 |

| | | | | | | | | | | | | | | | | |
|---|---|---|---|---|---|---|---|---|---|---|---|---|---|---|---|---|
| 3 | 48972892 | rs1134591 | 0.665 | -0.765 | 0.396 | -2.774 | 0.034 | -0.086 | -0.086 | 0 | 0.73 | 0.55 | 0.76 | 0.95 | 0.67 | 0.92 |
| 3 | 48982732 | rs9864243 | -0.594 | -0.766 | 0.419 | -2.774 | -0.024 | 0.026 | 0.026 | 0 | 0.82 | 0.9 | 0.86 | 1 | 0.78 | 0.96 |
| 3 | 48999323 | rs4974082 | 0.961 | -0.776 | 0.418 | -2.776 | 0.02 | -0.085 | -0.085 | 0 | 0.7 | 0.44 | 0.74 | 0.95 | 0.66 | 0.92 |
| 3 | 49040462 | rs6772452 | -0.043 | -0.766 | 0.239 | -2.754 | 0.034 | -0.118 | -0.118 | 0 | 0.79 | 0.53 | 0.83 | 0.98 | 0.78 | 0.96 |
| 3 | 49044767 | rs6781790 | 0.67 | -0.744 | 0.219 | -2.766 | 0.034 | -0.086 | -0.086 | 0 | 0.73 | 0.55 | 0.76 | 0.95 | 0.66 | 0.92 |
| 3 | 49053219 | rs7100 | -0.61 | -0.623 | 0.22 | -2.765 | -0.023 | 0.033 | 0.033 | 0 | 0.82 | 0.9 | 0.86 | 1 | 0.79 | 0.96 |
| 3 | 49054692 | rs3087866 | -0.61 | -0.623 | 0.22 | -2.765 | -0.023 | 0.033 | 0.033 | 0 | 0.82 | 0.9 | 0.86 | 1 | 0.79 | 0.96 |
| 3 | 49064110 | rs11706052 | 0.567 | -0.651 | 0.22 | -2.742 | -0.033 | -0.015 | -0.015 | 1.515 | 0.06 | 0.01 | 0.06 | 0.05 | 0.12 | 0.04 |
| 3 | 49113668 | rs35283189 | 0.573 | -0.642 | 0.22 | -2.746 | -0.033 | -0.015 | -0.015 | 0 | 0.06 | 0.01 | 0.06 | 0.05 | 0.12 | 0.04 |
| 3 | 49113713 | rs34580506 | 0.573 | -0.642 | 0.22 | -2.746 | -0.033 | -0.015 | -0.015 | 0 | 0.06 | 0.01 | 0.06 | 0.05 | 0.12 | 0.04 |
| 3 | 49133310 | rs9840050 | 0.637 | -0.654 | 0.19 | -2.766 | 0.034 | -0.084 | -0.084 | 0 | 0.73 | 0.55 | 0.76 | 0.95 | 0.66 | 0.92 |
| 3 | 49139406 | rs35673421 | 0.583 | -0.714 | 0.19 | -2.762 | -0.033 | -0.015 | -0.015 | 0.781 | 0.06 | 0.01 | 0.06 | 0.05 | 0.12 | 0.04 |
| 3 | 49141116 | rs5030795 | 0.583 | -0.714 | 0.19 | -2.762 | -0.033 | -0.015 | -0.015 | 0.024 | 0.06 | 0.01 | 0.06 | 0.05 | 0.12 | 0.04 |
| 3 | 49145304 | rs4955411 | -0.212 | -0.609 | 0.198 | -2.765 | 0.026 | -0.124 | -0.124 | 0 | 0.8 | 0.65 | 0.84 | 1 | 0.8 | 0.96 |
| 3 | 49145741 | rs3212 | 0.633 | -0.61 | 0.198 | -2.765 | 0.031 | -0.084 | -0.084 | 0.224 | 0.73 | 0.55 | 0.77 | 0.95 | 0.68 | 0.92 |
| 3 | 49148890 | rs4292260 | -0.176 | -0.628 | 0.191 | -2.773 | 0.025 | -0.124 | -0.124 | 0.916 | 0.8 | 0.65 | 0.84 | 1 | 0.8 | 0.96 |
| 3 | 49156473 | rs11552724 | 0.638 | -0.642 | 0.229 | -2.778 | -0.033 | -0.016 | -0.016 | 1.34 | 0.06 | 0.01 | 0.06 | 0.05 | 0.12 | 0.05 |
| 3 | 49156676 | rs4279134 | 0.721 | -0.647 | 0.237 | -2.779 | 0.029 | -0.091 | -0.091 | 6.345 | 0.72 | 0.53 | 0.77 | 0.95 | 0.68 | 0.92 |
| 3 | 49157771 | rs6779394 | 0.62 | -0.617 | 0.231 | -2.791 | 0.025 | -0.081 | -0.081 | 0.54 | 0.73 | 0.62 | 0.77 | 0.95 | 0.68 | 0.92 |
| 3 | 49162284 | rs34759087 | 0.554 | -0.66 | 0.023 | -2.961 | 0.013 | 0.107 | 0.107 | 0 | 0.17 | 0.01 | 0.04 | 0 | 0.11 | 0.14 |
| 3 | 49194331 | rs11715581 | 0.967 | -0.074 | 0.12 | -2.973 | 0.012 | -0.109 | -0.109 | 0.068 | 0.66 | 0.49 | 0.71 | 0.94 | 0.64 | 0.9 |
| 3 | 49196132 | rs12490393 | 0.931 | -0.074 | 0.121 | -2.973 | 0.013 | -0.112 | -0.112 | 0.085 | 0.66 | 0.53 | 0.72 | 0.94 | 0.65 | 0.9 |
| 3 | 49200137 | rs1134043 | -1.584 | -0.08 | 0.143 | -2.979 | -0.038 | -0.091 | -0.091 | 0 | 0.22 | 0.54 | 0.17 | 0.04 | 0.15 | 0.06 |
| 3 | 49200627 | rs4955418 | 0.503 | 0.104 | 0.264 | -2.844 | 0.014 | -0.116 | -0.116 | 0 | 0.67 | 0.57 | 0.72 | 0.94 | 0.65 | 0.9 |
| 3 | 49210732 | rs7617480 | -0.185 | 0.347 | 0.281 | -2.74 | 0.001 | -0.121 | -0.121 | 3.081 | 0.75 | 0.65 | 0.81 | 0.99 | 0.79 | 0.95 |
| 3 | 49211989 | rs10865954 | 0.7 | 0.346 | 0.288 | -2.747 | 0.003 | -0.085 | -0.085 | 0 | 0.65 | 0.45 | 0.73 | 0.94 | 0.67 | 0.9 |
| 3 | 49213030 | rs11713297 | 0.297 | 0.337 | 0.243 | -2.737 | 0.003 | -0.003 | -0.003 | 0.405 | 0.08 | 0.08 | 0.07 | 0.05 | 0.11 | 0.04 |
| 3 | 49213637 | rs9586 | -0.151 | 0.453 | 0.3 | -2.742 | -0.008 | -0.128 | -0.128 | 0 | 0.74 | 0.63 | 0.81 | 0.99 | 0.79 | 0.95 |
| 3 | 49215583 | rs9837237 | 0.181 | 0.216 | 0.361 | -2.717 | 0.012 | -0.006 | -0.006 | 0 | 0.08 | 0.11 | 0.08 | 0.04 | 0.11 | 0.04 |
| 3 | 49215966 | rs9837625 | 0.319 | 0.24 | 0.257 | -2.705 | 0.004 | -0.003 | -0.003 | 0 | 0.08 | 0.08 | 0.07 | 0.04 | 0.11 | 0.04 |
| 3 | 49228946 | rs11130183 | -0.105 | 0.305 | 0.339 | -2.74 | 0.007 | -0.123 | -0.123 | 2.225 | 0.74 | 0.6 | 0.79 | 0.99 | 0.76 | 0.94 |
| 3 | 49237334 | rs12637576 | 0.593 | 0.32 | 0.342 | -2.744 | 0.018 | -0.113 | -0.113 | 0 | 0.66 | 0.48 | 0.72 | 0.94 | 0.64 | 0.9 |
| 3 | 49237394 | rs12629759 | 0.624 | 0.298 | 0.352 | -2.748 | 0.007 | -0.084 | -0.084 | 0 | 0.65 | 0.41 | 0.71 | 0.94 | 0.64 | 0.9 |
| 3 | 49245077 | rs34566463 | 0.306 | 0.306 | 0.306 | -2.712 | 0.007 | -0.002 | -0.002 | 0.118 | 0.08 | 0.14 | 0.08 | 0.05 | 0.11 | 0.04 |
| 3 | 49245201 | rs11719220 | 0.68 | 0.308 | 0.296 | -2.711 | 0.004 | -0.001 | -0.001 | 0 | 0.07 | 0.06 | 0.07 | 0.04 | 0.11 | 0.04 |
| 3 | 49273996 | rs12631989 | 0.704 | 0.299 | 0.32 | -2.741 | 0.009 | -0.1 | -0.1 | 0 | 0.65 | 0.43 | 0.71 | 0.94 | 0.64 | 0.9 |
| 3 | 49275828 | rs11130186 | -0.107 | 0.322 | 0.294 | -2.733 | 0.005 | -0.118 | -0.118 | 0 | 0.74 | 0.61 | 0.79 | 0.99 | 0.76 | 0.94 |
| 3 | 49276077 | rs10154992 | -0.107 | 0.322 | 0.294 | -2.733 | 0.005 | -0.118 | -0.118 | 0.008 | 0.74 | 0.61 | 0.79 | 0.99 | 0.76 | 0.94 |
| 3 | 49294250 | rs13068038 | 0.731 | 0.34 | 0.155 | -2.714 | 0.003 | -0.001 | -0.001 | 0 | 0.07 | 0.06 | 0.07 | 0.04 | 0.11 | 0.04 |
| 3 | 49298139 | rs7626076 | -0.446 | 0.319 | 0.206 | -2.736 | -0.024 | 0.064 | 0.064 | 0 | 0.76 | 0.86 | 0.82 | 0.99 | 0.76 | 0.94 |
| 3 | 49309684 | rs10049462 | 0.712 | 0.229 | 0.137 | -2.66 | 0.003 | -0.001 | -0.001 | 0 | 0.07 | 0.06 | 0.07 | 0.04 | 0.11 | 0.04 |
| 3 | 49313978 | rs13077498 | 0.712 | 0.229 | 0.137 | -2.66 | 0.003 | -0.001 | -0.001 | 0 | 0.07 | 0.06 | 0.07 | 0.04 | 0.11 | 0.04 |
| 3 | 49317338 | rs6778080 | -0.169 | 0.178 | 0.283 | -2.694 | -0.001 | -0.046 | -0.046 | 0.177 | 0.74 | 0.71 | 0.8 | 0.99 | 0.76 | 0.94 |
| 3 | 49326178 | rs9874474 | 0.007 | 0.738 | 0.067 | -2.814 | 0.029 | -0.02 | -0.02 | 0 | 0.12 | 0.26 | 0.11 | 0.05 | 0.2 | 0.05 |
| 3 | 49336232 | rs11720964 | 0.965 | 0.529 | -0.068 | -2.893 | 0.006 | 0.055 | 0.055 | 0 | 0.52 | 0.32 | 0.64 | 0.93 | 0.54 | 0.78 |
| 3 | 49361791 | rs13078949 | 1.376 | 0.48 | -0.366 | -2.846 | -0.037 | 0.173 | 0.173 | 0 | 0.41 | 0.13 | 0.6 | 0.93 | 0.44 | 0.74 |
| 3 | 49389842 | rs17080528 | -0.205 | 0.461 | -0.153 | -2.742 | 0.002 | 0.01 | 0.01 | 0 | 0.31 | 0.27 | 0.17 | 0.07 | 0.34 | 0.2 |
| 3 | 49390250 | rs17650792 | 1.247 | 0.483 | -0.055 | -2.838 | -0.006 | 0.05 | 0.05 | 0 | 0.42 | 0.27 | 0.61 | 0.91 | 0.43 | 0.74 |
| 3 | 49396751 | rs3448 | -0.517 | 0.441 | 0.104 | -2.843 | 0.002 | -0.068 | -0.068 | 0 | 0.76 | 0.72 | 0.8 | 0.99 | 0.77 | 0.94 |
| 3 | 49400482 | rs974495 | -0.48 | 0.436 | 0.251 | -2.843 | 0.012 | -0.121 | -0.121 | 0.099 | 0.76 | 0.6 | 0.79 | 0.98 | 0.77 | 0.94 |
| 3 | 49441091 | rs11706370 | -0.176 | 0.358 | 0.274 | -2.944 | 0.002 | 0.028 | 0.028 | 0 | 0.3 | 0.21 | 0.17 | 0.07 | 0.34 | 0.2 |
| 3 | 49453834 | rs6997 | -0.157 | 0.345 | 0.204 | -2.95 | 0.002 | 0.01 | 0.01 | 0 | 0.31 | 0.27 | 0.17 | 0.07 | 0.34 | 0.2 |
| 3 | 49454277 | rs10640 | -0.118 | 0.345 | 0.206 | -2.95 | 0.003 | 0.023 | 0.023 | 0 | 0.3 | 0.23 | 0.17 | 0.07 | 0.34 | 0.2 |
| 3 | 49455330 | rs11715915 | -0.172 | 0.345 | 0.205 | -2.95 | 0.002 | 0.028 | 0.028 | 0 | 0.3 | 0.21 | 0.17 | 0.07 | 0.34 | 0.2 |
| 3 | 49458266 | rs1464568 | 1.375 | 0.339 | 0.193 | -2.961 | -0.006 | 0.05 | 0.05 | 0 | 0.42 | 0.26 | 0.61 | 0.91 | 0.43 | 0.74 |
| 3 | 49459376 | rs1464566 | 1.253 | 0.339 | 0.192 | -2.961 | -0.001 | 0.059 | 0.059 | 0 | 0.45 | 0.31 | 0.62 | 0.91 | 0.43 | 0.74 |
| 3 | 49460350 | rs1464569 | -0.579 | 0.327 | 0.218 | -2.967 | 0.014 | -0.12 | -0.12 | 0.12 | 0.76 | 0.6 | 0.79 | 0.98 | 0.77 | 0.94 |
| 3 | 49499829 | rs10865955 | 1.508 | 0.291 | 0.264 | -2.983 | -0.014 | 0.081 | 0.081 | 0 | 0.41 | 0.21 | 0.61 | 0.91 | 0.43 | 0.74 |
| 3 | 49507847 | rs4855839 | 1.265 | 0.329 | 0.26 | -2.966 | 0 | 0.06 | 0.06 | 0 | 0.45 | 0.31 | 0.62 | 0.91 | 0.43 | 0.74 |
| 3 | 49538932 | rs6766131 | -0.176 | 0.299 | 0.137 | -2.884 | 0.002 | 0.01 | 0.01 | 0 | 0.31 | 0.27 | 0.17 | 0.07 | 0.33 | 0.2 |
| 3 | 49557857 | rs3870336 | 1.762 | -0.186 | -0.279 | -2.988 | 0.027 | -0.003 | -0.003 | 0 | 0.22 | 0.1 | 0.39 | 0.77 | 0.08 | 0.42 |
| 3 | 49570882 | rs1050088 | 0.767 | -0.092 | -0.262 | -3.097 | -0.01 | 0.095 | 0.095 | 0.034 | 0.43 | 0.21 | 0.61 | 0.91 | 0.43 | 0.74 |
| 3 | 49572140 | rs4625 | 0.2 | -0.121 | -0.249 | -3.183 | 0.002 | 0.01 | 0.01 | 0 | 0.31 | 0.27 | 0.17 | 0.07 | 0.33 | 0.2 |
| 3 | 49581559 | rs10865956 | -0.861 | -0.134 | -0.135 | -3.173 | 0.003 | -0.074 | -0.074 | 0 | 0.76 | 0.72 | 0.8 | 0.99 | 0.77 | 0.94 |
| 3 | 49590770 | rs3811697 | 0.229 | -0.151 | -0.149 | -3.169 | 0.002 | 0.008 | 0.008 | 0 | 0.3 | 0.27 | 0.18 | 0.07 | 0.33 | 0.2 |
| 3 | 49601255 | rs2312462 | -0.827 | -0.09 | -0.194 | -3.091 | 0.003 | -0.074 | -0.074 | 0 | 0.76 | 0.72 | 0.8 | 0.99 | 0.77 | 0.94 |
| 3 | 49656789 | rs11919311 | 0.699 | -0.021 | -0.186 | -2.835 | -0.004 | 0.016 | 0.016 | 0 | 0.48 | 0.31 | 0.67 | 0.93 | 0.5 | 0.76 |
| 3 | 49678651 | rs4283605 | 0.136 | -0.089 | -0.064 | -2.819 | 0.005 | 0.012 | 0.012 | 0 | 0.3 | 0.26 | 0.17 | 0.06 | 0.32 | 0.2 |
| 3 | 49680973 | rs10452032 | -1.062 | -0.092 | -0.044 | -2.823 | 0.007 | -0.086 | -0.086 | 0.01 | 0.8 | 0.71 | 0.86 | 0.99 | 0.83 | 0.96 |

| | | | | | | | | | | | | | | | | |
|---|---|---|---|---|---|---|---|---|---|---|---|---|---|---|---|---|
| 3 | 49685592 | rs1078341 | -1.062 | -0.092 | -0.044 | -2.823 | 0.007 | -0.086 | -0.086 | 0.053 | 0.8 | 0.71 | 0.86 | 0.99 | 0.83 | 0.96 |
| 3 | 49690199 | rs4855885 | -1.403 | -0.168 | 0.118 | -2.832 | -0.018 | 0.034 | 0.034 | 0 | 0.83 | 0.9 | 0.88 | 0.99 | 0.83 | 0.96 |
| 3 | 49701298 | rs2005557 | 0.538 | -0.159 | 0.11 | -2.828 | 0.002 | -0.003 | -0.003 | 0 | 0.5 | 0.4 | 0.68 | 0.93 | 0.51 | 0.76 |
| 3 | 49701983 | rs9858542 | 0.299 | -0.175 | 0.1 | -2.833 | 0.005 | 0.025 | 0.025 | 0 | 0.29 | 0.22 | 0.17 | 0.05 | 0.32 | 0.2 |
| 3 | 49708502 | rs1060962 | 0.706 | -0.173 | 0.101 | -2.83 | 0 | 0.021 | 0.021 | 0 | 0.48 | 0.32 | 0.68 | 0.93 | 0.51 | 0.76 |
| 3 | 49708769 | rs1060970 | 1.785 | -0.04 | -0.062 | -2.845 | 0.022 | 0.014 | 0.014 | 0 | 0.27 | 0.11 | 0.47 | 0.78 | 0.2 | 0.44 |
| 3 | 49712297 | rs11130213 | 0.105 | 0.015 | -0.125 | -2.78 | 0.006 | 0.021 | 0.021 | 0 | 0.3 | 0.24 | 0.17 | 0.05 | 0.32 | 0.2 |
| 3 | 49714225 | rs1131095 | 0.105 | 0.017 | -0.124 | -2.777 | 0.006 | 0.021 | 0.021 | 0 | 0.3 | 0.24 | 0.17 | 0.06 | 0.32 | 0.2 |
| 3 | 49715354 | rs4855882 | -1.417 | 0.046 | -0.137 | -2.758 | -0.015 | 0.024 | 0.024 | 0 | 0.83 | 0.88 | 0.88 | 0.99 | 0.83 | 0.96 |
| 3 | 49721532 | rs3197999 | 0.112 | 0.08 | -0.161 | -2.611 | 0.006 | 0.026 | 0.026 | 0 | 0.29 | 0.22 | 0.17 | 0.06 | 0.31 | 0.19 |
| 3 | 49731861 | rs9858213 | 0.247 | 0.098 | -0.16 | -2.527 | 0.002 | 0.018 | 0.018 | 0 | 0.28 | 0.22 | 0.17 | 0.05 | 0.32 | 0.2 |
| 3 | 49736172 | rs11130216 | 0.527 | 0.473 | 0.299 | -2.25 | 0.015 | -0.009 | -0.009 | 0 | 0.21 | 0.21 | 0.2 | 0.15 | 0.31 | 0.32 |
| 3 | 49737323 | rs11130217 | -0.8 | 0.409 | 0.39 | -2.333 | 0.006 | -0.085 | -0.085 | 0 | 0.8 | 0.71 | 0.86 | 0.99 | 0.83 | 0.96 |
| 3 | 49738945 | rs6804655 | 0.588 | 0.404 | 0.387 | -2.334 | 0.016 | -0.009 | -0.009 | 0 | 0.21 | 0.21 | 0.2 | 0.15 | 0.31 | 0.32 |
| 3 | 49751585 | rs2291542 | 0.386 | 0.395 | 0.387 | -2.337 | 0.011 | -0.01 | -0.01 | 0.036 | 0.22 | 0.25 | 0.21 | 0.15 | 0.31 | 0.32 |
| 3 | 49752372 | rs11130218 | 0.658 | 0.377 | 0.412 | -2.343 | -0.005 | 0.035 | 0.035 | 0 | 0.56 | 0.33 | 0.64 | 0.83 | 0.52 | 0.64 |
| 3 | 49753788 | rs11720705 | -0.697 | 0.376 | 0.417 | -2.343 | 0.005 | -0.097 | -0.097 | 0 | 0.8 | 0.68 | 0.86 | 0.99 | 0.83 | 0.96 |
| 3 | 49754970 | rs7628207 | -0.675 | 0.374 | 0.418 | -2.344 | 0.005 | -0.096 | -0.096 | 0 | 0.79 | 0.67 | 0.86 | 0.99 | 0.83 | 0.96 |
| 3 | 49762662 | rs7634902 | 0.554 | 0.342 | 0.437 | -2.336 | -0.002 | -0.002 | -0.002 | 0 | 0.57 | 0.42 | 0.65 | 0.83 | 0.53 | 0.64 |
| 3 | 49763338 | rs1049256 | 0.554 | 0.342 | 0.437 | -2.336 | -0.002 | -0.002 | -0.002 | 0 | 0.57 | 0.42 | 0.65 | 0.83 | 0.53 | 0.64 |
| 3 | 49776178 | rs11714957 | 0.55 | 0.368 | 0.438 | -2.325 | 0.015 | -0.009 | -0.009 | 0 | 0.21 | 0.21 | 0.2 | 0.15 | 0.31 | 0.32 |
| 3 | 49792023 | rs9872864 | 0.658 | 0.378 | 0.419 | -2.288 | -0.005 | 0.033 | 0.033 | 0 | 0.56 | 0.33 | 0.64 | 0.84 | 0.53 | 0.64 |
| 3 | 49834571 | rs6809879 | 0.731 | 0.371 | 0.422 | -2.343 | 0.011 | -0.073 | -0.073 | 0 | 0.18 | 0.27 | 0.13 | 0.02 | 0.16 | 0.03 |
| 3 | 49836707 | rs7629936 | 0.424 | 0.678 | 0.382 | -2.385 | 0.011 | -0.018 | -0.018 | 0.843 | 0.21 | 0.26 | 0.18 | 0.15 | 0.24 | 0.31 |
| 3 | 49842625 | rs2234391 | -0.682 | 0.593 | 0.362 | -2.371 | 0.001 | -0.024 | -0.024 | 0.172 | 0.42 | 0.54 | 0.35 | 0.17 | 0.47 | 0.38 |
| 3 | 49847623 | rs1799845 | 0.344 | 0.608 | 0.554 | -2.379 | -0.016 | 0.013 | 0.013 | 0.369 | 0.22 | 0.14 | 0.2 | 0.15 | 0.31 | 0.34 |
| 3 | 49849723 | rs1799843 | 0.147 | 0.612 | 0.54 | -2.377 | 0.019 | -0.022 | -0.022 | 0 | 0.23 | 0.26 | 0.21 | 0.15 | 0.31 | 0.34 |
| 3 | 49853073 | rs6775384 | -0.669 | 0.562 | 0.545 | -2.374 | 0 | -0.013 | -0.013 | 0 | 0.42 | 0.58 | 0.35 | 0.17 | 0.47 | 0.34 |
| 3 | 49860854 | rs6446298 | -1.396 | 0.478 | 0.451 | -2.441 | 0.051 | 0.078 | 0.078 | 0 | 0.55 | 0.73 | 0.46 | 0.22 | 0.66 | 0.46 |
| 3 | 49877585 | rs9858428 | 1.23 | 0.983 | 0.242 | -1.969 | 0.001 | -0.001 | -0.001 | 0 | 0.06 | 0.05 | 0.07 | 0.03 | 0.11 | 0.04 |
| 3 | 49878652 | rs1317140 | -0.227 | 0.732 | 0.227 | -1.806 | 0.02 | -0.025 | -0.025 | 0 | 0.23 | 0.26 | 0.21 | 0.14 | 0.3 | 0.25 |
| 3 | 49898273 | rs2681781 | -0.522 | -0.033 | 0.153 | -1.698 | -0.006 | 0.009 | 0.009 | 0 | 0.4 | 0.69 | 0.37 | 0.14 | 0.49 | 0.21 |
| 3 | 49914397 | rs11712056 | -0.5 | -0.379 | 0.245 | -1.926 | -0.005 | 0.023 | 0.023 | 0.037 | 0.44 | 0.43 | 0.35 | 0.12 | 0.44 | 0.21 |
| 3 | 49924424 | rs11713193 | -0.365 | 0.371 | 0.463 | -1.563 | 0.022 | -0.065 | -0.065 | 0 | 0.37 | 0.53 | 0.36 | 0.14 | 0.5 | 0.21 |
| 3 | 49924940 | rs1062633 | -0.826 | 0.337 | 0.458 | -1.579 | -0.015 | 0.073 | 0.073 | 0 | 0.42 | 0.74 | 0.38 | 0.14 | 0.5 | 0.21 |
| 3 | 49936102 | rs2230590 | -0.847 | 0.188 | 0.211 | -1.538 | -0.013 | 0.065 | 0.065 | 0 | 0.43 | 0.74 | 0.38 | 0.14 | 0.5 | 0.21 |
| 3 | 49941436 | rs2280406 | -0.475 | 0.542 | 0.067 | -1.39 | 0.019 | -0.056 | -0.056 | 0 | 0.37 | 0.55 | 0.36 | 0.14 | 0.49 | 0.21 |
| 3 | 49949071 | rs868891 | -0.102 | 0.588 | 0.237 | -1.185 | -0.029 | 0.036 | 0.036 | 0.029 | 0.32 | 0.25 | 0.3 | 0.12 | 0.42 | 0.16 |
| 3 | 49956628 | rs11709573 | -0.096 | 0.571 | 0.212 | -1.186 | -0.029 | 0.035 | 0.035 | 0.003 | 0.31 | 0.25 | 0.3 | 0.12 | 0.42 | 0.16 |
| 3 | 49961780 | rs6774354 | -0.236 | 0.555 | 0.273 | -1.192 | 0.021 | -0.054 | -0.054 | 0 | 0.33 | 0.41 | 0.32 | 0.12 | 0.42 | 0.16 |
| 3 | 50085153 | rs17304079 | 0.432 | 1.144 | 0.252 | -1.373 | 0.015 | -0.055 | -0.055 | 0 | 0.19 | 0.23 | 0.2 | 0.01 | 0.26 | 0.05 |
| 3 | 50103917 | rs2023953 | 1.168 | 1.352 | 0.203 | -1.444 | 0.001 | 0 | 0 | 1.091 | 0.06 | 0.05 | 0.13 | 0.05 | 0.11 | 0.05 |
| 3 | 50113034 | rs2240327 | 0.845 | 1.373 | 0.416 | -1.533 | -0.003 | 0.001 | 0.001 | 0 | 0.6 | 0.31 | 0.62 | 0.86 | 0.49 | 0.79 |
| 3 | 50123417 | rs10049087 | 0.934 | 1.322 | 0.447 | -1.555 | -0.011 | 0.056 | 0.056 | 0 | 0.59 | 0.26 | 0.62 | 0.86 | 0.49 | 0.79 |
| 3 | 50128386 | rs2240326 | 0.941 | 1.315 | 0.444 | -1.558 | -0.011 | 0.056 | 0.056 | 0 | 0.59 | 0.26 | 0.62 | 0.86 | 0.49 | 0.79 |
| 3 | 50129399 | rs2013208 | 0.571 | 1.366 | 0.45 | -1.535 | 0.006 | -0.03 | -0.03 | 1.52 | 0.63 | 0.6 | 0.65 | 0.86 | 0.49 | 0.79 |
| 3 | 50132131 | rs2624834 | -0.964 | 1.545 | 0.221 | -1.523 | -0.052 | 0 | 0 | 0 | 0.87 | 0.98 | 0.89 | 0.89 | 0.82 | 0.88 |
| 3 | 50144951 | rs1061474 | 0.051 | 1.308 | 0.274 | -1.479 | -0.031 | 0.035 | 0.035 | 0 | 0.69 | 0.75 | 0.69 | 0.88 | 0.57 | 0.83 |
| 3 | 50153356 | rs1138536 | 0.04 | 1.31 | 0.275 | -1.478 | -0.031 | 0.034 | 0.034 | 0.006 | 0.69 | 0.75 | 0.69 | 0.88 | 0.57 | 0.83 |
| 3 | 50172397 | rs2014830 | 1.208 | 0.547 | -0.293 | -1.431 | 0.066 | 0.088 | 0.088 | 0 | 0.44 | 0.1 | 0.46 | 0.71 | 0.31 | 0.58 |
| 3 | 50174197 | rs2624847 | 0.066 | 0.912 | 0.025 | -1.609 | -0.001 | 0.002 | 0.002 | 0 | 0.79 | 0.76 | 0.84 | 0.87 | 0.75 | 0.84 |
| 3 | 50174572 | rs13100903 | -0.088 | 1.038 | 0.244 | -1.626 | 0.076 | -0.133 | -0.133 | 0 | 0.25 | 0.46 | 0.26 | 0.03 | 0.33 | 0.11 |
| 3 | 50178011 | rs739983 | 0.822 | 0.258 | -0.406 | -1.594 | 0 | -0.021 | -0.021 | 2.059 | 0.56 | 0.42 | 0.37 | 0.15 | 0.62 | 0.52 |
| 3 | 50191191 | rs2526392 | -0.451 | 0.577 | -0.539 | -0.898 | -0.007 | -0.009 | -0.009 | 0 | 0.87 | 0.93 | 0.88 | 0.88 | 0.88 | 0.87 |
| 3 | 50201924 | rs2188151 | -0.148 | -0.088 | -0.591 | -1.457 | 0.003 | -0.102 | -0.102 | 0 | 0.36 | 0.54 | 0.28 | 0.03 | 0.4 | 0.16 |
| 3 | 50207075 | rs2518796 | -0.233 | -0.106 | -0.578 | -1.465 | -0.001 | -0.051 | -0.051 | 0 | 0.37 | 0.57 | 0.28 | 0.03 | 0.4 | 0.16 |
| 3 | 50222693 | rs7642692 | -0.532 | -0.306 | -0.29 | -1.323 | -0.013 | 0.001 | 0.001 | 0.277 | 0.19 | 0.36 | 0.14 | 0.14 | 0.12 | 0.14 |
| 3 | 50224225 | rs12632110 | -0.304 | -0.275 | -0.387 | -0.933 | 0.014 | 0.028 | 0.028 | 0 | 0.7 | 0.57 | 0.48 | 0.37 | 0.66 | 0.88 |
| 3 | 50225029 | rs11711407 | 0.715 | -0.214 | -0.643 | -0.93 | -0.024 | 0.15 | 0.15 | 0.82 | 0.52 | 0.22 | 0.34 | 0.23 | 0.55 | 0.73 |
| 3 | 50233215 | rs11919418 | -0.15 | -0.139 | -0.501 | -0.832 | -0.002 | 0.028 | 0.028 | 0 | 0.34 | 0.72 | 0.28 | 0.03 | 0.33 | 0.12 |
| 3 | 50242387 | rs1858828 | -0.818 | -0.205 | -0.64 | -0.753 | -0.036 | 0.136 | 0.136 | 0.549 | 0.47 | 0.78 | 0.65 | 0.76 | 0.44 | 0.24 |
| 3 | 50243510 | rs2236939 | 0.34 | -0.169 | -0.599 | -0.746 | -0.003 | -0.003 | -0.003 | 0.058 | 0.13 | 0.09 | 0.1 | 0.13 | 0.11 | 0.12 |
| 3 | 50247824 | rs2236941 | -0.012 | -0.189 | -0.354 | -0.746 | 0.003 | 0.016 | 0.016 | 0 | 0.33 | 0.56 | 0.54 | 0.63 | 0.32 | 0.14 |
| 3 | 50255305 | rs739431 | -0.918 | -0.889 | -0.297 | -0.614 | -0.025 | -0.029 | -0.029 | 0 | 0.29 | 0.59 | 0.18 | 0.13 | 0.15 | 0.18 |
| 3 | 50267837 | rs9883222 | -0.099 | -0.941 | -0.252 | -0.563 | -0.095 | 0.152 | 0.152 | 0.646 | 0.18 | 0.57 | 0.35 | 0.62 | 0.04 | 0.06 |
| 3 | 50280881 | rs1894324 | 0.383 | -1.153 | -0.546 | -0.972 | -0.1 | -0.126 | -0.126 | 0.089 | 0.17 | 0.52 | 0.07 | 0 | 0.03 | 0.05 |
| 3 | 50291785 | rs2282751 | -0.753 | -0.887 | -0.674 | -0.727 | 0.058 | -0.012 | -0.012 | 0 | 0.3 | 0.76 | 0.44 | 0.68 | 0.14 | 0.19 |
| 3 | 50304845 | rs11916258 | 0.369 | -0.954 | -0.356 | -0.78 | -0.1 | -0.126 | -0.126 | 0.12 | 0.16 | 0.45 | 0.06 | 0 | 0.03 | 0.05 |
| 3 | 50306249 | rs3806708 | 0.485 | -0.67 | -0.55 | -0.726 | -0.023 | 0.012 | 0.012 | 0 | 0.12 | 0.02 | 0.08 | 0.07 | 0.12 | 0.12 |

| | | | | | | | | | | | | | | | | |
|---|---|---|---|---|---|---|---|---|---|---|---|---|---|---|---|---|
| 3 | 50316119 | rs6776145 | -0.308 | -0.733 | -0.401 | -0.594 | 0.007 | 0.072 | 0.072 | 0 | 0.16 | 0.25 | 0.38 | 0.69 | 0.13 | 0.13 |
| 3 | 50324672 | rs12629572 | -0.19 | -0.743 | -0.419 | -0.586 | 0.01 | 0.072 | 0.072 | 0 | 0.15 | 0.23 | 0.37 | 0.69 | 0.13 | 0.13 |
| 3 | 50329826 | rs1076872 | -1.454 | -0.803 | -1.515 | -1.216 | -0.083 | -0.042 | -0.042 | 0 | 0.71 | 0.99 | 0.83 | 0.78 | 0.68 | 0.52 |
| 3 | 50332697 | rs13100173 | 0.836 | -1.19 | -1.822 | -1.566 | -0.119 | 0.138 | 0.138 | 0 | 0.29 | 0.13 | 0.35 | 0.09 | 0.46 | 0.26 |
| 3 | 50340996 | rs28365992 | 1.154 | -0.402 | -1.861 | -1.41 | -0.045 | 0.004 | 0.004 | 0 | 0.12 | 0.01 | 0.08 | 0.07 | 0.12 | 0.12 |
| 3 | 50352458 | rs9877046 | -1.639 | -0.369 | -1.883 | -1.42 | -0.061 | -0.043 | -0.043 | 1.544 | 0.71 | 0.99 | 0.84 | 0.8 | 0.7 | 0.53 |
| 3 | 50357869 | rs709210 | -1.161 | -0.135 | -1.705 | -1.21 | -0.014 | 0.036 | 0.036 | 0.702 | 0.62 | 0.93 | 0.82 | 0.79 | 0.65 | 0.45 |
| 3 | 50369546 | rs2073498 | 0.728 | -0.144 | -1.001 | -0.521 | -0.045 | 0.006 | 0.006 | 0 | 0.11 | 0.01 | 0.08 | 0.06 | 0.12 | 0.12 |
| 3 | 50388607 | rs2233474 | -0.43 | -0.12 | -0.69 | -0.486 | 0.014 | -0.003 | -0.003 | 0 | 0.31 | 0.64 | 0.43 | 0.69 | 0.16 | 0.18 |
| 3 | 50421081 | rs2236951 | -0.507 | 0.314 | -0.761 | -0.614 | -0.005 | 0.001 | 0.001 | 0 | 0.3 | 0.59 | 0.46 | 0.7 | 0.21 | 0.19 |
| 3 | 50431194 | rs2298955 | -0.663 | 0.506 | -0.355 | 0.204 | 0.009 | 0.111 | 0.111 | 0 | 0.16 | 0.32 | 0.39 | 0.49 | 0.12 | 0.13 |
| 3 | 50440552 | rs17606778 | 1.04 | 0.895 | -1.081 | -0.261 | -0.052 | 0.062 | 0.062 | 0 | 0.11 | 0.01 | 0.1 | 0 | 0.18 | 0.06 |
| 3 | 50463935 | rs1540293 | 1.764 | 0.606 | -1.29 | 0.189 | -0.03 | -0.037 | -0.037 | 0 | 0.06 | 0 | 0.13 | 0.25 | 0.11 | 0.05 |
| 3 | 50468491 | rs2282757 | 1.111 | 0.476 | -0.851 | 0.118 | 0.065 | 0.155 | 0.155 | 0 | 0.06 | 0.37 | 0.03 | 0.06 | 0.01 | 0 |
| 3 | 50470047 | rs2236969 | 1.111 | 0.476 | -0.851 | 0.118 | 0.065 | 0.155 | 0.155 | 0 | 0.06 | 0.37 | 0.03 | 0.06 | 0.01 | 0 |
| 3 | 50478487 | rs12488468 | 0.12 | 0.637 | -1.234 | -0.145 | -0.049 | 0.02 | 0.02 | 0.846 | 0.73 | 0.82 | 0.7 | 0.69 | 0.57 | 0.67 |
| 3 | 50480863 | rs762898 | 0.921 | 0.649 | -1.294 | 0.304 | -0.043 | -0.019 | -0.019 | 3.561 | 0.08 | 0.01 | 0.35 | 0.42 | 0.11 | 0.13 |
| 3 | 50483437 | rs2236975 | 0.326 | 0.596 | -1.272 | 0.253 | -0.007 | 0.022 | 0.022 | 0 | 0.11 | 0.1 | 0.36 | 0.41 | 0.11 | 0.12 |
| 3 | 50486057 | rs2236977 | 0.897 | 0.579 | -1.208 | -0.013 | 0.025 | 0.104 | 0.104 | 0.233 | 0.07 | 0.51 | 0.05 | 0.06 | 0.01 | 0 |
| 3 | 50503170 | rs929047 | 0.497 | 0.769 | -1.681 | 0.099 | -0.014 | 0.006 | 0.006 | 0 | 0.1 | 0.07 | 0.36 | 0.62 | 0.11 | 0.11 |
| 3 | 50504896 | rs2236988 | 0.873 | 0.854 | -1.767 | 0.138 | -0.043 | -0.023 | -0.023 | 0.926 | 0.08 | 0.01 | 0.36 | 0.62 | 0.11 | 0.11 |
| 3 | 50505395 | rs2236989 | 1.994 | 0.505 | -2.111 | -0.351 | -0.026 | -0.021 | -0.021 | 0.21 | 0.27 | 0.03 | 0.15 | 0.19 | 0.27 | 0.43 |
| 3 | 50513157 | rs916288 | -0.422 | 0.929 | -1.248 | 0.023 | -0.109 | 0.065 | 0.065 | 0 | 0.81 | 0.98 | 0.84 | 0.92 | 0.67 | 0.7 |
| 3 | 50522063 | rs4688718 | -0.336 | 0.016 | -0.382 | 0.602 | -0.011 | -0.02 | -0.02 | 0.061 | 0.45 | 0.54 | 0.55 | 0.79 | 0.36 | 0.41 |
| 3 | 50532006 | rs6807916 | -0.062 | 0.636 | -0.123 | 1.306 | -0.002 | 0.006 | 0.006 | 0 | 0.09 | 0.06 | 0.14 | 0.31 | 0.11 | 0.11 |
| 3 | 50534635 | rs9814874 | -0.195 | 0.282 | 0.33 | 1.345 | 0.037 | 0.49 | 0.49 | 0.019 | 0.06 | 0.37 | 0.24 | 0.32 | 0.01 | 0.01 |
| 3 | 50574213 | rs398703 | 1.406 | 0.574 | 0.081 | 1.188 | 0.037 | 0.286 | 0.286 | 0 | 0.85 | 0.54 | 0.6 | 0.57 | 0.88 | 0.88 |
| 3 | 50577415 | rs13091933 | -0.03 | 0.75 | -0.268 | 0.996 | -0.003 | -0.004 | -0.004 | 0 | 0.06 | 0.03 | 0.14 | 0.08 | 0.11 | 0.13 |
| 3 | 50580658 | rs420536 | 0.297 | 0.911 | -1.138 | 0.319 | -0.152 | 0.073 | 0.073 | 0 | 0.3 | 0.02 | 0.2 | 0.14 | 0.43 | 0.37 |
| 3 | 50584999 | rs13076850 | 0.229 | 1.007 | -0.574 | 0.66 | 0.019 | 0.005 | 0.005 | 0.332 | 0.06 | 0.05 | 0.13 | 0.06 | 0.11 | 0.09 |
| 3 | 50597092 | rs1034405 | 0.749 | 0.889 | -0.13 | 0.628 | 0.161 | 0.094 | 0.094 | 0.079 | 0.91 | 0.64 | 0.84 | 0.94 | 0.88 | 0.92 |
| 3 | 50606374 | rs3804627 | 1.524 | 0.727 | -0.165 | 0.972 | 0.059 | 0.34 | 0.34 | 0 | 0.86 | 0.59 | 0.61 | 0.59 | 0.88 | 0.89 |
| 3 | 50607064 | rs34584636 | 0.357 | 0.731 | -0.202 | 0.98 | 0 | -0.003 | -0.003 | 0 | 0.06 | 0.03 | 0.13 | 0.06 | 0.11 | 0.08 |
| 3 | 50642975 | rs201194 | 0.584 | 0.802 | 0.092 | 0.825 | 0.117 | 0.049 | 0.049 | 0.105 | 0.92 | 0.73 | 0.85 | 0.95 | 0.88 | 0.92 |
| 3 | 50644134 | rs175634 | 1.236 | 0.752 | 0.091 | 0.934 | 0.084 | 0.364 | 0.364 | 0 | 0.88 | 0.63 | 0.62 | 0.6 | 0.87 | 0.89 |
| 3 | 50645413 | rs2239752 | -0.188 | 0.676 | 0.049 | 0.953 | -0.019 | 0.206 | 0.206 | 0.277 | 0.05 | 0.13 | 0.23 | 0.34 | 0.01 | 0.03 |
| 3 | 50651751 | rs873985 | 0.967 | 1.071 | -0.394 | 0.517 | -0.051 | 0.083 | 0.083 | 0 | 0.14 | 0.01 | 0.09 | 0.01 | 0.16 | 0.16 |
| 3 | 50657826 | rs375544 | 1.487 | 1.216 | -0.184 | 0.555 | 0.062 | 0.357 | 0.357 | 0 | 0.87 | 0.59 | 0.62 | 0.6 | 0.88 | 0.91 |
| 3 | 50668532 | rs616689 | 0.885 | 1.201 | -0.186 | 0.559 | 0.016 | 0.461 | 0.461 | 1.181 | 0.92 | 0.62 | 0.74 | 0.66 | 0.99 | 0.97 |
| 3 | 50670078 | rs2040397 | 0.871 | 1.313 | -0.538 | 0.367 | -0.084 | -0.003 | -0.003 | 0 | 0.18 | 0.02 | 0.12 | 0.13 | 0.28 | 0.2 |
| 3 | 50685642 | rs9879397 | 0.807 | 1.617 | -0.53 | 0.339 | 0.016 | 0.461 | 0.461 | 1.181 | 0.92 | 0.62 | 0.74 | 0.65 | 0.99 | 0.98 |
| 3 | 50686517 | rs2170840 | 0.807 | 1.416 | -0.517 | 0.351 | 0.016 | 0.461 | 0.461 | 1.181 | 0.92 | 0.62 | 0.74 | 0.65 | 0.99 | 0.97 |
| 3 | 50692172 | rs7637429 | 1.767 | 0.865 | -1.262 | -0.409 | -0.025 | 0.094 | 0.094 | 0 | 0.15 | 0.02 | 0.09 | 0 | 0.12 | 0.06 |
| 3 | 50720909 | rs13094785 | -0.23 | 0.821 | -1.148 | -0.414 | 0.122 | 0.058 | 0.058 | 0.004 | 0.93 | 0.76 | 0.86 | 0.95 | 0.89 | 0.94 |
| 3 | 50772664 | rs9825535 | 0.882 | -0.258 | -1.24 | -1.041 | -0.007 | 0.195 | 0.195 | 0 | 0.8 | 0.57 | 0.59 | 0.6 | 0.86 | 0.89 |
| 3 | 50849368 | rs9869945 | 0.896 | -0.494 | -1.276 | -1.151 | -0.001 | 0.199 | 0.199 | 0 | 0.81 | 0.62 | 0.6 | 0.6 | 0.86 | 0.89 |
| 3 | 50897386 | rs11921930 | 2.26 | -0.837 | -1.631 | -1.188 | -0.032 | -0.072 | -0.072 | 0 | 0.2 | 0.03 | 0.12 | 0.21 | 0.2 | 0.36 |
| 3 | 50927009 | rs11928389 | -1.032 | -0.8 | -1.562 | -1.173 | -0.039 | 0.039 | 0.039 | 0 | 0.62 | 0.93 | 0.78 | 0.79 | 0.67 | 0.56 |
| 3 | 50932324 | rs11130263 | -0.403 | -0.612 | -1.156 | -1.006 | 0.129 | 0.062 | 0.062 | 0.006 | 0.08 | 0.29 | 0.14 | 0.05 | 0.11 | 0.06 |
| 3 | 50968897 | rs4078296 | 0.308 | -0.704 | -1.224 | -0.986 | -0.025 | 0.057 | 0.057 | 0 | 0.11 | 0.13 | 0.25 | 0.35 | 0.03 | 0.05 |
| 3 | 50998816 | rs9869826 | 2.186 | -0.659 | -1.055 | -1.019 | 0 | 0.011 | 0.011 | 0 | 0.07 | 0.1 | 0.14 | 0.24 | 0.08 | 0.09 |
| 3 | 51038375 | rs7614881 | 0.793 | -0.674 | -0.868 | -1.054 | 0.051 | -0.006 | -0.006 | 0 | 0.88 | 0.71 | 0.83 | 0.94 | 0.88 | 0.94 |
| 3 | 51062402 | rs11919195 | 1.493 | -0.726 | -0.949 | -1.132 | -0.025 | 0.093 | 0.093 | 0 | 0.15 | 0.02 | 0.1 | 0 | 0.13 | 0.06 |
| 3 | 51122750 | rs7653451 | -0.693 | -0.543 | -0.408 | -0.688 | 0 | -0.002 | -0.002 | 0 | 0.94 | 0.98 | 0.87 | 0.95 | 0.89 | 0.94 |
| 3 | 51137089 | rs4356848 | 1.624 | -0.592 | -0.394 | -0.661 | -0.004 | 0.2 | 0.2 | 0 | 0.8 | 0.61 | 0.59 | 0.6 | 0.86 | 0.88 |
| 3 | 51147807 | rs4267675 | 0.343 | -0.569 | -0.464 | -0.693 | -0.005 | 0.002 | 0.002 | 0 | 0.9 | 0.93 | 0.87 | 0.94 | 0.89 | 0.94 |
| 3 | 51216578 | rs4346580 | 0.612 | -0.936 | -0.443 | -0.581 | -0.02 | 0.098 | 0.098 | 0 | 0.08 | 0.09 | 0.25 | 0.34 | 0.03 | 0.05 |
| 3 | 51245578 | rs4367100 | 0.923 | -0.898 | -0.445 | -0.66 | -0.009 | 0.056 | 0.056 | 0 | 0.83 | 0.81 | 0.62 | 0.6 | 0.88 | 0.88 |
| 3 | 51248198 | rs4619816 | 0.147 | -0.954 | -0.532 | -0.758 | 0.007 | 0.008 | 0.008 | 0 | 0.5 | 0.53 | 0.45 | 0.39 | 0.65 | 0.47 |
| 3 | 51369574 | rs4974105 | -0.553 | -0.404 | -0.113 | -0.434 | 0.007 | 0.001 | 0.001 | 0 | 0.92 | 0.91 | 0.85 | 0.94 | 0.88 | 0.91 |
| 3 | 51372940 | rs7625006 | 0.363 | -0.449 | -0.032 | -0.488 | 0.02 | -0.013 | -0.013 | 0 | 0.88 | 0.75 | 0.84 | 0.94 | 0.88 | 0.91 |
| 3 | 51373707 | rs9819195 | 0.801 | -0.533 | -0.009 | -0.422 | -0.029 | 0.207 | 0.207 | 0 | 0.06 | 0.12 | 0.25 | 0.36 | 0.02 | 0.04 |
| 3 | 51383354 | rs4974104 | 1.052 | -0.421 | -0.255 | -0.429 | -0.024 | 0.232 | 0.232 | 0 | 0.81 | 0.61 | 0.59 | 0.58 | 0.86 | 0.87 |
| 3 | 51389976 | rs17051762 | 0.585 | -0.2 | -0.525 | -0.518 | 0.007 | -0.01 | -0.01 | 0 | 0.08 | 0.1 | 0.1 | 0.04 | 0.08 | 0.07 |
| 3 | 51398958 | rs2127655 | -0.28 | -0.246 | -0.453 | -0.505 | 0.003 | -0.008 | -0.008 | 0 | 0.88 | 0.88 | 0.85 | 0.94 | 0.88 | 0.91 |
| 3 | 51409600 | rs11715034 | 1.367 | 0.002 | -0.927 | -0.647 | -0.14 | -0.012 | -0.012 | 0.679 | 0.2 | 0.02 | 0.17 | 0.17 | 0.37 | 0.2 |
| 3 | 51422188 | rs1370124 | -0.213 | -1.443 | -0.423 | -0.497 | -0.038 | 0.104 | 0.104 | 0 | 0.09 | 0.15 | 0.25 | 0.35 | 0.02 | 0.04 |
| 3 | 51425481 | rs2304512 | 1.098 | -1.436 | -0.429 | -0.494 | -0.026 | 0.197 | 0.197 | 0 | 0.06 | 0.12 | 0.25 | 0.35 | 0.02 | 0.04 |
| 3 | 51432895 | rs4687569 | -1.199 | -1.423 | -0.172 | -0.564 | 0.014 | 0.002 | 0.002 | 0 | 0.92 | 0.94 | 0.86 | 0.94 | 0.88 | 0.91 |

| | | | | | | | | | | | | | | | | |
|---|---|---|---|---|---|---|---|---|---|---|---|---|---|---|---|---|
| 3 | 51434215 | rs12637997 | 1.988 | -1.828 | -0.55 | -0.841 | -0.031 | -0.058 | -0.058 | 0 | 0.2 | 0.03 | 0.12 | 0.21 | 0.21 | 0.37 |
| 3 | 51436456 | rs17051782 | 2.068 | -1.446 | -0.256 | -0.91 | -0.031 | -0.055 | -0.055 | 0.597 | 0.21 | 0.04 | 0.12 | 0.21 | 0.21 | 0.35 |
| 3 | 51441667 | rs10510758 | 1.045 | -1.387 | -0.179 | -0.86 | -0.026 | 0.191 | 0.191 | 0 | 0.06 | 0.11 | 0.25 | 0.35 | 0.02 | 0.04 |
| 3 | 51469107 | rs4611808 | -1.254 | -1.383 | -0.182 | -0.868 | -0.007 | -0.002 | -0.002 | 1.871 | 0.93 | 0.98 | 0.86 | 0.94 | 0.88 | 0.91 |
| 3 | 51474182 | rs17712228 | 1.035 | -1.384 | -0.198 | -0.862 | -0.026 | 0.197 | 0.197 | 0 | 0.06 | 0.12 | 0.24 | 0.35 | 0.02 | 0.04 |
| 3 | 51476387 | rs4687800 | 0.823 | -1.407 | -0.373 | -0.914 | -0.028 | 0.208 | 0.208 | 0 | 0.81 | 0.62 | 0.59 | 0.59 | 0.86 | 0.86 |
| 3 | 51500591 | rs7620081 | 1.573 | -1.456 | -0.313 | -1.01 | -0.001 | -0.006 | -0.006 | 0 | 0.07 | 0.13 | 0.14 | 0.03 | 0.08 | 0.09 |
| 3 | 51568137 | rs6773188 | 0.803 | -1.654 | -0.692 | -0.831 | 0.012 | 0.014 | 0.014 | 0 | 0.5 | 0.53 | 0.43 | 0.38 | 0.63 | 0.46 |
| 3 | 51570508 | rs1055429 | -0.877 | -1.644 | -0.908 | -0.735 | 0.014 | 0.001 | 0.001 | 0 | 0.92 | 0.93 | 0.85 | 0.95 | 0.88 | 0.91 |
| 3 | 51593827 | rs11719995 | 1.038 | -1.605 | -1.191 | -0.665 | -0.032 | 0.199 | 0.199 | 0 | 0.8 | 0.62 | 0.59 | 0.59 | 0.86 | 0.86 |
| 3 | 51598254 | rs4687721 | -1.032 | -1.534 | -1.318 | -0.623 | -0.039 | -0.02 | -0.02 | 0 | 0.92 | 1 | 0.86 | 0.95 | 0.88 | 0.91 |
| 3 | 51623547 | rs11928276 | 0.91 | -1.785 | -0.333 | 0.008 | -0.023 | -0.051 | -0.051 | 0 | 0.21 | 0.04 | 0.12 | 0.21 | 0.21 | 0.35 |
| 3 | 51655856 | rs4687592 | -1.601 | -1.171 | 0.837 | 0.32 | -0.029 | 0.303 | 0.303 | 0 | 0.08 | 0.25 | 0.26 | 0.35 | 0.02 | 0.05 |
| 3 | 51755065 | rs4687770 | -2.178 | -0.667 | 1.676 | 1.341 | -0.025 | 0.195 | 0.195 | 0 | 0.19 | 0.36 | 0.37 | 0.4 | 0.13 | 0.16 |
| 3 | 51755441 | rs6800949 | -0.311 | -0.317 | 0.598 | 0.967 | -0.006 | -0.002 | -0.002 | 0 | 0.07 | 0.03 | 0.11 | 0.05 | 0.11 | 0.11 |
| 3 | 51772347 | rs6445959 | -1.788 | -0.5 | 0.943 | 1.06 | -0.015 | 0.158 | 0.158 | 0 | 0.19 | 0.38 | 0.41 | 0.46 | 0.14 | 0.18 |
| 3 | 51802912 | rs12330652 | -1.51 | -0.905 | 1.289 | 1.004 | -0.062 | 0.349 | 0.349 | 0.632 | 0.12 | 0.47 | 0.31 | 0.39 | 0.03 | 0.05 |
| 3 | 51860817 | rs12639503 | -1.018 | -0.537 | 1.278 | -0.159 | -0.01 | 0.052 | 0.052 | 15.84 | 0.1 | 0.5 | 0.07 | 0.06 | 0.02 | 0.04 |
| 3 | 51863926 | rs6795223 | -0.691 | -0.543 | 1.568 | -0.16 | -0.025 | -0.013 | -0.013 | 0 | 0.08 | 0.25 | 0.05 | 0.04 | 0.02 | 0.02 |
| 3 | 51874275 | rs9853511 | -1.209 | -0.543 | 1.518 | -0.16 | -0.01 | 0.052 | 0.052 | 15.84 | 0.1 | 0.5 | 0.07 | 0.06 | 0.02 | 0.04 |
| 3 | 51881578 | rs9874046 | -1.538 | -0.663 | 1.047 | -0.624 | -0.027 | -0.114 | -0.114 | 0.511 | 0.25 | 0.5 | 0.17 | 0.06 | 0.17 | 0.19 |
| 3 | 51895260 | rs2007607 | 0.524 | -0.729 | 1.317 | -0.651 | -0.012 | 0.003 | 0.003 | 20.371 | 0.91 | 0.62 | 0.93 | 0.96 | 0.98 | 0.98 |
| 3 | 51895850 | rs986539 | 0.34 | -0.697 | 1.331 | -0.637 | -0.004 | 0.012 | 0.012 | 17.506 | 0.91 | 0.62 | 0.93 | 0.96 | 0.98 | 0.98 |
| 3 | 51899242 | rs975258 | -0.295 | -0.704 | 1.328 | -0.642 | 0.001 | -0.06 | -0.06 | 0 | 0.16 | 0.23 | 0.12 | 0.01 | 0.15 | 0.17 |
| 3 | 51903914 | rs4280629 | 0.996 | -0.65 | 0.878 | -0.619 | -0.019 | -0.071 | -0.071 | 0 | 0.75 | 0.39 | 0.81 | 0.94 | 0.83 | 0.8 |
| 3 | 51907736 | rs7635409 | 1.026 | -0.636 | 0.832 | -0.616 | -0.018 | -0.046 | -0.046 | 0.708 | 0.74 | 0.38 | 0.81 | 0.92 | 0.83 | 0.79 |
| 3 | 51917015 | rs1605070 | 0.86 | -0.551 | 0.357 | -0.863 | -0.006 | -0.063 | -0.063 | 1.278 | 0.61 | 0.4 | 0.72 | 0.94 | 0.68 | 0.75 |
| 3 | 51927598 | rs4234645 | 1.053 | -1.109 | 0.444 | -0.662 | -0.004 | -0.012 | -0.012 | 0 | 0.61 | 0.43 | 0.75 | 0.87 | 0.71 | 0.79 |
| 3 | 51933178 | rs1847677 | 1.052 | -1.356 | 0.609 | -0.725 | -0.028 | -0.053 | -0.053 | 0 | 0.64 | 0.51 | 0.78 | 0.87 | 0.75 | 0.83 |
| 3 | 51936193 | rs11130298 | 1.195 | -1.332 | 0.511 | -0.721 | -0.001 | -0.006 | -0.006 | 0 | 0.62 | 0.43 | 0.78 | 0.87 | 0.75 | 0.83 |
| 3 | 51941785 | rs4416372 | -1.027 | -1.412 | 0.556 | -0.746 | -0.007 | 0 | 0 | 0.774 | 0.2 | 0.21 | 0.05 | 0.08 | 0.06 | 0.06 |
| 3 | 51943333 | rs6775482 | 1.003 | -1.234 | 0.34 | -0.662 | -0.009 | -0.021 | -0.021 | 0.881 | 0.62 | 0.45 | 0.78 | 0.87 | 0.75 | 0.83 |
| 3 | 51962081 | rs323881 | -1.479 | -1.306 | 0.347 | -0.392 | -0.063 | -0.043 | -0.043 | 0 | 0.18 | 0.37 | 0.08 | 0.07 | 0.06 | 0.08 |
| 3 | 51975577 | rs929533 | -0.136 | -1.805 | 0.333 | 0.141 | -0.035 | 0.079 | 0.079 | 0.003 | 0.26 | 0.49 | 0.43 | 0.42 | 0.21 | 0.14 |
| 3 | 51979120 | rs4986825 | -0.412 | -1.323 | -0.089 | 0.523 | 0.022 | -0.026 | -0.026 | 0.464 | 0.06 | 0 | 0 | 0.06 | 0.01 | 0.03 |
| 3 | 51982251 | rs188953 | -1.683 | -1.327 | -0.093 | 0.907 | -0.052 | 0.189 | 0.189 | 0.116 | 0.11 | 0.34 | 0.23 | 0.28 | 0.02 | 0.05 |
| 3 | 52013883 | rs13099450 | 0.765 | -0.673 | -0.594 | 0.366 | -0.119 | -0.067 | -0.067 | 2.681 | 0.18 | 0.02 | 0.26 | 0.29 | 0.34 | 0.31 |
| 3 | 52021316 | rs323895 | -2.014 | -1.491 | 0.292 | 0.42 | -0.053 | 0.001 | 0.001 | 0.176 | 0.12 | 0.34 | 0.04 | 0.07 | 0.02 | 0.07 |
| 3 | 52047421 | rs6446166 | -1.916 | -1.423 | 0.066 | 0.382 | -0.043 | -0.039 | -0.039 | 0.095 | 0.17 | 0.34 | 0.05 | 0.07 | 0.07 | 0.11 |
| 3 | 52059636 | rs9873175 | -1.219 | -1.358 | -0.091 | 0.678 | -0.006 | 0.01 | 0.01 | 0.099 | 0.26 | 0.24 | 0.33 | 0.47 | 0.12 | 0.18 |
| 3 | 52064510 | rs4687798 | -0.067 | -1.402 | -0.283 | 0.691 | -0.011 | 0.043 | 0.043 | 0 | 0.15 | 0.14 | 0.31 | 0.41 | 0.09 | 0.12 |
| 3 | 52068736 | rs9817697 | -1.732 | -1.306 | -0.196 | 0.74 | -0.043 | -0.025 | -0.025 | 0.301 | 0.14 | 0.31 | 0.04 | 0.07 | 0.04 | 0.11 |
| 3 | 52070955 | rs9848727 | -1.805 | -1.371 | -0.163 | 0.851 | -0.077 | 0.114 | 0.114 | 0.075 | 0.25 | 0.42 | 0.34 | 0.46 | 0.09 | 0.22 |
| 3 | 52080112 | rs13316105 | -1.61 | -1.266 | -0.306 | 0.865 | -0.055 | 0.085 | 0.085 | 0.004 | 0.25 | 0.4 | 0.34 | 0.48 | 0.1 | 0.21 |
| 3 | 52081811 | rs12495536 | -1.83 | -1.168 | -0.388 | 0.88 | -0.027 | 0.037 | 0.037 | 0 | 0.29 | 0.39 | 0.35 | 0.46 | 0.16 | 0.25 |
| 3 | 52084040 | rs3821841 | 0.469 | -1.21 | -0.448 | 0.91 | 0.005 | -0.067 | -0.067 | 0 | 0.1 | 0.01 | 0.29 | 0.37 | 0.07 | 0.13 |
| 3 | 52088651 | rs4687607 | -1.895 | -1.224 | -0.446 | 0.877 | -0.033 | 0.045 | 0.045 | 0 | 0.29 | 0.4 | 0.35 | 0.47 | 0.16 | 0.26 |
| 3 | 52099441 | rs17051957 | -0.972 | -1.189 | -0.414 | 0.8 | -0.021 | -0.027 | -0.027 | 0 | 0.16 | 0.28 | 0.05 | 0.09 | 0.09 | 0.08 |
| 3 | 52101799 | rs17259762 | 0.134 | -0.97 | -0.564 | 0.717 | -0.028 | 0.034 | 0.034 | 0 | 0.07 | 0 | 0.09 | 0 | 0.09 | 0.04 |
| 3 | 52107909 | rs876782 | -1.466 | -1.172 | -0.217 | 0.991 | -0.031 | -0.025 | -0.025 | 0.505 | 0.17 | 0.38 | 0.07 | 0.1 | 0.09 | 0.12 |
| 3 | 52108372 | rs4687801 | 0.248 | -1.18 | -0.231 | 1.054 | 0.012 | -0.072 | -0.072 | 0 | 0.11 | 0.01 | 0.29 | 0.37 | 0.07 | 0.14 |
| 3 | 52109407 | rs747343 | -1.934 | -1.182 | -0.23 | 1.053 | -0.038 | 0.059 | 0.059 | 0 | 0.28 | 0.42 | 0.36 | 0.48 | 0.16 | 0.26 |
| 3 | 52109564 | rs893056 | -1.048 | -1.185 | -0.224 | 1.061 | -0.019 | -0.035 | -0.035 | 0 | 0.15 | 0.27 | 0.05 | 0.06 | 0.09 | 0.08 |
| 3 | 52109668 | rs9857878 | -1.123 | -1.182 | -0.282 | 1.062 | -0.011 | -0.019 | -0.019 | 0 | 0.15 | 0.21 | 0.05 | 0.06 | 0.09 | 0.08 |
| 3 | 52109841 | rs731684 | -1.911 | -1.178 | -0.411 | 1.064 | -0.038 | 0.054 | 0.054 | 0 | 0.28 | 0.42 | 0.36 | 0.45 | 0.16 | 0.26 |
| 3 | 52115398 | rs13314125 | -0.662 | -1.133 | -0.507 | 0.982 | 0 | 0 | 0 | 0 | 0.13 | 0.13 | 0.04 | 0.09 | 0.09 | 0.08 |
| 3 | 52116714 | rs7618573 | 0.24 | -0.978 | -0.755 | 1.035 | 0.01 | -0.068 | -0.068 | 0 | 0.1 | 0.01 | 0.29 | 0.37 | 0.07 | 0.14 |
| 3 | 52121297 | rs12489539 | 0.221 | -0.951 | -0.742 | 1.05 | 0.011 | -0.071 | -0.071 | 0.951 | 0.11 | 0.01 | 0.29 | 0.38 | 0.07 | 0.14 |
| 3 | 52210700 | rs352152 | -0.879 | -0.751 | -0.571 | 0.857 | -0.045 | 0.071 | 0.071 | 0 | 0.28 | 0.45 | 0.36 | 0.44 | 0.16 | 0.26 |
| 3 | 52217088 | rs353551 | -0.249 | 0.114 | -1.244 | 0.538 | 0.002 | 0.041 | 0.041 | 2.893 | 0.48 | 0.26 | 0.48 | 0.37 | 0.52 | 0.38 |
| 3 | 52220203 | rs614288 | -0.416 | -0.168 | -0.325 | 0.554 | 0.003 | 0.008 | 0.008 | 3.096 | 0.52 | 0.49 | 0.53 | 0.39 | 0.53 | 0.42 |
| 3 | 52236077 | rs352170 | -0.278 | -0.095 | -0.208 | 0.561 | 0.014 | 0.025 | 0.025 | 3.359 | 0.5 | 0.32 | 0.49 | 0.4 | 0.53 | 0.39 |
| 3 | 52236762 | rs352169 | -0.252 | -0.094 | -0.199 | 0.562 | 0.013 | 0.028 | 0.028 | 3.407 | 0.49 | 0.27 | 0.49 | 0.39 | 0.53 | 0.39 |
| 3 | 52237970 | rs352168 | -0.229 | -0.131 | -0.207 | 0.541 | 0.01 | 0.028 | 0.028 | 2.811 | 0.49 | 0.26 | 0.49 | 0.39 | 0.53 | 0.39 |
| 3 | 52247110 | rs352163 | -0.277 | -0.189 | 0.194 | 0.475 | 0.011 | 0.028 | 0.028 | 1.243 | 0.5 | 0.32 | 0.5 | 0.39 | 0.54 | 0.43 |
| 3 | 52256697 | rs352140 | -0.226 | -0.313 | 0.98 | 0.313 | 0.009 | 0.027 | 0.027 | 1.096 | 0.49 | 0.29 | 0.5 | 0.39 | 0.55 | 0.42 |
| 3 | 52261031 | rs187084 | 0.051 | -0.563 | 0.776 | -0.113 | 0.006 | -0.014 | -0.014 | 0 | 0.37 | 0.29 | 0.44 | 0.4 | 0.43 | 0.37 |
| 3 | 52264907 | rs352143 | -1.129 | -0.625 | 1.187 | -0.216 | -0.011 | -0.094 | -0.094 | 0 | 0.21 | 0.42 | 0.17 | 0.04 | 0.19 | 0.11 |
| 3 | 52268246 | rs11717574 | -0.352 | -0.628 | 1.217 | -0.214 | 0.01 | -0.08 | -0.08 | 0 | 0.14 | 0.37 | 0.1 | 0.01 | 0.13 | 0.09 |

| | | | | | | | | | | | | | | | | |
|---|---|---|---|---|---|---|---|---|---|---|---|---|---|---|---|---|
| 3 | 52268662 | rs11714313 | -0.352 | -0.615 | 1.217 | -0.214 | 0.006 | -0.08 | -0.08 | 0 | 0.14 | 0.36 | 0.1 | 0.01 | 0.13 | 0.09 |
| 3 | 52279594 | rs7618915 | 0.545 | -0.453 | 1.498 | -0.09 | 0.003 | -0.013 | -0.013 | 0.023 | 0.3 | 0.2 | 0.32 | 0.37 | 0.35 | 0.31 |
| 3 | 52280378 | rs11715391 | 0.124 | -0.465 | 1.985 | -0.12 | 0.034 | -0.073 | -0.073 | 0 | 0.12 | 0.39 | 0.1 | 0 | 0.1 | 0.05 |
| 3 | 52287468 | rs11717383 | -1.07 | -0.519 | 1.815 | -0.154 | 0.023 | 0.019 | 0.019 | 0.196 | 0.49 | 0.67 | 0.48 | 0.4 | 0.52 | 0.41 |
| 3 | 52288945 | rs1060330 | -1.07 | -0.519 | 1.815 | -0.154 | 0.023 | 0.019 | 0.019 | 0.135 | 0.49 | 0.67 | 0.48 | 0.4 | 0.52 | 0.41 |
| 3 | 52321976 | rs11707056 | 0.599 | -0.491 | 1.816 | -0.153 | -0.012 | 0.034 | 0.034 | 1.705 | 0.1 | 0.03 | 0.07 | 0 | 0.1 | 0.05 |
| 3 | 52334593 | rs17052053 | -0.21 | -0.494 | 1.925 | -0.154 | 0.019 | -0.08 | -0.08 | 0 | 0.13 | 0.38 | 0.1 | 0.01 | 0.1 | 0.08 |
| 3 | 52342529 | rs699469 | 1.764 | -0.316 | 1.113 | -0.226 | -0.018 | -0.028 | -0.028 | 0 | 0.17 | 0.04 | 0.14 | 0.17 | 0.14 | 0.25 |
| 3 | 52343679 | rs10460962 | 0.481 | -0.418 | 1.198 | -0.293 | 0.03 | -0.077 | -0.077 | 0 | 0.12 | 0.38 | 0.1 | 0 | 0.1 | 0.05 |
| 3 | 52346240 | rs17052061 | -0.485 | -0.419 | 1.191 | -0.296 | 0.009 | -0.099 | -0.099 | 0.932 | 0.17 | 0.38 | 0.16 | 0.03 | 0.16 | 0.1 |
| 3 | 52348364 | rs730050 | 0.718 | -0.271 | 0.883 | -0.644 | -0.069 | -0.05 | -0.05 | 0 | 0.39 | 0.13 | 0.44 | 0.58 | 0.39 | 0.53 |
| 3 | 52349412 | rs3852064 | 1.248 | -0.216 | 1.067 | -0.389 | 0.002 | -0.009 | -0.009 | 0 | 0.3 | 0.23 | 0.32 | 0.38 | 0.35 | 0.3 |
| 3 | 52350212 | rs6445386 | -0.449 | -0.182 | 0.979 | -0.4 | 0.009 | -0.099 | -0.099 | 0 | 0.17 | 0.38 | 0.16 | 0.03 | 0.16 | 0.1 |
| 3 | 52382822 | rs7633687 | 1.057 | -0.173 | 0.888 | -0.392 | 0 | -0.03 | -0.03 | 0 | 0.1 | 0.09 | 0.07 | 0 | 0.1 | 0.06 |
| 3 | 52391735 | rs11714402 | -0.337 | -0.184 | 0.755 | -0.402 | 0.003 | -0.077 | -0.077 | 0 | 0.16 | 0.31 | 0.15 | 0.02 | 0.17 | 0.1 |
| 3 | 52396944 | rs11706515 | 1.109 | -0.175 | 0.644 | -0.371 | -0.012 | 0.034 | 0.034 | 1.394 | 0.1 | 0.03 | 0.07 | 0 | 0.1 | 0.06 |
| 3 | 52407041 | rs1546737 | 1.37 | -0.291 | 0.412 | -0.047 | -0.058 | 0.019 | 0.019 | 0 | 0.28 | 0.01 | 0.27 | 0.18 | 0.35 | 0.28 |
| 3 | 52430526 | rs12163565 | 1.437 | 0.352 | 0.321 | 0.127 | -0.063 | -0.07 | -0.07 | 1.355 | 0.18 | 0.02 | 0.12 | 0.29 | 0.2 | 0.23 |
| 3 | 52432214 | rs11711974 | 0.625 | 0.091 | 0.956 | 0.044 | 0.019 | -0.067 | -0.067 | 0 | 0.11 | 0.29 | 0.09 | 0 | 0.1 | 0.06 |
| 3 | 52450043 | rs13094687 | 0.934 | 0.168 | 0.434 | 0.113 | -0.06 | -0.155 | -0.155 | 0.128 | 0.28 | 0.02 | 0.3 | 0.38 | 0.35 | 0.29 |
| 3 | 52456973 | rs2272088 | 0.85 | 0.18 | 0.468 | 0.117 | -0.044 | -0.142 | -0.142 | 0.056 | 0.29 | 0.05 | 0.3 | 0.38 | 0.35 | 0.3 |
| 3 | 52477080 | rs2016575 | 0.671 | 0.114 | 1.022 | -0.191 | 0.002 | -0.075 | -0.075 | 0 | 0.84 | 0.69 | 0.85 | 0.97 | 0.84 | 0.9 |
| 3 | 52477866 | rs11716487 | -0.895 | 0.115 | 0.194 | -0.18 | -0.013 | -0.077 | -0.077 | 0 | 0.16 | 0.69 | 0.13 | 0 | 0.11 | 0.07 |
| 3 | 52482851 | rs634382 | 1.101 | 0.241 | 0.129 | -0.12 | -0.012 | -0.071 | -0.071 | 0 | 0.79 | 0.34 | 0.82 | 0.97 | 0.83 | 0.87 |
| 3 | 52492707 | rs1541495 | 0.322 | 0.4 | -0.079 | 0.102 | 0.002 | 0.006 | 0.006 | 0 | 0.94 | 0.96 | 0.93 | 0.98 | 0.93 | 0.96 |
| 3 | 52506848 | rs1011063 | 0.847 | 0.635 | 0.57 | -0.133 | 0.008 | -0.076 | -0.076 | 0 | 0.82 | 0.68 | 0.84 | 0.98 | 0.82 | 0.9 |
| 3 | 52523405 | rs887515 | 1.317 | 0.416 | 0.614 | -0.194 | -0.009 | -0.062 | -0.062 | 0 | 0.79 | 0.31 | 0.81 | 0.98 | 0.82 | 0.88 |
| 3 | 52530787 | rs17052201 | -0.292 | 0.477 | -0.15 | 0.666 | -0.024 | -0.085 | -0.085 | 1.897 | 0.13 | 0.01 | 0.1 | 0.27 | 0.14 | 0.12 |
| 3 | 52536308 | rs758801 | 0.557 | 0.845 | 0.514 | 0.345 | 0.014 | -0.051 | -0.051 | 1.521 | 0.89 | 0.8 | 0.87 | 0.98 | 0.87 | 0.95 |
| 3 | 52540773 | rs1010553 | 0.452 | 0.942 | -0.553 | 0.3 | -0.049 | -0.064 | -0.064 | 0.61 | 0.6 | 0.86 | 0.43 | 0.54 | 0.47 | 0.76 |
| 3 | 52555316 | rs13081028 | -1.456 | 0.566 | 0.064 | 0.175 | -0.002 | 0.007 | 0.007 | 0.05 | 0.51 | 0.7 | 0.65 | 0.46 | 0.56 | 0.33 |
| 3 | 52555957 | rs9853056 | -1.604 | 0.423 | 0.297 | 0.101 | -0.015 | -0.011 | -0.011 | 0.08 | 0.52 | 0.75 | 0.66 | 0.46 | 0.56 | 0.35 |
| 3 | 52558008 | rs13303 | -1.532 | 0.399 | 0.198 | 0.21 | -0.007 | -0.004 | -0.004 | 0 | 0.52 | 0.71 | 0.65 | 0.45 | 0.57 | 0.33 |
| 3 | 52566914 | rs7614981 | -0.92 | 0.4 | 0.891 | 0.203 | 0 | 0 | 0 | 0.008 | 0.45 | 0.47 | 0.61 | 0.45 | 0.55 | 0.26 |
| 3 | 52572056 | rs7638808 | -0.789 | 0.521 | 1.034 | 0.173 | -0.004 | 0.001 | 0.001 | 2.066 | 0.46 | 0.56 | 0.55 | 0.43 | 0.48 | 0.31 |
| 3 | 52584787 | rs2251219 | -0.24 | 0.234 | 1.116 | 0.204 | 0.01 | -0.069 | -0.069 | 0 | 0.36 | 0.15 | 0.47 | 0.43 | 0.42 | 0.25 |
| 3 | 52593230 | rs3796353 | -0.774 | 0.029 | 1.071 | 0.236 | 0.008 | -0.009 | -0.009 | 0 | 0.42 | 0.37 | 0.5 | 0.43 | 0.43 | 0.28 |
| 3 | 52607288 | rs10510759 | -1.213 | 0.019 | 1.067 | 0.237 | -0.013 | -0.028 | -0.028 | 2.212 | 0.06 | 0.23 | 0.04 | 0 | 0.01 | 0.04 |
| 3 | 52622086 | rs2590838 | -1.395 | 0.065 | 1.07 | 0.237 | 0.014 | -0.001 | -0.001 | 0 | 0.49 | 0.69 | 0.62 | 0.45 | 0.52 | 0.3 |
| 3 | 52643685 | rs3755806 | -0.215 | 0.075 | 1.082 | 0.243 | 0.005 | -0.072 | -0.072 | 0 | 0.36 | 0.15 | 0.46 | 0.43 | 0.42 | 0.25 |
| 3 | 52650348 | rs10510760 | 0.38 | 0.105 | 1.064 | 0.247 | -0.06 | -0.196 | -0.196 | 0.061 | 0.29 | 0.02 | 0.42 | 0.43 | 0.36 | 0.2 |
| 3 | 52680823 | rs2028216 | -0.085 | 0.143 | 1.122 | 0.274 | 0.002 | -0.07 | -0.07 | 0 | 0.35 | 0.15 | 0.46 | 0.43 | 0.42 | 0.2 |
| 3 | 52719398 | rs1866268 | -0.87 | -0.072 | 1.265 | 0.387 | -0.011 | 0.006 | 0.006 | 1.433 | 0.43 | 0.56 | 0.51 | 0.43 | 0.43 | 0.28 |
| 3 | 52720080 | rs1108842 | -1.234 | -0.064 | 1.261 | 0.384 | -0.001 | -0.001 | -0.001 | 0.546 | 0.48 | 0.57 | 0.61 | 0.45 | 0.52 | 0.3 |
| 3 | 52721305 | rs11177 | -0.199 | -0.037 | 1.29 | 0.396 | 0.001 | -0.07 | -0.07 | 0 | 0.35 | 0.15 | 0.46 | 0.42 | 0.42 | 0.21 |
| 3 | 52727257 | rs2289247 | -0.911 | -0.037 | 1.29 | 0.396 | -0.011 | 0.006 | 0.006 | 1.433 | 0.43 | 0.56 | 0.51 | 0.43 | 0.43 | 0.28 |
| 3 | 52728804 | rs6976 | -0.188 | -0.035 | 1.243 | 0.397 | -0.004 | -0.084 | -0.084 | 0 | 0.35 | 0.12 | 0.45 | 0.43 | 0.42 | 0.21 |
| 3 | 52731483 | rs11130317 | -0.182 | -0.031 | 1.23 | 0.349 | 0.004 | -0.072 | -0.072 | 0.576 | 0.36 | 0.15 | 0.46 | 0.43 | 0.42 | 0.24 |
| 3 | 52738165 | rs12635140 | -0.926 | -0.025 | 1.272 | 0.353 | -0.003 | 0.001 | 0.001 | 1.444 | 0.43 | 0.57 | 0.52 | 0.43 | 0.43 | 0.28 |
| 3 | 52741160 | rs3755798 | -0.941 | -0.037 | 1.236 | 0.369 | -0.01 | 0.005 | 0.005 | 1.291 | 0.44 | 0.55 | 0.51 | 0.43 | 0.43 | 0.34 |
| 3 | 52748857 | rs11716747 | -0.939 | -0.248 | 1.294 | 0.36 | -0.01 | 0.005 | 0.005 | 1.287 | 0.44 | 0.55 | 0.51 | 0.43 | 0.43 | 0.34 |
| 3 | 52764624 | rs10780035 | -0.201 | -0.237 | 0.994 | 0.386 | 0.001 | -0.068 | -0.068 | 0 | 0.35 | 0.15 | 0.46 | 0.43 | 0.42 | 0.2 |
| 3 | 52800284 | rs2230535 | -0.22 | -0.233 | 1.031 | 0.418 | 0.006 | -0.066 | -0.066 | 0.614 | 0.36 | 0.15 | 0.46 | 0.43 | 0.42 | 0.3 |
| 3 | 52804487 | rs13071584 | -0.844 | -0.246 | 1.165 | 0.411 | 0 | -0.001 | -0.001 | 0 | 0.43 | 0.46 | 0.5 | 0.43 | 0.43 | 0.34 |
| 3 | 52808341 | rs1014969 | -0.88 | -0.185 | 1.231 | 0.427 | -0.015 | 0.008 | 0.008 | 1.246 | 0.43 | 0.51 | 0.51 | 0.43 | 0.42 | 0.35 |
| 3 | 52809193 | rs3755799 | -0.432 | -0.406 | 1.174 | 0.334 | -0.001 | -0.001 | -0.001 | 0 | 0.35 | 0.39 | 0.43 | 0.33 | 0.34 | 0.26 |
| 3 | 52817675 | rs3774354 | -0.285 | -0.204 | 0.844 | 0.339 | 0.01 | -0.054 | -0.054 | 0.13 | 0.34 | 0.14 | 0.45 | 0.37 | 0.4 | 0.2 |
| 3 | 52822921 | rs2300149 | -0.619 | -0.213 | 0.995 | 0.164 | 0.006 | -0.007 | -0.007 | 0 | 0.38 | 0.29 | 0.47 | 0.37 | 0.41 | 0.21 |
| 3 | 52825462 | rs1076425 | -0.548 | -0.168 | 1.058 | 0.193 | 0.012 | -0.022 | -0.022 | 0.023 | 0.35 | 0.22 | 0.44 | 0.37 | 0.39 | 0.2 |
| 3 | 52825528 | rs1075653 | -0.68 | -0.177 | 1.123 | 0.189 | -0.018 | 0.016 | 0.016 | 0.867 | 0.37 | 0.53 | 0.47 | 0.37 | 0.39 | 0.2 |
| 3 | 52825585 | rs9324 | -0.726 | -0.191 | 1.165 | 0.183 | -0.008 | 0.005 | 0.005 | 0.949 | 0.37 | 0.58 | 0.47 | 0.37 | 0.39 | 0.2 |
| 3 | 52826846 | rs2071508 | -0.743 | -0.163 | 0.807 | 0.157 | 0.001 | -0.004 | -0.004 | 0 | 0.35 | 0.28 | 0.44 | 0.37 | 0.39 | 0.2 |
| 3 | 52833805 | rs3617 | -1.596 | -0.182 | 0.629 | 0.212 | 0.027 | -0.02 | -0.02 | 0 | 0.46 | 0.77 | 0.61 | 0.41 | 0.48 | 0.39 |
| 3 | 52835303 | rs2710334 | -1.339 | -0.241 | 1.018 | 0.197 | 0.057 | 0.077 | 0.077 | 0.244 | 0.06 | 0.47 | 0.05 | 0.02 | 0 | 0.17 |
| 3 | 52835354 | rs736408 | -1.152 | -0.21 | 0.872 | 0.193 | 0.043 | -0.035 | -0.035 | 0 | 0.4 | 0.75 | 0.5 | 0.38 | 0.38 | 0.35 |
| 3 | 52837793 | rs4481150 | -1.552 | -0.251 | 0.983 | 0.185 | 0.009 | -0.005 | -0.005 | 0 | 0.45 | 0.62 | 0.6 | 0.43 | 0.5 | 0.39 |
| 3 | 52844534 | rs17331151 | 0.159 | -0.213 | 0.69 | 0.214 | -0.033 | -0.008 | -0.008 | 0 | 0.06 | 0.01 | 0.11 | 0.04 | 0.1 | 0.03 |
| 3 | 52845105 | rs2535627 | -1.502 | -0.239 | 0.736 | 0.215 | 0.02 | -0.011 | -0.011 | 2.159 | 0.45 | 0.71 | 0.6 | 0.43 | 0.5 | 0.39 |
| 3 | 52851258 | rs2535619 | -1.459 | -0.214 | 1.124 | 0.311 | 0.072 | 0.087 | 0.087 | 1.35 | 0.05 | 0.42 | 0.05 | 0.01 | 0 | 0.26 |

| | | | | | | | | | | | | | | | | |
|---|---|---|---|---|---|---|---|---|---|---|---|---|---|---|---|---|
| 3 | 52853401 | rs3821831 | 1.172 | -0.343 | 0.584 | -0.021 | -0.027 | 0.086 | 0.086 | 0 | 0.33 | 0.15 | 0.2 | 0.09 | 0.34 | 0.22 |
| 3 | 52853480 | rs2276814 | -1.217 | -0.381 | 0.891 | 0.174 | 0.072 | 0.087 | 0.087 | 1.35 | 0.05 | 0.42 | 0.05 | 0.01 | 0 | 0.26 |
| 3 | 52855865 | rs2256332 | -1.484 | -0.432 | 0.791 | 0.296 | 0.023 | -0.036 | -0.036 | 0 | 0.38 | 0.73 | 0.56 | 0.37 | 0.39 | 0.45 |
| 3 | 52860816 | rs2276816 | 0.92 | -0.617 | 0.809 | 0.42 | -0.031 | -0.014 | -0.014 | 0 | 0.07 | 0 | 0.1 | 0.06 | 0.13 | 0.1 |
| 3 | 52860936 | rs2276817 | 0.316 | -0.675 | 0.835 | 0.376 | 0.005 | -0.013 | -0.013 | 0 | 0.2 | 0.2 | 0.38 | 0.32 | 0.26 | 0.12 |
| 3 | 52864584 | rs2071042 | -1.72 | -0.673 | 0.843 | 0.37 | 0.016 | -0.03 | -0.03 | 0 | 0.38 | 0.7 | 0.56 | 0.37 | 0.4 | 0.46 |
| 3 | 52872874 | rs2336155 | 0.948 | -0.664 | 0.871 | 0.374 | 0.074 | 0.088 | 0.088 | 1.35 | 0.95 | 0.58 | 0.95 | 0.99 | 1 | 0.73 |
| 3 | 52873984 | rs11626 | -0.703 | -0.666 | 0.881 | 0.375 | -0.009 | 0.018 | 0.018 | 0 | 0.13 | 0.08 | 0.13 | 0.04 | 0.14 | 0.07 |
| 3 | 52874288 | rs6445538 | 0.37 | -0.655 | 0.77 | 0.378 | 0.006 | -0.021 | -0.021 | 2.034 | 0.2 | 0.19 | 0.38 | 0.32 | 0.27 | 0.12 |
| 3 | 52874296 | rs6445539 | 0.37 | -0.682 | 0.762 | 0.378 | 0.005 | -0.012 | -0.012 | 0 | 0.2 | 0.2 | 0.38 | 0.32 | 0.27 | 0.12 |
| 3 | 52874446 | rs6769789 | 1.63 | -0.724 | 0.734 | 0.387 | 0.028 | -0.045 | -0.045 | 0.065 | 0.61 | 0.21 | 0.44 | 0.63 | 0.59 | 0.54 |
| 3 | 52880974 | rs2001733 | 1.095 | -0.605 | 0.765 | 0.295 | 0.07 | 0.084 | 0.084 | 1.138 | 0.94 | 0.58 | 0.95 | 0.99 | 1 | 0.73 |
| 3 | 52896855 | rs11130329 | 0.626 | -0.492 | 0.396 | 0.301 | 0.005 | -0.026 | -0.026 | 0 | 0.87 | 0.82 | 0.85 | 0.96 | 0.87 | 0.93 |
| 3 | 52907372 | rs4687680 | -1.134 | -0.732 | 0.513 | 0.499 | 0.008 | -0.03 | -0.03 | 1.195 | 0.32 | 0.32 | 0.5 | 0.36 | 0.38 | 0.18 |
| 3 | 52913006 | rs3821873 | 1.35 | -0.758 | 0.504 | 0.493 | -0.013 | 0.029 | 0.029 | 0.443 | 0.65 | 0.51 | 0.47 | 0.63 | 0.62 | 0.56 |
| 3 | 52916916 | rs9876403 | 0.223 | -0.712 | 0.484 | 0.521 | 0.008 | -0.03 | -0.03 | 0.209 | 0.2 | 0.2 | 0.37 | 0.32 | 0.27 | 0.12 |
| 3 | 52936763 | rs11922975 | 1.358 | -0.726 | 0.416 | 0.498 | -0.017 | 0.032 | 0.032 | 0.452 | 0.65 | 0.47 | 0.47 | 0.61 | 0.62 | 0.56 |
| 3 | 52937774 | rs1134546 | 0.73 | -0.759 | 0.363 | 0.58 | -0.03 | -0.011 | -0.011 | 0 | 0.17 | 0.01 | 0.18 | 0.1 | 0.26 | 0.11 |
| 3 | 52948332 | rs4687687 | 1.808 | -0.883 | 0.477 | 0.234 | 0.038 | -0.035 | -0.035 | 0 | 0.6 | 0.23 | 0.44 | 0.61 | 0.61 | 0.56 |
| 3 | 52956918 | rs11130331 | 1.54 | -0.921 | 0.528 | 0.241 | 0.003 | -0.003 | -0.003 | 0 | 0.62 | 0.31 | 0.45 | 0.61 | 0.62 | 0.56 |
| 3 | 52960859 | rs2581810 | 0.371 | -0.58 | 0.609 | 0.555 | -0.035 | 0.094 | 0.094 | 0.022 | 0.75 | 0.42 | 0.6 | 0.66 | 0.74 | 0.64 |
| 3 | 52968726 | rs2257216 | 0.998 | -0.568 | 0.531 | 0.557 | -0.016 | 0.078 | 0.078 | 0.597 | 0.71 | 0.48 | 0.51 | 0.6 | 0.66 | 0.57 |
| 3 | 52982296 | rs2952830 | 1.698 | -0.501 | 0.166 | 0.395 | -0.063 | -0.069 | -0.069 | 0 | 0.32 | 0.06 | 0.28 | 0.48 | 0.29 | 0.36 |
| 3 | 53003179 | rs11717619 | -0.967 | -0.424 | 0.354 | 0.521 | 0 | -0.001 | -0.001 | 0.095 | 0.07 | 0.15 | 0.12 | 0.06 | 0.08 | 0.07 |
| 3 | 53007995 | rs9846976 | 0.135 | -0.423 | 0.384 | 0.546 | -0.009 | 0.07 | 0.07 | 0 | 0.2 | 0.37 | 0.31 | 0.32 | 0.26 | 0.12 |
| 3 | 53015839 | rs11710394 | -0.036 | -0.36 | 0.155 | 0.541 | 0.003 | 0.002 | 0.002 | 0 | 0.06 | 0.1 | 0.11 | 0.06 | 0.08 | 0.07 |
| 3 | 53060637 | rs2255107 | -0.219 | -0.473 | 0.045 | 0.551 | -0.015 | 0.06 | 0.06 | 0 | 0.79 | 0.66 | 0.7 | 0.68 | 0.76 | 0.87 |
| 3 | 53061377 | rs12633064 | 0.906 | -0.207 | -0.002 | 0.45 | -0.02 | 0.068 | 0.068 | 0 | 0.33 | 0.14 | 0.18 | 0.12 | 0.32 | 0.2 |
| 3 | 53062661 | rs9847710 | 0.218 | -0.179 | -0.058 | 0.518 | -0.001 | -0.003 | -0.003 | 0 | 0.47 | 0.54 | 0.51 | 0.62 | 0.41 | 0.67 |
| 3 | 53070755 | rs17304995 | 0.858 | -0.122 | -0.313 | 0.673 | -0.03 | -0.109 | -0.109 | 0.002 | 0.17 | 0.02 | 0.28 | 0.27 | 0.26 | 0.13 |
| 3 | 53077372 | rs1346719 | -0.999 | -0.187 | 0.162 | 0.63 | 0.033 | 0.093 | 0.093 | 0.467 | 0.09 | 0.34 | 0.2 | 0.12 | 0.08 | 0.31 |
| 3 | 53084326 | rs11707780 | -0.53 | -0.176 | 0.111 | 0.634 | 0.002 | 0.021 | 0.021 | 0.245 | 0.07 | 0.14 | 0.18 | 0.12 | 0.08 | 0.31 |
| 3 | 53092375 | rs9870898 | 0.411 | -0.277 | 0.195 | 0.661 | 0.002 | -0.004 | -0.004 | 0 | 0.2 | 0.23 | 0.3 | 0.23 | 0.26 | 0.13 |
| 3 | 53094730 | rs2564926 | 0.163 | -0.285 | 0.275 | 0.657 | 0.007 | 0.035 | 0.035 | 0.161 | 0.93 | 0.86 | 0.82 | 0.88 | 0.92 | 0.69 |
| 3 | 53096394 | rs6794389 | 0.292 | -0.287 | 0.117 | 0.654 | -0.001 | 0.001 | 0.001 | 0 | 0.21 | 0.27 | 0.3 | 0.23 | 0.26 | 0.13 |
| 3 | 53098775 | rs1811351 | 0.893 | -0.183 | 0.026 | 0.281 | -0.023 | 0.04 | 0.04 | 0 | 0.32 | 0.14 | 0.19 | 0.15 | 0.33 | 0.2 |
| 3 | 53108776 | rs2581799 | 0.661 | -0.205 | -0.136 | 0.195 | -0.097 | 0.047 | 0.047 | 0.845 | 0.48 | 0.07 | 0.56 | 0.72 | 0.55 | 0.47 |
| 3 | 53118204 | rs2564929 | -0.849 | -0.047 | -0.392 | 0.343 | -0.01 | 0.027 | 0.027 | 0 | 0.68 | 0.78 | 0.81 | 0.87 | 0.67 | 0.8 |
| 3 | 53119703 | rs2253685 | 0.376 | -0.04 | -0.745 | 0.37 | -0.004 | 0.059 | 0.059 | 0.307 | 0.55 | 0.17 | 0.59 | 0.75 | 0.6 | 0.48 |
| 3 | 53123486 | rs1078968 | 0.376 | -0.038 | -0.744 | 0.371 | -0.004 | 0.059 | 0.059 | 0.198 | 0.55 | 0.17 | 0.59 | 0.75 | 0.6 | 0.48 |
| 3 | 53125922 | rs11242 | 0.376 | -0.038 | -0.744 | 0.371 | -0.004 | 0.059 | 0.059 | 0.198 | 0.55 | 0.17 | 0.59 | 0.75 | 0.6 | 0.48 |
| 3 | 53134098 | rs2581830 | 0.268 | 0.054 | -0.787 | 0.419 | 0.01 | 0.038 | 0.038 | 1.145 | 0.57 | 0.22 | 0.6 | 0.75 | 0.62 | 0.54 |
| 3 | 53138254 | rs1560333 | 0.67 | 0.328 | -1.047 | 0.577 | 0.004 | 0.062 | 0.062 | 0.665 | 0.89 | 0.54 | 0.79 | 0.88 | 0.91 | 0.74 |
| 3 | 53147886 | rs13433700 | 0.43 | 0.764 | -2.251 | 0.462 | -0.009 | 0.045 | 0.045 | 0 | 0.31 | 0.16 | 0.19 | 0.13 | 0.31 | 0.19 |
| 3 | 53155158 | rs9757079 | 0.438 | 0.749 | -2.248 | 0.466 | -0.011 | 0.044 | 0.044 | 0 | 0.31 | 0.16 | 0.19 | 0.13 | 0.31 | 0.19 |
| 3 | 53160360 | rs11713763 | -0.238 | 0.897 | -2.174 | 1.008 | -0.076 | -0.14 | -0.14 | 0 | 0.21 | 0.03 | 0.39 | 0.34 | 0.35 | 0.21 |
| 3 | 53175557 | rs7615501 | 0.243 | 0.911 | -2.528 | 0.431 | -0.103 | 0.009 | 0.009 | 0 | 0.66 | 0.95 | 0.81 | 0.77 | 0.62 | 0.7 |
| 3 | 53186704 | rs13085294 | 0.773 | -0.283 | -3.023 | -0.201 | 0.049 | 0.069 | 0.069 | 1.715 | 0.65 | 0.15 | 0.67 | 0.78 | 0.76 | 0.63 |
| 3 | 53199014 | rs1483185 | 0.562 | -0.148 | -1.671 | -0.072 | 0.078 | 0.112 | 0.112 | 2.052 | 0.86 | 0.56 | 0.75 | 0.77 | 0.85 | 0.88 |
| 3 | 53199080 | rs1483186 | 0.303 | -0.246 | -1.387 | -0.082 | 0 | 0.005 | 0.005 | 0.149 | 0.89 | 0.9 | 0.79 | 0.76 | 0.87 | 0.75 |
| 3 | 53200876 | rs3773732 | 1.19 | -0.633 | -1.892 | -0.181 | -0.001 | 0.003 | 0.003 | 0.133 | 0.67 | 0.38 | 0.68 | 0.66 | 0.72 | 0.63 |
| 3 | 53207401 | rs6764111 | 0.652 | -0.557 | -1.655 | -0.431 | -0.02 | -0.066 | -0.066 | 0 | 0.76 | 0.43 | 0.85 | 0.96 | 0.83 | 0.69 |
| 3 | 53208782 | rs6778964 | 1.172 | -0.9 | -1.848 | -0.607 | 0.02 | 0.044 | 0.044 | 1.05 | 0.59 | 0.34 | 0.64 | 0.74 | 0.67 | 0.56 |
| 3 | 53212716 | rs17052826 | -0.008 | -0.785 | -2.063 | -0.583 | 0.011 | 0.045 | 0.045 | 0 | 0.17 | 0.08 | 0.09 | 0.01 | 0.14 | 0.14 |
| 3 | 53213657 | rs2230493 | 0.249 | -0.758 | -2.18 | -0.571 | -0.003 | 0.065 | 0.065 | 0 | 0.16 | 0.04 | 0.08 | 0.02 | 0.14 | 0.14 |
| 3 | 53215875 | rs11130350 | 1.205 | -0.886 | -2.26 | -0.447 | 0.051 | 0.082 | 0.082 | 2.184 | 0.57 | 0.14 | 0.62 | 0.72 | 0.67 | 0.56 |
| 3 | 53217413 | rs2306572 | -0.418 | -1.022 | -2.179 | -0.569 | -0.003 | 0.032 | 0.032 | 0.245 | 0.29 | 0.08 | 0.27 | 0.21 | 0.27 | 0.24 |
| 3 | 53228919 | rs12490645 | -0.326 | -1.106 | -1.961 | -0.625 | -0.024 | 0.03 | 0.03 | 0.689 | 0.28 | 0.03 | 0.25 | 0.2 | 0.26 | 0.23 |
| 3 | 53234994 | rs11919522 | 0.431 | -0.802 | -1.782 | -0.592 | -0.012 | 0.008 | 0.008 | 1.192 | 0.42 | 0.26 | 0.38 | 0.61 | 0.39 | 0.46 |
| 3 | 53240854 | rs4328795 | -0.278 | -0.993 | -1.563 | -0.69 | -0.059 | -0.011 | -0.011 | 0 | 0.85 | 0.52 | 0.92 | 0.93 | 0.96 | 0.95 |
| 3 | 53241485 | rs4687712 | 0.301 | -0.696 | -1.686 | -0.617 | -0.039 | 0.082 | 0.082 | 0 | 0.42 | 0.22 | 0.53 | 0.28 | 0.56 | 0.35 |
| 3 | 53244603 | rs6772648 | 1.065 | -0.218 | -1.819 | -0.057 | 0.008 | -0.05 | -0.05 | 0 | 0.28 | 0.2 | 0.29 | 0.57 | 0.22 | 0.35 |
| 3 | 53252338 | rs11130358 | 0.359 | -0.061 | -2.231 | -0.275 | -0.101 | -0.071 | -0.071 | 0 | 0.43 | 0.06 | 0.39 | 0.6 | 0.44 | 0.48 |
| 3 | 53259608 | rs3736151 | 0.172 | -0.305 | -1.986 | -0.768 | 0.007 | 0.015 | 0.015 | 0 | 0.21 | 0.12 | 0.14 | 0.16 | 0.17 | 0.31 |
| 3 | 53259850 | rs3163 | 0.355 | -0.33 | -2.002 | -0.773 | -0.005 | 0.023 | 0.023 | 0 | 0.19 | 0.05 | 0.14 | 0.11 | 0.17 | 0.17 |
| 3 | 53259996 | rs3816767 | 0.415 | -0.37 | -1.787 | -0.787 | -0.007 | 0.045 | 0.045 | 1.247 | 0.49 | 0.21 | 0.42 | 0.6 | 0.46 | 0.55 |
| 3 | 53263279 | rs3736156 | -0.996 | -0.358 | -1.759 | -0.798 | -0.006 | 0.007 | 0.007 | 2.276 | 0.7 | 0.29 | 0.56 | 0.75 | 0.63 | 0.86 |
| 3 | 53265766 | rs11130362 | -0.34 | -0.61 | -1.277 | -0.943 | 0.001 | 0.009 | 0.009 | 0.002 | 0.26 | 0.23 | 0.18 | 0.18 | 0.22 | 0.38 |
| 3 | 53269046 | rs1051485 | 0.094 | -0.515 | -1.38 | -0.711 | 0.007 | 0.015 | 0.015 | 0 | 0.21 | 0.12 | 0.14 | 0.16 | 0.17 | 0.31 |

| | | | | | | | | | | | | | | | | |
|---|---|---|---|---|---|---|---|---|---|---|---|---|---|---|---|---|
| 3 | 53275777 | rs9864057 | -0.643 | -0.574 | -1.355 | -0.913 | 0 | 0.07 | 0.07 | 0 | 0.76 | 0.88 | 0.86 | 0.94 | 0.75 | 0.88 |
| 3 | 53278241 | rs12493802 | 0.091 | -0.486 | -1.017 | -0.903 | -0.021 | 0.028 | 0.028 | 0.329 | 0.45 | 0.29 | 0.63 | 0.37 | 0.59 | 0.26 |
| 3 | 53278653 | rs7646760 | -1.075 | -0.563 | -0.246 | -0.934 | -0.006 | 0.027 | 0.027 | 0.111 | 0.69 | 0.62 | 0.75 | 0.43 | 0.77 | 0.42 |
| 3 | 53282188 | rs4687717 | -0.567 | -0.586 | -0.036 | -0.919 | -0.008 | -0.015 | -0.015 | 1.519 | 0.69 | 0.28 | 0.57 | 0.63 | 0.62 | 0.64 |
| 3 | 53282303 | rs4687718 | 0.191 | -0.837 | 0.339 | -1.212 | -0.028 | -0.081 | -0.081 | 0.993 | 0.74 | 0.39 | 0.87 | 0.96 | 0.86 | 0.87 |
| 3 | 53286688 | rs9850440 | 0.348 | -0.782 | 0.514 | -1.23 | -0.021 | -0.104 | -0.104 | 0 | 0.29 | 0.38 | 0.17 | 0.07 | 0.25 | 0.14 |
| 3 | 53289527 | rs7633966 | -0.695 | -0.64 | 0.573 | -1.176 | 0.012 | 0.013 | 0.013 | 0 | 0.49 | 0.54 | 0.29 | 0.13 | 0.42 | 0.3 |
| 3 | 53293373 | rs11130367 | 1.177 | -0.657 | 0.679 | -1.089 | 0.006 | 0.006 | 0.006 | 0 | 0.25 | 0.3 | 0.5 | 0.28 | 0.35 | 0.19 |
| 3 | 53293986 | rs12490667 | 1.681 | -0.646 | 0.521 | -1.084 | -0.053 | -0.052 | -0.052 | 0.18 | 0.22 | 0.11 | 0.48 | 0.28 | 0.35 | 0.19 |
| 3 | 53294215 | rs17052957 | -1.462 | -0.658 | 0.584 | -1.062 | 0.001 | 0.021 | 0.021 | 0.375 | 0.73 | 0.7 | 0.77 | 0.46 | 0.77 | 0.67 |
| 3 | 53312915 | rs6798771 | 1.654 | -0.604 | 0.448 | -1.041 | -0.064 | -0.063 | -0.063 | 0 | 0.22 | 0.08 | 0.4 | 0.28 | 0.35 | 0.19 |
| 3 | 53318868 | rs11551405 | 0.937 | -0.724 | 0.439 | -1.129 | -0.006 | 0.131 | 0.131 | 0 | 0.25 | 0.04 | 0.14 | 0.07 | 0.25 | 0.13 |
| 3 | 53321002 | rs6792874 | -0.659 | -0.73 | 0.624 | -1.126 | -0.002 | -0.026 | -0.026 | 0.001 | 0.49 | 0.48 | 0.35 | 0.18 | 0.41 | 0.5 |
| 3 | 53321631 | rs9628 | -0.65 | -0.73 | 0.624 | -1.126 | -0.002 | -0.026 | -0.026 | 0.348 | 0.49 | 0.48 | 0.35 | 0.18 | 0.42 | 0.5 |
| 3 | 53323914 | rs2279319 | -0.591 | -0.722 | 0.676 | -1.122 | 0.011 | 0.021 | 0.021 | 0 | 0.48 | 0.4 | 0.34 | 0.18 | 0.41 | 0.49 |
| 3 | 53332363 | rs10461029 | -0.159 | -0.735 | 0.666 | -1.114 | 0.011 | 0.179 | 0.179 | 0 | 0.43 | 0.17 | 0.25 | 0.13 | 0.41 | 0.29 |
| 3 | 53362310 | rs6445576 | -1.42 | -0.635 | 0.539 | -1.056 | 0.002 | 0.039 | 0.039 | 0.248 | 0.72 | 0.65 | 0.76 | 0.48 | 0.76 | 0.68 |
| 3 | 53385418 | rs4687576 | 1.649 | -0.526 | 0.114 | -0.981 | -0.064 | -0.063 | -0.063 | 0 | 0.22 | 0.08 | 0.4 | 0.28 | 0.35 | 0.19 |
| 3 | 53397244 | rs2884525 | -1.134 | -0.466 | 0.182 | -0.912 | 0 | 0.062 | 0.062 | 0 | 0.71 | 0.57 | 0.75 | 0.47 | 0.76 | 0.68 |
| 3 | 53397481 | rs10510763 | 0.554 | -0.399 | -0.154 | -0.951 | -0.002 | 0.016 | 0.016 | 0 | 0.17 | 0.13 | 0.11 | 0.07 | 0.17 | 0.16 |
| 3 | 53415468 | rs12053905 | 0.25 | -0.586 | 0.027 | -0.963 | -0.004 | -0.047 | -0.047 | 0 | 0.31 | 0.36 | 0.17 | 0.07 | 0.25 | 0.13 |
| 3 | 53424056 | rs6783810 | 0.515 | -0.526 | -0.22 | -0.859 | -0.1 | -0.047 | -0.047 | 0 | 0.38 | 0.1 | 0.5 | 0.36 | 0.51 | 0.35 |
| 3 | 53434612 | rs4687728 | -0.973 | -0.654 | 0.026 | -0.293 | 0.002 | 0.033 | 0.033 | 0.187 | 0.72 | 0.6 | 0.76 | 0.6 | 0.76 | 0.67 |
| 3 | 53441376 | rs10452033 | 0.073 | -0.737 | -0.365 | -0.494 | -0.018 | -0.021 | -0.021 | 0 | 0.27 | 0.37 | 0.21 | 0.2 | 0.22 | 0.28 |
| 3 | 53445957 | rs674628 | -0.844 | -0.631 | -0.458 | -0.58 | 0.003 | 0.063 | 0.063 | 0 | 0.54 | 0.71 | 0.39 | 0.3 | 0.47 | 0.53 |
| 3 | 53454680 | rs660746 | 1.683 | -0.477 | -0.649 | -0.495 | -0.044 | -0.103 | -0.103 | 0.355 | 0.22 | 0.05 | 0.22 | 0.42 | 0.22 | 0.31 |
| 3 | 53456549 | rs13314396 | 0.247 | -0.726 | -0.486 | -0.681 | 0.027 | 0.155 | 0.155 | 0 | 0.41 | 0.19 | 0.21 | 0.09 | 0.36 | 0.27 |
| 3 | 53458934 | rs681751 | -0.059 | -0.726 | -0.221 | -0.743 | 0.026 | 0 | 0 | 0.056 | 0.88 | 0.63 | 0.91 | 0.92 | 0.89 | 0.93 |
| 3 | 53466077 | rs667217 | 1.614 | -0.8 | -0.457 | -0.724 | -0.001 | 0.002 | 0.002 | 0 | 0.24 | 0.21 | 0.23 | 0.4 | 0.22 | 0.3 |
| 3 | 53467506 | rs9880156 | 0.094 | -0.815 | -0.6 | -0.79 | 0.033 | 0.146 | 0.146 | 0 | 0.42 | 0.2 | 0.3 | 0.2 | 0.38 | 0.46 |
| 3 | 53470863 | rs511963 | 0.668 | -1.059 | -0.713 | -0.865 | 0.016 | -0.018 | -0.018 | 0.055 | 0.37 | 0.23 | 0.27 | 0.42 | 0.26 | 0.39 |
| 3 | 53481714 | rs551551 | -0.401 | -1.049 | -0.734 | -0.882 | 0.002 | 0.007 | 0.007 | 0 | 0.65 | 0.39 | 0.43 | 0.46 | 0.59 | 0.56 |
| 3 | 53481831 | rs9876191 | 0.82 | -0.768 | -0.91 | -0.813 | -0.022 | 0.089 | 0.089 | 0 | 0.29 | 0.15 | 0.17 | 0.05 | 0.32 | 0.19 |
| 3 | 53487279 | rs2930176 | 1.117 | -0.889 | -0.355 | -0.299 | 0 | -0.007 | -0.007 | 0 | 0.47 | 0.22 | 0.44 | 0.56 | 0.53 | 0.66 |
| 3 | 53489090 | rs9830114 | 0.495 | -0.427 | -0.519 | -0.257 | -0.006 | -0.021 | -0.021 | 0.135 | 0.07 | 0.12 | 0.03 | 0.04 | 0.05 | 0.04 |
| 3 | 53495847 | rs555480 | 0.987 | -0.256 | -0.333 | 0.079 | 0.015 | 0.025 | 0.025 | 0.698 | 0.51 | 0.23 | 0.38 | 0.43 | 0.56 | 0.49 |
| 3 | 53514416 | rs312476 | 0.198 | 0.247 | -0.731 | -0.242 | 0 | -0.006 | -0.006 | 0 | 0.24 | 0.23 | 0.37 | 0.32 | 0.25 | 0.18 |
| 3 | 53524051 | rs312483 | 0.715 | 0.46 | -0.801 | 0.007 | -0.091 | -0.119 | -0.119 | 0.507 | 0.2 | 0.03 | 0.36 | 0.32 | 0.28 | 0.17 |
| 3 | 53529140 | rs312480 | 0.146 | 0.649 | -0.171 | -0.046 | 0.113 | 0.084 | 0.084 | 0 | 0.93 | 0.64 | 0.91 | 0.96 | 0.91 | 0.96 |
| 3 | 53537879 | rs6763768 | -0.212 | 0.344 | -1.092 | -0.089 | 0.006 | 0 | 0 | 0 | 0.36 | 0.25 | 0.44 | 0.33 | 0.33 | 0.25 |
| 3 | 53545622 | rs312487 | 0.127 | 0.384 | -1.702 | -0.42 | -0.068 | -0.084 | -0.084 | 0 | 0.43 | 0.12 | 0.56 | 0.57 | 0.49 | 0.49 |
| 3 | 53551010 | rs3774421 | 0.625 | 0.596 | -1.924 | -0.655 | -0.026 | 0.032 | 0.032 | 0 | 0.22 | 0.08 | 0.17 | 0.13 | 0.28 | 0.11 |
| 3 | 53551902 | rs312481 | -1.164 | 0.385 | -1.652 | -0.694 | -0.024 | -0.007 | -0.007 | 0 | 0.81 | 0.88 | 0.83 | 0.86 | 0.77 | 0.83 |
| 3 | 53556223 | rs979220 | 0.09 | -0.108 | -1.869 | -1.138 | -0.009 | -0.03 | -0.03 | 0 | 0.35 | 0.63 | 0.23 | 0.12 | 0.24 | 0.22 |
| 3 | 53556658 | rs3774424 | 0.746 | 0.246 | -1.689 | -0.865 | -0.039 | 0.03 | 0.03 | 0 | 0.21 | 0.07 | 0.17 | 0.13 | 0.28 | 0.11 |
| 3 | 53565829 | rs3774428 | 0.586 | -0.104 | -1.265 | -0.807 | -0.001 | -0.012 | -0.012 | 0.744 | 0.25 | 0.21 | 0.16 | 0.13 | 0.24 | 0.1 |
| 3 | 53573517 | rs13078747 | 0.202 | -0.269 | -1.457 | -0.914 | -0.021 | 0.116 | 0.116 | 0.193 | 0.44 | 0.18 | 0.36 | 0.19 | 0.51 | 0.39 |
| 3 | 53583468 | rs6764908 | 0.988 | -0.205 | -1.117 | -0.915 | -0.003 | 0.012 | 0.012 | 0 | 0.23 | 0.17 | 0.15 | 0.13 | 0.26 | 0.11 |
| 3 | 53583724 | rs2109182 | -0.152 | -0.246 | -0.854 | -0.867 | -0.023 | 0.128 | 0.128 | 1.8 | 0.07 | 0.32 | 0.05 | 0.13 | 0.01 | 0.22 |
| 3 | 53585307 | rs3821846 | 0.198 | -0.202 | -0.678 | -0.996 | -0.004 | -0.051 | -0.051 | 0 | 0.34 | 0.44 | 0.22 | 0.18 | 0.32 | 0.33 |
| 3 | 53587089 | rs3774443 | 0.007 | -0.125 | -0.239 | -0.315 | 0.002 | 0.094 | 0.094 | 1.889 | 0.06 | 0.3 | 0.04 | 0.08 | 0 | 0.01 |
| 3 | 53587519 | rs991818 | 0.303 | 0.448 | -0.605 | -0.261 | 0.028 | -0.043 | -0.043 | 0.121 | 0.33 | 0.5 | 0.26 | 0.18 | 0.36 | 0.26 |
| 3 | 53592864 | rs2359453 | -0.003 | 0.709 | -1.224 | 0.706 | -0.018 | -0.13 | -0.13 | 0 | 0.78 | 0.94 | 0.58 | 0.45 | 0.77 | 0.71 |
| 3 | 53600137 | rs17053186 | -0.94 | 0.924 | -0.165 | 0.284 | 0.031 | -0.005 | -0.005 | 0 | 0.17 | 0.44 | 0.16 | 0.17 | 0.17 | 0.18 |
| 3 | 53605845 | rs10490772 | -0.265 | 0.619 | -0.821 | -0.113 | -0.018 | 0.079 | 0.079 | 0.024 | 0.16 | 0.01 | 0.12 | 0.02 | 0.15 | 0.1 |
| 3 | 53609610 | rs709317 | 0.35 | 0.815 | -0.689 | 0.369 | -0.001 | 0 | 0 | 0 | 0.87 | 0.86 | 0.85 | 0.8 | 0.87 | 0.85 |
| 3 | 53614484 | rs709323 | 0.823 | 0.487 | 0.412 | 0.031 | -0.002 | 0.007 | 0.007 | 0 | 0.79 | 0.72 | 0.81 | 0.77 | 0.8 | 0.81 |
| 3 | 53616338 | rs807193 | 0.604 | 0.545 | -0.123 | 0.677 | -0.007 | -0.049 | -0.049 | 0 | 0.82 | 0.86 | 0.81 | 0.65 | 0.82 | 0.72 |
| 3 | 53617868 | rs3774458 | -0.967 | 0.578 | 0.062 | 0.32 | 0.001 | 0.005 | 0.005 | 0.722 | 0.29 | 0.29 | 0.25 | 0.19 | 0.29 | 0.24 |
| 3 | 53629706 | rs3774468 | -0.911 | 0.804 | -0.395 | 0.154 | -0.011 | 0.052 | 0.052 | 0 | 0.34 | 0.21 | 0.3 | 0.21 | 0.38 | 0.35 |
| 3 | 53630679 | rs6445588 | -1.281 | 0.403 | 0.112 | 0.393 | 0.008 | 0.112 | 0.112 | 0.13 | 0.14 | 0.39 | 0.2 | 0.28 | 0.14 | 0.26 |
| 3 | 53634426 | rs936450 | -0.977 | -0.497 | -0.121 | 0.112 | 0.048 | -0.012 | -0.012 | 0 | 0.46 | 0.76 | 0.4 | 0.4 | 0.39 | 0.4 |
| 3 | 53637695 | rs7610071 | -0.709 | -0.614 | 0.51 | -0.118 | -0.02 | -0.004 | -0.004 | 0 | 0.38 | 0.51 | 0.31 | 0.28 | 0.34 | 0.26 |
| 3 | 53638711 | rs17309406 | -0.062 | -0.661 | 0.743 | -0.256 | 0.017 | 0.004 | 0.004 | 0 | 0.28 | 0.18 | 0.15 | 0 | 0.24 | 0.12 |
| 3 | 53638851 | rs3774473 | -0.036 | -0.481 | 0.821 | -0.029 | -0.002 | 0.008 | 0.008 | 0 | 0.07 | 0.12 | 0.14 | 0.17 | 0.09 | 0.14 |
| 3 | 53647812 | rs1460115 | -0.316 | -0.259 | 0.719 | -0.299 | 0.009 | 0.01 | 0.01 | 0 | 0.14 | 0.24 | 0.17 | 0.17 | 0.18 | 0.15 |
| 3 | 53649677 | rs13063116 | -0.425 | -0.373 | 0.838 | -0.091 | -0.007 | 0.015 | 0.015 | 0 | 0.08 | 0.13 | 0.07 | 0.13 | 0.08 | 0.12 |
| 3 | 53654893 | rs3774480 | 0.772 | -0.095 | -0.26 | -0.014 | -0.051 | -0.058 | -0.058 | 5.02 | 0.27 | 0.02 | 0.3 | 0.34 | 0.23 | 0.4 |
| 3 | 53656804 | rs9850569 | -0.317 | -0.252 | 0.073 | 0.055 | 0.002 | 0.003 | 0.003 | 0 | 0.41 | 0.55 | 0.38 | 0.36 | 0.46 | 0.36 |

| | | | | | | | | | | | | | | | | |
|---|---|---|---|---|---|---|---|---|---|---|---|---|---|---|---|---|
| 3 | 53662194 | rs3774487 | -0.615 | 0.079 | 0.596 | -0.119 | 0.028 | 0.119 | 0.119 | 0 | 0.08 | 0.26 | 0.06 | 0.13 | 0.08 | 0.09 |
| 3 | 53662752 | rs7630805 | -0.896 | -0.26 | 0.201 | -0.093 | -0.092 | 0.199 | 0.199 | 2.018 | 0.57 | 0.93 | 0.59 | 0.81 | 0.5 | 0.7 |
| 3 | 53663648 | rs3774490 | 0.231 | -0.249 | 0.314 | -0.053 | 0.021 | 0.094 | 0.094 | 0 | 0.07 | 0.17 | 0.05 | 0.14 | 0.08 | 0.09 |
| 3 | 53668708 | rs11707976 | 0.879 | -0.52 | 0.271 | -0.287 | -0.012 | -0.073 | -0.073 | 0 | 0.26 | 0.31 | 0.14 | 0 | 0.2 | 0.1 |
| 3 | 53669904 | rs11709171 | 1.356 | -0.499 | -0.091 | -0.276 | -0.004 | 0.142 | 0.142 | 0 | 0.22 | 0.03 | 0.12 | 0 | 0.2 | 0.1 |
| 3 | 53675256 | rs12486415 | 0.78 | -0.374 | 0.038 | 0.116 | -0.007 | -0.043 | -0.043 | 0.097 | 0.17 | 0.15 | 0.34 | 0.4 | 0.2 | 0.45 |
| 3 | 53685039 | rs2276836 | 0.78 | -0.373 | -0.011 | 0.128 | -0.008 | -0.046 | -0.046 | 4.33 | 0.17 | 0.15 | 0.35 | 0.58 | 0.2 | 0.46 |
| 3 | 53688958 | rs11712998 | 0.942 | -0.426 | 0.052 | -0.178 | -0.009 | -0.069 | -0.069 | 0 | 0.26 | 0.3 | 0.14 | 0 | 0.2 | 0.1 |
| 3 | 53696955 | rs1401494 | -1.161 | -0.081 | -0.411 | -0.191 | -0.127 | 0.199 | 0.199 | 1.742 | 0.59 | 0.93 | 0.61 | 0.84 | 0.53 | 0.72 |
| 3 | 53700550 | rs2250736 | -1.129 | -0.069 | -0.405 | -0.15 | -0.125 | 0.041 | 0.041 | 1.553 | 0.58 | 0.93 | 0.61 | 0.65 | 0.53 | 0.71 |
| 3 | 53700704 | rs3821856 | 1.24 | -0.032 | -0.47 | -0.256 | -0.047 | 0.081 | 0.081 | 0.019 | 0.2 | 0.02 | 0.14 | 0.11 | 0.19 | 0.07 |
| 3 | 53705003 | rs3774527 | 2.497 | 0.218 | -0.448 | -0.259 | -0.01 | 0.028 | 0.028 | 0.034 | 0.08 | 0.02 | 0.06 | 0.01 | 0.08 | 0.03 |
| 3 | 53709851 | rs1460118 | 0.828 | 0.497 | -0.214 | -0.12 | 0.001 | -0.004 | -0.004 | 0 | 0.32 | 0.34 | 0.31 | 0.26 | 0.34 | 0.19 |
| 3 | 53712910 | rs6809164 | 0.902 | 0.591 | -0.2 | -0.122 | -0.003 | -0.018 | -0.018 | 0 | 0.31 | 0.33 | 0.28 | 0.13 | 0.29 | 0.2 |
| 3 | 53715973 | rs2680672 | -0.498 | 0.906 | 0.214 | 0.079 | -0.005 | -0.005 | -0.005 | 1.051 | 0.64 | 0.67 | 0.54 | 0.61 | 0.59 | 0.42 |
| 3 | 53716912 | rs898411 | 0.844 | 0.87 | 0.24 | 0.078 | -0.004 | -0.006 | -0.006 | 0.317 | 0.36 | 0.36 | 0.32 | 0.22 | 0.29 | 0.24 |
| 3 | 53717282 | rs898422 | -0.737 | 0.924 | -0.071 | 0 | -0.027 | 0.118 | 0.118 | 0.076 | 0.61 | 0.77 | 0.74 | 0.89 | 0.6 | 0.8 |
| 3 | 53722874 | rs11130377 | 1.005 | 0.867 | -0.117 | -0.016 | 0.014 | -0.042 | -0.042 | 0 | 0.33 | 0.08 | 0.3 | 0.36 | 0.29 | 0.26 |
| 3 | 53724370 | rs1380608 | 0.768 | 0.768 | 0.09 | 0.164 | 0.011 | 0.028 | 0.028 | 0 | 0.06 | 0.09 | 0.06 | 0.18 | 0.11 | 0.08 |
| 3 | 53729391 | rs898417 | -1.781 | 0.769 | -0.077 | 0.094 | -0.043 | 0.085 | 0.085 | 0.17 | 0.83 | 0.97 | 0.9 | 1 | 0.79 | 0.87 |
| 3 | 53730735 | rs9841978 | 0.042 | 0.318 | 0.672 | 0.584 | -0.016 | -0.019 | -0.019 | 0 | 0.37 | 0.46 | 0.23 | 0.27 | 0.32 | 0.23 |
| 3 | 53734443 | rs7340705 | -0.134 | 0.422 | 0.64 | 0.636 | 0.024 | -0.011 | -0.011 | 0 | 0.4 | 0.67 | 0.26 | 0.39 | 0.32 | 0.26 |
| 3 | 53734531 | rs6445597 | -0.042 | 0.452 | 0.408 | 0.52 | -0.015 | -0.011 | -0.011 | 0 | 0.38 | 0.45 | 0.24 | 0.25 | 0.32 | 0.26 |
| 3 | 53735766 | rs9830632 | 0.226 | 0.448 | 0.106 | 0.017 | -0.024 | -0.062 | -0.062 | 3.318 | 0.36 | 0.49 | 0.23 | 0.15 | 0.3 | 0.19 |
| 3 | 53737575 | rs3774531 | 0.23 | 0.923 | -0.2 | 0.154 | 0.001 | 0.012 | 0.012 | 1.128 | 0.52 | 0.49 | 0.55 | 0.66 | 0.51 | 0.65 |
| 3 | 53738424 | rs2633731 | -0.237 | 1.135 | -1.006 | 0.323 | -0.029 | 0.093 | 0.093 | 0.467 | 0.63 | 0.82 | 0.63 | 0.87 | 0.62 | 0.76 |
| 3 | 53740699 | rs2633730 | 0.12 | 1.018 | -0.841 | 0.816 | -0.007 | -0.001 | -0.001 | 0 | 0.56 | 0.63 | 0.48 | 0.43 | 0.49 | 0.47 |
| 3 | 53742148 | rs2680661 | -0.331 | 0.227 | -0.495 | 0.823 | -0.012 | 0.038 | 0.038 | 0 | 0.25 | 0.32 | 0.34 | 0.47 | 0.23 | 0.23 |
| 3 | 53742331 | rs2612015 | -0.578 | 0.278 | -1.117 | -0.436 | 0.022 | 0.098 | 0.098 | 0 | 0.57 | 0.74 | 0.75 | 0.88 | 0.64 | 0.74 |
| 3 | 53743112 | rs13074345 | 0.253 | -0.172 | -0.944 | -0.07 | -0.038 | 0.031 | 0.031 | 0 | 0.35 | 0.23 | 0.47 | 0.76 | 0.41 | 0.48 |
| 3 | 53749498 | rs11709505 | 1.299 | -0.088 | -0.718 | -0.879 | 0.016 | -0.075 | -0.075 | 0 | 0.2 | 0.29 | 0.16 | 0.01 | 0.24 | 0.05 |
| 3 | 53758323 | rs7638857 | -0.14 | -0.917 | -0.947 | -0.953 | -0.007 | -0.004 | -0.004 | 0 | 0.36 | 0.37 | 0.42 | 0.73 | 0.29 | 0.57 |
| 3 | 53759389 | rs17053417 | 1.16 | -1.373 | -1.038 | -1.165 | 0.004 | -0.033 | -0.033 | 0 | 0.25 | 0.35 | 0.18 | 0.1 | 0.26 | 0.08 |
| 3 | 53766284 | rs11926741 | 1.407 | -1.648 | -1.271 | -0.83 | 0.024 | 0.047 | 0.047 | 0 | 0.15 | 0.02 | 0.06 | 0.08 | 0.09 | 0.06 |
| 3 | 53771914 | rs11915963 | 1.258 | -1.508 | -1.279 | -0.833 | 0.002 | 0.048 | 0.048 | 0 | 0.16 | 0.02 | 0.08 | 0.09 | 0.12 | 0.07 |
| 3 | 53773165 | rs2680652 | -0.931 | -1.878 | -1.088 | -0.938 | -0.002 | -0.012 | -0.012 | 0 | 0.81 | 0.81 | 0.9 | 0.91 | 0.84 | 0.85 |
| 3 | 53776678 | rs2612027 | 0.028 | -2.115 | -0.978 | -0.827 | -0.007 | 0.013 | 0.013 | 0 | 0.66 | 0.41 | 0.62 | 0.49 | 0.67 | 0.39 |
| 3 | 53776995 | rs2612026 | -0.7 | -2.229 | -0.716 | -0.912 | -0.02 | -0.159 | -0.159 | 1.555 | 0.79 | 0.53 | 0.88 | 1 | 0.83 | 0.88 |
| 3 | 53777176 | rs2680648 | 1.905 | -2.246 | -1.567 | -1.135 | -0.09 | -0.102 | -0.102 | 0.505 | 0.24 | 0.04 | 0.3 | 0.3 | 0.24 | 0.11 |
| 3 | 53778875 | rs2253795 | 2.029 | -2.257 | -1.573 | -1.204 | -0.092 | 0.03 | 0.03 | 0 | 0.23 | 0.04 | 0.28 | 0.13 | 0.24 | 0.1 |
| 3 | 53779776 | rs17053453 | 1.255 | -2.554 | -1.525 | -1.322 | 0.026 | 0.022 | 0.022 | 0 | 0.16 | 0.08 | 0.06 | 0.09 | 0.08 | 0.04 |
| 3 | 53779825 | rs3821860 | 0.922 | -2.508 | -1.445 | -1.301 | -0.031 | 0.044 | 0.044 | 0 | 0.26 | 0.15 | 0.25 | 0.15 | 0.35 | 0.3 |
| 3 | 53780994 | rs17053459 | 1.419 | -2.452 | -1.337 | -1.185 | -0.02 | 0.005 | 0.005 | 1.558 | 0.06 | 0.09 | 0.03 | 0.1 | 0.02 | 0.04 |
| 3 | 53788732 | rs17053472 | 0.189 | -2.403 | -1.323 | -1.245 | -0.022 | 0.057 | 0.057 | 0 | 0.37 | 0.26 | 0.32 | 0.09 | 0.44 | 0.26 |
| 3 | 53792890 | rs3796347 | -0.685 | -2.576 | -1.422 | -1.198 | -0.009 | 0.117 | 0.117 | 0 | 0.54 | 0.73 | 0.42 | 0.2 | 0.54 | 0.35 |
| 3 | 53799960 | rs3774573 | 0.886 | -2.468 | -0.641 | -0.725 | -0.001 | -0.007 | -0.007 | 0 | 0.31 | 0.3 | 0.33 | 0.18 | 0.25 | 0.12 |
| 3 | 53800143 | rs3774574 | 1.537 | -2.432 | -0.719 | -0.824 | -0.085 | -0.013 | -0.013 | 0 | 0.25 | 0.03 | 0.29 | 0.17 | 0.24 | 0.1 |
| 3 | 53800222 | rs1109959 | 1.398 | -1.855 | -0.199 | -0.483 | -0.019 | 0.039 | 0.039 | 1.319 | 0.05 | 0.14 | 0.03 | 0.14 | 0.02 | 0.04 |
| 3 | 53802748 | rs3774581 | 0.189 | -1.987 | -0.373 | -0.444 | 0.01 | 0.002 | 0.002 | 0 | 0.45 | 0.39 | 0.39 | 0.39 | 0.32 | 0.15 |
| 3 | 53804489 | rs17053501 | 0.733 | -1.833 | -0.852 | -0.451 | 0.006 | 0.005 | 0.005 | 0 | 0.1 | 0.07 | 0.04 | 0.01 | 0.03 | 0.02 |
| 3 | 53805023 | rs9881910 | -1.047 | -1.522 | -1.111 | -0.387 | 0 | 0.006 | 0.006 | 0 | 0.73 | 0.73 | 0.63 | 0.38 | 0.68 | 0.4 |
| 3 | 53806023 | rs11918583 | 0.871 | -1.473 | -1.309 | -0.64 | -0.071 | 0.088 | 0.088 | 0 | 0.22 | 0.1 | 0.21 | 0.05 | 0.34 | 0.21 |
| 3 | 53807148 | rs2278516 | -0.57 | -1.152 | -1.043 | -0.29 | -0.002 | -0.01 | -0.01 | 0 | 0.69 | 0.55 | 0.59 | 0.33 | 0.66 | 0.33 |
| 3 | 53813232 | rs7617553 | -0.636 | -0.964 | -1.167 | -0.384 | 0.002 | 0.025 | 0.025 | 0.077 | 0.72 | 0.64 | 0.61 | 0.45 | 0.67 | 0.34 |
| 3 | 53822893 | rs3774601 | 0.528 | -1.13 | -1.335 | 0.085 | -0.144 | -0.034 | -0.034 | 0 | 0.31 | 0.04 | 0.42 | 0.25 | 0.36 | 0.14 |
| 3 | 53832712 | rs3774608 | -0.792 | -0.718 | -0.946 | 0.334 | -0.001 | 0.042 | 0.042 | 0.191 | 0.53 | 0.87 | 0.54 | 0.41 | 0.4 | 0.2 |
| 3 | 53833509 | rs3796349 | -1.374 | -0.273 | -0.169 | 0.803 | -0.001 | 0.135 | 0.135 | 0.039 | 0.09 | 0.29 | 0.27 | 0.25 | 0.05 | 0.05 |
| 3 | 53840665 | rs3774611 | 0.707 | -0.116 | -0.992 | 0.796 | -0.106 | -0.074 | -0.074 | 0 | 0.41 | 0.1 | 0.42 | 0.42 | 0.51 | 0.63 |
| 3 | 53845295 | rs10154841 | -0.899 | -0.432 | -0.514 | 0.38 | -0.021 | -0.014 | -0.014 | 0 | 0.17 | 0.65 | 0.14 | 0.15 | 0.11 | 0.1 |
| 3 | 53846924 | rs877484 | 0.846 | -0.29 | -0.671 | 0.541 | -0.082 | 0.11 | 0.11 | 0.022 | 0.43 | 0.11 | 0.44 | 0.69 | 0.52 | 0.7 |
| 3 | 53850814 | rs4563403 | -1.414 | -0.544 | 0.293 | 0.497 | 0.019 | 0.009 | 0.009 | 0 | 0.16 | 0.48 | 0.28 | 0.18 | 0.14 | 0.1 |
| 3 | 53852835 | rs2289209 | -0.696 | -0.543 | -0.07 | 0.496 | -0.008 | 0.014 | 0.014 | 0 | 0.08 | 0.18 | 0.21 | 0.12 | 0.05 | 0.05 |
| 3 | 53855083 | rs9836592 | -0.586 | -0.541 | -0.833 | 0.359 | -0.01 | 0.154 | 0.154 | 0 | 0.63 | 0.89 | 0.78 | 0.85 | 0.67 | 0.8 |
| 3 | 53856050 | rs6445606 | -1.035 | -0.506 | -0.874 | 0.3 | -0.033 | 0.173 | 0.173 | 0 | 0.68 | 0.9 | 0.8 | 1 | 0.69 | 0.88 |
| 3 | 53857158 | rs2241807 | 0.987 | -0.566 | -0.924 | 0.593 | -0.116 | -0.038 | -0.038 | 0 | 0.43 | 0.08 | 0.43 | 0.57 | 0.53 | 0.63 |
| 3 | 53860103 | rs9835128 | -1.825 | -0.477 | -0.027 | 0.79 | 0.002 | -0.003 | -0.003 | 0 | 0.25 | 0.73 | 0.38 | 0.41 | 0.15 | 0.17 |
| 3 | 53870840 | rs13317328 | -2.319 | -0.32 | 0.247 | 1.137 | 0.018 | 0.183 | 0.183 | 0.514 | 0.09 | 0.25 | 0.29 | 0.38 | 0.08 | 0.12 |
| 3 | 53873578 | rs930367 | -1.864 | -0.421 | 0.266 | 1.075 | -0.011 | 0.104 | 0.104 | 0.725 | 0.07 | 0.16 | 0.24 | 0.3 | 0.05 | 0.1 |
| 3 | 53876218 | rs6801605 | 0.08 | -0.676 | -0.15 | 0.467 | 0.005 | 0.025 | 0.025 | 1.22 | 0.59 | 0.63 | 0.52 | 0.7 | 0.63 | 0.78 |

| | | | | | | | | | | | | | | | | |
|---|---|---|---|---|---|---|---|---|---|---|---|---|---|---|---|---|
| 3 | 53878616 | rs2289205 | 0.141 | -0.828 | -0.42 | 0.235 | 0.021 | 0.139 | 0.139 | 0.189 | 0.38 | 0.19 | 0.23 | 0.07 | 0.31 | 0.18 |
| 3 | 53883910 | rs4687751 | 0.105 | -0.885 | 0.137 | 0.631 | 0.016 | -0.001 | -0.001 | 0 | 0.54 | 0.37 | 0.49 | 0.47 | 0.64 | 0.63 |
| 3 | 53889981 | rs999515 | 0.009 | -1.028 | 0.503 | 1.151 | 0.002 | 0.013 | 0.013 | 0 | 0.27 | 0.31 | 0.28 | 0.46 | 0.33 | 0.34 |
| 3 | 53893840 | rs3821869 | -0.462 | -1.373 | 0.893 | 1.045 | 0.001 | 0.013 | 0.013 | 0 | 0.33 | 0.37 | 0.3 | 0.46 | 0.35 | 0.4 |
| 3 | 53894412 | rs6766099 | -0.682 | -1.859 | 1.049 | 0.996 | -0.038 | 0.052 | 0.052 | 0.532 | 0.16 | 0.26 | 0.29 | 0.34 | 0.1 | 0.23 |
| 3 | 53899276 | rs1043261 | 0.228 | -1.957 | 1.16 | 1.062 | -0.002 | 0.032 | 0.032 | 0 | 0.07 | 0.09 | 0.11 | 0.21 | 0.07 | 0.2 |
| 3 | 53899565 | rs709324 | -0.442 | -1.962 | 1.203 | 1.034 | -0.001 | -0.011 | -0.011 | 0.565 | 0.35 | 0.49 | 0.33 | 0.6 | 0.35 | 0.43 |
| 3 | 53899580 | rs3017 | -0.29 | -1.959 | 1.179 | 0.987 | 0.001 | 0.021 | 0.021 | 0 | 0.34 | 0.44 | 0.3 | 0.46 | 0.35 | 0.4 |
| 3 | 53899601 | rs3774626 | -0.312 | -1.981 | 1.126 | 0.915 | -0.03 | 0.001 | 0.001 | 1.026 | 0.08 | 0.13 | 0.17 | 0.13 | 0.03 | 0.07 |
| 3 | 53899798 | rs6798958 | -0.483 | -2.068 | 1.176 | 0.957 | -0.025 | 0.022 | 0.022 | 0 | 0.07 | 0.21 | 0.05 | 0.09 | 0.01 | 0.13 |
| 3 | 53905308 | rs1046677 | -1.351 | -1.847 | 1.245 | 0.966 | 0.036 | 0.159 | 0.159 | 0.75 | 0.5 | 0.72 | 0.62 | 0.94 | 0.45 | 0.7 |
| 3 | 53906334 | rs2289204 | -1.092 | -1.858 | 1.296 | 1.013 | -0.003 | 0.094 | 0.094 | 0 | 0.07 | 0.12 | 0.11 | 0.25 | 0.05 | 0.21 |
| 3 | 53906513 | rs4687757 | -1.374 | -1.851 | 1.261 | 0.981 | 0.036 | 0.154 | 0.154 | 0.728 | 0.5 | 0.71 | 0.62 | 0.94 | 0.45 | 0.7 |
| 3 | 53909731 | rs893367 | -1.481 | -1.88 | 1.293 | 0.968 | 0.031 | 0.205 | 0.205 | 0.941 | 0.51 | 0.79 | 0.63 | 0.94 | 0.45 | 0.7 |
| 3 | 53914093 | rs3733082 | -0.199 | -1.842 | 1.258 | 1.022 | -0.027 | 0.005 | 0.005 | 0.699 | 0.08 | 0.13 | 0.18 | 0.13 | 0.03 | 0.07 |
| 3 | 53919720 | rs34310576 | 0.964 | -1.843 | 1.237 | 1.012 | 0.001 | 0.03 | 0.03 | 0 | 0.06 | 0.02 | 0.02 | 0 | 0.04 | 0.05 |
| 3 | 53922257 | rs2304576 | 0.153 | -1.832 | 1.312 | 1.072 | 0.005 | 0.018 | 0.018 | 0 | 0.3 | 0.43 | 0.31 | 0.56 | 0.33 | 0.36 |
| 3 | 53928996 | rs726028 | -0.357 | -1.829 | 1.206 | 1.096 | -0.028 | 0.044 | 0.044 | 0.861 | 0.08 | 0.13 | 0.2 | 0.24 | 0.03 | 0.11 |
| 3 | 53929806 | rs6445612 | 0.424 | -1.824 | 1.135 | 1.079 | 0.008 | 0.041 | 0.041 | 0 | 0.28 | 0.36 | 0.28 | 0.46 | 0.32 | 0.33 |
| 3 | 53948316 | rs7624521 | -2.46 | -2.043 | 1.768 | 1.646 | -0.05 | 0.148 | 0.148 | 0 | 0.17 | 0.32 | 0.32 | 0.48 | 0.09 | 0.32 |
| 3 | 53971360 | rs7611432 | -1.121 | -1.756 | 1.549 | 1.539 | -0.021 | 0.044 | 0.044 | 0.037 | 0.1 | 0.4 | 0.2 | 0.12 | 0.04 | 0.06 |
| 3 | 53985919 | rs13064140 | -1.035 | -1.702 | 1.208 | 1.61 | -0.001 | 0.025 | 0.025 | 0 | 0.11 | 0.13 | 0.14 | 0.25 | 0.09 | 0.26 |
| 3 | 53992218 | rs7650055 | -1.121 | -1.943 | 0.924 | 1.271 | -0.039 | -0.002 | -0.002 | 0 | 0.1 | 0.19 | 0.19 | 0.13 | 0.04 | 0.06 |
| 3 | 53996479 | rs7639932 | -1.895 | -1.988 | 0.635 | 1.386 | -0.062 | 0.085 | 0.085 | 0.717 | 0.24 | 0.57 | 0.35 | 0.56 | 0.1 | 0.43 |
| 3 | 53997009 | rs4582049 | 0.873 | -1.919 | -0.672 | 1.023 | -0.082 | 0.041 | 0.041 | 0.323 | 0.38 | 0.08 | 0.37 | 0.28 | 0.54 | 0.42 |
| 3 | 54010039 | rs9876934 | -0.429 | -1.741 | 0.076 | 0.59 | -0.004 | -0.058 | -0.058 | 0 | 0.47 | 0.52 | 0.35 | 0.06 | 0.37 | 0.19 |
| 3 | 54017818 | rs4687765 | -0.349 | -1.755 | 0.09 | 0.587 | 0.002 | -0.048 | -0.048 | 0.099 | 0.46 | 0.48 | 0.29 | 0.03 | 0.37 | 0.19 |
| 3 | 54021866 | rs9817109 | -0.3 | -1.758 | 0.16 | 0.579 | 0.013 | 0.119 | 0.119 | 0 | 0.45 | 0.36 | 0.33 | 0.06 | 0.37 | 0.19 |
| 3 | 54036605 | rs6763067 | -1.497 | -1.679 | 0.25 | 1.096 | -0.016 | 0.257 | 0.257 | 0.206 | 0.11 | 0.5 | 0.15 | 0.4 | 0.06 | 0.28 |
| 3 | 54053981 | rs4927992 | 0.159 | -1.64 | -0.014 | 1.044 | -0.031 | 0.026 | 0.026 | 0.634 | 0.06 | 0.19 | 0.15 | 0.05 | 0.01 | 0.05 |
| 3 | 54063193 | rs9810229 | -0.308 | -1.76 | -0.181 | 1.05 | 0.005 | -0.004 | -0.004 | 0 | 0.42 | 0.67 | 0.34 | 0.46 | 0.37 | 0.37 |
| 3 | 54063875 | rs12492799 | 0.075 | -1.881 | -0.239 | 1 | 0 | -0.004 | -0.004 | 0 | 0.37 | 0.59 | 0.32 | 0.45 | 0.35 | 0.33 |
| 3 | 54065571 | rs9879164 | -1.387 | -1.896 | -0.335 | 0.998 | -0.088 | -0.106 | -0.106 | 0.888 | 0.56 | 0.95 | 0.52 | 0.52 | 0.44 | 0.47 |
| 3 | 54067037 | rs17053673 | -0.127 | -1.646 | -0.187 | 1.143 | -0.001 | -0.016 | -0.016 | 2.477 | 0.06 | 0.02 | 0.08 | 0.21 | 0.07 | 0.25 |
| 3 | 54068119 | rs9846278 | -1.993 | -1.756 | 0.141 | 0.747 | -0.06 | -0.004 | -0.004 | 1.471 | 0.23 | 0.6 | 0.28 | 0.2 | 0.08 | 0.28 |
| 3 | 54073008 | rs13088277 | -0.401 | -1.109 | -0.631 | 0.684 | 0 | 0.006 | 0.006 | 0 | 0.38 | 0.57 | 0.26 | 0.26 | 0.33 | 0.13 |
| 3 | 54073300 | rs9873127 | -0.736 | -0.673 | -0.439 | 0.882 | -0.029 | 0.035 | 0.035 | 0.589 | 0.06 | 0.17 | 0.15 | 0.1 | 0.01 | 0.05 |
| 3 | 54074534 | rs13095224 | -0.921 | -0.065 | -0.701 | 0.54 | -0.003 | 0.035 | 0.035 | 0 | 0.07 | 0.02 | 0.03 | 0 | 0.05 | 0.05 |
| 3 | 54077726 | rs1992965 | -1.147 | 0.188 | -0.391 | 0.73 | -0.006 | -0.008 | -0.008 | 0 | 0.17 | 0.31 | 0.21 | 0.14 | 0.16 | 0.19 |
| 3 | 54084917 | rs9862917 | -0.779 | 0.717 | -0.075 | 1.126 | 0.053 | 0.097 | 0.097 | 0.775 | 0.27 | 0.52 | 0.47 | 0.51 | 0.39 | 0.46 |
| 3 | 54092440 | rs1449339 | -0.363 | 0.847 | 0.317 | 1.473 | -0.003 | -0.003 | -0.003 | 0 | 0.19 | 0.17 | 0.32 | 0.28 | 0.29 | 0.3 |
| 3 | 54093419 | rs17744447 | -1.157 | 0.838 | 0.671 | 1.513 | 0.02 | 0.042 | 0.042 | 0.255 | 0.24 | 0.33 | 0.36 | 0.47 | 0.35 | 0.45 |
| 3 | 54093664 | rs4928044 | -0.638 | 0.69 | 0.731 | 1.513 | -0.001 | 0.034 | 0.034 | 0.009 | 0.05 | 0.07 | 0.08 | 0.2 | 0.06 | 0.15 |
| 3 | 54093832 | rs4928045 | -1.137 | 0.675 | 0.667 | 1.443 | 0.022 | 0.032 | 0.032 | 0.552 | 0.25 | 0.33 | 0.37 | 0.47 | 0.4 | 0.45 |
| 3 | 54101402 | rs1868514 | -0.352 | 0.638 | 1.178 | 1.592 | -0.003 | 0 | 0 | 0 | 0.19 | 0.18 | 0.32 | 0.19 | 0.32 | 0.29 |
| 3 | 54101939 | rs2359655 | -0.445 | 0.603 | 1.634 | 1.619 | 0.044 | -0.018 | -0.018 | 0.005 | 0.19 | 0.26 | 0.32 | 0.17 | 0.31 | 0.29 |
| 3 | 54104178 | rs12629593 | 0.193 | -0.921 | 0.699 | 1.384 | 0.036 | -0.033 | -0.033 | 0.153 | 0.23 | 0.14 | 0.15 | 0.38 | 0.13 | 0.12 |
| 3 | 54108916 | rs2359656 | -0.634 | -0.78 | 0.754 | 1.275 | 0.068 | 0.009 | 0.009 | 0.319 | 0.25 | 0.46 | 0.38 | 0.3 | 0.35 | 0.33 |
| 3 | 54113502 | rs729502 | -1.493 | -0.127 | 0.409 | 0.898 | 0.008 | 0.035 | 0.035 | 0 | 0.55 | 0.66 | 0.62 | 0.89 | 0.63 | 0.66 |
| 3 | 54117849 | rs1449334 | -1.584 | -0.14 | 0.403 | 0.895 | 0.009 | 0.073 | 0.073 | 0 | 0.55 | 0.67 | 0.62 | 0.89 | 0.62 | 0.66 |
| 3 | 54118798 | rs9835631 | -0.102 | 0.078 | 0.3 | 0.785 | 0.018 | -0.008 | -0.008 | 0 | 0.22 | 0.22 | 0.34 | 0.17 | 0.43 | 0.31 |
| 3 | 54121188 | rs6804875 | -0.109 | -0.428 | 0.341 | 1.352 | -0.013 | 0.004 | 0.004 | 0 | 0.22 | 0.23 | 0.18 | 0.28 | 0.15 | 0.14 |
| 3 | 54123011 | rs4472075 | 0.709 | -0.87 | -0.266 | 0.602 | -0.045 | 0.081 | 0.081 | 0 | 0.35 | 0.18 | 0.4 | 0.17 | 0.43 | 0.42 |
| 3 | 54128575 | rs17801638 | 0.67 | -0.964 | -0.268 | 0.604 | -0.053 | 0.065 | 0.065 | 0 | 0.36 | 0.18 | 0.41 | 0.22 | 0.45 | 0.41 |
| 3 | 54129143 | rs6768219 | -0.738 | -0.987 | -0.153 | 0.662 | 0.011 | 0.014 | 0.014 | 0 | 0.63 | 0.5 | 0.61 | 0.58 | 0.61 | 0.57 |
| 3 | 54129904 | rs10510767 | 0.975 | -0.978 | -0.246 | 0.369 | 0.024 | -0.024 | -0.024 | 0 | 0.18 | 0.06 | 0.14 | 0.28 | 0.12 | 0.1 |
| 3 | 54148378 | rs764664 | 0.226 | -1.128 | -0.08 | 0.103 | 0.023 | -0.005 | -0.005 | 0 | 0.18 | 0.44 | 0.18 | 0.12 | 0.18 | 0.13 |
| 3 | 54148716 | rs2029119 | -1.015 | -1.069 | -0.373 | 0.089 | -0.033 | -0.023 | -0.023 | 3.835 | 0.62 | 0.74 | 0.51 | 0.4 | 0.5 | 0.47 |
| 3 | 54148770 | rs2029118 | 1.502 | -0.757 | -0.224 | 0.054 | -0.015 | 0.001 | 0.001 | 0 | 0.11 | 0.17 | 0.12 | 0.09 | 0.06 | 0.05 |
| 3 | 54149801 | rs4320084 | -0.028 | -0.35 | -0.74 | 0.236 | 0.017 | 0.043 | 0.043 | 0 | 0.37 | 0.23 | 0.3 | 0.19 | 0.29 | 0.26 |
| 3 | 54152819 | rs11130396 | -0.128 | -0.499 | -0.751 | 0.356 | -0.006 | 0.063 | 0.063 | 0 | 0.38 | 0.26 | 0.31 | 0.19 | 0.35 | 0.25 |
| 3 | 54159417 | rs9881544 | 0.126 | 0.331 | -0.637 | 1.096 | 0.011 | 0.02 | 0.02 | 0 | 0.3 | 0.33 | 0.29 | 0.49 | 0.38 | 0.53 |
| 3 | 54161174 | rs1449338 | 0.164 | 0.361 | -0.66 | 1.116 | 0.003 | 0.005 | 0.005 | 0 | 0.3 | 0.3 | 0.28 | 0.62 | 0.38 | 0.54 |
| 3 | 54164974 | rs1375526 | -0.337 | 0 | -0.763 | 1.345 | -0.017 | -0.019 | -0.019 | 0 | 0.19 | 0.07 | 0.18 | 0.19 | 0.22 | 0.14 |
| 3 | 54167806 | rs7618672 | -1.144 | -0.547 | 0.166 | 0.96 | -0.041 | 0.053 | 0.053 | 2.468 | 0.14 | 0.33 | 0.07 | 0.23 | 0.08 | 0.16 |
| 3 | 54168246 | rs9825725 | -1.42 | -0.67 | 0.041 | 0.862 | -0.039 | 0.014 | 0.014 | 2.06 | 0.18 | 0.59 | 0.09 | 0.25 | 0.09 | 0.17 |
| 3 | 54173258 | rs6765381 | -0.035 | -0.815 | -1.095 | 0.473 | 0.012 | 0.02 | 0.02 | 0.024 | 0.69 | 0.87 | 0.59 | 0.75 | 0.75 | 0.72 |
| 3 | 54173568 | rs9852858 | 0.528 | -0.07 | -0.844 | 0.143 | 0.01 | -0.014 | -0.014 | 1.648 | 0.21 | 0.25 | 0.17 | 0.17 | 0.29 | 0.25 |
| 3 | 54178199 | rs9880023 | -0.35 | -0.344 | -0.784 | -0.572 | 0 | -0.053 | -0.053 | 1.564 | 0.48 | 0.49 | 0.62 | 0.96 | 0.59 | 0.6 |

| | | | | | | | | | | | | | | | | |
|---|---|---|---|---|---|---|---|---|---|---|---|---|---|---|---|---|
| 3 | 54179586 | rs725992 | 0.799 | -0.375 | -1.22 | -0.607 | 0.007 | 0.057 | 0.057 | 0 | 0.14 | 0 | 0.11 | 0.02 | 0.13 | 0.05 |
| 3 | 54187839 | rs6793459 | -0.123 | -0.672 | -0.568 | 0.117 | -0.02 | 0.02 | 0.02 | 0.012 | 0.36 | 0.4 | 0.43 | 0.43 | 0.28 | 0.35 |
| 3 | 54192836 | rs4927999 | -0.404 | -0.577 | -0.562 | 0.101 | 0.002 | 0.018 | 0.018 | 0 | 0.52 | 0.55 | 0.55 | 0.61 | 0.49 | 0.59 |
| 3 | 54195096 | rs11926147 | 0.773 | -0.195 | -0.491 | 0.201 | 0.009 | 0.018 | 0.018 | 2.606 | 0.15 | 0.22 | 0.2 | 0.35 | 0.25 | 0.21 |
| 3 | 54196466 | rs1530734 | -0.302 | -0.604 | -0.651 | 0.13 | -0.02 | 0.036 | 0.036 | 0 | 0.56 | 0.7 | 0.53 | 0.41 | 0.49 | 0.59 |
| 3 | 54202453 | rs4928005 | -0.266 | -0.688 | 0.007 | 0.087 | 0 | 0.001 | 0.001 | 0 | 0.53 | 0.39 | 0.5 | 0.41 | 0.49 | 0.59 |
| 3 | 54205850 | rs4928006 | -1.017 | -0.678 | -0.268 | -0.237 | -0.011 | -0.002 | -0.002 | 0.244 | 0.72 | 0.64 | 0.76 | 0.8 | 0.79 | 0.9 |
| 3 | 54209340 | rs9829714 | -1.255 | -0.594 | -0.562 | -0.366 | 0.016 | 0.014 | 0.014 | 0.035 | 0.74 | 0.81 | 0.82 | 1 | 0.79 | 0.9 |
| 3 | 54212182 | rs13077924 | 1.763 | -0.503 | -0.887 | -0.421 | 0.015 | 0.082 | 0.082 | 1.261 | 0.14 | 0.01 | 0.08 | 0 | 0.08 | 0.18 |
| 3 | 54216117 | rs9809064 | -0.514 | -0.509 | -0.624 | -0.365 | -0.018 | -0.023 | -0.023 | 0.458 | 0.61 | 0.69 | 0.5 | 0.45 | 0.54 | 0.62 |
| 3 | 54217223 | rs6445631 | -1.362 | -0.643 | -0.792 | -0.509 | 0.013 | 0.046 | 0.046 | 0 | 0.75 | 0.87 | 0.83 | 1 | 0.79 | 0.91 |
| 3 | 54221080 | rs4927987 | -0.592 | -1.008 | -0.832 | -0.56 | 0.038 | 0.082 | 0.082 | 1.172 | 0.63 | 0.81 | 0.75 | 1 | 0.71 | 0.73 |
| 3 | 54222149 | rs7653234 | 1.689 | -1.006 | -0.83 | -0.544 | 0.012 | 0.008 | 0.008 | 1.07 | 0.13 | 0.06 | 0.08 | 0 | 0.08 | 0.18 |
| 3 | 54222459 | rs6445632 | 0.171 | -0.688 | -0.58 | -0.38 | 0.007 | 0.021 | 0.021 | 0 | 0.2 | 0.17 | 0.19 | 0.06 | 0.18 | 0.15 |
| 3 | 54223872 | rs9880752 | -0.095 | -0.699 | -0.698 | -0.299 | 0.019 | 0.038 | 0.038 | 2.178 | 0.46 | 0.67 | 0.54 | 0.74 | 0.59 | 0.58 |
| 3 | 54224820 | rs4076708 | -0.464 | -0.348 | -0.541 | 0.494 | 0.008 | -0.032 | -0.032 | 0.527 | 0.2 | 0.14 | 0.23 | 0.25 | 0.18 | 0.16 |
| 3 | 54226000 | rs4077974 | 0.25 | -0.29 | -0.407 | 0.186 | 0.005 | 0.014 | 0.014 | 0 | 0.16 | 0.14 | 0.16 | 0.05 | 0.12 | 0.14 |
| 3 | 54234722 | rs4485754 | -0.64 | 0.638 | -0.013 | 0.754 | 0.085 | 0.225 | 0.225 | 0.716 | 0.11 | 0.33 | 0.18 | 0.38 | 0.18 | 0.2 |
| 3 | 54243646 | rs4643737 | -0.019 | 0.535 | -1.167 | 0.772 | -0.071 | -0.017 | -0.017 | 0 | 0.32 | 0.1 | 0.45 | 0.32 | 0.45 | 0.43 |
| 3 | 54246298 | rs7432602 | -0.864 | -0.315 | -1.038 | 0.418 | -0.026 | 0.002 | 0.002 | 3.334 | 0.27 | 0.67 | 0.4 | 0.71 | 0.17 | 0.32 |
| 3 | 54248525 | rs12631896 | 1.344 | -0.394 | -1.202 | 0.903 | -0.005 | -0.038 | -0.038 | 0 | 0.07 | 0 | 0.11 | 0.23 | 0.05 | 0.1 |
| 3 | 54249088 | rs12632508 | 0.211 | -0.429 | -0.629 | 1.189 | 0.02 | 0.176 | 0.176 | 0 | 0.06 | 0.19 | 0.14 | 0.27 | 0.08 | 0.08 |
| 3 | 54249650 | rs11130399 | 0.556 | -0.423 | -0.769 | 1.2 | -0.056 | 0.036 | 0.036 | 0.583 | 0.31 | 0.13 | 0.38 | 0.23 | 0.44 | 0.43 |
| 3 | 54254129 | rs4505744 | -1.402 | -0.304 | -0.064 | 1.393 | -0.07 | 0.178 | 0.178 | 2.483 | 0.2 | 0.53 | 0.27 | 0.43 | 0.12 | 0.23 |
| 3 | 54256559 | rs11711040 | 0.321 | -0.006 | -0.519 | 1.339 | 0.003 | 0.153 | 0.153 | 0 | 0.05 | 0.07 | 0.13 | 0.39 | 0.08 | 0.1 |
| 3 | 54258938 | rs11924650 | -0.333 | 0.182 | -0.924 | 1.021 | 0 | -0.018 | -0.018 | 0 | 0.26 | 0.21 | 0.42 | 0.45 | 0.3 | 0.41 |
| 3 | 54262746 | rs9880354 | -1.066 | -0.281 | -0.782 | 0.095 | -0.001 | -0.068 | -0.068 | 0 | 0.63 | 0.67 | 0.76 | 1 | 0.57 | 0.87 |
| 3 | 54264197 | rs9830760 | -1.252 | -0.408 | -0.579 | 1.241 | -0.034 | 0.078 | 0.078 | 0.389 | 0.19 | 0.39 | 0.25 | 0.62 | 0.14 | 0.19 |
| 3 | 54266231 | rs13073263 | -0.92 | 0.167 | -1.444 | 1.446 | -0.027 | -0.126 | -0.126 | 0 | 0.16 | 0.02 | 0.26 | 0.5 | 0.18 | 0.23 |
| 3 | 54267473 | rs12631566 | -0.334 | 0.09 | -1.317 | 1.546 | 0.004 | -0.02 | -0.02 | 0.18 | 0.08 | 0.03 | 0.14 | 0.24 | 0.1 | 0.14 |
| 3 | 54267624 | rs4928016 | -0.973 | 0.485 | -1.33 | 0.947 | -0.007 | -0.007 | -0.007 | 0 | 0.39 | 0.69 | 0.51 | 0.63 | 0.38 | 0.38 |
| 3 | 54267797 | rs9878916 | -1.038 | 0.581 | -1.128 | 0.991 | -0.001 | 0.003 | 0.003 | 0 | 0.34 | 0.63 | 0.48 | 0.62 | 0.33 | 0.39 |
| 3 | 54269786 | rs4420909 | 0.834 | 0.785 | -1.156 | 0.75 | -0.003 | -0.001 | -0.001 | 0 | 0.47 | 0.43 | 0.55 | 0.49 | 0.51 | 0.59 |
| 3 | 54270646 | rs11130405 | 0.485 | 0.432 | -1.446 | 0.536 | 0.03 | 0.116 | 0.116 | 0 | 0.26 | 0.07 | 0.18 | 0.06 | 0.2 | 0.16 |
| 3 | 54273364 | rs4928020 | -0.646 | 0.677 | -1.66 | 0.475 | -0.014 | 0.038 | 0.038 | 0 | 0.59 | 0.72 | 0.6 | 0.34 | 0.54 | 0.39 |
| 3 | 54280137 | rs4432683 | -0.686 | 0.03 | -1.412 | 0.879 | -0.007 | -0.026 | -0.026 | 0 | 0.92 | 0.96 | 0.89 | 0.73 | 0.9 | 0.92 |
| 3 | 54285016 | rs11130406 | -0.808 | 0.422 | -0.856 | 0.285 | -0.001 | 0.023 | 0.023 | 0 | 0.44 | 0.4 | 0.36 | 0.23 | 0.42 | 0.42 |
| 3 | 54288475 | rs9828485 | -0.15 | 0.695 | -0.59 | 0.525 | 0.039 | 0.009 | 0.009 | 0.175 | 0.18 | 0.36 | 0.18 | 0.22 | 0.21 | 0.26 |
| 3 | 54292014 | rs7340675 | 0.041 | -0.116 | -0.66 | 0.187 | -0.022 | -0.037 | -0.037 | 0 | 0.3 | 0.43 | 0.27 | 0.16 | 0.21 | 0.18 |
| 3 | 54293095 | rs11715046 | 0.933 | -0.142 | -0.819 | 0.145 | -0.009 | 0.074 | 0.074 | 0 | 0.23 | 0.06 | 0.23 | 0.08 | 0.2 | 0.18 |
| 3 | 54293298 | rs9854514 | 1.077 | -0.065 | -0.742 | 0.54 | -0.004 | 0.026 | 0.026 | 0.172 | 0.43 | 0.28 | 0.36 | 0.24 | 0.45 | 0.37 |
| 3 | 54293345 | rs11707360 | -0.093 | -0.241 | -0.612 | 0.497 | -0.015 | -0.029 | -0.029 | 0 | 0.3 | 0.4 | 0.27 | 0.15 | 0.21 | 0.18 |
| 3 | 54298050 | rs9852538 | -0.056 | 0.165 | -1.034 | 0.816 | -0.077 | -0.084 | -0.084 | 0 | 0.73 | 0.86 | 0.65 | 0.6 | 0.66 | 0.64 |
| 3 | 54301338 | rs11130409 | 0.341 | -0.085 | -0.467 | 1.055 | -0.009 | 0.151 | 0.151 | 0 | 0.09 | 0.21 | 0.14 | 0.36 | 0.08 | 0.11 |
| 3 | 54305670 | rs7615792 | -1.222 | -0.558 | -0.41 | 0.397 | 0.006 | 0.026 | 0.026 | 0.001 | 0.57 | 0.65 | 0.71 | 0.84 | 0.57 | 0.64 |
| 3 | 54308102 | rs11720066 | 1.334 | -0.417 | -0.569 | 0.429 | -0.024 | 0.013 | 0.013 | 0.688 | 0.15 | 0.04 | 0.22 | 0.08 | 0.18 | 0.18 |
| 3 | 54314698 | rs12374075 | 0.809 | -0.125 | -0.556 | 0.7 | -0.018 | -0.019 | -0.019 | 1.305 | 0.13 | 0.14 | 0.23 | 0.28 | 0.21 | 0.24 |
| 3 | 54315380 | rs7618850 | 1.379 | -0.528 | -0.398 | 0.729 | -0.008 | 0.041 | 0.041 | 0 | 0.15 | 0.21 | 0.12 | 0.3 | 0.1 | 0.14 |
| 3 | 54316176 | rs7619590 | -0.841 | -0.672 | -0.186 | 0.487 | 0.005 | 0.098 | 0.098 | 0 | 0.49 | 0.7 | 0.63 | 0.7 | 0.48 | 0.64 |
| 3 | 54316431 | rs7643501 | -0.587 | -0.725 | -0.076 | 0.491 | 0 | -0.003 | -0.003 | 0 | 0.44 | 0.56 | 0.62 | 0.64 | 0.47 | 0.62 |
| 3 | 54326631 | rs6804595 | -0.589 | -0.743 | -0.398 | 0.548 | -0.028 | -0.083 | -0.083 | 0 | 0.29 | 0.19 | 0.32 | 0.55 | 0.33 | 0.36 |
| 3 | 54331072 | rs6445643 | 0.326 | -0.654 | -0.424 | 0.5 | -0.021 | -0.026 | -0.026 | 0 | 0.2 | 0.18 | 0.23 | 0.32 | 0.25 | 0.37 |
| 3 | 54334723 | rs9851393 | 0.889 | -0.536 | -0.447 | 0.332 | -0.014 | 0.084 | 0.084 | 0.455 | 0.45 | 0.31 | 0.31 | 0.11 | 0.51 | 0.37 |
| 3 | 54338651 | rs6414573 | -1.821 | -0.378 | -0.312 | 0.515 | -0.001 | 0 | 0 | 0 | 0.85 | 0.89 | 0.77 | 0.88 | 0.84 | 0.82 |
| 3 | 54339912 | rs7431577 | -1.167 | -0.317 | -0.2 | 0.366 | 0 | 0.002 | 0.002 | 0 | 0.71 | 0.73 | 0.65 | 0.66 | 0.74 | 0.69 |
| 3 | 54341281 | rs4072628 | 0.369 | -0.357 | -0.168 | 0.351 | 0.001 | 0 | 0 | 0 | 0.5 | 0.49 | 0.36 | 0.3 | 0.48 | 0.26 |
| 3 | 54343019 | rs6445646 | 0.653 | -0.154 | -0.536 | 0.543 | -0.001 | 0.009 | 0.009 | 0 | 0.15 | 0.18 | 0.24 | 0.33 | 0.14 | 0.26 |
| 3 | 54354510 | rs9879885 | -1.018 | -0.176 | -0.755 | 0.139 | -0.001 | 0.022 | 0.022 | 0 | 0.7 | 0.63 | 0.61 | 0.6 | 0.73 | 0.68 |
| 3 | 54370763 | rs9855109 | 1.507 | -0.202 | -1.517 | -0.405 | -0.009 | 0.018 | 0.018 | 0.158 | 0.22 | 0.17 | 0.16 | 0.07 | 0.27 | 0.1 |
| 3 | 54371145 | rs9818348 | -0.17 | -0.877 | -1.642 | -0.418 | 0.025 | 0.004 | 0.004 | 0.113 | 0.5 | 0.43 | 0.47 | 0.55 | 0.61 | 0.53 |
| 3 | 54371791 | rs11708620 | 1.425 | -0.549 | -1.861 | -0.299 | 0.018 | 0.017 | 0.017 | 0 | 0.15 | 0.07 | 0.08 | 0.01 | 0.12 | 0.09 |
| 3 | 54372609 | rs7615581 | 0.533 | 0.221 | -1.767 | 0.301 | -0.006 | -0.048 | -0.048 | 0 | 0.11 | 0.01 | 0.09 | 0.2 | 0.1 | 0.06 |
| 3 | 54375609 | rs9683195 | -0.463 | 0.3 | -1.337 | 0.579 | -0.005 | 0.038 | 0.038 | 0 | 0.69 | 0.58 | 0.56 | 0.57 | 0.73 | 0.62 |
| 3 | 54378600 | rs11711956 | -0.238 | -0.436 | -1.216 | 0.445 | 0.004 | 0.001 | 0.001 | 0 | 0.41 | 0.6 | 0.29 | 0.36 | 0.28 | 0.37 |
| 3 | 54379733 | rs6784395 | -0.225 | -0.189 | -1.73 | 0.339 | 0.082 | -0.031 | -0.031 | 2.377 | 0.52 | 0.89 | 0.36 | 0.58 | 0.32 | 0.51 |
| 3 | 54379802 | rs9811393 | -0.383 | 0.233 | -0.809 | 0.163 | 0.009 | -0.012 | -0.012 | 0.072 | 0.54 | 0.71 | 0.39 | 0.53 | 0.39 | 0.52 |
| 3 | 54381071 | rs9816771 | -0.714 | 0.144 | 0.054 | 0.078 | -0.003 | -0.01 | -0.01 | 0 | 0.11 | 0.18 | 0.07 | 0.09 | 0.11 | 0.12 |
| 3 | 54389115 | rs1530733 | -0.179 | 0.34 | -0.222 | 0.042 | -0.043 | -0.024 | -0.024 | 0 | 0.12 | 0.01 | 0.24 | 0.15 | 0.14 | 0.07 |
| 3 | 54399608 | rs7427395 | -0.823 | 0.445 | 0.1 | 0.027 | 0 | -0.001 | -0.001 | 0 | 0.87 | 0.85 | 0.87 | 0.86 | 0.87 | 0.81 |

| | | | | | | | | | | | | | | | | |
|---|---|---|---|---|---|---|---|---|---|---|---|---|---|---|---|---|
| 3 | 54407040 | rs9311529 | -0.928 | 0.257 | 0.39 | -0.04 | -0.006 | -0.014 | -0.014 | 0.023 | 0.21 | 0.26 | 0.19 | 0.12 | 0.16 | 0.13 |
| 3 | 54407766 | rs11711662 | -1.772 | 0.472 | 0.776 | 0.113 | -0.001 | -0.01 | -0.01 | 5.255 | 0.34 | 0.5 | 0.34 | 0.36 | 0.31 | 0.25 |
| 3 | 54408995 | rs1473348 | -1.829 | 0.468 | 0.761 | 0.077 | 0.001 | -0.001 | -0.001 | 0 | 0.35 | 0.67 | 0.44 | 0.42 | 0.31 | 0.25 |
| 3 | 54417322 | rs7645545 | -1.509 | 0.402 | 0.628 | 0.047 | 0.001 | -0.006 | -0.006 | 0.555 | 0.34 | 0.39 | 0.44 | 0.34 | 0.32 | 0.27 |
| 3 | 54435563 | rs6445653 | -1.543 | 0.405 | 0.661 | 0.014 | 0.001 | 0.003 | 0.003 | 0.647 | 0.34 | 0.41 | 0.43 | 0.43 | 0.32 | 0.28 |
| 3 | 54447789 | rs7638153 | -0.789 | 0.741 | 1.192 | 0.468 | 0.002 | 0.003 | 0.003 | 0 | 0.85 | 0.87 | 0.86 | 0.87 | 0.86 | 0.81 |
| 3 | 54450193 | rs6803199 | 1.694 | 0.899 | 0.978 | 0.503 | 0.007 | -0.023 | -0.023 | 0.512 | 0.54 | 0.23 | 0.43 | 0.45 | 0.58 | 0.57 |
| 3 | 54451457 | rs4974370 | -1.202 | 0.831 | 0.993 | 0.015 | -0.001 | 0.002 | 0.002 | 0.096 | 0.2 | 0.17 | 0.31 | 0.18 | 0.21 | 0.19 |
| 3 | 54459688 | rs4131315 | -1.562 | 0.765 | 1.053 | 0.048 | 0.004 | -0.005 | -0.005 | 0 | 0.24 | 0.34 | 0.35 | 0.27 | 0.24 | 0.21 |
| 3 | 54463904 | rs9882299 | -1.287 | 0.803 | 0.824 | 0 | 0 | -0.001 | -0.001 | 1.633 | 0.22 | 0.23 | 0.33 | 0.25 | 0.24 | 0.2 |
| 3 | 54469592 | rs7644514 | 1.614 | 1.046 | 0.446 | -0.047 | -0.012 | 0.044 | 0.044 | 0 | 0.54 | 0.29 | 0.48 | 0.58 | 0.56 | 0.58 |
| 3 | 54471358 | rs7428295 | 0.693 | 1.025 | 0.7 | 0.281 | -0.014 | -0.009 | -0.009 | 0 | 0.84 | 0.88 | 0.71 | 0.74 | 0.78 | 0.85 |
| 3 | 54472975 | rs1375517 | -0.306 | 0.375 | 0.803 | 0.301 | -0.04 | -0.044 | -0.044 | 0 | 0.22 | 0.51 | 0.17 | 0.13 | 0.14 | 0.2 |
| 3 | 54473565 | rs1868500 | -0.354 | 0.457 | 0.744 | 0.273 | 0.002 | 0 | 0 | 0 | 0.11 | 0.1 | 0.27 | 0.15 | 0.14 | 0.08 |
| 3 | 54476640 | rs1375515 | 1.383 | 0.431 | 0.436 | 0.277 | 0.013 | 0.005 | 0.005 | 0 | 0.63 | 0.35 | 0.54 | 0.61 | 0.69 | 0.65 |
| 3 | 54480798 | rs1530732 | -0.066 | 0.863 | 0.356 | 0.431 | -0.028 | -0.033 | -0.033 | 0 | 0.09 | 0.04 | 0.27 | 0.23 | 0.14 | 0.07 |
| 3 | 54483667 | rs7624727 | 0.841 | 0.826 | 0.479 | 0.421 | 0.03 | 0.05 | 0.05 | 0.045 | 0.87 | 0.75 | 0.71 | 0.77 | 0.86 | 0.91 |
| 3 | 54484118 | rs17054035 | -0.159 | 0.972 | -0.198 | 0.196 | -0.024 | -0.031 | -0.031 | 0 | 0.19 | 0.33 | 0.16 | 0.11 | 0.13 | 0.19 |
| 3 | 54485016 | rs4974360 | -0.545 | 1.264 | -0.899 | 0.478 | -0.041 | -0.03 | -0.03 | 0 | 0.09 | 0.02 | 0.26 | 0.16 | 0.14 | 0.06 |
| 3 | 54486470 | rs9654001 | 0.356 | 1.048 | -0.949 | 0.336 | -0.003 | 0.045 | 0.045 | 0.175 | 0.53 | 0.25 | 0.49 | 0.6 | 0.56 | 0.53 |
| 3 | 54486790 | rs6789236 | -1.328 | 1.135 | 0.113 | 0.348 | 0.077 | 0.004 | 0.004 | 0.437 | 0.11 | 0.37 | 0.08 | 0.07 | 0.11 | 0.07 |
| 3 | 54508988 | rs6777286 | -1.197 | 0.467 | -0.167 | 0.246 | -0.026 | -0.026 | -0.026 | 0.003 | 0.06 | 0.13 | 0.03 | 0.01 | 0.01 | 0.01 |
| 3 | 54516884 | rs6773944 | -0.325 | 0.102 | -0.447 | 0.136 | -0.003 | -0.011 | -0.011 | 0 | 0.09 | 0 | 0.11 | 0.07 | 0.05 | 0.05 |
| 3 | 54529944 | rs7614229 | 0.21 | -0.032 | -0.108 | 0.401 | 0.001 | -0.003 | -0.003 | 0 | 0.45 | 0.48 | 0.5 | 0.45 | 0.57 | 0.58 |
| 3 | 54537372 | rs7372750 | 0.031 | -0.044 | -0.016 | 0.33 | 0.001 | 0.002 | 0.002 | 0 | 0.17 | 0.15 | 0.16 | 0.12 | 0.13 | 0.12 |
| 3 | 54537866 | rs7430102 | 1.196 | -0.182 | -0.372 | 0.112 | -0.021 | -0.073 | -0.073 | 0 | 0.17 | 0.04 | 0.16 | 0.27 | 0.14 | 0.19 |
| 3 | 54538880 | rs9879189 | -0.711 | -0.214 | -0.179 | 0.059 | -0.003 | -0.011 | -0.011 | 0.698 | 0.51 | 0.25 | 0.43 | 0.42 | 0.43 | 0.38 |
| 3 | 54542225 | rs6783107 | 0.836 | -0.312 | -0.007 | 0.152 | -0.013 | 0.047 | 0.047 | 2.102 | 0.91 | 0.57 | 0.9 | 0.88 | 0.98 | 0.95 |
| 3 | 54542933 | rs7372541 | -0.087 | -0.22 | -0.231 | 0.26 | -0.014 | -0.05 | -0.05 | 0 | 0.36 | 0.2 | 0.35 | 0.47 | 0.32 | 0.32 |
| 3 | 54555732 | rs7432845 | 0.158 | -0.193 | -0.184 | 0.338 | -0.001 | -0.001 | -0.001 | 0.015 | 0.62 | 0.6 | 0.55 | 0.4 | 0.66 | 0.65 |
| 3 | 54562556 | rs6798952 | 0.017 | -0.186 | -0.302 | 0.268 | -0.003 | -0.025 | -0.025 | 0 | 0.63 | 0.69 | 0.61 | 0.54 | 0.68 | 0.68 |
| 3 | 54574534 | rs6445691 | 1.317 | -0.009 | -0.813 | 0.003 | 0 | -0.005 | -0.005 | 0 | 0.26 | 0.21 | 0.21 | 0.09 | 0.31 | 0.28 |
| 3 | 54575770 | rs6445693 | 0.034 | -0.854 | -0.643 | 0.342 | -0.004 | 0.001 | 0.001 | 0.864 | 0.38 | 0.35 | 0.31 | 0.38 | 0.25 | 0.29 |
| 3 | 54578805 | rs11130424 | 0.815 | -0.62 | -0.577 | 0.322 | 0 | -0.007 | -0.007 | 5.88 | 0.44 | 0.36 | 0.5 | 0.45 | 0.61 | 0.58 |
| 3 | 54579051 | rs7428777 | -0.651 | -0.783 | -0.8 | 0.209 | 0.007 | 0.004 | 0.004 | 0.655 | 0.43 | 0.36 | 0.34 | 0.35 | 0.28 | 0.31 |
| 3 | 54585416 | rs7627415 | -0.67 | -0.889 | -1.138 | 0.365 | 0.03 | -0.005 | -0.005 | 0.114 | 0.27 | 0.19 | 0.22 | 0.27 | 0.16 | 0.13 |
| 3 | 54591243 | rs6764520 | 0.42 | -0.568 | -0.863 | 0.118 | 0.005 | -0.013 | -0.013 | 0 | 0.67 | 0.62 | 0.78 | 0.77 | 0.74 | 0.78 |
| 3 | 54591730 | rs7371994 | 0.761 | -0.483 | -1.032 | 0.107 | 0 | 0.002 | 0.002 | 0 | 0.27 | 0.24 | 0.28 | 0.27 | 0.31 | 0.16 |
| 3 | 54597829 | rs2139683 | 0.456 | -0.577 | -1.29 | -0.12 | 0 | 0.062 | 0.062 | 0.044 | 0.31 | 0.14 | 0.26 | 0.18 | 0.33 | 0.13 |
| 3 | 54599904 | rs4955897 | -0.118 | -0.859 | -1.198 | -0.158 | -0.004 | 0.011 | 0.011 | 0 | 0.25 | 0.14 | 0.15 | 0.2 | 0.18 | 0.15 |
| 3 | 54603516 | rs6794883 | 1.008 | -0.855 | -1.263 | -0.339 | -0.018 | 0.056 | 0.056 | 0 | 0.3 | 0.16 | 0.27 | 0.18 | 0.34 | 0.13 |
| 3 | 54604188 | rs2251788 | -0.735 | -0.704 | -1.19 | -0.264 | 0.003 | -0.009 | -0.009 | 0.235 | 0.62 | 0.4 | 0.53 | 0.45 | 0.66 | 0.61 |
| 3 | 54605919 | rs13318684 | 0.793 | -0.748 | -1.44 | -0.299 | -0.018 | 0.048 | 0.048 | 0 | 0.34 | 0.16 | 0.3 | 0.22 | 0.35 | 0.19 |
| 3 | 54609024 | rs1543208 | 0.101 | -0.623 | -1.342 | -0.225 | -0.024 | 0.002 | 0.002 | 0 | 0.24 | 0.07 | 0.15 | 0.19 | 0.18 | 0.14 |
| 3 | 54610463 | rs4955856 | 0.863 | -0.593 | -1.307 | -0.311 | -0.037 | 0.012 | 0.012 | 0 | 0.33 | 0.15 | 0.35 | 0.25 | 0.45 | 0.44 |
| 3 | 54613359 | rs6445700 | -0.138 | -0.959 | -1.27 | -0.244 | -0.014 | 0.037 | 0.037 | 0.837 | 0.53 | 0.74 | 0.63 | 0.36 | 0.54 | 0.52 |
| 3 | 54615971 | rs6782115 | 0.414 | -0.793 | -0.824 | -0.091 | -0.001 | -0.006 | -0.006 | 0.022 | 0.39 | 0.43 | 0.47 | 0.28 | 0.38 | 0.21 |
| 3 | 54621046 | rs1607754 | -0.019 | -0.845 | -1.357 | -0.162 | -0.02 | 0.143 | 0.143 | 0 | 0.36 | 0.12 | 0.22 | 0.07 | 0.34 | 0.42 |
| 3 | 54622000 | rs1851046 | 1.09 | -0.592 | -1.666 | -0.055 | -0.011 | 0.029 | 0.029 | 0 | 0.31 | 0.24 | 0.28 | 0.19 | 0.35 | 0.15 |
| 3 | 54622159 | rs11130428 | 0.725 | -0.405 | -1.425 | 0.207 | 0.001 | 0.011 | 0.011 | 0.017 | 0.37 | 0.34 | 0.47 | 0.3 | 0.37 | 0.22 |
| 3 | 54626332 | rs6445703 | -0.489 | 0.164 | -1.289 | 0.642 | -0.007 | -0.023 | -0.023 | 0.001 | 0.77 | 0.84 | 0.79 | 0.55 | 0.73 | 0.66 |
| 3 | 54630705 | rs11716867 | 0.136 | 0.109 | -1.001 | 0.624 | -0.063 | -0.045 | -0.045 | 0 | 0.23 | 0.07 | 0.43 | 0.35 | 0.31 | 0.31 |
| 3 | 54631902 | rs12495467 | 0.148 | 0.13 | -0.92 | 0.644 | 0.014 | 0.004 | 0.004 | 0 | 0.25 | 0.24 | 0.45 | 0.35 | 0.34 | 0.32 |
| 3 | 54633739 | rs1851043 | -0.423 | -0.069 | -1.121 | 0.608 | 0.004 | -0.008 | -0.008 | 0 | 0.45 | 0.49 | 0.54 | 0.4 | 0.39 | 0.61 |
| 3 | 54638547 | rs545517 | 0.365 | -0.665 | -1.581 | 0.804 | 0.056 | -0.041 | -0.041 | 0.191 | 0.72 | 0.9 | 0.85 | 0.71 | 0.81 | 0.68 |
| 3 | 54639209 | rs616788 | -0.614 | -1.066 | -0.833 | 0.871 | -0.062 | 0.054 | 0.054 | 0 | 0.29 | 0.54 | 0.41 | 0.4 | 0.18 | 0.22 |
| 3 | 54639268 | rs6445704 | -0.629 | -0.072 | -0.551 | 0.76 | 0.004 | 0.002 | 0.002 | 0.215 | 0.42 | 0.54 | 0.48 | 0.4 | 0.36 | 0.39 |
| 3 | 54639970 | rs603059 | 0.9 | -0.009 | 0.062 | 1.319 | -0.02 | 0.176 | 0.176 | 1.558 | 0.79 | 0.49 | 0.83 | 0.57 | 0.83 | 0.72 |
| 3 | 54641568 | rs2884808 | -0.485 | 0.41 | 0.449 | 0.438 | 0.04 | -0.136 | -0.136 | 0 | 0.2 | 0.48 | 0.15 | 0.01 | 0.24 | 0.17 |
| 3 | 54645869 | rs573279 | 0.331 | 0.143 | -0.363 | 0.375 | 0.012 | -0.04 | -0.04 | 0 | 0.27 | 0.21 | 0.3 | 0.53 | 0.2 | 0.44 |
| 3 | 54646035 | rs605284 | -1.093 | 0.01 | -0.435 | 0.312 | -0.018 | 0.005 | 0.005 | 0.803 | 0.52 | 0.8 | 0.47 | 0.56 | 0.46 | 0.62 |
| 3 | 54651208 | rs17054227 | -0.034 | 0.062 | 0.302 | 0.193 | 0.011 | -0.111 | -0.111 | 0 | 0.22 | 0.36 | 0.14 | 0.01 | 0.24 | 0.17 |
| 3 | 54653316 | rs513324 | -0.355 | -0.025 | -0.114 | 0.671 | 0 | 0.007 | 0.007 | 0.06 | 0.93 | 0.94 | 0.96 | 0.83 | 0.94 | 0.87 |
| 3 | 54659057 | rs4955988 | -0.661 | -0.088 | 0.009 | 0.683 | -0.006 | 0.008 | 0.008 | 0 | 0.77 | 0.71 | 0.71 | 0.73 | 0.81 | 0.6 |
| 3 | 54662474 | rs13084347 | 1.391 | -0.123 | -0.23 | 0.5 | 0.006 | 0.02 | 0.02 | 0.02 | 0.18 | 0.11 | 0.24 | 0.11 | 0.14 | 0.29 |
| 3 | 54663005 | rs496625 | 0.807 | -0.034 | -0.231 | 0.548 | -0.001 | 0.005 | 0.005 | 1.656 | 0.51 | 0.33 | 0.57 | 0.48 | 0.54 | 0.4 |
| 3 | 54675974 | rs17081825 | -0.757 | -0.241 | 0.046 | 0.438 | 0 | -0.004 | -0.004 | 0 | 0.08 | 0.48 | 0.04 | 0 | 0 | 0 |
| 3 | 54676642 | rs2359861 | 0.824 | -0.371 | -0.521 | 0.839 | 0.002 | -0.031 | -0.031 | 0.62 | 0.2 | 0.09 | 0.26 | 0.27 | 0.16 | 0.27 |
| 3 | 54679831 | rs1965072 | 0.816 | -0.453 | -0.971 | 0.23 | 0.016 | -0.024 | -0.024 | 0.958 | 0.72 | 0.32 | 0.81 | 0.58 | 0.8 | 0.75 |

| | | | | | | | | | | | | | | | | |
|---|---|---|---|---|---|---|---|---|---|---|---|---|---|---|---|---|
| 3 | 54688260 | rs530868 | 0.826 | -0.477 | -0.967 | 0.248 | -0.015 | 0.049 | 0.049 | 2.227 | 0.83 | 0.35 | 0.86 | 0.6 | 0.9 | 0.84 |
| 3 | 54688515 | rs533524 | -1.195 | -0.391 | -0.964 | 0.293 | -0.001 | 0.117 | 0.117 | 0.489 | 0.94 | 0.94 | 0.95 | 0.61 | 0.95 | 0.86 |
| 3 | 54690359 | rs6800366 | 0.363 | -0.41 | -0.958 | 0.154 | -0.007 | -0.047 | -0.047 | 0 | 0.17 | 0.19 | 0.1 | 0.01 | 0.15 | 0.1 |
| 3 | 54693752 | rs17054246 | 1.276 | -0.009 | -1.187 | 0.087 | -0.03 | 0.05 | 0.05 | 5.605 | 0.16 | 0.08 | 0.1 | 0.01 | 0.21 | 0.14 |
| 3 | 54695763 | rs646929 | 1.344 | -0.027 | -1.581 | 0.104 | -0.119 | -0.157 | -0.157 | 0 | 0.3 | 0.02 | 0.42 | 0.43 | 0.36 | 0.27 |
| 3 | 54696985 | rs4955876 | 1.689 | -0.227 | -1.679 | 0.235 | -0.013 | -0.079 | -0.079 | 0 | 0.13 | 0 | 0.25 | 0.27 | 0.12 | 0.11 |
| 3 | 54709162 | rs510605 | 0.104 | -0.248 | -1.357 | 0.267 | -0.023 | 0.245 | 0.245 | 1.482 | 0.86 | 0.51 | 0.9 | 0.6 | 0.92 | 0.83 |
| 3 | 54712786 | rs2099019 | 2.554 | -0.208 | -1.615 | 0.111 | -0.01 | 0.032 | 0.032 | 0.16 | 0.17 | 0.07 | 0.23 | 0.1 | 0.14 | 0.3 |
| 3 | 54718721 | rs515920 | 0.043 | -0.227 | -1.804 | -0.054 | -0.027 | -0.034 | -0.034 | 0 | 0.5 | 0.17 | 0.53 | 0.48 | 0.54 | 0.39 |
| 3 | 54721055 | rs10510771 | 0.864 | 0.189 | -1.38 | 0.059 | 0.01 | -0.064 | -0.064 | 0.542 | 0.2 | 0.25 | 0.15 | 0.01 | 0.27 | 0.15 |
| 3 | 54739910 | rs551114 | 1.617 | 0.394 | -1.626 | 0.104 | -0.066 | -0.035 | -0.035 | 0 | 0.17 | 0.01 | 0.16 | 0.14 | 0.21 | 0.16 |
| 3 | 54741995 | rs566884 | 0.296 | 0.263 | -1.315 | 0.007 | 0 | -0.008 | -0.008 | 0 | 0.68 | 0.36 | 0.77 | 0.6 | 0.68 | 0.71 |
| 3 | 54747244 | rs589281 | -0.045 | 0.201 | -1.539 | -0.111 | -0.059 | -0.079 | -0.079 | 0 | 0.4 | 0.14 | 0.47 | 0.48 | 0.46 | 0.36 |
| 3 | 54748368 | rs13095178 | 0.719 | 0.16 | -1.396 | -0.189 | -0.001 | -0.002 | -0.002 | 0 | 0.46 | 0.44 | 0.45 | 0.12 | 0.46 | 0.47 |
| 3 | 54765332 | rs11130431 | 0.6 | 0.202 | -1.453 | -0.108 | -0.018 | -0.043 | -0.043 | 0 | 0.27 | 0.16 | 0.41 | 0.41 | 0.33 | 0.27 |
| 3 | 54769696 | rs3906509 | 1.973 | -0.096 | -1.285 | -0.206 | 0.007 | 0.042 | 0.042 | 0.64 | 0.25 | 0.17 | 0.3 | 0.1 | 0.21 | 0.31 |
| 3 | 54772441 | rs13068008 | 1.997 | -0.074 | -1.282 | -0.199 | 0.008 | 0.038 | 0.038 | 0.432 | 0.25 | 0.21 | 0.29 | 0.1 | 0.2 | 0.31 |
| 3 | 54777529 | rs1008187 | 0.817 | 0.374 | -1.469 | -0.006 | -0.023 | -0.029 | -0.029 | 0.565 | 0.21 | 0.12 | 0.33 | 0.66 | 0.24 | 0.31 |
| 3 | 54780349 | rs12488046 | 1.68 | -0.075 | -0.95 | 0.023 | 0 | 0.049 | 0.049 | 0.399 | 0.24 | 0.14 | 0.3 | 0.1 | 0.21 | 0.31 |
| 3 | 54781631 | rs11914780 | 1.204 | -0.066 | -1.048 | 0.029 | -0.001 | -0.049 | -0.049 | 0 | 0.17 | 0.21 | 0.1 | 0.01 | 0.19 | 0.13 |
| 3 | 54790379 | rs934083 | 0.006 | 0.093 | -1.555 | 0.294 | -0.032 | -0.045 | -0.045 | 0.205 | 0.44 | 0.18 | 0.51 | 0.48 | 0.52 | 0.39 |
| 3 | 54793362 | rs7633762 | -1.002 | 0.091 | -1.303 | 0.612 | 0 | 0.091 | 0.091 | 1.076 | 0.94 | 0.93 | 0.95 | 0.62 | 0.95 | 0.86 |
| 3 | 54802351 | rs4955893 | 0.998 | 0.243 | -1.592 | 0.818 | -0.021 | -0.029 | -0.029 | 1.669 | 0.07 | 0 | 0.21 | 0.3 | 0.07 | 0.12 |
| 3 | 54807240 | rs1997463 | -0.044 | 0.008 | -1.287 | 0.437 | -0.008 | 0.018 | 0.018 | 2.331 | 0.29 | 0.6 | 0.18 | 0.41 | 0.27 | 0.28 |
| 3 | 54811259 | rs10510773 | 0.786 | -0.076 | -0.893 | -0.06 | 0 | -0.006 | -0.006 | 0 | 0.25 | 0.25 | 0.3 | 0.1 | 0.21 | 0.3 |
| 3 | 54819371 | rs6793171 | -0.558 | 0.22 | -0.643 | 0.638 | -0.015 | 0.131 | 0.131 | 0.015 | 0.23 | 0.41 | 0.3 | 0.51 | 0.19 | 0.23 |
| 3 | 54819686 | rs13081825 | 0.075 | 0.406 | -1.635 | -0.008 | -0.074 | 0.056 | 0.056 | 0 | 0.36 | 0.14 | 0.33 | 0.19 | 0.53 | 0.3 |
| 3 | 54825311 | rs7642224 | -0.005 | 0.304 | -0.996 | -0.071 | -0.008 | -0.008 | -0.008 | 0 | 0.28 | 0.39 | 0.25 | 0.18 | 0.21 | 0.21 |
| 3 | 54834603 | rs13093404 | 0.78 | 0.2 | -1.724 | 0.123 | -0.014 | -0.027 | -0.027 | 0.086 | 0.09 | 0.02 | 0.04 | 0.11 | 0.09 | 0.15 |
| 3 | 54836275 | rs1168693 | -1.47 | 0.189 | -2.224 | 0.222 | -0.031 | -0.075 | -0.075 | 0 | 0.76 | 0.94 | 0.7 | 0.62 | 0.72 | 0.66 |
| 3 | 54846329 | rs1465010 | 0.802 | -0.035 | -2.434 | 0.133 | -0.003 | -0.079 | -0.079 | 0 | 0.13 | 0.01 | 0.21 | 0.29 | 0.14 | 0.11 |
| 3 | 54848641 | rs7644892 | 1.567 | -0.313 | -1.772 | 0.013 | 0.004 | 0.004 | 0.004 | 0 | 0.16 | 0.12 | 0.17 | 0.16 | 0.17 | 0.17 |
| 3 | 54861274 | rs9860460 | 0.616 | -0.374 | -1.937 | -0.227 | -0.03 | 0.009 | 0.009 | 0 | 0.31 | 0.2 | 0.38 | 0.3 | 0.4 | 0.24 |
| 3 | 54862361 | rs11130434 | 1.459 | -0.48 | -1.739 | -0.162 | 0 | 0.002 | 0.002 | 0 | 0.1 | 0.09 | 0.06 | 0.15 | 0.11 | 0.18 |
| 3 | 54863260 | rs9832945 | -1.877 | -0.442 | -1.87 | -0.155 | -0.021 | -0.076 | -0.076 | 0.295 | 0.77 | 0.95 | 0.72 | 0.63 | 0.74 | 0.67 |
| 3 | 54867313 | rs11130435 | 0.638 | -0.067 | -1.43 | -0.03 | -0.01 | -0.004 | -0.004 | 0.005 | 0.16 | 0.25 | 0.09 | 0.16 | 0.13 | 0.21 |
| 3 | 54871826 | rs3773605 | 0.073 | 0.729 | -1.777 | 0.217 | -0.023 | -0.016 | -0.016 | 0.131 | 0.3 | 0.19 | 0.35 | 0.35 | 0.36 | 0.24 |
| 3 | 54880461 | rs3773603 | 0.403 | 0.974 | -1.284 | 0.192 | -0.002 | -0.002 | -0.002 | 0 | 0.17 | 0.11 | 0.19 | 0.18 | 0.19 | 0.19 |
| 3 | 54885173 | rs9849795 | -0.107 | 0.677 | -1.548 | -0.643 | -0.032 | 0.092 | 0.092 | 0 | 0.32 | 0.17 | 0.22 | 0.03 | 0.35 | 0.22 |
| 3 | 54886083 | rs2970533 | 0.446 | 0.312 | -1.635 | -0.11 | -0.018 | -0.027 | -0.027 | 0 | 0.46 | 0.73 | 0.55 | 0.48 | 0.45 | 0.34 |
| 3 | 54889044 | rs4955898 | 0.42 | -0.096 | -1.359 | -0.255 | -0.007 | 0.105 | 0.105 | 0.007 | 0.47 | 0.24 | 0.66 | 0.77 | 0.5 | 0.52 |
| 3 | 54891599 | rs6810058 | 0.672 | 0.149 | -1.436 | -0.342 | -0.046 | 0.114 | 0.114 | 0.011 | 0.25 | 0.11 | 0.15 | 0.04 | 0.28 | 0.2 |
| 3 | 54894304 | rs12634786 | 0.663 | -0.052 | -1.389 | -0.061 | -0.019 | 0.065 | 0.065 | 0 | 0.12 | 0.2 | 0.05 | 0.3 | 0.07 | 0.18 |
| 3 | 54894545 | rs1526596 | -0.628 | -0.108 | -0.755 | -0.117 | -0.001 | -0.004 | -0.004 | 0 | 0.42 | 0.52 | 0.39 | 0.36 | 0.4 | 0.4 |
| 3 | 54895922 | rs1179911 | 1.104 | 0.047 | -0.918 | -0.161 | -0.002 | 0.007 | 0.007 | 0 | 0.4 | 0.45 | 0.61 | 0.47 | 0.4 | 0.33 |
| 3 | 54896342 | rs7642067 | 0.234 | 0.528 | -1.546 | -0.548 | -0.032 | 0.121 | 0.121 | 0.131 | 0.27 | 0.12 | 0.17 | 0.03 | 0.28 | 0.21 |
| 3 | 54897396 | rs1168704 | -0.568 | 0.543 | -1.094 | 0.169 | -0.06 | -0.033 | -0.033 | 1.002 | 0.83 | 0.99 | 0.89 | 0.81 | 0.78 | 0.72 |
| 3 | 54901360 | rs1184882 | -1.256 | -0.019 | -0.209 | 0.246 | -0.006 | 0.081 | 0.081 | 0.363 | 0.08 | 0.22 | 0.03 | 0.1 | 0.03 | 0.15 |
| 3 | 54903615 | rs2703029 | 0.031 | -0.133 | -0.877 | -0.534 | -0.013 | 0.107 | 0.107 | 0 | 0.59 | 0.73 | 0.76 | 0.86 | 0.6 | 0.68 |
| 3 | 54903917 | rs2694125 | 0.095 | -0.094 | -1.069 | -0.661 | 0.019 | 0.179 | 0.179 | 0 | 0.55 | 0.77 | 0.68 | 0.86 | 0.45 | 0.66 |
| 3 | 54907358 | rs1168709 | -1.181 | -0.601 | -1.034 | -0.239 | -0.026 | 0.027 | 0.027 | 1.536 | 0.17 | 0.73 | 0.1 | 0.24 | 0.09 | 0.2 |
| 3 | 54909196 | rs4955903 | 1.187 | -0.34 | -1.28 | -0.523 | -0.009 | 0.006 | 0.006 | 0 | 0.37 | 0.04 | 0.57 | 0.68 | 0.34 | 0.47 |
| 3 | 54912798 | rs1568983 | -1.241 | -0.403 | 0.068 | -0.579 | -0.042 | 0.006 | 0.006 | 0.291 | 0.13 | 0.5 | 0.06 | 0.08 | 0.06 | 0.17 |
| 3 | 54918438 | rs2703022 | 0.476 | 0.235 | -0.033 | -0.866 | -0.028 | 0.025 | 0.025 | 0 | 0.7 | 0.78 | 0.77 | 0.87 | 0.57 | 0.73 |
| 3 | 54919185 | rs11716356 | 0.035 | -0.161 | 0.389 | -1.099 | -0.028 | -0.045 | -0.045 | 3.777 | 0.2 | 0.42 | 0.14 | 0.06 | 0.13 | 0.18 |
| 3 | 54919351 | rs10510774 | -0.085 | 0.271 | 0.198 | -1.163 | -0.018 | 0.015 | 0.015 | 0 | 0.08 | 0.04 | 0.07 | 0 | 0.14 | 0.04 |
| 3 | 54920515 | rs2694116 | -0.703 | -0.013 | 0.034 | -1.25 | -0.086 | 0.036 | 0.036 | 1.481 | 0.8 | 0.97 | 0.85 | 0.88 | 0.72 | 0.8 |
| 3 | 54922235 | rs2360659 | 0.721 | 0.02 | 0.383 | -1.235 | 0.009 | 0.011 | 0.011 | 0 | 0.16 | 0.14 | 0.09 | 0.06 | 0.1 | 0.16 |
| 3 | 54928174 | rs10510775 | -0.051 | 0.012 | 0.881 | -1.247 | 0.032 | -0.059 | -0.059 | 0 | 0.75 | 0.61 | 0.83 | 0.87 | 0.71 | 0.76 |
| 3 | 54928327 | rs11130438 | 1.686 | -0.007 | 0.891 | -1.3 | 0.036 | 0.09 | 0.09 | 0 | 0.47 | 0.24 | 0.62 | 0.79 | 0.44 | 0.54 |
| 3 | 54929590 | rs9848771 | 0.287 | 0.246 | 0.948 | -1.129 | -0.011 | -0.029 | -0.029 | 0 | 0.18 | 0.25 | 0.12 | 0.06 | 0.13 | 0.16 |
| 3 | 54932940 | rs17054525 | -0.569 | 0.235 | 0.896 | -1.16 | 0.002 | -0.019 | -0.019 | 0 | 0.23 | 0.32 | 0.15 | 0.06 | 0.22 | 0.21 |
| 3 | 54939122 | rs6772996 | 0.547 | 0.244 | 0.88 | -1.148 | -0.008 | -0.012 | -0.012 | 0.014 | 0.17 | 0.18 | 0.1 | 0.06 | 0.11 | 0.17 |
| 3 | 54939511 | rs10514715 | -0.694 | 0.318 | 0.818 | -1.143 | 0.033 | -0.026 | -0.026 | 0.043 | 0.07 | 0.15 | 0.06 | 0.01 | 0.11 | 0.04 |
| 3 | 54946239 | rs7637262 | 0.994 | 0.112 | 0.894 | -0.973 | -0.001 | -0.005 | -0.005 | 0 | 0.54 | 0.54 | 0.7 | 0.75 | 0.5 | 0.54 |
| 3 | 54946578 | rs1526589 | 0.95 | 0.105 | 0.926 | -0.974 | -0.003 | -0.016 | -0.016 | 0 | 0.55 | 0.49 | 0.7 | 0.76 | 0.5 | 0.54 |
| 3 | 54948002 | rs1125025 | 0.195 | 0.098 | 0.92 | -1.001 | -0.039 | 0.032 | 0.032 | 0 | 0.66 | 0.84 | 0.76 | 0.78 | 0.62 | 0.63 |
| 3 | 54948869 | rs4468946 | 0.737 | 0.072 | 0.907 | -1.003 | 0.003 | 0.002 | 0.002 | 0 | 0.16 | 0.15 | 0.09 | 0.09 | 0.1 | 0.16 |
| 3 | 54948936 | rs4245919 | 0.517 | 0.096 | 0.918 | -1.001 | -0.017 | 0.011 | 0.011 | 0 | 0.72 | 0.75 | 0.79 | 0.85 | 0.6 | 0.7 |

| | | | | | | | | | | | | | | | | |
|---|---|---|---|---|---|---|---|---|---|---|---|---|---|---|---|---|
| 3 | 54951752 | rs1880827 | 1.58 | 0.037 | 1.006 | -1.044 | 0.02 | 0.037 | 0.037 | 0 | 0.5 | 0.35 | 0.65 | 0.75 | 0.47 | 0.54 |
| 3 | 54953998 | rs2360661 | 0.802 | 0.064 | 0.835 | -0.992 | -0.003 | 0.008 | 0.008 | 0 | 0.59 | 0.6 | 0.72 | 0.76 | 0.59 | 0.57 |
| 3 | 54961022 | rs3806632 | -0.265 | 0.187 | 0.609 | -0.944 | -0.017 | 0.02 | 0.02 | 0.797 | 0.78 | 0.83 | 0.84 | 0.85 | 0.72 | 0.74 |
| 3 | 54978020 | rs1850141 | 0.303 | 0.174 | 0.481 | -0.97 | -0.022 | 0.04 | 0.04 | 2.086 | 0.64 | 0.81 | 0.76 | 0.78 | 0.62 | 0.63 |
| 3 | 54980729 | rs1515956 | -0.534 | 0.241 | 1.05 | -0.953 | 0.033 | -0.038 | -0.038 | 0 | 0.24 | 0.36 | 0.18 | 0.09 | 0.24 | 0.2 |
| 3 | 54981651 | rs17054588 | 0.167 | -0.305 | 0.871 | -0.905 | -0.008 | -0.008 | -0.008 | 0 | 0.19 | 0.26 | 0.12 | 0.09 | 0.12 | 0.17 |
| 3 | 54984224 | rs17054602 | -0.85 | -0.219 | 0.803 | -0.739 | -0.028 | -0.026 | -0.026 | 6.151 | 0.07 | 0.25 | 0.02 | 0.02 | 0.01 | 0.05 |
| 3 | 54987923 | rs3821659 | 0.845 | -0.02 | 0.349 | -0.678 | 0.055 | 0.124 | 0.124 | 0.771 | 0.49 | 0.18 | 0.62 | 0.74 | 0.47 | 0.53 |
| 3 | 54990157 | rs2048809 | 0.626 | -0.104 | -0.058 | -0.389 | -0.003 | 0 | 0 | 0 | 0.49 | 0.17 | 0.69 | 0.58 | 0.61 | 0.59 |
| 3 | 54990517 | rs6794724 | -0.435 | -0.718 | -0.001 | -0.405 | -0.003 | 0 | 0 | 0 | 0.48 | 0.48 | 0.25 | 0.26 | 0.26 | 0.25 |
| 3 | 54999412 | rs1356181 | 0.601 | -0.556 | -0.202 | 0.066 | -0.074 | -0.082 | -0.082 | 0 | 0.4 | 0.13 | 0.54 | 0.57 | 0.48 | 0.57 |
| 3 | 55002066 | rs12487923 | 0.553 | -0.536 | -0.174 | 0.069 | -0.04 | -0.043 | -0.043 | 0 | 0.41 | 0.18 | 0.55 | 0.57 | 0.48 | 0.57 |
| 3 | 55006182 | rs1492001 | -0.306 | -0.576 | -0.289 | 0.169 | -0.005 | 0.013 | 0.013 | 0.002 | 0.55 | 0.61 | 0.63 | 0.75 | 0.53 | 0.64 |
| 3 | 55006308 | rs17054609 | -1.019 | -0.751 | -0.209 | 0.375 | -0.044 | 0.004 | 0.004 | 0 | 0.17 | 0.24 | 0.09 | 0.19 | 0.07 | 0.1 |
| 3 | 55007718 | rs3821656 | -0.995 | -0.734 | -0.228 | 0.118 | -0.042 | -0.012 | -0.012 | 0 | 0.17 | 0.24 | 0.09 | 0.14 | 0.07 | 0.1 |
| 3 | 55017749 | rs3773582 | -0.742 | 0.437 | -0.964 | 0.712 | -0.006 | -0.037 | -0.037 | 0.196 | 0.16 | 0.1 | 0.29 | 0.32 | 0.18 | 0.26 |
| 3 | 55019386 | rs1491998 | -0.134 | 0.525 | -1.379 | -0.251 | -0.021 | 0.062 | 0.062 | 0.066 | 0.66 | 0.77 | 0.75 | 0.9 | 0.64 | 0.79 |
| 3 | 55022475 | rs6794253 | -0.859 | 0.555 | -0.767 | -0.77 | 0.01 | -0.104 | -0.104 | 0 | 0.23 | 0.35 | 0.14 | 0.01 | 0.22 | 0.12 |
| 3 | 55028941 | rs987505 | -0.424 | 0.35 | -0.431 | -0.865 | 0.004 | -0.067 | -0.067 | 12.335 | 0.14 | 0.21 | 0.07 | 0.01 | 0.11 | 0.06 |
| 3 | 55030204 | rs9851733 | -0.713 | -0.25 | -0.246 | -1.244 | -0.01 | -0.042 | -0.042 | 0.794 | 0.76 | 0.65 | 0.84 | 0.92 | 0.82 | 0.84 |
| 3 | 55032038 | rs4423737 | -0.825 | -0.163 | -0.276 | -1.273 | -0.017 | -0.043 | -0.043 | 0.086 | 0.78 | 0.65 | 0.85 | 0.92 | 0.84 | 0.84 |
| 3 | 55033713 | rs3773573 | 1.627 | -0.26 | -0.532 | -1.115 | -0.048 | -0.012 | -0.012 | 0 | 0.42 | 0.13 | 0.44 | 0.58 | 0.48 | 0.53 |
| 3 | 55035874 | rs4955826 | -0.967 | -0.359 | -0.108 | -1.11 | 0.009 | 0.004 | 0.004 | 0.099 | 0.31 | 0.58 | 0.37 | 0.35 | 0.33 | 0.29 |
| 3 | 55042361 | rs3912834 | -0.489 | -0.539 | -0.47 | -1.631 | -0.002 | -0.06 | -0.06 | 3.575 | 0.21 | 0.22 | 0.11 | 0.01 | 0.16 | 0.15 |
| 3 | 55043070 | rs2048806 | 0.229 | -0.637 | -1.013 | -1.507 | 0.007 | 0.024 | 0.024 | 0 | 0.51 | 0.4 | 0.47 | 0.16 | 0.54 | 0.45 |
| 3 | 55044876 | rs3773569 | 0.613 | -0.542 | -1.364 | -1.2 | -0.033 | -0.039 | -0.039 | 0 | 0.16 | 0.06 | 0.23 | 0.21 | 0.22 | 0.2 |
| 3 | 55045176 | rs3773568 | 1.914 | -0.113 | -1.37 | -1.196 | -0.032 | 0.044 | 0.044 | 0.042 | 0.3 | 0.13 | 0.34 | 0.14 | 0.37 | 0.29 |
| 3 | 55046102 | rs4955917 | -0.594 | -0.204 | -1.273 | -1.151 | 0.011 | -0.025 | -0.025 | 9.49 | 0.66 | 0.73 | 0.68 | 0.36 | 0.7 | 0.55 |
| 3 | 55047412 | rs3821653 | 0.662 | -0.113 | -1.463 | -1.017 | -0.045 | -0.06 | -0.06 | 0.088 | 0.13 | 0.01 | 0.21 | 0.2 | 0.17 | 0.15 |
| 3 | 55050678 | rs7649117 | 1.877 | -0.054 | -1.383 | -1.202 | -0.028 | 0.042 | 0.042 | 0.056 | 0.3 | 0.14 | 0.33 | 0.14 | 0.36 | 0.29 |
| 3 | 55050765 | rs358028 | -1.529 | -0.374 | -1.433 | -1.222 | -0.021 | 0.098 | 0.098 | 1.173 | 0.54 | 0.81 | 0.54 | 0.78 | 0.46 | 0.56 |
| 3 | 55050796 | rs358029 | -0.617 | -0.411 | -0.805 | -1.246 | 0.006 | 0.01 | 0.01 | 0.17 | 0.38 | 0.36 | 0.44 | 0.76 | 0.36 | 0.49 |
| 3 | 55055091 | rs358033 | -0.847 | -0.293 | -0.839 | -1.271 | -0.003 | 0.006 | 0.006 | 0.208 | 0.33 | 0.71 | 0.4 | 0.31 | 0.35 | 0.28 |
| 3 | 55057249 | rs358045 | -0.907 | -0.304 | -0.825 | -1.22 | -0.005 | 0.008 | 0.008 | 0.573 | 0.34 | 0.74 | 0.41 | 0.31 | 0.35 | 0.27 |
| 3 | 55058561 | rs358046 | -0.787 | -0.385 | -1.472 | -1.703 | -0.007 | 0.001 | 0.001 | 0.064 | 0.71 | 0.75 | 0.64 | 0.8 | 0.62 | 0.68 |
| 3 | 55059102 | rs2125543 | -0.193 | -0.05 | -0.805 | -1.658 | 0.028 | -0.013 | -0.013 | 0 | 0.25 | 0.32 | 0.3 | 0.16 | 0.32 | 0.32 |
| 3 | 55063628 | rs358056 | -0.543 | 0.018 | -0.424 | -1.573 | -0.006 | -0.032 | -0.032 | 0.421 | 0.28 | 0.59 | 0.16 | 0.11 | 0.25 | 0.19 |
| 3 | 55064184 | rs3773557 | 0.547 | 0.021 | -0.723 | -1.539 | -0.002 | -0.025 | -0.025 | 0 | 0.15 | 0.08 | 0.27 | 0.26 | 0.18 | 0.19 |
| 3 | 55064740 | rs1982862 | 1.123 | 0.746 | -0.422 | -1.282 | 0.032 | 0.005 | 0.005 | 0 | 0.09 | 0.15 | 0.22 | 0.08 | 0.17 | 0.16 |
| 3 | 55065684 | rs3773555 | 0.929 | 0.732 | -0.426 | -1.288 | 0.025 | 0.002 | 0.002 | 0 | 0.1 | 0.16 | 0.23 | 0.08 | 0.17 | 0.16 |
| 3 | 55065735 | rs3773554 | -1.193 | 0.429 | -0.274 | -1.312 | -0.001 | 0 | 0 | 0.226 | 0.45 | 0.55 | 0.64 | 0.45 | 0.59 | 0.54 |
| 3 | 55068255 | rs4955920 | -0.818 | 0.269 | -0.241 | -1.131 | 0.015 | -0.013 | -0.013 | 0 | 0.61 | 0.65 | 0.74 | 0.45 | 0.72 | 0.59 |
| 3 | 55072187 | rs358493 | -1.454 | 0.721 | 0.146 | -0.653 | 0.001 | -0.015 | -0.015 | 0.948 | 0.29 | 0.75 | 0.37 | 0.35 | 0.33 | 0.35 |
| 3 | 55073917 | rs6777055 | -0.519 | 0.764 | 0.459 | -1.014 | 0.006 | -0.01 | -0.01 | 0.72 | 0.15 | 0.21 | 0.1 | 0.1 | 0.18 | 0.17 |
| 3 | 55076797 | rs358498 | -1.225 | 0.678 | 0.383 | -1.005 | -0.013 | -0.031 | -0.031 | 0.771 | 0.19 | 0.62 | 0.12 | 0.09 | 0.16 | 0.17 |
| 3 | 55089374 | rs11130448 | -0.505 | 0.956 | -0.104 | -0.876 | 0.004 | -0.006 | -0.006 | 0.001 | 0.14 | 0.17 | 0.09 | 0.1 | 0.17 | 0.17 |
| 3 | 55091151 | rs358063 | 0.213 | 1.153 | 0.121 | -1.413 | 0.01 | -0.064 | -0.064 | 0 | 0.32 | 0.33 | 0.25 | 0.05 | 0.41 | 0.26 |
| 3 | 55091518 | rs358064 | -0.048 | 0.631 | 0.279 | -1.399 | 0.007 | 0.012 | 0.012 | 0.27 | 0.36 | 0.66 | 0.26 | 0.06 | 0.3 | 0.22 |
| 3 | 55096928 | rs3773540 | -1.221 | 0.588 | 0.666 | -0.251 | -0.005 | 0.317 | 0.317 | 2.046 | 0.08 | 0.38 | 0.35 | 0.25 | 0.04 | 0.18 |
| 3 | 55100492 | rs358067 | 0.424 | 0.603 | 0.305 | -0.376 | -0.028 | 0.028 | 0.028 | 0.909 | 0.72 | 0.49 | 0.51 | 0.7 | 0.78 | 0.72 |
| 3 | 55104615 | rs165440 | 0.077 | 0.631 | -0.584 | -0.756 | 0.014 | 0.06 | 0.06 | 0.156 | 0.76 | 0.83 | 0.87 | 0.94 | 0.83 | 0.87 |
| 3 | 55107944 | rs17323097 | 0.422 | 1.268 | -0.498 | -0.447 | 0.006 | -0.013 | -0.013 | 0 | 0.06 | 0.07 | 0.08 | 0 | 0.15 | 0.05 |
| 3 | 55108358 | rs358077 | 0.471 | 1.264 | -0.569 | -0.45 | -0.006 | 0 | 0 | 0 | 0.06 | 0.04 | 0.07 | 0 | 0.15 | 0.05 |
| 3 | 55108625 | rs17323187 | 0.165 | 1.117 | -0.527 | -0.382 | -0.001 | -0.002 | -0.002 | 0 | 0.06 | 0.06 | 0.03 | 0.01 | 0.07 | 0.05 |
| 3 | 55112415 | rs11130451 | -0.376 | 1.226 | -0.51 | -0.101 | -0.004 | 0 | 0 | 0 | 0.1 | 0.07 | 0.3 | 0.18 | 0.13 | 0.23 |
| 3 | 55112923 | rs358081 | 0.607 | 0.822 | -0.239 | -0.5 | 0 | 0 | 0 | 0 | 0.69 | 0.65 | 0.57 | 0.79 | 0.73 | 0.61 |
| 3 | 55118984 | rs10865992 | 0.38 | 0.652 | 0.132 | -0.783 | -0.013 | -0.13 | -0.13 | 0 | 0.68 | 0.47 | 0.78 | 0.96 | 0.71 | 0.78 |
| 3 | 55120948 | rs9865871 | 0.645 | 0.832 | -0.005 | -0.653 | -0.003 | 0.007 | 0.007 | 0.263 | 0.62 | 0.68 | 0.5 | 0.79 | 0.58 | 0.55 |
| 3 | 55122872 | rs358007 | -0.716 | 0.31 | -0.954 | -1.227 | 0.047 | 0.126 | 0.126 | 0 | 0.52 | 0.76 | 0.49 | 0.22 | 0.35 | 0.54 |
| 3 | 55125641 | rs358012 | 0.576 | 0.183 | -0.464 | -0.975 | 0.001 | 0.016 | 0.016 | 0 | 0.15 | 0.15 | 0.37 | 0.21 | 0.16 | 0.28 |
| 3 | 55127512 | rs9843442 | 0.376 | -0.304 | -0.115 | -1.336 | -0.026 | -0.002 | -0.002 | 0 | 0.68 | 0.53 | 0.53 | 0.75 | 0.75 | 0.63 |
| 3 | 55130269 | rs13061131 | 0.738 | -0.58 | -0.434 | -1.701 | 0.011 | 0 | 0 | 0 | 0.13 | 0.1 | 0.04 | 0 | 0.09 | 0.06 |
| 3 | 55132626 | rs11130454 | 1.043 | -0.635 | -0.407 | -1.729 | 0.02 | 0.018 | 0.018 | 2.054 | 0.54 | 0.4 | 0.48 | 0.75 | 0.65 | 0.51 |
| 3 | 55136997 | rs1458977 | 0.974 | -0.489 | -0.944 | -1.87 | 0.004 | 0.004 | 0.004 | 0.5 | 0.3 | 0.33 | 0.29 | 0.66 | 0.38 | 0.4 |
| 3 | 55138063 | rs1458978 | 1.797 | -0.125 | -1.144 | -1.552 | -0.049 | 0.088 | 0.088 | 2.31 | 0.25 | 0.07 | 0.19 | 0.11 | 0.27 | 0.19 |
| 3 | 55144055 | rs697928 | -1.63 | -0.19 | -0.995 | -1.407 | -0.001 | 0.005 | 0.005 | 0 | 0.87 | 0.9 | 0.72 | 0.81 | 0.89 | 0.75 |
| 3 | 55146011 | rs697929 | 1.196 | -0.043 | -1.132 | -1.498 | -0.002 | 0.061 | 0.061 | 0 | 0.15 | 0.06 | 0.08 | 0.03 | 0.11 | 0.14 |
| 3 | 55155778 | rs697931 | -0.331 | 0.038 | -0.599 | -1.241 | 0 | -0.006 | -0.006 | 0.01 | 0.58 | 0.5 | 0.55 | 0.68 | 0.59 | 0.56 |
| 3 | 55157606 | rs11708743 | 1.273 | 0.246 | -0.806 | -1.247 | -0.01 | 0.053 | 0.053 | 1.46 | 0.25 | 0.12 | 0.17 | 0.1 | 0.25 | 0.19 |

| | | | | | | | | | | | | | | | | |
|---|---|---|---|---|---|---|---|---|---|---|---|---|---|---|---|---|
| 3 | 55161652 | rs2122562 | -1.136 | 0.027 | -0.432 | -1.037 | -0.006 | 0.01 | 0.01 | 0 | 0.83 | 0.81 | 0.73 | 0.77 | 0.86 | 0.73 |
| 3 | 55161838 | rs7633147 | 0.741 | 0.887 | -0.452 | -0.471 | 0.029 | 0.1 | 0.1 | 0 | 0.12 | 0.19 | 0.25 | 0.46 | 0.26 | 0.2 |
| 3 | 55162101 | rs12493791 | 1.564 | 0.615 | -0.968 | -1.153 | -0.007 | 0.01 | 0.01 | 2.011 | 0.18 | 0.15 | 0.19 | 0.14 | 0.2 | 0.15 |
| 3 | 55164934 | rs17054861 | 1.591 | 0.516 | -0.96 | -1.174 | -0.001 | 0.008 | 0.008 | 1.974 | 0.18 | 0.15 | 0.14 | 0.13 | 0.18 | 0.13 |
| 3 | 55165237 | rs17054864 | 1.288 | 0.445 | -0.74 | -0.931 | -0.001 | 0.109 | 0.109 | 0 | 0.05 | 0.15 | 0.15 | 0.22 | 0.06 | 0.19 |
| 3 | 55166092 | rs1458980 | 1.488 | 0.405 | -1.123 | -1.303 | -0.006 | 0.038 | 0.038 | 0.029 | 0.13 | 0.06 | 0.13 | 0.03 | 0.13 | 0.13 |
| 3 | 55170034 | rs1388634 | -0.915 | 0.139 | -1.028 | -1.481 | 0.004 | -0.001 | -0.001 | 0 | 0.62 | 0.61 | 0.56 | 0.41 | 0.55 | 0.58 |
| 3 | 55177886 | rs1688334 | -0.121 | -0.161 | -0.958 | -1.353 | -0.002 | -0.003 | -0.003 | 0 | 0.33 | 0.37 | 0.37 | 0.27 | 0.29 | 0.43 |
| 3 | 55179111 | rs1674535 | -0.504 | 0.253 | -0.823 | -1.071 | 0.001 | 0.001 | 0.001 | 0 | 0.44 | 0.44 | 0.42 | 0.27 | 0.38 | 0.49 |
| 3 | 55179991 | rs4955830 | 0.817 | 0.536 | -0.659 | -0.384 | -0.004 | 0.105 | 0.105 | 0 | 0.06 | 0.16 | 0.16 | 0.22 | 0.06 | 0.19 |
| 3 | 55180636 | rs11130458 | 0.415 | 0.991 | -1.065 | -1.143 | -0.009 | 0.015 | 0.015 | 0.004 | 0.17 | 0.13 | 0.16 | 0.06 | 0.21 | 0.16 |
| 3 | 55181375 | rs4955932 | -0.375 | 0.026 | -0.944 | -1.218 | 0.031 | 0.061 | 0.061 | 0 | 0.48 | 0.69 | 0.44 | 0.33 | 0.4 | 0.57 |
| 3 | 55182993 | rs4955933 | 0.949 | 0.309 | -0.737 | -1.632 | -0.036 | 0.09 | 0.09 | 1.128 | 0.42 | 0.14 | 0.57 | 0.75 | 0.44 | 0.41 |
| 3 | 55183520 | rs9870828 | -0.815 | 0.694 | 0.178 | -1.547 | 0.009 | -0.048 | -0.048 | 1.321 | 0.38 | 0.48 | 0.28 | 0.16 | 0.4 | 0.4 |
| 3 | 55187140 | rs13433743 | 0.515 | 0.647 | 0.357 | -1.547 | -0.001 | -0.006 | -0.006 | 0 | 0.16 | 0.19 | 0.14 | 0.11 | 0.17 | 0.11 |
| 3 | 55188228 | rs11713980 | 0.098 | 0.684 | 0.284 | -1.528 | -0.001 | -0.028 | -0.028 | 0 | 0.2 | 0.32 | 0.12 | 0.06 | 0.19 | 0.17 |
| 3 | 55188514 | rs4955935 | -1.084 | 0.604 | 0.369 | -1.597 | 0.002 | -0.013 | -0.013 | 0 | 0.88 | 0.84 | 0.94 | 0.91 | 0.88 | 0.88 |
| 3 | 55189959 | rs9879126 | -1.039 | 0.532 | 0.333 | -1.545 | -0.001 | -0.03 | -0.03 | 0.717 | 0.37 | 0.57 | 0.27 | 0.17 | 0.37 | 0.28 |
| 3 | 55190515 | rs9859995 | 0.112 | 0.616 | -0.172 | -1.232 | 0.003 | -0.021 | -0.021 | 0.132 | 0.16 | 0.23 | 0.14 | 0.11 | 0.17 | 0.11 |
| 3 | 55193060 | rs12638400 | -0.645 | 0.867 | -0.583 | -1.208 | -0.009 | 0.17 | 0.17 | 0.167 | 0.48 | 0.16 | 0.35 | 0.2 | 0.5 | 0.47 |
| 3 | 55194016 | rs9880768 | 0.436 | 0.9 | -0.468 | -1.347 | -0.002 | -0.024 | -0.024 | 0 | 0.07 | 0.11 | 0.05 | 0 | 0.06 | 0.1 |
| 3 | 55194254 | rs4955937 | 0.982 | 0.82 | -0.372 | -1.235 | -0.024 | 0.082 | 0.082 | 0 | 0.45 | 0.19 | 0.57 | 0.72 | 0.49 | 0.41 |
| 3 | 55194450 | rs11917809 | -0.39 | 0.941 | -0.67 | -1.341 | -0.024 | 0.107 | 0.107 | 0.151 | 0.47 | 0.16 | 0.33 | 0.27 | 0.48 | 0.56 |
| 3 | 55194956 | rs753294 | -0.822 | 1.132 | -0.76 | -1.157 | 0.001 | -0.015 | -0.015 | 0 | 0.11 | 0.11 | 0.14 | 0.03 | 0.15 | 0.11 |
| 3 | 55196913 | rs10510776 | 0.741 | 0.901 | -0.93 | -1.135 | -0.009 | 0.136 | 0.136 | 0.615 | 0.49 | 0.11 | 0.51 | 0.68 | 0.58 | 0.41 |
| 3 | 55198849 | rs896109 | -0.693 | 0.949 | -0.816 | -1.124 | 0.014 | 0.025 | 0.025 | 0.048 | 0.7 | 0.82 | 0.83 | 0.79 | 0.77 | 0.65 |
| 3 | 55199267 | rs17054896 | 0.949 | 0.806 | -0.662 | -1.033 | -0.002 | 0.001 | 0.001 | 0 | 0.05 | 0.05 | 0.13 | 0.07 | 0.05 | 0.04 |
| 3 | 55204544 | rs1450643 | 1.033 | 0.874 | -0.739 | -1.097 | -0.012 | 0.026 | 0.026 | 5.103 | 0.1 | 0.04 | 0.07 | 0.01 | 0.13 | 0.05 |
| 3 | 55207133 | rs2036487 | 0.77 | 0.888 | -0.748 | -1.124 | 0.006 | 0.033 | 0.033 | 0 | 0.47 | 0.67 | 0.52 | 0.68 | 0.47 | 0.4 |
| 3 | 55208424 | rs6445716 | 0.151 | 0.776 | -0.526 | -1.207 | 0.042 | -0.007 | -0.007 | 0 | 0.08 | 0.48 | 0.07 | 0 | 0.03 | 0.04 |
| 3 | 55210070 | rs12632515 | 0.592 | 1.026 | -1.389 | -0.965 | 0.004 | -0.003 | -0.003 | 0 | 0.06 | 0.09 | 0.14 | 0.05 | 0.07 | 0.05 |
| 3 | 55212014 | rs716384 | 0.533 | 1.076 | -1.647 | -0.972 | -0.023 | 0.099 | 0.099 | 0 | 0.47 | 0.2 | 0.54 | 0.71 | 0.5 | 0.43 |
| 3 | 55212226 | rs1823499 | 0.947 | 1.109 | -1.703 | -0.832 | -0.011 | 0.007 | 0.007 | 0 | 0.4 | 0.17 | 0.44 | 0.64 | 0.41 | 0.35 |
| 3 | 55212253 | rs12491562 | 0.434 | 0.937 | -1.673 | -0.829 | -0.028 | -0.007 | -0.007 | 0 | 0.07 | 0.01 | 0.1 | 0.05 | 0.09 | 0.08 |
| 3 | 55216445 | rs13081750 | 0.611 | 0.517 | -1.496 | -0.88 | -0.023 | 0.033 | 0.033 | 0.917 | 0.1 | 0.03 | 0.07 | 0.01 | 0.13 | 0.04 |
| 3 | 55221542 | rs6770519 | 0.001 | 0.443 | -1.535 | -0.932 | 0.037 | 0.094 | 0.094 | 0 | 0.59 | 0.19 | 0.63 | 0.74 | 0.67 | 0.48 |
| 3 | 55222983 | rs4955834 | -0.594 | 0.776 | -0.718 | -0.526 | -0.001 | -0.008 | -0.008 | 0 | 0.17 | 0.18 | 0.2 | 0.06 | 0.17 | 0.09 |
| 3 | 55224080 | rs1461796 | 0.352 | 1.063 | -0.222 | -0.848 | 0.026 | -0.149 | -0.149 | 0 | 0.78 | 0.49 | 0.74 | 0.96 | 0.79 | 0.88 |
| 3 | 55224638 | rs6765362 | -0.224 | 0.873 | -0.313 | 0.07 | -0.001 | 0.05 | 0.05 | 0 | 0.19 | 0.25 | 0.25 | 0.57 | 0.2 | 0.44 |
| 3 | 55226941 | rs11130462 | -0.198 | 0.773 | -0.269 | -0.245 | 0 | 0.002 | 0.002 | 0 | 0.52 | 0.58 | 0.5 | 0.71 | 0.55 | 0.77 |
| 3 | 55228586 | rs9818965 | 0.067 | 0.516 | -0.088 | -0.174 | -0.023 | 0.016 | 0.016 | 0 | 0.44 | 0.24 | 0.36 | 0.65 | 0.54 | 0.73 |
| 3 | 55228684 | rs9878241 | 1.012 | 0.195 | -0.121 | -0.728 | -0.001 | -0.035 | -0.035 | 0 | 0.26 | 0.21 | 0.19 | 0 | 0.24 | 0.09 |
| 3 | 55229775 | rs6782687 | -0.545 | 0.291 | 0.088 | -0.583 | -0.033 | -0.037 | -0.037 | 0 | 0.8 | 0.55 | 0.79 | 0.94 | 0.87 | 0.9 |
| 3 | 55230058 | rs9311553 | -0.306 | 0.279 | -0.155 | -0.594 | -0.036 | -0.043 | -0.043 | 0.348 | 0.78 | 0.45 | 0.78 | 0.94 | 0.87 | 0.91 |
| 3 | 55231799 | rs1381389 | -0.003 | 0.721 | -0.222 | -0.518 | 0.011 | 0.017 | 0.017 | 0.218 | 0.75 | 0.57 | 0.64 | 0.7 | 0.75 | 0.84 |
| 3 | 55234425 | rs1563976 | -0.288 | 0.633 | -0.127 | -0.484 | -0.029 | -0.047 | -0.047 | 0 | 0.78 | 0.51 | 0.79 | 0.94 | 0.85 | 0.89 |
| 3 | 55236699 | rs12491607 | 0.673 | 0.587 | -0.49 | -0.67 | -0.027 | 0 | 0 | 0 | 0.42 | 0.23 | 0.33 | 0.63 | 0.49 | 0.71 |
| 3 | 55237738 | rs10510777 | 0.439 | 0.679 | -0.482 | -0.708 | -0.007 | 0.074 | 0.074 | 0 | 0.47 | 0.28 | 0.43 | 0.7 | 0.5 | 0.74 |
| 3 | 55241137 | rs6802697 | 0.222 | 0.725 | -0.586 | -0.66 | -0.002 | 0.014 | 0.014 | 0.477 | 0.37 | 0.32 | 0.46 | 0.29 | 0.36 | 0.19 |
| 3 | 55256432 | rs374563 | 0.434 | 0.812 | -1.058 | -0.404 | -0.068 | -0.112 | -0.112 | 1.166 | 0.37 | 0.11 | 0.29 | 0.55 | 0.42 | 0.67 |
| 3 | 55259031 | rs414538 | 0.198 | 0.432 | -0.99 | -0.573 | 0.001 | 0 | 0 | 0 | 0.6 | 0.41 | 0.67 | 0.88 | 0.66 | 0.66 |
| 3 | 55259743 | rs422712 | -1.56 | 0.489 | -0.497 | -0.602 | 0.003 | -0.089 | -0.089 | 0 | 0.85 | 0.69 | 0.89 | 1 | 0.86 | 0.94 |
| 3 | 55261956 | rs443840 | 0.289 | 0.608 | -0.661 | -0.71 | 0.004 | 0.023 | 0.023 | 0 | 0.46 | 0.36 | 0.38 | 0.16 | 0.41 | 0.24 |
| 3 | 55262928 | rs17709697 | 1.66 | 0.728 | -0.705 | -0.555 | -0.016 | -0.017 | -0.017 | 3.114 | 0.07 | 0.04 | 0.17 | 0.16 | 0.1 | 0.08 |
| 3 | 55263106 | rs373532 | 0.481 | 0.644 | -0.715 | -0.618 | 0.001 | 0.009 | 0.009 | 0 | 0.42 | 0.34 | 0.28 | 0.1 | 0.4 | 0.21 |
| 3 | 55263442 | rs535157 | -0.129 | 0.724 | -0.722 | -0.402 | 0.011 | 0.028 | 0.028 | 0.578 | 0.5 | 0.39 | 0.54 | 0.43 | 0.53 | 0.3 |
| 3 | 55263471 | rs535219 | -2.294 | 0.544 | -0.611 | -0.531 | 0.032 | -0.044 | -0.044 | 0 | 0.92 | 0.8 | 0.93 | 1 | 0.88 | 0.97 |
| 3 | 55263626 | rs6445725 | 0.071 | 0.554 | -0.731 | -0.498 | 0.003 | 0.046 | 0.046 | 0.474 | 0.49 | 0.39 | 0.46 | 0.27 | 0.5 | 0.29 |
| 3 | 55264003 | rs403286 | -0.198 | 0.773 | -0.706 | -0.397 | 0 | 0.008 | 0.008 | 1.535 | 0.53 | 0.41 | 0.54 | 0.32 | 0.51 | 0.32 |
| 3 | 55265321 | rs6801136 | 0.995 | 0.659 | -0.936 | -0.406 | 0.001 | 0.002 | 0.002 | 0 | 0.37 | 0.3 | 0.34 | 0.16 | 0.32 | 0.16 |
| 3 | 55265674 | rs393836 | -0.012 | 0.562 | -1.625 | -0.255 | -0.068 | -0.101 | -0.101 | 0 | 0.43 | 0.11 | 0.31 | 0.55 | 0.45 | 0.69 |
| 3 | 55267391 | rs381062 | -1.424 | 0.451 | -0.821 | -0.158 | 0.047 | -0.044 | -0.044 | 0.276 | 0.91 | 0.75 | 0.92 | 1 | 0.87 | 0.98 |
| 3 | 55267443 | rs716588 | 0.02 | 0.344 | -1.951 | 0.094 | 0.057 | 0.033 | 0.033 | 2.536 | 0.53 | 0.15 | 0.53 | 0.44 | 0.6 | 0.38 |
| 3 | 55268392 | rs392699 | 0.53 | 0.641 | -1.204 | 0.201 | 0.006 | -0.011 | -0.011 | 0 | 0.58 | 0.49 | 0.39 | 0.3 | 0.55 | 0.57 |
| 3 | 55268476 | rs11716583 | -0.947 | 0.432 | -1.123 | 0.01 | -0.007 | 0.021 | 0.021 | 0 | 0.44 | 0.37 | 0.61 | 0.78 | 0.45 | 0.5 |
| 3 | 55274426 | rs17760821 | 0.018 | 0.446 | -1.377 | 0.173 | -0.004 | 0.003 | 0.003 | 0 | 0.29 | 0.28 | 0.44 | 0.23 | 0.36 | 0.22 |
| 3 | 55274566 | rs11719604 | -0.164 | 0.304 | -1.96 | 0.095 | -0.072 | -0.041 | -0.041 | 0 | 0.27 | 0.02 | 0.13 | 0.25 | 0.28 | 0.24 |
| 3 | 55276518 | rs6772225 | 1.312 | -0.122 | -1.405 | 0.22 | -0.016 | 0.117 | 0.117 | 0 | 0.13 | 0.22 | 0.15 | 0.4 | 0.07 | 0.28 |
| 3 | 55276601 | rs17055018 | 1.579 | -0.022 | -1.62 | 0.187 | -0.005 | 0.003 | 0.003 | 0 | 0.15 | 0.01 | 0.06 | 0.08 | 0.13 | 0.14 |

| | | | | | | | | | | | | | | | | |
|---|---|---|---|---|---|---|---|---|---|---|---|---|---|---|---|---|
| 3 | 55281622 | rs6764543 | 0.282 | -0.045 | -0.971 | 0.136 | 0.003 | -0.001 | -0.001 | 0.02 | 0.51 | 0.55 | 0.46 | 0.48 | 0.48 | 0.62 |
| 3 | 55284390 | rs11715075 | 0.497 | -0.029 | -0.973 | 0.143 | 0.013 | -0.002 | -0.002 | 0 | 0.48 | 0.55 | 0.45 | 0.47 | 0.41 | 0.55 |
| 3 | 55290746 | rs7613572 | -0.207 | -0.046 | -1.274 | 0.189 | -0.021 | -0.038 | -0.038 | 0 | 0.45 | 0.2 | 0.51 | 0.48 | 0.52 | 0.36 |
| 3 | 55300666 | rs17761091 | 1.141 | -0.126 | -1.063 | 0.058 | 0.002 | 0.025 | 0.025 | 0 | 0.32 | 0.21 | 0.28 | 0.07 | 0.33 | 0.28 |
| 3 | 55302625 | rs4435604 | 0.714 | 0.161 | -1.727 | 0.466 | -0.077 | -0.088 | -0.088 | 0 | 0.14 | 0.01 | 0.33 | 0.26 | 0.18 | 0.07 |
| 3 | 55306395 | rs458308 | 0.258 | 0.098 | -1.859 | 0.487 | -0.031 | -0.003 | -0.003 | 0 | 0.29 | 0.13 | 0.18 | 0.3 | 0.35 | 0.31 |
| 3 | 55310644 | rs6801238 | -0.19 | 0.215 | -1.022 | 1.084 | 0.004 | 0.004 | 0.004 | 0 | 0.12 | 0.13 | 0.14 | 0.38 | 0.07 | 0.28 |
| 3 | 55311650 | rs17055089 | 0.804 | 0.23 | -1.101 | 0.663 | -0.004 | 0.008 | 0.008 | 0 | 0.11 | 0.06 | 0.08 | 0.08 | 0.1 | 0.13 |
| 3 | 55312099 | rs6805013 | 0.08 | 0.283 | -1.101 | 0.747 | 0.003 | 0.025 | 0.025 | 0 | 0.11 | 0.12 | 0.2 | 0.41 | 0.07 | 0.22 |
| 3 | 55313400 | rs358806 | -0.839 | 0.495 | -1.202 | 0.614 | -0.01 | -0.007 | -0.007 | 1.41 | 0.87 | 0.89 | 0.81 | 0.81 | 0.78 | 0.92 |
| 3 | 55314975 | rs358805 | 0.994 | 0.343 | -1.179 | 0.55 | 0 | -0.003 | -0.003 | 0 | 0.59 | 0.4 | 0.5 | 0.34 | 0.61 | 0.57 |
| 3 | 55318709 | rs358803 | 0.137 | 0.534 | -1.819 | 0.149 | 0.01 | -0.012 | -0.012 | 0 | 0.74 | 0.35 | 0.7 | 0.81 | 0.69 | 0.88 |
| 3 | 55322453 | rs13083875 | 0.592 | -0.153 | -2.033 | 0.454 | 0.001 | 0.005 | 0.005 | 0.06 | 0.1 | 0.07 | 0.08 | 0.09 | 0.09 | 0.19 |
| 3 | 55325605 | rs4955840 | -0.377 | -0.18 | -1.786 | 0.864 | -0.001 | 0.072 | 0.072 | 0 | 0.06 | 0.1 | 0.15 | 0.27 | 0.05 | 0.21 |
| 3 | 55332449 | rs749152 | 0.613 | -0.442 | -2.21 | 0.657 | -0.039 | 0.048 | 0.048 | 0.137 | 0.3 | 0.14 | 0.23 | 0.15 | 0.36 | 0.32 |
| 3 | 55333080 | rs2004214 | 0.436 | -0.413 | -2.24 | 0.934 | -0.014 | 0.085 | 0.085 | 0 | 0.43 | 0.69 | 0.33 | 0.15 | 0.44 | 0.35 |
| 3 | 55333096 | rs2004213 | 0.59 | -0.652 | -2.277 | 0.894 | 0.002 | 0.003 | 0.003 | 1.345 | 0.06 | 0.07 | 0.1 | 0.08 | 0.06 | 0.02 |
| 3 | 55335095 | rs358783 | -0.526 | -0.598 | -2.178 | 0.97 | 0.016 | -0.007 | -0.007 | 0 | 0.84 | 0.91 | 0.92 | 0.83 | 0.9 | 0.89 |
| 3 | 55341143 | rs358817 | -0.183 | 0.715 | -2.234 | 0.455 | -0.088 | -0.052 | -0.052 | 0 | 0.38 | 0.1 | 0.47 | 0.42 | 0.45 | 0.35 |
| 3 | 55354717 | rs4955978 | -0.136 | 0.09 | -1.38 | -0.108 | -0.018 | -0.021 | -0.021 | 0 | 0.42 | 0.49 | 0.53 | 0.73 | 0.31 | 0.36 |
| 3 | 55356725 | rs1000408 | 0.258 | -0.5 | -1.073 | -0.384 | -0.001 | 0 | 0 | 0 | 0.24 | 0.21 | 0.43 | 0.71 | 0.21 | 0.31 |
| 3 | 55356854 | rs9818883 | 0.33 | -1.048 | -0.983 | -0.968 | -0.022 | -0.026 | -0.026 | 0 | 0.13 | 0.23 | 0.13 | 0.09 | 0.09 | 0.31 |
| 3 | 55371976 | rs9842885 | 1.086 | -1.211 | -1.04 | -1.129 | 0.014 | 0.063 | 0.063 | 0.372 | 0.13 | 0.02 | 0.05 | 0 | 0.09 | 0.05 |
| 3 | 55372745 | rs7616961 | -1.279 | -1.801 | -1.396 | -1.452 | -0.063 | 0.147 | 0.147 | 3.556 | 0.58 | 0.93 | 0.68 | 0.85 | 0.41 | 0.71 |
| 3 | 55373924 | rs9873066 | 0.086 | -1.776 | -1.014 | -1.397 | -0.037 | -0.026 | -0.026 | 0 | 0.16 | 0.31 | 0.12 | 0.1 | 0.09 | 0.31 |
| 3 | 55377935 | rs9839808 | -1.493 | -0.335 | -1.928 | -1.924 | -0.114 | 0.025 | 0.025 | 0 | 0.83 | 0.99 | 0.85 | 0.93 | 0.7 | 0.85 |
| 3 | 55389356 | rs12492556 | 1.63 | -0.775 | -1.659 | -1.883 | -0.025 | -0.089 | -0.089 | 0 | 0.21 | 0.03 | 0.44 | 0.7 | 0.22 | 0.33 |
| 3 | 55390886 | rs11925054 | 1.09 | -0.669 | -1.98 | -2.289 | -0.021 | 0.064 | 0.064 | 0 | 0.13 | 0.03 | 0.1 | 0 | 0.12 | 0.05 |
| 3 | 55393851 | rs4580514 | 1.034 | -0.039 | -1.958 | -2.494 | -0.022 | 0.052 | 0.052 | 0 | 0.22 | 0.12 | 0.11 | 0.01 | 0.23 | 0.12 |
| 3 | 55406255 | rs9683125 | 0.691 | -0.058 | -1.505 | -2.519 | 0.005 | -0.052 | -0.052 | 0 | 0.24 | 0.28 | 0.17 | 0.03 | 0.25 | 0.13 |
| 3 | 55406634 | rs7629260 | 0.732 | -0.057 | -1.505 | -2.518 | 0.006 | -0.053 | -0.053 | 0.437 | 0.24 | 0.28 | 0.17 | 0.03 | 0.25 | 0.13 |
| 3 | 55408967 | rs7635898 | 1.001 | -0.052 | -1.477 | -2.516 | -0.004 | -0.015 | -0.015 | 0 | 0.22 | 0.19 | 0.13 | 0.01 | 0.24 | 0.12 |
| 3 | 55411011 | rs11130471 | 1.748 | -0.226 | -0.91 | -2.588 | 0.006 | -0.014 | -0.014 | 0.127 | 0.24 | 0.17 | 0.45 | 0.73 | 0.23 | 0.34 |
| 3 | 55412020 | rs7633876 | 0.892 | -0.408 | -0.534 | -2.649 | -0.028 | 0.053 | 0.053 | 0 | 0.09 | 0.3 | 0.09 | 0.09 | 0.01 | 0.26 |
| 3 | 55430471 | rs1879382 | 0.635 | -0.347 | -0.603 | -2.694 | -0.004 | 0.026 | 0.026 | 0.266 | 0.46 | 0.36 | 0.58 | 0.74 | 0.47 | 0.46 |
| 3 | 55430599 | rs9843897 | 1.219 | -0.312 | -0.724 | -2.698 | -0.014 | 0.019 | 0.019 | 0.036 | 0.22 | 0.16 | 0.13 | 0.02 | 0.24 | 0.12 |
| 3 | 55431615 | rs7618735 | -0.203 | -0.361 | -0.777 | -2.612 | -0.101 | 0.153 | 0.153 | 2.47 | 0.58 | 0.88 | 0.67 | 0.85 | 0.49 | 0.72 |
| 3 | 55438194 | rs1829555 | 0.815 | -0.457 | -0.41 | -2.505 | -0.014 | 0.055 | 0.055 | 0 | 0.09 | 0.34 | 0.09 | 0.07 | 0.01 | 0.25 |
| 3 | 55441527 | rs3846037 | -0.36 | -0.056 | -0.906 | -2.731 | -0.044 | 0.111 | 0.111 | 0.797 | 0.62 | 0.89 | 0.77 | 0.85 | 0.56 | 0.76 |
| 3 | 55445851 | rs11716743 | 1.147 | -0.158 | -0.897 | -2.775 | -0.002 | -0.021 | -0.021 | 0 | 0.2 | 0.19 | 0.11 | 0.01 | 0.19 | 0.11 |
| 3 | 55452900 | rs4955992 | 1.842 | -0.362 | -0.74 | -2.64 | -0.017 | -0.05 | -0.05 | 0.173 | 0.25 | 0.08 | 0.46 | 0.73 | 0.26 | 0.34 |
| 3 | 55453536 | rs9872536 | 0.423 | -0.335 | -0.336 | -2.54 | -0.034 | 0.058 | 0.058 | 0.018 | 0.1 | 0.46 | 0.1 | 0.09 | 0.01 | 0.25 |
| 3 | 55461980 | rs7640326 | 0.868 | -0.055 | -0.5 | -2.704 | 0.017 | -0.08 | -0.08 | 0.727 | 0.19 | 0.3 | 0.14 | 0.01 | 0.23 | 0.09 |
| 3 | 55463424 | rs1373634 | 0.362 | -0.516 | -0.384 | -2.941 | -0.049 | -0.017 | -0.017 | 0 | 0.14 | 0.26 | 0.09 | 0.1 | 0.04 | 0.28 |
| 3 | 55468209 | rs2885453 | 1.314 | -0.371 | -0.422 | -3.047 | 0.01 | -0.054 | -0.054 | 0.043 | 0.18 | 0.22 | 0.13 | 0.01 | 0.23 | 0.08 |
| 3 | 55469813 | rs1520703 | 0.815 | -0.821 | -0.75 | -2.948 | -0.01 | 0.012 | 0.012 | 0 | 0.42 | 0.37 | 0.6 | 0.68 | 0.49 | 0.42 |
| 3 | 55470190 | rs12488984 | 1.894 | -0.734 | -0.934 | -2.969 | -0.016 | -0.039 | -0.039 | 0.148 | 0.24 | 0.14 | 0.46 | 0.67 | 0.26 | 0.34 |
| 3 | 55472156 | rs865235 | -0.866 | -1.022 | -0.534 | -2.872 | -0.001 | 0.007 | 0.007 | 0 | 0.7 | 0.74 | 0.83 | 0.79 | 0.74 | 0.81 |
| 3 | 55474073 | rs1392224 | 1.277 | -1.007 | -0.661 | -2.856 | -0.029 | 0.024 | 0.024 | 0 | 0.41 | 0.31 | 0.59 | 0.69 | 0.5 | 0.42 |
| 3 | 55475273 | rs815533 | -2.316 | -1.059 | -0.926 | -2.854 | -0.048 | -0.047 | -0.047 | 0 | 0.84 | 0.99 | 0.91 | 0.85 | 0.85 | 0.89 |
| 3 | 55487428 | rs751194 | 1.256 | -0.959 | -0.539 | -2.328 | -0.016 | -0.006 | -0.006 | 0 | 0.39 | 0.31 | 0.58 | 0.63 | 0.45 | 0.42 |
| 3 | 55488281 | rs6445746 | 0.855 | -1.024 | -0.27 | -2.292 | -0.048 | -0.005 | -0.005 | 0 | 0.12 | 0.25 | 0.09 | 0.1 | 0.03 | 0.27 |
| 3 | 55491013 | rs629537 | 0.013 | -0.94 | -0.429 | -2.318 | 0.01 | -0.037 | -0.037 | 0.647 | 0.14 | 0.26 | 0.12 | 0.07 | 0.13 | 0.1 |
| 3 | 55491436 | rs503022 | 0.065 | -0.932 | -0.424 | -2.315 | 0.012 | -0.036 | -0.036 | 0 | 0.13 | 0.25 | 0.12 | 0.07 | 0.13 | 0.1 |
| 3 | 55493514 | rs476986 | -1.247 | -0.862 | -1.318 | -2.285 | -0.042 | -0.033 | -0.033 | 1.782 | 0.75 | 0.88 | 0.85 | 0.79 | 0.77 | 0.84 |
| 3 | 55499664 | rs12497254 | 0.84 | -0.532 | -0.72 | -2.219 | -0.01 | 0.003 | 0.003 | 0 | 0.37 | 0.34 | 0.59 | 0.7 | 0.45 | 0.38 |
| 3 | 55501175 | rs1829556 | 0.855 | -0.515 | -0.855 | -2.206 | -0.023 | 0.006 | 0.006 | 0 | 0.36 | 0.3 | 0.59 | 0.7 | 0.45 | 0.38 |
| 3 | 55511675 | rs472631 | -0.78 | -0.153 | -1.043 | -2.173 | -0.005 | 0.003 | 0.003 | 0 | 0.59 | 0.63 | 0.37 | 0.31 | 0.5 | 0.59 |
| 3 | 55519857 | rs648872 | -0.275 | -0.372 | -1.308 | -1.507 | -0.005 | -0.063 | -0.063 | 0 | 0.18 | 0.31 | 0.13 | 0.05 | 0.18 | 0.17 |
| 3 | 55520778 | rs566926 | -1.677 | 0.039 | -2.371 | -1.386 | -0.061 | -0.121 | -0.121 | 1.705 | 0.76 | 0.98 | 0.54 | 0.32 | 0.73 | 0.69 |
| 3 | 55531267 | rs557077 | -1.084 | -0.592 | -2.225 | -1.72 | 0.023 | 0.028 | 0.028 | 4.677 | 0.29 | 0.75 | 0.3 | 0.13 | 0.22 | 0.31 |
| 3 | 55531959 | rs1160047 | -0.544 | -0.222 | -2.325 | -2.596 | -0.008 | 0.069 | 0.069 | 0.034 | 0.44 | 0.3 | 0.32 | 0.22 | 0.49 | 0.34 |
| 3 | 55535274 | rs7624718 | 0.138 | 0.114 | -2.253 | -2.479 | -0.002 | 0.012 | 0.012 | 0 | 0.15 | 0.11 | 0.15 | 0.05 | 0.2 | 0.11 |
| 3 | 55536802 | rs17711589 | 1.425 | 0.161 | -2.447 | -2.543 | -0.078 | -0.035 | -0.035 | 0 | 0.2 | 0.01 | 0.1 | 0.16 | 0.24 | 0.15 |
| 3 | 55539560 | rs1795651 | 0.742 | 0.109 | -2.229 | -2.528 | -0.002 | -0.044 | -0.044 | 0 | 0.3 | 0.6 | 0.27 | 0.1 | 0.25 | 0.3 |
| 3 | 55546868 | rs1499886 | -0.847 | -0.275 | -2.638 | -2.528 | -0.009 | -0.015 | -0.015 | 0 | 0.67 | 0.89 | 0.57 | 0.28 | 0.69 | 0.57 |
| 3 | 55551064 | rs570374 | -0.604 | -1.083 | -2.818 | -2.67 | -0.024 | 0.308 | 0.308 | 0.632 | 0.5 | 0.9 | 0.4 | 0.13 | 0.43 | 0.48 |
| 3 | 55552815 | rs511843 | 0.652 | -1.377 | -2.071 | -2.773 | -0.004 | -0.049 | -0.049 | 0 | 0.31 | 0.57 | 0.27 | 0.08 | 0.25 | 0.35 |
| 3 | 55552848 | rs1392220 | 0.219 | -0.944 | -1.543 | -2.291 | -0.001 | 0.004 | 0.004 | 0 | 0.11 | 0.09 | 0.11 | 0.05 | 0.14 | 0.1 |

| | | | | | | | | | | | | | | | | |
|---|---|---|---|---|---|---|---|---|---|---|---|---|---|---|---|---|
| 3 | 55556129 | rs528574 | -0.794 | -1.143 | -1.71 | -1.882 | -0.07 | -0.085 | -0.085 | 0 | 0.69 | 0.91 | 0.61 | 0.35 | 0.62 | 0.61 |
| 3 | 55570952 | rs6767040 | 0.437 | -1.648 | -0.424 | -1.545 | 0.047 | 0.054 | 0.054 | 0.017 | 0.05 | 0.28 | 0.03 | 0.06 | 0.03 | 0.12 |
| 3 | 55571760 | rs1795648 | -0.425 | -1.897 | -0.783 | -1.74 | -0.001 | -0.013 | -0.013 | 0.212 | 0.72 | 0.41 | 0.63 | 0.86 | 0.75 | 0.7 |
| 3 | 55574938 | rs1167241 | 1.519 | -1.765 | -1.152 | -1.751 | -0.118 | -0.047 | -0.047 | 0.1 | 0.24 | 0.05 | 0.49 | 0.74 | 0.43 | 0.37 |
| 3 | 55575785 | rs1167242 | -0.422 | -1.939 | -0.812 | -1.825 | 0.016 | -0.015 | -0.015 | 0.12 | 0.7 | 0.55 | 0.62 | 0.8 | 0.7 | 0.57 |
| 3 | 55577935 | rs1167244 | 1.065 | -0.981 | -1.499 | -2.116 | -0.116 | -0.046 | -0.046 | 0.011 | 0.25 | 0.06 | 0.49 | 0.74 | 0.42 | 0.37 |
| 3 | 55586557 | rs11720559 | -0.406 | -1.924 | -0.86 | -1.791 | 0.008 | 0.073 | 0.073 | 2.318 | 0.33 | 0.06 | 0.14 | 0.13 | 0.28 | 0.24 |
| 3 | 55586738 | rs2047566 | -0.95 | -1.966 | -0.458 | -1.795 | 0.012 | 0.015 | 0.015 | 0.036 | 0.43 | 0.34 | 0.22 | 0.19 | 0.33 | 0.46 |
| 3 | 55591995 | rs11716980 | -1.128 | -1.649 | -0.704 | -1.647 | 0.002 | -0.003 | -0.003 | 0 | 0.51 | 0.52 | 0.29 | 0.2 | 0.41 | 0.5 |
| 3 | 55596916 | rs2173623 | -1.409 | -1.617 | -0.694 | -1.613 | -0.003 | -0.013 | -0.013 | 0.457 | 0.54 | 0.53 | 0.3 | 0.2 | 0.42 | 0.54 |
| 3 | 55602828 | rs11711663 | -1.206 | -1.458 | -0.646 | -1.062 | 0.025 | 0.032 | 0.032 | 0 | 0.45 | 0.39 | 0.25 | 0.2 | 0.35 | 0.53 |
| 3 | 55607737 | rs6772485 | -0.72 | -0.584 | -0.859 | -0.269 | 0.012 | -0.007 | -0.007 | 0 | 0.21 | 0.06 | 0.1 | 0.13 | 0.18 | 0.14 |
| 3 | 55610406 | rs11709710 | -0.483 | -0.617 | -0.721 | -0.218 | 0.024 | 0.007 | 0.007 | 0.009 | 0.19 | 0.07 | 0.09 | 0.11 | 0.16 | 0.12 |
| 3 | 55615260 | rs6794010 | -0.737 | -0.625 | -0.455 | -0.267 | -0.027 | -0.026 | -0.026 | 0 | 0.26 | 0.32 | 0.12 | 0.11 | 0.19 | 0.23 |
| 3 | 55626065 | rs1795656 | 0.38 | -0.576 | -0.582 | -0.825 | 0.005 | 0.024 | 0.024 | 0.05 | 0.68 | 0.79 | 0.7 | 0.83 | 0.72 | 0.67 |
| 3 | 55626915 | rs1875108 | -0.479 | -0.404 | -1.091 | -1.514 | -0.016 | 0.048 | 0.048 | 0.183 | 0.65 | 0.8 | 0.65 | 0.24 | 0.66 | 0.74 |
| 3 | 55631684 | rs4955814 | 0.511 | -0.417 | -0.577 | -1.321 | 0.008 | -0.038 | -0.038 | 0 | 0.28 | 0.39 | 0.53 | 0.15 | 0.32 | 0.46 |
| 3 | 55633742 | rs7627969 | -0.134 | -0.672 | -0.948 | -1.474 | -0.019 | 0.083 | 0.083 | 0 | 0.33 | 0.17 | 0.24 | 0.16 | 0.34 | 0.21 |
| 3 | 55637763 | rs1123080 | -0.686 | -0.518 | -0.53 | -1.156 | 0 | 0 | 0 | 0 | 0.2 | 0.24 | 0.33 | 0.2 | 0.19 | 0.31 |
| 3 | 55653812 | rs17055518 | -0.733 | -0.392 | -0.954 | -0.412 | -0.015 | -0.053 | -0.053 | 0.022 | 0.27 | 0.17 | 0.34 | 0.62 | 0.26 | 0.34 |
| 3 | 55663593 | rs358917 | -0.591 | -0.461 | -0.404 | -0.655 | -0.005 | -0.033 | -0.033 | 0 | 0.94 | 0.87 | 0.97 | 1 | 0.94 | 0.9 |
| 3 | 55665383 | rs2067745 | 0.366 | -0.259 | -0.402 | -0.175 | -0.038 | -0.067 | -0.067 | 0.667 | 0.16 | 0.05 | 0.19 | 0.41 | 0.19 | 0.22 |
| 3 | 55668470 | rs815460 | 1.542 | -0.743 | 0.257 | -0.356 | 0.006 | -0.015 | -0.015 | 0 | 0.6 | 0.33 | 0.61 | 0.4 | 0.65 | 0.52 |
| 3 | 55669764 | rs815457 | -0.874 | -0.694 | 0.26 | -0.583 | -0.004 | -0.03 | -0.03 | 0 | 0.94 | 0.89 | 0.97 | 1 | 0.94 | 0.89 |
| 3 | 55671759 | rs815466 | 1.035 | -0.769 | 0.764 | -0.603 | -0.041 | -0.018 | -0.018 | 0 | 0.78 | 0.53 | 0.82 | 0.82 | 0.84 | 0.76 |
| 3 | 55682726 | rs709336 | -0.778 | -0.147 | 0.086 | -0.714 | -0.001 | -0.012 | -0.012 | 0 | 0.94 | 0.93 | 0.97 | 1 | 0.94 | 0.89 |
| 3 | 55687722 | rs1126230 | 1.266 | 0.101 | -0.185 | -0.811 | -0.019 | 0.065 | 0.065 | 0 | 0.54 | 0.28 | 0.59 | 0.56 | 0.54 | 0.55 |
| 3 | 55696202 | rs2702139 | 0.086 | -0.329 | 0.151 | -0.773 | -0.001 | -0.001 | -0.001 | 0 | 0.76 | 0.72 | 0.83 | 0.64 | 0.76 | 0.67 |
| 3 | 55697980 | rs815471 | -0.335 | -0.542 | 0.384 | -0.897 | -0.051 | -0.098 | -0.098 | 0 | 0.21 | 0.47 | 0.18 | 0.07 | 0.11 | 0.09 |
| 3 | 55704079 | rs1495357 | 1.133 | -0.489 | 0.229 | -0.859 | -0.014 | -0.031 | -0.031 | 0 | 0.58 | 0.56 | 0.72 | 0.65 | 0.67 | 0.61 |
| 3 | 55711370 | rs815415 | -0.243 | -0.386 | -0.2 | -1.164 | -0.018 | -0.085 | -0.085 | 0 | 0.36 | 0.5 | 0.28 | 0.13 | 0.3 | 0.23 |
| 3 | 55716651 | rs863631 | 0.04 | -0.233 | -1.045 | -1.132 | 0.009 | 0.06 | 0.06 | 2.199 | 0.33 | 0.19 | 0.25 | 0.12 | 0.3 | 0.22 |
| 3 | 55717050 | rs815456 | -0.924 | -0.1 | -0.851 | -0.442 | -0.003 | -0.019 | -0.019 | 4.512 | 0.93 | 0.93 | 0.97 | 1 | 0.94 | 0.9 |
| 3 | 55719552 | rs1994978 | -0.751 | -0.097 | -0.864 | -0.433 | -0.011 | -0.029 | -0.029 | 1.762 | 0.92 | 0.89 | 0.96 | 1 | 0.94 | 0.91 |
| 3 | 55724442 | rs6805592 | 0.618 | 0.258 | -1.297 | -0.202 | 0.046 | 0.092 | 0.092 | 0 | 0.55 | 0.22 | 0.69 | 0.64 | 0.66 | 0.6 |
| 3 | 55732594 | rs17055764 | -0.117 | 0.689 | -0.861 | 0.13 | -0.003 | -0.018 | -0.018 | 0.874 | 0.05 | 0.06 | 0.03 | 0 | 0.04 | 0.01 |
| 3 | 55733390 | rs4955865 | -0.021 | 0.722 | -1.162 | 0.551 | 0.007 | -0.024 | -0.024 | 0 | 0.37 | 0.26 | 0.44 | 0.37 | 0.37 | 0.35 |
| 3 | 55734673 | rs1495371 | -1.226 | 0.369 | 0.312 | 0.259 | 0.021 | -0.017 | -0.017 | 0 | 0.1 | 0.51 | 0.08 | 0.05 | 0.07 | 0.03 |
| 3 | 55734681 | rs815444 | -0.119 | -0.133 | -0.499 | -0.561 | -0.022 | 0.11 | 0.11 | 0 | 0.31 | 0.13 | 0.17 | 0.05 | 0.3 | 0.18 |
| 3 | 55734822 | rs815443 | -0.361 | -0.156 | 0.122 | -0.126 | -0.007 | -0.06 | -0.06 | 0.006 | 0.29 | 0.48 | 0.19 | 0.14 | 0.24 | 0.17 |
| 3 | 55736867 | rs970930 | -0.937 | -1.195 | 0.773 | 0.372 | -0.063 | -0.028 | -0.028 | 0 | 0.27 | 0.63 | 0.41 | 0.23 | 0.17 | 0.14 |
| 3 | 55744160 | rs937311 | 1.001 | -0.379 | 0.873 | 1.046 | 0.004 | 0.011 | 0.011 | 0 | 0.6 | 0.48 | 0.41 | 0.55 | 0.54 | 0.71 |
| 3 | 55747996 | rs7610277 | 0.123 | -0.401 | 1.086 | 1.065 | -0.044 | -0.005 | -0.005 | 0 | 0.21 | 0.35 | 0.36 | 0.22 | 0.17 | 0.16 |
| 3 | 55749194 | rs815406 | 0.712 | -0.384 | 1.106 | 1.056 | 0.014 | 0.03 | 0.03 | 0 | 0.65 | 0.57 | 0.45 | 0.6 | 0.58 | 0.72 |
| 3 | 55750910 | rs815409 | 0.722 | -0.384 | 1.094 | 1.056 | 0.019 | 0.042 | 0.042 | 0 | 0.65 | 0.54 | 0.44 | 0.6 | 0.58 | 0.72 |
| 3 | 55755590 | rs1874311 | 1.207 | -0.426 | 1.217 | 1.037 | 0 | 0.001 | 0.001 | 0 | 0.6 | 0.47 | 0.42 | 0.55 | 0.54 | 0.71 |
| 3 | 55760165 | rs1495360 | -0.923 | -0.393 | 1.063 | 1.059 | -0.057 | -0.056 | -0.056 | 0.638 | 0.88 | 0.97 | 0.82 | 0.81 | 0.75 | 0.9 |
| 3 | 55762116 | rs1472309 | 0.869 | -0.428 | 1.196 | 1.058 | 0.005 | 0.013 | 0.013 | 0 | 0.64 | 0.4 | 0.43 | 0.6 | 0.58 | 0.73 |
| 3 | 55767988 | rs815453 | -0.872 | -0.396 | 1.159 | 1.073 | -0.034 | -0.033 | -0.033 | 0 | 0.88 | 0.92 | 0.81 | 0.79 | 0.75 | 0.9 |
| 3 | 55769985 | rs815448 | 0.98 | -0.412 | 1.053 | 1.054 | -0.004 | -0.011 | -0.011 | 0.716 | 0.62 | 0.34 | 0.43 | 0.54 | 0.58 | 0.72 |
| 3 | 55781842 | rs4955874 | 0.258 | -0.415 | 0.76 | 0.99 | 0.002 | 0 | 0 | 0 | 0.2 | 0.16 | 0.34 | 0.22 | 0.17 | 0.17 |
| 3 | 55789724 | rs9868399 | 0.804 | -0.411 | 0.887 | 1.099 | 0.058 | 0.087 | 0.087 | 0 | 0.78 | 0.43 | 0.54 | 0.73 | 0.71 | 0.85 |
| 3 | 55791368 | rs1495363 | 0.143 | 0.159 | 0.138 | 1.092 | -0.024 | -0.004 | -0.004 | 0 | 0.2 | 0.24 | 0.35 | 0.22 | 0.17 | 0.19 |
| 3 | 55796723 | rs9848037 | -0.31 | 0.316 | -0.636 | 0.922 | -0.081 | -0.09 | -0.09 | 0 | 0.83 | 0.96 | 0.79 | 0.72 | 0.74 | 0.8 |
| 3 | 55801004 | rs9311578 | -0.407 | 0.023 | -0.29 | 0.337 | -0.001 | 0.004 | 0.004 | 0 | 0.49 | 0.45 | 0.68 | 0.77 | 0.51 | 0.51 |
| 3 | 55802668 | rs1495365 | -0.822 | -0.628 | -0.036 | 0.367 | -0.042 | 0.006 | 0.006 | 0.022 | 0.26 | 0.49 | 0.33 | 0.3 | 0.15 | 0.33 |
| 3 | 55803332 | rs9852746 | -0.986 | -0.769 | -0.148 | 0.118 | -0.028 | -0.02 | -0.02 | 0 | 0.31 | 0.5 | 0.29 | 0.27 | 0.2 | 0.27 |
| 3 | 55807071 | rs2362852 | 1.403 | -0.94 | -0.9 | -0.51 | 0.014 | 0.084 | 0.084 | 0.004 | 0.47 | 0.09 | 0.5 | 0.63 | 0.62 | 0.45 |
| 3 | 55808202 | rs1495368 | -0.307 | -1.062 | -0.038 | -0.319 | -0.001 | -0.001 | -0.001 | 0.227 | 0.18 | 0.16 | 0.22 | 0.23 | 0.16 | 0.09 |
| 3 | 55811863 | rs9834489 | 0.115 | -1.059 | 0.228 | -0.379 | -0.005 | -0.025 | -0.025 | 0.475 | 0.07 | 0.13 | 0.03 | 0.01 | 0.04 | 0.06 |
| 3 | 55816899 | rs1495370 | -0.227 | -1.246 | 0.549 | -0.452 | -0.038 | -0.047 | -0.047 | 0 | 0.12 | 0.22 | 0.06 | 0.03 | 0.04 | 0.2 |
| 3 | 55821111 | rs7641474 | -1.043 | -1.323 | 0.68 | -0.439 | -0.005 | -0.003 | -0.003 | 0 | 0.24 | 0.33 | 0.3 | 0.25 | 0.19 | 0.12 |
| 3 | 55824412 | rs4572738 | -1.934 | -1.534 | 0.619 | -0.46 | 0.001 | 0.001 | 0.001 | 0 | 0.39 | 0.61 | 0.35 | 0.3 | 0.24 | 0.33 |
| 3 | 55826780 | rs4299453 | -0.056 | -0.727 | 0.582 | -0.479 | -0.016 | -0.012 | -0.012 | 2.058 | 0.17 | 0.13 | 0.27 | 0.23 | 0.22 | 0.07 |
| 3 | 55829678 | rs9873405 | 0.028 | -1.174 | 0.639 | -0.462 | 0.001 | -0.004 | -0.004 | 0 | 0.07 | 0.1 | 0.02 | 0.01 | 0.04 | 0.06 |
| 3 | 55834549 | rs12487362 | 0.053 | -1.197 | 0.639 | -0.471 | -0.001 | -0.011 | -0.011 | 2.751 | 0.16 | 0.12 | 0.25 | 0.23 | 0.15 | 0.07 |
| 3 | 55834871 | rs12488085 | -1.647 | -1.207 | 0.694 | -0.473 | -0.019 | -0.02 | -0.02 | 0 | 0.36 | 0.59 | 0.34 | 0.27 | 0.23 | 0.31 |
| 3 | 55850971 | rs9854247 | -1.306 | -1.157 | 1.03 | -0.458 | -0.037 | -0.023 | -0.023 | 0.94 | 0.29 | 0.5 | 0.31 | 0.27 | 0.19 | 0.26 |
| 3 | 55855820 | rs2315942 | -1.939 | -1.23 | 1.016 | -0.321 | 0 | 0 | 0 | 0 | 0.39 | 0.68 | 0.38 | 0.34 | 0.24 | 0.4 |

| | | | | | | | | | | | | | | | | |
|---|---|---|---|---|---|---|---|---|---|---|---|---|---|---|---|---|
| 3 | 55860526 | rs9874792 | -1.246 | -1.224 | 1.031 | -0.319 | -0.029 | -0.019 | -0.019 | 0 | 0.29 | 0.46 | 0.31 | 0.26 | 0.19 | 0.26 |
| 3 | 55874765 | rs9850057 | -0.765 | -1.177 | 0.924 | -0.301 | 0.009 | 0.002 | 0.002 | 0 | 0.09 | 0.08 | 0.06 | 0.08 | 0.05 | 0.15 |
| 3 | 55881520 | rs6782906 | -1.529 | -1.231 | 1.062 | -0.327 | -0.045 | -0.045 | -0.045 | 0.31 | 0.21 | 0.41 | 0.12 | 0.11 | 0.09 | 0.33 |
| 3 | 55883396 | rs7631846 | -0.525 | -1.11 | 1.126 | -0.227 | -0.025 | -0.04 | -0.04 | 0.552 | 0.11 | 0.31 | 0.05 | 0.03 | 0.04 | 0.16 |
| 3 | 55894951 | rs6766607 | -0.348 | -1.638 | 0.948 | -0.537 | 0.005 | -0.001 | -0.001 | 0 | 0.16 | 0.13 | 0.07 | 0.03 | 0.07 | 0.19 |
| 3 | 55895409 | rs6769408 | -0.529 | -1.654 | 1.023 | -0.545 | -0.013 | -0.028 | -0.028 | 0.151 | 0.17 | 0.2 | 0.08 | 0.03 | 0.07 | 0.19 |
| 3 | 55902452 | rs1993539 | -0.257 | -2.004 | 1.441 | -0.555 | -0.011 | 0.004 | 0.004 | 0.026 | 0.2 | 0.3 | 0.16 | 0.26 | 0.15 | 0.13 |
| 3 | 55911137 | rs7625888 | -1.716 | -1.836 | 1.317 | -0.477 | 0.067 | 0.024 | 0.024 | 0.255 | 0.4 | 0.8 | 0.4 | 0.35 | 0.23 | 0.37 |
| 3 | 55912786 | rs17056081 | -0.252 | -1.76 | 1.159 | -0.483 | 0.004 | -0.004 | -0.004 | 0 | 0.17 | 0.15 | 0.07 | 0.03 | 0.07 | 0.18 |
| 3 | 55918352 | rs6763408 | -0.333 | -1.756 | 1.149 | -0.481 | -0.025 | -0.045 | -0.045 | 0 | 0.17 | 0.24 | 0.08 | 0.03 | 0.07 | 0.18 |
| 3 | 55924967 | rs922002 | -1.382 | -1.932 | 0.972 | -0.124 | -0.026 | 0.004 | 0.004 | 0 | 0.38 | 0.57 | 0.38 | 0.41 | 0.23 | 0.37 |
| 3 | 55927979 | rs9881523 | -0.388 | -1.779 | 0.243 | -0.078 | 0 | -0.002 | -0.002 | 0 | 0.18 | 0.22 | 0.3 | 0.31 | 0.14 | 0.15 |
| 3 | 55929044 | rs9811620 | -0.892 | -1.639 | -0.214 | -0.037 | 0.003 | 0 | 0 | 0 | 0.36 | 0.67 | 0.39 | 0.36 | 0.22 | 0.34 |
| 3 | 55929848 | rs1156513 | 0.935 | -0.696 | -0.981 | 0.314 | -0.047 | -0.066 | -0.066 | 0 | 0.15 | 0.01 | 0.2 | 0.21 | 0.25 | 0.19 |
| 3 | 55934301 | rs13322687 | -0.571 | -0.886 | -0.676 | 0.424 | 0.002 | -0.002 | -0.002 | 0 | 0.35 | 0.67 | 0.4 | 0.43 | 0.24 | 0.35 |
| 3 | 55934939 | rs11708779 | 0.338 | -0.317 | -1.848 | 0.118 | -0.131 | -0.05 | -0.05 | 0.343 | 0.29 | 0.03 | 0.22 | 0.22 | 0.41 | 0.35 |
| 3 | 55935878 | rs6774873 | -0.17 | -0.874 | -1.323 | -0.064 | 0.007 | -0.021 | -0.021 | 0.573 | 0.8 | 0.88 | 0.91 | 0.99 | 0.88 | 0.79 |
| 3 | 55936417 | rs11926232 | 0.335 | -0.813 | -0.966 | 0.488 | 0.007 | 0.069 | 0.069 | 0 | 0.05 | 0.13 | 0.21 | 0.23 | 0.05 | 0.03 |
| 3 | 55943591 | rs13072547 | 0.701 | -1.155 | -0.828 | 0.296 | 0.009 | 0.002 | 0.002 | 1.794 | 0.55 | 0.4 | 0.57 | 0.45 | 0.66 | 0.57 |
| 3 | 55945666 | rs1110225 | 0.203 | -1.254 | -0.696 | 0.353 | -0.002 | 0 | 0 | 0 | 0.65 | 0.68 | 0.8 | 0.69 | 0.71 | 0.63 |
| 3 | 55952782 | rs11917617 | 0.144 | -1.252 | -0.703 | 0.341 | -0.003 | 0 | 0 | 0.973 | 0.66 | 0.68 | 0.8 | 0.69 | 0.71 | 0.61 |
| 3 | 55958416 | rs13076088 | 0.566 | -1.281 | -0.395 | 0.301 | 0.002 | -0.004 | -0.004 | 0 | 0.6 | 0.43 | 0.58 | 0.46 | 0.66 | 0.57 |
| 3 | 55969858 | rs4974153 | 1.486 | -1.14 | -0.108 | 0.502 | 0.008 | -0.006 | -0.006 | 0 | 0.2 | 0.2 | 0.26 | 0.2 | 0.27 | 0.19 |
| 3 | 55976133 | rs12635403 | 1.741 | -1.348 | -0.471 | 0.392 | -0.036 | 0.003 | 0.003 | 0 | 0.34 | 0.09 | 0.29 | 0.24 | 0.38 | 0.34 |
| 3 | 55979964 | rs11715867 | -0.211 | -1.463 | -0.24 | 0.431 | 0.004 | 0.009 | 0.009 | 0 | 0.72 | 0.7 | 0.82 | 0.7 | 0.73 | 0.61 |
| 3 | 55987823 | rs7641315 | -0.574 | -1.399 | -0.059 | 0.418 | -0.007 | 0.049 | 0.049 | 0 | 0.09 | 0.2 | 0.22 | 0.24 | 0.06 | 0.05 |
| 3 | 55990726 | rs9812727 | -1.075 | -1.406 | 0.017 | 0.334 | -0.003 | -0.043 | -0.043 | 0.003 | 0.86 | 0.86 | 0.92 | 1 | 0.9 | 0.8 |
| 3 | 55993656 | rs12495172 | 0.158 | -1.35 | -0.101 | 0.416 | 0.003 | 0.014 | 0.014 | 0 | 0.33 | 0.46 | 0.49 | 0.45 | 0.34 | 0.26 |
| 3 | 56012978 | rs4974155 | 0.918 | -1.155 | -0.44 | 0.45 | -0.023 | -0.042 | -0.042 | 0.112 | 0.42 | 0.2 | 0.54 | 0.48 | 0.47 | 0.37 |
| 3 | 56015670 | rs4974157 | -0.428 | -1.019 | -0.201 | 0.59 | 0.01 | -0.008 | -0.008 | 0 | 0.45 | 0.65 | 0.59 | 0.57 | 0.38 | 0.53 |
| 3 | 56015925 | rs1458063 | 1.011 | -1.244 | 0.139 | 0.285 | -0.02 | -0.008 | -0.008 | 0 | 0.67 | 0.41 | 0.65 | 0.7 | 0.82 | 0.7 |
| 3 | 56016020 | rs1458062 | 1.054 | -1.186 | 0.066 | 0.41 | 0.02 | 0.022 | 0.022 | 0 | 0.5 | 0.23 | 0.41 | 0.49 | 0.58 | 0.47 |
| 3 | 56017598 | rs9878600 | 1.721 | -1.227 | 0.107 | 0.561 | -0.011 | 0.112 | 0.112 | 0 | 0.38 | 0.16 | 0.25 | 0.16 | 0.41 | 0.29 |
| 3 | 56018046 | rs6797314 | 1.3 | -1.146 | 0.147 | 0.573 | 0.002 | 0.033 | 0.033 | 0 | 0.13 | 0.11 | 0.16 | 0.34 | 0.17 | 0.18 |
| 3 | 56020159 | rs7613818 | 0.625 | -1.218 | 0.183 | 0.613 | 0.007 | 0.076 | 0.076 | 0 | 0.53 | 0.31 | 0.44 | 0.67 | 0.58 | 0.53 |
| 3 | 56023131 | rs7650821 | -0.461 | -1.327 | 0.659 | 0.579 | 0.013 | 0.052 | 0.052 | 0 | 0.32 | 0.41 | 0.44 | 0.46 | 0.3 | 0.41 |
| 3 | 56025951 | rs4974159 | 0.282 | -1.345 | 0.681 | 0.544 | -0.007 | -0.043 | -0.043 | 1.065 | 0.18 | 0.28 | 0.27 | 0.11 | 0.13 | 0.24 |
| 3 | 56028793 | rs9809605 | 0.91 | -1.387 | 0.648 | 0.552 | 0.037 | 0.028 | 0.028 | 0 | 0.48 | 0.34 | 0.31 | 0.36 | 0.42 | 0.37 |
| 3 | 56035242 | rs1007155 | 0.293 | -1.388 | 0.679 | 0.547 | -0.006 | -0.032 | -0.032 | 0.016 | 0.19 | 0.22 | 0.28 | 0.11 | 0.13 | 0.24 |
| 3 | 56035991 | rs12631548 | 0.902 | -1.473 | 0.754 | 0.652 | 0.015 | 0.078 | 0.078 | 0.003 | 0.14 | 0.19 | 0.16 | 0.34 | 0.17 | 0.17 |
| 3 | 56042228 | rs9851203 | 1.426 | -1.397 | 0.083 | 0.484 | 0.01 | 0.018 | 0.018 | 0 | 0.41 | 0.2 | 0.28 | 0.33 | 0.42 | 0.35 |
| 3 | 56051038 | rs9850937 | -1.132 | -1.335 | 0.126 | 0.035 | 0.009 | 0.014 | 0.014 | 0.032 | 0.4 | 0.59 | 0.37 | 0.29 | 0.29 | 0.4 |
| 3 | 56051779 | rs1023880 | 1.693 | -1.177 | -0.265 | 0.124 | -0.005 | -0.046 | -0.046 | 0.229 | 0.36 | 0.14 | 0.32 | 0.51 | 0.39 | 0.39 |
| 3 | 56052060 | rs17825124 | 1.459 | -1.202 | -0.147 | 0.086 | 0.009 | -0.035 | -0.035 | 0 | 0.12 | 0.1 | 0.08 | 0 | 0.17 | 0.07 |
| 3 | 56052839 | rs3816575 | 1.532 | -1.154 | -0.344 | 0.244 | -0.004 | -0.045 | -0.045 | 0 | 0.36 | 0.13 | 0.32 | 0.51 | 0.39 | 0.39 |
| 3 | 56054077 | rs13061634 | 0.755 | -0.96 | 0.154 | 0.212 | 0.005 | -0.003 | -0.003 | 0.017 | 0.22 | 0.25 | 0.29 | 0.21 | 0.29 | 0.22 |
| 3 | 56054925 | rs6805331 | 0.524 | -1.139 | 0.488 | 0.186 | 0.009 | 0.008 | 0.008 | 0 | 0.16 | 0.21 | 0.11 | 0.16 | 0.17 | 0.16 |
| 3 | 56058004 | rs2222248 | 1.722 | -1.29 | 0.33 | 0.103 | 0.009 | -0.017 | -0.017 | 0.892 | 0.37 | 0.18 | 0.33 | 0.53 | 0.4 | 0.39 |
| 3 | 56058518 | rs17031174 | -0.701 | -1.318 | 0.546 | 0.087 | -0.018 | -0.039 | -0.039 | 0 | 0.34 | 0.47 | 0.34 | 0.26 | 0.27 | 0.39 |
| 3 | 56070758 | rs9862661 | 0.927 | -1.135 | 0.242 | 0.132 | -0.035 | 0.011 | 0.011 | 0 | 0.22 | 0.13 | 0.29 | 0.21 | 0.3 | 0.21 |
| 3 | 56098410 | rs9819904 | 1.511 | -1.089 | 0.332 | 0.124 | 0.012 | 0.004 | 0.004 | 0 | 0.4 | 0.26 | 0.36 | 0.54 | 0.42 | 0.41 |
| 3 | 56099091 | rs1401404 | 0.888 | -1.118 | 0.446 | 0.035 | -0.001 | -0.008 | -0.008 | 0 | 0.58 | 0.4 | 0.62 | 0.75 | 0.7 | 0.61 |
| 3 | 56104964 | rs9818750 | 1.838 | -1.088 | 0.176 | 0.089 | 0.01 | -0.032 | -0.032 | 0 | 0.37 | 0.15 | 0.32 | 0.53 | 0.41 | 0.4 |
| 3 | 56105349 | rs1485673 | 1.154 | -0.965 | 0.1 | 0.096 | -0.081 | -0.001 | -0.001 | 0 | 0.19 | 0.06 | 0.28 | 0.2 | 0.3 | 0.21 |
| 3 | 56114348 | rs17056416 | 0.838 | -1.086 | 0.437 | -0.136 | -0.008 | -0.021 | -0.021 | 0 | 0.15 | 0.17 | 0.23 | 0.08 | 0.1 | 0.24 |
| 3 | 56119289 | rs6779576 | -1.041 | -1.161 | 0.646 | -0.218 | -0.016 | -0.039 | -0.039 | 0 | 0.93 | 0.82 | 0.97 | 1 | 0.97 | 1 |
| 3 | 56121074 | rs9810354 | -0.138 | -1.217 | 0.736 | -0.217 | -0.001 | -0.002 | -0.002 | 0 | 0.2 | 0.22 | 0.12 | 0.17 | 0.18 | 0.16 |
| 3 | 56130392 | rs9850438 | -0.157 | -1.232 | 0.735 | -0.2 | -0.001 | -0.002 | -0.002 | 1.912 | 0.2 | 0.22 | 0.12 | 0.18 | 0.18 | 0.16 |
| 3 | 56130514 | rs17825189 | 1.07 | -1.245 | 0.737 | -0.211 | -0.004 | -0.002 | -0.002 | 0 | 0.14 | 0.11 | 0.1 | 0.14 | 0.16 | 0.14 |
| 3 | 56132088 | rs17235284 | 2.393 | -1.346 | 0.477 | 0.009 | -0.055 | 0.08 | 0.08 | 0 | 0.33 | 0.02 | 0.21 | 0.15 | 0.35 | 0.24 |
| 3 | 56136175 | rs7651042 | 0.584 | -1.423 | 0.814 | 0.24 | -0.003 | -0.005 | -0.005 | 0 | 0.16 | 0.17 | 0.25 | 0.12 | 0.15 | 0.28 |
| 3 | 56140451 | rs9855836 | -0.16 | -1.441 | 0.742 | 0.259 | -0.003 | -0.001 | -0.001 | 0 | 0.2 | 0.24 | 0.11 | 0.17 | 0.17 | 0.16 |
| 3 | 56147139 | rs12493083 | -0.025 | -1.421 | 0.891 | 0.27 | -0.025 | -0.032 | -0.032 | 0 | 0.19 | 0.38 | 0.29 | 0.13 | 0.15 | 0.28 |
| 3 | 56150778 | rs1385690 | 1.476 | -1.354 | 0.553 | 0.296 | 0.049 | 0.064 | 0.064 | 0 | 0.55 | 0.22 | 0.59 | 0.71 | 0.65 | 0.56 |
| 3 | 56151970 | rs2200432 | 1.294 | -1.297 | 0.572 | -0.083 | 0.029 | 0.109 | 0.109 | 0 | 0.57 | 0.24 | 0.61 | 0.85 | 0.64 | 0.63 |
| 3 | 56154017 | rs7629769 | 0.338 | -1.08 | 0.85 | 0.044 | 0.008 | -0.06 | -0.06 | 3.697 | 0.16 | 0.32 | 0.09 | 0.02 | 0.16 | 0.07 |
| 3 | 56178417 | rs4974169 | 1.279 | -0.487 | 0.469 | 0.338 | -0.034 | -0.017 | -0.017 | 0.691 | 0.15 | 0.05 | 0.24 | 0.21 | 0.19 | 0.14 |
| 3 | 56189053 | rs9873343 | -0.31 | -1.186 | 0.508 | 0.131 | 0.007 | 0.007 | 0.007 | 0 | 0.11 | 0.09 | 0.03 | 0.03 | 0.03 | 0.05 |
| 3 | 56191328 | rs2171073 | 0.329 | -1.293 | 0.535 | 0.042 | -0.031 | -0.05 | -0.05 | 0 | 0.82 | 0.74 | 0.93 | 0.98 | 0.93 | 0.96 |

| | | | | | | | | | | | | | | | | |
|---|---|---|---|---|---|---|---|---|---|---|---|---|---|---|---|---|
| 3 | 56193500 | rs1564907 | 0.079 | -1.252 | 0.755 | 0.186 | -0.018 | 0.02 | 0.02 | 0 | 0.85 | 0.58 | 0.76 | 0.86 | 0.92 | 0.96 |
| 3 | 56194236 | rs11917679 | -0.326 | -1.124 | 0.631 | 0.185 | -0.004 | 0.002 | 0.002 | 0 | 0.4 | 0.38 | 0.49 | 0.68 | 0.47 | 0.65 |
| 3 | 56207634 | rs6764295 | 0.121 | -1.013 | 0.744 | 0.247 | 0.021 | 0.074 | 0.074 | 0 | 0.07 | 0.19 | 0.2 | 0.14 | 0.04 | 0.05 |
| 3 | 56220993 | rs4974120 | 0.671 | -1.013 | 0.772 | 0.217 | 0.075 | 0.136 | 0.136 | 0 | 0.05 | 0.26 | 0.22 | 0.11 | 0.04 | 0.03 |
| 3 | 56223384 | rs7628255 | 1.151 | -1.214 | 0.771 | 0.11 | -0.037 | -0.027 | -0.027 | 0 | 0.75 | 0.39 | 0.73 | 0.86 | 0.9 | 0.9 |
| 3 | 56223717 | rs1466178 | 0.478 | -1.219 | 0.692 | 0.072 | -0.068 | -0.085 | -0.085 | 0 | 0.8 | 0.59 | 0.94 | 0.97 | 0.94 | 0.94 |
| 3 | 56234932 | rs991335 | 1.817 | -1.136 | 0.397 | 0.014 | -0.053 | 0.134 | 0.134 | 0 | 0.37 | 0.06 | 0.23 | 0.16 | 0.43 | 0.24 |
| 3 | 56237810 | rs17634512 | 2.049 | -1.135 | 0.375 | 0.038 | -0.072 | 0.104 | 0.104 | 0 | 0.35 | 0.04 | 0.23 | 0.16 | 0.43 | 0.22 |
| 3 | 56242724 | rs6800915 | -0.763 | -1.174 | 0.572 | 0.016 | -0.035 | -0.066 | -0.066 | 0 | 0.89 | 0.66 | 0.96 | 1 | 0.97 | 0.99 |
| 3 | 56243087 | rs13084765 | -0.249 | -1.199 | 0.6 | 0.048 | -0.022 | 0.015 | 0.015 | 0 | 0.39 | 0.32 | 0.49 | 0.71 | 0.47 | 0.64 |
| 3 | 56257202 | rs1452217 | 0.504 | -1.192 | 0.639 | 0.037 | -0.042 | -0.035 | -0.035 | 0.003 | 0.82 | 0.47 | 0.76 | 0.88 | 0.92 | 0.94 |
| 3 | 56258327 | rs1452220 | 1.629 | -1.034 | 0.408 | 0.109 | -0.065 | -0.017 | -0.017 | 1.112 | 0.16 | 0.06 | 0.27 | 0.22 | 0.27 | 0.16 |
| 3 | 56260841 | rs1452208 | 2.22 | -1.085 | 0.385 | 0.085 | -0.071 | 0.107 | 0.107 | 0 | 0.36 | 0.04 | 0.23 | 0.16 | 0.43 | 0.26 |
| 3 | 56263534 | rs4974186 | -0.972 | -0.944 | 0.569 | 0.169 | -0.002 | -0.029 | -0.029 | 0 | 0.44 | 0.5 | 0.69 | 0.82 | 0.51 | 0.67 |
| 3 | 56270338 | rs9873480 | 0.223 | -0.882 | 0.387 | 0.225 | -0.032 | 0.009 | 0.009 | 0 | 0.25 | 0.15 | 0.29 | 0.25 | 0.3 | 0.21 |
| 3 | 56271889 | rs2123184 | 0.165 | -0.771 | 0.48 | 0.422 | -0.004 | 0.017 | 0.017 | 0.224 | 0.23 | 0.26 | 0.22 | 0.5 | 0.19 | 0.53 |
| 3 | 56279415 | rs7653453 | -0.42 | -0.728 | 0.369 | 0.315 | -0.022 | 0.034 | 0.034 | 0 | 0.28 | 0.56 | 0.44 | 0.6 | 0.22 | 0.52 |
| 3 | 56284305 | rs17235466 | 1.753 | -0.731 | -0.069 | -0.049 | -0.023 | 0.097 | 0.097 | 0 | 0.36 | 0.12 | 0.24 | 0.15 | 0.45 | 0.22 |
| 3 | 56288250 | rs13081402 | 1.115 | -0.873 | 0.405 | 0.099 | -0.001 | -0.012 | -0.012 | 0 | 0.44 | 0.46 | 0.47 | 0.28 | 0.49 | 0.29 |
| 3 | 56292829 | rs13068081 | 0.448 | -0.905 | 0.366 | 0.099 | -0.01 | 0.001 | 0.001 | 0 | 0.63 | 0.57 | 0.74 | 0.45 | 0.76 | 0.44 |
| 3 | 56302401 | rs1470802 | 0.704 | -0.936 | 0.4 | 0.082 | -0.032 | -0.003 | -0.003 | 0.44 | 0.59 | 0.48 | 0.73 | 0.44 | 0.76 | 0.45 |
| 3 | 56307703 | rs6763275 | 1.77 | -0.901 | 0.269 | 0.004 | -0.008 | 0.086 | 0.086 | 0 | 0.38 | 0.2 | 0.25 | 0.14 | 0.44 | 0.22 |
| 3 | 56329264 | rs11712446 | 1.067 | -0.386 | 0.804 | 0.066 | -0.064 | 0.029 | 0.029 | 0 | 0.16 | 0.06 | 0.26 | 0.15 | 0.26 | 0.16 |
| 3 | 56343544 | rs2873303 | -1 | -0.552 | 0.801 | 0 | -0.004 | 0.002 | 0.002 | 0 | 0.31 | 0.32 | 0.32 | 0.18 | 0.31 | 0.25 |
| 3 | 56350395 | rs6780890 | -0.76 | -0.642 | 0.979 | 0.082 | -0.031 | -0.037 | -0.037 | 0 | 0.12 | 0.22 | 0.05 | 0.02 | 0.04 | 0.07 |
| 3 | 56374916 | rs2123183 | 1.345 | -0.636 | 0.856 | 0.301 | 0 | -0.017 | -0.017 | 0 | 0.61 | 0.49 | 0.64 | 0.68 | 0.67 | 0.71 |
| 3 | 56378263 | rs4974135 | -0.594 | -0.611 | 0.863 | 0.333 | -0.028 | 0.02 | 0.02 | 0 | 0.9 | 0.8 | 0.96 | 0.86 | 0.97 | 0.96 |
| 3 | 56383404 | rs6445782 | -0.098 | -0.475 | 0.637 | -0.036 | -0.012 | -0.005 | -0.005 | 0 | 0.09 | 0.12 | 0.04 | 0.02 | 0.02 | 0.07 |
| 3 | 56386290 | rs7610959 | 0.714 | -0.404 | 0.564 | -0.035 | -0.016 | 0.011 | 0.011 | 0 | 0.18 | 0.11 | 0.28 | 0.16 | 0.27 | 0.17 |
| 3 | 56398307 | rs4974136 | -0.803 | -0.512 | 0.557 | -0.035 | 0.014 | -0.03 | -0.03 | 0.063 | 0.29 | 0.35 | 0.32 | 0.17 | 0.31 | 0.24 |
| 3 | 56409134 | rs12494857 | 0.506 | -0.383 | 0.565 | 0.083 | -0.003 | -0.003 | -0.003 | 0 | 0.06 | 0.04 | 0.2 | 0.15 | 0.05 | 0.04 |
| 3 | 56411332 | rs11923975 | 0.827 | -0.421 | 0.525 | -0.005 | -0.002 | 0.002 | 0.002 | 0 | 0.17 | 0.16 | 0.27 | 0.15 | 0.28 | 0.17 |
| 3 | 56430513 | rs969242 | -0.024 | -0.632 | 0.183 | -0.017 | -0.03 | -0.082 | -0.082 | 0.037 | 0.87 | 0.72 | 0.95 | 1 | 0.95 | 0.99 |
| 3 | 56431988 | rs4974139 | 0.776 | -0.098 | -0.769 | 0.024 | -0.048 | 0.027 | 0.027 | 0 | 0.17 | 0.08 | 0.28 | 0.15 | 0.28 | 0.15 |
| 3 | 56432160 | rs10212434 | 1.171 | -0.356 | -0.896 | 0.146 | -0.029 | 0.045 | 0.045 | 0 | 0.46 | 0.18 | 0.39 | 0.63 | 0.53 | 0.63 |
| 3 | 56434215 | rs2130881 | 0.768 | -0.725 | -0.405 | 0.047 | -0.019 | -0.026 | -0.026 | 2.489 | 0.87 | 0.84 | 0.96 | 0.97 | 0.96 | 0.88 |
| 3 | 56434744 | rs6776772 | -0.858 | -0.37 | -0.309 | 0.011 | 0.004 | -0.004 | -0.004 | 0 | 0.37 | 0.38 | 0.57 | 0.34 | 0.5 | 0.43 |
| 3 | 56435395 | rs1546060 | -1.108 | -0.645 | -0.144 | -0.22 | 0 | -0.001 | -0.001 | 0.127 | 0.37 | 0.62 | 0.56 | 0.29 | 0.36 | 0.38 |
| 3 | 56437072 | rs9824429 | -1.517 | -1.158 | -0.251 | 0.459 | -0.002 | 0.026 | 0.026 | 0 | 0.46 | 0.67 | 0.66 | 0.68 | 0.41 | 0.55 |
| 3 | 56440966 | rs11130519 | 0.048 | -1.063 | -0.068 | 0.408 | 0.04 | -0.003 | -0.003 | 0 | 0.26 | 0.35 | 0.51 | 0.27 | 0.34 | 0.3 |
| 3 | 56441795 | rs9880091 | -0.577 | -1.043 | 0.029 | 0.446 | 0.047 | -0.019 | -0.019 | 0 | 0.29 | 0.46 | 0.52 | 0.27 | 0.33 | 0.31 |
| 3 | 56450723 | rs7625995 | 1.7 | -1.236 | -0.287 | 0.022 | -0.001 | 0.001 | 0.001 | 0.647 | 0.38 | 0.36 | 0.24 | 0.24 | 0.43 | 0.18 |
| 3 | 56453197 | rs9815821 | 0.908 | -1.145 | -0.248 | 0.105 | -0.031 | -0.009 | -0.009 | 0 | 0.06 | 0.15 | 0.03 | 0.02 | 0.01 | 0.07 |
| 3 | 56464460 | rs4974213 | 1.336 | -0.862 | -0.253 | 0.313 | 0.053 | 0.054 | 0.054 | 0 | 0.15 | 0.24 | 0.46 | 0.34 | 0.26 | 0.29 |
| 3 | 56467691 | rs11719646 | 1.866 | -0.906 | -0.133 | 0.503 | 0.001 | -0.007 | -0.007 | 0.002 | 0.35 | 0.38 | 0.25 | 0.24 | 0.41 | 0.13 |
| 3 | 56475423 | rs7646515 | -0.578 | -1.032 | -0.347 | 0.415 | -0.03 | 0.042 | 0.042 | 0 | 0.86 | 0.93 | 0.9 | 0.94 | 0.8 | 0.9 |
| 3 | 56477496 | rs1878267 | 0.882 | -1.147 | -0.344 | 0.421 | -0.005 | -0.004 | -0.004 | 0.057 | 0.3 | 0.25 | 0.52 | 0.38 | 0.33 | 0.45 |
| 3 | 56483680 | rs4974142 | 1.089 | -1.193 | -0.347 | 0.44 | 0.025 | -0.035 | -0.035 | 0 | 0.1 | 0.05 | 0.11 | 0.3 | 0.04 | 0.18 |
| 3 | 56484132 | rs7645578 | 0.36 | -1.108 | -0.317 | 0.378 | 0.001 | 0.015 | 0.015 | 0.892 | 0.56 | 0.67 | 0.38 | 0.56 | 0.48 | 0.44 |
| 3 | 56493678 | rs9819137 | -0.278 | -1.069 | -0.306 | 0.39 | 0.002 | -0.004 | -0.004 | 0.046 | 0.4 | 0.45 | 0.55 | 0.4 | 0.37 | 0.57 |
| 3 | 56497897 | rs9812701 | -0.334 | -1.061 | -0.416 | 0.639 | -0.028 | 0.025 | 0.025 | 0 | 0.86 | 0.92 | 0.9 | 0.93 | 0.8 | 0.9 |
| 3 | 56511089 | rs9853704 | 0.21 | -1.086 | -0.287 | 0.731 | 0.009 | 0.017 | 0.017 | 0.851 | 0.34 | 0.36 | 0.52 | 0.46 | 0.36 | 0.5 |
| 3 | 56511815 | rs6800117 | 1.812 | -1.157 | -0.213 | 0.493 | 0 | 0.02 | 0.02 | 0.547 | 0.36 | 0.31 | 0.24 | 0.14 | 0.39 | 0.13 |
| 3 | 56518184 | rs9882932 | -0.124 | -0.344 | -0.365 | 0.584 | -0.005 | 0.002 | 0.002 | 0 | 0.87 | 0.88 | 0.9 | 0.94 | 0.8 | 0.91 |
| 3 | 56520399 | rs7622465 | -0.292 | -0.567 | -0.492 | 0.968 | -0.003 | -0.001 | -0.001 | 1.044 | 0.32 | 0.28 | 0.53 | 0.38 | 0.36 | 0.46 |
| 3 | 56535237 | rs17216615 | -0.272 | -1.226 | -0.673 | 1.2 | -0.007 | 0.014 | 0.014 | 0 | 0.06 | 0.13 | 0.04 | 0.09 | 0.04 | 0.04 |
| 3 | 56541253 | rs1602456 | -0.665 | -1.464 | -0.842 | 1.64 | 0.003 | 0.038 | 0.038 | 0 | 0.08 | 0.07 | 0.11 | 0.35 | 0.03 | 0.21 |
| 3 | 56556790 | rs1491161 | -1.234 | -1.686 | -0.784 | 1.519 | -0.03 | 0.032 | 0.032 | 0 | 0.17 | 0.24 | 0.14 | 0.34 | 0.07 | 0.23 |
| 3 | 56590069 | rs9834064 | 1.338 | -2.006 | -0.9 | 1.27 | -0.018 | 0.013 | 0.013 | 0 | 0.7 | 0.61 | 0.79 | 0.54 | 0.85 | 0.53 |
| 3 | 56593680 | rs11130525 | 0.728 | -1.922 | -0.902 | 1.246 | 0.027 | -0.032 | -0.032 | 0 | 0.77 | 0.82 | 0.83 | 0.62 | 0.89 | 0.62 |
| 3 | 56594121 | rs17235684 | 0.809 | -1.935 | -0.915 | 1.241 | 0.008 | 0.031 | 0.031 | 0.123 | 0.11 | 0 | 0.03 | 0.04 | 0.05 | 0.15 |
| 3 | 56597954 | rs6769055 | 1.347 | -1.899 | -0.896 | 1.251 | -0.019 | 0.014 | 0.014 | 0 | 0.7 | 0.6 | 0.79 | 0.54 | 0.85 | 0.53 |
| 3 | 56608033 | rs13089169 | -0.357 | -2.09 | -1.038 | 1.155 | 0.002 | 0.038 | 0.038 | 0 | 0.11 | 0.02 | 0.05 | 0.04 | 0.1 | 0.11 |
| 3 | 56627048 | rs1388256 | 0.914 | -2.064 | -0.891 | 1.305 | -0.004 | -0.002 | -0.002 | 0 | 0.75 | 0.74 | 0.83 | 0.63 | 0.9 | 0.62 |
| 3 | 56647799 | rs17216685 | -1.956 | -2.107 | -0.727 | 1.277 | -0.012 | 0 | 0 | 0 | 0.36 | 0.37 | 0.2 | 0.41 | 0.16 | 0.46 |
| 3 | 56654322 | rs6556 | -1.714 | -2.106 | -0.737 | 1.281 | -0.054 | 0.026 | 0.026 | 0 | 0.29 | 0.41 | 0.18 | 0.37 | 0.12 | 0.42 |
| 3 | 56654703 | rs17288852 | 0.422 | -2.109 | -0.758 | 1.288 | 0.024 | -0.046 | -0.046 | 0.206 | 0.09 | 0 | 0.12 | 0.34 | 0.05 | 0.25 |
| 3 | 56656047 | rs3172355 | -1.228 | -1.94 | -0.66 | 1.247 | -0.036 | 0.016 | 0.016 | 0 | 0.29 | 0.35 | 0.18 | 0.38 | 0.12 | 0.39 |
| 3 | 56657197 | rs3732506 | 0.328 | -1.948 | -0.665 | 1.242 | 0.011 | 0.017 | 0.017 | 0 | 0.07 | 0.02 | 0.02 | 0.03 | 0.04 | 0.07 |

| | | | | | | | | | | | | | | | | |
|---|---|---|---|---|---|---|---|---|---|---|---|---|---|---|---|---|
| 3 | 56658871 | rs2291498 | -0.778 | -1.734 | -0.899 | 1.328 | -0.021 | 0.011 | 0.011 | 0 | 0.26 | 0.33 | 0.17 | 0.38 | 0.11 | 0.44 |
| 3 | 56684765 | rs9871265 | -0.705 | -1.931 | -0.93 | 1.39 | -0.002 | -0.004 | -0.004 | 0 | 0.25 | 0.27 | 0.17 | 0.38 | 0.11 | 0.38 |
| 3 | 56698266 | rs6794450 | -0.226 | -1.87 | -0.97 | 1.418 | 0.058 | -0.109 | -0.109 | 2.329 | 0.21 | 0.04 | 0.15 | 0.38 | 0.1 | 0.38 |
| 3 | 56740101 | rs6445815 | -1.656 | -1.167 | -1.279 | 0.89 | 0.041 | 0.181 | 0.181 | 0 | 0.53 | 0.84 | 0.65 | 0.85 | 0.41 | 0.75 |
| 3 | 56757717 | rs1489948 | -0.567 | -1.125 | -1.154 | 1.577 | 0.03 | -0.008 | -0.008 | 0.012 | 0.32 | 0.25 | 0.11 | 0.34 | 0.17 | 0.43 |
| 3 | 56761650 | rs808 | 0.524 | -0.943 | -1.113 | 1.745 | -0.013 | -0.014 | -0.014 | 0 | 0.08 | 0.07 | 0.29 | 0.26 | 0.14 | 0.21 |
| 3 | 56761715 | rs6978 | -0.71 | -1.627 | -1.062 | 1.664 | 0.077 | -0.025 | -0.025 | 0 | 0.31 | 0.16 | 0.11 | 0.34 | 0.17 | 0.43 |
| 3 | 56762245 | rs3821414 | -0.811 | -1.628 | -1.061 | 1.675 | 0.032 | -0.011 | -0.011 | 0 | 0.32 | 0.25 | 0.11 | 0.35 | 0.17 | 0.43 |
| 3 | 56763228 | rs1009119 | -0.71 | -1.628 | -1.062 | 1.624 | 0.077 | -0.025 | -0.025 | 0.018 | 0.31 | 0.17 | 0.11 | 0.34 | 0.16 | 0.43 |
| 3 | 56763619 | rs6765444 | -0.763 | -1.596 | -1.105 | 1.638 | 0.032 | -0.008 | -0.008 | 0 | 0.32 | 0.25 | 0.11 | 0.34 | 0.17 | 0.43 |
| 3 | 56768374 | rs7431479 | -0.747 | -1.839 | -1.141 | 1.439 | 0.045 | -0.037 | -0.037 | 0 | 0.32 | 0.24 | 0.16 | 0.44 | 0.14 | 0.48 |
| 3 | 56796180 | rs9825368 | -1.124 | -1.266 | -0.428 | 1.792 | 0.029 | 0.222 | 0.222 | 0.547 | 0.13 | 0.5 | 0.19 | 0.38 | 0.11 | 0.14 |
| 3 | 56798129 | rs10510788 | 0.861 | -1.071 | -1.191 | 1.355 | 0.018 | 0.031 | 0.031 | 1.495 | 0.46 | 0.13 | 0.31 | 0.42 | 0.45 | 0.25 |
| 3 | 56803175 | rs9871950 | -0.203 | -0.971 | -0.958 | 1.09 | 0.114 | 0.085 | 0.085 | 4.201 | 0.09 | 0.51 | 0.17 | 0.15 | 0.11 | 0.11 |
| 3 | 56806203 | rs6794153 | -0.879 | -1.103 | -0.993 | 1.029 | 0.036 | -0.002 | -0.002 | 0 | 0.13 | 0.61 | 0.19 | 0.15 | 0.15 | 0.11 |
| 3 | 56809628 | rs7646054 | -0.278 | -1.313 | -1.478 | 1.14 | -0.03 | 0.035 | 0.035 | 0.146 | 0.66 | 0.78 | 0.53 | 0.83 | 0.62 | 0.6 |
| 3 | 56815994 | rs4681859 | -0.651 | -1.23 | -1.813 | 0.808 | -0.078 | 0.117 | 0.117 | 0.423 | 0.69 | 0.88 | 0.61 | 0.93 | 0.62 | 0.66 |
| 3 | 56822377 | rs7611544 | 0.473 | -1.469 | -1.668 | 1.198 | 0.02 | 0.128 | 0.128 | 1.434 | 0.46 | 0.13 | 0.31 | 0.32 | 0.44 | 0.24 |
| 3 | 56826324 | rs1110866 | 0.352 | -1.314 | -1.831 | 1.161 | -0.02 | -0.097 | -0.097 | 0.07 | 0.41 | 0.13 | 0.29 | 0.5 | 0.41 | 0.23 |
| 3 | 56829661 | rs12635549 | 0.602 | -1.504 | -0.567 | 1.403 | -0.04 | 0.067 | 0.067 | 0 | 0.78 | 0.5 | 0.81 | 0.64 | 0.85 | 0.82 |
| 3 | 56830087 | rs4681888 | 1.017 | -2.267 | -0.194 | 1.602 | -0.043 | 0.064 | 0.064 | 0 | 0.74 | 0.64 | 0.84 | 0.58 | 0.83 | 0.66 |
| 3 | 56830579 | rs6800401 | 1.086 | -2.189 | -0.649 | 0.893 | -0.001 | 0.021 | 0.021 | 0 | 0.22 | 0.16 | 0.14 | 0.08 | 0.2 | 0.09 |
| 3 | 56831548 | rs6803697 | -0.461 | -2.371 | -1.045 | 1.275 | 0.053 | 0.028 | 0.028 | 0 | 0.45 | 0.73 | 0.42 | 0.69 | 0.3 | 0.41 |
| 3 | 56833979 | rs1392702 | 0.631 | -1.718 | -1.441 | 1.022 | -0.068 | 0.11 | 0.11 | 0 | 0.4 | 0.09 | 0.49 | 0.19 | 0.57 | 0.48 |
| 3 | 56836740 | rs4681898 | 0.098 | -1.351 | -1.151 | 1.295 | 0.047 | 0.076 | 0.076 | 0 | 0.51 | 0.37 | 0.56 | 0.2 | 0.66 | 0.52 |
| 3 | 56843028 | rs12638861 | -0.554 | -0.446 | -1.126 | 1.95 | -0.006 | -0.033 | -0.033 | 0 | 0.07 | 0.01 | 0.06 | 0.29 | 0.05 | 0.24 |
| 3 | 56849749 | rs1354034 | 0.408 | 0.089 | -1.726 | 1.419 | 0.046 | -0.014 | -0.014 | 0.058 | 0.51 | 0.21 | 0.44 | 0.57 | 0.6 | 0.48 |
| 3 | 56851387 | rs17288922 | 0.039 | -0.148 | -1.638 | 1.363 | 0.03 | -0.01 | -0.01 | 1.576 | 0.24 | 0.14 | 0.19 | 0.24 | 0.18 | 0.33 |
| 3 | 56852389 | rs17288929 | 0.373 | -0.159 | -1.627 | 1.367 | 0.026 | -0.022 | -0.022 | 0 | 0.19 | 0.12 | 0.16 | 0.24 | 0.12 | 0.31 |
| 3 | 56857433 | rs1344142 | 1.085 | -0.252 | -0.936 | 1.165 | 0 | -0.009 | -0.009 | 0 | 0.55 | 0.48 | 0.39 | 0.32 | 0.56 | 0.41 |
| 3 | 56858494 | rs10866002 | -0.193 | -0.121 | -0.944 | 1.277 | 0.04 | 0.102 | 0.102 | 0 | 0.11 | 0.24 | 0.28 | 0.36 | 0.15 | 0.07 |
| 3 | 56858686 | rs13062174 | 1.318 | -0.131 | -0.956 | 1.176 | 0.001 | -0.009 | -0.009 | 0 | 0.55 | 0.48 | 0.39 | 0.32 | 0.56 | 0.43 |
| 3 | 56859335 | rs1500712 | 0.215 | -0.472 | -1.18 | 0.815 | 0.01 | 0.011 | 0.011 | 0 | 0.22 | 0.16 | 0.12 | 0.15 | 0.18 | 0.26 |
| 3 | 56862006 | rs7640115 | 0.191 | -0.238 | -1.213 | 0.514 | -0.002 | -0.001 | -0.001 | 2.334 | 0.88 | 0.87 | 0.86 | 0.91 | 0.89 | 0.82 |
| 3 | 56865752 | rs4681908 | 0.303 | -0.116 | -1.268 | 0.393 | -0.008 | 0.002 | 0.002 | 0.038 | 0.08 | 0.04 | 0.12 | 0.04 | 0.1 | 0.13 |
| 3 | 56866376 | rs7630830 | 0.168 | 0.123 | -1.57 | -0.171 | 0.019 | 0.014 | 0.014 | 0 | 0.53 | 0.63 | 0.38 | 0.6 | 0.42 | 0.58 |
| 3 | 56870534 | rs9881237 | -0.015 | -0.391 | -1.764 | -0.206 | -0.005 | 0 | 0 | 0 | 0.86 | 0.81 | 0.79 | 0.89 | 0.88 | 0.86 |
| 3 | 56874376 | rs11717281 | 0.926 | -0.064 | -1.94 | -0.548 | -0.019 | -0.018 | -0.018 | 1.972 | 0.08 | 0.05 | 0.2 | 0.13 | 0.11 | 0.05 |
| 3 | 56874729 | rs1039381 | 1.764 | -0.248 | -2.342 | -0.686 | -0.03 | -0.169 | -0.169 | 0 | 0.24 | 0.01 | 0.3 | 0.54 | 0.23 | 0.31 |
| 3 | 56877788 | rs10510789 | 0.717 | 0.025 | -1.942 | -0.718 | 0.014 | 0.062 | 0.062 | 0.053 | 0.2 | 0.06 | 0.09 | 0.07 | 0.15 | 0.26 |
| 3 | 56880444 | rs11130549 | 0.924 | 0.106 | -1.794 | -0.548 | -0.057 | 0.051 | 0.051 | 0 | 0.27 | 0.15 | 0.2 | 0.16 | 0.37 | 0.2 |
| 3 | 56891446 | rs6445829 | 0.099 | 0.435 | -1.137 | -0.19 | 0.01 | 0.014 | 0.014 | 0 | 0.46 | 0.62 | 0.53 | 0.64 | 0.42 | 0.53 |
| 3 | 56896385 | rs7630738 | -0.347 | 0.346 | -1.049 | -0.349 | -0.002 | 0.045 | 0.045 | 0 | 0.54 | 0.64 | 0.72 | 0.77 | 0.53 | 0.58 |
| 3 | 56902096 | rs13317313 | -0.025 | 0.391 | -0.41 | -0.142 | 0.007 | -0.025 | -0.025 | 0 | 0.11 | 0.19 | 0.13 | 0.03 | 0.11 | 0.12 |
| 3 | 56902938 | rs11714628 | -0.276 | -0.403 | -0.439 | 0.115 | -0.006 | 0.008 | 0.008 | 0 | 0.11 | 0.13 | 0.19 | 0.28 | 0.07 | 0.2 |
| 3 | 56906501 | rs12485693 | -0.139 | -0.396 | -0.431 | 0.123 | -0.011 | 0.017 | 0.017 | 0 | 0.1 | 0.13 | 0.19 | 0.28 | 0.06 | 0.19 |
| 3 | 56909193 | rs6445833 | -0.219 | -0.54 | -0.674 | -0.452 | -0.002 | 0.001 | 0.001 | 0 | 0.59 | 0.6 | 0.54 | 0.46 | 0.61 | 0.58 |
| 3 | 56914154 | rs1500711 | 0.452 | -0.247 | -1.115 | -0.772 | -0.1 | -0.124 | -0.124 | 0 | 0.37 | 0.06 | 0.4 | 0.52 | 0.43 | 0.33 |
| 3 | 56915719 | rs6445834 | 0.017 | 0.017 | -1.092 | -0.838 | 0.022 | 0.01 | 0.01 | 0 | 0.56 | 0.67 | 0.58 | 0.35 | 0.68 | 0.49 |
| 3 | 56920909 | rs950154 | 1.184 | -0.032 | -0.693 | -0.914 | -0.016 | 0.047 | 0.047 | 0 | 0.37 | 0.26 | 0.26 | 0.17 | 0.41 | 0.34 |
| 3 | 56924711 | rs11916112 | 0.155 | 0.192 | -0.731 | -0.833 | -0.028 | 0.032 | 0.032 | 0 | 0.17 | 0.06 | 0.12 | 0.11 | 0.25 | 0.19 |
| 3 | 56926166 | rs4681928 | 0.914 | 0.072 | -0.313 | -0.751 | -0.019 | 0.012 | 0.012 | 0 | 0.28 | 0.41 | 0.47 | 0.7 | 0.22 | 0.35 |
| 3 | 56927023 | rs4681929 | 1.916 | 0.086 | -1.4 | -0.667 | -0.001 | -0.159 | -0.159 | 0.011 | 0.2 | 0.01 | 0.26 | 0.52 | 0.16 | 0.31 |
| 3 | 56929066 | rs874404 | 1.49 | 0.724 | -1.212 | -0.765 | -0.093 | 0.024 | 0.024 | 0 | 0.18 | 0.03 | 0.18 | 0.11 | 0.28 | 0.1 |
| 3 | 56942002 | rs13065124 | -0.029 | 0.629 | -0.556 | -0.466 | -0.001 | 0.068 | 0.068 | 0 | 0.88 | 0.8 | 0.79 | 0.68 | 0.9 | 0.79 |
| 3 | 56942973 | rs13070800 | 0.47 | 0.631 | -0.66 | -0.828 | 0.009 | 0.034 | 0.034 | 0.382 | 0.56 | 0.26 | 0.56 | 0.66 | 0.61 | 0.59 |
| 3 | 56949816 | rs17057445 | 0.48 | 0.571 | -0.367 | -0.687 | 0.003 | -0.029 | -0.029 | 0 | 0.11 | 0.12 | 0.1 | 0 | 0.15 | 0.02 |
| 3 | 56950055 | rs17825630 | 0.259 | 0.277 | -0.297 | -0.499 | 0.003 | 0.024 | 0.024 | 0 | 0.17 | 0.09 | 0.08 | 0.06 | 0.16 | 0.25 |
| 3 | 56958358 | rs6792170 | -0.52 | -0.499 | -0.416 | -0.84 | -0.005 | -0.006 | -0.006 | 0 | 0.56 | 0.55 | 0.7 | 0.7 | 0.61 | 0.58 |
| 3 | 56974739 | rs2291977 | 1.418 | -0.237 | -0.274 | -0.483 | -0.002 | -0.031 | -0.031 | 1.672 | 0.09 | 0.04 | 0.23 | 0.26 | 0.1 | 0.07 |
| 3 | 56989917 | rs13079459 | 1.18 | -0.043 | -0.487 | -0.685 | -0.053 | 0.078 | 0.078 | 0 | 0.13 | 0.01 | 0.08 | 0 | 0.16 | 0.11 |
| 3 | 56990504 | rs11130558 | 0.682 | -0.213 | -0.609 | -0.75 | -0.113 | -0.134 | -0.134 | 0.177 | 0.22 | 0.03 | 0.31 | 0.41 | 0.34 | 0.33 |
| 3 | 56996795 | rs9311634 | 0.04 | -0.242 | -0.583 | -0.771 | -0.183 | -0.124 | -0.124 | 0 | 0.34 | 0.04 | 0.32 | 0.4 | 0.46 | 0.41 |
| 3 | 57003081 | rs7371892 | -0.695 | -0.642 | -0.141 | -0.932 | -0.035 | 0.069 | 0.069 | 0 | 0.78 | 0.52 | 0.82 | 0.71 | 0.86 | 0.68 |
| 3 | 57006140 | rs12637209 | 1.51 | -0.585 | 0.047 | -1.052 | -0.004 | 0.016 | 0.016 | 0 | 0.1 | 0.11 | 0.24 | 0.26 | 0.1 | 0.08 |
| 3 | 57009097 | rs9834815 | 0.604 | -0.605 | 0.02 | -1.107 | 0.002 | -0.011 | -0.011 | 0 | 0.47 | 0.34 | 0.36 | 0.05 | 0.46 | 0.3 |
| 3 | 57011234 | rs13080107 | 0.134 | -0.522 | -0.197 | -1.059 | 0 | -0.011 | -0.011 | 0.364 | 0.56 | 0.47 | 0.61 | 0.31 | 0.57 | 0.37 |
| 3 | 57014104 | rs4464419 | 1.208 | -0.371 | -0.616 | -1.116 | -0.063 | 0.092 | 0.092 | 0 | 0.15 | 0.01 | 0.11 | 0 | 0.2 | 0.12 |
| 3 | 57015131 | rs4075857 | 1.523 | -0.579 | -0.257 | -1.132 | 0.005 | 0.002 | 0.002 | 0.064 | 0.29 | 0.24 | 0.23 | 0.07 | 0.25 | 0.18 |

| | | | | | | | | | | | | | | | | |
|---|---|---|---|---|---|---|---|---|---|---|---|---|---|---|---|---|
| 3 | 57015470 | rs4264698 | 1.024 | -0.31 | -0.387 | -0.904 | -0.106 | -0.131 | -0.131 | 0 | 0.21 | 0.03 | 0.22 | 0.42 | 0.29 | 0.31 |
| 3 | 57018178 | rs6809587 | -0.552 | -0.703 | 0.063 | -0.715 | -0.027 | 0.054 | 0.054 | 0 | 0.78 | 0.59 | 0.82 | 0.71 | 0.85 | 0.68 |
| 3 | 57020736 | rs3924042 | -0.434 | -0.393 | -0.437 | -1.219 | -0.005 | -0.021 | -0.021 | 0 | 0.53 | 0.44 | 0.5 | 0.71 | 0.55 | 0.73 |
| 3 | 57021793 | rs13091176 | 0.576 | -0.137 | -0.483 | -0.519 | -0.066 | 0.086 | 0.086 | 0.043 | 0.14 | 0.01 | 0.12 | 0 | 0.21 | 0.12 |
| 3 | 57026693 | rs9846324 | 0.564 | 0.055 | -0.203 | -0.266 | -0.007 | -0.011 | -0.011 | 0.331 | 0.13 | 0.07 | 0.24 | 0.22 | 0.19 | 0.14 |
| 3 | 57031469 | rs6801499 | 1.028 | 0.043 | -0.166 | -0.186 | -0.078 | -0.035 | -0.035 | 0 | 0.37 | 0.09 | 0.28 | 0.39 | 0.4 | 0.35 |
| 3 | 57032690 | rs17235827 | 0.793 | 0.017 | -0.159 | -0.275 | -0.012 | 0.01 | 0.01 | 0.004 | 0.16 | 0.1 | 0.13 | 0 | 0.2 | 0.14 |
| 3 | 57039080 | rs6784399 | 0.154 | -0.097 | 0.021 | -0.278 | 0.055 | 0.042 | 0.042 | 0 | 0.53 | 0.23 | 0.41 | 0.39 | 0.6 | 0.49 |
| 3 | 57042314 | rs2054857 | -0.945 | -0.101 | 0.363 | -0.134 | 0 | -0.003 | -0.003 | 0 | 0.58 | 0.48 | 0.68 | 0.61 | 0.6 | 0.65 |
| 3 | 57053921 | rs6445844 | -0.023 | 0.182 | 0.357 | -0.467 | -0.015 | -0.022 | -0.022 | 0.001 | 0.31 | 0.56 | 0.31 | 0.26 | 0.3 | 0.2 |
| 3 | 57059370 | rs11922193 | -0.699 | 0.197 | 0.281 | -0.487 | 0.001 | 0.009 | 0.009 | 0 | 0.46 | 0.57 | 0.4 | 0.26 | 0.47 | 0.34 |
| 3 | 57060589 | rs11130564 | -0.541 | 0.414 | -0.644 | -0.392 | -0.017 | 0.075 | 0.075 | 0 | 0.42 | 0.23 | 0.37 | 0.26 | 0.47 | 0.34 |
| 3 | 57071029 | rs752466 | 0.315 | 0.487 | -0.754 | -0.345 | 0.002 | 0.009 | 0.009 | 0 | 0.27 | 0.15 | 0.27 | 0.26 | 0.3 | 0.2 |
| 3 | 57071127 | rs2873611 | -0.458 | 0.577 | -0.691 | -0.156 | 0 | 0.029 | 0.029 | 0 | 0.45 | 0.39 | 0.38 | 0.26 | 0.47 | 0.34 |
| 3 | 57073070 | rs4681944 | -0.389 | 0.55 | -0.686 | 0.147 | 0 | -0.001 | -0.001 | 0 | 0.45 | 0.49 | 0.47 | 0.6 | 0.46 | 0.48 |
| 3 | 57074119 | rs11130567 | 0.167 | 0.423 | -0.606 | -0.848 | -0.01 | -0.102 | -0.102 | 1.017 | 0.63 | 0.51 | 0.71 | 0.95 | 0.67 | 0.78 |
| 3 | 57074218 | rs9682671 | 0.551 | 0.068 | -1.251 | -1.188 | 0.031 | 0.096 | 0.096 | 0.107 | 0.37 | 0.18 | 0.22 | 0.2 | 0.31 | 0.26 |
| 3 | 57076196 | rs6445846 | -0.406 | 0.135 | -0.824 | -1.489 | -0.006 | -0.043 | -0.043 | 1.03 | 0.79 | 0.65 | 0.8 | 0.96 | 0.8 | 0.87 |
| 3 | 57081514 | rs2318008 | 1.238 | -0.026 | -0.543 | -1.396 | 0.034 | 0.025 | 0.025 | 1.545 | 0.95 | 0.6 | 0.96 | 1 | 0.99 | 1 |
| 3 | 57084818 | rs4681947 | 0.214 | 0.413 | -0.761 | -1.347 | -0.036 | -0.043 | -0.043 | 1.23 | 0.19 | 0.11 | 0.39 | 0.63 | 0.25 | 0.38 |
| 3 | 57085010 | rs7622066 | 0.622 | 0.144 | -1.289 | -1.786 | 0.005 | -0.045 | -0.045 | 0 | 0.66 | 0.49 | 0.5 | 0.18 | 0.65 | 0.56 |
| 3 | 57086123 | rs11918968 | -0.581 | 0.047 | -1.092 | -1.578 | -0.016 | 0.016 | 0.016 | 7.226 | 0.86 | 0.7 | 0.91 | 0.81 | 0.89 | 0.95 |
| 3 | 57087830 | rs11706697 | 1.635 | 0.235 | -1.277 | -1.177 | -0.024 | -0.052 | -0.052 | 0 | 0.08 | 0 | 0.06 | 0.29 | 0.09 | 0.05 |
| 3 | 57088114 | rs11710867 | 1.613 | 0.235 | -1.277 | -1.177 | -0.024 | -0.053 | -0.053 | 2.966 | 0.08 | 0.01 | 0.06 | 0.29 | 0.09 | 0.05 |
| 3 | 57091575 | rs9830791 | -0.064 | 0.074 | -0.877 | -1.657 | 0.001 | -0.002 | -0.002 | 0 | 0.43 | 0.48 | 0.6 | 0.61 | 0.47 | 0.47 |
| 3 | 57104428 | rs9871274 | -0.018 | 0.222 | -0.072 | -1.032 | 0.011 | 0.118 | 0.118 | 0.114 | 0.22 | 0.44 | 0.4 | 0.48 | 0.18 | 0.22 |
| 3 | 57104606 | rs7427326 | -0.682 | 0.131 | -0.149 | -1.427 | 0.001 | 0.008 | 0.008 | 0.084 | 0.34 | 0.69 | 0.45 | 0.83 | 0.28 | 0.28 |
| 3 | 57110913 | rs7612293 | 0.456 | -0.076 | -0.149 | -1.92 | 0.018 | 0.147 | 0.147 | 0.196 | 0.5 | 0.25 | 0.36 | 0.1 | 0.56 | 0.51 |
| 3 | 57121557 | rs1872947 | -0.571 | 0.077 | 0.034 | -1.108 | -0.006 | 0.053 | 0.053 | 4.154 | 0.72 | 0.58 | 0.79 | 0.48 | 0.75 | 0.78 |
| 3 | 57125424 | rs17289035 | 0.897 | 0.093 | -0.084 | -0.953 | 0 | 0.001 | 0.001 | 0.112 | 0.21 | 0.17 | 0.37 | 0.38 | 0.19 | 0.27 |
| 3 | 57127991 | rs2035653 | -1.678 | 0.039 | 0.046 | -0.921 | -0.005 | 0.188 | 0.188 | 3.635 | 0.87 | 0.74 | 0.89 | 0.52 | 0.9 | 0.96 |
| 3 | 57128211 | rs11920551 | 1.365 | -0.029 | -0.275 | -1.228 | -0.055 | 0.13 | 0.13 | 0 | 0.22 | 0.03 | 0.17 | 0.04 | 0.22 | 0.36 |
| 3 | 57132200 | rs2291462 | -0.922 | -0.039 | -0.173 | -0.86 | -0.004 | 0.235 | 0.235 | 1.555 | 0.83 | 0.61 | 0.85 | 0.51 | 0.86 | 0.92 |
| 3 | 57133138 | rs6809523 | 0.255 | -0.082 | -0.299 | -0.987 | -0.012 | -0.1 | -0.1 | 0 | 0.66 | 0.57 | 0.52 | 0.13 | 0.71 | 0.69 |
| 3 | 57134230 | rs6807055 | -1.783 | -0.079 | -0.224 | -0.958 | -0.007 | 0.155 | 0.155 | 3.635 | 0.87 | 0.77 | 0.9 | 0.52 | 0.9 | 0.96 |
| 3 | 57136585 | rs17057718 | 1.646 | -0.031 | -0.86 | -0.952 | -0.011 | -0.092 | -0.092 | 0 | 0.17 | 0.04 | 0.33 | 0.38 | 0.15 | 0.23 |
| 3 | 57138419 | rs6780995 | -0.011 | 0.109 | -0.793 | -1.402 | -0.011 | -0.101 | -0.101 | 0 | 0.66 | 0.57 | 0.52 | 0.14 | 0.71 | 0.69 |
| 3 | 57142094 | rs6445854 | 1.758 | 0.115 | -1.362 | -1.674 | 0 | -0.05 | -0.05 | 0 | 0.22 | 0.3 | 0.15 | 0.07 | 0.22 | 0.22 |
| 3 | 57149659 | rs12487790 | -0.49 | 0.134 | -1.222 | -1.31 | -0.015 | -0.079 | -0.079 | 0.689 | 0.68 | 0.52 | 0.56 | 0.14 | 0.75 | 0.73 |
| 3 | 57157716 | rs6810042 | 1.643 | 0.29 | -1.667 | -1.323 | -0.031 | 0.083 | 0.083 | 0.422 | 0.21 | 0.07 | 0.18 | 0.02 | 0.27 | 0.12 |
| 3 | 57158685 | rs4596115 | 0.778 | 0.29 | -1.495 | -1.575 | -0.004 | -0.004 | -0.004 | 0 | 0.45 | 0.42 | 0.34 | 0.11 | 0.46 | 0.57 |
| 3 | 57160312 | rs12495640 | -0.242 | 0.171 | -1.534 | -0.925 | 0.004 | -0.009 | -0.009 | 0 | 0.42 | 0.54 | 0.52 | 0.41 | 0.39 | 0.35 |
| 3 | 57163751 | rs6787279 | 0.156 | 0.291 | -1.676 | -0.665 | -0.019 | 0.065 | 0.065 | 0.34 | 0.24 | 0.48 | 0.4 | 0.55 | 0.18 | 0.28 |
| 3 | 57165310 | rs4681957 | 1.573 | 0.336 | -2.311 | -0.623 | -0.003 | -0.066 | -0.066 | 0 | 0.16 | 0.06 | 0.31 | 0.38 | 0.14 | 0.23 |
| 3 | 57173915 | rs7374667 | -1.864 | 0.406 | -2.581 | -0.894 | -0.051 | -0.054 | -0.054 | 0.029 | 0.73 | 0.94 | 0.82 | 0.61 | 0.69 | 0.75 |
| 3 | 57174274 | rs4681960 | 0.704 | 0.361 | -2.015 | -0.691 | -0.067 | 0.142 | 0.142 | 0 | 0.23 | 0.03 | 0.2 | 0.04 | 0.26 | 0.33 |
| 3 | 57186580 | rs17235841 | 0.871 | 0.483 | -2.003 | -0.438 | -0.053 | 0.002 | 0.002 | 0 | 0.26 | 0.01 | 0.2 | 0.19 | 0.23 | 0.29 |
| 3 | 57187078 | rs3968279 | 0.276 | 0.317 | -1.276 | -0.151 | -0.032 | 0.182 | 0.182 | 0.307 | 0.19 | 0.47 | 0.39 | 0.38 | 0.12 | 0.27 |
| 3 | 57192294 | rs11710921 | 0.493 | 0.499 | -2.153 | -0.345 | -0.074 | 0.089 | 0.089 | 0.132 | 0.36 | 0.04 | 0.26 | 0.18 | 0.37 | 0.37 |
| 3 | 57192927 | rs12496746 | 1.659 | 0.505 | -1.801 | -0.098 | 0.008 | -0.051 | -0.051 | 0 | 0.11 | 0.03 | 0.32 | 0.37 | 0.08 | 0.21 |
| 3 | 57200758 | rs4283545 | -0.99 | 0.793 | -1.485 | -0.772 | -0.001 | -0.01 | -0.01 | 0.59 | 0.6 | 0.76 | 0.72 | 0.54 | 0.62 | 0.71 |
| 3 | 57204419 | rs11130575 | 1.127 | 0.844 | -1.983 | -1.565 | -0.087 | 0.18 | 0.18 | 0 | 0.28 | 0.01 | 0.18 | 0.02 | 0.32 | 0.17 |
| 3 | 57209197 | rs6445860 | -0.866 | 1.098 | -1.89 | -1.481 | -0.111 | 0.307 | 0.307 | 1.623 | 0.56 | 0.95 | 0.45 | 0.08 | 0.53 | 0.48 |
| 3 | 57219344 | rs6445861 | -0.558 | 1.015 | -1.53 | -1.469 | 0.017 | 0.207 | 0.207 | 0.639 | 0.52 | 0.79 | 0.44 | 0.09 | 0.56 | 0.48 |
| 3 | 57226591 | rs4325895 | -0.69 | 1.041 | -1.457 | -1.458 | 0.001 | 0.252 | 0.252 | 0.386 | 0.53 | 0.84 | 0.45 | 0.09 | 0.56 | 0.46 |
| 3 | 57235585 | rs3755639 | 0.685 | 0.95 | -1.221 | -1.466 | -0.011 | 0.251 | 0.251 | 0.473 | 0.44 | 0.11 | 0.52 | 0.91 | 0.4 | 0.49 |
| 3 | 57237221 | rs11710045 | 0.644 | 0.978 | -1.262 | -1.611 | -0.064 | 0.194 | 0.194 | 0 | 0.35 | 0.05 | 0.21 | 0.08 | 0.37 | 0.31 |
| 3 | 57238604 | rs975418 | 0.671 | 1.022 | -1.097 | -1.339 | -0.011 | 0.251 | 0.251 | 0.473 | 0.44 | 0.11 | 0.52 | 0.91 | 0.4 | 0.49 |
| 3 | 57280735 | rs1553992 | 0.72 | 0.986 | -0.563 | -1.159 | -0.01 | 0.221 | 0.221 | 0.183 | 0.45 | 0.11 | 0.52 | 0.84 | 0.41 | 0.5 |
| 3 | 57306319 | rs3087684 | -0.472 | 1.153 | -0.209 | -1.49 | 0.001 | 0.094 | 0.094 | 0 | 0.61 | 0.77 | 0.75 | 0.92 | 0.61 | 0.64 |
| 3 | 57306611 | rs17791685 | 0.502 | 1.141 | -0.081 | -1.499 | 0.109 | -0.033 | -0.033 | 0 | 0.08 | 0.29 | 0.16 | 0 | 0.17 | 0.1 |
| 3 | 57306961 | rs1913302 | -0.508 | 1.12 | -0.092 | -1.507 | 0.001 | 0.092 | 0.092 | 0 | 0.61 | 0.77 | 0.75 | 0.92 | 0.61 | 0.64 |
| 3 | 57308203 | rs17735309 | 0.459 | 1.244 | -0.379 | -1.388 | 0.111 | -0.032 | -0.032 | 0 | 0.08 | 0.29 | 0.16 | 0 | 0.17 | 0.1 |
| 3 | 57315557 | rs1355797 | 1.796 | 0.724 | -1.083 | -0.8 | 0.008 | -0.06 | -0.06 | 0 | 0.12 | 0.03 | 0.24 | 0.4 | 0.09 | 0.23 |
| 3 | 57317586 | rs11130579 | 0.602 | 0.838 | -1.027 | -1.046 | -0.005 | 0.268 | 0.268 | 0.202 | 0.46 | 0.11 | 0.53 | 0.91 | 0.4 | 0.5 |
| 3 | 57347878 | rs571879 | 0.456 | 0.75 | -1.034 | -1.138 | 0.059 | 0.26 | 0.26 | 0.277 | 0.45 | 0.1 | 0.48 | 0.91 | 0.33 | 0.54 |
| 3 | 57355028 | rs7642234 | 0.105 | 0.317 | -0.551 | -1.186 | -0.009 | -0.001 | -0.001 | 0 | 0.43 | 0.55 | 0.35 | 0.4 | 0.27 | 0.43 |
| 3 | 57369292 | rs6765079 | 0.352 | 0.299 | -0.438 | -1.171 | -0.042 | 0.004 | 0.004 | 0 | 0.4 | 0.48 | 0.32 | 0.4 | 0.23 | 0.42 |
| 3 | 57388346 | rs4681977 | 0.386 | 0.284 | -0.309 | -1.186 | -0.02 | 0.001 | 0.001 | 0 | 0.39 | 0.45 | 0.34 | 0.41 | 0.23 | 0.47 |

| | | | | | | | | | | | | | | | | |
|---|---|---|---|---|---|---|---|---|---|---|---|---|---|---|---|---|
| 3 | 57393368 | rs9311647 | -0.546 | 0.991 | -0.144 | -1.304 | 0.062 | -0.079 | -0.079 | 0.309 | 0.14 | 0.68 | 0.21 | 0 | 0.2 | 0.11 |
| 3 | 57412804 | rs9829584 | 0.525 | 0.673 | -0.153 | -1.41 | -0.016 | 0.002 | 0.002 | 0 | 0.39 | 0.45 | 0.34 | 0.42 | 0.24 | 0.47 |
| 3 | 57414330 | rs11915740 | 0.855 | 0.574 | -0.45 | -1.64 | -0.078 | 0.151 | 0.151 | 0 | 0.35 | 0.1 | 0.26 | 0.09 | 0.44 | 0.33 |
| 3 | 57414434 | rs4462937 | 1.229 | 0.535 | -0.43 | -1.627 | 0.057 | -0.055 | -0.055 | 0 | 0.36 | 0.12 | 0.31 | 0.42 | 0.24 | 0.47 |
| 3 | 57416701 | rs9850744 | 0.164 | 0.578 | 0.006 | -1.236 | 0.026 | 0.114 | 0.114 | 0 | 0.78 | 0.65 | 0.64 | 0.5 | 0.72 | 0.81 |
| 3 | 57417434 | rs13084781 | 0.656 | 0.579 | -0.034 | -1.235 | 0.026 | 0.109 | 0.109 | 0 | 0.74 | 0.56 | 0.61 | 0.5 | 0.68 | 0.79 |
| 3 | 57420002 | rs13070406 | 1.396 | 0.588 | -0.25 | -1.552 | 0.003 | 0.002 | 0.002 | 0 | 0.61 | 0.45 | 0.49 | 0.42 | 0.55 | 0.57 |
| 3 | 57421771 | rs17058156 | 0.127 | 0.603 | -0.711 | -1.514 | 0.001 | 0.003 | 0.003 | 0.003 | 0.13 | 0.12 | 0.12 | 0.08 | 0.13 | 0.22 |
| 3 | 57431985 | rs2121743 | 1.354 | 0.622 | -0.839 | -1.532 | -0.018 | -0.033 | -0.033 | 0 | 0.61 | 0.15 | 0.49 | 0.42 | 0.59 | 0.58 |
| 3 | 57443817 | rs6768661 | 0.232 | 0.596 | -0.501 | -1.547 | 0.025 | 0.111 | 0.111 | 0 | 0.77 | 0.61 | 0.62 | 0.5 | 0.71 | 0.74 |
| 3 | 57450988 | rs7431380 | -0.335 | 0.452 | -0.727 | -1.816 | 0 | 0.012 | 0.012 | 1.631 | 0.46 | 0.6 | 0.38 | 0.12 | 0.46 | 0.54 |
| 3 | 57473750 | rs10510793 | 0.553 | 0.769 | -0.53 | -1.22 | -0.015 | 0.012 | 0.012 | 0 | 0.09 | 0.03 | 0.08 | 0.08 | 0.12 | 0.15 |
| 3 | 57482888 | rs2084358 | 1.796 | 0.741 | -0.557 | -1.228 | -0.013 | -0.012 | -0.012 | 0 | 0.58 | 0.2 | 0.47 | 0.42 | 0.55 | 0.57 |
| 3 | 57489591 | rs4681992 | 1.284 | 0.802 | -0.46 | -1.179 | -0.01 | -0.018 | -0.018 | 0 | 0.64 | 0.29 | 0.51 | 0.42 | 0.59 | 0.59 |
| 3 | 57494138 | rs6445902 | -1.097 | 0.839 | -0.173 | -1.211 | 0 | -0.084 | -0.084 | 0.059 | 0.19 | 0.49 | 0.16 | 0.08 | 0.16 | 0.17 |
| 3 | 57499769 | rs6445905 | 1.097 | 0.819 | -0.286 | -1.16 | -0.015 | 0.027 | 0.027 | 0.231 | 0.68 | 0.45 | 0.64 | 0.43 | 0.71 | 0.65 |
| 3 | 57510407 | rs3926468 | 1.574 | 0.954 | -0.813 | -1.186 | -0.031 | -0.015 | -0.015 | 0 | 0.58 | 0.14 | 0.46 | 0.43 | 0.54 | 0.55 |
| 3 | 57520145 | rs11707555 | -0.158 | 1.233 | -0.608 | -1.028 | -0.018 | 0.006 | 0.006 | 3.127 | 0.07 | 0.02 | 0.1 | 0.06 | 0.08 | 0.13 |
| 3 | 57528503 | rs9311651 | -0.896 | 1.35 | -0.158 | -1.054 | 0.089 | -0.039 | -0.039 | 0 | 0.09 | 0.55 | 0.17 | 0 | 0.15 | 0.09 |
| 3 | 57532024 | rs11707037 | -0.474 | 1.381 | -0.233 | -1.106 | -0.002 | 0.005 | 0.005 | 1.844 | 0.12 | 0.12 | 0.11 | 0.08 | 0.13 | 0.23 |
| 3 | 57540733 | rs10049412 | -1.705 | 1.522 | -0.206 | -1.101 | 0.019 | -0.048 | -0.048 | 0 | 0.21 | 0.67 | 0.28 | 0.09 | 0.28 | 0.33 |
| 3 | 57543614 | rs2278755 | -0.596 | 2.04 | -0.514 | -1.229 | -0.009 | -0.004 | -0.004 | 0 | 0.86 | 0.92 | 0.77 | 0.52 | 0.8 | 0.88 |
| 3 | 57545154 | rs3732546 | -0.308 | 2.039 | -0.49 | -1.234 | -0.002 | 0.006 | 0.006 | 1.893 | 0.12 | 0.12 | 0.11 | 0.09 | 0.13 | 0.23 |
| 3 | 57545993 | rs7618684 | -1.554 | 2.038 | -0.551 | -1.233 | 0.019 | -0.05 | -0.05 | 0 | 0.21 | 0.67 | 0.28 | 0.09 | 0.28 | 0.32 |
| 3 | 57555793 | rs7635607 | -0.831 | 2.014 | -0.504 | -1.198 | 0.095 | -0.047 | -0.047 | 0 | 0.1 | 0.6 | 0.17 | 0 | 0.15 | 0.09 |
| 3 | 57559004 | rs4681737 | -0.785 | 2.016 | -0.505 | -1.197 | 0.098 | -0.045 | -0.045 | 0 | 0.1 | 0.57 | 0.17 | 0 | 0.15 | 0.09 |
| 3 | 57568583 | rs11130591 | 0.473 | 2.024 | -0.539 | -1.196 | -0.041 | -0.083 | -0.083 | 0 | 0.13 | 0.02 | 0.22 | 0.48 | 0.2 | 0.12 |
| 3 | 57573846 | rs9850535 | -0.46 | 1.977 | 0.028 | -1.208 | 0.132 | -0.012 | -0.012 | 0.011 | 0.94 | 0.73 | 0.84 | 1 | 0.85 | 0.91 |
| 3 | 57578276 | rs6767123 | 0.849 | 1.965 | 0.021 | -1.213 | 0.097 | -0.047 | -0.047 | 0 | 0.9 | 0.43 | 0.83 | 1 | 0.85 | 0.91 |
| 3 | 57585406 | rs4638933 | 0.481 | 1.873 | -0.115 | -1.18 | -0.048 | -0.093 | -0.093 | 0 | 0.13 | 0.02 | 0.22 | 0.48 | 0.2 | 0.12 |
| 3 | 57586825 | rs12488539 | -1.413 | 1.863 | -0.119 | -1.269 | -0.021 | -0.041 | -0.041 | 1.231 | 0.29 | 0.2 | 0.36 | 0.57 | 0.38 | 0.37 |
| 3 | 57601582 | rs11713689 | 0.503 | 1.862 | -0.124 | -1.256 | -0.048 | -0.096 | -0.096 | 0.027 | 0.13 | 0.02 | 0.22 | 0.48 | 0.2 | 0.12 |
| 3 | 57612623 | rs6762914 | -1.734 | 1.847 | -0.073 | -1.261 | 0.012 | 0.043 | 0.043 | 0 | 0.32 | 0.48 | 0.38 | 0.57 | 0.38 | 0.37 |
| 3 | 57630018 | rs12152351 | 0.671 | 1.9 | -0.29 | -1.242 | 0.079 | -0.053 | -0.053 | 1.526 | 0.9 | 0.42 | 0.83 | 1 | 0.85 | 0.91 |
| 3 | 57637294 | rs11708662 | 0.452 | 1.827 | -0.497 | -1.153 | -0.048 | -0.096 | -0.096 | 0.018 | 0.13 | 0.02 | 0.22 | 0.48 | 0.2 | 0.12 |
| 3 | 57683838 | rs11706044 | -0.72 | 1.655 | -0.425 | -1.215 | -0.007 | 0.024 | 0.024 | 0.052 | 0.15 | 0.1 | 0.13 | 0.09 | 0.17 | 0.25 |
| 3 | 57695201 | rs12715516 | 1.441 | 1.323 | -1.081 | -1.837 | 0.017 | 0.086 | 0.086 | 0 | 0.27 | 0.1 | 0.12 | 0.01 | 0.21 | 0.2 |
| 3 | 57702947 | rs7646913 | -0.93 | 1.382 | -0.94 | -1.719 | -0.001 | 0 | 0 | 0.376 | 0.57 | 0.4 | 0.49 | 0.57 | 0.59 | 0.56 |
| 3 | 57706177 | rs6802572 | 0.686 | 1.352 | -1.084 | -1.651 | -0.019 | -0.041 | -0.041 | 0 | 0.14 | 0.09 | 0.23 | 0.47 | 0.2 | 0.12 |
| 3 | 57721147 | rs13070936 | 1.173 | 1.184 | -1.258 | -1.806 | 0.016 | 0.094 | 0.094 | 0 | 0.29 | 0.11 | 0.14 | 0.01 | 0.24 | 0.18 |
| 3 | 57723143 | rs7624585 | 0.233 | 1.08 | -1.257 | -1.82 | 0.012 | 0 | 0 | 0 | 0.51 | 0.58 | 0.48 | 0.52 | 0.4 | 0.61 |
| 3 | 57724602 | rs4530498 | -0.681 | 1.047 | -1.158 | -1.941 | 0.005 | -0.098 | -0.098 | 0 | 0.65 | 0.6 | 0.7 | 0.99 | 0.6 | 0.73 |
| 3 | 57731725 | rs11714051 | 0.026 | 1.043 | -1.197 | -1.944 | -0.006 | 0.023 | 0.023 | 2.289 | 0.13 | 0.08 | 0.11 | 0.09 | 0.13 | 0.25 |
| 3 | 57741346 | rs6445927 | -0.661 | 1.042 | -1.199 | -1.949 | 0.004 | -0.09 | -0.09 | 0.081 | 0.65 | 0.61 | 0.7 | 0.99 | 0.6 | 0.75 |
| 3 | 57743246 | rs1053711 | 1.441 | 1.036 | -1.436 | -1.987 | -0.089 | 0.181 | 0.181 | 0.125 | 0.27 | 0.01 | 0.17 | 0.01 | 0.31 | 0.18 |
| 3 | 57774559 | rs17058538 | -0.24 | 0.792 | -1.158 | -1.834 | -0.001 | 0.007 | 0.007 | 2.924 | 0.13 | 0.12 | 0.11 | 0.09 | 0.14 | 0.25 |
| 3 | 57779836 | rs6768930 | 0.395 | 0.769 | -1.232 | -1.658 | 0.01 | -0.016 | -0.016 | 0 | 0.48 | 0.16 | 0.45 | 0.56 | 0.4 | 0.61 |
| 3 | 57808883 | rs11130601 | 0.684 | 1.064 | -0.796 | -1.135 | -0.046 | -0.091 | -0.091 | 0 | 0.13 | 0.02 | 0.22 | 0.43 | 0.2 | 0.13 |
| 3 | 57822950 | rs9848092 | 1.141 | 0.963 | -0.61 | -1.11 | -0.009 | 0.011 | 0.011 | 0.139 | 0.37 | 0.36 | 0.37 | 0.46 | 0.27 | 0.37 |
| 3 | 57826107 | rs1540580 | 0.455 | 1.012 | -0.617 | -1.137 | 0.001 | 0 | 0 | 1.188 | 0.47 | 0.52 | 0.46 | 0.55 | 0.4 | 0.53 |
| 3 | 57826854 | rs12492247 | -0.798 | 1.049 | -0.686 | -1.223 | -0.014 | -0.036 | -0.036 | 0 | 0.24 | 0.21 | 0.32 | 0.52 | 0.33 | 0.3 |
| 3 | 57826884 | rs17554850 | 0.34 | 1.02 | -0.613 | -1.141 | -0.005 | 0.005 | 0.005 | 0.003 | 0.09 | 0.07 | 0.09 | 0.09 | 0.13 | 0.16 |
| 3 | 57849296 | rs9844865 | 0.827 | 0.482 | -0.464 | -0.948 | -0.023 | 0.025 | 0.025 | 0.185 | 0.38 | 0.46 | 0.38 | 0.46 | 0.27 | 0.37 |
| 3 | 57864762 | rs6445930 | 0.853 | 0.483 | -0.463 | -0.948 | -0.023 | 0.026 | 0.026 | 0.181 | 0.38 | 0.46 | 0.38 | 0.46 | 0.27 | 0.37 |
| 3 | 57882601 | rs17058639 | 1.173 | 0.482 | -0.605 | -0.827 | 0.021 | -0.064 | -0.064 | 0 | 0.35 | 0.12 | 0.34 | 0.48 | 0.27 | 0.36 |
| 3 | 57913714 | rs1057719 | 1.014 | 0.564 | -0.769 | -0.711 | -0.01 | 0.012 | 0.012 | 0.065 | 0.37 | 0.35 | 0.36 | 0.46 | 0.27 | 0.37 |
| 3 | 57914178 | rs1043045 | 0.962 | 0.583 | -0.779 | -0.702 | -0.012 | 0.014 | 0.014 | 0.087 | 0.37 | 0.36 | 0.37 | 0.46 | 0.27 | 0.37 |
| 3 | 57945243 | rs1675382 | -0.143 | 0.827 | -1.145 | -0.378 | -0.016 | 0.071 | 0.071 | 0 | 0.56 | 0.64 | 0.57 | 0.78 | 0.47 | 0.59 |
| 3 | 57945526 | rs1658356 | -0.305 | 1.144 | -1.006 | 0.587 | -0.045 | -0.101 | -0.101 | 1.097 | 0.14 | 0.03 | 0.22 | 0.47 | 0.2 | 0.15 |
| 3 | 57961370 | rs11720285 | 0.164 | 0.82 | -0.924 | -0.295 | -0.006 | 0.118 | 0.118 | 0 | 0.28 | 0.12 | 0.16 | 0.01 | 0.28 | 0.19 |
| 3 | 57965749 | rs2577322 | 0.666 | 0.996 | -0.255 | 0.384 | 0 | 0.005 | 0.005 | 0 | 0.54 | 0.38 | 0.51 | 0.43 | 0.46 | 0.64 |
| 3 | 57971523 | rs1866164 | 0.031 | 0.866 | 0.093 | 0.791 | -0.033 | 0.214 | 0.214 | 0.052 | 0.09 | 0.29 | 0.14 | 0.32 | 0.03 | 0.1 |
| 3 | 57974822 | rs6445938 | -0.051 | 1.307 | -0.372 | 0.703 | 0.009 | 0.013 | 0.013 | 0.008 | 0.91 | 0.92 | 0.8 | 0.81 | 0.77 | 0.93 |
| 3 | 57977432 | rs839248 | 0.711 | 1.008 | -0.327 | 0.108 | 0.021 | 0.006 | 0.006 | 0 | 0.5 | 0.33 | 0.39 | 0.47 | 0.39 | 0.55 |
| 3 | 57978556 | rs839250 | 0.073 | 0.924 | -0.527 | -0.49 | -0.064 | 0.103 | 0.103 | 0 | 0.65 | 0.79 | 0.69 | 0.95 | 0.5 | 0.74 |
| 3 | 57989169 | rs11130605 | 0.675 | 0.579 | -0.187 | -0.59 | 0.009 | 0.05 | 0.05 | 0 | 0.42 | 0.28 | 0.26 | 0.01 | 0.4 | 0.25 |
| 3 | 57998986 | rs9828717 | 0.246 | 0.456 | 0.648 | -0.242 | -0.014 | 0.008 | 0.008 | 0.567 | 0.39 | 0.51 | 0.34 | 0.48 | 0.25 | 0.49 |
| 3 | 58006600 | rs839241 | -0.157 | 0.512 | 0.713 | -0.325 | 0.018 | 0.02 | 0.02 | 0.443 | 0.44 | 0.57 | 0.51 | 0.95 | 0.29 | 0.59 |
| 3 | 58010300 | rs1718459 | -0.624 | 0.864 | 0.627 | 0.007 | -0.062 | 0.023 | 0.023 | 0.766 | 0.88 | 0.98 | 0.78 | 0.95 | 0.69 | 0.87 |

| | | | | | | | | | | | | | | | | |
|---|---|---|---|---|---|---|---|---|---|---|---|---|---|---|---|---|
| 3 | 58012348 | rs1718456 | 0.018 | 0.508 | 0.907 | 0.123 | -0.002 | 0.366 | 0.366 | 0.016 | 0.09 | 0.36 | 0.21 | 0.46 | 0.04 | 0.1 |
| 3 | 58018879 | rs865726 | -0.552 | 0.794 | 0.355 | -0.216 | -0.037 | 0.022 | 0.022 | 0.816 | 0.87 | 0.93 | 0.78 | 0.95 | 0.68 | 0.85 |
| 3 | 58025451 | rs9856128 | 0.962 | 0.042 | 0.706 | -0.179 | 0.006 | -0.019 | -0.019 | 0 | 0.07 | 0.08 | 0.08 | 0.01 | 0.07 | 0.05 |
| 3 | 58025903 | rs1718481 | -0.615 | -0.197 | 0.794 | -0.171 | -0.019 | -0.018 | -0.018 | 0 | 0.58 | 0.76 | 0.52 | 0.51 | 0.44 | 0.4 |
| 3 | 58027197 | rs1623879 | -0.798 | -0.198 | 1.039 | -0.012 | -0.006 | 0.252 | 0.252 | 0 | 0.16 | 0.42 | 0.28 | 0.5 | 0.11 | 0.15 |
| 3 | 58036792 | rs839230 | -0.056 | -0.229 | 0.618 | -0.043 | 0.032 | 0.002 | 0.002 | 0 | 0.51 | 0.67 | 0.44 | 0.5 | 0.37 | 0.35 |
| 3 | 58045538 | rs1249968 | -0.621 | 0.122 | 0.727 | 0.635 | 0.01 | 0.135 | 0.135 | 0 | 0.14 | 0.28 | 0.31 | 0.53 | 0.13 | 0.18 |
| 3 | 58048699 | rs6445942 | -1.595 | -0.139 | 0.692 | 0.282 | -0.007 | 0.044 | 0.044 | 4.759 | 0.07 | 0.62 | 0.05 | 0.06 | 0.01 | 0.02 |
| 3 | 58049391 | rs1658347 | -0.763 | -0.352 | -0.039 | 0.374 | -0.012 | 0.009 | 0.009 | 0.963 | 0.22 | 0.65 | 0.35 | 0.61 | 0.13 | 0.16 |
| 3 | 58050265 | rs9809315 | 0.67 | -0.449 | -0.862 | -0.363 | 0.015 | 0.045 | 0.045 | 0 | 0.36 | 0.28 | 0.21 | 0.01 | 0.27 | 0.18 |
| 3 | 58053032 | rs1718480 | -0.363 | 0.04 | -1.271 | 0.22 | -0.124 | -0.037 | -0.037 | 0.974 | 0.59 | 0.96 | 0.53 | 0.6 | 0.4 | 0.37 |
| 3 | 58057684 | rs9822918 | -0.24 | 0.378 | -1.101 | -0.095 | -0.066 | -0.053 | -0.053 | 1.564 | 0.63 | 0.78 | 0.52 | 0.37 | 0.47 | 0.38 |
| 3 | 58069429 | rs2049239 | -0.349 | 0.482 | -1.238 | -1.366 | -0.038 | 0.151 | 0.151 | 1.319 | 0.72 | 0.93 | 0.79 | 1 | 0.65 | 0.83 |
| 3 | 58072471 | rs13315037 | 0.009 | 0.491 | -1.102 | -1.415 | 0 | -0.009 | -0.009 | 0.616 | 0.08 | 0.09 | 0.11 | 0 | 0.07 | 0.08 |
| 3 | 58088939 | rs11719557 | 0.848 | 0.324 | -1.11 | -1.855 | -0.01 | 0.065 | 0.065 | 0.254 | 0.33 | 0.18 | 0.19 | 0.02 | 0.29 | 0.13 |
| 3 | 58089516 | rs6445945 | -0.971 | 0.177 | -1.262 | -1.992 | -0.033 | 0.145 | 0.145 | 1.254 | 0.72 | 0.92 | 0.78 | 1 | 0.66 | 0.83 |
| 3 | 58089851 | rs1522385 | 0.771 | 0.216 | -0.787 | -1.473 | -0.073 | 0.185 | 0.185 | 0.372 | 0.31 | 0.04 | 0.18 | 0.02 | 0.29 | 0.13 |
| 3 | 58092346 | rs2177153 | 0.326 | 0.159 | -0.334 | -1.482 | -0.001 | -0.048 | -0.048 | 0.22 | 0.35 | 0.32 | 0.2 | 0.02 | 0.29 | 0.14 |
| 3 | 58123249 | rs2001972 | 0.126 | 0.364 | -0.484 | -1.317 | -0.005 | 0.03 | 0.03 | 0.196 | 0.37 | 0.27 | 0.27 | 0.03 | 0.34 | 0.17 |
| 3 | 58129450 | rs9826147 | 0.142 | 0.406 | -0.692 | -1.288 | -0.01 | 0.076 | 0.076 | 0.323 | 0.37 | 0.23 | 0.27 | 0.08 | 0.34 | 0.19 |
| 3 | 58138528 | rs9884098 | 0.647 | 0.676 | -0.95 | -1.298 | -0.029 | 0.113 | 0.113 | 1.388 | 0.31 | 0.05 | 0.16 | 0.07 | 0.25 | 0.14 |
| 3 | 58156176 | rs4681660 | -0.438 | 0.455 | -1.136 | -1.776 | -0.03 | 0.188 | 0.188 | 0.175 | 0.66 | 0.93 | 0.75 | 1 | 0.6 | 0.78 |
| 3 | 58163901 | rs6445953 | -0.339 | 0.585 | -0.927 | -1.855 | -0.004 | 0.137 | 0.137 | 0.162 | 0.65 | 0.85 | 0.75 | 1 | 0.63 | 0.79 |
| 3 | 58165233 | rs10470697 | 0.01 | 0.409 | -0.491 | -1.368 | 0.019 | 0 | 0 | 0 | 0.35 | 0.46 | 0.48 | 0.64 | 0.39 | 0.63 |
| 3 | 58169472 | rs7373012 | -1.38 | -0.031 | -0.962 | -1.762 | -0.029 | 0.12 | 0.12 | 0 | 0.74 | 0.94 | 0.84 | 1 | 0.68 | 0.89 |
| 3 | 58175360 | rs9845458 | -0.595 | -0.456 | -0.264 | -1.728 | -0.071 | 0.048 | 0.048 | 0.077 | 0.21 | 0.49 | 0.21 | 0.28 | 0.14 | 0.2 |
| 3 | 58176843 | rs9855113 | -1.382 | -0.55 | -0.517 | -2.01 | -0.024 | 0.155 | 0.155 | 0.184 | 0.68 | 0.91 | 0.81 | 1 | 0.65 | 0.85 |
| 3 | 58179028 | rs3732630 | 1.007 | -0.396 | -0.416 | -2.029 | 0.005 | 0.003 | 0.003 | 0.596 | 0.22 | 0.2 | 0.42 | 0.64 | 0.33 | 0.55 |
| 3 | 58185595 | rs12494813 | -0.162 | -0.39 | -0.383 | -1.743 | 0.009 | -0.001 | -0.001 | 0 | 0.43 | 0.61 | 0.39 | 0.43 | 0.35 | 0.29 |
| 3 | 58186719 | rs3772985 | 0.237 | -0.49 | 0 | -2.037 | -0.025 | 0.057 | 0.057 | 0 | 0.17 | 0.29 | 0.18 | 0.29 | 0.12 | 0.19 |
| 3 | 58186911 | rs4234389 | -1.285 | -0.08 | 0.035 | -1.878 | -0.027 | 0.164 | 0.164 | 0.535 | 0.68 | 0.92 | 0.82 | 1 | 0.65 | 0.84 |
| 3 | 58189680 | rs12490897 | 1.028 | 0.078 | 0.328 | -1.914 | 0.004 | 0.018 | 0.018 | 0 | 0.2 | 0.24 | 0.3 | 0.64 | 0.22 | 0.53 |
| 3 | 58192390 | rs17058991 | 0.51 | 0.109 | 0.645 | -1.94 | 0.04 | -0.024 | -0.024 | 0 | 0.07 | 0.24 | 0.12 | 0 | 0.07 | 0.04 |
| 3 | 58192585 | rs9815775 | -0.653 | 0.042 | 0.325 | -2.164 | -0.002 | -0.034 | -0.034 | 0 | 0.61 | 0.63 | 0.7 | 1 | 0.59 | 0.8 |
| 3 | 58194051 | rs3732631 | 0.774 | -0.052 | -0.153 | -1.955 | 0.004 | -0.01 | -0.01 | 0.125 | 0.35 | 0.15 | 0.35 | 0.35 | 0.32 | 0.25 |
| 3 | 58195039 | rs3772989 | -0.019 | 0.07 | -0.369 | -1.76 | -0.002 | -0.003 | -0.003 | 0.09 | 0.44 | 0.56 | 0.48 | 0.36 | 0.39 | 0.29 |
| 3 | 58209423 | rs12629451 | 1.014 | 0.58 | -0.596 | -1.333 | 0 | 0.006 | 0.006 | 0 | 0.18 | 0.16 | 0.31 | 0.63 | 0.21 | 0.53 |
| 3 | 58211083 | rs7636513 | 0.755 | 0.639 | -0.691 | -1.449 | -0.005 | -0.01 | -0.01 | 0.06 | 0.2 | 0.15 | 0.32 | 0.66 | 0.21 | 0.52 |
| 3 | 58217037 | rs6802482 | -1.623 | 0.68 | -1.154 | -2.38 | -0.053 | 0.093 | 0.093 | 0 | 0.78 | 0.95 | 0.82 | 1 | 0.65 | 0.85 |
| 3 | 58221822 | rs7640259 | -0.429 | 0.468 | -1.111 | -2.312 | 0.011 | 0.105 | 0.105 | 0 | 0.55 | 0.78 | 0.47 | 0.24 | 0.4 | 0.31 |
| 3 | 58227948 | rs11130619 | 0.754 | 0.5 | -0.702 | -2.384 | -0.005 | -0.043 | -0.043 | 1.005 | 0.17 | 0.17 | 0.05 | 0 | 0.08 | 0.08 |
| 3 | 58228082 | rs11130620 | -0.365 | 0.557 | -0.48 | -2.46 | 0.004 | 0.026 | 0.026 | 1.355 | 0.38 | 0.32 | 0.33 | 0.77 | 0.29 | 0.58 |
| 3 | 58232485 | rs4681829 | -1.63 | 0.302 | -0.834 | -2.804 | -0.043 | 0.155 | 0.155 | 0.015 | 0.7 | 0.94 | 0.77 | 1 | 0.62 | 0.84 |
| 3 | 58232659 | rs6445961 | -0.037 | 0.251 | -0.539 | -2.803 | 0.002 | 0.018 | 0.018 | 1.726 | 0.36 | 0.3 | 0.34 | 0.79 | 0.29 | 0.57 |
| 3 | 58247600 | rs6805249 | 0.737 | 0.009 | -0.097 | -2.691 | 0.035 | -0.025 | -0.025 | 0 | 0.06 | 0.22 | 0.11 | 0 | 0.07 | 0.05 |
| 3 | 58259999 | rs6769100 | -0.167 | -0.092 | 0.057 | -2.736 | -0.001 | -0.01 | -0.01 | 0.111 | 0.36 | 0.64 | 0.44 | 0.22 | 0.34 | 0.28 |
| 3 | 58267846 | rs13077539 | 0.822 | -0.205 | 0.012 | -2.796 | 0 | -0.032 | -0.032 | 0 | 0.17 | 0.13 | 0.05 | 0 | 0.08 | 0.08 |
| 3 | 58269712 | rs3773008 | -1.177 | -0.232 | -0.122 | -2.831 | -0.013 | 0.179 | 0.179 | 0 | 0.53 | 0.89 | 0.48 | 0.22 | 0.42 | 0.35 |
| 3 | 58269751 | rs3773010 | 0.664 | -0.307 | 0.257 | -2.847 | -0.008 | -0.019 | -0.019 | 0 | 0.27 | 0.36 | 0.32 | 0.22 | 0.26 | 0.21 |
| 3 | 58278244 | rs7626140 | -1.679 | 0.074 | -0.361 | -2.902 | 0.01 | 0.106 | 0.106 | 0 | 0.7 | 0.92 | 0.81 | 1 | 0.7 | 0.84 |
| 3 | 58280406 | rs9986 | 1.715 | 0.337 | -0.375 | -2.7 | -0.091 | -0.048 | -0.048 | 0 | 0.18 | 0.03 | 0.32 | 0.78 | 0.28 | 0.49 |
| 3 | 58290143 | rs13094898 | 0.935 | 0.191 | -0.248 | -2.793 | 0.018 | 0.045 | 0.045 | 2.212 | 0.16 | 0.06 | 0.04 | 0 | 0.08 | 0.08 |
| 3 | 58305033 | rs14927 | 0.046 | 0.242 | -0.206 | -2.808 | -0.049 | 0.086 | 0.086 | 2.188 | 0.35 | 0.14 | 0.36 | 0.81 | 0.36 | 0.58 |
| 3 | 58305235 | rs10687 | 1.764 | 0.249 | -0.262 | -2.803 | -0.101 | -0.045 | -0.045 | 0.414 | 0.18 | 0.02 | 0.32 | 0.81 | 0.28 | 0.49 |
| 3 | 58312297 | rs7622074 | -0.102 | 0.215 | -0.062 | -2.793 | -0.019 | 0.068 | 0.068 | 1.011 | 0.37 | 0.26 | 0.37 | 0.81 | 0.36 | 0.58 |
| 3 | 58316657 | rs3796215 | 0.113 | 0.214 | -0.075 | -2.794 | -0.089 | 0.088 | 0.088 | 2.302 | 0.35 | 0.07 | 0.36 | 0.81 | 0.36 | 0.58 |
| 3 | 58317692 | rs12637163 | -0.118 | 0.225 | -0.063 | -2.789 | -0.019 | 0.07 | 0.07 | 1.024 | 0.37 | 0.26 | 0.37 | 0.81 | 0.36 | 0.58 |
| 3 | 58320984 | rs11713310 | -1.236 | 0.063 | 0.213 | -2.947 | -0.001 | -0.085 | -0.085 | 0.044 | 0.65 | 0.61 | 0.7 | 1 | 0.62 | 0.79 |
| 3 | 58331186 | rs2176082 | -0.213 | 0.371 | 0.69 | -2.734 | 0.008 | -0.025 | -0.025 | 0 | 0.24 | 0.54 | 0.35 | 0.19 | 0.26 | 0.25 |
| 3 | 58332068 | rs4681679 | -2.093 | 0.394 | 0.6 | -2.764 | 0.012 | 0.122 | 0.122 | 0.48 | 0.63 | 0.86 | 0.72 | 1 | 0.62 | 0.83 |
| 3 | 58335573 | rs9817356 | 0.748 | 0.409 | 0.701 | -2.746 | -0.002 | -0.001 | -0.001 | 0 | 0.18 | 0.28 | 0.27 | 0.18 | 0.19 | 0.2 |
| 3 | 58336846 | rs9864325 | 0.615 | 0.382 | 0.564 | -2.767 | 0.016 | 0.083 | 0.083 | 0.717 | 0.18 | 0.04 | 0.04 | 0 | 0.09 | 0.09 |
| 3 | 58337723 | rs7625655 | -0.576 | 0.381 | 0.595 | -2.77 | -0.012 | -0.121 | -0.121 | 0.046 | 0.25 | 0.46 | 0.13 | 0 | 0.16 | 0.14 |
| 3 | 58339924 | rs6771642 | -0.429 | 0.41 | 0.598 | -2.731 | 0.068 | -0.014 | -0.014 | 0.182 | 0.07 | 0.35 | 0.08 | 0 | 0.07 | 0.05 |
| 3 | 58370177 | rs6445975 | 0.088 | 0.458 | 0.553 | -2.688 | 0.006 | -0.016 | -0.016 | 0.274 | 0.75 | 0.43 | 0.65 | 0.8 | 0.74 | 0.75 |
| 3 | 58375286 | rs4076852 | -0.03 | 0.376 | 0.57 | -2.676 | 0.015 | -0.016 | -0.016 | 0 | 0.76 | 0.49 | 0.65 | 0.8 | 0.74 | 0.75 |
| 3 | 58377039 | rs6809011 | 0.273 | 0.575 | 0.46 | -2.517 | 0.035 | -0.023 | -0.023 | 0 | 0.05 | 0.19 | 0.08 | 0 | 0.07 | 0.05 |
| 3 | 58377159 | rs11915480 | 1.486 | 0.509 | 0.007 | -2.467 | 0.019 | 0.09 | 0.09 | 0.649 | 0.38 | 0.2 | 0.28 | 0 | 0.38 | 0.17 |
| 3 | 58382846 | rs3191903 | 1.486 | 0.52 | -0.226 | -2.462 | 0.019 | 0.09 | 0.09 | 0.649 | 0.38 | 0.2 | 0.28 | 0 | 0.38 | 0.17 |

| | | | | | | | | | | | | | | | | |
|---|---|---|---|---|---|---|---|---|---|---|---|---|---|---|---|---|
| 3 | 58405947 | rs11130638 | 1.449 | 0.545 | -0.449 | -2.451 | 0.019 | 0.074 | 0.074 | 0.726 | 0.38 | 0.21 | 0.28 | 0 | 0.38 | 0.17 |
| 3 | 58408353 | rs12497256 | 1.289 | 0.567 | -0.472 | -2.456 | -0.065 | -0.028 | -0.028 | 2.639 | 0.17 | 0.04 | 0.32 | 0.8 | 0.28 | 0.49 |
| 3 | 58413518 | rs1126722 | 1.292 | 0.08 | -0.347 | -2.455 | 0.021 | 0.09 | 0.09 | 0.708 | 0.38 | 0.2 | 0.28 | 0 | 0.37 | 0.17 |
| 3 | 58423288 | rs11130643 | 1.248 | 0.103 | -0.348 | -2.445 | 0.022 | 0.076 | 0.076 | 1.004 | 0.39 | 0.21 | 0.28 | 0 | 0.38 | 0.17 |
| 3 | 58477280 | rs4681686 | 0.995 | 0.451 | -0.265 | -1.856 | 0.001 | 0.019 | 0.019 | 0 | 0.13 | 0.15 | 0.16 | 0.31 | 0.14 | 0.16 |
| 3 | 58478970 | rs11130651 | 1.139 | 0.362 | -0.589 | -2.072 | -0.093 | -0.096 | -0.096 | 0 | 0.19 | 0.01 | 0.35 | 0.69 | 0.29 | 0.5 |
| 3 | 58481891 | rs4681688 | -0.433 | -0.23 | -0.178 | -2.244 | 0.005 | -0.044 | -0.044 | 1.011 | 0.25 | 0.32 | 0.23 | 0.1 | 0.24 | 0.19 |
| 3 | 58483630 | rs7625947 | -1.044 | -0.261 | -0.08 | -2.34 | -0.013 | -0.024 | -0.024 | 1.128 | 0.41 | 0.47 | 0.34 | 0.32 | 0.32 | 0.3 |
| 3 | 58484521 | rs9853873 | 0.106 | -0.139 | -0.292 | -2.215 | 0.014 | -0.001 | -0.001 | 0.861 | 0.2 | 0.12 | 0.11 | 0.22 | 0.09 | 0.12 |
| 3 | 58497779 | rs6795193 | 1.078 | -0.066 | -0.683 | -2.258 | 0.019 | 0.04 | 0.04 | 1.337 | 0.36 | 0.26 | 0.23 | 0 | 0.29 | 0.17 |
| 3 | 58502708 | rs4681689 | 0.617 | -0.034 | -0.787 | -1.671 | 0.001 | -0.005 | -0.005 | 0.406 | 0.15 | 0.15 | 0.17 | 0.1 | 0.17 | 0.17 |
| 3 | 58512237 | rs1127745 | -1.172 | -0.891 | -0.379 | -1.236 | -0.023 | -0.005 | -0.005 | 0 | 0.29 | 0.7 | 0.16 | 0.23 | 0.08 | 0.08 |
| 3 | 58515566 | rs4681690 | 0.812 | -0.538 | -0.32 | -1.193 | -0.022 | -0.009 | -0.009 | 0.039 | 0.7 | 0.37 | 0.79 | 0.76 | 0.86 | 0.87 |
| 3 | 58516383 | rs4387990 | 1.158 | 0.291 | -0.686 | -1.301 | -0.14 | 0.219 | 0.219 | 0 | 0.38 | 0.02 | 0.34 | 0.1 | 0.43 | 0.32 |
| 3 | 58520859 | rs4681862 | 0.902 | -0.522 | 0.383 | -0.749 | -0.024 | -0.086 | -0.086 | 5.798 | 0.87 | 0.6 | 0.93 | 1 | 0.95 | 0.98 |
| 3 | 58543512 | rs12488633 | -0.894 | -0.203 | -0.489 | -0.388 | 0.056 | 0.106 | 0.106 | 0.629 | 0.49 | 0.69 | 0.44 | 0.83 | 0.35 | 0.53 |
| 3 | 58549297 | rs1982759 | -0.069 | -0.16 | -0.541 | 0.282 | -0.024 | -0.061 | -0.061 | 0 | 0.29 | 0.12 | 0.38 | 0.57 | 0.31 | 0.52 |
| 3 | 58549647 | rs1982758 | -0.769 | -0.23 | -0.387 | 0.029 | 0.014 | 0.058 | 0.058 | 0 | 0.44 | 0.69 | 0.47 | 0.79 | 0.35 | 0.56 |
| 3 | 58551800 | rs2120577 | 0.819 | -0.46 | 0.371 | 0.698 | -0.066 | 0.135 | 0.135 | 1.226 | 0.85 | 0.59 | 0.75 | 0.78 | 0.96 | 0.97 |
| 3 | 58558769 | rs6798348 | 0.93 | 0.233 | 0.494 | -0.066 | 0.012 | -0.123 | -0.123 | 0 | 0.82 | 0.42 | 0.81 | 1 | 0.83 | 0.88 |
| 3 | 58560043 | rs6776877 | 1.304 | 0.031 | -0.225 | 0.405 | 0.005 | -0.026 | -0.026 | 0 | 0.56 | 0.27 | 0.43 | 0.5 | 0.59 | 0.4 |
| 3 | 58562525 | rs13088795 | -0.111 | 0.081 | 0.175 | 0.353 | 0.026 | -0.032 | -0.032 | 0 | 0.07 | 0.14 | 0.11 | 0 | 0.14 | 0.04 |
| 3 | 58563151 | rs2306671 | 1.495 | -0.247 | -0.313 | -0.011 | -0.006 | -0.004 | -0.004 | 0.04 | 0.48 | 0.28 | 0.4 | 0.51 | 0.53 | 0.27 |
| 3 | 58565992 | rs4681868 | 1.032 | -0.237 | -0.244 | -0.216 | -0.004 | -0.102 | -0.102 | 0 | 0.72 | 0.37 | 0.77 | 1 | 0.74 | 0.73 |
| 3 | 58573534 | rs4681869 | 1.535 | -0.062 | -0.619 | 0.196 | -0.059 | 0.089 | 0.089 | 0.063 | 0.44 | 0.15 | 0.35 | 0.33 | 0.51 | 0.21 |
| 3 | 58584220 | rs4681871 | 0.512 | 0.048 | -0.745 | 0.631 | -0.004 | -0.098 | -0.098 | 0 | 0.2 | 0.07 | 0.34 | 0.46 | 0.15 | 0.46 |
| 3 | 58588019 | rs13076420 | -0.364 | 0.065 | -0.594 | 0.461 | -0.02 | 0.029 | 0.029 | 0 | 0.11 | 0.03 | 0.14 | 0 | 0.17 | 0.13 |
| 3 | 58590284 | rs4511910 | -1.296 | -0.038 | 0.435 | 0.699 | -0.005 | -0.006 | -0.006 | 0 | 0.4 | 0.56 | 0.49 | 0.63 | 0.32 | 0.66 |
| 3 | 58591151 | rs9860663 | 0.04 | -0.024 | 0.823 | 0.705 | 0.03 | -0.019 | -0.019 | 2.112 | 0.07 | 0.31 | 0.05 | 0 | 0.07 | 0.03 |
| 3 | 58593754 | rs4558779 | -0.883 | 0.065 | 0.158 | 0.862 | 0.005 | -0.004 | -0.004 | 0.102 | 0.33 | 0.22 | 0.43 | 0.63 | 0.24 | 0.57 |
| 3 | 58594418 | rs717323 | 0.418 | -0.021 | 0.175 | 0.976 | 0 | 0.022 | 0.022 | 0 | 0.1 | 0.11 | 0.06 | 0.16 | 0.07 | 0.1 |
| 3 | 58594827 | rs2306670 | -0.649 | -0.019 | 0.046 | 1.012 | 0.017 | -0.03 | -0.03 | 0 | 0.27 | 0.17 | 0.2 | 0.59 | 0.17 | 0.58 |
| 3 | 58595823 | rs4681875 | 0.028 | 0.276 | 0.156 | 0.814 | 0.001 | 0.007 | 0.007 | 0 | 0.88 | 0.87 | 0.91 | 0.86 | 0.89 | 0.89 |
| 3 | 58601047 | rs13080308 | -0.436 | 0.046 | -0.816 | 0.571 | -0.025 | -0.003 | -0.003 | 0 | 0.5 | 0.18 | 0.57 | 0.54 | 0.48 | 0.64 |
| 3 | 58603203 | rs17059498 | 0.112 | 0.02 | -0.654 | 0.26 | 0.035 | 0.021 | 0.021 | 1.62 | 0.45 | 0.26 | 0.49 | 0.43 | 0.38 | 0.56 |
| 3 | 58609492 | rs728969 | -0.13 | 0.433 | -0.467 | 0.3 | -0.003 | 0.006 | 0.006 | 1.357 | 0.47 | 0.3 | 0.56 | 0.44 | 0.48 | 0.61 |
| 3 | 58613329 | rs4681879 | -0.33 | 0.41 | -0.071 | 0.216 | -0.004 | 0.02 | 0.02 | 0.274 | 0.49 | 0.29 | 0.59 | 0.44 | 0.5 | 0.66 |
| 3 | 58614016 | rs11710358 | -0.894 | 0.487 | -0.146 | 0.179 | 0.009 | -0.024 | -0.024 | 0.038 | 0.1 | 0.12 | 0.17 | 0 | 0.2 | 0.1 |
| 3 | 58615670 | rs11130660 | 0.191 | 0.601 | 0.092 | 0.157 | 0.027 | -0.026 | -0.026 | 0 | 0.42 | 0.2 | 0.45 | 0.44 | 0.38 | 0.56 |
| 3 | 58616627 | rs9862897 | 0.186 | 0.482 | 0.134 | 0.175 | 0.05 | -0.016 | -0.016 | 0 | 0.94 | 0.79 | 0.91 | 1 | 0.9 | 0.95 |
| 3 | 58617544 | rs4681881 | -0.54 | 1.272 | -0.36 | 0.143 | 0.011 | 0.022 | 0.022 | 0 | 0.55 | 0.24 | 0.65 | 0.6 | 0.59 | 0.68 |
| 3 | 58619522 | rs3860562 | -0.136 | 1.318 | -0.392 | 0.287 | -0.038 | -0.029 | -0.029 | 0 | 0.12 | 0.04 | 0.15 | 0.15 | 0.18 | 0.08 |
| 3 | 58622620 | rs11130665 | 1.147 | 1.054 | -0.063 | 0.121 | -0.004 | 0.017 | 0.017 | 1.158 | 0.32 | 0.41 | 0.24 | 0.4 | 0.28 | 0.2 |
| 3 | 58624770 | rs731608 | 0.16 | 1.061 | -0.229 | 0.061 | -0.022 | -0.014 | -0.014 | 1.738 | 0.45 | 0.25 | 0.56 | 0.46 | 0.51 | 0.6 |
| 3 | 58626075 | rs6762192 | 0.934 | 0.812 | -1.104 | 0.197 | -0.051 | -0.05 | -0.05 | 0 | 0.18 | 0.01 | 0.1 | 0.23 | 0.17 | 0.15 |
| 3 | 58634551 | rs7620807 | -0.428 | 0.554 | -0.141 | -0.2 | 0.037 | -0.019 | -0.019 | 0.278 | 0.56 | 0.24 | 0.68 | 0.46 | 0.67 | 0.63 |
| 3 | 58642452 | rs17059602 | -1.164 | 0.595 | 0.533 | -0.036 | 0.011 | 0.063 | 0.063 | 0 | 0.11 | 0.29 | 0.2 | 0.15 | 0.12 | 0.11 |
| 3 | 58644623 | rs17059619 | 0.338 | 0.677 | 0.468 | -0.194 | -0.008 | 0.005 | 0.005 | 0 | 0.1 | 0.07 | 0.1 | 0 | 0.15 | 0.06 |
| 3 | 58645595 | rs1026952 | 0.588 | 0.618 | 0.622 | -0.356 | 0.021 | -0.011 | -0.011 | 0 | 0.48 | 0.21 | 0.46 | 0.48 | 0.46 | 0.56 |
| 3 | 58649117 | rs2099960 | 1.133 | 0.566 | 0.973 | -0.354 | 0.015 | 0.07 | 0.07 | 0.152 | 0.87 | 0.54 | 0.79 | 0.86 | 0.88 | 0.84 |
| 3 | 58649994 | rs12489751 | 1.205 | 0.58 | 0.82 | -0.354 | -0.002 | -0.014 | -0.014 | 0 | 0.76 | 0.34 | 0.68 | 0.85 | 0.74 | 0.78 |
| 3 | 58654095 | rs2364306 | -0.77 | 0.505 | 0.923 | -0.217 | 0.009 | 0.011 | 0.011 | 0 | 0.68 | 0.58 | 0.74 | 0.61 | 0.7 | 0.69 |
| 3 | 58655038 | rs13075889 | 0.858 | 0.51 | 0.837 | -0.152 | -0.011 | -0.013 | -0.013 | 0 | 0.29 | 0.25 | 0.22 | 0.38 | 0.28 | 0.26 |
| 3 | 58656618 | rs11130669 | 0.077 | 0.299 | 0.72 | -0.235 | 0.003 | 0.004 | 0.004 | 0 | 0.4 | 0.45 | 0.43 | 0.59 | 0.36 | 0.35 |
| 3 | 58658454 | rs11130670 | -0.844 | -0.19 | 1.011 | -0.757 | 0.034 | -0.018 | -0.018 | 0.018 | 0.7 | 0.62 | 0.62 | 0.73 | 0.71 | 0.74 |
| 3 | 58658866 | rs1373403 | 0.328 | -0.2 | 0.179 | -0.841 | 0.015 | 0.019 | 0.019 | 0 | 0.51 | 0.35 | 0.36 | 0.56 | 0.53 | 0.61 |
| 3 | 58673232 | rs7616579 | 0.852 | -0.767 | 0.184 | -1.098 | -0.005 | -0.011 | -0.011 | 0 | 0.77 | 0.37 | 0.69 | 0.83 | 0.79 | 0.85 |
| 3 | 58685959 | rs923034 | 0.625 | -0.909 | -0.849 | -1.81 | -0.03 | 0.136 | 0.136 | 0 | 0.46 | 0.21 | 0.45 | 0.71 | 0.54 | 0.6 |
| 3 | 58687200 | rs7633829 | -0.004 | -0.662 | -0.703 | -1.652 | 0.011 | 0.055 | 0.055 | 0 | 0.58 | 0.33 | 0.56 | 0.72 | 0.66 | 0.67 |
| 3 | 58691075 | rs936739 | -0.728 | -0.301 | -0.823 | -1.438 | 0.016 | -0.036 | -0.036 | 0 | 0.7 | 0.6 | 0.62 | 0.72 | 0.74 | 0.78 |
| 3 | 58691637 | rs900876 | 0.637 | 0.048 | -1.333 | -1.254 | -0.061 | 0.075 | 0.075 | 0 | 0.36 | 0.17 | 0.39 | 0.72 | 0.46 | 0.53 |
| 3 | 58691737 | rs9843676 | 0.002 | 0.508 | -0.691 | -0.264 | -0.001 | -0.02 | -0.02 | 0 | 0.64 | 0.42 | 0.58 | 0.72 | 0.68 | 0.73 |
| 3 | 58705874 | rs4681901 | -0.408 | 0.07 | -0.798 | -0.147 | 0.004 | 0.005 | 0.005 | 2.192 | 0.36 | 0.68 | 0.4 | 0.23 | 0.3 | 0.2 |
| 3 | 58711094 | rs9814023 | -1.167 | -0.31 | -0.141 | 0.872 | -0.014 | 0.232 | 0.232 | 0 | 0.07 | 0.25 | 0.15 | 0.38 | 0.04 | 0.13 |
| 3 | 58744141 | rs7624292 | -0.698 | -0.194 | -0.281 | -0.349 | -0.019 | -0.012 | -0.012 | 1.12 | 0.54 | 0.81 | 0.49 | 0.43 | 0.5 | 0.41 |
| 3 | 58779226 | rs7634348 | -0.84 | 0.033 | -0.434 | -0.535 | -0.086 | -0.101 | -0.101 | 0.42 | 0.58 | 0.96 | 0.52 | 0.42 | 0.54 | 0.41 |
| 3 | 58782604 | rs9818318 | -0.226 | -0.023 | -0.343 | -0.474 | -0.013 | 0.216 | 0.216 | 0 | 0.07 | 0.23 | 0.15 | 0.37 | 0.04 | 0.13 |
| 3 | 58785643 | rs17059849 | -0.138 | -0.024 | -0.3 | -0.474 | -0.013 | 0.174 | 0.174 | 0 | 0.07 | 0.19 | 0.14 | 0.37 | 0.04 | 0.13 |
| 3 | 58799602 | rs12637980 | -0.493 | -0.047 | -0.194 | -0.481 | -0.017 | 0.179 | 0.179 | 0 | 0.07 | 0.2 | 0.15 | 0.37 | 0.04 | 0.13 |

| | | | | | | | | | | | | | | | | |
|---|---|---|---|---|---|---|---|---|---|---|---|---|---|---|---|---|
| 3 | 58799884 | rs12635827 | -0.493 | -0.047 | -0.194 | -0.481 | -0.017 | 0.179 | 0.179 | 0 | 0.07 | 0.2 | 0.15 | 0.37 | 0.04 | 0.13 |
| 3 | 58819748 | rs7627981 | -1.247 | 0.153 | 0.853 | -0.575 | 0.024 | 0.371 | 0.371 | 1.031 | 0.1 | 0.47 | 0.17 | 0.37 | 0.06 | 0.14 |
| 3 | 58823639 | rs10510798 | 0.498 | 0.147 | 0.886 | -0.583 | 0.006 | 0.074 | 0.074 | 0 | 0.05 | 0.05 | 0.12 | 0.37 | 0.04 | 0.13 |
| 3 | 58825127 | rs2037165 | 0.564 | 0.173 | 0.513 | -1.09 | 0.002 | -0.046 | -0.046 | 3.719 | 0.27 | 0.29 | 0.25 | 0.05 | 0.27 | 0.21 |
| 3 | 58834824 | rs1554909 | -0.501 | 0.193 | 0.532 | -0.992 | -0.017 | 0.198 | 0.198 | 0 | 0.07 | 0.23 | 0.15 | 0.38 | 0.04 | 0.12 |
| 3 | 58834890 | rs6797698 | -1.178 | 0.193 | 0.527 | -1 | 0.024 | 0.362 | 0.362 | 1.306 | 0.1 | 0.48 | 0.17 | 0.37 | 0.06 | 0.13 |
| 3 | 58870148 | rs9826422 | 0.615 | 0.118 | 0.631 | -1.183 | -0.006 | 0.018 | 0.018 | 0.792 | 0.26 | 0.15 | 0.24 | 0.05 | 0.27 | 0.21 |
| 3 | 58910094 | rs1273551 | 1.771 | -0.043 | -0.259 | -1.478 | -0.105 | -0.18 | -0.18 | 0.033 | 0.34 | 0.02 | 0.42 | 0.55 | 0.38 | 0.52 |
| 3 | 58985078 | rs13082751 | 1.111 | 0.357 | 0.009 | -1.57 | -0.029 | 0.085 | 0.085 | 0.53 | 0.16 | 0.02 | 0.08 | 0 | 0.18 | 0.06 |
| 3 | 59000763 | rs1900621 | 1.214 | 0.544 | 0.232 | -1.655 | 0.022 | -0.056 | -0.056 | 0.503 | 0.13 | 0.2 | 0.15 | 0 | 0.2 | 0.11 |
| 3 | 59029189 | rs7649388 | 0.945 | 0.533 | 0.258 | -1.66 | 0.03 | -0.073 | -0.073 | 0 | 0.14 | 0.26 | 0.16 | 0 | 0.2 | 0.11 |
| 3 | 59048414 | rs9818329 | 1.136 | 0.418 | 0.225 | -1.718 | 0.003 | 0.015 | 0.015 | 0 | 0.11 | 0.03 | 0.08 | 0.05 | 0.07 | 0.1 |
| 3 | 59058905 | rs11717883 | 1.005 | 0.509 | 0.479 | -1.606 | -0.002 | -0.018 | -0.018 | 0 | 0.13 | 0.12 | 0.15 | 0 | 0.2 | 0.11 |
| 3 | 59071899 | rs9833252 | 0.867 | 0.501 | 0.536 | -1.61 | 0.011 | -0.049 | -0.049 | 0.065 | 0.14 | 0.16 | 0.15 | 0 | 0.2 | 0.11 |
| 3 | 59097593 | rs6446009 | -0.634 | 0.292 | 1.131 | -1.615 | -0.061 | -0.071 | -0.071 | 0 | 0.11 | 0.57 | 0.07 | 0 | 0.01 | 0.09 |
| 3 | 59112888 | rs6414591 | -2.311 | 0.016 | -0.194 | -1.834 | -0.033 | -0.185 | -0.185 | 2.329 | 0.76 | 0.99 | 0.83 | 0.46 | 0.82 | 0.68 |
| 3 | 59114369 | rs6797499 | -0.055 | -0.198 | 0.792 | -1.656 | -0.002 | -0.143 | -0.143 | 0 | 0.23 | 0.5 | 0.36 | 0 | 0.23 | 0.18 |
| 3 | 59125631 | rs12489749 | 0.926 | -0.195 | 0.722 | -1.646 | -0.02 | 0.043 | 0.043 | 0 | 0.17 | 0.07 | 0.3 | 0 | 0.22 | 0.16 |
| 3 | 59126777 | rs9824833 | -0.333 | -0.207 | 0.549 | -1.349 | -0.017 | -0.059 | -0.059 | 0 | 0.27 | 0.59 | 0.37 | 0.09 | 0.25 | 0.24 |
| 3 | 59153978 | rs6762031 | -1.678 | -0.238 | -0.286 | -1.243 | -0.022 | 0.058 | 0.058 | 0.757 | 0.54 | 0.77 | 0.64 | 0.3 | 0.51 | 0.45 |
| 3 | 59169333 | rs1512821 | -0.397 | -0.417 | -0.146 | -0.443 | -0.001 | -0.142 | -0.142 | 0 | 0.36 | 0.6 | 0.45 | 0.04 | 0.36 | 0.31 |
| 3 | 59170215 | rs1567728 | -1.714 | -0.45 | -0.127 | -0.265 | -0.013 | 0.027 | 0.027 | 0.145 | 0.6 | 0.82 | 0.67 | 0.31 | 0.6 | 0.47 |
| 3 | 59178030 | rs1512828 | -0.405 | -0.493 | 0.614 | 0.085 | 0.008 | -0.169 | -0.169 | 0 | 0.3 | 0.59 | 0.4 | 0 | 0.3 | 0.14 |
| 3 | 59185912 | rs1588678 | -2.206 | -1.067 | 1.293 | 2.041 | 0.007 | -0.013 | -0.013 | 0 | 0.22 | 0.74 | 0.2 | 0.36 | 0.11 | 0.26 |
| 3 | 59191320 | rs17060183 | -1.187 | -0.933 | 1.668 | 2.102 | -0.028 | 0.073 | 0.073 | 0 | 0.11 | 0.2 | 0.05 | 0.2 | 0.06 | 0.16 |
| 3 | 59224244 | rs6799967 | -1.095 | -0.946 | 1.537 | 2.374 | -0.014 | 0.052 | 0.052 | 0 | 0.13 | 0.18 | 0.19 | 0.31 | 0.12 | 0.19 |
| 3 | 59228745 | rs9880478 | 2.048 | -0.935 | 1.368 | 2.22 | -0.07 | 0.101 | 0.101 | 0 | 0.75 | 0.49 | 0.75 | 0.5 | 0.83 | 0.67 |
| 3 | 59232354 | rs6446016 | 1.279 | -0.664 | 1.392 | 2.154 | -0.041 | 0.102 | 0.102 | 0 | 0.88 | 0.7 | 0.93 | 0.8 | 0.95 | 0.86 |
| 3 | 59237106 | rs9311710 | 1.083 | -0.369 | 0.595 | 1.843 | -0.078 | 0.164 | 0.164 | 0 | 0.46 | 0.1 | 0.26 | 0.28 | 0.44 | 0.49 |
| 3 | 59247165 | rs1443936 | 1.523 | -0.374 | 1.202 | 1.871 | -0.002 | 0.036 | 0.036 | 3.845 | 0.85 | 0.77 | 0.72 | 0.31 | 0.8 | 0.66 |
| 3 | 59253941 | rs7375089 | -0.176 | -0.438 | 1.122 | 1.737 | 0 | 0.014 | 0.014 | 2.074 | 0.37 | 0.69 | 0.46 | 0.02 | 0.37 | 0.17 |
| 3 | 59259043 | rs6789380 | 1.342 | -0.545 | 1.022 | 1.74 | -0.079 | 0.154 | 0.154 | 0 | 0.45 | 0.13 | 0.26 | 0.27 | 0.43 | 0.43 |
| 3 | 59259536 | rs6446023 | 1.119 | -0.529 | 1.151 | 2.525 | -0.004 | 0.152 | 0.152 | 0.752 | 0.94 | 0.88 | 0.88 | 0.66 | 0.94 | 0.9 |
| 3 | 59265315 | rs9838009 | -0.267 | -0.493 | 1.797 | 2.295 | 0.028 | -0.074 | -0.074 | 0.019 | 0.12 | 0.32 | 0.1 | 0 | 0.14 | 0.07 |
| 3 | 59269042 | rs10510805 | -0.835 | -0.48 | 1.481 | 2.589 | 0.001 | -0.029 | -0.029 | 0 | 0.1 | 0.03 | 0.09 | 0.24 | 0.14 | 0.23 |
| 3 | 59269090 | rs17326080 | -0.353 | -0.499 | 1.534 | 2.012 | 0.001 | 0.026 | 0.026 | 2.352 | 0.37 | 0.66 | 0.46 | 0.04 | 0.37 | 0.17 |
| 3 | 59269320 | rs12054008 | -1.908 | -0.45 | 1.317 | 2.137 | 0.01 | -0.056 | -0.056 | 1.154 | 0.17 | 0.07 | 0.2 | 0.58 | 0.19 | 0.33 |
| 3 | 59288215 | rs7652341 | -0.19 | -0.342 | 1.765 | 1.545 | 0.001 | -0.101 | -0.101 | 0.02 | 0.36 | 0.43 | 0.44 | 0.04 | 0.37 | 0.17 |
| 3 | 59289140 | rs9854717 | -2.025 | -0.541 | 2.031 | 1.848 | -0.012 | 0.102 | 0.102 | 1.112 | 0.18 | 0.42 | 0.23 | 0.65 | 0.19 | 0.31 |
| 3 | 59290258 | rs11917063 | -0.957 | -0.467 | 1.91 | 2.106 | 0 | 0 | 0 | 0.683 | 0.47 | 0.58 | 0.56 | 0.47 | 0.52 | 0.4 |
| 3 | 59292559 | rs6807244 | -2.279 | -0.549 | 2.034 | 2.196 | -0.002 | 0.334 | 0.334 | 0.02 | 0.1 | 0.41 | 0.19 | 0.3 | 0.05 | 0.16 |
| 3 | 59295141 | rs13321256 | -1.732 | -0.491 | 2.05 | 2.035 | -0.013 | 0.111 | 0.111 | 0.964 | 0.14 | 0.19 | 0.18 | 0.53 | 0.14 | 0.29 |
| 3 | 59295659 | rs7646580 | 0.189 | -0.483 | 2.042 | 1.59 | -0.004 | -0.047 | -0.047 | 2.603 | 0.28 | 0.28 | 0.38 | 0.04 | 0.25 | 0.11 |
| 3 | 59300274 | rs1443933 | -0.357 | -0.339 | 1.762 | 1.024 | 0 | -0.05 | -0.05 | 0 | 0.36 | 0.39 | 0.47 | 0.04 | 0.42 | 0.18 |
| 3 | 59304765 | rs17060306 | -0.843 | -0.139 | 1.928 | 1.125 | 0.044 | -0.073 | -0.073 | 0 | 0.12 | 0.31 | 0.1 | 0 | 0.18 | 0.07 |
| 3 | 59306653 | rs9842553 | 0.301 | -0.167 | 2.006 | 1.138 | 0.027 | -0.044 | -0.044 | 0.121 | 0.92 | 0.79 | 0.92 | 1 | 0.89 | 0.95 |
| 3 | 59307311 | rs6446029 | 0.181 | -0.303 | 2.183 | 1.178 | 0.001 | -0.047 | -0.047 | 0.496 | 0.67 | 0.61 | 0.5 | 0.89 | 0.63 | 0.77 |
| 3 | 59307725 | rs6792486 | 0.576 | -0.299 | 1.759 | 1.19 | -0.009 | 0.086 | 0.086 | 0 | 0.24 | 0.07 | 0.37 | 0.04 | 0.25 | 0.11 |
| 3 | 59317981 | rs4679616 | 0.46 | -0.37 | 1.827 | 1.197 | 0 | 0.026 | 0.026 | 0 | 0.25 | 0.12 | 0.43 | 0.12 | 0.25 | 0.17 |
| 3 | 59318138 | rs6808731 | -0.815 | -0.345 | 1.797 | 1.094 | 0 | 0.025 | 0.025 | 0 | 0.42 | 0.28 | 0.63 | 0.85 | 0.43 | 0.55 |
| 3 | 59321520 | rs1373295 | -0.666 | -0.332 | 1.466 | 1.838 | -0.031 | -0.083 | -0.083 | 0.078 | 0.13 | 0.01 | 0.1 | 0.28 | 0.15 | 0.22 |
| 3 | 59321911 | rs6782301 | -0.95 | -0.369 | 1.628 | 1.582 | 0.003 | -0.013 | -0.013 | 0 | 0.46 | 0.53 | 0.66 | 0.84 | 0.44 | 0.55 |
| 3 | 59323387 | rs6414596 | 0.65 | -0.516 | 1.645 | 1.653 | 0.002 | -0.074 | -0.074 | 0.075 | 0.88 | 0.65 | 0.91 | 1 | 0.89 | 0.94 |
| 3 | 59324609 | rs6765646 | -1.87 | -0.611 | 1.2 | 1.821 | -0.009 | 0 | 0 | 1.911 | 0.21 | 0.36 | 0.22 | 0.72 | 0.18 | 0.38 |
| 3 | 59326134 | rs6778032 | 1.631 | -0.423 | 0.392 | 1.76 | -0.133 | 0.21 | 0.21 | 0.693 | 0.41 | 0.05 | 0.27 | 0.19 | 0.47 | 0.39 |
| 3 | 59327461 | rs1023793 | -0.604 | -0.199 | 0.471 | 1.926 | -0.039 | -0.086 | -0.086 | 0 | 0.12 | 0.01 | 0.1 | 0.26 | 0.14 | 0.21 |
| 3 | 59328217 | rs1443941 | 0.052 | -0.182 | 0.775 | 1.266 | -0.011 | -0.06 | -0.06 | 3.562 | 0.64 | 0.44 | 0.49 | 0.91 | 0.7 | 0.85 |
| 3 | 59331595 | rs1443934 | -0.92 | 0.004 | 0.802 | 1.626 | 0.003 | 0.04 | 0.04 | 2.516 | 0.16 | 0.13 | 0.13 | 0.5 | 0.19 | 0.3 |
| 3 | 59351946 | rs4679644 | 0.982 | 0.036 | 0.22 | 1.546 | 0.003 | 0.046 | 0.046 | 1.384 | 0.73 | 0.4 | 0.78 | 0.51 | 0.71 | 0.59 |
| 3 | 59353090 | rs4679519 | 0.438 | -0.005 | -0.536 | 1.025 | 0.016 | 0.052 | 0.052 | 0 | 0.26 | 0.12 | 0.4 | 0.09 | 0.21 | 0.12 |
| 3 | 59353452 | rs13092118 | 1.134 | 0.111 | -0.841 | 1.166 | -0.149 | 0.015 | 0.015 | 0.565 | 0.41 | 0.03 | 0.27 | 0.3 | 0.46 | 0.42 |
| 3 | 59357436 | rs2366434 | 0.309 | 0.209 | 0.158 | 1.566 | 0 | -0.003 | -0.003 | 0 | 0.61 | 0.73 | 0.65 | 0.52 | 0.64 | 0.61 |
| 3 | 59357910 | rs11130702 | 1.353 | 0.076 | 0.349 | 1.573 | -0.057 | 0.122 | 0.122 | 0.368 | 0.45 | 0.11 | 0.48 | 0.29 | 0.51 | 0.47 |
| 3 | 59358190 | rs11130703 | -0.049 | 0.128 | 0.54 | 2.074 | 0.013 | 0.031 | 0.031 | 0 | 0.33 | 0.15 | 0.31 | 0.21 | 0.3 | 0.33 |
| 3 | 59368074 | rs2034127 | 0.989 | 0.142 | 1.144 | 2.042 | 0.021 | 0.212 | 0.212 | 0.888 | 0.88 | 0.75 | 0.8 | 0.57 | 0.86 | 0.66 |
| 3 | 59372700 | rs717463 | -1.026 | 0.138 | 1.083 | 2.093 | 0.049 | 0.237 | 0.237 | 0.408 | 0.12 | 0.26 | 0.15 | 0.37 | 0.13 | 0.3 |
| 3 | 59389870 | rs4679547 | 0.357 | 0.15 | 0.409 | 1.622 | 0.021 | 0.102 | 0.102 | 0.125 | 0.31 | 0.1 | 0.23 | 0 | 0.25 | 0.11 |
| 3 | 59398634 | rs7610781 | -0.149 | 0.333 | 0.453 | 1.288 | 0.006 | -0.008 | -0.008 | 0.332 | 0.08 | 0.14 | 0.05 | 0.06 | 0.1 | 0.09 |
| 3 | 59400715 | rs9850077 | -1.443 | 0.347 | 0.445 | 1.39 | 0.001 | 0.017 | 0.017 | 2.801 | 0.19 | 0.18 | 0.24 | 0.48 | 0.25 | 0.39 |

| | | | | | | | | | | | | | | | | |
|---|---|---|---|---|---|---|---|---|---|---|---|---|---|---|---|---|
| 3 | 59404816 | rs12054029 | -0.301 | 0.401 | 0.009 | 1.506 | -0.027 | -0.015 | -0.015 | 0 | 0.1 | 0.02 | 0.18 | 0.46 | 0.14 | 0.3 |
| 3 | 59405125 | rs4679485 | -1.492 | 0.295 | 0.007 | 1.345 | -0.07 | 0.03 | 0.03 | 0 | 0.61 | 0.89 | 0.54 | 0.65 | 0.53 | 0.56 |
| 3 | 59405936 | rs2279811 | -0.679 | -0.039 | 0.711 | 0.796 | 0.026 | 0.205 | 0.205 | 0.833 | 0.07 | 0.33 | 0.05 | 0.17 | 0.03 | 0.06 |
| 3 | 59407908 | rs9841867 | -0.466 | 0.222 | -0.382 | 0.82 | 0.004 | -0.007 | -0.007 | 0 | 0.08 | 0.16 | 0.06 | 0.05 | 0.1 | 0.09 |
| 3 | 59408242 | rs10510808 | -0.303 | -0.04 | -0.345 | 0.748 | 0.034 | 0.046 | 0.046 | 0.514 | 0.36 | 0.7 | 0.28 | 0.18 | 0.28 | 0.16 |
| 3 | 59409404 | rs7628355 | -1.433 | 0.786 | -0.305 | 0.992 | 0.007 | 0.003 | 0.003 | 0.07 | 0.09 | 0.14 | 0.1 | 0.11 | 0.13 | 0.05 |
| 3 | 59409719 | rs7650098 | -0.753 | 0.724 | -0.897 | 0.634 | 0.015 | 0.011 | 0.011 | 0.525 | 0.42 | 0.66 | 0.36 | 0.36 | 0.34 | 0.37 |
| 3 | 59410065 | rs4073471 | -0.959 | 0.495 | -0.86 | 0.402 | 0.004 | 0.007 | 0.007 | 1.468 | 0.52 | 0.74 | 0.71 | 0.36 | 0.55 | 0.47 |
| 3 | 59413111 | rs9849137 | -0.267 | -0.508 | -0.634 | 0.066 | -0.069 | 0.024 | 0.024 | 3.466 | 0.16 | 0.28 | 0.1 | 0.23 | 0.05 | 0.1 |
| 3 | 59413618 | rs9869521 | -1.34 | -0.628 | -0.859 | -0.128 | -0.026 | 0.012 | 0.012 | 0.147 | 0.32 | 0.39 | 0.34 | 0.36 | 0.27 | 0.29 |
| 3 | 59417186 | rs962791 | -0.722 | -0.25 | -1.079 | -0.352 | 0.001 | -0.025 | -0.025 | 0 | 0.44 | 0.49 | 0.53 | 0.83 | 0.43 | 0.48 |
| 3 | 59422456 | rs4679561 | -0.927 | -0.103 | -0.818 | -0.201 | 0.001 | -0.002 | -0.002 | 0 | 0.52 | 0.58 | 0.71 | 0.73 | 0.62 | 0.61 |
| 3 | 59424638 | rs6810383 | 0.16 | -0.022 | -0.858 | 0.083 | -0.015 | -0.018 | -0.018 | 1.75 | 0.67 | 0.69 | 0.73 | 0.42 | 0.58 | 0.67 |
| 3 | 59424696 | rs17331297 | 0.353 | -0.496 | -0.769 | -0.083 | 0.003 | -0.006 | -0.006 | 0 | 0.14 | 0.1 | 0.11 | 0 | 0.16 | 0.08 |
| 3 | 59424791 | rs12628984 | 0.602 | -0.867 | -0.303 | 0.326 | 0.01 | 0.025 | 0.025 | 0 | 0.41 | 0.38 | 0.46 | 0.21 | 0.34 | 0.43 |
| 3 | 59424841 | rs17060691 | -1.128 | -0.989 | 0.527 | 1.192 | -0.032 | 0.199 | 0.199 | 0 | 0.11 | 0.42 | 0.16 | 0.27 | 0.04 | 0.2 |
| 3 | 59427797 | rs6783485 | 0.359 | -0.693 | -0.17 | 0.864 | -0.034 | -0.008 | -0.008 | 0 | 0.07 | 0.11 | 0.06 | 0.03 | 0.02 | 0.02 |
| 3 | 59430783 | rs12629522 | 0.681 | -0.655 | -0.262 | 0.856 | -0.028 | -0.009 | -0.009 | 0 | 0.06 | 0.1 | 0.05 | 0.02 | 0.01 | 0.02 |
| 3 | 59431342 | rs9813383 | -0.899 | -0.58 | -0.095 | 1.272 | -0.038 | 0.171 | 0.171 | 0 | 0.12 | 0.32 | 0.2 | 0.35 | 0.06 | 0.28 |
| 3 | 59434420 | rs1910236 | -1.028 | -0.122 | -1.532 | 0.296 | -0.042 | 0.282 | 0.282 | 1.306 | 0.52 | 0.92 | 0.61 | 0.89 | 0.45 | 0.48 |
| 3 | 59434498 | rs11926264 | 0.07 | -0.17 | -1.392 | 1.077 | -0.018 | -0.033 | -0.033 | 0.322 | 0.21 | 0.17 | 0.31 | 0.39 | 0.26 | 0.16 |
| 3 | 59435291 | rs4679563 | -1.048 | -0.196 | -1.396 | 1.036 | -0.034 | 0.017 | 0.017 | 0 | 0.34 | 0.28 | 0.55 | 0.68 | 0.44 | 0.32 |
| 3 | 59436205 | rs12490634 | -0.178 | -0.241 | -1.399 | 1.127 | 0 | -0.012 | -0.012 | 0 | 0.1 | 0.09 | 0.17 | 0.27 | 0.12 | 0.12 |
| 3 | 59436405 | rs11130709 | -1.172 | -0.199 | -1.393 | 1.129 | -0.035 | 0.032 | 0.032 | 0 | 0.35 | 0.28 | 0.55 | 0.69 | 0.44 | 0.32 |
| 3 | 59439053 | rs10510809 | -0.351 | -0.165 | -1.121 | 1.233 | 0 | -0.023 | -0.023 | 0 | 0.11 | 0.09 | 0.17 | 0.27 | 0.12 | 0.12 |
| 3 | 59443356 | rs6778006 | 0.062 | -0.427 | -0.625 | 1.102 | -0.016 | 0.119 | 0.119 | 0.395 | 0.1 | 0.47 | 0.09 | 0.2 | 0.03 | 0.17 |
| 3 | 59448694 | rs6762062 | -1.476 | -0.544 | -0.956 | 0.894 | -0.006 | 0.045 | 0.045 | 1.909 | 0.44 | 0.45 | 0.6 | 0.72 | 0.46 | 0.32 |
| 3 | 59449029 | rs2366696 | -0.033 | -0.432 | -1.22 | 0.946 | -0.022 | -0.035 | -0.035 | 0.423 | 0.23 | 0.2 | 0.38 | 0.42 | 0.31 | 0.2 |
| 3 | 59451699 | rs6805741 | -2.026 | -0.475 | -1.265 | 0.832 | -0.099 | 0.3 | 0.3 | 3.851 | 0.55 | 0.95 | 0.7 | 0.91 | 0.5 | 0.49 |
| 3 | 59452272 | rs6790861 | -0.738 | -0.308 | -1.168 | 1.094 | -0.03 | -0.016 | -0.016 | 0 | 0.3 | 0.26 | 0.52 | 0.57 | 0.4 | 0.26 |
| 3 | 59460320 | rs9883195 | -0.188 | -0.217 | -1.046 | 0.953 | -0.001 | 0.005 | 0.005 | 4.026 | 0.08 | 0.11 | 0.14 | 0.14 | 0.1 | 0.07 |
| 3 | 59461873 | rs9855380 | -1.741 | -0.314 | -1.262 | 0.479 | -0.084 | 0.286 | 0.286 | 3.602 | 0.55 | 0.93 | 0.69 | 0.92 | 0.5 | 0.5 |
| 3 | 59463037 | rs10454231 | -0.704 | -0.624 | -0.725 | 0.816 | -0.04 | 0.003 | 0.003 | 1.648 | 0.25 | 0.61 | 0.17 | 0.32 | 0.1 | 0.22 |
| 3 | 59465209 | rs17273249 | 0.867 | -0.495 | -1.39 | 0.036 | 0.006 | 0.094 | 0.094 | 0 | 0.3 | 0.04 | 0.27 | 0.12 | 0.24 | 0.14 |
| 3 | 59469187 | rs6775298 | -0.747 | -0.323 | -0.587 | 0.309 | 0.005 | -0.006 | -0.006 | 0 | 0.65 | 0.7 | 0.63 | 0.58 | 0.67 | 0.5 |
| 3 | 59470380 | rs9870523 | 0.083 | 0.222 | -0.01 | 0.593 | -0.004 | 0.016 | 0.016 | 0 | 0.52 | 0.38 | 0.53 | 0.42 | 0.49 | 0.34 |
| 3 | 59470554 | rs9833420 | 0.52 | 0.305 | 0.251 | 0.593 | 0.01 | 0.007 | 0.007 | 0 | 0.12 | 0.13 | 0.09 | 0.17 | 0.18 | 0.16 |
| 3 | 59470670 | rs4679575 | -0.339 | 0.047 | 0.375 | 0.632 | 0.002 | -0.001 | -0.001 | 0 | 0.59 | 0.41 | 0.57 | 0.41 | 0.59 | 0.4 |
| 3 | 59474911 | rs7626018 | -0.339 | 0.03 | 0.38 | 0.628 | 0.002 | -0.005 | -0.005 | 0 | 0.59 | 0.41 | 0.57 | 0.42 | 0.59 | 0.4 |
| 3 | 59481280 | rs9856979 | -1 | -0.014 | 0.608 | 0.626 | 0.004 | -0.021 | -0.021 | 0 | 0.67 | 0.81 | 0.66 | 0.57 | 0.68 | 0.5 |
| 3 | 59481588 | rs10514716 | 1.356 | 0.067 | 0.425 | 0.649 | 0.03 | 0.068 | 0.068 | 0 | 0.05 | 0.13 | 0.05 | 0.15 | 0.08 | 0.09 |
| 3 | 59481606 | rs10510812 | 1.394 | 0.036 | 0.105 | 0.581 | 0.032 | 0.105 | 0.105 | 0 | 0.31 | 0.07 | 0.29 | 0.11 | 0.24 | 0.15 |
| 3 | 59481821 | rs11130713 | 1.449 | 0.037 | 0.105 | 0.562 | 0.031 | 0.081 | 0.081 | 0 | 0.31 | 0.07 | 0.29 | 0.13 | 0.24 | 0.15 |
| 3 | 59488503 | rs4679582 | -0.299 | 0.012 | 0.333 | 0.556 | 0.011 | 0.002 | 0.002 | 0.121 | 0.27 | 0.33 | 0.29 | 0.28 | 0.35 | 0.28 |
| 3 | 59491762 | rs9866404 | -0.82 | -0.239 | 0.249 | 0.646 | 0.002 | -0.005 | -0.005 | 0 | 0.65 | 0.67 | 0.64 | 0.57 | 0.67 | 0.52 |
| 3 | 59492652 | rs11130714 | 1.09 | 0.138 | -0.458 | 0.572 | -0.008 | 0.103 | 0.103 | 0 | 0.33 | 0.08 | 0.4 | 0.13 | 0.31 | 0.2 |
| 3 | 59494579 | rs17274833 | -0.178 | 0.219 | -0.387 | 0.531 | 0.015 | 0.007 | 0.007 | 0 | 0.53 | 0.3 | 0.55 | 0.4 | 0.57 | 0.41 |
| 3 | 59501351 | rs12629410 | -0.342 | 0.74 | 0 | 0.762 | -0.005 | -0.005 | -0.005 | 1.036 | 0.19 | 0.23 | 0.14 | 0.27 | 0.25 | 0.22 |
| 3 | 59504585 | rs9865717 | 0.093 | 0.984 | 0.324 | 0.784 | 0.037 | 0.05 | 0.05 | 0 | 0.07 | 0.23 | 0.07 | 0.14 | 0.1 | 0.1 |
| 3 | 59504831 | rs6788477 | 0.543 | 0.861 | 0.389 | 0.158 | -0.02 | -0.06 | -0.06 | 0.016 | 0.25 | 0.33 | 0.15 | 0.13 | 0.21 | 0.29 |
| 3 | 59504894 | rs9832635 | -0.42 | 0.876 | 0.428 | 0.139 | 0.025 | 0.019 | 0.019 | 0 | 0.09 | 0.28 | 0.07 | 0.09 | 0.1 | 0.1 |
| 3 | 59505488 | rs17275189 | 1.15 | 0.647 | -0.235 | 0.011 | -0.021 | 0.12 | 0.12 | 0.033 | 0.35 | 0.04 | 0.42 | 0.13 | 0.32 | 0.19 |
| 3 | 59507994 | rs9840238 | -0.353 | 0.748 | 0.24 | -0.017 | 0.015 | -0.003 | -0.003 | 3.509 | 0.12 | 0.31 | 0.09 | 0.1 | 0.12 | 0.12 |
| 3 | 59508024 | rs12487257 | 1.215 | 0.706 | 0.102 | -0.04 | -0.004 | 0.121 | 0.121 | 0 | 0.36 | 0.07 | 0.42 | 0.13 | 0.32 | 0.19 |
| 3 | 59509607 | rs17060879 | 0.924 | 0.783 | 0.142 | -0.005 | 0.009 | 0.014 | 0.014 | 0 | 0.06 | 0.11 | 0.04 | 0.09 | 0.08 | 0.09 |
| 3 | 59511396 | rs11719324 | -0.067 | 0.789 | 0.247 | 0.286 | 0.001 | 0.001 | 0.001 | 0 | 0.42 | 0.4 | 0.28 | 0.43 | 0.47 | 0.49 |
| 3 | 59519764 | rs9855650 | 0.49 | 0.809 | 0.078 | 0.08 | 0 | -0.002 | -0.002 | 0 | 0.48 | 0.45 | 0.51 | 0.23 | 0.44 | 0.31 |
| 3 | 59527111 | rs4679491 | 0.543 | 1.128 | -0.107 | 0.648 | -0.008 | -0.016 | -0.016 | 0 | 0.75 | 0.77 | 0.66 | 0.53 | 0.68 | 0.66 |
| 3 | 59527196 | rs4679492 | -0.084 | 1.117 | -0.083 | 0.879 | -0.036 | -0.078 | -0.078 | 2.419 | 0.83 | 0.93 | 0.71 | 0.53 | 0.76 | 0.69 |
| 3 | 59527322 | rs4679493 | 0.104 | 1.13 | -0.046 | 0.846 | -0.013 | -0.033 | -0.033 | 0 | 0.82 | 0.88 | 0.72 | 0.53 | 0.76 | 0.69 |
| 3 | 59529264 | rs1995416 | 0.275 | 1.089 | 0.124 | 0.856 | 0 | 0.018 | 0.018 | 0.12 | 0.81 | 0.77 | 0.7 | 0.53 | 0.76 | 0.69 |
| 3 | 59529885 | rs13097743 | 1.061 | 1.046 | 0.115 | 0.546 | 0.012 | -0.005 | -0.005 | 0 | 0.61 | 0.46 | 0.43 | 0.36 | 0.51 | 0.5 |
| 3 | 59535997 | rs10510814 | 0.433 | 1.307 | -0.094 | 0.766 | -0.044 | -0.042 | -0.042 | 0.603 | 0.07 | 0.01 | 0.16 | 0.23 | 0.18 | 0.13 |
| 3 | 59536406 | rs17060928 | 0.841 | 1.066 | 0.108 | 0.207 | 0 | -0.001 | -0.001 | 0.016 | 0.16 | 0.11 | 0.13 | 0.16 | 0.18 | 0.16 |
| 3 | 59541974 | rs9841772 | 0.161 | 1.165 | -0.232 | -0.084 | 0.001 | 0.013 | 0.013 | 0 | 0.5 | 0.53 | 0.41 | 0.22 | 0.44 | 0.32 |
| 3 | 59542556 | rs12491058 | 0.418 | 0.875 | -0.134 | 0.155 | -0.007 | 0.004 | 0.004 | 0 | 0.18 | 0.11 | 0.24 | 0.16 | 0.24 | 0.19 |
| 3 | 59550000 | rs7619681 | 0.561 | 0.941 | -0.32 | 0.267 | 0 | -0.001 | -0.001 | 0 | 0.3 | 0.28 | 0.18 | 0.31 | 0.26 | 0.34 |
| 3 | 59554896 | rs734184 | -0.312 | 1.217 | -0.699 | 0.443 | 0.008 | -0.002 | -0.002 | 3.414 | 0.51 | 0.66 | 0.45 | 0.48 | 0.5 | 0.55 |
| 3 | 59559489 | rs6765225 | 0.134 | 1.303 | -1.093 | 0.444 | -0.027 | -0.011 | -0.011 | 1.384 | 0.22 | 0.1 | 0.16 | 0.26 | 0.25 | 0.28 |

| | | | | | | | | | | | | | | | | |
|---|---|---|---|---|---|---|---|---|---|---|---|---|---|---|---|---|
| 3 | 59562365 | rs4679495 | -0.432 | 0.975 | -0.669 | -0.004 | -0.001 | -0.04 | -0.04 | 0 | 0.61 | 0.45 | 0.58 | 0.85 | 0.69 | 0.73 |
| 3 | 59567691 | rs7619446 | -0.168 | 1.144 | -0.437 | 0.42 | 0.003 | 0.012 | 0.012 | 0 | 0.07 | 0.11 | 0.22 | 0.13 | 0.08 | 0.14 |
| 3 | 59570968 | rs7615380 | -0.288 | 0.688 | -0.126 | 1.028 | -0.012 | 0.109 | 0.109 | 0 | 0.07 | 0.17 | 0.08 | 0.29 | 0.06 | 0.11 |
| 3 | 59576007 | rs6791284 | 0.113 | 0.79 | -0.088 | 0.813 | -0.008 | -0.024 | -0.024 | 0 | 0.33 | 0.22 | 0.38 | 0.43 | 0.35 | 0.43 |
| 3 | 59579562 | rs6796064 | 0.687 | 0.675 | -0.095 | 0.569 | -0.001 | 0.003 | 0.003 | 0 | 0.11 | 0.13 | 0.2 | 0.2 | 0.13 | 0.21 |
| 3 | 59580150 | rs4679496 | -0.887 | 0.539 | 0.208 | 0.288 | 0.014 | 0.037 | 0.037 | 0 | 0.52 | 0.58 | 0.66 | 0.7 | 0.56 | 0.71 |
| 3 | 59587934 | rs9863719 | -0.084 | 0.619 | 0.376 | 0.566 | -0.018 | 0.1 | 0.1 | 0.052 | 0.09 | 0.26 | 0.1 | 0.29 | 0.06 | 0.12 |
| 3 | 59588996 | rs12634709 | -0.466 | 0.669 | 0.473 | 0.509 | 0.015 | 0.015 | 0.015 | 0.411 | 0.17 | 0.3 | 0.28 | 0.26 | 0.21 | 0.28 |
| 3 | 59589215 | rs4679497 | -1.144 | 0.716 | 0.258 | 0.345 | -0.001 | 0.122 | 0.122 | 4.741 | 0.26 | 0.58 | 0.36 | 0.54 | 0.27 | 0.4 |
| 3 | 59592180 | rs9810248 | -0.836 | 0.449 | -0.058 | -0.334 | 0.003 | 0.013 | 0.013 | 0 | 0.51 | 0.57 | 0.69 | 0.94 | 0.53 | 0.75 |
| 3 | 59593494 | rs13325549 | 0.529 | 0.47 | -0.479 | -0.09 | -0.025 | 0.029 | 0.029 | 0 | 0.29 | 0.15 | 0.22 | 0.2 | 0.37 | 0.3 |
| 3 | 59594892 | rs11720729 | 0.493 | 0.467 | -0.48 | -0.049 | -0.025 | 0.023 | 0.023 | 0 | 0.3 | 0.15 | 0.22 | 0.22 | 0.36 | 0.3 |
| 3 | 59598180 | rs1504912 | 0.576 | 0.533 | -0.122 | 0.424 | 0.021 | 0.137 | 0.137 | 0 | 0.06 | 0.11 | 0.19 | 0.27 | 0.05 | 0.12 |
| 3 | 59620429 | rs1281447 | -1.586 | 0.267 | -0.268 | -0.385 | -0.016 | 0.053 | 0.053 | 0.069 | 0.64 | 0.77 | 0.77 | 1 | 0.65 | 0.85 |
| 3 | 59620816 | rs4679500 | -0.177 | 0.029 | 0.123 | -0.18 | -0.006 | 0.03 | 0.03 | 0.184 | 0.12 | 0.15 | 0.14 | 0.29 | 0.17 | 0.2 |
| 3 | 59621041 | rs1281449 | -0.126 | -0.174 | 0.189 | -0.216 | -0.001 | 0.089 | 0.089 | 0 | 0.21 | 0.33 | 0.41 | 0.46 | 0.18 | 0.39 |
| 3 | 59622981 | rs2292130 | 0.717 | -0.133 | -0.008 | -0.35 | -0.008 | 0.016 | 0.016 | 1.366 | 0.29 | 0.2 | 0.21 | 0.23 | 0.32 | 0.26 |
| 3 | 59625877 | rs3821473 | 1.405 | 0.287 | -0.077 | -0.382 | 0.009 | 0.002 | 0.002 | 2.441 | 0.24 | 0.15 | 0.13 | 0.23 | 0.24 | 0.23 |
| 3 | 59626208 | rs1281459 | -1.779 | 0.304 | -0.076 | -0.55 | -0.012 | 0.04 | 0.04 | 0.103 | 0.64 | 0.77 | 0.77 | 1 | 0.66 | 0.85 |
| 3 | 59631424 | rs11914539 | 0.221 | 0.535 | 0.042 | 0.028 | -0.008 | -0.018 | -0.018 | 0 | 0.08 | 0.14 | 0.08 | 0.05 | 0.11 | 0.1 |
| 3 | 59635215 | rs9831300 | 0.1 | 0.462 | -0.369 | -0.046 | -0.001 | 0.016 | 0.016 | 1.289 | 0.68 | 0.63 | 0.52 | 0.54 | 0.72 | 0.55 |
| 3 | 59641369 | rs9872658 | -0.187 | 0.614 | -0.401 | 0.511 | -0.005 | -0.005 | -0.005 | 0 | 0.1 | 0.1 | 0.14 | 0.2 | 0.14 | 0.15 |
| 3 | 59643631 | rs17061008 | 0.258 | 0.473 | -1.041 | 0.12 | -0.043 | -0.05 | -0.05 | 0 | 0.4 | 0.09 | 0.3 | 0.47 | 0.43 | 0.42 |
| 3 | 59644233 | rs2121794 | -0.114 | 0.813 | 0.161 | 0.611 | 0.006 | 0.04 | 0.04 | 0 | 0.71 | 0.65 | 0.52 | 0.54 | 0.74 | 0.59 |
| 3 | 59649521 | rs6763159 | 0.606 | 0.704 | 0.151 | 0.522 | 0.004 | -0.005 | -0.005 | 0.031 | 0.59 | 0.44 | 0.37 | 0.34 | 0.6 | 0.44 |
| 3 | 59649815 | rs13066895 | 1.216 | 0.762 | -0.259 | 0.467 | 0.008 | 0.126 | 0.126 | 0.878 | 0.5 | 0.22 | 0.28 | 0.27 | 0.47 | 0.34 |
| 3 | 59658825 | rs9311717 | -1.46 | 0.65 | 0.559 | 0.93 | 0.005 | 0.009 | 0.009 | 1.404 | 0.25 | 0.52 | 0.43 | 0.34 | 0.32 | 0.4 |
| 3 | 59664687 | rs1281465 | -1.564 | 0.77 | 0.629 | 1.034 | -0.005 | -0.008 | -0.008 | 0 | 0.44 | 0.59 | 0.65 | 0.88 | 0.45 | 0.64 |
| 3 | 59675336 | rs7634315 | -1.665 | 0.632 | 0.602 | 1.85 | -0.012 | 0.028 | 0.028 | 3.088 | 0.18 | 0.68 | 0.24 | 0.3 | 0.1 | 0.26 |
| 3 | 59676546 | rs1447820 | -0.277 | 0.703 | 0.442 | 2.247 | -0.007 | 0.002 | 0.002 | 0 | 0.26 | 0.25 | 0.37 | 0.26 | 0.27 | 0.34 |
| 3 | 59678303 | rs7635981 | -2.144 | 0.693 | 0.038 | 2.198 | -0.072 | -0.071 | -0.071 | 0 | 0.44 | 0.93 | 0.61 | 0.56 | 0.37 | 0.59 |
| 3 | 59685402 | rs11927857 | -0.586 | 0.62 | 0.693 | 1.757 | -0.001 | 0.003 | 0.003 | 0.476 | 0.26 | 0.21 | 0.21 | 0.16 | 0.22 | 0.25 |
| 3 | 59686286 | rs17292696 | -0.141 | 0.76 | 0.244 | 1.794 | -0.039 | 0.028 | 0.028 | 0 | 0.22 | 0.05 | 0.2 | 0.15 | 0.22 | 0.25 |
| 3 | 59688158 | rs17061076 | 0.564 | 0.624 | 0.257 | 1.864 | 0.025 | -0.013 | -0.013 | 0.024 | 0.17 | 0.03 | 0.09 | 0.14 | 0.12 | 0.2 |
| 3 | 59692543 | rs7432640 | -0.667 | 0.788 | 1.392 | 2.131 | -0.017 | 0.021 | 0.021 | 0.822 | 0.25 | 0.42 | 0.33 | 0.31 | 0.21 | 0.45 |
| 3 | 59695594 | rs4679599 | -1.531 | 0.97 | 1.119 | 2.294 | 0.001 | 0.08 | 0.08 | 0 | 0.12 | 0.19 | 0.17 | 0.56 | 0.1 | 0.17 |
| 3 | 59696535 | rs4679600 | -1.034 | 1.106 | 1.191 | 1.753 | 0.033 | -0.005 | -0.005 | 0.024 | 0.08 | 0.17 | 0.12 | 0.04 | 0.09 | 0.15 |
| 3 | 59700603 | rs17061096 | -1.27 | 0.33 | 1.322 | 2.101 | 0.007 | 0.089 | 0.089 | 0.109 | 0.1 | 0.34 | 0.13 | 0.15 | 0.04 | 0.14 |
| 3 | 59700722 | rs6446079 | -0.344 | 0.381 | 0.64 | 2.096 | -0.012 | 0.038 | 0.038 | 0 | 0.22 | 0.2 | 0.15 | 0.53 | 0.19 | 0.27 |
| 3 | 59701430 | rs1868190 | -0.468 | 0.41 | 0.618 | 1.866 | -0.007 | 0.018 | 0.018 | 0 | 0.23 | 0.2 | 0.29 | 0.6 | 0.2 | 0.28 |
| 3 | 59708761 | rs6446081 | -0.146 | 0.337 | 0.314 | 1.348 | 0.066 | -0.067 | -0.067 | 0.319 | 0.61 | 0.89 | 0.82 | 0.46 | 0.73 | 0.65 |
| 3 | 59709050 | rs17359485 | 1.417 | 0.379 | 0.261 | 1.192 | -0.071 | 0.055 | 0.055 | 0.39 | 0.36 | 0.15 | 0.42 | 0.24 | 0.49 | 0.37 |
| 3 | 59711017 | rs7373248 | -0.26 | 0.685 | 0.667 | 1.256 | 0.144 | 0.032 | 0.032 | 0.002 | 0.07 | 0.31 | 0.25 | 0.06 | 0.13 | 0.1 |
| 3 | 59716384 | rs1947152 | 0.361 | 0.584 | 0.516 | 1.244 | 0.001 | 0 | 0 | 0 | 0.09 | 0.14 | 0.08 | 0.05 | 0.09 | 0.1 |
| 3 | 59716457 | rs1456155 | 0.633 | 0.491 | 0.614 | 1.248 | 0.002 | 0.038 | 0.038 | 1.636 | 0.46 | 0.32 | 0.5 | 0.31 | 0.57 | 0.46 |
| 3 | 59718817 | rs9311725 | 0.143 | 0.951 | 0.768 | 0.883 | -0.001 | -0.001 | -0.001 | 0 | 0.08 | 0.11 | 0.07 | 0.05 | 0.09 | 0.1 |
| 3 | 59719172 | rs17359709 | -2.104 | 0.914 | 0.713 | 0.898 | -0.003 | 0 | 0 | 0 | 0.25 | 0.77 | 0.41 | 0.23 | 0.25 | 0.29 |
| 3 | 59727802 | rs1155530 | -0.39 | 0.351 | 0.109 | 0.675 | 0.002 | 0.102 | 0.102 | 0.421 | 0.71 | 0.94 | 0.84 | 0.9 | 0.76 | 0.73 |
| 3 | 59731151 | rs780373 | -2.044 | 0.236 | 0.694 | 0.828 | 0.002 | 0.007 | 0.007 | 2.93 | 0.25 | 0.66 | 0.35 | 0.3 | 0.21 | 0.27 |
| 3 | 59733262 | rs1628424 | -1.002 | 0.277 | 0.867 | 0.59 | -0.006 | -0.019 | -0.019 | 0 | 0.16 | 0.25 | 0.11 | 0.07 | 0.12 | 0.16 |
| 3 | 59734372 | rs293602 | -0.844 | 0.289 | 0.572 | 0.585 | 0.102 | 0.048 | 0.048 | 0.322 | 0.16 | 0.49 | 0.29 | 0.19 | 0.18 | 0.19 |
| 3 | 59736299 | rs812965 | -0.384 | 0.217 | -0.664 | 0.49 | 0.031 | 0.105 | 0.105 | 2.052 | 0.72 | 0.93 | 0.85 | 0.92 | 0.78 | 0.72 |
| 3 | 59736584 | rs2077774 | 0.959 | 0.4 | -0.665 | 0.734 | -0.046 | 0.134 | 0.134 | 0.269 | 0.47 | 0.12 | 0.5 | 0.73 | 0.57 | 0.5 |
| 3 | 59739919 | rs6803403 | -0.975 | 0.386 | 0.653 | 1.21 | -0.01 | 0.086 | 0.086 | 0.124 | 0.25 | 0.39 | 0.18 | 0.43 | 0.16 | 0.5 |
| 3 | 59741515 | rs1385715 | -0.922 | 0.259 | 0.852 | 0.919 | 0.019 | 0.015 | 0.015 | 0 | 0.45 | 0.55 | 0.27 | 0.37 | 0.27 | 0.56 |
| 3 | 59741758 | rs994931 | 0.051 | 0.59 | 0.905 | 0.686 | 0.079 | -0.018 | -0.018 | 0 | 0.06 | 0.27 | 0.23 | 0 | 0.08 | 0.06 |
| 3 | 59745585 | rs993000 | 0.159 | 0.39 | 1.292 | 0.908 | -0.014 | -0.023 | -0.023 | 0 | 0.31 | 0.32 | 0.25 | 0.1 | 0.14 | 0.32 |
| 3 | 59749703 | rs1683346 | 0.331 | 0.451 | 1.188 | 0.974 | -0.02 | -0.032 | -0.032 | 0 | 0.61 | 0.6 | 0.7 | 0.87 | 0.81 | 0.6 |
| 3 | 59750791 | rs1126268 | 1.044 | 0.546 | 1.063 | 1.176 | 0.028 | 0.114 | 0.114 | 0 | 0.51 | 0.2 | 0.39 | 0.75 | 0.65 | 0.4 |
| 3 | 59754022 | rs167140 | -0.212 | 0.772 | 0.734 | 1.121 | -0.015 | -0.012 | -0.012 | 0 | 0.08 | 0.04 | 0.07 | 0.12 | 0.14 | 0.08 |
| 3 | 59754178 | rs17061173 | 0.73 | 0.575 | 0.881 | 0.902 | 0.036 | 0.045 | 0.045 | 0.994 | 0.25 | 0.09 | 0.21 | 0.09 | 0.11 | 0.27 |
| 3 | 59755570 | rs17295070 | 0.13 | 0.608 | 0.914 | 0.91 | 0 | -0.005 | -0.005 | 0 | 0.3 | 0.28 | 0.43 | 0.1 | 0.2 | 0.33 |
| 3 | 59755841 | rs17061176 | -0.237 | 0.613 | 1.032 | 0.898 | 0.01 | 0.024 | 0.024 | 0 | 0.05 | 0.08 | 0.08 | 0.12 | 0.07 | 0.15 |
| 3 | 59756364 | rs4679606 | 0.229 | 0.6 | 0.894 | 0.771 | -0.001 | -0.007 | -0.007 | 0 | 0.3 | 0.3 | 0.43 | 0.11 | 0.2 | 0.33 |
| 3 | 59756690 | rs17061180 | 1.424 | 0.928 | 0.242 | 0.665 | -0.008 | -0.078 | -0.078 | 0 | 0.4 | 0.15 | 0.26 | 0.53 | 0.43 | 0.29 |
| 3 | 59757484 | rs49413 | -0.346 | 0.724 | 0.43 | 0.878 | 0.012 | -0.032 | -0.032 | 0 | 0.7 | 0.8 | 0.67 | 0.6 | 0.79 | 0.56 |
| 3 | 59757575 | rs49411 | 0.61 | 0.845 | 0.362 | 0.606 | -0.005 | 0.002 | 0.002 | 0 | 0.46 | 0.36 | 0.32 | 0.61 | 0.57 | 0.39 |
| 3 | 59759572 | rs780362 | -0.324 | 1.276 | 0.393 | 0.334 | 0.003 | -0.017 | -0.017 | 0 | 0.07 | 0.06 | 0.09 | 0 | 0.15 | 0.07 |
| 3 | 59761158 | rs492836 | 0.468 | 1.182 | 0.5 | 0.487 | 0.009 | 0.045 | 0.045 | 0.017 | 0.5 | 0.29 | 0.36 | 0.42 | 0.57 | 0.32 |

| | | | | | | | | | | | | | | | | |
|---|---|---|---|---|---|---|---|---|---|---|---|---|---|---|---|---|
| 3 | 59761789 | rs780365 | -0.15 | 1.153 | 1.398 | 0.367 | -0.004 | -0.002 | -0.002 | 0 | 0.59 | 0.56 | 0.59 | 0.42 | 0.69 | 0.4 |
| 3 | 59762008 | rs517711 | -0.08 | 1.195 | 1.254 | 0.348 | -0.006 | -0.003 | -0.003 | 0 | 0.1 | 0.07 | 0.11 | 0 | 0.17 | 0.1 |
| 3 | 59762033 | rs517644 | 0.396 | 1.169 | 1.389 | 0.389 | 0 | 0.033 | 0.033 | 0.003 | 0.52 | 0.32 | 0.36 | 0.42 | 0.57 | 0.35 |
| 3 | 59762116 | rs780366 | 0.476 | 1.225 | 1.297 | 0.44 | 0.109 | -0.017 | -0.017 | 0 | 0.93 | 0.67 | 0.76 | 1 | 0.88 | 0.94 |
| 3 | 59768262 | rs4679610 | 1.252 | 1.169 | 0.831 | 0.666 | -0.004 | -0.027 | -0.027 | 0 | 0.4 | 0.25 | 0.28 | 0.5 | 0.41 | 0.27 |
| 3 | 59771515 | rs17361653 | 0.851 | 1.215 | 0.751 | 0.59 | -0.003 | 0.004 | 0.004 | 0 | 0.44 | 0.4 | 0.47 | 0.41 | 0.51 | 0.28 |
| 3 | 59772328 | rs17361780 | -0.193 | 1.662 | 0.264 | 0.414 | 0.099 | -0.02 | -0.02 | 0 | 0.06 | 0.21 | 0.23 | 0 | 0.12 | 0.07 |
| 3 | 59772342 | rs13092167 | 0.177 | 1.343 | -0.167 | 0.402 | 0.005 | -0.047 | -0.047 | 0 | 0.33 | 0.14 | 0.33 | 0.41 | 0.28 | 0.47 |
| 3 | 59772460 | rs780354 | -1.196 | 1.727 | 0.248 | 0.155 | 0.025 | -0.071 | -0.071 | 0 | 0.17 | 0.25 | 0.33 | 0.01 | 0.28 | 0.17 |
| 3 | 59781217 | rs1905864 | 0.974 | 1.448 | 0.353 | 0.765 | 0.043 | 0.053 | 0.053 | 2.341 | 0.74 | 0.5 | 0.73 | 0.67 | 0.67 | 0.68 |
| 3 | 59785980 | rs6781354 | 0.697 | 1.528 | 0.794 | 0.746 | 0.005 | 0.021 | 0.021 | 4.25 | 0.84 | 0.79 | 0.82 | 0.66 | 0.8 | 0.77 |
| 3 | 59787574 | rs670592 | 1.386 | 1.147 | 1.459 | 0.69 | 0.034 | 0.232 | 0.232 | 1.156 | 0.84 | 0.55 | 0.84 | 0.61 | 0.84 | 0.77 |
| 3 | 59788967 | rs589792 | -1.797 | 0.986 | 1.804 | 0.478 | 0.053 | -0.043 | -0.043 | 0 | 0.1 | 0.39 | 0.11 | 0 | 0.13 | 0.09 |
| 3 | 59789058 | rs12634148 | 2.545 | 0.715 | 0.731 | 0.237 | -0.07 | -0.116 | -0.116 | 0 | 0.55 | 0.05 | 0.6 | 0.53 | 0.55 | 0.44 |
| 3 | 59789614 | rs534185 | 0.908 | 0.525 | 1.501 | 0.595 | 0.041 | 0.266 | 0.266 | 0.446 | 0.88 | 0.67 | 0.85 | 0.56 | 0.86 | 0.76 |
| 3 | 59795925 | rs2251216 | 0.996 | 0.537 | 1.329 | 0.613 | 0.022 | 0.133 | 0.133 | 1.841 | 0.87 | 0.79 | 0.85 | 0.57 | 0.83 | 0.73 |
| 3 | 59802813 | rs6778206 | -1.422 | 0.329 | 0.964 | 0.185 | -0.01 | -0.039 | -0.039 | 0.207 | 0.14 | 0.14 | 0.1 | 0 | 0.09 | 0.18 |
| 3 | 59810153 | rs11130736 | 1.077 | 0.752 | 0.225 | 0.607 | -0.023 | 0.048 | 0.048 | 0.257 | 0.53 | 0.16 | 0.56 | 0.42 | 0.43 | 0.43 |
| 3 | 59810867 | rs4679614 | 1.115 | 0.856 | 0.881 | 0.485 | 0.005 | -0.091 | -0.091 | 0 | 0.84 | 0.4 | 0.83 | 0.97 | 0.81 | 0.9 |
| 3 | 59828477 | rs3112497 | 0.85 | 0.366 | 0.936 | -0.069 | 0.01 | -0.009 | -0.009 | 0 | 0.59 | 0.21 | 0.69 | 0.44 | 0.63 | 0.52 |
| 3 | 59842644 | rs684584 | -1.244 | 0.164 | 1.042 | -0.28 | 0.019 | 0.023 | 0.023 | 0.388 | 0.43 | 0.81 | 0.33 | 0.59 | 0.41 | 0.53 |
| 3 | 59846067 | rs12487204 | 1.585 | 0.071 | 0.87 | -0.353 | -0.032 | 0.044 | 0.044 | 0.373 | 0.49 | 0.09 | 0.58 | 0.41 | 0.49 | 0.45 |
| 3 | 59846186 | rs678981 | -0.974 | 0.096 | 1.15 | -0.313 | 0.002 | -0.029 | -0.029 | 0 | 0.07 | 0.13 | 0.08 | 0 | 0.12 | 0.08 |
| 3 | 59848463 | rs6446085 | -0.488 | 0.137 | 1.45 | -0.353 | -0.013 | 0.006 | 0.006 | 0 | 0.33 | 0.47 | 0.21 | 0.59 | 0.27 | 0.45 |
| 3 | 59849088 | rs646372 | -1.501 | 0.156 | 1.215 | -0.336 | 0.005 | 0.05 | 0.05 | 0.831 | 0.48 | 0.78 | 0.37 | 0.58 | 0.49 | 0.54 |
| 3 | 59859953 | rs662910 | -1.709 | 0.177 | 1.597 | -1.101 | 0.006 | -0.04 | -0.04 | 0 | 0.11 | 0.2 | 0.13 | 0 | 0.2 | 0.08 |
| 3 | 59871049 | rs602197 | 0.792 | -1.386 | 1.597 | -1.764 | 0.027 | -0.009 | -0.009 | 0 | 0.64 | 0.26 | 0.77 | 0.99 | 0.81 | 0.87 |
| 3 | 59875155 | rs17061352 | -1.798 | -0.298 | 1.559 | -1.136 | -0.025 | -0.04 | -0.04 | 0 | 0.09 | 0.22 | 0.04 | 0.01 | 0.04 | 0.09 |
| 3 | 59875996 | rs11924342 | -1.338 | -0.29 | 1.532 | -1.132 | -0.021 | -0.034 | -0.034 | 1.523 | 0.08 | 0.16 | 0.04 | 0.01 | 0.04 | 0.08 |
| 3 | 59877988 | rs17061377 | 0.971 | -0.246 | 1.298 | -1.29 | -0.012 | 0.052 | 0.052 | 5.236 | 0.11 | 0.03 | 0.03 | 0 | 0.1 | 0.13 |
| 3 | 59880142 | rs17299801 | -2.269 | -0.38 | 1.971 | -0.749 | -0.023 | -0.071 | -0.071 | 0.519 | 0.14 | 0.24 | 0.09 | 0.01 | 0.11 | 0.09 |
| 3 | 59884212 | rs9311739 | -2.399 | -0.294 | 1.753 | -0.699 | -0.03 | -0.114 | -0.114 | 0 | 0.18 | 0.6 | 0.17 | 0.01 | 0.13 | 0.09 |
| 3 | 59884453 | rs9883913 | -1.268 | -0.147 | 1.284 | -0.603 | 0.021 | -0.057 | -0.057 | 0.327 | 0.1 | 0.3 | 0.1 | 0 | 0.09 | 0.02 |
| 3 | 59889113 | rs9844365 | -1.56 | -0.201 | 0.274 | -0.597 | -0.029 | -0.091 | -0.091 | 0 | 0.17 | 0.66 | 0.17 | 0.01 | 0.11 | 0.1 |
| 3 | 59890976 | rs643629 | -0.256 | -0.109 | -0.211 | -0.267 | 0 | -0.004 | -0.004 | 0 | 0.1 | 0.09 | 0.05 | 0.11 | 0.14 | 0.11 |
| 3 | 59891010 | rs7641743 | 0.528 | 0.159 | -0.67 | -1.054 | 0.013 | 0.099 | 0.099 | 0 | 0.49 | 0.32 | 0.62 | 0.81 | 0.46 | 0.61 |
| 3 | 59891227 | rs11715245 | 0.714 | 0.321 | -0.495 | -1.225 | -0.036 | 0.099 | 0.099 | 0 | 0.3 | 0.11 | 0.19 | 0.03 | 0.34 | 0.23 |
| 3 | 59892473 | rs1026989 | 0.604 | 0.352 | -1.032 | -0.629 | 0.003 | -0.035 | -0.035 | 0 | 0.25 | 0.17 | 0.2 | 0.42 | 0.22 | 0.22 |
| 3 | 59893532 | rs9810994 | 0.805 | 0.194 | -0.873 | -0.552 | -0.024 | 0.158 | 0.158 | 0.194 | 0.1 | 0.24 | 0.08 | 0.39 | 0.04 | 0.11 |
| 3 | 59894061 | rs1485727 | 1.962 | 0.493 | -1.322 | -0.722 | -0.056 | -0.031 | -0.031 | 0 | 0.33 | 0.09 | 0.47 | 0.37 | 0.34 | 0.41 |
| 3 | 59898057 | rs294459 | -0.833 | 0.412 | -0.922 | -0.774 | -0.012 | 0.053 | 0.053 | 1.183 | 0.24 | 0.37 | 0.19 | 0.51 | 0.24 | 0.22 |
| 3 | 59898962 | rs4679622 | 2.443 | 0.749 | -1.097 | -0.814 | -0.049 | -0.058 | -0.058 | 0 | 0.3 | 0.08 | 0.46 | 0.39 | 0.32 | 0.41 |
| 3 | 59899215 | rs294457 | 0.717 | 0.565 | -0.558 | -0.923 | -0.029 | 0.089 | 0.089 | 0.004 | 0.75 | 0.51 | 0.76 | 0.56 | 0.83 | 0.78 |
| 3 | 59899683 | rs17366548 | 0.647 | 0.817 | -0.637 | -0.765 | -0.048 | 0.101 | 0.101 | 0 | 0.3 | 0.12 | 0.17 | 0.03 | 0.35 | 0.24 |
| 3 | 59900220 | rs7349527 | 1.573 | 0.865 | -0.521 | -0.689 | -0.011 | -0.001 | -0.001 | 6.021 | 0.15 | 0.1 | 0.08 | 0 | 0.19 | 0.13 |
| 3 | 59900251 | rs6787894 | 0.699 | 0.745 | 0.186 | -0.375 | 0.02 | 0.049 | 0.049 | 0.032 | 0.06 | 0.16 | 0.06 | 0.05 | 0.02 | 0.03 |
| 3 | 59900739 | rs294454 | 0.08 | 0.756 | -0.565 | -0.392 | -0.017 | 0.092 | 0.092 | 0.183 | 0.43 | 0.23 | 0.25 | 0.15 | 0.48 | 0.36 |
| 3 | 59900929 | rs294453 | 0.168 | 0.589 | -0.609 | -0.397 | -0.006 | 0.102 | 0.102 | 0.209 | 0.42 | 0.19 | 0.23 | 0.14 | 0.44 | 0.35 |
| 3 | 59902252 | rs7612797 | 0.727 | 0.801 | -0.626 | -0.406 | -0.067 | 0.016 | 0.016 | 0 | 0.42 | 0.17 | 0.58 | 0.43 | 0.48 | 0.54 |
| 3 | 59902801 | rs1825630 | -0.443 | 0.963 | -0.594 | -0.365 | 0.004 | 0.005 | 0.005 | 0 | 0.5 | 0.43 | 0.34 | 0.52 | 0.49 | 0.42 |
| 3 | 59904539 | rs1973780 | 0.513 | 0.619 | -0.953 | -1.05 | 0.04 | 0.183 | 0.183 | 0 | 0.5 | 0.3 | 0.63 | 0.92 | 0.62 | 0.68 |
| 3 | 59904811 | rs9825594 | -0.556 | 1.153 | -0.915 | -0.734 | -0.004 | 0.002 | 0.002 | 0 | 0.42 | 0.56 | 0.52 | 0.35 | 0.5 | 0.41 |
| 3 | 59905777 | rs1385815 | -0.48 | 1.279 | -0.781 | -0.934 | -0.007 | 0.021 | 0.021 | 0 | 0.21 | 0.18 | 0.27 | 0.01 | 0.24 | 0.22 |
| 3 | 59906598 | rs2293212 | -1.629 | 1.173 | -0.499 | -0.902 | -0.001 | 0.001 | 0.001 | 0 | 0.4 | 0.59 | 0.58 | 0.43 | 0.38 | 0.45 |
| 3 | 59907329 | rs9311741 | -1.962 | 1.234 | -0.471 | -0.742 | 0 | -0.006 | -0.006 | 0 | 0.59 | 0.65 | 0.71 | 0.43 | 0.6 | 0.57 |
| 3 | 59908126 | rs1385816 | -0.567 | 1.367 | -0.529 | -0.484 | -0.013 | -0.001 | -0.001 | 0.675 | 0.09 | 0.07 | 0.34 | 0.33 | 0.2 | 0.26 |
| 3 | 59911566 | rs12489018 | -1.363 | 1.116 | 0.056 | -0.625 | 0 | -0.029 | -0.029 | 0.043 | 0.53 | 0.46 | 0.54 | 0.1 | 0.54 | 0.39 |
| 3 | 59916314 | rs2366968 | 0.968 | 1.03 | -0.01 | -0.969 | 0.003 | -0.045 | -0.045 | 0 | 0.5 | 0.49 | 0.52 | 0.98 | 0.49 | 0.62 |
| 3 | 59920004 | rs1385812 | 0.834 | 0.963 | 0.247 | -0.981 | 0.011 | -0.035 | -0.035 | 0 | 0.81 | 0.62 | 0.76 | 0.9 | 0.82 | 0.79 |
| 3 | 59920697 | rs11719244 | 0.994 | 1.02 | 0.053 | -1.119 | -0.005 | 0.014 | 0.014 | 0.005 | 0.21 | 0.14 | 0.2 | 0.08 | 0.22 | 0.12 |
| 3 | 59920912 | rs17061494 | 0.497 | 0.881 | -0.001 | -1.128 | -0.002 | -0.023 | -0.023 | 4.629 | 0.08 | 0.1 | 0.11 | 0.01 | 0.11 | 0.13 |
| 3 | 59921786 | rs17374239 | 0.766 | 0.856 | -0.229 | -1.185 | -0.018 | 0.019 | 0.019 | 0 | 0.06 | 0.01 | 0.15 | 0.01 | 0.11 | 0.03 |
| 3 | 59922017 | rs3772448 | 1.283 | 1.113 | -0.073 | -1.369 | 0 | -0.013 | -0.013 | 0.134 | 0.51 | 0.52 | 0.47 | 0.9 | 0.46 | 0.6 |
| 3 | 59923459 | rs17374441 | 0.603 | 1.1 | -0.596 | -1.309 | 0.003 | 0.075 | 0.075 | 0 | 0.14 | 0.04 | 0.05 | 0.01 | 0.08 | 0.13 |
| 3 | 59923514 | rs3772453 | 0.501 | 1.509 | -0.726 | -0.947 | -0.045 | -0.073 | -0.073 | 0 | 0.11 | 0.01 | 0.17 | 0.34 | 0.11 | 0.22 |
| 3 | 59924068 | rs6802153 | 0.869 | 1.414 | -0.371 | -0.968 | -0.003 | 0.003 | 0.003 | 0 | 0.22 | 0.2 | 0.2 | 0.08 | 0.25 | 0.14 |
| 3 | 59924843 | rs17061517 | 1.195 | 1.057 | -0.858 | -0.833 | -0.023 | -0.086 | -0.086 | 0 | 0.14 | 0.03 | 0.16 | 0.51 | 0.15 | 0.25 |
| 3 | 59925591 | rs9877140 | -0.039 | 0.983 | -0.25 | -1.213 | 0 | -0.027 | -0.027 | 0 | 0.21 | 0.24 | 0.21 | 0.02 | 0.17 | 0.15 |
| 3 | 59926256 | rs931322 | 0.602 | 1.114 | -0.218 | -0.849 | -0.003 | 0.095 | 0.095 | 0 | 0.46 | 0.28 | 0.4 | 0.89 | 0.46 | 0.58 |

| | | | | | | | | | | | | | | | | |
|---|---|---|---|---|---|---|---|---|---|---|---|---|---|---|---|---|
| 3 | 59930672 | rs2366964 | -0.614 | 0.545 | 0.334 | 0.509 | 0 | 0.188 | 0.188 | 0 | 0.11 | 0.31 | 0.18 | 0.33 | 0.08 | 0.11 |
| 3 | 59933791 | rs9819930 | -0.701 | 0.644 | 0.309 | 0.42 | 0.002 | 0.082 | 0.082 | 0 | 0.14 | 0.28 | 0.18 | 0.34 | 0.1 | 0.14 |
| 3 | 59935451 | rs3821476 | 1.722 | 1.013 | -0.145 | -0.052 | -0.051 | -0.059 | -0.059 | 2.667 | 0.67 | 0.18 | 0.46 | 0.59 | 0.7 | 0.67 |
| 3 | 59939093 | rs12487051 | 0.733 | 0.898 | -0.041 | -0.269 | 0.005 | -0.035 | -0.035 | 0 | 0.08 | 0.14 | 0.13 | 0 | 0.12 | 0.15 |
| 3 | 59940252 | rs9862003 | 0.573 | 1.063 | 0.679 | -0.365 | -0.002 | -0.046 | -0.046 | 0 | 0.8 | 0.69 | 0.71 | 0.93 | 0.77 | 0.73 |
| 3 | 59940778 | rs13081936 | 2.219 | 0.2 | -0.067 | -0.443 | -0.072 | -0.043 | -0.043 | 0.41 | 0.37 | 0.07 | 0.24 | 0.64 | 0.45 | 0.4 |
| 3 | 59941389 | rs7629424 | 0.272 | 0.023 | 0.089 | -0.477 | 0.002 | 0.014 | 0.014 | 0 | 0.16 | 0.29 | 0.19 | 0.22 | 0.11 | 0.13 |
| 3 | 59943532 | rs3772460 | 1.272 | -0.007 | 0.014 | -0.663 | 0.052 | 0.055 | 0.055 | 0 | 0.18 | 0.06 | 0.08 | 0.05 | 0.08 | 0.16 |
| 3 | 59946354 | rs11715559 | 1.705 | -0.13 | -0.005 | -0.746 | 0.014 | -0.002 | -0.002 | 0 | 0.15 | 0.13 | 0.08 | 0 | 0.07 | 0.11 |
| 3 | 59946851 | rs3772464 | 2.227 | -0.115 | -0.5 | -0.776 | -0.071 | -0.129 | -0.129 | 0 | 0.19 | 0.03 | 0.15 | 0.49 | 0.26 | 0.28 |
| 3 | 59950763 | rs1485904 | 0.282 | -0.47 | 0.615 | -0.126 | -0.014 | -0.022 | -0.022 | 0.073 | 0.84 | 0.76 | 0.82 | 0.92 | 0.88 | 0.84 |
| 3 | 59951847 | rs13096138 | 2.972 | -0.466 | 0.501 | -0.132 | -0.081 | -0.078 | -0.078 | 0.623 | 0.37 | 0.07 | 0.24 | 0.58 | 0.45 | 0.4 |
| 3 | 59953207 | rs9865672 | -1.389 | -0.437 | 0.604 | -0.457 | -0.011 | -0.001 | -0.001 | 0 | 0.4 | 0.57 | 0.4 | 0.37 | 0.31 | 0.35 |
| 3 | 59957116 | rs6446102 | 2.061 | -0.466 | 0.474 | -0.405 | -0.017 | 0.016 | 0.016 | 2.179 | 0.49 | 0.1 | 0.46 | 0.61 | 0.61 | 0.64 |
| 3 | 59958388 | rs3772479 | 0.436 | -0.398 | 1.009 | -0.368 | 0.108 | -0.028 | -0.028 | 0 | 0.91 | 0.67 | 0.81 | 0.99 | 0.88 | 0.97 |
| 3 | 59959014 | rs13064036 | 2.033 | -0.472 | 0.953 | -0.407 | -0.03 | -0.007 | -0.007 | 0 | 0.61 | 0.26 | 0.44 | 0.62 | 0.57 | 0.62 |
| 3 | 59961536 | rs212034 | -0.295 | -0.472 | 1.075 | -0.393 | 0 | 0.054 | 0.054 | 0 | 0.16 | 0.23 | 0.19 | 0.37 | 0.11 | 0.12 |
| 3 | 59963131 | rs212032 | 1.623 | -0.492 | 1.093 | -0.424 | 0.003 | 0.005 | 0.005 | 1.444 | 0.55 | 0.44 | 0.45 | 0.9 | 0.57 | 0.54 |
| 3 | 59963156 | rs3772482 | 1.924 | -0.567 | 1.036 | -0.391 | -0.031 | -0.012 | -0.012 | 0.08 | 0.6 | 0.26 | 0.43 | 0.62 | 0.57 | 0.63 |
| 3 | 59964699 | rs10490808 | 2.692 | -0.4 | 0.626 | -0.281 | -0.074 | -0.114 | -0.114 | 0 | 0.38 | 0.1 | 0.25 | 0.53 | 0.45 | 0.41 |
| 3 | 59964805 | rs2736794 | -1.152 | -0.192 | 0.831 | -0.112 | 0.021 | 0.041 | 0.041 | 0 | 0.22 | 0.32 | 0.36 | 0.38 | 0.23 | 0.19 |
| 3 | 59968353 | rs802775 | -1.102 | -0.161 | 0.734 | -0.105 | 0.009 | 0.018 | 0.018 | 0 | 0.2 | 0.26 | 0.34 | 0.37 | 0.23 | 0.15 |
| 3 | 59973302 | rs3772497 | 2.163 | -0.352 | 0.687 | -0.156 | -0.007 | 0.01 | 0.01 | 0.185 | 0.46 | 0.3 | 0.31 | 0.48 | 0.52 | 0.41 |
| 3 | 59973677 | rs2630170 | 0.889 | -0.597 | 0.73 | -0.112 | -0.015 | 0.012 | 0.012 | 0 | 0.69 | 0.67 | 0.7 | 0.57 | 0.83 | 0.74 |
| 3 | 59976176 | rs3772498 | 0.102 | -0.582 | 0.511 | -0.085 | 0.034 | -0.062 | -0.062 | 0 | 0.2 | 0.12 | 0.21 | 0.41 | 0.09 | 0.18 |
| 3 | 59978703 | rs3772501 | -0.358 | -0.56 | 0.518 | -0.004 | 0.047 | -0.064 | -0.064 | 0.052 | 0.23 | 0.13 | 0.22 | 0.37 | 0.1 | 0.22 |
| 3 | 59979237 | rs12152244 | -0.437 | -0.494 | 0.566 | -0.123 | -0.001 | -0.033 | -0.033 | 0.774 | 0.16 | 0.12 | 0.18 | 0 | 0.2 | 0.25 |
| 3 | 59980424 | rs3772503 | -0.874 | -0.436 | 0.605 | -0.099 | -0.001 | -0.026 | -0.026 | 0 | 0.18 | 0.14 | 0.2 | 0.02 | 0.2 | 0.3 |
| 3 | 59981525 | rs212016 | -1.972 | -0.661 | 0.825 | -0.074 | -0.01 | 0.039 | 0.039 | 0 | 0.3 | 0.49 | 0.35 | 0.37 | 0.24 | 0.48 |
| 3 | 59981976 | rs17061708 | -0.487 | -0.644 | 0.84 | -0.12 | -0.005 | -0.037 | -0.037 | 0 | 0.14 | 0.21 | 0.16 | 0.02 | 0.14 | 0.3 |
| 3 | 59982993 | rs6784057 | 2.742 | -0.602 | 0.372 | -0.166 | -0.126 | -0.146 | -0.146 | 0 | 0.39 | 0.04 | 0.29 | 0.54 | 0.47 | 0.35 |
| 3 | 59984355 | rs3772507 | 0.248 | -0.468 | 0.511 | -0.484 | -0.014 | 0.003 | 0.003 | 0.157 | 0.11 | 0.02 | 0.18 | 0.07 | 0.16 | 0.27 |
| 3 | 59984501 | rs212017 | 0.329 | -0.524 | 0.675 | -0.443 | 0.01 | -0.054 | -0.054 | 0 | 0.11 | 0.23 | 0.05 | 0.01 | 0.08 | 0.05 |
| 3 | 59986929 | rs12487545 | 0.999 | -0.588 | 0.741 | -0.356 | -0.042 | 0.038 | 0.038 | 0.023 | 0.68 | 0.47 | 0.72 | 0.61 | 0.83 | 0.78 |
| 3 | 59988905 | rs212020 | -0.958 | -0.866 | 0.095 | -0.373 | 0.032 | 0.024 | 0.024 | 0 | 0.84 | 0.91 | 0.93 | 1 | 0.93 | 0.93 |
| 3 | 59991608 | rs809710 | -1.436 | -0.571 | 0.02 | -0.204 | 0.009 | 0.004 | 0.004 | 0 | 0.41 | 0.52 | 0.61 | 0.5 | 0.45 | 0.52 |
| 3 | 59995658 | rs3915503 | 1.122 | -0.087 | 0.223 | 0.314 | 0.007 | 0.033 | 0.033 | 0.717 | 0.52 | 0.34 | 0.46 | 0.38 | 0.62 | 0.58 |
| 3 | 59996969 | rs4679627 | 1.508 | -0.048 | 0.148 | 0.095 | -0.001 | 0 | 0 | 0.87 | 0.48 | 0.45 | 0.38 | 0.46 | 0.53 | 0.49 |
| 3 | 59998294 | rs2736792 | -0.692 | 0.141 | -0.104 | 0.36 | -0.003 | 0 | 0 | 1.213 | 0.36 | 0.33 | 0.32 | 0.1 | 0.34 | 0.2 |
| 3 | 59998714 | rs3772509 | -0.935 | -0.235 | -0.103 | 1.158 | 0.005 | 0.036 | 0.036 | 0 | 0.09 | 0.08 | 0.16 | 0.46 | 0.03 | 0.15 |
| 3 | 59999965 | rs17061789 | -1.377 | 0.018 | -0.041 | 0.318 | -0.004 | -0.007 | -0.007 | 1.108 | 0.11 | 0.1 | 0.07 | 0.01 | 0.1 | 0.19 |
| 3 | 60000052 | rs17061792 | -1.261 | -0.211 | 0.498 | 0.095 | -0.001 | -0.072 | -0.072 | 0 | 0.17 | 0.25 | 0.2 | 0.01 | 0.14 | 0.15 |
| 3 | 60000536 | rs3845970 | -0.947 | -0.244 | 1.097 | 0.836 | 0.003 | 0.036 | 0.036 | 0 | 0.23 | 0.35 | 0.23 | 0.46 | 0.26 | 0.31 |
| 3 | 60001825 | rs9311745 | -1.494 | -0.732 | 1.192 | 0.96 | -0.01 | 0.017 | 0.017 | 0 | 0.17 | 0.15 | 0.16 | 0.3 | 0.05 | 0.17 |
| 3 | 60002455 | rs3912492 | -2.235 | -0.736 | 1.414 | 0.884 | -0.04 | 0.04 | 0.04 | 0 | 0.23 | 0.29 | 0.21 | 0.31 | 0.11 | 0.25 |
| 3 | 60004908 | rs9861525 | -0.442 | -0.703 | 1.366 | 0.853 | -0.003 | -0.004 | -0.004 | 0 | 0.2 | 0.22 | 0.21 | 0.46 | 0.25 | 0.31 |
| 3 | 60007121 | rs3912493 | -2.28 | -0.696 | 1.545 | 0.61 | 0.021 | 0.04 | 0.04 | 0 | 0.44 | 0.66 | 0.45 | 0.77 | 0.36 | 0.59 |
| 3 | 60009173 | rs3856659 | -1.942 | -0.685 | 1.815 | 0.672 | 0.011 | 0.019 | 0.019 | 0 | 0.4 | 0.65 | 0.46 | 0.73 | 0.36 | 0.55 |
| 3 | 60010550 | rs6794968 | -2.228 | -0.661 | 1.588 | 0.646 | 0.044 | 0.103 | 0.103 | 0.51 | 0.46 | 0.78 | 0.48 | 0.74 | 0.39 | 0.58 |
| 3 | 60010769 | rs6803449 | -1.234 | -0.747 | 1.655 | 0.82 | -0.031 | 0.05 | 0.05 | 1.312 | 0.16 | 0.22 | 0.19 | 0.27 | 0.09 | 0.14 |
| 3 | 60011243 | rs6782804 | -2.07 | -0.609 | 1.254 | 0.859 | 0.037 | 0.084 | 0.084 | 0.131 | 0.44 | 0.77 | 0.46 | 0.73 | 0.38 | 0.58 |
| 3 | 60013145 | rs12636081 | -1.734 | 0.166 | 1.313 | 1.365 | -0.008 | 0.043 | 0.043 | 0.056 | 0.12 | 0.11 | 0.17 | 0.28 | 0.11 | 0.13 |
| 3 | 60016166 | rs9816955 | -1.134 | 0.708 | 0.665 | 1.1 | 0.014 | 0.077 | 0.077 | 2.278 | 0.22 | 0.35 | 0.26 | 0.47 | 0.28 | 0.39 |
| 3 | 60018326 | rs17061880 | -1.716 | 0.635 | 0.906 | 0.373 | 0.002 | -0.027 | -0.027 | 0 | 0.17 | 0.45 | 0.16 | 0.07 | 0.18 | 0.17 |
| 3 | 60020000 | rs182418 | 0.554 | 0.468 | 1.318 | 0.32 | -0.016 | 0.027 | 0.027 | 0 | 0.78 | 0.67 | 0.56 | 0.3 | 0.82 | 0.61 |
| 3 | 60020954 | rs17061897 | -1.189 | 0.402 | 1.2 | 0.119 | 0.015 | 0.006 | 0.006 | 0 | 0.1 | 0.06 | 0.04 | 0.05 | 0.06 | 0.16 |
| 3 | 60022161 | rs212063 | -2.456 | 0.941 | 1.79 | 0.009 | 0.062 | 0.017 | 0.017 | 3.23 | 0.09 | 0.37 | 0.09 | 0.07 | 0.08 | 0.2 |
| 3 | 60024883 | rs212059 | 0.088 | 0.882 | 1.642 | 0.14 | -0.003 | 0.009 | 0.009 | 0 | 0.8 | 0.83 | 0.58 | 0.33 | 0.82 | 0.63 |
| 3 | 60025038 | rs17257241 | -1.903 | 0.93 | 1.486 | 0.107 | -0.004 | 0.012 | 0.012 | 0 | 0.15 | 0.12 | 0.11 | 0.05 | 0.19 | 0.17 |
| 3 | 60028556 | rs3856662 | -0.122 | 0.889 | 1.515 | 0.095 | 0.017 | -0.025 | -0.025 | 0 | 0.05 | 0.1 | 0.06 | 0 | 0.12 | 0 |
| 3 | 60034194 | rs212052 | -2.642 | 1.38 | 1.873 | -0.03 | 0.002 | -0.007 | -0.007 | 0 | 0.18 | 0.3 | 0.13 | 0.12 | 0.17 | 0.37 |
| 3 | 60034455 | rs212051 | 1.297 | 1.335 | 1.993 | -0.107 | 0.017 | 0.017 | 0.017 | 0.179 | 0.68 | 0.71 | 0.49 | 0.25 | 0.72 | 0.45 |
| 3 | 60035070 | rs212050 | 1.817 | 1.348 | 2.083 | -0.118 | -0.007 | -0.032 | -0.032 | 0 | 0.6 | 0.51 | 0.41 | 0.25 | 0.58 | 0.41 |
| 3 | 60036213 | rs212046 | -0.878 | 1.3 | 1.871 | -0.134 | 0.005 | -0.001 | -0.001 | 0 | 0.08 | 0.07 | 0.09 | 0.05 | 0.18 | 0.09 |
| 3 | 60037145 | rs4083381 | 0.736 | 1.142 | 2.149 | -0.113 | 0 | -0.004 | -0.004 | 0 | 0.74 | 0.77 | 0.51 | 0.27 | 0.75 | 0.43 |
| 3 | 60037467 | rs169841 | 1.881 | 1.143 | 2.325 | -0.203 | 0.001 | -0.018 | -0.018 | 0.021 | 0.6 | 0.42 | 0.39 | 0.23 | 0.56 | 0.51 |
| 3 | 60038000 | rs212043 | 0.913 | 1.091 | 2.366 | -0.206 | -0.023 | -0.007 | -0.007 | 0 | 0.72 | 0.62 | 0.51 | 0.23 | 0.78 | 0.53 |
| 3 | 60038372 | rs212040 | -1.462 | 1.157 | 2.578 | -0.228 | 0.029 | -0.036 | -0.036 | 0 | 0.07 | 0.16 | 0.06 | 0 | 0.07 | 0.12 |
| 3 | 60038632 | rs212038 | -2.688 | 1.169 | 2.624 | -0.247 | 0.103 | -0.074 | -0.074 | 2.381 | 0.16 | 0.42 | 0.16 | 0.02 | 0.24 | 0.23 |

| | | | | | | | | | | | | | | | | |
|---|---|---|---|---|---|---|---|---|---|---|---|---|---|---|---|---|
| 3 | 60039488 | rs3845974 | -1.771 | 1.148 | 2.386 | -0.203 | 0.067 | -0.03 | -0.03 | 0 | 0.11 | 0.35 | 0.12 | 0.02 | 0.2 | 0.11 |
| 3 | 60040105 | rs9860107 | -1.693 | 1.014 | 2.013 | -0.171 | -0.005 | -0.007 | -0.007 | 0 | 0.13 | 0.15 | 0.06 | 0.05 | 0.08 | 0.18 |
| 3 | 60049738 | rs212006 | -2.658 | 1.201 | 2.21 | -0.143 | 0.007 | -0.021 | -0.021 | 1.485 | 0.21 | 0.72 | 0.21 | 0.06 | 0.27 | 0.28 |
| 3 | 60051071 | rs191927 | -2.659 | 1.192 | 2.238 | -0.162 | 0.014 | -0.032 | -0.032 | 0.623 | 0.21 | 0.67 | 0.21 | 0.06 | 0.27 | 0.27 |
| 3 | 60051227 | rs3856668 | -2.735 | 1.224 | 2.296 | -0.161 | 0.017 | -0.027 | -0.027 | 0 | 0.18 | 0.62 | 0.17 | 0.07 | 0.22 | 0.17 |
| 3 | 60051461 | rs3856669 | -2.339 | 1.199 | 2.37 | -0.174 | 0.053 | -0.046 | -0.046 | 0.143 | 0.15 | 0.57 | 0.15 | 0.05 | 0.21 | 0.08 |
| 3 | 60053331 | rs2630173 | -2.503 | 1.174 | 2.37 | -0.279 | 0.027 | -0.053 | -0.053 | 0 | 0.21 | 0.6 | 0.2 | 0.07 | 0.28 | 0.29 |
| 3 | 60055347 | rs2630181 | -2.42 | 1.175 | 2.371 | -0.278 | 0.031 | -0.051 | -0.051 | 0.075 | 0.2 | 0.6 | 0.2 | 0.08 | 0.28 | 0.29 |
| 3 | 60056853 | rs2630196 | -2.533 | 1.254 | 2.41 | -0.275 | 0.053 | -0.054 | -0.054 | 11.739 | 0.19 | 0.53 | 0.19 | 0.08 | 0.28 | 0.29 |
| 3 | 60059041 | rs2630207 | 0.983 | 1.41 | 2.124 | -0.269 | -0.003 | -0.003 | -0.003 | 1.145 | 0.71 | 0.62 | 0.49 | 0.25 | 0.75 | 0.57 |
| 3 | 60062400 | rs2736767 | 1.055 | 1.411 | 2.102 | -0.259 | -0.004 | -0.003 | -0.003 | 1.533 | 0.7 | 0.61 | 0.48 | 0.26 | 0.75 | 0.58 |
| 3 | 60062654 | rs17062101 | -0.26 | 1.475 | 2.074 | -0.311 | 0.04 | -0.03 | -0.03 | 0.061 | 0.06 | 0.15 | 0.06 | 0 | 0.12 | 0 |
| 3 | 60065257 | rs7615432 | -0.951 | 1.377 | 2.037 | 0.213 | -0.005 | -0.009 | -0.009 | 0 | 0.24 | 0.31 | 0.24 | 0.2 | 0.19 | 0.22 |
| 3 | 60069828 | rs3845977 | -0.7 | 1.367 | 2.169 | 0.273 | 0.003 | 0.006 | 0.006 | 0 | 0.22 | 0.22 | 0.24 | 0.2 | 0.19 | 0.21 |
| 3 | 60070931 | rs4145506 | 1.262 | 1.198 | 0.96 | -0.718 | -0.062 | 0.011 | 0.011 | 0.04 | 0.27 | 0.06 | 0.19 | 0.2 | 0.3 | 0.24 |
| 3 | 60073652 | rs6801817 | 1.498 | 1.859 | 1.121 | -0.881 | -0.009 | 0.019 | 0.019 | 0.586 | 0.57 | 0.3 | 0.37 | 0.26 | 0.53 | 0.4 |
| 3 | 60074752 | rs986015 | 2.572 | 1.699 | 0.973 | -0.943 | 0.037 | 0.127 | 0.127 | 5.093 | 0.23 | 0.02 | 0.07 | 0 | 0.16 | 0.03 |
| 3 | 60075192 | rs1040336 | -0.638 | 1.932 | 1.302 | -0.629 | 0.093 | 0.265 | 0.265 | 0.288 | 0.09 | 0.23 | 0.3 | 0.53 | 0.17 | 0.25 |
| 3 | 60076491 | rs9817116 | 0.525 | 1.221 | 1.097 | -0.821 | -0.004 | -0.012 | -0.012 | 0.586 | 0.71 | 0.62 | 0.71 | 0.79 | 0.75 | 0.68 |
| 3 | 60080887 | rs17376767 | 0.071 | 1.138 | 0.967 | -0.803 | 0.002 | -0.045 | -0.045 | 0 | 0.14 | 0.14 | 0.12 | 0 | 0.18 | 0.06 |
| 3 | 60084924 | rs7642268 | 1.45 | 1.01 | 1.294 | -0.994 | 0.002 | -0.123 | -0.123 | 0.37 | 0.71 | 0.45 | 0.75 | 0.98 | 0.67 | 0.78 |
| 3 | 60084971 | rs7609578 | 1.255 | 1.009 | 1.426 | -0.994 | 0.01 | -0.112 | -0.112 | 0 | 0.73 | 0.57 | 0.76 | 0.98 | 0.67 | 0.79 |
| 3 | 60086281 | rs6804216 | 1.396 | 1.054 | 1.489 | -0.926 | 0.008 | -0.075 | -0.075 | 0.009 | 0.84 | 0.52 | 0.86 | 0.98 | 0.85 | 0.85 |
| 3 | 60089286 | rs9871368 | -1.094 | 0.984 | 1.374 | -0.797 | 0.022 | 0.031 | 0.031 | 1.422 | 0.61 | 0.7 | 0.7 | 0.73 | 0.66 | 0.75 |
| 3 | 60089314 | rs10510825 | 1.683 | 0.973 | 1.333 | -0.797 | 0.031 | 0.031 | 0.031 | 2.718 | 0.33 | 0.14 | 0.24 | 0.26 | 0.27 | 0.14 |
| 3 | 60089378 | rs10510826 | -1.582 | 0.959 | 1.369 | -0.794 | 0.015 | -0.05 | -0.05 | 2.41 | 0.1 | 0.43 | 0.1 | 0 | 0.08 | 0.03 |
| 3 | 60089571 | rs10510827 | 0.249 | 0.96 | 1.114 | -0.71 | -0.005 | 0.031 | 0.031 | 0 | 0.17 | 0.08 | 0.13 | 0.08 | 0.2 | 0.19 |
| 3 | 60090013 | rs6766789 | -1.337 | 0.919 | 1.198 | -0.424 | 0.01 | 0.358 | 0.358 | 0 | 0.11 | 0.37 | 0.3 | 0.5 | 0.07 | 0.26 |
| 3 | 60090116 | rs12330253 | 1.093 | 0.953 | 0.91 | -0.475 | -0.04 | 0.04 | 0.04 | 0.326 | 0.3 | 0.12 | 0.24 | 0.15 | 0.35 | 0.26 |
| 3 | 60090683 | rs11917733 | 1.342 | 0.464 | 0.78 | -0.467 | 0.057 | 0.039 | 0.039 | 3.502 | 0.36 | 0.19 | 0.24 | 0.26 | 0.26 | 0.15 |
| 3 | 60091257 | rs2205361 | -1.121 | 0.781 | 0.47 | -0.251 | -0.006 | -0.074 | -0.074 | 0.134 | 0.28 | 0.16 | 0.42 | 0.55 | 0.31 | 0.48 |
| 3 | 60093362 | rs996174 | -0.804 | 0.681 | 0.746 | -0.177 | 0.03 | 0.03 | 0.03 | 2.61 | 0.56 | 0.36 | 0.67 | 0.73 | 0.67 | 0.72 |
| 3 | 60093775 | rs4679512 | -0.364 | 0.688 | 0.782 | 0.247 | -0.004 | 0.194 | 0.194 | 0 | 0.05 | 0.09 | 0.24 | 0.48 | 0.03 | 0.24 |
| 3 | 60094767 | rs13317306 | -0.27 | 0.854 | 0.745 | -0.05 | 0.004 | -0.045 | -0.045 | 0.001 | 0.08 | 0.22 | 0.04 | 0 | 0.04 | 0.03 |
| 3 | 60095070 | rs2142301 | -0.445 | 0.696 | 0.636 | -0.163 | 0.018 | 0.001 | 0.001 | 0 | 0.36 | 0.44 | 0.48 | 0.63 | 0.42 | 0.52 |
| 3 | 60095604 | rs9311749 | -1.499 | 0.343 | 0.415 | -0.296 | 0.006 | 0.005 | 0.005 | 0.768 | 0.58 | 0.63 | 0.49 | 0.33 | 0.61 | 0.57 |
| 3 | 60095853 | rs926141 | -0.856 | 0.319 | 0.379 | -0.491 | 0.018 | 0.061 | 0.061 | 0 | 0.49 | 0.6 | 0.38 | 0.23 | 0.52 | 0.49 |
| 3 | 60098968 | rs760316 | -2.057 | 0.295 | 0.405 | -0.441 | 0.005 | 0.059 | 0.059 | 1.777 | 0.65 | 0.82 | 0.75 | 0.81 | 0.65 | 0.82 |
| 3 | 60102668 | rs7634772 | -0.104 | 0.667 | 0.493 | -0.508 | -0.041 | -0.044 | -0.044 | 0 | 0.2 | 0.35 | 0.16 | 0.08 | 0.14 | 0.23 |
| 3 | 60105329 | rs9815054 | -0.729 | 0.72 | 0.482 | -0.502 | -0.001 | 0.007 | 0.007 | 0.019 | 0.45 | 0.53 | 0.34 | 0.25 | 0.48 | 0.44 |
| 3 | 60106852 | rs9882623 | 0.449 | 0.761 | 0.407 | -0.491 | -0.002 | 0.001 | 0.001 | 0 | 0.26 | 0.22 | 0.19 | 0.15 | 0.33 | 0.25 |
| 3 | 60108517 | rs35345892 | -0.333 | 0.63 | 0.166 | -0.232 | -0.001 | 0.024 | 0.024 | 0 | 0.46 | 0.58 | 0.35 | 0.22 | 0.48 | 0.44 |
| 3 | 60110529 | rs9311751 | -0.315 | 0.606 | -0.132 | -0.26 | -0.001 | -0.01 | -0.01 | 0 | 0.46 | 0.49 | 0.34 | 0.26 | 0.48 | 0.43 |
| 3 | 60116055 | rs2064016 | -1.046 | -0.364 | -0.321 | -0.069 | -0.012 | -0.003 | -0.003 | 0.325 | 0.2 | 0.23 | 0.34 | 0.15 | 0.17 | 0.25 |
| 3 | 60118879 | rs9682976 | -0.713 | -0.312 | -0.108 | -0.119 | -0.008 | -0.008 | -0.008 | 0 | 0.09 | 0.15 | 0.11 | 0.05 | 0.06 | 0.09 |
| 3 | 60120243 | rs10510828 | 1.142 | -0.389 | -0.406 | -0.164 | -0.028 | 0.06 | 0.06 | 0.167 | 0.44 | 0.2 | 0.44 | 0.22 | 0.51 | 0.39 |
| 3 | 60122156 | rs17062321 | -1.02 | -0.491 | 0.183 | -0.219 | -0.013 | -0.012 | -0.012 | 0 | 0.12 | 0.4 | 0.13 | 0.06 | 0.06 | 0.1 |
| 3 | 60122472 | rs12629469 | 1.524 | -0.55 | -0.061 | -0.281 | -0.062 | 0.051 | 0.051 | 0.17 | 0.41 | 0.12 | 0.41 | 0.21 | 0.49 | 0.37 |
| 3 | 60123522 | rs10510829 | 0.643 | -0.737 | -0.345 | -0.073 | -0.055 | 0.026 | 0.026 | 0.4 | 0.24 | 0.05 | 0.12 | 0.13 | 0.22 | 0.26 |
| 3 | 60124230 | rs7623479 | 1.459 | -0.687 | -0.101 | 0.07 | -0.01 | 0.058 | 0.058 | 0 | 0.46 | 0.27 | 0.5 | 0.23 | 0.57 | 0.4 |
| 3 | 60124388 | rs2361340 | 0.498 | -0.604 | -0.291 | 0.086 | -0.018 | 0.152 | 0.152 | 0 | 0.8 | 0.62 | 0.74 | 0.41 | 0.84 | 0.74 |
| 3 | 60124580 | rs6788821 | -0.233 | -0.613 | -0.312 | 0.118 | -0.009 | 0.044 | 0.044 | 1.041 | 0.37 | 0.28 | 0.27 | 0.18 | 0.34 | 0.42 |
| 3 | 60127027 | rs6787664 | -0.858 | -0.947 | 0.148 | 0.736 | -0.043 | 0.277 | 0.277 | 0.009 | 0.16 | 0.44 | 0.23 | 0.6 | 0.08 | 0.19 |
| 3 | 60128846 | rs17062350 | 0.308 | -1.371 | -0.634 | 0.004 | -0.004 | -0.009 | -0.009 | 0 | 0.43 | 0.59 | 0.28 | 0.21 | 0.31 | 0.27 |
| 3 | 60129997 | rs5014208 | -0.032 | -1.554 | -0.975 | 0.012 | -0.034 | -0.062 | -0.062 | 0 | 0.7 | 0.47 | 0.75 | 0.84 | 0.75 | 0.82 |
| 3 | 60130210 | rs7630823 | 1.179 | -1.646 | -0.823 | 0.033 | 0.015 | 0.027 | 0.027 | 0 | 0.32 | 0.07 | 0.17 | 0.2 | 0.22 | 0.25 |
| 3 | 60130721 | rs17062363 | -0.669 | -1.413 | -0.48 | 0.084 | -0.006 | -0.057 | -0.057 | 0 | 0.1 | 0.37 | 0.09 | 0 | 0.08 | 0.02 |
| 3 | 60134996 | rs4679640 | -0.305 | -1.076 | -0.963 | 0.222 | 0.005 | 0.007 | 0.007 | 0 | 0.92 | 0.95 | 0.79 | 0.99 | 0.94 | 0.94 |
| 3 | 60136927 | rs6788677 | 0.299 | -1.172 | -0.98 | 0.747 | -0.033 | 0.185 | 0.185 | 0.941 | 0.79 | 0.51 | 0.61 | 0.42 | 0.84 | 0.75 |
| 3 | 60138099 | rs2064020 | -0.087 | -0.458 | -0.921 | 0.42 | 0.004 | 0.009 | 0.009 | 0 | 0.44 | 0.39 | 0.43 | 0.19 | 0.61 | 0.5 |
| 3 | 60144588 | rs4679642 | 0.41 | 0.491 | -1.045 | 1.205 | -0.008 | -0.016 | -0.016 | 0 | 0.82 | 0.88 | 0.66 | 0.4 | 0.81 | 0.56 |
| 3 | 60145321 | rs12638728 | -0.348 | 0.962 | -0.21 | 1.303 | 0.072 | 0.109 | 0.109 | 0 | 0.06 | 0.23 | 0.13 | 0.08 | 0.09 | 0.1 |
| 3 | 60145726 | rs4679516 | -0.24 | 0.471 | -0.535 | 0.185 | -0.001 | 0.011 | 0.011 | 0 | 0.39 | 0.51 | 0.43 | 0.39 | 0.42 | 0.3 |
| 3 | 60145812 | rs4679643 | 1.636 | 0.168 | -1.055 | -0.093 | -0.075 | -0.116 | -0.116 | 0.321 | 0.17 | 0.01 | 0.23 | 0.31 | 0.25 | 0.13 |
| 3 | 60147355 | rs10510831 | -0.288 | -0.508 | -1.386 | -0.392 | -0.088 | -0.095 | -0.095 | 0.372 | 0.33 | 0.03 | 0.35 | 0.32 | 0.4 | 0.23 |
| 3 | 60148685 | rs6446115 | 0.285 | -0.561 | -1.022 | -0.131 | 0.009 | -0.031 | -0.031 | 0 | 0.82 | 0.94 | 0.69 | 0.62 | 0.86 | 0.75 |
| 3 | 60149936 | rs7613687 | -0.881 | -0.805 | -0.488 | -0.085 | -0.028 | 0.174 | 0.174 | 0.232 | 0.1 | 0.37 | 0.11 | 0.37 | 0.03 | 0.16 |
| 3 | 60150819 | rs17062405 | -1.177 | -0.751 | -0.394 | -0.043 | 0.003 | 0.277 | 0.277 | 2.57 | 0.07 | 0.39 | 0.1 | 0.36 | 0.01 | 0.11 |
| 3 | 60152188 | rs4411839 | 1.762 | -0.759 | -1.108 | -0.609 | -0.046 | 0.134 | 0.134 | 0 | 0.3 | 0.14 | 0.18 | 0.02 | 0.33 | 0.17 |

| | | | | | | | | | | | | | | | | |
|---|---|---|---|---|---|---|---|---|---|---|---|---|---|---|---|---|
| 3 | 60152583 | rs17062413 | 0.458 | -0.809 | -1.024 | -0.487 | -0.003 | 0.057 | 0.057 | 0 | 0.08 | 0.23 | 0.08 | 0.07 | 0.05 | 0.11 |
| 3 | 60159463 | rs177138 | 0.866 | -0.746 | -1.19 | -0.44 | -0.001 | -0.027 | -0.027 | 0 | 0.4 | 0.42 | 0.25 | 0.09 | 0.39 | 0.32 |
| 3 | 60167219 | rs9882944 | 1.067 | -0.631 | -1.033 | 0.114 | -0.007 | 0.055 | 0.055 | 0.663 | 0.09 | 0.09 | 0.2 | 0.44 | 0.02 | 0.18 |
| 3 | 60170083 | rs6797203 | -0.303 | -0.452 | -1.052 | -1.126 | -0.001 | -0.001 | -0.001 | 0.348 | 0.47 | 0.59 | 0.65 | 0.92 | 0.48 | 0.55 |
| 3 | 60186728 | rs213299 | 0.312 | -0.475 | -1.032 | -1.046 | 0.009 | 0.033 | 0.033 | 0.338 | 0.5 | 0.4 | 0.62 | 0.83 | 0.53 | 0.6 |
| 3 | 60188605 | rs213307 | -0.329 | -0.868 | -1.155 | -1.264 | 0.018 | 0 | 0 | 0 | 0.81 | 0.86 | 0.92 | 1 | 0.9 | 0.94 |
| 3 | 60189471 | rs1496650 | 0.531 | -0.704 | -0.899 | -1.492 | -0.001 | -0.048 | -0.048 | 0 | 0.29 | 0.3 | 0.24 | 0.11 | 0.28 | 0.32 |
| 3 | 60193900 | rs213319 | 1.455 | -0.274 | -1.141 | -1.663 | -0.015 | 0.069 | 0.069 | 0.034 | 0.24 | 0.12 | 0.3 | 0.04 | 0.26 | 0.2 |
| 3 | 60194059 | rs213322 | -0.151 | -0.359 | -0.966 | -1.044 | 0 | 0 | 0 | 0 | 0.47 | 0.48 | 0.55 | 0.29 | 0.49 | 0.43 |
| 3 | 60194570 | rs213323 | 1.735 | -0.355 | -0.875 | -1.078 | -0.004 | 0.021 | 0.021 | 0.014 | 0.25 | 0.19 | 0.3 | 0.04 | 0.26 | 0.2 |
| 3 | 60196537 | rs11130760 | 1.1 | -0.277 | -0.813 | -0.885 | -0.004 | 0.019 | 0.019 | 0 | 0.08 | 0.12 | 0.09 | 0.12 | 0.1 | 0.17 |
| 3 | 60197485 | rs802933 | -1.137 | -0.271 | -0.726 | -0.979 | 0.011 | -0.029 | -0.029 | 0 | 0.66 | 0.65 | 0.68 | 0.82 | 0.57 | 0.64 |
| 3 | 60210356 | rs2173056 | 0.613 | -0.275 | -0.759 | -0.876 | -0.001 | -0.086 | -0.086 | 0.056 | 0.19 | 0.07 | 0.15 | 0.59 | 0.14 | 0.25 |
| 3 | 60210372 | rs213412 | 1.166 | -0.52 | -0.646 | -0.998 | -0.001 | -0.008 | -0.008 | 0 | 0.32 | 0.36 | 0.37 | 0.14 | 0.32 | 0.37 |
| 3 | 60212448 | rs9880343 | -0.831 | -0.408 | -0.98 | -1.144 | -0.102 | -0.012 | -0.012 | 0.042 | 0.58 | 0.91 | 0.5 | 0.36 | 0.51 | 0.54 |
| 3 | 60212916 | rs213415 | -1.147 | -0.056 | -0.724 | -1.108 | -0.057 | 0.048 | 0.048 | 0.109 | 0.86 | 0.98 | 0.86 | 0.97 | 0.79 | 0.96 |
| 3 | 60213301 | rs9827643 | 0.165 | -0.331 | -0.122 | -0.814 | 0.024 | 0.114 | 0.114 | 0.001 | 0.09 | 0.19 | 0.19 | 0.25 | 0.16 | 0.19 |
| 3 | 60213805 | rs9832192 | -1.218 | -0.437 | -0.19 | -0.866 | -0.028 | 0.033 | 0.033 | 0.014 | 0.26 | 0.57 | 0.27 | 0.25 | 0.23 | 0.23 |
| 3 | 60215950 | rs6764230 | -0.01 | 0.048 | -0.571 | -1.026 | 0 | -0.048 | -0.048 | 0.042 | 0.32 | 0.31 | 0.34 | 0.03 | 0.27 | 0.17 |
| 3 | 60216788 | rs17062553 | 1.244 | 0.16 | -0.662 | -0.963 | 0 | 0.013 | 0.013 | 1.311 | 0.19 | 0.14 | 0.25 | 0.01 | 0.17 | 0.12 |
| 3 | 60217330 | rs9831415 | 0.34 | 0.24 | -0.5 | -0.958 | 0.016 | 0.077 | 0.077 | 0.037 | 0.11 | 0.18 | 0.15 | 0.24 | 0.17 | 0.2 |
| 3 | 60218337 | rs11717164 | 1.454 | 0.256 | -0.817 | -0.932 | -0.002 | 0.01 | 0.01 | 1.893 | 0.18 | 0.13 | 0.25 | 0.04 | 0.17 | 0.12 |
| 3 | 60220486 | rs7629068 | -0.731 | 0.246 | -0.924 | -0.755 | -0.001 | -0.002 | -0.002 | 0 | 0.45 | 0.55 | 0.5 | 0.29 | 0.43 | 0.37 |
| 3 | 60228660 | rs17338297 | -0.077 | 0.294 | -1.235 | -1.041 | -0.022 | 0.009 | 0.009 | 0 | 0.11 | 0.01 | 0.08 | 0.07 | 0.14 | 0.19 |
| 3 | 60236298 | rs1496652 | -0.106 | 0.478 | -0.679 | -1.191 | -0.003 | 0.003 | 0.003 | 0 | 0.24 | 0.21 | 0.34 | 0.19 | 0.27 | 0.36 |
| 3 | 60236847 | rs213392 | 0.5 | -0.374 | -0.628 | -1.796 | -0.037 | -0.08 | -0.08 | 0 | 0.19 | 0.29 | 0.07 | 0.02 | 0.11 | 0.24 |
| 3 | 60237710 | rs213396 | -0.979 | -0.474 | -0.683 | -1.672 | -0.049 | -0.043 | -0.043 | 0 | 0.32 | 0.58 | 0.19 | 0.19 | 0.23 | 0.49 |
| 3 | 60238326 | rs213398 | -1.001 | -0.289 | -0.763 | -1.941 | 0.003 | 0 | 0 | 0 | 0.47 | 0.6 | 0.45 | 0.2 | 0.41 | 0.61 |
| 3 | 60238818 | rs213400 | 0.951 | -0.624 | -0.715 | -1.78 | -0.021 | -0.056 | -0.056 | 0.21 | 0.14 | 0.22 | 0.06 | 0.03 | 0.1 | 0.24 |
| 3 | 60240638 | rs169913 | -0.463 | -0.434 | -1.305 | -2.244 | 0 | -0.048 | -0.048 | 0 | 0.55 | 0.58 | 0.49 | 0.13 | 0.51 | 0.52 |
| 3 | 60241445 | rs169915 | -0.508 | -0.594 | -1.216 | -2.226 | 0.014 | 0.028 | 0.028 | 0 | 0.61 | 0.7 | 0.56 | 0.19 | 0.69 | 0.68 |
| 3 | 60242432 | rs12487617 | 0.745 | -0.916 | -1.317 | -2.395 | -0.006 | -0.015 | -0.015 | 0.813 | 0.32 | 0.34 | 0.39 | 0.84 | 0.24 | 0.27 |
| 3 | 60246190 | rs143018 | 0.412 | -0.543 | -1.789 | -2.59 | 0.017 | 0.177 | 0.177 | 0 | 0.6 | 0.8 | 0.77 | 0.92 | 0.63 | 0.73 |
| 3 | 60248418 | rs213406 | -0.131 | -0.542 | -1.882 | -2.632 | 0.023 | 0.152 | 0.152 | 0 | 0.67 | 0.88 | 0.81 | 0.99 | 0.71 | 0.84 |
| 3 | 60252827 | rs213333 | 0.45 | -0.429 | -1.692 | -2.757 | 0.015 | 0.06 | 0.06 | 0 | 0.75 | 0.87 | 0.83 | 0.91 | 0.73 | 0.8 |
| 3 | 60254987 | rs7625421 | 1.097 | -0.416 | -1.669 | -2.866 | -0.003 | -0.003 | -0.003 | 0.991 | 0.08 | 0.08 | 0.06 | 0 | 0.11 | 0.04 |
| 3 | 60256485 | rs213339 | 0.416 | -0.799 | -1.454 | -2.997 | 0.001 | -0.007 | -0.007 | 0 | 0.48 | 0.52 | 0.49 | 0.26 | 0.51 | 0.64 |
| 3 | 60257587 | rs213341 | -0.489 | -0.5 | -1.338 | -2.822 | -0.001 | -0.012 | -0.012 | 0 | 0.87 | 0.9 | 0.9 | 0.99 | 0.84 | 0.93 |
| 3 | 60260223 | rs213347 | 0.277 | -0.404 | -1.307 | -2.812 | 0.01 | 0.009 | 0.009 | 0 | 0.68 | 0.75 | 0.81 | 0.92 | 0.71 | 0.79 |
| 3 | 60260504 | rs213348 | 1.329 | -0.058 | -1.284 | -2.836 | -0.001 | -0.001 | -0.001 | 0.439 | 0.21 | 0.21 | 0.14 | 0.17 | 0.18 | 0.36 |
| 3 | 60263721 | rs213354 | 0.313 | -0.071 | -1.316 | -2.831 | -0.004 | 0.051 | 0.051 | 0.008 | 0.42 | 0.3 | 0.47 | 0.25 | 0.5 | 0.64 |
| 3 | 60267024 | rs213366 | -0.578 | 0.8 | -1.372 | -2.412 | -0.014 | 0.01 | 0.01 | 0 | 0.75 | 0.76 | 0.62 | 0.92 | 0.63 | 0.69 |
| 3 | 60267052 | rs12487741 | 1.075 | 1.035 | -1.615 | -2.134 | -0.001 | -0.001 | -0.001 | 3.097 | 0.26 | 0.21 | 0.33 | 0.7 | 0.22 | 0.16 |
| 3 | 60270554 | rs9849856 | 0.292 | 0.88 | -1.456 | -2.077 | -0.013 | 0.045 | 0.045 | 0 | 0.42 | 0.3 | 0.64 | 0.76 | 0.51 | 0.38 |
| 3 | 60272317 | rs9879860 | 0.182 | 1.345 | -1.176 | -1.382 | -0.016 | 0.014 | 0.014 | 0 | 0.08 | 0.03 | 0.08 | 0.05 | 0.11 | 0.12 |
| 3 | 60274049 | rs213376 | 0.311 | 1.561 | -1.213 | -1.887 | -0.037 | 0.055 | 0.055 | 0.357 | 0.37 | 0.24 | 0.28 | 0.05 | 0.48 | 0.43 |
| 3 | 60276722 | rs732380 | 0.317 | 0.966 | -0.717 | -1.199 | -0.003 | -0.002 | -0.002 | 0 | 0.38 | 0.57 | 0.5 | 0.73 | 0.27 | 0.29 |
| 3 | 60277771 | rs2049847 | -0.529 | 1.252 | -1.024 | -1.365 | -0.036 | 0.114 | 0.114 | 0 | 0.58 | 0.77 | 0.68 | 0.93 | 0.48 | 0.48 |
| 3 | 60283702 | rs13317933 | 0.568 | 1.186 | -1.071 | -1.216 | -0.017 | -0.007 | -0.007 | 0 | 0.35 | 0.35 | 0.47 | 0.73 | 0.23 | 0.3 |
| 3 | 60283972 | rs9311756 | -0.786 | 1.272 | -1.174 | -1.292 | 0.006 | -0.094 | -0.094 | 1.407 | 0.59 | 0.61 | 0.66 | 0.93 | 0.48 | 0.54 |
| 3 | 60286486 | rs6768755 | -0.752 | 1.286 | -1.141 | -1.29 | -0.003 | -0.061 | -0.061 | 0.985 | 0.6 | 0.67 | 0.69 | 0.95 | 0.48 | 0.6 |
| 3 | 60288428 | rs10510836 | 0.68 | 1.402 | -1.177 | -1.154 | -0.013 | -0.014 | -0.014 | 0.446 | 0.18 | 0.09 | 0.09 | 0.14 | 0.14 | 0.15 |
| 3 | 60289842 | rs10510837 | -0.668 | 1.403 | -1.091 | -1.009 | -0.006 | -0.004 | -0.004 | 1.183 | 0.07 | 0.07 | 0.09 | 0.08 | 0.12 | 0.1 |
| 3 | 60292606 | rs4679524 | -0.097 | 1.37 | -1.124 | -1.059 | 0.001 | -0.002 | -0.002 | 0.007 | 0.42 | 0.61 | 0.6 | 0.81 | 0.35 | 0.45 |
| 3 | 60295081 | rs6776005 | -0.447 | 1.91 | -0.72 | -1.139 | 0.005 | -0.01 | -0.01 | 0 | 0.05 | 0.16 | 0.06 | 0.03 | 0.05 | 0.14 |
| 3 | 60298229 | rs7644138 | -0.63 | 1.887 | -0.905 | -1.108 | 0.024 | 0.007 | 0.007 | 0 | 0.07 | 0.13 | 0.12 | 0.11 | 0.14 | 0.17 |
| 3 | 60299260 | rs4679531 | -0.116 | 1.115 | -1.1 | -1.013 | -0.004 | -0.036 | -0.036 | 0 | 0.56 | 0.53 | 0.59 | 0.8 | 0.39 | 0.48 |
| 3 | 60305117 | rs186154 | -0.129 | 1.07 | -1.536 | -0.646 | -0.008 | 0.028 | 0.028 | 0.006 | 0.33 | 0.21 | 0.49 | 0.8 | 0.33 | 0.39 |
| 3 | 60306891 | rs213382 | 0.136 | 1.052 | -1.167 | -0.997 | 0.003 | -0.031 | -0.031 | 0 | 0.43 | 0.4 | 0.51 | 0.16 | 0.51 | 0.6 |
| 3 | 60310376 | rs1882898 | -0.529 | 0.913 | -1.401 | -1.133 | -0.012 | 0.039 | 0.039 | 0 | 0.62 | 0.75 | 0.67 | 0.2 | 0.6 | 0.72 |
| 3 | 60310790 | rs10510838 | 0.668 | 0.86 | -1.63 | -1.117 | -0.033 | 0.053 | 0.053 | 0.13 | 0.16 | 0.01 | 0.11 | 0.03 | 0.16 | 0.12 |
| 3 | 60315708 | rs6762886 | 0.371 | 0.892 | -1.499 | -1.104 | -0.034 | 0.108 | 0.108 | 0.448 | 0.41 | 0.09 | 0.53 | 0.2 | 0.43 | 0.6 |
| 3 | 60321082 | rs13074132 | 0.416 | 1.084 | -1.459 | -1.17 | 0.032 | 0.133 | 0.133 | 0 | 0.39 | 0.17 | 0.44 | 0.15 | 0.34 | 0.5 |
| 3 | 60321271 | rs9878437 | -0.003 | 0.941 | -1.198 | -1.201 | -0.009 | -0.061 | -0.061 | 0 | 0.18 | 0.32 | 0.1 | 0.03 | 0.16 | 0.13 |
| 3 | 60327391 | rs17599845 | 0.489 | 0.858 | -1.228 | -1.197 | -0.02 | 0.1 | 0.1 | 0.186 | 0.4 | 0.17 | 0.52 | 0.22 | 0.41 | 0.64 |
| 3 | 60328600 | rs6800393 | 0.244 | 0.915 | -1.22 | -1.286 | 0.001 | -0.004 | -0.004 | 0 | 0.17 | 0.18 | 0.1 | 0.03 | 0.16 | 0.14 |
| 3 | 60330356 | rs11708562 | 0.309 | 0.923 | -1.216 | -1.283 | 0.002 | 0.011 | 0.011 | 0 | 0.17 | 0.14 | 0.1 | 0.03 | 0.16 | 0.14 |
| 3 | 60330390 | rs11716358 | 0.291 | 0.957 | -1.216 | -1.284 | 0.001 | 0.011 | 0.011 | 0 | 0.17 | 0.14 | 0.1 | 0.03 | 0.17 | 0.14 |
| 3 | 60332301 | rs6771886 | -0.344 | 0.973 | -1.268 | -1.226 | -0.003 | 0.017 | 0.017 | 0.559 | 0.57 | 0.4 | 0.46 | 0.13 | 0.48 | 0.61 |

| | | | | | | | | | | | | | | | | |
|---|---|---|---|---|---|---|---|---|---|---|---|---|---|---|---|---|
| 3 | 60333449 | rs11920482 | -0.564 | 0.911 | -1.273 | -1.261 | -0.037 | 0.095 | 0.095 | 1.939 | 0.61 | 0.79 | 0.58 | 0.17 | 0.5 | 0.61 |
| 3 | 60337651 | rs6764498 | -0.729 | 1.147 | -1.778 | -0.695 | 0.002 | 0.014 | 0.014 | 1.672 | 0.55 | 0.4 | 0.65 | 0.38 | 0.56 | 0.74 |
| 3 | 60340650 | rs17606882 | 0.525 | 0.645 | -2.407 | -0.776 | -0.039 | 0.039 | 0.039 | 0 | 0.13 | 0.01 | 0.08 | 0.03 | 0.16 | 0.12 |
| 3 | 60341773 | rs6778312 | -0.033 | 0.414 | -1.921 | -0.406 | -0.006 | 0.013 | 0.013 | 0.063 | 0.34 | 0.08 | 0.47 | 0.3 | 0.31 | 0.45 |
| 3 | 60348943 | rs6784469 | -0.238 | 0.637 | -1.817 | -0.361 | -0.007 | 0.129 | 0.129 | 0.48 | 0.45 | 0.84 | 0.46 | 0.21 | 0.44 | 0.34 |
| 3 | 60348977 | rs6773303 | -0.263 | 0.714 | -2.089 | -0.852 | -0.013 | 0.208 | 0.208 | 0.049 | 0.55 | 0.84 | 0.56 | 0.07 | 0.54 | 0.4 |
| 3 | 60349109 | rs6773500 | 0.145 | 0.838 | -2.299 | -1.838 | -0.016 | 0.13 | 0.13 | 0.359 | 0.47 | 0.29 | 0.31 | 0.06 | 0.46 | 0.35 |
| 3 | 60349304 | rs965033 | -0.05 | 0.959 | -1.904 | -1.111 | -0.001 | 0 | 0 | 2.506 | 0.53 | 0.49 | 0.53 | 0.5 | 0.41 | 0.39 |
| 3 | 60349462 | rs970045 | 0.085 | 0.979 | -3.029 | -1.132 | -0.129 | 0.016 | 0.016 | 0 | 0.66 | 0.94 | 0.69 | 0.74 | 0.49 | 0.75 |
| 3 | 60350709 | rs6782631 | -0.234 | -2.08 | -1.023 | -0.51 | -0.004 | 0.001 | 0.001 | 0 | 0.43 | 0.47 | 0.42 | 0.45 | 0.58 | 0.51 |
| 3 | 60351746 | rs6786100 | -1.075 | -2.28 | -1.144 | -0.402 | 0.02 | 0.054 | 0.054 | 0.044 | 0.62 | 0.87 | 0.55 | 0.71 | 0.64 | 0.67 |
| 3 | 60353622 | rs12489534 | 0.128 | -2.123 | -0.927 | -0.571 | -0.017 | 0.008 | 0.008 | 3.207 | 0.35 | 0.29 | 0.34 | 0.27 | 0.44 | 0.27 |
| 3 | 60357382 | rs17062850 | 0.724 | -2.181 | -0.783 | -0.49 | 0.006 | -0.03 | -0.03 | 0 | 0.25 | 0.19 | 0.16 | 0.41 | 0.2 | 0.41 |
| 3 | 60358100 | rs3732551 | -1.036 | -2.265 | -0.64 | -0.549 | -0.005 | -0.006 | -0.006 | 0 | 0.6 | 0.46 | 0.53 | 0.69 | 0.63 | 0.67 |
| 3 | 60358517 | rs7636619 | 0.688 | -2.06 | -0.677 | -0.634 | -0.046 | -0.005 | -0.005 | 0 | 0.3 | 0.19 | 0.34 | 0.28 | 0.4 | 0.26 |
| 3 | 60360747 | rs6807084 | 0.881 | -2.488 | -0.489 | -0.509 | -0.029 | 0.018 | 0.018 | 2.914 | 0.2 | 0.25 | 0.14 | 0.27 | 0.06 | 0.17 |
| 3 | 60370362 | rs9847855 | 0.288 | -1.702 | -0.313 | -0.48 | -0.007 | 0.011 | 0.011 | 1.333 | 0.35 | 0.34 | 0.28 | 0.48 | 0.26 | 0.39 |
| 3 | 60372036 | rs2687169 | -0.432 | -1.663 | -0.411 | -0.595 | 0.019 | 0.002 | 0.002 | 0.108 | 0.51 | 0.39 | 0.5 | 0.48 | 0.63 | 0.54 |
| 3 | 60374028 | rs9868281 | 0.303 | -1.397 | -0.814 | -0.766 | -0.063 | 0.056 | 0.056 | 0 | 0.23 | 0.12 | 0.27 | 0 | 0.38 | 0.19 |
| 3 | 60379253 | rs17746541 | 1.197 | -0.877 | -0.82 | -0.55 | -0.048 | 0.027 | 0.027 | 0 | 0.16 | 0.07 | 0.25 | 0 | 0.37 | 0.15 |
| 3 | 60382023 | rs7646666 | 1.161 | -1.104 | -0.861 | -0.303 | -0.035 | -0.143 | -0.143 | 0.418 | 0.24 | 0.03 | 0.23 | 0.46 | 0.22 | 0.36 |
| 3 | 60388322 | rs7618479 | 1.157 | -1.15 | -0.394 | -0.37 | 0 | 0.036 | 0.036 | 0.541 | 0.26 | 0.31 | 0.26 | 0.46 | 0.24 | 0.38 |
| 3 | 60388743 | rs2734361 | -0.694 | -2.458 | 0.029 | -0.023 | 0.01 | 0.011 | 0.011 | 0 | 0.68 | 0.73 | 0.82 | 0.71 | 0.69 | 0.76 |
| 3 | 60396850 | rs242197 | 0.628 | -1.052 | -0.748 | 0.414 | 0.046 | -0.018 | -0.018 | 0 | 0.45 | 0.19 | 0.49 | 0.49 | 0.61 | 0.55 |
| 3 | 60405004 | rs13059601 | -0.479 | -0.832 | -0.493 | 0.908 | -0.042 | 0.029 | 0.029 | 0.064 | 0.17 | 0.24 | 0.26 | 0.26 | 0.05 | 0.25 |
| 3 | 60407864 | rs151743 | -0.309 | -0.45 | -0.822 | 0.468 | 0 | -0.005 | -0.005 | 0.466 | 0.42 | 0.5 | 0.46 | 0.44 | 0.42 | 0.38 |
| 3 | 60408404 | rs241693 | 0.344 | -3.191 | -1.178 | -0.024 | -0.115 | -0.122 | -0.122 | 0 | 0.29 | 0.08 | 0.31 | 0.46 | 0.55 | 0.37 |
| 3 | 60410706 | rs17063058 | 0.371 | -3.636 | -1.181 | -0.07 | -0.023 | -0.092 | -0.092 | 0.073 | 0.14 | 0.02 | 0.15 | 0.37 | 0.11 | 0.33 |
| 3 | 60416547 | rs1996828 | -0.272 | -4.355 | -0.69 | -0.426 | 0.006 | -0.005 | -0.005 | 0 | 0.66 | 0.7 | 0.52 | 0.46 | 0.73 | 0.54 |
| 3 | 60418721 | rs7621114 | 1.863 | -4.376 | -0.481 | -0.424 | 0.015 | -0.005 | -0.005 | 0 | 0.17 | 0.1 | 0.29 | 0.14 | 0.16 | 0.11 |
| 3 | 60420417 | rs409484 | 1.888 | -4.48 | -0.581 | -0.42 | -0.159 | -0.065 | -0.065 | 3.451 | 0.2 | 0.03 | 0.24 | 0.22 | 0.48 | 0.23 |
| 3 | 60421119 | rs2594250 | -0.111 | -4.165 | 0.392 | -0.107 | -0.007 | 0.005 | 0.005 | 0 | 0.67 | 0.63 | 0.52 | 0.47 | 0.73 | 0.55 |
| 3 | 60423009 | rs2687189 | -0.189 | -4.1 | 0.399 | -0.154 | -0.021 | 0.082 | 0.082 | 0.714 | 0.87 | 0.81 | 0.83 | 0.59 | 0.94 | 0.67 |
| 3 | 60434122 | rs381450 | -0.764 | -4.249 | 0.803 | -0.355 | 0.083 | 0.114 | 0.114 | 0.335 | 0.45 | 0.8 | 0.47 | 0.19 | 0.33 | 0.19 |
| 3 | 60438026 | rs410351 | -0.325 | -3.946 | 0.966 | -0.305 | -0.071 | -0.143 | -0.143 | 0.19 | 0.21 | 0.47 | 0.11 | 0.01 | 0.09 | 0.04 |
| 3 | 60439043 | rs17063156 | 0.518 | -3.734 | 0.987 | -0.147 | -0.02 | 0.085 | 0.085 | 0.081 | 0.12 | 0.18 | 0.14 | 0.36 | 0.05 | 0.3 |
| 3 | 60441079 | rs1947161 | -1.27 | -3.776 | 1.276 | -0.173 | 0.071 | 0.029 | 0.029 | 0.137 | 0.53 | 0.85 | 0.59 | 0.58 | 0.38 | 0.51 |
| 3 | 60445253 | rs391844 | -0.244 | -3.688 | 1.323 | -0.15 | -0.076 | -0.144 | -0.144 | 0 | 0.2 | 0.48 | 0.11 | 0.01 | 0.09 | 0.04 |
| 3 | 60447369 | rs445057 | -1.083 | -3.691 | 1.289 | -0.127 | -0.027 | 0.007 | 0.007 | 2.153 | 0.33 | 0.64 | 0.27 | 0.41 | 0.15 | 0.36 |
| 3 | 60448261 | rs424675 | -0.484 | -3.667 | 1.166 | -0.11 | -0.03 | 0.065 | 0.065 | 0 | 0.19 | 0.26 | 0.2 | 0.4 | 0.09 | 0.32 |
| 3 | 60448671 | rs9809480 | 1.116 | -3.387 | 0.375 | -0.132 | 0 | -0.007 | -0.007 | 0.229 | 0.2 | 0.13 | 0.31 | 0.17 | 0.23 | 0.15 |
| 3 | 60453227 | rs1597104 | 1.122 | -3.375 | 0.369 | -0.127 | 0.001 | -0.003 | -0.003 | 0.002 | 0.2 | 0.15 | 0.32 | 0.16 | 0.23 | 0.15 |
| 3 | 60458976 | rs3856654 | -0.387 | -3.203 | 0.203 | -0.215 | 0.015 | 0.007 | 0.007 | 0.132 | 0.46 | 0.66 | 0.55 | 0.57 | 0.35 | 0.51 |
| 3 | 60459291 | rs2734390 | -0.968 | -3.31 | 0.164 | -0.241 | 0.058 | 0.033 | 0.033 | 0.023 | 0.53 | 0.77 | 0.59 | 0.57 | 0.38 | 0.51 |
| 3 | 60461416 | rs7644852 | 0.243 | -3.4 | 0.155 | -0.202 | -0.008 | 0.007 | 0.007 | 0 | 0.32 | 0.41 | 0.47 | 0.53 | 0.28 | 0.46 |
| 3 | 60461994 | rs9868373 | 0.379 | -3.391 | -0.012 | -0.28 | 0.006 | -0.011 | -0.011 | 0 | 0.31 | 0.33 | 0.45 | 0.52 | 0.27 | 0.46 |
| 3 | 60464275 | rs453951 | -0.17 | -2.919 | 0.096 | -0.078 | 0.023 | 0.001 | 0.001 | 0 | 0.46 | 0.4 | 0.51 | 0.52 | 0.31 | 0.51 |
| 3 | 60465279 | rs1447971 | -0.218 | -1.916 | -0.475 | 0.455 | 0.047 | 0.013 | 0.013 | 0.141 | 0.53 | 0.72 | 0.57 | 0.55 | 0.35 | 0.55 |
| 3 | 60472107 | rs9818349 | -1.194 | -1.067 | -0.173 | 1.025 | -0.018 | 0.06 | 0.06 | 0.11 | 0.17 | 0.25 | 0.26 | 0.48 | 0.12 | 0.34 |
| 3 | 60472209 | rs9856634 | -0.167 | -0.988 | -0.583 | 1.078 | 0.009 | -0.035 | -0.035 | 0 | 0.11 | 0.07 | 0.18 | 0.38 | 0.07 | 0.24 |
| 3 | 60472710 | rs410072 | -0.879 | -0.715 | -0.492 | 1.03 | -0.035 | 0.054 | 0.054 | 0 | 0.24 | 0.36 | 0.31 | 0.48 | 0.15 | 0.34 |
| 3 | 60479380 | rs242219 | -0.267 | 0.071 | -1.016 | 0.647 | 0.006 | -0.005 | -0.005 | 0.977 | 0.47 | 0.31 | 0.5 | 0.43 | 0.51 | 0.49 |
| 3 | 60481459 | rs2197751 | -0.153 | -0.065 | -0.991 | -0.136 | 0.001 | -0.009 | -0.009 | 0.457 | 0.51 | 0.41 | 0.5 | 0.81 | 0.43 | 0.58 |
| 3 | 60484433 | rs6793324 | -0.34 | -0.24 | -1.196 | -0.186 | -0.045 | 0.031 | 0.031 | 1.67 | 0.63 | 0.71 | 0.5 | 0.73 | 0.55 | 0.59 |
| 3 | 60487074 | rs17661314 | -0.029 | 0.039 | -0.818 | 0.367 | 0.011 | 0.029 | 0.029 | 0 | 0.07 | 0.12 | 0.05 | 0.11 | 0.1 | 0.08 |
| 3 | 60487247 | rs9881736 | 0.338 | -0.454 | -0.939 | 0.324 | 0 | 0.045 | 0.045 | 3.274 | 0.19 | 0.08 | 0.22 | 0.08 | 0.24 | 0.06 |
| 3 | 60487313 | rs2121865 | -0.876 | -1.198 | -0.931 | 0.51 | 0.001 | 0.001 | 0.001 | 0 | 0.48 | 0.42 | 0.41 | 0.62 | 0.49 | 0.55 |
| 3 | 60487578 | rs17609699 | 0.8 | -1.272 | -1.011 | 0.678 | -0.009 | 0.019 | 0.019 | 0 | 0.15 | 0.03 | 0.08 | 0.12 | 0.13 | 0.22 |
| 3 | 60491103 | rs6789538 | -0.131 | -1.532 | -0.485 | 0.556 | -0.009 | -0.023 | -0.023 | 0 | 0.23 | 0.29 | 0.23 | 0.15 | 0.15 | 0.25 |
| 3 | 60491407 | rs6783772 | -0.34 | -1.626 | -0.384 | 0.513 | -0.022 | -0.04 | -0.04 | 2.134 | 0.25 | 0.4 | 0.25 | 0.15 | 0.15 | 0.26 |
| 3 | 60491503 | rs6783802 | -1.213 | -1.665 | -0.361 | 0.396 | -0.001 | -0.004 | -0.004 | 0 | 0.49 | 0.51 | 0.4 | 0.26 | 0.48 | 0.37 |
| 3 | 60492136 | rs10510846 | 0.325 | -1.597 | -0.426 | 0.396 | -0.019 | 0.041 | 0.041 | 1.517 | 0.23 | 0.11 | 0.15 | 0.11 | 0.33 | 0.12 |
| 3 | 60496077 | rs1447974 | -0.683 | -0.613 | -0.929 | 0.287 | 0 | -0.014 | -0.014 | 0 | 0.57 | 0.53 | 0.61 | 0.31 | 0.58 | 0.45 |
| 3 | 60496302 | rs9863826 | -0.825 | -0.554 | -0.834 | 0.509 | -0.003 | 0.007 | 0.007 | 0.064 | 0.58 | 0.54 | 0.62 | 0.47 | 0.63 | 0.52 |
| 3 | 60498130 | rs6771732 | -0.935 | -0.748 | -0.567 | 0.784 | -0.035 | 0.001 | 0.001 | 0.058 | 0.61 | 0.75 | 0.64 | 0.63 | 0.55 | 0.63 |
| 3 | 60498172 | rs13101027 | 0.821 | 0.063 | -0.602 | 0.665 | -0.015 | -0.001 | -0.001 | 0 | 0.19 | 0.29 | 0.21 | 0.2 | 0.11 | 0.25 |
| 3 | 60498520 | rs9828971 | 0.181 | 0.079 | -0.638 | 0.637 | 0.002 | -0.004 | -0.004 | 0 | 0.33 | 0.46 | 0.46 | 0.32 | 0.29 | 0.39 |
| 3 | 60498732 | rs7634921 | 1.518 | 0.116 | -0.854 | 0.563 | 0.015 | 0.092 | 0.092 | 5.954 | 0.21 | 0.07 | 0.12 | 0 | 0.23 | 0.04 |
| 3 | 60500549 | rs2017941 | 1.964 | 0.121 | -0.739 | 0.669 | 0.01 | 0.004 | 0.004 | 0 | 0.13 | 0.18 | 0.25 | 0.16 | 0.18 | 0.15 |

| | | | | | | | | | | | | | | | | |
|---|---|---|---|---|---|---|---|---|---|---|---|---|---|---|---|---|
| 3 | 60501861 | rs9829514 | -0.108 | -0.559 | -0.655 | 0.742 | -0.008 | -0.02 | -0.02 | 0 | 0.41 | 0.45 | 0.33 | 0.21 | 0.35 | 0.29 |
| 3 | 60501884 | rs9829628 | 1.32 | -0.524 | -1.028 | 0.583 | 0.016 | 0.087 | 0.087 | 6.051 | 0.2 | 0.07 | 0.1 | 0.04 | 0.18 | 0.04 |
| 3 | 60503108 | rs9311767 | -0.701 | -0.187 | -0.787 | 0.951 | 0.023 | 0.036 | 0.036 | 3.495 | 0.54 | 0.69 | 0.43 | 0.62 | 0.47 | 0.56 |
| 3 | 60506349 | rs17063425 | 0.593 | -0.063 | -0.351 | 1.364 | -0.03 | 0.155 | 0.155 | 0 | 0.07 | 0.11 | 0.05 | 0.3 | 0.03 | 0.2 |
| 3 | 60511133 | rs241706 | -0.418 | 0.161 | -0.673 | 0.773 | -0.01 | 0.008 | 0.008 | 0 | 0.41 | 0.43 | 0.45 | 0.54 | 0.35 | 0.53 |
| 3 | 60512505 | rs241705 | -1.003 | -0.364 | -1.183 | -0.184 | 0.017 | 0.123 | 0.123 | 1.849 | 0.59 | 0.8 | 0.77 | 0.92 | 0.58 | 0.79 |
| 3 | 60512522 | rs17063443 | 1.962 | -0.481 | -1.2 | 0.184 | -0.016 | -0.049 | -0.049 | 0 | 0.13 | 0.05 | 0.2 | 0.23 | 0.1 | 0.21 |
| 3 | 60512889 | rs241704 | -0.121 | -0.562 | -0.801 | 0.25 | 0.004 | 0.004 | 0.004 | 4.117 | 0.52 | 0.62 | 0.64 | 0.74 | 0.56 | 0.67 |
| 3 | 60513123 | rs4679545 | 1.524 | -0.572 | -0.791 | 0.326 | -0.008 | -0.005 | -0.005 | 0 | 0.26 | 0.23 | 0.23 | 0.18 | 0.21 | 0.14 |
| 3 | 60515778 | rs717821 | 1.082 | -0.47 | -0.424 | 0.23 | 0.051 | 0.032 | 0.032 | 0 | 0.07 | 0.26 | 0.05 | 0.08 | 0.09 | 0.07 |
| 3 | 60516256 | rs241700 | 0.421 | 0.472 | -0.986 | 0.324 | 0.002 | 0.019 | 0.019 | 0.143 | 0.32 | 0.44 | 0.22 | 0.42 | 0.3 | 0.39 |
| 3 | 60518998 | rs17063477 | 1.359 | 0.822 | -1.175 | 0.266 | -0.018 | -0.02 | -0.02 | 0 | 0.14 | 0.08 | 0.22 | 0.23 | 0.14 | 0.21 |
| 3 | 60519576 | rs241698 | -0.348 | 0.83 | -1.126 | 0.294 | 0.001 | 0.004 | 0.004 | 1.474 | 0.43 | 0.43 | 0.46 | 0.59 | 0.55 | 0.54 |
| 3 | 60521596 | rs17668453 | 1.35 | 1.093 | -1.126 | 0.274 | -0.035 | -0.012 | -0.012 | 0 | 0.13 | 0.07 | 0.24 | 0.17 | 0.26 | 0.14 |
| 3 | 60523330 | rs2594257 | -1.418 | 0.877 | -1.208 | 0.096 | 0.005 | 0.036 | 0.036 | 1.346 | 0.67 | 0.84 | 0.69 | 0.77 | 0.7 | 0.77 |
| 3 | 60531242 | rs6807877 | 2.221 | 0.855 | -1.291 | -0.033 | 0.006 | 0.024 | 0.024 | 0.212 | 0.11 | 0.1 | 0.07 | 0 | 0.07 | 0.1 |
| 3 | 60536420 | rs2006807 | 0.174 | 0.748 | -1.068 | 0.168 | 0.029 | -0.01 | -0.01 | 0 | 0.3 | 0.45 | 0.46 | 0.29 | 0.35 | 0.35 |
| 3 | 60536552 | rs107342 | 1.458 | 1.801 | -1.408 | 0.38 | -0.038 | -0.024 | -0.024 | 0.021 | 0.07 | 0.01 | 0.07 | 0.07 | 0.1 | 0.04 |
| 3 | 60540665 | rs241669 | -1.006 | 1.866 | -1.02 | 0.244 | -0.033 | -0.02 | -0.02 | 0 | 0.91 | 0.98 | 0.94 | 0.92 | 0.85 | 0.95 |
| 3 | 60542151 | rs241671 | 0.155 | 1.805 | -0.84 | 0.202 | -0.016 | -0.012 | -0.012 | 0 | 0.55 | 0.76 | 0.51 | 0.47 | 0.49 | 0.62 |
| 3 | 60542525 | rs241672 | 0.149 | 2.107 | -0.661 | 0.398 | -0.004 | -0.019 | -0.019 | 0 | 0.61 | 0.76 | 0.54 | 0.49 | 0.55 | 0.63 |
| 3 | 60543675 | rs17031618 | 2.146 | 2.102 | -0.695 | 0.282 | -0.021 | 0.074 | 0.074 | 7.499 | 0.16 | 0.03 | 0.1 | 0 | 0.18 | 0.02 |
| 3 | 60544132 | rs6805296 | 0.339 | 2.112 | -0.683 | 0.331 | 0 | 0 | 0 | 0 | 0.31 | 0.21 | 0.4 | 0.46 | 0.31 | 0.32 |
| 3 | 60545602 | rs180998 | 1.14 | 2.533 | -0.845 | 0.217 | -0.065 | 0.039 | 0.039 | 0 | 0.17 | 0.11 | 0.29 | 0.11 | 0.27 | 0.17 |
| 3 | 60546396 | rs12487370 | 0.995 | 2.462 | -0.769 | 0.311 | -0.001 | 0.027 | 0.027 | 0 | 0.14 | 0.1 | 0.3 | 0.39 | 0.12 | 0.31 |
| 3 | 60546807 | rs181002 | 0.808 | 2.363 | -0.55 | 0.203 | 0.011 | -0.008 | -0.008 | 0 | 0.43 | 0.39 | 0.26 | 0.5 | 0.36 | 0.5 |
| 3 | 60546856 | rs181003 | 0.792 | 2.595 | -0.338 | 0.296 | 0.001 | 0 | 0 | 0 | 0.59 | 0.46 | 0.33 | 0.5 | 0.47 | 0.52 |
| 3 | 60547602 | rs1447958 | -1.337 | 3.054 | -0.672 | 0.315 | -0.055 | -0.005 | -0.005 | 0.201 | 0.91 | 0.98 | 0.88 | 0.93 | 0.79 | 0.94 |
| 3 | 60548728 | rs1447959 | -0.342 | 3.122 | -0.545 | 0.437 | 0.001 | 0.007 | 0.007 | 0 | 0.32 | 0.39 | 0.53 | 0.45 | 0.31 | 0.46 |
| 3 | 60550891 | rs7634073 | 0.569 | 3.216 | -0.563 | 0.522 | 0.011 | -0.038 | -0.038 | 0.11 | 0.19 | 0.34 | 0.24 | 0.11 | 0.2 | 0.16 |
| 3 | 60554521 | rs2594259 | -1.221 | 3.006 | -1.262 | 0.589 | -0.055 | -0.042 | -0.042 | 0 | 0.91 | 0.98 | 0.88 | 0.89 | 0.79 | 0.94 |
| 3 | 60557685 | rs2301156 | 0.563 | 3.125 | -1.227 | 0.732 | 0.002 | 0.042 | 0.042 | 0.534 | 0.15 | 0.1 | 0.38 | 0.5 | 0.17 | 0.31 |
| 3 | 60557852 | rs2255772 | -0.201 | 3.008 | -1.232 | 0.68 | -0.026 | -0.03 | -0.03 | 0 | 0.3 | 0.22 | 0.61 | 0.62 | 0.42 | 0.46 |
| 3 | 60559047 | rs2856017 | 1.476 | 1.64 | -1.296 | 0.438 | -0.028 | -0.015 | -0.015 | 0.155 | 0.1 | 0.02 | 0.06 | 0.07 | 0.1 | 0.12 |
| 3 | 60562113 | rs12495226 | 0.812 | 1.638 | -0.707 | 0.487 | -0.013 | -0.042 | -0.042 | 1.386 | 0.11 | 0.02 | 0.31 | 0.34 | 0.14 | 0.3 |
| 3 | 60563119 | rs11130776 | 0.874 | 1.632 | -0.792 | 0.622 | 0.005 | -0.038 | -0.038 | 0 | 0.11 | 0.02 | 0.3 | 0.34 | 0.1 | 0.29 |
| 3 | 60563160 | rs10510849 | 0.246 | 1.585 | -0.801 | 0.532 | 0.015 | -0.012 | -0.012 | 0 | 0.26 | 0.13 | 0.36 | 0.34 | 0.2 | 0.31 |
| 3 | 60563632 | rs3937353 | -1.002 | 1.754 | -0.944 | 0.492 | -0.005 | 0.025 | 0.025 | 0 | 0.49 | 0.58 | 0.69 | 0.46 | 0.53 | 0.48 |
| 3 | 60566161 | rs17669257 | 0.333 | 1.785 | -0.937 | 0.271 | -0.067 | 0.039 | 0.039 | 0 | 0.19 | 0.13 | 0.3 | 0.13 | 0.34 | 0.18 |
| 3 | 60566492 | rs10451993 | -0.937 | 1.928 | -0.991 | 0.401 | 0.005 | 0.005 | 0.005 | 0 | 0.48 | 0.59 | 0.69 | 0.52 | 0.55 | 0.48 |
| 3 | 60567102 | rs10510850 | 0.669 | 1.51 | -1.607 | 0.831 | 0.002 | -0.023 | -0.023 | 0 | 0.1 | 0.02 | 0.29 | 0.4 | 0.1 | 0.29 |
| 3 | 60567427 | rs2734367 | -0.206 | 1.943 | -1.227 | 0.827 | -0.012 | -0.007 | -0.007 | 0 | 0.91 | 0.95 | 0.94 | 0.92 | 0.85 | 0.94 |
| 3 | 60568253 | rs9833533 | 0.998 | 1.916 | -1.421 | 0.762 | 0.009 | -0.007 | -0.007 | 0.19 | 0.15 | 0.09 | 0.06 | 0 | 0.09 | 0.01 |
| 3 | 60571772 | rs2682963 | -0.133 | 1.906 | -1.379 | 0.27 | -0.044 | -0.044 | -0.044 | 0 | 0.9 | 0.98 | 0.93 | 0.88 | 0.84 | 0.94 |
| 3 | 60572073 | rs7651468 | 0.69 | 1.642 | -0.825 | 0.509 | 0.007 | -0.002 | -0.002 | 0 | 0.16 | 0.09 | 0.07 | 0 | 0.11 | 0.02 |
| 3 | 60573681 | rs13079016 | 0.139 | 2.045 | -1.266 | 0.658 | -0.051 | 0.012 | 0.012 | 0 | 0.17 | 0.14 | 0.27 | 0.11 | 0.3 | 0.15 |
| 3 | 60573769 | rs9826800 | -0.56 | 0.639 | -0.47 | 1.207 | 0 | 0.001 | 0.001 | 4.476 | 0.24 | 0.19 | 0.36 | 0.46 | 0.18 | 0.22 |
| 3 | 60574069 | rs6785709 | -1.748 | 1.175 | -0.4 | 0.678 | 0.003 | -0.018 | -0.018 | 0 | 0.09 | 0.18 | 0.06 | 0.06 | 0.08 | 0.06 |
| 3 | 60574546 | rs9865739 | -1.095 | 1.163 | -0.545 | 0.803 | 0.005 | -0.008 | -0.008 | 0.886 | 0.07 | 0.19 | 0.06 | 0.06 | 0.07 | 0.05 |
| 3 | 60576324 | rs2049799 | -0.808 | 1.546 | -0.967 | 1.256 | -0.013 | -0.028 | -0.028 | 0 | 0.23 | 0.18 | 0.27 | 0.45 | 0.26 | 0.36 |
| 3 | 60576769 | rs2734370 | -0.357 | 1.471 | -1 | 0.206 | 0 | 0.022 | 0.022 | 1.509 | 0.51 | 0.42 | 0.45 | 0.17 | 0.54 | 0.31 |
| 3 | 60581806 | rs12638023 | 0.113 | 1.474 | -0.733 | 0.292 | 0.007 | 0.016 | 0.016 | 0 | 0.21 | 0.21 | 0.3 | 0.6 | 0.25 | 0.51 |
| 3 | 60585933 | rs2856025 | -1.297 | 0.914 | -0.619 | 0.027 | 0.001 | 0.002 | 0.002 | 0.68 | 0.77 | 0.74 | 0.9 | 0.83 | 0.89 | 0.89 |
| 3 | 60587580 | rs11712757 | 0.504 | 0.033 | -1.057 | -0.506 | -0.034 | 0.147 | 0.147 | 1.261 | 0.45 | 0.08 | 0.51 | 0.23 | 0.59 | 0.36 |
| 3 | 60588627 | rs2856028 | -2.029 | -0.294 | -1.121 | -0.558 | 0.027 | -0.006 | -0.006 | 0 | 0.81 | 0.97 | 0.93 | 0.81 | 0.88 | 0.88 |
| 3 | 60589410 | rs2856029 | -1.474 | -0.339 | -1.03 | -0.431 | 0.043 | -0.004 | -0.004 | 0 | 0.78 | 0.85 | 0.92 | 0.78 | 0.87 | 0.88 |
| 3 | 60589986 | rs1158425 | -1.405 | 0.932 | -0.923 | -0.689 | -0.016 | 0.005 | 0.005 | 0 | 0.76 | 0.85 | 0.83 | 0.77 | 0.7 | 0.84 |
| 3 | 60590063 | rs724025 | 0.572 | 1.035 | -0.809 | -0.784 | -0.016 | 0.111 | 0.111 | 0.012 | 0.4 | 0.1 | 0.44 | 0.22 | 0.42 | 0.31 |
| 3 | 60591506 | rs2682971 | 0.697 | 0.592 | -0.232 | -0.748 | -0.024 | -0.046 | -0.046 | 0 | 0.09 | 0.13 | 0.07 | 0 | 0.05 | 0.02 |
| 3 | 60592348 | rs11720214 | 0.288 | 0.875 | -0.378 | -0.743 | -0.02 | 0.147 | 0.147 | 1.143 | 0.46 | 0.12 | 0.5 | 0.23 | 0.58 | 0.36 |
| 3 | 60592751 | rs13100376 | 1.283 | 1.398 | -0.632 | -0.641 | -0.038 | 0.024 | 0.024 | 0.379 | 0.1 | 0.01 | 0.12 | 0.04 | 0.11 | 0.08 |
| 3 | 60594367 | rs2856034 | -0.334 | 0.866 | -1.135 | -1.047 | 0.019 | 0.092 | 0.092 | 0 | 0.29 | 0.1 | 0.18 | 0.07 | 0.27 | 0.11 |
| 3 | 60595403 | rs939496 | -1.32 | 0.441 | -1.173 | -1.225 | 0.059 | 0.087 | 0.087 | 0 | 0.6 | 0.75 | 0.73 | 0.81 | 0.75 | 0.74 |
| 3 | 60597001 | rs17670198 | 0.433 | 1.24 | -0.926 | -0.648 | 0.076 | 0.04 | 0.04 | 0 | 0.18 | 0.3 | 0.2 | 0.25 | 0.29 | 0.15 |
| 3 | 60599282 | rs715381 | 0.662 | 1.255 | -1.025 | -0.649 | 0.013 | 0.003 | 0.003 | 0 | 0.16 | 0.21 | 0.19 | 0.21 | 0.29 | 0.15 |
| 3 | 60600915 | rs2682979 | -2.873 | 0.681 | -1.028 | -0.767 | 0.017 | 0.012 | 0.012 | 0.236 | 0.76 | 0.75 | 0.89 | 0.82 | 0.88 | 0.87 |
| 3 | 60602816 | rs2246924 | 1.341 | 0.698 | -1.099 | -0.93 | 0.005 | 0.022 | 0.022 | 0.364 | 0.13 | 0.06 | 0.12 | 0 | 0.15 | 0.02 |
| 3 | 60603621 | rs7632868 | -0.661 | 0.688 | -1.112 | -0.806 | -0.015 | 0.079 | 0.079 | 0 | 0.45 | 0.59 | 0.49 | 0.81 | 0.47 | 0.7 |
| 3 | 60604489 | rs13093294 | 1.453 | 0.953 | -1.133 | -0.592 | -0.018 | 0.018 | 0.018 | 0 | 0.13 | 0.02 | 0.14 | 0.05 | 0.14 | 0.1 |

| | | | | | | | | | | | | | | | | |
|---|---|---|---|---|---|---|---|---|---|---|---|---|---|---|---|---|
| 3 | 60607896 | rs721207 | -0.528 | 0.786 | -0.779 | -0.34 | 0.001 | -0.007 | -0.007 | 0 | 0.42 | 0.52 | 0.54 | 0.47 | 0.45 | 0.57 |
| 3 | 60611613 | rs2363670 | -0.1 | 1.181 | -0.914 | -0.597 | -0.068 | 0.049 | 0.049 | 0 | 0.28 | 0.13 | 0.3 | 0.16 | 0.37 | 0.27 |
| 3 | 60613659 | rs7633853 | -0.171 | 1.139 | -0.698 | -0.662 | 0.01 | 0.003 | 0.003 | 0 | 0.32 | 0.42 | 0.44 | 0.64 | 0.34 | 0.59 |
| 3 | 60614899 | rs12152491 | -1.553 | -0.098 | -0.535 | -0.489 | 0.008 | 0 | 0 | 0.076 | 0.62 | 0.6 | 0.76 | 0.86 | 0.71 | 0.86 |
| 3 | 60615310 | rs11926787 | -0.14 | 0.262 | -0.502 | -0.471 | 0.028 | -0.006 | -0.006 | 0 | 0.35 | 0.49 | 0.5 | 0.64 | 0.45 | 0.61 |
| 3 | 60616153 | rs6762641 | -0.083 | 0.336 | -0.548 | -0.474 | -0.006 | 0.023 | 0.023 | 0 | 0.25 | 0.13 | 0.26 | 0.18 | 0.25 | 0.22 |
| 3 | 60616790 | rs10510852 | -0.034 | 0.477 | -0.502 | -0.402 | -0.002 | 0.005 | 0.005 | 0 | 0.24 | 0.17 | 0.25 | 0.18 | 0.24 | 0.22 |
| 3 | 60620284 | rs12637393 | -0.534 | 0.76 | -0.35 | 0.096 | 0.036 | -0.014 | -0.014 | 0 | 0.36 | 0.46 | 0.49 | 0.64 | 0.45 | 0.61 |
| 3 | 60625598 | rs2856058 | 0.049 | 0.245 | 0.111 | 0.467 | -0.003 | 0.013 | 0.013 | 0 | 0.82 | 0.74 | 0.77 | 0.83 | 0.83 | 0.85 |
| 3 | 60626006 | rs9843949 | -0.299 | 0.003 | 0.194 | 0.72 | -0.005 | 0.029 | 0.029 | 0 | 0.24 | 0.31 | 0.19 | 0.51 | 0.23 | 0.38 |
| 3 | 60631936 | rs2682931 | 0.277 | -0.103 | 0.406 | -0.063 | -0.009 | -0.009 | -0.009 | 0.573 | 0.77 | 0.66 | 0.74 | 0.83 | 0.79 | 0.79 |
| 3 | 60636061 | rs2856076 | 1.544 | -0.2 | 0.026 | -0.327 | 0.036 | 0.182 | 0.182 | 0 | 0.49 | 0.24 | 0.53 | 0.11 | 0.54 | 0.31 |
| 3 | 60639504 | rs7649096 | 0.059 | -0.336 | 0.263 | -0.15 | 0.002 | -0.004 | -0.004 | 0 | 0.22 | 0.21 | 0.17 | 0.74 | 0.21 | 0.42 |
| 3 | 60640906 | rs9311770 | -0.278 | -0.335 | 0.244 | -0.145 | -0.01 | 0.011 | 0.011 | 0 | 0.23 | 0.35 | 0.18 | 0.74 | 0.2 | 0.43 |
| 3 | 60643045 | rs2213215 | 0.355 | 0.337 | -0.19 | -0.343 | -0.049 | 0.004 | 0.004 | 0 | 0.13 | 0.05 | 0.26 | 0.09 | 0.21 | 0.12 |
| 3 | 60647540 | rs1735448 | 0.489 | -0.093 | 0.379 | -0.188 | -0.026 | -0.039 | -0.039 | 0 | 0.74 | 0.54 | 0.71 | 0.85 | 0.8 | 0.76 |
| 3 | 60657068 | rs17063778 | 0.582 | -0.248 | 0.387 | -0.081 | -0.015 | -0.035 | -0.035 | 0 | 0.07 | 0.12 | 0.05 | 0.01 | 0.04 | 0.1 |
| 3 | 60657213 | rs1716709 | 0.199 | -0.316 | 0.384 | -0.094 | -0.025 | 0.003 | 0.003 | 0.006 | 0.83 | 0.77 | 0.79 | 0.85 | 0.9 | 0.81 |
| 3 | 60658871 | rs1629179 | 1.205 | -0.413 | 0.326 | -0.238 | 0.041 | 0.067 | 0.067 | 0 | 0.58 | 0.31 | 0.61 | 0.13 | 0.69 | 0.39 |
| 3 | 60660179 | rs1716714 | -0.19 | -0.158 | 0.586 | 0.224 | -0.006 | 0.021 | 0.021 | 0 | 0.92 | 0.89 | 0.85 | 0.87 | 0.94 | 0.9 |
| 3 | 60661146 | rs17063781 | 0.543 | -0.201 | 0.707 | 0.196 | -0.019 | -0.032 | -0.032 | 0.294 | 0.07 | 0.12 | 0.05 | 0.01 | 0.03 | 0.1 |
| 3 | 60662282 | rs1735443 | 0.925 | -0.221 | 0.696 | 0.208 | -0.059 | 0.03 | 0.03 | 0.085 | 0.83 | 0.55 | 0.79 | 0.85 | 0.91 | 0.8 |
| 3 | 60664075 | rs1735444 | 1.144 | -0.27 | 0.775 | 0.275 | -0.061 | 0.027 | 0.027 | 0.324 | 0.82 | 0.51 | 0.78 | 0.81 | 0.91 | 0.8 |
| 3 | 60664643 | rs12633258 | 0.419 | -0.249 | 0.415 | 0.664 | 0.003 | -0.028 | -0.028 | 0 | 0.08 | 0 | 0.04 | 0.47 | 0.05 | 0.22 |
| 3 | 60681361 | rs1716722 | 1.401 | 0.036 | 0.745 | -0.224 | -0.063 | -0.016 | -0.016 | 4.77 | 0.8 | 0.59 | 0.77 | 0.88 | 0.88 | 0.8 |
| 3 | 60683737 | rs1735457 | -0.145 | 0.056 | 0.628 | -0.207 | 0.003 | -0.005 | -0.005 | 0.279 | 0.92 | 0.92 | 0.85 | 0.87 | 0.94 | 0.91 |
| 3 | 60689476 | rs9861551 | -0.566 | -0.015 | 0.795 | -0.305 | -0.02 | -0.045 | -0.045 | 0 | 0.1 | 0.36 | 0.06 | 0 | 0.03 | 0.1 |
| 3 | 60692133 | rs9838669 | -0.182 | 0.025 | 0.848 | -0.286 | -0.013 | -0.037 | -0.037 | 0 | 0.09 | 0.34 | 0.05 | 0 | 0.03 | 0.1 |
| 3 | 60693253 | rs1716725 | -0.502 | 0.647 | 0.487 | -0.328 | -0.008 | 0.003 | 0.003 | 0 | 0.11 | 0.07 | 0.22 | 0.08 | 0.2 | 0.13 |
| 3 | 60703616 | rs1735468 | 1.097 | 0.771 | 0.503 | -0.432 | -0.022 | -0.056 | -0.056 | 0 | 0.74 | 0.53 | 0.7 | 0.89 | 0.77 | 0.75 |
| 3 | 60708889 | rs1716748 | -1.393 | 0.804 | 0.308 | -0.421 | 0.04 | -0.027 | -0.027 | 0 | 0.3 | 0.49 | 0.4 | 0.21 | 0.38 | 0.34 |
| 3 | 60711659 | rs7643144 | 1.735 | 0.577 | 0.127 | -0.658 | -0.008 | 0.05 | 0.05 | 0.009 | 0.57 | 0.35 | 0.47 | 0.8 | 0.55 | 0.53 |
| 3 | 60712075 | rs1716753 | 1.427 | 0.773 | 0.074 | -0.687 | 0.003 | 0.004 | 0.004 | 0 | 0.61 | 0.42 | 0.62 | 0.88 | 0.64 | 0.63 |
| 3 | 60714931 | rs7636357 | 1.167 | 0.996 | -0.27 | -0.439 | 0.011 | -0.018 | -0.018 | 0.063 | 0.66 | 0.6 | 0.52 | 0.81 | 0.59 | 0.63 |
| 3 | 60720891 | rs17627738 | -0.022 | 1.365 | -0.702 | -0.156 | -0.027 | -0.026 | -0.026 | 0 | 0.07 | 0.02 | 0.1 | 0.11 | 0.14 | 0.08 |
| 3 | 60721185 | rs1735477 | -0.37 | 1.154 | -0.774 | -0.343 | -0.01 | 0.055 | 0.055 | 0 | 0.89 | 0.98 | 0.87 | 1 | 0.92 | 0.94 |
| 3 | 60723910 | rs741891 | 0.25 | 1.365 | -0.798 | -0.161 | -0.001 | -0.001 | -0.001 | 0.406 | 0.81 | 0.81 | 0.78 | 0.77 | 0.84 | 0.82 |
| 3 | 60727326 | rs17031633 | -0.142 | 0.493 | -0.211 | -0.523 | -0.044 | -0.082 | -0.082 | 0 | 0.13 | 0.34 | 0.08 | 0 | 0.08 | 0.13 |
| 3 | 60727499 | rs2205359 | 1.69 | 0.224 | -0.959 | -1.117 | -0.002 | -0.001 | -0.001 | 0 | 0.37 | 0.37 | 0.32 | 0.61 | 0.32 | 0.46 |
| 3 | 60727958 | rs760314 | -0.104 | 0.562 | -1.303 | -1.22 | -0.016 | 0.069 | 0.069 | 0 | 0.69 | 0.86 | 0.69 | 0.88 | 0.64 | 0.83 |
| 3 | 60729115 | rs11914567 | 0.865 | 0.281 | -1.579 | -1.461 | 0.046 | 0.075 | 0.075 | 0 | 0.39 | 0.23 | 0.26 | 0.03 | 0.28 | 0.27 |
| 3 | 60730525 | rs17063899 | 0.503 | 0.848 | -1.33 | -0.716 | -0.008 | -0.009 | -0.009 | 0 | 0.1 | 0.05 | 0.06 | 0.15 | 0.08 | 0.14 |
| 3 | 60731999 | rs6783059 | -0.63 | 0.876 | -1.235 | -0.766 | 0.009 | -0.013 | -0.013 | 0 | 0.41 | 0.51 | 0.49 | 0.31 | 0.44 | 0.36 |
| 3 | 60733152 | rs3920477 | 0.413 | 0.911 | -1.438 | -0.098 | -0.018 | -0.021 | -0.021 | 0 | 0.55 | 0.68 | 0.5 | 0.54 | 0.52 | 0.65 |
| 3 | 60735438 | rs17628405 | 1.164 | 0.874 | -1.775 | -0.435 | -0.042 | 0.026 | 0.026 | 0 | 0.19 | 0.07 | 0.26 | 0.17 | 0.22 | 0.25 |
| 3 | 60737133 | rs6446152 | -1.038 | 0.725 | -1.76 | -0.532 | -0.018 | -0.015 | -0.015 | 0.012 | 0.94 | 0.99 | 0.94 | 0.91 | 0.91 | 0.94 |
| 3 | 60740480 | rs9311779 | 0.125 | 0.464 | -1.746 | -1.084 | 0.005 | 0.023 | 0.023 | 0 | 0.53 | 0.57 | 0.4 | 0.7 | 0.5 | 0.58 |
| 3 | 60741689 | rs9850579 | -0.195 | 0.934 | -1.679 | -0.956 | 0.001 | -0.013 | -0.013 | 0.145 | 0.79 | 0.82 | 0.84 | 0.71 | 0.8 | 0.85 |
| 3 | 60742058 | rs13063937 | 1.213 | 1.335 | -1.767 | -1.35 | -0.055 | 0.05 | 0.05 | 0 | 0.15 | 0.06 | 0.27 | 0.01 | 0.26 | 0.22 |
| 3 | 60743058 | rs4974255 | 0.381 | 0.937 | -1.658 | -1.366 | -0.055 | 0.099 | 0.099 | 2.074 | 0.31 | 0.11 | 0.36 | 0.02 | 0.4 | 0.24 |
| 3 | 60744333 | rs7612949 | 0.2 | 0.444 | -1.08 | -0.72 | -0.002 | 0 | 0 | 0 | 0.44 | 0.51 | 0.46 | 0.58 | 0.39 | 0.56 |
| 3 | 60750062 | rs1569346 | -1.056 | 0.641 | -1.233 | -1.413 | -0.033 | 0.123 | 0.123 | 0 | 0.42 | 0.9 | 0.38 | 0.23 | 0.39 | 0.34 |
| 3 | 60752059 | rs11130781 | -1.206 | 0.909 | -1.37 | -1.496 | -0.072 | 0.177 | 0.177 | 0.514 | 0.48 | 0.96 | 0.49 | 0.23 | 0.53 | 0.36 |
| 3 | 60753126 | rs13098466 | -0.802 | 0.865 | -0.667 | -1.684 | 0.014 | -0.008 | -0.008 | 0 | 0.28 | 0.36 | 0.37 | 0.21 | 0.43 | 0.26 |
| 3 | 60759462 | rs11710207 | -0.929 | 0.83 | -0.68 | -1.699 | 0.007 | -0.005 | -0.005 | 0 | 0.29 | 0.35 | 0.37 | 0.21 | 0.43 | 0.26 |
| 3 | 60761015 | rs11130783 | 0.174 | 0.95 | -0.158 | -1.479 | -0.019 | -0.065 | -0.065 | 0.932 | 0.15 | 0.22 | 0.08 | 0 | 0.1 | 0.03 |
| 3 | 60767452 | rs12492901 | -0.011 | 1.489 | 0.069 | -1.211 | 0.099 | 0.067 | 0.067 | 0 | 0.06 | 0.21 | 0.13 | 0.1 | 0.11 | 0.1 |
| 3 | 60767700 | rs11709918 | 0.05 | 1.302 | 0.012 | -1.475 | 0.012 | 0.035 | 0.035 | 1.094 | 0.31 | 0.24 | 0.25 | 0.06 | 0.31 | 0.13 |
| 3 | 60770570 | rs6801955 | -0.398 | 1.426 | 0.458 | -1.478 | 0.014 | -0.109 | -0.109 | 0 | 0.35 | 0.46 | 0.32 | 0.1 | 0.39 | 0.17 |
| 3 | 60771914 | rs17063990 | -0.212 | 1.975 | 0.498 | -1.122 | 0.098 | 0.098 | 0.098 | 0 | 0.06 | 0.28 | 0.12 | 0.11 | 0.07 | 0.08 |
| 3 | 60775460 | rs4974241 | -1.046 | 2.042 | 0.561 | -1.12 | 0.225 | 0.12 | 0.12 | 0.005 | 0.1 | 0.48 | 0.18 | 0.14 | 0.15 | 0.12 |
| 3 | 60780913 | rs4974230 | -1.13 | 1.561 | -0.393 | -1.302 | -0.04 | 0.151 | 0.151 | 1.316 | 0.56 | 0.93 | 0.51 | 0.23 | 0.56 | 0.41 |
| 3 | 60793653 | rs2736736 | -1.016 | 1.341 | -0.516 | -1.31 | -0.032 | 0.187 | 0.187 | 1.649 | 0.54 | 0.88 | 0.49 | 0.22 | 0.54 | 0.42 |
| 3 | 60797810 | rs1900669 | -0.134 | 0.84 | -0.431 | -1.507 | 0.049 | 0.139 | 0.139 | 0.133 | 0.4 | 0.2 | 0.28 | 0.06 | 0.38 | 0.2 |
| 3 | 60802554 | rs6782022 | -0.352 | 0.846 | -0.432 | -1.461 | 0.055 | 0.131 | 0.131 | 0.42 | 0.42 | 0.25 | 0.29 | 0.09 | 0.4 | 0.2 |
| 3 | 60806310 | rs2736744 | -0.749 | 1.044 | -0.302 | -1.4 | 0.003 | 0.005 | 0.005 | 0.917 | 0.48 | 0.57 | 0.37 | 0.15 | 0.47 | 0.33 |
| 3 | 60806430 | rs2736745 | -0.5 | 1.534 | 0.289 | -0.44 | 0.174 | 0.041 | 0.041 | 0.018 | 0.08 | 0.4 | 0.09 | 0.05 | 0.09 | 0.14 |
| 3 | 60808285 | rs2594141 | -0.575 | 0.964 | -0.747 | -0.613 | 0.007 | 0.048 | 0.048 | 1.044 | 0.25 | 0.18 | 0.16 | 0.13 | 0.24 | 0.14 |
| 3 | 60808405 | rs2594140 | -1.217 | 1.651 | -0.26 | 0.141 | 0.027 | 0.029 | 0.029 | 2.812 | 0.2 | 0.57 | 0.25 | 0.25 | 0.19 | 0.2 |

| | | | | | | | | | | | | | | | | |
|---|---|---|---|---|---|---|---|---|---|---|---|---|---|---|---|---|
| 3 | 60808825 | rs2594139 | -0.02 | 1.005 | -0.463 | -0.382 | 0 | -0.002 | -0.002 | 0.431 | 0.64 | 0.59 | 0.48 | 0.25 | 0.64 | 0.3 |
| 3 | 60813191 | rs17064044 | 0.394 | 0.37 | -0.609 | -0.621 | 0.034 | 0.035 | 0.035 | 0.101 | 0.1 | 0.02 | 0.03 | 0 | 0.06 | 0.1 |
| 3 | 60817248 | rs2736751 | -0.638 | 0.42 | -0.531 | -1.289 | 0.002 | -0.052 | -0.052 | 1.317 | 0.83 | 0.84 | 0.85 | 1 | 0.77 | 0.93 |
| 3 | 60818874 | rs717292 | 0.887 | 0.401 | -0.689 | -1.269 | 0.038 | 0.047 | 0.047 | 0.041 | 0.09 | 0.01 | 0.03 | 0 | 0.06 | 0.09 |
| 3 | 60830545 | rs2594129 | 0.596 | 0.15 | 0.044 | -1.382 | -0.031 | -0.161 | -0.161 | 0 | 0.7 | 0.4 | 0.78 | 1 | 0.71 | 0.84 |
| 3 | 60838518 | rs11130789 | 0.754 | 0.118 | 0.423 | -1.436 | 0.025 | 0.057 | 0.057 | 1.044 | 0.69 | 0.17 | 0.75 | 1 | 0.7 | 0.84 |
| 3 | 60841656 | rs10510855 | -0.071 | 0.184 | 0.615 | -1.433 | -0.001 | -0.036 | -0.036 | 0.008 | 0.8 | 0.21 | 0.78 | 1 | 0.76 | 0.93 |
| 3 | 60841817 | rs4974235 | -0.08 | 0.178 | 0.612 | -1.435 | -0.001 | -0.038 | -0.038 | 0.417 | 0.8 | 0.21 | 0.78 | 1 | 0.76 | 0.93 |
| 3 | 60853686 | rs2734350 | -0.22 | 0.576 | -0.104 | -1.318 | -0.019 | 0.057 | 0.057 | 0.045 | 0.19 | 0.02 | 0.17 | 0.08 | 0.24 | 0.15 |
| 3 | 60858696 | rs9827591 | 0.426 | 0.459 | 0.668 | -1.381 | 0.037 | -0.131 | -0.131 | 0 | 0.18 | 0.34 | 0.17 | 0 | 0.24 | 0.06 |
| 3 | 60860868 | rs17064099 | 0.505 | 1.133 | 0.131 | -0.693 | -0.049 | -0.041 | -0.041 | 0 | 0.08 | 0.01 | 0.18 | 0.22 | 0.11 | 0.13 |
| 3 | 60871882 | rs9849318 | 0.65 | 0.888 | 0.499 | -0.986 | 0.042 | -0.13 | -0.13 | 0 | 0.17 | 0.38 | 0.18 | 0 | 0.24 | 0.06 |
| 3 | 60874077 | rs11130791 | 0.521 | 0.623 | 0.238 | -1.191 | 0.029 | 0.154 | 0.154 | 0.31 | 0.67 | 0.13 | 0.74 | 1 | 0.68 | 0.84 |
| 3 | 60876648 | rs6803394 | 0.634 | 0.67 | 0.597 | -1.161 | 0.033 | -0.133 | -0.133 | 0 | 0.18 | 0.48 | 0.19 | 0 | 0.24 | 0.06 |
| 3 | 60877992 | rs6446159 | 0.451 | 0.088 | 0.274 | -1.38 | -0.009 | -0.098 | -0.098 | 0 | 0.28 | 0.41 | 0.26 | 0.1 | 0.29 | 0.14 |
| 3 | 60879694 | rs1562521 | 0.189 | -0.047 | 0.018 | -1.473 | 0.081 | 0.239 | 0.239 | 0 | 0.54 | 0.15 | 0.68 | 0.91 | 0.63 | 0.75 |
| 3 | 60880607 | rs6768425 | -0.702 | 0.095 | -0.004 | -1.366 | 0.01 | 0.043 | 0.043 | 0 | 0.51 | 0.53 | 0.43 | 0.17 | 0.53 | 0.28 |
| 3 | 60888974 | rs17683604 | 1.239 | -0.292 | -0.265 | -1.712 | 0.036 | 0.049 | 0.049 | 0 | 0.12 | 0.02 | 0.04 | 0 | 0.07 | 0.09 |
| 3 | 60892741 | rs6765028 | 0.658 | -0.338 | -0.089 | -1.774 | 0 | -0.012 | -0.012 | 0.165 | 0.28 | 0.26 | 0.24 | 0.09 | 0.3 | 0.15 |
| 3 | 60895579 | rs17632681 | 1.283 | -0.282 | -0.279 | -1.698 | 0.036 | 0.058 | 0.058 | 0 | 0.11 | 0.01 | 0.04 | 0 | 0.07 | 0.09 |
| 3 | 60898109 | rs9821525 | -0.053 | -0.474 | -0.694 | -1.969 | -0.105 | 0.073 | 0.073 | 0 | 0.26 | 0.08 | 0.51 | 0.86 | 0.37 | 0.55 |
| 3 | 60898271 | rs6793737 | -0.825 | -0.601 | -0.414 | -2.051 | 0.008 | -0.029 | -0.029 | 0 | 0.58 | 0.67 | 0.78 | 0.95 | 0.67 | 0.72 |
| 3 | 60900460 | rs9311788 | -0.813 | -0.595 | -0.411 | -2.047 | 0.009 | -0.028 | -0.028 | 1.054 | 0.58 | 0.67 | 0.78 | 0.95 | 0.67 | 0.71 |
| 3 | 60903111 | rs4974249 | 1.009 | -0.527 | -0.473 | -1.695 | -0.041 | -0.087 | -0.087 | 0 | 0.17 | 0.06 | 0.32 | 0.64 | 0.24 | 0.41 |
| 3 | 60907899 | rs2121845 | 1.332 | -0.351 | -0.403 | -1.694 | -0.091 | 0.146 | 0.146 | 0 | 0.31 | 0.08 | 0.19 | 0.05 | 0.36 | 0.25 |
| 3 | 60908013 | rs2044613 | -0.445 | 0.2 | -0.12 | -1.271 | 0.001 | 0.042 | 0.042 | 0 | 0.5 | 0.66 | 0.6 | 0.73 | 0.55 | 0.58 |
| 3 | 60918609 | rs9852553 | -0.464 | 0.18 | -0.339 | -1.331 | 0.003 | 0.06 | 0.06 | 0 | 0.5 | 0.67 | 0.6 | 0.71 | 0.54 | 0.58 |
| 3 | 60918957 | rs9876711 | -0.451 | 0.184 | -0.339 | -1.33 | 0.002 | 0.061 | 0.061 | 0 | 0.5 | 0.68 | 0.6 | 0.71 | 0.54 | 0.58 |
| 3 | 60921419 | rs11926529 | 0.684 | -0.066 | -0.39 | -1.564 | -0.003 | -0.061 | -0.061 | 0 | 0.27 | 0.27 | 0.23 | 0.06 | 0.28 | 0.15 |
| 3 | 60931990 | rs1562519 | 0.48 | 0.11 | -1.26 | -1.173 | 0.04 | 0.118 | 0.118 | 0 | 0.41 | 0.19 | 0.21 | 0.09 | 0.33 | 0.28 |
| 3 | 60933886 | rs1123706 | -0.404 | 0.655 | -1.315 | -0.467 | -0.049 | -0.066 | -0.066 | 0 | 0.16 | 0.05 | 0.21 | 0.25 | 0.15 | 0.16 |
| 3 | 60937330 | rs9811973 | -0.058 | 0.245 | -1.019 | -0.962 | -0.012 | -0.056 | -0.056 | 0.149 | 0.13 | 0.24 | 0.09 | 0.01 | 0.09 | 0.09 |
| 3 | 60939763 | rs13070093 | 0.517 | 0.244 | -1.4 | -1.545 | -0.001 | 0.062 | 0.062 | 0 | 0.24 | 0.09 | 0.19 | 0 | 0.29 | 0.08 |
| 3 | 60952418 | rs6414610 | -0.702 | 0.487 | -1.548 | -1.207 | 0.004 | 0.046 | 0.046 | 0 | 0.88 | 0.92 | 0.92 | 1 | 0.9 | 0.91 |
| 3 | 60957450 | rs11130802 | 0.605 | 1.251 | -1.626 | -0.7 | -0.034 | -0.041 | -0.041 | 0 | 0.14 | 0.08 | 0.33 | 0.63 | 0.24 | 0.39 |
| 3 | 60957562 | rs12053964 | 0.775 | 0.655 | -1.492 | -0.717 | -0.037 | -0.041 | -0.041 | 0 | 0.1 | 0.02 | 0.18 | 0.23 | 0.11 | 0.17 |
| 3 | 60969373 | rs6446172 | -0.201 | -0.352 | -1.101 | -1.227 | -0.002 | -0.019 | -0.019 | 0 | 0.53 | 0.58 | 0.73 | 0.83 | 0.67 | 0.67 |
| 3 | 60975380 | rs7632682 | -0.254 | -0.332 | -1.051 | -1.263 | 0.005 | -0.016 | -0.016 | 0 | 0.56 | 0.62 | 0.75 | 0.89 | 0.69 | 0.71 |
| 3 | 60977517 | rs6797228 | 0.624 | -0.54 | -1.115 | -0.68 | 0.008 | 0.039 | 0.039 | 0.85 | 0.46 | 0.29 | 0.52 | 0.66 | 0.6 | 0.58 |
| 3 | 60983267 | rs13087545 | 1.127 | -0.419 | -1.42 | -1.094 | -0.141 | 0.142 | 0.142 | 0.412 | 0.36 | 0.03 | 0.3 | 0.12 | 0.51 | 0.32 |
| 3 | 60992932 | rs7374239 | -0.023 | -0.966 | -1.052 | -1.307 | 0 | -0.003 | -0.003 | 0.18 | 0.23 | 0.22 | 0.26 | 0.23 | 0.19 | 0.15 |
| 3 | 60993915 | rs4129352 | 1.264 | -1.196 | -0.7 | -1.364 | -0.02 | -0.015 | -0.015 | 0.008 | 0.05 | 0.1 | 0.03 | 0.02 | 0.03 | 0.07 |
| 3 | 60995350 | rs6773775 | -0.695 | -1.275 | -0.141 | -1.333 | -0.02 | -0.021 | -0.021 | 2.073 | 0.31 | 0.53 | 0.3 | 0.25 | 0.21 | 0.22 |
| 3 | 61004199 | rs6786392 | -1.174 | -1.28 | -0.091 | -1.444 | 0 | 0.004 | 0.004 | 0 | 0.86 | 0.86 | 0.88 | 0.98 | 0.87 | 0.9 |
| 3 | 61004306 | rs9832173 | -0.551 | -1.36 | -0.1 | -1.485 | -0.015 | -0.015 | -0.015 | 0 | 0.29 | 0.46 | 0.3 | 0.24 | 0.22 | 0.22 |
| 3 | 61004740 | rs11130806 | -0.949 | -1.262 | 0.047 | -1.459 | -0.003 | -0.052 | -0.052 | 0.087 | 0.84 | 0.75 | 0.87 | 0.98 | 0.87 | 0.88 |
| 3 | 61012013 | rs17064318 | -0.743 | -1.342 | -0.284 | -1.366 | -0.017 | -0.051 | -0.051 | 5.16 | 0.29 | 0.48 | 0.18 | 0.18 | 0.21 | 0.21 |
| 3 | 61014080 | rs4688142 | 0.802 | -1.086 | -0.235 | -1.28 | -0.004 | -0.017 | -0.017 | 0 | 0.57 | 0.47 | 0.57 | 0.73 | 0.64 | 0.71 |
| 3 | 61015134 | rs17064323 | 1.145 | -0.743 | -0.961 | -1.098 | 0.038 | 0.048 | 0.048 | 0 | 0.06 | 0.12 | 0.06 | 0.17 | 0.09 | 0.1 |
| 3 | 61016039 | rs11130807 | 0.937 | -0.895 | -1.219 | -1.007 | 0.008 | 0.116 | 0.116 | 4.215 | 0.5 | 0.07 | 0.51 | 0.7 | 0.63 | 0.64 |
| 3 | 61021188 | rs17064329 | -0.009 | -0.887 | -0.8 | -1.003 | -0.005 | -0.013 | -0.013 | 0 | 0.18 | 0.23 | 0.14 | 0.12 | 0.13 | 0.22 |
| 3 | 61021239 | rs2365045 | -0.38 | -1.078 | -0.883 | -1.244 | -0.026 | -0.137 | -0.137 | 1.604 | 0.71 | 0.45 | 0.82 | 0.98 | 0.79 | 0.88 |
| 3 | 61021526 | rs1898749 | 0.465 | -0.49 | -1.555 | -0.846 | -0.053 | -0.085 | -0.085 | 4.355 | 0.12 | 0.01 | 0.22 | 0.63 | 0.13 | 0.34 |
| 3 | 61023355 | rs6446179 | 0.559 | -1.139 | -1.29 | -1.135 | 0.064 | 0.183 | 0.183 | 2.519 | 0.49 | 0.17 | 0.55 | 0.79 | 0.65 | 0.65 |
| 3 | 61036905 | rs6791496 | 0.163 | -0.966 | -1.321 | -1.064 | 0.04 | 0.143 | 0.143 | 1.634 | 0.54 | 0.27 | 0.7 | 0.86 | 0.67 | 0.72 |
| 3 | 61040284 | rs2365052 | 0.475 | -0.726 | -1.733 | -0.923 | -0.032 | 0.176 | 0.176 | 0.527 | 0.45 | 0.1 | 0.54 | 0.84 | 0.63 | 0.61 |
| 3 | 61040705 | rs10510858 | 0.655 | -1.061 | -0.898 | -1.161 | 0.008 | -0.021 | -0.021 | 0 | 0.41 | 0.42 | 0.43 | 0.22 | 0.55 | 0.29 |
| 3 | 61040860 | rs10510859 | 0.014 | -1.788 | -0.365 | -1.565 | -0.048 | -0.152 | -0.152 | 0 | 0.38 | 0.45 | 0.17 | 0.01 | 0.16 | 0.26 |
| 3 | 61042317 | rs12108196 | 0.634 | -2.175 | -0.271 | -1.575 | -0.018 | -0.061 | -0.061 | 0.064 | 0.28 | 0.33 | 0.14 | 0.01 | 0.12 | 0.17 |
| 3 | 61046212 | rs6809101 | 0.979 | -1.728 | -0.237 | -1.309 | 0.001 | 0.027 | 0.027 | 2.705 | 0.42 | 0.28 | 0.45 | 0.21 | 0.64 | 0.33 |
| 3 | 61046863 | rs7616675 | 0.708 | -1.756 | -0.402 | -1.322 | -0.004 | -0.031 | -0.031 | 0.561 | 0.45 | 0.44 | 0.46 | 0.21 | 0.64 | 0.33 |
| 3 | 61048258 | rs9869954 | 0.755 | -1.722 | -0.554 | -1.306 | -0.001 | -0.007 | -0.007 | 0 | 0.44 | 0.41 | 0.45 | 0.22 | 0.64 | 0.33 |
| 3 | 61051714 | rs7622067 | -0.757 | -1.679 | -0.691 | -0.813 | -0.029 | -0.016 | -0.016 | 0 | 0.93 | 0.99 | 0.72 | 0.32 | 0.92 | 0.66 |
| 3 | 61052347 | rs6763300 | -0.877 | -1.612 | -0.474 | -1.033 | 0.046 | 0.162 | 0.162 | 0.074 | 0.06 | 0.35 | 0.05 | 0.08 | 0.01 | 0.02 |
| 3 | 61052481 | rs13087596 | 1.175 | -1.544 | -0.622 | -1.039 | -0.125 | 0.162 | 0.162 | 3.956 | 0.37 | 0.04 | 0.41 | 0.13 | 0.62 | 0.32 |
| 3 | 61054494 | rs13099398 | 0.7 | -1.341 | -0.483 | -0.871 | -0.002 | -0.016 | -0.016 | 0.043 | 0.43 | 0.43 | 0.45 | 0.25 | 0.63 | 0.34 |
| 3 | 61061362 | rs7636568 | 0.259 | -1.342 | -0.561 | -0.706 | -0.008 | -0.029 | -0.029 | 0 | 0.11 | 0.13 | 0.04 | 0 | 0.05 | 0.05 |
| 3 | 61061861 | rs1439006 | 0.206 | -0.715 | -0.767 | -0.936 | 0.002 | -0.007 | -0.007 | 0.003 | 0.74 | 0.71 | 0.52 | 0.15 | 0.7 | 0.66 |
| 3 | 61066925 | rs1439007 | 1.698 | -1.234 | -1.535 | -1.329 | -0.174 | 0.088 | 0.088 | 1.995 | 0.36 | 0.04 | 0.33 | 0.17 | 0.53 | 0.38 |

| | | | | | | | | | | | | | | | | |
|---|---|---|---|---|---|---|---|---|---|---|---|---|---|---|---|---|
| 3 | 61069814 | rs2365054 | -0.896 | -0.216 | -0.932 | 0.181 | -0.041 | -0.063 | -0.063 | 0 | 0.9 | 1 | 0.73 | 0.56 | 0.86 | 0.75 |
| 3 | 61077258 | rs17687542 | 0.879 | -0.287 | -0.807 | -0.428 | 0.005 | 0.035 | 0.035 | 0 | 0.14 | 0.04 | 0.12 | 0.02 | 0.13 | 0.22 |
| 3 | 61081348 | rs1530092 | 1.761 | -0.527 | -0.394 | -0.588 | 0.009 | 0.178 | 0.178 | 3.811 | 0.49 | 0.23 | 0.36 | 0.17 | 0.54 | 0.44 |
| 3 | 61081536 | rs9820971 | 0.066 | -0.541 | -0.313 | -0.612 | -0.014 | 0.011 | 0.011 | 2.231 | 0.75 | 0.82 | 0.53 | 0.19 | 0.7 | 0.67 |
| 3 | 61081768 | rs11921290 | -0.114 | -0.535 | -0.228 | -0.612 | 0.012 | 0.005 | 0.005 | 0 | 0.21 | 0.17 | 0.13 | 0.02 | 0.16 | 0.24 |
| 3 | 61082153 | rs9825823 | 1.66 | -0.479 | -0.113 | -0.558 | 0.011 | 0.113 | 0.113 | 3.637 | 0.5 | 0.36 | 0.37 | 0.17 | 0.55 | 0.44 |
| 3 | 61096082 | rs7642143 | 0.103 | -0.415 | -0.161 | -0.459 | -0.012 | -0.029 | -0.029 | 0 | 0.05 | 0.17 | 0.02 | 0 | 0.02 | 0 |
| 3 | 61098641 | rs17687956 | 0.871 | -0.403 | -0.152 | -0.46 | 0.014 | 0.025 | 0.025 | 0 | 0.15 | 0.06 | 0.11 | 0.02 | 0.12 | 0.22 |
| 3 | 61101006 | rs9850400 | 0.872 | 0.578 | 0.224 | -0.014 | -0.002 | 0.007 | 0.007 | 1.397 | 0.7 | 0.51 | 0.57 | 0.28 | 0.74 | 0.73 |
| 3 | 61101760 | rs17064482 | 0.014 | 0.732 | -0.471 | -0.234 | 0.031 | 0.033 | 0.033 | 0 | 0.05 | 0.17 | 0.1 | 0.1 | 0.11 | 0.05 |
| 3 | 61103609 | rs6806415 | 0.374 | 0.393 | -0.662 | -0.142 | 0.004 | -0.009 | -0.009 | 0 | 0.75 | 0.81 | 0.62 | 0.28 | 0.79 | 0.74 |
| 3 | 61119923 | rs12487188 | 0.59 | 0.439 | -0.724 | -0.137 | -0.003 | -0.011 | -0.011 | 0.777 | 0.73 | 0.77 | 0.6 | 0.31 | 0.74 | 0.72 |
| 3 | 61122318 | rs9830950 | -1.111 | 0.258 | -0.947 | -0.549 | 0.009 | 0.178 | 0.178 | 1.198 | 0.46 | 0.79 | 0.67 | 0.85 | 0.41 | 0.53 |
| 3 | 61126821 | rs2886261 | -0.599 | 0.732 | -1.231 | -0.427 | 0.024 | -0.02 | -0.02 | 0.589 | 0.1 | 0.12 | 0.09 | 0.02 | 0.16 | 0.04 |
| 3 | 61127205 | rs9877396 | -0.406 | 0.69 | -1.205 | 0.053 | 0.035 | 0.108 | 0.108 | 0 | 0.15 | 0.23 | 0.34 | 0.4 | 0.23 | 0.27 |
| 3 | 61132698 | rs4309721 | 1.349 | 0.321 | -1.333 | -0.392 | -0.002 | -0.055 | -0.055 | 0.104 | 0.54 | 0.53 | 0.42 | 0.14 | 0.54 | 0.41 |
| 3 | 61136894 | rs17637774 | -0.189 | 0.135 | -1.262 | -0.285 | 0.035 | 0.025 | 0.025 | 0.013 | 0.27 | 0.19 | 0.19 | 0.15 | 0.17 | 0.3 |
| 3 | 61141434 | rs9876185 | -0.196 | 0.218 | -1.252 | -0.149 | 0.005 | 0.025 | 0.025 | 0 | 0.16 | 0.17 | 0.34 | 0.4 | 0.27 | 0.26 |
| 3 | 61142441 | rs17637930 | 0.705 | -0.641 | -1.29 | -0.168 | 0.003 | 0.004 | 0.004 | 0 | 0.11 | 0.06 | 0.1 | 0.07 | 0.11 | 0.24 |
| 3 | 61145050 | rs12633494 | 1.487 | -0.522 | -1.276 | -0.076 | 0 | -0.034 | -0.034 | 0.799 | 0.09 | 0 | 0.22 | 0.38 | 0.06 | 0.24 |
| 3 | 61152287 | rs1978865 | -0.409 | -0.693 | -1.54 | -0.284 | 0.043 | 0.035 | 0.035 | 0.796 | 0.29 | 0.2 | 0.22 | 0.15 | 0.18 | 0.31 |
| 3 | 61153578 | rs1966136 | -1.598 | -0.606 | -1.48 | -0.195 | -0.003 | -0.03 | -0.03 | 0.004 | 0.49 | 0.47 | 0.6 | 0.83 | 0.42 | 0.6 |
| 3 | 61166821 | rs11917930 | 0.093 | -0.666 | -1.373 | -0.225 | 0.021 | 0.066 | 0.066 | 0 | 0.16 | 0.27 | 0.34 | 0.71 | 0.16 | 0.26 |
| 3 | 61168215 | rs4688332 | 0.88 | -0.744 | -1.382 | -0.239 | 0.076 | 0.008 | 0.008 | 1.546 | 0.16 | 0.01 | 0.09 | 0.07 | 0.06 | 0.06 |
| 3 | 61176788 | rs9831393 | 0.468 | -0.47 | -1.835 | -0.592 | -0.008 | 0.009 | 0.009 | 0 | 0.35 | 0.28 | 0.52 | 0.78 | 0.42 | 0.45 |
| 3 | 61177577 | rs7617424 | 0.423 | -1.073 | -1.816 | -0.564 | -0.004 | 0.007 | 0.007 | 0 | 0.35 | 0.28 | 0.5 | 0.78 | 0.38 | 0.45 |
| 3 | 61177751 | rs9819588 | -1.036 | -0.764 | -1.583 | -0.47 | 0.031 | -0.127 | -0.127 | 0 | 0.66 | 0.46 | 0.73 | 0.94 | 0.59 | 0.76 |
| 3 | 61184321 | rs1470035 | -1.314 | -1.581 | -1.065 | 0.252 | 0.006 | 0.004 | 0.004 | 0.058 | 0.48 | 0.41 | 0.54 | 0.56 | 0.38 | 0.57 |
| 3 | 61190233 | rs9856186 | 0.569 | -1.465 | -1.182 | -0.078 | 0.007 | -0.027 | -0.027 | 0.091 | 0.12 | 0.17 | 0.11 | 0.02 | 0.14 | 0.04 |
| 3 | 61190420 | rs6772523 | 1.058 | -1.631 | -1.327 | 0.011 | -0.001 | 0 | 0 | 5.057 | 0.12 | 0.1 | 0.1 | 0.08 | 0.06 | 0.06 |
| 3 | 61205924 | rs17668266 | 0.678 | -1.611 | -1.387 | 0.03 | 0.009 | 0.022 | 0.022 | 0 | 0.13 | 0.03 | 0.1 | 0.07 | 0.14 | 0.25 |
| 3 | 61211945 | rs12497257 | 0.951 | -1.638 | -1.371 | 0.175 | 0.014 | 0.001 | 0.001 | 0 | 0.09 | 0.04 | 0.22 | 0.37 | 0.05 | 0.22 |
| 3 | 61215497 | rs10866046 | -1.026 | -1.645 | -1.398 | 0.047 | 0.042 | -0.016 | -0.016 | 0 | 0.35 | 0.19 | 0.43 | 0.54 | 0.28 | 0.54 |
| 3 | 61216475 | rs7640496 | 0.133 | -1.709 | -1.222 | -0.208 | -0.007 | -0.108 | -0.108 | 0 | 0.15 | 0.55 | 0.15 | 0 | 0.11 | 0.03 |
| 3 | 61217770 | rs13099397 | 2.026 | -1.733 | -1.752 | -0.35 | -0.151 | 0.139 | 0.139 | 0 | 0.36 | 0.02 | 0.32 | 0.16 | 0.5 | 0.35 |
| 3 | 61232197 | rs2365491 | 0.253 | -1.743 | -1.389 | -0.252 | -0.032 | 0.004 | 0.004 | 1.756 | 0.17 | 0.49 | 0.15 | 0.1 | 0.09 | 0.06 |
| 3 | 61236328 | rs6791206 | -1.161 | -1.8 | -1.982 | -0.247 | 0.079 | 0.142 | 0.142 | 0.987 | 0.4 | 0.83 | 0.54 | 0.79 | 0.25 | 0.36 |
| 3 | 61241178 | rs12632761 | 1.061 | -1.644 | -1.862 | -0.042 | -0.003 | 0.025 | 0.025 | 1.236 | 0.11 | 0.16 | 0.15 | 0.31 | 0.1 | 0.05 |
| 3 | 61252469 | rs815719 | 0.383 | -1.277 | -1.677 | 0.118 | -0.013 | 0.029 | 0.029 | 0.54 | 0.24 | 0.35 | 0.25 | 0.41 | 0.18 | 0.1 |
| 3 | 61253045 | rs10510861 | 0.736 | -1.326 | -2.342 | -0.217 | 0.007 | 0.017 | 0.017 | 0 | 0.13 | 0.03 | 0.1 | 0.08 | 0.15 | 0.25 |
| 3 | 61253495 | rs1093194 | -0.351 | -1.331 | -2.333 | -0.218 | 0.031 | 0.046 | 0.046 | 0.576 | 0.37 | 0.72 | 0.52 | 0.79 | 0.24 | 0.33 |
| 3 | 61262149 | rs815710 | -0.76 | -0.951 | -2.192 | -0.351 | 0.09 | 0.236 | 0.236 | 0 | 0.51 | 0.82 | 0.63 | 0.88 | 0.4 | 0.6 |
| 3 | 61267416 | rs745830 | 0.613 | -0.776 | -2.169 | -0.421 | 0.037 | 0.042 | 0.042 | 0.791 | 0.22 | 0.15 | 0.15 | 0.07 | 0.13 | 0.08 |
| 3 | 61271104 | rs815699 | 0.694 | -0.678 | -2.122 | -0.376 | 0.034 | 0.039 | 0.039 | 0.813 | 0.21 | 0.16 | 0.15 | 0.07 | 0.13 | 0.07 |
| 3 | 61272046 | rs815701 | -0.469 | 0.324 | -1.369 | 0.294 | -0.013 | 0.055 | 0.055 | 0 | 0.89 | 0.98 | 0.95 | 1 | 0.89 | 0.92 |
| 3 | 61273564 | rs1349799 | -0.815 | 0.475 | -0.756 | 0.943 | -0.007 | 0.015 | 0.015 | 0 | 0.28 | 0.64 | 0.42 | 0.5 | 0.26 | 0.49 |
| 3 | 61280155 | rs11706338 | -0.394 | 2.09 | -1.503 | 0.636 | -0.112 | -0.024 | -0.024 | 0.122 | 0.17 | 0.02 | 0.18 | 0.16 | 0.29 | 0.29 |
| 3 | 61281037 | rs7626375 | -0.366 | 1.475 | -2.01 | 0.407 | 0.005 | 0.027 | 0.027 | 0.388 | 0.73 | 0.82 | 0.81 | 0.79 | 0.77 | 0.83 |
| 3 | 61281841 | rs7651321 | -0.22 | 0.515 | -1.487 | 0.299 | -0.048 | -0.016 | -0.016 | 0 | 0.84 | 0.57 | 0.79 | 0.92 | 0.91 | 0.85 |
| 3 | 61282140 | rs9877709 | 0.067 | 0.86 | -1.085 | 0.249 | -0.011 | -0.028 | -0.028 | 0 | 0.83 | 0.46 | 0.76 | 0.92 | 0.86 | 0.8 |
| 3 | 61287140 | rs7652939 | 0.501 | 0.739 | -0.637 | 0.878 | 0.036 | -0.013 | -0.013 | 0 | 0.52 | 0.2 | 0.47 | 0.49 | 0.64 | 0.58 |
| 3 | 61289533 | rs9812140 | 0.757 | 0.753 | -0.609 | 0.908 | -0.003 | -0.005 | -0.005 | 0.237 | 0.27 | 0.24 | 0.19 | 0.2 | 0.19 | 0.16 |
| 3 | 61291738 | rs12634258 | 0.917 | 0.357 | -1.052 | 0.844 | -0.004 | 0.01 | 0.01 | 0.699 | 0.8 | 0.25 | 0.66 | 0.69 | 0.87 | 0.77 |
| 3 | 61298955 | rs1453750 | -0.535 | 0.916 | -1.4 | 0.803 | -0.165 | -0.19 | -0.19 | 0 | 0.29 | 0.02 | 0.29 | 0.47 | 0.37 | 0.45 |
| 3 | 61299630 | rs6784508 | 0.859 | 0.671 | -1.191 | 0.689 | 0.03 | 0.018 | 0.018 | 0 | 0.4 | 0.3 | 0.38 | 0.34 | 0.4 | 0.3 |
| 3 | 61304491 | rs2365740 | 0.808 | 0.285 | -1.109 | 0.774 | 0.014 | -0.016 | -0.016 | 0 | 0.25 | 0.01 | 0.24 | 0.2 | 0.18 | 0.15 |
| 3 | 61310328 | rs7625594 | -1.511 | 1.069 | -0.35 | 1.091 | 0.002 | -0.011 | -0.011 | 0 | 0.16 | 0.1 | 0.13 | 0.18 | 0.13 | 0.22 |
| 3 | 61315350 | rs6445204 | 0.881 | 1.114 | 0.115 | 1.282 | 0.043 | 0.121 | 0.121 | 3.786 | 0.93 | 0.37 | 0.85 | 0.87 | 0.95 | 0.91 |
| 3 | 61318721 | rs1901892 | 1.965 | 1.098 | 0.016 | 1.147 | -0.001 | 0.005 | 0.005 | 0.043 | 0.76 | 0.3 | 0.7 | 0.69 | 0.83 | 0.68 |
| 3 | 61326783 | rs17064788 | -1.068 | 0.819 | -0.006 | 0.329 | 0.007 | -0.024 | -0.024 | 0.715 | 0.14 | 0.02 | 0.1 | 0.09 | 0.09 | 0.19 |
| 3 | 61331555 | rs4688444 | 1.84 | 0.799 | 0.15 | 0.28 | -0.008 | -0.021 | -0.021 | 0 | 0.78 | 0.32 | 0.74 | 0.91 | 0.85 | 0.74 |
| 3 | 61332071 | rs11714700 | -1.139 | 0.813 | -0.048 | 0.35 | 0 | -0.021 | -0.021 | 0.069 | 0.14 | 0.02 | 0.1 | 0.09 | 0.1 | 0.2 |
| 3 | 61348592 | rs9826581 | 1.178 | 0.947 | 0.276 | 0.434 | 0.002 | -0.009 | -0.009 | 1.622 | 0.84 | 0.36 | 0.78 | 0.9 | 0.86 | 0.82 |
| 3 | 61356250 | rs13061149 | -0.577 | 1.751 | -0.55 | 0.267 | -0.088 | 0.037 | 0.037 | 0 | 0.15 | 0.01 | 0.22 | 0.07 | 0.26 | 0.19 |
| 3 | 61357417 | rs13067383 | -0.618 | 1.717 | -0.563 | 0.522 | -0.069 | -0.044 | -0.044 | 0.005 | 0.15 | 0.01 | 0.19 | 0.19 | 0.24 | 0.17 |
| 3 | 61375686 | rs9822542 | -0.975 | 1.216 | -0.072 | 0.456 | 0 | 0 | 0 | 1.874 | 0.11 | 0.09 | 0.08 | 0.07 | 0.09 | 0.17 |
| 3 | 61389476 | rs9810775 | -1.212 | 1.152 | 0.208 | 0.445 | -0.012 | -0.004 | -0.004 | 0 | 0.12 | 0.15 | 0.09 | 0.07 | 0.09 | 0.17 |
| 3 | 61395236 | rs1563135 | -1.587 | 1.215 | 0.841 | 0.539 | 0.034 | 0.067 | 0.067 | 0.931 | 0.08 | 0.28 | 0.08 | 0.07 | 0.06 | 0.16 |
| 3 | 61399078 | rs9835007 | 0.34 | 0.961 | 0.358 | 0.336 | -0.01 | 0.014 | 0.014 | 2.969 | 0.53 | 0.28 | 0.43 | 0.55 | 0.46 | 0.56 |

| | | | | | | | | | | | | | | | | |
|---|---|---|---|---|---|---|---|---|---|---|---|---|---|---|---|---|
| 3 | 61402988 | rs11130841 | 0.212 | 0.871 | 0.383 | 0.274 | -0.03 | -0.001 | -0.001 | 0.439 | 0.35 | 0.15 | 0.38 | 0.36 | 0.44 | 0.27 |
| 3 | 61406345 | rs1540554 | -0.052 | 0.964 | 0.589 | 0.315 | -0.003 | 0.01 | 0.01 | 0.214 | 0.38 | 0.3 | 0.39 | 0.36 | 0.44 | 0.26 |
| 3 | 61416514 | rs6781195 | 0.815 | 0.843 | 1.151 | 0.203 | 0.055 | 0.059 | 0.059 | 0.011 | 0.89 | 0.56 | 0.89 | 0.9 | 0.91 | 0.83 |
| 3 | 61418941 | rs10433636 | 0.696 | 0.899 | 0.515 | 0.06 | -0.007 | 0.061 | 0.061 | 0.235 | 0.47 | 0.14 | 0.4 | 0.64 | 0.43 | 0.58 |
| 3 | 61429785 | rs1453754 | -0.865 | 0.833 | 1.17 | 0.171 | 0.047 | 0.056 | 0.056 | 0 | 0.11 | 0.34 | 0.1 | 0.1 | 0.08 | 0.17 |
| 3 | 61438210 | rs17320598 | 0.79 | 0.601 | 0.049 | 0.354 | -0.008 | -0.058 | -0.058 | 0 | 0.35 | 0.16 | 0.35 | 0.43 | 0.34 | 0.46 |
| 3 | 61444443 | rs17254891 | 0.46 | 0.228 | -0.121 | -0.193 | 0.012 | 0.03 | 0.03 | 1.128 | 0.22 | 0.13 | 0.12 | 0.04 | 0.18 | 0.14 |
| 3 | 61445321 | rs9311811 | 0.835 | 0.399 | 0.106 | -0.316 | -0.002 | -0.003 | -0.003 | 0.798 | 0.49 | 0.34 | 0.53 | 0.51 | 0.48 | 0.66 |
| 3 | 61446281 | rs1869405 | 0.813 | 0.793 | 0.07 | -0.344 | 0.023 | 0.002 | 0.002 | 0.005 | 0.5 | 0.35 | 0.6 | 0.52 | 0.6 | 0.67 |
| 3 | 61447968 | rs967329 | 0.567 | 0.238 | 0.277 | -0.486 | -0.042 | -0.082 | -0.082 | 0.495 | 0.8 | 0.54 | 0.88 | 0.93 | 0.86 | 0.83 |
| 3 | 61447981 | rs17652380 | 0.607 | 0.28 | 0.002 | -0.466 | -0.016 | 0.026 | 0.026 | 0.038 | 0.12 | 0.01 | 0.08 | 0.05 | 0.12 | 0.13 |
| 3 | 61452146 | rs4432637 | 1.595 | 0.384 | 0.159 | -0.47 | 0.007 | -0.006 | -0.006 | 0 | 0.44 | 0.19 | 0.38 | 0.43 | 0.41 | 0.47 |
| 3 | 61453255 | rs1453758 | 1.056 | 0.353 | 0.213 | -0.534 | 0.008 | 0.016 | 0.016 | 1.623 | 0.67 | 0.35 | 0.69 | 0.85 | 0.73 | 0.63 |
| 3 | 61454724 | rs11716449 | 0.666 | 0.385 | 0.183 | -0.502 | -0.048 | -0.07 | -0.07 | 0 | 0.23 | 0.16 | 0.3 | 0.42 | 0.32 | 0.16 |
| 3 | 61455771 | rs7638621 | 2.096 | 0.35 | 0.23 | -0.473 | 0 | -0.029 | -0.029 | 0 | 0.36 | 0.23 | 0.35 | 0.44 | 0.36 | 0.47 |
| 3 | 61459331 | rs6807238 | -0.239 | 0.257 | 0.474 | -0.744 | -0.008 | -0.044 | -0.044 | 0 | 0.24 | 0.24 | 0.13 | 0.04 | 0.18 | 0.14 |
| 3 | 61465329 | rs1023859 | 1.347 | 0.178 | 0.303 | -0.447 | 0.045 | 0.097 | 0.097 | 1.226 | 0.65 | 0.22 | 0.68 | 0.85 | 0.74 | 0.64 |
| 3 | 61466361 | rs1975729 | 0.629 | 0.144 | 0.479 | -0.381 | -0.035 | -0.058 | -0.058 | 0 | 0.79 | 0.43 | 0.87 | 0.93 | 0.86 | 0.85 |
| 3 | 61472886 | rs4544606 | 1.537 | 0.613 | -0.185 | -0.023 | 0.013 | -0.057 | -0.057 | 0 | 0.43 | 0.13 | 0.37 | 0.5 | 0.37 | 0.46 |
| 3 | 61474575 | rs4234688 | -0.449 | 0.979 | 0.443 | -0.087 | 0.008 | -0.004 | -0.004 | 0.001 | 0.19 | 0.33 | 0.25 | 0.14 | 0.16 | 0.24 |
| 3 | 61474877 | rs3885648 | 0.552 | 1.146 | 0.366 | -0.222 | 0.026 | -0.055 | -0.055 | 0 | 0.86 | 0.74 | 0.84 | 0.97 | 0.79 | 0.87 |
| 3 | 61480657 | rs10866048 | -1.029 | 1.154 | -0.135 | -0.049 | -0.006 | 0.002 | 0.002 | 0 | 0.29 | 0.36 | 0.35 | 0.3 | 0.43 | 0.26 |
| 3 | 61490771 | rs11130845 | -0.542 | 0.811 | -0.798 | -0.221 | -0.049 | -0.015 | -0.015 | 0 | 0.25 | 0.12 | 0.21 | 0.29 | 0.27 | 0.21 |
| 3 | 61501001 | rs2886524 | -1.107 | 1.018 | -0.505 | -0.074 | -0.028 | -0.007 | -0.007 | 0.052 | 0.24 | 0.16 | 0.34 | 0.31 | 0.41 | 0.22 |
| 3 | 61504316 | rs7616413 | 1.282 | 0.386 | -0.827 | -0.358 | 0.004 | 0.041 | 0.041 | 0 | 0.53 | 0.3 | 0.53 | 0.63 | 0.61 | 0.61 |
| 3 | 61506098 | rs6809350 | 1.6 | 0.161 | -0.895 | -0.292 | 0.035 | 0.054 | 0.054 | 0 | 0.48 | 0.19 | 0.43 | 0.63 | 0.45 | 0.57 |
| 3 | 61509137 | rs7427981 | -1.096 | 0.548 | -0.023 | -0.215 | 0.048 | -0.019 | -0.019 | 0.468 | 0.28 | 0.39 | 0.37 | 0.22 | 0.41 | 0.25 |
| 3 | 61512473 | rs2886522 | 0.969 | 1.097 | -0.185 | 0.021 | -0.042 | -0.042 | -0.042 | 0 | 0.07 | 0.01 | 0.21 | 0.19 | 0.2 | 0.08 |
| 3 | 61512919 | rs12108098 | -0.454 | 0.614 | 0.429 | -0.162 | -0.023 | -0.034 | -0.034 | 0.142 | 0.17 | 0.27 | 0.08 | 0.06 | 0.11 | 0.18 |
| 3 | 61513495 | rs6794009 | 1.95 | 0.683 | 0.455 | -0.526 | 0.053 | 0.068 | 0.068 | 0 | 0.49 | 0.27 | 0.43 | 0.65 | 0.43 | 0.47 |
| 3 | 61515679 | rs9819373 | 0.663 | 0.825 | 0.834 | -0.496 | 0.147 | 0.061 | 0.061 | 0 | 0.83 | 0.61 | 0.59 | 0.74 | 0.65 | 0.86 |
| 3 | 61516927 | rs2365953 | 0.073 | 0.832 | 0.708 | -0.52 | 0.023 | 0.004 | 0.004 | 0 | 0.11 | 0.13 | 0.26 | 0.18 | 0.25 | 0.1 |
| 3 | 61518116 | rs1320173 | 1.404 | 0.165 | 0.601 | -0.414 | 0.001 | 0.007 | 0.007 | 0 | 0.54 | 0.43 | 0.48 | 0.65 | 0.48 | 0.5 |
| 3 | 61528270 | rs2365956 | -0.553 | 0.079 | 0.633 | -0.152 | 0.031 | 0.009 | 0.009 | 0 | 0.27 | 0.37 | 0.37 | 0.27 | 0.31 | 0.32 |
| 3 | 61534766 | rs7627124 | 1.619 | 0.103 | 0 | -0.293 | 0.041 | 0.02 | 0.02 | 0.055 | 0.47 | 0.33 | 0.38 | 0.64 | 0.35 | 0.46 |
| 3 | 61549295 | rs9845678 | 0.374 | 0.35 | -0.121 | -0.48 | 0.021 | -0.061 | -0.061 | 0 | 0.74 | 0.53 | 0.73 | 0.87 | 0.7 | 0.82 |
| 3 | 61549568 | rs9808938 | 0.433 | 0.347 | -0.122 | -0.482 | 0.021 | -0.061 | -0.061 | 0 | 0.74 | 0.53 | 0.73 | 0.87 | 0.7 | 0.82 |
| 3 | 61555637 | rs4688636 | -0.659 | 0.056 | 0.001 | -0.336 | 0.066 | 0.035 | 0.035 | 0 | 0.93 | 0.7 | 0.84 | 0.91 | 0.87 | 0.95 |
| 3 | 61557718 | rs7621604 | -0.384 | 0.021 | -0.427 | -0.302 | -0.082 | 0.004 | 0.004 | 0.067 | 0.38 | 0.15 | 0.44 | 0.3 | 0.46 | 0.42 |
| 3 | 61557787 | rs11709537 | 1.392 | -0.864 | -0.433 | -0.363 | 0.044 | -0.023 | -0.023 | 0 | 0.4 | 0.12 | 0.31 | 0.6 | 0.34 | 0.45 |
| 3 | 61559771 | rs938959 | -0.664 | -0.45 | -0.577 | -0.219 | 0.009 | 0.001 | 0.001 | 0.136 | 0.44 | 0.5 | 0.59 | 0.41 | 0.57 | 0.45 |
| 3 | 61562014 | rs13061182 | 0.399 | -0.203 | -1.247 | -0.29 | -0.012 | -0.019 | -0.019 | 0 | 0.26 | 0.25 | 0.21 | 0.15 | 0.19 | 0.18 |
| 3 | 61567988 | rs7643514 | -0.685 | 0.265 | -1.94 | -0.096 | -0.013 | -0.098 | -0.098 | 0.012 | 0.67 | 0.95 | 0.78 | 0.45 | 0.65 | 0.57 |
| 3 | 61569213 | rs1317639 | -0.935 | 0.709 | -0.901 | -0.808 | -0.004 | -0.079 | -0.079 | 0.047 | 0.34 | 0.41 | 0.17 | 0.04 | 0.25 | 0.32 |
| 3 | 61569753 | rs1320172 | -0.689 | 1.37 | -0.867 | -0.143 | -0.031 | -0.026 | -0.026 | 0.243 | 0.37 | 0.18 | 0.38 | 0.39 | 0.35 | 0.31 |
| 3 | 61577341 | rs9863064 | 0.233 | 1.412 | -0.122 | -0.264 | -0.022 | -0.061 | -0.061 | 0 | 0.14 | 0.3 | 0.08 | 0.02 | 0.1 | 0.06 |
| 3 | 61584646 | rs9828680 | -0.348 | 1.487 | -0.001 | -0.23 | 0.034 | 0.065 | 0.065 | 0 | 0.18 | 0.35 | 0.17 | 0.25 | 0.21 | 0.13 |
| 3 | 61586124 | rs12715572 | -0.585 | 1.53 | 0.129 | -0.165 | 0.008 | 0.019 | 0.019 | 0 | 0.2 | 0.25 | 0.24 | 0.32 | 0.24 | 0.14 |
| 3 | 61587570 | rs9862251 | 0.589 | 1.352 | 0.115 | -0.283 | -0.006 | -0.026 | -0.026 | 0 | 0.14 | 0.16 | 0.08 | 0.02 | 0.11 | 0.1 |
| 3 | 61589713 | rs4688639 | 1.09 | 1.5 | -0.085 | -0.29 | 0.013 | -0.017 | -0.017 | 0 | 0.54 | 0.27 | 0.48 | 0.53 | 0.51 | 0.63 |
| 3 | 61590181 | rs4390919 | 1.486 | 1.514 | 0.298 | -0.483 | 0.04 | 0.002 | 0.002 | 0.108 | 0.5 | 0.23 | 0.46 | 0.53 | 0.48 | 0.56 |
| 3 | 61590517 | rs9832251 | 0.284 | 0.841 | 0.804 | -0.263 | -0.017 | -0.063 | -0.063 | 0 | 0.15 | 0.36 | 0.1 | 0.02 | 0.1 | 0.06 |
| 3 | 61594769 | rs4688640 | 0.844 | 0.952 | 0.599 | -0.228 | 0 | 0 | 0 | 0 | 0.75 | 0.36 | 0.76 | 0.75 | 0.73 | 0.85 |
| 3 | 61600528 | rs9837256 | -0.162 | 0.939 | 0.746 | -0.24 | 0.049 | -0.002 | -0.002 | 0 | 0.07 | 0.3 | 0.07 | 0 | 0.07 | 0.04 |
| 3 | 61606059 | rs11130850 | 0.753 | 1.002 | 0.6 | -0.073 | 0 | 0.023 | 0.023 | 0 | 0.12 | 0.07 | 0.07 | 0.02 | 0.11 | 0.08 |
| 3 | 61609594 | rs12635919 | -0.861 | 0.854 | 0.811 | -0.139 | -0.001 | 0.004 | 0.004 | 0 | 0.38 | 0.67 | 0.46 | 0.32 | 0.36 | 0.36 |
| 3 | 61615565 | rs1320234 | -1.14 | 0.959 | 0.442 | 0.136 | 0.02 | -0.002 | -0.002 | 0 | 0.55 | 0.66 | 0.55 | 0.54 | 0.52 | 0.43 |
| 3 | 61618935 | rs12497685 | -0.466 | 0.988 | -0.241 | 0.073 | -0.075 | 0.081 | 0.081 | 0.065 | 0.43 | 0.1 | 0.36 | 0.26 | 0.39 | 0.36 |
| 3 | 61620466 | rs17624623 | -0.424 | 1.032 | -0.092 | 0.084 | -0.012 | 0.063 | 0.063 | 0.722 | 0.45 | 0.2 | 0.37 | 0.27 | 0.4 | 0.36 |
| 3 | 61628388 | rs9826684 | -0.938 | 1.604 | 0.018 | 0.299 | 0.073 | 0.149 | 0.149 | 1.648 | 0.13 | 0.56 | 0.22 | 0.27 | 0.16 | 0.1 |
| 3 | 61632170 | rs4688649 | -0.529 | 1.32 | -0.544 | 0.168 | -0.019 | 0.006 | 0.006 | 1.308 | 0.33 | 0.71 | 0.4 | 0.28 | 0.37 | 0.3 |
| 3 | 61633850 | rs2365929 | 0.274 | 0.379 | -0.395 | 0.556 | -0.007 | 0.006 | 0.006 | 3.517 | 0.75 | 0.71 | 0.76 | 0.77 | 0.79 | 0.81 |
| 3 | 61635594 | rs6807692 | -0.152 | 0.991 | -0.717 | 0.536 | -0.012 | -0.02 | -0.02 | 0.326 | 0.1 | 0.06 | 0.14 | 0.18 | 0.13 | 0.04 |
| 3 | 61638592 | rs13064689 | -0.087 | 0.771 | -0.712 | 0.641 | -0.01 | -0.042 | -0.042 | 1.822 | 0.22 | 0.12 | 0.28 | 0.35 | 0.23 | 0.26 |
| 3 | 61639764 | rs17670940 | 0.881 | 0.868 | -0.93 | 0.696 | -0.068 | -0.048 | -0.048 | 0 | 0.35 | 0.09 | 0.39 | 0.42 | 0.42 | 0.38 |
| 3 | 61645378 | rs2365933 | 1.029 | 0.653 | 0.517 | 0.797 | 0.001 | 0.067 | 0.067 | 0.655 | 0.81 | 0.61 | 0.81 | 0.76 | 0.84 | 0.9 |
| 3 | 61661311 | rs1320232 | 0.858 | 1.079 | 0.268 | 0.092 | 0.037 | -0.068 | -0.068 | 0.412 | 0.8 | 0.6 | 0.82 | 0.9 | 0.79 | 0.85 |
| 3 | 61662996 | rs13065463 | -0.418 | 1.117 | -0.237 | 0.104 | -0.006 | 0.004 | 0.004 | 0 | 0.07 | 0.03 | 0.07 | 0 | 0.11 | 0.03 |
| 3 | 61663778 | rs6784338 | -0.067 | 0.966 | -0.519 | 0.367 | -0.002 | -0.018 | -0.018 | 0.143 | 0.06 | 0.04 | 0.04 | 0.1 | 0.06 | 0.1 |

| | | | | | | | | | | | | | | | | |
|---|---|---|---|---|---|---|---|---|---|---|---|---|---|---|---|---|
| 3 | 61664946 | rs6787964 | 1.154 | 1.117 | -0.296 | 0.447 | -0.003 | 0.001 | 0.001 | 0.352 | 0.65 | 0.35 | 0.67 | 0.66 | 0.63 | 0.7 |
| 3 | 61666047 | rs7636402 | -0.86 | 0.372 | -0.807 | 0.155 | -0.003 | -0.01 | -0.01 | 0 | 0.4 | 0.5 | 0.31 | 0.22 | 0.31 | 0.26 |
| 3 | 61668923 | rs10446491 | 0.162 | 0.799 | -2.063 | 0.344 | -0.165 | -0.088 | -0.088 | 0 | 0.35 | 0.04 | 0.42 | 0.6 | 0.49 | 0.23 |
| 3 | 61670137 | rs4688268 | -0.239 | 0.574 | -1.86 | 0.009 | 0.046 | 0.196 | 0.196 | 0.084 | 0.48 | 0.19 | 0.57 | 0.79 | 0.62 | 0.38 |
| 3 | 61685759 | rs17065198 | 0.122 | 0.585 | -1.105 | 0.212 | -0.007 | -0.029 | -0.029 | 0 | 0.05 | 0.16 | 0.02 | 0 | 0.02 | 0.04 |
| 3 | 61688122 | rs2204128 | -0.222 | 0.349 | -1.441 | 0.021 | -0.005 | 0 | 0 | 0 | 0.92 | 0.99 | 0.93 | 0.93 | 0.95 | 0.78 |
| 3 | 61688641 | rs1403197 | -0.488 | 0.627 | -1.607 | 0.113 | -0.06 | -0.045 | -0.045 | 0 | 0.11 | 0.01 | 0.13 | 0.15 | 0.18 | 0.11 |
| 3 | 61693826 | rs11130851 | -0.228 | 0.634 | -1.588 | 0.168 | -0.047 | -0.048 | -0.048 | 0 | 0.09 | 0.01 | 0.12 | 0.16 | 0.15 | 0.1 |
| 3 | 61699969 | rs1915992 | -0.349 | 0.52 | -1.264 | -0.01 | -0.006 | -0.038 | -0.038 | 0.179 | 0.07 | 0.21 | 0.04 | 0 | 0.03 | 0.04 |
| 3 | 61704297 | rs9856705 | -0.35 | 0.554 | -1.227 | -0.015 | -0.005 | -0.039 | -0.039 | 0.314 | 0.07 | 0.21 | 0.05 | 0.01 | 0.04 | 0.07 |
| 3 | 61706745 | rs11718714 | -1.307 | 0.754 | -0.836 | 0.401 | 0.004 | 0.008 | 0.008 | 0 | 0.2 | 0.2 | 0.33 | 0.7 | 0.24 | 0.26 |
| 3 | 61708608 | rs7637558 | -1.213 | 0.711 | -0.847 | 0.419 | 0.005 | 0.007 | 0.007 | 0 | 0.2 | 0.2 | 0.33 | 0.71 | 0.26 | 0.27 |
| 3 | 61720229 | rs17065240 | -0.255 | 0.634 | -0.759 | 0.817 | -0.003 | 0.126 | 0.126 | 0 | 0.1 | 0.16 | 0.18 | 0.48 | 0.09 | 0.13 |
| 3 | 61722135 | rs2680238 | 0.651 | 0.489 | -0.214 | 0.667 | -0.003 | -0.004 | -0.004 | 0 | 0.81 | 0.82 | 0.77 | 0.73 | 0.8 | 0.66 |
| 3 | 61722196 | rs4688661 | -0.714 | 0.352 | -0.181 | 0.672 | -0.009 | 0.129 | 0.129 | 0.304 | 0.15 | 0.24 | 0.22 | 0.53 | 0.12 | 0.2 |
| 3 | 61727718 | rs648546 | 0.725 | 0.372 | -0.317 | 0.596 | -0.004 | -0.007 | -0.007 | 0.292 | 0.81 | 0.85 | 0.78 | 0.72 | 0.8 | 0.67 |
| 3 | 61728831 | rs585388 | 0.851 | 0.328 | -0.225 | 0.583 | 0.001 | 0.003 | 0.003 | 0 | 0.79 | 0.74 | 0.76 | 0.71 | 0.8 | 0.64 |
| 3 | 61729061 | rs632091 | 1.903 | 0.305 | -0.195 | 0.592 | 0 | -0.023 | -0.023 | 0 | 0.66 | 0.55 | 0.56 | 0.19 | 0.69 | 0.47 |
| 3 | 61729301 | rs9311830 | -0.002 | 0.489 | -0.398 | 0.625 | -0.038 | -0.038 | -0.038 | 0 | 0.09 | 0.02 | 0.14 | 0.2 | 0.15 | 0.11 |
| 3 | 61734539 | rs3821880 | 0.703 | 0.075 | -0.341 | -0.139 | -0.001 | 0.002 | 0.002 | 0 | 0.59 | 0.55 | 0.54 | 0.73 | 0.57 | 0.59 |
| 3 | 61739784 | rs636624 | 0.645 | -0.327 | -0.288 | -0.3 | -0.008 | 0.036 | 0.036 | 0.526 | 0.62 | 0.7 | 0.56 | 0.74 | 0.57 | 0.59 |
| 3 | 61740126 | rs625163 | 1.836 | -0.525 | -0.065 | -0.261 | -0.004 | 0.012 | 0.012 | 2.766 | 0.43 | 0.35 | 0.32 | 0.22 | 0.45 | 0.37 |
| 3 | 61741051 | rs4688663 | 0.169 | -0.492 | -0.042 | -0.143 | -0.005 | 0.077 | 0.077 | 0.397 | 0.14 | 0.17 | 0.2 | 0.51 | 0.11 | 0.2 |
| 3 | 61742742 | rs11708000 | 0.492 | -0.488 | -0.205 | -0.176 | -0.067 | -0.059 | -0.059 | 1.032 | 0.13 | 0.02 | 0.17 | 0.21 | 0.25 | 0.12 |
| 3 | 61746338 | rs2037501 | -1.441 | -0.54 | -0.135 | -0.162 | 0.002 | 0.001 | 0.001 | 0 | 0.32 | 0.37 | 0.4 | 0.68 | 0.36 | 0.34 |
| 3 | 61747737 | rs17065308 | 0.116 | -0.506 | -0.12 | -0.116 | -0.01 | 0.1 | 0.1 | 0.219 | 0.14 | 0.2 | 0.2 | 0.51 | 0.1 | 0.22 |
| 3 | 61748765 | rs9879444 | -0.624 | -0.568 | -0.171 | 0.008 | -0.006 | -0.007 | -0.007 | 0 | 0.2 | 0.16 | 0.19 | 0.29 | 0.25 | 0.12 |
| 3 | 61749709 | rs12491760 | -1.617 | -0.582 | -0.144 | -0.056 | -0.001 | 0 | 0 | 0 | 0.33 | 0.31 | 0.4 | 0.79 | 0.36 | 0.31 |
| 3 | 61751127 | rs7648560 | 2.045 | -0.726 | -0.033 | -0.196 | -0.004 | 0.054 | 0.054 | 3.305 | 0.46 | 0.34 | 0.45 | 0.13 | 0.52 | 0.44 |
| 3 | 61755838 | rs665974 | -0.402 | -0.826 | -0.035 | -0.147 | -0.008 | -0.026 | -0.026 | 0 | 0.81 | 0.76 | 0.86 | 0.94 | 0.88 | 0.77 |
| 3 | 61757206 | rs2680237 | 1.026 | -0.843 | -0.029 | 0.154 | -0.024 | 0.035 | 0.035 | 5.306 | 0.67 | 0.54 | 0.69 | 0.54 | 0.79 | 0.61 |
| 3 | 61762630 | rs660323 | -0.481 | -0.804 | -0.074 | 0.1 | -0.012 | -0.034 | -0.034 | 0.513 | 0.82 | 0.75 | 0.87 | 0.95 | 0.89 | 0.75 |
| 3 | 61771711 | rs17065393 | 0.892 | -0.611 | 0.038 | 0.277 | -0.001 | 0.091 | 0.091 | 0 | 0.08 | 0.09 | 0.16 | 0.4 | 0.07 | 0.1 |
| 3 | 61772727 | rs17627732 | 2.051 | -0.261 | -0.308 | -0.688 | 0.004 | -0.016 | -0.016 | 0.357 | 0.18 | 0.15 | 0.11 | 0.07 | 0.21 | 0.09 |
| 3 | 61773374 | rs11130852 | -0.275 | -0.178 | -0.357 | -0.332 | -0.031 | 0.178 | 0.178 | 0 | 0.17 | 0.38 | 0.2 | 0.5 | 0.1 | 0.15 |
| 3 | 61775863 | rs10433639 | 0.75 | 0.336 | -0.943 | -0.326 | -0.072 | -0.072 | -0.072 | 0.982 | 0.12 | 0.01 | 0.19 | 0.27 | 0.24 | 0.12 |
| 3 | 61776260 | rs654033 | -0.777 | -0.295 | -0.7 | 0.01 | -0.015 | 0.024 | 0.024 | 4.288 | 0.33 | 0.4 | 0.32 | 0.51 | 0.23 | 0.44 |
| 3 | 61783170 | rs657813 | 0.406 | 0.878 | -1.154 | 0.156 | -0.024 | 0.03 | 0.03 | 0 | 0.65 | 0.85 | 0.61 | 0.67 | 0.58 | 0.47 |
| 3 | 61788620 | rs1980283 | 0.946 | 0.723 | -0.921 | 0.553 | 0.008 | 0.016 | 0.016 | 0.088 | 0.44 | 0.63 | 0.3 | 0.37 | 0.42 | 0.59 |
| 3 | 61789749 | rs1980281 | 2.094 | 0.261 | -0.524 | 0.608 | -0.004 | -0.003 | -0.003 | 0 | 0.34 | 0.28 | 0.2 | 0.07 | 0.36 | 0.29 |
| 3 | 61792133 | rs636636 | -1.788 | -0.601 | -0.116 | 1.329 | 0.062 | 0.038 | 0.038 | 2.485 | 0.47 | 0.71 | 0.42 | 0.66 | 0.31 | 0.52 |
| 3 | 61793618 | rs17065421 | -0.765 | -0.649 | 0.077 | 1.346 | -0.016 | 0.077 | 0.077 | 0 | 0.13 | 0.33 | 0.13 | 0.3 | 0.11 | 0.35 |
| 3 | 61794054 | rs652889 | -1.787 | -0.576 | 0.015 | 1.532 | 0.048 | -0.007 | -0.007 | 0 | 0.46 | 0.69 | 0.37 | 0.5 | 0.31 | 0.51 |
| 3 | 61803428 | rs615854 | -1.278 | -0.653 | -0.128 | 1.314 | 0.015 | 0 | 0 | 0 | 0.39 | 0.35 | 0.36 | 0.56 | 0.27 | 0.34 |
| 3 | 61812690 | rs1522549 | -0.281 | -0.491 | -0.318 | 0.913 | -0.009 | -0.048 | -0.048 | 0 | 0.21 | 0.11 | 0.28 | 0.29 | 0.21 | 0.13 |
| 3 | 61814338 | rs11130857 | -0.65 | -0.424 | -0.201 | 0.944 | -0.01 | -0.029 | -0.029 | 0 | 0.3 | 0.14 | 0.32 | 0.28 | 0.28 | 0.18 |
| 3 | 61817359 | rs2687147 | -0.472 | -0.626 | 0.779 | 0.669 | 0.046 | 0.078 | 0.078 | 1.162 | 0.08 | 0.48 | 0.06 | 0.07 | 0.02 | 0.01 |
| 3 | 61830799 | rs652936 | -0.131 | -0.71 | 0.489 | 0.566 | -0.005 | -0.013 | -0.013 | 0 | 0.24 | 0.25 | 0.14 | 0.1 | 0.15 | 0.13 |
| 3 | 61832151 | rs669005 | -0.02 | -0.714 | 0.502 | 0.563 | 0.007 | 0.008 | 0.008 | 1.157 | 0.24 | 0.19 | 0.14 | 0.1 | 0.15 | 0.13 |
| 3 | 61834315 | rs6802156 | -1.518 | -0.95 | 0.91 | 0.843 | 0.022 | 0.01 | 0.01 | 0.621 | 0.46 | 0.64 | 0.38 | 0.59 | 0.34 | 0.5 |
| 3 | 61834942 | rs6766943 | -1.097 | -0.941 | 0.955 | 0.864 | -0.013 | 0.093 | 0.093 | 0 | 0.22 | 0.4 | 0.25 | 0.49 | 0.18 | 0.37 |
| 3 | 61838890 | rs6792064 | -0.625 | -0.951 | 1 | 0.874 | 0.02 | -0.015 | -0.015 | 0 | 0.08 | 0.3 | 0.1 | 0.03 | 0.06 | 0.02 |
| 3 | 61839880 | rs689236 | -1.434 | -0.967 | 0.946 | 0.867 | 0 | 0 | 0 | 0 | 0.45 | 0.59 | 0.38 | 0.59 | 0.33 | 0.49 |
| 3 | 61842176 | rs596904 | -1.53 | -0.965 | 0.984 | 0.815 | 0.023 | 0.014 | 0.014 | 0.38 | 0.46 | 0.65 | 0.38 | 0.61 | 0.34 | 0.5 |
| 3 | 61847695 | rs591988 | -0.81 | -1.001 | 0.94 | 0.833 | 0.029 | -0.021 | -0.021 | 0.087 | 0.36 | 0.23 | 0.27 | 0.57 | 0.27 | 0.48 |
| 3 | 61853342 | rs13076126 | 0.072 | -0.957 | 0.927 | 0.797 | 0.016 | -0.007 | -0.007 | 0 | 0.07 | 0.22 | 0.09 | 0.03 | 0.06 | 0.02 |
| 3 | 61853529 | rs630806 | -0.753 | -0.978 | 0.669 | 0.805 | -0.021 | -0.044 | -0.044 | 0 | 0.33 | 0.57 | 0.25 | 0.13 | 0.21 | 0.15 |
| 3 | 61859265 | rs2687149 | -0.708 | -0.946 | 0.592 | 0.809 | -0.018 | -0.039 | -0.039 | 1.183 | 0.33 | 0.57 | 0.25 | 0.13 | 0.21 | 0.15 |
| 3 | 61868242 | rs12637323 | -0.608 | -0.357 | 0.347 | 1.463 | 0.007 | -0.053 | -0.053 | 0 | 0.14 | 0.08 | 0.14 | 0.48 | 0.12 | 0.35 |
| 3 | 61869264 | rs1534809 | -1.014 | -0.114 | 0.048 | 1.322 | -0.02 | -0.048 | -0.048 | 0 | 0.88 | 0.96 | 0.7 | 0.68 | 0.82 | 0.86 |
| 3 | 61887743 | rs7625335 | -1.791 | -0.097 | 0.574 | 1.35 | -0.018 | 0.108 | 0.108 | 0.225 | 0.24 | 0.6 | 0.25 | 0.51 | 0.19 | 0.37 |
| 3 | 61899499 | rs17065607 | 0.024 | 0.031 | -0.119 | 1.306 | 0.006 | -0.053 | -0.053 | 0 | 0.09 | 0.01 | 0.05 | 0.21 | 0.07 | 0.14 |
| 3 | 61901792 | rs7630364 | -0.876 | 0.287 | 0.463 | 0.916 | 0.093 | 0.05 | 0.05 | 0.03 | 0.06 | 0.34 | 0.09 | 0.03 | 0.06 | 0.01 |
| 3 | 61904945 | rs4234692 | -0.852 | -0.158 | -0.435 | 1.23 | -0.021 | -0.049 | -0.049 | 0 | 0.88 | 0.96 | 0.75 | 0.7 | 0.83 | 0.88 |
| 3 | 61907965 | rs2366461 | 0.003 | -0.623 | -0.668 | 0.714 | -0.006 | -0.009 | -0.009 | 0 | 0.7 | 0.82 | 0.6 | 0.65 | 0.65 | 0.78 |
| 3 | 61910192 | rs6797184 | -0.495 | -1.008 | -0.418 | 0.506 | 0 | -0.001 | -0.001 | 0.09 | 0.43 | 0.56 | 0.33 | 0.51 | 0.29 | 0.52 |
| 3 | 61914978 | rs28695326 | -0.058 | -1.037 | -0.516 | 0.664 | -0.014 | 0.067 | 0.067 | 0 | 0.15 | 0.2 | 0.13 | 0.41 | 0.09 | 0.31 |
| 3 | 61916364 | rs11706742 | 1.002 | -0.976 | -0.72 | 0.703 | -0.001 | 0.047 | 0.047 | 0 | 0.07 | 0.04 | 0.09 | 0.41 | 0.07 | 0.26 |
| 3 | 61924199 | rs1479401 | -0.398 | -0.994 | -0.651 | 0.316 | -0.026 | 0.008 | 0.008 | 0 | 0.4 | 0.55 | 0.31 | 0.52 | 0.24 | 0.46 |

| | | | | | | | | | | | | | | | | |
|---|---|---|---|---|---|---|---|---|---|---|---|---|---|---|---|---|
| 3 | 61936606 | rs4688668 | -1.115 | -1.087 | -0.735 | 0.45 | -0.003 | -0.009 | -0.009 | 0 | 0.81 | 0.85 | 0.68 | 0.66 | 0.76 | 0.74 |
| 3 | 61936843 | rs4688276 | -0.564 | -1.095 | -0.755 | 0.68 | -0.02 | 0.004 | 0.004 | 0 | 0.41 | 0.58 | 0.29 | 0.52 | 0.23 | 0.56 |
| 3 | 61938693 | rs10514719 | -0.428 | -0.145 | -1.08 | 1.153 | -0.022 | 0.094 | 0.094 | 0 | 0.16 | 0.27 | 0.14 | 0.42 | 0.09 | 0.3 |
| 3 | 61940856 | rs1006588 | 0.358 | 0.668 | -1.739 | 0.544 | -0.085 | 0.058 | 0.058 | 0.607 | 0.28 | 0.1 | 0.26 | 0.13 | 0.4 | 0.14 |
| 3 | 61942066 | rs40494 | -0.835 | 0.166 | -1.547 | 1.171 | 0.008 | -0.046 | -0.046 | 0 | 0.11 | 0.03 | 0.16 | 0.22 | 0.12 | 0.24 |
| 3 | 61942816 | rs1479397 | -0.333 | -0.239 | -1.322 | 1.08 | -0.003 | 0 | 0 | 0 | 0.18 | 0.15 | 0.17 | 0.26 | 0.14 | 0.12 |
| 3 | 61950168 | rs1479394 | 0.201 | 0.464 | -0.841 | 0.89 | 0.048 | 0.022 | 0.022 | 0 | 0.2 | 0.39 | 0.19 | 0.21 | 0.2 | 0.14 |
| 3 | 61951683 | rs33080 | -1.184 | 0.265 | -1.707 | 0.474 | -0.059 | 0.064 | 0.064 | 0 | 0.76 | 0.96 | 0.77 | 0.9 | 0.73 | 0.66 |
| 3 | 61951903 | rs9836440 | 0.256 | 0.373 | -0.947 | 0.93 | 0.047 | 0.008 | 0.008 | 0 | 0.21 | 0.4 | 0.19 | 0.21 | 0.21 | 0.14 |
| 3 | 61952379 | rs9817604 | -0.963 | 0.404 | -1.122 | 1.179 | -0.01 | -0.004 | -0.004 | 0 | 0.42 | 0.71 | 0.49 | 0.66 | 0.36 | 0.38 |
| 3 | 61953101 | rs1029508 | -0.587 | 0.046 | -0.822 | 1.32 | -0.007 | 0.041 | 0.041 | 0 | 0.2 | 0.21 | 0.29 | 0.47 | 0.16 | 0.24 |
| 3 | 61956588 | rs251039 | -0.693 | 0.128 | -0.896 | 1.243 | -0.004 | 0.028 | 0.028 | 0 | 0.23 | 0.25 | 0.33 | 0.51 | 0.2 | 0.25 |
| 3 | 61958023 | rs809764 | -0.511 | 0.248 | -0.546 | 0.604 | -0.002 | -0.007 | -0.007 | 0 | 0.35 | 0.6 | 0.27 | 0.28 | 0.31 | 0.21 |
| 3 | 61959582 | rs4141291 | -0.835 | 0.466 | -0.484 | 0.313 | -0.002 | 0.018 | 0.018 | 0 | 0.53 | 0.74 | 0.41 | 0.4 | 0.5 | 0.43 |
| 3 | 61960632 | rs13085295 | -0.791 | 0.46 | -0.219 | 0.281 | -0.001 | 0.008 | 0.008 | 0 | 0.52 | 0.65 | 0.41 | 0.41 | 0.5 | 0.44 |
| 3 | 61964956 | rs4447756 | -0.456 | 0.435 | -0.339 | 0.065 | 0.003 | -0.003 | -0.003 | 0 | 0.41 | 0.68 | 0.34 | 0.43 | 0.39 | 0.43 |
| 3 | 61967036 | rs7647959 | 0.743 | 0.569 | -0.401 | -0.083 | -0.04 | 0.069 | 0.069 | 0 | 0.12 | 0.02 | 0.07 | 0 | 0.12 | 0.11 |
| 3 | 61968016 | rs13320692 | -0.341 | 0.354 | -0.035 | 0.039 | -0.005 | -0.044 | -0.044 | 0.065 | 0.38 | 0.51 | 0.3 | 0.2 | 0.36 | 0.27 |
| 3 | 61981197 | rs17334158 | 0.251 | 0.561 | -0.189 | -0.041 | -0.025 | 0.118 | 0.118 | 0.032 | 0.21 | 0.04 | 0.1 | 0 | 0.16 | 0.14 |
| 3 | 61983459 | rs6788568 | 0.344 | 1.001 | -0.02 | 0.08 | -0.024 | -0.012 | -0.012 | 0 | 0.14 | 0.09 | 0.19 | 0.18 | 0.23 | 0.12 |
| 3 | 61983786 | rs804418 | -0.801 | 0.956 | 0.614 | -0.025 | 0.019 | -0.009 | -0.009 | 0 | 0.25 | 0.39 | 0.27 | 0.22 | 0.3 | 0.18 |
| 3 | 61986342 | rs804415 | -1.452 | 0.915 | 0.734 | -0.021 | 0.014 | 0.019 | 0.019 | 1.747 | 0.31 | 0.63 | 0.31 | 0.38 | 0.37 | 0.35 |
| 3 | 61987503 | rs6445245 | -1.036 | 0.57 | 0.797 | -0.054 | 0.036 | 0.148 | 0.148 | 0 | 0.06 | 0.27 | 0.04 | 0.14 | 0.04 | 0.17 |
| 3 | 61989945 | rs6785475 | 0.48 | 0.572 | 0.723 | -0.21 | 0.009 | -0.02 | -0.02 | 0 | 0.11 | 0.26 | 0.08 | 0.04 | 0.07 | 0.06 |
| 3 | 61990795 | rs9847300 | -0.759 | 0.806 | 0.671 | -0.097 | -0.016 | 0.006 | 0.006 | 0.105 | 0.26 | 0.4 | 0.3 | 0.27 | 0.21 | 0.16 |
| 3 | 61991787 | rs11706999 | -1.663 | 1.442 | 0.214 | -0.376 | -0.001 | -0.002 | -0.002 | 0.266 | 0.52 | 0.51 | 0.42 | 0.36 | 0.46 | 0.46 |
| 3 | 61993536 | rs2107561 | -0.881 | 1.569 | 0.149 | -0.465 | -0.026 | 0.098 | 0.098 | 0 | 0.8 | 0.93 | 0.9 | 1 | 0.83 | 0.88 |
| 3 | 61993666 | rs9311833 | -0.949 | 1.514 | 0.285 | -0.431 | 0.001 | -0.006 | -0.006 | 0 | 0.31 | 0.43 | 0.34 | 0.28 | 0.3 | 0.28 |
| 3 | 61997367 | rs6794405 | -2.039 | 1.543 | 0.297 | -0.533 | -0.001 | 0.014 | 0.014 | 0 | 0.55 | 0.68 | 0.46 | 0.35 | 0.49 | 0.52 |
| 3 | 62000317 | rs739984 | -0.62 | 1.504 | 0.343 | -0.569 | 0.044 | 0.015 | 0.015 | 0 | 0.17 | 0.39 | 0.12 | 0.13 | 0.18 | 0.24 |
| 3 | 62000344 | rs11710174 | 1.601 | 1.535 | 0.163 | -0.521 | -0.023 | -0.008 | -0.008 | 0 | 0.07 | 0 | 0.04 | 0.04 | 0.07 | 0.06 |
| 3 | 62000476 | rs6804425 | -0.451 | 1.532 | 0.171 | -0.533 | 0.013 | 0.052 | 0.052 | 0 | 0.35 | 0.19 | 0.31 | 0.22 | 0.29 | 0.21 |
| 3 | 62004450 | rs11714561 | -0.866 | 1.532 | 0.457 | -0.48 | 0 | -0.005 | -0.005 | 0.152 | 0.26 | 0.29 | 0.3 | 0.38 | 0.25 | 0.28 |
| 3 | 62004585 | rs11718787 | -0.866 | 1.532 | 0.459 | -0.484 | 0.001 | -0.009 | -0.009 | 1.06 | 0.26 | 0.28 | 0.3 | 0.38 | 0.25 | 0.28 |
| 3 | 62005962 | rs6445251 | -0.694 | 1.508 | 0.467 | -0.478 | -0.001 | 0.002 | 0.002 | 0 | 0.24 | 0.29 | 0.28 | 0.35 | 0.22 | 0.21 |
| 3 | 62007036 | rs11717816 | -0.149 | 1.513 | 0.423 | -0.479 | 0.005 | -0.02 | -0.02 | 0 | 0.22 | 0.14 | 0.27 | 0.26 | 0.2 | 0.15 |
| 3 | 62007307 | rs1544420 | -0.449 | 1.585 | 0.494 | -0.525 | 0.074 | -0.023 | -0.023 | 0 | 0.09 | 0.34 | 0.07 | 0.02 | 0.09 | 0.14 |
| 3 | 62011294 | rs10490779 | 1.318 | 1.568 | 0.36 | -0.506 | -0.005 | 0 | 0 | 0 | 0.07 | 0.05 | 0.04 | 0 | 0.07 | 0.07 |
| 3 | 62012687 | rs7434122 | -2.088 | 1.487 | 0.606 | -0.556 | -0.001 | -0.002 | -0.002 | 0 | 0.54 | 0.76 | 0.47 | 0.4 | 0.49 | 0.53 |
| 3 | 62015120 | rs986613 | -1.341 | 1.496 | 0.634 | -0.57 | -0.004 | 0.012 | 0.012 | 0 | 0.39 | 0.63 | 0.24 | 0.15 | 0.34 | 0.45 |
| 3 | 62015268 | rs10490778 | -0.541 | 1.468 | 0.571 | -0.537 | 0.013 | -0.121 | -0.121 | 0 | 0.3 | 0.44 | 0.18 | 0.04 | 0.26 | 0.32 |
| 3 | 62016422 | rs7642746 | 0.533 | 1.397 | 0.321 | -0.351 | 0.008 | -0.022 | -0.022 | 0 | 0.15 | 0.11 | 0.22 | 0.23 | 0.14 | 0.08 |
| 3 | 62023467 | rs7637925 | -1.083 | 1.491 | 0.437 | -0.507 | 0 | 0.006 | 0.006 | 0 | 0.41 | 0.45 | 0.36 | 0.24 | 0.35 | 0.21 |
| 3 | 62027634 | rs17767097 | 0.642 | 1.347 | 0.268 | -0.527 | -0.003 | 0.069 | 0.069 | 0 | 0.19 | 0.07 | 0.09 | 0 | 0.14 | 0.09 |
| 3 | 62028246 | rs7619444 | 0.06 | 0.713 | 0.951 | -0.143 | -0.042 | 0.155 | 0.155 | 0 | 0.86 | 0.77 | 0.79 | 0.42 | 0.93 | 0.83 |
| 3 | 62030649 | rs4688121 | -0.017 | 0.72 | 0.758 | -0.208 | -0.031 | 0.131 | 0.131 | 0 | 0.86 | 0.79 | 0.79 | 0.42 | 0.92 | 0.86 |
| 3 | 62030718 | rs12489693 | 1.01 | 0.652 | 0.625 | -0.31 | 0.003 | -0.041 | -0.041 | 0 | 0.12 | 0.04 | 0.2 | 0.19 | 0.12 | 0.05 |
| 3 | 62030767 | rs10490777 | 0.931 | 0.836 | 0.624 | -0.395 | -0.007 | 0.002 | 0.002 | 0 | 0.08 | 0.04 | 0.06 | 0.05 | 0.08 | 0.1 |
| 3 | 62030849 | rs9311834 | 0.1 | 0.834 | 0.699 | -0.419 | -0.033 | 0.132 | 0.132 | 0.04 | 0.84 | 0.74 | 0.76 | 0.39 | 0.9 | 0.72 |
| 3 | 62032616 | rs7638067 | 1.092 | 0.816 | 0.715 | -0.435 | -0.021 | 0.056 | 0.056 | 0.875 | 0.75 | 0.62 | 0.7 | 0.35 | 0.82 | 0.65 |
| 3 | 62033353 | rs7640737 | 0.368 | 0.844 | 0.689 | -0.445 | 0.003 | -0.002 | -0.002 | 0.108 | 0.1 | 0.11 | 0.09 | 0.08 | 0.11 | 0.24 |
| 3 | 62034833 | rs1125465 | -0.44 | 0.849 | 0.675 | -0.457 | -0.002 | -0.015 | -0.015 | 0 | 0.23 | 0.17 | 0.29 | 0.27 | 0.22 | 0.29 |
| 3 | 62035492 | rs7647357 | 1.414 | 0.809 | 0.826 | -0.494 | -0.02 | -0.053 | -0.053 | 0 | 0.62 | 0.52 | 0.49 | 0.15 | 0.7 | 0.57 |
| 3 | 62037499 | rs6774478 | 1.448 | 1.141 | 0.854 | -0.511 | 0.007 | -0.042 | -0.042 | 0 | 0.61 | 0.57 | 0.43 | 0.16 | 0.57 | 0.57 |
| 3 | 62040007 | rs7615943 | -0.183 | 1.168 | 0.797 | -0.44 | 0.021 | 0.124 | 0.124 | 0 | 0.84 | 0.77 | 0.72 | 0.42 | 0.79 | 0.89 |
| 3 | 62042825 | rs17065828 | -0.237 | 1.26 | 0.618 | -0.457 | 0.001 | -0.006 | -0.006 | 0 | 0.21 | 0.2 | 0.26 | 0.25 | 0.2 | 0.16 |
| 3 | 62043111 | rs17767976 | 0.801 | 1.285 | 0.568 | -0.449 | -0.002 | 0.001 | 0.001 | 0 | 0.08 | 0.07 | 0.06 | 0.05 | 0.08 | 0.1 |
| 3 | 62044326 | rs12715576 | 1.668 | 1.215 | 0.731 | -0.648 | -0.001 | 0.091 | 0.091 | 3.358 | 0.49 | 0.35 | 0.36 | 0.12 | 0.51 | 0.49 |
| 3 | 62044608 | rs9311835 | 0.14 | 1.599 | 0.998 | -0.228 | 0.096 | 0.328 | 0.328 | 0.397 | 0.89 | 0.64 | 0.74 | 0.4 | 0.83 | 0.95 |
| 3 | 62047076 | rs9311837 | -0.205 | 1.78 | 0.534 | -0.915 | 0.061 | -0.039 | -0.039 | 0 | 0.07 | 0.21 | 0.05 | 0 | 0.09 | 0.05 |
| 3 | 62048629 | rs17768204 | 0.437 | 1.371 | 0.373 | -0.676 | -0.007 | 0.002 | 0.002 | 0.041 | 0.08 | 0.05 | 0.06 | 0.05 | 0.08 | 0.1 |
| 3 | 62048657 | rs6445254 | 0.311 | 1.224 | 0.485 | -0.51 | 0.025 | 0.009 | 0.009 | 0.192 | 0.77 | 0.54 | 0.54 | 0.21 | 0.72 | 0.9 |
| 3 | 62050151 | rs7610872 | 0.413 | 1.098 | -0.117 | -1.149 | 0.012 | 0.024 | 0.024 | 7.355 | 0.49 | 0.44 | 0.31 | 0.13 | 0.44 | 0.46 |
| 3 | 62050279 | rs6445255 | 0.383 | 0.751 | -0.039 | -1.018 | 0.001 | -0.041 | -0.041 | 0 | 0.36 | 0.37 | 0.29 | 0.1 | 0.44 | 0.28 |
| 3 | 62052829 | rs168371 | 0.4 | 0.143 | 0.981 | -0.062 | -0.009 | -0.013 | -0.013 | 0 | 0.83 | 0.65 | 0.91 | 0.88 | 0.9 | 0.85 |
| 3 | 62052909 | rs13094307 | -0.938 | 0.188 | 1.057 | 0.311 | 0.01 | 0.039 | 0.039 | 1.661 | 0.11 | 0.27 | 0.17 | 0.26 | 0.1 | 0.2 |
| 3 | 62053586 | rs2107557 | 1.264 | 0.146 | 0.913 | 0.272 | -0.003 | 0.016 | 0.016 | 1.24 | 0.73 | 0.43 | 0.75 | 0.62 | 0.81 | 0.66 |
| 3 | 62056869 | rs9879067 | 1.192 | 0.13 | 0.631 | 0.329 | -0.001 | 0.055 | 0.055 | 0.31 | 0.53 | 0.36 | 0.42 | 0.16 | 0.51 | 0.24 |
| 3 | 62056891 | rs2526431 | 0.476 | 0.009 | 0.443 | 0.429 | -0.023 | -0.035 | -0.035 | 0 | 0.65 | 0.78 | 0.61 | 0.47 | 0.6 | 0.45 |

| | | | | | | | | | | | | | | | | |
|---|---|---|---|---|---|---|---|---|---|---|---|---|---|---|---|---|
| 3 | 62064925 | rs11715438 | -1.073 | -0.295 | 0.82 | 0.457 | -0.029 | -0.005 | -0.005 | 0.552 | 0.19 | 0.06 | 0.12 | 0.17 | 0.17 | 0.29 |
| 3 | 62069750 | rs2526424 | -1.861 | -0.391 | 1.362 | 0.424 | -0.004 | -0.006 | -0.006 | 0 | 0.27 | 0.65 | 0.2 | 0.26 | 0.2 | 0.32 |
| 3 | 62073316 | rs9870965 | -1.745 | -0.445 | 1.484 | 0.439 | -0.001 | -0.009 | -0.009 | 0.059 | 0.25 | 0.5 | 0.18 | 0.27 | 0.2 | 0.32 |
| 3 | 62078867 | rs2661884 | -1.186 | -0.159 | 0.398 | 0.502 | 0 | -0.002 | -0.002 | 0.741 | 0.21 | 0.2 | 0.14 | 0.21 | 0.17 | 0.3 |
| 3 | 62079016 | rs191558 | -1.186 | -0.159 | 0.383 | 0.502 | 0 | -0.002 | -0.002 | 0 | 0.21 | 0.2 | 0.14 | 0.21 | 0.17 | 0.3 |
| 3 | 62081966 | rs886562 | -1.236 | -0.359 | 0.409 | 1.098 | -0.016 | 0.186 | 0.186 | 1.396 | 0.19 | 0.41 | 0.19 | 0.58 | 0.16 | 0.28 |
| 3 | 62086558 | rs6772567 | 0.06 | -0.149 | -0.17 | 0.748 | -0.013 | -0.003 | -0.003 | 1.044 | 0.1 | 0.04 | 0.09 | 0.12 | 0.1 | 0.14 |
| 3 | 62089804 | rs10510870 | -0.862 | -0.338 | -0.373 | 0.54 | -0.007 | 0.034 | 0.034 | 0 | 0.78 | 0.57 | 0.67 | 0.71 | 0.78 | 0.66 |
| 3 | 62095691 | rs17632018 | 0.851 | -0.066 | -0.802 | 0.457 | -0.016 | 0.021 | 0.021 | 1.105 | 0.08 | 0.03 | 0.04 | 0 | 0.08 | 0.08 |
| 3 | 62098678 | rs6784506 | 0.139 | -0.261 | -0.579 | 0.674 | 0 | -0.002 | -0.002 | 0 | 0.11 | 0.11 | 0.1 | 0.13 | 0.1 | 0.12 |
| 3 | 62108081 | rs880417 | 0.689 | -0.108 | -1.208 | 0.344 | -0.012 | -0.041 | -0.041 | 0 | 0.32 | 0.17 | 0.35 | 0.48 | 0.38 | 0.28 |
| 3 | 62108226 | rs880416 | -0.43 | -0.389 | -1.128 | -0.248 | -0.003 | 0.028 | 0.028 | 0.929 | 0.33 | 0.24 | 0.21 | 0.11 | 0.3 | 0.24 |
| 3 | 62114109 | rs17066154 | -0.828 | -1.406 | -1.316 | 0.32 | 0.037 | -0.036 | -0.036 | 0 | 0.2 | 0.08 | 0.07 | 0.36 | 0.08 | 0.2 |
| 3 | 62114428 | rs4515044 | -1.037 | -1.459 | -1.216 | 0.009 | 0.054 | -0.045 | -0.045 | 0 | 0.26 | 0.07 | 0.1 | 0.36 | 0.13 | 0.21 |
| 3 | 62117037 | rs6797646 | 0.118 | -1.326 | -1.946 | -0.825 | -0.048 | 0.111 | 0.111 | 0 | 0.25 | 0.07 | 0.18 | 0.05 | 0.3 | 0.24 |
| 3 | 62117185 | rs6764198 | -1.229 | -2.03 | -1.193 | -1.149 | 0.002 | -0.002 | -0.002 | 0.034 | 0.58 | 0.37 | 0.31 | 0.42 | 0.45 | 0.45 |
| 3 | 62131691 | rs4440115 | -1.589 | -2.776 | -0.608 | -1.243 | 0.024 | 0.013 | 0.013 | 5.035 | 0.48 | 0.57 | 0.25 | 0.41 | 0.31 | 0.35 |
| 3 | 62133323 | rs9881025 | 0.163 | -2.959 | -0.372 | -1.387 | -0.019 | -0.033 | -0.033 | 0.061 | 0.1 | 0.39 | 0.06 | 0.01 | 0.03 | 0.05 |
| 3 | 62133492 | rs17066205 | 0.101 | -3.143 | -0.406 | -1.484 | 0.02 | 0.005 | 0.005 | 0 | 0.16 | 0.08 | 0.06 | 0 | 0.06 | 0.08 |
| 3 | 62134802 | rs7632091 | -1.564 | -3.145 | -0.453 | -1.494 | -0.048 | -0.153 | -0.153 | 0.36 | 0.3 | 0.49 | 0.16 | 0.02 | 0.2 | 0.19 |
| 3 | 62137083 | rs9826645 | -1.641 | -2.58 | -0.222 | -1.169 | -0.029 | -0.002 | -0.002 | 0 | 0.37 | 0.53 | 0.18 | 0.38 | 0.18 | 0.3 |
| 3 | 62151054 | rs4505694 | -0.342 | -2.155 | -0.313 | -1.026 | 0.068 | -0.021 | -0.021 | 0 | 0.07 | 0.27 | 0.08 | 0.01 | 0.11 | 0.05 |
| 3 | 62151448 | rs7612628 | -0.445 | -2.193 | -0.316 | -1.034 | 0.033 | 0.01 | 0.01 | 2.161 | 0.06 | 0.3 | 0.03 | 0 | 0.01 | 0 |
| 3 | 62157497 | rs11130879 | -1.591 | -2.281 | -0.433 | -0.998 | -0.011 | 0.011 | 0.011 | 1.103 | 0.3 | 0.38 | 0.16 | 0.38 | 0.22 | 0.21 |
| 3 | 62159990 | rs17066236 | 0.19 | -2.245 | -0.5 | -0.971 | -0.009 | -0.101 | -0.101 | 0 | 0.14 | 0.01 | 0.07 | 0.37 | 0.1 | 0.15 |
| 3 | 62161615 | rs7627359 | -1.578 | -2.286 | -0.399 | -1.013 | -0.052 | 0 | 0 | 0 | 0.36 | 0.49 | 0.17 | 0.38 | 0.19 | 0.29 |
| 3 | 62166837 | rs4586801 | -0.433 | -1.81 | -0.262 | -0.723 | 0.057 | -0.025 | -0.025 | 0 | 0.07 | 0.25 | 0.08 | 0.01 | 0.11 | 0.04 |
| 3 | 62167361 | rs6791264 | -0.623 | -1.82 | -0.321 | -0.694 | 0.04 | -0.058 | -0.058 | 0 | 0.22 | 0.08 | 0.07 | 0.37 | 0.11 | 0.15 |
| 3 | 62171253 | rs9857089 | -1.36 | -1.858 | -0.339 | -0.618 | -0.019 | 0.023 | 0.023 | 0.468 | 0.31 | 0.47 | 0.17 | 0.39 | 0.22 | 0.2 |
| 3 | 62174532 | rs7641268 | -0.56 | -1.478 | -0.423 | -0.414 | -0.003 | 0.002 | 0.002 | 0 | 0.08 | 0.1 | 0.09 | 0.03 | 0.15 | 0.12 |
| 3 | 62184407 | rs6764475 | -0.864 | -2.156 | -0.286 | -0.336 | -0.088 | 0.093 | 0.093 | 6.527 | 0.21 | 0.45 | 0.11 | 0.36 | 0.07 | 0.14 |
| 3 | 62188720 | rs17066275 | 0.378 | -1.821 | -0.648 | -0.115 | 0.007 | 0.015 | 0.015 | 0.72 | 0.09 | 0.05 | 0.05 | 0.37 | 0.05 | 0.12 |
| 3 | 62189189 | rs2292245 | -0.934 | -2.316 | -1.093 | -0.612 | 0.049 | -0.056 | -0.056 | 2.288 | 0.25 | 0.15 | 0.11 | 0.36 | 0.16 | 0.29 |
| 3 | 62198335 | rs17685418 | -0.745 | -2.59 | -1.041 | -0.914 | 0.004 | 0.015 | 0.015 | 0.008 | 0.17 | 0.17 | 0.12 | 0.08 | 0.15 | 0.17 |
| 3 | 62199471 | rs1007555 | -1.092 | -3.252 | -0.986 | -1.702 | 0.01 | 0.01 | 0.01 | 0.122 | 0.47 | 0.56 | 0.33 | 0.39 | 0.35 | 0.33 |
| 3 | 62206239 | rs10510873 | 1.464 | -2.155 | -1.142 | -1.962 | 0 | 0.01 | 0.01 | 0 | 0.08 | 0.05 | 0.09 | 0.01 | 0.09 | 0.2 |
| 3 | 62221041 | rs6785764 | -1.067 | -2.763 | -1.301 | -1.894 | -0.049 | 0.008 | 0.008 | 6.072 | 0.33 | 0.54 | 0.16 | 0.39 | 0.14 | 0.21 |
| 3 | 62226704 | rs1388615 | -0.788 | -2.872 | -0.829 | -1.966 | -0.031 | 0.018 | 0.018 | 0.048 | 0.24 | 0.36 | 0.14 | 0.33 | 0.15 | 0.25 |
| 3 | 62231319 | rs1352883 | 0.966 | -2.881 | -0.365 | -1.972 | 0.001 | 0.001 | 0.001 | 0 | 0.08 | 0.12 | 0.06 | 0.09 | 0.08 | 0.05 |
| 3 | 62233769 | rs1491963 | 1.464 | -3.02 | -0.511 | -2.103 | -0.002 | 0.048 | 0.048 | 0 | 0.17 | 0.06 | 0.12 | 0.03 | 0.16 | 0.1 |
| 3 | 62234885 | rs17066305 | -1.363 | -3.048 | -0.432 | -2.089 | -0.029 | -0.005 | -0.005 | 0 | 0.37 | 0.5 | 0.19 | 0.37 | 0.22 | 0.25 |
| 3 | 62239596 | rs11709670 | 1.715 | -2.851 | -0.461 | -2 | -0.023 | 0.026 | 0.026 | 0.012 | 0.05 | 0.03 | 0.04 | 0.01 | 0.09 | 0.12 |
| 3 | 62244444 | rs1388612 | -0.358 | -3.05 | -0.15 | -1.998 | -0.023 | 0.005 | 0.005 | 0.483 | 0.25 | 0.33 | 0.14 | 0.31 | 0.16 | 0.25 |
| 3 | 62244570 | rs1388613 | -1.332 | -2.489 | -0.161 | -2.085 | -0.03 | 0.002 | 0.002 | 0.237 | 0.3 | 0.47 | 0.18 | 0.33 | 0.21 | 0.29 |
| 3 | 62246422 | rs875423 | -2.15 | -2.715 | -0.326 | -2.186 | -0.024 | -0.006 | -0.006 | 0 | 0.62 | 0.73 | 0.39 | 0.44 | 0.47 | 0.58 |
| 3 | 62263047 | rs2291540 | 0.212 | -2.544 | 0.113 | -2.051 | -0.021 | -0.033 | -0.033 | 0 | 0.16 | 0.36 | 0.09 | 0.09 | 0.1 | 0.06 |
| 3 | 62272627 | rs11712908 | -0.658 | -2.362 | -0.345 | -1.93 | 0 | -0.004 | -0.004 | 0 | 0.32 | 0.26 | 0.21 | 0.37 | 0.31 | 0.32 |
| 3 | 62275613 | rs7621740 | -1.374 | -2.519 | -0.411 | -2.09 | 0.004 | 0.007 | 0.007 | 0.584 | 0.5 | 0.55 | 0.35 | 0.39 | 0.41 | 0.34 |
| 3 | 62281436 | rs2242352 | -1.134 | -2.275 | -0.37 | -1.962 | -0.009 | -0.003 | -0.003 | 0 | 0.37 | 0.53 | 0.24 | 0.39 | 0.31 | 0.31 |
| 3 | 62309701 | rs1881269 | -1.188 | -2.227 | -0.499 | -1.737 | 0.009 | 0.011 | 0.011 | 0.083 | 0.52 | 0.59 | 0.4 | 0.4 | 0.39 | 0.37 |
| 3 | 62315312 | rs813064 | 1.447 | -2.408 | -0.748 | -1.464 | -0.036 | -0.002 | -0.002 | 0.053 | 0.4 | 0.23 | 0.5 | 0.56 | 0.51 | 0.42 |
| 3 | 62316394 | rs3774700 | 0.543 | -2.538 | -1.085 | -1.311 | 0.002 | -0.004 | -0.004 | 0.573 | 0.15 | 0.1 | 0.09 | 0.31 | 0.06 | 0.13 |
| 3 | 62324103 | rs6445263 | -0.849 | -2.51 | -0.86 | -1.392 | 0.001 | 0.012 | 0.012 | 0 | 0.54 | 0.64 | 0.47 | 0.42 | 0.44 | 0.58 |
| 3 | 62327705 | rs13062701 | 0.797 | -2.306 | -1.264 | -1.215 | -0.007 | 0.079 | 0.079 | 0 | 0.16 | 0.01 | 0.18 | 0.03 | 0.14 | 0.27 |
| 3 | 62331478 | rs17031696 | -0.06 | -2 | -0.804 | -1.018 | 0 | 0 | 0 | 0 | 0.26 | 0.28 | 0.15 | 0.17 | 0.2 | 0.25 |
| 3 | 62341014 | rs11130880 | -0.716 | -1.867 | -0.811 | -0.758 | -0.031 | -0.034 | -0.034 | 0 | 0.7 | 0.45 | 0.67 | 0.8 | 0.79 | 0.63 |
| 3 | 62341044 | rs11707184 | 0.078 | -1.204 | -0.915 | -0.45 | -0.002 | 0.016 | 0.016 | 0 | 0.23 | 0.19 | 0.14 | 0.17 | 0.21 | 0.23 |
| 3 | 62344960 | rs6788223 | -0.114 | -0.939 | -0.845 | -0.258 | 0.002 | 0.02 | 0.02 | 0 | 0.29 | 0.25 | 0.18 | 0.19 | 0.26 | 0.23 |
| 3 | 62352659 | rs1528192 | 0.169 | 0.084 | -1.939 | 0.497 | -0.092 | -0.085 | -0.085 | 0 | 0.35 | 0.07 | 0.51 | 0.57 | 0.59 | 0.4 |
| 3 | 62354425 | rs6445264 | -0.026 | -0.431 | -1.307 | 0.584 | 0.061 | 0.003 | 0.003 | 0.002 | 0.51 | 0.33 | 0.67 | 0.38 | 0.66 | 0.62 |
| 3 | 62369592 | rs17066359 | -0.529 | -0.531 | -0.64 | 0.482 | -0.02 | -0.022 | -0.022 | 0 | 0.14 | 0.27 | 0.08 | 0.05 | 0.11 | 0.24 |
| 3 | 62371558 | rs1979673 | -0.405 | -1.015 | -0.648 | 0.664 | -0.083 | 0.02 | 0.02 | 3.105 | 0.22 | 0.47 | 0.15 | 0.27 | 0.12 | 0.17 |
| 3 | 62376645 | rs6777784 | 0.22 | -0.813 | -1.38 | 0.574 | -0.009 | 0.002 | 0.002 | 0.098 | 0.55 | 0.53 | 0.7 | 0.52 | 0.62 | 0.66 |
| 3 | 62376859 | rs17687440 | 0.524 | 0.12 | -2.045 | 0.427 | -0.082 | -0.082 | -0.082 | 0.046 | 0.3 | 0.08 | 0.51 | 0.39 | 0.37 | 0.41 |
| 3 | 62379483 | rs17687518 | -0.037 | -0.111 | -2.039 | 0.618 | -0.025 | -0.035 | -0.035 | 0 | 0.16 | 0.01 | 0.11 | 0.14 | 0.18 | 0.07 |
| 3 | 62390589 | rs10866054 | 0.177 | 0.105 | -1.738 | 0.416 | 0.013 | 0.044 | 0.044 | 0.448 | 0.73 | 0.61 | 0.78 | 0.57 | 0.7 | 0.75 |
| 3 | 62394268 | rs4688128 | 1.199 | 0.598 | -2.415 | 0.697 | -0.018 | -0.031 | -0.031 | 0 | 0.19 | 0.12 | 0.35 | 0.35 | 0.25 | 0.45 |
| 3 | 62394559 | rs12630374 | -0.31 | 0.146 | -2.641 | -0.027 | -0.009 | -0.03 | -0.03 | 0.052 | 0.29 | 0.05 | 0.32 | 0.2 | 0.28 | 0.15 |
| 3 | 62398776 | rs13066791 | -0.117 | 0.265 | -2.578 | -0.326 | -0.039 | 0.044 | 0.044 | 0 | 0.22 | 0.06 | 0.16 | 0.13 | 0.27 | 0.17 |

| | | | | | | | | | | | | | | | | |
|---|---|---|---|---|---|---|---|---|---|---|---|---|---|---|---|---|
| 3 | 62399206 | rs13090161 | 0.373 | -0.003 | -2.191 | -0.166 | 0.009 | 0.006 | 0.006 | 0 | 0.1 | 0.05 | 0.06 | 0.06 | 0.12 | 0.04 |
| 3 | 62400329 | rs11716318 | 0.027 | -0.036 | -2.454 | -0.245 | -0.036 | -0.014 | -0.014 | 0 | 0.21 | 0.01 | 0.2 | 0.16 | 0.22 | 0.13 |
| 3 | 62402895 | rs7611626 | 0.58 | 0.325 | -1.857 | -0.006 | 0.002 | 0.001 | 0.001 | 0 | 0.09 | 0.08 | 0.06 | 0.06 | 0.12 | 0.04 |
| 3 | 62404892 | rs1528197 | -0.412 | 0.366 | -1.303 | -0.169 | 0.009 | 0.016 | 0.016 | 0 | 0.51 | 0.49 | 0.44 | 0.25 | 0.46 | 0.28 |
| 3 | 62414134 | rs3344 | -0.278 | 1.113 | -0.952 | -0.321 | 0.007 | -0.019 | -0.019 | 1.213 | 0.72 | 0.71 | 0.68 | 0.84 | 0.55 | 0.54 |
| 3 | 62419093 | rs2037816 | -0.954 | 0.803 | -0.76 | 0.534 | 0.002 | 0.01 | 0.01 | 0 | 0.22 | 0.29 | 0.45 | 0.38 | 0.26 | 0.26 |
| 3 | 62421292 | rs4688129 | -0.905 | 0.552 | -0.631 | 1.002 | -0.006 | -0.025 | -0.025 | 0 | 0.13 | 0.1 | 0.21 | 0.24 | 0.13 | 0.11 |
| 3 | 62423076 | rs1881267 | 0.469 | -0.809 | -0.348 | 0.186 | -0.005 | -0.074 | -0.074 | 0 | 0.36 | 0.36 | 0.2 | 0.09 | 0.31 | 0.31 |
| 3 | 62426240 | rs13081552 | 2.161 | -1.175 | -0.796 | 0.379 | -0.093 | -0.126 | -0.126 | 0.052 | 0.26 | 0.05 | 0.33 | 0.48 | 0.37 | 0.37 |
| 3 | 62428968 | rs1534723 | 1.048 | -1.519 | -0.169 | 0.319 | -0.007 | 0.04 | 0.04 | 0 | 0.75 | 0.56 | 0.6 | 0.62 | 0.77 | 0.75 |
| 3 | 62429584 | rs1534724 | 1.302 | -1.565 | -0.331 | 0.237 | -0.087 | -0.105 | -0.105 | 1.203 | 0.34 | 0.09 | 0.46 | 0.51 | 0.44 | 0.42 |
| 3 | 62430908 | rs6445270 | 0.102 | -1.391 | -0.127 | 0.358 | -0.016 | 0.005 | 0.005 | 0 | 0.85 | 0.75 | 0.79 | 0.83 | 0.88 | 0.82 |
| 3 | 62433040 | rs7611540 | 0.721 | -1.339 | -0.557 | 0.265 | -0.002 | -0.035 | -0.035 | 0.247 | 0.38 | 0.32 | 0.21 | 0.09 | 0.31 | 0.31 |
| 3 | 62433496 | rs6766993 | 1.451 | -1.244 | -0.557 | 0.392 | 0.005 | 0.063 | 0.063 | 0.248 | 0.71 | 0.4 | 0.56 | 0.59 | 0.7 | 0.72 |
| 3 | 62434127 | rs12639116 | 0.643 | -1.119 | -0.903 | 0.123 | -0.016 | -0.015 | -0.015 | 0 | 0.07 | 0.01 | 0.13 | 0.06 | 0.1 | 0.15 |
| 3 | 62434256 | rs9311839 | 0.561 | -0.998 | -0.584 | -0.018 | 0.007 | 0.087 | 0.087 | 0.109 | 0.33 | 0.12 | 0.18 | 0.1 | 0.28 | 0.31 |
| 3 | 62439586 | rs1997366 | -0.776 | -0.96 | -0.751 | 0.052 | -0.006 | 0.004 | 0.004 | 1.534 | 0.74 | 0.91 | 0.84 | 0.78 | 0.77 | 0.77 |
| 3 | 62442622 | rs6792816 | -0.107 | -0.925 | -0.09 | -0.138 | 0.01 | -0.035 | -0.035 | 0.057 | 0.11 | 0.26 | 0.17 | 0.02 | 0.11 | 0.1 |
| 3 | 62447365 | rs6445272 | -0.65 | -1.379 | 0.499 | -0.304 | -0.011 | -0.073 | -0.073 | 0.021 | 0.15 | 0.47 | 0.09 | 0.02 | 0.07 | 0.05 |
| 3 | 62450760 | rs4688131 | 0.909 | -1.798 | 0.48 | -0.343 | 0.067 | 0.121 | 0.121 | 3.027 | 0.5 | 0.17 | 0.7 | 0.74 | 0.67 | 0.69 |
| 3 | 62451935 | rs6445273 | -0.973 | -1.65 | 0.734 | -0.051 | -0.041 | -0.093 | -0.093 | 0.023 | 0.17 | 0.58 | 0.08 | 0.02 | 0.07 | 0.04 |
| 3 | 62458083 | rs17688430 | 0.156 | -1.083 | 0.641 | 0.551 | -0.033 | -0.051 | -0.051 | 0 | 0.1 | 0.02 | 0.16 | 0.26 | 0.18 | 0.16 |
| 3 | 62460883 | rs4688298 | -0.362 | -1.263 | 0.663 | 0.051 | 0.014 | 0.004 | 0.004 | 0 | 0.13 | 0.09 | 0.12 | 0 | 0.08 | 0.08 |
| 3 | 62462818 | rs9874646 | 0.388 | -1.121 | 0.407 | 0.168 | 0.022 | 0.055 | 0.055 | 0 | 0.61 | 0.9 | 0.75 | 0.74 | 0.67 | 0.66 |
| 3 | 62470510 | rs1349594 | -1.363 | -0.908 | 1.189 | 0.477 | 0.081 | 0.016 | 0.016 | 0.508 | 0.07 | 0.33 | 0.18 | 0.02 | 0.1 | 0.02 |
| 3 | 62471086 | rs6774533 | 0.309 | -0.879 | 0.764 | 0.479 | 0.027 | 0.019 | 0.019 | 0.398 | 0.64 | 0.78 | 0.74 | 0.73 | 0.68 | 0.65 |
| 3 | 62471282 | rs9968060 | 1.137 | -0.915 | 0.747 | 0.432 | 0.072 | 0.071 | 0.071 | 0.033 | 0.54 | 0.12 | 0.66 | 0.7 | 0.63 | 0.61 |
| 3 | 62476790 | rs2367089 | -2.058 | -0.442 | 1.931 | 0.648 | 0.099 | 0.014 | 0.014 | 0.228 | 0.09 | 0.45 | 0.2 | 0.02 | 0.11 | 0.03 |
| 3 | 62477699 | rs9871763 | -2.318 | -0.461 | 1.982 | 0.954 | 0.096 | 0.183 | 0.183 | 0.195 | 0.11 | 0.54 | 0.16 | 0.24 | 0.15 | 0.12 |
| 3 | 62478194 | rs13325751 | -1.089 | -0.562 | 1.929 | 0.857 | 0.011 | 0.014 | 0.014 | 0 | 0.1 | 0.22 | 0.24 | 0.05 | 0.11 | 0.07 |
| 3 | 62481063 | rs1452075 | 0.489 | -0.823 | 1.826 | 0.894 | 0.008 | 0.004 | 0.004 | 0 | 0.71 | 0.77 | 0.76 | 0.73 | 0.72 | 0.64 |
| 3 | 62482552 | rs2049760 | 0.886 | -0.804 | 2.051 | 0.872 | -0.001 | 0 | 0 | 0 | 0.66 | 0.57 | 0.6 | 0.64 | 0.62 | 0.58 |
| 3 | 62482636 | rs9839020 | 1.543 | -0.965 | 1.995 | 0.772 | 0.002 | 0.008 | 0.008 | 0 | 0.57 | 0.55 | 0.57 | 0.63 | 0.57 | 0.54 |
| 3 | 62487267 | rs11925875 | 1.748 | -1.037 | 1.931 | 0.739 | 0.038 | 0.017 | 0.017 | 0 | 0.52 | 0.29 | 0.5 | 0.57 | 0.58 | 0.52 |
| 3 | 62491047 | rs7650561 | 2.537 | -1.173 | 1.678 | 0.744 | -0.052 | -0.081 | -0.081 | 0.675 | 0.44 | 0.09 | 0.4 | 0.5 | 0.48 | 0.46 |
| 3 | 62494608 | rs7646021 | 1.576 | -0.789 | 2.428 | 1.27 | 0.003 | 0.187 | 0.187 | 0 | 0.87 | 0.71 | 0.72 | 0.55 | 0.87 | 0.85 |
| 3 | 62501634 | rs9863956 | -1.224 | -0.803 | 2.647 | 1.213 | -0.014 | -0.065 | -0.065 | 0.002 | 0.11 | 0.31 | 0.05 | 0.01 | 0.06 | 0.07 |
| 3 | 62506533 | rs13065590 | 2.302 | -0.826 | 2.56 | 1.252 | -0.027 | 0.152 | 0.152 | 0 | 0.81 | 0.61 | 0.72 | 0.6 | 0.86 | 0.85 |
| 3 | 62509407 | rs1355543 | 2 | -0.817 | 2.441 | 1.245 | 0.055 | 0.337 | 0.337 | 0.224 | 0.9 | 0.63 | 0.74 | 0.57 | 0.88 | 0.87 |
| 3 | 62512846 | rs9843019 | 2.345 | -0.827 | 2.398 | 1.245 | -0.032 | 0.119 | 0.119 | 0 | 0.8 | 0.6 | 0.71 | 0.59 | 0.86 | 0.83 |
| 3 | 62523546 | rs2068187 | 2.613 | -0.806 | 2.33 | 1.072 | 0.013 | 0.134 | 0.134 | 0.329 | 0.8 | 0.61 | 0.67 | 0.5 | 0.78 | 0.73 |
| 3 | 62525450 | rs855586 | -3.053 | -0.841 | 2.437 | -0.036 | -0.014 | 0.002 | 0.002 | 0.271 | 0.22 | 0.64 | 0.18 | 0.22 | 0.18 | 0.22 |
| 3 | 62526657 | rs833673 | -0.806 | -0.787 | 2.329 | -0.008 | -0.006 | -0.006 | -0.006 | 0 | 0.06 | 0.06 | 0.04 | 0.17 | 0.1 | 0.12 |
| 3 | 62530059 | rs9857515 | -1.924 | -0.82 | 2.154 | -0.088 | -0.026 | -0.09 | -0.09 | 0 | 0.19 | 0.25 | 0.08 | 0.01 | 0.12 | 0.14 |
| 3 | 62534469 | rs755358 | -0.726 | -0.697 | 1.939 | -0.085 | 0.007 | 0.005 | 0.005 | 0 | 0.18 | 0.12 | 0.07 | 0 | 0.14 | 0.05 |
| 3 | 62539736 | rs2251423 | 1.895 | -0.72 | 2.083 | 0.003 | 0 | 0 | 0 | 0 | 0.7 | 0.33 | 0.79 | 0.72 | 0.77 | 0.74 |
| 3 | 62541049 | rs10470701 | -1.135 | -0.716 | 2.267 | -0.095 | -0.025 | -0.017 | -0.017 | 2.271 | 0.22 | 0.26 | 0.16 | 0.12 | 0.13 | 0.06 |
| 3 | 62543436 | rs2600854 | 2.183 | -0.694 | 2.044 | -0.121 | 0.018 | 0.02 | 0.02 | 0 | 0.62 | 0.32 | 0.75 | 0.69 | 0.7 | 0.64 |
| 3 | 62544639 | rs2034610 | -1.454 | -0.62 | 1.849 | -0.385 | -0.018 | -0.088 | -0.088 | 0 | 0.24 | 0.33 | 0.12 | 0.02 | 0.18 | 0.14 |
| 3 | 62548081 | rs11708441 | -0.223 | -0.621 | 1.235 | -0.277 | 0.029 | 0.057 | 0.057 | 0.003 | 0.16 | 0.04 | 0.05 | 0 | 0.09 | 0.04 |
| 3 | 62549168 | rs9311843 | -1.605 | -0.363 | 1.847 | 0.038 | 0.09 | 0.205 | 0.205 | 0 | 0.08 | 0.23 | 0.12 | 0.28 | 0.13 | 0.19 |
| 3 | 62551775 | rs833657 | 0.414 | -0.64 | 1.89 | -0.206 | -0.026 | 0.013 | 0.013 | 0 | 0.8 | 0.61 | 0.85 | 0.73 | 0.84 | 0.8 |
| 3 | 62556868 | rs833664 | 1.079 | -0.745 | 1.962 | -0.406 | 0.044 | 0.035 | 0.035 | 0 | 0.94 | 0.67 | 0.95 | 0.98 | 0.95 | 0.89 |
| 3 | 62561035 | rs7620572 | -0.885 | -0.847 | 1.353 | -0.731 | 0.021 | 0.087 | 0.087 | 0 | 0.22 | 0.1 | 0.13 | 0.04 | 0.2 | 0.32 |
| 3 | 62562791 | rs1512019 | 1.65 | -0.819 | 1.561 | -0.785 | 0.038 | 0.052 | 0.052 | 0.498 | 0.87 | 0.37 | 0.83 | 0.78 | 0.87 | 0.84 |
| 3 | 62564515 | rs833649 | 2.07 | -0.826 | 1.266 | -0.832 | 0.013 | 0.125 | 0.125 | 2.953 | 0.58 | 0.12 | 0.65 | 0.72 | 0.59 | 0.45 |
| 3 | 62565383 | rs6445279 | -1.234 | -0.628 | 1.509 | -0.567 | 0.073 | 0.135 | 0.135 | 0 | 0.07 | 0.39 | 0.1 | 0.2 | 0.08 | 0.09 |
| 3 | 62571037 | rs2659467 | 0.272 | -0.496 | 0.736 | -0.577 | -0.009 | -0.002 | -0.002 | 0 | 0.19 | 0.04 | 0.27 | 0.13 | 0.18 | 0.08 |
| 3 | 62573943 | rs866248 | 0.841 | -0.403 | 1.179 | -0.583 | -0.026 | -0.07 | -0.07 | 0 | 0.82 | 0.3 | 0.83 | 0.94 | 0.86 | 0.75 |
| 3 | 62574188 | rs3411 | -1.376 | -0.112 | 0.43 | -0.426 | 0.034 | 0.095 | 0.095 | 0 | 0.09 | 0.3 | 0.11 | 0.27 | 0.1 | 0.13 |
| 3 | 62574621 | rs861303 | 1.18 | -0.368 | -0.535 | -0.999 | -0.023 | -0.106 | -0.106 | 0.535 | 0.7 | 0.43 | 0.76 | 0.93 | 0.69 | 0.64 |
| 3 | 62578336 | rs17066673 | -1.217 | 0.304 | -1.19 | -0.147 | 0 | 0 | 0 | 0 | 0.06 | 0.08 | 0.05 | 0.07 | 0.05 | 0.07 |
| 3 | 62588619 | rs833635 | 0.038 | 0.37 | -1.491 | -0.595 | 0.059 | 0.227 | 0.227 | 4.556 | 0.49 | 0.17 | 0.47 | 0.13 | 0.6 | 0.38 |
| 3 | 62604010 | rs1567092 | -0.064 | 0.367 | -1.822 | -0.924 | -0.043 | 0.146 | 0.146 | 0.201 | 0.45 | 0.17 | 0.49 | 0.78 | 0.5 | 0.68 |
| 3 | 62605941 | rs6772352 | -0.223 | -0.003 | -2.027 | -0.869 | 0.024 | -0.05 | -0.05 | 3.332 | 0.34 | 0.19 | 0.35 | 0.52 | 0.29 | 0.49 |
| 3 | 62610057 | rs1398095 | 0.298 | 0.884 | -1.034 | -1.162 | 0.003 | -0.003 | -0.003 | 0 | 0.22 | 0.25 | 0.27 | 0.19 | 0.27 | 0.16 |
| 3 | 62611998 | rs2600853 | 0.719 | -0.414 | -0.776 | -2.066 | 0.01 | 0.061 | 0.061 | 0.64 | 0.22 | 0.08 | 0.15 | 0.09 | 0.17 | 0.03 |
| 3 | 62612342 | rs2248023 | -0.9 | -0.482 | -0.571 | -2.04 | 0.042 | 0.014 | 0.014 | 0 | 0.95 | 0.71 | 0.96 | 1 | 0.97 | 0.97 |
| 3 | 62615298 | rs833668 | 0.678 | -0.548 | -0.911 | -2.157 | 0.013 | 0.067 | 0.067 | 0.157 | 0.22 | 0.08 | 0.16 | 0.09 | 0.18 | 0.14 |

| | | | | | | | | | | | | | | | | |
|---|---|---|---|---|---|---|---|---|---|---|---|---|---|---|---|---|
| 3 | 62617576 | rs4688300 | 0.952 | -0.281 | -1.169 | -2.075 | -0.144 | -0.025 | -0.025 | 0 | 0.23 | 0.04 | 0.3 | 0.21 | 0.36 | 0.35 |
| 3 | 62619642 | rs7645811 | -1.136 | -0.382 | -0.693 | -2.173 | -0.011 | -0.048 | -0.048 | 0 | 0.8 | 0.73 | 0.82 | 0.91 | 0.84 | 0.86 |
| 3 | 62620091 | rs12489036 | 0.883 | -0.403 | -0.875 | -2.165 | -0.103 | -0.036 | -0.036 | 0 | 0.14 | 0.02 | 0.27 | 0.18 | 0.32 | 0.19 |
| 3 | 62620131 | rs10510880 | -0.111 | -0.434 | -0.612 | -2.172 | -0.09 | 0.003 | 0.003 | 0 | 0.24 | 0.11 | 0.31 | 0.22 | 0.36 | 0.35 |
| 3 | 62620324 | rs10510881 | -0.098 | -0.431 | -0.611 | -2.17 | -0.09 | 0.002 | 0.002 | 0 | 0.24 | 0.11 | 0.31 | 0.22 | 0.36 | 0.35 |
| 3 | 62620763 | rs9855708 | 1.515 | -0.557 | -0.477 | -2.117 | -0.002 | 0 | 0 | 0 | 0.08 | 0.12 | 0.06 | 0.08 | 0.05 | 0.07 |
| 3 | 62623923 | rs1237030 | 0.403 | -0.53 | -0.541 | -2.084 | -0.001 | -0.001 | -0.001 | 0 | 0.6 | 0.38 | 0.59 | 0.78 | 0.53 | 0.62 |
| 3 | 62631553 | rs4588354 | 0.672 | -0.696 | -0.562 | -2.063 | 0.014 | 0.004 | 0.004 | 0.112 | 0.65 | 0.34 | 0.77 | 0.92 | 0.75 | 0.74 |
| 3 | 62640221 | rs7633623 | -0.134 | -0.792 | -0.032 | -1.338 | -0.07 | -0.039 | -0.039 | 0 | 0.84 | 0.48 | 0.88 | 0.92 | 0.94 | 0.84 |
| 3 | 62642970 | rs9821411 | -0.175 | 0.157 | -0.605 | -0.62 | 0.001 | 0.003 | 0.003 | 0 | 0.12 | 0.13 | 0.12 | 0.08 | 0.14 | 0.06 |
| 3 | 62645585 | rs9833297 | -0.575 | 0.164 | -0.575 | -1.452 | 0.002 | 0.007 | 0.007 | 0 | 0.55 | 0.61 | 0.63 | 0.86 | 0.49 | 0.79 |
| 3 | 62645954 | rs10510882 | 0.733 | 0.686 | -0.976 | -1.616 | -0.029 | -0.002 | -0.002 | 0 | 0.06 | 0.01 | 0.1 | 0.04 | 0.09 | 0.04 |
| 3 | 62647115 | rs12491112 | 0.911 | 0.197 | -1.249 | -1.406 | 0.007 | -0.06 | -0.06 | 0 | 0.28 | 0.06 | 0.39 | 0.69 | 0.27 | 0.61 |
| 3 | 62647174 | rs6794531 | -0.124 | 0.276 | -0.233 | -1.225 | -0.013 | -0.009 | -0.009 | 0 | 0.2 | 0.31 | 0.15 | 0.15 | 0.18 | 0.1 |
| 3 | 62649666 | rs2135582 | -0.885 | 0.848 | -0.235 | -1.73 | 0.004 | -0.017 | -0.017 | 0 | 0.76 | 0.71 | 0.81 | 0.89 | 0.74 | 0.83 |
| 3 | 62656891 | rs4688305 | -0.625 | 1.234 | -0.133 | -1.819 | 0.008 | -0.034 | -0.034 | 0 | 0.74 | 0.63 | 0.8 | 0.89 | 0.73 | 0.81 |
| 3 | 62657017 | rs17355939 | 0.514 | 0.745 | -0.25 | -1.756 | 0.002 | 0.001 | 0.001 | 0 | 0.17 | 0.14 | 0.16 | 0.14 | 0.18 | 0.1 |
| 3 | 62657357 | rs11706594 | 1.386 | 0.813 | -0.39 | -1.708 | 0.004 | 0.017 | 0.017 | 0.365 | 0.19 | 0.16 | 0.16 | 0.11 | 0.16 | 0.08 |
| 3 | 62658151 | rs6445281 | -0.13 | 0.92 | -0.031 | -1.398 | -0.011 | -0.01 | -0.01 | 0 | 0.19 | 0.29 | 0.16 | 0.14 | 0.18 | 0.1 |
| 3 | 62659658 | rs1858385 | 0.233 | 1.527 | -0.484 | -1.407 | -0.037 | 0.03 | 0.03 | 0.261 | 0.06 | 0.01 | 0.23 | 0.01 | 0.15 | 0.06 |
| 3 | 62660566 | rs17066793 | 0.055 | 1.225 | -0.48 | -1.557 | -0.004 | 0 | 0 | 0 | 0.1 | 0.09 | 0.1 | 0.1 | 0.06 | 0.07 |
| 3 | 62661945 | rs10510884 | 1.154 | 1.076 | -0.561 | -2.035 | 0.008 | 0.014 | 0.014 | 0 | 0.21 | 0.16 | 0.11 | 0.02 | 0.13 | 0.19 |
| 3 | 62662213 | rs2175527 | -0.49 | 1.342 | -0.48 | -2.145 | -0.003 | 0.007 | 0.007 | 0 | 0.73 | 0.77 | 0.84 | 0.96 | 0.76 | 0.74 |
| 3 | 62663519 | rs978879 | 0.45 | 1.281 | -0.35 | -2.077 | 0.002 | -0.007 | -0.007 | 1.042 | 0.83 | 0.82 | 0.85 | 0.88 | 0.83 | 0.85 |
| 3 | 62665975 | rs1876253 | -0.482 | 1.194 | -0.499 | -2.09 | -0.006 | 0.018 | 0.018 | 0 | 0.71 | 0.76 | 0.82 | 0.95 | 0.71 | 0.72 |
| 3 | 62672786 | rs17066808 | 0.487 | 1.15 | -0.547 | -1.963 | 0.003 | -0.003 | -0.003 | 0 | 0.1 | 0.05 | 0.1 | 0.1 | 0.06 | 0.08 |
| 3 | 62674434 | rs4688308 | -0.109 | 1.273 | -0.554 | -2.035 | -0.015 | 0.011 | 0.011 | 0.611 | 0.09 | 0.05 | 0.25 | 0.08 | 0.15 | 0.16 |
| 3 | 62676116 | rs2367318 | 0.376 | 1.006 | 0.089 | -2.329 | 0.019 | -0.109 | -0.109 | 0.653 | 0.79 | 0.45 | 0.82 | 0.97 | 0.74 | 0.89 |
| 3 | 62676188 | rs2135581 | -0.182 | 1.024 | 0.122 | -2.304 | 0.025 | -0.021 | -0.021 | 0.009 | 0.92 | 0.8 | 0.93 | 0.97 | 0.89 | 0.92 |
| 3 | 62678071 | rs1948926 | -0.537 | 1.003 | -0.016 | -2.397 | -0.007 | 0.017 | 0.017 | 0 | 0.7 | 0.74 | 0.8 | 0.95 | 0.7 | 0.7 |
| 3 | 62678151 | rs9868884 | 1.411 | 1.01 | -0.044 | -2.394 | -0.002 | 0.073 | 0.073 | 0 | 0.22 | 0.09 | 0.13 | 0.02 | 0.18 | 0.22 |
| 3 | 62678836 | rs6781363 | 0.253 | 1.004 | 0.02 | -2.417 | -0.002 | 0.036 | 0.036 | 0 | 0.57 | 0.36 | 0.69 | 0.95 | 0.56 | 0.67 |
| 3 | 62679656 | rs13070975 | 1.184 | 0.989 | -0.13 | -2.381 | -0.035 | 0.004 | 0.004 | 0.204 | 0.06 | 0.01 | 0.05 | 0.03 | 0.11 | 0.08 |
| 3 | 62680329 | rs6763320 | 0.457 | 0.923 | -0.034 | -2.4 | 0 | -0.102 | -0.102 | 0 | 0.14 | 0.39 | 0.12 | 0 | 0.15 | 0.03 |
| 3 | 62682335 | rs7634365 | 0.213 | 0.994 | -0.202 | -2.392 | -0.003 | 0.047 | 0.047 | 0 | 0.56 | 0.36 | 0.69 | 0.95 | 0.55 | 0.67 |
| 3 | 62686004 | rs2367315 | 0.846 | 1.018 | -0.225 | -2.351 | -0.009 | 0.08 | 0.08 | 0.007 | 0.47 | 0.31 | 0.6 | 0.85 | 0.49 | 0.59 |
| 3 | 62686403 | rs13090570 | 0.379 | 0.947 | -0.202 | -2.315 | 0.028 | 0.09 | 0.09 | 0 | 0.53 | 0.31 | 0.64 | 0.88 | 0.6 | 0.67 |
| 3 | 62691870 | rs6445290 | -0.34 | 1.065 | -0.371 | -1.886 | -0.016 | 0.049 | 0.049 | 0 | 0.64 | 0.71 | 0.72 | 0.84 | 0.64 | 0.51 |
| 3 | 62693396 | rs4688139 | -0.819 | 1.038 | -0.273 | -1.902 | -0.006 | -0.044 | -0.044 | 0 | 0.7 | 0.58 | 0.76 | 0.86 | 0.75 | 0.6 |
| 3 | 62695867 | rs17066966 | 0.369 | 0.996 | -0.243 | -1.901 | 0 | -0.003 | -0.003 | 0.061 | 0.08 | 0.13 | 0.09 | 0.11 | 0.06 | 0.19 |
| 3 | 62696567 | rs17357323 | 1.106 | 0.93 | -0.263 | -1.937 | 0.007 | -0.005 | -0.005 | 2.958 | 0.09 | 0.03 | 0.1 | 0.07 | 0.06 | 0.05 |
| 3 | 62698382 | rs9881771 | 0.85 | 0.897 | -0.204 | -2.033 | 0.014 | -0.079 | -0.079 | 0.223 | 0.12 | 0.36 | 0.11 | 0 | 0.14 | 0.03 |
| 3 | 62698892 | rs952040 | 2.269 | 1.072 | -0.43 | -2.129 | -0.043 | -0.088 | -0.088 | 0 | 0.33 | 0.01 | 0.26 | 0.67 | 0.31 | 0.25 |
| 3 | 62699831 | rs6768519 | 1.198 | 0.885 | -0.244 | -2.037 | -0.03 | 0.048 | 0.048 | 0.39 | 0.41 | 0.12 | 0.5 | 0.73 | 0.44 | 0.4 |
| 3 | 62700227 | rs304202 | -0.91 | 0.892 | -0.218 | -2.057 | -0.009 | -0.013 | -0.013 | 2.615 | 0.24 | 0.23 | 0.24 | 0.24 | 0.24 | 0.34 |
| 3 | 62701312 | rs9822224 | 0.642 | 0.971 | -0.179 | -1.972 | 0.016 | -0.087 | -0.087 | 0 | 0.13 | 0.43 | 0.12 | 0 | 0.14 | 0.03 |
| 3 | 62703006 | rs6445293 | 0.335 | 1.01 | -0.372 | -1.893 | 0 | -0.011 | -0.011 | 0 | 0.53 | 0.47 | 0.61 | 0.73 | 0.58 | 0.44 |
| 3 | 62704180 | rs1506746 | -0.177 | 0.83 | -0.305 | -1.825 | -0.011 | -0.026 | -0.026 | 0 | 0.6 | 0.48 | 0.66 | 0.74 | 0.69 | 0.52 |
| 3 | 62709377 | rs9836677 | 0.744 | 0.682 | -0.002 | -1.719 | 0.004 | -0.072 | -0.072 | 0 | 0.11 | 0.31 | 0.11 | 0 | 0.12 | 0.03 |
| 3 | 62710444 | rs1395440 | 0.273 | 0.629 | 0.11 | -1.733 | 0.005 | 0.041 | 0.041 | 0.048 | 0.08 | 0.19 | 0.1 | 0.11 | 0.06 | 0.18 |
| 3 | 62711374 | rs1658789 | -0.761 | 0.737 | -0.205 | -1.999 | -0.051 | 0.197 | 0.197 | 0 | 0.5 | 0.88 | 0.4 | 0.22 | 0.49 | 0.54 |
| 3 | 62713280 | rs557993 | 1.293 | 0.81 | -0.229 | -1.969 | -0.008 | -0.062 | -0.062 | 0 | 0.24 | 0.34 | 0.15 | 0.09 | 0.2 | 0.25 |
| 3 | 62714620 | rs481319 | -1.705 | 0.853 | -0.455 | -1.676 | -0.123 | -0.062 | -0.062 | 0 | 0.62 | 0.97 | 0.53 | 0.34 | 0.59 | 0.7 |
| 3 | 62714737 | rs523452 | 0.422 | 0.832 | -0.035 | -1.492 | -0.009 | -0.035 | -0.035 | 0 | 0.32 | 0.54 | 0.25 | 0.19 | 0.26 | 0.43 |
| 3 | 62715646 | rs494186 | 0.471 | 0.848 | -0.066 | -1.485 | -0.009 | -0.038 | -0.038 | 0 | 0.32 | 0.52 | 0.25 | 0.19 | 0.26 | 0.43 |
| 3 | 62728523 | rs504739 | 0.118 | 0.639 | -0.39 | -1.672 | 0 | -0.014 | -0.014 | 0.036 | 0.34 | 0.34 | 0.19 | 0.14 | 0.31 | 0.47 |
| 3 | 62729720 | rs472365 | 1.283 | 0.632 | -0.577 | -1.61 | 0 | 0.001 | 0.001 | 0 | 0.23 | 0.21 | 0.15 | 0.09 | 0.2 | 0.24 |
| 3 | 62731267 | rs531700 | -0.268 | 0.609 | -0.766 | -1.602 | 0.016 | 0.169 | 0.169 | 0 | 0.47 | 0.76 | 0.32 | 0.14 | 0.44 | 0.53 |
| 3 | 62731592 | rs523320 | 0.196 | 0.848 | -0.809 | -1.416 | -0.014 | 0.003 | 0.003 | 0 | 0.87 | 0.93 | 0.68 | 0.93 | 0.82 | 0.82 |
| 3 | 62733032 | rs2971927 | -0.099 | 0.784 | -0.71 | -1.321 | 0.001 | -0.002 | -0.002 | 0.068 | 0.92 | 0.9 | 0.79 | 0.9 | 0.91 | 0.81 |
| 3 | 62733212 | rs3843397 | 0.316 | 0.676 | -0.572 | -1.763 | -0.003 | 0.11 | 0.11 | 0 | 0.5 | 0.27 | 0.48 | 0.82 | 0.49 | 0.47 |
| 3 | 62735090 | rs4423760 | 1.244 | 0.643 | -0.691 | -1.858 | -0.031 | 0.099 | 0.099 | 0 | 0.39 | 0.15 | 0.34 | 0.81 | 0.39 | 0.34 |
| 3 | 62738577 | rs504093 | -0.127 | 0.702 | -0.34 | -1.655 | -0.003 | 0.003 | 0.003 | 0 | 0.91 | 0.89 | 0.79 | 0.92 | 0.91 | 0.82 |
| 3 | 62738603 | rs504024 | 0.465 | 0.805 | -0.051 | -1.789 | 0.053 | -0.083 | -0.083 | 0.05 | 0.13 | 0.4 | 0.13 | 0 | 0.17 | 0.05 |
| 3 | 62741041 | rs13313979 | 1.212 | 0.418 | -0.56 | -1.568 | -0.001 | 0.021 | 0.021 | 0 | 0.24 | 0.17 | 0.15 | 0.06 | 0.23 | 0.24 |
| 3 | 62745160 | rs9830865 | 1.117 | 0.407 | -0.59 | -1.567 | -0.001 | 0.031 | 0.031 | 0 | 0.25 | 0.17 | 0.16 | 0.06 | 0.23 | 0.24 |
| 3 | 62750973 | rs17067161 | 1.212 | 0.409 | -0.552 | -0.601 | 0.003 | 0.086 | 0.086 | 0.295 | 0.07 | 0.13 | 0.13 | 0.26 | 0.08 | 0.08 |
| 3 | 62753439 | rs304196 | 1.115 | 0.262 | -0.085 | -1.376 | 0.001 | -0.117 | -0.117 | 0 | 0.25 | 0.49 | 0.2 | 0.06 | 0.24 | 0.24 |
| 3 | 62755758 | rs2367221 | 0.495 | 0.158 | -0.754 | -1.467 | 0.014 | 0.023 | 0.023 | 0 | 0.17 | 0.12 | 0.15 | 0.05 | 0.12 | 0.19 |

| | | | | | | | | | | | | | | | | |
|---|---|---|---|---|---|---|---|---|---|---|---|---|---|---|---|---|
| 3 | 62756449 | rs304191 | -0.735 | -0.03 | -0.904 | -1.511 | 0.053 | 0.191 | 0.191 | 0 | 0.49 | 0.77 | 0.41 | 0.22 | 0.41 | 0.52 |
| 3 | 62758410 | rs7633796 | -0.078 | -0.075 | -1.012 | -1.549 | 0.018 | 0.097 | 0.097 | 0 | 0.61 | 0.27 | 0.6 | 0.82 | 0.65 | 0.63 |
| 3 | 62764872 | rs12639117 | 0.714 | 0.047 | -1.212 | -1.661 | -0.003 | 0.209 | 0.209 | 0 | 0.47 | 0.12 | 0.46 | 0.79 | 0.53 | 0.55 |
| 3 | 62769867 | rs216059 | -1.422 | -0.05 | -0.849 | -1.788 | -0.006 | 0.044 | 0.044 | 1.045 | 0.75 | 0.84 | 0.71 | 0.95 | 0.71 | 0.74 |
| 3 | 62771450 | rs304205 | -1.477 | -0.005 | -0.852 | -1.793 | -0.008 | 0.041 | 0.041 | 1.007 | 0.75 | 0.84 | 0.71 | 0.95 | 0.71 | 0.74 |
| 3 | 62773179 | rs1913228 | -0.097 | -0.117 | -0.819 | -1.828 | 0.019 | 0.112 | 0.112 | 0.045 | 0.63 | 0.2 | 0.61 | 0.87 | 0.66 | 0.67 |
| 3 | 62773860 | rs12636581 | -0.079 | -0.118 | -0.821 | -1.829 | 0.02 | 0.117 | 0.117 | 0.108 | 0.63 | 0.19 | 0.61 | 0.88 | 0.66 | 0.68 |
| 3 | 62778742 | rs1513143 | 0.927 | 0.024 | -1.129 | -1.882 | -0.064 | 0.184 | 0.184 | 0 | 0.42 | 0.1 | 0.38 | 0.83 | 0.45 | 0.48 |
| 3 | 62782468 | rs304172 | 0.162 | -0.12 | -0.032 | -1.657 | -0.013 | 0.001 | 0.001 | 0.039 | 0.08 | 0.18 | 0.06 | 0.1 | 0.05 | 0.06 |
| 3 | 62786855 | rs2063621 | 0.862 | -0.189 | -0.264 | -1.76 | 0.027 | 0.055 | 0.055 | 0.424 | 0.72 | 0.21 | 0.67 | 0.87 | 0.78 | 0.83 |
| 3 | 62788035 | rs13067730 | 0.546 | -0.199 | -0.435 | -1.798 | 0.032 | 0.264 | 0.264 | 0 | 0.5 | 0.12 | 0.51 | 0.81 | 0.54 | 0.58 |
| 3 | 62790064 | rs2221910 | 2.018 | -0.09 | -0.424 | -1.705 | -0.025 | 0.089 | 0.089 | 0.771 | 0.21 | 0.05 | 0.16 | 0.06 | 0.23 | 0.24 |
| 3 | 62790623 | rs487321 | -0.537 | -0.001 | -0.236 | -1.555 | 0 | -0.001 | -0.001 | 1.476 | 0.92 | 0.95 | 0.85 | 0.97 | 0.91 | 0.93 |
| 3 | 62796691 | rs304225 | 0.285 | -0.023 | 0.19 | -1.53 | 0.025 | 0.018 | 0.018 | 0 | 0.91 | 0.69 | 0.82 | 0.9 | 0.92 | 0.97 |
| 3 | 62800243 | rs1866257 | 0.471 | -0.442 | -0.652 | -2.11 | -0.066 | 0.195 | 0.195 | 0.73 | 0.42 | 0.1 | 0.47 | 0.81 | 0.49 | 0.56 |
| 3 | 62801073 | rs304230 | 0.735 | -0.349 | -0.277 | -2.026 | -0.006 | -0.049 | -0.049 | 0 | 0.72 | 0.35 | 0.63 | 0.88 | 0.72 | 0.82 |
| 3 | 62802226 | rs13077848 | 0.739 | 0.112 | -0.609 | -2.06 | -0.03 | -0.019 | -0.019 | 0 | 0.06 | 0.01 | 0.07 | 0.06 | 0.09 | 0.15 |
| 3 | 62803545 | rs7610255 | 0.49 | 0.105 | -0.6 | -2.356 | -0.003 | -0.069 | -0.069 | 0 | 0.23 | 0.26 | 0.16 | 0 | 0.21 | 0.1 |
| 3 | 62805758 | rs6769778 | -0.265 | -0.381 | -0.331 | -2.008 | -0.011 | -0.142 | -0.142 | 0.044 | 0.8 | 0.51 | 0.72 | 1 | 0.82 | 0.89 |
| 3 | 62808969 | rs11707381 | 1.496 | -0.591 | -1.143 | -2.007 | 0.021 | 0.119 | 0.119 | 3.746 | 0.27 | 0.07 | 0.15 | 0.06 | 0.23 | 0.26 |
| 3 | 62810786 | rs2118535 | 1.117 | -0.372 | -0.901 | -1.797 | 0.005 | -0.002 | -0.002 | 0 | 0.13 | 0.14 | 0.1 | 0 | 0.12 | 0.1 |
| 3 | 62814528 | rs2164881 | 0.832 | -0.109 | -1.062 | -1.252 | -0.04 | -0.053 | -0.053 | 4.186 | 0.29 | 0.12 | 0.29 | 0.47 | 0.4 | 0.33 |
| 3 | 62818566 | rs304181 | -0.894 | 0.073 | -0.998 | -1.129 | 0.001 | 0.026 | 0.026 | 0.011 | 0.84 | 0.84 | 0.7 | 0.61 | 0.82 | 0.81 |
| 3 | 62819885 | rs304180 | -0.284 | -0.06 | -1.312 | -1.296 | -0.05 | 0.11 | 0.11 | 0 | 0.59 | 0.87 | 0.45 | 0.24 | 0.46 | 0.52 |
| 3 | 62826497 | rs304185 | -0.876 | -0.19 | -0.727 | -1.044 | 0.002 | 0.034 | 0.034 | 0.302 | 0.85 | 0.83 | 0.72 | 0.68 | 0.84 | 0.82 |
| 3 | 62826934 | rs11925708 | 0.329 | -0.293 | -0.5 | -1.175 | 0.032 | 0.04 | 0.04 | 0.551 | 0.49 | 0.41 | 0.32 | 0.13 | 0.36 | 0.47 |
| 3 | 62830773 | rs549249 | 0.497 | -0.111 | -0.656 | -1.008 | -0.007 | -0.007 | -0.007 | 0 | 0.37 | 0.34 | 0.39 | 0.54 | 0.48 | 0.39 |
| 3 | 62832230 | rs514606 | -0.303 | 0.173 | -0.658 | -0.528 | 0.001 | 0.045 | 0.045 | 0.149 | 0.86 | 0.82 | 0.73 | 0.67 | 0.86 | 0.81 |
| 3 | 62848326 | rs11711506 | 0.735 | 0.02 | -1.662 | -1.333 | 0.022 | 0.066 | 0.066 | 0.69 | 0.25 | 0.11 | 0.16 | 0.08 | 0.2 | 0.27 |
| 3 | 62861192 | rs968808 | 0.135 | 0.461 | -1.544 | -0.894 | 0 | 0.099 | 0.099 | 0 | 0.47 | 0.72 | 0.35 | 0.23 | 0.43 | 0.46 |
| 3 | 62862981 | rs1398623 | 0.034 | 0.363 | -1.277 | -0.72 | 0.004 | 0.056 | 0.056 | 0 | 0.52 | 0.65 | 0.4 | 0.23 | 0.51 | 0.47 |
| 3 | 62863469 | rs304234 | 0.371 | 0.506 | -1.281 | -0.622 | 0.019 | 0.065 | 0.065 | 0 | 0.48 | 0.6 | 0.37 | 0.17 | 0.41 | 0.42 |
| 3 | 62871502 | rs526163 | -0.316 | 0.491 | -1.679 | -0.126 | -0.041 | 0 | 0 | 0.058 | 0.66 | 0.85 | 0.47 | 0.32 | 0.58 | 0.62 |
| 3 | 62881272 | rs7634212 | 0.058 | 0.78 | -1.503 | -0.033 | -0.004 | 0.051 | 0.051 | 0 | 0.56 | 0.71 | 0.42 | 0.25 | 0.52 | 0.5 |
| 3 | 62883050 | rs17358923 | -0.429 | 0.945 | -1.679 | 0.297 | -0.042 | 0.036 | 0.036 | 0 | 0.4 | 0.21 | 0.3 | 0.26 | 0.46 | 0.37 |
| 3 | 62883380 | rs1992192 | -0.428 | 0.404 | -0.784 | 0.687 | -0.007 | 0.03 | 0.03 | 0 | 0.32 | 0.52 | 0.37 | 0.39 | 0.25 | 0.3 |
| 3 | 62889109 | rs6796067 | 0.815 | 0.053 | 0.289 | 1.028 | 0.02 | 0.211 | 0.211 | 0.693 | 0.92 | 0.6 | 0.95 | 0.8 | 0.98 | 0.88 |
| 3 | 62910807 | rs2367412 | 0.956 | -0.193 | 0.434 | 0.967 | -0.013 | 0.176 | 0.176 | 5.067 | 0.91 | 0.57 | 0.95 | 0.81 | 0.99 | 0.91 |
| 3 | 62923347 | rs9842130 | 0.338 | -0.792 | 0.1 | 0.428 | -0.015 | -0.032 | -0.032 | 0.057 | 0.12 | 0.16 | 0.06 | 0.02 | 0.05 | 0.05 |
| 3 | 62933597 | rs12631887 | 0.064 | 0.511 | -0.947 | 0.912 | -0.125 | -0.14 | -0.14 | 0 | 0.27 | 0.03 | 0.4 | 0.47 | 0.39 | 0.33 |
| 3 | 62938561 | rs12631661 | 0.406 | 0.299 | -0.481 | 0.504 | 0.014 | 0.053 | 0.053 | 0.012 | 0.51 | 0.27 | 0.6 | 0.59 | 0.59 | 0.43 |
| 3 | 62939437 | rs10866061 | 0.044 | 0.353 | -0.101 | -0.048 | -0.001 | 0.028 | 0.028 | 0.388 | 0.35 | 0.29 | 0.28 | 0.22 | 0.34 | 0.48 |
| 3 | 62941248 | rs9311853 | -0.171 | 0.5 | 0.167 | 0.226 | -0.011 | 0.067 | 0.067 | 0.197 | 0.43 | 0.32 | 0.37 | 0.22 | 0.44 | 0.47 |
| 3 | 62948246 | rs17360047 | 0.454 | 0.2 | 0.416 | 0.706 | -0.005 | -0.025 | -0.025 | 0.019 | 0.07 | 0 | 0.07 | 0.07 | 0.06 | 0.08 |
| 3 | 62958569 | rs6762469 | 0.838 | 0.384 | 1.155 | 0.649 | 0.066 | 0.074 | 0.074 | 4.242 | 0.94 | 0.52 | 0.94 | 0.98 | 0.99 | 0.97 |
| 3 | 62968507 | rs4583651 | -0.297 | 1.293 | 0.607 | 1.225 | -0.058 | -0.055 | -0.055 | 0 | 0.1 | 0.02 | 0.23 | 0.3 | 0.21 | 0.12 |
| 3 | 62984179 | rs9311855 | -0.771 | 0.952 | 0.403 | 0.849 | 0.009 | 0.011 | 0.011 | 0 | 0.4 | 0.52 | 0.46 | 0.4 | 0.43 | 0.44 |
| 3 | 62987175 | rs4147406 | -0.84 | 1.447 | 0.386 | 1.453 | -0.037 | -0.036 | -0.036 | 0 | 0.12 | 0.06 | 0.25 | 0.31 | 0.22 | 0.13 |
| 3 | 62993083 | rs11130909 | -0.505 | 1.464 | 0.417 | 1.39 | -0.01 | 0.006 | 0.006 | 0.048 | 0.1 | 0.11 | 0.24 | 0.3 | 0.2 | 0.12 |
| 3 | 63008579 | rs6777724 | 0.916 | 1.117 | 0.733 | 1.368 | 0.032 | 0.258 | 0.258 | 0.729 | 0.87 | 0.48 | 0.73 | 0.7 | 0.89 | 0.91 |
| 3 | 63020787 | rs17361695 | -0.095 | 1.278 | -0.256 | 0.746 | -0.081 | 0.094 | 0.094 | 0 | 0.17 | 0 | 0.18 | 0.05 | 0.23 | 0.17 |
| 3 | 63022365 | rs6789592 | -0.742 | 1.237 | 0.34 | 1.416 | 0.045 | 0.248 | 0.248 | 0 | 0.11 | 0.36 | 0.26 | 0.32 | 0.11 | 0.1 |
| 3 | 63024368 | rs6445307 | 0.665 | 0.982 | -0.298 | 0.858 | 0.003 | 0.02 | 0.02 | 0 | 0.92 | 0.97 | 0.97 | 0.95 | 0.93 | 0.94 |
| 3 | 63028072 | rs1031246 | 0.207 | 0.992 | 0.19 | 0.881 | 0.003 | -0.011 | -0.011 | 0 | 0.94 | 0.85 | 0.98 | 0.96 | 0.95 | 0.96 |
| 3 | 63030602 | rs6445309 | 0.081 | 0.972 | 0.193 | 0.882 | -0.004 | -0.012 | -0.012 | 0 | 0.94 | 0.9 | 0.98 | 0.96 | 0.96 | 0.96 |
| 3 | 63031308 | rs11715184 | -0.107 | 0.747 | 0.051 | 1.14 | 0.002 | -0.008 | -0.008 | 1.251 | 0.11 | 0.08 | 0.17 | 0.2 | 0.14 | 0.29 |
| 3 | 63035696 | rs2121671 | -0.716 | 0.79 | -0.076 | 1.425 | -0.062 | -0.034 | -0.034 | 0 | 0.35 | 0.16 | 0.57 | 0.58 | 0.48 | 0.56 |
| 3 | 63037605 | rs873306 | -0.275 | 0.996 | -0.139 | 1.457 | -0.045 | 0.017 | 0.017 | 0.612 | 0.27 | 0.08 | 0.34 | 0.3 | 0.37 | 0.46 |
| 3 | 63037948 | rs749244 | 0.942 | 0.886 | 0.319 | 1.41 | -0.001 | 0.002 | 0.002 | 0.028 | 0.63 | 0.61 | 0.56 | 0.66 | 0.56 | 0.48 |
| 3 | 63044186 | rs1447436 | 0.914 | 0.81 | 0.31 | 1.405 | -0.035 | 0.014 | 0.014 | 0 | 0.06 | 0.01 | 0.22 | 0.3 | 0.1 | 0.09 |
| 3 | 63051342 | rs2367598 | 1.47 | 0.707 | 0.61 | 1.462 | -0.001 | -0.009 | -0.009 | 0.118 | 0.54 | 0.48 | 0.36 | 0.32 | 0.45 | 0.38 |
| 3 | 63055960 | rs6445312 | 0.832 | 0.724 | 0.616 | 1.493 | 0.015 | 0.118 | 0.118 | 0 | 0.86 | 0.76 | 0.75 | 0.67 | 0.85 | 0.87 |
| 3 | 63063240 | rs1868065 | -0.637 | 0.699 | 0.595 | 1.4 | -0.002 | -0.011 | -0.011 | 0 | 0.94 | 0.88 | 0.98 | 0.96 | 0.96 | 0.96 |
| 3 | 63065094 | rs7629251 | 1.67 | 0.335 | 0.954 | 1.212 | 0.003 | 0.009 | 0.009 | 0.103 | 0.46 | 0.48 | 0.35 | 0.27 | 0.39 | 0.29 |
| 3 | 63071000 | rs2367600 | -1.116 | 0.397 | 0.951 | 1.19 | 0 | -0.005 | -0.005 | 0 | 0.47 | 0.5 | 0.63 | 0.68 | 0.53 | 0.66 |
| 3 | 63073556 | rs2053758 | -0.91 | 0.263 | 0.911 | 1.155 | -0.002 | -0.006 | -0.006 | 1.088 | 0.94 | 0.94 | 0.99 | 0.96 | 0.96 | 0.97 |
| 3 | 63074219 | rs4688370 | -0.437 | 0.291 | 1.082 | 1.398 | 0.007 | 0.134 | 0.134 | 0 | 0.18 | 0.31 | 0.27 | 0.33 | 0.15 | 0.19 |
| 3 | 63075267 | rs1374679 | 0.38 | 0.266 | 1.042 | 1.381 | -0.032 | 0.008 | 0.008 | 0.424 | 0.65 | 0.83 | 0.63 | 0.61 | 0.56 | 0.48 |
| 3 | 63078242 | rs2034318 | 0.346 | 0.315 | 1.096 | 1.427 | 0.022 | 0.138 | 0.138 | 0 | 0.08 | 0.14 | 0.22 | 0.28 | 0.1 | 0.1 |

| | | | | | | | | | | | | | | | | |
|---|---|---|---|---|---|---|---|---|---|---|---|---|---|---|---|---|
| 3 | 63087173 | rs7611505 | 1.787 | 0.262 | 1.078 | 1.601 | 0.008 | 0.035 | 0.035 | 0.014 | 0.44 | 0.36 | 0.29 | 0.18 | 0.39 | 0.28 |
| 3 | 63088315 | rs982678 | -0.978 | 0.183 | 1.052 | 1.611 | 0 | -0.002 | -0.002 | 0 | 0.41 | 0.39 | 0.65 | 0.73 | 0.52 | 0.6 |
| 3 | 63088792 | rs921160 | -1.521 | 0.143 | 1.01 | 1.621 | 0.009 | 0.035 | 0.035 | 0 | 0.51 | 0.57 | 0.69 | 0.79 | 0.57 | 0.69 |
| 3 | 63090351 | rs1900585 | 1.35 | 0.171 | 0.979 | 1.44 | 0.001 | 0.019 | 0.019 | 0 | 0.5 | 0.39 | 0.54 | 0.54 | 0.49 | 0.38 |
| 3 | 63091430 | rs921162 | 0.787 | -0.022 | 0.789 | 1.38 | -0.014 | -0.007 | -0.007 | 0.233 | 0.1 | 0.14 | 0.03 | 0.05 | 0.04 | 0.09 |
| 3 | 63093250 | rs12487147 | 0.013 | 0.302 | 0.607 | 1.605 | -0.063 | -0.027 | -0.027 | 0 | 0.31 | 0.12 | 0.37 | 0.36 | 0.43 | 0.47 |
| 3 | 63094473 | rs4688376 | -1.377 | 0.268 | 0.836 | 1.654 | 0.009 | 0.017 | 0.017 | 0 | 0.52 | 0.56 | 0.66 | 0.68 | 0.6 | 0.69 |
| 3 | 63103449 | rs1447449 | -1.262 | 0.155 | 0.932 | 1.701 | -0.001 | -0.002 | -0.002 | 0 | 0.51 | 0.49 | 0.65 | 0.64 | 0.59 | 0.66 |
| 3 | 63106623 | rs7374870 | -1.538 | 0.105 | 1.026 | 1.647 | 0.017 | 0.039 | 0.039 | 0 | 0.54 | 0.68 | 0.68 | 0.69 | 0.6 | 0.67 |
| 3 | 63108164 | rs1447437 | 0.613 | 0.103 | 0.871 | 1.938 | 0.015 | 0.084 | 0.084 | 0 | 0.07 | 0.12 | 0.22 | 0.28 | 0.12 | 0.11 |
| 3 | 63109093 | rs6445317 | -0.218 | 0.105 | 0.728 | 1.821 | -0.073 | 0.021 | 0.021 | 0.456 | 0.67 | 0.89 | 0.64 | 0.72 | 0.55 | 0.54 |
| 3 | 63116512 | rs939093 | 1.582 | 0.118 | 0.776 | 1.798 | -0.003 | -0.006 | -0.006 | 0 | 0.4 | 0.38 | 0.32 | 0.28 | 0.34 | 0.27 |
| 3 | 63118161 | rs9848161 | 0.219 | 0.16 | 0.686 | 1.623 | -0.009 | -0.004 | -0.004 | 0.004 | 0.11 | 0.14 | 0.05 | 0.05 | 0.04 | 0.1 |
| 3 | 63130869 | rs9813741 | 0.745 | 0.225 | 0.66 | 1.652 | -0.022 | -0.008 | -0.008 | 0 | 0.1 | 0.18 | 0.04 | 0.05 | 0.04 | 0.09 |
| 3 | 63133304 | rs6445318 | -0.015 | 0.27 | 0.434 | 1.659 | -0.078 | 0.032 | 0.032 | 0.454 | 0.65 | 0.87 | 0.63 | 0.71 | 0.54 | 0.52 |
| 3 | 63136580 | rs9841232 | -0.297 | 0.291 | 0.968 | 1.823 | -0.036 | 0.024 | 0.024 | 0.708 | 0.12 | 0.35 | 0.06 | 0.11 | 0.04 | 0.1 |
| 3 | 63143707 | rs9854179 | -0.937 | 0.403 | 0.597 | 1.7 | -0.019 | 0.12 | 0.12 | 0 | 0.23 | 0.5 | 0.31 | 0.35 | 0.18 | 0.2 |
| 3 | 63152052 | rs1447441 | -0.124 | 0.66 | -0.176 | 1.593 | -0.043 | -0.108 | -0.108 | 0 | 0.83 | 0.97 | 0.82 | 0.49 | 0.81 | 0.64 |
| 3 | 63152907 | rs6763613 | -0.184 | 1.43 | 0.274 | 0.86 | 0.033 | -0.059 | -0.059 | 0 | 0.1 | 0.2 | 0.12 | 0 | 0.18 | 0.05 |
| 3 | 63153020 | rs6763709 | 0.156 | 1.962 | -0.199 | 0.175 | -0.07 | -0.038 | -0.038 | 0.206 | 0.65 | 0.78 | 0.5 | 0.38 | 0.53 | 0.5 |
| 3 | 63156589 | rs6776815 | -0.122 | 2.012 | -0.081 | 0.147 | 0.078 | -0.072 | -0.072 | 0 | 0.11 | 0.32 | 0.12 | 0 | 0.18 | 0.04 |
| 3 | 63162753 | rs2887073 | -0.902 | 2.093 | -0.363 | 0.297 | -0.001 | -0.008 | -0.008 | 0 | 0.85 | 0.89 | 0.82 | 0.44 | 0.84 | 0.67 |
| 3 | 63163127 | rs1470543 | 0.26 | 2.038 | -0.345 | 0.178 | -0.072 | -0.044 | -0.044 | 0 | 0.66 | 0.8 | 0.51 | 0.39 | 0.53 | 0.49 |
| 3 | 63165718 | rs1900577 | -0.879 | 1.704 | -0.099 | 0.221 | -0.022 | 0.135 | 0.135 | 0.047 | 0.85 | 0.76 | 0.85 | 0.44 | 0.89 | 0.67 |
| 3 | 63167851 | rs1838990 | 1.278 | 1.033 | -0.414 | -0.509 | 0.056 | 0.016 | 0.016 | 0 | 0.4 | 0.18 | 0.32 | 0.35 | 0.29 | 0.37 |
| 3 | 63173302 | rs7649454 | -1.12 | 0.662 | -0.002 | -0.461 | -0.004 | 0.112 | 0.112 | 0.853 | 0.83 | 0.74 | 0.83 | 0.44 | 0.86 | 0.65 |
| 3 | 63181128 | rs6770731 | -1.139 | 0.757 | 0.05 | -0.449 | 0.003 | 0.1 | 0.1 | 0.846 | 0.83 | 0.74 | 0.82 | 0.43 | 0.85 | 0.66 |
| 3 | 63191817 | rs7635111 | -0.759 | 0.634 | -0.087 | -0.592 | 0 | -0.032 | -0.032 | 0 | 0.32 | 0.58 | 0.29 | 0.16 | 0.35 | 0.23 |
| 3 | 63194035 | rs7613378 | -0.359 | 0.579 | 0.051 | -0.708 | 0.073 | -0.103 | -0.103 | 0 | 0.14 | 0.43 | 0.17 | 0 | 0.21 | 0.05 |
| 3 | 63206756 | rs3856941 | -1.066 | 0.591 | -0.063 | -0.607 | 0.01 | 0.099 | 0.099 | 0.741 | 0.81 | 0.67 | 0.8 | 0.38 | 0.84 | 0.65 |
| 3 | 63207284 | rs4688380 | 1.881 | 0.103 | -0.667 | -1.092 | -0.046 | 0.053 | 0.053 | 0 | 0.25 | 0.13 | 0.34 | 0.1 | 0.29 | 0.19 |
| 3 | 63213635 | rs2367763 | -1.674 | -0.114 | -0.612 | -1.055 | 0.005 | 0.105 | 0.105 | 0.64 | 0.83 | 0.76 | 0.82 | 0.43 | 0.85 | 0.66 |
| 3 | 63213913 | rs17300166 | 2.024 | -0.291 | -0.72 | -1.174 | -0.009 | 0.072 | 0.072 | 0 | 0.12 | 0.01 | 0.05 | 0 | 0.1 | 0.09 |
| 3 | 63214900 | rs9311860 | -0.902 | -0.1 | -0.619 | -1.114 | -0.004 | 0.004 | 0.004 | 0.86 | 0.58 | 0.61 | 0.48 | 0.33 | 0.55 | 0.46 |
| 3 | 63220615 | rs9857872 | -1.542 | -0.024 | -0.453 | -0.939 | 0.005 | 0.128 | 0.128 | 0.539 | 0.83 | 0.73 | 0.83 | 0.42 | 0.86 | 0.7 |
| 3 | 63226290 | rs11710912 | 0.728 | -0.027 | -0.218 | -1.043 | 0.031 | -0.051 | -0.051 | 0 | 0.1 | 0.15 | 0.14 | 0 | 0.2 | 0.04 |
| 3 | 63228414 | rs2367764 | -1.697 | -0.173 | -0.199 | -0.889 | -0.001 | 0.018 | 0.018 | 0.83 | 0.84 | 0.84 | 0.83 | 0.42 | 0.86 | 0.72 |
| 3 | 63229893 | rs7631923 | 0.408 | -0.345 | -0.537 | -1.07 | 0 | -0.058 | -0.058 | 0 | 0.49 | 0.51 | 0.54 | 0.05 | 0.59 | 0.34 |
| 3 | 63233311 | rs11130919 | 1.454 | -0.31 | -0.59 | -0.982 | 0.028 | 0.028 | 0.028 | 0 | 0.28 | 0.13 | 0.21 | 0.22 | 0.23 | 0.31 |
| 3 | 63247928 | rs2367773 | -1.157 | -0.426 | -0.389 | -0.689 | -0.005 | -0.055 | -0.055 | 0.282 | 0.82 | 0.93 | 0.84 | 0.42 | 0.86 | 0.69 |
| 3 | 63249608 | rs6445336 | -0.846 | -0.262 | -0.218 | -0.639 | 0.007 | 0.111 | 0.111 | 0.497 | 0.81 | 0.64 | 0.81 | 0.37 | 0.85 | 0.71 |
| 3 | 63251670 | rs9829040 | 1.391 | -0.542 | -0.522 | -0.99 | 0.016 | 0.001 | 0.001 | 0 | 0.11 | 0.02 | 0.07 | 0.12 | 0.07 | 0.14 |
| 3 | 63254501 | rs17373870 | 1.984 | -0.111 | -1.003 | -1.213 | -0.053 | 0.105 | 0.105 | 0 | 0.26 | 0.06 | 0.33 | 0.05 | 0.33 | 0.22 |
| 3 | 63255742 | rs11714284 | 0.528 | -0.147 | -0.79 | -1.137 | -0.004 | -0.007 | -0.007 | 0 | 0.09 | 0.07 | 0.1 | 0 | 0.16 | 0.03 |
| 3 | 63257892 | rs7647554 | 0.118 | -0.37 | -0.784 | -0.804 | 0.054 | 0.115 | 0.115 | 0.022 | 0.74 | 0.55 | 0.7 | 0.52 | 0.69 | 0.71 |
| 3 | 63258941 | rs2367779 | -0.112 | -0.69 | -0.799 | -0.84 | 0.059 | -0.022 | -0.022 | 0 | 0.39 | 0.22 | 0.32 | 0.41 | 0.3 | 0.47 |
| 3 | 63261972 | rs7433582 | -0.714 | -0.415 | -0.534 | -0.59 | 0.017 | 0.181 | 0.181 | 0.174 | 0.81 | 0.64 | 0.79 | 0.51 | 0.83 | 0.73 |
| 3 | 63265740 | rs4688384 | 0.308 | -0.504 | -0.58 | -0.618 | 0.018 | 0.076 | 0.076 | 0.008 | 0.71 | 0.51 | 0.69 | 0.52 | 0.7 | 0.7 |
| 3 | 63266007 | rs4688387 | -0.172 | -0.603 | -0.771 | -0.77 | 0.035 | -0.008 | -0.008 | 2.571 | 0.4 | 0.3 | 0.34 | 0.4 | 0.32 | 0.46 |
| 3 | 63266941 | rs12491231 | 2.174 | -0.607 | -0.761 | -0.776 | -0.084 | 0.111 | 0.111 | 0 | 0.3 | 0.07 | 0.34 | 0.12 | 0.38 | 0.23 |
| 3 | 63269193 | rs9846545 | 1.093 | -1.046 | -0.8 | -0.766 | 0.009 | 0 | 0 | 0 | 0.11 | 0.1 | 0.05 | 0.06 | 0.06 | 0.08 |
| 3 | 63269800 | rs4688165 | 0.134 | -0.854 | -0.783 | -0.686 | 0.009 | -0.009 | -0.009 | 0 | 0.29 | 0.19 | 0.3 | 0.34 | 0.26 | 0.38 |
| 3 | 63276836 | rs6767034 | -0.495 | -0.891 | -0.417 | -0.423 | -0.018 | 0.186 | 0.186 | 0.139 | 0.79 | 0.58 | 0.78 | 0.51 | 0.83 | 0.71 |
| 3 | 63291970 | rs1874927 | -0.423 | -0.885 | -0.584 | -0.421 | -0.02 | 0.175 | 0.175 | 0.17 | 0.78 | 0.54 | 0.78 | 0.51 | 0.83 | 0.7 |
| 3 | 63298187 | rs36093116 | -1.084 | -0.862 | -1.081 | -0.401 | 0.003 | -0.08 | -0.08 | 0 | 0.83 | 0.94 | 0.82 | 0.51 | 0.85 | 0.72 |
| 3 | 63309890 | rs17311528 | 0.751 | -0.994 | -1.326 | -0.524 | -0.031 | -0.012 | -0.012 | 0 | 0.21 | 0.05 | 0.18 | 0.22 | 0.21 | 0.26 |
| 3 | 63310713 | rs938926 | -1.085 | -1.038 | -1.146 | -0.436 | 0.006 | -0.037 | -0.037 | 0.248 | 0.82 | 0.87 | 0.82 | 0.51 | 0.84 | 0.7 |
| 3 | 63315687 | rs17068247 | 1.142 | -1.059 | -1.541 | -0.511 | 0.044 | -0.016 | -0.016 | 0 | 0.11 | 0.01 | 0.04 | 0.06 | 0.06 | 0.08 |
| 3 | 63329842 | rs1523439 | 1.089 | -0.87 | -1.236 | -0.86 | 0.001 | 0.022 | 0.022 | 0 | 0.45 | 0.35 | 0.45 | 0.13 | 0.54 | 0.25 |
| 3 | 63331331 | rs13060806 | 0.053 | -0.757 | -1.608 | -0.574 | -0.014 | -0.047 | -0.047 | 0.038 | 0.28 | 0.07 | 0.28 | 0.32 | 0.24 | 0.38 |
| 3 | 63333375 | rs11712437 | 0.145 | -0.354 | -1.368 | -0.86 | 0.01 | -0.051 | -0.051 | 0 | 0.14 | 0.17 | 0.13 | 0 | 0.19 | 0.09 |
| 3 | 63337860 | rs17068288 | 0.243 | -0.436 | -1.47 | -0.848 | 0.002 | -0.031 | -0.031 | 0 | 0.13 | 0.15 | 0.12 | 0 | 0.18 | 0.09 |
| 3 | 63337921 | rs13093115 | 0.33 | -0.38 | -1.773 | -0.583 | -0.011 | 0.034 | 0.034 | 0.631 | 0.22 | 0.1 | 0.27 | 0.14 | 0.23 | 0.24 |
| 3 | 63339909 | rs9829816 | -0.702 | -0.363 | -1.587 | -0.542 | 0.037 | 0.103 | 0.103 | 0 | 0.43 | 0.77 | 0.38 | 0.71 | 0.36 | 0.52 |
| 3 | 63340627 | rs7648383 | -0.723 | -0.232 | -1.013 | -0.45 | 0 | -0.006 | -0.006 | 0.003 | 0.81 | 0.8 | 0.83 | 0.89 | 0.8 | 0.86 |
| 3 | 63342866 | rs935786 | 0.817 | -0.604 | -0.766 | -0.405 | -0.003 | 0.036 | 0.036 | 0 | 0.58 | 0.3 | 0.64 | 0.75 | 0.59 | 0.62 |
| 3 | 63344264 | rs6764201 | 0.739 | -0.586 | -0.721 | -0.425 | -0.002 | 0.032 | 0.032 | 0 | 0.59 | 0.3 | 0.65 | 0.75 | 0.61 | 0.62 |
| 3 | 63344664 | rs9876045 | 0.512 | -0.569 | -0.752 | -0.554 | -0.002 | -0.004 | -0.004 | 0 | 0.65 | 0.32 | 0.67 | 0.89 | 0.63 | 0.77 |
| 3 | 63346006 | rs6445339 | -1.144 | -0.398 | -0.412 | -0.459 | -0.009 | 0.014 | 0.014 | 0 | 0.84 | 0.84 | 0.82 | 0.98 | 0.81 | 0.86 |

| | | | | | | | | | | | | | | | | |
|---|---|---|---|---|---|---|---|---|---|---|---|---|---|---|---|---|
| 3 | 63348134 | rs6762466 | 1.229 | -1.219 | -0.531 | -0.33 | 0.014 | 0.052 | 0.052 | 0 | 0.33 | 0.2 | 0.31 | 0.2 | 0.3 | 0.31 |
| 3 | 63349451 | rs13062956 | 0.436 | -1.341 | 0.152 | -0.361 | 0.01 | 0.056 | 0.056 | 0 | 0.51 | 0.39 | 0.43 | 0.2 | 0.5 | 0.4 |
| 3 | 63350070 | rs1376479 | -1.402 | -1.275 | -0.281 | -0.442 | -0.081 | 0.098 | 0.098 | 0.323 | 0.79 | 0.94 | 0.69 | 0.98 | 0.68 | 0.78 |
| 3 | 63350194 | rs9311874 | -0.263 | -1.633 | 0.312 | 0.142 | 0.013 | 0.151 | 0.151 | 0 | 0.09 | 0.34 | 0.07 | 0.2 | 0.04 | 0.14 |
| 3 | 63350469 | rs13068980 | 0.711 | -1.468 | -0.516 | 0.209 | -0.018 | -0.092 | -0.092 | 0 | 0.14 | 0.01 | 0.16 | 0.57 | 0.14 | 0.24 |
| 3 | 63363166 | rs17068400 | 1.239 | -1.474 | -0.374 | -0.179 | 0.02 | 0.065 | 0.065 | 0 | 0.31 | 0.13 | 0.28 | 0.1 | 0.26 | 0.22 |
| 3 | 63369620 | rs17068421 | 0.367 | -1.182 | -0.486 | 0.378 | 0.006 | -0.116 | -0.116 | 0 | 0.16 | 0.01 | 0.16 | 0.59 | 0.11 | 0.25 |
| 3 | 63373421 | rs6445341 | -1.188 | -1.022 | -0.487 | 0.513 | -0.092 | -0.017 | -0.017 | 0 | 0.74 | 0.93 | 0.65 | 0.81 | 0.62 | 0.61 |
| 3 | 63376482 | rs6790302 | -0.632 | -1.238 | 0.204 | 0.572 | -0.014 | 0.034 | 0.034 | 0 | 0.07 | 0.21 | 0.03 | 0.04 | 0.02 | 0.01 |
| 3 | 63376887 | rs4643705 | 0.88 | -1.504 | 0.568 | 0.428 | 0.014 | 0.035 | 0.035 | 0 | 0.45 | 0.37 | 0.37 | 0.13 | 0.41 | 0.25 |
| 3 | 63378710 | rs7641300 | -0.461 | -1.366 | 0.608 | 0.645 | -0.04 | 0.134 | 0.134 | 0 | 0.21 | 0.42 | 0.16 | 0.56 | 0.13 | 0.24 |
| 3 | 63379696 | rs6800579 | -1.05 | -1.3 | 0.617 | 0.67 | -0.031 | 0.048 | 0.048 | 0 | 0.27 | 0.43 | 0.25 | 0.7 | 0.2 | 0.35 |
| 3 | 63380099 | rs11709894 | 0.641 | -1.279 | 0.61 | 0.526 | -0.001 | -0.018 | -0.018 | 1.671 | 0.48 | 0.48 | 0.39 | 0.13 | 0.42 | 0.25 |
| 3 | 63380775 | rs1996918 | -0.403 | -1.35 | 0.314 | 0.447 | -0.006 | -0.009 | -0.009 | 0 | 0.23 | 0.26 | 0.2 | 0.15 | 0.21 | 0.15 |
| 3 | 63380863 | rs13066814 | 0.143 | -0.521 | 0.819 | 1.26 | 0.001 | -0.09 | -0.09 | 0 | 0.15 | 0.01 | 0.1 | 0.44 | 0.11 | 0.23 |
| 3 | 63382986 | rs17068484 | -0.349 | -0.675 | 1.316 | 0.435 | 0.011 | 0.084 | 0.084 | 0 | 0.07 | 0.17 | 0.11 | 0.15 | 0.08 | 0.11 |
| 3 | 63385835 | rs17031723 | 0.446 | -0.769 | 1.004 | 0.311 | 0.006 | 0.043 | 0.043 | 0 | 0.47 | 0.31 | 0.44 | 0.28 | 0.49 | 0.36 |
| 3 | 63390741 | rs1901304 | -0.17 | -0.821 | 0.596 | 0.221 | -0.031 | 0.001 | 0.001 | 1.258 | 0.54 | 0.87 | 0.51 | 0.38 | 0.5 | 0.36 |
| 3 | 63392409 | rs9311875 | 0.564 | -0.587 | 0.267 | 0.279 | -0.006 | -0.022 | -0.022 | 0 | 0.46 | 0.51 | 0.42 | 0.22 | 0.4 | 0.25 |
| 3 | 63396026 | rs11710162 | 0.385 | -0.683 | -0.35 | 0.341 | 0.007 | 0.082 | 0.082 | 0.098 | 0.44 | 0.28 | 0.31 | 0.22 | 0.48 | 0.31 |
| 3 | 63396603 | rs2887097 | 0.149 | 0.284 | -0.249 | 0.728 | -0.003 | 0 | 0 | 2.923 | 0.57 | 0.43 | 0.48 | 0.38 | 0.52 | 0.4 |
| 3 | 63400081 | rs13316595 | -1.163 | 0.442 | -0.071 | 1.086 | 0.03 | 0.103 | 0.103 | 0 | 0.08 | 0.22 | 0.07 | 0.15 | 0.1 | 0.12 |
| 3 | 63405972 | rs727777 | -0.425 | 0.19 | -0.753 | 0.769 | 0.01 | -0.005 | -0.005 | 0.103 | 0.36 | 0.28 | 0.26 | 0.61 | 0.29 | 0.49 |
| 3 | 63410966 | rs17068548 | 0.374 | 0.075 | -0.665 | 0.173 | -0.004 | -0.051 | -0.051 | 0.34 | 0.13 | 0.15 | 0.12 | 0.01 | 0.13 | 0.18 |
| 3 | 63412640 | rs2122798 | 0.44 | -0.422 | -0.936 | 0.66 | 0.026 | -0.015 | -0.015 | 0 | 0.11 | 0.03 | 0.06 | 0.44 | 0.03 | 0.22 |
| 3 | 63415195 | rs1403700 | -0.93 | 0.141 | -1.439 | -0.327 | -0.065 | 0.079 | 0.079 | 0.071 | 0.78 | 0.88 | 0.65 | 0.96 | 0.69 | 0.79 |
| 3 | 63415387 | rs17394121 | 0.046 | -0.341 | -1.005 | -0.363 | -0.005 | 0.003 | 0.003 | 0 | 0.18 | 0.16 | 0.15 | 0.15 | 0.19 | 0.2 |
| 3 | 63417340 | rs1545458 | -0.216 | -0.712 | -1.114 | -0.412 | -0.001 | 0.016 | 0.016 | 0.779 | 0.45 | 0.68 | 0.38 | 0.44 | 0.42 | 0.45 |
| 3 | 63417713 | rs6773908 | -0.523 | -1.149 | -0.845 | -0.588 | -0.011 | 0.002 | 0.002 | 0 | 0.26 | 0.39 | 0.21 | 0.25 | 0.17 | 0.29 |
| 3 | 63420897 | rs6445345 | -0.599 | -1.094 | -1.404 | -0.199 | 0.006 | 0.013 | 0.013 | 4.248 | 0.53 | 0.76 | 0.6 | 0.58 | 0.45 | 0.46 |
| 3 | 63422149 | rs2367834 | 0.413 | -0.8 | -1.466 | 0.151 | -0.018 | -0.059 | -0.059 | 0.016 | 0.13 | 0.04 | 0.12 | 0.34 | 0.12 | 0.21 |
| 3 | 63423459 | rs13085353 | 0.644 | -0.789 | -1.384 | 0.093 | 0.002 | 0.011 | 0.011 | 0 | 0.06 | 0.05 | 0.29 | 0.16 | 0.1 | 0.12 |
| 3 | 63426277 | rs9810713 | 0.93 | -0.372 | -2.014 | -0.232 | -0.005 | -0.085 | -0.085 | 0 | 0.19 | 0.08 | 0.16 | 0.41 | 0.19 | 0.39 |
| 3 | 63427243 | rs11707206 | 0.483 | -0.237 | -1.463 | -0.155 | 0 | -0.021 | -0.021 | 0 | 0.27 | 0.38 | 0.44 | 0.18 | 0.29 | 0.16 |
| 3 | 63428111 | rs1530856 | 0.43 | 0.41 | -2.607 | -0.298 | -0.063 | -0.016 | -0.016 | 0.176 | 0.66 | 0.86 | 0.67 | 0.64 | 0.56 | 0.7 |
| 3 | 63430538 | rs12374033 | 1.382 | -0.289 | -2.305 | -0.23 | -0.026 | 0.029 | 0.029 | 0 | 0.15 | 0.07 | 0.18 | 0.02 | 0.27 | 0.06 |
| 3 | 63432152 | rs935794 | -0.408 | -0.458 | -2.327 | -0.346 | 0.001 | -0.004 | -0.004 | 0 | 0.7 | 0.61 | 0.52 | 0.74 | 0.69 | 0.75 |
| 3 | 63432680 | rs935793 | -1.476 | -0.776 | -1.947 | -0.036 | 0.001 | -0.01 | -0.01 | 0 | 0.82 | 0.79 | 0.82 | 0.9 | 0.8 | 0.88 |
| 3 | 63434206 | rs13099553 | 0.283 | -1.332 | -1.376 | 0.53 | -0.013 | 0.006 | 0.006 | 0.135 | 0.17 | 0.11 | 0.18 | 0.02 | 0.27 | 0.06 |
| 3 | 63435226 | rs13082843 | 1.161 | -1.2 | -1.365 | 0.603 | -0.006 | 0.005 | 0.005 | 0 | 0.13 | 0.1 | 0.12 | 0.02 | 0.19 | 0.05 |
| 3 | 63436469 | rs4688398 | -0.984 | -1.321 | -1.316 | 0.725 | 0.039 | 0.043 | 0.043 | 0.809 | 0.52 | 0.57 | 0.38 | 0.71 | 0.4 | 0.62 |
| 3 | 63436581 | rs6790210 | 0.215 | -0.902 | -1.111 | 0.634 | -0.123 | 0.054 | 0.054 | 0 | 0.24 | 0.43 | 0.1 | 0.32 | 0.09 | 0.38 |
| 3 | 63437107 | rs6766009 | -0.354 | 0.938 | -1.48 | 0.623 | -0.014 | -0.007 | -0.007 | 0 | 0.23 | 0.11 | 0.24 | 0.25 | 0.31 | 0.12 |
| 3 | 63437858 | rs6806334 | -0.942 | 1.495 | -0.969 | 0.68 | -0.011 | 0.005 | 0.005 | 0 | 0.24 | 0.21 | 0.22 | 0.17 | 0.33 | 0.19 |
| 3 | 63438258 | rs17068597 | -0.78 | 1.074 | -0.855 | 0.865 | 0.007 | -0.009 | -0.009 | 0.807 | 0.19 | 0.13 | 0.08 | 0.29 | 0.16 | 0.26 |
| 3 | 63438754 | rs10866066 | -1.153 | 1.662 | -0.207 | 0.656 | 0 | -0.001 | -0.001 | 0.041 | 0.38 | 0.36 | 0.29 | 0.33 | 0.4 | 0.38 |
| 3 | 63438886 | rs1975649 | -2.252 | 1.395 | -0.827 | 0.94 | -0.014 | -0.052 | -0.052 | 0.753 | 0.66 | 0.96 | 0.7 | 0.66 | 0.65 | 0.68 |
| 3 | 63439209 | rs1562859 | -0.371 | 1.127 | -0.308 | 0.999 | -0.005 | 0.035 | 0.035 | 0 | 0.26 | 0.57 | 0.39 | 0.31 | 0.24 | 0.31 |
| 3 | 63439281 | rs1975650 | -0.334 | 1.118 | -0.22 | 0.995 | -0.005 | 0.036 | 0.036 | 0 | 0.25 | 0.54 | 0.39 | 0.31 | 0.24 | 0.31 |
| 3 | 63439334 | rs9823776 | 0.085 | 1.021 | 0.34 | 1.059 | -0.002 | 0 | 0 | 0 | 0.18 | 0.18 | 0.23 | 0.11 | 0.22 | 0.15 |
| 3 | 63439414 | rs4688169 | -0.816 | 0.94 | 0.493 | 1.163 | 0.018 | 0.275 | 0.275 | 0.468 | 0.07 | 0.36 | 0.15 | 0.21 | 0.02 | 0.15 |
| 3 | 63439733 | rs4688399 | -0.837 | 1.136 | 0.622 | 1.002 | 0 | 0.106 | 0.106 | 0.951 | 0.24 | 0.45 | 0.24 | 0.39 | 0.22 | 0.37 |
| 3 | 63439824 | rs4688400 | 0.151 | 1.173 | 0.916 | 1.005 | 0.014 | 0.043 | 0.043 | 0.086 | 0.16 | 0.24 | 0.2 | 0.28 | 0.18 | 0.23 |
| 3 | 63441925 | rs13319407 | -0.085 | 0.75 | 0.998 | 0.821 | 0.005 | -0.056 | -0.056 | 0 | 0.2 | 0.27 | 0.21 | 0.04 | 0.19 | 0.07 |
| 3 | 63445205 | rs9855522 | -1.051 | 0.86 | 1.075 | 0.777 | -0.007 | -0.069 | -0.069 | 1.135 | 0.38 | 0.51 | 0.41 | 0.13 | 0.35 | 0.23 |
| 3 | 63449206 | rs2122797 | 0.157 | 1.034 | 0.835 | 0.829 | 0.006 | -0.05 | -0.05 | 1.96 | 0.18 | 0.26 | 0.21 | 0.04 | 0.19 | 0.07 |
| 3 | 63449344 | rs1839733 | -2.15 | 1.112 | 0.512 | 0.974 | -0.009 | -0.084 | -0.084 | 0 | 0.62 | 0.88 | 0.66 | 0.53 | 0.61 | 0.56 |
| 3 | 63458063 | rs6790592 | -0.481 | 1.494 | 1.191 | 1.283 | 0.04 | 0.105 | 0.105 | 0.061 | 0.2 | 0.34 | 0.25 | 0.41 | 0.24 | 0.31 |
| 3 | 63463754 | rs935789 | -1.215 | 1.038 | 1.076 | 1.203 | 0.007 | 0.008 | 0.008 | 0 | 0.34 | 0.39 | 0.44 | 0.47 | 0.42 | 0.37 |
| 3 | 63468136 | rs12630244 | 0.067 | 1.144 | 0.929 | 1.356 | -0.014 | -0.016 | -0.016 | 0 | 0.16 | 0.15 | 0.26 | 0.29 | 0.24 | 0.27 |
| 3 | 63468228 | rs17068745 | -0.1 | 0.978 | 1.05 | 1.213 | -0.002 | -0.006 | -0.006 | 0 | 0.18 | 0.18 | 0.19 | 0.08 | 0.16 | 0.17 |
| 3 | 63469983 | rs6783555 | -1.106 | 0.978 | 1.105 | 1.258 | 0.005 | 0.003 | 0.003 | 0 | 0.33 | 0.36 | 0.43 | 0.35 | 0.41 | 0.33 |
| 3 | 63470203 | rs6799191 | -1.236 | 0.966 | 1.153 | 1.253 | 0.011 | 0.018 | 0.018 | 0 | 0.35 | 0.46 | 0.45 | 0.48 | 0.41 | 0.37 |
| 3 | 63470819 | rs764924 | 0.171 | 0.977 | 1.155 | 1.262 | -0.014 | -0.017 | -0.017 | 4.087 | 0.16 | 0.16 | 0.23 | 0.25 | 0.22 | 0.25 |
| 3 | 63475570 | rs11130928 | -0.237 | 0.968 | 1.264 | 1.249 | -0.014 | -0.031 | -0.031 | 0 | 0.19 | 0.25 | 0.19 | 0.08 | 0.16 | 0.17 |
| 3 | 63476147 | rs1812207 | -2.658 | 0.931 | 0.854 | 1.237 | -0.038 | 0.02 | 0.02 | 0 | 0.59 | 0.92 | 0.67 | 0.65 | 0.61 | 0.65 |
| 3 | 63477074 | rs6765108 | -2.157 | 0.94 | 0.777 | 1.233 | 0.004 | 0.006 | 0.006 | 0 | 0.53 | 0.64 | 0.63 | 0.56 | 0.57 | 0.54 |
| 3 | 63477970 | rs11130935 | -0.187 | 0.954 | 0.755 | 1.23 | -0.002 | -0.003 | -0.003 | 4.316 | 0.18 | 0.21 | 0.17 | 0.18 | 0.17 | 0.1 |
| 3 | 63478720 | rs6796188 | 0.543 | 0.992 | 0.695 | 1.19 | -0.009 | 0.019 | 0.019 | 0 | 0.14 | 0.08 | 0.15 | 0.06 | 0.17 | 0.06 |

| | | | | | | | | | | | | | | | | |
|---|---|---|---|---|---|---|---|---|---|---|---|---|---|---|---|---|
| 3 | 63484444 | rs4688410 | -0.862 | 1.129 | 0.585 | 1.287 | 0.004 | 0.014 | 0.014 | 0 | 0.31 | 0.39 | 0.42 | 0.45 | 0.36 | 0.4 |
| 3 | 63484923 | rs7642916 | -0.61 | 0.863 | 0.449 | 1.212 | -0.027 | -0.047 | -0.047 | 0 | 0.27 | 0.49 | 0.21 | 0.16 | 0.22 | 0.22 |
| 3 | 63488841 | rs9866272 | -0.905 | 0.953 | 0.474 | 1.233 | 0.011 | 0.014 | 0.014 | 0 | 0.3 | 0.36 | 0.46 | 0.38 | 0.39 | 0.4 |
| 3 | 63489319 | rs6766569 | -0.103 | 0.946 | 0.542 | 1.211 | -0.012 | -0.051 | -0.051 | 0 | 0.22 | 0.31 | 0.17 | 0.06 | 0.18 | 0.11 |
| 3 | 63491051 | rs17068857 | 0.241 | 0.834 | 0.338 | 1.233 | 0.002 | 0 | 0 | 0 | 0.14 | 0.24 | 0.23 | 0.13 | 0.17 | 0.15 |
| 3 | 63491141 | rs11715745 | -1.597 | 0.848 | -0.337 | 1.241 | 0.01 | 0.011 | 0.011 | 0 | 0.55 | 0.73 | 0.64 | 0.56 | 0.58 | 0.55 |
| 3 | 63491925 | rs4688412 | -0.706 | 0.869 | -1.342 | 1.063 | -0.015 | -0.002 | -0.002 | 1.403 | 0.37 | 0.26 | 0.4 | 0.32 | 0.4 | 0.38 |
| 3 | 63498703 | rs9311876 | -0.995 | 1.095 | -1.724 | 0.962 | 0.008 | 0.028 | 0.028 | 0 | 0.61 | 0.92 | 0.68 | 0.66 | 0.66 | 0.7 |
| 3 | 63498876 | rs13086978 | -0.733 | 1.567 | -0.779 | 1.109 | -0.022 | -0.006 | -0.006 | 0 | 0.15 | 0.08 | 0.24 | 0.16 | 0.23 | 0.25 |
| 3 | 63500435 | rs9877681 | -1.006 | 0.375 | -0.414 | 1.291 | -0.021 | 0.026 | 0.026 | 2.356 | 0.17 | 0.21 | 0.15 | 0.28 | 0.1 | 0.13 |
| 3 | 63504450 | rs6805951 | -1.404 | 0.645 | -0.104 | 0.953 | -0.017 | 0.006 | 0.006 | 2.103 | 0.28 | 0.38 | 0.23 | 0.29 | 0.21 | 0.23 |
| 3 | 63504600 | rs11708840 | -1.357 | 1.11 | -0.466 | 0.922 | -0.004 | -0.002 | -0.002 | 0 | 0.29 | 0.22 | 0.25 | 0.28 | 0.28 | 0.25 |
| 3 | 63505191 | rs11918416 | -1.254 | 1.376 | 0.32 | 1.504 | 0.048 | 0.186 | 0.186 | 0 | 0.13 | 0.33 | 0.35 | 0.52 | 0.18 | 0.24 |
| 3 | 63507244 | rs7614676 | -1.093 | 1.053 | -0.121 | 0.848 | -0.002 | -0.004 | -0.004 | 0 | 0.18 | 0.16 | 0.28 | 0.31 | 0.2 | 0.18 |
| 3 | 63507428 | rs1394898 | 1.142 | 0.625 | 0.007 | -0.072 | 0 | 0 | 0 | 0 | 0.54 | 0.5 | 0.53 | 0.68 | 0.57 | 0.74 |
| 3 | 63508820 | rs11708252 | 0.082 | 0.558 | 0.043 | 0.294 | -0.004 | -0.036 | -0.036 | 0 | 0.08 | 0.04 | 0.16 | 0.17 | 0.08 | 0.08 |
| 3 | 63510487 | rs6802712 | 0.222 | 0.456 | 0.101 | 0.186 | 0.017 | 0.032 | 0.032 | 0 | 0.19 | 0.1 | 0.11 | 0 | 0.15 | 0.03 |
| 3 | 63510844 | rs17068905 | -0.805 | 0.472 | 0.162 | 0.223 | 0 | 0.009 | 0.009 | 0 | 0.19 | 0.21 | 0.29 | 0.31 | 0.2 | 0.18 |
| 3 | 63515234 | rs6801282 | -1.169 | 0.431 | 0.504 | 0.167 | 0 | -0.005 | -0.005 | 0 | 0.36 | 0.34 | 0.39 | 0.19 | 0.34 | 0.21 |
| 3 | 63515740 | rs6766087 | -0.575 | 0.438 | 0.394 | 0.026 | 0.002 | 0.041 | 0.041 | 0 | 0.19 | 0.28 | 0.28 | 0.31 | 0.2 | 0.19 |
| 3 | 63516421 | rs11924708 | -1.341 | 0.433 | 0.335 | 0.025 | -0.012 | -0.006 | -0.006 | 0 | 0.39 | 0.48 | 0.4 | 0.32 | 0.35 | 0.25 |
| 3 | 63526806 | rs1505597 | -0.273 | 0.513 | -0.445 | -0.394 | 0.011 | -0.002 | -0.002 | 0 | 0.19 | 0.09 | 0.11 | 0 | 0.14 | 0.03 |
| 3 | 63532532 | rs4688173 | 0.385 | 1.032 | -0.396 | -0.761 | 0.001 | -0.051 | -0.051 | 0 | 0.7 | 0.65 | 0.77 | 0.98 | 0.67 | 0.91 |
| 3 | 63532999 | rs6445357 | -0.464 | 1.252 | -1.133 | -0.469 | -0.021 | -0.063 | -0.063 | 0.82 | 0.62 | 0.84 | 0.46 | 0.5 | 0.62 | 0.59 |
| 3 | 63533576 | rs9856025 | 0.624 | 1.337 | -0.496 | -0.753 | -0.006 | 0.004 | 0.004 | 0.047 | 0.3 | 0.33 | 0.19 | 0.37 | 0.28 | 0.39 |
| 3 | 63535676 | rs17399296 | 1.071 | 1.842 | -0.57 | -0.723 | -0.03 | 0.026 | 0.026 | 0 | 0.07 | 0.02 | 0.12 | 0.01 | 0.13 | 0.08 |
| 3 | 63543254 | rs949553 | 0.116 | 1.586 | -0.319 | -0.787 | 0.033 | -0.011 | -0.011 | 0.016 | 0.34 | 0.19 | 0.21 | 0.36 | 0.31 | 0.4 |
| 3 | 63543345 | rs13097916 | -0.286 | 1.61 | 0.41 | -0.722 | 0.019 | -0.011 | -0.011 | 0 | 0.13 | 0.18 | 0.16 | 0.06 | 0.21 | 0.16 |
| 3 | 63548238 | rs6414558 | 0.038 | 1.509 | 1.167 | -0.71 | -0.009 | 0.019 | 0.019 | 0 | 0.3 | 0.6 | 0.19 | 0.47 | 0.27 | 0.46 |
| 3 | 63554424 | rs11711276 | 1.41 | 1.47 | 0.918 | -0.505 | -0.001 | -0.062 | -0.062 | 0 | 0.18 | 0.02 | 0.12 | 0.27 | 0.19 | 0.35 |
| 3 | 63555337 | rs5000576 | -0.776 | 1.925 | 1.02 | -0.662 | 0.05 | -0.026 | -0.026 | 0 | 0.95 | 0.89 | 0.92 | 1 | 0.89 | 0.97 |
| 3 | 63560769 | rs11712941 | 0.453 | 1.518 | 1.097 | -0.519 | -0.011 | 0.066 | 0.066 | 0 | 0.27 | 0.38 | 0.17 | 0.41 | 0.27 | 0.46 |
| 3 | 63563282 | rs6809477 | -1.102 | 1.349 | 1.015 | -0.579 | -0.004 | -0.009 | -0.009 | 0 | 0.49 | 0.72 | 0.37 | 0.49 | 0.51 | 0.57 |
| 3 | 63564385 | rs13316611 | 0.014 | 1.311 | 1.164 | -0.601 | 0.032 | -0.091 | -0.091 | 0 | 0.22 | 0.46 | 0.22 | 0.07 | 0.26 | 0.1 |
| 3 | 63567881 | rs11923419 | 0.57 | 1.305 | 1.258 | -0.591 | 0.002 | -0.009 | -0.009 | 0.054 | 0.26 | 0.25 | 0.15 | 0.41 | 0.25 | 0.47 |
| 3 | 63569526 | rs11917016 | -1.164 | 1.336 | 0.886 | -0.585 | -0.002 | -0.002 | -0.002 | 0 | 0.49 | 0.71 | 0.37 | 0.49 | 0.51 | 0.57 |
| 3 | 63572197 | rs7642139 | 0.757 | 0.883 | 0.813 | -0.507 | -0.017 | -0.049 | -0.049 | 0 | 0.17 | 0.35 | 0.1 | 0.06 | 0.13 | 0.05 |
| 3 | 63580540 | rs11709292 | 0.424 | 0.969 | 0.74 | -0.454 | -0.007 | 0.03 | 0.03 | 0 | 0.27 | 0.3 | 0.14 | 0.41 | 0.25 | 0.47 |
| 3 | 63583133 | rs6445365 | -0.577 | 1.126 | 0.091 | -0.369 | 0.018 | -0.01 | -0.01 | 0 | 0.44 | 0.66 | 0.25 | 0.47 | 0.38 | 0.51 |
| 3 | 63590814 | rs1876092 | -0.081 | 1.456 | -0.836 | 0.293 | -0.05 | -0.071 | -0.071 | 0 | 0.68 | 0.93 | 0.62 | 0.66 | 0.63 | 0.66 |
| 3 | 63594207 | rs11710939 | 0.614 | 1.016 | -0.282 | 0.09 | 0.004 | 0.004 | 0.004 | 0 | 0.5 | 0.4 | 0.4 | 0.46 | 0.5 | 0.48 |
| 3 | 63597115 | rs6796563 | 0.606 | 1.015 | -0.25 | 0.09 | 0.004 | 0.004 | 0.004 | 0 | 0.5 | 0.39 | 0.4 | 0.46 | 0.5 | 0.48 |
| 3 | 63598974 | rs7646965 | 0.954 | 1.059 | -1.021 | 0.348 | 0.019 | 0.036 | 0.036 | 0 | 0.39 | 0.26 | 0.23 | 0.19 | 0.34 | 0.38 |
| 3 | 63599888 | rs4688416 | 0.997 | 1.102 | -1.684 | 0.305 | -0.081 | -0.075 | -0.075 | 0 | 0.18 | 0.03 | 0.25 | 0.19 | 0.27 | 0.21 |
| 3 | 63600501 | rs6785323 | 1.491 | 1.176 | -1.572 | 0.353 | -0.004 | -0.007 | -0.007 | 0 | 0.08 | 0.08 | 0.06 | 0 | 0.12 | 0.06 |
| 3 | 63603040 | rs6445370 | -2.68 | 1.118 | -1.588 | 0.48 | 0.002 | 0.005 | 0.005 | 0.001 | 0.88 | 0.96 | 0.9 | 0.9 | 0.89 | 0.91 |
| 3 | 63603510 | rs905941 | 0.909 | 0.823 | -1.317 | 0.443 | -0.013 | 0.002 | 0.002 | 0 | 0.41 | 0.46 | 0.34 | 0.4 | 0.37 | 0.4 |
| 3 | 63611223 | rs12492432 | -1.099 | 0.944 | -1.452 | 0.742 | -0.002 | 0.003 | 0.003 | 0 | 0.77 | 0.78 | 0.69 | 0.77 | 0.78 | 0.82 |
| 3 | 63612013 | rs6445375 | 0.15 | 0.831 | -1.629 | 0.544 | -0.006 | 0.016 | 0.016 | 0 | 0.52 | 0.68 | 0.46 | 0.66 | 0.51 | 0.55 |
| 3 | 63612431 | rs7646765 | 0.144 | 0.845 | -1.622 | 0.487 | -0.006 | -0.01 | -0.01 | 0 | 0.52 | 0.68 | 0.46 | 0.58 | 0.51 | 0.53 |
| 3 | 63612877 | rs1505593 | 0.18 | 0.855 | -1.584 | 0.51 | -0.003 | 0.017 | 0.017 | 0 | 0.52 | 0.67 | 0.46 | 0.47 | 0.51 | 0.52 |
| 3 | 63615084 | rs903747 | 0.659 | 0.936 | -1.996 | 0.415 | -0.119 | -0.038 | -0.038 | 0 | 0.24 | 0.03 | 0.33 | 0.16 | 0.37 | 0.22 |
| 3 | 63615622 | rs751191 | 0.002 | 0.757 | -1.644 | 0.266 | 0.024 | -0.003 | -0.003 | 0 | 0.5 | 0.72 | 0.43 | 0.59 | 0.44 | 0.52 |
| 3 | 63615879 | rs751188 | 0.488 | 0.345 | -1.387 | 0.234 | -0.018 | -0.006 | -0.006 | 0.088 | 0.42 | 0.54 | 0.26 | 0.47 | 0.33 | 0.46 |
| 3 | 63616643 | rs964902 | 0.013 | 0.683 | -1.357 | 0.312 | 0.002 | 0.001 | 0.001 | 0 | 0.56 | 0.58 | 0.42 | 0.53 | 0.54 | 0.63 |
| 3 | 63616874 | rs13076701 | 0.003 | 1.077 | -0.995 | 0.229 | -0.002 | 0 | 0 | 0.04 | 0.26 | 0.21 | 0.38 | 0.21 | 0.35 | 0.25 |
| 3 | 63617158 | rs6782591 | -0.082 | 0.6 | -1.24 | 0.272 | 0.007 | 0.032 | 0.032 | 0 | 0.23 | 0.53 | 0.24 | 0.29 | 0.24 | 0.22 |
| 3 | 63618827 | rs7609960 | -0.644 | 0.653 | -1.499 | -0.398 | 0.008 | -0.02 | -0.02 | 0 | 0.61 | 0.62 | 0.53 | 0.74 | 0.58 | 0.68 |
| 3 | 63623896 | rs13086496 | -1.032 | 0.906 | -1.323 | -0.232 | 0.034 | -0.016 | -0.016 | 0 | 0.7 | 0.62 | 0.57 | 0.76 | 0.66 | 0.78 |
| 3 | 63624266 | rs2135220 | -0.649 | 0.76 | -1.403 | -0.16 | 0.017 | -0.026 | -0.026 | 0 | 0.61 | 0.51 | 0.49 | 0.76 | 0.55 | 0.75 |
| 3 | 63628019 | rs6799742 | 0.454 | 0.859 | -1.473 | -0.127 | 0 | -0.001 | -0.001 | 0 | 0.17 | 0.23 | 0.13 | 0.12 | 0.18 | 0.2 |
| 3 | 63637090 | rs17069190 | 0.501 | 0.617 | -1.975 | 0.039 | -0.013 | -0.011 | -0.011 | 0 | 0.22 | 0.03 | 0.12 | 0.19 | 0.2 | 0.28 |
| 3 | 63639624 | rs1158143 | -1.043 | 0.624 | -1.389 | -0.176 | -0.013 | 0.048 | 0.048 | 0 | 0.59 | 0.33 | 0.48 | 0.77 | 0.57 | 0.77 |
| 3 | 63640601 | rs1565983 | 1.419 | 0.599 | -1.45 | -0.16 | -0.001 | -0.011 | -0.011 | 0 | 0.16 | 0.23 | 0.14 | 0.15 | 0.16 | 0.22 |
| 3 | 63643367 | rs17069221 | 1.646 | 0.575 | -1.677 | 0.112 | -0.035 | -0.032 | -0.032 | 0 | 0.08 | 0.01 | 0.1 | 0.11 | 0.09 | 0.18 |
| 3 | 63644694 | rs924067 | -2.207 | 1.037 | -1.562 | -0.129 | -0.007 | -0.02 | -0.02 | 0 | 0.75 | 0.74 | 0.79 | 0.86 | 0.78 | 0.83 |
| 3 | 63648088 | rs17069229 | 1.784 | 1.071 | -1.817 | -0.054 | -0.035 | -0.032 | -0.032 | 0.791 | 0.09 | 0.01 | 0.1 | 0.11 | 0.09 | 0.18 |
| 3 | 63649001 | rs7631976 | -0.423 | 1.083 | -1.825 | -0.068 | -0.018 | 0.042 | 0.042 | 0 | 0.43 | 0.1 | 0.34 | 0.62 | 0.41 | 0.55 |
| 3 | 63654362 | rs11718760 | 1.822 | 1.001 | -1.876 | -0.118 | -0.028 | -0.042 | -0.042 | 0.184 | 0.17 | 0.01 | 0.1 | 0.19 | 0.16 | 0.24 |

| | | | | | | | | | | | | | | | | |
|---|---|---|---|---|---|---|---|---|---|---|---|---|---|---|---|---|
| 3 | 63655742 | rs7626892 | 0.506 | 1.235 | -1.664 | -0.092 | 0.02 | -0.076 | -0.076 | 0 | 0.34 | 0.1 | 0.25 | 0.51 | 0.31 | 0.37 |
| 3 | 63657263 | rs11130955 | -0.823 | 1.303 | -1.287 | -0.131 | 0.015 | -0.04 | -0.04 | 0 | 0.66 | 0.52 | 0.56 | 0.77 | 0.63 | 0.76 |
| 3 | 63660258 | rs6784922 | -0.804 | 1.307 | -1.289 | -0.127 | 0.013 | -0.038 | -0.038 | 0 | 0.66 | 0.52 | 0.56 | 0.76 | 0.63 | 0.75 |
| 3 | 63662859 | rs2201582 | -0.408 | 1.392 | -1.408 | 0.223 | 0.008 | 0.053 | 0.053 | 0.187 | 0.49 | 0.32 | 0.43 | 0.64 | 0.47 | 0.55 |
| 3 | 63663215 | rs1390442 | 0.126 | 0.67 | -2.202 | 0.256 | -0.012 | 0.098 | 0.098 | 0 | 0.45 | 0.24 | 0.54 | 0.71 | 0.51 | 0.58 |
| 3 | 63665132 | rs6764278 | -0.122 | 0.841 | -1.773 | 0.647 | 0.025 | 0.095 | 0.095 | 0.301 | 0.55 | 0.25 | 0.62 | 0.72 | 0.6 | 0.62 |
| 3 | 63670140 | rs7629447 | -0.411 | 0.951 | -1.093 | 0.514 | -0.001 | 0.021 | 0.021 | 0 | 0.49 | 0.59 | 0.51 | 0.7 | 0.5 | 0.6 |
| 3 | 63671448 | rs13083429 | 0.09 | 0.886 | -1.518 | 0.124 | 0.004 | 0.057 | 0.057 | 0 | 0.17 | 0.07 | 0.12 | 0.03 | 0.17 | 0.16 |
| 3 | 63672506 | rs6785901 | -0.797 | 1.183 | -1.369 | 0.094 | -0.013 | 0.041 | 0.041 | 0 | 0.34 | 0.3 | 0.39 | 0.17 | 0.41 | 0.33 |
| 3 | 63672752 | rs6797570 | -1.018 | 1.673 | -1.985 | 0.125 | -0.086 | -0.065 | -0.065 | 0 | 0.87 | 0.99 | 0.84 | 0.8 | 0.74 | 0.87 |
| 3 | 63673795 | rs12108177 | 0.117 | 1.263 | -1.829 | 0.161 | 0.004 | 0.04 | 0.04 | 0 | 0.2 | 0.14 | 0.13 | 0.1 | 0.19 | 0.19 |
| 3 | 63674343 | rs6414560 | -0.978 | 0.842 | -1.659 | -0.059 | -0.026 | 0.106 | 0.106 | 0.501 | 0.57 | 0.87 | 0.61 | 0.75 | 0.53 | 0.68 |
| 3 | 63694931 | rs17069292 | 0.547 | 0.44 | -0.757 | 0.84 | -0.005 | 0.032 | 0.032 | 0 | 0.17 | 0.27 | 0.14 | 0.22 | 0.1 | 0.3 |
| 3 | 63728571 | rs7635168 | -0.423 | 0.505 | 0.068 | 1.055 | 0.005 | 0.002 | 0.002 | 0.697 | 0.37 | 0.41 | 0.45 | 0.42 | 0.41 | 0.37 |
| 3 | 63738745 | rs971575 | -1.221 | 0.635 | -0.088 | 0.882 | 0.016 | 0.018 | 0.018 | 0 | 0.57 | 0.67 | 0.64 | 0.72 | 0.65 | 0.5 |
| 3 | 63744733 | rs1221474 | -1.056 | 0.573 | -0.311 | 1.073 | -0.056 | -0.042 | -0.042 | 0 | 0.87 | 0.99 | 0.78 | 0.88 | 0.86 | 0.85 |
| 3 | 63749405 | rs13098181 | 0.883 | 0.919 | -0.003 | 1.418 | -0.001 | 0.051 | 0.051 | 0.211 | 0.17 | 0.27 | 0.22 | 0.28 | 0.17 | 0.15 |
| 3 | 63749444 | rs1472337 | -1.624 | 0.931 | -0.065 | 1.465 | 0.03 | 0.06 | 0.06 | 0.073 | 0.56 | 0.74 | 0.64 | 0.75 | 0.64 | 0.5 |
| 3 | 63752269 | rs17699988 | 0.966 | 0.966 | -0.358 | 1.4 | -0.031 | 0.029 | 0.029 | 0 | 0.15 | 0.08 | 0.22 | 0.11 | 0.23 | 0.17 |
| 3 | 63755029 | rs17069353 | 0.721 | 1.122 | -0.319 | 1.648 | -0.01 | -0.004 | -0.004 | 0 | 0.16 | 0.1 | 0.17 | 0.19 | 0.22 | 0.1 |
| 3 | 63762994 | rs10510908 | 0.625 | 1.015 | -0.366 | 1.52 | -0.001 | -0.003 | -0.003 | 0.364 | 0.14 | 0.11 | 0.21 | 0.28 | 0.17 | 0.15 |
| 3 | 63770138 | rs6804524 | -0.38 | 0.983 | -0.345 | 1.512 | -0.001 | 0.002 | 0.002 | 0.02 | 0.21 | 0.22 | 0.23 | 0.32 | 0.23 | 0.22 |
| 3 | 63772196 | rs17654983 | -0.515 | 1.221 | -0.364 | 1.49 | -0.002 | -0.002 | -0.002 | 0 | 0.22 | 0.21 | 0.27 | 0.32 | 0.27 | 0.23 |
| 3 | 63775595 | rs7650014 | 0.372 | 1.413 | -0.371 | 1.524 | -0.003 | -0.002 | -0.002 | 0 | 0.16 | 0.14 | 0.26 | 0.27 | 0.26 | 0.17 |
| 3 | 63781484 | rs2201103 | -0.324 | 1.566 | -0.323 | 1.679 | 0.003 | 0.022 | 0.022 | 0 | 0.84 | 0.77 | 0.88 | 0.64 | 0.79 | 0.87 |
| 3 | 63788094 | rs4688424 | -0.245 | 1.638 | -0.339 | 1.766 | 0.004 | 0.024 | 0.024 | 0 | 0.84 | 0.77 | 0.88 | 0.63 | 0.79 | 0.87 |
| 3 | 63792326 | rs6804141 | -1.086 | 1.16 | 0.101 | 1.464 | -0.026 | -0.033 | -0.033 | 0 | 0.07 | 0.28 | 0.05 | 0 | 0.02 | 0.03 |
| 3 | 63813654 | rs7614311 | 0.295 | 1.466 | -0.171 | 1.692 | -0.046 | -0.026 | -0.026 | 0 | 0.15 | 0.07 | 0.21 | 0.15 | 0.2 | 0.13 |
| 3 | 63820493 | rs2087704 | -0.637 | 1.378 | 0.045 | 1.675 | 0.006 | -0.001 | -0.001 | 1.255 | 0.19 | 0.29 | 0.24 | 0.15 | 0.22 | 0.16 |
| 3 | 63821632 | rs2637983 | -0.701 | 1.421 | 0.06 | 1.654 | 0.006 | -0.004 | -0.004 | 0 | 0.19 | 0.29 | 0.24 | 0.15 | 0.22 | 0.16 |
| 3 | 63827381 | rs13059603 | 0.173 | 1.723 | -0.078 | 1.701 | -0.006 | -0.006 | -0.006 | 0.42 | 0.8 | 0.77 | 0.86 | 0.64 | 0.74 | 0.86 |
| 3 | 63841395 | rs832188 | -0.492 | 1.567 | -0.033 | 1.768 | 0.004 | 0.022 | 0.022 | 0 | 0.84 | 0.77 | 0.89 | 0.64 | 0.78 | 0.87 |
| 3 | 63854445 | rs832193 | -0.577 | 1.656 | -0.124 | 1.861 | 0.001 | 0.023 | 0.023 | 0 | 0.85 | 0.82 | 0.92 | 0.64 | 0.83 | 0.87 |
| 3 | 63857071 | rs832194 | -0.539 | 1.656 | -0.128 | 1.861 | 0.003 | 0.035 | 0.035 | 0 | 0.85 | 0.77 | 0.91 | 0.64 | 0.83 | 0.85 |
| 3 | 63881549 | rs3774708 | -0.268 | 1.699 | -0.109 | 1.855 | -0.001 | 0 | 0 | 0 | 0.17 | 0.18 | 0.21 | 0.14 | 0.15 | 0.13 |
| 3 | 63893477 | rs6806547 | -1.616 | 1.998 | -0.593 | 1.931 | -0.075 | -0.068 | -0.068 | 0.354 | 0.86 | 0.99 | 0.81 | 0.74 | 0.77 | 0.8 |
| 3 | 63897215 | rs2292662 | -1.369 | 1.175 | 0.024 | 1.65 | 0.021 | 0.15 | 0.15 | 0 | 0.18 | 0.44 | 0.11 | 0.38 | 0.17 | 0.17 |
| 3 | 63924956 | rs17069507 | 0.668 | 1.317 | -0.267 | 1.697 | -0.034 | 0.006 | 0.006 | 0 | 0.06 | 0.01 | 0.06 | 0.27 | 0.13 | 0.09 |
| 3 | 63927739 | rs6796317 | -0.051 | 1.164 | -0.225 | 1.339 | -0.034 | -0.038 | -0.038 | 0.454 | 0.15 | 0.01 | 0.18 | 0.15 | 0.15 | 0.13 |
| 3 | 63976145 | rs2241821 | 0.654 | 0.967 | -0.386 | 1.333 | -0.01 | -0.046 | -0.046 | 0 | 0.09 | 0.01 | 0.15 | 0.14 | 0.09 | 0.1 |
| 3 | 63982082 | rs3774729 | -1.914 | 1.016 | -0.143 | 1.456 | 0 | 0.008 | 0.008 | 0.096 | 0.35 | 0.64 | 0.34 | 0.46 | 0.33 | 0.32 |
| 3 | 63986170 | rs13272 | -2.454 | 0.837 | -0.145 | 1.424 | 0.004 | 0.002 | 0.002 | 0.342 | 0.39 | 0.74 | 0.37 | 0.56 | 0.37 | 0.39 |
| 3 | 63990807 | rs11922435 | -1.974 | 1.032 | -0.044 | 1.018 | 0.002 | -0.005 | -0.005 | 0 | 0.35 | 0.72 | 0.34 | 0.47 | 0.31 | 0.31 |
| 3 | 63994505 | rs9865811 | -2 | 0.986 | -0.087 | 1.015 | -0.001 | 0.023 | 0.023 | 0.176 | 0.35 | 0.59 | 0.33 | 0.42 | 0.33 | 0.31 |
| 3 | 63997491 | rs2280162 | -0.645 | 1.05 | -0.084 | 1.01 | -0.001 | -0.002 | -0.002 | 0.123 | 0.18 | 0.22 | 0.24 | 0.14 | 0.17 | 0.16 |
| 3 | 63999322 | rs3816157 | -1.153 | 1.12 | 0.003 | 1.089 | 0.027 | 0.137 | 0.137 | 0 | 0.16 | 0.44 | 0.12 | 0.26 | 0.16 | 0.18 |
| 3 | 64009508 | rs40610 | -0.804 | 1.449 | -0.286 | 0.9 | 0.001 | 0.01 | 0.01 | 0 | 0.19 | 0.25 | 0.13 | 0.41 | 0.25 | 0.23 |
| 3 | 64012756 | rs13087186 | 1.164 | 1.003 | -0.789 | 0.165 | -0.001 | 0.081 | 0.081 | 0.181 | 0.22 | 0.09 | 0.16 | 0.02 | 0.24 | 0.11 |
| 3 | 64013249 | rs35837 | -0.409 | 1.065 | -0.696 | 0.25 | 0.001 | 0.018 | 0.018 | 0 | 0.2 | 0.28 | 0.12 | 0.38 | 0.2 | 0.22 |
| 3 | 64014239 | rs17344570 | 0.496 | 0.963 | -0.846 | 0.188 | -0.001 | 0 | 0 | 0.194 | 0.16 | 0.12 | 0.2 | 0.13 | 0.16 | 0.15 |
| 3 | 64020447 | rs6764606 | -0.259 | 0.89 | -0.719 | 0.087 | 0.001 | -0.006 | -0.006 | 1.202 | 0.44 | 0.43 | 0.41 | 0.15 | 0.44 | 0.27 |
| 3 | 64021228 | rs166229 | 1.323 | 0.743 | -0.687 | 0.161 | -0.003 | -0.01 | -0.01 | 0 | 0.38 | 0.28 | 0.47 | 0.54 | 0.36 | 0.51 |
| 3 | 64032223 | rs17069779 | 0.517 | 0.39 | -0.584 | 0.091 | -0.008 | 0.003 | 0.003 | 0 | 0.16 | 0.24 | 0.18 | 0.13 | 0.12 | 0.16 |
| 3 | 64033043 | rs42456 | 0.449 | 0.236 | -0.721 | 0.024 | -0.007 | -0.006 | -0.006 | 0 | 0.64 | 0.62 | 0.68 | 0.48 | 0.58 | 0.62 |
| 3 | 64043631 | rs704375 | 0.972 | 0.288 | -0.928 | 0.496 | 0.002 | -0.005 | -0.005 | 0 | 0.36 | 0.27 | 0.46 | 0.45 | 0.3 | 0.49 |
| 3 | 64066754 | rs35835 | 0.747 | 0.288 | -0.811 | 0.603 | 0.001 | -0.009 | -0.009 | 0.485 | 0.39 | 0.27 | 0.47 | 0.47 | 0.37 | 0.51 |
| 3 | 64069310 | rs26935 | 0.69 | 0.289 | -0.771 | 0.624 | 0.007 | -0.017 | -0.017 | 0 | 0.39 | 0.2 | 0.46 | 0.47 | 0.36 | 0.55 |
| 3 | 64072378 | rs1870919 | 0.462 | 0.261 | -1.077 | 0.64 | -0.038 | -0.032 | -0.032 | 1.819 | 0.19 | 0.06 | 0.22 | 0.2 | 0.22 | 0.15 |
| 3 | 64072872 | rs254855 | 1.328 | 0.328 | -1.118 | 0.619 | 0.013 | 0 | 0 | 5.674 | 0.37 | 0.2 | 0.44 | 0.38 | 0.32 | 0.51 |
| 3 | 64074794 | rs1007234 | -0.034 | -0.072 | -0.676 | 0.422 | -0.053 | 0 | 0 | 0 | 0.35 | 0.16 | 0.26 | 0.3 | 0.43 | 0.31 |
| 3 | 64078009 | rs153734 | 0.768 | 0.171 | -0.417 | 0.381 | 0.066 | 0.159 | 0.159 | 0.458 | 0.81 | 0.52 | 0.83 | 0.59 | 0.77 | 0.78 |
| 3 | 64080489 | rs14056 | 0.876 | 0.071 | -0.478 | 0.289 | -0.015 | 0.124 | 0.124 | 0 | 0.22 | 0.03 | 0.15 | 0.02 | 0.23 | 0.13 |
| 3 | 64082003 | rs12494731 | 0.922 | 0.065 | -0.451 | 0.294 | 0.004 | 0.122 | 0.122 | 0 | 0.21 | 0.04 | 0.13 | 0.01 | 0.19 | 0.13 |
| 3 | 64082964 | rs26939 | 1.489 | 0.464 | -0.269 | 0.499 | 0.002 | -0.016 | -0.016 | 0.021 | 0.36 | 0.2 | 0.45 | 0.45 | 0.34 | 0.51 |
| 3 | 64083417 | rs17069879 | -0.102 | 0.597 | -0.331 | 0.583 | 0.01 | 0.02 | 0.02 | 0 | 0.13 | 0.16 | 0.12 | 0.28 | 0.19 | 0.16 |
| 3 | 64084032 | rs26937 | 0.611 | 0.46 | -0.328 | 0.157 | 0.021 | 0.039 | 0.039 | 0.073 | 0.49 | 0.36 | 0.57 | 0.81 | 0.54 | 0.69 |
| 3 | 64092217 | rs7621907 | -0.259 | 1.079 | -0.413 | 0.785 | -0.065 | -0.065 | -0.065 | 0.393 | 0.1 | 0.01 | 0.14 | 0.3 | 0.17 | 0.15 |
| 3 | 64092863 | rs697263 | 1.299 | 0.444 | 0.18 | 0.243 | 0.01 | 0.063 | 0.063 | 6.016 | 0.93 | 0.36 | 0.94 | 0.91 | 0.99 | 1 |
| 3 | 64094417 | rs40158 | 0.5 | 0.461 | 0.01 | 0.07 | -0.007 | -0.017 | -0.017 | 0 | 0.69 | 0.39 | 0.69 | 0.78 | 0.7 | 0.81 |

| | | | | | | | | | | | | | | | | |
|---|---|---|---|---|---|---|---|---|---|---|---|---|---|---|---|---|
| 3 | 64095658 | rs36239 | 0.252 | 0.173 | -0.26 | 0.119 | -0.012 | -0.015 | -0.015 | 0.525 | 0.72 | 0.64 | 0.72 | 0.8 | 0.73 | 0.82 |
| 3 | 64097705 | rs153716 | 0.437 | 0.199 | -0.295 | 0.134 | -0.008 | -0.013 | -0.013 | 0 | 0.68 | 0.63 | 0.69 | 0.79 | 0.69 | 0.77 |
| 3 | 64097856 | rs11130957 | -0.717 | 0.17 | -1.018 | -0.24 | 0.005 | 0.226 | 0.226 | 0 | 0.47 | 0.13 | 0.41 | 0.14 | 0.49 | 0.34 |
| 3 | 64099197 | rs830354 | 0.83 | 0.485 | 0.004 | 0.392 | 0.087 | 0.234 | 0.234 | 0.148 | 0.87 | 0.49 | 0.85 | 0.69 | 0.87 | 0.87 |
| 3 | 64103135 | rs1096171 | 0.876 | 0.537 | -0.103 | 0.274 | 0.062 | 0.135 | 0.135 | 0.496 | 0.82 | 0.46 | 0.81 | 0.69 | 0.8 | 0.85 |
| 3 | 64104388 | rs1096170 | 1.201 | 0.185 | -0.368 | 0.215 | 0 | 0 | 0 | 0 | 0.54 | 0.4 | 0.57 | 0.57 | 0.55 | 0.63 |
| 3 | 64107568 | rs704384 | -0.509 | 0.394 | -0.291 | 0.227 | -0.007 | -0.002 | -0.002 | 0 | 0.89 | 0.91 | 0.9 | 1 | 0.83 | 0.97 |
| 3 | 64108273 | rs704387 | 0.403 | 0.358 | 0.017 | 0.234 | 0.024 | -0.041 | -0.041 | 0.087 | 0.86 | 0.73 | 0.89 | 0.97 | 0.83 | 0.96 |
| 3 | 64108737 | rs830010 | 1.092 | 0.083 | 0.205 | 0.1 | 0.007 | 0.018 | 0.018 | 0 | 0.65 | 0.49 | 0.61 | 0.59 | 0.61 | 0.71 |
| 3 | 64109244 | rs6791571 | 0.264 | 0.123 | 0.032 | 0.097 | -0.002 | 0.03 | 0.03 | 0 | 0.2 | 0.05 | 0.22 | 0.11 | 0.21 | 0.17 |
| 3 | 64110135 | rs1069977 | -0.559 | 0.217 | 0.097 | 0.089 | -0.001 | -0.001 | -0.001 | 0 | 0.89 | 0.89 | 0.9 | 0.98 | 0.83 | 0.96 |
| 3 | 64111719 | rs696009 | 1.034 | 0.026 | 0.04 | 0.081 | 0 | -0.002 | -0.002 | 0 | 0.58 | 0.56 | 0.6 | 0.58 | 0.57 | 0.67 |
| 3 | 64111897 | rs696010 | 1.273 | 0.082 | -0.066 | 0.119 | 0.003 | 0.003 | 0.003 | 0 | 0.53 | 0.35 | 0.56 | 0.56 | 0.54 | 0.64 |
| 3 | 64113047 | rs986150 | 1.122 | -0.618 | -0.728 | -0.925 | 0.012 | 0.02 | 0.02 | 0.601 | 0.18 | 0.1 | 0.09 | 0.04 | 0.13 | 0.14 |
| 3 | 64113795 | rs152295 | 0.092 | -0.249 | -0.834 | -0.772 | 0.016 | 0.069 | 0.069 | 0.01 | 0.58 | 0.71 | 0.77 | 0.94 | 0.64 | 0.77 |
| 3 | 64115156 | rs161658 | 0.772 | 0.133 | -0.821 | -0.274 | 0.017 | -0.055 | -0.055 | 0 | 0.38 | 0.51 | 0.53 | 0.84 | 0.41 | 0.57 |
| 3 | 64119198 | rs9873216 | -0.638 | -0.175 | -1.769 | 0.201 | -0.039 | -0.054 | -0.054 | 0 | 0.57 | 0.86 | 0.46 | 0.59 | 0.45 | 0.51 |
| 3 | 64120052 | rs9878228 | 1.581 | 0.005 | -1.388 | 0.057 | -0.002 | 0 | 0 | 0.152 | 0.26 | 0.23 | 0.29 | 0.46 | 0.33 | 0.33 |
| 3 | 64121290 | rs1835898 | -0.639 | -0.107 | -1.127 | -0.234 | 0.01 | 0.019 | 0.019 | 0 | 0.48 | 0.32 | 0.39 | 0.43 | 0.43 | 0.47 |
| 3 | 64123647 | rs4688433 | 0.635 | -0.176 | -1.536 | -0.205 | 0.003 | -0.004 | -0.004 | 0.026 | 0.17 | 0.13 | 0.16 | 0.29 | 0.09 | 0.15 |
| 3 | 64125528 | rs696012 | 1.079 | -0.25 | -1.427 | -0.351 | -0.005 | 0.021 | 0.021 | 0 | 0.47 | 0.34 | 0.48 | 0.59 | 0.5 | 0.52 |
| 3 | 64126596 | rs697267 | 0.336 | -0.14 | -1.761 | -0.436 | 0.008 | -0.065 | -0.065 | 0 | 0.63 | 0.45 | 0.62 | 0.85 | 0.56 | 0.67 |
| 3 | 64127088 | rs696014 | 1.068 | 0.359 | -2.519 | 0.005 | 0.001 | 0.014 | 0.014 | 0.959 | 0.16 | 0.05 | 0.19 | 0.12 | 0.16 | 0.14 |
| 3 | 64130066 | rs697269 | 0.253 | 0.463 | -2.481 | 0.044 | 0.03 | 0.042 | 0.042 | 0 | 0.53 | 0.21 | 0.6 | 0.59 | 0.62 | 0.58 |
| 3 | 64130521 | rs697270 | -0.147 | 0.77 | -1.9 | 0.152 | -0.002 | 0.035 | 0.035 | 0.356 | 0.24 | 0.15 | 0.34 | 0.12 | 0.27 | 0.21 |
| 3 | 64134120 | rs11714038 | 0.206 | 0.976 | -1.702 | 0.56 | -0.038 | -0.038 | -0.038 | 0.049 | 0.08 | 0.01 | 0.1 | 0.11 | 0.13 | 0.09 |
| 3 | 64135500 | rs6445392 | 0.937 | 0.76 | -1.845 | 0.499 | -0.061 | -0.087 | -0.087 | 0 | 0.32 | 0.12 | 0.46 | 0.48 | 0.38 | 0.37 |
| 3 | 64136428 | rs1078398 | -0.466 | 0.471 | -1.099 | 0.563 | -0.001 | 0 | 0 | 0 | 0.36 | 0.34 | 0.36 | 0.39 | 0.34 | 0.33 |
| 3 | 64137371 | rs753562 | 0.685 | 0.502 | -0.715 | 0.699 | 0.034 | 0.006 | 0.006 | 0 | 0.51 | 0.3 | 0.53 | 0.52 | 0.54 | 0.54 |
| 3 | 64137693 | rs929701 | 0.213 | 0.337 | -0.707 | 0.262 | -0.001 | 0.003 | 0.003 | 0 | 0.07 | 0.07 | 0.09 | 0.03 | 0.12 | 0.07 |
| 3 | 64139759 | rs11706882 | 0.387 | 0.05 | -0.849 | 0.316 | 0.003 | -0.008 | -0.008 | 0.377 | 0.16 | 0.03 | 0.09 | 0.22 | 0.14 | 0.14 |
| 3 | 64142859 | rs17720698 | 0.316 | 0.522 | -0.677 | -0.076 | 0.055 | -0.072 | -0.072 | 0 | 0.16 | 0.3 | 0.13 | 0.01 | 0.25 | 0.19 |
| 3 | 64146322 | rs17664202 | -0.712 | 0.517 | -0.519 | -0.108 | -0.008 | 0.007 | 0.007 | 0.021 | 0.35 | 0.33 | 0.22 | 0.24 | 0.41 | 0.33 |
| 3 | 64147086 | rs11707013 | -0.396 | 0.603 | -0.512 | -0.111 | -0.003 | 0.002 | 0.002 | 0.024 | 0.31 | 0.3 | 0.2 | 0.23 | 0.37 | 0.31 |
| 3 | 64150206 | rs184452 | 0.823 | 0.458 | 0.133 | -0.117 | -0.008 | 0.008 | 0.008 | 0.228 | 0.62 | 0.36 | 0.69 | 0.71 | 0.56 | 0.62 |
| 3 | 64154268 | rs39687 | 0.146 | -0.024 | -0.025 | 0.147 | -0.026 | 0.033 | 0.033 | 0 | 0.24 | 0.26 | 0.22 | 0.43 | 0.14 | 0.25 |
| 3 | 64154710 | rs13080962 | 1.447 | -0.018 | 0.007 | 0.144 | -0.06 | 0.057 | 0.057 | 0 | 0.42 | 0.15 | 0.49 | 0.3 | 0.44 | 0.38 |
| 3 | 64157031 | rs167787 | -1.251 | 0.013 | 0.21 | 0.16 | 0 | -0.001 | -0.001 | 0 | 0.53 | 0.52 | 0.4 | 0.65 | 0.5 | 0.56 |
| 3 | 64157235 | rs7637364 | 1.702 | -0.002 | 0.178 | 0.163 | -0.051 | 0.042 | 0.042 | 0.005 | 0.39 | 0.11 | 0.46 | 0.28 | 0.41 | 0.37 |
| 3 | 64157720 | rs11714978 | 1.179 | 0.023 | 0.171 | 0.199 | -0.026 | -0.039 | -0.039 | 0 | 0.14 | 0.01 | 0.09 | 0.24 | 0.14 | 0.13 |
| 3 | 64160053 | rs153740 | 0.261 | 0.023 | 0.255 | 0.194 | -0.006 | 0.007 | 0.007 | 0 | 0.23 | 0.23 | 0.21 | 0.42 | 0.14 | 0.21 |
| 3 | 64165625 | rs13068250 | -0.491 | 0.186 | 0.29 | -0.159 | -0.007 | 0.008 | 0.008 | 0.242 | 0.33 | 0.32 | 0.21 | 0.24 | 0.39 | 0.32 |
| 3 | 64165770 | rs17665101 | 0.351 | 0.275 | 0.389 | -0.091 | -0.012 | -0.002 | -0.002 | 0 | 0.18 | 0.18 | 0.12 | 0.01 | 0.24 | 0.19 |
| 3 | 64171754 | rs11917643 | 0.239 | 0.259 | 0.35 | -0.055 | 0.031 | -0.052 | -0.052 | 2.815 | 0.18 | 0.24 | 0.13 | 0.02 | 0.26 | 0.18 |
| 3 | 64171993 | rs153717 | -0.982 | 0.079 | 0.545 | 0.116 | 0 | 0 | 0 | 0.02 | 0.41 | 0.57 | 0.36 | 0.49 | 0.4 | 0.42 |
| 3 | 64174577 | rs103323 | 0.202 | 0.227 | 0.397 | -0.038 | 0.058 | -0.062 | -0.062 | 0 | 0.77 | 0.44 | 0.78 | 0.94 | 0.69 | 0.78 |
| 3 | 64177029 | rs161666 | -0.062 | -0.249 | 0.404 | 0.249 | -0.042 | 0.065 | 0.065 | 0 | 0.22 | 0.31 | 0.22 | 0.48 | 0.14 | 0.24 |
| 3 | 64184343 | rs696016 | 0.83 | -0.321 | 0.223 | 0.423 | 0.016 | -0.037 | -0.037 | 0.432 | 0.13 | 0.06 | 0.19 | 0.39 | 0.11 | 0.19 |
| 3 | 64184956 | rs696018 | -0.78 | -0.355 | 0.411 | 0.248 | -0.011 | -0.003 | -0.003 | 0 | 0.37 | 0.56 | 0.32 | 0.67 | 0.28 | 0.35 |
| 3 | 64191981 | rs695938 | 0.105 | -0.219 | 0.564 | 0.268 | -0.023 | 0.085 | 0.085 | 0 | 0.87 | 0.56 | 0.9 | 0.89 | 0.92 | 0.91 |
| 3 | 64194547 | rs10510915 | -0.076 | -0.106 | 0.06 | 0.005 | -0.019 | 0.04 | 0.04 | 0.39 | 0.32 | 0.25 | 0.21 | 0.23 | 0.35 | 0.31 |
| 3 | 64199364 | rs7610343 | 1.785 | -0.298 | -0.113 | 0.002 | -0.116 | 0.058 | 0.058 | 0 | 0.38 | 0.06 | 0.48 | 0.25 | 0.47 | 0.4 |
| 3 | 64200864 | rs2008299 | 1.783 | -0.286 | -0.133 | 0 | -0.118 | 0.046 | 0.046 | 0.348 | 0.38 | 0.04 | 0.47 | 0.26 | 0.45 | 0.4 |
| 3 | 64201702 | rs888405 | 0.546 | -0.312 | -0.049 | 0.014 | 0.011 | -0.012 | -0.012 | 0 | 0.7 | 0.3 | 0.69 | 0.49 | 0.81 | 0.73 |
| 3 | 64202707 | rs696023 | 0.871 | -0.289 | -0.016 | 0.058 | -0.021 | 0.051 | 0.051 | 1.981 | 0.53 | 0.17 | 0.67 | 0.64 | 0.55 | 0.6 |
| 3 | 64205099 | rs697286 | 0.917 | -0.294 | -0.023 | 0.058 | -0.027 | 0.056 | 0.056 | 1.996 | 0.53 | 0.13 | 0.67 | 0.65 | 0.56 | 0.6 |
| 3 | 64205735 | rs9877584 | 1.296 | -0.339 | 0.031 | 0.05 | 0.004 | -0.007 | -0.007 | 0 | 0.15 | 0.1 | 0.1 | 0.23 | 0.15 | 0.15 |
| 3 | 64206296 | rs696231 | 1.28 | -0.403 | -0.066 | 0.026 | 0.011 | -0.023 | -0.023 | 0.351 | 0.13 | 0.08 | 0.19 | 0.38 | 0.11 | 0.19 |
| 3 | 64206604 | rs696232 | 0.483 | -0.323 | 0.058 | 0.039 | -0.006 | 0.002 | 0.002 | 0.445 | 0.71 | 0.35 | 0.79 | 0.66 | 0.77 | 0.79 |
| 3 | 64208418 | rs17666116 | 0.391 | -0.173 | 0.007 | -0.129 | -0.025 | 0.026 | 0.026 | 0 | 0.22 | 0.16 | 0.17 | 0 | 0.28 | 0.24 |
| 3 | 64211102 | rs697287 | 0.298 | -0.478 | 0.22 | 0.008 | -0.032 | 0.075 | 0.075 | 0.513 | 0.18 | 0.31 | 0.21 | 0.38 | 0.11 | 0.2 |
| 3 | 64212196 | rs697291 | -0.069 | -0.432 | 0.222 | -0.209 | 0.005 | 0.05 | 0.05 | 0 | 0.87 | 0.57 | 0.89 | 0.9 | 0.88 | 0.92 |
| 3 | 64213414 | rs1564115 | 0.718 | -0.421 | 0.1 | -0.202 | 0.017 | -0.017 | -0.017 | 0.346 | 0.68 | 0.29 | 0.67 | 0.5 | 0.76 | 0.72 |
| 3 | 64215913 | rs697293 | 0.893 | -0.439 | -0.068 | -0.166 | 0.027 | -0.058 | -0.058 | 0 | 0.16 | 0.07 | 0.19 | 0.4 | 0.11 | 0.2 |
| 3 | 64225376 | rs7641821 | 0.873 | -0.394 | -0.016 | -0.207 | 0.079 | 0.099 | 0.099 | 0 | 0.54 | 0.23 | 0.59 | 0.27 | 0.65 | 0.57 |
| 3 | 64225898 | rs221991 | -0.354 | -0.44 | 0.083 | -0.229 | -0.012 | -0.01 | -0.01 | 1.773 | 0.18 | 0.43 | 0.16 | 0.1 | 0.17 | 0.12 |
| 3 | 64226281 | rs6801029 | 1.275 | -0.395 | -0.012 | -0.206 | 0.063 | 0.129 | 0.129 | 0 | 0.49 | 0.23 | 0.53 | 0.27 | 0.57 | 0.53 |
| 3 | 64227748 | rs3911777 | 0.822 | -0.33 | -0.074 | -0.161 | -0.007 | 0 | 0 | 0 | 0.18 | 0.23 | 0.11 | 0.23 | 0.15 | 0.16 |
| 3 | 64230433 | rs3911778 | 2.296 | -0.101 | -0.708 | -0.171 | -0.04 | 0.032 | 0.032 | 0 | 0.3 | 0.1 | 0.4 | 0.24 | 0.32 | 0.34 |

| | | | | | | | | | | | | | | | | |
|---|---|---|---|---|---|---|---|---|---|---|---|---|---|---|---|---|
| 3 | 64232901 | rs186417 | -0.944 | -0.23 | -0.671 | 0.148 | 0.023 | -0.064 | -0.064 | 0.021 | 0.85 | 0.94 | 0.81 | 0.58 | 0.89 | 0.8 |
| 3 | 64242106 | rs704404 | -0.172 | -0.188 | -0.471 | 0.046 | -0.008 | -0.011 | -0.011 | 0.072 | 0.23 | 0.33 | 0.17 | 0.24 | 0.21 | 0.18 |
| 3 | 64244118 | rs864089 | -1.186 | -0.099 | -0.291 | 0.242 | -0.01 | 0.02 | 0.02 | 0 | 0.16 | 0.37 | 0.16 | 0.13 | 0.17 | 0.12 |
| 3 | 64246321 | rs704408 | -0.487 | -0.322 | -0.321 | 0.441 | -0.013 | 0.082 | 0.082 | 0 | 0.12 | 0.35 | 0.09 | 0.14 | 0.08 | 0.09 |
| 3 | 64246829 | rs704409 | 0.696 | 0.13 | -1.192 | 0.496 | -0.008 | 0.03 | 0.03 | 0 | 0.44 | 0.35 | 0.5 | 0.28 | 0.45 | 0.4 |
| 3 | 64247327 | rs704410 | 0.595 | -0.258 | -1.122 | 1.264 | 0.007 | -0.006 | -0.006 | 0 | 0.84 | 0.82 | 0.77 | 0.54 | 0.88 | 0.79 |
| 3 | 64251390 | rs704416 | 0.594 | -0.06 | -1.376 | 1.231 | 0.006 | -0.017 | -0.017 | 0 | 0.85 | 0.9 | 0.78 | 0.53 | 0.87 | 0.79 |
| 3 | 64252424 | rs704417 | -0.269 | 0.019 | -2.033 | 0.102 | -0.045 | 0.244 | 0.244 | 1.986 | 0.52 | 0.91 | 0.57 | 0.88 | 0.49 | 0.59 |
| 3 | 64252803 | rs704418 | 1.084 | -0.57 | -1.383 | 0.349 | 0.014 | 0.066 | 0.066 | 0 | 0.78 | 0.93 | 0.86 | 0.98 | 0.81 | 0.84 |
| 3 | 64272923 | rs6803116 | -1.501 | -0.636 | 0.24 | 2 | -0.018 | 0.231 | 0.231 | 0.831 | 0.16 | 0.41 | 0.27 | 0.47 | 0.1 | 0.22 |
| 3 | 64276852 | rs10490792 | -1.188 | -0.591 | 0.468 | 1.738 | 0.025 | 0.043 | 0.043 | 0.329 | 0.09 | 0.32 | 0.09 | 0.08 | 0.07 | 0.1 |
| 3 | 64279458 | rs17404898 | -0.453 | -0.636 | 0.201 | 1.504 | -0.005 | 0.021 | 0.021 | 0 | 0.09 | 0.06 | 0.09 | 0.01 | 0.1 | 0.09 |
| 3 | 64290819 | rs1381807 | 1.297 | -0.788 | 1.359 | 1.753 | 0.003 | 0.001 | 0.001 | 0.638 | 0.05 | 0.11 | 0.15 | 0.04 | 0.04 | 0.05 |
| 3 | 64292545 | rs741015 | -1.83 | -1.124 | 1.608 | 1.785 | -0.008 | 0.021 | 0.021 | 0 | 0.17 | 0.19 | 0.26 | 0.27 | 0.12 | 0.24 |
| 3 | 64294923 | rs6445395 | -1.929 | -1.078 | 1.72 | 1.836 | -0.012 | 0.115 | 0.115 | 0.021 | 0.17 | 0.37 | 0.26 | 0.28 | 0.11 | 0.21 |
| 3 | 64314552 | rs853260 | -1.737 | -1.273 | 1.695 | 1.636 | -0.011 | -0.041 | -0.041 | 0.112 | 0.23 | 0.59 | 0.31 | 0.11 | 0.21 | 0.26 |
| 3 | 64321102 | rs853254 | -1.749 | -1.241 | 1.949 | 1.757 | 0.075 | 0.035 | 0.035 | 2.534 | 0.08 | 0.48 | 0.18 | 0.03 | 0.06 | 0.09 |
| 3 | 64326232 | rs6762476 | -1.335 | -1.303 | 1.63 | 1.886 | 0.014 | 0.146 | 0.146 | 0 | 0.06 | 0.21 | 0.2 | 0.21 | 0.04 | 0.07 |
| 3 | 64328025 | rs6791551 | -1.237 | -1.391 | 1.26 | 1.803 | 0.008 | 0.039 | 0.039 | 1.229 | 0.19 | 0.11 | 0.14 | 0.08 | 0.17 | 0.18 |
| 3 | 64329989 | rs17070461 | -1.396 | -1.377 | 1.202 | 1.817 | -0.016 | -0.047 | -0.047 | 0 | 0.21 | 0.08 | 0.33 | 0.28 | 0.22 | 0.26 |
| 3 | 64335910 | rs6445398 | -1.379 | -1.195 | 2.189 | 2.513 | -0.007 | 0.337 | 0.337 | 0.426 | 0.11 | 0.39 | 0.19 | 0.39 | 0.07 | 0.15 |
| 3 | 64339256 | rs7623833 | -2.02 | -1.383 | 2.197 | 2.422 | -0.015 | 0.144 | 0.144 | 0.004 | 0.14 | 0.42 | 0.37 | 0.3 | 0.1 | 0.16 |
| 3 | 64340527 | rs17070488 | -1.565 | -1.293 | 2.04 | 2.755 | 0.004 | 0.315 | 0.315 | 0.334 | 0.07 | 0.3 | 0.06 | 0.34 | 0.02 | 0.12 |
| 3 | 64341070 | rs9874741 | -1.13 | -1.337 | 1.907 | 2.125 | 0.011 | 0.025 | 0.025 | 0 | 0.19 | 0.36 | 0.2 | 0.23 | 0.21 | 0.24 |
| 3 | 64341819 | rs17070497 | -1.972 | -1.691 | 1.241 | 1.926 | 0.016 | 0.006 | 0.006 | 0 | 0.33 | 0.68 | 0.37 | 0.24 | 0.28 | 0.31 |
| 3 | 64344544 | rs733316 | -0.995 | -1.308 | 1.054 | 1.913 | 0 | -0.01 | -0.01 | 0.712 | 0.19 | 0.27 | 0.19 | 0.1 | 0.19 | 0.24 |
| 3 | 64348954 | rs1860819 | -1.155 | -1.474 | 0.104 | 2.024 | 0.007 | -0.009 | -0.009 | 0 | 0.14 | 0.12 | 0.24 | 0.71 | 0.12 | 0.21 |
| 3 | 64355896 | rs2052150 | -0.552 | -1.137 | 0.231 | 2.184 | -0.003 | 0.013 | 0.013 | 0 | 0.12 | 0.15 | 0.07 | 0.22 | 0.1 | 0.11 |
| 3 | 64356331 | rs929555 | -1.417 | -1.088 | -1.066 | 1.21 | -0.031 | 0.21 | 0.21 | 0.084 | 0.54 | 0.95 | 0.55 | 0.84 | 0.38 | 0.55 |
| 3 | 64372233 | rs17070648 | -0.746 | -1.134 | 0.075 | 1.702 | -0.023 | 0.041 | 0.041 | 0 | 0.28 | 0.52 | 0.21 | 0.43 | 0.22 | 0.25 |
| 3 | 64381219 | rs4688188 | -0.542 | -1.151 | 0.841 | 2.089 | -0.002 | 0.171 | 0.171 | 0 | 0.11 | 0.23 | 0.26 | 0.58 | 0.11 | 0.15 |
| 3 | 64385407 | rs1992565 | 0.226 | -0.759 | 0.858 | 1.494 | 0.003 | -0.054 | -0.054 | 1.047 | 0.79 | 0.67 | 0.85 | 0.92 | 0.8 | 0.79 |
| 3 | 64391051 | rs1867302 | 1.168 | -0.969 | 0.591 | 1.523 | 0.009 | -0.026 | -0.026 | 0 | 0.69 | 0.27 | 0.6 | 0.38 | 0.71 | 0.69 |
| 3 | 64393476 | rs1443518 | -0.03 | -0.98 | 0.919 | 1.045 | 0.023 | -0.136 | -0.136 | 0 | 0.81 | 0.57 | 0.73 | 0.99 | 0.8 | 0.79 |
| 3 | 64394241 | rs4688448 | -1.738 | -0.349 | 0.29 | 0.943 | -0.031 | -0.057 | -0.057 | 2.195 | 0.17 | 0.18 | 0.4 | 0.65 | 0.18 | 0.3 |
| 3 | 64396264 | rs9830368 | -0.282 | -0.231 | -0.173 | 0.183 | -0.055 | 0.054 | 0.054 | 0 | 0.16 | 0.08 | 0.12 | 0.04 | 0.19 | 0.13 |
| 3 | 64396723 | rs13087994 | -1.266 | -0.297 | 0.257 | 0.235 | 0 | 0 | 0 | 1.391 | 0.26 | 0.27 | 0.22 | 0.23 | 0.29 | 0.26 |
| 3 | 64396762 | rs6445401 | 0.376 | -0.59 | 0.308 | 0.151 | -0.026 | 0.069 | 0.069 | 0 | 0.75 | 0.52 | 0.55 | 0.38 | 0.79 | 0.75 |
| 3 | 64405996 | rs4688454 | -0.107 | -0.644 | 0.819 | -0.001 | 0.047 | -0.135 | -0.135 | 0 | 0.72 | 0.55 | 0.68 | 0.98 | 0.68 | 0.8 |
| 3 | 64407943 | rs12490122 | 1.511 | -0.769 | 0.631 | -0.111 | 0.003 | 0.083 | 0.083 | 0 | 0.51 | 0.37 | 0.36 | 0.15 | 0.51 | 0.46 |
| 3 | 64407972 | rs1822440 | -0.739 | -0.602 | 0.395 | 0.156 | 0.025 | -0.053 | -0.053 | 0 | 0.17 | 0.03 | 0.29 | 0.79 | 0.15 | 0.23 |
| 3 | 64422803 | rs6445403 | -1.074 | -0.361 | 0.795 | 0.168 | 0.025 | 0.062 | 0.062 | 3.352 | 0.07 | 0.32 | 0.08 | 0.07 | 0.04 | 0.16 |
| 3 | 64424271 | rs1443513 | -0.299 | -0.101 | 0.504 | -0.299 | -0.002 | -0.038 | -0.038 | 0.056 | 0.69 | 0.77 | 0.65 | 0.99 | 0.65 | 0.85 |
| 3 | 64427102 | rs894741 | 0.587 | -0.051 | 0.491 | -0.179 | -0.001 | -0.006 | -0.006 | 0 | 0.57 | 0.61 | 0.57 | 0.85 | 0.52 | 0.62 |
| 3 | 64435236 | rs6445404 | -0.168 | 0.139 | 0.354 | -0.078 | 0.013 | -0.07 | -0.07 | 0.364 | 0.68 | 0.71 | 0.64 | 0.99 | 0.62 | 0.81 |
| 3 | 64442663 | rs1443523 | 0.694 | 0.164 | -0.172 | 0.229 | 0 | 0.032 | 0.032 | 0 | 0.49 | 0.38 | 0.34 | 0.34 | 0.55 | 0.66 |
| 3 | 64451058 | rs11917164 | 0.115 | 0.488 | -0.407 | 0.685 | -0.014 | -0.012 | -0.012 | 0 | 0.3 | 0.25 | 0.26 | 0.62 | 0.4 | 0.31 |
| 3 | 64455564 | rs4327386 | -0.915 | 0.083 | -0.241 | 0.424 | -0.034 | -0.064 | -0.064 | 1.06 | 0.87 | 0.81 | 0.97 | 1 | 0.97 | 0.93 |
| 3 | 64457067 | rs11130964 | 0.728 | 0.231 | -0.321 | 0.779 | -0.059 | -0.003 | -0.003 | 0 | 0.23 | 0.09 | 0.22 | 0.17 | 0.34 | 0.29 |
| 3 | 64457663 | rs13061549 | -0.122 | 0.695 | -0.61 | 0.174 | 0.004 | -0.028 | -0.028 | 0.162 | 0.79 | 0.77 | 0.87 | 0.91 | 0.79 | 0.83 |
| 3 | 64458447 | rs17667547 | 0.928 | 0.777 | -0.89 | 0.32 | -0.093 | 0.042 | 0.042 | 0 | 0.22 | 0.05 | 0.21 | 0.09 | 0.33 | 0.24 |
| 3 | 64459007 | rs17070793 | 0.404 | 0.951 | -0.546 | 0.339 | 0.016 | -0.013 | -0.013 | 0 | 0.11 | 0.09 | 0.21 | 0.17 | 0.03 | 0.11 |
| 3 | 64468933 | rs4688466 | 0.185 | 1.872 | -0.353 | -0.01 | -0.001 | -0.012 | -0.012 | 0.8 | 0.38 | 0.42 | 0.42 | 0.14 | 0.38 | 0.4 |
| 3 | 64472243 | rs6787443 | 0.003 | 1.304 | -1.432 | 0.321 | -0.005 | 0.028 | 0.028 | 0 | 0.45 | 0.38 | 0.5 | 0.22 | 0.54 | 0.49 |
| 3 | 64475164 | rs4591504 | -0.657 | 1.926 | -1.764 | 0.936 | -0.076 | -0.032 | -0.032 | 0 | 0.2 | 0.06 | 0.25 | 0.3 | 0.44 | 0.19 |
| 3 | 64477004 | rs2371357 | -0.22 | 0.63 | -1.857 | 0.823 | 0.032 | 0.003 | 0.003 | 0.322 | 0.49 | 0.19 | 0.6 | 0.47 | 0.59 | 0.56 |
| 3 | 64478133 | rs2371358 | -0.072 | 0.603 | -2.006 | 0.699 | -0.034 | 0.08 | 0.08 | 0.017 | 0.57 | 0.81 | 0.67 | 0.73 | 0.48 | 0.74 |
| 3 | 64480785 | rs9871177 | -0.372 | 1.102 | -2.085 | 1.326 | -0.037 | -0.025 | -0.025 | 0 | 0.18 | 0.06 | 0.15 | 0.23 | 0.24 | 0.3 |
| 3 | 64480906 | rs17070862 | -0.261 | 0.954 | -2.274 | 1.777 | -0.001 | -0.025 | -0.025 | 0 | 0.14 | 0.12 | 0.2 | 0.32 | 0.18 | 0.29 |
| 3 | 64481661 | rs17070866 | -0.153 | 0.83 | -2.143 | 1.817 | 0.002 | -0.004 | -0.004 | 0 | 0.13 | 0.15 | 0.18 | 0.27 | 0.13 | 0.25 |
| 3 | 64482709 | rs4488822 | -0.475 | 0.954 | -1.95 | 1.527 | -0.023 | 0.011 | 0.011 | 0 | 0.25 | 0.14 | 0.17 | 0.2 | 0.31 | 0.28 |
| 3 | 64483115 | rs6768681 | -0.725 | 0.921 | -1.368 | 1.586 | -0.006 | 0.012 | 0.012 | 1.497 | 0.31 | 0.68 | 0.32 | 0.38 | 0.3 | 0.43 |
| 3 | 64485040 | rs13088425 | -0.185 | 1.026 | -1.392 | 1.742 | 0 | -0.009 | -0.009 | 0 | 0.14 | 0.13 | 0.2 | 0.31 | 0.17 | 0.26 |
| 3 | 64486041 | rs17724728 | 0.537 | 1.411 | -1.484 | 1.075 | -0.109 | 0.133 | 0.133 | 0 | 0.21 | 0.01 | 0.17 | 0 | 0.29 | 0.11 |
| 3 | 64499343 | rs2888015 | 1.165 | 0.4 | -0.614 | 2.234 | 0.014 | -0.05 | -0.05 | 0.707 | 0.77 | 0.22 | 0.75 | 0.61 | 0.84 | 0.77 |
| 3 | 64505542 | rs11713906 | 1.126 | 0.378 | -0.633 | 2.477 | 0.012 | -0.053 | -0.053 | 0.362 | 0.78 | 0.22 | 0.75 | 0.51 | 0.84 | 0.76 |
| 3 | 64509267 | rs13095481 | 1.151 | 0.377 | -0.419 | 2.427 | 0.004 | -0.015 | -0.015 | 0 | 0.78 | 0.27 | 0.84 | 0.57 | 0.84 | 0.78 |
| 3 | 64535687 | rs6445409 | 0.403 | 0.238 | 0.164 | 1.778 | -0.03 | -0.048 | -0.048 | 0.048 | 0.88 | 0.86 | 0.97 | 1 | 0.97 | 0.99 |
| 3 | 64539775 | rs9311896 | -1.241 | 0.974 | 0.499 | 2.32 | 0.107 | 0.297 | 0.297 | 0.349 | 0.09 | 0.32 | 0.09 | 0.3 | 0.13 | 0.17 |

| | | | | | | | | | | | | | | | | |
|---|---|---|---|---|---|---|---|---|---|---|---|---|---|---|---|---|
| 3 | 64541669 | rs6782835 | -0.762 | 0.962 | -0.492 | 1.646 | -0.089 | -0.006 | -0.006 | 0 | 0.34 | 0.17 | 0.42 | 0.37 | 0.47 | 0.22 |
| 3 | 64543563 | rs1036920 | 0.509 | 0.872 | 0.293 | 1.958 | 0.063 | 0.046 | 0.046 | 0 | 0.85 | 0.72 | 0.69 | 0.78 | 0.74 | 0.93 |
| 3 | 64543615 | rs1014640 | -0.89 | 0.368 | 0.217 | 2.205 | 0.021 | -0.049 | -0.049 | 0 | 0.28 | 0.19 | 0.27 | 0.55 | 0.22 | 0.46 |
| 3 | 64550958 | rs17071079 | -0.601 | 0.259 | 0.103 | 2.56 | -0.008 | -0.05 | -0.05 | 2.191 | 0.14 | 0.06 | 0.09 | 0.35 | 0.15 | 0.26 |
| 3 | 64553217 | rs17726803 | -1.542 | 0.059 | 0.315 | 2.3 | -0.036 | -0.002 | -0.002 | 0 | 0.3 | 0.17 | 0.34 | 0.32 | 0.4 | 0.38 |
| 3 | 64553452 | rs17726893 | -0.356 | 0.115 | 0.254 | 2.227 | -0.034 | 0.026 | 0.026 | 0 | 0.14 | 0.05 | 0.16 | 0 | 0.25 | 0.07 |
| 3 | 64556698 | rs6809120 | -0.444 | -0.125 | 0.939 | 2.283 | -0.045 | 0.271 | 0.271 | 0.02 | 0.15 | 0.38 | 0.33 | 0.5 | 0.06 | 0.15 |
| 3 | 64557634 | rs9833903 | -2.211 | -0.146 | 0.936 | 2.093 | -0.001 | 0.016 | 0.016 | 0.872 | 0.46 | 0.56 | 0.58 | 0.77 | 0.46 | 0.52 |
| 3 | 64560013 | rs9820942 | -1.271 | 0.07 | 1.045 | 2.113 | -0.016 | 0.007 | 0.007 | 0 | 0.26 | 0.16 | 0.34 | 0.26 | 0.32 | 0.21 |
| 3 | 64562105 | rs13318141 | -0.693 | -0.554 | 1.348 | 2.333 | -0.044 | 0.15 | 0.15 | 0 | 0.15 | 0.36 | 0.28 | 0.32 | 0.06 | 0.18 |
| 3 | 64565501 | rs7648540 | -0.696 | -0.55 | 1.12 | 2.297 | -0.018 | 0.013 | 0.013 | 0.054 | 0.14 | 0.2 | 0.18 | 0.26 | 0.06 | 0.15 |
| 3 | 64565516 | rs7638248 | -0.671 | -0.536 | 1.126 | 2.428 | -0.018 | 0.051 | 0.051 | 0 | 0.14 | 0.2 | 0.27 | 0.37 | 0.07 | 0.19 |
| 3 | 64575629 | rs4405909 | -1.201 | -0.642 | 1.396 | 2.16 | -0.024 | 0.036 | 0.036 | 1.925 | 0.21 | 0.63 | 0.24 | 0.31 | 0.14 | 0.29 |
| 3 | 64580169 | rs13320542 | -1.732 | -1.131 | 1.04 | 1.73 | 0.026 | 0.026 | 0.026 | 0.3 | 0.39 | 0.68 | 0.55 | 0.81 | 0.22 | 0.53 |
| 3 | 64594603 | rs4688489 | -1.248 | -1.079 | 1.423 | 1.597 | -0.038 | 0.029 | 0.029 | 0.076 | 0.31 | 0.58 | 0.25 | 0.57 | 0.16 | 0.48 |
| 3 | 64595816 | rs4356814 | -1.898 | -0.642 | 1.266 | 1.406 | -0.001 | -0.001 | -0.001 | 0.074 | 0.44 | 0.6 | 0.42 | 0.53 | 0.45 | 0.52 |
| 3 | 64599279 | rs6806842 | -0.296 | -0.399 | 0.591 | 1.353 | 0.024 | -0.023 | -0.023 | 0 | 0.16 | 0.2 | 0.18 | 0.04 | 0.3 | 0.13 |
| 3 | 64603654 | rs4340697 | 0.416 | -0.89 | 0.318 | 1.623 | 0.005 | -0.064 | -0.064 | 0.795 | 0.11 | 0 | 0.16 | 0.27 | 0.08 | 0.19 |
| 3 | 64621779 | rs7652820 | -1.351 | -1.066 | 0.683 | 1.489 | 0 | -0.003 | -0.003 | 0.045 | 0.35 | 0.67 | 0.36 | 0.53 | 0.17 | 0.45 |
| 3 | 64627274 | rs9831846 | -1.32 | -0.98 | 1.089 | 1.514 | -0.058 | 0.01 | 0.01 | 0.102 | 0.35 | 0.54 | 0.36 | 0.5 | 0.18 | 0.45 |
| 3 | 64627752 | rs7632802 | -0.98 | -1.004 | 1.233 | 1.332 | -0.07 | 0.037 | 0.037 | 0.064 | 0.22 | 0.55 | 0.33 | 0.28 | 0.11 | 0.25 |
| 3 | 64629698 | rs11916325 | -0.576 | 0.24 | 0.84 | 0.877 | 0.047 | 0.028 | 0.028 | 0 | 0.15 | 0.28 | 0.27 | 0.15 | 0.25 | 0.12 |
| 3 | 64632256 | rs9830450 | -0.936 | 0.137 | 0.305 | 0.232 | -0.001 | -0.009 | -0.009 | 0 | 0.52 | 0.46 | 0.65 | 0.86 | 0.51 | 0.63 |
| 3 | 64632414 | rs9868005 | -0.796 | 0.068 | 0.29 | 0.314 | 0.002 | 0.109 | 0.109 | 0.013 | 0.51 | 0.32 | 0.64 | 0.82 | 0.51 | 0.6 |
| 3 | 64634340 | rs13095235 | 1.269 | -0.144 | -0.271 | -0.541 | -0.009 | 0.024 | 0.024 | 0.795 | 0.29 | 0.23 | 0.23 | 0.04 | 0.35 | 0.28 |
| 3 | 64634411 | rs9861153 | -0.002 | -0.018 | -0.817 | -0.454 | 0.028 | 0.023 | 0.023 | 0 | 0.19 | 0.3 | 0.3 | 0.22 | 0.3 | 0.22 |
| 3 | 64634578 | rs4371513 | 0.856 | 0.072 | -1.112 | -0.378 | 0.004 | -0.127 | -0.127 | 0.293 | 0.32 | 0.03 | 0.35 | 0.61 | 0.21 | 0.38 |
| 3 | 64634736 | rs4605539 | -0.954 | 0.407 | -0.254 | -0.15 | -0.015 | -0.091 | -0.091 | 0 | 0.82 | 0.55 | 0.88 | 0.91 | 0.86 | 0.91 |
| 3 | 64637823 | rs9853285 | -1.091 | 1.251 | 0.217 | 0.402 | 0.042 | 0.027 | 0.027 | 0 | 0.22 | 0.41 | 0.31 | 0.22 | 0.32 | 0.24 |
| 3 | 64641698 | rs7615771 | -0.36 | 1.568 | -0.006 | -0.244 | 0.006 | -0.066 | -0.066 | 0 | 0.25 | 0.33 | 0.2 | 0.08 | 0.3 | 0.17 |
| 3 | 64641906 | rs7615977 | 0.311 | 2.114 | 0.034 | -0.657 | 0.003 | -0.049 | -0.049 | 0.545 | 0.81 | 0.78 | 0.84 | 1 | 0.76 | 0.89 |
| 3 | 64642056 | rs9855230 | 0.564 | 1.878 | -0.315 | -0.432 | 0.017 | -0.033 | -0.033 | 0 | 0.64 | 0.41 | 0.63 | 0.88 | 0.6 | 0.73 |
| 3 | 64642313 | rs7646362 | 0.248 | 1.618 | -0.031 | -0.035 | 0.002 | -0.025 | -0.025 | 0 | 0.19 | 0.21 | 0.19 | 0.07 | 0.26 | 0.12 |
| 3 | 64642342 | rs7646449 | 0.292 | 1.625 | -0.029 | -0.032 | 0.002 | -0.025 | -0.025 | 0 | 0.19 | 0.21 | 0.18 | 0.07 | 0.26 | 0.12 |
| 3 | 64649194 | rs7635505 | -1.045 | 1.48 | 0.075 | 0.65 | 0 | 0 | 0 | 0 | 0.47 | 0.56 | 0.56 | 0.41 | 0.63 | 0.57 |
| 3 | 64650587 | rs13085838 | 0 | 2.198 | 0.096 | 0.861 | -0.042 | -0.031 | -0.031 | 0 | 0.16 | 0.1 | 0.32 | 0.27 | 0.31 | 0.26 |
| 3 | 64650996 | rs9836710 | -0.085 | 1.734 | 0.635 | 1.402 | 0 | 0.037 | 0.037 | 0.037 | 0.13 | 0.3 | 0.08 | 0.12 | 0.09 | 0.17 |
| 3 | 64655357 | rs6793277 | -0.065 | 1.662 | 0.589 | 1.236 | -0.013 | 0.025 | 0.025 | 0 | 0.22 | 0.52 | 0.23 | 0.2 | 0.23 | 0.21 |
| 3 | 64662383 | rs4505693 | -1.136 | 1.476 | -0.141 | 1.034 | 0.013 | 0.012 | 0.012 | 0.463 | 0.47 | 0.6 | 0.75 | 0.57 | 0.64 | 0.6 |
| 3 | 64664314 | rs12490804 | -0.574 | 2.256 | -0.573 | 1.203 | -0.09 | -0.053 | -0.053 | 0 | 0.2 | 0.09 | 0.33 | 0.3 | 0.34 | 0.28 |
| 3 | 64673815 | rs6789298 | 0.438 | 2.212 | -0.504 | 1.003 | -0.013 | 0.002 | 0.002 | 0 | 0.08 | 0.04 | 0.11 | 0.03 | 0.16 | 0.08 |
| 3 | 64674346 | rs4392413 | 0.398 | 3.401 | -0.458 | 0.711 | 0.022 | -0.016 | -0.016 | 0 | 0.87 | 0.9 | 0.87 | 1 | 0.67 | 0.96 |
| 3 | 64675254 | rs4234677 | -0.282 | 1.648 | -0.636 | 0.62 | -0.009 | -0.001 | -0.001 | 0.883 | 0.39 | 0.62 | 0.48 | 0.5 | 0.18 | 0.52 |
| 3 | 64676293 | rs4688500 | -0.992 | 1.637 | -0.612 | 0.603 | 0.086 | 0.059 | 0.059 | 0 | 0.51 | 0.77 | 0.61 | 0.53 | 0.37 | 0.61 |
| 3 | 64677505 | rs11929329 | 0.069 | 0.702 | -0.167 | 0.715 | -0.006 | 0.091 | 0.091 | 0.001 | 0.07 | 0.24 | 0.19 | 0.07 | 0.03 | 0.06 |
| 3 | 64677737 | rs10470707 | -0.307 | 0.973 | -0.932 | 0.547 | 0.086 | -0.022 | -0.022 | 0.143 | 0.4 | 0.75 | 0.5 | 0.5 | 0.17 | 0.52 |
| 3 | 64679365 | rs4234678 | 0.982 | 1.122 | -0.838 | 0.66 | 0.035 | 0.001 | 0.001 | 0.677 | 0.39 | 0.17 | 0.27 | 0.46 | 0.31 | 0.32 |
| 3 | 64679496 | rs4234679 | 0.981 | 1.184 | -1.425 | 0.684 | 0.036 | -0.015 | -0.015 | 0.254 | 0.39 | 0.13 | 0.26 | 0.47 | 0.32 | 0.35 |
| 3 | 64683150 | rs9859317 | 1.021 | 1.256 | -1.666 | 0.695 | 0.005 | -0.001 | -0.001 | 0 | 0.36 | 0.15 | 0.25 | 0.39 | 0.33 | 0.4 |
| 3 | 64684378 | rs9865029 | -1.003 | 1.069 | -1.665 | 0.59 | -0.041 | 0.021 | 0.021 | 0.108 | 0.63 | 0.84 | 0.71 | 0.62 | 0.59 | 0.59 |
| 3 | 64688063 | rs12638651 | -0.168 | 0.505 | -1.443 | 1.048 | 0.037 | -0.083 | -0.083 | 0 | 0.2 | 0.11 | 0.26 | 0.49 | 0.1 | 0.23 |
| 3 | 64692746 | rs12488554 | -0.257 | 1.49 | -1.021 | 0.88 | 0.008 | 0.022 | 0.022 | 0 | 0.09 | 0.1 | 0.25 | 0.16 | 0.22 | 0.13 |
| 3 | 64693100 | rs9828674 | -0.437 | 1.438 | -0.791 | 0.853 | 0.041 | 0.072 | 0.072 | 0 | 0.11 | 0.23 | 0.26 | 0.16 | 0.24 | 0.13 |
| 3 | 64697088 | rs9850029 | -0.61 | 0.993 | -0.976 | 0.359 | 0.006 | 0.048 | 0.048 | 0 | 0.5 | 0.34 | 0.42 | 0.23 | 0.54 | 0.41 |
| 3 | 64700425 | rs6772129 | 0.048 | 0.429 | -1.276 | 0.337 | 0.029 | 0.073 | 0.073 | 0 | 0.64 | 0.85 | 0.67 | 0.75 | 0.3 | 0.75 |
| 3 | 64703665 | rs4411878 | 1.026 | -0.2 | -1.369 | 0.437 | 0.119 | 0.139 | 0.139 | 0 | 0.46 | 0.21 | 0.2 | 0.22 | 0.22 | 0.48 |
| 3 | 64708114 | rs4132228 | 0.137 | 0.476 | -0.693 | 0.75 | 0.001 | -0.002 | -0.002 | 0 | 0.63 | 0.69 | 0.5 | 0.6 | 0.28 | 0.69 |
| 3 | 64710253 | rs918466 | -0.36 | 0.9 | -1.267 | 0.772 | -0.085 | -0.077 | -0.077 | 0.868 | 0.7 | 0.87 | 0.56 | 0.6 | 0.42 | 0.69 |
| 3 | 64714182 | rs17727064 | 0.171 | 0.956 | -1.261 | 0.206 | -0.092 | 0.041 | 0.041 | 0 | 0.29 | 0.13 | 0.28 | 0.25 | 0.56 | 0.22 |
| 3 | 64733440 | rs17727363 | -0.365 | 0.604 | -0.97 | 0.253 | -0.017 | 0.063 | 0.063 | 0.007 | 0.4 | 0.25 | 0.29 | 0.24 | 0.54 | 0.22 |
| 3 | 64735531 | rs2371766 | -0.491 | 0.671 | -0.239 | 0.346 | -0.001 | 0.037 | 0.037 | 0 | 0.49 | 0.59 | 0.67 | 0.82 | 0.51 | 0.49 |
| 3 | 64744052 | rs2194093 | 1.154 | -0.059 | -0.228 | 0.537 | 0 | -0.016 | -0.016 | 1.11 | 0.31 | 0.18 | 0.37 | 0.25 | 0.34 | 0.28 |
| 3 | 64750562 | rs17676681 | 0.316 | 0.061 | 0.621 | 0.727 | -0.012 | 0.022 | 0.022 | 0 | 0.35 | 0.25 | 0.23 | 0.12 | 0.43 | 0.26 |
| 3 | 64750744 | rs10510927 | 0.233 | 0.082 | 0.652 | 0.805 | -0.004 | 0 | 0 | 0 | 0.36 | 0.29 | 0.42 | 0.27 | 0.45 | 0.29 |
| 3 | 64751742 | rs17071395 | 2.446 | -0.011 | 0.522 | 0.916 | 0.011 | -0.019 | -0.019 | 0 | 0.08 | 0.01 | 0.03 | 0.09 | 0.05 | 0.03 |
| 3 | 64755337 | rs318725 | -0.83 | -0.004 | 0.834 | 0.882 | 0.029 | 0.021 | 0.021 | 0 | 0.53 | 0.71 | 0.55 | 0.67 | 0.57 | 0.45 |
| 3 | 64756158 | rs318726 | -0.644 | 0.024 | 0.714 | 0.895 | 0.008 | 0.006 | 0.006 | 0 | 0.51 | 0.47 | 0.52 | 0.67 | 0.57 | 0.44 |
| 3 | 64761884 | rs318738 | -0.734 | 0.045 | 0.617 | 0.729 | 0.001 | -0.013 | -0.013 | 0 | 0.52 | 0.45 | 0.7 | 0.83 | 0.58 | 0.49 |
| 3 | 64789111 | rs13075154 | -0.168 | 0.082 | 0.059 | 0.975 | -0.014 | 0.016 | 0.016 | 0 | 0.05 | 0.04 | 0.05 | 0 | 0.06 | 0.08 |

| | | | | | | | | | | | | | | | | |
|---|---|---|---|---|---|---|---|---|---|---|---|---|---|---|---|---|
| 3 | 64792082 | rs446047 | -0.197 | -0.081 | 0.251 | 0.84 | 0.006 | 0.015 | 0.015 | 0 | 0.4 | 0.55 | 0.46 | 0.61 | 0.45 | 0.41 |
| 3 | 64798527 | rs9881486 | -0.197 | 0.123 | -0.161 | 0.719 | -0.001 | -0.001 | -0.001 | 0 | 0.43 | 0.66 | 0.48 | 0.43 | 0.46 | 0.46 |
| 3 | 64801187 | rs998394 | -0.534 | 0.202 | -0.136 | 0.8 | 0.007 | 0.173 | 0.173 | 0 | 0.5 | 0.76 | 0.74 | 0.84 | 0.54 | 0.6 |
| 3 | 64820003 | rs13080734 | 1.725 | -0.101 | -0.517 | 0.647 | -0.01 | 0.015 | 0.015 | 0.33 | 0.08 | 0 | 0.03 | 0.05 | 0.06 | 0.02 |
| 3 | 64829480 | rs9811632 | -0.293 | 0.105 | -0.069 | 0.697 | -0.016 | -0.011 | -0.011 | 0 | 0.22 | 0.44 | 0.18 | 0.26 | 0.21 | 0.29 |
| 3 | 64830316 | rs318783 | 0.809 | 0.422 | -0.474 | 1.294 | 0.002 | 0.069 | 0.069 | 0 | 0.92 | 0.89 | 0.74 | 0.39 | 0.88 | 0.86 |
| 3 | 64832519 | rs1851671 | 0.109 | -0.086 | -0.698 | 1.33 | -0.016 | 0.013 | 0.013 | 0 | 0.09 | 0.04 | 0.22 | 0.47 | 0.1 | 0.09 |
| 3 | 64835861 | rs9847789 | 0.705 | -0.291 | -1.488 | 0.256 | -0.036 | 0.125 | 0.125 | 0 | 0.28 | 0.07 | 0.17 | 0.07 | 0.27 | 0.11 |
| 3 | 64842029 | rs11923135 | 0.262 | -0.961 | -1.128 | -0.051 | 0.006 | 0.013 | 0.013 | 0 | 0.45 | 0.37 | 0.39 | 0.23 | 0.34 | 0.34 |
| 3 | 64852147 | rs17071651 | 0.039 | -0.883 | -0.962 | -0.011 | -0.001 | -0.002 | -0.002 | 0 | 0.48 | 0.46 | 0.4 | 0.34 | 0.36 | 0.34 |
| 3 | 64857528 | rs2197116 | 0.961 | -0.569 | -1.558 | -0.899 | -0.069 | 0.092 | 0.092 | 0 | 0.28 | 0.11 | 0.24 | 0.04 | 0.49 | 0.31 |
| 3 | 64858993 | rs9830285 | 0.242 | -0.507 | -1.13 | -0.339 | 0.001 | 0.017 | 0.017 | 0 | 0.35 | 0.24 | 0.22 | 0.2 | 0.3 | 0.12 |
| 3 | 64861470 | rs793299 | 1.166 | -0.284 | -1.159 | 0.283 | -0.015 | -0.039 | -0.039 | 2.176 | 0.08 | 0.01 | 0.08 | 0.1 | 0.06 | 0.2 |
| 3 | 64867534 | rs704436 | 0.609 | -0.218 | -1.275 | 0.341 | -0.03 | -0.022 | -0.022 | 0 | 0.11 | 0.03 | 0.07 | 0.09 | 0.1 | 0.11 |
| 3 | 64867724 | rs6784720 | 0.26 | -1.084 | -0.432 | 0.756 | -0.026 | 0.078 | 0.078 | 0 | 0.15 | 0.3 | 0.27 | 0.28 | 0.09 | 0.15 |
| 3 | 64869857 | rs6791760 | -0.214 | -1.793 | -0.498 | 0.777 | -0.013 | 0.005 | 0.005 | 0 | 0.27 | 0.24 | 0.41 | 0.57 | 0.13 | 0.32 |
| 3 | 64874125 | rs697297 | -0.466 | -0.689 | -0.44 | 0.275 | -0.001 | 0 | 0 | 0.868 | 0.43 | 0.4 | 0.49 | 0.51 | 0.51 | 0.45 |
| 3 | 64878444 | rs704442 | -0.982 | -0.458 | -0.432 | 0.206 | 0.081 | -0.005 | -0.005 | 0 | 0.51 | 0.79 | 0.61 | 0.59 | 0.62 | 0.47 |
| 3 | 64885634 | rs11926307 | 0.097 | 0.303 | 0.047 | 0.026 | -0.013 | -0.038 | -0.038 | 0 | 0.07 | 0.21 | 0.05 | 0 | 0.02 | 0 |
| 3 | 64886335 | rs1036796 | -0.62 | 0.233 | -0.445 | 0.025 | 0.003 | 0.043 | 0.043 | 0.155 | 0.42 | 0.73 | 0.37 | 0.28 | 0.56 | 0.32 |
| 3 | 64891359 | rs704448 | -0.308 | 0.312 | -0.605 | -0.008 | -0.003 | 0.001 | 0.001 | 0 | 0.35 | 0.34 | 0.27 | 0.2 | 0.45 | 0.3 |
| 3 | 64900159 | rs793292 | -0.799 | -0.006 | -0.663 | 0.418 | 0.011 | 0.006 | 0.006 | 0 | 0.79 | 0.75 | 0.82 | 0.74 | 0.71 | 0.69 |
| 3 | 64902050 | rs7620511 | 0.164 | -0.657 | -1.078 | 0.394 | 0.026 | -0.03 | -0.03 | 4.555 | 0.32 | 0.2 | 0.35 | 0.41 | 0.25 | 0.28 |
| 3 | 64902828 | rs11710147 | 1.254 | -0.886 | -1.359 | -0.126 | -0.105 | 0.092 | 0.092 | 0.016 | 0.22 | 0.05 | 0.18 | 0.09 | 0.35 | 0.16 |
| 3 | 64905426 | rs1442023 | 0.424 | -1.057 | -1.308 | 0.169 | 0.014 | -0.01 | -0.01 | 0 | 0.28 | 0.21 | 0.34 | 0.32 | 0.24 | 0.28 |
| 3 | 64907284 | rs2280585 | -0.096 | -1.302 | -0.983 | 0.143 | -0.025 | -0.007 | -0.007 | 0 | 0.36 | 0.51 | 0.51 | 0.6 | 0.26 | 0.38 |
| 3 | 64907649 | rs793291 | -0.486 | -0.92 | -0.704 | 0.39 | -0.003 | -0.017 | -0.017 | 0.109 | 0.63 | 0.56 | 0.71 | 0.7 | 0.66 | 0.58 |
| 3 | 64907956 | rs7622698 | 0.592 | -0.744 | -0.78 | 0.906 | -0.003 | -0.022 | -0.022 | 1.291 | 0.11 | 0.06 | 0.07 | 0.1 | 0.11 | 0.12 |
| 3 | 64910843 | rs1851676 | 0.752 | -0.699 | -0.677 | 0.996 | -0.003 | 0.065 | 0.065 | 1.898 | 0.07 | 0.06 | 0.15 | 0.28 | 0.04 | 0.1 |
| 3 | 64916597 | rs728270 | -0.111 | -0.379 | -0.537 | 0.522 | 0.011 | 0.063 | 0.063 | 0 | 0.13 | 0.3 | 0.12 | 0.18 | 0.13 | 0.1 |
| 3 | 64930075 | rs704457 | 0 | -0.046 | -1.028 | 0.375 | -0.005 | 0.003 | 0.003 | 0 | 0.76 | 0.77 | 0.82 | 0.74 | 0.72 | 0.72 |
| 3 | 64936775 | rs34348747 | -0.259 | -1.06 | -0.967 | 0.621 | 0.009 | -0.014 | -0.014 | 0 | 0.18 | 0.13 | 0.27 | 0.52 | 0.12 | 0.12 |
| 3 | 64941064 | rs938826 | 1.033 | -1.47 | -0.478 | 0.193 | 0.028 | 0.007 | 0.007 | 0.77 | 0.62 | 0.25 | 0.48 | 0.39 | 0.74 | 0.78 |
| 3 | 64941146 | rs17071877 | 0.14 | -1.394 | -0.629 | 0.506 | 0.007 | -0.004 | -0.004 | 0 | 0.17 | 0.12 | 0.27 | 0.42 | 0.11 | 0.11 |
| 3 | 64941603 | rs1077953 | 0.764 | -1.439 | -0.641 | 0.943 | 0 | -0.001 | -0.001 | 0 | 0.09 | 0.04 | 0.1 | 0.16 | 0.08 | 0.02 |
| 3 | 64944033 | rs4688219 | 1.043 | -1.67 | -1.382 | -0.25 | -0.129 | -0.056 | -0.056 | 0 | 0.26 | 0.05 | 0.29 | 0.35 | 0.52 | 0.39 |
| 3 | 64945813 | rs1523450 | 1.138 | -1.788 | -0.187 | -0.928 | 0.061 | 0.001 | 0.001 | 0 | 0.08 | 0.23 | 0.21 | 0.08 | 0.13 | 0.03 |
| 3 | 64945871 | rs2244503 | -0.656 | -2.26 | -0.297 | -0.949 | -0.004 | -0.053 | -0.053 | 0 | 0.62 | 0.6 | 0.82 | 0.97 | 0.79 | 0.62 |
| 3 | 64947998 | rs12054188 | 0.76 | -2.031 | -0.611 | -0.575 | 0.017 | -0.017 | -0.017 | 12.586 | 0.07 | 0.02 | 0.02 | 0.12 | 0.02 | 0.09 |
| 3 | 64948958 | rs11924641 | 0.504 | -2.464 | -0.4 | -0.987 | 0.025 | -0.004 | -0.004 | 1.378 | 0.37 | 0.27 | 0.2 | 0.03 | 0.19 | 0.37 |
| 3 | 64952753 | rs1720545 | -0.281 | -2.646 | -0.188 | -1.023 | 0.018 | 0.015 | 0.015 | 0 | 0.46 | 0.41 | 0.37 | 0.2 | 0.27 | 0.46 |
| 3 | 64955186 | rs7623134 | 0.881 | -2.502 | -0.607 | -0.948 | 0.008 | 0.002 | 0.002 | 0 | 0.08 | 0.07 | 0.02 | 0.08 | 0.04 | 0.1 |
| 3 | 64958608 | rs1916629 | -0.767 | -2.624 | -0.306 | -1.042 | -0.068 | -0.147 | -0.147 | 0 | 0.67 | 0.53 | 0.83 | 0.97 | 0.86 | 0.67 |
| 3 | 64958963 | rs1829891 | 0.406 | -2.467 | -0.518 | -0.931 | 0.093 | 0.121 | 0.121 | 0 | 0.5 | 0.26 | 0.59 | 0.79 | 0.72 | 0.53 |
| 3 | 64961027 | rs1523460 | 0.904 | -2.424 | -0.484 | -0.897 | 0.002 | 0 | 0 | 0 | 0.08 | 0.08 | 0.03 | 0.09 | 0.04 | 0.1 |
| 3 | 64962522 | rs1851675 | -0.075 | -2.37 | -0.452 | -0.92 | 0.022 | 0.015 | 0.015 | 0 | 0.58 | 0.34 | 0.62 | 0.87 | 0.76 | 0.63 |
| 3 | 64962621 | rs1878633 | -0.075 | -2.37 | -0.452 | -0.92 | 0.022 | 0.015 | 0.015 | 0 | 0.58 | 0.34 | 0.62 | 0.87 | 0.76 | 0.63 |
| 3 | 64969450 | rs4688519 | -0.259 | -1.819 | -0.548 | -0.428 | 0.003 | 0.001 | 0.001 | 0 | 0.15 | 0.14 | 0.2 | 0.17 | 0.13 | 0.12 |
| 3 | 64970601 | rs10514722 | 0.512 | -1.809 | -0.568 | -0.391 | 0.006 | 0.001 | 0.001 | 0 | 0.08 | 0.06 | 0.02 | 0.09 | 0.04 | 0.1 |
| 3 | 64971331 | rs2678321 | 0.545 | -3.402 | -0.672 | -1.173 | 0.013 | 0.015 | 0.015 | 0.118 | 0.26 | 0.2 | 0.23 | 0.11 | 0.1 | 0.17 |
| 3 | 64985326 | rs9827112 | -0.578 | -3.029 | 0.006 | -1.119 | -0.001 | -0.035 | -0.035 | 0 | 0.32 | 0.46 | 0.26 | 0.29 | 0.34 | 0.39 |
| 3 | 64990398 | rs9825783 | -0.364 | -3.495 | 0.156 | -1.219 | -0.016 | -0.018 | -0.018 | 0 | 0.26 | 0.56 | 0.22 | 0.29 | 0.23 | 0.37 |
| 3 | 64992289 | rs938823 | 0.397 | -3.739 | 0.089 | -1.347 | 0.024 | 0.025 | 0.025 | 0.393 | 0.36 | 0.29 | 0.31 | 0.11 | 0.22 | 0.18 |
| 3 | 64993316 | rs9841542 | -0.411 | -3.466 | 0.249 | -1.491 | -0.006 | -0.022 | -0.022 | 0 | 0.27 | 0.43 | 0.23 | 0.29 | 0.28 | 0.39 |
| 3 | 64993427 | rs4688533 | -1.663 | -3.901 | -0.214 | -0.781 | 0.018 | -0.051 | -0.051 | 0 | 0.89 | 0.98 | 0.92 | 0.67 | 0.91 | 0.71 |
| 3 | 64993562 | rs9841977 | 0.226 | -3.874 | 0.002 | -0.96 | 0.037 | -0.01 | -0.01 | 0 | 0.19 | 0.49 | 0.22 | 0.21 | 0.24 | 0.3 |
| 3 | 64994295 | rs1983357 | 0.62 | -3.849 | -0.1 | -0.978 | 0.022 | 0.021 | 0.021 | 0.412 | 0.36 | 0.29 | 0.33 | 0.11 | 0.22 | 0.19 |
| 3 | 64995557 | rs9311929 | -1.065 | -3.633 | -0.242 | -0.696 | -0.014 | 0.003 | 0.003 | 0.033 | 0.63 | 0.74 | 0.53 | 0.42 | 0.49 | 0.56 |
| 3 | 65004205 | rs17072075 | 0.907 | -3.286 | -0.194 | -0.592 | 0 | -0.018 | -0.018 | 0 | 0.12 | 0.13 | 0.08 | 0.01 | 0.1 | 0.19 |
| 3 | 65005951 | rs13078878 | 1.227 | -3.354 | -0.146 | -0.589 | -0.002 | -0.005 | -0.005 | 0 | 0.1 | 0.09 | 0.08 | 0 | 0.14 | 0.02 |
| 3 | 65007135 | rs1517928 | -0.283 | -3.903 | -0.194 | -0.78 | 0.004 | 0.002 | 0.002 | 0.702 | 0.37 | 0.66 | 0.44 | 0.38 | 0.36 | 0.27 |
| 3 | 65010837 | rs6785581 | 0.428 | -3.902 | -0.05 | -0.849 | -0.046 | -0.094 | -0.094 | 0.159 | 0.3 | 0.46 | 0.31 | 0.11 | 0.21 | 0.16 |
| 3 | 65014778 | rs9863931 | 0.3 | -3.933 | -0.071 | -0.807 | -0.068 | -0.076 | -0.076 | 0 | 0.32 | 0.52 | 0.3 | 0.12 | 0.13 | 0.32 |
| 3 | 65018403 | rs1517931 | -0.549 | -3.654 | -0.603 | -0.743 | 0.003 | -0.015 | -0.015 | 0.192 | 0.32 | 0.47 | 0.26 | 0.3 | 0.36 | 0.28 |
| 3 | 65019427 | rs11715854 | 1.121 | -3.297 | -0.957 | -0.578 | -0.047 | -0.035 | -0.035 | 0 | 0.12 | 0.01 | 0.14 | 0.16 | 0.15 | 0.08 |
| 3 | 65019442 | rs13071619 | 1.417 | -3.84 | -1.382 | -0.661 | -0.151 | -0.067 | -0.067 | 0.099 | 0.3 | 0.03 | 0.25 | 0.33 | 0.47 | 0.44 |
| 3 | 65020796 | rs11717899 | -0.029 | -3.259 | -0.637 | -0.656 | 0.003 | 0.002 | 0.002 | 0 | 0.19 | 0.24 | 0.19 | 0.27 | 0.27 | 0.12 |
| 3 | 65021946 | rs9873203 | -0.157 | -3.139 | -0.937 | -0.609 | -0.007 | -0.006 | -0.006 | 2.166 | 0.39 | 0.56 | 0.42 | 0.38 | 0.33 | 0.27 |
| 3 | 65027043 | rs12054271 | -0.87 | -2.435 | -1.121 | 0.067 | 0.055 | 0.007 | 0.007 | 0 | 0.46 | 0.82 | 0.32 | 0.49 | 0.36 | 0.41 |

| | | | | | | | | | | | | | | | | |
|---|---|---|---|---|---|---|---|---|---|---|---|---|---|---|---|---|
| 3 | 65033653 | rs12631821 | 0.627 | -2.43 | -1.189 | 0.067 | -0.04 | -0.048 | -0.048 | 0 | 0.15 | 0.05 | 0.17 | 0.31 | 0.21 | 0.15 |
| 3 | 65034421 | rs9834596 | 0.264 | -2.891 | -0.683 | -0.219 | 0.05 | 0.038 | 0.038 | 0 | 0.69 | 0.2 | 0.84 | 0.78 | 0.85 | 0.75 |
| 3 | 65038985 | rs7433057 | 0.41 | -2.51 | 0.041 | 0.012 | 0.001 | 0.12 | 0.12 | 0 | 0.91 | 0.67 | 0.95 | 0.79 | 0.95 | 0.92 |
| 3 | 65047603 | rs9875970 | -0.013 | -1.92 | -0.115 | -0.138 | -0.029 | 0.016 | 0.016 | 0 | 0.83 | 0.65 | 0.91 | 0.79 | 0.9 | 0.88 |
| 3 | 65048345 | rs6445448 | -0.047 | -1.92 | -0.114 | -0.137 | -0.028 | 0.016 | 0.016 | 0 | 0.83 | 0.64 | 0.91 | 0.79 | 0.9 | 0.87 |
| 3 | 65048510 | rs6445451 | 0.241 | -1.865 | -0.457 | -0.429 | -0.059 | -0.109 | -0.109 | 0 | 0.23 | 0.43 | 0.12 | 0.02 | 0.12 | 0.21 |
| 3 | 65054400 | rs6772379 | -0.482 | -1.483 | -1.101 | 0.375 | 0.007 | -0.065 | -0.065 | 0 | 0.24 | 0.13 | 0.15 | 0.57 | 0.22 | 0.31 |
| 3 | 65057623 | rs1482599 | -1.011 | -2.215 | -0.528 | 0.027 | 0.068 | 0.05 | 0.05 | 0 | 0.48 | 0.73 | 0.44 | 0.63 | 0.32 | 0.49 |
| 3 | 65064116 | rs17072226 | 1.128 | -1.944 | -0.588 | 0.742 | -0.011 | -0.025 | -0.025 | 0 | 0.08 | 0.06 | 0.08 | 0.31 | 0.13 | 0.12 |
| 3 | 65065491 | rs12636227 | 0.375 | -2.066 | -0.228 | 0.793 | -0.002 | -0.002 | -0.002 | 0 | 0.14 | 0.15 | 0.07 | 0.1 | 0.1 | 0.16 |
| 3 | 65067975 | rs1482597 | -1.349 | -2.628 | -1.054 | 0.281 | -0.077 | 0.012 | 0.012 | 0 | 0.7 | 0.91 | 0.53 | 0.73 | 0.44 | 0.69 |
| 3 | 65068148 | rs1482596 | -0.495 | -2.8 | -0.524 | -0.042 | 0.049 | 0.069 | 0.069 | 0.086 | 0.58 | 0.72 | 0.42 | 0.26 | 0.32 | 0.53 |
| 3 | 65072366 | rs17342237 | 1.905 | -1.979 | -0.477 | -0.055 | -0.001 | 0.003 | 0.003 | 0 | 0.08 | 0.05 | 0.06 | 0.06 | 0.08 | 0.01 |
| 3 | 65077476 | rs6797402 | 0.737 | -1.714 | -0.034 | -0.214 | 0.002 | -0.037 | -0.037 | 0 | 0.08 | 0.18 | 0.05 | 0 | 0.09 | 0.07 |
| 3 | 65077900 | rs6773617 | 0.774 | -2.309 | -0.033 | -0.165 | -0.033 | 0.014 | 0.014 | 0 | 0.25 | 0.33 | 0.3 | 0.24 | 0.14 | 0.19 |
| 3 | 65082924 | rs871630 | 0.46 | -2.076 | -0.491 | -0.494 | 0.047 | 0.049 | 0.049 | 0.818 | 0.38 | 0.25 | 0.17 | 0.09 | 0.2 | 0.33 |
| 3 | 65086196 | rs6809503 | -0.266 | -0.462 | -0.307 | -0.104 | 0.017 | -0.029 | -0.029 | 0.013 | 0.22 | 0.31 | 0.16 | 0.14 | 0.28 | 0.11 |
| 3 | 65090427 | rs9842777 | -0.401 | -0.461 | -0.301 | 0.158 | 0.017 | 0.062 | 0.062 | 0 | 0.23 | 0.33 | 0.16 | 0.33 | 0.29 | 0.15 |
| 3 | 65094803 | rs11706125 | 0.022 | 0.21 | -0.553 | -0.189 | -0.009 | 0.004 | 0.004 | 0 | 0.37 | 0.34 | 0.3 | 0.27 | 0.47 | 0.27 |
| 3 | 65099997 | rs2128406 | -0.165 | -0.281 | -0.741 | -0.405 | 0.013 | 0.028 | 0.028 | 0.354 | 0.51 | 0.37 | 0.36 | 0.29 | 0.54 | 0.39 |
| 3 | 65103850 | rs9790242 | 0.844 | -0.504 | -0.433 | -0.392 | -0.003 | 0.043 | 0.043 | 0.067 | 0.1 | 0.25 | 0.04 | 0.15 | 0.09 | 0.04 |
| 3 | 65103930 | rs9311933 | -0.577 | -0.67 | -1.008 | -0.496 | -0.094 | 0.043 | 0.043 | 0.849 | 0.59 | 0.87 | 0.35 | 0.35 | 0.51 | 0.5 |
| 3 | 65104040 | rs17403629 | 2.373 | -0.823 | -1.006 | -0.516 | 0.036 | 0.044 | 0.044 | 0.926 | 0.14 | 0.04 | 0.06 | 0.02 | 0.08 | 0.13 |
| 3 | 65104234 | rs9831020 | -0.079 | -0.26 | -0.654 | -0.211 | -0.037 | 0.046 | 0.046 | 0.915 | 0.18 | 0.56 | 0.1 | 0.25 | 0.13 | 0.16 |
| 3 | 65112055 | rs17403915 | 1.662 | -0.004 | -1.051 | -0.019 | -0.014 | -0.011 | -0.011 | 0.012 | 0.1 | 0.07 | 0.1 | 0.18 | 0.17 | 0.05 |
| 3 | 65116677 | rs9833943 | -1.151 | -0.085 | -1.136 | -0.116 | -0.061 | -0.116 | -0.116 | 0 | 0.69 | 0.94 | 0.45 | 0.54 | 0.67 | 0.55 |
| 3 | 65121265 | rs6445464 | -1.793 | 0.012 | -1.328 | 0.077 | -0.018 | -0.054 | -0.054 | 0.931 | 0.86 | 0.98 | 0.75 | 0.8 | 0.86 | 0.75 |
| 3 | 65121760 | rs1482611 | -0.525 | -0.119 | -0.905 | 0.024 | -0.003 | 0.04 | 0.04 | 1.685 | 0.67 | 0.8 | 0.44 | 0.2 | 0.66 | 0.51 |
| 3 | 65122313 | rs1482613 | 1.064 | 0.106 | -0.466 | -0.243 | 0.015 | 0.008 | 0.008 | 2.158 | 0.46 | 0.36 | 0.26 | 0.09 | 0.34 | 0.32 |
| 3 | 65123689 | rs13097168 | 1.534 | 0.285 | -0.495 | -0.043 | 0.095 | 0.043 | 0.043 | 0 | 0.07 | 0.12 | 0.11 | 0.16 | 0.19 | 0.06 |
| 3 | 65124098 | rs6792509 | 2.071 | 0.279 | -0.406 | -0.105 | 0.023 | -0.008 | -0.008 | 0 | 0.06 | 0.07 | 0.1 | 0.01 | 0.19 | 0.05 |
| 3 | 65128809 | rs9917782 | 1.235 | -0.179 | -0.352 | -0.118 | -0.007 | -0.001 | -0.001 | 0 | 0.39 | 0.36 | 0.18 | 0.44 | 0.2 | 0.35 |
| 3 | 65131072 | rs6767794 | -0.053 | -0.1 | -0.304 | -0.145 | -0.012 | 0.003 | 0.003 | 0 | 0.7 | 0.6 | 0.66 | 0.7 | 0.72 | 0.61 |
| 3 | 65133040 | rs9883021 | -0.134 | -0.068 | -0.303 | -0.037 | -0.014 | 0.013 | 0.013 | 2.293 | 0.62 | 0.74 | 0.34 | 0.35 | 0.47 | 0.5 |
| 3 | 65133697 | rs6807775 | 0.181 | -0.23 | -0.34 | -0.08 | 0.092 | -0.004 | -0.004 | 0 | 0.54 | 0.75 | 0.25 | 0.38 | 0.32 | 0.47 |
| 3 | 65137329 | rs1482588 | 1.117 | -0.096 | 0.251 | 0.096 | -0.016 | 0.043 | 0.043 | 0 | 0.08 | 0.31 | 0.05 | 0.1 | 0.05 | 0.12 |
| 3 | 65139561 | rs7647787 | 0.203 | 0.142 | 0.006 | 0.086 | 0.03 | 0.019 | 0.019 | 0 | 0.28 | 0.37 | 0.4 | 0.41 | 0.34 | 0.23 |
| 3 | 65148324 | rs6650907 | 0.544 | -0.11 | 0.106 | 0.236 | 0.049 | -0.003 | -0.003 | 0 | 0.15 | 0.21 | 0.16 | 0.15 | 0.22 | 0.07 |
| 3 | 65148368 | rs6650908 | -0.125 | -0.729 | -0.238 | 0.219 | 0.013 | 0.002 | 0.002 | 0 | 0.4 | 0.41 | 0.36 | 0.59 | 0.43 | 0.29 |
| 3 | 65148431 | rs959940 | -1.025 | -0.666 | -0.653 | 0.128 | -0.061 | 0.003 | 0.003 | 0 | 0.75 | 0.9 | 0.66 | 0.8 | 0.63 | 0.7 |
| 3 | 65148588 | rs959942 | 1.311 | -2.05 | -0.187 | 0.327 | -0.022 | 0.055 | 0.055 | 0 | 0.07 | 0.27 | 0.03 | 0.11 | 0.02 | 0.11 |
| 3 | 65148621 | rs10510933 | 0.881 | -1.643 | -0.925 | 0.292 | -0.001 | 0.019 | 0.019 | 0.351 | 0.23 | 0.1 | 0.19 | 0.25 | 0.2 | 0.19 |
| 3 | 65153035 | rs931233 | -0.266 | -1.037 | -0.697 | 0.113 | -0.002 | -0.001 | -0.001 | 0 | 0.66 | 0.69 | 0.46 | 0.64 | 0.44 | 0.73 |
| 3 | 65153699 | rs9311935 | 0.088 | -1.517 | -0.942 | -0.017 | 0.06 | 0.033 | 0.033 | 0.074 | 0.48 | 0.64 | 0.28 | 0.6 | 0.31 | 0.47 |
| 3 | 65154680 | rs1466506 | 0.662 | -2.077 | -0.951 | -0.697 | -0.015 | -0.038 | -0.038 | 0.016 | 0.28 | 0.31 | 0.13 | 0.06 | 0.18 | 0.26 |
| 3 | 65154788 | rs11922361 | 1.015 | -2.735 | -0.49 | -0.762 | -0.04 | 0.022 | 0.022 | 0 | 0.1 | 0.25 | 0.04 | 0.13 | 0.03 | 0.09 |
| 3 | 65156233 | rs17072493 | 0.915 | -2.817 | -0.434 | -0.825 | -0.01 | -0.03 | -0.03 | 0 | 0.28 | 0.31 | 0.13 | 0.05 | 0.18 | 0.26 |
| 3 | 65159808 | rs11914286 | 0.061 | -3.161 | -0.63 | -0.771 | -0.056 | 0.022 | 0.022 | 0.094 | 0.16 | 0.25 | 0.05 | 0.24 | 0.05 | 0.15 |
| 3 | 65160691 | rs9851742 | 0.969 | -3.071 | -0.687 | -0.768 | -0.021 | -0.047 | -0.047 | 0 | 0.28 | 0.33 | 0.13 | 0.05 | 0.18 | 0.26 |
| 3 | 65161935 | rs9877359 | 1.085 | -2.063 | -0.926 | -0.915 | -0.06 | -0.067 | -0.067 | 0 | 0.29 | 0.11 | 0.48 | 0.54 | 0.35 | 0.43 |
| 3 | 65162485 | rs7644210 | 1.014 | -1.743 | -0.899 | -0.832 | -0.014 | -0.016 | -0.016 | 0.019 | 0.29 | 0.32 | 0.17 | 0.11 | 0.19 | 0.26 |
| 3 | 65162793 | rs7632720 | -2.096 | -1.132 | -1.147 | -1.004 | -0.082 | 0.079 | 0.079 | 0 | 0.82 | 0.99 | 0.87 | 0.94 | 0.76 | 0.87 |
| 3 | 65163618 | rs9826120 | 0.969 | -1.036 | -0.917 | -0.836 | -0.007 | -0.01 | -0.01 | 0 | 0.29 | 0.31 | 0.16 | 0.11 | 0.19 | 0.26 |
| 3 | 65165334 | rs12330417 | 0.363 | -0.967 | -0.865 | -0.903 | -0.023 | -0.039 | -0.039 | 0 | 0.35 | 0.41 | 0.19 | 0.16 | 0.27 | 0.3 |
| 3 | 65171587 | rs9311936 | -1.062 | -0.502 | -0.632 | -0.766 | -0.055 | 0.05 | 0.05 | 0 | 0.76 | 0.86 | 0.76 | 0.85 | 0.66 | 0.81 |
| 3 | 65172403 | rs1586070 | -0.24 | -0.606 | -0.629 | -0.791 | -0.025 | 0.029 | 0.029 | 0 | 0.69 | 0.76 | 0.7 | 0.81 | 0.57 | 0.77 |
| 3 | 65172519 | rs9828000 | 0.342 | -0.918 | -0.724 | -0.935 | 0.008 | -0.029 | -0.029 | 0.048 | 0.36 | 0.45 | 0.57 | 0.76 | 0.39 | 0.54 |
| 3 | 65173235 | rs12489371 | 0.069 | -0.835 | -0.663 | -0.658 | -0.02 | -0.028 | -0.028 | 0 | 0.19 | 0.16 | 0.31 | 0.45 | 0.29 | 0.35 |
| 3 | 65173555 | rs9851004 | 2.553 | -1.088 | -0.642 | -0.731 | -0.013 | -0.03 | -0.03 | 0 | 0.17 | 0.2 | 0.08 | 0.04 | 0.11 | 0.11 |
| 3 | 65173884 | rs6771469 | -0.223 | -0.152 | -0.577 | -0.227 | 0.008 | -0.029 | -0.029 | 0 | 0.36 | 0.45 | 0.57 | 0.76 | 0.39 | 0.53 |
| 3 | 65177996 | rs9822235 | 1.053 | -0.001 | -0.81 | -0.146 | -0.039 | -0.048 | -0.048 | 3.059 | 0.13 | 0.1 | 0.25 | 0.4 | 0.21 | 0.31 |
| 3 | 65180001 | rs9815340 | -2.252 | 0.016 | -0.91 | -0.199 | -0.046 | 0.027 | 0.027 | 0 | 0.83 | 0.98 | 0.91 | 0.9 | 0.83 | 0.85 |
| 3 | 65184899 | rs9822094 | -0.624 | 0.022 | -0.901 | -0.18 | -0.116 | -0.139 | -0.139 | 0 | 0.71 | 0.91 | 0.68 | 0.5 | 0.63 | 0.56 |
| 3 | 65185849 | rs1994111 | 0.767 | -0.416 | -0.627 | -0.188 | -0.015 | 0.015 | 0.015 | 0 | 0.23 | 0.26 | 0.31 | 0.3 | 0.18 | 0.21 |
| 3 | 65193246 | rs2062133 | 0.073 | -0.577 | -0.459 | -0.456 | -0.001 | 0 | 0 | 0.977 | 0.37 | 0.36 | 0.59 | 0.7 | 0.43 | 0.55 |
| 3 | 65196123 | rs9852846 | 0.231 | -0.556 | -0.488 | -0.452 | -0.004 | 0.003 | 0.003 | 0.863 | 0.36 | 0.32 | 0.56 | 0.71 | 0.41 | 0.53 |
| 3 | 65201343 | rs9871283 | 0.443 | -1.042 | -0.398 | -0.507 | -0.024 | -0.065 | -0.065 | 0.044 | 0.36 | 0.39 | 0.14 | 0.05 | 0.2 | 0.25 |
| 3 | 65206583 | rs6768701 | 0.51 | -0.757 | -0.936 | -0.657 | 0.006 | -0.015 | -0.015 | 0.437 | 0.34 | 0.3 | 0.14 | 0.05 | 0.2 | 0.23 |
| 3 | 65211420 | rs17072569 | -0.007 | 0.641 | -0.988 | -0.322 | 0.037 | -0.011 | -0.011 | 0.017 | 0.09 | 0.17 | 0.18 | 0.08 | 0.22 | 0.04 |

| | | | | | | | | | | | | | | | | |
|---|---|---|---|---|---|---|---|---|---|---|---|---|---|---|---|---|
| 3 | 65212596 | rs781291 | -0.39 | 0.561 | -1.348 | -0.437 | -0.036 | 0.027 | 0.027 | 0 | 0.75 | 0.81 | 0.74 | 0.84 | 0.63 | 0.8 |
| 3 | 65214533 | rs9837180 | 0.431 | -1.256 | -1.455 | -0.071 | 0.031 | -0.04 | -0.04 | 0 | 0.25 | 0.18 | 0.16 | 0.43 | 0.13 | 0.37 |
| 3 | 65215804 | rs10510935 | 1.328 | -0.569 | -1.351 | -0.208 | -0.007 | 0 | 0 | 0 | 0.13 | 0.05 | 0.31 | 0.14 | 0.13 | 0.12 |
| 3 | 65219095 | rs1482590 | 0.617 | -1.121 | -1.001 | -0.335 | -0.053 | 0.019 | 0.019 | 0 | 0.37 | 0.47 | 0.25 | 0.41 | 0.2 | 0.4 |
| 3 | 65220481 | rs1109195 | -0.972 | -0.985 | -0.984 | -0.348 | 0.003 | 0.001 | 0.001 | 0 | 0.67 | 0.73 | 0.82 | 0.7 | 0.74 | 0.69 |
| 3 | 65222650 | rs2624863 | -0.097 | -0.649 | -0.966 | -0.383 | 0.012 | 0 | 0 | 0 | 0.51 | 0.6 | 0.5 | 0.48 | 0.55 | 0.54 |
| 3 | 65222847 | rs17348975 | 0.588 | -0.37 | -0.888 | -0.267 | -0.002 | -0.001 | -0.001 | 0 | 0.06 | 0.06 | 0.05 | 0.05 | 0.06 | 0.03 |
| 3 | 65225209 | rs17072610 | -0.466 | -0.428 | -0.856 | -0.217 | 0.001 | 0.039 | 0.039 | 0 | 0.09 | 0.19 | 0.08 | 0.19 | 0.09 | 0.07 |
| 3 | 65226990 | rs17349171 | -0.285 | 0.277 | -1.083 | -0.365 | -0.096 | 0.043 | 0.043 | 0 | 0.13 | 0.02 | 0.25 | 0.04 | 0.4 | 0.15 |
| 3 | 65227760 | rs1471504 | -0.287 | -0.177 | -1.337 | -0.491 | -0.119 | 0 | 0 | 0 | 0.72 | 0.94 | 0.64 | 0.79 | 0.41 | 0.71 |
| 3 | 65228519 | rs1383067 | 0.549 | -0.572 | -1.261 | -0.593 | -0.003 | -0.001 | -0.001 | 0 | 0.3 | 0.32 | 0.35 | 0.32 | 0.2 | 0.28 |
| 3 | 65228793 | rs1479955 | 1.986 | -0.742 | -1.335 | -0.683 | 0.022 | 0.018 | 0.018 | 0 | 0.16 | 0.09 | 0.05 | 0.01 | 0.07 | 0.13 |
| 3 | 65229598 | rs4688549 | 1.337 | -0.719 | -1.331 | -0.669 | 0.001 | 0.017 | 0.017 | 0 | 0.15 | 0.21 | 0.3 | 0.31 | 0.14 | 0.15 |
| 3 | 65231027 | rs17349380 | 2.207 | -0.675 | -1.338 | -0.668 | 0.042 | 0.057 | 0.057 | 0 | 0.14 | 0.04 | 0.05 | 0.01 | 0.06 | 0.13 |
| 3 | 65232324 | rs1479950 | 1.636 | -1.372 | -0.853 | -0.905 | -0.037 | -0.048 | -0.048 | 1.525 | 0.14 | 0.24 | 0.11 | 0.02 | 0.05 | 0.11 |
| 3 | 65232772 | rs4557104 | -0.757 | -1.255 | -0.873 | -0.885 | -0.071 | 0.013 | 0.013 | 0 | 0.72 | 0.82 | 0.66 | 0.77 | 0.48 | 0.72 |
| 3 | 65234047 | rs922577 | -0.004 | -1.373 | -0.793 | -0.877 | -0.016 | -0.01 | -0.01 | 1.317 | 0.44 | 0.53 | 0.46 | 0.34 | 0.26 | 0.38 |
| 3 | 65234067 | rs9866792 | 0.336 | -1.354 | -0.779 | -0.866 | -0.031 | 0.004 | 0.004 | 0 | 0.29 | 0.47 | 0.41 | 0.33 | 0.19 | 0.26 |
| 3 | 65234145 | rs7653116 | 0.296 | -1.187 | -0.789 | -0.781 | 0.02 | 0.027 | 0.027 | 0 | 0.53 | 0.6 | 0.5 | 0.71 | 0.35 | 0.57 |
| 3 | 65235610 | rs7612552 | 2.097 | -1.156 | -0.859 | -0.647 | -0.024 | 0.054 | 0.054 | 0.04 | 0.2 | 0.27 | 0.11 | 0.38 | 0.13 | 0.28 |
| 3 | 65237119 | rs781308 | -2.414 | -0.543 | -1.029 | -0.806 | -0.005 | -0.005 | -0.005 | 0 | 0.84 | 0.89 | 0.89 | 0.84 | 0.81 | 0.86 |
| 3 | 65237921 | rs781304 | 1.319 | -0.46 | -0.793 | -0.857 | -0.016 | -0.023 | -0.023 | 0.623 | 0.13 | 0.15 | 0.1 | 0.02 | 0.05 | 0.11 |
| 3 | 65241732 | rs1479948 | 0.398 | 0.173 | -0.863 | -0.704 | 0.006 | 0.005 | 0.005 | 0 | 0.09 | 0.08 | 0.21 | 0.06 | 0.28 | 0.12 |
| 3 | 65248757 | rs4688554 | 1.097 | -0.087 | -0.957 | -0.7 | -0.021 | -0.004 | -0.004 | 1.044 | 0.14 | 0.05 | 0.32 | 0.18 | 0.18 | 0.12 |
| 3 | 65249142 | rs6770043 | 0.496 | -0.529 | -0.991 | -0.934 | 0.005 | 0.049 | 0.049 | 0 | 0.3 | 0.12 | 0.39 | 0.18 | 0.26 | 0.26 |
| 3 | 65249702 | rs2620505 | -2.323 | -0.458 | -1.057 | -0.821 | -0.008 | -0.01 | -0.01 | 0.66 | 0.84 | 0.91 | 0.89 | 0.83 | 0.82 | 0.86 |
| 3 | 65250003 | rs2620504 | 0.07 | -0.379 | -0.663 | -0.805 | 0.027 | -0.044 | -0.044 | 0 | 0.24 | 0.28 | 0.33 | 0.06 | 0.4 | 0.25 |
| 3 | 65251311 | rs17415670 | 0.84 | -0.415 | -0.955 | -0.709 | -0.002 | 0.004 | 0.004 | 0 | 0.05 | 0.05 | 0.04 | 0.03 | 0.06 | 0.03 |
| 3 | 65253750 | rs936611 | 0.615 | -0.293 | -0.956 | -0.711 | -0.025 | 0.027 | 0.027 | 0 | 0.42 | 0.13 | 0.46 | 0.34 | 0.38 | 0.41 |
| 3 | 65254272 | rs1479945 | 0.37 | -0.311 | -0.87 | -0.611 | -0.036 | 0 | 0 | 0 | 0.36 | 0.57 | 0.23 | 0.64 | 0.24 | 0.37 |
| 3 | 65258935 | rs12638637 | 0.807 | 0.158 | -1.296 | -0.415 | -0.019 | 0.008 | 0.008 | 0 | 0.16 | 0.05 | 0.33 | 0.14 | 0.16 | 0.14 |
| 3 | 65262507 | rs1531642 | -0.062 | 0.467 | -1.207 | -0.467 | -0.028 | 0.017 | 0.017 | 0 | 0.22 | 0.16 | 0.33 | 0.02 | 0.43 | 0.24 |
| 3 | 65266072 | rs10510938 | -0.509 | -0.517 | -1.078 | 0.184 | 0.009 | -0.006 | -0.006 | 0.016 | 0.13 | 0.06 | 0.09 | 0.15 | 0.1 | 0.09 |
| 3 | 65271225 | rs9875900 | 0.123 | -1.149 | -0.784 | 0.134 | 0.006 | 0.002 | 0.002 | 0 | 0.6 | 0.61 | 0.72 | 0.37 | 0.68 | 0.48 |
| 3 | 65271254 | rs9875914 | -0.441 | -0.575 | -0.954 | -0.285 | 0.002 | 0.044 | 0.044 | 0 | 0.72 | 0.84 | 0.82 | 0.86 | 0.74 | 0.71 |
| 3 | 65271571 | rs9876518 | 0.202 | -0.229 | -0.457 | 0.205 | -0.029 | -0.034 | -0.034 | 0.937 | 0.1 | 0.22 | 0.09 | 0.01 | 0.05 | 0.11 |
| 3 | 65273299 | rs9848932 | -0.014 | 0.498 | -0.829 | 0.491 | 0 | 0.002 | 0.002 | 0 | 0.51 | 0.46 | 0.42 | 0.57 | 0.52 | 0.58 |
| 3 | 65274417 | rs17416597 | 0.716 | 0.82 | -1.252 | 0.21 | -0.06 | 0.028 | 0.028 | 0 | 0.24 | 0.09 | 0.3 | 0.16 | 0.4 | 0.19 |
| 3 | 65275536 | rs1158602 | 0.777 | 0.552 | -1.341 | 0.564 | -0.076 | -0.027 | -0.027 | 0.328 | 0.27 | 0.08 | 0.37 | 0.34 | 0.44 | 0.23 |
| 3 | 65276107 | rs1479944 | -0.409 | -0.465 | -1.001 | 0.024 | -0.015 | -0.022 | -0.022 | 0.151 | 0.65 | 0.59 | 0.63 | 0.79 | 0.75 | 0.6 |
| 3 | 65279590 | rs9809428 | -0.11 | -0.467 | -0.615 | 0.271 | 0.065 | 0.098 | 0.098 | 0 | 0.06 | 0.28 | 0.06 | 0.12 | 0.09 | 0.05 |
| 3 | 65280299 | rs13084546 | 0.157 | -1.279 | -0.773 | 0.058 | 0.054 | 0.041 | 0.041 | 0 | 0.39 | 0.32 | 0.15 | 0.21 | 0.19 | 0.25 |
| 3 | 65289469 | rs7638581 | 1.357 | -1.134 | -0.894 | 0.136 | -0.011 | -0.052 | -0.052 | 0 | 0.09 | 0.01 | 0.06 | 0.21 | 0.09 | 0.06 |
| 3 | 65289530 | rs1479942 | -0.66 | -1.26 | -0.882 | -0.047 | -0.001 | 0 | 0 | 0 | 0.45 | 0.61 | 0.28 | 0.39 | 0.37 | 0.44 |
| 3 | 65292489 | rs4688557 | -0.632 | -0.88 | -0.777 | 0.19 | 0.034 | 0.193 | 0.193 | 0.179 | 0.93 | 0.73 | 0.75 | 0.79 | 0.96 | 0.77 |
| 3 | 65295327 | rs13100540 | -0.706 | -0.54 | -0.789 | 0.076 | 0.071 | -0.056 | -0.056 | 0 | 0.19 | 0.41 | 0.21 | 0.11 | 0.28 | 0.17 |
| 3 | 65296020 | rs4688558 | 0.163 | -0.516 | -0.872 | -0.006 | 0.01 | -0.072 | -0.072 | 3.179 | 0.81 | 0.78 | 0.82 | 0.99 | 0.77 | 0.81 |
| 3 | 65296128 | rs13086188 | 0.796 | -0.441 | -1.448 | 0.31 | -0.086 | -0.102 | -0.102 | 0 | 0.35 | 0.05 | 0.45 | 0.42 | 0.58 | 0.34 |
| 3 | 65302322 | rs1479958 | -0.668 | -0.257 | -1.121 | 0.505 | -0.005 | 0.004 | 0.004 | 1.17 | 0.36 | 0.62 | 0.24 | 0.4 | 0.29 | 0.4 |
| 3 | 65303595 | rs1383072 | 0.553 | -0.021 | -1.932 | 0.394 | -0.064 | -0.064 | -0.064 | 0 | 0.4 | 0.05 | 0.48 | 0.42 | 0.58 | 0.34 |
| 3 | 65306966 | rs2371951 | 1.018 | 0.499 | -1.864 | 0.398 | -0.154 | -0.108 | -0.108 | 0 | 0.34 | 0.04 | 0.6 | 0.42 | 0.52 | 0.29 |
| 3 | 65311745 | rs13095497 | 0.888 | 0.572 | -1.813 | 0.45 | -0.147 | -0.102 | -0.102 | 0.741 | 0.36 | 0.04 | 0.61 | 0.42 | 0.52 | 0.32 |
| 3 | 65313459 | rs9847783 | 0.102 | 0.747 | -1.433 | 0.481 | 0.038 | 0.048 | 0.048 | 0 | 0.05 | 0.15 | 0.03 | 0.1 | 0.07 | 0.08 |
| 3 | 65315323 | rs4422246 | -0.566 | 0.566 | -1.357 | 0.449 | -0.011 | -0.006 | -0.006 | 0 | 0.65 | 0.53 | 0.6 | 0.65 | 0.71 | 0.65 |
| 3 | 65317569 | rs9832005 | -0.408 | 1.372 | -1.356 | 0.811 | 0.131 | 0.03 | 0.03 | 0.407 | 0.1 | 0.32 | 0.12 | 0.11 | 0.18 | 0.15 |
| 3 | 65322335 | rs17031792 | 0.473 | 1.03 | -2.555 | 1.028 | 0.015 | -0.032 | -0.032 | 0 | 0.08 | 0.01 | 0.04 | 0.23 | 0.05 | 0.06 |
| 3 | 65322448 | rs1842307 | 0.633 | 1.26 | -2.439 | 0.767 | -0.084 | -0.095 | -0.095 | 1.478 | 0.31 | 0.04 | 0.39 | 0.42 | 0.41 | 0.25 |
| 3 | 65326802 | rs6800292 | 0.543 | 0.768 | -2.388 | 0.763 | -0.113 | -0.099 | -0.099 | 0 | 0.32 | 0.04 | 0.39 | 0.42 | 0.43 | 0.27 |
| 3 | 65326882 | rs6764388 | 0.415 | 0.538 | -2.251 | 0.829 | -0.139 | -0.118 | -0.118 | 0 | 0.36 | 0.04 | 0.43 | 0.47 | 0.49 | 0.31 |
| 3 | 65328946 | rs12633114 | 0.183 | -0.107 | -1.892 | 0.511 | 0 | 0.025 | 0.025 | 0.507 | 0.54 | 0.05 | 0.69 | 0.68 | 0.61 | 0.73 |
| 3 | 65345028 | rs11708083 | 0.255 | -0.543 | -1.068 | 0.157 | 0.008 | 0.079 | 0.079 | 1.931 | 0.13 | 0.03 | 0.04 | 0 | 0.06 | 0.02 |
| 3 | 65362393 | rs9854780 | -1.065 | -0.471 | -0.146 | 1.545 | -0.004 | 0.401 | 0.401 | 0.057 | 0.07 | 0.3 | 0.27 | 0.45 | 0.03 | 0.25 |
| 3 | 65362797 | rs2371948 | -1.014 | -0.481 | -0.049 | 1.534 | 0.021 | 0.422 | 0.422 | 1.629 | 0.06 | 0.32 | 0.26 | 0.41 | 0.03 | 0.24 |
| 3 | 65367088 | rs3772193 | 0.115 | -0.48 | -1.569 | 0.572 | -0.124 | -0.141 | -0.141 | 0 | 0.37 | 0.02 | 0.32 | 0.58 | 0.43 | 0.38 |
| 3 | 65379294 | rs1016553 | 0.871 | -0.37 | -0.429 | 0.819 | 0.007 | 0.184 | 0.184 | 0 | 0.92 | 0.76 | 0.78 | 0.65 | 0.96 | 0.76 |
| 3 | 65383903 | rs13086093 | -0.238 | -1.056 | -0.83 | -0.012 | -0.012 | -0.051 | -0.051 | 0 | 0.35 | 0.41 | 0.25 | 0.13 | 0.28 | 0.41 |
| 3 | 65389993 | rs12486728 | -0.162 | -1.161 | -1.554 | 0.246 | 0.025 | -0.02 | -0.02 | 0 | 0.41 | 0.09 | 0.35 | 0.4 | 0.37 | 0.43 |
| 3 | 65394171 | rs9876177 | -0.542 | -1.62 | -1.505 | 0.131 | 0.023 | 0.028 | 0.028 | 0.987 | 0.47 | 0.26 | 0.43 | 0.54 | 0.43 | 0.55 |
| 3 | 65396401 | rs11131018 | -0.174 | -1.239 | -1.607 | -0.003 | -0.004 | -0.039 | -0.039 | 0 | 0.41 | 0.07 | 0.35 | 0.4 | 0.39 | 0.44 |

| | | | | | | | | | | | | | | | | |
|---|---|---|---|---|---|---|---|---|---|---|---|---|---|---|---|---|
| 3 | 65396481 | rs9869670 | -0.174 | -1.239 | -1.597 | -0.003 | 0.007 | -0.026 | -0.026 | 0.808 | 0.41 | 0.09 | 0.35 | 0.4 | 0.39 | 0.44 |
| 3 | 65399949 | rs11712543 | -0.468 | -1.309 | -1.231 | -0.07 | 0.033 | 0.004 | 0.004 | 0.868 | 0.47 | 0.24 | 0.36 | 0.44 | 0.42 | 0.49 |
| 3 | 65401156 | rs11922267 | 0.225 | -1.044 | -1.117 | -0.678 | 0.019 | 0.021 | 0.021 | 0 | 0.15 | 0.06 | 0.14 | 0 | 0.12 | 0.14 |
| 3 | 65402094 | rs7615097 | 0.003 | -1.151 | -0.312 | -0.672 | -0.005 | -0.029 | -0.029 | 0 | 0.13 | 0.36 | 0.11 | 0.05 | 0.1 | 0.1 |
| 3 | 65403341 | rs9878560 | -0.456 | -1.368 | -0.368 | -0.792 | -0.044 | -0.072 | -0.072 | 0 | 0.23 | 0.33 | 0.22 | 0.06 | 0.15 | 0.24 |
| 3 | 65404262 | rs13078213 | 0.848 | 0.316 | -1.38 | -1.095 | -0.084 | 0.131 | 0.131 | 0 | 0.28 | 0.07 | 0.21 | 0 | 0.43 | 0.12 |
| 3 | 65404615 | rs4688233 | -0.527 | 0.326 | -1.26 | -1.027 | -0.018 | 0.017 | 0.017 | 0 | 0.67 | 0.73 | 0.75 | 0.94 | 0.52 | 0.84 |
| 3 | 65405625 | rs12637665 | -0.372 | 0.074 | -1.423 | -0.657 | 0.004 | -0.019 | -0.019 | 0 | 0.2 | 0.14 | 0.39 | 0.69 | 0.19 | 0.52 |
| 3 | 65406021 | rs6776155 | -0.695 | 0.163 | -1.58 | -0.804 | 0.104 | 0.105 | 0.105 | 0.773 | 0.55 | 0.15 | 0.68 | 0.73 | 0.73 | 0.66 |
| 3 | 65406253 | rs6778911 | 0.768 | 0.288 | -1.299 | -0.692 | 0.05 | 0.047 | 0.047 | 0 | 0.93 | 0.8 | 0.94 | 0.92 | 0.9 | 0.9 |
| 3 | 65408072 | rs12493497 | -0.22 | 0.258 | -1.493 | -0.772 | 0.023 | 0.177 | 0.177 | 0 | 0.48 | 0.2 | 0.42 | 0.07 | 0.58 | 0.27 |
| 3 | 65408234 | rs13093148 | -0.545 | 0.599 | -1.294 | -0.216 | -0.003 | 0.039 | 0.039 | 0 | 0.2 | 0.09 | 0.2 | 0.05 | 0.16 | 0.19 |
| 3 | 65409511 | rs7428806 | -0.308 | 0.319 | -1.206 | 0.197 | 0.025 | -0.061 | -0.061 | 0 | 0.16 | 0.05 | 0.35 | 0.67 | 0.12 | 0.49 |
| 3 | 65414051 | rs4467382 | -0.054 | 1.092 | -0.759 | -0.996 | 0.001 | -0.021 | -0.021 | 0.681 | 0.64 | 0.61 | 0.62 | 0.91 | 0.47 | 0.68 |
| 3 | 65414719 | rs9837844 | 0.192 | 0.295 | -0.688 | -1.062 | -0.001 | 0.014 | 0.014 | 0 | 0.33 | 0.29 | 0.32 | 0.24 | 0.28 | 0.31 |
| 3 | 65419616 | rs4308214 | -0.223 | 0.499 | -0.289 | -1.402 | -0.003 | -0.011 | -0.011 | 0 | 0.55 | 0.46 | 0.68 | 0.77 | 0.62 | 0.58 |
| 3 | 65426299 | rs6798518 | -0.819 | 0.945 | -0.274 | -1.631 | -0.007 | -0.104 | -0.104 | 0.227 | 0.66 | 0.56 | 0.68 | 0.94 | 0.69 | 0.78 |
| 3 | 65430402 | rs6763617 | 0.393 | 1.078 | -0.438 | -1.472 | -0.037 | -0.04 | -0.04 | 1.117 | 0.21 | 0.1 | 0.36 | 0.7 | 0.31 | 0.37 |
| 3 | 65432841 | rs12635983 | 0.42 | 1.059 | -0.476 | -1.499 | -0.057 | -0.058 | -0.058 | 0 | 0.2 | 0.08 | 0.36 | 0.71 | 0.31 | 0.35 |
| 3 | 65439175 | rs9824280 | 0.903 | 0.884 | -0.17 | -1.764 | -0.006 | -0.063 | -0.063 | 0 | 0.31 | 0.33 | 0.25 | 0.03 | 0.25 | 0.21 |
| 3 | 65442578 | rs4618171 | -0.138 | 0.778 | -0.215 | -2.093 | 0.017 | 0.147 | 0.147 | 0 | 0.51 | 0.31 | 0.65 | 0.95 | 0.56 | 0.69 |
| 3 | 65461535 | rs7614228 | 0.765 | 0.8 | -0.192 | -2.059 | 0.005 | 0.025 | 0.025 | 0 | 0.32 | 0.23 | 0.26 | 0.19 | 0.24 | 0.32 |
| 3 | 65461675 | rs4688568 | -0.375 | 0.877 | -0.178 | -2.098 | 0.03 | 0.125 | 0.125 | 0 | 0.53 | 0.3 | 0.65 | 0.94 | 0.59 | 0.69 |
| 3 | 65467238 | rs1917519 | 1.514 | 0.799 | 0.051 | -2.099 | -0.001 | -0.037 | -0.037 | 0 | 0.29 | 0.24 | 0.23 | 0.03 | 0.24 | 0.21 |
| 3 | 65476044 | rs6775267 | 0.089 | 0.756 | 0.387 | -2.078 | -0.001 | -0.005 | -0.005 | 0 | 0.51 | 0.47 | 0.63 | 0.78 | 0.59 | 0.55 |
| 3 | 65483906 | rs2222584 | 0.858 | 0.879 | 0.069 | -1.947 | -0.071 | -0.066 | -0.066 | 0 | 0.2 | 0.05 | 0.37 | 0.75 | 0.33 | 0.31 |
| 3 | 65484875 | rs17430548 | 0.703 | 0.741 | 0.375 | -1.848 | 0.017 | -0.006 | -0.006 | 0 | 0.05 | 0.12 | 0.09 | 0.02 | 0.07 | 0.05 |
| 3 | 65489584 | rs17430709 | 0.564 | 0.629 | 0.358 | -1.846 | 0.011 | -0.007 | -0.007 | 0 | 0.06 | 0.12 | 0.09 | 0.02 | 0.06 | 0.05 |
| 3 | 65493125 | rs9851493 | 0.828 | 0.222 | 0.435 | -1.966 | -0.017 | -0.019 | -0.019 | 0 | 0.29 | 0.36 | 0.2 | 0.18 | 0.18 | 0.3 |
| 3 | 65501649 | rs1554664 | -0.648 | 0.453 | 0.134 | -1.782 | -0.002 | 0.003 | 0.003 | 0 | 0.85 | 0.85 | 0.91 | 0.97 | 0.84 | 0.87 |
| 3 | 65506761 | rs7612930 | 0.729 | 0.449 | 0.013 | -1.784 | -0.019 | 0.041 | 0.041 | 0 | 0.13 | 0.05 | 0.08 | 0.03 | 0.16 | 0.12 |
| 3 | 65510396 | rs1524972 | 0.865 | 0.227 | -0.161 | -1.884 | 0.022 | 0.051 | 0.051 | 0 | 0.29 | 0.1 | 0.29 | 0.18 | 0.2 | 0.28 |
| 3 | 65514229 | rs17353211 | 0.951 | 0.145 | 0.035 | -1.901 | 0.016 | 0.097 | 0.097 | 0.021 | 0.28 | 0.08 | 0.19 | 0.1 | 0.2 | 0.25 |
| 3 | 65514406 | rs1357830 | 0.325 | 0.198 | 0.253 | -1.779 | 0.003 | 0.01 | 0.01 | 0 | 0.45 | 0.4 | 0.38 | 0.18 | 0.43 | 0.29 |
| 3 | 65515613 | rs11711657 | 1.134 | 0.196 | 0.076 | -1.784 | -0.002 | -0.026 | -0.026 | 0 | 0.32 | 0.28 | 0.17 | 0.08 | 0.27 | 0.22 |
| 3 | 65521587 | rs7620620 | 0.734 | 0.375 | -0.524 | -1.697 | -0.017 | -0.058 | -0.058 | 0.363 | 0.37 | 0.5 | 0.26 | 0.12 | 0.3 | 0.26 |
| 3 | 65522392 | rs4688240 | 0.468 | 0.695 | -0.946 | -1.105 | -0.039 | -0.02 | -0.02 | 0 | 0.12 | 0.03 | 0.19 | 0.11 | 0.16 | 0.07 |
| 3 | 65530706 | rs11721065 | -0.084 | 0.824 | -0.899 | -1.276 | -0.046 | -0.016 | -0.016 | 0 | 0.25 | 0.22 | 0.37 | 0.72 | 0.41 | 0.42 |
| 3 | 65530833 | rs1524966 | -0.82 | -1.099 | -0.569 | -1.765 | -0.06 | -0.088 | -0.088 | 0.012 | 0.77 | 0.7 | 0.93 | 0.95 | 0.89 | 0.81 |
| 3 | 65535262 | rs6809997 | 0.475 | -0.798 | 0.358 | -1.567 | -0.009 | 0.029 | 0.029 | 1.019 | 0.08 | 0.21 | 0.15 | 0.11 | 0.06 | 0.05 |
| 3 | 65536289 | rs11916056 | 0.005 | -1.131 | 0.249 | -1.933 | -0.03 | -0.02 | -0.02 | 0 | 0.35 | 0.47 | 0.42 | 0.29 | 0.25 | 0.33 |
| 3 | 65543313 | rs1880524 | 0.315 | -0.512 | -0.029 | -2.047 | -0.045 | 0.004 | 0.004 | 0 | 0.3 | 0.25 | 0.46 | 0.7 | 0.45 | 0.47 |
| 3 | 65544252 | rs6787045 | -1.008 | -0.64 | 0.081 | -2.155 | -0.001 | -0.021 | -0.021 | 0 | 0.44 | 0.57 | 0.66 | 0.81 | 0.57 | 0.54 |
| 3 | 65547455 | rs4688577 | -1.004 | -0.642 | 0.049 | -2.088 | -0.005 | -0.077 | -0.077 | 0.359 | 0.61 | 0.62 | 0.79 | 0.95 | 0.65 | 0.74 |
| 3 | 65560167 | rs1917524 | 0.716 | -0.359 | -0.197 | -1.705 | -0.079 | -0.054 | -0.054 | 1.819 | 0.26 | 0.13 | 0.38 | 0.62 | 0.43 | 0.4 |
| 3 | 65562398 | rs12634999 | 1.301 | -0.506 | 0.036 | -1.641 | -0.006 | 0.042 | 0.042 | 0 | 0.06 | 0.14 | 0.16 | 0.15 | 0.06 | 0.06 |
| 3 | 65566806 | rs9821646 | 0.857 | -0.639 | -0.09 | -1.721 | -0.004 | 0.002 | 0.002 | 0 | 0.08 | 0.05 | 0.15 | 0.14 | 0.06 | 0.08 |
| 3 | 65567543 | rs13326823 | -0.808 | -0.889 | -0.037 | -1.997 | -0.002 | -0.042 | -0.042 | 0 | 0.56 | 0.52 | 0.78 | 0.93 | 0.66 | 0.7 |
| 3 | 65568609 | rs1880521 | -0.466 | -0.873 | -0.313 | -1.986 | 0.061 | 0.167 | 0.167 | 0 | 0.51 | 0.27 | 0.69 | 0.9 | 0.64 | 0.65 |
| 3 | 65572048 | rs1404544 | -1.649 | -0.875 | -0.148 | -1.996 | 0.005 | 0.019 | 0.019 | 0 | 0.7 | 0.8 | 0.84 | 0.95 | 0.73 | 0.77 |
| 3 | 65580498 | rs1917527 | 0.65 | -0.354 | -0.49 | -1.263 | -0.071 | -0.082 | -0.082 | 0 | 0.24 | 0.14 | 0.35 | 0.6 | 0.39 | 0.4 |
| 3 | 65581480 | rs4408825 | 0.999 | -0.552 | -0.294 | -1.519 | 0.002 | 0.011 | 0.011 | 0 | 0.21 | 0.19 | 0.19 | 0.13 | 0.17 | 0.18 |
| 3 | 65582041 | rs13326145 | 0.916 | -0.474 | -0.168 | -1.576 | -0.007 | 0.002 | 0.002 | 0.108 | 0.08 | 0.06 | 0.15 | 0.11 | 0.06 | 0.08 |
| 3 | 65582137 | rs12638335 | -1.01 | -0.325 | 0.089 | -1.715 | 0.008 | 0.016 | 0.016 | 0 | 0.59 | 0.68 | 0.79 | 0.9 | 0.66 | 0.71 |
| 3 | 65584681 | rs1524962 | 0.293 | 0.79 | -0.035 | -1.255 | 0.007 | -0.032 | -0.032 | 0 | 0.14 | 0.19 | 0.14 | 0 | 0.25 | 0.05 |
| 3 | 65587758 | rs7629574 | -0.382 | 0.797 | 0.04 | -1.23 | 0.036 | -0.055 | -0.055 | 0.093 | 0.17 | 0.31 | 0.2 | 0.02 | 0.27 | 0.1 |
| 3 | 65595897 | rs4450738 | 0.144 | -0.007 | -0.004 | -0.525 | -0.007 | 0.04 | 0.04 | 0 | 0.77 | 0.61 | 0.53 | 0.35 | 0.78 | 0.59 |
| 3 | 65597953 | rs4407359 | 0.145 | -0.206 | -0.229 | -0.869 | -0.014 | 0.036 | 0.036 | 0.921 | 0.4 | 0.34 | 0.33 | 0.26 | 0.5 | 0.34 |
| 3 | 65604551 | rs9880851 | 0.895 | -0.221 | -0.483 | -0.924 | 0.013 | 0.006 | 0.006 | 0.099 | 0.37 | 0.27 | 0.2 | 0.08 | 0.28 | 0.25 |
| 3 | 65608736 | rs7431819 | -0.536 | 0.02 | -0.343 | -0.37 | -0.001 | -0.002 | -0.002 | 0 | 0.82 | 0.87 | 0.64 | 0.36 | 0.82 | 0.62 |
| 3 | 65609303 | rs17073207 | 1.4 | -0.063 | -0.274 | -0.491 | -0.017 | -0.031 | -0.031 | 0 | 0.06 | 0.09 | 0.03 | 0 | 0.02 | 0.01 |
| 3 | 65611029 | rs11918964 | -0.284 | -0.053 | -0.444 | -0.391 | 0 | 0.002 | 0.002 | 0 | 0.24 | 0.18 | 0.39 | 0.61 | 0.25 | 0.41 |
| 3 | 65611042 | rs11926742 | -0.799 | -0.359 | -0.268 | -1.093 | -0.026 | -0.088 | -0.088 | 0 | 0.59 | 0.5 | 0.71 | 0.92 | 0.69 | 0.73 |
| 3 | 65611384 | rs13084817 | 0.914 | -0.293 | -0.378 | -1.455 | 0.01 | 0.002 | 0.002 | 0.066 | 0.38 | 0.22 | 0.22 | 0 | 0.35 | 0.26 |
| 3 | 65613051 | rs2140755 | 0.6 | 0.773 | -0.398 | -0.536 | -0.025 | -0.056 | -0.056 | 0 | 0.13 | 0.05 | 0.36 | 0.62 | 0.18 | 0.34 |
| 3 | 65620088 | rs9311947 | -0.496 | 1.091 | 0.015 | -1.135 | 0.042 | -0.104 | -0.104 | 0 | 0.17 | 0.4 | 0.17 | 0 | 0.24 | 0.06 |
| 3 | 65624729 | rs9859634 | -0.259 | 0.626 | -0.077 | -1.099 | -0.008 | -0.093 | -0.093 | 0.257 | 0.18 | 0.33 | 0.12 | 0 | 0.14 | 0.07 |
| 3 | 65636485 | rs7633294 | -0.265 | 0.002 | -0.477 | -0.938 | 0.001 | -0.001 | -0.001 | 0.097 | 0.67 | 0.75 | 0.52 | 0.36 | 0.71 | 0.45 |
| 3 | 65638248 | rs11923504 | 0.193 | 0.223 | -0.241 | -0.561 | -0.008 | 0.087 | 0.087 | 0 | 0.2 | 0.31 | 0.38 | 0.57 | 0.17 | 0.34 |

| | | | | | | | | | | | | | | | | |
|---|---|---|---|---|---|---|---|---|---|---|---|---|---|---|---|---|
| 3 | 65643473 | rs6794189 | 0.55 | 0.733 | 0.116 | -0.474 | 0.049 | -0.028 | -0.028 | 0 | 0.07 | 0.09 | 0.12 | 0 | 0.23 | 0.04 |
| 3 | 65643558 | rs6794294 | 0.641 | 0.413 | -0.186 | -0.829 | -0.005 | 0.008 | 0.008 | 0.052 | 0.41 | 0.21 | 0.61 | 0.63 | 0.6 | 0.56 |
| 3 | 65645149 | rs13065777 | 1.047 | 0.191 | -0.409 | -0.731 | 0.002 | -0.031 | -0.031 | 0 | 0.36 | 0.26 | 0.48 | 0.57 | 0.35 | 0.45 |
| 3 | 65652373 | rs7628120 | -0.208 | -0.08 | -0.45 | -0.959 | -0.018 | -0.093 | -0.093 | 0.362 | 0.6 | 0.6 | 0.73 | 0.97 | 0.69 | 0.7 |
| 3 | 65652762 | rs884067 | -0.396 | 0.311 | -1.081 | -0.714 | -0.006 | -0.006 | -0.006 | 0 | 0.25 | 0.27 | 0.23 | 0.35 | 0.32 | 0.26 |
| 3 | 65658302 | rs1499503 | 0.994 | 0.354 | -0.806 | -0.289 | 0.003 | -0.019 | -0.019 | 0 | 0.35 | 0.26 | 0.48 | 0.57 | 0.35 | 0.45 |
| 3 | 65663133 | rs17434189 | -0.205 | 0.559 | -0.313 | 0.093 | -0.012 | 0.002 | 0.002 | 0 | 0.12 | 0.06 | 0.09 | 0 | 0.2 | 0.09 |
| 3 | 65674722 | rs12635081 | -0.959 | -0.037 | 0.137 | 0.806 | -0.034 | 0.109 | 0.109 | 6.014 | 0.17 | 0.31 | 0.11 | 0.44 | 0.06 | 0.26 |
| 3 | 65687741 | rs6783171 | -0.726 | 0.266 | 0.172 | 0.86 | -0.008 | 0.021 | 0.021 | 0 | 0.17 | 0.18 | 0.09 | 0.25 | 0.11 | 0.23 |
| 3 | 65688382 | rs9311951 | -0.728 | 1.101 | 0.585 | 0.691 | 0.051 | 0.002 | 0.002 | 0 | 0.2 | 0.37 | 0.22 | 0.24 | 0.27 | 0.18 |
| 3 | 65690647 | rs4437178 | -0.651 | 0.623 | 0.75 | 0.652 | -0.019 | 0.062 | 0.062 | 0.004 | 0.18 | 0.34 | 0.1 | 0.26 | 0.12 | 0.19 |
| 3 | 65694905 | rs11131035 | -1.107 | 1.45 | 0.663 | 0.556 | 0.057 | 0.017 | 0.017 | 0 | 0.25 | 0.43 | 0.3 | 0.31 | 0.42 | 0.17 |
| 3 | 65699405 | rs11714649 | -1.077 | 1.007 | 0.66 | 0.354 | 0.006 | -0.002 | -0.002 | 0 | 0.41 | 0.65 | 0.45 | 0.62 | 0.34 | 0.47 |
| 3 | 65702885 | rs9815082 | -1.622 | 1.643 | 0.986 | 0.041 | 0.05 | -0.104 | -0.104 | 11.325 | 0.26 | 0.39 | 0.25 | 0 | 0.43 | 0.13 |
| 3 | 65704484 | rs9859419 | 0.091 | 1.853 | 1.064 | 0.154 | 0.065 | -0.061 | -0.061 | 0 | 0.11 | 0.27 | 0.12 | 0 | 0.2 | 0.05 |
| 3 | 65706257 | rs7610241 | -1.515 | 1.328 | 1.084 | 0.21 | 0.101 | -0.007 | -0.007 | 0 | 0.27 | 0.53 | 0.29 | 0.24 | 0.42 | 0.17 |
| 3 | 65708515 | rs9814511 | -1.103 | 1.183 | 1.164 | -0.461 | 0.084 | -0.106 | -0.106 | 0.438 | 0.19 | 0.39 | 0.2 | 0 | 0.32 | 0.09 |
| 3 | 65711119 | rs1499498 | -1.197 | 1.234 | 0.813 | -0.439 | 0.102 | -0.152 | -0.152 | 0.183 | 0.21 | 0.53 | 0.21 | 0 | 0.32 | 0.09 |
| 3 | 65717427 | rs6782678 | -1.157 | 1.269 | 0.234 | -0.455 | 0.052 | -0.119 | -0.119 | 6.89 | 0.24 | 0.59 | 0.22 | 0 | 0.35 | 0.11 |
| 3 | 65718229 | rs13092999 | 0.131 | 0.981 | -0.129 | -0.125 | -0.002 | -0.029 | -0.029 | 0.151 | 0.09 | 0.02 | 0.04 | 0.24 | 0.08 | 0.21 |
| 3 | 65718426 | rs9877160 | -0.848 | 1.3 | 0.122 | -0.692 | 0.06 | -0.142 | -0.142 | 0 | 0.19 | 0.47 | 0.16 | 0 | 0.24 | 0.06 |
| 3 | 65721857 | rs2062475 | -0.096 | 0.716 | -0.661 | -0.841 | 0 | 0.015 | 0.015 | 0 | 0.52 | 0.6 | 0.53 | 0.92 | 0.44 | 0.76 |
| 3 | 65725215 | rs4688597 | 0.046 | 0.63 | -1.121 | 0.174 | 0.01 | 0.006 | 0.006 | 0.623 | 0.42 | 0.18 | 0.29 | 0.33 | 0.4 | 0.3 |
| 3 | 65726434 | rs11131036 | -0.039 | 1.607 | -1.564 | 0.822 | -0.071 | -0.088 | -0.088 | 0.142 | 0.12 | 0.02 | 0.39 | 0.38 | 0.24 | 0.19 |
| 3 | 65729827 | rs17073625 | -0.012 | 1.822 | -0.844 | 0.246 | 0.091 | -0.02 | -0.02 | 0 | 0.05 | 0.18 | 0.06 | 0 | 0.11 | 0.02 |
| 3 | 65730801 | rs9861877 | 0.388 | 0.469 | -2.084 | -0.178 | 0.068 | 0.044 | 0.044 | 0.162 | 0.3 | 0.18 | 0.16 | 0.19 | 0.13 | 0.24 |
| 3 | 65732674 | rs4362751 | 0.773 | 0.429 | -2.221 | -0.148 | 0.072 | 0.033 | 0.033 | 0 | 0.26 | 0.13 | 0.14 | 0.17 | 0.12 | 0.2 |
| 3 | 65733351 | rs11707106 | -0.006 | 1.459 | -2.213 | 0.775 | -0.061 | -0.103 | -0.103 | 0 | 0.23 | 0.06 | 0.44 | 0.43 | 0.33 | 0.25 |
| 3 | 65733841 | rs869557 | 0.101 | 0.985 | -1.505 | 0.807 | 0.019 | -0.038 | -0.038 | 0 | 0.24 | 0.1 | 0.14 | 0.37 | 0.18 | 0.45 |
| 3 | 65745435 | rs264096 | 0.714 | 2.522 | -1.307 | 0.659 | -0.027 | -0.017 | -0.017 | 0 | 0.92 | 0.98 | 0.88 | 0.89 | 0.81 | 0.92 |
| 3 | 65745558 | rs17073710 | -0.296 | 2.519 | -1.394 | 1.1 | -0.001 | -0.018 | -0.018 | 0 | 0.17 | 0.16 | 0.39 | 0.47 | 0.17 | 0.18 |
| 3 | 65746365 | rs264083 | 1.226 | 1.421 | -0.594 | 0.567 | 0.004 | 0.004 | 0.004 | 1.242 | 0.64 | 0.37 | 0.43 | 0.43 | 0.61 | 0.74 |
| 3 | 65750652 | rs2372192 | -1.118 | 0.692 | 0.055 | 0.494 | -0.031 | 0.057 | 0.057 | 0.025 | 0.32 | 0.54 | 0.52 | 0.5 | 0.29 | 0.2 |
| 3 | 65753176 | rs704463 | 0.986 | 0.148 | -0.754 | -0.12 | 0.004 | -0.06 | -0.06 | 0.25 | 0.23 | 0.07 | 0.16 | 0.32 | 0.17 | 0.42 |
| 3 | 65753414 | rs814756 | -1.641 | 0.516 | -1.111 | -0.541 | -0.036 | -0.106 | -0.106 | 0.2 | 0.68 | 0.57 | 0.82 | 0.92 | 0.78 | 0.71 |
| 3 | 65757246 | rs17073748 | 0.664 | 0.524 | -1.044 | -0.594 | 0 | 0.004 | 0.004 | 0.54 | 0.17 | 0.19 | 0.44 | 0.51 | 0.21 | 0.17 |
| 3 | 65760472 | rs17073750 | 1.572 | 0.181 | -0.518 | -0.68 | 0.001 | -0.02 | -0.02 | 0 | 0.31 | 0.3 | 0.17 | 0.08 | 0.23 | 0.29 |
| 3 | 65762252 | rs984038 | -1.034 | 0.488 | -0.299 | -0.439 | 0.002 | 0.002 | 0.002 | 0 | 0.46 | 0.55 | 0.67 | 0.6 | 0.61 | 0.3 |
| 3 | 65762547 | rs1495453 | -0.489 | 0.535 | -0.402 | -0.38 | -0.015 | -0.006 | -0.006 | 0.102 | 0.37 | 0.33 | 0.61 | 0.6 | 0.53 | 0.29 |
| 3 | 65773011 | rs1390241 | 0.568 | 0.619 | -0.234 | -0.371 | 0.023 | 0.011 | 0.011 | 0.113 | 0.49 | 0.39 | 0.6 | 0.56 | 0.46 | 0.48 |
| 3 | 65774634 | rs2643738 | 0.79 | 0.799 | -0.887 | -0.481 | 0 | -0.05 | -0.05 | 0.929 | 0.26 | 0.1 | 0.17 | 0.32 | 0.2 | 0.38 |
| 3 | 65774722 | rs13063316 | 0.358 | 1.305 | -0.437 | -0.612 | 0.031 | -0.011 | -0.011 | 0 | 0.17 | 0.21 | 0.18 | 0.12 | 0.28 | 0.12 |
| 3 | 65775478 | rs9814172 | -0.812 | 0.809 | -0.059 | -0.405 | -0.011 | -0.056 | -0.056 | 0 | 0.12 | 0.19 | 0.08 | 0.01 | 0.08 | 0.03 |
| 3 | 65780102 | rs2649188 | -0.656 | 0.708 | -0.907 | -0.227 | -0.007 | 0.088 | 0.088 | 0 | 0.46 | 0.29 | 0.62 | 0.75 | 0.49 | 0.55 |
| 3 | 65780869 | rs4688602 | -0.977 | 0.985 | -0.718 | -0.172 | -0.008 | -0.013 | -0.013 | 3.372 | 0.24 | 0.22 | 0.48 | 0.43 | 0.32 | 0.22 |
| 3 | 65787275 | rs12631008 | -0.892 | 1.082 | -0.869 | -0.134 | -0.022 | -0.04 | -0.04 | 0 | 0.21 | 0.19 | 0.31 | 0.44 | 0.28 | 0.22 |
| 3 | 65793390 | rs4688605 | -0.863 | 1.03 | -0.337 | -0.083 | 0.052 | 0.165 | 0.165 | 0.325 | 0.18 | 0.34 | 0.27 | 0.42 | 0.25 | 0.19 |
| 3 | 65796628 | rs2643723 | -0.659 | 0.565 | -0.814 | -0.9 | -0.004 | 0.004 | 0.004 | 0.977 | 0.43 | 0.36 | 0.47 | 0.87 | 0.41 | 0.58 |
| 3 | 65797540 | rs1909492 | -0.554 | 0.315 | -0.683 | -0.788 | 0 | -0.004 | -0.004 | 3.067 | 0.52 | 0.55 | 0.65 | 0.72 | 0.58 | 0.55 |
| 3 | 65800232 | rs1473550 | -1.167 | -0.2 | -0.735 | -0.776 | 0.002 | -0.005 | -0.005 | 0.397 | 0.58 | 0.67 | 0.72 | 0.93 | 0.6 | 0.82 |
| 3 | 65801606 | rs17295791 | 1.049 | -0.151 | -1.107 | -0.634 | -0.002 | 0.015 | 0.015 | 0 | 0.07 | 0.12 | 0.23 | 0.05 | 0.07 | 0.03 |
| 3 | 65804583 | rs1108718 | -0.286 | -0.363 | -1.223 | -0.914 | -0.019 | 0.112 | 0.112 | 1.219 | 0.45 | 0.23 | 0.57 | 0.75 | 0.53 | 0.52 |
| 3 | 65806838 | rs17361892 | 0.876 | -0.356 | -1.32 | -0.732 | -0.018 | -0.077 | -0.077 | 0 | 0.17 | 0.01 | 0.08 | 0.22 | 0.15 | 0.23 |
| 3 | 65809842 | rs1874320 | -0.24 | -0.361 | -0.54 | -1.334 | -0.002 | -0.082 | -0.082 | 0.127 | 0.53 | 0.53 | 0.67 | 0.92 | 0.58 | 0.82 |
| 3 | 65813373 | rs12490463 | -0.253 | -0.159 | -0.538 | -1.263 | 0.001 | -0.034 | -0.034 | 1.001 | 0.56 | 0.62 | 0.75 | 0.94 | 0.59 | 0.83 |
| 3 | 65817691 | rs10510946 | -0.728 | 0.148 | -0.869 | -0.869 | -0.012 | 0.073 | 0.073 | 0.056 | 0.07 | 0.21 | 0.05 | 0.17 | 0.03 | 0.26 |
| 3 | 65821701 | rs17073937 | -0.475 | 0.332 | -1.388 | -0.833 | 0 | -0.111 | -0.111 | 0 | 0.59 | 0.53 | 0.71 | 0.94 | 0.62 | 0.88 |
| 3 | 65822747 | rs7638342 | -1.23 | 0.35 | -1.442 | 0.567 | -0.002 | 0.002 | 0.002 | 0 | 0.09 | 0.08 | 0.06 | 0.17 | 0.05 | 0.31 |
| 3 | 65846859 | rs2132254 | -0.068 | 0.657 | -2.355 | 0.086 | -0.115 | 0.072 | 0.072 | 0 | 0.62 | 0.98 | 0.76 | 0.83 | 0.58 | 0.77 |
| 3 | 65847080 | rs12491013 | 0.096 | 0.461 | -2.295 | -0.477 | -0.001 | 0.003 | 0.003 | 0 | 0.13 | 0.08 | 0.09 | 0.01 | 0.15 | 0.04 |
| 3 | 65848944 | rs17073988 | -0.525 | 0.284 | -1.129 | 0.392 | 0.035 | 0.248 | 0.248 | 0.274 | 0.1 | 0.31 | 0.35 | 0.32 | 0.11 | 0.32 |
| 3 | 65849896 | rs924022 | -0.704 | 0.48 | -0.878 | 0.346 | 0.04 | 0.204 | 0.204 | 0.006 | 0.1 | 0.25 | 0.34 | 0.32 | 0.13 | 0.32 |
| 3 | 65851664 | rs7617634 | -0.707 | 0.313 | -0.859 | 0.52 | 0.031 | 0.247 | 0.247 | 0.001 | 0.09 | 0.25 | 0.33 | 0.37 | 0.11 | 0.35 |
| 3 | 65852328 | rs6793201 | -0.707 | 0.313 | -0.859 | 0.479 | 0.031 | 0.223 | 0.223 | 0.001 | 0.09 | 0.25 | 0.34 | 0.33 | 0.11 | 0.34 |
| 3 | 65854178 | rs2643727 | -1.153 | 0.006 | -0.801 | -0.066 | 0.018 | 0.021 | 0.021 | 0.698 | 0.38 | 0.77 | 0.49 | 0.34 | 0.35 | 0.51 |
| 3 | 65854650 | rs2372154 | -0.667 | -0.263 | -0.117 | -0.013 | -0.011 | 0.076 | 0.076 | 0 | 0.21 | 0.4 | 0.38 | 0.33 | 0.2 | 0.41 |
| 3 | 65868822 | rs11923705 | -0.73 | -0.588 | 0.681 | -0.341 | 0.008 | 0.017 | 0.017 | 0 | 0.25 | 0.17 | 0.17 | 0.11 | 0.23 | 0.19 |
| 3 | 65869363 | rs264709 | -1.111 | -0.676 | 0.866 | -0.371 | -0.031 | -0.073 | -0.073 | 0 | 0.31 | 0.6 | 0.2 | 0.11 | 0.23 | 0.19 |
| 3 | 65873106 | rs9876471 | -0.323 | -0.64 | 0.969 | -0.219 | -0.014 | -0.032 | -0.032 | 0.814 | 0.13 | 0.17 | 0.1 | 0.08 | 0.09 | 0.14 |

| | | | | | | | | | | | | | | | | |
|---|---|---|---|---|---|---|---|---|---|---|---|---|---|---|---|---|
| 3 | 65879245 | rs264668 | -1.44 | -0.72 | 0.801 | -0.314 | -0.05 | -0.104 | -0.104 | 0 | 0.22 | 0.58 | 0.17 | 0.1 | 0.14 | 0.16 |
| 3 | 65881273 | rs9864600 | -0.621 | -0.436 | -0.125 | -0.204 | 0 | 0.016 | 0.016 | 0.011 | 0.56 | 0.54 | 0.41 | 0.66 | 0.51 | 0.71 |
| 3 | 65886519 | rs845312 | -0.246 | -1.543 | -0.588 | -0.555 | 0.002 | 0.002 | 0.002 | 0 | 0.36 | 0.33 | 0.34 | 0.32 | 0.25 | 0.28 |
| 3 | 65886745 | rs264677 | -0.216 | -1.558 | -0.533 | -0.608 | 0.032 | 0.015 | 0.015 | 0 | 0.34 | 0.26 | 0.33 | 0.29 | 0.22 | 0.26 |
| 3 | 65888984 | rs174652 | -1.163 | -0.845 | -0.859 | -0.901 | -0.108 | 0.118 | 0.118 | 0.02 | 0.75 | 0.9 | 0.69 | 0.99 | 0.65 | 0.9 |
| 3 | 65890657 | rs1552966 | 0.578 | -1.35 | -0.809 | -0.885 | -0.006 | 0.005 | 0.005 | 0.018 | 0.18 | 0.17 | 0.28 | 0.24 | 0.15 | 0.15 |
| 3 | 65891022 | rs3850570 | 1.005 | -1.354 | -1.036 | -0.858 | 0.015 | -0.014 | -0.014 | 0 | 0.16 | 0.1 | 0.26 | 0.23 | 0.12 | 0.12 |
| 3 | 65891956 | rs4688610 | 0.996 | -1.339 | -1.049 | -0.84 | 0.033 | -0.036 | -0.036 | 0 | 0.16 | 0.05 | 0.25 | 0.23 | 0.12 | 0.12 |
| 3 | 65893577 | rs1390245 | -0.486 | -1.1 | -1.564 | -0.784 | -0.045 | 0.049 | 0.049 | 0.433 | 0.59 | 0.75 | 0.45 | 0.74 | 0.54 | 0.77 |
| 3 | 65904425 | rs264692 | -1.166 | -0.86 | -1.514 | -0.831 | -0.05 | 0.056 | 0.056 | 0.105 | 0.77 | 0.9 | 0.72 | 0.94 | 0.68 | 0.91 |
| 3 | 65905920 | rs264695 | 0.365 | -1.502 | -1.458 | -0.986 | -0.024 | 0.021 | 0.021 | 0.037 | 0.22 | 0.29 | 0.3 | 0.26 | 0.18 | 0.21 |
| 3 | 65908894 | rs264686 | -1.19 | -1.274 | -1.777 | -1.049 | -0.139 | 0.161 | 0.161 | 0.261 | 0.76 | 0.97 | 0.71 | 0.99 | 0.66 | 0.91 |
| 3 | 65917408 | rs17441629 | 0.789 | -1.474 | -1.251 | -0.616 | 0.047 | 0.055 | 0.055 | 0 | 0.15 | 0.02 | 0.05 | 0 | 0.06 | 0.1 |
| 3 | 65937976 | rs10866075 | 0.071 | -1.262 | -0.781 | -0.647 | -0.003 | -0.015 | -0.015 | 0.245 | 0.46 | 0.48 | 0.39 | 0.76 | 0.45 | 0.73 |
| 3 | 65940092 | rs6784567 | -0.797 | -1.165 | -0.872 | -0.615 | -0.028 | 0.042 | 0.042 | 0.484 | 0.63 | 0.71 | 0.47 | 0.81 | 0.53 | 0.79 |
| 3 | 65944066 | rs9880270 | -0.242 | -1.239 | -0.667 | -0.084 | 0.001 | -0.001 | -0.001 | 0 | 0.44 | 0.55 | 0.27 | 0.52 | 0.41 | 0.67 |
| 3 | 65950836 | rs17382966 | 0.078 | -1.353 | -0.663 | 0.107 | 0.007 | -0.002 | -0.002 | 0 | 0.39 | 0.26 | 0.22 | 0.4 | 0.39 | 0.61 |
| 3 | 65951752 | rs1430416 | 0.906 | -1.296 | -0.606 | 0.028 | -0.016 | -0.007 | -0.007 | 0.696 | 0.42 | 0.3 | 0.65 | 0.42 | 0.49 | 0.25 |
| 3 | 65952898 | rs7637355 | -0.132 | -0.749 | -0.75 | 0.186 | -0.002 | 0.017 | 0.017 | 0.772 | 0.37 | 0.27 | 0.23 | 0.3 | 0.4 | 0.35 |
| 3 | 65963727 | rs6795228 | -0.329 | -0.423 | -0.703 | 0.455 | -0.004 | -0.003 | -0.003 | 0 | 0.11 | 0.11 | 0.07 | 0.27 | 0.15 | 0.1 |
| 3 | 65964941 | rs17074213 | 0.1 | -0.719 | -0.6 | -0.111 | -0.004 | -0.028 | -0.028 | 2.456 | 0.1 | 0.13 | 0.06 | 0 | 0.09 | 0.12 |
| 3 | 65965134 | rs4688621 | 1.229 | -0.711 | -1.737 | -1.019 | -0.116 | 0.228 | 0.228 | 1.572 | 0.32 | 0.01 | 0.36 | 0.01 | 0.34 | 0.09 |
| 3 | 65965346 | rs4688255 | 0.81 | -1.276 | -0.693 | -0.477 | -0.008 | 0.019 | 0.019 | 0 | 0.74 | 0.72 | 0.85 | 0.62 | 0.75 | 0.75 |
| 3 | 65965366 | rs10510949 | 0.645 | -1.329 | -0.688 | -0.365 | 0.003 | -0.007 | -0.007 | 0 | 0.1 | 0.06 | 0.05 | 0.1 | 0.11 | 0.16 |
| 3 | 65965425 | rs17074219 | -0.92 | -1.34 | -0.588 | -0.374 | 0.012 | -0.035 | -0.035 | 0 | 0.23 | 0.15 | 0.13 | 0.38 | 0.24 | 0.24 |
| 3 | 65966296 | rs6445523 | -0.431 | -1.225 | -0.497 | -0.319 | -0.006 | 0.024 | 0.024 | 0 | 0.86 | 0.87 | 0.92 | 0.73 | 0.87 | 0.92 |
| 3 | 65969410 | rs7624164 | -1.017 | -1.224 | -0.029 | -0.733 | -0.001 | -0.052 | -0.052 | 0 | 0.77 | 0.65 | 0.86 | 0.95 | 0.78 | 0.63 |
| 3 | 65971019 | rs3733137 | 2.02 | -1.25 | 0.018 | -0.735 | -0.001 | 0.003 | 0.003 | 0 | 0.06 | 0.06 | 0.04 | 0.11 | 0.06 | 0.07 |
| 3 | 65971499 | rs7620106 | 0.694 | -1.416 | 0.302 | -0.945 | 0.005 | 0.204 | 0.204 | 0.95 | 0.55 | 0.1 | 0.71 | 0.84 | 0.59 | 0.41 |
| 3 | 65972904 | rs6807782 | 0.054 | -1.393 | 0.398 | -0.92 | 0.009 | 0.097 | 0.097 | 0.262 | 0.65 | 0.13 | 0.77 | 0.84 | 0.7 | 0.55 |
| 3 | 65977869 | rs17821707 | -0.523 | -1.515 | 0.452 | -0.885 | 0.006 | -0.013 | -0.013 | 0 | 0.25 | 0.14 | 0.15 | 0.26 | 0.25 | 0.23 |
| 3 | 65979643 | rs6445524 | -1.298 | -1.428 | 0.601 | -0.918 | -0.004 | -0.078 | -0.078 | 0 | 0.78 | 0.62 | 0.87 | 0.96 | 0.79 | 0.64 |
| 3 | 65986180 | rs4688258 | 0.29 | -1.17 | 0.506 | -0.682 | 0.003 | 0.012 | 0.012 | 0.075 | 0.54 | 0.5 | 0.72 | 0.7 | 0.54 | 0.42 |
| 3 | 65987793 | rs4688629 | 1.241 | -1.195 | 0.507 | -0.659 | -0.031 | -0.002 | -0.002 | 0.38 | 0.44 | 0.14 | 0.66 | 0.62 | 0.47 | 0.35 |
| 3 | 65987866 | rs4688630 | 0.517 | -0.997 | 0.318 | -0.72 | 0.001 | 0.007 | 0.007 | 0.055 | 0.52 | 0.44 | 0.71 | 0.7 | 0.53 | 0.42 |
| 3 | 65990790 | rs11919056 | 0.014 | -0.777 | 0.464 | -1.092 | 0.012 | -0.072 | -0.072 | 0 | 0.12 | 0.28 | 0.1 | 0 | 0.13 | 0.13 |
| 3 | 65991856 | rs1541855 | 0.425 | -1.356 | 0.035 | -1.132 | 0.02 | 0.053 | 0.053 | 0 | 0.51 | 0.57 | 0.71 | 0.73 | 0.6 | 0.42 |
| 3 | 65992303 | rs10510951 | 0.543 | -1.57 | -0.4 | -1.225 | 0.006 | -0.018 | -0.018 | 0 | 0.15 | 0.01 | 0.07 | 0.14 | 0.11 | 0.09 |
| 3 | 65994050 | rs4688631 | 1.138 | -1.795 | -0.548 | -1.731 | -0.082 | 0.032 | 0.032 | 0 | 0.31 | 0.13 | 0.58 | 0.73 | 0.47 | 0.38 |
| 3 | 65994746 | rs6809559 | 1.72 | -1.805 | -0.056 | -1.796 | -0.036 | -0.12 | -0.12 | 0 | 0.19 | 0.34 | 0.11 | 0 | 0.12 | 0.03 |
| 3 | 65999532 | rs2060023 | 1.367 | -1.675 | -0.425 | -1.513 | -0.022 | -0.009 | -0.009 | 0 | 0.23 | 0.14 | 0.55 | 0.66 | 0.42 | 0.32 |
| 3 | 65999755 | rs2060022 | 0.689 | -1.782 | -0.447 | -1.578 | -0.01 | 0.006 | 0.006 | 0 | 0.29 | 0.24 | 0.6 | 0.73 | 0.48 | 0.37 |
| 3 | 66000105 | rs6767345 | -0.021 | -1.827 | -0.265 | -1.633 | -0.002 | 0.004 | 0.004 | 0 | 0.4 | 0.37 | 0.66 | 0.73 | 0.54 | 0.46 |
| 3 | 66003182 | rs9810565 | 1.344 | -1.628 | -0.212 | -1.517 | -0.025 | -0.008 | -0.008 | 0 | 0.26 | 0.18 | 0.56 | 0.65 | 0.42 | 0.33 |
| 3 | 66003745 | rs6762527 | -0.254 | -1.412 | 0.114 | -1.669 | 0.003 | -0.016 | -0.016 | 0 | 0.56 | 0.44 | 0.75 | 0.84 | 0.67 | 0.57 |
| 3 | 66009385 | rs28496178 | -0.352 | -1.122 | -0.206 | -1.506 | 0.031 | 0.117 | 0.117 | 0 | 0.52 | 0.71 | 0.32 | 0.16 | 0.41 | 0.5 |
| 3 | 66023592 | rs28696109 | -0.346 | -0.812 | 0.134 | -0.895 | -0.02 | -0.027 | -0.027 | 5.607 | 0.74 | 0.69 | 0.84 | 0.84 | 0.83 | 0.6 |
| 3 | 66029293 | rs28705165 | -0.414 | -0.149 | 0.185 | -0.493 | 0.023 | 0.002 | 0.002 | 0.059 | 0.12 | 0.27 | 0.06 | 0.1 | 0.13 | 0.09 |
| 3 | 66039580 | rs28758977 | -0.678 | 0.913 | 0.083 | -0.364 | 0.033 | -0.022 | -0.022 | 0 | 0.8 | 0.65 | 0.7 | 0.87 | 0.78 | 0.87 |
| 3 | 66050568 | rs28561801 | -0.381 | 1.039 | -0.617 | -0.339 | -0.007 | 0.037 | 0.037 | 0 | 0.46 | 0.34 | 0.42 | 0.64 | 0.48 | 0.55 |
| 3 | 66061727 | rs28433140 | 0.211 | 1.227 | -0.619 | -0.019 | 0.011 | -0.024 | -0.024 | 2.07 | 0.39 | 0.2 | 0.38 | 0.53 | 0.37 | 0.47 |
| 3 | 66069898 | rs28569266 | -1.123 | 1.479 | 0.173 | 0.05 | 0.074 | 0.01 | 0.01 | 0 | 0.85 | 0.68 | 0.72 | 0.88 | 0.81 | 0.89 |
| 3 | 66079814 | rs28594923 | -0.76 | 1.41 | 0.066 | 0.069 | 0.079 | -0.045 | -0.045 | 0.322 | 0.8 | 0.6 | 0.68 | 0.91 | 0.75 | 0.89 |
| 3 | 66101934 | rs28421915 | 0.13 | 0.592 | 0.023 | 0.433 | -0.028 | -0.034 | -0.034 | 0 | 0.33 | 0.55 | 0.16 | 0.19 | 0.23 | 0.18 |
| 3 | 66110121 | rs28528235 | -0.442 | 1.001 | -0.237 | 0.642 | 0.02 | -0.001 | -0.001 | 0 | 0.48 | 0.65 | 0.31 | 0.54 | 0.41 | 0.44 |
| 3 | 66119382 | rs28602282 | -0.362 | 0.769 | -0.131 | 0.721 | 0.017 | -0.002 | -0.002 | 0 | 0.45 | 0.62 | 0.3 | 0.48 | 0.37 | 0.4 |
| 3 | 66122047 | rs28438945 | -0.375 | 0.781 | -0.114 | 0.73 | 0.015 | -0.004 | -0.004 | 0 | 0.45 | 0.62 | 0.3 | 0.5 | 0.37 | 0.4 |
| 3 | 66139937 | rs28534383 | -0.983 | 2.646 | 1.391 | 1.087 | 0.007 | 0.016 | 0.016 | 0.668 | 0.29 | 0.44 | 0.23 | 0.39 | 0.3 | 0.36 |
| 3 | 66293480 | rs2311298 | -2.322 | 2.363 | 2.304 | 1.061 | 0 | 0.279 | 0.279 | 0.831 | 0.07 | 0.3 | 0.14 | 0.27 | 0.06 | 0.06 |
| 3 | 66293688 | rs3772197 | -0.657 | 2.64 | 1.915 | 1.027 | -0.008 | -0.004 | -0.004 | 0 | 0.05 | 0.03 | 0.05 | 0.04 | 0.07 | 0.1 |
| 3 | 66300547 | rs1680394 | -1.561 | 3.126 | 2.244 | 1.007 | 0.006 | 0.014 | 0.014 | 0 | 0.27 | 0.61 | 0.53 | 0.45 | 0.4 | 0.18 |
| 3 | 66310451 | rs6777571 | -1.059 | 3.02 | 2.163 | 0.251 | 0.056 | -0.062 | -0.062 | 0 | 0.22 | 0.44 | 0.42 | 0.15 | 0.34 | 0.12 |
| 3 | 66311773 | rs2306499 | 0.081 | 3.121 | 1.992 | 0.193 | -0.056 | 0.012 | 0.012 | 0 | 0.14 | 0.03 | 0.13 | 0.13 | 0.22 | 0.07 |
| 3 | 66319062 | rs13088067 | -0.91 | 3.119 | 2.159 | 0.189 | -0.009 | 0.007 | 0.007 | 0 | 0.2 | 0.17 | 0.38 | 0.13 | 0.33 | 0.11 |
| 3 | 66326302 | rs782728 | -2.127 | 3.069 | 2.171 | 0.173 | -0.042 | 0.056 | 0.056 | 0.364 | 0.33 | 0.79 | 0.55 | 0.2 | 0.47 | 0.25 |
| 3 | 66329205 | rs782733 | -2.473 | 2.524 | 2.569 | 0.169 | 0.024 | -0.05 | -0.05 | 0 | 0.16 | 0.55 | 0.38 | 0.04 | 0.24 | 0.18 |
| 3 | 66335732 | rs1091407 | -2.349 | 2.618 | 2.395 | 0.211 | 0.048 | -0.043 | -0.043 | 0 | 0.16 | 0.48 | 0.37 | 0.07 | 0.24 | 0.19 |
| 3 | 66378999 | rs10510953 | -0.484 | 2.667 | 2.162 | -0.094 | -0.006 | -0.008 | -0.008 | 0 | 0.21 | 0.16 | 0.42 | 0.68 | 0.29 | 0.42 |
| 3 | 66415349 | rs9827734 | -1.512 | 2.442 | 2.047 | -0.212 | -0.01 | 0.001 | 0.001 | 0.682 | 0.33 | 0.24 | 0.48 | 0.69 | 0.4 | 0.59 |

| | | | | | | | | | | | | | | | | |
|---|---|---|---|---|---|---|---|---|---|---|---|---|---|---|---|---|
| 3 | 66421147 | rs332378 | -0.524 | 2.424 | 2.196 | -0.856 | 0.053 | -0.047 | -0.047 | 0 | 0.19 | 0.4 | 0.2 | 0.1 | 0.24 | 0.13 |
| 3 | 66426912 | rs7627907 | -0.859 | 2.547 | 1.959 | -0.904 | -0.049 | -0.01 | -0.01 | 0.012 | 0.26 | 0.1 | 0.27 | 0.71 | 0.36 | 0.5 |
| 3 | 66428282 | rs1046844 | -0.816 | 2.406 | 2.326 | -0.969 | 0.021 | -0.1 | -0.1 | 0 | 0.23 | 0.56 | 0.36 | 0.05 | 0.23 | 0.17 |
| 3 | 66430811 | rs2279290 | 0.51 | 2.409 | 2.247 | -0.967 | -0.03 | 0.033 | 0.033 | 0 | 0.15 | 0.08 | 0.11 | 0.06 | 0.2 | 0.1 |
| 3 | 66433010 | rs7642319 | -1.238 | 2.475 | 2.147 | -0.969 | 0 | -0.048 | -0.048 | 0 | 0.3 | 0.66 | 0.42 | 0.07 | 0.3 | 0.26 |
| 3 | 66433676 | rs900171 | -2.282 | 2.309 | 2.128 | -0.982 | 0.033 | 0.104 | 0.104 | 0 | 0.53 | 0.72 | 0.64 | 0.8 | 0.6 | 0.68 |
| 3 | 66434643 | rs2306272 | -0.604 | 2.364 | 1.985 | -0.941 | -0.019 | -0.007 | -0.007 | 0 | 0.23 | 0.15 | 0.24 | 0.73 | 0.29 | 0.42 |
| 3 | 66437086 | rs17831815 | -1.133 | 2.355 | 1.994 | -0.933 | -0.038 | 0.009 | 0.009 | 0 | 0.29 | 0.15 | 0.29 | 0.75 | 0.36 | 0.49 |
| 3 | 66441661 | rs1283532 | -1.971 | 1.708 | 2.36 | -0.569 | -0.003 | 0.095 | 0.095 | 0.024 | 0.47 | 0.69 | 0.64 | 0.8 | 0.55 | 0.66 |
| 3 | 66442545 | rs9840088 | -0.309 | 1.708 | 1.938 | -0.458 | -0.003 | -0.01 | -0.01 | 0.895 | 0.18 | 0.15 | 0.2 | 0.71 | 0.2 | 0.38 |
| 3 | 66455797 | rs3796148 | -0.814 | 1.716 | 1.649 | -0.256 | 0 | -0.009 | -0.009 | 3.797 | 0.21 | 0.23 | 0.35 | 0.09 | 0.24 | 0.19 |
| 3 | 66466444 | rs11925003 | -0.695 | 1.309 | 1.496 | -0.265 | -0.001 | 0.003 | 0.003 | 0 | 0.2 | 0.18 | 0.31 | 0.1 | 0.2 | 0.18 |
| 3 | 66474487 | rs13073932 | 0.023 | 1.401 | 1.467 | -0.23 | 0 | 0 | 0 | 0 | 0.15 | 0.16 | 0.3 | 0.07 | 0.2 | 0.11 |
| 3 | 66487542 | rs10049330 | -0.685 | 1.257 | 1.844 | -0.35 | -0.029 | -0.087 | -0.087 | 0.526 | 0.15 | 0.45 | 0.09 | 0.04 | 0.09 | 0.17 |
| 3 | 66488134 | rs13096446 | 0.031 | 1.335 | 1.776 | -0.324 | 0.001 | -0.003 | -0.003 | 1.948 | 0.15 | 0.15 | 0.3 | 0.05 | 0.2 | 0.1 |
| 3 | 66490125 | rs3845895 | 1.31 | 1.248 | 1.391 | -0.31 | 0.013 | 0.082 | 0.082 | 0 | 0.67 | 0.2 | 0.53 | 0.88 | 0.7 | 0.71 |
| 3 | 66496592 | rs9849546 | -1.068 | 1.223 | 1.332 | -0.247 | -0.002 | -0.053 | -0.053 | 0.207 | 0.29 | 0.58 | 0.38 | 0.1 | 0.27 | 0.26 |
| 3 | 66504064 | rs3845899 | -0.266 | 1.11 | 0.567 | 0.465 | -0.032 | -0.038 | -0.038 | 0.001 | 0.16 | 0.11 | 0.21 | 0.69 | 0.25 | 0.34 |
| 3 | 66507442 | rs6549120 | 0.739 | 0.898 | 0.854 | 0.437 | 0.005 | -0.01 | -0.01 | 0 | 0.67 | 0.43 | 0.65 | 0.26 | 0.63 | 0.48 |
| 3 | 66513947 | rs4856941 | 1.693 | 0.183 | 1.79 | 0.77 | -0.01 | 0.078 | 0.078 | 0.478 | 0.91 | 0.61 | 0.7 | 0.87 | 0.95 | 0.94 |
| 3 | 66514637 | rs10510956 | 2.025 | 0.083 | 2.124 | 0.726 | -0.045 | 0.029 | 0.029 | 2.26 | 0.89 | 0.4 | 0.68 | 0.87 | 0.95 | 0.95 |
| 3 | 66525963 | rs3845905 | 1.927 | 0.072 | 2.127 | 0.72 | -0.044 | 0.018 | 0.018 | 3.98 | 0.9 | 0.41 | 0.72 | 0.86 | 0.96 | 0.95 |
| 3 | 66526602 | rs3845906 | 1.76 | 0.081 | 2.122 | 0.722 | -0.009 | 0.079 | 0.079 | 1.032 | 0.91 | 0.59 | 0.72 | 0.85 | 0.96 | 0.95 |
| 3 | 66531096 | rs12637858 | -0.054 | 0.283 | 0.852 | 1.322 | -0.042 | -0.072 | -0.072 | 0 | 0.11 | 0.01 | 0.19 | 0.48 | 0.15 | 0.28 |
| 3 | 66539628 | rs11709533 | 1.878 | -0.075 | 1.319 | 0.851 | -0.034 | 0 | 0 | 10.773 | 0.9 | 0.18 | 0.87 | 0.86 | 0.97 | 0.96 |
| 3 | 66543696 | rs7646811 | 1.545 | 0.061 | 1.672 | 0.814 | 0.025 | 0.087 | 0.087 | 5.349 | 0.93 | 0.5 | 0.91 | 0.94 | 0.98 | 0.96 |
| 3 | 66558036 | rs7630018 | 1.116 | 0.108 | 1.677 | 0.83 | 0.033 | 0.084 | 0.084 | 1.546 | 0.95 | 0.69 | 0.93 | 0.94 | 0.99 | 0.97 |
| 3 | 66562776 | rs4402870 | -0.022 | 0.223 | 0.675 | 0.344 | 0.003 | 0.032 | 0.032 | 0 | 0.53 | 0.64 | 0.62 | 0.41 | 0.48 | 0.56 |
| 3 | 66568741 | rs6789596 | 0.102 | 0.274 | 0.427 | 0.286 | -0.016 | 0.029 | 0.029 | 0.247 | 0.45 | 0.08 | 0.38 | 0.52 | 0.54 | 0.42 |
| 3 | 66569114 | rs11707571 | -0.135 | 0.355 | 0.666 | 0.261 | -0.009 | 0.001 | 0.001 | 0 | 0.11 | 0.07 | 0.08 | 0.13 | 0.11 | 0.18 |
| 3 | 66570008 | rs1523295 | 1.27 | 0.087 | 1.069 | 0.212 | -0.025 | 0.081 | 0.081 | 0 | 0.91 | 0.66 | 0.85 | 0.8 | 0.96 | 0.88 |
| 3 | 66577919 | rs1403624 | 0.953 | 0.538 | 1.217 | 0.158 | 0.041 | 0.006 | 0.006 | 0 | 0.87 | 0.66 | 0.9 | 0.84 | 0.88 | 0.8 |
| 3 | 66580551 | rs4241442 | 0.65 | 0.457 | 0.73 | 0.028 | 0 | -0.013 | -0.013 | 0 | 0.71 | 0.41 | 0.81 | 0.81 | 0.71 | 0.78 |
| 3 | 66586374 | rs6783434 | -0.566 | -0.547 | 0.39 | -0.11 | -0.016 | -0.001 | -0.001 | 0.419 | 0.12 | 0.16 | 0.1 | 0.1 | 0.08 | 0.14 |
| 3 | 66589531 | rs13079535 | -0.41 | -0.393 | 0.418 | -0.166 | 0 | -0.015 | -0.015 | 0 | 0.07 | 0.11 | 0.15 | 0.03 | 0.06 | 0.09 |
| 3 | 66600918 | rs6549201 | -0.863 | -0.027 | 0.723 | 0.297 | 0 | -0.002 | -0.002 | 0 | 0.32 | 0.61 | 0.25 | 0.28 | 0.31 | 0.41 |
| 3 | 66613245 | rs9881490 | -1.079 | -0.189 | 0.67 | 0.173 | -0.002 | -0.005 | -0.005 | 0 | 0.34 | 0.6 | 0.25 | 0.27 | 0.3 | 0.41 |
| 3 | 66623998 | rs17777649 | 0.145 | -0.164 | 0.695 | -0.014 | -0.082 | 0.101 | 0.101 | 0 | 0.2 | 0.03 | 0.13 | 0.03 | 0.22 | 0.25 |
| 3 | 66627521 | rs4547637 | 0.015 | -0.962 | 0.427 | -0.163 | 0.021 | 0.016 | 0.016 | 0 | 0.14 | 0.07 | 0.1 | 0.06 | 0.09 | 0.14 |
| 3 | 66635873 | rs4856952 | -1.886 | -1.139 | 0.872 | 0.077 | -0.053 | 0.01 | 0.01 | 0.045 | 0.29 | 0.44 | 0.3 | 0.34 | 0.17 | 0.29 |
| 3 | 66636185 | rs4856953 | -0.109 | -0.877 | 0.653 | 0.083 | -0.002 | 0.005 | 0.005 | 0 | 0.07 | 0.06 | 0.17 | 0.07 | 0.07 | 0.1 |
| 3 | 66637645 | rs6802463 | 0.425 | -0.847 | 0.199 | -0.239 | 0.004 | 0.073 | 0.073 | 0 | 0.32 | 0.13 | 0.23 | 0.16 | 0.34 | 0.18 |
| 3 | 66638886 | rs6805929 | 0.4 | -0.677 | 0.141 | -0.246 | -0.043 | 0.049 | 0.049 | 0 | 0.31 | 0.14 | 0.33 | 0.2 | 0.33 | 0.32 |
| 3 | 66648162 | rs7612441 | 0.472 | -0.574 | -0.065 | -0.211 | 0.003 | 0.125 | 0.125 | 0 | 0.3 | 0.1 | 0.2 | 0.04 | 0.3 | 0.14 |
| 3 | 66654179 | rs1456916 | 0.776 | -0.699 | -0.292 | 0.701 | -0.026 | -0.034 | -0.034 | 0.181 | 0.54 | 0.17 | 0.5 | 0.5 | 0.62 | 0.54 |
| 3 | 66656277 | rs1379195 | 0.768 | -0.418 | -0.08 | 1.179 | -0.004 | -0.001 | -0.001 | 0 | 0.32 | 0.35 | 0.27 | 0.27 | 0.4 | 0.22 |
| 3 | 66659759 | rs1379194 | 1.282 | -0.108 | 0.202 | 1.496 | -0.025 | -0.04 | -0.04 | 1.242 | 0.58 | 0.17 | 0.53 | 0.53 | 0.66 | 0.59 |
| 3 | 66668365 | rs11128158 | 1.397 | -0.19 | 0.769 | 1.174 | 0.005 | -0.026 | -0.026 | 0 | 0.67 | 0.21 | 0.75 | 0.69 | 0.73 | 0.7 |
| 3 | 66677341 | rs6769418 | -1.119 | -0.328 | 0.807 | 1.288 | -0.001 | 0.003 | 0.003 | 0 | 0.25 | 0.27 | 0.21 | 0.23 | 0.22 | 0.2 |
| 3 | 66689366 | rs1870033 | 1.594 | -0.171 | 0.661 | 0.783 | 0.013 | 0.004 | 0.004 | 0 | 0.67 | 0.27 | 0.75 | 0.74 | 0.75 | 0.62 |
| 3 | 66692191 | rs12629940 | 0.867 | -0.449 | 0.998 | 1.048 | -0.023 | 0.052 | 0.052 | 0 | 0.89 | 0.66 | 0.95 | 0.92 | 0.94 | 0.82 |
| 3 | 66692943 | rs10048978 | 0.835 | -0.408 | 0.257 | 1.032 | -0.054 | 0.055 | 0.055 | 0 | 0.85 | 0.45 | 0.88 | 0.85 | 0.93 | 0.8 |
| 3 | 66697961 | rs11711551 | 0.899 | -0.427 | 0.223 | 1.086 | -0.065 | 0.081 | 0.081 | 0 | 0.84 | 0.46 | 0.88 | 0.8 | 0.93 | 0.8 |
| 3 | 66699668 | rs1001763 | -0.545 | -0.199 | -0.19 | 0.701 | 0 | 0 | 0 | 0 | 0.22 | 0.2 | 0.36 | 0.25 | 0.2 | 0.33 |
| 3 | 66700816 | rs6795912 | -1.067 | -0.524 | 0.566 | 0.891 | 0.003 | 0.063 | 0.063 | 0 | 0.06 | 0.25 | 0.04 | 0.11 | 0.03 | 0.04 |
| 3 | 66700959 | rs11128172 | 0.455 | -0.002 | 0.053 | 0.627 | -0.047 | 0.007 | 0.007 | 2.156 | 0.27 | 0.17 | 0.5 | 0.25 | 0.47 | 0.29 |
| 3 | 66703685 | rs9862866 | -0.079 | 0.18 | -0.178 | 0.613 | -0.031 | 0.002 | 0.002 | 0 | 0.34 | 0.22 | 0.56 | 0.32 | 0.52 | 0.39 |
| 3 | 66704744 | rs9846070 | -0.057 | 0.214 | 0.01 | 0.657 | 0.016 | 0.012 | 0.012 | 0 | 0.33 | 0.42 | 0.55 | 0.43 | 0.51 | 0.35 |
| 3 | 66706240 | rs9870520 | -0.076 | -1.287 | 0.042 | 0.218 | 0.057 | -0.038 | -0.038 | 0 | 0.62 | 0.79 | 0.77 | 0.56 | 0.83 | 0.64 |
| 3 | 66707908 | rs9813839 | 1.338 | -1.667 | 0.184 | 0.076 | -0.002 | -0.001 | -0.001 | 0 | 0.3 | 0.3 | 0.55 | 0.32 | 0.46 | 0.32 |
| 3 | 66709318 | rs1563541 | -0.336 | -1.44 | -0.033 | -0.082 | 0.012 | -0.034 | -0.034 | 1.498 | 0.61 | 0.81 | 0.7 | 0.46 | 0.71 | 0.58 |
| 3 | 66712564 | rs9839845 | 1.794 | -1.404 | -0.085 | -0.056 | -0.016 | 0.003 | 0.003 | 0 | 0.28 | 0.2 | 0.52 | 0.25 | 0.44 | 0.26 |
| 3 | 66712640 | rs6801170 | -0.047 | -1.403 | 0.029 | -0.086 | 0.013 | -0.003 | -0.003 | 0 | 0.55 | 0.7 | 0.66 | 0.46 | 0.68 | 0.51 |
| 3 | 66713714 | rs17835506 | 1.417 | -1.364 | -0.215 | -0.096 | -0.035 | 0.037 | 0.037 | 0 | 0.08 | 0 | 0.04 | 0.01 | 0.13 | 0.07 |
| 3 | 66714410 | rs11705970 | 0.335 | -1.536 | 0.14 | -0.071 | 0.005 | 0.007 | 0.007 | 0 | 0.47 | 0.41 | 0.63 | 0.34 | 0.62 | 0.46 |
| 3 | 66721682 | rs6782747 | 0.41 | -1.737 | -0.085 | 0.442 | -0.002 | -0.014 | -0.014 | 0 | 0.26 | 0.19 | 0.23 | 0.48 | 0.24 | 0.3 |
| 3 | 66726579 | rs6781616 | -0.444 | -1.903 | 0.523 | 0.652 | 0.002 | -0.005 | -0.005 | 0 | 0.44 | 0.44 | 0.28 | 0.68 | 0.43 | 0.5 |
| 3 | 66728124 | rs2886375 | 0.233 | -1.891 | 0.28 | 0.634 | -0.014 | -0.013 | -0.013 | 0 | 0.37 | 0.23 | 0.21 | 0.49 | 0.38 | 0.37 |
| 3 | 66729080 | rs1598705 | 1.121 | -1.826 | 0.289 | 0.677 | -0.063 | 0.108 | 0.108 | 0.459 | 0.38 | 0.19 | 0.6 | 0.13 | 0.54 | 0.39 |

| | | | | | | | | | | | | | | | | |
|---|---|---|---|---|---|---|---|---|---|---|---|---|---|---|---|---|
| 3 | 66734825 | rs4437080 | -1.183 | -1.791 | -0.14 | 0.647 | 0.013 | -0.009 | -0.009 | 0.07 | 0.59 | 0.47 | 0.36 | 0.69 | 0.45 | 0.57 |
| 3 | 66735566 | rs7638955 | -0.95 | -1.978 | -0.022 | 0.743 | -0.013 | 0.021 | 0.021 | 0 | 0.17 | 0.23 | 0.15 | 0.32 | 0.07 | 0.23 |
| 3 | 66740282 | rs6803411 | -1.13 | -1.9 | 0.076 | 0.877 | -0.063 | 0.131 | 0.131 | 0 | 0.15 | 0.35 | 0.06 | 0.32 | 0.05 | 0.17 |
| 3 | 66743157 | rs11717234 | 0.713 | -1.975 | -0.437 | 0.741 | 0.002 | -0.005 | -0.005 | 0 | 0.73 | 0.72 | 0.8 | 0.64 | 0.9 | 0.72 |
| 3 | 66744271 | rs1456919 | -0.665 | -1.987 | -1.029 | 0.387 | 0.064 | 0.023 | 0.023 | 1.316 | 0.54 | 0.77 | 0.37 | 0.64 | 0.36 | 0.3 |
| 3 | 66747371 | rs6549359 | -0.16 | -2.125 | -0.534 | 0.345 | 0.048 | 0.02 | 0.02 | 0 | 0.46 | 0.65 | 0.4 | 0.64 | 0.35 | 0.3 |
| 3 | 66753674 | rs11710109 | -0.325 | -2.157 | -0.303 | 0.338 | 0.041 | 0.018 | 0.018 | 0 | 0.48 | 0.68 | 0.28 | 0.55 | 0.41 | 0.26 |
| 3 | 66755179 | rs17045018 | 1.302 | -2.663 | -0.475 | 0.857 | 0.029 | -0.014 | -0.014 | 1.531 | 0.06 | 0.02 | 0.05 | 0.19 | 0.02 | 0.04 |
| 3 | 66756087 | rs11713957 | -0.239 | -2.791 | 0.365 | 0.654 | 0.017 | -0.003 | -0.003 | 0 | 0.41 | 0.34 | 0.32 | 0.56 | 0.38 | 0.24 |
| 3 | 66757136 | rs6774173 | 0.564 | -2.936 | 0.339 | 0.385 | 0.007 | 0.015 | 0.015 | 0 | 0.29 | 0.15 | 0.24 | 0.3 | 0.31 | 0.16 |
| 3 | 66757323 | rs11919842 | 1.367 | -2.977 | 0.375 | 0.241 | -0.061 | -0.063 | -0.063 | 0.265 | 0.12 | 0.23 | 0.05 | 0 | 0 | 0 |
| 3 | 66759545 | rs9873886 | -1.228 | -2.8 | 0.411 | 0.487 | 0.027 | 0.002 | 0.002 | 1.323 | 0.54 | 0.67 | 0.36 | 0.53 | 0.38 | 0.25 |
| 3 | 66761125 | rs7637555 | -2.107 | -2.563 | -0.363 | 0.291 | -0.131 | -0.116 | -0.116 | 0 | 0.69 | 0.91 | 0.36 | 0.57 | 0.51 | 0.41 |
| 3 | 66764725 | rs1379189 | -0.353 | -2.975 | -0.252 | 0.219 | 0 | -0.002 | -0.002 | 0 | 0.18 | 0.14 | 0.13 | 0.23 | 0.12 | 0.23 |
| 3 | 66765251 | rs7622744 | 0.052 | -3.272 | 0.005 | 0.028 | -0.02 | -0.062 | -0.062 | 0 | 0.37 | 0.54 | 0.21 | 0.14 | 0.29 | 0.1 |
| 3 | 66768059 | rs11128202 | 0.048 | -2.892 | 0.112 | -0.148 | 0.014 | 0.03 | 0.03 | 2.145 | 0.37 | 0.25 | 0.33 | 0.13 | 0.34 | 0.12 |
| 3 | 66768164 | rs1350249 | -0.519 | -2.916 | 0.054 | -0.133 | 0.01 | 0.028 | 0.028 | 0 | 0.47 | 0.32 | 0.37 | 0.31 | 0.42 | 0.25 |
| 3 | 66770644 | rs13081684 | -2.662 | -2.814 | -0.367 | -0.02 | -0.112 | -0.109 | -0.109 | 0 | 0.73 | 0.9 | 0.51 | 0.58 | 0.59 | 0.44 |
| 3 | 66772425 | rs13079502 | 0.937 | -2.828 | -0.359 | -0.03 | -0.009 | 0.024 | 0.024 | 0.299 | 0.13 | 0.06 | 0.04 | 0.05 | 0.1 | 0.14 |
| 3 | 66773595 | rs9879990 | 0.913 | -2.828 | -0.249 | -0.03 | -0.001 | 0.011 | 0.011 | 0.261 | 0.13 | 0.11 | 0.04 | 0.05 | 0.1 | 0.14 |
| 3 | 66774036 | rs7647735 | -1.624 | -2.849 | -0.176 | -0.018 | -0.011 | -0.025 | -0.025 | 0.154 | 0.58 | 0.66 | 0.47 | 0.51 | 0.49 | 0.3 |
| 3 | 66774407 | rs7643123 | 0.817 | -2.842 | -0.168 | -0.009 | 0 | 0.007 | 0.007 | 0.834 | 0.1 | 0.08 | 0.04 | 0.18 | 0.08 | 0.13 |
| 3 | 66777695 | rs7613063 | -1.635 | -2.846 | -0.168 | -0.015 | -0.01 | -0.023 | -0.023 | 0 | 0.58 | 0.65 | 0.47 | 0.51 | 0.49 | 0.3 |
| 3 | 66781296 | rs12493832 | 0.699 | -2.428 | -0.36 | -0.32 | -0.001 | 0.004 | 0.004 | 0.856 | 0.1 | 0.09 | 0.13 | 0.18 | 0.08 | 0.14 |
| 3 | 66783668 | rs7646301 | -1.1 | -2.487 | -0.415 | -0.553 | -0.011 | 0.112 | 0.112 | 0.873 | 0.55 | 0.69 | 0.41 | 0.18 | 0.47 | 0.27 |
| 3 | 66786091 | rs13095384 | 1.265 | -2.562 | -0.899 | -0.59 | -0.093 | 0.047 | 0.047 | 0.214 | 0.21 | 0.01 | 0.19 | 0.14 | 0.32 | 0.11 |
| 3 | 66789030 | rs754820 | 0.461 | -2.571 | -0.748 | -0.597 | 0.047 | 0.085 | 0.085 | 0.399 | 0.25 | 0.08 | 0.09 | 0.05 | 0.11 | 0.15 |
| 3 | 66791147 | rs2364884 | -2.388 | -2.344 | -0.878 | -0.261 | -0.097 | -0.111 | -0.111 | 0.159 | 0.73 | 0.9 | 0.51 | 0.47 | 0.59 | 0.42 |
| 3 | 66791685 | rs11705804 | 1.488 | -2.142 | -0.889 | -0.163 | -0.046 | -0.021 | -0.021 | 0 | 0.07 | 0.01 | 0.21 | 0.14 | 0.18 | 0.06 |
| 3 | 66793227 | rs17045047 | -1.362 | -2.392 | -0.492 | -0.26 | 0.069 | 0.055 | 0.055 | 0.051 | 0.49 | 0.72 | 0.33 | 0.31 | 0.27 | 0.32 |
| 3 | 66794904 | rs9819324 | -0.422 | -2.007 | -0.329 | -0.443 | -0.029 | -0.076 | -0.076 | 0 | 0.31 | 0.4 | 0.13 | 0.05 | 0.16 | 0.15 |
| 3 | 66798575 | rs13068339 | -0.761 | -1.921 | -0.234 | 0.024 | 0.003 | 0.002 | 0.002 | 2.946 | 0.47 | 0.54 | 0.35 | 0.41 | 0.42 | 0.21 |
| 3 | 66799169 | rs17045066 | 0.838 | -2.174 | -0.386 | -0.12 | 0.032 | 0.02 | 0.02 | 5.943 | 0.18 | 0.07 | 0.08 | 0.1 | 0.08 | 0.06 |
| 3 | 66801080 | rs9866506 | -0.574 | -2.145 | 0.356 | 0.254 | -0.044 | 0.032 | 0.032 | 0 | 0.23 | 0.5 | 0.22 | 0.23 | 0.13 | 0.16 |
| 3 | 66802514 | rs6783763 | -0.495 | -2.167 | 0.399 | 0.244 | -0.037 | 0.03 | 0.03 | 0 | 0.23 | 0.45 | 0.14 | 0.23 | 0.13 | 0.18 |
| 3 | 66802559 | rs6798342 | -0.495 | -2.167 | 0.399 | 0.244 | -0.037 | 0.03 | 0.03 | 0 | 0.23 | 0.45 | 0.22 | 0.23 | 0.13 | 0.18 |
| 3 | 66804200 | rs13080378 | 1.306 | -2.101 | 0.249 | 0.178 | -0.053 | -0.044 | -0.044 | 0 | 0.09 | 0.01 | 0.2 | 0.2 | 0.18 | 0.06 |
| 3 | 66804504 | rs9843859 | 0.828 | -2.161 | 0.371 | 0.124 | 0.01 | -0.033 | -0.033 | 0 | 0.08 | 0.23 | 0.03 | 0 | 0.03 | 0.04 |
| 3 | 66806549 | rs6796945 | -2.389 | -2.059 | -0.203 | 0.134 | -0.055 | -0.055 | -0.055 | 0 | 0.68 | 0.82 | 0.5 | 0.53 | 0.57 | 0.37 |
| 3 | 66807053 | rs17781617 | 0.29 | -2.119 | -0.153 | 0.106 | 0.054 | 0.101 | 0.101 | 2.361 | 0.35 | 0.12 | 0.14 | 0.11 | 0.25 | 0.13 |
| 3 | 66807194 | rs6796957 | -1.246 | -2.018 | -0.141 | 0.181 | 0.007 | 0.001 | 0.001 | 0.77 | 0.58 | 0.63 | 0.29 | 0.34 | 0.39 | 0.31 |
| 3 | 66809671 | rs9870722 | -2.147 | -2 | -0.163 | 0.166 | -0.025 | -0.025 | -0.025 | 0 | 0.67 | 0.75 | 0.5 | 0.53 | 0.57 | 0.35 |
| 3 | 66811847 | rs11128217 | -0.428 | -1.948 | -0.768 | 0.125 | 0.013 | 0.058 | 0.058 | 0.931 | 0.44 | 0.26 | 0.28 | 0.31 | 0.44 | 0.18 |
| 3 | 66823157 | rs11709625 | 0.463 | -1.061 | -1.333 | 0.522 | -0.005 | -0.005 | -0.005 | 0.918 | 0.1 | 0.09 | 0.13 | 0.2 | 0.18 | 0.07 |
| 3 | 66823237 | rs11128218 | -1.211 | -1.571 | -1.414 | 0.599 | -0.073 | -0.063 | -0.063 | 0 | 0.68 | 0.86 | 0.5 | 0.57 | 0.57 | 0.36 |
| 3 | 66833437 | rs6768767 | -0.79 | -2.051 | -0.84 | 0.708 | 0.071 | -0.042 | -0.042 | 2.125 | 0.37 | 0.78 | 0.35 | 0.47 | 0.18 | 0.32 |
| 3 | 66834901 | rs7635161 | 0.233 | -2.411 | -0.461 | 0.435 | 0.01 | -0.004 | -0.004 | 0 | 0.17 | 0.14 | 0.04 | 0.02 | 0.06 | 0.06 |
| 3 | 66835382 | rs7635053 | -1.438 | -2.045 | -0.791 | 0.428 | 0.063 | 0.085 | 0.085 | 0.438 | 0.51 | 0.76 | 0.5 | 0.39 | 0.35 | 0.34 |
| 3 | 66843365 | rs1001689 | 0.802 | -1.633 | -0.822 | 0.531 | -0.016 | -0.01 | -0.01 | 1.426 | 0.06 | 0.02 | 0.06 | 0.1 | 0.14 | 0.07 |
| 3 | 66845554 | rs4487196 | -1.253 | -1.903 | -0.048 | 0.376 | 0 | 0.004 | 0.004 | 0.591 | 0.43 | 0.63 | 0.56 | 0.64 | 0.44 | 0.41 |
| 3 | 66848809 | rs13096599 | 1.214 | -1.702 | -0.289 | 0.465 | -0.066 | 0.048 | 0.048 | 0.142 | 0.14 | 0.01 | 0.17 | 0.07 | 0.18 | 0.07 |
| 3 | 66852927 | rs12714778 | -1.605 | -1.72 | -0.169 | 0.401 | -0.028 | 0.024 | 0.024 | 0 | 0.56 | 0.71 | 0.58 | 0.65 | 0.5 | 0.48 |
| 3 | 66854987 | rs989343 | -1.539 | -1.611 | 0.29 | 0.464 | -0.001 | -0.001 | -0.001 | 0 | 0.53 | 0.57 | 0.57 | 0.66 | 0.5 | 0.48 |
| 3 | 66860401 | rs725161 | -0.68 | -1.582 | 0.343 | 0.514 | -0.013 | 0.024 | 0.024 | 0 | 0.34 | 0.49 | 0.49 | 0.57 | 0.3 | 0.31 |
| 3 | 66861656 | rs12495879 | 1.525 | -1.579 | -0.056 | 0.601 | -0.064 | -0.015 | -0.015 | 0 | 0.13 | 0.01 | 0.16 | 0.14 | 0.18 | 0.08 |
| 3 | 66862644 | rs9813067 | -0.738 | -1.661 | -0.045 | 0.645 | -0.013 | 0.044 | 0.044 | 0 | 0.33 | 0.53 | 0.47 | 0.54 | 0.31 | 0.33 |
| 3 | 66866536 | rs6779112 | 0.871 | -2.311 | 0.589 | 0.659 | -0.019 | 0.037 | 0.037 | 0 | 0.05 | 0.14 | 0.03 | 0.12 | 0.01 | 0.11 |
| 3 | 66866621 | rs17045211 | 0.735 | -2.302 | 0.554 | 0.628 | 0.017 | -0.008 | -0.008 | 0.048 | 0.07 | 0.04 | 0.19 | 0.08 | 0.04 | 0.03 |
| 3 | 66866903 | rs17794276 | 1.359 | -2.438 | 0.231 | 0.513 | -0.004 | -0.006 | -0.006 | 0.041 | 0.18 | 0.01 | 0.09 | 0.14 | 0.18 | 0.16 |
| 3 | 66868609 | rs10049466 | -2.154 | -1.647 | 1.372 | 0.677 | 0.014 | -0.009 | -0.009 | 0 | 0.35 | 0.72 | 0.39 | 0.35 | 0.3 | 0.25 |
| 3 | 66868827 | rs10049470 | -0.826 | -1.553 | 1.403 | 0.689 | 0.006 | -0.042 | -0.042 | 0.01 | 0.2 | 0.3 | 0.12 | 0.08 | 0.19 | 0.13 |
| 3 | 66869791 | rs3884663 | -0.574 | -1.507 | 1.348 | 0.71 | 0.002 | -0.014 | -0.014 | 0 | 0.19 | 0.2 | 0.12 | 0.08 | 0.18 | 0.13 |
| 3 | 66873213 | rs13434181 | 0.204 | -0.896 | 1.229 | 1.173 | 0.006 | 0.008 | 0.008 | 0 | 0.05 | 0.04 | 0.05 | 0.19 | 0.07 | 0.09 |
| 3 | 66877773 | rs7643146 | -0.636 | -0.811 | 1.342 | 0.94 | 0.069 | -0.024 | -0.024 | 0 | 0.13 | 0.35 | 0.12 | 0.07 | 0.15 | 0.13 |
| 3 | 66879115 | rs2218406 | -1.907 | -0.997 | 0.479 | 0.967 | 0.027 | 0.025 | 0.025 | 1.241 | 0.31 | 0.75 | 0.39 | 0.24 | 0.25 | 0.31 |
| 3 | 66879884 | rs11916711 | -2.281 | -1.023 | 0.364 | 1.022 | 0.049 | -0.11 | -0.11 | 0 | 0.34 | 0.89 | 0.41 | 0.47 | 0.27 | 0.35 |
| 3 | 66880001 | rs12630651 | -0.157 | -0.814 | 0.44 | 1.2 | -0.022 | -0.002 | -0.002 | 0.536 | 0.1 | 0.17 | 0.07 | 0.12 | 0.07 | 0.13 |
| 3 | 66880124 | rs17045315 | -0.052 | -0.802 | 0.418 | 1.192 | 0.02 | -0.003 | -0.003 | 0.267 | 0.07 | 0.04 | 0.19 | 0.06 | 0.04 | 0.03 |
| 3 | 66882799 | rs7627693 | -0.482 | -0.692 | 0.47 | 1.217 | -0.039 | 0.068 | 0.068 | 0 | 0.18 | 0.34 | 0.28 | 0.4 | 0.12 | 0.22 |

| | | | | | | | | | | | | | | | | |
|---|---|---|---|---|---|---|---|---|---|---|---|---|---|---|---|---|
| 3 | 66892025 | rs7635135 | -1.658 | -0.614 | 0.08 | 1.175 | 0.029 | -0.035 | -0.035 | 0.599 | 0.34 | 0.78 | 0.41 | 0.46 | 0.28 | 0.39 |
| 3 | 66900101 | rs6772906 | -0.857 | -0.798 | -0.295 | 1.404 | -0.016 | 0.015 | 0.015 | 0 | 0.19 | 0.24 | 0.27 | 0.35 | 0.12 | 0.19 |
| 3 | 66905775 | rs9862849 | -0.947 | 0.822 | -0.138 | 1.003 | 0.021 | -0.002 | -0.002 | 3.201 | 0.09 | 0.17 | 0.12 | 0.07 | 0.08 | 0.11 |
| 3 | 66906996 | rs9875865 | -1.81 | 0.35 | 0.513 | 1.336 | 0.007 | 0.093 | 0.093 | 0 | 0.14 | 0.38 | 0.15 | 0.26 | 0.11 | 0.18 |
| 3 | 66912679 | rs4411828 | -0.249 | 0.082 | 0.089 | 0.418 | -0.019 | 0.092 | 0.092 | 3.202 | 0.76 | 0.95 | 0.74 | 0.94 | 0.79 | 0.9 |
| 3 | 66914581 | rs1470354 | -0.886 | -0.183 | 0.891 | 0.028 | 0.021 | -0.058 | -0.058 | 0.057 | 0.22 | 0.44 | 0.19 | 0.1 | 0.23 | 0.19 |
| 3 | 66914689 | rs7648743 | 1.219 | -0.234 | 0.938 | -0.025 | 0.003 | 0.06 | 0.06 | 0.258 | 0.52 | 0.35 | 0.53 | 0.84 | 0.55 | 0.72 |
| 3 | 66917635 | rs7650991 | 0.181 | -0.224 | 1.189 | -0.138 | -0.002 | -0.043 | -0.043 | 0.001 | 0.14 | 0.22 | 0.07 | 0.03 | 0.13 | 0.07 |
| 3 | 66928844 | rs9861183 | -0.469 | -0.432 | 0.736 | -0.625 | -0.003 | 0.006 | 0.006 | 0.774 | 0.18 | 0.19 | 0.14 | 0.08 | 0.2 | 0.25 |
| 3 | 66929752 | rs6790110 | 1.568 | -0.591 | 0.611 | -0.656 | -0.09 | 0.054 | 0.054 | 3.093 | 0.41 | 0.13 | 0.51 | 0.72 | 0.48 | 0.56 |
| 3 | 66930228 | rs7634770 | -1.045 | -0.557 | 0.702 | -0.694 | -0.002 | -0.061 | -0.061 | 1.21 | 0.31 | 0.42 | 0.2 | 0.11 | 0.3 | 0.31 |
| 3 | 66931371 | rs9876289 | 0.59 | -0.409 | 0.833 | -0.567 | 0.004 | 0.01 | 0.01 | 3.952 | 0.11 | 0.08 | 0.08 | 0.06 | 0.1 | 0.12 |
| 3 | 66940414 | rs9310249 | -0.323 | -0.424 | 0.973 | -0.553 | 0.002 | -0.021 | -0.021 | 0 | 0.18 | 0.23 | 0.15 | 0.08 | 0.2 | 0.25 |
| 3 | 66955944 | rs6776236 | -0.084 | -0.476 | 1.096 | -0.594 | -0.003 | -0.062 | -0.062 | 0 | 0.6 | 0.54 | 0.68 | 0.97 | 0.69 | 0.84 |
| 3 | 66959472 | rs1456217 | -0.566 | -0.717 | 0.862 | -0.82 | -0.017 | -0.13 | -0.13 | 0 | 0.68 | 0.53 | 0.85 | 0.97 | 0.75 | 0.91 |
| 3 | 66961432 | rs1378849 | 1.214 | -0.692 | 0.561 | -0.757 | 0.009 | 0.025 | 0.025 | 0.156 | 0.08 | 0.04 | 0.17 | 0 | 0.06 | 0.07 |
| 3 | 66980276 | rs4856832 | 1.428 | -0.651 | 0.57 | -0.621 | -0.109 | 0.021 | 0.021 | 2.929 | 0.41 | 0.08 | 0.51 | 0.71 | 0.48 | 0.58 |
| 3 | 66990123 | rs936207 | 0.311 | -0.672 | 0.951 | -0.657 | -0.022 | -0.066 | -0.066 | 0.081 | 0.11 | 0.34 | 0.2 | 0 | 0.07 | 0.07 |
| 3 | 66990499 | rs2045308 | -0.405 | -0.685 | 0.977 | -0.713 | 0.008 | -0.003 | -0.003 | 0.482 | 0.67 | 0.34 | 0.85 | 0.97 | 0.76 | 0.91 |
| 3 | 66995885 | rs9843667 | -0.316 | -0.597 | 0.998 | -0.598 | 0.008 | -0.025 | -0.025 | 0 | 0.19 | 0.27 | 0.16 | 0.09 | 0.22 | 0.24 |
| 3 | 66996179 | rs1463409 | -0.494 | -0.655 | 0.922 | -0.717 | -0.013 | -0.093 | -0.093 | 0.902 | 0.67 | 0.49 | 0.85 | 0.97 | 0.76 | 0.91 |
| 3 | 67007232 | rs2198795 | -1.279 | -0.64 | 1.26 | -0.649 | -0.008 | -0.006 | -0.006 | 0 | 0.3 | 0.67 | 0.36 | 0.26 | 0.29 | 0.33 |
| 3 | 67010311 | rs6800348 | -0.348 | -0.735 | 1.271 | -0.669 | 0.007 | 0.03 | 0.03 | 1.197 | 0.21 | 0.35 | 0.17 | 0.26 | 0.22 | 0.25 |
| 3 | 67011039 | rs726083 | -1.276 | -0.771 | 1.186 | -0.682 | -0.009 | -0.005 | -0.005 | 0 | 0.31 | 0.63 | 0.37 | 0.27 | 0.29 | 0.33 |
| 3 | 67013258 | rs9874651 | -0.91 | -0.769 | 1.184 | -0.68 | -0.001 | -0.02 | -0.02 | 0 | 0.28 | 0.32 | 0.33 | 0.09 | 0.28 | 0.3 |
| 3 | 67015910 | rs6807497 | -0.817 | -0.763 | 0.956 | -1.04 | -0.009 | -0.08 | -0.08 | 0 | 0.71 | 0.69 | 0.87 | 0.97 | 0.77 | 0.92 |
| 3 | 67023150 | rs1869900 | 1.391 | -0.082 | 0.522 | -1.026 | -0.074 | 0.179 | 0.179 | 3.043 | 0.39 | 0.09 | 0.49 | 0.89 | 0.45 | 0.61 |
| 3 | 67033323 | rs1009032 | 0.929 | -0.083 | 0.532 | -0.985 | 0.006 | 0.032 | 0.032 | 0 | 0.09 | 0.04 | 0.17 | 0 | 0.08 | 0.07 |
| 3 | 67039109 | rs9877428 | 0.019 | -0.243 | 0.657 | -1.112 | -0.005 | 0.01 | 0.01 | 1.328 | 0.18 | 0.18 | 0.15 | 0.09 | 0.21 | 0.24 |
| 3 | 67040146 | rs12486952 | 1.129 | -0.296 | 1.2 | -1.236 | -0.012 | 0.052 | 0.052 | 2.173 | 0.44 | 0.31 | 0.55 | 0.89 | 0.53 | 0.63 |
| 3 | 67046187 | rs9875436 | -0.308 | -0.359 | 1.287 | -1.182 | 0.004 | -0.045 | -0.045 | 2.204 | 0.21 | 0.34 | 0.17 | 0.09 | 0.22 | 0.26 |
| 3 | 67047977 | rs9833720 | -0.992 | -0.287 | 1.068 | -1.136 | 0 | -0.029 | -0.029 | 0 | 0.72 | 0.79 | 0.87 | 0.98 | 0.76 | 0.92 |
| 3 | 67054649 | rs13096789 | 0.693 | 0.776 | 0.63 | -0.976 | -0.026 | 0.042 | 0.042 | 0 | 0.11 | 0.05 | 0.23 | 0.02 | 0.14 | 0.09 |
| 3 | 67055144 | rs6549565 | -1.124 | 0.764 | 0.834 | -1.035 | 0.019 | -0.09 | -0.09 | 0.094 | 0.31 | 0.58 | 0.42 | 0.1 | 0.34 | 0.33 |
| 3 | 67059553 | rs13078632 | 0.389 | 0.784 | 0.395 | -0.905 | -0.021 | 0.019 | 0.019 | 0 | 0.07 | 0.05 | 0.23 | 0.02 | 0.13 | 0.1 |
| 3 | 67061960 | rs6549575 | -0.442 | 0.865 | 0.098 | -1.183 | -0.004 | 0.099 | 0.099 | 0 | 0.51 | 0.68 | 0.75 | 0.85 | 0.56 | 0.63 |
| 3 | 67073768 | rs11128375 | 0.004 | 0.334 | -0.923 | -0.295 | -0.003 | 0.026 | 0.026 | 0.29 | 0.48 | 0.27 | 0.44 | 0.57 | 0.49 | 0.67 |
| 3 | 67078548 | rs13086666 | 0.648 | 0.177 | -1.734 | -0.271 | -0.066 | -0.11 | -0.11 | 0 | 0.34 | 0.08 | 0.36 | 0.51 | 0.34 | 0.54 |
| 3 | 67079104 | rs7653502 | 0.375 | 0.209 | -1.031 | 0.138 | -0.006 | -0.01 | -0.01 | 0 | 0.12 | 0.11 | 0.2 | 0.4 | 0.15 | 0.11 |
| 3 | 67081996 | rs4241443 | -1.636 | 0.475 | -1.472 | -0.2 | -0.044 | 0.197 | 0.197 | 0 | 0.57 | 0.86 | 0.77 | 0.89 | 0.56 | 0.61 |
| 3 | 67087513 | rs4856981 | -1.279 | 0.274 | -0.659 | -0.203 | -0.001 | -0.002 | -0.002 | 0 | 0.41 | 0.51 | 0.58 | 0.46 | 0.42 | 0.37 |
| 3 | 67089778 | rs7427956 | 0.074 | 0.31 | -0.462 | -0.2 | 0.001 | 0.002 | 0.002 | 0 | 0.19 | 0.19 | 0.36 | 0.29 | 0.19 | 0.13 |
| 3 | 67094975 | rs9814525 | -1.279 | 0.269 | -0.531 | -0.241 | 0.002 | -0.008 | -0.008 | 0.373 | 0.36 | 0.46 | 0.55 | 0.37 | 0.37 | 0.26 |
| 3 | 67095678 | rs6805111 | 0.495 | 0.451 | -0.419 | -0.188 | -0.006 | 0.02 | 0.02 | 2.659 | 0.15 | 0.11 | 0.17 | 0.07 | 0.18 | 0.09 |
| 3 | 67102024 | rs6549588 | -1.929 | 0.386 | -0.28 | -0.098 | -0.002 | -0.017 | -0.017 | 0 | 0.54 | 0.57 | 0.74 | 0.78 | 0.56 | 0.62 |
| 3 | 67104628 | rs6786681 | -1.263 | 0.519 | 0.83 | 0.378 | -0.003 | -0.024 | -0.024 | 0 | 0.41 | 0.57 | 0.57 | 0.77 | 0.38 | 0.56 |
| 3 | 67112672 | rs11128388 | 0.411 | 0.656 | 0.542 | 0.482 | -0.018 | -0.038 | -0.038 | 0 | 0.13 | 0.1 | 0.2 | 0.31 | 0.15 | 0.11 |
| 3 | 67113777 | rs4462892 | -0.318 | 0.659 | 0.551 | 0.487 | -0.012 | -0.019 | -0.019 | 0 | 0.17 | 0.14 | 0.35 | 0.3 | 0.19 | 0.15 |
| 3 | 67127674 | rs4561768 | -0.255 | 0.677 | 0.761 | 0.695 | -0.001 | 0.003 | 0.003 | 0 | 0.14 | 0.16 | 0.34 | 0.33 | 0.16 | 0.11 |
| 3 | 67147194 | rs6804724 | -1.308 | 0.659 | 0.846 | 0.687 | -0.001 | 0.03 | 0.03 | 0 | 0.21 | 0.34 | 0.38 | 0.36 | 0.21 | 0.24 |
| 3 | 67154946 | rs7645834 | -0.266 | 0.736 | 0.583 | 0.68 | -0.006 | -0.013 | -0.013 | 0 | 0.14 | 0.13 | 0.34 | 0.3 | 0.16 | 0.12 |
| 3 | 67160633 | rs9867793 | 0.736 | 0.752 | 0.518 | 0.721 | -0.01 | -0.022 | -0.022 | 1.207 | 0.1 | 0.1 | 0.17 | 0.32 | 0.12 | 0.09 |
| 3 | 67162104 | rs7632236 | 0.478 | 0.222 | 0.927 | 0.613 | -0.004 | 0.093 | 0.093 | 0.069 | 0.18 | 0.3 | 0.21 | 0.35 | 0.14 | 0.28 |
| 3 | 67163881 | rs6765905 | -0.584 | 0.222 | 0.755 | 0.022 | -0.02 | 0.058 | 0.058 | 1.086 | 0.29 | 0.57 | 0.38 | 0.48 | 0.23 | 0.45 |
| 3 | 67164944 | rs7616284 | 0.571 | 0.174 | 1.059 | -0.886 | 0.001 | 0.002 | 0.002 | 0 | 0.28 | 0.34 | 0.37 | 0.71 | 0.29 | 0.34 |
| 3 | 67165103 | rs7616394 | -0.63 | 0.19 | 0.741 | -1.044 | 0.025 | 0.159 | 0.159 | 2.496 | 0.48 | 0.84 | 0.67 | 0.81 | 0.42 | 0.56 |
| 3 | 67177419 | rs4602330 | -0.041 | -0.071 | 1.398 | -0.843 | -0.019 | 0.075 | 0.075 | 2.451 | 0.34 | 0.14 | 0.34 | 0.15 | 0.41 | 0.4 |
| 3 | 67180677 | rs6774674 | -0.659 | -0.083 | 1.716 | -0.857 | -0.001 | -0.003 | -0.003 | 0 | 0.42 | 0.43 | 0.47 | 0.2 | 0.46 | 0.4 |
| 3 | 67195060 | rs7609636 | -0.736 | -0.03 | 1.91 | -0.834 | -0.001 | -0.032 | -0.032 | 0 | 0.43 | 0.5 | 0.47 | 0.2 | 0.46 | 0.4 |
| 3 | 67195990 | rs6549715 | 0.095 | -0.016 | 1.83 | -0.827 | -0.007 | 0.049 | 0.049 | 0.929 | 0.33 | 0.23 | 0.21 | 0.15 | 0.39 | 0.37 |
| 3 | 67211709 | rs11128434 | -0.334 | -0.19 | 1.892 | -0.775 | -0.004 | 0 | 0 | 0 | 0.94 | 0.94 | 0.97 | 0.95 | 0.95 | 0.85 |
| 3 | 67222606 | rs9829523 | 0.481 | -0.219 | 2.281 | -0.784 | 0.006 | 0.02 | 0.02 | 0 | 0.93 | 0.79 | 0.96 | 0.95 | 0.95 | 0.85 |
| 3 | 67226757 | rs11128459 | 1.372 | -0.256 | 2.315 | -0.803 | -0.017 | -0.008 | -0.008 | 0 | 0.89 | 0.7 | 0.8 | 0.93 | 0.92 | 0.83 |
| 3 | 67231601 | rs4347959 | 1.68 | -0.278 | 2.361 | -0.823 | -0.024 | -0.05 | -0.05 | 0 | 0.83 | 0.55 | 0.7 | 0.89 | 0.88 | 0.82 |
| 3 | 67245703 | rs6549776 | 0.257 | -0.269 | 2.292 | -0.819 | -0.001 | 0.007 | 0.007 | 0 | 0.92 | 0.82 | 0.96 | 0.95 | 0.95 | 0.85 |
| 3 | 67250048 | rs4857002 | 0.671 | -0.244 | 2.244 | -0.852 | -0.001 | -0.008 | -0.008 | 0 | 0.54 | 0.5 | 0.64 | 0.73 | 0.52 | 0.46 |
| 3 | 67261017 | rs9844493 | 1.202 | -0.246 | 2.217 | -0.86 | 0.007 | 0.092 | 0.092 | 1.278 | 0.49 | 0.34 | 0.48 | 0.71 | 0.49 | 0.43 |
| 3 | 67261856 | rs6769831 | 0.277 | -0.13 | 2.239 | -0.85 | -0.032 | 0.068 | 0.068 | 2.078 | 0.32 | 0.11 | 0.21 | 0.18 | 0.39 | 0.39 |
| 3 | 67263028 | rs4368439 | 0.065 | -0.198 | 2.277 | -0.87 | 0.006 | 0.015 | 0.015 | 0 | 0.93 | 0.8 | 0.96 | 0.95 | 0.95 | 0.85 |

| | | | | | | | | | | | | | | | | |
|---|---|---|---|---|---|---|---|---|---|---|---|---|---|---|---|---|
| 3 | 67263927 | rs13067571 | 1.176 | -0.183 | 2.245 | -0.874 | 0.008 | 0.14 | 0.14 | 1.462 | 0.49 | 0.29 | 0.49 | 0.72 | 0.49 | 0.43 |
| 3 | 67264139 | rs4578968 | 0.785 | -0.119 | 1.701 | -0.844 | 0.002 | 0.021 | 0.021 | 1.017 | 0.53 | 0.46 | 0.64 | 0.73 | 0.52 | 0.46 |
| 3 | 67266117 | rs4443108 | 0.362 | -0.11 | 1.577 | -0.834 | -0.002 | -0.021 | -0.021 | 0.014 | 0.6 | 0.62 | 0.74 | 0.77 | 0.56 | 0.47 |
| 3 | 67270120 | rs4411829 | 0.872 | -0.18 | 1.835 | -0.381 | 0.004 | 0.015 | 0.015 | 0 | 0.91 | 0.73 | 0.96 | 0.95 | 0.94 | 0.85 |
| 3 | 67271303 | rs9825439 | 0.632 | -0.033 | 1.463 | -0.315 | 0.011 | 0.033 | 0.033 | 0.355 | 0.56 | 0.43 | 0.52 | 0.72 | 0.6 | 0.47 |
| 3 | 67290851 | rs6809990 | -1.287 | 0.115 | 2.04 | 0.41 | 0.04 | -0.009 | -0.009 | 0 | 0.08 | 0.32 | 0.12 | 0.04 | 0.04 | 0.01 |
| 3 | 67290959 | rs17045959 | -0.669 | 0.064 | 2.044 | 0.409 | 0.003 | -0.025 | -0.025 | 0.015 | 0.07 | 0.15 | 0.11 | 0.04 | 0.04 | 0.01 |
| 3 | 67291478 | rs6549843 | 1.46 | 0.049 | 1.669 | 0.422 | -0.012 | 0.029 | 0.029 | 5.939 | 0.89 | 0.46 | 0.86 | 0.86 | 0.96 | 0.86 |
| 3 | 67297481 | rs9874171 | -1.167 | 0.367 | 0.377 | 0.283 | 0.019 | -0.013 | -0.013 | 0 | 0.08 | 0.36 | 0.06 | 0.04 | 0.04 | 0.01 |
| 3 | 67299754 | rs7612076 | 0.177 | 0.463 | -0.952 | 0.431 | 0.04 | 0.041 | 0.041 | 0 | 0.48 | 0.19 | 0.46 | 0.55 | 0.6 | 0.65 |
| 3 | 67300242 | rs6548418 | 0.175 | 0.829 | -0.785 | 0.745 | 0.01 | 0.008 | 0.008 | 1.648 | 0.48 | 0.4 | 0.66 | 0.39 | 0.57 | 0.61 |
| 3 | 67300424 | rs13084679 | -0.449 | 0.668 | -0.198 | 0.503 | -0.005 | 0.01 | 0.01 | 0 | 0.06 | 0.16 | 0.03 | 0.06 | 0.03 | 0.04 |
| 3 | 67302250 | rs11712613 | 0.374 | 1.365 | -0.321 | 0.709 | -0.003 | 0.084 | 0.084 | 0.445 | 0.45 | 0.27 | 0.52 | 0.29 | 0.45 | 0.56 |
| 3 | 67308939 | rs4638986 | 0.302 | 1.431 | -0.159 | 0.806 | -0.054 | 0.085 | 0.085 | 0.927 | 0.44 | 0.14 | 0.52 | 0.32 | 0.45 | 0.57 |
| 3 | 67317284 | rs1490244 | -0.102 | 1.168 | -0.396 | 0.411 | 0.013 | 0.012 | 0.012 | 1.096 | 0.3 | 0.47 | 0.31 | 0.62 | 0.32 | 0.34 |
| 3 | 67320692 | rs4626110 | -0.455 | 1.302 | -0.408 | 0.389 | 0 | 0.036 | 0.036 | 0 | 0.76 | 0.82 | 0.84 | 0.94 | 0.75 | 0.91 |
| 3 | 67321882 | rs12485939 | 1.005 | 1.266 | -0.476 | 0.352 | 0.008 | 0.072 | 0.072 | 0.661 | 0.45 | 0.29 | 0.53 | 0.32 | 0.44 | 0.57 |
| 3 | 67323670 | rs1490235 | -0.361 | 1.377 | -0.692 | 0.619 | -0.024 | 0.047 | 0.047 | 0 | 0.75 | 0.86 | 0.79 | 0.87 | 0.7 | 0.89 |
| 3 | 67328279 | rs13095630 | -0.228 | 1.291 | -0.196 | 0.542 | 0.002 | 0.044 | 0.044 | 0.33 | 0.29 | 0.4 | 0.26 | 0.56 | 0.26 | 0.33 |
| 3 | 67336952 | rs7611269 | 0.377 | 1.11 | -0.163 | -0.144 | -0.002 | -0.019 | -0.019 | 0 | 0.59 | 0.33 | 0.61 | 0.95 | 0.55 | 0.86 |
| 3 | 67340025 | rs1490270 | -1.04 | 1.711 | -0.501 | -0.013 | -0.042 | 0.033 | 0.033 | 0 | 0.19 | 0.11 | 0.39 | 0.08 | 0.28 | 0.22 |
| 3 | 67342414 | rs1490273 | -0.401 | 1.3 | -0.717 | -0.194 | 0 | 0.001 | 0.001 | 4.749 | 0.87 | 0.88 | 0.86 | 0.86 | 0.84 | 0.87 |
| 3 | 67345223 | rs1490271 | 0.33 | -0.403 | -0.844 | -0.613 | -0.085 | 0.008 | 0.008 | 1.336 | 0.3 | 0.39 | 0.12 | 0.29 | 0.16 | 0.15 |
| 3 | 67348332 | rs902323 | 0.742 | -0.444 | -0.828 | -0.622 | -0.051 | -0.029 | -0.029 | 1.553 | 0.27 | 0.29 | 0.09 | 0.15 | 0.12 | 0.05 |
| 3 | 67348748 | rs2885873 | 1.731 | -0.322 | -0.528 | -0.71 | -0.016 | -0.002 | -0.002 | 0 | 0.4 | 0.35 | 0.41 | 0.63 | 0.45 | 0.6 |
| 3 | 67351039 | rs2054956 | 0.051 | -0.509 | -0.513 | -0.656 | -0.034 | -0.019 | -0.019 | 0 | 0.6 | 0.59 | 0.77 | 0.71 | 0.72 | 0.82 |
| 3 | 67356977 | rs4856846 | 0.333 | -0.34 | -0.485 | -0.677 | -0.04 | -0.041 | -0.041 | 0.86 | 0.31 | 0.33 | 0.35 | 0.15 | 0.19 | 0.12 |
| 3 | 67366936 | rs17805868 | 0.711 | -0.489 | -0.422 | -0.697 | 0.08 | 0.048 | 0.048 | 1.72 | 0.27 | 0.08 | 0.08 | 0.15 | 0.13 | 0.08 |
| 3 | 67377843 | rs1552621 | 1.862 | -0.524 | -0.57 | -0.984 | -0.047 | -0.023 | -0.023 | 0.48 | 0.35 | 0.26 | 0.37 | 0.63 | 0.47 | 0.52 |
| 3 | 67379361 | rs9876253 | 0.144 | -0.648 | -0.478 | -1.034 | -0.013 | -0.016 | -0.016 | 0 | 0.57 | 0.62 | 0.51 | 0.73 | 0.65 | 0.79 |
| 3 | 67397253 | rs7636311 | 1.866 | -0.437 | -0.6 | -0.9 | -0.037 | -0.023 | -0.023 | 0.436 | 0.34 | 0.26 | 0.36 | 0.63 | 0.43 | 0.5 |
| 3 | 67400029 | rs1552616 | 0.073 | -0.261 | -0.672 | -0.979 | -0.009 | 0.011 | 0.011 | 0 | 0.73 | 0.74 | 0.76 | 0.9 | 0.67 | 0.77 |
| 3 | 67405492 | rs7629435 | 0.1 | -0.508 | -0.497 | -0.766 | 0.022 | 0.023 | 0.023 | 0.35 | 0.39 | 0.28 | 0.2 | 0.23 | 0.26 | 0.1 |
| 3 | 67415330 | rs1120653 | 0.208 | -0.821 | -0.482 | -0.832 | -0.017 | -0.014 | -0.014 | 0 | 0.77 | 0.62 | 0.66 | 0.79 | 0.81 | 0.82 |
| 3 | 67416322 | rs17806888 | 1.336 | -0.825 | -0.799 | -0.861 | 0.055 | 0.08 | 0.08 | 0.154 | 0.21 | 0.04 | 0.07 | 0.04 | 0.13 | 0.03 |
| 3 | 67418124 | rs12637020 | 0.234 | -0.156 | -0.39 | 0.097 | 0.039 | 0.045 | 0.045 | 0 | 0.13 | 0.19 | 0.21 | 0.24 | 0.24 | 0.19 |
| 3 | 67421818 | rs3952322 | 0.744 | -0.02 | -0.417 | 0.051 | 0.034 | 0.04 | 0.04 | 0 | 0.1 | 0.17 | 0.17 | 0.15 | 0.16 | 0.1 |
| 3 | 67422015 | rs1979639 | 1.039 | -0.348 | -0.821 | -0.127 | -0.001 | -0.001 | -0.001 | 0.249 | 0.6 | 0.44 | 0.45 | 0.55 | 0.57 | 0.62 |
| 3 | 67427443 | rs11923377 | -0.067 | 0.345 | -0.805 | 0.403 | 0.003 | -0.009 | -0.009 | 0 | 0.07 | 0.05 | 0.03 | 0.07 | 0.06 | 0.18 |
| 3 | 67430315 | rs9816677 | 0.085 | 0.413 | -0.804 | -0.015 | 0.003 | 0.033 | 0.033 | 3.219 | 0.84 | 0.9 | 0.72 | 0.97 | 0.85 | 0.88 |
| 3 | 67430355 | rs2101566 | -0.026 | 0.328 | -0.527 | 0.472 | -0.012 | -0.003 | -0.003 | 0.01 | 0.62 | 0.59 | 0.63 | 0.61 | 0.71 | 0.69 |
| 3 | 67431071 | rs13064490 | 0.92 | 1.63 | 0.034 | 0.972 | 0.026 | 0.029 | 0.029 | 0 | 0.7 | 0.61 | 0.64 | 0.64 | 0.62 | 0.71 |
| 3 | 67433189 | rs1352658 | 0.657 | 2.212 | -0.041 | 1.042 | -0.027 | -0.015 | -0.015 | 0 | 0.74 | 0.78 | 0.68 | 0.61 | 0.59 | 0.58 |
| 3 | 67437836 | rs6791324 | -1.323 | 2.353 | 0.446 | 1.29 | 0.063 | 0.101 | 0.101 | 0 | 0.1 | 0.33 | 0.34 | 0.15 | 0.13 | 0.13 |
| 3 | 67438396 | rs17194629 | 0.388 | 2.393 | 0.47 | 1.071 | 0.002 | -0.012 | -0.012 | 0 | 0.11 | 0.11 | 0.09 | 0.06 | 0.15 | 0.1 |
| 3 | 67439555 | rs17194748 | 1.322 | 2.09 | 0.212 | 1.029 | -0.007 | -0.017 | -0.017 | 0 | 0.1 | 0.01 | 0.1 | 0.09 | 0.08 | 0.02 |
| 3 | 67439784 | rs7647085 | 1.944 | 2.011 | 0.29 | 0.999 | 0.088 | 0.059 | 0.059 | 1.019 | 0.48 | 0.28 | 0.3 | 0.32 | 0.3 | 0.41 |
| 3 | 67441915 | rs17194986 | -0.533 | 2.126 | 0.373 | 1.332 | -0.003 | -0.003 | -0.003 | 0 | 0.06 | 0.06 | 0.14 | 0 | 0.07 | 0.08 |
| 3 | 67449926 | rs7652000 | 0.757 | 2.083 | 0.391 | 1.39 | 0.01 | 0.002 | 0.002 | 1.995 | 0.11 | 0.06 | 0.1 | 0.09 | 0.08 | 0.02 |
| 3 | 67450230 | rs1552626 | -0.513 | 2.467 | 0.41 | 1.536 | 0.002 | 0 | 0 | 0.196 | 0.85 | 0.86 | 0.89 | 0.74 | 0.78 | 0.78 |
| 3 | 67452891 | rs1352656 | -1.553 | 1.765 | 0.575 | 1.251 | 0.035 | 0.056 | 0.056 | 0 | 0.14 | 0.4 | 0.36 | 0.16 | 0.16 | 0.16 |
| 3 | 67453018 | rs1352657 | 1.882 | 1.846 | 0.018 | 1.264 | 0.009 | 0.018 | 0.018 | 0 | 0.56 | 0.31 | 0.38 | 0.37 | 0.46 | 0.5 |
| 3 | 67453084 | rs1387867 | -0.757 | 2.495 | -0.464 | 1.436 | -0.003 | -0.007 | -0.007 | 0 | 0.08 | 0.08 | 0.15 | 0 | 0.1 | 0.08 |
| 3 | 67454918 | rs4450832 | -1.081 | 2.44 | -0.464 | 1.475 | 0.003 | 0.006 | 0.006 | 0 | 0.12 | 0.19 | 0.35 | 0.16 | 0.16 | 0.16 |
| 3 | 67455803 | rs4132748 | 0.532 | 2.207 | -0.73 | 1.231 | -0.011 | -0.01 | -0.01 | 0 | 0.79 | 0.83 | 0.84 | 0.62 | 0.7 | 0.64 |
| 3 | 67458381 | rs4577484 | 0.182 | 1.052 | -0.549 | 0.86 | -0.006 | -0.008 | -0.008 | 0 | 0.35 | 0.44 | 0.37 | 0.37 | 0.33 | 0.42 |
| 3 | 67458562 | rs12497038 | 0.641 | 1.047 | -1.018 | 0.901 | 0 | -0.022 | -0.022 | 0 | 0.31 | 0.21 | 0.3 | 0.31 | 0.28 | 0.3 |
| 3 | 67459050 | rs4320072 | -0.891 | 1.269 | -1.119 | 0.745 | -0.015 | 0.003 | 0.003 | 0 | 0.84 | 0.82 | 0.9 | 0.87 | 0.77 | 0.8 |
| 3 | 67464789 | rs13076634 | 0.914 | 1.163 | -0.218 | 0.947 | -0.001 | -0.003 | -0.003 | 0 | 0.38 | 0.38 | 0.37 | 0.37 | 0.35 | 0.42 |
| 3 | 67468422 | rs4856784 | -0.173 | 1.011 | 0.219 | 1.325 | 0.012 | -0.054 | -0.054 | 1.557 | 0.18 | 0.07 | 0.33 | 0.38 | 0.17 | 0.18 |
| 3 | 67470690 | rs4856863 | 1.782 | 1.005 | 0.222 | 1.263 | -0.011 | 0.029 | 0.029 | 0 | 0.55 | 0.34 | 0.49 | 0.37 | 0.51 | 0.48 |
| 3 | 67474874 | rs4377498 | -0.572 | 1.085 | 0.564 | 1.58 | -0.023 | -0.003 | -0.003 | 0 | 0.84 | 0.87 | 0.91 | 0.85 | 0.78 | 0.8 |
| 3 | 67475097 | rs12489223 | -0.27 | 1.072 | 0.621 | 1.561 | -0.002 | 0 | 0 | 0 | 0.83 | 0.74 | 0.9 | 0.85 | 0.78 | 0.8 |
| 3 | 67476165 | rs4130105 | -1.12 | 0.967 | 0.679 | 1.624 | 0.047 | 0.058 | 0.058 | 0 | 0.07 | 0.18 | 0.08 | 0.11 | 0.1 | 0.12 |
| 3 | 67506870 | rs6810055 | 0.733 | 0.748 | 0.594 | 1.687 | 0.038 | 0.081 | 0.081 | 0 | 0.76 | 0.54 | 0.72 | 0.49 | 0.7 | 0.6 |
| 3 | 67511157 | rs4645146 | 1.642 | 0.682 | 0.628 | 1.588 | 0.007 | 0.005 | 0.005 | 0 | 0.67 | 0.39 | 0.67 | 0.39 | 0.66 | 0.5 |
| 3 | 67512114 | rs4423787 | -0.286 | 0.96 | 0.385 | 1.718 | -0.031 | -0.005 | -0.005 | 0 | 0.84 | 0.86 | 0.86 | 0.85 | 0.74 | 0.81 |
| 3 | 67513491 | rs9824365 | -1.695 | 0.845 | 0.494 | 1.625 | 0.021 | 0.064 | 0.064 | 0.924 | 0.08 | 0.28 | 0.06 | 0.07 | 0.05 | 0.1 |
| 3 | 67515392 | rs9851908 | -1.783 | 0.841 | 0.469 | 1.65 | 0.001 | 0.083 | 0.083 | 0.069 | 0.11 | 0.28 | 0.09 | 0.23 | 0.08 | 0.15 |

| | | | | | | | | | | | | | | | | |
|---|---|---|---|---|---|---|---|---|---|---|---|---|---|---|---|---|
| 3 | 67522017 | rs9809635 | -0.707 | 1.162 | 0.152 | 1.743 | 0.011 | 0.055 | 0.055 | 0 | 0.06 | 0.15 | 0.05 | 0.17 | 0.05 | 0.14 |
| 3 | 67527259 | rs6788383 | -0.474 | 1.201 | 0.067 | 1.781 | 0.007 | 0.043 | 0.043 | 0 | 0.05 | 0.07 | 0.05 | 0.18 | 0.05 | 0.14 |
| 3 | 67531388 | rs6792584 | 0.089 | 1.136 | 0.053 | 1.918 | -0.027 | -0.018 | -0.018 | 0.19 | 0.83 | 0.86 | 0.85 | 0.77 | 0.74 | 0.78 |
| 3 | 67536688 | rs6548593 | -0.082 | 1.092 | 0.062 | 1.955 | -0.019 | -0.016 | -0.016 | 0 | 0.84 | 0.83 | 0.87 | 0.77 | 0.75 | 0.79 |
| 3 | 67554942 | rs9872341 | 0.391 | 0.938 | 0.786 | 2.018 | 0.052 | 0.066 | 0.066 | 0 | 0.78 | 0.67 | 0.8 | 0.63 | 0.69 | 0.75 |
| 3 | 67564351 | rs9860008 | -1.244 | 0.207 | 1.514 | 1.901 | -0.017 | 0.179 | 0.179 | 0.021 | 0.11 | 0.31 | 0.13 | 0.22 | 0.04 | 0.17 |
| 3 | 67567254 | rs4334654 | -1.488 | -0.261 | 1.354 | 1.919 | -0.004 | 0.004 | 0.004 | 0 | 0.29 | 0.64 | 0.3 | 0.49 | 0.19 | 0.44 |
| 3 | 67569703 | rs7624455 | 0.652 | 0.003 | 1.661 | 1.868 | 0.043 | 0.086 | 0.086 | 0 | 0.77 | 0.69 | 0.77 | 0.47 | 0.7 | 0.63 |
| 3 | 67574830 | rs17028544 | -0.487 | 0.02 | 1.773 | 1.922 | -0.007 | 0.147 | 0.147 | 0 | 0.07 | 0.14 | 0.12 | 0.25 | 0.04 | 0.17 |
| 3 | 67577666 | rs11709337 | 1.744 | -0.022 | 1.703 | 1.93 | 0.006 | -0.007 | -0.007 | 0 | 0.66 | 0.43 | 0.66 | 0.27 | 0.64 | 0.49 |
| 3 | 67584363 | rs11919650 | -0.625 | -0.007 | 1.667 | 1.948 | 0.006 | 0.055 | 0.055 | 0 | 0.07 | 0.12 | 0.09 | 0.21 | 0.05 | 0.14 |
| 3 | 67585721 | rs2772480 | -0.933 | -0.112 | 1.664 | 1.909 | 0.01 | 0.194 | 0.194 | 0 | 0.08 | 0.23 | 0.12 | 0.22 | 0.04 | 0.17 |
| 3 | 67587313 | rs12496418 | 1.661 | -0.076 | 1.583 | 1.87 | -0.003 | 0.002 | 0.002 | 0 | 0.66 | 0.42 | 0.67 | 0.27 | 0.64 | 0.49 |
| 3 | 67596107 | rs2772473 | -0.806 | -0.027 | 1.797 | 1.986 | 0.002 | 0.167 | 0.167 | 0.095 | 0.07 | 0.18 | 0.11 | 0.21 | 0.04 | 0.2 |
| 3 | 67597756 | rs1610008 | -1.967 | -0.069 | 1.832 | 1.84 | -0.028 | 0.286 | 0.286 | 0.277 | 0.14 | 0.41 | 0.18 | 0.4 | 0.06 | 0.32 |
| 3 | 67598859 | rs2772471 | -3.039 | -0.096 | 1.724 | 1.815 | -0.039 | 0.098 | 0.098 | 0.383 | 0.19 | 0.61 | 0.23 | 0.42 | 0.09 | 0.36 |
| 3 | 67603407 | rs1590874 | -2.889 | -0.1 | 1.8 | 1.812 | -0.039 | 0.121 | 0.121 | 0.122 | 0.18 | 0.61 | 0.23 | 0.42 | 0.09 | 0.36 |
| 3 | 67608177 | rs2772450 | -2.187 | -0.095 | 1.684 | 1.78 | -0.044 | 0.285 | 0.285 | 0.901 | 0.16 | 0.52 | 0.22 | 0.46 | 0.07 | 0.37 |
| 3 | 67611271 | rs2634741 | -0.514 | -0.214 | 1.457 | 1.683 | -0.034 | -0.025 | -0.025 | 0 | 0.82 | 0.92 | 0.89 | 0.74 | 0.75 | 0.8 |
| 3 | 67611478 | rs2634742 | -2.693 | -0.267 | 1.507 | 1.701 | -0.047 | 0.211 | 0.211 | 0.091 | 0.19 | 0.54 | 0.23 | 0.47 | 0.09 | 0.37 |
| 3 | 67615531 | rs1028950 | 1.061 | -0.136 | 1.39 | 1.215 | -0.015 | 0.018 | 0.018 | 1.053 | 0.6 | 0.68 | 0.63 | 0.31 | 0.54 | 0.43 |
| 3 | 67622948 | rs17046620 | -0.742 | -0.046 | 1.501 | 1.327 | -0.003 | 0.026 | 0.026 | 0 | 0.2 | 0.21 | 0.24 | 0.39 | 0.18 | 0.35 |
| 3 | 67628201 | rs7621788 | -0.607 | -0.132 | 1.658 | 1.325 | -0.008 | 0.129 | 0.129 | 0 | 0.19 | 0.42 | 0.26 | 0.38 | 0.17 | 0.33 |
| 3 | 67628283 | rs2250952 | 1.074 | -0.086 | 1.028 | 1.34 | -0.004 | 0.006 | 0.006 | 0 | 0.73 | 0.33 | 0.71 | 0.58 | 0.8 | 0.62 |
| 3 | 67645798 | rs2772465 | -0.434 | -0.067 | 1.274 | 1.33 | 0.001 | 0.001 | 0.001 | 0 | 0.37 | 0.57 | 0.25 | 0.49 | 0.42 | 0.34 |
| 3 | 67657428 | rs9849270 | -0.005 | -0.05 | 1.679 | 1.229 | 0.002 | 0.001 | 0.001 | 0 | 0.32 | 0.32 | 0.21 | 0.41 | 0.4 | 0.26 |
| 3 | 67657497 | rs6796037 | -0.626 | -0.092 | 1.609 | 1.217 | -0.027 | -0.009 | -0.009 | 0.032 | 0.39 | 0.73 | 0.27 | 0.47 | 0.43 | 0.29 |
| 3 | 67660908 | rs9814167 | -0.528 | -0.056 | 1.649 | 1.223 | -0.015 | -0.004 | -0.004 | 0 | 0.38 | 0.68 | 0.26 | 0.44 | 0.43 | 0.3 |
| 3 | 67663925 | rs9877255 | -0.535 | -0.064 | 1.646 | 1.22 | -0.015 | -0.003 | -0.003 | 1.118 | 0.38 | 0.68 | 0.26 | 0.44 | 0.43 | 0.3 |
| 3 | 67667800 | rs1868674 | -0.52 | -0.108 | 1.854 | 1.227 | 0.003 | 0 | 0 | 0 | 0.38 | 0.59 | 0.26 | 0.42 | 0.43 | 0.29 |
| 3 | 67671703 | rs7643466 | 0.047 | -0.093 | 2.043 | 1.212 | 0.01 | 0.003 | 0.003 | 0 | 0.32 | 0.33 | 0.21 | 0.37 | 0.4 | 0.24 |
| 3 | 67672171 | rs7646094 | -0.764 | -0.238 | 1.486 | 1.095 | 0.003 | 0.004 | 0.004 | 0.026 | 0.49 | 0.54 | 0.45 | 0.56 | 0.58 | 0.44 |
| 3 | 67678536 | rs11717561 | 0.668 | -0.31 | 1.501 | 0.855 | -0.023 | -0.002 | -0.002 | 0.129 | 0.27 | 0.2 | 0.17 | 0.28 | 0.36 | 0.16 |
| 3 | 67678753 | rs6808106 | 0.338 | -0.202 | 1.603 | 0.847 | -0.004 | 0 | 0 | 0 | 0.29 | 0.28 | 0.18 | 0.28 | 0.36 | 0.16 |
| 3 | 67681107 | rs2167180 | -1.045 | 0.056 | 1.368 | 0.746 | 0.009 | 0.026 | 0.026 | 0.006 | 0.81 | 0.8 | 0.52 | 0.75 | 0.73 | 0.67 |
| 3 | 67685999 | rs1376030 | 0.435 | 0.075 | 1.222 | -0.189 | -0.036 | -0.007 | -0.007 | 3.724 | 0.27 | 0.16 | 0.17 | 0.28 | 0.35 | 0.16 |
| 3 | 67699371 | rs935734 | 0.058 | 0.017 | 1.509 | -0.267 | 0.021 | 0.003 | 0.003 | 0 | 0.3 | 0.4 | 0.2 | 0.34 | 0.36 | 0.16 |
| 3 | 67704102 | rs2198015 | 0.177 | 0.028 | 1.485 | -0.292 | 0.015 | 0.001 | 0.001 | 0 | 0.29 | 0.37 | 0.2 | 0.33 | 0.36 | 0.17 |
| 3 | 67709737 | rs6548690 | -0.338 | -0.215 | 1.376 | -0.539 | -0.018 | -0.049 | -0.049 | 0 | 0.84 | 0.72 | 0.9 | 0.9 | 0.89 | 0.82 |
| 3 | 67711742 | rs13069251 | 0.101 | 0.636 | 0.836 | -0.176 | 0.003 | -0.002 | -0.002 | 3.172 | 0.29 | 0.32 | 0.2 | 0.28 | 0.35 | 0.16 |
| 3 | 67720488 | rs7640220 | 0.025 | 0.613 | 0.732 | -0.408 | -0.065 | 0.027 | 0.027 | 0 | 0.31 | 0.14 | 0.2 | 0.3 | 0.4 | 0.25 |
| 3 | 67730333 | rs9824834 | -0.385 | 0.609 | 0.81 | -0.402 | 0.021 | -0.015 | -0.015 | 0 | 0.34 | 0.43 | 0.23 | 0.35 | 0.4 | 0.26 |
| 3 | 67732386 | rs10510966 | -0.103 | 0.647 | 0.131 | -0.388 | -0.046 | 0.029 | 0.029 | 0 | 0.32 | 0.19 | 0.21 | 0.3 | 0.39 | 0.25 |
| 3 | 67737120 | rs6787099 | 0.016 | 0.347 | 0.253 | -0.299 | -0.021 | 0.025 | 0.025 | 0.082 | 0.41 | 0.34 | 0.25 | 0.44 | 0.46 | 0.36 |
| 3 | 67737700 | rs13087619 | -0.096 | 0.35 | 0.103 | -0.443 | -0.029 | 0.08 | 0.08 | 0.403 | 0.43 | 0.2 | 0.26 | 0.38 | 0.49 | 0.37 |
| 3 | 67740112 | rs6802783 | -0.162 | 0.438 | 0.141 | -0.395 | 0.001 | 0.039 | 0.039 | 0.079 | 0.46 | 0.42 | 0.28 | 0.38 | 0.49 | 0.37 |
| 3 | 67740402 | rs6779419 | -0.032 | 0.606 | 0.136 | -0.164 | -0.016 | 0.022 | 0.022 | 0.15 | 0.38 | 0.39 | 0.24 | 0.38 | 0.42 | 0.35 |
| 3 | 67755257 | rs11715476 | -0.719 | 0.23 | 0.469 | -0.376 | 0 | -0.032 | -0.032 | 0.054 | 0.13 | 0.16 | 0.09 | 0 | 0.12 | 0.03 |
| 3 | 67756224 | rs11925909 | -0.742 | -0.081 | 0.63 | -0.455 | -0.011 | -0.053 | -0.053 | 0.708 | 0.14 | 0.21 | 0.11 | 0 | 0.12 | 0.02 |
| 3 | 67761916 | rs17046864 | -1.025 | -0.138 | 1.227 | -0.48 | -0.025 | -0.092 | -0.092 | 0 | 0.15 | 0.34 | 0.13 | 0 | 0.12 | 0.02 |
| 3 | 67767250 | rs1459914 | 1.305 | -0.361 | 1.205 | -0.596 | -0.04 | -0.127 | -0.127 | 0 | 0.78 | 0.46 | 0.84 | 0.94 | 0.83 | 0.88 |
| 3 | 67772319 | rs4856797 | 0.499 | -0.332 | 1.149 | -0.583 | -0.001 | -0.029 | -0.029 | 0 | 0.86 | 0.86 | 0.89 | 1 | 0.88 | 0.98 |
| 3 | 67777268 | rs7614322 | 0.608 | -0.303 | 1.156 | -0.583 | -0.011 | -0.097 | -0.097 | 0 | 0.85 | 0.64 | 0.87 | 1 | 0.87 | 0.98 |
| 3 | 67784974 | rs7630256 | 1.059 | 0.035 | 1.589 | -0.145 | 0.007 | -0.017 | -0.017 | 0 | 0.92 | 0.7 | 0.95 | 0.94 | 0.95 | 0.9 |
| 3 | 67789621 | rs2362440 | 1.108 | 0.352 | 1.019 | -0.101 | 0.008 | -0.042 | -0.042 | 0 | 0.72 | 0.53 | 0.86 | 0.77 | 0.73 | 0.77 |
| 3 | 67794965 | rs10510967 | -0.054 | 0.637 | 1.153 | 0.079 | -0.009 | -0.011 | -0.011 | 0 | 0.06 | 0.16 | 0.04 | 0.06 | 0.04 | 0.11 |
| 3 | 67799690 | rs2125812 | 0.344 | 0.652 | 0.67 | 0.121 | 0.022 | -0.024 | -0.024 | 0 | 0.77 | 0.5 | 0.87 | 0.78 | 0.74 | 0.8 |
| 3 | 67802866 | rs6548756 | -0.259 | 0.57 | 0.364 | 0.753 | 0 | -0.009 | -0.009 | 0 | 0.18 | 0.11 | 0.26 | 0.24 | 0.21 | 0.42 |
| 3 | 67833749 | rs4331688 | 0.079 | 0.637 | -0.769 | 0.592 | 0.011 | 0.002 | 0.002 | 0 | 0.49 | 0.24 | 0.67 | 0.43 | 0.54 | 0.5 |
| 3 | 67841137 | rs9309877 | 1.103 | 0.637 | 1.146 | 0.447 | 0.207 | 0.108 | 0.108 | 0 | 0.95 | 0.59 | 0.95 | 0.99 | 0.95 | 0.98 |
| 3 | 67856341 | rs1585383 | 0.861 | 0.248 | 0.567 | 0.235 | -0.004 | -0.034 | -0.034 | 0.003 | 0.74 | 0.68 | 0.81 | 0.85 | 0.76 | 0.93 |
| 3 | 67857576 | rs9865535 | -0.471 | 0.242 | 0.548 | 0.175 | -0.005 | 0.021 | 0.021 | 0 | 0.2 | 0.1 | 0.16 | 0.14 | 0.21 | 0.05 |
| 3 | 67859042 | rs1459920 | 1.663 | -0.184 | 0.213 | 0.196 | -0.008 | -0.011 | -0.011 | 1.873 | 0.62 | 0.62 | 0.62 | 0.75 | 0.69 | 0.67 |
| 3 | 67861686 | rs6789499 | 1.703 | -0.184 | 0.368 | 0.175 | -0.012 | -0.024 | -0.024 | 0 | 0.61 | 0.53 | 0.6 | 0.76 | 0.69 | 0.67 |
| 3 | 67866836 | rs9814396 | 0.31 | -0.171 | 0.347 | 0.165 | -0.006 | -0.042 | -0.042 | 0 | 0.78 | 0.7 | 0.83 | 0.88 | 0.79 | 0.95 |
| 3 | 67875807 | rs9810776 | 1.639 | -0.163 | 0.214 | 0.19 | -0.012 | -0.023 | -0.023 | 0 | 0.61 | 0.56 | 0.6 | 0.74 | 0.69 | 0.67 |
| 3 | 67884900 | rs7626153 | -0.59 | -0.158 | 0.34 | 0.413 | 0.01 | -0.033 | -0.033 | 0 | 0.11 | 0.01 | 0.2 | 0.1 | 0.07 | 0.26 |
| 3 | 67894662 | rs4586824 | 1.28 | -0.174 | 0.553 | 0.389 | -0.038 | -0.018 | -0.018 | 4.39 | 0.8 | 0.6 | 0.76 | 0.89 | 0.9 | 0.73 |
| 3 | 67905314 | rs6787449 | 1.269 | 0.085 | 1.268 | 0.794 | 0.001 | 0.038 | 0.038 | 0 | 0.87 | 0.75 | 0.88 | 0.83 | 0.89 | 0.75 |

| | | | | | | | | | | | | | | | | |
|---|---|---|---|---|---|---|---|---|---|---|---|---|---|---|---|---|
| 3 | 67908896 | rs7642515 | 1.278 | -0.173 | 1.404 | 0.682 | -0.011 | 0.027 | 0.027 | 0 | 0.85 | 0.66 | 0.87 | 0.83 | 0.88 | 0.72 |
| 3 | 67918792 | rs6801973 | 1.678 | -0.019 | 1.075 | 0.723 | -0.008 | 0.033 | 0.033 | 0 | 0.75 | 0.57 | 0.79 | 0.69 | 0.75 | 0.69 |
| 3 | 68065679 | rs9818541 | 0.17 | -0.777 | 0.435 | -0.069 | 0.005 | 0.01 | 0.01 | 0 | 0.66 | 0.69 | 0.77 | 0.8 | 0.71 | 0.8 |
| 3 | 68066484 | rs6800897 | 0.522 | -0.963 | 0.851 | 0.324 | -0.012 | -0.012 | -0.012 | 0 | 0.6 | 0.57 | 0.73 | 0.7 | 0.69 | 0.79 |
| 3 | 68069643 | rs11920675 | -1.544 | -0.974 | 0.889 | 0.277 | 0.001 | 0.144 | 0.144 | 0.151 | 0.12 | 0.37 | 0.08 | 0.19 | 0.06 | 0.03 |
| 3 | 68071041 | rs1349005 | -0.503 | -0.997 | 0.773 | 0.225 | -0.011 | 0.139 | 0.139 | 0 | 0.72 | 0.96 | 0.81 | 0.89 | 0.75 | 0.81 |
| 3 | 68087612 | rs9866037 | 0.335 | -1.402 | 0.65 | 0.048 | 0.001 | 0.001 | 0.001 | 0 | 0.1 | 0.14 | 0.04 | 0.03 | 0.05 | 0.17 |
| 3 | 68090341 | rs9881594 | 0.592 | -1.446 | 0.862 | 0.072 | 0.006 | 0.008 | 0.008 | 0 | 0.56 | 0.57 | 0.73 | 0.76 | 0.66 | 0.64 |
| 3 | 68090811 | rs7373042 | 0.9 | -1.543 | 1.407 | 0.052 | 0.017 | 0.016 | 0.016 | 0 | 0.51 | 0.45 | 0.69 | 0.66 | 0.63 | 0.62 |
| 3 | 68092835 | rs9869498 | 0.974 | -1.56 | 1.348 | 0.044 | 0.043 | 0.041 | 0.041 | 0 | 0.5 | 0.38 | 0.69 | 0.66 | 0.63 | 0.63 |
| 3 | 68093710 | rs7429717 | -0.133 | -1.357 | 1.117 | 0.063 | 0 | -0.002 | -0.002 | 0 | 0.67 | 0.71 | 0.77 | 0.79 | 0.71 | 0.81 |
| 3 | 68106746 | rs9822658 | -0.248 | -1.336 | 0.784 | 0.34 | 0.01 | 0.005 | 0.005 | 0 | 0.53 | 0.39 | 0.34 | 0.57 | 0.42 | 0.32 |
| 3 | 68121791 | rs6548962 | 0.463 | -1.561 | 1.135 | 0.296 | -0.002 | 0.001 | 0.001 | 0 | 0.41 | 0.41 | 0.62 | 0.33 | 0.56 | 0.66 |
| 3 | 68122404 | rs6548964 | -0.679 | -1.541 | 1.21 | 0.306 | 0.004 | 0.015 | 0.015 | 0.483 | 0.95 | 0.89 | 0.96 | 0.91 | 0.96 | 0.98 |
| 3 | 68123731 | rs7610023 | 0.224 | -1.701 | 1.479 | 0.333 | 0.056 | 0.017 | 0.017 | 0 | 0.48 | 0.25 | 0.3 | 0.48 | 0.38 | 0.31 |
| 3 | 68138729 | rs7650405 | 0.192 | -1.701 | 1.512 | 0.353 | 0.052 | 0.02 | 0.02 | 0 | 0.49 | 0.29 | 0.3 | 0.48 | 0.38 | 0.31 |
| 3 | 68150331 | rs9850969 | 0.392 | -1.764 | 1.404 | 0.039 | -0.009 | 0.045 | 0.045 | 0 | 0.89 | 0.81 | 0.94 | 0.81 | 0.93 | 0.97 |
| 3 | 68151655 | rs7636462 | 0.145 | -1.73 | 1.441 | 0.02 | -0.002 | -0.001 | -0.001 | 0 | 0.49 | 0.47 | 0.31 | 0.48 | 0.37 | 0.31 |
| 3 | 68171164 | rs6774687 | 0.44 | -1.761 | 1.056 | -0.06 | 0.007 | 0.013 | 0.013 | 0 | 0.42 | 0.38 | 0.65 | 0.33 | 0.61 | 0.66 |
| 3 | 68174413 | rs9309968 | 1.194 | -2.227 | 0.635 | -0.124 | 0.025 | -0.047 | -0.047 | 1.061 | 0.2 | 0.13 | 0.11 | 0.35 | 0.12 | 0.1 |
| 3 | 68179836 | rs7624036 | 1.141 | -2.187 | 0.358 | -0.439 | 0.015 | 0.005 | 0.005 | 2.373 | 0.19 | 0.1 | 0.08 | 0.17 | 0.11 | 0.12 |
| 3 | 68180657 | rs9868432 | 0.919 | -2.211 | 0.14 | -0.476 | 0.006 | 0.014 | 0.014 | 0 | 0.41 | 0.33 | 0.65 | 0.33 | 0.61 | 0.66 |
| 3 | 68187910 | rs6776172 | 0.469 | -2.117 | -0.143 | -0.489 | 0.031 | -0.026 | -0.026 | 0 | 0.32 | 0.2 | 0.21 | 0.41 | 0.23 | 0.2 |
| 3 | 68197774 | rs2218053 | 1.47 | -1.592 | -0.063 | -0.817 | 0.019 | 0.011 | 0.011 | 1.001 | 0.14 | 0.02 | 0.05 | 0.12 | 0.09 | 0.08 |
| 3 | 68202602 | rs6549021 | 1.003 | -0.546 | -0.346 | -1.038 | -0.035 | 0.057 | 0.057 | 0.696 | 0.19 | 0.02 | 0.1 | 0.11 | 0.2 | 0.17 |
| 3 | 68204352 | rs17047218 | 0.871 | -0.617 | -0.5 | -1.043 | 0.001 | -0.007 | -0.007 | 0.953 | 0.07 | 0.04 | 0.05 | 0.08 | 0.05 | 0.02 |
| 3 | 68207278 | rs7638281 | 0.466 | -0.538 | -0.453 | -0.99 | -0.013 | 0.021 | 0.021 | 0.683 | 0.27 | 0.06 | 0.15 | 0.19 | 0.26 | 0.19 |
| 3 | 68207759 | rs1007583 | 0.206 | -0.623 | -0.079 | -1.014 | -0.01 | -0.016 | -0.016 | 0.122 | 0.65 | 0.53 | 0.8 | 0.71 | 0.7 | 0.79 |
| 3 | 68208180 | rs961915 | 0.886 | -0.535 | 0.075 | -0.944 | 0.002 | 0.014 | 0.014 | 0 | 0.05 | 0.13 | 0.03 | 0.1 | 0.04 | 0.02 |
| 3 | 68211926 | rs1595489 | -0.065 | -0.449 | -0.182 | -0.943 | 0.066 | 0.09 | 0.09 | 1.652 | 0.92 | 0.61 | 0.95 | 0.9 | 0.96 | 0.98 |
| 3 | 68233947 | rs11716986 | 0.605 | -0.138 | -0.957 | -1.087 | -0.024 | 0.069 | 0.069 | 0 | 0.19 | 0.06 | 0.11 | 0.03 | 0.2 | 0.16 |
| 3 | 68238985 | rs4446218 | 0.053 | -0.217 | -0.624 | -0.928 | -0.005 | -0.007 | -0.007 | 0 | 0.64 | 0.48 | 0.79 | 0.7 | 0.7 | 0.79 |
| 3 | 68252679 | rs7650009 | 0.104 | 0.094 | -0.666 | -0.789 | -0.01 | -0.02 | -0.02 | 0 | 0.6 | 0.55 | 0.75 | 0.72 | 0.65 | 0.77 |
| 3 | 68256979 | rs11711058 | 0.672 | 1.616 | -1.07 | -0.83 | -0.067 | 0.15 | 0.15 | 0 | 0.24 | 0.04 | 0.14 | 0.02 | 0.28 | 0.13 |
| 3 | 68257048 | rs1388495 | 1.055 | 1.797 | -0.851 | -0.696 | 0.064 | 0.02 | 0.02 | 2.148 | 0.44 | 0.3 | 0.36 | 0.61 | 0.27 | 0.49 |
| 3 | 68263389 | rs4311217 | 0.535 | 2.06 | -0.616 | -0.944 | -0.001 | -0.003 | -0.003 | 0.009 | 0.56 | 0.59 | 0.63 | 0.69 | 0.4 | 0.59 |
| 3 | 68270017 | rs12487860 | -0.042 | 2.151 | -0.679 | -1.007 | -0.025 | 0.056 | 0.056 | 0.001 | 0.33 | 0.21 | 0.4 | 0.02 | 0.41 | 0.2 |
| 3 | 68277282 | rs12490102 | 1.714 | 2.191 | -0.846 | -0.817 | -0.002 | -0.009 | -0.009 | 0.831 | 0.07 | 0.08 | 0.26 | 0 | 0.13 | 0.05 |
| 3 | 68283146 | rs6549072 | -0.478 | 2.739 | -0.944 | -0.604 | -0.05 | -0.001 | -0.001 | 0 | 0.9 | 0.98 | 0.88 | 0.9 | 0.79 | 0.78 |
| 3 | 68283196 | rs6771678 | 0.677 | 2.324 | -1.018 | -0.664 | -0.067 | 0.15 | 0.15 | 0 | 0.24 | 0.05 | 0.14 | 0.02 | 0.28 | 0.14 |
| 3 | 68284661 | rs17047387 | -0.097 | 2.567 | 0.006 | 0.322 | 0.042 | 0.05 | 0.05 | 0 | 0.1 | 0.21 | 0.11 | 0.17 | 0.12 | 0.09 |
| 3 | 68286023 | rs1491722 | 0.395 | 2.204 | -0.3 | 0.296 | 0.132 | 0.052 | 0.052 | 0.954 | 0.79 | 0.55 | 0.77 | 0.71 | 0.68 | 0.71 |
| 3 | 68288626 | rs9863981 | 0.027 | 1.002 | -0.948 | 0.385 | -0.017 | -0.002 | -0.002 | 0 | 0.28 | 0.18 | 0.38 | 0.28 | 0.32 | 0.2 |
| 3 | 68289861 | rs11706310 | -0.161 | 0.492 | -0.583 | 0.124 | -0.001 | -0.008 | -0.008 | 0 | 0.24 | 0.28 | 0.3 | 0.19 | 0.24 | 0.17 |
| 3 | 68299637 | rs6549090 | -0.019 | 0.087 | -0.745 | 0.286 | -0.009 | 0.036 | 0.036 | 0 | 0.36 | 0.46 | 0.58 | 0.53 | 0.32 | 0.49 |
| 3 | 68305892 | rs10440086 | -0.742 | 0.017 | -1.127 | 0.021 | -0.005 | -0.004 | -0.004 | 3.444 | 0.43 | 0.72 | 0.59 | 0.5 | 0.39 | 0.41 |
| 3 | 68306724 | rs6764336 | -1.078 | 0.057 | -1.248 | -0.21 | -0.039 | 0.079 | 0.079 | 0.479 | 0.69 | 0.82 | 0.8 | 0.98 | 0.59 | 0.74 |
| 3 | 68308222 | rs1352851 | -0.676 | -0.034 | -0.596 | 1.041 | -0.003 | 0.009 | 0.009 | 0 | 0.3 | 0.31 | 0.2 | 0.37 | 0.26 | 0.21 |
| 3 | 68317975 | rs1032376 | -0.179 | 0.151 | -1.089 | 1.113 | -0.004 | -0.03 | -0.03 | 2.953 | 0.22 | 0.15 | 0.17 | 0.35 | 0.21 | 0.13 |
| 3 | 68320679 | rs9826468 | -0.481 | -0.075 | -0.779 | 0.738 | 0.008 | 0 | 0 | 0 | 0.1 | 0.07 | 0.05 | 0.11 | 0.05 | 0.1 |
| 3 | 68322450 | rs2313273 | 0.386 | 0.242 | -0.562 | 0.807 | -0.001 | 0.001 | 0.001 | 0 | 0.66 | 0.6 | 0.61 | 0.68 | 0.66 | 0.65 |
| 3 | 68322897 | rs1388489 | 0.24 | 0.599 | -0.732 | 1.015 | -0.015 | -0.022 | -0.022 | 0 | 0.22 | 0.17 | 0.29 | 0.36 | 0.29 | 0.33 |
| 3 | 68323280 | rs6549098 | -1.086 | 0.435 | -0.82 | 0.79 | -0.007 | -0.004 | -0.004 | 0 | 0.68 | 0.85 | 0.6 | 0.73 | 0.64 | 0.82 |
| 3 | 68324960 | rs17047423 | 0.526 | 0.29 | -0.828 | 0.671 | -0.02 | -0.044 | -0.044 | 0 | 0.25 | 0.19 | 0.32 | 0.46 | 0.28 | 0.29 |
| 3 | 68328378 | rs4428185 | -1.481 | 0.181 | -1.003 | -0.378 | -0.016 | 0.062 | 0.062 | 4.423 | 0.76 | 0.87 | 0.66 | 0.93 | 0.69 | 0.93 |
| 3 | 68328441 | rs11708702 | 1.227 | -0.055 | -1.355 | -0.476 | -0.01 | -0.005 | -0.005 | 0.275 | 0.18 | 0.02 | 0.12 | 0.13 | 0.14 | 0.08 |
| 3 | 68328674 | rs11709646 | 0.963 | 0.627 | -0.661 | -0.075 | 0.04 | 0.063 | 0.063 | 0 | 0.34 | 0.17 | 0.19 | 0.15 | 0.21 | 0.22 |
| 3 | 68328854 | rs17047430 | 0.349 | 0.611 | -0.559 | -0.062 | 0.02 | -0.007 | -0.007 | 1.108 | 0.29 | 0.4 | 0.36 | 0.73 | 0.34 | 0.51 |
| 3 | 68329788 | rs9829202 | 0.912 | 0.67 | -1.396 | -0.02 | 0.036 | 0.057 | 0.057 | 0 | 0.34 | 0.18 | 0.2 | 0.14 | 0.22 | 0.22 |
| 3 | 68330052 | rs1027238 | -0.825 | 1.109 | -1.861 | -0.039 | -0.073 | 0.088 | 0.088 | 4.213 | 0.75 | 0.98 | 0.66 | 0.93 | 0.69 | 0.93 |
| 3 | 68332557 | rs17231705 | -0.226 | 1.444 | -0.812 | 0.582 | 0.004 | -0.012 | -0.012 | 0.625 | 0.1 | 0.11 | 0.09 | 0.05 | 0.13 | 0.21 |
| 3 | 68334875 | rs17043602 | -0.499 | 1.396 | -0.908 | 1.026 | 0.006 | 0.015 | 0.015 | 0 | 0.28 | 0.31 | 0.31 | 0.47 | 0.34 | 0.44 |
| 3 | 68336119 | rs9837363 | -0.043 | 1.067 | -1.641 | 0.624 | -0.009 | 0.069 | 0.069 | 0.309 | 0.53 | 0.76 | 0.63 | 0.67 | 0.54 | 0.48 |
| 3 | 68336930 | rs17047443 | -0.523 | -0.001 | -1.648 | 1.1 | 0.013 | -0.02 | -0.02 | 0 | 0.19 | 0.1 | 0.18 | 0.62 | 0.12 | 0.35 |
| 3 | 68336960 | rs17047444 | -0.964 | -0.175 | -2.06 | 0.708 | 0.03 | -0.04 | -0.04 | 0.549 | 0.26 | 0.12 | 0.24 | 0.64 | 0.23 | 0.39 |
| 3 | 68337202 | rs902720 | -0.311 | 0.276 | -1.578 | -0.431 | 0.022 | -0.152 | -0.152 | 0 | 0.69 | 0.56 | 0.64 | 0.98 | 0.65 | 0.85 |
| 3 | 68337634 | rs9823980 | 0.385 | -0.052 | -1.523 | -0.495 | 0.002 | -0.002 | -0.002 | 0.017 | 0.26 | 0.21 | 0.23 | 0.74 | 0.16 | 0.43 |
| 3 | 68337818 | rs11708892 | 0.483 | -0.344 | -1.582 | -0.635 | 0.022 | -0.096 | -0.096 | 0 | 0.33 | 0.43 | 0.38 | 0.06 | 0.37 | 0.17 |
| 3 | 68339327 | rs4241390 | -1.421 | -0.411 | -1.85 | -0.932 | -0.047 | 0.059 | 0.059 | 0.343 | 0.78 | 0.92 | 0.75 | 0.91 | 0.68 | 0.74 |

| | | | | | | | | | | | | | | | | |
|---|---|---|---|---|---|---|---|---|---|---|---|---|---|---|---|---|
| 3 | 68341414 | rs7618050 | -0.501 | -0.422 | -1.855 | -0.98 | -0.024 | 0.102 | 0.102 | 0 | 0.66 | 0.84 | 0.69 | 0.89 | 0.58 | 0.65 |
| 3 | 68342451 | rs2172219 | 0.184 | -0.565 | -1.376 | -0.55 | -0.027 | -0.025 | -0.025 | 0.302 | 0.29 | 0.41 | 0.29 | 0.83 | 0.19 | 0.45 |
| 3 | 68357852 | rs9822608 | -1.577 | -0.086 | -0.669 | -0.189 | 0.029 | 0.027 | 0.027 | 0.06 | 0.85 | 0.73 | 0.75 | 0.78 | 0.79 | 0.71 |
| 3 | 68364358 | rs17047489 | 1.007 | -0.647 | -0.894 | 0.459 | 0.033 | -0.108 | -0.108 | 0 | 0.24 | 0.01 | 0.17 | 0.35 | 0.15 | 0.23 |
| 3 | 68365162 | rs1155324 | 1.007 | -0.647 | -0.859 | 0.459 | 0.044 | -0.094 | -0.094 | 0 | 0.24 | 0.04 | 0.17 | 0.35 | 0.15 | 0.23 |
| 3 | 68365879 | rs1491758 | 0.398 | -0.793 | -0.216 | 0.484 | -0.043 | 0.226 | 0.226 | 4.1 | 0.16 | 0.36 | 0.12 | 0.45 | 0.1 | 0.23 |
| 3 | 68368649 | rs10155009 | -1.921 | -0.402 | -0.611 | 0.257 | -0.03 | 0.019 | 0.019 | 0.006 | 0.86 | 0.93 | 0.74 | 0.91 | 0.77 | 0.75 |
| 3 | 68369288 | rs1491756 | -1.961 | -0.4 | -0.628 | 0.256 | -0.032 | 0.016 | 0.016 | 0 | 0.86 | 0.93 | 0.75 | 0.91 | 0.77 | 0.74 |
| 3 | 68373086 | rs1388505 | 0.361 | -0.582 | -0.646 | 0.284 | 0.018 | -0.019 | -0.019 | 1.131 | 0.19 | 0.1 | 0.12 | 0.28 | 0.12 | 0.21 |
| 3 | 68373756 | rs6549116 | -1.976 | -0.531 | -0.419 | 0.115 | 0.002 | -0.005 | -0.005 | 0.055 | 0.84 | 0.8 | 0.75 | 0.92 | 0.78 | 0.74 |
| 3 | 68374110 | rs6770136 | -1.905 | -0.529 | -0.354 | 0.116 | 0.013 | -0.02 | -0.02 | 0 | 0.84 | 0.73 | 0.74 | 0.92 | 0.78 | 0.74 |
| 3 | 68374432 | rs9832486 | 2.311 | -0.45 | -0.358 | 0.117 | 0.003 | -0.026 | -0.026 | 0.009 | 0.05 | 0.08 | 0.03 | 0 | 0.04 | 0.02 |
| 3 | 68376852 | rs983963 | 1.13 | -0.693 | -0.487 | 0.102 | 0.058 | -0.065 | -0.065 | 0 | 0.26 | 0.09 | 0.18 | 0.37 | 0.17 | 0.33 |
| 3 | 68378613 | rs17047523 | 0.947 | -0.073 | 0.267 | 0.168 | 0.004 | -0.038 | -0.038 | 0 | 0.07 | 0.18 | 0.04 | 0 | 0.04 | 0.1 |
| 3 | 68379929 | rs1158994 | -0.993 | 0.077 | 0.228 | 0.125 | 0.026 | -0.128 | -0.128 | 0.06 | 0.69 | 0.62 | 0.63 | 1 | 0.57 | 0.89 |
| 3 | 68385527 | rs6792971 | 0.9 | 0.157 | -0.015 | 0.604 | 0.006 | 0.07 | 0.07 | 0.012 | 0.07 | 0.09 | 0.07 | 0.26 | 0.07 | 0.09 |
| 3 | 68387992 | rs1491748 | 0.753 | -0.102 | -0.201 | 0.41 | 0.017 | -0.007 | -0.007 | 0 | 0.45 | 0.2 | 0.43 | 0.46 | 0.41 | 0.53 |
| 3 | 68391629 | rs262210 | 0.084 | 0.132 | -0.073 | 0.379 | -0.014 | 0.092 | 0.092 | 0 | 0.53 | 0.31 | 0.51 | 0.72 | 0.49 | 0.69 |
| 3 | 68393781 | rs262215 | -0.004 | -0.051 | 0.551 | 0.377 | 0.002 | -0.014 | -0.014 | 0 | 0.54 | 0.53 | 0.52 | 0.72 | 0.49 | 0.69 |
| 3 | 68397737 | rs2665523 | 0.911 | -0.094 | 0.547 | 0.398 | 0.014 | 0.004 | 0.004 | 0.075 | 0.45 | 0.36 | 0.45 | 0.45 | 0.41 | 0.53 |
| 3 | 68402933 | rs1491753 | 0.235 | -0.193 | 0.452 | 0.356 | 0.017 | -0.032 | -0.032 | 0 | 0.13 | 0.06 | 0.1 | 0.31 | 0.09 | 0.2 |
| 3 | 68404612 | rs262249 | 1.034 | -0.192 | 0.607 | 0.364 | 0.004 | -0.019 | -0.019 | 0 | 0.46 | 0.17 | 0.45 | 0.45 | 0.43 | 0.53 |
| 3 | 68405259 | rs262248 | -0.044 | -0.176 | 0.688 | 0.32 | 0.008 | -0.023 | -0.023 | 0 | 0.58 | 0.55 | 0.53 | 0.71 | 0.51 | 0.7 |
| 3 | 68405363 | rs4855463 | 0.784 | 0.231 | 1.036 | 0.092 | -0.051 | 0.062 | 0.062 | 0.374 | 0.3 | 0.18 | 0.36 | 0 | 0.42 | 0.11 |
| 3 | 68409374 | rs262234 | 0.172 | 0.226 | 0.912 | 0.155 | -0.003 | 0.007 | 0.007 | 0.434 | 0.58 | 0.4 | 0.55 | 0.74 | 0.5 | 0.73 |
| 3 | 68413184 | rs262227 | 0.169 | 0.244 | 0.96 | 0.148 | 0.002 | -0.007 | -0.007 | 0 | 0.58 | 0.44 | 0.54 | 0.75 | 0.49 | 0.73 |
| 3 | 68416626 | rs17047606 | 0.291 | 0.078 | 0.853 | 0.194 | 0.005 | 0.078 | 0.078 | 0.033 | 0.08 | 0.19 | 0.08 | 0.23 | 0.07 | 0.15 |
| 3 | 68417939 | rs262180 | -0.782 | 0.352 | 0.768 | -0.034 | 0.014 | -0.106 | -0.106 | 0.096 | 0.7 | 0.66 | 0.63 | 1 | 0.61 | 0.89 |
| 3 | 68426920 | rs171893 | -0.507 | 0.384 | 0.667 | -0.051 | 0.021 | -0.116 | -0.116 | 0 | 0.7 | 0.65 | 0.64 | 1 | 0.61 | 0.9 |
| 3 | 68436430 | rs7610318 | -0.589 | 0.129 | 0.663 | 0.078 | -0.006 | 0.018 | 0.018 | 0.712 | 0.43 | 0.59 | 0.44 | 0.22 | 0.48 | 0.27 |
| 3 | 68442332 | rs7642781 | -0.087 | 0.134 | 0.75 | 0.082 | 0.027 | -0.024 | -0.024 | 0 | 0.38 | 0.46 | 0.43 | 0.22 | 0.47 | 0.18 |
| 3 | 68449637 | rs10222670 | -0.855 | -0.241 | 0.528 | 0.526 | 0.013 | 0.014 | 0.014 | 0.218 | 0.47 | 0.72 | 0.54 | 0.39 | 0.52 | 0.33 |
| 3 | 68450201 | rs17047630 | -0.168 | 0.883 | 1.223 | 0.644 | 0.114 | 0.085 | 0.085 | 0.078 | 0.14 | 0.4 | 0.16 | 0.18 | 0.19 | 0.09 |
| 3 | 68452616 | rs2136715 | -0.131 | 0.729 | 1.207 | 0.816 | 0.023 | -0.024 | -0.024 | 0 | 0.32 | 0.39 | 0.4 | 0.19 | 0.44 | 0.16 |
| 3 | 68453623 | rs4855467 | -0.372 | 0.698 | 1.028 | 0.794 | 0.012 | 0.006 | 0.006 | 0.056 | 0.36 | 0.55 | 0.45 | 0.37 | 0.47 | 0.19 |
| 3 | 68457521 | rs9833308 | -0.909 | 0.625 | 0.938 | 0.815 | -0.013 | -0.011 | -0.011 | 1.183 | 0.4 | 0.63 | 0.49 | 0.46 | 0.48 | 0.26 |
| 3 | 68457880 | rs12489559 | -0.308 | 0.645 | 0.949 | 0.841 | 0.004 | 0.004 | 0.004 | 0.162 | 0.35 | 0.58 | 0.44 | 0.38 | 0.44 | 0.22 |
| 3 | 68458440 | rs4855469 | -0.742 | 0.607 | 0.933 | 0.783 | -0.027 | -0.012 | -0.012 | 0.016 | 0.38 | 0.75 | 0.5 | 0.39 | 0.47 | 0.27 |
| 3 | 68459113 | rs9822547 | -0.819 | 0.656 | 0.789 | 0.779 | -0.011 | 0.012 | 0.012 | 0.026 | 0.4 | 0.65 | 0.47 | 0.31 | 0.46 | 0.23 |
| 3 | 68462634 | rs4277695 | 0.922 | 0.163 | 0.003 | 0.658 | -0.003 | -0.114 | -0.114 | 0 | 0.36 | 0.02 | 0.25 | 0.41 | 0.23 | 0.38 |
| 3 | 68464413 | rs17047650 | 0.964 | 0.176 | 0.007 | 0.636 | 0.032 | -0.048 | -0.048 | 0 | 0.35 | 0.02 | 0.21 | 0.33 | 0.2 | 0.37 |
| 3 | 68468152 | rs9850187 | 0.233 | 0.165 | 0.313 | 0.599 | 0.024 | -0.052 | -0.052 | 0.882 | 0.33 | 0.41 | 0.4 | 0.13 | 0.45 | 0.12 |
| 3 | 68470326 | rs2136716 | 1.56 | 0.098 | 0.148 | 0.639 | 0.019 | -0.067 | -0.067 | 0.048 | 0.31 | 0.02 | 0.2 | 0.32 | 0.18 | 0.34 |
| 3 | 68472602 | rs1397253 | -2.095 | 0.144 | 0.094 | 0.554 | -0.039 | 0.021 | 0.021 | 0 | 0.81 | 0.96 | 0.76 | 0.87 | 0.75 | 0.71 |
| 3 | 68474875 | rs17047692 | 0.969 | 0.079 | 0.213 | 0.596 | 0.063 | -0.033 | -0.033 | 0 | 0.36 | 0.15 | 0.22 | 0.42 | 0.2 | 0.41 |
| 3 | 68475128 | rs1039878 | -0.148 | 0.592 | 0.509 | 0.769 | 0.03 | -0.018 | -0.018 | 0 | 0.36 | 0.43 | 0.46 | 0.26 | 0.49 | 0.17 |
| 3 | 68477210 | rs6767029 | -0.119 | 0.607 | 0.512 | 0.698 | 0.035 | -0.022 | -0.022 | 0 | 0.35 | 0.42 | 0.46 | 0.24 | 0.49 | 0.17 |
| 3 | 68480290 | rs6782797 | -1.448 | 0.299 | 0.009 | 0.37 | -0.032 | 0.081 | 0.081 | 0 | 0.73 | 0.96 | 0.73 | 0.88 | 0.71 | 0.67 |
| 3 | 68484388 | rs4855475 | -0.788 | 0.56 | 0.15 | 0.293 | -0.023 | 0.023 | 0.023 | 0 | 0.66 | 0.84 | 0.68 | 0.72 | 0.6 | 0.59 |
| 3 | 68484997 | rs6549127 | 1.076 | 0.472 | 0.169 | 0.282 | 0.009 | -0.01 | -0.01 | 0.03 | 0.29 | 0.27 | 0.23 | 0.61 | 0.18 | 0.41 |
| 3 | 68485146 | rs7622452 | 1.913 | 0.553 | 0.302 | 0.139 | -0.002 | 0.006 | 0.006 | 1.735 | 0.18 | 0.16 | 0.27 | 0.1 | 0.21 | 0.08 |
| 3 | 68485443 | rs7634101 | 1.777 | 0.4 | 0.079 | 0.353 | 0.002 | -0.109 | -0.109 | 0 | 0.23 | 0.01 | 0.17 | 0.36 | 0.15 | 0.3 |
| 3 | 68492075 | rs1510363 | 0.378 | 0.283 | 0.214 | -0.041 | 0.016 | 0.003 | 0.003 | 0.934 | 0.36 | 0.29 | 0.27 | 0.67 | 0.24 | 0.47 |
| 3 | 68494698 | rs1039879 | -1.409 | 0.313 | 0.224 | -0.068 | -0.011 | 0.02 | 0.02 | 0 | 0.72 | 0.84 | 0.71 | 0.81 | 0.67 | 0.65 |
| 3 | 68496610 | rs9310080 | 0.027 | 0.358 | 0.149 | -0.095 | 0.004 | -0.035 | -0.035 | 0.814 | 0.37 | 0.56 | 0.44 | 0.11 | 0.42 | 0.17 |
| 3 | 68497840 | rs6766498 | 0.968 | 0.042 | -0.242 | 0.074 | 0.025 | -0.083 | -0.083 | 0.358 | 0.3 | 0.08 | 0.22 | 0.43 | 0.2 | 0.39 |
| 3 | 68503271 | rs1510352 | -0.564 | 0.163 | 0.062 | 0.31 | 0.03 | 0.026 | 0.026 | 0 | 0.65 | 0.83 | 0.71 | 0.71 | 0.71 | 0.57 |
| 3 | 68505398 | rs13081659 | -0.079 | 0.189 | 0.244 | 0.234 | 0.025 | -0.026 | -0.026 | 0 | 0.31 | 0.4 | 0.43 | 0.13 | 0.45 | 0.17 |
| 3 | 68513822 | rs907081 | -1.111 | 0.095 | 0.472 | 0.48 | 0.06 | 0.532 | 0.532 | 0.896 | 0.05 | 0.37 | 0.07 | 0.31 | 0.01 | 0.1 |
| 3 | 68531655 | rs6801913 | 0.82 | 0.232 | 0.258 | -0.037 | -0.008 | -0.003 | -0.003 | 0 | 0.35 | 0.52 | 0.33 | 0.66 | 0.27 | 0.37 |
| 3 | 68531742 | rs9836240 | -0.56 | 0.373 | 0.314 | -0.088 | -0.008 | 0.004 | 0.004 | 0 | 0.37 | 0.38 | 0.41 | 0.14 | 0.47 | 0.22 |
| 3 | 68534078 | rs1877001 | -0.445 | 0.369 | 0.302 | -0.095 | -0.003 | -0.006 | -0.006 | 0 | 0.35 | 0.38 | 0.41 | 0.14 | 0.47 | 0.21 |
| 3 | 68535918 | rs10510970 | -0.225 | 0.501 | 0.48 | -0.054 | 0.005 | -0.027 | -0.027 | 0 | 0.19 | 0.24 | 0.14 | 0.07 | 0.21 | 0.1 |
| 3 | 68542647 | rs7653709 | -0.171 | 0.472 | 0.67 | -0.066 | 0.001 | -0.01 | -0.01 | 1.068 | 0.19 | 0.21 | 0.14 | 0.09 | 0.21 | 0.1 |
| 3 | 68544398 | rs13061250 | -0.348 | 0.469 | 0.759 | -0.067 | 0.015 | -0.049 | -0.049 | 0 | 0.2 | 0.3 | 0.14 | 0.09 | 0.21 | 0.1 |
| 3 | 68547301 | rs1511894 | -0.532 | 0.31 | 0.71 | -0.158 | 0 | -0.013 | -0.013 | 0 | 0.23 | 0.24 | 0.15 | 0.08 | 0.24 | 0.1 |
| 3 | 68551172 | rs17184043 | 0.606 | 0.284 | 0.621 | -0.188 | -0.059 | 0.081 | 0.081 | 0 | 0.14 | 0.02 | 0.11 | 0 | 0.2 | 0.05 |
| 3 | 68554434 | rs1397249 | -1.532 | 0.261 | 0.522 | -0.177 | 0.005 | 0.03 | 0.03 | 0 | 0.76 | 0.92 | 0.76 | 0.82 | 0.78 | 0.6 |
| 3 | 68560669 | rs2665549 | 0.169 | 0.236 | 0.315 | 0.109 | 0.003 | -0.047 | -0.047 | 0 | 0.22 | 0.26 | 0.15 | 0.05 | 0.24 | 0.11 |

| | | | | | | | | | | | | | | | | |
|---|---|---|---|---|---|---|---|---|---|---|---|---|---|---|---|---|
| 3 | 68561209 | rs2665551 | 0.27 | 0.26 | 0.096 | 0.108 | -0.003 | 0.001 | 0.001 | 0 | 0.22 | 0.21 | 0.15 | 0.08 | 0.24 | 0.11 |
| 3 | 68563209 | rs4855488 | 1.022 | 0.325 | -0.224 | -0.004 | -0.007 | -0.011 | -0.011 | 1.962 | 0.35 | 0.35 | 0.31 | 0.72 | 0.27 | 0.4 |
| 3 | 68565190 | rs12489309 | 0.142 | 0.626 | -0.82 | -0.227 | 0.02 | 0.025 | 0.025 | 0 | 0.49 | 0.32 | 0.56 | 0.56 | 0.54 | 0.47 |
| 3 | 68565824 | rs9846667 | 1.394 | 0.667 | -1.275 | -0.304 | 0.006 | -0.008 | -0.008 | 1.224 | 0.15 | 0.17 | 0.26 | 0.05 | 0.23 | 0.09 |
| 3 | 68566475 | rs9838165 | 0.766 | 0.654 | -1.286 | -0.287 | 0.001 | -0.001 | -0.001 | 0.019 | 0.19 | 0.19 | 0.34 | 0.14 | 0.28 | 0.14 |
| 3 | 68567423 | rs9647394 | -0.381 | 0.633 | -0.972 | -0.308 | -0.014 | 0.044 | 0.044 | 0 | 0.79 | 0.87 | 0.85 | 0.92 | 0.76 | 0.89 |
| 3 | 68569440 | rs4855489 | 0.932 | 0.589 | -1.08 | 0.063 | -0.067 | -0.19 | -0.19 | 0.462 | 0.3 | 0.01 | 0.28 | 0.48 | 0.27 | 0.34 |
| 3 | 68571521 | rs13094511 | 0.191 | 1.043 | 0.123 | -0.263 | 0.026 | -0.03 | -0.03 | 0.195 | 0.08 | 0.19 | 0.21 | 0.02 | 0.08 | 0.07 |
| 3 | 68583643 | rs17047811 | -0.692 | 0.864 | -0.091 | -0.287 | -0.015 | -0.077 | -0.077 | 0 | 0.24 | 0.39 | 0.15 | 0.1 | 0.21 | 0.1 |
| 3 | 68587884 | rs3828366 | -0.274 | 1.832 | -0.449 | -0.281 | -0.008 | 0.005 | 0.005 | 0.17 | 0.19 | 0.17 | 0.36 | 0.12 | 0.33 | 0.13 |
| 3 | 68589664 | rs1511906 | 0.233 | 1.573 | -0.908 | -1.088 | -0.012 | 0.098 | 0.098 | 0 | 0.46 | 0.34 | 0.45 | 0.12 | 0.5 | 0.47 |
| 3 | 68594889 | rs2137182 | 0.99 | 1.606 | -0.466 | -0.412 | -0.001 | -0.001 | -0.001 | 0 | 0.31 | 0.3 | 0.26 | 0.27 | 0.3 | 0.4 |
| 3 | 68595634 | rs7431217 | -0.131 | 1.366 | 0.098 | -0.159 | -0.004 | 0.001 | 0.001 | 0 | 0.45 | 0.57 | 0.44 | 0.39 | 0.39 | 0.48 |
| 3 | 68603563 | rs872114 | 0.879 | 1.362 | 0.474 | 0.029 | -0.004 | 0.023 | 0.023 | 0 | 0.06 | 0.19 | 0.04 | 0.05 | 0.02 | 0.03 |
| 3 | 68603582 | rs872113 | -0.037 | 1.399 | 0.615 | -0.211 | 0.005 | 0.021 | 0.021 | 0.038 | 0.46 | 0.69 | 0.43 | 0.33 | 0.4 | 0.45 |
| 3 | 68604623 | rs7646567 | 0.082 | 1.518 | 0.629 | -0.342 | 0 | -0.03 | -0.03 | 0 | 0.08 | 0.13 | 0.05 | 0 | 0.07 | 0.04 |
| 3 | 68610847 | rs9835386 | 1.456 | 1.605 | 0.431 | -0.359 | -0.053 | 0.156 | 0.156 | 0 | 0.28 | 0.05 | 0.2 | 0.05 | 0.28 | 0.34 |
| 3 | 68642732 | rs5014131 | 0.487 | 1.49 | 0.28 | -0.539 | -0.001 | -0.007 | -0.007 | 0.097 | 0.68 | 0.37 | 0.63 | 0.88 | 0.66 | 0.85 |
| 3 | 68645662 | rs2221393 | 0.714 | 1.467 | 0.124 | -0.522 | 0.014 | -0.045 | -0.045 | 0.106 | 0.37 | 0.18 | 0.29 | 0.6 | 0.34 | 0.47 |
| 3 | 68650788 | rs11721243 | 1.63 | 1.497 | 0.101 | -0.533 | -0.075 | 0.164 | 0.164 | 0.018 | 0.29 | 0.05 | 0.3 | 0.05 | 0.31 | 0.35 |
| 3 | 68652335 | rs4855496 | 0.334 | 1.491 | 0.536 | -0.536 | 0.008 | -0.05 | -0.05 | 0 | 0.69 | 0.6 | 0.64 | 0.88 | 0.67 | 0.86 |
| 3 | 68667357 | rs4305427 | 1.229 | 1.494 | 0.494 | -0.48 | 0.002 | -0.002 | -0.002 | 0.315 | 0.32 | 0.4 | 0.35 | 0.28 | 0.32 | 0.39 |
| 3 | 68680923 | rs1511909 | -0.897 | 1.528 | 0.453 | -0.559 | 0.013 | -0.013 | -0.013 | 0 | 0.81 | 0.77 | 0.7 | 0.87 | 0.77 | 0.9 |
| 3 | 68684050 | rs9867648 | 1.17 | 1.495 | 0.067 | -0.431 | -0.075 | 0.178 | 0.178 | 0.001 | 0.32 | 0.06 | 0.31 | 0.06 | 0.35 | 0.36 |
| 3 | 68686991 | rs9825378 | 1.162 | 1.489 | 0.193 | -0.302 | 0.002 | -0.003 | -0.003 | 0.018 | 0.31 | 0.36 | 0.34 | 0.27 | 0.32 | 0.39 |
| 3 | 68700769 | rs9831891 | -0.495 | 1.116 | -0.033 | -0.208 | 0.035 | -0.034 | -0.034 | 0 | 0.63 | 0.79 | 0.72 | 0.4 | 0.69 | 0.54 |
| 3 | 68707980 | rs9837173 | 1.422 | 1.292 | -0.387 | -0.335 | -0.071 | 0.148 | 0.148 | 0 | 0.29 | 0.06 | 0.46 | 0.06 | 0.33 | 0.35 |
| 3 | 68708150 | rs1394251 | 0.396 | 1.275 | -0.211 | -0.212 | 0.014 | -0.031 | -0.031 | 0.003 | 0.37 | 0.21 | 0.28 | 0.63 | 0.32 | 0.48 |
| 3 | 68719119 | rs1875829 | -0.025 | 1.228 | 0.212 | -0.203 | 0.003 | -0.082 | -0.082 | 0.1 | 0.68 | 0.53 | 0.79 | 0.91 | 0.67 | 0.86 |
| 3 | 68719218 | rs9813609 | 1.201 | 1.2 | -0.577 | -0.446 | -0.127 | 0.211 | 0.211 | 0 | 0.35 | 0.06 | 0.51 | 0.05 | 0.41 | 0.38 |
| 3 | 68720455 | rs6549138 | 1.007 | 1.184 | -0.403 | -0.279 | -0.034 | 0.012 | 0.012 | 0 | 0.37 | 0.19 | 0.54 | 0.27 | 0.41 | 0.36 |
| 3 | 68720845 | rs7626112 | 0.418 | 0.835 | -0.24 | -0.069 | 0.018 | -0.047 | -0.047 | 0.75 | 0.29 | 0.14 | 0.22 | 0.64 | 0.24 | 0.47 |
| 3 | 68723014 | rs11718279 | 0.375 | 0.774 | -0.228 | 0.17 | 0.017 | -0.05 | -0.05 | 0 | 0.29 | 0.19 | 0.22 | 0.61 | 0.25 | 0.47 |
| 3 | 68728679 | rs17247167 | -0.453 | 0.946 | -0.349 | -0.202 | 0.007 | 0.015 | 0.015 | 0.276 | 0.53 | 0.25 | 0.36 | 0.65 | 0.48 | 0.56 |
| 3 | 68728821 | rs13068024 | -0.23 | 1.128 | 0.049 | -0.29 | 0.044 | -0.005 | -0.005 | 4.534 | 0.85 | 0.64 | 0.7 | 0.91 | 0.84 | 0.88 |
| 3 | 68739025 | rs11713512 | 1.017 | 1.194 | -0.089 | -0.334 | -0.117 | 0.22 | 0.22 | 0.099 | 0.37 | 0.07 | 0.24 | 0.05 | 0.42 | 0.32 |
| 3 | 68748667 | rs7614844 | 0.049 | 0.935 | 0.061 | -0.37 | 0.007 | -0.062 | -0.062 | 0 | 0.68 | 0.6 | 0.61 | 0.92 | 0.66 | 0.87 |
| 3 | 68752713 | rs11128087 | 0.117 | 0.896 | 0.055 | -0.388 | 0.005 | -0.059 | -0.059 | 0 | 0.67 | 0.61 | 0.61 | 0.91 | 0.66 | 0.87 |
| 3 | 68766552 | rs4855499 | -0.346 | 0.643 | 0.384 | -0.125 | 0.008 | -0.092 | -0.092 | 0 | 0.68 | 0.49 | 0.61 | 0.91 | 0.66 | 0.87 |
| 3 | 68775668 | rs6772391 | 0.776 | 0.597 | 0.301 | 0.051 | -0.07 | 0.028 | 0.028 | 0 | 0.39 | 0.18 | 0.38 | 0.27 | 0.45 | 0.37 |
| 3 | 68776817 | rs9856483 | -0.221 | 0.535 | 0.284 | 0.179 | -0.001 | -0.062 | -0.062 | 0 | 0.65 | 0.61 | 0.61 | 0.9 | 0.65 | 0.85 |
| 3 | 68777626 | rs867310 | -0.148 | 0.53 | 0.255 | 0.192 | 0 | -0.038 | -0.038 | 0.733 | 0.65 | 0.42 | 0.59 | 0.83 | 0.64 | 0.84 |
| 3 | 68779816 | rs9853012 | 1.72 | 0.611 | 0.175 | 0.15 | -0.102 | 0.167 | 0.167 | 0.288 | 0.31 | 0.06 | 0.24 | 0.03 | 0.37 | 0.31 |
| 3 | 68780703 | rs12630480 | -0.062 | 1.005 | 0.718 | 0.519 | -0.015 | 0.006 | 0.006 | 0 | 0.1 | 0.2 | 0.1 | 0.12 | 0.06 | 0.07 |
| 3 | 68781025 | rs1504299 | 0.123 | 1.065 | 0.783 | 0.33 | 0 | -0.022 | -0.022 | 2.379 | 0.64 | 0.43 | 0.57 | 0.8 | 0.63 | 0.84 |
| 3 | 68785278 | rs11927926 | 0.051 | 0.986 | 0.692 | 0.37 | -0.009 | 0.002 | 0.002 | 0 | 0.1 | 0.16 | 0.05 | 0.13 | 0.07 | 0.07 |
| 3 | 68786217 | rs1394255 | -0.613 | 1.065 | 0.508 | 0.115 | 0.003 | -0.079 | -0.079 | 0.023 | 0.69 | 0.66 | 0.62 | 0.97 | 0.66 | 0.88 |
| 3 | 68786540 | rs983964 | 0.055 | 0.974 | 0.07 | 0.188 | 0.018 | -0.035 | -0.035 | 0 | 0.31 | 0.16 | 0.25 | 0.64 | 0.26 | 0.51 |
| 3 | 68788363 | rs1504301 | 0.054 | 0.37 | 0.387 | 0.303 | 0 | -0.056 | -0.056 | 0 | 0.64 | 0.46 | 0.59 | 0.9 | 0.65 | 0.84 |
| 3 | 68793833 | rs9683308 | 0.151 | 0.105 | 0.38 | 0.708 | 0.016 | -0.023 | -0.023 | 0 | 0.27 | 0.16 | 0.21 | 0.64 | 0.2 | 0.47 |
| 3 | 68807365 | rs1387729 | 1.114 | 0.42 | 0.087 | 0.407 | -0.043 | 0.157 | 0.157 | 0.296 | 0.36 | 0.13 | 0.27 | 0.07 | 0.38 | 0.31 |
| 3 | 68808225 | rs13070250 | -0.14 | 0.337 | 0.332 | 0.373 | -0.014 | -0.015 | -0.015 | 0.032 | 0.3 | 0.34 | 0.29 | 0.85 | 0.2 | 0.5 |
| 3 | 68813388 | rs13433766 | 0.527 | 0.358 | 0.692 | 0.353 | 0.003 | 0 | 0 | 0 | 0.44 | 0.55 | 0.42 | 0.37 | 0.47 | 0.41 |
| 3 | 68823301 | rs7623995 | 0.083 | 0.354 | 0.678 | 0.36 | 0.039 | -0.063 | -0.063 | 0 | 0.29 | 0.13 | 0.19 | 0.61 | 0.19 | 0.48 |
| 3 | 68823432 | rs4855508 | 0.19 | 0.581 | 1.07 | 0.528 | 0.01 | 0.19 | 0.19 | 0 | 0.08 | 0.27 | 0.15 | 0.3 | 0.08 | 0.1 |
| 3 | 68836710 | rs902247 | -0.58 | 0.601 | 0.802 | 0.511 | 0.012 | 0.002 | 0.002 | 0 | 0.72 | 0.68 | 0.53 | 0.75 | 0.63 | 0.85 |
| 3 | 68872978 | rs2171869 | 0.001 | 0.499 | 0.798 | 0.406 | 0.035 | 0.049 | 0.049 | 0 | 0.1 | 0.34 | 0.07 | 0.07 | 0.09 | 0.07 |
| 3 | 68873857 | rs1032304 | 0.464 | 0.504 | 0.435 | 0.258 | -0.015 | 0.085 | 0.085 | 0 | 0.46 | 0.33 | 0.33 | 0.17 | 0.47 | 0.39 |
| 3 | 68884600 | rs6781395 | 1.045 | 0.499 | 0.18 | 0.246 | -0.028 | 0.145 | 0.145 | 0.316 | 0.36 | 0.13 | 0.26 | 0.1 | 0.36 | 0.32 |
| 3 | 68884857 | rs6784291 | 0.946 | 0.495 | 0.222 | 0.319 | -0.009 | 0.093 | 0.093 | 0.185 | 0.37 | 0.24 | 0.27 | 0.14 | 0.36 | 0.32 |
| 3 | 68900822 | rs17048061 | 0.093 | 0.584 | 0.602 | 0.453 | 0.037 | 0.033 | 0.033 | 0 | 0.1 | 0.33 | 0.07 | 0.07 | 0.1 | 0.07 |
| 3 | 68901094 | rs1014405 | 0.144 | 0.584 | 0.628 | 0.453 | 0.043 | 0.038 | 0.038 | 0 | 0.1 | 0.34 | 0.07 | 0.07 | 0.1 | 0.07 |
| 3 | 68902950 | rs902251 | -0.227 | 0.399 | 0.757 | 0.494 | 0.02 | 0.089 | 0.089 | 0.008 | 0.11 | 0.48 | 0.07 | 0.11 | 0.09 | 0.07 |
| 3 | 68903387 | rs6780607 | 0.193 | 0.416 | 0.618 | 0.482 | 0 | 0.006 | 0.006 | 0 | 0.1 | 0.23 | 0.05 | 0.06 | 0.09 | 0.07 |
| 3 | 68905094 | rs2171871 | 0.356 | 0.438 | 0.161 | 0.379 | -0.002 | 0.005 | 0.005 | 0 | 0.46 | 0.51 | 0.37 | 0.2 | 0.46 | 0.39 |
| 3 | 68910803 | rs4855530 | 0.119 | 0.28 | -0.165 | 0.727 | 0.011 | -0.039 | -0.039 | 0 | 0.24 | 0.14 | 0.2 | 0.62 | 0.19 | 0.46 |
| 3 | 68913396 | rs1490000 | 1.227 | 0.339 | -0.74 | 0.467 | -0.054 | 0.104 | 0.104 | 0.424 | 0.37 | 0.07 | 0.26 | 0.18 | 0.35 | 0.29 |
| 3 | 68915507 | rs4104180 | -0.395 | 0.786 | -0.159 | 0.852 | -0.002 | 0.001 | 0.001 | 0 | 0.72 | 0.75 | 0.79 | 0.84 | 0.63 | 0.87 |
| 3 | 68934434 | rs4855535 | 0.025 | 0.548 | -0.667 | 1.066 | -0.02 | -0.08 | -0.08 | 0 | 0.15 | 0.06 | 0.35 | 0.52 | 0.15 | 0.39 |

| | | | | | | | | | | | | | | | | |
|---|---|---|---|---|---|---|---|---|---|---|---|---|---|---|---|---|
| 3 | 68944949 | rs902249 | -0.009 | 0.501 | -0.264 | 0.73 | 0.004 | -0.013 | -0.013 | 0 | 0.61 | 0.46 | 0.71 | 0.75 | 0.53 | 0.73 |
| 3 | 68945048 | rs902250 | -0.009 | 0.489 | -0.264 | 0.73 | 0.004 | -0.013 | -0.013 | 0 | 0.61 | 0.46 | 0.71 | 0.75 | 0.54 | 0.73 |
| 3 | 68945812 | rs17048128 | 0.031 | 0.485 | -0.225 | 0.975 | -0.004 | 0.107 | 0.107 | 0 | 0.15 | 0.21 | 0.35 | 0.5 | 0.14 | 0.38 |
| 3 | 68955591 | rs12714748 | -0.298 | 0.378 | -0.354 | 0.62 | -0.002 | 0.008 | 0.008 | 0 | 0.41 | 0.61 | 0.28 | 0.3 | 0.42 | 0.26 |
| 3 | 68961010 | rs6770678 | 0.16 | 0.47 | -0.4 | 0.715 | -0.022 | -0.067 | -0.067 | 0 | 0.75 | 0.88 | 0.48 | 0.34 | 0.69 | 0.47 |
| 3 | 68963500 | rs3925004 | -0.163 | 0.307 | -0.769 | 0.893 | -0.028 | -0.138 | -0.138 | 0 | 0.78 | 0.94 | 0.53 | 0.47 | 0.76 | 0.54 |
| 3 | 68964333 | rs7643035 | 0.753 | 0.231 | -0.667 | 0.621 | -0.009 | 0.012 | 0.012 | 0 | 0.23 | 0.18 | 0.14 | 0.19 | 0.24 | 0.19 |
| 3 | 68965552 | rs7609906 | 0.712 | 0.226 | -0.663 | 0.619 | -0.001 | 0.001 | 0.001 | 0 | 0.23 | 0.26 | 0.15 | 0.19 | 0.24 | 0.18 |
| 3 | 68971860 | rs9871910 | 0.134 | 0.245 | -0.798 | 0.875 | -0.004 | -0.094 | -0.094 | 0 | 0.77 | 0.9 | 0.52 | 0.42 | 0.75 | 0.51 |
| 3 | 68977647 | rs4541414 | -0.583 | 0.309 | -0.859 | 0.898 | -0.039 | -0.11 | -0.11 | 0 | 0.83 | 0.99 | 0.69 | 0.61 | 0.82 | 0.62 |
| 3 | 68995937 | rs6787753 | -0.404 | 0.143 | -0.041 | 0.875 | 0.002 | 0.171 | 0.171 | 0 | 0.14 | 0.35 | 0.17 | 0.37 | 0.14 | 0.24 |
| 3 | 68996638 | rs6803374 | -0.33 | 0.13 | -0.048 | 0.869 | -0.003 | 0.119 | 0.119 | 0 | 0.13 | 0.27 | 0.16 | 0.37 | 0.14 | 0.24 |
| 3 | 69000370 | rs13094584 | 1.076 | 0.173 | -0.226 | 0.385 | 0 | -0.042 | -0.042 | 0.696 | 0.15 | 0.21 | 0.1 | 0.02 | 0.15 | 0.05 |
| 3 | 69007855 | rs1948936 | -0.394 | 0.095 | -0.592 | 0.388 | -0.052 | -0.13 | -0.13 | 0 | 0.7 | 0.94 | 0.57 | 0.51 | 0.71 | 0.57 |
| 3 | 69008447 | rs6549161 | 0.901 | 0.068 | -0.435 | 0.33 | 0 | -0.021 | -0.021 | 0.007 | 0.55 | 0.42 | 0.38 | 0.1 | 0.58 | 0.33 |
| 3 | 69019978 | rs7628996 | -0.754 | 0.045 | 0.13 | 0.765 | -0.017 | 0.235 | 0.235 | 0 | 0.16 | 0.54 | 0.2 | 0.44 | 0.11 | 0.23 |
| 3 | 69030301 | rs7637369 | -0.127 | 0.052 | -0.127 | 0.482 | -0.008 | 0.052 | 0.052 | 0.129 | 0.33 | 0.51 | 0.34 | 0.52 | 0.3 | 0.48 |
| 3 | 69032484 | rs9813487 | -0.463 | -0.112 | -0.387 | 0.242 | -0.002 | 0.013 | 0.013 | 0 | 0.76 | 0.73 | 0.85 | 0.58 | 0.79 | 0.73 |
| 3 | 69033283 | rs10049354 | 0.37 | -0.535 | -0.048 | 0.098 | 0 | 0.001 | 0.001 | 0 | 0.3 | 0.32 | 0.4 | 0.53 | 0.3 | 0.45 |
| 3 | 69034315 | rs9818973 | -0.105 | -0.532 | 0.299 | 0.297 | -0.001 | -0.001 | -0.001 | 0.293 | 0.66 | 0.63 | 0.59 | 0.33 | 0.7 | 0.53 |
| 3 | 69034385 | rs2033773 | 0.197 | -0.478 | 0.627 | 0.598 | -0.011 | 0.064 | 0.064 | 0 | 0.29 | 0.42 | 0.29 | 0.49 | 0.31 | 0.42 |
| 3 | 69038217 | rs4855543 | 0.747 | -0.485 | 0.579 | 0.192 | -0.002 | -0.004 | -0.004 | 0 | 0.45 | 0.43 | 0.44 | 0.11 | 0.47 | 0.31 |
| 3 | 69038615 | rs5019847 | 0.269 | -0.378 | 0.419 | 0.326 | -0.001 | 0 | 0 | 0 | 0.33 | 0.35 | 0.42 | 0.55 | 0.33 | 0.46 |
| 3 | 69047141 | rs4855544 | 0.648 | -0.412 | 0.22 | 0.217 | 0.011 | -0.062 | -0.062 | 0.059 | 0.28 | 0.16 | 0.36 | 0.51 | 0.28 | 0.47 |
| 3 | 69047612 | rs11128107 | -0.916 | -0.572 | 0.244 | 0.254 | -0.004 | -0.009 | -0.009 | 0 | 0.8 | 0.84 | 0.87 | 0.66 | 0.8 | 0.77 |
| 3 | 69059023 | rs9812253 | 1.037 | -0.577 | -0.06 | -0.349 | -0.006 | -0.039 | -0.039 | 0 | 0.24 | 0.22 | 0.12 | 0.06 | 0.17 | 0.1 |
| 3 | 69063936 | rs12488463 | 1.521 | -0.287 | -0.425 | -0.087 | 0 | -0.002 | -0.002 | 0 | 0.25 | 0.25 | 0.29 | 0.52 | 0.26 | 0.38 |
| 3 | 69070457 | rs1745 | 1.538 | -0.292 | -0.512 | -0.124 | -0.002 | -0.062 | -0.062 | 0.112 | 0.24 | 0.14 | 0.28 | 0.53 | 0.26 | 0.38 |
| 3 | 69082657 | rs937859 | 1.11 | 0.063 | -0.727 | -0.241 | -0.035 | -0.062 | -0.062 | 0 | 0.33 | 0.14 | 0.3 | 0.43 | 0.36 | 0.38 |
| 3 | 69087412 | rs9809889 | -0.395 | 0.117 | -0.562 | -0.297 | -0.001 | 0.012 | 0.012 | 0 | 0.52 | 0.45 | 0.62 | 0.21 | 0.53 | 0.37 |
| 3 | 69093598 | rs2271118 | -1.223 | 0.135 | -0.621 | -0.219 | -0.006 | -0.012 | -0.012 | 9.714 | 0.78 | 0.87 | 0.88 | 0.68 | 0.8 | 0.75 |
| 3 | 69096614 | rs2292199 | -1.234 | 0.164 | -0.597 | -0.209 | 0.002 | 0.005 | 0.005 | 0 | 0.78 | 0.79 | 0.87 | 0.68 | 0.8 | 0.75 |
| 3 | 69103000 | rs12489922 | 0.54 | 0.09 | -0.805 | -0.299 | -0.061 | -0.058 | -0.058 | 0.162 | 0.41 | 0.13 | 0.43 | 0.43 | 0.45 | 0.46 |
| 3 | 69113153 | rs13090606 | 1.254 | 0.134 | -0.702 | -0.215 | 0 | 0.005 | 0.005 | 0 | 0.21 | 0.2 | 0.14 | 0.34 | 0.2 | 0.25 |
| 3 | 69114005 | rs2118539 | -0.219 | 0.166 | -0.706 | -0.305 | 0 | -0.001 | -0.001 | 0 | 0.55 | 0.46 | 0.51 | 0.8 | 0.55 | 0.71 |
| 3 | 69114370 | rs6549181 | 0.823 | 0.189 | -0.573 | -0.213 | 0 | 0.012 | 0.012 | 0 | 0.23 | 0.28 | 0.15 | 0.37 | 0.22 | 0.27 |
| 3 | 69121641 | rs2009602 | 0.713 | 0.188 | -0.511 | -0.207 | 0.009 | 0.028 | 0.028 | 0 | 0.24 | 0.36 | 0.16 | 0.38 | 0.23 | 0.27 |
| 3 | 69125881 | rs3853154 | -0.085 | 0.17 | -0.624 | -0.266 | -0.002 | -0.006 | -0.006 | 0.017 | 0.58 | 0.59 | 0.47 | 0.8 | 0.6 | 0.65 |
| 3 | 69126495 | rs3853156 | -0.314 | 0.212 | -0.562 | -0.247 | -0.007 | -0.026 | -0.026 | 0.596 | 0.61 | 0.55 | 0.46 | 0.8 | 0.6 | 0.68 |
| 3 | 69128682 | rs3755718 | 0.538 | 0.203 | -0.402 | -0.122 | -0.023 | -0.026 | -0.026 | 0 | 0.4 | 0.25 | 0.43 | 0.43 | 0.44 | 0.47 |
| 3 | 69132554 | rs6787569 | 0.542 | -0.087 | -0.296 | -0.105 | -0.021 | 0.124 | 0.124 | 0.124 | 0.11 | 0.27 | 0.06 | 0.33 | 0.06 | 0.2 |
| 3 | 69137219 | rs3853158 | 0.025 | -0.185 | -0.724 | -0.148 | 0 | 0.013 | 0.013 | 0 | 0.4 | 0.45 | 0.29 | 0.5 | 0.4 | 0.51 |
| 3 | 69139300 | rs12714750 | -0.404 | -0.337 | -0.851 | -0.192 | 0 | -0.004 | -0.004 | 0 | 0.42 | 0.42 | 0.54 | 0.27 | 0.49 | 0.42 |
| 3 | 69145632 | rs11128111 | -0.499 | -0.998 | 0.064 | 0.7 | 0 | 0.004 | 0.004 | 0.458 | 0.5 | 0.53 | 0.57 | 0.56 | 0.51 | 0.62 |
| 3 | 69153854 | rs7038 | -0.417 | -1.647 | 0.034 | 1.062 | 0.027 | 0.014 | 0.014 | 0 | 0.47 | 0.7 | 0.31 | 0.61 | 0.41 | 0.52 |
| 3 | 69177524 | rs4615083 | 0.194 | -0.788 | 1.206 | 0.649 | -0.03 | -0.043 | -0.043 | 0 | 0.08 | 0.22 | 0.02 | 0 | 0.01 | 0.01 |
| 3 | 69177913 | rs4241395 | -1.616 | -0.935 | 1.732 | 0.769 | -0.068 | 0.152 | 0.152 | 7.903 | 0.14 | 0.57 | 0.15 | 0.32 | 0.04 | 0.19 |
| 3 | 69183004 | rs1995664 | -2.168 | -1.006 | 1.638 | 0.606 | -0.016 | 0.006 | 0.006 | 1.291 | 0.3 | 0.64 | 0.23 | 0.4 | 0.19 | 0.32 |
| 3 | 69185740 | rs4855550 | -1.225 | -0.858 | 1.741 | 0.683 | -0.009 | 0.009 | 0.009 | 0 | 0.2 | 0.35 | 0.11 | 0.21 | 0.13 | 0.24 |
| 3 | 69190768 | rs9824792 | 1.602 | -1.07 | 1.626 | 0.529 | 0.029 | -0.003 | -0.003 | 0 | 0.57 | 0.17 | 0.67 | 0.57 | 0.66 | 0.62 |
| 3 | 69191221 | rs4407408 | 1.347 | -0.507 | 2.425 | 1.071 | 0.081 | 0.517 | 0.517 | 4.231 | 0.95 | 0.58 | 0.89 | 0.68 | 0.98 | 0.84 |
| 3 | 69192246 | rs4621333 | 1.459 | -0.508 | 2.353 | 1.072 | 0.058 | 0.467 | 0.467 | 4.151 | 0.95 | 0.63 | 0.9 | 0.68 | 0.98 | 0.83 |
| 3 | 69197507 | rs4635707 | 1.632 | -0.449 | 1.631 | 1.045 | 0.02 | -0.011 | -0.011 | 0.095 | 0.68 | 0.24 | 0.75 | 0.59 | 0.77 | 0.62 |
| 3 | 69206654 | rs9840077 | 1.356 | -0.831 | 1.737 | 1.565 | -0.064 | 0.114 | 0.114 | 11.393 | 0.87 | 0.35 | 0.87 | 0.67 | 0.98 | 0.83 |
| 3 | 69222863 | rs4473559 | 1.482 | -0.545 | 1.762 | 1.827 | 0.015 | 0.368 | 0.368 | 1.156 | 0.94 | 0.65 | 0.96 | 0.7 | 0.99 | 0.82 |
| 3 | 69230801 | rs9836305 | 1.98 | -0.624 | 1.944 | 1.934 | -0.001 | 0.41 | 0.41 | 1.112 | 0.93 | 0.61 | 0.88 | 0.64 | 0.99 | 0.81 |
| 3 | 69246697 | rs9847983 | 1.966 | -0.772 | 2.245 | 1.738 | -0.042 | 0.246 | 0.246 | 0 | 0.87 | 0.54 | 0.92 | 0.66 | 0.94 | 0.79 |
| 3 | 69246903 | rs7647940 | -1.286 | -0.769 | 2.17 | 1.737 | 0 | 0.008 | 0.008 | 0 | 0.09 | 0.1 | 0.06 | 0.24 | 0.05 | 0.19 |
| 3 | 69248608 | rs9875511 | 2.511 | -0.84 | 2.457 | 1.688 | -0.055 | 0.144 | 0.144 | 0 | 0.85 | 0.4 | 0.91 | 0.66 | 0.94 | 0.79 |
| 3 | 69249770 | rs9868056 | -1.549 | -0.226 | 2.322 | 1.159 | 0.044 | -0.072 | -0.072 | 0.685 | 0.14 | 0.42 | 0.08 | 0.02 | 0.11 | 0.06 |
| 3 | 69252606 | rs12494107 | -0.597 | 0.178 | 1.806 | 1.615 | -0.001 | 0.056 | 0.056 | 0 | 0.07 | 0.11 | 0.1 | 0.34 | 0.08 | 0.18 |
| 3 | 69252899 | rs6806528 | -0.381 | -0.008 | 1.766 | 1.493 | 0.004 | -0.011 | -0.011 | 0.033 | 0.09 | 0.08 | 0.05 | 0.02 | 0.1 | 0.03 |
| 3 | 69253005 | rs4855364 | 1.849 | -0.104 | 2.051 | 1.795 | 0.025 | 0.237 | 0.237 | 0.003 | 0.81 | 0.5 | 0.75 | 0.49 | 0.81 | 0.78 |
| 3 | 69253318 | rs6784519 | -1.581 | -0.101 | 2.012 | 1.795 | 0.006 | 0.025 | 0.025 | 0 | 0.16 | 0.22 | 0.15 | 0.35 | 0.18 | 0.21 |
| 3 | 69256663 | rs13088684 | 1.837 | -0.189 | 2.004 | 1.756 | 0.009 | 0.137 | 0.137 | 0 | 0.76 | 0.46 | 0.73 | 0.51 | 0.77 | 0.75 |
| 3 | 69257817 | rs6775076 | -0.426 | -0.192 | 1.77 | 1.739 | -0.001 | -0.002 | -0.002 | 0 | 0.11 | 0.12 | 0.12 | 0.34 | 0.11 | 0.21 |
| 3 | 69259117 | rs9809129 | 2.112 | -0.085 | 1.795 | 1.512 | -0.001 | 0.023 | 0.023 | 0.824 | 0.68 | 0.37 | 0.76 | 0.51 | 0.73 | 0.7 |
| 3 | 69263857 | rs12636686 | 1.487 | -0.179 | 1.519 | 1.462 | 0.007 | 0.14 | 0.14 | 0 | 0.76 | 0.52 | 0.76 | 0.52 | 0.78 | 0.77 |
| 3 | 69264796 | rs7640502 | -0.365 | -0.098 | 1.358 | 1.099 | 0.013 | -0.053 | -0.053 | 0 | 0.11 | 0.16 | 0.06 | 0.01 | 0.11 | 0.03 |

| | | | | | | | | | | | | | | | | |
|---|---|---|---|---|---|---|---|---|---|---|---|---|---|---|---|---|
| 3 | 69273190 | rs9848178 | 1.186 | -0.223 | 1.306 | 1.096 | -0.013 | -0.014 | -0.014 | 0.132 | 0.81 | 0.63 | 0.83 | 0.91 | 0.87 | 0.95 |
| 3 | 69274066 | rs9849591 | 0.393 | -0.259 | 1.017 | 1.201 | -0.032 | 0.031 | 0.031 | 0.826 | 0.92 | 0.79 | 0.9 | 0.92 | 0.99 | 0.99 |
| 3 | 69274314 | rs9840469 | 0.62 | -0.025 | 0.368 | 0.656 | 0.02 | 0.017 | 0.017 | 0 | 0.51 | 0.33 | 0.48 | 0.56 | 0.6 | 0.58 |
| 3 | 69280041 | rs4530530 | 0.871 | 0.149 | 0.506 | 0.209 | -0.017 | -0.069 | -0.069 | 0.785 | 0.7 | 0.49 | 0.79 | 0.82 | 0.74 | 0.88 |
| 3 | 69286048 | rs4343629 | 1.17 | -0.071 | 0.672 | 0.169 | -0.001 | -0.01 | -0.01 | 1.178 | 0.81 | 0.79 | 0.88 | 0.92 | 0.85 | 0.91 |
| 3 | 69290119 | rs4855302 | 1.271 | -0.205 | 1.24 | 0.156 | -0.021 | -0.03 | -0.03 | 0 | 0.88 | 0.61 | 0.93 | 0.92 | 0.97 | 0.93 |
| 3 | 69295821 | rs7629818 | -1.932 | 0.793 | 1.598 | 0.454 | 0.064 | -0.029 | -0.029 | 5.575 | 0.2 | 0.51 | 0.17 | 0.18 | 0.24 | 0.17 |
| 3 | 69300337 | rs13322901 | -1.54 | 0.402 | 1.48 | 1.172 | 0.007 | 0.115 | 0.115 | 0 | 0.07 | 0.17 | 0.13 | 0.2 | 0.04 | 0.1 |
| 3 | 69304150 | rs9826784 | -0.184 | -0.354 | 1.017 | 1.168 | 0.02 | -0.028 | -0.028 | 0.025 | 0.31 | 0.24 | 0.34 | 0.5 | 0.22 | 0.39 |
| 3 | 69307885 | rs7623989 | -0.319 | -0.38 | 1.1 | 1.158 | -0.001 | 0 | 0 | 0.172 | 0.33 | 0.31 | 0.34 | 0.5 | 0.22 | 0.4 |
| 3 | 69319503 | rs2046374 | 0.364 | -0.447 | 0.974 | 1.098 | -0.038 | 0.045 | 0.045 | 0 | 0.66 | 0.47 | 0.64 | 0.54 | 0.77 | 0.6 |
| 3 | 69320101 | rs2019265 | 0.129 | -0.363 | 0.522 | 1.096 | 0 | -0.078 | -0.078 | 0.185 | 0.24 | 0.06 | 0.23 | 0.36 | 0.2 | 0.32 |
| 3 | 69328971 | rs9847661 | 0.723 | -0.117 | 0.795 | 0.61 | 0.041 | -0.055 | -0.055 | 0 | 0.84 | 0.63 | 0.87 | 0.95 | 0.75 | 0.85 |
| 3 | 69333045 | rs13100429 | 0.357 | -0.254 | 0.817 | 1.001 | 0.001 | -0.076 | -0.076 | 0 | 0.25 | 0.07 | 0.24 | 0.35 | 0.2 | 0.3 |
| 3 | 69336740 | rs4855381 | 1.141 | -0.195 | 0.844 | 1.008 | 0 | -0.001 | -0.001 | 0 | 0.56 | 0.41 | 0.59 | 0.5 | 0.58 | 0.49 |
| 3 | 69337702 | rs11720505 | -0.584 | 0.118 | 0.438 | 0.889 | 0.013 | -0.009 | -0.009 | 0 | 0.1 | 0.11 | 0.11 | 0.05 | 0.19 | 0.13 |
| 3 | 69343566 | rs1384934 | 1.103 | 0.067 | 0.416 | 0.837 | 0.001 | -0.014 | -0.014 | 0 | 0.79 | 0.68 | 0.82 | 0.85 | 0.76 | 0.77 |
| 3 | 69344290 | rs9813130 | 1.327 | -0.045 | 0.091 | 0.812 | -0.001 | 0.002 | 0.002 | 0 | 0.76 | 0.75 | 0.81 | 0.85 | 0.73 | 0.75 |
| 3 | 69346680 | rs9863228 | 0.655 | -0.288 | 0.278 | 0.905 | -0.002 | 0 | 0 | 0 | 0.57 | 0.55 | 0.59 | 0.51 | 0.6 | 0.49 |
| 3 | 69347542 | rs4855305 | 0.351 | -0.293 | 0.03 | 0.798 | 0.015 | 0.024 | 0.024 | 0 | 0.51 | 0.25 | 0.54 | 0.45 | 0.53 | 0.5 |
| 3 | 69347922 | rs7640861 | 0.097 | -0.034 | -0.799 | 0.571 | -0.05 | -0.03 | -0.03 | 0 | 0.39 | 0.2 | 0.47 | 0.56 | 0.44 | 0.52 |
| 3 | 69350874 | rs4855384 | -0.054 | -0.04 | -0.366 | -0.366 | 0.006 | -0.001 | -0.001 | 0.95 | 0.51 | 0.48 | 0.55 | 0.19 | 0.51 | 0.39 |
| 3 | 69355917 | rs9844501 | -0.255 | 0.073 | -0.048 | -0.286 | 0.041 | 0.006 | 0.006 | 0 | 0.11 | 0.32 | 0.16 | 0.17 | 0.12 | 0.1 |
| 3 | 69357338 | rs11707515 | -1.122 | -0.448 | 0.062 | -0.22 | -0.005 | 0.024 | 0.024 | 0 | 0.26 | 0.46 | 0.29 | 0.33 | 0.24 | 0.29 |
| 3 | 69362706 | rs1483895 | 1.091 | -0.587 | 0.575 | -0.139 | -0.011 | 0.01 | 0.01 | 0.013 | 0.74 | 0.34 | 0.73 | 0.59 | 0.8 | 0.71 |
| 3 | 69365528 | rs1872152 | -1.195 | -0.499 | 0.712 | -0.17 | -0.019 | 0.037 | 0.037 | 0 | 0.26 | 0.58 | 0.28 | 0.41 | 0.22 | 0.29 |
| 3 | 69370396 | rs12152436 | 1.113 | -0.346 | 0.06 | -0.698 | -0.046 | 0.164 | 0.164 | 0 | 0.3 | 0.08 | 0.19 | 0.03 | 0.32 | 0.15 |
| 3 | 69373074 | rs6549200 | 0.244 | -0.122 | 0.383 | -0.566 | -0.024 | 0.017 | 0.017 | 0.558 | 0.13 | 0.33 | 0.13 | 0.26 | 0.09 | 0.08 |
| 3 | 69375655 | rs4855388 | 0.397 | -0.071 | 0.284 | -0.459 | -0.001 | 0.002 | 0.002 | 0 | 0.1 | 0.11 | 0.13 | 0.15 | 0.13 | 0.19 |
| 3 | 69383013 | rs2170800 | -0.527 | -0.308 | -0.012 | -0.568 | 0.003 | -0.017 | -0.017 | 0 | 0.58 | 0.9 | 0.7 | 0.44 | 0.63 | 0.57 |
| 3 | 69385650 | rs11128123 | 0.451 | -0.327 | 0.378 | -0.536 | 0 | 0.012 | 0.012 | 0 | 0.46 | 0.4 | 0.54 | 0.31 | 0.5 | 0.38 |
| 3 | 69389842 | rs10865649 | -0.509 | -0.306 | 0.328 | -0.573 | 0.032 | 0.011 | 0.011 | 0 | 0.6 | 0.77 | 0.69 | 0.44 | 0.66 | 0.57 |
| 3 | 69393419 | rs6774290 | 0.321 | -0.094 | 0.254 | -0.545 | 0.029 | 0.017 | 0.017 | 0.2 | 0.13 | 0.31 | 0.16 | 0.26 | 0.14 | 0.1 |
| 3 | 69401248 | rs4428187 | -0.676 | -0.333 | -0.032 | -0.589 | 0.026 | -0.007 | -0.007 | 0 | 0.63 | 0.86 | 0.71 | 0.43 | 0.69 | 0.59 |
| 3 | 69402086 | rs767285 | 1.308 | -0.332 | -0.047 | -0.674 | -0.072 | 0.159 | 0.159 | 0 | 0.36 | 0.13 | 0.42 | 0.05 | 0.41 | 0.31 |
| 3 | 69402489 | rs1564756 | 0.998 | -0.356 | 0.141 | -0.495 | 0.012 | -0.013 | -0.013 | 0 | 0.71 | 0.18 | 0.7 | 0.62 | 0.72 | 0.73 |
| 3 | 69403918 | rs1843050 | -0.953 | -0.333 | 0.157 | -0.485 | 0.005 | -0.007 | -0.007 | 0.07 | 0.28 | 0.77 | 0.29 | 0.38 | 0.28 | 0.28 |
| 3 | 69406469 | rs2008239 | 0.662 | -0.346 | 0.367 | -0.413 | -0.002 | 0.025 | 0.025 | 0.008 | 0.11 | 0.25 | 0.13 | 0.26 | 0.1 | 0.08 |
| 3 | 69406856 | rs881031 | 0.401 | -0.334 | 0.517 | -0.418 | 0.002 | 0.009 | 0.009 | 0 | 0.14 | 0.24 | 0.14 | 0.13 | 0.18 | 0.19 |
| 3 | 69411188 | rs12629224 | -0.955 | -0.403 | 0.372 | -0.486 | 0.002 | -0.002 | -0.002 | 0 | 0.29 | 0.69 | 0.27 | 0.38 | 0.27 | 0.27 |
| 3 | 69412134 | rs6549208 | -0.964 | -0.413 | 0.368 | -0.49 | 0.002 | -0.002 | -0.002 | 0.584 | 0.29 | 0.69 | 0.27 | 0.38 | 0.28 | 0.27 |
| 3 | 69413694 | rs9868990 | -0.901 | -0.452 | 0.096 | -0.484 | 0.021 | -0.049 | -0.049 | 0 | 0.66 | 0.86 | 0.71 | 0.44 | 0.71 | 0.59 |
| 3 | 69418612 | rs4855394 | 0.128 | -0.215 | 0.125 | -0.245 | 0 | -0.011 | -0.011 | 0.792 | 0.14 | 0.28 | 0.14 | 0.09 | 0.16 | 0.2 |
| 3 | 69421495 | rs7635662 | -0.2 | -0.203 | -0.105 | 0.295 | 0.002 | -0.003 | -0.003 | 0 | 0.54 | 0.38 | 0.58 | 0.48 | 0.58 | 0.69 |
| 3 | 69421910 | rs6549213 | 0.236 | -0.045 | 0.004 | 0.436 | 0.011 | 0.051 | 0.051 | 0.053 | 0.15 | 0.48 | 0.17 | 0.28 | 0.13 | 0.1 |
| 3 | 69422051 | rs4279104 | -0.202 | 0.004 | -0.454 | 0.39 | 0.009 | 0.012 | 0.012 | 0.147 | 0.52 | 0.33 | 0.62 | 0.67 | 0.55 | 0.59 |
| 3 | 69422396 | rs9985329 | -0.432 | 0.604 | -0.41 | 0.036 | 0 | -0.024 | -0.024 | 0 | 0.84 | 0.82 | 0.72 | 0.97 | 0.85 | 0.8 |
| 3 | 69426612 | rs2872805 | -0.287 | 0.352 | 0.004 | 0.579 | 0.003 | 0.004 | 0.004 | 0 | 0.25 | 0.37 | 0.25 | 0.32 | 0.26 | 0.32 |
| 3 | 69427512 | rs2128972 | 0.045 | 0.439 | -0.456 | 0.435 | -0.001 | -0.004 | -0.004 | 0.65 | 0.93 | 0.92 | 0.94 | 0.97 | 0.92 | 0.94 |
| 3 | 69430007 | rs2200247 | -0.536 | 0.301 | -0.282 | 0.711 | -0.012 | 0.034 | 0.034 | 0 | 0.28 | 0.47 | 0.28 | 0.38 | 0.26 | 0.39 |
| 3 | 69430610 | rs7610842 | 0.013 | 0.176 | -0.649 | 1.116 | 0.003 | -0.007 | -0.007 | 2.792 | 0.2 | 0.18 | 0.1 | 0.39 | 0.14 | 0.25 |
| 3 | 69432555 | rs12330712 | -0.555 | 0.356 | 0.429 | 0.717 | 0.026 | -0.014 | -0.014 | 0 | 0.15 | 0.41 | 0.28 | 0.1 | 0.15 | 0.19 |
| 3 | 69434867 | rs17005789 | 0.058 | 0.376 | 0.219 | 1.009 | -0.005 | -0.024 | -0.024 | 0 | 0.06 | 0.02 | 0.15 | 0.13 | 0.07 | 0.21 |
| 3 | 69435706 | rs6769509 | -0.375 | 0.236 | -0.503 | 0.561 | 0.001 | 0.005 | 0.005 | 0 | 0.65 | 0.54 | 0.67 | 0.62 | 0.69 | 0.6 |
| 3 | 69437171 | rs981054 | 0.739 | 0.144 | -0.453 | 0.521 | -0.002 | -0.009 | -0.009 | 0 | 0.12 | 0.09 | 0.25 | 0.02 | 0.14 | 0.18 |
| 3 | 69437399 | rs986204 | 0.466 | 0.217 | -0.437 | 0.57 | -0.006 | 0.023 | 0.023 | 0.037 | 0.53 | 0.39 | 0.42 | 0.6 | 0.54 | 0.42 |
| 3 | 69444935 | rs1967034 | -0.328 | 0.056 | 0.045 | 0.723 | 0.001 | -0.007 | -0.007 | 0.036 | 0.66 | 0.65 | 0.7 | 0.7 | 0.69 | 0.62 |
| 3 | 69445700 | rs2314981 | -0.156 | 0.006 | 0.175 | 0.718 | -0.004 | -0.019 | -0.019 | 0 | 0.64 | 0.53 | 0.68 | 0.7 | 0.69 | 0.62 |
| 3 | 69460531 | rs6549217 | 0.553 | 0.094 | -0.191 | 0.897 | 0.001 | -0.016 | -0.016 | 0 | 0.35 | 0.3 | 0.22 | 0.42 | 0.32 | 0.32 |
| 3 | 69463560 | rs4855403 | 0.42 | 0.11 | -0.061 | 0.863 | 0.001 | 0.016 | 0.016 | 0 | 0.33 | 0.5 | 0.31 | 0.42 | 0.3 | 0.37 |
| 3 | 69472504 | rs10451956 | 0.289 | 0.183 | -0.018 | 0.797 | 0.002 | -0.025 | -0.025 | 0 | 0.35 | 0.48 | 0.3 | 0.29 | 0.3 | 0.34 |
| 3 | 69475108 | rs7640488 | -0.795 | 0.063 | -0.23 | 0.759 | -0.037 | 0.12 | 0.12 | 0 | 0.7 | 0.91 | 0.57 | 0.91 | 0.61 | 0.72 |
| 3 | 69477220 | rs7625432 | 0.419 | 0.156 | -0.11 | 0.803 | 0.003 | -0.023 | -0.023 | 0.068 | 0.34 | 0.41 | 0.3 | 0.29 | 0.3 | 0.34 |
| 3 | 69483752 | rs6797697 | 0.292 | 0.135 | -0.08 | 0.832 | 0 | 0 | 0 | 0.646 | 0.36 | 0.59 | 0.31 | 0.3 | 0.3 | 0.35 |
| 3 | 69483946 | rs6795389 | 0.18 | 0.157 | -0.166 | 0.913 | 0.001 | -0.002 | -0.002 | 0.841 | 0.37 | 0.66 | 0.33 | 0.39 | 0.31 | 0.35 |
| 3 | 69490081 | rs6762603 | 0.958 | 0.081 | -0.299 | 0.985 | -0.034 | -0.088 | -0.088 | 5.448 | 0.3 | 0.12 | 0.19 | 0.43 | 0.31 | 0.3 |
| 3 | 69500831 | rs4303865 | 0.522 | 0.143 | -0.158 | 0.767 | -0.001 | -0.011 | -0.011 | 0 | 0.33 | 0.39 | 0.25 | 0.31 | 0.29 | 0.36 |
| 3 | 69506162 | rs6549219 | 0.443 | 0.191 | -0.195 | 0.77 | -0.001 | -0.016 | -0.016 | 0 | 0.33 | 0.47 | 0.26 | 0.31 | 0.29 | 0.36 |
| 3 | 69508222 | rs9868115 | -0.456 | 0.294 | -0.502 | 0.676 | -0.011 | 0.04 | 0.04 | 0 | 0.66 | 0.82 | 0.52 | 0.84 | 0.6 | 0.68 |

| | | | | | | | | | | | | | | | | |
|---|---|---|---|---|---|---|---|---|---|---|---|---|---|---|---|---|
| 3 | 69512024 | rs1122023 | -0.031 | 0.293 | -0.116 | 0.754 | 0.001 | 0.007 | 0.007 | 0 | 0.33 | 0.25 | 0.31 | 0.33 | 0.32 | 0.44 |
| 3 | 69513028 | rs17354320 | 0.572 | 0.138 | 0.258 | 0.164 | 0 | -0.039 | -0.039 | 0 | 0.26 | 0.25 | 0.16 | 0.03 | 0.3 | 0.16 |
| 3 | 69513198 | rs9843168 | 0.334 | 0.077 | 0.71 | 0.133 | 0.011 | -0.136 | -0.136 | 0 | 0.27 | 0.41 | 0.17 | 0.03 | 0.3 | 0.16 |
| 3 | 69513547 | rs17367084 | 0.605 | 0.105 | 0.008 | 0.147 | -0.024 | 0.062 | 0.062 | 0 | 0.25 | 0.12 | 0.15 | 0.03 | 0.29 | 0.16 |
| 3 | 69516320 | rs1001498 | -0.934 | 0.319 | -0.129 | 0.759 | -0.054 | -0.064 | -0.064 | 0 | 0.62 | 0.84 | 0.44 | 0.49 | 0.54 | 0.54 |
| 3 | 69516937 | rs1498839 | -0.268 | 0.232 | 0.142 | 0.976 | 0.021 | -0.024 | -0.024 | 0.582 | 0.36 | 0.69 | 0.29 | 0.54 | 0.25 | 0.38 |
| 3 | 69518013 | rs6792118 | -1.16 | 0.505 | -0.27 | 0.816 | -0.041 | 0.002 | 0.002 | 0.093 | 0.6 | 0.88 | 0.6 | 0.66 | 0.53 | 0.68 |
| 3 | 69518133 | rs1472464 | -0.519 | 0.327 | -0.104 | 0.725 | -0.001 | -0.001 | -0.001 | 0 | 0.39 | 0.39 | 0.5 | 0.47 | 0.41 | 0.45 |
| 3 | 69519682 | rs6809340 | -1.444 | 0.336 | -0.188 | 0.718 | -0.043 | 0.003 | 0.003 | 0.113 | 0.59 | 0.86 | 0.6 | 0.68 | 0.53 | 0.68 |
| 3 | 69520888 | rs10510980 | -0.395 | 0.387 | -0.158 | 0.686 | -0.003 | 0.013 | 0.013 | 0 | 0.19 | 0.23 | 0.37 | 0.38 | 0.16 | 0.3 |
| 3 | 69523979 | rs9869412 | 0.707 | 0.285 | -0.092 | 0.56 | -0.033 | -0.038 | -0.038 | 0.673 | 0.2 | 0.47 | 0.1 | 0.17 | 0.12 | 0.23 |
| 3 | 69524160 | rs13089266 | 1.773 | 0.462 | -0.311 | 0.436 | 0.002 | -0.026 | -0.026 | 0 | 0.18 | 0.16 | 0.11 | 0.03 | 0.23 | 0.14 |
| 3 | 69531183 | rs10510979 | 0.662 | 0.297 | -0.438 | 0.398 | -0.013 | 0.044 | 0.044 | 0 | 0.11 | 0.01 | 0.25 | 0.03 | 0.07 | 0.15 |
| 3 | 69532105 | rs1498861 | -0.73 | 0.303 | -0.091 | 0.503 | -0.01 | -0.011 | -0.011 | 1.213 | 0.42 | 0.7 | 0.53 | 0.59 | 0.42 | 0.47 |
| 3 | 69533040 | rs10510977 | 1.041 | 0.255 | -0.075 | -0.203 | 0.008 | 0.04 | 0.04 | 0 | 0.17 | 0.06 | 0.07 | 0.07 | 0.12 | 0.22 |
| 3 | 69537330 | rs13092108 | 0.38 | 0.293 | 0.071 | 0.017 | -0.012 | 0.037 | 0.037 | 0 | 0.36 | 0.25 | 0.18 | 0.1 | 0.36 | 0.37 |
| 3 | 69540495 | rs7647186 | 0.037 | 0.488 | -0.364 | 0.255 | -0.001 | 0.017 | 0.017 | 0 | 0.46 | 0.59 | 0.26 | 0.27 | 0.41 | 0.39 |
| 3 | 69541799 | rs9809725 | -0.157 | 0.8 | -1.106 | 0.511 | -0.063 | 0.026 | 0.026 | 1.941 | 0.55 | 0.9 | 0.41 | 0.37 | 0.46 | 0.43 |
| 3 | 69546846 | rs1995177 | -1.312 | 1.385 | -0.849 | 0.067 | -0.035 | 0.145 | 0.145 | 0 | 0.76 | 0.98 | 0.84 | 0.99 | 0.75 | 0.79 |
| 3 | 69547173 | rs1995178 | -0.373 | 1.293 | -0.369 | 0.769 | -0.002 | -0.012 | -0.012 | 2.067 | 0.55 | 0.73 | 0.44 | 0.49 | 0.51 | 0.47 |
| 3 | 69557771 | rs904419 | -0.293 | 1.318 | -0.702 | 0.779 | -0.042 | -0.063 | -0.063 | 0 | 0.56 | 0.91 | 0.45 | 0.5 | 0.51 | 0.48 |
| 3 | 69566662 | rs11924415 | 0.184 | 1.356 | -0.083 | 0.775 | -0.009 | 0.006 | 0.006 | 0 | 0.25 | 0.3 | 0.26 | 0.25 | 0.2 | 0.29 |
| 3 | 69575680 | rs11915680 | 0.507 | 1.395 | -0.205 | 0.233 | 0.003 | 0.008 | 0.008 | 0 | 0.22 | 0.15 | 0.15 | 0.14 | 0.18 | 0.26 |
| 3 | 69588295 | rs6798211 | 0.986 | 1.282 | -0.021 | 0.164 | 0.004 | -0.013 | -0.013 | 0 | 0.19 | 0.3 | 0.1 | 0.14 | 0.17 | 0.16 |
| 3 | 69597148 | rs6549238 | -0.457 | 1.009 | 0.278 | 0.157 | -0.007 | 0.02 | 0.02 | 0 | 0.47 | 0.63 | 0.34 | 0.31 | 0.49 | 0.45 |
| 3 | 69597700 | rs1126301 | -1.727 | 0.933 | -0.075 | -0.292 | 0.002 | -0.01 | -0.01 | 0 | 0.72 | 0.79 | 0.68 | 0.99 | 0.68 | 0.81 |
| 3 | 69597914 | rs12629703 | 0.728 | 0.736 | -0.493 | -0.223 | -0.021 | 0.052 | 0.052 | 0.036 | 0.24 | 0.09 | 0.19 | 0.1 | 0.23 | 0.3 |
| 3 | 69598341 | rs9818139 | -0.032 | 0.755 | -0.422 | -0.228 | 0.014 | 0.009 | 0.009 | 0 | 0.41 | 0.29 | 0.26 | 0.32 | 0.34 | 0.42 |
| 3 | 69599093 | rs12633166 | 0.677 | 0.779 | -0.563 | -0.191 | -0.026 | 0.055 | 0.055 | 0.056 | 0.24 | 0.06 | 0.18 | 0.1 | 0.23 | 0.31 |
| 3 | 69600026 | rs10510984 | 0.142 | 0.455 | -0.035 | 1.144 | 0.033 | -0.042 | -0.042 | 0.058 | 0.15 | 0.1 | 0.17 | 0.35 | 0.06 | 0.19 |
| 3 | 69600854 | rs4855413 | -0.433 | 0.596 | 0.037 | 0.369 | 0.006 | -0.046 | -0.046 | 0 | 0.27 | 0.15 | 0.35 | 0.67 | 0.23 | 0.37 |
| 3 | 69600987 | rs7653611 | 0.461 | 0.617 | 0.794 | 0.851 | -0.008 | 0.009 | 0.009 | 0.853 | 0.17 | 0.2 | 0.07 | 0.21 | 0.1 | 0.11 |
| 3 | 69607472 | rs7653574 | -0.71 | 1.337 | 0.377 | 0.7 | -0.008 | -0.001 | -0.001 | 0 | 0.36 | 0.29 | 0.28 | 0.3 | 0.36 | 0.41 |
| 3 | 69614895 | rs1021667 | 1.016 | 1.454 | 0.357 | 0.765 | -0.001 | 0.006 | 0.006 | 0.043 | 0.12 | 0.16 | 0.04 | 0.09 | 0.08 | 0.08 |
| 3 | 69616385 | rs6774198 | 0.123 | 1.632 | 0.232 | 1.334 | -0.006 | -0.002 | -0.002 | 1.63 | 0.08 | 0.1 | 0.38 | 0.26 | 0.14 | 0.19 |
| 3 | 69616814 | rs904418 | -0.39 | 1.577 | 0.269 | 1.338 | -0.001 | 0.022 | 0.022 | 0.22 | 0.1 | 0.14 | 0.42 | 0.33 | 0.18 | 0.2 |
| 3 | 69621800 | rs1121758 | -1.421 | 1.461 | 0.277 | 1.09 | -0.002 | 0.006 | 0.006 | 0 | 0.44 | 0.39 | 0.66 | 0.55 | 0.49 | 0.56 |
| 3 | 69631756 | rs1498858 | -1.576 | 0.997 | 0.437 | 1.057 | 0.012 | 0.05 | 0.05 | 0 | 0.51 | 0.42 | 0.67 | 0.83 | 0.52 | 0.77 |
| 3 | 69637329 | rs7630350 | -0.727 | 1.155 | 0.379 | 1.414 | 0 | -0.004 | -0.004 | 0 | 0.31 | 0.29 | 0.2 | 0.55 | 0.27 | 0.38 |
| 3 | 69640802 | rs7623679 | 0.655 | 1.057 | 0.224 | 1.571 | -0.001 | 0.129 | 0.129 | 0.333 | 0.81 | 0.6 | 0.56 | 0.47 | 0.79 | 0.72 |
| 3 | 69642255 | rs7619319 | -0.85 | 1.39 | 0.003 | 1.578 | -0.012 | -0.064 | -0.064 | 0.003 | 0.21 | 0.12 | 0.31 | 0.45 | 0.21 | 0.33 |
| 3 | 69656569 | rs746947 | -0.903 | 1.194 | 0.029 | 1.014 | -0.014 | 0.133 | 0.133 | 0 | 0.46 | 0.32 | 0.52 | 0.87 | 0.52 | 0.55 |
| 3 | 69663871 | rs10510985 | -0.843 | 1.199 | -0.181 | 1.057 | -0.037 | 0.137 | 0.137 | 0 | 0.43 | 0.24 | 0.62 | 0.82 | 0.44 | 0.54 |
| 3 | 69667120 | rs6773506 | -1.042 | 1.152 | 0.087 | 1.036 | -0.008 | 0.067 | 0.067 | 0 | 0.44 | 0.39 | 0.49 | 0.82 | 0.44 | 0.54 |
| 3 | 69668385 | rs13097605 | -0.495 | 1.281 | 0.005 | 1.189 | -0.029 | -0.063 | -0.063 | 0 | 0.21 | 0.12 | 0.34 | 0.54 | 0.27 | 0.26 |
| 3 | 69669591 | rs11705949 | 0.74 | 0.755 | 0.106 | 1.295 | 0.004 | -0.051 | -0.051 | 0 | 0.22 | 0.1 | 0.13 | 0.28 | 0.17 | 0.28 |
| 3 | 69694906 | rs17006293 | 0.4 | 0.768 | -0.065 | 1.613 | -0.011 | -0.009 | -0.009 | 0.041 | 0.05 | 0.01 | 0.13 | 0.22 | 0.05 | 0.07 |
| 3 | 69696134 | rs7619198 | 0.182 | 1.025 | 0.047 | 1.247 | -0.002 | -0.01 | -0.01 | 0 | 0.27 | 0.23 | 0.19 | 0.3 | 0.24 | 0.35 |
| 3 | 69697398 | rs17686341 | 0.371 | 1.034 | -0.283 | 1.246 | -0.08 | -0.091 | -0.091 | 0.111 | 0.24 | 0.02 | 0.18 | 0.28 | 0.24 | 0.35 |
| 3 | 69699700 | rs6792167 | 1.514 | 1.019 | -0.165 | 0.778 | -0.016 | 0.189 | 0.189 | 0 | 0.4 | 0.13 | 0.23 | 0.06 | 0.4 | 0.3 |
| 3 | 69701806 | rs17635394 | 0.558 | 1.132 | -0.41 | 1.009 | -0.042 | -0.042 | -0.042 | 0 | 0.25 | 0.07 | 0.17 | 0.28 | 0.23 | 0.35 |
| 3 | 69707585 | rs13090187 | -0.423 | 0.933 | -0.327 | 1.136 | -0.019 | -0.086 | -0.086 | 0.644 | 0.26 | 0.1 | 0.36 | 0.54 | 0.3 | 0.24 |
| 3 | 69717329 | rs4855433 | -0.898 | 0.98 | 0.078 | 0.584 | 0.003 | 0.027 | 0.027 | 0 | 0.51 | 0.44 | 0.5 | 0.83 | 0.47 | 0.56 |
| 3 | 69723217 | rs12631428 | 0.774 | 0.726 | 0.086 | 0.824 | -0.003 | 0.004 | 0.004 | 0.287 | 0.67 | 0.35 | 0.54 | 0.54 | 0.68 | 0.58 |
| 3 | 69728231 | rs6796248 | 0.859 | 0.961 | 0.27 | 1.011 | 0.011 | 0.159 | 0.159 | 0 | 0.78 | 0.51 | 0.66 | 0.46 | 0.76 | 0.68 |
| 3 | 69732280 | rs2114659 | 1.014 | 0.829 | 0.059 | 0.771 | 0.01 | -0.003 | -0.003 | 0 | 0.23 | 0.09 | 0.12 | 0.22 | 0.16 | 0.28 |
| 3 | 69733322 | rs11128137 | 0.056 | 0.846 | 0.256 | 1.054 | -0.006 | 0.004 | 0.004 | 0 | 0.2 | 0.31 | 0.18 | 0.18 | 0.19 | 0.14 |
| 3 | 69734210 | rs9682528 | -0.759 | 0.942 | -0.066 | 1.23 | -0.008 | 0.004 | 0.004 | 0 | 0.14 | 0.13 | 0.23 | 0.51 | 0.17 | 0.23 |
| 3 | 69744135 | rs4855320 | 0.046 | 0.543 | 0.117 | 0.533 | -0.024 | -0.055 | -0.055 | 0.007 | 0.9 | 0.73 | 0.94 | 1 | 0.93 | 0.88 |
| 3 | 69748374 | rs1427434 | 1.086 | 0.805 | -0.358 | 0.322 | -0.041 | 0.139 | 0.139 | 0 | 0.41 | 0.21 | 0.33 | 0.14 | 0.5 | 0.29 |
| 3 | 69749122 | rs6549246 | 0.061 | 0.793 | -0.34 | 0.666 | 0.013 | -0.03 | -0.03 | 1.1 | 0.41 | 0.29 | 0.43 | 0.5 | 0.36 | 0.5 |
| 3 | 69755567 | rs1427430 | 0.387 | 0.824 | -0.357 | 0.766 | 0.005 | 0.019 | 0.019 | 0.058 | 0.06 | 0.16 | 0.07 | 0.08 | 0.05 | 0.12 |
| 3 | 69755743 | rs9284823 | 1.272 | 0.7 | -0.511 | 0.856 | 0.007 | -0.022 | -0.022 | 0 | 0.22 | 0.05 | 0.11 | 0.25 | 0.15 | 0.28 |
| 3 | 69756983 | rs10514724 | 0.182 | 0.735 | -0.338 | 0.873 | -0.001 | 0.012 | 0.012 | 0 | 0.19 | 0.23 | 0.31 | 0.26 | 0.17 | 0.22 |
| 3 | 69758910 | rs1030439 | 0.609 | 0.708 | -0.289 | 0.864 | 0 | -0.003 | -0.003 | 0 | 0.16 | 0.17 | 0.14 | 0.17 | 0.16 | 0.12 |
| 3 | 69762000 | rs9869666 | 0.362 | 0.623 | -0.2 | 0.863 | -0.002 | 0.005 | 0.005 | 0 | 0.17 | 0.26 | 0.14 | 0.17 | 0.16 | 0.12 |
| 3 | 69762267 | rs9856864 | -0.452 | 0.557 | -0.212 | 0.848 | -0.023 | -0.021 | -0.021 | 0 | 0.69 | 0.66 | 0.83 | 0.75 | 0.78 | 0.62 |
| 3 | 69763731 | rs7617168 | 0.216 | 0.655 | -0.23 | 0.832 | -0.002 | 0.001 | 0.001 | 0 | 0.6 | 0.48 | 0.65 | 0.4 | 0.67 | 0.51 |
| 3 | 69764792 | rs6762835 | -0.403 | 0.673 | -0.095 | 0.84 | -0.015 | -0.048 | -0.048 | 0 | 0.91 | 0.71 | 0.94 | 1 | 0.93 | 0.9 |

| | | | | | | | | | | | | | | | | |
|---|---|---|---|---|---|---|---|---|---|---|---|---|---|---|---|---|
| 3 | 69765938 | rs17006384 | 0.489 | 0.71 | -0.161 | 0.879 | 0.015 | 0.088 | 0.088 | 0 | 0.08 | 0.11 | 0.18 | 0.35 | 0.12 | 0.1 |
| 3 | 69768996 | rs4855446 | 0.563 | 0.688 | -0.144 | 0.865 | 0.013 | 0.119 | 0.119 | 0 | 0.82 | 0.49 | 0.76 | 0.64 | 0.82 | 0.79 |
| 3 | 69769329 | rs1946427 | 0.321 | 0.848 | -0.43 | 0.988 | 0.03 | 0.129 | 0.129 | 0 | 0.08 | 0.16 | 0.19 | 0.35 | 0.12 | 0.1 |
| 3 | 69774171 | rs6780999 | 0.261 | 0.876 | -0.568 | 0.809 | -0.006 | -0.016 | -0.016 | 0 | 0.31 | 0.23 | 0.3 | 0.61 | 0.28 | 0.41 |
| 3 | 69774391 | rs17006409 | 0.441 | 0.836 | -0.393 | 0.73 | 0 | -0.001 | -0.001 | 0 | 0.16 | 0.17 | 0.14 | 0.17 | 0.16 | 0.15 |
| 3 | 69778439 | rs990922 | -0.125 | 0.382 | -0.506 | 0.59 | 0.001 | 0.001 | 0.001 | 0 | 0.6 | 0.44 | 0.61 | 0.66 | 0.71 | 0.48 |
| 3 | 69782597 | rs12632773 | 0.614 | 0.338 | -0.718 | 0.067 | -0.002 | 0.001 | 0.001 | 0 | 0.14 | 0.16 | 0.12 | 0.16 | 0.1 | 0.1 |
| 3 | 69783567 | rs6794315 | 0.891 | 0.801 | -0.767 | -0.207 | -0.1 | 0.195 | 0.195 | 0 | 0.36 | 0.1 | 0.28 | 0.05 | 0.47 | 0.16 |
| 3 | 69784265 | rs1427422 | 0.552 | 0.518 | 0.258 | 0.719 | -0.014 | -0.057 | -0.057 | 0 | 0.89 | 0.68 | 0.94 | 0.99 | 0.92 | 0.91 |
| 3 | 69787714 | rs1427421 | 0.518 | 0.611 | 0.108 | 0.76 | -0.01 | -0.05 | -0.05 | 0.038 | 0.9 | 0.72 | 0.95 | 0.99 | 0.93 | 0.91 |
| 3 | 69791738 | rs9869047 | -0.865 | 1.432 | -0.892 | 1.379 | -0.004 | -0.005 | -0.005 | 0.014 | 0.19 | 0.15 | 0.26 | 0.55 | 0.25 | 0.28 |
| 3 | 69797326 | rs1121878 | 0.67 | 1.104 | -1.934 | -0.051 | -0.081 | 0.168 | 0.168 | 0 | 0.33 | 0.1 | 0.18 | 0.02 | 0.4 | 0.2 |
| 3 | 69803328 | rs4855447 | 0.068 | 1.059 | -2.116 | 0.773 | -0.055 | -0.116 | -0.116 | 0.207 | 0.17 | 0.05 | 0.23 | 0.45 | 0.22 | 0.18 |
| 3 | 69808638 | rs13073599 | 0.573 | 0.769 | -1.917 | 0.754 | -0.088 | 0.068 | 0.068 | 0 | 0.34 | 0.12 | 0.19 | 0.14 | 0.43 | 0.28 |
| 3 | 69825970 | rs17043851 | -0.47 | 0.543 | -1.251 | 0.904 | -0.005 | -0.032 | -0.032 | 0 | 0.21 | 0.16 | 0.25 | 0.47 | 0.24 | 0.2 |
| 3 | 69834454 | rs10865653 | -1.335 | -0.293 | -0.995 | 0.498 | -0.038 | -0.043 | -0.043 | 0.271 | 0.67 | 0.57 | 0.61 | 0.73 | 0.81 | 0.66 |
| 3 | 69834555 | rs11128143 | -0.212 | 0.262 | -1.421 | 0.341 | 0.027 | -0.02 | -0.02 | 0 | 0.47 | 0.2 | 0.37 | 0.48 | 0.58 | 0.38 |
| 3 | 69843971 | rs6777363 | 1.4 | -0.234 | -1.413 | -0.174 | -0.046 | 0.068 | 0.068 | 0 | 0.31 | 0.18 | 0.18 | 0.05 | 0.41 | 0.21 |
| 3 | 69846272 | rs6787022 | -0.05 | -0.249 | -1.4 | 0.371 | 0.078 | 0.137 | 0.137 | 0 | 0.5 | 0.24 | 0.34 | 0.23 | 0.63 | 0.3 |
| 3 | 69870595 | rs1529585 | 1.366 | -0.151 | -1.49 | 0.315 | -0.097 | 0.158 | 0.158 | 0 | 0.31 | 0.1 | 0.19 | 0.03 | 0.41 | 0.12 |
| 3 | 69887656 | rs1367370 | 0.088 | -0.141 | -1.422 | 0.412 | -0.015 | -0.014 | -0.014 | 1.34 | 0.17 | 0.09 | 0.15 | 0.19 | 0.21 | 0.17 |
| 3 | 69900137 | rs1544978 | 1.356 | -0.031 | -1.362 | 0.258 | -0.089 | 0.135 | 0.135 | 0 | 0.3 | 0.12 | 0.19 | 0.03 | 0.41 | 0.11 |
| 3 | 69902514 | rs13071701 | 1.378 | -0.038 | -1.375 | 0.255 | -0.099 | 0.146 | 0.146 | 0 | 0.3 | 0.1 | 0.19 | 0.03 | 0.41 | 0.11 |
| 3 | 69909094 | rs7623486 | 0.176 | 0.196 | -0.918 | 0.699 | 0.007 | 0.008 | 0.008 | 0.04 | 0.15 | 0.21 | 0.14 | 0.16 | 0.19 | 0.17 |
| 3 | 69916637 | rs2320172 | 2 | -0.43 | -0.562 | 0.41 | 0.01 | -0.001 | -0.001 | 1.576 | 0.09 | 0.14 | 0.22 | 0.08 | 0.08 | 0.06 |
| 3 | 69934529 | rs17006579 | -1.137 | -0.996 | -0.891 | 0.37 | -0.006 | 0.004 | 0.004 | 0.439 | 0.39 | 0.5 | 0.42 | 0.38 | 0.37 | 0.45 |
| 3 | 69978466 | rs7430957 | -0.203 | -1.218 | -0.934 | 0.405 | -0.037 | 0.071 | 0.071 | 0.264 | 0.27 | 0.5 | 0.28 | 0.38 | 0.2 | 0.32 |
| 3 | 70005281 | rs7623610 | -1.739 | -1.088 | -1.875 | -0.067 | -0.116 | -0.034 | -0.034 | 0.28 | 0.57 | 0.96 | 0.67 | 0.54 | 0.52 | 0.61 |
| 3 | 70016467 | rs576 | -0.331 | -0.658 | -1.188 | -0.461 | -0.008 | 0.002 | 0.002 | 0.48 | 0.32 | 0.45 | 0.32 | 0.29 | 0.26 | 0.34 |
| 3 | 70016904 | rs704246 | -0.071 | -0.534 | -1.36 | -0.404 | -0.001 | 0.003 | 0.003 | 0.04 | 0.28 | 0.35 | 0.3 | 0.27 | 0.24 | 0.3 |
| 3 | 70017701 | rs1491687 | 0.004 | -0.521 | -1.353 | -0.396 | -0.001 | 0.002 | 0.002 | 0.007 | 0.72 | 0.65 | 0.7 | 0.73 | 0.75 | 0.7 |
| 3 | 70027374 | rs11128156 | -2.317 | 0.328 | -1.801 | -0.48 | -0.127 | 0.15 | 0.15 | 0 | 0.78 | 0.99 | 0.82 | 1 | 0.65 | 0.93 |
| 3 | 70039781 | rs9310179 | -0.646 | -0.695 | -1.296 | -0.029 | 0.019 | 0.05 | 0.05 | 0 | 0.57 | 0.79 | 0.56 | 0.71 | 0.43 | 0.65 |
| 3 | 70044334 | rs6809108 | 0.083 | -0.782 | -0.804 | -0.033 | -0.001 | 0.011 | 0.011 | 0 | 0.07 | 0.17 | 0.06 | 0.1 | 0.09 | 0.14 |
| 3 | 70069907 | rs17785152 | 0.018 | -1.198 | -0.416 | -0.098 | -0.011 | -0.002 | -0.002 | 0 | 0.43 | 0.46 | 0.47 | 0.62 | 0.32 | 0.49 |
| 3 | 70099299 | rs9815027 | -0.18 | -1.228 | -0.626 | -0.103 | 0 | 0 | 0 | 0 | 0.45 | 0.54 | 0.48 | 0.62 | 0.33 | 0.48 |
| 3 | 70110627 | rs1873628 | 0.034 | -1.638 | -0.249 | -0.168 | -0.007 | -0.036 | -0.036 | 0 | 0.15 | 0.23 | 0.07 | 0.04 | 0.1 | 0.05 |
| 3 | 70115600 | rs1027234 | -0.728 | -1.419 | -0.283 | -0.044 | 0.02 | 0.027 | 0.027 | 0 | 0.55 | 0.66 | 0.53 | 0.66 | 0.41 | 0.55 |
| 3 | 70121452 | rs7634340 | -0.645 | -1.278 | -0.557 | -0.657 | 0.025 | 0.126 | 0.126 | 0 | 0.56 | 0.83 | 0.56 | 0.77 | 0.43 | 0.66 |
| 3 | 70129459 | rs729038 | 0.53 | -0.605 | -0.288 | -0.287 | -0.033 | -0.039 | -0.039 | 0 | 0.07 | 0.01 | 0.09 | 0.13 | 0.08 | 0.09 |
| 3 | 70134474 | rs6549268 | -1.032 | -0.93 | -0.313 | -0.551 | -0.008 | -0.003 | -0.003 | 0 | 0.32 | 0.51 | 0.39 | 0.31 | 0.24 | 0.35 |
| 3 | 70141750 | rs6762229 | 1.065 | -1.12 | -0.45 | -1.159 | -0.005 | 0.03 | 0.03 | 0 | 0.39 | 0.31 | 0.53 | 0.07 | 0.56 | 0.22 |
| 3 | 70143095 | rs755353 | 1.726 | -1.106 | -0.511 | -1.202 | -0.186 | 0.24 | 0.24 | 0 | 0.33 | 0.02 | 0.3 | 0 | 0.48 | 0.14 |
| 3 | 70143459 | rs902710 | 0.475 | -1.046 | -0.436 | -1.158 | 0.082 | 0.032 | 0.032 | 0 | 0.06 | 0.26 | 0.22 | 0.06 | 0.08 | 0.08 |
| 3 | 70149696 | rs6782608 | -0.15 | -1.413 | -0.501 | -1.027 | -0.017 | -0.01 | -0.01 | 0.116 | 0.4 | 0.51 | 0.33 | 0.7 | 0.3 | 0.53 |
| 3 | 70155835 | rs7632699 | -0.295 | -1.4 | -0.784 | -1.234 | -0.013 | -0.024 | -0.024 | 0 | 0.45 | 0.49 | 0.4 | 0.75 | 0.34 | 0.6 |
| 3 | 70165358 | rs9855111 | -0.863 | -1.479 | -0.601 | -1.169 | -0.001 | 0.002 | 0.002 | 0 | 0.3 | 0.31 | 0.23 | 0.43 | 0.2 | 0.42 |
| 3 | 70173049 | rs9857007 | 0.972 | -1.525 | -0.658 | -1.129 | -0.042 | 0.067 | 0.067 | 0 | 0.26 | 0.35 | 0.21 | 0.5 | 0.18 | 0.35 |
| 3 | 70182149 | rs9824741 | 0.105 | -1.471 | -0.645 | -1.122 | -0.038 | 0.002 | 0.002 | 0 | 0.34 | 0.44 | 0.27 | 0.7 | 0.27 | 0.53 |
| 3 | 70186696 | rs7633462 | 0.457 | -1.43 | -0.655 | -1.147 | 0.001 | -0.007 | -0.007 | 0 | 0.15 | 0.12 | 0.15 | 0.22 | 0.1 | 0.23 |
| 3 | 70199824 | rs11128165 | 0.296 | -1.096 | -0.52 | -1.548 | 0.011 | -0.009 | -0.009 | 0 | 0.51 | 0.48 | 0.58 | 0.08 | 0.64 | 0.28 |
| 3 | 70207638 | rs9847642 | -0.081 | -1.106 | -0.505 | -1.53 | 0.002 | 0 | 0 | 0.181 | 0.46 | 0.42 | 0.4 | 0.72 | 0.34 | 0.59 |
| 3 | 70209796 | rs7614990 | 0.967 | -0.993 | -0.549 | -1.546 | -0.001 | -0.037 | -0.037 | 0 | 0.41 | 0.4 | 0.54 | 0.05 | 0.56 | 0.22 |
| 3 | 70223684 | rs13069349 | 0.443 | -0.872 | -0.507 | -1.475 | 0.018 | 0.009 | 0.009 | 0 | 0.49 | 0.46 | 0.58 | 0.1 | 0.64 | 0.26 |
| 3 | 70224184 | rs7642356 | 2.035 | -0.855 | -0.553 | -1.459 | -0.019 | -0.008 | -0.008 | 0 | 0.06 | 0.01 | 0.04 | 0.03 | 0.08 | 0.04 |
| 3 | 70224323 | rs6549288 | 0.865 | -0.832 | -0.441 | -1.525 | 0 | -0.021 | -0.021 | 0.043 | 0.41 | 0.38 | 0.54 | 0.05 | 0.56 | 0.22 |
| 3 | 70229761 | rs4564939 | 0.456 | -0.906 | -0.804 | -1.451 | -0.001 | -0.042 | -0.042 | 0.119 | 0.51 | 0.53 | 0.64 | 0.12 | 0.64 | 0.37 |
| 3 | 70232626 | rs6549292 | -1.862 | -1.404 | -0.845 | -1.428 | 0.044 | -0.065 | -0.065 | 0.135 | 0.82 | 0.92 | 0.84 | 0.46 | 0.88 | 0.69 |
| 3 | 70242154 | rs6549297 | 0.659 | -1.269 | -0.844 | -1.593 | 0.001 | -0.038 | -0.038 | 0 | 0.49 | 0.48 | 0.58 | 0.12 | 0.6 | 0.36 |
| 3 | 70245870 | rs12637141 | 1.855 | -1.321 | -0.821 | -1.505 | 0.05 | -0.113 | -0.113 | 0 | 0.17 | 0.03 | 0.16 | 0.56 | 0.12 | 0.29 |
| 3 | 70246225 | rs7431876 | 0.149 | -1.35 | -0.814 | -1.524 | -0.03 | -0.015 | -0.015 | 1.066 | 0.62 | 0.58 | 0.72 | 0.3 | 0.79 | 0.47 |
| 3 | 70247251 | rs13064328 | 1.852 | -1.25 | -0.834 | -1.475 | 0.038 | -0.117 | -0.117 | 0 | 0.18 | 0.03 | 0.16 | 0.55 | 0.15 | 0.29 |
| 3 | 70248282 | rs7623693 | -0.167 | -0.983 | -0.672 | -1.378 | -0.032 | 0.012 | 0.012 | 0.004 | 0.82 | 0.64 | 0.88 | 0.86 | 0.92 | 0.77 |
| 3 | 70249543 | rs7427775 | -0.18 | -0.977 | -0.664 | -1.372 | -0.039 | 0.003 | 0.003 | 0.009 | 0.82 | 0.62 | 0.88 | 0.87 | 0.92 | 0.77 |
| 3 | 70253714 | rs17006834 | 1.252 | -0.917 | -1.081 | -1.348 | -0.004 | -0.017 | -0.017 | 0 | 0.09 | 0.25 | 0.04 | 0.03 | 0.04 | 0.13 |
| 3 | 70258677 | rs7430257 | 0.555 | -0.792 | -1.395 | -1.563 | -0.015 | 0.167 | 0.167 | 0 | 0.46 | 0.23 | 0.43 | 0.11 | 0.51 | 0.34 |
| 3 | 70258762 | rs7430272 | 0.75 | -0.823 | -1.389 | -1.61 | 0.001 | 0.005 | 0.005 | 0 | 0.62 | 0.32 | 0.7 | 0.68 | 0.67 | 0.64 |
| 3 | 70263077 | rs7432941 | 0.768 | -0.15 | -1.266 | -1.077 | -0.01 | 0.154 | 0.154 | 0 | 0.46 | 0.24 | 0.44 | 0.11 | 0.51 | 0.34 |
| 3 | 70268860 | rs17787105 | 1.294 | -0.006 | -1.652 | -1.014 | -0.06 | 0.014 | 0.014 | 0 | 0.11 | 0.01 | 0.14 | 0.03 | 0.19 | 0.07 |

| | | | | | | | | | | | | | | | | |
|---|---|---|---|---|---|---|---|---|---|---|---|---|---|---|---|---|
| 3 | 70270499 | rs11712898 | 0.228 | 0.247 | -1.147 | -0.874 | -0.096 | 0.214 | 0.214 | 0.192 | 0.44 | 0.07 | 0.43 | 0.11 | 0.5 | 0.34 |
| 3 | 70274229 | rs1403069 | 0.287 | 0.27 | -0.882 | -0.483 | -0.005 | -0.01 | -0.01 | 2.291 | 0.39 | 0.44 | 0.5 | 0.71 | 0.41 | 0.46 |
| 3 | 70283132 | rs8180009 | -0.354 | 0.688 | 0.058 | 0.247 | -0.002 | 0.079 | 0.079 | 0 | 0.09 | 0.31 | 0.04 | 0.18 | 0.04 | 0.19 |
| 3 | 70285244 | rs6790613 | -0.828 | 1.176 | -0.725 | -0.024 | 0.04 | 0.064 | 0.064 | 0 | 0.07 | 0.25 | 0.18 | 0.12 | 0.04 | 0.11 |
| 3 | 70286279 | rs1546734 | -0.412 | 1.375 | -1.791 | -0.097 | -0.085 | 0.109 | 0.109 | 0 | 0.17 | 0.01 | 0.16 | 0 | 0.27 | 0.06 |
| 3 | 70300475 | rs12486303 | -0.646 | 0.352 | -1.528 | 0.795 | 0.003 | -0.037 | -0.037 | 0 | 0.06 | 0 | 0.07 | 0.14 | 0.04 | 0.09 |
| 3 | 70306417 | rs1522344 | 0.297 | -0.626 | -1.169 | 1.64 | -0.019 | 0.054 | 0.054 | 0.366 | 0.8 | 0.71 | 0.87 | 0.54 | 0.88 | 0.69 |
| 3 | 70307605 | rs7616565 | 1.319 | -0.219 | -0.957 | 1.79 | 0.01 | 0.006 | 0.006 | 0 | 0.68 | 0.68 | 0.72 | 0.27 | 0.72 | 0.56 |
| 3 | 70307767 | rs12497885 | -0.648 | -0.251 | -0.902 | 2.569 | -0.002 | 0.067 | 0.067 | 0 | 0.08 | 0.08 | 0.09 | 0.35 | 0.07 | 0.15 |
| 3 | 70309725 | rs13089371 | 1.345 | -0.467 | -1.061 | 1.263 | -0.042 | 0.097 | 0.097 | 0.053 | 0.4 | 0.24 | 0.38 | 0.02 | 0.48 | 0.16 |
| 3 | 70311639 | rs12489265 | 0.601 | -0.717 | -1.372 | 1.398 | -0.007 | -0.02 | -0.02 | 0 | 0.09 | 0 | 0.08 | 0.07 | 0.07 | 0.13 |
| 3 | 70312134 | rs1608098 | 0.75 | -0.737 | -1.211 | 1.463 | 0.011 | -0.004 | -0.004 | 0 | 0.17 | 0.13 | 0.11 | 0.24 | 0.09 | 0.24 |
| 3 | 70313844 | rs2030527 | 0.26 | -0.821 | -0.306 | 1.449 | -0.005 | -0.006 | -0.006 | 0 | 0.22 | 0.14 | 0.31 | 0.3 | 0.29 | 0.3 |
| 3 | 70319708 | rs9846850 | -1.296 | -0.945 | 0.014 | 1.345 | 0 | -0.001 | -0.001 | 2.74 | 0.42 | 0.57 | 0.52 | 0.73 | 0.42 | 0.58 |
| 3 | 70320864 | rs6549322 | -0.465 | -0.799 | 0.205 | 1.53 | -0.02 | 0.004 | 0.004 | 0.019 | 0.81 | 0.72 | 0.88 | 0.73 | 0.91 | 0.73 |
| 3 | 70321857 | rs1015097 | -1.178 | -0.559 | 0.321 | 1.857 | 0.007 | 0.001 | 0.001 | 0.782 | 0.34 | 0.43 | 0.43 | 0.6 | 0.39 | 0.48 |
| 3 | 70322156 | rs1522337 | -0.636 | -0.416 | 0.461 | 1.912 | 0.03 | 0.124 | 0.124 | 1.616 | 0.24 | 0.42 | 0.35 | 0.5 | 0.32 | 0.35 |
| 3 | 70323379 | rs9681907 | -1.072 | -0.716 | 0.439 | 2.507 | -0.001 | 0.149 | 0.149 | 0 | 0.09 | 0.13 | 0.27 | 0.46 | 0.14 | 0.14 |
| 3 | 70326261 | rs9855222 | 0.793 | -1.333 | 0.668 | 2.257 | -0.051 | 0.052 | 0.052 | 0.019 | 0.77 | 0.44 | 0.74 | 0.39 | 0.83 | 0.64 |
| 3 | 70326288 | rs9868241 | 0.354 | -1.403 | 0.639 | 1.952 | 0 | -0.005 | -0.005 | 0.225 | 0.87 | 0.89 | 0.87 | 0.98 | 0.91 | 0.88 |
| 3 | 70327250 | rs2320474 | 1.162 | -1.35 | 0.322 | 1.939 | 0 | 0 | 0 | 0 | 0.64 | 0.39 | 0.6 | 0.33 | 0.73 | 0.49 |
| 3 | 70328120 | rs1357270 | 0.524 | -0.726 | -0.042 | 1.691 | -0.018 | -0.013 | -0.013 | 0.007 | 0.06 | 0.02 | 0.04 | 0.07 | 0.09 | 0.05 |
| 3 | 70328225 | rs7651174 | 1.289 | -1.14 | 0.091 | 1.449 | -0.007 | -0.022 | -0.022 | 0 | 0.54 | 0.52 | 0.54 | 0.21 | 0.59 | 0.42 |
| 3 | 70330440 | rs1105032 | -0.689 | -0.813 | 0.283 | 1.748 | 0.006 | 0.002 | 0.002 | 0 | 0.22 | 0.31 | 0.22 | 0.25 | 0.28 | 0.31 |
| 3 | 70331707 | rs4498002 | -1.534 | -0.695 | 0.334 | 1.781 | 0 | -0.036 | -0.036 | 0 | 0.37 | 0.43 | 0.43 | 0.79 | 0.39 | 0.53 |
| 3 | 70341051 | rs6549329 | -1.793 | -0.738 | 0.382 | 1.755 | -0.01 | 0.013 | 0.013 | 0.033 | 0.4 | 0.48 | 0.34 | 0.5 | 0.34 | 0.59 |
| 3 | 70347392 | rs1403081 | -0.37 | -0.697 | 0.322 | 1.585 | 0.004 | -0.056 | -0.056 | 0 | 0.19 | 0.26 | 0.13 | 0.07 | 0.21 | 0.27 |
| 3 | 70369162 | rs17007071 | -0.917 | -0.804 | 0.231 | 1.702 | 0.01 | 0.004 | 0.004 | 0 | 0.13 | 0.09 | 0.06 | 0.08 | 0.1 | 0.18 |
| 3 | 70370965 | rs1533815 | 1.512 | -0.842 | 0.287 | 1.78 | -0.003 | 0.006 | 0.006 | 0 | 0.66 | 0.65 | 0.67 | 0.4 | 0.68 | 0.5 |
| 3 | 70373502 | rs1403072 | 0.841 | -0.766 | 0.353 | 1.832 | -0.005 | 0.05 | 0.05 | 0 | 0.75 | 0.68 | 0.75 | 0.59 | 0.78 | 0.61 |
| 3 | 70374190 | rs9870891 | 0.349 | -0.768 | 0.364 | 1.85 | -0.001 | 0.07 | 0.07 | 0 | 0.08 | 0.09 | 0.16 | 0.35 | 0.06 | 0.15 |
| 3 | 70374871 | rs1522334 | 0.074 | -0.768 | 0.368 | 1.789 | 0 | -0.028 | -0.028 | 0 | 0.82 | 0.81 | 0.89 | 0.93 | 0.82 | 0.75 |
| 3 | 70375780 | rs1522335 | -0.043 | -0.789 | 0.35 | 1.809 | -0.002 | 0.095 | 0.095 | 0 | 0.09 | 0.1 | 0.19 | 0.45 | 0.06 | 0.16 |
| 3 | 70382857 | rs13093307 | 1.81 | -0.752 | 0.289 | 1.832 | -0.006 | -0.001 | -0.001 | 0 | 0.61 | 0.56 | 0.65 | 0.39 | 0.67 | 0.42 |
| 3 | 70384008 | rs6763116 | 1.784 | -0.777 | 0.285 | 1.79 | -0.007 | 0.015 | 0.015 | 0 | 0.62 | 0.56 | 0.67 | 0.51 | 0.68 | 0.42 |
| 3 | 70384363 | rs4246676 | 0.809 | -0.75 | 0.288 | 1.803 | -0.003 | 0.035 | 0.035 | 0 | 0.75 | 0.71 | 0.74 | 0.59 | 0.78 | 0.62 |
| 3 | 70391895 | rs805468 | 0.002 | -0.773 | 0.216 | 1.35 | 0 | -0.011 | -0.011 | 0.33 | 0.83 | 0.85 | 0.91 | 0.93 | 0.83 | 0.76 |
| 3 | 70391905 | rs805469 | -1.156 | -0.776 | 0.084 | 1.296 | 0.004 | 0.007 | 0.007 | 0 | 0.14 | 0.13 | 0.08 | 0.08 | 0.12 | 0.2 |
| 3 | 70397985 | rs12490035 | 1.531 | -0.753 | 0.067 | 1.339 | -0.008 | -0.004 | -0.004 | 0.031 | 0.61 | 0.53 | 0.61 | 0.39 | 0.65 | 0.41 |
| 3 | 70399711 | rs813397 | -0.583 | -0.533 | -0.19 | 1.344 | -0.013 | -0.018 | -0.018 | 0 | 0.07 | 0.03 | 0.04 | 0.07 | 0.09 | 0.1 |
| 3 | 70403938 | rs805490 | 0.61 | -0.648 | 0.022 | 1.445 | -0.003 | 0.074 | 0.074 | 0 | 0.75 | 0.68 | 0.72 | 0.48 | 0.78 | 0.61 |
| 3 | 70406133 | rs805492 | -1.021 | -0.443 | -0.093 | 0.995 | -0.003 | -0.004 | -0.004 | 0.215 | 0.13 | 0.17 | 0.07 | 0.08 | 0.11 | 0.17 |
| 3 | 70410209 | rs810912 | -0.247 | -0.404 | -0.053 | 1.049 | 0.006 | 0.013 | 0.013 | 0 | 0.06 | 0.11 | 0.06 | 0.09 | 0.09 | 0.06 |
| 3 | 70417993 | rs13314351 | -0.738 | -0.016 | -0.402 | 0.507 | 0.004 | 0.028 | 0.028 | 0 | 0.35 | 0.24 | 0.22 | 0.14 | 0.37 | 0.33 |
| 3 | 70425261 | rs2058890 | -2.002 | 0.027 | -0.069 | 0.38 | -0.003 | 0.012 | 0.012 | 0 | 0.58 | 0.61 | 0.6 | 0.88 | 0.6 | 0.67 |
| 3 | 70427067 | rs805478 | -0.419 | 0.146 | -0.091 | 0.375 | -0.006 | 0.035 | 0.035 | 0 | 0.37 | 0.24 | 0.25 | 0.14 | 0.41 | 0.35 |
| 3 | 70427085 | rs805479 | -1.822 | 0.083 | -0.083 | 0.949 | -0.006 | 0.004 | 0.004 | 0 | 0.54 | 0.59 | 0.48 | 0.59 | 0.52 | 0.61 |
| 3 | 70430309 | rs805486 | -1.846 | 0.086 | -0.109 | 0.95 | -0.006 | 0.004 | 0.004 | 0 | 0.55 | 0.59 | 0.48 | 0.59 | 0.52 | 0.61 |
| 3 | 70436904 | rs17007189 | 0.757 | -0.334 | 0.19 | 1.207 | -0.016 | -0.041 | -0.041 | 0.653 | 0.08 | 0.17 | 0.08 | 0.04 | 0.06 | 0.1 |
| 3 | 70438588 | rs17007192 | 1.047 | -0.333 | 0.155 | 1.538 | 0 | 0.064 | 0.064 | 0 | 0.07 | 0.09 | 0.14 | 0.32 | 0.05 | 0.11 |
| 3 | 70442904 | rs9310191 | -0.943 | -0.341 | 0.448 | 1.447 | -0.014 | 0.173 | 0.173 | 0 | 0.16 | 0.4 | 0.23 | 0.45 | 0.16 | 0.21 |
| 3 | 70444705 | rs9831386 | -0.102 | 0.101 | -0.078 | 1.439 | 0.009 | 0.215 | 0.215 | 0.07 | 0.07 | 0.18 | 0.16 | 0.41 | 0.1 | 0.11 |
| 3 | 70445283 | rs4974287 | 0.041 | 0.126 | -0.685 | 1.003 | 0.002 | -0.017 | -0.017 | 0 | 0.73 | 0.8 | 0.75 | 0.65 | 0.75 | 0.73 |
| 3 | 70446896 | rs9825323 | 0.396 | 0.329 | -0.668 | 0.909 | 0.002 | -0.006 | -0.006 | 0 | 0.61 | 0.68 | 0.67 | 0.59 | 0.65 | 0.59 |
| 3 | 70450505 | rs7649557 | -0.33 | 0.619 | -0.622 | 0.584 | 0.007 | 0.005 | 0.005 | 0.318 | 0.51 | 0.44 | 0.59 | 0.38 | 0.6 | 0.59 |
| 3 | 70452730 | rs2216367 | -0.084 | 1.529 | -0.727 | 0.605 | -0.009 | -0.007 | -0.007 | 0.726 | 0.06 | 0.04 | 0.13 | 0.1 | 0.12 | 0.06 |
| 3 | 70453579 | rs9833791 | -0.213 | 1.902 | -0.889 | -0.009 | 0 | -0.045 | -0.045 | 0.197 | 0.29 | 0.26 | 0.15 | 0.04 | 0.22 | 0.32 |
| 3 | 70454534 | rs13070827 | -0.201 | 1.997 | -0.965 | 0.193 | -0.05 | -0.037 | -0.037 | 0 | 0.26 | 0.15 | 0.47 | 0.33 | 0.42 | 0.32 |
| 3 | 70466110 | rs918190 | 0.426 | 1.58 | -0.631 | 0.335 | -0.001 | 0.005 | 0.005 | 0 | 0.24 | 0.2 | 0.29 | 0.33 | 0.22 | 0.27 |
| 3 | 70475237 | rs11927774 | 0.46 | 1.599 | -0.631 | 0.438 | -0.001 | 0 | 0 | 0.06 | 0.23 | 0.19 | 0.24 | 0.21 | 0.22 | 0.27 |
| 3 | 70479629 | rs9840539 | -0.037 | 1.426 | -0.074 | 0.225 | 0.003 | -0.028 | -0.028 | 0 | 0.22 | 0.34 | 0.17 | 0.15 | 0.18 | 0.15 |
| 3 | 70482080 | rs7432792 | -0.697 | 1.366 | -0.091 | -0.031 | 0.002 | 0.005 | 0.005 | 0 | 0.45 | 0.53 | 0.41 | 0.35 | 0.4 | 0.42 |
| 3 | 70508727 | rs717669 | -0.443 | 1.41 | 0.038 | -0.062 | -0.02 | 0.021 | 0.021 | 0 | 0.24 | 0.19 | 0.31 | 0.03 | 0.29 | 0.16 |
| 3 | 70511919 | rs4974291 | 0.525 | 1.28 | 0.181 | -0.033 | -0.004 | 0.013 | 0.013 | 0 | 0.25 | 0.26 | 0.29 | 0.32 | 0.22 | 0.28 |
| 3 | 70524412 | rs4491863 | -1.577 | 1.148 | 0.082 | 0.176 | 0 | -0.01 | -0.01 | 1.16 | 0.63 | 0.51 | 0.76 | 0.68 | 0.62 | 0.71 |
| 3 | 70528833 | rs4638930 | -1.617 | 1.144 | 0.056 | 0.16 | -0.001 | 0 | 0 | 0 | 0.64 | 0.64 | 0.76 | 0.67 | 0.62 | 0.7 |
| 3 | 70539559 | rs11915371 | -0.212 | 0.754 | 0.268 | 0.257 | -0.002 | -0.047 | -0.047 | 0 | 0.23 | 0.24 | 0.32 | 0.04 | 0.19 | 0.15 |
| 3 | 70543096 | rs9830720 | -1.492 | 0.768 | 0.292 | 0.303 | -0.012 | 0.008 | 0.008 | 0 | 0.61 | 0.67 | 0.71 | 0.68 | 0.52 | 0.69 |
| 3 | 70546864 | rs12487810 | 0.152 | 0.807 | 0.072 | 0.251 | -0.006 | 0.036 | 0.036 | 0 | 0.21 | 0.1 | 0.3 | 0.03 | 0.19 | 0.14 |

| | | | | | | | | | | | | | | | | |
|---|---|---|---|---|---|---|---|---|---|---|---|---|---|---|---|---|
| 3 | 70554524 | rs1491610 | -0.017 | 0.865 | 0.109 | 0.409 | 0.012 | -0.031 | -0.031 | 0 | 0.36 | 0.23 | 0.39 | 0.53 | 0.35 | 0.54 |
| 3 | 70560258 | rs3732730 | 0.591 | 0.81 | 0.081 | 0.423 | 0.012 | -0.021 | -0.021 | 0 | 0.13 | 0.11 | 0.1 | 0.2 | 0.11 | 0.27 |
| 3 | 70560553 | rs12492328 | 0.214 | 0.927 | 0.13 | 0.105 | -0.015 | 0.061 | 0.061 | 0 | 0.2 | 0.08 | 0.3 | 0.03 | 0.18 | 0.14 |
| 3 | 70565820 | rs17790790 | -1.461 | 0.972 | 0.198 | 0.156 | 0.01 | -0.015 | -0.015 | 0.214 | 0.6 | 0.52 | 0.7 | 0.67 | 0.53 | 0.7 |
| 3 | 70575898 | rs7627684 | -0.177 | 0.834 | -0.302 | 0.182 | 0.021 | 0.005 | 0.005 | 0 | 0.38 | 0.27 | 0.39 | 0.64 | 0.33 | 0.55 |
| 3 | 70582231 | rs9871358 | -0.919 | 1.115 | -0.633 | 0.349 | -0.014 | -0.002 | -0.002 | 1.986 | 0.56 | 0.33 | 0.61 | 0.56 | 0.51 | 0.67 |
| 3 | 70592215 | rs11128180 | 0.697 | 1.16 | -0.728 | -0.228 | -0.004 | -0.027 | -0.027 | 0 | 0.22 | 0.1 | 0.21 | 0.26 | 0.22 | 0.27 |
| 3 | 70593081 | rs11708540 | 0.398 | 0.822 | -0.953 | -0.303 | -0.004 | -0.003 | -0.003 | 0 | 0.12 | 0.09 | 0.28 | 0.02 | 0.14 | 0.12 |
| 3 | 70595276 | rs931473 | -1.078 | 0.643 | -1.333 | -0.263 | -0.012 | -0.09 | -0.09 | 0 | 0.68 | 0.53 | 0.74 | 0.89 | 0.67 | 0.89 |
| 3 | 70595849 | rs6762974 | -0.488 | 0.556 | -1.887 | 0.568 | -0.003 | -0.064 | -0.064 | 1.863 | 0.32 | 0.16 | 0.25 | 0.59 | 0.28 | 0.49 |
| 3 | 70606921 | rs11927332 | -0.432 | 0.107 | -1.874 | 0.028 | 0.017 | 0.065 | 0.065 | 0.367 | 0.27 | 0.18 | 0.26 | 0.11 | 0.21 | 0.14 |
| 3 | 70607208 | rs9880540 | 0.105 | -0.542 | -1.823 | -0.097 | 0 | -0.009 | -0.009 | 0 | 0.45 | 0.52 | 0.47 | 0.32 | 0.46 | 0.36 |
| 3 | 70608205 | rs11128181 | -0.531 | -0.416 | -2.401 | 0.115 | -0.053 | 0.157 | 0.157 | 0 | 0.46 | 0.19 | 0.59 | 0.26 | 0.46 | 0.42 |
| 3 | 70613911 | rs11708678 | 0.56 | -0.735 | -2.166 | 0.129 | -0.005 | 0.055 | 0.055 | 0 | 0.09 | 0.11 | 0.09 | 0.35 | 0.04 | 0.19 |
| 3 | 70616737 | rs6790542 | 1.126 | -0.739 | -2.16 | 0.142 | 0.003 | 0.015 | 0.015 | 0 | 0.07 | 0.06 | 0.09 | 0.28 | 0.04 | 0.19 |
| 3 | 70621521 | rs1491603 | 1.231 | -0.81 | -1.515 | 0.098 | 0.019 | 0.077 | 0.077 | 0 | 0.19 | 0.3 | 0.15 | 0.36 | 0.23 | 0.22 |
| 3 | 70624868 | rs10511003 | 0.102 | -0.915 | -1.535 | 0.012 | -0.007 | 0.01 | 0.01 | 0 | 0.35 | 0.38 | 0.2 | 0.46 | 0.29 | 0.35 |
| 3 | 70624977 | rs9817104 | 1.931 | -0.912 | -1.407 | -0.28 | -0.003 | -0.078 | -0.078 | 0 | 0.24 | 0.24 | 0.15 | 0 | 0.26 | 0.11 |
| 3 | 70629840 | rs1565553 | 0.082 | -0.791 | -1.215 | 0.235 | 0.013 | -0.003 | -0.003 | 0 | 0.27 | 0.2 | 0.27 | 0.27 | 0.22 | 0.29 |
| 3 | 70631018 | rs10511002 | 1.603 | -1.088 | -1.273 | -0.024 | 0.017 | 0.083 | 0.083 | 0 | 0.32 | 0.16 | 0.17 | 0.08 | 0.3 | 0.21 |
| 3 | 70638311 | rs1491599 | -0.648 | -1.074 | -1.162 | 0.138 | 0.004 | 0.007 | 0.007 | 0.024 | 0.76 | 0.74 | 0.74 | 0.9 | 0.8 | 0.83 |
| 3 | 70638565 | rs1491598 | 1.008 | -0.905 | -1.414 | 0.296 | -0.088 | -0.137 | -0.137 | 0 | 0.19 | 0.02 | 0.27 | 0.58 | 0.24 | 0.43 |
| 3 | 70641810 | rs7628382 | -0.528 | -0.935 | -1.341 | 0.168 | -0.002 | 0.151 | 0.151 | 0 | 0.44 | 0.75 | 0.48 | 0.15 | 0.43 | 0.24 |
| 3 | 70645255 | rs9832620 | 0.809 | -0.64 | -0.905 | 0.328 | -0.007 | -0.011 | -0.011 | 0 | 0.38 | 0.34 | 0.26 | 0.27 | 0.34 | 0.34 |
| 3 | 70655368 | rs7611165 | -1.149 | -0.32 | -0.424 | 0.211 | -0.001 | -0.024 | -0.024 | 0 | 0.86 | 0.87 | 0.87 | 1 | 0.79 | 0.98 |
| 3 | 70657915 | rs1491590 | 0.08 | -0.602 | -0.686 | 0.49 | 0.006 | 0.005 | 0.005 | 0 | 0.36 | 0.18 | 0.38 | 0.36 | 0.31 | 0.44 |
| 3 | 70663233 | rs11128184 | 0.114 | -0.619 | -0.712 | 0.473 | 0.006 | 0.005 | 0.005 | 0 | 0.36 | 0.16 | 0.37 | 0.35 | 0.3 | 0.44 |
| 3 | 70664189 | rs4594588 | 1.233 | -0.747 | -1.16 | 0.746 | -0.017 | -0.053 | -0.053 | 0.039 | 0.34 | 0.05 | 0.24 | 0.29 | 0.31 | 0.29 |
| 3 | 70668698 | rs2130996 | 0.743 | -0.725 | -0.719 | 0.817 | -0.005 | -0.001 | -0.001 | 0.003 | 0.28 | 0.16 | 0.27 | 0.29 | 0.26 | 0.34 |
| 3 | 70671427 | rs725609 | 1.293 | -0.837 | -0.43 | 0.803 | 0.015 | 0.002 | 0.002 | 0 | 0.34 | 0.21 | 0.25 | 0.3 | 0.31 | 0.3 |
| 3 | 70687749 | rs9855970 | 0.907 | -0.837 | -0.167 | 0.899 | -0.011 | 0.015 | 0.015 | 0 | 0.27 | 0.12 | 0.26 | 0.25 | 0.26 | 0.31 |
| 3 | 70688041 | rs6801496 | 0.524 | -0.914 | 0.035 | 0.703 | -0.007 | -0.01 | -0.01 | 0 | 0.59 | 0.31 | 0.48 | 0.54 | 0.54 | 0.64 |
| 3 | 70697033 | rs10511006 | 1.098 | -1.157 | 0.141 | 1.164 | 0.001 | -0.035 | -0.035 | 0 | 0.07 | 0 | 0.1 | 0.09 | 0.04 | 0.1 |
| 3 | 70700679 | rs7632714 | 1.07 | -1.127 | 0.195 | 1.083 | -0.016 | 0.011 | 0.011 | 0 | 0.26 | 0.08 | 0.26 | 0.25 | 0.26 | 0.31 |
| 3 | 70703747 | rs1565552 | 0.389 | -1.032 | 0.367 | 0.961 | -0.028 | 0.043 | 0.043 | 0 | 0.19 | 0.1 | 0.22 | 0.07 | 0.25 | 0.13 |
| 3 | 70705510 | rs745675 | 0.949 | -1.12 | 0.787 | 1.099 | -0.007 | -0.006 | -0.006 | 2.294 | 0.38 | 0.52 | 0.27 | 0.3 | 0.32 | 0.36 |
| 3 | 70708646 | rs2320879 | -0.676 | -0.528 | 0.95 | 1.456 | -0.008 | -0.018 | -0.018 | 0 | 0.84 | 0.83 | 0.85 | 0.75 | 0.77 | 0.93 |
| 3 | 70734546 | rs6798132 | -1.808 | -0.67 | 1.167 | 1.195 | 0.071 | 0.191 | 0.191 | 2.951 | 0.05 | 0.44 | 0.03 | 0.11 | 0 | 0.12 |
| 3 | 70752697 | rs2060501 | 1.079 | -0.735 | 0.141 | 1.132 | 0.068 | 0.053 | 0.053 | 1.132 | 0.48 | 0.14 | 0.31 | 0.33 | 0.39 | 0.41 |
| 3 | 70757099 | rs1344819 | -0.011 | -0.062 | 0.593 | 1.096 | 0.029 | 0.018 | 0.018 | 0.183 | 0.88 | 0.76 | 0.86 | 0.83 | 0.85 | 0.83 |
| 3 | 70764744 | rs17737336 | 0.334 | -0.092 | 0.316 | 1.07 | -0.08 | -0.022 | -0.022 | 0 | 0.18 | 0.01 | 0.33 | 0.18 | 0.22 | 0.22 |
| 3 | 70768688 | rs6781443 | -0.391 | -0.506 | 1.177 | 0.804 | 0.088 | 0.039 | 0.039 | 0 | 0.12 | 0.34 | 0.14 | 0.13 | 0.14 | 0.16 |
| 3 | 70783491 | rs6766000 | -0.641 | -0.686 | 1.094 | 0.819 | 0.056 | 0.046 | 0.046 | 0.492 | 0.13 | 0.39 | 0.24 | 0.14 | 0.12 | 0.13 |
| 3 | 70791173 | rs7635109 | -0.75 | -0.717 | 0.95 | 0.811 | 0.035 | 0.025 | 0.025 | 0.227 | 0.15 | 0.46 | 0.26 | 0.14 | 0.14 | 0.14 |
| 3 | 70791669 | rs7633132 | 0.786 | -0.562 | 0.758 | 0.33 | 0.018 | 0.027 | 0.027 | 1.093 | 0.56 | 0.36 | 0.27 | 0.32 | 0.41 | 0.52 |
| 3 | 70836444 | rs6804071 | 0.582 | -0.641 | 0.099 | -0.367 | 0.052 | 0.192 | 0.192 | 4.439 | 0.56 | 0.79 | 0.29 | 0.03 | 0.4 | 0.29 |
| 3 | 70857460 | rs10865656 | 0.445 | -0.715 | 0.47 | 0.275 | 0.036 | 0.033 | 0.033 | 4.625 | 0.57 | 0.8 | 0.29 | 0.41 | 0.42 | 0.42 |
| 3 | 70873003 | rs17801787 | 0.981 | -0.886 | 0.457 | 0.237 | 0.057 | 0.069 | 0.069 | 2.141 | 0.51 | 0.31 | 0.26 | 0.36 | 0.39 | 0.41 |
| 3 | 70878343 | rs6796438 | -1.028 | -1.31 | -0.404 | 0.096 | -0.027 | 0.025 | 0.025 | 2.203 | 0.15 | 0.71 | 0.08 | 0.35 | 0.07 | 0.26 |
| 3 | 70883177 | rs2573174 | -1.788 | -1.253 | -0.934 | -0.223 | 0.059 | -0.038 | -0.038 | 0 | 0.33 | 0.87 | 0.2 | 0.44 | 0.21 | 0.38 |
| 3 | 70888070 | rs11128198 | 0.251 | -1.364 | -0.825 | -0.179 | 0.048 | 0.014 | 0.014 | 0.757 | 0.38 | 0.77 | 0.24 | 0.4 | 0.31 | 0.37 |
| 3 | 70889227 | rs17007912 | 0.5 | -1.365 | -0.655 | -0.639 | 0.017 | -0.006 | -0.006 | 0 | 0.35 | 0.67 | 0.23 | 0.02 | 0.28 | 0.24 |
| 3 | 70900385 | rs1403201 | -0.11 | -1.133 | -0.827 | 0.358 | 0.024 | -0.097 | -0.097 | 0.809 | 0.2 | 0.09 | 0.19 | 0.44 | 0.15 | 0.51 |
| 3 | 70905581 | rs2687195 | -0.921 | -0.881 | 0.248 | 0.407 | -0.005 | -0.054 | -0.054 | 0 | 0.7 | 0.52 | 0.78 | 0.87 | 0.7 | 0.82 |
| 3 | 70907252 | rs1522552 | -0.34 | -1.103 | 0.253 | 0.363 | 0.023 | -0.068 | -0.068 | 0 | 0.65 | 0.23 | 0.77 | 0.45 | 0.69 | 0.69 |
| 3 | 70918127 | rs1018340 | -0.406 | -1.148 | 0.894 | 0.387 | -0.006 | 0.014 | 0.014 | 0.531 | 0.66 | 0.47 | 0.76 | 0.48 | 0.69 | 0.69 |
| 3 | 70927234 | rs11716788 | 1.585 | -0.555 | 0.131 | -0.03 | -0.085 | 0.184 | 0.184 | 0.001 | 0.27 | 0.03 | 0.18 | 0 | 0.33 | 0.07 |
| 3 | 70928930 | rs2687201 | 0.019 | -0.268 | 1.094 | 0.949 | 0.016 | -0.043 | -0.043 | 0.279 | 0.66 | 0.25 | 0.72 | 0.51 | 0.68 | 0.71 |
| 3 | 70929441 | rs9825238 | -1.214 | -0.201 | 1.073 | 1.112 | -0.024 | 0.113 | 0.113 | 0 | 0.07 | 0.15 | 0.01 | 0.38 | 0.02 | 0.13 |
| 3 | 70930264 | rs9839462 | -1.043 | -0.065 | 1.127 | 1.169 | -0.013 | 0.231 | 0.231 | 0 | 0.05 | 0.2 | 0.02 | 0.38 | 0.02 | 0.11 |
| 3 | 70959438 | rs9837992 | -0.317 | -0.204 | 1.305 | 1.054 | 0.018 | -0.032 | -0.032 | 0 | 0.36 | 0.7 | 0.3 | 0.53 | 0.32 | 0.32 |
| 3 | 70965044 | rs13065564 | -0.217 | -0.974 | 1.451 | 1.112 | 0.01 | -0.008 | -0.008 | 0 | 0.35 | 0.64 | 0.24 | 0.41 | 0.32 | 0.3 |
| 3 | 70969345 | rs9822779 | -1.89 | -1.162 | 1.842 | 1.173 | -0.002 | 0.498 | 0.498 | 0.012 | 0.09 | 0.44 | 0.07 | 0.45 | 0.03 | 0.17 |
| 3 | 70973305 | rs17802967 | 0.38 | -1.106 | 1.375 | 0.96 | -0.001 | -0.072 | -0.072 | 1.186 | 0.29 | 0.29 | 0.22 | 0 | 0.29 | 0.14 |
| 3 | 70976765 | rs12636225 | -1.386 | -0.947 | 1.116 | 1.389 | -0.025 | 0.261 | 0.261 | 0 | 0.07 | 0.2 | 0.02 | 0.38 | 0.02 | 0.17 |
| 3 | 71003125 | rs7640237 | -1.694 | -0.98 | 0.691 | 1.349 | -0.035 | 0.35 | 0.35 | 0.676 | 0.1 | 0.56 | 0.04 | 0.39 | 0.02 | 0.15 |
| 3 | 71012580 | rs3773494 | 0.33 | -0.662 | -0.845 | 1.57 | -0.022 | 0.03 | 0.03 | 0 | 0.05 | 0.01 | 0.09 | 0.31 | 0.06 | 0.32 |
| 3 | 71015644 | rs3773497 | 0.552 | -0.695 | -0.998 | 0.445 | -0.019 | 0.062 | 0.062 | 0 | 0.24 | 0.09 | 0.18 | 0.1 | 0.26 | 0.18 |
| 3 | 71019750 | rs9874207 | -1.452 | -0.809 | -1.501 | -0.023 | -0.134 | 0.334 | 0.334 | 0 | 0.58 | 0.97 | 0.47 | 1 | 0.49 | 0.78 |

| | | | | | | | | | | | | | | | | |
|---|---|---|---|---|---|---|---|---|---|---|---|---|---|---|---|---|
| 3 | 71020900 | rs3755733 | -0.58 | -0.76 | -1.119 | 0.851 | -0.021 | -0.052 | -0.052 | 0 | 0.09 | 0.01 | 0.07 | 0.32 | 0.09 | 0.15 |
| 3 | 71024676 | rs8179901 | -0.635 | -1.162 | -0.956 | 0.585 | 0.067 | -0.05 | -0.05 | 0.064 | 0.34 | 0.87 | 0.3 | 0.43 | 0.25 | 0.28 |
| 3 | 71042518 | rs9841227 | -0.759 | -0.98 | -0.08 | 0.754 | -0.01 | 0.188 | 0.188 | 0 | 0.08 | 0.23 | 0.04 | 0.27 | 0.04 | 0.09 |
| 3 | 71043496 | rs1474305 | -0.543 | -1.018 | -0.257 | 0.633 | -0.001 | -0.007 | -0.007 | 0 | 0.17 | 0.15 | 0.12 | 0.52 | 0.15 | 0.44 |
| 3 | 71045000 | rs17008114 | -0.967 | -1.111 | 0.085 | 0.487 | -0.004 | 0.041 | 0.041 | 0 | 0.2 | 0.27 | 0.13 | 0.57 | 0.16 | 0.49 |
| 3 | 71047711 | rs17697822 | 1.272 | -1.205 | -0.413 | 0.36 | -0.012 | 0.041 | 0.041 | 0 | 0.19 | 0.03 | 0.13 | 0.1 | 0.19 | 0.16 |
| 3 | 71048229 | rs6768948 | -0.444 | -1.035 | -0.088 | 0.404 | 0.018 | 0.261 | 0.261 | 0 | 0.07 | 0.27 | 0.14 | 0.32 | 0.07 | 0.1 |
| 3 | 71052376 | rs17008127 | 0.022 | -1.068 | -0.327 | -0.433 | -0.034 | -0.048 | -0.048 | 0.65 | 0.12 | 0.27 | 0.05 | 0.05 | 0.05 | 0.08 |
| 3 | 71060640 | rs11720703 | 1.232 | -0.614 | -1.367 | -0.46 | -0.125 | 0.325 | 0.325 | 0 | 0.43 | 0.06 | 0.51 | 0.01 | 0.51 | 0.2 |
| 3 | 71074194 | rs2177291 | -0.685 | -0.483 | -1.18 | 0.214 | -0.051 | 0.009 | 0.009 | 0 | 0.77 | 0.99 | 0.83 | 0.86 | 0.76 | 0.82 |
| 3 | 71081818 | rs9861314 | -1.25 | -0.858 | -0.474 | 0.53 | -0.001 | 0.002 | 0.002 | 2.796 | 0.12 | 0.85 | 0.17 | 0.1 | 0.05 | 0.1 |
| 3 | 71127834 | rs11717053 | 0.051 | -0.865 | -1.154 | 0.96 | -0.043 | -0.155 | -0.155 | 0 | 0.21 | 0.01 | 0.13 | 0.45 | 0.18 | 0.41 |
| 3 | 71128028 | rs1288974 | 0.985 | -1.237 | -1.021 | 0.717 | 0.05 | -0.05 | -0.05 | 0 | 0.64 | 0.15 | 0.69 | 0.45 | 0.76 | 0.5 |
| 3 | 71151219 | rs831436 | 1.331 | -1.67 | -1.234 | 0.255 | 0.086 | 0.233 | 0.233 | 0.186 | 0.53 | 0.06 | 0.63 | 0.18 | 0.69 | 0.38 |
| 3 | 71153110 | rs6549377 | 0.688 | -2.052 | -0.444 | 0.353 | 0.012 | -0.02 | -0.02 | 0.803 | 0.67 | 0.74 | 0.8 | 0.51 | 0.76 | 0.52 |
| 3 | 71154796 | rs831438 | 0.469 | -1.936 | -0.853 | 0.417 | 0.014 | 0.017 | 0.017 | 1.801 | 0.93 | 0.95 | 0.98 | 0.98 | 0.96 | 0.96 |
| 3 | 71154818 | rs831439 | 0.4 | -2.109 | -0.997 | 0.641 | -0.003 | -0.144 | -0.144 | 1.167 | 0.7 | 0.93 | 0.83 | 0.44 | 0.76 | 0.51 |
| 3 | 71155250 | rs3846028 | 0.137 | -2.1 | -1.065 | 0.631 | -0.031 | -0.162 | -0.162 | 0.199 | 0.24 | 0.01 | 0.11 | 0.56 | 0.18 | 0.44 |
| 3 | 71156340 | rs831440 | 0.959 | -2.024 | -0.806 | 0.619 | 0.059 | -0.021 | -0.021 | 0 | 0.62 | 0.2 | 0.78 | 0.38 | 0.75 | 0.51 |
| 3 | 71158354 | rs831444 | 0.652 | -2.092 | -0.738 | 0.546 | 0.063 | -0.054 | -0.054 | 6.162 | 0.65 | 0.22 | 0.84 | 0.5 | 0.8 | 0.54 |
| 3 | 71162001 | rs3843363 | 0.214 | -1.514 | -0.893 | 0.277 | 0.023 | 0.004 | 0.004 | 0 | 0.62 | 0.33 | 0.65 | 0.33 | 0.7 | 0.46 |
| 3 | 71174475 | rs2121783 | 0.271 | -0.627 | -0.47 | 0.373 | -0.005 | 0.027 | 0.027 | 0.358 | 0.58 | 0.39 | 0.54 | 0.32 | 0.58 | 0.42 |
| 3 | 71174872 | rs2593852 | -0.091 | -0.777 | 0.121 | 0.549 | 0.004 | -0.013 | -0.013 | 0 | 0.37 | 0.2 | 0.29 | 0.6 | 0.32 | 0.53 |
| 3 | 71177701 | rs1474273 | -0.201 | -0.026 | 0.49 | 0.629 | 0.018 | 0.03 | 0.03 | 0 | 0.22 | 0.34 | 0.3 | 0.29 | 0.21 | 0.27 |
| 3 | 71186626 | rs9881149 | 1.321 | -0.286 | 0.757 | 0.515 | 0.05 | 0.063 | 0.063 | 4.548 | 0.95 | 0.45 | 0.94 | 0.96 | 0.99 | 0.99 |
| 3 | 71192476 | rs9833452 | -2.17 | -0.331 | 0.73 | 0.887 | 0.007 | 0.143 | 0.143 | 5.584 | 0.06 | 0.6 | 0.07 | 0.13 | 0.01 | 0.07 |
| 3 | 71193197 | rs1568555 | 0.767 | -0.447 | 0.037 | 0.865 | 0.009 | -0.05 | -0.05 | 0 | 0.71 | 0.12 | 0.66 | 0.65 | 0.77 | 0.68 |
| 3 | 71193929 | rs13070108 | 0.428 | -0.875 | 0.459 | 0.481 | -0.035 | -0.051 | -0.051 | 1.25 | 0.73 | 0.39 | 0.77 | 0.95 | 0.87 | 0.84 |
| 3 | 71199024 | rs1522177 | -0.374 | -0.252 | 0.163 | 0.366 | -0.009 | 0.012 | 0.012 | 0.606 | 0.39 | 0.4 | 0.44 | 0.88 | 0.45 | 0.69 |
| 3 | 71199723 | rs10865657 | 0.247 | -0.227 | 0.101 | 0.545 | -0.014 | -0.013 | -0.013 | 0.064 | 0.33 | 0.23 | 0.35 | 0.63 | 0.37 | 0.59 |
| 3 | 71199965 | rs12633797 | 0.149 | -0.247 | 0.151 | 0.565 | -0.008 | -0.007 | -0.007 | 0.977 | 0.33 | 0.28 | 0.35 | 0.63 | 0.37 | 0.6 |
| 3 | 71200157 | rs7647909 | 0.245 | -0.255 | 0.052 | 0.808 | -0.011 | 0.006 | 0.006 | 0.501 | 0.67 | 0.4 | 0.66 | 0.36 | 0.75 | 0.63 |
| 3 | 71200872 | rs9829152 | 0.559 | -0.2 | -0.052 | 0 | 0.05 | 0.1 | 0.1 | 0 | 0.05 | 0.19 | 0.09 | 0.17 | 0.08 | 0.16 |
| 3 | 71201897 | rs7632470 | 0.264 | -0.644 | -0.124 | 0.035 | -0.016 | -0.006 | -0.006 | 0 | 0.31 | 0.24 | 0.38 | 0.71 | 0.38 | 0.48 |
| 3 | 71203369 | rs1402994 | 0.574 | -0.521 | -0.254 | 0.024 | -0.015 | 0.057 | 0.057 | 0 | 0.44 | 0.36 | 0.42 | 0.2 | 0.49 | 0.34 |
| 3 | 71206903 | rs4334611 | 0.639 | -0.615 | 0.089 | 0.527 | 0.005 | -0.001 | -0.001 | 0.095 | 0.59 | 0.45 | 0.53 | 0.41 | 0.66 | 0.5 |
| 3 | 71216984 | rs1554536 | 0.54 | -1.452 | -0.204 | -0.175 | -0.029 | -0.028 | -0.028 | 0 | 0.72 | 0.66 | 0.8 | 0.91 | 0.86 | 0.87 |
| 3 | 71224429 | rs6763814 | -0.521 | -1.278 | -0.279 | 0.067 | 0.002 | 0.001 | 0.001 | 0.087 | 0.27 | 0.24 | 0.26 | 0.25 | 0.17 | 0.17 |
| 3 | 71225021 | rs7623157 | 0.985 | -1.185 | -0.52 | -0.49 | 0.004 | 0.009 | 0.009 | 0 | 0.29 | 0.24 | 0.24 | 0.08 | 0.26 | 0.11 |
| 3 | 71228953 | rs2056467 | -0.397 | -1.344 | 0.197 | -0.587 | -0.038 | 0.016 | 0.016 | 0.437 | 0.3 | 0.49 | 0.32 | 0.33 | 0.21 | 0.2 |
| 3 | 71230143 | rs4677029 | -0.885 | -1.445 | 0.058 | -0.631 | 0.001 | 0.012 | 0.012 | 0.037 | 0.59 | 0.64 | 0.51 | 0.33 | 0.43 | 0.3 |
| 3 | 71230554 | rs4677031 | -0.977 | -0.932 | 0.525 | 0.22 | -0.026 | -0.038 | -0.038 | 0.019 | 0.29 | 0.36 | 0.23 | 0.09 | 0.18 | 0.15 |
| 3 | 71236180 | rs9825176 | -0.696 | -0.671 | -0.179 | 0.267 | -0.001 | 0 | 0 | 0 | 0.27 | 0.23 | 0.3 | 0.33 | 0.22 | 0.19 |
| 3 | 71247304 | rs2037474 | -1.117 | -0.565 | 0.042 | 0.977 | -0.005 | 0.139 | 0.139 | 0 | 0.16 | 0.27 | 0.34 | 0.44 | 0.14 | 0.25 |
| 3 | 71250901 | rs11128208 | -0.629 | -1.04 | 0.082 | 0.994 | 0.011 | -0.015 | -0.015 | 0.308 | 0.17 | 0.15 | 0.14 | 0.33 | 0.09 | 0.15 |
| 3 | 71254193 | rs1474272 | -0.388 | -1.56 | 0.783 | 1.152 | -0.017 | 0.075 | 0.075 | 0.075 | 0.12 | 0.39 | 0.08 | 0.18 | 0.08 | 0.28 |
| 3 | 71256462 | rs6549381 | -0.338 | -0.61 | 0.436 | 1.239 | 0.056 | 0.02 | 0.02 | 0 | 0.9 | 0.8 | 0.93 | 0.86 | 0.83 | 0.9 |
| 3 | 71257098 | rs17008343 | -0.026 | -0.66 | 0.268 | 1.257 | 0.013 | -0.146 | -0.146 | 0 | 0.17 | 0.01 | 0.15 | 0.49 | 0.11 | 0.37 |
| 3 | 71257157 | rs11928137 | 1.751 | -0.957 | -0.262 | 0.138 | 0.042 | 0.094 | 0.094 | 0 | 0.29 | 0.12 | 0.17 | 0 | 0.2 | 0.08 |
| 3 | 71261183 | rs4677597 | 1.424 | 0.168 | -0.445 | 0.308 | -0.06 | 0.153 | 0.153 | 0 | 0.35 | 0.13 | 0.37 | 0.04 | 0.4 | 0.2 |
| 3 | 71263218 | rs6786408 | -0.875 | 0.103 | -0.116 | 0.284 | 0.014 | 0.049 | 0.049 | 0 | 0.54 | 0.65 | 0.55 | 0.83 | 0.43 | 0.71 |
| 3 | 71266885 | rs6549383 | 0.914 | 0.042 | -0.015 | 0.447 | -0.011 | 0.059 | 0.059 | 0 | 0.45 | 0.34 | 0.48 | 0.22 | 0.48 | 0.49 |
| 3 | 71267832 | rs1522175 | -0.256 | 0.575 | 0.276 | 0.76 | 0.028 | 0.009 | 0.009 | 0 | 0.89 | 0.85 | 0.92 | 0.85 | 0.82 | 0.91 |
| 3 | 71270788 | rs7649879 | -0.269 | 0.614 | 0.071 | 0.636 | 0.014 | -0.003 | -0.003 | 0 | 0.9 | 0.88 | 0.92 | 0.93 | 0.83 | 0.91 |
| 3 | 71278815 | rs17008381 | 0.569 | 0.478 | 0.006 | 0.955 | 0.005 | 0.04 | 0.04 | 0.233 | 0.07 | 0.12 | 0.06 | 0.16 | 0.06 | 0.03 |
| 3 | 71279122 | rs880543 | 0.097 | 0.177 | 0.105 | 0.847 | 0.002 | 0.015 | 0.015 | 0 | 0.28 | 0.38 | 0.38 | 0.35 | 0.3 | 0.29 |
| 3 | 71281430 | rs1288814 | 0.111 | 0.238 | 0.058 | 0.918 | 0.004 | -0.008 | -0.008 | 0 | 0.33 | 0.32 | 0.42 | 0.46 | 0.31 | 0.35 |
| 3 | 71287528 | rs9310212 | 0.923 | 0.634 | -0.045 | 0.406 | 0.007 | 0.039 | 0.039 | 0.064 | 0.06 | 0.11 | 0.06 | 0.15 | 0.06 | 0.04 |
| 3 | 71288864 | rs2568844 | -0.595 | 0.719 | -0.15 | 0.261 | -0.014 | 0.012 | 0.012 | 0 | 0.43 | 0.69 | 0.5 | 0.67 | 0.46 | 0.57 |
| 3 | 71292039 | rs4055824 | -0.494 | 1.832 | 0.249 | 0.567 | -0.017 | 0.011 | 0.011 | 0.427 | 0.77 | 0.9 | 0.66 | 0.83 | 0.75 | 0.72 |
| 3 | 71302616 | rs17008404 | -1.141 | 1.834 | 0.503 | 0.694 | -0.003 | 0 | 0 | 0.379 | 0.19 | 0.17 | 0.14 | 0.19 | 0.23 | 0.11 |
| 3 | 71318993 | rs2280518 | -0.147 | 1.331 | 1.07 | 1.139 | -0.018 | 0.377 | 0.377 | 0 | 0.1 | 0.37 | 0.14 | 0.46 | 0.06 | 0.29 |
| 3 | 71323222 | rs7647520 | 0.577 | 2.036 | 0.751 | 0.789 | -0.024 | 0.056 | 0.056 | 0 | 0.26 | 0.11 | 0.33 | 0.13 | 0.29 | 0.21 |
| 3 | 71324328 | rs1541903 | -0.341 | 2.164 | 0.789 | 0.781 | -0.007 | -0.012 | -0.012 | 0 | 0.11 | 0.06 | 0.06 | 0.25 | 0.12 | 0.13 |
| 3 | 71324580 | rs1288694 | -1.34 | 2.074 | 1.018 | 0.704 | 0.049 | 0.006 | 0.006 | 0.012 | 0.49 | 0.71 | 0.63 | 0.47 | 0.6 | 0.49 |
| 3 | 71324696 | rs1288696 | -1.143 | 2.126 | 0.863 | 0.588 | 0.105 | 0.114 | 0.114 | 0 | 0.15 | 0.32 | 0.23 | 0.36 | 0.27 | 0.25 |
| 3 | 71327283 | rs1529669 | 0.055 | 2.115 | 0.561 | 0.005 | 0 | 0.006 | 0.006 | 0.035 | 0.31 | 0.32 | 0.37 | 0.11 | 0.31 | 0.23 |
| 3 | 71330583 | rs1288821 | -0.325 | 2.176 | 0.196 | -0.027 | 0.038 | -0.047 | -0.047 | 0 | 0.35 | 0.52 | 0.39 | 0.22 | 0.44 | 0.3 |
| 3 | 71334477 | rs1288829 | -1.585 | 0.286 | -0.679 | 0.101 | -0.081 | 0.237 | 0.237 | 0 | 0.58 | 0.9 | 0.56 | 0.94 | 0.51 | 0.75 |

| | | | | | | | | | | | | | | | | |
|---|---|---|---|---|---|---|---|---|---|---|---|---|---|---|---|---|
| 3 | 71337856 | rs12495693 | -0.336 | 1.156 | -0.186 | 0.471 | -0.013 | 0.014 | 0.014 | 0 | 0.27 | 0.14 | 0.23 | 0.24 | 0.29 | 0.22 |
| 3 | 71342132 | rs12485826 | -0.766 | 1.377 | -0.992 | 0.506 | -0.009 | -0.008 | -0.008 | 0.014 | 0.08 | 0.04 | 0.04 | 0.09 | 0.1 | 0.09 |
| 3 | 71346348 | rs17008473 | -0.373 | 0.93 | -0.955 | 0.576 | -0.024 | -0.005 | -0.005 | 1.343 | 0.26 | 0.12 | 0.31 | 0.26 | 0.29 | 0.22 |
| 3 | 71347176 | rs17655393 | -0.379 | 0.933 | -0.946 | 0.033 | -0.014 | 0.027 | 0.027 | 0 | 0.26 | 0.15 | 0.32 | 0.19 | 0.29 | 0.21 |
| 3 | 71349375 | rs17008474 | -1.332 | 0.628 | -0.369 | 0.037 | -0.008 | 0.015 | 0.015 | 0 | 0.05 | 0.11 | 0.02 | 0.08 | 0.04 | 0.07 |
| 3 | 71352844 | rs1733535 | -1.161 | 0.308 | -0.245 | -0.209 | -0.015 | -0.007 | -0.007 | 0.018 | 0.1 | 0.1 | 0.06 | 0.06 | 0.08 | 0.15 |
| 3 | 71354427 | rs11718210 | -0.623 | 0.753 | -0.584 | -0.625 | -0.029 | 0.041 | 0.041 | 0 | 0.09 | 0.03 | 0.08 | 0 | 0.15 | 0.09 |
| 3 | 71360708 | rs7633896 | -0.446 | 1.103 | 0.689 | -1.461 | 0.029 | -0.071 | -0.071 | 1.005 | 0.15 | 0.32 | 0.14 | 0.01 | 0.2 | 0.14 |
| 3 | 71365371 | rs17655524 | -0.653 | 1.38 | 0.677 | -1.579 | -0.016 | 0.011 | 0.011 | 0 | 0.12 | 0.07 | 0.1 | 0 | 0.18 | 0.13 |
| 3 | 71369174 | rs9875477 | -0.255 | 1.419 | 0.879 | -1.584 | 0.002 | -0.027 | -0.027 | 0 | 0.14 | 0.18 | 0.13 | 0.01 | 0.21 | 0.15 |
| 3 | 71371734 | rs4677035 | 1.56 | 1.23 | 0.84 | -1.818 | -0.001 | -0.006 | -0.006 | 0.01 | 0.73 | 0.36 | 0.72 | 0.91 | 0.72 | 0.71 |
| 3 | 71379546 | rs11712688 | 2.398 | 1.539 | 0.696 | -1.867 | -0.021 | 0.298 | 0.298 | 1.752 | 0.51 | 0.09 | 0.54 | 0.9 | 0.43 | 0.52 |
| 3 | 71382085 | rs7644066 | 2.041 | 1.556 | 0.83 | -1.873 | 0.002 | 0.087 | 0.087 | 1.449 | 0.66 | 0.25 | 0.67 | 0.91 | 0.66 | 0.66 |
| 3 | 71389142 | rs1300170 | -0.996 | 1.659 | 1.024 | -1.655 | 0.002 | 0.016 | 0.016 | 0 | 0.07 | 0.09 | 0.13 | 0.1 | 0.09 | 0.15 |
| 3 | 71389498 | rs1288701 | 1.687 | 1.153 | 0.921 | -1.644 | -0.006 | 0.012 | 0.012 | 0.781 | 0.61 | 0.35 | 0.74 | 0.83 | 0.59 | 0.73 |
| 3 | 71396478 | rs9859586 | 1.714 | 1.183 | 0.781 | -1.64 | -0.024 | -0.023 | -0.023 | 0.813 | 0.71 | 0.52 | 0.73 | 0.79 | 0.73 | 0.7 |
| 3 | 71398494 | rs1997479 | 2.296 | 1.106 | 0.186 | -1.615 | -0.07 | 0.144 | 0.144 | 2.803 | 0.53 | 0.1 | 0.59 | 0.73 | 0.5 | 0.57 |
| 3 | 71401928 | rs6774982 | -0.801 | 1.226 | 0.31 | -1.582 | -0.006 | -0.011 | -0.011 | 0.001 | 0.2 | 0.2 | 0.23 | 0.07 | 0.3 | 0.17 |
| 3 | 71403136 | rs1288691 | -0.642 | 1.241 | -0.089 | -1.574 | -0.074 | 0.083 | 0.083 | 0 | 0.19 | 0.09 | 0.23 | 0.07 | 0.29 | 0.17 |
| 3 | 71408319 | rs17008544 | -1.03 | 1.08 | -0.025 | -1.431 | 0.009 | 0.017 | 0.017 | 0 | 0.18 | 0.1 | 0.1 | 0.12 | 0.17 | 0.12 |
| 3 | 71410012 | rs10511017 | -0.663 | 1.176 | -0.031 | -1.515 | -0.019 | 0.012 | 0.012 | 0 | 0.12 | 0.1 | 0.12 | 0 | 0.19 | 0.11 |
| 3 | 71412654 | rs1288688 | 0.819 | 1.18 | 0.422 | -1.414 | -0.01 | 0.026 | 0.026 | 0.045 | 0.79 | 0.65 | 0.87 | 0.8 | 0.81 | 0.74 |
| 3 | 71413077 | rs1288685 | 0.648 | 1.15 | 0.293 | -1.422 | -0.022 | 0.027 | 0.027 | 0.105 | 0.8 | 0.68 | 0.9 | 0.8 | 0.84 | 0.74 |
| 3 | 71414045 | rs13072512 | 1.608 | 1.168 | -0.911 | -1.454 | -0.085 | 0.124 | 0.124 | 2.337 | 0.55 | 0.1 | 0.6 | 0.75 | 0.5 | 0.58 |
| 3 | 71417200 | rs17717911 | -0.17 | 1.358 | -0.668 | -1.316 | -0.056 | 0.02 | 0.02 | 0 | 0.11 | 0.01 | 0.07 | 0.06 | 0.16 | 0.1 |
| 3 | 71421814 | rs13065253 | 1 | 0.906 | -0.807 | -2.149 | 0.02 | -0.002 | -0.002 | 0 | 0.4 | 0.26 | 0.23 | 0.01 | 0.35 | 0.19 |
| 3 | 71427707 | rs831072 | -0.337 | 1.184 | -0.945 | -2.13 | -0.026 | 0.138 | 0.138 | 0 | 0.36 | 0.13 | 0.48 | 0.89 | 0.34 | 0.46 |
| 3 | 71430917 | rs831077 | 0.163 | 1.337 | -1.142 | -1.997 | -0.043 | -0.003 | -0.003 | 0 | 0.23 | 0.05 | 0.43 | 0.77 | 0.23 | 0.37 |
| 3 | 71432139 | rs6549387 | -0.166 | 1.081 | -0.692 | -1.912 | -0.012 | -0.001 | -0.001 | 0 | 0.12 | 0.15 | 0.03 | 0.11 | 0.09 | 0.07 |
| 3 | 71432595 | rs12494939 | -0.51 | 1.517 | -0.635 | -2.177 | -0.009 | 0.017 | 0.017 | 0.075 | 0.15 | 0.09 | 0.17 | 0.1 | 0.18 | 0.22 |
| 3 | 71432608 | rs831081 | 1.056 | 1.794 | 0.222 | -2.318 | 0.043 | -0.097 | -0.097 | 2.176 | 0.79 | 0.43 | 0.75 | 0.9 | 0.75 | 0.75 |
| 3 | 71434538 | rs831084 | 0.612 | 1.925 | -0.462 | -2.458 | -0.04 | -0.042 | -0.042 | 0 | 0.17 | 0.02 | 0.38 | 0.77 | 0.16 | 0.34 |
| 3 | 71435374 | rs10470554 | 0.929 | 1.853 | -0.235 | -3.38 | 0 | -0.106 | -0.106 | 0 | 0.45 | 0.53 | 0.34 | 0.01 | 0.51 | 0.24 |
| 3 | 71437594 | rs9310213 | -1.496 | 2.313 | 0.003 | -3.369 | 0.109 | -0.05 | -0.05 | 3.386 | 0.1 | 0.42 | 0.12 | 0 | 0.15 | 0.06 |
| 3 | 71439472 | rs9815974 | 0.327 | 1.886 | -0.177 | -3.443 | 0.003 | 0.01 | 0.01 | 0 | 0.58 | 0.7 | 0.51 | 0.09 | 0.66 | 0.54 |
| 3 | 71449057 | rs11711425 | 0.055 | 1.757 | -0.157 | -3.422 | 0.008 | -0.001 | -0.001 | 0 | 0.62 | 0.7 | 0.53 | 0.09 | 0.73 | 0.58 |
| 3 | 71454340 | rs12494677 | 0.319 | 1.86 | -0.262 | -3.457 | -0.023 | 0.026 | 0.026 | 0.032 | 0.15 | 0.04 | 0.18 | 0.08 | 0.19 | 0.24 |
| 3 | 71455775 | rs7633837 | 0.097 | 1.731 | -0.224 | -3.407 | -0.006 | -0.049 | -0.049 | 2.69 | 0.61 | 0.61 | 0.53 | 0.09 | 0.73 | 0.58 |
| 3 | 71469845 | rs1864896 | 1.33 | 1.622 | -0.81 | -3.34 | -0.028 | 0.057 | 0.057 | 0 | 0.42 | 0.25 | 0.32 | 0.01 | 0.56 | 0.23 |
| 3 | 71474696 | rs10511018 | -0.252 | 1.552 | -0.971 | -2.659 | -0.013 | 0.02 | 0.02 | 0 | 0.16 | 0.05 | 0.18 | 0.09 | 0.17 | 0.25 |
| 3 | 71480566 | rs9809818 | 0.074 | 0.912 | -1.202 | -2.23 | 0.016 | 0.006 | 0.006 | 0 | 0.82 | 0.98 | 0.65 | 0.17 | 0.91 | 0.59 |
| 3 | 71496008 | rs6771130 | -0.287 | 1.025 | -1.09 | -3.126 | 0.003 | -0.061 | -0.061 | 0.814 | 0.32 | 0.41 | 0.28 | 0.07 | 0.4 | 0.25 |
| 3 | 71501077 | rs6803690 | 1.365 | 1.501 | -1.032 | -3.566 | 0.017 | -0.038 | -0.038 | 0 | 0.11 | 0.17 | 0.15 | 0 | 0.19 | 0.12 |
| 3 | 71501895 | rs2135548 | 0.158 | 0.678 | -1.12 | -3.499 | -0.008 | -0.12 | -0.12 | 2.072 | 0.62 | 0.59 | 0.51 | 0.04 | 0.74 | 0.39 |
| 3 | 71505650 | rs11720980 | 0.364 | 0.958 | -1.018 | -3.493 | -0.004 | -0.062 | -0.062 | 0 | 0.29 | 0.34 | 0.25 | 0.06 | 0.32 | 0.22 |
| 3 | 71517643 | rs11706279 | 1.263 | 0.946 | -0.833 | -3.631 | -0.021 | -0.006 | -0.006 | 0 | 0.37 | 0.38 | 0.28 | 0 | 0.5 | 0.31 |
| 3 | 71521494 | rs7617596 | -0.338 | 0.58 | -0.507 | -3.765 | -0.006 | -0.084 | -0.084 | 0.986 | 0.69 | 0.69 | 0.47 | 0.04 | 0.81 | 0.5 |
| 3 | 71523110 | rs9819066 | 1.238 | 1.036 | -0.328 | -3.443 | -0.006 | -0.033 | -0.033 | 0 | 0.35 | 0.39 | 0.25 | 0 | 0.42 | 0.3 |
| 3 | 71529915 | rs6549391 | 1.787 | 1.006 | -0.705 | -3.408 | -0.013 | 0.051 | 0.051 | 0 | 0.31 | 0.24 | 0.23 | 0 | 0.36 | 0.17 |
| 3 | 71546744 | rs6789751 | 1.961 | 1.017 | -0.821 | -3.415 | -0.051 | 0.161 | 0.161 | 0 | 0.29 | 0.08 | 0.22 | 0 | 0.35 | 0.17 |
| 3 | 71552366 | rs9837383 | 1.501 | 0.231 | -0.565 | -3.316 | -0.018 | -0.003 | -0.003 | 0 | 0.33 | 0.3 | 0.27 | 0 | 0.44 | 0.18 |
| 3 | 71555673 | rs13067120 | 1.082 | 0.037 | -1.242 | -3.217 | -0.017 | 0.049 | 0.049 | 0 | 0.22 | 0.01 | 0.1 | 0.14 | 0.12 | 0.11 |
| 3 | 71568501 | rs9866496 | -0.145 | 0.26 | -0.452 | -3.072 | 0 | -0.148 | -0.148 | 0 | 0.25 | 0.46 | 0.15 | 0.03 | 0.27 | 0.25 |
| 3 | 71573135 | rs17008723 | 0.778 | 0.177 | -0.55 | -3.043 | 0.011 | 0.062 | 0.062 | 0 | 0.18 | 0.08 | 0.11 | 0.06 | 0.19 | 0.12 |
| 3 | 71580024 | rs1079409 | -0.273 | 0.106 | -0.475 | -3.002 | 0.011 | -0.027 | -0.027 | 0 | 0.55 | 0.66 | 0.41 | 0.03 | 0.7 | 0.42 |
| 3 | 71590347 | rs830604 | -0.604 | 0.217 | -0.316 | -2.95 | 0.024 | 0.076 | 0.076 | 0 | 0.58 | 0.8 | 0.45 | 0.03 | 0.71 | 0.41 |
| 3 | 71592594 | rs6790644 | 1.484 | 0.499 | -0.637 | -2.865 | -0.062 | 0.151 | 0.151 | 0 | 0.3 | 0.12 | 0.24 | 0 | 0.41 | 0.17 |
| 3 | 71612149 | rs6773658 | 0.002 | -0.006 | -0.262 | -2.414 | 0.004 | -0.061 | -0.061 | 0 | 0.55 | 0.59 | 0.43 | 0.03 | 0.72 | 0.41 |
| 3 | 71615018 | rs13316357 | -0.082 | 0.029 | -0.238 | -2.398 | 0.014 | -0.01 | -0.01 | 0 | 0.57 | 0.69 | 0.45 | 0.03 | 0.72 | 0.41 |
| 3 | 71626123 | rs9828619 | 0.079 | 0.195 | -0.503 | -2.236 | -0.006 | -0.108 | -0.108 | 0 | 0.53 | 0.51 | 0.4 | 0.03 | 0.68 | 0.4 |
| 3 | 71647281 | rs830652 | 0.008 | 0.468 | -0.588 | -1.256 | -0.032 | 0.176 | 0.176 | 0 | 0.45 | 0.79 | 0.51 | 0.17 | 0.53 | 0.23 |
| 3 | 71647908 | rs830653 | -0.529 | 0.698 | -0.727 | -0.691 | -0.046 | -0.076 | -0.076 | 0 | 0.89 | 0.99 | 0.8 | 0.39 | 0.89 | 0.74 |
| 3 | 71652888 | rs7643734 | -0.381 | 0.391 | 0.122 | -1.129 | 0.005 | -0.016 | -0.016 | 0 | 0.08 | 0.14 | 0.21 | 0.04 | 0.09 | 0.03 |
| 3 | 71666535 | rs864380 | -0.543 | 0.315 | 0.678 | -1.198 | -0.001 | -0.058 | -0.058 | 0 | 0.45 | 0.51 | 0.48 | 0.07 | 0.6 | 0.35 |
| 3 | 71668564 | rs830640 | 0.482 | 0.383 | 1.252 | -1.254 | 0.012 | -0.126 | -0.126 | 0 | 0.33 | 0.42 | 0.26 | 0.01 | 0.43 | 0.16 |
| 3 | 71670465 | rs830637 | -0.144 | 0.405 | 1.369 | -1.231 | -0.002 | -0.045 | -0.045 | 0 | 0.42 | 0.61 | 0.35 | 0.05 | 0.5 | 0.33 |
| 3 | 71672306 | rs10460943 | -1.312 | 0.511 | 1.387 | -1.074 | 0.032 | 0 | 0 | 1.106 | 0.14 | 0.27 | 0.25 | 0.14 | 0.17 | 0.2 |
| 3 | 71674676 | rs830628 | -0.275 | 0.488 | 1.22 | -1.115 | 0.004 | -0.094 | -0.094 | 0 | 0.44 | 0.44 | 0.42 | 0.05 | 0.52 | 0.19 |
| 3 | 71677817 | rs830622 | -1.446 | 0.657 | 1.118 | -0.635 | 0.054 | -0.041 | -0.041 | 0.332 | 0.14 | 0.5 | 0.25 | 0.06 | 0.17 | 0.2 |

| | | | | | | | | | | | | | | | |
|---|---|---|---|---|---|---|---|---|---|---|---|---|---|---|---|
| 3 | 71681950 | rs830617 | -1.181 | 0.363 | 0.494 | -0.431 | -0.005 | 0.01 | 0.01 | 0.374 | 0.21 | 0.36 | 0.25 | 0.27 | 0.21 | 0.15 |
| 3 | 71684845 | rs6782714 | 0.447 | 0.502 | 0.273 | -0.313 | -0.051 | -0.011 | -0.011 | 3.178 | 0.29 | 0.14 | 0.26 | 0.28 | 0.42 | 0.35 |
| 3 | 71684904 | rs830609 | -1.171 | 0.42 | 0.368 | -0.451 | 0 | -0.065 | -0.065 | 0.056 | 0.15 | 0.43 | 0.2 | 0.05 | 0.1 | 0.06 |
| 3 | 71686082 | rs1288124 | -0.572 | 0.43 | -0.375 | -0.338 | -0.028 | 0.026 | 0.026 | 0 | 0.46 | 0.75 | 0.45 | 0.36 | 0.53 | 0.46 |
| 3 | 71687308 | rs7612407 | -0.022 | 0.908 | -0.033 | -1.193 | 0.029 | -0.018 | -0.018 | 0 | 0.06 | 0.16 | 0.08 | 0.01 | 0.08 | 0.03 |
| 3 | 71687381 | rs17663544 | -0.189 | 1.183 | -0.658 | -1.448 | 0.002 | -0.069 | -0.069 | 0 | 0.26 | 0.31 | 0.18 | 0 | 0.31 | 0.17 |
| 3 | 71689923 | rs17663591 | 0.346 | 1.206 | -1.076 | -1.281 | -0.08 | 0.146 | 0.146 | 2.763 | 0.22 | 0.05 | 0.17 | 0 | 0.31 | 0.11 |
| 3 | 71697761 | rs9872923 | -0.104 | 0.901 | -0.725 | -0.913 | 0 | 0.038 | 0.038 | 0.394 | 0.47 | 0.66 | 0.45 | 0.26 | 0.5 | 0.42 |
| 3 | 71699114 | rs704259 | -0.263 | 1.067 | -1.213 | -1.343 | 0 | -0.016 | -0.016 | 0.002 | 0.57 | 0.58 | 0.54 | 0.05 | 0.59 | 0.43 |
| 3 | 71700507 | rs704264 | -0.17 | 0.982 | -1.216 | -1.226 | -0.001 | 0.012 | 0.012 | 0.112 | 0.55 | 0.57 | 0.54 | 0.18 | 0.58 | 0.42 |
| 3 | 71704119 | rs11710870 | -0.472 | 1.086 | -1.26 | -0.779 | 0.029 | -0.077 | -0.077 | 0 | 0.15 | 0.31 | 0.17 | 0 | 0.18 | 0.06 |
| 3 | 71708412 | rs704275 | -0.081 | 0.127 | -2.369 | -0.426 | 0.044 | -0.035 | -0.035 | 0 | 0.72 | 0.97 | 0.77 | 0.2 | 0.79 | 0.36 |
| 3 | 71708437 | rs13077861 | -0.279 | 1.335 | -1.621 | -0.737 | -0.029 | 0.009 | 0.009 | 0 | 0.28 | 0.17 | 0.22 | 0 | 0.39 | 0.09 |
| 3 | 71720566 | rs704285 | -0.21 | 0.867 | -1.758 | 0.281 | -0.031 | -0.043 | -0.043 | 0 | 0.55 | 0.86 | 0.63 | 0.52 | 0.46 | 0.6 |
| 3 | 71726259 | rs704292 | -0.968 | 0.591 | -0.595 | 1.055 | -0.009 | 0.06 | 0.06 | 0.675 | 0.19 | 0.29 | 0.25 | 0.6 | 0.16 | 0.26 |
| 3 | 71737107 | rs3796226 | -0.966 | 0.622 | -0.552 | 1.083 | -0.007 | 0.02 | 0.02 | 0.553 | 0.19 | 0.28 | 0.48 | 0.69 | 0.17 | 0.28 |
| 3 | 71739564 | rs9864677 | -0.556 | 0.653 | -0.524 | 1.102 | -0.005 | -0.008 | -0.008 | 0.08 | 0.16 | 0.23 | 0.1 | 0.12 | 0.12 | 0.11 |
| 3 | 71741068 | rs6767160 | -0.904 | 0.452 | 0.16 | 1.43 | 0.011 | 0.055 | 0.055 | 0.3 | 0.06 | 0.26 | 0.04 | 0.14 | 0.05 | 0.18 |
| 3 | 71743681 | rs4423732 | 0.895 | 0.095 | 0.279 | 1.471 | -0.043 | 0.116 | 0.116 | 2.965 | 0.84 | 0.65 | 0.58 | 0.34 | 0.91 | 0.74 |
| 3 | 71744801 | rs9310219 | -1.578 | 1.119 | 1.063 | 1.511 | 0.086 | 0.03 | 0.03 | 0.202 | 0.12 | 0.34 | 0.13 | 0.13 | 0.18 | 0.15 |
| 3 | 71747670 | rs9859923 | -1.396 | 1.179 | 1.352 | 1.498 | 0.029 | -0.029 | -0.029 | 2.916 | 0.07 | 0.2 | 0.03 | 0.01 | 0.06 | 0.08 |
| 3 | 71748073 | rs4995840 | -2.526 | 1.55 | 1.48 | 1.453 | 0.07 | 0.02 | 0.02 | 0.097 | 0.11 | 0.31 | 0.13 | 0.14 | 0.18 | 0.12 |
| 3 | 71752407 | rs1447904 | -2.586 | 1.631 | 1.605 | 2.331 | 0.045 | 0.135 | 0.135 | 0.845 | 0.24 | 0.49 | 0.29 | 0.5 | 0.31 | 0.3 |
| 3 | 71755488 | rs17008958 | -0.801 | 1.608 | 1.65 | 2.366 | 0 | 0.015 | 0.015 | 0 | 0.13 | 0.13 | 0.09 | 0.37 | 0.13 | 0.18 |
| 3 | 71756272 | rs17008972 | -2.05 | 1.458 | 2.262 | 2.076 | 0.082 | 0.063 | 0.063 | 0.262 | 0.08 | 0.4 | 0.12 | 0.13 | 0.13 | 0.05 |
| 3 | 71756765 | rs4677039 | 1.273 | 1.446 | 0.955 | 1.312 | 0.027 | 0.123 | 0.123 | 0.057 | 0.4 | 0.06 | 0.27 | 0.2 | 0.33 | 0.39 |
| 3 | 71757417 | rs1447899 | -1.017 | 1.448 | 0.989 | 1.304 | 0.024 | 0.15 | 0.15 | 0 | 0.21 | 0.51 | 0.22 | 0.49 | 0.26 | 0.23 |
| 3 | 71757540 | rs13085841 | 0.733 | 1.687 | 0.96 | 1.268 | 0.018 | 0.066 | 0.066 | 0 | 0.47 | 0.29 | 0.3 | 0.21 | 0.4 | 0.48 |
| 3 | 71758015 | rs6804667 | -1.066 | 1.66 | 0.793 | 1.298 | -0.018 | -0.003 | -0.003 | 0.678 | 0.69 | 0.85 | 0.52 | 0.7 | 0.64 | 0.72 |
| 3 | 71767413 | rs7648953 | -1.706 | 1.946 | 1.112 | 1.418 | 0.021 | 0.065 | 0.065 | 0.154 | 0.28 | 0.55 | 0.23 | 0.46 | 0.31 | 0.32 |
| 3 | 71768770 | rs13073428 | -1.671 | 1.942 | 1.123 | 1.42 | 0.02 | 0.065 | 0.065 | 0.088 | 0.27 | 0.57 | 0.25 | 0.47 | 0.31 | 0.33 |
| 3 | 71778737 | rs2044609 | -0.974 | 2.396 | 1.149 | 1.67 | 0.007 | 0.049 | 0.049 | 0 | 0.12 | 0.18 | 0.14 | 0.32 | 0.16 | 0.14 |
| 3 | 71795743 | rs6801173 | -0.973 | 2.409 | 1.212 | 1.669 | 0.009 | 0.056 | 0.056 | 0 | 0.12 | 0.21 | 0.15 | 0.32 | 0.16 | 0.13 |
| 3 | 71808566 | rs2654439 | 1.457 | 2.451 | 1.647 | 1.672 | 0.037 | 0.104 | 0.104 | 0 | 0.84 | 0.65 | 0.8 | 0.64 | 0.76 | 0.83 |
| 3 | 71809397 | rs11716884 | -0.408 | 1.954 | 1.202 | 1.364 | 0.025 | -0.025 | -0.025 | 0 | 0.2 | 0.12 | 0.14 | 0.29 | 0.13 | 0.35 |
| 3 | 71811875 | rs1374912 | 1.49 | 2.39 | 1.252 | 0.924 | -0.009 | -0.002 | -0.002 | 0 | 0.62 | 0.49 | 0.44 | 0.36 | 0.68 | 0.35 |
| 3 | 71821490 | rs6782813 | 1.021 | 2.327 | 1.046 | 0.803 | 0.063 | -0.015 | -0.015 | 0 | 0.92 | 0.49 | 0.94 | 0.99 | 0.95 | 0.91 |
| 3 | 71823582 | rs3796224 | -0.222 | 2.289 | 0.766 | 0.839 | 0.025 | -0.015 | -0.015 | 0.116 | 0.2 | 0.12 | 0.14 | 0.27 | 0.12 | 0.35 |
| 3 | 71825873 | rs2322143 | 0.164 | 2.438 | 0.742 | 0.831 | -0.023 | 0.009 | 0.009 | 0 | 0.78 | 0.67 | 0.83 | 0.71 | 0.88 | 0.61 |
| 3 | 71828420 | rs1320015 | -1.452 | 2.674 | -0.174 | 0.861 | -0.032 | -0.008 | -0.008 | 0 | 0.24 | 0.2 | 0.57 | 0.71 | 0.39 | 0.37 |
| 3 | 71829467 | rs7634474 | 0.723 | 2.167 | 0.085 | 1.058 | -0.009 | -0.004 | -0.004 | 0 | 0.73 | 0.37 | 0.8 | 0.72 | 0.82 | 0.56 |
| 3 | 71829914 | rs10779992 | -0.601 | 2.421 | 0.066 | 1.18 | -0.003 | 0.011 | 0.011 | 0 | 0.11 | 0.14 | 0.15 | 0.31 | 0.16 | 0.21 |
| 3 | 71840892 | rs4484160 | 0.193 | 2.042 | 0.029 | 1.123 | -0.001 | 0.019 | 0.019 | 0 | 0.07 | 0.05 | 0.07 | 0.19 | 0.1 | 0.15 |
| 3 | 71847123 | rs7629632 | -0.96 | 2.263 | 0.056 | 0.709 | 0.001 | 0 | 0 | 0 | 0.27 | 0.37 | 0.42 | 0.29 | 0.31 | 0.38 |
| 3 | 71850562 | rs9873991 | -1.095 | 1.704 | 0.118 | 1.447 | -0.007 | 0.079 | 0.079 | 0 | 0.18 | 0.25 | 0.21 | 0.54 | 0.17 | 0.38 |
| 3 | 71851723 | rs4677048 | -0.718 | 2.016 | 0.13 | 1.236 | 0.031 | -0.007 | -0.007 | 0 | 0.18 | 0.23 | 0.17 | 0.18 | 0.32 | 0.2 |
| 3 | 71854868 | rs1343953 | -1.499 | 0.689 | 0.166 | 1.705 | -0.009 | 0.04 | 0.04 | 0.024 | 0.21 | 0.27 | 0.21 | 0.55 | 0.17 | 0.39 |
| 3 | 71855936 | rs17721983 | -0.385 | 1.152 | -0.352 | 1.05 | -0.005 | -0.031 | -0.031 | 0 | 0.11 | 0.03 | 0.07 | 0.16 | 0.11 | 0.16 |
| 3 | 71868169 | rs10514728 | -0.984 | 1.121 | 0.158 | 0.84 | 0.036 | 0.111 | 0.111 | 0 | 0.06 | 0.2 | 0.07 | 0.17 | 0.06 | 0.08 |
| 3 | 71869001 | rs4677051 | 0.594 | 1.153 | -0.399 | -0.084 | 0.008 | 0.097 | 0.097 | 1.22 | 0.51 | 0.3 | 0.34 | 0.15 | 0.52 | 0.25 |
| 3 | 71872014 | rs9881946 | 0.282 | 1.387 | -0.596 | 0.403 | -0.002 | -0.007 | -0.007 | 0 | 0.7 | 0.71 | 0.56 | 0.62 | 0.66 | 0.61 |
| 3 | 71872388 | rs11721051 | -0.49 | 1.747 | -0.837 | -0.259 | -0.024 | 0.024 | 0.024 | 0 | 0.18 | 0.11 | 0.13 | 0.01 | 0.26 | 0.16 |
| 3 | 71873579 | rs6770667 | -0.002 | 1.327 | -0.605 | 0.024 | 0 | -0.001 | -0.001 | 0 | 0.77 | 0.76 | 0.63 | 0.84 | 0.72 | 0.7 |
| 3 | 71879031 | rs2322157 | 0.368 | -0.084 | -0.113 | -0.107 | -0.047 | -0.012 | -0.012 | 0 | 0.71 | 0.49 | 0.57 | 0.76 | 0.85 | 0.71 |
| 3 | 71879327 | rs9812198 | 0.128 | 0.119 | -0.62 | -0.283 | 0.004 | -0.008 | -0.008 | 0 | 0.35 | 0.37 | 0.32 | 0.25 | 0.4 | 0.17 |
| 3 | 71880578 | rs7622817 | -0.576 | -0.388 | -0.189 | -0.142 | -0.013 | 0.053 | 0.053 | 6.402 | 0.05 | 0.21 | 0.03 | 0.05 | 0 | 0.22 |
| 3 | 71880648 | rs6549411 | -0.102 | -0.577 | -0.021 | -0.355 | -0.038 | -0.033 | -0.033 | 0 | 0.66 | 0.55 | 0.56 | 0.78 | 0.82 | 0.7 |
| 3 | 71881411 | rs9878896 | 0.657 | -0.143 | -0.223 | -0.413 | 0.034 | 0.032 | 0.032 | 0.192 | 0.57 | 0.3 | 0.46 | 0.71 | 0.74 | 0.45 |
| 3 | 71882256 | rs11929696 | 1.555 | -0.49 | 0.158 | -0.263 | -0.003 | -0.005 | -0.005 | 0.267 | 0.08 | 0.09 | 0.04 | 0.04 | 0.04 | 0.06 |
| 3 | 71883565 | rs4676844 | 0.638 | -0.345 | -0.029 | -0.238 | -0.01 | 0.012 | 0.012 | 0 | 0.24 | 0.19 | 0.2 | 0.19 | 0.27 | 0.11 |
| 3 | 71885614 | rs9310222 | -0.919 | -0.419 | 0.381 | -0.224 | -0.011 | -0.002 | -0.002 | 0 | 0.82 | 0.8 | 0.62 | 0.84 | 0.9 | 0.78 |
| 3 | 71897255 | rs937661 | -0.603 | -0.577 | 0.48 | -0.129 | -0.019 | 0.003 | 0.003 | 0 | 0.78 | 0.74 | 0.61 | 0.75 | 0.89 | 0.77 |
| 3 | 71898645 | rs9861616 | 0.837 | -0.456 | 0.92 | -0.277 | -0.006 | 0 | 0 | 0 | 0.4 | 0.38 | 0.37 | 0.64 | 0.54 | 0.41 |
| 3 | 71899694 | rs7640019 | 1.297 | -0.508 | 0.973 | -0.462 | -0.017 | -0.031 | -0.031 | 0 | 0.1 | 0.15 | 0.05 | 0.02 | 0.04 | 0.15 |
| 3 | 71905876 | rs6780193 | 0.973 | -0.593 | 0.925 | -0.6 | -0.02 | 0.056 | 0.056 | 0 | 0.25 | 0.13 | 0.16 | 0.01 | 0.28 | 0.08 |
| 3 | 71906516 | rs9845463 | -1.589 | -0.693 | 1.041 | -0.382 | -0.022 | 0.062 | 0.062 | 0 | 0.82 | 0.71 | 0.57 | 0.67 | 0.9 | 0.77 |
| 3 | 71911170 | rs11718674 | 0.724 | -0.892 | 0.737 | -0.454 | 0.066 | -0.035 | -0.035 | 0 | 0.61 | 0.26 | 0.46 | 0.47 | 0.79 | 0.49 |
| 3 | 71913071 | rs7631578 | -0.556 | -0.596 | 0.697 | -0.422 | 0 | -0.005 | -0.005 | 0 | 0.44 | 0.47 | 0.27 | 0.12 | 0.4 | 0.38 |
| 3 | 71919973 | rs13085641 | 1.389 | -0.941 | 0.171 | -0.268 | -0.166 | -0.14 | -0.14 | 0.868 | 0.34 | 0.05 | 0.29 | 0.44 | 0.51 | 0.37 |

| | | | | | | | | | | | | | | | | |
|---|---|---|---|---|---|---|---|---|---|---|---|---|---|---|---|---|
| 3 | 71922551 | rs6549415 | -0.335 | -0.869 | 0.528 | -0.21 | -0.016 | 0.003 | 0.003 | 0 | 0.72 | 0.66 | 0.54 | 0.67 | 0.85 | 0.72 |
| 3 | 71930263 | rs11921763 | -0.916 | 0.271 | 0.749 | -0.066 | 0.147 | 0.004 | 0.004 | 0.486 | 0.06 | 0.27 | 0.06 | 0 | 0.12 | 0.03 |
| 3 | 71930936 | rs12714781 | -1.374 | -0.091 | 0.365 | -0.216 | 0.03 | 0.026 | 0.026 | 0 | 0.65 | 0.74 | 0.53 | 0.7 | 0.78 | 0.73 |
| 3 | 71934042 | rs7648714 | 1.014 | -0.159 | 0.001 | -0.188 | -0.139 | -0.125 | -0.125 | 1.655 | 0.33 | 0.06 | 0.34 | 0.48 | 0.51 | 0.38 |
| 3 | 71935481 | rs17009066 | 0.037 | -0.28 | 0.246 | -0.251 | -0.004 | 0 | 0 | 0 | 0.12 | 0.15 | 0.07 | 0.1 | 0.09 | 0.24 |
| 3 | 71937853 | rs1319094 | -1.311 | -0.1 | 0.395 | -0.317 | -0.007 | -0.003 | -0.003 | 0 | 0.63 | 0.6 | 0.5 | 0.61 | 0.77 | 0.7 |
| 3 | 71939885 | rs17009073 | -0.304 | -0.008 | 0.244 | -0.364 | 0 | -0.003 | -0.003 | 0 | 0.22 | 0.22 | 0.1 | 0.09 | 0.18 | 0.26 |
| 3 | 71948799 | rs1995453 | 0.683 | 0.226 | -0.368 | -0.291 | -0.088 | -0.073 | -0.073 | 0 | 0.35 | 0.09 | 0.36 | 0.51 | 0.57 | 0.43 |
| 3 | 71951781 | rs7616034 | -1.017 | -0.049 | 0.112 | -0.253 | -0.014 | -0.007 | -0.007 | 0 | 0.61 | 0.55 | 0.49 | 0.6 | 0.76 | 0.72 |
| 3 | 71955461 | rs9310225 | 1.321 | -0.054 | -0.139 | -0.31 | 0.034 | 0.029 | 0.029 | 0.033 | 0.34 | 0.17 | 0.4 | 0.29 | 0.22 | 0.25 |
| 3 | 71956442 | rs2874710 | 0.163 | 0.263 | -0.318 | -0.464 | -0.056 | 0.036 | 0.036 | 0.029 | 0.42 | 0.14 | 0.45 | 0.59 | 0.61 | 0.63 |
| 3 | 71959310 | rs6790891 | 0.497 | 0.524 | 0.438 | -0.453 | 0 | -0.003 | -0.003 | 0 | 0.14 | 0.11 | 0.07 | 0.11 | 0.14 | 0.08 |
| 3 | 71961277 | rs12636756 | 0.637 | 0.499 | 0.021 | -0.443 | -0.01 | 0.038 | 0.038 | 0.405 | 0.14 | 0.03 | 0.06 | 0.11 | 0.13 | 0.08 |
| 3 | 71961393 | rs4234146 | -1.82 | 0.744 | -0.03 | -0.521 | 0.002 | 0.034 | 0.034 | 0.727 | 0.41 | 0.65 | 0.22 | 0.25 | 0.43 | 0.46 |
| 3 | 71962547 | rs9832640 | 0.424 | 0.766 | -0.058 | -0.514 | 0 | -0.001 | -0.001 | 0 | 0.16 | 0.14 | 0.07 | 0.13 | 0.15 | 0.09 |
| 3 | 71963209 | rs6804315 | -0.671 | 0.79 | -0.305 | -0.437 | -0.014 | 0.01 | 0.01 | 0.003 | 0.22 | 0.15 | 0.13 | 0.13 | 0.29 | 0.38 |
| 3 | 71969532 | rs7648163 | -0.388 | 0.783 | -0.088 | -0.353 | 0.011 | -0.095 | -0.095 | 0 | 0.28 | 0.44 | 0.16 | 0.15 | 0.33 | 0.2 |
| 3 | 71976489 | rs9874131 | 1.162 | 0.389 | -1.199 | -0.09 | 0.006 | 0.054 | 0.054 | 0.099 | 0.37 | 0.16 | 0.4 | 0.26 | 0.26 | 0.27 |
| 3 | 71980438 | rs4677062 | -0.503 | 0.044 | -0.542 | 0.415 | -0.007 | 0.007 | 0.007 | 0.011 | 0.12 | 0.25 | 0.08 | 0.09 | 0.1 | 0.26 |
| 3 | 71980643 | rs7631897 | -0.176 | 0.484 | -1.121 | 0.588 | -0.057 | -0.142 | -0.142 | 1.623 | 0.23 | 0.03 | 0.34 | 0.53 | 0.3 | 0.29 |
| 3 | 71982824 | rs6796359 | 0.005 | 0.154 | -0.118 | 0.038 | -0.035 | -0.122 | -0.122 | 0.04 | 0.25 | 0.39 | 0.14 | 0.06 | 0.21 | 0.13 |
| 3 | 71983098 | rs11706411 | 1.134 | 0.021 | -0.026 | 0.081 | 0.053 | 0.088 | 0.088 | 0.003 | 0.53 | 0.28 | 0.49 | 0.25 | 0.57 | 0.42 |
| 3 | 71983334 | rs6549419 | 0.619 | 0.111 | 0.05 | 0.182 | -0.006 | 0.014 | 0.014 | 0.055 | 0.79 | 0.73 | 0.63 | 0.32 | 0.78 | 0.56 |
| 3 | 71984747 | rs6549420 | 0.506 | 0.077 | 0.094 | 0.581 | -0.008 | 0.066 | 0.066 | 0 | 0.81 | 0.75 | 0.68 | 0.48 | 0.83 | 0.64 |
| 3 | 71986893 | rs6809759 | 1.061 | 0.049 | 0.293 | 0.556 | 0.022 | 0.027 | 0.027 | 0 | 0.56 | 0.36 | 0.5 | 0.26 | 0.6 | 0.45 |
| 3 | 71987258 | rs9880789 | 0.724 | 0.365 | 0.49 | 0.184 | -0.003 | 0.032 | 0.032 | 0 | 0.89 | 0.79 | 0.88 | 0.83 | 0.88 | 0.82 |
| 3 | 71987961 | rs11706033 | -0.055 | 0.342 | 0.613 | -0.046 | -0.01 | -0.055 | -0.055 | 0 | 0.24 | 0.24 | 0.12 | 0.06 | 0.21 | 0.11 |
| 3 | 71992763 | rs6801209 | -0.321 | 0.531 | 0.326 | 0.211 | -0.008 | -0.038 | -0.038 | 0 | 0.08 | 0.26 | 0.03 | 0.01 | 0.02 | 0.1 |
| 3 | 71993530 | rs6802036 | -0.733 | 0.619 | 0.048 | 0.234 | 0.003 | 0.04 | 0.04 | 0 | 0.35 | 0.31 | 0.23 | 0.21 | 0.34 | 0.29 |
| 3 | 72000714 | rs7642104 | -0.024 | 0.523 | -0.443 | 0.563 | 0.001 | 0.047 | 0.047 | 0 | 0.25 | 0.14 | 0.18 | 0.19 | 0.28 | 0.19 |
| 3 | 72003089 | rs4234152 | 1.159 | 0.688 | 0.427 | 1.162 | -0.001 | 0.054 | 0.054 | 0 | 0.93 | 0.79 | 0.91 | 0.86 | 0.97 | 0.91 |
| 3 | 72005435 | rs1109569 | 0.621 | 0.98 | 0.205 | 0.721 | 0.001 | -0.013 | -0.013 | 1.579 | 0.6 | 0.56 | 0.57 | 0.27 | 0.59 | 0.46 |
| 3 | 72012377 | rs9856074 | 0.921 | 1.181 | 0.662 | 1.3 | 0.038 | 0.083 | 0.083 | 0.534 | 0.81 | 0.55 | 0.79 | 0.69 | 0.81 | 0.67 |
| 3 | 72019020 | rs870691 | 0.662 | 1.225 | 1.018 | 1.558 | 0.054 | 0.133 | 0.133 | 0 | 0.95 | 0.8 | 0.87 | 0.84 | 0.92 | 0.96 |
| 3 | 72021775 | rs4365591 | -0.643 | 1.299 | 0.696 | 2.05 | 0.01 | 0.05 | 0.05 | 0 | 0.05 | 0.08 | 0.05 | 0.21 | 0.08 | 0.13 |
| 3 | 72022752 | rs12491110 | -0.204 | 1.364 | 0.635 | 2.622 | 0.001 | -0.014 | -0.014 | 0 | 0.09 | 0.04 | 0.21 | 0.27 | 0.09 | 0.18 |
| 3 | 72024544 | rs4365590 | -1.279 | 1.42 | 0.562 | 1.826 | -0.008 | -0.026 | -0.026 | 0 | 0.34 | 0.25 | 0.36 | 0.49 | 0.35 | 0.47 |
| 3 | 72026826 | rs2102313 | -1.288 | 1.41 | 0.526 | 1.802 | -0.013 | -0.043 | -0.043 | 3.095 | 0.34 | 0.21 | 0.36 | 0.49 | 0.35 | 0.47 |
| 3 | 72033639 | rs13090166 | -0.201 | 1.453 | 0.776 | 1.671 | 0.051 | 0.063 | 0.063 | 1.69 | 0.07 | 0.34 | 0.06 | 0.08 | 0.04 | 0.07 |
| 3 | 72035934 | rs7619697 | 0.318 | 1.198 | 0.112 | 1.449 | 0.002 | 0.049 | 0.049 | 1.279 | 0.84 | 0.42 | 0.78 | 0.76 | 0.86 | 0.89 |
| 3 | 72036558 | rs1505972 | -0.952 | 1.206 | 0.017 | 1.503 | 0.001 | 0.051 | 0.051 | 0.684 | 0.17 | 0.21 | 0.19 | 0.46 | 0.18 | 0.36 |
| 3 | 72037324 | rs9829101 | 1.797 | 1.105 | -0.091 | 1.52 | -0.067 | 0.099 | 0.099 | 0.113 | 0.49 | 0.1 | 0.5 | 0.3 | 0.5 | 0.39 |
| 3 | 72041932 | rs11720188 | 0.538 | 0.744 | 0.658 | 1.616 | 0.009 | 0.047 | 0.047 | 1.107 | 0.84 | 0.37 | 0.77 | 0.76 | 0.85 | 0.89 |
| 3 | 72048692 | rs9879563 | 0.457 | 0.706 | -0.637 | 1.661 | -0.004 | 0.043 | 0.043 | 2.07 | 0.84 | 0.35 | 0.78 | 0.75 | 0.87 | 0.89 |
| 3 | 72049020 | rs9864349 | 0.773 | 0.756 | -0.8 | 1.19 | 0.003 | -0.008 | -0.008 | 0 | 0.73 | 0.3 | 0.58 | 0.43 | 0.78 | 0.72 |
| 3 | 72050088 | rs2174546 | 0.66 | 0.736 | -1.79 | 0.729 | -0.022 | 0.034 | 0.034 | 0 | 0.42 | 0.1 | 0.23 | 0.31 | 0.39 | 0.39 |
| 3 | 72058945 | rs11711611 | 0.582 | 0.603 | -1.992 | 0.653 | -0.018 | 0.049 | 0.049 | 0.024 | 0.46 | 0.15 | 0.43 | 0.31 | 0.54 | 0.31 |
| 3 | 72065348 | rs6810056 | -0.439 | 0.788 | -1.374 | 1.043 | -0.004 | -0.002 | -0.002 | 0.386 | 0.75 | 0.77 | 0.61 | 0.72 | 0.72 | 0.69 |
| 3 | 72067302 | rs1553574 | 0.261 | 0.486 | -1.299 | 1.265 | 0.002 | -0.04 | -0.04 | 0.044 | 0.14 | 0.08 | 0.14 | 0.3 | 0.16 | 0.23 |
| 3 | 72071821 | rs9874496 | -0.631 | 0.481 | -1.233 | 1.209 | -0.006 | -0.003 | -0.003 | 0.055 | 0.76 | 0.79 | 0.61 | 0.69 | 0.71 | 0.7 |
| 3 | 72081057 | rs9823122 | -0.832 | 0.035 | -0.967 | 0.931 | -0.039 | 0.051 | 0.051 | 0 | 0.27 | 0.47 | 0.23 | 0.37 | 0.21 | 0.3 |
| 3 | 72082185 | rs17009307 | 0.42 | -0.033 | -0.552 | 0.804 | -0.007 | -0.005 | -0.005 | 0 | 0.12 | 0.14 | 0.09 | 0.09 | 0.09 | 0.11 |
| 3 | 72090145 | rs10511021 | 0.596 | 0.152 | -0.857 | 0.796 | -0.02 | -0.037 | -0.037 | 0.858 | 0.13 | 0.08 | 0.14 | 0.32 | 0.17 | 0.23 |
| 3 | 72095871 | rs6549433 | -0.684 | 0.058 | -0.97 | 0.802 | -0.016 | 0.009 | 0.009 | 0 | 0.31 | 0.53 | 0.41 | 0.33 | 0.26 | 0.34 |
| 3 | 72097527 | rs9682418 | -0.674 | 0.071 | -1.066 | 0.81 | -0.013 | 0.034 | 0.034 | 0 | 0.31 | 0.56 | 0.46 | 0.41 | 0.27 | 0.34 |
| 3 | 72108144 | rs13326050 | 1.017 | 0.164 | -1.216 | 0.293 | 0.005 | 0.037 | 0.037 | 0 | 0.4 | 0.3 | 0.35 | 0.14 | 0.41 | 0.27 |
| 3 | 72109408 | rs1505971 | 0.507 | 0.204 | -1.812 | 0.328 | -0.045 | 0.027 | 0.027 | 0 | 0.49 | 0.12 | 0.44 | 0.58 | 0.52 | 0.46 |
| 3 | 72110578 | rs2322269 | -1.097 | 0.022 | -1.833 | -0.203 | -0.009 | 0.02 | 0.02 | 0 | 0.85 | 0.92 | 0.9 | 1 | 0.85 | 0.8 |
| 3 | 72111169 | rs9814799 | -0.628 | -0.012 | -1.096 | -0.107 | 0.017 | 0.036 | 0.036 | 0 | 0.28 | 0.41 | 0.4 | 0.41 | 0.31 | 0.27 |
| 3 | 72114276 | rs879526 | -0.041 | 0.289 | -0.891 | -0.245 | 0.055 | 0.039 | 0.039 | 0 | 0.13 | 0.36 | 0.08 | 0.16 | 0.14 | 0.11 |
| 3 | 72118981 | rs7649841 | 0.446 | -0.126 | -1.282 | -0.386 | -0.043 | -0.059 | -0.059 | 0 | 0.26 | 0.52 | 0.4 | 0.14 | 0.17 | 0.11 |
| 3 | 72126377 | rs924961 | -1.105 | -0.163 | -1.871 | -0.565 | -0.022 | 0.062 | 0.062 | 0.008 | 0.58 | 0.67 | 0.43 | 0.81 | 0.48 | 0.55 |
| 3 | 72126604 | rs924962 | 0.829 | -0.185 | -1.842 | -0.883 | -0.031 | -0.114 | -0.114 | 0 | 0.26 | 0.56 | 0.19 | 0.02 | 0.17 | 0.08 |
| 3 | 72126831 | rs10746538 | -1.437 | -0.178 | -2.003 | -0.974 | 0.005 | -0.045 | -0.045 | 0 | 0.86 | 0.84 | 0.89 | 1 | 0.84 | 0.85 |
| 3 | 72128884 | rs1547458 | -1.174 | -0.216 | -1.944 | -0.869 | -0.03 | 0.075 | 0.075 | 0.003 | 0.59 | 0.69 | 0.45 | 0.84 | 0.49 | 0.57 |
| 3 | 72129064 | rs1546389 | 1.279 | -0.083 | -2.08 | -0.493 | -0.023 | -0.021 | -0.021 | 0 | 0.06 | 0.02 | 0.04 | 0.12 | 0.13 | 0.1 |
| 3 | 72131974 | rs2322276 | -0.182 | -0.111 | -2.045 | -1.075 | -0.036 | 0.058 | 0.058 | 0 | 0.32 | 0.08 | 0.24 | 0.81 | 0.32 | 0.49 |
| 3 | 72141723 | rs11707080 | -0.154 | 0.005 | -1.947 | -1.001 | -0.01 | 0.047 | 0.047 | 0 | 0.33 | 0.13 | 0.24 | 0.8 | 0.31 | 0.44 |
| 3 | 72146520 | rs6549436 | 0.735 | -0.17 | -2 | -0.93 | -0.027 | 0.013 | 0.013 | 1.041 | 0.36 | 0.26 | 0.49 | 0.31 | 0.5 | 0.47 |

| | | | | | | | | | | | | | | | | |
|---|---|---|---|---|---|---|---|---|---|---|---|---|---|---|---|---|
| 3 | 72150676 | rs4677083 | 0.82 | -0.219 | -2.097 | -0.979 | -0.029 | 0.01 | 0.01 | 1.177 | 0.36 | 0.24 | 0.49 | 0.32 | 0.5 | 0.46 |
| 3 | 72151732 | rs6782778 | -0.315 | -0.414 | -2.118 | -1.12 | 0.062 | 0.055 | 0.055 | 2.962 | 0.52 | 0.63 | 0.39 | 0.67 | 0.33 | 0.4 |
| 3 | 72152533 | rs11128246 | 0.767 | -0.355 | -2.225 | -1.031 | -0.013 | -0.154 | -0.154 | 1.888 | 0.24 | 0.01 | 0.16 | 0.54 | 0.19 | 0.31 |
| 3 | 72156041 | rs4677085 | -1.677 | -0.201 | -1.934 | -1.003 | -0.002 | -0.005 | -0.005 | 0 | 0.89 | 0.9 | 0.89 | 1 | 0.85 | 0.87 |
| 3 | 72158910 | rs9832693 | -0.123 | -0.291 | -1.922 | -1.016 | 0.05 | 0.021 | 0.021 | 3.358 | 0.49 | 0.61 | 0.37 | 0.57 | 0.33 | 0.4 |
| 3 | 72160602 | rs2055644 | -0.132 | -0.478 | -2.123 | -1.16 | 0.034 | 0.035 | 0.035 | 0.193 | 0.59 | 0.35 | 0.68 | 0.97 | 0.7 | 0.76 |
| 3 | 72161259 | rs6549439 | -0.66 | -0.209 | -2.269 | -1.181 | -0.027 | 0.088 | 0.088 | 0.74 | 0.57 | 0.71 | 0.45 | 0.82 | 0.46 | 0.55 |
| 3 | 72163161 | rs7646485 | -1.957 | -0.201 | -2.321 | -1.178 | -0.04 | 0.096 | 0.096 | 0 | 0.85 | 0.99 | 0.89 | 1 | 0.84 | 0.83 |
| 3 | 72163948 | rs4677092 | 1.495 | -0.388 | -2.262 | -1.147 | -0.032 | 0.027 | 0.027 | 0 | 0.27 | 0.17 | 0.43 | 0.17 | 0.38 | 0.29 |
| 3 | 72166172 | rs9863207 | 0.502 | -0.148 | -2.216 | -1.021 | -0.01 | -0.056 | -0.056 | 0.965 | 0.28 | 0.62 | 0.22 | 0.02 | 0.2 | 0.09 |
| 3 | 72167619 | rs11718202 | 0.606 | 0.308 | -2.71 | -0.535 | -0.029 | -0.129 | -0.129 | 0 | 0.23 | 0.02 | 0.21 | 0.62 | 0.2 | 0.33 |
| 3 | 72170887 | rs17009411 | 0.751 | 0.334 | -2.686 | -0.576 | -0.028 | -0.15 | -0.15 | 1.175 | 0.22 | 0.01 | 0.16 | 0.55 | 0.19 | 0.31 |
| 3 | 72171883 | rs17009412 | -0.053 | 0.512 | -1.914 | -0.511 | 0.002 | -0.007 | -0.007 | 0.013 | 0.3 | 0.38 | 0.25 | 0.31 | 0.27 | 0.22 |
| 3 | 72176615 | rs2322272 | -0.083 | 1.45 | -1.135 | -0.573 | -0.001 | -0.032 | -0.032 | 1.265 | 0.34 | 0.59 | 0.44 | 0.15 | 0.35 | 0.3 |
| 3 | 72179475 | rs7629471 | -0.692 | 1.938 | -0.372 | -0.115 | 0.156 | -0.027 | -0.027 | 0 | 0.08 | 0.28 | 0.07 | 0.01 | 0.14 | 0.09 |
| 3 | 72180203 | rs4677096 | -0.774 | 1.165 | -1.342 | 0.238 | -0.02 | -0.078 | -0.078 | 0 | 0.24 | 0.06 | 0.2 | 0.65 | 0.27 | 0.36 |
| 3 | 72180966 | rs6779450 | 0.057 | 1.408 | -1.374 | -0.03 | -0.01 | 0.009 | 0.009 | 0 | 0.8 | 0.86 | 0.64 | 0.87 | 0.76 | 0.78 |
| 3 | 72181219 | rs6779548 | 0.766 | 0.995 | -0.265 | -0.143 | -0.018 | -0.009 | -0.009 | 0 | 0.72 | 0.43 | 0.6 | 0.81 | 0.74 | 0.71 |
| 3 | 72181924 | rs4677097 | 1.324 | 1.063 | -0.295 | -0.619 | 0.025 | 0.117 | 0.117 | 0 | 0.51 | 0.24 | 0.39 | 0.19 | 0.44 | 0.38 |
| 3 | 72186063 | rs4417833 | 1.591 | 0.714 | 1.079 | -0.248 | 0.009 | -0.06 | -0.06 | 0 | 0.81 | 0.5 | 0.86 | 0.92 | 0.83 | 0.91 |
| 3 | 72187269 | rs6802792 | -2.224 | 0.466 | 1.582 | -0.13 | 0.061 | 0.153 | 0.153 | 3.845 | 0.06 | 0.53 | 0.1 | 0.06 | 0.01 | 0.06 |
| 3 | 72193422 | rs4676861 | 0.973 | 0.468 | 1.34 | -0.379 | -0.026 | -0.007 | -0.007 | 3.932 | 0.89 | 0.74 | 0.94 | 0.92 | 0.94 | 0.92 |
| 3 | 72200540 | rs7617884 | 1.986 | 0.537 | 0.944 | -0.44 | -0.005 | -0.021 | -0.021 | 0 | 0.76 | 0.31 | 0.83 | 0.92 | 0.8 | 0.85 |
| 3 | 72207045 | rs6549445 | 1.109 | 0.699 | 0.712 | -0.278 | 0 | -0.015 | -0.015 | 0.521 | 0.84 | 0.78 | 0.9 | 0.92 | 0.84 | 0.91 |
| 3 | 72209538 | rs9865222 | -1.096 | 0.505 | 0.453 | -0.424 | -0.01 | -0.012 | -0.012 | 0 | 0.27 | 0.33 | 0.22 | 0.18 | 0.25 | 0.24 |
| 3 | 72209597 | rs4074259 | -0.672 | 1.247 | 0.314 | -0.187 | 0.036 | 0.032 | 0.032 | 0 | 0.28 | 0.53 | 0.4 | 0.64 | 0.35 | 0.4 |
| 3 | 72210920 | rs9847728 | -0.56 | 0.735 | 0.168 | -0.276 | 0.004 | -0.038 | -0.038 | 0 | 0.17 | 0.22 | 0.18 | 0.08 | 0.2 | 0.12 |
| 3 | 72213025 | rs4677105 | 0.481 | 0.492 | -0.166 | 0.176 | -0.008 | -0.098 | -0.098 | 0.327 | 0.2 | 0.06 | 0.27 | 0.56 | 0.2 | 0.36 |
| 3 | 72216448 | rs7643591 | -2.245 | 0.621 | 0.135 | -0.103 | -0.016 | 0.165 | 0.165 | 0.102 | 0.57 | 0.87 | 0.57 | 0.85 | 0.57 | 0.62 |
| 3 | 72217210 | rs7646549 | -1.441 | 0.634 | 0.465 | -0.098 | 0.003 | -0.019 | -0.019 | 0 | 0.43 | 0.56 | 0.45 | 0.79 | 0.49 | 0.51 |
| 3 | 72222004 | rs6793065 | -2.132 | 0.798 | 0.289 | 0.021 | -0.008 | 0.068 | 0.068 | 0.112 | 0.55 | 0.85 | 0.56 | 0.72 | 0.56 | 0.61 |
| 3 | 72222082 | rs6784559 | -0.635 | 0.846 | 0.412 | 0.011 | 0.027 | -0.03 | -0.03 | 0.031 | 0.15 | 0.23 | 0.14 | 0.07 | 0.22 | 0.04 |
| 3 | 72222650 | rs6793710 | -1.212 | 0.872 | 0.172 | 0.055 | 0.037 | -0.073 | -0.073 | 0 | 0.25 | 0.48 | 0.19 | 0.13 | 0.29 | 0.14 |
| 3 | 72222689 | rs10470587 | -0.66 | 0.986 | 0.152 | 0.531 | -0.007 | 0.018 | 0.018 | 0 | 0.3 | 0.37 | 0.37 | 0.58 | 0.27 | 0.46 |
| 3 | 72224748 | rs11707718 | -0.638 | 1.176 | -0.149 | 0.537 | -0.009 | -0.011 | -0.011 | 0 | 0.16 | 0.28 | 0.12 | 0.12 | 0.13 | 0.19 |
| 3 | 72225773 | rs13065721 | -1.161 | 1.74 | -0.385 | 0.845 | 0.004 | 0 | 0 | 0 | 0.08 | 0.1 | 0.11 | 0.1 | 0.18 | 0.05 |
| 3 | 72227587 | rs9829428 | -1.253 | 1.143 | 0.198 | 0.641 | 0.041 | 0.034 | 0.034 | 0 | 0.1 | 0.29 | 0.05 | 0.15 | 0.08 | 0.12 |
| 3 | 72227868 | rs7644006 | -0.812 | 0.605 | -0.234 | 0.21 | -0.003 | 0.013 | 0.013 | 0 | 0.55 | 0.69 | 0.53 | 0.72 | 0.51 | 0.63 |
| 3 | 72228716 | rs7632787 | -0.994 | 0.419 | -0.01 | 0.068 | -0.043 | 0.056 | 0.056 | 0.002 | 0.61 | 0.86 | 0.54 | 0.72 | 0.53 | 0.63 |
| 3 | 72231202 | rs4677107 | 0.408 | -0.62 | 0.019 | 0.235 | 0.044 | -0.002 | -0.002 | 0.048 | 0.25 | 0.14 | 0.33 | 0.24 | 0.16 | 0.1 |
| 3 | 72236434 | rs4677109 | -0.641 | -0.643 | 0.043 | 0.1 | 0.008 | 0.031 | 0.031 | 0 | 0.48 | 0.41 | 0.5 | 0.56 | 0.48 | 0.54 |
| 3 | 72239901 | rs6767307 | -1.352 | -0.463 | -0.244 | 0.12 | -0.053 | 0.008 | 0.008 | 0.08 | 0.6 | 0.85 | 0.56 | 0.64 | 0.53 | 0.62 |
| 3 | 72248600 | rs11720697 | 0.199 | -0.193 | -0.363 | 0.295 | -0.007 | -0.009 | -0.009 | 0 | 0.08 | 0.21 | 0.05 | 0.07 | 0.04 | 0.07 |
| 3 | 72250823 | rs13071822 | -0.109 | -0.078 | -1.008 | 0.372 | 0 | 0.002 | 0.002 | 0 | 0.22 | 0.26 | 0.15 | 0.33 | 0.3 | 0.43 |
| 3 | 72251412 | rs6789965 | 0.932 | -0.125 | -0.858 | 0.604 | 0.003 | -0.03 | -0.03 | 0 | 0.06 | 0 | 0.26 | 0.24 | 0.05 | 0.02 |
| 3 | 72254890 | rs9790178 | 0.229 | 0.314 | -0.633 | 0.001 | 0.018 | -0.012 | -0.012 | 0 | 0.39 | 0.23 | 0.46 | 0.55 | 0.4 | 0.48 |
| 3 | 72256315 | rs4131907 | -0.722 | 0.951 | -0.401 | 0.235 | 0.027 | -0.04 | -0.04 | 0 | 0.08 | 0.24 | 0.04 | 0 | 0.06 | 0.05 |
| 3 | 72263880 | rs7432938 | -0.281 | 0.795 | -1.187 | 0.367 | -0.082 | 0.013 | 0.013 | 0.37 | 0.62 | 0.87 | 0.59 | 0.7 | 0.52 | 0.66 |
| 3 | 72264836 | rs4676864 | 0.044 | 1.021 | -0.6 | 0.257 | 0.003 | -0.009 | -0.009 | 0.04 | 0.71 | 0.64 | 0.75 | 0.75 | 0.67 | 0.71 |
| 3 | 72266311 | rs4677113 | -0.408 | 1.153 | -0.725 | 0.854 | -0.028 | -0.068 | -0.068 | 0.011 | 0.17 | 0.04 | 0.22 | 0.58 | 0.24 | 0.36 |
| 3 | 72273331 | rs4676866 | -1.462 | 0.959 | -0.348 | 0.619 | -0.001 | -0.005 | -0.005 | 0.476 | 0.26 | 0.23 | 0.26 | 0.58 | 0.28 | 0.4 |
| 3 | 72277024 | rs4303838 | 0.039 | 0.989 | -0.356 | 0.559 | -0.003 | -0.018 | -0.018 | 0.647 | 0.05 | 0.05 | 0.02 | 0 | 0.03 | 0.01 |
| 3 | 72278809 | rs7615690 | -1.549 | 1.023 | 0.191 | 0.476 | 0.005 | -0.055 | -0.055 | 0.075 | 0.13 | 0.4 | 0.16 | 0.04 | 0.08 | 0.07 |
| 3 | 72279564 | rs4677116 | -0.255 | 1.135 | -0.529 | 0.833 | -0.028 | -0.069 | -0.069 | 0 | 0.17 | 0.04 | 0.22 | 0.57 | 0.24 | 0.36 |
| 3 | 72282932 | rs11713031 | -1.465 | 0.849 | -0.108 | 0.409 | -0.001 | -0.038 | -0.038 | 0.164 | 0.11 | 0.25 | 0.15 | 0.04 | 0.08 | 0.07 |
| 3 | 72285612 | rs11128254 | -0.141 | 0.684 | -0.808 | 0.191 | 0.014 | 0.052 | 0.052 | 0 | 0.54 | 0.31 | 0.59 | 0.73 | 0.6 | 0.68 |
| 3 | 72288745 | rs12632003 | -0.579 | 0.385 | -0.317 | 0.067 | -0.001 | 0 | 0 | 0 | 0.63 | 0.68 | 0.59 | 0.74 | 0.67 | 0.67 |
| 3 | 72289425 | rs4078475 | 0.172 | 0.002 | -1.358 | -0.215 | -0.074 | 0.024 | 0.024 | 0 | 0.39 | 0.13 | 0.46 | 0.71 | 0.51 | 0.56 |
| 3 | 72294743 | rs4525828 | 0.081 | 0.444 | -1.828 | -0.04 | -0.032 | 0.023 | 0.023 | 0 | 0.42 | 0.22 | 0.48 | 0.69 | 0.51 | 0.53 |
| 3 | 72295961 | rs12635882 | 0.9 | 1.009 | -1.995 | 0.131 | -0.06 | -0.087 | -0.087 | 0 | 0.14 | 0.03 | 0.27 | 0.47 | 0.27 | 0.28 |
| 3 | 72300561 | rs4677122 | -0.601 | 0.796 | -1.912 | -0.398 | 0.015 | 0.094 | 0.094 | 0 | 0.6 | 0.79 | 0.7 | 0.85 | 0.66 | 0.7 |
| 3 | 72302917 | rs4676868 | -0.767 | 0.703 | -1.977 | -0.457 | 0.009 | 0.091 | 0.091 | 0 | 0.63 | 0.8 | 0.67 | 0.85 | 0.68 | 0.7 |
| 3 | 72305546 | rs9878609 | 0.23 | 0.83 | -2.182 | -0.254 | -0.004 | -0.032 | -0.032 | 0 | 0.46 | 0.14 | 0.46 | 0.55 | 0.57 | 0.49 |
| 3 | 72310765 | rs6549449 | 0.349 | 1.033 | -1.72 | -0.054 | -0.012 | 0.016 | 0.016 | 1.995 | 0.35 | 0.49 | 0.34 | 0.45 | 0.32 | 0.33 |
| 3 | 72315956 | rs4532099 | -1.083 | 1.171 | -1.661 | -0.285 | -0.022 | 0.085 | 0.085 | 0.151 | 0.71 | 0.95 | 0.74 | 0.84 | 0.75 | 0.73 |
| 3 | 72316559 | rs6796081 | 0.703 | 1.409 | -1.649 | -0.406 | 0 | -0.075 | -0.075 | 1.227 | 0.22 | 0.26 | 0.19 | 0.02 | 0.25 | 0.18 |
| 3 | 72316954 | rs9830272 | 0.353 | 1.796 | -1.357 | 0.622 | 0 | -0.012 | -0.012 | 0 | 0.18 | 0.14 | 0.21 | 0.22 | 0.17 | 0.23 |
| 3 | 72317470 | rs9815397 | -0.702 | 1.855 | -1.299 | 0.425 | -0.015 | 0.067 | 0.067 | 0 | 0.45 | 0.32 | 0.51 | 0.71 | 0.5 | 0.55 |
| 3 | 72322710 | rs11928668 | -0.143 | 1.682 | -1.079 | 0.341 | 0.006 | 0.047 | 0.047 | 0.079 | 0.17 | 0.27 | 0.21 | 0.32 | 0.17 | 0.27 |

| | | | | | | | | | | | | | | | | |
|---|---|---|---|---|---|---|---|---|---|---|---|---|---|---|---|---|
| 3 | 72326691 | rs9858942 | 0.059 | 1.743 | -1.051 | 0.546 | -0.014 | 0.049 | 0.049 | 2.251 | 0.3 | 0.44 | 0.3 | 0.55 | 0.24 | 0.29 |
| 3 | 72329337 | rs7642076 | -0.727 | 2.805 | -1.427 | 0.13 | -0.042 | 0.034 | 0.034 | 0.09 | 0.17 | 0.07 | 0.23 | 0.01 | 0.33 | 0.17 |
| 3 | 72334772 | rs7429249 | -0.865 | 3.141 | -1.548 | 0.965 | -0.052 | -0.048 | -0.048 | 0.136 | 0.14 | 0.04 | 0.2 | 0.28 | 0.27 | 0.19 |
| 3 | 72336095 | rs4677136 | 0.003 | 2.643 | -1.766 | 0.479 | 0.002 | 0.011 | 0.011 | 0 | 0.83 | 0.82 | 0.86 | 0.92 | 0.82 | 0.93 |
| 3 | 72344601 | rs11709621 | -0.217 | 2.004 | -1.949 | 0.811 | -0.006 | -0.051 | -0.051 | 1.493 | 0.22 | 0.11 | 0.28 | 0.38 | 0.22 | 0.27 |
| 3 | 72344971 | rs9847707 | 1.061 | 1.472 | -1.298 | 0.249 | 0 | 0.024 | 0.024 | 0.868 | 0.5 | 0.39 | 0.47 | 0.27 | 0.51 | 0.41 |
| 3 | 72347479 | rs4677138 | 0.312 | 0.317 | -1.426 | -0.634 | -0.01 | 0.006 | 0.006 | 0.568 | 0.26 | 0.52 | 0.26 | 0.28 | 0.19 | 0.29 |
| 3 | 72351680 | rs11715508 | 0.397 | 0.174 | -0.632 | -0.462 | -0.017 | 0.064 | 0.064 | 0.269 | 0.28 | 0.43 | 0.35 | 0.44 | 0.24 | 0.32 |
| 3 | 72353415 | rs7431828 | 1.981 | -0.079 | -0.593 | -0.577 | 0.007 | -0.012 | -0.012 | 0.033 | 0.13 | 0.08 | 0.18 | 0.22 | 0.08 | 0.26 |
| 3 | 72353932 | rs9809213 | 0.887 | -0.178 | -0.288 | -0.567 | 0 | 0.001 | 0.001 | 0.079 | 0.23 | 0.18 | 0.32 | 0.42 | 0.22 | 0.26 |
| 3 | 72354511 | rs7643143 | -1.028 | -0.276 | -0.39 | -0.766 | -0.011 | -0.12 | -0.12 | 0.623 | 0.63 | 0.53 | 0.72 | 0.98 | 0.65 | 0.79 |
| 3 | 72355054 | rs11706917 | 1.639 | 0.001 | -0.428 | -0.55 | -0.025 | -0.021 | -0.021 | 0.754 | 0.11 | 0.04 | 0.15 | 0.32 | 0.21 | 0.17 |
| 3 | 72356680 | rs4677143 | 0.088 | -0.106 | -0.344 | -0.645 | -0.02 | -0.014 | -0.014 | 0 | 0.3 | 0.24 | 0.37 | 0.58 | 0.37 | 0.45 |
| 3 | 72357857 | rs4677144 | -1.104 | -0.129 | -0.334 | -0.593 | -0.021 | -0.026 | -0.026 | 0.184 | 0.95 | 0.8 | 0.99 | 1 | 0.99 | 0.95 |
| 3 | 72361466 | rs4073919 | -0.15 | -0.214 | -0.365 | -0.65 | -0.046 | -0.091 | -0.091 | 0.154 | 0.86 | 0.65 | 0.97 | 1 | 0.95 | 0.91 |
| 3 | 72363674 | rs9820604 | 0.911 | -0.533 | -1.048 | -0.75 | 0.021 | -0.068 | -0.068 | 0.479 | 0.25 | 0.1 | 0.31 | 0.4 | 0.19 | 0.38 |
| 3 | 72364338 | rs11708327 | 1.406 | -0.047 | -0.856 | -0.15 | -0.023 | -0.023 | -0.023 | 0 | 0.1 | 0.05 | 0.15 | 0.27 | 0.21 | 0.14 |
| 3 | 72364412 | rs3934820 | -0.483 | -0.409 | -1.016 | -0.708 | -0.026 | 0.092 | 0.092 | 1 | 0.71 | 0.88 | 0.8 | 0.98 | 0.66 | 0.85 |
| 3 | 72365204 | rs4676870 | -1.212 | -0.311 | -0.946 | -0.585 | -0.037 | 0.069 | 0.069 | 0.167 | 0.78 | 0.92 | 0.81 | 0.98 | 0.69 | 0.9 |
| 3 | 72371023 | rs9826269 | 0.232 | -0.435 | -0.29 | 0.054 | 0.002 | -0.009 | -0.009 | 0 | 0.57 | 0.48 | 0.41 | 0.4 | 0.45 | 0.49 |
| 3 | 72371722 | rs9848266 | 0.521 | -0.589 | -0.161 | -0.152 | -0.037 | -0.065 | -0.065 | 1.109 | 0.11 | 0.26 | 0.02 | 0 | 0.03 | 0.04 |
| 3 | 72375717 | rs17726037 | 1.538 | -0.298 | -0.581 | 0.164 | -0.021 | -0.02 | -0.02 | 0 | 0.1 | 0.04 | 0.17 | 0.27 | 0.19 | 0.11 |
| 3 | 72376808 | rs6790069 | 0.411 | -0.469 | -0.385 | 0.084 | 0.003 | 0.005 | 0.005 | 0 | 0.16 | 0.16 | 0.18 | 0.27 | 0.2 | 0.13 |
| 3 | 72377190 | rs12486275 | 1.033 | -0.663 | -1.005 | 0.06 | -0.063 | -0.071 | -0.071 | 0.033 | 0.39 | 0.07 | 0.34 | 0.39 | 0.36 | 0.46 |
| 3 | 72378733 | rs8179972 | -0.09 | -0.572 | 0.194 | 0.035 | -0.041 | -0.078 | -0.078 | 0 | 0.87 | 0.76 | 0.94 | 1 | 0.93 | 0.94 |
| 3 | 72378950 | rs11920844 | -0.216 | -0.084 | 0.152 | 0.897 | 0.024 | 0.052 | 0.052 | 0 | 0.17 | 0.22 | 0.24 | 0.37 | 0.27 | 0.15 |
| 3 | 72380757 | rs6798001 | -0.713 | -0.194 | 0.177 | 0.858 | 0.02 | 0.134 | 0.134 | 0.112 | 0.18 | 0.48 | 0.22 | 0.35 | 0.2 | 0.13 |
| 3 | 72383495 | rs9883587 | 0.495 | -0.179 | -0.106 | 0.176 | -0.026 | -0.073 | -0.073 | 2.236 | 0.73 | 0.57 | 0.78 | 0.88 | 0.75 | 0.83 |
| 3 | 72385884 | rs11918732 | -0.601 | 0.062 | -0.445 | 0.545 | -0.003 | 0 | 0 | 0 | 0.47 | 0.51 | 0.39 | 0.49 | 0.39 | 0.43 |
| 3 | 72393312 | rs9815886 | -1.033 | 0.276 | -0.557 | 0.453 | 0.003 | 0.041 | 0.041 | 0 | 0.54 | 0.73 | 0.61 | 0.67 | 0.52 | 0.55 |
| 3 | 72398604 | rs9870681 | -0.102 | 0.222 | -0.07 | 0.71 | 0.005 | 0.087 | 0.087 | 0.281 | 0.23 | 0.37 | 0.33 | 0.51 | 0.22 | 0.2 |
| 3 | 72400695 | rs12497693 | 1.34 | 0.268 | -0.768 | 0.308 | -0.025 | 0.003 | 0.003 | 0.575 | 0.38 | 0.1 | 0.36 | 0.32 | 0.42 | 0.47 |
| 3 | 72403467 | rs9841241 | -0.633 | 0.022 | -0.042 | -0.406 | 0 | -0.01 | -0.01 | 0 | 0.42 | 0.6 | 0.31 | 0.18 | 0.36 | 0.36 |
| 3 | 72412943 | rs4677152 | 0.197 | 0.096 | -0.814 | -0.223 | -0.014 | -0.029 | -0.029 | 0 | 0.24 | 0.05 | 0.24 | 0.26 | 0.22 | 0.28 |
| 3 | 72420314 | rs12631820 | 1.244 | 0.067 | -0.952 | -0.487 | -0.011 | -0.03 | -0.03 | 0.173 | 0.41 | 0.17 | 0.4 | 0.44 | 0.37 | 0.44 |
| 3 | 72422131 | rs2272078 | 0.716 | 0.311 | 0.189 | -0.444 | -0.001 | 0.008 | 0.008 | 1.419 | 0.67 | 0.4 | 0.55 | 0.55 | 0.65 | 0.48 |
| 3 | 72432888 | rs7628338 | 0.719 | 0.458 | 0.57 | -0.407 | 0 | 0 | 0 | 0.034 | 0.64 | 0.4 | 0.63 | 0.69 | 0.65 | 0.65 |
| 3 | 72437413 | rs9863706 | 0.607 | 0.4 | 1.122 | -0.425 | 0.005 | 0.015 | 0.015 | 0 | 0.17 | 0.24 | 0.25 | 0.26 | 0.21 | 0.19 |
| 3 | 72455797 | rs11128265 | 0.178 | 0.366 | 1.354 | -0.435 | 0.047 | 0.107 | 0.107 | 0.131 | 0.19 | 0.41 | 0.26 | 0.27 | 0.21 | 0.19 |
| 3 | 72459756 | rs6797249 | 0.475 | 0.318 | 0.932 | -0.46 | 0.005 | 0.01 | 0.01 | 0 | 0.14 | 0.08 | 0.1 | 0.01 | 0.13 | 0.14 |
| 3 | 72469651 | rs4676875 | 0.773 | 0.45 | 0.995 | -0.463 | 0.008 | 0.023 | 0.023 | 0.138 | 0.69 | 0.4 | 0.56 | 0.55 | 0.6 | 0.59 |
| 3 | 72477819 | rs13086339 | 1.843 | 0.335 | 0.126 | -0.609 | -0.08 | -0.134 | -0.134 | 0.041 | 0.42 | 0.03 | 0.37 | 0.44 | 0.32 | 0.37 |
| 3 | 72478611 | rs17009723 | 0.174 | 0.61 | 0.798 | -0.366 | -0.005 | -0.005 | -0.005 | 0.178 | 0.11 | 0.09 | 0.19 | 0.2 | 0.21 | 0.22 |
| 3 | 72505316 | rs9870697 | -0.082 | 0.027 | 0.661 | -0.364 | 0.047 | 0.038 | 0.038 | 0.094 | 0.26 | 0.47 | 0.33 | 0.31 | 0.38 | 0.22 |
| 3 | 72510771 | rs6796880 | -0.976 | -0.156 | 0.46 | -0.648 | -0.008 | 0.016 | 0.016 | 0 | 0.42 | 0.64 | 0.4 | 0.33 | 0.48 | 0.32 |
| 3 | 72511359 | rs6549457 | 0.438 | -0.2 | 0.5 | -0.627 | -0.008 | -0.01 | -0.01 | 0.014 | 0.67 | 0.33 | 0.54 | 0.54 | 0.57 | 0.58 |
| 3 | 72512244 | rs17009757 | 0.464 | -0.076 | 1.11 | -1.003 | 0 | 0.001 | 0.001 | 0 | 0.1 | 0.11 | 0.15 | 0.17 | 0.13 | 0.19 |
| 3 | 72512380 | rs6790808 | 0.64 | -0.271 | 0.876 | -1.274 | -0.001 | -0.032 | -0.032 | 0 | 0.6 | 0.48 | 0.62 | 0.72 | 0.59 | 0.67 |
| 3 | 72512902 | rs2028241 | 0.228 | -0.483 | 0.368 | -1.077 | -0.003 | -0.022 | -0.022 | 0 | 0.93 | 0.81 | 0.96 | 0.95 | 0.92 | 0.94 |
| 3 | 72515841 | rs3943699 | 0.972 | -0.35 | 0.143 | -1.093 | -0.021 | 0.117 | 0.117 | 0.711 | 0.47 | 0.21 | 0.63 | 0.74 | 0.51 | 0.62 |
| 3 | 72515960 | rs11128275 | 0.985 | -0.373 | -0.086 | -1.107 | -0.025 | 0.142 | 0.142 | 0.572 | 0.47 | 0.16 | 0.62 | 0.73 | 0.5 | 0.6 |
| 3 | 72516056 | rs6549458 | -0.107 | 0.153 | 0.279 | -0.812 | -0.002 | -0.026 | -0.026 | 0 | 0.2 | 0.19 | 0.09 | 0.05 | 0.15 | 0.15 |
| 3 | 72516206 | rs934841 | 0.393 | 0.191 | -0.026 | -0.359 | -0.012 | 0.127 | 0.127 | 0 | 0.47 | 0.2 | 0.62 | 0.73 | 0.48 | 0.58 |
| 3 | 72530482 | rs2043813 | 0.945 | -0.68 | 0.027 | 1.001 | -0.001 | 0 | 0 | 0 | 0.67 | 0.33 | 0.7 | 0.61 | 0.76 | 0.7 |
| 3 | 72533768 | rs4677169 | 0.866 | -0.711 | 0.818 | 0.918 | -0.027 | 0.002 | 0.002 | 0 | 0.69 | 0.45 | 0.71 | 0.65 | 0.76 | 0.73 |
| 3 | 72534713 | rs1438318 | 0.733 | -0.854 | 1.594 | 0.66 | -0.036 | -0.069 | -0.069 | 0 | 0.78 | 0.65 | 0.87 | 0.89 | 0.85 | 0.89 |
| 3 | 72541705 | rs9831827 | 0.46 | -1.113 | 1.852 | 0.679 | -0.005 | -0.006 | -0.006 | 0 | 0.84 | 0.79 | 0.94 | 0.93 | 0.92 | 0.89 |
| 3 | 72544196 | rs10511022 | 1.856 | -1.771 | 1.839 | 0.552 | 0.058 | 0.068 | 0.068 | 0 | 0.64 | 0.21 | 0.74 | 0.76 | 0.78 | 0.76 |
| 3 | 72549515 | rs6794917 | 1.325 | -1.42 | 2.158 | 0.677 | 0.002 | 0.008 | 0.008 | 0 | 0.6 | 0.44 | 0.54 | 0.51 | 0.55 | 0.57 |
| 3 | 72550115 | rs1898634 | 0.922 | -1.445 | 1.964 | 0.649 | -0.007 | 0.009 | 0.009 | 0.091 | 0.67 | 0.53 | 0.69 | 0.67 | 0.64 | 0.65 |
| 3 | 72553512 | rs6768310 | 0.54 | -1.403 | 1.199 | 0.932 | 0.008 | 0.025 | 0.025 | 0 | 0.07 | 0.08 | 0.14 | 0.22 | 0.09 | 0.08 |
| 3 | 72554528 | rs9849164 | 1.95 | -1.452 | 1.048 | 0.906 | 0.017 | 0.024 | 0.024 | 0 | 0.5 | 0.15 | 0.49 | 0.44 | 0.51 | 0.53 |
| 3 | 72556982 | rs6549465 | 0.7 | -1.014 | 0.836 | 0.706 | -0.008 | -0.034 | -0.034 | 0.005 | 0.72 | 0.81 | 0.73 | 0.68 | 0.68 | 0.67 |
| 3 | 72558999 | rs4074154 | 0.124 | -1.683 | 1.056 | 0.63 | -0.002 | -0.002 | -0.002 | 0 | 0.07 | 0.03 | 0.14 | 0.15 | 0.09 | 0.09 |
| 3 | 72564523 | rs6776824 | 0.898 | -2.273 | 1.393 | 0.446 | -0.022 | -0.045 | -0.045 | 0 | 0.9 | 0.79 | 0.97 | 0.99 | 0.94 | 0.95 |
| 3 | 72568038 | rs9811139 | -1.684 | -2.45 | 1.615 | 0.373 | -0.031 | -0.073 | -0.073 | 0 | 0.15 | 0.29 | 0.08 | 0.01 | 0.1 | 0.07 |
| 3 | 72568369 | rs13061465 | -1.251 | -2.671 | 1.662 | 0.871 | -0.022 | 0.105 | 0.105 | 0 | 0.12 | 0.19 | 0.16 | 0.32 | 0.06 | 0.23 |
| 3 | 72570502 | rs6790541 | -1.483 | -3.053 | 1.858 | 0.367 | -0.089 | -0.115 | -0.115 | 2.352 | 0.2 | 0.42 | 0.08 | 0.01 | 0.08 | 0.09 |
| 3 | 72571732 | rs4677179 | 2.078 | -3.155 | 1.368 | 0.427 | 0.095 | -0.003 | -0.003 | 0 | 0.65 | 0.18 | 0.74 | 0.69 | 0.82 | 0.68 |

| | | | | | | | | | | | | | | | | |
|---|---|---|---|---|---|---|---|---|---|---|---|---|---|---|---|---|
| 3 | 72580844 | rs13080756 | 1.707 | -3.059 | 1.915 | 0.762 | -0.056 | 0.066 | 0.066 | 0 | 0.75 | 0.44 | 0.82 | 0.62 | 0.86 | 0.77 |
| 3 | 72582045 | rs6801930 | -1.049 | -3.096 | 1.951 | 0.659 | -0.023 | -0.031 | -0.031 | 0 | 0.1 | 0.25 | 0.05 | 0.04 | 0.05 | 0.08 |
| 3 | 72584333 | rs7610309 | 2.355 | -3.555 | 1.667 | 0.517 | 0.133 | -0.067 | -0.067 | 0 | 0.63 | 0.13 | 0.77 | 0.61 | 0.82 | 0.71 |
| 3 | 72586140 | rs11922009 | -1.392 | -3.538 | 2.234 | 0.567 | -0.013 | 0.217 | 0.217 | 0.007 | 0.12 | 0.34 | 0.12 | 0.39 | 0.07 | 0.21 |
| 3 | 72588855 | rs1472064 | 1.945 | -4.231 | 2.266 | 0.083 | -0.03 | 0.008 | 0.008 | 0.424 | 0.66 | 0.38 | 0.78 | 0.62 | 0.82 | 0.76 |
| 3 | 72590900 | rs13066103 | 2.462 | -4.269 | 1.892 | 0.045 | 0.17 | -0.061 | -0.061 | 0.351 | 0.54 | 0.06 | 0.72 | 0.6 | 0.78 | 0.7 |
| 3 | 72596598 | rs6786199 | 0.706 | -4.186 | 2.138 | -0.333 | -0.009 | 0.025 | 0.025 | 0 | 0.88 | 0.77 | 0.94 | 0.8 | 0.93 | 0.82 |
| 3 | 72597927 | rs7642544 | 1.741 | -4.291 | 2.357 | -0.359 | -0.051 | -0.004 | -0.004 | 0.031 | 0.73 | 0.59 | 0.86 | 0.75 | 0.84 | 0.76 |
| 3 | 72599203 | rs17009945 | 0.242 | -4.02 | 2.178 | -0.383 | -0.013 | -0.035 | -0.035 | 0 | 0.1 | 0.16 | 0.04 | 0 | 0.03 | 0 |
| 3 | 72615908 | rs6775154 | -1.071 | -4.114 | 2.534 | -0.381 | -0.036 | 0.036 | 0.036 | 0 | 0.21 | 0.36 | 0.1 | 0.28 | 0.12 | 0.23 |
| 3 | 72618072 | rs4677187 | 2.234 | -4.22 | 2.3 | -0.473 | 0.11 | 0.065 | 0.065 | 0 | 0.57 | 0.22 | 0.83 | 0.7 | 0.81 | 0.73 |
| 3 | 72622040 | rs6785907 | -0.9 | -4.185 | 2.258 | -0.497 | -0.001 | 0 | 0 | 0 | 0.2 | 0.22 | 0.09 | 0.28 | 0.13 | 0.24 |
| 3 | 72622061 | rs2029507 | 2.666 | -4.265 | 2.166 | -0.558 | 0.181 | 0.159 | 0.159 | 0.115 | 0.5 | 0.09 | 0.78 | 0.71 | 0.76 | 0.71 |
| 3 | 72622532 | rs11924047 | 0.41 | -4.192 | 2.355 | -0.535 | -0.027 | -0.055 | -0.055 | 0 | 0.1 | 0.25 | 0.04 | 0 | 0.03 | 0.01 |
| 3 | 72628207 | rs11711356 | 2.25 | -4.093 | 1.31 | -0.473 | 0.172 | 0.155 | 0.155 | 0.064 | 0.51 | 0.11 | 0.77 | 0.7 | 0.76 | 0.69 |
| 3 | 72629854 | rs2172011 | -0.949 | -3.662 | 1.306 | -0.206 | -0.002 | 0.002 | 0.002 | 0 | 0.19 | 0.21 | 0.09 | 0.31 | 0.13 | 0.26 |
| 3 | 72635104 | rs6778908 | -1.189 | -3.336 | 0.879 | -0.161 | -0.053 | -0.028 | -0.028 | 0 | 0.31 | 0.5 | 0.13 | 0.26 | 0.17 | 0.3 |
| 3 | 72638322 | rs2322483 | -1.482 | -3.208 | 0.086 | -0.22 | 0.145 | 0.142 | 0.142 | 0.117 | 0.49 | 0.8 | 0.23 | 0.28 | 0.25 | 0.31 |
| 3 | 72638400 | rs2874805 | -1.111 | -2.382 | 0.789 | 0.398 | -0.046 | 0.024 | 0.024 | 0 | 0.2 | 0.38 | 0.1 | 0.26 | 0.09 | 0.23 |
| 3 | 72639229 | rs6810278 | -1.096 | -1.77 | 0.65 | 0.773 | 0.013 | 0.183 | 0.183 | 0.58 | 0.15 | 0.44 | 0.12 | 0.34 | 0.13 | 0.18 |
| 3 | 72639769 | rs17677643 | -0.508 | -1.515 | -0.27 | -0.319 | 0.006 | 0.011 | 0.011 | 0.668 | 0.31 | 0.23 | 0.17 | 0.19 | 0.27 | 0.35 |
| 3 | 72640585 | rs9837318 | -0.695 | -1.667 | -0.356 | -0.561 | -0.003 | -0.042 | -0.042 | 0 | 0.29 | 0.32 | 0.23 | 0.05 | 0.23 | 0.24 |
| 3 | 72645502 | rs12486939 | -0.721 | -0.27 | -0.003 | -0.166 | 0.042 | -0.011 | -0.011 | 0.094 | 0.06 | 0.16 | 0.08 | 0.04 | 0.15 | 0.04 |
| 3 | 72648980 | rs10049135 | -0.579 | -0.482 | -0.171 | 0.221 | 0.005 | -0.006 | -0.006 | 0.344 | 0.21 | 0.2 | 0.21 | 0.35 | 0.13 | 0.27 |
| 3 | 72649326 | rs9310243 | -1.645 | -1.124 | 0.295 | 0.646 | -0.012 | 0.028 | 0.028 | 0.562 | 0.2 | 0.5 | 0.23 | 0.25 | 0.19 | 0.09 |
| 3 | 72649530 | rs9844133 | 1.328 | -1.074 | 0.202 | 0.481 | -0.009 | -0.012 | -0.012 | 0.395 | 0.05 | 0.05 | 0.02 | 0 | 0.03 | 0.02 |
| 3 | 72650801 | rs1873351 | -1.486 | -1.05 | 0.32 | 0.607 | -0.01 | 0.009 | 0.009 | 0.058 | 0.35 | 0.62 | 0.34 | 0.61 | 0.29 | 0.35 |
| 3 | 72655715 | rs13076653 | 0.608 | -0.731 | -0.614 | -0.06 | 0.038 | 0.143 | 0.143 | 0 | 0.3 | 0.08 | 0.12 | 0.05 | 0.24 | 0.28 |
| 3 | 72660579 | rs4676882 | -0.507 | -0.483 | -0.17 | 0.26 | 0.034 | 0.055 | 0.055 | 0 | 0.51 | 0.34 | 0.41 | 0.33 | 0.42 | 0.54 |
| 3 | 72663045 | rs8179915 | -0.149 | -0.343 | -0.083 | 0.337 | -0.001 | -0.016 | -0.016 | 0 | 0.19 | 0.14 | 0.3 | 0.28 | 0.18 | 0.27 |
| 3 | 72667611 | rs4677197 | -1.14 | -0.301 | 0.163 | -0.005 | 0.007 | -0.039 | -0.039 | 0 | 0.92 | 0.77 | 0.96 | 1 | 0.95 | 0.98 |
| 3 | 72668254 | rs902421 | -0.666 | 0.037 | 0.103 | 0.245 | 0.005 | 0.004 | 0.004 | 0 | 0.6 | 0.5 | 0.56 | 0.64 | 0.59 | 0.63 |
| 3 | 72675146 | rs7647258 | 0.384 | -0.566 | -0.024 | 0.421 | -0.01 | -0.004 | -0.004 | 0.017 | 0.43 | 0.47 | 0.25 | 0.37 | 0.29 | 0.36 |
| 3 | 72677091 | rs923576 | -0.006 | -0.33 | -0.228 | 0.606 | -0.002 | -0.028 | -0.028 | 0 | 0.18 | 0.1 | 0.29 | 0.26 | 0.15 | 0.26 |
| 3 | 72677572 | rs9809964 | 1.672 | 0.236 | -0.094 | 0.574 | -0.001 | -0.004 | -0.004 | 0 | 0.05 | 0.05 | 0.03 | 0 | 0.04 | 0.01 |
| 3 | 72682269 | rs7644890 | 0.136 | 0.164 | 0.033 | 0.825 | -0.007 | 0.059 | 0.059 | 0 | 0.05 | 0.1 | 0.06 | 0.27 | 0.04 | 0.08 |
| 3 | 72682319 | rs4677201 | -0.68 | 0.436 | -0.023 | 0.713 | 0.003 | 0.003 | 0.003 | 0 | 0.55 | 0.61 | 0.58 | 0.63 | 0.47 | 0.53 |
| 3 | 72683651 | rs10049353 | 0.521 | 0.534 | -0.535 | 0.354 | 0.02 | 0.03 | 0.03 | 0 | 0.35 | 0.29 | 0.26 | 0.08 | 0.3 | 0.28 |
| 3 | 72686090 | rs4676888 | -0.065 | 0.485 | -0.553 | 1.362 | 0.001 | 0.006 | 0.006 | 0 | 0.12 | 0.12 | 0.17 | 0.34 | 0.11 | 0.18 |
| 3 | 72696228 | rs13085034 | 0.739 | 0.632 | -0.375 | 1.065 | 0.01 | 0.098 | 0.098 | 0 | 0.76 | 0.48 | 0.69 | 0.37 | 0.77 | 0.75 |
| 3 | 72696627 | rs923573 | 0.342 | 1.262 | -0.798 | -0.036 | -0.087 | 0.163 | 0.163 | 0 | 0.36 | 0.15 | 0.26 | 0.01 | 0.52 | 0.24 |
| 3 | 72697011 | rs923572 | 0.52 | 1.295 | 0.182 | 1.34 | 0.063 | 0.166 | 0.166 | 0 | 0.92 | 0.68 | 0.88 | 0.75 | 0.91 | 0.89 |
| 3 | 72701660 | rs2046869 | -0.676 | 1.52 | -0.593 | 1.196 | 0 | 0 | 0 | 0.11 | 0.26 | 0.3 | 0.3 | 0.3 | 0.34 | 0.21 |
| 3 | 72704079 | rs4676891 | 0.355 | 0.552 | -1.148 | 1.125 | 0.04 | -0.044 | -0.044 | 1.489 | 0.25 | 0.07 | 0.21 | 0.34 | 0.12 | 0.35 |
| 3 | 72704836 | rs11708505 | -1.162 | 0.714 | -1.041 | 1.077 | 0 | 0.014 | 0.014 | 4.979 | 0.53 | 0.48 | 0.54 | 0.86 | 0.43 | 0.61 |
| 3 | 72705741 | rs7642116 | -1.358 | 0.971 | -1.047 | 0.726 | -0.005 | -0.078 | -0.078 | 1.847 | 0.59 | 0.59 | 0.67 | 0.96 | 0.61 | 0.63 |
| 3 | 72709390 | rs2322573 | -1.158 | 0.833 | -0.876 | 1.002 | -0.001 | -0.01 | -0.01 | 0.051 | 0.49 | 0.56 | 0.6 | 0.91 | 0.5 | 0.58 |
| 3 | 72709402 | rs2322574 | -0.645 | 1.198 | -1.204 | 1.384 | -0.026 | -0.011 | -0.011 | 0.024 | 0.38 | 0.17 | 0.43 | 0.59 | 0.36 | 0.49 |
| 3 | 72709792 | rs13084927 | -0.013 | 1.291 | -1.045 | 1.287 | -0.039 | -0.038 | -0.038 | 0 | 0.1 | 0.05 | 0.13 | 0.18 | 0.18 | 0.12 |
| 3 | 72723252 | rs1878770 | -0.41 | -0.029 | -0.641 | 1.106 | 0.013 | -0.01 | -0.01 | 0 | 0.37 | 0.62 | 0.16 | 0.55 | 0.22 | 0.46 |
| 3 | 72724329 | rs11926619 | -0.488 | -0.113 | -0.501 | 1.208 | 0.042 | -0.009 | -0.009 | 0 | 0.38 | 0.68 | 0.17 | 0.67 | 0.22 | 0.46 |
| 3 | 72725554 | rs9822901 | 0.098 | -0.35 | 0.529 | 1.719 | 0.049 | -0.006 | -0.006 | 0 | 0.29 | 0.13 | 0.08 | 0.3 | 0.09 | 0.37 |
| 3 | 72731997 | rs9878934 | -0.127 | -0.38 | 0.931 | 1.794 | 0.012 | -0.007 | -0.007 | 1.701 | 0.31 | 0.24 | 0.1 | 0.69 | 0.1 | 0.44 |
| 3 | 72733739 | rs6771281 | -0.477 | -0.482 | 1.072 | 1.641 | -0.04 | 0.034 | 0.034 | 0 | 0.35 | 0.48 | 0.15 | 0.55 | 0.22 | 0.47 |
| 3 | 72734318 | rs9814581 | -0.705 | -0.437 | 0.881 | 1.264 | 0.039 | 0.013 | 0.013 | 1.035 | 0.38 | 0.68 | 0.16 | 0.32 | 0.22 | 0.39 |
| 3 | 72736592 | rs1565471 | 0.165 | 0.283 | 0.284 | 0.997 | -0.054 | -0.065 | -0.065 | 0 | 0.13 | 0.04 | 0.43 | 0.37 | 0.26 | 0.16 |
| 3 | 72739192 | rs11705733 | -0.338 | -0.538 | 0.784 | 0.479 | -0.04 | 0.007 | 0.007 | 0 | 0.33 | 0.45 | 0.2 | 0.34 | 0.22 | 0.4 |
| 3 | 72747253 | rs9985487 | -0.191 | -0.627 | -0.134 | 0.192 | 0.04 | 0.009 | 0.009 | 0 | 0.34 | 0.12 | 0.2 | 0.28 | 0.24 | 0.33 |
| 3 | 72766902 | rs1490508 | -1.045 | -1.613 | 0.386 | -0.396 | -0.018 | -0.044 | -0.044 | 2.85 | 0.24 | 0.72 | 0.06 | 0 | 0.04 | 0.06 |
| 3 | 72781075 | rs2046867 | 0.183 | -0.598 | 1.144 | -0.548 | 0.023 | -0.038 | -0.038 | 0.031 | 0.17 | 0.26 | 0.16 | 0.04 | 0.25 | 0.16 |
| 3 | 72809591 | rs9811830 | -0.926 | -0.846 | 1.893 | -0.653 | -0.001 | -0.068 | -0.068 | 0 | 0.26 | 0.64 | 0.21 | 0.04 | 0.25 | 0.17 |
| 3 | 72812746 | rs7632726 | -1.574 | -0.766 | 1.697 | -0.187 | -0.03 | 0.009 | 0.009 | 2.597 | 0.32 | 0.92 | 0.29 | 0.26 | 0.29 | 0.19 |
| 3 | 72833889 | rs17010142 | 0.394 | -0.713 | 2.206 | -0.839 | -0.034 | -0.036 | -0.036 | 0 | 0.07 | 0.19 | 0.02 | 0 | 0.01 | 0 |
| 3 | 72863504 | rs3821561 | -1.299 | -0.681 | 1.506 | -1.103 | -0.103 | -0.046 | -0.046 | 0.076 | 0.67 | 0.98 | 0.77 | 0.7 | 0.62 | 0.56 |
| 3 | 72865126 | rs1502758 | 0.917 | -0.652 | 1.808 | -1.132 | 0.019 | -0.032 | -0.032 | 0 | 0.15 | 0.21 | 0.15 | 0.04 | 0.23 | 0.13 |
| 3 | 72868028 | rs11924329 | 0.811 | -0.656 | 1.857 | -1.134 | -0.036 | -0.038 | -0.038 | 0 | 0.07 | 0.18 | 0.02 | 0 | 0.01 | 0 |
| 3 | 72868234 | rs11929144 | -0.756 | -0.691 | 2.007 | -1.149 | 0.006 | 0.025 | 0.025 | 3.924 | 0.27 | 0.77 | 0.21 | 0.04 | 0.24 | 0.13 |
| 3 | 72879765 | rs17010207 | 0.81 | -0.647 | 2.15 | -1.035 | -0.035 | -0.037 | -0.037 | 0.846 | 0.07 | 0.18 | 0.03 | 0 | 0.01 | 0.01 |
| 3 | 72880470 | rs17043990 | 1.264 | -0.639 | 1.967 | -1.032 | -0.031 | -0.032 | -0.032 | 0 | 0.06 | 0.11 | 0.02 | 0 | 0.01 | 0 |

| | | | | | | | | | | | | | | | | |
|---|---|---|---|---|---|---|---|---|---|---|---|---|---|---|---|---|
| 3 | 72892988 | rs4676894 | 0.316 | -0.627 | 1.771 | -1.033 | 0.006 | 0.014 | 0.014 | 0.47 | 0.42 | 0.33 | 0.56 | 0.66 | 0.38 | 0.41 |
| 3 | 72898844 | rs1875624 | -0.201 | -0.072 | 1.749 | -0.744 | 0.038 | 0.048 | 0.048 | 0.161 | 0.52 | 0.73 | 0.6 | 0.65 | 0.38 | 0.42 |
| 3 | 72899742 | rs3755715 | 0.371 | -0.068 | 1.93 | -0.744 | 0.011 | 0.008 | 0.008 | 0.209 | 0.44 | 0.37 | 0.54 | 0.65 | 0.34 | 0.38 |
| 3 | 72915315 | rs7617996 | -1.489 | 0.559 | 2.784 | -0.697 | -0.006 | 0.006 | 0.006 | 0.139 | 0.25 | 0.77 | 0.22 | 0.05 | 0.28 | 0.18 |
| 3 | 72922181 | rs13079651 | -0.816 | 0.614 | 2.459 | -0.764 | 0.06 | -0.118 | -0.118 | 1.242 | 0.23 | 0.46 | 0.26 | 0.04 | 0.33 | 0.18 |
| 3 | 72925997 | rs7640629 | -0.592 | 0.633 | 1.829 | -0.755 | -0.008 | 0 | 0 | 0 | 0.22 | 0.19 | 0.23 | 0.04 | 0.33 | 0.18 |
| 3 | 72933767 | rs6805260 | -0.405 | 0.762 | 1.448 | -0.869 | -0.04 | 0.047 | 0.047 | 0 | 0.2 | 0.11 | 0.25 | 0.01 | 0.33 | 0.12 |
| 3 | 72939767 | rs6807627 | -0.928 | 0.754 | 0.666 | -0.797 | 0.03 | -0.129 | -0.129 | 0 | 0.27 | 0.59 | 0.31 | 0.01 | 0.36 | 0.15 |
| 3 | 72956898 | rs7643654 | -1.306 | 0.171 | 0.871 | -0.358 | 0.002 | -0.075 | -0.075 | 0 | 0.13 | 0.32 | 0.15 | 0 | 0.11 | 0.11 |
| 3 | 72956987 | rs7613392 | -1.644 | 0.035 | 1.571 | -0.418 | 0 | -0.105 | -0.105 | 0 | 0.15 | 0.49 | 0.17 | 0 | 0.11 | 0.11 |
| 3 | 72963769 | rs7616069 | -2.059 | -0.064 | 1.374 | -0.461 | 0.009 | 0.012 | 0.012 | 1.665 | 0.21 | 0.79 | 0.2 | 0.02 | 0.14 | 0.13 |
| 3 | 72973732 | rs7640661 | -2.094 | 0.024 | 1.73 | 0.609 | -0.006 | 0.016 | 0.016 | 0.997 | 0.18 | 0.76 | 0.31 | 0.29 | 0.13 | 0.19 |
| 3 | 72982575 | rs2322693 | 0.602 | -0.551 | 1.429 | 0.304 | 0.046 | 0.006 | 0.006 | 0 | 0.52 | 0.29 | 0.65 | 0.57 | 0.6 | 0.54 |
| 3 | 72983866 | rs884766 | 0.648 | -0.599 | 1.346 | 0.282 | 0.043 | 0.009 | 0.009 | 0.096 | 0.51 | 0.31 | 0.65 | 0.57 | 0.6 | 0.53 |
| 3 | 72994544 | rs4677212 | 0.545 | -0.838 | 2.262 | 0.307 | 0.001 | 0 | 0 | 0 | 0.52 | 0.46 | 0.68 | 0.57 | 0.62 | 0.53 |
| 3 | 72994594 | rs4677213 | -0.543 | -1.335 | 1.925 | 0.324 | 0.012 | 0.043 | 0.043 | 0 | 0.74 | 0.77 | 0.87 | 0.92 | 0.76 | 0.89 |
| 3 | 72998456 | rs7617057 | 1.201 | -2.143 | 1.186 | -0.03 | 0.021 | -0.035 | -0.035 | 4.001 | 0.16 | 0.09 | 0.27 | 0.42 | 0.06 | 0.13 |
| 3 | 73000499 | rs6785758 | -0.263 | -1.983 | 1.581 | 0.069 | -0.019 | 0.064 | 0.064 | 0.656 | 0.23 | 0.35 | 0.21 | 0.35 | 0.17 | 0.36 |
| 3 | 73000767 | rs6762394 | -0.021 | -2.017 | 1.713 | 0.267 | -0.03 | 0.062 | 0.062 | 0.418 | 0.19 | 0.29 | 0.09 | 0.3 | 0.1 | 0.28 |
| 3 | 73004057 | rs4455270 | 0.151 | -2.351 | 1.556 | 0.143 | -0.011 | 0.001 | 0.001 | 0.008 | 0.56 | 0.44 | 0.59 | 0.51 | 0.67 | 0.75 |
| 3 | 73008718 | rs13099810 | 1.237 | -2.353 | 1.082 | 0.12 | -0.029 | 0.093 | 0.093 | 0.064 | 0.39 | 0.12 | 0.52 | 0.2 | 0.61 | 0.47 |
| 3 | 73010237 | rs13080133 | 1.955 | -2.406 | 1.021 | 0.096 | -0.151 | 0.07 | 0.07 | 0.227 | 0.32 | 0.04 | 0.4 | 0.14 | 0.54 | 0.37 |
| 3 | 73010544 | rs4635643 | -1.237 | -2.375 | 1.014 | 0.094 | -0.052 | 0.111 | 0.111 | 0 | 0.74 | 0.98 | 0.86 | 0.92 | 0.73 | 0.88 |
| 3 | 73011532 | rs4585135 | 1.21 | -2.007 | 1.311 | 0.238 | 0.009 | -0.007 | -0.007 | 0 | 0.13 | 0.09 | 0.23 | 0.42 | 0.06 | 0.13 |
| 3 | 73012077 | rs4677214 | -0.224 | -1.974 | 1.384 | 0.259 | 0.005 | 0.002 | 0.002 | 0 | 0.56 | 0.57 | 0.78 | 0.61 | 0.67 | 0.61 |
| 3 | 73016974 | rs3821379 | -0.116 | -1.962 | 1.277 | 0.269 | -0.008 | -0.003 | -0.003 | 0 | 0.55 | 0.48 | 0.78 | 0.61 | 0.66 | 0.6 |
| 3 | 73022475 | rs9829843 | 0.458 | -1.922 | 0.792 | 0.381 | -0.056 | 0.189 | 0.189 | 1.256 | 0.17 | 0.48 | 0.27 | 0.38 | 0.06 | 0.12 |
| 3 | 73024350 | rs1052278 | 0.064 | -1.859 | 0.748 | 0.31 | 0.004 | 0.001 | 0.001 | 0.067 | 0.52 | 0.54 | 0.67 | 0.53 | 0.59 | 0.49 |
| 3 | 73036622 | rs6789471 | -0.703 | -1.651 | 0.918 | 0.617 | -0.035 | 0.11 | 0.11 | 0.236 | 0.21 | 0.54 | 0.38 | 0.44 | 0.13 | 0.24 |
| 3 | 73050334 | rs17010333 | -0.387 | -2.063 | 0.368 | 0.332 | 0.023 | -0.019 | -0.019 | 0 | 0.26 | 0.17 | 0.09 | 0.35 | 0.11 | 0.33 |
| 3 | 73052765 | rs4677225 | 0.391 | -1.49 | 0.362 | -0.343 | 0.052 | -0.089 | -0.089 | 0 | 0.29 | 0.42 | 0.27 | 0.1 | 0.39 | 0.29 |
| 3 | 73088864 | rs9870408 | -1.188 | -1.548 | 0.796 | -0.326 | -0.002 | -0.001 | -0.001 | 0.501 | 0.54 | 0.62 | 0.36 | 0.45 | 0.5 | 0.59 |
| 3 | 73090462 | rs4677234 | -1.192 | -1.554 | 0.797 | -0.329 | -0.001 | 0 | 0 | 0 | 0.54 | 0.62 | 0.36 | 0.45 | 0.5 | 0.59 |
| 3 | 73091447 | rs12638019 | 0.207 | -1.502 | 0.801 | -0.381 | 0.027 | -0.024 | -0.024 | 0 | 0.26 | 0.15 | 0.09 | 0.35 | 0.12 | 0.33 |
| 3 | 73091837 | rs9869215 | -0.372 | -1.75 | 0.688 | -0.387 | 0.002 | -0.006 | -0.006 | 0 | 0.45 | 0.46 | 0.31 | 0.38 | 0.45 | 0.57 |
| 3 | 73092941 | rs7614412 | -1.554 | -1.707 | 0.267 | -0.47 | -0.015 | -0.026 | -0.026 | 0 | 0.57 | 0.76 | 0.39 | 0.45 | 0.51 | 0.63 |
| 3 | 73102013 | rs17010449 | -0.019 | -1.717 | 0.35 | -0.463 | 0.036 | -0.034 | -0.034 | 0 | 0.24 | 0.12 | 0.08 | 0.35 | 0.11 | 0.33 |
| 3 | 73102725 | rs9824246 | 0.104 | -1.766 | 0.391 | -0.53 | -0.012 | -0.03 | -0.03 | 0 | 0.09 | 0.16 | 0.1 | 0.02 | 0.05 | 0.05 |
| 3 | 73108092 | rs4113632 | -0.101 | -1.767 | 0.215 | -0.519 | 0.032 | -0.035 | -0.035 | 0 | 0.25 | 0.12 | 0.09 | 0.35 | 0.12 | 0.32 |
| 3 | 73109891 | rs2291527 | 0.674 | -1.722 | 0.305 | -0.465 | -0.034 | -0.004 | -0.004 | 0 | 0.1 | 0.19 | 0.05 | 0.08 | 0.06 | 0.04 |
| 3 | 73111973 | rs2231928 | -1.437 | -1.457 | 0.228 | -0.258 | -0.057 | -0.092 | -0.092 | 0 | 0.59 | 0.89 | 0.4 | 0.49 | 0.51 | 0.63 |
| 3 | 73111998 | rs1060584 | -0.923 | -1.413 | 0.211 | -0.288 | 0.004 | -0.002 | -0.002 | 0 | 0.55 | 0.62 | 0.37 | 0.45 | 0.51 | 0.61 |
| 3 | 73131351 | rs902652 | -0.481 | -1.236 | 0.21 | -0.024 | 0.008 | 0.046 | 0.046 | 0 | 0.15 | 0.25 | 0.32 | 0.31 | 0.15 | 0.2 |
| 3 | 73137972 | rs6792265 | -0.959 | -1.433 | -0.047 | -0.194 | 0.01 | 0.01 | 0.01 | 0 | 0.72 | 0.66 | 0.61 | 0.68 | 0.66 | 0.73 |
| 3 | 73139477 | rs2839770 | 0.194 | -0.897 | -0.006 | -0.341 | 0.021 | -0.105 | -0.105 | 0 | 0.34 | 0.49 | 0.27 | 0.09 | 0.4 | 0.27 |
| 3 | 73144090 | rs2630615 | -0.791 | -0.893 | 0.492 | -0.04 | -0.026 | 0.018 | 0.018 | 0 | 0.46 | 0.83 | 0.58 | 0.42 | 0.5 | 0.47 |
| 3 | 73146038 | rs2637748 | 0.106 | -0.743 | 0.574 | 0.004 | -0.004 | 0.002 | 0.002 | 0 | 0.13 | 0.11 | 0.29 | 0.33 | 0.15 | 0.2 |
| 3 | 73151050 | rs7636827 | 0.823 | -0.749 | 0.455 | -0.118 | -0.004 | -0.008 | -0.008 | 0.39 | 0.26 | 0.22 | 0.18 | 0.07 | 0.35 | 0.24 |
| 3 | 73152124 | rs880547 | 1.068 | -0.986 | 0.241 | -0.024 | 0.027 | -0.021 | -0.021 | 0.636 | 0.27 | 0.16 | 0.12 | 0.35 | 0.14 | 0.29 |
| 3 | 73156334 | rs6809020 | 0.768 | -0.611 | 0.172 | -0.042 | -0.011 | 0.016 | 0.016 | 0 | 0.26 | 0.21 | 0.24 | 0.16 | 0.34 | 0.25 |
| 3 | 73157190 | rs172526 | -1.161 | -0.469 | 0.104 | 0.084 | -0.028 | 0.005 | 0.005 | 0.167 | 0.71 | 0.8 | 0.65 | 0.76 | 0.63 | 0.76 |
| 3 | 73157288 | rs6766673 | -0.065 | -0.465 | 0.184 | 0.185 | 0 | 0 | 0 | 0.24 | 0.55 | 0.48 | 0.36 | 0.49 | 0.47 | 0.56 |
| 3 | 73159908 | rs301539 | -0.182 | -0.524 | 0.259 | -0.071 | -0.009 | -0.009 | -0.009 | 0 | 0.38 | 0.47 | 0.46 | 0.67 | 0.3 | 0.52 |
| 3 | 73167031 | rs1552688 | 1.015 | -0.351 | 0.13 | 0.13 | 0.017 | -0.045 | -0.045 | 0 | 0.25 | 0.18 | 0.25 | 0.49 | 0.14 | 0.33 |
| 3 | 73168351 | rs11916160 | 0.546 | -0.067 | 0.29 | -0.095 | 0.013 | -0.008 | -0.008 | 0 | 0.1 | 0.17 | 0.13 | 0.09 | 0.2 | 0.06 |
| 3 | 73182547 | rs9647401 | -0.096 | -0.403 | 0.426 | -0.144 | 0.009 | 0 | 0 | 0.261 | 0.77 | 0.47 | 0.64 | 0.76 | 0.78 | 0.81 |
| 3 | 73183673 | rs6549508 | 0.632 | -0.375 | 0.383 | -0.197 | -0.003 | -0.09 | -0.09 | 0 | 0.18 | 0.36 | 0.19 | 0.05 | 0.17 | 0.16 |
| 3 | 73185026 | rs2003425 | 1.142 | -0.6 | -0.498 | -0.339 | 0.011 | 0.041 | 0.041 | 0 | 0.16 | 0.04 | 0.1 | 0.08 | 0.13 | 0.25 |
| 3 | 73187203 | rs6774835 | -0.275 | -0.319 | -0.308 | -0.149 | -0.006 | 0.03 | 0.03 | 0 | 0.4 | 0.67 | 0.42 | 0.21 | 0.48 | 0.47 |
| 3 | 73191526 | rs7617496 | 1.675 | -0.247 | -0.935 | -0.252 | -0.112 | -0.065 | -0.065 | 0 | 0.26 | 0.03 | 0.3 | 0.29 | 0.35 | 0.18 |
| 3 | 73193620 | rs17010557 | 1.634 | -0.208 | -0.889 | -0.191 | 0.056 | -0.08 | -0.08 | 0 | 0.22 | 0.06 | 0.08 | 0.32 | 0.09 | 0.32 |
| 3 | 73195688 | rs13074421 | 1.629 | -0.205 | -0.884 | -0.19 | 0.054 | -0.07 | -0.07 | 0 | 0.22 | 0.07 | 0.11 | 0.32 | 0.1 | 0.32 |
| 3 | 73209521 | rs2322847 | 1.62 | -0.148 | -0.846 | -0.993 | -0.115 | -0.067 | -0.067 | 0 | 0.26 | 0.03 | 0.3 | 0.28 | 0.35 | 0.19 |
| 3 | 73220728 | rs6775461 | -0.584 | 0.753 | -0.255 | -0.607 | 0.004 | -0.035 | -0.035 | 0.075 | 0.39 | 0.45 | 0.4 | 0.21 | 0.49 | 0.46 |
| 3 | 73222581 | rs9846109 | 1.012 | 0.555 | -0.46 | -0.515 | 0.025 | -0.067 | -0.067 | 0 | 0.25 | 0.14 | 0.24 | 0.51 | 0.14 | 0.36 |
| 3 | 73229019 | rs17694568 | 0.048 | 0.583 | -0.505 | -0.704 | 0 | 0.04 | 0.04 | 0 | 0.3 | 0.2 | 0.27 | 0.11 | 0.32 | 0.38 |
| 3 | 73232449 | rs9870859 | 0.887 | 0.313 | -0.619 | -0.465 | 0.005 | -0.007 | -0.007 | 0 | 0.26 | 0.24 | 0.27 | 0.54 | 0.15 | 0.42 |
| 3 | 73235473 | rs6772476 | -1.126 | 1.124 | -0.442 | -0.659 | 0.008 | -0.07 | -0.07 | 0 | 0.4 | 0.52 | 0.39 | 0.19 | 0.49 | 0.41 |
| 3 | 73238693 | rs11719409 | -0.71 | 0.785 | -0.779 | 0.135 | -0.001 | -0.01 | -0.01 | 0.047 | 0.13 | 0.11 | 0.15 | 0.03 | 0.14 | 0.14 |

| | | | | | | | | | | | | | | | | |
|---|---|---|---|---|---|---|---|---|---|---|---|---|---|---|---|---|
| 3 | 73241653 | rs6775039 | -0.098 | 0.962 | -0.78 | 0.356 | -0.001 | 0.002 | 0.002 | 0.239 | 0.51 | 0.5 | 0.62 | 0.64 | 0.47 | 0.59 |
| 3 | 73242344 | rs1488380 | 0.551 | 1.596 | -0.507 | 0.569 | 0.01 | 0.005 | 0.005 | 0 | 0.9 | 0.84 | 0.81 | 0.9 | 0.82 | 0.85 |
| 3 | 73247430 | rs13074807 | 0.399 | 1.614 | -0.374 | 0.474 | -0.008 | 0.011 | 0.011 | 0.031 | 0.54 | 0.33 | 0.5 | 0.6 | 0.47 | 0.54 |
| 3 | 73248855 | rs11128316 | 0.322 | 1.581 | 0.502 | 0.412 | -0.002 | 0.001 | 0.001 | 0.007 | 0.55 | 0.56 | 0.52 | 0.6 | 0.47 | 0.53 |
| 3 | 73252662 | rs9826629 | -0.288 | 1.423 | 0.799 | 0.377 | -0.039 | -0.003 | -0.003 | 0.123 | 0.41 | 0.21 | 0.43 | 0.36 | 0.52 | 0.44 |
| 3 | 73255057 | rs983775 | 0.564 | 1.461 | 0.732 | 0.599 | 0 | 0 | 0 | 0 | 0.53 | 0.54 | 0.5 | 0.55 | 0.45 | 0.49 |
| 3 | 73258312 | rs2035580 | 0.697 | 2.113 | 0.802 | 0.454 | 0.002 | 0.001 | 0.001 | 0.281 | 0.51 | 0.5 | 0.47 | 0.58 | 0.37 | 0.5 |
| 3 | 73261224 | rs17751397 | -0.466 | 2.176 | 0.357 | 0.249 | -0.028 | 0.014 | 0.014 | 1.645 | 0.44 | 0.25 | 0.42 | 0.36 | 0.51 | 0.49 |
| 3 | 73262379 | rs17010608 | -0.314 | 1.83 | 0.417 | 0.333 | 0.031 | -0.013 | -0.013 | 0.022 | 0.42 | 0.22 | 0.34 | 0.39 | 0.34 | 0.47 |
| 3 | 73264785 | rs7618585 | -0.713 | 1.611 | -0.196 | 0.341 | -0.037 | -0.032 | -0.032 | 1.48 | 0.59 | 0.22 | 0.43 | 0.5 | 0.52 | 0.59 |
| 3 | 73274994 | rs4676908 | 0.592 | 1.74 | 0.383 | 0.513 | 0.01 | 0.005 | 0.005 | 2.349 | 0.42 | 0.47 | 0.52 | 0.44 | 0.49 | 0.42 |
| 3 | 73276952 | rs12486903 | 1.084 | 1.673 | 0.577 | 0.658 | -0.034 | -0.037 | -0.037 | 0 | 0.36 | 0.16 | 0.47 | 0.43 | 0.43 | 0.39 |
| 3 | 73279050 | rs9829217 | 0.863 | 1.427 | 0.17 | 0.777 | -0.054 | 0.007 | 0.007 | 0.53 | 0.31 | 0.02 | 0.26 | 0.23 | 0.31 | 0.25 |
| 3 | 73281120 | rs7633357 | 0.373 | 1.275 | 0.419 | 0.778 | -0.019 | 0.018 | 0.018 | 0 | 0.73 | 0.63 | 0.78 | 0.66 | 0.79 | 0.65 |
| 3 | 73291914 | rs616508 | 1.033 | 1.314 | 0.438 | 0.942 | -0.004 | 0.13 | 0.13 | 0 | 0.91 | 0.69 | 0.88 | 0.78 | 0.94 | 0.76 |
| 3 | 73293512 | rs9880113 | 0.586 | 1.507 | 0.018 | 0.918 | -0.064 | -0.006 | -0.006 | 0 | 0.3 | 0.02 | 0.25 | 0.23 | 0.31 | 0.22 |
| 3 | 73297035 | rs649822 | 1.143 | 1.488 | 0.278 | 0.813 | -0.017 | 0.109 | 0.109 | 0.761 | 0.9 | 0.67 | 0.88 | 0.79 | 0.94 | 0.77 |
| 3 | 73298314 | rs12635520 | -0.694 | 1.646 | -0.25 | 0.857 | 0.004 | 0.024 | 0.024 | 0 | 0.08 | 0.1 | 0.11 | 0.16 | 0.09 | 0.26 |
| 3 | 73298666 | rs1873009 | 0.433 | 1.35 | -0.456 | 0.848 | -0.065 | 0.03 | 0.03 | 0.388 | 0.3 | 0.02 | 0.25 | 0.17 | 0.3 | 0.22 |
| 3 | 73303979 | rs12633844 | -0.198 | 1.165 | 0.101 | 1.07 | -0.005 | 0.064 | 0.064 | 0 | 0.05 | 0.13 | 0.07 | 0.12 | 0.02 | 0.22 |
| 3 | 73304713 | rs11713919 | 0.459 | 1.295 | -0.491 | 1.067 | -0.093 | -0.043 | -0.043 | 3.002 | 0.27 | 0.03 | 0.29 | 0.17 | 0.31 | 0.23 |
| 3 | 73304827 | rs11711247 | 0.404 | 1.262 | -0.506 | 1.207 | -0.092 | -0.027 | -0.027 | 0 | 0.27 | 0.02 | 0.28 | 0.16 | 0.31 | 0.23 |
| 3 | 73305809 | rs574715 | -0.924 | 0.763 | -0.771 | 1.184 | -0.003 | 0.017 | 0.017 | 3.491 | 0.56 | 0.73 | 0.48 | 0.51 | 0.52 | 0.58 |
| 3 | 73308948 | rs671986 | -1.15 | 0.733 | -0.903 | 1.112 | -0.049 | 0.01 | 0.01 | 5.615 | 0.58 | 0.89 | 0.5 | 0.55 | 0.52 | 0.56 |
| 3 | 73309708 | rs814232 | -0.439 | 0.495 | 0.417 | 0.959 | -0.014 | -0.06 | -0.06 | 0.601 | 0.24 | 0.43 | 0.14 | 0.08 | 0.19 | 0.14 |
| 3 | 73314679 | rs506713 | -0.22 | -0.077 | 0.967 | 1.094 | -0.027 | -0.076 | -0.076 | 0 | 0.19 | 0.38 | 0.1 | 0.05 | 0.12 | 0.07 |
| 3 | 73326446 | rs6784750 | -0.114 | 0.051 | 0.915 | 1.424 | -0.016 | 0.062 | 0.062 | 1.346 | 0.2 | 0.08 | 0.18 | 0.06 | 0.2 | 0.08 |
| 3 | 73327602 | rs1482176 | -0.784 | 0.173 | 0.948 | 1.47 | -0.014 | 0.042 | 0.042 | 4.806 | 0.42 | 0.27 | 0.43 | 0.25 | 0.4 | 0.29 |
| 3 | 73331002 | rs791443 | -0.989 | 0.112 | 1.309 | 1.496 | -0.005 | -0.005 | -0.005 | 0 | 0.47 | 0.46 | 0.52 | 0.47 | 0.48 | 0.45 |
| 3 | 73331242 | rs616708 | -1.781 | 0.103 | 1.31 | 1.505 | 0.013 | 0.129 | 0.129 | 0.059 | 0.07 | 0.31 | 0.11 | 0.22 | 0.08 | 0.16 |
| 3 | 73332590 | rs590203 | -0.793 | -0.194 | 1.113 | 1.447 | -0.004 | 0.037 | 0.037 | 5.134 | 0.45 | 0.33 | 0.45 | 0.29 | 0.43 | 0.29 |
| 3 | 73333240 | rs1482172 | 0.895 | -0.114 | 0.737 | 1.389 | -0.016 | -0.014 | -0.014 | 1.937 | 0.21 | 0.09 | 0.23 | 0.18 | 0.19 | 0.2 |
| 3 | 73335374 | rs664786 | -0.838 | -0.13 | 0.986 | 1.475 | -0.009 | -0.007 | -0.007 | 0.006 | 0.45 | 0.35 | 0.51 | 0.45 | 0.47 | 0.45 |
| 3 | 73336315 | rs650141 | -1.187 | -0.157 | 1.295 | 1.45 | -0.001 | -0.001 | -0.001 | 0.307 | 0.49 | 0.61 | 0.52 | 0.46 | 0.47 | 0.46 |
| 3 | 73337297 | rs2170474 | -0.217 | -0.248 | 0.775 | 1.418 | 0.007 | 0.054 | 0.054 | 0 | 0.76 | 0.66 | 0.59 | 0.6 | 0.75 | 0.63 |
| 3 | 73341692 | rs922788 | 0.857 | -0.159 | 0.819 | 1.437 | -0.008 | -0.007 | -0.007 | 0 | 0.54 | 0.64 | 0.41 | 0.51 | 0.53 | 0.55 |
| 3 | 73344631 | rs6770138 | 0.922 | -0.146 | 0.857 | 1.265 | -0.005 | 0.014 | 0.014 | 3.122 | 0.54 | 0.62 | 0.41 | 0.41 | 0.53 | 0.53 |
| 3 | 73346745 | rs10212501 | 0.175 | -0.256 | 0.77 | 1.275 | 0 | 0.008 | 0.008 | 0 | 0.23 | 0.21 | 0.36 | 0.48 | 0.21 | 0.37 |
| 3 | 73347207 | rs17701256 | 0.641 | -0.236 | 0.734 | 1.287 | -0.044 | -0.041 | -0.041 | 0 | 0.21 | 0.02 | 0.28 | 0.19 | 0.19 | 0.24 |
| 3 | 73347416 | rs17752247 | 0.672 | -0.224 | 0.735 | 1.285 | -0.048 | -0.016 | -0.016 | 0 | 0.2 | 0.02 | 0.2 | 0.15 | 0.19 | 0.24 |
| 3 | 73348014 | rs6549526 | 0.16 | -0.262 | 0.919 | 1.173 | -0.009 | -0.07 | -0.07 | 0 | 0.75 | 0.61 | 0.77 | 0.92 | 0.76 | 0.91 |
| 3 | 73351986 | rs7612743 | 0.105 | -0.182 | 1.127 | 0.585 | -0.004 | 0.074 | 0.074 | 0 | 0.73 | 0.59 | 0.6 | 0.49 | 0.76 | 0.63 |
| 3 | 73362826 | rs7630253 | -0.106 | -0.012 | 1.07 | 0.575 | 0.003 | -0.065 | -0.065 | 0.09 | 0.2 | 0.31 | 0.2 | 0.05 | 0.2 | 0.06 |
| 3 | 73363151 | rs7619189 | -0.22 | -0.11 | 0.796 | 0.449 | 0 | 0.059 | 0.059 | 0 | 0.29 | 0.44 | 0.39 | 0.48 | 0.26 | 0.4 |
| 3 | 73363208 | rs1482171 | 0.846 | -0.044 | 0.644 | 0.622 | -0.005 | 0.031 | 0.031 | 2.678 | 0.49 | 0.33 | 0.4 | 0.41 | 0.52 | 0.52 |
| 3 | 73373092 | rs1905091 | -0.341 | 0.265 | 0.024 | 0.78 | -0.009 | 0.012 | 0.012 | 0.193 | 0.41 | 0.28 | 0.53 | 0.57 | 0.41 | 0.48 |
| 3 | 73393147 | rs10865676 | 0.215 | 0.405 | 0.505 | 0.961 | -0.01 | 0.005 | 0.005 | 0 | 0.61 | 0.53 | 0.55 | 0.44 | 0.65 | 0.6 |
| 3 | 73394611 | rs3901920 | 0.551 | 0.696 | -0.24 | 0.587 | 0.015 | 0.141 | 0.141 | 0.033 | 0.53 | 0.24 | 0.4 | 0.21 | 0.56 | 0.42 |
| 3 | 73398644 | rs13319277 | -1.228 | 0.575 | 0.432 | 1.199 | 0.01 | 0.243 | 0.243 | 0.947 | 0.1 | 0.29 | 0.25 | 0.35 | 0.07 | 0.16 |
| 3 | 73398839 | rs11710982 | 0.122 | 0.731 | 0.248 | 0.867 | -0.011 | -0.001 | -0.001 | 0 | 0.43 | 0.48 | 0.37 | 0.45 | 0.37 | 0.42 |
| 3 | 73398901 | rs11711792 | -0.667 | 0.801 | 0.188 | 0.874 | 0.01 | 0.151 | 0.151 | 0.928 | 0.54 | 0.77 | 0.62 | 0.8 | 0.45 | 0.58 |
| 3 | 73399384 | rs13070556 | 0.093 | 0.842 | 0.338 | 0.915 | -0.004 | -0.001 | -0.001 | 0 | 0.44 | 0.44 | 0.36 | 0.45 | 0.36 | 0.42 |
| 3 | 73400018 | rs13319638 | -1.046 | 0.871 | 0.06 | 0.915 | 0.015 | 0.28 | 0.28 | 2.087 | 0.1 | 0.29 | 0.25 | 0.37 | 0.08 | 0.16 |
| 3 | 73405263 | rs9866906 | -0.507 | 1.103 | -0.759 | 0.792 | -0.003 | 0.153 | 0.153 | 1.015 | 0.55 | 0.82 | 0.63 | 0.79 | 0.44 | 0.57 |
| 3 | 73411670 | rs11128329 | -0.397 | 1.115 | -0.896 | 0.775 | 0.011 | 0.074 | 0.074 | 0 | 0.07 | 0.08 | 0.19 | 0.34 | 0.15 | 0.16 |
| 3 | 73411687 | rs3912601 | 0.235 | 0.942 | -0.876 | 0.554 | -0.016 | 0.004 | 0.004 | 0.67 | 0.43 | 0.4 | 0.34 | 0.53 | 0.32 | 0.45 |
| 3 | 73412599 | rs3905755 | -0.169 | 0.956 | -1.859 | -0.347 | -0.088 | 0.241 | 0.241 | 1.69 | 0.55 | 0.9 | 0.68 | 0.89 | 0.5 | 0.62 |
| 3 | 73417094 | rs3843347 | 0.185 | 0.746 | -1.532 | -0.213 | 0.004 | 0.033 | 0.033 | 0.194 | 0.72 | 0.81 | 0.8 | 0.82 | 0.72 | 0.57 |
| 3 | 73425846 | rs6777921 | -0.189 | -0.685 | -1.043 | 0.059 | -0.084 | 0.048 | 0.048 | 0.407 | 0.18 | 0.5 | 0.15 | 0.21 | 0.06 | 0.14 |
| 3 | 73427609 | rs3845868 | -0.201 | -0.522 | -1.689 | -0.112 | 0.007 | 0.042 | 0.042 | 0 | 0.66 | 0.79 | 0.7 | 0.75 | 0.66 | 0.54 |
| 3 | 73430237 | rs11128332 | 0.612 | 0.108 | -2.465 | -0.201 | -0.092 | -0.083 | -0.083 | 0.145 | 0.41 | 0.07 | 0.46 | 0.57 | 0.47 | 0.29 |
| 3 | 73431830 | rs7064 | -0.238 | 0.048 | -1.312 | -0.252 | 0.001 | -0.01 | -0.01 | 0.928 | 0.54 | 0.5 | 0.61 | 0.67 | 0.5 | 0.43 |
| 3 | 73434086 | rs6787588 | -1.321 | 0.764 | -1.096 | -0.334 | 0.017 | 0.002 | 0.002 | 0.627 | 0.33 | 0.43 | 0.31 | 0.31 | 0.31 | 0.32 |
| 3 | 73435210 | rs3845869 | -1.051 | 0.891 | -1.047 | -0.615 | 0.003 | 0 | 0 | 5.843 | 0.21 | 0.73 | 0.23 | 0.19 | 0.18 | 0.24 |
| 3 | 73437112 | rs6549532 | -1.09 | 1.027 | -1.285 | -1.17 | -0.058 | 0.073 | 0.073 | 0 | 0.75 | 0.93 | 0.74 | 0.86 | 0.67 | 0.64 |
| 3 | 73437875 | rs7647488 | -1.02 | 1.006 | -1.092 | -1.194 | -0.034 | 0.077 | 0.077 | 0 | 0.74 | 0.9 | 0.74 | 0.86 | 0.67 | 0.64 |
| 3 | 73438059 | rs11128333 | 0.063 | 0.899 | -0.591 | -1.418 | 0 | -0.013 | -0.013 | 0 | 0.61 | 0.39 | 0.67 | 0.85 | 0.61 | 0.56 |
| 3 | 73438465 | rs11128334 | 0.971 | 1.068 | -0.692 | -1.443 | 0.021 | 0.035 | 0.035 | 0 | 0.43 | 0.33 | 0.52 | 0.68 | 0.36 | 0.33 |
| 3 | 73444439 | rs9868179 | -0.896 | 1.122 | -0.455 | -1.361 | -0.005 | 0.001 | 0.001 | 1.375 | 0.19 | 0.16 | 0.18 | 0.17 | 0.27 | 0.22 |

| | | | | | | | | | | | | | | | | |
|---|---|---|---|---|---|---|---|---|---|---|---|---|---|---|---|---|
| 3 | 73447595 | rs7648118 | -0.046 | 0.791 | -0.317 | -1.417 | 0.001 | -0.014 | -0.014 | 0 | 0.59 | 0.41 | 0.63 | 0.85 | 0.6 | 0.57 |
| 3 | 73449019 | rs3845876 | -1.205 | 0.758 | -0.411 | -1.331 | -0.02 | 0.058 | 0.058 | 0.068 | 0.73 | 0.9 | 0.73 | 0.84 | 0.68 | 0.64 |
| 3 | 73450198 | rs3816831 | 0.102 | 0.856 | -0.398 | -1.397 | -0.02 | -0.002 | -0.002 | 1.998 | 0.14 | 0.04 | 0.1 | 0.13 | 0.14 | 0.17 |
| 3 | 73453325 | rs2291463 | 0.172 | 0.524 | -0.134 | -1.315 | -0.002 | -0.059 | -0.059 | 0 | 0.57 | 0.48 | 0.63 | 0.84 | 0.56 | 0.55 |
| 3 | 73459773 | rs9816862 | -0.016 | 0.36 | -0.175 | -1.279 | -0.01 | -0.048 | -0.048 | 0 | 0.09 | 0.19 | 0.04 | 0 | 0.05 | 0.04 |
| 3 | 73461096 | rs11128337 | 0.58 | 0.566 | -0.216 | -1.297 | 0.011 | 0.047 | 0.047 | 0 | 0.5 | 0.39 | 0.6 | 0.72 | 0.54 | 0.46 |
| 3 | 73461683 | rs6780143 | -0.405 | 0.676 | -0.09 | -1.134 | -0.015 | -0.083 | -0.083 | 1.961 | 0.68 | 0.5 | 0.7 | 0.86 | 0.69 | 0.63 |
| 3 | 73468527 | rs4054221 | -0.417 | 0.654 | -0.891 | -1.065 | -0.012 | 0.035 | 0.035 | 0.999 | 0.68 | 0.75 | 0.69 | 0.86 | 0.62 | 0.6 |
| 3 | 73471572 | rs4677281 | -0.336 | 0.461 | -1.392 | -1.349 | -0.09 | 0.164 | 0.164 | 0 | 0.66 | 0.88 | 0.68 | 0.88 | 0.58 | 0.65 |
| 3 | 73475521 | rs6797484 | 0.433 | -0.083 | -1.07 | -1.32 | -0.001 | -0.002 | -0.002 | 0 | 0.34 | 0.33 | 0.51 | 0.71 | 0.31 | 0.46 |
| 3 | 73476107 | rs1872956 | -0.311 | -0.5 | -1.44 | -1.617 | -0.029 | 0.18 | 0.18 | 1.667 | 0.56 | 0.81 | 0.63 | 0.82 | 0.48 | 0.56 |
| 3 | 73476665 | rs13063715 | 0.666 | -0.648 | -1.39 | -1.606 | 0.009 | -0.002 | -0.002 | 0 | 0.13 | 0.03 | 0.07 | 0.1 | 0.11 | 0.1 |
| 3 | 73479377 | rs4054220 | -0.973 | -0.88 | -1.5 | -1.932 | -0.073 | 0.189 | 0.189 | 0.437 | 0.63 | 0.86 | 0.67 | 0.96 | 0.55 | 0.61 |
| 3 | 73479416 | rs953582 | 0.814 | -1.241 | -0.893 | -2.06 | 0.017 | -0.034 | -0.034 | 0 | 0.33 | 0.45 | 0.5 | 0.76 | 0.37 | 0.5 |
| 3 | 73480230 | rs17011123 | 0.069 | -0.783 | -0.708 | -1.745 | 0.047 | 0.056 | 0.056 | 0 | 0.06 | 0.22 | 0.06 | 0.1 | 0.07 | 0.02 |
| 3 | 73480931 | rs6771685 | 0.311 | -0.871 | -0.83 | -1.727 | 0.001 | 0.005 | 0.005 | 0 | 0.46 | 0.56 | 0.53 | 0.75 | 0.44 | 0.58 |
| 3 | 73485170 | rs3856581 | 0.07 | -0.687 | -0.789 | -1.753 | -0.039 | -0.022 | -0.022 | 0.731 | 0.16 | 0.66 | 0.11 | 0.09 | 0.06 | 0.09 |
| 3 | 73487467 | rs17754369 | 0.338 | -0.467 | -1.535 | -1.693 | -0.077 | 0.237 | 0.237 | 0.402 | 0.46 | 0.09 | 0.56 | 0.85 | 0.48 | 0.58 |
| 3 | 73494803 | rs4677284 | -0.074 | -0.827 | -0.722 | -1.593 | -0.015 | -0.03 | -0.03 | 0 | 0.75 | 0.74 | 0.85 | 0.92 | 0.81 | 0.83 |
| 3 | 73497023 | rs6800289 | 0.836 | -0.894 | -0.802 | -1.588 | -0.02 | -0.003 | -0.003 | 0 | 0.1 | 0.14 | 0.05 | 0.08 | 0.03 | 0.08 |
| 3 | 73498998 | rs3863973 | -0.171 | -0.619 | -0.537 | -1.595 | -0.012 | -0.04 | -0.04 | 0.034 | 0.75 | 0.68 | 0.83 | 0.92 | 0.79 | 0.82 |
| 3 | 73506785 | rs7614618 | -0.6 | -0.059 | -0.418 | -1.608 | 0.041 | -0.086 | -0.086 | 0.025 | 0.74 | 0.5 | 0.74 | 0.9 | 0.69 | 0.79 |
| 3 | 73510597 | rs4677286 | -0.017 | -0.395 | -0.623 | -1.721 | 0.005 | -0.023 | -0.023 | 0.101 | 0.6 | 0.56 | 0.65 | 0.75 | 0.56 | 0.76 |
| 3 | 73516119 | rs4677287 | 1.714 | -0.971 | -1.427 | -0.894 | 0.024 | -0.111 | -0.111 | 0.523 | 0.17 | 0.03 | 0.23 | 0.31 | 0.11 | 0.19 |
| 3 | 73523329 | rs1565176 | 1.056 | -0.89 | -1.598 | -1.878 | -0.078 | -0.029 | -0.029 | 0 | 0.32 | 0.04 | 0.46 | 0.68 | 0.32 | 0.57 |
| 3 | 73523496 | rs6549544 | 0.77 | -1.04 | -0.891 | -1.778 | -0.03 | 0.003 | 0.003 | 0.004 | 0.13 | 0.48 | 0.1 | 0.09 | 0.04 | 0.09 |
| 3 | 73526484 | rs1872961 | -0.586 | -0.916 | -1.1 | -1.833 | -0.003 | -0.034 | -0.034 | 0 | 0.84 | 0.75 | 0.85 | 0.93 | 0.85 | 0.96 |
| 3 | 73527288 | rs17703851 | 1 | -0.689 | -1.263 | -1.878 | -0.028 | 0.029 | 0.029 | 0.146 | 0.16 | 0.06 | 0.14 | 0.1 | 0.16 | 0.18 |
| 3 | 73527986 | rs3855577 | 0.287 | -1.181 | -0.789 | -2.15 | -0.016 | -0.075 | -0.075 | 0 | 0.67 | 0.39 | 0.77 | 0.92 | 0.69 | 0.88 |
| 3 | 73528482 | rs9861503 | -0.17 | -0.718 | -0.393 | -1.773 | -0.016 | 0.012 | 0.012 | 0 | 0.31 | 0.27 | 0.32 | 0.16 | 0.44 | 0.24 |
| 3 | 73530323 | rs4263231 | 0.296 | -1.269 | -0.287 | -1.549 | -0.006 | -0.003 | -0.003 | 0 | 0.66 | 0.68 | 0.65 | 0.85 | 0.56 | 0.76 |
| 3 | 73532069 | rs7627890 | 0.668 | -1.356 | -0.408 | -1.522 | -0.002 | -0.004 | -0.004 | 0 | 0.58 | 0.59 | 0.63 | 0.75 | 0.55 | 0.61 |
| 3 | 73536586 | rs6783760 | 0.029 | -1.129 | -0.947 | -1.162 | 0.016 | 0.015 | 0.015 | 6.444 | 0.14 | 0.03 | 0.07 | 0.11 | 0.08 | 0.22 |
| 3 | 73538049 | rs4234171 | -0.046 | -0.689 | -0.642 | -1.043 | -0.044 | 0.047 | 0.047 | 0 | 0.74 | 0.8 | 0.72 | 0.85 | 0.62 | 0.76 |
| 3 | 73540081 | rs3213822 | -0.066 | -0.625 | -0.469 | -0.947 | 0.003 | -0.004 | -0.004 | 0 | 0.73 | 0.68 | 0.71 | 0.82 | 0.61 | 0.76 |
| 3 | 73546789 | rs4677292 | 0.794 | -0.963 | -0.823 | -0.841 | -0.055 | 0.052 | 0.052 | 1.107 | 0.41 | 0.14 | 0.55 | 0.66 | 0.48 | 0.53 |
| 3 | 73550817 | rs13081823 | 0.361 | -0.612 | -0.743 | -0.942 | -0.107 | 0.127 | 0.127 | 0 | 0.28 | 0.04 | 0.26 | 0.11 | 0.38 | 0.24 |
| 3 | 73559480 | rs5029197 | -0.513 | -0.396 | 0.175 | -0.845 | -0.003 | -0.078 | -0.078 | 0 | 0.64 | 0.56 | 0.73 | 0.89 | 0.66 | 0.78 |
| 3 | 73566832 | rs11128343 | -0.513 | -0.397 | -0.035 | -0.797 | -0.003 | -0.047 | -0.047 | 0 | 0.63 | 0.61 | 0.74 | 0.88 | 0.67 | 0.78 |
| 3 | 73572544 | rs13069228 | 0.427 | -0.374 | -0.915 | -0.695 | -0.007 | 0.122 | 0.122 | 0.585 | 0.36 | 0.15 | 0.25 | 0.15 | 0.35 | 0.23 |
| 3 | 73578778 | rs11128344 | -0.883 | -0.75 | -0.998 | -0.386 | -0.003 | -0.004 | -0.004 | 0 | 0.63 | 0.69 | 0.46 | 0.47 | 0.6 | 0.54 |
| 3 | 73588223 | rs4535167 | 0.72 | -0.062 | -1.676 | -0.021 | -0.035 | 0.011 | 0.011 | 1.079 | 0.44 | 0.2 | 0.58 | 0.54 | 0.49 | 0.48 |
| 3 | 73594445 | rs4677299 | 0.111 | 0.566 | -1.676 | -0.134 | -0.049 | 0.094 | 0.094 | 3.001 | 0.65 | 0.89 | 0.77 | 0.79 | 0.66 | 0.73 |
| 3 | 73601210 | rs9842067 | 0.181 | 0.683 | -1.546 | -0.15 | -0.018 | 0.085 | 0.085 | 2.305 | 0.65 | 0.85 | 0.78 | 0.8 | 0.67 | 0.73 |
| 3 | 73612949 | rs6802500 | -0.212 | 0.81 | -1.247 | 0.333 | -0.005 | -0.007 | -0.007 | 0 | 0.37 | 0.3 | 0.42 | 0.38 | 0.36 | 0.37 |
| 3 | 73614083 | rs3889642 | -0.028 | -0.01 | -2.073 | 0.001 | 0.052 | 0.049 | 0.049 | 0.47 | 0.68 | 0.18 | 0.79 | 0.78 | 0.77 | 0.83 |
| 3 | 73617708 | rs4677301 | -0.203 | -0.525 | -0.286 | -0.239 | -0.002 | -0.006 | -0.006 | 0 | 0.86 | 0.83 | 0.88 | 0.9 | 0.86 | 0.9 |
| 3 | 73620180 | rs4342050 | 0.058 | -0.515 | -0.268 | -0.234 | -0.004 | -0.044 | -0.044 | 0 | 0.84 | 0.53 | 0.86 | 0.9 | 0.86 | 0.89 |
| 3 | 73624645 | rs9866965 | -0.677 | -0.866 | -0.166 | -0.177 | -0.012 | -0.001 | -0.001 | 0 | 0.27 | 0.32 | 0.28 | 0.25 | 0.19 | 0.15 |
| 3 | 73626561 | rs13060646 | 0.119 | -0.275 | -0.216 | -0.16 | 0.01 | 0.004 | 0.004 | 0.533 | 0.41 | 0.35 | 0.24 | 0.37 | 0.28 | 0.34 |
| 3 | 73626847 | rs4677304 | -1.044 | -0.262 | -0.337 | -0.194 | -0.029 | -0.025 | -0.025 | 0 | 0.93 | 0.99 | 0.91 | 0.9 | 0.88 | 0.91 |
| 3 | 73627014 | rs12496312 | -0.276 | -0.312 | -0.308 | -0.22 | 0 | -0.001 | -0.001 | 0 | 0.9 | 0.9 | 0.88 | 0.9 | 0.88 | 0.91 |
| 3 | 73629487 | rs4440068 | 0.341 | -0.345 | -0.381 | -0.237 | 0.002 | 0.001 | 0.001 | 2.696 | 0.49 | 0.54 | 0.64 | 0.53 | 0.59 | 0.56 |
| 3 | 73629968 | rs13079376 | 1.611 | -0.3 | -0.887 | -0.229 | -0.027 | -0.029 | -0.029 | 0 | 0.25 | 0.06 | 0.18 | 0.26 | 0.21 | 0.28 |
| 3 | 73632188 | rs4677306 | 1.433 | -0.217 | -0.875 | -0.24 | -0.172 | -0.13 | -0.13 | 0.184 | 0.32 | 0.03 | 0.38 | 0.38 | 0.46 | 0.45 |
| 3 | 73636020 | rs9830956 | -0.704 | -0.25 | 0.045 | -0.56 | -0.02 | -0.018 | -0.018 | 0 | 0.27 | 0.33 | 0.25 | 0.2 | 0.19 | 0.14 |
| 3 | 73639587 | rs4596089 | -0.429 | -0.382 | -0.081 | -0.377 | 0.001 | 0.003 | 0.003 | 1.273 | 0.61 | 0.36 | 0.67 | 0.71 | 0.66 | 0.62 |
| 3 | 73640279 | rs11128348 | 1.559 | -0.375 | -0.316 | -0.367 | -0.044 | -0.036 | -0.036 | 0 | 0.26 | 0.01 | 0.16 | 0.21 | 0.21 | 0.27 |
| 3 | 73642690 | rs4677310 | 0.762 | -0.376 | -0.211 | -0.341 | 0.004 | -0.009 | -0.009 | 0 | 0.13 | 0.07 | 0.21 | 0.16 | 0.12 | 0.08 |
| 3 | 73643655 | rs4234183 | -0.656 | -0.347 | -0.084 | -0.338 | -0.004 | 0 | 0 | 2.093 | 0.92 | 0.95 | 0.88 | 0.92 | 0.87 | 0.89 |
| 3 | 73647454 | rs4677313 | -0.656 | -0.347 | -0.084 | -0.338 | -0.004 | 0 | 0 | 2.093 | 0.92 | 0.95 | 0.88 | 0.92 | 0.87 | 0.89 |
| 3 | 73649851 | rs13319740 | 0.388 | -0.294 | 0.091 | -0.3 | -0.004 | 0.003 | 0.003 | 0 | 0.14 | 0.16 | 0.21 | 0.16 | 0.12 | 0.11 |
| 3 | 73651469 | rs6797804 | -0.887 | -0.339 | 0.287 | -0.268 | -0.01 | 0.009 | 0.009 | 1.691 | 0.64 | 0.85 | 0.72 | 0.7 | 0.66 | 0.62 |
| 3 | 73652833 | rs9879918 | -0.161 | 0.741 | 0.008 | -0.515 | 0.002 | -0.001 | -0.001 | 0.137 | 0.19 | 0.18 | 0.29 | 0.16 | 0.23 | 0.16 |
| 3 | 73653773 | rs9822882 | -0.014 | 0.068 | 0.261 | -0.442 | -0.035 | -0.047 | -0.047 | 0 | 0.14 | 0.28 | 0.05 | 0.04 | 0.07 | 0.05 |
| 3 | 73660741 | rs4677315 | 0.758 | 0.254 | 0.051 | -0.255 | -0.058 | -0.06 | -0.06 | 0 | 0.41 | 0.16 | 0.43 | 0.51 | 0.51 | 0.48 |
| 3 | 73666816 | rs4676940 | -0.658 | 0.174 | 0.272 | -0.381 | 0.026 | 0.04 | 0.04 | 1.187 | 0.64 | 0.8 | 0.73 | 0.74 | 0.69 | 0.62 |
| 3 | 73667917 | rs9828164 | -1.247 | 0.286 | 0.833 | -0.485 | 0.039 | 0.012 | 0.012 | 1.708 | 0.09 | 0.51 | 0.06 | 0.04 | 0.06 | 0.06 |
| 3 | 73682075 | rs6549556 | -0.658 | 0.249 | 0.433 | -0.446 | 0.03 | 0.046 | 0.046 | 3.25 | 0.64 | 0.79 | 0.74 | 0.75 | 0.69 | 0.6 |

| | | | | | | | | | | | | | | | | |
|---|---|---|---|---|---|---|---|---|---|---|---|---|---|---|---|---|
| 3 | 73687477 | rs4677319 | -0.615 | 0.289 | 0.46 | -0.424 | 0.024 | 0.034 | 0.034 | 3.009 | 0.64 | 0.75 | 0.74 | 0.75 | 0.7 | 0.6 |
| 3 | 73690263 | rs7653458 | -0.271 | 0.514 | 0.524 | -0.578 | 0.001 | 0.005 | 0.005 | 0.368 | 0.6 | 0.57 | 0.69 | 0.75 | 0.61 | 0.58 |
| 3 | 73695465 | rs11707332 | 0.521 | 0.059 | 0.418 | -0.459 | 0.028 | 0.023 | 0.023 | 1.374 | 0.36 | 0.25 | 0.24 | 0.27 | 0.27 | 0.39 |
| 3 | 73699523 | rs12486279 | 1.038 | -0.014 | 0.038 | -0.523 | 0.029 | 0.037 | 0.037 | 0.203 | 0.29 | 0.02 | 0.14 | 0.18 | 0.18 | 0.28 |
| 3 | 73704141 | rs9310276 | -0.933 | 0.11 | 0.438 | -0.408 | 0.004 | 0.008 | 0.008 | 0.389 | 0.21 | 0.27 | 0.34 | 0.26 | 0.22 | 0.25 |
| 3 | 73705032 | rs1022171 | 0.743 | 0.081 | 0.389 | -0.364 | 0.003 | -0.021 | -0.021 | 0 | 0.1 | 0.05 | 0.21 | 0.17 | 0.09 | 0.11 |
| 3 | 73710945 | rs9310279 | -0.621 | 0.056 | 0.468 | -0.438 | 0.007 | 0.008 | 0.008 | 0.272 | 0.54 | 0.39 | 0.5 | 0.43 | 0.41 | 0.53 |
| 3 | 73711340 | rs9823789 | -1.017 | 0.191 | 0.552 | -0.359 | 0.011 | -0.006 | -0.006 | 0 | 0.11 | 0.23 | 0.09 | 0.08 | 0.1 | 0.09 |
| 3 | 73716503 | rs2875192 | 0.555 | 0.101 | 0.718 | -0.376 | -0.014 | 0.034 | 0.034 | 2.196 | 0.81 | 0.42 | 0.69 | 0.76 | 0.84 | 0.86 |
| 3 | 73718136 | rs9828758 | 0.677 | -0.009 | 0.509 | -0.478 | 0.03 | 0.061 | 0.061 | 0 | 0.36 | 0.23 | 0.23 | 0.19 | 0.27 | 0.35 |
| 3 | 73719148 | rs12493615 | -0.552 | 0.034 | 0.864 | -0.46 | 0 | 0.031 | 0.031 | 0.679 | 0.17 | 0.32 | 0.3 | 0.24 | 0.18 | 0.15 |
| 3 | 73721258 | rs4599238 | -0.508 | -0.003 | 0.86 | -0.496 | -0.026 | -0.049 | -0.049 | 0 | 0.61 | 0.47 | 0.74 | 0.8 | 0.71 | 0.65 |
| 3 | 73724034 | rs9821729 | -0.265 | 0.013 | 0.929 | -0.483 | 0.015 | -0.011 | -0.011 | 0 | 0.06 | 0.17 | 0.03 | 0.02 | 0.05 | 0.03 |
| 3 | 73724526 | rs4676943 | 1.45 | 0 | 0.883 | -0.488 | 0.055 | 0.016 | 0.016 | 3.085 | 0.91 | 0.48 | 0.93 | 0.97 | 0.94 | 0.97 |
| 3 | 73727760 | rs11714828 | 1.251 | -0.098 | 0.688 | -0.526 | 0.023 | 0.067 | 0.067 | 0.327 | 0.31 | 0.06 | 0.2 | 0.17 | 0.23 | 0.34 |
| 3 | 73733491 | rs868709 | 1.054 | 0.081 | 0.862 | -0.446 | -0.004 | -0.001 | -0.001 | 0 | 0.78 | 0.25 | 0.7 | 0.8 | 0.84 | 0.88 |
| 3 | 73734443 | rs6789737 | -0.91 | 0.103 | 0.89 | -0.435 | 0.032 | 0.022 | 0.022 | 0 | 0.06 | 0.22 | 0.08 | 0.06 | 0.08 | 0.06 |
| 3 | 73735674 | rs1511532 | 0.263 | 0.039 | 0.679 | -0.464 | 0.056 | 0.031 | 0.031 | 2.905 | 0.43 | 0.11 | 0.4 | 0.35 | 0.32 | 0.39 |
| 3 | 73742625 | rs9852827 | -0.414 | 0.179 | 0.872 | -0.382 | 0 | -0.013 | -0.013 | 0 | 0.06 | 0.13 | 0.03 | 0.02 | 0.04 | 0.03 |
| 3 | 73743436 | rs907341 | 1.595 | 0.139 | 0.895 | -0.402 | 0.009 | -0.003 | -0.003 | 0 | 0.74 | 0.17 | 0.64 | 0.74 | 0.77 | 0.81 |
| 3 | 73744048 | rs972750 | -0.733 | 0.193 | 0.965 | -0.357 | -0.003 | 0.019 | 0.019 | 0.427 | 0.17 | 0.28 | 0.29 | 0.23 | 0.17 | 0.16 |
| 3 | 73750993 | rs2137064 | -0.407 | 0.109 | 0.085 | -0.405 | 0.008 | 0.017 | 0.017 | 0.107 | 0.52 | 0.18 | 0.5 | 0.42 | 0.44 | 0.48 |
| 3 | 73753986 | rs13327091 | 0.553 | 0.091 | 0.417 | -0.402 | 0.041 | 0.057 | 0.057 | 0.02 | 0.37 | 0.21 | 0.23 | 0.19 | 0.27 | 0.31 |
| 3 | 73756857 | rs1949120 | -0.855 | 0.036 | 0.359 | -0.441 | -0.018 | -0.004 | -0.004 | 0 | 0.09 | 0.02 | 0.09 | 0.08 | 0.11 | 0.09 |
| 3 | 73756870 | rs13060404 | 1.099 | -0.033 | 0.411 | -0.482 | 0.032 | 0.038 | 0.038 | 0.83 | 0.32 | 0.2 | 0.2 | 0.18 | 0.22 | 0.28 |
| 3 | 73769764 | rs925431 | 0.406 | 0.25 | -0.225 | -0.238 | -0.004 | -0.021 | -0.021 | 0 | 0.44 | 0.17 | 0.46 | 0.58 | 0.57 | 0.52 |
| 3 | 73783030 | rs3845881 | -0.283 | 0.523 | -0.589 | -0.391 | 0.052 | 0.114 | 0.114 | 0 | 0.54 | 0.24 | 0.69 | 0.8 | 0.64 | 0.68 |
| 3 | 73785176 | rs4676952 | 0.33 | 1.611 | -0.873 | -0.586 | -0.098 | 0.201 | 0.201 | 2.108 | 0.45 | 0.05 | 0.37 | 0.16 | 0.45 | 0.42 |
| 3 | 73785389 | rs12633549 | 0.254 | 1.609 | -0.839 | -0.584 | -0.012 | 0.095 | 0.095 | 0.06 | 0.44 | 0.17 | 0.54 | 0.74 | 0.46 | 0.55 |
| 3 | 73786057 | rs17756459 | -0.102 | 1.652 | -0.784 | -0.555 | -0.012 | 0.009 | 0.009 | 2.019 | 0.09 | 0.07 | 0.16 | 0.06 | 0.19 | 0.13 |
| 3 | 73788184 | rs12496240 | 0.702 | 1.232 | -0.633 | -0.4 | 0.009 | 0.012 | 0.012 | 5.634 | 0.9 | 0.28 | 0.91 | 0.9 | 0.9 | 0.97 |
| 3 | 73788701 | rs3856583 | 0.71 | 1.232 | -0.636 | -0.4 | 0.008 | 0.011 | 0.011 | 5.546 | 0.9 | 0.28 | 0.91 | 0.9 | 0.9 | 0.97 |
| 3 | 73789292 | rs11716081 | -0.013 | 1.179 | -0.561 | -0.238 | 0.004 | 0.075 | 0.075 | 0.068 | 0.45 | 0.25 | 0.54 | 0.74 | 0.46 | 0.54 |
| 3 | 73797420 | rs3911952 | 0.768 | 1.119 | -0.779 | -0.198 | -0.049 | -0.12 | -0.12 | 0 | 0.33 | 0.02 | 0.35 | 0.6 | 0.32 | 0.43 |
| 3 | 73797559 | rs3911953 | -0.237 | 1.081 | -0.683 | -0.226 | 0.013 | 0.089 | 0.089 | 0.013 | 0.45 | 0.24 | 0.57 | 0.76 | 0.43 | 0.54 |
| 3 | 73802248 | rs11709003 | 0.589 | 0.961 | -1.257 | -0.037 | -0.016 | -0.012 | -0.012 | 0 | 0.29 | 0.16 | 0.37 | 0.68 | 0.34 | 0.46 |
| 3 | 73803981 | rs881521 | 0.538 | 0.982 | -1.248 | 0.049 | -0.013 | -0.04 | -0.04 | 0 | 0.31 | 0.17 | 0.38 | 0.59 | 0.34 | 0.45 |
| 3 | 73805967 | rs4622849 | 0.414 | 0.934 | -1.376 | 0.035 | -0.006 | -0.01 | -0.01 | 0 | 0.09 | 0.04 | 0.05 | 0.09 | 0.09 | 0.1 |
| 3 | 73806887 | rs3856584 | 0.296 | 1.028 | -1.815 | 0.046 | -0.112 | -0.112 | -0.112 | 1.301 | 0.34 | 0.02 | 0.51 | 0.59 | 0.39 | 0.44 |
| 3 | 73807858 | rs291477 | -0.291 | 0.721 | -1.915 | -0.463 | -0.004 | 0.056 | 0.056 | 2.499 | 0.32 | 0.17 | 0.52 | 0.8 | 0.33 | 0.44 |
| 3 | 73808557 | rs12633728 | -0.402 | 0.726 | -1.58 | -0.298 | 0.002 | 0.025 | 0.025 | 0.105 | 0.05 | 0.11 | 0.02 | 0.15 | 0.04 | 0.04 |
| 3 | 73808594 | rs291478 | -1.585 | 1.044 | -1.154 | -0.563 | 0.006 | -0.004 | -0.004 | 0.001 | 0.55 | 0.44 | 0.73 | 0.85 | 0.66 | 0.71 |
| 3 | 73811569 | rs291482 | -0.395 | 0.812 | -1.108 | -0.519 | -0.024 | 0.047 | 0.047 | 0 | 0.28 | 0.18 | 0.22 | 0.1 | 0.33 | 0.33 |
| 3 | 73812978 | rs17011531 | -0.826 | 0.726 | -1.054 | -0.369 | 0.01 | 0.049 | 0.049 | 1.342 | 0.06 | 0.19 | 0.03 | 0.15 | 0.03 | 0.04 |
| 3 | 73816737 | rs291545 | -1.353 | 0.576 | -0.623 | -0.347 | 0.017 | 0.05 | 0.05 | 0.089 | 0.48 | 0.67 | 0.68 | 0.76 | 0.54 | 0.55 |
| 3 | 73817526 | rs17011553 | 0.908 | 0.304 | -0.782 | 0.059 | -0.065 | -0.119 | -0.119 | 0 | 0.15 | 0.01 | 0.42 | 0.49 | 0.18 | 0.24 |
| 3 | 73818249 | rs3845887 | -0.778 | 0.249 | -0.448 | 0.043 | -0.008 | 0.118 | 0.118 | 2.769 | 0.06 | 0.25 | 0.03 | 0.16 | 0.01 | 0.04 |
| 3 | 73822711 | rs291538 | -1.057 | 0.282 | -0.438 | -0.297 | -0.001 | 0 | 0 | 1.261 | 0.28 | 0.34 | 0.5 | 0.71 | 0.26 | 0.37 |
| 3 | 73829024 | rs291501 | 0.191 | 0.312 | -0.542 | -0.286 | -0.004 | 0.012 | 0.012 | 1.153 | 0.08 | 0.02 | 0.06 | 0.08 | 0.08 | 0.11 |
| 3 | 73835569 | rs9865572 | -0.171 | 0.303 | -0.368 | -0.252 | -0.003 | 0.035 | 0.035 | 1.596 | 0.22 | 0.36 | 0.45 | 0.61 | 0.19 | 0.25 |
| 3 | 73836712 | rs487986 | 0.111 | 0.46 | -0.722 | -0.299 | 0.003 | 0.03 | 0.03 | 0.328 | 0.11 | 0.02 | 0.05 | 0.1 | 0.09 | 0.11 |
| 3 | 73837087 | rs484332 | -1.739 | 0.615 | -0.808 | -0.708 | -0.004 | 0.056 | 0.056 | 0 | 0.57 | 0.73 | 0.69 | 0.78 | 0.51 | 0.57 |
| 3 | 73837141 | rs17706958 | 0.279 | 0.855 | -0.542 | -0.618 | 0.005 | -0.038 | -0.038 | 0 | 0.19 | 0.24 | 0.15 | 0.04 | 0.22 | 0.19 |
| 3 | 73840367 | rs291529 | -0.122 | 0.741 | -0.657 | -0.376 | 0.049 | 0.047 | 0.047 | 0.134 | 0.86 | 0.62 | 0.83 | 0.83 | 0.85 | 0.84 |
| 3 | 73844649 | rs291517 | 0.112 | 0.377 | -1.471 | -0.887 | 0.031 | 0.096 | 0.096 | 0 | 0.24 | 0.07 | 0.12 | 0.03 | 0.17 | 0.2 |
| 3 | 73844739 | rs291516 | -0.413 | 0.292 | -1.813 | -1.335 | -0.005 | 0.19 | 0.19 | 0 | 0.47 | 0.23 | 0.26 | 0.06 | 0.44 | 0.4 |
| 3 | 73845507 | rs291515 | 0.527 | 0.046 | -1.658 | -1.027 | 0.026 | 0.128 | 0.128 | 0.018 | 0.23 | 0.04 | 0.12 | 0.03 | 0.16 | 0.19 |
| 3 | 73851885 | rs4392360 | 0.945 | 0.36 | -1.054 | 0.012 | 0.014 | 0.174 | 0.174 | 0.909 | 0.47 | 0.15 | 0.26 | 0.2 | 0.45 | 0.42 |
| 3 | 73852036 | rs6770929 | -0.018 | 0.651 | -1.009 | 1.119 | -0.058 | -0.049 | -0.049 | 0.89 | 0.32 | 0.14 | 0.37 | 0.39 | 0.39 | 0.38 |
| 3 | 73853704 | rs13070417 | -0.181 | 0.764 | -0.934 | 0.447 | 0.002 | 0 | 0 | 0.057 | 0.34 | 0.4 | 0.56 | 0.63 | 0.4 | 0.41 |
| 3 | 73860264 | rs291535 | 0.047 | 0.654 | -0.86 | 0.591 | 0 | 0.003 | 0.003 | 0.611 | 0.57 | 0.54 | 0.69 | 0.53 | 0.6 | 0.55 |
| 3 | 73861164 | rs13081760 | 1.528 | -0.045 | -1.317 | 0.493 | 0.049 | -0.036 | -0.036 | 7.357 | 0.17 | 0.04 | 0.24 | 0.21 | 0.11 | 0.16 |
| 3 | 73861226 | rs7631627 | 0.545 | -0.492 | -1.021 | 0.229 | -0.007 | 0 | 0 | 0 | 0.25 | 0.26 | 0.27 | 0.21 | 0.13 | 0.26 |
| 3 | 73861878 | rs291533 | -0.119 | -0.024 | -1.174 | 0.012 | 0.03 | 0.06 | 0.06 | 0.532 | 0.42 | 0.29 | 0.4 | 0.23 | 0.37 | 0.34 |
| 3 | 73867480 | rs6796430 | 0.021 | 0.388 | -1.315 | -0.043 | 0.063 | 0.026 | 0.026 | 0 | 0.72 | 0.49 | 0.61 | 0.38 | 0.62 | 0.69 |
| 3 | 73869793 | rs9848344 | -0.017 | 0.208 | -0.776 | -0.141 | 0.005 | 0.003 | 0.003 | 0 | 0.57 | 0.51 | 0.68 | 0.59 | 0.54 | 0.58 |
| 3 | 73871925 | rs4627704 | 1.41 | 0.17 | -1.153 | -0.191 | 0.002 | 0.033 | 0.033 | 0 | 0.29 | 0.18 | 0.21 | 0.14 | 0.26 | 0.32 |
| 3 | 73872065 | rs3899618 | 1.424 | 0.455 | -1.038 | 0.381 | -0.026 | -0.06 | -0.06 | 0 | 0.14 | 0.05 | 0.12 | 0.25 | 0.15 | 0.23 |
| 3 | 73872496 | rs7649822 | -0.515 | 0.393 | -0.903 | 0.367 | -0.001 | -0.02 | -0.02 | 0 | 0.7 | 0.36 | 0.77 | 0.85 | 0.7 | 0.81 |

| | | | | | | | | | | | | | | | | |
|---|---|---|---|---|---|---|---|---|---|---|---|---|---|---|---|---|
| 3 | 73874248 | rs7611630 | 1.359 | 0.437 | -1.295 | 0.372 | -0.026 | -0.06 | -0.06 | 0 | 0.14 | 0.05 | 0.12 | 0.25 | 0.15 | 0.23 |
| 3 | 73874699 | rs167096 | 0.892 | 0.164 | -0.961 | -0.063 | 0 | -0.078 | -0.078 | 0 | 0.25 | 0.33 | 0.16 | 0.03 | 0.26 | 0.18 |
| 3 | 73874777 | rs291503 | 1.095 | 0.295 | -1.762 | -0.001 | -0.047 | 0.128 | 0.128 | 0 | 0.23 | 0.03 | 0.15 | 0.03 | 0.26 | 0.18 |
| 3 | 73876725 | rs1107768 | 1.149 | 0.328 | -1.774 | 0.004 | -0.048 | 0.13 | 0.13 | 0 | 0.22 | 0.03 | 0.14 | 0.03 | 0.26 | 0.18 |
| 3 | 73877100 | rs907336 | 0.159 | 0.604 | -1.55 | 0.394 | -0.039 | -0.056 | -0.056 | 0 | 0.25 | 0.16 | 0.33 | 0.49 | 0.33 | 0.28 |
| 3 | 73877349 | rs291484 | 0.262 | 0.014 | -1.575 | 0.056 | -0.015 | 0.078 | 0.078 | 1.137 | 0.39 | 0.04 | 0.38 | 0.25 | 0.37 | 0.36 |
| 3 | 73878667 | rs291485 | -0.191 | -0.11 | -0.845 | -0.048 | -0.001 | -0.002 | -0.002 | 0 | 0.46 | 0.45 | 0.33 | 0.52 | 0.45 | 0.54 |
| 3 | 73881814 | rs9310281 | 0.977 | 0.143 | -0.573 | -0.542 | 0.011 | -0.044 | -0.044 | 0 | 0.17 | 0.18 | 0.16 | 0.01 | 0.26 | 0.21 |
| 3 | 73883107 | rs291490 | 0.531 | 0.14 | -0.233 | -0.53 | 0.003 | 0.009 | 0.009 | 0 | 0.28 | 0.25 | 0.49 | 0.4 | 0.34 | 0.34 |
| 3 | 73887678 | rs6767612 | 1.143 | 0.239 | -0.459 | -0.591 | 0.015 | -0.051 | -0.051 | 0 | 0.18 | 0.19 | 0.16 | 0.02 | 0.26 | 0.21 |
| 3 | 73897047 | rs1441447 | -0.333 | -0.308 | -0.442 | -0.345 | -0.007 | -0.027 | -0.027 | 0 | 0.41 | 0.49 | 0.54 | 0.71 | 0.35 | 0.36 |
| 3 | 73897704 | rs991270 | 0.552 | -0.105 | -1.22 | -0.146 | -0.104 | -0.135 | -0.135 | 0.054 | 0.27 | 0.04 | 0.48 | 0.4 | 0.34 | 0.34 |
| 3 | 73898068 | rs17707903 | 0.998 | -0.094 | -1.055 | -0.385 | 0.001 | 0.067 | 0.067 | 0.713 | 0.15 | 0.06 | 0.05 | 0.02 | 0.13 | 0.26 |
| 3 | 73898514 | rs4677353 | -1.418 | -0.282 | -0.639 | -0.233 | -0.037 | 0.068 | 0.068 | 11.779 | 0.89 | 0.82 | 0.97 | 0.68 | 0.99 | 0.98 |
| 3 | 73898855 | rs7629754 | -1.43 | -0.286 | -0.584 | -0.268 | -0.046 | 0.089 | 0.089 | 11.782 | 0.89 | 0.8 | 0.97 | 0.69 | 0.99 | 0.98 |
| 3 | 73899934 | rs4677360 | 0.65 | -0.228 | -0.285 | -0.584 | -0.019 | 0.006 | 0.006 | 2.385 | 0.62 | 0.41 | 0.73 | 0.58 | 0.77 | 0.58 |
| 3 | 73900052 | rs12485807 | 1.629 | 0.325 | -0.936 | -1.048 | -0.043 | 0.051 | 0.051 | 0.869 | 0.16 | 0.07 | 0.13 | 0.01 | 0.25 | 0.19 |
| 3 | 73901630 | rs9858596 | 1.708 | -0.102 | -0.879 | -1.056 | 0.01 | 0.05 | 0.05 | 0 | 0.27 | 0.11 | 0.19 | 0.17 | 0.26 | 0.18 |
| 3 | 73903631 | rs7645267 | -0.213 | -0.354 | -0.15 | -0.619 | -0.019 | -0.021 | -0.021 | 0.305 | 0.69 | 0.44 | 0.55 | 0.28 | 0.75 | 0.78 |
| 3 | 73906871 | rs973538 | 0.506 | -0.202 | -0.231 | -0.61 | -0.002 | -0.023 | -0.023 | 0 | 0.61 | 0.45 | 0.42 | 0.21 | 0.65 | 0.63 |
| 3 | 73913622 | rs17759315 | 0.508 | 0.703 | -0.464 | -0.471 | -0.029 | 0.019 | 0.019 | 0 | 0.09 | 0.04 | 0.14 | 0.08 | 0.1 | 0.15 |
| 3 | 73914516 | rs2197054 | 0.57 | 0.199 | 0.083 | -0.126 | 0.006 | 0.112 | 0.112 | 0 | 0.2 | 0.34 | 0.41 | 0.4 | 0.24 | 0.2 |
| 3 | 73915047 | rs1866929 | 0.278 | 0.139 | -0.361 | -0.641 | 0 | -0.022 | -0.022 | 0 | 0.16 | 0.18 | 0.06 | 0.02 | 0.12 | 0.26 |
| 3 | 73916849 | rs6776662 | 0.495 | 0.437 | -0.154 | -0.625 | 0.008 | -0.081 | -0.081 | 0.363 | 0.24 | 0.4 | 0.14 | 0.09 | 0.22 | 0.24 |
| 3 | 73919008 | rs6799987 | -0.235 | 0.164 | -0.344 | -1.006 | 0.002 | -0.02 | -0.02 | 0.009 | 0.44 | 0.41 | 0.24 | 0.1 | 0.35 | 0.49 |
| 3 | 73919119 | rs7432145 | 0.512 | 0.267 | -0.685 | -0.205 | 0.002 | -0.049 | -0.049 | 0 | 0.22 | 0.11 | 0.39 | 0.4 | 0.29 | 0.15 |
| 3 | 73923210 | rs1441443 | 0.266 | -0.047 | 0.011 | -0.078 | -0.039 | 0 | 0 | 0 | 0.25 | 0.43 | 0.45 | 0.74 | 0.25 | 0.21 |
| 3 | 73928496 | rs4676962 | -0.593 | 0.452 | 0.168 | 0.382 | 0.075 | 0.389 | 0.389 | 0 | 0.05 | 0.26 | 0.12 | 0.37 | 0.08 | 0.05 |
| 3 | 73930583 | rs4383445 | 1.104 | 0.379 | 0.149 | 0.412 | 0.01 | -0.037 | -0.037 | 0.009 | 0.19 | 0.1 | 0.34 | 0.41 | 0.21 | 0.16 |
| 3 | 73931460 | rs1992419 | 0.523 | 0.64 | -0.31 | -0.165 | -0.004 | 0.015 | 0.015 | 0.256 | 0.23 | 0.22 | 0.18 | 0.07 | 0.24 | 0.26 |
| 3 | 73936006 | rs13077887 | -0.338 | 0.616 | -0.153 | -0.166 | 0.005 | -0.006 | -0.006 | 0 | 0.44 | 0.42 | 0.54 | 0.47 | 0.46 | 0.42 |
| 3 | 73940255 | rs2119609 | -0.188 | 0.698 | -0.452 | -0.37 | 0.016 | -0.011 | -0.011 | 0 | 0.44 | 0.4 | 0.53 | 0.49 | 0.44 | 0.58 |
| 3 | 73943974 | rs6808996 | 1.126 | 0.595 | -0.212 | -0.484 | -0.031 | 0.14 | 0.14 | 1.117 | 0.41 | 0.18 | 0.35 | 0.13 | 0.42 | 0.38 |
| 3 | 73944436 | rs6549566 | 0.527 | 0.691 | -0.123 | -0.462 | -0.004 | -0.001 | -0.001 | 0 | 0.47 | 0.52 | 0.42 | 0.5 | 0.46 | 0.4 |
| 3 | 73945032 | rs9284824 | -1.777 | 0.623 | -0.234 | -0.28 | 0.016 | -0.085 | -0.085 | 0.281 | 0.8 | 0.93 | 0.7 | 0.6 | 0.79 | 0.85 |
| 3 | 73946281 | rs10460948 | 0.311 | 1.061 | -0.091 | -0.448 | -0.025 | -0.057 | -0.057 | 0 | 0.22 | 0.11 | 0.25 | 0.47 | 0.23 | 0.15 |
| 3 | 73951152 | rs2323206 | -0.757 | 0.961 | -0.026 | -0.735 | 0.015 | -0.067 | -0.067 | 0.983 | 0.67 | 0.59 | 0.77 | 0.88 | 0.62 | 0.73 |
| 3 | 73952351 | rs9869897 | 1.024 | 0.602 | -0.011 | -0.966 | -0.014 | 0.044 | 0.044 | 0 | 0.29 | 0.16 | 0.2 | 0.05 | 0.33 | 0.27 |
| 3 | 73954634 | rs1405398 | -0.372 | 0.801 | -0.227 | -0.652 | 0.001 | -0.062 | -0.062 | 1.739 | 0.55 | 0.55 | 0.63 | 0.87 | 0.58 | 0.62 |
| 3 | 73958239 | rs1852378 | -0.09 | 0.924 | -0.177 | -0.15 | 0.025 | 0.049 | 0.049 | 0.393 | 0.79 | 0.47 | 0.83 | 0.73 | 0.77 | 0.74 |
| 3 | 73958469 | rs5012667 | 1.05 | 0.513 | -0.624 | -0.116 | 0.014 | -0.046 | -0.046 | 3.997 | 0.19 | 0.09 | 0.35 | 0.48 | 0.19 | 0.13 |
| 3 | 73959971 | rs1405389 | -0.065 | 0.631 | -0.509 | -0.395 | 0.002 | -0.007 | -0.007 | 0 | 0.64 | 0.69 | 0.49 | 0.52 | 0.62 | 0.65 |
| 3 | 73960394 | rs1405392 | -0.988 | 0.636 | -0.582 | -0.478 | 0.005 | -0.075 | -0.075 | 0 | 0.81 | 0.7 | 0.83 | 0.97 | 0.82 | 0.77 |
| 3 | 73961525 | rs6785650 | 0.44 | 0.392 | -0.871 | -0.533 | 0 | -0.003 | -0.003 | 0 | 0.56 | 0.38 | 0.35 | 0.15 | 0.52 | 0.57 |
| 3 | 73961594 | rs17709406 | 0.857 | 0.457 | -2.155 | -0.628 | -0.079 | 0.064 | 0.064 | 1.07 | 0.17 | 0.01 | 0.13 | 0.08 | 0.17 | 0.29 |
| 3 | 73964281 | rs1526705 | 1.382 | 0.021 | -1.709 | -0.416 | 0.017 | -0.056 | -0.056 | 0.093 | 0.17 | 0.07 | 0.34 | 0.45 | 0.19 | 0.12 |
| 3 | 73965145 | rs17709608 | 0.505 | 0.133 | -1.547 | -1.076 | -0.005 | 0.154 | 0.154 | 2.371 | 0.36 | 0.14 | 0.21 | 0.08 | 0.35 | 0.28 |
| 3 | 73965973 | rs17709673 | -0.057 | -0.221 | -0.882 | -0.579 | -0.001 | 0.001 | 0.001 | 0 | 0.33 | 0.31 | 0.25 | 0.31 | 0.37 | 0.34 |
| 3 | 73967610 | rs6795429 | 0.208 | -1.445 | -0.405 | -0.085 | -0.044 | 0.162 | 0.162 | 0 | 0.2 | 0.36 | 0.4 | 0.44 | 0.1 | 0.17 |
| 3 | 73968051 | rs1526709 | -1.021 | -1.408 | -0.869 | -0.511 | -0.015 | 0.027 | 0.027 | 0 | 0.61 | 0.66 | 0.68 | 0.82 | 0.52 | 0.57 |
| 3 | 73971723 | rs13098784 | -0.246 | -0.573 | -1.25 | -0.496 | -0.042 | 0.037 | 0.037 | 0.114 | 0.34 | 0.1 | 0.3 | 0.31 | 0.41 | 0.39 |
| 3 | 73974415 | rs13089605 | 1.281 | -0.885 | -1.219 | -0.642 | -0.051 | 0.076 | 0.076 | 0 | 0.16 | 0.01 | 0.13 | 0.07 | 0.18 | 0.15 |
| 3 | 73977773 | rs13075070 | 0.127 | -0.808 | -1.022 | -0.466 | -0.027 | 0.031 | 0.031 | 0 | 0.33 | 0.1 | 0.29 | 0.31 | 0.38 | 0.37 |
| 3 | 73979983 | rs6549571 | 0.358 | -0.617 | -0.378 | -0.371 | 0.041 | -0.008 | -0.008 | 0.046 | 0.06 | 0.21 | 0.11 | 0 | 0.05 | 0.03 |
| 3 | 73982358 | rs6549573 | 0.082 | -0.615 | -0.386 | -0.291 | 0.064 | 0.107 | 0.107 | 0 | 0.06 | 0.27 | 0.12 | 0.07 | 0.06 | 0.03 |
| 3 | 73984721 | rs9861737 | -0.066 | -0.891 | -1.38 | -0.414 | 0.015 | 0.064 | 0.064 | 0.003 | 0.39 | 0.14 | 0.29 | 0.32 | 0.4 | 0.45 |
| 3 | 73986513 | rs1526724 | 0.357 | -1.389 | -0.505 | -0.043 | -0.057 | 0.109 | 0.109 | 0 | 0.18 | 0.27 | 0.28 | 0.44 | 0.05 | 0.15 |
| 3 | 73986720 | rs9880722 | 0.339 | -1.291 | -0.889 | -1.13 | 0.035 | 0.106 | 0.106 | 0 | 0.27 | 0.09 | 0.16 | 0.08 | 0.23 | 0.26 |
| 3 | 73986975 | rs6775849 | -0.144 | -1.212 | -0.313 | -1.127 | -0.035 | -0.037 | -0.037 | 0 | 0.39 | 0.46 | 0.54 | 0.74 | 0.24 | 0.42 |
| 3 | 73987684 | rs11706107 | -0.564 | -0.818 | 0.136 | -1.064 | 0.033 | 0.043 | 0.043 | 0.084 | 0.52 | 0.7 | 0.43 | 0.31 | 0.63 | 0.48 |
| 3 | 73987905 | rs11718192 | -0.004 | -0.91 | 0.277 | -1.42 | -0.026 | -0.025 | -0.025 | 0 | 0.14 | 0.23 | 0.12 | 0.09 | 0.09 | 0.12 |
| 3 | 73988264 | rs17760827 | 0.399 | -0.248 | 0.123 | -1.591 | -0.084 | 0.135 | 0.135 | 0 | 0.34 | 0.2 | 0.25 | 0.08 | 0.45 | 0.3 |
| 3 | 73994728 | rs7618992 | 0.837 | -0.544 | 0.659 | -1.455 | 0.012 | 0.045 | 0.045 | 0.595 | 0.42 | 0.3 | 0.48 | 0.79 | 0.38 | 0.53 |
| 3 | 73996892 | rs6792155 | -0.911 | -0.548 | 0.736 | -1.456 | -0.009 | -0.045 | -0.045 | 0.057 | 0.58 | 0.56 | 0.5 | 0.21 | 0.62 | 0.47 |
| 3 | 74011687 | rs4416330 | 0.481 | -0.524 | 0.188 | -1.458 | 0.016 | 0.064 | 0.064 | 0.528 | 0.45 | 0.33 | 0.5 | 0.79 | 0.4 | 0.54 |
| 3 | 74019380 | rs9852485 | 0.742 | 0.321 | -0.937 | -1.612 | -0.019 | 0.016 | 0.016 | 0 | 0.2 | 0.21 | 0.29 | 0.08 | 0.34 | 0.14 |
| 3 | 74028783 | rs9872846 | 0.5 | 0.321 | -0.955 | -1.611 | -0.009 | 0.005 | 0.005 | 0.026 | 0.22 | 0.24 | 0.29 | 0.08 | 0.34 | 0.14 |
| 3 | 74030761 | rs17761103 | -0.309 | 0.504 | -1.34 | -1.338 | -0.002 | 0.068 | 0.068 | 0 | 0.31 | 0.16 | 0.19 | 0.13 | 0.28 | 0.33 |
| 3 | 74032848 | rs17011909 | -0.182 | -0.111 | -1.073 | -0.9 | 0 | 0.001 | 0.001 | 0 | 0.14 | 0.13 | 0.1 | 0.13 | 0.1 | 0.14 |

| | | | | | | | | | | | | | | | | |
|---|---|---|---|---|---|---|---|---|---|---|---|---|---|---|---|---|
| 3 | 74033442 | rs1526722 | -0.603 | 0.271 | -1.22 | -0.34 | 0.05 | -0.007 | -0.007 | 0 | 0.6 | 0.79 | 0.55 | 0.41 | 0.68 | 0.54 |
| 3 | 74034341 | rs6771004 | -0.432 | 0.244 | -1.127 | -0.817 | 0.026 | 0.098 | 0.098 | 0.004 | 0.55 | 0.72 | 0.51 | 0.23 | 0.61 | 0.45 |
| 3 | 74055676 | rs17011927 | -0.185 | 0.104 | -1.091 | -0.398 | 0.015 | -0.017 | -0.017 | 0.248 | 0.1 | 0 | 0.05 | 0.14 | 0.08 | 0.12 |
| 3 | 74059816 | rs11716019 | 0.154 | 0.409 | 0.14 | -0.837 | 0.015 | -0.024 | -0.024 | 0 | 0.14 | 0.3 | 0.1 | 0.13 | 0.16 | 0.14 |
| 3 | 74060217 | rs9990420 | 0.477 | 0.565 | 0.07 | -1.336 | -0.002 | -0.026 | -0.026 | 0 | 0.42 | 0.5 | 0.66 | 0.78 | 0.39 | 0.56 |
| 3 | 74065520 | rs17710919 | 1.147 | 0.541 | -0.016 | -1.641 | 0.002 | 0.002 | 0.002 | 0 | 0.17 | 0.1 | 0.09 | 0.16 | 0.17 | 0.27 |
| 3 | 74066222 | rs13077605 | 0.509 | 0.441 | 0.179 | -1.686 | 0.021 | 0.132 | 0.132 | 0.155 | 0.49 | 0.3 | 0.71 | 0.76 | 0.51 | 0.59 |
| 3 | 74066457 | rs6793236 | -0.779 | 0.403 | 0.279 | -1.705 | 0.003 | -0.063 | -0.063 | 0.061 | 0.74 | 0.63 | 0.82 | 0.94 | 0.7 | 0.87 |
| 3 | 74071505 | rs6800733 | 0.692 | 0.555 | 0.242 | -1.696 | -0.037 | -0.015 | -0.015 | 0 | 0.21 | 0.3 | 0.1 | 0.18 | 0.16 | 0.28 |
| 3 | 74072243 | rs2030875 | -1.74 | 0.841 | 0.181 | -1.891 | -0.012 | -0.015 | -0.015 | 0 | 0.71 | 0.85 | 0.55 | 0.39 | 0.69 | 0.64 |
| 3 | 74073182 | rs1949931 | 0.732 | 0.852 | 0.191 | -1.859 | -0.025 | -0.019 | -0.019 | 0.296 | 0.21 | 0.33 | 0.11 | 0.18 | 0.18 | 0.29 |
| 3 | 74080776 | rs2323205 | 0.027 | 0.921 | 0.194 | -1.809 | 0.003 | -0.013 | -0.013 | 0.037 | 0.14 | 0.24 | 0.22 | 0.13 | 0.12 | 0.15 |
| 3 | 74081555 | rs9790237 | 1.338 | 1.13 | -0.173 | -1.509 | -0.02 | -0.019 | -0.019 | 0 | 0.29 | 0.12 | 0.46 | 0.62 | 0.31 | 0.41 |
| 3 | 74085057 | rs12639599 | 1.383 | 0.88 | -0.685 | -1.23 | -0.043 | -0.114 | -0.114 | 0.476 | 0.29 | 0.03 | 0.37 | 0.57 | 0.28 | 0.41 |
| 3 | 74086175 | rs2222688 | 0.624 | 0.956 | -0.645 | -1.203 | -0.027 | -0.009 | -0.009 | 0 | 0.39 | 0.19 | 0.44 | 0.56 | 0.41 | 0.43 |
| 3 | 74095314 | rs1526727 | 0.431 | 1.237 | -0.596 | -0.983 | 0.003 | -0.06 | -0.06 | 0 | 0.62 | 0.61 | 0.75 | 0.92 | 0.59 | 0.81 |
| 3 | 74095484 | rs1526728 | 0.312 | 1.305 | -0.701 | -0.954 | 0 | -0.033 | -0.033 | 3.984 | 0.62 | 0.66 | 0.76 | 0.92 | 0.59 | 0.81 |
| 3 | 74096374 | rs1569000 | -0.38 | 1.137 | -1.056 | -0.49 | -0.002 | 0.028 | 0.028 | 0 | 0.27 | 0.22 | 0.19 | 0.19 | 0.3 | 0.23 |
| 3 | 74097623 | rs4677373 | 0.109 | 1.917 | -1.291 | 0.901 | -0.074 | -0.075 | -0.075 | 0.123 | 0.11 | 0.01 | 0.44 | 0.41 | 0.23 | 0.23 |
| 3 | 74098901 | rs9990068 | -0.594 | 1.155 | -0.923 | -0.025 | -0.021 | -0.075 | -0.075 | 0 | 0.29 | 0.6 | 0.17 | 0.08 | 0.22 | 0.12 |
| 3 | 74105648 | rs1526710 | 0.111 | 1.368 | -1.461 | 0.256 | -0.03 | -0.038 | -0.038 | 0.073 | 0.08 | 0 | 0.07 | 0.1 | 0.09 | 0.15 |
| 3 | 74109475 | rs7645365 | 0.469 | 0.79 | -0.796 | 0.342 | -0.004 | -0.008 | -0.008 | 1.015 | 0.68 | 0.68 | 0.55 | 0.58 | 0.61 | 0.64 |
| 3 | 74124566 | rs7427597 | 0.471 | -0.52 | -0.502 | -0.386 | -0.005 | 0.007 | 0.007 | 0 | 0.37 | 0.34 | 0.64 | 0.68 | 0.51 | 0.32 |
| 3 | 74128102 | rs6549582 | 0.698 | -0.58 | -0.4 | -0.383 | -0.002 | 0 | 0 | 0.234 | 0.34 | 0.34 | 0.63 | 0.66 | 0.49 | 0.32 |
| 3 | 74140019 | rs13098451 | -0.11 | -1.121 | 0.468 | -0.792 | -0.001 | -0.059 | -0.059 | 0 | 0.48 | 0.48 | 0.28 | 0.06 | 0.41 | 0.51 |
| 3 | 74143047 | rs1447676 | -0.182 | -1.145 | 0.498 | -0.798 | -0.008 | -0.078 | -0.078 | 0 | 0.54 | 0.5 | 0.73 | 0.94 | 0.61 | 0.49 |
| 3 | 74146788 | rs6549584 | -0.698 | -1.415 | 0.535 | -0.613 | 0.012 | 0.072 | 0.072 | 0.241 | 0.06 | 0.21 | 0.05 | 0.14 | 0.04 | 0.05 |
| 3 | 74151374 | rs9861321 | -0.49 | -1.775 | 0.674 | -0.946 | -0.007 | -0.052 | -0.052 | 0.399 | 0.72 | 0.64 | 0.81 | 0.83 | 0.75 | 0.52 |
| 3 | 74155133 | rs278481 | -0.682 | -1.635 | 0.865 | -0.956 | 0.017 | 0.052 | 0.052 | 0.001 | 0.68 | 0.85 | 0.84 | 0.97 | 0.75 | 0.56 |
| 3 | 74159454 | rs1440625 | 0.848 | -1.896 | 0.778 | -1.035 | 0.011 | 0.025 | 0.025 | 0 | 0.15 | 0.07 | 0.1 | 0.02 | 0.14 | 0.07 |
| 3 | 74160426 | rs6779811 | 1.044 | -2.015 | 0.935 | -0.823 | -0.007 | 0.062 | 0.062 | 0.073 | 0.42 | 0.26 | 0.52 | 0.73 | 0.54 | 0.44 |
| 3 | 74163712 | rs1440627 | -0.196 | -1.947 | 1.043 | -0.788 | -0.019 | -0.051 | -0.051 | 0 | 0.63 | 0.59 | 0.79 | 0.79 | 0.72 | 0.52 |
| 3 | 74168617 | rs1440629 | -0.601 | -1.896 | 1.011 | -0.813 | -0.006 | -0.026 | -0.026 | 0 | 0.67 | 0.68 | 0.81 | 0.83 | 0.74 | 0.52 |
| 3 | 74174264 | rs7631363 | 1.272 | -1.185 | 0.846 | -0.244 | -0.118 | -0.055 | -0.055 | 0.424 | 0.33 | 0.08 | 0.47 | 0.6 | 0.47 | 0.27 |
| 3 | 74175307 | rs2875149 | -1.394 | -0.824 | 1.492 | 0.108 | -0.025 | 0.034 | 0.034 | 0.072 | 0.14 | 0.26 | 0.22 | 0.21 | 0.11 | 0.18 |
| 3 | 74177841 | rs172142 | -1.036 | -0.734 | 1.646 | -0.061 | -0.002 | 0.056 | 0.056 | 0 | 0.1 | 0.34 | 0.08 | 0.16 | 0.08 | 0.18 |
| 3 | 74184872 | rs649661 | -2.202 | -0.954 | 1.851 | -0.109 | -0.001 | -0.001 | -0.001 | 0 | 0.2 | 0.8 | 0.15 | 0.28 | 0.13 | 0.24 |
| 3 | 74191690 | rs278405 | -2.157 | -0.962 | 1.999 | 0.011 | -0.013 | 0.007 | 0.007 | 2.132 | 0.2 | 0.76 | 0.16 | 0.3 | 0.13 | 0.24 |
| 3 | 74194608 | rs278424 | -1.684 | -0.973 | 2.663 | 0.013 | -0.024 | 0.085 | 0.085 | 0.056 | 0.16 | 0.55 | 0.13 | 0.28 | 0.13 | 0.16 |
| 3 | 74206048 | rs13072365 | 1.286 | -1.134 | 2.039 | -0.33 | -0.162 | -0.086 | -0.086 | 0 | 0.34 | 0.03 | 0.6 | 0.62 | 0.49 | 0.26 |
| 3 | 74212519 | rs278459 | -0.427 | -1.107 | 2.154 | -0.337 | 0.05 | 0.178 | 0.178 | 0 | 0.54 | 0.79 | 0.76 | 0.92 | 0.62 | 0.5 |
| 3 | 74216920 | rs278463 | -0.54 | -1.141 | 2.197 | -0.301 | -0.013 | 0.021 | 0.021 | 0.047 | 0.1 | 0.26 | 0.07 | 0.13 | 0.08 | 0.13 |
| 3 | 74227419 | rs188077 | -0.376 | -1.065 | 1.939 | -0.362 | 0.065 | 0.173 | 0.173 | 0 | 0.53 | 0.78 | 0.72 | 0.92 | 0.64 | 0.52 |
| 3 | 74253010 | rs17582032 | 1.339 | -0.859 | 1.378 | -0.173 | -0.155 | -0.096 | -0.096 | 0 | 0.34 | 0.04 | 0.53 | 0.6 | 0.49 | 0.27 |
| 3 | 74253678 | rs36003 | -0.093 | -0.896 | 1.632 | -0.26 | 0.055 | 0.217 | 0.217 | 0.018 | 0.51 | 0.75 | 0.67 | 0.91 | 0.6 | 0.52 |
| 3 | 74263465 | rs6780663 | -0.221 | -1.068 | 1.658 | -0.276 | 0.02 | 0.028 | 0.028 | 0 | 0.53 | 0.56 | 0.7 | 0.86 | 0.64 | 0.54 |
| 3 | 74263840 | rs6765892 | -0.337 | -0.95 | 1.222 | -0.159 | -0.003 | 0.065 | 0.065 | 0.06 | 0.09 | 0.16 | 0.08 | 0.26 | 0.09 | 0.18 |
| 3 | 74263894 | rs6789681 | 0.809 | -0.824 | 0.951 | -0.152 | -0.061 | -0.025 | -0.025 | 1.748 | 0.38 | 0.09 | 0.56 | 0.61 | 0.53 | 0.27 |
| 3 | 74266309 | rs4234186 | 1.615 | -0.941 | 1.326 | -0.602 | -0.02 | -0.051 | -0.051 | 0 | 0.87 | 0.42 | 0.88 | 0.97 | 0.91 | 0.93 |
| 3 | 74269761 | rs35989 | 0.322 | -0.615 | 1.238 | -0.428 | -0.017 | 0.046 | 0.046 | 0 | 0.43 | 0.69 | 0.52 | 0.69 | 0.48 | 0.35 |
| 3 | 74277232 | rs3845989 | 1.201 | -0.563 | 0.654 | -0.38 | -0.066 | -0.017 | -0.017 | 0 | 0.34 | 0.16 | 0.44 | 0.63 | 0.45 | 0.28 |
| 3 | 74280205 | rs17012210 | 0.044 | -0.666 | 0.519 | -0.295 | -0.007 | 0.007 | 0.007 | 0 | 0.1 | 0.17 | 0.07 | 0.14 | 0.09 | 0.16 |
| 3 | 74282522 | rs35975 | -0.053 | -0.599 | 0.073 | -0.32 | 0.037 | 0.182 | 0.182 | 0 | 0.51 | 0.82 | 0.59 | 0.82 | 0.57 | 0.5 |
| 3 | 74289885 | rs35961 | -0.79 | -0.956 | -0.099 | -0.034 | 0.081 | 0.009 | 0.009 | 0.345 | 0.4 | 0.86 | 0.4 | 0.33 | 0.3 | 0.35 |
| 3 | 74307352 | rs7651173 | -0.537 | -0.341 | -0.593 | -0.164 | -0.011 | -0.006 | -0.006 | 0.022 | 0.34 | 0.41 | 0.42 | 0.28 | 0.29 | 0.28 |
| 3 | 74321141 | rs541653 | 0.192 | 0.491 | -1.358 | -0.059 | 0.001 | 0.141 | 0.141 | 0 | 0.49 | 0.73 | 0.53 | 0.8 | 0.46 | 0.53 |
| 3 | 74329839 | rs12492560 | -0.418 | -0.05 | -1.621 | 0.27 | 0.042 | -0.035 | -0.035 | 0 | 0.36 | 0.22 | 0.4 | 0.5 | 0.25 | 0.55 |
| 3 | 74332837 | rs17650372 | -0.319 | 1.458 | -1.944 | 0.267 | -0.041 | -0.025 | -0.025 | 0 | 0.07 | 0.01 | 0.12 | 0.1 | 0.23 | 0.08 |
| 3 | 74335764 | rs862825 | -0.007 | 0.874 | -1.942 | 0.183 | -0.001 | 0 | 0 | 0 | 0.11 | 0.12 | 0.09 | 0.16 | 0.12 | 0.11 |
| 3 | 74343804 | rs6779469 | -0.374 | -0.549 | -1.44 | -0.473 | 0.019 | 0.006 | 0.006 | 0 | 0.53 | 0.41 | 0.45 | 0.54 | 0.32 | 0.64 |
| 3 | 74344356 | rs10490832 | 0.449 | -1.11 | -1.908 | -0.711 | -0.014 | 0.073 | 0.073 | 0 | 0.16 | 0.01 | 0.07 | 0.03 | 0.14 | 0.22 |
| 3 | 74345173 | rs9867036 | -0.708 | -0.64 | -1.294 | -0.694 | 0.003 | 0.001 | 0.001 | 0 | 0.56 | 0.42 | 0.5 | 0.54 | 0.42 | 0.65 |
| 3 | 74348587 | rs614023 | -1.21 | -1.289 | -0.661 | -0.369 | 0.028 | 0.027 | 0.027 | 1.156 | 0.43 | 0.39 | 0.4 | 0.7 | 0.25 | 0.6 |
| 3 | 74349062 | rs626578 | 0.584 | -1.299 | -0.783 | -0.16 | -0.003 | 0.024 | 0.024 | 0 | 0.09 | 0.12 | 0.07 | 0.23 | 0.07 | 0.1 |
| 3 | 74350297 | rs558917 | -1.193 | -1.469 | -0.163 | -0.248 | 0.076 | 0.07 | 0.07 | 1.248 | 0.46 | 0.73 | 0.44 | 0.7 | 0.25 | 0.59 |
| 3 | 74352261 | rs1545384 | 0.184 | -1.562 | -0.581 | -0.332 | 0.002 | -0.167 | -0.167 | 0.696 | 0.21 | 0.01 | 0.28 | 0.47 | 0.16 | 0.5 |
| 3 | 74358189 | rs507833 | -1.05 | -1.08 | -0.429 | -0.132 | -0.028 | -0.026 | -0.026 | 0 | 0.44 | 0.53 | 0.41 | 0.7 | 0.25 | 0.59 |
| 3 | 74365556 | rs667986 | -0.064 | 0.208 | -0.581 | 0.404 | -0.004 | 0.022 | 0.022 | 0 | 0.08 | 0.12 | 0.07 | 0.2 | 0.06 | 0.08 |
| 3 | 74371936 | rs677171 | -0.275 | 0.227 | -0.573 | 0.37 | -0.003 | 0.02 | 0.02 | 0 | 0.09 | 0.12 | 0.07 | 0.23 | 0.07 | 0.11 |

| | | | | | | | | | | | | | | | | |
|---|---|---|---|---|---|---|---|---|---|---|---|---|---|---|---|---|
| 3 | 74378567 | rs540365 | 0.797 | 0.67 | -0.419 | 0.219 | 0.027 | -0.02 | -0.02 | 0 | 0.71 | 0.52 | 0.72 | 0.74 | 0.7 | 0.73 |
| 3 | 74380566 | rs479652 | 0.413 | 0.812 | -0.94 | 0.319 | -0.118 | -0.047 | -0.047 | 0 | 0.77 | 0.96 | 0.74 | 0.73 | 0.66 | 0.72 |
| 3 | 74391718 | rs491492 | -0.09 | 0.715 | -0.295 | 0.134 | 0.026 | 0.018 | 0.018 | 1.35 | 0.06 | 0.28 | 0.03 | 0 | 0.01 | 0.01 |
| 3 | 74391757 | rs2044594 | 0.475 | 0.693 | -1.107 | 0.019 | 0.011 | 0.016 | 0.016 | 0.557 | 0.45 | 0.44 | 0.51 | 0.68 | 0.39 | 0.62 |
| 3 | 74392917 | rs589344 | -0.221 | 0.064 | -1.205 | 0.05 | -0.011 | -0.035 | -0.035 | 0.489 | 0.41 | 0.48 | 0.37 | 0.17 | 0.34 | 0.23 |
| 3 | 74398369 | rs549560 | -0.234 | -0.226 | -2.192 | 0.591 | 0.069 | -0.043 | -0.043 | 0 | 0.38 | 0.87 | 0.33 | 0.36 | 0.32 | 0.29 |
| 3 | 74401129 | rs551595 | 0.103 | 0.226 | -2.349 | 0.35 | 0.007 | 0.087 | 0.087 | 0.974 | 0.58 | 0.84 | 0.65 | 0.79 | 0.59 | 0.73 |
| 3 | 74403730 | rs17012416 | -0.018 | -0.332 | -2.25 | 0.805 | 0.021 | -0.026 | -0.026 | 0 | 0.08 | 0.03 | 0.1 | 0.24 | 0.04 | 0.19 |
| 3 | 74405726 | rs4677384 | 0.415 | -0.234 | -2.608 | 0.825 | -0.018 | -0.028 | -0.028 | 0 | 0.25 | 0.05 | 0.28 | 0.17 | 0.26 | 0.09 |
| 3 | 74409330 | rs17012429 | -0.318 | -0.407 | -2.601 | 1.122 | 0.035 | -0.053 | -0.053 | 0 | 0.11 | 0.03 | 0.11 | 0.31 | 0.06 | 0.25 |
| 3 | 74409801 | rs11715775 | 0.518 | -0.551 | -2.481 | 0.963 | 0.007 | -0.021 | -0.021 | 0 | 0.22 | 0.12 | 0.24 | 0.21 | 0.22 | 0.14 |
| 3 | 74413676 | rs6549590 | -0.933 | 0.052 | -2.193 | 0.378 | 0.003 | 0.018 | 0.018 | 1.375 | 0.52 | 0.31 | 0.52 | 0.47 | 0.56 | 0.46 |
| 3 | 74432477 | rs11713787 | 0.392 | 0.062 | -1.851 | 0.195 | -0.078 | 0.022 | 0.022 | 2.048 | 0.27 | 0.1 | 0.33 | 0.14 | 0.39 | 0.15 |
| 3 | 74435355 | rs10865681 | 0.884 | 0.002 | -2.596 | 0.158 | -0.069 | -0.144 | -0.144 | 0 | 0.21 | 0.01 | 0.18 | 0.38 | 0.24 | 0.35 |
| 3 | 74444271 | rs12494066 | 0.613 | -0.881 | -1.801 | -0.457 | -0.045 | 0 | 0 | 0 | 0.36 | 0.24 | 0.37 | 0.66 | 0.43 | 0.56 |
| 3 | 74447090 | rs6780510 | -0.124 | -1.201 | -1.919 | -0.935 | 0.06 | 0.08 | 0.08 | 0 | 0.71 | 0.94 | 0.76 | 0.81 | 0.81 | 0.67 |
| 3 | 74452081 | rs10865682 | 0.511 | -0.868 | -1.277 | -0.759 | -0.002 | 0.001 | 0.001 | 2.497 | 0.37 | 0.35 | 0.4 | 0.69 | 0.41 | 0.52 |
| 3 | 74458518 | rs4677391 | -0.36 | -1.433 | -1.384 | -0.977 | 0.041 | 0.071 | 0.071 | 0 | 0.74 | 0.89 | 0.85 | 0.83 | 0.83 | 0.69 |
| 3 | 74460896 | rs11128385 | 0.252 | -1.435 | -1.001 | -1.036 | 0.023 | 0.051 | 0.051 | 0 | 0.66 | 0.77 | 0.79 | 0.81 | 0.73 | 0.61 |
| 3 | 74461792 | rs7433180 | 0.701 | -1.556 | -1.09 | -1.126 | 0.066 | 0.125 | 0.125 | 0.263 | 0.57 | 0.21 | 0.71 | 0.8 | 0.71 | 0.6 |
| 3 | 74465691 | rs4073942 | 0.69 | -0.974 | -1.275 | -0.874 | -0.069 | 0.003 | 0.003 | 0.918 | 0.41 | 0.11 | 0.5 | 0.64 | 0.5 | 0.49 |
| 3 | 74479445 | rs13098874 | 0.423 | -0.672 | -0.469 | -0.553 | -0.009 | -0.004 | -0.004 | 0 | 0.37 | 0.29 | 0.4 | 0.59 | 0.42 | 0.38 |
| 3 | 74485139 | rs13064307 | 0.549 | -0.635 | -0.51 | -0.537 | -0.027 | -0.014 | -0.014 | 0 | 0.37 | 0.24 | 0.39 | 0.59 | 0.42 | 0.38 |
| 3 | 74490422 | rs6549595 | -0.999 | -0.622 | -0.913 | -0.582 | -0.026 | 0.101 | 0.101 | 0.125 | 0.71 | 0.93 | 0.84 | 0.83 | 0.72 | 0.61 |
| 3 | 74497115 | rs6764156 | 1.459 | -0.276 | -0.208 | -0.313 | 0.01 | 0.008 | 0.008 | 1.958 | 0.18 | 0.28 | 0.31 | 0.14 | 0.22 | 0.06 |
| 3 | 74498707 | rs7432763 | 0.328 | -1.456 | -0.162 | 0.22 | -0.011 | 0.138 | 0.138 | 0.991 | 0.16 | 0.24 | 0.17 | 0.47 | 0.13 | 0.18 |
| 3 | 74508426 | rs7428834 | 1.152 | -1.555 | -0.194 | 0.064 | 0.001 | -0.005 | -0.005 | 0.464 | 0.41 | 0.54 | 0.58 | 0.29 | 0.52 | 0.3 |
| 3 | 74513576 | rs7430250 | -0.009 | -1.403 | -0.289 | 0.147 | 0.03 | 0.138 | 0.138 | 0.057 | 0.58 | 0.81 | 0.74 | 0.76 | 0.65 | 0.47 |
| 3 | 74515252 | rs7432669 | 1.321 | -1.392 | -0.182 | 0.157 | 0.004 | -0.012 | -0.012 | 0 | 0.41 | 0.44 | 0.56 | 0.29 | 0.52 | 0.3 |
| 3 | 74517138 | rs7429592 | 0.06 | -1.463 | -0.412 | 0.182 | 0.036 | 0.143 | 0.143 | 0.12 | 0.58 | 0.8 | 0.74 | 0.76 | 0.65 | 0.46 |
| 3 | 74517413 | rs7433810 | 1.216 | -1.329 | -0.662 | 0.24 | -0.003 | 0.007 | 0.007 | 0.506 | 0.41 | 0.4 | 0.57 | 0.29 | 0.54 | 0.31 |
| 3 | 74523135 | rs7635421 | 0.002 | -0.492 | -1.025 | 0.976 | -0.006 | 0.092 | 0.092 | 2.109 | 0.13 | 0.15 | 0.14 | 0.46 | 0.11 | 0.15 |
| 3 | 74526216 | rs11918154 | -0.327 | -0.496 | -1.449 | 0.224 | -0.024 | 0.077 | 0.077 | 0 | 0.28 | 0.06 | 0.16 | 0.18 | 0.27 | 0.36 |
| 3 | 74527165 | rs7429855 | 0.533 | -0.672 | -0.551 | 0.949 | -0.018 | 0.049 | 0.049 | 0 | 0.71 | 0.6 | 0.75 | 0.59 | 0.75 | 0.76 |
| 3 | 74529226 | rs7430041 | 0.001 | -0.69 | -0.671 | 0.415 | 0.002 | 0.002 | 0.002 | 0.019 | 0.15 | 0.1 | 0.07 | 0.08 | 0.14 | 0.11 |
| 3 | 74529592 | rs7431887 | 0.001 | -0.69 | -0.671 | 0.415 | 0.002 | 0.002 | 0.002 | 0.019 | 0.15 | 0.1 | 0.07 | 0.08 | 0.14 | 0.11 |
| 3 | 74538638 | rs9830820 | 0.611 | -0.563 | -0.757 | 0.529 | -0.023 | 0.058 | 0.058 | 0 | 0.44 | 0.27 | 0.44 | 0.32 | 0.51 | 0.44 |
| 3 | 74541725 | rs7639846 | 0.768 | -0.609 | -0.237 | 0.309 | 0.031 | 0.133 | 0.133 | 2.041 | 0.5 | 0.26 | 0.34 | 0.13 | 0.57 | 0.33 |
| 3 | 74543312 | rs7427025 | 0.329 | -0.865 | 0.486 | 1.269 | -0.055 | 0.046 | 0.046 | 0.262 | 0.84 | 0.67 | 0.83 | 0.74 | 0.94 | 0.87 |
| 3 | 74545777 | rs6549604 | 0.72 | -0.782 | 0.38 | 1.31 | -0.058 | 0.099 | 0.099 | 0.992 | 0.8 | 0.53 | 0.81 | 0.71 | 0.91 | 0.8 |
| 3 | 74548611 | rs6765009 | 1.081 | -0.142 | 0.639 | 2.018 | 0.032 | 0.005 | 0.005 | 0.132 | 0.73 | 0.6 | 0.5 | 0.31 | 0.71 | 0.63 |
| 3 | 74550084 | rs6549607 | 1.158 | -0.185 | 0.821 | 2.042 | 0.029 | 0.235 | 0.235 | 0 | 0.91 | 0.69 | 0.76 | 0.7 | 0.93 | 0.9 |
| 3 | 74551469 | rs7428981 | 1.009 | -0.175 | 0.855 | 2.046 | 0.014 | 0.167 | 0.167 | 0 | 0.91 | 0.8 | 0.77 | 0.7 | 0.93 | 0.89 |
| 3 | 74557169 | rs6549614 | 0.979 | -0.215 | 0.848 | 2.067 | 0.014 | 0.07 | 0.07 | 0.024 | 0.78 | 0.68 | 0.53 | 0.43 | 0.78 | 0.68 |
| 3 | 74563488 | rs7430259 | 1.175 | -0.257 | 0.809 | 2.127 | 0.027 | 0.08 | 0.08 | 0.028 | 0.78 | 0.72 | 0.66 | 0.51 | 0.75 | 0.61 |
| 3 | 74571050 | rs4677408 | 1.894 | -0.362 | 0.915 | 1.719 | 0.009 | 0.006 | 0.006 | 0 | 0.89 | 0.73 | 0.86 | 0.91 | 0.92 | 0.91 |
| 3 | 74577921 | rs6767471 | 2.135 | -0.476 | 1.034 | 1.625 | 0.023 | -0.041 | -0.041 | 0 | 0.91 | 0.71 | 0.93 | 1 | 0.94 | 0.94 |
| 3 | 74584605 | rs9831609 | 2.341 | -0.493 | 1.025 | 1.631 | 0.004 | -0.018 | -0.018 | 0.009 | 0.88 | 0.67 | 0.85 | 0.94 | 0.92 | 0.91 |
| 3 | 74607020 | rs4677414 | 1.799 | -0.48 | 0.853 | 1.638 | 0 | -0.001 | -0.001 | 0 | 0.92 | 0.89 | 0.89 | 0.94 | 0.92 | 0.91 |
| 3 | 74613171 | rs1447830 | 0.468 | -0.53 | 0.028 | 1.002 | -0.042 | 0.021 | 0.021 | 0 | 0.15 | 0.02 | 0.08 | 0.09 | 0.18 | 0.11 |
| 3 | 74616253 | rs11928037 | 1.069 | -0.963 | -0.065 | 0.754 | -0.021 | 0.037 | 0.037 | 0 | 0.75 | 0.59 | 0.7 | 0.61 | 0.78 | 0.72 |
| 3 | 74618288 | rs733144 | 0.781 | -0.881 | -0.336 | 0.719 | 0.006 | -0.027 | -0.027 | 0 | 0.12 | 0.07 | 0.15 | 0.35 | 0.13 | 0.17 |
| 3 | 74621026 | rs7429759 | 0.158 | -0.816 | -0.272 | 0.724 | -0.001 | -0.029 | -0.029 | 0 | 0.8 | 0.84 | 0.75 | 0.62 | 0.78 | 0.74 |
| 3 | 74621972 | rs9814962 | 0.094 | -0.816 | -0.282 | 0.724 | -0.005 | -0.047 | -0.047 | 0.086 | 0.81 | 0.87 | 0.75 | 0.62 | 0.78 | 0.74 |
| 3 | 74625123 | rs7630534 | 0.177 | -0.84 | -0.255 | 0.744 | 0 | -0.005 | -0.005 | 0 | 0.8 | 0.8 | 0.74 | 0.62 | 0.79 | 0.74 |
| 3 | 74627703 | rs13073838 | 1.747 | -1.395 | -0.943 | -0.076 | -0.077 | 0.178 | 0.178 | 0.013 | 0.32 | 0.07 | 0.2 | 0 | 0.38 | 0.2 |
| 3 | 74628590 | rs1579905 | 0.655 | -1.548 | -0.445 | 0.33 | 0.002 | -0.001 | -0.001 | 0 | 0.61 | 0.63 | 0.64 | 0.54 | 0.62 | 0.62 |
| 3 | 74629235 | rs6549631 | 0.188 | -1.774 | 0.458 | 0.817 | -0.016 | 0.135 | 0.135 | 0 | 0.79 | 0.62 | 0.62 | 0.47 | 0.81 | 0.7 |
| 3 | 74632701 | rs7429609 | 0.198 | -1.84 | 0.967 | 0.674 | -0.044 | 0.124 | 0.124 | 0 | 0.88 | 0.54 | 0.82 | 0.82 | 0.97 | 0.87 |
| 3 | 74642677 | rs13314386 | -1.601 | -1.975 | 0.809 | 0.662 | -0.045 | 0.031 | 0.031 | 0.249 | 0.21 | 0.57 | 0.27 | 0.3 | 0.09 | 0.32 |
| 3 | 74650559 | rs9808933 | -1.257 | -1.972 | 0.646 | 0.709 | -0.049 | 0.064 | 0.064 | 0.042 | 0.18 | 0.53 | 0.24 | 0.3 | 0.06 | 0.29 |
| 3 | 74654727 | rs2121798 | -1.59 | -2.031 | 0.384 | 0.688 | -0.041 | 0.017 | 0.017 | 0.294 | 0.22 | 0.62 | 0.24 | 0.29 | 0.08 | 0.31 |
| 3 | 74669078 | rs10511037 | 0.184 | -2.121 | -0.094 | 0.377 | 0.028 | -0.042 | -0.042 | 0 | 0.16 | 0.06 | 0.14 | 0.24 | 0.15 | 0.14 |
| 3 | 74670150 | rs17711625 | 0.032 | -2.436 | 0.041 | -0.321 | -0.002 | -0.018 | -0.018 | 0 | 0.56 | 0.5 | 0.43 | 0.35 | 0.6 | 0.53 |
| 3 | 74672571 | rs17012697 | -0.871 | -2.402 | 0.034 | -0.233 | -0.008 | 0.002 | 0.002 | 0 | 0.26 | 0.28 | 0.2 | 0.35 | 0.2 | 0.31 |
| 3 | 74675911 | rs11715812 | 0.001 | -2.427 | -0.504 | -0.113 | 0.04 | -0.056 | -0.056 | 0 | 0.18 | 0.06 | 0.14 | 0.25 | 0.16 | 0.14 |
| 3 | 74676957 | rs1374866 | 0.756 | -2.273 | -0.588 | -0.688 | 0.001 | -0.047 | -0.047 | 0 | 0.37 | 0.57 | 0.28 | 0.11 | 0.41 | 0.4 |
| 3 | 74688458 | rs9310301 | -0.342 | -2.162 | -0.847 | -0.385 | -0.056 | -0.012 | -0.012 | 0 | 0.63 | 0.8 | 0.47 | 0.41 | 0.57 | 0.57 |
| 3 | 74703689 | rs6775796 | 0.306 | -2.161 | -0.6 | -0.28 | 0.013 | -0.014 | -0.014 | 0 | 0.23 | 0.21 | 0.2 | 0.36 | 0.13 | 0.16 |

| | | | | | | | | | | | | | | | | |
|---|---|---|---|---|---|---|---|---|---|---|---|---|---|---|---|---|
| 3 | 74710143 | rs9853168 | 1.607 | -2.23 | -1.194 | -1.323 | -0.089 | 0.197 | 0.197 | 0.548 | 0.34 | 0.05 | 0.27 | 0.01 | 0.45 | 0.26 |
| 3 | 74710819 | rs935523 | -1.552 | -1.973 | -1.362 | -1.259 | -0.053 | 0.033 | 0.033 | 0 | 0.86 | 0.97 | 0.89 | 0.95 | 0.81 | 0.88 |
| 3 | 74718651 | rs17012751 | 1.156 | -1.76 | -0.914 | -1.156 | 0.004 | -0.001 | -0.001 | 0 | 0.2 | 0.12 | 0.2 | 0.24 | 0.12 | 0.16 |
| 3 | 74733670 | rs1880966 | -0.882 | -1.793 | 0.017 | -1.232 | -0.004 | 0.011 | 0.011 | 0 | 0.8 | 0.82 | 0.89 | 0.93 | 0.81 | 0.85 |
| 3 | 74734628 | rs6549636 | 0.79 | -1.856 | 0.275 | -1.178 | 0.003 | -0.016 | -0.016 | 0.553 | 0.58 | 0.4 | 0.5 | 0.28 | 0.62 | 0.47 |
| 3 | 74737022 | rs12714810 | 0.839 | -1.862 | 0.263 | -1.203 | -0.007 | 0.033 | 0.033 | 0.635 | 0.57 | 0.3 | 0.49 | 0.27 | 0.62 | 0.47 |
| 3 | 74740625 | rs12494320 | 0.63 | -1.597 | 0.392 | -1.044 | 0.007 | 0.039 | 0.039 | 0 | 0.22 | 0.37 | 0.38 | 0.64 | 0.19 | 0.38 |
| 3 | 74742523 | rs6808687 | 1.997 | -1.578 | 0.052 | -1.169 | -0.078 | 0.174 | 0.174 | 0.179 | 0.34 | 0.09 | 0.27 | 0 | 0.45 | 0.26 |
| 3 | 74743990 | rs4234187 | 0.642 | -1.505 | 0.314 | -0.989 | -0.002 | -0.006 | -0.006 | 0.134 | 0.22 | 0.23 | 0.41 | 0.65 | 0.19 | 0.4 |
| 3 | 74762379 | rs6803546 | 0.917 | -1.181 | 0.355 | -1.018 | 0.008 | -0.052 | -0.052 | 0 | 0.4 | 0.49 | 0.35 | 0.1 | 0.51 | 0.38 |
| 3 | 74770492 | rs9837543 | 0.933 | -1.156 | 0.426 | -1.06 | 0.016 | -0.105 | -0.105 | 0 | 0.39 | 0.42 | 0.31 | 0.03 | 0.51 | 0.3 |
| 3 | 74781833 | rs13067637 | 1.469 | -1.142 | 0.653 | -1.083 | -0.019 | 0.005 | 0.005 | 0.169 | 0.35 | 0.22 | 0.29 | 0 | 0.47 | 0.26 |
| 3 | 74783321 | rs7635628 | 1.297 | -1.147 | 0.501 | -1.079 | 0.002 | -0.083 | -0.083 | 0 | 0.36 | 0.32 | 0.29 | 0.01 | 0.47 | 0.26 |
| 3 | 74787022 | rs13061773 | 1.144 | -1.142 | 0.428 | -0.987 | 0.009 | -0.077 | -0.077 | 0.447 | 0.38 | 0.4 | 0.4 | 0.1 | 0.48 | 0.34 |
| 3 | 74796435 | rs11712042 | 2.777 | 0.149 | -0.377 | -1.249 | -0.085 | 0.076 | 0.076 | 0.039 | 0.14 | 0.02 | 0.14 | 0 | 0.3 | 0.07 |
| 3 | 74805509 | rs9846280 | 1.285 | -0.772 | 0.073 | -1.204 | -0.048 | 0.101 | 0.101 | 0.559 | 0.36 | 0.19 | 0.39 | 0.09 | 0.51 | 0.34 |
| 3 | 74812287 | rs1918398 | 1.289 | -0.783 | 0.037 | -1.209 | -0.065 | 0.133 | 0.133 | 0.552 | 0.36 | 0.16 | 0.38 | 0.09 | 0.51 | 0.35 |
| 3 | 74813794 | rs2141178 | -1.895 | -0.957 | -0.262 | -1.039 | -0.039 | -0.053 | -0.053 | 0.008 | 0.87 | 0.99 | 0.94 | 0.81 | 0.87 | 0.88 |
| 3 | 74821600 | rs2049623 | 1.057 | -0.94 | 0.366 | -1.093 | 0.001 | -0.019 | -0.019 | 0 | 0.41 | 0.52 | 0.42 | 0.14 | 0.53 | 0.37 |
| 3 | 74831395 | rs7621878 | 1.128 | -0.989 | 0.448 | -1.093 | 0.016 | -0.059 | -0.059 | 0.188 | 0.4 | 0.47 | 0.42 | 0.14 | 0.53 | 0.37 |
| 3 | 74836374 | rs7651319 | 1.069 | -0.968 | -0.065 | -1.083 | 0.015 | -0.058 | -0.058 | 0 | 0.41 | 0.46 | 0.41 | 0.14 | 0.54 | 0.37 |
| 3 | 74853611 | rs7627286 | -0.244 | -0.998 | -0.877 | -1.175 | 0.027 | 0.068 | 0.068 | 0 | 0.45 | 0.67 | 0.52 | 0.72 | 0.35 | 0.53 |
| 3 | 74861269 | rs6549659 | -0.596 | -0.573 | -0.848 | -0.775 | 0.003 | -0.002 | -0.002 | 0.022 | 0.81 | 0.8 | 0.9 | 0.79 | 0.87 | 0.88 |
| 3 | 74865631 | rs6549660 | 0.137 | -0.873 | -1.011 | -1.236 | -0.006 | 0.029 | 0.029 | 0.103 | 0.63 | 0.66 | 0.52 | 0.14 | 0.68 | 0.54 |
| 3 | 74867417 | rs1918400 | 0.924 | -1.93 | -1.292 | -1.162 | 0.053 | 0.099 | 0.099 | 0.046 | 0.19 | 0.01 | 0.07 | 0.05 | 0.13 | 0.15 |
| 3 | 74880857 | rs1918392 | -1.48 | -1.629 | -1.37 | -0.766 | -0.019 | -0.062 | -0.062 | 0 | 0.81 | 0.98 | 0.86 | 0.73 | 0.84 | 0.85 |
| 3 | 74885751 | rs13073856 | 0.885 | -1.583 | -1.338 | -0.855 | -0.003 | 0.014 | 0.014 | 0 | 0.08 | 0.03 | 0.03 | 0.03 | 0.09 | 0.1 |
| 3 | 74887781 | rs941304 | -0.106 | -2.037 | -0.394 | -0.839 | -0.038 | 0.001 | 0.001 | 0.525 | 0.71 | 0.47 | 0.8 | 0.66 | 0.81 | 0.83 |
| 3 | 74908822 | rs9310304 | -1.02 | -2.086 | 0.306 | -0.848 | 0.014 | -0.01 | -0.01 | 0 | 0.77 | 0.85 | 0.85 | 0.67 | 0.83 | 0.8 |
| 3 | 74915001 | rs9817859 | -0.224 | -2.072 | 0.309 | -0.784 | -0.044 | 0.009 | 0.009 | 0.445 | 0.72 | 0.56 | 0.81 | 0.65 | 0.81 | 0.8 |
| 3 | 74916063 | rs4417905 | 0.747 | -2.337 | 0.092 | -1.118 | 0.025 | 0.104 | 0.104 | 0 | 0.52 | 0.31 | 0.44 | 0.13 | 0.62 | 0.49 |
| 3 | 74916429 | rs4261917 | -0.146 | -2.415 | 0.269 | -1.143 | -0.036 | -0.004 | -0.004 | 0 | 0.7 | 0.43 | 0.8 | 0.66 | 0.81 | 0.8 |
| 3 | 74978829 | rs4395431 | 2.934 | -1.746 | -0.284 | -1.186 | -0.098 | 0.101 | 0.101 | 0.118 | 0.17 | 0.01 | 0.18 | 0 | 0.32 | 0.09 |
| 3 | 74985519 | rs7629942 | 1.531 | -2.44 | 0.053 | -0.736 | -0.009 | -0.021 | -0.021 | 0 | 0.17 | 0.11 | 0.4 | 0.6 | 0.19 | 0.38 |
| 3 | 75004381 | rs9874613 | -0.245 | -2.596 | 0.498 | -0.697 | 0.005 | 0.068 | 0.068 | 0.727 | 0.76 | 0.47 | 0.54 | 0.4 | 0.78 | 0.62 |
| 3 | 75009019 | rs11712680 | 1.267 | -2.722 | 0.38 | -0.731 | -0.027 | -0.034 | -0.034 | 0 | 0.21 | 0.01 | 0.1 | 0.22 | 0.18 | 0.17 |
| 3 | 75012674 | rs9880427 | -0.114 | -2.535 | 0.773 | -0.409 | 0.006 | 0.062 | 0.062 | 0.732 | 0.76 | 0.49 | 0.54 | 0.39 | 0.78 | 0.62 |
| 3 | 75051562 | rs9882666 | 0.954 | -2.52 | 0.768 | -0.39 | -0.003 | -0.005 | -0.005 | 4.397 | 0.18 | 0.15 | 0.4 | 0.61 | 0.19 | 0.37 |
| 3 | 75063638 | rs4428227 | -0.303 | -2.537 | 0.814 | -0.39 | 0.035 | 0.042 | 0.042 | 1.291 | 0.94 | 0.67 | 0.95 | 0.98 | 0.98 | 1 |
| 3 | 75082450 | rs6784085 | 0.076 | -2.534 | 0.817 | -0.388 | 0.003 | 0.036 | 0.036 | 0.4 | 0.23 | 0.33 | 0.44 | 0.59 | 0.21 | 0.38 |
| 3 | 75099324 | rs9850353 | 0.885 | -2.514 | 0.814 | -0.405 | 0.001 | 0.012 | 0.012 | 2.236 | 0.18 | 0.16 | 0.4 | 0.6 | 0.19 | 0.38 |
| 3 | 75100364 | rs9868696 | 0.112 | -2.494 | 0.799 | -0.391 | 0.005 | 0.041 | 0.041 | 0.401 | 0.23 | 0.36 | 0.44 | 0.6 | 0.21 | 0.38 |
| 3 | 75120432 | rs13091717 | 1.173 | -2.532 | 0.636 | -0.506 | 0.064 | 0.243 | 0.243 | 0 | 0.48 | 0.12 | 0.45 | 0.12 | 0.66 | 0.45 |
| 3 | 75179763 | rs7432248 | 0.436 | -2.747 | 0.749 | -0.213 | -0.001 | -0.01 | -0.01 | 0 | 0.21 | 0.19 | 0.43 | 0.6 | 0.22 | 0.39 |
| 3 | 75186326 | rs12636203 | -0.026 | -2.75 | 0.789 | -0.252 | 0.003 | -0.013 | -0.013 | 0.147 | 0.24 | 0.2 | 0.44 | 0.61 | 0.22 | 0.39 |
| 3 | 75198904 | rs13094233 | 1.279 | -2.789 | 0.715 | -0.367 | 0.064 | 0.255 | 0.255 | 0 | 0.48 | 0.12 | 0.34 | 0.06 | 0.65 | 0.4 |
| 3 | 75202372 | rs9877751 | -0.258 | -2.022 | 0.658 | 0.371 | 0.003 | -0.017 | -0.017 | 0.888 | 0.23 | 0.18 | 0.44 | 0.61 | 0.22 | 0.39 |
| 3 | 75209574 | rs9824395 | -0.402 | -2.02 | 0.639 | 0.372 | -0.001 | 0.001 | 0.001 | 0.034 | 0.24 | 0.26 | 0.44 | 0.61 | 0.22 | 0.39 |
| 3 | 75234791 | rs12488229 | 0.209 | -1.985 | 0.624 | 0.411 | 0.001 | 0.016 | 0.016 | 0.603 | 0.2 | 0.21 | 0.4 | 0.6 | 0.21 | 0.39 |
| 3 | 75249701 | rs6809268 | -1.092 | -2.171 | 0.301 | 0.187 | 0.004 | 0.004 | 0.004 | 0.052 | 0.4 | 0.63 | 0.61 | 0.73 | 0.31 | 0.6 |
| 3 | 75253207 | rs7622916 | -0.77 | -2.172 | 0.364 | 0.25 | -0.003 | -0.001 | -0.001 | 0 | 0.36 | 0.39 | 0.48 | 0.65 | 0.3 | 0.53 |
| 3 | 75276627 | rs9856720 | -0.862 | -2.03 | 0.5 | 1.153 | -0.009 | 0.289 | 0.289 | 0 | 0.06 | 0.27 | 0.16 | 0.27 | 0.01 | 0.07 |
| 3 | 75277620 | rs4615102 | -0.85 | -2.284 | 0.078 | 0.955 | -0.013 | -0.025 | -0.025 | 0 | 0.78 | 0.99 | 0.94 | 0.81 | 0.86 | 0.79 |
| 3 | 75278358 | rs12491189 | -1.143 | -1.827 | 1.516 | 2.012 | 0.013 | -0.044 | -0.044 | 0 | 0.36 | 0.13 | 0.47 | 0.45 | 0.31 | 0.52 |
| 3 | 75279989 | rs9310319 | -1.322 | -1.857 | 1.469 | 1.995 | -0.01 | 0.284 | 0.284 | 0 | 0.06 | 0.28 | 0.16 | 0.27 | 0.01 | 0.07 |
| 3 | 75283711 | rs9814680 | -1.322 | -1.857 | 1.469 | 1.995 | -0.01 | 0.284 | 0.284 | 0 | 0.06 | 0.27 | 0.16 | 0.26 | 0.01 | 0.07 |
| 3 | 75285488 | rs7429544 | -0.431 | -1.854 | 1.238 | 1.99 | -0.011 | -0.082 | -0.082 | 0 | 0.22 | 0.07 | 0.38 | 0.44 | 0.21 | 0.43 |
| 3 | 75311829 | rs4536851 | -1.929 | -1.923 | 1.473 | 1.996 | 0.009 | 0.01 | 0.01 | 0 | 0.48 | 0.55 | 0.63 | 0.74 | 0.33 | 0.63 |
| 3 | 75313621 | rs12497509 | -1.437 | -1.869 | 1.327 | 2.024 | 0.033 | -0.023 | -0.023 | 0 | 0.41 | 0.21 | 0.47 | 0.48 | 0.32 | 0.56 |
| 3 | 75321960 | rs9843420 | -0.626 | -2.051 | 2.113 | 2.275 | 0.014 | 0.019 | 0.019 | 0 | 0.17 | 0.12 | 0.1 | 0.07 | 0.1 | 0.12 |
| 3 | 75322243 | rs9843930 | -1.891 | -2.08 | 1.988 | 2.474 | -0.059 | 0.043 | 0.043 | 0 | 0.29 | 0.4 | 0.3 | 0.42 | 0.11 | 0.3 |
| 3 | 75344973 | rs4677489 | -1.751 | -1.941 | 1.615 | 1.456 | -0.088 | -0.061 | -0.061 | 0.515 | 0.29 | 0.51 | 0.24 | 0.18 | 0.11 | 0.24 |
| 3 | 75353238 | rs9849051 | -1.16 | -1.825 | 1.546 | 1.481 | -0.021 | -0.002 | -0.002 | 0.088 | 0.1 | 0.12 | 0.13 | 0.11 | 0.01 | 0.12 |
| 3 | 75361506 | rs2007444 | -1.518 | -2.282 | 1.65 | 1.35 | -0.087 | -0.066 | -0.066 | 1.835 | 0.3 | 0.5 | 0.24 | 0.17 | 0.11 | 0.24 |
| 3 | 75372071 | rs9831266 | -0.837 | -2.269 | 0.358 | 1.221 | -0.039 | -0.007 | -0.007 | 0 | 0.1 | 0.16 | 0.09 | 0.06 | 0.01 | 0.08 |
| 3 | 75374095 | rs6768580 | -1.027 | -2.047 | -0.318 | 1.34 | 0.026 | -0.015 | -0.015 | 0 | 0.32 | 0.74 | 0.37 | 0.42 | 0.1 | 0.32 |
| 3 | 75379085 | rs12490683 | 1.144 | -0.611 | -0.478 | 1.103 | 0.036 | -0.05 | -0.05 | 0.052 | 0.58 | 0.13 | 0.45 | 0.44 | 0.62 | 0.52 |
| 3 | 75381599 | rs4677498 | -0.998 | -0.774 | -0.059 | 0.98 | 0.013 | -0.019 | -0.019 | 0 | 0.36 | 0.71 | 0.52 | 0.56 | 0.32 | 0.45 |
| 3 | 75387457 | rs4677503 | 1.104 | -0.897 | -0.091 | 0.951 | 0.027 | 0.004 | 0.004 | 0.378 | 0.62 | 0.24 | 0.72 | 0.55 | 0.67 | 0.59 |

| | | | | | | | | | | | | | | | | |
|---|---|---|---|---|---|---|---|---|---|---|---|---|---|---|---|---|
| 3 | 75388161 | rs6765044 | 1.242 | -0.889 | -0.134 | 0.892 | 0.039 | -0.051 | -0.051 | 0 | 0.6 | 0.12 | 0.48 | 0.46 | 0.64 | 0.55 |
| 3 | 75392157 | rs6549726 | 0.57 | -0.731 | 0.295 | 0.119 | -0.049 | -0.107 | -0.107 | 0 | 0.75 | 0.45 | 0.85 | 0.94 | 0.82 | 0.76 |
| 3 | 75394787 | rs6767960 | -1.099 | 0.267 | 0.156 | 0.763 | 0.017 | 0.042 | 0.042 | 0.005 | 0.12 | 0.24 | 0.13 | 0.24 | 0.16 | 0.17 |
| 3 | 75407145 | rs7638598 | -0.982 | 0.276 | -0.194 | 0.052 | -0.007 | -0.048 | -0.048 | 0 | 0.25 | 0.43 | 0.23 | 0.11 | 0.21 | 0.21 |
| 3 | 75428675 | rs4677005 | -0.838 | 0.872 | -0.578 | -0.282 | -0.013 | 0.029 | 0.029 | 0 | 0.35 | 0.27 | 0.38 | 0.13 | 0.39 | 0.25 |
| 3 | 75453100 | rs6793001 | 0.697 | -0.059 | 0.157 | -0.311 | -0.045 | -0.009 | -0.009 | 0 | 0.83 | 0.68 | 0.87 | 0.86 | 0.91 | 0.84 |
| 3 | 75469493 | rs11707608 | -1.04 | 0.934 | 0.212 | 0.009 | 0.03 | -0.048 | -0.048 | 0 | 0.13 | 0.23 | 0.16 | 0.02 | 0.16 | 0.06 |
| 3 | 75605157 | rs9830748 | 0.558 | 1.019 | 0.214 | 0.128 | 0.005 | 0.01 | 0.01 | 0 | 0.62 | 0.49 | 0.61 | 0.63 | 0.56 | 0.66 |
| 3 | 75803471 | rs3887345 | -1.367 | 0.228 | -0.451 | 0.182 | 0.091 | -0.019 | -0.019 | 0 | 0.52 | 0.9 | 0.65 | 0.4 | 0.65 | 0.45 |
| 3 | 75836858 | rs7427663 | -0.891 | 0.554 | 0.165 | 0.245 | 0.012 | 0.013 | 0.013 | 0 | 0.11 | 0.17 | 0.15 | 0.09 | 0.16 | 0.14 |
| 3 | 75861696 | rs7633390 | -1.435 | 0.228 | 0.716 | 0.068 | -0.036 | -0.045 | -0.045 | 0 | 0.28 | 0.54 | 0.27 | 0.11 | 0.2 | 0.21 |
| 3 | 75875239 | rs7627370 | -0.68 | -0.191 | 1.139 | 0.009 | 0.082 | 0.077 | 0.077 | 5.224 | 0.05 | 0.43 | 0.04 | 0 | 0 | 0 |
| 3 | 75923055 | rs9834641 | -2.073 | -0.42 | 0.539 | -0.055 | 0.122 | 0.026 | 0.026 | 0 | 0.54 | 0.84 | 0.64 | 0.39 | 0.68 | 0.47 |
| 3 | 75926961 | rs2078354 | 1.894 | -0.472 | 0.5 | -0.074 | 0.129 | 0.018 | 0.018 | 1.143 | 0.47 | 0.1 | 0.35 | 0.61 | 0.32 | 0.52 |
| 3 | 75930359 | rs2324363 | -0.124 | -0.222 | 0.597 | 0.014 | -0.039 | 0.04 | 0.04 | 3.109 | 0.34 | 0.17 | 0.46 | 0.19 | 0.53 | 0.31 |
| 3 | 75942670 | rs7650372 | 2.045 | -0.468 | 0.697 | -0.026 | 0.116 | 0.013 | 0.013 | 0 | 0.44 | 0.16 | 0.35 | 0.63 | 0.3 | 0.5 |
| 3 | 75953505 | rs2203217 | -0.197 | 0.283 | -0.07 | 0.093 | -0.041 | 0.051 | 0.051 | 3.427 | 0.34 | 0.15 | 0.45 | 0.17 | 0.54 | 0.32 |
| 3 | 75961218 | rs9310350 | 1.016 | 0.957 | 1.065 | 0.594 | -0.007 | 0.005 | 0.005 | 0 | 0.89 | 0.89 | 0.86 | 0.94 | 0.84 | 0.85 |
| 3 | 75963401 | rs9310351 | 1.465 | 0.996 | 1.057 | 0.609 | 0.01 | -0.013 | -0.013 | 0 | 0.88 | 0.79 | 0.85 | 0.94 | 0.84 | 0.84 |
| 3 | 75963686 | rs4856024 | 1.262 | 1.091 | 1.12 | 0.666 | 0.014 | -0.012 | -0.012 | 0.895 | 0.89 | 0.79 | 0.85 | 0.94 | 0.84 | 0.85 |
| 3 | 75974251 | rs7611923 | 1.462 | 0.992 | 0.492 | 0.711 | -0.003 | -0.002 | -0.002 | 0 | 0.84 | 0.28 | 0.82 | 0.81 | 0.87 | 0.86 |
| 3 | 75991451 | rs4095546 | -0.92 | 1.157 | 0.426 | 1.314 | 0.021 | 0.022 | 0.022 | 0.766 | 0.33 | 0.58 | 0.52 | 0.44 | 0.47 | 0.32 |
| 3 | 76000442 | rs6766044 | -0.152 | 0.858 | 0.735 | 1.023 | -0.012 | -0.014 | -0.014 | 0 | 0.05 | 0.23 | 0.02 | 0 | 0 | 0 |
| 3 | 76005683 | rs12488213 | 1.221 | 0.75 | 0.612 | 1.275 | -0.029 | -0.03 | -0.03 | 0.17 | 0.6 | 0.18 | 0.46 | 0.43 | 0.53 | 0.65 |
| 3 | 76011920 | rs643476 | 1.855 | 0.622 | 0.134 | 1.154 | -0.019 | 0.174 | 0.174 | 1.566 | 0.42 | 0.06 | 0.35 | 0.21 | 0.37 | 0.49 |
| 3 | 76013002 | rs7642330 | -0.57 | 1.311 | 1.024 | 1.553 | -0.022 | 0.057 | 0.057 | 0.352 | 0.06 | 0.22 | 0.01 | 0.12 | 0 | 0.01 |
| 3 | 76024339 | rs13073504 | 0.147 | 2.145 | 0.959 | 1.414 | -0.001 | 0.002 | 0.002 | 0.063 | 0.07 | 0.05 | 0.1 | 0.03 | 0.14 | 0.08 |
| 3 | 76029306 | rs616166 | 1.17 | 2.084 | 1.44 | 1.366 | 0.08 | -0.051 | -0.051 | 0.125 | 0.86 | 0.54 | 0.85 | 0.97 | 0.85 | 0.9 |
| 3 | 76043296 | rs3892905 | 0.075 | 2.147 | 1.412 | 1.602 | -0.037 | -0.03 | -0.03 | 0 | 0.21 | 0.15 | 0.37 | 0.38 | 0.33 | 0.22 |
| 3 | 76047655 | rs648535 | 1.126 | 1.861 | 1.849 | 1.398 | 0.06 | -0.046 | -0.046 | 2.111 | 0.86 | 0.63 | 0.85 | 0.97 | 0.85 | 0.9 |
| 3 | 76051624 | rs686591 | 1.558 | 1.86 | 1.614 | 1.432 | 0.013 | 0.024 | 0.024 | 0.369 | 0.82 | 0.42 | 0.84 | 0.81 | 0.85 | 0.89 |
| 3 | 76055157 | rs662919 | 1.241 | 1.868 | 1.517 | 1.437 | 0.078 | 0.102 | 0.102 | 0.877 | 0.86 | 0.55 | 0.85 | 0.81 | 0.85 | 0.89 |
| 3 | 76077143 | rs679219 | 0.143 | 1.89 | 1.35 | 1.355 | -0.006 | -0.018 | -0.018 | 0 | 0.93 | 0.71 | 0.95 | 1 | 0.98 | 0.98 |
| 3 | 76077808 | rs618677 | 1.283 | 1.922 | 0.59 | 1.406 | 0 | 0.01 | 0.01 | 2.081 | 0.81 | 0.33 | 0.83 | 0.8 | 0.85 | 0.89 |
| 3 | 76082929 | rs1352505 | -1.611 | 1.922 | 0.59 | 1.406 | 0 | 0.01 | 0.01 | 2.081 | 0.19 | 0.67 | 0.17 | 0.2 | 0.15 | 0.11 |
| 3 | 76083846 | rs2130094 | -0.4 | 1.968 | 0.309 | 1.245 | -0.012 | -0.014 | -0.014 | 0.509 | 0.05 | 0.22 | 0.02 | 0 | 0 | 0 |
| 3 | 76084525 | rs1489101 | -0.519 | 2.436 | -0.54 | 1.415 | -0.056 | -0.042 | -0.042 | 0 | 0.25 | 0.15 | 0.39 | 0.38 | 0.41 | 0.23 |
| 3 | 76094221 | rs4856002 | 0.36 | 2.276 | -0.569 | 1.293 | -0.039 | 0.018 | 0.018 | 0.922 | 0.06 | 0.01 | 0.25 | 0.45 | 0.16 | 0.19 |
| 3 | 76097894 | rs4856003 | 1.528 | 2.065 | -0.58 | 0.855 | 0.022 | 0.091 | 0.091 | 1.543 | 0.56 | 0.23 | 0.42 | 0.25 | 0.46 | 0.6 |
| 3 | 76097994 | rs4856004 | 0.283 | 2.107 | -0.586 | 0.859 | -0.108 | -0.092 | -0.092 | 0 | 0.78 | 0.91 | 0.56 | 0.48 | 0.58 | 0.7 |
| 3 | 76101402 | rs9880437 | 0.52 | 1.755 | -0.285 | 0.728 | 0 | -0.013 | -0.013 | 0.017 | 0.08 | 0.06 | 0.09 | 0 | 0.14 | 0.04 |
| 3 | 76103687 | rs9868043 | -1.218 | 1.722 | -0.043 | 0.733 | -0.01 | 0.001 | 0.001 | 0.582 | 0.2 | 0.57 | 0.15 | 0.22 | 0.16 | 0.07 |
| 3 | 76110123 | rs6778360 | 0.032 | 1.835 | -0.526 | 0.494 | -0.046 | -0.061 | -0.061 | 0 | 0.1 | 0.27 | 0.04 | 0 | 0.01 | 0 |
| 3 | 76114576 | rs9833153 | 1.04 | 2.948 | -1.391 | 1.002 | -0.021 | -0.01 | -0.01 | 0.165 | 0.74 | 0.77 | 0.55 | 0.36 | 0.61 | 0.66 |
| 3 | 76118545 | rs7620001 | 0.81 | 2.982 | -1.384 | 1.022 | -0.076 | -0.047 | -0.047 | 0 | 0.76 | 0.87 | 0.56 | 0.38 | 0.61 | 0.7 |
| 3 | 76121506 | rs1489092 | 0.968 | 2.923 | -1.404 | 1.006 | -0.018 | -0.009 | -0.009 | 1.244 | 0.75 | 0.77 | 0.56 | 0.36 | 0.62 | 0.67 |
| 3 | 76126167 | rs7630937 | 1.814 | 2.771 | -1.518 | 0.898 | -0.003 | 0.005 | 0.005 | 3.729 | 0.63 | 0.69 | 0.47 | 0.36 | 0.47 | 0.62 |
| 3 | 76131251 | rs9863209 | 1.014 | 3.01 | -2.223 | 0.76 | -0.099 | -0.088 | -0.088 | 0 | 0.74 | 0.9 | 0.57 | 0.47 | 0.62 | 0.71 |
| 3 | 76135368 | rs9847106 | 0.028 | 3.487 | -1.995 | 0.88 | -0.029 | 0.003 | 0.003 | 0.056 | 0.85 | 0.92 | 0.83 | 0.92 | 0.79 | 0.89 |
| 3 | 76144252 | rs9815763 | -0.085 | 2.372 | -1.886 | 0.411 | -0.001 | -0.013 | -0.013 | 0.078 | 0.08 | 0.06 | 0.09 | 0 | 0.14 | 0.04 |
| 3 | 76154084 | rs7643807 | 0.566 | 1.917 | -1.841 | 0.802 | -0.095 | -0.082 | -0.082 | 0 | 0.76 | 0.9 | 0.56 | 0.47 | 0.62 | 0.7 |
| 3 | 76158661 | rs6549833 | 0.175 | 0.249 | -1.915 | -0.279 | 0.054 | 0.126 | 0.126 | 0 | 0.63 | 0.24 | 0.81 | 0.92 | 0.76 | 0.75 |
| 3 | 76158947 | rs7632601 | -0.455 | 0.465 | -1.986 | -0.261 | 0.002 | 0.109 | 0.109 | 0 | 0.43 | 0.71 | 0.38 | 0.17 | 0.35 | 0.28 |
| 3 | 76161487 | rs4856008 | -1.343 | -0.24 | -1.886 | 1.012 | 0.013 | 0 | 0 | 0 | 0.08 | 0.04 | 0.15 | 0.38 | 0.06 | 0.1 |
| 3 | 76163422 | rs11128503 | -0.795 | -0.288 | -1.452 | 1.091 | -0.006 | 0.013 | 0.013 | 0.333 | 0.43 | 0.77 | 0.5 | 0.61 | 0.35 | 0.35 |
| 3 | 76164330 | rs7639460 | -0.807 | -0.512 | -0.346 | 1.028 | -0.022 | 0.113 | 0.113 | 0.185 | 0.07 | 0.31 | 0.14 | 0.14 | 0.01 | 0.06 |
| 3 | 76168569 | rs4124239 | -0.653 | -0.492 | -0.1 | 1.032 | -0.019 | 0.107 | 0.107 | 0 | 0.07 | 0.26 | 0.14 | 0.14 | 0.01 | 0.06 |
| 3 | 76169642 | rs4393913 | 0.44 | -0.509 | 0.368 | 0.96 | 0 | -0.004 | -0.004 | 0 | 0.62 | 0.37 | 0.67 | 0.76 | 0.71 | 0.75 |
| 3 | 76172232 | rs1400237 | 2.083 | -0.525 | 0.839 | 1.038 | -0.064 | 0.161 | 0.161 | 0.567 | 0.83 | 0.38 | 0.68 | 0.47 | 0.93 | 0.85 |
| 3 | 76178350 | rs17013853 | -1.641 | -0.281 | 0.892 | 1.11 | -0.002 | 0.306 | 0.306 | 0 | 0.09 | 0.27 | 0.3 | 0.53 | 0.07 | 0.14 |
| 3 | 76178771 | rs17577554 | 0.018 | -0.172 | 0.362 | 1.018 | -0.036 | 0.061 | 0.061 | 0 | 0.28 | 0.14 | 0.23 | 0.09 | 0.29 | 0.19 |
| 3 | 76184968 | rs1516468 | -0.467 | -0.006 | 0.212 | 0.9 | 0.005 | 0.008 | 0.008 | 0 | 0.35 | 0.35 | 0.45 | 0.47 | 0.36 | 0.31 |
| 3 | 76197852 | rs11128504 | -1.483 | -0.223 | 0.21 | 0.73 | -0.001 | -0.014 | -0.014 | 0.026 | 0.06 | 0.07 | 0.04 | 0 | 0.03 | 0.05 |
| 3 | 76204716 | rs2875360 | -0.15 | -0.184 | 0.88 | 1.19 | -0.029 | 0.058 | 0.058 | 0.011 | 0.34 | 0.2 | 0.22 | 0.18 | 0.36 | 0.3 |
| 3 | 76205107 | rs4490371 | 0.245 | -0.265 | 0.638 | 1.115 | -0.104 | 0.045 | 0.045 | 0.048 | 0.27 | 0.05 | 0.25 | 0.18 | 0.33 | 0.26 |
| 3 | 76209314 | rs13096638 | 0.166 | -0.243 | 0.618 | 1.091 | -0.117 | 0.064 | 0.064 | 0.013 | 0.27 | 0.05 | 0.21 | 0.17 | 0.34 | 0.26 |
| 3 | 76210082 | rs12497612 | -0.996 | -0.193 | 1.205 | 1.132 | 0.012 | -0.024 | -0.024 | 0 | 0.06 | 0.17 | 0.04 | 0.01 | 0.04 | 0.04 |
| 3 | 76210743 | rs12496209 | -0.84 | -0.191 | 0.926 | 1.133 | -0.002 | -0.016 | -0.016 | 0 | 0.06 | 0.07 | 0.04 | 0.01 | 0.04 | 0.05 |
| 3 | 76221679 | rs7636341 | -1.6 | -0.578 | 0.9 | 1.289 | -0.015 | 0.06 | 0.06 | 0 | 0.54 | 0.86 | 0.67 | 0.63 | 0.52 | 0.5 |

| | | | | | | | | | | | | | | | | |
|---|---|---|---|---|---|---|---|---|---|---|---|---|---|---|---|---|
| 3 | 76222174 | rs12497502 | -1.455 | -0.43 | 1.376 | 1.395 | -0.014 | 0.07 | 0.07 | 0 | 0.09 | 0.2 | 0.17 | 0.24 | 0.06 | 0.1 |
| 3 | 76225040 | rs2037190 | -0.206 | -1.255 | 0.416 | 0.625 | -0.028 | 0.167 | 0.167 | 0.427 | 0.46 | 0.21 | 0.32 | 0.17 | 0.43 | 0.39 |
| 3 | 76225482 | rs4856011 | -1.953 | -1.381 | 0.099 | 0.697 | -0.043 | -0.008 | -0.008 | 0 | 0.68 | 0.86 | 0.78 | 0.64 | 0.58 | 0.65 |
| 3 | 76404546 | rs7609927 | 1.019 | -1.248 | 1.322 | 0.641 | 0.024 | -0.05 | -0.05 | 0 | 0.88 | 0.61 | 0.8 | 0.96 | 0.89 | 0.9 |
| 3 | 76412145 | rs7638155 | -0.692 | -1.243 | 1.304 | 0.641 | 0.013 | -0.054 | -0.054 | 0.009 | 0.09 | 0.24 | 0.09 | 0 | 0.11 | 0.08 |
| 3 | 76413209 | rs11920211 | 0.006 | -1.315 | 1.326 | 1.383 | -0.011 | 0.177 | 0.177 | 0 | 0.05 | 0.08 | 0.21 | 0.4 | 0.03 | 0.07 |
| 3 | 76413820 | rs7611506 | 0.006 | -1.315 | 1.326 | 1.383 | -0.011 | 0.177 | 0.177 | 0 | 0.05 | 0.08 | 0.21 | 0.4 | 0.03 | 0.07 |
| 3 | 76418463 | rs7627309 | -2.038 | -1.44 | 1.657 | 1.146 | -0.029 | -0.094 | -0.094 | 0 | 0.18 | 0.64 | 0.24 | 0.04 | 0.12 | 0.1 |
| 3 | 76425559 | rs3843874 | -1.557 | -1.469 | 0.903 | 1.065 | 0.027 | 0.02 | 0.02 | 3.16 | 0.57 | 0.81 | 0.74 | 0.64 | 0.61 | 0.66 |
| 3 | 76425822 | rs9830053 | -1.385 | -1.262 | 0.88 | 0.683 | -0.024 | -0.112 | -0.112 | 0 | 0.21 | 0.62 | 0.25 | 0.04 | 0.15 | 0.11 |
| 3 | 76439750 | rs3913581 | 0.951 | -1.051 | -0.118 | 0.827 | -0.167 | -0.081 | -0.081 | 0 | 0.24 | 0.02 | 0.24 | 0.25 | 0.41 | 0.33 |
| 3 | 76447667 | rs9815217 | -1.302 | -1.25 | 0.426 | 0.608 | -0.018 | -0.13 | -0.13 | 0 | 0.2 | 0.43 | 0.16 | 0 | 0.16 | 0.13 |
| 3 | 76453297 | rs17791350 | 0.621 | -1.255 | 0.347 | 0.6 | -0.014 | 0.04 | 0.04 | 0 | 0.08 | 0.01 | 0.03 | 0 | 0.07 | 0.03 |
| 3 | 76453596 | rs6762968 | -0.898 | -1.265 | 0.524 | 0.599 | -0.004 | -0.092 | -0.092 | 0 | 0.17 | 0.33 | 0.14 | 0 | 0.15 | 0.08 |
| 3 | 76456239 | rs3849483 | -2.022 | -1.463 | 0.159 | 0.847 | 0.016 | 0.037 | 0.037 | 0.629 | 0.61 | 0.86 | 0.76 | 0.7 | 0.63 | 0.69 |
| 3 | 76456574 | rs7630091 | -2.025 | -1.48 | 0.2 | 0.806 | 0.021 | 0.035 | 0.035 | 0.589 | 0.61 | 0.83 | 0.76 | 0.71 | 0.64 | 0.69 |
| 3 | 76458784 | rs6762174 | -1.761 | -1.337 | 0.457 | 1.073 | -0.026 | 0.056 | 0.056 | 0 | 0.25 | 0.56 | 0.38 | 0.41 | 0.19 | 0.2 |
| 3 | 76467191 | rs7647603 | -0.134 | -1.522 | 0.317 | 1.052 | 0.027 | 0.024 | 0.024 | 0 | 0.37 | 0.48 | 0.5 | 0.52 | 0.46 | 0.37 |
| 3 | 76469582 | rs13091632 | 0.552 | -1.629 | 0.017 | 1.063 | -0.184 | -0.04 | -0.04 | 0 | 0.31 | 0.03 | 0.33 | 0.24 | 0.46 | 0.44 |
| 3 | 76470473 | rs6796472 | -1.851 | -1.771 | 0.245 | 0.999 | -0.019 | 0.004 | 0.004 | 5.404 | 0.2 | 0.73 | 0.35 | 0.33 | 0.12 | 0.14 |
| 3 | 76471166 | rs7621495 | -1.63 | -1.796 | 0.151 | 0.982 | 0 | -0.037 | -0.037 | 2.04 | 0.55 | 0.84 | 0.68 | 0.55 | 0.58 | 0.65 |
| 3 | 76473163 | rs9817359 | 0.321 | -1.829 | 0.676 | 0.568 | -0.001 | -0.01 | -0.01 | 1.008 | 0.07 | 0.4 | 0.05 | 0 | 0.01 | 0 |
| 3 | 76477010 | rs6548429 | -1.807 | -1.864 | -0.007 | 0.88 | -0.025 | 0.034 | 0.034 | 0 | 0.61 | 0.84 | 0.71 | 0.67 | 0.56 | 0.62 |
| 3 | 76481056 | rs4856032 | 0.153 | -1.674 | -0.347 | 0.651 | -0.138 | -0.085 | -0.085 | 3.126 | 0.35 | 0.05 | 0.4 | 0.33 | 0.46 | 0.48 |
| 3 | 76483391 | rs10514733 | -0.625 | -1.522 | 0.026 | 0.485 | -0.006 | 0.001 | 0.001 | 0 | 0.06 | 0.03 | 0.07 | 0.02 | 0.01 | 0.12 |
| 3 | 76488608 | rs9882550 | -0.315 | -1.331 | 0.496 | 0.515 | -0.04 | -0.095 | -0.095 | 0.488 | 0.14 | 0.4 | 0.05 | 0 | 0.05 | 0.04 |
| 3 | 76489848 | rs7612860 | -0.92 | -1.118 | 0.267 | 0.628 | -0.002 | -0.001 | -0.001 | 0 | 0.46 | 0.58 | 0.37 | 0.36 | 0.47 | 0.35 |
| 3 | 76490030 | rs13063821 | 0.123 | -0.204 | -0.103 | 1.138 | -0.177 | -0.17 | -0.17 | 0 | 0.27 | 0.02 | 0.29 | 0.34 | 0.41 | 0.31 |
| 3 | 76494066 | rs13087864 | -0.874 | 0.303 | 0.306 | 1.031 | 0.014 | 0.057 | 0.057 | 0 | 0.29 | 0.41 | 0.28 | 0.4 | 0.34 | 0.3 |
| 3 | 76500144 | rs6769256 | -0.532 | 0.812 | 0.304 | 0.776 | 0.021 | 0.018 | 0.018 | 0 | 0.29 | 0.36 | 0.3 | 0.32 | 0.38 | 0.32 |
| 3 | 76504796 | rs6786214 | 1.508 | 0.585 | 0.342 | 0.697 | 0.046 | 0.076 | 0.076 | 0 | 0.49 | 0.24 | 0.29 | 0.37 | 0.39 | 0.46 |
| 3 | 76515427 | rs3913578 | -0.316 | 0.703 | 0.771 | 0.965 | 0 | 0.001 | 0.001 | 0 | 0.18 | 0.2 | 0.33 | 0.23 | 0.19 | 0.18 |
| 3 | 76520342 | rs12107824 | -0.21 | 0.554 | 0.866 | 0.876 | 0.012 | 0.043 | 0.043 | 0 | 0.3 | 0.4 | 0.27 | 0.42 | 0.35 | 0.3 |
| 3 | 76524177 | rs9866083 | -0.868 | 0.31 | 1.157 | 0.722 | -0.024 | 0.076 | 0.076 | 0.416 | 0.2 | 0.5 | 0.34 | 0.29 | 0.16 | 0.17 |
| 3 | 76525546 | rs4645140 | -0.477 | 0.536 | 1.116 | 0.697 | -0.023 | -0.016 | -0.016 | 0.47 | 0.34 | 0.23 | 0.53 | 0.59 | 0.41 | 0.37 |
| 3 | 76533907 | rs3899654 | -1.531 | 0.422 | 1.193 | 0.655 | 0.004 | 0.01 | 0.01 | 0 | 0.5 | 0.6 | 0.66 | 0.6 | 0.57 | 0.52 |
| 3 | 76538694 | rs6788691 | -0.031 | 0.579 | 1.045 | 0.684 | -0.024 | -0.037 | -0.037 | 0 | 0.29 | 0.18 | 0.33 | 0.39 | 0.39 | 0.33 |
| 3 | 76540493 | rs17014275 | -0.999 | 0.543 | 1.095 | 0.67 | -0.018 | -0.007 | -0.007 | 3.313 | 0.22 | 0.53 | 0.35 | 0.23 | 0.18 | 0.19 |
| 3 | 76540573 | rs4856036 | -1.727 | 0.514 | 0.58 | 0.697 | 0.016 | 0.07 | 0.07 | 0 | 0.51 | 0.78 | 0.68 | 0.63 | 0.57 | 0.52 |
| 3 | 76548812 | rs17014282 | 0.298 | 0.612 | 0.273 | 0.709 | -0.002 | -0.015 | -0.015 | 0 | 0.07 | 0.07 | 0.25 | 0.21 | 0.09 | 0.12 |
| 3 | 76551403 | rs7431017 | -0.862 | 0.611 | 0.499 | 0.573 | 0.003 | -0.002 | -0.002 | 0.05 | 0.4 | 0.58 | 0.61 | 0.58 | 0.48 | 0.48 |
| 3 | 76559964 | rs9849425 | -1.617 | 0.543 | 0.314 | 0.484 | 0.012 | 0.046 | 0.046 | 0 | 0.53 | 0.72 | 0.66 | 0.61 | 0.6 | 0.58 |
| 3 | 76575853 | rs3913574 | -0.597 | 0.577 | -0.132 | 0.486 | -0.076 | -0.069 | -0.069 | 0 | 0.42 | 0.1 | 0.35 | 0.41 | 0.51 | 0.38 |
| 3 | 76578176 | rs196532 | -2.684 | 0.322 | -0.678 | 0.382 | -0.002 | -0.06 | -0.06 | 0 | 0.89 | 0.99 | 0.91 | 0.82 | 0.88 | 0.96 |
| 3 | 76579227 | rs774217 | -0.956 | 1.164 | 0.282 | 0.859 | 0.004 | 0 | 0 | 0 | 0.41 | 0.57 | 0.63 | 0.57 | 0.5 | 0.44 |
| 3 | 76579933 | rs774216 | -0.848 | 1.113 | 0.756 | 0.75 | 0.077 | 0.151 | 0.151 | 0.3 | 0.1 | 0.35 | 0.31 | 0.25 | 0.11 | 0.16 |
| 3 | 76602394 | rs1562499 | -1.237 | 1.183 | -0.109 | 1.017 | 0.002 | 0 | 0 | 2.017 | 0.5 | 0.56 | 0.63 | 0.54 | 0.57 | 0.5 |
| 3 | 76608904 | rs1562498 | 1.189 | 1.174 | -0.349 | 0.974 | 0.013 | 0.055 | 0.055 | 0 | 0.49 | 0.33 | 0.34 | 0.33 | 0.41 | 0.49 |
| 3 | 76633507 | rs1868189 | -0.028 | 1.291 | -0.316 | 1.321 | -0.096 | -0.087 | -0.087 | 0 | 0.24 | 0.02 | 0.27 | 0.29 | 0.34 | 0.27 |
| 3 | 76636887 | rs6799201 | 0.635 | 1.062 | 0.289 | 1.636 | -0.001 | -0.004 | -0.004 | 0 | 0.58 | 0.37 | 0.48 | 0.44 | 0.52 | 0.4 |
| 3 | 76639128 | rs17014431 | 0.093 | 1.598 | -0.405 | 1.6 | 0.001 | 0.046 | 0.046 | 0 | 0.06 | 0.08 | 0.23 | 0.28 | 0.08 | 0.13 |
| 3 | 76646013 | rs13093006 | 0.407 | 1.11 | -1.29 | 1.631 | -0.092 | -0.17 | -0.17 | 0.116 | 0.26 | 0.02 | 0.21 | 0.41 | 0.33 | 0.36 |
| 3 | 76664559 | rs10511051 | 0.436 | -0.164 | -1.178 | 1.385 | -0.028 | -0.059 | -0.059 | 0 | 0.43 | 0.09 | 0.38 | 0.42 | 0.39 | 0.43 |
| 3 | 76672692 | rs7623289 | 0.514 | -0.358 | -0.222 | 1.289 | 0 | 0.001 | 0.001 | 0 | 0.35 | 0.37 | 0.45 | 0.48 | 0.39 | 0.5 |
| 3 | 76674553 | rs2875404 | 0.409 | -0.396 | 0.295 | 0.809 | -0.021 | -0.054 | -0.054 | 0.989 | 0.78 | 0.62 | 0.84 | 0.9 | 0.76 | 0.91 |
| 3 | 76676537 | rs12639248 | -0.752 | -0.493 | 0.779 | 1.076 | 0.017 | 0.05 | 0.05 | 0 | 0.54 | 0.63 | 0.6 | 0.72 | 0.62 | 0.5 |
| 3 | 76681495 | rs7634503 | -0.732 | -0.505 | 0.77 | 1.109 | 0.019 | 0.066 | 0.066 | 0 | 0.55 | 0.75 | 0.6 | 0.69 | 0.62 | 0.51 |
| 3 | 76682623 | rs1975573 | 0.682 | -0.441 | 0.786 | 1.144 | -0.003 | -0.002 | -0.002 | 0 | 0.33 | 0.3 | 0.45 | 0.62 | 0.38 | 0.42 |
| 3 | 76684657 | rs13061512 | -0.272 | -0.124 | 1.023 | 0.843 | -0.003 | 0.004 | 0.004 | 0 | 0.14 | 0.11 | 0.12 | 0.09 | 0.24 | 0.06 |
| 3 | 76689925 | rs9867514 | -1.707 | -0.2 | 1.708 | 0.812 | -0.007 | -0.033 | -0.033 | 0 | 0.24 | 0.62 | 0.18 | 0.11 | 0.28 | 0.17 |
| 3 | 76693749 | rs6764287 | -0.982 | -0.253 | 1.807 | 1.135 | 0.012 | -0.002 | -0.002 | 0.698 | 0.53 | 0.6 | 0.54 | 0.48 | 0.65 | 0.53 |
| 3 | 76702186 | rs2121791 | 0.302 | -0.161 | 1.538 | 1.095 | 0.009 | 0.002 | 0.002 | 0 | 0.36 | 0.55 | 0.42 | 0.38 | 0.39 | 0.53 |
| 3 | 76713783 | rs1816955 | -0.077 | -0.064 | 0.956 | 1.04 | -0.005 | 0 | 0 | 0 | 0.43 | 0.63 | 0.51 | 0.43 | 0.45 | 0.59 |
| 3 | 76720428 | rs2593865 | 0.545 | -0.449 | 0.615 | 0.85 | -0.02 | -0.043 | -0.043 | 0 | 0.81 | 0.7 | 0.87 | 0.91 | 0.78 | 0.93 |
| 3 | 76721413 | rs1349008 | 0.845 | -0.477 | 0.42 | 0.882 | -0.023 | -0.044 | -0.044 | 0 | 0.41 | 0.13 | 0.42 | 0.54 | 0.38 | 0.38 |
| 3 | 76722454 | rs197097 | 0.212 | -0.226 | 0.368 | 1.038 | 0.01 | -0.007 | -0.007 | 0 | 0.39 | 0.54 | 0.46 | 0.37 | 0.4 | 0.53 |
| 3 | 76725768 | rs264533 | 0.572 | -0.409 | 0.047 | 0.853 | -0.047 | 0.103 | 0.103 | 0.053 | 0.42 | 0.18 | 0.35 | 0.22 | 0.54 | 0.41 |
| 3 | 76725903 | rs10511052 | -0.833 | -0.764 | 0.706 | 1.015 | -0.02 | 0.043 | 0.043 | 0 | 0.09 | 0.29 | 0.09 | 0.16 | 0.04 | 0.17 |
| 3 | 76730474 | rs9851014 | -0.521 | -0.667 | 0.475 | 1.053 | -0.007 | 0.03 | 0.03 | 0 | 0.56 | 0.6 | 0.55 | 0.77 | 0.49 | 0.55 |

| | | | | | | | | | | | | | | | | |
|---|---|---|---|---|---|---|---|---|---|---|---|---|---|---|---|---|
| 3 | 76733323 | rs7632207 | -0.531 | -0.668 | 0.499 | 1.054 | -0.01 | 0.044 | 0.044 | 0 | 0.56 | 0.62 | 0.54 | 0.77 | 0.49 | 0.55 |
| 3 | 76735823 | rs978586 | -0.98 | -0.656 | 0.494 | 1.047 | -0.043 | 0.069 | 0.069 | 0.032 | 0.59 | 0.79 | 0.57 | 0.78 | 0.49 | 0.61 |
| 3 | 76739295 | rs3849508 | -0.42 | -1.253 | 0.356 | 1.015 | -0.081 | -0.002 | -0.002 | 0.009 | 0.23 | 0.56 | 0.19 | 0.22 | 0.08 | 0.2 |
| 3 | 76739333 | rs155473 | 0.296 | -1.122 | 0.114 | 1.058 | -0.044 | -0.064 | -0.064 | 0 | 0.35 | 0.12 | 0.37 | 0.57 | 0.41 | 0.4 |
| 3 | 76749882 | rs2593875 | 0.177 | -0.998 | -0.038 | 1.095 | -0.056 | -0.088 | -0.088 | 0.015 | 0.36 | 0.14 | 0.38 | 0.56 | 0.4 | 0.38 |
| 3 | 76753766 | rs2608141 | 0.888 | -0.596 | 0.315 | 0.913 | -0.028 | 0.039 | 0.039 | 0.103 | 0.38 | 0.23 | 0.38 | 0.18 | 0.44 | 0.38 |
| 3 | 76754936 | rs182040 | 0.602 | -0.642 | 0.632 | 0.89 | -0.004 | 0.005 | 0.005 | 0 | 0.42 | 0.36 | 0.4 | 0.18 | 0.44 | 0.41 |
| 3 | 76758806 | rs267164 | 0.514 | -0.687 | 0.449 | 1.057 | -0.006 | 0.014 | 0.014 | 0.007 | 0.43 | 0.32 | 0.49 | 0.29 | 0.49 | 0.5 |
| 3 | 76761924 | rs166439 | 0.857 | -0.72 | 0.432 | 1.06 | -0.023 | 0.032 | 0.032 | 0.093 | 0.38 | 0.22 | 0.39 | 0.18 | 0.44 | 0.4 |
| 3 | 76763490 | rs264543 | 0.965 | -0.731 | 0.246 | 1.034 | -0.053 | 0.061 | 0.061 | 0.181 | 0.37 | 0.15 | 0.37 | 0.18 | 0.44 | 0.38 |
| 3 | 76771797 | rs166130 | 0.5 | -0.803 | 0.166 | 1.038 | 0.022 | 0.026 | 0.026 | 1.103 | 0.46 | 0.35 | 0.57 | 0.3 | 0.62 | 0.57 |
| 3 | 76772527 | rs17014579 | -0.193 | -1.248 | -0.117 | 1.199 | -0.06 | 0.02 | 0.02 | 0 | 0.32 | 0.54 | 0.28 | 0.42 | 0.2 | 0.28 |
| 3 | 76780491 | rs6808931 | 0.137 | -0.615 | -0.367 | 1.131 | -0.015 | -0.02 | -0.02 | 0.016 | 0.7 | 0.76 | 0.67 | 0.58 | 0.62 | 0.64 |
| 3 | 76780594 | rs774610 | -0.323 | -0.431 | -0.381 | 1.171 | 0.002 | -0.001 | -0.001 | 0 | 0.1 | 0.12 | 0.2 | 0.12 | 0.22 | 0.2 |
| 3 | 76782340 | rs774601 | -0.33 | -0.483 | -0.381 | 1.17 | 0 | -0.001 | -0.001 | 0 | 0.1 | 0.14 | 0.2 | 0.12 | 0.21 | 0.2 |
| 3 | 76783510 | rs810184 | -1.032 | -0.553 | -0.387 | 1.157 | 0.008 | -0.021 | -0.021 | 0 | 0.79 | 0.9 | 0.87 | 0.7 | 0.83 | 0.83 |
| 3 | 76787711 | rs7630234 | -0.181 | -0.719 | -0.523 | 1.112 | -0.01 | 0 | 0 | 0 | 0.34 | 0.63 | 0.26 | 0.39 | 0.21 | 0.28 |
| 3 | 76792468 | rs1375963 | 0.014 | -0.705 | -0.576 | 1.091 | -0.006 | -0.01 | -0.01 | 0 | 0.69 | 0.75 | 0.67 | 0.58 | 0.62 | 0.63 |
| 3 | 76802905 | rs3884314 | -0.111 | -0.713 | -0.512 | 1.071 | -0.005 | -0.007 | -0.007 | 0 | 0.7 | 0.74 | 0.67 | 0.59 | 0.62 | 0.63 |
| 3 | 76806018 | rs7609557 | -0.927 | -0.734 | -0.584 | 1.07 | 0.024 | -0.016 | -0.016 | 1.416 | 0.44 | 0.72 | 0.45 | 0.51 | 0.43 | 0.49 |
| 3 | 76816266 | rs197867 | -1.214 | -0.759 | -0.704 | 1.085 | 0.002 | -0.004 | -0.004 | 0 | 0.8 | 0.87 | 0.87 | 0.7 | 0.83 | 0.85 |
| 3 | 76817443 | rs3884404 | -0.081 | -0.739 | -0.795 | 1.103 | -0.035 | 0.005 | 0.005 | 0 | 0.33 | 0.57 | 0.26 | 0.4 | 0.21 | 0.27 |
| 3 | 76819666 | rs267131 | 0.15 | -0.921 | -0.586 | 1.128 | -0.014 | 0.005 | 0.005 | 1.135 | 0.6 | 0.49 | 0.69 | 0.5 | 0.75 | 0.68 |
| 3 | 76842705 | rs976603 | 1.316 | -0.535 | -1.305 | 1.009 | -0.155 | 0.024 | 0.024 | 0.435 | 0.34 | 0.04 | 0.4 | 0.19 | 0.4 | 0.33 |
| 3 | 76850916 | rs1167035 | -1.182 | -0.419 | -0.573 | 1.069 | 0.022 | -0.014 | -0.014 | 0 | 0.44 | 0.73 | 0.46 | 0.52 | 0.43 | 0.52 |
| 3 | 76853328 | rs17014723 | -0.188 | -0.146 | -0.558 | 0.953 | -0.009 | 0.004 | 0.004 | 0 | 0.14 | 0.25 | 0.09 | 0.19 | 0.13 | 0.1 |
| 3 | 76857965 | rs267152 | -0.599 | -0.073 | -1.189 | 1.038 | -0.012 | -0.04 | -0.04 | 0 | 0.16 | 0.01 | 0.08 | 0.16 | 0.16 | 0.07 |
| 3 | 76859234 | rs1146006 | -0.994 | 0.037 | -1.056 | 1.053 | 0.005 | -0.008 | -0.008 | 0 | 0.24 | 0.23 | 0.18 | 0.31 | 0.24 | 0.35 |
| 3 | 76863912 | rs1146007 | 0.932 | 0.411 | -1.964 | 0.904 | -0.099 | 0.007 | 0.007 | 0 | 0.28 | 0.03 | 0.28 | 0.19 | 0.34 | 0.17 |
| 3 | 76866452 | rs7618823 | -0.33 | 0.009 | -0.613 | 1.281 | -0.022 | -0.027 | -0.027 | 0 | 0.79 | 0.62 | 0.77 | 0.84 | 0.84 | 0.77 |
| 3 | 76869831 | rs1146013 | -0.069 | -0.059 | -0.688 | 1.527 | 0.003 | 0.015 | 0.015 | 0.65 | 0.09 | 0.17 | 0.13 | 0.24 | 0.12 | 0.2 |
| 3 | 76879809 | rs1146017 | 0.055 | -0.066 | -0.665 | 1.394 | 0 | 0.002 | 0.002 | 0.288 | 0.09 | 0.13 | 0.12 | 0.22 | 0.12 | 0.2 |
| 3 | 76881893 | rs1146021 | 1.572 | -0.138 | -0.171 | 1.456 | 0.025 | 0.068 | 0.068 | 0.765 | 0.49 | 0.29 | 0.5 | 0.29 | 0.55 | 0.38 |
| 3 | 76882785 | rs1166940 | -1.106 | -0.077 | -0.17 | 1.381 | 0.002 | -0.008 | -0.008 | 0 | 0.26 | 0.3 | 0.18 | 0.46 | 0.25 | 0.37 |
| 3 | 76885786 | rs7645328 | 0.302 | -0.052 | -0.24 | 1.37 | 0.014 | -0.042 | -0.042 | 0 | 0.16 | 0.06 | 0.05 | 0.23 | 0.13 | 0.16 |
| 3 | 76890421 | rs6775462 | 0.068 | -0.011 | -0.22 | 1.372 | 0.016 | -0.021 | -0.021 | 0.431 | 0.17 | 0.12 | 0.06 | 0.24 | 0.13 | 0.17 |
| 3 | 76895761 | rs1166957 | 0.507 | 0.032 | -0.194 | 1.366 | -0.001 | 0 | 0 | 0.216 | 0.07 | 0.09 | 0.12 | 0.1 | 0.12 | 0.14 |
| 3 | 76899560 | rs1166958 | -1.024 | 0.019 | -0.035 | 1.41 | -0.003 | 0.01 | 0.01 | 0 | 0.27 | 0.34 | 0.18 | 0.46 | 0.25 | 0.36 |
| 3 | 76910935 | rs3911526 | -0.373 | 0.075 | 0.092 | 1.491 | -0.005 | 0.012 | 0.012 | 0 | 0.19 | 0.22 | 0.07 | 0.36 | 0.13 | 0.22 |
| 3 | 76917321 | rs7428044 | 0.057 | 0.742 | 0.422 | 0.54 | -0.015 | -0.028 | -0.028 | 0.529 | 0.8 | 0.7 | 0.78 | 0.85 | 0.84 | 0.77 |
| 3 | 76917552 | rs7433861 | -0.054 | 1.361 | 0.275 | 1.036 | 0.019 | 0.04 | 0.04 | 1.397 | 0.91 | 0.8 | 0.82 | 0.84 | 0.9 | 0.79 |
| 3 | 76917601 | rs1166972 | -1.079 | 1.374 | -0.42 | 0.798 | 0.017 | 0.026 | 0.026 | 0.758 | 0.1 | 0.22 | 0.13 | 0.21 | 0.12 | 0.18 |
| 3 | 76921632 | rs1166978 | -0.876 | 1.782 | -1.212 | 0.463 | -0.067 | 0.04 | 0.04 | 0 | 0.18 | 0.06 | 0.18 | 0.11 | 0.25 | 0.18 |
| 3 | 76924329 | rs10446372 | 0.179 | 1.327 | -1.113 | 0.499 | 0.004 | 0.002 | 0.002 | 0.204 | 0.52 | 0.39 | 0.57 | 0.63 | 0.43 | 0.53 |
| 3 | 76925298 | rs17818793 | 1.154 | 0.888 | -1.547 | 0.484 | 0 | -0.006 | -0.006 | 0.4 | 0.18 | 0.09 | 0.08 | 0.17 | 0.19 | 0.24 |
| 3 | 76926040 | rs10446375 | 1.15 | 0.872 | -1.55 | 0.5 | -0.002 | -0.01 | -0.01 | 0 | 0.18 | 0.07 | 0.08 | 0.17 | 0.19 | 0.24 |
| 3 | 76928990 | rs1166980 | 1.069 | 1.009 | -1.127 | -0.006 | -0.009 | -0.044 | -0.044 | 0 | 0.2 | 0.26 | 0.11 | 0.07 | 0.19 | 0.14 |
| 3 | 76931014 | rs1166981 | -0.48 | 1.399 | -0.815 | 0.048 | 0.029 | 0.038 | 0.038 | 0.547 | 0.72 | 0.61 | 0.71 | 0.68 | 0.68 | 0.67 |
| 3 | 76933219 | rs11127568 | 1.08 | 0.91 | -0.868 | -0.201 | -0.004 | -0.016 | -0.016 | 0 | 0.13 | 0.15 | 0.12 | 0.08 | 0.08 | 0.17 |
| 3 | 76940455 | rs3911529 | 1.09 | 1.442 | -0.653 | 0.101 | 0.007 | 0.02 | 0.02 | 0.702 | 0.16 | 0.2 | 0.17 | 0.18 | 0.17 | 0.17 |
| 3 | 76943398 | rs3936484 | 1.081 | 0.583 | -0.887 | 0.055 | -0.008 | 0.014 | 0.014 | 0.029 | 0.33 | 0.14 | 0.34 | 0.33 | 0.36 | 0.32 |
| 3 | 76943884 | rs1177498 | 0.755 | 0.49 | -0.564 | 0.004 | -0.008 | 0.002 | 0.002 | 0 | 0.26 | 0.3 | 0.25 | 0.26 | 0.26 | 0.17 |
| 3 | 76947105 | rs12630815 | 1.693 | 0.553 | -0.6 | -0.19 | -0.016 | -0.003 | -0.003 | 0 | 0.08 | 0 | 0.1 | 0.08 | 0.06 | 0.17 |
| 3 | 76949295 | rs13085650 | 1.548 | 0.459 | -0.536 | -0.123 | -0.001 | -0.001 | -0.001 | 0 | 0.2 | 0.13 | 0.12 | 0.19 | 0.24 | 0.22 |
| 3 | 76954084 | rs1166992 | -0.565 | 0.637 | -0.403 | 0.077 | 0.004 | 0.014 | 0.014 | 0 | 0.76 | 0.73 | 0.62 | 0.62 | 0.7 | 0.69 |
| 3 | 76955007 | rs3884325 | 0.68 | 0.423 | -0.614 | -0.212 | 0.011 | 0.022 | 0.022 | 0 | 0.25 | 0.11 | 0.3 | 0.29 | 0.22 | 0.31 |
| 3 | 76955188 | rs1166995 | 0.642 | 0.432 | -0.304 | -0.288 | -0.001 | 0 | 0 | 0 | 0.29 | 0.27 | 0.21 | 0.16 | 0.32 | 0.15 |
| 3 | 76956434 | rs1166996 | 1.544 | 0.337 | -0.656 | -0.343 | -0.052 | 0.091 | 0.091 | 0 | 0.18 | 0.01 | 0.13 | 0.05 | 0.24 | 0.11 |
| 3 | 76960792 | rs17014895 | 0.016 | -0.49 | -0.173 | -0.162 | -0.01 | -0.003 | -0.003 | 0 | 0.43 | 0.49 | 0.38 | 0.49 | 0.33 | 0.36 |
| 3 | 76961685 | rs13088432 | 0.453 | -0.723 | 0.083 | -0.01 | -0.023 | -0.002 | -0.002 | 2.837 | 0.29 | 0.48 | 0.16 | 0.28 | 0.24 | 0.26 |
| 3 | 76967285 | rs1166998 | 0.206 | -0.509 | 0.088 | 0.268 | -0.001 | 0.003 | 0.003 | 0 | 0.43 | 0.46 | 0.38 | 0.47 | 0.39 | 0.5 |
| 3 | 76972804 | rs9827948 | 0.125 | -0.471 | 0.046 | 0.276 | 0 | -0.001 | -0.001 | 1.42 | 0.43 | 0.6 | 0.41 | 0.45 | 0.39 | 0.5 |
| 3 | 76976439 | rs13091953 | 0.935 | -0.539 | 0.086 | 0.338 | 0 | 0.005 | 0.005 | 0 | 0.22 | 0.21 | 0.28 | 0.27 | 0.22 | 0.33 |
| 3 | 76982025 | rs17819911 | 2.056 | -0.552 | -0.059 | 0.328 | -0.001 | -0.02 | -0.02 | 0 | 0.17 | 0.05 | 0.09 | 0.18 | 0.17 | 0.16 |
| 3 | 76982992 | rs11127569 | 1.918 | -0.539 | -0.072 | 0.319 | -0.016 | -0.021 | -0.021 | 0 | 0.18 | 0.06 | 0.09 | 0.18 | 0.21 | 0.19 |
| 3 | 76986200 | rs9841520 | 0.125 | -0.532 | 0.195 | 0.332 | -0.002 | 0.003 | 0.003 | 0 | 0.42 | 0.57 | 0.4 | 0.45 | 0.39 | 0.5 |
| 3 | 76986296 | rs9859240 | 0.162 | -0.532 | 0.172 | 0.332 | -0.001 | 0.003 | 0.003 | 0 | 0.42 | 0.43 | 0.39 | 0.45 | 0.39 | 0.49 |
| 3 | 76986961 | rs10155005 | 0.07 | -0.535 | -0.307 | 0.311 | 0 | -0.001 | -0.001 | 0 | 0.43 | 0.54 | 0.4 | 0.45 | 0.39 | 0.49 |
| 3 | 76992952 | rs9858491 | -0.015 | -0.532 | -0.452 | 0.317 | 0.009 | -0.014 | -0.014 | 0 | 0.44 | 0.72 | 0.41 | 0.45 | 0.38 | 0.5 |

| | | | | | | | | | | | | | | | | |
|---|---|---|---|---|---|---|---|---|---|---|---|---|---|---|---|---|
| 3 | 76996273 | rs13058966 | 2 | -0.447 | -0.581 | 0.323 | -0.034 | -0.042 | -0.042 | 2.25 | 0.17 | 0.02 | 0.09 | 0.18 | 0.21 | 0.2 |
| 3 | 77002317 | rs4855995 | 0.892 | -0.288 | -0.374 | 0.378 | -0.003 | -0.009 | -0.009 | 0 | 0.22 | 0.16 | 0.29 | 0.27 | 0.21 | 0.35 |
| 3 | 77007830 | rs12496653 | 0.937 | -0.329 | -0.759 | 0.431 | -0.001 | -0.006 | -0.006 | 0 | 0.22 | 0.16 | 0.28 | 0.27 | 0.18 | 0.32 |
| 3 | 77008129 | rs6766804 | -1.783 | -0.11 | -1.205 | 0.282 | -0.077 | 0.03 | 0.03 | 0 | 0.8 | 0.97 | 0.83 | 0.91 | 0.72 | 0.83 |
| 3 | 77014498 | rs4855997 | 0.86 | 0.252 | -0.743 | 0.512 | -0.001 | 0.005 | 0.005 | 0 | 0.23 | 0.2 | 0.29 | 0.27 | 0.17 | 0.32 |
| 3 | 77022269 | rs6764089 | 0.405 | 0.603 | -1.937 | 0.404 | 0.006 | -0.012 | -0.012 | 0 | 0.35 | 0.61 | 0.38 | 0.38 | 0.27 | 0.44 |
| 3 | 77026208 | rs4585239 | 0.441 | 0.904 | -1.727 | 0.234 | -0.083 | -0.002 | -0.002 | 0 | 0.18 | 0.02 | 0.29 | 0.21 | 0.22 | 0.13 |
| 3 | 77026968 | rs6548470 | 0.083 | 0.59 | -0.57 | 0.684 | 0 | 0.013 | 0.013 | 0 | 0.7 | 0.77 | 0.76 | 0.76 | 0.67 | 0.73 |
| 3 | 77037626 | rs12630195 | -0.644 | 0.242 | -0.217 | 1.139 | 0.009 | 0 | 0 | 0.276 | 0.33 | 0.24 | 0.43 | 0.63 | 0.24 | 0.38 |
| 3 | 77038193 | rs7617945 | -0.674 | 0.666 | 0.379 | 0.906 | 0.021 | -0.006 | -0.006 | 0 | 0.13 | 0.23 | 0.12 | 0.18 | 0.19 | 0.19 |
| 3 | 77041173 | rs7427534 | -1.208 | 0.543 | 0.846 | 0.837 | 0.03 | 0.078 | 0.078 | 0 | 0.46 | 0.69 | 0.57 | 0.79 | 0.42 | 0.52 |
| 3 | 77046740 | rs6786589 | -1.143 | 0.533 | 0.881 | 0.833 | 0.016 | 0.041 | 0.041 | 0 | 0.45 | 0.62 | 0.57 | 0.79 | 0.42 | 0.52 |
| 3 | 77058627 | rs9835363 | 0.287 | 0.599 | 1.14 | 0.889 | -0.019 | 0 | 0 | 0 | 0.67 | 0.58 | 0.53 | 0.38 | 0.75 | 0.63 |
| 3 | 77060574 | rs6773575 | -1.465 | 0.585 | 0.792 | 0.413 | -0.002 | -0.124 | -0.124 | 1.036 | 0.16 | 0.48 | 0.08 | 0.01 | 0.11 | 0.08 |
| 3 | 77069799 | rs6762489 | 0.408 | 0.567 | 0.87 | 1.218 | -0.002 | 0.014 | 0.014 | 1.494 | 0.11 | 0.12 | 0.08 | 0.18 | 0.1 | 0.15 |
| 3 | 77070110 | rs6762635 | -0.721 | 0.565 | 1.193 | 1.599 | 0.025 | 0.078 | 0.078 | 0 | 0.08 | 0.19 | 0.08 | 0.26 | 0.11 | 0.15 |
| 3 | 77073964 | rs9812289 | -0.986 | 0.664 | 0.838 | 1.316 | 0.043 | 0.038 | 0.038 | 1.535 | 0.11 | 0.56 | 0.1 | 0.21 | 0.11 | 0.15 |
| 3 | 77075657 | rs4558796 | -1.994 | 0.703 | 0.871 | 1.301 | 0.004 | -0.028 | -0.028 | 0.364 | 0.17 | 0.64 | 0.14 | 0.21 | 0.18 | 0.17 |
| 3 | 77084262 | rs6795561 | -2.354 | 0.697 | 0.726 | 1.667 | 0.001 | 0.002 | 0.002 | 4.391 | 0.23 | 0.73 | 0.24 | 0.37 | 0.25 | 0.31 |
| 3 | 77086844 | rs11127575 | -2.046 | 0.781 | 0.722 | 1.714 | 0.004 | 0.011 | 0.011 | 0.025 | 0.15 | 0.18 | 0.31 | 0.21 | 0.17 | 0.19 |
| 3 | 77089395 | rs3923745 | -1.644 | 0.776 | 0.409 | 1.668 | 0.016 | 0.017 | 0.017 | 0.506 | 0.39 | 0.76 | 0.44 | 0.4 | 0.34 | 0.39 |
| 3 | 77089699 | rs3923744 | -2.274 | 1.765 | 0.598 | 2.235 | -0.01 | -0.015 | -0.015 | 0 | 0.14 | 0.1 | 0.29 | 0.22 | 0.18 | 0.19 |
| 3 | 77094799 | rs3934881 | -2.048 | 1.86 | 0.603 | 2.309 | -0.01 | -0.019 | -0.019 | 0 | 0.13 | 0.07 | 0.28 | 0.25 | 0.18 | 0.21 |
| 3 | 77099218 | rs3884526 | -2.002 | 1.54 | 1.267 | 2.411 | 0.016 | 0.121 | 0.121 | 0 | 0.08 | 0.22 | 0.15 | 0.23 | 0.06 | 0.08 |
| 3 | 77102870 | rs11712939 | -0.921 | 2.544 | 0.702 | 1.85 | -0.091 | -0.042 | -0.042 | 0 | 0.14 | 0.01 | 0.1 | 0.12 | 0.23 | 0.24 |
| 3 | 77104596 | rs12714456 | 1.23 | 1.344 | 1.875 | 1.775 | 0.101 | 0.174 | 0.174 | 0 | 0.94 | 0.61 | 0.93 | 0.79 | 0.92 | 0.89 |
| 3 | 77133304 | rs9856171 | -1.009 | 1.242 | 0.629 | 1.65 | -0.001 | 0.025 | 0.025 | 0 | 0.05 | 0.04 | 0.11 | 0.12 | 0.05 | 0.08 |
| 3 | 77133873 | rs7628843 | -1.375 | 1.612 | 0.812 | 1.921 | 0.019 | 0.092 | 0.092 | 0.031 | 0.17 | 0.32 | 0.29 | 0.56 | 0.24 | 0.43 |
| 3 | 77138278 | rs9816491 | -1.117 | 1.594 | 0.25 | 1.923 | 0 | 0.044 | 0.044 | 0 | 0.06 | 0.09 | 0.12 | 0.13 | 0.05 | 0.08 |
| 3 | 77147039 | rs9873691 | -1.259 | 1.69 | -0.128 | 1.807 | -0.017 | -0.027 | -0.027 | 0.002 | 0.17 | 0.16 | 0.27 | 0.57 | 0.23 | 0.41 |
| 3 | 77151787 | rs4624600 | 0.879 | 0.851 | 0 | 0.992 | -0.013 | -0.009 | -0.009 | 0 | 0.63 | 0.47 | 0.54 | 0.31 | 0.71 | 0.43 |
| 3 | 77154433 | rs9859970 | -0.949 | 1.749 | -0.754 | 0.125 | -0.081 | 0.09 | 0.09 | 0 | 0.15 | 0.03 | 0.14 | 0.03 | 0.23 | 0.09 |
| 3 | 77155824 | rs13322517 | -1.639 | 1.357 | 0.039 | -0.143 | 0.009 | -0.076 | -0.076 | 0 | 0.21 | 0.3 | 0.16 | 0.03 | 0.24 | 0.11 |
| 3 | 77165296 | rs12185957 | -1.87 | 1.107 | 0.621 | -0.016 | -0.012 | -0.038 | -0.038 | 0 | 0.16 | 0.55 | 0.13 | 0.13 | 0.14 | 0.14 |
| 3 | 77167475 | rs6796664 | -1.609 | 0.95 | 0.847 | 0.055 | -0.012 | -0.059 | -0.059 | 0.092 | 0.12 | 0.49 | 0.09 | 0.03 | 0.07 | 0.11 |
| 3 | 77168168 | rs6769992 | -1.005 | 0.55 | -0.222 | 0.019 | 0.004 | -0.002 | -0.002 | 0 | 0.56 | 0.67 | 0.45 | 0.51 | 0.54 | 0.51 |
| 3 | 77169971 | rs6785747 | -1.195 | 0.1 | 0.145 | 0.398 | -0.044 | 0.121 | 0.121 | 0 | 0.15 | 0.49 | 0.11 | 0.26 | 0.08 | 0.22 |
| 3 | 77180639 | rs7648530 | -0.634 | 0.584 | -0.146 | 0.222 | -0.002 | -0.002 | -0.002 | 0.865 | 0.63 | 0.65 | 0.57 | 0.5 | 0.57 | 0.53 |
| 3 | 77188041 | rs4077419 | 0.252 | 0.417 | -0.558 | -0.264 | -0.006 | 0.198 | 0.198 | 0.112 | 0.44 | 0.15 | 0.31 | 0.14 | 0.45 | 0.28 |
| 3 | 77196171 | rs7615545 | -0.63 | 0.192 | 0.173 | -0.025 | 0 | 0.136 | 0.136 | 0 | 0.12 | 0.28 | 0.16 | 0.3 | 0.06 | 0.17 |
| 3 | 77197856 | rs12486426 | 0.729 | 0.222 | -0.182 | -0.059 | -0.02 | -0.016 | -0.016 | 0.581 | 0.06 | 0 | 0.12 | 0.08 | 0.04 | 0.04 |
| 3 | 77207087 | rs7432300 | -0.227 | 0.201 | 0.267 | -0.211 | 0.003 | 0.062 | 0.062 | 0.255 | 0.52 | 0.31 | 0.43 | 0.22 | 0.49 | 0.34 |
| 3 | 77219008 | rs7428284 | -0.361 | 0.219 | 0.324 | -0.195 | 0.028 | 0.008 | 0.008 | 0.086 | 0.09 | 0.23 | 0.13 | 0.08 | 0.05 | 0.05 |
| 3 | 77224268 | rs4683993 | -0.643 | 0.176 | 0.396 | -0.076 | 0.005 | -0.005 | -0.005 | 0.134 | 0.57 | 0.48 | 0.48 | 0.44 | 0.51 | 0.46 |
| 3 | 77235898 | rs6548485 | -0.209 | 0.071 | 0.383 | -0.28 | 0.001 | 0.009 | 0.009 | 0.481 | 0.53 | 0.41 | 0.42 | 0.17 | 0.47 | 0.36 |
| 3 | 77240101 | rs4683981 | 0.831 | -0.354 | -0.339 | -0.724 | -0.007 | -0.056 | -0.056 | 0 | 0.27 | 0.34 | 0.23 | 0.08 | 0.26 | 0.21 |
| 3 | 77244307 | rs17820386 | -0.003 | -0.256 | -0.69 | -0.645 | -0.034 | 0.099 | 0.099 | 0 | 0.27 | 0.04 | 0.22 | 0.12 | 0.22 | 0.15 |
| 3 | 77249737 | rs9812479 | 0.248 | -0.242 | -0.79 | -0.617 | -0.039 | 0.065 | 0.065 | 0 | 0.24 | 0.01 | 0.19 | 0.12 | 0.19 | 0.14 |
| 3 | 77253192 | rs11713960 | -1.547 | -0.129 | -0.241 | -0.353 | -0.004 | -0.004 | -0.004 | 0 | 0.64 | 0.71 | 0.55 | 0.52 | 0.54 | 0.56 |
| 3 | 77288105 | rs6768395 | 0.165 | -0.803 | -0.588 | -1.224 | -0.04 | -0.048 | -0.048 | 0 | 0.31 | 0.54 | 0.39 | 0.79 | 0.24 | 0.47 |
| 3 | 77290845 | rs975808 | 1.474 | -0.427 | -0.884 | -1.154 | -0.002 | 0.103 | 0.103 | 0 | 0.26 | 0.1 | 0.22 | 0.08 | 0.26 | 0.19 |
| 3 | 77295661 | rs6764328 | 1.329 | -0.303 | -1.081 | -1.229 | -0.161 | 0.042 | 0.042 | 1.634 | 0.22 | 0.02 | 0.28 | 0.13 | 0.34 | 0.24 |
| 3 | 77295675 | rs6788710 | 0.062 | -0.538 | -0.607 | -1.171 | 0.039 | 0.065 | 0.065 | 0 | 0.51 | 0.4 | 0.52 | 0.2 | 0.6 | 0.43 |
| 3 | 77297960 | rs7618355 | -0.196 | -0.591 | -0.096 | -0.71 | -0.023 | -0.043 | -0.043 | 0 | 0.66 | 0.55 | 0.72 | 0.79 | 0.7 | 0.66 |
| 3 | 77299434 | rs4072806 | -0.571 | -0.36 | 0.022 | -0.769 | -0.019 | -0.114 | -0.114 | 0.057 | 0.7 | 0.57 | 0.82 | 0.95 | 0.73 | 0.78 |
| 3 | 77307051 | rs1441951 | -1.538 | 0.503 | -0.444 | -0.887 | -0.032 | 0.031 | 0.031 | 0.426 | 0.85 | 0.94 | 0.84 | 0.96 | 0.74 | 0.79 |
| 3 | 77309262 | rs1441954 | -1.529 | 0.5 | -0.446 | -0.881 | -0.032 | 0.031 | 0.031 | 0.499 | 0.84 | 0.94 | 0.84 | 0.96 | 0.74 | 0.8 |
| 3 | 77317940 | rs1372425 | 0.758 | 0.127 | -0.224 | -0.324 | -0.004 | 0.068 | 0.068 | 0 | 0.12 | 0.15 | 0.28 | 0.63 | 0.12 | 0.14 |
| 3 | 77319487 | rs17015112 | 0.615 | 0.097 | -0.043 | -0.337 | -0.007 | 0.119 | 0.119 | 0 | 0.12 | 0.2 | 0.28 | 0.63 | 0.12 | 0.13 |
| 3 | 77320482 | rs13064976 | 1.127 | -0.208 | -0.198 | -0.876 | 0.009 | 0.013 | 0.013 | 0 | 0.29 | 0.18 | 0.15 | 0.01 | 0.27 | 0.28 |
| 3 | 77320740 | rs17822079 | 0.707 | -0.343 | -0.359 | -0.807 | -0.016 | 0.011 | 0.011 | 0 | 0.37 | 0.12 | 0.35 | 0.3 | 0.31 | 0.37 |
| 3 | 77322334 | rs9882239 | 0.597 | -0.247 | -0.044 | -0.322 | 0.002 | 0.013 | 0.013 | 0 | 0.53 | 0.45 | 0.41 | 0.27 | 0.45 | 0.37 |
| 3 | 77323280 | rs6776288 | -0.623 | -0.329 | -0.134 | -0.383 | 0.009 | 0.011 | 0.011 | 0 | 0.27 | 0.75 | 0.3 | 0.09 | 0.16 | 0.26 |
| 3 | 77325417 | rs11718389 | 1.194 | 0.045 | -0.185 | -0.179 | -0.002 | -0.053 | -0.053 | 0.526 | 0.15 | 0.33 | 0.17 | 0.03 | 0.1 | 0.17 |
| 3 | 77325741 | rs9309764 | -0.203 | 0.463 | -0.667 | 0.235 | -0.054 | -0.042 | -0.042 | 0 | 0.73 | 0.89 | 0.65 | 0.36 | 0.6 | 0.61 |
| 3 | 77333765 | rs7433256 | -1.43 | -0.225 | -0.002 | 0.084 | 0.026 | 0.014 | 0.014 | 0.656 | 0.95 | 0.75 | 0.97 | 1 | 0.99 | 1 |
| 3 | 77334841 | rs7629183 | 0.115 | 0.256 | -0.502 | -0.022 | -0.032 | 0.039 | 0.039 | 2.482 | 0.28 | 0.23 | 0.24 | 0.13 | 0.37 | 0.27 |
| 3 | 77334932 | rs9309765 | 0.388 | 0.383 | -0.337 | 1.061 | 0.032 | 0.194 | 0.194 | 0 | 0.09 | 0.21 | 0.11 | 0.35 | 0.12 | 0.15 |
| 3 | 77336156 | rs2875436 | 0.75 | 0.382 | -0.286 | 1.065 | -0.002 | 0.038 | 0.038 | 0 | 0.08 | 0.06 | 0.09 | 0.34 | 0.12 | 0.15 |

| | | | | | | | | | | | | | | | | |
|---|---|---|---|---|---|---|---|---|---|---|---|---|---|---|---|---|
| 3 | 77353410 | rs1899373 | 1.197 | -0.105 | -1.21 | 0.807 | -0.033 | -0.027 | -0.027 | 0 | 0.41 | 0.03 | 0.34 | 0.36 | 0.4 | 0.35 |
| 3 | 77358376 | rs12637117 | 0.773 | -0.129 | -0.685 | 1.363 | -0.011 | 0.002 | 0.002 | 0 | 0.08 | 0.05 | 0.09 | 0.33 | 0.12 | 0.16 |
| 3 | 77365899 | rs9820365 | 0.735 | -0.131 | -0.69 | 1.362 | -0.014 | -0.009 | -0.009 | 0 | 0.08 | 0.03 | 0.09 | 0.34 | 0.12 | 0.16 |
| 3 | 77369911 | rs1978940 | 0.005 | 0.043 | -0.434 | 1.085 | 0.034 | 0.063 | 0.063 | 0.431 | 0.16 | 0.42 | 0.2 | 0.24 | 0.16 | 0.24 |
| 3 | 77374207 | rs11921508 | 1.046 | -0.16 | -0.686 | 1.088 | -0.002 | -0.001 | -0.001 | 2.061 | 0.17 | 0.21 | 0.32 | 0.14 | 0.11 | 0.22 |
| 3 | 77383456 | rs2028512 | 0.427 | -0.117 | -0.758 | 1.269 | 0.009 | -0.023 | -0.023 | 1.671 | 0.62 | 0.27 | 0.47 | 0.57 | 0.72 | 0.54 |
| 3 | 77383950 | rs9868442 | 0.836 | 0.144 | -0.803 | 1.456 | -0.005 | -0.002 | -0.002 | 3.315 | 0.07 | 0.05 | 0.09 | 0.31 | 0.12 | 0.15 |
| 3 | 77385970 | rs17015201 | 0.901 | -0.659 | -1.171 | 0.752 | -0.022 | -0.036 | -0.036 | 0.61 | 0.18 | 0.05 | 0.14 | 0.25 | 0.19 | 0.13 |
| 3 | 77386855 | rs9818075 | 0.098 | -0.914 | -0.898 | 0.541 | 0 | 0.003 | 0.003 | 0.068 | 0.57 | 0.4 | 0.39 | 0.32 | 0.59 | 0.4 |
| 3 | 77389366 | rs6548499 | 1.021 | -0.958 | -0.949 | 0.172 | 0.017 | 0.02 | 0.02 | 0 | 0.2 | 0.12 | 0.14 | 0.13 | 0.13 | 0.17 |
| 3 | 77394540 | rs4449296 | 0.042 | -1.031 | -0.864 | 0.189 | -0.046 | 0.022 | 0.022 | 0 | 0.37 | 0.24 | 0.51 | 0.72 | 0.48 | 0.44 |
| 3 | 77402615 | rs7431317 | 1.092 | -1.092 | -0.797 | 0.065 | 0 | -0.035 | -0.035 | 0.33 | 0.37 | 0.35 | 0.24 | 0.06 | 0.38 | 0.27 |
| 3 | 77406994 | rs7429743 | -1.307 | -1.102 | -0.683 | 0.085 | 0.009 | 0.009 | 0.009 | 0 | 0.8 | 0.84 | 0.86 | 0.87 | 0.87 | 0.83 |
| 3 | 77409190 | rs7642312 | 0.997 | -1.123 | -0.65 | 0.058 | -0.003 | 0.02 | 0.02 | 0 | 0.36 | 0.26 | 0.23 | 0.06 | 0.38 | 0.27 |
| 3 | 77412233 | rs9869100 | 0.083 | -0.993 | -0.705 | 0.064 | -0.022 | 0.006 | 0.006 | 0 | 0.34 | 0.26 | 0.49 | 0.73 | 0.44 | 0.44 |
| 3 | 77415062 | rs9809640 | 0.829 | -0.97 | -0.742 | 0.205 | -0.01 | -0.015 | -0.015 | 0.338 | 0.2 | 0.21 | 0.39 | 0.54 | 0.29 | 0.33 |
| 3 | 77418423 | rs9876951 | 1.158 | -1.03 | -0.713 | 0.075 | -0.008 | 0.064 | 0.064 | 0 | 0.36 | 0.21 | 0.23 | 0.05 | 0.39 | 0.26 |
| 3 | 77418859 | rs4683973 | -0.831 | -0.804 | -0.531 | 0.247 | 0.001 | -0.041 | -0.041 | 0 | 0.77 | 0.76 | 0.9 | 1 | 0.83 | 0.92 |
| 3 | 77420922 | rs4684000 | 0.006 | -0.799 | -0.516 | 0.306 | -0.017 | 0.186 | 0.186 | 0 | 0.07 | 0.27 | 0.19 | 0.24 | 0.01 | 0.2 |
| 3 | 77424989 | rs9872242 | -0.271 | -0.6 | -0.624 | 0.377 | -0.04 | -0.019 | -0.019 | 0 | 0.69 | 0.47 | 0.71 | 0.76 | 0.82 | 0.71 |
| 3 | 77428877 | rs12488430 | 1.405 | -0.306 | -1.089 | 0.092 | -0.018 | -0.011 | -0.011 | 0.254 | 0.13 | 0.03 | 0.09 | 0.17 | 0.15 | 0.12 |
| 3 | 77429959 | rs9870610 | 0.641 | -0.565 | -1.19 | -0.243 | 0.003 | -0.051 | -0.051 | 0 | 0.29 | 0.21 | 0.26 | 0.45 | 0.28 | 0.33 |
| 3 | 77430630 | rs6768736 | -0.014 | -0.421 | -0.659 | -0.277 | -0.001 | -0.001 | -0.001 | 0 | 0.43 | 0.56 | 0.6 | 0.54 | 0.48 | 0.55 |
| 3 | 77435623 | rs9873887 | 1.419 | -0.416 | -1.29 | -0.31 | -0.093 | 0.067 | 0.067 | 1.934 | 0.2 | 0.03 | 0.16 | 0.05 | 0.31 | 0.22 |
| 3 | 77437181 | rs4684004 | -0.705 | -0.392 | -0.713 | -0.025 | -0.025 | -0.147 | -0.147 | 0.51 | 0.76 | 0.6 | 0.89 | 1 | 0.84 | 0.91 |
| 3 | 77439564 | rs6548506 | -0.522 | 0.168 | -1.042 | 1.023 | -0.001 | -0.037 | -0.037 | 0 | 0.35 | 0.25 | 0.31 | 0.47 | 0.35 | 0.36 |
| 3 | 77444784 | rs9809056 | -0.136 | 0.45 | -0.835 | 1.105 | -0.001 | 0.007 | 0.007 | 0 | 0.12 | 0.1 | 0.18 | 0.26 | 0.15 | 0.22 |
| 3 | 77450363 | rs7432690 | -1.282 | 0.344 | -0.433 | 1.022 | 0.008 | 0.042 | 0.042 | 0 | 0.32 | 0.44 | 0.35 | 0.46 | 0.36 | 0.5 |
| 3 | 77461355 | rs4327364 | 0.398 | 0.355 | -0.563 | 1.039 | 0.017 | 0.015 | 0.015 | 0 | 0.1 | 0.23 | 0.1 | 0.11 | 0.11 | 0.11 |
| 3 | 77466531 | rs12497293 | 1.026 | 0.097 | -1.655 | 0.67 | -0.023 | 0.042 | 0.042 | 0.402 | 0.28 | 0.01 | 0.15 | 0.23 | 0.27 | 0.2 |
| 3 | 77468883 | rs9654017 | -1.915 | -0.21 | -1.765 | 0.52 | -0.12 | -0.082 | -0.082 | 0.265 | 0.69 | 0.92 | 0.56 | 0.7 | 0.65 | 0.72 |
| 3 | 77473461 | rs4459877 | 1.739 | -0.161 | -1.494 | 0.491 | 0.002 | -0.021 | -0.021 | 0 | 0.07 | 0.12 | 0.04 | 0.01 | 0.07 | 0.1 |
| 3 | 77476152 | rs4458348 | 0.956 | -0.185 | -1.498 | 0.565 | -0.014 | -0.035 | -0.035 | 0.488 | 0.11 | 0.04 | 0.17 | 0.28 | 0.15 | 0.21 |
| 3 | 77481335 | rs9817088 | -0.654 | -0.208 | -1.109 | 0.36 | 0.009 | 0.099 | 0.099 | 0 | 0.25 | 0.36 | 0.3 | 0.47 | 0.28 | 0.41 |
| 3 | 77486138 | rs12634690 | 0.498 | -0.243 | -1.184 | 0.316 | -0.007 | 0.093 | 0.093 | 0.036 | 0.35 | 0.13 | 0.19 | 0.25 | 0.36 | 0.3 |
| 3 | 77498624 | rs6777631 | -0.939 | -0.249 | -1.121 | 0.477 | -0.01 | 0.021 | 0.021 | 0.199 | 0.3 | 0.6 | 0.3 | 0.39 | 0.24 | 0.4 |
| 3 | 77503498 | rs9840819 | 0.695 | 0.44 | -0.748 | 0.814 | 0.043 | 0.019 | 0.019 | 0 | 0.09 | 0.28 | 0.1 | 0.09 | 0.11 | 0.1 |
| 3 | 77504146 | rs13096330 | -0.308 | 0.496 | -1.23 | 1.057 | 0 | -0.018 | -0.018 | 0 | 0.15 | 0.09 | 0.19 | 0.36 | 0.16 | 0.29 |
| 3 | 77505705 | rs9872416 | 0.051 | 0.684 | -1.211 | 0.761 | 0.003 | 0.011 | 0.011 | 0.398 | 0.48 | 0.55 | 0.32 | 0.32 | 0.47 | 0.4 |
| 3 | 77507524 | rs7651943 | -0.26 | 0.855 | -1.446 | 1.365 | -0.02 | -0.05 | -0.05 | 0.327 | 0.1 | 0.02 | 0.16 | 0.27 | 0.13 | 0.2 |
| 3 | 77509454 | rs2166802 | -0.393 | 1.235 | -1.697 | 0.706 | -0.039 | 0.033 | 0.033 | 0.303 | 0.29 | 0.13 | 0.44 | 0.29 | 0.37 | 0.31 |
| 3 | 77517611 | rs4684011 | -1.012 | 1.337 | -2.01 | 0.858 | -0.04 | -0.031 | -0.031 | 0.003 | 0.26 | 0.12 | 0.29 | 0.36 | 0.34 | 0.4 |
| 3 | 77521306 | rs7614439 | 0.571 | 1.274 | -1.794 | 0.776 | 0.004 | -0.018 | -0.018 | 0.061 | 0.36 | 0.26 | 0.44 | 0.45 | 0.35 | 0.32 |
| 3 | 77525133 | rs1345179 | 0.52 | 1.211 | -1.406 | 0.947 | -0.002 | 0.004 | 0.004 | 0 | 0.36 | 0.37 | 0.47 | 0.42 | 0.35 | 0.31 |
| 3 | 77526031 | rs2166801 | -0.214 | 1.234 | -1.519 | 0.965 | 0 | 0.048 | 0.048 | 0 | 0.48 | 0.37 | 0.64 | 0.62 | 0.55 | 0.57 |
| 3 | 77526065 | rs12634489 | 0.612 | 1.132 | -1.313 | 0.916 | -0.005 | -0.005 | -0.005 | 0 | 0.12 | 0.09 | 0.09 | 0.18 | 0.14 | 0.14 |
| 3 | 77531926 | rs1470569 | -0.199 | 1.113 | -1.189 | 0.883 | 0.001 | 0.025 | 0.025 | 0 | 0.49 | 0.41 | 0.65 | 0.62 | 0.55 | 0.57 |
| 3 | 77536346 | rs7610881 | 0.198 | 1.228 | -1.429 | 0.945 | -0.042 | -0.062 | -0.062 | 0.099 | 0.11 | 0.02 | 0.16 | 0.18 | 0.15 | 0.22 |
| 3 | 77547882 | rs12489432 | 0.435 | 1.028 | -1.204 | 0.983 | -0.004 | -0.012 | -0.012 | 0 | 0.12 | 0.09 | 0.09 | 0.21 | 0.15 | 0.14 |
| 3 | 77558385 | rs13077150 | 0.428 | 1.091 | -1.148 | 0.924 | -0.009 | -0.017 | -0.017 | 0 | 0.36 | 0.25 | 0.48 | 0.42 | 0.4 | 0.35 |
| 3 | 77565186 | rs7641269 | -0.952 | 0.611 | -1.745 | 0.34 | -0.088 | 0.039 | 0.039 | 0.004 | 0.72 | 0.97 | 0.85 | 0.82 | 0.74 | 0.77 |
| 3 | 77571639 | rs775779 | -0.966 | 0.471 | -1.168 | 0.981 | -0.001 | 0 | 0 | 0 | 0.63 | 0.64 | 0.61 | 0.66 | 0.61 | 0.68 |
| 3 | 77577912 | rs775715 | -0.985 | 0.545 | -0.538 | 0.775 | -0.017 | 0.041 | 0.041 | 0.023 | 0.71 | 0.86 | 0.84 | 0.82 | 0.74 | 0.77 |
| 3 | 77588489 | rs775762 | 0.614 | 0.197 | 0.081 | 1.059 | -0.002 | 0.002 | 0.002 | 0 | 0.4 | 0.4 | 0.44 | 0.43 | 0.4 | 0.44 |
| 3 | 77591572 | rs775767 | -0.328 | 0.208 | -0.023 | 0.943 | -0.002 | 0.002 | 0.002 | 0.064 | 0.56 | 0.61 | 0.71 | 0.69 | 0.59 | 0.65 |
| 3 | 77591607 | rs775768 | 0.241 | 0.174 | -0.045 | 0.936 | 0.007 | 0.012 | 0.012 | 0 | 0.48 | 0.59 | 0.55 | 0.58 | 0.45 | 0.49 |
| 3 | 77598943 | rs1666130 | 0.063 | 0.061 | 0.079 | 0.896 | 0 | -0.001 | -0.001 | 1.703 | 0.57 | 0.58 | 0.57 | 0.56 | 0.56 | 0.57 |
| 3 | 77604242 | rs1523759 | -1.111 | 0.06 | 0.084 | 0.905 | 0.012 | 0.067 | 0.067 | 0 | 0.17 | 0.41 | 0.28 | 0.25 | 0.19 | 0.23 |
| 3 | 77604690 | rs10514734 | 0.757 | 0.105 | -0.102 | 0.917 | -0.034 | -0.036 | -0.036 | 0 | 0.08 | 0.01 | 0.16 | 0.13 | 0.15 | 0.16 |
| 3 | 77604987 | rs6797719 | 0.77 | 0.024 | -0.078 | 0.943 | -0.038 | 0.017 | 0.017 | 0 | 0.15 | 0.01 | 0.1 | 0.13 | 0.14 | 0.14 |
| 3 | 77609776 | rs808966 | -1.2 | 0.034 | 0.196 | 1.002 | -0.011 | 0.016 | 0.016 | 0.369 | 0.73 | 0.84 | 0.84 | 0.82 | 0.75 | 0.8 |
| 3 | 77633425 | rs1523765 | -0.566 | 0.065 | 0.096 | 0.956 | 0 | -0.011 | -0.011 | 0 | 0.1 | 0.07 | 0.16 | 0.13 | 0.13 | 0.12 |
| 3 | 77643174 | rs775723 | -0.109 | 0.02 | 0.223 | 0.935 | -0.001 | -0.001 | -0.001 | 0 | 0.6 | 0.6 | 0.59 | 0.57 | 0.6 | 0.64 |
| 3 | 77650799 | rs1666125 | 0.131 | -0.176 | 0.172 | 0.849 | 0.003 | 0.001 | 0.001 | 0.007 | 0.58 | 0.49 | 0.56 | 0.57 | 0.6 | 0.63 |
| 3 | 77651695 | rs3732916 | -0.834 | -0.2 | 0.186 | 0.898 | 0 | -0.001 | -0.001 | 0 | 0.1 | 0.11 | 0.18 | 0.13 | 0.13 | 0.1 |
| 3 | 77651985 | rs9852724 | -0.847 | -0.202 | 0.185 | 0.883 | 0 | -0.001 | -0.001 | 0 | 0.1 | 0.11 | 0.18 | 0.13 | 0.13 | 0.12 |
| 3 | 77668346 | rs9848106 | -0.939 | -0.229 | 0.331 | 1.134 | 0.003 | 0.083 | 0.083 | 1.027 | 0.09 | 0.25 | 0.15 | 0.21 | 0.06 | 0.07 |
| 3 | 77673908 | rs4684016 | -0.058 | -0.185 | 0.334 | 1.122 | 0.001 | 0.006 | 0.006 | 0.163 | 0.08 | 0.11 | 0.19 | 0.15 | 0.13 | 0.12 |
| 3 | 77675638 | rs876675 | 0.63 | -0.268 | 0.172 | 0.851 | 0.001 | 0 | 0 | 0 | 0.52 | 0.39 | 0.48 | 0.51 | 0.5 | 0.62 |

| | | | | | | | | | | | | | | | | |
|---|---|---|---|---|---|---|---|---|---|---|---|---|---|---|---|---|
| 3 | 77677857 | rs4684017 | -1.134 | -0.233 | 0.243 | 0.713 | -0.007 | 0.018 | 0.018 | 1.811 | 0.7 | 0.78 | 0.83 | 0.87 | 0.69 | 0.81 |
| 3 | 77678306 | rs10779982 | -0.062 | -0.236 | 0.067 | 0.655 | 0.004 | -0.007 | -0.007 | 7.031 | 0.6 | 0.5 | 0.67 | 0.66 | 0.63 | 0.74 |
| 3 | 77693125 | rs1839795 | 0.343 | 0.007 | 0.052 | 0.698 | -0.008 | 0.015 | 0.015 | 0.646 | 0.36 | 0.53 | 0.52 | 0.43 | 0.4 | 0.4 |
| 3 | 77696422 | rs1031377 | -0.492 | 0.012 | -0.158 | 0.513 | 0.018 | 0.019 | 0.019 | 0 | 0.62 | 0.56 | 0.65 | 0.56 | 0.58 | 0.57 |
| 3 | 77698442 | rs1163750 | -0.292 | -0.041 | -0.15 | 0.467 | -0.011 | -0.008 | -0.008 | 0 | 0.58 | 0.62 | 0.63 | 0.56 | 0.55 | 0.56 |
| 3 | 77701041 | rs809603 | -0.321 | 0.005 | -0.188 | 0.246 | 0.001 | -0.006 | -0.006 | 0.199 | 0.58 | 0.54 | 0.64 | 0.76 | 0.57 | 0.62 |
| 3 | 77706807 | rs776144 | 0.878 | -0.233 | -0.476 | 0.283 | 0.015 | 0.046 | 0.046 | 0 | 0.19 | 0.05 | 0.09 | 0.13 | 0.13 | 0.15 |
| 3 | 77708605 | rs1178778 | -0.348 | -0.217 | -0.298 | 0.372 | -0.003 | 0.012 | 0.012 | 0.08 | 0.57 | 0.72 | 0.48 | 0.65 | 0.46 | 0.48 |
| 3 | 77709064 | rs12629236 | 0.866 | -0.299 | -0.167 | 0.455 | 0 | 0 | 0 | 0 | 0.31 | 0.37 | 0.35 | 0.48 | 0.28 | 0.33 |
| 3 | 77712583 | rs1163740 | 0.326 | -0.157 | -0.347 | 0.446 | 0.012 | 0.038 | 0.038 | 0 | 0.24 | 0.11 | 0.11 | 0.16 | 0.17 | 0.16 |
| 3 | 77714832 | rs9842169 | 0.836 | 0.293 | -0.538 | 0.423 | -0.026 | -0.034 | -0.034 | 0 | 0.23 | 0.12 | 0.24 | 0.25 | 0.27 | 0.14 |
| 3 | 77716749 | rs810270 | -0.805 | -0.061 | -0.314 | 0.17 | -0.013 | 0.021 | 0.021 | 0.68 | 0.79 | 0.85 | 0.7 | 0.88 | 0.73 | 0.78 |
| 3 | 77717398 | rs9828866 | -0.196 | -0.102 | -0.288 | 0.176 | 0.017 | 0.059 | 0.059 | 6.327 | 0.49 | 0.22 | 0.34 | 0.37 | 0.4 | 0.43 |
| 3 | 77720924 | rs9883373 | -0.253 | -0.117 | -0.006 | 0.156 | 0.021 | 0.048 | 0.048 | 5.78 | 0.5 | 0.32 | 0.35 | 0.38 | 0.4 | 0.43 |
| 3 | 77722292 | rs1451627 | -0.698 | -0.13 | 0.552 | 0.022 | 0.031 | -0.07 | -0.07 | 0 | 0.78 | 0.66 | 0.69 | 0.88 | 0.73 | 0.78 |
| 3 | 77724063 | rs1721179 | -0.814 | -0.146 | 0.217 | -0.023 | -0.013 | 0.023 | 0.023 | 0.75 | 0.79 | 0.85 | 0.71 | 0.89 | 0.74 | 0.8 |
| 3 | 77724642 | rs10511059 | 1.719 | -0.216 | 0.145 | 0.368 | 0 | 0 | 0 | 0 | 0.25 | 0.29 | 0.32 | 0.26 | 0.32 | 0.3 |
| 3 | 77731974 | rs1523752 | 0.556 | -0.217 | -0.377 | 0.483 | 0.019 | -0.009 | -0.009 | 0 | 0.32 | 0.06 | 0.14 | 0.3 | 0.2 | 0.36 |
| 3 | 77739202 | rs1851740 | 1.642 | -0.102 | -0.269 | 0.663 | -0.01 | -0.071 | -0.071 | 0 | 0.18 | 0.04 | 0.19 | 0.27 | 0.17 | 0.11 |
| 3 | 77740954 | rs1997421 | 1.07 | -0.163 | -0.1 | 0.418 | -0.003 | -0.023 | -0.023 | 0.081 | 0.21 | 0.17 | 0.23 | 0.33 | 0.22 | 0.13 |
| 3 | 77742421 | rs11922844 | 1.779 | -0.134 | -0.133 | 0.265 | 0.004 | 0.004 | 0.004 | 1.321 | 0.24 | 0.31 | 0.32 | 0.26 | 0.31 | 0.3 |
| 3 | 77742551 | rs776106 | -0.728 | -0.108 | -0.121 | 0.24 | -0.003 | -0.001 | -0.001 | 0 | 0.79 | 0.79 | 0.7 | 0.88 | 0.74 | 0.79 |
| 3 | 77745383 | rs13091812 | -0.153 | -0.496 | -0.255 | 0.509 | 0 | -0.001 | -0.001 | 0.314 | 0.55 | 0.49 | 0.45 | 0.55 | 0.5 | 0.62 |
| 3 | 77745533 | rs7642693 | 1.269 | -0.043 | -0.32 | 0.695 | 0.003 | 0.005 | 0.005 | 1.603 | 0.23 | 0.31 | 0.31 | 0.25 | 0.31 | 0.26 |
| 3 | 77746082 | rs1163761 | 0.269 | 0.053 | -1.119 | 0.786 | -0.004 | -0.042 | -0.042 | 0 | 0.28 | 0.13 | 0.23 | 0.4 | 0.27 | 0.31 |
| 3 | 77746164 | rs12487085 | -0.28 | -0.166 | -0.853 | 0.733 | 0.002 | 0.003 | 0.003 | 0 | 0.57 | 0.53 | 0.51 | 0.58 | 0.57 | 0.59 |
| 3 | 77749428 | rs1669215 | -0.151 | -0.382 | -1.274 | 0.877 | 0.002 | -0.056 | -0.056 | 0 | 0.24 | 0.13 | 0.16 | 0.39 | 0.23 | 0.31 |
| 3 | 77752758 | rs776122 | -0.459 | 0.25 | -0.887 | 0.203 | 0.024 | -0.054 | -0.054 | 0 | 0.84 | 0.67 | 0.89 | 0.93 | 0.85 | 0.78 |
| 3 | 77754333 | rs2044890 | 0.359 | 0.225 | -0.919 | 0.295 | 0.009 | -0.018 | -0.018 | 0 | 0.71 | 0.56 | 0.68 | 0.77 | 0.72 | 0.68 |
| 3 | 77755170 | rs776119 | -0.259 | -0.176 | -1.073 | 0.775 | 0 | -0.017 | -0.017 | 0 | 0.06 | 0.08 | 0.04 | 0 | 0.04 | 0.01 |
| 3 | 77756847 | rs12634677 | -0.365 | -0.248 | -1.245 | 1.119 | -0.003 | -0.077 | -0.077 | 0 | 0.22 | 0.08 | 0.31 | 0.52 | 0.22 | 0.29 |
| 3 | 77763258 | rs9881868 | -0.815 | -0.135 | -0.6 | 0.405 | -0.015 | -0.049 | -0.049 | 0 | 0.09 | 0.25 | 0.06 | 0 | 0.04 | 0.01 |
| 3 | 77764243 | rs9849387 | 0.185 | -0.506 | -2.035 | 0.424 | -0.056 | 0.002 | 0.002 | 0.437 | 0.44 | 0.06 | 0.55 | 0.3 | 0.6 | 0.53 |
| 3 | 77764672 | rs776112 | -0.021 | -0.679 | -1.752 | 1.053 | 0 | 0.008 | 0.008 | 0.668 | 0.81 | 0.85 | 0.81 | 0.67 | 0.79 | 0.69 |
| 3 | 77765100 | rs12635472 | -0.087 | -0.957 | -0.817 | 1.472 | -0.006 | 0.342 | 0.342 | 1.424 | 0.08 | 0.28 | 0.09 | 0.48 | 0.06 | 0.19 |
| 3 | 77767063 | rs1523755 | 1.207 | -1.235 | -1.28 | 0.7 | -0.142 | 0.073 | 0.073 | 0.109 | 0.36 | 0.07 | 0.47 | 0.18 | 0.48 | 0.3 |
| 3 | 77769689 | rs10511058 | 0.014 | -1.196 | -1.016 | 0.835 | 0.028 | 0.387 | 0.387 | 2.443 | 0.09 | 0.4 | 0.09 | 0.39 | 0.06 | 0.19 |
| 3 | 77778813 | rs17274650 | 0.395 | -1.273 | -1.384 | 0.749 | -0.041 | -0.081 | -0.081 | 0 | 0.13 | 0.01 | 0.17 | 0.31 | 0.17 | 0.27 |
| 3 | 77790068 | rs9810156 | 1.324 | -1.206 | -1.362 | 0.742 | -0.143 | 0.073 | 0.073 | 0.176 | 0.35 | 0.07 | 0.47 | 0.18 | 0.48 | 0.3 |
| 3 | 77801683 | rs17274847 | -0.396 | -1.385 | -1.032 | 0.939 | 0.004 | -0.053 | -0.053 | 0 | 0.23 | 0.07 | 0.22 | 0.31 | 0.23 | 0.34 |
| 3 | 77804193 | rs6794930 | -0.187 | -1.22 | -0.634 | 1.06 | 0.003 | 0.284 | 0.284 | 1.45 | 0.08 | 0.29 | 0.09 | 0.39 | 0.06 | 0.19 |
| 3 | 77836956 | rs6798119 | 0.015 | -1.184 | -0.675 | 1.064 | 0.004 | 0.287 | 0.287 | 1.079 | 0.08 | 0.28 | 0.09 | 0.4 | 0.06 | 0.19 |
| 3 | 77862410 | rs3852019 | -0.932 | -1.387 | -0.554 | 0.619 | -0.007 | -0.003 | -0.003 | 0 | 0.32 | 0.35 | 0.3 | 0.7 | 0.29 | 0.53 |
| 3 | 77868589 | rs3852021 | -0.904 | -2.009 | -0.335 | 0.065 | 0.022 | 0.055 | 0.055 | 0 | 0.75 | 0.87 | 0.84 | 0.88 | 0.82 | 0.84 |
| 3 | 77887303 | rs7629921 | -0.556 | -2.218 | -0.076 | -0.014 | 0.002 | 0.001 | 0.001 | 0 | 0.31 | 0.28 | 0.3 | 0.7 | 0.29 | 0.53 |
| 3 | 77909964 | rs6793440 | 1.157 | -2.19 | 0.043 | -0.026 | 0 | 0.049 | 0.049 | 0 | 0.43 | 0.36 | 0.53 | 0.19 | 0.53 | 0.31 |
| 3 | 77916461 | rs7615325 | -0.988 | -2.084 | 0.075 | 0.063 | 0.019 | 0.053 | 0.053 | 0 | 0.76 | 0.87 | 0.85 | 0.88 | 0.82 | 0.85 |
| 3 | 77920575 | rs1529990 | 1.677 | -1.895 | 0.038 | 0.085 | -0.137 | 0.069 | 0.069 | 0.197 | 0.35 | 0.08 | 0.49 | 0.18 | 0.5 | 0.31 |
| 3 | 77935049 | rs6548526 | -0.82 | -1.762 | 0.345 | 0.43 | -0.022 | -0.005 | -0.005 | 0.11 | 0.33 | 0.45 | 0.32 | 0.69 | 0.29 | 0.53 |
| 3 | 77950703 | rs1437528 | 0.72 | -1.76 | 0.394 | 0.408 | -0.017 | -0.006 | -0.006 | 0 | 0.67 | 0.43 | 0.68 | 0.31 | 0.71 | 0.48 |
| 3 | 77959159 | rs17015624 | -0.192 | -1.769 | 0.372 | 0.757 | -0.015 | 0.173 | 0.173 | 1.048 | 0.09 | 0.19 | 0.08 | 0.39 | 0.06 | 0.16 |
| 3 | 77960033 | rs17015627 | 0.602 | -1.958 | 0.226 | 0.489 | 0.025 | 0.026 | 0.026 | 0 | 0.11 | 0.07 | 0.06 | 0.01 | 0.06 | 0.11 |
| 3 | 77968488 | rs9880985 | 0.481 | -2.401 | 0.25 | 0.194 | -0.018 | -0.043 | -0.043 | 0.32 | 0.28 | 0.32 | 0.15 | 0.12 | 0.16 | 0.16 |
| 3 | 77974318 | rs7615248 | 0.398 | -0.996 | -0.191 | -0.022 | 0.014 | 0.034 | 0.034 | 0 | 0.65 | 0.51 | 0.7 | 0.51 | 0.57 | 0.63 |
| 3 | 77976587 | rs986449 | -0.506 | -0.976 | -0.221 | -0.012 | 0.045 | 0.079 | 0.079 | 0 | 0.77 | 0.64 | 0.76 | 0.52 | 0.63 | 0.74 |
| 3 | 77980641 | rs6548534 | -0.457 | -0.976 | -0.209 | -0.051 | -0.013 | -0.043 | -0.043 | 0 | 0.86 | 0.78 | 0.91 | 0.99 | 0.91 | 0.89 |
| 3 | 78001054 | rs17015683 | 1.145 | -0.983 | -0.208 | -0.045 | 0.015 | 0.011 | 0.011 | 0 | 0.11 | 0.09 | 0.06 | 0.01 | 0.06 | 0.11 |
| 3 | 78002349 | rs1018067 | 0.269 | -0.929 | -0.216 | -0.003 | 0.008 | 0.019 | 0.019 | 0 | 0.66 | 0.57 | 0.7 | 0.51 | 0.57 | 0.63 |
| 3 | 78012759 | rs17775669 | 1.6 | -0.99 | -0.537 | -0.166 | -0.059 | 0.031 | 0.031 | 0.034 | 0.23 | 0.02 | 0.38 | 0.09 | 0.24 | 0.19 |
| 3 | 78056120 | rs17832529 | 1.575 | -0.902 | -0.575 | -0.166 | -0.034 | 0.046 | 0.046 | 0.03 | 0.23 | 0.05 | 0.38 | 0.09 | 0.24 | 0.19 |
| 3 | 78064944 | rs11127613 | 0.791 | -0.861 | -0.484 | 0.222 | 0.011 | 0.022 | 0.022 | 0 | 0.2 | 0.21 | 0.2 | 0.45 | 0.33 | 0.25 |
| 3 | 78069352 | rs870564 | -0.315 | -0.818 | -0.52 | 0.223 | 0.011 | -0.003 | -0.003 | 0 | 0.57 | 0.65 | 0.69 | 0.55 | 0.66 | 0.56 |
| 3 | 78101577 | rs6781390 | 1.037 | -0.469 | -0.762 | -0.099 | -0.039 | 0.047 | 0.047 | 0.181 | 0.24 | 0.1 | 0.41 | 0.09 | 0.31 | 0.21 |
| 3 | 78106208 | rs1495598 | 0.337 | 0.452 | -0.674 | 0.132 | -0.004 | -0.011 | -0.011 | 0.161 | 0.22 | 0.21 | 0.2 | 0.41 | 0.29 | 0.28 |
| 3 | 78107128 | rs2872373 | -0.033 | 0.496 | -0.623 | -0.027 | 0.025 | -0.008 | -0.008 | 0.004 | 0.46 | 0.35 | 0.63 | 0.5 | 0.64 | 0.47 |
| 3 | 78110954 | rs7639226 | 0.68 | 0.368 | -0.479 | 0.076 | 0.003 | 0.01 | 0.01 | 0.109 | 0.28 | 0.31 | 0.46 | 0.39 | 0.34 | 0.28 |
| 3 | 78132623 | rs7614879 | -1.385 | 0.325 | -1.038 | 0.227 | -0.013 | -0.054 | -0.054 | 0.638 | 0.91 | 1 | 0.9 | 0.73 | 0.91 | 0.79 |
| 3 | 78144437 | rs1344075 | -0.348 | 0.015 | -0.8 | -0.28 | -0.019 | -0.033 | -0.033 | 7.398 | 0.73 | 0.8 | 0.72 | 0.47 | 0.64 | 0.51 |
| 3 | 78161505 | rs9868401 | 1.63 | -0.261 | -0.51 | -0.198 | -0.003 | -0.015 | -0.015 | 0 | 0.08 | 0.07 | 0.05 | 0 | 0.06 | 0.03 |

| | | | | | | | | | | | | | | | | |
|---|---|---|---|---|---|---|---|---|---|---|---|---|---|---|---|---|
| 3 | 78166986 | rs1917091 | -0.237 | -0.207 | 0.013 | 0.098 | 0.066 | 0.342 | 0.342 | 3.504 | 0.06 | 0.51 | 0.07 | 0.24 | 0.04 | 0.1 |
| 3 | 78168724 | rs4680974 | 1.671 | -0.023 | -0.769 | -0.028 | -0.074 | -0.019 | -0.019 | 0 | 0.2 | 0.03 | 0.38 | 0.14 | 0.27 | 0.17 |
| 3 | 78182203 | rs1917083 | -0.146 | -0.307 | -0.866 | -0.187 | 0.007 | -0.002 | -0.002 | 0 | 0.39 | 0.68 | 0.53 | 0.39 | 0.39 | 0.32 |
| 3 | 78192927 | rs10511131 | 0.532 | 0.141 | -1.156 | -0.816 | 0.004 | -0.02 | -0.02 | 0 | 0.3 | 0.4 | 0.47 | 0.14 | 0.35 | 0.22 |
| 3 | 78207187 | rs1917072 | 1.496 | 0.392 | -1.37 | -0.567 | -0.04 | -0.004 | -0.004 | 0.283 | 0.19 | 0.1 | 0.39 | 0.14 | 0.26 | 0.17 |
| 3 | 78208877 | rs17833444 | 0.393 | 0.507 | -1.524 | -0.797 | -0.003 | 0.055 | 0.055 | 0 | 0.47 | 0.32 | 0.57 | 0.26 | 0.49 | 0.32 |
| 3 | 78234456 | rs9882906 | 0.254 | 0.507 | -1.485 | -0.672 | 0.012 | 0.046 | 0.046 | 0.404 | 0.36 | 0.21 | 0.24 | 0.12 | 0.29 | 0.16 |
| 3 | 78275119 | rs2140646 | 0.814 | 0.658 | -1.782 | -0.267 | -0.012 | 0.094 | 0.094 | 0 | 0.33 | 0.11 | 0.53 | 0.14 | 0.35 | 0.23 |
| 3 | 78276135 | rs7433616 | -0.354 | 1.309 | -1.451 | 0.396 | -0.06 | -0.061 | -0.061 | 0.762 | 0.67 | 0.77 | 0.73 | 0.54 | 0.49 | 0.6 |
| 3 | 78279461 | rs7645322 | 0.262 | 0.853 | -1.089 | -0.488 | 0.012 | 0.05 | 0.05 | 0.051 | 0.51 | 0.36 | 0.5 | 0.1 | 0.41 | 0.34 |
| 3 | 78280658 | rs9836421 | -0.587 | 0.539 | -0.767 | 0.171 | -0.003 | 0.014 | 0.014 | 0 | 0.38 | 0.54 | 0.48 | 0.49 | 0.35 | 0.37 |
| 3 | 78287935 | rs11713404 | 1.197 | 0.36 | -1.396 | -0.207 | -0.156 | 0.226 | 0.226 | 0.014 | 0.45 | 0.02 | 0.41 | 0.16 | 0.51 | 0.3 |
| 3 | 78288555 | rs1880363 | 0.441 | 0.122 | -0.626 | 0.156 | -0.009 | 0.227 | 0.227 | 1.121 | 0.13 | 0.39 | 0.11 | 0.34 | 0.08 | 0.18 |
| 3 | 78289887 | rs7616235 | -1.208 | 0.135 | -0.293 | -0.044 | -0.007 | -0.05 | -0.05 | 0 | 0.9 | 0.84 | 0.94 | 0.99 | 0.91 | 0.92 |
| 3 | 78292249 | rs6766104 | -0.46 | 0.312 | -0.224 | -0.095 | -0.008 | 0.024 | 0.024 | 0 | 0.29 | 0.24 | 0.37 | 0.15 | 0.29 | 0.22 |
| 3 | 78295664 | rs2872375 | 0.932 | 0.244 | 0.252 | -0.13 | -0.009 | -0.052 | -0.052 | 0 | 0.63 | 0.47 | 0.57 | 0.85 | 0.66 | 0.73 |
| 3 | 78300528 | rs1917100 | 0.917 | 0.244 | 0.402 | -0.129 | -0.009 | -0.052 | -0.052 | 0 | 0.63 | 0.48 | 0.57 | 0.85 | 0.66 | 0.73 |
| 3 | 78304225 | rs1917098 | -0.521 | 0.327 | 0.382 | -0.085 | 0.001 | -0.011 | -0.011 | 0 | 0.29 | 0.38 | 0.38 | 0.15 | 0.29 | 0.22 |
| 3 | 78307578 | rs2140645 | 0.525 | 0.043 | 0.202 | -0.088 | -0.001 | -0.006 | -0.006 | 0.004 | 0.16 | 0.15 | 0.29 | 0.14 | 0.18 | 0.12 |
| 3 | 78307694 | rs9309793 | 0.574 | 0.092 | 0.323 | 0.035 | 0 | 0.154 | 0.154 | 0.605 | 0.12 | 0.28 | 0.11 | 0.33 | 0.11 | 0.18 |
| 3 | 78310110 | rs9309796 | -0.885 | 0.078 | 0.304 | 0.025 | -0.008 | 0.02 | 0.02 | 0.009 | 0.34 | 0.43 | 0.43 | 0.47 | 0.32 | 0.35 |
| 3 | 78322422 | rs9881821 | 0.035 | 0.032 | 0.125 | 0.038 | 0.029 | 0.157 | 0.157 | 1.428 | 0.15 | 0.35 | 0.14 | 0.34 | 0.14 | 0.21 |
| 3 | 78324964 | rs1512487 | -1.031 | 0.031 | 0.08 | 0.036 | -0.001 | 0.009 | 0.009 | 0.176 | 0.37 | 0.6 | 0.47 | 0.48 | 0.35 | 0.39 |
| 3 | 78329438 | rs1512474 | -1.281 | 0.037 | -0.072 | -0.049 | -0.029 | -0.056 | -0.056 | 3.49 | 0.9 | 0.83 | 0.95 | 1 | 0.94 | 0.94 |
| 3 | 78331851 | rs12631515 | -0.031 | 0.3 | -0.447 | 0.088 | 0.002 | 0.005 | 0.005 | 0.99 | 0.21 | 0.16 | 0.32 | 0.14 | 0.21 | 0.18 |
| 3 | 78337174 | rs9845829 | -0.737 | 0.428 | -0.506 | 0.12 | 0.005 | -0.014 | -0.014 | 0 | 0.4 | 0.67 | 0.48 | 0.46 | 0.36 | 0.44 |
| 3 | 78341723 | rs11927217 | -0.457 | 0.334 | -0.331 | -0.069 | 0.004 | 0.023 | 0.023 | 0.117 | 0.29 | 0.37 | 0.41 | 0.43 | 0.34 | 0.32 |
| 3 | 78351238 | rs12486667 | 0.723 | 0.277 | -1.074 | -0.34 | -0.065 | -0.04 | -0.04 | 0 | 0.2 | 0.01 | 0.21 | 0.23 | 0.22 | 0.36 |
| 3 | 78357272 | rs1355625 | 0.757 | -0.278 | -0.82 | -0.855 | 0.001 | 0.015 | 0.015 | 0.234 | 0.49 | 0.46 | 0.36 | 0.3 | 0.48 | 0.28 |
| 3 | 78371577 | rs1028024 | 0.865 | 0.008 | -1.012 | -0.565 | -0.035 | -0.065 | -0.065 | 1.354 | 0.15 | 0.06 | 0.34 | 0.46 | 0.21 | 0.3 |
| 3 | 78381212 | rs11127617 | -0.415 | -0.245 | -0.542 | -0.79 | 0.001 | 0.006 | 0.006 | 1.462 | 0.34 | 0.68 | 0.29 | 0.28 | 0.29 | 0.41 |
| 3 | 78386422 | rs1512483 | 0.989 | -0.206 | -0.213 | -0.769 | -0.006 | -0.002 | -0.002 | 0 | 0.19 | 0.12 | 0.19 | 0.22 | 0.2 | 0.34 |
| 3 | 78387430 | rs6765243 | 0.754 | -0.268 | 0.364 | -0.796 | -0.019 | -0.017 | -0.017 | 0.006 | 0.09 | 0.32 | 0.04 | 0.05 | 0.04 | 0.03 |
| 3 | 78396840 | rs1512495 | 0.933 | -0.258 | 0.234 | -0.782 | -0.004 | -0.002 | -0.002 | 0 | 0.19 | 0.13 | 0.19 | 0.23 | 0.2 | 0.34 |
| 3 | 78399328 | rs9309797 | 0.543 | -0.334 | -0.308 | -0.869 | -0.053 | -0.109 | -0.109 | 0.613 | 0.26 | 0.08 | 0.41 | 0.45 | 0.34 | 0.34 |
| 3 | 78399857 | rs9840973 | -1.155 | -0.614 | -0.163 | -1.114 | 0.028 | 0.081 | 0.081 | 4.758 | 0.59 | 0.82 | 0.69 | 0.73 | 0.63 | 0.74 |
| 3 | 78400215 | rs11921066 | 0.214 | -0.392 | 0.09 | -0.94 | -0.022 | -0.021 | -0.021 | 1.453 | 0.09 | 0.27 | 0.04 | 0.05 | 0.04 | 0.03 |
| 3 | 78417968 | rs11915895 | 1.168 | -0.602 | -0.512 | -1.214 | 0.011 | 0.035 | 0.035 | 0 | 0.33 | 0.2 | 0.22 | 0.24 | 0.31 | 0.25 |
| 3 | 78423185 | rs6548554 | 0.544 | -0.786 | -0.58 | -1.205 | -0.08 | -0.133 | -0.133 | 0 | 0.19 | 0.02 | 0.41 | 0.52 | 0.24 | 0.33 |
| 3 | 78434843 | rs11920974 | 1.317 | -0.67 | -0.453 | -1.206 | 0.001 | 0.038 | 0.038 | 0 | 0.35 | 0.22 | 0.24 | 0.24 | 0.34 | 0.26 |
| 3 | 78435488 | rs997705 | 1.181 | -0.147 | -0.375 | -1.034 | -0.063 | -0.041 | -0.041 | 1.457 | 0.15 | 0.01 | 0.17 | 0.19 | 0.19 | 0.31 |
| 3 | 78439894 | rs4680983 | -0.106 | -0.235 | 0.024 | -1.119 | 0.012 | -0.041 | -0.041 | 0 | 0.72 | 0.64 | 0.75 | 0.81 | 0.68 | 0.67 |
| 3 | 78444051 | rs10865564 | -1.455 | -0.29 | 0.08 | -1.105 | 0.001 | -0.012 | -0.012 | 0 | 0.88 | 0.91 | 0.93 | 1 | 0.86 | 0.98 |
| 3 | 78455571 | rs10865566 | 1.186 | -0.279 | 0.113 | -1.111 | 0.002 | -0.001 | -0.001 | 0 | 0.16 | 0.16 | 0.17 | 0.19 | 0.2 | 0.31 |
| 3 | 78455767 | rs7627884 | 0.989 | -0.374 | 0.236 | -1.096 | -0.006 | -0.01 | -0.01 | 0.603 | 0.08 | 0.13 | 0.03 | 0.05 | 0.04 | 0.05 |
| 3 | 78459918 | rs17016040 | 0.62 | -0.385 | 0.283 | -1.101 | -0.013 | -0.023 | -0.023 | 0 | 0.09 | 0.19 | 0.04 | 0.05 | 0.04 | 0.04 |
| 3 | 78460675 | rs9819574 | -1.14 | -0.012 | 0.042 | -0.867 | -0.003 | -0.074 | -0.074 | 0 | 0.87 | 0.8 | 0.92 | 1 | 0.87 | 0.98 |
| 3 | 78468807 | rs7647185 | 0.666 | 0.091 | -0.425 | -0.759 | -0.078 | -0.113 | -0.113 | 0.734 | 0.16 | 0.02 | 0.41 | 0.52 | 0.23 | 0.32 |
| 3 | 78469764 | rs9839172 | 0.139 | -0.115 | 0.126 | -0.885 | 0.012 | -0.056 | -0.056 | 0.546 | 0.71 | 0.61 | 0.74 | 0.81 | 0.66 | 0.67 |
| 3 | 78473138 | rs7626026 | 0.399 | -0.4 | 0.06 | -0.881 | -0.011 | -0.066 | -0.066 | 0.797 | 0.15 | 0.17 | 0.08 | 0 | 0.14 | 0.06 |
| 3 | 78474658 | rs2314126 | 0.601 | -0.213 | -0.642 | -0.85 | -0.091 | -0.143 | -0.143 | 0 | 0.22 | 0.02 | 0.43 | 0.51 | 0.29 | 0.38 |
| 3 | 78475747 | rs17016095 | 1.805 | -0.921 | -0.465 | -1.239 | -0.027 | -0.048 | -0.048 | 2.723 | 0.1 | 0.12 | 0.05 | 0 | 0.07 | 0 |
| 3 | 78477614 | rs1512493 | -0.538 | -1.131 | -0.205 | -1.344 | -0.013 | -0.025 | -0.025 | 0.206 | 0.38 | 0.42 | 0.27 | 0.29 | 0.34 | 0.34 |
| 3 | 78480021 | rs7620963 | 0.221 | -1.177 | -0.185 | -1.379 | -0.041 | -0.13 | -0.13 | 0 | 0.25 | 0.34 | 0.13 | 0 | 0.21 | 0.09 |
| 3 | 78481969 | rs2068704 | -1.983 | -1.272 | 0.109 | -1.497 | -0.002 | 0.018 | 0.018 | 0.017 | 0.68 | 0.75 | 0.75 | 0.85 | 0.69 | 0.76 |
| 3 | 78485890 | rs12233386 | 1.707 | -1.183 | -0.073 | -1.436 | -0.033 | -0.057 | -0.057 | 2.544 | 0.12 | 0.03 | 0.14 | 0.29 | 0.13 | 0.25 |
| 3 | 78489549 | rs3923142 | 0.403 | -1.242 | -0.257 | -1.475 | -0.123 | -0.148 | -0.148 | 0 | 0.33 | 0.03 | 0.48 | 0.51 | 0.41 | 0.44 |
| 3 | 78499137 | rs13087593 | 0.422 | -1.059 | 0.064 | -1.414 | -0.038 | -0.128 | -0.128 | 0.016 | 0.25 | 0.34 | 0.13 | 0 | 0.21 | 0.09 |
| 3 | 78503357 | rs7650723 | -0.756 | -0.998 | 0.016 | -1.415 | 0.015 | 0.035 | 0.035 | 0.894 | 0.46 | 0.62 | 0.31 | 0.34 | 0.4 | 0.38 |
| 3 | 78505090 | rs12631698 | -1.724 | -1.056 | -0.022 | -1.695 | 0 | -0.012 | -0.012 | 0 | 0.68 | 0.64 | 0.74 | 0.85 | 0.68 | 0.76 |
| 3 | 78505213 | rs9866490 | -0.436 | -0.998 | -0.027 | -1.65 | -0.005 | 0.013 | 0.013 | 0.46 | 0.43 | 0.42 | 0.63 | 0.85 | 0.48 | 0.68 |
| 3 | 78511515 | rs4680987 | 0.263 | -1.146 | -0.257 | -1.402 | -0.024 | 0.025 | 0.025 | 1.134 | 0.34 | 0.12 | 0.44 | 0.76 | 0.39 | 0.6 |
| 3 | 78519523 | rs4266157 | 0 | -1.268 | 0.088 | -1.477 | -0.019 | -0.107 | -0.107 | 0 | 0.31 | 0.39 | 0.27 | 0.05 | 0.26 | 0.15 |
| 3 | 78524001 | rs4330230 | 0.01 | -1.263 | 0.091 | -1.475 | -0.019 | -0.107 | -0.107 | 2.177 | 0.3 | 0.39 | 0.27 | 0.05 | 0.26 | 0.15 |
| 3 | 78527213 | rs4446181 | 1.513 | -1.204 | -0.113 | -1.344 | -0.013 | -0.044 | -0.044 | 1.696 | 0.13 | 0.01 | 0.14 | 0.21 | 0.14 | 0.24 |
| 3 | 78535374 | rs13066384 | -0.566 | -1.16 | 0.551 | -1.549 | -0.011 | -0.074 | -0.074 | 0.265 | 0.32 | 0.66 | 0.29 | 0.05 | 0.26 | 0.17 |
| 3 | 78551628 | rs4680992 | -1.466 | -1.15 | 0.301 | -1.535 | -0.026 | 0.03 | 0.03 | 0.013 | 0.67 | 0.83 | 0.75 | 0.78 | 0.67 | 0.75 |
| 3 | 78551907 | rs17016200 | 0.212 | -0.664 | 0.79 | -1.265 | -0.035 | -0.115 | -0.115 | 1.306 | 0.23 | 0.26 | 0.11 | 0 | 0.17 | 0.11 |
| 3 | 78552232 | rs13089920 | -0.542 | -0.664 | 0.829 | -1.266 | -0.03 | -0.139 | -0.139 | 0.497 | 0.31 | 0.47 | 0.27 | 0.05 | 0.26 | 0.17 |

| | | | | | | | | | | | | | | | | |
|---|---|---|---|---|---|---|---|---|---|---|---|---|---|---|---|---|
| 3 | 78559012 | rs1444483 | 0.31 | -0.668 | 0.399 | -1.282 | -0.001 | 0.103 | 0.103 | 0 | 0.5 | 0.36 | 0.63 | 0.78 | 0.51 | 0.69 |
| 3 | 78580228 | rs6763358 | -0.837 | -0.38 | -0.063 | -1.101 | 0.031 | -0.015 | -0.015 | 0 | 0.72 | 0.59 | 0.73 | 0.78 | 0.68 | 0.77 |
| 3 | 78580690 | rs2120705 | -1.146 | -0.32 | -0.653 | -1.075 | -0.023 | 0.004 | 0.004 | 0 | 0.74 | 0.85 | 0.75 | 0.78 | 0.68 | 0.77 |
| 3 | 78590131 | rs6789918 | 0.479 | -0.224 | -0.693 | -0.778 | -0.029 | 0.075 | 0.075 | 0 | 0.4 | 0.08 | 0.45 | 0.73 | 0.42 | 0.64 |
| 3 | 78594819 | rs9876636 | -0.072 | -0.124 | 0.704 | -0.297 | -0.014 | -0.081 | -0.081 | 3.755 | 0.84 | 0.64 | 0.8 | 0.95 | 0.87 | 0.94 |
| 3 | 78595335 | rs4401330 | 0.503 | -0.123 | 0.073 | -0.296 | -0.03 | -0.085 | -0.085 | 0 | 0.8 | 0.31 | 0.76 | 0.95 | 0.87 | 0.94 |
| 3 | 78609963 | rs9682532 | -0.124 | 0.453 | -0.377 | 0.764 | -0.012 | -0.027 | -0.027 | 0 | 0.17 | 0.11 | 0.2 | 0.4 | 0.24 | 0.27 |
| 3 | 78610831 | rs1507413 | -0.872 | 0.572 | 0.233 | 0.685 | 0.006 | 0.035 | 0.035 | 0 | 0.27 | 0.3 | 0.28 | 0.47 | 0.31 | 0.32 |
| 3 | 78614378 | rs11716795 | -0.016 | 0.557 | 0.234 | 0.849 | -0.009 | -0.019 | -0.019 | 0 | 0.18 | 0.13 | 0.21 | 0.35 | 0.24 | 0.27 |
| 3 | 78615037 | rs6801989 | 0.329 | 0.325 | 0.588 | 0.546 | -0.013 | -0.05 | -0.05 | 0 | 0.86 | 0.68 | 0.91 | 0.92 | 0.91 | 0.95 |
| 3 | 78617646 | rs17016341 | 0.999 | 0.306 | 0.572 | 0.551 | -0.003 | -0.018 | -0.018 | 0 | 0.09 | 0.16 | 0.07 | 0.07 | 0.08 | 0.04 |
| 3 | 78620239 | rs11127627 | -0.099 | 0.267 | 0.2 | 0.246 | 0.001 | 0.067 | 0.067 | 0 | 0.51 | 0.3 | 0.44 | 0.71 | 0.48 | 0.53 |
| 3 | 78626222 | rs1507424 | -0.064 | 0.253 | 0.262 | 0.234 | -0.01 | -0.058 | -0.058 | 1.56 | 0.86 | 0.69 | 0.9 | 0.93 | 0.9 | 0.92 |
| 3 | 78629833 | rs6548589 | -0.152 | 0.245 | 0.108 | 0.478 | 0.001 | -0.02 | -0.02 | 0 | 0.79 | 0.84 | 0.79 | 0.59 | 0.78 | 0.73 |
| 3 | 78640492 | rs6807667 | -0.157 | 0.21 | 0.433 | 0.428 | -0.001 | 0.016 | 0.016 | 0 | 0.79 | 0.8 | 0.78 | 0.58 | 0.79 | 0.73 |
| 3 | 78643050 | rs11706918 | 0.652 | 0.222 | 0.259 | 0.437 | -0.002 | -0.008 | -0.008 | 0 | 0.17 | 0.13 | 0.2 | 0.41 | 0.2 | 0.27 |
| 3 | 78646942 | rs7626143 | 0.748 | 0.233 | 0.141 | 0.428 | -0.049 | -0.123 | -0.123 | 0.083 | 0.17 | 0.02 | 0.19 | 0.41 | 0.2 | 0.27 |
| 3 | 78648532 | rs1444480 | 0.836 | 0.229 | 0.185 | 0.381 | 0.001 | 0.006 | 0.006 | 0 | 0.34 | 0.25 | 0.25 | 0.33 | 0.28 | 0.27 |
| 3 | 78652831 | rs12496962 | 1.225 | 0.269 | -0.009 | 0.432 | -0.034 | 0.04 | 0.04 | 0 | 0.33 | 0.13 | 0.43 | 0.19 | 0.41 | 0.33 |
| 3 | 78654924 | rs3773190 | 0.221 | 0.092 | 0.356 | 0.275 | 0.013 | 0.037 | 0.037 | 0 | 0.52 | 0.37 | 0.64 | 0.62 | 0.62 | 0.65 |
| 3 | 78657774 | rs723766 | -0.289 | -0.065 | 0.45 | 0.348 | -0.012 | -0.013 | -0.013 | 0.103 | 0.67 | 0.58 | 0.76 | 0.69 | 0.73 | 0.77 |
| 3 | 78659692 | rs9811063 | 1.191 | -0.354 | 0.87 | 0.186 | -0.033 | -0.036 | -0.036 | 1.202 | 0.87 | 0.7 | 0.94 | 0.95 | 0.93 | 0.84 |
| 3 | 78660289 | rs1507419 | 0.895 | -0.304 | 0.799 | 0.208 | 0.009 | 0.007 | 0.007 | 0.816 | 0.95 | 0.82 | 0.97 | 0.96 | 0.97 | 0.91 |
| 3 | 78662193 | rs3773194 | 0.895 | -0.304 | 0.804 | 0.207 | 0.011 | 0.006 | 0.006 | 0.816 | 0.95 | 0.82 | 0.97 | 0.98 | 0.97 | 0.91 |
| 3 | 78663956 | rs1027832 | 1.403 | -0.374 | 0.52 | 0.345 | -0.03 | 0.002 | 0.002 | 0.845 | 0.41 | 0.17 | 0.5 | 0.33 | 0.43 | 0.44 |
| 3 | 78666765 | rs9839790 | 1.299 | -0.493 | 0.685 | 0.517 | -0.008 | 0.017 | 0.017 | 1.629 | 0.67 | 0.45 | 0.64 | 0.58 | 0.69 | 0.59 |
| 3 | 78668453 | rs3773201 | 0.464 | -0.14 | 0.474 | 0.704 | 0.011 | 0.061 | 0.061 | 0.991 | 0.78 | 0.66 | 0.69 | 0.59 | 0.73 | 0.66 |
| 3 | 78670654 | rs17016413 | 0.362 | 0.001 | -0.282 | 0.77 | -0.022 | -0.045 | -0.045 | 1.143 | 0.16 | 0.09 | 0.27 | 0.37 | 0.23 | 0.25 |
| 3 | 78676405 | rs7625555 | -0.845 | -0.118 | 0.026 | 0.487 | 0 | 0.032 | 0.032 | 0 | 0.51 | 0.66 | 0.5 | 0.7 | 0.52 | 0.54 |
| 3 | 78676422 | rs7614084 | 1.172 | -0.191 | -0.005 | 0.447 | -0.006 | 0.033 | 0.033 | 1.823 | 0.42 | 0.24 | 0.45 | 0.26 | 0.42 | 0.38 |
| 3 | 78678196 | rs4681006 | -0.652 | -0.198 | 0.188 | 0.497 | -0.001 | 0.008 | 0.008 | 0 | 0.48 | 0.54 | 0.5 | 0.7 | 0.52 | 0.49 |
| 3 | 78700901 | rs6795556 | -0.029 | -0.364 | 0.034 | 0.362 | 0 | 0.019 | 0.019 | 0 | 0.28 | 0.17 | 0.22 | 0.22 | 0.26 | 0.24 |
| 3 | 78702439 | rs9851228 | 1.002 | -0.003 | -0.191 | 0.713 | 0.021 | -0.003 | -0.003 | 0 | 0.59 | 0.27 | 0.68 | 0.59 | 0.67 | 0.66 |
| 3 | 78704821 | rs997274 | -0.117 | 0.283 | 0.609 | 0.566 | 0.02 | -0.011 | -0.011 | 0 | 0.23 | 0.32 | 0.23 | 0.23 | 0.25 | 0.22 |
| 3 | 78705219 | rs7610686 | -0.029 | 0.279 | 0.667 | 0.565 | 0.012 | -0.007 | -0.007 | 0 | 0.22 | 0.24 | 0.22 | 0.22 | 0.24 | 0.22 |
| 3 | 78714118 | rs7619742 | 1.38 | 0.352 | 0.663 | 0.677 | -0.003 | 0.086 | 0.086 | 0.922 | 0.52 | 0.29 | 0.41 | 0.23 | 0.5 | 0.43 |
| 3 | 78717508 | rs2255164 | 1.205 | 0.349 | 0.915 | 0.66 | -0.016 | 0.066 | 0.066 | 0.142 | 0.55 | 0.34 | 0.42 | 0.23 | 0.51 | 0.43 |
| 3 | 78733721 | rs2043661 | 0.064 | 0.338 | 0.882 | 0.766 | -0.013 | -0.022 | -0.022 | 4.155 | 0.14 | 0.08 | 0.29 | 0.49 | 0.2 | 0.28 |
| 3 | 78744206 | rs9856624 | 1.582 | 0.31 | 0.864 | 0.729 | -0.012 | 0.122 | 0.122 | 0.557 | 0.52 | 0.19 | 0.4 | 0.23 | 0.5 | 0.44 |
| 3 | 78754631 | rs767182 | -0.067 | 0.445 | 1.086 | 0.71 | 0.023 | -0.02 | -0.02 | 0.885 | 0.24 | 0.38 | 0.24 | 0.22 | 0.25 | 0.21 |
| 3 | 78759564 | rs9808950 | -0.649 | 0.374 | 0.983 | 0.677 | 0.003 | -0.035 | -0.035 | 0 | 0.28 | 0.51 | 0.26 | 0.22 | 0.26 | 0.22 |
| 3 | 78774034 | rs7626082 | -0.047 | 0.493 | 1.01 | 0.931 | -0.036 | -0.075 | -0.075 | 0 | 0.14 | 0.06 | 0.21 | 0.45 | 0.2 | 0.28 |
| 3 | 78774255 | rs3773232 | -0.47 | 0.479 | 1.194 | 0.925 | 0.022 | 0.148 | 0.148 | 0 | 0.17 | 0.29 | 0.23 | 0.45 | 0.21 | 0.28 |
| 3 | 78775567 | rs9837496 | 1.764 | 0.447 | 1.118 | 0.827 | 0.005 | 0.089 | 0.089 | 0.06 | 0.51 | 0.26 | 0.47 | 0.27 | 0.49 | 0.47 |
| 3 | 78777511 | rs3773234 | 0.113 | 0.521 | 0.997 | 0.857 | -0.002 | 0 | 0 | 0 | 0.22 | 0.21 | 0.23 | 0.24 | 0.25 | 0.21 |
| 3 | 78778901 | rs3773236 | -0.113 | 0.601 | 0.898 | 0.971 | -0.028 | -0.06 | -0.06 | 0.71 | 0.14 | 0.07 | 0.21 | 0.45 | 0.2 | 0.28 |
| 3 | 78789362 | rs2608021 | 1.278 | 0.621 | 1.048 | 0.856 | 0.002 | 0.118 | 0.118 | 3.067 | 0.76 | 0.52 | 0.74 | 0.47 | 0.77 | 0.72 |
| 3 | 78796078 | rs2304503 | 1.389 | 0.69 | 1.103 | 0.691 | -0.005 | 0.024 | 0.024 | 0 | 0.55 | 0.32 | 0.52 | 0.25 | 0.52 | 0.55 |
| 3 | 78797159 | rs2125123 | -1.456 | 0.728 | 0.862 | 0.637 | -0.019 | -0.024 | -0.024 | 0 | 0.07 | 0.25 | 0.05 | 0.02 | 0.02 | 0.02 |
| 3 | 78803784 | rs1865863 | 0.258 | 0.962 | 0.689 | 0.664 | 0.006 | -0.002 | -0.002 | 0 | 0.21 | 0.27 | 0.23 | 0.23 | 0.26 | 0.19 |
| 3 | 78803850 | rs1865862 | -1.24 | 1.118 | 0.725 | 0.77 | 0.005 | 0.058 | 0.058 | 2.537 | 0.23 | 0.34 | 0.25 | 0.51 | 0.25 | 0.26 |
| 3 | 78803935 | rs510006 | 1.879 | 1.012 | 0.304 | 0.709 | -0.069 | 0.077 | 0.077 | 1.369 | 0.47 | 0.04 | 0.46 | 0.24 | 0.45 | 0.52 |
| 3 | 78808850 | rs333491 | -1.503 | 1.103 | 0.42 | 0.648 | 0.001 | 0.005 | 0.005 | 0 | 0.49 | 0.54 | 0.48 | 0.72 | 0.55 | 0.46 |
| 3 | 78824912 | rs333472 | -1.073 | 1.156 | 0.414 | 0.735 | -0.003 | -0.016 | -0.016 | 0.096 | 0.27 | 0.27 | 0.28 | 0.5 | 0.31 | 0.29 |
| 3 | 78844274 | rs7650002 | 0.087 | 1.365 | 0.542 | 0.781 | 0.032 | -0.021 | -0.021 | 0 | 0.24 | 0.4 | 0.25 | 0.23 | 0.29 | 0.21 |
| 3 | 78853284 | rs333483 | -0.116 | 1.352 | 0.553 | 0.772 | 0.012 | -0.019 | -0.019 | 0 | 0.75 | 0.43 | 0.74 | 0.77 | 0.72 | 0.8 |
| 3 | 78857802 | rs11127635 | -0.17 | 1.37 | 0.518 | 0.756 | 0.013 | 0.073 | 0.073 | 2.952 | 0.16 | 0.26 | 0.24 | 0.48 | 0.23 | 0.25 |
| 3 | 78873936 | rs3856695 | -0.116 | 1.366 | 0.372 | 0.754 | 0.014 | 0.075 | 0.075 | 3.037 | 0.16 | 0.26 | 0.23 | 0.48 | 0.23 | 0.26 |
| 3 | 78877217 | rs162870 | -2.157 | 1.313 | 0.223 | 0.69 | 0.034 | 0.071 | 0.071 | 2.8 | 0.55 | 0.88 | 0.57 | 0.76 | 0.64 | 0.52 |
| 3 | 78877509 | rs162871 | 0.404 | 1.343 | 0.226 | 0.613 | 0.001 | 0.037 | 0.037 | 0 | 0.09 | 0.02 | 0.04 | 0.03 | 0.08 | 0.03 |
| 3 | 78890598 | rs17016554 | -1.018 | 1.422 | 0.333 | 0.756 | 0.003 | 0.023 | 0.023 | 3.037 | 0.22 | 0.28 | 0.33 | 0.52 | 0.27 | 0.29 |
| 3 | 78890766 | rs167944 | -0.475 | 1.294 | 0.404 | 0.631 | -0.012 | -0.022 | -0.022 | 0 | 0.94 | 0.81 | 0.95 | 0.98 | 0.98 | 0.99 |
| 3 | 78892352 | rs331158 | -0.215 | 1.313 | 0.325 | 0.606 | 0.007 | -0.007 | -0.007 | 0 | 0.28 | 0.27 | 0.27 | 0.25 | 0.37 | 0.22 |
| 3 | 78896598 | rs9835958 | 2.102 | 1.171 | 0.047 | 0.96 | -0.02 | -0.084 | -0.084 | 0.725 | 0.43 | 0.03 | 0.35 | 0.35 | 0.34 | 0.52 |
| 3 | 78902252 | rs9863758 | 1.692 | 1.359 | 0.22 | 0.978 | 0.047 | 0.035 | 0.035 | 0.264 | 0.48 | 0.25 | 0.38 | 0.38 | 0.36 | 0.56 |
| 3 | 78908171 | rs1383407 | 2.052 | 0.994 | 0.009 | 1.004 | -0.07 | -0.108 | -0.108 | 0 | 0.43 | 0.02 | 0.39 | 0.41 | 0.42 | 0.56 |
| 3 | 78913203 | rs1031932 | 1.963 | 1.032 | 0.06 | 0.786 | -0.08 | -0.099 | -0.099 | 0 | 0.44 | 0.06 | 0.41 | 0.44 | 0.44 | 0.57 |
| 3 | 78917012 | rs331147 | 0.029 | 0.881 | 0.202 | 0.851 | -0.003 | 0.012 | 0.012 | 4.171 | 0.84 | 0.81 | 0.69 | 0.55 | 0.75 | 0.74 |
| 3 | 78922852 | rs11127636 | 1.579 | 0.488 | 0.246 | 0.623 | 0.025 | 0.003 | 0.003 | 0.02 | 0.49 | 0.28 | 0.42 | 0.45 | 0.43 | 0.57 |

| | | | | | | | | | | | | | | | | |
|---|---|---|---|---|---|---|---|---|---|---|---|---|---|---|---|---|
| 3 | 78939727 | rs329819 | -1.323 | 0.192 | 0.17 | 0.215 | 0.002 | 0.108 | 0.108 | 0.586 | 0.42 | 0.81 | 0.34 | 0.25 | 0.38 | 0.3 |
| 3 | 78956204 | rs162263 | -0.257 | -0.023 | 0.308 | 0.195 | 0.003 | -0.011 | -0.011 | 2.214 | 0.3 | 0.32 | 0.26 | 0.19 | 0.34 | 0.26 |
| 3 | 78957109 | rs329814 | -0.646 | -0.012 | 0.311 | 0.199 | 0.005 | -0.035 | -0.035 | 4.63 | 0.33 | 0.52 | 0.29 | 0.21 | 0.35 | 0.26 |
| 3 | 78962148 | rs162268 | -0.887 | -0.039 | 0.278 | 0.208 | -0.001 | -0.015 | -0.015 | 0 | 0.37 | 0.57 | 0.31 | 0.21 | 0.36 | 0.29 |
| 3 | 78968841 | rs498805 | -0.944 | -0.035 | 0.278 | 0.169 | -0.007 | -0.029 | -0.029 | 0 | 0.42 | 0.51 | 0.31 | 0.19 | 0.37 | 0.31 |
| 3 | 78972040 | rs328047 | -1.411 | -0.041 | 0.176 | 0.153 | 0.061 | 0.161 | 0.161 | 0.003 | 0.49 | 0.75 | 0.35 | 0.25 | 0.39 | 0.29 |
| 3 | 78983923 | rs9839572 | -0.805 | 0.212 | -0.107 | 0.208 | 0.002 | 0.003 | 0.003 | 2.638 | 0.09 | 0.43 | 0.06 | 0.05 | 0.02 | 0.02 |
| 3 | 78985805 | rs328049 | 1.529 | 0.303 | -0.26 | 0.263 | -0.104 | 0.024 | 0.024 | 1.863 | 0.44 | 0.07 | 0.41 | 0.32 | 0.46 | 0.46 |
| 3 | 78995040 | rs13325291 | -1.071 | 0.639 | 0.809 | 0 | 0.005 | 0.006 | 0.006 | 1.744 | 0.08 | 0.37 | 0.06 | 0.05 | 0.02 | 0.02 |
| 3 | 79005935 | rs1031933 | -1.178 | 0.697 | 0.756 | -0.001 | -0.024 | -0.086 | -0.086 | 0 | 0.14 | 0.42 | 0.11 | 0.03 | 0.09 | 0.04 |
| 3 | 79013878 | rs17313732 | -0.488 | 0.807 | 0.488 | 0.026 | 0.011 | -0.015 | -0.015 | 0.035 | 0.24 | 0.33 | 0.24 | 0.21 | 0.3 | 0.23 |
| 3 | 79015464 | rs9832632 | -1.533 | 0.305 | 0.677 | 0.23 | -0.023 | -0.079 | -0.079 | 0.719 | 0.14 | 0.36 | 0.11 | 0.03 | 0.09 | 0.04 |
| 3 | 79024541 | rs17379690 | -1.11 | 0.404 | 0.155 | 0.28 | -0.012 | -0.05 | -0.05 | 0.093 | 0.11 | 0.18 | 0.07 | 0 | 0.08 | 0.04 |
| 3 | 79025569 | rs4680919 | -0.42 | 0.515 | -0.313 | 1.199 | -0.034 | -0.066 | -0.066 | 0 | 0.11 | 0.02 | 0.28 | 0.42 | 0.16 | 0.14 |
| 3 | 79027354 | rs1447834 | 0.846 | 0.919 | 0.154 | 0.221 | 0.003 | -0.019 | -0.019 | 0.057 | 0.67 | 0.61 | 0.74 | 0.75 | 0.65 | 0.68 |
| 3 | 79061220 | rs6790493 | -0.737 | 1.329 | 0.279 | 0.253 | -0.021 | 0.009 | 0.009 | 0.744 | 0.26 | 0.16 | 0.2 | 0.25 | 0.33 | 0.32 |
| 3 | 79077887 | rs6766522 | -0.512 | 1.006 | 0.367 | 0.609 | 0.017 | 0.025 | 0.025 | 0.12 | 0.06 | 0.09 | 0.18 | 0.16 | 0.15 | 0.09 |
| 3 | 79137230 | rs1455824 | -0.558 | 0.599 | -0.116 | 0.436 | -0.021 | -0.001 | -0.001 | 0 | 0.23 | 0.16 | 0.2 | 0.26 | 0.28 | 0.33 |
| 3 | 79157354 | rs9866364 | 1.244 | 0.328 | -0.058 | 0.287 | -0.007 | 0.019 | 0.019 | 0.933 | 0.57 | 0.25 | 0.56 | 0.59 | 0.52 | 0.52 |
| 3 | 79175867 | rs1455833 | 0.071 | 0.214 | 0.164 | 0.241 | 0.001 | -0.001 | -0.001 | 0 | 0.78 | 0.76 | 0.79 | 0.8 | 0.71 | 0.69 |
| 3 | 79176116 | rs1455832 | 0.015 | 0.204 | 0.116 | 0.236 | -0.015 | 0.004 | 0.004 | 0 | 0.78 | 0.85 | 0.8 | 0.8 | 0.71 | 0.68 |
| 3 | 79190911 | rs7631687 | 0.116 | 0.005 | -0.078 | 0.069 | -0.029 | 0.009 | 0.009 | 0 | 0.77 | 0.8 | 0.75 | 0.79 | 0.67 | 0.62 |
| 3 | 79192921 | rs9856034 | 1.411 | -0.23 | 0.456 | 0.02 | -0.019 | 0.017 | 0.017 | 0 | 0.58 | 0.26 | 0.54 | 0.6 | 0.5 | 0.48 |
| 3 | 79197710 | rs12486635 | 0.189 | 0.017 | 0.267 | -0.063 | -0.004 | 0.001 | 0.001 | 0 | 0.77 | 0.76 | 0.75 | 0.8 | 0.67 | 0.62 |
| 3 | 79205480 | rs17016763 | -1.109 | -0.227 | 0.687 | -0.218 | -0.027 | -0.083 | -0.083 | 0.303 | 0.13 | 0.31 | 0.1 | 0 | 0.08 | 0.04 |
| 3 | 79212886 | rs7611697 | -0.287 | -0.225 | 0.695 | -0.212 | -0.01 | -0.034 | -0.034 | 0.041 | 0.08 | 0.09 | 0.04 | 0 | 0.05 | 0.03 |
| 3 | 79227552 | rs11925923 | -0.781 | -0.273 | 0.75 | -0.233 | -0.018 | -0.056 | -0.056 | 1.545 | 0.1 | 0.23 | 0.06 | 0 | 0.05 | 0.03 |
| 3 | 79286801 | rs11924816 | -1.436 | -0.254 | 0.743 | -0.246 | -0.011 | -0.075 | -0.075 | 0 | 0.12 | 0.34 | 0.1 | 0 | 0.08 | 0.04 |
| 3 | 79318988 | rs9880736 | -1.603 | -0.868 | -0.375 | -0.382 | -0.089 | 0.007 | 0.007 | 2.575 | 0.53 | 0.96 | 0.43 | 0.37 | 0.55 | 0.45 |
| 3 | 79319977 | rs12629261 | -0.356 | -1.073 | 0.016 | -0.48 | -0.015 | -0.012 | -0.012 | 0 | 0.17 | 0.29 | 0.12 | 0.13 | 0.15 | 0.06 |
| 3 | 79329159 | rs1489850 | 0.569 | -1.064 | -0.08 | -0.486 | 0.013 | -0.008 | -0.008 | 0.179 | 0.12 | 0.06 | 0.06 | 0.13 | 0.11 | 0.05 |
| 3 | 79339590 | rs7618349 | 0.121 | -0.97 | -0.029 | -0.484 | 0.005 | -0.037 | -0.037 | 0 | 0.07 | 0.13 | 0.05 | 0.01 | 0.07 | 0.03 |
| 3 | 79354187 | rs9832396 | -1.748 | -1.068 | 0.063 | -0.529 | 0.066 | 0.101 | 0.101 | 3.719 | 0.33 | 0.86 | 0.22 | 0.14 | 0.24 | 0.11 |
| 3 | 79393357 | rs4362686 | -0.925 | -1.465 | -0.083 | -0.712 | -0.043 | -0.092 | -0.092 | 0 | 0.21 | 0.59 | 0.12 | 0.03 | 0.12 | 0.13 |
| 3 | 79399480 | rs13073607 | 0.631 | -1.127 | -0.596 | -0.655 | -0.008 | -0.008 | -0.008 | 0 | 0.13 | 0.08 | 0.08 | 0.13 | 0.15 | 0.05 |
| 3 | 79399575 | rs6548616 | -1.539 | -1.074 | -0.596 | -0.63 | 0.044 | 0.139 | 0.139 | 2.669 | 0.39 | 0.92 | 0.26 | 0.16 | 0.3 | 0.2 |
| 3 | 79404693 | rs7622139 | 0.054 | -1.071 | -0.374 | -0.613 | -0.006 | -0.038 | -0.038 | 0 | 0.11 | 0.1 | 0.05 | 0.01 | 0.09 | 0.03 |
| 3 | 79405038 | rs1489848 | -2.141 | -1.395 | -1.234 | -0.284 | -0.114 | -0.14 | -0.14 | 0 | 0.67 | 0.99 | 0.69 | 0.59 | 0.68 | 0.62 |
| 3 | 79406519 | rs9842630 | -0.722 | -1.264 | -0.689 | -0.361 | 0.015 | 0.015 | 0.015 | 1.875 | 0.31 | 0.75 | 0.2 | 0.16 | 0.22 | 0.17 |
| 3 | 79409145 | rs1387667 | 0.786 | -1.109 | -0.765 | 0.178 | -0.034 | -0.041 | -0.041 | 0.014 | 0.2 | 0.1 | 0.22 | 0.37 | 0.32 | 0.27 |
| 3 | 79412192 | rs6784911 | -1.235 | -1.384 | -1.123 | -0.052 | -0.085 | -0.104 | -0.104 | 0 | 0.57 | 0.94 | 0.43 | 0.51 | 0.58 | 0.57 |
| 3 | 79429811 | rs1387665 | -0.652 | -1.296 | -0.422 | -0.223 | -0.001 | 0.006 | 0.006 | 0 | 0.46 | 0.56 | 0.38 | 0.36 | 0.54 | 0.52 |
| 3 | 79461529 | rs4680962 | 1.288 | -1.354 | -0.327 | -0.237 | 0.008 | -0.046 | -0.046 | 0 | 0.42 | 0.19 | 0.55 | 0.5 | 0.37 | 0.42 |
| 3 | 79469022 | rs4130991 | 1.39 | -1.376 | 0.014 | -0.248 | -0.018 | -0.083 | -0.083 | 4.812 | 0.42 | 0.11 | 0.54 | 0.5 | 0.37 | 0.42 |
| 3 | 79489971 | rs4535189 | 0.595 | -1.393 | 0.177 | -0.266 | 0 | -0.003 | -0.003 | 0 | 0.55 | 0.55 | 0.62 | 0.63 | 0.46 | 0.48 |
| 3 | 79522372 | rs7628757 | -1.529 | -1.339 | -0.276 | -0.195 | -0.036 | -0.105 | -0.105 | 0.263 | 0.59 | 0.93 | 0.46 | 0.51 | 0.63 | 0.59 |
| 3 | 79533095 | rs4564923 | 0.774 | -1.039 | -0.067 | -0.144 | 0.022 | 0.019 | 0.019 | 0 | 0.22 | 0.21 | 0.24 | 0.33 | 0.35 | 0.35 |
| 3 | 79550373 | rs6548621 | -1.47 | -1.438 | -0.221 | 0.086 | -0.009 | -0.019 | -0.019 | 3.333 | 0.55 | 0.87 | 0.65 | 0.52 | 0.61 | 0.62 |
| 3 | 79557927 | rs7622444 | -0.605 | -1.43 | 0.159 | -0.016 | -0.014 | -0.088 | -0.088 | 0.203 | 0.28 | 0.39 | 0.17 | 0 | 0.22 | 0.21 |
| 3 | 79559914 | rs9832405 | 0.242 | -1.342 | -0.095 | 0.07 | -0.002 | -0.001 | -0.001 | 0 | 0.29 | 0.24 | 0.28 | 0.46 | 0.43 | 0.38 |
| 3 | 79560604 | rs7637338 | 0.693 | -1.399 | -0.21 | -0.057 | -0.014 | 0.055 | 0.055 | 0 | 0.13 | 0.05 | 0.08 | 0 | 0.15 | 0.13 |
| 3 | 79562067 | rs9880584 | 0.843 | -1.182 | -0.033 | 0.179 | 0.024 | 0.017 | 0.017 | 0 | 0.23 | 0.22 | 0.24 | 0.32 | 0.35 | 0.33 |
| 3 | 79567629 | rs7614913 | -1.956 | -1.689 | -0.578 | 0.184 | -0.076 | -0.028 | -0.028 | 0.151 | 0.63 | 0.97 | 0.67 | 0.66 | 0.66 | 0.56 |
| 3 | 79569558 | rs6548628 | -0.943 | -1.147 | 0.152 | -0.224 | -0.002 | -0.007 | -0.007 | 0 | 0.48 | 0.5 | 0.38 | 0.33 | 0.54 | 0.44 |
| 3 | 79580070 | rs9882489 | 1.151 | -1.138 | 0.106 | -0.225 | -0.013 | -0.011 | -0.011 | 0 | 0.21 | 0.17 | 0.23 | 0.32 | 0.35 | 0.25 |
| 3 | 79580982 | rs4479556 | 1.756 | -1.174 | 0.01 | -0.075 | -0.01 | -0.042 | -0.042 | 0 | 0.07 | 0.01 | 0.04 | 0.14 | 0.08 | 0.03 |
| 3 | 79585158 | rs9853895 | -1.36 | -1.147 | 0.129 | -0.094 | -0.001 | 0 | 0 | 0.114 | 0.56 | 0.57 | 0.42 | 0.48 | 0.62 | 0.47 |
| 3 | 79604064 | rs7617400 | -0.898 | -0.656 | 0.489 | -0.002 | -0.025 | -0.034 | -0.034 | 0 | 0.26 | 0.38 | 0.14 | 0.14 | 0.21 | 0.17 |
| 3 | 79610625 | rs11924965 | -0.938 | -0.657 | 0.498 | -0.002 | -0.027 | -0.036 | -0.036 | 2.662 | 0.26 | 0.39 | 0.14 | 0.14 | 0.21 | 0.17 |
| 3 | 79629094 | rs9863111 | 0.279 | -0.479 | 0.279 | 0.02 | -0.006 | -0.042 | -0.042 | 0 | 0.15 | 0.18 | 0.08 | 0.01 | 0.12 | 0.13 |
| 3 | 79634066 | rs9872108 | -0.929 | -0.525 | -0.094 | 0.195 | 0.012 | 0.021 | 0.021 | 0 | 0.5 | 0.65 | 0.35 | 0.39 | 0.53 | 0.43 |
| 3 | 79646157 | rs7644918 | -1.046 | -0.565 | 0.378 | 0.068 | -0.035 | -0.037 | -0.037 | 0 | 0.27 | 0.59 | 0.15 | 0.13 | 0.18 | 0.17 |
| 3 | 79648663 | rs9837046 | -0.165 | -0.462 | 0.788 | 0.06 | -0.033 | -0.095 | -0.095 | 0 | 0.17 | 0.33 | 0.08 | 0 | 0.1 | 0.12 |
| 3 | 79652575 | rs13071987 | -0.227 | 0.092 | 0.667 | 0.293 | 0.002 | 0.02 | 0.02 | 0 | 0.84 | 0.83 | 0.84 | 0.73 | 0.84 | 0.87 |
| 3 | 79652654 | rs13072324 | -0.226 | 0.079 | 0.643 | 0.303 | -0.012 | -0.033 | -0.033 | 0 | 0.86 | 0.94 | 0.84 | 0.73 | 0.84 | 0.87 |
| 3 | 79652716 | rs13072475 | -0.162 | 0.583 | 0.868 | -0.016 | 0.006 | -0.101 | -0.101 | 0.233 | 0.19 | 0.39 | 0.11 | 0.01 | 0.17 | 0.15 |
| 3 | 79662973 | rs4680966 | -0.352 | 0.538 | 0.605 | 0.091 | 0.002 | 0.006 | 0.006 | 0 | 0.86 | 0.91 | 0.88 | 0.89 | 0.87 | 0.88 |
| 3 | 79663077 | rs4680967 | 1.391 | 0.831 | 1.04 | -0.075 | -0.004 | 0.039 | 0.039 | 0 | 0.63 | 0.3 | 0.72 | 0.87 | 0.63 | 0.71 |
| 3 | 79674538 | rs9881556 | -0.593 | 0.835 | 1.242 | -0.077 | 0.026 | -0.144 | -0.144 | 0.142 | 0.22 | 0.5 | 0.14 | 0.02 | 0.24 | 0.17 |

| | | | | | | | | | | | | | | | | |
|---|---|---|---|---|---|---|---|---|---|---|---|---|---|---|---|---|
| 3 | 79678609 | rs7429525 | -0.402 | 0.639 | 1.102 | 0.043 | 0.001 | 0.005 | 0.005 | 0 | 0.86 | 0.92 | 0.88 | 0.89 | 0.87 | 0.88 |
| 3 | 79685538 | rs7433360 | 0.564 | 0.503 | 0.523 | 0.059 | -0.003 | -0.007 | -0.007 | 0.015 | 0.59 | 0.62 | 0.52 | 0.51 | 0.53 | 0.66 |
| 3 | 79704090 | rs4856420 | -1.587 | 0.445 | 0.331 | 0.069 | -0.015 | -0.017 | -0.017 | 0 | 0.9 | 0.99 | 0.91 | 0.9 | 0.89 | 0.89 |
| 3 | 79710734 | rs11127653 | 0.776 | 0.444 | 0.48 | 0.108 | -0.005 | -0.006 | -0.006 | 0 | 0.23 | 0.2 | 0.34 | 0.27 | 0.3 | 0.2 |
| 3 | 79712822 | rs9869899 | 0.172 | 0.4 | 0.54 | 0.111 | -0.013 | -0.033 | -0.033 | 0 | 0.63 | 0.76 | 0.56 | 0.54 | 0.56 | 0.67 |
| 3 | 79724179 | rs6548646 | -0.155 | 0.572 | 0.682 | -0.06 | 0.015 | -0.124 | -0.124 | 2.724 | 0.22 | 0.46 | 0.16 | 0.03 | 0.24 | 0.16 |
| 3 | 79724489 | rs12488868 | 1.998 | 0.426 | 0.4 | -0.106 | -0.058 | -0.182 | -0.182 | 0.019 | 0.37 | 0.01 | 0.38 | 0.53 | 0.32 | 0.5 |
| 3 | 79730175 | rs9829565 | -0.386 | 0.563 | 0.628 | -0.114 | 0.018 | -0.148 | -0.148 | 0 | 0.24 | 0.51 | 0.17 | 0.03 | 0.25 | 0.16 |
| 3 | 79737657 | rs7427034 | 0.089 | 0.528 | 0.648 | -0.075 | 0.015 | -0.087 | -0.087 | 0.13 | 0.21 | 0.37 | 0.16 | 0.03 | 0.24 | 0.16 |
| 3 | 79757107 | rs7433871 | 0.11 | 0.501 | 0.389 | -0.066 | 0.002 | 0.003 | 0.003 | 0 | 0.69 | 0.68 | 0.61 | 0.6 | 0.63 | 0.69 |
| 3 | 79778355 | rs7613917 | 0.011 | 0.548 | 0.387 | 0 | 0.023 | 0.027 | 0.027 | 0 | 0.24 | 0.32 | 0.33 | 0.31 | 0.3 | 0.21 |
| 3 | 79780667 | rs6798360 | 1.7 | 0.517 | -0.109 | -0.039 | 0.002 | -0.064 | -0.064 | 0.004 | 0.4 | 0.1 | 0.4 | 0.58 | 0.32 | 0.49 |
| 3 | 79782273 | rs11706934 | 0.433 | 0.497 | 0.076 | 0.138 | -0.009 | 0.009 | 0.009 | 0 | 0.07 | 0.02 | 0.04 | 0.05 | 0.12 | 0.04 |
| 3 | 79782922 | rs7627096 | 1.146 | 0.42 | 0.51 | 0.175 | 0.006 | 0.011 | 0.011 | 0 | 0.61 | 0.47 | 0.54 | 0.57 | 0.57 | 0.68 |
| 3 | 79784416 | rs6548651 | -1.139 | 0.442 | 0.321 | 0.169 | 0 | -0.001 | -0.001 | 1.534 | 0.89 | 0.89 | 0.9 | 0.89 | 0.89 | 0.89 |
| 3 | 79785218 | rs7431063 | 1.007 | 0.435 | 0.443 | 0.198 | -0.017 | -0.021 | -0.021 | 0.226 | 0.63 | 0.74 | 0.66 | 0.57 | 0.58 | 0.67 |
| 3 | 79789314 | rs9847684 | -0.447 | 0.392 | 0.384 | 0.183 | 0.008 | 0.003 | 0.003 | 0 | 0.85 | 0.89 | 0.9 | 0.89 | 0.88 | 0.87 |
| 3 | 79803193 | rs1502311 | 2.03 | 0.255 | 0.235 | 0.423 | -0.053 | 0.189 | 0.189 | 0 | 0.44 | 0.06 | 0.38 | 0.24 | 0.39 | 0.47 |
| 3 | 79803516 | rs11127664 | 2.004 | 0.255 | 0.235 | 0.423 | -0.051 | 0.192 | 0.192 | 0 | 0.44 | 0.06 | 0.38 | 0.24 | 0.39 | 0.47 |
| 3 | 79805396 | rs7428483 | -0.481 | 0.291 | 0.38 | 0.56 | -0.008 | -0.009 | -0.009 | 0 | 0.3 | 0.24 | 0.28 | 0.36 | 0.3 | 0.29 |
| 3 | 79808437 | rs1502288 | -0.554 | 0.395 | 0.491 | 0.626 | -0.001 | -0.001 | -0.001 | 0 | 0.33 | 0.32 | 0.29 | 0.36 | 0.3 | 0.29 |
| 3 | 79818831 | rs12632490 | 0.828 | 0.469 | 0.626 | 0.788 | 0.012 | 0.086 | 0.086 | 0 | 0.77 | 0.43 | 0.67 | 0.68 | 0.7 | 0.7 |
| 3 | 79819713 | rs9821785 | -0.126 | 0.527 | 0.843 | 0.859 | 0.021 | 0.119 | 0.119 | 0 | 0.15 | 0.34 | 0.27 | 0.25 | 0.17 | 0.25 |
| 3 | 79824232 | rs1827435 | 0.601 | 0.565 | 0.886 | 0.88 | 0.011 | 0.083 | 0.083 | 0 | 0.06 | 0.06 | 0.22 | 0.24 | 0.12 | 0.14 |
| 3 | 79835258 | rs13067870 | 1.402 | 0.419 | 0.84 | 0.725 | -0.015 | 0.05 | 0.05 | 0 | 0.56 | 0.31 | 0.48 | 0.36 | 0.52 | 0.53 |
| 3 | 79842200 | rs17017186 | 1.051 | 0.609 | 0.699 | 0.67 | 0.001 | -0.001 | -0.001 | 0 | 0.11 | 0.15 | 0.1 | 0.1 | 0.13 | 0.12 |
| 3 | 79847532 | rs2055452 | 0.244 | 0.606 | 0.759 | 0.695 | 0.001 | 0.013 | 0.013 | 0 | 0.21 | 0.24 | 0.19 | 0.33 | 0.19 | 0.18 |
| 3 | 79848057 | rs1393358 | 1.262 | 0.661 | 0.376 | 0.682 | -0.016 | 0.055 | 0.055 | 0 | 0.56 | 0.3 | 0.48 | 0.36 | 0.52 | 0.53 |
| 3 | 79848404 | rs7429136 | 1.123 | 0.671 | 0.35 | 0.686 | -0.002 | 0 | 0 | 0 | 0.11 | 0.11 | 0.09 | 0.1 | 0.13 | 0.12 |
| 3 | 79851205 | rs13100461 | 1.344 | 0.633 | 0.466 | 0.814 | -0.011 | 0.039 | 0.039 | 0 | 0.57 | 0.35 | 0.49 | 0.35 | 0.52 | 0.53 |
| 3 | 79855423 | rs7612833 | 0.64 | 0.679 | 0.222 | 0.878 | -0.003 | -0.005 | -0.005 | 0 | 0.18 | 0.15 | 0.17 | 0.32 | 0.18 | 0.15 |
| 3 | 79860607 | rs10511119 | 1.211 | 0.622 | 0.139 | 0.646 | -0.023 | 0.071 | 0.071 | 0 | 0.56 | 0.24 | 0.48 | 0.35 | 0.52 | 0.53 |
| 3 | 79881449 | rs9812051 | -0.045 | 0.506 | 0.35 | 0.203 | 0.014 | -0.068 | -0.068 | 0.066 | 0.17 | 0.28 | 0.13 | 0.01 | 0.23 | 0.15 |
| 3 | 79884675 | rs9848056 | 0.609 | 0.255 | 0.348 | 0.344 | -0.001 | -0.002 | -0.002 | 0 | 0.57 | 0.43 | 0.6 | 0.3 | 0.54 | 0.57 |
| 3 | 79888401 | rs4856360 | -0.387 | 0.182 | 0.52 | 0.324 | 0.002 | -0.005 | -0.005 | 0 | 0.38 | 0.38 | 0.38 | 0.7 | 0.45 | 0.39 |
| 3 | 79892470 | rs953223 | 1.626 | 0.213 | 0.132 | 0.309 | -0.022 | 0.044 | 0.044 | 0 | 0.08 | 0 | 0.07 | 0 | 0.11 | 0.02 |
| 3 | 79897447 | rs13095041 | 1.606 | 0.101 | 0.122 | 0.29 | 0.002 | 0.076 | 0.076 | 0.104 | 0.39 | 0.06 | 0.46 | 0.28 | 0.31 | 0.42 |
| 3 | 79907219 | rs9835250 | -0.375 | 0.161 | 0.428 | 0.635 | 0.021 | 0.042 | 0.042 | 0 | 0.22 | 0.36 | 0.22 | 0.26 | 0.3 | 0.28 |
| 3 | 79914090 | rs6796632 | 0.482 | 0.067 | 0.611 | 0.673 | 0 | 0.006 | 0.006 | 0 | 0.62 | 0.44 | 0.69 | 0.54 | 0.6 | 0.71 |
| 3 | 79919894 | rs11719555 | 1.073 | 0.088 | 0.64 | 0.636 | -0.003 | 0.008 | 0.008 | 0 | 0.13 | 0.1 | 0.07 | 0.07 | 0.17 | 0.04 |
| 3 | 79927812 | rs4856212 | 0.747 | 0.094 | 0.617 | 0.609 | 0 | 0.004 | 0.004 | 0 | 0.61 | 0.43 | 0.68 | 0.55 | 0.6 | 0.71 |
| 3 | 79951653 | rs2653395 | 0.453 | 0.207 | 0.636 | 0.643 | 0.005 | 0.019 | 0.019 | 0.926 | 0.2 | 0.21 | 0.22 | 0.4 | 0.22 | 0.22 |
| 3 | 79954627 | rs7630950 | 0.607 | 0.158 | 0.795 | 0.664 | 0.001 | 0.007 | 0.007 | 0 | 0.63 | 0.55 | 0.67 | 0.5 | 0.61 | 0.7 |
| 3 | 79955295 | rs2625548 | 0.392 | 0.214 | 0.535 | 0.569 | -0.001 | 0.002 | 0.002 | 0.978 | 0.21 | 0.22 | 0.22 | 0.4 | 0.22 | 0.23 |
| 3 | 79960599 | rs13317859 | 1.137 | 0.212 | 0.469 | 0.522 | -0.011 | 0.024 | 0.024 | 0.006 | 0.12 | 0.05 | 0.06 | 0.06 | 0.17 | 0.04 |
| 3 | 79987748 | rs2374752 | 0.508 | 0.174 | 1.145 | 0.555 | 0.009 | 0.013 | 0.013 | 0 | 0.66 | 0.42 | 0.68 | 0.55 | 0.61 | 0.73 |
| 3 | 79988304 | rs922918 | -0.654 | 0.231 | 1.231 | 0.561 | 0.017 | 0.042 | 0.042 | 0 | 0.81 | 0.69 | 0.76 | 0.61 | 0.78 | 0.77 |
| 3 | 79999534 | rs9811926 | 0.258 | 0.09 | 0.893 | 0.328 | -0.006 | 0.031 | 0.031 | 0 | 0.17 | 0.09 | 0.12 | 0.01 | 0.22 | 0.16 |
| 3 | 80010209 | rs17449605 | 2.27 | -0.023 | 0.568 | 0.164 | -0.053 | 0.232 | 0.232 | 1.693 | 0.35 | 0.01 | 0.4 | 0.06 | 0.27 | 0.43 |
| 3 | 80013246 | rs9810312 | -0.751 | 0.341 | 1.01 | 0.431 | 0.002 | 0.011 | 0.011 | 0 | 0.81 | 0.77 | 0.77 | 0.61 | 0.79 | 0.77 |
| 3 | 80024037 | rs4496441 | -0.511 | 0.352 | 1.113 | 0.482 | 0.036 | 0.09 | 0.09 | 0.437 | 0.8 | 0.56 | 0.75 | 0.61 | 0.78 | 0.77 |
| 3 | 80035078 | rs1384762 | -0.181 | 0.355 | 0.479 | 0.504 | 0.009 | 0.035 | 0.035 | 2.035 | 0.78 | 0.33 | 0.73 | 0.61 | 0.78 | 0.77 |
| 3 | 80044904 | rs6548668 | -0.87 | 0.404 | 0.312 | 0.526 | -0.001 | 0.003 | 0.003 | 0 | 0.81 | 0.81 | 0.78 | 0.61 | 0.78 | 0.77 |
| 3 | 80054997 | rs9825395 | 0.419 | 0.276 | -0.225 | 0.698 | -0.031 | -0.033 | -0.033 | 1.506 | 0.61 | 0.24 | 0.63 | 0.4 | 0.57 | 0.72 |
| 3 | 80072578 | rs7630918 | 0.427 | 0.321 | -0.24 | 0.471 | -0.02 | 0.001 | 0.001 | 0.325 | 0.59 | 0.29 | 0.61 | 0.57 | 0.5 | 0.73 |
| 3 | 80082660 | rs3856690 | -0.143 | 0.295 | -0.096 | 0.491 | 0.007 | -0.002 | -0.002 | 0 | 0.69 | 0.39 | 0.74 | 0.67 | 0.65 | 0.83 |
| 3 | 80091406 | rs9833473 | 0.192 | 0.408 | -0.342 | 0.345 | -0.001 | -0.018 | -0.018 | 0.164 | 0.17 | 0.16 | 0.12 | 0.02 | 0.23 | 0.16 |
| 3 | 80103068 | rs2341389 | 0.873 | 0.203 | -1.113 | 0.422 | -0.068 | 0.004 | 0.004 | 0 | 0.29 | 0.05 | 0.42 | 0.24 | 0.27 | 0.4 |
| 3 | 80118232 | rs12636593 | 0.88 | 0.205 | -1.113 | 0.423 | -0.069 | 0.003 | 0.003 | 0 | 0.29 | 0.05 | 0.42 | 0.24 | 0.27 | 0.4 |
| 3 | 80124685 | rs9823341 | 0.215 | -0.014 | -0.81 | 1.022 | -0.001 | -0.011 | -0.011 | 0.253 | 0.21 | 0.19 | 0.23 | 0.22 | 0.27 | 0.27 |
| 3 | 80130323 | rs17017524 | -0.592 | -0.322 | -0.01 | 0.793 | -0.008 | 0.035 | 0.035 | 0 | 0.3 | 0.65 | 0.34 | 0.47 | 0.34 | 0.34 |
| 3 | 80131214 | rs2197383 | 1.183 | -0.344 | 0.346 | 0.965 | 0.003 | 0.177 | 0.177 | 0 | 0.05 | 0.12 | 0.1 | 0.34 | 0.06 | 0.04 |
| 3 | 80137908 | rs12489982 | -0.965 | -0.35 | 0.423 | 0.95 | 0.021 | 0.455 | 0.455 | 0.116 | 0.12 | 0.54 | 0.22 | 0.46 | 0.1 | 0.16 |
| 3 | 80143818 | rs13317946 | 0.399 | -0.355 | 0.313 | 0.841 | -0.001 | 0.013 | 0.013 | 0 | 0.21 | 0.19 | 0.21 | 0.13 | 0.27 | 0.27 |
| 3 | 80147995 | rs6778068 | -0.91 | -0.412 | 0.531 | 0.811 | -0.002 | 0 | 0 | 0 | 0.34 | 0.44 | 0.36 | 0.63 | 0.37 | 0.37 |
| 3 | 80158184 | rs9811785 | -1.195 | -0.44 | 0.328 | 0.587 | -0.007 | 0.12 | 0.12 | 0.463 | 0.49 | 0.82 | 0.46 | 0.67 | 0.55 | 0.41 |
| 3 | 80158934 | rs9816658 | -0.211 | -0.428 | 0.198 | 0.601 | -0.003 | -0.007 | -0.007 | 0.043 | 0.3 | 0.29 | 0.35 | 0.63 | 0.37 | 0.34 |
| 3 | 80171401 | rs10511117 | 1.634 | -0.392 | 0.126 | 0.595 | -0.01 | 0.026 | 0.026 | 0 | 0.13 | 0.04 | 0.06 | 0.05 | 0.18 | 0.04 |
| 3 | 80173567 | rs7622781 | -1.121 | -0.596 | 0.427 | 0.828 | -0.018 | 0.026 | 0.026 | 0 | 0.74 | 0.91 | 0.84 | 0.76 | 0.78 | 0.78 |

| | | | | | | | | | | | | | | | | |
|---|---|---|---|---|---|---|---|---|---|---|---|---|---|---|---|---|
| 3 | 80194428 | rs1584112 | 0.489 | -0.53 | 0.645 | 0.953 | -0.012 | 0.06 | 0.06 | 0 | 0.34 | 0.2 | 0.27 | 0.18 | 0.45 | 0.34 |
| 3 | 80206970 | rs17017593 | 1.225 | -0.544 | 0.785 | 1.073 | 0.006 | 0.205 | 0.205 | 0 | 0.05 | 0.15 | 0.1 | 0.34 | 0.06 | 0.04 |
| 3 | 80212927 | rs9823914 | -0.486 | -0.572 | 1.161 | 0.955 | -0.014 | 0.086 | 0.086 | 0.12 | 0.44 | 0.86 | 0.43 | 0.68 | 0.51 | 0.41 |
| 3 | 80218598 | rs9827022 | 0.3 | -0.527 | 1.146 | 0.95 | -0.001 | -0.018 | -0.018 | 0.239 | 0.37 | 0.4 | 0.29 | 0.18 | 0.45 | 0.35 |
| 3 | 80229232 | rs6805096 | -0.79 | -0.541 | 1.066 | 0.92 | -0.009 | 0.067 | 0.067 | 0 | 0.68 | 0.89 | 0.8 | 0.76 | 0.74 | 0.78 |
| 3 | 80233532 | rs1509278 | -0.468 | -0.417 | 1.362 | 1.313 | -0.001 | 0.001 | 0.001 | 0 | 0.41 | 0.43 | 0.4 | 0.68 | 0.51 | 0.41 |
| 3 | 80235308 | rs9867579 | -0.654 | -0.416 | 1.279 | 1.313 | 0 | -0.002 | -0.002 | 0.76 | 0.43 | 0.58 | 0.41 | 0.68 | 0.51 | 0.41 |
| 3 | 80242147 | rs12493467 | 0.272 | -0.351 | 0.722 | 1.364 | 0.01 | 0.274 | 0.274 | 0 | 0.06 | 0.2 | 0.13 | 0.5 | 0.06 | 0.07 |
| 3 | 80244121 | rs4856565 | -0.392 | -0.331 | 0.649 | 0.856 | -0.003 | 0.024 | 0.024 | 0 | 0.41 | 0.38 | 0.39 | 0.68 | 0.51 | 0.41 |
| 3 | 80261661 | rs6788230 | 0.066 | -0.339 | 0.328 | 1.072 | -0.018 | -0.073 | -0.073 | 0 | 0.34 | 0.1 | 0.28 | 0.4 | 0.44 | 0.36 |
| 3 | 80264806 | rs7642828 | 0.864 | -0.286 | 0.618 | 0.733 | 0.025 | 0.042 | 0.042 | 0 | 0.05 | 0.18 | 0.09 | 0.11 | 0.06 | 0.04 |
| 3 | 80265933 | rs9861664 | 0.661 | -0.342 | 0.383 | 0.715 | -0.054 | 0.129 | 0.129 | 0 | 0.28 | 0.05 | 0.18 | 0.05 | 0.4 | 0.24 |
| 3 | 80269036 | rs11920819 | -1.315 | -0.582 | 1.315 | 1.201 | -0.014 | 0.142 | 0.142 | 0.648 | 0.12 | 0.52 | 0.15 | 0.28 | 0.07 | 0.09 |
| 3 | 80288675 | rs2202862 | 0.432 | -0.414 | 0.413 | 0.577 | -0.047 | 0.136 | 0.136 | 0.301 | 0.36 | 0.12 | 0.22 | 0.06 | 0.48 | 0.24 |
| 3 | 80294041 | rs6795725 | -0.984 | -0.584 | 0.727 | 0.946 | -0.005 | 0.167 | 0.167 | 1.83 | 0.07 | 0.37 | 0.05 | 0.22 | 0.01 | 0.05 |
| 3 | 80303968 | rs9866516 | 0.475 | -0.423 | 0.243 | 0.446 | -0.064 | 0.163 | 0.163 | 0.8 | 0.35 | 0.09 | 0.22 | 0.06 | 0.48 | 0.24 |
| 3 | 80316867 | rs13064754 | 1.127 | -0.569 | 0.094 | 0.209 | -0.004 | 0.029 | 0.029 | 1.177 | 0.3 | 0.26 | 0.41 | 0.08 | 0.26 | 0.37 |
| 3 | 80320561 | rs4131900 | 0.755 | -0.541 | -0.057 | 0.096 | -0.05 | 0.136 | 0.136 | 0.37 | 0.35 | 0.11 | 0.23 | 0.06 | 0.48 | 0.23 |
| 3 | 80347270 | rs951357 | 1.071 | -0.681 | -0.224 | 0.407 | -0.053 | -0.065 | -0.065 | 0 | 0.14 | 0.01 | 0.12 | 0.21 | 0.15 | 0.19 |
| 3 | 80350894 | rs12493163 | -0.431 | -0.738 | 0.276 | 0.487 | -0.001 | -0.006 | -0.006 | 0.729 | 0.8 | 0.69 | 0.74 | 0.21 | 0.82 | 0.65 |
| 3 | 80355773 | rs1876089 | 1.074 | -0.468 | -0.239 | -0.129 | -0.002 | 0.018 | 0.018 | 0.661 | 0.27 | 0.24 | 0.38 | 0.07 | 0.23 | 0.36 |
| 3 | 80374602 | rs6548685 | 0.45 | -0.553 | -0.161 | 0.124 | 0.008 | 0.01 | 0.01 | 0 | 0.65 | 0.33 | 0.63 | 0.12 | 0.74 | 0.61 |
| 3 | 80393506 | rs11127684 | -0.132 | -0.179 | 0.133 | 0.453 | 0.019 | -0.008 | -0.008 | 0 | 0.75 | 0.63 | 0.65 | 0.67 | 0.73 | 0.77 |
| 3 | 80399400 | rs6783799 | -0.648 | -0.22 | 0.225 | 0.708 | -0.026 | 0.197 | 0.197 | 3.008 | 0.1 | 0.29 | 0.22 | 0.53 | 0.06 | 0.17 |
| 3 | 80403787 | rs6798031 | -0.235 | -0.219 | 0.163 | 0.708 | 0.004 | 0.045 | 0.045 | 0 | 0.08 | 0.07 | 0.19 | 0.54 | 0.06 | 0.17 |
| 3 | 80416552 | rs1028089 | 1.345 | -0.085 | -0.406 | 0.193 | -0.035 | 0.058 | 0.058 | 3.109 | 0.26 | 0.09 | 0.21 | 0.12 | 0.25 | 0.35 |
| 3 | 80420033 | rs1028087 | -0.848 | -0.039 | -0.004 | 0.348 | -0.024 | 0.252 | 0.252 | 4.154 | 0.11 | 0.35 | 0.22 | 0.54 | 0.06 | 0.17 |
| 3 | 80446450 | rs6794396 | 1.398 | -0.152 | -0.407 | -0.085 | -0.034 | 0.111 | 0.111 | 0.054 | 0.27 | 0.08 | 0.18 | 0.02 | 0.35 | 0.21 |
| 3 | 80450254 | rs9848696 | 0.637 | -0.369 | -0.488 | -0.279 | 0 | -0.003 | -0.003 | 0 | 0.37 | 0.36 | 0.44 | 0.66 | 0.36 | 0.52 |
| 3 | 80456863 | rs1028082 | -0.228 | -0.324 | -0.442 | -0.289 | -0.003 | -0.029 | -0.029 | 0.001 | 0.68 | 0.48 | 0.64 | 0.68 | 0.71 | 0.76 |
| 3 | 80458878 | rs13060155 | 1.56 | -0.411 | -0.577 | 0.565 | -0.032 | -0.044 | -0.044 | 0 | 0.26 | 0.08 | 0.21 | 0.26 | 0.31 | 0.2 |
| 3 | 80461424 | rs6809002 | 0.394 | -0.363 | -0.475 | 0.764 | -0.004 | -0.034 | -0.034 | 0 | 0.41 | 0.25 | 0.43 | 0.48 | 0.4 | 0.57 |
| 3 | 80463682 | rs7614859 | 0.449 | -0.347 | -0.485 | 0.459 | -0.006 | 0.005 | 0.005 | 0.797 | 0.4 | 0.25 | 0.42 | 0.36 | 0.4 | 0.57 |
| 3 | 80472420 | rs1514555 | -0.148 | -0.363 | -0.429 | 0.445 | -0.007 | -0.035 | -0.035 | 0 | 0.67 | 0.48 | 0.64 | 0.68 | 0.71 | 0.76 |
| 3 | 80483952 | rs13084486 | 1.436 | -0.244 | -0.685 | 0.468 | -0.03 | -0.073 | -0.073 | 0 | 0.26 | 0.08 | 0.21 | 0.32 | 0.31 | 0.2 |
| 3 | 80487479 | rs4279055 | -1.185 | -0.203 | -0.68 | 0.587 | -0.035 | 0.1 | 0.1 | 0.221 | 0.67 | 0.91 | 0.76 | 0.87 | 0.71 | 0.61 |
| 3 | 80515770 | rs13097229 | 0.808 | -0.131 | -0.977 | 0.542 | 0.015 | -0.011 | -0.011 | 0 | 0.21 | 0.04 | 0.09 | 0.23 | 0.14 | 0.18 |
| 3 | 80517889 | rs7624315 | -0.591 | -0.349 | -0.343 | 0.563 | -0.003 | 0.031 | 0.031 | 0 | 0.35 | 0.48 | 0.49 | 0.54 | 0.36 | 0.36 |
| 3 | 80518843 | rs7627861 | 0.663 | -0.351 | -0.425 | 0.492 | -0.084 | -0.014 | -0.014 | 0 | 0.23 | 0.02 | 0.27 | 0.24 | 0.26 | 0.21 |
| 3 | 80525044 | rs6548699 | -0.515 | -0.312 | -0.307 | 0.501 | -0.004 | 0.023 | 0.023 | 0 | 0.35 | 0.5 | 0.49 | 0.54 | 0.36 | 0.35 |
| 3 | 80536788 | rs12630978 | -1.385 | -0.246 | -0.442 | 0.32 | -0.016 | 0.083 | 0.083 | 1.244 | 0.68 | 0.85 | 0.75 | 0.87 | 0.72 | 0.64 |
| 3 | 80575042 | rs7631866 | -1.261 | -0.186 | -0.469 | 0.025 | 0 | 0.011 | 0.011 | 1.187 | 0.67 | 0.7 | 0.74 | 0.87 | 0.72 | 0.64 |
| 3 | 80595022 | rs1406781 | 0.295 | -0.22 | -0.364 | 0.081 | -0.016 | 0.204 | 0.204 | 0.015 | 0.11 | 0.42 | 0.22 | 0.31 | 0.11 | 0.15 |
| 3 | 80599812 | rs7431461 | -1.325 | -0.167 | -0.484 | 0.013 | -0.013 | 0.079 | 0.079 | 1.286 | 0.67 | 0.82 | 0.75 | 0.87 | 0.72 | 0.64 |
| 3 | 80603691 | rs12487535 | 1.771 | -0.351 | -0.668 | -0.149 | -0.024 | 0.014 | 0.014 | 0 | 0.21 | 0.01 | 0.17 | 0.07 | 0.15 | 0.24 |
| 3 | 80611818 | rs4416346 | 1.781 | -0.242 | -0.337 | 0.146 | -0.057 | -0.098 | -0.098 | 0 | 0.25 | 0.05 | 0.2 | 0.33 | 0.33 | 0.22 |
| 3 | 80654883 | rs2138110 | -1.491 | -0.317 | -0.306 | 0.145 | -0.026 | 0.092 | 0.092 | 1.349 | 0.67 | 0.89 | 0.75 | 0.87 | 0.72 | 0.64 |
| 3 | 80662463 | rs2056163 | -0.371 | -0.311 | -0.232 | 0.124 | -0.007 | 0.029 | 0.029 | 0 | 0.34 | 0.48 | 0.48 | 0.54 | 0.34 | 0.35 |
| 3 | 80665977 | rs11127705 | 1.522 | -0.294 | -0.206 | 0.136 | -0.009 | -0.015 | -0.015 | 0 | 0.26 | 0.17 | 0.22 | 0.33 | 0.35 | 0.23 |
| 3 | 80687359 | rs485190 | -0.375 | -0.3 | -0.185 | 0.096 | -0.007 | 0.029 | 0.029 | 0 | 0.34 | 0.48 | 0.48 | 0.54 | 0.34 | 0.35 |
| 3 | 80696606 | rs6775791 | 0.181 | -0.301 | -0.262 | -0.018 | -0.015 | -0.034 | -0.034 | 2.225 | 0.06 | 0.16 | 0.04 | 0 | 0.03 | 0.06 |
| 3 | 80713938 | rs658551 | -1.41 | -0.304 | -0.495 | -0.047 | -0.054 | 0.096 | 0.096 | 1.311 | 0.67 | 0.94 | 0.75 | 0.87 | 0.72 | 0.63 |
| 3 | 80714827 | rs10511108 | 1.543 | -0.266 | -0.481 | 0.079 | -0.055 | -0.081 | -0.081 | 0 | 0.26 | 0.08 | 0.21 | 0.34 | 0.34 | 0.23 |
| 3 | 80721103 | rs548608 | -0.391 | -0.296 | -0.138 | 0.112 | -0.007 | 0.027 | 0.027 | 0 | 0.34 | 0.56 | 0.49 | 0.54 | 0.35 | 0.35 |
| 3 | 80721289 | rs7634914 | 0.691 | -0.339 | 0.164 | 0.254 | -0.024 | 0.062 | 0.062 | 0 | 0.1 | 0.29 | 0.19 | 0.18 | 0.11 | 0.13 |
| 3 | 80785415 | rs13078019 | 0.555 | -0.261 | -0.145 | 0.184 | -0.014 | 0.022 | 0.022 | 0 | 0.24 | 0.16 | 0.26 | 0.22 | 0.24 | 0.2 |
| 3 | 80792633 | rs17391287 | 1.328 | -0.258 | -0.176 | 0.226 | -0.008 | -0.016 | -0.016 | 0 | 0.27 | 0.22 | 0.23 | 0.34 | 0.34 | 0.23 |
| 3 | 80799801 | rs9872779 | -0.677 | -0.314 | -0.047 | 0.254 | -0.009 | 0.226 | 0.226 | 0 | 0.89 | 0.59 | 0.77 | 0.69 | 0.89 | 0.85 |
| 3 | 80809059 | rs9828864 | -0.427 | -0.256 | -0.571 | 0.198 | 0.007 | 0 | 0 | 0 | 0.56 | 0.4 | 0.52 | 0.56 | 0.61 | 0.49 |
| 3 | 80839513 | rs13086426 | 1.532 | -0.138 | -0.793 | 0.212 | -0.056 | -0.096 | -0.096 | 0 | 0.26 | 0.06 | 0.21 | 0.34 | 0.34 | 0.26 |
| 3 | 80859757 | rs7642458 | -0.2 | -0.186 | -0.791 | 0.121 | 0.001 | 0.039 | 0.039 | 0.716 | 0.6 | 0.19 | 0.52 | 0.34 | 0.55 | 0.59 |
| 3 | 80864748 | rs11719015 | 0.578 | -0.202 | -0.81 | 0.114 | -0.041 | 0.019 | 0.019 | 0 | 0.24 | 0.06 | 0.25 | 0.22 | 0.24 | 0.2 |
| 3 | 80912002 | rs9881131 | 0.85 | -0.413 | -0.754 | 0.128 | -0.02 | 0.136 | 0.136 | 0.701 | 0.37 | 0.13 | 0.26 | 0.12 | 0.3 | 0.38 |
| 3 | 80918205 | rs6777275 | -0.262 | -0.389 | -0.451 | 0.366 | -0.031 | 0.166 | 0.166 | 0.113 | 0.88 | 0.43 | 0.76 | 0.69 | 0.89 | 0.86 |
| 3 | 80984958 | rs4298011 | -1.042 | -0.458 | -0.777 | 0.273 | 0.022 | -0.065 | -0.065 | 0 | 0.41 | 0.85 | 0.55 | 0.53 | 0.38 | 0.4 |
| 3 | 81007665 | rs2372736 | 1.428 | -0.424 | -0.245 | 0.332 | -0.023 | -0.009 | -0.009 | 0 | 0.07 | 0.12 | 0.06 | 0.06 | 0.04 | 0.02 |
| 3 | 81033150 | rs11716250 | 1.241 | -0.32 | -0.461 | 0.291 | -0.065 | -0.09 | -0.09 | 0 | 0.27 | 0.08 | 0.22 | 0.38 | 0.36 | 0.33 |
| 3 | 81033263 | rs6773716 | 0.558 | -0.515 | -0.222 | 0.321 | -0.028 | 0.034 | 0.034 | 0 | 0.1 | 0.18 | 0.19 | 0.18 | 0.11 | 0.14 |
| 3 | 81033323 | rs2639222 | -0.281 | -0.512 | -0.487 | 0.306 | -0.033 | 0.177 | 0.177 | 0.31 | 0.88 | 0.48 | 0.77 | 0.7 | 0.89 | 0.84 |

| | | | | | | | | | | | | | | | | |
|---|---|---|---|---|---|---|---|---|---|---|---|---|---|---|---|---|
| 3 | 81036085 | rs2578309 | 1.038 | -0.585 | -0.575 | 0.197 | 0.005 | 0.029 | 0.029 | 0.568 | 0.36 | 0.32 | 0.27 | 0.13 | 0.27 | 0.32 |
| 3 | 81037906 | rs12635918 | 0.409 | -0.679 | -0.715 | 0.18 | -0.046 | 0.01 | 0.01 | 0 | 0.24 | 0.06 | 0.26 | 0.23 | 0.24 | 0.21 |
| 3 | 81038666 | rs9862577 | 0.366 | -0.625 | -0.68 | 0.203 | -0.002 | 0.004 | 0.004 | 0 | 0.64 | 0.43 | 0.5 | 0.47 | 0.64 | 0.64 |
| 3 | 81052987 | rs2639256 | 0.321 | -0.478 | -0.802 | 0.057 | -0.009 | 0.065 | 0.065 | 0.44 | 0.55 | 0.15 | 0.5 | 0.35 | 0.52 | 0.53 |
| 3 | 81062618 | rs1438564 | 0.732 | -0.153 | -0.234 | 0.063 | -0.024 | -0.018 | -0.018 | 0 | 0.8 | 0.5 | 0.73 | 0.81 | 0.79 | 0.77 |
| 3 | 81069531 | rs2639234 | 0.242 | -0.122 | -0.122 | 0.139 | -0.007 | 0.047 | 0.047 | 0.218 | 0.86 | 0.55 | 0.77 | 0.81 | 0.84 | 0.84 |
| 3 | 81071684 | rs2578295 | 1.078 | -0.227 | -0.725 | -0.162 | -0.012 | 0.056 | 0.056 | 1.653 | 0.29 | 0.15 | 0.23 | 0.12 | 0.25 | 0.31 |
| 3 | 81081489 | rs2578349 | -0.167 | -0.419 | -0.25 | -0.023 | 0.027 | 0.077 | 0.077 | 0 | 0.77 | 0.5 | 0.69 | 0.63 | 0.77 | 0.76 |
| 3 | 81083861 | rs1370894 | 0.599 | -0.145 | -0.786 | -0.231 | -0.001 | 0.15 | 0.15 | 3.073 | 0.47 | 0.18 | 0.45 | 0.22 | 0.55 | 0.54 |
| 3 | 81096365 | rs2639280 | 0.628 | -0.151 | -0.789 | -0.233 | -0.002 | 0.148 | 0.148 | 3.235 | 0.47 | 0.18 | 0.46 | 0.22 | 0.55 | 0.54 |
| 3 | 81113949 | rs2639210 | 0.363 | -0.525 | -0.466 | -0.235 | 0.026 | 0.002 | 0.002 | 1.741 | 0.56 | 0.36 | 0.7 | 0.61 | 0.69 | 0.66 |
| 3 | 81120509 | rs13072710 | 0.272 | -0.844 | -0.704 | -0.414 | -0.014 | 0.208 | 0.208 | 1.053 | 0.29 | 0.02 | 0.13 | 0 | 0.21 | 0.19 |
| 3 | 81121118 | rs6548746 | 0.821 | -0.876 | -0.222 | -0.403 | -0.032 | 0.165 | 0.165 | 0.572 | 0.13 | 0.25 | 0.15 | 0.39 | 0.1 | 0.15 |
| 3 | 81125424 | rs2639229 | -0.295 | -0.813 | -0.005 | -0.495 | 0.006 | -0.039 | -0.039 | 0 | 0.69 | 0.74 | 0.85 | 1 | 0.79 | 0.81 |
| 3 | 81126562 | rs2639227 | -0.284 | -0.813 | -0.002 | -0.495 | -0.012 | -0.103 | -0.103 | 0 | 0.69 | 0.63 | 0.84 | 1 | 0.79 | 0.81 |
| 3 | 81127652 | rs9830652 | -1.552 | -0.686 | 0.125 | -0.434 | -0.007 | 0.019 | 0.019 | 2.851 | 0.6 | 0.66 | 0.6 | 0.89 | 0.6 | 0.66 |
| 3 | 81129944 | rs17018482 | 0.078 | -0.588 | 0.83 | 0.528 | 0.006 | 0.025 | 0.025 | 0 | 0.14 | 0.19 | 0.19 | 0.35 | 0.18 | 0.26 |
| 3 | 81135775 | rs11127721 | -0.912 | -0.645 | 0.808 | 0.5 | 0.001 | -0.009 | -0.009 | 0 | 0.27 | 0.33 | 0.32 | 0.74 | 0.28 | 0.38 |
| 3 | 81146297 | rs7643316 | -0.782 | -0.544 | 0.759 | 0.585 | 0.004 | 0.048 | 0.048 | 0 | 0.25 | 0.39 | 0.3 | 0.64 | 0.26 | 0.35 |
| 3 | 81148588 | rs17018517 | -0.132 | 0.242 | 0.307 | 0.294 | 0.062 | 0.077 | 0.077 | 0 | 0.13 | 0.32 | 0.17 | 0.24 | 0.16 | 0.22 |
| 3 | 81157287 | rs2578317 | -0.16 | 0.155 | -1.089 | 0.001 | -0.066 | 0.016 | 0.016 | 2.645 | 0.29 | 0.07 | 0.3 | 0.22 | 0.35 | 0.19 |
| 3 | 81165761 | rs2372752 | 0.828 | 0.015 | -0.789 | 0.085 | 0.001 | 0.014 | 0.014 | 0 | 0.52 | 0.28 | 0.46 | 0.52 | 0.49 | 0.55 |
| 3 | 81171255 | rs1477694 | 0.235 | 0.141 | -1.825 | 0.053 | -0.071 | 0.015 | 0.015 | 0.171 | 0.26 | 0.07 | 0.31 | 0.2 | 0.34 | 0.22 |
| 3 | 81177939 | rs11533825 | 0.092 | 0.102 | -1.658 | 0.067 | -0.001 | -0.003 | -0.003 | 0 | 0.18 | 0.19 | 0.18 | 0.28 | 0.16 | 0.23 |
| 3 | 81181268 | rs4856330 | -0.04 | 0.622 | -1.864 | -0.271 | -0.071 | 0.015 | 0.015 | 0.171 | 0.74 | 0.93 | 0.69 | 0.8 | 0.66 | 0.78 |
| 3 | 81187867 | rs6806220 | -0.494 | 0.021 | -1.481 | 0.023 | 0.04 | 0.045 | 0.045 | 0 | 0.42 | 0.71 | 0.43 | 0.71 | 0.36 | 0.53 |
| 3 | 81191942 | rs10511104 | 0.905 | 0.451 | -1.799 | -0.041 | -0.068 | 0.064 | 0.064 | 0 | 0.39 | 0.08 | 0.28 | 0.2 | 0.44 | 0.34 |
| 3 | 81191984 | rs11923166 | 0.615 | 0.131 | -1.215 | 0.333 | -0.007 | 0.034 | 0.034 | 0 | 0.18 | 0.24 | 0.29 | 0.39 | 0.18 | 0.2 |
| 3 | 81194949 | rs4856333 | -0.455 | 0.262 | -1.281 | 0.161 | -0.01 | 0.065 | 0.065 | 0 | 0.8 | 0.93 | 0.84 | 0.95 | 0.81 | 0.8 |
| 3 | 81201878 | rs11127727 | 1.06 | 0.164 | -1.206 | 0.18 | 0 | 0.008 | 0.008 | 0 | 0.16 | 0.13 | 0.28 | 0.39 | 0.18 | 0.19 |
| 3 | 81205014 | rs899062 | 0.827 | 0.075 | -0.278 | 0.099 | -0.002 | 0.001 | 0.001 | 0 | 0.62 | 0.47 | 0.54 | 0.55 | 0.64 | 0.61 |
| 3 | 81210101 | rs11914499 | -0.188 | 0.106 | -0.327 | 0.114 | 0.001 | 0.024 | 0.024 | 0.028 | 0.23 | 0.38 | 0.25 | 0.36 | 0.2 | 0.27 |
| 3 | 81211514 | rs10865593 | -0.394 | 0.09 | 0.276 | 0.06 | -0.001 | -0.061 | -0.061 | 0 | 0.8 | 0.76 | 0.83 | 0.95 | 0.81 | 0.8 |
| 3 | 81212815 | rs6764658 | -1.152 | 0.294 | 0.59 | 0.36 | -0.004 | -0.018 | -0.018 | 0 | 0.39 | 0.54 | 0.54 | 0.76 | 0.37 | 0.46 |
| 3 | 81217102 | rs6803251 | -0.387 | 0.306 | 0.578 | 0.394 | 0.001 | 0.021 | 0.021 | 0.039 | 0.23 | 0.37 | 0.25 | 0.36 | 0.2 | 0.27 |
| 3 | 81220591 | rs969152 | 0.644 | 0.317 | 0.432 | 0.525 | 0 | 0.013 | 0.013 | 0 | 0.16 | 0.14 | 0.29 | 0.4 | 0.17 | 0.19 |
| 3 | 81221960 | rs12634100 | 0.591 | 0.535 | 0.837 | 0.314 | 0.029 | 0.038 | 0.038 | 0 | 0.74 | 0.46 | 0.77 | 0.62 | 0.73 | 0.74 |
| 3 | 81222552 | rs2372757 | 0.011 | 0.296 | 0.718 | 0.191 | -0.02 | 0.125 | 0.125 | 0 | 0.21 | 0.45 | 0.45 | 0.41 | 0.23 | 0.28 |
| 3 | 81224298 | rs4856202 | 0.256 | 0.29 | 0.644 | 0.2 | -0.011 | -0.016 | -0.016 | 0 | 0.77 | 0.81 | 0.8 | 0.61 | 0.73 | 0.73 |
| 3 | 81227042 | rs6802413 | 1.041 | 0.286 | 0.583 | 0.056 | -0.002 | 0.016 | 0.016 | 0 | 0.55 | 0.42 | 0.35 | 0.2 | 0.49 | 0.44 |
| 3 | 81234859 | rs1477676 | 0.333 | 0.366 | 0.689 | -0.002 | 0.014 | 0.062 | 0.062 | 0 | 0.78 | 0.67 | 0.81 | 0.6 | 0.77 | 0.74 |
| 3 | 81245962 | rs7628131 | -0.091 | 0.19 | 0.403 | 0.048 | -0.016 | -0.085 | -0.085 | 0 | 0.17 | 0.03 | 0.14 | 0.41 | 0.17 | 0.2 |
| 3 | 81246367 | rs12632640 | 1.74 | 0.445 | -0.101 | -0.351 | -0.062 | 0.126 | 0.126 | 0 | 0.33 | 0.05 | 0.18 | 0.11 | 0.32 | 0.31 |
| 3 | 81247480 | rs6801771 | 0.069 | 0.058 | 1.342 | 0.284 | -0.02 | 0.093 | 0.093 | 0 | 0.77 | 0.56 | 0.55 | 0.61 | 0.77 | 0.69 |
| 3 | 81251347 | rs6548752 | -0.314 | -0.038 | 1.483 | 0.345 | -0.01 | 0.091 | 0.091 | 0 | 0.8 | 0.63 | 0.57 | 0.61 | 0.79 | 0.66 |
| 3 | 81257186 | rs1545746 | -0.164 | -0.062 | 1.384 | 0.331 | -0.017 | 0.073 | 0.073 | 0 | 0.79 | 0.47 | 0.55 | 0.61 | 0.8 | 0.66 |
| 3 | 81279099 | rs17043256 | -1.335 | -0.002 | 1.146 | 0.371 | -0.007 | 0.027 | 0.027 | 0 | 0.33 | 0.39 | 0.25 | 0.47 | 0.3 | 0.25 |
| 3 | 81283362 | rs727824 | 0.252 | -0.134 | 1.029 | 0.425 | -0.003 | -0.001 | -0.001 | 0 | 0.08 | 0.06 | 0.06 | 0.05 | 0.08 | 0.03 |
| 3 | 81308774 | rs9856313 | -1.555 | -0.02 | 1.393 | 0.372 | 0.001 | -0.014 | -0.014 | 0 | 0.35 | 0.64 | 0.28 | 0.49 | 0.32 | 0.27 |
| 3 | 81309923 | rs13076541 | -0.316 | -0.069 | 1.271 | 0.329 | -0.023 | -0.084 | -0.084 | 0.01 | 0.18 | 0.05 | 0.16 | 0.43 | 0.19 | 0.22 |
| 3 | 81312096 | rs17280690 | 0.589 | -0.039 | 1.196 | 0.392 | -0.034 | -0.028 | -0.028 | 0 | 0.07 | 0.01 | 0.05 | 0.08 | 0.08 | 0.03 |
| 3 | 81317402 | rs12107250 | 0.271 | -0.041 | 1.262 | 0.391 | 0 | 0 | 0 | 0 | 0.08 | 0.08 | 0.06 | 0.08 | 0.08 | 0.03 |
| 3 | 81331222 | rs13097418 | -0.623 | -0.075 | 1.414 | 0.35 | 0.013 | 0.091 | 0.091 | 0 | 0.2 | 0.23 | 0.17 | 0.44 | 0.2 | 0.23 |
| 3 | 81331298 | rs763855 | 1.865 | -0.158 | 0.939 | 0.013 | -0.123 | 0.193 | 0.193 | 0 | 0.41 | 0.06 | 0.27 | 0.13 | 0.48 | 0.39 |
| 3 | 81344873 | rs7612863 | -0.274 | 0.336 | 1.964 | 0.445 | 0.026 | 0.002 | 0.002 | 0.218 | 0.06 | 0.3 | 0.06 | 0 | 0.05 | 0.02 |
| 3 | 81348262 | rs13093389 | -0.243 | 0.436 | 1.828 | 0.707 | 0.002 | 0.009 | 0.009 | 0 | 0.14 | 0.11 | 0.13 | 0.29 | 0.19 | 0.09 |
| 3 | 81349843 | rs11714085 | -0.947 | 0.45 | 1.856 | 0.765 | -0.001 | -0.005 | -0.005 | 0.011 | 0.11 | 0.11 | 0.09 | 0.23 | 0.1 | 0.16 |
| 3 | 81356647 | rs13094279 | -1.606 | 0.457 | 1.718 | 0.753 | -0.018 | -0.039 | -0.039 | 1.311 | 0.24 | 0.14 | 0.2 | 0.5 | 0.28 | 0.24 |
| 3 | 81373557 | rs2129522 | -0.259 | 0.53 | 1.586 | 0.768 | 0.001 | 0.006 | 0.006 | 0 | 0.14 | 0.11 | 0.13 | 0.29 | 0.19 | 0.09 |
| 3 | 81376268 | rs7650907 | -1.97 | 0.554 | 1.207 | 0.773 | -0.001 | -0.001 | -0.001 | 0 | 0.32 | 0.67 | 0.28 | 0.5 | 0.33 | 0.24 |
| 3 | 81394501 | rs6784564 | -0.434 | 0.651 | 0.733 | 0.82 | -0.024 | -0.047 | -0.047 | 0 | 0.09 | 0.02 | 0.08 | 0.21 | 0.09 | 0.11 |
| 3 | 81398154 | rs1461612 | -0.434 | 0.651 | 0.706 | 0.831 | -0.032 | -0.056 | -0.056 | 0 | 0.09 | 0.01 | 0.08 | 0.21 | 0.09 | 0.11 |
| 3 | 81404812 | rs7645184 | -0.427 | 0.561 | 1.158 | 0.505 | -0.003 | -0.027 | -0.027 | 0.395 | 0.06 | 0.26 | 0.05 | 0 | 0.05 | 0.02 |
| 3 | 81406155 | rs9877021 | -1.245 | 0.621 | 0.325 | 0.544 | 0.004 | 0.01 | 0.01 | 1.296 | 0.24 | 0.66 | 0.19 | 0.28 | 0.24 | 0.14 |
| 3 | 81413725 | rs275365 | 0.826 | 0.284 | 0.192 | 0.461 | -0.004 | 0.012 | 0.012 | 1.627 | 0.71 | 0.29 | 0.75 | 0.61 | 0.78 | 0.8 |
| 3 | 81414530 | rs275366 | -0.075 | 0.287 | 0.442 | 0.533 | 0.012 | 0.008 | 0.008 | 0 | 0.8 | 0.65 | 0.85 | 0.78 | 0.8 | 0.88 |
| 3 | 81414970 | rs6781602 | -0.651 | 0.091 | 0.659 | 0.954 | -0.028 | 0.076 | 0.076 | 0.263 | 0.1 | 0.34 | 0.11 | 0.18 | 0.03 | 0.08 |
| 3 | 81420196 | rs1461616 | -0.404 | 0.745 | -0.525 | 0.845 | 0 | 0.006 | 0.006 | 0.005 | 0.26 | 0.18 | 0.48 | 0.49 | 0.33 | 0.33 |
| 3 | 81427818 | rs275360 | 0.609 | -0.023 | -0.449 | 0.267 | -0.038 | -0.044 | -0.044 | 0 | 0.66 | 0.51 | 0.77 | 0.76 | 0.79 | 0.67 |

| | | | | | | | | | | | | | | | | |
|---|---|---|---|---|---|---|---|---|---|---|---|---|---|---|---|---|
| 3 | 81428048 | rs6778246 | 0.261 | 0.172 | -0.61 | -0.023 | 0 | 0.01 | 0.01 | 0 | 0.52 | 0.57 | 0.7 | 0.62 | 0.55 | 0.56 |
| 3 | 81428061 | rs6802544 | -0.608 | 0.491 | -0.877 | 0.016 | 0.025 | 0.064 | 0.064 | 2.6 | 0.39 | 0.14 | 0.2 | 0.25 | 0.31 | 0.19 |
| 3 | 81429555 | rs4856343 | -1.415 | 1.496 | 0.481 | 1.213 | 0.062 | 0.211 | 0.211 | 0.282 | 0.18 | 0.47 | 0.32 | 0.47 | 0.26 | 0.22 |
| 3 | 81439098 | rs17018881 | -0.918 | 1.39 | 0.22 | 1.137 | 0.004 | -0.005 | -0.005 | 0.404 | 0.13 | 0.15 | 0.12 | 0.09 | 0.19 | 0.14 |
| 3 | 81440354 | rs6786066 | -0.599 | 1.279 | 0.161 | 1.206 | 0.001 | 0.002 | 0.002 | 0 | 0.1 | 0.1 | 0.19 | 0.25 | 0.09 | 0.09 |
| 3 | 81441511 | rs13318351 | -0.73 | 1.344 | 0.058 | 1.163 | 0.002 | -0.003 | -0.003 | 0 | 0.13 | 0.14 | 0.12 | 0.09 | 0.19 | 0.14 |
| 3 | 81443206 | rs17018889 | 0.227 | 1.64 | -0.135 | 1.284 | -0.026 | -0.017 | -0.017 | 0 | 0.05 | 0.01 | 0.07 | 0.2 | 0.1 | 0.1 |
| 3 | 81448115 | rs17018894 | 0.101 | 1.503 | -0.084 | 1.355 | -0.001 | 0 | 0 | 0 | 0.06 | 0.03 | 0.17 | 0.23 | 0.08 | 0.08 |
| 3 | 81456652 | rs9856502 | -1.728 | 1.011 | 0.68 | 1.019 | -0.016 | 0.079 | 0.079 | 0.443 | 0.25 | 0.46 | 0.29 | 0.43 | 0.19 | 0.21 |
| 3 | 81459535 | rs13087520 | -0.181 | 1.318 | 0.573 | 0.987 | -0.002 | 0.003 | 0.003 | 0 | 0.07 | 0.07 | 0.09 | 0.21 | 0.15 | 0.11 |
| 3 | 81464918 | rs7629444 | -0.405 | 1.264 | 1.24 | 1.289 | -0.025 | -0.012 | -0.012 | 0.164 | 0.11 | 0.2 | 0.08 | 0.06 | 0.03 | 0.05 |
| 3 | 81465962 | rs1478289 | -0.11 | 1.303 | 1.209 | 1.29 | -0.002 | 0.003 | 0.003 | 0 | 0.07 | 0.07 | 0.09 | 0.21 | 0.16 | 0.11 |
| 3 | 81468139 | rs968778 | 0.8 | 1.866 | 1.468 | 1.135 | 0.081 | 0.236 | 0.236 | 0 | 0.81 | 0.53 | 0.71 | 0.55 | 0.72 | 0.82 |
| 3 | 81476841 | rs275356 | -0.342 | 1.745 | 1.296 | 1.483 | -0.001 | 0.01 | 0.01 | 0 | 0.93 | 0.92 | 0.82 | 0.82 | 0.92 | 0.91 |
| 3 | 81481169 | rs9853372 | -1.102 | 1.702 | 1.416 | 1.364 | 0.003 | 0.011 | 0.011 | 0.132 | 0.18 | 0.24 | 0.18 | 0.26 | 0.18 | 0.16 |
| 3 | 81496341 | rs9309871 | -0.309 | 1.753 | 1.464 | 1.374 | -0.001 | 0.002 | 0.002 | 0.01 | 0.12 | 0.12 | 0.15 | 0.07 | 0.24 | 0.12 |
| 3 | 81502116 | rs17018940 | -1.548 | 1.737 | 1.674 | 1.605 | 0.06 | 0.163 | 0.163 | 0 | 0.21 | 0.41 | 0.3 | 0.41 | 0.3 | 0.17 |
| 3 | 81502655 | rs4856346 | 1.514 | 1.958 | 1.74 | 1.501 | 0.021 | 0.103 | 0.103 | 0.177 | 0.75 | 0.53 | 0.62 | 0.5 | 0.72 | 0.72 |
| 3 | 81512525 | rs973689 | -0.44 | 1.963 | 2.082 | 1.507 | 0.02 | -0.019 | -0.019 | 0 | 0.13 | 0.17 | 0.14 | 0.05 | 0.24 | 0.08 |
| 3 | 81513408 | rs898763 | 1.581 | 1.75 | 2.223 | 1.66 | 0.023 | 0.107 | 0.107 | 0.175 | 0.75 | 0.53 | 0.63 | 0.5 | 0.72 | 0.72 |
| 3 | 81534944 | rs6783132 | -0.513 | 1.755 | 2.291 | 1.681 | 0.016 | 0.332 | 0.332 | 1.061 | 0.07 | 0.24 | 0.14 | 0.36 | 0.05 | 0.07 |
| 3 | 81539406 | rs9820089 | -0.107 | 1.67 | 2.466 | 1.664 | -0.015 | -0.011 | -0.011 | 0 | 0.1 | 0.14 | 0.08 | 0.06 | 0.03 | 0.04 |
| 3 | 81547264 | rs13082940 | -0.88 | 1.723 | 2.379 | 1.725 | -0.018 | -0.018 | -0.018 | 0 | 0.14 | 0.1 | 0.28 | 0.44 | 0.25 | 0.23 |
| 3 | 81550315 | rs7622741 | -1.82 | 1.893 | 2.333 | 1.671 | 0.054 | 0.228 | 0.228 | 0.014 | 0.18 | 0.41 | 0.26 | 0.41 | 0.24 | 0.15 |
| 3 | 81557686 | rs10440071 | -0.561 | 1.886 | 2.228 | 1.637 | 0.026 | -0.013 | -0.013 | 0.199 | 0.1 | 0.13 | 0.11 | 0.05 | 0.18 | 0.08 |
| 3 | 81566655 | rs6789701 | 0.418 | 1.891 | 2.127 | 1.703 | -0.007 | -0.008 | -0.008 | 0 | 0.07 | 0.03 | 0.18 | 0.24 | 0.1 | 0.09 |
| 3 | 81586439 | rs2307058 | -1.778 | 1.745 | 2.325 | 1.479 | 0.048 | 0.22 | 0.22 | 0.014 | 0.18 | 0.37 | 0.26 | 0.42 | 0.24 | 0.15 |
| 3 | 81613942 | rs7610703 | -1.834 | 1.683 | 2.33 | 1.448 | 0.01 | 0.081 | 0.081 | 0 | 0.27 | 0.52 | 0.37 | 0.48 | 0.28 | 0.29 |
| 3 | 81614473 | rs9845467 | -1.589 | 1.685 | 2.35 | 1.449 | 0.013 | 0.089 | 0.089 | 0 | 0.27 | 0.49 | 0.37 | 0.48 | 0.28 | 0.28 |
| 3 | 81639454 | rs13092113 | -0.743 | 1.682 | 2.353 | 1.485 | 0.015 | 0.346 | 0.346 | 1.675 | 0.08 | 0.25 | 0.15 | 0.36 | 0.06 | 0.07 |
| 3 | 81642131 | rs13082609 | -0.47 | 1.698 | 2.423 | 1.483 | 0.012 | 0.039 | 0.039 | 0 | 0.15 | 0.16 | 0.28 | 0.43 | 0.25 | 0.24 |
| 3 | 81643583 | rs9877327 | -2.415 | 1.665 | 2.22 | 1.394 | 0.002 | 0.002 | 0.002 | 0.003 | 0.36 | 0.64 | 0.49 | 0.55 | 0.47 | 0.35 |
| 3 | 81644911 | rs13324454 | -0.398 | 1.41 | 2.202 | 1.469 | 0.003 | 0 | 0 | 0 | 0.09 | 0.11 | 0.12 | 0.06 | 0.19 | 0.09 |
| 3 | 81668050 | rs1031265 | -1.438 | 1.239 | 2.268 | 1.567 | 0.013 | 0.381 | 0.381 | 3.636 | 0.09 | 0.34 | 0.16 | 0.36 | 0.06 | 0.08 |
| 3 | 81675281 | rs2291593 | -0.646 | 1.063 | 2.068 | 1.122 | -0.033 | -0.016 | -0.016 | 0 | 0.12 | 0.26 | 0.09 | 0.06 | 0.03 | 0.05 |
| 3 | 81698130 | rs2229519 | -0.964 | 1.461 | 2.462 | 1.45 | 0.032 | 0.072 | 0.072 | 0 | 0.2 | 0.26 | 0.3 | 0.44 | 0.31 | 0.31 |
| 3 | 81720076 | rs13320194 | -0.226 | 1.375 | 2.431 | 1.405 | -0.027 | -0.005 | -0.005 | 0.119 | 0.1 | 0.26 | 0.09 | 0.07 | 0.02 | 0.03 |
| 3 | 81723009 | rs3821550 | -1.796 | 1.405 | 2.581 | 1.314 | 0.013 | -0.023 | -0.023 | 0 | 0.21 | 0.49 | 0.22 | 0.14 | 0.18 | 0.11 |
| 3 | 81723365 | rs3772902 | -2.418 | 1.448 | 2.571 | 1.33 | -0.045 | -0.014 | -0.014 | 0 | 0.41 | 0.77 | 0.53 | 0.58 | 0.49 | 0.41 |
| 3 | 81762787 | rs7620240 | -1.62 | 1.459 | 2.632 | 1.326 | 0.022 | 0.088 | 0.088 | 0 | 0.29 | 0.5 | 0.4 | 0.51 | 0.33 | 0.34 |
| 3 | 81763410 | rs7623056 | -0.455 | 1.455 | 2.619 | 1.308 | -0.009 | 0.002 | 0.002 | 0 | 0.1 | 0.06 | 0.11 | 0.07 | 0.16 | 0.07 |
| 3 | 81789795 | rs9811796 | -0.05 | 1.439 | 2.766 | 1.308 | -0.021 | -0.006 | -0.006 | 3.446 | 0.1 | 0.2 | 0.09 | 0.07 | 0.02 | 0.03 |
| 3 | 81811224 | rs3755713 | -0.681 | 1.643 | 2.783 | 1.426 | 0.033 | 0.064 | 0.064 | 0 | 0.19 | 0.23 | 0.31 | 0.43 | 0.31 | 0.31 |
| 3 | 81823806 | rs13323851 | -0.441 | 1.624 | 2.678 | 1.378 | -0.012 | 0.003 | 0.003 | 0.031 | 0.09 | 0.06 | 0.11 | 0.07 | 0.16 | 0.06 |
| 3 | 81851714 | rs9813371 | -0.657 | 1.58 | 2.703 | 1.388 | 0.021 | -0.007 | -0.007 | 0.022 | 0.11 | 0.18 | 0.11 | 0.06 | 0.15 | 0.07 |
| 3 | 81854425 | rs2315966 | -2.055 | 1.59 | 2.687 | 1.379 | 0.016 | -0.004 | -0.004 | 1.258 | 0.19 | 0.45 | 0.2 | 0.13 | 0.16 | 0.1 |
| 3 | 81858615 | rs2691081 | 1.706 | 1.59 | 2.647 | 1.379 | 0.023 | -0.001 | -0.001 | 0 | 0.81 | 0.49 | 0.79 | 0.87 | 0.84 | 0.9 |
| 3 | 81873269 | rs6804453 | -0.471 | 1.598 | 2.342 | 1.302 | 0.001 | 0 | 0 | 0 | 0.09 | 0.07 | 0.1 | 0.06 | 0.14 | 0.06 |
| 3 | 81885274 | rs2691093 | 2.143 | 1.588 | 2.327 | 1.323 | -0.058 | -0.051 | -0.051 | 0 | 0.61 | 0.17 | 0.46 | 0.5 | 0.5 | 0.59 |
| 3 | 81892031 | rs1524557 | -0.584 | 1.638 | 2.562 | 1.388 | 0.072 | 0.126 | 0.126 | 0 | 0.18 | 0.32 | 0.36 | 0.43 | 0.32 | 0.31 |
| 3 | 81896410 | rs6798822 | -0.969 | 1.647 | 2.624 | 1.368 | 0.005 | -0.008 | -0.008 | 0.048 | 0.12 | 0.14 | 0.12 | 0.05 | 0.16 | 0.06 |
| 3 | 81898999 | rs17019353 | -0.251 | 1.644 | 2.569 | 1.4 | -0.014 | -0.005 | -0.005 | 0 | 0.08 | 0.27 | 0.05 | 0.04 | 0.02 | 0.04 |
| 3 | 81905441 | rs6798680 | -1.131 | 1.596 | 2.089 | 1.331 | 0.032 | 0.073 | 0.073 | 0.032 | 0.27 | 0.58 | 0.42 | 0.46 | 0.34 | 0.35 |
| 3 | 81944233 | rs7650373 | -1.975 | 1.566 | 2.132 | 1.519 | 0.06 | 0.293 | 0.293 | 0 | 0.14 | 0.4 | 0.27 | 0.41 | 0.19 | 0.11 |
| 3 | 81949335 | rs17019375 | -0.41 | 1.6 | 1.914 | 1.593 | -0.011 | 0.153 | 0.153 | 0 | 0.07 | 0.14 | 0.14 | 0.34 | 0.07 | 0.08 |
| 3 | 81950302 | rs10511101 | -0.966 | 1.511 | 2.015 | 1.513 | 0.001 | 0.01 | 0.01 | 0 | 0.28 | 0.3 | 0.37 | 0.48 | 0.33 | 0.33 |
| 3 | 81951005 | rs9881991 | -0.389 | 1.463 | 1.986 | 1.628 | 0.036 | 0.015 | 0.015 | 0 | 0.08 | 0.12 | 0.12 | 0.07 | 0.16 | 0.08 |
| 3 | 81985189 | rs9865893 | -0.42 | 1.652 | 0.849 | 1.232 | -0.011 | -0.001 | -0.001 | 0 | 0.08 | 0.05 | 0.11 | 0.06 | 0.16 | 0.07 |
| 3 | 81993902 | rs9829799 | -0.278 | 1.545 | 0.777 | 1.225 | -0.024 | -0.009 | -0.009 | 0.907 | 0.08 | 0.03 | 0.08 | 0.06 | 0.12 | 0.08 |
| 3 | 81994008 | rs9847586 | -0.33 | 1.544 | 0.853 | 1.225 | 0.006 | 0 | 0 | 0 | 0.08 | 0.09 | 0.09 | 0.06 | 0.12 | 0.08 |
| 3 | 81994093 | rs9810554 | -0.33 | 1.544 | 0.853 | 1.225 | 0.006 | 0 | 0 | 0 | 0.08 | 0.09 | 0.09 | 0.06 | 0.12 | 0.08 |
| 3 | 82010368 | rs9968070 | -1.091 | 1.259 | 0.693 | 1.262 | 0.002 | 0.005 | 0.005 | 0 | 0.31 | 0.31 | 0.4 | 0.46 | 0.39 | 0.31 |
| 3 | 82019603 | rs7431326 | -1.094 | 0.634 | 0.462 | 0.964 | 0 | -0.003 | -0.003 | 0.736 | 0.36 | 0.38 | 0.38 | 0.43 | 0.37 | 0.44 |
| 3 | 82024432 | rs11929553 | -1.22 | 0.357 | 1.195 | 1.024 | 0.004 | 0.048 | 0.048 | 2.782 | 0.11 | 0.54 | 0.09 | 0.11 | 0.06 | 0.03 |
| 3 | 82031499 | rs9309884 | 1.083 | -0.086 | 0.201 | 0.607 | -0.013 | -0.008 | -0.008 | 0 | 0.42 | 0.3 | 0.62 | 0.49 | 0.47 | 0.38 |
| 3 | 82034178 | rs6776655 | -0.58 | -0.126 | 0.612 | 0.594 | 0.061 | 0.111 | 0.111 | 1.719 | 0.08 | 0.37 | 0.08 | 0.11 | 0.06 | 0.03 |
| 3 | 82046207 | rs9816690 | -0.491 | -0.14 | 1.073 | 0.596 | 0.046 | 0.091 | 0.091 | 1.402 | 0.08 | 0.34 | 0.08 | 0.11 | 0.06 | 0.03 |
| 3 | 82049826 | rs1529028 | -1.053 | -0.191 | 0.641 | 0.684 | -0.027 | -0.061 | -0.061 | 0 | 0.88 | 0.99 | 0.94 | 0.85 | 0.91 | 0.83 |
| 3 | 82050284 | rs9875387 | -1.647 | -0.233 | 0.697 | 0.514 | -0.012 | -0.017 | -0.017 | 0 | 0.23 | 0.62 | 0.21 | 0.16 | 0.18 | 0.14 |

| | | | | | | | | | | | | | | | | |
|---|---|---|---|---|---|---|---|---|---|---|---|---|---|---|---|---|
| 3 | 82074024 | rs17784061 | 0.416 | -0.235 | 0.615 | 0.5 | -0.029 | 0.005 | 0.005 | 0 | 0.11 | 0.01 | 0.11 | 0.05 | 0.12 | 0.1 |
| 3 | 82098814 | rs4856375 | -0.306 | -0.203 | 0.624 | 0.51 | -0.022 | -0.048 | -0.048 | 0 | 0.64 | 0.93 | 0.82 | 0.64 | 0.65 | 0.53 |
| 3 | 82103629 | rs7612651 | 1.545 | -0.206 | 0.329 | -0.074 | -0.107 | 0.093 | 0.093 | 0 | 0.38 | 0.07 | 0.35 | 0.21 | 0.44 | 0.36 |
| 3 | 82113892 | rs9869419 | -0.944 | -0.191 | 0.781 | 0.047 | 0.02 | 0.058 | 0.058 | 2.234 | 0.11 | 0.5 | 0.1 | 0.11 | 0.06 | 0.03 |
| 3 | 82120191 | rs6769852 | -1.181 | -0.196 | 0.602 | 0.045 | -0.01 | 0.019 | 0.019 | 2.967 | 0.11 | 0.62 | 0.1 | 0.11 | 0.06 | 0.03 |
| 3 | 82127643 | rs7639020 | -0.412 | -0.033 | 0.46 | 0.073 | 0.016 | 0.012 | 0.012 | 0 | 0.07 | 0.14 | 0.09 | 0.08 | 0.1 | 0.04 |
| 3 | 82127705 | rs7639102 | -0.473 | -0.034 | 0.478 | 0.078 | 0.024 | 0.021 | 0.021 | 0 | 0.08 | 0.18 | 0.09 | 0.08 | 0.1 | 0.04 |
| 3 | 82145477 | rs9813907 | -1.178 | -0.049 | 0.597 | -0.032 | 0.024 | -0.024 | -0.024 | 0 | 0.17 | 0.44 | 0.19 | 0.11 | 0.18 | 0.12 |
| 3 | 82148325 | rs10514737 | 0.717 | 0.013 | 0.444 | 0.018 | -0.023 | 0.023 | 0.023 | 0 | 0.08 | 0.01 | 0.07 | 0.03 | 0.08 | 0.08 |
| 3 | 82163235 | rs12638629 | 1.473 | 0.096 | 0.251 | -0.247 | -0.02 | 0.071 | 0.071 | 0 | 0.36 | 0.16 | 0.32 | 0.21 | 0.38 | 0.36 |
| 3 | 82183188 | rs6548795 | -1.101 | 0.188 | 0.158 | -0.103 | 0.006 | -0.026 | -0.026 | 0 | 0.18 | 0.57 | 0.18 | 0.11 | 0.14 | 0.12 |
| 3 | 82194083 | rs6797285 | -1.104 | 0.207 | -0.06 | -0.118 | 0.006 | -0.03 | -0.03 | 0 | 0.18 | 0.54 | 0.18 | 0.11 | 0.14 | 0.12 |
| 3 | 82212239 | rs1406547 | 1.146 | -0.019 | -0.276 | -0.231 | -0.046 | 0.083 | 0.083 | 0.058 | 0.36 | 0.07 | 0.31 | 0.21 | 0.34 | 0.32 |
| 3 | 82212402 | rs1406549 | -0.124 | -0.07 | -0.189 | -0.243 | 0.011 | 0.044 | 0.044 | 0 | 0.49 | 0.62 | 0.42 | 0.31 | 0.41 | 0.37 |
| 3 | 82213185 | rs4856210 | 0.588 | 0.209 | -0.28 | -0.166 | 0.01 | 0.06 | 0.06 | 0 | 0.4 | 0.28 | 0.33 | 0.24 | 0.35 | 0.33 |
| 3 | 82214617 | rs1534322 | 0.589 | 0.288 | -0.435 | -0.185 | -0.111 | 0.091 | 0.091 | 0.009 | 0.4 | 0.03 | 0.34 | 0.21 | 0.41 | 0.32 |
| 3 | 82223946 | rs6779875 | 0.943 | 0.264 | -0.364 | -0.169 | -0.005 | 0.076 | 0.076 | 0 | 0.37 | 0.19 | 0.31 | 0.21 | 0.34 | 0.33 |
| 3 | 82225560 | rs12490125 | -0.334 | 0.262 | -0.185 | -0.115 | 0 | -0.013 | -0.013 | 0.003 | 0.36 | 0.57 | 0.37 | 0.32 | 0.32 | 0.35 |
| 3 | 82235386 | rs7624419 | 0.71 | 0.289 | -0.322 | -0.122 | 0.001 | 0.006 | 0.006 | 0 | 0.57 | 0.39 | 0.58 | 0.69 | 0.6 | 0.66 |
| 3 | 82237565 | rs13088543 | 0.137 | 0.3 | -0.255 | -0.118 | 0.02 | 0.032 | 0.032 | 0 | 0.91 | 0.74 | 0.82 | 0.9 | 0.93 | 0.96 |
| 3 | 82239399 | rs4561854 | 0.011 | 0.297 | -0.302 | -0.122 | -0.011 | 0.054 | 0.054 | 0 | 0.34 | 0.18 | 0.22 | 0.21 | 0.33 | 0.3 |
| 3 | 82249474 | rs4384977 | 0.503 | 0.548 | 0.142 | 0.325 | 0 | -0.006 | -0.006 | 0.907 | 0.64 | 0.4 | 0.62 | 0.65 | 0.68 | 0.66 |
| 3 | 82258737 | rs13066341 | 0.881 | 0.497 | 0.201 | 0.36 | 0.047 | 0.062 | 0.062 | 0.106 | 0.91 | 0.65 | 0.81 | 0.86 | 0.93 | 0.96 |
| 3 | 82280224 | rs13087528 | 0.934 | 0.445 | 0.016 | 0.428 | -0.001 | 0 | 0 | 0.173 | 0.51 | 0.48 | 0.56 | 0.49 | 0.54 | 0.49 |
| 3 | 82281924 | rs12633958 | 0.41 | 0.48 | -0.026 | 0.422 | -0.007 | 0.014 | 0.014 | 0 | 0.26 | 0.17 | 0.17 | 0.21 | 0.25 | 0.29 |
| 3 | 82288497 | rs7614064 | -0.215 | 0.483 | -0.028 | 0.409 | -0.005 | -0.012 | -0.012 | 0 | 0.81 | 0.87 | 0.76 | 0.77 | 0.8 | 0.79 |
| 3 | 82311711 | rs11127766 | -0.19 | 0.521 | -0.185 | 0.411 | 0.005 | 0.014 | 0.014 | 0 | 0.8 | 0.75 | 0.76 | 0.76 | 0.79 | 0.79 |
| 3 | 82316849 | rs4632567 | 1.162 | 0.398 | -0.183 | 0.391 | -0.003 | -0.001 | -0.001 | 1.605 | 0.52 | 0.38 | 0.55 | 0.49 | 0.54 | 0.49 |
| 3 | 82335965 | rs9309888 | -0.1 | 0.574 | -0.549 | 0.476 | 0 | 0.002 | 0.002 | 0 | 0.2 | 0.2 | 0.24 | 0.24 | 0.21 | 0.21 |
| 3 | 82359035 | rs1449862 | -0.088 | 0.594 | -0.684 | 0.488 | 0.009 | 0.027 | 0.027 | 0 | 0.2 | 0.27 | 0.25 | 0.24 | 0.21 | 0.21 |
| 3 | 82363196 | rs7649805 | 0.324 | 0.674 | -1.074 | 0.501 | -0.028 | 0.007 | 0.007 | 0 | 0.26 | 0.08 | 0.17 | 0.2 | 0.25 | 0.29 |
| 3 | 82389573 | rs7625954 | -1.213 | 0.856 | -0.935 | 0.636 | 0.001 | 0.003 | 0.003 | 0 | 0.44 | 0.53 | 0.42 | 0.5 | 0.42 | 0.5 |
| 3 | 82441812 | rs4856387 | -0.871 | 0.907 | -0.251 | 0.831 | 0.008 | 0.039 | 0.039 | 0 | 0.23 | 0.44 | 0.28 | 0.31 | 0.23 | 0.22 |
| 3 | 82448567 | rs1992983 | -0.529 | 1.53 | -1.035 | 1.001 | -0.081 | -0.085 | -0.085 | 0.005 | 0.22 | 0.09 | 0.44 | 0.41 | 0.38 | 0.26 |
| 3 | 82448822 | rs1992984 | -1.181 | 1.013 | -0.358 | 0.778 | 0.008 | 0.006 | 0.006 | 0.008 | 0.36 | 0.36 | 0.51 | 0.54 | 0.49 | 0.39 |
| 3 | 82453932 | rs1375805 | 0.604 | 0.27 | -0.845 | 0.238 | 0.023 | -0.046 | -0.046 | 0.536 | 0.26 | 0.12 | 0.23 | 0.39 | 0.19 | 0.36 |
| 3 | 82455946 | rs13066021 | 0.495 | 0.074 | -0.602 | 0.239 | -0.019 | -0.019 | -0.019 | 0 | 0.14 | 0.26 | 0.07 | 0.13 | 0.11 | 0.13 |
| 3 | 82458130 | rs17019771 | 0.768 | 1.008 | -1.237 | 0.439 | -0.058 | -0.082 | -0.082 | 0.755 | 0.13 | 0.03 | 0.23 | 0.34 | 0.2 | 0.17 |
| 3 | 82461279 | rs12494049 | 1.219 | 0.436 | -0.936 | -0.094 | -0.006 | 0.163 | 0.163 | 0 | 0.34 | 0.1 | 0.22 | 0.07 | 0.31 | 0.25 |
| 3 | 82469264 | rs1375799 | -0.376 | -0.041 | -0.34 | 0.28 | -0.054 | -0.029 | -0.029 | 0 | 0.68 | 0.44 | 0.86 | 0.74 | 0.86 | 0.66 |
| 3 | 82469376 | rs13095290 | 0.341 | 0.037 | -0.304 | 0.638 | 0.008 | 0.071 | 0.071 | 0 | 0.09 | 0.26 | 0.2 | 0.21 | 0.12 | 0.11 |
| 3 | 82472857 | rs6805523 | 0.286 | -0.003 | -0.322 | 0.807 | 0.001 | 0.128 | 0.128 | 0 | 0.1 | 0.26 | 0.29 | 0.26 | 0.12 | 0.12 |
| 3 | 82477805 | rs9875783 | 0.223 | 0.139 | -0.679 | 0.559 | 0.11 | -0.046 | -0.046 | 1.592 | 0.58 | 0.18 | 0.57 | 0.49 | 0.74 | 0.56 |
| 3 | 82480713 | rs4856216 | -0.659 | 0.187 | -0.798 | 0.287 | 0.037 | 0.094 | 0.094 | 0 | 0.53 | 0.27 | 0.52 | 0.7 | 0.54 | 0.73 |
| 3 | 82480866 | rs13325641 | 0.028 | 0.193 | -0.83 | 0.351 | -0.138 | -0.083 | -0.083 | 0 | 0.28 | 0.07 | 0.33 | 0.35 | 0.42 | 0.39 |
| 3 | 82488068 | rs10460926 | 1.237 | 0.634 | -1.032 | 0.404 | -0.06 | -0.065 | -0.065 | 0 | 0.1 | 0.01 | 0.1 | 0.29 | 0.17 | 0.18 |
| 3 | 82488254 | rs9309903 | -1.648 | -0.157 | -0.608 | 0.267 | 0.009 | 0.063 | 0.063 | 0 | 0.69 | 0.84 | 0.76 | 0.9 | 0.69 | 0.83 |
| 3 | 82491482 | rs7432307 | 0.195 | -0.069 | -1.094 | 0.326 | 0.048 | 0.151 | 0.151 | 0 | 0.49 | 0.84 | 0.48 | 0.3 | 0.46 | 0.28 |
| 3 | 82492458 | rs9831660 | 0.718 | -0.116 | -0.982 | 0.559 | 0.037 | -0.055 | -0.055 | 0 | 0.24 | 0.09 | 0.2 | 0.34 | 0.15 | 0.35 |
| 3 | 82495988 | rs7644123 | 1.45 | 0.152 | -1.228 | 0.437 | -0.02 | 0.113 | 0.113 | 0 | 0.28 | 0.08 | 0.27 | 0.1 | 0.3 | 0.17 |
| 3 | 82498272 | rs6548818 | -0.049 | 1.083 | -0.312 | 0.963 | -0.012 | -0.008 | -0.008 | 1.318 | 0.7 | 0.73 | 0.73 | 0.65 | 0.6 | 0.66 |
| 3 | 82499607 | rs7651342 | -0.706 | 0.608 | 0.146 | 0.849 | -0.009 | 0.012 | 0.012 | 0 | 0.16 | 0.38 | 0.25 | 0.21 | 0.15 | 0.14 |
| 3 | 82507158 | rs4856388 | -0.552 | 1.247 | -0.006 | 1.257 | 0.03 | 0.069 | 0.069 | 0 | 0.14 | 0.24 | 0.17 | 0.38 | 0.18 | 0.26 |
| 3 | 82512579 | rs9817863 | -0.807 | 1.007 | -0.057 | 0.843 | 0.003 | 0.005 | 0.005 | 0 | 0.4 | 0.55 | 0.4 | 0.54 | 0.41 | 0.4 |
| 3 | 82517313 | rs4856390 | -1.103 | 1.156 | -0.793 | 0.896 | -0.043 | -0.064 | -0.064 | 0 | 0.85 | 0.99 | 0.79 | 0.83 | 0.82 | 0.81 |
| 3 | 82541304 | rs13080791 | 0.242 | 1.406 | -0.466 | 0.949 | 0.016 | 0.013 | 0.013 | 0 | 0.07 | 0.13 | 0.13 | 0.14 | 0.11 | 0.09 |
| 3 | 82541380 | rs9850201 | 0.146 | 1.282 | -0.074 | 0.828 | 0.021 | -0.004 | -0.004 | 0.458 | 0.32 | 0.48 | 0.27 | 0.38 | 0.35 | 0.3 |
| 3 | 82542364 | rs7429328 | -0.933 | 1.292 | -0.034 | 0.881 | -0.005 | -0.011 | -0.011 | 0 | 0.46 | 0.76 | 0.54 | 0.66 | 0.53 | 0.5 |
| 3 | 82545990 | rs12636819 | 0.388 | 1.255 | -0.047 | 0.887 | 0.017 | 0.014 | 0.014 | 0 | 0.07 | 0.13 | 0.12 | 0.14 | 0.1 | 0.09 |
| 3 | 82550856 | rs7612815 | 0.828 | 1.072 | -0.029 | 0.853 | 0 | 0 | 0 | 0 | 0.08 | 0.07 | 0.13 | 0.14 | 0.07 | 0.11 |
| 3 | 82558381 | rs6548823 | -0.526 | 1.158 | -0.013 | 0.869 | -0.004 | -0.004 | -0.004 | 0 | 0.39 | 0.62 | 0.41 | 0.52 | 0.42 | 0.41 |
| 3 | 82559498 | rs7623845 | 0.999 | 1.138 | -0.021 | 0.873 | -0.005 | 0.015 | 0.015 | 0.07 | 0.07 | 0.12 | 0.13 | 0.14 | 0.07 | 0.11 |
| 3 | 82562418 | rs6788150 | 0.999 | 1.138 | -0.021 | 0.873 | -0.005 | 0.015 | 0.015 | 0.07 | 0.07 | 0.12 | 0.13 | 0.14 | 0.07 | 0.11 |
| 3 | 82563862 | rs9856109 | 0.115 | 1.167 | -0.053 | 0.861 | 0.018 | -0.004 | -0.004 | 0 | 0.32 | 0.5 | 0.28 | 0.38 | 0.35 | 0.3 |
| 3 | 82564293 | rs7427791 | 1.571 | 1.191 | 0.058 | 0.824 | -0.001 | -0.005 | -0.005 | 0 | 0.16 | 0.14 | 0.09 | 0 | 0.17 | 0.03 |
| 3 | 82565903 | rs4856219 | -1.218 | 1.147 | -0.122 | 0.654 | -0.027 | -0.049 | -0.049 | 2.193 | 0.51 | 0.91 | 0.58 | 0.66 | 0.56 | 0.51 |
| 3 | 82566012 | rs4856220 | 0.686 | 1.459 | 0.189 | 0.903 | 0.052 | 0.116 | 0.116 | 0 | 0.15 | 0.31 | 0.18 | 0.37 | 0.18 | 0.27 |
| 3 | 82566370 | rs4856222 | 1.163 | 1.622 | 0.137 | 0.89 | 0 | 0.018 | 0.018 | 0 | 0.1 | 0.08 | 0.13 | 0.36 | 0.13 | 0.2 |
| 3 | 82570656 | rs6797568 | -0.608 | 1.489 | -0.029 | 0.913 | 0.026 | 0.125 | 0.125 | 0.368 | 0.24 | 0.44 | 0.33 | 0.53 | 0.26 | 0.37 |

| | | | | | | | | | | | | | | | | |
|---|---|---|---|---|---|---|---|---|---|---|---|---|---|---|---|---|
| 3 | 82582827 | rs12497630 | 0.438 | 1.344 | -0.061 | 0.879 | -0.008 | 0.005 | 0.005 | 0.517 | 0.1 | 0.14 | 0.15 | 0.13 | 0.08 | 0.11 |
| 3 | 82585128 | rs12630116 | 0.164 | 1.487 | -0.077 | 0.836 | 0.015 | 0.001 | 0.001 | 0 | 0.08 | 0.13 | 0.14 | 0.11 | 0.13 | 0.09 |
| 3 | 82596134 | rs1463336 | -0.72 | 1.312 | -0.045 | 0.954 | 0.019 | 0.104 | 0.104 | 0.424 | 0.25 | 0.45 | 0.34 | 0.52 | 0.25 | 0.37 |
| 3 | 82602965 | rs4856401 | 0.419 | 1.378 | -0.172 | 1.025 | 0.056 | 0.147 | 0.147 | 0 | 0.14 | 0.27 | 0.18 | 0.4 | 0.18 | 0.26 |
| 3 | 82614903 | rs13079811 | -0.232 | 1.15 | -0.269 | 0.677 | 0.005 | 0.019 | 0.019 | 0 | 0.69 | 0.71 | 0.61 | 0.43 | 0.65 | 0.52 |
| 3 | 82616037 | rs13097360 | 0.958 | 0.802 | -0.843 | 0.447 | 0.01 | 0.013 | 0.013 | 0 | 0.31 | 0.23 | 0.23 | 0.2 | 0.28 | 0.24 |
| 3 | 82618784 | rs1454262 | -0.145 | 0.831 | -0.483 | 0.542 | 0.005 | 0.008 | 0.008 | 0 | 0.69 | 0.71 | 0.64 | 0.4 | 0.67 | 0.52 |
| 3 | 82621024 | rs9813850 | 0.787 | 0.486 | -0.418 | 0.038 | 0.009 | -0.007 | -0.007 | 0 | 0.23 | 0.22 | 0.1 | 0 | 0.17 | 0.08 |
| 3 | 82628053 | rs10865598 | -0.375 | 0.872 | -0.334 | 0.415 | 0.01 | 0.014 | 0.014 | 0 | 0.7 | 0.71 | 0.66 | 0.45 | 0.64 | 0.56 |
| 3 | 82629021 | rs2317203 | 0.658 | 0.456 | 0.081 | 0.134 | -0.002 | 0.004 | 0.004 | 0 | 0.36 | 0.36 | 0.48 | 0.25 | 0.46 | 0.28 |
| 3 | 82642734 | rs7647242 | 0.642 | 0.445 | 0.398 | 0.11 | 0.003 | -0.029 | -0.029 | 0 | 0.36 | 0.44 | 0.46 | 0.19 | 0.41 | 0.28 |
| 3 | 82652995 | rs834895 | 0.451 | 0.366 | 0.443 | 0.075 | 0 | -0.042 | -0.042 | 0 | 0.38 | 0.51 | 0.47 | 0.19 | 0.41 | 0.29 |
| 3 | 82658735 | rs1598118 | 0.869 | 0.298 | 0.549 | 0.022 | -0.011 | 0.028 | 0.028 | 0 | 0.33 | 0.22 | 0.44 | 0.19 | 0.41 | 0.28 |
| 3 | 82663150 | rs834864 | -0.103 | 0.442 | 0.757 | 0.474 | 0.013 | 0.011 | 0.011 | 0 | 0.51 | 0.56 | 0.62 | 0.65 | 0.59 | 0.53 |
| 3 | 82665045 | rs17019957 | 1.316 | 0.726 | 0.682 | 0.574 | -0.012 | -0.01 | -0.01 | 0.019 | 0.05 | 0.02 | 0.08 | 0.08 | 0.08 | 0.12 |
| 3 | 82668363 | rs17019962 | 0.767 | 0.559 | 0.69 | 0.485 | 0.014 | 0.061 | 0.061 | 0 | 0.15 | 0.15 | 0.16 | 0.46 | 0.18 | 0.23 |
| 3 | 82669493 | rs17731998 | 1.832 | 0.816 | 0.46 | 0.317 | -0.052 | 0.071 | 0.071 | 0 | 0.25 | 0.05 | 0.18 | 0.09 | 0.28 | 0.18 |
| 3 | 82675810 | rs17019967 | 0.791 | 0.794 | 0.62 | 0.316 | 0.019 | -0.002 | -0.002 | 0.041 | 0.07 | 0.09 | 0.26 | 0.11 | 0.13 | 0.1 |
| 3 | 82678168 | rs10433516 | 0.606 | 0.775 | 0.672 | 0.316 | -0.001 | 0.001 | 0.001 | 0 | 0.36 | 0.38 | 0.46 | 0.19 | 0.41 | 0.3 |
| 3 | 82691819 | rs11127782 | 1.036 | 0.455 | 0.657 | 0.296 | -0.021 | 0.012 | 0.012 | 0 | 0.11 | 0.17 | 0.09 | 0.14 | 0.04 | 0.13 |
| 3 | 82694000 | rs1598120 | -0.199 | 0.688 | 0.83 | 0.414 | 0.024 | 0.021 | 0.021 | 0 | 0.5 | 0.6 | 0.63 | 0.65 | 0.59 | 0.53 |
| 3 | 82694503 | rs11711911 | 1.705 | 0.731 | 0.596 | 0.407 | -0.015 | 0.069 | 0.069 | 0 | 0.26 | 0.09 | 0.18 | 0.09 | 0.28 | 0.18 |
| 3 | 82697984 | rs2873392 | 0.922 | 0.587 | 0.429 | 0.315 | 0.036 | 0.063 | 0.063 | 0 | 0.33 | 0.18 | 0.16 | 0.14 | 0.22 | 0.29 |
| 3 | 82701950 | rs936035 | -1.597 | 0.671 | 0.17 | 0.383 | -0.052 | -0.084 | -0.084 | 0 | 0.86 | 0.99 | 0.8 | 0.79 | 0.81 | 0.82 |
| 3 | 82719412 | rs12636480 | 1.047 | 0.743 | 0.355 | 0.256 | -0.013 | 0.033 | 0.033 | 0 | 0.3 | 0.19 | 0.42 | 0.19 | 0.37 | 0.21 |
| 3 | 82720444 | rs7632237 | -0.628 | 0.779 | 0.609 | 0.353 | 0.025 | 0.085 | 0.085 | 0 | 0.83 | 0.73 | 0.81 | 0.54 | 0.79 | 0.74 |
| 3 | 82734786 | rs4856414 | 0.512 | 0.645 | 0.366 | -0.027 | 0.043 | 0.02 | 0.02 | 0 | 0.06 | 0.13 | 0.24 | 0.11 | 0.1 | 0.08 |
| 3 | 82737773 | rs2169642 | 0.463 | 0.856 | 0.336 | -0.211 | -0.004 | 0.034 | 0.034 | 0 | 0.35 | 0.3 | 0.44 | 0.15 | 0.39 | 0.27 |
| 3 | 82738464 | rs1979373 | 0.485 | 0.872 | 0.122 | -0.404 | -0.008 | -0.032 | -0.032 | 0 | 0.29 | 0.36 | 0.39 | 0.16 | 0.27 | 0.21 |
| 3 | 82739213 | rs1351043 | 0.518 | 0.916 | 0.301 | -0.302 | -0.005 | -0.003 | -0.003 | 0 | 0.41 | 0.42 | 0.24 | 0.29 | 0.34 | 0.32 |
| 3 | 82740425 | rs7639667 | -0.162 | 0.95 | 0.421 | -0.141 | -0.01 | -0.013 | -0.013 | 0 | 0.68 | 0.7 | 0.62 | 0.45 | 0.6 | 0.54 |
| 3 | 82747257 | rs1382066 | 0.158 | 0.896 | 0.496 | -0.162 | 0.023 | 0.035 | 0.035 | 0 | 0.65 | 0.43 | 0.61 | 0.45 | 0.6 | 0.53 |
| 3 | 82749782 | rs12494363 | 0.17 | 0.933 | 0.503 | -0.166 | 0.005 | 0.008 | 0.008 | 0 | 0.66 | 0.36 | 0.6 | 0.44 | 0.6 | 0.54 |
| 3 | 82752602 | rs11712003 | 1.792 | 0.794 | 0.097 | -0.339 | 0.024 | 0.012 | 0.012 | 0 | 0.19 | 0.01 | 0.09 | 0.09 | 0.1 | 0.12 |
| 3 | 82758743 | rs6548843 | -0.059 | 1.243 | 0.771 | -0.167 | 0.011 | 0.016 | 0.016 | 0 | 0.68 | 0.63 | 0.62 | 0.44 | 0.6 | 0.54 |
| 3 | 82765927 | rs12714541 | 0.02 | 1.273 | 0.846 | -0.222 | 0.003 | 0.008 | 0.008 | 0 | 0.72 | 0.67 | 0.64 | 0.43 | 0.67 | 0.57 |
| 3 | 82788885 | rs7619008 | -0.054 | 1.345 | 0.523 | -0.251 | -0.011 | -0.016 | -0.016 | 0 | 0.73 | 0.72 | 0.64 | 0.43 | 0.67 | 0.58 |
| 3 | 82789651 | rs7621875 | 1.696 | 0.899 | 0.944 | -0.592 | -0.01 | -0.05 | -0.05 | 0.341 | 0.89 | 0.28 | 0.69 | 0.93 | 0.9 | 0.91 |
| 3 | 82818242 | rs9870034 | -0.692 | 1.02 | 0.936 | -0.539 | 0.111 | 0.034 | 0.034 | 3.853 | 0.07 | 0.41 | 0.26 | 0.07 | 0.09 | 0.08 |
| 3 | 82832243 | rs9858734 | -1.633 | 1.245 | 0.633 | -0.352 | -0.05 | -0.048 | -0.048 | 0.996 | 0.91 | 0.99 | 0.82 | 0.88 | 0.87 | 0.85 |
| 3 | 82835605 | rs7639777 | 0.526 | 0.334 | 0.815 | -1.013 | 0.013 | 0 | 0 | 0 | 0.32 | 0.28 | 0.14 | 0 | 0.29 | 0.23 |
| 3 | 82850096 | rs9880341 | -0.375 | 0.331 | 0.748 | -1.017 | 0.02 | 0.062 | 0.062 | 0.303 | 0.4 | 0.73 | 0.4 | 0.08 | 0.38 | 0.3 |
| 3 | 82851186 | rs13324550 | 1.117 | 0.362 | 0.762 | -1.037 | 0.017 | 0.043 | 0.043 | 0 | 0.27 | 0.15 | 0.12 | 0 | 0.26 | 0.16 |
| 3 | 82851367 | rs9309910 | 1.189 | 0.369 | 0.721 | -1.034 | 0.015 | 0.101 | 0.101 | 0 | 0.27 | 0.09 | 0.11 | 0 | 0.26 | 0.16 |
| 3 | 82853590 | rs1032554 | -0.445 | 0.369 | 0.931 | -1.031 | 0.05 | -0.021 | -0.021 | 0 | 0.08 | 0.21 | 0.24 | 0.07 | 0.1 | 0.13 |
| 3 | 82856192 | rs35868010 | 1.076 | 0.147 | 0.54 | -0.92 | 0.018 | 0.068 | 0.068 | 0 | 0.27 | 0.14 | 0.12 | 0 | 0.26 | 0.15 |
| 3 | 82867892 | rs9824524 | -0.671 | 0.326 | 0.809 | -0.367 | 0.038 | 0.029 | 0.029 | 3.8 | 0.05 | 0.38 | 0.04 | 0 | 0.02 | 0 |
| 3 | 82903854 | rs9875881 | -0.5 | 0.35 | 0.367 | -0.5 | 0.009 | 0.049 | 0.049 | 0.343 | 0.44 | 0.74 | 0.44 | 0.28 | 0.44 | 0.37 |
| 3 | 82913445 | rs1995065 | -1.307 | 0.256 | -0.452 | -0.681 | -0.046 | -0.061 | -0.061 | 0 | 0.67 | 0.86 | 0.59 | 0.41 | 0.6 | 0.64 |
| 3 | 82920077 | rs9878786 | 1.442 | 0.054 | -0.378 | -0.85 | 0.009 | 0.101 | 0.101 | 0 | 0.27 | 0.1 | 0.12 | 0 | 0.27 | 0.15 |
| 3 | 82922156 | rs13325164 | 0.808 | 0.131 | -0.181 | -0.828 | 0.021 | 0.082 | 0.082 | 0 | 0.33 | 0.18 | 0.15 | 0 | 0.32 | 0.21 |
| 3 | 82922579 | rs17020233 | 0.671 | 0.241 | -0.069 | -0.731 | 0.019 | 0.087 | 0.087 | 0.082 | 0.06 | 0.05 | 0.27 | 0.28 | 0.11 | 0.16 |
| 3 | 82925647 | rs6791124 | -0.245 | 0.136 | 0.071 | -0.889 | 0.009 | 0.073 | 0.073 | 0.514 | 0.44 | 0.74 | 0.43 | 0.25 | 0.44 | 0.37 |
| 3 | 82929459 | rs7619844 | -1.02 | 0.077 | -0.047 | -0.967 | -0.052 | 0.01 | 0.01 | 0 | 0.62 | 0.8 | 0.51 | 0.33 | 0.52 | 0.48 |
| 3 | 82953061 | rs982064 | -1.076 | 0.178 | -0.057 | -1.077 | 0.013 | 0.029 | 0.029 | 0 | 0.66 | 0.77 | 0.83 | 1 | 0.66 | 0.78 |
| 3 | 82969840 | rs1532428 | 0.869 | 0.114 | -0.166 | -0.952 | -0.056 | 0.02 | 0.02 | 0 | 0.39 | 0.19 | 0.61 | 0.71 | 0.49 | 0.52 |
| 3 | 82972073 | rs13099031 | 1.371 | 0.023 | -0.119 | -1.042 | 0.018 | 0.062 | 0.062 | 0 | 0.3 | 0.14 | 0.12 | 0 | 0.29 | 0.2 |
| 3 | 82973130 | rs9869425 | -0.105 | 0.026 | 0.075 | -1.04 | 0.002 | -0.014 | -0.014 | 0.127 | 0.73 | 0.79 | 0.76 | 0.73 | 0.77 | 0.72 |
| 3 | 82974977 | rs9309920 | 1.346 | -0.042 | 0.058 | -1.098 | 0.016 | 0.041 | 0.041 | 0 | 0.3 | 0.16 | 0.12 | 0 | 0.29 | 0.2 |
| 3 | 82975102 | rs9309921 | -0.169 | 0.184 | -0.14 | -0.993 | 0.041 | 0.082 | 0.082 | 0 | 0.53 | 0.63 | 0.22 | 0.08 | 0.37 | 0.32 |
| 3 | 82988570 | rs6782805 | -1.074 | 0.062 | -0.226 | -0.924 | -0.007 | -0.006 | -0.006 | 0 | 0.84 | 0.83 | 0.8 | 0.82 | 0.76 | 0.82 |
| 3 | 83004185 | rs4856423 | 0.295 | -0.682 | -0.24 | -1.183 | 0.103 | 0.158 | 0.158 | 0 | 0.54 | 0.23 | 0.79 | 0.92 | 0.71 | 0.7 |
| 3 | 83009526 | rs9868657 | -1.058 | -0.726 | -0.291 | -1.138 | 0.028 | 0.027 | 0.027 | 0 | 0.82 | 0.87 | 0.93 | 0.92 | 0.91 | 0.88 |
| 3 | 83012312 | rs1482617 | -1.058 | -0.726 | -0.291 | -1.138 | 0.028 | 0.027 | 0.027 | 0 | 0.82 | 0.87 | 0.92 | 0.92 | 0.91 | 0.88 |
| 3 | 83030152 | rs11127801 | 1.284 | -0.8 | -0.061 | -1.14 | 0.028 | 0.017 | 0.017 | 0 | 0.14 | 0.08 | 0.04 | 0.08 | 0.05 | 0.11 |
| 3 | 83039277 | rs9827152 | 0.799 | -0.192 | -0.292 | -0.943 | 0.024 | 0.058 | 0.058 | 0 | 0.23 | 0.11 | 0.1 | 0 | 0.19 | 0.18 |
| 3 | 83049086 | rs17423261 | 0.668 | 0.223 | -0.408 | -0.896 | -0.02 | -0.011 | -0.011 | 0 | 0.19 | 0.11 | 0.21 | 0.21 | 0.26 | 0.32 |
| 3 | 83053687 | rs2873498 | -0.516 | 0.188 | -0.321 | -0.79 | -0.019 | 0.052 | 0.052 | 0 | 0.42 | 0.26 | 0.31 | 0.21 | 0.46 | 0.5 |
| 3 | 83071349 | rs9878800 | 0.603 | -0.053 | -0.04 | -0.954 | -0.002 | -0.052 | -0.052 | 0 | 0.24 | 0.22 | 0.11 | 0 | 0.2 | 0.18 |

| | | | | | | | | | | | | | | | | |
|---|---|---|---|---|---|---|---|---|---|---|---|---|---|---|---|---|
| 3 | 83089596 | rs7618878 | 1.063 | -0.023 | -0.05 | -0.838 | -0.001 | -0.001 | -0.001 | 0 | 0.43 | 0.42 | 0.61 | 0.71 | 0.46 | 0.38 |
| 3 | 83107505 | rs1080728 | 0.941 | 0.07 | -0.163 | -0.823 | -0.032 | -0.022 | -0.022 | 0 | 0.18 | 0.08 | 0.19 | 0.21 | 0.25 | 0.3 |
| 3 | 83115092 | rs9857267 | -0.253 | -0.092 | -0.122 | -0.901 | 0.001 | 0.001 | 0.001 | 0 | 0.87 | 0.84 | 0.93 | 0.92 | 0.92 | 0.87 |
| 3 | 83124229 | rs9877278 | 1.887 | 0.178 | -0.291 | -0.5 | -0.001 | -0.001 | -0.001 | 0 | 0.3 | 0.3 | 0.46 | 0.26 | 0.29 | 0.28 |
| 3 | 83130081 | rs11127807 | 0.924 | 0.131 | -0.326 | -0.606 | 0 | -0.001 | -0.001 | 0 | 0.43 | 0.42 | 0.61 | 0.71 | 0.46 | 0.4 |
| 3 | 83132865 | rs6806032 | 1.434 | 0.256 | -0.491 | -0.514 | -0.032 | -0.022 | -0.022 | 0.895 | 0.18 | 0.08 | 0.19 | 0.21 | 0.25 | 0.29 |
| 3 | 83166899 | rs17020566 | 1.151 | 0.089 | -0.532 | -0.484 | 0.003 | -0.023 | -0.023 | 0 | 0.08 | 0 | 0.02 | 0.09 | 0.05 | 0.12 |
| 3 | 83169976 | rs1473624 | 1.562 | 0.177 | -0.41 | -0.345 | -0.013 | -0.018 | -0.018 | 0.049 | 0.13 | 0.1 | 0.16 | 0.44 | 0.18 | 0.1 |
| 3 | 83171687 | rs11920785 | 0.235 | 0.455 | -0.406 | -0.54 | -0.001 | -0.011 | -0.011 | 0 | 0.06 | 0.08 | 0.06 | 0 | 0.08 | 0.01 |
| 3 | 83207518 | rs7624990 | 0.851 | 0.413 | -0.468 | -0.648 | -0.021 | -0.008 | -0.008 | 0 | 0.21 | 0.14 | 0.21 | 0.2 | 0.26 | 0.37 |
| 3 | 83212404 | rs7624009 | 0.928 | 0.325 | -0.463 | -0.631 | -0.01 | -0.026 | -0.026 | 0.083 | 0.05 | 0.11 | 0.05 | 0 | 0.04 | 0.01 |
| 3 | 83241703 | rs11127809 | 0.632 | 0.459 | -0.369 | -0.567 | 0.001 | 0.021 | 0.021 | 0 | 0.5 | 0.44 | 0.51 | 0.75 | 0.48 | 0.5 |
| 3 | 83261027 | rs9828520 | 1.176 | -0.03 | -0.227 | -0.781 | -0.005 | 0.129 | 0.129 | 0 | 0.22 | 0.05 | 0.09 | 0 | 0.19 | 0.12 |
| 3 | 83305448 | rs4856450 | 0.385 | 0.169 | -0.157 | -0.676 | 0.002 | 0.013 | 0.013 | 0 | 0.54 | 0.44 | 0.69 | 0.8 | 0.57 | 0.57 |
| 3 | 83329109 | rs365527 | -2.266 | -0.03 | -0.461 | -0.657 | -0.061 | -0.005 | -0.005 | 0 | 0.71 | 0.96 | 0.56 | 0.79 | 0.72 | 0.65 |
| 3 | 83412009 | rs823452 | -0.532 | 0.155 | 0.122 | -0.79 | -0.001 | -0.022 | -0.022 | 0 | 0.44 | 0.46 | 0.31 | 0.2 | 0.44 | 0.44 |
| 3 | 83419162 | rs2100340 | 1.581 | 0.26 | 0.15 | -0.806 | -0.002 | -0.001 | -0.001 | 0 | 0.16 | 0.15 | 0.16 | 0.14 | 0.18 | 0.14 |
| 3 | 83426152 | rs2198281 | 1.34 | 0.294 | 0.451 | -0.759 | -0.002 | -0.047 | -0.047 | 0 | 0.2 | 0.2 | 0.09 | 0 | 0.19 | 0.12 |
| 3 | 83501948 | rs1376967 | 1.516 | 0.469 | -0.231 | -0.093 | -0.061 | -0.058 | -0.058 | 0 | 0.12 | 0.01 | 0.15 | 0.19 | 0.2 | 0.1 |
| 3 | 83512237 | rs1470801 | -0.805 | 0.13 | 0.201 | -0.308 | -0.011 | -0.095 | -0.095 | 0.19 | 0.77 | 0.72 | 0.89 | 1 | 0.8 | 0.81 |
| 3 | 83532973 | rs2060956 | -1.123 | 0.177 | 0.199 | 0.232 | -0.002 | -0.013 | -0.013 | 0 | 0.61 | 0.58 | 0.45 | 0.75 | 0.61 | 0.55 |
| 3 | 83536528 | rs12491148 | 0.655 | -0.436 | 0.371 | 0.518 | 0.011 | 0.025 | 0.025 | 1.823 | 0.55 | 0.26 | 0.67 | 0.66 | 0.6 | 0.53 |
| 3 | 83546019 | rs4374540 | 0.531 | -0.477 | 0.788 | 0.652 | 0.001 | -0.001 | -0.001 | 1.655 | 0.57 | 0.4 | 0.69 | 0.44 | 0.6 | 0.53 |
| 3 | 83547249 | rs8180047 | -1.595 | -0.596 | 0.695 | 0.442 | -0.011 | -0.077 | -0.077 | 0.61 | 0.83 | 0.78 | 0.92 | 1 | 0.86 | 0.89 |
| 3 | 83550973 | rs4060642 | 0.905 | -0.585 | 0.664 | 0.491 | 0.011 | -0.003 | -0.003 | 0 | 0.55 | 0.25 | 0.67 | 0.44 | 0.6 | 0.53 |
| 3 | 83558368 | rs11127817 | 0.208 | -0.543 | 0.741 | 0.667 | -0.011 | 0.022 | 0.022 | 0 | 0.07 | 0.17 | 0.04 | 0.12 | 0.03 | 0.06 |
| 3 | 83567210 | rs964243 | -0.502 | -0.586 | 0.723 | 0.712 | 0 | 0.009 | 0.009 | 0 | 0.27 | 0.31 | 0.23 | 0.57 | 0.27 | 0.36 |
| 3 | 83576033 | rs3214002 | 1.254 | -0.765 | 0.313 | 0.157 | 0.023 | 0.036 | 0.036 | 0.443 | 0.48 | 0.21 | 0.65 | 0.65 | 0.58 | 0.44 |
| 3 | 83609319 | rs4856461 | 1.28 | -0.604 | -0.228 | 0.184 | -0.009 | 0.037 | 0.037 | 0.919 | 0.48 | 0.12 | 0.65 | 0.65 | 0.58 | 0.44 |
| 3 | 83625938 | rs1453724 | 0.977 | -0.544 | -0.2 | 0.161 | -0.016 | -0.021 | -0.021 | 0 | 0.15 | 0.08 | 0.11 | 0.17 | 0.18 | 0.18 |
| 3 | 83626697 | rs7641552 | -1.603 | -0.614 | 0.183 | 0.121 | 0.01 | -0.006 | -0.006 | 0 | 0.35 | 0.71 | 0.28 | 0.35 | 0.28 | 0.46 |
| 3 | 83629693 | rs17020896 | -0.333 | -0.654 | 0.494 | 0.105 | -0.017 | 0.023 | 0.023 | 0 | 0.08 | 0.17 | 0.05 | 0.12 | 0.03 | 0.07 |
| 3 | 83645386 | rs1598035 | -1.751 | -0.543 | 0.255 | 0.07 | -0.001 | -0.042 | -0.042 | 0.559 | 0.84 | 0.83 | 0.93 | 1 | 0.86 | 0.9 |
| 3 | 83651289 | rs6779777 | -1.86 | -0.567 | 0.24 | 0.07 | 0.003 | -0.007 | -0.007 | 0.013 | 0.84 | 0.88 | 0.93 | 1 | 0.86 | 0.9 |
| 3 | 83662948 | rs1453733 | 0.021 | -0.286 | 0.122 | 0.044 | 0 | -0.019 | -0.019 | 0 | 0.64 | 0.6 | 0.72 | 0.81 | 0.63 | 0.78 |
| 3 | 83663733 | rs9853891 | -0.845 | -0.218 | 0.347 | 0.158 | -0.003 | 0.005 | 0.005 | 0.262 | 0.32 | 0.37 | 0.21 | 0.35 | 0.28 | 0.47 |
| 3 | 83668614 | rs2045078 | 1.244 | -0.265 | 0.191 | 0.184 | 0.03 | 0.058 | 0.058 | 1.087 | 0.5 | 0.26 | 0.7 | 0.65 | 0.58 | 0.43 |
| 3 | 83699033 | rs1880463 | -1.633 | -0.147 | 0.338 | 0.171 | -0.012 | -0.085 | -0.085 | 2.256 | 0.82 | 0.76 | 0.92 | 1 | 0.86 | 0.9 |
| 3 | 83716074 | rs6548900 | -1.178 | -0.179 | 0.42 | 0.156 | -0.024 | -0.108 | -0.108 | 1.501 | 0.79 | 0.69 | 0.91 | 1 | 0.86 | 0.9 |
| 3 | 83723656 | rs6548901 | -1.459 | -0.159 | 0.348 | 0.165 | -0.014 | -0.085 | -0.085 | 0.012 | 0.81 | 0.76 | 0.92 | 1 | 0.86 | 0.9 |
| 3 | 83758828 | rs33983617 | 2.278 | -0.181 | -0.195 | 0.244 | -0.046 | -0.091 | -0.091 | 0 | 0.33 | 0.08 | 0.51 | 0.46 | 0.36 | 0.27 |
| 3 | 83761790 | rs1917376 | -0.285 | -0.136 | -0.14 | -0.061 | 0.018 | 0.004 | 0.004 | 0.016 | 0.26 | 0.46 | 0.23 | 0.22 | 0.3 | 0.38 |
| 3 | 83763541 | rs10511093 | 0.707 | -0.076 | -0.384 | -0.046 | -0.008 | -0.042 | -0.042 | 0 | 0.09 | 0.15 | 0.06 | 0 | 0.06 | 0.09 |
| 3 | 83768568 | rs17021032 | 2.439 | -0.033 | -0.581 | 0.052 | -0.098 | -0.175 | -0.175 | 0 | 0.33 | 0.01 | 0.5 | 0.48 | 0.35 | 0.28 |
| 3 | 83770825 | rs7651607 | 1.757 | -0.195 | -0.369 | -0.149 | -0.108 | 0.029 | 0.029 | 1.748 | 0.45 | 0.05 | 0.64 | 0.67 | 0.53 | 0.39 |
| 3 | 83775078 | rs1357797 | -0.129 | -0.217 | 0.259 | -0.226 | -0.031 | -0.087 | -0.087 | 0 | 0.72 | 0.45 | 0.86 | 0.9 | 0.83 | 0.74 |
| 3 | 83791220 | rs11715563 | 0.196 | 0.122 | -0.047 | -0.107 | -0.001 | -0.019 | -0.019 | 0.971 | 0.62 | 0.6 | 0.76 | 0.82 | 0.62 | 0.64 |
| 3 | 83793414 | rs1917372 | -1.91 | 0.12 | -0.333 | -0.137 | 0.002 | 0.017 | 0.017 | 0 | 0.85 | 0.93 | 0.94 | 1 | 0.87 | 0.9 |
| 3 | 83808458 | rs6768298 | 0.288 | 0.114 | -0.378 | -0.067 | -0.024 | -0.023 | -0.023 | 0 | 0.19 | 0.07 | 0.14 | 0.18 | 0.21 | 0.25 |
| 3 | 83832320 | rs6548904 | -1.519 | 0.114 | -0.013 | -0.212 | -0.005 | -0.079 | -0.079 | 5.732 | 0.82 | 0.7 | 0.91 | 1 | 0.85 | 0.9 |
| 3 | 83834878 | rs6780231 | 1.829 | -0.246 | -0.356 | -0.449 | -0.052 | 0.039 | 0.039 | 0.97 | 0.4 | 0.16 | 0.62 | 0.7 | 0.5 | 0.38 |
| 3 | 83838977 | rs4856472 | -0.318 | -0.299 | -0.234 | -0.541 | -0.001 | -0.053 | -0.053 | 0 | 0.69 | 0.32 | 0.82 | 0.99 | 0.75 | 0.7 |
| 3 | 83842624 | rs11127824 | 2.529 | -0.163 | -0.434 | -0.24 | -0.015 | -0.07 | -0.07 | 0 | 0.29 | 0.12 | 0.5 | 0.5 | 0.33 | 0.28 |
| 3 | 83843079 | rs17021121 | 0.202 | -0.146 | -0.318 | -0.166 | 0.01 | 0.01 | 0.01 | 0 | 0.13 | 0.1 | 0.05 | 0.11 | 0.07 | 0.18 |
| 3 | 83849641 | rs7617600 | 0.936 | 0.107 | -0.336 | -0.128 | -0.012 | -0.022 | -0.022 | 0 | 0.1 | 0.05 | 0.09 | 0.16 | 0.1 | 0.13 |
| 3 | 83849658 | rs7617742 | 1.319 | 0.226 | -0.267 | -0.061 | -0.016 | -0.025 | -0.025 | 0 | 0.16 | 0.05 | 0.12 | 0.16 | 0.18 | 0.14 |
| 3 | 83856348 | rs4330264 | -1.966 | 0.115 | 0.15 | -0.115 | 0 | -0.102 | -0.102 | 1.387 | 0.84 | 0.72 | 0.92 | 1 | 0.86 | 0.9 |
| 3 | 83860129 | rs4383539 | 1.508 | 0.004 | 0.039 | -0.145 | -0.011 | -0.01 | -0.01 | 0 | 0.4 | 0.23 | 0.56 | 0.61 | 0.4 | 0.43 |
| 3 | 83863312 | rs4856473 | -2.161 | 0.103 | 0.113 | -0.362 | -0.001 | -0.096 | -0.096 | 0.266 | 0.85 | 0.74 | 0.93 | 1 | 0.86 | 0.9 |
| 3 | 83870864 | rs4613466 | -0.429 | 0.125 | 0.114 | -0.369 | 0.008 | -0.016 | -0.016 | 0 | 0.71 | 0.7 | 0.77 | 0.78 | 0.66 | 0.79 |
| 3 | 83879054 | rs10470566 | 0.913 | 0.001 | 0.199 | 0.174 | -0.016 | 0.028 | 0.028 | 0 | 0.2 | 0.25 | 0.13 | 0.35 | 0.16 | 0.17 |
| 3 | 83884502 | rs4547715 | 1.1 | 0.233 | -0.03 | -0.014 | -0.013 | -0.023 | -0.023 | 1.237 | 0.12 | 0.07 | 0.12 | 0.22 | 0.16 | 0.09 |
| 3 | 83903609 | rs4241420 | -1.103 | 0.146 | 0.151 | -0.185 | 0.007 | -0.089 | -0.089 | 0.754 | 0.79 | 0.79 | 0.87 | 1 | 0.79 | 0.82 |
| 3 | 83904413 | rs4589949 | -0.497 | 0.181 | 0.108 | -0.192 | 0.005 | -0.067 | -0.067 | 1.096 | 0.74 | 0.73 | 0.75 | 0.98 | 0.74 | 0.71 |
| 3 | 83907715 | rs9841953 | 1.333 | 0.22 | 0.084 | -0.13 | -0.002 | -0.001 | -0.001 | 0 | 0.11 | 0.09 | 0.13 | 0.21 | 0.16 | 0.09 |
| 3 | 83907981 | rs7628076 | 0.936 | 0.375 | -0.003 | 0.04 | 0.003 | -0.008 | -0.008 | 0 | 0.21 | 0.22 | 0.19 | 0.42 | 0.2 | 0.2 |
| 3 | 83909132 | rs4401373 | -0.399 | 0.549 | -0.155 | -0.02 | 0.01 | -0.107 | -0.107 | 0 | 0.73 | 0.6 | 0.74 | 0.98 | 0.74 | 0.71 |
| 3 | 83920089 | rs4270515 | 2.314 | 0.485 | -1.022 | 0.124 | -0.086 | -0.053 | -0.053 | 0.146 | 0.25 | 0.01 | 0.33 | 0.23 | 0.29 | 0.21 |
| 3 | 83924470 | rs4314181 | -0.527 | 0.566 | -0.815 | 0.163 | 0.025 | -0.032 | -0.032 | 0 | 0.38 | 0.21 | 0.38 | 0.57 | 0.34 | 0.52 |

| | | | | | | | | | | | | | | | | |
|---|---|---|---|---|---|---|---|---|---|---|---|---|---|---|---|---|
| 3 | 83931100 | rs6802899 | -0.11 | 0.655 | -0.481 | -0.413 | -0.028 | 0.146 | 0.146 | 0 | 0.53 | 0.73 | 0.55 | 0.86 | 0.53 | 0.52 |
| 3 | 83931598 | rs12490245 | 2.205 | 0.447 | -0.324 | -0.334 | -0.007 | -0.078 | -0.078 | 0 | 0.26 | 0.1 | 0.33 | 0.54 | 0.27 | 0.21 |
| 3 | 83938414 | rs6548921 | -1.233 | 0.458 | -0.39 | -0.462 | 0 | -0.012 | -0.012 | 0.37 | 0.95 | 0.96 | 0.95 | 1 | 0.95 | 0.95 |
| 3 | 83940448 | rs4128523 | -0.184 | 0.402 | -0.187 | -0.341 | -0.004 | 0.08 | 0.08 | 0 | 0.53 | 0.62 | 0.55 | 0.86 | 0.55 | 0.52 |
| 3 | 83943579 | rs6767363 | 0.217 | 0.476 | -0.509 | -0.259 | 0 | 0 | 0 | 0 | 0.13 | 0.14 | 0.07 | 0.12 | 0.1 | 0.19 |
| 3 | 83945154 | rs9990164 | 0.733 | 0.474 | -0.218 | -0.261 | 0.002 | -0.009 | -0.009 | 0.136 | 0.4 | 0.42 | 0.48 | 0.74 | 0.45 | 0.34 |
| 3 | 83946126 | rs9857160 | -1.042 | 0.836 | 0.085 | -0.447 | 0.007 | -0.004 | -0.004 | 0.02 | 0.28 | 0.44 | 0.23 | 0.31 | 0.3 | 0.31 |
| 3 | 83951956 | rs12629641 | -0.283 | 0.84 | -0.161 | -0.426 | 0 | 0 | 0 | 1.196 | 0.17 | 0.23 | 0.24 | 0.21 | 0.19 | 0.2 |
| 3 | 83952437 | rs4568144 | 1.345 | 0.751 | -0.476 | -0.889 | 0.003 | 0.104 | 0.104 | 1.314 | 0.47 | 0.19 | 0.53 | 0.68 | 0.47 | 0.47 |
| 3 | 83970229 | rs9853129 | 1.433 | 0.578 | -0.007 | -0.628 | -0.001 | -0.003 | -0.003 | 0 | 0.64 | 0.35 | 0.6 | 0.62 | 0.64 | 0.56 |
| 3 | 83976987 | rs6778107 | 0.704 | 0.638 | 0.027 | -0.566 | -0.005 | -0.006 | -0.006 | 0 | 0.12 | 0.09 | 0.13 | 0.17 | 0.16 | 0.12 |
| 3 | 84004695 | rs9878647 | 1.343 | 0.532 | 0.256 | -0.665 | 0 | -0.001 | -0.001 | 0 | 0.65 | 0.5 | 0.61 | 0.64 | 0.64 | 0.56 |
| 3 | 84006037 | rs7632535 | 0.541 | 0.499 | 0.343 | -0.672 | 0.067 | 0.125 | 0.125 | 0 | 0.92 | 0.66 | 0.83 | 0.92 | 0.94 | 0.86 |
| 3 | 84017630 | rs4856477 | 0.675 | 0.508 | 0.34 | -0.678 | 0.088 | 0.11 | 0.11 | 0 | 0.91 | 0.67 | 0.83 | 0.92 | 0.92 | 0.86 |
| 3 | 84032058 | rs1997254 | 0.898 | 0.315 | 0.217 | -0.701 | -0.008 | 0.005 | 0.005 | 0 | 0.79 | 0.57 | 0.75 | 0.81 | 0.82 | 0.68 |
| 3 | 84037979 | rs9840698 | 1.236 | 0.764 | -0.26 | -0.896 | -0.003 | -0.02 | -0.02 | 0.134 | 0.14 | 0.15 | 0.07 | 0 | 0.17 | 0.1 |
| 3 | 84045211 | rs4856480 | -0.642 | 0.329 | -0.664 | -0.798 | -0.046 | -0.051 | -0.051 | 0 | 0.22 | 0.02 | 0.18 | 0.27 | 0.24 | 0.28 |
| 3 | 84063290 | rs6798585 | 1.056 | 0.216 | 0.395 | -0.896 | 0.002 | 0.003 | 0.003 | 0 | 0.73 | 0.7 | 0.69 | 0.69 | 0.72 | 0.62 |
| 3 | 84072534 | rs17793314 | -0.137 | 0.207 | 0.261 | -0.899 | -0.06 | -0.05 | -0.05 | 3.676 | 0.18 | 0.02 | 0.16 | 0.24 | 0.22 | 0.22 |
| 3 | 84083262 | rs2222745 | 0.422 | 0.193 | 0.205 | -1.353 | 0.011 | -0.035 | -0.035 | 0 | 0.07 | 0.16 | 0.08 | 0 | 0.08 | 0.06 |
| 3 | 84094818 | rs4856241 | 0.95 | 0.196 | 0.254 | -1.346 | 0.002 | 0.004 | 0.004 | 0 | 0.72 | 0.58 | 0.68 | 0.69 | 0.72 | 0.62 |
| 3 | 84108706 | rs12639222 | 1.216 | 0.036 | 0.541 | -1.339 | 0.009 | -0.01 | -0.01 | 0 | 0.66 | 0.55 | 0.61 | 0.69 | 0.64 | 0.56 |
| 3 | 84113708 | rs4501122 | 0.919 | 0.098 | 0.262 | -1.357 | -0.005 | 0.01 | 0.01 | 0 | 0.2 | 0.15 | 0.08 | 0 | 0.19 | 0.16 |
| 3 | 84143107 | rs4605562 | 1.591 | 0.034 | 0.146 | -1.321 | -0.008 | 0.071 | 0.071 | 5.547 | 0.43 | 0.12 | 0.5 | 0.69 | 0.45 | 0.4 |
| 3 | 84147330 | rs6778602 | 1.177 | -0.072 | 0.405 | -1.207 | -0.002 | -0.001 | -0.001 | 0 | 0.67 | 0.35 | 0.62 | 0.69 | 0.69 | 0.6 |
| 3 | 84157105 | rs6767775 | -1.738 | 0.032 | 0.005 | -1.198 | -0.077 | 0.049 | 0.049 | 2.902 | 0.54 | 0.94 | 0.48 | 0.32 | 0.51 | 0.57 |
| 3 | 84182782 | rs9852182 | -0.589 | 0.04 | 0.297 | -1.189 | 0.003 | -0.006 | -0.006 | 0 | 0.25 | 0.39 | 0.25 | 0.25 | 0.26 | 0.26 |
| 3 | 84200533 | rs6767473 | -1.609 | 0.016 | -0.073 | -1.14 | -0.009 | 0.045 | 0.045 | 0.806 | 0.55 | 0.74 | 0.55 | 0.32 | 0.53 | 0.59 |
| 3 | 84202116 | rs9822210 | 0.97 | -0.015 | -0.123 | -1.243 | -0.015 | 0.035 | 0.035 | 0 | 0.2 | 0.07 | 0.09 | 0 | 0.2 | 0.15 |
| 3 | 84211841 | rs9878280 | -1.325 | -0.079 | 0.14 | -1.097 | 0.007 | -0.006 | -0.006 | 0.613 | 0.36 | 0.83 | 0.48 | 0.32 | 0.33 | 0.43 |
| 3 | 84223725 | rs4362730 | 1.59 | -0.287 | -0.224 | -1.161 | 0.007 | 0.014 | 0.014 | 0 | 0.11 | 0.01 | 0.05 | 0.08 | 0.1 | 0.1 |
| 3 | 84229385 | rs7639920 | -0.143 | 0.249 | 0.571 | -0.944 | 0.027 | -0.056 | -0.056 | 0 | 0.1 | 0.29 | 0.11 | 0 | 0.09 | 0.05 |
| 3 | 84244787 | rs4321538 | -0.124 | 0.156 | 0.262 | -0.834 | -0.048 | -0.097 | -0.097 | 0 | 0.18 | 0.02 | 0.22 | 0.27 | 0.19 | 0.27 |
| 3 | 84245741 | rs9985384 | -0.314 | 0.254 | 0.704 | -0.724 | 0.04 | 0.054 | 0.054 | 0 | 0.06 | 0.19 | 0.15 | 0.07 | 0.07 | 0.13 |
| 3 | 84261595 | rs9832716 | -1.246 | 0.219 | 0.48 | -0.74 | 0.071 | 0.09 | 0.09 | 0 | 0.08 | 0.43 | 0.16 | 0.07 | 0.07 | 0.14 |
| 3 | 84292356 | rs9837479 | -0.264 | 0.246 | 0.572 | -0.708 | 0.03 | 0.052 | 0.052 | 0 | 0.06 | 0.2 | 0.15 | 0.07 | 0.06 | 0.14 |
| 3 | 84296569 | rs7617594 | -1.705 | 0.166 | 0.39 | -0.829 | -0.043 | -0.025 | -0.025 | 1.832 | 0.39 | 0.86 | 0.52 | 0.34 | 0.39 | 0.49 |
| 3 | 84304934 | rs4518142 | -1.181 | 0.03 | 0.537 | -0.814 | 0.015 | -0.016 | -0.016 | 0.567 | 0.3 | 0.54 | 0.36 | 0.23 | 0.31 | 0.29 |
| 3 | 84306139 | rs12636524 | -1.954 | 0.133 | 0.247 | -0.941 | -0.058 | 0.005 | 0.005 | 2.07 | 0.43 | 0.87 | 0.54 | 0.34 | 0.45 | 0.5 |
| 3 | 84325987 | rs4856501 | -0.074 | 0.175 | 0.364 | -0.919 | 0.013 | 0.085 | 0.085 | 0 | 0.83 | 0.56 | 0.8 | 0.76 | 0.85 | 0.8 |
| 3 | 84336945 | rs7638444 | -0.009 | 0.198 | 0.329 | -0.965 | 0.03 | -0.087 | -0.087 | 0 | 0.14 | 0.31 | 0.14 | 0 | 0.15 | 0.09 |
| 3 | 84339246 | rs4536842 | 0.186 | 0.215 | 0.114 | -0.913 | -0.008 | -0.004 | -0.004 | 2.082 | 0.11 | 0.04 | 0.18 | 0.11 | 0.14 | 0.2 |
| 3 | 84342855 | rs17352297 | 1.849 | 0.189 | -0.081 | -1.03 | -0.013 | -0.015 | -0.015 | 0 | 0.41 | 0.11 | 0.41 | 0.65 | 0.4 | 0.38 |
| 3 | 84357635 | rs4856502 | -0.185 | 0.339 | 0.292 | -0.924 | 0.004 | 0.032 | 0.032 | 0 | 0.95 | 0.86 | 0.89 | 0.92 | 0.95 | 0.85 |
| 3 | 84377718 | rs12636052 | -0.272 | 0.373 | 0.303 | -0.921 | 0.011 | 0.037 | 0.037 | 0 | 0.95 | 0.86 | 0.89 | 0.92 | 0.95 | 0.85 |
| 3 | 84385686 | rs7630474 | 1.071 | 0.328 | -0.107 | -0.827 | -0.001 | 0.012 | 0.012 | 0.233 | 0.7 | 0.38 | 0.6 | 0.63 | 0.69 | 0.57 |
| 3 | 84387950 | rs12714592 | 0.054 | 0.505 | -0.29 | -0.872 | -0.022 | 0.067 | 0.067 | 0 | 0.26 | 0.1 | 0.24 | 0.11 | 0.29 | 0.32 |
| 3 | 84404086 | rs17250522 | 0.34 | 0.607 | -0.071 | -0.761 | 0.04 | -0.06 | -0.06 | 0 | 0.11 | 0.25 | 0.13 | 0 | 0.13 | 0.06 |
| 3 | 84407703 | rs4572785 | -0.778 | 0.551 | -0.154 | -0.735 | 0.024 | -0.002 | -0.002 | 0 | 0.25 | 0.41 | 0.22 | 0.19 | 0.27 | 0.15 |
| 3 | 84427337 | rs9844018 | 0.331 | 0.545 | -0.44 | -0.793 | 0.017 | 0.002 | 0.002 | 1.53 | 0.76 | 0.64 | 0.8 | 0.81 | 0.75 | 0.85 |
| 3 | 84443114 | rs12714597 | 0.954 | 0.645 | -0.476 | -0.775 | -0.039 | 0.103 | 0.103 | 0 | 0.16 | 0.01 | 0.06 | 0 | 0.16 | 0.12 |
| 3 | 84451512 | rs6794880 | -0.15 | 0.748 | -0.208 | -0.585 | 0.049 | -0.053 | -0.053 | 0.213 | 0.89 | 0.76 | 0.87 | 0.98 | 0.86 | 0.9 |
| 3 | 84481068 | rs10511089 | 0.418 | 0.145 | -0.771 | -0.651 | -0.004 | -0.063 | -0.063 | 0 | 0.17 | 0.02 | 0.08 | 0.19 | 0.12 | 0.09 |
| 3 | 84494692 | rs9881708 | -1.106 | 0.157 | -0.466 | -0.926 | -0.001 | -0.037 | -0.037 | 0 | 0.83 | 0.86 | 0.92 | 1 | 0.84 | 0.89 |
| 3 | 84495097 | rs4348007 | 0.073 | 0.134 | 0.007 | -1.055 | 0.02 | -0.169 | -0.169 | 0 | 0.72 | 0.54 | 0.8 | 1 | 0.72 | 0.82 |
| 3 | 84509525 | rs4362733 | -0.052 | 0.129 | 0.049 | -1.058 | 0.012 | -0.116 | -0.116 | 2.265 | 0.73 | 0.64 | 0.81 | 1 | 0.72 | 0.83 |
| 3 | 84542267 | rs12486306 | 1.741 | -0.041 | -0.253 | -0.979 | -0.037 | -0.078 | -0.078 | 0 | 0.37 | 0.1 | 0.39 | 0.61 | 0.4 | 0.38 |
| 3 | 84556332 | rs7631338 | -0.705 | -0.023 | -0.002 | -0.96 | -0.013 | 0.018 | 0.018 | 0.028 | 0.35 | 0.54 | 0.42 | 0.4 | 0.3 | 0.44 |
| 3 | 84569395 | rs9877868 | -0.417 | 0.001 | -0.025 | -0.986 | 0.009 | -0.099 | -0.099 | 0 | 0.73 | 0.7 | 0.82 | 1 | 0.72 | 0.82 |
| 3 | 84572818 | rs17021585 | 0.101 | -0.049 | -0.187 | -0.911 | -0.005 | -0.01 | -0.01 | 0.102 | 0.15 | 0.09 | 0.3 | 0.18 | 0.18 | 0.38 |
| 3 | 84585678 | rs1485584 | 1.497 | -0.053 | -0.269 | -0.863 | -0.034 | -0.066 | -0.066 | 0 | 0.39 | 0.13 | 0.4 | 0.6 | 0.43 | 0.38 |
| 3 | 84623711 | rs12629186 | 0.936 | -0.093 | -0.156 | -0.851 | 0.014 | -0.047 | -0.047 | 0 | 0.17 | 0.04 | 0.08 | 0.22 | 0.11 | 0.07 |
| 3 | 84630305 | rs9872666 | 0.732 | -0.106 | 0.067 | -0.857 | -0.002 | 0.001 | 0.001 | 0.209 | 0.18 | 0.16 | 0.09 | 0.22 | 0.12 | 0.07 |
| 3 | 84644186 | rs6810190 | 1.511 | -0.063 | -0.13 | -0.889 | -0.029 | -0.056 | -0.056 | 0 | 0.38 | 0.15 | 0.4 | 0.6 | 0.43 | 0.38 |
| 3 | 84665102 | rs2326036 | -0.641 | -0.096 | 0.098 | -0.937 | -0.022 | 0.027 | 0.027 | 0.046 | 0.34 | 0.51 | 0.42 | 0.4 | 0.29 | 0.45 |
| 3 | 84673260 | rs6782828 | -0.278 | 0.046 | 0.165 | -1.005 | 0.009 | -0.099 | -0.099 | 0 | 0.73 | 0.68 | 0.82 | 1 | 0.72 | 0.82 |
| 3 | 84679083 | rs7650286 | 0.916 | 0.099 | 0.125 | -0.993 | 0.011 | -0.032 | -0.032 | 0 | 0.09 | 0.14 | 0.1 | 0 | 0.12 | 0.06 |
| 3 | 84688173 | rs901134 | 1.378 | -0.148 | -0.093 | -0.65 | -0.041 | -0.082 | -0.082 | 0.816 | 0.4 | 0.1 | 0.4 | 0.58 | 0.42 | 0.4 |
| 3 | 84696539 | rs9860330 | -0.991 | -0.064 | 0.202 | -0.492 | 0 | -0.004 | -0.004 | 0.176 | 0.42 | 0.65 | 0.5 | 0.42 | 0.39 | 0.48 |

| | | | | | | | | | | | | | | | | |
|---|---|---|---|---|---|---|---|---|---|---|---|---|---|---|---|---|
| 3 | 84707940 | rs9809799 | 0.552 | -0.239 | 0.539 | -0.379 | -0.029 | 0.055 | 0.055 | 0.714 | 0.18 | 0.29 | 0.17 | 0.3 | 0.09 | 0.1 |
| 3 | 84709148 | rs6806546 | -0.345 | 0.163 | 0.778 | -0.69 | 0.012 | -0.076 | -0.076 | 0 | 0.24 | 0.34 | 0.23 | 0.04 | 0.31 | 0.31 |
| 3 | 84716208 | rs4374542 | -0.588 | -0.303 | 0.216 | -0.557 | 0.052 | 0.011 | 0.011 | 0 | 0.57 | 0.84 | 0.81 | 0.67 | 0.66 | 0.71 |
| 3 | 84730413 | rs4582072 | -0.295 | -0.203 | 0.138 | -0.613 | -0.016 | 0.032 | 0.032 | 0 | 0.77 | 0.83 | 0.79 | 0.96 | 0.69 | 0.69 |
| 3 | 84742570 | rs9835484 | 1.719 | -0.135 | -0.218 | -0.724 | -0.051 | 0.076 | 0.076 | 0 | 0.13 | 0.01 | 0.06 | 0 | 0.14 | 0.12 |
| 3 | 84743057 | rs4426692 | 0.386 | -0.105 | 0.265 | -0.414 | 0.009 | -0.003 | -0.003 | 0 | 0.46 | 0.4 | 0.3 | 0.42 | 0.36 | 0.33 |
| 3 | 84746287 | rs4572783 | 0.279 | -0.082 | 0.216 | -0.416 | 0.016 | -0.087 | -0.087 | 0.834 | 0.72 | 0.44 | 0.59 | 0.89 | 0.67 | 0.66 |
| 3 | 84747008 | rs3890731 | 1.069 | 0.184 | 0.258 | -0.05 | -0.004 | -0.007 | -0.007 | 0 | 0.28 | 0.25 | 0.31 | 0.48 | 0.31 | 0.33 |
| 3 | 84779079 | rs9820177 | -0.715 | 0.273 | 0.235 | 0.191 | -0.005 | -0.022 | -0.022 | 0 | 0.31 | 0.73 | 0.47 | 0.49 | 0.36 | 0.43 |
| 3 | 84785549 | rs16878 | 1.706 | 0.334 | -0.203 | -0.081 | -0.009 | 0.053 | 0.053 | 0.629 | 0.23 | 0.15 | 0.12 | 0.1 | 0.23 | 0.21 |
| 3 | 84790582 | rs7647725 | 1.479 | 0.296 | 0.088 | -0.098 | -0.001 | -0.02 | -0.02 | 0 | 0.24 | 0.28 | 0.13 | 0.1 | 0.23 | 0.21 |
| 3 | 84806803 | rs17501615 | 0.066 | 0.289 | 0.268 | -0.209 | -0.009 | 0.006 | 0.006 | 0 | 0.25 | 0.21 | 0.2 | 0.05 | 0.32 | 0.34 |
| 3 | 84822412 | rs9839064 | 1.004 | 0.255 | 0.176 | 0.037 | -0.002 | 0.086 | 0.086 | 0.866 | 0.28 | 0.47 | 0.35 | 0.51 | 0.26 | 0.3 |
| 3 | 84832334 | rs1584859 | -1.054 | 0.312 | -1.627 | -0.073 | 0.011 | 0.01 | 0.01 | 0 | 0.62 | 0.87 | 0.79 | 0.72 | 0.66 | 0.76 |
| 3 | 84833354 | rs1454094 | 1.205 | 0.516 | -1.106 | 0.166 | -0.028 | 0.07 | 0.07 | 0.841 | 0.22 | 0.06 | 0.12 | 0.1 | 0.23 | 0.22 |
| 3 | 84835951 | rs7618782 | -0.174 | 0.911 | -1.075 | 0.36 | -0.005 | -0.017 | -0.017 | 0 | 0.41 | 0.74 | 0.57 | 0.56 | 0.45 | 0.56 |
| 3 | 84836382 | rs9876866 | 0.309 | 0.869 | -1.062 | 0.765 | -0.023 | -0.048 | -0.048 | 0 | 0.86 | 0.92 | 0.7 | 0.47 | 0.78 | 0.66 |
| 3 | 84838494 | rs13082591 | 1.219 | 1.02 | -1.489 | 0.178 | -0.035 | 0.064 | 0.064 | 0 | 0.13 | 0.03 | 0.05 | 0.01 | 0.1 | 0.15 |
| 3 | 84838638 | rs980332 | -0.127 | 1.112 | -1.185 | 0.152 | 0.001 | 0.02 | 0.02 | 0 | 0.84 | 0.83 | 0.67 | 0.46 | 0.77 | 0.64 |
| 3 | 84838697 | rs980331 | 0.703 | 1.053 | -0.899 | 0.138 | -0.008 | -0.009 | -0.009 | 2.361 | 0.71 | 0.72 | 0.61 | 0.46 | 0.67 | 0.5 |
| 3 | 84852101 | rs1454089 | -0.515 | 1.144 | -1.072 | 0.689 | -0.047 | -0.088 | -0.088 | 1.593 | 0.88 | 0.99 | 0.92 | 0.64 | 0.82 | 0.74 |
| 3 | 84856220 | rs2647716 | -0.071 | 1.069 | -1.056 | 0.482 | -0.024 | -0.059 | -0.059 | 1.14 | 0.84 | 0.92 | 0.68 | 0.47 | 0.77 | 0.65 |
| 3 | 84857927 | rs13092059 | 1.081 | 1.032 | -0.29 | 0.372 | 0.01 | -0.066 | -0.066 | 0.085 | 0.14 | 0.24 | 0.07 | 0.01 | 0.1 | 0.14 |
| 3 | 84865117 | rs2326199 | 1.241 | 0.117 | -0.299 | -0.213 | 0.009 | 0.038 | 0.038 | 0 | 0.55 | 0.62 | 0.4 | 0.26 | 0.59 | 0.36 |
| 3 | 84868834 | rs395807 | -0.049 | 0.017 | -0.319 | -0.25 | 0.005 | 0.005 | 0.005 | 0 | 0.29 | 0.29 | 0.28 | 0.21 | 0.18 | 0.29 |
| 3 | 84870750 | rs955644 | 1.219 | 0.154 | 0.016 | -0.388 | -0.001 | -0.005 | -0.005 | 0 | 0.55 | 0.53 | 0.39 | 0.26 | 0.59 | 0.37 |
| 3 | 84872804 | rs10440084 | 1.444 | 0.204 | -0.028 | -0.68 | 0 | -0.001 | -0.001 | 0.059 | 0.27 | 0.23 | 0.23 | 0.25 | 0.31 | 0.16 |
| 3 | 84873993 | rs1146762 | 0.202 | 0.193 | 0.025 | -0.71 | -0.014 | -0.043 | -0.043 | 0.036 | 0.28 | 0.37 | 0.25 | 0.16 | 0.18 | 0.25 |
| 3 | 84876250 | rs9877971 | 2.11 | 0.173 | 0.284 | -0.732 | -0.025 | 0.066 | 0.066 | 0 | 0.28 | 0.14 | 0.15 | 0.01 | 0.28 | 0.21 |
| 3 | 84876486 | rs1146760 | 0.192 | 0.16 | 0.329 | -0.731 | 0 | -0.007 | -0.007 | 0 | 0.29 | 0.31 | 0.23 | 0.16 | 0.18 | 0.24 |
| 3 | 84879722 | rs9823132 | 0.229 | 0.172 | 0.281 | -0.73 | -0.002 | 0.004 | 0.004 | 0 | 0.57 | 0.6 | 0.41 | 0.16 | 0.46 | 0.46 |
| 3 | 84879887 | rs13074892 | 2.184 | 0.231 | 0.168 | -0.7 | -0.077 | 0.165 | 0.165 | 0.003 | 0.27 | 0.06 | 0.15 | 0.01 | 0.28 | 0.21 |
| 3 | 84887157 | rs1146755 | 0.944 | 0.022 | 0.296 | -0.828 | 0.006 | -0.018 | -0.018 | 0.059 | 0.08 | 0.12 | 0.15 | 0.05 | 0.09 | 0.11 |
| 3 | 84889597 | rs1146751 | -0.967 | -0.026 | 0.227 | -0.83 | -0.011 | 0.016 | 0.016 | 0 | 0.56 | 0.67 | 0.49 | 0.41 | 0.5 | 0.41 |
| 3 | 84891973 | rs401242 | -0.888 | -0.009 | 0.019 | -0.781 | -0.016 | -0.046 | -0.046 | 0 | 0.84 | 0.9 | 0.64 | 0.41 | 0.77 | 0.62 |
| 3 | 84893668 | rs1603201 | 2.01 | -0.132 | -0.147 | -0.95 | -0.001 | 0.01 | 0.01 | 0 | 0.17 | 0.16 | 0.15 | 0.29 | 0.17 | 0.13 |
| 3 | 84897368 | rs6802379 | 0.993 | -0.138 | -0.135 | -0.986 | -0.005 | -0.009 | -0.009 | 0 | 0.2 | 0.21 | 0.1 | 0.11 | 0.1 | 0.14 |
| 3 | 84911212 | rs1032506 | 1.061 | -0.144 | -0.199 | -0.988 | 0.028 | 0.038 | 0.038 | 0 | 0.19 | 0.12 | 0.08 | 0.11 | 0.09 | 0.14 |
| 3 | 84920368 | rs7642644 | 2.451 | -0.129 | -0.183 | -1.015 | -0.001 | -0.003 | -0.003 | 0 | 0.1 | 0.08 | 0.08 | 0 | 0.15 | 0.03 |
| 3 | 84926472 | rs12495388 | 0.965 | -0.166 | -0.114 | -0.99 | -0.004 | -0.008 | -0.008 | 0 | 0.2 | 0.21 | 0.1 | 0.11 | 0.09 | 0.14 |
| 3 | 84927805 | rs12493342 | 1.638 | -0.188 | -0.301 | -0.935 | 0.007 | 0.003 | 0.003 | 0.09 | 0.11 | 0.12 | 0.06 | 0.11 | 0.05 | 0.09 |
| 3 | 84930139 | rs9847448 | 1.34 | -0.225 | -0.228 | -0.915 | 0.004 | -0.006 | -0.006 | 0 | 0.25 | 0.2 | 0.18 | 0.29 | 0.21 | 0.17 |
| 3 | 84931731 | rs7649296 | 1.427 | -0.165 | -0.212 | -1.035 | -0.001 | 0.007 | 0.007 | 0.18 | 0.39 | 0.35 | 0.22 | 0.12 | 0.33 | 0.3 |
| 3 | 84932956 | rs1248816 | 0.794 | -0.153 | -0.122 | -0.931 | 0.038 | -0.024 | -0.024 | 0 | 0.09 | 0.26 | 0.16 | 0.05 | 0.08 | 0.11 |
| 3 | 84940927 | rs452496 | 0.251 | -0.147 | -0.293 | -0.894 | -0.014 | 0.007 | 0.007 | 0 | 0.34 | 0.47 | 0.34 | 0.33 | 0.29 | 0.28 |
| 3 | 84951716 | rs1694933 | -1.209 | -0.122 | -0.369 | -0.926 | -0.007 | -0.003 | -0.003 | 0.032 | 0.56 | 0.67 | 0.47 | 0.45 | 0.49 | 0.4 |
| 3 | 84954203 | rs1691483 | -0.578 | -0.149 | -0.363 | -0.888 | 0.009 | 0 | 0 | 0 | 0.45 | 0.59 | 0.4 | 0.45 | 0.34 | 0.37 |
| 3 | 84961750 | rs1614844 | 0.795 | -0.19 | -0.55 | -0.911 | -0.003 | -0.004 | -0.004 | 0 | 0.6 | 0.51 | 0.49 | 0.34 | 0.57 | 0.48 |
| 3 | 84963982 | rs7633333 | -0.237 | -0.339 | -0.404 | -1.142 | -0.014 | 0.016 | 0.016 | 0 | 0.27 | 0.55 | 0.39 | 0.28 | 0.25 | 0.22 |
| 3 | 84964399 | rs7625608 | 0.997 | -0.354 | -0.337 | -1.148 | -0.019 | -0.006 | -0.006 | 0.122 | 0.13 | 0.38 | 0.07 | 0.11 | 0.05 | 0.09 |
| 3 | 84965355 | rs9849768 | 2.169 | -0.185 | -0.706 | -1.233 | -0.098 | 0.186 | 0.186 | 0 | 0.3 | 0.04 | 0.17 | 0.01 | 0.34 | 0.24 |
| 3 | 84966018 | rs382210 | -0.649 | -0.221 | -0.633 | -1.187 | 0.013 | -0.011 | -0.011 | 0 | 0.65 | 0.9 | 0.73 | 0.34 | 0.68 | 0.56 |
| 3 | 84970988 | rs12496982 | -0.34 | -0.016 | -0.329 | -0.811 | -0.014 | 0.018 | 0.018 | 0 | 0.27 | 0.55 | 0.39 | 0.28 | 0.25 | 0.22 |
| 3 | 84977191 | rs9683309 | -0.34 | -0.016 | -0.329 | -0.833 | -0.014 | 0.016 | 0.016 | 0 | 0.27 | 0.55 | 0.39 | 0.28 | 0.25 | 0.22 |
| 3 | 84978010 | rs11127866 | -0.36 | -0.012 | -0.351 | -0.818 | -0.015 | 0.01 | 0.01 | 0 | 0.27 | 0.56 | 0.39 | 0.27 | 0.25 | 0.21 |
| 3 | 84983294 | rs9683133 | -0.395 | -0.018 | -0.41 | -0.841 | -0.016 | 0.012 | 0.012 | 0 | 0.27 | 0.55 | 0.39 | 0.28 | 0.25 | 0.24 |
| 3 | 84987790 | rs12489914 | -0.29 | 0.316 | -0.2 | -0.638 | 0.006 | 0.056 | 0.056 | 0 | 0.21 | 0.55 | 0.38 | 0.28 | 0.23 | 0.23 |
| 3 | 84993411 | rs1248825 | -0.944 | 0.205 | -0.651 | -0.722 | -0.007 | -0.061 | -0.061 | 0 | 0.67 | 0.93 | 0.77 | 0.42 | 0.68 | 0.61 |
| 3 | 84995928 | rs1248869 | -0.314 | 0.293 | -0.289 | -0.675 | -0.002 | 0.004 | 0.004 | 0 | 0.1 | 0.08 | 0.17 | 0.05 | 0.08 | 0.12 |
| 3 | 85003523 | rs1691473 | -0.417 | 0.298 | -0.232 | -0.797 | 0.001 | 0.002 | 0.002 | 0 | 0.1 | 0.09 | 0.17 | 0.05 | 0.07 | 0.11 |
| 3 | 85006021 | rs450495 | -1.425 | 0.265 | -0.033 | -0.87 | -0.001 | -0.009 | -0.009 | 2.368 | 0.13 | 0.6 | 0.11 | 0.12 | 0.09 | 0.13 |
| 3 | 85009817 | rs9870384 | 1 | 0.26 | -0.472 | -1.083 | -0.068 | 0.163 | 0.163 | 0 | 0.36 | 0.12 | 0.22 | 0.14 | 0.39 | 0.32 |
| 3 | 85016560 | rs17022126 | 0.252 | 0.382 | -0.313 | -0.909 | 0.003 | 0.008 | 0.008 | 0 | 0.07 | 0.05 | 0.13 | 0.03 | 0.04 | 0.06 |
| 3 | 85022821 | rs4856554 | 0.766 | 0.514 | 0.306 | -0.967 | -0.006 | 0.017 | 0.017 | 0 | 0.4 | 0.3 | 0.32 | 0.06 | 0.39 | 0.32 |
| 3 | 85023651 | rs1248853 | -0.044 | 0.536 | 0.404 | -0.931 | 0.084 | 0.028 | 0.028 | 0.355 | 0.05 | 0.3 | 0.05 | 0 | 0.05 | 0.06 |
| 3 | 85040284 | rs9871198 | 0.304 | 0.48 | 0.187 | -0.897 | 0.001 | -0.044 | -0.044 | 0 | 0.46 | 0.5 | 0.38 | 0.17 | 0.43 | 0.39 |
| 3 | 85049088 | rs1872552 | 0.826 | 0.469 | 0.055 | -0.819 | -0.001 | -0.002 | -0.002 | 0 | 0.4 | 0.4 | 0.44 | 0.2 | 0.39 | 0.38 |
| 3 | 85057281 | rs4301023 | -0.08 | 0.514 | -0.2 | -0.774 | 0.004 | 0.034 | 0.034 | 0 | 0.51 | 0.63 | 0.62 | 0.34 | 0.47 | 0.51 |
| 3 | 85072584 | rs2200472 | -0.436 | 0.674 | -0.287 | -0.56 | 0 | 0.002 | 0.002 | 0 | 0.1 | 0.14 | 0.17 | 0.14 | 0.07 | 0.12 |

| | | | | | | | | | | | | | | | | |
|---|---|---|---|---|---|---|---|---|---|---|---|---|---|---|---|---|
| 3 | 85082588 | rs1016306 | -0.644 | 0.573 | -0.813 | -0.93 | -0.038 | 0.005 | 0.005 | 0.015 | 0.62 | 0.91 | 0.68 | 0.33 | 0.55 | 0.56 |
| 3 | 85084508 | rs2171143 | 1.53 | 0.456 | -0.8 | -0.957 | -0.1 | 0.202 | 0.202 | 0 | 0.32 | 0.06 | 0.18 | 0.03 | 0.37 | 0.26 |
| 3 | 85086403 | rs6549009 | -0.891 | 0.462 | -0.4 | -0.399 | -0.039 | -0.102 | -0.102 | 0 | 0.74 | 0.96 | 0.78 | 0.41 | 0.71 | 0.6 |
| 3 | 85090935 | rs6801271 | -0.128 | 0.469 | 0.083 | -0.413 | -0.008 | -0.004 | -0.004 | 2.161 | 0.65 | 0.59 | 0.63 | 0.38 | 0.66 | 0.54 |
| 3 | 85093629 | rs11914646 | -0.996 | 0.507 | 0.513 | -0.518 | 0.021 | -0.013 | -0.013 | 0.41 | 0.11 | 0.37 | 0.18 | 0.06 | 0.05 | 0.09 |
| 3 | 85096492 | rs6549011 | 0.086 | 0.459 | 0.541 | -0.7 | -0.002 | -0.012 | -0.012 | 0 | 0.62 | 0.62 | 0.41 | 0.21 | 0.68 | 0.46 |
| 3 | 85138172 | rs9820587 | 0.071 | 0.379 | 0.28 | -0.725 | -0.002 | -0.01 | -0.01 | 0 | 0.62 | 0.61 | 0.42 | 0.21 | 0.68 | 0.46 |
| 3 | 85140944 | rs11127878 | -0.237 | 0.467 | 0.213 | -0.047 | -0.005 | 0.093 | 0.093 | 0 | 0.07 | 0.13 | 0.35 | 0.24 | 0.03 | 0.15 |
| 3 | 85147735 | rs4856561 | -0.187 | 0.475 | 0.212 | -0.046 | -0.001 | 0.09 | 0.09 | 0 | 0.07 | 0.13 | 0.36 | 0.24 | 0.04 | 0.15 |
| 3 | 85163167 | rs6773806 | -0.751 | 0.48 | 0.453 | -0.12 | 0.081 | 0.045 | 0.045 | 0.229 | 0.06 | 0.32 | 0.05 | 0.04 | 0.04 | 0.05 |
| 3 | 85200179 | rs17022545 | 1.467 | 0.372 | -0.398 | 0.253 | 0.005 | -0.053 | -0.053 | 0 | 0.17 | 0.02 | 0.11 | 0.21 | 0.13 | 0.12 |
| 3 | 85248477 | rs9810830 | -1.063 | 0.272 | -0.32 | -0.017 | -0.013 | 0.01 | 0.01 | 0.017 | 0.17 | 0.58 | 0.1 | 0.19 | 0.1 | 0.14 |
| 3 | 85263778 | rs7630177 | -1.063 | 0.328 | -0.314 | -0.016 | -0.01 | 0.012 | 0.012 | 0.017 | 0.17 | 0.58 | 0.1 | 0.19 | 0.1 | 0.14 |
| 3 | 85304062 | rs1551213 | 1.143 | 0.554 | -0.674 | -0.309 | -0.068 | 0.099 | 0.099 | 0 | 0.44 | 0.14 | 0.61 | 0.26 | 0.54 | 0.38 |
| 3 | 85336559 | rs7637876 | -0.767 | 0.72 | -0.491 | -0.229 | -0.014 | 0.017 | 0.017 | 0 | 0.16 | 0.56 | 0.1 | 0.18 | 0.1 | 0.14 |
| 3 | 85349089 | rs1452122 | 0.292 | 0.649 | -0.454 | -0.25 | 0 | 0.024 | 0.024 | 0 | 0.1 | 0.19 | 0.04 | 0.14 | 0.07 | 0.09 |
| 3 | 85352957 | rs11926317 | -0.418 | 0.647 | -0.577 | -0.295 | -0.025 | -0.045 | -0.045 | 0.01 | 0.63 | 0.85 | 0.73 | 0.44 | 0.64 | 0.53 |
| 3 | 85358043 | rs6780644 | -1.046 | 0.864 | -0.272 | 0.035 | -0.012 | 0.018 | 0.018 | 0 | 0.17 | 0.59 | 0.11 | 0.19 | 0.11 | 0.14 |
| 3 | 85362101 | rs2290338 | -1.196 | 1.047 | -0.788 | 0.128 | -0.067 | -0.106 | -0.106 | 0 | 0.83 | 0.99 | 0.86 | 0.66 | 0.79 | 0.63 |
| 3 | 85371400 | rs818215 | -0.554 | 0.802 | -0.413 | -0.282 | 0.002 | 0.03 | 0.03 | 0 | 0.5 | 0.78 | 0.64 | 0.45 | 0.47 | 0.47 |
| 3 | 85372814 | rs818217 | -0.264 | 0.962 | -0.409 | -0.634 | 0 | 0.028 | 0.028 | 0 | 0.09 | 0.19 | 0.04 | 0.14 | 0.07 | 0.09 |
| 3 | 85391672 | rs9811546 | 0.68 | 0.884 | -0.817 | -1.01 | -0.019 | 0.094 | 0.094 | 0 | 0.29 | 0.12 | 0.35 | 0.1 | 0.35 | 0.29 |
| 3 | 85402469 | rs9861237 | 0.144 | 0.951 | -0.261 | -1.104 | 0.021 | -0.11 | -0.11 | 0.007 | 0.33 | 0.45 | 0.4 | 0.12 | 0.39 | 0.37 |
| 3 | 85404783 | rs2117152 | 0.285 | 0.389 | -0.292 | -1.275 | -0.002 | -0.083 | -0.083 | 0.256 | 0.62 | 0.43 | 0.59 | 0.83 | 0.64 | 0.53 |
| 3 | 85405501 | rs9822731 | 1.261 | -0.102 | -0.126 | -1.093 | -0.047 | 0.13 | 0.13 | 0 | 0.21 | 0.02 | 0.15 | 0.03 | 0.23 | 0.18 |
| 3 | 85406367 | rs12498010 | 0.258 | -0.087 | 0.212 | -1.084 | -0.004 | -0.035 | -0.035 | 0 | 0.29 | 0.37 | 0.32 | 0.12 | 0.24 | 0.34 |
| 3 | 85414291 | rs7627971 | 0.111 | -0.074 | 0.193 | -1.083 | -0.017 | -0.089 | -0.089 | 0.036 | 0.67 | 0.44 | 0.64 | 0.83 | 0.71 | 0.57 |
| 3 | 85428963 | rs1433711 | 0.119 | -0.159 | 0.132 | -1.16 | 0.007 | 0.016 | 0.016 | 0 | 0.3 | 0.3 | 0.32 | 0.12 | 0.24 | 0.34 |
| 3 | 85430839 | rs9824840 | -0.147 | 0.019 | 0.262 | -1.28 | -0.004 | -0.085 | -0.085 | 0.305 | 0.83 | 0.47 | 0.88 | 0.95 | 0.87 | 0.82 |
| 3 | 85460131 | rs1549979 | 0.529 | 0.011 | -0.014 | -1.199 | 0 | 0.262 | 0.262 | 0.183 | 0.55 | 0.08 | 0.58 | 0.83 | 0.62 | 0.48 |
| 3 | 85477017 | rs2326309 | 0.527 | 0.005 | 0.413 | -1.218 | 0.002 | -0.012 | -0.012 | 0 | 0.57 | 0.37 | 0.6 | 0.83 | 0.62 | 0.48 |
| 3 | 85482661 | rs35827242 | 0.525 | 0.005 | 0.334 | -1.218 | 0.019 | 0.221 | 0.221 | 0.172 | 0.55 | 0.12 | 0.58 | 0.83 | 0.62 | 0.48 |
| 3 | 85496758 | rs2033526 | 0.525 | 0.005 | 0.334 | -1.218 | 0.019 | 0.221 | 0.221 | 0.172 | 0.55 | 0.12 | 0.58 | 0.83 | 0.62 | 0.48 |
| 3 | 85499333 | rs993136 | 0.622 | 0.018 | 0.409 | -1.204 | 0.017 | 0.118 | 0.118 | 0.005 | 0.56 | 0.23 | 0.59 | 0.83 | 0.62 | 0.48 |
| 3 | 85504982 | rs1433713 | -0.711 | 0.015 | 0.434 | -1.198 | 0.001 | -0.028 | -0.028 | 0 | 0.87 | 0.78 | 0.91 | 0.95 | 0.87 | 0.82 |
| 3 | 85515031 | rs12485813 | -0.208 | -0.067 | 0.396 | -1.172 | -0.01 | 0.001 | 0.001 | 0 | 0.07 | 0.22 | 0.14 | 0.08 | 0.02 | 0.15 |
| 3 | 85518580 | rs2875907 | 0.034 | -0.079 | 0.375 | -1.236 | 0.003 | -0.011 | -0.011 | 0 | 0.63 | 0.38 | 0.73 | 0.93 | 0.65 | 0.64 |
| 3 | 85519957 | rs892366 | 1.549 | -0.219 | -0.162 | -0.931 | -0.047 | -0.072 | -0.072 | 0 | 0.28 | 0.04 | 0.26 | 0.33 | 0.26 | 0.26 |
| 3 | 85525323 | rs12637791 | -0.308 | -0.198 | -0.114 | -0.923 | -0.003 | -0.075 | -0.075 | 0.084 | 0.63 | 0.44 | 0.74 | 0.92 | 0.65 | 0.64 |
| 3 | 85533359 | rs7637879 | 0.788 | -0.208 | -0.597 | -0.925 | -0.037 | -0.07 | -0.07 | 0.246 | 0.28 | 0.12 | 0.33 | 0.52 | 0.37 | 0.25 |
| 3 | 85551820 | rs12638798 | -0.281 | -0.201 | -0.551 | -0.972 | 0.003 | -0.017 | -0.017 | 0.808 | 0.63 | 0.38 | 0.73 | 0.94 | 0.65 | 0.64 |
| 3 | 85570561 | rs10511084 | 0.742 | -0.136 | -1.175 | -0.893 | -0.037 | -0.065 | -0.065 | 0 | 0.28 | 0.12 | 0.33 | 0.54 | 0.37 | 0.25 |
| 3 | 85577752 | rs10511079 | 1.508 | -0.129 | -1.195 | -0.88 | -0.073 | -0.095 | -0.095 | 0.022 | 0.27 | 0.01 | 0.26 | 0.32 | 0.26 | 0.25 |
| 3 | 85591467 | rs1449403 | 1.052 | -0.023 | -0.704 | -1.195 | -0.029 | 0.035 | 0.035 | 0 | 0.11 | 0 | 0.06 | 0.05 | 0.11 | 0.18 |
| 3 | 85594695 | rs10865610 | 0.468 | 0.045 | -0.63 | -1.163 | 0.014 | 0.062 | 0.062 | 0 | 0.57 | 0.3 | 0.6 | 0.86 | 0.63 | 0.49 |
| 3 | 85603104 | rs1449409 | 0.619 | -0.037 | -1.033 | -1.056 | 0.023 | 0.042 | 0.042 | 0.227 | 0.76 | 0.17 | 0.75 | 0.87 | 0.86 | 0.68 |
| 3 | 85608137 | rs1375547 | 1.268 | 0.059 | -1.217 | -1.122 | -0.043 | 0.126 | 0.126 | 0 | 0.21 | 0.03 | 0.15 | 0.01 | 0.22 | 0.18 |
| 3 | 85617450 | rs7623947 | -0.717 | 0.093 | -0.652 | -0.905 | -0.001 | -0.019 | -0.019 | 0 | 0.87 | 0.82 | 0.92 | 0.95 | 0.88 | 0.82 |
| 3 | 85653433 | rs10511074 | 1.277 | -0.162 | -1.124 | -0.909 | -0.054 | 0.066 | 0.066 | 0.006 | 0.28 | 0.1 | 0.24 | 0.23 | 0.28 | 0.25 |
| 3 | 85656222 | rs9812845 | 1.103 | -0.38 | -1.261 | -1.141 | -0.025 | 0.092 | 0.092 | 0 | 0.23 | 0.08 | 0.27 | 0.09 | 0.25 | 0.26 |
| 3 | 85657609 | rs4856599 | -0.591 | -0.582 | -0.724 | -0.96 | -0.039 | -0.019 | -0.019 | 0.276 | 0.83 | 0.41 | 0.9 | 0.9 | 0.9 | 0.84 |
| 3 | 85658202 | rs1375562 | 0.042 | -0.652 | -0.722 | -1.004 | -0.006 | -0.004 | -0.004 | 0 | 0.8 | 0.27 | 0.89 | 0.9 | 0.89 | 0.8 |
| 3 | 85661265 | rs11928368 | 0.381 | -0.736 | -0.834 | -1.048 | -0.004 | 0 | 0 | 0 | 0.31 | 0.22 | 0.37 | 0.6 | 0.34 | 0.31 |
| 3 | 85663849 | rs6779752 | 0.363 | -0.721 | -0.787 | -0.982 | 0.038 | 0.151 | 0.151 | 0 | 0.57 | 0.2 | 0.62 | 0.82 | 0.64 | 0.55 |
| 3 | 85667683 | rs1449395 | 1.229 | -0.674 | -0.796 | -0.951 | -0.032 | 0.096 | 0.096 | 0 | 0.23 | 0.06 | 0.27 | 0.09 | 0.25 | 0.26 |
| 3 | 85669226 | rs9816536 | -0.264 | -0.663 | -0.805 | -0.943 | 0.016 | 0.053 | 0.053 | 0 | 0.4 | 0.68 | 0.38 | 0.18 | 0.35 | 0.42 |
| 3 | 85676752 | rs9867437 | -0.69 | -0.603 | -0.465 | -0.844 | 0.016 | 0.067 | 0.067 | 0 | 0.48 | 0.63 | 0.47 | 0.69 | 0.47 | 0.49 |
| 3 | 85690143 | rs4507269 | 0.781 | -0.422 | -0.806 | -0.569 | -0.008 | -0.071 | -0.071 | 0 | 0.22 | 0.12 | 0.24 | 0.52 | 0.22 | 0.14 |
| 3 | 85697614 | rs9810211 | 0.194 | -0.448 | -0.883 | -0.703 | -0.066 | -0.012 | -0.012 | 0 | 0.42 | 0.16 | 0.39 | 0.52 | 0.47 | 0.3 |
| 3 | 85710336 | rs11127905 | -1.56 | -0.537 | -0.355 | -0.785 | 0 | -0.015 | -0.015 | 0 | 0.69 | 0.68 | 0.73 | 0.78 | 0.72 | 0.76 |
| 3 | 85712974 | rs7629690 | -0.412 | -0.494 | -0.389 | -0.755 | -0.029 | -0.001 | -0.001 | 1.86 | 0.27 | 0.52 | 0.34 | 0.26 | 0.24 | 0.46 |
| 3 | 85719890 | rs9844720 | -1.762 | -0.395 | -0.525 | -0.806 | -0.086 | 0.048 | 0.048 | 0.03 | 0.73 | 0.97 | 0.76 | 0.78 | 0.72 | 0.76 |
| 3 | 85720888 | rs7644873 | 0.59 | -0.385 | -0.042 | -0.771 | 0.005 | 0.003 | 0.003 | 0 | 0.1 | 0.23 | 0.14 | 0.08 | 0.14 | 0.17 |
| 3 | 85725069 | rs9871618 | -0.332 | -0.685 | -0.235 | -0.058 | -0.002 | -0.033 | -0.033 | 0.255 | 0.72 | 0.79 | 0.66 | 0.55 | 0.68 | 0.76 |
| 3 | 85729502 | rs7612090 | 0.233 | -0.82 | -0.136 | -0.086 | 0.012 | 0.061 | 0.061 | 0 | 0.48 | 0.65 | 0.43 | 0.36 | 0.43 | 0.54 |
| 3 | 85763107 | rs1037231 | 0.665 | -0.752 | -0.012 | 0.003 | 0 | -0.011 | -0.011 | 0 | 0.41 | 0.45 | 0.4 | 0.37 | 0.42 | 0.54 |
| 3 | 85778718 | rs6793697 | -0.065 | -0.578 | 0.187 | 0.297 | -0.038 | 0.102 | 0.102 | 0.381 | 0.11 | 0.29 | 0.11 | 0.19 | 0.02 | 0.07 |
| 3 | 85792137 | rs7355953 | 1.3 | -0.277 | -0.159 | -0.004 | -0.035 | 0.091 | 0.091 | 0 | 0.2 | 0.04 | 0.14 | 0 | 0.22 | 0.13 |
| 3 | 85792456 | rs7620948 | 0.491 | -0.282 | 0.105 | 0.109 | 0.015 | 0.015 | 0.015 | 0 | 0.1 | 0.25 | 0.14 | 0.09 | 0.15 | 0.17 |

| | | | | | | | | | | | | | | | | |
|---|---|---|---|---|---|---|---|---|---|---|---|---|---|---|---|---|
| 3 | 85793588 | rs3158 | -0.482 | -0.361 | 0.047 | 0.21 | 0.003 | -0.012 | -0.012 | 0 | 0.29 | 0.26 | 0.39 | 0.62 | 0.28 | 0.29 |
| 3 | 85797093 | rs1448610 | -0.481 | -0.398 | 0.264 | 0.241 | 0 | -0.002 | -0.002 | 0.013 | 0.29 | 0.28 | 0.4 | 0.59 | 0.28 | 0.3 |
| 3 | 85799507 | rs9309989 | 0.81 | -0.38 | 0.508 | 0.259 | 0.019 | 0.058 | 0.058 | 0 | 0.08 | 0.22 | 0.14 | 0.11 | 0.12 | 0.14 |
| 3 | 85803785 | rs7616936 | -0.339 | -0.45 | 0.536 | 0.243 | -0.002 | 0.058 | 0.058 | 0 | 0.12 | 0.32 | 0.14 | 0.16 | 0.13 | 0.29 |
| 3 | 85804321 | rs7653685 | 1.316 | -0.408 | 0.298 | 0.228 | -0.001 | 0.001 | 0.001 | 0 | 0.07 | 0.08 | 0.12 | 0.1 | 0.12 | 0.14 |
| 3 | 85804585 | rs11925878 | 0.188 | -0.479 | 0.235 | 0.476 | -0.005 | 0.031 | 0.031 | 0 | 0.06 | 0.07 | 0.08 | 0.19 | 0.01 | 0.11 |
| 3 | 85811233 | rs2325035 | 0.088 | -0.469 | 0.102 | 0.404 | -0.037 | -0.129 | -0.129 | 1.737 | 0.24 | 0.08 | 0.3 | 0.46 | 0.27 | 0.31 |
| 3 | 85826736 | rs9852127 | 1.412 | -0.318 | 0.068 | 0.08 | -0.03 | 0.078 | 0.078 | 0 | 0.18 | 0.05 | 0.13 | 0 | 0.21 | 0.09 |
| 3 | 85835907 | rs6549055 | -0.849 | -0.312 | 0.223 | 0.099 | 0.001 | 0.012 | 0.012 | 1.098 | 0.37 | 0.35 | 0.52 | 0.76 | 0.41 | 0.56 |
| 3 | 85850041 | rs7640660 | 0.55 | -0.329 | 0.367 | 0.081 | 0.005 | -0.156 | -0.156 | 0 | 0.23 | 0.43 | 0.16 | 0 | 0.22 | 0.11 |
| 3 | 85870303 | rs1003985 | -0.222 | -0.475 | 0.222 | 0.46 | -0.014 | 0.018 | 0.018 | 0 | 0.18 | 0.24 | 0.29 | 0.27 | 0.16 | 0.35 |
| 3 | 85876799 | rs10511067 | -0.356 | -0.487 | 0.484 | 0.094 | 0.001 | 0.017 | 0.017 | 0.404 | 0.33 | 0.41 | 0.46 | 0.57 | 0.39 | 0.45 |
| 3 | 85881725 | rs11915638 | 0.427 | -0.578 | 0.393 | -0.013 | 0.001 | 0.003 | 0.003 | 0 | 0.24 | 0.15 | 0.21 | 0.19 | 0.22 | 0.2 |
| 3 | 85884150 | rs13078807 | 1.298 | -0.573 | 0.29 | -0.052 | -0.035 | 0.087 | 0.087 | 0 | 0.18 | 0.04 | 0.13 | 0 | 0.21 | 0.09 |
| 3 | 85895494 | rs7629375 | -0.522 | -0.544 | 0.258 | 0.031 | -0.001 | 0.001 | 0.001 | 0.04 | 0.36 | 0.63 | 0.47 | 0.57 | 0.4 | 0.48 |
| 3 | 85903031 | rs3901898 | -1.604 | -0.574 | -0.151 | -0.011 | -0.116 | 0.085 | 0.085 | 0.276 | 0.62 | 0.98 | 0.7 | 0.75 | 0.62 | 0.68 |
| 3 | 85906928 | rs3911063 | 0.466 | -0.519 | 0.136 | 0.092 | -0.005 | -0.013 | -0.013 | 0.083 | 0.24 | 0.21 | 0.37 | 0.45 | 0.34 | 0.31 |
| 3 | 85986720 | rs9812103 | -0.293 | -0.731 | 0.837 | 0.18 | 0.022 | 0.014 | 0.014 | 0 | 0.17 | 0.36 | 0.17 | 0.12 | 0.17 | 0.15 |
| 3 | 85989000 | rs9861174 | -1.345 | -0.846 | 0.821 | 0.172 | -0.021 | 0.011 | 0.011 | 0 | 0.24 | 0.5 | 0.23 | 0.23 | 0.19 | 0.28 |
| 3 | 85990690 | rs6765959 | 0.006 | -0.766 | 0.682 | 0.18 | 0 | 0 | 0 | 0 | 0.15 | 0.14 | 0.16 | 0.12 | 0.17 | 0.15 |
| 3 | 86006210 | rs2875492 | 0.194 | -1.005 | 0.574 | 0.111 | 0.019 | -0.003 | -0.003 | 0 | 0.06 | 0.27 | 0.03 | 0 | 0.02 | 0.05 |
| 3 | 86029127 | rs12636273 | -0.902 | -1.052 | 0.267 | 0.182 | -0.003 | 0.037 | 0.037 | 0.001 | 0.56 | 0.68 | 0.67 | 0.68 | 0.6 | 0.61 |
| 3 | 86037162 | rs12486982 | 0.796 | -1.011 | 0.037 | 0.248 | -0.109 | -0.141 | -0.141 | 0.754 | 0.32 | 0.05 | 0.42 | 0.43 | 0.41 | 0.31 |
| 3 | 86050795 | rs7619493 | 0.048 | -0.922 | 0.351 | 0.348 | -0.001 | 0.003 | 0.003 | 0 | 0.15 | 0.17 | 0.16 | 0.21 | 0.16 | 0.26 |
| 3 | 86063190 | rs35215849 | 0.081 | -0.923 | 0.353 | 0.348 | -0.001 | 0.005 | 0.005 | 0 | 0.15 | 0.18 | 0.16 | 0.21 | 0.16 | 0.26 |
| 3 | 86073977 | rs6804410 | -0.395 | -1.006 | 0.543 | 0.288 | 0.007 | 0.039 | 0.039 | 0 | 0.08 | 0.33 | 0.05 | 0.06 | 0.03 | 0.08 |
| 3 | 86082200 | rs11922904 | -0.376 | -0.948 | 0.449 | 0.342 | -0.008 | 0.081 | 0.081 | 0 | 0.18 | 0.34 | 0.2 | 0.3 | 0.17 | 0.3 |
| 3 | 86112234 | rs9310001 | -0.78 | -0.968 | 0.398 | 0.319 | -0.02 | 0.038 | 0.038 | 0 | 0.22 | 0.34 | 0.22 | 0.3 | 0.18 | 0.3 |
| 3 | 86117669 | rs7630752 | -1.61 | -1.049 | 0.074 | 0.294 | -0.128 | 0.018 | 0.018 | 0.126 | 0.64 | 0.98 | 0.69 | 0.72 | 0.61 | 0.68 |
| 3 | 86118206 | rs17024876 | -1.204 | -1.328 | 0.624 | 0.373 | -0.042 | 0.062 | 0.062 | 0 | 0.24 | 0.49 | 0.23 | 0.3 | 0.18 | 0.3 |
| 3 | 86142120 | rs6783870 | -0.323 | -1.079 | 0.332 | 0.041 | -0.04 | -0.063 | -0.063 | 0 | 0.11 | 0.26 | 0.05 | 0 | 0.03 | 0.07 |
| 3 | 86167873 | rs13093396 | 0.864 | -0.704 | -0.253 | -0.006 | -0.174 | -0.126 | -0.126 | 0 | 0.31 | 0.05 | 0.46 | 0.44 | 0.48 | 0.35 |
| 3 | 86173480 | rs9853144 | 1.006 | -0.613 | -0.014 | -0.1 | -0.01 | 0.003 | 0.003 | 0.039 | 0.09 | 0.06 | 0.07 | 0 | 0.14 | 0.06 |
| 3 | 86173613 | rs4327387 | 1.871 | -0.817 | -0.404 | -0.115 | -0.034 | -0.059 | -0.059 | 0.01 | 0.24 | 0.01 | 0.2 | 0.27 | 0.22 | 0.24 |
| 3 | 86177581 | rs2325001 | -0.557 | -0.939 | 0.093 | 0.006 | -0.015 | 0.035 | 0.035 | 0 | 0.82 | 0.7 | 0.82 | 0.76 | 0.87 | 0.75 |
| 3 | 86189861 | rs12494510 | 0.695 | -0.705 | -0.331 | 0.096 | 0.004 | 0.002 | 0.002 | 0.059 | 0.66 | 0.29 | 0.7 | 0.71 | 0.7 | 0.61 |
| 3 | 86196341 | rs6549077 | 0.481 | -1.195 | -0.317 | -0.065 | -0.026 | -0.001 | -0.001 | 0 | 0.32 | 0.35 | 0.23 | 0.32 | 0.16 | 0.2 |
| 3 | 86199626 | rs7428773 | 0.21 | -1.415 | -0.411 | -0.14 | 0.028 | 0.009 | 0.009 | 0 | 0.69 | 0.27 | 0.75 | 0.77 | 0.84 | 0.69 |
| 3 | 86203809 | rs9833726 | 1.092 | -1.075 | -0.574 | -0.222 | -0.03 | 0.035 | 0.035 | 0.05 | 0.09 | 0.02 | 0.07 | 0 | 0.14 | 0.06 |
| 3 | 86204867 | rs13093222 | 0.876 | -1.046 | -0.573 | -0.104 | -0.102 | -0.128 | -0.128 | 0.002 | 0.38 | 0.08 | 0.5 | 0.5 | 0.57 | 0.44 |
| 3 | 86206285 | rs12485540 | 0.82 | -1.053 | -0.198 | -0.107 | -0.022 | -0.036 | -0.036 | 1.525 | 0.39 | 0.24 | 0.51 | 0.5 | 0.57 | 0.44 |
| 3 | 86222710 | rs7340540 | -0.697 | -0.859 | -0.117 | -0.124 | -0.008 | -0.005 | -0.005 | 0 | 0.8 | 0.74 | 0.83 | 0.85 | 0.85 | 0.7 |
| 3 | 86222845 | rs9814386 | 1.174 | -0.865 | -0.367 | -0.097 | -0.044 | -0.031 | -0.031 | 0 | 0.28 | 0.03 | 0.23 | 0.27 | 0.27 | 0.25 |
| 3 | 86224663 | rs7340745 | 1.222 | -0.898 | -0.214 | -0.103 | -0.047 | -0.011 | -0.011 | 0 | 0.11 | 0.37 | 0.07 | 0.04 | 0.02 | 0.01 |
| 3 | 86229700 | rs7651338 | 0.175 | -0.846 | -0.314 | -0.084 | -0.022 | -0.009 | -0.009 | 0 | 0.39 | 0.44 | 0.32 | 0.35 | 0.29 | 0.26 |
| 3 | 86239049 | rs6772266 | 1.28 | -0.769 | -0.239 | -0.019 | 0.018 | 0.003 | 0.003 | 1.331 | 0.1 | 0.07 | 0.06 | 0.08 | 0.02 | 0.01 |
| 3 | 86242930 | rs992803 | -0.333 | -0.595 | -0.2 | 0.008 | -0.017 | -0.013 | -0.013 | 0.929 | 0.78 | 0.59 | 0.82 | 0.85 | 0.85 | 0.7 |
| 3 | 86246160 | rs11707021 | 0.807 | -0.556 | -0.929 | -0.032 | -0.137 | -0.134 | -0.134 | 2.913 | 0.35 | 0.09 | 0.49 | 0.5 | 0.54 | 0.41 |
| 3 | 86251633 | rs6549085 | -0.518 | -1.019 | -0.193 | 0.367 | -0.018 | 0.009 | 0.009 | 0 | 0.81 | 0.73 | 0.84 | 0.79 | 0.89 | 0.68 |
| 3 | 86255672 | rs7428796 | 0.7 | -1.25 | -0.199 | 0.187 | 0.028 | 0.003 | 0.003 | 0 | 0.45 | 0.28 | 0.53 | 0.46 | 0.6 | 0.43 |
| 3 | 86261481 | rs9833236 | -0.906 | -1.166 | -0.147 | 0.239 | 0.006 | -0.005 | -0.005 | 0 | 0.81 | 0.81 | 0.85 | 0.79 | 0.89 | 0.68 |
| 3 | 86264283 | rs2175671 | -0.2 | -1.361 | 0.16 | 0.197 | -0.025 | 0.01 | 0.01 | 0.602 | 0.6 | 0.52 | 0.64 | 0.5 | 0.76 | 0.5 |
| 3 | 86266042 | rs1567737 | -0.922 | -1.43 | 0.207 | 0.215 | -0.006 | 0.002 | 0.002 | 0 | 0.81 | 0.79 | 0.84 | 0.79 | 0.9 | 0.68 |
| 3 | 86267150 | rs1512926 | 1.273 | -1.416 | 0.204 | 0.181 | 0 | 0 | 0 | 0.569 | 0.24 | 0.22 | 0.25 | 0.29 | 0.27 | 0.24 |
| 3 | 86268067 | rs2048590 | -0.893 | -1.431 | 0.22 | 0.181 | -0.022 | 0.01 | 0.01 | 0 | 0.8 | 0.72 | 0.84 | 0.79 | 0.9 | 0.68 |
| 3 | 86268100 | rs2048589 | 0.022 | -1.424 | 0.213 | 0.181 | -0.008 | 0.006 | 0.006 | 1.23 | 0.57 | 0.5 | 0.59 | 0.5 | 0.62 | 0.43 |
| 3 | 86271772 | rs1913188 | 1.203 | -1.311 | 0.014 | 0.224 | -0.001 | -0.001 | -0.001 | 1.978 | 0.24 | 0.21 | 0.25 | 0.29 | 0.27 | 0.24 |
| 3 | 86278951 | rs13063456 | 0.163 | -1.592 | 0.063 | 0.13 | -0.072 | 0 | 0 | 0 | 0.68 | 0.47 | 0.79 | 0.74 | 0.87 | 0.67 |
| 3 | 86282294 | rs17026843 | 0.761 | -0.952 | -0.316 | 0.135 | -0.034 | -0.031 | -0.031 | 0 | 0.42 | 0.12 | 0.49 | 0.46 | 0.54 | 0.43 |
| 3 | 86282586 | rs7645670 | -0.231 | -0.576 | -0.43 | 0.295 | 0.01 | -0.022 | -0.022 | 0 | 0.58 | 0.75 | 0.56 | 0.53 | 0.55 | 0.42 |
| 3 | 86286280 | rs7625346 | 0.394 | -1.27 | 0.096 | 0.286 | -0.041 | 0.007 | 0.007 | 0.868 | 0.09 | 0.28 | 0.04 | 0.06 | 0.01 | 0.07 |
| 3 | 86286959 | rs4263293 | 0.183 | -1.369 | 0.247 | 0.179 | 0.016 | -0.002 | -0.002 | 0 | 0.52 | 0.32 | 0.54 | 0.52 | 0.57 | 0.42 |
| 3 | 86289456 | rs9860570 | -0.429 | -1.389 | 0.164 | 0.198 | -0.034 | 0.006 | 0.006 | 0 | 0.8 | 0.66 | 0.84 | 0.83 | 0.9 | 0.71 |
| 3 | 86290724 | rs1512935 | 0.982 | -1.469 | 0.038 | 0.168 | -0.038 | -0.038 | -0.038 | 0.63 | 0.42 | 0.11 | 0.49 | 0.48 | 0.54 | 0.41 |
| 3 | 86294252 | rs1567734 | -1.918 | -1.82 | 0.081 | 0.231 | 0.011 | -0.027 | -0.027 | 0.006 | 0.89 | 0.94 | 0.88 | 0.89 | 0.91 | 0.78 |
| 3 | 86312396 | rs6549088 | -0.446 | -1.742 | 0.415 | 0.226 | -0.072 | 0.041 | 0.041 | 0 | 0.78 | 0.53 | 0.8 | 0.79 | 0.89 | 0.71 |
| 3 | 86318521 | rs9858836 | 1.331 | -1.583 | 0.238 | 0.214 | -0.001 | -0.002 | -0.002 | 0 | 0.25 | 0.22 | 0.26 | 0.3 | 0.29 | 0.28 |
| 3 | 86331094 | rs6781294 | -0.741 | -1.632 | 0.026 | 0.205 | -0.068 | 0.025 | 0.025 | 0 | 0.79 | 0.53 | 0.82 | 0.83 | 0.9 | 0.71 |
| 3 | 86345140 | rs1355686 | 0.854 | -1.701 | -0.16 | 0.272 | 0.014 | 0.003 | 0.003 | 0 | 0.42 | 0.15 | 0.52 | 0.45 | 0.58 | 0.42 |

| | | | | | | | | | | | | | | | | |
|---|---|---|---|---|---|---|---|---|---|---|---|---|---|---|---|---|
| 3 | 86356130 | rs11926519 | 1.562 | -1.586 | -0.198 | 0.375 | -0.03 | -0.023 | -0.023 | 0 | 0.26 | 0.08 | 0.25 | 0.28 | 0.29 | 0.28 |
| 3 | 86360829 | rs2644217 | -0.075 | -1.625 | -0.084 | 0.342 | -0.049 | -0.013 | -0.013 | 0 | 0.71 | 0.62 | 0.81 | 0.84 | 0.87 | 0.7 |
| 3 | 86362913 | rs12488883 | 0.712 | -1.476 | -0.221 | 0.403 | 0.004 | 0.001 | 0.001 | 0.653 | 0.42 | 0.17 | 0.51 | 0.46 | 0.57 | 0.42 |
| 3 | 86365826 | rs1353422 | -0.536 | -1.251 | -0.066 | 0.5 | -0.075 | 0.031 | 0.031 | 0 | 0.79 | 0.52 | 0.83 | 0.83 | 0.9 | 0.71 |
| 3 | 86381883 | rs1170214 | -0.228 | -1.185 | -0.346 | 0.527 | 0.009 | -0.003 | -0.003 | 0 | 0.54 | 0.56 | 0.6 | 0.53 | 0.62 | 0.44 |
| 3 | 86384224 | rs9848250 | 1.153 | -0.763 | -0.807 | 0.654 | -0.047 | -0.061 | -0.061 | 0 | 0.24 | 0.05 | 0.25 | 0.3 | 0.28 | 0.25 |
| 3 | 86411485 | rs13059931 | 0.651 | -1.194 | -0.731 | 0.461 | -0.018 | -0.01 | -0.01 | 0.798 | 0.39 | 0.09 | 0.5 | 0.42 | 0.59 | 0.43 |
| 3 | 86413934 | rs9879023 | 0.041 | -1.646 | 0.09 | 0.211 | -0.032 | -0.045 | -0.045 | 0 | 0.19 | 0.47 | 0.12 | 0.11 | 0.15 | 0.06 |
| 3 | 86421805 | rs13089170 | 1.948 | -1.004 | -0.419 | 0.083 | -0.086 | -0.044 | -0.044 | 0 | 0.2 | 0.02 | 0.26 | 0.27 | 0.31 | 0.23 |
| 3 | 86426224 | rs9867082 | 0.085 | -2.196 | 0.117 | 0.34 | 0.025 | 0.025 | 0.025 | 0 | 0.07 | 0.17 | 0.08 | 0.11 | 0.13 | 0.06 |
| 3 | 86426324 | rs9870892 | 0.145 | -2.153 | -0.112 | 0.361 | -0.025 | -0.035 | -0.035 | 0.516 | 0.18 | 0.39 | 0.12 | 0.11 | 0.15 | 0.06 |
| 3 | 86435209 | rs9859509 | -0.887 | -2.376 | -0.153 | 0.466 | 0.057 | 0.016 | 0.016 | 0 | 0.39 | 0.77 | 0.28 | 0.28 | 0.25 | 0.37 |
| 3 | 86457535 | rs9820995 | -0.22 | -2.225 | -0.209 | 0.514 | -0.004 | 0 | 0 | 0 | 0.58 | 0.54 | 0.63 | 0.54 | 0.74 | 0.49 |
| 3 | 86466286 | rs6800203 | -0.111 | -2.135 | -0.85 | 0.568 | -0.024 | 0.005 | 0.005 | 0 | 0.57 | 0.46 | 0.62 | 0.54 | 0.74 | 0.49 |
| 3 | 86485937 | rs291963 | 0.284 | -1.962 | -1.008 | 0.664 | -0.021 | -0.003 | -0.003 | 0 | 0.26 | 0.45 | 0.21 | 0.18 | 0.22 | 0.34 |
| 3 | 86486239 | rs12488703 | 0.837 | -1.964 | -1.082 | 0.675 | -0.016 | 0.005 | 0.005 | 0 | 0.41 | 0.11 | 0.51 | 0.43 | 0.59 | 0.43 |
| 3 | 86490729 | rs7647614 | 0.179 | -1.617 | -0.943 | 0.697 | -0.003 | 0 | 0 | 0.127 | 0.54 | 0.5 | 0.59 | 0.54 | 0.63 | 0.46 |
| 3 | 86500544 | rs291956 | -1.324 | -1.081 | -1.28 | 0.934 | -0.002 | -0.072 | -0.072 | 0.361 | 0.8 | 0.96 | 0.8 | 0.72 | 0.85 | 0.8 |
| 3 | 86508755 | rs291960 | -0.628 | -1.082 | -0.827 | 0.801 | -0.005 | -0.001 | -0.001 | 0.029 | 0.37 | 0.6 | 0.45 | 0.36 | 0.25 | 0.36 |
| 3 | 86511758 | rs291961 | 0.133 | -1.19 | -0.725 | 0.706 | -0.036 | 0.052 | 0.052 | 0.743 | 0.26 | 0.49 | 0.4 | 0.34 | 0.17 | 0.28 |
| 3 | 86526422 | rs1603320 | -1.042 | -1.082 | -0.966 | 0.738 | -0.015 | 0 | 0 | 0.94 | 0.75 | 0.8 | 0.69 | 0.72 | 0.55 | 0.69 |
| 3 | 86529127 | rs9825899 | 1.235 | -1.318 | -0.764 | 0.516 | -0.006 | -0.066 | -0.066 | 0.387 | 0.35 | 0.08 | 0.23 | 0.36 | 0.3 | 0.33 |
| 3 | 86529994 | rs9848467 | 1.132 | -1.326 | -0.728 | 0.512 | 0.018 | -0.019 | -0.019 | 0 | 0.36 | 0.21 | 0.24 | 0.36 | 0.3 | 0.33 |
| 3 | 86554319 | rs13061855 | 1.229 | -1.19 | -0.772 | 0.579 | -0.016 | -0.048 | -0.048 | 0 | 0.36 | 0.07 | 0.23 | 0.35 | 0.31 | 0.34 |
| 3 | 86555969 | rs2035985 | 0.46 | -1.285 | -0.644 | 0.603 | -0.045 | -0.095 | -0.095 | 0 | 0.24 | 0.47 | 0.13 | 0.07 | 0.16 | 0.15 |
| 3 | 86556132 | rs10511062 | 1.606 | -1.305 | -0.681 | 0.413 | 0 | -0.004 | -0.004 | 0 | 0.05 | 0.07 | 0.04 | 0.01 | 0.04 | 0.06 |
| 3 | 86557037 | rs1552883 | 1.284 | -1.297 | -0.644 | 0.419 | 0.008 | -0.022 | -0.022 | 2.363 | 0.36 | 0.11 | 0.23 | 0.35 | 0.31 | 0.34 |
| 3 | 86558884 | rs1948483 | -0.201 | -1.286 | -0.607 | 0.402 | -0.008 | -0.041 | -0.041 | 3.031 | 0.3 | 0.64 | 0.17 | 0.05 | 0.19 | 0.2 |
| 3 | 86571808 | rs9866050 | 1.259 | -1.236 | -0.792 | 0.487 | 0.006 | -0.022 | -0.022 | 0 | 0.35 | 0.15 | 0.23 | 0.36 | 0.31 | 0.35 |
| 3 | 86573780 | rs7616310 | 0.777 | -1.073 | -0.713 | 0.563 | 0.02 | 0.009 | 0.009 | 0.013 | 0.41 | 0.23 | 0.28 | 0.37 | 0.34 | 0.39 |
| 3 | 86602453 | rs6549099 | -0.517 | -1.124 | -0.795 | 0.56 | -0.05 | -0.031 | -0.031 | 0.12 | 0.67 | 0.76 | 0.4 | 0.4 | 0.5 | 0.54 |
| 3 | 86602789 | rs7623440 | 0.471 | -1.196 | -0.722 | 0.572 | -0.005 | -0.035 | -0.035 | 0 | 0.08 | 0.21 | 0.05 | 0.01 | 0.04 | 0.05 |
| 3 | 86603154 | rs725579 | 0.513 | -1.195 | -0.803 | 0.573 | -0.012 | -0.036 | -0.036 | 0.005 | 0.07 | 0.17 | 0.05 | 0.01 | 0.04 | 0.05 |
| 3 | 86609267 | rs9829809 | 1.145 | -1.029 | -0.931 | 0.675 | -0.037 | -0.065 | -0.065 | 0 | 0.35 | 0.06 | 0.22 | 0.35 | 0.31 | 0.36 |
| 3 | 86629088 | rs7617821 | -0.617 | -1.205 | -0.81 | 0.414 | 0.088 | 0.106 | 0.106 | 0.042 | 0.37 | 0.84 | 0.33 | 0.18 | 0.21 | 0.21 |
| 3 | 86635618 | rs6795064 | -0.006 | -1.235 | -0.558 | 0.277 | -0.042 | -0.112 | -0.112 | 0.042 | 0.31 | 0.53 | 0.17 | 0.05 | 0.19 | 0.16 |
| 3 | 86654296 | rs4293705 | -0.343 | -1.237 | -0.715 | 0.286 | 0.089 | 0.143 | 0.143 | 0.167 | 0.36 | 0.83 | 0.19 | 0.05 | 0.19 | 0.16 |
| 3 | 86654664 | rs9810220 | 1.402 | -1.213 | -0.97 | 0.618 | -0.07 | -0.083 | -0.083 | 0 | 0.33 | 0.01 | 0.2 | 0.34 | 0.3 | 0.35 |
| 3 | 86665408 | rs4333087 | 0.259 | -0.897 | -0.765 | 0.34 | 0.101 | 0.183 | 0.183 | 0.514 | 0.64 | 0.15 | 0.8 | 0.95 | 0.81 | 0.84 |
| 3 | 86673923 | rs6804348 | 1.016 | -0.8 | -0.711 | 0.567 | -0.023 | -0.064 | -0.064 | 0.174 | 0.33 | 0.13 | 0.37 | 0.55 | 0.31 | 0.41 |
| 3 | 86693504 | rs9829630 | 1.007 | -0.677 | -0.605 | 0.624 | 0.002 | -0.04 | -0.04 | 0 | 0.35 | 0.16 | 0.37 | 0.55 | 0.3 | 0.42 |
| 3 | 86703247 | rs4855157 | 1.056 | -0.658 | -0.612 | 0.637 | -0.028 | -0.08 | -0.08 | 0.041 | 0.34 | 0.11 | 0.37 | 0.55 | 0.3 | 0.42 |
| 3 | 86714839 | rs9821289 | -0.742 | -0.686 | -0.638 | 0.544 | -0.09 | -0.04 | -0.04 | 0 | 0.68 | 0.86 | 0.55 | 0.6 | 0.49 | 0.58 |
| 3 | 86720613 | rs7652288 | 1.363 | -0.638 | -0.716 | 0.667 | -0.039 | -0.121 | -0.121 | 0 | 0.32 | 0.04 | 0.22 | 0.42 | 0.28 | 0.36 |
| 3 | 86720689 | rs7652534 | 1.363 | -0.644 | -0.716 | 0.667 | -0.043 | -0.121 | -0.121 | 0.024 | 0.32 | 0.04 | 0.22 | 0.42 | 0.28 | 0.36 |
| 3 | 86722823 | rs6780199 | 0.492 | -0.758 | -0.019 | 0.188 | -0.001 | -0.001 | -0.001 | 0 | 0.44 | 0.44 | 0.42 | 0.56 | 0.34 | 0.42 |
| 3 | 86723625 | rs4522768 | 1.122 | -0.812 | -0.015 | 0.099 | -0.01 | -0.057 | -0.057 | 0 | 0.1 | 0.28 | 0.05 | 0 | 0.03 | 0.01 |
| 3 | 86730354 | rs7622550 | 0.054 | -0.344 | -0.304 | 0.461 | -0.009 | -0.029 | -0.029 | 0 | 0.22 | 0.22 | 0.12 | 0.05 | 0.16 | 0.15 |
| 3 | 86738113 | rs4855161 | 0.165 | 0.832 | -0.302 | 1.156 | -0.043 | -0.014 | -0.014 | 0 | 0.67 | 0.76 | 0.54 | 0.6 | 0.49 | 0.56 |
| 3 | 86749427 | rs1370209 | -0.681 | 1.316 | 0.252 | 1.601 | 0.027 | 0.048 | 0.048 | 0 | 0.29 | 0.42 | 0.54 | 0.55 | 0.41 | 0.43 |
| 3 | 86752414 | rs6549110 | -0.358 | 1.377 | -0.094 | 1.297 | -0.037 | 0.036 | 0.036 | 0 | 0.25 | 0.17 | 0.29 | 0.18 | 0.39 | 0.34 |
| 3 | 86759022 | rs1437050 | -0.358 | 1.372 | -0.001 | 1.456 | -0.015 | -0.001 | -0.001 | 0.211 | 0.26 | 0.23 | 0.3 | 0.33 | 0.39 | 0.39 |
| 3 | 86760197 | rs5007299 | -0.3 | 1.389 | -0.092 | 1.354 | -0.054 | 0.045 | 0.045 | 0 | 0.24 | 0.12 | 0.29 | 0.18 | 0.39 | 0.34 |
| 3 | 86767029 | rs4855143 | -1.45 | 1.172 | 0.869 | 1.912 | 0.072 | 0.451 | 0.451 | 1.491 | 0.05 | 0.35 | 0.27 | 0.32 | 0.02 | 0.06 |
| 3 | 86782973 | rs17027938 | -1.611 | 1.168 | 1.041 | 1.964 | 0.038 | 0.472 | 0.472 | 0.365 | 0.05 | 0.26 | 0.28 | 0.47 | 0.03 | 0.11 |
| 3 | 86790149 | rs9827403 | -0.208 | 1.224 | 0.987 | 1.773 | 0.011 | -0.016 | -0.016 | 0 | 0.25 | 0.28 | 0.3 | 0.19 | 0.39 | 0.34 |
| 3 | 86796801 | rs1982153 | -0.751 | 1.207 | 1.093 | 1.948 | 0.016 | 0.014 | 0.014 | 0 | 0.3 | 0.61 | 0.38 | 0.46 | 0.41 | 0.42 |
| 3 | 86824662 | rs4435642 | -0.164 | 0.608 | 0.234 | 1.167 | -0.01 | -0.073 | -0.073 | 0 | 0.71 | 0.64 | 0.78 | 0.95 | 0.75 | 0.83 |
| 3 | 86853485 | rs1583578 | 0.366 | 0.86 | 0.605 | 1.417 | 0.004 | -0.008 | -0.008 | 0.107 | 0.25 | 0.27 | 0.3 | 0.18 | 0.39 | 0.33 |
| 3 | 86860115 | rs9881535 | 0.123 | 0.857 | 0.597 | 1.429 | 0.014 | -0.008 | -0.008 | 0 | 0.27 | 0.33 | 0.33 | 0.28 | 0.39 | 0.35 |
| 3 | 86890846 | rs12714668 | 0.559 | 0.636 | 0.272 | 1.143 | -0.007 | 0.002 | 0.002 | 0.033 | 0.6 | 0.37 | 0.34 | 0.32 | 0.47 | 0.49 |
| 3 | 86900344 | rs9683034 | 1.414 | 0.631 | -0.127 | 1.122 | -0.044 | -0.071 | -0.071 | 0.062 | 0.37 | 0.02 | 0.2 | 0.31 | 0.32 | 0.4 |
| 3 | 86901096 | rs9310040 | -0.4 | 0.687 | 0.299 | 1.144 | 0.003 | -0.008 | -0.008 | 0 | 0.75 | 0.73 | 0.7 | 0.82 | 0.61 | 0.67 |
| 3 | 86906208 | rs9310042 | -0.4 | 0.687 | 0.297 | 1.159 | 0.002 | -0.006 | -0.006 | 0 | 0.75 | 0.74 | 0.7 | 0.82 | 0.61 | 0.67 |
| 3 | 86965294 | rs7644356 | 0.062 | 0.747 | 0.476 | 1.134 | 0.024 | -0.009 | -0.009 | 0 | 0.72 | 0.63 | 0.67 | 0.7 | 0.61 | 0.63 |
| 3 | 86975258 | rs1370034 | 0.169 | 0.712 | 1.06 | 1.053 | 0.04 | -0.017 | -0.017 | 0 | 0.72 | 0.54 | 0.68 | 0.7 | 0.61 | 0.63 |
| 3 | 86975853 | rs1370032 | 0.267 | 0.699 | 1.062 | 1.049 | 0.016 | -0.005 | -0.005 | 0.054 | 0.69 | 0.39 | 0.64 | 0.67 | 0.6 | 0.6 |
| 3 | 86980246 | rs11127967 | 0.233 | 0.637 | 0.794 | 1.055 | 0.052 | -0.023 | -0.023 | 0.016 | 0.71 | 0.48 | 0.67 | 0.7 | 0.61 | 0.63 |
| 3 | 86986783 | rs2028088 | -0.243 | 0.203 | 0.09 | 0.919 | 0.044 | 0.067 | 0.067 | 0.046 | 0.35 | 0.17 | 0.26 | 0.15 | 0.27 | 0.29 |

| | | | | | | | | | | | | | | | | |
|---|---|---|---|---|---|---|---|---|---|---|---|---|---|---|---|---|
| 3 | 86986820 | rs9863011 | -0.556 | 0.588 | 0.661 | 1.33 | 0.01 | 0.006 | 0.006 | 0 | 0.21 | 0.25 | 0.46 | 0.36 | 0.34 | 0.31 |
| 3 | 86989334 | rs13074432 | 0.45 | 0.208 | 0.67 | 1.404 | -0.009 | -0.002 | -0.002 | 0.052 | 0.7 | 0.71 | 0.48 | 0.63 | 0.61 | 0.63 |
| 3 | 86995124 | rs1436639 | 0.576 | 0.193 | 0.844 | 1.426 | 0.034 | 0.035 | 0.035 | 0 | 0.68 | 0.5 | 0.46 | 0.53 | 0.6 | 0.6 |
| 3 | 86999330 | rs1898270 | 0.556 | -0.223 | 0.77 | 1.174 | -0.001 | 0.002 | 0.002 | 0 | 0.73 | 0.32 | 0.7 | 0.67 | 0.74 | 0.69 |
| 3 | 87001049 | rs13087787 | -0.177 | -0.144 | 0.489 | 1.136 | 0.007 | -0.01 | -0.01 | 0 | 0.78 | 0.69 | 0.73 | 0.82 | 0.75 | 0.75 |
| 3 | 87021374 | rs1025631 | -0.033 | -0.12 | 0.542 | 1.146 | 0.023 | -0.005 | -0.005 | 0 | 0.77 | 0.58 | 0.7 | 0.77 | 0.74 | 0.71 |
| 3 | 87038154 | rs2043663 | -0.068 | -0.119 | 0.53 | 1.168 | 0.018 | 0.014 | 0.014 | 0 | 0.77 | 0.63 | 0.69 | 0.74 | 0.74 | 0.71 |
| 3 | 87042798 | rs2575775 | 0.08 | -0.151 | 0.548 | 1.197 | 0.01 | 0.008 | 0.008 | 0 | 0.75 | 0.55 | 0.69 | 0.74 | 0.74 | 0.72 |
| 3 | 87050997 | rs1898274 | 0.465 | -0.184 | 0.277 | 1.132 | 0.005 | -0.017 | -0.017 | 1.169 | 0.71 | 0.23 | 0.66 | 0.64 | 0.74 | 0.69 |
| 3 | 87055061 | rs2196523 | 0.964 | -0.464 | 0.083 | 1.296 | 0.011 | -0.126 | -0.126 | 0 | 0.66 | 0.09 | 0.65 | 0.5 | 0.73 | 0.59 |
| 3 | 87063357 | rs2118082 | 0.022 | -0.194 | 0.476 | 0.917 | 0.011 | 0.005 | 0.005 | 0 | 0.75 | 0.52 | 0.69 | 0.74 | 0.75 | 0.72 |
| 3 | 87064799 | rs2660785 | -0.229 | -0.185 | 0.456 | 0.906 | 0.006 | -0.006 | -0.006 | 0 | 0.77 | 0.64 | 0.71 | 0.79 | 0.76 | 0.75 |
| 3 | 87079981 | rs2660768 | -0.043 | -0.156 | -0.824 | 0.519 | -0.049 | 0.103 | 0.103 | 0 | 0.25 | 0.01 | 0.18 | 0.08 | 0.25 | 0.16 |
| 3 | 87087385 | rs6763673 | -1.308 | -0.437 | 0.063 | 0.988 | 0.004 | 0.134 | 0.134 | 1.364 | 0.08 | 0.53 | 0.08 | 0.15 | 0.03 | 0.06 |
| 3 | 87087905 | rs2660775 | -0.169 | -0.156 | -0.242 | 0.852 | 0.004 | -0.001 | -0.001 | 0 | 0.79 | 0.73 | 0.73 | 0.78 | 0.76 | 0.78 |
| 3 | 87090544 | rs13078933 | -0.77 | -0.815 | -0.121 | 1.022 | -0.034 | 0.016 | 0.016 | 0 | 0.27 | 0.4 | 0.17 | 0.32 | 0.15 | 0.23 |
| 3 | 87095589 | rs973557 | 0.857 | -0.949 | 0.439 | 0.961 | -0.024 | 0.007 | 0.007 | 0 | 0.71 | 0.39 | 0.81 | 0.68 | 0.84 | 0.77 |
| 3 | 87097872 | rs9836203 | -0.572 | -1.023 | 0.603 | 0.929 | -0.016 | 0.021 | 0.021 | 0 | 0.23 | 0.31 | 0.14 | 0.32 | 0.15 | 0.23 |
| 3 | 87101747 | rs2660745 | 0.62 | -0.995 | 0.619 | 0.978 | -0.042 | 0.041 | 0.041 | 0 | 0.75 | 0.5 | 0.83 | 0.7 | 0.84 | 0.77 |
| 3 | 87104899 | rs17023854 | 0.538 | -1.149 | -0.01 | 0.756 | 0.003 | -0.002 | -0.002 | 0 | 0.31 | 0.32 | 0.35 | 0.29 | 0.38 | 0.35 |
| 3 | 87110674 | rs2660753 | 0.461 | -1.307 | -0.135 | 0.702 | -0.059 | 0.018 | 0.018 | 0 | 0.73 | 0.41 | 0.84 | 0.73 | 0.89 | 0.79 |
| 3 | 87113522 | rs2575758 | 1.406 | -1.327 | -0.19 | 0.881 | -0.16 | -0.036 | -0.036 | 0.788 | 0.39 | 0.03 | 0.45 | 0.36 | 0.47 | 0.41 |
| 3 | 87119543 | rs1865866 | 0.801 | -1.171 | 0.502 | 1.108 | -0.058 | 0.019 | 0.019 | 0 | 0.73 | 0.41 | 0.84 | 0.73 | 0.89 | 0.79 |
| 3 | 87123762 | rs9872223 | -0.623 | -1.07 | 0.32 | 1.128 | -0.003 | 0.006 | 0.006 | 0 | 0.23 | 0.26 | 0.15 | 0.32 | 0.15 | 0.23 |
| 3 | 87126120 | rs2660761 | 1.02 | -1.174 | 0.411 | 1.244 | -0.005 | 0.11 | 0.11 | 0.241 | 0.88 | 0.55 | 0.92 | 0.87 | 0.95 | 0.91 |
| 3 | 87127455 | rs2575762 | 1.234 | -1.296 | 0.34 | 1.21 | -0.057 | 0.015 | 0.015 | 0 | 0.73 | 0.47 | 0.86 | 0.76 | 0.87 | 0.83 |
| 3 | 87134800 | rs17023900 | 0.364 | -1.29 | 0.249 | 1.201 | 0.038 | -0.003 | -0.003 | 0 | 0.15 | 0.05 | 0.06 | 0.12 | 0.08 | 0.07 |
| 3 | 87138182 | rs2660757 | 1.503 | -1.16 | 0.301 | 1.187 | -0.016 | 0.002 | 0.002 | 0 | 0.71 | 0.35 | 0.83 | 0.76 | 0.84 | 0.82 |
| 3 | 87158116 | rs2340382 | -1.168 | -1.1 | 0.387 | 1.206 | -0.009 | 0.094 | 0.094 | 0.266 | 0.12 | 0.43 | 0.07 | 0.12 | 0.05 | 0.09 |
| 3 | 87162301 | rs17181079 | 0.672 | -1.027 | -0.392 | 1.128 | -0.007 | -0.003 | -0.003 | 0 | 0.46 | 0.32 | 0.48 | 0.52 | 0.53 | 0.49 |
| 3 | 87172005 | rs9836404 | 0.372 | -1.092 | -0.421 | 0.99 | -0.016 | -0.004 | -0.004 | 0 | 0.15 | 0.16 | 0.06 | 0.11 | 0.07 | 0.07 |
| 3 | 87173324 | rs17181170 | 1.027 | -1.391 | -0.902 | 1.075 | -0.076 | 0.053 | 0.053 | 0.04 | 0.4 | 0.15 | 0.38 | 0.28 | 0.5 | 0.35 |
| 3 | 87173731 | rs7639047 | 0.195 | -1.368 | -0.849 | 1.078 | -0.007 | 0.027 | 0.027 | 0.382 | 0.53 | 0.2 | 0.44 | 0.39 | 0.57 | 0.42 |
| 3 | 87175298 | rs9852738 | -0.974 | -1.589 | -0.464 | 1.189 | -0.033 | 0.032 | 0.032 | 2.335 | 0.15 | 0.6 | 0.09 | 0.13 | 0.06 | 0.09 |
| 3 | 87189459 | rs6766573 | -0.97 | -1.587 | -0.5 | 1.19 | -0.032 | 0.026 | 0.026 | 2.229 | 0.15 | 0.63 | 0.1 | 0.13 | 0.06 | 0.09 |
| 3 | 87204024 | rs7639237 | -0.541 | -1.83 | -0.503 | 1.215 | -0.015 | -0.024 | -0.024 | 0 | 0.66 | 0.75 | 0.52 | 0.51 | 0.62 | 0.51 |
| 3 | 87205079 | rs6788616 | 0.303 | -1.842 | -0.443 | 1.177 | 0.015 | 0.001 | 0.001 | 0 | 0.52 | 0.7 | 0.47 | 0.4 | 0.55 | 0.44 |
| 3 | 87219009 | rs1825896 | 0.058 | -1.775 | -0.547 | 1.321 | -0.007 | 0.024 | 0.024 | 0 | 0.09 | 0.16 | 0.08 | 0.23 | 0.07 | 0.16 |
| 3 | 87223646 | rs963622 | -1.624 | -1.839 | -0.727 | 1.255 | -0.096 | -0.059 | -0.059 | 0 | 0.76 | 0.97 | 0.61 | 0.73 | 0.68 | 0.63 |
| 3 | 87223963 | rs1812519 | -1.306 | -1.841 | -0.576 | 1.245 | -0.016 | -0.003 | -0.003 | 0.006 | 0.74 | 0.79 | 0.58 | 0.72 | 0.66 | 0.62 |
| 3 | 87225147 | rs12638524 | -0.449 | -1.993 | -0.642 | 1.22 | 0.03 | -0.015 | -0.015 | 0.165 | 0.24 | 0.17 | 0.13 | 0.33 | 0.13 | 0.22 |
| 3 | 87228472 | rs6805890 | 1.416 | -2.05 | -0.727 | 1.21 | -0.166 | 0.019 | 0.019 | 0.702 | 0.38 | 0.03 | 0.37 | 0.28 | 0.49 | 0.32 |
| 3 | 87234066 | rs9858014 | -1.481 | -2.007 | -0.638 | 1.078 | 0.067 | -0.034 | -0.034 | 2.871 | 0.36 | 0.76 | 0.21 | 0.44 | 0.17 | 0.3 |
| 3 | 87236541 | rs9310049 | 0.1 | -1.918 | -0.089 | 1.024 | 0.007 | 0.16 | 0.16 | 0 | 0.08 | 0.32 | 0.06 | 0.12 | 0.03 | 0.08 |
| 3 | 87236790 | rs4859044 | 1.754 | -2.035 | -0.468 | 0.977 | -0.159 | 0.031 | 0.031 | 0.277 | 0.33 | 0.03 | 0.34 | 0.23 | 0.47 | 0.3 |
| 3 | 87239338 | rs1825901 | 0.145 | -1.707 | -0.366 | 1.048 | 0.001 | -0.002 | -0.002 | 0.19 | 0.86 | 0.84 | 0.9 | 0.77 | 0.89 | 0.83 |
| 3 | 87249364 | rs1488367 | 0.925 | -1.779 | -0.297 | 1.047 | -0.011 | -0.005 | -0.005 | 0 | 0.71 | 0.62 | 0.8 | 0.68 | 0.79 | 0.73 |
| 3 | 87253551 | rs12107900 | 0.861 | -1.807 | -0.298 | 1.082 | -0.009 | 0.012 | 0.012 | 0.893 | 0.45 | 0.59 | 0.41 | 0.33 | 0.51 | 0.37 |
| 3 | 87257423 | rs17024019 | 0.95 | -1.724 | -0.272 | 1.155 | -0.002 | 0 | 0 | 0 | 0.13 | 0.14 | 0.08 | 0.12 | 0.07 | 0.08 |
| 3 | 87262998 | rs1994487 | 0.696 | -1.369 | -0.199 | 1.182 | 0.046 | 0.154 | 0.154 | 0.111 | 0.06 | 0.29 | 0.05 | 0.11 | 0.04 | 0.07 |
| 3 | 87263497 | rs17024046 | 0.301 | -1.429 | -0.801 | 1.238 | -0.008 | -0.08 | -0.08 | 0.371 | 0.39 | 0.13 | 0.44 | 0.44 | 0.35 | 0.43 |
| 3 | 87274437 | rs904115 | 0.52 | -1.358 | -0.699 | 1.198 | -0.004 | 0.003 | 0.003 | 0.095 | 0.84 | 0.75 | 0.87 | 0.76 | 0.86 | 0.8 |
| 3 | 87276699 | rs2279720 | 0.752 | -1.438 | -0.666 | 1.181 | 0.014 | -0.012 | -0.012 | 0.015 | 0.14 | 0.11 | 0.07 | 0.19 | 0.07 | 0.08 |
| 3 | 87277062 | rs1002765 | -0.701 | -1.376 | -0.591 | 1.233 | -0.015 | 0.021 | 0.021 | 0.108 | 0.24 | 0.63 | 0.13 | 0.3 | 0.13 | 0.15 |
| 3 | 87280088 | rs13061754 | 1.724 | -1.412 | -0.777 | 1.166 | -0.129 | 0.094 | 0.094 | 0.239 | 0.34 | 0.02 | 0.34 | 0.19 | 0.45 | 0.3 |
| 3 | 87284401 | rs10511133 | 0.059 | -1.371 | -0.742 | 1.17 | -0.005 | -0.028 | -0.028 | 0 | 0.41 | 0.28 | 0.5 | 0.5 | 0.39 | 0.54 |
| 3 | 87293542 | rs2879985 | 1.499 | -1.42 | -0.775 | 1.177 | -0.049 | -0.069 | -0.069 | 0 | 0.26 | 0.06 | 0.38 | 0.33 | 0.28 | 0.35 |
| 3 | 87299075 | rs1044499 | 0.918 | -1.408 | -0.765 | 1.186 | 0.014 | -0.011 | -0.011 | 0 | 0.14 | 0.11 | 0.07 | 0.19 | 0.07 | 0.08 |
| 3 | 87301395 | rs2171525 | -0.777 | -1.056 | -0.372 | 1.186 | -0.003 | -0.002 | -0.002 | 0.433 | 0.15 | 0.22 | 0.12 | 0.17 | 0.11 | 0.19 |
| 3 | 87307547 | rs7428816 | 0.205 | -0.95 | -0.216 | 1.251 | 0.044 | 0.147 | 0.147 | 2.622 | 0.9 | 0.52 | 0.94 | 0.89 | 0.94 | 0.93 |
| 3 | 87321757 | rs17024118 | 1.16 | -0.994 | -0.467 | 1.196 | -0.058 | 0.043 | 0.043 | 0 | 0.45 | 0.12 | 0.4 | 0.38 | 0.52 | 0.39 |
| 3 | 87341538 | rs300992 | -0.088 | -0.823 | -0.528 | 1.251 | -0.021 | -0.015 | -0.015 | 0 | 0.1 | 0.02 | 0.09 | 0.13 | 0.11 | 0.1 |
| 3 | 87346707 | rs13082131 | -0.296 | -0.897 | -0.934 | 1.218 | -0.017 | -0.026 | -0.026 | 0 | 0.4 | 0.31 | 0.53 | 0.54 | 0.43 | 0.53 |
| 3 | 87367401 | rs2642347 | -0.296 | -0.928 | -0.957 | 1.243 | -0.02 | -0.031 | -0.031 | 0.206 | 0.4 | 0.28 | 0.53 | 0.54 | 0.43 | 0.53 |
| 3 | 87367945 | rs2646237 | -0.296 | -0.928 | -0.957 | 1.243 | -0.02 | -0.031 | -0.031 | 0 | 0.4 | 0.28 | 0.53 | 0.54 | 0.43 | 0.53 |
| 3 | 87370782 | rs2642350 | -0.149 | -0.909 | -0.966 | 1.227 | -0.027 | -0.046 | -0.046 | 0.161 | 0.38 | 0.24 | 0.5 | 0.53 | 0.4 | 0.52 |
| 3 | 87371282 | rs6551471 | -0.461 | -1.027 | -0.994 | 0.76 | -0.005 | -0.02 | -0.02 | 0.047 | 0.42 | 0.3 | 0.44 | 0.51 | 0.42 | 0.54 |
| 3 | 87371474 | rs1826637 | 1.15 | -1.099 | -1.159 | 0.755 | -0.076 | -0.128 | -0.128 | 0 | 0.26 | 0.03 | 0.3 | 0.34 | 0.27 | 0.37 |
| 3 | 87371723 | rs7637935 | -0.544 | -0.972 | -1.059 | 0.775 | -0.023 | -0.008 | -0.008 | 0 | 0.44 | 0.31 | 0.45 | 0.53 | 0.47 | 0.62 |

| | | | | | | | | | | | | | | | | |
|---|---|---|---|---|---|---|---|---|---|---|---|---|---|---|---|---|
| 3 | 87373894 | rs4859055 | 1.537 | -0.881 | -0.872 | 0.53 | 0.025 | 0.083 | 0.083 | 2.283 | 0.71 | 0.15 | 0.79 | 0.84 | 0.77 | 0.84 |
| 3 | 87374853 | rs11713655 | -0.073 | -0.828 | -0.763 | 0.558 | -0.022 | -0.074 | -0.074 | 0 | 0.86 | 0.43 | 0.92 | 0.98 | 0.93 | 0.95 |
| 3 | 87386352 | rs903295 | 0.923 | -0.794 | -1.362 | 0.649 | -0.069 | -0.079 | -0.079 | 0 | 0.43 | 0.11 | 0.47 | 0.49 | 0.49 | 0.45 |
| 3 | 87401121 | rs1011609 | 0.261 | -0.613 | -1.058 | 0.797 | -0.001 | 0 | 0 | 0 | 0.57 | 0.43 | 0.6 | 0.62 | 0.61 | 0.55 |
| 3 | 87402474 | rs11711267 | 0.14 | -0.614 | -0.993 | 0.679 | -0.009 | -0.075 | -0.075 | 0 | 0.84 | 0.45 | 0.91 | 0.97 | 0.89 | 0.93 |
| 3 | 87403560 | rs9841254 | -0.182 | -0.69 | -0.914 | 0.725 | -0.011 | -0.064 | -0.064 | 0 | 0.87 | 0.53 | 0.94 | 0.98 | 0.94 | 0.95 |
| 3 | 87406236 | rs6789301 | -0.324 | -0.376 | -1.468 | 1.079 | -0.012 | -0.041 | -0.041 | 0.056 | 0.37 | 0.25 | 0.42 | 0.49 | 0.38 | 0.5 |
| 3 | 87407246 | rs9861650 | -1.149 | -0.698 | -0.668 | 0.845 | -0.003 | 0.004 | 0.004 | 0 | 0.1 | 0.22 | 0.11 | 0.13 | 0.1 | 0.07 |
| 3 | 87417191 | rs9866824 | 0.779 | -0.948 | -1.548 | 0.817 | 0.101 | -0.041 | -0.041 | 0 | 0.56 | 0.07 | 0.71 | 0.62 | 0.69 | 0.63 |
| 3 | 87426052 | rs6801412 | 0.859 | -1.329 | -1.269 | 0.813 | 0.028 | -0.019 | -0.019 | 0 | 0.77 | 0.19 | 0.83 | 0.69 | 0.88 | 0.81 |
| 3 | 87428732 | rs2342172 | -0.69 | -1.539 | -0.184 | 0.764 | 0.075 | 0.065 | 0.065 | 2.808 | 0.05 | 0.47 | 0.05 | 0 | 0.01 | 0.01 |
| 3 | 87438524 | rs6776212 | 1.248 | -1.583 | -0.035 | 0.769 | 0.024 | -0.019 | -0.019 | 3.449 | 0.77 | 0.2 | 0.86 | 0.69 | 0.92 | 0.81 |
| 3 | 87447834 | rs6777295 | 0.149 | -1.625 | 0.355 | 0.81 | -0.031 | -0.002 | -0.002 | 0.016 | 0.07 | 0.12 | 0.04 | 0.05 | 0.02 | 0.12 |
| 3 | 87447949 | rs17024472 | -0.162 | -1.647 | 0.384 | 0.835 | -0.027 | 0.089 | 0.089 | 0 | 0.09 | 0.18 | 0.04 | 0.26 | 0.05 | 0.06 |
| 3 | 87449493 | rs7640232 | -0.67 | -1.644 | 0.361 | 0.824 | 0.071 | 0.067 | 0.067 | 2.529 | 0.05 | 0.43 | 0.05 | 0 | 0.01 | 0.01 |
| 3 | 87452064 | rs7636251 | -1.77 | -1.625 | 0.059 | 0.835 | 0.021 | -0.02 | -0.02 | 0 | 0.22 | 0.8 | 0.14 | 0.31 | 0.08 | 0.19 |
| 3 | 87473426 | rs4858960 | 1.043 | -0.928 | 0.008 | 1.076 | 0.002 | 0.093 | 0.093 | 0.357 | 0.83 | 0.33 | 0.87 | 0.65 | 0.81 | 0.83 |
| 3 | 87485329 | rs6551482 | 0.118 | -0.936 | 0.336 | 1.085 | -0.01 | 0.092 | 0.092 | 0 | 0.89 | 0.77 | 0.94 | 0.73 | 0.89 | 0.94 |
| 3 | 87489249 | rs9846815 | 1.114 | -1.069 | 0.128 | 0.909 | 0.02 | -0.025 | -0.025 | 0 | 0.62 | 0.19 | 0.74 | 0.6 | 0.63 | 0.68 |
| 3 | 87494596 | rs9857625 | -0.518 | -1.045 | 0.136 | 0.912 | 0.144 | 0.132 | 0.132 | 1.825 | 0.06 | 0.43 | 0.07 | 0.09 | 0.08 | 0.09 |
| 3 | 87495550 | rs9834692 | -0.333 | -1.044 | 0.165 | 0.912 | 0.126 | 0.115 | 0.115 | 1.126 | 0.05 | 0.34 | 0.06 | 0.09 | 0.08 | 0.09 |
| 3 | 87496499 | rs6769150 | 1.467 | -1.094 | -0.112 | 0.881 | 0.017 | -0.122 | -0.122 | 0.405 | 0.55 | 0.06 | 0.7 | 0.55 | 0.61 | 0.56 |
| 3 | 87499338 | rs6782359 | 1.777 | -0.782 | -0.8 | 0.751 | -0.133 | -0.11 | -0.11 | 0 | 0.22 | 0.02 | 0.42 | 0.36 | 0.32 | 0.36 |
| 3 | 87520900 | rs6796548 | 0.96 | -0.835 | -0.583 | 0.591 | 0.012 | 0.029 | 0.029 | 0 | 0.21 | 0.14 | 0.13 | 0.05 | 0.18 | 0.15 |
| 3 | 87525449 | rs9883474 | -0.171 | -0.833 | -0.455 | 0.6 | 0.135 | 0.124 | 0.124 | 1.825 | 0.06 | 0.41 | 0.07 | 0.09 | 0.08 | 0.09 |
| 3 | 87525540 | rs7614061 | 0.777 | -0.873 | -0.536 | 0.626 | 0.023 | -0.023 | -0.023 | 0.032 | 0.62 | 0.19 | 0.74 | 0.6 | 0.63 | 0.68 |
| 3 | 87527204 | rs7647651 | -0.523 | -0.9 | -0.533 | 0.602 | 0.007 | 0.012 | 0.012 | 0 | 0.33 | 0.66 | 0.24 | 0.18 | 0.29 | 0.36 |
| 3 | 87528312 | rs9823976 | 0.883 | -0.966 | -0.531 | 0.587 | -0.006 | 0.056 | 0.056 | 0.286 | 0.81 | 0.29 | 0.87 | 0.64 | 0.81 | 0.83 |
| 3 | 87528969 | rs17465479 | 0.995 | -0.93 | -0.52 | 0.591 | 0.015 | 0.063 | 0.063 | 0 | 0.2 | 0.09 | 0.13 | 0.05 | 0.18 | 0.15 |
| 3 | 87529297 | rs6551495 | -0.106 | -0.933 | -0.495 | 0.61 | -0.014 | 0.058 | 0.058 | 0 | 0.88 | 0.8 | 0.94 | 0.73 | 0.9 | 0.92 |
| 3 | 87529807 | rs12714688 | 0.587 | -0.811 | -0.608 | 0.629 | -0.03 | 0.003 | 0.003 | 0 | 0.62 | 0.52 | 0.77 | 0.64 | 0.69 | 0.65 |
| 3 | 87547508 | rs873293 | 1.148 | -0.607 | -0.966 | 0.733 | -0.068 | -0.014 | -0.014 | 0 | 0.34 | 0.02 | 0.28 | 0.25 | 0.32 | 0.25 |
| 3 | 87552073 | rs6790377 | -0.673 | -0.842 | 0.019 | 0.73 | -0.027 | -0.051 | -0.051 | 0 | 0.33 | 0.5 | 0.22 | 0.18 | 0.28 | 0.34 |
| 3 | 87552760 | rs9820876 | -0.205 | -0.721 | -0.043 | 0.779 | 0.012 | 0.033 | 0.033 | 2.717 | 0.12 | 0.34 | 0.09 | 0.13 | 0.09 | 0.17 |
| 3 | 87555607 | rs9828066 | -0.088 | -0.718 | -0.193 | 0.781 | 0.008 | 0.027 | 0.027 | 2.044 | 0.11 | 0.3 | 0.08 | 0.13 | 0.09 | 0.17 |
| 3 | 87565136 | rs4859077 | 1.39 | -0.77 | -0.566 | 0.896 | 0.029 | -0.107 | -0.107 | 0.018 | 0.56 | 0.08 | 0.7 | 0.55 | 0.62 | 0.6 |
| 3 | 87567430 | rs966978 | -0.823 | -0.733 | -0.284 | 0.884 | -0.015 | -0.008 | -0.008 | 0 | 0.14 | 0.63 | 0.11 | 0.13 | 0.1 | 0.19 |
| 3 | 87574738 | rs2102480 | -0.141 | -0.683 | -0.526 | 0.953 | -0.003 | 0.062 | 0.062 | 0 | 0.9 | 0.84 | 0.94 | 0.73 | 0.89 | 0.92 |
| 3 | 87575896 | rs1912247 | 0.084 | -0.526 | -0.831 | 0.871 | -0.018 | 0.004 | 0.004 | 0 | 0.66 | 0.43 | 0.77 | 0.68 | 0.69 | 0.8 |
| 3 | 87586061 | rs1509771 | -0.928 | -0.048 | 0.074 | 0.86 | 0.011 | 0.022 | 0.022 | 1.924 | 0.12 | 0.36 | 0.09 | 0.13 | 0.09 | 0.18 |
| 3 | 87591630 | rs7618312 | -0.136 | -0.086 | -0.237 | 1.007 | -0.007 | 0.033 | 0.033 | 0 | 0.23 | 0.36 | 0.18 | 0.31 | 0.18 | 0.17 |
| 3 | 87594454 | rs4858963 | 0.752 | -0.085 | -0.212 | 0.755 | -0.007 | -0.006 | -0.006 | 0 | 0.69 | 0.42 | 0.73 | 0.75 | 0.69 | 0.69 |
| 3 | 87600636 | rs9859851 | -0.26 | -0.213 | -0.121 | 0.721 | -0.021 | -0.013 | -0.013 | 0 | 0.12 | 0.18 | 0.09 | 0.06 | 0.09 | 0.06 |
| 3 | 87601624 | rs9865089 | 1.32 | -0.189 | 0.358 | 0.767 | -0.006 | 0 | 0 | 0.168 | 0.7 | 0.54 | 0.75 | 0.73 | 0.69 | 0.68 |
| 3 | 87607867 | rs6793602 | 1.392 | -0.222 | 0.4 | 0.696 | -0.005 | -0.005 | -0.005 | 1.265 | 0.68 | 0.41 | 0.73 | 0.73 | 0.68 | 0.68 |
| 3 | 87609221 | rs2048284 | 0.623 | -0.181 | 0.492 | 0.709 | 0.029 | 0.02 | 0.02 | 0 | 0.79 | 0.5 | 0.8 | 0.79 | 0.77 | 0.74 |
| 3 | 87626088 | rs1567405 | 0.031 | -0.222 | 0.448 | 0.742 | 0 | 0.001 | 0.001 | 0 | 0.84 | 0.86 | 0.85 | 0.78 | 0.79 | 0.74 |
| 3 | 87630433 | rs6767721 | -0.128 | 0.057 | 0.49 | 0.742 | -0.004 | -0.005 | -0.005 | 0 | 0.11 | 0.13 | 0.08 | 0.06 | 0.12 | 0.06 |
| 3 | 87635697 | rs2221729 | 0.134 | 0.107 | 0.409 | 0.869 | 0.008 | -0.005 | -0.005 | 0 | 0.09 | 0.03 | 0.07 | 0.08 | 0.11 | 0.07 |
| 3 | 87643214 | rs7631474 | -1.384 | 0.154 | 0.674 | 0.948 | 0.003 | 0.037 | 0.037 | 0 | 0.29 | 0.45 | 0.25 | 0.42 | 0.28 | 0.24 |
| 3 | 87645884 | rs9877171 | 2.187 | -0.208 | -0.328 | 0.414 | -0.052 | 0.011 | 0.011 | 0 | 0.34 | 0.04 | 0.41 | 0.28 | 0.31 | 0.4 |
| 3 | 87649520 | rs4499631 | 1.611 | -0.362 | -0.294 | 0.216 | -0.001 | -0.001 | -0.001 | 0.058 | 0.11 | 0.12 | 0.08 | 0.08 | 0.07 | 0.06 |
| 3 | 87657432 | rs7646773 | -1.55 | -0.366 | -0.264 | 0.222 | 0.026 | -0.039 | -0.039 | 1.316 | 0.43 | 0.82 | 0.36 | 0.5 | 0.35 | 0.31 |
| 3 | 87670474 | rs4367091 | -0.322 | -0.368 | -0.185 | 0.225 | 0.004 | 0.046 | 0.046 | 0 | 0.28 | 0.39 | 0.23 | 0.44 | 0.24 | 0.25 |
| 3 | 87673665 | rs4449347 | 0.995 | -0.371 | -0.213 | 0.227 | 0.011 | 0.01 | 0.01 | 0 | 0.11 | 0.09 | 0.08 | 0.06 | 0.11 | 0.07 |
| 3 | 87682031 | rs6798732 | -1.471 | -0.291 | -0.2 | 0.237 | 0.008 | -0.009 | -0.009 | 0 | 0.43 | 0.7 | 0.36 | 0.5 | 0.37 | 0.32 |
| 3 | 87691029 | rs4569679 | 2.594 | -0.518 | -0.297 | 0.241 | -0.067 | 0.024 | 0.024 | 0 | 0.33 | 0.03 | 0.38 | 0.27 | 0.3 | 0.38 |
| 3 | 87695226 | rs12486756 | -1.405 | -0.208 | 0.174 | 0.705 | -0.004 | 0.011 | 0.011 | 0 | 0.39 | 0.47 | 0.39 | 0.5 | 0.37 | 0.31 |
| 3 | 87695705 | rs6809812 | 1.393 | -0.171 | 0.059 | 0.73 | 0.004 | -0.021 | -0.021 | 0 | 0.1 | 0.02 | 0.09 | 0.09 | 0.08 | 0.07 |
| 3 | 87702109 | rs9808901 | -1.611 | -0.123 | 0.093 | 0.733 | 0.001 | -0.003 | -0.003 | 0 | 0.41 | 0.66 | 0.41 | 0.51 | 0.37 | 0.31 |
| 3 | 87714516 | rs4621351 | 1.162 | -0.035 | -0.097 | 0.67 | -0.05 | -0.07 | -0.07 | 0 | 0.11 | 0.01 | 0.15 | 0.36 | 0.14 | 0.15 |
| 3 | 87722254 | rs12632280 | -0.85 | -0.078 | 0.018 | 0.6 | 0.004 | 0.021 | 0.021 | 0 | 0.26 | 0.32 | 0.27 | 0.42 | 0.29 | 0.24 |
| 3 | 87733256 | rs9843580 | -1.883 | -0.095 | -0.068 | 0.602 | 0.027 | -0.025 | -0.025 | 0.677 | 0.44 | 0.79 | 0.41 | 0.48 | 0.37 | 0.32 |
| 3 | 87739286 | rs4549297 | -0.833 | 0.351 | 0.408 | 1.006 | -0.037 | -0.039 | -0.039 | 0 | 0.91 | 0.98 | 0.91 | 0.82 | 0.82 | 0.84 |
| 3 | 87766543 | rs6769025 | -0.804 | -0.514 | 0.754 | 1.273 | 0.009 | 0.083 | 0.083 | 0.21 | 0.06 | 0.15 | 0.06 | 0.18 | 0.06 | 0.04 |
| 3 | 87766735 | rs4547727 | -0.427 | -0.46 | 0.664 | 1.247 | 0.027 | 0.03 | 0.03 | 0 | 0.17 | 0.34 | 0.21 | 0.25 | 0.19 | 0.17 |
| 3 | 87783976 | rs9860340 | -1.765 | -0.432 | 0.08 | 1.218 | 0 | 0.007 | 0.007 | 0.371 | 0.27 | 0.74 | 0.29 | 0.44 | 0.25 | 0.15 |
| 3 | 87816137 | rs4546169 | -0.397 | -0.275 | -0.459 | 1.123 | -0.002 | -0.002 | -0.002 | 0 | 0.15 | 0.18 | 0.2 | 0.2 | 0.17 | 0.11 |
| 3 | 87833897 | rs2880695 | -1.135 | -0.379 | -0.546 | 1.174 | -0.012 | -0.044 | -0.044 | 0 | 0.28 | 0.22 | 0.27 | 0.42 | 0.31 | 0.19 |

| | | | | | | | | | | | | | | | | |
|---|---|---|---|---|---|---|---|---|---|---|---|---|---|---|---|---|
| 3 | 87842202 | rs1873210 | -1.262 | -0.44 | -0.523 | 1.178 | -0.001 | -0.028 | -0.028 | 0 | 0.29 | 0.27 | 0.26 | 0.42 | 0.3 | 0.2 |
| 3 | 87854277 | rs959132 | 1.982 | -0.592 | -1.323 | 0.996 | -0.118 | 0.044 | 0.044 | 0 | 0.35 | 0.02 | 0.43 | 0.26 | 0.4 | 0.53 |
| 3 | 87858977 | rs2932270 | -1.909 | -1.046 | -0.168 | 1.501 | 0.031 | -0.052 | -0.052 | 0 | 0.34 | 0.8 | 0.29 | 0.45 | 0.25 | 0.2 |
| 3 | 87859947 | rs1812620 | -1.127 | -1.072 | 0.284 | 1.487 | 0.034 | 0.025 | 0.025 | 1.951 | 0.07 | 0.54 | 0.06 | 0.01 | 0.01 | 0 |
| 3 | 87863280 | rs1489777 | -2.027 | -1.034 | 0.18 | 1.492 | 0.032 | -0.051 | -0.051 | 0.635 | 0.37 | 0.79 | 0.31 | 0.45 | 0.29 | 0.2 |
| 3 | 87866257 | rs6799017 | 2.532 | -1.05 | 0.004 | 1.532 | -0.032 | 0.103 | 0.103 | 0 | 0.53 | 0.03 | 0.62 | 0.34 | 0.6 | 0.67 |
| 3 | 87875813 | rs17025015 | -1.436 | -0.957 | 0.175 | 1.524 | 0.006 | -0.046 | -0.046 | 0 | 0.3 | 0.21 | 0.25 | 0.44 | 0.28 | 0.2 |
| 3 | 87884244 | rs2932306 | -2.231 | -0.912 | 0.773 | 1.628 | 0.025 | -0.036 | -0.036 | 0 | 0.27 | 0.82 | 0.25 | 0.38 | 0.2 | 0.15 |
| 3 | 87892012 | rs2938279 | -0.961 | -0.874 | 0.737 | 1.75 | -0.012 | 0.034 | 0.034 | 0 | 0.14 | 0.28 | 0.13 | 0.22 | 0.12 | 0.07 |
| 3 | 87909882 | rs17025082 | -1.279 | -0.85 | 0.788 | 1.747 | -0.003 | 0.054 | 0.054 | 0 | 0.09 | 0.21 | 0.07 | 0.16 | 0.07 | 0.07 |
| 3 | 87910931 | rs2932286 | -0.634 | -0.803 | 0.563 | 1.859 | -0.033 | -0.048 | -0.048 | 0 | 0.92 | 0.99 | 0.95 | 0.87 | 0.89 | 0.9 |
| 3 | 87927515 | rs2938265 | -2.446 | -0.838 | 0.611 | 1.733 | 0.035 | -0.056 | -0.056 | 3.106 | 0.27 | 0.86 | 0.25 | 0.39 | 0.2 | 0.15 |
| 3 | 87939047 | rs6764864 | -0.993 | -0.675 | 0.799 | 1.204 | -0.005 | 0.085 | 0.085 | 0 | 0.14 | 0.41 | 0.14 | 0.22 | 0.12 | 0.07 |
| 3 | 87945282 | rs2932275 | -2.035 | -0.726 | 0.735 | 1.134 | 0.004 | -0.032 | -0.032 | 0 | 0.12 | 0.5 | 0.1 | 0.04 | 0.07 | 0.08 |
| 3 | 87950229 | rs1465648 | -2.453 | -0.679 | 0.72 | 1.187 | 0.017 | 0.001 | 0.001 | 0.409 | 0.26 | 0.86 | 0.26 | 0.26 | 0.23 | 0.15 |
| 3 | 87977722 | rs923474 | -2.102 | -0.739 | 0.675 | 1.127 | -0.012 | -0.017 | -0.017 | 0 | 0.37 | 0.5 | 0.33 | 0.31 | 0.37 | 0.27 |
| 3 | 87983547 | rs6770080 | -0.665 | -0.714 | 0.69 | 0.574 | 0.018 | 0.066 | 0.066 | 0 | 0.14 | 0.34 | 0.16 | 0.21 | 0.15 | 0.07 |
| 3 | 87989436 | rs12487603 | 0.283 | -0.694 | 0.486 | 0.388 | -0.014 | 0.03 | 0.03 | 0 | 0.15 | 0.01 | 0.11 | 0.06 | 0.14 | 0.11 |
| 3 | 87998114 | rs17025185 | -0.941 | -0.664 | 0.562 | 0.404 | -0.001 | -0.006 | -0.006 | 0.102 | 0.08 | 0.11 | 0.09 | 0.04 | 0.07 | 0.1 |
| 3 | 87998942 | rs1960034 | -2.368 | -0.769 | 0.691 | 0.403 | 0.008 | 0.013 | 0.013 | 0 | 0.4 | 0.64 | 0.35 | 0.3 | 0.38 | 0.27 |
| 3 | 88001713 | rs17025198 | -1.699 | -0.464 | 0.832 | 0.54 | 0.001 | 0.006 | 0.006 | 0 | 0.21 | 0.41 | 0.24 | 0.24 | 0.23 | 0.17 |
| 3 | 88022535 | rs10865624 | 1.792 | -0.274 | 0.903 | 0.58 | 0.004 | -0.032 | -0.032 | 0 | 0.88 | 0.66 | 0.88 | 0.95 | 0.91 | 0.89 |
| 3 | 88022858 | rs12489305 | -1.516 | -0.267 | 0.837 | 0.582 | -0.001 | -0.004 | -0.004 | 0.027 | 0.09 | 0.12 | 0.08 | 0.04 | 0.08 | 0.11 |
| 3 | 88031698 | rs6770013 | 1.975 | -0.369 | 0.858 | 0.548 | 0.002 | -0.022 | -0.022 | 0 | 0.62 | 0.49 | 0.68 | 0.7 | 0.61 | 0.75 |
| 3 | 88032327 | rs12714696 | 1.942 | -0.372 | 0.87 | 0.546 | 0.003 | -0.023 | -0.023 | 0 | 0.62 | 0.51 | 0.69 | 0.7 | 0.61 | 0.75 |
| 3 | 88033939 | rs9863076 | 1.97 | -0.365 | 0.874 | 0.55 | 0.003 | -0.023 | -0.023 | 0.576 | 0.62 | 0.5 | 0.69 | 0.7 | 0.61 | 0.75 |
| 3 | 88037200 | rs1503433 | 1.944 | -0.363 | 0.87 | 0.546 | 0.003 | -0.023 | -0.023 | 0 | 0.62 | 0.51 | 0.69 | 0.7 | 0.61 | 0.75 |
| 3 | 88041650 | rs2016224 | 2.012 | -0.363 | 0.918 | 0.546 | 0 | -0.01 | -0.01 | 0.201 | 0.62 | 0.43 | 0.68 | 0.7 | 0.61 | 0.75 |
| 3 | 88044780 | rs7629565 | 2.012 | -0.363 | 0.918 | 0.546 | 0 | -0.01 | -0.01 | 0 | 0.62 | 0.43 | 0.68 | 0.7 | 0.61 | 0.75 |
| 3 | 88053396 | rs17025241 | -0.839 | -0.356 | 0.874 | 0.544 | -0.059 | -0.005 | -0.005 | 0 | 0.27 | 0.09 | 0.2 | 0.26 | 0.33 | 0.15 |
| 3 | 88064242 | rs4858930 | 1.815 | -0.342 | 1.005 | 0.287 | -0.012 | -0.011 | -0.011 | 0 | 0.89 | 0.57 | 0.89 | 0.95 | 0.95 | 0.9 |
| 3 | 88082883 | rs11915058 | -1.735 | -0.137 | 1.063 | 0.448 | -0.011 | 0.022 | 0.022 | 0 | 0.33 | 0.25 | 0.29 | 0.27 | 0.37 | 0.2 |
| 3 | 88085633 | rs6785333 | 1.825 | -0.186 | 1.112 | 0.447 | 0.003 | -0.02 | -0.02 | 0 | 0.64 | 0.41 | 0.68 | 0.73 | 0.62 | 0.8 |
| 3 | 88091543 | rs6551264 | 2.039 | -0.068 | 1.457 | 0.627 | 0.003 | 0.026 | 0.026 | 0 | 0.65 | 0.27 | 0.71 | 0.73 | 0.66 | 0.81 |
| 3 | 88092679 | rs2033389 | 2.107 | 0.397 | 1.951 | 0.743 | 0.018 | 0.009 | 0.009 | 0.541 | 0.9 | 0.5 | 0.88 | 0.95 | 0.95 | 0.96 |
| 3 | 88097372 | rs6551266 | 3.028 | 0.365 | 1.911 | 0.698 | -0.003 | -0.039 | -0.039 | 0 | 0.78 | 0.3 | 0.8 | 0.92 | 0.82 | 0.92 |
| 3 | 88101530 | rs1051436 | 2.694 | 0.36 | 1.964 | 0.97 | 0.016 | 0.025 | 0.025 | 0.955 | 0.82 | 0.43 | 0.84 | 0.83 | 0.84 | 0.91 |
| 3 | 88101551 | rs1063603 | 2.265 | 0.364 | 1.971 | 0.972 | 0.048 | 0.058 | 0.058 | 0 | 0.85 | 0.57 | 0.85 | 0.84 | 0.84 | 0.91 |
| 3 | 88102293 | rs11128020 | -0.601 | 0.364 | 2.015 | 0.972 | -0.002 | -0.003 | -0.003 | 0 | 0.06 | 0.08 | 0.07 | 0.04 | 0.05 | 0.05 |
| 3 | 88103602 | rs2116149 | 2.277 | 0.367 | 1.867 | 0.973 | 0.048 | 0.058 | 0.058 | 0 | 0.85 | 0.57 | 0.85 | 0.84 | 0.84 | 0.91 |
| 3 | 88103932 | rs1063597 | 2.277 | 0.367 | 1.867 | 0.973 | 0.048 | 0.058 | 0.058 | 0 | 0.85 | 0.57 | 0.85 | 0.84 | 0.84 | 0.91 |
| 3 | 88104411 | rs1006896 | -0.154 | 0.379 | 1.788 | 0.963 | -0.032 | 0 | 0 | 0 | 0.05 | 0.01 | 0.04 | 0.02 | 0.11 | 0.03 |
| 3 | 88139016 | rs11128021 | -2.163 | 0.368 | 1.854 | 0.96 | 0.017 | -0.014 | -0.014 | 0 | 0.12 | 0.16 | 0.12 | 0.07 | 0.15 | 0.08 |
| 3 | 88145070 | rs9310074 | 3.189 | 0.351 | 1.909 | 0.965 | -0.007 | -0.005 | -0.005 | 1.91 | 0.78 | 0.26 | 0.8 | 0.83 | 0.84 | 0.91 |
| 3 | 88171027 | rs13063511 | 3.189 | 0.358 | 1.757 | 0.968 | -0.007 | -0.005 | -0.005 | 0.011 | 0.78 | 0.26 | 0.8 | 0.83 | 0.84 | 0.91 |
| 3 | 88176189 | rs6551273 | 3.189 | 0.358 | 1.757 | 0.968 | -0.007 | -0.005 | -0.005 | 0.164 | 0.78 | 0.26 | 0.8 | 0.83 | 0.84 | 0.91 |
| 3 | 88176770 | rs4432672 | 2.454 | 0.715 | 1.215 | 1.146 | 0.048 | 0.058 | 0.058 | 0 | 0.85 | 0.57 | 0.85 | 0.84 | 0.84 | 0.91 |
| 3 | 88188420 | rs9813894 | 2.454 | 0.715 | 1.215 | 1.146 | 0.048 | 0.058 | 0.058 | 0 | 0.85 | 0.57 | 0.85 | 0.84 | 0.84 | 0.91 |
| 3 | 88190768 | rs7432826 | 0.303 | 0.72 | 1.193 | 1.143 | -0.032 | 0 | 0 | 0 | 0.95 | 0.99 | 0.96 | 0.98 | 0.89 | 0.97 |
| 3 | 88190809 | rs7432838 | 1.909 | 0.704 | 1.224 | 1.145 | 0.018 | -0.014 | -0.014 | 0.212 | 0.88 | 0.84 | 0.88 | 0.93 | 0.85 | 0.92 |
| 3 | 88197331 | rs7617691 | 2.197 | 0.713 | 1.054 | 1.126 | 0.048 | 0.058 | 0.058 | 0 | 0.85 | 0.57 | 0.85 | 0.84 | 0.84 | 0.91 |
| 3 | 88199179 | rs6551277 | 2.278 | 0.713 | 1.055 | 1.126 | 0.056 | 0.066 | 0.066 | 0 | 0.86 | 0.6 | 0.86 | 0.84 | 0.84 | 0.91 |
| 3 | 88199298 | rs6551278 | 2.168 | 0.717 | 1.024 | 1.128 | 0.048 | 0.058 | 0.058 | 0 | 0.85 | 0.57 | 0.85 | 0.84 | 0.84 | 0.91 |
| 3 | 88258450 | rs7427047 | 2.124 | 0.691 | 0.986 | 1.185 | 0.046 | 0.068 | 0.068 | 0 | 0.84 | 0.54 | 0.85 | 0.84 | 0.84 | 0.9 |
| 3 | 88273071 | rs9284814 | 1.166 | 0.872 | 0.754 | 1.222 | 0.066 | 0.084 | 0.084 | 0 | 0.93 | 0.76 | 0.93 | 0.87 | 0.89 | 0.96 |
| 3 | 88277712 | rs7634064 | -0.017 | 1.126 | -0.415 | 1.341 | -0.063 | -0.085 | -0.085 | 0.153 | 0.22 | 0.02 | 0.15 | 0.26 | 0.27 | 0.25 |
| 3 | 88287719 | rs6419870 | 0.847 | 1.265 | -0.336 | 1.088 | -0.037 | 0.003 | 0.003 | 0 | 0.92 | 0.98 | 0.93 | 0.94 | 0.85 | 0.96 |
| 3 | 88296807 | rs6769326 | 0.783 | 1.267 | -0.037 | 1.089 | 0.025 | -0.004 | -0.004 | 0 | 0.93 | 0.92 | 0.93 | 0.94 | 0.85 | 0.96 |
| 3 | 88316277 | rs2011414 | 1.394 | 1.295 | -0.356 | 1.351 | -0.107 | -0.025 | -0.025 | 4.286 | 0.56 | 0.06 | 0.58 | 0.39 | 0.52 | 0.64 |
| 3 | 88327426 | rs936361 | -0.149 | 1.349 | 0.568 | 1.598 | -0.008 | 0.026 | 0.026 | 0 | 0.26 | 0.35 | 0.21 | 0.4 | 0.26 | 0.27 |
| 3 | 88334311 | rs6785442 | 1.272 | 1.454 | 1.33 | 1.517 | 0.082 | 0.047 | 0.047 | 0.979 | 0.83 | 0.42 | 0.79 | 0.79 | 0.79 | 0.91 |
| 3 | 88347562 | rs2199016 | 1.215 | 1.464 | 1.253 | 1.52 | 0.114 | 0.073 | 0.073 | 0.581 | 0.84 | 0.48 | 0.79 | 0.79 | 0.79 | 0.91 |
| 3 | 88377822 | rs6796650 | 0.903 | 0.642 | -0.398 | 1.378 | -0.029 | 0.187 | 0.187 | 0 | 0.53 | 0.1 | 0.31 | 0.23 | 0.53 | 0.56 |
| 3 | 88378348 | rs9858909 | 0.565 | 0.956 | -0.553 | 1.802 | 0.028 | -0.014 | -0.014 | 0.236 | 0.61 | 0.24 | 0.69 | 0.41 | 0.66 | 0.6 |
| 3 | 88379459 | rs9826824 | 1.142 | 0.94 | -0.545 | 1.775 | -0.003 | 0.065 | 0.065 | 0 | 0.55 | 0.23 | 0.32 | 0.3 | 0.54 | 0.57 |
| 3 | 88383142 | rs4859013 | -0.728 | 0.744 | 0.493 | 2.48 | 0.008 | 0.035 | 0.035 | 0 | 0.3 | 0.38 | 0.56 | 0.43 | 0.34 | 0.25 |
| 3 | 88387623 | rs9861542 | 1.48 | 0.742 | 0.451 | 2.412 | 0 | 0.06 | 0.06 | 0 | 0.56 | 0.23 | 0.33 | 0.3 | 0.54 | 0.59 |
| 3 | 88420443 | rs9830719 | 1.526 | 0.565 | 1.139 | 2.75 | 0 | 0.155 | 0.155 | 0 | 0.54 | 0.13 | 0.32 | 0.23 | 0.54 | 0.58 |
| 3 | 88425671 | rs1458012 | 0.222 | 0.574 | 1.258 | 2.896 | -0.01 | -0.009 | -0.009 | 0.459 | 0.21 | 0.24 | 0.14 | 0.18 | 0.19 | 0.19 |

| | | | | | | | | | | | | | | | | |
|---|---|---|---|---|---|---|---|---|---|---|---|---|---|---|---|---|
| 3 | 88431461 | rs1458001 | -0.175 | 0.593 | 1.136 | 2.968 | -0.036 | -0.036 | -0.036 | 0.12 | 0.06 | 0.01 | 0.36 | 0.13 | 0.12 | 0.03 |
| 3 | 88435576 | rs6551321 | -0.327 | 0.572 | 1.178 | 2.865 | -0.004 | -0.009 | -0.009 | 0 | 0.27 | 0.22 | 0.5 | 0.31 | 0.31 | 0.22 |
| 3 | 88445108 | rs9828049 | 0.354 | 0.65 | 1.237 | 2.679 | -0.012 | 0.08 | 0.08 | 2.287 | 0.86 | 0.61 | 0.88 | 0.73 | 0.88 | 0.85 |
| 3 | 88452746 | rs2344502 | 1.739 | 0.385 | 0.501 | 2.369 | -0.12 | 0.107 | 0.107 | 0.364 | 0.46 | 0.02 | 0.29 | 0.26 | 0.51 | 0.5 |
| 3 | 88454307 | rs1458015 | 0.225 | 0.414 | 0.502 | 2.398 | 0.002 | -0.035 | -0.035 | 0 | 0.11 | 0.05 | 0.08 | 0.24 | 0.11 | 0.19 |
| 3 | 88455499 | rs1463473 | -1.802 | 0.394 | 0.721 | 2.391 | 0 | 0.097 | 0.097 | 0 | 0.52 | 0.75 | 0.68 | 0.71 | 0.49 | 0.51 |
| 3 | 88461825 | rs1458006 | -0.912 | 0.343 | 0.853 | 2.327 | -0.003 | 0.008 | 0.008 | 0 | 0.38 | 0.53 | 0.58 | 0.51 | 0.37 | 0.31 |
| 3 | 88467206 | rs7645228 | -1.65 | 0.319 | 0.66 | 2.119 | 0.011 | 0.052 | 0.052 | 0 | 0.5 | 0.55 | 0.66 | 0.75 | 0.48 | 0.51 |
| 3 | 88471321 | rs11922350 | -0.301 | 0.348 | 0.387 | 2.224 | 0.003 | 0 | 0 | 0 | 0.13 | 0.01 | 0.08 | 0.1 | 0.11 | 0.16 |
| 3 | 88472522 | rs3911553 | -0.42 | 0.403 | 0.742 | 2.466 | 0.01 | 0.004 | 0.004 | 0.092 | 0.29 | 0.37 | 0.26 | 0.25 | 0.32 | 0.21 |
| 3 | 88476595 | rs6789349 | 1.148 | 0.439 | 0.711 | 2.602 | -0.005 | -0.042 | -0.042 | 0.508 | 0.58 | 0.22 | 0.6 | 0.57 | 0.58 | 0.6 |
| 3 | 88478666 | rs4859028 | 1.074 | -0.158 | 0.107 | 2.189 | -0.007 | 0.01 | 0.01 | 0 | 0.14 | 0.01 | 0.32 | 0.08 | 0.09 | 0.07 |
| 3 | 88479786 | rs4563418 | 0.81 | -0.09 | 0.303 | 2.257 | -0.02 | 0.011 | 0.011 | 0.061 | 0.21 | 0.31 | 0.15 | 0.19 | 0.18 | 0.19 |
| 3 | 88481090 | rs6795028 | 0.951 | -0.115 | 0.298 | 2.246 | 0.063 | -0.001 | -0.001 | 0 | 0.51 | 0.32 | 0.39 | 0.54 | 0.62 | 0.55 |
| 3 | 88484194 | rs4470525 | -0.379 | -0.073 | 0.353 | 1.912 | -0.018 | 0.066 | 0.066 | 0 | 0.86 | 0.63 | 0.86 | 0.82 | 0.89 | 0.81 |
| 3 | 88492824 | rs4292237 | 0.153 | -0.022 | 0.325 | 2.041 | -0.03 | 0.091 | 0.091 | 0 | 0.26 | 0.35 | 0.21 | 0.49 | 0.2 | 0.28 |
| 3 | 88498034 | rs11128045 | -0.924 | 0.201 | 0.406 | 1.903 | 0 | -0.001 | -0.001 | 0 | 0.49 | 0.51 | 0.61 | 0.64 | 0.42 | 0.38 |
| 3 | 88502687 | rs13089642 | -0.487 | -0.137 | 0.485 | 1.766 | -0.022 | 0.096 | 0.096 | 0 | 0.06 | 0.22 | 0.03 | 0.22 | 0.02 | 0.09 |
| 3 | 88507849 | rs13101122 | -0.602 | -0.075 | 0.418 | 1.805 | 0.007 | 0.007 | 0.007 | 0.746 | 0.43 | 0.39 | 0.58 | 0.68 | 0.33 | 0.4 |
| 3 | 88518323 | rs4859033 | 1.432 | -0.149 | 0.291 | 1.728 | 0.008 | 0.005 | 0.005 | 0.337 | 0.72 | 0.18 | 0.8 | 0.7 | 0.77 | 0.71 |
| 3 | 88522589 | rs4543020 | 0.327 | -0.117 | 0.441 | 1.741 | 0.027 | -0.059 | -0.059 | 0 | 0.8 | 0.55 | 0.85 | 0.92 | 0.79 | 0.8 |
| 3 | 88524175 | rs2880961 | 0.117 | -0.115 | 0.427 | 1.701 | 0.029 | -0.04 | -0.04 | 0 | 0.36 | 0.17 | 0.55 | 0.46 | 0.3 | 0.31 |
| 3 | 88524999 | rs9851482 | -0.546 | -0.093 | 0.456 | 1.692 | -0.008 | 0.142 | 0.142 | 0 | 0.07 | 0.28 | 0.04 | 0.22 | 0.02 | 0.09 |
| 3 | 88526986 | rs2344815 | 0.22 | -0.064 | 0.435 | 1.645 | 0.002 | -0.01 | -0.01 | 0.02 | 0.07 | 0.07 | 0.07 | 0 | 0.13 | 0.02 |
| 3 | 88532455 | rs4859036 | 0.188 | -0.12 | 0.377 | 1.677 | 0.027 | -0.066 | -0.066 | 0 | 0.8 | 0.55 | 0.84 | 0.93 | 0.79 | 0.8 |
| 3 | 88532530 | rs6776423 | 1.798 | -0.081 | -0.022 | 1.692 | -0.091 | 0.075 | 0.075 | 1.268 | 0.37 | 0.11 | 0.26 | 0.25 | 0.47 | 0.4 |
| 3 | 88536336 | rs2344956 | 1.359 | 0.042 | 0.267 | 1.718 | 0.001 | -0.009 | -0.009 | 0.128 | 0.71 | 0.24 | 0.77 | 0.55 | 0.74 | 0.67 |
| 3 | 88544569 | rs13077366 | -1.162 | -0.085 | 0.445 | 1.802 | -0.002 | 0.241 | 0.241 | 0 | 0.09 | 0.31 | 0.07 | 0.36 | 0.07 | 0.15 |
| 3 | 88558883 | rs13098649 | -1.132 | -0.091 | 0.515 | 1.872 | -0.009 | 0.197 | 0.197 | 0 | 0.09 | 0.29 | 0.07 | 0.36 | 0.07 | 0.15 |
| 3 | 88576606 | rs7372143 | 0.081 | -0.177 | 0.152 | 1.802 | 0.004 | -0.042 | -0.042 | 0 | 0.34 | 0.06 | 0.51 | 0.29 | 0.28 | 0.27 |
| 3 | 88580067 | rs7430972 | 1.577 | 0.08 | 0.561 | 1.778 | 0.014 | 0.149 | 0.149 | 0.118 | 0.82 | 0.51 | 0.85 | 0.64 | 0.81 | 0.84 |
| 3 | 88585081 | rs6551338 | 0.852 | 0.109 | 0.159 | 1.686 | -0.003 | 0.01 | 0.01 | 0.074 | 0.89 | 0.94 | 0.93 | 0.91 | 0.87 | 0.91 |
| 3 | 88604818 | rs6551343 | -0.365 | -0.015 | 0.132 | 1.639 | -0.002 | -0.009 | -0.009 | 0.137 | 0.28 | 0.31 | 0.22 | 0.17 | 0.28 | 0.18 |
| 3 | 88609467 | rs6419877 | 0.141 | -0.003 | 0.125 | 1.727 | 0.007 | -0.005 | -0.005 | 0 | 0.92 | 0.94 | 0.94 | 0.76 | 0.94 | 0.88 |
| 3 | 88616805 | rs7613601 | 0.06 | 0.002 | 0.149 | 1.678 | 0 | -0.002 | -0.002 | 0.659 | 0.25 | 0.24 | 0.2 | 0.17 | 0.28 | 0.18 |
| 3 | 88627781 | rs7642570 | 1.151 | 0.042 | 0.118 | 1.68 | 0 | 0.016 | 0.016 | 0 | 0.84 | 0.8 | 0.87 | 0.76 | 0.83 | 0.87 |
| 3 | 88633850 | rs9868943 | 0.924 | 0.06 | 0.089 | 1.692 | 0.001 | 0.011 | 0.011 | 0.019 | 0.87 | 0.85 | 0.9 | 0.76 | 0.84 | 0.85 |
| 3 | 88636898 | rs6773719 | 0.826 | -0.058 | 0.14 | 1.633 | -0.006 | 0.01 | 0.01 | 0 | 0.63 | 0.55 | 0.71 | 0.48 | 0.66 | 0.67 |
| 3 | 88643544 | rs17025980 | -0.033 | 0.022 | 0.345 | 1.873 | -0.003 | 0.024 | 0.024 | 0 | 0.93 | 0.91 | 0.94 | 0.8 | 0.94 | 0.88 |
| 3 | 88648609 | rs6551355 | 0.37 | -0.001 | 0.401 | 1.863 | -0.011 | 0.059 | 0.059 | 0 | 0.92 | 0.82 | 0.94 | 0.8 | 0.94 | 0.88 |
| 3 | 88659709 | rs963736 | -0.314 | -0.098 | 0.359 | 1.837 | 0.003 | -0.006 | -0.006 | 0 | 0.28 | 0.28 | 0.21 | 0.51 | 0.23 | 0.3 |
| 3 | 88665467 | rs6777020 | 0.648 | -0.095 | 0.316 | 1.841 | 0.012 | -0.046 | -0.046 | 0 | 0.2 | 0.11 | 0.15 | 0.31 | 0.17 | 0.19 |
| 3 | 88668147 | rs7646260 | -1.569 | -0.317 | 0.33 | 1.469 | 0.009 | 0.061 | 0.061 | 0.044 | 0.51 | 0.63 | 0.67 | 0.65 | 0.45 | 0.43 |
| 3 | 88670359 | rs7629888 | -0.024 | -0.28 | 0.088 | 1.504 | 0.016 | -0.038 | -0.038 | 0 | 0.27 | 0.19 | 0.2 | 0.51 | 0.23 | 0.3 |
| 3 | 88670387 | rs10511138 | 0.981 | -0.283 | 0.073 | 1.5 | 0.008 | -0.015 | -0.015 | 0.251 | 0.19 | 0.08 | 0.15 | 0.17 | 0.17 | 0.16 |
| 3 | 88672567 | rs9832715 | -0.143 | -0.256 | 0.11 | 1.505 | -0.008 | 0.016 | 0.016 | 0 | 0.28 | 0.3 | 0.22 | 0.5 | 0.23 | 0.31 |
| 3 | 88675576 | rs1495838 | 1.161 | -0.213 | 0.044 | 1.439 | -0.038 | -0.046 | -0.046 | 0 | 0.14 | 0.01 | 0.36 | 0.15 | 0.11 | 0.11 |
| 3 | 88689456 | rs2344816 | 0.331 | -0.244 | 0.155 | 1.496 | 0.009 | -0.01 | -0.01 | 0 | 0.21 | 0.27 | 0.44 | 0.15 | 0.21 | 0.12 |
| 3 | 88700486 | rs7652836 | 0.529 | -0.366 | -0.125 | 1.415 | 0.009 | 0 | 0 | 0 | 0.22 | 0.16 | 0.15 | 0.17 | 0.17 | 0.16 |
| 3 | 88708334 | rs7638288 | 0.753 | 0.678 | -0.334 | 0.85 | -0.008 | 0.031 | 0.031 | 0.299 | 0.62 | 0.29 | 0.4 | 0.78 | 0.63 | 0.69 |
| 3 | 88715097 | rs2047253 | -0.137 | 0.87 | -0.532 | 0.671 | 0.013 | -0.051 | -0.051 | 1.786 | 0.83 | 0.59 | 0.86 | 0.92 | 0.84 | 0.83 |
| 3 | 88720805 | rs1390390 | 1.009 | 1.035 | -1.351 | 0.855 | -0.107 | 0.031 | 0.031 | 0.586 | 0.35 | 0.02 | 0.21 | 0.26 | 0.4 | 0.39 |
| 3 | 88731112 | rs9814076 | 1.225 | 1.295 | -1.587 | 0.792 | -0.075 | -0.021 | -0.021 | 0.146 | 0.28 | 0.02 | 0.19 | 0.27 | 0.34 | 0.32 |
| 3 | 88733658 | rs1495824 | -0.603 | 1.29 | -0.698 | 0.57 | 0.006 | -0.012 | -0.012 | 0 | 0.62 | 0.55 | 0.7 | 0.65 | 0.52 | 0.55 |
| 3 | 88753628 | rs17026135 | 0.561 | 0.972 | -1.371 | 0.824 | -0.047 | -0.069 | -0.069 | 0 | 0.27 | 0.03 | 0.15 | 0.29 | 0.23 | 0.25 |
| 3 | 88753647 | rs10511140 | 1.561 | 0.843 | -1.702 | 0.987 | -0.024 | -0.073 | -0.073 | 2.883 | 0.13 | 0.01 | 0.36 | 0.19 | 0.1 | 0.12 |
| 3 | 88756407 | rs6419880 | -1.147 | 0.769 | -1.566 | 0.973 | -0.009 | 0.01 | 0.01 | 0.018 | 0.57 | 0.62 | 0.66 | 0.61 | 0.46 | 0.47 |
| 3 | 88761581 | rs1603312 | -0.623 | 0.909 | -1.11 | 0.933 | 0.029 | 0.005 | 0.005 | 0 | 0.93 | 0.77 | 0.93 | 0.93 | 0.93 | 0.94 |
| 3 | 88763832 | rs1587152 | -1.723 | 0.79 | -1.075 | 0.961 | -0.002 | 0.001 | 0.001 | 0.105 | 0.62 | 0.62 | 0.7 | 0.65 | 0.52 | 0.55 |
| 3 | 88765697 | rs1845754 | -0.511 | 0.77 | -0.748 | 0.998 | -0.008 | -0.007 | -0.007 | 0.003 | 0.42 | 0.5 | 0.28 | 0.39 | 0.36 | 0.33 |
| 3 | 88767061 | rs11128059 | -1.786 | 0.774 | -0.805 | 0.964 | -0.011 | 0.011 | 0.011 | 0.109 | 0.63 | 0.68 | 0.7 | 0.65 | 0.53 | 0.55 |
| 3 | 88773079 | rs970485 | -1.786 | 0.774 | -0.805 | 0.964 | -0.011 | 0.011 | 0.011 | 0.109 | 0.63 | 0.68 | 0.7 | 0.65 | 0.52 | 0.55 |
| 3 | 88777721 | rs11914552 | 0.412 | 0.749 | -0.29 | 0.903 | -0.002 | -0.013 | -0.013 | 0 | 0.13 | 0.23 | 0.11 | 0.1 | 0.13 | 0.08 |
| 3 | 88782993 | rs12638255 | 0.82 | 0.767 | -0.834 | 1.064 | -0.055 | 0.017 | 0.017 | 0.902 | 0.27 | 0.01 | 0.15 | 0.14 | 0.22 | 0.22 |
| 3 | 88788435 | rs9877297 | 0.507 | 1.163 | -0.516 | 1.051 | 0.024 | 0.012 | 0.012 | 0 | 0.07 | 0.11 | 0.06 | 0.08 | 0.09 | 0.11 |
| 3 | 88790309 | rs9846491 | 0.507 | 1.163 | -0.516 | 1.051 | 0.024 | 0.012 | 0.012 | 0 | 0.07 | 0.11 | 0.06 | 0.08 | 0.09 | 0.11 |
| 3 | 88793337 | rs9828389 | 0.361 | 1.161 | -0.507 | 1.05 | 0.024 | 0.012 | 0.012 | 0 | 0.07 | 0.12 | 0.06 | 0.08 | 0.09 | 0.11 |
| 3 | 88793424 | rs11914750 | -0.374 | 0.882 | -0.549 | 1.057 | -0.004 | -0.002 | -0.002 | 0 | 0.42 | 0.55 | 0.28 | 0.39 | 0.34 | 0.32 |
| 3 | 88793982 | rs2046327 | 1.273 | 0.952 | -0.842 | 1.135 | -0.026 | -0.073 | -0.073 | 0.349 | 0.13 | 0.01 | 0.36 | 0.19 | 0.1 | 0.12 |

| | | | | | | | | | | | | | | | | |
|---|---|---|---|---|---|---|---|---|---|---|---|---|---|---|---|---|
| 3 | 88795331 | rs13061110 | -1.001 | 0.928 | -0.468 | 0.809 | -0.001 | 0 | 0 | 1.874 | 0.58 | 0.6 | 0.65 | 0.59 | 0.45 | 0.44 |
| 3 | 88795626 | rs6808681 | 1.003 | 0.576 | -0.764 | 0.595 | 0.068 | 0.071 | 0.071 | 4.797 | 0.5 | 0.21 | 0.4 | 0.39 | 0.64 | 0.6 |
| 3 | 88795783 | rs2170716 | 1.449 | 1.322 | -0.87 | 0.532 | 0 | 0.033 | 0.033 | 0 | 0.4 | 0.23 | 0.28 | 0.36 | 0.45 | 0.43 |
| 3 | 88803526 | rs2345428 | 1.391 | 1.395 | -0.782 | 0.414 | -0.002 | 0.037 | 0.037 | 0 | 0.4 | 0.28 | 0.28 | 0.35 | 0.46 | 0.46 |
| 3 | 88815135 | rs12633508 | -0.125 | 2.162 | -0.397 | 0.424 | 0.023 | 0.002 | 0.002 | 0 | 0.13 | 0.18 | 0.16 | 0.17 | 0.25 | 0.07 |
| 3 | 88853354 | rs13093533 | -0.594 | 1.378 | -0.061 | 0.494 | -0.008 | -0.025 | -0.025 | 0.22 | 0.33 | 0.29 | 0.53 | 0.43 | 0.34 | 0.31 |
| 3 | 88857169 | rs1483480 | -0.94 | 1.346 | -0.223 | 0.467 | -0.043 | -0.006 | -0.006 | 0 | 0.78 | 0.99 | 0.86 | 0.85 | 0.8 | 0.77 |
| 3 | 88858018 | rs9853806 | 2.506 | 1.562 | -0.384 | 0.419 | -0.037 | -0.016 | -0.016 | 0.052 | 0.34 | 0.03 | 0.19 | 0.32 | 0.35 | 0.39 |
| 3 | 88859041 | rs13093031 | 0.605 | 0.922 | -0.178 | 0.527 | 0.017 | 0.01 | 0.01 | 0 | 0.11 | 0.17 | 0.12 | 0.17 | 0.16 | 0.05 |
| 3 | 88859625 | rs11712997 | 1.94 | 0.924 | -0.115 | 0.53 | -0.01 | 0.028 | 0.028 | 0 | 0.4 | 0.23 | 0.28 | 0.38 | 0.46 | 0.46 |
| 3 | 88859724 | rs4855810 | 0.542 | 0.826 | -0.392 | 0.609 | 0 | -0.073 | -0.073 | 1.216 | 0.22 | 0.07 | 0.41 | 0.26 | 0.17 | 0.26 |
| 3 | 88863054 | rs10511142 | 1.899 | 0.975 | -0.368 | 0.757 | -0.023 | -0.074 | -0.074 | 0 | 0.13 | 0.01 | 0.35 | 0.2 | 0.08 | 0.12 |
| 3 | 88863594 | rs9827227 | -0.742 | 1.025 | -0.226 | 0.728 | -0.032 | 0.001 | 0.001 | 0 | 0.78 | 0.97 | 0.86 | 0.85 | 0.81 | 0.77 |
| 3 | 88865890 | rs4855811 | -0.742 | 1.025 | -0.226 | 0.728 | -0.032 | 0.001 | 0.001 | 0 | 0.78 | 0.97 | 0.86 | 0.85 | 0.81 | 0.77 |
| 3 | 88911119 | rs12053900 | 1.861 | 1.024 | -0.298 | 0.844 | -0.042 | -0.075 | -0.075 | 0 | 0.12 | 0.01 | 0.36 | 0.2 | 0.1 | 0.12 |
| 3 | 88911404 | rs4462970 | 0.612 | 0.989 | -0.288 | 0.855 | -0.017 | -0.066 | -0.066 | 0 | 0.18 | 0.07 | 0.41 | 0.26 | 0.17 | 0.25 |
| 3 | 88933154 | rs6774956 | 0.927 | 0.971 | -0.155 | 0.818 | 0.001 | -0.003 | -0.003 | 0 | 0.06 | 0.06 | 0.05 | 0.06 | 0.07 | 0.13 |
| 3 | 88936216 | rs9847347 | 1.524 | 1.055 | 0.094 | 0.698 | 0.01 | 0.025 | 0.025 | 0 | 0.44 | 0.35 | 0.3 | 0.38 | 0.46 | 0.46 |
| 3 | 88949759 | rs9817483 | 0.759 | 1.059 | -0.028 | 0.719 | -0.002 | 0.001 | 0.001 | 0 | 0.56 | 0.55 | 0.41 | 0.54 | 0.64 | 0.52 |
| 3 | 88963322 | rs13100058 | 0.227 | 1.204 | -0.406 | 0.792 | -0.017 | -0.011 | -0.011 | 0.103 | 0.1 | 0.06 | 0.11 | 0.17 | 0.17 | 0.06 |
| 3 | 88973311 | rs12053823 | 0.15 | 1.054 | -0.539 | 0.822 | -0.057 | -0.104 | -0.104 | 0.099 | 0.23 | 0.07 | 0.42 | 0.42 | 0.25 | 0.19 |
| 3 | 88974863 | rs4552364 | 1.618 | 0.652 | -0.24 | 0.64 | -0.001 | 0.007 | 0.007 | 0 | 0.41 | 0.37 | 0.34 | 0.34 | 0.49 | 0.45 |
| 3 | 88978059 | rs4368505 | 0.062 | 0.602 | -0.055 | 0.649 | -0.024 | -0.042 | -0.042 | 0.036 | 0.71 | 0.6 | 0.83 | 0.85 | 0.8 | 0.76 |
| 3 | 88978538 | rs7649188 | 1.269 | 0.602 | -0.082 | 0.658 | -0.001 | 0 | 0 | 0 | 0.47 | 0.46 | 0.39 | 0.37 | 0.54 | 0.57 |
| 3 | 88985151 | rs9864351 | -0.122 | 0.463 | -0.13 | 0.643 | 0.006 | 0.012 | 0.012 | 0.941 | 0.73 | 0.76 | 0.84 | 0.85 | 0.8 | 0.76 |
| 3 | 88992369 | rs4311220 | 0.517 | 0.599 | -0.002 | 0.848 | 0.022 | 0.008 | 0.008 | 0.449 | 0.54 | 0.64 | 0.5 | 0.64 | 0.69 | 0.62 |
| 3 | 88993275 | rs4270512 | 1.799 | 0.76 | 0.019 | 0.934 | -0.003 | -0.027 | -0.027 | 0 | 0.37 | 0.44 | 0.32 | 0.33 | 0.42 | 0.42 |
| 3 | 88994672 | rs958144 | 1.114 | 0.446 | -0.196 | 0.905 | -0.003 | -0.068 | -0.068 | 0.455 | 0.23 | 0.02 | 0.33 | 0.21 | 0.13 | 0.15 |
| 3 | 88999499 | rs17026486 | -0.946 | -0.096 | -0.268 | 0.57 | -0.007 | -0.029 | -0.029 | 0.647 | 0.18 | 0.05 | 0.1 | 0.22 | 0.15 | 0.06 |
| 3 | 89004916 | rs11924091 | -0.431 | 0.769 | 0.349 | 0.922 | 0.003 | 0.004 | 0.004 | 0 | 0.1 | 0.22 | 0.09 | 0.08 | 0.12 | 0.14 |
| 3 | 89012893 | rs1496522 | -1.165 | 0.816 | 0.372 | 0.891 | -0.001 | -0.001 | -0.001 | 0 | 0.56 | 0.47 | 0.58 | 0.45 | 0.43 | 0.52 |
| 3 | 89013058 | rs1496523 | -0.568 | 1.057 | 0.174 | 1.179 | 0 | 0.008 | 0.008 | 0 | 0.93 | 0.93 | 0.9 | 0.78 | 0.85 | 0.94 |
| 3 | 89034040 | rs1496515 | -1.331 | 0.061 | 0.686 | 1.332 | -0.005 | -0.004 | -0.004 | 0 | 0.55 | 0.38 | 0.64 | 0.46 | 0.45 | 0.54 |
| 3 | 89040057 | rs12495112 | -1.95 | 0.08 | 0.847 | 1.332 | -0.026 | 0.05 | 0.05 | 0.108 | 0.62 | 0.73 | 0.73 | 0.73 | 0.51 | 0.59 |
| 3 | 89044238 | rs2881281 | -0.208 | -0.02 | 0.675 | 1.411 | 0 | -0.031 | -0.031 | 0 | 0.42 | 0.07 | 0.47 | 0.33 | 0.3 | 0.34 |
| 3 | 89048023 | rs17026569 | -0.401 | -0.02 | 0.763 | 1.403 | 0.011 | 0.015 | 0.015 | 0 | 0.11 | 0.21 | 0.16 | 0.12 | 0.14 | 0.19 |
| 3 | 89048700 | rs7636945 | 2.035 | 0.002 | 0.179 | 1.378 | -0.051 | 0.078 | 0.078 | 0.1 | 0.37 | 0.16 | 0.26 | 0.28 | 0.49 | 0.41 |
| 3 | 89051472 | rs875005 | 0.772 | -0.092 | -0.108 | 1.415 | -0.026 | 0.02 | 0.02 | 0 | 0.19 | 0.01 | 0.13 | 0.13 | 0.17 | 0.2 |
| 3 | 89054190 | rs35631553 | 2.187 | -0.033 | -0.063 | 1.424 | -0.128 | 0.077 | 0.077 | 0.078 | 0.37 | 0.06 | 0.25 | 0.28 | 0.49 | 0.41 |
| 3 | 89066588 | rs4571247 | 1.097 | -0.084 | -0.035 | 1.37 | -0.006 | -0.068 | -0.068 | 0 | 0.22 | 0.03 | 0.34 | 0.2 | 0.14 | 0.14 |
| 3 | 89074355 | rs9836220 | 1.451 | 0.025 | 0.482 | 1.393 | -0.005 | 0 | 0 | 0 | 0.43 | 0.35 | 0.31 | 0.49 | 0.54 | 0.45 |
| 3 | 89075886 | rs4555526 | -2.005 | -0.205 | 0.894 | 0.97 | 0.002 | 0.067 | 0.067 | 0.155 | 0.23 | 0.63 | 0.27 | 0.39 | 0.23 | 0.25 |
| 3 | 89087160 | rs9825532 | -0.452 | -0.192 | 0.663 | 0.413 | 0.005 | 0.024 | 0.024 | 0 | 0.28 | 0.2 | 0.21 | 0.22 | 0.28 | 0.31 |
| 3 | 89088459 | rs2088304 | -0.425 | -0.191 | 0.602 | 0.414 | 0.003 | 0.028 | 0.028 | 0 | 0.28 | 0.16 | 0.21 | 0.22 | 0.28 | 0.31 |
| 3 | 89097831 | rs1496502 | -1.373 | -0.176 | 1.344 | 0.812 | 0.02 | 0.141 | 0.141 | 0.505 | 0.11 | 0.31 | 0.14 | 0.22 | 0.09 | 0.15 |
| 3 | 89103302 | rs1496504 | -1.38 | -0.327 | 1.468 | 0.73 | -0.003 | 0.009 | 0.009 | 0 | 0.37 | 0.58 | 0.35 | 0.48 | 0.34 | 0.43 |
| 3 | 89110767 | rs10511144 | -0.236 | -0.327 | 1.377 | 0.731 | 0 | 0 | 0 | 0.108 | 0.26 | 0.26 | 0.21 | 0.25 | 0.25 | 0.28 |
| 3 | 89121921 | rs9942009 | -1.304 | -0.33 | 1.447 | 0.67 | -0.006 | 0.041 | 0.041 | 0 | 0.35 | 0.48 | 0.34 | 0.48 | 0.34 | 0.43 |
| 3 | 89132282 | rs9850867 | -0.741 | -0.317 | 1.349 | 0.673 | -0.001 | 0.008 | 0.008 | 0 | 0.3 | 0.31 | 0.25 | 0.46 | 0.3 | 0.34 |
| 3 | 89156519 | rs13074291 | 0.346 | -0.446 | 1.299 | 0.435 | -0.055 | -0.069 | -0.069 | 0 | 0.26 | 0.41 | 0.14 | 0.07 | 0.13 | 0.15 |
| 3 | 89164099 | rs9860797 | -0.101 | -0.496 | 1.106 | 0.41 | 0.012 | 0.014 | 0.014 | 0 | 0.27 | 0.21 | 0.2 | 0.19 | 0.22 | 0.3 |
| 3 | 89178317 | rs9883387 | -0.152 | -0.485 | 1.117 | 0.417 | 0.01 | 0.01 | 0.01 | 0 | 0.28 | 0.23 | 0.19 | 0.2 | 0.22 | 0.3 |
| 3 | 89187123 | rs7428598 | 1.476 | -0.502 | 1.111 | 0.447 | 0.012 | -0.024 | -0.024 | 0 | 0.41 | 0.32 | 0.65 | 0.53 | 0.6 | 0.49 |
| 3 | 89211284 | rs9868686 | 1.153 | -0.584 | 1.288 | 0.452 | -0.007 | 0.001 | 0.001 | 0.356 | 0.44 | 0.48 | 0.67 | 0.55 | 0.61 | 0.49 |
| 3 | 89212941 | rs12633482 | 1.29 | -0.576 | 1.289 | 0.445 | -0.003 | 0.001 | 0.001 | 0.293 | 0.43 | 0.46 | 0.66 | 0.53 | 0.61 | 0.49 |
| 3 | 89213871 | rs13058761 | 0.827 | -0.592 | 1.096 | 0.422 | 0.008 | 0.005 | 0.005 | 0 | 0.23 | 0.17 | 0.1 | 0.05 | 0.12 | 0.15 |
| 3 | 89237732 | rs4128782 | 1.913 | -0.393 | 0.803 | 0.53 | -0.151 | -0.123 | -0.123 | 0.018 | 0.35 | 0.03 | 0.5 | 0.44 | 0.5 | 0.41 |
| 3 | 89250573 | rs7374904 | -0.069 | -0.025 | 1.629 | 0.683 | 0.079 | 0.089 | 0.089 | 0 | 0.07 | 0.34 | 0.16 | 0.08 | 0.11 | 0.08 |
| 3 | 89251079 | rs7432113 | 0.589 | -0.104 | 1.747 | 0.671 | -0.003 | 0.001 | 0.001 | 0 | 0.64 | 0.38 | 0.7 | 0.65 | 0.72 | 0.72 |
| 3 | 89262358 | rs9852594 | 0.248 | -0.079 | 1.778 | 0.511 | 0.02 | 0.026 | 0.026 | 0 | 0.06 | 0.18 | 0.15 | 0.08 | 0.11 | 0.07 |
| 3 | 89265366 | rs7432948 | 0.764 | -0.104 | 1.833 | 0.509 | 0.025 | -0.004 | -0.004 | 0 | 0.63 | 0.27 | 0.68 | 0.67 | 0.72 | 0.72 |
| 3 | 89268714 | rs6797343 | 0.178 | -0.09 | 1.844 | 0.518 | -0.041 | -0.004 | -0.004 | 1.212 | 0.7 | 0.51 | 0.84 | 0.76 | 0.83 | 0.79 |
| 3 | 89270605 | rs6551407 | 0.856 | 0.147 | 1.847 | 0.648 | 0.023 | 0.107 | 0.107 | 0 | 0.9 | 0.75 | 0.83 | 0.72 | 0.85 | 0.87 |
| 3 | 89271087 | rs6804377 | 0.814 | -0.158 | 1.788 | 0.615 | 0.029 | 0.151 | 0.151 | 0 | 0.94 | 0.82 | 0.95 | 0.8 | 0.94 | 0.95 |
| 3 | 89299059 | rs6773815 | -0.111 | 0.681 | 1.612 | 0.366 | 0.196 | 0.053 | 0.053 | 0 | 0.08 | 0.32 | 0.2 | 0.09 | 0.26 | 0.09 |
| 3 | 89309406 | rs11916939 | -0.2 | 0.464 | 1.604 | 0.306 | -0.043 | -0.022 | -0.022 | 0 | 0.32 | 0.43 | 0.18 | 0.19 | 0.18 | 0.21 |
| 3 | 89318462 | rs994129 | -0.996 | 0.491 | 1.431 | 0.305 | -0.018 | 0.016 | 0.016 | 0 | 0.4 | 0.75 | 0.38 | 0.28 | 0.44 | 0.29 |
| 3 | 89319577 | rs9866618 | 0.292 | 0.544 | 1.313 | 0.319 | 0.069 | 0.007 | 0.007 | 0.105 | 0.07 | 0.13 | 0.19 | 0.08 | 0.26 | 0.08 |
| 3 | 89322805 | rs9863058 | 0.213 | 0.508 | 1.316 | 0.319 | 0.083 | 0.009 | 0.009 | 0.048 | 0.07 | 0.15 | 0.19 | 0.08 | 0.27 | 0.08 |

| | | | | | | | | | | | | | | | | |
|---|---|---|---|---|---|---|---|---|---|---|---|---|---|---|---|---|
| 3 | 89329632 | rs1040017 | -0.826 | 0.378 | 1.081 | 0.32 | -0.016 | 0.012 | 0.012 | 0 | 0.39 | 0.7 | 0.38 | 0.28 | 0.44 | 0.29 |
| 3 | 89370107 | rs7644070 | -0.774 | 0.308 | 1.265 | 0.171 | 0.007 | -0.015 | -0.015 | 0 | 0.39 | 0.55 | 0.37 | 0.26 | 0.44 | 0.3 |
| 3 | 89379935 | rs9866959 | -0.821 | 0.296 | 1.42 | 0.166 | 0 | -0.003 | -0.003 | 0 | 0.4 | 0.58 | 0.38 | 0.27 | 0.44 | 0.3 |
| 3 | 89384566 | rs9817553 | -0.762 | 0.313 | 1.382 | 0.174 | 0 | -0.002 | -0.002 | 0 | 0.4 | 0.59 | 0.37 | 0.27 | 0.44 | 0.3 |
| 3 | 89402447 | rs2063589 | -0.088 | 0.768 | 1.428 | 0.105 | -0.028 | -0.01 | -0.01 | 0.861 | 0.8 | 0.99 | 0.88 | 0.8 | 0.85 | 0.82 |
| 3 | 89405754 | rs9836340 | -0.874 | 0.91 | 2.031 | 0.154 | 0.182 | 0.008 | 0.008 | 0.171 | 0.12 | 0.5 | 0.24 | 0.1 | 0.29 | 0.09 |
| 3 | 89407256 | rs9842599 | -0.563 | 0.935 | 2.027 | 0.165 | 0.18 | 0.017 | 0.017 | 0.013 | 0.11 | 0.33 | 0.22 | 0.1 | 0.29 | 0.09 |
| 3 | 89414555 | rs17801001 | -0.362 | 0.97 | 1.819 | 0.151 | 0.116 | 0.001 | 0.001 | 0.055 | 0.1 | 0.2 | 0.21 | 0.09 | 0.28 | 0.09 |
| 3 | 89417064 | rs870898 | -0.427 | 0.97 | 1.693 | 0.151 | 0.161 | 0.012 | 0.012 | 0.03 | 0.1 | 0.27 | 0.21 | 0.09 | 0.28 | 0.09 |
| 3 | 89417171 | rs870899 | -0.363 | 0.976 | 1.231 | 0.153 | 0.07 | -0.005 | -0.005 | 0 | 0.1 | 0.16 | 0.2 | 0.09 | 0.28 | 0.09 |
| 3 | 94213473 | rs13327303 | -1.34 | 1.177 | 0.581 | 0.936 | 0.003 | 0.008 | 0.008 | 0.431 | 0.31 | 0.37 | 0.44 | 0.46 | 0.41 | 0.43 |
| 3 | 94219431 | rs1352463 | -1.34 | 1.177 | 0.581 | 0.936 | 0.003 | 0.008 | 0.008 | 0.431 | 0.31 | 0.37 | 0.44 | 0.46 | 0.41 | 0.43 |
| 3 | 94229960 | rs9290284 | -0.765 | 1.219 | 0.582 | 0.926 | -0.005 | 0.026 | 0.026 | 0.095 | 0.23 | 0.29 | 0.22 | 0.44 | 0.21 | 0.24 |
| 3 | 94234304 | rs1948183 | -1.608 | 1.189 | 0.503 | 0.923 | 0.005 | 0.016 | 0.016 | 0.12 | 0.33 | 0.57 | 0.46 | 0.48 | 0.41 | 0.44 |
| 3 | 94235153 | rs9290298 | -1.495 | 1.333 | 0.04 | 0.919 | 0.025 | 0.061 | 0.061 | 0.337 | 0.32 | 0.45 | 0.45 | 0.46 | 0.41 | 0.43 |
| 3 | 94255612 | rs1387122 | 0.855 | 0.82 | -0.214 | 0.218 | -0.018 | -0.041 | -0.041 | 0 | 0.8 | 0.7 | 0.78 | 0.89 | 0.85 | 0.75 |
| 3 | 94256654 | rs6763149 | 0.94 | 1.013 | -0.824 | -0.112 | 0.006 | 0.039 | 0.039 | 0 | 0.59 | 0.72 | 0.48 | 0.7 | 0.57 | 0.47 |
| 3 | 94258967 | rs13098825 | -0.975 | 0.566 | -1.497 | -0.033 | -0.023 | -0.063 | -0.063 | 0.42 | 0.19 | 0.01 | 0.14 | 0.23 | 0.16 | 0.25 |
| 3 | 94260037 | rs12488331 | -0.305 | 0.725 | -1.698 | 0.127 | -0.033 | 0.057 | 0.057 | 0 | 0.44 | 0.19 | 0.61 | 0.35 | 0.53 | 0.53 |
| 3 | 94268157 | rs957303 | -1.777 | 0.333 | -0.766 | 0.074 | 0.002 | 0.031 | 0.031 | 0 | 0.68 | 0.45 | 0.78 | 0.61 | 0.7 | 0.78 |
| 3 | 94275549 | rs12494596 | -1.777 | 0.333 | -0.766 | 0.074 | 0.002 | 0.031 | 0.031 | 0 | 0.68 | 0.45 | 0.78 | 0.6 | 0.69 | 0.78 |
| 3 | 94289683 | rs13082399 | -0.119 | 0.909 | -0.983 | -0.134 | -0.055 | 0.02 | 0.02 | 0 | 0.4 | 0.17 | 0.6 | 0.38 | 0.54 | 0.5 |
| 3 | 94303542 | rs901922 | 1.765 | 0.702 | -0.4 | 0.002 | -0.002 | -0.008 | -0.008 | 0.994 | 0.32 | 0.28 | 0.2 | 0.37 | 0.31 | 0.27 |
| 3 | 94321390 | rs10936849 | -1.815 | 0.692 | -0.396 | 0.061 | 0 | 0.013 | 0.013 | 0 | 0.67 | 0.58 | 0.79 | 0.64 | 0.69 | 0.73 |
| 3 | 94328364 | rs4857458 | -0.031 | 0.584 | -0.52 | 0.013 | -0.056 | 0.01 | 0.01 | 0 | 0.39 | 0.16 | 0.59 | 0.38 | 0.51 | 0.48 |
| 3 | 94334170 | rs12493833 | 0.498 | 1.09 | -0.705 | -0.21 | -0.113 | -0.083 | -0.083 | 0 | 0.31 | 0.01 | 0.38 | 0.34 | 0.4 | 0.34 |
| 3 | 94335368 | rs6782460 | 1.939 | 1.187 | -0.652 | -0.262 | -0.001 | 0.013 | 0.013 | 0 | 0.33 | 0.43 | 0.21 | 0.37 | 0.29 | 0.27 |
| 3 | 94342986 | rs7629048 | -0.164 | 1.033 | -0.624 | -0.252 | -0.056 | 0.005 | 0.005 | 0 | 0.39 | 0.15 | 0.58 | 0.38 | 0.5 | 0.48 |
| 3 | 94346384 | rs2885053 | -0.718 | 0.848 | -0.452 | -0.253 | 0.02 | 0.038 | 0.038 | 0 | 0.49 | 0.29 | 0.63 | 0.42 | 0.6 | 0.58 |
| 3 | 94377411 | rs9844990 | 1.836 | 0.188 | 0.165 | -0.405 | -0.004 | -0.009 | -0.009 | 0 | 0.31 | 0.24 | 0.34 | 0.38 | 0.3 | 0.26 |
| 3 | 94415048 | rs7634511 | -1.617 | 0.032 | 0.312 | -0.503 | 0.001 | 0.014 | 0.014 | 0 | 0.66 | 0.46 | 0.63 | 0.62 | 0.69 | 0.74 |
| 3 | 94437412 | rs2166805 | -1.807 | -0.113 | 0.243 | -0.567 | -0.002 | -0.006 | -0.006 | 0 | 0.67 | 0.73 | 0.66 | 0.62 | 0.7 | 0.74 |
| 3 | 94452002 | rs7630912 | -1.02 | -0.037 | -0.005 | -0.719 | 0 | -0.001 | -0.001 | 0.123 | 0.45 | 0.48 | 0.42 | 0.56 | 0.5 | 0.53 |
| 3 | 94455418 | rs13088303 | -0.781 | 0.084 | 0.002 | -0.656 | -0.003 | -0.015 | -0.015 | 0.496 | 0.52 | 0.21 | 0.55 | 0.54 | 0.59 | 0.65 |
| 3 | 94470447 | rs9856524 | 1.847 | 0.272 | 0.047 | -0.566 | -0.005 | -0.012 | -0.012 | 0.144 | 0.31 | 0.24 | 0.34 | 0.38 | 0.3 | 0.26 |
| 3 | 94479849 | rs1447853 | 1.713 | 0.176 | 0.061 | -0.546 | 0.002 | 0.002 | 0.002 | 0 | 0.32 | 0.3 | 0.38 | 0.38 | 0.32 | 0.27 |
| 3 | 94495036 | rs13081582 | -1.033 | -0.278 | -0.048 | -0.711 | 0 | -0.004 | -0.004 | 0.004 | 0.61 | 0.39 | 0.53 | 0.75 | 0.61 | 0.74 |
| 3 | 94500476 | rs13321334 | 0.337 | -0.182 | 0.452 | -0.543 | -0.015 | -0.052 | -0.052 | 0 | 0.1 | 0.3 | 0.07 | 0 | 0.07 | 0.06 |
| 3 | 94523335 | rs13066018 | 0.088 | -0.656 | -0.138 | -0.551 | -0.09 | -0.121 | -0.121 | 0 | 0.24 | 0.07 | 0.25 | 0.48 | 0.4 | 0.51 |
| 3 | 94531485 | rs13063373 | -1.231 | -0.885 | 0.125 | -1.218 | -0.006 | 0.101 | 0.101 | 0 | 0.55 | 0.24 | 0.5 | 0.75 | 0.56 | 0.67 |
| 3 | 94534163 | rs4493432 | 1.116 | -1.063 | 0.162 | -1.314 | -0.011 | 0.095 | 0.095 | 0 | 0.47 | 0.21 | 0.45 | 0.29 | 0.47 | 0.36 |
| 3 | 94548034 | rs9846361 | 1.913 | -0.548 | 0.404 | -1.031 | -0.017 | 0.02 | 0.02 | 0 | 0.33 | 0.2 | 0.41 | 0.24 | 0.37 | 0.28 |
| 3 | 94549686 | rs4411883 | 1.241 | -0.546 | 0.185 | -0.98 | -0.018 | -0.029 | -0.029 | 0 | 0.08 | 0 | 0.06 | 0.1 | 0.08 | 0.19 |
| 3 | 94552269 | rs9848675 | 1.906 | -0.354 | 0.462 | -1.22 | -0.053 | 0.032 | 0.032 | 0 | 0.32 | 0.11 | 0.4 | 0.24 | 0.37 | 0.28 |
| 3 | 94561214 | rs9829425 | 1.447 | -0.366 | 0.644 | -1.225 | 0.009 | -0.028 | -0.028 | 0 | 0.36 | 0.42 | 0.43 | 0.25 | 0.37 | 0.28 |
| 3 | 94562501 | rs9874001 | 1.857 | -0.315 | 0.287 | -1.203 | -0.025 | 0.024 | 0.024 | 0 | 0.32 | 0.18 | 0.41 | 0.24 | 0.37 | 0.28 |
| 3 | 94596072 | rs7613024 | -0.634 | -0.172 | 0.65 | -0.964 | -0.058 | -0.1 | -0.1 | 0.535 | 0.15 | 0.6 | 0.11 | 0 | 0.07 | 0.06 |
| 3 | 94601976 | rs9859333 | 0.328 | -0.158 | -0.301 | -1.025 | -0.019 | 0.026 | 0.026 | 1.469 | 0.38 | 0.17 | 0.37 | 0.67 | 0.43 | 0.58 |
| 3 | 94640761 | rs380053 | 1.7 | -0.348 | 0.094 | -0.544 | -0.093 | -0.022 | -0.022 | 1.057 | 0.27 | 0.07 | 0.39 | 0.28 | 0.36 | 0.27 |
| 3 | 94641757 | rs300089 | 0.302 | -0.474 | 0.295 | -0.615 | -0.011 | 0.04 | 0.04 | 0.189 | 0.42 | 0.72 | 0.5 | 0.29 | 0.43 | 0.34 |
| 3 | 94643907 | rs11718065 | 1.872 | -0.111 | 0.063 | -0.571 | -0.077 | 0.119 | 0.119 | 0 | 0.24 | 0.03 | 0.28 | 0.03 | 0.27 | 0.2 |
| 3 | 94651998 | rs1991752 | 1.597 | -0.909 | 0.278 | -0.591 | -0.041 | 0 | 0 | 0.865 | 0.28 | 0.19 | 0.4 | 0.28 | 0.36 | 0.27 |
| 3 | 94658656 | rs9714169 | 1.281 | -0.971 | 0.281 | -0.588 | -0.037 | 0.01 | 0.01 | 0 | 0.31 | 0.15 | 0.33 | 0.27 | 0.37 | 0.08 |
| 3 | 94675697 | rs9790140 | 0.963 | -0.966 | -0.22 | -0.52 | 0.016 | 0.035 | 0.035 | 0 | 0.14 | 0.01 | 0.06 | 0.06 | 0.09 | 0.07 |
| 3 | 94679788 | rs11926470 | 0.6 | -0.253 | 0.652 | -0.114 | -0.025 | -0.068 | -0.068 | 0.634 | 0.11 | 0.37 | 0.09 | 0 | 0.07 | 0.06 |
| 3 | 94682972 | rs1897462 | -0.009 | -0.241 | -0.095 | -0.108 | -0.045 | -0.078 | -0.078 | 0 | 0.86 | 0.32 | 0.89 | 0.99 | 0.93 | 0.93 |
| 3 | 94690130 | rs9880324 | 1.34 | -0.126 | -0.328 | -0.07 | -0.068 | -0.011 | -0.011 | 4.11 | 0.26 | 0.1 | 0.4 | 0.28 | 0.36 | 0.26 |
| 3 | 94714475 | rs4527380 | -0.08 | -0.08 | -0.33 | -0.048 | -0.052 | -0.09 | -0.09 | 0 | 0.14 | 0.61 | 0.11 | 0.01 | 0.07 | 0.06 |
| 3 | 94721501 | rs49698 | 0.998 | 0.84 | -0.913 | 0.527 | -0.079 | -0.016 | -0.016 | 0 | 0.26 | 0.08 | 0.4 | 0.28 | 0.36 | 0.26 |
| 3 | 94759504 | rs8180036 | -0.042 | 0.414 | -0.585 | 0.409 | 0.001 | -0.003 | -0.003 | 0.466 | 0.44 | 0.51 | 0.52 | 0.64 | 0.48 | 0.58 |
| 3 | 94772205 | rs3905886 | -0.197 | -0.529 | -0.916 | 0.364 | 0.045 | -0.035 | -0.035 | 0 | 0.33 | 0.17 | 0.33 | 0.59 | 0.24 | 0.49 |
| 3 | 94774673 | rs13067978 | 0.803 | -1.267 | -0.582 | -0.269 | -0.052 | 0.073 | 0.073 | 0 | 0.27 | 0.08 | 0.16 | 0.15 | 0.34 | 0.18 |
| 3 | 94784932 | rs9832221 | 0.011 | -1.312 | -0.587 | -0.042 | 0.043 | -0.085 | -0.085 | 0.592 | 0.33 | 0.17 | 0.28 | 0.46 | 0.23 | 0.47 |
| 3 | 94786646 | rs9862168 | 2.108 | -1.505 | -0.319 | -0.148 | -0.055 | -0.049 | -0.049 | 0 | 0.27 | 0.11 | 0.45 | 0.41 | 0.37 | 0.29 |
| 3 | 94803053 | rs10933879 | 0.991 | -1.537 | -0.383 | 0.098 | -0.058 | 0.033 | 0.033 | 0 | 0.26 | 0.08 | 0.18 | 0.17 | 0.35 | 0.18 |
| 3 | 94807394 | rs7612444 | 0.714 | -1.611 | 0.099 | 0.109 | -0.043 | -0.085 | -0.085 | 1.243 | 0.13 | 0.4 | 0.09 | 0 | 0.06 | 0.06 |
| 3 | 94832290 | rs3849435 | -0.068 | -1.739 | 0.265 | 0.183 | 0.042 | -0.037 | -0.037 | 0 | 0.34 | 0.21 | 0.29 | 0.4 | 0.23 | 0.45 |
| 3 | 94836467 | rs3849437 | 1.201 | -1.505 | 0.443 | 0.22 | -0.015 | -0.002 | -0.002 | 0 | 0.36 | 0.33 | 0.51 | 0.38 | 0.44 | 0.38 |
| 3 | 94840443 | rs3849439 | 0.001 | -1.991 | 0.175 | 0.034 | 0.006 | -0.01 | -0.01 | 0 | 0.2 | 0.2 | 0.1 | 0.33 | 0.14 | 0.43 |

| | | | | | | | | | | | | | | | | |
|---|---|---|---|---|---|---|---|---|---|---|---|---|---|---|---|---|
| 3 | 94840605 | rs3843866 | -1.4 | -2.066 | -0.027 | -0.135 | -0.011 | 0.016 | 0.016 | 0 | 0.78 | 0.84 | 0.86 | 0.9 | 0.71 | 0.84 |
| 3 | 94840854 | rs9790095 | 1.585 | -2.184 | -0.15 | 0.122 | -0.027 | -0.063 | -0.063 | 0 | 0.1 | 0 | 0.06 | 0.23 | 0.08 | 0.14 |
| 3 | 94863471 | rs4407407 | 1.586 | -2.224 | 0.005 | 0.045 | -0.022 | -0.06 | -0.06 | 0 | 0.1 | 0.27 | 0.07 | 0 | 0.07 | 0.06 |
| 3 | 94867122 | rs13080340 | 1.294 | -2.308 | 0.027 | -0.005 | -0.077 | -0.085 | -0.085 | 0.063 | 0.41 | 0.13 | 0.64 | 0.48 | 0.47 | 0.35 |
| 3 | 94868812 | rs17497698 | 3.498 | -1.924 | -0.411 | -0.326 | -0.061 | 0.048 | 0.048 | 0 | 0.13 | 0.02 | 0.28 | 0.03 | 0.25 | 0.1 |
| 3 | 94869032 | rs3849442 | -1.664 | -2.306 | -0.174 | -0.266 | -0.016 | 0.024 | 0.024 | 0 | 0.78 | 0.86 | 0.87 | 0.9 | 0.71 | 0.84 |
| 3 | 94869371 | rs9821870 | 1.892 | -2.322 | -0.151 | -0.244 | -0.017 | -0.052 | -0.052 | 0.509 | 0.1 | 0.21 | 0.06 | 0 | 0.07 | 0.06 |
| 3 | 94869452 | rs2310773 | -0.414 | -2.314 | -0.103 | -0.226 | -0.014 | -0.007 | -0.007 | 0 | 0.63 | 0.74 | 0.78 | 0.58 | 0.59 | 0.5 |
| 3 | 94876659 | rs3913809 | 1.592 | -2.285 | -0.142 | -0.155 | -0.025 | -0.034 | -0.034 | 0 | 0.3 | 0.21 | 0.48 | 0.5 | 0.4 | 0.27 |
| 3 | 94896920 | rs9814404 | 0.394 | -2.394 | -0.085 | -0.211 | -0.048 | -0.057 | -0.057 | 0 | 0.28 | 0.38 | 0.28 | 0.08 | 0.18 | 0.17 |
| 3 | 94909396 | rs13091391 | -0.116 | -2.403 | -0.087 | -0.21 | -0.005 | 0.006 | 0.006 | 0 | 0.42 | 0.35 | 0.23 | 0.42 | 0.42 | 0.56 |
| 3 | 94912049 | rs12632730 | -1.451 | -2.404 | -0.142 | -0.222 | -0.033 | -0.022 | -0.022 | 0.226 | 0.82 | 0.77 | 0.78 | 0.92 | 0.89 | 0.89 |
| 3 | 94912737 | rs12632888 | 2.448 | -2.389 | -0.18 | -0.202 | -0.09 | -0.086 | -0.086 | 3.035 | 0.24 | 0.07 | 0.44 | 0.4 | 0.36 | 0.25 |
| 3 | 94913312 | rs6802456 | 0.521 | -2.153 | -0.224 | -0.309 | -0.002 | -0.045 | -0.045 | 0 | 0.32 | 0.43 | 0.2 | 0.1 | 0.33 | 0.22 |
| 3 | 94916211 | rs13081907 | 2.368 | -2.272 | -0.638 | -0.47 | -0.044 | 0.014 | 0.014 | 0 | 0.11 | 0.03 | 0.09 | 0.05 | 0.21 | 0.09 |
| 3 | 94918524 | rs17383280 | 0.965 | -2.223 | -0.584 | -0.44 | -0.016 | 0.038 | 0.038 | 0 | 0.22 | 0.14 | 0.12 | 0.09 | 0.26 | 0.15 |
| 3 | 94925544 | rs6800736 | 1.246 | -1.801 | -0.37 | -0.186 | -0.016 | 0.005 | 0.005 | 0 | 0.33 | 0.25 | 0.5 | 0.73 | 0.42 | 0.37 |
| 3 | 94943493 | rs2871670 | 0.062 | -1.657 | -0.424 | -0.146 | 0.006 | -0.032 | -0.032 | 0 | 0.2 | 0.13 | 0.1 | 0.32 | 0.16 | 0.4 |
| 3 | 94952474 | rs10511150 | 0.433 | -1.652 | -0.374 | -0.228 | 0.003 | -0.039 | -0.039 | 0 | 0.2 | 0.13 | 0.13 | 0.42 | 0.18 | 0.38 |
| 3 | 94986449 | rs11715081 | -0.334 | -1.287 | 0.221 | -0.245 | -0.039 | -0.09 | -0.09 | 3.236 | 0.86 | 0.58 | 0.9 | 0.99 | 0.92 | 0.87 |
| 3 | 94997266 | rs9809394 | -0.473 | -1.334 | 0 | -0.274 | 0.016 | 0.002 | 0.002 | 0.007 | 0.62 | 0.51 | 0.45 | 0.6 | 0.56 | 0.63 |
| 3 | 95006992 | rs9824774 | -1.744 | -1.32 | 0.045 | -0.267 | -0.026 | -0.024 | -0.024 | 0 | 0.76 | 0.83 | 0.55 | 0.6 | 0.63 | 0.75 |
| 3 | 95012506 | rs9834958 | 0.445 | -1.337 | 0.105 | -0.28 | -0.026 | -0.063 | -0.063 | 0.256 | 0.13 | 0.26 | 0.08 | 0 | 0.08 | 0.12 |
| 3 | 95015652 | rs11928664 | -1.748 | -1.145 | 0.112 | -0.168 | -0.042 | -0.037 | -0.037 | 0 | 0.76 | 0.87 | 0.55 | 0.61 | 0.64 | 0.75 |
| 3 | 95017585 | rs7627614 | -1.634 | -1.135 | 0.108 | -0.166 | 0.005 | 0.004 | 0.004 | 0 | 0.75 | 0.77 | 0.53 | 0.6 | 0.63 | 0.75 |
| 3 | 95026777 | rs9853603 | -0.846 | -0.822 | 0.13 | -0.076 | 0.01 | -0.001 | -0.001 | 0 | 0.66 | 0.57 | 0.5 | 0.63 | 0.59 | 0.69 |
| 3 | 95028066 | rs13097088 | 1.744 | -0.791 | -0.161 | -0.171 | -0.045 | 0.028 | 0.028 | 0 | 0.11 | 0.03 | 0.1 | 0.05 | 0.22 | 0.09 |
| 3 | 95033388 | rs1607052 | -0.666 | -1.011 | 0.222 | -0.007 | 0.008 | -0.003 | -0.003 | 0 | 0.65 | 0.45 | 0.5 | 0.63 | 0.59 | 0.69 |
| 3 | 95034056 | rs9875634 | 0.548 | -0.995 | 0.27 | -0.006 | -0.013 | -0.068 | -0.068 | 0 | 0.11 | 0.39 | 0.08 | 0 | 0.07 | 0.07 |
| 3 | 95045970 | rs1607490 | 0.23 | -1.45 | -0.378 | -0.077 | -0.008 | -0.011 | -0.011 | 0 | 0.48 | 0.23 | 0.28 | 0.53 | 0.48 | 0.53 |
| 3 | 95049205 | rs9847492 | 0.366 | -1.408 | 0.106 | 0.094 | -0.02 | -0.056 | -0.056 | 0 | 0.1 | 0.26 | 0.07 | 0 | 0.06 | 0.07 |
| 3 | 95058076 | rs7642437 | 0.34 | -1.408 | 0.027 | 0.094 | -0.02 | -0.051 | -0.051 | 0 | 0.1 | 0.24 | 0.07 | 0 | 0.07 | 0.06 |
| 3 | 95062190 | rs498006 | -0.437 | -1.541 | 0.514 | 0.717 | -0.064 | 0.055 | 0.055 | 0.229 | 0.15 | 0.4 | 0.1 | 0.17 | 0.07 | 0.19 |
| 3 | 95062486 | rs520258 | -1.185 | -1.835 | 0.338 | 0.724 | -0.037 | 0.003 | 0.003 | 0 | 0.26 | 0.62 | 0.17 | 0.27 | 0.13 | 0.22 |
| 3 | 95068715 | rs547833 | -0.209 | -1.788 | 0.492 | 0.687 | -0.058 | 0.031 | 0.031 | 0.428 | 0.15 | 0.36 | 0.09 | 0.16 | 0.07 | 0.19 |
| 3 | 95070749 | rs578737 | -0.217 | -1.766 | 0.62 | 0.699 | -0.049 | 0.026 | 0.026 | 0.36 | 0.14 | 0.29 | 0.08 | 0.16 | 0.07 | 0.19 |
| 3 | 95078219 | rs474509 | -0.737 | -1.036 | -0.092 | 0.854 | 0.004 | 0.019 | 0.019 | 0 | 0.37 | 0.52 | 0.36 | 0.48 | 0.42 | 0.53 |
| 3 | 95080377 | rs523450 | -1.091 | -1.136 | -0.955 | 0.78 | 0.029 | -0.013 | -0.013 | 0 | 0.48 | 0.79 | 0.4 | 0.54 | 0.46 | 0.57 |
| 3 | 95083343 | rs10933994 | -0.039 | -0.348 | -0.893 | 0.794 | -0.019 | -0.011 | -0.011 | 0 | 0.23 | 0.17 | 0.23 | 0.28 | 0.33 | 0.35 |
| 3 | 95086836 | rs9853362 | 0.159 | -1.08 | -1.121 | 0.836 | 0.024 | -0.012 | -0.012 | 0 | 0.18 | 0.07 | 0.13 | 0.19 | 0.11 | 0.15 |
| 3 | 95104671 | rs555809 | -0.498 | -1.745 | -0.851 | 0.634 | 0.006 | 0.008 | 0.008 | 0 | 0.37 | 0.82 | 0.22 | 0.35 | 0.36 | 0.5 |
| 3 | 95105974 | rs489085 | -1.805 | -1.855 | -1.479 | 0.571 | -0.127 | -0.038 | -0.038 | 0 | 0.67 | 0.93 | 0.42 | 0.69 | 0.57 | 0.68 |
| 3 | 95108147 | rs537146 | -0.907 | -1.771 | -1.357 | 0.569 | 0.008 | -0.04 | -0.04 | 0 | 0.49 | 0.89 | 0.31 | 0.52 | 0.47 | 0.53 |
| 3 | 95115832 | rs16857046 | 1.104 | -1.774 | -1.319 | 0.606 | 0.008 | 0.004 | 0.004 | 0 | 0.12 | 0.07 | 0.08 | 0.12 | 0.1 | 0.04 |
| 3 | 95116949 | rs501118 | -0.102 | -1.698 | -1.297 | 0.522 | 0.003 | -0.018 | -0.018 | 3.196 | 0.36 | 0.82 | 0.22 | 0.4 | 0.36 | 0.49 |
| 3 | 95144399 | rs9860786 | 0.566 | -2.114 | 0.251 | 0.321 | 0.005 | -0.008 | -0.008 | 0 | 0.07 | 0.36 | 0.04 | 0 | 0.01 | 0 |
| 3 | 95146114 | rs7620512 | -0.284 | -2.331 | 0.268 | 0.277 | -0.078 | -0.061 | -0.061 | 0 | 0.16 | 0.62 | 0.08 | 0.06 | 0.04 | 0.13 |
| 3 | 95148926 | rs10049290 | 0.61 | -2.213 | 0.196 | 0.297 | 0.003 | -0.01 | -0.01 | 0 | 0.07 | 0.35 | 0.04 | 0 | 0.01 | 0 |
| 3 | 95150253 | rs11925662 | 1.04 | -2.316 | -0.031 | 0.219 | 0.011 | 0.035 | 0.035 | 0 | 0.14 | 0.07 | 0.09 | 0.06 | 0.13 | 0.04 |
| 3 | 95154073 | rs6783439 | -0.773 | -2.165 | -0.08 | 0.256 | 0.001 | 0.001 | 0.001 | 0 | 0.45 | 0.57 | 0.28 | 0.46 | 0.45 | 0.55 |
| 3 | 95156135 | rs12493110 | -0.535 | -2.183 | -0.049 | 0.3 | -0.001 | -0.001 | -0.001 | 0 | 0.43 | 0.4 | 0.26 | 0.46 | 0.46 | 0.55 |
| 3 | 95159211 | rs9847422 | 0.898 | -2.092 | -0.124 | 0.298 | -0.003 | -0.015 | -0.015 | 0 | 0.07 | 0.32 | 0.03 | 0 | 0.01 | 0 |
| 3 | 95165174 | rs4478076 | -1.188 | -2.092 | -0.477 | 0.341 | -0.006 | -0.002 | -0.002 | 0.064 | 0.51 | 0.89 | 0.32 | 0.47 | 0.47 | 0.56 |
| 3 | 95176888 | rs4342104 | -2.011 | -2.336 | -0.673 | 0.358 | -0.128 | -0.063 | -0.063 | 0 | 0.7 | 0.93 | 0.42 | 0.67 | 0.57 | 0.69 |
| 3 | 95177386 | rs7633906 | 0.969 | -2.008 | 0.007 | 0.265 | -0.001 | 0 | 0 | 0.259 | 0.18 | 0.18 | 0.12 | 0.32 | 0.27 | 0.37 |
| 3 | 95186258 | rs6437879 | 0.333 | -2.09 | 0.109 | 0.096 | -0.009 | -0.022 | -0.022 | 0 | 0.08 | 0.37 | 0.04 | 0 | 0.01 | 0 |
| 3 | 95225762 | rs7631830 | -1.442 | -2.181 | 0.041 | 0.104 | 0.012 | 0.018 | 0.018 | 0 | 0.51 | 0.84 | 0.32 | 0.47 | 0.46 | 0.55 |
| 3 | 95227299 | rs4438655 | -1.535 | -2.182 | 0.024 | 0.104 | -0.014 | -0.01 | -0.01 | 1.442 | 0.51 | 0.9 | 0.32 | 0.47 | 0.47 | 0.55 |
| 3 | 95235091 | rs6804576 | 0.148 | -2.116 | 0.244 | 0.173 | 0.001 | -0.013 | -0.013 | 0.896 | 0.08 | 0.38 | 0.04 | 0 | 0.01 | 0 |
| 3 | 95237562 | rs10439950 | 0.836 | -2.106 | 0.356 | 0.2 | -0.017 | -0.023 | -0.023 | 0 | 0.06 | 0.22 | 0.03 | 0 | 0.01 | 0 |
| 3 | 95257786 | rs4507254 | 1.833 | -2.144 | 0.125 | 0.234 | 0.026 | -0.02 | -0.02 | 0 | 0.06 | 0 | 0.01 | 0.07 | 0.02 | 0.14 |
| 3 | 95270483 | rs6773466 | 0.198 | -2.089 | 0.313 | 0.214 | 0.042 | 0.061 | 0.061 | 0 | 0.23 | 0.38 | 0.17 | 0.39 | 0.31 | 0.38 |
| 3 | 95281021 | rs11918851 | -0.06 | -2.112 | 0.07 | 0.145 | -0.027 | -0.01 | -0.01 | 0 | 0.34 | 0.2 | 0.23 | 0.38 | 0.46 | 0.43 |
| 3 | 95291185 | rs13091676 | -0.06 | -2.112 | 0.07 | 0.144 | -0.027 | -0.009 | -0.009 | 0 | 0.34 | 0.21 | 0.23 | 0.38 | 0.46 | 0.43 |
| 3 | 95298427 | rs4073617 | -0.672 | -2.084 | 0.166 | 0.157 | 0 | -0.002 | -0.002 | 0.101 | 0.42 | 0.56 | 0.27 | 0.38 | 0.47 | 0.44 |
| 3 | 95310843 | rs11708806 | 1.264 | -2.108 | -0.229 | 0.15 | -0.055 | -0.038 | -0.038 | 0 | 0.46 | 0.1 | 0.68 | 0.54 | 0.51 | 0.41 |
| 3 | 95315128 | rs13325836 | 1.596 | -2.044 | -0.142 | 0.182 | -0.006 | -0.001 | -0.001 | 0 | 0.07 | 0.06 | 0.02 | 0.08 | 0.01 | 0.14 |
| 3 | 95334138 | rs6808351 | -0.081 | -2.035 | -0.247 | 0.18 | -0.028 | -0.006 | -0.006 | 0 | 0.35 | 0.2 | 0.22 | 0.38 | 0.46 | 0.44 |
| 3 | 95334933 | rs6803348 | 0.313 | -1.709 | -0.882 | 0.201 | -0.109 | -0.093 | -0.093 | 0 | 0.29 | 0.01 | 0.18 | 0.32 | 0.37 | 0.41 |

| | | | | | | | | | | | | | | | | |
|---|---|---|---|---|---|---|---|---|---|---|---|---|---|---|---|---|
| 3 | 95338953 | rs6414250 | -0.912 | -1.455 | -0.451 | 0.253 | 0.006 | 0.011 | 0.011 | 0 | 0.52 | 0.66 | 0.3 | 0.46 | 0.49 | 0.59 |
| 3 | 95341528 | rs11708935 | 0.881 | -1.374 | -1.343 | 0.282 | -0.06 | -0.039 | -0.039 | 0 | 0.46 | 0.1 | 0.68 | 0.53 | 0.51 | 0.41 |
| 3 | 95353065 | rs4857166 | 0.892 | -1.427 | -1.356 | 0.269 | -0.058 | -0.041 | -0.041 | 1.142 | 0.46 | 0.1 | 0.68 | 0.53 | 0.51 | 0.41 |
| 3 | 95359287 | rs13097583 | -0.998 | -1.338 | -1.272 | 0.563 | 0.02 | -0.103 | -0.103 | 0 | 0.41 | 0.89 | 0.25 | 0.52 | 0.34 | 0.61 |
| 3 | 95368663 | rs7620600 | -0.435 | -1.53 | -0.286 | 0.589 | 0.008 | 0.029 | 0.029 | 0 | 0.27 | 0.65 | 0.21 | 0.43 | 0.32 | 0.44 |
| 3 | 95373610 | rs13066397 | -0.289 | -1.756 | 0.184 | 0.339 | -0.003 | -0.004 | -0.004 | 0.592 | 0.26 | 0.25 | 0.18 | 0.43 | 0.34 | 0.45 |
| 3 | 95374182 | rs6767658 | -0.125 | -1.688 | 0.094 | 0.491 | 0.009 | 0.003 | 0.003 | 0 | 0.22 | 0.25 | 0.17 | 0.26 | 0.34 | 0.28 |
| 3 | 95382552 | rs7631170 | 0.252 | -1.363 | 0.502 | 0.311 | -0.039 | -0.043 | -0.043 | 0 | 0.21 | 0.14 | 0.16 | 0.43 | 0.37 | 0.44 |
| 3 | 95383494 | rs12233539 | -0.775 | -1.449 | 0.522 | 0.333 | 0.017 | 0.034 | 0.034 | 0 | 0.32 | 0.59 | 0.23 | 0.52 | 0.4 | 0.6 |
| 3 | 95385129 | rs7618693 | 0.108 | -1.424 | 0.899 | 0.357 | 0.046 | 0.063 | 0.063 | 0 | 0.22 | 0.35 | 0.18 | 0.43 | 0.37 | 0.44 |
| 3 | 95390622 | rs7612111 | -0.208 | -1.444 | 1.019 | 0.368 | 0.075 | 0.094 | 0.094 | 0 | 0.26 | 0.5 | 0.2 | 0.43 | 0.38 | 0.45 |
| 3 | 95392545 | rs7624415 | -0.772 | -1.416 | 0.939 | 0.361 | 0.01 | 0.022 | 0.022 | 0 | 0.33 | 0.6 | 0.23 | 0.51 | 0.4 | 0.6 |
| 3 | 95427920 | rs9856079 | 1.049 | -1.425 | 1.074 | 0.333 | -0.028 | -0.03 | -0.03 | 0.534 | 0.06 | 0.14 | 0.02 | 0 | 0 | 0.02 |
| 3 | 95441803 | rs13316538 | 1.539 | -1.449 | 1.146 | 0.353 | -0.018 | 0.005 | 0.005 | 0 | 0.06 | 0.08 | 0.03 | 0.08 | 0.02 | 0.14 |
| 3 | 95455361 | rs1397977 | 1.754 | -1.976 | 0.467 | -0.053 | -0.107 | 0.193 | 0.193 | 0.005 | 0.38 | 0.07 | 0.43 | 0.07 | 0.45 | 0.21 |
| 3 | 95456045 | rs6414256 | -0.975 | -1.812 | 0.839 | 0.202 | -0.001 | -0.006 | -0.006 | 0 | 0.39 | 0.63 | 0.24 | 0.5 | 0.41 | 0.62 |
| 3 | 95471052 | rs6764619 | -1.093 | -1.803 | 0.91 | 0.209 | -0.001 | -0.016 | -0.016 | 0 | 0.4 | 0.68 | 0.24 | 0.5 | 0.41 | 0.62 |
| 3 | 95483745 | rs7623417 | -0.882 | -1.821 | 0.887 | 0.348 | -0.002 | -0.007 | -0.007 | 0 | 0.38 | 0.62 | 0.23 | 0.53 | 0.41 | 0.63 |
| 3 | 95514239 | rs7610595 | -0.106 | -1.82 | 0.942 | 0.265 | -0.006 | -0.006 | -0.006 | 0 | 0.29 | 0.27 | 0.18 | 0.42 | 0.39 | 0.47 |
| 3 | 95518055 | rs7625250 | -0.694 | -1.808 | 1.037 | 0.264 | 0.006 | 0.017 | 0.017 | 0 | 0.37 | 0.43 | 0.21 | 0.5 | 0.41 | 0.61 |
| 3 | 95519536 | rs6780568 | -1.064 | -1.815 | 0.826 | 0.261 | 0 | 0 | 0 | 0 | 0.4 | 0.59 | 0.24 | 0.5 | 0.42 | 0.62 |
| 3 | 95531331 | rs34437438 | 0.657 | -1.866 | 0.289 | 0.513 | 0.027 | -0.017 | -0.017 | 8.718 | 0.17 | 0.07 | 0.1 | 0.2 | 0.09 | 0.1 |
| 3 | 95534825 | rs13080677 | 1.013 | -1.812 | 0.313 | 0.564 | 0.026 | -0.025 | -0.025 | 0 | 0.16 | 0.06 | 0.09 | 0.2 | 0.08 | 0.09 |
| 3 | 95548755 | rs13081814 | -0.303 | -1.83 | 0.387 | 0.5 | 0.023 | -0.006 | -0.006 | 0 | 0.27 | 0.17 | 0.12 | 0.28 | 0.11 | 0.25 |
| 3 | 95549039 | rs1368515 | -1.531 | -1.818 | 0.538 | 0.49 | 0.001 | -0.012 | -0.012 | 1.814 | 0.54 | 0.51 | 0.32 | 0.7 | 0.51 | 0.71 |
| 3 | 95550475 | rs2312155 | 0.426 | -1.81 | 0.545 | 0.491 | 0.001 | 0.001 | 0.001 | 0 | 0.26 | 0.28 | 0.19 | 0.42 | 0.39 | 0.45 |
| 3 | 95557371 | rs10804512 | -1.436 | -1.701 | 0.49 | 0.488 | 0.001 | -0.012 | -0.012 | 0.127 | 0.54 | 0.51 | 0.32 | 0.7 | 0.51 | 0.72 |
| 3 | 95599457 | rs9826974 | -1.561 | -1.506 | 0.474 | 0.378 | -0.021 | 0.067 | 0.067 | 0 | 0.56 | 0.78 | 0.39 | 0.72 | 0.53 | 0.72 |
| 3 | 95600254 | rs7624435 | -0.066 | -1.523 | 0.039 | 0.424 | 0.001 | 0.001 | 0.001 | 0 | 0.31 | 0.33 | 0.2 | 0.42 | 0.4 | 0.47 |
| 3 | 95600730 | rs7624833 | 0.261 | -1.359 | -0.247 | 0.441 | 0.032 | -0.048 | -0.048 | 0 | 0.22 | 0.07 | 0.15 | 0.3 | 0.13 | 0.24 |
| 3 | 95617191 | rs980856 | -1.513 | -1.356 | -0.293 | 0.297 | -0.061 | 0.14 | 0.14 | 0.043 | 0.56 | 0.88 | 0.39 | 0.75 | 0.53 | 0.72 |
| 3 | 95625994 | rs13071548 | 0.389 | -1.42 | 0.219 | 0.373 | -0.012 | -0.014 | -0.014 | 0 | 0.24 | 0.2 | 0.18 | 0.47 | 0.39 | 0.45 |
| 3 | 95633649 | rs13095700 | -0.563 | -1.703 | 0.09 | 0.155 | -0.024 | 0.046 | 0.046 | 0 | 0.42 | 0.27 | 0.31 | 0.69 | 0.51 | 0.55 |
| 3 | 95638744 | rs892256 | -2.675 | -1.42 | -0.332 | -0.269 | -0.033 | 0.03 | 0.03 | 0 | 0.88 | 0.95 | 0.85 | 1 | 0.81 | 0.91 |
| 3 | 95639149 | rs9861559 | -0.464 | -1.863 | 0.315 | 0.148 | -0.073 | -0.116 | -0.116 | 0 | 0.15 | 0.48 | 0.09 | 0 | 0.06 | 0.07 |
| 3 | 95649705 | rs6803023 | -2.037 | -1.846 | 0.296 | 0.167 | -0.039 | 0.07 | 0.07 | 0 | 0.64 | 0.79 | 0.43 | 0.77 | 0.59 | 0.77 |
| 3 | 95654017 | rs9824704 | -0.82 | -1.831 | 0.358 | 0.203 | -0.102 | -0.073 | -0.073 | 0 | 0.21 | 0.5 | 0.11 | 0.08 | 0.08 | 0.22 |
| 3 | 95688843 | rs9814870 | -0.707 | -1.318 | -0.298 | 0.484 | 0.037 | 0.021 | 0.021 | 0 | 0.28 | 0.1 | 0.14 | 0.19 | 0.23 | 0.32 |
| 3 | 95697725 | rs9851532 | 1.012 | -1.14 | -0.211 | 0.528 | 0.011 | -0.03 | -0.03 | 0 | 0.15 | 0.07 | 0.14 | 0.22 | 0.11 | 0.09 |
| 3 | 95702105 | rs4589952 | 0.083 | -1.28 | -0.285 | 0.456 | 0.012 | 0.029 | 0.029 | 0 | 0.18 | 0.07 | 0.08 | 0.1 | 0.16 | 0.15 |
| 3 | 95702596 | rs13323585 | 0.822 | -1.199 | -0.058 | 0.517 | 0.001 | -0.001 | -0.001 | 0 | 0.16 | 0.12 | 0.14 | 0.22 | 0.11 | 0.09 |
| 3 | 95711037 | rs6773783 | -0.414 | -1.199 | 0.53 | 0.459 | -0.024 | -0.04 | -0.04 | 0.258 | 0.2 | 0.33 | 0.1 | 0.1 | 0.17 | 0.13 |
| 3 | 95712310 | rs9863217 | -0.575 | -1.142 | 0.467 | 0.479 | -0.019 | -0.033 | -0.033 | 0 | 0.78 | 0.85 | 0.84 | 0.63 | 0.71 | 0.65 |
| 3 | 95716185 | rs12495594 | 2.043 | -1.263 | 0.456 | 0.407 | -0.059 | 0.074 | 0.074 | 0 | 0.36 | 0.16 | 0.56 | 0.24 | 0.42 | 0.25 |
| 3 | 95723140 | rs4642148 | 2.114 | -1.273 | 0.387 | 0.403 | -0.088 | 0.073 | 0.073 | 0.048 | 0.35 | 0.11 | 0.55 | 0.23 | 0.42 | 0.25 |
| 3 | 95741376 | rs10934322 | 1.11 | -1.242 | 0.632 | 0.485 | 0 | 0.001 | 0.001 | 0 | 0.14 | 0.12 | 0.14 | 0.22 | 0.11 | 0.1 |
| 3 | 95745701 | rs11713687 | 0.659 | -1.307 | 0.87 | 0.428 | 0 | -0.001 | -0.001 | 2.922 | 0.6 | 0.44 | 0.68 | 0.41 | 0.6 | 0.56 |
| 3 | 95760926 | rs9990104 | -0.09 | -1.341 | 0.891 | 0.324 | 0.018 | 0.01 | 0.01 | 0 | 0.23 | 0.14 | 0.11 | 0.17 | 0.18 | 0.3 |
| 3 | 95765590 | rs4459918 | 0.063 | -1.306 | 1.09 | 0.544 | -0.003 | -0.007 | -0.007 | 0 | 0.67 | 0.68 | 0.73 | 0.46 | 0.65 | 0.57 |
| 3 | 95768172 | rs13090997 | 1.045 | -1.25 | 1.104 | 0.576 | -0.003 | 0.007 | 0.007 | 0 | 0.15 | 0.15 | 0.15 | 0.22 | 0.11 | 0.1 |
| 3 | 95773241 | rs7615218 | 0.06 | -1.197 | 1.157 | 0.542 | 0.005 | 0.002 | 0.002 | 4.033 | 0.07 | 0.28 | 0.06 | 0.04 | 0.05 | 0.02 |
| 3 | 95778099 | rs4462982 | 0.941 | -0.9 | 1.115 | 0.717 | -0.001 | -0.001 | -0.001 | 0.131 | 0.54 | 0.52 | 0.67 | 0.52 | 0.52 | 0.35 |
| 3 | 95783063 | rs9813567 | -0.488 | -1.114 | 0.903 | 0.699 | -0.011 | -0.011 | -0.011 | 0 | 0.76 | 0.82 | 0.84 | 0.67 | 0.69 | 0.62 |
| 3 | 95784872 | rs4393926 | 1.105 | -1.123 | 1.107 | 0.704 | 0.001 | 0 | 0 | 0 | 0.53 | 0.43 | 0.72 | 0.45 | 0.51 | 0.35 |
| 3 | 95787760 | rs9824978 | 0.253 | -1.146 | 1.043 | 0.238 | 0.005 | 0.01 | 0.01 | 0 | 0.17 | 0.16 | 0.09 | 0.1 | 0.16 | 0.14 |
| 3 | 95793742 | rs12487224 | 0.924 | -1.133 | 1.02 | 0.332 | -0.009 | 0 | 0 | 0.028 | 0.16 | 0.18 | 0.15 | 0.14 | 0.1 | 0.1 |
| 3 | 95796421 | rs4383543 | -1.998 | -1.121 | 0.755 | 0.335 | -0.027 | 0.054 | 0.054 | 0 | 0.63 | 0.76 | 0.43 | 0.77 | 0.59 | 0.75 |
| 3 | 95798225 | rs6781601 | 0.295 | -1.054 | 0.732 | 0.251 | -0.004 | -0.007 | -0.007 | 1.546 | 0.24 | 0.21 | 0.16 | 0.4 | 0.29 | 0.36 |
| 3 | 95803758 | rs6438285 | -2.004 | -0.888 | 0.341 | 0.27 | -0.041 | 0.037 | 0.037 | 0 | 0.63 | 0.81 | 0.44 | 0.72 | 0.59 | 0.74 |
| 3 | 95825479 | rs13085875 | 0.537 | -0.492 | 0.788 | 0.345 | -0.016 | 0.007 | 0.007 | 0.126 | 0.16 | 0.27 | 0.16 | 0.14 | 0.11 | 0.1 |
| 3 | 95825679 | rs7647183 | -2.003 | -0.537 | 0.521 | 0.286 | -0.028 | 0.057 | 0.057 | 0 | 0.59 | 0.81 | 0.44 | 0.72 | 0.59 | 0.74 |
| 3 | 95838373 | rs1453624 | -0.202 | -0.545 | 0.778 | 0.422 | -0.01 | -0.019 | -0.019 | 0.257 | 0.2 | 0.3 | 0.12 | 0.12 | 0.19 | 0.25 |
| 3 | 95838707 | rs1453626 | -0.506 | -0.551 | 0.693 | 0.43 | -0.021 | -0.006 | -0.006 | 0 | 0.22 | 0.45 | 0.13 | 0.17 | 0.19 | 0.26 |
| 3 | 95848906 | rs9846401 | 0.229 | -0.373 | 0.317 | 0.422 | -0.017 | -0.03 | -0.03 | 0.016 | 0.22 | 0.15 | 0.15 | 0.4 | 0.28 | 0.37 |
| 3 | 95850094 | rs12492664 | -1.084 | -0.303 | 0.606 | 0.442 | -0.004 | -0.001 | -0.001 | 0 | 0.35 | 0.56 | 0.26 | 0.31 | 0.29 | 0.36 |
| 3 | 95851156 | rs6438313 | -0.34 | -0.184 | 1.227 | 0.625 | -0.021 | 0.057 | 0.057 | 0.164 | 0.2 | 0.37 | 0.18 | 0.29 | 0.13 | 0.25 |
| 3 | 95861229 | rs1453621 | 0.466 | -0.121 | 0.846 | 0.629 | -0.025 | 0.02 | 0.02 | 1.597 | 0.76 | 0.42 | 0.81 | 0.72 | 0.87 | 0.75 |
| 3 | 95873964 | rs4857176 | 0.496 | 0.065 | 0.436 | 0.578 | -0.021 | -0.02 | -0.02 | 0 | 0.63 | 0.31 | 0.7 | 0.53 | 0.51 | 0.45 |
| 3 | 95886313 | rs9855928 | -0.371 | 0.081 | 0.791 | 0.589 | 0.035 | 0.018 | 0.018 | 0 | 0.33 | 0.41 | 0.29 | 0.42 | 0.49 | 0.54 |

| | | | | | | | | | | | | | | | | |
|---|---|---|---|---|---|---|---|---|---|---|---|---|---|---|---|---|
| 3 | 95889438 | rs1901874 | 1.208 | -0.021 | 0.653 | 0.656 | 0.001 | -0.002 | -0.002 | 4.094 | 0.06 | 0.05 | 0.03 | 0.08 | 0.02 | 0.12 |
| 3 | 95896470 | rs6785076 | -0.131 | -0.055 | 0.711 | 0.629 | -0.099 | -0.069 | -0.069 | 0.005 | 0.31 | 0.11 | 0.25 | 0.42 | 0.49 | 0.54 |
| 3 | 95901617 | rs9289089 | -0.256 | -0.053 | 1.023 | 0.643 | -0.002 | -0.002 | -0.002 | 0.188 | 0.32 | 0.31 | 0.27 | 0.42 | 0.49 | 0.54 |
| 3 | 95905989 | rs9827296 | -0.135 | -0.044 | 0.917 | 0.628 | -0.047 | -0.031 | -0.031 | 0.62 | 0.31 | 0.18 | 0.26 | 0.42 | 0.49 | 0.54 |
| 3 | 95908799 | rs7636345 | -0.121 | -0.044 | 0.927 | 0.628 | -0.081 | -0.055 | -0.055 | 0.016 | 0.31 | 0.13 | 0.25 | 0.42 | 0.49 | 0.54 |
| 3 | 95912089 | rs4857185 | -0.516 | -0.316 | 0.855 | 0.606 | -0.024 | 0.103 | 0.103 | 2.08 | 0.87 | 0.61 | 0.88 | 0.78 | 0.94 | 0.91 |
| 3 | 95912978 | rs9860584 | 0.022 | -0.256 | 0.281 | 0.568 | -0.043 | -0.028 | -0.028 | 0 | 0.31 | 0.18 | 0.26 | 0.42 | 0.49 | 0.52 |
| 3 | 95917311 | rs10934423 | 1.115 | -0.213 | -0.817 | 0.304 | -0.02 | 0.121 | 0.121 | 0 | 0.48 | 0.15 | 0.57 | 0.28 | 0.42 | 0.24 |
| 3 | 95941123 | rs12695362 | -0.16 | 0.469 | -1.001 | 0.517 | -0.044 | -0.091 | -0.091 | 0 | 0.26 | 0.11 | 0.29 | 0.51 | 0.39 | 0.48 |
| 3 | 95951596 | rs9879845 | -1.597 | 0.35 | -1.166 | 0.1 | 0.004 | 0.161 | 0.161 | 0 | 0.52 | 0.77 | 0.58 | 0.79 | 0.56 | 0.72 |
| 3 | 95957943 | rs11709188 | 2.169 | 0.42 | -1.803 | -0.446 | -0.049 | 0.112 | 0.112 | 0 | 0.17 | 0.01 | 0.13 | 0 | 0.18 | 0.13 |
| 3 | 95967949 | rs16829342 | 0.843 | 0.573 | -1.249 | 0.223 | -0.03 | -0.034 | -0.034 | 0 | 0.16 | 0.01 | 0.11 | 0.15 | 0.13 | 0.1 |
| 3 | 95980193 | rs7612359 | -0.471 | 0.553 | -1.027 | 0.318 | -0.007 | 0.007 | 0.007 | 0 | 0.26 | 0.6 | 0.18 | 0.28 | 0.16 | 0.22 |
| 3 | 95991618 | rs9881321 | -0.197 | 0.647 | -1.175 | 0.252 | -0.067 | -0.123 | -0.123 | 0 | 0.29 | 0.07 | 0.43 | 0.53 | 0.43 | 0.51 |
| 3 | 95995099 | rs1020612 | -0.605 | 0.637 | -1.154 | 0.108 | -0.042 | -0.018 | -0.018 | 0 | 0.33 | 0.16 | 0.45 | 0.66 | 0.45 | 0.64 |
| 3 | 95998146 | rs34937040 | 0.113 | 0.663 | -1.282 | 0.208 | -0.009 | -0.084 | -0.084 | 0 | 0.22 | 0.13 | 0.23 | 0.5 | 0.24 | 0.45 |
| 3 | 96003204 | rs7641736 | 0.068 | 0.662 | -1.121 | 0.219 | -0.001 | -0.023 | -0.023 | 0 | 0.22 | 0.2 | 0.24 | 0.5 | 0.24 | 0.45 |
| 3 | 96010677 | rs7641316 | 0.081 | 0.666 | -1.231 | 0.221 | -0.002 | -0.041 | -0.041 | 1.097 | 0.22 | 0.17 | 0.23 | 0.5 | 0.24 | 0.45 |
| 3 | 96028986 | rs10511153 | 0.401 | 0.675 | -1.258 | 0.225 | -0.032 | -0.122 | -0.122 | 0.087 | 0.2 | 0.03 | 0.22 | 0.5 | 0.24 | 0.45 |
| 3 | 96040392 | rs9843398 | -0.73 | 0.785 | -1.228 | 0.222 | 0 | 0.001 | 0.001 | 0.048 | 0.34 | 0.62 | 0.39 | 0.32 | 0.32 | 0.29 |
| 3 | 96043488 | rs9863581 | -0.748 | 0.783 | -1.409 | 0.221 | 0 | 0.003 | 0.003 | 0.075 | 0.34 | 0.66 | 0.39 | 0.31 | 0.32 | 0.28 |
| 3 | 96045751 | rs4857192 | 0.014 | 0.916 | -1.125 | -0.13 | 0.004 | -0.014 | -0.014 | 0 | 0.27 | 0.27 | 0.35 | 0.2 | 0.3 | 0.17 |
| 3 | 96049190 | rs1473914 | -1.001 | 1.033 | -1.807 | -0.333 | -0.017 | 0.136 | 0.136 | 0 | 0.56 | 0.77 | 0.62 | 0.81 | 0.57 | 0.72 |
| 3 | 96056259 | rs7613763 | -0.793 | 0.798 | -1.526 | -0.005 | 0.032 | 0.125 | 0.125 | 2.426 | 0.44 | 0.75 | 0.53 | 0.8 | 0.35 | 0.69 |
| 3 | 96066172 | rs9821537 | 0.085 | 0.949 | -0.956 | 0.716 | 0.004 | 0.004 | 0.004 | 0 | 0.62 | 0.54 | 0.64 | 0.55 | 0.6 | 0.58 |
| 3 | 96073585 | rs12490647 | 0.053 | 0.871 | -1.439 | 0.674 | -0.1 | -0.057 | -0.057 | 0 | 0.34 | 0.06 | 0.33 | 0.4 | 0.41 | 0.42 |
| 3 | 96078864 | rs1514422 | 0.356 | 1.119 | -1.474 | 0.913 | -0.015 | -0.044 | -0.044 | 0 | 0.08 | 0 | 0.06 | 0.26 | 0.07 | 0.16 |
| 3 | 96094190 | rs6800770 | 0.756 | 1.026 | -1.461 | 0.742 | 0.012 | 0.102 | 0.102 | 0.935 | 0.4 | 0.11 | 0.34 | 0.22 | 0.34 | 0.22 |
| 3 | 96117228 | rs16831515 | 0.749 | 1.021 | -1.363 | 0.895 | 0.013 | 0.053 | 0.053 | 0.953 | 0.4 | 0.1 | 0.34 | 0.28 | 0.35 | 0.22 |
| 3 | 96125199 | rs12497820 | -0.206 | 1.268 | -0.834 | 1.171 | -0.062 | -0.073 | -0.073 | 0 | 0.12 | 0.03 | 0.25 | 0.26 | 0.22 | 0.21 |
| 3 | 96140650 | rs980031 | -0.011 | 1.142 | -0.366 | 0.792 | 0.009 | 0.015 | 0.015 | 0 | 0.94 | 0.9 | 0.93 | 0.95 | 0.94 | 0.88 |
| 3 | 96144029 | rs7614294 | 0.041 | 0.846 | -0.924 | 0.499 | -0.079 | -0.074 | -0.074 | 0 | 0.43 | 0.13 | 0.47 | 0.42 | 0.48 | 0.48 |
| 3 | 96144237 | rs10934513 | 0.501 | 0.601 | -0.142 | 0.462 | -0.006 | 0.009 | 0.009 | 0 | 0.44 | 0.26 | 0.34 | 0.41 | 0.44 | 0.44 |
| 3 | 96150160 | rs10934524 | 0.177 | 0.613 | -0.47 | 0.467 | -0.026 | -0.026 | -0.026 | 0.452 | 0.43 | 0.21 | 0.48 | 0.42 | 0.47 | 0.48 |
| 3 | 96152471 | rs326305 | 0.811 | 0.729 | -0.726 | 0.387 | -0.064 | -0.039 | -0.039 | 0 | 0.4 | 0.06 | 0.21 | 0.38 | 0.4 | 0.39 |
| 3 | 96165411 | rs326251 | 0.611 | 0.85 | -0.531 | 0.42 | -0.051 | 0.002 | 0.002 | 0 | 0.45 | 0.08 | 0.33 | 0.42 | 0.45 | 0.46 |
| 3 | 96167389 | rs9842685 | 0.722 | 0.848 | -0.66 | 0.398 | -0.089 | -0.07 | -0.07 | 0.039 | 0.34 | 0.04 | 0.38 | 0.36 | 0.42 | 0.34 |
| 3 | 96171522 | rs1156953 | -0.02 | 0.688 | 0.059 | -0.053 | 0.003 | -0.006 | -0.006 | 0 | 0.08 | 0.07 | 0.14 | 0.19 | 0.08 | 0.06 |
| 3 | 96198737 | rs9831202 | -0.081 | 0.951 | 0.807 | -0.24 | -0.025 | 0.058 | 0.058 | 0 | 0.14 | 0.07 | 0.13 | 0 | 0.2 | 0.08 |
| 3 | 96226112 | rs9289152 | -1.68 | 0.936 | 1.24 | -0.205 | 0.005 | 0.001 | 0.001 | 0 | 0.28 | 0.64 | 0.37 | 0.25 | 0.29 | 0.26 |
| 3 | 96233924 | rs9835413 | -1.587 | 0.967 | 1.218 | -0.215 | 0.01 | 0.003 | 0.003 | 0 | 0.27 | 0.58 | 0.35 | 0.26 | 0.27 | 0.26 |
| 3 | 96244775 | rs690994 | -0.915 | 1.117 | 0.697 | -0.499 | -0.029 | 0.097 | 0.097 | 0.096 | 0.67 | 0.99 | 0.84 | 0.88 | 0.73 | 0.77 |
| 3 | 96257497 | rs1389030 | -0.649 | 1.818 | 2.054 | 0.306 | -0.023 | 0.051 | 0.051 | 0 | 0.13 | 0.06 | 0.1 | 0 | 0.17 | 0.08 |
| 3 | 96297380 | rs610112 | -1.81 | 1.711 | 2.005 | 0.572 | 0.02 | 0.186 | 0.186 | 0 | 0.12 | 0.37 | 0.23 | 0.28 | 0.09 | 0.15 |
| 3 | 96301741 | rs9849572 | -0.455 | 1.9 | 1.987 | 0.498 | -0.028 | 0.031 | 0.031 | 0 | 0.12 | 0.08 | 0.1 | 0 | 0.17 | 0.08 |
| 3 | 96303888 | rs1487735 | 1.669 | 1.866 | 2.313 | 0.518 | -0.005 | 0.221 | 0.221 | 0 | 0.92 | 0.46 | 0.83 | 0.76 | 0.96 | 0.93 |
| 3 | 96308055 | rs7631577 | 1.917 | 1.862 | 2.321 | 0.51 | 0.008 | 0.008 | 0.008 | 0 | 0.76 | 0.33 | 0.66 | 0.72 | 0.74 | 0.77 |
| 3 | 96317764 | rs1906310 | 1.284 | 1.327 | 2.091 | 0.596 | 0.031 | 0.298 | 0.298 | 0 | 0.93 | 0.64 | 0.84 | 0.76 | 0.96 | 0.92 |
| 3 | 96333159 | rs6794290 | 1.314 | 1.467 | 1.572 | 0.476 | 0.029 | 0.064 | 0.064 | 0 | 0.8 | 0.51 | 0.74 | 0.74 | 0.78 | 0.85 |
| 3 | 96346543 | rs6784955 | -1.006 | 1.425 | 1.842 | 0.503 | -0.003 | 0.171 | 0.171 | 0 | 0.06 | 0.22 | 0.14 | 0.23 | 0.04 | 0.07 |
| 3 | 96387449 | rs3884655 | 0.658 | 1.482 | 1.786 | 0.404 | 0.034 | -0.052 | -0.052 | 1.276 | 0.86 | 0.69 | 0.87 | 0.97 | 0.82 | 0.92 |
| 3 | 96406364 | rs9819410 | -0.465 | 1.484 | 1.668 | 0.401 | -0.024 | 0.023 | 0.023 | 0 | 0.12 | 0.07 | 0.09 | 0 | 0.17 | 0.08 |
| 3 | 96408475 | rs1507382 | 1.929 | 1.435 | 2.022 | 0.523 | 0.003 | 0.024 | 0.024 | 0 | 0.75 | 0.34 | 0.7 | 0.64 | 0.75 | 0.78 |
| 3 | 96411154 | rs13096088 | 0.857 | 1.444 | 1.909 | 0.108 | 0.05 | -0.068 | -0.068 | 0 | 0.85 | 0.58 | 0.87 | 0.97 | 0.82 | 0.92 |
| 3 | 96442435 | rs1395769 | -2.611 | 0.694 | 2.026 | 0.453 | -0.04 | 0.173 | 0.173 | 1.025 | 0.12 | 0.58 | 0.32 | 0.26 | 0.06 | 0.1 |
| 3 | 96464374 | rs10511157 | -0.897 | 0.73 | 1.826 | 0.54 | -0.007 | 0.073 | 0.073 | 0 | 0.05 | 0.09 | 0.23 | 0.23 | 0.03 | 0.09 |
| 3 | 96472739 | rs937878 | 2.554 | 0.728 | 1.431 | 0.165 | 0.008 | -0.07 | -0.07 | 0 | 0.73 | 0.06 | 0.55 | 0.66 | 0.82 | 0.75 |
| 3 | 96479016 | rs2318166 | -2.74 | 0.64 | 2.237 | 0.308 | -0.045 | 0.139 | 0.139 | 3.405 | 0.11 | 0.65 | 0.3 | 0.23 | 0.03 | 0.1 |
| 3 | 96492922 | rs4857230 | -0.867 | 0.656 | 2.218 | 0.316 | -0.013 | 0.086 | 0.086 | 0 | 0.05 | 0.13 | 0.23 | 0.2 | 0.02 | 0.09 |
| 3 | 96514665 | rs6763562 | -2.156 | 0.605 | 2.29 | 0.279 | -0.011 | 0.021 | 0.021 | 0 | 0.13 | 0.19 | 0.27 | 0.26 | 0.08 | 0.17 |
| 3 | 96516039 | rs1507383 | -1.47 | 0.61 | 2.295 | 0.283 | -0.019 | 0.092 | 0.092 | 0.197 | 0.07 | 0.18 | 0.24 | 0.19 | 0.03 | 0.09 |
| 3 | 96547856 | rs9826636 | -1.768 | 0.585 | 2.464 | 0.241 | 0.01 | 0.03 | 0.03 | 0 | 0.09 | 0.34 | 0.09 | 0.11 | 0.06 | 0.09 |
| 3 | 96559488 | rs2318154 | -0.407 | 0.449 | 2.317 | 0.281 | 0.009 | 0.01 | 0.01 | 0 | 0.06 | 0.01 | 0.03 | 0.06 | 0.03 | 0.08 |
| 3 | 96560537 | rs9846289 | -1.359 | 0.447 | 2.506 | 0.236 | 0.005 | 0.221 | 0.221 | 0.004 | 0.07 | 0.25 | 0.25 | 0.23 | 0.02 | 0.1 |
| 3 | 96584999 | rs7639342 | -2.069 | 0.429 | 2.509 | 0.225 | -0.061 | 0.113 | 0.113 | 0.108 | 0.15 | 0.46 | 0.3 | 0.28 | 0.06 | 0.18 |
| 3 | 96669046 | rs6438843 | -2.3 | 0.438 | 1.621 | 0.23 | -0.049 | 0.034 | 0.034 | 0.836 | 0.17 | 0.71 | 0.33 | 0.29 | 0.07 | 0.18 |
| 3 | 96681220 | rs9824190 | -1.378 | 0.543 | 1.569 | -0.327 | -0.041 | -0.048 | -0.048 | 0 | 0.16 | 0.4 | 0.11 | 0.09 | 0.08 | 0.1 |
| 3 | 96681559 | rs4621347 | -0.924 | 0.53 | 1.837 | -0.267 | -0.008 | 0.025 | 0.025 | 0 | 0.05 | 0.12 | 0.22 | 0.13 | 0.03 | 0.05 |
| 3 | 96710451 | rs1547399 | -0.918 | 1.279 | 1.605 | -0.759 | 0.046 | -0.095 | -0.095 | 0 | 0.2 | 0.52 | 0.19 | 0.04 | 0.23 | 0.1 |

| | | | | | | | | | | | | | | | | |
|---|---|---|---|---|---|---|---|---|---|---|---|---|---|---|---|---|
| 3 | 96732700 | rs17509071 | -0.456 | 1.304 | 1.696 | -0.748 | -0.03 | 0.038 | 0.038 | 0 | 0.17 | 0.09 | 0.13 | 0 | 0.23 | 0.1 |
| 3 | 96738205 | rs1029282 | -2.153 | 1.152 | 1.991 | -0.82 | 0.005 | 0.092 | 0.092 | 0.217 | 0.37 | 0.82 | 0.48 | 0.2 | 0.33 | 0.22 |
| 3 | 96739595 | rs2001023 | -1.642 | 1.16 | 2.318 | -0.833 | 0.004 | -0.117 | -0.117 | 0 | 0.31 | 0.5 | 0.24 | 0.08 | 0.3 | 0.18 |
| 3 | 96740888 | rs2213254 | -0.56 | 1.171 | 2.106 | -0.827 | 0.001 | -0.011 | -0.011 | 0 | 0.18 | 0.17 | 0.17 | 0.05 | 0.23 | 0.1 |
| 3 | 96767743 | rs2856478 | -1.76 | 1.165 | 2.154 | -0.831 | -0.001 | -0.046 | -0.046 | 0.117 | 0.32 | 0.65 | 0.25 | 0.08 | 0.3 | 0.17 |
| 3 | 96771625 | rs10470430 | -0.5 | 1.218 | 1.732 | -0.808 | -0.023 | 0.025 | 0.025 | 0 | 0.17 | 0.11 | 0.13 | 0 | 0.23 | 0.1 |
| 3 | 96776008 | rs2612269 | -0.187 | 1.36 | 1.604 | -0.748 | 0.009 | 0.009 | 0.009 | 0 | 0.1 | 0.05 | 0.08 | 0.07 | 0.06 | 0.08 |
| 3 | 96777760 | rs9877403 | -1.236 | 1.936 | 1.475 | -1.05 | -0.026 | 0.03 | 0.03 | 0 | 0.24 | 0.16 | 0.38 | 0.14 | 0.31 | 0.14 |
| 3 | 96778204 | rs762746 | -0.491 | 1.677 | 2.113 | -0.92 | -0.014 | -0.058 | -0.058 | 0 | 0.11 | 0.33 | 0.07 | 0.03 | 0.07 | 0.08 |
| 3 | 96785335 | rs6767738 | -2.247 | 1.805 | 2.251 | -0.975 | 0.002 | 0.072 | 0.072 | 0.412 | 0.34 | 0.8 | 0.44 | 0.17 | 0.31 | 0.19 |
| 3 | 96789865 | rs7640007 | -0.383 | 1.756 | 2.424 | -0.973 | -0.001 | -0.048 | -0.048 | 0 | 0.11 | 0.39 | 0.09 | 0.04 | 0.07 | 0.08 |
| 3 | 96793051 | rs9838310 | -1.254 | 1.821 | 2.237 | -0.951 | -0.033 | -0.069 | -0.069 | 0.037 | 0.14 | 0.64 | 0.12 | 0.04 | 0.08 | 0.08 |
| 3 | 96801401 | rs9872845 | -0.39 | 1.825 | 2.031 | -0.973 | -0.017 | 0.016 | 0.016 | 0 | 0.15 | 0.09 | 0.12 | 0 | 0.2 | 0.07 |
| 3 | 96801930 | rs7629006 | -2.21 | 1.436 | 2.014 | -0.863 | 0.041 | 0.039 | 0.039 | 2.841 | 0.22 | 0.86 | 0.35 | 0.17 | 0.11 | 0.12 |
| 3 | 96811128 | rs9875126 | -0.226 | 1.534 | 2.01 | -0.834 | -0.002 | -0.01 | -0.01 | 0 | 0.1 | 0.12 | 0.06 | 0.04 | 0.07 | 0.08 |
| 3 | 96816161 | rs4857238 | -1.078 | 1.452 | 2.06 | -0.841 | -0.01 | 0.012 | 0.012 | 0 | 0.06 | 0.14 | 0.22 | 0.14 | 0.03 | 0.05 |
| 3 | 96818570 | rs9861668 | -1.618 | 1.48 | 2.296 | -0.882 | -0.035 | -0.043 | -0.043 | 0.04 | 0.18 | 0.47 | 0.3 | 0.17 | 0.11 | 0.13 |
| 3 | 96825782 | rs9832209 | -1.853 | 1.445 | 2.485 | -0.897 | -0.044 | -0.05 | -0.05 | 0.033 | 0.2 | 0.63 | 0.32 | 0.17 | 0.11 | 0.13 |
| 3 | 96838282 | rs4857240 | -2.206 | 1.602 | 2.336 | -1.007 | 0.014 | 0.11 | 0.11 | 0.934 | 0.34 | 0.85 | 0.42 | 0.17 | 0.27 | 0.19 |
| 3 | 96838442 | rs13434261 | -0.249 | 1.599 | 2.356 | -1.021 | -0.016 | 0.025 | 0.025 | 0 | 0.14 | 0.07 | 0.09 | 0 | 0.16 | 0.06 |
| 3 | 96849665 | rs7614446 | -1.356 | 2.288 | 2.226 | -1.017 | 0.016 | 0.007 | 0.007 | 0 | 0.07 | 0.16 | 0.27 | 0.15 | 0.12 | 0.08 |
| 3 | 96854119 | rs7620542 | -1.738 | 2.065 | 2.186 | -1.007 | -0.05 | -0.054 | -0.054 | 0 | 0.2 | 0.6 | 0.33 | 0.17 | 0.11 | 0.15 |
| 3 | 96874360 | rs2221205 | -0.625 | 2.027 | 1.786 | -0.982 | -0.022 | -0.057 | -0.057 | 0 | 0.13 | 0.21 | 0.07 | 0.03 | 0.08 | 0.09 |
| 3 | 96885697 | rs9818654 | -1.879 | 2.107 | 1.396 | -0.959 | 0.016 | 0.12 | 0.12 | 1.139 | 0.34 | 0.87 | 0.43 | 0.18 | 0.27 | 0.22 |
| 3 | 96907996 | rs9853340 | -0.46 | 2.373 | 1.447 | -0.676 | -0.031 | 0.04 | 0.04 | 0 | 0.16 | 0.07 | 0.1 | 0 | 0.2 | 0.07 |
| 3 | 96922673 | rs9825558 | -0.741 | 2.423 | 1.67 | -0.423 | 0.015 | -0.056 | -0.056 | 0.817 | 0.17 | 0.21 | 0.12 | 0 | 0.24 | 0.07 |
| 3 | 97014070 | rs6796268 | -3.11 | 1.025 | 2.762 | 0.113 | -0.019 | 0.006 | 0.006 | 4.893 | 0.09 | 0.73 | 0.32 | 0.11 | 0.03 | 0.09 |
| 3 | 97035796 | rs4857057 | -2.737 | 1.031 | 2.867 | 0.116 | 0.027 | 0.06 | 0.06 | 2.664 | 0.07 | 0.61 | 0.31 | 0.11 | 0.03 | 0.09 |
| 3 | 97062536 | rs9848435 | -3.017 | 1.038 | 3.085 | 0.092 | -0.012 | 0.007 | 0.007 | 5.225 | 0.09 | 0.71 | 0.32 | 0.1 | 0.03 | 0.09 |
| 3 | 97086184 | rs6770349 | -2.831 | 1.045 | 3.081 | 0.093 | 0.035 | 0.037 | 0.037 | 3.272 | 0.07 | 0.55 | 0.08 | 0.01 | 0.01 | 0.01 |
| 3 | 97105552 | rs1598915 | 2.734 | 1.069 | 2.707 | 0.006 | -0.014 | -0.018 | -0.018 | 0.925 | 0.86 | 0.2 | 0.65 | 0.88 | 0.91 | 0.87 |
| 3 | 97140890 | rs11925955 | -2.774 | 0.971 | 2.922 | 0.149 | 0.049 | 0.069 | 0.069 | 4.395 | 0.06 | 0.49 | 0.07 | 0.02 | 0.01 | 0.01 |
| 3 | 97146650 | rs9289363 | -2.743 | 0.977 | 2.924 | 0.145 | 0.052 | 0.065 | 0.065 | 4.51 | 0.06 | 0.49 | 0.07 | 0.02 | 0.01 | 0.01 |
| 3 | 97188153 | rs6784670 | -2.78 | 1.563 | 3.155 | 0.141 | 0.198 | 0.101 | 0.101 | 3.689 | 0.07 | 0.44 | 0.27 | 0.1 | 0.09 | 0.11 |
| 3 | 97188583 | rs4857258 | -2.808 | 1.562 | 3.127 | 0.141 | 0.191 | 0.095 | 0.095 | 3.674 | 0.07 | 0.43 | 0.27 | 0.1 | 0.09 | 0.11 |
| 3 | 97191339 | rs6801975 | -2.262 | 1.576 | 3.125 | 0.147 | 0.133 | 0.058 | 0.058 | 0.934 | 0.06 | 0.26 | 0.25 | 0.1 | 0.09 | 0.11 |
| 3 | 97211634 | rs9882320 | -2.651 | 1.432 | 2.83 | 0.125 | 0.041 | 0.054 | 0.054 | 3.416 | 0.05 | 0.43 | 0.06 | 0.01 | 0.01 | 0.01 |
| 3 | 97252553 | rs2118350 | -2.579 | 1.583 | 2.366 | 0.133 | 0.082 | 0.013 | 0.013 | 3.75 | 0.1 | 0.63 | 0.31 | 0.11 | 0.15 | 0.14 |
| 3 | 97275606 | rs1370454 | -2.854 | 1.546 | 2.356 | 0.121 | 0.044 | -0.012 | -0.012 | 2.568 | 0.11 | 0.69 | 0.32 | 0.11 | 0.14 | 0.14 |
| 3 | 97279087 | rs9878433 | -2.58 | 1.545 | 2.435 | 0.121 | 0.077 | 0.008 | 0.008 | 3.831 | 0.1 | 0.62 | 0.31 | 0.11 | 0.14 | 0.13 |
| 3 | 97282841 | rs4857274 | -2.973 | 1.541 | 2.41 | 0.119 | 0.038 | -0.015 | -0.015 | 0 | 0.11 | 0.71 | 0.33 | 0.11 | 0.14 | 0.14 |
| 3 | 97284992 | rs6810360 | -2.741 | 1.545 | 2.387 | 0.12 | 0.051 | -0.007 | -0.007 | 0 | 0.11 | 0.68 | 0.31 | 0.11 | 0.14 | 0.14 |
| 3 | 97286565 | rs9868829 | -2.815 | 1.543 | 2.479 | 0.12 | 0.06 | -0.003 | -0.003 | 0.839 | 0.11 | 0.66 | 0.31 | 0.11 | 0.14 | 0.14 |
| 3 | 97289022 | rs6775629 | -2.68 | 1.545 | 2.5 | 0.12 | 0.105 | 0.025 | 0.025 | 2.496 | 0.1 | 0.57 | 0.31 | 0.11 | 0.14 | 0.13 |
| 3 | 97292036 | rs7614909 | -2.687 | 1.527 | 2.211 | 0.113 | 0.049 | -0.008 | -0.008 | 0 | 0.11 | 0.68 | 0.31 | 0.11 | 0.14 | 0.14 |
| 3 | 97304955 | rs6439265 | -2.686 | 1.518 | 2.216 | 0.128 | 0.044 | -0.012 | -0.012 | 4.871 | 0.11 | 0.69 | 0.31 | 0.12 | 0.14 | 0.14 |
| 3 | 97315448 | rs1545127 | -0.034 | 1.469 | 0.717 | 0.206 | -0.047 | -0.019 | -0.019 | 0 | 0.21 | 0.01 | 0.11 | 0.13 | 0.17 | 0.21 |
| 3 | 97339649 | rs2317211 | -1.416 | 1.54 | 1.595 | 0.998 | 0.173 | 0.064 | 0.064 | 0.034 | 0.06 | 0.29 | 0.28 | 0.1 | 0.14 | 0.13 |
| 3 | 97349304 | rs6809865 | -1.773 | 1.493 | 1.259 | 0.999 | 0.187 | 0.069 | 0.069 | 1.747 | 0.08 | 0.48 | 0.29 | 0.11 | 0.14 | 0.13 |
| 3 | 97359037 | rs301937 | -0.522 | 1.143 | -0.58 | 0.677 | -0.02 | -0.004 | -0.004 | 0 | 0.39 | 0.26 | 0.48 | 0.43 | 0.46 | 0.56 |
| 3 | 97365038 | rs301948 | 0.193 | 1.034 | -0.61 | 0.442 | -0.002 | 0.004 | 0.004 | 0 | 0.6 | 0.68 | 0.57 | 0.7 | 0.58 | 0.67 |
| 3 | 97376223 | rs301957 | 0.147 | 1.018 | -0.643 | 0.408 | -0.005 | 0.011 | 0.011 | 0 | 0.61 | 0.73 | 0.58 | 0.71 | 0.58 | 0.68 |
| 3 | 97378912 | rs4857282 | 0.397 | 1.61 | -0.807 | 0.664 | -0.005 | -0.004 | -0.004 | 0 | 0.1 | 0.08 | 0.17 | 0.23 | 0.18 | 0.25 |
| 3 | 97381406 | rs301960 | -0.557 | 1.823 | -0.937 | 0.655 | 0.02 | 0.057 | 0.057 | 0 | 0.94 | 0.89 | 0.93 | 0.87 | 0.91 | 0.95 |
| 3 | 97386798 | rs1009521 | 0.191 | 1.561 | -1.336 | 0.423 | -0.013 | 0.085 | 0.085 | 0 | 0.33 | 0.1 | 0.35 | 0.16 | 0.33 | 0.27 |
| 3 | 97395137 | rs13061095 | 0.615 | 1.456 | -1.896 | 0.117 | -0.032 | 0.044 | 0.044 | 0 | 0.26 | 0.06 | 0.32 | 0.15 | 0.28 | 0.2 |
| 3 | 97404612 | rs719278 | 0.629 | 0.856 | -1.66 | 0.236 | 0.006 | 0.094 | 0.094 | 0 | 0.43 | 0.16 | 0.36 | 0.24 | 0.38 | 0.43 |
| 3 | 97411617 | rs301923 | 0.576 | 0.436 | -0.874 | 0.555 | -0.006 | -0.063 | -0.063 | 0.46 | 0.28 | 0.14 | 0.32 | 0.4 | 0.3 | 0.2 |
| 3 | 97440269 | rs4857288 | 0.077 | 0.53 | -0.494 | 0.384 | -0.006 | -0.004 | -0.004 | 0 | 0.62 | 0.54 | 0.64 | 0.63 | 0.64 | 0.77 |
| 3 | 97443050 | rs4257593 | 0.453 | 0.213 | -0.292 | 0.535 | -0.017 | 0.019 | 0.019 | 0 | 0.27 | 0.36 | 0.25 | 0.37 | 0.24 | 0.17 |
| 3 | 97452146 | rs2873394 | 0.667 | 0.948 | -0.12 | 0.404 | -0.003 | 0.035 | 0.035 | 0 | 0.46 | 0.36 | 0.45 | 0.27 | 0.47 | 0.45 |
| 3 | 97453438 | rs7634548 | 0.286 | 1.393 | -0.596 | 0.669 | 0.01 | 0 | 0 | 0 | 0.12 | 0.13 | 0.2 | 0.16 | 0.19 | 0.21 |
| 3 | 97455560 | rs2087070 | 1.07 | 1.199 | -0.569 | 0.621 | 0.027 | -0.072 | -0.072 | 0 | 0.28 | 0.06 | 0.23 | 0.36 | 0.24 | 0.19 |
| 3 | 97455908 | rs9864542 | -0.485 | 1.149 | -0.39 | 0.414 | 0.004 | 0.085 | 0.085 | 0 | 0.5 | 0.33 | 0.51 | 0.72 | 0.52 | 0.53 |
| 3 | 97462795 | rs9822683 | 0.539 | 1.118 | -0.39 | 0.413 | 0.002 | 0.05 | 0.05 | 0 | 0.48 | 0.62 | 0.48 | 0.27 | 0.47 | 0.45 |
| 3 | 97474630 | rs9881515 | -0.727 | 1.077 | -0.518 | 0.477 | -0.013 | -0.012 | -0.012 | 0 | 0.22 | 0.19 | 0.27 | 0.31 | 0.26 | 0.29 |
| 3 | 97477083 | rs4857065 | -1.177 | 1.039 | -0.49 | 0.428 | 0.002 | 0.002 | 0.002 | 0 | 0.26 | 0.3 | 0.31 | 0.34 | 0.32 | 0.33 |
| 3 | 97512420 | rs2279896 | -0.405 | 1.056 | -0.718 | 0.315 | -0.044 | 0.057 | 0.057 | 0.151 | 0.44 | 0.19 | 0.49 | 0.66 | 0.47 | 0.49 |
| 3 | 97514266 | rs10511163 | 1.532 | 1.08 | -0.765 | 0.358 | -0.02 | -0.133 | -0.133 | 2.944 | 0.23 | 0.01 | 0.22 | 0.35 | 0.21 | 0.19 |

| | | | | | | | | | | | | | | | | |
|---|---|---|---|---|---|---|---|---|---|---|---|---|---|---|---|---|
| 3 | 97526324 | rs12491333 | 0.781 | 1.221 | -0.766 | 0.192 | -0.041 | -0.039 | -0.039 | 0 | 0.08 | 0 | 0.05 | 0.1 | 0.09 | 0.08 |
| 3 | 97527233 | rs9853330 | -1.422 | 1.133 | -0.709 | -0.22 | -0.012 | -0.019 | -0.019 | 1.005 | 0.67 | 0.41 | 0.8 | 0.83 | 0.77 | 0.71 |
| 3 | 97529147 | rs2873322 | -1.831 | 1.036 | -0.27 | -0.292 | 0.013 | 0.033 | 0.033 | 0 | 0.72 | 0.82 | 0.86 | 0.86 | 0.79 | 0.73 |
| 3 | 97550849 | rs3907775 | -1.392 | 1.162 | -0.327 | -0.237 | 0.002 | 0.001 | 0.001 | 0.045 | 0.67 | 0.36 | 0.8 | 0.83 | 0.77 | 0.71 |
| 3 | 97570446 | rs4857296 | 0.092 | 1.204 | -0.386 | -0.203 | -0.034 | 0.082 | 0.082 | 0.163 | 0.46 | 0.2 | 0.5 | 0.68 | 0.51 | 0.51 |
| 3 | 97593380 | rs2279898 | 1.054 | 1.255 | -0.372 | -0.151 | -0.02 | -0.006 | -0.006 | 0 | 0.1 | 0.06 | 0.17 | 0.18 | 0.17 | 0.15 |
| 3 | 97593485 | rs2279897 | 0.159 | 1.224 | -0.319 | -0.219 | -0.017 | 0.032 | 0.032 | 0 | 0.22 | 0.11 | 0.32 | 0.16 | 0.28 | 0.21 |
| 3 | 97595379 | rs6786354 | 0.072 | 1.207 | -0.308 | -0.226 | -0.023 | 0.072 | 0.072 | 0.125 | 0.46 | 0.26 | 0.5 | 0.67 | 0.51 | 0.51 |
| 3 | 97602737 | rs10935130 | 0.167 | 1.21 | -0.322 | -0.254 | -0.017 | 0.032 | 0.032 | 0 | 0.22 | 0.11 | 0.32 | 0.16 | 0.28 | 0.21 |
| 3 | 97606672 | rs12637414 | 0.828 | 1.213 | -0.354 | -0.246 | -0.045 | -0.039 | -0.039 | 1.171 | 0.11 | 0.01 | 0.06 | 0.11 | 0.13 | 0.1 |
| 3 | 97609951 | rs831899 | -1.465 | 1.08 | 0.186 | -0.409 | -0.026 | -0.047 | -0.047 | 0 | 0.69 | 0.47 | 0.83 | 0.86 | 0.79 | 0.73 |
| 3 | 97620528 | rs1498652 | 0.405 | 1.079 | -0.008 | -0.41 | -0.023 | 0.078 | 0.078 | 0.125 | 0.46 | 0.25 | 0.5 | 0.68 | 0.51 | 0.51 |
| 3 | 97632716 | rs831878 | -1.391 | 1.027 | -0.04 | -0.427 | -0.024 | -0.049 | -0.049 | 0 | 0.71 | 0.6 | 0.85 | 0.86 | 0.79 | 0.73 |
| 3 | 97633737 | rs1391665 | 0.416 | 1.068 | -0.207 | -0.392 | -0.033 | 0.084 | 0.084 | 0.149 | 0.46 | 0.2 | 0.5 | 0.68 | 0.51 | 0.51 |
| 3 | 97650715 | rs17302656 | 0.976 | 1.249 | -0.282 | -0.357 | -0.002 | -0.004 | -0.004 | 0 | 0.07 | 0.04 | 0.23 | 0.15 | 0.17 | 0.04 |
| 3 | 97657476 | rs2172258 | 1.244 | 0.963 | -0.266 | -0.314 | -0.016 | -0.064 | -0.064 | 0 | 0.34 | 0.08 | 0.32 | 0.5 | 0.34 | 0.35 |
| 3 | 97660084 | rs11544446 | -0.393 | 0.346 | -0.038 | -0.38 | -0.017 | 0.034 | 0.034 | 0 | 0.22 | 0.11 | 0.31 | 0.15 | 0.28 | 0.21 |
| 3 | 97660106 | rs4857302 | 0.67 | 0.381 | 0.039 | -0.368 | -0.013 | 0.067 | 0.067 | 0.149 | 0.46 | 0.26 | 0.51 | 0.69 | 0.51 | 0.51 |
| 3 | 97660198 | rs832089 | -1.247 | 0.379 | 0.113 | -0.374 | -0.011 | -0.023 | -0.023 | 0 | 0.71 | 0.66 | 0.85 | 0.86 | 0.8 | 0.73 |
| 3 | 97664725 | rs2172257 | 0.503 | 0.599 | -0.716 | -0.151 | -0.015 | 0.056 | 0.056 | 0.133 | 0.46 | 0.26 | 0.51 | 0.67 | 0.51 | 0.51 |
| 3 | 97684754 | rs9877544 | 0.252 | 0.638 | -0.726 | -0.249 | -0.008 | 0.115 | 0.115 | 0.072 | 0.48 | 0.25 | 0.54 | 0.76 | 0.53 | 0.53 |
| 3 | 97696446 | rs832072 | -0.194 | 0.957 | -0.757 | -0.242 | -0.035 | 0.039 | 0.039 | 0 | 0.83 | 0.93 | 0.74 | 0.9 | 0.75 | 0.82 |
| 3 | 97697528 | rs1492030 | 0.327 | 0.704 | -0.818 | -0.216 | -0.001 | 0.103 | 0.103 | 0.086 | 0.49 | 0.21 | 0.55 | 0.71 | 0.54 | 0.54 |
| 3 | 97701082 | rs9818084 | -0.539 | 0.887 | -0.182 | -0.227 | 0.031 | -0.063 | -0.063 | 0 | 0.2 | 0.31 | 0.28 | 0.1 | 0.25 | 0.17 |
| 3 | 97702896 | rs1492038 | 1.038 | 0.946 | -0.892 | 0.066 | -0.028 | -0.074 | -0.074 | 0.925 | 0.36 | 0.21 | 0.34 | 0.43 | 0.39 | 0.37 |
| 3 | 97704434 | rs6780855 | -0.169 | 1.039 | -0.647 | -0.013 | 0.006 | -0.018 | -0.018 | 0.025 | 0.18 | 0.21 | 0.27 | 0.1 | 0.25 | 0.17 |
| 3 | 97708955 | rs832100 | 1.093 | 0.593 | -1.165 | 0.078 | -0.003 | -0.01 | -0.01 | 0 | 0.26 | 0.04 | 0.14 | 0.19 | 0.21 | 0.29 |
| 3 | 97711997 | rs937130 | 0.918 | 1.078 | -1.483 | -0.07 | -0.055 | -0.07 | -0.07 | 0 | 0.37 | 0.19 | 0.41 | 0.47 | 0.46 | 0.35 |
| 3 | 97712289 | rs1874864 | 0.215 | 1.497 | -1.571 | 0.6 | -0.09 | -0.088 | -0.088 | 0 | 0.4 | 0.11 | 0.38 | 0.48 | 0.5 | 0.39 |
| 3 | 97713502 | rs7638369 | -1.027 | 1.406 | -0.678 | 0.12 | 0.01 | -0.014 | -0.014 | 1.711 | 0.26 | 0.61 | 0.32 | 0.21 | 0.32 | 0.21 |
| 3 | 97713804 | rs1844934 | 0.712 | 0.936 | -0.667 | 0.455 | -0.012 | -0.104 | -0.104 | 0.248 | 0.36 | 0.17 | 0.35 | 0.44 | 0.37 | 0.4 |
| 3 | 97721299 | rs10511164 | -0.162 | 0.744 | -0.633 | 0.164 | 0.001 | -0.002 | -0.002 | 0.421 | 0.17 | 0.14 | 0.2 | 0.19 | 0.15 | 0.17 |
| 3 | 97721745 | rs1688377 | -0.098 | 0.856 | -0.516 | -0.222 | 0.019 | 0.089 | 0.089 | 0.047 | 0.5 | 0.35 | 0.42 | 0.22 | 0.51 | 0.6 |
| 3 | 97727555 | rs832031 | -0.118 | 0.029 | 0.007 | -0.501 | -0.042 | -0.151 | -0.151 | 0.035 | 0.7 | 0.54 | 0.81 | 0.99 | 0.81 | 0.83 |
| 3 | 97728248 | rs16839096 | -0.192 | 0.154 | -0.543 | -0.073 | -0.026 | -0.111 | -0.111 | 0.992 | 0.25 | 0.1 | 0.26 | 0.44 | 0.29 | 0.33 |
| 3 | 97729328 | rs832027 | -0.598 | 0.102 | -0.186 | -0.181 | -0.03 | -0.122 | -0.122 | 0.266 | 0.71 | 0.65 | 0.82 | 0.99 | 0.81 | 0.83 |
| 3 | 97739149 | rs17376145 | 0.976 | 0.135 | -0.296 | 0.043 | -0.027 | -0.047 | -0.047 | 0 | 0.11 | 0.01 | 0.08 | 0.12 | 0.14 | 0.08 |
| 3 | 97739493 | rs832035 | -0.77 | 0.04 | -0.068 | -0.03 | -0.014 | -0.08 | -0.08 | 0.01 | 0.72 | 0.72 | 0.82 | 0.99 | 0.81 | 0.83 |
| 3 | 97741821 | rs832038 | 1.069 | 0.054 | -0.185 | 0.044 | -0.002 | -0.004 | -0.004 | 0 | 0.44 | 0.45 | 0.56 | 0.55 | 0.51 | 0.49 |
| 3 | 97744611 | rs832041 | -0.813 | 0.561 | -0.27 | 0.129 | -0.044 | -0.081 | -0.081 | 0 | 0.26 | 0.12 | 0.28 | 0.44 | 0.34 | 0.34 |
| 3 | 97749522 | rs1498646 | 0.991 | 0.191 | 0.103 | 0.076 | -0.001 | -0.003 | -0.003 | 0.139 | 0.44 | 0.45 | 0.55 | 0.54 | 0.51 | 0.49 |
| 3 | 97754170 | rs832067 | -0.773 | 0.138 | 0.077 | -0.085 | 0.005 | -0.015 | -0.015 | 0 | 0.68 | 0.78 | 0.8 | 0.98 | 0.72 | 0.77 |
| 3 | 97755270 | rs11711963 | 1.199 | 0.203 | -0.34 | 0.039 | 0.001 | -0.04 | -0.04 | 0.027 | 0.1 | 0.01 | 0.07 | 0.1 | 0.1 | 0.08 |
| 3 | 97766185 | rs832052 | -0.409 | 0.142 | -0.165 | -0.15 | -0.021 | -0.16 | -0.16 | 0.013 | 0.65 | 0.57 | 0.77 | 0.98 | 0.72 | 0.78 |
| 3 | 97767182 | rs832049 | -0.338 | 0.242 | -0.435 | -0.019 | -0.013 | -0.055 | -0.055 | 0 | 0.21 | 0.11 | 0.24 | 0.47 | 0.26 | 0.28 |
| 3 | 97769777 | rs9289510 | 1.029 | 0.316 | -0.614 | 0.043 | -0.014 | -0.015 | -0.015 | 0 | 0.43 | 0.34 | 0.52 | 0.52 | 0.46 | 0.49 |
| 3 | 97771603 | rs7643404 | -0.604 | 0.491 | -1.181 | -0.248 | 0.003 | 0.138 | 0.138 | 0 | 0.69 | 0.93 | 0.8 | 0.98 | 0.72 | 0.78 |
| 3 | 97783335 | rs1388646 | -0.587 | 1.024 | -0.897 | 0.215 | -0.022 | 0.001 | 0.001 | 0 | 0.87 | 0.94 | 0.78 | 0.9 | 0.8 | 0.9 |
| 3 | 97784175 | rs1352891 | 0.628 | 0.751 | -0.808 | 0.187 | 0.006 | 0.004 | 0.004 | 0.104 | 0.47 | 0.56 | 0.51 | 0.45 | 0.41 | 0.41 |
| 3 | 97784559 | rs6764595 | -0.673 | 0.384 | -0.651 | 0.49 | 0 | 0 | 0 | 0 | 0.34 | 0.38 | 0.33 | 0.53 | 0.35 | 0.38 |
| 3 | 97786424 | rs1492042 | 0.246 | 0.401 | -0.73 | 0.405 | -0.082 | -0.13 | -0.13 | 0 | 0.64 | 0.95 | 0.66 | 0.46 | 0.58 | 0.65 |
| 3 | 97787260 | rs1826351 | 0.584 | 0.384 | -0.747 | 0.694 | -0.027 | -0.026 | -0.026 | 0 | 0.25 | 0.03 | 0.26 | 0.22 | 0.26 | 0.13 |
| 3 | 97791188 | rs1826352 | 0.176 | 0.32 | -0.378 | 0.498 | 0 | 0.01 | 0.01 | 0 | 0.6 | 0.66 | 0.57 | 0.37 | 0.56 | 0.6 |
| 3 | 97791448 | rs7617121 | 0.109 | 0.199 | 0.052 | 0.552 | 0.004 | 0.054 | 0.054 | 0.021 | 0.11 | 0.31 | 0.21 | 0.17 | 0.06 | 0.14 |
| 3 | 97794808 | rs1602584 | 0.114 | 0.202 | 0.014 | 0.552 | 0 | 0 | 0 | 0.004 | 0.58 | 0.52 | 0.55 | 0.37 | 0.54 | 0.59 |
| 3 | 97799532 | rs12054459 | 0.033 | 0.274 | -0.261 | 0.583 | -0.001 | 0.009 | 0.009 | 0 | 0.59 | 0.62 | 0.55 | 0.37 | 0.54 | 0.59 |
| 3 | 97799624 | rs12054461 | 0.439 | 0.444 | -0.627 | 0.472 | 0 | 0.034 | 0.034 | 0.028 | 0.09 | 0.17 | 0.19 | 0.17 | 0.06 | 0.14 |
| 3 | 97800433 | rs10935192 | 0.132 | 0.629 | -1.101 | 0.473 | -0.022 | -0.036 | -0.036 | 0 | 0.63 | 0.75 | 0.63 | 0.45 | 0.55 | 0.63 |
| 3 | 97806616 | rs4518168 | 0.397 | 0.663 | -0.843 | 0.454 | -0.001 | -0.002 | -0.002 | 0 | 0.56 | 0.54 | 0.55 | 0.45 | 0.51 | 0.58 |
| 3 | 97807234 | rs4857314 | -0.13 | 0.199 | -1.041 | -0.63 | 0.039 | 0.134 | 0.134 | 0 | 0.48 | 0.3 | 0.69 | 0.84 | 0.52 | 0.61 |
| 3 | 97816039 | rs4317139 | -0.425 | 0.264 | -1.503 | -0.331 | -0.068 | -0.066 | -0.066 | 0.122 | 0.53 | 0.9 | 0.59 | 0.53 | 0.54 | 0.57 |
| 3 | 97824577 | rs9837490 | -0.407 | 0.252 | -1.427 | -0.337 | -0.049 | -0.048 | -0.048 | 0 | 0.53 | 0.86 | 0.59 | 0.52 | 0.53 | 0.56 |
| 3 | 97853159 | rs4519741 | 0.256 | -0.224 | -1.748 | -0.742 | -0.068 | -0.091 | -0.091 | 0 | 0.28 | 0.08 | 0.46 | 0.39 | 0.37 | 0.35 |
| 3 | 97854166 | rs4469014 | 1.137 | -0.356 | -0.647 | -0.808 | -0.014 | -0.045 | -0.045 | 0 | 0.21 | 0.46 | 0.09 | 0.13 | 0.16 | 0.22 |
| 3 | 97854334 | rs4449360 | -0.601 | -0.175 | 0.014 | -0.367 | -0.03 | -0.029 | -0.029 | 0 | 0.52 | 0.84 | 0.58 | 0.52 | 0.53 | 0.56 |
| 3 | 97854425 | rs4271952 | -1.704 | -0.248 | -0.19 | -0.237 | -0.092 | -0.072 | -0.072 | 0 | 0.82 | 0.97 | 0.82 | 0.75 | 0.75 | 0.77 |
| 3 | 97856333 | rs4376074 | -0.83 | -0.087 | 0.316 | -0.149 | -0.03 | -0.029 | -0.029 | 0.348 | 0.52 | 0.84 | 0.58 | 0.52 | 0.53 | 0.56 |
| 3 | 97858246 | rs9289534 | 1.293 | -0.077 | 0.691 | -0.285 | 0.004 | 0.031 | 0.031 | 0 | 0.28 | 0.19 | 0.2 | 0.01 | 0.35 | 0.23 |
| 3 | 97861768 | rs6793365 | 0.437 | -0.076 | 0.707 | -0.262 | -0.006 | -0.063 | -0.063 | 0 | 0.37 | 0.47 | 0.25 | 0.14 | 0.4 | 0.34 |

| | | | | | | | | | | | | | | | | |
|---|---|---|---|---|---|---|---|---|---|---|---|---|---|---|---|---|
| 3 | 97862651 | rs4271951 | -0.852 | -0.077 | 0.82 | -0.256 | -0.006 | -0.006 | -0.006 | 0 | 0.51 | 0.71 | 0.57 | 0.52 | 0.53 | 0.56 |
| 3 | 97866177 | rs4857072 | 0.446 | -0.07 | 0.846 | -0.25 | -0.01 | -0.014 | -0.014 | 0 | 0.12 | 0.12 | 0.31 | 0.39 | 0.12 | 0.22 |
| 3 | 97867846 | rs4302408 | -0.814 | -0.074 | 0.859 | -0.256 | -0.004 | -0.004 | -0.004 | 0.336 | 0.51 | 0.71 | 0.56 | 0.52 | 0.53 | 0.56 |
| 3 | 97875767 | rs4550856 | 1.284 | -0.058 | 0.822 | -0.262 | 0.004 | 0.031 | 0.031 | 0 | 0.28 | 0.19 | 0.2 | 0.01 | 0.35 | 0.23 |
| 3 | 97879569 | rs9810825 | 0.416 | -0.035 | 0.729 | -0.245 | -0.007 | -0.064 | -0.064 | 0.232 | 0.37 | 0.47 | 0.25 | 0.14 | 0.4 | 0.35 |
| 3 | 97880870 | rs13098247 | 0.541 | -0.036 | 0.504 | -0.202 | -0.019 | -0.033 | -0.033 | 0 | 0.12 | 0.07 | 0.31 | 0.39 | 0.12 | 0.22 |
| 3 | 97883653 | rs6772614 | -0.906 | -0.117 | 0.528 | -0.23 | -0.021 | -0.021 | -0.021 | 0 | 0.52 | 0.82 | 0.58 | 0.51 | 0.54 | 0.57 |
| 3 | 97885241 | rs4857079 | -0.722 | -0.109 | 0.715 | -0.238 | 0.008 | 0.048 | 0.048 | 0 | 0.19 | 0.25 | 0.34 | 0.48 | 0.18 | 0.33 |
| 3 | 97888197 | rs7623031 | -0.79 | -0.092 | 0.487 | -0.09 | -0.035 | -0.031 | -0.031 | 0 | 0.53 | 0.85 | 0.58 | 0.51 | 0.54 | 0.57 |
| 3 | 97889348 | rs6804026 | -0.819 | -0.087 | 0.489 | -0.088 | -0.04 | -0.036 | -0.036 | 0 | 0.53 | 0.87 | 0.59 | 0.51 | 0.54 | 0.57 |
| 3 | 97901263 | rs7615262 | -0.793 | -0.083 | 0.489 | -0.086 | -0.035 | -0.031 | -0.031 | 1.123 | 0.53 | 0.85 | 0.58 | 0.51 | 0.54 | 0.57 |
| 3 | 97902287 | rs13092516 | 0.453 | -0.066 | 0.5 | -0.076 | -0.021 | -0.036 | -0.036 | 0.119 | 0.12 | 0.07 | 0.31 | 0.38 | 0.12 | 0.22 |
| 3 | 97903374 | rs4358331 | -0.417 | -0.09 | 0.622 | 0.066 | -0.035 | -0.031 | -0.031 | 0 | 0.53 | 0.85 | 0.58 | 0.51 | 0.54 | 0.57 |
| 3 | 97906208 | rs11716786 | -0.423 | -0.131 | 0.451 | 0.177 | -0.016 | -0.014 | -0.014 | 1.931 | 0.54 | 0.79 | 0.51 | 0.49 | 0.55 | 0.56 |
| 3 | 97909666 | rs9859733 | -0.415 | -0.133 | 0.45 | 0.176 | -0.016 | -0.014 | -0.014 | 0.101 | 0.54 | 0.79 | 0.5 | 0.49 | 0.55 | 0.56 |
| 3 | 97917256 | rs2047391 | -0.308 | -0.149 | 0.469 | 0.169 | -0.008 | -0.006 | -0.006 | 0 | 0.53 | 0.77 | 0.5 | 0.49 | 0.55 | 0.56 |
| 3 | 97919363 | rs12695682 | -0.182 | -0.146 | 0.841 | 0.172 | 0.001 | 0.002 | 0.002 | 0 | 0.52 | 0.66 | 0.49 | 0.49 | 0.55 | 0.56 |
| 3 | 97937035 | rs5000373 | -0.418 | -0.154 | 0.755 | 0.169 | -0.056 | -0.052 | -0.052 | 0 | 0.56 | 0.88 | 0.52 | 0.49 | 0.55 | 0.56 |
| 3 | 97939961 | rs16839427 | 0.636 | -0.127 | 0.912 | 0.153 | 0.003 | 0.06 | 0.06 | 0 | 0.09 | 0.12 | 0.22 | 0.4 | 0.09 | 0.17 |
| 3 | 97940745 | rs16839441 | 0.359 | -0.255 | 1.058 | 0.005 | -0.002 | -0.073 | -0.073 | 0 | 0.43 | 0.47 | 0.27 | 0.1 | 0.46 | 0.4 |
| 3 | 97942333 | rs2316298 | 0.63 | -0.245 | 0.903 | -0.237 | -0.005 | -0.14 | -0.14 | 0 | 0.38 | 0.44 | 0.26 | 0.04 | 0.42 | 0.31 |
| 3 | 97958253 | rs9847708 | -0.417 | -0.165 | 0.031 | 0.024 | -0.019 | -0.019 | -0.019 | 0 | 0.54 | 0.77 | 0.51 | 0.5 | 0.55 | 0.56 |
| 3 | 97958635 | rs13321188 | -0.195 | -0.13 | -0.079 | 0.048 | -0.001 | 0.002 | 0.002 | 0 | 0.52 | 0.68 | 0.5 | 0.49 | 0.54 | 0.56 |
| 3 | 97960399 | rs9875700 | -0.439 | -0.254 | -0.223 | 0.931 | -0.041 | -0.059 | -0.059 | 0.058 | 0.69 | 0.87 | 0.63 | 0.6 | 0.69 | 0.63 |
| 3 | 97961301 | rs12639163 | -0.875 | -0.076 | 0.142 | 1.204 | -0.004 | -0.002 | -0.002 | 0 | 0.15 | 0.15 | 0.25 | 0.44 | 0.12 | 0.26 |
| 3 | 97962137 | rs4857355 | -0.002 | -0.018 | -0.054 | 1.346 | -0.017 | -0.019 | -0.019 | 0 | 0.09 | 0.06 | 0.22 | 0.42 | 0.08 | 0.16 |
| 3 | 97983942 | rs9853906 | -0.06 | -0.28 | 0.245 | 0.816 | -0.007 | -0.014 | -0.014 | 0 | 0.66 | 0.72 | 0.62 | 0.62 | 0.69 | 0.64 |
| 3 | 97984556 | rs17195199 | 0.926 | -0.369 | 0.846 | 0.778 | 0.003 | 0.016 | 0.016 | 0 | 0.49 | 0.37 | 0.35 | 0.13 | 0.56 | 0.38 |
| 3 | 97984838 | rs11927378 | -0.761 | -0.594 | 0.94 | 0.915 | 0.002 | 0.063 | 0.063 | 0 | 0.15 | 0.19 | 0.25 | 0.49 | 0.12 | 0.25 |
| 3 | 97988441 | rs13064749 | 0.28 | -0.582 | 0.82 | 0.924 | -0.026 | -0.024 | -0.024 | 0 | 0.09 | 0.03 | 0.22 | 0.47 | 0.08 | 0.16 |
| 3 | 97992964 | rs1566169 | -0.879 | -0.794 | 0.72 | 0.906 | 0.005 | 0.005 | 0.005 | 0 | 0.35 | 0.34 | 0.42 | 0.75 | 0.23 | 0.39 |
| 3 | 97994268 | rs1011205 | 0.28 | -0.833 | 0.839 | 0.929 | -0.044 | 0.014 | 0.014 | 0 | 0.22 | 0.37 | 0.18 | 0.26 | 0.11 | 0.13 |
| 3 | 97999365 | rs4857359 | -0.372 | -0.859 | 0.846 | 0.93 | 0.027 | 0.01 | 0.01 | 0 | 0.29 | 0.2 | 0.39 | 0.73 | 0.18 | 0.28 |
| 3 | 98001332 | rs2316276 | 0.896 | -0.852 | 0.786 | 0.938 | 0.045 | -0.04 | -0.04 | 0.667 | 0.19 | 0.08 | 0.16 | 0.26 | 0.1 | 0.12 |
| 3 | 98002587 | rs16839611 | -1.12 | -0.947 | 0.811 | 0.873 | 0.006 | 0.006 | 0.006 | 0 | 0.39 | 0.66 | 0.45 | 0.75 | 0.23 | 0.38 |
| 3 | 98019629 | rs13100068 | 0.153 | -0.692 | 0.862 | 1.451 | 0.003 | 0.084 | 0.084 | 0 | 0.1 | 0.13 | 0.22 | 0.44 | 0.09 | 0.16 |
| 3 | 98027888 | rs2132895 | -0.982 | -0.733 | 0.965 | 1.422 | -0.023 | -0.021 | -0.021 | 0 | 0.35 | 0.59 | 0.43 | 0.73 | 0.21 | 0.34 |
| 3 | 98037111 | rs13080412 | 0.241 | -0.673 | 0.731 | 1.483 | 0.004 | 0.092 | 0.092 | 0 | 0.1 | 0.14 | 0.23 | 0.44 | 0.09 | 0.16 |
| 3 | 98049409 | rs4857374 | 0.675 | -0.791 | 0.701 | 1.402 | 0.039 | -0.023 | -0.023 | 0 | 0.2 | 0.15 | 0.17 | 0.27 | 0.1 | 0.13 |
| 3 | 98060332 | rs11920195 | 0.042 | -0.522 | 0.963 | 1.512 | 0.012 | 0.142 | 0.142 | 0 | 0.1 | 0.18 | 0.23 | 0.44 | 0.08 | 0.15 |
| 3 | 98063570 | rs1948608 | 0.365 | -0.439 | 0.487 | 1.548 | -0.032 | -0.024 | -0.024 | 0 | 0.09 | 0.02 | 0.21 | 0.44 | 0.08 | 0.15 |
| 3 | 98073313 | rs9822460 | -0.311 | -0.065 | 0.859 | 1.074 | 0.027 | -0.009 | -0.009 | 0 | 0.18 | 0.25 | 0.23 | 0.17 | 0.22 | 0.29 |
| 3 | 98075543 | rs11708509 | -0.393 | -0.087 | 0.849 | 1.064 | 0.024 | -0.009 | -0.009 | 0 | 0.19 | 0.28 | 0.23 | 0.17 | 0.22 | 0.29 |
| 3 | 98082519 | rs4630981 | -0.307 | -0.078 | 0.853 | 1.068 | 0.027 | -0.009 | -0.009 | 0 | 0.18 | 0.25 | 0.23 | 0.17 | 0.22 | 0.29 |
| 3 | 98087516 | rs9821488 | 0.846 | -0.202 | 1.264 | 1.028 | -0.004 | 0.177 | 0.177 | 0 | 0.51 | 0.22 | 0.37 | 0.13 | 0.6 | 0.41 |
| 3 | 98093705 | rs13093620 | 0.247 | -0.024 | 1.353 | 1.126 | 0.058 | 0.009 | 0.009 | 0 | 0.11 | 0.2 | 0.2 | 0.14 | 0.17 | 0.23 |
| 3 | 98100691 | rs9862953 | 0.116 | -0.048 | 1.403 | 1.124 | 0.05 | 0.005 | 0.005 | 0 | 0.12 | 0.21 | 0.19 | 0.14 | 0.17 | 0.22 |
| 3 | 98101101 | rs9289584 | -0.572 | -0.216 | 1.439 | 1.272 | 0.037 | -0.044 | -0.044 | 0 | 0.69 | 0.8 | 0.64 | 0.43 | 0.82 | 0.78 |
| 3 | 98102684 | rs6439822 | -0.37 | -0.165 | 1.494 | 1.197 | 0.021 | -0.019 | -0.019 | 0 | 0.67 | 0.73 | 0.59 | 0.43 | 0.79 | 0.78 |
| 3 | 98102801 | rs6439823 | -0.148 | -0.131 | 1.644 | 1.088 | 0.069 | 0.007 | 0.007 | 0 | 0.13 | 0.31 | 0.21 | 0.14 | 0.17 | 0.22 |
| 3 | 98106152 | rs9858863 | 0.603 | -0.177 | 1.645 | 1.038 | 0.005 | 0.073 | 0.073 | 0 | 0.54 | 0.28 | 0.38 | 0.2 | 0.63 | 0.46 |
| 3 | 98107127 | rs9844815 | -1.51 | 0.195 | 2.111 | 1.778 | 0.073 | 0.115 | 0.115 | 0.273 | 0.18 | 0.37 | 0.24 | 0.38 | 0.22 | 0.35 |
| 3 | 98109489 | rs13069360 | -1.062 | 0.379 | 1.652 | 1.86 | 0.084 | 0.066 | 0.066 | 0 | 0.15 | 0.29 | 0.14 | 0.26 | 0.2 | 0.26 |
| 3 | 98109640 | rs13068323 | -0.004 | 0.5 | 1.446 | 1.918 | 0.033 | 0.035 | 0.035 | 0 | 0.1 | 0.13 | 0.12 | 0.22 | 0.16 | 0.19 |
| 3 | 98120608 | rs9847827 | -0.889 | 0.578 | 1.513 | 1.982 | 0.108 | 0.191 | 0.191 | 0.068 | 0.11 | 0.33 | 0.23 | 0.34 | 0.2 | 0.26 |
| 3 | 98147260 | rs4241475 | 0.298 | -1.078 | 0.903 | 1.252 | 0.07 | -0.005 | -0.005 | 0 | 0.48 | 0.22 | 0.41 | 0.46 | 0.34 | 0.41 |
| 3 | 98148355 | rs13091483 | 0.152 | -1.135 | 1.3 | 1.226 | 0.005 | 0 | 0 | 0.31 | 0.5 | 0.52 | 0.44 | 0.46 | 0.34 | 0.41 |
| 3 | 98153308 | rs4303908 | 0.051 | -1.138 | 1.325 | 1.22 | 0.033 | 0 | 0 | 0.163 | 0.51 | 0.65 | 0.45 | 0.46 | 0.34 | 0.41 |
| 3 | 98156712 | rs1112393 | -0.059 | -1.227 | 1.294 | 1.22 | 0.001 | -0.003 | -0.003 | 0 | 0.57 | 0.58 | 0.47 | 0.48 | 0.38 | 0.47 |
| 3 | 98157847 | rs9857087 | 0.425 | -1.138 | 1.347 | 1.196 | -0.016 | -0.003 | -0.003 | 1.925 | 0.37 | 0.2 | 0.39 | 0.4 | 0.58 | 0.43 |
| 3 | 98170450 | rs9809157 | -0.025 | -1.136 | 1.205 | 1.191 | 0.008 | 0.022 | 0.022 | 0 | 0.57 | 0.23 | 0.55 | 0.6 | 0.41 | 0.58 |
| 3 | 98174233 | rs9843090 | -2.102 | -0.855 | 1.243 | 1.122 | 0.01 | 0.151 | 0.151 | 3.319 | 0.08 | 0.51 | 0.14 | 0.12 | 0.03 | 0.09 |
| 3 | 98174258 | rs7614425 | -1.638 | -1.087 | 1.166 | 1.129 | -0.056 | 0.056 | 0.056 | 0.53 | 0.15 | 0.46 | 0.19 | 0.25 | 0.1 | 0.28 |
| 3 | 98174864 | rs1406559 | 0.192 | -1.128 | 0.685 | 1.015 | -0.036 | -0.012 | -0.012 | 0 | 0.38 | 0.12 | 0.37 | 0.37 | 0.52 | 0.39 |
| 3 | 98175411 | rs9882767 | -0.643 | -0.882 | 0.903 | 1.146 | 0.014 | 0.003 | 0.003 | 0 | 0.65 | 0.58 | 0.61 | 0.63 | 0.49 | 0.65 |
| 3 | 98175458 | rs13073282 | 0.009 | -0.688 | 1.09 | 1.37 | 0.015 | -0.004 | -0.004 | 0 | 0.52 | 0.31 | 0.49 | 0.54 | 0.44 | 0.55 |
| 3 | 98191245 | rs11928315 | 0.169 | -0.585 | 0.473 | 1.543 | 0.015 | 0 | 0 | 0 | 0.5 | 0.51 | 0.43 | 0.45 | 0.39 | 0.44 |
| 3 | 98204198 | rs6798905 | 0.513 | -0.583 | 0.214 | 1.512 | 0.022 | 0.023 | 0.023 | 0 | 0.46 | 0.31 | 0.4 | 0.34 | 0.37 | 0.35 |
| 3 | 98205140 | rs10935397 | -0.266 | -0.233 | -0.766 | 1.568 | -0.021 | 0.001 | 0.001 | 0 | 0.61 | 0.78 | 0.5 | 0.6 | 0.48 | 0.62 |

| | | | | | | | | | | | | | | | | |
|---|---|---|---|---|---|---|---|---|---|---|---|---|---|---|---|---|
| 3 | 98208873 | rs10935405 | -0.276 | -0.213 | -1.102 | 1.311 | -0.036 | -0.044 | -0.044 | 0 | 0.61 | 0.82 | 0.5 | 0.57 | 0.48 | 0.6 |
| 3 | 98212517 | rs10511167 | -0.822 | 0.117 | -1.202 | 1.232 | -0.006 | -0.031 | -0.031 | 0.091 | 0.08 | 0.01 | 0.05 | 0.15 | 0.08 | 0.18 |
| 3 | 98216252 | rs12639369 | -0.466 | 0.368 | -1.646 | 1.461 | -0.023 | -0.092 | -0.092 | 0.472 | 0.21 | 0.07 | 0.2 | 0.4 | 0.19 | 0.15 |
| 3 | 98221749 | rs6772735 | -1.093 | 0.541 | -2.279 | 0.931 | -0.028 | 0.141 | 0.141 | 0.599 | 0.55 | 0.86 | 0.68 | 0.76 | 0.5 | 0.57 |
| 3 | 98224945 | rs1675517 | -0.334 | 0.087 | -1.051 | 0.843 | -0.04 | 0.045 | 0.045 | 0 | 0.3 | 0.51 | 0.3 | 0.43 | 0.19 | 0.24 |
| 3 | 98225441 | rs1132750 | -1.722 | 0.151 | -1.076 | 0.838 | -0.027 | 0.141 | 0.141 | 0.506 | 0.55 | 0.87 | 0.68 | 0.76 | 0.51 | 0.59 |
| 3 | 98225660 | rs3762660 | -0.028 | 0.601 | -1.002 | 0.97 | -0.002 | -0.03 | -0.03 | 0 | 0.14 | 0.07 | 0.26 | 0.27 | 0.11 | 0.22 |
| 3 | 98230489 | rs2316131 | 0.524 | 0.654 | -0.929 | 0.702 | 0.047 | -0.006 | -0.006 | 0 | 0.09 | 0.18 | 0.13 | 0.06 | 0.2 | 0.11 |
| 3 | 98231270 | rs1529045 | -0.02 | 0.179 | -0.353 | 0.775 | -0.03 | -0.03 | -0.03 | 1.747 | 0.18 | 0.33 | 0.1 | 0.1 | 0.13 | 0.1 |
| 3 | 98249665 | rs1381087 | -1.662 | 0.438 | -1.21 | 0.764 | -0.021 | 0.146 | 0.146 | 0.454 | 0.54 | 0.85 | 0.68 | 0.76 | 0.5 | 0.57 |
| 3 | 98250215 | rs4857399 | -0.472 | 0.536 | -1.162 | 1.403 | -0.021 | -0.05 | -0.05 | 0 | 0.15 | 0.07 | 0.24 | 0.26 | 0.15 | 0.34 |
| 3 | 98250986 | rs2230344 | -0.24 | 0.186 | -1.575 | 1.61 | 0.011 | -0.067 | -0.067 | 0 | 0.19 | 0.04 | 0.29 | 0.36 | 0.14 | 0.2 |
| 3 | 98252027 | rs1529047 | -1.548 | 0.242 | -1.344 | 1.354 | -0.008 | 0.092 | 0.092 | 0 | 0.47 | 0.29 | 0.61 | 0.78 | 0.47 | 0.58 |
| 3 | 98252220 | rs1881672 | -1.926 | 0.029 | -1.249 | 0.454 | -0.01 | -0.093 | -0.093 | 0.045 | 0.69 | 0.62 | 0.78 | 0.95 | 0.7 | 0.7 |
| 3 | 98255392 | rs1350790 | -0.951 | 0.019 | -1.235 | 0.755 | 0.01 | 0.021 | 0.021 | 0.067 | 0.6 | 0.35 | 0.73 | 0.83 | 0.63 | 0.61 |
| 3 | 98261658 | rs9784358 | -0.36 | 0.249 | -1.118 | 1.074 | -0.018 | -0.009 | -0.009 | 0 | 0.22 | 0.17 | 0.25 | 0.26 | 0.3 | 0.23 |
| 3 | 98262890 | rs7430346 | -0.841 | 0.114 | -1.044 | 1.018 | 0.003 | 0.004 | 0.004 | 0 | 0.5 | 0.49 | 0.43 | 0.56 | 0.56 | 0.49 |
| 3 | 98267289 | rs9289628 | -0.676 | 0.296 | -0.937 | 0.828 | -0.001 | 0.008 | 0.008 | 0 | 0.52 | 0.43 | 0.58 | 0.59 | 0.49 | 0.51 |
| 3 | 98271961 | rs4857401 | 0.992 | 0.52 | -0.903 | 0.845 | 0.001 | 0.001 | 0.001 | 0 | 0.22 | 0.2 | 0.19 | 0.17 | 0.19 | 0.15 |
| 3 | 98272751 | rs13069468 | -0.151 | 0.451 | -1.052 | 0.85 | -0.045 | -0.05 | -0.05 | 0 | 0.11 | 0.02 | 0.17 | 0.2 | 0.15 | 0.16 |
| 3 | 98273979 | rs9827866 | -0.44 | -0.019 | -0.826 | 0.781 | -0.007 | -0.001 | -0.001 | 1.245 | 0.32 | 0.21 | 0.27 | 0.34 | 0.31 | 0.36 |
| 3 | 98277697 | rs16839940 | -0.349 | 0.4 | -0.494 | 0.983 | 0.014 | -0.001 | -0.001 | 0 | 0.41 | 0.32 | 0.45 | 0.4 | 0.33 | 0.46 |
| 3 | 98277721 | rs1350789 | -1.569 | -0.02 | -0.101 | 0.922 | 0.005 | 0.009 | 0.009 | 0 | 0.62 | 0.73 | 0.67 | 0.67 | 0.65 | 0.63 |
| 3 | 98281078 | rs6797035 | -0.679 | 0.177 | -0.412 | 0.904 | -0.004 | 0.004 | 0.004 | 0 | 0.48 | 0.4 | 0.52 | 0.46 | 0.49 | 0.52 |
| 3 | 98282957 | rs6778616 | -1.737 | 0.018 | -0.314 | 0.886 | 0.003 | 0.006 | 0.006 | 0 | 0.63 | 0.68 | 0.66 | 0.7 | 0.65 | 0.64 |
| 3 | 98283375 | rs1729964 | 0.843 | 0.368 | -0.589 | 0.929 | 0.023 | -0.004 | -0.004 | 0 | 0.48 | 0.6 | 0.42 | 0.46 | 0.41 | 0.45 |
| 3 | 98284330 | rs1729976 | -1.115 | 0.644 | -0.475 | 1.163 | -0.003 | -0.033 | -0.033 | 0.846 | 0.9 | 0.99 | 0.93 | 0.92 | 0.91 | 0.84 |
| 3 | 98284443 | rs1569234 | 0.948 | 0.609 | -0.422 | 1.142 | 0.03 | 0.033 | 0.033 | 0 | 0.52 | 0.65 | 0.38 | 0.37 | 0.42 | 0.41 |
| 3 | 98285043 | rs1675505 | 1.023 | 0.787 | -0.628 | 1.149 | 0.003 | 0.009 | 0.009 | 0 | 0.71 | 0.81 | 0.68 | 0.71 | 0.77 | 0.67 |
| 3 | 98286775 | rs4857097 | 0.216 | 0.89 | -0.797 | 1.215 | -0.023 | -0.009 | -0.009 | 0 | 0.76 | 0.92 | 0.7 | 0.75 | 0.78 | 0.71 |
| 3 | 98289131 | rs16839969 | 0.289 | 0.908 | -0.676 | 1.237 | 0.036 | 0.045 | 0.045 | 0 | 0.26 | 0.42 | 0.32 | 0.35 | 0.35 | 0.26 |
| 3 | 98290200 | rs9853063 | 0.649 | 0.931 | -0.721 | 1.093 | -0.025 | -0.003 | -0.003 | 0.494 | 0.13 | 0.04 | 0.23 | 0.2 | 0.14 | 0.16 |
| 3 | 98292683 | rs1675511 | 0.699 | 0.23 | -0.466 | 0.99 | 0.011 | 0.012 | 0.012 | 0 | 0.46 | 0.38 | 0.36 | 0.36 | 0.42 | 0.43 |
| 3 | 98297057 | rs1729963 | 0.401 | 0.517 | -0.621 | 0.669 | 0.008 | 0.013 | 0.013 | 0 | 0.5 | 0.45 | 0.39 | 0.4 | 0.47 | 0.46 |
| 3 | 98297937 | rs9283596 | -0.317 | 0.641 | -1.339 | 0.745 | -0.003 | 0.004 | 0.004 | 0.606 | 0.15 | 0.17 | 0.12 | 0.16 | 0.12 | 0.08 |
| 3 | 98298600 | rs7103 | 0.472 | 0.663 | -1.82 | 0.577 | 0.026 | 0.017 | 0.017 | 0 | 0.4 | 0.16 | 0.32 | 0.37 | 0.37 | 0.41 |
| 3 | 98303182 | rs3804622 | 0.412 | 0.916 | -1.816 | 0.565 | -0.02 | -0.029 | -0.029 | 0 | 0.54 | 0.18 | 0.49 | 0.5 | 0.54 | 0.54 |
| 3 | 98308915 | rs4558798 | 0.163 | 1.076 | -1.719 | 0.592 | -0.02 | 0.008 | 0.008 | 0 | 0.7 | 0.83 | 0.67 | 0.73 | 0.7 | 0.69 |
| 3 | 98317836 | rs1729987 | -0.859 | 1.095 | -1.957 | 0.575 | -0.015 | -0.051 | -0.051 | 0 | 0.88 | 0.99 | 0.92 | 0.88 | 0.87 | 0.91 |
| 3 | 98319101 | rs1675526 | 0.795 | 1.045 | -1.916 | 0.526 | -0.016 | 0.011 | 0.011 | 0 | 0.69 | 0.84 | 0.68 | 0.74 | 0.7 | 0.69 |
| 3 | 98323790 | rs1824548 | -0.229 | 1.62 | -2.101 | 0.204 | -0.05 | 0.041 | 0.041 | 0 | 0.16 | 0.03 | 0.24 | 0.1 | 0.18 | 0.18 |
| 3 | 98337517 | rs7630058 | -0.662 | 0.981 | -1.637 | 0.089 | 0 | -0.006 | -0.006 | 0 | 0.26 | 0.62 | 0.22 | 0.25 | 0.25 | 0.26 |
| 3 | 98342907 | rs1000003 | 0.096 | 0.701 | -1.113 | 0.116 | -0.028 | 0.02 | 0.02 | 0 | 0.19 | 0.53 | 0.25 | 0.24 | 0.17 | 0.2 |
| 3 | 98343242 | rs1531377 | -0.703 | 0.46 | -1.285 | -0.069 | 0.038 | 0.01 | 0.01 | 0.414 | 0.25 | 0.88 | 0.29 | 0.26 | 0.2 | 0.22 |
| 3 | 98346506 | rs7653204 | 0.017 | 0.439 | -0.986 | -0.075 | -0.025 | 0.009 | 0.009 | 0.029 | 0.19 | 0.62 | 0.26 | 0.24 | 0.17 | 0.2 |
| 3 | 98348504 | rs1841541 | -0.849 | 0.443 | -1.077 | -0.075 | 0.03 | 0.01 | 0.01 | 0.066 | 0.25 | 0.84 | 0.29 | 0.26 | 0.19 | 0.22 |
| 3 | 98367525 | rs6797163 | 0.274 | 0.911 | -1.405 | 0.195 | -0.11 | -0.099 | -0.099 | 0 | 0.72 | 0.97 | 0.74 | 0.71 | 0.68 | 0.71 |
| 3 | 98377357 | rs16840059 | -0.17 | 0.769 | -1.102 | 0.224 | 0.026 | -0.005 | -0.005 | 0 | 0.06 | 0.29 | 0.03 | 0.02 | 0.02 | 0.02 |
| 3 | 98385816 | rs9825798 | -0.734 | 0.645 | -0.545 | 0.347 | 0.036 | -0.006 | -0.006 | 0 | 0.25 | 0.88 | 0.28 | 0.28 | 0.2 | 0.22 |
| 3 | 98386666 | rs11918855 | -1.015 | 0.629 | -0.428 | 0.342 | -0.003 | -0.039 | -0.039 | 0 | 0.1 | 0.52 | 0.07 | 0.02 | 0.04 | 0.05 |
| 3 | 98391040 | rs4857412 | 0.632 | 0.735 | -0.402 | 0.441 | -0.007 | 0.058 | 0.058 | 0 | 0.15 | 0.35 | 0.21 | 0.26 | 0.16 | 0.17 |
| 3 | 98416900 | rs10935473 | 1.411 | 0.758 | -1.215 | 0.246 | -0.063 | -0.045 | -0.045 | 0.069 | 0.43 | 0.09 | 0.44 | 0.43 | 0.45 | 0.48 |
| 3 | 98432559 | rs4857414 | 1.428 | 0.144 | -1.446 | 0.088 | -0.076 | -0.093 | -0.093 | 0 | 0.39 | 0.13 | 0.43 | 0.52 | 0.46 | 0.45 |
| 3 | 98433611 | rs828588 | 0.062 | -0.167 | -0.802 | -0.044 | 0 | 0 | 0 | 0 | 0.19 | 0.18 | 0.12 | 0.12 | 0.17 | 0.15 |
| 3 | 98439637 | rs278364 | -0.108 | -0.015 | -0.424 | -0.132 | -0.008 | -0.02 | -0.02 | 0 | 0.21 | 0.26 | 0.11 | 0.07 | 0.19 | 0.15 |
| 3 | 98442701 | rs7628381 | -0.227 | -0.073 | -0.432 | 0.078 | -0.001 | -0.001 | -0.001 | 0 | 0.35 | 0.38 | 0.47 | 0.38 | 0.26 | 0.4 |
| 3 | 98443648 | rs844159 | -1.664 | -0.098 | -0.392 | 0.067 | -0.007 | -0.011 | -0.011 | 0.225 | 0.57 | 0.69 | 0.6 | 0.45 | 0.45 | 0.55 |
| 3 | 98445324 | rs865474 | -1.647 | -0.09 | -0.383 | 0.071 | -0.006 | -0.01 | -0.01 | 0 | 0.56 | 0.69 | 0.59 | 0.45 | 0.45 | 0.55 |
| 3 | 98449742 | rs3755569 | -0.662 | -0.06 | -0.372 | 0.073 | -0.003 | 0 | 0 | 0 | 0.35 | 0.38 | 0.48 | 0.39 | 0.26 | 0.4 |
| 3 | 98489011 | rs3755573 | -0.846 | 0.284 | -0.241 | 0.021 | -0.003 | 0.004 | 0.004 | 0 | 0.44 | 0.63 | 0.56 | 0.41 | 0.4 | 0.51 |
| 3 | 98489410 | rs3772097 | -0.148 | 0.148 | 0.21 | 0.044 | -0.012 | 0.016 | 0.016 | 0.187 | 0.26 | 0.32 | 0.33 | 0.36 | 0.15 | 0.3 |
| 3 | 98493644 | rs7431448 | -0.77 | 0.383 | 0.154 | -0.624 | -0.004 | 0.019 | 0.019 | 0 | 0.4 | 0.38 | 0.34 | 0.2 | 0.4 | 0.33 |
| 3 | 98509587 | rs278390 | 1.713 | 0.227 | -0.016 | -0.801 | -0.031 | 0.035 | 0.035 | 0 | 0.36 | 0.08 | 0.51 | 0.7 | 0.38 | 0.43 |
| 3 | 98515161 | rs704586 | 1.668 | 0.202 | 0.235 | -0.812 | -0.032 | 0.026 | 0.026 | 0 | 0.37 | 0.08 | 0.51 | 0.7 | 0.38 | 0.43 |
| 3 | 98515569 | rs8833 | 1.709 | 0.202 | 0.235 | -0.812 | -0.03 | 0.024 | 0.024 | 0.003 | 0.36 | 0.08 | 0.51 | 0.7 | 0.38 | 0.43 |
| 3 | 98517013 | rs1062196 | -0.664 | 0.188 | 0.385 | -0.847 | 0 | -0.01 | -0.01 | 0 | 0.42 | 0.46 | 0.37 | 0.23 | 0.41 | 0.33 |
| 3 | 98518072 | rs17270986 | 1.393 | 0.132 | 0.267 | -0.982 | -0.012 | -0.037 | -0.037 | 3.99 | 0.19 | 0.01 | 0.2 | 0.19 | 0.12 | 0.18 |
| 3 | 98519481 | rs1822017 | 0.832 | 0.542 | 0.802 | -0.516 | 0.004 | -0.056 | -0.056 | 0.866 | 0.17 | 0.25 | 0.1 | 0.05 | 0.16 | 0.19 |
| 3 | 98523960 | rs1866677 | 0.258 | 0.529 | 0.977 | -0.524 | 0.002 | 0.03 | 0.03 | 0 | 0.38 | 0.3 | 0.32 | 0.23 | 0.34 | 0.32 |

| | | | | | | | | | | | | | | | | |
|---|---|---|---|---|---|---|---|---|---|---|---|---|---|---|---|---|
| 3 | 98535909 | rs278387 | 1.372 | 0.552 | 0.781 | -0.4 | -0.034 | -0.005 | -0.005 | 2.354 | 0.36 | 0.08 | 0.51 | 0.68 | 0.38 | 0.42 |
| 3 | 98537341 | rs11916285 | 0.846 | 0.553 | 0.799 | -0.402 | 0.004 | -0.056 | -0.056 | 0 | 0.17 | 0.25 | 0.09 | 0.05 | 0.17 | 0.12 |
| 3 | 98541116 | rs828616 | 1.359 | 0.545 | 0.724 | -0.382 | -0.034 | -0.005 | -0.005 | 0.452 | 0.36 | 0.08 | 0.51 | 0.68 | 0.38 | 0.42 |
| 3 | 98541633 | rs828618 | 2.122 | 0.58 | 0.599 | -0.068 | -0.086 | -0.122 | -0.122 | 10.663 | 0.29 | 0.01 | 0.3 | 0.42 | 0.32 | 0.31 |
| 3 | 98587977 | rs7615856 | 1.144 | 0.891 | 1.06 | -1.091 | 0.001 | 0.04 | 0.04 | 0.072 | 0.4 | 0.18 | 0.53 | 0.71 | 0.42 | 0.47 |
| 3 | 98613773 | rs2439222 | 0.157 | 0.86 | 1.072 | -1.105 | -0.006 | 0.066 | 0.066 | 0 | 0.37 | 0.22 | 0.32 | 0.23 | 0.35 | 0.33 |
| 3 | 98615611 | rs10935518 | 0.686 | 0.9 | 0.977 | -1.08 | 0.003 | -0.043 | -0.043 | 0 | 0.17 | 0.22 | 0.09 | 0.05 | 0.17 | 0.19 |
| 3 | 98634704 | rs17278278 | 0.741 | 0.822 | 0.839 | -1.063 | -0.025 | 0.059 | 0.059 | 0 | 0.18 | 0.07 | 0.1 | 0.04 | 0.18 | 0.21 |
| 3 | 98637768 | rs4577404 | 1.502 | 0.812 | 0.776 | -1.073 | -0.011 | 0.006 | 0.006 | 0 | 0.36 | 0.11 | 0.52 | 0.68 | 0.37 | 0.42 |
| 3 | 98668768 | rs11489075 | -0.202 | 0.852 | 0.93 | -1.04 | 0.016 | 0.002 | 0.002 | 0 | 0.06 | 0.12 | 0.06 | 0.05 | 0.11 | 0.07 |
| 3 | 98683264 | rs996966 | -0.466 | 0.859 | 0.693 | -1.047 | 0.013 | -0.051 | -0.051 | 0 | 0.25 | 0.34 | 0.17 | 0.07 | 0.29 | 0.27 |
| 3 | 98697389 | rs2062059 | -0.204 | 0.747 | 0.482 | -0.992 | 0.008 | 0.031 | 0.031 | 0 | 0.41 | 0.31 | 0.33 | 0.25 | 0.38 | 0.36 |
| 3 | 98722598 | rs2449027 | 0.014 | 0.763 | 0.405 | -0.982 | 0.009 | 0.025 | 0.025 | 0 | 0.59 | 0.66 | 0.67 | 0.74 | 0.63 | 0.64 |
| 3 | 98730871 | rs2250022 | 0.346 | 0.807 | 0.432 | -0.991 | 0.003 | 0.025 | 0.025 | 0 | 0.56 | 0.59 | 0.64 | 0.73 | 0.57 | 0.64 |
| 3 | 98732578 | rs2462221 | 0.016 | 0.791 | 0.345 | -0.979 | 0.009 | 0.025 | 0.025 | 0 | 0.59 | 0.66 | 0.67 | 0.74 | 0.63 | 0.64 |
| 3 | 98734914 | rs2462227 | 0.07 | 0.671 | 0.469 | -0.878 | 0.008 | 0.017 | 0.017 | 0 | 0.6 | 0.68 | 0.7 | 0.73 | 0.65 | 0.64 |
| 3 | 98740816 | rs9849048 | 0.208 | 0.655 | 0.187 | -0.888 | 0.007 | 0.056 | 0.056 | 0 | 0.35 | 0.18 | 0.29 | 0.23 | 0.34 | 0.31 |
| 3 | 98766326 | rs11705895 | 0.648 | 0.942 | -0.098 | -0.886 | -0.019 | 0.058 | 0.058 | 0.134 | 0.17 | 0.07 | 0.1 | 0.04 | 0.16 | 0.21 |
| 3 | 98768784 | rs1403436 | -1.485 | 0.845 | -0.193 | -0.748 | -0.011 | 0.006 | 0.006 | 0 | 0.64 | 0.82 | 0.47 | 0.33 | 0.62 | 0.62 |
| 3 | 98769721 | rs2462215 | -0.644 | 0.088 | 0.093 | -0.178 | -0.005 | -0.01 | -0.01 | 0 | 0.4 | 0.39 | 0.31 | 0.27 | 0.35 | 0.36 |
| 3 | 98786601 | rs1038273 | -0.621 | -0.074 | -0.592 | -0.336 | 0.006 | 0.006 | 0.006 | 0 | 0.41 | 0.36 | 0.33 | 0.18 | 0.34 | 0.35 |
| 3 | 98790767 | rs1404100 | -0.375 | -0.315 | -0.518 | -0.228 | -0.018 | -0.015 | -0.015 | 0 | 0.32 | 0.33 | 0.27 | 0.19 | 0.2 | 0.29 |
| 3 | 98791847 | rs13082809 | 0.765 | 0.606 | -1.123 | -0.32 | -0.079 | 0.095 | 0.095 | 0.023 | 0.3 | 0.07 | 0.23 | 0.13 | 0.37 | 0.11 |
| 3 | 98793597 | rs7634836 | -0.841 | 0.341 | -0.864 | 0.381 | -0.006 | -0.024 | -0.024 | 0 | 0.33 | 0.87 | 0.35 | 0.32 | 0.32 | 0.42 |
| 3 | 98796057 | rs16840545 | 0.317 | 0.438 | -0.986 | 0.549 | -0.027 | -0.037 | -0.037 | 0.548 | 0.06 | 0.01 | 0.06 | 0.2 | 0.09 | 0.08 |
| 3 | 98798173 | rs2177275 | -0.76 | 0.065 | 0.534 | 0.139 | 0.024 | 0.029 | 0.029 | 1.278 | 0.24 | 0.51 | 0.29 | 0.25 | 0.27 | 0.27 |
| 3 | 98799599 | rs4928194 | -0.044 | 0.02 | 1.008 | 0.174 | -0.003 | 0.065 | 0.065 | 0 | 0.06 | 0.19 | 0.12 | 0.11 | 0.04 | 0.12 |
| 3 | 98804408 | rs12493380 | -0.107 | 0.012 | 1.32 | 0.159 | 0.028 | -0.03 | -0.03 | 2.854 | 0.18 | 0.32 | 0.17 | 0.13 | 0.22 | 0.15 |
| 3 | 98805315 | rs2140280 | -1.033 | 0.048 | 1.382 | 0.161 | 0.011 | 0.027 | 0.027 | 1.129 | 0.24 | 0.47 | 0.27 | 0.25 | 0.24 | 0.26 |
| 3 | 98806212 | rs13324173 | -0.074 | 0.004 | 1.346 | 0.172 | -0.005 | 0.041 | 0.041 | 0 | 0.06 | 0.16 | 0.11 | 0.11 | 0.04 | 0.12 |
| 3 | 98807198 | rs4928156 | -0.074 | 0.004 | 1.346 | 0.172 | -0.005 | 0.041 | 0.041 | 0 | 0.06 | 0.15 | 0.11 | 0.11 | 0.04 | 0.12 |
| 3 | 98810191 | rs9833346 | -1.012 | 0.011 | 1.284 | 0.16 | 0.024 | 0.027 | 0.027 | 1.2 | 0.24 | 0.51 | 0.29 | 0.25 | 0.27 | 0.26 |
| 3 | 98814043 | rs1404092 | -1.528 | 0.028 | 1.221 | 0.147 | -0.001 | -0.017 | -0.017 | 0 | 0.32 | 0.83 | 0.34 | 0.33 | 0.32 | 0.43 |
| 3 | 98835121 | rs4928196 | -1.513 | -0.172 | 1.677 | 0.12 | -0.034 | 0.096 | 0.096 | 0.07 | 0.16 | 0.54 | 0.17 | 0.19 | 0.11 | 0.28 |
| 3 | 98843989 | rs7620833 | -1.261 | -0.178 | 1.826 | 0.13 | -0.032 | 0.064 | 0.064 | 0.174 | 0.14 | 0.39 | 0.16 | 0.13 | 0.09 | 0.25 |
| 3 | 98850962 | rs704592 | 1.212 | -0.149 | 1.908 | 0.078 | -0.001 | 0.007 | 0.007 | 0.192 | 0.71 | 0.51 | 0.69 | 0.73 | 0.7 | 0.61 |
| 3 | 98862025 | rs2687686 | 1.599 | -0.163 | 1.547 | 0.07 | 0.006 | -0.013 | -0.013 | 0.076 | 0.67 | 0.15 | 0.65 | 0.67 | 0.68 | 0.57 |
| 3 | 98862083 | rs774950 | 1.279 | -0.169 | 1.732 | 0.07 | -0.001 | 0.006 | 0.006 | 0.292 | 0.71 | 0.41 | 0.68 | 0.73 | 0.69 | 0.61 |
| 3 | 98862644 | rs774947 | 1.316 | -0.196 | 1.485 | 0.09 | -0.015 | 0.013 | 0.013 | 0.247 | 0.83 | 0.25 | 0.81 | 0.81 | 0.89 | 0.72 |
| 3 | 98869960 | rs774731 | 1.46 | -0.056 | 1.419 | -0.636 | -0.009 | -0.014 | -0.014 | 0.392 | 0.83 | 0.22 | 0.9 | 0.92 | 0.89 | 0.74 |
| 3 | 98872593 | rs774736 | 1.512 | 0.008 | 1.364 | -0.64 | 0 | 0.047 | 0.047 | 0.47 | 0.67 | 0.15 | 0.74 | 0.79 | 0.68 | 0.59 |
| 3 | 98894246 | rs2687670 | 0.962 | 0.153 | 1.801 | -0.559 | 0.074 | 0.027 | 0.027 | 0.482 | 0.91 | 0.52 | 0.94 | 0.94 | 0.94 | 0.88 |
| 3 | 98895124 | rs1851066 | 1.088 | 0.127 | 1.983 | -0.56 | 0.043 | 0.021 | 0.021 | 0.082 | 0.91 | 0.52 | 0.94 | 0.94 | 0.94 | 0.88 |
| 3 | 98895650 | rs2682410 | -0.183 | 0.147 | 1.722 | -0.535 | 0.008 | -0.008 | -0.008 | 0.051 | 0.84 | 0.75 | 0.84 | 0.85 | 0.78 | 0.84 |
| 3 | 98899033 | rs4928154 | -0.308 | 0.187 | 1.739 | -0.529 | 0.01 | -0.007 | -0.007 | 3.318 | 0.84 | 0.75 | 0.84 | 0.85 | 0.78 | 0.84 |
| 3 | 98902911 | rs774972 | -0.308 | 0.187 | 1.739 | -0.529 | 0.01 | -0.007 | -0.007 | 0.192 | 0.84 | 0.75 | 0.84 | 0.85 | 0.78 | 0.84 |
| 3 | 98919134 | rs774951 | 0.792 | 0.121 | 1.823 | -0.576 | 0.015 | -0.052 | -0.052 | 0 | 0.77 | 0.52 | 0.8 | 0.84 | 0.74 | 0.76 |
| 3 | 98930445 | rs9681261 | 1.204 | 0.012 | 1.39 | -0.604 | -0.005 | -0.019 | -0.019 | 0 | 0.88 | 0.3 | 0.91 | 0.92 | 0.91 | 0.85 |
| 3 | 98943479 | rs13076500 | 1.33 | -0.643 | 0.49 | -1.051 | -0.058 | 0.084 | 0.084 | 0 | 0.42 | 0.1 | 0.51 | 0.71 | 0.51 | 0.43 |
| 3 | 98957110 | rs12490980 | 0.652 | -0.623 | 0.51 | -1.044 | 0.061 | 0.137 | 0.137 | 0 | 0.27 | 0.03 | 0.22 | 0.05 | 0.15 | 0.13 |
| 3 | 98960340 | rs2139700 | 0.87 | 0.183 | 1.232 | -0.561 | -0.003 | 0.004 | 0.004 | 1.047 | 0.75 | 0.25 | 0.77 | 0.77 | 0.7 | 0.68 |
| 3 | 98966922 | rs4928175 | 1.037 | 0.196 | 1.005 | -0.549 | -0.012 | 0.023 | 0.023 | 1.695 | 0.73 | 0.15 | 0.77 | 0.77 | 0.69 | 0.68 |
| 3 | 98970607 | rs7624383 | 0.646 | 0.348 | 1.034 | -0.273 | 0.076 | 0.033 | 0.033 | 0.348 | 0.92 | 0.61 | 0.94 | 0.94 | 0.94 | 0.88 |
| 3 | 98972835 | rs4928176 | 0.494 | -0.001 | -0.51 | -0.504 | -0.042 | 0.095 | 0.095 | 0.01 | 0.4 | 0.14 | 0.55 | 0.73 | 0.57 | 0.38 |
| 3 | 98987607 | rs6800453 | 0.154 | -0.424 | 0.65 | -0.743 | 0.001 | -0.002 | -0.002 | 0 | 0.48 | 0.56 | 0.59 | 0.75 | 0.6 | 0.49 |
| 3 | 98988161 | rs1402268 | -0.02 | -0.405 | 0.645 | -0.825 | 0.064 | 0.146 | 0.146 | 0.771 | 0.52 | 0.75 | 0.76 | 0.75 | 0.64 | 0.53 |
| 3 | 99003734 | rs12494255 | -0.454 | -0.873 | 0.244 | -1.226 | 0.012 | 0.023 | 0.023 | 0 | 0.74 | 0.8 | 0.84 | 0.82 | 0.78 | 0.73 |
| 3 | 99006939 | rs975276 | 0.782 | -1.098 | 0.759 | -1.353 | 0.006 | 0.002 | 0.002 | 0.379 | 0.51 | 0.46 | 0.74 | 0.75 | 0.65 | 0.48 |
| 3 | 99007270 | rs4539971 | 0.03 | -1.038 | 0.775 | -1.281 | -0.024 | -0.064 | -0.064 | 0.036 | 0.24 | 0.33 | 0.1 | 0.06 | 0.12 | 0.19 |
| 3 | 99015616 | rs6809700 | 0.833 | -0.868 | 0.997 | -1.117 | 0.03 | 0.139 | 0.139 | 1.866 | 0.95 | 0.69 | 0.92 | 0.85 | 0.98 | 0.82 |
| 3 | 99034331 | rs16840977 | 0.338 | -0.804 | 0.812 | -1.115 | 0.007 | 0.006 | 0.006 | 0 | 0.25 | 0.21 | 0.11 | 0.06 | 0.21 | 0.21 |
| 3 | 99037859 | rs9818711 | -0.253 | -0.812 | 0.421 | -1.117 | 0.002 | -0.004 | -0.004 | 0 | 0.74 | 0.75 | 0.89 | 0.89 | 0.78 | 0.67 |
| 3 | 99038331 | rs9871288 | 1.615 | 0.002 | -0.333 | -0.687 | -0.031 | -0.004 | -0.004 | 0 | 0.26 | 0.15 | 0.21 | 0.3 | 0.35 | 0.2 |
| 3 | 99038644 | rs1383843 | 0.392 | -0.214 | -0.258 | -0.576 | 0.008 | 0.007 | 0.007 | 0 | 0.25 | 0.2 | 0.1 | 0.05 | 0.21 | 0.22 |
| 3 | 99043024 | rs7615723 | 0.275 | -0.253 | -0.383 | -0.634 | 0.022 | 0.064 | 0.064 | 0 | 0.25 | 0.12 | 0.1 | 0.03 | 0.21 | 0.16 |
| 3 | 99045218 | rs985633 | -0.483 | 0.024 | 0.13 | -0.104 | 0.005 | 0.009 | 0.009 | 0 | 0.27 | 0.69 | 0.35 | 0.21 | 0.18 | 0.33 |
| 3 | 99056448 | rs9835559 | 1.647 | 0.153 | 1.059 | 0.922 | -0.012 | 0.123 | 0.123 | 0 | 0.86 | 0.71 | 0.86 | 0.56 | 0.9 | 0.66 |
| 3 | 99056994 | rs6798863 | -1.058 | 1.289 | 1.372 | 0.555 | 0.132 | -0.048 | -0.048 | 0 | 0.1 | 0.19 | 0.25 | 0.02 | 0.21 | 0.11 |
| 3 | 99058195 | rs6762194 | -0.983 | 1.283 | 1.378 | 0.552 | 0.062 | -0.034 | -0.034 | 0 | 0.09 | 0.1 | 0.23 | 0.02 | 0.21 | 0.11 |

| | | | | | | | | | | | | | | | | |
|---|---|---|---|---|---|---|---|---|---|---|---|---|---|---|---|---|
| 3 | 99060359 | rs16841061 | -2.042 | 0.381 | 1.738 | 0.713 | -0.02 | 0.202 | 0.202 | 0 | 0.13 | 0.51 | 0.09 | 0.31 | 0.07 | 0.14 |
| 3 | 99060454 | rs7652907 | -2.562 | 0.312 | 1.561 | 1.043 | -0.04 | 0.115 | 0.115 | 0.096 | 0.16 | 0.64 | 0.13 | 0.44 | 0.09 | 0.35 |
| 3 | 99060484 | rs4133236 | -2.748 | 0.321 | 1.622 | 0.796 | -0.019 | 0.189 | 0.189 | 0.987 | 0.09 | 0.62 | 0.09 | 0.28 | 0.03 | 0.24 |
| 3 | 99071647 | rs2470691 | -2.911 | 0.226 | 1.495 | 0.794 | -0.039 | 0.129 | 0.129 | 6.029 | 0.1 | 0.68 | 0.09 | 0.26 | 0.02 | 0.09 |
| 3 | 99084862 | rs2929443 | -2.811 | 0.265 | 2.076 | 0.843 | -0.013 | 0.224 | 0.224 | 2.836 | 0.08 | 0.38 | 0.07 | 0.28 | 0.02 | 0.14 |
| 3 | 99085806 | rs2700667 | -2.69 | 0.527 | 1.909 | 0.674 | 0.032 | 0.022 | 0.022 | 1.664 | 0.21 | 0.62 | 0.32 | 0.33 | 0.24 | 0.38 |
| 3 | 99093509 | rs1383847 | -2.608 | 0.643 | 1.973 | 0.667 | 0.067 | 0.047 | 0.047 | 1.288 | 0.2 | 0.52 | 0.31 | 0.3 | 0.24 | 0.26 |
| 3 | 99095112 | rs12497126 | -0.759 | 0.578 | 1.432 | 0.644 | 0.003 | 0.009 | 0.009 | 0 | 0.08 | 0.03 | 0.07 | 0.05 | 0.05 | 0.1 |
| 3 | 99097665 | rs11918189 | -1.271 | 0.685 | 1.579 | 0.694 | 0.07 | 0.175 | 0.175 | 0 | 0.05 | 0.25 | 0.05 | 0.19 | 0.04 | 0.18 |
| 3 | 99105577 | rs13325971 | -1.145 | 0.674 | 1.64 | 0.672 | 0.036 | -0.04 | -0.04 | 0 | 0.11 | 0.16 | 0.22 | 0.02 | 0.19 | 0.11 |
| 3 | 99107683 | rs2700648 | -3 | 0.664 | 1.662 | 0.684 | 0.093 | 0.074 | 0.074 | 0.198 | 0.19 | 0.45 | 0.3 | 0.33 | 0.24 | 0.38 |
| 3 | 99113755 | rs1481542 | -1.855 | 0.663 | 1.566 | 0.657 | 0.033 | -0.013 | -0.013 | 0 | 0.13 | 0.2 | 0.25 | 0.12 | 0.2 | 0.15 |
| 3 | 99115940 | rs7631745 | -1.143 | 0.66 | 1.578 | 0.655 | 0.017 | -0.021 | -0.021 | 0 | 0.1 | 0.12 | 0.22 | 0.02 | 0.19 | 0.11 |
| 3 | 99116207 | rs1383852 | -2.119 | 0.656 | 1.806 | 0.655 | 0.083 | -0.014 | -0.014 | 0.31 | 0.16 | 0.41 | 0.27 | 0.16 | 0.2 | 0.33 |
| 3 | 99126094 | rs969987 | -1.87 | 0.664 | 1.725 | 0.658 | 0.062 | -0.013 | -0.013 | 0 | 0.15 | 0.31 | 0.26 | 0.14 | 0.2 | 0.2 |
| 3 | 99131534 | rs2623324 | -1.685 | 0.667 | 1.692 | 0.655 | 0.042 | -0.019 | -0.019 | 0 | 0.14 | 0.24 | 0.25 | 0.12 | 0.2 | 0.15 |
| 3 | 99136488 | rs2700605 | -1.856 | -0.04 | 0.632 | 0.758 | -0.142 | -0.082 | -0.082 | 0 | 0.56 | 0.98 | 0.58 | 0.58 | 0.5 | 0.77 |
| 3 | 99137028 | rs2448969 | -1.577 | 0.524 | 1.617 | 0.756 | 0.042 | -0.019 | -0.019 | 0 | 0.14 | 0.25 | 0.25 | 0.12 | 0.2 | 0.15 |
| 3 | 99137350 | rs2623348 | -1.577 | 0.524 | 1.617 | 0.756 | 0.042 | -0.019 | -0.019 | 0 | 0.14 | 0.25 | 0.25 | 0.12 | 0.2 | 0.15 |
| 3 | 99138140 | rs980842 | -1.577 | 0.524 | 1.617 | 0.76 | 0.042 | -0.017 | -0.017 | 0 | 0.14 | 0.25 | 0.25 | 0.12 | 0.2 | 0.15 |
| 3 | 99144245 | rs1997405 | -2.401 | 0.509 | 1.596 | 0.784 | 0.059 | 0.047 | 0.047 | 0.402 | 0.2 | 0.55 | 0.31 | 0.33 | 0.24 | 0.38 |
| 3 | 99147683 | rs9833999 | -2.617 | 0.468 | 1.295 | 0.534 | 0.027 | -0.039 | -0.039 | 0.413 | 0.24 | 0.42 | 0.35 | 0.22 | 0.25 | 0.42 |
| 3 | 99148533 | rs4536866 | -1.244 | 0.323 | 0.707 | 0.621 | -0.009 | -0.008 | -0.008 | 0 | 0.09 | 0.12 | 0.08 | 0.08 | 0.05 | 0.22 |
| 3 | 99158618 | rs2700650 | -2.069 | 0.383 | 1.031 | 0.564 | -0.001 | 0.001 | 0.001 | 0.355 | 0.33 | 0.67 | 0.42 | 0.36 | 0.32 | 0.44 |
| 3 | 99161022 | rs1997404 | -1.617 | 0.396 | 1.594 | 0.553 | 0.006 | -0.046 | -0.046 | 0 | 0.27 | 0.35 | 0.24 | 0.13 | 0.28 | 0.31 |
| 3 | 99162536 | rs2670338 | -2.387 | 0.404 | 1.563 | 0.555 | -0.005 | 0.006 | 0.006 | 0.355 | 0.33 | 0.68 | 0.43 | 0.36 | 0.33 | 0.44 |
| 3 | 99165693 | rs6807474 | -1.836 | 0.426 | 1.631 | 0.561 | 0.017 | -0.083 | -0.083 | 0 | 0.28 | 0.49 | 0.38 | 0.19 | 0.29 | 0.4 |
| 3 | 99178757 | rs11708005 | -1.269 | 0.87 | 1.442 | 0.646 | 0.027 | -0.044 | -0.044 | 0 | 0.13 | 0.21 | 0.13 | 0.03 | 0.22 | 0.08 |
| 3 | 99184544 | rs2700628 | -2.108 | 0.308 | 1.195 | 0.986 | 0.009 | 0.096 | 0.096 | 0 | 0.11 | 0.22 | 0.26 | 0.32 | 0.08 | 0.2 |
| 3 | 99189574 | rs9289822 | -2.333 | 0.293 | 1.15 | 1.008 | 0.003 | 0.055 | 0.055 | 0 | 0.13 | 0.2 | 0.28 | 0.32 | 0.12 | 0.22 |
| 3 | 99190963 | rs1355762 | -0.979 | 0.2 | 1.878 | 0.741 | 0.06 | -0.002 | -0.002 | 0 | 0.07 | 0.33 | 0.04 | 0.01 | 0.06 | 0.03 |
| 3 | 99199528 | rs6805837 | -0.478 | 0.197 | 2.13 | 0.748 | 0.02 | -0.026 | -0.026 | 0.144 | 0.06 | 0.17 | 0.03 | 0 | 0.05 | 0.03 |
| 3 | 99205316 | rs2670320 | -1.675 | -0.334 | 1.241 | 0.721 | -0.092 | -0.112 | -0.112 | 0 | 0.57 | 0.98 | 0.49 | 0.53 | 0.42 | 0.67 |
| 3 | 99207253 | rs6798177 | -0.271 | 0.186 | 2.006 | 1.163 | 0.02 | -0.026 | -0.026 | 0 | 0.06 | 0.17 | 0.03 | 0 | 0.05 | 0.03 |
| 3 | 99219235 | rs13101237 | -1.589 | 0.189 | 2.163 | 1.26 | 0.032 | 0.055 | 0.055 | 0 | 0.09 | 0.23 | 0.06 | 0.17 | 0.1 | 0.08 |
| 3 | 99221871 | rs2670339 | -3.058 | -0.124 | 2.089 | 1.119 | 0.013 | -0.028 | -0.028 | 0.207 | 0.35 | 0.86 | 0.38 | 0.46 | 0.27 | 0.35 |
| 3 | 99235720 | rs17775122 | -1.112 | -0.113 | 2.427 | 1.053 | 0.015 | -0.019 | -0.019 | 0.092 | 0.09 | 0.2 | 0.17 | 0.06 | 0.09 | 0.15 |
| 3 | 99235739 | rs4605585 | -2.816 | -0.147 | 2.494 | 1.046 | -0.012 | -0.015 | -0.015 | 0 | 0.3 | 0.62 | 0.33 | 0.28 | 0.22 | 0.31 |
| 3 | 99240824 | rs12487789 | 2.077 | -0.114 | 2.523 | 1.075 | 0.005 | -0.026 | -0.026 | 0 | 0.83 | 0.39 | 0.77 | 0.84 | 0.86 | 0.82 |
| 3 | 99242097 | rs12490079 | -2.462 | -0.114 | 2.616 | 1.137 | -0.005 | 0.024 | 0.024 | 0 | 0.26 | 0.38 | 0.31 | 0.39 | 0.21 | 0.32 |
| 3 | 99248498 | rs2063638 | -0.617 | -0.051 | 2.524 | 1.138 | 0.007 | 0.003 | 0.003 | 0 | 0.13 | 0.07 | 0.1 | 0.12 | 0.08 | 0.13 |
| 3 | 99254625 | rs10935836 | -0.617 | -0.11 | 2.504 | 1.146 | 0.008 | 0.003 | 0.003 | 0 | 0.13 | 0.07 | 0.1 | 0.12 | 0.08 | 0.13 |
| 3 | 99256035 | rs16841326 | 3.303 | -0.108 | 2.331 | 1.146 | 0.011 | -0.023 | -0.023 | 0 | 0.66 | 0.24 | 0.63 | 0.51 | 0.73 | 0.65 |
| 3 | 99262240 | rs6804164 | 2.443 | -0.097 | 2.365 | 1.019 | -0.031 | -0.078 | -0.078 | 0 | 0.72 | 0.52 | 0.82 | 0.87 | 0.81 | 0.76 |
| 3 | 99262827 | rs6440679 | 0.81 | -0.208 | 2.429 | 1.072 | -0.028 | -0.011 | -0.011 | 0.005 | 0.86 | 0.78 | 0.89 | 0.88 | 0.92 | 0.87 |
| 3 | 99264583 | rs7646873 | 1.484 | -0.209 | 2.057 | 0.856 | -0.042 | -0.062 | -0.062 | 0.946 | 0.79 | 0.59 | 0.86 | 0.88 | 0.87 | 0.84 |
| 3 | 99269750 | rs1513304 | -2.081 | 0.166 | 1.431 | 1.388 | 0.008 | 0.102 | 0.102 | 0.189 | 0.25 | 0.45 | 0.39 | 0.5 | 0.25 | 0.29 |
| 3 | 99271993 | rs2089983 | 2.408 | 0.251 | 1.229 | 0.718 | -0.016 | -0.008 | -0.008 | 0 | 0.65 | 0.17 | 0.61 | 0.58 | 0.67 | 0.61 |
| 3 | 99274206 | rs3914296 | -0.855 | 0.206 | 1.494 | 0.712 | -0.007 | 0.003 | 0.003 | 0 | 0.12 | 0.15 | 0.11 | 0.14 | 0.08 | 0.16 |
| 3 | 99281234 | rs2175746 | -0.861 | 0.206 | 1.493 | 0.712 | -0.007 | 0.002 | 0.002 | 0 | 0.12 | 0.15 | 0.11 | 0.14 | 0.08 | 0.16 |
| 3 | 99281436 | rs4928138 | 1.421 | 0.227 | 1.877 | 0.698 | 0.062 | 0.089 | 0.089 | 0.407 | 0.87 | 0.6 | 0.82 | 0.74 | 0.84 | 0.86 |
| 3 | 99284056 | rs4367103 | 2.345 | 0.218 | 1.948 | 0.693 | -0.001 | 0.001 | 0.001 | 0 | 0.74 | 0.3 | 0.69 | 0.59 | 0.75 | 0.7 |
| 3 | 99294991 | rs3914294 | 2.871 | 0.192 | 1.832 | 0.743 | -0.046 | -0.092 | -0.092 | 2.202 | 0.64 | 0.09 | 0.59 | 0.51 | 0.67 | 0.61 |
| 3 | 99295844 | rs7647722 | -1.344 | 0.163 | 1.973 | 0.744 | -0.026 | 0.014 | 0.014 | 0 | 0.13 | 0.3 | 0.13 | 0.14 | 0.08 | 0.16 |
| 3 | 99317673 | rs7633643 | 0.919 | 0.214 | 1.843 | 0.753 | 0.054 | 0.184 | 0.184 | 0.21 | 0.94 | 0.68 | 0.96 | 0.82 | 0.95 | 0.97 |
| 3 | 99326501 | rs10511176 | -0.54 | 0.125 | 1.834 | 0.724 | -0.007 | 0.008 | 0.008 | 0 | 0.08 | 0.11 | 0.1 | 0.13 | 0.06 | 0.12 |
| 3 | 99328145 | rs1849496 | -0.234 | 0.102 | 1.86 | 0.714 | 0.007 | 0.033 | 0.033 | 0.388 | 0.07 | 0.15 | 0.13 | 0.14 | 0.07 | 0.09 |
| 3 | 99328500 | rs6440719 | -1.599 | 0.08 | 1.866 | 0.677 | 0.021 | -0.017 | -0.017 | 0 | 0.16 | 0.36 | 0.22 | 0.15 | 0.15 | 0.18 |
| 3 | 99330847 | rs9872048 | 2.093 | 0.079 | 1.869 | 0.679 | -0.001 | 0.009 | 0.009 | 0.388 | 0.78 | 0.32 | 0.74 | 0.67 | 0.8 | 0.79 |
| 3 | 99331072 | rs12486836 | -0.402 | 0.085 | 1.849 | 0.671 | 0.003 | -0.025 | -0.025 | 0.162 | 0.09 | 0.11 | 0.08 | 0.01 | 0.08 | 0.09 |
| 3 | 99331973 | rs6792073 | 0.846 | 0.094 | 1.696 | 0.681 | 0.054 | 0.184 | 0.184 | 0.21 | 0.94 | 0.68 | 0.96 | 0.82 | 0.95 | 0.97 |
| 3 | 99341249 | rs9882045 | -2.228 | 0.081 | 1.672 | 0.696 | -0.009 | -0.008 | -0.008 | 0.431 | 0.25 | 0.55 | 0.32 | 0.27 | 0.2 | 0.31 |
| 3 | 99345689 | rs561391 | 1.136 | 0.041 | 1.467 | 0.673 | 0.05 | 0.184 | 0.184 | 0.414 | 0.93 | 0.6 | 0.96 | 0.82 | 0.96 | 0.97 |
| 3 | 99347057 | rs1618515 | 2.406 | 0.054 | 0.684 | 0.652 | -0.006 | -0.114 | -0.114 | 0 | 0.67 | 0.05 | 0.62 | 0.53 | 0.75 | 0.67 |
| 3 | 99358539 | rs2280505 | -1.74 | 0.536 | 0.858 | 0.772 | 0.032 | 0.046 | 0.046 | 0 | 0.22 | 0.47 | 0.26 | 0.34 | 0.24 | 0.26 |
| 3 | 99363985 | rs7645305 | -0.318 | 0.563 | -0.171 | 0.1 | -0.003 | 0.027 | 0.027 | 0 | 0.22 | 0.11 | 0.21 | 0.13 | 0.2 | 0.18 |
| 3 | 99364270 | rs16841568 | -0.912 | 0.598 | -0.002 | 0.56 | 0.023 | 0.016 | 0.016 | 0 | 0.08 | 0.15 | 0.14 | 0.1 | 0.12 | 0.14 |
| 3 | 99375069 | rs647882 | 0.71 | 0.001 | -0.967 | 0.267 | 0.017 | 0.016 | 0.016 | 0 | 0.63 | 0.25 | 0.56 | 0.69 | 0.69 | 0.51 |
| 3 | 99375151 | rs648006 | 1.039 | 0.008 | -1.874 | 0.098 | -0.102 | -0.127 | -0.127 | 0 | 0.37 | 0.07 | 0.42 | 0.46 | 0.47 | 0.36 |

| | | | | | | | | | | | | | | | | |
|---|---|---|---|---|---|---|---|---|---|---|---|---|---|---|---|---|
| 3 | 99383411 | rs1965397 | 0.527 | 0.055 | -2.029 | -0.275 | -0.026 | 0.046 | 0.046 | 0 | 0.11 | 0.04 | 0.07 | 0.01 | 0.12 | 0.08 |
| 3 | 99391933 | rs1711560 | 0.028 | -0.642 | -1.32 | -0.466 | -0.001 | -0.039 | -0.039 | 0.033 | 0.61 | 0.55 | 0.58 | 0.77 | 0.65 | 0.52 |
| 3 | 99394092 | rs9872469 | 1.24 | -0.596 | -1.437 | -0.426 | -0.021 | 0.011 | 0.011 | 0.022 | 0.05 | 0.08 | 0.03 | 0.07 | 0.02 | 0.01 |
| 3 | 99398156 | rs13059876 | 1.123 | -0.598 | -1.844 | -0.544 | 0.003 | 0.013 | 0.013 | 0 | 0.1 | 0.04 | 0.08 | 0.07 | 0.11 | 0.09 |
| 3 | 99401638 | rs1717608 | 1.733 | -0.534 | -1.826 | -0.358 | -0.103 | -0.15 | -0.15 | 3.302 | 0.3 | 0.05 | 0.36 | 0.47 | 0.39 | 0.3 |
| 3 | 99410162 | rs1111615 | 0.13 | -0.497 | -1.857 | -0.446 | -0.001 | 0.004 | 0.004 | 0 | 0.16 | 0.08 | 0.12 | 0.1 | 0.13 | 0.21 |
| 3 | 99410293 | rs9836385 | 0.203 | -0.715 | -1.711 | -0.438 | 0.004 | 0 | 0 | 0 | 0.14 | 0.09 | 0.09 | 0.1 | 0.1 | 0.19 |
| 3 | 99410661 | rs621061 | 1.237 | -0.848 | -1.812 | -0.538 | -0.053 | -0.06 | -0.06 | 0 | 0.4 | 0.17 | 0.45 | 0.48 | 0.51 | 0.35 |
| 3 | 99414479 | rs9858145 | -0.138 | -0.636 | -1.339 | -0.528 | -0.02 | -0.011 | -0.011 | 3.39 | 0.16 | 0.22 | 0.1 | 0.1 | 0.09 | 0.18 |
| 3 | 99418950 | rs518138 | -0.274 | -0.595 | -1.266 | -0.626 | -0.002 | -0.057 | -0.057 | 0 | 0.7 | 0.53 | 0.64 | 0.81 | 0.73 | 0.65 |
| 3 | 99422587 | rs1711564 | 1.005 | -0.396 | -1.225 | -0.685 | 0.01 | 0.018 | 0.018 | 0 | 0.54 | 0.42 | 0.53 | 0.62 | 0.59 | 0.49 |
| 3 | 99425669 | rs1711557 | -0.177 | -0.879 | -1.321 | -0.845 | 0.005 | 0.003 | 0.003 | 0 | 0.78 | 0.82 | 0.75 | 0.79 | 0.79 | 0.73 |
| 3 | 99427774 | rs1607993 | 0.473 | -0.967 | -1.315 | -0.94 | 0.009 | 0.008 | 0.008 | 0 | 0.68 | 0.72 | 0.69 | 0.74 | 0.7 | 0.65 |
| 3 | 99435102 | rs17778253 | 1.005 | -0.946 | -1.254 | -0.905 | -0.011 | -0.018 | -0.018 | 0 | 0.11 | 0.2 | 0.07 | 0.05 | 0.1 | 0.09 |
| 3 | 99436424 | rs668668 | 0.353 | -0.949 | -1.366 | -1.024 | -0.012 | 0.074 | 0.074 | 0 | 0.67 | 0.9 | 0.65 | 0.76 | 0.67 | 0.66 |
| 3 | 99442745 | rs4928219 | -0.068 | -0.962 | -1.246 | -1.036 | -0.006 | -0.007 | -0.007 | 0 | 0.19 | 0.23 | 0.12 | 0.14 | 0.16 | 0.22 |
| 3 | 99442809 | rs652026 | 1.798 | -0.964 | -1.227 | -1.035 | -0.001 | 0.003 | 0.003 | 0 | 0.46 | 0.49 | 0.51 | 0.62 | 0.51 | 0.44 |
| 3 | 99450469 | rs1971381 | -0.31 | -0.961 | -1.22 | -1.021 | -0.013 | -0.014 | -0.014 | 0.424 | 0.2 | 0.38 | 0.2 | 0.14 | 0.16 | 0.22 |
| 3 | 99462647 | rs792829 | 0.37 | -0.937 | -1.311 | -0.998 | -0.001 | 0.066 | 0.066 | 0 | 0.67 | 0.88 | 0.64 | 0.76 | 0.66 | 0.66 |
| 3 | 99463810 | rs13080329 | -0.114 | -0.889 | -1.014 | -0.883 | -0.016 | -0.01 | -0.01 | 0.921 | 0.18 | 0.27 | 0.09 | 0.14 | 0.14 | 0.21 |
| 3 | 99471251 | rs792840 | 0.734 | -0.842 | -1.287 | -0.935 | 0.005 | 0.075 | 0.075 | 0.011 | 0.6 | 0.83 | 0.59 | 0.74 | 0.6 | 0.61 |
| 3 | 99471308 | rs792841 | 0.734 | -0.842 | -1.287 | -0.935 | 0.005 | 0.075 | 0.075 | 0.011 | 0.6 | 0.83 | 0.59 | 0.74 | 0.6 | 0.61 |
| 3 | 99476295 | rs792835 | 1.501 | -0.835 | -1.04 | -0.937 | 0.003 | 0.01 | 0.01 | 0 | 0.5 | 0.53 | 0.52 | 0.65 | 0.54 | 0.47 |
| 3 | 99481539 | rs13081855 | 1.361 | -0.849 | -1.051 | -0.945 | 0.004 | 0.016 | 0.016 | 0 | 0.1 | 0.03 | 0.05 | 0.05 | 0.1 | 0.1 |
| 3 | 99483773 | rs704574 | 1.371 | -0.732 | -0.982 | -0.874 | 0.003 | 0.01 | 0.01 | 0 | 0.5 | 0.53 | 0.52 | 0.65 | 0.54 | 0.47 |
| 3 | 99494215 | rs35535760 | 1.115 | -0.357 | -0.963 | -0.807 | -0.046 | 0.049 | 0.049 | 0 | 0.11 | 0.01 | 0.1 | 0.03 | 0.12 | 0.06 |
| 3 | 99503728 | rs1038294 | -0.675 | -0.772 | -0.974 | -0.813 | 0.016 | 0.025 | 0.025 | 0 | 0.79 | 0.94 | 0.76 | 0.84 | 0.8 | 0.76 |
| 3 | 99505610 | rs793497 | 1.021 | -0.631 | -0.921 | -0.888 | -0.001 | -0.003 | -0.003 | 0 | 0.59 | 0.45 | 0.57 | 0.71 | 0.63 | 0.56 |
| 3 | 99524417 | rs1072820 | 1.248 | -0.527 | -0.953 | -0.852 | -0.016 | 0.044 | 0.044 | 0 | 0.1 | 0.03 | 0.09 | 0.03 | 0.09 | 0.06 |
| 3 | 99573547 | rs793443 | 0.954 | -0.549 | -0.914 | -0.91 | -0.003 | -0.024 | -0.024 | 0 | 0.6 | 0.52 | 0.57 | 0.72 | 0.63 | 0.55 |
| 3 | 99602555 | rs3921767 | -0.093 | -0.547 | -0.878 | -0.911 | 0.096 | 0.108 | 0.108 | 0.198 | 0.07 | 0.3 | 0.07 | 0.09 | 0.06 | 0.13 |
| 3 | 99614626 | rs17338680 | 1.201 | -0.55 | -0.928 | -0.924 | -0.029 | 0.047 | 0.047 | 0 | 0.1 | 0.02 | 0.1 | 0.03 | 0.1 | 0.06 |
| 3 | 99637783 | rs793439 | -1.622 | -0.632 | -0.947 | -0.973 | 0.009 | 0.037 | 0.037 | 0 | 0.48 | 0.59 | 0.48 | 0.32 | 0.45 | 0.53 |
| 3 | 99640716 | rs6791889 | 1.986 | -0.606 | -1.241 | -0.962 | 0.001 | 0.002 | 0.002 | 0 | 0.07 | 0.05 | 0.05 | 0.04 | 0.07 | 0.09 |
| 3 | 99643176 | rs793440 | -0.481 | -0.521 | -1.322 | -0.915 | 0.004 | 0.081 | 0.081 | 0.073 | 0.33 | 0.16 | 0.35 | 0.2 | 0.31 | 0.31 |
| 3 | 99643693 | rs704583 | -0.323 | -0.522 | -1.323 | -0.917 | -0.001 | 0.082 | 0.082 | 0.062 | 0.31 | 0.08 | 0.34 | 0.2 | 0.3 | 0.31 |
| 3 | 99647850 | rs1688766 | 0.553 | -0.522 | -1.319 | -0.913 | 0.016 | 0.027 | 0.027 | 0 | 0.21 | 0.06 | 0.24 | 0.16 | 0.2 | 0.25 |
| 3 | 99674093 | rs10936005 | 0.326 | -0.521 | -1.319 | -0.914 | 0.004 | 0.081 | 0.081 | 0.073 | 0.67 | 0.84 | 0.65 | 0.8 | 0.69 | 0.69 |
| 3 | 99679461 | rs4244712 | 1.886 | -0.791 | -1.677 | -0.879 | -0.152 | -0.187 | -0.187 | 0.019 | 0.33 | 0.02 | 0.41 | 0.47 | 0.43 | 0.33 |
| 3 | 99680086 | rs4075038 | -0.668 | -0.439 | -1.068 | -0.673 | 0.016 | 0.027 | 0.027 | 0 | 0.79 | 0.94 | 0.76 | 0.84 | 0.8 | 0.75 |
| 3 | 99701134 | rs7616988 | -0.458 | -0.442 | -1.058 | -0.664 | 0.013 | 0.033 | 0.033 | 0 | 0.78 | 0.85 | 0.74 | 0.85 | 0.79 | 0.75 |
| 3 | 99722396 | rs6440967 | 0.885 | -0.436 | -0.938 | -0.703 | 0 | 0.02 | 0.02 | 0 | 0.61 | 0.64 | 0.59 | 0.72 | 0.63 | 0.56 |
| 3 | 99724873 | rs10511177 | 0.312 | -0.428 | -1.008 | -0.688 | 0.013 | 0.029 | 0.029 | 0 | 0.22 | 0.14 | 0.26 | 0.16 | 0.21 | 0.25 |
| 3 | 99754498 | rs17392948 | 0.838 | -0.459 | -1.032 | -0.707 | 0.002 | 0.079 | 0.079 | 0 | 0.18 | 0.04 | 0.23 | 0.03 | 0.18 | 0.21 |
| 3 | 99758465 | rs6440991 | -0.622 | -0.434 | -0.948 | -0.711 | -0.016 | 0.007 | 0.007 | 0 | 0.85 | 0.92 | 0.86 | 0.88 | 0.84 | 0.81 |
| 3 | 99760864 | rs6807176 | 0.091 | -0.504 | -1.006 | -0.753 | 0.017 | 0.035 | 0.035 | 0 | 0.23 | 0.14 | 0.26 | 0.16 | 0.22 | 0.25 |
| 3 | 99767541 | rs13064704 | 0.478 | -0.44 | -0.954 | -0.704 | 0.013 | 0.026 | 0.026 | 0 | 0.21 | 0.06 | 0.24 | 0.16 | 0.2 | 0.25 |
| 3 | 99786289 | rs1021341 | 0.269 | -0.439 | -0.952 | -0.705 | 0.013 | 0.028 | 0.028 | 0 | 0.22 | 0.14 | 0.26 | 0.16 | 0.21 | 0.25 |
| 3 | 99795507 | rs9823908 | 0.481 | -0.437 | -0.993 | -0.697 | 0.011 | 0.025 | 0.025 | 0 | 0.21 | 0.04 | 0.24 | 0.16 | 0.2 | 0.25 |
| 3 | 99822536 | rs12494994 | 0.108 | -0.354 | -0.867 | -0.688 | 0.01 | -0.008 | -0.008 | 0 | 0.16 | 0.21 | 0.15 | 0.12 | 0.16 | 0.19 |
| 3 | 99835254 | rs12488245 | -0.919 | -0.332 | -0.813 | -0.335 | -0.001 | 0.001 | 0.001 | 0 | 0.39 | 0.42 | 0.43 | 0.34 | 0.37 | 0.45 |
| 3 | 99844293 | rs9811920 | -0.994 | -0.386 | -0.593 | -0.217 | -0.008 | -0.014 | -0.014 | 0 | 0.4 | 0.49 | 0.43 | 0.35 | 0.36 | 0.46 |
| 3 | 99848340 | rs17315194 | 1.896 | -0.54 | -0.978 | -0.223 | -0.015 | -0.038 | -0.038 | 0.047 | 0.15 | 0.02 | 0.08 | 0.17 | 0.12 | 0.12 |
| 3 | 99863183 | rs6767441 | -1.124 | -0.558 | -0.485 | -0.269 | 0.002 | 0.006 | 0.006 | 0 | 0.39 | 0.4 | 0.42 | 0.35 | 0.36 | 0.47 |
| 3 | 99873317 | rs2130369 | -1.232 | -0.561 | -0.452 | -0.269 | -0.008 | -0.014 | -0.014 | 0 | 0.4 | 0.48 | 0.43 | 0.35 | 0.36 | 0.46 |
| 3 | 99883598 | rs7641782 | -0.109 | -0.626 | -0.486 | -0.294 | 0.01 | -0.002 | -0.002 | 0 | 0.16 | 0.18 | 0.16 | 0.15 | 0.16 | 0.2 |
| 3 | 99884509 | rs9875878 | 0.938 | -0.621 | -0.598 | -0.343 | 0.007 | 0.087 | 0.087 | 0.089 | 0.18 | 0.02 | 0.22 | 0.04 | 0.16 | 0.21 |
| 3 | 99886662 | rs11537816 | 0.888 | -0.619 | -0.594 | -0.342 | 0.01 | 0.086 | 0.086 | 0.059 | 0.18 | 0.02 | 0.22 | 0.04 | 0.16 | 0.21 |
| 3 | 99893111 | rs12487143 | -0.735 | -0.572 | -0.526 | -0.302 | 0.01 | 0.044 | 0.044 | 0 | 0.2 | 0.31 | 0.18 | 0.31 | 0.18 | 0.24 |
| 3 | 99900096 | rs2316263 | -1.06 | -0.461 | -0.617 | -0.232 | 0 | 0 | 0 | 0 | 0.46 | 0.53 | 0.47 | 0.39 | 0.42 | 0.55 |
| 3 | 99905803 | rs1017967 | -1.072 | -0.486 | -0.63 | -0.093 | 0.001 | 0.001 | 0.001 | 0 | 0.46 | 0.55 | 0.48 | 0.47 | 0.43 | 0.55 |
| 3 | 99906993 | rs9290003 | -1.091 | -0.441 | -0.485 | -0.067 | 0.005 | 0.005 | 0.005 | 0 | 0.47 | 0.57 | 0.48 | 0.47 | 0.43 | 0.54 |
| 3 | 99909642 | rs13096852 | 1.851 | -0.318 | -0.62 | -0.093 | 0 | -0.001 | -0.001 | 0 | 0.07 | 0.05 | 0.05 | 0.04 | 0.08 | 0.09 |
| 3 | 99914139 | rs9868170 | -1.144 | -0.451 | -0.583 | -0.084 | 0.013 | 0.018 | 0.018 | 0 | 0.49 | 0.59 | 0.49 | 0.47 | 0.44 | 0.56 |
| 3 | 99917359 | rs9860514 | -1.167 | 0.215 | -0.489 | 0.189 | 0.01 | 0.023 | 0.023 | 0 | 0.49 | 0.61 | 0.5 | 0.44 | 0.45 | 0.57 |
| 3 | 99920884 | rs11719857 | -0.714 | 0.474 | -0.479 | -0.032 | 0.005 | -0.011 | -0.011 | 0 | 0.17 | 0.13 | 0.18 | 0.09 | 0.19 | 0.22 |
| 3 | 99924774 | rs6800438 | 0.555 | 0.556 | -0.72 | -0.008 | -0.036 | 0.024 | 0.024 | 0.137 | 0.07 | 0 | 0.07 | 0.05 | 0.1 | 0.18 |
| 3 | 99929024 | rs7625101 | 0.929 | 0.654 | -1.193 | 0.028 | -0.001 | -0.104 | -0.104 | 0.482 | 0.25 | 0.1 | 0.32 | 0.51 | 0.27 | 0.33 |
| 3 | 99937577 | rs17279479 | 1.187 | 0.85 | -1.51 | -0.344 | -0.09 | 0.109 | 0.109 | 0 | 0.19 | 0.01 | 0.13 | 0.04 | 0.19 | 0.2 |

| | | | | | | | | | | | | | | | | |
|---|---|---|---|---|---|---|---|---|---|---|---|---|---|---|---|---|
| 3 | 99944381 | rs17213695 | 2.622 | 0.883 | -1.495 | -0.33 | -0.034 | 0.074 | 0.074 | 0 | 0.12 | 0.01 | 0.05 | 0 | 0.09 | 0.03 |
| 3 | 99946822 | rs12492610 | 0.414 | 0.545 | -1.092 | -0.261 | -0.006 | 0.002 | 0.002 | 0 | 0.3 | 0.21 | 0.25 | 0.09 | 0.28 | 0.24 |
| 3 | 99954721 | rs6802765 | -1.03 | 0.464 | -1.015 | -0.26 | 0 | 0 | 0 | 0.245 | 0.56 | 0.4 | 0.58 | 0.6 | 0.56 | 0.58 |
| 3 | 99962992 | rs12490140 | 0.408 | 0.512 | -0.992 | -0.24 | -0.003 | -0.008 | -0.008 | 0 | 0.3 | 0.23 | 0.25 | 0.09 | 0.28 | 0.24 |
| 3 | 99970995 | rs6772774 | 0.976 | 0.549 | -1.177 | -0.191 | 0.004 | -0.084 | -0.084 | 0.167 | 0.27 | 0.15 | 0.33 | 0.51 | 0.28 | 0.34 |
| 3 | 99976007 | rs10936147 | 0.912 | 0.548 | -1.174 | -0.191 | 0.007 | -0.062 | -0.062 | 0.118 | 0.27 | 0.17 | 0.33 | 0.51 | 0.28 | 0.34 |
| 3 | 99980281 | rs3772701 | 1.146 | 0.56 | -1.244 | -0.187 | 0.003 | -0.098 | -0.098 | 0.014 | 0.25 | 0.12 | 0.33 | 0.52 | 0.27 | 0.34 |
| 3 | 99981711 | rs7637157 | 1.266 | 0.387 | -1.234 | -0.324 | -0.02 | 0.174 | 0.174 | 0 | 0.37 | 0.14 | 0.36 | 0.05 | 0.41 | 0.34 |
| 3 | 99982607 | rs9863738 | 1.35 | 0.367 | -1.245 | -0.333 | -0.047 | 0.223 | 0.223 | 0 | 0.36 | 0.07 | 0.35 | 0.05 | 0.41 | 0.33 |
| 3 | 100011853 | rs4263320 | 1.212 | 0.422 | -1.773 | -0.249 | 0.003 | -0.101 | -0.101 | 0.245 | 0.25 | 0.12 | 0.33 | 0.51 | 0.27 | 0.34 |
| 3 | 100014606 | rs17395659 | 1.344 | 0.413 | -1.791 | -0.268 | -0.025 | -0.006 | -0.006 | 0.317 | 0.08 | 0.01 | 0.1 | 0.05 | 0.09 | 0.04 |
| 3 | 100041703 | rs10511178 | 0.983 | 0.385 | -1.682 | -0.262 | 0.004 | -0.083 | -0.083 | 2.785 | 0.27 | 0.16 | 0.33 | 0.51 | 0.28 | 0.34 |
| 3 | 100044952 | rs1214373 | -0.868 | 0.206 | -1.56 | -0.761 | -0.019 | 0.17 | 0.17 | 0 | 0.63 | 0.85 | 0.64 | 0.95 | 0.59 | 0.66 |
| 3 | 100055125 | rs11714045 | 0.933 | -0.018 | -1.326 | 0.005 | -0.055 | -0.018 | -0.018 | 0 | 0.33 | 0.04 | 0.25 | 0.22 | 0.35 | 0.4 |
| 3 | 100058205 | rs392066 | -0.033 | -0.034 | -0.51 | 0.199 | 0.074 | 0.209 | 0.209 | 1.376 | 0.07 | 0.33 | 0.19 | 0.2 | 0.04 | 0.07 |
| 3 | 100064902 | rs2289506 | 1.059 | -0.047 | -0.711 | 0.11 | -0.08 | -0.037 | -0.037 | 0.034 | 0.32 | 0.02 | 0.23 | 0.22 | 0.35 | 0.4 |
| 3 | 100065632 | rs1214375 | 0.265 | -0.164 | -0.486 | 0.342 | -0.003 | -0.027 | -0.027 | 0.024 | 0.4 | 0.4 | 0.44 | 0.74 | 0.39 | 0.48 |
| 3 | 100072332 | rs9873033 | 0.726 | -0.196 | -0.555 | 0.064 | 0.045 | -0.052 | -0.052 | 0.961 | 0.26 | 0.45 | 0.18 | 0.19 | 0.3 | 0.23 |
| 3 | 100086477 | rs2316701 | 0.087 | -0.255 | -0.453 | 0.101 | -0.02 | 0.001 | 0.001 | 0.125 | 0.6 | 0.55 | 0.43 | 0.42 | 0.64 | 0.65 |
| 3 | 100086861 | rs277640 | -0.671 | -0.265 | -0.53 | 0.099 | 0.026 | 0.169 | 0.169 | 0.11 | 0.67 | 0.92 | 0.63 | 0.94 | 0.68 | 0.72 |
| 3 | 100096419 | rs277633 | -0.07 | -0.254 | -0.291 | 0.13 | 0.055 | 0.539 | 0.539 | 0.57 | 0.07 | 0.31 | 0.2 | 0.51 | 0.04 | 0.07 |
| 3 | 100104058 | rs3772692 | -0.807 | -0.233 | -0.47 | 0.116 | 0.053 | 0.521 | 0.521 | 0.551 | 0.09 | 0.41 | 0.2 | 0.51 | 0.05 | 0.08 |
| 3 | 100107095 | rs10936180 | -0.241 | -0.23 | -0.476 | 0.124 | 0.053 | 0.555 | 0.555 | 1.184 | 0.08 | 0.37 | 0.2 | 0.51 | 0.04 | 0.07 |
| 3 | 100112168 | rs11711112 | -0.517 | -0.193 | -0.783 | -0.237 | 0.026 | 0.169 | 0.169 | 0.11 | 0.67 | 0.92 | 0.63 | 0.94 | 0.68 | 0.71 |
| 3 | 100133977 | rs10936193 | 1.172 | 0.159 | -0.564 | 0.025 | -0.101 | 0.12 | 0.12 | 0 | 0.2 | 0.01 | 0.11 | 0.02 | 0.24 | 0.15 |
| 3 | 100154117 | rs9290057 | -0.465 | 0.027 | -0.188 | -0.514 | 0.017 | 0.005 | 0.005 | 1.654 | 0.94 | 0.73 | 0.96 | 0.93 | 0.98 | 0.97 |
| 3 | 100170825 | rs6797397 | 0.393 | -0.043 | -0.757 | -0.691 | -0.001 | -0.032 | -0.032 | 0 | 0.46 | 0.52 | 0.45 | 0.72 | 0.42 | 0.48 |
| 3 | 100174722 | rs1132022 | 0.302 | -0.032 | -1.213 | -0.686 | -0.002 | 0.046 | 0.046 | 0 | 0.45 | 0.38 | 0.44 | 0.72 | 0.42 | 0.48 |
| 3 | 100176360 | rs7643442 | -0.306 | 0.06 | -1.692 | -0.549 | 0.005 | 0.119 | 0.119 | 0 | 0.74 | 0.94 | 0.64 | 0.92 | 0.71 | 0.74 |
| 3 | 100181868 | rs9290061 | -0.276 | 0.012 | -1.796 | -0.528 | 0.003 | 0.135 | 0.135 | 0 | 0.73 | 0.96 | 0.64 | 0.92 | 0.71 | 0.69 |
| 3 | 100186962 | rs9857793 | -0.511 | 0.286 | -0.85 | 0.339 | -0.004 | 0.011 | 0.011 | 0 | 0.15 | 0.18 | 0.24 | 0.21 | 0.11 | 0.13 |
| 3 | 100209602 | rs11707581 | 0.56 | 0.431 | -1.424 | -0.42 | -0.052 | 0.139 | 0.139 | 0 | 0.26 | 0.04 | 0.13 | 0.01 | 0.28 | 0.22 |
| 3 | 100211294 | rs7616839 | -1.034 | -0.173 | -1.684 | -0.442 | -0.028 | 0.193 | 0.193 | 0.459 | 0.52 | 0.88 | 0.46 | 0.74 | 0.46 | 0.41 |
| 3 | 100213778 | rs4928097 | -0.521 | -0.669 | -0.564 | 0.077 | -0.01 | 0.025 | 0.025 | 0 | 0.2 | 0.29 | 0.28 | 0.37 | 0.17 | 0.18 |
| 3 | 100225478 | rs12639465 | -1.035 | -0.695 | -0.141 | -0.053 | -0.015 | 0.007 | 0.007 | 0.456 | 0.25 | 0.7 | 0.32 | 0.28 | 0.17 | 0.17 |
| 3 | 100238060 | rs7429392 | -1.491 | -1.101 | -0.53 | -0.343 | 0.058 | 0.005 | 0.005 | 0 | 0.36 | 0.78 | 0.58 | 0.33 | 0.26 | 0.41 |
| 3 | 100240509 | rs6784997 | -0.125 | -1.113 | 0.364 | -0.398 | -0.033 | -0.051 | -0.051 | 0 | 0.12 | 0.26 | 0.24 | 0.05 | 0.06 | 0.24 |
| 3 | 100242358 | rs6441308 | -1.442 | -1.1 | 0.388 | -0.311 | -0.021 | 0.004 | 0.004 | 0 | 0.33 | 0.61 | 0.54 | 0.37 | 0.23 | 0.41 |
| 3 | 100247142 | rs6777810 | -1.111 | -0.342 | 0.273 | -0.45 | -0.049 | 0.02 | 0.02 | 0 | 0.58 | 0.82 | 0.67 | 0.37 | 0.49 | 0.49 |
| 3 | 100247165 | rs3924054 | 0.333 | -0.296 | 0.507 | -0.294 | 0.004 | 0.002 | 0.002 | 0.188 | 0.11 | 0.12 | 0.18 | 0.26 | 0.06 | 0.09 |
| 3 | 100249015 | rs12636452 | -0.495 | -0.327 | 0.58 | -0.33 | 0.001 | -0.001 | -0.001 | 0 | 0.17 | 0.21 | 0.26 | 0.31 | 0.12 | 0.16 |
| 3 | 100253841 | rs6441316 | -0.576 | -0.14 | 0.697 | -0.242 | 0.013 | 0.035 | 0.035 | 0.362 | 0.49 | 0.62 | 0.42 | 0.32 | 0.44 | 0.25 |
| 3 | 100257128 | rs11719592 | -0.722 | -0.138 | 0.531 | -0.242 | 0.01 | 0.097 | 0.097 | 0.54 | 0.51 | 0.84 | 0.44 | 0.32 | 0.44 | 0.26 |
| 3 | 100263559 | rs4928112 | -0.223 | -0.087 | 0.518 | -0.219 | -0.011 | 0.056 | 0.056 | 0.377 | 0.44 | 0.22 | 0.38 | 0.32 | 0.44 | 0.26 |
| 3 | 100267619 | rs9843498 | -0.185 | -0.074 | 0.598 | -0.213 | 0.001 | 0.007 | 0.007 | 0.066 | 0.46 | 0.42 | 0.39 | 0.32 | 0.44 | 0.25 |
| 3 | 100280637 | rs7647499 | -1.108 | -0.175 | 0.583 | -0.075 | -0.025 | 0.102 | 0.102 | 1.087 | 0.19 | 0.52 | 0.27 | 0.32 | 0.11 | 0.14 |
| 3 | 100283838 | rs9864288 | 0.412 | 0.033 | 0.515 | -0.284 | -0.009 | -0.109 | -0.109 | 0.969 | 0.36 | 0.48 | 0.18 | 0.01 | 0.32 | 0.12 |
| 3 | 100284007 | rs7429915 | -0.718 | 0.013 | 0.528 | -0.24 | 0.003 | 0.078 | 0.078 | 1.156 | 0.52 | 0.8 | 0.43 | 0.33 | 0.44 | 0.25 |
| 3 | 100308233 | rs9829641 | -0.828 | -0.155 | 0.738 | -0.05 | -0.001 | 0.072 | 0.072 | 1.217 | 0.17 | 0.38 | 0.25 | 0.29 | 0.12 | 0.16 |
| 3 | 100322043 | rs9856338 | 0.369 | 0.09 | 0.451 | -0.241 | 0.003 | 0.111 | 0.111 | 1.35 | 0.49 | 0.28 | 0.54 | 0.71 | 0.51 | 0.71 |
| 3 | 100324522 | rs1718272 | 0.924 | -0.088 | 0.419 | -0.325 | -0.032 | 0.149 | 0.149 | 0.717 | 0.3 | 0.07 | 0.18 | 0.01 | 0.36 | 0.13 |
| 3 | 100334256 | rs1676345 | 0.508 | -0.119 | 0.913 | -0.337 | -0.001 | -0.072 | -0.072 | 2.308 | 0.33 | 0.31 | 0.19 | 0.02 | 0.37 | 0.13 |
| 3 | 100354524 | rs1144122 | 1.026 | -0.287 | 0.935 | -0.407 | -0.019 | 0.074 | 0.074 | 0.325 | 0.28 | 0.11 | 0.18 | 0.02 | 0.36 | 0.12 |
| 3 | 100354683 | rs1520652 | -0.637 | -0.136 | 1.119 | -0.339 | -0.007 | -0.125 | -0.125 | 0.565 | 0.66 | 0.53 | 0.79 | 0.98 | 0.64 | 0.87 |
| 3 | 100357671 | rs6793682 | 0.356 | -0.077 | 0.533 | -0.567 | -0.076 | 0.106 | 0.106 | 0.935 | 0.32 | 0.17 | 0.43 | 0.1 | 0.36 | 0.51 |
| 3 | 100367257 | rs1145341 | 0.568 | -0.076 | 0.706 | -0.481 | 0.089 | 0.084 | 0.084 | 3.296 | 0.95 | 0.57 | 0.96 | 1 | 1 | 1 |
| 3 | 100374740 | rs9866111 | -0.357 | -0.161 | 0.538 | 0.046 | 0.043 | 0.068 | 0.068 | 2.407 | 0.53 | 0.22 | 0.61 | 0.43 | 0.57 | 0.78 |
| 3 | 100375572 | rs619574 | 0.829 | -0.094 | 0.994 | 0.386 | 0.069 | 0.638 | 0.638 | 3.834 | 0.95 | 0.58 | 0.96 | 0.69 | 1 | 0.99 |
| 3 | 100379856 | rs4928072 | 0.83 | -0.006 | 1.01 | 0.203 | -0.006 | 0.025 | 0.025 | 0 | 0.23 | 0.29 | 0.21 | 0.65 | 0.22 | 0.28 |
| 3 | 100387709 | rs1144110 | 0.978 | 0.101 | 1.419 | 0.24 | 0.072 | 0.671 | 0.671 | 2.668 | 0.95 | 0.56 | 0.96 | 0.68 | 1 | 0.99 |
| 3 | 100396915 | rs7629279 | -0.187 | 0.07 | 0.884 | -0.084 | -0.065 | 0.12 | 0.12 | 1.132 | 0.34 | 0.19 | 0.42 | 0.1 | 0.37 | 0.52 |
| 3 | 100410724 | rs4928080 | 0.747 | -0.064 | 0.928 | -0.048 | 0.021 | -0.053 | -0.053 | 0 | 0.3 | 0.15 | 0.35 | 0.56 | 0.25 | 0.34 |
| 3 | 100411334 | rs7614943 | -0.111 | -0.011 | 1.022 | -0.143 | -0.037 | 0.069 | 0.069 | 0.646 | 0.35 | 0.26 | 0.43 | 0.09 | 0.37 | 0.53 |
| 3 | 100411361 | rs4928081 | -1.177 | 0.012 | 1.111 | -0.15 | -0.01 | 0.022 | 0.022 | 0.525 | 0.66 | 0.62 | 0.8 | 0.65 | 0.64 | 0.87 |
| 3 | 100427937 | rs6810101 | -0.003 | 0.053 | 1.002 | -0.095 | -0.066 | 0.086 | 0.086 | 0.493 | 0.34 | 0.2 | 0.43 | 0.12 | 0.37 | 0.54 |
| 3 | 100437892 | rs6775161 | -0.234 | 0.049 | 0.965 | -0.096 | 0.014 | -0.039 | -0.039 | 1.107 | 0.37 | 0.5 | 0.45 | 0.12 | 0.39 | 0.54 |
| 3 | 100468500 | rs1048364 | 0.858 | 0.005 | 0.642 | -0.052 | 0.023 | -0.039 | -0.039 | 0 | 0.3 | 0.14 | 0.35 | 0.53 | 0.25 | 0.33 |
| 3 | 100473505 | rs10936352 | -0.059 | 0.139 | 0.608 | -0.148 | -0.117 | 0.097 | 0.097 | 0.628 | 0.33 | 0.12 | 0.42 | 0.12 | 0.37 | 0.53 |
| 3 | 100479384 | rs7642041 | -0.403 | 0.252 | 1.099 | -0.139 | 0.008 | -0.038 | -0.038 | 0.127 | 0.37 | 0.49 | 0.45 | 0.12 | 0.37 | 0.54 |

| | | | | | | | | | | | | | | | | |
|---|---|---|---|---|---|---|---|---|---|---|---|---|---|---|---|---|
| 3 | 100480812 | rs4928087 | -1.272 | 0.188 | 0.998 | -0.071 | -0.001 | 0 | 0 | 0.505 | 0.68 | 0.72 | 0.81 | 0.65 | 0.64 | 0.87 |
| 3 | 100485451 | rs544500 | -0.096 | 0.565 | 1.017 | -0.127 | 0 | 0.002 | 0.002 | 0 | 0.78 | 0.76 | 0.77 | 0.67 | 0.8 | 0.53 |
| 3 | 100489286 | rs7642942 | -0.491 | 0.449 | 1.079 | -0.051 | -0.01 | 0.174 | 0.174 | 0 | 0.1 | 0.3 | 0.14 | 0.29 | 0.05 | 0.2 |
| 3 | 100493422 | rs9846590 | 0.769 | 0.47 | 1.239 | -0.209 | 0 | -0.044 | -0.044 | 0.024 | 0.13 | 0.16 | 0.1 | 0 | 0.15 | 0.07 |
| 3 | 100499321 | rs7613610 | -0.698 | 0.442 | 1.186 | -0.209 | -0.01 | 0.027 | 0.027 | 0 | 0.24 | 0.37 | 0.22 | 0.3 | 0.19 | 0.26 |
| 3 | 100500860 | rs9842968 | -1.297 | 0.601 | 1.318 | -0.246 | -0.001 | 0 | 0 | 0 | 0.29 | 0.71 | 0.27 | 0.29 | 0.23 | 0.27 |
| 3 | 100501854 | rs7648469 | -0.124 | 0.583 | 1.091 | -0.253 | 0.004 | -0.011 | -0.011 | 0 | 0.09 | 0.32 | 0.14 | 0.05 | 0.05 | 0.19 |
| 3 | 100507827 | rs7628455 | -1.09 | 0.558 | 0.235 | -0.442 | -0.099 | 0.261 | 0.261 | 0 | 0.58 | 0.93 | 0.62 | 0.94 | 0.48 | 0.6 |
| 3 | 100514481 | rs12696195 | -0.123 | 0.862 | 0.873 | 0.34 | -0.002 | 0.006 | 0.006 | 0 | 0.17 | 0.19 | 0.1 | 0.24 | 0.15 | 0.08 |
| 3 | 100523359 | rs9290224 | -0.24 | 0.876 | 0.877 | 0.336 | -0.003 | -0.011 | -0.011 | 5.805 | 0.08 | 0.21 | 0.13 | 0.04 | 0.05 | 0.18 |
| 3 | 100523771 | rs10511184 | 0.383 | 1.019 | 0.625 | 0.295 | -0.086 | 0.021 | 0.021 | 0.081 | 0.14 | 0.01 | 0.1 | 0.05 | 0.17 | 0.28 |
| 3 | 100529700 | rs7609779 | -1.291 | 1.026 | 0.844 | 0.297 | -0.009 | 0.007 | 0.007 | 0.161 | 0.28 | 0.69 | 0.27 | 0.28 | 0.22 | 0.27 |
| 3 | 100530856 | rs764684 | -1.865 | 0.988 | 0.579 | 0.342 | -0.016 | 0.006 | 0.006 | 0.962 | 0.43 | 0.77 | 0.37 | 0.34 | 0.37 | 0.55 |
| 3 | 100536284 | rs12632235 | -0.498 | 0.868 | 0.589 | 0.571 | 0.013 | 0.015 | 0.015 | 0 | 0.05 | 0.19 | 0.11 | 0.04 | 0.03 | 0.15 |
| 3 | 100539887 | rs7615467 | -0.598 | 0.834 | -0.03 | 0.818 | -0.005 | 0.035 | 0.035 | 0 | 0.15 | 0.24 | 0.21 | 0.24 | 0.14 | 0.08 |
| 3 | 100550136 | rs6808242 | -0.42 | 0.905 | -0.072 | 0.855 | 0.002 | 0.033 | 0.033 | 0 | 0.14 | 0.19 | 0.2 | 0.24 | 0.15 | 0.08 |
| 3 | 100550650 | rs9840549 | -1.038 | 1.082 | -0.164 | 0.549 | -0.018 | -0.009 | -0.009 | 1.043 | 0.36 | 0.71 | 0.45 | 0.34 | 0.35 | 0.51 |
| 3 | 100558416 | rs9841585 | -0.558 | 0.364 | -0.328 | 0.322 | 0.008 | 0.003 | 0.003 | 0 | 0.27 | 0.23 | 0.29 | 0.28 | 0.2 | 0.24 |
| 3 | 100558905 | rs1354374 | 0.167 | 0.291 | -0.457 | 0.302 | 0.005 | -0.001 | -0.001 | 0.878 | 0.11 | 0.06 | 0.19 | 0.16 | 0.11 | 0.08 |
| 3 | 100561198 | rs1503840 | -1.37 | 0.49 | -0.999 | 0.221 | -0.026 | -0.103 | -0.103 | 0.054 | 0.65 | 0.88 | 0.68 | 0.5 | 0.64 | 0.6 |
| 3 | 100561439 | rs17284369 | 0.898 | 0.284 | -0.9 | -0.072 | -0.049 | 0.065 | 0.065 | 1.228 | 0.22 | 0.04 | 0.25 | 0.06 | 0.28 | 0.34 |
| 3 | 100564866 | rs16842881 | 0.424 | 0.209 | -0.614 | 0.266 | 0.002 | 0.002 | 0.002 | 0 | 0.19 | 0.17 | 0.11 | 0.14 | 0.1 | 0.17 |
| 3 | 100566571 | rs3806656 | 0.158 | 0.172 | -0.397 | 0.167 | -0.019 | -0.012 | -0.012 | 0.974 | 0.2 | 0.25 | 0.11 | 0.12 | 0.1 | 0.17 |
| 3 | 100571493 | rs4928089 | 0.487 | 0.986 | -0.219 | 0.425 | 0.026 | 0.043 | 0.043 | 0 | 0.37 | 0.18 | 0.32 | 0.32 | 0.28 | 0.24 |
| 3 | 100575753 | rs6780804 | 0.109 | 1.07 | -0.019 | 0.491 | 0.008 | 0.006 | 0.006 | 2.237 | 0.27 | 0.68 | 0.36 | 0.24 | 0.33 | 0.34 |
| 3 | 100576353 | rs7631720 | 0.169 | 1.066 | 0.377 | 0.517 | 0.053 | -0.069 | -0.069 | 0 | 0.25 | 0.5 | 0.34 | 0.09 | 0.33 | 0.33 |
| 3 | 100587759 | rs2033708 | -0.757 | 0.867 | -0.305 | 0.406 | -0.002 | 0.006 | 0.006 | 0 | 0.65 | 0.62 | 0.67 | 0.61 | 0.66 | 0.67 |
| 3 | 100590925 | rs9833094 | 0.337 | 0.984 | -0.685 | 0.385 | 0.007 | 0.029 | 0.029 | 0 | 0.39 | 0.31 | 0.37 | 0.33 | 0.34 | 0.24 |
| 3 | 100591006 | rs973309 | 0.025 | 1.017 | -0.532 | 0.426 | 0 | 0.001 | 0.001 | 0 | 0.62 | 0.66 | 0.65 | 0.57 | 0.61 | 0.57 |
| 3 | 100593266 | rs9863850 | -0.467 | 0.977 | -0.595 | -0.124 | -0.001 | -0.005 | -0.005 | 0 | 0.36 | 0.38 | 0.32 | 0.17 | 0.33 | 0.37 |
| 3 | 100594640 | rs2118170 | 0.601 | 0.751 | -0.328 | 0.117 | 0.001 | 0.002 | 0.002 | 0 | 0.5 | 0.45 | 0.65 | 0.46 | 0.62 | 0.53 |
| 3 | 100594684 | rs2118171 | 0.83 | 0.938 | -0.14 | -0.046 | -0.035 | -0.022 | -0.022 | 1.503 | 0.44 | 0.11 | 0.55 | 0.46 | 0.48 | 0.5 |
| 3 | 100596371 | rs6778625 | -0.101 | 0.749 | -0.174 | -0.23 | -0.017 | 0.059 | 0.059 | 0 | 0.69 | 0.5 | 0.74 | 0.5 | 0.77 | 0.61 |
| 3 | 100597517 | rs1821794 | -0.118 | 0.744 | -0.178 | -0.232 | -0.017 | 0.055 | 0.055 | 0 | 0.7 | 0.51 | 0.75 | 0.5 | 0.77 | 0.61 |
| 3 | 100598403 | rs1318859 | -0.728 | 0.559 | 0.537 | -0.003 | 0.009 | 0.034 | 0.034 | 0 | 0.3 | 0.43 | 0.23 | 0.31 | 0.31 | 0.16 |
| 3 | 100598675 | rs9880062 | 0.306 | 0.321 | 0.595 | 0.101 | -0.01 | 0.248 | 0.248 | 0.01 | 0.06 | 0.23 | 0.06 | 0.27 | 0.02 | 0.04 |
| 3 | 100601171 | rs2118172 | 0.193 | 0.708 | 0.567 | -0.462 | -0.007 | -0.033 | -0.033 | 0 | 0.73 | 0.42 | 0.76 | 0.77 | 0.77 | 0.65 |
| 3 | 100601751 | rs4928093 | -0.075 | 0.627 | 0.56 | -0.44 | -0.005 | -0.016 | -0.016 | 0 | 0.75 | 0.6 | 0.79 | 0.77 | 0.79 | 0.65 |
| 3 | 100605206 | rs1987384 | -0.317 | 0.357 | 0.191 | -0.473 | -0.037 | -0.038 | -0.038 | 3.459 | 0.79 | 0.47 | 0.81 | 0.82 | 0.85 | 0.79 |
| 3 | 100608425 | rs1436939 | -0.297 | 0.76 | -0.384 | -0.32 | -0.002 | 0.016 | 0.016 | 0 | 0.2 | 0.1 | 0.09 | 0.04 | 0.15 | 0.08 |
| 3 | 100613293 | rs12330346 | 0.107 | 0.724 | -0.474 | -0.284 | 0.007 | 0.033 | 0.033 | 0.255 | 0.16 | 0.05 | 0.06 | 0.03 | 0.09 | 0.06 |
| 3 | 100613996 | rs2576384 | -0.472 | 0.661 | -0.724 | -0.854 | -0.015 | 0.108 | 0.108 | 0 | 0.43 | 0.28 | 0.33 | 0.07 | 0.46 | 0.26 |
| 3 | 100617680 | rs2245370 | 0.232 | 0.727 | -0.593 | -0.911 | -0.022 | 0.11 | 0.11 | 0.021 | 0.58 | 0.75 | 0.67 | 0.93 | 0.52 | 0.73 |
| 3 | 100643807 | rs1529955 | 1.337 | 0.909 | -0.492 | -0.51 | -0.001 | -0.016 | -0.016 | 0.482 | 0.25 | 0.16 | 0.41 | 0.45 | 0.27 | 0.37 |
| 3 | 100668763 | rs2576389 | -0.336 | 0.737 | -0.359 | -0.699 | -0.013 | 0.053 | 0.053 | 0 | 0.6 | 0.67 | 0.67 | 0.94 | 0.55 | 0.74 |
| 3 | 100672975 | rs2576390 | -0.06 | 0.529 | 0.061 | -0.816 | -0.005 | 0.011 | 0.011 | 0 | 0.56 | 0.59 | 0.64 | 0.91 | 0.52 | 0.72 |
| 3 | 100677661 | rs7641631 | 1.667 | 0.578 | -0.15 | -0.954 | -0.021 | 0.066 | 0.066 | 0.13 | 0.19 | 0.01 | 0.14 | 0.03 | 0.16 | 0.13 |
| 3 | 100688890 | rs2576368 | 0.239 | 0.624 | 0.055 | -0.943 | -0.005 | 0.021 | 0.021 | 0 | 0.31 | 0.38 | 0.22 | 0.46 | 0.25 | 0.35 |
| 3 | 100689310 | rs9844950 | 1.019 | 0.593 | -0.089 | -0.967 | -0.019 | 0.052 | 0.052 | 0 | 0.17 | 0.07 | 0.08 | 0.03 | 0.14 | 0.09 |
| 3 | 100696838 | rs1463189 | 1.494 | 0.522 | -0.241 | -0.491 | -0.043 | 0.014 | 0.014 | 0.01 | 0.28 | 0.07 | 0.38 | 0.29 | 0.31 | 0.46 |
| 3 | 100697944 | rs7643050 | 1.344 | 0.672 | -0.232 | -0.734 | 0 | 0.067 | 0.067 | 0 | 0.23 | 0.1 | 0.16 | 0.04 | 0.18 | 0.15 |
| 3 | 100698454 | rs2595899 | 0.105 | 0.575 | -0.124 | -0.739 | -0.007 | 0.025 | 0.025 | 0 | 0.56 | 0.6 | 0.65 | 0.91 | 0.52 | 0.71 |
| 3 | 100699827 | rs7647099 | 0.519 | 0.517 | -0.198 | -0.696 | -0.002 | 0.007 | 0.007 | 0 | 0.2 | 0.17 | 0.11 | 0.05 | 0.17 | 0.1 |
| 3 | 100703124 | rs1449311 | 1.292 | 0.554 | -0.326 | -0.677 | -0.002 | -0.031 | -0.031 | 0 | 0.3 | 0.17 | 0.43 | 0.52 | 0.31 | 0.5 |
| 3 | 100708127 | rs1530731 | -1.675 | 0.366 | -0.417 | -0.601 | 0.013 | 0.052 | 0.052 | 0 | 0.8 | 0.89 | 0.85 | 0.96 | 0.84 | 0.87 |
| 3 | 100708996 | rs1449308 | 0.189 | 0.282 | -0.213 | -0.546 | 0.007 | -0.025 | -0.025 | 0 | 0.23 | 0.23 | 0.17 | 0.34 | 0.16 | 0.13 |
| 3 | 100712059 | rs3732896 | -0.057 | 0.424 | -0.424 | -0.613 | 0 | 0.009 | 0.009 | 0.213 | 0.2 | 0.22 | 0.15 | 0.41 | 0.13 | 0.24 |
| 3 | 100712249 | rs3732895 | -0.057 | 0.387 | -0.436 | -0.613 | 0 | 0.009 | 0.009 | 0.213 | 0.2 | 0.24 | 0.15 | 0.41 | 0.13 | 0.24 |
| 3 | 100715576 | rs4928102 | -0.048 | 0.385 | -0.436 | -0.605 | 0 | 0.009 | 0.009 | 0.224 | 0.2 | 0.24 | 0.15 | 0.41 | 0.13 | 0.24 |
| 3 | 100730716 | rs16843153 | 1.633 | 0.344 | -0.428 | -0.722 | 0.008 | 0.018 | 0.018 | 0 | 0.2 | 0.14 | 0.15 | 0.03 | 0.16 | 0.13 |
| 3 | 100735017 | rs13320885 | 1.2 | 0.265 | -0.134 | -0.758 | -0.002 | -0.039 | -0.039 | 0 | 0.1 | 0.16 | 0.07 | 0.02 | 0.09 | 0.05 |
| 3 | 100736117 | rs13060756 | 1.345 | 0.292 | -0.306 | -0.691 | 0.008 | -0.025 | -0.025 | 1.872 | 0.09 | 0.02 | 0.09 | 0.13 | 0.04 | 0.15 |
| 3 | 100737036 | rs13085823 | 1.221 | 0.438 | -0.278 | -0.785 | -0.009 | 0.024 | 0.024 | 0 | 0.23 | 0.17 | 0.24 | 0.03 | 0.31 | 0.19 |
| 3 | 100739808 | rs2595917 | -0.005 | 0.391 | -0.132 | -0.582 | -0.004 | 0.045 | 0.045 | 0.26 | 0.79 | 0.7 | 0.83 | 0.58 | 0.85 | 0.76 |
| 3 | 100740159 | rs2713773 | -0.063 | 0.382 | -0.134 | -0.58 | -0.005 | 0.046 | 0.046 | 0.216 | 0.8 | 0.72 | 0.85 | 0.58 | 0.87 | 0.76 |
| 3 | 100745823 | rs9811022 | 1.116 | 0.426 | -0.048 | -0.64 | 0.004 | -0.046 | -0.046 | 0 | 0.1 | 0.17 | 0.08 | 0.02 | 0.09 | 0.05 |
| 3 | 100753024 | rs1868496 | -0.324 | 0.427 | -0.023 | -0.764 | 0.01 | 0.028 | 0.028 | 0 | 0.51 | 0.38 | 0.31 | 0.13 | 0.4 | 0.36 |
| 3 | 100761838 | rs1449317 | -0.094 | 0.43 | 0.189 | -0.524 | 0.009 | 0.075 | 0.075 | 0 | 0.21 | 0.32 | 0.14 | 0.32 | 0.19 | 0.15 |
| 3 | 100763611 | rs4928106 | -0.555 | 0.442 | -0.046 | -0.497 | 0 | 0.015 | 0.015 | 0 | 0.82 | 0.8 | 0.87 | 0.68 | 0.83 | 0.87 |

| | | | | | | | | | | | | | | | | |
|---|---|---|---|---|---|---|---|---|---|---|---|---|---|---|---|---|
| 3 | 100764309 | rs2246096 | 0.259 | 0.251 | -0.321 | -0.56 | 0 | 0.003 | 0.003 | 0 | 0.21 | 0.21 | 0.11 | 0.14 | 0.17 | 0.15 |
| 3 | 100764857 | rs1868493 | 0.001 | 0.241 | -0.394 | -0.769 | 0.027 | 0.135 | 0.135 | 0.225 | 0.4 | 0.22 | 0.24 | 0.1 | 0.31 | 0.29 |
| 3 | 100768698 | rs16843185 | 0.937 | 0.282 | -0.145 | -0.614 | 0 | -0.013 | -0.013 | 0 | 0.22 | 0.18 | 0.14 | 0.03 | 0.17 | 0.15 |
| 3 | 100769942 | rs13088814 | 0.328 | 0.261 | 0.232 | -0.371 | -0.001 | -0.001 | -0.001 | 0.864 | 0.4 | 0.42 | 0.62 | 0.54 | 0.51 | 0.59 |
| 3 | 100771624 | rs4928108 | 1.53 | 0.292 | 0.028 | -0.425 | -0.011 | -0.042 | -0.042 | 0 | 0.24 | 0.14 | 0.44 | 0.4 | 0.3 | 0.33 |
| 3 | 100787768 | rs7430518 | -0.542 | 0.452 | 0.3 | -0.469 | -0.002 | -0.005 | -0.005 | 0.256 | 0.6 | 0.59 | 0.38 | 0.48 | 0.49 | 0.42 |
| 3 | 100788425 | rs12492516 | 0.495 | 0.441 | 0.321 | -0.474 | -0.009 | -0.018 | -0.018 | 0.067 | 0.39 | 0.34 | 0.61 | 0.52 | 0.51 | 0.58 |
| 3 | 100792449 | rs9835849 | -0.528 | 0.363 | 0.041 | -0.656 | -0.001 | -0.045 | -0.045 | 0 | 0.63 | 0.58 | 0.76 | 0.89 | 0.69 | 0.73 |
| 3 | 100793911 | rs6781428 | 0.84 | 0.346 | -0.308 | -0.563 | 0.001 | 0.003 | 0.003 | 0.078 | 0.38 | 0.38 | 0.62 | 0.53 | 0.5 | 0.57 |
| 3 | 100796642 | rs2317338 | -0.349 | 0.463 | -0.713 | -1.004 | -0.001 | 0.021 | 0.021 | 0 | 0.65 | 0.69 | 0.77 | 0.88 | 0.7 | 0.73 |
| 3 | 100799213 | rs9881220 | -0.329 | 0.419 | -0.741 | -1.005 | -0.004 | 0.082 | 0.082 | 0 | 0.64 | 0.77 | 0.77 | 0.88 | 0.7 | 0.73 |
| 3 | 100809267 | rs1349202 | -0.431 | 0.264 | -1.123 | -0.352 | 0.001 | 0.078 | 0.078 | 0 | 0.26 | 0.14 | 0.15 | 0.07 | 0.2 | 0.16 |
| 3 | 100818827 | rs1848252 | 0.258 | -0.348 | -0.817 | 0.422 | 0.021 | -0.012 | -0.012 | 0.89 | 0.31 | 0.29 | 0.4 | 0.39 | 0.21 | 0.34 |
| 3 | 100819495 | rs4518100 | -0.609 | -0.087 | -0.303 | 0.786 | 0 | 0.097 | 0.097 | 0.036 | 0.18 | 0.5 | 0.25 | 0.3 | 0.22 | 0.1 |
| 3 | 100820506 | rs10048990 | -0.886 | 0.002 | 0.137 | 0.342 | 0.076 | -0.03 | -0.03 | 0 | 0.09 | 0.43 | 0.12 | 0 | 0.11 | 0.05 |
| 3 | 100825306 | rs13071772 | -0.586 | 0.109 | 0.172 | 0.388 | 0.098 | -0.009 | -0.009 | 0 | 0.08 | 0.44 | 0.12 | 0 | 0.11 | 0.05 |
| 3 | 100825521 | rs9872728 | -0.617 | 0.104 | 0.169 | 0.386 | 0.096 | -0.011 | -0.011 | 0 | 0.08 | 0.44 | 0.12 | 0 | 0.11 | 0.05 |
| 3 | 100834385 | rs7615517 | -0.388 | -0.15 | -0.159 | 0.457 | 0.017 | 0.006 | 0.006 | 0 | 0.37 | 0.7 | 0.51 | 0.37 | 0.3 | 0.35 |
| 3 | 100834729 | rs2036650 | 0.235 | -0.175 | 0.251 | 0.42 | 0.01 | -0.005 | -0.005 | 0 | 0.3 | 0.3 | 0.4 | 0.37 | 0.2 | 0.31 |
| 3 | 100834876 | rs4234488 | -0.41 | -0.146 | -0.025 | 0.429 | 0.019 | 0.009 | 0.009 | 0 | 0.38 | 0.7 | 0.51 | 0.37 | 0.3 | 0.37 |
| 3 | 100839483 | rs1507434 | -0.437 | -0.198 | -0.092 | 0.432 | 0.03 | 0.015 | 0.015 | 0 | 0.38 | 0.75 | 0.51 | 0.37 | 0.3 | 0.39 |
| 3 | 100841396 | rs10470492 | -0.427 | 0.069 | 0.597 | 0.341 | 0.094 | 0.026 | 0.026 | 0 | 0.06 | 0.36 | 0.08 | 0 | 0.07 | 0.03 |
| 3 | 100843147 | rs9290311 | -0.446 | -0.119 | 0.22 | 0.37 | -0.013 | -0.011 | -0.011 | 0 | 0.44 | 0.51 | 0.59 | 0.65 | 0.4 | 0.45 |
| 3 | 100843711 | rs12696270 | 0.479 | 0.746 | -0.291 | 0.301 | -0.038 | -0.036 | -0.036 | 0 | 0.13 | 0.04 | 0.07 | 0.23 | 0.18 | 0.23 |
| 3 | 100844145 | rs9869642 | 0.354 | 0.096 | -0.161 | 0.544 | -0.028 | 0.01 | 0.01 | 0.658 | 0.31 | 0.46 | 0.44 | 0.36 | 0.22 | 0.33 |
| 3 | 100844373 | rs6784790 | 0.898 | 0.683 | -0.086 | 0.945 | 0.045 | 0.1 | 0.1 | 0.054 | 0.07 | 0.13 | 0.18 | 0.23 | 0.21 | 0.08 |
| 3 | 100850928 | rs6790241 | 1.066 | 0.691 | -0.104 | 0.956 | 0.039 | 0.094 | 0.094 | 0.067 | 0.07 | 0.12 | 0.18 | 0.24 | 0.21 | 0.08 |
| 3 | 100852656 | rs1395787 | -1.01 | 0.668 | -0.462 | 0.48 | -0.015 | 0.182 | 0.182 | 0 | 0.58 | 0.82 | 0.73 | 0.89 | 0.62 | 0.64 |
| 3 | 100854588 | rs6441582 | -0.481 | 0.579 | 0.144 | 0.615 | 0.025 | 0.096 | 0.096 | 0.547 | 0.25 | 0.39 | 0.25 | 0.49 | 0.37 | 0.28 |
| 3 | 100861030 | rs9847830 | 0.595 | 0.369 | 0.104 | 0.618 | -0.008 | -0.03 | -0.03 | 0.051 | 0.16 | 0.05 | 0.06 | 0.26 | 0.16 | 0.2 |
| 3 | 100864107 | rs1004115 | 0.317 | 0.517 | 0.316 | 0.626 | -0.013 | 0.024 | 0.024 | 0.611 | 0.31 | 0.43 | 0.47 | 0.39 | 0.24 | 0.36 |
| 3 | 100871443 | rs6441583 | 0.641 | 0.154 | 0.346 | 0.044 | -0.015 | -0.07 | -0.07 | 0 | 0.18 | 0.26 | 0.14 | 0.03 | 0.14 | 0.14 |
| 3 | 100872536 | rs13073439 | 0.747 | 0.246 | -0.372 | 0 | -0.01 | 0.019 | 0.019 | 0 | 0.24 | 0.15 | 0.14 | 0.19 | 0.21 | 0.2 |
| 3 | 100878496 | rs13099833 | -0.19 | 0.278 | -0.218 | 0.136 | 0 | -0.001 | -0.001 | 0 | 0.5 | 0.55 | 0.62 | 0.65 | 0.6 | 0.48 |
| 3 | 100879020 | rs10936540 | -0.821 | 0.142 | -0.339 | 0.079 | -0.002 | 0.076 | 0.076 | 0 | 0.66 | 0.93 | 0.81 | 0.76 | 0.71 | 0.61 |
| 3 | 100885183 | rs9815994 | -0.837 | 0.148 | -0.337 | 0.08 | -0.003 | 0.061 | 0.061 | 0 | 0.66 | 0.93 | 0.81 | 0.73 | 0.71 | 0.6 |
| 3 | 100886622 | rs6771745 | -0.675 | 0.138 | -0.214 | 0.077 | 0.006 | 0.017 | 0.017 | 0 | 0.64 | 0.75 | 0.8 | 0.73 | 0.71 | 0.6 |
| 3 | 100891071 | rs1905561 | 0.58 | 0.353 | -0.336 | 0.195 | 0.012 | 0.037 | 0.037 | 0 | 0.36 | 0.16 | 0.26 | 0.3 | 0.3 | 0.36 |
| 3 | 100892054 | rs1484231 | 0.58 | 0.353 | -0.336 | 0.195 | 0.012 | 0.037 | 0.037 | 0 | 0.36 | 0.16 | 0.26 | 0.3 | 0.3 | 0.36 |
| 3 | 100892487 | rs7630546 | -0.497 | 0.548 | -0.318 | 0.249 | -0.015 | 0.046 | 0.046 | 0 | 0.6 | 0.71 | 0.63 | 0.79 | 0.61 | 0.63 |
| 3 | 100895753 | rs4683862 | -1.037 | 0.631 | -0.599 | 0.174 | -0.025 | 0.054 | 0.054 | 0 | 0.73 | 0.85 | 0.81 | 0.88 | 0.73 | 0.79 |
| 3 | 100897035 | rs1354867 | -0.344 | 0.54 | -0.132 | 0.352 | 0 | -0.003 | -0.003 | 0 | 0.37 | 0.6 | 0.55 | 0.64 | 0.43 | 0.43 |
| 3 | 100905910 | rs906380 | 0.126 | 0.431 | -0.384 | 0.209 | -0.02 | -0.02 | -0.02 | 0.096 | 0.37 | 0.24 | 0.48 | 0.59 | 0.47 | 0.44 |
| 3 | 100908066 | rs348862 | -0.94 | 0.345 | -0.62 | 0.041 | -0.033 | 0.069 | 0.069 | 0 | 0.71 | 0.85 | 0.77 | 0.86 | 0.71 | 0.77 |
| 3 | 100912701 | rs4270483 | -1.214 | 0.592 | -0.45 | -0.003 | -0.001 | 0.016 | 0.016 | 0 | 0.74 | 0.79 | 0.83 | 0.86 | 0.81 | 0.79 |
| 3 | 100915962 | rs1471738 | 0.18 | 0.418 | 0.009 | -0.02 | 0.015 | -0.001 | -0.001 | 0 | 0.4 | 0.46 | 0.55 | 0.59 | 0.57 | 0.45 |
| 3 | 100923230 | rs6762738 | 0.636 | 0.378 | 0.121 | -0.014 | 0.011 | 0.006 | 0.006 | 0 | 0.33 | 0.28 | 0.28 | 0.27 | 0.24 | 0.34 |
| 3 | 100924203 | rs17400307 | 1.745 | 0.325 | 0.098 | -0.052 | 0 | 0 | 0 | 0 | 0.13 | 0.13 | 0.17 | 0.1 | 0.12 | 0.15 |
| 3 | 100939516 | rs10460874 | 1.45 | 0.45 | -0.006 | 0.662 | -0.004 | -0.005 | -0.005 | 0 | 0.14 | 0.13 | 0.3 | 0.32 | 0.17 | 0.2 |
| 3 | 100949842 | rs348867 | -1.27 | 0.408 | -0.115 | 0.808 | -0.001 | 0.02 | 0.02 | 0 | 0.73 | 0.83 | 0.82 | 0.82 | 0.81 | 0.78 |
| 3 | 100953212 | rs513154 | -0.588 | 0.326 | 0.201 | 0.819 | -0.008 | -0.011 | -0.011 | 0.004 | 0.54 | 0.49 | 0.71 | 0.66 | 0.68 | 0.6 |
| 3 | 100963154 | rs571391 | -0.548 | 0.3 | -0.027 | 0.807 | 0.023 | 0.03 | 0.03 | 0 | 0.53 | 0.62 | 0.72 | 0.65 | 0.69 | 0.59 |
| 3 | 100970240 | rs9870114 | 1 | 0.072 | -0.46 | 0.625 | -0.169 | -0.106 | -0.106 | 0 | 0.28 | 0.02 | 0.3 | 0.34 | 0.5 | 0.28 |
| 3 | 100970920 | rs348881 | -0.52 | 0.304 | -0.072 | 0.768 | 0.024 | 0.041 | 0.041 | 0 | 0.54 | 0.68 | 0.72 | 0.68 | 0.69 | 0.6 |
| 3 | 100978288 | rs348877 | -1.273 | 0.316 | -0.191 | 0.756 | -0.002 | 0.028 | 0.028 | 0 | 0.73 | 0.85 | 0.83 | 0.85 | 0.8 | 0.78 |
| 3 | 100980849 | rs35902489 | 1.637 | 0.291 | -0.279 | 0.812 | 0.023 | 0.014 | 0.014 | 0 | 0.19 | 0.11 | 0.1 | 0.17 | 0.12 | 0.17 |
| 3 | 100991546 | rs3773920 | 1.412 | 0.453 | -0.072 | 0.985 | 0 | 0.019 | 0.019 | 0 | 0.14 | 0.17 | 0.3 | 0.33 | 0.17 | 0.2 |
| 3 | 100994497 | rs533852 | 0.396 | 0.368 | 0.098 | 0.809 | -0.047 | 0.006 | 0.006 | 0 | 0.38 | 0.19 | 0.4 | 0.35 | 0.51 | 0.4 |
| 3 | 101005268 | rs482436 | -0.763 | 0.119 | 0.358 | 0.564 | -0.031 | -0.077 | -0.077 | 0.874 | 0.68 | 0.57 | 0.83 | 0.83 | 0.8 | 0.79 |
| 3 | 101022241 | rs475521 | -0.75 | 0.124 | 0.453 | 0.565 | -0.033 | -0.079 | -0.079 | 0 | 0.68 | 0.56 | 0.84 | 0.83 | 0.8 | 0.79 |
| 3 | 101042594 | rs7431086 | 0.429 | 0.501 | 0.34 | 0.582 | -0.005 | -0.011 | -0.011 | 0 | 0.34 | 0.35 | 0.55 | 0.48 | 0.41 | 0.4 |
| 3 | 101043565 | rs497953 | 1.065 | 0.505 | 0.344 | 0.574 | -0.017 | -0.006 | -0.006 | 0 | 0.34 | 0.18 | 0.28 | 0.34 | 0.39 | 0.36 |
| 3 | 101058030 | rs2553422 | -1.914 | 0.474 | 0.558 | 0.522 | 0.012 | -0.065 | -0.065 | 0 | 0.88 | 0.82 | 0.94 | 1 | 0.92 | 0.96 |
| 3 | 101068620 | rs2553419 | 0.291 | 0.455 | 0.526 | 0.519 | -0.005 | -0.004 | -0.004 | 0 | 0.54 | 0.32 | 0.38 | 0.52 | 0.5 | 0.56 |
| 3 | 101071429 | rs9870022 | 1.66 | 0.523 | 0.535 | 0.569 | -0.004 | -0.007 | -0.007 | 0 | 0.15 | 0.15 | 0.38 | 0.33 | 0.17 | 0.2 |
| 3 | 101080694 | rs7616677 | 0.467 | 0.563 | 0.551 | 0.667 | -0.005 | -0.01 | -0.01 | 0.004 | 0.33 | 0.35 | 0.55 | 0.49 | 0.41 | 0.4 |
| 3 | 101085413 | rs939443 | 0.467 | 0.563 | 0.551 | 0.629 | -0.005 | -0.011 | -0.011 | 0 | 0.33 | 0.35 | 0.55 | 0.48 | 0.41 | 0.4 |
| 3 | 101106527 | rs2553427 | 1.134 | 0.554 | 0.568 | 0.624 | -0.015 | -0.004 | -0.004 | 0 | 0.34 | 0.18 | 0.28 | 0.34 | 0.39 | 0.36 |
| 3 | 101120030 | rs11710016 | 1.965 | 0.559 | 0.562 | 0.626 | -0.066 | -0.036 | -0.036 | 0.14 | 0.17 | 0.02 | 0.16 | 0.15 | 0.24 | 0.2 |

| | | | | | | | | | | | | | | | | |
|---|---|---|---|---|---|---|---|---|---|---|---|---|---|---|---|---|
| 3 | 101160339 | rs10936621 | 0.454 | 0.581 | 0.579 | 0.578 | -0.005 | -0.011 | -0.011 | 0 | 0.33 | 0.35 | 0.55 | 0.48 | 0.41 | 0.4 |
| 3 | 101220400 | rs3846092 | 1.299 | 0.52 | 0.422 | 0.46 | 0.015 | -0.087 | -0.087 | 0 | 0.13 | 0.3 | 0.07 | 0 | 0.09 | 0.04 |
| 3 | 101231731 | rs11708616 | 1.622 | 0.654 | 0.29 | 0.691 | -0.002 | 0 | 0 | 1.224 | 0.19 | 0.19 | 0.25 | 0.18 | 0.24 | 0.2 |
| 3 | 101231921 | rs9682313 | 0.408 | 0.645 | 0.31 | 0.595 | -0.009 | -0.013 | -0.013 | 0 | 0.34 | 0.34 | 0.62 | 0.52 | 0.41 | 0.4 |
| 3 | 101232736 | rs6772703 | -1.168 | 0.702 | 0.141 | 0.54 | -0.012 | 0.011 | 0.011 | 0 | 0.66 | 0.81 | 0.79 | 0.7 | 0.61 | 0.64 |
| 3 | 101237890 | rs7623537 | 2.08 | 0.802 | -0.047 | 0.747 | -0.01 | -0.008 | -0.008 | 0 | 0.12 | 0.08 | 0.24 | 0.11 | 0.12 | 0.15 |
| 3 | 101279006 | rs7641261 | 1.467 | 0.858 | 0.33 | 1.096 | -0.019 | -0.022 | -0.022 | 0.895 | 0.18 | 0.15 | 0.25 | 0.24 | 0.24 | 0.19 |
| 3 | 101280710 | rs4683930 | -0.982 | 0.721 | 0.801 | 0.934 | 0.032 | -0.105 | -0.105 | 0 | 0.86 | 0.53 | 0.93 | 1 | 0.92 | 0.96 |
| 3 | 101281534 | rs4618204 | 0.136 | 0.688 | 0.828 | 0.873 | -0.001 | -0.002 | -0.002 | 0 | 0.41 | 0.38 | 0.48 | 0.59 | 0.44 | 0.48 |
| 3 | 101281665 | rs13072301 | 0.867 | 0.697 | 0.787 | 0.884 | -0.009 | 0 | 0 | 0.569 | 0.46 | 0.21 | 0.44 | 0.41 | 0.48 | 0.49 |
| 3 | 101285420 | rs1976944 | -0.278 | 0.674 | 0.803 | 0.854 | -0.017 | -0.081 | -0.081 | 2.79 | 0.67 | 0.48 | 0.82 | 0.88 | 0.77 | 0.75 |
| 3 | 101289589 | rs6788864 | -1.218 | 0.745 | 0.785 | 0.874 | 0.018 | -0.088 | -0.088 | 0.325 | 0.87 | 0.68 | 0.93 | 1 | 0.92 | 0.96 |
| 3 | 101294572 | rs3804778 | 1.811 | 0.788 | 0.713 | 1.05 | -0.074 | -0.103 | -0.103 | 0 | 0.17 | 0.01 | 0.23 | 0.27 | 0.24 | 0.19 |
| 3 | 101308776 | rs6777292 | -0.813 | 0.86 | 0.939 | 1.021 | 0.045 | 0.027 | 0.027 | 0 | 0.84 | 0.56 | 0.8 | 0.8 | 0.87 | 0.89 |
| 3 | 101333441 | rs12696325 | -1.311 | 0.863 | 0.672 | 0.94 | 0.011 | -0.075 | -0.075 | 0 | 0.87 | 0.74 | 0.94 | 1 | 0.92 | 0.96 |
| 3 | 101334609 | rs12638134 | 0.258 | 0.89 | 0.427 | 1.114 | 0.001 | -0.002 | -0.002 | 0 | 0.39 | 0.36 | 0.36 | 0.39 | 0.39 | 0.4 |
| 3 | 101349644 | rs6441620 | -0.105 | 0.811 | 0.353 | 1.146 | 0.003 | -0.005 | -0.005 | 0 | 0.45 | 0.54 | 0.53 | 0.62 | 0.51 | 0.53 |
| 3 | 101373305 | rs13069443 | 1.113 | 0.812 | 0.365 | 1.351 | 0.002 | 0.004 | 0.004 | 0.406 | 0.23 | 0.26 | 0.29 | 0.29 | 0.33 | 0.21 |
| 3 | 101378476 | rs2288272 | 1.195 | 0.805 | 0.408 | 1.348 | -0.015 | -0.017 | -0.017 | 0 | 0.22 | 0.22 | 0.29 | 0.29 | 0.33 | 0.21 |
| 3 | 101383321 | rs13081846 | 1.123 | 0.925 | 0.409 | 1.401 | 0.002 | 0.004 | 0.004 | 0.4 | 0.23 | 0.26 | 0.29 | 0.29 | 0.33 | 0.21 |
| 3 | 101399126 | rs17411983 | 1.123 | 0.925 | 0.409 | 1.142 | 0.002 | 0.004 | 0.004 | 0.4 | 0.23 | 0.26 | 0.29 | 0.29 | 0.33 | 0.21 |
| 3 | 101404813 | rs2303473 | 0.647 | 0.74 | 0.424 | 0.96 | 0.027 | 0.03 | 0.03 | 0.53 | 0.43 | 0.18 | 0.39 | 0.37 | 0.4 | 0.42 |
| 3 | 101409665 | rs6794860 | -1.759 | 0.682 | 0.611 | 0.794 | 0.005 | -0.044 | -0.044 | 0.166 | 0.9 | 0.85 | 0.94 | 1 | 0.92 | 0.96 |
| 3 | 101445118 | rs2554962 | 1.257 | 0.694 | 0.623 | 0.913 | 0.005 | -0.029 | -0.029 | 0 | 0.22 | 0.16 | 0.24 | 0.48 | 0.19 | 0.34 |
| 3 | 101453858 | rs11914318 | 1.292 | 0.717 | 0.742 | 0.882 | 0.018 | 0.028 | 0.028 | 0.349 | 0.23 | 0.31 | 0.3 | 0.29 | 0.34 | 0.21 |
| 3 | 101457391 | rs2466369 | -1.071 | 0.528 | 0.615 | 0.711 | 0 | -0.011 | -0.011 | 0.27 | 0.86 | 0.88 | 0.93 | 0.99 | 0.9 | 0.83 |
| 3 | 101463895 | rs1031710 | 1.402 | 0.533 | 0.617 | 0.714 | 0.018 | 0.028 | 0.028 | 0.349 | 0.23 | 0.31 | 0.3 | 0.29 | 0.34 | 0.21 |
| 3 | 101467239 | rs9845802 | 0.908 | 0.489 | 0.552 | 0.701 | 0.019 | 0.101 | 0.101 | 0.618 | 0.4 | 0.17 | 0.38 | 0.22 | 0.37 | 0.29 |
| 3 | 101484335 | rs2625282 | -0.952 | 0.593 | 0.632 | 0.707 | 0 | -0.01 | -0.01 | 0 | 0.86 | 0.88 | 0.93 | 0.99 | 0.9 | 0.83 |
| 3 | 101486050 | rs6764700 | 1.626 | 0.621 | 0.641 | 0.713 | 0.001 | -0.026 | -0.026 | 0 | 0.1 | 0.1 | 0.05 | 0 | 0.08 | 0.04 |
| 3 | 101489629 | rs13099674 | 1.038 | 0.396 | 0.625 | 0.801 | 0.058 | 0.085 | 0.085 | 0.649 | 0.24 | 0.5 | 0.31 | 0.29 | 0.34 | 0.21 |
| 3 | 101497569 | rs1460097 | 1.297 | 0.439 | 0.504 | 0.809 | 0.017 | 0.02 | 0.02 | 0.649 | 0.23 | 0.3 | 0.29 | 0.27 | 0.33 | 0.21 |
| 3 | 101497925 | rs6774950 | 1.647 | 0.547 | 0.394 | 0.763 | -0.003 | -0.035 | -0.035 | 7.435 | 0.09 | 0.12 | 0.05 | 0 | 0.06 | 0.04 |
| 3 | 101501827 | rs2466367 | -0.159 | 0.237 | 0.103 | 0.081 | 0.003 | 0.001 | 0.001 | 0 | 0.51 | 0.52 | 0.59 | 0.76 | 0.62 | 0.55 |
| 3 | 101503765 | rs4342086 | 1.69 | 0.305 | -0.013 | 0.111 | -0.002 | -0.001 | -0.001 | 0 | 0.22 | 0.25 | 0.28 | 0.27 | 0.33 | 0.2 |
| 3 | 101504624 | rs2625289 | -1.245 | 0.29 | -0.297 | 0.101 | 0.001 | -0.02 | -0.02 | 0.018 | 0.86 | 0.85 | 0.94 | 1 | 0.91 | 0.83 |
| 3 | 101504674 | rs17347156 | 1.597 | 0.532 | -0.65 | 0.338 | -0.024 | -0.027 | -0.027 | 0 | 0.21 | 0.19 | 0.28 | 0.27 | 0.33 | 0.2 |
| 3 | 101507015 | rs11916199 | 0.715 | 0.227 | -0.771 | 0.482 | 0.007 | 0.044 | 0.044 | 0 | 0.33 | 0.15 | 0.33 | 0.24 | 0.3 | 0.29 |
| 3 | 101545319 | rs2625288 | 0.496 | 0.606 | -1.135 | 1.313 | 0.003 | -0.037 | -0.037 | 0 | 0.21 | 0.13 | 0.25 | 0.48 | 0.2 | 0.35 |
| 3 | 101552553 | rs11718446 | 0.646 | 0.871 | -1.101 | 0.929 | 0.002 | 0.006 | 0.006 | 0 | 0.23 | 0.23 | 0.25 | 0.27 | 0.28 | 0.2 |
| 3 | 101556546 | rs7644388 | -0.12 | 0.723 | -1.495 | 0.402 | -0.036 | 0.024 | 0.024 | 0 | 0.2 | 0.09 | 0.15 | 0.13 | 0.26 | 0.19 |
| 3 | 101569726 | rs616597 | -0.416 | 0.017 | -1.821 | 0.596 | 0.004 | -0.115 | -0.115 | 0.117 | 0.74 | 0.95 | 0.8 | 0.6 | 0.78 | 0.83 |
| 3 | 101574503 | rs677011 | 0.152 | -0.186 | -0.575 | 0.368 | -0.014 | -0.002 | -0.002 | 0 | 0.56 | 0.49 | 0.69 | 0.44 | 0.66 | 0.49 |
| 3 | 101576175 | rs14134 | 0.05 | -0.127 | -0.605 | 0.326 | -0.006 | -0.002 | -0.002 | 0 | 0.55 | 0.5 | 0.68 | 0.44 | 0.63 | 0.48 |
| 3 | 101584469 | rs683403 | 0.405 | -0.025 | -0.85 | 0.202 | 0 | 0.003 | 0.003 | 0 | 0.29 | 0.24 | 0.47 | 0.3 | 0.28 | 0.29 |
| 3 | 101584983 | rs9844218 | 1.515 | 0.008 | -0.888 | 0.152 | -0.044 | 0.015 | 0.015 | 0 | 0.22 | 0.11 | 0.19 | 0.13 | 0.32 | 0.19 |
| 3 | 101585467 | rs688058 | 0.169 | 0.186 | -0.721 | -0.209 | 0.03 | 0.002 | 0.002 | 0 | 0.51 | 0.34 | 0.66 | 0.47 | 0.6 | 0.48 |
| 3 | 101587566 | rs641432 | 0.649 | 0.151 | -0.637 | -0.175 | -0.028 | -0.016 | -0.016 | 5.381 | 0.36 | 0.22 | 0.28 | 0.37 | 0.43 | 0.35 |
| 3 | 101592286 | rs617688 | 0.593 | 0.242 | -0.23 | -0.17 | 0.01 | 0.009 | 0.009 | 0 | 0.36 | 0.5 | 0.29 | 0.38 | 0.41 | 0.34 |
| 3 | 101594402 | rs650567 | 0.761 | 0.277 | -0.318 | -0.172 | -0.001 | -0.001 | -0.001 | 0 | 0.35 | 0.35 | 0.27 | 0.37 | 0.41 | 0.34 |
| 3 | 101600028 | rs665211 | 0.271 | 0.177 | -0.23 | -0.628 | -0.001 | -0.001 | -0.001 | 0 | 0.4 | 0.37 | 0.31 | 0.47 | 0.46 | 0.42 |
| 3 | 101600724 | rs651907 | 0.316 | 0.188 | -0.263 | -0.623 | 0.005 | 0.004 | 0.004 | 0 | 0.41 | 0.53 | 0.31 | 0.47 | 0.46 | 0.43 |
| 3 | 101601159 | rs3846094 | -0.121 | 0.011 | -0.33 | -0.886 | 0.009 | 0.044 | 0.044 | 0 | 0.44 | 0.34 | 0.47 | 0.27 | 0.35 | 0.51 |
| 3 | 101602954 | rs10936815 | 0.402 | 0.25 | -0.191 | -0.676 | -0.035 | 0.074 | 0.074 | 0 | 0.21 | 0.08 | 0.4 | 0.08 | 0.24 | 0.14 |
| 3 | 101603126 | rs6794896 | 0.195 | 0.284 | -0.085 | -0.636 | 0.078 | 0.08 | 0.08 | 0 | 0.05 | 0.3 | 0.05 | 0.03 | 0.05 | 0.08 |
| 3 | 101607164 | rs17842780 | 0.267 | 0.29 | -0.42 | -0.655 | 0.009 | -0.021 | -0.021 | 0 | 0.21 | 0.24 | 0.42 | 0.1 | 0.25 | 0.21 |
| 3 | 101620682 | rs771574 | 1.47 | 0.272 | -0.598 | -0.561 | -0.016 | -0.048 | -0.048 | 1.243 | 0.26 | 0.19 | 0.19 | 0.43 | 0.32 | 0.31 |
| 3 | 101630232 | rs10936822 | 0.515 | -0.02 | -0.624 | -0.496 | -0.018 | 0.058 | 0.058 | 0 | 0.19 | 0.07 | 0.34 | 0.08 | 0.21 | 0.13 |
| 3 | 101632783 | rs1672382 | -1.211 | -0.071 | -0.468 | -0.471 | -0.002 | 0.003 | 0.003 | 0.519 | 0.85 | 0.85 | 0.8 | 0.74 | 0.86 | 0.94 |
| 3 | 101635645 | rs1473857 | 0.874 | -0.105 | -0.43 | -0.537 | 0.017 | 0.054 | 0.054 | 0 | 0.3 | 0.19 | 0.17 | 0.12 | 0.19 | 0.35 |
| 3 | 101636054 | rs1672376 | -1.255 | -0.087 | -0.401 | -0.517 | 0 | 0 | 0 | 0.519 | 0.85 | 0.83 | 0.8 | 0.74 | 0.86 | 0.94 |
| 3 | 101645933 | rs1707606 | -0.064 | -0.076 | -0.299 | -0.536 | -0.009 | -0.01 | -0.01 | 0 | 0.54 | 0.48 | 0.62 | 0.6 | 0.67 | 0.59 |
| 3 | 101646079 | rs1616007 | -0.027 | 0.015 | -0.564 | -0.496 | 0.026 | 0.027 | 0.027 | 0 | 0.55 | 0.64 | 0.63 | 0.6 | 0.67 | 0.59 |
| 3 | 101654298 | rs2951443 | 0.714 | 0.222 | -1.088 | -0.297 | -0.036 | -0.038 | -0.038 | 0.842 | 0.34 | 0.21 | 0.27 | 0.52 | 0.44 | 0.39 |
| 3 | 101658671 | rs2943301 | -0.905 | 0.189 | -1.099 | -0.4 | 0.006 | -0.034 | -0.034 | 0 | 0.8 | 0.87 | 0.76 | 0.65 | 0.78 | 0.86 |
| 3 | 101658945 | rs1456200 | -0.185 | 0.217 | -1.091 | -0.414 | 0.005 | 0.083 | 0.083 | 0 | 0.51 | 0.61 | 0.53 | 0.22 | 0.42 | 0.54 |
| 3 | 101662803 | rs9857771 | 0.233 | 0.444 | -0.975 | -0.126 | -0.009 | -0.035 | -0.035 | 0 | 0.35 | 0.39 | 0.2 | 0.17 | 0.24 | 0.42 |
| 3 | 101663317 | rs9881690 | 0.03 | 1.108 | -1.246 | 0.539 | -0.016 | -0.035 | -0.035 | 0 | 0.29 | 0.22 | 0.22 | 0.43 | 0.37 | 0.31 |
| 3 | 101668885 | rs13059489 | -0.085 | 0.944 | -1.138 | 0.384 | -0.002 | 0.001 | 0.001 | 0 | 0.27 | 0.24 | 0.23 | 0.26 | 0.3 | 0.4 |

| | | | | | | | | | | | | | | | | |
|---|---|---|---|---|---|---|---|---|---|---|---|---|---|---|---|---|
| 3 | 101670259 | rs1828669 | -0.107 | 0.973 | -1.592 | 0.428 | -0.019 | -0.004 | -0.004 | 0 | 0.64 | 0.84 | 0.68 | 0.67 | 0.62 | 0.65 |
| 3 | 101671755 | rs1707574 | 0.497 | 1.064 | -0.815 | -0.133 | 0.008 | -0.01 | -0.01 | 0.293 | 0.81 | 0.73 | 0.67 | 0.82 | 0.76 | 0.75 |
| 3 | 101673386 | rs9848728 | 0.42 | 0.639 | -0.715 | -0.768 | -0.014 | -0.043 | -0.043 | 0 | 0.4 | 0.47 | 0.28 | 0.12 | 0.35 | 0.22 |
| 3 | 101679080 | rs1606462 | 0.125 | 0.927 | -0.95 | -0.476 | 0.009 | -0.001 | -0.001 | 0 | 0.73 | 0.82 | 0.75 | 0.71 | 0.71 | 0.61 |
| 3 | 101681481 | rs7626110 | 0.99 | 0.584 | -0.826 | -0.802 | 0.008 | 0.024 | 0.024 | 0 | 0.27 | 0.17 | 0.22 | 0.11 | 0.27 | 0.16 |
| 3 | 101691814 | rs771829 | -0.825 | 1.044 | -0.18 | -0.574 | 0.001 | -0.006 | -0.006 | 0 | 0.12 | 0.14 | 0.13 | 0.06 | 0.15 | 0.14 |
| 3 | 101692131 | rs771828 | -0.786 | 1.043 | -0.181 | -0.558 | 0.001 | -0.004 | -0.004 | 0 | 0.11 | 0.13 | 0.13 | 0.07 | 0.15 | 0.14 |
| 3 | 101697208 | rs1514471 | 1.347 | 1.15 | -0.218 | -0.795 | 0.033 | 0.032 | 0.032 | 2.927 | 0.44 | 0.28 | 0.25 | 0.35 | 0.34 | 0.26 |
| 3 | 101697679 | rs1513120 | -0.133 | 1.128 | 0.108 | -0.893 | -0.01 | -0.01 | -0.01 | 0 | 0.32 | 0.27 | 0.48 | 0.59 | 0.41 | 0.39 |
| 3 | 101705550 | rs708233 | -1.005 | 1.095 | 0.322 | -0.929 | -0.001 | -0.009 | -0.009 | 0.347 | 0.45 | 0.52 | 0.61 | 0.63 | 0.56 | 0.55 |
| 3 | 101705763 | rs13077965 | 0.988 | 1.184 | 0.348 | -0.898 | -0.006 | 0.003 | 0.003 | 0 | 0.05 | 0.03 | 0.04 | 0 | 0.07 | 0.02 |
| 3 | 101705809 | rs771785 | -0.781 | 1.261 | 0.335 | -0.936 | 0.008 | -0.003 | -0.003 | 0.095 | 0.39 | 0.41 | 0.57 | 0.63 | 0.5 | 0.51 |
| 3 | 101712056 | rs17349394 | 0.805 | 1.235 | 0.242 | -0.931 | -0.001 | -0.05 | -0.05 | 0.589 | 0.11 | 0.18 | 0.13 | 0.01 | 0.1 | 0.21 |
| 3 | 101712255 | rs771788 | 0.323 | 1.235 | 0.226 | -0.911 | 0 | -0.007 | -0.007 | 0 | 0.06 | 0.15 | 0.09 | 0.04 | 0.08 | 0.12 |
| 3 | 101712739 | rs12639221 | 0.143 | 1.26 | 0.182 | -0.916 | -0.002 | -0.002 | -0.002 | 0 | 0.28 | 0.27 | 0.45 | 0.59 | 0.35 | 0.37 |
| 3 | 101714180 | rs771790 | 1.282 | 1.197 | 0.188 | -0.86 | 0.042 | 0.048 | 0.048 | 3.206 | 0.5 | 0.33 | 0.34 | 0.39 | 0.42 | 0.35 |
| 3 | 101719377 | rs771798 | -0.681 | 1.284 | 0.219 | -0.901 | 0.044 | 0.127 | 0.127 | 0.855 | 0.78 | 0.54 | 0.56 | 0.44 | 0.69 | 0.67 |
| 3 | 101720536 | rs1081617 | 0.39 | 1.291 | 0.186 | -0.92 | 0.005 | 0.009 | 0.009 | 2.078 | 0.65 | 0.46 | 0.43 | 0.43 | 0.59 | 0.47 |
| 3 | 101721005 | rs9848014 | -0.656 | 1.252 | 0.048 | -1.047 | -0.049 | 0.063 | 0.063 | 0 | 0.14 | 0.04 | 0.09 | 0 | 0.17 | 0.11 |
| 3 | 101723923 | rs2943306 | -0.357 | 1.23 | 0.027 | -1.061 | 0.005 | 0.032 | 0.032 | 0 | 0.17 | 0.11 | 0.21 | 0.04 | 0.17 | 0.32 |
| 3 | 101735647 | rs771757 | 0.865 | 1.236 | 0.11 | -1.051 | 0.034 | -0.096 | -0.096 | 0 | 0.83 | 0.66 | 0.89 | 1 | 0.83 | 0.89 |
| 3 | 101748638 | rs771767 | 1.29 | 1.215 | 0.061 | -1.048 | 0.012 | -0.119 | -0.119 | 0 | 0.71 | 0.59 | 0.76 | 0.99 | 0.73 | 0.69 |
| 3 | 101755738 | rs1398607 | 1.189 | 1.127 | 0.234 | -0.852 | 0.004 | -0.113 | -0.113 | 0 | 0.74 | 0.55 | 0.73 | 0.96 | 0.76 | 0.65 |
| 3 | 101756221 | rs10804863 | 1.189 | 1.127 | 0.234 | -0.852 | 0.004 | -0.113 | -0.113 | 0.005 | 0.74 | 0.55 | 0.73 | 0.96 | 0.76 | 0.65 |
| 3 | 101765067 | rs1513121 | -0.519 | 1.159 | 0.089 | -0.843 | -0.005 | -0.004 | -0.004 | 0 | 0.18 | 0.14 | 0.17 | 0.01 | 0.16 | 0.23 |
| 3 | 101765100 | rs11717579 | 1.109 | 1.186 | -0.004 | -0.764 | 0.001 | 0.043 | 0.043 | 0.316 | 0.53 | 0.29 | 0.3 | 0.39 | 0.45 | 0.32 |
| 3 | 101766211 | rs12696408 | 0.858 | 0.687 | 0.109 | -0.554 | 0.003 | -0.007 | -0.007 | 0 | 0.59 | 0.53 | 0.36 | 0.39 | 0.52 | 0.34 |
| 3 | 101773841 | rs812220 | 0.103 | 0.669 | 0.641 | -0.114 | 0.018 | 0.077 | 0.077 | 0.319 | 0.76 | 0.47 | 0.57 | 0.4 | 0.73 | 0.67 |
| 3 | 101774820 | rs771809 | -0.884 | 0.62 | 0.731 | -0.174 | -0.016 | 0.049 | 0.049 | 0 | 0.08 | 0.19 | 0.18 | 0.12 | 0.07 | 0.27 |
| 3 | 101787222 | rs6791238 | 0.9 | 0.681 | 0.459 | -0.254 | -0.015 | -0.02 | -0.02 | 0 | 0.8 | 0.54 | 0.74 | 0.88 | 0.82 | 0.67 |
| 3 | 101795138 | rs974059 | 0.168 | 0.726 | 0.093 | -0.131 | -0.012 | -0.025 | -0.025 | 1.854 | 0.21 | 0.16 | 0.4 | 0.62 | 0.26 | 0.34 |
| 3 | 101797959 | rs6770164 | -0.563 | 1.025 | -0.192 | -0.176 | 0.001 | -0.014 | -0.014 | 0 | 0.15 | 0.3 | 0.18 | 0.07 | 0.13 | 0.22 |
| 3 | 101805624 | rs2175927 | 0.56 | 0.928 | -1.381 | -0.439 | 0.006 | 0.072 | 0.072 | 0.026 | 0.57 | 0.3 | 0.38 | 0.28 | 0.59 | 0.43 |
| 3 | 101807592 | rs1877818 | 0.515 | 1.01 | -1.566 | -0.403 | 0.004 | 0.072 | 0.072 | 0.082 | 0.57 | 0.29 | 0.38 | 0.28 | 0.58 | 0.43 |
| 3 | 101810454 | rs7645815 | -0.424 | 1.113 | -1.827 | -0.496 | -0.025 | 0.089 | 0.089 | 0 | 0.36 | 0.19 | 0.48 | 0.84 | 0.4 | 0.45 |
| 3 | 101816344 | rs12486534 | 0.136 | 0.975 | -2.287 | -0.949 | -0.034 | 0.138 | 0.138 | 0.142 | 0.41 | 0.13 | 0.5 | 0.83 | 0.39 | 0.52 |
| 3 | 101823824 | rs6774331 | 0.15 | 0.85 | -1.498 | -1.017 | -0.019 | 0.125 | 0.125 | 0.266 | 0.41 | 0.15 | 0.5 | 0.82 | 0.39 | 0.52 |
| 3 | 101831122 | rs16844748 | 0.409 | 0.316 | -1.552 | -1.501 | 0.013 | 0.01 | 0.01 | 0 | 0.15 | 0.14 | 0.07 | 0.01 | 0.08 | 0.16 |
| 3 | 101834050 | rs6796074 | -0.937 | 0.582 | -0.976 | -1.871 | 0.021 | -0.083 | -0.083 | 0.011 | 0.84 | 0.6 | 0.87 | 0.98 | 0.79 | 0.91 |
| 3 | 101835194 | rs4634092 | 1.544 | 0.609 | -1.41 | -1.322 | -0.019 | -0.051 | -0.051 | 0 | 0.15 | 0.06 | 0.16 | 0.41 | 0.2 | 0.24 |
| 3 | 101835236 | rs17417560 | 1.314 | 0.82 | -1.364 | -1.689 | 0.009 | -0.008 | -0.008 | 0 | 0.19 | 0.25 | 0.24 | 0.15 | 0.22 | 0.14 |
| 3 | 101836229 | rs6441647 | 0.029 | 0.77 | -1.362 | -1.659 | -0.001 | -0.02 | -0.02 | 0 | 0.35 | 0.41 | 0.31 | 0.16 | 0.3 | 0.36 |
| 3 | 101841161 | rs6810085 | 1.713 | 0.854 | -1.287 | -1.467 | -0.003 | -0.004 | -0.004 | 0.226 | 0.08 | 0.05 | 0.07 | 0 | 0.1 | 0.04 |
| 3 | 101855907 | rs9844461 | 1.363 | 0.824 | -1.147 | -1.467 | 0.002 | -0.001 | -0.001 | 0.253 | 0.17 | 0.2 | 0.22 | 0.15 | 0.21 | 0.15 |
| 3 | 101860560 | rs938988 | -0.249 | 0.846 | -1.532 | -1.602 | -0.014 | -0.133 | -0.133 | 0 | 0.68 | 0.54 | 0.81 | 0.96 | 0.7 | 0.69 |
| 3 | 101860798 | rs938987 | 0.881 | 0.864 | -1.541 | -1.608 | -0.01 | 0.133 | 0.133 | 0 | 0.42 | 0.1 | 0.5 | 0.81 | 0.39 | 0.5 |
| 3 | 101863476 | rs2619298 | 0.488 | 0.98 | -1.271 | -1.844 | -0.001 | -0.06 | -0.06 | 0 | 0.6 | 0.46 | 0.75 | 0.96 | 0.61 | 0.65 |
| 3 | 101882411 | rs2619284 | -0.469 | 0.959 | -1.086 | -1.586 | 0.011 | 0.03 | 0.03 | 0 | 0.68 | 0.63 | 0.7 | 0.55 | 0.56 | 0.65 |
| 3 | 101890895 | rs2619283 | 0.322 | 0.937 | -1.096 | -1.26 | 0.002 | -0.001 | -0.001 | 0 | 0.45 | 0.43 | 0.58 | 0.54 | 0.38 | 0.4 |
| 3 | 101900270 | rs6801429 | -0.317 | 0.816 | -0.841 | -0.988 | -0.001 | 0 | 0 | 0.015 | 0.47 | 0.51 | 0.36 | 0.45 | 0.52 | 0.56 |
| 3 | 101901465 | rs4683874 | -0.225 | 0.586 | -0.512 | -1.288 | -0.011 | 0.045 | 0.045 | 0 | 0.43 | 0.3 | 0.35 | 0.15 | 0.5 | 0.45 |
| 3 | 101911058 | rs11927186 | -1.49 | 0.074 | 0.404 | -1.123 | -0.016 | -0.106 | -0.106 | 1.447 | 0.17 | 0.65 | 0.13 | 0 | 0.14 | 0.06 |
| 3 | 101922655 | rs11922137 | -1.486 | 0.171 | 0.807 | -1.092 | 0.042 | -0.056 | -0.056 | 0 | 0.11 | 0.48 | 0.09 | 0 | 0.1 | 0.03 |
| 3 | 101925787 | rs7628422 | -0.997 | 0.12 | 1.096 | -1.086 | 0.023 | -0.027 | -0.027 | 1.386 | 0.06 | 0.15 | 0.04 | 0 | 0.08 | 0.04 |
| 3 | 101943151 | rs13082870 | -2.054 | 0.123 | 0.992 | -1.085 | 0.035 | -0.042 | -0.042 | 0 | 0.1 | 0.44 | 0.07 | 0 | 0.08 | 0.04 |
| 3 | 101968917 | rs11720464 | -1.502 | -0.582 | 0.487 | -1.503 | -0.036 | -0.099 | -0.099 | 0 | 0.21 | 0.39 | 0.13 | 0.01 | 0.16 | 0.08 |
| 3 | 101988001 | rs7644557 | -0.719 | -0.679 | -0.631 | -2.234 | 0.063 | 0.2 | 0.2 | 0.482 | 0.51 | 0.76 | 0.35 | 0.05 | 0.37 | 0.24 |
| 3 | 101999972 | rs1600329 | -0.772 | -0.641 | -0.462 | -2.305 | 0.071 | 0.195 | 0.195 | 0.121 | 0.51 | 0.76 | 0.35 | 0.05 | 0.37 | 0.24 |
| 3 | 102000567 | rs1585772 | -0.735 | -0.637 | -0.516 | -2.303 | 0.052 | 0.103 | 0.103 | 0.14 | 0.5 | 0.67 | 0.34 | 0.05 | 0.37 | 0.24 |
| 3 | 102010586 | rs6441657 | -0.97 | -0.351 | -1.001 | -2.335 | 0.004 | 0.25 | 0.25 | 0.192 | 0.55 | 0.87 | 0.39 | 0.05 | 0.4 | 0.29 |
| 3 | 102036275 | rs9862862 | -1.494 | -0.245 | -0.114 | -2.201 | 0.028 | -0.023 | -0.023 | 0 | 0.07 | 0.22 | 0.06 | 0 | 0.05 | 0.05 |
| 3 | 102038560 | rs9815746 | -1.647 | -0.265 | 0.251 | -2.209 | 0.051 | -0.01 | -0.01 | 0 | 0.07 | 0.29 | 0.07 | 0 | 0.05 | 0.05 |
| 3 | 102058887 | rs7625204 | -0.423 | -0.937 | -0.257 | -2.802 | -0.023 | 0.129 | 0.129 | 0 | 0.22 | 0.04 | 0.12 | 0.01 | 0.2 | 0.14 |
| 3 | 102064584 | rs1435209 | -1.363 | -0.92 | -0.234 | -2.791 | 0.05 | -0.006 | -0.006 | 0 | 0.07 | 0.29 | 0.06 | 0 | 0.05 | 0.05 |
| 3 | 102067254 | rs985374 | -1.45 | -1.033 | 0.246 | -2.73 | -0.016 | -0.128 | -0.128 | 0.055 | 0.3 | 0.54 | 0.2 | 0.04 | 0.25 | 0.19 |
| 3 | 102093452 | rs10511198 | 0.919 | -1.467 | 0.255 | -2.89 | 0.027 | 0.068 | 0.068 | 1.523 | 0.22 | 0.09 | 0.17 | 0.07 | 0.15 | 0.1 |
| 3 | 102093492 | rs10511197 | 0.802 | -1.475 | 0.306 | -2.893 | 0.028 | 0.057 | 0.057 | 1.326 | 0.23 | 0.16 | 0.17 | 0.07 | 0.15 | 0.1 |
| 3 | 102098086 | rs6804554 | -0.963 | -1.48 | 0.432 | -2.923 | 0.038 | 0.091 | 0.091 | 0.14 | 0.46 | 0.38 | 0.32 | 0.08 | 0.36 | 0.24 |
| 3 | 102100781 | rs6766223 | -0.436 | -1.628 | 0.486 | -2.976 | 0.003 | -0.01 | -0.01 | 0.286 | 0.35 | 0.41 | 0.29 | 0.08 | 0.25 | 0.19 |

| | | | | | | | | | | | | | | | | |
|---|---|---|---|---|---|---|---|---|---|---|---|---|---|---|---|---|
| 3 | 102112443 | rs16845008 | 1.535 | -0.905 | -0.636 | -2.411 | -0.094 | -0.141 | -0.141 | 0 | 0.24 | 0.01 | 0.29 | 0.39 | 0.33 | 0.32 |
| 3 | 102116525 | rs9811444 | 0.902 | -1.183 | -0.547 | -2.576 | -0.007 | -0.048 | -0.048 | 0 | 0.26 | 0.29 | 0.21 | 0.04 | 0.16 | 0.13 |
| 3 | 102124339 | rs9870734 | -0.997 | -1.167 | -0.455 | -2.619 | 0.018 | -0.005 | -0.005 | 0 | 0.5 | 0.62 | 0.36 | 0.05 | 0.4 | 0.26 |
| 3 | 102141451 | rs3891857 | -1.054 | -1.157 | -0.296 | -2.629 | 0.049 | 0.15 | 0.15 | 0.375 | 0.51 | 0.78 | 0.36 | 0.05 | 0.38 | 0.25 |
| 3 | 102151310 | rs10937024 | 1.474 | -1.173 | -0.371 | -2.624 | 0.028 | 0.081 | 0.081 | 1.002 | 0.21 | 0.09 | 0.16 | 0.04 | 0.14 | 0.08 |
| 3 | 102157417 | rs6784389 | 1.465 | -1.185 | -0.376 | -2.629 | 0.028 | 0.081 | 0.081 | 0.961 | 0.21 | 0.09 | 0.16 | 0.04 | 0.14 | 0.08 |
| 3 | 102174089 | rs1473269 | -0.989 | -1.201 | -0.491 | -2.637 | 0.004 | -0.052 | -0.052 | 0 | 0.49 | 0.55 | 0.34 | 0.05 | 0.39 | 0.25 |
| 3 | 102176125 | rs1533934 | -0.132 | -1.182 | -1.249 | -2.597 | -0.036 | 0.128 | 0.128 | 0 | 0.2 | 0.02 | 0.12 | 0.01 | 0.21 | 0.12 |
| 3 | 102181131 | rs12054046 | 1.415 | -1.159 | -1.169 | -2.528 | 0.028 | 0.081 | 0.081 | 0.961 | 0.21 | 0.09 | 0.16 | 0.04 | 0.14 | 0.08 |
| 3 | 102185938 | rs16845047 | 1.075 | -0.307 | -1.488 | -1.608 | -0.036 | -0.077 | -0.077 | 0 | 0.22 | 0.03 | 0.22 | 0.31 | 0.2 | 0.38 |
| 3 | 102189169 | rs17822656 | 0.47 | -0.426 | -1.199 | -1.493 | -0.036 | 0.033 | 0.033 | 0 | 0.13 | 0.01 | 0.05 | 0.04 | 0.15 | 0.09 |
| 3 | 102194775 | rs2688700 | -0.721 | -0.443 | -1.53 | -1.709 | -0.014 | 0.101 | 0.101 | 0 | 0.29 | 0.09 | 0.14 | 0.04 | 0.27 | 0.2 |
| 3 | 102195680 | rs7627117 | 0.045 | -0.542 | -1.039 | -1.703 | 0.01 | 0.008 | 0.008 | 0 | 0.35 | 0.73 | 0.32 | 0.29 | 0.25 | 0.33 |
| 3 | 102198169 | rs7647414 | -0.773 | -0.129 | -1.071 | -1.544 | -0.025 | -0.031 | -0.031 | 1.143 | 0.93 | 0.98 | 0.85 | 0.8 | 0.89 | 0.77 |
| 3 | 102221520 | rs4683838 | 1.994 | -0.126 | -1.188 | -1.242 | -0.096 | -0.144 | -0.144 | 0.45 | 0.22 | 0.02 | 0.31 | 0.42 | 0.33 | 0.31 |
| 3 | 102221592 | rs1151318 | 1.421 | -0.279 | -0.071 | -1.401 | -0.015 | 0.063 | 0.063 | 0 | 0.46 | 0.25 | 0.66 | 0.75 | 0.55 | 0.56 |
| 3 | 102228007 | rs9861714 | -0.543 | -0.259 | 0.499 | -1.411 | 0.01 | 0.037 | 0.037 | 0 | 0.37 | 0.25 | 0.17 | 0.06 | 0.3 | 0.19 |
| 3 | 102240202 | rs1620217 | 1.278 | -0.01 | 0.694 | -1.235 | 0.003 | 0.021 | 0.021 | 0 | 0.5 | 0.46 | 0.68 | 0.74 | 0.5 | 0.57 |
| 3 | 102240277 | rs1299752 | -0.331 | -0.432 | 0.526 | -0.803 | -0.025 | -0.034 | -0.034 | 0 | 0.92 | 0.98 | 0.9 | 0.79 | 0.88 | 0.77 |
| 3 | 102242724 | rs1151324 | -0.202 | -0.625 | 0.424 | -0.983 | 0.031 | 0.139 | 0.139 | 0.073 | 0.25 | 0.05 | 0.09 | 0.02 | 0.17 | 0.12 |
| 3 | 102253323 | rs1284718 | 2.274 | -0.578 | 0.846 | -0.623 | -0.016 | 0.032 | 0.032 | 0 | 0.47 | 0.21 | 0.68 | 0.65 | 0.56 | 0.57 |
| 3 | 102259425 | rs9854367 | -0.274 | -0.525 | 1.401 | -0.503 | 0.012 | -0.01 | -0.01 | 0 | 0.17 | 0.28 | 0.11 | 0.11 | 0.16 | 0.09 |
| 3 | 102267396 | rs1284757 | -0.052 | -0.295 | 1.746 | -0.358 | 0.045 | 0.12 | 0.12 | 0.018 | 0.9 | 0.7 | 0.88 | 0.77 | 0.88 | 0.78 |
| 3 | 102269072 | rs16845116 | -0.289 | -0.408 | 1.068 | -0.598 | 0.002 | -0.009 | -0.009 | 0 | 0.19 | 0.16 | 0.07 | 0.02 | 0.17 | 0.1 |
| 3 | 102275521 | rs1284767 | 1.582 | -0.228 | 1.019 | -0.492 | 0.042 | 0.068 | 0.068 | 0 | 0.5 | 0.29 | 0.71 | 0.66 | 0.6 | 0.61 |
| 3 | 102293262 | rs1284726 | 1.906 | -0.413 | 1.104 | -0.567 | -0.045 | 0.01 | 0.01 | 0 | 0.46 | 0.12 | 0.65 | 0.64 | 0.54 | 0.59 |
| 3 | 102297540 | rs1715530 | 1.747 | -0.417 | 1.171 | -0.569 | -0.001 | 0.027 | 0.027 | 0 | 0.48 | 0.29 | 0.68 | 0.64 | 0.54 | 0.59 |
| 3 | 102317512 | rs6802319 | 0.013 | -0.497 | 1.173 | -0.647 | -0.055 | -0.12 | -0.12 | 0 | 0.21 | 0.54 | 0.12 | 0.03 | 0.11 | 0.07 |
| 3 | 102328237 | rs17773589 | 1.142 | -0.432 | 0.908 | -0.618 | 0.013 | 0.059 | 0.059 | 0 | 0.15 | 0.04 | 0.08 | 0.03 | 0.11 | 0.06 |
| 3 | 102329365 | rs6441667 | 0.14 | -0.44 | 0.836 | -0.663 | -0.035 | 0.086 | 0.086 | 0 | 0.18 | 0.01 | 0.08 | 0.03 | 0.18 | 0.08 |
| 3 | 102335916 | rs1284749 | 2.008 | -0.743 | 1.322 | -0.652 | -0.05 | 0.013 | 0.013 | 0 | 0.46 | 0.12 | 0.65 | 0.64 | 0.54 | 0.58 |
| 3 | 102345638 | rs1356106 | 0.673 | -0.608 | 0.874 | -0.58 | -0.027 | -0.023 | -0.023 | 0.325 | 0.68 | 0.85 | 0.8 | 0.67 | 0.66 | 0.65 |
| 3 | 102349532 | rs1701679 | 1.343 | -0.313 | 1.258 | -0.238 | 0.019 | -0.005 | -0.005 | 0.129 | 0.52 | 0.22 | 0.67 | 0.57 | 0.58 | 0.6 |
| 3 | 102354380 | rs13083534 | 1.344 | -0.315 | 0.726 | -0.238 | 0.014 | -0.014 | -0.014 | 0 | 0.52 | 0.17 | 0.66 | 0.57 | 0.58 | 0.6 |
| 3 | 102355402 | rs17774042 | 0.133 | 0.077 | 0.091 | -0.082 | 0.038 | 0.123 | 0.123 | 0 | 0.24 | 0.03 | 0.11 | 0.05 | 0.17 | 0.1 |
| 3 | 102361006 | rs9985428 | -1.251 | 0.495 | -0.875 | 0.105 | -0.003 | -0.013 | -0.013 | 0 | 0.42 | 0.29 | 0.28 | 0.43 | 0.42 | 0.4 |
| 3 | 102364297 | rs9875063 | -0.722 | 1.538 | -0.619 | 1.034 | -0.009 | -0.005 | -0.005 | 0 | 0.17 | 0.12 | 0.16 | 0.38 | 0.25 | 0.3 |
| 3 | 102372037 | rs6807772 | -0.28 | 0.952 | -0.353 | 0.349 | 0.002 | -0.032 | -0.032 | 0 | 0.06 | 0.15 | 0.04 | 0 | 0.04 | 0.04 |
| 3 | 102386156 | rs1506436 | -0.153 | 0.646 | -0.916 | -0.025 | 0.007 | 0.036 | 0.036 | 0 | 0.8 | 0.91 | 0.89 | 0.84 | 0.83 | 0.91 |
| 3 | 102413432 | rs1581304 | 0.512 | 0.066 | 0.294 | -0.284 | -0.044 | -0.071 | -0.071 | 0.324 | 0.75 | 0.56 | 0.84 | 0.84 | 0.83 | 0.91 |
| 3 | 102415127 | rs11720453 | 1.106 | 0.001 | 0.23 | -0.313 | -0.018 | -0.048 | -0.048 | 0.094 | 0.68 | 0.38 | 0.8 | 0.84 | 0.78 | 0.87 |
| 3 | 102424262 | rs13434070 | 0.415 | 0.456 | -0.459 | -0.175 | -0.041 | -0.042 | -0.042 | 0 | 0.09 | 0.01 | 0.13 | 0.09 | 0.13 | 0.14 |
| 3 | 102434725 | rs1391419 | 1.699 | 0.772 | -1.114 | -0.611 | -0.095 | -0.038 | -0.038 | 0.793 | 0.21 | 0.01 | 0.26 | 0.24 | 0.28 | 0.23 |
| 3 | 102455600 | rs10470520 | 0.02 | 1.287 | -0.499 | 0.041 | -0.036 | -0.059 | -0.059 | 0 | 0.15 | 0.08 | 0.3 | 0.47 | 0.23 | 0.38 |
| 3 | 102457118 | rs6768201 | 0.406 | 0.875 | -0.224 | -0.234 | 0.012 | 0.042 | 0.042 | 0 | 0.21 | 0.13 | 0.11 | 0.14 | 0.17 | 0.1 |
| 3 | 102467051 | rs9860789 | 0.918 | 0.819 | -0.083 | -0.254 | 0 | -0.025 | -0.025 | 0 | 0.06 | 0.1 | 0.03 | 0 | 0.04 | 0.04 |
| 3 | 102467369 | rs931652 | 0.663 | 0.683 | 0.719 | -0.333 | -0.004 | -0.022 | -0.022 | 0.302 | 0.69 | 0.31 | 0.8 | 0.87 | 0.78 | 0.86 |
| 3 | 102468142 | rs9883402 | -0.335 | 0.684 | 0.506 | -0.331 | 0.014 | 0.052 | 0.052 | 0 | 0.26 | 0.17 | 0.13 | 0.13 | 0.21 | 0.14 |
| 3 | 102477716 | rs6801256 | 0.348 | 0.959 | 1.277 | -0.254 | -0.004 | -0.081 | -0.081 | 3.959 | 0.74 | 0.52 | 0.83 | 0.87 | 0.78 | 0.85 |
| 3 | 102496824 | rs2398777 | 0.113 | 0.967 | 1.489 | -0.25 | 0 | -0.002 | -0.002 | 0 | 0.75 | 0.73 | 0.85 | 0.87 | 0.78 | 0.85 |
| 3 | 102499522 | rs1498083 | 0.601 | 0.953 | 1.45 | -0.257 | -0.009 | -0.065 | -0.065 | 0 | 0.72 | 0.44 | 0.82 | 0.87 | 0.77 | 0.85 |
| 3 | 102502059 | rs825353 | -0.631 | 1.073 | 0.951 | -0.206 | -0.005 | 0.047 | 0.047 | 0 | 0.8 | 0.93 | 0.88 | 0.87 | 0.81 | 0.89 |
| 3 | 102506798 | rs9872254 | 0.154 | 1.074 | 0.997 | -0.187 | -0.002 | -0.033 | -0.033 | 0 | 0.74 | 0.63 | 0.84 | 0.87 | 0.78 | 0.85 |
| 3 | 102521272 | rs6795619 | 0.201 | 1.376 | 1.034 | -0.081 | 0.053 | 0.073 | 0.073 | 0 | 0.07 | 0.17 | 0.12 | 0.14 | 0.1 | 0.07 |
| 3 | 102529319 | rs7634477 | -0.344 | 1.125 | 1.025 | -0.326 | 0.006 | -0.003 | -0.003 | 0 | 0.17 | 0.24 | 0.14 | 0.13 | 0.14 | 0.13 |
| 3 | 102533446 | rs1498075 | 1.309 | 1.121 | 1.686 | -0.41 | -0.002 | 0.002 | 0.002 | 0 | 0.66 | 0.3 | 0.7 | 0.74 | 0.67 | 0.77 |
| 3 | 102581948 | rs1151307 | -0.1 | 1.124 | 1.864 | -0.387 | 0.07 | 0.093 | 0.093 | 0 | 0.07 | 0.28 | 0.13 | 0.13 | 0.11 | 0.07 |
| 3 | 102583687 | rs6441682 | 1.143 | 1.093 | 1.996 | -0.406 | -0.018 | -0.059 | -0.059 | 0 | 0.11 | 0.21 | 0.03 | 0 | 0.03 | 0.06 |
| 3 | 102585782 | rs1546208 | 0.458 | 1.192 | 2.064 | -0.405 | 0.045 | 0.016 | 0.016 | 0 | 0.78 | 0.59 | 0.75 | 0.74 | 0.7 | 0.83 |
| 3 | 102598113 | rs1241287 | -0.629 | 1.3 | 1.742 | -0.403 | -0.004 | -0.004 | -0.004 | 0 | 0.22 | 0.2 | 0.4 | 0.6 | 0.34 | 0.46 |
| 3 | 102599596 | rs4856107 | 1.141 | 0.791 | 1.971 | -0.464 | 0.014 | -0.019 | -0.019 | 1.539 | 0.67 | 0.43 | 0.71 | 0.74 | 0.67 | 0.77 |
| 3 | 102602750 | rs16845761 | 1.37 | 0.787 | 1.706 | -0.467 | -0.004 | -0.027 | -0.027 | 0.054 | 0.1 | 0.14 | 0.03 | 0 | 0.03 | 0.05 |
| 3 | 102605470 | rs1151303 | 0.3 | 0.786 | 1.698 | -0.466 | -0.001 | -0.01 | -0.01 | 0.653 | 0.75 | 0.73 | 0.85 | 0.88 | 0.78 | 0.85 |
| 3 | 102607793 | rs1705894 | -0.449 | 0.663 | 1.566 | -0.381 | 0.023 | 0 | 0 | 0 | 0.18 | 0.47 | 0.16 | 0.13 | 0.14 | 0.13 |
| 3 | 102608943 | rs11919331 | 0.524 | 0.74 | 0.953 | -0.347 | -0.001 | 0.002 | 0.002 | 0.039 | 0.13 | 0.1 | 0.07 | 0.13 | 0.13 | 0.09 |
| 3 | 102609928 | rs11919584 | 0.682 | 0.752 | 0.833 | -0.341 | -0.006 | 0.005 | 0.005 | 0.011 | 0.12 | 0.06 | 0.07 | 0.13 | 0.13 | 0.09 |
| 3 | 102635852 | rs17778288 | 0.23 | 0.769 | 1.054 | -0.349 | -0.001 | 0.002 | 0.002 | 0.02 | 0.14 | 0.13 | 0.14 | 0.14 | 0.16 | 0.1 |
| 3 | 102649293 | rs708287 | 0.096 | 0.625 | 1.35 | -0.449 | 0.018 | 0.008 | 0.008 | 0.383 | 0.15 | 0.27 | 0.13 | 0.14 | 0.13 | 0.16 |
| 3 | 102649650 | rs16845818 | 1.649 | 0.626 | 1.236 | -0.458 | -0.002 | -0.02 | -0.02 | 0 | 0.09 | 0.07 | 0.02 | 0 | 0.03 | 0.08 |

| | | | | | | | | | | | | | | | | |
|---|---|---|---|---|---|---|---|---|---|---|---|---|---|---|---|---|
| 3 | 102651023 | rs3849511 | 0.527 | 0.665 | 0.814 | -0.388 | -0.03 | -0.095 | -0.095 | 0.015 | 0.13 | 0.01 | 0.2 | 0.43 | 0.13 | 0.37 |
| 3 | 102651365 | rs16845833 | 0.095 | 0.48 | 1.321 | -0.739 | -0.002 | -0.002 | -0.002 | 0 | 0.15 | 0.18 | 0.1 | 0.15 | 0.16 | 0.1 |
| 3 | 102673948 | rs344675 | 1.645 | 0.322 | 1.312 | -0.789 | -0.001 | 0.018 | 0.018 | 0.081 | 0.66 | 0.19 | 0.72 | 0.71 | 0.7 | 0.73 |
| 3 | 102676267 | rs1470146 | 0.422 | 0.354 | 1.271 | -0.769 | 0.042 | 0.066 | 0.066 | 0 | 0.06 | 0.12 | 0.11 | 0.14 | 0.1 | 0.08 |
| 3 | 102690992 | rs17779020 | 1.132 | -0.263 | 0.323 | -1.283 | 0.024 | 0.099 | 0.099 | 0 | 0.17 | 0.03 | 0.08 | 0.01 | 0.09 | 0.12 |
| 3 | 102724770 | rs17175072 | 0.263 | 0.046 | 0.494 | -1.138 | -0.003 | 0.008 | 0.008 | 0.037 | 0.15 | 0.07 | 0.1 | 0.13 | 0.17 | 0.1 |
| 3 | 102751470 | rs346879 | 1.462 | 0.035 | 0.375 | -1.224 | 0.008 | 0.114 | 0.114 | 0 | 0.53 | 0.15 | 0.64 | 0.71 | 0.62 | 0.64 |
| 3 | 102769976 | rs7634719 | 0.016 | 0.679 | 0.968 | -0.81 | 0.004 | -0.006 | -0.006 | 0 | 0.15 | 0.2 | 0.1 | 0.14 | 0.17 | 0.09 |
| 3 | 102772946 | rs346890 | 0.131 | 0.592 | 1.042 | -0.814 | 0.026 | -0.002 | -0.002 | 0.165 | 0.82 | 0.43 | 0.83 | 0.87 | 0.86 | 0.84 |
| 3 | 102782838 | rs3849512 | 0.256 | 0.58 | 1.046 | -0.817 | 0.022 | -0.008 | -0.008 | 0.224 | 0.82 | 0.44 | 0.82 | 0.87 | 0.85 | 0.83 |
| 3 | 102787456 | rs12634503 | -0.109 | 0.616 | 0.905 | -0.7 | 0.007 | -0.005 | -0.005 | 0 | 0.14 | 0.23 | 0.1 | 0.14 | 0.16 | 0.08 |
| 3 | 102806604 | rs6767265 | -0.181 | 0.71 | 0.527 | -0.735 | -0.006 | 0.004 | 0.004 | 0 | 0.15 | 0.15 | 0.13 | 0.13 | 0.21 | 0.09 |
| 3 | 102816124 | rs6782902 | -0.812 | 0.175 | 0.127 | -0.969 | -0.014 | 0.085 | 0.085 | 0 | 0.28 | 0.12 | 0.14 | 0.13 | 0.26 | 0.18 |
| 3 | 102827950 | rs1399781 | 1.11 | 0.394 | 1.059 | -0.927 | 0.016 | -0.036 | -0.036 | 0 | 0.64 | 0.48 | 0.7 | 0.75 | 0.63 | 0.71 |
| 3 | 102833131 | rs4643721 | 1.569 | 0.307 | 1.087 | -0.945 | -0.026 | 0.077 | 0.077 | 0.51 | 0.6 | 0.18 | 0.67 | 0.75 | 0.63 | 0.71 |
| 3 | 102837063 | rs1120426 | 0.177 | 0.301 | 1.276 | -0.72 | 0.136 | 0.141 | 0.141 | 0 | 0.94 | 0.69 | 0.87 | 0.88 | 0.9 | 0.92 |
| 3 | 102837863 | rs13071487 | 0.537 | 0.269 | 0.63 | -0.737 | -0.021 | -0.027 | -0.027 | 0 | 0.78 | 0.37 | 0.82 | 0.88 | 0.87 | 0.86 |
| 3 | 102840677 | rs3849514 | -0.304 | 0.662 | -0.293 | -0.44 | -0.047 | 0.015 | 0.015 | 1.165 | 0.73 | 0.87 | 0.8 | 0.79 | 0.67 | 0.8 |
| 3 | 102842671 | rs10937285 | 1.203 | -0.092 | -0.642 | -0.468 | -0.05 | -0.019 | -0.019 | 0 | 0.43 | 0.15 | 0.51 | 0.54 | 0.51 | 0.62 |
| 3 | 102843523 | rs16845978 | -0.126 | -0.416 | 0.181 | -0.714 | 0 | 0.001 | 0.001 | 0.024 | 0.33 | 0.33 | 0.34 | 0.46 | 0.34 | 0.28 |
| 3 | 102846337 | rs10511205 | 0.299 | -0.509 | 0.209 | -0.71 | -0.026 | -0.03 | -0.03 | 0 | 0.28 | 0.19 | 0.29 | 0.38 | 0.33 | 0.26 |
| 3 | 102853034 | rs6788651 | 0.114 | -0.544 | 0.877 | -0.733 | -0.038 | -0.157 | -0.157 | 0 | 0.77 | 0.54 | 0.85 | 1 | 0.85 | 0.9 |
| 3 | 102854945 | rs3914494 | 1.382 | -0.494 | 0.503 | -0.683 | -0.014 | 0.002 | 0.002 | 0 | 0.44 | 0.28 | 0.52 | 0.54 | 0.51 | 0.62 |
| 3 | 102862268 | rs1515526 | -0.667 | -0.727 | 0.655 | -0.697 | 0.007 | 0 | 0 | 0 | 0.39 | 0.43 | 0.41 | 0.38 | 0.45 | 0.3 |
| 3 | 102864676 | rs10514747 | 0.854 | -0.57 | 0.113 | -0.763 | -0.071 | -0.004 | -0.004 | 0 | 0.25 | 0.01 | 0.14 | 0.19 | 0.24 | 0.21 |
| 3 | 102872264 | rs16846002 | -0.174 | -0.484 | 0.538 | -0.617 | 0.01 | -0.051 | -0.051 | 0 | 0.1 | 0.19 | 0.11 | 0 | 0.12 | 0.03 |
| 3 | 102883817 | rs11707000 | 0.222 | -0.452 | 0.3 | -0.603 | -0.002 | -0.013 | -0.013 | 0 | 0.09 | 0.08 | 0.11 | 0 | 0.12 | 0.03 |
| 3 | 102892138 | rs1849925 | -1.313 | -0.561 | 0.184 | -0.6 | -0.065 | -0.033 | -0.033 | 1.203 | 0.59 | 0.86 | 0.5 | 0.45 | 0.49 | 0.39 |
| 3 | 102904391 | rs1379497 | 0.7 | -0.425 | 0.289 | -0.417 | -0.025 | -0.007 | -0.007 | 0 | 0.05 | 0.09 | 0.04 | 0.08 | 0.01 | 0.02 |
| 3 | 102912069 | rs4399903 | -0.383 | -0.166 | 0.136 | -0.426 | -0.002 | -0.012 | -0.012 | 0 | 0.92 | 0.95 | 0.88 | 1 | 0.86 | 0.96 |
| 3 | 102927717 | rs6786935 | 0.985 | -0.407 | 0.157 | -0.517 | -0.017 | -0.007 | -0.007 | 0 | 0.4 | 0.29 | 0.52 | 0.56 | 0.51 | 0.62 |
| 3 | 102946216 | rs13063401 | 0.407 | -0.414 | 0.066 | -0.413 | 0.006 | -0.007 | -0.007 | 8.135 | 0.37 | 0.28 | 0.3 | 0.37 | 0.31 | 0.27 |
| 3 | 102948192 | rs3926576 | -0.241 | -0.167 | 0.203 | -0.458 | 0.049 | -0.054 | -0.054 | 0 | 0.91 | 0.77 | 0.87 | 1 | 0.86 | 0.96 |
| 3 | 102948373 | rs9825113 | 0.685 | -0.379 | 0.324 | -0.549 | -0.025 | 0.009 | 0.009 | 0 | 0.45 | 0.27 | 0.52 | 0.56 | 0.54 | 0.67 |
| 3 | 102950695 | rs2398838 | 0.043 | -0.175 | 0.386 | -0.554 | 0.049 | -0.054 | -0.054 | 0 | 0.91 | 0.77 | 0.87 | 1 | 0.86 | 0.96 |
| 3 | 102965023 | rs1492873 | -0.492 | -0.28 | 0.039 | -0.52 | 0.001 | -0.03 | -0.03 | 0 | 0.42 | 0.48 | 0.56 | 0.3 | 0.48 | 0.38 |
| 3 | 102966862 | rs6777050 | -0.753 | -0.116 | -0.386 | -0.289 | -0.039 | -0.027 | -0.027 | 0 | 0.82 | 0.89 | 0.8 | 0.72 | 0.75 | 0.82 |
| 3 | 102980444 | rs1492872 | 1.425 | -0.029 | -0.437 | -0.74 | -0.051 | -0.06 | -0.06 | 0 | 0.13 | 0.05 | 0.32 | 0.29 | 0.23 | 0.08 |
| 3 | 102980751 | rs11919883 | -0.251 | -0.024 | 0.417 | -0.658 | -0.005 | -0.059 | -0.059 | 0 | 0.11 | 0.39 | 0.05 | 0 | 0.07 | 0.05 |
| 3 | 102985465 | rs6441713 | -1.005 | -0.032 | 0.395 | -0.647 | -0.014 | -0.113 | -0.113 | 0 | 0.2 | 0.45 | 0.2 | 0.02 | 0.18 | 0.26 |
| 3 | 102993405 | rs4269090 | 1.255 | -0.264 | 0.403 | -0.518 | 0.029 | -0.045 | -0.045 | 0 | 0.33 | 0.22 | 0.28 | 0.46 | 0.24 | 0.44 |
| 3 | 103001990 | rs13098370 | 0.801 | -0.09 | 0.46 | -0.468 | -0.004 | -0.039 | -0.039 | 0 | 0.08 | 0.2 | 0.04 | 0 | 0.07 | 0.05 |
| 3 | 103009576 | rs9840550 | 0.738 | 0.104 | -0.392 | -0.388 | -0.004 | 0.007 | 0.007 | 0 | 0.08 | 0.06 | 0.15 | 0.02 | 0.12 | 0.2 |
| 3 | 103011287 | rs1492885 | -0.387 | 0.132 | -0.46 | -0.372 | -0.007 | -0.084 | -0.084 | 0.047 | 0.18 | 0.3 | 0.18 | 0.02 | 0.18 | 0.26 |
| 3 | 103016226 | rs1353025 | -0.6 | 0.429 | -1.008 | 0.016 | -0.039 | 0 | 0 | 0 | 0.8 | 0.87 | 0.87 | 0.8 | 0.74 | 0.84 |
| 3 | 103017948 | rs2029659 | 0.762 | 0.394 | -1.186 | 0.073 | -0.014 | -0.017 | -0.017 | 1.192 | 0.27 | 0.15 | 0.38 | 0.32 | 0.31 | 0.14 |
| 3 | 103023075 | rs7639547 | 0.295 | 0.499 | -1.027 | 0.359 | -0.002 | 0.001 | 0.001 | 1.09 | 0.43 | 0.53 | 0.46 | 0.48 | 0.36 | 0.65 |
| 3 | 103026304 | rs1994818 | -0.432 | 1.326 | -1.235 | 0.402 | -0.054 | -0.001 | -0.001 | 0.004 | 0.18 | 0.08 | 0.12 | 0.16 | 0.25 | 0.15 |
| 3 | 103035266 | rs9876832 | -0.358 | -0.119 | -1.127 | 0.313 | 0.019 | 0.02 | 0.02 | 2.285 | 0.29 | 0.76 | 0.31 | 0.2 | 0.17 | 0.26 |
| 3 | 103037822 | rs6766007 | -0.254 | -0.164 | -0.268 | 0.353 | -0.011 | -0.02 | -0.02 | 0 | 0.28 | 0.62 | 0.29 | 0.16 | 0.17 | 0.26 |
| 3 | 103052036 | rs8179908 | -0.394 | -0.225 | 0.503 | 0.301 | 0.019 | 0.02 | 0.02 | 2.24 | 0.29 | 0.76 | 0.31 | 0.19 | 0.17 | 0.26 |
| 3 | 103056815 | rs1492869 | -0.65 | -0.847 | 1.051 | 0.112 | -0.025 | 0.042 | 0.042 | 1.09 | 0.11 | 0.48 | 0.13 | 0.08 | 0.02 | 0.11 |
| 3 | 103057272 | rs9862353 | -1.283 | -1.307 | 0.253 | 0.467 | -0.022 | -0.068 | -0.068 | 0.021 | 0.57 | 0.92 | 0.54 | 0.6 | 0.32 | 0.67 |
| 3 | 103062026 | rs1492865 | -0.461 | -1.148 | 0.584 | 0.254 | -0.026 | 0.041 | 0.041 | 1.399 | 0.11 | 0.49 | 0.13 | 0.08 | 0.02 | 0.11 |
| 3 | 103079255 | rs6788478 | -0.639 | -1.148 | 0.397 | 0.254 | 0.04 | 0.036 | 0.036 | 4.983 | 0.38 | 0.71 | 0.31 | 0.21 | 0.19 | 0.37 |
| 3 | 103087583 | rs4306866 | 0.025 | -0.9 | 0.432 | 0.223 | -0.009 | -0.025 | -0.025 | 0 | 0.08 | 0.08 | 0.04 | 0 | 0.05 | 0.06 |
| 3 | 103088982 | rs11706808 | -0.792 | -1.02 | 1.015 | 0.62 | 0.007 | 0.108 | 0.108 | 0 | 0.08 | 0.24 | 0.11 | 0.25 | 0.05 | 0.18 |
| 3 | 103089194 | rs6806931 | -1.498 | -0.874 | -0.034 | 0.571 | -0.038 | -0.011 | -0.011 | 0.493 | 0.48 | 0.86 | 0.57 | 0.6 | 0.51 | 0.59 |
| 3 | 103089635 | rs6792532 | 0.726 | -0.918 | 0.25 | 0.557 | -0.021 | -0.044 | -0.044 | 0.062 | 0.08 | 0.15 | 0.04 | 0 | 0.05 | 0.06 |
| 3 | 103089936 | rs9819225 | 0.861 | -0.603 | 0.07 | 0.523 | 0.002 | -0.003 | -0.003 | 0.847 | 0.06 | 0.07 | 0.09 | 0.03 | 0.1 | 0.11 |
| 3 | 103090087 | rs9290901 | -1.637 | -0.639 | -0.004 | 0.535 | -0.022 | -0.007 | -0.007 | 0 | 0.49 | 0.86 | 0.57 | 0.6 | 0.53 | 0.59 |
| 3 | 103090241 | rs7653853 | 0.813 | -0.771 | -0.019 | 0.733 | -0.05 | -0.007 | -0.007 | 0 | 0.15 | 0.08 | 0.27 | 0.17 | 0.31 | 0.2 |
| 3 | 103093137 | rs12630356 | 1.484 | -1.086 | -0.492 | 0.556 | -0.01 | -0.081 | -0.081 | 0.127 | 0.15 | 0.01 | 0.2 | 0.22 | 0.13 | 0.15 |
| 3 | 103094740 | rs4521238 | -1.907 | -0.982 | -0.405 | 0.407 | -0.028 | 0.11 | 0.11 | 0.164 | 0.64 | 0.92 | 0.78 | 0.82 | 0.66 | 0.74 |
| 3 | 103097277 | rs6441729 | -2.167 | -1.41 | -0.468 | 0.321 | -0.042 | 0.081 | 0.081 | 0.239 | 0.67 | 0.92 | 0.77 | 0.8 | 0.67 | 0.75 |
| 3 | 103099956 | rs4317112 | -0.014 | -0.848 | 1.124 | 0.867 | 0.002 | 0.002 | 0.002 | 0 | 0.23 | 0.21 | 0.22 | 0.3 | 0.33 | 0.27 |
| 3 | 103104977 | rs10804921 | -0.865 | -0.89 | 0.824 | 0.868 | 0.018 | -0.001 | -0.001 | 0 | 0.36 | 0.5 | 0.34 | 0.33 | 0.43 | 0.38 |
| 3 | 103115144 | rs4328810 | -0.367 | -1.013 | 1.401 | 1.296 | 0.049 | 0.041 | 0.041 | 0 | 0.23 | 0.44 | 0.24 | 0.3 | 0.32 | 0.27 |
| 3 | 103119876 | rs11919013 | -0.367 | -1.013 | 1.401 | 1.296 | 0.049 | 0.041 | 0.041 | 0 | 0.23 | 0.44 | 0.24 | 0.3 | 0.32 | 0.27 |

| | | | | | | | | | | | | | | | | |
|---|---|---|---|---|---|---|---|---|---|---|---|---|---|---|---|---|
| 3 | 103120384 | rs6778525 | -1.07 | -1.128 | 1.409 | 1.315 | -0.017 | 0.005 | 0.005 | 0 | 0.41 | 0.72 | 0.53 | 0.54 | 0.45 | 0.42 |
| 3 | 103126594 | rs12636260 | -1.071 | -1.128 | 1.389 | 1.315 | -0.017 | 0.005 | 0.005 | 0 | 0.41 | 0.73 | 0.53 | 0.54 | 0.45 | 0.41 |
| 3 | 103140700 | rs4314176 | -1.146 | -1.094 | 0.951 | 1.33 | -0.026 | 0.005 | 0.005 | 0 | 0.41 | 0.77 | 0.53 | 0.54 | 0.45 | 0.41 |
| 3 | 103144212 | rs4585205 | 0.86 | -0.983 | 0.868 | 1.368 | 0.007 | -0.042 | -0.042 | 0 | 0.15 | 0.05 | 0.28 | 0.24 | 0.14 | 0.14 |
| 3 | 103156593 | rs4507267 | -1.129 | -0.966 | 0.791 | 1.378 | -0.016 | 0.001 | 0.001 | 0 | 0.4 | 0.69 | 0.53 | 0.54 | 0.45 | 0.41 |
| 3 | 103158005 | rs12637597 | 0 | -1.618 | 0.671 | 1.417 | -0.007 | 0.023 | 0.023 | 0 | 0.16 | 0.18 | 0.29 | 0.24 | 0.14 | 0.14 |
| 3 | 103173919 | rs9851044 | -0.575 | -1.531 | 0.787 | 0.788 | 0.026 | 0.041 | 0.041 | 0 | 0.5 | 0.43 | 0.49 | 0.29 | 0.36 | 0.42 |
| 3 | 103174143 | rs9851376 | 0.264 | -1.952 | 0.86 | 0.68 | 0.021 | 0.005 | 0.005 | 0 | 0.27 | 0.23 | 0.28 | 0.26 | 0.14 | 0.29 |
| 3 | 103180397 | rs11919124 | 0.209 | -2.018 | 1.123 | 0.725 | -0.011 | -0.001 | -0.001 | 0 | 0.27 | 0.29 | 0.29 | 0.27 | 0.14 | 0.29 |
| 3 | 103186883 | rs7640570 | 0.308 | -2.063 | 1.412 | 0.721 | -0.028 | -0.022 | -0.022 | 0 | 0.25 | 0.28 | 0.16 | 0.17 | 0.13 | 0.24 |
| 3 | 103203766 | rs1584440 | 0.302 | -2.095 | 1.559 | 0.651 | 0.003 | 0.01 | 0.01 | 0 | 0.24 | 0.2 | 0.2 | 0.03 | 0.2 | 0.13 |
| 3 | 103206133 | rs2122131 | -1.139 | -2.142 | 1.929 | 0.659 | 0.053 | 0.079 | 0.079 | 0 | 0.52 | 0.72 | 0.45 | 0.3 | 0.34 | 0.46 |
| 3 | 103209899 | rs4856154 | -1.309 | -2.221 | 2.108 | 1.213 | 0.054 | -0.04 | -0.04 | 0.774 | 0.54 | 0.87 | 0.49 | 0.56 | 0.34 | 0.49 |
| 3 | 103214808 | rs12488469 | 0.431 | -2.285 | 2.285 | 1.193 | -0.039 | -0.011 | -0.011 | 1.299 | 0.24 | 0.31 | 0.19 | 0.18 | 0.09 | 0.22 |
| 3 | 103215012 | rs16846300 | 0.82 | -2.28 | 2.181 | 1.168 | -0.024 | -0.022 | -0.022 | 0 | 0.21 | 0.23 | 0.11 | 0.08 | 0.08 | 0.12 |
| 3 | 103227532 | rs9839927 | -1.35 | -2.242 | 2.033 | 1.173 | 0.047 | -0.054 | -0.054 | 0 | 0.54 | 0.89 | 0.5 | 0.56 | 0.34 | 0.49 |
| 3 | 103245092 | rs1397924 | -1.112 | -2.475 | 2.031 | 1.248 | 0.077 | 0.09 | 0.09 | 0 | 0.51 | 0.73 | 0.45 | 0.29 | 0.31 | 0.45 |
| 3 | 103252139 | rs7626713 | 0.005 | -2.172 | 2.161 | 1.397 | -0.054 | 0.048 | 0.048 | 0 | 0.25 | 0.33 | 0.22 | 0.45 | 0.09 | 0.25 |
| 3 | 103271449 | rs721778 | -0.142 | -1.984 | 2.046 | 1.128 | -0.007 | -0.076 | -0.076 | 0 | 0.25 | 0.43 | 0.27 | 0.11 | 0.24 | 0.3 |
| 3 | 103274074 | rs10937480 | 0.141 | -1.986 | 2.107 | 1.135 | -0.008 | -0.055 | -0.055 | 0 | 0.25 | 0.33 | 0.26 | 0.1 | 0.24 | 0.31 |
| 3 | 103291237 | rs9883994 | -1.481 | -2.024 | 2.448 | 1.313 | 0.023 | 0.412 | 0.412 | 0.004 | 0.06 | 0.3 | 0.15 | 0.39 | 0.03 | 0.23 |
| 3 | 103292226 | rs4856162 | -0.944 | -2.319 | 2.464 | 1.193 | 0.059 | -0.004 | -0.004 | 0 | 0.48 | 0.68 | 0.46 | 0.49 | 0.32 | 0.48 |
| 3 | 103295384 | rs9853928 | -0.194 | -2.281 | 2.435 | 1.204 | -0.019 | 0.044 | 0.044 | 0 | 0.28 | 0.55 | 0.36 | 0.39 | 0.25 | 0.35 |
| 3 | 103298212 | rs9817638 | -0.158 | -2.289 | 2.443 | 1.2 | -0.018 | 0.05 | 0.05 | 0 | 0.27 | 0.53 | 0.36 | 0.39 | 0.25 | 0.35 |
| 3 | 103307642 | rs7612581 | -1.383 | -2.373 | 2.288 | 1.196 | 0.091 | -0.03 | -0.03 | 0.01 | 0.53 | 0.91 | 0.51 | 0.56 | 0.33 | 0.57 |
| 3 | 103323653 | rs7629407 | 0.832 | -2.306 | 2.102 | 1.209 | 0.03 | 0.028 | 0.028 | 0.188 | 0.21 | 0.17 | 0.1 | 0.1 | 0.08 | 0.12 |
| 3 | 103331483 | rs9820695 | -0.952 | -2.276 | 1.919 | 1.228 | 0.059 | -0.004 | -0.004 | 0 | 0.48 | 0.69 | 0.47 | 0.49 | 0.32 | 0.48 |
| 3 | 103333897 | rs2068840 | 0.739 | -2.222 | 1.594 | 1.252 | 0 | 0.005 | 0.005 | 0 | 0.21 | 0.19 | 0.21 | 0.01 | 0.22 | 0.13 |
| 3 | 103345439 | rs9821642 | -0.732 | -2.176 | 1.278 | 1.237 | -0.015 | 0.001 | 0.001 | 0 | 0.47 | 0.5 | 0.45 | 0.49 | 0.32 | 0.48 |
| 3 | 103348699 | rs2063485 | 0.845 | -1.538 | 0.086 | 1.358 | -0.035 | -0.036 | -0.036 | 0 | 0.06 | 0 | 0.05 | 0.15 | 0.12 | 0.1 |
| 3 | 103354281 | rs4856184 | 0.602 | -2.063 | 0.42 | 1.082 | 0.066 | 0.049 | 0.049 | 0.273 | 0.2 | 0.06 | 0.09 | 0.1 | 0.07 | 0.12 |
| 3 | 103359218 | rs10937540 | -0.954 | -1.929 | 0.72 | 1.095 | 0.013 | 0.004 | 0.004 | 0 | 0.52 | 0.46 | 0.5 | 0.59 | 0.34 | 0.6 |
| 3 | 103363061 | rs10937544 | -0.15 | -1.855 | 0.674 | 1.284 | -0.003 | 0.003 | 0.003 | 0 | 0.28 | 0.34 | 0.39 | 0.33 | 0.27 | 0.43 |
| 3 | 103379682 | rs6772983 | -0.266 | -2.067 | 0.86 | 1.254 | -0.059 | 0.038 | 0.038 | 0 | 0.31 | 0.38 | 0.27 | 0.56 | 0.12 | 0.47 |
| 3 | 103380852 | rs6775909 | -1.288 | -2.099 | 0.927 | 1.15 | 0.031 | 0.005 | 0.005 | 0 | 0.54 | 0.68 | 0.51 | 0.59 | 0.35 | 0.6 |
| 3 | 103387440 | rs2398880 | -0.206 | -2.117 | 1.052 | 1.208 | -0.045 | 0.026 | 0.026 | 0 | 0.31 | 0.38 | 0.27 | 0.54 | 0.12 | 0.47 |
| 3 | 103388364 | rs16846592 | -0.196 | -2.107 | 1.173 | 1.187 | 0.011 | -0.016 | -0.016 | 0 | 0.3 | 0.26 | 0.25 | 0.53 | 0.12 | 0.47 |
| 3 | 103394499 | rs6803099 | -0.271 | -2.119 | 1.196 | 1.182 | -0.04 | 0.023 | 0.023 | 0 | 0.31 | 0.36 | 0.26 | 0.53 | 0.12 | 0.47 |
| 3 | 103397769 | rs12485744 | -0.413 | -2.184 | 1.278 | 1.27 | -0.015 | 0.005 | 0.005 | 0 | 0.3 | 0.49 | 0.39 | 0.31 | 0.27 | 0.43 |
| 3 | 103400082 | rs7628767 | -1.431 | -2.198 | 1.287 | 1.271 | 0.04 | 0.004 | 0.004 | 0 | 0.55 | 0.7 | 0.5 | 0.57 | 0.35 | 0.6 |
| 3 | 103403514 | rs10511217 | -0.406 | -2.198 | 1.278 | 1.273 | -0.015 | 0.005 | 0.005 | 0 | 0.3 | 0.49 | 0.39 | 0.31 | 0.27 | 0.43 |
| 3 | 103404111 | rs16846607 | 0.625 | -2.186 | 1.187 | 1.285 | 0.065 | -0.018 | -0.018 | 0 | 0.23 | 0.08 | 0.1 | 0.26 | 0.08 | 0.17 |
| 3 | 103408208 | rs16846615 | -0.719 | -2.124 | 1.067 | 1.296 | -0.007 | 0.136 | 0.136 | 0 | 0.07 | 0.21 | 0.15 | 0.27 | 0.04 | 0.31 |
| 3 | 103421621 | rs10937564 | 0.45 | -2.128 | 0.853 | 1.197 | -0.004 | -0.055 | -0.055 | 0.048 | 0.23 | 0.34 | 0.24 | 0.04 | 0.23 | 0.13 |
| 3 | 103422112 | rs4370037 | -0.964 | -2.17 | 0.912 | 1.273 | 0.045 | 0.035 | 0.035 | 0 | 0.52 | 0.65 | 0.39 | 0.35 | 0.32 | 0.39 |
| 3 | 103430322 | rs9291039 | 0.045 | -2.068 | 0.129 | 1.315 | -0.016 | -0.093 | -0.093 | 0.006 | 0.28 | 0.54 | 0.29 | 0.1 | 0.24 | 0.23 |
| 3 | 103433922 | rs4856085 | 0.521 | -1.985 | -0.229 | 1.245 | 0.055 | -0.012 | -0.012 | 0 | 0.24 | 0.1 | 0.1 | 0.26 | 0.08 | 0.16 |
| 3 | 103438874 | rs4493441 | -1.192 | -1.978 | -0.234 | 1.302 | 0.043 | -0.003 | -0.003 | 0 | 0.55 | 0.75 | 0.51 | 0.56 | 0.35 | 0.6 |
| 3 | 103439060 | rs4493442 | -0.104 | -1.731 | -0.541 | 1.361 | 0.05 | -0.01 | -0.01 | 0.018 | 0.24 | 0.11 | 0.1 | 0.26 | 0.08 | 0.16 |
| 3 | 103464814 | rs2100653 | 0.401 | -0.796 | -0.553 | 1.527 | 0.005 | -0.023 | -0.023 | 0.001 | 0.59 | 0.32 | 0.5 | 0.45 | 0.63 | 0.35 |
| 3 | 103465052 | rs2085534 | -0.726 | -0.937 | -0.338 | 1.531 | -0.031 | 0.027 | 0.027 | 1.945 | 0.77 | 0.89 | 0.75 | 0.84 | 0.75 | 0.5 |
| 3 | 103465219 | rs2608432 | 0.068 | -0.979 | 0.048 | 1.346 | 0.003 | 0.021 | 0.021 | 1.028 | 0.7 | 0.41 | 0.55 | 0.54 | 0.7 | 0.4 |
| 3 | 103466610 | rs2608430 | -0.299 | -0.988 | 0.757 | 1.341 | 0.018 | 0.065 | 0.065 | 1.673 | 0.73 | 0.6 | 0.58 | 0.54 | 0.7 | 0.4 |
| 3 | 103470236 | rs9825206 | 0.694 | -0.996 | 0.587 | 1.742 | 0.004 | 0.018 | 0.018 | 0 | 0.59 | 0.34 | 0.48 | 0.32 | 0.62 | 0.32 |
| 3 | 103482015 | rs12635765 | 0.504 | -1.203 | 0.452 | 1.803 | 0.004 | -0.005 | -0.005 | 0.004 | 0.11 | 0.08 | 0.07 | 0.22 | 0.07 | 0.06 |
| 3 | 103485884 | rs16849322 | 0.573 | -1.201 | 0.43 | 1.804 | 0.011 | -0.016 | -0.016 | 0 | 0.11 | 0.07 | 0.07 | 0.22 | 0.08 | 0.06 |
| 3 | 103497868 | rs2677251 | -0.213 | -0.198 | 0.43 | 1.344 | 0.004 | 0.003 | 0.003 | 0 | 0.67 | 0.69 | 0.47 | 0.52 | 0.54 | 0.36 |
| 3 | 103504420 | rs16849343 | -0.197 | -1.338 | 0.051 | 0.875 | -0.051 | 0.005 | 0.005 | 0 | 0.36 | 0.51 | 0.22 | 0.32 | 0.22 | 0.17 |
| 3 | 103506909 | rs16849346 | -0.217 | -1.359 | 0.095 | 0.33 | -0.023 | -0.009 | -0.009 | 10.942 | 0.25 | 0.37 | 0.14 | 0.17 | 0.16 | 0.08 |
| 3 | 103512912 | rs1457835 | 0.057 | -1.183 | 0.035 | 0.407 | 0 | 0.094 | 0.094 | 0 | 0.39 | 0.15 | 0.3 | 0.22 | 0.41 | 0.22 |
| 3 | 103518852 | rs6441763 | 0.082 | -1.123 | 0.303 | 0.381 | -0.009 | 0.013 | 0.013 | 0.017 | 0.06 | 0.2 | 0.06 | 0.07 | 0.06 | 0.05 |
| 3 | 103525796 | rs1902834 | 1.122 | -1.285 | 0.094 | 0.247 | -0.044 | 0.066 | 0.066 | 0 | 0.24 | 0.05 | 0.22 | 0.13 | 0.27 | 0.15 |
| 3 | 103532130 | rs1673869 | 0.585 | -1.255 | 0.391 | 0.604 | -0.02 | 0.085 | 0.085 | 0 | 0.35 | 0.11 | 0.25 | 0.18 | 0.36 | 0.18 |
| 3 | 103538656 | rs1673865 | -0.045 | -1.093 | 0.447 | 0.585 | 0.009 | 0.007 | 0.007 | 0 | 0.45 | 0.64 | 0.36 | 0.39 | 0.44 | 0.25 |
| 3 | 103556995 | rs1993527 | 0.226 | -0.775 | 0.692 | 0.808 | 0.016 | 0.108 | 0.108 | 0 | 0.27 | 0.54 | 0.37 | 0.51 | 0.33 | 0.31 |
| 3 | 103558678 | rs1733213 | 0.648 | -0.903 | 0.498 | 0.637 | -0.022 | 0.14 | 0.14 | 0 | 0.4 | 0.1 | 0.28 | 0.15 | 0.4 | 0.19 |
| 3 | 103559931 | rs9826326 | -0.777 | -0.931 | 0.614 | 0.68 | -0.017 | 0.04 | 0.04 | 0 | 0.71 | 0.87 | 0.68 | 0.84 | 0.74 | 0.53 |
| 3 | 103568109 | rs12054076 | 0.992 | -0.918 | 1.102 | 0.743 | 0.01 | 0.035 | 0.035 | 0 | 0.21 | 0.29 | 0.31 | 0.45 | 0.3 | 0.26 |
| 3 | 103568533 | rs6799792 | 0.667 | -0.92 | 1.143 | 0.728 | 0.028 | 0.136 | 0.136 | 0 | 0.23 | 0.5 | 0.34 | 0.51 | 0.3 | 0.28 |

| | | | | | | | | | | | | | | | | |
|---|---|---|---|---|---|---|---|---|---|---|---|---|---|---|---|---|
| 3 | 103581156 | rs9824527 | 0.064 | -0.916 | 0.982 | 0.753 | -0.001 | 0 | 0 | 0.762 | 0.29 | 0.73 | 0.39 | 0.51 | 0.32 | 0.29 |
| 3 | 103586985 | rs1673874 | -0.818 | -0.927 | 0.862 | 0.726 | -0.012 | -0.034 | -0.034 | 0 | 0.69 | 0.84 | 0.67 | 0.67 | 0.72 | 0.49 |
| 3 | 103588426 | rs2061345 | -0.747 | -0.931 | 0.853 | 0.719 | -0.016 | 0.043 | 0.043 | 0 | 0.7 | 0.88 | 0.68 | 0.84 | 0.74 | 0.52 |
| 3 | 103595704 | rs17531418 | 0.544 | -0.959 | 1.29 | 0.708 | 0.003 | 0.084 | 0.084 | 0 | 0.42 | 0.16 | 0.34 | 0.23 | 0.4 | 0.22 |
| 3 | 103595761 | rs7611877 | 1.16 | -0.98 | 1.534 | 0.709 | -0.011 | -0.024 | -0.024 | 0 | 0.21 | 0.13 | 0.3 | 0.45 | 0.3 | 0.26 |
| 3 | 103599833 | rs2608486 | 0.641 | -1.085 | 1.318 | 0.736 | -0.009 | 0.085 | 0.085 | 0 | 0.4 | 0.12 | 0.34 | 0.23 | 0.4 | 0.22 |
| 3 | 103601325 | rs1540565 | 0.49 | -1.086 | 1.479 | 0.736 | 0.005 | 0.059 | 0.059 | 0 | 0.42 | 0.2 | 0.35 | 0.23 | 0.4 | 0.22 |
| 3 | 103601778 | rs1673887 | -0.703 | -0.967 | 1.499 | 0.712 | -0.001 | -0.016 | -0.016 | 0 | 0.69 | 0.78 | 0.71 | 0.67 | 0.73 | 0.5 |
| 3 | 103615361 | rs6437472 | -0.84 | -0.946 | 1.395 | 0.621 | -0.005 | 0.045 | 0.045 | 0 | 0.69 | 0.85 | 0.72 | 0.85 | 0.75 | 0.53 |
| 3 | 103629938 | rs13076727 | 0.399 | -1.453 | 1.476 | 0.742 | 0 | 0.064 | 0.064 | 0 | 0.4 | 0.22 | 0.34 | 0.22 | 0.4 | 0.22 |
| 3 | 103632129 | rs10511223 | 0.274 | -1.45 | 1.555 | 0.757 | 0 | 0.005 | 0.005 | 0 | 0.41 | 0.35 | 0.37 | 0.35 | 0.42 | 0.25 |
| 3 | 103635435 | rs4132806 | -0.566 | -1.489 | 1.318 | 0.639 | -0.003 | 0.036 | 0.036 | 0 | 0.44 | 0.78 | 0.43 | 0.69 | 0.43 | 0.37 |
| 3 | 103636914 | rs9833794 | 0.881 | -1.447 | 1.379 | 0.66 | -0.012 | -0.014 | -0.014 | 0 | 0.28 | 0.23 | 0.34 | 0.46 | 0.39 | 0.28 |
| 3 | 103637945 | rs9856810 | 0.566 | -1.447 | 1.433 | 0.657 | 0.032 | 0.058 | 0.058 | 0 | 0.31 | 0.5 | 0.38 | 0.5 | 0.39 | 0.28 |
| 3 | 103639646 | rs9845713 | 0.614 | -1.439 | 1.316 | 0.661 | 0.02 | 0.035 | 0.035 | 0 | 0.3 | 0.36 | 0.36 | 0.5 | 0.39 | 0.28 |
| 3 | 103641158 | rs4856091 | 0.871 | -1.389 | 1.195 | 0.664 | -0.012 | -0.014 | -0.014 | 1.449 | 0.28 | 0.23 | 0.34 | 0.45 | 0.39 | 0.28 |
| 3 | 103642313 | rs16849459 | 0.843 | -1.375 | 1.024 | 0.641 | -0.015 | 0.003 | 0.003 | 0 | 0.1 | 0.13 | 0.03 | 0.13 | 0.04 | 0.05 |
| 3 | 103647989 | rs9288769 | -0.292 | -1.401 | 1.204 | 0.63 | -0.001 | 0.005 | 0.005 | 0 | 0.42 | 0.67 | 0.41 | 0.68 | 0.43 | 0.37 |
| 3 | 103648320 | rs6765661 | 1.073 | -1.518 | 1.079 | 0.538 | -0.084 | 0.056 | 0.056 | 0 | 0.29 | 0.06 | 0.31 | 0.16 | 0.39 | 0.17 |
| 3 | 103655069 | rs7617988 | 0.802 | -1.143 | 1.418 | 0.797 | 0.012 | -0.007 | -0.007 | 0 | 0.1 | 0.04 | 0.02 | 0.13 | 0.04 | 0.05 |
| 3 | 103655090 | rs9815880 | 0.504 | -1.15 | 1.571 | 0.819 | -0.003 | -0.004 | -0.004 | 0 | 0.29 | 0.31 | 0.35 | 0.5 | 0.39 | 0.29 |
| 3 | 103659255 | rs7653181 | -0.807 | -1.152 | 1.454 | 0.797 | 0 | 0.033 | 0.033 | 0 | 0.44 | 0.77 | 0.43 | 0.69 | 0.43 | 0.37 |
| 3 | 103665933 | rs9288771 | 0.093 | -1.053 | 1.416 | 0.81 | 0.018 | 0.038 | 0.038 | 0 | 0.33 | 0.62 | 0.4 | 0.56 | 0.39 | 0.31 |
| 3 | 103678133 | rs7634768 | 0.199 | -1.047 | 1.275 | 0.802 | 0.017 | 0.038 | 0.038 | 0 | 0.32 | 0.49 | 0.38 | 0.54 | 0.39 | 0.31 |
| 3 | 103697819 | rs10048962 | 0.62 | -0.996 | 0.321 | 0.775 | -0.023 | -0.026 | -0.026 | 0 | 0.28 | 0.22 | 0.34 | 0.44 | 0.39 | 0.28 |
| 3 | 103698105 | rs1900678 | -0.447 | -0.729 | -0.32 | 0.557 | -0.001 | 0.033 | 0.033 | 0 | 0.44 | 0.77 | 0.43 | 0.68 | 0.43 | 0.37 |
| 3 | 103701756 | rs2166826 | -0.009 | -0.369 | -0.05 | 0.224 | -0.005 | -0.029 | -0.029 | 0 | 0.58 | 0.5 | 0.66 | 0.74 | 0.65 | 0.78 |
| 3 | 103704244 | rs12054174 | -0.74 | -1.049 | 0.381 | 0.977 | -0.02 | 0.057 | 0.057 | 0 | 0.29 | 0.38 | 0.4 | 0.49 | 0.24 | 0.5 |
| 3 | 103716354 | rs7628112 | -1.521 | -1.075 | 0.1 | 0.933 | 0.061 | 0.089 | 0.089 | 0.614 | 0.49 | 0.84 | 0.57 | 0.63 | 0.43 | 0.66 |
| 3 | 103719786 | rs2398945 | -0.06 | -0.939 | 0.132 | 1.103 | 0.005 | 0.082 | 0.082 | 0 | 0.22 | 0.32 | 0.38 | 0.44 | 0.23 | 0.49 |
| 3 | 103722343 | rs6808622 | -0.033 | -0.586 | -0.228 | 0.905 | -0.045 | 0.053 | 0.053 | 0 | 0.16 | 0.47 | 0.09 | 0.17 | 0.1 | 0.14 |
| 3 | 103732994 | rs1900677 | -0.518 | 0.274 | -0.888 | 0.997 | 0.003 | -0.008 | -0.008 | 0 | 0.34 | 0.42 | 0.49 | 0.46 | 0.33 | 0.52 |
| 3 | 103746185 | rs2029019 | 0.034 | 0.595 | -1.078 | 0.847 | 0.013 | 0.001 | 0.001 | 0 | 0.51 | 0.29 | 0.44 | 0.57 | 0.5 | 0.48 |
| 3 | 103747624 | rs9858370 | -0.573 | 0.545 | -0.737 | 0.896 | 0.008 | -0.001 | -0.001 | 0.831 | 0.39 | 0.49 | 0.49 | 0.35 | 0.44 | 0.59 |
| 3 | 103748729 | rs1165925 | -0.32 | 0.746 | -0.496 | 0.83 | 0.007 | 0.01 | 0.01 | 0 | 0.3 | 0.35 | 0.35 | 0.3 | 0.35 | 0.51 |
| 3 | 103750905 | rs1165930 | -1.061 | 0.814 | -0.846 | 0.849 | -0.02 | -0.056 | -0.056 | 0 | 0.79 | 0.89 | 0.74 | 0.67 | 0.73 | 0.73 |
| 3 | 103752461 | rs1165931 | -1.029 | 0.797 | -0.853 | 0.842 | -0.02 | -0.057 | -0.057 | 0 | 0.79 | 0.89 | 0.74 | 0.67 | 0.73 | 0.72 |
| 3 | 103754387 | rs6804331 | 0.766 | 0.541 | -0.599 | 0.661 | 0.015 | 0.073 | 0.073 | 0.029 | 0.46 | 0.23 | 0.36 | 0.37 | 0.38 | 0.23 |
| 3 | 103759995 | rs4420904 | -0.03 | 0.292 | -0.067 | 0.636 | -0.035 | -0.001 | -0.001 | 0 | 0.16 | 0.28 | 0.17 | 0.09 | 0.14 | 0.11 |
| 3 | 103760176 | rs4420906 | -0.512 | 0.232 | 0.197 | 0.645 | -0.068 | -0.039 | -0.039 | 0 | 0.19 | 0.49 | 0.18 | 0.08 | 0.13 | 0.1 |
| 3 | 103761138 | rs1920329 | -0.934 | 0.421 | 0.164 | 0.517 | -0.016 | -0.04 | -0.04 | 0.756 | 0.23 | 0.37 | 0.33 | 0.13 | 0.25 | 0.18 |
| 3 | 103762454 | rs10154899 | -1.274 | 0.583 | 0.064 | 0.75 | -0.011 | -0.009 | -0.009 | 0 | 0.28 | 0.54 | 0.4 | 0.24 | 0.3 | 0.29 |
| 3 | 103765618 | rs1165919 | -1.411 | 1.339 | -0.314 | 1.236 | -0.037 | -0.035 | -0.035 | 0 | 0.85 | 0.97 | 0.87 | 0.88 | 0.83 | 0.75 |
| 3 | 103766321 | rs16849564 | 0.775 | 1.264 | -0.199 | 1.144 | -0.002 | -0.002 | -0.002 | 0.104 | 0.07 | 0.06 | 0.16 | 0.15 | 0.12 | 0.08 |
| 3 | 103777910 | rs1447963 | 0.017 | 1.187 | -0.123 | 1.101 | 0 | -0.001 | -0.001 | 0.192 | 0.28 | 0.21 | 0.27 | 0.24 | 0.3 | 0.37 |
| 3 | 103777955 | rs1447964 | -0.75 | 0.954 | -0.07 | 1.036 | 0.001 | 0 | 0 | 0.056 | 0.4 | 0.57 | 0.5 | 0.4 | 0.47 | 0.55 |
| 3 | 103778214 | rs1447966 | -0.7 | 0.978 | -0.214 | 1.047 | -0.002 | -0.002 | -0.002 | 0 | 0.39 | 0.38 | 0.49 | 0.4 | 0.47 | 0.55 |
| 3 | 103795249 | rs2700429 | 1.31 | 0.971 | -0.626 | 0.362 | 0.007 | 0.062 | 0.062 | 0 | 0.32 | 0.18 | 0.22 | 0.09 | 0.28 | 0.12 |
| 3 | 103801180 | rs1731649 | -0.09 | 0.79 | -0.333 | 0.65 | -0.002 | 0.056 | 0.056 | 0.081 | 0.1 | 0.32 | 0.04 | 0.14 | 0.06 | 0.04 |
| 3 | 103806619 | rs1666559 | -0.489 | 0.466 | -1.69 | 0.612 | -0.064 | 0.003 | 0.003 | 0 | 0.54 | 0.86 | 0.56 | 0.65 | 0.46 | 0.52 |
| 3 | 103811702 | rs1345189 | 0.912 | 0.576 | -1.23 | 0.388 | -0.017 | 0.012 | 0.012 | 0 | 0.29 | 0.22 | 0.3 | 0.21 | 0.36 | 0.33 |
| 3 | 103811750 | rs1731642 | 0.816 | 0.086 | -0.821 | 0.47 | -0.025 | 0.018 | 0.018 | 0 | 0.2 | 0.42 | 0.16 | 0.26 | 0.14 | 0.13 |
| 3 | 103812426 | rs17533293 | 1.759 | -0.033 | -0.962 | 0.428 | 0.011 | 0.042 | 0.042 | 0.268 | 0.3 | 0.18 | 0.23 | 0.13 | 0.21 | 0.46 |
| 3 | 103813326 | rs13077637 | 1.515 | 0.108 | -0.995 | 0.502 | -0.017 | 0.013 | 0.013 | 0 | 0.29 | 0.22 | 0.3 | 0.21 | 0.36 | 0.33 |
| 3 | 103817461 | rs1165907 | -0.812 | 0.054 | -0.987 | 0.791 | -0.028 | -0.036 | -0.036 | 0 | 0.87 | 0.91 | 0.72 | 0.73 | 0.75 | 0.94 |
| 3 | 103820843 | rs6808059 | 0.981 | -0.087 | -0.772 | 0.524 | -0.026 | 0.018 | 0.018 | 0.055 | 0.09 | 0.12 | 0.02 | 0.14 | 0.03 | 0.02 |
| 3 | 103825912 | rs9855732 | 0.906 | 0.129 | -0.845 | 0.435 | -0.005 | -0.003 | -0.003 | 0 | 0.06 | 0.08 | 0.06 | 0 | 0.1 | 0.02 |
| 3 | 103829117 | rs1447982 | 1.634 | -0.034 | -0.914 | 0.276 | 0.009 | 0.035 | 0.035 | 0.334 | 0.29 | 0.19 | 0.24 | 0.13 | 0.22 | 0.46 |
| 3 | 103830499 | rs2044616 | 0.258 | -0.091 | -0.89 | 0.324 | -0.008 | 0.004 | 0.004 | 1.467 | 0.43 | 0.39 | 0.38 | 0.36 | 0.49 | 0.36 |
| 3 | 103830771 | rs7632326 | 1.756 | -0.143 | -0.846 | 0.324 | -0.047 | 0.016 | 0.016 | 0 | 0.28 | 0.14 | 0.29 | 0.21 | 0.36 | 0.31 |
| 3 | 103831303 | rs10212614 | 1.188 | -0.137 | -0.86 | 0.334 | -0.018 | 0.01 | 0.01 | 0.065 | 0.09 | 0.1 | 0.02 | 0.14 | 0.03 | 0.02 |
| 3 | 103831895 | rs9812361 | 0.031 | -0.133 | -0.86 | 0.332 | -0.001 | 0 | 0 | 0 | 0.15 | 0.18 | 0.08 | 0.14 | 0.13 | 0.04 |
| 3 | 103832345 | rs13321574 | 1.204 | -0.057 | -0.869 | 0.318 | -0.004 | -0.003 | -0.003 | 0 | 0.06 | 0.08 | 0.05 | 0 | 0.1 | 0.02 |
| 3 | 103835791 | rs10933738 | 1.703 | -0.157 | -0.873 | 0.337 | 0.011 | 0.041 | 0.041 | 0.29 | 0.3 | 0.18 | 0.23 | 0.13 | 0.21 | 0.46 |
| 3 | 103843936 | rs7620137 | -0.156 | -0.2 | -0.578 | 0.495 | -0.011 | 0.073 | 0.073 | 0 | 0.72 | 0.58 | 0.61 | 0.48 | 0.7 | 0.82 |
| 3 | 103855920 | rs9855313 | 1.271 | -0.191 | -0.684 | 0.528 | -0.014 | 0.013 | 0.013 | 0 | 0.08 | 0.09 | 0.02 | 0.14 | 0.03 | 0.03 |
| 3 | 103860285 | rs13093575 | 1.584 | -0.102 | -0.786 | 0.422 | -0.049 | 0.02 | 0.02 | 0 | 0.28 | 0.13 | 0.27 | 0.21 | 0.36 | 0.31 |
| 3 | 103864780 | rs7650653 | 1.499 | -0.099 | -0.768 | 0.424 | -0.034 | 0.02 | 0.02 | 0 | 0.28 | 0.15 | 0.27 | 0.21 | 0.36 | 0.31 |
| 3 | 103868553 | rs12630106 | 0.376 | -0.067 | -0.764 | 0.475 | -0.001 | -0.01 | -0.01 | 2.315 | 0.43 | 0.35 | 0.35 | 0.48 | 0.42 | 0.37 |

| | | | | | | | | | | | | | | | | |
|---|---|---|---|---|---|---|---|---|---|---|---|---|---|---|---|---|
| 3 | 103869519 | rs2141617 | -0.313 | -0.079 | -0.432 | 0.816 | -0.003 | 0.079 | 0.079 | 0 | 0.77 | 0.56 | 0.65 | 0.62 | 0.73 | 0.84 |
| 3 | 103870946 | rs7622522 | 0.003 | -0.036 | -0.703 | 0.586 | -0.003 | 0 | 0 | 0 | 0.13 | 0.13 | 0.12 | 0.13 | 0.13 | 0.05 |
| 3 | 103874997 | rs1920328 | 0.096 | -0.492 | -0.411 | 0.59 | -0.064 | 0.078 | 0.078 | 0.086 | 0.87 | 0.49 | 0.93 | 0.85 | 0.95 | 0.97 |
| 3 | 103879036 | rs4894836 | 0.809 | -0.443 | -0.449 | 0.557 | -0.021 | 0 | 0 | 0.689 | 0.81 | 0.41 | 0.87 | 0.85 | 0.85 | 0.95 |
| 3 | 103882496 | rs6783554 | 0.837 | -0.535 | -0.49 | 0.49 | -0.028 | -0.011 | -0.011 | 0 | 0.8 | 0.41 | 0.87 | 0.86 | 0.85 | 0.96 |
| 3 | 103888802 | rs6437492 | 0.629 | -0.441 | -0.359 | 0.488 | -0.003 | -0.005 | -0.005 | 0.094 | 0.06 | 0.07 | 0.06 | 0 | 0.09 | 0.02 |
| 3 | 103890240 | rs11719905 | 0.903 | 0.005 | -1.132 | 0.321 | -0.113 | -0.118 | -0.118 | 0 | 0.3 | 0.04 | 0.34 | 0.43 | 0.4 | 0.2 |
| 3 | 103890979 | rs1104964 | 1.244 | -0.038 | -1.159 | 0.124 | 0.011 | -0.04 | -0.04 | 0 | 0.54 | 0.12 | 0.57 | 0.55 | 0.56 | 0.63 |
| 3 | 103897067 | rs13060667 | 0.523 | 0.085 | -0.969 | 0.106 | -0.026 | -0.009 | -0.009 | 0.967 | 0.8 | 0.4 | 0.87 | 0.86 | 0.85 | 0.96 |
| 3 | 103900668 | rs4419405 | 1.018 | -0.011 | -1.017 | 0.101 | -0.001 | 0 | 0 | 0 | 0.66 | 0.38 | 0.66 | 0.68 | 0.64 | 0.67 |
| 3 | 103903449 | rs12639084 | 0.048 | -0.212 | -0.766 | 0.238 | -0.071 | 0.049 | 0.049 | 0 | 0.86 | 0.46 | 0.93 | 0.86 | 0.94 | 0.97 |
| 3 | 103912131 | rs1920331 | -0.039 | -0.013 | -0.799 | 0.276 | -0.04 | 0.012 | 0.012 | 0 | 0.78 | 0.53 | 0.84 | 0.75 | 0.84 | 0.87 |
| 3 | 103915789 | rs2141621 | 2.288 | 0.131 | -1.203 | 0.058 | 0.003 | 0.036 | 0.036 | 1.744 | 0.26 | 0.12 | 0.3 | 0.13 | 0.22 | 0.47 |
| 3 | 103921999 | rs1882006 | 0.893 | -0.058 | -0.889 | 0.496 | -0.001 | 0.005 | 0.005 | 0.343 | 0.52 | 0.35 | 0.6 | 0.46 | 0.51 | 0.71 |
| 3 | 103922216 | rs35695636 | 1.34 | 0.042 | -0.951 | 0.48 | -0.024 | -0.052 | -0.052 | 0.004 | 0.2 | 0.1 | 0.17 | 0.29 | 0.24 | 0.14 |
| 3 | 103931234 | rs1920346 | 1.34 | 0.042 | -0.951 | 0.48 | -0.024 | -0.052 | -0.052 | 0 | 0.2 | 0.1 | 0.17 | 0.29 | 0.24 | 0.14 |
| 3 | 103933915 | rs7616188 | 1.306 | 0.038 | -0.945 | 0.478 | -0.02 | -0.048 | -0.048 | 0.753 | 0.2 | 0.12 | 0.18 | 0.3 | 0.24 | 0.14 |
| 3 | 103943698 | rs2895264 | 0.266 | 0.044 | -0.937 | 0.438 | -0.007 | -0.02 | -0.02 | 0 | 0.72 | 0.47 | 0.78 | 0.75 | 0.75 | 0.84 |
| 3 | 103949117 | rs11925785 | 1.001 | 0.042 | -0.94 | 0.436 | -0.003 | 0.006 | 0.006 | 0.284 | 0.51 | 0.35 | 0.6 | 0.45 | 0.51 | 0.7 |
| 3 | 103955623 | rs13325125 | 1.426 | 0.055 | -1.049 | 0.411 | -0.051 | -0.063 | -0.063 | 0 | 0.19 | 0.06 | 0.19 | 0.29 | 0.26 | 0.13 |
| 3 | 103958244 | rs11717324 | 1.252 | 0.059 | -1.045 | 0.413 | -0.036 | -0.048 | -0.048 | 0.584 | 0.2 | 0.12 | 0.2 | 0.29 | 0.27 | 0.14 |
| 3 | 103960764 | rs2895261 | 1.753 | -0.732 | -1.18 | 0.338 | -0.07 | 0.116 | 0.116 | 0.543 | 0.37 | 0.09 | 0.42 | 0.18 | 0.4 | 0.64 |
| 3 | 103964141 | rs11716008 | 1.153 | -0.716 | -1.005 | 0.427 | -0.005 | 0.033 | 0.033 | 0.857 | 0.45 | 0.32 | 0.53 | 0.36 | 0.44 | 0.67 |
| 3 | 103975791 | rs4501143 | -1.424 | -0.719 | -1.049 | 0.402 | -0.011 | 0.036 | 0.036 | 0 | 0.87 | 0.98 | 0.91 | 0.92 | 0.89 | 0.88 |
| 3 | 103976415 | rs6437500 | 0.486 | -0.763 | -0.82 | 0.467 | -0.004 | -0.001 | -0.001 | 0 | 0.65 | 0.44 | 0.73 | 0.65 | 0.68 | 0.8 |
| 3 | 103979493 | rs13099947 | 1.267 | -0.767 | -0.871 | 0.465 | -0.017 | -0.048 | -0.048 | 0 | 0.2 | 0.11 | 0.2 | 0.3 | 0.25 | 0.14 |
| 3 | 103980782 | rs11706663 | 1.267 | -0.767 | -0.871 | 0.465 | -0.017 | -0.048 | -0.048 | 0 | 0.2 | 0.11 | 0.2 | 0.3 | 0.25 | 0.14 |
| 3 | 103981414 | rs11707610 | 1.504 | -0.748 | -0.928 | 0.475 | -0.041 | -0.077 | -0.077 | 0 | 0.19 | 0.04 | 0.18 | 0.3 | 0.24 | 0.13 |
| 3 | 103985553 | rs4895044 | 0.597 | -0.619 | -0.847 | 0.375 | -0.003 | 0 | 0 | 0 | 0.65 | 0.45 | 0.73 | 0.66 | 0.68 | 0.8 |
| 3 | 103990584 | rs34053484 | 1.228 | -0.546 | -0.917 | 0.191 | -0.054 | 0.107 | 0.107 | 0.257 | 0.37 | 0.1 | 0.42 | 0.18 | 0.4 | 0.64 |
| 3 | 104007457 | rs6437504 | 0.157 | -0.353 | -0.192 | 0.589 | -0.008 | 0.013 | 0.013 | 0 | 0.1 | 0.1 | 0.07 | 0.14 | 0.07 | 0.03 |
| 3 | 104017607 | rs9833635 | 0.148 | -0.34 | 0.223 | 0.549 | -0.006 | 0.144 | 0.144 | 1.359 | 0.95 | 0.72 | 0.88 | 0.8 | 0.98 | 0.97 |
| 3 | 104017741 | rs17434624 | 1.149 | -0.491 | 0.107 | 0.279 | -0.031 | 0.079 | 0.079 | 0.25 | 0.41 | 0.19 | 0.45 | 0.21 | 0.44 | 0.66 |
| 3 | 104019345 | rs2895281 | 0.878 | -0.435 | 0.087 | 0.317 | -0.039 | 0.129 | 0.129 | 0.721 | 0.44 | 0.15 | 0.46 | 0.21 | 0.45 | 0.71 |
| 3 | 104022366 | rs9879700 | -0.665 | -0.177 | 0.018 | 0.26 | 0 | 0.004 | 0.004 | 0.065 | 0.5 | 0.53 | 0.42 | 0.59 | 0.53 | 0.26 |
| 3 | 104031239 | rs2398993 | -0.043 | -0.331 | 0.19 | 0.131 | -0.012 | -0.03 | -0.03 | 0 | 0.23 | 0.29 | 0.13 | 0.05 | 0.16 | 0.06 |
| 3 | 104032299 | rs2398996 | 0.442 | -0.07 | 0.054 | 0.363 | 0.001 | 0.015 | 0.015 | 0.061 | 0.5 | 0.44 | 0.58 | 0.41 | 0.47 | 0.74 |
| 3 | 104057483 | rs7620614 | -1.438 | 0.023 | 0.39 | 1.089 | -0.027 | 0 | 0 | 0.532 | 0.38 | 0.29 | 0.53 | 0.6 | 0.41 | 0.46 |
| 3 | 104060810 | rs2399009 | 0.617 | -0.432 | 0.763 | 1.5 | -0.043 | 0.125 | 0.125 | 7.345 | 0.9 | 0.3 | 0.82 | 0.71 | 0.96 | 0.94 |
| 3 | 104061922 | rs6437511 | -0.655 | 0.095 | 0.335 | 1.935 | -0.02 | -0.055 | -0.055 | 0.141 | 0.14 | 0.04 | 0.26 | 0.34 | 0.16 | 0.23 |
| 3 | 104078908 | rs3749315 | -0.687 | -0.025 | 0.356 | 1.195 | 0.037 | 0.11 | 0.11 | 0 | 0.06 | 0.29 | 0.15 | 0.2 | 0.07 | 0.09 |
| 3 | 104079543 | rs12491237 | -0.738 | 0.011 | -0.692 | 1.138 | -0.016 | -0.05 | -0.05 | 0 | 0.14 | 0.04 | 0.25 | 0.32 | 0.15 | 0.25 |
| 3 | 104101105 | rs9828652 | -1.023 | -0.036 | -0.71 | 0.544 | -0.004 | 0.018 | 0.018 | 0 | 0.34 | 0.21 | 0.25 | 0.24 | 0.28 | 0.35 |
| 3 | 104103488 | rs17369659 | -0.444 | -0.043 | -0.749 | 0.566 | 0.004 | -0.016 | -0.016 | 1.453 | 0.26 | 0.14 | 0.26 | 0.28 | 0.2 | 0.31 |
| 3 | 104106742 | rs17435554 | 0.182 | 0.177 | -1.031 | 0.078 | -0.006 | -0.02 | -0.02 | 0 | 0.06 | 0 | 0.17 | 0.09 | 0.03 | 0.26 |
| 3 | 104107435 | rs11720877 | 1.208 | 0.148 | -1.713 | -0.444 | -0.135 | 0.306 | 0.306 | 0 | 0.43 | 0.03 | 0.26 | 0.09 | 0.51 | 0.36 |
| 3 | 104110851 | rs9823032 | -0.431 | 0.421 | -0.605 | 0.582 | 0.008 | 0.038 | 0.038 | 0 | 0.05 | 0.12 | 0.12 | 0.17 | 0.1 | 0.1 |
| 3 | 104111719 | rs9848147 | -0.727 | 0.379 | -0.868 | 0.323 | -0.002 | -0.011 | -0.011 | 0 | 0.09 | 0.08 | 0.07 | 0 | 0.07 | 0.05 |
| 3 | 104121004 | rs9852058 | 0.4 | -0.177 | -1.035 | 0.484 | 0.042 | 0.038 | 0.038 | 0 | 0.53 | 0.3 | 0.5 | 0.34 | 0.64 | 0.4 |
| 3 | 104127819 | rs2398963 | 0.96 | -0.44 | -1.173 | 0.415 | 0.019 | 0.207 | 0.207 | 0.02 | 0.45 | 0.15 | 0.43 | 0.16 | 0.59 | 0.33 |
| 3 | 104128989 | rs13061353 | 1.001 | -0.488 | -1.244 | 0.342 | -0.018 | 0.2 | 0.2 | 0.31 | 0.43 | 0.12 | 0.42 | 0.13 | 0.55 | 0.29 |
| 3 | 104135255 | rs6806731 | 0.339 | -1.059 | -0.569 | 0.656 | -0.03 | 0.017 | 0.017 | 0 | 0.67 | 0.57 | 0.78 | 0.68 | 0.76 | 0.79 |
| 3 | 104135554 | rs9871458 | 0.199 | -1.02 | -0.654 | 0.79 | 0.011 | -0.021 | -0.021 | 0 | 0.06 | 0.02 | 0.05 | 0.18 | 0.04 | 0.08 |
| 3 | 104157047 | rs2048784 | 0.053 | -1.029 | -0.536 | 0.934 | -0.002 | 0.022 | 0.022 | 0 | 0.19 | 0.19 | 0.32 | 0.48 | 0.16 | 0.44 |
| 3 | 104175286 | rs2673460 | 0.705 | -1.341 | -0.573 | 0.79 | 0.03 | -0.027 | -0.027 | 0 | 0.61 | 0.76 | 0.59 | 0.44 | 0.68 | 0.51 |
| 3 | 104175604 | rs1877997 | 1.935 | -1.463 | -0.578 | 0.594 | 0.02 | 0.182 | 0.182 | 0.16 | 0.43 | 0.15 | 0.45 | 0.15 | 0.57 | 0.28 |
| 3 | 104178878 | rs2673468 | -0.658 | -1.532 | -0.595 | 0.627 | 0.002 | 0.049 | 0.049 | 0 | 0.81 | 0.96 | 0.89 | 0.93 | 0.81 | 0.93 |
| 3 | 104179183 | rs10049306 | -0.909 | -1.603 | 0.244 | 0.701 | -0.048 | 0.099 | 0.099 | 0.463 | 0.12 | 0.31 | 0.08 | 0.25 | 0.05 | 0.2 |
| 3 | 104187485 | rs7653621 | 1.845 | -1.603 | 0.446 | 0.636 | 0.061 | 0.154 | 0.154 | 0 | 0.47 | 0.23 | 0.39 | 0.11 | 0.6 | 0.29 |
| 3 | 104201533 | rs9827573 | 0.542 | -1.544 | 0.491 | 0.77 | 0.028 | 0.103 | 0.103 | 0 | 0.05 | 0.28 | 0.05 | 0.08 | 0.02 | 0.06 |
| 3 | 104213388 | rs6790921 | -0.07 | -1.258 | 0.236 | 0.588 | -0.005 | -0.022 | -0.022 | 0 | 0.23 | 0.17 | 0.44 | 0.59 | 0.23 | 0.48 |
| 3 | 104214903 | rs3911814 | 1.031 | -1.207 | 0.341 | 0.624 | -0.042 | -0.002 | -0.002 | 0 | 0.67 | 0.47 | 0.79 | 0.69 | 0.76 | 0.72 |
| 3 | 104219900 | rs4131120 | 0.85 | -1.186 | 0.279 | 0.812 | -0.052 | 0.096 | 0.096 | 0.338 | 0.88 | 0.69 | 0.94 | 0.76 | 0.97 | 0.8 |
| 3 | 104225600 | rs1599760 | 1.129 | -1.229 | 0.316 | 0.8 | -0.069 | 0.096 | 0.096 | 2.127 | 0.84 | 0.65 | 0.93 | 0.73 | 0.96 | 0.76 |
| 3 | 104229825 | rs2961353 | 0.627 | -0.671 | -0.039 | 0.426 | -0.035 | 0.072 | 0.072 | 0 | 0.15 | 0.04 | 0.11 | 0 | 0.18 | 0.06 |
| 3 | 104232663 | rs4895081 | 0.841 | -0.714 | -0.414 | 1.082 | -0.004 | -0.053 | -0.053 | 5.155 | 0.11 | 0 | 0.18 | 0.22 | 0.1 | 0.18 |
| 3 | 104235998 | rs1606834 | 1.988 | -0.977 | -0.656 | 0.543 | 0.013 | 0.042 | 0.042 | 0 | 0.5 | 0.42 | 0.4 | 0.13 | 0.6 | 0.25 |
| 3 | 104238470 | rs2961193 | 0.119 | -1.014 | -0.613 | 0.745 | -0.047 | 0.122 | 0.122 | 0.637 | 0.88 | 0.7 | 0.93 | 0.76 | 0.97 | 0.8 |
| 3 | 104240760 | rs2961321 | 1.702 | -0.99 | -0.766 | 0.539 | 0.004 | 0.016 | 0.016 | 0 | 0.54 | 0.42 | 0.39 | 0.15 | 0.61 | 0.29 |

| | | | | | | | | | | | | | | | | |
|---|---|---|---|---|---|---|---|---|---|---|---|---|---|---|---|---|
| 3 | 104245490 | rs16850051 | 0.735 | -1.007 | -1.057 | 0.574 | -0.031 | -0.041 | -0.041 | 0.233 | 0.09 | 0.01 | 0.25 | 0.35 | 0.14 | 0.25 |
| 3 | 104249703 | rs9860685 | 2.861 | -1.084 | -0.996 | -0.281 | -0.029 | 0.072 | 0.072 | 0 | 0.33 | 0.16 | 0.33 | 0.16 | 0.4 | 0.2 |
| 3 | 104250596 | rs7635661 | 0.71 | -1.052 | -1.056 | -0.266 | -0.039 | 0.046 | 0.046 | 0 | 0.13 | 0.03 | 0.1 | 0 | 0.14 | 0.07 |
| 3 | 104255356 | rs9844288 | 1.553 | -1.239 | -0.888 | -0.19 | 0.017 | 0.07 | 0.07 | 0 | 0.54 | 0.36 | 0.53 | 0.17 | 0.64 | 0.32 |
| 3 | 104256753 | rs2961341 | 1.477 | -1.242 | -0.813 | -0.191 | 0.009 | 0.035 | 0.035 | 0 | 0.54 | 0.4 | 0.53 | 0.17 | 0.64 | 0.32 |
| 3 | 104259387 | rs1610247 | 0.949 | -1.279 | -0.783 | -0.161 | -0.005 | 0.06 | 0.06 | 0 | 0.62 | 0.81 | 0.6 | 0.24 | 0.66 | 0.38 |
| 3 | 104263821 | rs7643516 | -0.602 | -1.293 | -0.774 | -0.118 | -0.013 | -0.009 | -0.009 | 0 | 0.32 | 0.19 | 0.39 | 0.69 | 0.33 | 0.49 |
| 3 | 104265023 | rs2943735 | -0.873 | -1.293 | -0.778 | -0.141 | 0.021 | -0.037 | -0.037 | 0 | 0.94 | 1 | 0.99 | 0.88 | 0.99 | 0.87 |
| 3 | 104265382 | rs2943733 | 1.601 | -1.306 | -0.762 | -0.151 | 0.016 | 0.075 | 0.075 | 0 | 0.54 | 0.37 | 0.53 | 0.13 | 0.64 | 0.32 |
| 3 | 104266905 | rs11720334 | 0.57 | -1.296 | -0.737 | -0.306 | -0.056 | 0.09 | 0.09 | 0.791 | 0.87 | 0.61 | 0.92 | 0.82 | 0.97 | 0.81 |
| 3 | 104270022 | rs2048790 | -0.378 | -1.146 | -1.237 | -0.195 | 0.001 | -0.002 | -0.002 | 4.96 | 0.93 | 0.95 | 0.98 | 0.88 | 0.99 | 0.87 |
| 3 | 104274366 | rs2961223 | 0.718 | -1.036 | -1.258 | -0.272 | -0.043 | 0.059 | 0.059 | 0 | 0.14 | 0.05 | 0.19 | 0 | 0.22 | 0.09 |
| 3 | 104280035 | rs6802846 | -0.89 | -1.147 | -1.118 | -0.132 | -0.005 | 0.003 | 0.003 | 0 | 0.33 | 0.25 | 0.43 | 0.72 | 0.33 | 0.49 |
| 3 | 104289464 | rs6437523 | 0.279 | -1.083 | -1.472 | -0.76 | -0.017 | 0.052 | 0.052 | 0 | 0.18 | 0.04 | 0.19 | 0 | 0.22 | 0.15 |
| 3 | 104290380 | rs9870668 | 1.769 | -0.814 | -1.486 | -0.672 | -0.062 | 0.118 | 0.118 | 0 | 0.33 | 0.1 | 0.34 | 0.08 | 0.4 | 0.21 |
| 3 | 104305891 | rs6414228 | -1.846 | -0.755 | -1.577 | -0.626 | -0.058 | 0.118 | 0.118 | 0 | 0.66 | 0.89 | 0.66 | 0.91 | 0.6 | 0.79 |
| 3 | 104311987 | rs7428418 | -1.642 | -0.9 | -1.765 | -0.726 | -0.095 | 0.115 | 0.115 | 0.093 | 0.64 | 0.91 | 0.64 | 0.86 | 0.56 | 0.72 |
| 3 | 104315714 | rs4342125 | -0.337 | -1.443 | -1.269 | -0.857 | 0.002 | 0.002 | 0.002 | 0 | 0.36 | 0.29 | 0.25 | 0.21 | 0.3 | 0.31 |
| 3 | 104326725 | rs7648727 | -1.79 | -1.602 | -1.274 | -0.728 | 0.041 | 0.18 | 0.18 | 0.447 | 0.45 | 0.85 | 0.46 | 0.86 | 0.34 | 0.58 |
| 3 | 104341720 | rs12488015 | -0.18 | -1.726 | -0.971 | -0.424 | 0.001 | 0.012 | 0.012 | 0 | 0.18 | 0.21 | 0.34 | 0.6 | 0.22 | 0.39 |
| 3 | 104341975 | rs6785752 | 0.621 | -1.824 | -1.041 | -0.486 | 0.023 | -0.047 | -0.047 | 1.04 | 0.14 | 0.01 | 0.05 | 0.2 | 0.08 | 0.16 |
| 3 | 104343243 | rs718007 | -0.185 | -1.809 | -0.966 | -0.429 | 0.001 | 0.012 | 0.012 | 0 | 0.18 | 0.21 | 0.34 | 0.6 | 0.22 | 0.39 |
| 3 | 104348709 | rs16850153 | -0.851 | -1.734 | -0.959 | -0.382 | -0.01 | -0.019 | -0.019 | 0 | 0.23 | 0.23 | 0.32 | 0.59 | 0.25 | 0.41 |
| 3 | 104351297 | rs777942 | 0.671 | -1.837 | -1.066 | -0.494 | 0.024 | -0.052 | -0.052 | 0.584 | 0.13 | 0.02 | 0.05 | 0.19 | 0.08 | 0.16 |
| 3 | 104355010 | rs1083411 | -0.367 | -1.338 | -0.889 | -0.723 | -0.06 | -0.067 | -0.067 | 0 | 0.19 | 0.06 | 0.3 | 0.64 | 0.24 | 0.36 |
| 3 | 104363856 | rs796589 | -1.191 | -1.424 | -1.031 | -1.1 | -0.045 | 0.03 | 0.03 | 0 | 0.41 | 0.21 | 0.49 | 0.64 | 0.46 | 0.51 |
| 3 | 104369646 | rs2171524 | 1.356 | -1.642 | -0.477 | -1.129 | 0.01 | 0.076 | 0.076 | 0 | 0.5 | 0.3 | 0.46 | 0.35 | 0.51 | 0.45 |
| 3 | 104380356 | rs2947772 | -0.977 | -1.65 | -0.064 | -1.13 | -0.021 | 0.023 | 0.023 | 0 | 0.42 | 0.31 | 0.5 | 0.64 | 0.45 | 0.51 |
| 3 | 104381193 | rs9848965 | 1.354 | -1.642 | -0.131 | -1.123 | 0.01 | 0.076 | 0.076 | 0 | 0.5 | 0.3 | 0.46 | 0.35 | 0.51 | 0.45 |
| 3 | 104383689 | rs777875 | -0.772 | -1.576 | -0.253 | -1.098 | -0.075 | 0.017 | 0.017 | 0 | 0.39 | 0.12 | 0.48 | 0.64 | 0.45 | 0.51 |
| 3 | 104389814 | rs777937 | -0.024 | -1.536 | 0.175 | -1.098 | -0.012 | -0.168 | -0.168 | 0 | 0.72 | 0.55 | 0.78 | 0.99 | 0.74 | 0.83 |
| 3 | 104396611 | rs777929 | -1.462 | -1.671 | -0.313 | -1.356 | -0.021 | 0.018 | 0.018 | 0 | 0.73 | 0.85 | 0.68 | 0.85 | 0.65 | 0.76 |
| 3 | 104404772 | rs777917 | -1.211 | -1.684 | -0.275 | -1.365 | 0.034 | -0.044 | -0.044 | 0 | 0.72 | 0.67 | 0.66 | 0.85 | 0.65 | 0.76 |
| 3 | 104414871 | rs11711077 | 0.855 | -1.721 | -0.359 | -1.382 | -0.008 | 0.081 | 0.081 | 0 | 0.26 | 0.07 | 0.19 | 0.01 | 0.25 | 0.17 |
| 3 | 104418573 | rs2685056 | -0.313 | -1.728 | -0.425 | -1.362 | 0.048 | 0.106 | 0.106 | 0 | 0.48 | 0.78 | 0.48 | 0.84 | 0.4 | 0.59 |
| 3 | 104421626 | rs1521615 | 0.019 | -1.709 | -0.547 | -1.361 | 0.02 | 0.047 | 0.047 | 0 | 0.44 | 0.39 | 0.45 | 0.84 | 0.39 | 0.57 |
| 3 | 104422636 | rs2633650 | -0.431 | -1.705 | -0.539 | -1.368 | -0.008 | -0.098 | -0.098 | 1.882 | 0.7 | 0.43 | 0.76 | 0.99 | 0.74 | 0.81 |
| 3 | 104424997 | rs12633386 | 1.518 | -1.694 | -0.535 | -1.292 | -0.118 | 0.042 | 0.042 | 0 | 0.26 | 0.03 | 0.31 | 0.16 | 0.35 | 0.25 |
| 3 | 104427141 | rs9813174 | 1.174 | -1.758 | -0.587 | -1.338 | -0.124 | 0.061 | 0.061 | 0.159 | 0.29 | 0.03 | 0.32 | 0.16 | 0.36 | 0.25 |
| 3 | 104427677 | rs9851845 | 1.488 | -1.714 | -0.557 | -1.317 | -0.118 | 0.045 | 0.045 | 0 | 0.26 | 0.03 | 0.31 | 0.16 | 0.35 | 0.25 |
| 3 | 104428281 | rs2607960 | -0.494 | -1.646 | -0.487 | -1.358 | -0.003 | -0.09 | -0.09 | 0 | 0.69 | 0.42 | 0.74 | 0.99 | 0.72 | 0.81 |
| 3 | 104431587 | rs777897 | -0.485 | -1.651 | -0.478 | -1.36 | -0.003 | -0.09 | -0.09 | 0.126 | 0.69 | 0.42 | 0.74 | 0.99 | 0.72 | 0.81 |
| 3 | 104432140 | rs777896 | -0.537 | -1.637 | -0.461 | -1.354 | -0.005 | -0.148 | -0.148 | 0 | 0.7 | 0.51 | 0.75 | 0.99 | 0.72 | 0.81 |
| 3 | 104439114 | rs2947786 | 1.027 | -1.653 | -0.562 | -0.798 | -0.036 | -0.038 | -0.038 | 1.178 | 0.11 | 0.04 | 0.2 | 0.5 | 0.16 | 0.31 |
| 3 | 104452654 | rs13325883 | 0.339 | -1.608 | -0.756 | -0.867 | 0.089 | 0.194 | 0.194 | 0 | 0.52 | 0.1 | 0.52 | 0.18 | 0.63 | 0.42 |
| 3 | 104472192 | rs9847245 | 1.084 | -1.828 | -0.378 | -0.804 | 0.002 | -0.002 | -0.002 | 1.102 | 0.26 | 0.28 | 0.2 | 0.32 | 0.19 | 0.25 |
| 3 | 104473149 | rs777967 | -0.702 | -1.815 | -0.543 | -0.798 | 0.056 | 0.005 | 0.005 | 0 | 0.69 | 0.21 | 0.75 | 0.68 | 0.8 | 0.75 |
| 3 | 104487853 | rs796590 | 0.449 | -1.717 | -0.821 | -1.076 | -0.022 | 0.058 | 0.058 | 0 | 0.42 | 0.12 | 0.52 | 0.67 | 0.52 | 0.56 |
| 3 | 104493257 | rs6809018 | -0.6 | -2.095 | -0.589 | -1.117 | 0.078 | 0.032 | 0.032 | 0 | 0.68 | 0.17 | 0.72 | 0.75 | 0.8 | 0.73 |
| 3 | 104500264 | rs7640427 | 0.679 | -2.052 | -0.49 | -1.098 | -0.063 | -0.019 | -0.019 | 0 | 0.29 | 0.49 | 0.22 | 0.25 | 0.19 | 0.25 |
| 3 | 104513265 | rs6437539 | -0.157 | -1.784 | -0.651 | -1.1 | 0 | 0 | 0 | 0 | 0.64 | 0.66 | 0.61 | 0.43 | 0.63 | 0.55 |
| 3 | 104515782 | rs1436326 | -0.223 | -1.73 | -0.948 | -1.059 | -0.002 | -0.003 | -0.003 | 0 | 0.65 | 0.68 | 0.61 | 0.43 | 0.63 | 0.55 |
| 3 | 104521570 | rs9844843 | 1.124 | -1.833 | -1.073 | -1.024 | -0.029 | -0.007 | -0.007 | 0.042 | 0.25 | 0.36 | 0.21 | 0.24 | 0.18 | 0.22 |
| 3 | 104528615 | rs13066998 | 1.132 | -1.478 | -1.567 | -1.11 | -0.125 | 0.02 | 0.02 | 0 | 0.29 | 0.03 | 0.32 | 0.16 | 0.35 | 0.3 |
| 3 | 104530166 | rs3996202 | -0.109 | -1.613 | -1.465 | -1.259 | 0.019 | 0.111 | 0.111 | 0 | 0.53 | 0.73 | 0.4 | 0.26 | 0.5 | 0.38 |
| 3 | 104530538 | rs3996204 | 1.516 | -1.6 | -1.523 | -1.204 | -0.13 | 0.027 | 0.027 | 0 | 0.26 | 0.03 | 0.33 | 0.12 | 0.35 | 0.28 |
| 3 | 104533206 | rs9879873 | -0.597 | -1.568 | -1.393 | -1.143 | 0.001 | -0.01 | -0.01 | 0 | 0.81 | 0.81 | 0.76 | 0.37 | 0.86 | 0.69 |
| 3 | 104536448 | rs10446267 | 0.499 | -1.922 | -0.842 | -1.047 | -0.059 | -0.074 | -0.074 | 0.808 | 0.27 | 0.46 | 0.14 | 0.12 | 0.19 | 0.15 |
| 3 | 104539742 | rs6768068 | -0.196 | -1.872 | -0.936 | -1.106 | 0.055 | 0.154 | 0.154 | 0 | 0.46 | 0.74 | 0.33 | 0.12 | 0.39 | 0.3 |
| 3 | 104542237 | rs1836862 | 1.4 | -1.565 | -1.012 | -0.627 | -0.126 | -0.141 | -0.141 | 0 | 0.28 | 0.03 | 0.38 | 0.49 | 0.38 | 0.42 |
| 3 | 104551878 | rs11710356 | 0.606 | -1.921 | 0.084 | -0.647 | 0.01 | 0.013 | 0.013 | 0.024 | 0.1 | 0.05 | 0.03 | 0.03 | 0.06 | 0.02 |
| 3 | 104556687 | rs7646681 | 0.963 | -1.924 | -0.197 | -0.649 | -0.036 | -0.038 | -0.038 | 0 | 0.16 | 0.34 | 0.1 | 0.08 | 0.13 | 0.13 |
| 3 | 104562324 | rs1975269 | 0.342 | -1.844 | -0.476 | -0.679 | -0.064 | -0.081 | -0.081 | 0 | 0.71 | 0.48 | 0.83 | 0.89 | 0.8 | 0.84 |
| 3 | 104570060 | rs9851350 | 0.85 | -1.802 | -1.054 | -0.632 | 0.084 | 0.043 | 0.043 | 0.187 | 0.64 | 0.17 | 0.73 | 0.78 | 0.76 | 0.77 |
| 3 | 104572458 | rs10933793 | -0.471 | -0.907 | -0.855 | -0.073 | -0.053 | -0.06 | -0.06 | 0 | 0.82 | 0.6 | 0.87 | 0.92 | 0.87 | 0.87 |
| 3 | 104575864 | rs12635480 | -0.181 | 0.002 | -1.798 | 0.4 | -0.005 | -0.022 | -0.022 | 0.095 | 0.05 | 0.01 | 0.07 | 0.11 | 0.03 | 0.07 |
| 3 | 104580647 | rs1946726 | 0.022 | -0.349 | -2.199 | -0.607 | 0.068 | 0.221 | 0.221 | 0.018 | 0.59 | 0.13 | 0.66 | 0.92 | 0.67 | 0.69 |
| 3 | 104581866 | rs13061652 | -0.426 | 0.308 | -1.837 | -0.239 | -0.007 | -0.004 | -0.004 | 3.428 | 0.38 | 0.3 | 0.36 | 0.39 | 0.4 | 0.35 |
| 3 | 104586195 | rs9880353 | 0.83 | 0.23 | -1.565 | -0.215 | 0.017 | -0.009 | -0.009 | 0 | 0.26 | 0.31 | 0.29 | 0.26 | 0.3 | 0.42 |

| | | | | | | | | | | | | | | | | |
|---|---|---|---|---|---|---|---|---|---|---|---|---|---|---|---|---|
| 3 | 104586688 | rs9825604 | 1.215 | 0.253 | -1.586 | -0.559 | 0 | -0.032 | -0.032 | 0 | 0.16 | 0.15 | 0.16 | 0 | 0.15 | 0.1 |
| 3 | 104587776 | rs7649811 | 1.54 | -0.31 | -1.792 | -0.527 | 0.018 | 0.034 | 0.034 | 0 | 0.18 | 0.1 | 0.1 | 0.11 | 0.16 | 0.15 |
| 3 | 104594467 | rs16850638 | 2.381 | -0.367 | -1.791 | -0.466 | -0.005 | -0.057 | -0.057 | 0 | 0.09 | 0.01 | 0.19 | 0.21 | 0.09 | 0.06 |
| 3 | 104595269 | rs1436339 | 1.423 | 0.095 | -1.74 | -0.512 | -0.04 | -0.02 | -0.02 | 1.607 | 0.08 | 0 | 0.06 | 0.06 | 0.11 | 0.08 |
| 3 | 104595338 | rs11713106 | 1.216 | 0.025 | -1.492 | -0.517 | -0.009 | -0.015 | -0.015 | 0 | 0.19 | 0.29 | 0.11 | 0.12 | 0.17 | 0.16 |
| 3 | 104599330 | rs11926023 | 0.878 | 0.52 | -1.31 | -0.192 | 0.017 | -0.009 | -0.009 | 0 | 0.26 | 0.31 | 0.29 | 0.26 | 0.3 | 0.42 |
| 3 | 104604488 | rs1344860 | 0.285 | 0.023 | -1.692 | -0.483 | -0.002 | 0.04 | 0.04 | 0 | 0.27 | 0.13 | 0.17 | 0.2 | 0.3 | 0.24 |
| 3 | 104605525 | rs1946725 | 0.799 | 0.295 | -1.538 | -0.823 | 0 | -0.04 | -0.04 | 0.315 | 0.29 | 0.36 | 0.23 | 0.22 | 0.29 | 0.36 |
| 3 | 104607077 | rs1436345 | -0.278 | -0.037 | -1.549 | -0.682 | -0.002 | -0.001 | -0.001 | 0 | 0.72 | 0.69 | 0.63 | 0.7 | 0.68 | 0.69 |
| 3 | 104607231 | rs10511230 | 1.446 | 0.188 | -1.686 | -0.75 | -0.043 | -0.071 | -0.071 | 1.685 | 0.12 | 0.03 | 0.17 | 0.29 | 0.14 | 0.14 |
| 3 | 104615905 | rs1436349 | -1.135 | 0.144 | -1.596 | -0.808 | 0.026 | -0.066 | -0.066 | 0 | 0.85 | 0.72 | 0.81 | 0.99 | 0.85 | 0.83 |
| 3 | 104616256 | rs1436350 | -1.135 | 0.144 | -1.547 | -0.808 | 0.02 | -0.059 | -0.059 | 0 | 0.85 | 0.74 | 0.82 | 0.99 | 0.85 | 0.83 |
| 3 | 104617973 | rs1436351 | -0.674 | 0.023 | -1.61 | -0.823 | -0.015 | 0.012 | 0.012 | 0 | 0.7 | 0.51 | 0.59 | 0.71 | 0.76 | 0.75 |
| 3 | 104618616 | rs11914438 | 2.139 | 0.022 | -1.619 | -0.816 | -0.022 | 0.027 | 0.027 | 0 | 0.16 | 0.2 | 0.23 | 0.28 | 0.09 | 0.08 |
| 3 | 104620103 | rs9874718 | 0.878 | 0.3 | -1.213 | -0.751 | 0.002 | 0.031 | 0.031 | 0 | 0.28 | 0.16 | 0.17 | 0.2 | 0.3 | 0.24 |
| 3 | 104623906 | rs12630650 | 1.175 | 0.31 | -1.398 | -0.724 | -0.05 | -0.078 | -0.078 | 0 | 0.14 | 0.03 | 0.18 | 0.29 | 0.17 | 0.15 |
| 3 | 104625385 | rs9873656 | 1.216 | 0.139 | -1.038 | -1.351 | 0.025 | 0.021 | 0.021 | 0 | 0.2 | 0.13 | 0.11 | 0.18 | 0.19 | 0.16 |
| 3 | 104628143 | rs12486139 | 0.97 | 0.025 | -0.633 | -1.216 | -0.001 | 0.001 | 0.001 | 0 | 0.25 | 0.23 | 0.21 | 0.24 | 0.29 | 0.36 |
| 3 | 104631603 | rs9830592 | -0.294 | 0.205 | -0.641 | -1.244 | 0.001 | -0.008 | -0.008 | 0 | 0.56 | 0.46 | 0.4 | 0.42 | 0.59 | 0.6 |
| 3 | 104642165 | rs1865784 | -0.042 | 0.207 | -0.671 | -1.158 | 0.003 | 0.014 | 0.014 | 0 | 0.28 | 0.2 | 0.18 | 0.2 | 0.31 | 0.24 |
| 3 | 104646784 | rs4895117 | 0.161 | 0.248 | -0.679 | -1.066 | 0.002 | 0.019 | 0.019 | 0 | 0.87 | 0.93 | 0.89 | 0.95 | 0.92 | 0.89 |
| 3 | 104651404 | rs999704 | 0.617 | 0.033 | -0.094 | -1.162 | -0.01 | -0.037 | -0.037 | 1.229 | 0.86 | 0.78 | 0.88 | 0.95 | 0.92 | 0.86 |
| 3 | 104652489 | rs7432288 | 1.089 | 0.491 | 0.345 | -1.615 | 0.023 | -0.037 | -0.037 | 0 | 0.14 | 0.21 | 0.08 | 0.03 | 0.14 | 0.06 |
| 3 | 104653154 | rs993462 | -0.236 | 0.481 | 0.348 | -1.612 | 0.003 | 0.008 | 0.008 | 0 | 0.87 | 0.92 | 0.89 | 0.95 | 0.92 | 0.86 |
| 3 | 104658328 | rs6783013 | 0.827 | 0.226 | -0.155 | -2.377 | 0.006 | 0.023 | 0.023 | 0 | 0.63 | 0.69 | 0.78 | 0.92 | 0.72 | 0.74 |
| 3 | 104660707 | rs11921789 | 1.603 | 0.301 | -0.14 | -2.329 | 0.018 | -0.031 | -0.031 | 0 | 0.14 | 0.19 | 0.08 | 0.03 | 0.14 | 0.06 |
| 3 | 104663005 | rs9860129 | -0.165 | 0.293 | -0.133 | -2.303 | 0.005 | -0.035 | -0.035 | 4.029 | 0.74 | 0.72 | 0.81 | 0.92 | 0.77 | 0.8 |
| 3 | 104669238 | rs1436337 | 1.498 | -0.226 | -0.099 | -2.504 | -0.004 | 0.008 | 0.008 | 0 | 0.37 | 0.36 | 0.53 | 0.85 | 0.49 | 0.63 |
| 3 | 104693252 | rs13319705 | -0.158 | -0.39 | -0.101 | -2.29 | 0.01 | 0.199 | 0.199 | 0 | 0.36 | 0.1 | 0.31 | 0.06 | 0.29 | 0.19 |
| 3 | 104704485 | rs6771790 | 1.41 | -0.384 | 0.121 | -2.343 | 0.016 | -0.027 | -0.027 | 1.044 | 0.15 | 0.21 | 0.09 | 0.08 | 0.15 | 0.07 |
| 3 | 104714210 | rs9869633 | 0.222 | -0.482 | 0.349 | -2.406 | -0.004 | 0.078 | 0.078 | 0 | 0.26 | 0.14 | 0.17 | 0.02 | 0.22 | 0.15 |
| 3 | 104715038 | rs13324580 | 0.552 | -0.475 | 0.361 | -2.403 | -0.015 | 0.099 | 0.099 | 0 | 0.24 | 0.1 | 0.16 | 0.02 | 0.21 | 0.14 |
| 3 | 104715856 | rs6769796 | -1.308 | -0.559 | 0.665 | -2.425 | -0.01 | 0.048 | 0.048 | 0.116 | 0.55 | 0.64 | 0.43 | 0.15 | 0.44 | 0.28 |
| 3 | 104722799 | rs6797921 | 0.234 | -0.484 | 0.262 | -2.411 | -0.004 | 0.078 | 0.078 | 0 | 0.26 | 0.14 | 0.17 | 0.03 | 0.22 | 0.14 |
| 3 | 104722931 | rs17452316 | 1.554 | -0.487 | 0.307 | -2.421 | 0.014 | -0.028 | -0.028 | 0.148 | 0.14 | 0.19 | 0.08 | 0.04 | 0.15 | 0.06 |
| 3 | 104724634 | rs971551 | 0.127 | -0.435 | 0.293 | -2.375 | 0.002 | 0.027 | 0.027 | 0 | 0.26 | 0.22 | 0.17 | 0.03 | 0.22 | 0.14 |
| 3 | 104730056 | rs9682035 | -0.826 | -0.522 | 0.382 | -2.4 | 0.001 | -0.017 | -0.017 | 0.548 | 0.49 | 0.49 | 0.4 | 0.15 | 0.42 | 0.25 |
| 3 | 104730581 | rs9288796 | 0.578 | -0.422 | 0.121 | -2.316 | 0.001 | 0.032 | 0.032 | 0 | 0.21 | 0.17 | 0.16 | 0.03 | 0.19 | 0.11 |
| 3 | 104736772 | rs17387070 | 0.617 | -0.411 | 0.289 | -2.356 | 0.003 | -0.002 | -0.002 | 0 | 0.51 | 0.42 | 0.55 | 0.89 | 0.59 | 0.75 |
| 3 | 104736837 | rs16850772 | 0.269 | -0.427 | 0.217 | -2.429 | -0.001 | -0.07 | -0.07 | 0 | 0.63 | 0.42 | 0.7 | 0.92 | 0.68 | 0.82 |
| 3 | 104736846 | rs6781572 | 0.258 | -0.493 | 0.293 | -2.461 | -0.001 | -0.031 | -0.031 | 0.044 | 0.2 | 0.25 | 0.16 | 0.03 | 0.18 | 0.11 |
| 3 | 104737039 | rs17452768 | 2.267 | -0.33 | 0.139 | -2.368 | 0.009 | 0.059 | 0.059 | 0 | 0.12 | 0 | 0.15 | 0.03 | 0.09 | 0.07 |
| 3 | 104740516 | rs1503081 | -1.56 | -0.283 | 0.221 | -2.406 | 0.004 | -0.05 | -0.05 | 0.67 | 0.83 | 0.67 | 0.87 | 0.95 | 0.87 | 0.91 |
| 3 | 104742187 | rs7626601 | 0.45 | -0.417 | 0.337 | -2.453 | 0.007 | -0.089 | -0.089 | 1.598 | 0.19 | 0.27 | 0.16 | 0.02 | 0.17 | 0.07 |
| 3 | 104746619 | rs2399031 | 1.966 | -0.495 | 0.098 | -2.306 | 0.013 | 0.074 | 0.074 | 0 | 0.12 | 0 | 0.07 | 0.01 | 0.09 | 0.07 |
| 3 | 104748323 | rs9288798 | -0.642 | -0.466 | 0.331 | -2.325 | -0.018 | -0.092 | -0.092 | 0.963 | 0.69 | 0.56 | 0.78 | 0.94 | 0.76 | 0.82 |
| 3 | 104754616 | rs9819891 | 0.661 | -0.153 | 0.137 | -2.43 | 0.006 | -0.039 | -0.039 | 0 | 0.08 | 0.15 | 0.04 | 0 | 0.07 | 0.05 |
| 3 | 104755481 | rs13325696 | 0.274 | -0.272 | 0.781 | -2.683 | 0.013 | -0.137 | -0.137 | 0 | 0.2 | 0.48 | 0.17 | 0.01 | 0.17 | 0.1 |
| 3 | 104759032 | rs1503079 | 0.854 | -0.254 | 0.487 | -2.76 | -0.04 | 0.187 | 0.187 | 0.347 | 0.41 | 0.17 | 0.62 | 0.93 | 0.51 | 0.7 |
| 3 | 104763677 | rs4894824 | 0.198 | -0.331 | 0.849 | -2.827 | 0.008 | -0.116 | -0.116 | 0.782 | 0.35 | 0.42 | 0.23 | 0.06 | 0.32 | 0.16 |
| 3 | 104768678 | rs9875475 | 0.162 | -0.322 | 0.84 | -2.835 | 0.008 | -0.115 | -0.115 | 0 | 0.35 | 0.42 | 0.23 | 0.06 | 0.32 | 0.16 |
| 3 | 104773065 | rs9863924 | -0.392 | -0.317 | 0.916 | -2.636 | 0.018 | -0.025 | -0.025 | 0 | 0.95 | 0.8 | 0.95 | 1 | 0.95 | 0.95 |
| 3 | 104783637 | rs9814542 | 0.174 | -0.301 | 0.854 | -2.601 | 0.011 | -0.126 | -0.126 | 1.043 | 0.35 | 0.44 | 0.23 | 0.06 | 0.32 | 0.16 |
| 3 | 104786133 | rs1587707 | 0.843 | -0.319 | 0.783 | -2.59 | -0.018 | 0.13 | 0.13 | 0.168 | 0.42 | 0.23 | 0.62 | 0.93 | 0.51 | 0.7 |
| 3 | 104790415 | rs2895295 | 0.843 | -0.319 | 0.783 | -2.596 | -0.018 | 0.13 | 0.13 | 0.168 | 0.42 | 0.23 | 0.62 | 0.93 | 0.51 | 0.7 |
| 3 | 104793035 | rs2221332 | -0.43 | -0.327 | 0.725 | -2.573 | 0.001 | -0.04 | -0.04 | 0 | 0.76 | 0.71 | 0.86 | 0.99 | 0.84 | 0.86 |
| 3 | 104800350 | rs10933809 | 0.831 | -0.297 | 0.344 | -2.574 | -0.033 | 0.189 | 0.189 | 0.205 | 0.42 | 0.19 | 0.62 | 0.93 | 0.51 | 0.7 |
| 3 | 104805417 | rs1503089 | -0.071 | 0.115 | 0.446 | -2.255 | 0.005 | -0.087 | -0.087 | 0 | 0.35 | 0.4 | 0.24 | 0.06 | 0.34 | 0.18 |
| 3 | 104815999 | rs6437566 | -0.268 | 0.143 | -0.564 | -2.243 | -0.029 | 0.102 | 0.102 | 0 | 0.67 | 0.84 | 0.79 | 0.95 | 0.68 | 0.84 |
| 3 | 104821917 | rs9881967 | 0.6 | 0.238 | -0.7 | -2.196 | -0.012 | 0.016 | 0.016 | 0 | 0.14 | 0.1 | 0.13 | 0.01 | 0.16 | 0.06 |
| 3 | 104822019 | rs9881866 | 0.384 | 0.009 | -0.686 | -2.129 | -0.001 | -0.032 | -0.032 | 0 | 0.47 | 0.54 | 0.68 | 0.9 | 0.54 | 0.72 |
| 3 | 104833842 | rs4894825 | -0.23 | 0.281 | -0.556 | -0.854 | 0.002 | 0.017 | 0.017 | 0 | 0.81 | 0.72 | 0.77 | 0.76 | 0.78 | 0.79 |
| 3 | 104839517 | rs17389434 | -0.204 | 0.26 | -0.716 | -1.043 | 0.001 | -0.003 | -0.003 | 0 | 0.61 | 0.61 | 0.63 | 0.74 | 0.59 | 0.64 |
| 3 | 104850687 | rs12630126 | 1.14 | 0.157 | -1.231 | -0.482 | 0.012 | -0.038 | -0.038 | 0.45 | 0.13 | 0.07 | 0.2 | 0.34 | 0.14 | 0.14 |
| 3 | 104858070 | rs4894905 | -0.345 | 0.403 | -1.177 | -0.489 | 0.006 | -0.012 | -0.012 | 1.087 | 0.58 | 0.45 | 0.56 | 0.72 | 0.55 | 0.61 |
| 3 | 104859722 | rs3843985 | 1.178 | 1.104 | -0.984 | 0.741 | -0.03 | -0.019 | -0.019 | 3.24 | 0.14 | 0.04 | 0.16 | 0.14 | 0.18 | 0.15 |
| 3 | 104864415 | rs9880276 | 1.176 | 1.173 | -0.918 | 0.77 | -0.003 | 0.002 | 0.002 | 0 | 0.36 | 0.26 | 0.41 | 0.39 | 0.37 | 0.28 |
| 3 | 104869523 | rs9861329 | 1.093 | 0.436 | -0.119 | 0.812 | 0.02 | -0.001 | -0.001 | 0 | 0.5 | 0.29 | 0.58 | 0.53 | 0.55 | 0.43 |
| 3 | 104869888 | rs16850895 | 0.322 | 0.744 | -1.159 | 1.124 | 0.006 | -0.024 | -0.024 | 0.312 | 0.17 | 0.07 | 0.11 | 0.24 | 0.16 | 0.25 |

| | | | | | | | | | | | | | | | | |
|---|---|---|---|---|---|---|---|---|---|---|---|---|---|---|---|---|
| 3 | 104870677 | rs16850901 | -0.456 | 0.953 | -0.647 | 0.931 | 0.002 | 0 | 0 | 0.049 | 0.05 | 0.07 | 0.08 | 0.08 | 0.1 | 0.07 |
| 3 | 104878270 | rs9880693 | 0.46 | 0.863 | -0.691 | 0.882 | -0.001 | -0.003 | -0.003 | 0 | 0.72 | 0.65 | 0.69 | 0.78 | 0.72 | 0.72 |
| 3 | 104881936 | rs6770943 | -0.734 | 1.115 | -0.82 | 1.471 | -0.006 | 0.002 | 0.002 | 0 | 0.13 | 0.11 | 0.18 | 0.32 | 0.22 | 0.09 |
| 3 | 104887557 | rs12695107 | -0.198 | 1.223 | -1.05 | 0.913 | -0.003 | 0.004 | 0.004 | 0 | 0.76 | 0.79 | 0.64 | 0.87 | 0.77 | 0.74 |
| 3 | 104890715 | rs1393784 | 1.182 | 0.282 | -1.401 | 0.086 | -0.039 | 0.143 | 0.143 | 0 | 0.35 | 0.12 | 0.28 | 0.11 | 0.39 | 0.34 |
| 3 | 104892984 | rs16850936 | -0.28 | -0.138 | -0.505 | 0.472 | -0.008 | 0.019 | 0.019 | 2.431 | 0.3 | 0.34 | 0.25 | 0.41 | 0.28 | 0.21 |
| 3 | 104895565 | rs13094039 | 0.84 | -0.158 | -0.515 | 0.476 | -0.002 | 0.003 | 0.003 | 0 | 0.46 | 0.45 | 0.4 | 0.44 | 0.5 | 0.54 |
| 3 | 104907697 | rs7626224 | -0.885 | 0.023 | -0.262 | 0.473 | -0.002 | 0.002 | 0.002 | 0 | 0.75 | 0.78 | 0.82 | 0.88 | 0.77 | 0.73 |
| 3 | 104907931 | rs7628453 | -0.141 | -0.196 | -0.061 | 0.455 | -0.003 | 0.012 | 0.012 | 0 | 0.23 | 0.24 | 0.23 | 0.32 | 0.24 | 0.15 |
| 3 | 104915169 | rs10511238 | 1.076 | -0.11 | -0.133 | 0.355 | -0.004 | 0.009 | 0.009 | 1.86 | 0.05 | 0.09 | 0.04 | 0.08 | 0.04 | 0.06 |
| 3 | 104916079 | rs1525851 | -0.792 | -0.13 | 0.01 | 0.256 | 0.008 | -0.019 | -0.019 | 0 | 0.75 | 0.71 | 0.82 | 0.87 | 0.77 | 0.77 |
| 3 | 104917227 | rs1568421 | 0.327 | -0.373 | 0.34 | 0.272 | -0.025 | 0.099 | 0.099 | 1.106 | 0.89 | 0.62 | 0.74 | 0.89 | 0.92 | 0.87 |
| 3 | 104917414 | rs9874124 | 0.914 | -0.527 | 0.315 | 0.365 | -0.025 | 0.04 | 0.04 | 0.957 | 0.68 | 0.54 | 0.54 | 0.51 | 0.7 | 0.75 |
| 3 | 104928448 | rs9846852 | 0.923 | -0.508 | 0.374 | 0.374 | -0.029 | 0.047 | 0.047 | 1.76 | 0.68 | 0.5 | 0.53 | 0.51 | 0.7 | 0.74 |
| 3 | 104935343 | rs9818420 | 1.16 | -0.778 | 0.447 | -0.399 | -0.018 | 0.151 | 0.151 | 0 | 0.36 | 0.13 | 0.29 | 0.12 | 0.37 | 0.38 |
| 3 | 104959195 | rs1716127 | -0.498 | -0.74 | 0.94 | -0.289 | -0.009 | 0.112 | 0.112 | 0.838 | 0.11 | 0.47 | 0.28 | 0.11 | 0.09 | 0.11 |
| 3 | 104962935 | rs806658 | -1.011 | -0.585 | 0.144 | -0.677 | -0.01 | 0.152 | 0.152 | 0 | 0.63 | 0.84 | 0.7 | 0.89 | 0.63 | 0.62 |
| 3 | 104966715 | rs11708894 | -0.417 | -0.489 | 0.18 | -0.646 | 0.006 | -0.021 | -0.021 | 0.368 | 0.52 | 0.47 | 0.63 | 0.87 | 0.62 | 0.53 |
| 3 | 104973471 | rs2971376 | 0.455 | -0.819 | 0.001 | -0.499 | 0.001 | -0.01 | -0.01 | 0 | 0.5 | 0.43 | 0.4 | 0.16 | 0.6 | 0.47 |
| 3 | 104978553 | rs9879334 | 0.868 | -0.809 | -0.105 | -0.53 | -0.001 | 0.107 | 0.107 | 0.824 | 0.46 | 0.23 | 0.37 | 0.2 | 0.55 | 0.41 |
| 3 | 104979686 | rs7624664 | 0.778 | -0.881 | -0.2 | -0.565 | -0.029 | 0.148 | 0.148 | 1.703 | 0.44 | 0.13 | 0.36 | 0.2 | 0.54 | 0.41 |
| 3 | 104981251 | rs952365 | 0.399 | -1.403 | -0.643 | 0.275 | 0.005 | 0.015 | 0.015 | 0 | 0.14 | 0.11 | 0.3 | 0.51 | 0.09 | 0.22 |
| 3 | 104986735 | rs9683082 | -0.331 | -0.989 | -0.464 | -0.197 | 0.003 | -0.054 | -0.054 | 0 | 0.24 | 0.33 | 0.15 | 0.11 | 0.23 | 0.23 |
| 3 | 104998924 | rs2700344 | 0.468 | -0.45 | -0.821 | 0.111 | -0.007 | -0.029 | -0.029 | 0 | 0.55 | 0.47 | 0.43 | 0.2 | 0.61 | 0.48 |
| 3 | 105001491 | rs4144965 | -0.124 | 0.858 | -1.723 | 0.75 | -0.11 | -0.154 | -0.154 | 2.051 | 0.23 | 0.02 | 0.28 | 0.52 | 0.3 | 0.35 |
| 3 | 105002637 | rs13322578 | 0.207 | 0.739 | -1.558 | 0.742 | -0.07 | -0.097 | -0.097 | 0.628 | 0.21 | 0.07 | 0.29 | 0.55 | 0.3 | 0.37 |
| 3 | 105004853 | rs1795306 | 0.391 | 0.687 | -1.441 | 0.498 | 0 | 0.001 | 0.001 | 2.963 | 0.14 | 0.13 | 0.19 | 0.15 | 0.15 | 0.2 |
| 3 | 105021645 | rs6437577 | 1.105 | -0.506 | -0.865 | -0.177 | 0.004 | -0.024 | -0.024 | 0 | 0.58 | 0.51 | 0.49 | 0.3 | 0.57 | 0.44 |
| 3 | 105024155 | rs4361309 | -0.854 | -0.464 | -0.846 | 0.41 | -0.001 | 0.012 | 0.012 | 2.951 | 0.86 | 0.87 | 0.73 | 0.75 | 0.85 | 0.8 |
| 3 | 105034818 | rs4894918 | 0.671 | -1.3 | -0.686 | -0.387 | 0.005 | 0.099 | 0.099 | 0.568 | 0.45 | 0.29 | 0.23 | 0.12 | 0.42 | 0.35 |
| 3 | 105036097 | rs1920201 | 1.512 | -1.377 | -0.76 | -0.334 | -0.064 | -0.102 | -0.102 | 0.375 | 0.19 | 0.07 | 0.2 | 0.46 | 0.27 | 0.33 |
| 3 | 105041300 | rs9288800 | 1.285 | -1.444 | -0.654 | -0.372 | -0.007 | 0.088 | 0.088 | 0.02 | 0.42 | 0.26 | 0.22 | 0.12 | 0.41 | 0.34 |
| 3 | 105052776 | rs1438544 | -0.471 | -1.473 | -0.464 | -0.386 | -0.009 | 0.022 | 0.022 | 0.01 | 0.69 | 0.65 | 0.46 | 0.58 | 0.69 | 0.69 |
| 3 | 105062595 | rs1438552 | 0.885 | -1.442 | -0.437 | -0.381 | 0.009 | 0.032 | 0.032 | 0 | 0.22 | 0.26 | 0.23 | 0.46 | 0.27 | 0.33 |
| 3 | 105070234 | rs9880342 | -0.004 | -1.482 | -0.42 | -0.398 | -0.022 | 0.018 | 0.018 | 0 | 0.66 | 0.54 | 0.45 | 0.58 | 0.68 | 0.69 |
| 3 | 105070826 | rs4894921 | 1.563 | -1.011 | -0.841 | -1.264 | -0.009 | 0.026 | 0.026 | 0.018 | 0.23 | 0.16 | 0.12 | 0.09 | 0.27 | 0.22 |
| 3 | 105071369 | rs5012306 | 1.987 | -0.211 | -0.945 | -0.826 | -0.017 | -0.064 | -0.064 | 0 | 0.15 | 0.07 | 0.29 | 0.39 | 0.17 | 0.19 |
| 3 | 105072927 | rs16851083 | 0.643 | -0.571 | -0.941 | -0.945 | -0.004 | -0.001 | -0.001 | 0 | 0.46 | 0.34 | 0.34 | 0.5 | 0.47 | 0.48 |
| 3 | 105073420 | rs9879039 | 0.561 | -0.478 | -1.059 | -0.883 | 0.02 | 0.02 | 0.02 | 0 | 0.1 | 0.18 | 0.07 | 0.12 | 0.11 | 0.14 |
| 3 | 105075089 | rs13095274 | -0.144 | -0.846 | -0.951 | -1.017 | 0.021 | 0.017 | 0.017 | 0 | 0.47 | 0.36 | 0.59 | 0.38 | 0.42 | 0.38 |
| 3 | 105082472 | rs12497260 | 0.495 | -0.752 | -1.131 | -0.918 | 0.002 | 0.008 | 0.008 | 0 | 0.37 | 0.25 | 0.57 | 0.36 | 0.4 | 0.36 |
| 3 | 105097427 | rs6778460 | 1.603 | -0.349 | -0.97 | -1.025 | -0.02 | 0.036 | 0.036 | 0 | 0.17 | 0.09 | 0.1 | 0.04 | 0.19 | 0.16 |
| 3 | 105099947 | rs1438549 | 0.903 | -0.359 | -0.847 | -0.984 | 0.019 | -0.023 | -0.023 | 0 | 0.28 | 0.18 | 0.5 | 0.35 | 0.27 | 0.28 |
| 3 | 105100176 | rs16851098 | 1.882 | -0.197 | -0.981 | -0.832 | 0.01 | -0.082 | -0.082 | 3.491 | 0.13 | 0.01 | 0.23 | 0.23 | 0.11 | 0.18 |
| 3 | 105122214 | rs12638646 | 1.163 | -0.134 | -0.854 | -0.906 | -0.007 | -0.012 | -0.012 | 0.368 | 0.24 | 0.22 | 0.2 | 0.52 | 0.25 | 0.33 |
| 3 | 105122452 | rs13093663 | -1.117 | -0.276 | -0.748 | -1.089 | 0.016 | -0.034 | -0.034 | 0 | 0.74 | 0.68 | 0.84 | 0.91 | 0.7 | 0.78 |
| 3 | 105129061 | rs12488994 | -0.756 | -0.303 | -0.735 | -1.177 | 0.01 | -0.062 | -0.062 | 0 | 0.71 | 0.67 | 0.83 | 0.93 | 0.68 | 0.75 |
| 3 | 105132008 | rs9288803 | 0.386 | -0.427 | -0.695 | -1.247 | 0 | -0.007 | -0.007 | 0 | 0.43 | 0.49 | 0.48 | 0.65 | 0.41 | 0.43 |
| 3 | 105133395 | rs13085657 | 1.003 | -0.256 | -0.594 | -1.205 | 0.025 | -0.017 | -0.017 | 0 | 0.31 | 0.23 | 0.5 | 0.36 | 0.27 | 0.28 |
| 3 | 105134490 | rs6780238 | -0.613 | -0.114 | -0.659 | -1.146 | 0.009 | -0.057 | -0.057 | 0 | 0.71 | 0.67 | 0.83 | 0.93 | 0.68 | 0.75 |
| 3 | 105136914 | rs1438540 | 2.081 | 0.024 | -0.926 | -0.938 | -0.032 | -0.066 | -0.066 | 0.129 | 0.17 | 0.07 | 0.14 | 0.39 | 0.18 | 0.18 |
| 3 | 105137166 | rs2007681 | 2.081 | 0.024 | -0.926 | -0.938 | -0.032 | -0.066 | -0.066 | 2.375 | 0.17 | 0.07 | 0.14 | 0.39 | 0.18 | 0.18 |
| 3 | 105150752 | rs6804574 | 1.9 | 0.026 | -0.865 | -1.079 | -0.02 | 0.024 | 0.024 | 0 | 0.15 | 0.09 | 0.1 | 0.03 | 0.19 | 0.16 |
| 3 | 105152186 | rs9876906 | 2.434 | -0.101 | -0.858 | -1.043 | -0.008 | 0.015 | 0.015 | 0 | 0.1 | 0.03 | 0.05 | 0.04 | 0.11 | 0.06 |
| 3 | 105164872 | rs1955135 | 1.453 | -0.194 | -0.731 | -1.055 | -0.001 | -0.006 | -0.006 | 0.022 | 0.14 | 0.18 | 0.13 | 0.05 | 0.16 | 0.14 |
| 3 | 105182533 | rs10511244 | 1.925 | -0.118 | -0.751 | -0.913 | -0.02 | -0.046 | -0.046 | 0 | 0.18 | 0.11 | 0.16 | 0.39 | 0.19 | 0.19 |
| 3 | 105189978 | rs2399046 | 0.762 | -0.676 | -0.671 | -0.977 | 0.01 | -0.009 | -0.009 | 0 | 0.31 | 0.27 | 0.51 | 0.36 | 0.27 | 0.28 |
| 3 | 105198374 | rs7431531 | 1.199 | -0.274 | -0.832 | -1.01 | -0.046 | 0.062 | 0.062 | 0 | 0.24 | 0.1 | 0.14 | 0.07 | 0.32 | 0.23 |
| 3 | 105203665 | rs9857961 | -0.928 | -0.489 | -0.633 | -0.903 | -0.005 | -0.002 | -0.002 | 0.857 | 0.58 | 0.67 | 0.69 | 0.54 | 0.51 | 0.58 |
| 3 | 105205066 | rs6769329 | -0.664 | -0.46 | -0.594 | -0.869 | 0.003 | 0.003 | 0.003 | 0 | 0.55 | 0.67 | 0.68 | 0.54 | 0.49 | 0.56 |
| 3 | 105217018 | rs9848760 | 1.471 | -0.455 | -0.429 | -0.777 | 0.012 | 0.034 | 0.034 | 0.021 | 0.07 | 0.14 | 0.06 | 0.12 | 0.07 | 0.15 |
| 3 | 105217648 | rs2399047 | -0.305 | -0.614 | -0.522 | -0.875 | -0.007 | 0.001 | 0.001 | 0 | 0.42 | 0.57 | 0.58 | 0.49 | 0.34 | 0.44 |
| 3 | 105219511 | rs2291378 | 1.482 | -0.744 | -0.753 | -0.858 | 0.024 | -0.038 | -0.038 | 0 | 0.16 | 0.07 | 0.23 | 0.23 | 0.11 | 0.18 |
| 3 | 105221416 | rs1017915 | 0.362 | -0.567 | -0.581 | -0.872 | -0.022 | 0.001 | 0.001 | 0 | 0.27 | 0.49 | 0.35 | 0.26 | 0.23 | 0.26 |
| 3 | 105229549 | rs978436 | 1.542 | -0.389 | -1.034 | -0.948 | -0.001 | 0.013 | 0.013 | 0.623 | 0.13 | 0.09 | 0.11 | 0.06 | 0.15 | 0.13 |
| 3 | 105232816 | rs9846152 | -0.55 | -0.001 | -0.992 | -1.025 | 0.006 | -0.053 | -0.053 | 0 | 0.65 | 0.44 | 0.62 | 0.85 | 0.64 | 0.74 |
| 3 | 105234139 | rs7624777 | 1.349 | 0.201 | -1.009 | -0.73 | -0.027 | -0.092 | -0.092 | 0 | 0.26 | 0.08 | 0.2 | 0.45 | 0.28 | 0.26 |
| 3 | 105234486 | rs7613684 | -0.168 | 0.236 | -0.968 | -0.799 | 0.009 | 0.085 | 0.085 | 0 | 0.49 | 0.35 | 0.5 | 0.79 | 0.47 | 0.59 |
| 3 | 105237959 | rs1017914 | -0.817 | 0.084 | -0.824 | -0.894 | -0.006 | -0.018 | -0.018 | 0 | 0.66 | 0.66 | 0.78 | 0.92 | 0.62 | 0.7 |

| | | | | | | | | | | | | | | | | |
|---|---|---|---|---|---|---|---|---|---|---|---|---|---|---|---|---|
| 3 | 105238615 | rs1017913 | -0.611 | 0.198 | -0.892 | -0.867 | 0.003 | -0.1 | -0.1 | 1.09 | 0.64 | 0.53 | 0.77 | 0.92 | 0.64 | 0.7 |
| 3 | 105239394 | rs2171147 | 0.042 | 0.151 | -1.051 | -0.488 | -0.007 | 0.026 | 0.026 | 0 | 0.23 | 0.26 | 0.3 | 0.35 | 0.18 | 0.34 |
| 3 | 105248969 | rs3821492 | 0.507 | -0.069 | -1.043 | -0.443 | -0.02 | 0.043 | 0.043 | 0 | 0.19 | 0.27 | 0.28 | 0.29 | 0.13 | 0.23 |
| 3 | 105257485 | rs3772554 | 1.021 | 0.024 | -1.107 | -0.584 | -0.006 | 0.007 | 0.007 | 0 | 0.14 | 0.1 | 0.27 | 0.12 | 0.16 | 0.1 |
| 3 | 105258861 | rs1044240 | 2.127 | 0.116 | -1.362 | -0.715 | -0.068 | 0.057 | 0.057 | 0 | 0.14 | 0.01 | 0.07 | 0.03 | 0.18 | 0.16 |
| 3 | 105259312 | rs2279389 | 0.017 | -0.017 | -1.426 | -0.82 | 0.003 | 0.063 | 0.063 | 0 | 0.5 | 0.41 | 0.53 | 0.74 | 0.49 | 0.62 |
| 3 | 105266274 | rs16851279 | 1.347 | -0.175 | -1.23 | -0.91 | 0.023 | -0.02 | -0.02 | 0.089 | 0.15 | 0.07 | 0.23 | 0.17 | 0.1 | 0.18 |
| 3 | 105266331 | rs9855810 | 2.394 | -0.203 | -1.302 | -1.026 | -0.01 | 0.015 | 0.015 | 0 | 0.09 | 0.01 | 0.05 | 0.04 | 0.11 | 0.06 |
| 3 | 105267500 | rs495691 | -1.687 | -0.256 | -1.224 | -1.087 | -0.018 | -0.039 | -0.039 | 0 | 0.76 | 0.49 | 0.84 | 0.82 | 0.8 | 0.79 |
| 3 | 105270550 | rs13073764 | 2.283 | -0.091 | -1.847 | -0.926 | 0.006 | 0.009 | 0.009 | 0 | 0.07 | 0.11 | 0.03 | 0.04 | 0.07 | 0.08 |
| 3 | 105273679 | rs9990253 | 2.444 | -0.084 | -1.901 | -0.93 | -0.01 | 0.015 | 0.015 | 0 | 0.09 | 0.01 | 0.05 | 0.04 | 0.11 | 0.06 |
| 3 | 105276595 | rs2030515 | -0.112 | -0.077 | -1.939 | -0.943 | -0.001 | 0.07 | 0.07 | 0.109 | 0.55 | 0.22 | 0.53 | 0.7 | 0.58 | 0.61 |
| 3 | 105277521 | rs6792801 | 0.278 | -0.075 | -1.936 | -0.941 | -0.031 | 0.047 | 0.047 | 0 | 0.46 | 0.21 | 0.48 | 0.66 | 0.47 | 0.55 |
| 3 | 105285497 | rs2049217 | 1.861 | 0.601 | -2.932 | -0.67 | -0.055 | 0.061 | 0.061 | 0 | 0.14 | 0.02 | 0.07 | 0.03 | 0.18 | 0.16 |
| 3 | 105285975 | rs610047 | 0.903 | 0.594 | -2.926 | -0.666 | -0.001 | 0.001 | 0.001 | 0 | 0.14 | 0.13 | 0.27 | 0.09 | 0.15 | 0.09 |
| 3 | 105289655 | rs11705950 | 0.757 | 0.603 | -2.921 | -0.639 | -0.073 | -0.088 | -0.088 | 0.396 | 0.31 | 0.13 | 0.25 | 0.49 | 0.38 | 0.38 |
| 3 | 105292234 | rs627925 | -0.081 | 0.567 | -3.002 | -0.546 | 0.005 | 0.008 | 0.008 | 0 | 0.22 | 0.31 | 0.38 | 0.18 | 0.23 | 0.18 |
| 3 | 105295461 | rs1157 | 1.163 | 0.605 | -2.949 | -0.182 | 0.007 | 0.008 | 0.008 | 0 | 0.22 | 0.01 | 0.21 | 0.15 | 0.18 | 0.22 |
| 3 | 105298379 | rs13081408 | 0.806 | 1.569 | -1.97 | 0.612 | 0.007 | 0.057 | 0.057 | 0 | 0.07 | 0.16 | 0.1 | 0.1 | 0.09 | 0.09 |
| 3 | 105300908 | rs620165 | 1.35 | 1.291 | -1.851 | 0.665 | 0.001 | -0.002 | -0.002 | 0.762 | 0.08 | 0.07 | 0.23 | 0.08 | 0.1 | 0.02 |
| 3 | 105301006 | rs6778668 | 0.101 | 1.2 | -1.997 | 0.866 | -0.011 | -0.001 | -0.001 | 0.08 | 0.21 | 0.17 | 0.17 | 0.12 | 0.26 | 0.25 |
| 3 | 105304484 | rs2053627 | -0.143 | 0.607 | -1.772 | 1.205 | -0.021 | -0.074 | -0.074 | 0.507 | 0.18 | 0.09 | 0.18 | 0.47 | 0.21 | 0.2 |
| 3 | 105328255 | rs9822180 | -0.117 | -0.362 | -1.861 | 0.458 | 0.083 | 0.068 | 0.068 | 0.082 | 0.38 | 0.15 | 0.19 | 0.18 | 0.28 | 0.21 |
| 3 | 105332960 | rs736652 | -0.262 | 0.81 | -1.665 | 0.971 | -0.058 | -0.07 | -0.07 | 0.287 | 0.75 | 0.86 | 0.47 | 0.45 | 0.65 | 0.59 |
| 3 | 105333715 | rs7621204 | -0.075 | 0.668 | -1.182 | 0.91 | -0.016 | -0.021 | -0.021 | 0 | 0.29 | 0.24 | 0.54 | 0.56 | 0.36 | 0.44 |
| 3 | 105337398 | rs1867427 | -0.072 | 0.57 | -1.274 | 0.88 | -0.055 | -0.074 | -0.074 | 0.713 | 0.35 | 0.16 | 0.55 | 0.49 | 0.4 | 0.47 |
| 3 | 105338346 | rs894932 | -0.09 | -0.018 | -0.197 | 1.039 | -0.028 | 0.11 | 0.11 | 0.748 | 0.83 | 0.67 | 0.91 | 0.63 | 0.87 | 0.9 |
| 3 | 105341214 | rs1978973 | -0.784 | 0.23 | -0.171 | 1.178 | 0 | 0.044 | 0.044 | 0 | 0.09 | 0.13 | 0.04 | 0.34 | 0.09 | 0.17 |
| 3 | 105343015 | rs4894936 | 0.162 | 0.487 | -0.197 | 0.476 | -0.009 | 0.066 | 0.066 | 0 | 0.46 | 0.31 | 0.33 | 0.18 | 0.5 | 0.32 |
| 3 | 105353145 | rs930567 | -0.353 | 0.387 | -0.107 | 0.661 | 0.001 | 0.004 | 0.004 | 0 | 0.53 | 0.6 | 0.39 | 0.4 | 0.6 | 0.4 |
| 3 | 105356236 | rs7647969 | 0.625 | 1.685 | -0.15 | 0.476 | 0.008 | -0.014 | -0.014 | 0 | 0.19 | 0.2 | 0.15 | 0.07 | 0.27 | 0.12 |
| 3 | 105362282 | rs9843391 | 0.102 | 1.386 | -0.476 | 0.712 | -0.003 | 0.001 | 0.001 | 0.314 | 0.58 | 0.37 | 0.7 | 0.64 | 0.49 | 0.65 |
| 3 | 105369607 | rs7650120 | 0.366 | 1.708 | -0.781 | 0.386 | 0.011 | -0.037 | -0.037 | 0 | 0.1 | 0.11 | 0.15 | 0 | 0.13 | 0.12 |
| 3 | 105370343 | rs9857831 | 0.57 | 1.178 | -1.124 | 0.727 | 0.054 | -0.034 | -0.034 | 0 | 0.46 | 0.16 | 0.53 | 0.49 | 0.36 | 0.47 |
| 3 | 105372196 | rs11706098 | -0.041 | 1.501 | -0.57 | 0.512 | 0.009 | -0.032 | -0.032 | 0 | 0.31 | 0.43 | 0.22 | 0.14 | 0.36 | 0.26 |
| 3 | 105375746 | rs3732555 | -0.719 | 0.839 | -0.754 | 1.917 | 0.003 | -0.002 | -0.002 | 0 | 0.18 | 0.19 | 0.29 | 0.55 | 0.14 | 0.27 |
| 3 | 105377515 | rs1042852 | -0.122 | 1.896 | -0.426 | 1.715 | 0.04 | 0.008 | 0.008 | 0 | 0.17 | 0.23 | 0.31 | 0.19 | 0.27 | 0.16 |
| 3 | 105398047 | rs1550712 | -0.216 | 1.832 | -0.074 | 1.725 | 0.003 | -0.001 | -0.001 | 0 | 0.18 | 0.19 | 0.29 | 0.54 | 0.15 | 0.27 |
| 3 | 105409321 | rs1546733 | 1.174 | 1.772 | 0.138 | 1.706 | 0.019 | -0.015 | -0.015 | 0.685 | 0.64 | 0.48 | 0.38 | 0.28 | 0.58 | 0.56 |
| 3 | 105415038 | rs2301044 | 1.264 | 2.067 | 0.295 | 1.679 | 0.014 | -0.014 | -0.014 | 0 | 0.65 | 0.54 | 0.4 | 0.28 | 0.62 | 0.6 |
| 3 | 105422844 | rs2305037 | 0.227 | 2.208 | 0.502 | 1.936 | 0.052 | 0.013 | 0.013 | 0 | 0.82 | 0.73 | 0.68 | 0.78 | 0.72 | 0.83 |
| 3 | 105430251 | rs7615317 | -0.008 | 2.238 | 0.557 | 1.924 | 0.003 | -0.001 | -0.001 | 0 | 0.15 | 0.13 | 0.29 | 0.18 | 0.22 | 0.13 |
| 3 | 105438320 | rs726443 | 1.398 | 2.151 | 0.711 | 1.916 | 0.012 | -0.013 | -0.013 | 0 | 0.65 | 0.44 | 0.39 | 0.24 | 0.62 | 0.59 |
| 3 | 105438806 | rs7624400 | -0.319 | 2.094 | 0.76 | 2.121 | -0.001 | 0.018 | 0.018 | 0 | 0.85 | 0.85 | 0.9 | 0.59 | 0.9 | 0.8 |
| 3 | 105438957 | rs2305036 | -0.076 | 2.086 | 0.743 | 2.113 | 0.024 | 0.008 | 0.008 | 0 | 0.16 | 0.19 | 0.3 | 0.22 | 0.24 | 0.13 |
| 3 | 105439026 | rs2305035 | -0.037 | 2.088 | 0.744 | 2.128 | 0.025 | 0.008 | 0.008 | 0 | 0.16 | 0.19 | 0.3 | 0.22 | 0.24 | 0.13 |
| 3 | 105439522 | rs9288817 | 1.359 | 1.76 | 0.176 | 1.43 | 0.005 | 0.009 | 0.009 | 0 | 0.23 | 0.08 | 0.12 | 0.16 | 0.18 | 0.29 |
| 3 | 105441133 | rs1075739 | 0.54 | 1.914 | 0.328 | 1.733 | 0.008 | -0.036 | -0.036 | 0 | 0.15 | 0.09 | 0.1 | 0.41 | 0.1 | 0.2 |
| 3 | 105442573 | rs13082334 | 0.484 | 2.236 | 0.258 | 1.674 | -0.03 | -0.034 | -0.034 | 0 | 0.08 | 0.02 | 0.14 | 0.17 | 0.12 | 0.07 |
| 3 | 105444536 | rs17280845 | 1.572 | 1.868 | 0.306 | 1.252 | -0.003 | 0.067 | 0.067 | 0 | 0.24 | 0.08 | 0.16 | 0.09 | 0.28 | 0.14 |
| 3 | 105449793 | rs9876767 | -0.168 | 1.916 | 0.354 | 1.324 | 0.025 | 0.005 | 0.005 | 0 | 0.17 | 0.2 | 0.31 | 0.22 | 0.28 | 0.17 |
| 3 | 105455366 | rs9859953 | -0.634 | 1.573 | 0.425 | 1.352 | 0.041 | 0.048 | 0.048 | 0 | 0.2 | 0.33 | 0.5 | 0.35 | 0.28 | 0.21 |
| 3 | 105455955 | rs2289746 | 0.585 | 1.497 | 0.337 | 1.245 | 0.025 | 0.025 | 0.025 | 0 | 0.69 | 0.57 | 0.59 | 0.37 | 0.66 | 0.67 |
| 3 | 105500218 | rs6784447 | 1.227 | 1.117 | 0.367 | 1.345 | -0.025 | 0.077 | 0.077 | 0 | 0.26 | 0.08 | 0.18 | 0.08 | 0.32 | 0.17 |
| 3 | 105505307 | rs6437616 | -1.156 | 0.994 | 1.274 | 1.971 | 0.056 | 0.073 | 0.073 | 0 | 0.08 | 0.24 | 0.34 | 0.18 | 0.14 | 0.18 |
| 3 | 105518948 | rs12486371 | 0.091 | 0.995 | 1.276 | 1.96 | 0.057 | -0.016 | -0.016 | 0 | 0.05 | 0.14 | 0.15 | 0.01 | 0.08 | 0.08 |
| 3 | 105523704 | rs12632980 | 1.248 | 0.988 | 1.375 | 1.969 | 0.083 | 0.105 | 0.105 | 0 | 0.91 | 0.66 | 0.65 | 0.82 | 0.86 | 0.82 |
| 3 | 105526724 | rs1561922 | -0.023 | 0.935 | 1.317 | 2.081 | 0.011 | -0.03 | -0.03 | 0 | 0.15 | 0.09 | 0.1 | 0.41 | 0.1 | 0.18 |
| 3 | 105560638 | rs6767627 | -0.901 | 0.87 | 1.375 | 2.073 | 0.002 | 0.082 | 0.082 | 0 | 0.21 | 0.31 | 0.25 | 0.47 | 0.19 | 0.25 |
| 3 | 105568602 | rs2089091 | -0.81 | 0.882 | 1.372 | 2.081 | 0.002 | 0.041 | 0.041 | 0 | 0.2 | 0.24 | 0.24 | 0.42 | 0.18 | 0.25 |
| 3 | 105576911 | rs3996189 | 1.071 | 0.888 | 1.247 | 1.857 | -0.003 | 0.079 | 0.079 | 0 | 0.75 | 0.47 | 0.55 | 0.4 | 0.76 | 0.64 |
| 3 | 105643849 | rs6765145 | 1.425 | 0.983 | 0.876 | 1.714 | -0.011 | 0.066 | 0.066 | 0 | 0.7 | 0.47 | 0.53 | 0.48 | 0.75 | 0.65 |
| 3 | 105651647 | rs2174233 | -0.842 | 1.026 | 1.058 | 1.505 | -0.004 | 0.105 | 0.105 | 0 | 0.15 | 0.32 | 0.16 | 0.33 | 0.11 | 0.23 |
| 3 | 105661508 | rs2455882 | 0.035 | 0.814 | 1.136 | 1.493 | 0.014 | -0.049 | -0.049 | 0 | 0.91 | 0.81 | 0.86 | 1 | 0.93 | 0.95 |
| 3 | 105662637 | rs1566804 | -0.781 | 0.858 | 1.153 | 1.5 | -0.014 | 0.107 | 0.107 | 0 | 0.14 | 0.32 | 0.16 | 0.33 | 0.1 | 0.21 |
| 3 | 105663687 | rs2455881 | 0.343 | 1.08 | 0.724 | 1.01 | 0.015 | -0.051 | -0.051 | 0 | 0.91 | 0.81 | 0.86 | 1 | 0.93 | 0.95 |
| 3 | 105671329 | rs2470709 | 0.477 | 1.149 | 0.654 | 1.492 | 0.025 | 0.045 | 0.045 | 0 | 0.82 | 0.72 | 0.68 | 0.72 | 0.77 | 0.9 |
| 3 | 105673976 | rs12629811 | -0.745 | 0.554 | 0.795 | 1.574 | 0.011 | 0.037 | 0.037 | 0.408 | 0.21 | 0.29 | 0.16 | 0.34 | 0.27 | 0.14 |
| 3 | 105675360 | rs2470696 | 1.398 | 0.194 | 0.591 | 1.505 | 0.002 | 0 | 0 | 3.196 | 0.67 | 0.32 | 0.57 | 0.39 | 0.67 | 0.6 |

| | | | | | | | | | | | | | | | | |
|---|---|---|---|---|---|---|---|---|---|---|---|---|---|---|---|---|
| 3 | 105684092 | rs1503920 | 1.374 | -0.067 | 0.138 | 1.16 | 0.014 | 0.038 | 0.038 | 0.19 | 0.54 | 0.33 | 0.49 | 0.39 | 0.55 | 0.55 |
| 3 | 105687634 | rs16851846 | -0.968 | -0.068 | 0.286 | 1.138 | -0.001 | -0.006 | -0.006 | 0 | 0.33 | 0.37 | 0.22 | 0.27 | 0.32 | 0.16 |
| 3 | 105688886 | rs6437632 | 0.495 | -0.07 | 0.331 | 1.129 | 0.005 | -0.001 | -0.001 | 0 | 0.15 | 0.15 | 0.09 | 0 | 0.12 | 0.05 |
| 3 | 105689194 | rs2655045 | 0.352 | -0.053 | 0.09 | 1.437 | -0.022 | 0.136 | 0.136 | 0 | 0.95 | 0.84 | 0.84 | 0.84 | 0.98 | 0.88 |
| 3 | 105691565 | rs13062596 | 0.694 | 0.003 | -0.131 | 1.411 | -0.006 | 0.008 | 0.008 | 0 | 0.62 | 0.44 | 0.63 | 0.56 | 0.67 | 0.72 |
| 3 | 105693298 | rs6781481 | -0.479 | 0.458 | -0.379 | 0.643 | -0.003 | -0.008 | -0.008 | 0.011 | 0.33 | 0.42 | 0.35 | 0.28 | 0.32 | 0.16 |
| 3 | 105695304 | rs2618316 | 0.11 | -0.122 | -1.006 | 0.03 | 0.024 | 0.043 | 0.043 | 0 | 0.73 | 0.83 | 0.88 | 0.91 | 0.81 | 0.87 |
| 3 | 105695575 | rs2655033 | 0.425 | 0.303 | -0.75 | 0.367 | -0.001 | -0.001 | -0.001 | 1.01 | 0.56 | 0.42 | 0.6 | 0.46 | 0.51 | 0.47 |
| 3 | 105695700 | rs2655032 | 0.115 | 0.422 | -0.528 | 1.036 | 0 | 0.017 | 0.017 | 1.33 | 0.9 | 0.88 | 0.89 | 0.74 | 0.89 | 0.83 |
| 3 | 105697533 | rs16851901 | -0.129 | 0.55 | -0.131 | 0.899 | 0.016 | 0.064 | 0.064 | 0 | 0.28 | 0.47 | 0.38 | 0.36 | 0.26 | 0.4 |
| 3 | 105698149 | rs2455879 | 0.114 | 0.515 | 0.157 | 1.133 | -0.007 | -0.012 | -0.012 | 0 | 0.24 | 0.22 | 0.22 | 0.29 | 0.25 | 0.38 |
| 3 | 105698868 | rs9863266 | 0.855 | 0.115 | 0.309 | 0.654 | -0.014 | -0.033 | -0.033 | 0.582 | 0.6 | 0.17 | 0.44 | 0.64 | 0.68 | 0.55 |
| 3 | 105699113 | rs2618321 | 0.419 | 0.14 | 0.286 | 0.675 | -0.009 | -0.015 | -0.015 | 1.675 | 0.24 | 0.22 | 0.25 | 0.29 | 0.25 | 0.37 |
| 3 | 105699882 | rs2655023 | 0.263 | 0.074 | 0.358 | 0.642 | 0.012 | 0.053 | 0.053 | 0 | 0.26 | 0.4 | 0.37 | 0.36 | 0.25 | 0.41 |
| 3 | 105700120 | rs9868510 | 1.203 | -0.347 | 0.304 | 0.112 | -0.03 | 0.006 | 0.006 | 0 | 0.48 | 0.19 | 0.38 | 0.63 | 0.59 | 0.48 |
| 3 | 105703940 | rs3898308 | 0.176 | -0.32 | 0.822 | 0.14 | 0.006 | -0.043 | -0.043 | 0 | 0.17 | 0.42 | 0.21 | 0.07 | 0.12 | 0.1 |
| 3 | 105717509 | rs340060 | 1.321 | -0.192 | 0.634 | 0.254 | -0.031 | -0.053 | -0.053 | 0 | 0.2 | 0.1 | 0.22 | 0.29 | 0.23 | 0.37 |
| 3 | 105722581 | rs17194003 | 1.828 | -0.545 | 0.073 | -0.424 | -0.035 | 0.041 | 0.041 | 0 | 0.2 | 0.02 | 0.19 | 0.14 | 0.23 | 0.13 |
| 3 | 105723046 | rs340058 | 1.176 | -0.498 | 0.796 | -0.002 | 0.037 | 0.066 | 0.066 | 0.173 | 0.69 | 0.25 | 0.61 | 0.93 | 0.86 | 0.85 |
| 3 | 105727133 | rs6802262 | -1.411 | -0.457 | 0.97 | 0.057 | 0.038 | 0.098 | 0.098 | 0.582 | 0.07 | 0.41 | 0.15 | 0.07 | 0.02 | 0.03 |
| 3 | 105731843 | rs7642650 | 0.13 | -0.455 | 0.767 | 0.012 | -0.018 | -0.082 | -0.082 | 0 | 0.16 | 0.23 | 0.09 | 0 | 0.1 | 0.07 |
| 3 | 105736306 | rs9846331 | -1.423 | -0.477 | 0.471 | 0.007 | 0.033 | 0.107 | 0.107 | 0.343 | 0.31 | 0.81 | 0.39 | 0.07 | 0.17 | 0.17 |
| 3 | 105741844 | rs6437639 | -1.381 | -0.465 | 0.569 | 0.011 | 0.033 | 0.091 | 0.091 | 0.284 | 0.31 | 0.78 | 0.38 | 0.07 | 0.17 | 0.17 |
| 3 | 105749376 | rs12485870 | -0.268 | -0.477 | 0.382 | 0.027 | 0 | -0.039 | -0.039 | 0.963 | 0.2 | 0.15 | 0.19 | 0 | 0.11 | 0.1 |
| 3 | 105759149 | rs1159290 | 1.648 | -0.436 | -0.199 | 0.177 | -0.054 | 0.121 | 0.121 | 0 | 0.41 | 0.09 | 0.31 | 0.25 | 0.48 | 0.33 |
| 3 | 105762360 | rs2399060 | -0.894 | -0.596 | 0.744 | 0.317 | 0.002 | 0.001 | 0.001 | 0.051 | 0.23 | 0.24 | 0.32 | 0.07 | 0.12 | 0.13 |
| 3 | 105765123 | rs9840375 | -1.702 | -0.65 | 0.469 | 0.465 | 0.034 | 0.054 | 0.054 | 0.275 | 0.31 | 0.8 | 0.38 | 0.17 | 0.17 | 0.22 |
| 3 | 105765517 | rs7632765 | 0.147 | -0.577 | 0.984 | 0.032 | -0.014 | -0.07 | -0.07 | 0 | 0.13 | 0.17 | 0.05 | 0 | 0.08 | 0.06 |
| 3 | 105769227 | rs2127089 | 0.835 | -0.554 | 0.707 | 0.043 | 0.001 | -0.016 | -0.016 | 0 | 0.11 | 0.08 | 0.04 | 0 | 0.08 | 0.06 |
| 3 | 105777569 | rs10511248 | -1.835 | -0.541 | 0.886 | 0.323 | -0.002 | 0.011 | 0.011 | 0.701 | 0.19 | 0.6 | 0.23 | 0.2 | 0.17 | 0.14 |
| 3 | 105783456 | rs9833579 | 0.011 | -0.727 | 0.88 | 0.215 | 0.007 | -0.003 | -0.003 | 0 | 0.85 | 0.84 | 0.94 | 1 | 0.92 | 0.92 |
| 3 | 105790348 | rs6788447 | -0.495 | -0.512 | 1.291 | 0.293 | -0.004 | -0.107 | -0.107 | 0.192 | 0.17 | 0.31 | 0.18 | 0 | 0.1 | 0.1 |
| 3 | 105807324 | rs6793499 | -1.247 | -0.497 | 1.144 | 0.306 | -0.016 | -0.103 | -0.103 | 1.566 | 0.22 | 0.55 | 0.22 | 0 | 0.14 | 0.14 |
| 3 | 105807773 | rs767152 | 0.024 | -0.472 | 0.584 | 0.402 | 0.002 | -0.006 | -0.006 | 0 | 0.08 | 0.07 | 0.04 | 0.2 | 0.09 | 0.06 |
| 3 | 105809730 | rs2035254 | 1.784 | -0.486 | 0.589 | 0.401 | 0.026 | 0.054 | 0.054 | 0.104 | 0.67 | 0.14 | 0.71 | 0.8 | 0.76 | 0.78 |
| 3 | 105841011 | rs16852060 | -0.465 | -0.123 | 0.191 | 0.938 | 0.003 | -0.005 | -0.005 | 0 | 0.08 | 0.08 | 0.05 | 0.16 | 0.08 | 0.06 |
| 3 | 105847367 | rs4894982 | 0.739 | 0.132 | 0.462 | 0.616 | 0.01 | -0.025 | -0.025 | 0 | 0.67 | 0.55 | 0.61 | 0.77 | 0.69 | 0.56 |
| 3 | 105851489 | rs2169865 | 0.388 | -0.099 | 0.392 | 0.513 | 0.005 | -0.011 | -0.011 | 0 | 0.87 | 0.84 | 0.96 | 1 | 0.95 | 0.93 |
| 3 | 105851672 | rs4894843 | 0.801 | -0.077 | 0.47 | 0.518 | -0.009 | 0.027 | 0.027 | 0.04 | 0.69 | 0.75 | 0.75 | 0.79 | 0.72 | 0.59 |
| 3 | 105857527 | rs6437646 | 0.467 | -0.163 | 0.542 | 0.437 | -0.011 | -0.032 | -0.032 | 0 | 0.87 | 0.8 | 0.95 | 1 | 0.95 | 0.94 |
| 3 | 105894098 | rs7628390 | 1.26 | -0.325 | 0.54 | 0.75 | 0.004 | -0.006 | -0.006 | 0 | 0.63 | 0.49 | 0.63 | 0.7 | 0.68 | 0.41 |
| 3 | 105895352 | rs9838807 | -0.528 | -0.35 | 0.577 | 0.743 | 0.002 | 0.001 | 0.001 | 0.033 | 0.26 | 0.28 | 0.32 | 0.3 | 0.28 | 0.55 |
| 3 | 105897477 | rs1871272 | 0.249 | -0.425 | 0.474 | 0.705 | -0.064 | -0.043 | -0.043 | 0 | 0.21 | 0.04 | 0.26 | 0.22 | 0.23 | 0.45 |
| 3 | 105897816 | rs6793744 | 0.182 | -0.498 | 0.499 | 0.661 | -0.027 | -0.054 | -0.054 | 0 | 0.88 | 0.78 | 0.95 | 1 | 0.96 | 0.95 |
| 3 | 105911052 | rs9790068 | -0.311 | -0.299 | 0.201 | 0.817 | -0.005 | -0.002 | -0.002 | 0 | 0.24 | 0.24 | 0.31 | 0.22 | 0.25 | 0.46 |
| 3 | 105912130 | rs12487066 | -0.317 | -0.374 | 0.204 | 0.82 | -0.003 | -0.001 | -0.001 | 0 | 0.23 | 0.24 | 0.31 | 0.23 | 0.24 | 0.46 |
| 3 | 105916460 | rs12488982 | -0.355 | -0.48 | 0.149 | 0.825 | -0.002 | 0 | 0 | 0 | 0.27 | 0.21 | 0.3 | 0.24 | 0.24 | 0.42 |
| 3 | 105924836 | rs6414232 | 0.897 | -0.613 | 0.76 | 0.761 | -0.011 | 0.009 | 0.009 | 0.739 | 0.88 | 0.55 | 0.93 | 0.92 | 0.94 | 0.88 |
| 3 | 105925276 | rs12489296 | -0.157 | -0.625 | 0.726 | 1.015 | 0.001 | 0.001 | 0.001 | 0 | 0.25 | 0.25 | 0.3 | 0.25 | 0.23 | 0.36 |
| 3 | 105927328 | rs1838554 | -0.035 | -0.604 | 0.725 | 1.042 | -0.001 | 0.001 | 0.001 | 0 | 0.24 | 0.24 | 0.26 | 0.25 | 0.2 | 0.35 |
| 3 | 105943369 | rs7617081 | 1.086 | -0.617 | 0.65 | 1.041 | 0.005 | 0 | 0 | 1.027 | 0.63 | 0.31 | 0.63 | 0.67 | 0.71 | 0.52 |
| 3 | 105960464 | rs12185995 | 0.389 | -0.653 | 0.429 | 1.059 | -0.052 | -0.047 | -0.047 | 0 | 0.2 | 0.01 | 0.19 | 0.2 | 0.19 | 0.32 |
| 3 | 105965913 | rs9879608 | 0.975 | -0.74 | 0.677 | 0.982 | -0.022 | 0.021 | 0.021 | 0.642 | 0.87 | 0.55 | 0.93 | 0.92 | 0.95 | 0.83 |
| 3 | 105965996 | rs13096639 | -0.266 | -0.656 | 0.101 | 1.28 | 0.003 | -0.035 | -0.035 | 0 | 0.25 | 0.14 | 0.33 | 0.34 | 0.21 | 0.36 |
| 3 | 105967242 | rs13060369 | -0.126 | -0.627 | -0.05 | 1.294 | -0.006 | -0.066 | -0.066 | 0 | 0.24 | 0.09 | 0.33 | 0.34 | 0.21 | 0.36 |
| 3 | 105988605 | rs9848627 | -0.617 | -1.068 | -0.203 | 1.245 | -0.004 | -0.027 | -0.027 | 0 | 0.39 | 0.1 | 0.26 | 0.43 | 0.33 | 0.55 |
| 3 | 105991356 | rs10222671 | 1.748 | -1.45 | -0.352 | 1.248 | -0.04 | 0.024 | 0.024 | 0 | 0.42 | 0.09 | 0.47 | 0.35 | 0.6 | 0.21 |
| 3 | 105992865 | rs2077909 | -1.74 | -1.127 | 0.526 | 1.4 | -0.005 | 0.166 | 0.166 | 0.418 | 0.13 | 0.57 | 0.25 | 0.25 | 0.1 | 0.23 |
| 3 | 106009677 | rs13071798 | -1.059 | -1.317 | 1.017 | 1.285 | -0.021 | 0.122 | 0.122 | 0.928 | 0.12 | 0.34 | 0.15 | 0.23 | 0.07 | 0.19 |
| 3 | 106014926 | rs9880416 | -0.39 | -1.277 | 0.348 | 0.897 | -0.03 | 0.088 | 0.088 | 0.76 | 0.48 | 0.1 | 0.48 | 0.37 | 0.44 | 0.55 |
| 3 | 106016673 | rs4429664 | 1.543 | -1.286 | 0.468 | 0.853 | -0.002 | -0.001 | -0.001 | 0.551 | 0.37 | 0.38 | 0.35 | 0.35 | 0.48 | 0.22 |
| 3 | 106021500 | rs7614488 | 1.208 | -1.154 | 0.726 | 0.775 | -0.04 | 0.145 | 0.145 | 1.809 | 0.86 | 0.48 | 0.85 | 0.77 | 0.93 | 0.81 |
| 3 | 106033336 | rs9853623 | 1.056 | -1.076 | 0.714 | 0.726 | -0.029 | 0.12 | 0.12 | 1.281 | 0.87 | 0.57 | 0.92 | 0.78 | 0.93 | 0.81 |
| 3 | 106036014 | rs9878197 | -0.108 | -1.039 | 0.316 | 0.304 | 0.001 | -0.01 | -0.01 | 1.784 | 0.36 | 0.36 | 0.51 | 0.23 | 0.38 | 0.36 |
| 3 | 106048760 | rs13094374 | 0.959 | -0.999 | -0.036 | 0.212 | -0.075 | 0.008 | 0.008 | 0 | 0.22 | 0.01 | 0.31 | 0.17 | 0.22 | 0.27 |
| 3 | 106050024 | rs2895309 | -0.951 | -1.11 | 0.399 | 0.461 | -0.001 | 0.001 | 0.001 | 0 | 0.5 | 0.51 | 0.53 | 0.55 | 0.48 | 0.59 |
| 3 | 106051778 | rs12630324 | -0.758 | -1.022 | 0.157 | 0.454 | -0.002 | 0.014 | 0.014 | 0 | 0.17 | 0.17 | 0.09 | 0.36 | 0.15 | 0.3 |
| 3 | 106053688 | rs7620758 | -0.984 | -0.979 | 0.317 | 0.577 | -0.006 | 0.004 | 0.004 | 0 | 0.52 | 0.6 | 0.58 | 0.58 | 0.49 | 0.64 |
| 3 | 106059186 | rs7627494 | -0.773 | -0.961 | 0.132 | 0.486 | -0.003 | -0.008 | -0.008 | 0 | 0.9 | 0.89 | 0.97 | 0.96 | 0.94 | 0.87 |

| | | | | | | | | | | | | | | | | |
|---|---|---|---|---|---|---|---|---|---|---|---|---|---|---|---|---|
| 3 | 106063963 | rs6437662 | -0.69 | -0.97 | 0.025 | 0.495 | 0.008 | 0.013 | 0.013 | 0 | 0.9 | 0.95 | 0.97 | 0.96 | 0.95 | 0.87 |
| 3 | 106073491 | rs4594651 | 1.609 | -0.983 | -0.135 | 0.536 | -0.024 | -0.021 | -0.021 | 0 | 0.35 | 0.23 | 0.37 | 0.38 | 0.44 | 0.22 |
| 3 | 106081419 | rs10933844 | -0.458 | -0.945 | -0.151 | 0.592 | -0.02 | 0.09 | 0.09 | 0 | 0.11 | 0.17 | 0.05 | 0.36 | 0.09 | 0.27 |
| 3 | 106089227 | rs9831569 | 1.543 | -0.933 | -0.373 | 0.467 | -0.035 | -0.036 | -0.036 | 0 | 0.35 | 0.22 | 0.39 | 0.4 | 0.46 | 0.22 |
| 3 | 106090339 | rs7630432 | -0.165 | -0.662 | -0.252 | 0.532 | -0.026 | 0.107 | 0.107 | 0.887 | 0.39 | 0.18 | 0.42 | 0.22 | 0.36 | 0.32 |
| 3 | 106091518 | rs10511256 | -1.146 | -0.687 | 0.128 | 0.897 | -0.029 | 0.084 | 0.084 | 0 | 0.13 | 0.2 | 0.06 | 0.31 | 0.1 | 0.27 |
| 3 | 106093730 | rs2399093 | 0.147 | -0.587 | -0.789 | 0.506 | 0 | -0.037 | -0.037 | 0.035 | 0.65 | 0.9 | 0.8 | 0.69 | 0.7 | 0.68 |
| 3 | 106098490 | rs4895004 | 0.358 | -0.598 | 0.088 | 0.165 | -0.004 | -0.083 | -0.083 | 0.443 | 0.23 | 0.37 | 0.31 | 0.07 | 0.2 | 0.24 |
| 3 | 106111946 | rs16852439 | 0.831 | -0.522 | 0.112 | 1.011 | -0.018 | 0.005 | 0.005 | 0 | 0.4 | 0.29 | 0.49 | 0.64 | 0.5 | 0.4 |
| 3 | 106119522 | rs6777538 | 1.251 | -0.509 | 0.279 | 0.969 | -0.026 | -0.017 | -0.017 | 0.144 | 0.36 | 0.21 | 0.34 | 0.38 | 0.45 | 0.25 |
| 3 | 106124739 | rs11710616 | 1.152 | -0.528 | 0.102 | 0.958 | -0.053 | -0.037 | -0.037 | 0 | 0.37 | 0.15 | 0.34 | 0.38 | 0.46 | 0.25 |
| 3 | 106127714 | rs7620623 | 0.873 | -0.643 | 0.216 | 0.898 | -0.032 | 0.015 | 0.015 | 0 | 0.41 | 0.19 | 0.51 | 0.67 | 0.56 | 0.44 |
| 3 | 106131151 | rs7616267 | 0.623 | -0.758 | 0.681 | 0.854 | 0 | -0.001 | -0.001 | 1.218 | 0.57 | 0.53 | 0.64 | 0.5 | 0.63 | 0.48 |
| 3 | 106132923 | rs9870951 | -1.461 | -0.877 | 0.643 | 0.814 | -0.043 | 0.216 | 0.216 | 0 | 0.19 | 0.45 | 0.21 | 0.46 | 0.11 | 0.34 |
| 3 | 106133779 | rs13065261 | -0.075 | -0.7 | 0.094 | 0.702 | -0.021 | 0.132 | 0.132 | 0 | 0.69 | 0.88 | 0.82 | 0.91 | 0.7 | 0.68 |
| 3 | 106135465 | rs7651003 | -0.641 | -0.726 | 0.172 | 0.813 | 0.014 | 0.015 | 0.015 | 0 | 0.91 | 0.93 | 0.97 | 0.95 | 0.97 | 0.86 |
| 3 | 106139801 | rs10933849 | -0.095 | -0.565 | 0.498 | 1.111 | -0.021 | 0.102 | 0.102 | 0 | 0.76 | 0.56 | 0.61 | 0.6 | 0.84 | 0.67 |
| 3 | 106142880 | rs4342128 | 0.399 | -0.443 | 0.033 | 1.006 | 0.001 | -0.004 | -0.004 | 0.97 | 0.2 | 0.15 | 0.36 | 0.25 | 0.16 | 0.29 |
| 3 | 106150812 | rs4572800 | 0.215 | -0.307 | 0.339 | 0.96 | -0.011 | 0.004 | 0.004 | 0 | 0.6 | 0.43 | 0.69 | 0.59 | 0.68 | 0.55 |
| 3 | 106156027 | rs7628506 | -0.185 | -0.219 | -0.533 | 0.823 | 0.005 | -0.01 | -0.01 | 0 | 0.37 | 0.28 | 0.34 | 0.52 | 0.33 | 0.53 |
| 3 | 106159948 | rs1445059 | -0.107 | 0.56 | -1.265 | 0.726 | 0 | 0.068 | 0.068 | 0 | 0.62 | 0.78 | 0.64 | 0.74 | 0.67 | 0.71 |
| 3 | 106175524 | rs722813 | -0.318 | 0.122 | -1.142 | 1.098 | -0.024 | -0.002 | -0.002 | 0.343 | 0.57 | 0.77 | 0.63 | 0.59 | 0.53 | 0.63 |
| 3 | 106175546 | rs722812 | 0.212 | -0.98 | -0.489 | 0.951 | 0 | 0.026 | 0.026 | 0 | 0.38 | 0.33 | 0.24 | 0.13 | 0.38 | 0.29 |
| 3 | 106184511 | rs9831914 | 0.935 | -0.842 | 0.12 | 1.646 | 0.02 | 0.352 | 0.352 | 0 | 0.94 | 0.83 | 0.79 | 0.49 | 0.94 | 0.76 |
| 3 | 106186920 | rs968072 | 0.27 | -0.951 | 0.545 | 1.224 | 0 | -0.001 | -0.001 | 0 | 0.63 | 0.41 | 0.55 | 0.32 | 0.66 | 0.51 |
| 3 | 106189100 | rs7431430 | -0.017 | -0.978 | 0.536 | 0.235 | -0.002 | 0.016 | 0.016 | 0 | 0.32 | 0.27 | 0.27 | 0.23 | 0.31 | 0.29 |
| 3 | 106191942 | rs1445076 | 0.698 | -0.713 | 0.79 | 0.262 | 0.013 | 0.091 | 0.091 | 0.061 | 0.42 | 0.32 | 0.24 | 0.18 | 0.37 | 0.33 |
| 3 | 106192908 | rs1955156 | -1.446 | -0.465 | 0.971 | 0.708 | 0.031 | 0.416 | 0.416 | 0 | 0.05 | 0.24 | 0.2 | 0.55 | 0.04 | 0.21 |
| 3 | 106197762 | rs9845337 | 0.157 | -0.48 | 0.963 | 0.679 | 0 | -0.015 | -0.015 | 0 | 0.23 | 0.25 | 0.21 | 0.12 | 0.22 | 0.2 |
| 3 | 106199490 | rs2921264 | -0.366 | -0.423 | 0.909 | 0.703 | -0.001 | 0.026 | 0.026 | 0 | 0.28 | 0.51 | 0.41 | 0.67 | 0.26 | 0.41 |
| 3 | 106199700 | rs2590208 | -0.356 | -0.422 | 0.879 | 0.703 | 0 | 0.03 | 0.03 | 0 | 0.28 | 0.49 | 0.4 | 0.67 | 0.26 | 0.41 |
| 3 | 106199956 | rs2712464 | -0.301 | -0.403 | 0.888 | 0.692 | 0.003 | 0.105 | 0.105 | 0 | 0.28 | 0.48 | 0.4 | 0.53 | 0.26 | 0.4 |
| 3 | 106208882 | rs2712475 | 0.083 | 0.193 | 0.753 | 0.973 | -0.002 | 0.018 | 0.018 | 0 | 0.19 | 0.3 | 0.19 | 0.25 | 0.19 | 0.21 |
| 3 | 106212498 | rs1445065 | -0.27 | 0.13 | 0.763 | 1.016 | 0.006 | 0.063 | 0.063 | 0 | 0.21 | 0.36 | 0.38 | 0.66 | 0.22 | 0.4 |
| 3 | 106217161 | rs2953042 | 0.739 | -0.541 | 0.846 | 0.544 | -0.008 | 0.011 | 0.011 | 0 | 0.38 | 0.35 | 0.36 | 0.25 | 0.47 | 0.39 |
| 3 | 106218553 | rs2712473 | 0.353 | -0.512 | 0.782 | 0.437 | -0.001 | 0.004 | 0.004 | 0 | 0.44 | 0.41 | 0.61 | 0.76 | 0.55 | 0.64 |
| 3 | 106219745 | rs17794180 | -0.098 | -0.635 | 0.819 | 0.324 | -0.006 | 0.016 | 0.016 | 0 | 0.11 | 0.16 | 0.14 | 0.15 | 0.1 | 0.05 |
| 3 | 106220130 | rs2921253 | -0.341 | -0.634 | 0.779 | 0.312 | 0.004 | -0.036 | -0.036 | 0 | 0.55 | 0.55 | 0.75 | 0.91 | 0.65 | 0.69 |
| 3 | 106220749 | rs17179448 | 0.257 | -0.57 | 0.603 | 0.325 | 0 | 0.021 | 0.021 | 0 | 0.09 | 0.05 | 0.13 | 0.02 | 0.1 | 0.05 |
| 3 | 106220816 | rs2953039 | 0.412 | -0.53 | 0.608 | 0.368 | -0.001 | 0.007 | 0.007 | 0 | 0.44 | 0.41 | 0.61 | 0.76 | 0.55 | 0.64 |
| 3 | 106228764 | rs2921259 | 0.604 | -0.443 | 0.58 | 0.582 | -0.001 | 0.015 | 0.015 | 0 | 0.85 | 0.88 | 0.63 | 0.47 | 0.81 | 0.69 |
| 3 | 106231571 | rs9859768 | 0.158 | -0.545 | 0.797 | 0.445 | 0.009 | 0.016 | 0.016 | 0 | 0.46 | 0.47 | 0.27 | 0.22 | 0.35 | 0.31 |
| 3 | 106233426 | rs12485867 | 0.128 | -0.789 | 0.743 | 0.695 | -0.027 | 0.056 | 0.056 | 0.134 | 0.05 | 0.01 | 0.24 | 0.51 | 0.08 | 0.27 |
| 3 | 106236128 | rs2712489 | -0.103 | -0.79 | 0.771 | 0.69 | 0.009 | 0.174 | 0.174 | 0.445 | 0.94 | 0.94 | 0.75 | 0.49 | 0.91 | 0.73 |
| 3 | 106247828 | rs7627016 | 0.09 | -0.827 | 0.709 | 0.607 | -0.015 | 0.027 | 0.027 | 0 | 0.09 | 0.02 | 0.12 | 0.01 | 0.09 | 0.05 |
| 3 | 106247932 | rs2590174 | 0.515 | -1.111 | 0.86 | 0.335 | 0.007 | 0.005 | 0.005 | 0 | 0.5 | 0.5 | 0.49 | 0.47 | 0.6 | 0.44 |
| 3 | 106258932 | rs9827829 | 1.812 | -1.098 | 1.044 | 0.268 | -0.002 | -0.008 | -0.008 | 0 | 0.71 | 0.62 | 0.52 | 0.25 | 0.71 | 0.55 |
| 3 | 106262422 | rs9845005 | 0.733 | -1.065 | 1.007 | 0.279 | 0.012 | -0.024 | -0.024 | 0.795 | 0.33 | 0.29 | 0.13 | 0 | 0.22 | 0.21 |
| 3 | 106263029 | rs2440834 | -0.791 | -1.061 | 0.993 | 0.297 | 0 | -0.002 | -0.002 | 0 | 0.51 | 0.55 | 0.46 | 0.74 | 0.41 | 0.6 |
| 3 | 106271726 | rs1445073 | 0.708 | -1.192 | 0.89 | 0.271 | 0.02 | -0.005 | -0.005 | 0 | 0.33 | 0.27 | 0.13 | 0 | 0.22 | 0.21 |
| 3 | 106271855 | rs4145136 | 0.319 | -1.145 | 0.767 | 0.363 | -0.028 | 0.055 | 0.055 | 0.207 | 0.05 | 0.01 | 0.24 | 0.52 | 0.09 | 0.29 |
| 3 | 106275857 | rs1867559 | -0.445 | -1.208 | 1.002 | 0.06 | -0.008 | -0.017 | -0.017 | 0.018 | 0.47 | 0.49 | 0.4 | 0.75 | 0.35 | 0.57 |
| 3 | 106275998 | rs10933858 | -0.848 | -1.122 | 1.01 | 0.118 | -0.013 | 0 | 0 | 0 | 0.07 | 0.07 | 0.02 | 0.09 | 0.04 | 0.06 |
| 3 | 106280197 | rs13058741 | -0.606 | -1.083 | 0.982 | 0.13 | 0.001 | 0.004 | 0.004 | 0.009 | 0.11 | 0.06 | 0.07 | 0.09 | 0.1 | 0.09 |
| 3 | 106280822 | rs2712494 | -0.37 | -1.097 | 0.861 | -0.012 | 0.003 | 0.005 | 0.005 | 0 | 0.89 | 0.93 | 0.87 | 1 | 0.9 | 0.94 |
| 3 | 106283820 | rs13077519 | -1.485 | -1.052 | 0.858 | 0.103 | -0.001 | -0.001 | -0.001 | 0 | 0.18 | 0.13 | 0.32 | 0.75 | 0.19 | 0.39 |
| 3 | 106288013 | rs920937 | -0.455 | -0.681 | 0.856 | 0.409 | -0.002 | -0.007 | -0.007 | 1.604 | 0.51 | 0.49 | 0.46 | 0.74 | 0.41 | 0.6 |
| 3 | 106299324 | rs2921262 | -2.473 | -0.319 | 1.45 | 1.659 | 0.01 | 0.05 | 0.05 | 0.559 | 0.12 | 0.41 | 0.16 | 0.15 | 0.08 | 0.07 |
| 3 | 106302060 | rs2590199 | 0.61 | -0.338 | 1.182 | 1.854 | -0.003 | -0.035 | -0.035 | 0 | 0.83 | 0.9 | 0.7 | 0.52 | 0.83 | 0.61 |
| 3 | 106312262 | rs2953303 | -1.735 | -0.306 | 1.387 | 1.757 | 0.026 | 0.292 | 0.292 | 1.133 | 0.07 | 0.4 | 0.04 | 0.23 | 0.02 | 0.03 |
| 3 | 106320733 | rs2921263 | 1.251 | 0.049 | 0.433 | 1.894 | -0.009 | -0.059 | -0.059 | 0 | 0.77 | 0.85 | 0.62 | 0.53 | 0.76 | 0.57 |
| 3 | 106328940 | rs747451 | 0.975 | -0.036 | 0.587 | 1.43 | 0 | 0.03 | 0.03 | 0.951 | 0.8 | 0.79 | 0.73 | 0.38 | 0.81 | 0.62 |
| 3 | 106335082 | rs7626568 | -1.505 | -0.154 | 0.664 | 0.818 | -0.009 | 0.054 | 0.054 | 0 | 0.12 | 0.42 | 0.07 | 0.14 | 0.09 | 0.09 |
| 3 | 106335897 | rs2895313 | 0.242 | -0.066 | 0.277 | 0.121 | 0 | 0.012 | 0.012 | 0 | 0.53 | 0.56 | 0.59 | 0.84 | 0.48 | 0.55 |
| 3 | 106337570 | rs2399106 | 0.804 | -0.81 | 0.802 | 0.171 | -0.024 | -0.043 | -0.043 | 0 | 0.8 | 0.43 | 0.88 | 0.88 | 0.88 | 0.84 |
| 3 | 106337852 | rs4895016 | 1.588 | -1.376 | 0.17 | 0.443 | 0.081 | 0.048 | 0.048 | 0 | 0.54 | 0.12 | 0.69 | 0.63 | 0.72 | 0.51 |
| 3 | 106343354 | rs13089498 | 1.577 | -1.3 | 0.178 | 0.822 | 0.071 | -0.034 | -0.034 | 0 | 0.54 | 0.13 | 0.69 | 0.56 | 0.72 | 0.57 |
| 3 | 106351642 | rs7644108 | 1.718 | -1.253 | 0.216 | 0.842 | 0.063 | -0.036 | -0.036 | 1.912 | 0.55 | 0.13 | 0.69 | 0.56 | 0.72 | 0.57 |
| 3 | 106352340 | rs10933863 | -0.602 | -1.279 | 0.26 | 1.255 | -0.01 | 0.027 | 0.027 | 0 | 0.25 | 0.3 | 0.19 | 0.4 | 0.16 | 0.28 |

| | | | | | | | | | | | | | | | | |
|---|---|---|---|---|---|---|---|---|---|---|---|---|---|---|---|---|
| 3 | 106355415 | rs7634049 | -0.453 | -1.129 | 0.747 | 0.889 | 0.001 | -0.05 | -0.05 | 0.233 | 0.89 | 0.81 | 0.93 | 1 | 0.92 | 0.91 |
| 3 | 106359385 | rs7636315 | 0.735 | -1.149 | 0.673 | 0.886 | -0.032 | -0.037 | -0.037 | 0 | 0.82 | 0.52 | 0.9 | 0.88 | 0.9 | 0.84 |
| 3 | 106360511 | rs6437684 | 0.77 | -1.146 | 0.688 | 0.888 | -0.033 | -0.038 | -0.038 | 0 | 0.81 | 0.46 | 0.89 | 0.88 | 0.9 | 0.84 |
| 3 | 106364831 | rs4456895 | 0.335 | -1.147 | 0.819 | 0.934 | 0.002 | 0.106 | 0.106 | 0 | 0.92 | 0.64 | 0.96 | 0.88 | 0.98 | 0.93 |
| 3 | 106367054 | rs7632792 | -0.028 | -1.166 | 0.481 | 0.942 | -0.002 | -0.019 | -0.019 | 0 | 0.13 | 0.05 | 0.16 | 0.32 | 0.11 | 0.19 |
| 3 | 106379000 | rs6774644 | -0.994 | -1.176 | 0.485 | 0.944 | -0.019 | 0.041 | 0.041 | 0 | 0.25 | 0.41 | 0.21 | 0.32 | 0.16 | 0.33 |
| 3 | 106380005 | rs12633892 | -0.982 | -1.172 | 0.487 | 0.946 | -0.019 | 0.042 | 0.042 | 0 | 0.25 | 0.41 | 0.21 | 0.32 | 0.16 | 0.33 |
| 3 | 106380604 | rs7613153 | -0.982 | -1.129 | 0.487 | 0.946 | -0.017 | 0.042 | 0.042 | 0 | 0.25 | 0.41 | 0.21 | 0.32 | 0.17 | 0.33 |
| 3 | 106392429 | rs7617430 | -0.513 | -1.137 | -0.002 | 0.972 | -0.002 | 0.012 | 0.012 | 0.009 | 0.2 | 0.2 | 0.19 | 0.32 | 0.13 | 0.25 |
| 3 | 106393843 | rs2895315 | 0.999 | -0.938 | -0.66 | 0.98 | 0.056 | -0.059 | -0.059 | 0.01 | 0.57 | 0.12 | 0.7 | 0.55 | 0.74 | 0.49 |
| 3 | 106401155 | rs6437690 | 0.523 | -0.711 | 0.091 | 1.233 | -0.035 | 0.166 | 0.166 | 0 | 0.81 | 0.62 | 0.81 | 0.66 | 0.86 | 0.66 |
| 3 | 106402233 | rs9829756 | 0.298 | 0.343 | -0.313 | 1.412 | -0.043 | -0.044 | -0.044 | 0.092 | 0.07 | 0.01 | 0.15 | 0.2 | 0.12 | 0.09 |
| 3 | 106404147 | rs11928487 | -0.321 | 0.04 | -0.414 | 0.675 | 0 | 0 | 0 | 0.233 | 0.53 | 0.51 | 0.58 | 0.49 | 0.53 | 0.4 |
| 3 | 106405877 | rs9853812 | 0.203 | 0.201 | -0.394 | 0.228 | 0.014 | -0.004 | -0.004 | 0 | 0.66 | 0.57 | 0.53 | 0.66 | 0.64 | 0.79 |
| 3 | 106410820 | rs9818810 | -0.391 | 0.249 | -0.736 | 0.143 | -0.068 | -0.091 | -0.091 | 0.011 | 0.34 | 0.11 | 0.45 | 0.47 | 0.42 | 0.24 |
| 3 | 106416700 | rs9860520 | 1.323 | 0.054 | -0.793 | 0.169 | -0.033 | -0.029 | -0.029 | 0 | 0.41 | 0.21 | 0.43 | 0.44 | 0.45 | 0.59 |
| 3 | 106416840 | rs7644333 | -0.832 | 0.023 | -0.585 | 0.167 | -0.003 | 0.001 | 0.001 | 0 | 0.49 | 0.57 | 0.45 | 0.47 | 0.46 | 0.35 |
| 3 | 106419208 | rs9837471 | 0.117 | 0.699 | -0.463 | 0.701 | -0.012 | -0.018 | -0.018 | 0 | 0.75 | 0.69 | 0.82 | 0.84 | 0.84 | 0.79 |
| 3 | 106420030 | rs12488048 | 0.475 | 0.764 | -1.308 | 0.485 | -0.011 | -0.01 | -0.01 | 0 | 0.6 | 0.28 | 0.55 | 0.42 | 0.6 | 0.62 |
| 3 | 106421617 | rs6774396 | -0.35 | 1.002 | -1.215 | 0.434 | -0.004 | -0.004 | -0.004 | 0 | 0.39 | 0.4 | 0.52 | 0.5 | 0.47 | 0.47 |
| 3 | 106421910 | rs2399114 | -0.155 | 1.047 | -1.705 | -0.345 | -0.03 | 0.106 | 0.106 | 1.43 | 0.41 | 0.21 | 0.36 | 0.19 | 0.47 | 0.29 |
| 3 | 106423903 | rs11713414 | 0.195 | 0.61 | -1.438 | -0.148 | 0.016 | 0.052 | 0.052 | 0 | 0.56 | 0.29 | 0.63 | 0.69 | 0.66 | 0.58 |
| 3 | 106434399 | rs4443209 | -1.168 | 0.768 | -0.149 | 0.303 | 0.02 | 0.09 | 0.09 | 0 | 0.21 | 0.45 | 0.23 | 0.41 | 0.19 | 0.26 |
| 3 | 106435440 | rs6437697 | -1.168 | 0.768 | -0.149 | 0.281 | 0.02 | 0.084 | 0.084 | 0 | 0.21 | 0.45 | 0.23 | 0.4 | 0.19 | 0.26 |
| 3 | 106458422 | rs1915327 | 0.24 | 0.732 | 0.064 | 0.355 | 0.023 | 0.224 | 0.224 | 0.75 | 0.07 | 0.24 | 0.15 | 0.28 | 0.04 | 0.21 |
| 3 | 106464513 | rs4895023 | 1.504 | 0.539 | 0.593 | 0.152 | -0.002 | 0.016 | 0.016 | 0 | 0.75 | 0.35 | 0.73 | 0.6 | 0.81 | 0.74 |
| 3 | 106471129 | rs6781971 | 1.313 | 0.691 | 0.347 | -0.033 | 0 | -0.001 | -0.001 | 0 | 0.67 | 0.34 | 0.66 | 0.59 | 0.68 | 0.64 |
| 3 | 106481150 | rs1356931 | 1.301 | 0.694 | 0.491 | -0.027 | -0.001 | -0.001 | -0.001 | 0 | 0.67 | 0.34 | 0.66 | 0.59 | 0.68 | 0.65 |
| 3 | 106490276 | rs9830289 | 1.322 | 0.688 | 0.487 | -0.03 | -0.002 | -0.002 | -0.002 | 0 | 0.67 | 0.34 | 0.66 | 0.59 | 0.68 | 0.64 |
| 3 | 106495290 | rs11713942 | 0.236 | 0.747 | 0.651 | -0.044 | 0.003 | -0.01 | -0.01 | 2.619 | 0.12 | 0.13 | 0.08 | 0.12 | 0.14 | 0.04 |
| 3 | 106495324 | rs1402304 | 0.653 | 0.937 | 0.62 | 0.067 | 0.008 | 0.075 | 0.075 | 0.564 | 0.79 | 0.49 | 0.74 | 0.71 | 0.82 | 0.69 |
| 3 | 106523235 | rs4895039 | 0.641 | 1.054 | 0.54 | 0.166 | 0.011 | 0.017 | 0.017 | 0.01 | 0.74 | 0.71 | 0.77 | 0.58 | 0.72 | 0.65 |
| 3 | 106529642 | rs939452 | 1 | 0.908 | 0.953 | 0.102 | 0.018 | 0.031 | 0.031 | 0 | 0.72 | 0.6 | 0.76 | 0.58 | 0.71 | 0.65 |
| 3 | 106530324 | rs16852987 | 0.233 | 0.886 | 0.709 | 0.17 | 0.008 | 0.055 | 0.055 | 0.277 | 0.14 | 0.22 | 0.15 | 0.32 | 0.14 | 0.3 |
| 3 | 106543224 | rs6787966 | 0.355 | 0.702 | 0.678 | 0.199 | -0.004 | 0.031 | 0.031 | 0.214 | 0.18 | 0.22 | 0.15 | 0.37 | 0.15 | 0.35 |
| 3 | 106545997 | rs954995 | 1.342 | 0.752 | 0.74 | -0.037 | -0.001 | -0.002 | -0.002 | 0.315 | 0.62 | 0.36 | 0.65 | 0.53 | 0.67 | 0.56 |
| 3 | 106562094 | rs17798007 | -0.188 | 0.871 | 0.309 | 0.009 | -0.034 | 0.037 | 0.037 | 0 | 0.15 | 0.03 | 0.09 | 0.1 | 0.15 | 0.07 |
| 3 | 106567549 | rs16853011 | 0.166 | 0.784 | 0.389 | 0.114 | -0.001 | 0.016 | 0.016 | 0 | 0.21 | 0.24 | 0.18 | 0.38 | 0.21 | 0.41 |
| 3 | 106589586 | rs1709942 | 0.541 | 0.799 | 0.537 | 0.059 | -0.003 | 0.025 | 0.025 | 0 | 0.71 | 0.42 | 0.7 | 0.61 | 0.75 | 0.54 |
| 3 | 106594233 | rs1473964 | 0.396 | 0.984 | 0.634 | -0.098 | 0.009 | -0.036 | -0.036 | 0 | 0.82 | 0.71 | 0.78 | 0.91 | 0.84 | 0.92 |
| 3 | 106607795 | rs773233 | 0.819 | 0.825 | 0.773 | 0.366 | 0.001 | 0.003 | 0.003 | 0 | 0.6 | 0.44 | 0.59 | 0.48 | 0.63 | 0.48 |
| 3 | 106613301 | rs16853065 | 0.248 | 0.835 | 0.779 | 0.483 | -0.012 | 0.004 | 0.004 | 0 | 0.13 | 0.1 | 0.2 | 0.15 | 0.14 | 0.05 |
| 3 | 106613436 | rs1709915 | 0.808 | 0.834 | 0.798 | 0.507 | 0.003 | 0.003 | 0.003 | 0 | 0.6 | 0.46 | 0.59 | 0.42 | 0.62 | 0.47 |
| 3 | 106615905 | rs7618427 | -0.015 | 0.896 | 0.475 | 0.201 | 0.007 | -0.008 | -0.008 | 0.405 | 0.86 | 0.78 | 0.79 | 0.85 | 0.86 | 0.95 |
| 3 | 106624294 | rs1517605 | 0.511 | 1.105 | -0.104 | -0.073 | 0.023 | -0.046 | -0.046 | 0 | 0.09 | 0.16 | 0.07 | 0 | 0.13 | 0.03 |
| 3 | 106640874 | rs7624717 | 1.673 | 0.697 | -1.027 | -0.021 | -0.016 | -0.051 | -0.051 | 0.42 | 0.17 | 0.02 | 0.24 | 0.23 | 0.17 | 0.17 |
| 3 | 106647916 | rs7374615 | 0.212 | 1.047 | -0.518 | 0.031 | -0.004 | 0.007 | 0.007 | 0 | 0.6 | 0.61 | 0.57 | 0.4 | 0.6 | 0.48 |
| 3 | 106658062 | rs1712566 | 0.572 | 0.901 | -0.565 | -0.523 | -0.004 | -0.009 | -0.009 | 0 | 0.4 | 0.41 | 0.38 | 0.31 | 0.41 | 0.34 |
| 3 | 106658486 | rs13088732 | 0.031 | 1.137 | -0.615 | -0.654 | -0.006 | -0.003 | -0.003 | 0 | 0.62 | 0.61 | 0.55 | 0.88 | 0.59 | 0.78 |
| 3 | 106659501 | rs1709965 | 0.669 | 1.17 | -0.598 | -0.669 | -0.001 | -0.004 | -0.004 | 0 | 0.42 | 0.43 | 0.4 | 0.32 | 0.45 | 0.37 |
| 3 | 106671971 | rs1709949 | 0.728 | 1.17 | -0.652 | -0.668 | -0.001 | -0.008 | -0.008 | 0.005 | 0.41 | 0.43 | 0.41 | 0.32 | 0.45 | 0.37 |
| 3 | 106675278 | rs2056115 | 1.218 | 1.181 | -0.657 | -0.601 | -0.002 | 0.013 | 0.013 | 0.486 | 0.23 | 0.21 | 0.18 | 0.56 | 0.18 | 0.44 |
| 3 | 106683507 | rs6437715 | 0.774 | 1.27 | -0.633 | -0.819 | -0.004 | 0.004 | 0.004 | 0 | 0.15 | 0.12 | 0.21 | 0.11 | 0.16 | 0.06 |
| 3 | 106689103 | rs1515395 | -1.501 | 1.221 | -0.664 | -0.9 | -0.001 | -0.019 | -0.019 | 0 | 0.79 | 0.8 | 0.8 | 0.99 | 0.79 | 0.8 |
| 3 | 106693307 | rs6437717 | 2.033 | 1.245 | -0.663 | -0.998 | 0 | -0.011 | -0.011 | 0 | 0.07 | 0.07 | 0.07 | 0 | 0.14 | 0.04 |
| 3 | 106700416 | rs13063911 | -0.066 | 1.219 | -0.617 | -0.869 | -0.005 | 0.011 | 0.011 | 0 | 0.64 | 0.68 | 0.59 | 0.88 | 0.63 | 0.74 |
| 3 | 106705307 | rs1709938 | -1.07 | 1.217 | -0.452 | -0.638 | -0.001 | -0.042 | -0.042 | 0.011 | 0.78 | 0.77 | 0.8 | 0.99 | 0.79 | 0.8 |
| 3 | 106706931 | rs7650236 | 0.779 | 1.014 | -0.455 | -0.228 | -0.003 | 0.019 | 0.019 | 0.531 | 0.22 | 0.24 | 0.19 | 0.57 | 0.18 | 0.39 |
| 3 | 106722004 | rs11914364 | 1.116 | 0.737 | -0.424 | -0.248 | -0.01 | 0.002 | 0.002 | 0 | 0.27 | 0.3 | 0.31 | 0.31 | 0.27 | 0.26 |
| 3 | 106722802 | rs7632114 | 0.66 | 1.034 | 0.096 | -0.287 | 0.001 | 0 | 0 | 0.013 | 0.08 | 0.09 | 0.17 | 0.12 | 0.11 | 0.05 |
| 3 | 106743798 | rs9828598 | -0.191 | 0.839 | 0.173 | -0.27 | 0.003 | -0.002 | -0.002 | 0.309 | 0.11 | 0.16 | 0.19 | 0.12 | 0.12 | 0.07 |
| 3 | 106743875 | rs9869656 | -1.232 | 0.854 | 0.109 | -0.159 | 0.003 | 0.022 | 0.022 | 0 | 0.5 | 0.58 | 0.52 | 0.3 | 0.46 | 0.4 |
| 3 | 106764743 | rs4395396 | -0.568 | 0.873 | 0.076 | -0.144 | -0.001 | -0.014 | -0.014 | 0.025 | 0.39 | 0.4 | 0.33 | 0.18 | 0.35 | 0.33 |
| 3 | 106764796 | rs4599303 | 0.281 | 0.819 | 0.096 | -0.171 | -0.005 | -0.014 | -0.014 | 0.006 | 0.22 | 0.18 | 0.21 | 0.01 | 0.23 | 0.22 |
| 3 | 106769904 | rs6792401 | 0.702 | 1.018 | -0.094 | -0.207 | -0.001 | 0.015 | 0.015 | 0 | 0.16 | 0.1 | 0.09 | 0 | 0.15 | 0.19 |
| 3 | 106772492 | rs6781540 | -1.192 | 0.831 | -0.077 | -0.083 | 0.002 | 0.007 | 0.007 | 0 | 0.5 | 0.55 | 0.52 | 0.28 | 0.46 | 0.4 |
| 3 | 106777359 | rs12493716 | -1.067 | 0.522 | 0.11 | 0.138 | 0.001 | -0.005 | -0.005 | 1.1 | 0.15 | 0.21 | 0.29 | 0.11 | 0.15 | 0.09 |
| 3 | 106794051 | rs9844646 | -0.222 | 0.71 | -0.142 | 0.135 | -0.008 | 0.001 | 0.001 | 0.337 | 0.07 | 0.08 | 0.05 | 0.01 | 0.09 | 0.06 |
| 3 | 106796923 | rs1528313 | 0.805 | 0.502 | 0.14 | 0.132 | -0.019 | -0.007 | -0.007 | 6.815 | 0.75 | 0.68 | 0.71 | 0.77 | 0.81 | 0.77 |

| | | | | | | | | | | | | | | | | |
|---|---|---|---|---|---|---|---|---|---|---|---|---|---|---|---|---|
| 3 | 106798418 | rs6785880 | 1.397 | 0.504 | 0.137 | 0.128 | -0.003 | -0.024 | -0.024 | 0 | 0.68 | 0.59 | 0.66 | 0.76 | 0.73 | 0.73 |
| 3 | 106799860 | rs4422306 | 0.119 | 0.468 | 0.235 | 0.109 | 0.008 | -0.004 | -0.004 | 0 | 0.15 | 0.09 | 0.08 | 0 | 0.12 | 0.16 |
| 3 | 106800570 | rs1528308 | 1.606 | 0.459 | 0.291 | 0.106 | -0.005 | -0.036 | -0.036 | 0 | 0.67 | 0.51 | 0.65 | 0.76 | 0.72 | 0.71 |
| 3 | 106805013 | rs7610448 | -0.372 | 0.479 | 0.245 | 0.113 | 0.008 | -0.028 | -0.028 | 0.398 | 0.08 | 0.17 | 0.06 | 0.01 | 0.09 | 0.06 |
| 3 | 106807621 | rs9288843 | -0.18 | 0.481 | 0.231 | 0.114 | 0 | -0.013 | -0.013 | 0.001 | 0.07 | 0.14 | 0.06 | 0.01 | 0.09 | 0.06 |
| 3 | 106809176 | rs1819051 | 1.517 | 0.465 | 0.236 | 0.132 | -0.005 | -0.034 | -0.034 | 2.59 | 0.67 | 0.5 | 0.65 | 0.75 | 0.72 | 0.72 |
| 3 | 106809635 | rs11917688 | -0.254 | -0.068 | 0.484 | 0.522 | 0.01 | 0.112 | 0.112 | 0 | 0.09 | 0.23 | 0.21 | 0.23 | 0.07 | 0.07 |
| 3 | 106826316 | rs13072544 | 1.091 | -0.116 | 0.459 | 0.523 | -0.032 | 0.01 | 0.01 | 0 | 0.73 | 0.47 | 0.68 | 0.74 | 0.79 | 0.89 |
| 3 | 106833272 | rs9876108 | -0.337 | 0.148 | -0.455 | 0.191 | 0.003 | -0.02 | -0.02 | 0 | 0.16 | 0.1 | 0.1 | 0 | 0.13 | 0.05 |
| 3 | 106836240 | rs9863794 | -1.168 | -0.067 | -0.216 | 0.595 | -0.016 | 0.01 | 0.01 | 0.003 | 0.23 | 0.36 | 0.18 | 0.29 | 0.22 | 0.13 |
| 3 | 106841915 | rs1919005 | -0.051 | -0.171 | -0.432 | 0.527 | -0.005 | -0.006 | -0.006 | 0 | 0.58 | 0.59 | 0.47 | 0.71 | 0.62 | 0.61 |
| 3 | 106849597 | rs9830097 | -1.068 | -0.211 | -0.541 | 0.498 | 0.006 | 0.005 | 0.005 | 0 | 0.7 | 0.8 | 0.56 | 0.75 | 0.72 | 0.64 |
| 3 | 106850421 | rs7640923 | 0.786 | -0.021 | -0.967 | 0.397 | 0.002 | -0.009 | -0.009 | 0.646 | 0.1 | 0.07 | 0.08 | 0 | 0.1 | 0.04 |
| 3 | 106860495 | rs13323477 | 0.461 | 0.305 | -1.035 | 0.744 | -0.03 | -0.027 | -0.027 | 0 | 0.08 | 0 | 0.06 | 0.13 | 0.12 | 0.05 |
| 3 | 106861626 | rs6798375 | -0.365 | 0.328 | -0.975 | 0.664 | -0.008 | -0.009 | -0.009 | 0 | 0.57 | 0.5 | 0.51 | 0.72 | 0.62 | 0.61 |
| 3 | 106887467 | rs7634774 | 1.301 | 0.764 | -1.243 | 0.654 | -0.008 | -0.026 | -0.026 | 2.19 | 0.27 | 0.3 | 0.19 | 0.16 | 0.23 | 0.26 |
| 3 | 106903210 | rs6791146 | 0.546 | 0.867 | -1.255 | 0.683 | -0.08 | -0.072 | -0.072 | 0 | 0.19 | 0.08 | 0.18 | 0.28 | 0.27 | 0.23 |
| 3 | 106907345 | rs9867707 | -0.05 | 0.423 | -0.689 | 0.839 | -0.006 | -0.002 | -0.002 | 0 | 0.38 | 0.41 | 0.28 | 0.42 | 0.35 | 0.35 |
| 3 | 106922747 | rs7430092 | 0.385 | 0.597 | -0.837 | 0.986 | 0 | -0.004 | -0.004 | 0.232 | 0.73 | 0.7 | 0.65 | 0.5 | 0.71 | 0.55 |
| 3 | 106925898 | rs1881309 | -0.37 | 1.226 | -0.978 | 0.733 | -0.019 | -0.01 | -0.01 | 2.883 | 0.19 | 0.11 | 0.26 | 0.22 | 0.25 | 0.26 |
| 3 | 106928929 | rs11915523 | -0.945 | 0.495 | -0.632 | 0.54 | -0.024 | -0.01 | -0.01 | 0 | 0.08 | 0.17 | 0.03 | 0.04 | 0.02 | 0.05 |
| 3 | 106935865 | rs11929184 | -0.457 | -0.025 | -0.572 | 0.32 | 0.003 | 0 | 0 | 0 | 0.52 | 0.56 | 0.39 | 0.52 | 0.35 | 0.5 |
| 3 | 106938007 | rs6772017 | 0.47 | 0.388 | -0.42 | 0.475 | -0.021 | 0.064 | 0.064 | 0 | 0.41 | 0.26 | 0.28 | 0.2 | 0.43 | 0.33 |
| 3 | 106938456 | rs7651766 | 0.054 | 0.203 | -0.112 | 0.247 | 0.006 | 0.004 | 0.004 | 0.767 | 0.48 | 0.59 | 0.51 | 0.51 | 0.57 | 0.42 |
| 3 | 106950962 | rs1283092 | -1.291 | -0.25 | -0.122 | 0 | -0.005 | -0.061 | -0.061 | 0.258 | 0.81 | 0.84 | 0.9 | 1 | 0.86 | 0.94 |
| 3 | 106958845 | rs1283109 | 0.448 | -0.363 | -0.299 | 0.281 | 0.092 | 0.012 | 0.012 | 0.082 | 0.48 | 0.23 | 0.5 | 0.51 | 0.65 | 0.44 |
| 3 | 106961498 | rs1283106 | -0.021 | 0.052 | -0.08 | 0.284 | 0 | 0.001 | 0.001 | 0 | 0.42 | 0.38 | 0.52 | 0.46 | 0.4 | 0.57 |
| 3 | 106963543 | rs1283101 | 0.004 | 0.028 | -0.055 | 0.267 | -0.004 | -0.004 | -0.004 | 0 | 0.42 | 0.49 | 0.53 | 0.46 | 0.39 | 0.56 |
| 3 | 106966689 | rs9862762 | 1.108 | -0.05 | -0.297 | -0.055 | -0.051 | 0.134 | 0.134 | 0.214 | 0.33 | 0.05 | 0.19 | 0.13 | 0.33 | 0.27 |
| 3 | 106967877 | rs1283098 | -0.431 | 0.284 | 0.324 | 0.238 | 0.002 | 0.01 | 0.01 | 0 | 0.49 | 0.51 | 0.68 | 0.78 | 0.54 | 0.65 |
| 3 | 106972315 | rs793618 | 0.241 | 0.116 | -0.308 | 0.579 | 0.006 | 0.004 | 0.004 | 0 | 0.52 | 0.48 | 0.29 | 0.3 | 0.42 | 0.36 |
| 3 | 106978360 | rs17806089 | 0.558 | 0.385 | -0.61 | 0.36 | -0.009 | -0.023 | -0.023 | 0 | 0.06 | 0.11 | 0.02 | 0 | 0.03 | 0 |
| 3 | 106978527 | rs13074296 | 0.349 | 0.8 | -0.869 | 0.727 | -0.042 | -0.044 | -0.044 | 0 | 0.25 | 0.14 | 0.28 | 0.35 | 0.34 | 0.23 |
| 3 | 106981727 | rs793610 | -0.675 | -0.015 | -0.601 | 0.145 | -0.003 | -0.026 | -0.026 | 2.511 | 0.61 | 0.62 | 0.79 | 0.79 | 0.65 | 0.77 |
| 3 | 106982317 | rs709567 | -0.536 | -0.437 | -0.093 | 0.743 | -0.019 | 0.014 | 0.014 | 0 | 0.22 | 0.31 | 0.37 | 0.3 | 0.16 | 0.51 |
| 3 | 106985680 | rs12489299 | -0.616 | -0.506 | -0.529 | 0.97 | 0.017 | -0.066 | -0.066 | 0 | 0.13 | 0.03 | 0.33 | 0.36 | 0.11 | 0.45 |
| 3 | 106987474 | rs7645377 | -1.432 | -0.839 | 0.175 | 0.698 | -0.009 | -0.002 | -0.002 | 0 | 0.37 | 0.49 | 0.48 | 0.6 | 0.3 | 0.68 |
| 3 | 106988439 | rs9867626 | -0.345 | -0.917 | 0.222 | 0.696 | -0.043 | 0.107 | 0.107 | 0.006 | 0.11 | 0.39 | 0.06 | 0.2 | 0.05 | 0.07 |
| 3 | 106990475 | rs2553942 | 0.746 | -0.555 | 0.195 | 0.214 | 0 | -0.012 | -0.012 | 0.19 | 0.14 | 0.12 | 0.16 | 0.02 | 0.21 | 0.04 |
| 3 | 107003691 | rs16853338 | -0.959 | -0.847 | 0.204 | 0.449 | 0.005 | -0.054 | -0.054 | 0 | 0.28 | 0.19 | 0.44 | 0.56 | 0.26 | 0.62 |
| 3 | 107007263 | rs12495060 | -0.735 | -0.864 | 0.084 | 0.43 | -0.042 | -0.145 | -0.145 | 2.002 | 0.27 | 0.05 | 0.41 | 0.39 | 0.26 | 0.61 |
| 3 | 107014573 | rs11918378 | 0.059 | -0.765 | 0.369 | 0.424 | -0.027 | -0.038 | -0.038 | 0 | 0.09 | 0.18 | 0.04 | 0.03 | 0.04 | 0.06 |
| 3 | 107019713 | rs809421 | -0.996 | -0.772 | 0.744 | 0.336 | -0.01 | 0.109 | 0.109 | 0 | 0.89 | 0.81 | 0.9 | 0.64 | 0.95 | 0.93 |
| 3 | 107035683 | rs7641355 | -1.03 | -0.804 | 0.693 | -0.602 | -0.013 | -0.026 | -0.026 | 0 | 0.38 | 0.53 | 0.54 | 0.75 | 0.31 | 0.68 |
| 3 | 107041230 | rs12629430 | -0.91 | -0.786 | 0.521 | -0.782 | -0.002 | -0.009 | -0.009 | 0 | 0.37 | 0.42 | 0.52 | 0.81 | 0.31 | 0.69 |
| 3 | 107047443 | rs2676364 | -1.705 | -0.556 | 0.212 | -1.288 | -0.003 | 0.113 | 0.113 | 0 | 0.53 | 0.73 | 0.67 | 0.98 | 0.45 | 0.77 |
| 3 | 107049360 | rs17194101 | -0.272 | -0.103 | -0.004 | -0.772 | 0.002 | -0.024 | -0.024 | 0 | 0.18 | 0.2 | 0.1 | 0.04 | 0.16 | 0.19 |
| 3 | 107060396 | rs9856825 | -0.285 | -0.103 | -0.008 | -0.772 | 0.004 | -0.032 | -0.032 | 0.981 | 0.18 | 0.23 | 0.1 | 0.04 | 0.16 | 0.19 |
| 3 | 107073105 | rs709543 | -0.377 | -0.162 | -0.106 | -0.421 | -0.001 | -0.025 | -0.025 | 0 | 0.11 | 0.09 | 0.08 | 0 | 0.12 | 0.05 |
| 3 | 107074910 | rs709547 | 0.19 | -0.194 | -0.078 | -0.283 | -0.003 | -0.001 | -0.001 | 0.034 | 0.63 | 0.63 | 0.53 | 0.3 | 0.68 | 0.43 |
| 3 | 107075797 | rs12695170 | 0.412 | -0.362 | -0.048 | -0.894 | -0.013 | 0.061 | 0.061 | 0 | 0.25 | 0.14 | 0.15 | 0 | 0.23 | 0.1 |
| 3 | 107078058 | rs709550 | -0.588 | -0.302 | 0.546 | -0.855 | -0.002 | 0.006 | 0.006 | 0 | 0.48 | 0.57 | 0.66 | 0.68 | 0.48 | 0.6 |
| 3 | 107078487 | rs697963 | 0.079 | -0.478 | 0.607 | -0.855 | -0.02 | 0.027 | 0.027 | 0 | 0.94 | 0.81 | 0.98 | 0.94 | 0.98 | 0.96 |
| 3 | 107087074 | rs17807525 | 1.493 | -0.189 | 0.09 | -1.051 | -0.041 | 0.143 | 0.143 | 0 | 0.34 | 0.08 | 0.22 | 0.07 | 0.37 | 0.23 |
| 3 | 107089332 | rs11706666 | 1.501 | -0.195 | 0.105 | -1.039 | -0.034 | 0.135 | 0.135 | 0 | 0.34 | 0.09 | 0.22 | 0.07 | 0.37 | 0.23 |
| 3 | 107091046 | rs6782559 | 0.599 | -0.225 | 0.336 | -1.071 | -0.006 | 0.054 | 0.054 | 0 | 0.48 | 0.43 | 0.32 | 0.11 | 0.5 | 0.36 |
| 3 | 107097346 | rs3811061 | 0.443 | 0.044 | 0.104 | -1.206 | -0.041 | 0.031 | 0.031 | 0 | 0.1 | 0.02 | 0.07 | 0.04 | 0.13 | 0.11 |
| 3 | 107110769 | rs793556 | -0.358 | -0.16 | 0.587 | -1.106 | -0.006 | 0.032 | 0.032 | 0 | 0.95 | 0.84 | 0.98 | 0.94 | 0.96 | 0.95 |
| 3 | 107121773 | rs796981 | 0.93 | -0.272 | 0.836 | -1.183 | -0.028 | 0 | 0 | 0.129 | 0.9 | 0.66 | 0.95 | 0.94 | 0.95 | 0.94 |
| 3 | 107122219 | rs809417 | 0.768 | -0.215 | 0.507 | -1.158 | -0.026 | -0.004 | -0.004 | 0 | 0.91 | 0.79 | 0.95 | 0.94 | 0.95 | 0.94 |
| 3 | 107127171 | rs871211 | 1.138 | -0.201 | 0.096 | -1.275 | 0 | 0.075 | 0.075 | 0 | 0.37 | 0.19 | 0.31 | 0.17 | 0.36 | 0.33 |
| 3 | 107135327 | rs793540 | -0.498 | -0.09 | 0.164 | -1.203 | -0.009 | 0.027 | 0.027 | 0 | 0.95 | 0.84 | 0.98 | 0.94 | 0.96 | 0.95 |
| 3 | 107140163 | rs7611467 | 0.974 | 0.012 | -1.393 | -1.426 | -0.006 | 0.092 | 0.092 | 0 | 0.37 | 0.13 | 0.31 | 0.16 | 0.36 | 0.33 |
| 3 | 107162649 | rs6809605 | 0.035 | 0.133 | -1.257 | -1.697 | -0.01 | 0.079 | 0.079 | 0 | 0.3 | 0.17 | 0.17 | 0.03 | 0.28 | 0.11 |
| 3 | 107175934 | rs4597706 | -0.186 | 0.23 | -1.573 | -1.962 | -0.03 | 0.099 | 0.099 | 0 | 0.35 | 0.2 | 0.2 | 0.04 | 0.33 | 0.22 |
| 3 | 107183065 | rs3914623 | 0.485 | 0.096 | -1.502 | -1.83 | -0.018 | 0.023 | 0.023 | 0 | 0.26 | 0.08 | 0.28 | 0.8 | 0.23 | 0.38 |
| 3 | 107185073 | rs1850344 | 0.489 | 0.56 | -2.097 | -2.466 | -0.008 | 0.135 | 0.135 | 0 | 0.41 | 0.13 | 0.3 | 0.17 | 0.4 | 0.36 |
| 3 | 107200353 | rs9880129 | 0.473 | 0.233 | -1.73 | -2.678 | -0.003 | -0.072 | -0.072 | 0 | 0.37 | 0.56 | 0.21 | 0.04 | 0.27 | 0.16 |
| 3 | 107210629 | rs1356915 | 0.508 | 0.489 | -1.73 | -2.924 | 0.014 | -0.064 | -0.064 | 0 | 0.69 | 0.62 | 0.6 | 0.96 | 0.6 | 0.67 |

| | | | | | | | | | | | | | | | | |
|---|---|---|---|---|---|---|---|---|---|---|---|---|---|---|---|---|
| 3 | 107214662 | rs686564 | -0.273 | 0.798 | -1.738 | -3.024 | 0 | -0.015 | -0.015 | 0 | 0.86 | 0.83 | 0.76 | 1 | 0.86 | 0.85 |
| 3 | 107216817 | rs674217 | -0.9 | 0.669 | -1.738 | -3.024 | -0.007 | -0.029 | -0.029 | 0 | 0.94 | 0.9 | 0.81 | 1 | 0.94 | 0.88 |
| 3 | 107222388 | rs6781000 | 0.826 | 0.593 | -1.827 | -3.056 | -0.009 | -0.034 | -0.034 | 0 | 0.38 | 0.5 | 0.35 | 0.83 | 0.28 | 0.44 |
| 3 | 107223736 | rs1426954 | 0.742 | 0.907 | -1.984 | -2.851 | 0.01 | -0.012 | -0.012 | 0 | 0.11 | 0.11 | 0.07 | 0.04 | 0.12 | 0.12 |
| 3 | 107233097 | rs10511260 | 1.119 | 0.769 | -2.116 | -2.717 | -0.019 | 0.117 | 0.117 | 0 | 0.32 | 0.03 | 0.24 | 0.12 | 0.32 | 0.23 |
| 3 | 107233327 | rs594635 | 0.022 | 1.043 | -1.41 | -2.61 | -0.004 | -0.013 | -0.013 | 0.205 | 0.13 | 0.15 | 0.13 | 0 | 0.22 | 0.08 |
| 3 | 107235614 | rs2174016 | -0.058 | 0.362 | -1.541 | -2.904 | 0.058 | 0.129 | 0.129 | 0.186 | 0.49 | 0.67 | 0.48 | 0.93 | 0.36 | 0.54 |
| 3 | 107244359 | rs623738 | 0.163 | 0.927 | -1.198 | -3.484 | 0.003 | -0.077 | -0.077 | 0 | 0.25 | 0.33 | 0.21 | 0.01 | 0.27 | 0.07 |
| 3 | 107266759 | rs9877739 | 1.463 | 0.893 | -1.443 | -3.507 | -0.055 | 0.088 | 0.088 | 0 | 0.14 | 0.01 | 0.1 | 0.01 | 0.17 | 0.03 |
| 3 | 107274256 | rs669706 | 0.882 | 0.874 | -1.395 | -3.501 | -0.008 | 0.008 | 0.008 | 0 | 0.07 | 0.07 | 0.09 | 0 | 0.1 | 0.02 |
| 3 | 107283393 | rs576698 | 0.234 | 0.798 | -1.415 | -3.533 | -0.005 | 0.007 | 0.007 | 0 | 0.08 | 0.11 | 0.09 | 0 | 0.1 | 0.02 |
| 3 | 107285743 | rs1648890 | 0.161 | 0.784 | -1.253 | -3.574 | -0.033 | 0.062 | 0.062 | 0 | 0.23 | 0.16 | 0.19 | 0.01 | 0.27 | 0.06 |
| 3 | 107293971 | rs583305 | 0.854 | 0.788 | -1.158 | -3.55 | -0.003 | -0.008 | -0.008 | 0 | 0.07 | 0.09 | 0.09 | 0 | 0.11 | 0.02 |
| 3 | 107295008 | rs9851055 | 0.655 | 0.406 | -1.022 | -4.085 | 0.004 | 0.068 | 0.068 | 0 | 0.53 | 0.32 | 0.51 | 0.1 | 0.54 | 0.48 |
| 3 | 107316938 | rs9288851 | 1.812 | 0.416 | -1.118 | -3.988 | -0.03 | 0.059 | 0.059 | 0 | 0.15 | 0.05 | 0.1 | 0.01 | 0.16 | 0.03 |
| 3 | 107320843 | rs526034 | -0.185 | 0.328 | -0.725 | -3.984 | 0.019 | -0.067 | -0.067 | 0.369 | 0.89 | 0.66 | 0.9 | 1 | 0.89 | 0.98 |
| 3 | 107343402 | rs609777 | 0.891 | 0.608 | -1.251 | -4.544 | 0.001 | -0.055 | -0.055 | 0 | 0.71 | 0.36 | 0.64 | 0.93 | 0.7 | 0.82 |
| 3 | 107345012 | rs9836754 | 1.585 | 0.847 | -1.412 | -4.553 | -0.095 | 0.274 | 0.274 | 0 | 0.37 | 0.03 | 0.26 | 0.04 | 0.36 | 0.33 |
| 3 | 107346018 | rs17204920 | 1.416 | 1.062 | -1.562 | -4.721 | -0.017 | 0.026 | 0.026 | 0 | 0.06 | 0.02 | 0.13 | 0 | 0.09 | 0.09 |
| 3 | 107354130 | rs643938 | 0.402 | 0.985 | -1.511 | -4.703 | -0.01 | 0.063 | 0.063 | 0 | 0.56 | 0.31 | 0.54 | 0.93 | 0.55 | 0.79 |
| 3 | 107354432 | rs16844855 | 1.496 | 0.896 | -1.218 | -4.574 | 0.034 | -0.024 | -0.024 | 0 | 0.06 | 0.24 | 0.11 | 0.01 | 0.08 | 0.04 |
| 3 | 107371482 | rs658595 | -1.296 | 1.068 | -1.6 | -4.582 | -0.035 | 0.24 | 0.24 | 0 | 0.63 | 0.91 | 0.8 | 0.97 | 0.65 | 0.74 |
| 3 | 107379837 | rs167924 | -0.013 | 1.096 | -1.361 | -4.169 | -0.007 | 0.047 | 0.047 | 0 | 0.65 | 0.23 | 0.67 | 0.98 | 0.64 | 0.87 |
| 3 | 107385910 | rs16853756 | -0.631 | 1.318 | -1.139 | -3.876 | 0.011 | -0.051 | -0.051 | 0 | 0.19 | 0.28 | 0.27 | 0.09 | 0.21 | 0.17 |
| 3 | 107393593 | rs9867809 | -0.098 | 1.702 | -1.421 | -4.199 | -0.044 | 0.048 | 0.048 | 0 | 0.1 | 0.02 | 0.16 | 0.02 | 0.14 | 0.14 |
| 3 | 107398611 | rs6779452 | 0.811 | 1.472 | -1.509 | -4.531 | -0.007 | 0.039 | 0.039 | 0 | 0.26 | 0.16 | 0.39 | 0.92 | 0.29 | 0.5 |
| 3 | 107401605 | rs1021504 | 0.006 | 1.326 | -1.351 | -4.418 | 0.016 | -0.059 | -0.059 | 0 | 0.12 | 0.32 | 0.15 | 0.04 | 0.12 | 0.11 |
| 3 | 107401883 | rs9848674 | 0.66 | 1.162 | -2.144 | -4.591 | -0.103 | 0.268 | 0.268 | 0 | 0.4 | 0.03 | 0.28 | 0.07 | 0.39 | 0.39 |
| 3 | 107424090 | rs9846246 | -0.38 | 1.059 | -2.064 | -5.213 | -0.018 | 0.165 | 0.165 | 0 | 0.58 | 0.83 | 0.52 | 0.04 | 0.56 | 0.29 |
| 3 | 107445366 | rs923952 | -0.458 | 0.869 | -1.257 | -4.786 | -0.028 | -0.122 | -0.122 | 0 | 0.17 | 0.53 | 0.17 | 0.01 | 0.12 | 0.05 |
| 3 | 107450331 | rs9862385 | 1.762 | 1.001 | -1.47 | -4.912 | -0.068 | 0.074 | 0.074 | 0 | 0.16 | 0.04 | 0.11 | 0.04 | 0.18 | 0.06 |
| 3 | 107464170 | rs11710737 | -0.953 | 0.913 | -1.528 | -5.016 | 0 | -0.029 | -0.029 | 0 | 0.55 | 0.57 | 0.43 | 0.07 | 0.45 | 0.45 |
| 3 | 107466411 | rs13066717 | -0.681 | 0.964 | -1.4 | -4.982 | 0.02 | -0.072 | -0.072 | 0 | 0.2 | 0.3 | 0.29 | 0.04 | 0.26 | 0.19 |
| 3 | 107471699 | rs2049339 | 1.473 | 0.861 | -1.589 | -4.874 | -0.017 | 0.185 | 0.185 | 0 | 0.26 | 0.03 | 0.1 | 0.03 | 0.2 | 0.25 |
| 3 | 107473614 | rs10804463 | 1.034 | 1.058 | -0.972 | -4.525 | -0.013 | 0.043 | 0.043 | 0 | 0.3 | 0.22 | 0.45 | 0.92 | 0.38 | 0.53 |
| 3 | 107481168 | rs16853820 | 0.183 | 1.05 | -0.422 | -3.993 | 0.049 | -0.044 | -0.044 | 0 | 0.08 | 0.22 | 0.13 | 0.01 | 0.12 | 0.05 |
| 3 | 107485897 | rs2204260 | -1.199 | 0.784 | -0.461 | -4.16 | 0.011 | -0.093 | -0.093 | 0.808 | 0.84 | 0.74 | 0.89 | 0.99 | 0.85 | 0.96 |
| 3 | 107489708 | rs9845070 | 1.637 | 0.777 | -0.582 | -4.163 | -0.046 | 0.072 | 0.072 | 0 | 0.13 | 0.03 | 0.09 | 0.01 | 0.15 | 0.04 |
| 3 | 107494345 | rs9871235 | -0.193 | 0.143 | -0.83 | -4.515 | 0.005 | -0.035 | -0.035 | 0 | 0.56 | 0.62 | 0.75 | 0.92 | 0.62 | 0.64 |
| 3 | 107507776 | rs9839178 | 1.023 | -0.215 | -1.007 | -4.75 | -0.033 | 0.04 | 0.04 | 0 | 0.33 | 0.24 | 0.53 | 0.92 | 0.46 | 0.55 |
| 3 | 107513782 | rs11714512 | 1.168 | 0.276 | -1.239 | -4.397 | -0.016 | 0.169 | 0.169 | 0 | 0.27 | 0.02 | 0.14 | 0.07 | 0.23 | 0.32 |
| 3 | 107513866 | rs12330842 | 1.006 | 1.169 | 0.066 | -3.572 | 0.002 | -0.044 | -0.044 | 0 | 0.15 | 0.17 | 0.09 | 0.01 | 0.15 | 0.04 |
| 3 | 107527596 | rs9831296 | 0.861 | 1.317 | -0.351 | -3.51 | -0.004 | -0.009 | -0.009 | 0.004 | 0.15 | 0.13 | 0.09 | 0.01 | 0.15 | 0.05 |
| 3 | 107538740 | rs6802043 | 0.289 | 1.342 | -0.326 | -3.531 | -0.022 | -0.002 | -0.002 | 0 | 0.21 | 0.17 | 0.25 | 0.01 | 0.34 | 0.12 |
| 3 | 107543491 | rs2172730 | -0.711 | 0.83 | -0.081 | -3.572 | 0.003 | -0.091 | -0.091 | 0 | 0.37 | 0.39 | 0.44 | 0.02 | 0.51 | 0.2 |
| 3 | 107548789 | rs9849845 | 0.507 | 0.855 | -0.885 | -3.508 | -0.044 | 0.116 | 0.116 | 0.475 | 0.63 | 0.78 | 0.54 | 0.97 | 0.48 | 0.77 |
| 3 | 107555466 | rs10933916 | 0.883 | 0.203 | -0.643 | -3.703 | 0.003 | -0.038 | -0.038 | 0.087 | 0.47 | 0.43 | 0.47 | 0.95 | 0.41 | 0.67 |
| 3 | 107559552 | rs4129549 | 1.472 | 0.24 | -1.031 | -3.805 | -0.045 | 0.101 | 0.101 | 0 | 0.17 | 0.02 | 0.17 | 0.01 | 0.18 | 0.08 |
| 3 | 107563796 | rs10460810 | 1.058 | 0.352 | -0.891 | -3.099 | -0.001 | -0.02 | -0.02 | 0 | 0.45 | 0.45 | 0.43 | 0.84 | 0.37 | 0.64 |
| 3 | 107567130 | rs10933917 | 0.959 | -0.238 | -0.657 | -3.161 | 0.004 | 0.034 | 0.034 | 0.191 | 0.5 | 0.44 | 0.49 | 0.86 | 0.48 | 0.66 |
| 3 | 107569755 | rs9849624 | 0.413 | -0.165 | -0.672 | -3.121 | 0.022 | 0.069 | 0.069 | 0 | 0.25 | 0.11 | 0.19 | 0.01 | 0.2 | 0.18 |
| 3 | 107569873 | rs17227439 | -0.003 | -0.145 | -0.482 | -3.139 | -0.015 | -0.116 | -0.116 | 0 | 0.29 | 0.35 | 0.21 | 0.01 | 0.23 | 0.19 |
| 3 | 107570882 | rs1897496 | -0.848 | -0.251 | -0.519 | -3.13 | 0 | -0.022 | -0.022 | 0 | 0.44 | 0.55 | 0.45 | 0.13 | 0.45 | 0.33 |
| 3 | 107575074 | rs9852257 | -0.706 | -0.004 | -0.97 | -3.006 | -0.007 | 0.001 | 0.001 | 0.088 | 0.67 | 0.65 | 0.6 | 0.11 | 0.64 | 0.47 |
| 3 | 107576916 | rs13097121 | -0.176 | 0.344 | -0.997 | -2.701 | 0.017 | -0.045 | -0.045 | 7.935 | 0.18 | 0.23 | 0.23 | 0.04 | 0.25 | 0.16 |
| 3 | 107585300 | rs2099308 | -1.062 | 0.65 | -1.046 | -2.567 | -0.004 | 0.002 | 0.002 | 0.807 | 0.9 | 0.92 | 0.77 | 0.9 | 0.81 | 0.89 |
| 3 | 107591015 | rs4404444 | 0.298 | 0.096 | -1.277 | -2.866 | 0 | -0.016 | -0.016 | 1.301 | 0.52 | 0.51 | 0.53 | 0.84 | 0.5 | 0.67 |
| 3 | 107592185 | rs11707950 | 0.468 | 0.636 | -1.997 | -2.394 | 0.004 | 0.001 | 0.001 | 0 | 0.27 | 0.18 | 0.2 | 0.03 | 0.25 | 0.11 |
| 3 | 107594886 | rs1991783 | 0.701 | 0.928 | -2.527 | -2.199 | -0.12 | 0.028 | 0.028 | 0 | 0.26 | 0.06 | 0.35 | 0.8 | 0.34 | 0.45 |
| 3 | 107596954 | rs6773129 | 0.363 | 0.485 | -2.245 | -2.278 | -0.017 | 0.121 | 0.121 | 0 | 0.25 | 0.05 | 0.19 | 0.03 | 0.25 | 0.1 |
| 3 | 107599953 | rs10933921 | -0.031 | 0.282 | -2.864 | -2.316 | -0.102 | 0.214 | 0.214 | 0 | 0.44 | 0.05 | 0.46 | 0.82 | 0.51 | 0.67 |
| 3 | 107603762 | rs16853945 | 1.073 | 0.311 | -3.045 | -1.536 | -0.046 | -0.126 | -0.126 | 0 | 0.21 | 0.04 | 0.21 | 0.64 | 0.22 | 0.41 |
| 3 | 107603830 | rs960470 | 0.346 | -0.02 | -1.959 | -2.593 | 0.014 | -0.042 | -0.042 | 0 | 0.17 | 0.3 | 0.16 | 0.07 | 0.14 | 0.09 |
| 3 | 107604090 | rs960471 | -0.02 | -0.162 | -0.982 | -2.644 | 0.015 | -0.068 | -0.068 | 0 | 0.19 | 0.6 | 0.2 | 0.07 | 0.15 | 0.09 |
| 3 | 107604277 | rs840286 | -1.364 | -0.353 | -0.927 | -2.903 | -0.015 | 0.068 | 0.068 | 0 | 0.75 | 0.92 | 0.72 | 0.89 | 0.74 | 0.86 |
| 3 | 107611941 | rs709521 | -0.218 | -0.637 | -0.761 | -3.093 | 0 | -0.051 | -0.051 | 0 | 0.6 | 0.65 | 0.64 | 0.97 | 0.62 | 0.77 |
| 3 | 107614584 | rs6778247 | 1.049 | -0.717 | -0.047 | -3.109 | -0.01 | -0.052 | -0.052 | 2.041 | 0.09 | 0.2 | 0.05 | 0 | 0.06 | 0.09 |
| 3 | 107615040 | rs709519 | 0.676 | -0.751 | -0.065 | -3.132 | -0.069 | 0.247 | 0.247 | 0 | 0.44 | 0.1 | 0.49 | 0.89 | 0.52 | 0.61 |
| 3 | 107619385 | rs709516 | 0.687 | -0.762 | -0.089 | -3.08 | -0.079 | 0.23 | 0.23 | 0 | 0.44 | 0.09 | 0.48 | 0.88 | 0.52 | 0.61 |

| | | | | | | | | | | | | | | | | |
|---|---|---|---|---|---|---|---|---|---|---|---|---|---|---|---|---|
| 3 | 107624530 | rs4247450 | -1.204 | -0.751 | -0.025 | -3.216 | -0.035 | 0.094 | 0.094 | 0 | 0.72 | 0.92 | 0.71 | 0.97 | 0.7 | 0.87 |
| 3 | 107629427 | rs1700456 | -0.375 | -0.619 | -0.043 | -3.103 | -0.001 | -0.12 | -0.12 | 0.123 | 0.59 | 0.49 | 0.63 | 0.97 | 0.62 | 0.77 |
| 3 | 107634915 | rs697954 | -0.354 | -0.736 | -0.039 | -2.903 | -0.002 | -0.034 | -0.034 | 0 | 0.52 | 0.6 | 0.53 | 0.96 | 0.49 | 0.67 |
| 3 | 107640533 | rs13086279 | 0.54 | -0.815 | -0.658 | -2.76 | -0.02 | 0.164 | 0.164 | 0 | 0.38 | 0.13 | 0.37 | 0.89 | 0.38 | 0.48 |
| 3 | 107645774 | rs10933925 | -0.095 | -1.033 | -0.23 | -2.62 | -0.028 | -0.015 | -0.015 | 0 | 0.19 | 0.21 | 0.15 | 0.08 | 0.07 | 0.1 |
| 3 | 107645827 | rs1437240 | 0.175 | -1.206 | 0.005 | -2.937 | 0.004 | -0.046 | -0.046 | 0 | 0.42 | 0.56 | 0.42 | 0.88 | 0.49 | 0.6 |
| 3 | 107646728 | rs4855779 | -0.335 | -1.407 | 0.11 | -2.995 | 0.001 | 0.015 | 0.015 | 0.489 | 0.5 | 0.4 | 0.48 | 0.96 | 0.5 | 0.62 |
| 3 | 107646807 | rs9837335 | 1.107 | -1.182 | 0.047 | -2.876 | -0.077 | 0.072 | 0.072 | 0 | 0.29 | 0.11 | 0.33 | 0.87 | 0.39 | 0.52 |
| 3 | 107650668 | rs6799066 | -0.955 | -1.277 | 0.319 | -2.902 | -0.014 | -0.005 | -0.005 | 0 | 0.61 | 0.72 | 0.53 | 0.97 | 0.54 | 0.65 |
| 3 | 107652635 | rs1437238 | -0.95 | -1.276 | 0.337 | -2.901 | -0.014 | 0.003 | 0.003 | 0.287 | 0.61 | 0.71 | 0.54 | 0.97 | 0.56 | 0.66 |
| 3 | 107654556 | rs12638978 | -1.042 | -1.278 | 0.337 | -2.902 | -0.022 | 0.032 | 0.032 | 0.598 | 0.61 | 0.78 | 0.54 | 0.97 | 0.56 | 0.65 |
| 3 | 107654661 | rs17228803 | 1.179 | -1.361 | 0.245 | -2.946 | -0.113 | 0.083 | 0.083 | 0 | 0.29 | 0.04 | 0.32 | 0.89 | 0.39 | 0.51 |
| 3 | 107661284 | rs1370285 | -0.698 | -1.449 | 0.522 | -2.989 | 0 | -0.009 | -0.009 | 0 | 0.32 | 0.73 | 0.23 | 0.07 | 0.17 | 0.14 |
| 3 | 107661951 | rs4855614 | -1.151 | -1.422 | 0.515 | -3.003 | -0.035 | 0.07 | 0.07 | 1.017 | 0.62 | 0.84 | 0.55 | 0.97 | 0.56 | 0.65 |
| 3 | 107662734 | rs7625167 | 0.681 | -1.367 | 0.568 | -2.967 | -0.001 | -0.012 | -0.012 | 0 | 0.34 | 0.33 | 0.34 | 0.9 | 0.41 | 0.52 |
| 3 | 107671945 | rs10933928 | -0.433 | -1.402 | 0.648 | -2.886 | -0.049 | -0.069 | -0.069 | 0.702 | 0.27 | 0.46 | 0.21 | 0.07 | 0.15 | 0.14 |
| 3 | 107676808 | rs13090662 | -0.273 | -1.401 | 0.564 | -2.888 | -0.009 | -0.018 | -0.018 | 0 | 0.26 | 0.33 | 0.19 | 0.07 | 0.14 | 0.14 |
| 3 | 107678877 | rs934674 | -0.285 | -1.412 | 0.473 | -2.893 | -0.009 | -0.018 | -0.018 | 0.001 | 0.26 | 0.34 | 0.19 | 0.07 | 0.14 | 0.14 |
| 3 | 107682964 | rs6790356 | 0.387 | -0.915 | 0.654 | -2.7 | 0.012 | -0.079 | -0.079 | 0 | 0.35 | 0.46 | 0.45 | 0.9 | 0.42 | 0.52 |
| 3 | 107688670 | rs6807905 | 0.535 | -0.943 | 0.385 | -2.625 | -0.012 | -0.053 | -0.053 | 0 | 0.09 | 0.26 | 0.05 | 0 | 0.07 | 0.07 |
| 3 | 107689210 | rs11708384 | 0.898 | -0.85 | -0.061 | -2.664 | -0.068 | 0.081 | 0.081 | 0 | 0.29 | 0.08 | 0.31 | 0.89 | 0.38 | 0.51 |
| 3 | 107691697 | rs840281 | -1.003 | -0.535 | 0.693 | -2.046 | -0.045 | -0.018 | -0.018 | 0 | 0.21 | 0.46 | 0.27 | 0.14 | 0.15 | 0.14 |
| 3 | 107692467 | rs709500 | -1.207 | -0.516 | 0.292 | -2.038 | -0.029 | -0.017 | -0.017 | 0 | 0.24 | 0.69 | 0.29 | 0.14 | 0.15 | 0.14 |
| 3 | 107694834 | rs709498 | -0.306 | -0.191 | -0.096 | -2.453 | -0.046 | 0.085 | 0.085 | 2.132 | 0.65 | 0.87 | 0.65 | 0.95 | 0.57 | 0.67 |
| 3 | 107697666 | rs709494 | -0.143 | -0.203 | -0.111 | -2.406 | -0.051 | 0.099 | 0.099 | 1.497 | 0.63 | 0.83 | 0.55 | 0.9 | 0.53 | 0.66 |
| 3 | 107700952 | rs709488 | 0.232 | -0.224 | 0.028 | -2.386 | -0.002 | -0.006 | -0.006 | 0.253 | 0.54 | 0.57 | 0.49 | 0.9 | 0.49 | 0.62 |
| 3 | 107707411 | rs709467 | 0.253 | -0.208 | -0.195 | -2.376 | -0.001 | 0.026 | 0.026 | 1.958 | 0.54 | 0.61 | 0.5 | 0.9 | 0.49 | 0.62 |
| 3 | 107707753 | rs697957 | -0.25 | 0.57 | -0.465 | -2.089 | -0.012 | 0.009 | 0.009 | 0 | 0.79 | 0.84 | 0.71 | 0.93 | 0.75 | 0.92 |
| 3 | 107708956 | rs9845309 | -0.579 | 0.064 | -0.281 | -1.949 | 0.005 | 0.016 | 0.016 | 4.334 | 0.49 | 0.57 | 0.55 | 0.15 | 0.45 | 0.47 |
| 3 | 107714075 | rs877859 | 0.551 | -0.378 | -0.932 | -1.812 | 0.015 | 0.163 | 0.163 | 2.846 | 0.59 | 0.07 | 0.46 | 0.83 | 0.68 | 0.66 |
| 3 | 107720888 | rs7649924 | -0.708 | -0.36 | -0.065 | -1.743 | 0.003 | 0.018 | 0.018 | 2.464 | 0.11 | 0.56 | 0.13 | 0.1 | 0.07 | 0.14 |
| 3 | 107721213 | rs9823579 | 0.816 | -0.449 | -1.006 | -2.119 | 0.007 | 0.225 | 0.225 | 1.568 | 0.56 | 0.07 | 0.44 | 0.85 | 0.63 | 0.6 |
| 3 | 107721902 | rs709448 | -0.688 | -0.514 | -0.082 | -1.992 | -0.057 | -0.114 | -0.114 | 0 | 0.16 | 0.46 | 0.08 | 0 | 0.07 | 0.07 |
| 3 | 107722014 | rs9828514 | 0.514 | 0.065 | -0.985 | -1.956 | 0 | -0.004 | -0.004 | 0 | 0.33 | 0.28 | 0.29 | 0.87 | 0.26 | 0.45 |
| 3 | 107725831 | rs1866001 | 0.291 | 0.869 | -0.982 | -1.763 | -0.009 | 0.014 | 0.014 | 0.055 | 0.18 | 0.11 | 0.24 | 0.03 | 0.21 | 0.1 |
| 3 | 107726194 | rs9821923 | 0.099 | 0.491 | -1.053 | -0.633 | 0.028 | 0.026 | 0.026 | 1.694 | 0.46 | 0.29 | 0.42 | 0.67 | 0.36 | 0.57 |
| 3 | 107731924 | rs9835822 | -0.129 | 0.556 | -1.304 | -0.923 | 0.039 | 0.101 | 0.101 | 4.873 | 0.45 | 0.3 | 0.49 | 0.82 | 0.37 | 0.59 |
| 3 | 107732083 | rs696375 | 0.087 | 0.821 | -0.887 | -1.418 | 0.002 | -0.138 | -0.138 | 0.015 | 0.74 | 0.49 | 0.7 | 0.95 | 0.72 | 0.74 |
| 3 | 107732977 | rs10804466 | 0.063 | 0.847 | -2.295 | -1.18 | -0.121 | 0.095 | 0.095 | 0 | 0.4 | 0.04 | 0.46 | 0.18 | 0.53 | 0.34 |
| 3 | 107733600 | rs9859911 | -0.882 | 0.164 | -1.836 | -1.413 | -0.038 | -0.072 | -0.072 | 0 | 0.85 | 0.44 | 0.8 | 0.97 | 0.88 | 0.81 |
| 3 | 107737386 | rs1344877 | 0.832 | 0.413 | -2.117 | -1.394 | -0.04 | 0.112 | 0.112 | 0 | 0.2 | 0.04 | 0.11 | 0.03 | 0.17 | 0.08 |
| 3 | 107739094 | rs6790456 | 0.468 | 0.187 | -2.103 | -1.419 | 0.012 | -0.002 | -0.002 | 0 | 0.25 | 0.21 | 0.21 | 0.75 | 0.18 | 0.38 |
| 3 | 107741880 | rs9860959 | -0.654 | 1.191 | -2.005 | -1.339 | 0.009 | 0.091 | 0.091 | 0 | 0.53 | 0.36 | 0.56 | 0.97 | 0.55 | 0.6 |
| 3 | 107744023 | rs7622560 | -0.051 | 0.51 | -1.769 | -1.07 | 0.022 | 0.081 | 0.081 | 0.007 | 0.45 | 0.32 | 0.33 | 0.78 | 0.35 | 0.46 |
| 3 | 107750920 | rs16854140 | -0.251 | 0.888 | -1.526 | 0.091 | -0.02 | -0.013 | -0.013 | 0.015 | 0.09 | 0.03 | 0.18 | 0.54 | 0.11 | 0.24 |
| 3 | 107757086 | rs1006044 | 0.348 | 0.707 | -1.423 | 0.295 | -0.002 | 0.089 | 0.089 | 0 | 0.06 | 0.06 | 0.15 | 0.51 | 0.05 | 0.26 |
| 3 | 107760800 | rs696365 | 0.07 | 0.456 | -1.276 | -0.993 | -0.018 | 0.064 | 0.064 | 0.965 | 0.41 | 0.25 | 0.37 | 0.14 | 0.44 | 0.3 |
| 3 | 107761411 | rs2116693 | 0.228 | 0.53 | -0.899 | -0.99 | 0.007 | 0.05 | 0.05 | 0 | 0.52 | 0.62 | 0.47 | 0.32 | 0.5 | 0.44 |
| 3 | 107763040 | rs2271054 | 1.326 | 0.265 | -1.311 | -1.421 | -0.036 | 0.1 | 0.1 | 0 | 0.19 | 0.04 | 0.11 | 0.03 | 0.18 | 0.09 |
| 3 | 107765392 | rs9879947 | 0.14 | 0.087 | -1.019 | -1.516 | -0.002 | -0.018 | -0.018 | 0.447 | 0.47 | 0.51 | 0.43 | 0.32 | 0.47 | 0.41 |
| 3 | 107776646 | rs12695175 | 1.324 | 0.209 | -1.221 | -1.353 | -0.042 | 0.113 | 0.113 | 0 | 0.19 | 0.02 | 0.1 | 0.03 | 0.18 | 0.09 |
| 3 | 107787525 | rs10933931 | 0.974 | 0.344 | -0.477 | -0.676 | -0.002 | 0.07 | 0.07 | 0 | 0.06 | 0.05 | 0.15 | 0.55 | 0.05 | 0.26 |
| 3 | 107794140 | rs17827795 | -0.001 | 0.29 | -0.362 | -0.903 | -0.018 | 0.049 | 0.049 | 1 | 0.41 | 0.27 | 0.37 | 0.14 | 0.45 | 0.3 |
| 3 | 107805334 | rs17827915 | 0.406 | 0.417 | -0.169 | -1.025 | -0.002 | 0.012 | 0.012 | 0.001 | 0.56 | 0.6 | 0.5 | 0.34 | 0.52 | 0.45 |
| 3 | 107811142 | rs326341 | -0.775 | 0.266 | 0.059 | -0.891 | 0.001 | -0.003 | -0.003 | 0.013 | 0.44 | 0.48 | 0.37 | 0.38 | 0.46 | 0.35 |
| 3 | 107811787 | rs3762681 | -0.015 | 1.025 | -0.274 | -1.048 | 0.014 | 0 | 0 | 0 | 0.11 | 0.16 | 0.2 | 0.13 | 0.18 | 0.1 |
| 3 | 107817299 | rs17231503 | 0.712 | 1.035 | -0.387 | -1.061 | -0.052 | 0.128 | 0.128 | 0.317 | 0.22 | 0.04 | 0.12 | 0 | 0.24 | 0.17 |
| 3 | 107818777 | rs7635063 | 0.44 | 0.865 | -0.368 | -1.088 | -0.066 | 0.095 | 0.095 | 0 | 0.24 | 0.08 | 0.14 | 0 | 0.33 | 0.2 |
| 3 | 107824061 | rs162064 | -0.091 | 0.631 | -0.132 | -0.952 | -0.017 | 0.029 | 0.029 | 0 | 0.68 | 0.84 | 0.81 | 0.74 | 0.64 | 0.74 |
| 3 | 107829766 | rs4855620 | 1.038 | 0.599 | 0.268 | -0.988 | 0.001 | -0.005 | -0.005 | 1.282 | 0.55 | 0.48 | 0.61 | 0.62 | 0.52 | 0.64 |
| 3 | 107833289 | rs7648181 | 0.136 | 0.62 | 0.182 | -1.017 | -0.001 | 0.005 | 0.005 | 0 | 0.65 | 0.72 | 0.79 | 0.74 | 0.63 | 0.73 |
| 3 | 107840830 | rs170534 | 0.106 | 0.65 | -0.063 | -1.003 | -0.015 | 0.038 | 0.038 | 0 | 0.66 | 0.84 | 0.81 | 0.74 | 0.63 | 0.74 |
| 3 | 107853364 | rs326352 | 1.179 | 0.659 | 0.077 | -1.032 | 0.004 | 0.02 | 0.02 | 0 | 0.48 | 0.37 | 0.51 | 0.11 | 0.49 | 0.37 |
| 3 | 107858443 | rs6769732 | -0.258 | 0.735 | -0.156 | -1.021 | -0.064 | 0.121 | 0.121 | 0 | 0.35 | 0.23 | 0.24 | 0.12 | 0.43 | 0.29 |
| 3 | 107864693 | rs326325 | 1.261 | 0.743 | -0.136 | -1.165 | -0.001 | -0.041 | -0.041 | 0.053 | 0.49 | 0.43 | 0.52 | 0.12 | 0.48 | 0.42 |
| 3 | 107864963 | rs13093976 | 0.234 | 0.947 | -0.474 | -1.101 | -0.076 | 0.154 | 0.154 | 0.006 | 0.28 | 0.07 | 0.16 | 0 | 0.34 | 0.2 |
| 3 | 107865932 | rs16854238 | 0.88 | 0.957 | -0.3 | -0.818 | 0.001 | 0.165 | 0.165 | 0 | 0.05 | 0.12 | 0.17 | 0.51 | 0.03 | 0.22 |
| 3 | 107880473 | rs17828926 | 0.566 | 0.888 | -0.329 | -0.895 | -0.002 | 0 | 0 | 0.669 | 0.07 | 0.09 | 0.09 | 0.11 | 0.11 | 0.09 |
| 3 | 107883768 | rs13318436 | 0.566 | 0.888 | -0.329 | -0.895 | -0.002 | 0 | 0 | 4.166 | 0.07 | 0.09 | 0.09 | 0.11 | 0.11 | 0.09 |

| | | | | | | | | | | | | | | | | |
|---|---|---|---|---|---|---|---|---|---|---|---|---|---|---|---|---|
| 3 | 107886925 | rs17232836 | 0.642 | 0.888 | -0.331 | -0.895 | -0.011 | -0.001 | -0.001 | 0.054 | 0.07 | 0.06 | 0.08 | 0.11 | 0.11 | 0.09 |
| 3 | 107897239 | rs11922835 | 0.407 | 0.88 | -0.333 | -0.887 | -0.003 | -0.001 | -0.001 | 0.092 | 0.08 | 0.09 | 0.09 | 0.11 | 0.11 | 0.1 |
| 3 | 107926340 | rs1289744 | 0.544 | 0.866 | -0.155 | -0.82 | 0.027 | 0.097 | 0.097 | 0 | 0.87 | 0.68 | 0.74 | 0.39 | 0.86 | 0.69 |
| 3 | 107936558 | rs1289764 | -0.446 | 0.85 | -0.151 | -0.802 | 0.014 | 0.282 | 0.282 | 0 | 0.94 | 0.75 | 0.82 | 0.5 | 0.97 | 0.78 |
| 3 | 107937942 | rs1289757 | 0.377 | 0.835 | -0.205 | -0.869 | 0.027 | 0.097 | 0.097 | 0 | 0.87 | 0.68 | 0.74 | 0.39 | 0.86 | 0.69 |
| 3 | 107945078 | rs2701063 | 1.02 | 0.481 | -0.183 | -1.153 | 0.002 | -0.017 | -0.017 | 0 | 0.48 | 0.44 | 0.52 | 0.12 | 0.49 | 0.41 |
| 3 | 107961441 | rs9827766 | 0.199 | 0.257 | -0.072 | -1.045 | -0.011 | 0.001 | 0.001 | 0 | 0.65 | 0.69 | 0.63 | 0.39 | 0.63 | 0.51 |
| 3 | 107970084 | rs1531191 | -0.082 | 0.293 | -0.112 | -1.116 | -0.003 | 0 | 0 | 2.144 | 0.16 | 0.18 | 0.22 | 0.76 | 0.09 | 0.3 |
| 3 | 107975332 | rs1870007 | -0.091 | 0.29 | -0.154 | -1.132 | -0.001 | -0.011 | -0.011 | 0.128 | 0.71 | 0.71 | 0.8 | 0.88 | 0.68 | 0.73 |
| 3 | 107998620 | rs2100581 | -0.082 | 0.291 | -0.161 | -1.132 | -0.004 | -0.003 | -0.003 | 0.107 | 0.71 | 0.72 | 0.8 | 0.88 | 0.68 | 0.73 |
| 3 | 108000330 | rs1563515 | 1.512 | 0.026 | -0.267 | -1.708 | 0.004 | 0.02 | 0.02 | 0.024 | 0.22 | 0.15 | 0.17 | 0.05 | 0.2 | 0.11 |
| 3 | 108015247 | rs768342 | 0.688 | 0.121 | -0.267 | -1.605 | 0 | -0.043 | -0.043 | 0.343 | 0.54 | 0.44 | 0.57 | 0.12 | 0.59 | 0.42 |
| 3 | 108018926 | rs9637379 | 0.636 | -0.043 | -0.455 | -1.117 | 0.003 | 0.207 | 0.207 | 0 | 0.06 | 0.16 | 0.19 | 0.51 | 0.04 | 0.23 |
| 3 | 108019111 | rs9637380 | -0.302 | -0.106 | -0.417 | -1.157 | -0.028 | 0.007 | 0.007 | 0.283 | 0.17 | 0.29 | 0.23 | 0.76 | 0.09 | 0.31 |
| 3 | 108021294 | rs10933936 | -0.166 | -0.082 | -0.528 | -1.146 | -0.014 | 0.003 | 0.003 | 0.119 | 0.17 | 0.23 | 0.22 | 0.76 | 0.09 | 0.3 |
| 3 | 108021634 | rs10933937 | 0.906 | 0.016 | -0.6 | -1.086 | 0 | 0.151 | 0.151 | 0 | 0.06 | 0.09 | 0.18 | 0.51 | 0.04 | 0.23 |
| 3 | 108023965 | rs2016469 | 1.002 | -0.036 | -0.58 | -1.236 | -0.004 | 0.004 | 0.004 | 0 | 0.3 | 0.23 | 0.4 | 0.07 | 0.38 | 0.32 |
| 3 | 108024901 | rs6768207 | -0.531 | -0.396 | -0.831 | -0.527 | 0.039 | -0.01 | -0.01 | 0 | 0.21 | 0.12 | 0.07 | 0.26 | 0.15 | 0.13 |
| 3 | 108030714 | rs16854433 | -0.68 | 0.028 | -0.761 | -0.365 | 0.006 | -0.018 | -0.018 | 0.103 | 0.16 | 0.13 | 0.22 | 0.51 | 0.15 | 0.26 |
| 3 | 108041345 | rs12634962 | -0.513 | 0.052 | -0.686 | -0.322 | -0.002 | 0.02 | 0.02 | 0 | 0.16 | 0.21 | 0.23 | 0.51 | 0.15 | 0.21 |
| 3 | 108046671 | rs2061245 | 0.525 | 0.118 | -0.552 | -1.284 | -0.004 | -0.062 | -0.062 | 6.839 | 0.59 | 0.49 | 0.53 | 0.21 | 0.62 | 0.56 |
| 3 | 108049819 | rs2951512 | -0.567 | -0.113 | -0.745 | -1.506 | 0.011 | 0 | 0 | 1.683 | 0.86 | 0.86 | 0.76 | 0.21 | 0.93 | 0.7 |
| 3 | 108051504 | rs17241659 | 1.409 | 0.005 | -1.282 | -2.061 | -0.057 | 0.106 | 0.106 | 0 | 0.19 | 0.04 | 0.11 | 0 | 0.25 | 0.22 |
| 3 | 108056020 | rs6771609 | 0.195 | -0.201 | -0.753 | -1.648 | -0.009 | 0.082 | 0.082 | 0 | 0.67 | 0.81 | 0.59 | 0.21 | 0.69 | 0.6 |
| 3 | 108062185 | rs7643006 | 0.254 | -0.225 | -0.652 | -1.659 | -0.001 | 0.054 | 0.054 | 0 | 0.67 | 0.77 | 0.6 | 0.21 | 0.68 | 0.6 |
| 3 | 108066808 | rs6788771 | 0.537 | -0.333 | -0.429 | -1.751 | -0.006 | -0.03 | -0.03 | 0 | 0.06 | 0.15 | 0.06 | 0 | 0.05 | 0.02 |
| 3 | 108072298 | rs6779254 | 0.177 | -0.325 | -0.479 | -1.747 | -0.002 | 0.056 | 0.056 | 0 | 0.67 | 0.79 | 0.6 | 0.21 | 0.68 | 0.62 |
| 3 | 108072704 | rs6779650 | 0.711 | -0.319 | -0.5 | -1.105 | -0.002 | -0.026 | -0.026 | 0 | 0.6 | 0.58 | 0.54 | 0.21 | 0.64 | 0.58 |
| 3 | 108081217 | rs3792332 | 0.562 | -0.456 | -0.197 | -1.302 | 0.032 | 0.06 | 0.06 | 0 | 0.05 | 0.17 | 0.07 | 0.14 | 0.04 | 0.08 |
| 3 | 108081247 | rs16854512 | 1.318 | -0.779 | -0.294 | -1.804 | 0.001 | -0.02 | -0.02 | 0 | 0.22 | 0.23 | 0.17 | 0.05 | 0.19 | 0.11 |
| 3 | 108081277 | rs6779094 | 0.917 | -0.492 | -0.367 | -1.609 | -0.02 | 0.028 | 0.028 | 0 | 0.18 | 0.08 | 0.16 | 0 | 0.23 | 0.1 |
| 3 | 108085758 | rs4855644 | 1.039 | -0.505 | -0.352 | -1.619 | 0.007 | -0.003 | -0.003 | 0 | 0.55 | 0.39 | 0.46 | 0.06 | 0.6 | 0.49 |
| 3 | 108096496 | rs12497469 | -0.006 | -0.059 | -0.065 | -0.977 | 0.043 | 0.08 | 0.08 | 0 | 0.05 | 0.19 | 0.07 | 0.14 | 0.04 | 0.08 |
| 3 | 108103427 | rs2290600 | -0.357 | -0.751 | -0.467 | -1.179 | 0.018 | 0.019 | 0.019 | 0.06 | 0.5 | 0.4 | 0.46 | 0.5 | 0.43 | 0.51 |
| 3 | 108106667 | rs2278980 | 0.937 | -0.7 | -0.341 | -1.116 | 0 | 0.008 | 0.008 | 0 | 0.14 | 0.2 | 0.18 | 0.21 | 0.12 | 0.22 |
| 3 | 108107704 | rs1463431 | -0.437 | -0.944 | -0.272 | -1.042 | -0.014 | 0.101 | 0.101 | 0.121 | 0.93 | 0.86 | 0.8 | 0.47 | 0.97 | 0.84 |
| 3 | 108107925 | rs7616341 | 1.529 | -0.761 | -0.132 | -1.557 | 0.002 | -0.056 | -0.056 | 0 | 0.26 | 0.24 | 0.2 | 0.01 | 0.31 | 0.25 |
| 3 | 108109031 | rs4855650 | 0.782 | -0.99 | -0.25 | -1.762 | 0.01 | 0.055 | 0.055 | 0.139 | 0.31 | 0.12 | 0.23 | 0.25 | 0.28 | 0.28 |
| 3 | 108114572 | rs10511271 | -0.448 | -1.045 | -0.126 | -1.715 | -0.007 | 0.037 | 0.037 | 0 | 0.76 | 0.61 | 0.66 | 0.5 | 0.78 | 0.77 |
| 3 | 108115036 | rs4855559 | 0.837 | -0.944 | -0.337 | -1.696 | -0.011 | 0.063 | 0.063 | 0.15 | 0.31 | 0.06 | 0.23 | 0.25 | 0.28 | 0.28 |
| 3 | 108117694 | rs13087507 | 1.699 | -0.804 | -0.122 | -1.736 | -0.003 | -0.038 | -0.038 | 0.343 | 0.26 | 0.23 | 0.19 | 0.01 | 0.31 | 0.25 |
| 3 | 108119071 | rs4273371 | 0.459 | -0.861 | -0.102 | -1.686 | 0 | -0.031 | -0.031 | 0 | 0.47 | 0.46 | 0.42 | 0.22 | 0.5 | 0.49 |
| 3 | 108123001 | rs7630352 | -0.607 | -1.013 | 0.054 | -1.615 | 0.031 | 0.054 | 0.054 | 0 | 0.48 | 0.66 | 0.51 | 0.75 | 0.38 | 0.47 |
| 3 | 108126291 | rs6795741 | -0.453 | -0.713 | 0.329 | -1.152 | -0.018 | 0.272 | 0.272 | 0.331 | 0.92 | 0.72 | 0.79 | 0.5 | 0.97 | 0.83 |
| 3 | 108126872 | rs3957543 | 0.123 | -0.518 | -0.439 | -1.396 | 0.026 | 0.125 | 0.125 | 0.019 | 0.52 | 0.23 | 0.42 | 0.28 | 0.58 | 0.51 |
| 3 | 108129348 | rs1078455 | 1.012 | -0.208 | -0.455 | -1.365 | -0.023 | 0.018 | 0.018 | 0 | 0.25 | 0.16 | 0.2 | 0.01 | 0.32 | 0.24 |
| 3 | 108132759 | rs9810676 | 0.227 | -0.524 | -0.32 | -1.558 | 0.001 | -0.024 | -0.024 | 0 | 0.46 | 0.58 | 0.52 | 0.76 | 0.46 | 0.48 |
| 3 | 108135680 | rs9873504 | 0.74 | -0.646 | -0.175 | -1.199 | -0.007 | 0.114 | 0.114 | 0 | 0.06 | 0.12 | 0.17 | 0.55 | 0.04 | 0.19 |
| 3 | 108145443 | rs9842751 | 0.183 | -0.848 | -0.106 | -1.331 | -0.002 | 0.024 | 0.024 | 0 | 0.86 | 0.85 | 0.79 | 0.48 | 0.87 | 0.78 |
| 3 | 108147295 | rs1350235 | 0.29 | -0.934 | 0.009 | -1.378 | -0.009 | 0.054 | 0.054 | 0 | 0.84 | 0.8 | 0.78 | 0.48 | 0.87 | 0.78 |
| 3 | 108147728 | rs3900940 | 1.482 | -1 | -0.111 | -1.631 | -0.01 | 0.015 | 0.015 | 0 | 0.27 | 0.12 | 0.2 | 0.01 | 0.31 | 0.22 |
| 3 | 108148532 | rs11716039 | 1.65 | -1.247 | 0.166 | -1.858 | -0.002 | 0 | 0 | 0 | 0.19 | 0.23 | 0.15 | 0.23 | 0.12 | 0.24 |
| 3 | 108153593 | rs6798256 | -1.343 | -1.156 | 0.141 | -1.825 | 0.003 | -0.002 | -0.002 | 0 | 0.85 | 0.88 | 0.79 | 0.78 | 0.87 | 0.75 |
| 3 | 108158574 | rs3996040 | -0.29 | -1.209 | 0.257 | -1.849 | -0.008 | -0.005 | -0.005 | 0 | 0.79 | 0.73 | 0.62 | 0.22 | 0.83 | 0.57 |
| 3 | 108159977 | rs12638212 | 1.429 | -1.171 | 0.145 | -1.906 | 0.003 | 0.056 | 0.056 | 0 | 0.05 | 0.04 | 0.16 | 0.56 | 0.04 | 0.18 |
| 3 | 108163303 | rs3996017 | -0.007 | -1.201 | 0.162 | -1.928 | 0 | -0.001 | -0.001 | 0 | 0.76 | 0.78 | 0.74 | 0.58 | 0.78 | 0.7 |
| 3 | 108167382 | rs6766564 | -0.18 | -1.202 | 0.157 | -1.925 | 0.005 | -0.012 | -0.012 | 0.717 | 0.77 | 0.8 | 0.75 | 0.55 | 0.79 | 0.7 |
| 3 | 108172466 | rs16854645 | 1.418 | -1.187 | 0.103 | -1.915 | 0.007 | -0.049 | -0.049 | 0 | 0.13 | 0.05 | 0.2 | 0.42 | 0.13 | 0.25 |
| 3 | 108175071 | rs2140338 | 0.067 | -1.191 | 0.121 | -1.92 | -0.003 | 0.006 | 0.006 | 0 | 0.75 | 0.75 | 0.74 | 0.55 | 0.78 | 0.7 |
| 3 | 108184858 | rs3957559 | -1.344 | -1.167 | 0.1 | -1.934 | 0.001 | 0.072 | 0.072 | 0 | 0.95 | 0.95 | 0.84 | 0.44 | 0.96 | 0.82 |
| 3 | 108188755 | rs11917965 | 0.811 | -1.219 | 0.034 | -2.264 | -0.007 | -0.013 | -0.013 | 0.01 | 0.34 | 0.22 | 0.25 | 0.01 | 0.41 | 0.26 |
| 3 | 108188993 | rs9868484 | 0.533 | -1.019 | -0.133 | -2.052 | -0.004 | -0.072 | -0.072 | 0 | 0.7 | 0.69 | 0.57 | 0.02 | 0.74 | 0.52 |
| 3 | 108194093 | rs12633296 | 1.223 | -0.945 | -0.533 | -1.721 | 0.002 | 0.062 | 0.062 | 0 | 0.05 | 0.04 | 0.22 | 0.57 | 0.04 | 0.17 |
| 3 | 108196878 | rs4855654 | 1.176 | -0.979 | -0.416 | -1.734 | 0.008 | -0.023 | -0.023 | 0 | 0.14 | 0.07 | 0.14 | 0.41 | 0.12 | 0.24 |
| 3 | 108219393 | rs2603133 | -0.706 | -0.534 | -0.352 | -0.845 | -0.004 | 0.1 | 0.1 | 0 | 0.95 | 0.93 | 0.77 | 0.44 | 0.96 | 0.81 |
| 3 | 108236829 | rs1454196 | 0.219 | -0.649 | -0.52 | -1.501 | -0.022 | 0.034 | 0.034 | 0 | 0.4 | 0.23 | 0.47 | 0.62 | 0.5 | 0.45 |
| 3 | 108238592 | rs1840274 | 1.022 | -0.399 | -0.468 | -1.702 | -0.014 | -0.05 | -0.05 | 0 | 0.14 | 0.05 | 0.17 | 0.38 | 0.16 | 0.22 |
| 3 | 108264948 | rs2060986 | 0.669 | -0.716 | 0.368 | -2.021 | -0.02 | 0.201 | 0.201 | 0 | 0.39 | 0.12 | 0.31 | 0.02 | 0.34 | 0.33 |
| 3 | 108278077 | rs13069780 | 0.926 | -0.571 | 0.227 | -1.942 | -0.033 | -0.017 | -0.017 | 0.863 | 0.34 | 0.19 | 0.37 | 0.53 | 0.46 | 0.33 |

| | | | | | | | | | | | | | | | | |
|---|---|---|---|---|---|---|---|---|---|---|---|---|---|---|---|---|
| 3 | 108285722 | rs9866406 | 0.185 | -0.746 | 0.83 | -1.921 | 0.001 | 0.034 | 0.034 | 0 | 0.21 | 0.3 | 0.3 | 0.46 | 0.22 | 0.34 |
| 3 | 108293368 | rs2924708 | -0.244 | -0.747 | 0.965 | -1.922 | 0.003 | 0.072 | 0.072 | 0.011 | 0.78 | 0.63 | 0.7 | 0.54 | 0.78 | 0.66 |
| 3 | 108298260 | rs2278911 | 0.606 | -0.705 | 0.894 | -2.035 | -0.001 | 0.045 | 0.045 | 0 | 0.1 | 0.1 | 0.17 | 0.51 | 0.11 | 0.14 |
| 3 | 108307670 | rs11709183 | 1.588 | -0.939 | 0.494 | -2.284 | 0.011 | 0.009 | 0.009 | 0 | 0.22 | 0.16 | 0.16 | 0.02 | 0.13 | 0.26 |
| 3 | 108309323 | rs10511272 | 1.95 | -0.858 | 0.501 | -2.271 | -0.035 | 0.067 | 0.067 | 0 | 0.26 | 0.09 | 0.2 | 0.01 | 0.33 | 0.19 |
| 3 | 108309709 | rs9816505 | -0.777 | -0.802 | 0.335 | -2.205 | -0.004 | 0.135 | 0.135 | 0 | 0.61 | 0.82 | 0.69 | 0.98 | 0.66 | 0.67 |
| 3 | 108364392 | rs4855660 | 0.417 | -0.764 | 0.284 | -2.156 | -0.003 | 0.018 | 0.018 | 0 | 0.2 | 0.24 | 0.29 | 0.46 | 0.19 | 0.33 |
| 3 | 108386811 | rs6437789 | 0.528 | -0.779 | 0.129 | -2.167 | 0.007 | 0.007 | 0.007 | 0 | 0.41 | 0.36 | 0.32 | 0.02 | 0.34 | 0.33 |
| 3 | 108396656 | rs4410447 | 0.679 | -0.764 | 0.105 | -2.15 | -0.023 | -0.073 | -0.073 | 0 | 0.19 | 0.08 | 0.27 | 0.46 | 0.22 | 0.34 |
| 3 | 108397958 | rs7636263 | 1.921 | -0.731 | 0.128 | -2.143 | -0.063 | 0.106 | 0.106 | 0 | 0.17 | 0.02 | 0.15 | 0 | 0.21 | 0.06 |
| 3 | 108414595 | rs9878553 | 0.568 | -0.657 | 0.354 | -1.995 | 0.006 | 0.003 | 0.003 | 0 | 0.41 | 0.36 | 0.32 | 0.02 | 0.34 | 0.33 |
| 3 | 108441807 | rs4561844 | 0.396 | -0.422 | 0.012 | -1.375 | 0 | 0.047 | 0.047 | 0 | 0.09 | 0.08 | 0.16 | 0.5 | 0.11 | 0.15 |
| 3 | 108447943 | rs12637723 | -0.632 | -0.364 | -0.018 | -1.696 | -0.004 | 0.014 | 0.014 | 0.064 | 0.3 | 0.2 | 0.45 | 0.97 | 0.33 | 0.5 |
| 3 | 108454021 | rs12494937 | 0.333 | -0.384 | -0.026 | -1.34 | 0.003 | 0.023 | 0.023 | 0 | 0.1 | 0.07 | 0.17 | 0.5 | 0.1 | 0.16 |
| 3 | 108456254 | rs6437794 | -0.151 | -0.653 | -0.079 | -1.797 | 0.004 | -0.001 | -0.001 | 0 | 0.42 | 0.65 | 0.42 | 0.52 | 0.44 | 0.34 |
| 3 | 108459960 | rs9830861 | -0.448 | -0.88 | 0.372 | -1.815 | -0.049 | 0.257 | 0.257 | 0.011 | 0.16 | 0.47 | 0.2 | 0.5 | 0.1 | 0.17 |
| 3 | 108461900 | rs34240828 | 0.718 | -0.904 | 0.071 | -1.901 | -0.006 | -0.001 | -0.001 | 0 | 0.09 | 0.05 | 0.22 | 0.45 | 0.1 | 0.2 |
| 3 | 108465457 | rs6804995 | 1.077 | -0.823 | -0.178 | -2.382 | -0.032 | 0.163 | 0.163 | 0.261 | 0.32 | 0.1 | 0.23 | 0.04 | 0.28 | 0.38 |
| 3 | 108466827 | rs1975569 | 1.704 | -0.916 | -0.101 | -2.446 | -0.019 | 0.018 | 0.018 | 0.117 | 0.26 | 0.16 | 0.24 | 0.01 | 0.35 | 0.2 |
| 3 | 108474524 | rs9878093 | 2.169 | -0.946 | 0.019 | -2.456 | -0.003 | -0.016 | -0.016 | 0 | 0.17 | 0.11 | 0.12 | 0 | 0.19 | 0.07 |
| 3 | 108475808 | rs1107337 | 1.731 | -0.954 | 0.003 | -2.481 | -0.019 | 0.018 | 0.018 | 0.117 | 0.26 | 0.16 | 0.24 | 0.01 | 0.35 | 0.2 |
| 3 | 108475974 | rs11708527 | 1.159 | -0.892 | -0.01 | -2.502 | -0.017 | 0.142 | 0.142 | 0.243 | 0.32 | 0.12 | 0.23 | 0.04 | 0.28 | 0.38 |
| 3 | 108481730 | rs3762793 | 0.252 | -0.909 | 0.328 | -2.511 | -0.005 | -0.133 | -0.133 | 0.013 | 0.62 | 0.5 | 0.5 | 0.05 | 0.63 | 0.58 |
| 3 | 108489714 | rs9809154 | 0.652 | -0.745 | 0.266 | -2.18 | -0.008 | 0.14 | 0.14 | 0.339 | 0.49 | 0.28 | 0.36 | 0.04 | 0.47 | 0.45 |
| 3 | 108495463 | rs17223219 | 1.473 | -0.238 | -0.178 | -2.369 | -0.001 | -0.016 | -0.016 | 0.035 | 0.17 | 0.15 | 0.14 | 0.01 | 0.18 | 0.14 |
| 3 | 108496727 | rs10933962 | 2.591 | -0.228 | -0.2 | -2.304 | -0.044 | 0 | 0 | 0.018 | 0.09 | 0.01 | 0.18 | 0.07 | 0.18 | 0.06 |
| 3 | 108497048 | rs4146282 | -0.604 | -0.376 | 0.091 | -2.103 | -0.026 | 0.163 | 0.163 | 0 | 0.87 | 0.7 | 0.84 | 0.44 | 0.92 | 0.84 |
| 3 | 108498699 | rs16844883 | 0.539 | -0.355 | 0.042 | -2.1 | -0.003 | 0.056 | 0.056 | 1.681 | 0.11 | 0.2 | 0.15 | 0.31 | 0.09 | 0.19 |
| 3 | 108499238 | rs6794031 | -0.091 | -0.37 | -0.029 | -2.137 | -0.023 | 0.024 | 0.024 | 0.42 | 0.7 | 0.53 | 0.69 | 0.42 | 0.75 | 0.7 |
| 3 | 108504265 | rs2100139 | 0.749 | -0.344 | -0.289 | -2.148 | 0.011 | 0.026 | 0.026 | 1.022 | 0.59 | 0.33 | 0.55 | 0.11 | 0.65 | 0.51 |
| 3 | 108516567 | rs2715705 | 0.228 | -0.272 | -0.059 | -1.897 | -0.03 | 0.125 | 0.125 | 0 | 0.14 | 0.29 | 0.17 | 0.55 | 0.08 | 0.17 |
| 3 | 108516898 | rs7635702 | 1.218 | -0.5 | -0.329 | -1.99 | 0.002 | -0.011 | -0.011 | 0 | 0.19 | 0.16 | 0.15 | 0.01 | 0.18 | 0.15 |
| 3 | 108527107 | rs11919162 | -1.006 | -0.426 | -0.456 | -1.997 | 0.007 | -0.001 | -0.001 | 0 | 0.43 | 0.61 | 0.46 | 0.89 | 0.36 | 0.52 |
| 3 | 108531959 | rs2715738 | -2.384 | -0.176 | -0.642 | -1.947 | -0.018 | 0.009 | 0.009 | 0 | 0.9 | 0.95 | 0.82 | 0.93 | 0.82 | 0.94 |
| 3 | 108537807 | rs7631962 | 1.654 | -0.068 | -0.93 | -2.135 | -0.022 | 0.04 | 0.04 | 0.291 | 0.12 | 0.05 | 0.08 | 0 | 0.14 | 0.11 |
| 3 | 108538384 | rs4507261 | 0.738 | -0.341 | -0.68 | -1.901 | 0.003 | 0.059 | 0.059 | 7.405 | 0.1 | 0.16 | 0.15 | 0.33 | 0.09 | 0.2 |
| 3 | 108540610 | rs2953339 | -0.853 | -0.775 | -0.871 | -1.883 | -0.007 | 0.048 | 0.048 | 0 | 0.54 | 0.64 | 0.54 | 0.88 | 0.5 | 0.63 |
| 3 | 108541143 | rs2715694 | -0.068 | -1.423 | -0.902 | -1.634 | -0.038 | 0.213 | 0.213 | 0 | 0.15 | 0.42 | 0.18 | 0.54 | 0.08 | 0.17 |
| 3 | 108541761 | rs9879812 | 0.742 | -0.62 | -1.807 | -1.972 | -0.015 | 0.131 | 0.131 | 0.011 | 0.34 | 0.09 | 0.27 | 0.04 | 0.32 | 0.29 |
| 3 | 108549720 | rs2290055 | 0.176 | -0.716 | -1.677 | -1.16 | 0.057 | -0.005 | -0.005 | 0.206 | 0.35 | 0.13 | 0.34 | 0.59 | 0.23 | 0.44 |
| 3 | 108552960 | rs2715725 | -0.334 | -0.297 | -0.861 | -1.33 | -0.013 | -0.038 | -0.038 | 0 | 0.91 | 0.84 | 0.96 | 1 | 0.96 | 0.94 |
| 3 | 108556425 | rs1374818 | 0.047 | -0.162 | -1.167 | -1.9 | 0.013 | 0.03 | 0.03 | 5.595 | 0.59 | 0.36 | 0.64 | 0.91 | 0.64 | 0.79 |
| 3 | 108559624 | rs9861244 | 1.354 | 0.048 | -1.18 | -1.929 | -0.007 | -0.01 | -0.01 | 0 | 0.13 | 0.08 | 0.12 | 0 | 0.19 | 0.07 |
| 3 | 108565407 | rs2715713 | -0.848 | -0.107 | -0.754 | -2.041 | 0.001 | -0.042 | -0.042 | 0.043 | 0.9 | 0.85 | 0.95 | 1 | 0.94 | 0.89 |
| 3 | 108567680 | rs7609997 | -0.235 | 0.346 | -0.684 | -2.542 | -0.005 | 0.032 | 0.032 | 2.939 | 0.39 | 0.29 | 0.36 | 0.09 | 0.41 | 0.19 |
| 3 | 108570300 | rs6796621 | 1.078 | -0.234 | -1.267 | -2.672 | 0.014 | 0.026 | 0.026 | 0 | 0.13 | 0.01 | 0.06 | 0.05 | 0.09 | 0.1 |
| 3 | 108570396 | rs2715730 | 0.261 | 0.123 | -1.008 | -2.492 | -0.001 | -0.032 | -0.032 | 1.722 | 0.52 | 0.56 | 0.59 | 0.91 | 0.53 | 0.71 |
| 3 | 108580778 | rs9879707 | 1.359 | 0.041 | -0.871 | -2.017 | 0.001 | -0.027 | -0.027 | 0 | 0.11 | 0.12 | 0.11 | 0 | 0.15 | 0.05 |
| 3 | 108583871 | rs7640727 | 0.284 | -1.126 | -0.591 | -2.043 | -0.069 | -0.089 | -0.089 | 0 | 0.7 | 0.46 | 0.84 | 0.9 | 0.84 | 0.85 |
| 3 | 108588395 | rs4358307 | 0.448 | -1.073 | -0.688 | -2.228 | 0.024 | 0.054 | 0.054 | 1.637 | 0.57 | 0.31 | 0.73 | 0.89 | 0.68 | 0.78 |
| 3 | 108590475 | rs6437802 | -0.102 | -1.627 | -1.118 | -2.732 | 0.049 | 0.086 | 0.086 | 0 | 0.71 | 0.85 | 0.88 | 0.96 | 0.83 | 0.79 |
| 3 | 108596911 | rs1962162 | 0.826 | -1.708 | -0.643 | -2.905 | 0 | -0.073 | -0.073 | 0 | 0.22 | 0.29 | 0.18 | 0.02 | 0.24 | 0.13 |
| 3 | 108606743 | rs6437804 | -0.293 | -1.716 | -0.704 | -2.916 | 0.044 | 0.08 | 0.08 | 0 | 0.72 | 0.86 | 0.89 | 0.96 | 0.83 | 0.78 |
| 3 | 108607015 | rs11715374 | 0.174 | -1.377 | -1.01 | -2.713 | -0.048 | 0.094 | 0.094 | 0 | 0.41 | 0.19 | 0.52 | 0.78 | 0.52 | 0.54 |
| 3 | 108608767 | rs13327115 | 1.587 | -1.338 | -0.729 | -2.667 | -0.014 | 0.009 | 0.009 | 0.026 | 0.24 | 0.17 | 0.32 | 0.18 | 0.31 | 0.23 |
| 3 | 108611057 | rs4522770 | 0.472 | -1.33 | -0.542 | -2.676 | -0.024 | -0.009 | -0.009 | 0.366 | 0.22 | 0.16 | 0.37 | 0.76 | 0.28 | 0.41 |
| 3 | 108614492 | rs4535234 | 0.164 | -1.352 | -0.502 | -2.679 | 0.008 | -0.014 | -0.014 | 0 | 0.47 | 0.43 | 0.69 | 0.94 | 0.59 | 0.66 |
| 3 | 108619120 | rs2399248 | -0.006 | -1.434 | -0.234 | -1.455 | -0.002 | -0.003 | -0.003 | 2.169 | 0.42 | 0.6 | 0.53 | 0.54 | 0.45 | 0.47 |
| 3 | 108619666 | rs2399250 | -0.03 | -1.124 | -0.516 | -1.2 | -0.01 | 0.119 | 0.119 | 0.002 | 0.18 | 0.51 | 0.22 | 0.36 | 0.14 | 0.24 |
| 3 | 108623388 | rs10933971 | 0.415 | -0.968 | -0.826 | -1.092 | 0 | 0.013 | 0.013 | 0 | 0.35 | 0.53 | 0.37 | 0.41 | 0.36 | 0.35 |
| 3 | 108631337 | rs4855675 | -0.131 | -0.524 | -1.124 | -0.745 | -0.017 | -0.02 | -0.02 | 0 | 0.9 | 0.83 | 0.95 | 0.97 | 0.96 | 0.96 |
| 3 | 108632495 | rs9875407 | -0.202 | -0.418 | -1.749 | -0.585 | -0.029 | -0.089 | -0.089 | 0 | 0.23 | 0.09 | 0.24 | 0.4 | 0.22 | 0.3 |
| 3 | 108633078 | rs16854924 | 1.177 | -0.415 | -1.778 | -0.57 | -0.034 | -0.049 | -0.049 | 0 | 0.09 | 0.01 | 0.18 | 0.37 | 0.11 | 0.18 |
| 3 | 108633112 | rs6437807 | -0.724 | -0.186 | -1.56 | -0.628 | -0.003 | -0.007 | -0.007 | 0.381 | 0.65 | 0.68 | 0.64 | 0.6 | 0.58 | 0.6 |
| 3 | 108634973 | rs10933973 | 0.297 | 0.456 | -1.952 | -1.064 | -0.01 | 0.02 | 0.02 | 0 | 0.27 | 0.25 | 0.31 | 0.18 | 0.3 | 0.22 |
| 3 | 108635099 | rs9845098 | -0.638 | 0.323 | -1.708 | -0.466 | 0.011 | 0.055 | 0.055 | 0 | 0.17 | 0.08 | 0.07 | 0.04 | 0.12 | 0.12 |
| 3 | 108637430 | rs2715689 | 0.259 | 0.916 | -1.89 | -0.518 | -0.036 | 0.101 | 0.101 | 0.019 | 0.62 | 0.8 | 0.6 | 0.84 | 0.57 | 0.63 |
| 3 | 108638602 | rs7633206 | -0.251 | 0.674 | -1.67 | -0.725 | 0 | -0.006 | -0.006 | 0 | 0.32 | 0.42 | 0.24 | 0.24 | 0.34 | 0.26 |
| 3 | 108639119 | rs2715688 | 0.44 | 0.837 | -1.797 | -1.35 | -0.03 | 0.092 | 0.092 | 0 | 0.34 | 0.13 | 0.37 | 0.13 | 0.36 | 0.36 |

| | | | | | | | | | | | | | | | | |
|---|---|---|---|---|---|---|---|---|---|---|---|---|---|---|---|---|
| 3 | 108639423 | rs2715687 | -0.875 | 0.009 | -1.656 | -1.755 | 0.005 | 0.026 | 0.026 | 0 | 0.77 | 0.71 | 0.63 | 0.38 | 0.72 | 0.65 |
| 3 | 108644054 | rs2593974 | 0.801 | -0.06 | -1.702 | -1.825 | -0.019 | 0.089 | 0.089 | 0 | 0.33 | 0.13 | 0.19 | 0.12 | 0.35 | 0.3 |
| 3 | 108654067 | rs2399260 | -0.917 | 0.038 | -1.524 | -1.818 | -0.002 | 0.014 | 0.014 | 0 | 0.78 | 0.73 | 0.66 | 0.35 | 0.79 | 0.65 |
| 3 | 108656519 | rs751299 | 0.982 | -0.184 | -1.056 | -1.865 | -0.019 | -0.037 | -0.037 | 0 | 0.13 | 0.12 | 0.04 | 0.01 | 0.02 | 0.04 |
| 3 | 108659823 | rs6799826 | 1.051 | -0.151 | -1.384 | -2.219 | 0.002 | 0.024 | 0.024 | 0 | 0.23 | 0.18 | 0.23 | 0.09 | 0.2 | 0.22 |
| 3 | 108665371 | rs6801753 | 1.44 | -0.211 | -1.056 | -2.267 | -0.002 | -0.014 | -0.014 | 0 | 0.21 | 0.19 | 0.14 | 0.02 | 0.19 | 0.22 |
| 3 | 108676951 | rs2593931 | 0.645 | -0.295 | -1.177 | -1.88 | -0.002 | -0.003 | -0.003 | 0.081 | 0.14 | 0.1 | 0.07 | 0.08 | 0.16 | 0.13 |
| 3 | 108677821 | rs2197737 | -0.288 | -0.534 | -1.192 | -1.987 | 0.008 | -0.009 | -0.009 | 0 | 0.47 | 0.39 | 0.24 | 0.11 | 0.37 | 0.37 |
| 3 | 108681891 | rs10490845 | 1.062 | -0.618 | -1.227 | -2.083 | -0.043 | 0.008 | 0.008 | 0 | 0.3 | 0.22 | 0.39 | 0.26 | 0.42 | 0.28 |
| 3 | 108690221 | rs2593943 | -0.118 | -0.843 | -1.251 | -2.306 | 0.001 | -0.022 | -0.022 | 0 | 0.51 | 0.43 | 0.28 | 0.11 | 0.43 | 0.38 |
| 3 | 108692989 | rs2593950 | -0.142 | -0.84 | -1.25 | -2.305 | 0.001 | -0.024 | -0.024 | 0 | 0.51 | 0.43 | 0.28 | 0.11 | 0.43 | 0.38 |
| 3 | 108693851 | rs921174 | 1.76 | -0.759 | -1.612 | -2.469 | -0.03 | 0.004 | 0.004 | 0 | 0.09 | 0.03 | 0.21 | 0.12 | 0.17 | 0.06 |
| 3 | 108703548 | rs2593959 | 0.081 | -0.849 | -1.58 | -2.513 | 0 | -0.036 | -0.036 | 0.177 | 0.52 | 0.5 | 0.29 | 0.11 | 0.43 | 0.39 |
| 3 | 108703587 | rs2290057 | 2.175 | -1.195 | -1.316 | -2.439 | -0.063 | -0.081 | -0.081 | 0.133 | 0.12 | 0.01 | 0.3 | 0.64 | 0.19 | 0.32 |
| 3 | 108707698 | rs3804705 | 2.103 | -1.238 | -1.343 | -2.499 | -0.061 | 0.139 | 0.139 | 0 | 0.25 | 0.05 | 0.16 | 0.02 | 0.26 | 0.24 |
| 3 | 108714045 | rs3804701 | 0.548 | -1.254 | -1.334 | -2.481 | -0.073 | -0.038 | -0.038 | 3.324 | 0.26 | 0.12 | 0.35 | 0.25 | 0.37 | 0.27 |
| 3 | 108739203 | rs6765616 | 1.112 | -1.257 | -0.996 | -2.529 | 0.015 | 0.042 | 0.042 | 0 | 0.36 | 0.25 | 0.2 | 0.03 | 0.27 | 0.25 |
| 3 | 108753936 | rs13094869 | 2.031 | -1.276 | -1.011 | -2.539 | -0.043 | -0.049 | -0.049 | 0 | 0.11 | 0.18 | 0.04 | 0 | 0.01 | 0.02 |
| 3 | 108754214 | rs3762697 | 1.102 | -1.176 | -0.893 | -2.497 | 0.009 | -0.045 | -0.045 | 0 | 0.15 | 0.23 | 0.09 | 0.08 | 0.16 | 0.13 |
| 3 | 108754260 | rs3762696 | 0.201 | -1.09 | -0.918 | -2.388 | 0 | -0.038 | -0.038 | 0 | 0.52 | 0.51 | 0.29 | 0.11 | 0.43 | 0.38 |
| 3 | 108764725 | rs3804681 | 0.22 | -1.078 | -0.914 | -2.391 | 0 | -0.04 | -0.04 | 0.644 | 0.51 | 0.51 | 0.29 | 0.12 | 0.43 | 0.38 |
| 3 | 108779344 | rs7627562 | 1.728 | -1.268 | -0.877 | -2.3 | -0.043 | -0.052 | -0.052 | 1.737 | 0.12 | 0.19 | 0.04 | 0 | 0.02 | 0.02 |
| 3 | 108780831 | rs7653651 | 1.898 | -1.229 | -1.071 | -2.434 | -0.036 | 0.101 | 0.101 | 0 | 0.25 | 0.09 | 0.16 | 0.02 | 0.26 | 0.23 |
| 3 | 108781952 | rs13060344 | 1.14 | -0.89 | -0.908 | -2.292 | -0.002 | -0.003 | -0.003 | 0 | 0.14 | 0.1 | 0.07 | 0.09 | 0.15 | 0.13 |
| 3 | 108788461 | rs2305238 | 0.675 | -0.965 | -1.064 | -2.376 | -0.11 | -0.009 | -0.009 | 0 | 0.28 | 0.08 | 0.4 | 0.7 | 0.38 | 0.54 |
| 3 | 108802057 | rs4855688 | -1.178 | -0.952 | -0.643 | -2.337 | 0.011 | -0.042 | -0.042 | 0.295 | 0.86 | 0.79 | 0.91 | 0.91 | 0.85 | 0.87 |
| 3 | 108810524 | rs4555525 | 0.361 | -0.948 | -0.851 | -2.427 | -0.05 | 0.1 | 0.1 | 0 | 0.32 | 0.21 | 0.61 | 0.86 | 0.45 | 0.54 |
| 3 | 108811036 | rs9873588 | 0.121 | -0.988 | -0.686 | -2.331 | 0.004 | 0.124 | 0.124 | 0 | 0.53 | 0.7 | 0.36 | 0.2 | 0.43 | 0.35 |
| 3 | 108824811 | rs7626269 | -0.032 | -1.063 | -0.406 | -2.403 | -0.03 | -0.124 | -0.124 | 0 | 0.76 | 0.58 | 0.87 | 0.91 | 0.83 | 0.84 |
| 3 | 108829362 | rs7641406 | 2.249 | -0.849 | -0.63 | -2.242 | -0.03 | 0.001 | 0.001 | 0 | 0.09 | 0.03 | 0.21 | 0.12 | 0.17 | 0.06 |
| 3 | 108840227 | rs9868354 | 1.403 | -0.982 | -0.403 | -2.306 | -0.039 | -0.048 | -0.048 | 0 | 0.09 | 0.31 | 0.05 | 0 | 0.01 | 0.03 |
| 3 | 108851200 | rs4241399 | 0.092 | -0.9 | -0.694 | -2.437 | -0.089 | 0.191 | 0.191 | 0 | 0.38 | 0.16 | 0.65 | 0.88 | 0.56 | 0.58 |
| 3 | 108851791 | rs4441659 | -0.172 | -1.291 | -0.319 | -2.251 | -0.032 | -0.102 | -0.102 | 0 | 0.74 | 0.62 | 0.87 | 0.9 | 0.84 | 0.84 |
| 3 | 108867128 | rs4855697 | 0.31 | -0.885 | -0.871 | -2.122 | -0.09 | 0.136 | 0.136 | 0.015 | 0.31 | 0.12 | 0.61 | 0.84 | 0.41 | 0.56 |
| 3 | 108873618 | rs4638978 | 0.393 | -1.24 | -0.411 | -2.065 | 0.002 | -0.021 | -0.021 | 0.009 | 0.65 | 0.44 | 0.83 | 0.86 | 0.8 | 0.82 |
| 3 | 108874110 | rs938397 | -0.068 | -1.035 | -0.025 | -1.919 | -0.025 | -0.07 | -0.07 | 0 | 0.75 | 0.69 | 0.88 | 0.86 | 0.84 | 0.85 |
| 3 | 108875579 | rs2673351 | -0.94 | -0.91 | -0.068 | -1.859 | 0 | -0.007 | -0.007 | 0 | 0.86 | 0.88 | 0.92 | 0.86 | 0.85 | 0.86 |
| 3 | 108884070 | rs1920028 | 1.046 | -1.033 | -0.174 | -1.752 | -0.002 | 0.003 | 0.003 | 0 | 0.17 | 0.26 | 0.16 | 0.12 | 0.18 | 0.2 |
| 3 | 108894652 | rs1163346 | 1.063 | -1.033 | -0.173 | -1.752 | -0.002 | 0.004 | 0.004 | 0 | 0.16 | 0.26 | 0.16 | 0.12 | 0.18 | 0.2 |
| 3 | 108896155 | rs34037947 | 2.224 | -1.248 | -0.434 | -1.877 | 0.018 | 0.015 | 0.015 | 2.564 | 0.1 | 0.06 | 0.03 | 0 | 0.01 | 0.01 |
| 3 | 108897541 | rs9288889 | 1.324 | -1.214 | -0.465 | -2.084 | -0.004 | 0.088 | 0.088 | 0 | 0.43 | 0.3 | 0.26 | 0.04 | 0.43 | 0.31 |
| 3 | 108901645 | rs13067620 | 1.147 | -1.294 | -0.318 | -1.932 | -0.023 | -0.027 | -0.027 | 0.367 | 0.14 | 0.18 | 0.21 | 0.07 | 0.05 | 0.05 |
| 3 | 108905508 | rs6774626 | 0.819 | -1.336 | -0.276 | -1.87 | -0.008 | 0.091 | 0.091 | 0.309 | 0.48 | 0.35 | 0.47 | 0.11 | 0.5 | 0.36 |
| 3 | 108906621 | rs1183171 | 0.063 | -1.175 | -0.602 | -1.626 | -0.002 | 0.014 | 0.014 | 0.415 | 0.64 | 0.75 | 0.61 | 0.24 | 0.65 | 0.55 |
| 3 | 108936698 | rs1163378 | -1.256 | -0.607 | -0.709 | -1.302 | -0.019 | -0.022 | -0.022 | 0 | 0.83 | 0.88 | 0.69 | 0.3 | 0.74 | 0.6 |
| 3 | 108943340 | rs9882896 | 1.291 | -0.944 | -1.117 | -1.521 | -0.11 | 0.158 | 0.158 | 0.054 | 0.38 | 0.07 | 0.47 | 0.17 | 0.46 | 0.29 |
| 3 | 108946466 | rs1163368 | 0.016 | -1.073 | -0.49 | -1.537 | -0.004 | -0.004 | -0.004 | 0 | 0.65 | 0.43 | 0.6 | 0.3 | 0.68 | 0.55 |
| 3 | 108946543 | rs1163367 | 0.842 | -1.228 | -0.418 | -1.625 | 0.001 | 0.006 | 0.006 | 0 | 0.24 | 0.22 | 0.12 | 0.11 | 0.22 | 0.26 |
| 3 | 108950809 | rs6807870 | 0.128 | -1.196 | -0.233 | -1.614 | -0.093 | -0.131 | -0.131 | 0 | 0.19 | 0.49 | 0.1 | 0 | 0.06 | 0.04 |
| 3 | 108950879 | rs6805141 | 0.128 | -1.196 | -0.233 | -1.614 | -0.093 | -0.131 | -0.131 | 0.097 | 0.19 | 0.49 | 0.1 | 0 | 0.06 | 0.04 |
| 3 | 108961555 | rs17305051 | 0.663 | -0.764 | -0.488 | -1.402 | 0.001 | 0.079 | 0.079 | 0.087 | 0.48 | 0.33 | 0.5 | 0.21 | 0.47 | 0.29 |
| 3 | 108962866 | rs1920031 | 0.178 | -0.555 | -0.823 | -1.283 | 0.007 | 0.012 | 0.012 | 0 | 0.33 | 0.25 | 0.17 | 0.09 | 0.28 | 0.34 |
| 3 | 108963256 | rs1515379 | -1.217 | -0.062 | -0.892 | -0.8 | -0.078 | -0.093 | -0.093 | 0 | 0.84 | 0.96 | 0.7 | 0.33 | 0.74 | 0.63 |
| 3 | 108969821 | rs4855584 | 1.633 | -0.265 | -0.816 | -0.845 | -0.047 | -0.074 | -0.074 | 0.252 | 0.11 | 0.01 | 0.25 | 0.59 | 0.15 | 0.35 |
| 3 | 108970719 | rs2699975 | -0.672 | -0.868 | 0.012 | -1.332 | -0.051 | -0.059 | -0.059 | 0.045 | 0.89 | 0.66 | 0.95 | 1 | 0.98 | 0.99 |
| 3 | 108971725 | rs2943701 | -0.976 | -0.865 | 0.04 | -1.331 | -0.042 | -0.048 | -0.048 | 1.324 | 0.91 | 0.79 | 0.97 | 1 | 0.98 | 0.99 |
| 3 | 108973465 | rs2699972 | -0.976 | -0.79 | -0.275 | -1.298 | -0.042 | -0.048 | -0.048 | 0 | 0.91 | 0.77 | 0.97 | 1 | 0.98 | 0.99 |
| 3 | 108976446 | rs2673391 | -0.682 | -0.102 | -1.189 | -1.435 | 0.016 | 0.104 | 0.104 | 0 | 0.42 | 0.21 | 0.27 | 0.09 | 0.41 | 0.37 |
| 3 | 108978877 | rs1163380 | -0.199 | -0.077 | -0.826 | -0.978 | 0.009 | 0.027 | 0.027 | 0 | 0.27 | 0.16 | 0.14 | 0.09 | 0.24 | 0.31 |
| 3 | 108982704 | rs1163391 | 0.205 | 0.748 | -0.401 | -0.548 | 0.028 | 0.052 | 0.052 | 0 | 0.16 | 0.32 | 0.32 | 0.23 | 0.21 | 0.16 |
| 3 | 108989277 | rs4855586 | -0.612 | 0.559 | -0.851 | -0.971 | -0.006 | 0.018 | 0.018 | 0.002 | 0.62 | 0.8 | 0.62 | 0.31 | 0.59 | 0.58 |
| 3 | 108991441 | rs7652543 | 0.798 | 0.479 | -0.823 | -1.686 | 0 | 0.001 | 0.001 | 0.477 | 0.25 | 0.17 | 0.16 | 0.03 | 0.24 | 0.23 |
| 3 | 108996328 | rs6777382 | 2.484 | 0.318 | -0.412 | -1.645 | -0.003 | -0.009 | -0.009 | 0 | 0.05 | 0.23 | 0.03 | 0 | 0.02 | 0 |
| 3 | 108997447 | rs11714349 | 0.268 | 0.455 | -0.619 | -1.809 | -0.015 | -0.11 | -0.11 | 0 | 0.2 | 0.37 | 0.12 | 0.01 | 0.16 | 0.22 |
| 3 | 109001501 | rs6808221 | 0.278 | 0.47 | -0.614 | -1.803 | -0.014 | -0.11 | -0.11 | 0 | 0.2 | 0.37 | 0.12 | 0.01 | 0.16 | 0.22 |
| 3 | 109002032 | rs9876733 | 0.367 | 0.276 | -0.517 | -1.999 | 0.021 | 0.071 | 0.071 | 0 | 0.53 | 0.29 | 0.71 | 0.96 | 0.6 | 0.56 |
| 3 | 109002114 | rs9857002 | -0.646 | 0.348 | -1.092 | -1.718 | -0.005 | -0.091 | -0.091 | 0 | 0.73 | 0.67 | 0.83 | 0.97 | 0.76 | 0.77 |
| 3 | 109013108 | rs9861358 | 1.131 | 0.572 | -1.185 | -0.814 | -0.073 | 0.036 | 0.036 | 0.488 | 0.56 | 0.06 | 0.51 | 0.72 | 0.61 | 0.61 |
| 3 | 109027157 | rs7611135 | 0.65 | 0.559 | -0.154 | -0.816 | 0.004 | -0.062 | -0.062 | 0 | 0.72 | 0.64 | 0.8 | 0.91 | 0.73 | 0.79 |

| | | | | | | | | | | | | | | | | |
|---|---|---|---|---|---|---|---|---|---|---|---|---|---|---|---|---|
| 3 | 109034120 | rs11712428 | 1.225 | 0.484 | -0.443 | -0.699 | -0.003 | 0.07 | 0.07 | 0 | 0.48 | 0.32 | 0.5 | 0.73 | 0.53 | 0.58 |
| 3 | 109036817 | rs11713133 | 1.209 | 0.4 | -0.018 | -0.755 | -0.007 | 0.1 | 0.1 | 0 | 0.51 | 0.21 | 0.54 | 0.74 | 0.59 | 0.59 |
| 3 | 109037859 | rs2699983 | -0.744 | 0.352 | 0.701 | -0.787 | -0.005 | 0.211 | 0.211 | 0 | 0.12 | 0.54 | 0.23 | 0.25 | 0.05 | 0.21 |
| 3 | 109044307 | rs12488676 | 1.815 | 0.441 | 0.126 | -0.936 | -0.103 | 0.043 | 0.043 | 0.012 | 0.45 | 0.06 | 0.46 | 0.75 | 0.48 | 0.58 |
| 3 | 109045055 | rs1044266 | 0.054 | 0.572 | 0.102 | -1.081 | -0.062 | 0.084 | 0.084 | 0.074 | 0.81 | 0.95 | 0.81 | 1 | 0.7 | 0.91 |
| 3 | 109047928 | rs1163439 | 0.674 | 0.383 | 0.769 | -0.966 | 0 | 0.065 | 0.065 | 0 | 0.09 | 0.22 | 0.21 | 0.14 | 0.05 | 0.15 |
| 3 | 109052732 | rs3762648 | 1.891 | 0.429 | 0.804 | -1.034 | -0.007 | 0.071 | 0.071 | 0 | 0.46 | 0.22 | 0.47 | 0.75 | 0.48 | 0.57 |
| 3 | 109057397 | rs1163448 | 0.427 | 0.323 | 1.15 | -0.911 | 0.011 | 0.092 | 0.092 | 0 | 0.09 | 0.26 | 0.21 | 0.14 | 0.05 | 0.15 |
| 3 | 109068774 | rs16844893 | -2.257 | 0.225 | 1.221 | -0.983 | 0.038 | 0.164 | 0.164 | 2.884 | 0.07 | 0.45 | 0.05 | 0.1 | 0.02 | 0.06 |
| 3 | 109072323 | rs13069030 | 0.046 | 0.659 | 0.492 | -0.964 | -0.059 | 0.123 | 0.123 | 1.801 | 0.25 | 0.03 | 0.16 | 0.04 | 0.28 | 0.13 |
| 3 | 109079886 | rs7649662 | -0.947 | 0.545 | 0.668 | -0.926 | -0.076 | 0.198 | 0.198 | 0 | 0.4 | 0.14 | 0.28 | 0.04 | 0.48 | 0.21 |
| 3 | 109086278 | rs1125903 | 0.397 | 0.962 | 0.936 | -0.858 | 0.037 | 0.06 | 0.06 | 0 | 0.09 | 0.19 | 0.23 | 0.14 | 0.11 | 0.16 |
| 3 | 109091698 | rs6437847 | -1.423 | 0.887 | 0.849 | -0.839 | 0.031 | 0.062 | 0.062 | 0 | 0.5 | 0.32 | 0.51 | 0.18 | 0.6 | 0.38 |
| 3 | 109091955 | rs1947963 | 1.331 | 0.243 | 0.068 | -0.583 | 0.04 | -0.041 | -0.041 | 0 | 0.07 | 0.03 | 0.13 | 0.19 | 0.01 | 0.1 |
| 3 | 109101783 | rs4855735 | 1.779 | 0.4 | 0.182 | -0.617 | 0.005 | 0.018 | 0.018 | 0.071 | 0.4 | 0.2 | 0.33 | 0.66 | 0.38 | 0.53 |
| 3 | 109106911 | rs17193050 | 0.072 | 0.776 | 0.762 | -0.647 | 0.041 | 0.003 | 0.003 | 0 | 0.12 | 0.14 | 0.24 | 0.12 | 0.18 | 0.16 |
| 3 | 109111132 | rs3887341 | 2.249 | 0.382 | 0.316 | -0.877 | -0.033 | -0.005 | -0.005 | 0 | 0.37 | 0.12 | 0.33 | 0.67 | 0.39 | 0.49 |
| 3 | 109116136 | rs769504 | 0.229 | -0.552 | 0.334 | -1.041 | -0.022 | 0.172 | 0.172 | 0.049 | 0.65 | 0.86 | 0.75 | 0.96 | 0.6 | 0.76 |
| 3 | 109121182 | rs769495 | 1.46 | -0.544 | 0.484 | -0.992 | 0.004 | 0.002 | 0.002 | 0.702 | 0.5 | 0.39 | 0.57 | 0.78 | 0.57 | 0.66 |
| 3 | 109136484 | rs6775559 | -0.819 | -0.272 | 0.602 | -0.996 | -0.059 | -0.061 | -0.061 | 0 | 0.24 | 0.47 | 0.25 | 0.1 | 0.13 | 0.1 |
| 3 | 109139316 | rs769507 | 1.108 | -0.101 | 0.759 | -0.91 | 0 | -0.017 | -0.017 | 0 | 0.57 | 0.45 | 0.59 | 0.87 | 0.59 | 0.78 |
| 3 | 109142424 | rs6778096 | 2.238 | 0.094 | 0.755 | -0.696 | -0.002 | 0.022 | 0.022 | 1.342 | 0.46 | 0.18 | 0.41 | 0.61 | 0.48 | 0.55 |
| 3 | 109151680 | rs12496499 | 0.78 | -0.042 | 0.821 | -0.682 | -0.022 | -0.022 | -0.022 | 0 | 0.08 | 0.02 | 0.2 | 0.19 | 0.13 | 0.13 |
| 3 | 109151709 | rs769520 | -0.351 | -0.046 | 0.885 | -0.696 | 0.071 | 0.102 | 0.102 | 0 | 0.14 | 0.26 | 0.23 | 0.26 | 0.19 | 0.24 |
| 3 | 109156199 | rs769518 | 0.227 | -0.122 | 0.755 | -0.769 | 0.003 | 0.122 | 0.122 | 1.625 | 0.67 | 0.84 | 0.69 | 0.9 | 0.7 | 0.83 |
| 3 | 109161766 | rs1905351 | 2.543 | -0.132 | 0.734 | -0.77 | -0.005 | 0.004 | 0.004 | 0.602 | 0.41 | 0.13 | 0.36 | 0.63 | 0.4 | 0.55 |
| 3 | 109182586 | rs769529 | 1.054 | -0.22 | 0.883 | -0.559 | 0.005 | -0.008 | -0.008 | 0.273 | 0.57 | 0.52 | 0.6 | 0.9 | 0.6 | 0.79 |
| 3 | 109195046 | rs769568 | 0.51 | -0.064 | 0.915 | -0.465 | 0.009 | 0.036 | 0.036 | 1.526 | 0.65 | 0.66 | 0.68 | 0.9 | 0.69 | 0.83 |
| 3 | 109196914 | rs769563 | 0.484 | -0.056 | 0.791 | -0.461 | 0.01 | 0.092 | 0.092 | 1.419 | 0.66 | 0.78 | 0.69 | 0.9 | 0.7 | 0.83 |
| 3 | 109202103 | rs769558 | -0.819 | 0.24 | 0.838 | -0.464 | -0.019 | 0.042 | 0.042 | 3.239 | 0.38 | 0.29 | 0.33 | 0.09 | 0.4 | 0.18 |
| 3 | 109203399 | rs769556 | -1.379 | 0.105 | 0.943 | 0.047 | 0.002 | 0.003 | 0.003 | 1.359 | 0.49 | 0.62 | 0.48 | 0.56 | 0.52 | 0.32 |
| 3 | 109210303 | rs769544 | -1.197 | 0.11 | 0.907 | 0.049 | -0.001 | 0.001 | 0.001 | 1.424 | 0.47 | 0.41 | 0.47 | 0.56 | 0.52 | 0.32 |
| 3 | 109224312 | rs13063502 | 0.885 | 0.177 | 0.753 | -0.038 | -0.062 | 0.067 | 0.067 | 0 | 0.11 | 0.01 | 0.07 | 0 | 0.14 | 0.04 |
| 3 | 109237931 | rs1403250 | -0.489 | -0.352 | 1.008 | 0.173 | -0.019 | 0.042 | 0.042 | 3.239 | 0.38 | 0.29 | 0.33 | 0.09 | 0.4 | 0.18 |
| 3 | 109249961 | rs10933993 | -0.81 | -0.038 | 0.868 | 0.189 | 0.001 | 0.002 | 0.002 | 0 | 0.43 | 0.58 | 0.46 | 0.56 | 0.48 | 0.3 |
| 3 | 109250510 | rs11717790 | -0.755 | -0.42 | 1.057 | 0.398 | 0 | 0.002 | 0.002 | 1.536 | 0.48 | 0.6 | 0.48 | 0.56 | 0.52 | 0.33 |
| 3 | 109254828 | rs6800450 | 0.165 | -0.448 | 1.073 | 0.399 | 0.015 | 0.061 | 0.061 | 0 | 0.9 | 0.87 | 0.76 | 0.72 | 0.85 | 0.76 |
| 3 | 109256410 | rs9832962 | 1.379 | -0.479 | 1.102 | 0.363 | 0.027 | 0.06 | 0.06 | 0.12 | 0.42 | 0.26 | 0.28 | 0.15 | 0.33 | 0.44 |
| 3 | 109265353 | rs1732220 | -0.767 | 0.396 | 1.001 | 0.955 | 0.023 | 0.084 | 0.084 | 0 | 0.11 | 0.17 | 0.24 | 0.28 | 0.15 | 0.24 |
| 3 | 109270100 | rs1729614 | -0.733 | 0.433 | 0.928 | 0.931 | 0.005 | 0.032 | 0.032 | 0 | 0.1 | 0.11 | 0.22 | 0.25 | 0.15 | 0.24 |
| 3 | 109284286 | rs17499967 | 0.415 | 0.378 | 0.865 | 1.007 | 0.001 | 0.085 | 0.085 | 0 | 0.07 | 0.04 | 0.14 | 0.5 | 0.11 | 0.14 |
| 3 | 109300041 | rs1732187 | -1.135 | 0.284 | 0.951 | 0.719 | 0.008 | -0.003 | -0.003 | 0 | 0.24 | 0.32 | 0.42 | 0.74 | 0.31 | 0.4 |
| 3 | 109316638 | rs10511276 | 1.51 | 0.413 | 0.684 | 0.519 | 0.007 | 0.005 | 0.005 | 0 | 0.43 | 0.33 | 0.27 | 0.2 | 0.31 | 0.46 |
| 3 | 109323640 | rs1729620 | -1.577 | 0.509 | 0.347 | 0.76 | -0.003 | -0.017 | -0.017 | 0 | 0.28 | 0.49 | 0.47 | 0.72 | 0.31 | 0.38 |
| 3 | 109324308 | rs1528905 | -0.858 | 0.63 | 0.122 | 0.898 | 0.004 | 0.027 | 0.027 | 0 | 0.11 | 0.13 | 0.18 | 0.23 | 0.14 | 0.23 |
| 3 | 109338852 | rs1612794 | -1.534 | 0.591 | 0.19 | 0.874 | 0.04 | 0.118 | 0.118 | 0.067 | 0.14 | 0.37 | 0.21 | 0.22 | 0.15 | 0.23 |
| 3 | 109350052 | rs1106682 | -0.239 | 0.604 | 0.199 | 0.64 | 0.026 | 0.029 | 0.029 | 0 | 0.09 | 0.15 | 0.17 | 0.15 | 0.15 | 0.16 |
| 3 | 109350523 | rs1106193 | -1.259 | 0.607 | 0.111 | 0.721 | 0.012 | 0.039 | 0.039 | 0.082 | 0.25 | 0.45 | 0.46 | 0.65 | 0.32 | 0.31 |
| 3 | 109359465 | rs12485442 | -0.597 | 0.614 | 0.124 | 0.562 | 0.049 | 0.057 | 0.057 | 0 | 0.1 | 0.28 | 0.19 | 0.15 | 0.15 | 0.17 |
| 3 | 109361115 | rs1534634 | 0.735 | 0.615 | 0.001 | 0.547 | 0.064 | 0.071 | 0.071 | 0 | 0.89 | 0.64 | 0.8 | 0.85 | 0.85 | 0.83 |
| 3 | 109363105 | rs1534630 | 0.107 | 0.576 | -0.113 | 0.584 | 0.01 | -0.021 | -0.021 | 0.001 | 0.83 | 0.87 | 0.73 | 0.43 | 0.83 | 0.79 |
| 3 | 109405414 | rs1950052 | -1.211 | 0.651 | -0.024 | 0.506 | 0.012 | 0.028 | 0.028 | 0.132 | 0.25 | 0.45 | 0.46 | 0.67 | 0.32 | 0.31 |
| 3 | 109421199 | rs1529186 | -0.028 | 0.676 | -0.676 | 0.633 | 0.006 | -0.012 | -0.012 | 0 | 0.14 | 0.1 | 0.27 | 0.53 | 0.17 | 0.15 |
| 3 | 109429313 | rs1184781 | -0.067 | 1.081 | -0.564 | -0.049 | -0.109 | 0.151 | 0.151 | 0.11 | 0.28 | 0.08 | 0.18 | 0.01 | 0.38 | 0.17 |
| 3 | 109451739 | rs2253444 | 0.726 | 0.863 | 0.221 | 0.627 | 0.022 | -0.022 | -0.022 | 0 | 0.64 | 0.46 | 0.55 | 0.74 | 0.6 | 0.56 |
| 3 | 109454370 | rs6797827 | -0.882 | 0.958 | 0.301 | 0.659 | 0.036 | 0.021 | 0.021 | 0 | 0.09 | 0.19 | 0.28 | 0.15 | 0.13 | 0.2 |
| 3 | 109454436 | rs6798011 | -0.621 | 0.958 | 0.297 | 0.66 | 0.009 | 0.001 | 0.001 | 0 | 0.08 | 0.11 | 0.2 | 0.09 | 0.13 | 0.19 |
| 3 | 109462400 | rs1164064 | 1.507 | 0.86 | 0.167 | 0.781 | 0.007 | 0.032 | 0.032 | 0 | 0.48 | 0.37 | 0.32 | 0.31 | 0.45 | 0.39 |
| 3 | 109470819 | rs1920083 | 0.709 | 0.934 | 0.206 | 0.707 | 0.023 | -0.022 | -0.022 | 2.092 | 0.64 | 0.47 | 0.56 | 0.74 | 0.6 | 0.57 |
| 3 | 109484302 | rs1733265 | -0.104 | 1.112 | -0.048 | 0.57 | 0 | -0.024 | -0.024 | 0 | 0.73 | 0.68 | 0.84 | 0.89 | 0.74 | 0.74 |
| 3 | 109486238 | rs13081911 | 0.418 | 1.105 | -0.302 | 0.569 | -0.03 | 0.084 | 0.084 | 0 | 0.23 | 0.07 | 0.13 | 0.02 | 0.25 | 0.21 |
| 3 | 109489534 | rs1182789 | 1.315 | 1.362 | -0.412 | 0.681 | 0.006 | 0.024 | 0.024 | 0 | 0.48 | 0.34 | 0.42 | 0.35 | 0.46 | 0.38 |
| 3 | 109496029 | rs1915337 | 0.307 | 1.63 | -1.35 | 0.841 | 0.011 | -0.093 | -0.093 | 0 | 0.16 | 0.05 | 0.21 | 0.44 | 0.15 | 0.16 |
| 3 | 109503273 | rs11719237 | -1.294 | 2.084 | -0.656 | 0.177 | 0.013 | 0.002 | 0.002 | 0 | 0.08 | 0.13 | 0.2 | 0.09 | 0.13 | 0.19 |
| 3 | 109510473 | rs694022 | -0.582 | 1.295 | -0.863 | 0.408 | -0.006 | 0.005 | 0.005 | 0 | 0.29 | 0.26 | 0.29 | 0.21 | 0.32 | 0.41 |
| 3 | 109536495 | rs605172 | -0.098 | 1.064 | -0.773 | 0.45 | -0.004 | 0.004 | 0.004 | 1.208 | 0.42 | 0.45 | 0.43 | 0.39 | 0.47 | 0.27 |
| 3 | 109543489 | rs600790 | 0.45 | 1.204 | -0.705 | 0.009 | -0.008 | -0.008 | -0.008 | 0 | 0.17 | 0.15 | 0.27 | 0.39 | 0.25 | 0.21 |
| 3 | 109547714 | rs677533 | 0.473 | 1.176 | -0.688 | 0.027 | -0.043 | -0.053 | -0.053 | 0.186 | 0.17 | 0.11 | 0.27 | 0.39 | 0.25 | 0.21 |
| 3 | 109550055 | rs591350 | 0.256 | 0.928 | -0.4 | 0.017 | 0.007 | 0.015 | 0.015 | 0.111 | 0.5 | 0.4 | 0.48 | 0.69 | 0.54 | 0.61 |

| | | | | | | | | | | | | | | | |
|---|---|---|---|---|---|---|---|---|---|---|---|---|---|---|---|
| 3 | 109555870 | rs1881949 | -0.796 | 0.71 | 0.122 | 0.101 | -0.003 | -0.008 | -0.008 | 0.16 | 0.77 | 0.78 | 0.64 | 0.71 | 0.76 | 0.7 |
| 3 | 109575764 | rs12489624 | 0.933 | 0.433 | 0.072 | 0.251 | -0.052 | -0.044 | -0.044 | 0.094 | 0.19 | 0.05 | 0.21 | 0.22 | 0.24 | 0.23 |
| 3 | 109582957 | rs7649120 | 1.352 | 0.403 | 0.129 | 0.183 | 0.011 | 0.01 | 0.01 | 0.23 | 0.32 | 0.2 | 0.2 | 0.3 | 0.3 | 0.4 |
| 3 | 109591203 | rs595694 | 0.333 | 0.399 | 0.232 | 0.175 | 0.009 | 0.026 | 0.026 | 0 | 0.51 | 0.4 | 0.48 | 0.69 | 0.54 | 0.61 |
| 3 | 109604806 | rs1164194 | -0.467 | 0.708 | 0.42 | 0.154 | -0.004 | -0.005 | -0.005 | 0 | 0.76 | 0.84 | 0.62 | 0.71 | 0.72 | 0.69 |
| 3 | 109607659 | rs7653550 | 0.893 | 0.197 | 0.21 | -0.018 | 0.014 | 0.015 | 0.015 | 0 | 0.38 | 0.28 | 0.23 | 0.3 | 0.32 | 0.41 |
| 3 | 109616585 | rs9875746 | 1.125 | 0.346 | 0.54 | -0.061 | -0.002 | -0.016 | -0.016 | 0.436 | 0.2 | 0.16 | 0.09 | 0.02 | 0.15 | 0.07 |
| 3 | 109618919 | rs652972 | 0.346 | 0.415 | 0.461 | -0.01 | 0 | 0.002 | 0.002 | 0 | 0.55 | 0.43 | 0.5 | 0.69 | 0.56 | 0.63 |
| 3 | 109621818 | rs652760 | 1.218 | 0.536 | 0.378 | 0.041 | -0.024 | -0.033 | -0.033 | 0 | 0.18 | 0.13 | 0.26 | 0.39 | 0.23 | 0.21 |
| 3 | 109635659 | rs605795 | 0.051 | 0.858 | 0.355 | -0.133 | -0.009 | -0.002 | -0.002 | 0 | 0.61 | 0.53 | 0.59 | 0.69 | 0.63 | 0.7 |
| 3 | 109642605 | rs688898 | 0.348 | 0.904 | 0.411 | -0.137 | 0.001 | -0.008 | -0.008 | 0 | 0.57 | 0.5 | 0.51 | 0.7 | 0.56 | 0.64 |
| 3 | 109642716 | rs678058 | 1.114 | 0.991 | 0.367 | -0.034 | -0.018 | -0.027 | -0.027 | 0 | 0.19 | 0.15 | 0.26 | 0.39 | 0.23 | 0.22 |
| 3 | 109646918 | rs618829 | 0.064 | 0.973 | 0.417 | -0.123 | -0.001 | 0 | 0 | 0.05 | 0.38 | 0.3 | 0.36 | 0.42 | 0.38 | 0.29 |
| 3 | 109651181 | rs593904 | -0.297 | 0.925 | 0.433 | -0.154 | 0 | 0.01 | 0.01 | 0.112 | 0.75 | 0.66 | 0.66 | 0.73 | 0.75 | 0.75 |
| 3 | 109655820 | rs3915173 | 1.056 | 0.925 | 0.512 | -0.16 | -0.001 | 0.002 | 0.002 | 1.279 | 0.37 | 0.33 | 0.31 | 0.31 | 0.37 | 0.46 |
| 3 | 109661559 | rs602763 | -0.178 | 0.948 | 0.47 | -0.165 | -0.002 | 0.001 | 0.001 | 0 | 0.42 | 0.38 | 0.44 | 0.42 | 0.44 | 0.33 |
| 3 | 109670321 | rs616996 | 0.142 | 0.944 | 0.419 | -0.144 | -0.001 | -0.001 | -0.001 | 0 | 0.38 | 0.29 | 0.35 | 0.42 | 0.38 | 0.28 |
| 3 | 109677974 | rs619878 | 0.02 | 0.581 | 0.721 | -0.067 | 0.001 | 0.022 | 0.022 | 0.023 | 0.76 | 0.63 | 0.66 | 0.73 | 0.75 | 0.75 |
| 3 | 109678542 | rs16856195 | 0.777 | 0.852 | 0.631 | -0.047 | 0.017 | 0.01 | 0.01 | 0.04 | 0.18 | 0.08 | 0.08 | 0 | 0.15 | 0.03 |
| 3 | 109686614 | rs9845253 | -0.148 | 1.034 | 1.665 | 0.323 | 0.006 | 0.014 | 0.014 | 0 | 0.19 | 0.43 | 0.25 | 0.12 | 0.19 | 0.05 |
| 3 | 109687808 | rs9850837 | 0.512 | 1.015 | 1.783 | 0.184 | -0.009 | -0.052 | -0.052 | 0 | 0.15 | 0.19 | 0.09 | 0 | 0.14 | 0.03 |
| 3 | 109689619 | rs9878737 | 0.509 | 1.016 | 1.725 | 0.184 | 0.012 | 0.015 | 0.015 | 0.191 | 0.15 | 0.08 | 0.08 | 0 | 0.14 | 0.03 |
| 3 | 109696602 | rs709581 | 1.323 | 1.172 | 1.891 | 0.007 | -0.019 | 0.045 | 0.045 | 0.468 | 0.68 | 0.22 | 0.7 | 0.99 | 0.64 | 0.77 |
| 3 | 109702825 | rs10934005 | 0.101 | 0.451 | 2.209 | 0.173 | -0.004 | -0.15 | -0.15 | 0 | 0.8 | 0.51 | 0.84 | 1 | 0.8 | 0.96 |
| 3 | 109708722 | rs9860143 | -1.952 | 0.545 | 2.125 | 0.463 | 0.074 | 0.121 | 0.121 | 0 | 0.06 | 0.24 | 0.21 | 0.15 | 0.09 | 0.08 |
| 3 | 109711329 | rs2693644 | -2.907 | 0.394 | 2.274 | 0.592 | 0.033 | 0.109 | 0.109 | 2.116 | 0.16 | 0.57 | 0.4 | 0.21 | 0.15 | 0.17 |
| 3 | 109714119 | rs2701578 | 1.89 | 0.508 | 2.282 | 0.377 | 0.005 | 0.01 | 0.01 | 0.631 | 0.9 | 0.68 | 0.8 | 0.94 | 0.94 | 0.89 |
| 3 | 109724461 | rs7616317 | -0.695 | 0.523 | 2.242 | 0.048 | 0.001 | -0.062 | -0.062 | 6.538 | 0.28 | 0.35 | 0.33 | 0.06 | 0.3 | 0.19 |
| 3 | 109727353 | rs775525 | -1.301 | 0.235 | 2.131 | 0.572 | 0 | 0.021 | 0.021 | 0 | 0.45 | 0.67 | 0.57 | 0.54 | 0.43 | 0.42 |
| 3 | 109733870 | rs775534 | 0.59 | 0.295 | 2.016 | 0.641 | -0.001 | 0.027 | 0.027 | 0 | 0.14 | 0.15 | 0.24 | 0.46 | 0.13 | 0.21 |
| 3 | 109756664 | rs11921495 | -0.585 | 0.309 | 1.937 | 0.535 | 0 | -0.066 | -0.066 | 0 | 0.28 | 0.37 | 0.34 | 0.06 | 0.3 | 0.19 |
| 3 | 109758198 | rs11919475 | 1.263 | 0.286 | 1.862 | 0.615 | 0.002 | 0.022 | 0.022 | 0 | 0.54 | 0.23 | 0.42 | 0.49 | 0.57 | 0.6 |
| 3 | 109761983 | rs4682543 | -0.437 | 0.185 | 1.931 | 0.539 | 0 | -0.008 | -0.008 | 0.01 | 0.16 | 0.17 | 0.23 | 0.06 | 0.15 | 0.16 |
| 3 | 109765649 | rs6437888 | 0.545 | 0.361 | 1.785 | 0.656 | -0.006 | 0.039 | 0.039 | 0 | 0.7 | 0.4 | 0.64 | 0.55 | 0.72 | 0.76 |
| 3 | 109767320 | rs6796053 | 0.766 | 0.336 | 1.937 | 0.211 | -0.002 | -0.048 | -0.048 | 0 | 0.12 | 0.18 | 0.11 | 0 | 0.15 | 0.03 |
| 3 | 109789570 | rs10934011 | 1.495 | 0.186 | 2.168 | 0.376 | 0.059 | 0.014 | 0.014 | 0.585 | 0.8 | 0.5 | 0.69 | 0.85 | 0.79 | 0.85 |
| 3 | 109792367 | rs11719423 | 1.796 | 0.306 | 2.181 | 0.312 | -0.008 | 0.017 | 0.017 | 0.027 | 0.65 | 0.28 | 0.56 | 0.85 | 0.63 | 0.75 |
| 3 | 109799304 | rs1358018 | 2.881 | 0.325 | 2.349 | 0.335 | 0.051 | 0.123 | 0.123 | 3.54 | 0.93 | 0.64 | 0.87 | 0.9 | 0.96 | 0.97 |
| 3 | 109825034 | rs2895365 | 1.502 | 0.213 | 2.321 | 0.322 | 0 | -0.025 | -0.025 | 0 | 0.67 | 0.34 | 0.67 | 0.93 | 0.67 | 0.79 |
| 3 | 109831919 | rs4469005 | -0.992 | 0.228 | 2.391 | 0.367 | 0.04 | -0.092 | -0.092 | 1.116 | 0.25 | 0.47 | 0.3 | 0.06 | 0.3 | 0.14 |
| 3 | 109873808 | rs10755067 | 1.732 | 0.194 | 2.111 | 0.423 | 0 | -0.002 | -0.002 | 0 | 0.66 | 0.32 | 0.6 | 0.84 | 0.67 | 0.76 |
| 3 | 109877966 | rs2169401 | 1.363 | 0.205 | 2.266 | 0.201 | 0.02 | -0.048 | -0.048 | 0 | 0.81 | 0.64 | 0.79 | 0.92 | 0.82 | 0.81 |
| 3 | 109878008 | rs17667034 | 0.229 | 0.255 | 2.167 | 0.245 | -0.006 | -0.055 | -0.055 | 0 | 0.14 | 0.21 | 0.12 | 0 | 0.16 | 0.03 |
| 3 | 109883531 | rs6414243 | 0.429 | 0.4 | 1.92 | 0.331 | 0.017 | -0.012 | -0.012 | 1.281 | 0.87 | 0.78 | 0.81 | 0.93 | 0.84 | 0.89 |
| 3 | 109894018 | rs1551796 | -1.013 | 0.265 | 1.823 | 0.48 | 0.014 | 0.005 | 0.005 | 0.657 | 0.15 | 0.44 | 0.21 | 0.12 | 0.12 | 0.09 |
| 3 | 109901937 | rs6789041 | 0.363 | 0.24 | 1.566 | 0.449 | -0.015 | 0.026 | 0.026 | 0 | 0.14 | 0.07 | 0.12 | 0.01 | 0.19 | 0.08 |
| 3 | 109904601 | rs6783875 | -1.676 | 0.137 | 1.581 | 0.408 | 0.001 | -0.075 | -0.075 | 0.847 | 0.33 | 0.53 | 0.35 | 0.12 | 0.33 | 0.22 |
| 3 | 109912624 | rs10511281 | 0.113 | 0.344 | 1.322 | 0.475 | -0.006 | 0.009 | 0.009 | 0 | 0.15 | 0.12 | 0.12 | 0.01 | 0.19 | 0.08 |
| 3 | 109918190 | rs7633843 | 0.303 | 0.351 | 1.225 | 0.478 | -0.017 | 0.034 | 0.034 | 0 | 0.14 | 0.06 | 0.12 | 0 | 0.2 | 0.08 |
| 3 | 109942107 | rs716534 | 1.459 | 0.39 | 0.852 | 0.767 | -0.069 | 0.074 | 0.074 | 0 | 0.41 | 0.11 | 0.4 | 0.31 | 0.51 | 0.5 |
| 3 | 109944669 | rs9812947 | -1.347 | 1.104 | 1.72 | 0.968 | -0.009 | -0.079 | -0.079 | 0 | 0.23 | 0.39 | 0.19 | 0.06 | 0.21 | 0.17 |
| 3 | 109956216 | rs7631884 | 0.667 | 1.156 | 1.123 | 1.279 | 0.024 | 0.047 | 0.047 | 0 | 0.78 | 0.56 | 0.72 | 0.7 | 0.75 | 0.83 |
| 3 | 109963054 | rs16856531 | -1.226 | 0.726 | 1.054 | 1.202 | 0.015 | 0.006 | 0.006 | 0 | 0.07 | 0.09 | 0.14 | 0.06 | 0.09 | 0.03 |
| 3 | 109965453 | rs2399301 | -1.586 | 0.598 | 1.43 | 1.422 | 0.004 | 0.121 | 0.121 | 0.426 | 0.06 | 0.25 | 0.1 | 0.19 | 0.04 | 0.07 |
| 3 | 109966769 | rs1381356 | 2.315 | 0.202 | 1.687 | 1.23 | -0.022 | 0.021 | 0.021 | 0.017 | 0.79 | 0.35 | 0.72 | 0.75 | 0.85 | 0.85 |
| 3 | 109976128 | rs9869506 | 1.624 | 0.66 | 1.525 | 0.876 | 0.01 | -0.03 | -0.03 | 0 | 0.91 | 0.89 | 0.87 | 1 | 0.85 | 0.93 |
| 3 | 109987098 | rs6790010 | 2.467 | 0.55 | 2.056 | 1.329 | 0.01 | 0.041 | 0.041 | 0 | 0.81 | 0.52 | 0.79 | 0.8 | 0.84 | 0.88 |
| 3 | 109988573 | rs2712996 | -1.924 | 0.615 | 2.002 | 1.279 | -0.028 | 0.104 | 0.104 | 0 | 0.13 | 0.34 | 0.1 | 0.27 | 0.06 | 0.09 |
| 3 | 109992184 | rs2614693 | 1.892 | 0.725 | 1.571 | 0.928 | 0.053 | -0.07 | -0.07 | 0 | 0.88 | 0.69 | 0.85 | 1 | 0.85 | 0.85 |
| 3 | 110000728 | rs2713001 | -2.096 | 0.216 | 1.631 | 1.311 | -0.036 | 0.053 | 0.053 | 0 | 0.19 | 0.7 | 0.14 | 0.25 | 0.1 | 0.2 |
| 3 | 110007205 | rs2927047 | -2.007 | 0.082 | 2.262 | 1.337 | -0.064 | 0.11 | 0.11 | 0 | 0.17 | 0.56 | 0.12 | 0.24 | 0.08 | 0.18 |
| 3 | 110015748 | rs6802410 | 1.849 | 0.026 | 3.003 | 1.348 | -0.054 | 0.052 | 0.052 | 0 | 0.82 | 0.31 | 0.83 | 0.76 | 0.92 | 0.81 |
| 3 | 110024368 | rs6790478 | 1.709 | 0.026 | 3.047 | 1.342 | -0.067 | 0.099 | 0.099 | 0 | 0.82 | 0.41 | 0.84 | 0.76 | 0.92 | 0.81 |
| 3 | 110026835 | rs11707827 | 2.122 | -0.147 | 3.465 | 1.895 | -0.01 | 0.008 | 0.008 | 0.015 | 0.8 | 0.21 | 0.74 | 0.66 | 0.92 | 0.78 |
| 3 | 110028041 | rs11717542 | 2.09 | -0.147 | 3.47 | 1.895 | -0.012 | 0.012 | 0.012 | 0 | 0.8 | 0.22 | 0.74 | 0.66 | 0.92 | 0.78 |
| 3 | 110029068 | rs6782955 | -1.758 | -0.148 | 3.547 | 1.896 | -0.044 | 0.145 | 0.145 | 0 | 0.14 | 0.43 | 0.15 | 0.23 | 0.06 | 0.11 |
| 3 | 110031662 | rs9865422 | 2.37 | -0.147 | 3.545 | 1.896 | 0.012 | 0.138 | 0.138 | 0 | 0.95 | 0.77 | 0.9 | 0.89 | 0.98 | 0.89 |
| 3 | 110034038 | rs6781863 | -1.61 | -0.02 | 3.638 | 1.891 | -0.028 | 0.125 | 0.125 | 0 | 0.13 | 0.34 | 0.14 | 0.23 | 0.06 | 0.11 |
| 3 | 110035949 | rs12636638 | 0.813 | -0.204 | 2.177 | 1.334 | -0.084 | 0.145 | 0.145 | 1.552 | 0.42 | 0.11 | 0.41 | 0.23 | 0.51 | 0.55 |

| | | | | | | | | | | | | | | | | |
|---|---|---|---|---|---|---|---|---|---|---|---|---|---|---|---|---|
| 3 | 110044601 | rs12185907 | 0.757 | -0.191 | 2.228 | 1.333 | -0.074 | -0.033 | -0.033 | 0 | 0.37 | 0.07 | 0.33 | 0.43 | 0.4 | 0.24 |
| 3 | 110065332 | rs13091887 | 0.708 | -0.285 | 2.258 | 1.291 | -0.074 | -0.031 | -0.031 | 0.252 | 0.38 | 0.07 | 0.33 | 0.43 | 0.4 | 0.25 |
| 3 | 110072231 | rs7636367 | -0.877 | -0.293 | 2.44 | 1.315 | -0.038 | 0.154 | 0.154 | 0 | 0.13 | 0.41 | 0.16 | 0.23 | 0.05 | 0.11 |
| 3 | 110072760 | rs6437907 | -1.135 | -0.308 | 2.433 | 1.308 | -0.047 | 0.149 | 0.149 | 0 | 0.14 | 0.59 | 0.17 | 0.23 | 0.06 | 0.11 |
| 3 | 110079847 | rs2399342 | 1.088 | -0.398 | 2.265 | 1.264 | -0.011 | 0.004 | 0.004 | 0 | 0.95 | 0.89 | 0.98 | 0.97 | 0.96 | 0.81 |
| 3 | 110079927 | rs2001554 | 1.752 | -0.409 | 2.307 | 1.302 | -0.007 | 0.125 | 0.125 | 0 | 0.93 | 0.8 | 0.9 | 0.86 | 0.96 | 0.78 |
| 3 | 110080893 | rs9818820 | -0.295 | -0.447 | 2.249 | 1.332 | -0.022 | 0.132 | 0.132 | 0 | 0.11 | 0.28 | 0.15 | 0.24 | 0.06 | 0.11 |
| 3 | 110083045 | rs12634736 | 0.751 | -0.582 | 2.12 | 1.24 | -0.066 | -0.026 | -0.026 | 0.136 | 0.39 | 0.08 | 0.34 | 0.44 | 0.4 | 0.25 |
| 3 | 110083511 | rs4682576 | -0.055 | -0.573 | 2.146 | 1.223 | 0.001 | 0.004 | 0.004 | 2.215 | 0.52 | 0.46 | 0.53 | 0.43 | 0.56 | 0.52 |
| 3 | 110084303 | rs12635627 | -0.362 | -0.559 | 1.954 | 1.241 | -0.015 | 0.001 | 0.001 | 2.946 | 0.56 | 0.78 | 0.64 | 0.54 | 0.56 | 0.56 |
| 3 | 110087264 | rs6800418 | 0.419 | -0.536 | 1.802 | 1.142 | -0.057 | 0.176 | 0.176 | 1.082 | 0.44 | 0.19 | 0.4 | 0.2 | 0.54 | 0.46 |
| 3 | 110088488 | rs9848886 | -1.096 | -0.51 | 1.895 | 1.285 | -0.029 | 0.107 | 0.107 | 0 | 0.14 | 0.32 | 0.16 | 0.25 | 0.09 | 0.2 |
| 3 | 110091627 | rs13099844 | 0.65 | -0.205 | 1.017 | 1.094 | -0.066 | -0.029 | -0.029 | 0 | 0.37 | 0.08 | 0.33 | 0.43 | 0.4 | 0.24 |
| 3 | 110100138 | rs1546538 | 0.839 | -0.783 | 1.048 | 0.411 | -0.001 | -0.001 | -0.001 | 0 | 0.32 | 0.29 | 0.45 | 0.35 | 0.36 | 0.4 |
| 3 | 110103126 | rs10511285 | 0.1 | -0.587 | 1.282 | 0.355 | 0.001 | 0.061 | 0.061 | 0 | 0.53 | 0.39 | 0.33 | 0.09 | 0.49 | 0.38 |
| 3 | 110104986 | rs953237 | 0.162 | -0.636 | 1.394 | 0.492 | -0.003 | -0.001 | -0.001 | 2.655 | 0.41 | 0.49 | 0.51 | 0.41 | 0.38 | 0.47 |
| 3 | 110110033 | rs17726211 | 1.343 | -0.71 | 1.34 | 0.508 | -0.03 | 0.062 | 0.062 | 0 | 0.3 | 0.16 | 0.33 | 0.16 | 0.34 | 0.34 |
| 3 | 110112637 | rs1606368 | 0.342 | -0.719 | 1.419 | 0.508 | 0.001 | 0.013 | 0.013 | 0 | 0.38 | 0.33 | 0.38 | 0.22 | 0.36 | 0.41 |
| 3 | 110115936 | rs2669090 | -0.693 | -0.696 | 1.443 | 0.594 | -0.032 | -0.004 | -0.004 | 0 | 0.09 | 0.21 | 0.06 | 0.06 | 0.02 | 0.07 |
| 3 | 110126563 | rs6772187 | -0.002 | -0.992 | 0.394 | 0.194 | -0.002 | -0.023 | -0.023 | 0.119 | 0.16 | 0.27 | 0.07 | 0.03 | 0.1 | 0.15 |
| 3 | 110141727 | rs2068253 | 1.232 | -0.774 | -0.108 | 0.374 | 0.016 | 0.011 | 0.011 | 0.027 | 0.66 | 0.24 | 0.62 | 0.28 | 0.73 | 0.57 |
| 3 | 110146216 | rs13081374 | 0.646 | -0.626 | -0.363 | -0.217 | -0.033 | 0.145 | 0.145 | 0.082 | 0.36 | 0.08 | 0.26 | 0.06 | 0.39 | 0.23 |
| 3 | 110148936 | rs11720997 | 0.427 | -0.934 | -0.642 | -0.732 | -0.065 | 0.185 | 0.185 | 0.017 | 0.34 | 0.12 | 0.23 | 0.02 | 0.39 | 0.2 |
| 3 | 110152882 | rs13080207 | -0.839 | -1.36 | -0.296 | -1.125 | -0.003 | 0 | 0 | 0 | 0.53 | 0.53 | 0.33 | 0.05 | 0.51 | 0.42 |
| 3 | 110155824 | rs13062229 | -0.544 | -1.379 | -0.176 | -0.97 | -0.041 | -0.095 | -0.095 | 0 | 0.2 | 0.41 | 0.1 | 0.04 | 0.12 | 0.22 |
| 3 | 110184714 | rs9834665 | 1.126 | -1.034 | -0.444 | -0.433 | -0.037 | -0.001 | -0.001 | 0 | 0.07 | 0.01 | 0.14 | 0.41 | 0.11 | 0.15 |
| 3 | 110186611 | rs1533213 | 0.514 | -1.348 | -0.339 | -0.856 | -0.063 | 0.181 | 0.181 | 0.209 | 0.34 | 0.12 | 0.23 | 0.02 | 0.38 | 0.2 |
| 3 | 110187027 | rs11715989 | 1.269 | -1.289 | -0.512 | -1.012 | -0.079 | 0.158 | 0.158 | 0.043 | 0.24 | 0.02 | 0.17 | 0 | 0.28 | 0.11 |
| 3 | 110209251 | rs9818091 | 1.481 | -1.735 | 0.612 | -0.688 | -0.063 | -0.017 | -0.017 | 0.339 | 0.38 | 0.16 | 0.51 | 0.6 | 0.47 | 0.51 |
| 3 | 110224585 | rs1028026 | -1.425 | -1.692 | 0.794 | -0.669 | -0.017 | -0.001 | -0.001 | 0 | 0.6 | 0.71 | 0.48 | 0.4 | 0.52 | 0.49 |
| 3 | 110238230 | rs10934044 | 0.548 | -1.616 | 0.373 | -0.713 | -0.062 | 0.182 | 0.182 | 0.069 | 0.34 | 0.12 | 0.23 | 0.02 | 0.38 | 0.2 |
| 3 | 110256056 | rs9834281 | 1.444 | -1.443 | 0.356 | -0.6 | -0.124 | -0.064 | -0.064 | 0.113 | 0.37 | 0.08 | 0.5 | 0.59 | 0.47 | 0.51 |
| 3 | 110257313 | rs9820235 | 1.287 | -1.427 | 0.571 | -0.592 | -0.046 | -0.011 | -0.011 | 0.014 | 0.39 | 0.21 | 0.51 | 0.59 | 0.47 | 0.52 |
| 3 | 110262639 | rs1512519 | 1.288 | -1.43 | 0.54 | -0.593 | -0.075 | -0.025 | -0.025 | 1.035 | 0.39 | 0.17 | 0.5 | 0.59 | 0.48 | 0.51 |
| 3 | 110264260 | rs9877235 | -0.079 | -1.496 | 0.752 | -0.628 | -0.003 | -0.004 | -0.004 | 1.617 | 0.57 | 0.57 | 0.6 | 0.63 | 0.6 | 0.72 |
| 3 | 110271671 | rs11918532 | -0.601 | -1.591 | 1.196 | -0.332 | -0.014 | -0.049 | -0.049 | 0 | 0.17 | 0.28 | 0.09 | 0.04 | 0.12 | 0.21 |
| 3 | 110271975 | rs1512516 | -1.089 | -1.597 | 1.109 | -0.326 | -0.061 | -0.098 | -0.098 | 0.631 | 0.2 | 0.48 | 0.13 | 0.09 | 0.12 | 0.23 |
| 3 | 110287410 | rs6774121 | -0.761 | -1.521 | 0.915 | -0.378 | -0.044 | -0.111 | -0.111 | 0.921 | 0.18 | 0.39 | 0.1 | 0.04 | 0.12 | 0.21 |
| 3 | 110290490 | rs1996038 | 0.195 | -1.319 | 0.424 | -0.278 | 0.001 | -0.003 | -0.003 | 0.037 | 0.1 | 0.15 | 0.06 | 0.03 | 0.1 | 0.15 |
| 3 | 110302513 | rs1913097 | 0 | -0.976 | 0.625 | -0.189 | -0.001 | -0.006 | -0.006 | 0 | 0.61 | 0.57 | 0.65 | 0.69 | 0.6 | 0.73 |
| 3 | 110304185 | rs1022206 | 0.008 | -0.98 | 0.623 | -0.191 | -0.001 | -0.006 | -0.006 | 0 | 0.61 | 0.57 | 0.65 | 0.69 | 0.6 | 0.73 |
| 3 | 110306535 | rs4682013 | 1.357 | -1.238 | 0.479 | -0.319 | -0.049 | 0.003 | 0.003 | 0.007 | 0.46 | 0.16 | 0.57 | 0.61 | 0.51 | 0.52 |
| 3 | 110308735 | rs1398356 | -0.894 | -1.246 | 0.771 | -0.339 | -0.019 | 0.071 | 0.071 | 0 | 0.67 | 0.79 | 0.79 | 0.98 | 0.63 | 0.79 |
| 3 | 110313934 | rs1588652 | -0.168 | -1.265 | 1.069 | -0.236 | 0.018 | 0.334 | 0.334 | 0.584 | 0.94 | 0.78 | 0.85 | 0.71 | 0.98 | 0.94 |
| 3 | 110314691 | rs7433936 | -0.537 | -1.297 | 1.005 | -0.587 | -0.002 | -0.015 | -0.015 | 0 | 0.43 | 0.62 | 0.27 | 0.1 | 0.45 | 0.39 |
| 3 | 110319702 | rs13097144 | -0.756 | -1.261 | 1.015 | -0.571 | 0.001 | 0.004 | 0.004 | 0 | 0.47 | 0.51 | 0.28 | 0.1 | 0.47 | 0.43 |
| 3 | 110330282 | rs11919628 | -0.83 | -1.325 | 1.11 | -0.489 | -0.035 | -0.045 | -0.045 | 0 | 0.15 | 0.37 | 0.1 | 0.09 | 0.08 | 0.2 |
| 3 | 110332375 | rs10511287 | 0.702 | -1.22 | 0.906 | -0.43 | -0.064 | 0.164 | 0.164 | 0 | 0.32 | 0.1 | 0.2 | 0.02 | 0.37 | 0.21 |
| 3 | 110343621 | rs907658 | 1.843 | -1.166 | 1.521 | -0.282 | 0.054 | 0.117 | 0.117 | 1.837 | 0.94 | 0.64 | 0.97 | 0.93 | 0.99 | 0.97 |
| 3 | 110349192 | rs1512522 | 1.403 | -1.287 | 1.558 | -0.161 | -0.007 | 0.003 | 0.003 | 1.405 | 0.79 | 0.33 | 0.75 | 0.63 | 0.92 | 0.76 |
| 3 | 110360918 | rs1473828 | 1.205 | -1.284 | 1.461 | -0.239 | 0.005 | 0.032 | 0.032 | 0 | 0.48 | 0.22 | 0.56 | 0.62 | 0.54 | 0.55 |
| 3 | 110382273 | rs6768474 | -0.947 | -0.838 | 1.607 | 0.034 | 0.013 | 0.435 | 0.435 | 1.88 | 0.08 | 0.37 | 0.17 | 0.35 | 0.02 | 0.06 |
| 3 | 110399313 | rs4234398 | 0.511 | -1.026 | 1.453 | -0.162 | 0.002 | -0.002 | -0.002 | 0.093 | 0.53 | 0.48 | 0.71 | 0.89 | 0.55 | 0.6 |
| 3 | 110407281 | rs13096690 | 0.532 | -1.048 | 1.457 | -0.18 | -0.022 | 0.111 | 0.111 | 0.368 | 0.35 | 0.19 | 0.21 | 0.02 | 0.4 | 0.21 |
| 3 | 110421654 | rs11915300 | -0.816 | -1.015 | 1.376 | -0.14 | 0.008 | 0.345 | 0.345 | 2.215 | 0.08 | 0.36 | 0.17 | 0.28 | 0.02 | 0.05 |
| 3 | 110427655 | rs6772250 | 0.364 | -1.025 | 1.367 | -0.155 | -0.004 | -0.021 | -0.021 | 0 | 0.1 | 0.14 | 0.07 | 0.02 | 0.05 | 0.17 |
| 3 | 110430895 | rs9837589 | 0.276 | -1.028 | 1.328 | -0.154 | 0 | 0.08 | 0.08 | 0.076 | 0.56 | 0.65 | 0.72 | 0.89 | 0.55 | 0.61 |
| 3 | 110436187 | rs1903678 | 1.157 | -1.001 | 1.115 | -0.137 | -0.009 | 0.026 | 0.026 | 0.124 | 0.47 | 0.21 | 0.55 | 0.61 | 0.52 | 0.57 |
| 3 | 110453270 | rs11925026 | -0.937 | -0.986 | 1.147 | -0.109 | -0.003 | 0.316 | 0.316 | 2.869 | 0.09 | 0.36 | 0.17 | 0.27 | 0.02 | 0.05 |
| 3 | 110455019 | rs7633936 | 0.297 | -0.976 | 0.842 | -0.133 | 0 | 0.06 | 0.06 | 0.192 | 0.56 | 0.64 | 0.72 | 0.89 | 0.55 | 0.61 |
| 3 | 110463319 | rs9848649 | 0.898 | -0.216 | 0.054 | 0.211 | -0.04 | -0.015 | -0.015 | 0.26 | 0.08 | 0.01 | 0.16 | 0.43 | 0.11 | 0.13 |
| 3 | 110467766 | rs1156777 | 0.946 | -0.569 | 0.394 | 0.033 | 0.007 | 0.03 | 0.03 | 0.01 | 0.48 | 0.34 | 0.55 | 0.61 | 0.53 | 0.57 |
| 3 | 110468120 | rs7636192 | 0.019 | -0.47 | 0.016 | 0.084 | -0.01 | -0.04 | -0.04 | 0 | 0.1 | 0.18 | 0.07 | 0.02 | 0.06 | 0.17 |
| 3 | 110468507 | rs1350929 | 0.462 | -0.152 | -0.362 | 0.375 | 0.009 | 0.017 | 0.017 | 0.121 | 0.5 | 0.47 | 0.57 | 0.61 | 0.53 | 0.57 |
| 3 | 110487084 | rs4610259 | 0.257 | 0.112 | -0.334 | 0.289 | 0.002 | 0.01 | 0.01 | 0 | 0.54 | 0.46 | 0.58 | 0.69 | 0.54 | 0.59 |
| 3 | 110493936 | rs16857218 | -0.513 | 0 | -0.08 | 0.967 | -0.001 | 0.16 | 0.16 | 0 | 0.06 | 0.17 | 0.13 | 0.3 | 0.06 | 0.1 |
| 3 | 110494308 | rs9873710 | 0.121 | 1.252 | -0.37 | 0.867 | -0.021 | -0.001 | -0.001 | 0.371 | 0.06 | 0.01 | 0.21 | 0.04 | 0.17 | 0.1 |
| 3 | 110496376 | rs13089846 | 0.231 | 0.859 | -0.942 | -0.042 | -0.019 | 0.108 | 0.108 | 0 | 0.3 | 0.12 | 0.23 | 0.02 | 0.33 | 0.13 |
| 3 | 110498167 | rs10934066 | 1.535 | 0.373 | -1.434 | -0.03 | 0.03 | 0.083 | 0.083 | 0.051 | 0.33 | 0.08 | 0.23 | 0.21 | 0.28 | 0.39 |

| | | | | | | | | | | | | | | | | |
|---|---|---|---|---|---|---|---|---|---|---|---|---|---|---|---|---|
| 3 | 110501848 | rs1462307 | -0.252 | -0.819 | 0.333 | 0.036 | -0.021 | -0.041 | -0.041 | 0 | 0.79 | 0.61 | 0.82 | 0.91 | 0.85 | 0.74 |
| 3 | 110504049 | rs7622941 | 0.644 | -0.81 | 0.012 | -0.034 | 0 | -0.037 | -0.037 | 0.025 | 0.26 | 0.3 | 0.2 | 0.02 | 0.25 | 0.11 |
| 3 | 110506842 | rs2399363 | 0.873 | -0.731 | -0.606 | 0.265 | -0.092 | -0.048 | -0.048 | 0 | 0.41 | 0.1 | 0.45 | 0.59 | 0.46 | 0.5 |
| 3 | 110508263 | rs10934067 | -0.699 | -1.132 | 0.229 | 0.324 | -0.006 | -0.014 | -0.014 | 0.2 | 0.18 | 0.26 | 0.16 | 0.09 | 0.15 | 0.26 |
| 3 | 110512771 | rs10511288 | -0.977 | -1.124 | -0.05 | 0.32 | -0.01 | -0.03 | -0.03 | 5.756 | 0.28 | 0.33 | 0.22 | 0.1 | 0.2 | 0.27 |
| 3 | 110520698 | rs9863003 | 0.972 | -1.554 | -0.613 | 0.956 | 0.153 | 0.006 | 0.006 | 1.658 | 0.56 | 0.08 | 0.6 | 0.35 | 0.73 | 0.59 |
| 3 | 110528076 | rs10755069 | 1.156 | -0.734 | 1.15 | 1.498 | -0.041 | 0.018 | 0.018 | 1.995 | 0.87 | 0.38 | 0.84 | 0.9 | 0.95 | 0.95 |
| 3 | 110549398 | rs6809948 | 1.388 | -0.777 | 1.504 | 1.498 | -0.048 | -0.005 | -0.005 | 0.109 | 0.86 | 0.26 | 0.83 | 0.9 | 0.95 | 0.94 |
| 3 | 110558980 | rs9881407 | 0.937 | -0.823 | 2.662 | 1.524 | -0.011 | 0.067 | 0.067 | 0.53 | 0.92 | 0.6 | 0.86 | 0.9 | 0.97 | 0.95 |
| 3 | 110593478 | rs1350981 | 1.322 | -0.966 | 2.992 | 2.28 | -0.005 | 0.092 | 0.092 | 2.169 | 0.9 | 0.46 | 0.85 | 0.9 | 0.97 | 0.95 |
| 3 | 110602726 | rs10514755 | -0.522 | -0.955 | 3.212 | 2.333 | -0.002 | 0.002 | 0.002 | 0 | 0.09 | 0.09 | 0.07 | 0.13 | 0.09 | 0.14 |
| 3 | 110603974 | rs6763050 | 1.219 | -0.721 | 3.513 | 2.392 | 0.009 | 0.103 | 0.103 | 1.54 | 0.92 | 0.55 | 0.86 | 0.9 | 0.97 | 0.95 |
| 3 | 110692374 | rs7637068 | -1.794 | -0.688 | 3.646 | 2.385 | -0.021 | 0.393 | 0.393 | 1.232 | 0.12 | 0.49 | 0.25 | 0.47 | 0.09 | 0.11 |
| 3 | 110817982 | rs1477848 | 1.426 | -0.706 | 2.541 | 2.468 | -0.016 | 0.03 | 0.03 | 1.522 | 0.88 | 0.42 | 0.83 | 0.9 | 0.93 | 0.95 |
| 3 | 110822496 | rs1477846 | 1.426 | -0.706 | 2.541 | 2.468 | -0.016 | 0.03 | 0.03 | 1.522 | 0.88 | 0.42 | 0.83 | 0.9 | 0.93 | 0.95 |
| 3 | 110830515 | rs4682032 | -0.099 | -0.793 | 2.326 | 2.34 | -0.013 | 0.045 | 0.045 | 0 | 0.08 | 0 | 0.05 | 0 | 0.08 | 0 |
| 3 | 110860188 | rs11929280 | -1.432 | -0.664 | 2.538 | 2.849 | -0.029 | 0.158 | 0.158 | 1.274 | 0.11 | 0.54 | 0.15 | 0.21 | 0.03 | 0.08 |
| 3 | 110877031 | rs1477850 | -1.207 | -0.67 | 2.535 | 2.801 | -0.019 | 0.067 | 0.067 | 0.61 | 0.1 | 0.48 | 0.14 | 0.1 | 0.03 | 0.04 |
| 3 | 110886586 | rs7610760 | -2.153 | -0.638 | 2.114 | 3.14 | -0.03 | 0.037 | 0.037 | 3.922 | 0.14 | 0.79 | 0.19 | 0.29 | 0.09 | 0.09 |
| 3 | 110889974 | rs2169643 | -1.221 | -0.732 | 2.005 | 2.85 | -0.024 | 0.321 | 0.321 | 1.105 | 0.11 | 0.53 | 0.22 | 0.33 | 0.04 | 0.06 |
| 3 | 110908395 | rs7620372 | -1.236 | -0.759 | 1.664 | 2.739 | -0.033 | 0.378 | 0.378 | 0.458 | 0.11 | 0.45 | 0.22 | 0.44 | 0.03 | 0.11 |
| 3 | 110913972 | rs7610754 | -1.131 | -0.478 | 0.904 | 1.985 | -0.029 | 0.332 | 0.332 | 0.532 | 0.1 | 0.45 | 0.22 | 0.36 | 0.03 | 0.06 |
| 3 | 110929245 | rs4682242 | 0.292 | -0.062 | -0.066 | 0.98 | 0.014 | 0.042 | 0.042 | 0 | 0.49 | 0.46 | 0.57 | 0.29 | 0.47 | 0.46 |
| 3 | 110930787 | rs7432694 | 0.563 | -0.269 | -1.049 | 0.921 | 0.054 | 0.035 | 0.035 | 0 | 0.51 | 0.11 | 0.44 | 0.43 | 0.51 | 0.53 |
| 3 | 110934048 | rs7617793 | 0.173 | -0.61 | -0.92 | 0.816 | 0.011 | 0.017 | 0.017 | 0 | 0.13 | 0.08 | 0.1 | 0.05 | 0.12 | 0.07 |
| 3 | 110934660 | rs6437957 | -0.802 | -0.676 | -0.201 | 1.367 | 0.007 | 0.001 | 0.001 | 0 | 0.34 | 0.67 | 0.37 | 0.33 | 0.28 | 0.24 |
| 3 | 110940193 | rs6792042 | -0.75 | -0.831 | 0.343 | 1.04 | 0.021 | 0.006 | 0.006 | 0.017 | 0.25 | 0.1 | 0.19 | 0.2 | 0.26 | 0.13 |
| 3 | 110940332 | rs6437962 | -0.485 | -0.583 | 0.295 | 0.946 | -0.025 | 0.024 | 0.024 | 0.07 | 0.95 | 0.99 | 0.93 | 1 | 0.89 | 0.93 |
| 3 | 110941158 | rs7372838 | 0.559 | -0.17 | 0.143 | 0.932 | -0.037 | -0.048 | -0.048 | 0.885 | 0.13 | 0.01 | 0.12 | 0.15 | 0.18 | 0.06 |
| 3 | 110942148 | rs7433376 | 1.929 | -0.275 | -0.323 | 0.428 | -0.024 | 0.177 | 0.177 | 0 | 0.5 | 0.14 | 0.53 | 0.78 | 0.43 | 0.69 |
| 3 | 110950540 | rs7430044 | -0.409 | -0.232 | -0.358 | 0.422 | -0.073 | 0 | 0 | 0 | 0.81 | 0.98 | 0.85 | 0.86 | 0.72 | 0.87 |
| 3 | 110956475 | rs17202536 | 1.348 | -0.287 | -0.369 | 0.394 | 0.015 | 0.014 | 0.014 | 0 | 0.09 | 0.01 | 0.03 | 0.05 | 0.07 | 0.03 |
| 3 | 110962617 | rs2958070 | 1.849 | -0.366 | 0.437 | 0.386 | 0 | -0.005 | -0.005 | 0 | 0.55 | 0.42 | 0.62 | 0.74 | 0.49 | 0.71 |
| 3 | 110965554 | rs11924769 | -0.672 | -0.34 | 0.639 | 0.394 | 0.046 | -0.046 | -0.046 | 0 | 0.14 | 0.43 | 0.18 | 0.07 | 0.16 | 0.12 |
| 3 | 110967648 | rs6437970 | 1.131 | -0.342 | 0.63 | 0.396 | 0.021 | -0.035 | -0.035 | 0 | 0.67 | 0.55 | 0.66 | 0.79 | 0.56 | 0.74 |
| 3 | 110973976 | rs2958067 | 1.479 | -0.364 | 0.898 | 0.239 | 0.004 | 0.014 | 0.014 | 0 | 0.61 | 0.37 | 0.71 | 0.8 | 0.58 | 0.8 |
| 3 | 110978898 | rs2932054 | -1.075 | -0.404 | 1.506 | 0.634 | -0.009 | 0.038 | 0.038 | 0 | 0.25 | 0.4 | 0.24 | 0.34 | 0.24 | 0.29 |
| 3 | 110983620 | rs2634539 | -0.781 | -0.335 | 1.54 | 0.642 | 0.006 | 0.037 | 0.037 | 0 | 0.15 | 0.23 | 0.2 | 0.29 | 0.18 | 0.25 |
| 3 | 110985281 | rs6806881 | 0.876 | -0.367 | 1.631 | 0.635 | -0.016 | -0.02 | -0.02 | 1.017 | 0.05 | 0.08 | 0.01 | 0 | 0.02 | 0.01 |
| 3 | 110987911 | rs2675580 | -0.835 | -0.341 | 1.593 | 0.649 | 0.018 | 0.091 | 0.091 | 0 | 0.15 | 0.31 | 0.21 | 0.29 | 0.17 | 0.25 |
| 3 | 110989903 | rs6437972 | -0.526 | -0.311 | 1.396 | 0.639 | -0.073 | -0.002 | -0.002 | 1.26 | 0.81 | 0.98 | 0.84 | 0.86 | 0.72 | 0.87 |
| 3 | 110996938 | rs13084169 | 2.361 | -0.328 | 1.414 | 0.651 | -0.037 | -0.039 | -0.039 | 1.007 | 0.46 | 0.14 | 0.51 | 0.51 | 0.41 | 0.55 |
| 3 | 110998545 | rs9875279 | -1.151 | -0.326 | 1.512 | 0.668 | -0.003 | 0.035 | 0.035 | 0 | 0.19 | 0.27 | 0.21 | 0.29 | 0.19 | 0.25 |
| 3 | 111002374 | rs1489024 | 1.523 | -0.311 | 1.199 | 0.43 | 0.015 | 0.049 | 0.049 | 0 | 0.59 | 0.27 | 0.7 | 0.8 | 0.58 | 0.8 |
| 3 | 111003863 | rs13083225 | -0.247 | -0.33 | 1.24 | 0.419 | -0.038 | -0.067 | -0.067 | 0 | 0.16 | 0.46 | 0.11 | 0.05 | 0.12 | 0.06 |
| 3 | 111008156 | rs7629508 | -0.592 | -0.114 | 0.943 | 0.538 | 0 | -0.009 | -0.009 | 0 | 0.25 | 0.26 | 0.19 | 0.14 | 0.3 | 0.14 |
| 3 | 111023831 | rs17281151 | -0.704 | -0.02 | 0.188 | 0.262 | 0.002 | -0.011 | -0.011 | 0 | 0.2 | 0.2 | 0.28 | 0.13 | 0.21 | 0.15 |
| 3 | 111025158 | rs2971355 | 2.383 | -0.15 | -0.194 | 0.383 | -0.023 | -0.039 | -0.039 | 0 | 0.4 | 0.16 | 0.36 | 0.41 | 0.31 | 0.42 |
| 3 | 111028725 | rs12637512 | 2.128 | -0.352 | 0.396 | 0.316 | 0.013 | 0.012 | 0.012 | 0 | 0.49 | 0.39 | 0.44 | 0.46 | 0.39 | 0.53 |
| 3 | 111033241 | rs2958083 | -1.304 | -0.301 | 0.761 | 0.657 | -0.007 | 0.074 | 0.074 | 0 | 0.21 | 0.32 | 0.3 | 0.43 | 0.23 | 0.33 |
| 3 | 111038212 | rs2932073 | -0.002 | -0.33 | 0.784 | 0.588 | 0.037 | 0.024 | 0.024 | 0 | 0.08 | 0.18 | 0.21 | 0.1 | 0.12 | 0.09 |
| 3 | 111039820 | rs10804486 | 1.393 | -0.323 | 0.852 | 0.532 | 0.017 | -0.018 | -0.018 | 0 | 0.66 | 0.62 | 0.56 | 0.8 | 0.55 | 0.79 |
| 3 | 111040163 | rs2958081 | -0.217 | -0.209 | 0.383 | 0.523 | -0.018 | 0.017 | 0.017 | 0 | 0.8 | 0.85 | 0.79 | 0.9 | 0.69 | 0.89 |
| 3 | 111042752 | rs1873105 | 2.497 | -0.167 | 0.039 | 0.64 | -0.072 | -0.006 | -0.006 | 0 | 0.39 | 0.07 | 0.35 | 0.35 | 0.35 | 0.45 |
| 3 | 111043193 | rs12106997 | 0.34 | -0.262 | 0.512 | 0.748 | -0.017 | 0.073 | 0.073 | 0 | 0.09 | 0.16 | 0.08 | 0.34 | 0.09 | 0.24 |
| 3 | 111050246 | rs2634553 | -0.124 | -0.312 | 1.249 | 0.725 | -0.037 | 0.197 | 0.197 | 0.056 | 0.11 | 0.32 | 0.09 | 0.34 | 0.09 | 0.24 |
| 3 | 111050709 | rs6783451 | 2.228 | -0.332 | 1.01 | 0.595 | 0.026 | -0.032 | -0.032 | 0 | 0.56 | 0.13 | 0.64 | 0.56 | 0.54 | 0.63 |
| 3 | 111051871 | rs2971365 | -1.68 | -0.536 | 1.075 | 0.615 | -0.096 | 0.078 | 0.078 | 0.749 | 0.22 | 0.55 | 0.14 | 0.34 | 0.15 | 0.26 |
| 3 | 111052059 | rs2971366 | -1.495 | -0.524 | 1.014 | 0.621 | -0.094 | 0.101 | 0.101 | 0.155 | 0.21 | 0.48 | 0.13 | 0.34 | 0.15 | 0.26 |
| 3 | 111054726 | rs1552540 | 2.697 | -0.276 | 0.358 | 0.647 | -0.098 | 0.003 | 0.003 | 0 | 0.37 | 0.05 | 0.33 | 0.31 | 0.34 | 0.45 |
| 3 | 111056828 | rs11920390 | -1.27 | -0.093 | 1.11 | 0.698 | -0.016 | 0.033 | 0.033 | 0 | 0.23 | 0.35 | 0.14 | 0.35 | 0.21 | 0.3 |
| 3 | 111059861 | rs2087337 | -1.686 | 0.007 | 0.579 | 0.685 | -0.013 | -0.008 | -0.008 | 1.002 | 0.89 | 0.96 | 0.89 | 0.86 | 0.85 | 0.97 |
| 3 | 111060392 | rs9878888 | -0.677 | -0.012 | 0.821 | 0.675 | 0.001 | 0.023 | 0.023 | 0 | 0.18 | 0.26 | 0.17 | 0.36 | 0.24 | 0.32 |
| 3 | 111066327 | rs1844237 | -1.634 | 0.188 | 0.452 | 0.786 | -0.008 | -0.005 | -0.005 | 0 | 0.9 | 0.92 | 0.89 | 0.86 | 0.85 | 0.97 |
| 3 | 111074604 | rs7610103 | 1.734 | 0.199 | -0.058 | 0.705 | 0.023 | -0.042 | -0.042 | 0 | 0.56 | 0.17 | 0.65 | 0.5 | 0.53 | 0.64 |
| 3 | 111080687 | rs973208 | 1.565 | -0.053 | 0.389 | 0.891 | 0.056 | -0.047 | -0.047 | 0.444 | 0.7 | 0.16 | 0.73 | 0.61 | 0.77 | 0.67 |
| 3 | 111087564 | rs7633973 | -1.45 | -0.211 | 2.057 | 0.815 | -0.038 | 0.055 | 0.055 | 0 | 0.25 | 0.56 | 0.25 | 0.39 | 0.21 | 0.33 |
| 3 | 111119350 | rs6778463 | 1.71 | -0.224 | 2.074 | 0.808 | 0.048 | -0.039 | -0.039 | 0.606 | 0.71 | 0.18 | 0.73 | 0.61 | 0.77 | 0.67 |
| 3 | 111121960 | rs1529068 | 2.58 | 0.433 | 1.482 | 0.684 | -0.031 | 0.054 | 0.054 | 0.036 | 0.4 | 0.03 | 0.23 | 0.26 | 0.29 | 0.47 |

| | | | | | | | | | | | | | | | | |
|---|---|---|---|---|---|---|---|---|---|---|---|---|---|---|---|---|
| 3 | 111124619 | rs7623819 | -1.016 | 0.736 | 2.75 | 0.889 | -0.017 | 0.091 | 0.091 | 0 | 0.19 | 0.29 | 0.22 | 0.39 | 0.19 | 0.31 |
| 3 | 111125434 | rs4682264 | 1.583 | 0.731 | 2.4 | 0.887 | -0.006 | 0.009 | 0.009 | 0 | 0.75 | 0.32 | 0.75 | 0.61 | 0.79 | 0.67 |
| 3 | 111159353 | rs2698370 | 1.012 | 1.006 | 1.784 | 0.886 | 0.037 | -0.021 | -0.021 | 0 | 0.94 | 0.85 | 0.96 | 0.99 | 0.92 | 0.94 |
| 3 | 111159403 | rs11706807 | 2.152 | 0.981 | 1.308 | 0.906 | 0.021 | -0.046 | -0.046 | 0 | 0.69 | 0.18 | 0.7 | 0.59 | 0.71 | 0.64 |
| 3 | 111167008 | rs9865917 | -0.139 | 0.942 | 1.19 | 0.984 | -0.013 | -0.067 | -0.067 | 0 | 0.14 | 0.03 | 0.19 | 0.39 | 0.18 | 0.28 |
| 3 | 111180433 | rs1553090 | 2.033 | 1.173 | 1.526 | 0.864 | 0.005 | -0.033 | -0.033 | 0 | 0.79 | 0.28 | 0.8 | 0.93 | 0.8 | 0.87 |
| 3 | 111181808 | rs16858101 | -0.188 | 1.068 | 1.324 | 0.94 | -0.016 | -0.088 | -0.088 | 0.435 | 0.14 | 0.03 | 0.19 | 0.44 | 0.17 | 0.29 |
| 3 | 111189285 | rs4522763 | 2.171 | 1.039 | 1.398 | 0.837 | -0.002 | 0.003 | 0.003 | 0.917 | 0.78 | 0.22 | 0.79 | 0.93 | 0.79 | 0.87 |
| 3 | 111193516 | rs17281269 | -1.565 | 1.051 | 1.867 | 0.859 | 0.027 | -0.014 | -0.014 | 0.268 | 0.12 | 0.19 | 0.14 | 0.07 | 0.18 | 0.1 |
| 3 | 111197213 | rs1113850 | 2.594 | 1.027 | 1.894 | 0.954 | -0.01 | -0.048 | -0.048 | 0 | 0.64 | 0.2 | 0.6 | 0.49 | 0.62 | 0.58 |
| 3 | 111197934 | rs12495894 | -0.203 | 1.147 | 1.618 | 1.041 | -0.016 | -0.088 | -0.088 | 0.28 | 0.14 | 0.03 | 0.19 | 0.44 | 0.17 | 0.29 |
| 3 | 111219929 | rs9829839 | 1.542 | 1.177 | 2.036 | 0.979 | 0.058 | -0.025 | -0.025 | 0.014 | 0.87 | 0.75 | 0.85 | 0.94 | 0.82 | 0.89 |
| 3 | 111220516 | rs9868270 | -0.632 | 1.149 | 2.474 | 1.01 | -0.004 | 0.06 | 0.06 | 0 | 0.16 | 0.18 | 0.21 | 0.44 | 0.18 | 0.3 |
| 3 | 111225977 | rs4682278 | -1.27 | 1.147 | 2.346 | 1.009 | 0.01 | -0.003 | -0.003 | 0.626 | 0.11 | 0.12 | 0.14 | 0.06 | 0.18 | 0.11 |
| 3 | 111226730 | rs9834853 | 2.085 | 1.191 | 2.479 | 1.022 | 0.038 | 0.075 | 0.075 | 0 | 0.71 | 0.51 | 0.63 | 0.49 | 0.64 | 0.59 |
| 3 | 111233364 | rs1609935 | -0.276 | 1.167 | 1.862 | 0.969 | -0.006 | -0.064 | -0.064 | 0 | 0.14 | 0.05 | 0.2 | 0.44 | 0.17 | 0.29 |
| 3 | 111240427 | rs4682048 | 2.553 | 1.333 | 2.062 | 0.971 | -0.025 | -0.054 | -0.054 | 0.558 | 0.65 | 0.2 | 0.54 | 0.49 | 0.62 | 0.62 |
| 3 | 111253819 | rs6773238 | -1.44 | 1.64 | 2.126 | 0.917 | -0.012 | 0.008 | 0.008 | 0 | 0.12 | 0.1 | 0.23 | 0.09 | 0.24 | 0.17 |
| 3 | 111254219 | rs2895385 | -2.072 | 1.622 | 2.476 | 1.038 | 0.068 | -0.001 | -0.001 | 0 | 0.15 | 0.3 | 0.25 | 0.14 | 0.26 | 0.19 |
| 3 | 111254393 | rs2221065 | 2.339 | 1.628 | 2.168 | 1.032 | -0.029 | -0.036 | -0.036 | 0.014 | 0.64 | 0.22 | 0.52 | 0.48 | 0.56 | 0.61 |
| 3 | 111257405 | rs1391361 | 1.64 | 1.397 | 2.35 | 0.967 | 0.087 | -0.071 | -0.071 | 0 | 0.82 | 0.59 | 0.79 | 0.95 | 0.72 | 0.82 |
| 3 | 111260186 | rs1391356 | 1.572 | 1.541 | 1.749 | 1.063 | -0.012 | -0.04 | -0.04 | 0.001 | 0.71 | 0.79 | 0.66 | 0.59 | 0.67 | 0.62 |
| 3 | 111260975 | rs2276873 | 2.273 | 1.518 | 1.784 | 0.999 | 0.02 | -0.063 | -0.063 | 0 | 0.62 | 0.17 | 0.53 | 0.48 | 0.65 | 0.57 |
| 3 | 111267564 | rs10511298 | 1.563 | 1.536 | 1.722 | 1.019 | -0.004 | -0.016 | -0.016 | 0 | 0.66 | 0.77 | 0.63 | 0.62 | 0.6 | 0.59 |
| 3 | 111269236 | rs2173318 | 2.554 | 1.511 | 0.903 | 0.853 | -0.102 | -0.069 | -0.069 | 0 | 0.46 | 0.07 | 0.45 | 0.45 | 0.47 | 0.52 |
| 3 | 111283350 | rs7619313 | 0.161 | 1.58 | 1.261 | 0.922 | 0.003 | -0.018 | -0.018 | 0 | 0.12 | 0.13 | 0.06 | 0.04 | 0.12 | 0.04 |
| 3 | 111285876 | rs1513322 | -1.042 | 1.714 | 1.681 | 1.046 | 0.037 | -0.073 | -0.073 | 0 | 0.21 | 0.32 | 0.18 | 0.04 | 0.25 | 0.14 |
| 3 | 111288539 | rs7635078 | -0.955 | 0.777 | 1.653 | 0.918 | 0.001 | -0.059 | -0.059 | 0 | 0.2 | 0.28 | 0.16 | 0.05 | 0.19 | 0.14 |
| 3 | 111291695 | rs7628039 | 1.141 | 0.507 | 1.433 | 1.02 | -0.003 | 0.016 | 0.016 | 0 | 0.7 | 0.64 | 0.58 | 0.48 | 0.66 | 0.66 |
| 3 | 111291946 | rs9865195 | 1.12 | 0.511 | 1.332 | 1.021 | -0.001 | 0.002 | 0.002 | 0 | 0.7 | 0.67 | 0.58 | 0.48 | 0.66 | 0.66 |
| 3 | 111300775 | rs12497656 | 1.505 | 0.531 | 1.316 | 0.571 | 0.039 | -0.048 | -0.048 | 0.39 | 0.77 | 0.52 | 0.67 | 0.84 | 0.72 | 0.75 |
| 3 | 111309224 | rs1464061 | 2.246 | 0.542 | 0.755 | 0.778 | -0.042 | -0.039 | -0.039 | 2.095 | 0.45 | 0.14 | 0.43 | 0.44 | 0.48 | 0.51 |
| 3 | 111313457 | rs10804491 | 0.647 | 0.51 | 1.768 | 0.8 | -0.038 | -0.009 | -0.009 | 2.15 | 0.84 | 0.36 | 0.83 | 0.84 | 0.92 | 0.93 |
| 3 | 111313854 | rs938213 | 0.153 | 0.635 | 1.457 | 0.613 | -0.024 | -0.058 | -0.058 | 0 | 0.88 | 0.72 | 0.93 | 0.96 | 0.93 | 0.94 |
| 3 | 111330922 | rs9858544 | -1.677 | 1.077 | 1.77 | 0.83 | 0.064 | 0.074 | 0.074 | 0.008 | 0.16 | 0.51 | 0.21 | 0.25 | 0.18 | 0.22 |
| 3 | 111338169 | rs9868266 | -0.134 | 1.108 | 1.379 | 1.083 | -0.028 | -0.086 | -0.086 | 0.792 | 0.14 | 0 | 0.21 | 0.29 | 0.17 | 0.16 |
| 3 | 111343756 | rs6780210 | -1.518 | 1.169 | 1.844 | 1.118 | 0.06 | 0.068 | 0.068 | 0 | 0.16 | 0.47 | 0.2 | 0.26 | 0.18 | 0.22 |
| 3 | 111368813 | rs3733176 | -1.518 | 1.169 | 1.839 | 1.118 | 0.061 | 0.069 | 0.069 | 0 | 0.16 | 0.47 | 0.2 | 0.26 | 0.18 | 0.22 |
| 3 | 111369473 | rs6437999 | 0.994 | 1.081 | 1.88 | 0.889 | 0.028 | 0.02 | 0.02 | 0 | 0.95 | 0.75 | 0.96 | 0.97 | 0.97 | 0.96 |
| 3 | 111385006 | rs13078891 | 2.428 | 0.897 | 0.899 | 0.824 | -0.035 | -0.057 | -0.057 | 0.124 | 0.5 | 0.07 | 0.49 | 0.44 | 0.53 | 0.51 |
| 3 | 111399209 | rs7651713 | -1.091 | 1.028 | 1.122 | 0.626 | -0.014 | 0.056 | 0.056 | 0 | 0.26 | 0.43 | 0.3 | 0.33 | 0.27 | 0.28 |
| 3 | 111400162 | rs17423232 | -0.524 | 1.113 | 1.137 | 0.618 | -0.002 | -0.008 | -0.008 | 0 | 0.08 | 0.06 | 0.08 | 0 | 0.14 | 0.01 |
| 3 | 111405407 | rs4481128 | -2.017 | 1.053 | 1.091 | 0.612 | -0.001 | -0.005 | -0.005 | 0 | 0.4 | 0.54 | 0.38 | 0.34 | 0.43 | 0.34 |
| 3 | 111411812 | rs6796920 | 0.267 | 0.593 | 0.678 | 0.742 | -0.029 | -0.037 | -0.037 | 0 | 0.1 | 0.15 | 0.03 | 0 | 0.03 | 0.01 |
| 3 | 111416310 | rs1355767 | 1.237 | 0.919 | 0.284 | 0.962 | 0.054 | -0.039 | -0.039 | 0.128 | 0.59 | 0.22 | 0.58 | 0.49 | 0.68 | 0.55 |
| 3 | 111419486 | rs1553971 | -1.937 | 1.875 | 0.981 | 1.239 | 0.01 | 0.025 | 0.025 | 0 | 0.11 | 0.22 | 0.12 | 0.17 | 0.16 | 0.23 |
| 3 | 111423342 | rs4325897 | -2.319 | 2.513 | 1.532 | 0.934 | 0.055 | -0.116 | -0.116 | 0 | 0.17 | 0.5 | 0.14 | 0 | 0.18 | 0.12 |
| 3 | 111427455 | rs1877575 | -0.974 | 2.943 | 1.104 | 1.367 | -0.047 | -0.079 | -0.079 | 0 | 0.13 | 0.02 | 0.25 | 0.32 | 0.21 | 0.16 |
| 3 | 111435957 | rs6796976 | -1.529 | 3.046 | 1.561 | 1.262 | 0.046 | -0.031 | -0.031 | 0 | 0.1 | 0.46 | 0.09 | 0 | 0.07 | 0.09 |
| 3 | 111440689 | rs9877859 | 2.95 | 3.323 | 1.435 | 1.273 | -0.09 | 0.044 | 0.044 | 2.451 | 0.72 | 0.07 | 0.69 | 0.82 | 0.62 | 0.68 |
| 3 | 111441672 | rs11929568 | -1.63 | 3.347 | 1.532 | 1.284 | 0.101 | 0.022 | 0.022 | 3.389 | 0.13 | 0.49 | 0.16 | 0.12 | 0.15 | 0.16 |
| 3 | 111442959 | rs17423838 | -1.193 | 3.289 | 1.535 | 1.316 | 0.024 | 0.003 | 0.003 | 0 | 0.06 | 0.16 | 0.06 | 0.05 | 0.12 | 0.13 |
| 3 | 111445546 | rs2292518 | -1.904 | 3.175 | 1.535 | 1.38 | -0.005 | -0.004 | -0.004 | 0 | 0.19 | 0.18 | 0.31 | 0.37 | 0.33 | 0.28 |
| 3 | 111447773 | rs1398751 | 2.004 | 3.159 | 1.598 | 1.378 | 0.04 | 0.071 | 0.071 | 0 | 0.78 | 0.63 | 0.65 | 0.62 | 0.65 | 0.7 |
| 3 | 111448944 | rs1105448 | 1.823 | 3.162 | 1.556 | 1.337 | 0.008 | 0.016 | 0.016 | 0 | 0.79 | 0.76 | 0.66 | 0.63 | 0.65 | 0.7 |
| 3 | 111456174 | rs13317985 | -1.192 | 3.437 | 1.669 | 1.274 | 0.031 | -0.011 | -0.011 | 0 | 0.09 | 0.18 | 0.18 | 0.07 | 0.19 | 0.13 |
| 3 | 111456859 | rs4557129 | -2.425 | 3.31 | 1.63 | 1.341 | 0.007 | 0.053 | 0.053 | 2.114 | 0.25 | 0.63 | 0.33 | 0.45 | 0.25 | 0.32 |
| 3 | 111462570 | rs7629156 | -2.122 | 3.364 | 1.804 | 1.367 | 0.033 | 0.132 | 0.132 | 0.676 | 0.23 | 0.48 | 0.32 | 0.45 | 0.25 | 0.32 |
| 3 | 111478337 | rs6784441 | -1.992 | 3.362 | 2.222 | 1.417 | 0.024 | 0.105 | 0.105 | 0.003 | 0.2 | 0.33 | 0.3 | 0.52 | 0.25 | 0.31 |
| 3 | 111480946 | rs9868898 | -1.562 | 3.432 | 2.567 | 0.931 | 0.1 | -0.009 | -0.009 | 0.032 | 0.11 | 0.3 | 0.21 | 0.08 | 0.22 | 0.15 |
| 3 | 111488413 | rs13095929 | 2.224 | 3.466 | 2.566 | 0.931 | 0.008 | -0.018 | -0.018 | 0 | 0.77 | 0.32 | 0.77 | 0.84 | 0.74 | 0.79 |
| 3 | 111488596 | rs6800922 | -1.222 | 3.452 | 2.577 | 0.905 | 0.065 | -0.007 | -0.007 | 0.132 | 0.09 | 0.23 | 0.19 | 0.07 | 0.19 | 0.13 |
| 3 | 111489638 | rs4682062 | 2.106 | 3.457 | 2.644 | 0.902 | 0.073 | -0.034 | -0.034 | 0.603 | 0.91 | 0.64 | 0.89 | 0.99 | 0.89 | 0.97 |
| 3 | 111491772 | rs9858228 | -1.236 | 3.399 | 2.382 | 0.92 | 0.065 | -0.007 | -0.007 | 0 | 0.09 | 0.23 | 0.19 | 0.07 | 0.19 | 0.13 |
| 3 | 111493868 | rs6789977 | -2.047 | 3.391 | 2.406 | 0.916 | 0.071 | -0.027 | -0.027 | 0 | 0.13 | 0.55 | 0.21 | 0.07 | 0.19 | 0.13 |
| 3 | 111500814 | rs4682313 | 2 | 3.415 | 2.253 | 0.91 | 0.053 | -0.055 | -0.055 | 0 | 0.84 | 0.39 | 0.72 | 0.93 | 0.73 | 0.86 |
| 3 | 111505613 | rs4682314 | 2.533 | 3.465 | 1.583 | 0.931 | -0.166 | 0.063 | 0.063 | 1.216 | 0.66 | 0.05 | 0.55 | 0.76 | 0.52 | 0.64 |
| 3 | 111508146 | rs4516581 | -1.774 | 2.919 | 1.986 | 1.087 | 0.115 | 0.008 | 0.008 | 0 | 0.1 | 0.35 | 0.2 | 0.07 | 0.19 | 0.13 |
| 3 | 111513865 | rs9857303 | -1.721 | 2.945 | 1.504 | 1.011 | 0.044 | -0.026 | -0.026 | 0 | 0.13 | 0.18 | 0.16 | 0.06 | 0.26 | 0.14 |

| | | | | | | | | | | | | | | | | |
|---|---|---|---|---|---|---|---|---|---|---|---|---|---|---|---|---|
| 3 | 111517048 | rs6438014 | -2.274 | 2.055 | 2.01 | 1.63 | 0.006 | 0.342 | 0.342 | 0.436 | 0.13 | 0.49 | 0.29 | 0.42 | 0.07 | 0.22 |
| 3 | 111521616 | rs13062508 | 1.887 | 2.064 | 2.282 | 1.261 | 0.022 | 0.028 | 0.028 | 0.095 | 0.93 | 0.73 | 0.88 | 0.98 | 0.97 | 0.93 |
| 3 | 111528539 | rs9876709 | -2.494 | 3.234 | 2.284 | 1.684 | 0.051 | 0.064 | 0.064 | 0.627 | 0.2 | 0.28 | 0.48 | 0.42 | 0.35 | 0.5 |
| 3 | 111532971 | rs6809414 | -2.66 | 3.223 | 2.981 | 1.736 | 0.089 | 0.179 | 0.179 | 0 | 0.18 | 0.44 | 0.44 | 0.41 | 0.29 | 0.32 |
| 3 | 111542578 | rs4411858 | -2.765 | 3.294 | 3.234 | 1.836 | 0.026 | 0.227 | 0.227 | 0 | 0.1 | 0.34 | 0.29 | 0.32 | 0.09 | 0.26 |
| 3 | 111543285 | rs9872438 | -1.155 | 3.368 | 3.091 | 1.829 | 0.009 | -0.001 | -0.001 | 0 | 0.08 | 0.07 | 0.15 | 0.09 | 0.2 | 0.07 |
| 3 | 111544295 | rs6769535 | -3.209 | 3.229 | 3.454 | 1.825 | 0.029 | 0.272 | 0.272 | 0.068 | 0.12 | 0.5 | 0.31 | 0.32 | 0.09 | 0.26 |
| 3 | 111547852 | rs12491776 | -2.962 | 3.038 | 3.52 | 1.817 | 0.051 | 0.303 | 0.303 | 0.53 | 0.06 | 0.34 | 0.19 | 0.26 | 0.03 | 0.19 |
| 3 | 111556141 | rs3925081 | 2.153 | 2.91 | 3.271 | 1.689 | 0.03 | -0.002 | -0.002 | 0 | 0.84 | 0.83 | 0.7 | 0.85 | 0.71 | 0.67 |
| 3 | 111562805 | rs10934114 | 2.302 | 2.592 | 3.162 | 1.546 | 0.041 | -0.011 | -0.011 | 0 | 0.81 | 0.77 | 0.69 | 0.85 | 0.7 | 0.67 |
| 3 | 111564839 | rs937557 | -0.417 | 2.643 | 2.909 | 1.547 | -0.034 | -0.012 | -0.012 | 0 | 0.05 | 0 | 0.1 | 0.05 | 0.11 | 0.2 |
| 3 | 111570330 | rs1282977 | -1.848 | 2.664 | 2.886 | 1.544 | 0.016 | -0.006 | -0.006 | 0.217 | 0.13 | 0.14 | 0.2 | 0.11 | 0.18 | 0.13 |
| 3 | 111574943 | rs10934116 | 2.58 | 2.637 | 2.64 | 1.513 | -0.001 | 0.076 | 0.076 | 0.059 | 0.79 | 0.4 | 0.58 | 0.6 | 0.8 | 0.69 |
| 3 | 111589962 | rs1913494 | 2.699 | 2.882 | 2.377 | 0.987 | 0.022 | -0.007 | -0.007 | 0.161 | 0.74 | 0.38 | 0.67 | 0.77 | 0.69 | 0.52 |
| 3 | 111591934 | rs13323152 | -0.768 | 2.842 | 1.952 | 1.003 | -0.013 | -0.006 | -0.006 | 0.636 | 0.08 | 0.04 | 0.1 | 0.1 | 0.1 | 0.07 |
| 3 | 111599363 | rs6781879 | 1.761 | 2.69 | 1.617 | 0.376 | -0.027 | 0.042 | 0.042 | 0 | 0.61 | 0.32 | 0.51 | 0.8 | 0.52 | 0.63 |
| 3 | 111606391 | rs10934118 | 2.073 | 2.563 | 1.58 | 0.37 | -0.035 | 0.041 | 0.041 | 0 | 0.58 | 0.26 | 0.47 | 0.7 | 0.51 | 0.43 |
| 3 | 111612191 | rs7640773 | 1.076 | 2.436 | 2.037 | 0.334 | 0.036 | -0.081 | -0.081 | 0 | 0.73 | 0.46 | 0.69 | 0.94 | 0.63 | 0.71 |
| 3 | 111618531 | rs9811953 | -1.525 | 2.452 | 2.439 | 0.319 | -0.001 | -0.002 | -0.002 | 0.063 | 0.15 | 0.21 | 0.24 | 0.15 | 0.17 | 0.09 |
| 3 | 111619171 | rs1587456 | -1.64 | 2.394 | 2.409 | 0.36 | 0.007 | -0.01 | -0.01 | 0 | 0.13 | 0.12 | 0.2 | 0.25 | 0.09 | 0.27 |
| 3 | 111625374 | rs11716038 | -0.97 | 2.576 | 1.715 | -0.335 | -0.006 | 0.016 | 0.016 | 0 | 0.29 | 0.24 | 0.33 | 0.16 | 0.34 | 0.17 |
| 3 | 111631564 | rs1282948 | 1.575 | 2.326 | 1.631 | -0.455 | -0.007 | 0.012 | 0.012 | 0 | 0.62 | 0.66 | 0.55 | 0.74 | 0.54 | 0.56 |
| 3 | 111634902 | rs12491508 | 2.629 | 1.795 | 1.25 | -0.737 | -0.042 | -0.028 | -0.028 | 0 | 0.39 | 0.05 | 0.39 | 0.67 | 0.35 | 0.33 |
| 3 | 111635605 | rs968825 | 2.117 | 1.924 | 1.637 | -0.753 | 0.021 | 0.076 | 0.076 | 0 | 0.47 | 0.28 | 0.44 | 0.74 | 0.39 | 0.48 |
| 3 | 111637904 | rs2399399 | 0.479 | 1.939 | 1.998 | -0.749 | 0.017 | 0.026 | 0.026 | 0.021 | 0.91 | 0.79 | 0.87 | 0.91 | 0.89 | 0.74 |
| 3 | 111640282 | rs10934125 | 2.067 | 1.938 | 2.08 | -0.76 | 0.024 | 0.074 | 0.074 | 0 | 0.48 | 0.32 | 0.44 | 0.74 | 0.39 | 0.48 |
| 3 | 111641393 | rs4234408 | -1.488 | 1.94 | 2.011 | -0.78 | 0.007 | -0.044 | -0.044 | 0 | 0.43 | 0.48 | 0.43 | 0.17 | 0.5 | 0.25 |
| 3 | 111642822 | rs1282965 | 0.724 | 1.473 | 1.346 | -1.005 | -0.002 | -0.002 | -0.002 | 0 | 0.88 | 0.85 | 0.87 | 0.91 | 0.87 | 0.72 |
| 3 | 111643561 | rs2137897 | 0.333 | 1.711 | 0.925 | -0.918 | -0.009 | 0.013 | 0.013 | 0.217 | 0.15 | 0.12 | 0.12 | 0.07 | 0.18 | 0.22 |
| 3 | 111646052 | rs13091880 | -1.434 | 1.698 | 0.988 | -0.934 | -0.004 | 0.052 | 0.052 | 0 | 0.45 | 0.35 | 0.45 | 0.23 | 0.53 | 0.4 |
| 3 | 111651285 | rs754914 | -0.035 | 1.757 | 0.914 | -0.879 | -0.033 | 0.026 | 0.026 | 0 | 0.23 | 0.09 | 0.25 | 0.16 | 0.25 | 0.16 |
| 3 | 111658514 | rs2272289 | 0.892 | 1.598 | 0.94 | -0.799 | -0.001 | -0.01 | -0.01 | 0 | 0.12 | 0.12 | 0.09 | 0.01 | 0.15 | 0.07 |
| 3 | 111669631 | rs6804475 | 1.075 | 1.546 | 0.684 | -0.817 | 0 | 0.005 | 0.005 | 0 | 0.73 | 0.29 | 0.75 | 0.91 | 0.73 | 0.65 |
| 3 | 111672825 | rs3749298 | 0.306 | 1.524 | 0.499 | -0.809 | 0.008 | -0.018 | -0.018 | 0.247 | 0.09 | 0.06 | 0.13 | 0.16 | 0.09 | 0.06 |
| 3 | 111673147 | rs3749299 | 1.953 | 1.606 | 0.536 | -1.018 | -0.01 | 0.23 | 0.23 | 0 | 0.49 | 0.12 | 0.55 | 0.84 | 0.44 | 0.39 |
| 3 | 111677526 | rs11713044 | 1.127 | 1.516 | 0.7 | -1.026 | 0 | 0.02 | 0.02 | 0 | 0.73 | 0.27 | 0.75 | 0.91 | 0.73 | 0.65 |
| 3 | 111678633 | rs13062225 | 0.89 | 1.584 | 0.594 | -1.003 | -0.024 | 0.041 | 0.041 | 0 | 0.11 | 0.05 | 0.09 | 0.01 | 0.15 | 0.08 |
| 3 | 111679072 | rs9816238 | -1.736 | 1.589 | 0.779 | -1.049 | -0.006 | 0.113 | 0.113 | 0 | 0.47 | 0.67 | 0.4 | 0.09 | 0.53 | 0.47 |
| 3 | 111680469 | rs4682068 | 0.251 | 1.478 | 0.951 | -0.926 | -0.005 | -0.018 | -0.018 | 0 | 0.21 | 0.15 | 0.17 | 0 | 0.26 | 0.11 |
| 3 | 111680599 | rs16858936 | -1.47 | 1.211 | 1.138 | -0.565 | 0.002 | -0.049 | -0.049 | 0 | 0.22 | 0.52 | 0.22 | 0.09 | 0.23 | 0.34 |
| 3 | 111682042 | rs6788215 | -0.534 | 1.181 | 1.308 | -0.555 | 0.033 | -0.068 | -0.068 | 0.448 | 0.12 | 0.38 | 0.11 | 0 | 0.12 | 0.08 |
| 3 | 111683753 | rs16858950 | -0.402 | 1.171 | 1.325 | -0.554 | 0.009 | -0.065 | -0.065 | 0.431 | 0.11 | 0.28 | 0.1 | 0.01 | 0.12 | 0.08 |
| 3 | 111685572 | rs774857 | -0.58 | 1.139 | 1.375 | -0.569 | 0.02 | -0.071 | -0.071 | 0.01 | 0.12 | 0.35 | 0.11 | 0 | 0.12 | 0.08 |
| 3 | 111687756 | rs10511302 | 1.717 | 0.802 | 0.735 | -0.566 | -0.02 | 0.017 | 0.017 | 0 | 0.46 | 0.25 | 0.45 | 0.6 | 0.48 | 0.33 |
| 3 | 111688578 | rs774854 | -0.503 | 0.731 | 0.957 | -0.617 | 0.011 | -0.047 | -0.047 | 0 | 0.16 | 0.29 | 0.13 | 0.07 | 0.16 | 0.22 |
| 3 | 111691725 | rs2399400 | 1.188 | 0.761 | 1.192 | -0.589 | 0.007 | 0.02 | 0.02 | 0 | 0.53 | 0.41 | 0.56 | 0.72 | 0.59 | 0.6 |
| 3 | 111692829 | rs774852 | 0.056 | 0.586 | 1.016 | -0.628 | 0.02 | -0.025 | -0.025 | 1.278 | 0.87 | 0.93 | 0.8 | 0.79 | 0.9 | 0.9 |
| 3 | 111695124 | rs3177104 | 1.977 | -0.126 | -0.332 | -0.899 | 0.008 | 0.02 | 0.02 | 0 | 0.26 | 0.2 | 0.22 | 0.12 | 0.22 | 0.12 |
| 3 | 111700305 | rs510549 | -0.281 | -0.201 | -0.279 | -0.768 | 0.007 | 0.03 | 0.03 | 0 | 0.62 | 0.72 | 0.51 | 0.19 | 0.64 | 0.51 |
| 3 | 111701010 | rs2289597 | 1.57 | -0.254 | -0.365 | -0.396 | 0.008 | -0.05 | -0.05 | 0.239 | 0.09 | 0.01 | 0.18 | 0.29 | 0.08 | 0.06 |
| 3 | 111704406 | rs1609796 | 0.893 | -0.185 | -0.119 | -0.591 | -0.019 | 0.022 | 0.022 | 0 | 0.25 | 0.19 | 0.19 | 0 | 0.3 | 0.18 |
| 3 | 111705992 | rs3804765 | 1.392 | -0.176 | -0.059 | -0.577 | -0.003 | -0.015 | -0.015 | 0 | 0.27 | 0.23 | 0.29 | 0.53 | 0.27 | 0.39 |
| 3 | 111706366 | rs7615972 | 2.249 | -0.199 | -0.071 | -0.619 | 0.007 | 0.015 | 0.015 | 0 | 0.27 | 0.22 | 0.22 | 0.11 | 0.22 | 0.12 |
| 3 | 111711594 | rs1127853 | 2.307 | -0.158 | -0.163 | -0.661 | 0.009 | 0.022 | 0.022 | 0 | 0.26 | 0.19 | 0.22 | 0.11 | 0.22 | 0.12 |
| 3 | 111713296 | rs4682327 | 1.483 | 0.177 | -0.058 | -0.53 | -0.004 | -0.004 | -0.004 | 0 | 0.2 | 0.16 | 0.13 | 0 | 0.19 | 0.11 |
| 3 | 111714066 | rs12638225 | -1.655 | 0.401 | -0.285 | 0.26 | -0.006 | 0.032 | 0.032 | 0 | 0.29 | 0.46 | 0.49 | 0.68 | 0.28 | 0.54 |
| 3 | 111714795 | rs7429121 | -0.674 | 0.767 | -1.116 | 0.221 | 0.003 | -0.001 | -0.001 | 0 | 0.13 | 0.13 | 0.27 | 0.13 | 0.24 | 0.16 |
| 3 | 111716195 | rs3811705 | 0.116 | 0.721 | -1.337 | 0.261 | 0.001 | 0.005 | 0.005 | 0 | 0.39 | 0.32 | 0.47 | 0.29 | 0.42 | 0.26 |
| 3 | 111719330 | rs7639791 | 0.753 | 0.762 | -1.605 | 0.354 | 0.007 | -0.027 | -0.027 | 0.146 | 0.33 | 0.18 | 0.38 | 0.46 | 0.28 | 0.24 |
| 3 | 111719493 | rs774764 | -1.236 | 0.885 | -1.566 | 0.279 | -0.001 | -0.001 | -0.001 | 0.891 | 0.53 | 0.56 | 0.68 | 0.57 | 0.54 | 0.46 |
| 3 | 111719879 | rs2292582 | 1.348 | 0.389 | -1.584 | 0.548 | 0.003 | 0.012 | 0.012 | 0 | 0.24 | 0.17 | 0.21 | 0.16 | 0.21 | 0.09 |
| 3 | 111720465 | rs4682328 | 0.888 | 0.709 | -1.614 | 0.655 | 0.002 | -0.024 | -0.024 | 0.092 | 0.09 | 0.06 | 0.16 | 0.3 | 0.06 | 0.16 |
| 3 | 111722092 | rs774759 | 0.744 | 0.789 | -1.844 | 0.25 | -0.01 | -0.013 | -0.013 | 0 | 0.31 | 0.43 | 0.4 | 0.25 | 0.25 | 0.17 |
| 3 | 111723535 | rs774757 | 0.596 | 0.667 | -2.225 | 0.09 | 0.007 | 0.037 | 0.037 | 0 | 0.46 | 0.41 | 0.29 | 0.16 | 0.39 | 0.21 |
| 3 | 111723770 | rs1282731 | -0.226 | 0.539 | -1.766 | 0.912 | -0.017 | 0.213 | 0.213 | 0 | 0.85 | 0.5 | 0.78 | 0.52 | 0.92 | 0.65 |
| 3 | 111723978 | rs814035 | -0.226 | 0.539 | -1.766 | 0.901 | -0.017 | 0.217 | 0.217 | 0 | 0.85 | 0.5 | 0.78 | 0.52 | 0.92 | 0.65 |
| 3 | 111724401 | rs13096825 | 0.009 | 0.633 | -2.141 | 0.232 | 0.052 | 0.025 | 0.025 | 0 | 0.62 | 0.2 | 0.5 | 0.27 | 0.77 | 0.49 |
| 3 | 111725406 | rs6773379 | 0.43 | 0.775 | -2.141 | 0.199 | -0.016 | 0.089 | 0.089 | 0 | 0.43 | 0.3 | 0.37 | 0.17 | 0.49 | 0.47 |
| 3 | 111725434 | rs774754 | 0.27 | 0.366 | -2.259 | 0.754 | 0.007 | -0.017 | -0.017 | 0.22 | 0.27 | 0.25 | 0.37 | 0.52 | 0.18 | 0.2 |

| | | | | | | | | | | | | | | | | |
|---|---|---|---|---|---|---|---|---|---|---|---|---|---|---|---|---|
| 3 | 111728514 | rs11719077 | -0.136 | 0.832 | -2.251 | 0.521 | -0.045 | -0.078 | -0.078 | 0.564 | 0.77 | 0.89 | 0.66 | 0.66 | 0.73 | 0.69 |
| 3 | 111730546 | rs3749310 | -0.446 | 0.708 | -2.601 | 0.244 | -0.061 | -0.052 | -0.052 | 0 | 0.56 | 0.12 | 0.53 | 0.39 | 0.57 | 0.45 |
| 3 | 111730591 | rs3749311 | -0.453 | 0.701 | -2.517 | 0.243 | -0.035 | -0.032 | -0.032 | 0 | 0.56 | 0.17 | 0.53 | 0.39 | 0.57 | 0.45 |
| 3 | 111732608 | rs8070 | 0.217 | 0.726 | -2.345 | 0.364 | -0.024 | -0.054 | -0.054 | 0 | 0.22 | 0.11 | 0.34 | 0.33 | 0.25 | 0.29 |
| 3 | 111734945 | rs1000326 | 0.644 | 0.411 | -2.086 | 0.327 | 0 | 0.039 | 0.039 | 0.251 | 0.06 | 0.09 | 0.21 | 0.14 | 0.04 | 0.12 |
| 3 | 111739614 | rs9828227 | 0.484 | 0.243 | -2.057 | 0.08 | -0.042 | 0.16 | 0.16 | 0.173 | 0.32 | 0.02 | 0.17 | 0.06 | 0.3 | 0.13 |
| 3 | 111741577 | rs9839073 | 0.12 | 0.332 | -1.92 | 0.114 | 0.004 | 0.072 | 0.072 | 0.227 | 0.38 | 0.11 | 0.38 | 0.2 | 0.35 | 0.25 |
| 3 | 111755066 | rs4386465 | 0.154 | 0.379 | -1.924 | 0.218 | 0.022 | 0.047 | 0.047 | 0 | 0.56 | 0.78 | 0.56 | 0.67 | 0.61 | 0.69 |
| 3 | 111758076 | rs2197205 | 0.046 | 0.381 | -2.239 | 0.266 | -0.017 | 0.019 | 0.019 | 0.428 | 0.39 | 0.07 | 0.4 | 0.3 | 0.35 | 0.25 |
| 3 | 111762518 | rs4682081 | 0.006 | 0.266 | -1.214 | 0.01 | -0.004 | -0.021 | -0.021 | 0 | 0.3 | 0.43 | 0.27 | 0.22 | 0.33 | 0.31 |
| 3 | 111768423 | rs7622025 | -0.033 | 0.166 | -1.141 | -0.081 | -0.004 | -0.027 | -0.027 | 0 | 0.31 | 0.36 | 0.26 | 0.17 | 0.33 | 0.32 |
| 3 | 111770948 | rs10934131 | -0.033 | 0.166 | -1.132 | -0.081 | -0.004 | -0.025 | -0.025 | 0 | 0.31 | 0.36 | 0.26 | 0.17 | 0.33 | 0.32 |
| 3 | 111772292 | rs9288935 | 0.393 | -0.148 | -0.933 | -0.303 | 0.016 | 0.117 | 0.117 | 0 | 0.62 | 0.17 | 0.68 | 0.83 | 0.63 | 0.69 |
| 3 | 111774361 | rs16859115 | 0.518 | -0.22 | -0.435 | -0.28 | -0.001 | 0.005 | 0.005 | 0 | 0.25 | 0.23 | 0.2 | 0.17 | 0.28 | 0.27 |
| 3 | 111777504 | rs4682352 | 0.514 | -0.221 | -0.462 | -0.28 | -0.003 | 0.023 | 0.023 | 0 | 0.25 | 0.18 | 0.19 | 0.17 | 0.28 | 0.27 |
| 3 | 111781799 | rs9864383 | -0.345 | -0.676 | -0.574 | 0.079 | 0.03 | -0.015 | -0.015 | 0 | 0.41 | 0.2 | 0.35 | 0.48 | 0.35 | 0.53 |
| 3 | 111784889 | rs340146 | -0.092 | 0.356 | -0.135 | 0.163 | 0.109 | 0.013 | 0.013 | 0.006 | 0.95 | 0.74 | 0.92 | 1 | 0.92 | 0.99 |
| 3 | 111793316 | rs191454 | 0.228 | 0.194 | -0.445 | 0.03 | 0.004 | -0.003 | -0.003 | 0.329 | 0.64 | 0.59 | 0.63 | 0.6 | 0.64 | 0.56 |
| 3 | 111793957 | rs10511304 | 0.947 | 0.104 | -1.009 | -0.245 | -0.039 | 0.106 | 0.106 | 0 | 0.17 | 0.02 | 0.11 | 0 | 0.19 | 0.05 |
| 3 | 111799277 | rs902854 | -0.727 | -0.161 | -0.451 | -0.054 | -0.002 | 0.003 | 0.003 | 0.506 | 0.36 | 0.41 | 0.4 | 0.44 | 0.32 | 0.41 |
| 3 | 111799327 | rs168150 | 0.942 | -0.484 | -0.069 | -0.054 | 0.003 | -0.001 | -0.001 | 0 | 0.53 | 0.49 | 0.55 | 0.48 | 0.59 | 0.47 |
| 3 | 111807719 | rs4682355 | 1.45 | -0.776 | 0.279 | -0.01 | 0.01 | -0.001 | -0.001 | 0 | 0.73 | 0.3 | 0.76 | 0.75 | 0.85 | 0.66 |
| 3 | 111809773 | rs9847052 | 0.069 | -0.869 | 0.798 | -0.021 | -0.016 | 0.013 | 0.013 | 0 | 0.07 | 0.15 | 0.05 | 0.06 | 0.04 | 0.11 |
| 3 | 111815278 | rs340157 | -0.439 | -0.883 | 1.121 | -0.027 | -0.004 | 0.046 | 0.046 | 0.257 | 0.08 | 0.28 | 0.06 | 0.06 | 0.04 | 0.11 |
| 3 | 111819064 | rs12172949 | -0.156 | -0.946 | 1.068 | -0.023 | -0.017 | -0.04 | -0.04 | 0 | 0.08 | 0.01 | 0.07 | 0.13 | 0.06 | 0.11 |
| 3 | 111821747 | rs16859190 | -0.368 | -0.932 | 1.437 | -0.056 | -0.002 | 0.049 | 0.049 | 0.045 | 0.08 | 0.28 | 0.06 | 0.06 | 0.04 | 0.11 |
| 3 | 111824279 | rs16859207 | -1.162 | -0.918 | 1.509 | -0.053 | -0.008 | 0.005 | 0.005 | 0 | 0.16 | 0.21 | 0.12 | 0.19 | 0.11 | 0.22 |
| 3 | 111836566 | rs11824 | 1.786 | -0.692 | 0.811 | -0.22 | 0.02 | 0.066 | 0.066 | 0.447 | 0.57 | 0.25 | 0.63 | 0.73 | 0.61 | 0.61 |
| 3 | 111840513 | rs7649379 | 0.522 | -0.317 | 0.576 | -0.104 | -0.011 | 0.006 | 0.006 | 0 | 0.06 | 0.09 | 0.04 | 0.06 | 0.04 | 0.11 |
| 3 | 111841303 | rs1492489 | 0.488 | -0.327 | 0.616 | -0.125 | -0.022 | 0.033 | 0.033 | 0 | 0.18 | 0.1 | 0.14 | 0.01 | 0.25 | 0.06 |
| 3 | 111841346 | rs1492488 | 0.445 | -0.412 | 0.606 | 0.063 | -0.011 | 0.007 | 0.007 | 0 | 0.06 | 0.09 | 0.04 | 0.06 | 0.04 | 0.11 |
| 3 | 111841535 | rs16859259 | 0.789 | -0.403 | 0.42 | 0.058 | 0.011 | -0.024 | -0.024 | 0.129 | 0.05 | 0 | 0.04 | 0.06 | 0.04 | 0.11 |
| 3 | 111860222 | rs1492480 | 1.1 | -0.017 | 0.376 | 0.028 | 0 | -0.004 | -0.004 | 0.033 | 0.41 | 0.35 | 0.44 | 0.42 | 0.41 | 0.47 |
| 3 | 111873996 | rs7622733 | 0.543 | 0.093 | 0.472 | -0.13 | -0.006 | 0.007 | 0.007 | 0.008 | 0.27 | 0.36 | 0.4 | 0.33 | 0.24 | 0.33 |
| 3 | 111897540 | rs4682090 | 0.315 | 0.086 | 0.482 | -0.171 | -0.003 | 0.02 | 0.02 | 0 | 0.53 | 0.37 | 0.45 | 0.65 | 0.49 | 0.6 |
| 3 | 111919002 | rs7637682 | -0.696 | 0.056 | 0.498 | -0.197 | 0.008 | -0.073 | -0.073 | 0.38 | 0.79 | 0.7 | 0.85 | 0.99 | 0.73 | 0.93 |
| 3 | 111921116 | rs9288938 | 0.583 | 0.05 | 0.535 | -0.194 | -0.008 | 0.009 | 0.009 | 0.008 | 0.27 | 0.37 | 0.4 | 0.34 | 0.24 | 0.34 |
| 3 | 111921225 | rs6781844 | -0.67 | 0.115 | 0.466 | -0.2 | 0.009 | -0.073 | -0.073 | 0 | 0.79 | 0.7 | 0.85 | 0.99 | 0.73 | 0.93 |
| 3 | 111923123 | rs4434123 | -0.675 | 0.108 | 0.469 | -0.197 | 0.008 | -0.073 | -0.073 | 0 | 0.79 | 0.7 | 0.85 | 0.99 | 0.73 | 0.93 |
| 3 | 111936961 | rs17446282 | 0.751 | 0.111 | 0.335 | -0.195 | -0.013 | 0.015 | 0.015 | 0 | 0.19 | 0.13 | 0.14 | 0.01 | 0.26 | 0.07 |
| 3 | 111951023 | rs6781344 | 0.706 | 0.052 | 0.376 | -0.113 | 0.003 | -0.034 | -0.034 | 0.13 | 0.63 | 0.48 | 0.71 | 0.77 | 0.6 | 0.72 |
| 3 | 111951184 | rs17446744 | -0.241 | 0.038 | 0.451 | -0.128 | 0 | -0.002 | -0.002 | 0 | 0.29 | 0.32 | 0.23 | 0.22 | 0.34 | 0.18 |
| 3 | 111962851 | rs6768523 | 0.68 | 0.064 | 0.071 | -0.056 | -0.005 | 0.007 | 0.007 | 0.007 | 0.26 | 0.34 | 0.4 | 0.33 | 0.24 | 0.33 |
| 3 | 111965246 | rs9822786 | 0.569 | 0.064 | 0.071 | -0.056 | 0.003 | -0.034 | -0.034 | 0.001 | 0.63 | 0.48 | 0.71 | 0.77 | 0.6 | 0.72 |
| 3 | 111966018 | rs6766122 | 0.606 | 0.075 | 0.103 | 0.004 | -0.005 | 0.007 | 0.007 | 0.007 | 0.26 | 0.34 | 0.4 | 0.33 | 0.24 | 0.33 |
| 3 | 111978848 | rs10934156 | 1.615 | 0.077 | -0.079 | -0.014 | -0.023 | -0.071 | -0.071 | 0 | 0.37 | 0.14 | 0.31 | 0.43 | 0.36 | 0.38 |
| 3 | 111981924 | rs9809404 | 0.574 | -0.406 | 0.076 | -0.054 | 0.003 | -0.034 | -0.034 | 0 | 0.64 | 0.48 | 0.71 | 0.77 | 0.6 | 0.72 |
| 3 | 111982526 | rs4234414 | 1.76 | -0.415 | 0.059 | -0.054 | -0.022 | -0.071 | -0.071 | 0.179 | 0.38 | 0.14 | 0.31 | 0.43 | 0.36 | 0.38 |
| 3 | 111985107 | rs9872691 | 0.667 | -0.402 | 0.075 | -0.05 | -0.005 | 0.007 | 0.007 | 0.007 | 0.26 | 0.34 | 0.4 | 0.34 | 0.24 | 0.34 |
| 3 | 111985428 | rs4264699 | 0.573 | -0.378 | 0.074 | -0.054 | 0.004 | -0.034 | -0.034 | 5.238 | 0.64 | 0.48 | 0.71 | 0.77 | 0.6 | 0.72 |
| 3 | 111990941 | rs13059342 | 1.725 | -0.312 | -0.011 | -0.041 | -0.017 | -0.071 | -0.071 | 1.124 | 0.38 | 0.14 | 0.31 | 0.43 | 0.36 | 0.38 |
| 3 | 111996554 | rs9828502 | 0.605 | -0.444 | 0.174 | 0.057 | -0.003 | 0.005 | 0.005 | 0 | 0.26 | 0.31 | 0.4 | 0.3 | 0.23 | 0.34 |
| 3 | 111996726 | rs6791182 | -0.678 | -0.206 | 0.281 | 0.004 | -0.003 | 0.012 | 0.012 | 0 | 0.16 | 0.19 | 0.14 | 0.21 | 0.14 | 0.23 |
| 3 | 112000812 | rs7626450 | 0.721 | -0.219 | 0.227 | -0.018 | -0.003 | -0.027 | -0.027 | 0.01 | 0.06 | 0.07 | 0.04 | 0 | 0.05 | 0.1 |
| 3 | 112004516 | rs10934163 | 0.057 | -0.212 | 0.255 | -0.145 | -0.001 | -0.078 | -0.078 | 0 | 0.69 | 0.68 | 0.76 | 1 | 0.66 | 0.78 |
| 3 | 112006172 | rs4682392 | 0.434 | -0.24 | 0.295 | 0.221 | 0.008 | 0.023 | 0.023 | 0.139 | 0.24 | 0.4 | 0.35 | 0.31 | 0.25 | 0.33 |
| 3 | 112008124 | rs11923778 | -0.295 | -0.215 | 0.278 | -0.067 | -0.002 | 0.033 | 0.033 | 0.069 | 0.08 | 0.1 | 0.08 | 0.21 | 0.08 | 0.1 |
| 3 | 112013546 | rs12491414 | -0.18 | -0.003 | 0.378 | -0.612 | 0.019 | -0.039 | -0.039 | 1.764 | 0.07 | 0.16 | 0.08 | 0 | 0.09 | 0.11 |
| 3 | 112015292 | rs7636107 | 0.291 | -0.14 | 0.382 | -0.911 | 0.004 | -0.095 | -0.095 | 0 | 0.69 | 0.62 | 0.75 | 1 | 0.66 | 0.78 |
| 3 | 112020317 | rs6795820 | -0.177 | -0.326 | 0.323 | -0.847 | -0.01 | -0.01 | -0.01 | 0.091 | 0.24 | 0.21 | 0.23 | 0 | 0.3 | 0.14 |
| 3 | 112022024 | rs1317295 | -0.008 | -0.328 | 0.284 | -0.848 | -0.029 | 0.04 | 0.04 | 0 | 0.23 | 0.14 | 0.22 | 0 | 0.3 | 0.14 |
| 3 | 112025858 | rs9818526 | -0.649 | -0.314 | 0.394 | -0.771 | 0.004 | -0.005 | -0.005 | 0.078 | 0.94 | 0.91 | 0.88 | 0.97 | 0.93 | 0.85 |
| 3 | 112026120 | rs6773206 | 1.331 | -0.729 | 0.763 | -0.375 | 0.001 | -0.001 | -0.001 | 2.529 | 0.48 | 0.4 | 0.38 | 0.51 | 0.46 | 0.44 |
| 3 | 112033979 | rs6771014 | -0.957 | -0.662 | 0.609 | -0.336 | -0.002 | 0 | 0 | 0 | 0.95 | 0.95 | 0.88 | 0.97 | 0.93 | 0.85 |
| 3 | 112043158 | rs2895395 | 0.518 | -0.79 | 0.839 | -0.25 | -0.005 | 0.018 | 0.018 | 0 | 0.22 | 0.27 | 0.26 | 0.44 | 0.16 | 0.26 |
| 3 | 112053246 | rs1317243 | 1.301 | -0.743 | 1.022 | -0.307 | 0.004 | -0.006 | -0.006 | 0 | 0.48 | 0.28 | 0.38 | 0.51 | 0.47 | 0.49 |
| 3 | 112053565 | rs1317244 | -0.609 | -1.029 | 0.899 | -0.372 | 0.004 | 0.005 | 0.005 | 0 | 0.32 | 0.34 | 0.31 | 0.36 | 0.34 | 0.49 |
| 3 | 112055793 | rs4682103 | -1.404 | -0.969 | 0.693 | -0.418 | -0.003 | 0.024 | 0.024 | 0 | 0.55 | 0.61 | 0.58 | 0.81 | 0.51 | 0.77 |
| 3 | 112059213 | rs2399416 | 2.02 | -0.997 | 0.424 | -0.41 | -0.04 | 0.155 | 0.155 | 0 | 0.35 | 0.1 | 0.27 | 0.15 | 0.39 | 0.14 |

| | | | | | | | | | | | | | | | | |
|---|---|---|---|---|---|---|---|---|---|---|---|---|---|---|---|---|
| 3 | 112060256 | rs16859476 | 0.85 | -0.904 | 0.718 | -0.124 | -0.004 | 0.067 | 0.067 | 0 | 0.11 | 0.17 | 0.05 | 0.26 | 0.07 | 0.29 |
| 3 | 112061731 | rs12106673 | 1.376 | -1.077 | 0.699 | -0.209 | -0.001 | 0.008 | 0.008 | 0 | 0.45 | 0.37 | 0.34 | 0.4 | 0.47 | 0.43 |
| 3 | 112063850 | rs2272022 | 2.175 | -1.029 | 0.469 | -0.204 | -0.042 | 0.137 | 0.137 | 0 | 0.34 | 0.12 | 0.27 | 0.14 | 0.39 | 0.14 |
| 3 | 112065737 | rs2280462 | -0.967 | -0.892 | 0.621 | -0.155 | -0.001 | 0.004 | 0.004 | 0 | 0.91 | 0.92 | 0.88 | 0.97 | 0.9 | 0.9 |
| 3 | 112066360 | rs7612748 | 0.562 | -0.919 | 0.677 | -0.217 | -0.017 | 0.003 | 0.003 | 0 | 0.19 | 0.14 | 0.21 | 0 | 0.27 | 0.15 |
| 3 | 112066562 | rs1050572 | 0.509 | -0.94 | 0.702 | -0.175 | -0.009 | 0.112 | 0.112 | 0.614 | 0.13 | 0.31 | 0.07 | 0.26 | 0.07 | 0.29 |
| 3 | 112069765 | rs2399420 | -0.854 | -0.879 | 0.466 | -0.267 | -0.001 | -0.016 | -0.016 | 0.993 | 0.9 | 0.95 | 0.83 | 0.86 | 0.9 | 0.85 |
| 3 | 112071934 | rs2399422 | 0.382 | -0.743 | 0.61 | -0.315 | 0.031 | -0.072 | -0.072 | 8.387 | 0.2 | 0.29 | 0.23 | 0.01 | 0.27 | 0.15 |
| 3 | 112072833 | rs13093633 | 0.995 | -0.468 | 0.709 | -0.168 | 0.014 | 0.017 | 0.017 | 0.813 | 0.54 | 0.41 | 0.5 | 0.09 | 0.66 | 0.29 |
| 3 | 112074608 | rs9990220 | -1.12 | -0.043 | 1.022 | 0.307 | 0.015 | 0.018 | 0.018 | 0 | 0.38 | 0.46 | 0.48 | 0.46 | 0.42 | 0.42 |
| 3 | 112074977 | rs9863371 | 0.017 | -0.253 | 0.773 | 0.632 | -0.007 | 0.027 | 0.027 | 0.215 | 0.21 | 0.23 | 0.27 | 0.45 | 0.16 | 0.28 |
| 3 | 112077880 | rs17447803 | 0.094 | 0.326 | 0.703 | 0.291 | -0.001 | -0.002 | -0.002 | 1.697 | 0.19 | 0.17 | 0.24 | 0.05 | 0.28 | 0.17 |
| 3 | 112079277 | rs9867544 | -0.978 | -0.084 | 0.735 | 0.376 | 0.003 | 0.013 | 0.013 | 0 | 0.48 | 0.62 | 0.5 | 0.57 | 0.43 | 0.49 |
| 3 | 112083859 | rs4682401 | -0.279 | -0.032 | 0.496 | 0.517 | 0.027 | -0.041 | -0.041 | 0 | 0.8 | 0.64 | 0.75 | 0.94 | 0.75 | 0.8 |
| 3 | 112090292 | rs10934170 | 0.424 | -0.133 | 0.423 | 0.527 | 0.013 | -0.016 | -0.016 | 0 | 0.23 | 0.17 | 0.18 | 0.39 | 0.14 | 0.23 |
| 3 | 112091696 | rs7641311 | 0.799 | 0.183 | 0.045 | 0.242 | 0.012 | 0.032 | 0.032 | 0 | 0.52 | 0.28 | 0.44 | 0.41 | 0.6 | 0.5 |
| 3 | 112093900 | rs1879512 | 0.899 | 0.24 | 0.069 | 0.266 | -0.005 | 0.041 | 0.041 | 0 | 0.5 | 0.27 | 0.39 | 0.41 | 0.53 | 0.46 |
| 3 | 112094289 | rs2399426 | 0.383 | 1.161 | 0.348 | -0.034 | 0.029 | -0.031 | -0.031 | 2.189 | 0.8 | 0.63 | 0.68 | 0.92 | 0.74 | 0.81 |
| 3 | 112103140 | rs6438069 | -1.527 | 0.45 | 0.532 | 0.204 | 0.002 | 0.011 | 0.011 | 0 | 0.08 | 0.22 | 0.11 | 0.08 | 0.03 | 0.04 |
| 3 | 112104875 | rs1032239 | 0.179 | 0.217 | -0.024 | -0.024 | 0.069 | 0.106 | 0.106 | 0 | 0.51 | 0.23 | 0.47 | 0.26 | 0.62 | 0.45 |
| 3 | 112105769 | rs1386897 | -0.754 | 0.161 | 0.26 | -0.193 | -0.003 | -0.023 | -0.023 | 0 | 0.94 | 0.93 | 0.98 | 1 | 0.96 | 0.99 |
| 3 | 112110022 | rs2683904 | -0.486 | 0.124 | 0.27 | -0.209 | -0.003 | -0.022 | -0.022 | 0 | 0.94 | 0.92 | 0.98 | 1 | 0.96 | 0.99 |
| 3 | 112112749 | rs2683900 | 0.953 | 0.76 | 0.464 | -0.535 | -0.038 | 0.051 | 0.051 | 0 | 0.41 | 0.16 | 0.35 | 0.27 | 0.46 | 0.3 |
| 3 | 112113334 | rs11720514 | -0.253 | 0.754 | 0.572 | -0.685 | 0.039 | -0.056 | -0.056 | 0 | 0.13 | 0.21 | 0.15 | 0 | 0.22 | 0.1 |
| 3 | 112116643 | rs4682404 | 0.742 | 0.26 | 0.803 | -0.603 | 0.06 | -0.048 | -0.048 | 0.063 | 0.31 | 0.13 | 0.34 | 0.64 | 0.22 | 0.49 |
| 3 | 112118529 | rs9878780 | 0.019 | 0.282 | 1.058 | -0.596 | -0.036 | -0.019 | -0.019 | 1.562 | 0.39 | 0.48 | 0.47 | 0.73 | 0.28 | 0.59 |
| 3 | 112127280 | rs999802 | 1.179 | 0.382 | 1.141 | -0.713 | 0.018 | -0.045 | -0.045 | 0 | 0.77 | 0.38 | 0.75 | 0.91 | 0.75 | 0.89 |
| 3 | 112127559 | rs11708711 | 1.104 | 0.04 | 0.415 | -0.278 | 0.019 | -0.138 | -0.138 | 0 | 0.24 | 0.04 | 0.18 | 0.49 | 0.18 | 0.35 |
| 3 | 112128143 | rs1872987 | -0.077 | 0.536 | 0.973 | -0.971 | -0.007 | -0.038 | -0.038 | 0 | 0.93 | 0.83 | 0.97 | 1 | 0.96 | 0.99 |
| 3 | 112144026 | rs9847446 | 1.087 | 0.634 | 1.048 | -0.98 | 0.023 | -0.041 | -0.041 | 0 | 0.84 | 0.71 | 0.81 | 0.95 | 0.82 | 0.93 |
| 3 | 112144152 | rs2633546 | -0.004 | 0.703 | 0.835 | -0.945 | 0.001 | -0.007 | -0.007 | 0 | 0.07 | 0.05 | 0.1 | 0 | 0.13 | 0.06 |
| 3 | 112156632 | rs2705537 | -0.448 | 0.56 | 0.736 | -0.864 | 0.014 | -0.037 | -0.037 | 0 | 0.91 | 0.87 | 0.96 | 0.99 | 0.91 | 0.98 |
| 3 | 112164105 | rs1488265 | 0.306 | 0.555 | 0.618 | -0.848 | 0.037 | -0.035 | -0.035 | 0 | 0.87 | 0.65 | 0.87 | 0.95 | 0.89 | 0.97 |
| 3 | 112164278 | rs1488266 | 0.362 | 0.552 | 0.404 | -0.849 | 0.044 | -0.041 | -0.041 | 0.164 | 0.86 | 0.6 | 0.86 | 0.95 | 0.88 | 0.97 |
| 3 | 112165339 | rs13090873 | 0.899 | 0.04 | -0.869 | -0.699 | -0.12 | 0.016 | 0.016 | 0 | 0.34 | 0.05 | 0.38 | 0.29 | 0.41 | 0.26 |
| 3 | 112166582 | rs2633611 | -0.337 | -0.831 | 0.035 | 0.181 | 0.022 | 0.009 | 0.009 | 0 | 0.37 | 0.27 | 0.32 | 0.66 | 0.25 | 0.57 |
| 3 | 112171438 | rs2633568 | -0.396 | -0.591 | -0.063 | 0.071 | -0.021 | -0.017 | -0.017 | 0 | 0.37 | 0.41 | 0.39 | 0.71 | 0.24 | 0.58 |
| 3 | 112182740 | rs1982809 | -0.364 | -0.624 | -0.416 | 0.043 | -0.001 | -0.002 | -0.002 | 0 | 0.35 | 0.31 | 0.43 | 0.79 | 0.23 | 0.41 |
| 3 | 112184927 | rs2171513 | 1.137 | -1.205 | -0.296 | 0.442 | -0.015 | -0.001 | -0.001 | 0 | 0.81 | 0.24 | 0.81 | 0.78 | 0.96 | 0.8 |
| 3 | 112185025 | rs9288952 | 1.33 | -1.334 | -0.313 | 0.647 | 0.044 | -0.019 | -0.019 | 0 | 0.8 | 0.18 | 0.75 | 0.72 | 0.96 | 0.8 |
| 3 | 112196075 | rs6438079 | 1.296 | -1.304 | 0.4 | 0.651 | -0.052 | 0.023 | 0.023 | 2.073 | 0.83 | 0.28 | 0.75 | 0.73 | 0.96 | 0.82 |
| 3 | 112196431 | rs2633562 | -1.087 | -1.325 | 1.517 | 0.625 | -0.044 | 0.082 | 0.082 | 0.011 | 0.12 | 0.41 | 0.16 | 0.2 | 0.04 | 0.12 |
| 3 | 112220781 | rs2633578 | -2.088 | -1.233 | 1.303 | 0.486 | -0.057 | 0.032 | 0.032 | 0.854 | 0.19 | 0.62 | 0.27 | 0.25 | 0.07 | 0.19 |
| 3 | 112225934 | rs923349 | 0.116 | -0.571 | 0.648 | 0.283 | -0.005 | 0.001 | 0.001 | 0 | 0.09 | 0.05 | 0.16 | 0.05 | 0.1 | 0.25 |
| 3 | 112235371 | rs4682413 | 1.627 | -0.937 | 0.654 | 0.308 | -0.031 | 0.003 | 0.003 | 1.441 | 0.78 | 0.29 | 0.73 | 0.74 | 0.92 | 0.81 |
| 3 | 112250419 | rs2131008 | -1.911 | -0.953 | 0.77 | 0.301 | -0.052 | 0.021 | 0.021 | 1.369 | 0.19 | 0.66 | 0.25 | 0.25 | 0.07 | 0.17 |
| 3 | 112251775 | rs9855848 | 0.807 | -0.613 | -0.098 | -0.047 | -0.065 | -0.157 | -0.157 | 0 | 0.28 | 0.03 | 0.31 | 0.55 | 0.32 | 0.25 |
| 3 | 112252390 | rs2638037 | -1.385 | -0.625 | 0.37 | 0.042 | -0.058 | 0.038 | 0.038 | 1.095 | 0.18 | 0.59 | 0.25 | 0.25 | 0.07 | 0.17 |
| 3 | 112256630 | rs2969896 | -0.603 | -0.653 | 0.535 | 0.054 | -0.056 | 0.112 | 0.112 | 0.587 | 0.12 | 0.48 | 0.12 | 0.2 | 0.03 | 0.11 |
| 3 | 112262957 | rs3736271 | -0.411 | -0.648 | 0.514 | 0.054 | 0.002 | -0.004 | -0.004 | 0.004 | 0.06 | 0.11 | 0.13 | 0.04 | 0.04 | 0.06 |
| 3 | 112264067 | rs13071492 | -0.411 | -0.648 | 0.514 | 0.054 | 0.002 | -0.004 | -0.004 | 0 | 0.06 | 0.11 | 0.13 | 0.04 | 0.04 | 0.06 |
| 3 | 112265776 | rs2638022 | -1.707 | -0.646 | 0.475 | 0.055 | -0.052 | 0.023 | 0.023 | 1.553 | 0.19 | 0.66 | 0.25 | 0.25 | 0.07 | 0.17 |
| 3 | 112289372 | rs9869579 | 1.555 | -0.659 | 0.452 | 0.091 | -0.032 | 0.01 | 0.01 | 1.408 | 0.79 | 0.28 | 0.74 | 0.74 | 0.92 | 0.82 |
| 3 | 112304526 | rs1994711 | 1.204 | -0.473 | 0.368 | 0.067 | 0.008 | -0.005 | -0.005 | 0 | 0.92 | 0.77 | 0.86 | 0.94 | 0.95 | 0.92 |
| 3 | 112329334 | rs9868112 | 0.549 | 0.138 | -0.264 | -0.126 | -0.1 | -0.142 | -0.142 | 0 | 0.32 | 0.06 | 0.34 | 0.55 | 0.38 | 0.25 |
| 3 | 112332629 | rs4340680 | 0.606 | 0.034 | 0.062 | -0.281 | 0.008 | 0.09 | 0.09 | 0 | 0.5 | 0.29 | 0.4 | 0.2 | 0.54 | 0.56 |
| 3 | 112335039 | rs4290780 | 1.757 | 0.314 | 0.77 | -0.17 | 0.028 | -0.017 | -0.017 | 0 | 0.87 | 0.76 | 0.84 | 0.92 | 0.88 | 0.89 |
| 3 | 112337467 | rs3817188 | 0.916 | -0.098 | 0.296 | -0.446 | 0.004 | 0.128 | 0.128 | 0 | 0.34 | 0.12 | 0.19 | 0.14 | 0.35 | 0.28 |
| 3 | 112338027 | rs13084615 | 0.401 | -0.139 | 0.762 | -0.173 | 0.026 | 0.068 | 0.068 | 0.419 | 0.84 | 0.54 | 0.73 | 0.8 | 0.87 | 0.69 |
| 3 | 112339845 | rs16859850 | -1.203 | -0.105 | 1.063 | -0.122 | -0.008 | 0.114 | 0.114 | 1.041 | 0.06 | 0.3 | 0.11 | 0.13 | 0.01 | 0.04 |
| 3 | 112341941 | rs9870432 | 0.622 | -0.219 | 0.921 | -0.071 | -0.057 | -0.026 | -0.026 | 0.024 | 0.41 | 0.17 | 0.4 | 0.56 | 0.49 | 0.3 |
| 3 | 112345995 | rs7349538 | 0.633 | -0.208 | 0.741 | -0.074 | -0.133 | -0.084 | -0.084 | 0 | 0.4 | 0.08 | 0.4 | 0.56 | 0.49 | 0.31 |
| 3 | 112347079 | rs10511316 | 1.163 | -0.231 | 0.796 | -0.103 | 0.001 | 0.131 | 0.131 | 0 | 0.34 | 0.1 | 0.19 | 0.13 | 0.35 | 0.3 |
| 3 | 112350156 | rs4682417 | 0.974 | -0.237 | 0.925 | -0.081 | 0.017 | 0.169 | 0.169 | 0.915 | 0.94 | 0.62 | 0.85 | 0.88 | 0.99 | 0.91 |
| 3 | 112350480 | rs7648759 | 1.146 | -0.191 | 0.782 | -0.117 | 0.001 | 0.131 | 0.131 | 0 | 0.34 | 0.1 | 0.19 | 0.13 | 0.35 | 0.3 |
| 3 | 112351948 | rs16859875 | 0.483 | -0.285 | 1.089 | 0.11 | 0.026 | -0.008 | -0.008 | 0 | 0.09 | 0.12 | 0.11 | 0.06 | 0.11 | 0.24 |
| 3 | 112353238 | rs981181 | 1.231 | -0.325 | 1.005 | 0.109 | -0.007 | 0.056 | 0.056 | 0.538 | 0.77 | 0.48 | 0.61 | 0.69 | 0.83 | 0.59 |
| 3 | 112355977 | rs6794305 | 0.856 | -0.553 | 0.573 | -0.043 | 0.002 | 0.085 | 0.085 | 0 | 0.31 | 0.1 | 0.26 | 0.14 | 0.33 | 0.21 |
| 3 | 112359926 | rs6807798 | 0.801 | -0.582 | 0.572 | -0.019 | 0.022 | 0.093 | 0.093 | 0 | 0.32 | 0.1 | 0.26 | 0.14 | 0.32 | 0.21 |

| | | | | | | | | | | | | | | | | |
|---|---|---|---|---|---|---|---|---|---|---|---|---|---|---|---|---|
| 3 | 112359986 | rs6438086 | -0.544 | -0.536 | 1.29 | -0.05 | 0.004 | 0.003 | 0.003 | 0 | 0.64 | 0.64 | 0.7 | 0.86 | 0.64 | 0.76 |
| 3 | 112365127 | rs9815819 | -0.135 | -0.496 | 1.73 | 0.879 | 0.003 | -0.003 | -0.003 | 0 | 0.59 | 0.37 | 0.68 | 0.86 | 0.62 | 0.71 |
| 3 | 112371578 | rs7620601 | 0.833 | -0.569 | 1.344 | 1.212 | -0.051 | 0.001 | 0.001 | 0.067 | 0.43 | 0.06 | 0.35 | 0.34 | 0.53 | 0.42 |
| 3 | 112372317 | rs9870933 | -0.014 | -0.491 | 1.983 | 1.004 | 0.015 | -0.005 | -0.005 | 0 | 0.58 | 0.35 | 0.57 | 0.52 | 0.59 | 0.72 |
| 3 | 112372965 | rs9683135 | -0.178 | -0.509 | 2.191 | 0.874 | -0.017 | -0.054 | -0.054 | 0 | 0.65 | 0.43 | 0.73 | 0.86 | 0.68 | 0.77 |
| 3 | 112375089 | rs9821100 | -0.319 | -0.415 | 2.406 | 0.948 | 0.011 | 0.023 | 0.023 | 0.072 | 0.39 | 0.7 | 0.31 | 0.12 | 0.36 | 0.24 |
| 3 | 112375450 | rs9821338 | 0.392 | -0.409 | 2.496 | 0.95 | 0.011 | 0.022 | 0.022 | 0 | 0.33 | 0.29 | 0.26 | 0.14 | 0.31 | 0.2 |
| 3 | 112375793 | rs9859566 | 0.248 | -0.399 | 2.489 | 1.036 | -0.011 | -0.028 | -0.028 | 0 | 0.34 | 0.37 | 0.27 | 0.16 | 0.31 | 0.21 |
| 3 | 112376308 | rs12634229 | -1.757 | -0.361 | 2.379 | 1.008 | 0.075 | 0.313 | 0.313 | 0.094 | 0.05 | 0.3 | 0.13 | 0.33 | 0.06 | 0.05 |
| 3 | 112394340 | rs9818848 | 0.518 | -0.449 | 1.281 | 0.883 | -0.009 | 0.147 | 0.147 | 0 | 0.32 | 0.12 | 0.26 | 0.06 | 0.34 | 0.22 |
| 3 | 112396247 | rs9876497 | 0.365 | -0.402 | 1.356 | 0.905 | 0.01 | 0.053 | 0.053 | 0 | 0.33 | 0.25 | 0.27 | 0.06 | 0.34 | 0.21 |
| 3 | 112400174 | rs2399449 | 0.177 | -0.258 | 0.565 | 1.037 | 0.027 | -0.028 | -0.028 | 0 | 0.58 | 0.11 | 0.59 | 0.56 | 0.62 | 0.72 |
| 3 | 112406563 | rs820544 | -0.247 | -0.28 | -0.031 | 1.048 | 0.032 | 0.001 | 0.001 | 0.335 | 0.41 | 0.79 | 0.39 | 0.44 | 0.36 | 0.28 |
| 3 | 112407697 | rs17457593 | 0.166 | -0.254 | -0.653 | 0.929 | 0.044 | 0.013 | 0.013 | 0.903 | 0.58 | 0.11 | 0.6 | 0.59 | 0.64 | 0.72 |
| 3 | 112434931 | rs7627205 | 1.328 | -0.797 | 0.113 | 1.393 | -0.062 | 0.129 | 0.129 | 0 | 0.83 | 0.54 | 0.93 | 0.61 | 0.92 | 0.88 |
| 3 | 112436847 | rs3846043 | 0.405 | -0.602 | -0.562 | 0.867 | -0.003 | 0.033 | 0.033 | 0.069 | 0.37 | 0.3 | 0.37 | 0.18 | 0.37 | 0.24 |
| 3 | 112440393 | rs9821598 | -0.289 | -0.466 | -0.601 | 0.96 | -0.018 | -0.028 | -0.028 | 0 | 0.95 | 0.82 | 0.98 | 1 | 0.98 | 0.95 |
| 3 | 112440948 | rs9863500 | 0.488 | -0.786 | -0.586 | 0.877 | -0.029 | 0.003 | 0.003 | 0 | 0.37 | 0.24 | 0.5 | 0.7 | 0.48 | 0.43 |
| 3 | 112441312 | rs9826586 | 0.099 | -0.871 | -0.502 | 0.931 | -0.002 | 0.007 | 0.007 | 0 | 0.53 | 0.52 | 0.45 | 0.3 | 0.47 | 0.51 |
| 3 | 112453129 | rs6438101 | 0.371 | -0.504 | -0.886 | 0.842 | -0.039 | -0.007 | -0.007 | 0 | 0.37 | 0.23 | 0.53 | 0.37 | 0.5 | 0.43 |
| 3 | 112457920 | rs6800139 | 0.228 | -1.074 | -1.212 | 1.024 | 0.048 | 0.039 | 0.039 | 0.133 | 0.52 | 0.23 | 0.61 | 0.38 | 0.65 | 0.47 |
| 3 | 112460195 | rs717706 | 0.659 | -0.911 | -1.153 | 0.915 | -0.059 | 0.073 | 0.073 | 0 | 0.44 | 0.12 | 0.47 | 0.28 | 0.54 | 0.39 |
| 3 | 112463791 | rs9850350 | 0.832 | -1.034 | -1.353 | 0.731 | -0.076 | 0.042 | 0.042 | 0 | 0.44 | 0.14 | 0.46 | 0.3 | 0.51 | 0.38 |
| 3 | 112467307 | rs1387020 | 0.441 | -1.352 | -0.507 | 1.213 | -0.001 | 0.004 | 0.004 | 0.133 | 0.36 | 0.46 | 0.33 | 0.36 | 0.3 | 0.4 |
| 3 | 112469992 | rs820554 | 0.049 | -1.347 | -0.54 | 1.245 | -0.006 | 0.038 | 0.038 | 0 | 0.93 | 0.9 | 0.98 | 0.76 | 0.95 | 0.88 |
| 3 | 112470586 | rs820553 | 1.313 | -1.679 | -0.349 | 1.32 | -0.015 | 0.015 | 0.015 | 0.697 | 0.24 | 0.16 | 0.28 | 0.16 | 0.22 | 0.13 |
| 3 | 112474891 | rs3912626 | -0.187 | -1.42 | -0.634 | 1.228 | 0.008 | 0.025 | 0.025 | 0 | 0.08 | 0.03 | 0.07 | 0.02 | 0.09 | 0.07 |
| 3 | 112478401 | rs6784132 | 1.215 | -1.446 | -0.655 | 1.231 | -0.076 | 0.046 | 0.046 | 0 | 0.44 | 0.14 | 0.46 | 0.3 | 0.51 | 0.37 |
| 3 | 112478591 | rs4682421 | -1.89 | -1.678 | -0.494 | 1.076 | -0.021 | -0.026 | -0.026 | 0 | 0.15 | 0.26 | 0.18 | 0.11 | 0.14 | 0.19 |
| 3 | 112479872 | rs712527 | -0.723 | -1.556 | -0.535 | 1.148 | 0 | 0.001 | 0.001 | 0 | 0.45 | 0.56 | 0.41 | 0.37 | 0.4 | 0.48 |
| 3 | 112486737 | rs1154278 | 0.12 | -1.324 | -0.246 | 1.122 | 0.003 | -0.005 | -0.005 | 0.1 | 0.36 | 0.39 | 0.35 | 0.31 | 0.34 | 0.39 |
| 3 | 112496539 | rs1249895 | -1.306 | -1.528 | -0.397 | 1.165 | -0.022 | 0.03 | 0.03 | 0 | 0.55 | 0.78 | 0.51 | 0.66 | 0.47 | 0.64 |
| 3 | 112501357 | rs4682426 | -0.449 | -1.533 | -0.187 | 1.222 | 0.01 | -0.005 | -0.005 | 0 | 0.46 | 0.64 | 0.33 | 0.55 | 0.4 | 0.51 |
| 3 | 112501790 | rs7648520 | -0.15 | -1.466 | -0.154 | 1.248 | -0.01 | 0.005 | 0.005 | 0 | 0.09 | 0.13 | 0.18 | 0.1 | 0.07 | 0.06 |
| 3 | 112505650 | rs6781211 | -0.926 | -1.425 | -0.18 | 1.343 | -0.029 | 0.092 | 0.092 | 0 | 0.1 | 0.27 | 0.04 | 0.26 | 0.06 | 0.16 |
| 3 | 112507801 | rs7611423 | 0.891 | -1.14 | -0.542 | 1.257 | -0.014 | 0.033 | 0.033 | 0 | 0.46 | 0.24 | 0.49 | 0.35 | 0.53 | 0.43 |
| 3 | 112511598 | rs9862107 | -0.256 | -0.849 | -0.081 | 1.113 | -0.021 | -0.001 | -0.001 | 0 | 0.39 | 0.43 | 0.26 | 0.31 | 0.3 | 0.37 |
| 3 | 112513358 | rs10804503 | 1.296 | -0.632 | -0.418 | 1.079 | -0.074 | 0.015 | 0.015 | 0 | 0.37 | 0.15 | 0.44 | 0.33 | 0.43 | 0.42 |
| 3 | 112513482 | rs10934187 | 0.345 | -0.633 | 0.057 | 1.182 | 0.005 | 0.01 | 0.01 | 0 | 0.49 | 0.42 | 0.5 | 0.6 | 0.51 | 0.64 |
| 3 | 112516328 | rs6789210 | 0.552 | -0.631 | -0.177 | 1.2 | -0.007 | 0.006 | 0.006 | 0 | 0.47 | 0.3 | 0.49 | 0.6 | 0.5 | 0.58 |
| 3 | 112519276 | rs1352425 | 0.758 | -0.558 | 0.234 | 1.435 | -0.046 | 0.136 | 0.136 | 0 | 0.76 | 0.52 | 0.86 | 0.54 | 0.82 | 0.7 |
| 3 | 112520828 | rs1994509 | 0.512 | -0.453 | -0.004 | 1.332 | 0.003 | -0.012 | -0.012 | 0 | 0.79 | 0.81 | 0.74 | 0.69 | 0.79 | 0.69 |
| 3 | 112521973 | rs4682429 | -0.447 | -0.522 | 0.085 | 1.047 | -0.009 | 0.081 | 0.081 | 0 | 0.44 | 0.28 | 0.34 | 0.1 | 0.44 | 0.38 |
| 3 | 112526784 | rs9941979 | -0.459 | -0.534 | 0.05 | 1.116 | -0.007 | 0.079 | 0.079 | 0 | 0.44 | 0.28 | 0.35 | 0.15 | 0.44 | 0.32 |
| 3 | 112528969 | rs1387015 | 1.13 | -0.78 | 0.429 | 1.174 | -0.001 | -0.001 | -0.001 | 0 | 0.73 | 0.29 | 0.69 | 0.68 | 0.78 | 0.56 |
| 3 | 112534069 | rs4682434 | 1.667 | -0.789 | 0.879 | 1.346 | 0.035 | -0.041 | -0.041 | 0 | 0.68 | 0.21 | 0.67 | 0.49 | 0.75 | 0.62 |
| 3 | 112535057 | rs11918522 | -0.663 | -0.735 | 1.253 | 1.163 | -0.032 | -0.061 | -0.061 | 1.602 | 0.2 | 0.4 | 0.25 | 0.12 | 0.16 | 0.16 |
| 3 | 112536584 | rs6773083 | 1.053 | -0.699 | 1.216 | 1.214 | -0.036 | 0.03 | 0.03 | 0 | 0.75 | 0.42 | 0.68 | 0.7 | 0.76 | 0.67 |
| 3 | 112544367 | rs16860127 | -0.81 | -0.649 | 1.381 | 0.968 | -0.025 | -0.071 | -0.071 | 0.075 | 0.2 | 0.41 | 0.28 | 0.1 | 0.18 | 0.28 |
| 3 | 112544750 | rs13094280 | 1.562 | -0.516 | 0.874 | 0.884 | -0.067 | 0.001 | 0.001 | 0 | 0.38 | 0.09 | 0.37 | 0.38 | 0.4 | 0.31 |
| 3 | 112547601 | rs1387024 | -0.747 | -0.465 | 0.912 | 0.868 | -0.015 | -0.017 | -0.017 | 0 | 0.68 | 0.79 | 0.69 | 0.7 | 0.61 | 0.71 |
| 3 | 112554871 | rs4682435 | -1.533 | -0.471 | 1.06 | 0.899 | 0.002 | 0.005 | 0.005 | 0 | 0.06 | 0.07 | 0.04 | 0.2 | 0.05 | 0.16 |
| 3 | 112556393 | rs16860145 | -1.314 | -0.471 | 1.062 | 0.914 | -0.001 | 0.006 | 0.006 | 0 | 0.1 | 0.14 | 0.06 | 0.2 | 0.08 | 0.19 |
| 3 | 112557954 | rs7642984 | -0.306 | -0.553 | 1.267 | 0.88 | -0.036 | -0.056 | -0.056 | 0 | 0.18 | 0.43 | 0.23 | 0.1 | 0.13 | 0.21 |
| 3 | 112562919 | rs1114404 | 0.915 | -0.549 | 1.175 | 0.922 | -0.013 | 0.011 | 0.011 | 0.007 | 0.47 | 0.28 | 0.41 | 0.59 | 0.44 | 0.37 |
| 3 | 112566491 | rs12489734 | 1.591 | -0.499 | 1.142 | 0.847 | -0.021 | -0.01 | -0.01 | 2.516 | 0.35 | 0.18 | 0.35 | 0.39 | 0.34 | 0.31 |
| 3 | 112567789 | rs12233479 | -0.339 | -0.479 | 1.141 | 0.863 | -0.002 | 0.025 | 0.025 | 0 | 0.07 | 0.1 | 0.02 | 0.2 | 0.04 | 0.06 |
| 3 | 112597706 | rs4682123 | 0.075 | -0.552 | 1.467 | 1.086 | -0.02 | 0.018 | 0.018 | 0 | 0.15 | 0.38 | 0.12 | 0.19 | 0.13 | 0.22 |
| 3 | 112610825 | rs7641119 | 1.248 | -0.505 | 0.964 | 1.081 | -0.015 | -0.02 | -0.02 | 0 | 0.38 | 0.21 | 0.35 | 0.42 | 0.35 | 0.39 |
| 3 | 112616037 | rs6798395 | 1.553 | -0.487 | 0.978 | 0.98 | -0.017 | 0.04 | 0.04 | 0 | 0.35 | 0.21 | 0.35 | 0.22 | 0.34 | 0.3 |
| 3 | 112628205 | rs9818564 | -0.092 | -0.466 | 0.643 | 1.034 | -0.005 | -0.017 | -0.017 | 0.012 | 0.58 | 0.74 | 0.63 | 0.5 | 0.53 | 0.59 |
| 3 | 112641046 | rs9879005 | 0.101 | -0.18 | 0.744 | 0.935 | -0.008 | 0.006 | 0.006 | 0 | 0.54 | 0.7 | 0.59 | 0.48 | 0.5 | 0.54 |
| 3 | 112642122 | rs16860233 | -0.901 | -0.14 | 0.711 | 1.022 | 0.001 | -0.004 | -0.004 | 0 | 0.06 | 0.02 | 0.02 | 0.23 | 0.03 | 0.12 |
| 3 | 112642209 | rs16860236 | -0.901 | -0.14 | 0.732 | 1.022 | 0.001 | 0 | 0 | 0.102 | 0.06 | 0.02 | 0.02 | 0.23 | 0.03 | 0.12 |
| 3 | 112643087 | rs16860242 | 0.777 | -0.249 | 1.126 | 0.985 | -0.018 | 0.022 | 0.022 | 0 | 0.12 | 0.21 | 0.07 | 0.21 | 0.09 | 0.17 |
| 3 | 112643560 | rs9870568 | -0.221 | -0.206 | 0.984 | 0.968 | -0.009 | 0.014 | 0.014 | 0 | 0.6 | 0.72 | 0.61 | 0.7 | 0.53 | 0.66 |
| 3 | 112661354 | rs9871985 | -0.293 | -0.216 | 0.928 | 0.963 | -0.004 | 0.016 | 0.016 | 0 | 0.6 | 0.73 | 0.61 | 0.71 | 0.55 | 0.7 |
| 3 | 112665551 | rs9847757 | 0.259 | -0.25 | 0.995 | 0.98 | -0.031 | 0.038 | 0.038 | 0.31 | 0.14 | 0.33 | 0.08 | 0.21 | 0.09 | 0.18 |
| 3 | 112665968 | rs6438120 | -0.488 | -0.19 | 0.291 | 0.886 | -0.024 | 0.003 | 0.003 | 0 | 0.63 | 0.81 | 0.63 | 0.71 | 0.55 | 0.72 |

| | | | | | | | | | | | | | | | | |
|---|---|---|---|---|---|---|---|---|---|---|---|---|---|---|---|---|
| 3 | 112666859 | rs2133946 | 1.48 | -0.087 | 0.259 | 1.036 | -0.124 | 0.022 | 0.022 | 0.393 | 0.34 | 0.02 | 0.32 | 0.22 | 0.36 | 0.32 |
| 3 | 112705874 | rs984706 | 0.228 | 0.473 | 0.743 | 1.064 | -0.002 | -0.008 | -0.008 | 0.112 | 0.83 | 0.81 | 0.78 | 0.72 | 0.78 | 0.84 |
| 3 | 112718063 | rs10460813 | -0.429 | 0.349 | 1.066 | 1.198 | 0 | -0.01 | -0.01 | 0 | 0.87 | 0.87 | 0.81 | 0.73 | 0.82 | 0.87 |
| 3 | 112726479 | rs9844028 | 1.593 | 0.308 | 0.357 | 0.911 | -0.103 | 0.055 | 0.055 | 1.148 | 0.34 | 0.02 | 0.32 | 0.2 | 0.34 | 0.32 |
| 3 | 112740145 | rs3872610 | 1.222 | 0.207 | 0.745 | 1.037 | -0.014 | 0.013 | 0.013 | 1.663 | 0.75 | 0.37 | 0.76 | 0.35 | 0.79 | 0.7 |
| 3 | 112754626 | rs7650483 | 0.843 | 0.97 | 0.426 | 0.915 | -0.007 | -0.008 | -0.008 | 0 | 0.79 | 0.8 | 0.76 | 0.7 | 0.68 | 0.73 |
| 3 | 112771505 | rs2399460 | 0.536 | 0.949 | 0.35 | 0.942 | -0.013 | -0.017 | -0.017 | 0 | 0.81 | 0.85 | 0.77 | 0.7 | 0.71 | 0.83 |
| 3 | 112782186 | rs3872612 | -0.295 | 0.897 | 0.106 | 0.397 | -0.029 | 0.097 | 0.097 | 2.01 | 0.32 | 0.14 | 0.4 | 0.14 | 0.34 | 0.25 |
| 3 | 112788968 | rs3872614 | 0.249 | 0.932 | 0.193 | 0.498 | 0.016 | 0.054 | 0.054 | 1.739 | 0.27 | 0.13 | 0.34 | 0.13 | 0.24 | 0.22 |
| 3 | 112789865 | rs11918021 | -0.283 | 0.881 | 0.73 | 0.645 | 0.005 | 0.04 | 0.04 | 0 | 0.08 | 0.21 | 0.03 | 0.14 | 0.06 | 0.07 |
| 3 | 112790685 | rs6438127 | 1.139 | 0.777 | 1.078 | 0.752 | 0.011 | 0.051 | 0.051 | 0.344 | 0.54 | 0.24 | 0.44 | 0.36 | 0.55 | 0.58 |
| 3 | 112796030 | rs1795172 | 1.07 | 0.805 | 1.256 | 0.788 | 0.067 | 0.291 | 0.291 | 1.074 | 0.9 | 0.65 | 0.81 | 0.62 | 0.84 | 0.88 |
| 3 | 112797260 | rs6784662 | 1.819 | 0.813 | 1.034 | 0.789 | 0.017 | 0.148 | 0.148 | 1.314 | 0.81 | 0.41 | 0.78 | 0.48 | 0.78 | 0.8 |
| 3 | 112797997 | rs3912609 | 0.569 | 0.842 | 0.557 | 0.814 | 0.062 | 0.256 | 0.256 | 0.017 | 0.92 | 0.77 | 0.82 | 0.62 | 0.84 | 0.82 |
| 3 | 112805149 | rs10934215 | 0.716 | 0.833 | -0.264 | 0.248 | 0.006 | 0.013 | 0.013 | 0.801 | 0.59 | 0.35 | 0.44 | 0.34 | 0.58 | 0.57 |
| 3 | 112807518 | rs2613958 | 0.165 | 0.692 | -0.785 | 0.204 | 0.006 | 0.016 | 0.016 | 1.605 | 0.27 | 0.21 | 0.37 | 0.13 | 0.25 | 0.21 |
| 3 | 112811208 | rs7647147 | 1.065 | 0.958 | -0.989 | 0.153 | -0.1 | 0.175 | 0.175 | 0.835 | 0.45 | 0.04 | 0.37 | 0.19 | 0.43 | 0.38 |
| 3 | 112814706 | rs1920082 | 0.525 | 1.254 | -0.202 | 0.855 | 0.046 | 0.171 | 0.171 | 2.592 | 0.81 | 0.49 | 0.77 | 0.45 | 0.77 | 0.68 |
| 3 | 112816192 | rs9816697 | -0.236 | 1.402 | -1.215 | 0.016 | -0.019 | 0.041 | 0.041 | 0 | 0.25 | 0.16 | 0.22 | 0.14 | 0.26 | 0.15 |
| 3 | 112816746 | rs12695286 | -0.951 | 0.817 | -1.577 | -0.171 | 0.007 | 0.076 | 0.076 | 0.377 | 0.53 | 0.68 | 0.59 | 0.68 | 0.49 | 0.55 |
| 3 | 112818765 | rs11717576 | -0.34 | 0.722 | -1.927 | 0.006 | -0.042 | -0.03 | -0.03 | 0 | 0.24 | 0.05 | 0.36 | 0.22 | 0.25 | 0.17 |
| 3 | 112823658 | rs4682465 | -0.001 | 0.656 | -2.052 | -0.115 | -0.058 | -0.031 | -0.031 | 2.088 | 0.2 | 0.01 | 0.3 | 0.18 | 0.17 | 0.19 |
| 3 | 112825573 | rs2613946 | 1.073 | 0.718 | -1.62 | -0.85 | -0.003 | -0.02 | -0.02 | 0 | 0.39 | 0.45 | 0.34 | 0.73 | 0.39 | 0.54 |
| 3 | 112826253 | rs1797626 | 1.314 | 0.678 | -1.647 | -0.491 | 0 | -0.004 | -0.004 | 0 | 0.36 | 0.43 | 0.33 | 0.63 | 0.35 | 0.49 |
| 3 | 112829996 | rs1574478 | -0.46 | 0.612 | -1.973 | -0.615 | 0.002 | -0.002 | -0.002 | 0.006 | 0.37 | 0.5 | 0.57 | 0.59 | 0.41 | 0.37 |
| 3 | 112831824 | rs1797620 | 2.172 | 0.507 | -1.589 | -0.74 | -0.004 | 0.028 | 0.028 | 0.018 | 0.31 | 0.11 | 0.19 | 0.28 | 0.28 | 0.39 |
| 3 | 112832434 | rs1795173 | 0.507 | 0.477 | -1.558 | -0.968 | -0.002 | -0.03 | -0.03 | 0 | 0.64 | 0.58 | 0.72 | 0.9 | 0.66 | 0.79 |
| 3 | 112836352 | rs1797617 | -0.459 | 0.461 | -1.542 | -0.947 | -0.025 | -0.04 | -0.04 | 0 | 0.78 | 0.64 | 0.83 | 0.9 | 0.82 | 0.86 |
| 3 | 112838945 | rs1461106 | 0.749 | 0.49 | -1.486 | -0.903 | 0.016 | -0.037 | -0.037 | 0.091 | 0.12 | 0.21 | 0.1 | 0.04 | 0.14 | 0.14 |
| 3 | 112840885 | rs2613922 | -1.044 | 0.505 | -1.589 | -0.876 | 0.018 | 0.047 | 0.047 | 0 | 0.48 | 0.63 | 0.66 | 0.62 | 0.55 | 0.49 |
| 3 | 112841569 | rs2700204 | -0.356 | 0.517 | -1.594 | -0.888 | 0.007 | -0.002 | -0.002 | 0 | 0.34 | 0.57 | 0.54 | 0.62 | 0.39 | 0.42 |
| 3 | 112841864 | rs6438130 | 0.875 | 0.49 | -1.539 | -0.836 | 0.022 | 0.057 | 0.057 | 0 | 0.51 | 0.35 | 0.34 | 0.38 | 0.43 | 0.51 |
| 3 | 112844086 | rs3852011 | -0.663 | 0.38 | -1.536 | -0.829 | -0.006 | 0.009 | 0.009 | 0.027 | 0.41 | 0.61 | 0.59 | 0.61 | 0.44 | 0.43 |
| 3 | 112845175 | rs9811344 | 2.351 | 0.284 | -1.671 | -0.883 | -0.065 | -0.013 | -0.013 | 0 | 0.29 | 0.02 | 0.17 | 0.28 | 0.27 | 0.39 |
| 3 | 112846786 | rs753693 | -0.151 | 0.249 | -1.194 | -0.871 | 0 | -0.01 | -0.01 | 2.045 | 0.36 | 0.37 | 0.46 | 0.25 | 0.35 | 0.32 |
| 3 | 112847562 | rs871595 | 0.973 | 0.034 | -1.324 | -0.917 | -0.008 | 0.027 | 0.027 | 0 | 0.44 | 0.36 | 0.4 | 0.76 | 0.52 | 0.6 |
| 3 | 112848851 | rs7617193 | 0.979 | -0.119 | -1.438 | -0.987 | -0.017 | 0.051 | 0.051 | 0 | 0.43 | 0.27 | 0.39 | 0.73 | 0.52 | 0.56 |
| 3 | 112854339 | rs2700230 | -0.439 | -0.295 | -1.027 | -1.116 | -0.019 | -0.115 | -0.115 | 0 | 0.37 | 0.55 | 0.29 | 0.04 | 0.28 | 0.23 |
| 3 | 112854675 | rs9822730 | 0.765 | -0.495 | -1.298 | -1.114 | 0.039 | 0.134 | 0.134 | 0 | 0.15 | 0.26 | 0.2 | 0.36 | 0.22 | 0.17 |
| 3 | 112855888 | rs1881943 | 1.259 | -0.599 | -1.345 | -1.196 | 0.007 | 0.091 | 0.091 | 0 | 0.43 | 0.25 | 0.33 | 0.3 | 0.39 | 0.5 |
| 3 | 112856961 | rs11705763 | 1.53 | -0.621 | -1.302 | -1.108 | -0.031 | 0.085 | 0.085 | 0.046 | 0.15 | 0.01 | 0.11 | 0.01 | 0.15 | 0.07 |
| 3 | 112858184 | rs12486305 | 1.064 | -0.64 | -1.232 | -1.128 | 0.003 | -0.024 | -0.024 | 1.494 | 0.18 | 0.2 | 0.15 | 0 | 0.13 | 0.12 |
| 3 | 112860780 | rs2613964 | 0.149 | -0.631 | -1.126 | -1.082 | 0.004 | 0.005 | 0.005 | 0.581 | 0.59 | 0.58 | 0.52 | 0.65 | 0.64 | 0.62 |
| 3 | 112863027 | rs3732812 | -1.484 | -0.399 | -1.085 | -0.928 | -0.001 | -0.001 | -0.001 | 0 | 0.8 | 0.81 | 0.68 | 0.77 | 0.8 | 0.82 |
| 3 | 112867035 | rs1881945 | 0.468 | -0.372 | -1.004 | -0.837 | 0.004 | 0.051 | 0.051 | 0 | 0.2 | 0.27 | 0.24 | 0.35 | 0.23 | 0.19 |
| 3 | 112869412 | rs9845807 | 0.428 | -0.347 | -0.864 | -0.832 | 0.002 | 0.044 | 0.044 | 0 | 0.2 | 0.25 | 0.25 | 0.37 | 0.23 | 0.22 |
| 3 | 112875949 | rs11923603 | 0.282 | -0.321 | -0.693 | -0.948 | -0.002 | 0.077 | 0.077 | 0 | 0.52 | 0.69 | 0.5 | 0.76 | 0.5 | 0.67 |
| 3 | 112878536 | rs10934223 | 0.06 | -0.333 | -0.538 | -0.762 | -0.002 | 0.001 | 0.001 | 0.734 | 0.25 | 0.24 | 0.28 | 0.29 | 0.23 | 0.24 |
| 3 | 112879711 | rs13086237 | 0.264 | 0.109 | -0.276 | -0.57 | -0.001 | -0.042 | -0.042 | 0 | 0.18 | 0.23 | 0.16 | 0.01 | 0.15 | 0.12 |
| 3 | 112881539 | rs9881633 | -1.215 | 0.222 | -0.102 | -0.264 | 0.06 | 0.533 | 0.533 | 1.083 | 0.1 | 0.42 | 0.17 | 0.42 | 0.11 | 0.17 |
| 3 | 112886066 | rs1499904 | -0.647 | 0.074 | -0.588 | -0.754 | 0.004 | -0.068 | -0.068 | 0.086 | 0.33 | 0.53 | 0.25 | 0.12 | 0.33 | 0.21 |
| 3 | 112893640 | rs6770031 | 0.531 | -0.195 | -0.688 | -0.143 | -0.022 | 0.057 | 0.057 | 0 | 0.73 | 0.38 | 0.67 | 0.61 | 0.77 | 0.69 |
| 3 | 112895679 | rs9682694 | 1.89 | -0.055 | -0.825 | -0.021 | -0.059 | -0.031 | -0.031 | 0 | 0.22 | 0.03 | 0.24 | 0.27 | 0.26 | 0.36 |
| 3 | 112896373 | rs12495201 | -0.231 | -0.06 | -0.27 | -0.072 | 0.013 | -0.105 | -0.105 | 0.659 | 0.2 | 0.39 | 0.2 | 0.02 | 0.17 | 0.12 |
| 3 | 112897695 | rs953201 | -1.13 | -0.169 | -0.199 | 0.081 | 0.005 | -0.007 | -0.007 | 0 | 0.4 | 0.6 | 0.36 | 0.39 | 0.36 | 0.35 |
| 3 | 112900595 | rs12492489 | -0.241 | -0.027 | 0.13 | 0.224 | 0.002 | -0.014 | -0.014 | 1.214 | 0.06 | 0.25 | 0.12 | 0.01 | 0.02 | 0.01 |
| 3 | 112901729 | rs9856514 | 0.349 | 0.097 | 0.063 | 0.464 | 0.07 | 0.333 | 0.333 | 0.12 | 0.05 | 0.21 | 0.1 | 0.24 | 0.07 | 0.1 |
| 3 | 112905415 | rs1499900 | -1.004 | 0.073 | -0.369 | 0.202 | 0 | 0.001 | 0.001 | 0 | 0.64 | 0.65 | 0.56 | 0.66 | 0.67 | 0.57 |
| 3 | 112906970 | rs11718413 | 1.058 | 0.034 | -0.479 | 0.417 | -0.014 | -0.018 | -0.018 | 0 | 0.28 | 0.18 | 0.32 | 0.33 | 0.33 | 0.27 |
| 3 | 112911260 | rs1499899 | -0.024 | -0.38 | -0.161 | 0.377 | -0.001 | -0.001 | -0.001 | 0.007 | 0.49 | 0.56 | 0.46 | 0.53 | 0.51 | 0.42 |
| 3 | 112911525 | rs1392228 | -0.481 | -0.815 | 0.025 | 0.271 | -0.024 | -0.027 | -0.027 | 0 | 0.85 | 0.83 | 0.94 | 0.97 | 0.93 | 0.9 |
| 3 | 112913309 | rs9288978 | -0.165 | -0.399 | 0.016 | 0.512 | -0.022 | 0.008 | 0.008 | 0 | 0.87 | 0.8 | 0.92 | 0.85 | 0.9 | 0.86 |
| 3 | 112914615 | rs972554 | 0.026 | -0.035 | -0.479 | 0.333 | 0.013 | 0.049 | 0.049 | 0.022 | 0.48 | 0.4 | 0.55 | 0.35 | 0.42 | 0.48 |
| 3 | 112919344 | rs9831167 | -0.536 | 0.017 | -0.668 | 0.299 | 0.021 | 0.018 | 0.018 | 0.264 | 0.65 | 0.74 | 0.66 | 0.7 | 0.71 | 0.54 |
| 3 | 112921268 | rs4682475 | -0.345 | 0.062 | -0.593 | 0.395 | 0.014 | 0.018 | 0.018 | 0.02 | 0.61 | 0.67 | 0.56 | 0.69 | 0.68 | 0.49 |
| 3 | 112923015 | rs11706932 | 0.835 | 0.247 | -0.503 | 0.152 | 0.001 | -0.023 | -0.023 | 0.113 | 0.18 | 0.23 | 0.09 | 0.03 | 0.15 | 0.08 |
| 3 | 112923072 | rs13321908 | 0.214 | 0.307 | -0.262 | 0.705 | 0.01 | 0.01 | 0.01 | 0.365 | 0.4 | 0.4 | 0.37 | 0.5 | 0.5 | 0.37 |
| 3 | 112924776 | rs12630707 | 0.304 | 0.786 | -0.46 | 0.942 | -0.001 | 0.006 | 0.006 | 0.52 | 0.45 | 0.39 | 0.49 | 0.42 | 0.44 | 0.47 |
| 3 | 112933975 | rs9288979 | -0.357 | 1.162 | -0.047 | 1.354 | 0.092 | 0.194 | 0.194 | 0.305 | 0.06 | 0.2 | 0.09 | 0.21 | 0.11 | 0.09 |

| | | | | | | | | | | | | | | | | |
|---|---|---|---|---|---|---|---|---|---|---|---|---|---|---|---|---|
| 3 | 112934752 | rs931702 | -0.742 | 0.552 | -0.48 | 0.85 | 0.003 | 0.009 | 0.009 | 0 | 0.05 | 0.14 | 0.01 | 0.09 | 0.05 | 0.03 |
| 3 | 112936609 | rs1553191 | 0.059 | 0.261 | -1.177 | 0.46 | 0.016 | 0.066 | 0.066 | 0 | 0.41 | 0.2 | 0.36 | 0.29 | 0.39 | 0.3 |
| 3 | 112936907 | rs6782058 | 0.067 | 0.24 | -1.179 | 0.457 | 0.018 | 0.066 | 0.066 | 0 | 0.41 | 0.2 | 0.35 | 0.29 | 0.39 | 0.3 |
| 3 | 112938646 | rs904627 | 0.366 | 0.086 | -1.365 | 0.286 | -0.002 | -0.001 | -0.001 | 1.063 | 0.24 | 0.18 | 0.25 | 0.3 | 0.24 | 0.25 |
| 3 | 112938683 | rs904626 | -0.488 | -0.447 | -0.995 | 0.462 | 0.025 | 0.011 | 0.011 | 0 | 0.79 | 0.84 | 0.77 | 0.88 | 0.85 | 0.85 |
| 3 | 112941557 | rs9874798 | -1.286 | -0.252 | -0.649 | 0.813 | -0.001 | 0 | 0 | 0.12 | 0.2 | 0.75 | 0.2 | 0.26 | 0.16 | 0.11 |
| 3 | 112941684 | rs16860670 | -0.198 | -0.237 | -0.11 | 0.808 | 0.002 | 0.01 | 0.01 | 0 | 0.07 | 0.13 | 0.05 | 0.1 | 0.09 | 0.03 |
| 3 | 112942269 | rs9859731 | -0.22 | 0.243 | -0.091 | 0.676 | -0.002 | -0.002 | -0.002 | 0.941 | 0.32 | 0.32 | 0.4 | 0.57 | 0.35 | 0.37 |
| 3 | 112947155 | rs1875113 | -0.985 | 0.172 | -0.698 | 0.352 | -0.008 | 0.202 | 0.202 | 0.005 | 0.48 | 0.83 | 0.57 | 0.78 | 0.45 | 0.44 |
| 3 | 112947495 | rs1875112 | -0.72 | 0.349 | -0.493 | 0.774 | -0.004 | 0.004 | 0.004 | 0 | 0.36 | 0.41 | 0.38 | 0.56 | 0.33 | 0.36 |
| 3 | 112951348 | rs6769798 | -0.574 | 0.716 | -1.046 | 0.811 | -0.06 | -0.071 | -0.071 | 0.283 | 0.18 | 0 | 0.12 | 0.21 | 0.15 | 0.14 |
| 3 | 112951608 | rs9842173 | -0.763 | 0.461 | -0.401 | 0.136 | -0.005 | 0.016 | 0.016 | 0 | 0.42 | 0.41 | 0.51 | 0.7 | 0.43 | 0.47 |
| 3 | 112951832 | rs9825012 | 0.456 | 0.347 | 0.12 | 0.081 | 0.007 | -0.03 | -0.03 | 0 | 0.06 | 0.17 | 0.03 | 0 | 0.03 | 0.01 |
| 3 | 112955448 | rs6783739 | -0.487 | 0.222 | 0.447 | 0.217 | -0.005 | 0.001 | 0.001 | 0.031 | 0.37 | 0.32 | 0.51 | 0.65 | 0.4 | 0.45 |
| 3 | 112966416 | rs12491930 | -0.917 | 0.298 | 0.483 | 0.301 | -0.002 | 0.009 | 0.009 | 0 | 0.33 | 0.39 | 0.46 | 0.54 | 0.29 | 0.37 |
| 3 | 112966904 | rs10511324 | 0.604 | 0.347 | 0.442 | 0.295 | -0.002 | -0.03 | -0.03 | 0 | 0.06 | 0.12 | 0.03 | 0 | 0.03 | 0.01 |
| 3 | 112969270 | rs4682131 | 0.357 | 0.334 | 0.402 | 0.579 | -0.005 | -0.076 | -0.076 | 0.521 | 0.22 | 0.12 | 0.41 | 0.44 | 0.21 | 0.31 |
| 3 | 112970622 | rs9849714 | -0.85 | 0.332 | 0.435 | 0.532 | -0.006 | -0.003 | -0.003 | 0.515 | 0.11 | 0.26 | 0.05 | 0.1 | 0.08 | 0.06 |
| 3 | 112977275 | rs9820139 | 0.219 | 0.371 | 0.246 | 0.549 | -0.004 | -0.006 | -0.006 | 0 | 0.08 | 0.05 | 0.03 | 0.1 | 0.08 | 0.06 |
| 3 | 112978764 | rs1875111 | -0.982 | 0.362 | 0.401 | 0.559 | -0.002 | 0.048 | 0.048 | 0.118 | 0.33 | 0.51 | 0.46 | 0.48 | 0.28 | 0.35 |
| 3 | 112980483 | rs999367 | -1.277 | 0.313 | 0.788 | 0.506 | 0.004 | -0.003 | -0.003 | 0 | 0.38 | 0.72 | 0.49 | 0.58 | 0.33 | 0.4 |
| 3 | 112988511 | rs2936725 | -2.114 | 0.118 | 0.698 | 0.346 | -0.038 | 0.15 | 0.15 | 0.253 | 0.53 | 0.84 | 0.57 | 0.74 | 0.49 | 0.51 |
| 3 | 112990308 | rs9843707 | -0.921 | -0.001 | 1.079 | 0.315 | -0.007 | -0.007 | -0.007 | 0 | 0.27 | 0.42 | 0.14 | 0.26 | 0.24 | 0.18 |
| 3 | 112991117 | rs12631733 | 0.687 | -0.091 | 0.961 | 0.45 | 0.001 | 0.029 | 0.029 | 0.09 | 0.18 | 0.22 | 0.38 | 0.41 | 0.19 | 0.28 |
| 3 | 112991959 | rs11710894 | 0.52 | -0.164 | 0.984 | 0.4 | 0.001 | -0.004 | -0.004 | 0 | 0.23 | 0.22 | 0.39 | 0.41 | 0.2 | 0.32 |
| 3 | 112993063 | rs13093350 | 0.536 | -0.176 | 1.059 | 0.36 | -0.001 | 0 | 0 | 0 | 0.16 | 0.12 | 0.09 | 0.16 | 0.16 | 0.12 |
| 3 | 112996316 | rs3846046 | 0.719 | -0.178 | 1.035 | 0.38 | 0 | -0.047 | -0.047 | 0 | 0.22 | 0.12 | 0.38 | 0.41 | 0.2 | 0.32 |
| 3 | 112997554 | rs775228 | -1.937 | -0.2 | 1.13 | 0.337 | -0.025 | 0.115 | 0.115 | 0.189 | 0.53 | 0.77 | 0.54 | 0.72 | 0.48 | 0.52 |
| 3 | 112998090 | rs3856720 | 0.756 | -0.207 | 1.145 | 0.357 | -0.003 | -0.069 | -0.069 | 0 | 0.22 | 0.09 | 0.38 | 0.41 | 0.2 | 0.32 |
| 3 | 112998265 | rs2649878 | -1.394 | -0.302 | 1.062 | 0.244 | 0.013 | 0.025 | 0.025 | 0.177 | 0.42 | 0.84 | 0.47 | 0.61 | 0.34 | 0.42 |
| 3 | 113001706 | rs775230 | -0.6 | -0.224 | 1.28 | 0.273 | 0.001 | 0.013 | 0.013 | 0 | 0.35 | 0.53 | 0.44 | 0.56 | 0.32 | 0.4 |
| 3 | 113002150 | rs808951 | -0.743 | -0.064 | 0.631 | 0.24 | -0.014 | -0.049 | -0.049 | 0 | 0.25 | 0.52 | 0.14 | 0.15 | 0.18 | 0.12 |
| 3 | 113007525 | rs11919134 | 0.244 | 0.066 | 0.02 | 0.477 | -0.012 | -0.039 | -0.039 | 0 | 0.22 | 0.14 | 0.39 | 0.44 | 0.24 | 0.32 |
| 3 | 113008606 | rs7652801 | 0.716 | -0.282 | 0.088 | 0.228 | 0 | 0 | 0 | 1.527 | 0.06 | 0.06 | 0.02 | 0 | 0.03 | 0.01 |
| 3 | 113010303 | rs17321330 | 0.948 | -0.269 | -0.355 | 0.129 | -0.084 | 0.054 | 0.054 | 1.289 | 0.47 | 0.08 | 0.56 | 0.3 | 0.54 | 0.53 |
| 3 | 113010950 | rs775214 | 0.671 | -0.265 | 0.403 | 0.268 | -0.007 | -0.009 | -0.009 | 0.43 | 0.71 | 0.48 | 0.83 | 0.71 | 0.75 | 0.79 |
| 3 | 113015660 | rs2270781 | 1.425 | -0.3 | 0.277 | 0.301 | -0.023 | -0.078 | -0.078 | 0.067 | 0.19 | 0.06 | 0.24 | 0.4 | 0.21 | 0.25 |
| 3 | 113022996 | rs2270782 | 0.719 | -0.376 | 0.801 | 0.211 | -0.011 | -0.012 | -0.012 | 1.315 | 0.71 | 0.55 | 0.84 | 0.72 | 0.75 | 0.79 |
| 3 | 113023509 | rs3773697 | 1.553 | -0.293 | 0.346 | 0.258 | -0.044 | -0.013 | -0.013 | 2.275 | 0.11 | 0.01 | 0.08 | 0.12 | 0.15 | 0.11 |
| 3 | 113028334 | rs1353903 | -0.2 | -0.43 | 0.845 | 0.302 | -0.005 | 0.019 | 0.019 | 0 | 0.34 | 0.38 | 0.32 | 0.56 | 0.3 | 0.36 |
| 3 | 113034316 | rs7617723 | -0.518 | -0.491 | 0.827 | 0.302 | -0.017 | 0.019 | 0.019 | 0 | 0.1 | 0.16 | 0.05 | 0.16 | 0.06 | 0.1 |
| 3 | 113046640 | rs13064411 | 1.598 | -0.453 | 0.736 | 0.259 | -0.044 | -0.006 | -0.006 | 0 | 0.11 | 0.01 | 0.07 | 0.12 | 0.14 | 0.1 |
| 3 | 113049873 | rs7429847 | -0.351 | -0.447 | 0.91 | 0.282 | -0.02 | 0.039 | 0.039 | 0 | 0.1 | 0.2 | 0.05 | 0.16 | 0.06 | 0.1 |
| 3 | 113062350 | rs12489411 | 1.605 | -0.767 | 0.165 | -0.243 | -0.007 | 0 | 0 | 0 | 0.16 | 0.04 | 0.27 | 0.12 | 0.18 | 0.28 |
| 3 | 113062986 | rs4682133 | 1.227 | -0.905 | 0.379 | -0.249 | -0.003 | -0.015 | -0.015 | 1.936 | 0.24 | 0.18 | 0.27 | 0.4 | 0.24 | 0.26 |
| 3 | 113064142 | rs9863193 | 1.241 | -0.905 | 0.37 | -0.249 | -0.009 | -0.038 | -0.038 | 0.214 | 0.24 | 0.14 | 0.27 | 0.4 | 0.24 | 0.26 |
| 3 | 113064394 | rs6438140 | -0.949 | -1.02 | 0.581 | -0.31 | -0.012 | 0.061 | 0.061 | 0.977 | 0.52 | 0.8 | 0.43 | 0.68 | 0.45 | 0.46 |
| 3 | 113076730 | rs4682485 | 1.912 | -0.895 | 0.511 | -0.228 | -0.018 | -0.069 | -0.069 | 0 | 0.19 | 0.06 | 0.24 | 0.4 | 0.21 | 0.25 |
| 3 | 113085434 | rs7650500 | 0.024 | -0.824 | 0.344 | -0.262 | -0.005 | 0.007 | 0.007 | 0 | 0.09 | 0.11 | 0.05 | 0.16 | 0.06 | 0.1 |
| 3 | 113117165 | rs6771972 | 1.785 | -0.668 | -0.006 | -0.388 | -0.074 | 0.135 | 0.135 | 0 | 0.28 | 0.02 | 0.27 | 0.04 | 0.32 | 0.18 |
| 3 | 113117829 | rs6438143 | -1.123 | -1.212 | 0.77 | -0.08 | -0.003 | 0.066 | 0.066 | 0.72 | 0.51 | 0.74 | 0.43 | 0.69 | 0.45 | 0.46 |
| 3 | 113117897 | rs7643843 | -0.127 | -1.242 | 1.037 | -0.063 | 0.006 | 0.017 | 0.017 | 0.496 | 0.38 | 0.51 | 0.37 | 0.52 | 0.38 | 0.36 |
| 3 | 113122524 | rs6808394 | 0.841 | -1.252 | 1.057 | -0.071 | 0.009 | -0.02 | -0.02 | 0 | 0.16 | 0.22 | 0.11 | 0.12 | 0.17 | 0.11 |
| 3 | 113127991 | rs16845107 | -0.141 | -1.256 | 0.998 | -0.064 | -0.001 | 0.003 | 0.003 | 0 | 0.09 | 0.09 | 0.05 | 0.16 | 0.06 | 0.1 |
| 3 | 113128051 | rs13059888 | 2.342 | -1.528 | 0.758 | -0.27 | 0.015 | 0.089 | 0.089 | 0.341 | 0.14 | 0.01 | 0.08 | 0 | 0.14 | 0.03 |
| 3 | 113135345 | rs12635786 | 0.028 | -0.449 | 1.625 | 0.629 | -0.006 | -0.081 | -0.081 | 0 | 0.28 | 0.09 | 0.29 | 0.57 | 0.27 | 0.34 |
| 3 | 113136418 | rs7623122 | -1.162 | -0.445 | 1.65 | 0.623 | 0 | 0.022 | 0.022 | 0.058 | 0.47 | 0.61 | 0.42 | 0.69 | 0.44 | 0.46 |
| 3 | 113138763 | rs2129434 | 0.365 | -0.364 | 1.534 | 0.635 | 0.019 | -0.03 | -0.03 | 0 | 0.17 | 0.26 | 0.11 | 0.12 | 0.17 | 0.11 |
| 3 | 113146697 | rs10934229 | 1.116 | -0.329 | 1.147 | 0.832 | -0.046 | -0.115 | -0.115 | 0.027 | 0.19 | 0.02 | 0.24 | 0.41 | 0.21 | 0.24 |
| 3 | 113161519 | rs1906141 | -0.508 | 0.129 | 1.657 | 0.994 | -0.001 | 0.044 | 0.044 | 0 | 0.06 | 0.14 | 0.03 | 0.11 | 0.05 | 0.07 |
| 3 | 113162095 | rs6775574 | -1.6 | -0.013 | 1.736 | 0.949 | -0.021 | 0.063 | 0.063 | 0 | 0.55 | 0.75 | 0.43 | 0.71 | 0.46 | 0.47 |
| 3 | 113162117 | rs6787026 | 0.416 | -0.035 | 1.86 | 0.946 | 0.001 | -0.001 | -0.001 | 0 | 0.16 | 0.16 | 0.1 | 0.13 | 0.17 | 0.11 |
| 3 | 113162570 | rs11926687 | -0.412 | -0.015 | 1.797 | 0.957 | -0.001 | 0.044 | 0.044 | 0 | 0.06 | 0.14 | 0.03 | 0.11 | 0.05 | 0.07 |
| 3 | 113162699 | rs11923128 | -0.412 | -0.015 | 1.797 | 0.957 | -0.001 | 0.044 | 0.044 | 0 | 0.06 | 0.14 | 0.03 | 0.11 | 0.05 | 0.07 |
| 3 | 113164109 | rs2054823 | -1.578 | -0.019 | 1.702 | 0.954 | -0.02 | 0.066 | 0.066 | 0 | 0.55 | 0.75 | 0.43 | 0.71 | 0.46 | 0.47 |
| 3 | 113169205 | rs11920725 | -0.2 | 0.011 | 1.725 | 0.969 | -0.003 | 0.036 | 0.036 | 0 | 0.05 | 0.12 | 0.03 | 0.11 | 0.05 | 0.07 |
| 3 | 113174063 | rs9866748 | -1.483 | 0.007 | 1.72 | 0.966 | -0.022 | 0.062 | 0.062 | 0 | 0.55 | 0.75 | 0.43 | 0.71 | 0.46 | 0.47 |
| 3 | 113176122 | rs7637618 | -1.42 | -0.19 | 1.948 | 0.969 | -0.007 | -0.001 | -0.001 | 0 | 0.3 | 0.58 | 0.18 | 0.3 | 0.25 | 0.21 |
| 3 | 113184186 | rs16861023 | -0.241 | -0.154 | 1.83 | 0.881 | -0.001 | 0.041 | 0.041 | 0 | 0.05 | 0.11 | 0.03 | 0.11 | 0.05 | 0.07 |

| | | | | | | | | | | | | | | | | |
|---|---|---|---|---|---|---|---|---|---|---|---|---|---|---|---|---|
| 3 | 113211513 | rs1486894 | -0.462 | -0.159 | 1.88 | 0.877 | 0.003 | -0.03 | -0.03 | 0.697 | 0.21 | 0.31 | 0.12 | 0.13 | 0.19 | 0.12 |
| 3 | 113212562 | rs4396864 | -1.037 | -0.156 | 1.877 | 0.879 | 0.002 | -0.006 | -0.006 | 0 | 0.25 | 0.37 | 0.14 | 0.24 | 0.23 | 0.19 |
| 3 | 113233188 | rs16861103 | -0.888 | -0.117 | 1.85 | 0.864 | 0.001 | -0.024 | -0.024 | 0.039 | 0.25 | 0.23 | 0.11 | 0.22 | 0.22 | 0.1 |
| 3 | 113240667 | rs9682108 | -1.144 | -0.108 | 1.73 | 0.901 | -0.009 | 0.064 | 0.064 | 0 | 0.09 | 0.23 | 0.05 | 0.16 | 0.06 | 0.1 |
| 3 | 113248748 | rs6799610 | -0.402 | -0.075 | 1.598 | 0.839 | 0.001 | -0.018 | -0.018 | 1.428 | 0.21 | 0.27 | 0.11 | 0.13 | 0.18 | 0.12 |
| 3 | 113255143 | rs4580515 | -1.3 | -0.045 | 1.574 | 0.924 | -0.005 | 0.03 | 0.03 | 0 | 0.13 | 0.3 | 0.07 | 0.17 | 0.09 | 0.1 |
| 3 | 113259413 | rs6769970 | -1.485 | -0.084 | 1.465 | 0.931 | 0.006 | 0.076 | 0.076 | 0 | 0.53 | 0.73 | 0.41 | 0.71 | 0.44 | 0.45 |
| 3 | 113264994 | rs2254267 | -2.103 | 0.064 | 1.634 | 0.836 | -0.04 | -0.001 | -0.001 | 0 | 0.18 | 0.45 | 0.09 | 0.17 | 0.1 | 0.1 |
| 3 | 113265727 | rs2669909 | -1.431 | 0.09 | 1.585 | 0.866 | -0.01 | -0.006 | -0.006 | 0.025 | 0.3 | 0.54 | 0.17 | 0.3 | 0.24 | 0.2 |
| 3 | 113268970 | rs16861165 | -1.202 | 0.213 | 1.292 | 0.985 | -0.009 | 0.023 | 0.023 | 0 | 0.12 | 0.21 | 0.05 | 0.17 | 0.08 | 0.08 |
| 3 | 113269201 | rs2614191 | -0.512 | 0.267 | 1.006 | 0.997 | 0 | 0 | 0 | 0.002 | 0.19 | 0.21 | 0.11 | 0.23 | 0.19 | 0.16 |
| 3 | 113270402 | rs7612448 | -0.295 | 0.161 | 0.703 | 0.98 | -0.005 | 0.002 | 0.002 | 0 | 0.07 | 0.13 | 0.03 | 0.07 | 0.03 | 0.02 |
| 3 | 113271240 | rs11922598 | -1.233 | 0.108 | 0.84 | 0.718 | -0.012 | -0.004 | -0.004 | 0.018 | 0.12 | 0.28 | 0.07 | 0.12 | 0.08 | 0.07 |
| 3 | 113271406 | rs2399488 | 0.282 | 0.175 | 0.562 | 0.815 | 0.003 | 0.006 | 0.006 | 0 | 0.61 | 0.71 | 0.68 | 0.36 | 0.63 | 0.6 |
| 3 | 113273634 | rs13074313 | 0.366 | 0.6 | -0.057 | 0.622 | -0.003 | 0.002 | 0.002 | 0 | 0.15 | 0.12 | 0.09 | 0.1 | 0.17 | 0.14 |
| 3 | 113275624 | rs7611694 | -0.912 | 0.56 | 0.073 | 0.705 | 0.003 | 0.029 | 0.029 | 0 | 0.43 | 0.32 | 0.34 | 0.75 | 0.42 | 0.46 |
| 3 | 113276812 | rs9810754 | 0.119 | 0.638 | -0.213 | 0.62 | -0.001 | -0.033 | -0.033 | 0 | 0.06 | 0.17 | 0.03 | 0 | 0.03 | 0.01 |
| 3 | 113290468 | rs13092825 | 0.342 | 0.815 | -0.238 | 0.786 | -0.005 | 0.006 | 0.006 | 0 | 0.15 | 0.12 | 0.09 | 0.1 | 0.17 | 0.14 |
| 3 | 113300863 | rs9868243 | -0.035 | 0.734 | -0.142 | 0.841 | -0.014 | 0.02 | 0.02 | 0 | 0.31 | 0.38 | 0.26 | 0.6 | 0.24 | 0.27 |
| 3 | 113302425 | rs2271493 | 0.168 | 0.829 | -0.672 | 0.884 | 0.004 | -0.01 | -0.01 | 0 | 0.28 | 0.26 | 0.26 | 0.59 | 0.23 | 0.27 |
| 3 | 113309149 | rs6808932 | -0.422 | 0.866 | -0.9 | 0.873 | 0.001 | 0.003 | 0.003 | 0 | 0.46 | 0.47 | 0.36 | 0.69 | 0.41 | 0.42 |
| 3 | 113313136 | rs2669910 | 0.053 | 1.282 | -1.555 | 1.411 | -0.034 | -0.067 | -0.067 | 0.006 | 0.6 | 0.86 | 0.57 | 0.53 | 0.51 | 0.59 |
| 3 | 113318748 | rs13070953 | -0.631 | 2.382 | -0.563 | 1.379 | 0.007 | 0.003 | 0.003 | 0.1 | 0.16 | 0.18 | 0.26 | 0.18 | 0.24 | 0.21 |
| 3 | 113321199 | rs3773688 | -0.871 | 2.313 | -0.579 | 0.772 | 0.007 | -0.026 | -0.026 | 0.148 | 0.16 | 0.19 | 0.19 | 0.04 | 0.21 | 0.15 |
| 3 | 113326669 | rs1567778 | -0.773 | 2.29 | -0.652 | 1.387 | 0.001 | 0.009 | 0.009 | 0 | 0.25 | 0.28 | 0.42 | 0.43 | 0.27 | 0.37 |
| 3 | 113328960 | rs6792799 | 0.796 | 2.066 | -1.251 | 0.985 | 0.003 | 0.028 | 0.028 | 0.084 | 0.53 | 0.63 | 0.48 | 0.4 | 0.4 | 0.47 |
| 3 | 113331289 | rs9842516 | 0.925 | 1.863 | -1.624 | 0.733 | 0.038 | 0.009 | 0.009 | 0 | 0.44 | 0.67 | 0.44 | 0.44 | 0.29 | 0.44 |
| 3 | 113332235 | rs2203251 | 1.051 | 1.862 | -1.554 | 0.668 | 0.021 | 0.006 | 0.006 | 0 | 0.42 | 0.62 | 0.34 | 0.42 | 0.28 | 0.42 |
| 3 | 113334233 | rs950357 | -0.104 | 2.212 | -1.827 | 1.154 | -0.152 | -0.025 | -0.025 | 0 | 0.3 | 0.02 | 0.35 | 0.28 | 0.35 | 0.28 |
| 3 | 113341007 | rs3773683 | -0.11 | 1.982 | -1.105 | 1.308 | -0.004 | -0.009 | -0.009 | 0 | 0.27 | 0.27 | 0.24 | 0.28 | 0.37 | 0.29 |
| 3 | 113343646 | rs7638449 | -0.207 | 1.946 | -1.057 | 1.287 | 0 | 0.002 | 0.002 | 4.385 | 0.27 | 0.33 | 0.3 | 0.31 | 0.36 | 0.3 |
| 3 | 113345565 | rs2895415 | -0.123 | 1.854 | -0.782 | 1.286 | -0.002 | -0.006 | -0.006 | 0 | 0.27 | 0.28 | 0.23 | 0.29 | 0.37 | 0.29 |
| 3 | 113349451 | rs9882193 | -0.383 | 1.666 | -0.315 | 1.226 | -0.002 | 0.038 | 0.038 | 7.203 | 0.3 | 0.49 | 0.32 | 0.32 | 0.37 | 0.3 |
| 3 | 113370010 | rs1026364 | -0.422 | 1.765 | -0.151 | 1.449 | 0.003 | 0.037 | 0.037 | 5.539 | 0.29 | 0.44 | 0.32 | 0.31 | 0.38 | 0.27 |
| 3 | 113373019 | rs12493635 | -0.439 | 1.771 | 0.255 | 1.431 | 0.007 | 0.057 | 0.057 | 5.593 | 0.29 | 0.5 | 0.32 | 0.31 | 0.38 | 0.27 |
| 3 | 113377361 | rs9813630 | -0.233 | 1.699 | 0.141 | 1.404 | 0.002 | -0.005 | -0.005 | 0 | 0.27 | 0.44 | 0.27 | 0.22 | 0.36 | 0.18 |
| 3 | 113379865 | rs9866806 | 0.07 | 1.669 | 0.172 | 1.352 | -0.015 | 0.004 | 0.004 | 4.206 | 0.24 | 0.15 | 0.18 | 0.2 | 0.35 | 0.16 |
| 3 | 113398889 | rs12632748 | -0.21 | 1.673 | 0.111 | 1.355 | 0 | -0.005 | -0.005 | 0 | 0.27 | 0.34 | 0.19 | 0.2 | 0.36 | 0.17 |
| 3 | 113414031 | rs9881563 | -0.231 | 1.66 | 0.105 | 1.566 | 0 | 0 | 0 | 6.364 | 0.27 | 0.34 | 0.19 | 0.28 | 0.36 | 0.17 |
| 3 | 113436847 | rs12695294 | 0.678 | 1.62 | -0.133 | 1.295 | -0.021 | 0.062 | 0.062 | 4.218 | 0.11 | 0.01 | 0.07 | 0 | 0.15 | 0.03 |
| 3 | 113460614 | rs2712348 | -1.093 | 1.491 | 0.385 | 1.375 | 0 | 0.028 | 0.028 | 5.888 | 0.29 | 0.39 | 0.27 | 0.39 | 0.38 | 0.26 |
| 3 | 113464859 | rs1048892 | -0.838 | 1.494 | 0.396 | 1.405 | 0.001 | 0.014 | 0.014 | 6.544 | 0.27 | 0.33 | 0.19 | 0.35 | 0.38 | 0.18 |
| 3 | 113474866 | rs12695295 | -0.989 | 1.494 | 0.394 | 1.398 | 0.001 | 0.029 | 0.029 | 6.477 | 0.28 | 0.43 | 0.27 | 0.36 | 0.38 | 0.19 |
| 3 | 113495648 | rs2712353 | -0.904 | 1.573 | 0.392 | 1.494 | 0.004 | 0.036 | 0.036 | 4.16 | 0.23 | 0.36 | 0.21 | 0.36 | 0.3 | 0.24 |
| 3 | 113518602 | rs9828449 | -1.009 | 1.411 | 0.119 | 1.328 | 0 | 0.003 | 0.003 | 4.983 | 0.28 | 0.32 | 0.2 | 0.35 | 0.39 | 0.16 |
| 3 | 113552722 | rs2247225 | 0.233 | 1.305 | -0.239 | 0.924 | -0.005 | 0.001 | 0.001 | 0 | 0.05 | 0.05 | 0.03 | 0.06 | 0.05 | 0.03 |
| 3 | 113560020 | rs2566980 | -0.959 | 0.944 | 0.017 | 0.582 | -0.02 | -0.046 | -0.046 | 0 | 0.15 | 0.32 | 0.12 | 0.06 | 0.09 | 0.14 |
| 3 | 113561504 | rs2399490 | -0.985 | 0.994 | -0.373 | 0.376 | 0.013 | -0.001 | -0.001 | 0 | 0.39 | 0.49 | 0.47 | 0.34 | 0.45 | 0.38 |
| 3 | 113582544 | rs4610201 | 0.162 | 0.927 | -0.028 | 0.345 | -0.11 | -0.069 | -0.069 | 0.085 | 0.25 | 0.1 | 0.34 | 0.28 | 0.36 | 0.26 |
| 3 | 113582767 | rs13082304 | 0.432 | 0.1 | 0.044 | 0.255 | -0.01 | 0.012 | 0.012 | 0 | 0.31 | 0.16 | 0.34 | 0.31 | 0.34 | 0.24 |
| 3 | 113585154 | rs1025400 | -0.207 | 0.588 | 0.407 | 0.306 | 0.052 | 0.025 | 0.025 | 0 | 0.73 | 0.62 | 0.6 | 0.72 | 0.63 | 0.71 |
| 3 | 113585633 | rs1025398 | 0.316 | 0.589 | 0.405 | 0.306 | -0.026 | -0.015 | -0.015 | 1.275 | 0.25 | 0.23 | 0.34 | 0.28 | 0.36 | 0.26 |
| 3 | 113586776 | rs4682506 | 0.713 | 0.562 | 0.273 | 0.293 | -0.002 | 0 | 0 | 0 | 0.6 | 0.37 | 0.53 | 0.66 | 0.54 | 0.61 |
| 3 | 113588445 | rs1469559 | -0.135 | 0.646 | -0.182 | 0.31 | -0.012 | -0.034 | -0.034 | 4.28 | 0.14 | 0.25 | 0.07 | 0.06 | 0.09 | 0.1 |
| 3 | 113594377 | rs13079383 | 0.307 | 0.877 | -0.486 | 0.502 | -0.14 | -0.093 | -0.093 | 0.609 | 0.24 | 0.07 | 0.33 | 0.27 | 0.36 | 0.26 |
| 3 | 113595909 | rs2632246 | 0.473 | -0.065 | -0.337 | 0.065 | 0.008 | 0.005 | 0.005 | 0.903 | 0.51 | 0.36 | 0.4 | 0.43 | 0.54 | 0.51 |
| 3 | 113596326 | rs2712355 | -0.571 | 0.804 | 0.283 | -0.038 | 0.005 | -0.025 | -0.025 | 0.282 | 0.41 | 0.48 | 0.42 | 0.26 | 0.46 | 0.41 |
| 3 | 113598223 | rs2632253 | 0.403 | 0.845 | 0.298 | -0.119 | -0.023 | 0.018 | 0.018 | 0 | 0.68 | 0.75 | 0.68 | 0.74 | 0.57 | 0.67 |
| 3 | 113604898 | rs3796244 | 1.176 | 0.853 | 0.291 | -0.107 | 0.001 | -0.01 | -0.01 | 0 | 0.21 | 0.2 | 0.16 | 0.12 | 0.24 | 0.16 |
| 3 | 113607019 | rs4682144 | 1.366 | 0.877 | 0.445 | -0.092 | 0.09 | 0.028 | 0.028 | 0 | 0.43 | 0.25 | 0.47 | 0.61 | 0.31 | 0.45 |
| 3 | 113608975 | rs6438172 | -0.18 | 0.784 | 0.375 | 0.103 | 0.012 | -0.012 | -0.012 | 0 | 0.83 | 0.78 | 0.87 | 0.86 | 0.76 | 0.87 |
| 3 | 113609403 | rs9862440 | 0.821 | 0.669 | -0.017 | 0.136 | 0 | -0.008 | -0.008 | 0 | 0.21 | 0.2 | 0.16 | 0.13 | 0.24 | 0.16 |
| 3 | 113617074 | rs974952 | 0.675 | 0.664 | 0.008 | 0.136 | 0 | -0.013 | -0.013 | 0.02 | 0.22 | 0.27 | 0.2 | 0.13 | 0.24 | 0.2 |
| 3 | 113620344 | rs2632245 | -1.12 | 0.684 | -0.395 | 0.074 | -0.02 | -0.031 | -0.031 | 0 | 0.92 | 0.86 | 0.96 | 1 | 0.96 | 0.93 |
| 3 | 113624931 | rs4682511 | 0.942 | 0.876 | -0.59 | 0.166 | -0.001 | -0.023 | -0.023 | 0.872 | 0.21 | 0.25 | 0.15 | 0.12 | 0.24 | 0.11 |
| 3 | 113631780 | rs6798319 | 0.309 | 1.236 | -0.566 | 0.25 | 0.054 | 0.013 | 0.013 | 0 | 0.09 | 0.22 | 0.11 | 0.15 | 0.2 | 0.07 |
| 3 | 113631865 | rs16861409 | 1.147 | 1.008 | -1.352 | 0.377 | -0.003 | -0.086 | -0.086 | 0 | 0.13 | 0.01 | 0.25 | 0.3 | 0.1 | 0.14 |
| 3 | 113631884 | rs2712341 | 0.013 | 1.213 | -1.296 | 0.26 | 0.009 | -0.012 | -0.012 | 0 | 0.36 | 0.36 | 0.31 | 0.27 | 0.39 | 0.33 |
| 3 | 113637424 | rs6794522 | 0.478 | 2.018 | -1.023 | 0.651 | -0.029 | 0.006 | 0.006 | 0 | 0.15 | 0.09 | 0.16 | 0.15 | 0.15 | 0.22 |

| | | | | | | | | | | | | | | | | |
|---|---|---|---|---|---|---|---|---|---|---|---|---|---|---|---|---|
| 3 | 113645858 | rs9870411 | -1.255 | 2.093 | -0.787 | 0.845 | 0.006 | 0.002 | 0.002 | 0 | 0.48 | 0.52 | 0.61 | 0.61 | 0.51 | 0.55 |
| 3 | 113646657 | rs4422272 | -1.382 | 2.295 | -1.015 | 1.402 | -0.038 | -0.058 | -0.058 | 0 | 0.21 | 0.13 | 0.35 | 0.44 | 0.3 | 0.21 |
| 3 | 113646751 | rs17666485 | -0.936 | 2.683 | -0.66 | 0.794 | -0.015 | -0.011 | -0.011 | 0 | 0.08 | 0.06 | 0.1 | 0.14 | 0.19 | 0.07 |
| 3 | 113652774 | rs9865782 | 0.408 | 2.311 | -0.97 | 0.699 | -0.023 | -0.03 | -0.03 | 0 | 0.6 | 0.13 | 0.5 | 0.46 | 0.62 | 0.5 |
| 3 | 113654705 | rs7626657 | 0.758 | 2.167 | -0.521 | 0.068 | 0.005 | 0.005 | 0.005 | 0 | 0.46 | 0.42 | 0.36 | 0.38 | 0.38 | 0.37 |
| 3 | 113659549 | rs2029227 | 0.817 | 2.12 | -0.243 | 0.038 | 0.033 | 0.025 | 0.025 | 0 | 0.45 | 0.29 | 0.35 | 0.38 | 0.36 | 0.37 |
| 3 | 113673125 | rs11921691 | 0.339 | 2.247 | -0.004 | 0.505 | -0.006 | 0.004 | 0.004 | 0 | 0.61 | 0.65 | 0.53 | 0.67 | 0.56 | 0.54 |
| 3 | 113674025 | rs6799683 | -0.012 | 2.472 | 0.061 | 0.903 | 0.009 | 0.067 | 0.067 | 0.359 | 0.12 | 0.13 | 0.21 | 0.39 | 0.18 | 0.19 |
| 3 | 113675662 | rs9866271 | 0.552 | 2.4 | 0.24 | 0.886 | 0.003 | -0.01 | -0.01 | 0 | 0.57 | 0.48 | 0.52 | 0.65 | 0.56 | 0.54 |
| 3 | 113679589 | rs12629415 | -0.507 | 2.183 | 0.443 | 1.038 | 0.026 | 0.118 | 0.118 | 0.098 | 0.13 | 0.3 | 0.21 | 0.37 | 0.17 | 0.19 |
| 3 | 113679725 | rs3732785 | 0.111 | 1.882 | -0.232 | 0.716 | -0.054 | -0.001 | -0.001 | 0 | 0.69 | 0.93 | 0.72 | 0.76 | 0.65 | 0.77 |
| 3 | 113680090 | rs9816344 | 0.692 | 1.293 | -0.124 | 0.926 | -0.001 | -0.001 | -0.001 | 0 | 0.6 | 0.34 | 0.59 | 0.4 | 0.59 | 0.57 |
| 3 | 113682207 | rs1487268 | 1.083 | 1.002 | -0.248 | 1.123 | -0.021 | 0.023 | 0.023 | 0.187 | 0.46 | 0.09 | 0.43 | 0.39 | 0.44 | 0.4 |
| 3 | 113690427 | rs13096290 | 1.817 | 0.985 | -0.28 | 1.025 | -0.011 | 0.028 | 0.028 | 0.42 | 0.36 | 0.04 | 0.26 | 0.29 | 0.29 | 0.3 |
| 3 | 113700726 | rs3903132 | 0.06 | 1.267 | 0.194 | 1.021 | -0.005 | -0.003 | -0.003 | 0 | 0.6 | 0.64 | 0.55 | 0.58 | 0.57 | 0.52 |
| 3 | 113744268 | rs3796249 | -0.241 | 0.905 | 0.111 | 1.088 | -0.008 | -0.01 | -0.01 | 0.061 | 0.07 | 0.04 | 0.18 | 0.16 | 0.14 | 0.08 |
| 3 | 113749765 | rs2632241 | 1.559 | 0.908 | 0.079 | 1.06 | 0.032 | 0.094 | 0.094 | 0.981 | 0.41 | 0.11 | 0.27 | 0.27 | 0.31 | 0.31 |
| 3 | 113752500 | rs16861467 | -0.109 | 0.883 | 0.147 | 1.098 | -0.004 | 0 | 0 | 0.114 | 0.07 | 0.04 | 0.06 | 0.04 | 0.08 | 0.06 |
| 3 | 113753571 | rs2129571 | -0.753 | 0.881 | 0.179 | 1.099 | 0.01 | 0.029 | 0.029 | 0 | 0.09 | 0.14 | 0.07 | 0.14 | 0.08 | 0.13 |
| 3 | 113769097 | rs9859901 | -0.789 | 0.826 | 0.217 | 1.096 | 0.01 | 0.031 | 0.031 | 0 | 0.09 | 0.16 | 0.07 | 0.14 | 0.08 | 0.13 |
| 3 | 113774271 | rs6438180 | 0.28 | 0.808 | 0.206 | 1.089 | 0.006 | 0.001 | 0.001 | 0 | 0.57 | 0.5 | 0.53 | 0.58 | 0.52 | 0.51 |
| 3 | 113781736 | rs2306730 | 0.93 | 0.878 | 0.095 | 1.099 | -0.105 | -0.062 | -0.062 | 0.334 | 0.25 | 0.02 | 0.26 | 0.25 | 0.33 | 0.22 |
| 3 | 113783883 | rs16861476 | -0.372 | 0.804 | 0.348 | 0.924 | -0.002 | 0 | 0 | 0 | 0.08 | 0.05 | 0.06 | 0.04 | 0.08 | 0.06 |
| 3 | 113794154 | rs6782373 | -2.059 | 0.791 | 0.362 | 0.968 | 0.038 | 0.119 | 0.119 | 0 | 0.17 | 0.37 | 0.25 | 0.31 | 0.22 | 0.21 |
| 3 | 113794985 | rs11715834 | -0.101 | 0.814 | 0.256 | 0.924 | -0.009 | -0.01 | -0.01 | 0.281 | 0.07 | 0.04 | 0.17 | 0.16 | 0.14 | 0.08 |
| 3 | 113797117 | rs13314266 | 0.075 | 0.795 | 0.274 | 0.88 | 0 | 0 | 0 | 0.621 | 0.06 | 0.05 | 0.05 | 0.04 | 0.07 | 0.04 |
| 3 | 113804859 | rs3732788 | -0.654 | 0.787 | 0.282 | 0.89 | 0.006 | 0.015 | 0.015 | 0 | 0.09 | 0.14 | 0.08 | 0.13 | 0.08 | 0.11 |
| 3 | 113804979 | rs1872575 | 0.247 | 0.412 | 0.047 | 0.734 | 0 | -0.003 | -0.003 | 0 | 0.54 | 0.48 | 0.54 | 0.58 | 0.53 | 0.5 |
| 3 | 113808427 | rs9853414 | 0.283 | 0.426 | 0.057 | 0.618 | -0.005 | -0.007 | -0.007 | 0 | 0.55 | 0.63 | 0.54 | 0.51 | 0.52 | 0.48 |
| 3 | 113811278 | rs7632864 | -0.975 | 0.422 | -0.104 | 0.558 | 0.002 | 0.012 | 0.012 | 0.169 | 0.55 | 0.6 | 0.64 | 0.68 | 0.6 | 0.66 |
| 3 | 113811688 | rs13318232 | -0.048 | 0.473 | -0.413 | 0.504 | -0.001 | 0 | 0 | 0 | 0.08 | 0.07 | 0.06 | 0.04 | 0.08 | 0.08 |
| 3 | 113812644 | rs12630513 | -0.925 | 0.455 | -0.487 | 0.543 | -0.002 | -0.004 | -0.004 | 0 | 0.16 | 0.15 | 0.32 | 0.24 | 0.22 | 0.28 |
| 3 | 113812918 | rs10934250 | -0.814 | 0.449 | -0.631 | 0.512 | -0.008 | -0.021 | -0.021 | 0 | 0.16 | 0.1 | 0.32 | 0.23 | 0.22 | 0.28 |
| 3 | 113815108 | rs9824256 | -0.874 | 0.445 | -0.609 | 0.504 | 0.003 | 0.076 | 0.076 | 0.312 | 0.56 | 0.83 | 0.66 | 0.7 | 0.61 | 0.68 |
| 3 | 113818348 | rs13094677 | 1.551 | 0.219 | -0.881 | 0.52 | 0.004 | 0.064 | 0.064 | 2.56 | 0.36 | 0.09 | 0.29 | 0.26 | 0.3 | 0.24 |
| 3 | 113826308 | rs10934251 | -0.384 | 0.167 | -0.473 | 0.781 | -0.002 | 0.011 | 0.011 | 0 | 0.06 | 0.08 | 0.17 | 0.17 | 0.11 | 0.08 |
| 3 | 113829346 | rs4682522 | 0.449 | 0.053 | -0.273 | 0.525 | 0.001 | 0.003 | 0.003 | 0 | 0.91 | 0.88 | 0.95 | 0.92 | 0.93 | 0.89 |
| 3 | 113830519 | rs7631540 | 0.606 | -0.224 | -0.677 | 0.512 | -0.033 | -0.002 | -0.002 | 0 | 0.5 | 0.11 | 0.53 | 0.42 | 0.53 | 0.45 |
| 3 | 113847108 | rs9817063 | -0.055 | -0.185 | -0.058 | 0.403 | 0.024 | 0.021 | 0.021 | 0 | 0.34 | 0.36 | 0.46 | 0.45 | 0.45 | 0.44 |
| 3 | 113859173 | rs9288993 | 0.233 | -0.212 | 0.01 | 0.211 | 0.012 | -0.016 | -0.016 | 0.342 | 0.06 | 0.24 | 0.02 | 0 | 0.02 | 0.04 |
| 3 | 113862887 | rs963468 | 0.805 | -0.084 | -0.461 | 0.326 | -0.112 | -0.109 | -0.109 | 0.882 | 0.24 | 0.05 | 0.35 | 0.39 | 0.35 | 0.31 |
| 3 | 113870078 | rs3773678 | 1.517 | -0.805 | 0.546 | 0.078 | -0.007 | -0.005 | -0.005 | 0 | 0.71 | 0.31 | 0.82 | 0.85 | 0.87 | 0.71 |
| 3 | 113873372 | rs2630349 | 0.565 | -0.833 | 0.391 | 0.117 | -0.022 | 0.019 | 0.019 | 0 | 0.89 | 0.65 | 0.88 | 0.89 | 0.94 | 0.87 |
| 3 | 113876275 | rs167771 | 1.983 | -0.892 | -0.013 | 0.018 | 0.085 | 0.089 | 0.089 | 0.509 | 0.65 | 0.16 | 0.73 | 0.83 | 0.8 | 0.6 |
| 3 | 113881623 | rs324029 | 1.938 | -0.629 | -0.032 | -0.144 | 0.011 | 0.01 | 0.01 | 0.161 | 0.65 | 0.26 | 0.6 | 0.72 | 0.7 | 0.66 |
| 3 | 113887298 | rs324022 | 1.839 | -0.642 | -0.034 | -0.147 | 0.016 | 0.012 | 0.012 | 0.317 | 0.64 | 0.18 | 0.59 | 0.71 | 0.7 | 0.61 |
| 3 | 113888532 | rs1486009 | -0.22 | -0.658 | 0.053 | -0.192 | 0.002 | 0.006 | 0.006 | 0 | 0.07 | 0.07 | 0.09 | 0.01 | 0.08 | 0.1 |
| 3 | 113889069 | rs6806217 | 0.233 | -0.692 | 0.357 | -0.212 | 0.014 | -0.013 | -0.013 | 0 | 0.05 | 0.23 | 0.03 | 0 | 0.03 | 0.04 |
| 3 | 113890310 | rs16822416 | -0.694 | -0.674 | 0.285 | -0.197 | -0.006 | -0.026 | -0.026 | 0 | 0.08 | 0.14 | 0.1 | 0.01 | 0.08 | 0.13 |
| 3 | 113890789 | rs3732783 | -0.729 | -0.674 | 0.284 | -0.196 | -0.006 | -0.024 | -0.024 | 0.004 | 0.08 | 0.14 | 0.1 | 0.01 | 0.08 | 0.15 |
| 3 | 113890815 | rs6280 | 2.034 | -0.651 | 0.112 | -0.183 | -0.003 | 0.008 | 0.008 | 0.428 | 0.63 | 0.18 | 0.57 | 0.69 | 0.66 | 0.58 |
| 3 | 113900220 | rs9825563 | -1.803 | -0.718 | 0.338 | -0.308 | -0.005 | -0.006 | -0.006 | 1.031 | 0.31 | 0.52 | 0.32 | 0.29 | 0.28 | 0.34 |
| 3 | 113902749 | rs6787134 | -0.769 | -0.717 | 0.128 | -0.313 | 0.026 | 0.042 | 0.042 | 0.642 | 0.21 | 0.11 | 0.07 | 0.06 | 0.12 | 0.2 |
| 3 | 113902835 | rs1587757 | 0.175 | -0.755 | 0.101 | -0.3 | -0.016 | -0.007 | -0.007 | 0 | 0.07 | 0.05 | 0.23 | 0.23 | 0.16 | 0.08 |
| 3 | 113903791 | rs9880494 | -2.212 | -0.795 | 0.114 | -0.324 | 0.008 | 0.021 | 0.021 | 0.2 | 0.37 | 0.75 | 0.42 | 0.3 | 0.34 | 0.39 |
| 3 | 113908226 | rs9813633 | -1.141 | -0.744 | 0.165 | -0.292 | 0.036 | 0.006 | 0.006 | 0 | 0.07 | 0.38 | 0.1 | 0.01 | 0.06 | 0.11 |
| 3 | 113909889 | rs1394016 | -2.318 | -0.937 | -0.365 | -0.274 | 0.036 | 0.055 | 0.055 | 0 | 0.48 | 0.91 | 0.47 | 0.36 | 0.38 | 0.5 |
| 3 | 113924143 | rs7611535 | -1.465 | -0.36 | 0.125 | 0.039 | -0.017 | -0.016 | -0.016 | 0 | 0.21 | 0.14 | 0.33 | 0.24 | 0.27 | 0.21 |
| 3 | 113928332 | rs7646185 | 0.481 | -0.327 | -0.089 | 0.055 | 0.003 | 0.004 | 0.004 | 4.44 | 0.1 | 0.07 | 0.03 | 0.04 | 0.03 | 0.15 |
| 3 | 113935873 | rs7610054 | -0.434 | -0.293 | -0.278 | 0.183 | -0.023 | -0.025 | -0.025 | 0.024 | 0.12 | 0.06 | 0.3 | 0.24 | 0.22 | 0.13 |
| 3 | 113937999 | rs7616367 | -1.52 | -0.467 | -0.441 | 0.207 | 0.025 | 0.018 | 0.018 | 0.194 | 0.36 | 0.73 | 0.4 | 0.28 | 0.3 | 0.34 |
| 3 | 113939244 | rs7653787 | -0.808 | -0.891 | -0.576 | -0.144 | -0.053 | -0.073 | -0.073 | 0 | 0.22 | 0.35 | 0.08 | 0.05 | 0.08 | 0.22 |
| 3 | 113947863 | rs1503673 | 0.092 | -0.55 | -1.571 | 0.143 | 0.014 | -0.003 | -0.003 | 0 | 0.22 | 0.05 | 0.17 | 0.2 | 0.18 | 0.12 |
| 3 | 113947975 | rs7620754 | -0.248 | -0.625 | -1.911 | 0.198 | 0.009 | 0.004 | 0.004 | 0 | 0.33 | 0.25 | 0.27 | 0.32 | 0.26 | 0.34 |
| 3 | 113950127 | rs2062834 | -0.528 | -0.016 | -2.153 | 0.17 | -0.059 | -0.062 | -0.062 | 2.243 | 0.73 | 0.84 | 0.55 | 0.55 | 0.59 | 0.66 |
| 3 | 113952161 | rs2279828 | 0.244 | 0.355 | -2.109 | 0.017 | 0.002 | 0.001 | 0.001 | 0 | 0.08 | 0.11 | 0.08 | 0.09 | 0.12 | 0.06 |
| 3 | 113955164 | rs3732782 | -0.989 | 0.516 | -2.281 | 0.171 | -0.031 | -0.052 | -0.052 | 0 | 0.78 | 0.85 | 0.6 | 0.55 | 0.68 | 0.71 |
| 3 | 113955187 | rs3732781 | 1.279 | -0.378 | -3.022 | -0.458 | -0.022 | 0.089 | 0.089 | 0.018 | 0.34 | 0.03 | 0.24 | 0.16 | 0.28 | 0.29 |
| 3 | 113955265 | rs6438190 | -1.488 | -0.215 | -2.965 | -0.348 | -0.031 | -0.052 | -0.052 | 0 | 0.78 | 0.85 | 0.6 | 0.54 | 0.68 | 0.71 |

| | | | | | | | | | | | | | | | | |
|---|---|---|---|---|---|---|---|---|---|---|---|---|---|---|---|---|
| 3 | 113955320 | rs6438191 | -1.488 | -0.215 | -2.965 | -0.348 | -0.031 | -0.052 | -0.052 | 0 | 0.78 | 0.85 | 0.6 | 0.54 | 0.68 | 0.71 |
| 3 | 113958599 | rs905569 | -0.21 | -0.486 | -3.05 | -0.529 | -0.002 | -0.028 | -0.028 | 0 | 0.24 | 0.17 | 0.23 | 0.37 | 0.24 | 0.25 |
| 3 | 113960449 | rs1354350 | 0.669 | -0.919 | -2.069 | -0.774 | -0.005 | -0.021 | -0.021 | 0 | 0.43 | 0.47 | 0.3 | 0.17 | 0.34 | 0.38 |
| 3 | 113961934 | rs1983079 | 0.66 | -0.402 | -2.299 | -0.464 | 0.001 | -0.039 | -0.039 | 0.008 | 0.23 | 0.14 | 0.32 | 0.35 | 0.2 | 0.28 |
| 3 | 113962275 | rs9809768 | -0.161 | -0.115 | -2.167 | -0.343 | 0.004 | 0.011 | 0.011 | 0 | 0.7 | 0.64 | 0.62 | 0.53 | 0.61 | 0.73 |
| 3 | 113963869 | rs9819579 | 0.96 | 0.004 | -2.335 | -0.482 | -0.006 | -0.03 | -0.03 | 0 | 0.21 | 0.15 | 0.32 | 0.37 | 0.19 | 0.27 |
| 3 | 113973702 | rs1503672 | -0.218 | -0.594 | -2.181 | -1.183 | -0.005 | 0.206 | 0.206 | 0 | 0.54 | 0.87 | 0.28 | 0.13 | 0.42 | 0.43 |
| 3 | 113986760 | rs2733424 | 0.132 | -0.546 | -1.301 | -1.173 | 0.037 | 0.208 | 0.208 | 0 | 0.49 | 0.8 | 0.26 | 0.13 | 0.4 | 0.43 |
| 3 | 113991889 | rs2683782 | 0.028 | -0.291 | -1.111 | -1.198 | 0.011 | -0.015 | -0.015 | 0 | 0.1 | 0.22 | 0.06 | 0.06 | 0.11 | 0.13 |
| 3 | 113993582 | rs2683779 | 0.138 | -0.042 | -1.616 | -0.917 | -0.012 | 0.195 | 0.195 | 0 | 0.52 | 0.83 | 0.29 | 0.19 | 0.44 | 0.47 |
| 3 | 114007076 | rs1521070 | 0.252 | -0.026 | -1.527 | -0.942 | 0.009 | 0.163 | 0.163 | 0 | 0.51 | 0.77 | 0.28 | 0.19 | 0.44 | 0.47 |
| 3 | 114012274 | rs1039524 | -0.303 | 0.025 | -1.453 | -1.217 | -0.049 | 0.169 | 0.169 | 0.439 | 0.61 | 0.83 | 0.3 | 0.09 | 0.45 | 0.47 |
| 3 | 114012467 | rs2693052 | -0.206 | 0.102 | -1.146 | -1.183 | -0.003 | 0.057 | 0.057 | 0 | 0.57 | 0.65 | 0.28 | 0.09 | 0.45 | 0.47 |
| 3 | 114016047 | rs2654758 | 0.184 | -0.075 | -1.014 | -1.155 | 0.014 | 0.003 | 0.003 | 0 | 0.46 | 0.52 | 0.19 | 0.09 | 0.33 | 0.37 |
| 3 | 114016610 | rs2683789 | 0.461 | -0.014 | -0.882 | -0.915 | -0.008 | -0.032 | -0.032 | 0 | 0.39 | 0.37 | 0.17 | 0.15 | 0.34 | 0.37 |
| 3 | 114017258 | rs2693051 | -1.054 | -0.032 | -1.249 | -1.283 | 0.023 | 0.054 | 0.054 | 0.141 | 0.69 | 0.86 | 0.88 | 0.89 | 0.79 | 0.73 |
| 3 | 114024063 | rs1316642 | 0.191 | -0.331 | -1.077 | -1.114 | -0.001 | -0.002 | -0.002 | 0 | 0.33 | 0.3 | 0.37 | 0.61 | 0.32 | 0.36 |
| 3 | 114024530 | rs10934259 | 1.278 | -0.548 | -1.157 | -2.059 | -0.053 | 0.184 | 0.184 | 0 | 0.35 | 0.1 | 0.32 | 0.07 | 0.36 | 0.37 |
| 3 | 114028633 | rs9825470 | -0.272 | -0.436 | -0.842 | -2.013 | -0.001 | -0.042 | -0.042 | 0 | 0.5 | 0.46 | 0.6 | 0.92 | 0.55 | 0.46 |
| 3 | 114034098 | rs4682534 | 0.142 | -0.456 | -0.268 | -1.271 | 0.05 | -0.016 | -0.016 | 0.032 | 0.93 | 0.75 | 0.95 | 1 | 0.94 | 0.91 |
| 3 | 114050607 | rs4682536 | -0.09 | -0.831 | -0.463 | -1.33 | -0.045 | 0.086 | 0.086 | 0 | 0.37 | 0.26 | 0.46 | 0.82 | 0.46 | 0.36 |
| 3 | 114054707 | rs9851950 | -0.247 | -0.811 | -0.304 | -1.327 | -0.029 | 0.089 | 0.089 | 0 | 0.41 | 0.31 | 0.54 | 0.85 | 0.49 | 0.37 |
| 3 | 114055355 | rs2693039 | -0.716 | -1.142 | 0.637 | -0.945 | -0.037 | -0.056 | -0.056 | 0.32 | 0.11 | 0.33 | 0.03 | 0.01 | 0.02 | 0.18 |
| 3 | 114067518 | rs2722015 | 0.735 | -0.711 | 0.332 | -0.854 | 0.001 | -0.015 | -0.015 | 0 | 0.41 | 0.39 | 0.37 | 0.12 | 0.41 | 0.46 |
| 3 | 114072839 | rs9871676 | -0.601 | -0.772 | 0.363 | -0.841 | 0.003 | -0.02 | -0.02 | 0 | 0.54 | 0.55 | 0.61 | 0.88 | 0.56 | 0.53 |
| 3 | 114078257 | rs17670847 | 1.079 | -0.965 | 0.359 | -0.834 | -0.006 | 0.066 | 0.066 | 0 | 0.37 | 0.23 | 0.34 | 0.12 | 0.4 | 0.4 |
| 3 | 114085053 | rs1830095 | 0.721 | -0.933 | 0.382 | -0.825 | 0.004 | -0.062 | -0.062 | 0.016 | 0.42 | 0.47 | 0.39 | 0.13 | 0.46 | 0.49 |
| 3 | 114097441 | rs2683792 | 0.833 | -0.925 | 0.192 | -0.431 | -0.001 | 0.003 | 0.003 | 0 | 0.41 | 0.41 | 0.39 | 0.29 | 0.44 | 0.43 |
| 3 | 114104227 | rs9881461 | -0.523 | -0.902 | 0.093 | -0.392 | -0.001 | -0.011 | -0.011 | 0.012 | 0.52 | 0.48 | 0.59 | 0.7 | 0.52 | 0.5 |
| 3 | 114110200 | rs17615035 | 1.561 | -1.865 | -0.435 | -0.214 | -0.107 | 0.064 | 0.064 | 0 | 0.34 | 0.03 | 0.3 | 0.21 | 0.37 | 0.41 |
| 3 | 114110546 | rs9859157 | 0.176 | -2.1 | -0.152 | -0.056 | -0.034 | -0.009 | -0.009 | 0 | 0.36 | 0.22 | 0.36 | 0.34 | 0.4 | 0.38 |
| 3 | 114111407 | rs1105821 | -0.346 | -2.074 | -0.03 | 0.009 | -0.015 | -0.022 | -0.022 | 0 | 0.9 | 0.71 | 0.93 | 0.97 | 0.94 | 0.99 |
| 3 | 114112056 | rs3957585 | -0.79 | -1.801 | 0.01 | 0.084 | 0 | -0.005 | -0.005 | 0 | 0.45 | 0.5 | 0.39 | 0.33 | 0.42 | 0.4 |
| 3 | 114115034 | rs10439990 | -0.425 | -1.605 | -0.008 | 0.187 | 0.001 | -0.001 | -0.001 | 0.005 | 0.4 | 0.43 | 0.44 | 0.35 | 0.41 | 0.38 |
| 3 | 114117327 | rs16822593 | -0.212 | -1.343 | 0.254 | -0.21 | -0.046 | -0.013 | -0.013 | 0 | 0.18 | 0.37 | 0.24 | 0.17 | 0.13 | 0.05 |
| 3 | 114119500 | rs9878858 | 0.512 | -1.341 | -0.107 | -0.343 | -0.004 | 0.008 | 0.008 | 0 | 0.11 | 0.09 | 0.09 | 0.03 | 0.12 | 0.15 |
| 3 | 114121019 | rs10511330 | -0.1 | -1.209 | 0.014 | -0.238 | -0.045 | -0.005 | -0.005 | 0 | 0.18 | 0.42 | 0.25 | 0.18 | 0.13 | 0.05 |
| 3 | 114121762 | rs2733403 | -0.311 | -1.548 | 0.091 | -0.396 | -0.026 | -0.097 | -0.097 | 0 | 0.24 | 0.4 | 0.14 | 0.05 | 0.19 | 0.25 |
| 3 | 114123929 | rs7428451 | -0.059 | -1.609 | -0.368 | -0.33 | -0.045 | -0.005 | -0.005 | 0.013 | 0.18 | 0.42 | 0.26 | 0.18 | 0.13 | 0.05 |
| 3 | 114131674 | rs2733399 | -1.451 | -1.108 | -1.687 | -0.026 | -0.097 | 0.062 | 0.062 | 0 | 0.64 | 0.95 | 0.71 | 0.76 | 0.61 | 0.58 |
| 3 | 114132129 | rs9843972 | 0.132 | -0.641 | -1.295 | 0.269 | -0.067 | -0.09 | -0.09 | 0 | 0.22 | 0.06 | 0.31 | 0.53 | 0.3 | 0.28 |
| 3 | 114132816 | rs4510355 | -0.712 | -1.818 | -0.447 | -0.201 | -0.009 | -0.027 | -0.027 | 0.581 | 0.36 | 0.6 | 0.31 | 0.14 | 0.29 | 0.23 |
| 3 | 114133443 | rs6787896 | 0.972 | -2.061 | -0.095 | -0.161 | -0.032 | -0.049 | -0.049 | 0 | 0.77 | 0.35 | 0.72 | 0.91 | 0.86 | 0.87 |
| 3 | 114133924 | rs2722013 | 1.23 | -2.613 | -0.23 | 0.324 | 0.085 | 0.026 | 0.026 | 0.012 | 0.55 | 0.21 | 0.58 | 0.63 | 0.74 | 0.58 |
| 3 | 114134268 | rs2722012 | -1.075 | -2.816 | 0.436 | -0.21 | -0.008 | -0.022 | -0.022 | 0 | 0.33 | 0.58 | 0.33 | 0.1 | 0.18 | 0.25 |
| 3 | 114140714 | rs2733393 | -0.73 | -2.799 | 0.834 | -0.167 | -0.02 | -0.019 | -0.019 | 0 | 0.3 | 0.33 | 0.38 | 0.25 | 0.17 | 0.29 |
| 3 | 114142624 | rs16822651 | -0.473 | -2.935 | 0.727 | -0.017 | 0.008 | -0.013 | -0.013 | 1.361 | 0.08 | 0.04 | 0.07 | 0.17 | 0.06 | 0.15 |
| 3 | 114144733 | rs2722019 | -1.459 | -2.981 | 0.609 | -0.035 | -0.007 | -0.001 | -0.001 | 0.103 | 0.39 | 0.36 | 0.38 | 0.41 | 0.23 | 0.44 |
| 3 | 114150334 | rs2722033 | 0.778 | -2.901 | 0.447 | -0.003 | -0.031 | 0.104 | 0.104 | 1.037 | 0.88 | 0.59 | 0.9 | 0.83 | 0.93 | 0.84 |
| 3 | 114155161 | rs2718430 | -0.744 | -2.863 | 0.061 | -0.056 | -0.003 | -0.003 | -0.003 | 0.602 | 0.3 | 0.25 | 0.3 | 0.24 | 0.17 | 0.29 |
| 3 | 114162723 | rs9882269 | 1.167 | -2.447 | -0.696 | -0.126 | 0.146 | 0.053 | 0.053 | 0.734 | 0.53 | 0.17 | 0.51 | 0.58 | 0.76 | 0.54 |
| 3 | 114171390 | rs9883949 | -1.126 | -2.328 | 0.176 | 0.041 | -0.09 | 0.009 | 0.009 | 0 | 0.23 | 0.44 | 0.31 | 0.25 | 0.09 | 0.21 |
| 3 | 114174552 | rs4131837 | 0.893 | -2.335 | 0.281 | 0.056 | -0.102 | 0.015 | 0.015 | 0 | 0.77 | 0.54 | 0.75 | 0.74 | 0.93 | 0.83 |
| 3 | 114174924 | rs9856597 | 1.771 | -2.282 | 0.491 | -0.129 | 0.018 | 0.004 | 0.004 | 1.147 | 0.66 | 0.26 | 0.64 | 0.69 | 0.84 | 0.66 |
| 3 | 114178479 | rs10934275 | 1.78 | -2.286 | 0.489 | -0.13 | 0.021 | 0.005 | 0.005 | 1.11 | 0.66 | 0.26 | 0.64 | 0.69 | 0.84 | 0.66 |
| 3 | 114183575 | rs11917631 | -1.135 | -1.95 | 0.782 | -0.143 | -0.065 | 0.026 | 0.026 | 0 | 0.21 | 0.54 | 0.19 | 0.25 | 0.1 | 0.15 |
| 3 | 114183612 | rs11921658 | -1.697 | -1.89 | 0.723 | -0.142 | -0.021 | 0 | 0 | 0 | 0.3 | 0.62 | 0.37 | 0.32 | 0.19 | 0.32 |
| 3 | 114191863 | rs10511334 | -0.97 | -1.873 | 0.34 | -0.022 | -0.026 | 0.007 | 0.007 | 0 | 0.21 | 0.24 | 0.19 | 0.25 | 0.09 | 0.16 |
| 3 | 114193935 | rs1609969 | -1.453 | -1.899 | 0.305 | -0.38 | -0.017 | -0.035 | -0.035 | 0 | 0.14 | 0.27 | 0.12 | 0.07 | 0.09 | 0.17 |
| 3 | 114196123 | rs2718424 | -1.587 | -2.712 | -0.004 | -0.738 | 0.045 | 0.05 | 0.05 | 1.062 | 0.27 | 0.17 | 0.17 | 0.11 | 0.13 | 0.08 |
| 3 | 114199151 | rs2669898 | 1.55 | -2.431 | 0.05 | -1.014 | 0.01 | -0.002 | -0.002 | 0 | 0.71 | 0.42 | 0.57 | 0.7 | 0.69 | 0.61 |
| 3 | 114199650 | rs9873460 | 2.335 | -2.453 | 0.125 | -0.996 | 0.057 | 0.078 | 0.078 | 0.682 | 0.51 | 0.15 | 0.45 | 0.62 | 0.64 | 0.55 |
| 3 | 114205687 | rs2718425 | 0.742 | -2.035 | -0.382 | -1.504 | 0.006 | -0.01 | -0.01 | 2.008 | 0.7 | 0.42 | 0.56 | 0.75 | 0.62 | 0.68 |
| 3 | 114208597 | rs2718423 | 0.323 | -1.916 | -0.118 | -1.324 | 0.017 | 0.028 | 0.028 | 0 | 0.82 | 0.44 | 0.69 | 0.76 | 0.81 | 0.7 |
| 3 | 114221822 | rs7643975 | -0.601 | -1.892 | 0.338 | -1.639 | 0.089 | -0.014 | -0.014 | 0 | 0.17 | 0.44 | 0.22 | 0.15 | 0.21 | 0.26 |
| 3 | 114233596 | rs7620649 | -1.701 | -2.21 | 0.073 | -1.972 | 0.009 | 0.024 | 0.024 | 0.086 | 0.37 | 0.63 | 0.29 | 0.17 | 0.28 | 0.32 |
| 3 | 114236979 | rs11920889 | -0.391 | -2.22 | -0.185 | -2.107 | 0.03 | 0.031 | 0.031 | 0 | 0.19 | 0.11 | 0.06 | 0.03 | 0.07 | 0.06 |
| 3 | 114237183 | rs9878302 | -0.352 | -1.737 | 0.131 | -1.973 | 0.02 | -0.011 | -0.011 | 0 | 0.12 | 0.17 | 0.15 | 0.1 | 0.17 | 0.18 |
| 3 | 114244796 | rs16822875 | -0.583 | -1.91 | -0.213 | -1.887 | 0.007 | 0.007 | 0.007 | 1.088 | 0.1 | 0.09 | 0.12 | 0.06 | 0.03 | 0.12 |

| | | | | | | | | | | | | | | | | |
|---|---|---|---|---|---|---|---|---|---|---|---|---|---|---|---|---|
| 3 | 114246048 | rs16822883 | -0.388 | -1.891 | -0.222 | -1.884 | 0.008 | 0.008 | 0.008 | 1.266 | 0.09 | 0.07 | 0.12 | 0.06 | 0.03 | 0.12 |
| 3 | 114250796 | rs2669885 | -0.25 | -1.774 | 0.075 | -2.133 | 0.001 | -0.076 | -0.076 | 0 | 0.22 | 0.4 | 0.2 | 0.08 | 0.24 | 0.17 |
| 3 | 114257298 | rs2684290 | 1.132 | -1.718 | -0.198 | -2.333 | -0.007 | -0.028 | -0.028 | 0 | 0.55 | 0.52 | 0.6 | 0.8 | 0.69 | 0.59 |
| 3 | 114274333 | rs7615798 | -0.925 | -1.069 | 0.307 | -2.204 | 0.001 | -0.04 | -0.04 | 0 | 0.16 | 0.24 | 0.15 | 0.09 | 0.12 | 0.29 |
| 3 | 114275543 | rs6772469 | -0.378 | -1.427 | -0.384 | -2.511 | -0.001 | 0.066 | 0.066 | 0 | 0.19 | 0.05 | 0.23 | 0.06 | 0.13 | 0.18 |
| 3 | 114278138 | rs13066213 | 0.203 | -1.244 | -0.338 | -2.736 | 0.001 | 0.008 | 0.008 | 2.825 | 0.19 | 0.14 | 0.19 | 0.07 | 0.22 | 0.1 |
| 3 | 114278322 | rs2718417 | -1.546 | -1.304 | -0.259 | -2.778 | -0.001 | -0.028 | -0.028 | 0 | 0.38 | 0.57 | 0.35 | 0.16 | 0.35 | 0.4 |
| 3 | 114280380 | rs1589435 | 1.53 | -1.625 | -0.836 | -2.44 | 0.015 | 0.218 | 0.218 | 0 | 0.58 | 0.1 | 0.57 | 0.85 | 0.74 | 0.46 |
| 3 | 114293296 | rs11914467 | -0.348 | -1.751 | 0.228 | -2.265 | 0.001 | -0.062 | -0.062 | 0 | 0.12 | 0.25 | 0.08 | 0.02 | 0.08 | 0.08 |
| 3 | 114300652 | rs4300983 | -0.464 | -1.714 | 0.145 | -2.482 | -0.007 | 0.059 | 0.059 | 0 | 0.19 | 0.06 | 0.22 | 0.04 | 0.14 | 0.2 |
| 3 | 114305702 | rs7637678 | 2.186 | -1.761 | 0.54 | -2.822 | 0.023 | 0.105 | 0.105 | 0.011 | 0.48 | 0.34 | 0.56 | 0.91 | 0.63 | 0.6 |
| 3 | 114315566 | rs6796509 | -0.263 | -1.838 | 0.688 | -2.814 | -0.003 | 0.035 | 0.035 | 0 | 0.19 | 0.12 | 0.23 | 0.04 | 0.14 | 0.18 |
| 3 | 114318110 | rs9869646 | -0.257 | -1.816 | 0.699 | -2.805 | -0.004 | 0.037 | 0.037 | 0 | 0.18 | 0.12 | 0.23 | 0.04 | 0.14 | 0.18 |
| 3 | 114319046 | rs950992 | 0.118 | -1.828 | 0.963 | -2.8 | -0.002 | -0.041 | -0.041 | 0 | 0.11 | 0.19 | 0.08 | 0.01 | 0.08 | 0.08 |
| 3 | 114325385 | rs2947483 | -0.439 | -1.843 | 1.06 | -2.813 | -0.024 | -0.076 | -0.076 | 0 | 0.81 | 0.75 | 0.88 | 0.98 | 0.85 | 0.97 |
| 3 | 114327437 | rs16823002 | 0.252 | -1.837 | 0.77 | -2.812 | -0.001 | -0.033 | -0.033 | 0 | 0.11 | 0.15 | 0.07 | 0.01 | 0.08 | 0.08 |
| 3 | 114329409 | rs1274270 | -1.692 | -1.865 | 0.783 | -2.822 | 0.083 | 0.073 | 0.073 | 1.991 | 0.05 | 0.4 | 0.03 | 0 | 0.01 | 0.01 |
| 3 | 114329961 | rs6438213 | -1.824 | -1.765 | 0.381 | -2.826 | 0.021 | 0.129 | 0.129 | 0.291 | 0.46 | 0.29 | 0.4 | 0.08 | 0.36 | 0.38 |
| 3 | 114330091 | rs6438214 | -1.445 | -1.868 | 0.442 | -2.771 | 0.001 | -0.009 | -0.009 | 0 | 0.31 | 0.32 | 0.31 | 0.05 | 0.21 | 0.34 |
| 3 | 114331913 | rs4682162 | -0.262 | -1.737 | 0.05 | -2.678 | -0.015 | 0.06 | 0.06 | 0 | 0.18 | 0.07 | 0.22 | 0.04 | 0.14 | 0.18 |
| 3 | 114334478 | rs9852045 | -0.533 | -1.717 | 0.09 | -2.669 | 0.002 | 0.026 | 0.026 | 0 | 0.2 | 0.17 | 0.25 | 0.05 | 0.14 | 0.27 |
| 3 | 114341380 | rs1818757 | 2.005 | -1.489 | -0.046 | -2.327 | -0.053 | 0.092 | 0.092 | 0 | 0.39 | 0.14 | 0.46 | 0.72 | 0.55 | 0.42 |
| 3 | 114347164 | rs11717204 | 0.112 | -1.557 | 0.439 | -2.028 | 0.052 | 0.075 | 0.075 | 7.91 | 0.16 | 0.01 | 0.12 | 0.03 | 0.06 | 0.01 |
| 3 | 114353140 | rs1290894 | -1.542 | -1.486 | 1.378 | -1.433 | 0.107 | 0.106 | 0.106 | 0 | 0.1 | 0.46 | 0.18 | 0.17 | 0.07 | 0.17 |
| 3 | 114353168 | rs9820958 | -0.885 | -1.459 | 1.001 | -1.414 | 0.028 | 0.038 | 0.038 | 0 | 0.08 | 0.16 | 0.17 | 0.18 | 0.07 | 0.17 |
| 3 | 114357203 | rs1274262 | -1.75 | -1.585 | 0.688 | -1.605 | 0.025 | 0.033 | 0.033 | 1.188 | 0.37 | 0.68 | 0.4 | 0.2 | 0.22 | 0.24 |
| 3 | 114379193 | rs17681207 | 0.747 | -1.28 | -0.068 | -1.427 | 0.038 | 0.046 | 0.046 | 2.886 | 0.11 | 0.02 | 0.11 | 0.03 | 0.04 | 0.01 |
| 3 | 114391780 | rs1921493 | 1.947 | -1.177 | -0.099 | -1.532 | 0.041 | 0.145 | 0.145 | 0.778 | 0.57 | 0.2 | 0.56 | 0.79 | 0.72 | 0.68 |
| 3 | 114399296 | rs17681451 | -0.25 | -1.591 | -0.195 | -1.18 | -0.031 | 0.039 | 0.039 | 0 | 0.1 | 0.01 | 0.05 | 0.01 | 0.07 | 0.07 |
| 3 | 114417343 | rs10934276 | 0.707 | -1.408 | 0.474 | -0.869 | -0.026 | -0.034 | -0.034 | 0.018 | 0.82 | 0.54 | 0.73 | 0.85 | 0.87 | 0.81 |
| 3 | 114424965 | rs13072722 | 0.973 | -1.474 | -0.093 | -0.895 | -0.01 | -0.014 | -0.014 | 1.306 | 0.69 | 0.31 | 0.57 | 0.83 | 0.84 | 0.8 |
| 3 | 114434158 | rs2178555 | -0.887 | -1.154 | -0.281 | -0.375 | 0.013 | -0.024 | -0.024 | 1.134 | 0.14 | 0.07 | 0.19 | 0.34 | 0.09 | 0.27 |
| 3 | 114440159 | rs2222855 | -0.507 | -1.587 | 0.531 | -0.613 | -0.049 | -0.073 | -0.073 | 0 | 0.19 | 0.36 | 0.18 | 0.06 | 0.08 | 0.12 |
| 3 | 114443668 | rs16823137 | -0.373 | -1.665 | 0.633 | -0.6 | -0.058 | -0.057 | -0.057 | 0 | 0.18 | 0.32 | 0.17 | 0.07 | 0.06 | 0.09 |
| 3 | 114444310 | rs1274226 | -1.959 | -1.642 | 0.625 | -0.949 | 0.045 | 0.038 | 0.038 | 0 | 0.43 | 0.65 | 0.44 | 0.43 | 0.22 | 0.46 |
| 3 | 114449264 | rs9873528 | -0.542 | -1.313 | 0.436 | -0.729 | 0.038 | -0.038 | -0.038 | 0 | 0.14 | 0.05 | 0.17 | 0.36 | 0.08 | 0.27 |
| 3 | 114453255 | rs1274224 | -0.27 | -1.341 | 1.083 | -0.725 | 0.023 | -0.049 | -0.049 | 0 | 0.11 | 0.26 | 0.08 | 0.01 | 0.08 | 0.09 |
| 3 | 114461208 | rs1882289 | 0.539 | -1.264 | 1.302 | -0.702 | -0.02 | 0.061 | 0.061 | 0 | 0.84 | 0.62 | 0.8 | 0.64 | 0.88 | 0.71 |
| 3 | 114464858 | rs9826148 | -0.568 | -1.251 | 0.746 | -0.674 | 0.016 | -0.009 | -0.009 | 1.1 | 0.15 | 0.06 | 0.18 | 0.26 | 0.11 | 0.29 |
| 3 | 114466100 | rs16823178 | -0.168 | -1.13 | 0.822 | -0.692 | -0.008 | 0.002 | 0.002 | 0.006 | 0.1 | 0.04 | 0.07 | 0.09 | 0.07 | 0.09 |
| 3 | 114467929 | rs16823186 | -1.178 | -1.148 | 1.542 | -0.829 | 0.019 | -0.02 | -0.02 | 0 | 0.14 | 0.26 | 0.1 | 0.1 | 0.11 | 0.1 |
| 3 | 114469199 | rs2951800 | -0.025 | -1.166 | 1.739 | -0.887 | 0.022 | -0.036 | -0.036 | 0 | 0.91 | 0.78 | 0.93 | 0.99 | 0.93 | 0.93 |
| 3 | 114471931 | rs2140345 | 1.275 | -1.195 | 2.043 | -0.899 | 0.04 | -0.087 | -0.087 | 0.512 | 0.85 | 0.53 | 0.89 | 0.99 | 0.89 | 0.9 |
| 3 | 114478473 | rs7648823 | -0.089 | -1.739 | 1.843 | -1.115 | -0.059 | -0.074 | -0.074 | 2.49 | 0.16 | 0.36 | 0.1 | 0.03 | 0.05 | 0.11 |
| 3 | 114483413 | rs3732481 | -0.219 | -1.669 | 1.831 | -1.125 | -0.042 | -0.078 | -0.078 | 0 | 0.17 | 0.37 | 0.11 | 0.04 | 0.06 | 0.13 |
| 3 | 114497903 | rs4626069 | 1.465 | -1.61 | 1.42 | -1.138 | 0.031 | 0.131 | 0.131 | 0.198 | 0.69 | 0.16 | 0.79 | 0.95 | 0.83 | 0.77 |
| 3 | 114499741 | rs9848625 | 0.184 | -1.503 | 1.662 | -1.123 | 0.041 | -0.034 | -0.034 | 0 | 0.9 | 0.64 | 0.93 | 0.99 | 0.93 | 0.92 |
| 3 | 114500255 | rs9853475 | 2.28 | -1.72 | 0.915 | -1.028 | 0.05 | 0.094 | 0.094 | 1.41 | 0.55 | 0.13 | 0.66 | 0.61 | 0.73 | 0.54 |
| 3 | 114503412 | rs11718235 | 0.481 | -1.522 | 1.036 | -1.09 | -0.009 | 0.016 | 0.016 | 0 | 0.08 | 0.03 | 0.05 | 0.01 | 0.06 | 0.07 |
| 3 | 114511356 | rs12490319 | 1.47 | -1.732 | 0.949 | -1.253 | 0.04 | 0.139 | 0.139 | 0.099 | 0.69 | 0.15 | 0.79 | 0.96 | 0.83 | 0.76 |
| 3 | 114524645 | rs7650201 | -0.06 | -1.764 | 1.526 | -1.272 | -0.008 | -0.047 | -0.047 | 0.166 | 0.16 | 0.22 | 0.12 | 0.02 | 0.09 | 0.13 |
| 3 | 114530679 | rs9289004 | 1.353 | -1.779 | 1.82 | -1.282 | 0.015 | 0.023 | 0.023 | 0.175 | 0.72 | 0.24 | 0.81 | 0.97 | 0.85 | 0.8 |
| 3 | 114553234 | rs2399510 | 0.862 | -1.597 | 1.908 | -1.181 | -0.002 | -0.063 | -0.063 | 0.003 | 0.87 | 0.47 | 0.92 | 0.99 | 0.94 | 0.93 |
| 3 | 114565949 | rs4146802 | 0.151 | -1.585 | 2.112 | -1.17 | 0.005 | -0.026 | -0.026 | 0 | 0.91 | 0.69 | 0.94 | 0.98 | 0.95 | 0.95 |
| 3 | 114566709 | rs1533931 | 1.024 | -2.141 | 1.063 | -0.858 | 0.063 | 0.111 | 0.111 | 0.489 | 0.56 | 0.15 | 0.62 | 0.79 | 0.7 | 0.73 |
| 3 | 114569567 | rs9845450 | 0.695 | -2.034 | 1.533 | -0.832 | -0.02 | -0.072 | -0.072 | 0.169 | 0.87 | 0.39 | 0.92 | 0.99 | 0.95 | 0.84 |
| 3 | 114576087 | rs6796834 | 1.613 | -2.141 | 1.636 | -0.923 | 0.003 | -0.019 | -0.019 | 0 | 0.8 | 0.27 | 0.86 | 0.99 | 0.91 | 0.77 |
| 3 | 114585004 | rs677754 | -0.718 | -2.151 | 1.715 | -0.928 | 0 | -0.039 | -0.039 | 0 | 0.92 | 0.84 | 0.96 | 1 | 0.96 | 0.95 |
| 3 | 114585716 | rs664244 | 0.31 | -2.141 | 1.719 | -0.923 | -0.001 | -0.064 | -0.064 | 1.447 | 0.88 | 0.65 | 0.95 | 0.99 | 0.95 | 0.84 |
| 3 | 114593970 | rs1354056 | 0.506 | -1.909 | 1.089 | -1.347 | -0.007 | 0.045 | 0.045 | 1.445 | 0.31 | 0.13 | 0.31 | 0.18 | 0.3 | 0.15 |
| 3 | 114599978 | rs2937259 | 0.524 | -1.862 | 1.125 | -1.418 | -0.018 | -0.099 | -0.099 | 0.052 | 0.84 | 0.63 | 0.9 | 0.99 | 0.92 | 0.78 |
| 3 | 114611829 | rs9817665 | -0.232 | -1.94 | 0.55 | -0.817 | 0.033 | 0.188 | 0.188 | 1.295 | 0.47 | 0.2 | 0.56 | 0.78 | 0.41 | 0.46 |
| 3 | 114622323 | rs4682165 | -0.478 | -1.915 | 0.088 | -0.803 | 0.034 | 0.187 | 0.187 | 1.232 | 0.5 | 0.23 | 0.61 | 0.78 | 0.44 | 0.52 |
| 3 | 114627784 | rs6766555 | 0.575 | -1.687 | 0.023 | 0.025 | 0.002 | -0.072 | -0.072 | 0 | 0.33 | 0.14 | 0.37 | 0.41 | 0.31 | 0.35 |
| 3 | 114652462 | rs16823404 | -0.689 | -0.536 | 0.769 | -0.411 | -0.011 | -0.026 | -0.026 | 0.006 | 0.14 | 0.3 | 0.21 | 0.05 | 0.08 | 0.13 |
| 3 | 114671715 | rs11923714 | -1.96 | -0.739 | 1.617 | -0.487 | 0.005 | -0.014 | -0.014 | 0 | 0.08 | 0.39 | 0.08 | 0.01 | 0.02 | 0.14 |
| 3 | 114711353 | rs16823443 | -0.119 | -1.339 | 1.307 | -0.527 | 0.022 | -0.039 | -0.039 | 0.297 | 0.18 | 0.1 | 0.16 | 0.57 | 0.11 | 0.3 |
| 3 | 114714244 | rs2055650 | 1.267 | -1.488 | 0.963 | -0.853 | 0.006 | 0.021 | 0.021 | 3.48 | 0.33 | 0.11 | 0.31 | 0.24 | 0.3 | 0.22 |
| 3 | 114730257 | rs9859987 | 2.331 | -1.42 | 0.728 | -1.071 | -0.148 | 0.097 | 0.097 | 2.954 | 0.23 | 0.04 | 0.25 | 0.12 | 0.47 | 0.21 |

| | | | | | | | | | | | | | | | | |
|---|---|---|---|---|---|---|---|---|---|---|---|---|---|---|---|---|
| 3 | 114743234 | rs9825670 | -1.034 | -1.385 | 0.956 | -0.99 | 0.031 | -0.004 | -0.004 | 0 | 0.32 | 0.18 | 0.33 | 0.64 | 0.21 | 0.42 |
| 3 | 114774962 | rs1506024 | 1.98 | -1.421 | 1.094 | -0.999 | 0.074 | 0.043 | 0.043 | 0.217 | 0.58 | 0.25 | 0.57 | 0.35 | 0.77 | 0.43 |
| 3 | 114793620 | rs7648400 | -1.835 | -1.343 | 1.401 | -0.981 | -0.028 | -0.07 | -0.07 | 0 | 0.12 | 0.43 | 0.1 | 0.01 | 0.04 | 0.19 |
| 3 | 114812777 | rs6768251 | -2.219 | -1.348 | 1.279 | -0.963 | -0.045 | 0.01 | 0.01 | 0 | 0.14 | 0.46 | 0.21 | 0.07 | 0.06 | 0.2 |
| 3 | 114820006 | rs9289008 | -2.232 | -0.876 | 1.125 | -1.107 | 0.061 | 0.059 | 0.059 | 0.324 | 0.43 | 0.75 | 0.44 | 0.65 | 0.27 | 0.6 |
| 3 | 114821299 | rs2177041 | -1.291 | -0.555 | 1.497 | -0.798 | -0.011 | 0.098 | 0.098 | 4.063 | 0.59 | 0.9 | 0.65 | 0.78 | 0.65 | 0.72 |
| 3 | 114843510 | rs6782656 | 1.756 | -0.531 | 1.491 | -0.785 | -0.082 | 0.076 | 0.076 | 0.784 | 0.26 | 0.15 | 0.26 | 0.13 | 0.46 | 0.21 |
| 3 | 114855021 | rs6793973 | -1.84 | -0.509 | 1.616 | -0.711 | -0.007 | -0.001 | -0.001 | 0 | 0.32 | 0.62 | 0.38 | 0.65 | 0.18 | 0.51 |
| 3 | 114889454 | rs6795297 | -0.691 | -0.501 | 1.677 | -0.704 | 0.098 | 0.044 | 0.044 | 0 | 0.05 | 0.34 | 0.03 | 0.01 | 0.03 | 0.12 |
| 3 | 114913508 | rs6438234 | -2.04 | 0.053 | 1.465 | -0.729 | 0 | 0 | 0 | 0 | 0.34 | 0.62 | 0.4 | 0.65 | 0.24 | 0.53 |
| 3 | 114926956 | rs6786991 | -1.623 | -0.199 | 1.594 | -0.627 | -0.017 | -0.001 | -0.001 | 4.885 | 0.3 | 0.61 | 0.37 | 0.64 | 0.19 | 0.49 |
| 3 | 114940210 | rs17755786 | 1.216 | -0.075 | 1.481 | -0.769 | -0.078 | 0.101 | 0.101 | 1.419 | 0.3 | 0.16 | 0.3 | 0.13 | 0.48 | 0.23 |
| 3 | 114948198 | rs1380362 | -2.277 | -0.147 | 1.559 | -0.742 | 0.039 | 0.06 | 0.06 | 0 | 0.11 | 0.45 | 0.2 | 0.08 | 0.07 | 0.2 |
| 3 | 114955447 | rs9867281 | -0.165 | -0.122 | 1.567 | -0.735 | 0.011 | -0.016 | -0.016 | 0 | 0.19 | 0.16 | 0.17 | 0.57 | 0.12 | 0.29 |
| 3 | 114955517 | rs13433861 | -0.744 | -0.09 | 1.585 | -0.754 | 0.102 | 0.047 | 0.047 | 0 | 0.05 | 0.35 | 0.04 | 0.01 | 0.03 | 0.12 |
| 3 | 114957135 | rs9859887 | -1.363 | -0.095 | 1.514 | -0.763 | 0.004 | 0.083 | 0.083 | 5.636 | 0.62 | 0.88 | 0.67 | 0.78 | 0.67 | 0.72 |
| 3 | 114965974 | rs2219522 | -1.702 | -0.586 | 1.867 | -0.345 | -0.017 | -0.002 | -0.002 | 0 | 0.3 | 0.61 | 0.37 | 0.65 | 0.19 | 0.49 |
| 3 | 114971759 | rs9831948 | -2.02 | -0.474 | 1.767 | -0.361 | 0.028 | 0.016 | 0.016 | 0.261 | 0.36 | 0.72 | 0.42 | 0.65 | 0.25 | 0.51 |
| 3 | 114983867 | rs9851621 | -1.458 | -0.553 | 1.73 | -0.284 | -0.008 | -0.001 | -0.001 | 0 | 0.3 | 0.32 | 0.38 | 0.69 | 0.21 | 0.4 |
| 3 | 114984256 | rs1459515 | -1.588 | -0.555 | 1.787 | -0.285 | -0.035 | -0.007 | -0.007 | 1.705 | 0.31 | 0.42 | 0.38 | 0.69 | 0.22 | 0.4 |
| 3 | 114984814 | rs9853369 | 0.67 | -0.657 | 1.732 | -0.311 | 0 | 0.041 | 0.041 | 0 | 0.11 | 0.1 | 0.11 | 0.56 | 0.07 | 0.26 |
| 3 | 114988271 | rs9882905 | -2.344 | -0.659 | 1.891 | -0.321 | -0.029 | -0.008 | -0.008 | 0 | 0.33 | 0.57 | 0.4 | 0.69 | 0.22 | 0.4 |
| 3 | 114994740 | rs4831217 | 0.171 | -0.615 | 1.507 | -0.302 | -0.003 | 0.005 | 0.005 | 0 | 0.08 | 0.12 | 0.16 | 0.07 | 0.07 | 0.03 |
| 3 | 115000005 | rs11720709 | 0.19 | -0.568 | 1.271 | -0.303 | 0.013 | 0.017 | 0.017 | 0 | 0.1 | 0.04 | 0.1 | 0.06 | 0.06 | 0.08 |
| 3 | 115025326 | rs2125700 | -1.055 | -0.589 | 1.438 | -0.27 | -0.026 | 0.051 | 0.051 | 0 | 0.2 | 0.36 | 0.29 | 0.63 | 0.15 | 0.31 |
| 3 | 115025477 | rs11717514 | -1.555 | -0.575 | 1.4 | -0.292 | -0.032 | -0.036 | -0.036 | 2.576 | 0.21 | 0.37 | 0.29 | 0.13 | 0.15 | 0.14 |
| 3 | 115055887 | rs6782096 | 0.281 | -0.84 | 1.2 | -0.075 | -0.009 | 0.075 | 0.075 | 0 | 0.11 | 0.13 | 0.1 | 0.55 | 0.07 | 0.26 |
| 3 | 115063672 | rs7645613 | -1.542 | -1.003 | 1.302 | -0.292 | -0.021 | 0.001 | 0.001 | 0.067 | 0.27 | 0.62 | 0.35 | 0.68 | 0.13 | 0.38 |
| 3 | 115069397 | rs6809037 | -0.805 | -1.016 | 1.506 | -0.276 | -0.026 | 0.015 | 0.015 | 0 | 0.22 | 0.24 | 0.27 | 0.62 | 0.11 | 0.31 |
| 3 | 115079383 | rs6438246 | 1.367 | -1.017 | 1.544 | -0.277 | 0.077 | 0.097 | 0.097 | 0 | 0.94 | 0.62 | 0.92 | 0.94 | 0.97 | 0.93 |
| 3 | 115082070 | rs9839774 | 1.284 | -0.987 | 1.047 | -0.29 | -0.025 | 0.001 | 0.001 | 0.113 | 0.73 | 0.39 | 0.65 | 0.33 | 0.87 | 0.62 |
| 3 | 115089790 | rs7431473 | 1.103 | -0.007 | -0.106 | -0.619 | -0.102 | 0.136 | 0.136 | 1.616 | 0.28 | 0.11 | 0.29 | 0.08 | 0.47 | 0.22 |
| 3 | 115099856 | rs9840925 | -1.221 | 0.016 | 0.309 | -0.437 | 0.016 | -0.005 | -0.005 | 0 | 0.07 | 0.14 | 0.06 | 0.06 | 0.06 | 0.18 |
| 3 | 115099894 | rs7426620 | -0.44 | -0.12 | 0.73 | -0.692 | 0 | -0.027 | -0.027 | 0 | 0.63 | 0.37 | 0.61 | 0.86 | 0.47 | 0.59 |
| 3 | 115108682 | rs12637405 | -0.31 | -0.242 | 0.455 | 0.099 | 0.032 | -0.042 | -0.042 | 1.205 | 0.12 | 0.04 | 0.16 | 0.22 | 0.05 | 0.04 |
| 3 | 115114203 | rs6782599 | -0.915 | -0.373 | 0.659 | 0.019 | -0.004 | 0.03 | 0.03 | 2.744 | 0.61 | 0.74 | 0.62 | 0.7 | 0.64 | 0.66 |
| 3 | 115116868 | rs7431323 | 0.858 | -0.368 | 0.502 | 0.021 | -0.029 | 0.035 | 0.035 | 3.667 | 0.39 | 0.14 | 0.37 | 0.3 | 0.36 | 0.34 |
| 3 | 115118900 | rs9821873 | -0.396 | -0.384 | 0.659 | -0.036 | 0.039 | -0.039 | -0.039 | 0 | 0.08 | 0.18 | 0.03 | 0.01 | 0.06 | 0.13 |
| 3 | 115119044 | rs7428561 | 1.071 | -0.357 | 0.2 | 0.018 | -0.07 | -0.001 | -0.001 | 0 | 0.37 | 0.08 | 0.35 | 0.3 | 0.36 | 0.34 |
| 3 | 115135009 | rs6438258 | -1.408 | -0.136 | 0.201 | 0.011 | -0.016 | 0.031 | 0.031 | 2.458 | 0.64 | 0.91 | 0.69 | 0.76 | 0.68 | 0.71 |
| 3 | 115138637 | rs6769092 | -0.714 | -0.315 | 0.787 | 0.049 | -0.007 | 0.125 | 0.125 | 0 | 0.2 | 0.47 | 0.15 | 0.44 | 0.14 | 0.36 |
| 3 | 115142939 | rs9817986 | -0.574 | -0.308 | 0.89 | 0.051 | 0.001 | 0.101 | 0.101 | 0 | 0.19 | 0.37 | 0.14 | 0.44 | 0.14 | 0.36 |
| 3 | 115144495 | rs11716698 | -0.265 | -0.15 | 0.871 | -0.085 | -0.021 | 0.054 | 0.054 | 0 | 0.11 | 0.18 | 0.18 | 0.22 | 0.05 | 0.05 |
| 3 | 115144685 | rs1392261 | 1.095 | 0.126 | 0.567 | -0.309 | -0.008 | -0.027 | -0.027 | 0 | 0.31 | 0.25 | 0.31 | 0.04 | 0.48 | 0.24 |
| 3 | 115151369 | rs9870703 | -1.493 | 0.478 | 0.409 | -0.208 | -0.031 | 0.049 | 0.049 | 0 | 0.74 | 0.87 | 0.73 | 0.9 | 0.63 | 0.8 |
| 3 | 115151530 | rs9833318 | -0.89 | 0.434 | 1.116 | -0.327 | 0.091 | -0.018 | -0.018 | 0.041 | 0.09 | 0.34 | 0.04 | 0.03 | 0.07 | 0.1 |
| 3 | 115151612 | rs9871040 | -1.63 | 0.434 | 1.03 | -0.327 | -0.049 | 0.077 | 0.077 | 0 | 0.74 | 0.91 | 0.73 | 0.9 | 0.63 | 0.8 |
| 3 | 115155648 | rs9826938 | -1.956 | 0.162 | 0.983 | -0.139 | -0.003 | 0.002 | 0.002 | 1.242 | 0.23 | 0.73 | 0.23 | 0.24 | 0.13 | 0.16 |
| 3 | 115157029 | rs1463869 | 0.292 | 0.215 | 0.887 | -0.139 | -0.036 | 0.042 | 0.042 | 0.413 | 0.52 | 0.14 | 0.5 | 0.66 | 0.5 | 0.62 |
| 3 | 115161656 | rs1499954 | -0.602 | 0.33 | 1.095 | -0.283 | -0.004 | -0.017 | -0.017 | 0.186 | 0.64 | 0.44 | 0.67 | 0.87 | 0.56 | 0.68 |
| 3 | 115162379 | rs9825881 | -0.414 | 0.083 | 1.024 | -0.05 | 0.059 | -0.028 | -0.028 | 0 | 0.08 | 0.17 | 0.02 | 0.02 | 0.07 | 0.1 |
| 3 | 115192910 | rs9830165 | -1.458 | 0.121 | 0.345 | -0.192 | -0.061 | 0.101 | 0.101 | 0.104 | 0.69 | 0.86 | 0.7 | 0.88 | 0.57 | 0.73 |
| 3 | 115195109 | rs1499951 | -1.552 | -0.065 | 0.717 | -0.252 | 0.012 | -0.067 | -0.067 | 0 | 0.13 | 0.58 | 0.07 | 0.02 | 0.08 | 0.12 |
| 3 | 115200732 | rs11712245 | -1.531 | 0 | 0.079 | -0.225 | -0.059 | 0.096 | 0.096 | 0.157 | 0.69 | 0.86 | 0.7 | 0.88 | 0.57 | 0.73 |
| 3 | 115205520 | rs1499946 | 0.722 | -0.487 | -0.002 | 0.249 | -0.033 | -0.059 | -0.059 | 0 | 0.1 | 0.01 | 0.11 | 0.41 | 0.14 | 0.28 |
| 3 | 115209138 | rs6806816 | -1.253 | -0.418 | 0.3 | -0.083 | -0.021 | -0.011 | -0.011 | 0.101 | 0.17 | 0.74 | 0.2 | 0.08 | 0.08 | 0.07 |
| 3 | 115219719 | rs1353908 | -0.703 | -0.273 | 0.365 | -0.206 | -0.015 | 0.024 | 0.024 | 0.238 | 0.62 | 0.69 | 0.68 | 0.87 | 0.51 | 0.63 |
| 3 | 115219766 | rs1499945 | 0.43 | -0.014 | -0.299 | 0.012 | 0.009 | 0.039 | 0.039 | 0.998 | 0.47 | 0.16 | 0.48 | 0.66 | 0.44 | 0.56 |
| 3 | 115223004 | rs1553208 | -0.832 | 0.073 | -0.042 | -0.065 | -0.019 | 0.026 | 0.026 | 0.232 | 0.63 | 0.65 | 0.69 | 0.79 | 0.54 | 0.65 |
| 3 | 115231715 | rs10446272 | 0.921 | -0.161 | -0.278 | -0.366 | -0.005 | -0.008 | -0.008 | 1.417 | 0.09 | 0.09 | 0.09 | 0 | 0.12 | 0.08 |
| 3 | 115232912 | rs10511343 | 0.228 | -0.31 | 0.137 | 0.252 | -0.003 | 0.017 | 0.017 | 0 | 0.51 | 0.38 | 0.56 | 0.57 | 0.53 | 0.6 |
| 3 | 115236973 | rs1499955 | 0.439 | -0.278 | -0.195 | 0.26 | 0.011 | 0.018 | 0.018 | 0.198 | 0.48 | 0.38 | 0.53 | 0.57 | 0.48 | 0.57 |
| 3 | 115237226 | rs937542 | -0.093 | 0.113 | -0.644 | 0.643 | -0.001 | -0.001 | -0.001 | 0 | 0.6 | 0.62 | 0.6 | 0.57 | 0.54 | 0.62 |
| 3 | 115239407 | rs16823817 | 1.078 | -0.305 | -0.221 | 0.944 | 0 | 0 | 0 | 0 | 0.72 | 0.28 | 0.63 | 0.74 | 0.73 | 0.67 |
| 3 | 115240189 | rs6785900 | -1.567 | -0.642 | 0.073 | 1.57 | 0.002 | 0.074 | 0.074 | 0.016 | 0.1 | 0.19 | 0.11 | 0.26 | 0.12 | 0.11 |
| 3 | 115240953 | rs1499942 | -0.918 | -1.425 | 0.432 | 1.593 | -0.019 | 0.19 | 0.19 | 0 | 0.05 | 0.18 | 0.17 | 0.22 | 0.02 | 0.17 |
| 3 | 115241785 | rs9844692 | -0.967 | -1.699 | 1.161 | 1.361 | -0.039 | 0.112 | 0.112 | 0 | 0.13 | 0.31 | 0.28 | 0.25 | 0.07 | 0.29 |
| 3 | 115243731 | rs1499940 | 0.167 | -1.755 | 0.89 | 0.982 | 0 | -0.016 | -0.016 | 0 | 0.67 | 0.47 | 0.64 | 0.72 | 0.67 | 0.79 |
| 3 | 115251858 | rs4425251 | -0.267 | -1.674 | 0.836 | 0.996 | 0.007 | -0.025 | -0.025 | 0.051 | 0.31 | 0.46 | 0.35 | 0.27 | 0.33 | 0.21 |

| | | | | | | | | | | | | | | | | |
|---|---|---|---|---|---|---|---|---|---|---|---|---|---|---|---|---|
| 3 | 115252026 | rs4608693 | 0.697 | -1.474 | 0.17 | 1.058 | -0.086 | -0.008 | -0.008 | 1.539 | 0.21 | 0.01 | 0.25 | 0.19 | 0.27 | 0.09 |
| 3 | 115259865 | rs2133644 | 0.199 | -0.839 | 0.786 | 0.86 | 0 | -0.009 | -0.009 | 0 | 0.95 | 0.93 | 0.96 | 1 | 0.94 | 0.97 |
| 3 | 115262701 | rs1553207 | 0.002 | -1.185 | 0.647 | 0.814 | -0.013 | 0.005 | 0.005 | 0 | 0.4 | 0.49 | 0.53 | 0.46 | 0.33 | 0.41 |
| 3 | 115265050 | rs9849343 | -0.291 | -1.39 | 0.595 | 0.717 | 0.012 | 0 | 0 | 0.016 | 0.46 | 0.59 | 0.58 | 0.54 | 0.38 | 0.47 |
| 3 | 115273456 | rs6766852 | 1.137 | -1.36 | 0.333 | 0.647 | -0.124 | -0.053 | -0.053 | 0.919 | 0.39 | 0.04 | 0.3 | 0.33 | 0.5 | 0.4 |
| 3 | 115274499 | rs16823850 | -0.942 | -1.197 | 1.773 | 0.646 | -0.017 | 0.036 | 0.036 | 0 | 0.06 | 0.16 | 0.05 | 0.09 | 0.05 | 0.06 |
| 3 | 115278682 | rs16823858 | -1.192 | -1.188 | 1.593 | 0.525 | -0.022 | 0.017 | 0.017 | 0 | 0.07 | 0.18 | 0.05 | 0.05 | 0.05 | 0.05 |
| 3 | 115278983 | rs9857602 | 1.978 | -1.168 | 1.714 | 0.534 | -0.01 | -0.008 | -0.008 | 0 | 0.77 | 0.31 | 0.81 | 0.88 | 0.84 | 0.85 |
| 3 | 115279547 | rs9820822 | 1.207 | -0.642 | 1.003 | 0.466 | -0.021 | -0.048 | -0.048 | 0 | 0.84 | 0.6 | 0.88 | 0.94 | 0.89 | 0.89 |
| 3 | 115287554 | rs11919820 | -1.739 | -0.672 | 0.936 | 0.667 | -0.024 | 0.016 | 0.016 | 0.109 | 0.13 | 0.58 | 0.16 | 0.11 | 0.06 | 0.19 |
| 3 | 115289876 | rs6438269 | -1.607 | -1.05 | 1.223 | 1.119 | -0.034 | 0.093 | 0.093 | 0 | 0.16 | 0.6 | 0.13 | 0.26 | 0.11 | 0.24 |
| 3 | 115290159 | rs9870687 | -0.425 | -1.112 | 0.94 | 1.039 | -0.001 | 0.004 | 0.004 | 0 | 0.24 | 0.23 | 0.29 | 0.17 | 0.27 | 0.23 |
| 3 | 115291490 | rs9818763 | 0.942 | -1.869 | 0.475 | 0.56 | 0.014 | 0 | 0 | 0 | 0.71 | 0.27 | 0.82 | 0.75 | 0.84 | 0.72 |
| 3 | 115293391 | rs7631630 | -0.303 | -1.633 | 0.884 | 0.657 | -0.015 | 0.027 | 0.027 | 0 | 0.07 | 0.18 | 0.03 | 0.08 | 0.05 | 0.16 |
| 3 | 115297011 | rs9859842 | 0.296 | -1.568 | 0.845 | 0.556 | -0.001 | 0.012 | 0.012 | 0 | 0.15 | 0.26 | 0.21 | 0.16 | 0.16 | 0.14 |
| 3 | 115297103 | rs7611310 | 1.086 | -1.234 | 0.24 | 0.928 | 0.057 | -0.058 | -0.058 | 0 | 0.48 | 0.09 | 0.46 | 0.51 | 0.56 | 0.5 |
| 3 | 115301990 | rs10934299 | -1.177 | -0.753 | 1.202 | 1.326 | 0.03 | 0.236 | 0.236 | 0 | 0.06 | 0.3 | 0.07 | 0.18 | 0.03 | 0.1 |
| 3 | 115306577 | rs736884 | -1.243 | -0.365 | 0.286 | 0.87 | -0.005 | 0.065 | 0.065 | 0 | 0.07 | 0.29 | 0.04 | 0.08 | 0.04 | 0.16 |
| 3 | 115319600 | rs2043820 | -0.577 | 0.55 | -0.781 | 1.129 | -0.057 | -0.097 | -0.097 | 0 | 0.24 | 0.08 | 0.32 | 0.58 | 0.3 | 0.46 |
| 3 | 115320434 | rs2028245 | 0.641 | 0.164 | -1.108 | 0.259 | 0.018 | 0.103 | 0.103 | 0 | 0.46 | 0.72 | 0.48 | 0.23 | 0.39 | 0.35 |
| 3 | 115320700 | rs2028246 | -0.826 | 0.113 | -1.341 | 0.141 | 0.005 | 0.063 | 0.063 | 1.525 | 0.78 | 0.91 | 0.87 | 0.95 | 0.78 | 0.89 |
| 3 | 115321358 | rs12495327 | 1.042 | -0.338 | -0.489 | -0.034 | 0 | 0 | 0 | 0 | 0.41 | 0.42 | 0.44 | 0.2 | 0.38 | 0.29 |
| 3 | 115323184 | rs4831114 | -0.108 | -0.276 | -0.085 | 0.17 | -0.004 | -0.017 | -0.017 | 0 | 0.74 | 0.78 | 0.84 | 0.92 | 0.77 | 0.84 |
| 3 | 115326354 | rs4831117 | -0.405 | -0.232 | -0.382 | 0.234 | 0.008 | 0.009 | 0.009 | 0.886 | 0.77 | 0.85 | 0.84 | 0.85 | 0.78 | 0.89 |
| 3 | 115329174 | rs7634702 | -0.111 | -0.1 | -0.25 | 0.082 | -0.001 | 0 | 0 | 0.767 | 0.78 | 0.81 | 0.87 | 0.95 | 0.76 | 0.79 |
| 3 | 115332362 | rs13074182 | -0.3 | -0.268 | -0.17 | -0.067 | -0.003 | -0.01 | -0.01 | 0 | 0.14 | 0.1 | 0.07 | 0.06 | 0.09 | 0.17 |
| 3 | 115335684 | rs16823934 | 0.422 | 0.821 | -0.739 | 0.445 | -0.031 | -0.084 | -0.084 | 0 | 0.2 | 0.05 | 0.22 | 0.56 | 0.23 | 0.33 |
| 3 | 115339747 | rs13433884 | -0.766 | 0.807 | -0.305 | 0.064 | 0.003 | 0.011 | 0.011 | 0 | 0.45 | 0.41 | 0.45 | 0.78 | 0.54 | 0.62 |
| 3 | 115340900 | rs3827496 | 0.527 | 0.942 | -0.188 | 0.03 | 0.003 | 0.013 | 0.013 | 0 | 0.55 | 0.7 | 0.6 | 0.33 | 0.47 | 0.3 |
| 3 | 115340985 | rs6806885 | 0.72 | 0.861 | -0.218 | 0.213 | 0.002 | 0.067 | 0.067 | 0 | 0.07 | 0.17 | 0.08 | 0.17 | 0.07 | 0.06 |
| 3 | 115343213 | rs1837205 | 0.258 | 0.991 | -0.597 | -0.205 | -0.004 | 0.01 | 0.01 | 0 | 0.71 | 0.72 | 0.79 | 0.84 | 0.68 | 0.72 |
| 3 | 115347556 | rs13059137 | 1.051 | 0.823 | -0.995 | 0.071 | 0.001 | -0.067 | -0.067 | 0 | 0.31 | 0.18 | 0.28 | 0.37 | 0.29 | 0.21 |
| 3 | 115347676 | rs11922043 | 0.172 | 1.067 | -0.898 | -0.269 | -0.006 | 0.021 | 0.021 | 0 | 0.35 | 0.27 | 0.23 | 0.1 | 0.36 | 0.31 |
| 3 | 115353819 | rs1837206 | 0.031 | 0.991 | -0.515 | 0.409 | -0.001 | 0.032 | 0.032 | 0 | 0.18 | 0.18 | 0.23 | 0.51 | 0.19 | 0.41 |
| 3 | 115355964 | rs6803101 | 0.934 | 0.6 | -1.357 | -0.428 | 0.018 | 0.141 | 0.141 | 0 | 0.37 | 0.12 | 0.2 | 0.07 | 0.33 | 0.28 |
| 3 | 115358155 | rs714697 | 0.067 | 0.292 | -0.753 | 0.014 | -0.005 | -0.007 | -0.007 | 0.017 | 0.56 | 0.41 | 0.44 | 0.58 | 0.52 | 0.69 |
| 3 | 115362647 | rs9860586 | 0.628 | 0.248 | -1.181 | 0.173 | -0.03 | 0.019 | 0.019 | 0 | 0.48 | 0.11 | 0.4 | 0.46 | 0.51 | 0.53 |
| 3 | 115366966 | rs2028248 | 0.393 | 0.459 | -0.641 | 0.407 | 0.001 | -0.01 | -0.01 | 0 | 0.61 | 0.31 | 0.64 | 0.53 | 0.63 | 0.65 |
| 3 | 115367801 | rs9879241 | -0.201 | 0.274 | -0.312 | 0.637 | -0.021 | 0.032 | 0.032 | 0 | 0.07 | 0.19 | 0.04 | 0.14 | 0.01 | 0.18 |
| 3 | 115367990 | rs4544611 | 0.09 | 0.471 | -0.991 | 0.552 | -0.002 | 0.016 | 0.016 | 0.008 | 0.71 | 0.3 | 0.7 | 0.55 | 0.73 | 0.68 |
| 3 | 115371533 | rs6794935 | 0.17 | 0.717 | -1.199 | 0.489 | -0.002 | -0.003 | -0.003 | 0.567 | 0.78 | 0.76 | 0.81 | 0.67 | 0.75 | 0.68 |
| 3 | 115378117 | rs6782517 | 0.884 | 0.448 | -1.753 | 0.089 | -0.042 | 0.047 | 0.047 | 0 | 0.22 | 0.03 | 0.14 | 0.11 | 0.2 | 0.24 |
| 3 | 115386571 | rs12488667 | 0.486 | 0.332 | -1.092 | 0.266 | 0 | 0.011 | 0.011 | 0 | 0.2 | 0.25 | 0.3 | 0.2 | 0.16 | 0.13 |
| 3 | 115392763 | rs2164930 | -0.477 | -0.139 | -1.123 | -0.078 | 0.007 | 0.005 | 0.005 | 0 | 0.55 | 0.6 | 0.76 | 0.87 | 0.67 | 0.56 |
| 3 | 115393626 | rs1438367 | 0.709 | 0.199 | -1.266 | 0.401 | -0.002 | -0.029 | -0.029 | 0 | 0.14 | 0.1 | 0.16 | 0.41 | 0.16 | 0.19 |
| 3 | 115393874 | rs1837208 | -1.352 | -0.165 | -1.168 | 0.074 | 0.005 | 0.048 | 0.048 | 0 | 0.69 | 0.83 | 0.8 | 0.87 | 0.7 | 0.6 |
| 3 | 115394293 | rs11926976 | -0.74 | -0.046 | 0.028 | 0.269 | 0.001 | 0.007 | 0.007 | 0 | 0.49 | 0.59 | 0.5 | 0.66 | 0.55 | 0.47 |
| 3 | 115396251 | rs3772917 | -0.749 | 0.658 | 0.157 | 0.614 | 0.051 | 0.057 | 0.057 | 0 | 0.95 | 0.8 | 0.96 | 0.93 | 0.93 | 0.92 |
| 3 | 115396269 | rs3772918 | 1.368 | 0.403 | -0.145 | 0.427 | -0.006 | 0.042 | 0.042 | 0.493 | 0.46 | 0.07 | 0.45 | 0.25 | 0.4 | 0.43 |
| 3 | 115396528 | rs9873303 | 0.25 | 0.554 | 0.084 | 0.195 | 0.001 | -0.008 | -0.008 | 0 | 0.15 | 0.14 | 0.09 | 0 | 0.16 | 0.11 |
| 3 | 115396581 | rs1370807 | 0.948 | 0.499 | 0.232 | 0.303 | -0.026 | -0.019 | -0.019 | 0 | 0.19 | 0.13 | 0.3 | 0.27 | 0.31 | 0.17 |
| 3 | 115397785 | rs1370808 | 1.532 | -0.013 | 1.245 | 0.521 | 0.007 | -0.021 | -0.021 | 0 | 0.71 | 0.33 | 0.8 | 0.5 | 0.74 | 0.68 |
| 3 | 115402742 | rs10934305 | 0.429 | 0.199 | 1.326 | 0.631 | -0.011 | 0.011 | 0.011 | 0 | 0.83 | 0.64 | 0.9 | 0.86 | 0.87 | 0.8 |
| 3 | 115403818 | rs2067703 | -0.173 | 0.29 | 1.291 | 0.701 | 0.021 | 0.133 | 0.133 | 0 | 0.1 | 0.23 | 0.1 | 0.35 | 0.14 | 0.11 |
| 3 | 115404991 | rs283371 | 0.334 | 0.367 | 1.212 | 0.042 | 0.029 | 0.098 | 0.098 | 0 | 0.93 | 0.79 | 0.94 | 0.86 | 0.92 | 0.89 |
| 3 | 115406086 | rs283374 | 1.153 | 0.081 | 0.602 | -0.625 | -0.011 | -0.025 | -0.025 | 0 | 0.93 | 0.84 | 0.95 | 0.98 | 0.97 | 0.91 |
| 3 | 115407784 | rs9858175 | -0.046 | 0.269 | 0.381 | -0.38 | -0.004 | -0.027 | -0.027 | 2.413 | 0.1 | 0.13 | 0.05 | 0 | 0.05 | 0.08 |
| 3 | 115417865 | rs3772934 | -0.304 | 1.079 | 0.275 | 0.013 | -0.06 | -0.005 | -0.005 | 2.679 | 0.24 | 0.07 | 0.33 | 0.2 | 0.31 | 0.16 |
| 3 | 115420104 | rs2972474 | 1.246 | 0.812 | 0.543 | -0.1 | -0.019 | 0.033 | 0.033 | 0.313 | 0.61 | 0.23 | 0.56 | 0.73 | 0.59 | 0.52 |
| 3 | 115424974 | rs824354 | -1.205 | 0.993 | 0.193 | -0.018 | 0.014 | -0.013 | -0.013 | 2.998 | 0.36 | 0.59 | 0.42 | 0.27 | 0.41 | 0.48 |
| 3 | 115437175 | rs2918208 | 1.137 | 1.097 | 0.243 | -0.104 | 0.006 | -0.009 | -0.009 | 0 | 0.63 | 0.36 | 0.57 | 0.76 | 0.58 | 0.52 |
| 3 | 115440007 | rs14360 | -0.183 | 1.379 | 0.074 | -0.053 | -0.043 | -0.007 | -0.007 | 0 | 0.24 | 0.06 | 0.31 | 0.21 | 0.27 | 0.18 |
| 3 | 115440430 | rs7651867 | -1.5 | 0.968 | 0.523 | 0.096 | 0.055 | -0.043 | -0.043 | 0 | 0.14 | 0.53 | 0.11 | 0.06 | 0.16 | 0.31 |
| 3 | 115441608 | rs7638530 | -1.057 | 1.211 | -0.768 | 0.203 | 0.009 | -0.016 | -0.016 | 0 | 0.12 | 0.19 | 0.09 | 0.06 | 0.16 | 0.3 |
| 3 | 115448121 | rs2972469 | -0.055 | 0.83 | -1.318 | 0.101 | 0.031 | 0.101 | 0.101 | 0.581 | 0.49 | 0.72 | 0.5 | 0.29 | 0.41 | 0.31 |
| 3 | 115461467 | rs12635328 | 0.19 | 0.854 | -1.411 | 0.108 | -0.014 | -0.023 | -0.023 | 0.541 | 0.05 | 0.03 | 0.18 | 0.1 | 0.09 | 0.06 |
| 3 | 115462762 | rs2972464 | 0.514 | 0.737 | -0.814 | -0.124 | 0.018 | -0.009 | -0.009 | 0 | 0.07 | 0.16 | 0.06 | 0.07 | 0.05 | 0.06 |
| 3 | 115478263 | rs2972506 | -0.928 | 0.661 | -0.682 | -0.157 | 0.009 | 0.008 | 0.008 | 0 | 0.13 | 0.22 | 0.25 | 0.17 | 0.14 | 0.12 |
| 3 | 115484045 | rs11929078 | 0.744 | 1.136 | -1.379 | 0.171 | -0.048 | 0.017 | 0.017 | 0.033 | 0.3 | 0.16 | 0.22 | 0.24 | 0.36 | 0.31 |

| | | | | | | | | | | | | | | | | |
|---|---|---|---|---|---|---|---|---|---|---|---|---|---|---|---|---|
| 3 | 115502481 | rs2972491 | -0.643 | 0.867 | -0.678 | 0.115 | 0.018 | 0.015 | 0.015 | 0 | 0.14 | 0.27 | 0.26 | 0.17 | 0.13 | 0.12 |
| 3 | 115504836 | rs2972490 | 0.423 | 0.542 | -0.432 | -0.201 | -0.005 | -0.018 | -0.018 | 0 | 0.35 | 0.54 | 0.28 | 0.18 | 0.26 | 0.2 |
| 3 | 115523299 | rs11719103 | 1.122 | 0.394 | -0.039 | -0.283 | -0.016 | -0.034 | -0.034 | 0 | 0.27 | 0.3 | 0.21 | 0.12 | 0.22 | 0.16 |
| 3 | 115528760 | rs2289271 | -0.558 | 0.346 | -0.551 | -0.295 | -0.029 | 0.036 | 0.036 | 0 | 0.55 | 0.85 | 0.52 | 0.35 | 0.43 | 0.41 |
| 3 | 115529696 | rs3772946 | 0.457 | 0.552 | 0.034 | -0.429 | 0 | -0.052 | -0.052 | 0 | 0.31 | 0.33 | 0.33 | 0.11 | 0.24 | 0.19 |
| 3 | 115536212 | rs12485597 | 1 | 0.665 | -0.423 | -0.427 | -0.019 | 0.077 | 0.077 | 0 | 0.25 | 0.05 | 0.26 | 0.08 | 0.22 | 0.16 |
| 3 | 115543891 | rs7646077 | -0.337 | 0.901 | 0.585 | -0.182 | 0.108 | -0.016 | -0.016 | 0 | 0.1 | 0.37 | 0.09 | 0.03 | 0.1 | 0.07 |
| 3 | 115544663 | rs1550305 | -0.335 | 0.914 | 0.586 | -0.176 | 0.115 | -0.006 | -0.006 | 0 | 0.1 | 0.37 | 0.09 | 0.03 | 0.1 | 0.07 |
| 3 | 115548018 | rs3772948 | 0.636 | 0.796 | -0.597 | 0.06 | -0.076 | -0.167 | -0.167 | 0.23 | 0.3 | 0.03 | 0.25 | 0.44 | 0.35 | 0.28 |
| 3 | 115551931 | rs1878219 | -0.552 | 0.49 | 0.325 | -0.305 | 0.003 | 0.018 | 0.018 | 0 | 0.43 | 0.64 | 0.54 | 0.3 | 0.38 | 0.34 |
| 3 | 115556209 | rs7634813 | 0.368 | 0.278 | 0.614 | -0.38 | 0.071 | -0.018 | -0.018 | 0.148 | 0.08 | 0.38 | 0.07 | 0.01 | 0.05 | 0.01 |
| 3 | 115557462 | rs3772958 | 0.98 | 0.172 | -0.15 | -0.282 | -0.02 | 0.081 | 0.081 | 0 | 0.23 | 0.08 | 0.14 | 0.07 | 0.21 | 0.16 |
| 3 | 115561556 | rs2944401 | -0.247 | 0.216 | -0.123 | -0.149 | 0.004 | -0.008 | -0.008 | 0 | 0.09 | 0.05 | 0.32 | 0.19 | 0.09 | 0.12 |
| 3 | 115565490 | rs2918216 | -0.247 | 0.138 | 0.267 | -0.245 | -0.014 | -0.024 | -0.024 | 0.303 | 0.35 | 0.44 | 0.49 | 0.25 | 0.3 | 0.31 |
| 3 | 115566027 | rs2972479 | -0.024 | 0.249 | -0.016 | -0.18 | 0 | 0.004 | 0.004 | 0 | 0.08 | 0.07 | 0.32 | 0.19 | 0.08 | 0.13 |
| 3 | 115568314 | rs3772969 | -0.273 | 0.298 | 0.033 | -0.142 | -0.017 | -0.015 | -0.015 | 0 | 0.34 | 0.49 | 0.5 | 0.28 | 0.29 | 0.31 |
| 3 | 115571364 | rs3821560 | -0.333 | 0.397 | -0.142 | 0.141 | 0.003 | -0.025 | -0.025 | 0 | 0.08 | 0.03 | 0.3 | 0.18 | 0.09 | 0.15 |
| 3 | 115575949 | rs938521 | -1.311 | 0.386 | 0.115 | 0.009 | -0.016 | -0.005 | -0.005 | 0 | 0.19 | 0.63 | 0.39 | 0.19 | 0.14 | 0.19 |
| 3 | 115582838 | rs6763869 | -0.248 | 0.681 | -0.017 | -0.124 | 0.004 | 0.024 | 0.024 | 0 | 0.13 | 0.33 | 0.34 | 0.17 | 0.13 | 0.16 |
| 3 | 115586884 | rs1516519 | -0.265 | 0.398 | -0.564 | -0.3 | -0.017 | -0.003 | -0.003 | 0 | 0.3 | 0.38 | 0.44 | 0.29 | 0.23 | 0.34 |
| 3 | 115594175 | rs10511351 | 0.6 | 1.238 | -0.802 | -0.342 | -0.051 | 0.057 | 0.057 | 0 | 0.1 | 0.01 | 0.11 | 0 | 0.18 | 0.06 |
| 3 | 115596317 | rs9830559 | 0.771 | 0.905 | -0.738 | 0.094 | 0.021 | 0.008 | 0.008 | 0 | 0.48 | 0.29 | 0.46 | 0.46 | 0.47 | 0.49 |
| 3 | 115598604 | rs2090360 | -0.15 | 0.499 | -0.138 | 0.425 | 0.012 | 0.02 | 0.02 | 0 | 0.74 | 0.72 | 0.64 | 0.56 | 0.68 | 0.71 |
| 3 | 115602492 | rs2724466 | -0.102 | 0.529 | -0.424 | 0.416 | 0.041 | 0.061 | 0.061 | 0 | 0.74 | 0.6 | 0.62 | 0.57 | 0.68 | 0.71 |
| 3 | 115603783 | rs16824243 | -0.217 | -0.032 | -0.26 | 0.182 | -0.041 | -0.02 | -0.02 | 0.128 | 0.2 | 0.31 | 0.11 | 0.11 | 0.1 | 0.19 |
| 3 | 115605955 | rs17712069 | -0.368 | 0.789 | -0.808 | 0.456 | -0.026 | -0.033 | -0.033 | 0 | 0.1 | 0 | 0.05 | 0.09 | 0.11 | 0.16 |
| 3 | 115608057 | rs13070724 | -0.303 | 1.29 | -0.52 | 0.666 | -0.009 | -0.01 | -0.01 | 0 | 0.18 | 0.13 | 0.19 | 0.34 | 0.24 | 0.21 |
| 3 | 115609894 | rs16824248 | -0.566 | 1.157 | -0.547 | 0.495 | 0.001 | 0.004 | 0.004 | 0 | 0.19 | 0.18 | 0.12 | 0.19 | 0.16 | 0.19 |
| 3 | 115623564 | rs7612435 | -1.452 | 0.973 | 0.026 | 0.415 | 0.003 | 0.001 | 0.001 | 0.012 | 0.39 | 0.55 | 0.33 | 0.53 | 0.4 | 0.4 |
| 3 | 115625346 | rs1516520 | -0.471 | 0.705 | 0.062 | 0.397 | 0.003 | 0.015 | 0.015 | 0 | 0.18 | 0.1 | 0.11 | 0.19 | 0.16 | 0.19 |
| 3 | 115635433 | rs11712467 | -0.264 | 0.891 | -0.315 | 0.478 | -0.01 | 0.011 | 0.011 | 0 | 0.16 | 0.05 | 0.11 | 0.19 | 0.16 | 0.19 |
| 3 | 115643277 | rs10511352 | 0.275 | 0.998 | -0.796 | 0.348 | -0.032 | 0.037 | 0.037 | 0 | 0.1 | 0.01 | 0.08 | 0.06 | 0.11 | 0.1 |
| 3 | 115644194 | rs4640557 | -0.631 | 0.978 | -0.564 | 0.526 | 0.005 | 0.007 | 0.007 | 0 | 0.38 | 0.5 | 0.31 | 0.53 | 0.38 | 0.4 |
| 3 | 115649466 | rs7650115 | 0.326 | 1.016 | -1.104 | 0.18 | -0.032 | 0.132 | 0.132 | 0 | 0.54 | 0.17 | 0.36 | 0.26 | 0.51 | 0.65 |
| 3 | 115649821 | rs7638681 | 0.452 | 1.169 | -1.172 | 0.101 | -0.002 | 0.165 | 0.165 | 0 | 0.52 | 0.17 | 0.33 | 0.26 | 0.42 | 0.64 |
| 3 | 115650448 | rs4378954 | 0.105 | 1.2 | -0.069 | -0.043 | 0.063 | -0.001 | -0.001 | 2.986 | 0.92 | 0.61 | 0.89 | 0.98 | 0.92 | 0.97 |
| 3 | 115654421 | rs4240161 | -0.4 | 1.14 | 0.471 | -0.062 | 0.001 | -0.007 | -0.007 | 3.072 | 0.42 | 0.4 | 0.58 | 0.72 | 0.49 | 0.33 |
| 3 | 115664207 | rs3884593 | 0.21 | 1.239 | 0.068 | 0.455 | -0.013 | -0.072 | -0.072 | 0 | 0.22 | 0.12 | 0.26 | 0.51 | 0.29 | 0.15 |
| 3 | 115672610 | rs6769601 | 0.364 | 1.232 | -0.289 | 0.418 | 0.004 | -0.054 | -0.054 | 0.087 | 0.89 | 0.37 | 0.88 | 0.98 | 0.91 | 0.95 |
| 3 | 115675328 | rs17697123 | -0.115 | 1.316 | -0.795 | 0.645 | -0.07 | -0.07 | -0.07 | 0 | 0.17 | 0.01 | 0.27 | 0.19 | 0.19 | 0.17 |
| 3 | 115677750 | rs7632246 | 0.658 | 0.902 | -0.595 | 0.517 | 0.02 | -0.021 | -0.021 | 0.405 | 0.57 | 0.31 | 0.58 | 0.47 | 0.58 | 0.77 |
| 3 | 115679228 | rs9849995 | 0.548 | 0.384 | -0.794 | 0.28 | 0.045 | 0.032 | 0.032 | 0 | 0.47 | 0.64 | 0.25 | 0.29 | 0.27 | 0.49 |
| 3 | 115679429 | rs4831211 | -0.119 | 1.235 | -0.353 | 0.214 | -0.001 | 0.001 | 0.001 | 0 | 0.2 | 0.23 | 0.19 | 0.12 | 0.23 | 0.09 |
| 3 | 115679691 | rs6806562 | -0.72 | 1.158 | -0.279 | 0.125 | -0.001 | -0.03 | -0.03 | 0 | 0.41 | 0.45 | 0.4 | 0.25 | 0.39 | 0.28 |
| 3 | 115680748 | rs13315620 | -0.308 | 1.721 | 0.245 | 0.465 | 0.002 | -0.001 | -0.001 | 0 | 0.08 | 0.07 | 0.16 | 0.12 | 0.17 | 0.09 |
| 3 | 115680799 | rs9856549 | -0.975 | 1.673 | 0.636 | 0.444 | 0.058 | 0.022 | 0.022 | 0.741 | 0.11 | 0.28 | 0.19 | 0.12 | 0.18 | 0.11 |
| 3 | 115685453 | rs4518123 | -0.148 | 1.707 | 0.598 | 0.456 | -0.01 | -0.008 | -0.008 | 0 | 0.08 | 0.04 | 0.15 | 0.12 | 0.16 | 0.07 |
| 3 | 115687191 | rs12494795 | -0.75 | 1.892 | 0.871 | 0.432 | 0.035 | -0.028 | -0.028 | 0 | 0.17 | 0.51 | 0.3 | 0.15 | 0.18 | 0.19 |
| 3 | 115704434 | rs4831218 | -1.691 | 1.868 | 0.852 | 0.416 | 0.007 | -0.002 | -0.002 | 0 | 0.26 | 0.7 | 0.46 | 0.27 | 0.34 | 0.27 |
| 3 | 115721099 | rs4401357 | -0.924 | 2.147 | 1.341 | 0.418 | 0.134 | 0 | 0 | 0.424 | 0.11 | 0.32 | 0.21 | 0.09 | 0.18 | 0.1 |
| 3 | 115728651 | rs4398428 | -1.595 | 2.025 | 1.389 | 0.39 | 0.057 | -0.005 | -0.005 | 0 | 0.25 | 0.41 | 0.43 | 0.26 | 0.35 | 0.26 |
| 3 | 115731643 | rs1920388 | -1.904 | 1.983 | 1.251 | 0.371 | -0.009 | 0.004 | 0.004 | 0.164 | 0.28 | 0.73 | 0.47 | 0.26 | 0.36 | 0.28 |
| 3 | 115746815 | rs9289038 | -0.628 | 2.064 | 1.586 | 0.378 | 0.02 | -0.018 | -0.018 | 0 | 0.16 | 0.28 | 0.3 | 0.14 | 0.21 | 0.2 |
| 3 | 115751126 | rs9821040 | -0.003 | 2.102 | 1.473 | 0.397 | 0.008 | 0.002 | 0.002 | 0.253 | 0.07 | 0.08 | 0.15 | 0.12 | 0.15 | 0.07 |
| 3 | 115764342 | rs1920383 | -1.956 | 2.027 | 1.623 | 0.39 | -0.001 | 0 | 0 | 0.218 | 0.28 | 0.73 | 0.47 | 0.26 | 0.35 | 0.27 |
| 3 | 115775346 | rs6778817 | -0.89 | 2.135 | 1.682 | 0.378 | 0.017 | -0.013 | -0.013 | 0 | 0.18 | 0.23 | 0.31 | 0.15 | 0.22 | 0.21 |
| 3 | 115778365 | rs9289040 | -2.003 | 2.016 | 2.055 | 0.392 | 0.01 | -0.004 | -0.004 | 0 | 0.28 | 0.72 | 0.49 | 0.27 | 0.36 | 0.27 |
| 3 | 115786741 | rs7610371 | -1.57 | 2.025 | 2.028 | 0.396 | 0.021 | -0.001 | -0.001 | 1.429 | 0.25 | 0.34 | 0.44 | 0.27 | 0.36 | 0.27 |
| 3 | 115789995 | rs9878286 | -2.175 | 1.868 | 2.09 | 0.148 | 0.031 | -0.029 | -0.029 | 0 | 0.33 | 0.48 | 0.47 | 0.28 | 0.4 | 0.28 |
| 3 | 115794679 | rs6772113 | -1.527 | 1.416 | 1.919 | 0.183 | 0.027 | -0.026 | -0.026 | 0.064 | 0.17 | 0.58 | 0.18 | 0.13 | 0.17 | 0.07 |
| 3 | 115818840 | rs1920379 | -1.593 | 1.486 | 1.87 | 0.151 | 0.046 | -0.032 | -0.032 | 0.432 | 0.18 | 0.49 | 0.19 | 0.12 | 0.19 | 0.08 |
| 3 | 115832162 | rs4831229 | -0.894 | 1.501 | 1.725 | 0.185 | 0 | 0 | 0 | 0.356 | 0.14 | 0.13 | 0.14 | 0.12 | 0.19 | 0.08 |
| 3 | 115849331 | rs9809229 | -0.689 | 1.446 | 0.883 | 0.371 | -0.049 | -0.051 | -0.051 | 0.032 | 0.13 | 0.02 | 0.12 | 0.18 | 0.17 | 0.1 |
| 3 | 115852764 | rs13078076 | 1.549 | 0.606 | 1.416 | -0.192 | -0.033 | -0.064 | -0.064 | 0.624 | 0.7 | 0.5 | 0.8 | 0.89 | 0.81 | 0.84 |
| 3 | 115853285 | rs10511354 | -0.183 | 0.504 | 1.603 | -0.237 | -0.001 | -0.02 | -0.02 | 0 | 0.15 | 0.1 | 0.05 | 0 | 0.05 | 0.08 |
| 3 | 115855921 | rs1920374 | 2.369 | 0.817 | 1.423 | -0.345 | 0.002 | 0.045 | 0.045 | 0 | 0.57 | 0.34 | 0.54 | 0.74 | 0.61 | 0.64 |
| 3 | 115856130 | rs1882012 | 2.333 | 0.826 | 1.428 | -0.37 | 0.001 | 0.041 | 0.041 | 0 | 0.57 | 0.35 | 0.54 | 0.74 | 0.61 | 0.64 |
| 3 | 115864422 | rs1920363 | 0.163 | 0.896 | 1.257 | -0.345 | -0.002 | 0.001 | 0.001 | 1.396 | 0.13 | 0.12 | 0.26 | 0.15 | 0.19 | 0.2 |
| 3 | 115869444 | rs1381065 | -1.271 | 0.846 | 1.386 | -0.36 | 0 | -0.065 | -0.065 | 0 | 0.3 | 0.33 | 0.33 | 0.15 | 0.26 | 0.28 |

| | | | | | | | | | | | | | | | | |
|---|---|---|---|---|---|---|---|---|---|---|---|---|---|---|---|---|
| 3 | 115883224 | rs9812548 | -0.484 | 1.029 | 1.096 | -0.184 | 0 | -0.005 | -0.005 | 1.447 | 0.12 | 0.12 | 0.12 | 0.1 | 0.12 | 0.09 |
| 3 | 115886270 | rs7641464 | 2.076 | 0.866 | 1.159 | -0.411 | 0.002 | 0.084 | 0.084 | 0 | 0.54 | 0.32 | 0.56 | 0.75 | 0.61 | 0.64 |
| 3 | 115891691 | rs9289046 | 2.168 | 0.735 | 1.023 | -0.483 | -0.021 | 0.129 | 0.129 | 0 | 0.54 | 0.14 | 0.54 | 0.74 | 0.61 | 0.64 |
| 3 | 115892561 | rs7644082 | 0.38 | 0.728 | 1.503 | -0.509 | 0 | 0.002 | 0.002 | 0 | 0.85 | 0.74 | 0.88 | 0.9 | 0.87 | 0.92 |
| 3 | 115901032 | rs4296590 | 2.323 | 0.705 | 1.585 | -0.516 | -0.019 | 0.132 | 0.132 | 0 | 0.53 | 0.14 | 0.54 | 0.74 | 0.61 | 0.64 |
| 3 | 115901347 | rs7651707 | 0.321 | 0.648 | 1.9 | -0.529 | -0.025 | -0.064 | -0.064 | 0 | 0.15 | 0.23 | 0.05 | 0.02 | 0.06 | 0.07 |
| 3 | 115902720 | rs9875609 | -0.033 | 0.628 | 1.905 | -0.551 | -0.013 | -0.015 | -0.015 | 0.253 | 0.92 | 0.95 | 0.95 | 0.9 | 0.91 | 0.93 |
| 3 | 115908480 | rs2199349 | 0.226 | 0.656 | 1.867 | -0.514 | -0.035 | -0.082 | -0.082 | 0.163 | 0.16 | 0.33 | 0.06 | 0.02 | 0.06 | 0.07 |
| 3 | 115913887 | rs7641198 | 1.112 | 0.863 | 1.732 | -0.301 | 0.003 | 0.002 | 0.002 | 0 | 0.83 | 0.54 | 0.86 | 0.9 | 0.87 | 0.9 |
| 3 | 115914897 | rs998807 | 0.783 | 0.909 | 1.638 | -0.273 | 0.004 | 0.003 | 0.003 | 0 | 0.84 | 0.6 | 0.87 | 0.9 | 0.87 | 0.91 |
| 3 | 115917134 | rs9834589 | 0.541 | 0.904 | 1.504 | -0.247 | 0.004 | 0.005 | 0.005 | 0 | 0.85 | 0.68 | 0.87 | 0.9 | 0.87 | 0.91 |
| 3 | 115945873 | rs4831124 | 1.334 | 0.783 | 1.153 | -0.291 | 0.022 | -0.022 | -0.022 | 0 | 0.7 | 0.46 | 0.6 | 0.77 | 0.67 | 0.7 |
| 3 | 115952799 | rs1870709 | 0.885 | 0.428 | 1.254 | -0.029 | -0.029 | -0.014 | -0.014 | 0.121 | 0.76 | 0.46 | 0.69 | 0.8 | 0.8 | 0.85 |
| 3 | 115965295 | rs4831126 | 0.832 | 0.382 | 1.326 | -0.054 | -0.023 | -0.013 | -0.013 | 0 | 0.76 | 0.43 | 0.68 | 0.8 | 0.8 | 0.85 |
| 3 | 115965648 | rs9874470 | 1.66 | 0.366 | 0.261 | -0.292 | -0.009 | 0.044 | 0.044 | 0 | 0.63 | 0.06 | 0.58 | 0.76 | 0.69 | 0.8 |
| 3 | 115969765 | rs4831128 | 0.482 | -0.033 | 0.457 | -0.273 | -0.041 | -0.014 | -0.014 | 0 | 0.76 | 0.61 | 0.7 | 0.8 | 0.81 | 0.83 |
| 3 | 115970908 | rs16845196 | 0.334 | -0.074 | 0.513 | -0.277 | 0.006 | 0.016 | 0.016 | 0 | 0.08 | 0.09 | 0.23 | 0.16 | 0.14 | 0.08 |
| 3 | 115971442 | rs9830749 | 0.557 | -0.071 | 0.516 | -0.282 | -0.034 | -0.01 | -0.01 | 0 | 0.76 | 0.64 | 0.73 | 0.8 | 0.81 | 0.84 |
| 3 | 115972028 | rs4831089 | 1.335 | -0.406 | -0.429 | -0.309 | -0.176 | -0.162 | -0.162 | 0.018 | 0.35 | 0.02 | 0.32 | 0.5 | 0.45 | 0.32 |
| 3 | 115974374 | rs2279109 | 0.676 | -0.222 | 0.865 | -0.222 | -0.032 | -0.01 | -0.01 | 0.144 | 0.76 | 0.65 | 0.7 | 0.79 | 0.81 | 0.83 |
| 3 | 115981443 | rs4831129 | 1.931 | -0.144 | 0.688 | -0.205 | 0.023 | 0.05 | 0.05 | 0 | 0.62 | 0.1 | 0.59 | 0.75 | 0.7 | 0.79 |
| 3 | 115984749 | rs12496133 | -0.813 | -0.933 | 0.855 | -0.179 | -0.009 | -0.002 | -0.002 | 0 | 0.22 | 0.22 | 0.28 | 0.2 | 0.18 | 0.17 |
| 3 | 115986343 | rs10934326 | -0.787 | -0.925 | 0.852 | -0.472 | -0.013 | -0.021 | -0.021 | 0 | 0.22 | 0.24 | 0.28 | 0.13 | 0.18 | 0.17 |
| 3 | 115994501 | rs13073491 | -1.222 | -0.959 | 0.814 | -0.451 | -0.053 | -0.049 | -0.049 | 0 | 0.27 | 0.65 | 0.14 | 0.08 | 0.11 | 0.11 |
| 3 | 116011091 | rs6809953 | -1.193 | -0.933 | 0.859 | -0.442 | -0.059 | -0.052 | -0.052 | 0 | 0.26 | 0.64 | 0.14 | 0.08 | 0.09 | 0.11 |
| 3 | 116017931 | rs957783 | -1.235 | -0.888 | 0.854 | -0.445 | -0.041 | -0.051 | -0.051 | 0.116 | 0.27 | 0.64 | 0.14 | 0.08 | 0.12 | 0.1 |
| 3 | 116021453 | rs1993733 | 1.435 | -0.967 | 0.783 | -0.582 | 0.006 | 0.001 | 0.001 | 0 | 0.67 | 0.32 | 0.82 | 0.91 | 0.83 | 0.84 |
| 3 | 116023990 | rs2100807 | 1.454 | -1.01 | 1.022 | -0.583 | -0.016 | -0.025 | -0.025 | 0 | 0.67 | 0.35 | 0.82 | 0.91 | 0.84 | 0.84 |
| 3 | 116024024 | rs17645807 | 0.441 | -1.096 | 1.222 | -0.615 | 0.012 | 0.012 | 0.012 | 0 | 0.14 | 0.1 | 0.05 | 0.03 | 0.04 | 0.08 |
| 3 | 116024083 | rs7611085 | 0.821 | -0.889 | 1.337 | -0.547 | -0.007 | -0.001 | -0.001 | 0 | 0.88 | 0.47 | 0.91 | 0.95 | 0.94 | 0.94 |
| 3 | 116030986 | rs17646052 | -0.927 | -0.747 | 0.717 | -0.528 | -0.007 | 0.039 | 0.039 | 0 | 0.14 | 0.03 | 0.08 | 0.04 | 0.12 | 0.09 |
| 3 | 116037190 | rs9845203 | -0.594 | -0.356 | 0.925 | -0.492 | 0.046 | -0.029 | -0.029 | 0.336 | 0.08 | 0.13 | 0.24 | 0.05 | 0.2 | 0.08 |
| 3 | 116039842 | rs4831130 | 0.582 | -0.593 | 0.844 | -0.279 | 0 | 0 | 0 | 0 | 0.77 | 0.84 | 0.68 | 0.8 | 0.73 | 0.85 |
| 3 | 116041338 | rs6810089 | 0.553 | -0.588 | 0.847 | -0.288 | 0 | -0.002 | -0.002 | 1.545 | 0.77 | 0.84 | 0.68 | 0.85 | 0.74 | 0.85 |
| 3 | 116047619 | rs6782605 | -0.148 | -0.684 | 1.501 | -0.374 | -0.01 | -0.049 | -0.049 | 0.015 | 0.09 | 0.26 | 0.07 | 0 | 0.06 | 0.02 |
| 3 | 116065133 | rs2033406 | 2.246 | -1.04 | 1.447 | -0.477 | 0.053 | 0.164 | 0.164 | 0 | 0.56 | 0.09 | 0.54 | 0.81 | 0.64 | 0.75 |
| 3 | 116068599 | rs16824799 | 0.559 | -1.061 | 1.987 | -0.474 | -0.009 | -0.003 | -0.003 | 0 | 0.08 | 0.15 | 0.04 | 0.04 | 0.03 | 0.08 |
| 3 | 116070163 | rs7634137 | -1.754 | -1.074 | 2.095 | -0.489 | 0.052 | 0.049 | 0.049 | 0 | 0.36 | 0.76 | 0.23 | 0.15 | 0.17 | 0.17 |
| 3 | 116093754 | rs1367770 | -1.694 | -1.067 | 2.177 | -0.486 | -0.006 | -0.005 | -0.005 | 0.138 | 0.35 | 0.69 | 0.22 | 0.15 | 0.16 | 0.16 |
| 3 | 116094044 | rs9856796 | 0.088 | -1.052 | 2.183 | -0.478 | 0.032 | 0.002 | 0.002 | 0 | 0.92 | 0.9 | 0.77 | 0.91 | 0.8 | 0.92 |
| 3 | 116103099 | rs11924642 | -1.703 | -1.057 | 2.283 | -0.542 | -0.014 | -0.012 | -0.012 | 3.268 | 0.35 | 0.68 | 0.22 | 0.15 | 0.16 | 0.16 |
| 3 | 116104521 | rs17647091 | -0.761 | -1.085 | 2.669 | -0.523 | -0.032 | -0.016 | -0.016 | 0 | 0.25 | 0.28 | 0.13 | 0.14 | 0.1 | 0.15 |
| 3 | 116108165 | rs7627104 | 0.022 | -1.052 | 2.593 | -0.521 | 0.034 | -0.022 | -0.022 | 0 | 0.92 | 0.9 | 0.77 | 0.95 | 0.81 | 0.93 |
| 3 | 116110635 | rs9860260 | -0.845 | -1.088 | 2.619 | -0.523 | -0.061 | -0.035 | -0.035 | 0 | 0.26 | 0.35 | 0.15 | 0.16 | 0.1 | 0.14 |
| 3 | 116119143 | rs13075678 | -0.133 | -1.132 | 2.528 | -0.54 | -0.042 | -0.022 | -0.022 | 0.095 | 0.19 | 0.25 | 0.12 | 0.12 | 0.07 | 0.06 |
| 3 | 116120738 | rs1920190 | 2.326 | -1.142 | 2.468 | -0.561 | 0.065 | 0.174 | 0.174 | 0 | 0.56 | 0.1 | 0.54 | 0.79 | 0.64 | 0.77 |
| 3 | 116128090 | rs4831136 | -1.041 | -1.257 | 2.528 | -0.451 | -0.071 | -0.043 | -0.043 | 0 | 0.27 | 0.43 | 0.16 | 0.16 | 0.1 | 0.14 |
| 3 | 116128405 | rs4831139 | -0.004 | -1.225 | 2.222 | -0.434 | -0.007 | 0 | 0 | 0 | 0.18 | 0.13 | 0.1 | 0.12 | 0.07 | 0.06 |
| 3 | 116131208 | rs1731596 | -1.692 | -1.22 | 2.134 | -0.436 | 0.001 | 0.003 | 0.003 | 0 | 0.36 | 0.66 | 0.22 | 0.16 | 0.16 | 0.16 |
| 3 | 116131936 | rs1712999 | -1.26 | -0.54 | 1.987 | -0.206 | -0.034 | -0.061 | -0.061 | 0 | 0.17 | 0.41 | 0.11 | 0.04 | 0.09 | 0.1 |
| 3 | 116141816 | rs1713001 | 0.817 | -0.263 | 1.772 | -0.099 | 0.054 | 0.015 | 0.015 | 0.428 | 0.94 | 0.83 | 0.8 | 0.93 | 0.86 | 0.94 |
| 3 | 116144192 | rs1731607 | 1.706 | -0.609 | 2.164 | -0.027 | 0.01 | 0.008 | 0.008 | 0 | 0.61 | 0.39 | 0.63 | 0.71 | 0.66 | 0.76 |
| 3 | 116150232 | rs4831097 | -0.791 | -0.648 | 2.296 | -0.036 | -0.016 | -0.006 | -0.006 | 0.322 | 0.27 | 0.28 | 0.14 | 0.18 | 0.15 | 0.17 |
| 3 | 116159418 | rs1713007 | 0.557 | -0.614 | 2.276 | -0.013 | 0.062 | 0.02 | 0.02 | 0.343 | 0.94 | 0.83 | 0.8 | 0.93 | 0.87 | 0.93 |
| 3 | 116159810 | rs16824905 | 0.838 | -0.591 | 2.131 | -0.005 | 0 | -0.001 | -0.001 | 0 | 0.05 | 0.06 | 0.03 | 0.04 | 0.05 | 0.01 |
| 3 | 116164892 | rs10513826 | -0.737 | -0.7 | 2.195 | -0.046 | -0.021 | -0.008 | -0.008 | 0 | 0.27 | 0.27 | 0.15 | 0.17 | 0.15 | 0.17 |
| 3 | 116166378 | rs1795263 | 0.72 | -0.633 | 2.168 | -0.044 | 0.073 | 0.026 | 0.026 | 0.542 | 0.94 | 0.77 | 0.8 | 0.92 | 0.88 | 0.94 |
| 3 | 116167240 | rs9881385 | -1.162 | -0.775 | 2.38 | -0.104 | -0.004 | -0.001 | -0.001 | 0 | 0.32 | 0.29 | 0.17 | 0.21 | 0.2 | 0.22 |
| 3 | 116167302 | rs1920192 | 1.735 | -0.68 | 2.41 | -0.114 | 0.018 | 0.018 | 0.018 | 0 | 0.6 | 0.31 | 0.62 | 0.72 | 0.67 | 0.71 |
| 3 | 116168669 | rs2902853 | 1.767 | -0.668 | 2.103 | -0.073 | 0.042 | -0.014 | -0.014 | 0 | 0.86 | 0.56 | 0.77 | 0.89 | 0.82 | 0.88 |
| 3 | 116168964 | rs1920193 | 1.276 | -0.643 | 2.078 | -0.05 | 0.093 | 0.032 | 0.032 | 0.408 | 0.92 | 0.6 | 0.79 | 0.92 | 0.87 | 0.94 |
| 3 | 116180805 | rs1920196 | 0.45 | -0.716 | 1.976 | -0.067 | 0.027 | -0.02 | -0.02 | 0 | 0.17 | 0.02 | 0.1 | 0.12 | 0.11 | 0.06 |
| 3 | 116183381 | rs9863373 | 1.314 | -0.56 | 2.074 | -0.034 | 0.086 | 0.027 | 0.027 | 0.669 | 0.92 | 0.64 | 0.79 | 0.92 | 0.88 | 0.93 |
| 3 | 116185514 | rs4831100 | 0.896 | -0.595 | 2.048 | -0.043 | 0.014 | -0.01 | -0.01 | 0 | 0.85 | 0.59 | 0.77 | 0.87 | 0.84 | 0.83 |
| 3 | 116191226 | rs1731604 | -1.057 | -0.68 | 2.129 | -0.005 | -0.009 | -0.008 | -0.008 | 0.649 | 0.27 | 0.35 | 0.32 | 0.23 | 0.27 | 0.16 |
| 3 | 116192091 | rs9826952 | -0.43 | -0.821 | 1.879 | 0.651 | 0.004 | -0.005 | -0.005 | 0.491 | 0.22 | 0.2 | 0.14 | 0.47 | 0.17 | 0.28 |
| 3 | 116233415 | rs12629401 | 0.829 | -1.003 | 1.794 | 0.593 | 0.003 | -0.002 | -0.002 | 0 | 0.17 | 0.13 | 0.11 | 0.41 | 0.13 | 0.18 |
| 3 | 116234806 | rs1628472 | -0.466 | -0.711 | 1.765 | 0.421 | -0.01 | -0.016 | -0.016 | 0 | 0.25 | 0.37 | 0.32 | 0.2 | 0.22 | 0.1 |
| 3 | 116236180 | rs17706338 | 0.745 | -0.693 | 1.523 | 0.434 | 0.016 | 0.002 | 0.002 | 1.154 | 0.19 | 0.04 | 0.13 | 0.14 | 0.13 | 0.07 |

| | | | | | | | | | | | | | | | | |
|---|---|---|---|---|---|---|---|---|---|---|---|---|---|---|---|---|
| 3 | 116247126 | rs10934350 | -0.674 | -0.77 | 1.535 | 0.454 | 0.015 | 0.009 | 0.009 | 0.493 | 0.3 | 0.13 | 0.33 | 0.25 | 0.26 | 0.21 |
| 3 | 116247826 | rs1712997 | -0.947 | -0.802 | 1.578 | 0.464 | -0.011 | 0.042 | 0.042 | 1.219 | 0.82 | 0.77 | 0.89 | 0.58 | 0.87 | 0.82 |
| 3 | 116249215 | rs10511358 | 1.179 | -0.774 | 1.501 | 0.446 | 0.023 | -0.001 | -0.001 | 0 | 0.08 | 0.02 | 0.03 | 0.05 | 0.04 | 0.1 |
| 3 | 116253395 | rs1518334 | -1.198 | -0.747 | 1.585 | 0.465 | 0.012 | 0.014 | 0.014 | 0.194 | 0.47 | 0.35 | 0.43 | 0.67 | 0.39 | 0.39 |
| 3 | 116269592 | rs1795145 | -1.282 | -0.751 | 1.431 | 0.462 | -0.004 | -0.005 | -0.005 | 2.134 | 0.48 | 0.44 | 0.45 | 0.67 | 0.39 | 0.39 |
| 3 | 116270151 | rs2010874 | 1.659 | -0.797 | 1.493 | 0.425 | 0.01 | 0.043 | 0.043 | 0 | 0.46 | 0.36 | 0.51 | 0.3 | 0.55 | 0.59 |
| 3 | 116275754 | rs1795156 | -1.396 | -0.79 | 0.994 | 0.469 | 0.001 | 0.001 | 0.001 | 0 | 0.5 | 0.46 | 0.46 | 0.73 | 0.39 | 0.39 |
| 3 | 116276315 | rs1599769 | -0.419 | -0.836 | 0.99 | 0.508 | 0.005 | -0.009 | -0.009 | 0 | 0.26 | 0.21 | 0.14 | 0.47 | 0.17 | 0.28 |
| 3 | 116279888 | rs1401069 | 0.453 | -0.871 | 1.075 | 0.475 | -0.024 | -0.029 | -0.029 | 0 | 0.83 | 0.45 | 0.9 | 0.92 | 0.89 | 0.88 |
| 3 | 116280518 | rs1619470 | -0.434 | -1.013 | 1.114 | 0.474 | 0.055 | 0.023 | 0.023 | 0 | 0.92 | 0.57 | 0.93 | 0.97 | 0.93 | 0.98 |
| 3 | 116281408 | rs4305418 | 1.196 | -1.096 | 1.037 | 0.405 | 0.024 | 0.021 | 0.021 | 0 | 0.59 | 0.3 | 0.57 | 0.66 | 0.67 | 0.77 |
| 3 | 116281707 | rs1401072 | 0.416 | -0.742 | 1.219 | 0.518 | -0.021 | 0.049 | 0.049 | 0.962 | 0.74 | 0.38 | 0.82 | 0.56 | 0.8 | 0.8 |
| 3 | 116282192 | rs2864817 | -0.15 | -0.745 | 1.17 | 0.527 | 0.001 | 0.002 | 0.002 | 0 | 0.24 | 0.16 | 0.32 | 0.26 | 0.23 | 0.11 |
| 3 | 116283355 | rs9861415 | 0.55 | -0.743 | 1.171 | 0.527 | -0.008 | 0.029 | 0.029 | 1.184 | 0.18 | 0.2 | 0.11 | 0.41 | 0.13 | 0.18 |
| 3 | 116283564 | rs9862002 | -0.945 | -0.736 | 1.135 | 1.003 | 0 | -0.001 | -0.001 | 0 | 0.41 | 0.35 | 0.43 | 0.59 | 0.35 | 0.28 |
| 3 | 116291220 | rs4089676 | 0.218 | -0.597 | 0.851 | 0.796 | -0.005 | 0.004 | 0.004 | 0.76 | 0.21 | 0.04 | 0.16 | 0.2 | 0.2 | 0.09 |
| 3 | 116296460 | rs4831145 | -0.098 | -0.762 | 0.875 | 0.835 | 0.043 | 0.018 | 0.018 | 0 | 0.92 | 0.58 | 0.93 | 0.97 | 0.93 | 0.98 |
| 3 | 116301834 | rs2030230 | -0.097 | -0.754 | 0.879 | 0.838 | 0.043 | 0.018 | 0.018 | 0 | 0.92 | 0.58 | 0.93 | 0.97 | 0.93 | 0.98 |
| 3 | 116311374 | rs1518321 | -0.149 | 0.347 | 0.492 | 0.89 | 0.002 | -0.002 | -0.002 | 0.152 | 0.25 | 0.23 | 0.37 | 0.26 | 0.33 | 0.13 |
| 3 | 116311891 | rs1996490 | -0.816 | 0.465 | -0.052 | 0.71 | 0.001 | -0.003 | -0.003 | 1.619 | 0.43 | 0.49 | 0.48 | 0.68 | 0.46 | 0.31 |
| 3 | 116315423 | rs4831149 | 0.154 | 0.811 | -0.536 | 0.853 | 0.023 | 0.042 | 0.042 | 0.016 | 0.51 | 0.28 | 0.54 | 0.61 | 0.61 | 0.55 |
| 3 | 116320333 | rs9863866 | 0.248 | 0.74 | -0.138 | 0.932 | 0.002 | -0.004 | -0.004 | 0 | 0.41 | 0.38 | 0.58 | 0.38 | 0.37 | 0.59 |
| 3 | 116320409 | rs4358296 | 1.163 | 1.104 | -0.151 | 0.874 | -0.003 | -0.016 | -0.016 | 0 | 0.24 | 0.24 | 0.16 | 0.09 | 0.2 | 0.29 |
| 3 | 116321494 | rs7621196 | 1.252 | 1.678 | -0.07 | 1.115 | 0.015 | 0.085 | 0.085 | 0 | 0.42 | 0.25 | 0.34 | 0.16 | 0.38 | 0.51 |
| 3 | 116322626 | rs1467214 | 0.504 | 1.679 | 0.627 | 1.23 | -0.003 | 0.008 | 0.008 | 0 | 0.55 | 0.57 | 0.59 | 0.33 | 0.51 | 0.71 |
| 3 | 116322918 | rs4405915 | 0.499 | 1.675 | 0.628 | 1.227 | -0.004 | 0.007 | 0.007 | 0 | 0.55 | 0.56 | 0.59 | 0.33 | 0.51 | 0.71 |
| 3 | 116325786 | rs11929035 | 0.471 | 1.675 | 0.626 | 1.233 | -0.003 | 0.007 | 0.007 | 0 | 0.55 | 0.56 | 0.59 | 0.33 | 0.51 | 0.71 |
| 3 | 116327809 | rs6771286 | 0.183 | 1.67 | 0.683 | 1.295 | 0.042 | 0.11 | 0.11 | 0 | 0.12 | 0.32 | 0.25 | 0.17 | 0.12 | 0.2 |
| 3 | 116328422 | rs13094939 | 0.987 | 1.915 | 0.385 | 1.777 | -0.019 | -0.035 | -0.035 | 2.3 | 0.18 | 0.11 | 0.16 | 0.33 | 0.24 | 0.07 |
| 3 | 116336036 | rs9834397 | 0.818 | 1.761 | 0.553 | 1.807 | 0.001 | -0.001 | -0.001 | 0 | 0.61 | 0.4 | 0.48 | 0.42 | 0.59 | 0.65 |
| 3 | 116337995 | rs938913 | -0.305 | 1.702 | 0.536 | 1.891 | 0.016 | 0.06 | 0.06 | 0 | 0.74 | 0.48 | 0.7 | 0.59 | 0.71 | 0.88 |
| 3 | 116342669 | rs10934356 | 0.992 | 1.729 | 0.491 | 1.874 | 0 | 0.03 | 0.03 | 0 | 0.52 | 0.32 | 0.54 | 0.32 | 0.49 | 0.71 |
| 3 | 116347790 | rs1518343 | 0.635 | 1.466 | 0.373 | 1.589 | 0.001 | -0.005 | -0.005 | 0.1 | 0.19 | 0.16 | 0.13 | 0.26 | 0.19 | 0.13 |
| 3 | 116349916 | rs10428159 | 0.722 | 1.381 | 0.448 | 1.526 | 0 | 0.089 | 0.089 | 0 | 0.43 | 0.24 | 0.35 | 0.17 | 0.41 | 0.52 |
| 3 | 116353532 | rs7649435 | -2.337 | 1.851 | 0.96 | 2.482 | -0.006 | 0.007 | 0.007 | 0 | 0.36 | 0.66 | 0.55 | 0.68 | 0.41 | 0.45 |
| 3 | 116356191 | rs12491897 | -0.682 | 2.119 | 0.665 | 2.955 | 0.015 | 0.074 | 0.074 | 0.02 | 0.15 | 0.29 | 0.27 | 0.36 | 0.19 | 0.24 |
| 3 | 116356970 | rs1401086 | -1.316 | 2.154 | 0.181 | 2.644 | 0.03 | 0.096 | 0.096 | 0 | 0.12 | 0.22 | 0.24 | 0.3 | 0.16 | 0.13 |
| 3 | 116364914 | rs13324635 | -0.55 | 2.143 | 0.16 | 2.662 | 0.01 | 0.053 | 0.053 | 0 | 0.06 | 0.19 | 0.09 | 0.08 | 0.05 | 0.1 |
| 3 | 116366738 | rs16825260 | -1.433 | 2.166 | 0.16 | 2.685 | 0.026 | 0.088 | 0.088 | 0 | 0.12 | 0.19 | 0.24 | 0.31 | 0.16 | 0.13 |
| 3 | 116371020 | rs9843317 | -0.748 | 1.972 | 0.125 | 2.347 | -0.021 | 0.045 | 0.045 | 0 | 0.1 | 0.23 | 0.1 | 0.19 | 0.06 | 0.2 |
| 3 | 116374090 | rs1401085 | -1.195 | 2.287 | -0.346 | 2.058 | -0.008 | -0.006 | -0.006 | 0.455 | 0.2 | 0.2 | 0.4 | 0.37 | 0.3 | 0.23 |
| 3 | 116378627 | rs2867833 | -1.206 | 1.459 | -0.074 | 1.636 | 0.009 | 0.09 | 0.09 | 1.431 | 0.28 | 0.54 | 0.39 | 0.47 | 0.3 | 0.39 |
| 3 | 116381899 | rs10934362 | 1.014 | 0.789 | 0.148 | 1.426 | -0.023 | 0.11 | 0.11 | 0 | 0.79 | 0.56 | 0.66 | 0.33 | 0.84 | 0.79 |
| 3 | 116382045 | rs11923597 | 0.993 | 0.779 | 0.197 | 1.825 | -0.021 | 0.192 | 0.192 | 0 | 0.82 | 0.57 | 0.77 | 0.6 | 0.89 | 0.81 |
| 3 | 116386909 | rs1543175 | 1.056 | 0.286 | -0.621 | 0.889 | -0.028 | 0.092 | 0.092 | 0 | 0.43 | 0.13 | 0.35 | 0.26 | 0.58 | 0.35 |
| 3 | 116389615 | rs16825296 | 0.538 | 0.282 | -0.47 | 0.797 | -0.001 | -0.016 | -0.016 | 0 | 0.08 | 0.04 | 0.02 | 0.01 | 0.05 | 0.09 |
| 3 | 116394204 | rs11711002 | 1.219 | 0.216 | -0.503 | 0.771 | -0.047 | 0.083 | 0.083 | 0.047 | 0.43 | 0.09 | 0.35 | 0.26 | 0.58 | 0.35 |
| 3 | 116395578 | rs9812193 | 0.791 | 0.085 | -0.037 | 0.7 | 0.004 | -0.001 | -0.001 | 0 | 0.51 | 0.58 | 0.52 | 0.52 | 0.67 | 0.38 |
| 3 | 116395779 | rs4429628 | 0.887 | 0.072 | -0.027 | 0.721 | -0.009 | 0.109 | 0.109 | 0 | 0.81 | 0.69 | 0.77 | 0.58 | 0.88 | 0.81 |
| 3 | 116396116 | rs13087427 | -0.847 | 0.111 | 0.024 | 0.717 | 0 | -0.017 | -0.017 | 0 | 0.17 | 0.09 | 0.2 | 0.42 | 0.12 | 0.19 |
| 3 | 116398764 | rs7635760 | -0.25 | 0.037 | 0.08 | 0.151 | -0.007 | -0.005 | -0.005 | 0.136 | 0.39 | 0.63 | 0.43 | 0.33 | 0.3 | 0.45 |
| 3 | 116404027 | rs2903681 | -1.169 | 0.118 | 0.024 | 0.306 | 0.015 | 0.02 | 0.02 | 0 | 0.51 | 0.43 | 0.46 | 0.46 | 0.38 | 0.63 |
| 3 | 116406199 | rs13066320 | -0.36 | 0.059 | -0.02 | 0.269 | 0.039 | 0.122 | 0.122 | 0 | 0.42 | 0.27 | 0.31 | 0.08 | 0.3 | 0.53 |
| 3 | 116420202 | rs938911 | -1.029 | 0.278 | -0.115 | 0.289 | 0.023 | -0.036 | -0.036 | 0 | 0.17 | 0.36 | 0.1 | 0.14 | 0.15 | 0.18 |
| 3 | 116422778 | rs6790547 | 1.034 | 0.042 | -0.331 | 0.206 | 0.064 | 0.111 | 0.111 | 0.368 | 0.5 | 0.06 | 0.52 | 0.67 | 0.63 | 0.47 |
| 3 | 116426757 | rs4831153 | 2.151 | 0.32 | -0.697 | -0.073 | -0.113 | 0.035 | 0.035 | 0 | 0.27 | 0.01 | 0.2 | 0.16 | 0.33 | 0.29 |
| 3 | 116427109 | rs1401073 | 0.513 | -0.466 | 0.127 | -0.258 | 0.063 | -0.007 | -0.007 | 0 | 0.93 | 0.41 | 0.93 | 1 | 0.93 | 1 |
| 3 | 116427251 | rs16825350 | 0.93 | -0.811 | -0.063 | -0.389 | -0.001 | 0.022 | 0.022 | 0 | 0.14 | 0 | 0.17 | 0.05 | 0.1 | 0.28 |
| 3 | 116427291 | rs1401070 | -0.436 | -0.389 | 0.276 | -0.094 | 0.065 | 0.049 | 0.049 | 0 | 0.41 | 0.17 | 0.39 | 0.34 | 0.3 | 0.53 |
| 3 | 116433126 | rs12488450 | -0.575 | -0.302 | -0.379 | -0.001 | -0.016 | 0.038 | 0.038 | 0 | 0.11 | 0.18 | 0.15 | 0.27 | 0.08 | 0.1 |
| 3 | 116440045 | rs7640361 | 1.222 | -0.226 | -0.764 | -0.159 | -0.047 | -0.025 | -0.025 | 0 | 0.45 | 0.05 | 0.49 | 0.54 | 0.59 | 0.39 |
| 3 | 116442326 | rs6773550 | -0.219 | -0.305 | -0.321 | -1.061 | 0.034 | -0.07 | -0.07 | 0.013 | 0.11 | 0.38 | 0.06 | 0.01 | 0.1 | 0.08 |
| 3 | 116444174 | rs4831156 | -0.303 | -0.289 | -0.641 | -0.498 | 0.017 | 0.119 | 0.119 | 0 | 0.3 | 0.05 | 0.25 | 0.1 | 0.22 | 0.42 |
| 3 | 116455442 | rs1347060 | 0.198 | 0.99 | -0.304 | 0.52 | -0.004 | -0.003 | -0.003 | 0 | 0.19 | 0.19 | 0.12 | 0.2 | 0.19 | 0.19 |
| 3 | 116455826 | rs9861262 | 0.1 | 0.527 | -0.26 | 0.413 | 0.003 | 0.003 | 0.003 | 0.121 | 0.23 | 0.22 | 0.19 | 0.13 | 0.19 | 0.25 |
| 3 | 116456208 | rs17659043 | -0.017 | 1.111 | -0.53 | 0.668 | -0.035 | -0.015 | -0.015 | 0 | 0.07 | 0.01 | 0.06 | 0.08 | 0.1 | 0.02 |
| 3 | 116457044 | rs12695333 | -0.565 | 0.69 | -0.089 | 0.614 | -0.007 | 0.052 | 0.052 | 0 | 0.76 | 0.7 | 0.67 | 0.5 | 0.77 | 0.79 |
| 3 | 116460965 | rs4541407 | 0.665 | 0.764 | -0.347 | 0.656 | -0.015 | -0.102 | -0.102 | 0.005 | 0.21 | 0.05 | 0.3 | 0.49 | 0.23 | 0.18 |
| 3 | 116470835 | rs9817566 | 0.957 | 1.02 | 0.445 | 0.483 | 0.054 | 0.047 | 0.047 | 0.291 | 0.93 | 0.73 | 0.81 | 0.93 | 0.9 | 0.9 |

| | | | | | | | | | | | | | | | | |
|---|---|---|---|---|---|---|---|---|---|---|---|---|---|---|---|---|
| 3 | 116472294 | rs6438332 | 0.846 | 1.006 | 0.447 | 0.102 | -0.003 | 0.005 | 0.005 | 0 | 0.3 | 0.24 | 0.18 | 0.21 | 0.33 | 0.29 |
| 3 | 116474034 | rs9837798 | 0.208 | 0.712 | 0.58 | -0.033 | 0.006 | 0.014 | 0.014 | 0 | 0.38 | 0.33 | 0.3 | 0.22 | 0.34 | 0.43 |
| 3 | 116475071 | rs2116574 | 0.126 | 0.649 | 0.746 | -0.049 | 0.038 | 0.06 | 0.06 | 0 | 0.56 | 0.26 | 0.53 | 0.74 | 0.65 | 0.49 |
| 3 | 116475786 | rs7618116 | 0.105 | 0.665 | 0.722 | -0.049 | -0.014 | -0.036 | -0.036 | 0 | 0.39 | 0.45 | 0.31 | 0.22 | 0.34 | 0.43 |
| 3 | 116478269 | rs7638734 | 1.08 | 0.612 | 0.567 | -0.043 | -0.033 | 0.027 | 0.027 | 0 | 0.3 | 0.11 | 0.18 | 0.21 | 0.33 | 0.31 |
| 3 | 116483229 | rs10511359 | 1.049 | 0.578 | 0.801 | -0.029 | -0.018 | 0.017 | 0.017 | 0 | 0.3 | 0.16 | 0.18 | 0.21 | 0.33 | 0.29 |
| 3 | 116485013 | rs9839736 | 0.845 | 0.821 | 0.841 | 0.014 | -0.04 | -0.078 | -0.078 | 2.398 | 0.27 | 0.13 | 0.37 | 0.52 | 0.35 | 0.21 |
| 3 | 116486794 | rs6766679 | -0.153 | 0.544 | 0.62 | -0.198 | 0.014 | 0.009 | 0.009 | 0.095 | 0.68 | 0.25 | 0.6 | 0.76 | 0.74 | 0.65 |
| 3 | 116489442 | rs1347063 | 0.318 | 0.553 | 0.593 | -0.202 | -0.045 | 0.074 | 0.074 | 0.655 | 0.43 | 0.17 | 0.28 | 0.27 | 0.47 | 0.47 |
| 3 | 116498380 | rs11709313 | 1.35 | 0.584 | 0.762 | 0.156 | -0.034 | -0.083 | -0.083 | 0 | 0.24 | 0.07 | 0.21 | 0.39 | 0.27 | 0.17 |
| 3 | 116498715 | rs10934369 | -0.483 | 0.535 | 0.774 | 0.076 | -0.01 | 0.007 | 0.007 | 0.2 | 0.85 | 0.85 | 0.85 | 0.91 | 0.78 | 0.82 |
| 3 | 116499373 | rs10511360 | -0.314 | 0.455 | 0.859 | 0.055 | -0.002 | 0.002 | 0.002 | 0.006 | 0.55 | 0.55 | 0.69 | 0.73 | 0.53 | 0.54 |
| 3 | 116499774 | rs1821194 | 0.862 | 0.419 | 0.717 | 0.06 | -0.018 | -0.011 | -0.011 | 4.219 | 0.26 | 0.29 | 0.19 | 0.17 | 0.17 | 0.26 |
| 3 | 116501764 | rs12631018 | -0.282 | 0.572 | 0.618 | 0.171 | -0.001 | -0.035 | -0.035 | 1.524 | 0.84 | 0.65 | 0.75 | 0.92 | 0.86 | 0.76 |
| 3 | 116501978 | rs12631538 | 0.471 | 0.467 | 0.638 | 0.176 | -0.012 | 0.008 | 0.008 | 0 | 0.55 | 0.32 | 0.37 | 0.38 | 0.51 | 0.55 |
| 3 | 116505544 | rs7614098 | 0.832 | 0.461 | 0.577 | 0.339 | 0.002 | 0.022 | 0.022 | 0 | 0.41 | 0.28 | 0.35 | 0.28 | 0.38 | 0.43 |
| 3 | 116506471 | rs6806089 | -0.164 | 0.098 | 0.293 | 0.573 | 0.011 | -0.004 | -0.004 | 0 | 0.41 | 0.28 | 0.47 | 0.62 | 0.39 | 0.39 |
| 3 | 116507763 | rs13090380 | -0.197 | 0.079 | 0.179 | 0.581 | 0.005 | -0.001 | -0.001 | 0 | 0.41 | 0.37 | 0.47 | 0.62 | 0.39 | 0.38 |
| 3 | 116508847 | rs1991745 | -0.262 | -0.126 | 0.151 | 0.438 | -0.022 | -0.003 | -0.003 | 0 | 0.2 | 0.27 | 0.27 | 0.21 | 0.13 | 0.21 |
| 3 | 116509824 | rs9847281 | 0.515 | 0.01 | 0.098 | 0.086 | -0.003 | -0.026 | -0.026 | 0 | 0.68 | 0.59 | 0.73 | 0.81 | 0.68 | 0.49 |
| 3 | 116510192 | rs1368141 | 0.499 | -0.115 | 0.578 | 0.086 | 0.017 | 0.014 | 0.014 | 0.793 | 0.92 | 0.69 | 0.94 | 0.95 | 0.93 | 0.78 |
| 3 | 116511825 | rs9820027 | -0.889 | 0.148 | 0.404 | 0.827 | -0.009 | -0.034 | -0.034 | 0 | 0.33 | 0.22 | 0.28 | 0.47 | 0.37 | 0.55 |
| 3 | 116515862 | rs11927819 | 0.939 | 0.235 | 0.209 | 0.408 | 0.027 | 0.008 | 0.008 | 0.201 | 0.49 | 0.34 | 0.4 | 0.42 | 0.44 | 0.29 |
| 3 | 116520381 | rs16825642 | 1.406 | -0.555 | 0.044 | -0.116 | -0.011 | 0.027 | 0.027 | 0 | 0.21 | 0.31 | 0.17 | 0.27 | 0.21 | 0.16 |
| 3 | 116525589 | rs2020375 | 0.324 | -0.572 | -0.136 | -0.203 | 0.009 | -0.004 | -0.004 | 0 | 0.48 | 0.36 | 0.6 | 0.53 | 0.51 | 0.33 |
| 3 | 116528670 | rs9816739 | 0.332 | -0.528 | -0.194 | -0.21 | 0.01 | -0.004 | -0.004 | 0 | 0.48 | 0.35 | 0.61 | 0.53 | 0.51 | 0.33 |
| 3 | 116532683 | rs6805241 | 0.78 | -0.576 | -0.42 | -0.248 | 0.002 | 0.021 | 0.021 | 0.01 | 0.21 | 0.1 | 0.15 | 0.15 | 0.21 | 0.32 |
| 3 | 116536464 | rs7627376 | 0.104 | -0.361 | -0.247 | -0.141 | 0.016 | -0.004 | -0.004 | 0 | 0.48 | 0.34 | 0.62 | 0.53 | 0.51 | 0.33 |
| 3 | 116550622 | rs7430526 | -0.027 | -0.339 | -0.251 | -0.176 | 0.003 | 0.001 | 0.001 | 0.011 | 0.5 | 0.47 | 0.6 | 0.52 | 0.51 | 0.35 |
| 3 | 116554502 | rs7433718 | 0.415 | -0.369 | -0.247 | -0.191 | 0.008 | -0.007 | -0.007 | 0 | 0.43 | 0.35 | 0.32 | 0.43 | 0.42 | 0.48 |
| 3 | 116564876 | rs6438349 | 0.049 | -0.264 | -0.555 | -0.215 | 0.014 | -0.003 | -0.003 | 0 | 0.49 | 0.35 | 0.59 | 0.54 | 0.51 | 0.33 |
| 3 | 116568065 | rs9840374 | -0.883 | 0.032 | -1.199 | -0.075 | 0.004 | -0.01 | -0.01 | 0 | 0.93 | 0.88 | 0.94 | 0.95 | 0.93 | 0.81 |
| 3 | 116573921 | rs7613406 | -1.303 | -0.143 | -1.427 | -0.305 | 0.016 | 0.026 | 0.026 | 0.815 | 0.84 | 0.91 | 0.91 | 1 | 0.9 | 0.97 |
| 3 | 116574642 | rs7614031 | 0.097 | 0.61 | -1.057 | 0.242 | -0.003 | -0.004 | -0.004 | 0.065 | 0.32 | 0.36 | 0.51 | 0.54 | 0.42 | 0.31 |
| 3 | 116584082 | rs7433623 | 0.302 | 0.463 | -1.288 | -0.084 | 0.002 | 0.039 | 0.039 | 0 | 0.71 | 0.77 | 0.78 | 0.85 | 0.69 | 0.54 |
| 3 | 116585281 | rs7643014 | -0.305 | 0.465 | -1.435 | -0.121 | 0.01 | 0.022 | 0.022 | 0 | 0.77 | 0.81 | 0.81 | 0.85 | 0.77 | 0.66 |
| 3 | 116586863 | rs12631842 | 0.585 | 0.667 | -1.704 | -0.095 | -0.031 | 0 | 0 | 0 | 0.17 | 0.09 | 0.15 | 0.22 | 0.2 | 0.11 |
| 3 | 116589267 | rs6438359 | 0.322 | 0.423 | -1.552 | -0.242 | 0.002 | 0.052 | 0.052 | 0 | 0.72 | 0.81 | 0.78 | 0.85 | 0.7 | 0.55 |
| 3 | 116591877 | rs4855961 | -0.864 | 0.476 | -1.535 | -0.209 | 0.002 | -0.022 | -0.022 | 0.316 | 0.94 | 0.92 | 0.97 | 1 | 0.94 | 0.89 |
| 3 | 116595111 | rs6777508 | 0.018 | 0.349 | -1.431 | -0.003 | 0.008 | 0.021 | 0.021 | 0 | 0.23 | 0.19 | 0.18 | 0.16 | 0.24 | 0.34 |
| 3 | 116599486 | rs6438364 | -0.322 | 0.577 | -1.035 | 0.224 | 0 | -0.013 | -0.013 | 0 | 0.83 | 0.75 | 0.87 | 0.9 | 0.86 | 0.88 |
| 3 | 116600332 | rs7427875 | 0.242 | 0.539 | -1.098 | 0.412 | -0.001 | 0.006 | 0.006 | 0 | 0.76 | 0.71 | 0.8 | 0.67 | 0.76 | 0.67 |
| 3 | 116603705 | rs6438366 | 0.406 | 0.405 | -1.191 | -0.254 | -0.001 | 0 | 0 | 0 | 0.47 | 0.49 | 0.61 | 0.65 | 0.51 | 0.57 |
| 3 | 116606168 | rs6438367 | 0.521 | 0.838 | -0.932 | 0.203 | -0.014 | -0.004 | -0.004 | 0 | 0.3 | 0.45 | 0.29 | 0.23 | 0.26 | 0.38 |
| 3 | 116607255 | rs7429880 | 1.055 | 1.313 | -1.001 | 0.563 | -0.005 | -0.013 | -0.013 | 0 | 0.14 | 0.07 | 0.22 | 0.28 | 0.19 | 0.16 |
| 3 | 116608306 | rs4855963 | 0.238 | 1.149 | -1.113 | 0.348 | -0.008 | 0.032 | 0.032 | 0 | 0.47 | 0.27 | 0.53 | 0.42 | 0.5 | 0.3 |
| 3 | 116613461 | rs9866579 | 1.304 | 0.798 | -0.807 | 0.117 | 0.034 | -0.044 | -0.044 | 0 | 0.1 | 0.34 | 0.07 | 0 | 0.07 | 0.13 |
| 3 | 116616793 | rs9874549 | -0.458 | 0.742 | -0.996 | 0.197 | -0.005 | 0.01 | 0.01 | 0 | 0.71 | 0.65 | 0.61 | 0.4 | 0.72 | 0.68 |
| 3 | 116624325 | rs6780710 | 1.072 | 0.763 | -1.095 | 0.171 | -0.027 | 0.218 | 0.218 | 0 | 0.42 | 0.12 | 0.27 | 0.11 | 0.4 | 0.32 |
| 3 | 116627531 | rs2102626 | 2.272 | 0.661 | -1.119 | 0.363 | -0.022 | -0.034 | -0.034 | 0.743 | 0.07 | 0 | 0.08 | 0.27 | 0.09 | 0.08 |
| 3 | 116630343 | rs6777242 | 0.313 | 0.667 | -1.101 | 0.351 | 0.017 | 0.002 | 0.002 | 0 | 0.61 | 0.26 | 0.53 | 0.4 | 0.65 | 0.53 |
| 3 | 116631443 | rs12631952 | 1.602 | 0.674 | -0.997 | 0.38 | -0.001 | 0.021 | 0.021 | 0 | 0.09 | 0.09 | 0.09 | 0.27 | 0.09 | 0.08 |
| 3 | 116634122 | rs1980081 | 1.038 | 0.829 | -1.031 | 0.295 | -0.021 | -0.019 | -0.019 | 0 | 0.18 | 0.09 | 0.25 | 0.29 | 0.27 | 0.21 |
| 3 | 116635262 | rs4525877 | -1.399 | 0.715 | -0.92 | 0.401 | 0.009 | 0.209 | 0.209 | 0 | 0.91 | 0.77 | 0.81 | 0.6 | 0.91 | 0.92 |
| 3 | 116637776 | rs4452335 | 1.328 | 0.742 | -0.987 | 0.379 | 0.006 | -0.026 | -0.026 | 0 | 0.2 | 0.12 | 0.2 | 0.22 | 0.19 | 0.25 |
| 3 | 116647795 | rs535342 | 1.28 | 0.753 | -0.907 | 0.307 | 0.026 | -0.049 | -0.049 | 1.044 | 0.1 | 0.4 | 0.07 | 0 | 0.07 | 0.13 |
| 3 | 116648690 | rs10934370 | 1.53 | 0.738 | -1.094 | 0.333 | -0.003 | -0.03 | -0.03 | 0 | 0.19 | 0.05 | 0.2 | 0.2 | 0.19 | 0.24 |
| 3 | 116650801 | rs1133604 | -0.612 | 0.783 | -0.969 | 0.333 | -0.007 | -0.008 | -0.008 | 0 | 0.48 | 0.65 | 0.53 | 0.49 | 0.51 | 0.6 |
| 3 | 116655223 | rs7651803 | 1.311 | 0.774 | -0.833 | 0.315 | 0.024 | -0.052 | -0.052 | 0 | 0.1 | 0.39 | 0.07 | 0 | 0.07 | 0.13 |
| 3 | 116655296 | rs9873638 | 0.115 | 0.789 | -0.872 | 0.333 | -0.012 | -0.021 | -0.021 | 0 | 0.37 | 0.28 | 0.45 | 0.49 | 0.43 | 0.44 |
| 3 | 116658169 | rs7616813 | 1.249 | 0.775 | -0.767 | 0.254 | 0.012 | -0.056 | -0.056 | 0 | 0.11 | 0.35 | 0.07 | 0 | 0.07 | 0.13 |
| 3 | 116658337 | rs9814086 | 0.987 | 0.821 | -0.791 | 0.301 | 0.006 | 0.008 | 0.008 | 0 | 0.18 | 0.17 | 0.25 | 0.29 | 0.25 | 0.21 |
| 3 | 116662669 | rs6790499 | -0.171 | 0.819 | -0.88 | 0.481 | 0.018 | 0.013 | 0.013 | 0 | 0.28 | 0.52 | 0.32 | 0.36 | 0.32 | 0.34 |
| 3 | 116669163 | rs11921574 | 0.618 | 0.81 | -1.11 | 0.409 | 0.004 | -0.015 | -0.015 | 0 | 0.22 | 0.16 | 0.19 | 0.29 | 0.22 | 0.33 |
| 3 | 116670390 | rs1523262 | 1.509 | 0.819 | -1.138 | 0.338 | 0.009 | -0.026 | -0.026 | 0 | 0.2 | 0.12 | 0.21 | 0.22 | 0.19 | 0.26 |
| 3 | 116671791 | rs2868477 | 1.902 | 0.88 | -1.05 | 0.038 | -0.023 | 0.125 | 0.125 | 0.296 | 0.28 | 0.1 | 0.19 | 0.01 | 0.25 | 0.07 |
| 3 | 116673914 | rs1009215 | -0.115 | 0.908 | -0.773 | 0.334 | 0.013 | 0.001 | 0.001 | 0.005 | 0.27 | 0.54 | 0.3 | 0.29 | 0.31 | 0.34 |
| 3 | 116680676 | rs480756 | 0.14 | 0.442 | -0.801 | 0.357 | -0.025 | 0.091 | 0.091 | 1.832 | 0.8 | 0.42 | 0.74 | 0.62 | 0.84 | 0.73 |
| 3 | 116682766 | rs2868482 | 0.419 | 0.244 | -0.992 | 0.165 | -0.031 | 0.04 | 0.04 | 0 | 0.75 | 0.42 | 0.72 | 0.62 | 0.82 | 0.72 |

| | | | | | | | | | | | | | | | | |
|---|---|---|---|---|---|---|---|---|---|---|---|---|---|---|---|---|
| 3 | 116693645 | rs500877 | 1.561 | 0.376 | -1.151 | -0.309 | 0.007 | 0.026 | 0.026 | 0 | 0.26 | 0.16 | 0.17 | 0 | 0.24 | 0.06 |
| 3 | 116695998 | rs7647072 | -0.257 | 0.354 | -0.82 | -0.176 | -0.016 | 0.024 | 0.024 | 0 | 0.36 | 0.54 | 0.33 | 0.51 | 0.29 | 0.35 |
| 3 | 116698728 | rs16825904 | 1.526 | 0.431 | -0.393 | -0.081 | 0.005 | 0.121 | 0.121 | 0.216 | 0.09 | 0.12 | 0.17 | 0.4 | 0.09 | 0.11 |
| 3 | 116699163 | rs16825910 | 1.344 | 0.423 | -0.401 | -0.084 | 0.001 | 0.087 | 0.087 | 0.047 | 0.09 | 0.1 | 0.17 | 0.4 | 0.09 | 0.11 |
| 3 | 116699869 | rs11926630 | -0.566 | 0.699 | -0.528 | 0.076 | -0.009 | -0.002 | -0.002 | 0 | 0.75 | 0.8 | 0.59 | 0.75 | 0.71 | 0.59 |
| 3 | 116708204 | rs2090584 | -0.156 | 0.229 | -0.445 | -0.049 | -0.017 | 0.023 | 0.023 | 0 | 0.37 | 0.47 | 0.32 | 0.51 | 0.31 | 0.37 |
| 3 | 116711857 | rs558329 | 2.184 | 0.311 | -0.745 | -0.363 | 0.002 | -0.023 | -0.023 | 0 | 0.28 | 0.25 | 0.2 | 0 | 0.26 | 0.13 |
| 3 | 116712193 | rs4855890 | 0.47 | 0.537 | -0.799 | -0.316 | 0.02 | 0.139 | 0.139 | 0 | 0.5 | 0.27 | 0.53 | 0.23 | 0.52 | 0.46 |
| 3 | 116714091 | rs1914812 | 1.842 | 0.576 | -0.755 | -0.264 | 0 | 0.023 | 0.023 | 0 | 0.12 | 0.11 | 0.09 | 0.26 | 0.17 | 0.14 |
| 3 | 116714180 | rs2068215 | 2.12 | 0.563 | -1.004 | -0.251 | 0.001 | 0.083 | 0.083 | 0 | 0.09 | 0.07 | 0.17 | 0.43 | 0.1 | 0.11 |
| 3 | 116714620 | rs1518904 | 2.576 | 0.479 | -1.173 | -0.679 | -0.006 | 0.036 | 0.036 | 0 | 0.25 | 0.15 | 0.18 | 0.01 | 0.26 | 0.09 |
| 3 | 116717099 | rs7642132 | 0.214 | 0.614 | -1.214 | -0.397 | -0.034 | -0.007 | -0.007 | 0 | 0.23 | 0.11 | 0.34 | 0.23 | 0.26 | 0.33 |
| 3 | 116729273 | rs35258147 | 0.338 | 0.139 | -0.775 | -0.167 | 0.01 | 0.088 | 0.088 | 0 | 0.26 | 0.13 | 0.17 | 0.08 | 0.2 | 0.25 |
| 3 | 116735107 | rs10511363 | 1.221 | 0.517 | -0.66 | 0.08 | -0.026 | 0.022 | 0.022 | 0 | 0.14 | 0.06 | 0.14 | 0.09 | 0.16 | 0.08 |
| 3 | 116735993 | rs693456 | 0.299 | 0.326 | -0.276 | 0.283 | -0.012 | 0.058 | 0.058 | 0 | 0.68 | 0.53 | 0.65 | 0.53 | 0.69 | 0.68 |
| 3 | 116741261 | rs2203817 | -0.4 | 0.715 | -0.016 | 0.521 | 0.023 | 0.103 | 0.103 | 0.245 | 0.87 | 0.71 | 0.87 | 0.72 | 0.84 | 0.85 |
| 3 | 116746735 | rs3914909 | -1.351 | 0.804 | -0.277 | 0.223 | -0.003 | 0.016 | 0.016 | 0 | 0.55 | 0.59 | 0.66 | 0.7 | 0.5 | 0.69 |
| 3 | 116749018 | rs2868486 | -0.107 | 0.672 | -0.139 | 0.231 | 0.015 | 0.018 | 0.018 | 0 | 0.34 | 0.29 | 0.43 | 0.25 | 0.32 | 0.45 |
| 3 | 116753688 | rs10934380 | 0.199 | 0.73 | -0.077 | 0.312 | 0.002 | 0.05 | 0.05 | 0 | 0.2 | 0.23 | 0.22 | 0.45 | 0.19 | 0.24 |
| 3 | 116755113 | rs869407 | 1.34 | 0.615 | -0.289 | 0.247 | -0.011 | 0.028 | 0.028 | 0 | 0.14 | 0.01 | 0.19 | 0.07 | 0.14 | 0.16 |
| 3 | 116755365 | rs7651083 | -0.157 | 0.926 | 0.114 | 0.483 | 0.013 | 0.111 | 0.111 | 0.31 | 0.17 | 0.39 | 0.24 | 0.66 | 0.19 | 0.26 |
| 3 | 116755540 | rs13318637 | 2.214 | 0.787 | 0.076 | 0.299 | -0.021 | 0.065 | 0.065 | 0 | 0.34 | 0.2 | 0.23 | 0.01 | 0.36 | 0.2 |
| 3 | 116755811 | rs938110 | -1.582 | 0.777 | 0.154 | 0.37 | 0.001 | -0.001 | -0.001 | 0 | 0.6 | 0.57 | 0.62 | 0.63 | 0.61 | 0.7 |
| 3 | 116758159 | rs1851247 | -0.796 | 0.861 | 0.435 | 0.865 | 0.002 | 0.01 | 0.01 | 0 | 0.82 | 0.83 | 0.82 | 0.63 | 0.74 | 0.79 |
| 3 | 116759424 | rs553110 | 0.456 | 0.851 | 0.036 | 0.903 | -0.008 | 0.354 | 0.354 | 0.19 | 0.9 | 0.51 | 0.84 | 0.63 | 0.94 | 0.79 |
| 3 | 116759983 | rs6788569 | 0.961 | 0.867 | -0.04 | 0.704 | 0.005 | -0.012 | -0.012 | 1.451 | 0.68 | 0.42 | 0.65 | 0.26 | 0.66 | 0.61 |
| 3 | 116771107 | rs6801409 | 0.564 | 0.569 | 0.304 | 0.997 | 0.002 | 0.384 | 0.384 | 0.053 | 0.9 | 0.57 | 0.84 | 0.63 | 0.94 | 0.79 |
| 3 | 116775274 | rs938115 | -0.987 | 1.024 | -0.052 | 0.733 | 0 | -0.006 | -0.006 | 0.473 | 0.56 | 0.57 | 0.62 | 0.75 | 0.57 | 0.71 |
| 3 | 116780022 | rs13088089 | 0.55 | 1.019 | -0.498 | 0.51 | -0.011 | 0.128 | 0.128 | 0.782 | 0.33 | 0.06 | 0.19 | 0.13 | 0.29 | 0.46 |
| 3 | 116782852 | rs6438377 | 0.632 | 1.255 | -0.076 | 0.874 | 0.018 | 0.087 | 0.087 | 0 | 0.76 | 0.6 | 0.6 | 0.56 | 0.7 | 0.67 |
| 3 | 116784852 | rs9820102 | 2.335 | 1.244 | -0.705 | 0.515 | -0.002 | 0.066 | 0.066 | 0.973 | 0.31 | 0.15 | 0.21 | 0.01 | 0.3 | 0.1 |
| 3 | 116788351 | rs1521412 | 0.361 | 0.973 | -0.515 | 0.879 | 0.01 | -0.015 | -0.015 | 0.014 | 0.57 | 0.69 | 0.63 | 0.46 | 0.61 | 0.47 |
| 3 | 116789451 | rs4855919 | -0.018 | 0.939 | -0.538 | 0.973 | -0.01 | 0.009 | 0.009 | 0.016 | 0.6 | 0.86 | 0.73 | 0.59 | 0.61 | 0.48 |
| 3 | 116791001 | rs2221847 | 0.204 | 0.941 | -0.604 | 0.974 | 0.009 | -0.012 | -0.012 | 0 | 0.57 | 0.69 | 0.63 | 0.47 | 0.61 | 0.49 |
| 3 | 116791023 | rs12634401 | 0.75 | 0.681 | -0.173 | 1.314 | 0.003 | 0.032 | 0.032 | 0 | 0.1 | 0.1 | 0.26 | 0.31 | 0.13 | 0.13 |
| 3 | 116795417 | rs4101568 | 0.804 | 0.786 | -1.839 | 1.042 | 0.005 | 0.038 | 0.038 | 0 | 0.18 | 0.07 | 0.08 | 0.09 | 0.14 | 0.23 |
| 3 | 116795694 | rs12493930 | 0.293 | 0.775 | -1.456 | 1.611 | 0.023 | 0.153 | 0.153 | 0 | 0.05 | 0.1 | 0.25 | 0.37 | 0.1 | 0.1 |
| 3 | 116795942 | rs1402625 | 0.555 | 0.643 | -1.46 | 1.239 | 0.016 | 0.008 | 0.008 | 0 | 0.17 | 0.09 | 0.19 | 0.15 | 0.14 | 0.24 |
| 3 | 116796116 | rs12494712 | -0.922 | 1.081 | -1.253 | 1.26 | -0.019 | -0.051 | -0.051 | 0 | 0.2 | 0.14 | 0.35 | 0.49 | 0.25 | 0.3 |
| 3 | 116799394 | rs9842731 | -0.425 | 0.836 | -0.83 | 0.947 | 0 | 0.028 | 0.028 | 0 | 0.1 | 0.11 | 0.26 | 0.31 | 0.12 | 0.13 |
| 3 | 116802560 | rs9878980 | -0.008 | 0.615 | -0.994 | -0.004 | -0.031 | -0.041 | -0.041 | 0.228 | 0.39 | 0.25 | 0.42 | 0.55 | 0.44 | 0.39 |
| 3 | 116803821 | rs9289067 | -1.099 | 0.419 | -0.652 | -0.303 | -0.014 | 0.076 | 0.076 | 0 | 0.74 | 0.52 | 0.66 | 0.62 | 0.8 | 0.66 |
| 3 | 116804591 | rs1554333 | -1.089 | 0.415 | -0.654 | -0.294 | -0.015 | 0.08 | 0.08 | 0 | 0.73 | 0.52 | 0.66 | 0.61 | 0.8 | 0.66 |
| 3 | 116811470 | rs17731594 | 1.133 | 0.476 | -1.002 | -0.565 | -0.034 | 0.025 | 0.025 | 0 | 0.2 | 0.12 | 0.12 | 0.01 | 0.24 | 0.09 |
| 3 | 116814285 | rs16826094 | 0.46 | -0.066 | -1.15 | -0.492 | 0 | -0.022 | -0.022 | 0 | 0.31 | 0.36 | 0.27 | 0.18 | 0.29 | 0.23 |
| 3 | 116814934 | rs9289071 | 0.266 | -0.233 | -0.716 | -0.492 | 0.001 | 0.01 | 0.01 | 0 | 0.44 | 0.38 | 0.56 | 0.75 | 0.44 | 0.56 |
| 3 | 116817560 | rs7653556 | -1.226 | -0.218 | -0.948 | -0.397 | 0.006 | 0.064 | 0.064 | 0 | 0.73 | 0.51 | 0.62 | 0.61 | 0.77 | 0.64 |
| 3 | 116824230 | rs9855076 | 1.228 | -0.191 | -1.047 | -0.427 | 0.002 | -0.072 | -0.072 | 0.047 | 0.17 | 0.03 | 0.2 | 0.37 | 0.2 | 0.23 |
| 3 | 116825894 | rs9836910 | 0.27 | -0.041 | -1 | -0.62 | 0.02 | -0.002 | -0.002 | 0 | 0.5 | 0.31 | 0.51 | 0.46 | 0.5 | 0.56 |
| 3 | 116826426 | rs9875598 | -0.22 | -0.553 | -1.015 | -0.4 | -0.001 | -0.005 | -0.005 | 0 | 0.75 | 0.78 | 0.81 | 0.93 | 0.73 | 0.78 |
| 3 | 116827967 | rs9810432 | 0.782 | -0.56 | -1 | -0.343 | -0.002 | -0.004 | -0.004 | 0 | 0.56 | 0.54 | 0.6 | 0.63 | 0.52 | 0.58 |
| 3 | 116828165 | rs9811011 | 0.218 | -0.399 | -1.346 | 0.19 | -0.006 | 0.023 | 0.023 | 0 | 0.18 | 0.22 | 0.21 | 0.29 | 0.2 | 0.2 |
| 3 | 116828715 | rs7637468 | 0.121 | -0.534 | -1.111 | -0.042 | 0.003 | -0.008 | -0.008 | 0 | 0.09 | 0.06 | 0.06 | 0 | 0.07 | 0.02 |
| 3 | 116830475 | rs7621475 | -0.328 | -0.422 | -1.175 | 0.296 | 0.006 | -0.006 | -0.006 | 0 | 0.35 | 0.31 | 0.32 | 0.38 | 0.33 | 0.38 |
| 3 | 116830612 | rs7633227 | -0.408 | -0.177 | -1.244 | 0.174 | 0.004 | -0.007 | -0.007 | 0.004 | 0.39 | 0.35 | 0.47 | 0.51 | 0.39 | 0.45 |
| 3 | 116838635 | rs6784605 | -0.273 | -0.593 | -1.69 | -0.626 | 0.055 | 0.138 | 0.138 | 0 | 0.3 | 0.11 | 0.16 | 0.03 | 0.25 | 0.17 |
| 3 | 116839004 | rs1818768 | 0.539 | -0.257 | -0.493 | -0.201 | 0.024 | 0.012 | 0.012 | 0.52 | 0.91 | 0.68 | 0.81 | 0.91 | 0.93 | 0.94 |
| 3 | 116839289 | rs1357103 | 0.188 | -0.093 | -1.239 | 0.224 | 0.012 | -0.012 | -0.012 | 0 | 0.62 | 0.39 | 0.59 | 0.61 | 0.74 | 0.71 |
| 3 | 116839982 | rs9846104 | -1.244 | 0.827 | -1.258 | 0.015 | 0 | -0.005 | -0.005 | 0 | 0.1 | 0.1 | 0.13 | 0.04 | 0.17 | 0.05 |
| 3 | 116840077 | rs13058767 | -0.617 | 0.038 | -1.893 | 0 | 0.002 | 0.009 | 0.009 | 0.103 | 0.3 | 0.06 | 0.32 | 0.31 | 0.32 | 0.27 |
| 3 | 116842805 | rs1554328 | -0.412 | 0.136 | -1.486 | 0.874 | -0.003 | 0.079 | 0.079 | 0 | 0.12 | 0.14 | 0.09 | 0.43 | 0.11 | 0.28 |
| 3 | 116844543 | rs16826176 | -0.656 | 0.131 | -1.502 | 0.838 | -0.004 | 0.073 | 0.073 | 0 | 0.13 | 0.15 | 0.1 | 0.44 | 0.13 | 0.27 |
| 3 | 116845163 | rs9844832 | -0.136 | 0.115 | -1.37 | 0.238 | 0.006 | 0.02 | 0.02 | 0 | 0.4 | 0.31 | 0.44 | 0.34 | 0.44 | 0.42 |
| 3 | 116847881 | rs938111 | -0.693 | 0.41 | -1.146 | 0.82 | -0.001 | 0.07 | 0.07 | 0 | 0.14 | 0.17 | 0.11 | 0.45 | 0.13 | 0.29 |
| 3 | 116852151 | rs6796212 | 0.297 | 0.569 | -1.134 | 0.056 | -0.002 | -0.015 | -0.015 | 0 | 0.87 | 0.88 | 0.89 | 0.99 | 0.84 | 0.82 |
| 3 | 116852469 | rs6808814 | -0.031 | 0.915 | -0.994 | 0.546 | 0.006 | 0.016 | 0.016 | 0 | 0.24 | 0.23 | 0.19 | 0.35 | 0.27 | 0.31 |
| 3 | 116854801 | rs9818327 | 0.049 | 0.899 | -1.018 | 0.551 | -0.013 | -0.021 | -0.021 | 0 | 0.23 | 0.14 | 0.18 | 0.35 | 0.27 | 0.31 |
| 3 | 116868163 | rs4145154 | 1.023 | 0.268 | -0.835 | 0.517 | 0.012 | 0.047 | 0.047 | 1.149 | 0.49 | 0.28 | 0.49 | 0.46 | 0.47 | 0.4 |
| 3 | 116879279 | rs4855926 | 1.215 | -0.341 | -1.303 | 0.197 | -0.06 | -0.057 | -0.057 | 0.04 | 0.41 | 0.07 | 0.41 | 0.46 | 0.44 | 0.37 |

| | | | | | | | | | | | | | | | | |
|---|---|---|---|---|---|---|---|---|---|---|---|---|---|---|---|---|
| 3 | 116880684 | rs6807401 | 0.267 | -0.36 | -0.932 | 0.315 | -0.015 | 0 | 0 | 0.484 | 0.16 | 0.59 | 0.22 | 0.15 | 0.12 | 0.11 |
| 3 | 116881714 | rs7649284 | 0.815 | -0.348 | -0.998 | 0.259 | -0.06 | -0.095 | -0.095 | 0.199 | 0.23 | 0.06 | 0.23 | 0.35 | 0.27 | 0.32 |
| 3 | 116883875 | rs16826215 | 1.172 | -0.449 | -0.755 | 0.154 | -0.028 | -0.037 | -0.037 | 0 | 0.07 | 0.13 | 0.03 | 0 | 0.03 | 0.03 |
| 3 | 116885358 | rs6806238 | -0.775 | -0.448 | -0.773 | 0.264 | -0.008 | -0.017 | -0.017 | 0 | 0.4 | 0.66 | 0.46 | 0.52 | 0.4 | 0.46 |
| 3 | 116886656 | rs10511366 | -0.433 | -0.486 | -0.245 | 0.268 | 0.002 | 0.071 | 0.071 | 0.382 | 0.15 | 0.39 | 0.23 | 0.21 | 0.13 | 0.14 |
| 3 | 116890209 | rs16826240 | 1.649 | -0.481 | -1.222 | 0.178 | -0.017 | -0.054 | -0.054 | 0.025 | 0.1 | 0.01 | 0.05 | 0.22 | 0.09 | 0.14 |
| 3 | 116890295 | rs11916895 | 0.19 | -0.22 | -1.027 | 0.187 | -0.045 | -0.098 | -0.098 | 0 | 0.15 | 0.38 | 0.11 | 0.01 | 0.09 | 0.12 |
| 3 | 116891094 | rs7651981 | 1.748 | -0.572 | -1.431 | 0.01 | -0.104 | -0.057 | -0.057 | 0.059 | 0.23 | 0.02 | 0.33 | 0.23 | 0.3 | 0.2 |
| 3 | 116896559 | rs16826255 | 0.219 | -0.559 | -0.576 | -0.3 | -0.048 | -0.102 | -0.102 | 0 | 0.14 | 0.37 | 0.09 | 0 | 0.07 | 0.09 |
| 3 | 116896629 | rs13077308 | -0.022 | -0.111 | -1.313 | 0.036 | -0.006 | -0.007 | -0.007 | 0.111 | 0.41 | 0.66 | 0.47 | 0.55 | 0.4 | 0.47 |
| 3 | 116897722 | rs9831095 | 0.151 | -0.079 | -0.999 | -0.419 | -0.003 | -0.052 | -0.052 | 0 | 0.83 | 0.76 | 0.87 | 1 | 0.83 | 0.87 |
| 3 | 116899786 | rs6788007 | -0.173 | 0.639 | -1.587 | -0.217 | -0.058 | -0.027 | -0.027 | 0 | 0.31 | 0.15 | 0.41 | 0.41 | 0.39 | 0.45 |
| 3 | 116900419 | rs868114 | -0.381 | 0.14 | -1.558 | -0.932 | 0.008 | 0.013 | 0.013 | 1.02 | 0.55 | 0.36 | 0.53 | 0.41 | 0.59 | 0.61 |
| 3 | 116910324 | rs9845043 | 0.639 | -0.075 | -1.47 | -0.724 | -0.031 | -0.047 | -0.047 | 0 | 0.3 | 0.39 | 0.25 | 0.11 | 0.22 | 0.22 |
| 3 | 116918892 | rs4855930 | 0.187 | 0.384 | -2.413 | -0.685 | -0.073 | -0.014 | -0.014 | 0 | 0.44 | 0.06 | 0.51 | 0.68 | 0.52 | 0.44 |
| 3 | 116922745 | rs2137213 | -0.363 | 0.815 | -1.985 | -0.685 | -0.111 | 0.041 | 0.041 | 0.405 | 0.75 | 0.92 | 0.66 | 0.79 | 0.56 | 0.68 |
| 3 | 116932264 | rs1599674 | -2.56 | 0.583 | -1.884 | -0.911 | -0.01 | 0.011 | 0.011 | 0.194 | 0.94 | 0.99 | 0.93 | 1 | 0.91 | 0.94 |
| 3 | 116933851 | rs9856565 | -0.253 | 0.732 | -2.034 | -0.999 | -0.113 | 0.109 | 0.109 | 0.765 | 0.73 | 0.92 | 0.65 | 0.92 | 0.53 | 0.71 |
| 3 | 116935744 | rs2903971 | 0.567 | 0.405 | -1.918 | -1.212 | 0 | 0.002 | 0.002 | 0 | 0.21 | 0.19 | 0.14 | 0.19 | 0.19 | 0.27 |
| 3 | 116943010 | rs1462845 | 0.788 | 0.289 | -1.503 | -1.272 | 0.006 | 0.006 | 0.006 | 1.086 | 0.54 | 0.56 | 0.5 | 0.74 | 0.37 | 0.44 |
| 3 | 116945297 | rs7426831 | -0.88 | 0.287 | -1.572 | -1.305 | -0.127 | 0.102 | 0.102 | 0.24 | 0.77 | 0.95 | 0.67 | 0.93 | 0.56 | 0.72 |
| 3 | 116946218 | rs1462844 | 0.729 | 0.192 | -1.55 | -1.311 | 0 | 0.005 | 0.005 | 0 | 0.2 | 0.12 | 0.13 | 0.18 | 0.19 | 0.27 |
| 3 | 116946253 | rs1462843 | -0.378 | 0.406 | -1.383 | -1.227 | 0.026 | -0.041 | -0.041 | 0.592 | 0.74 | 0.7 | 0.63 | 0.92 | 0.55 | 0.71 |
| 3 | 116962328 | rs4522775 | 1.738 | -0.277 | -1.708 | -1.382 | 0.011 | 0.071 | 0.071 | 0.003 | 0.14 | 0.01 | 0.07 | 0.01 | 0.09 | 0.05 |
| 3 | 116964048 | rs11712223 | 0.699 | -0.146 | -1.772 | -1.617 | -0.071 | 0.07 | 0.07 | 0 | 0.24 | 0.1 | 0.3 | 0.07 | 0.38 | 0.24 |
| 3 | 116968649 | rs3845227 | -1.374 | -0.333 | -1.748 | -1.662 | -0.012 | 0.009 | 0.009 | 0.369 | 0.79 | 0.86 | 0.81 | 0.8 | 0.73 | 0.73 |
| 3 | 116970253 | rs3900989 | 0.861 | -0.194 | -1.896 | -1.613 | -0.122 | 0.115 | 0.115 | 0 | 0.23 | 0.07 | 0.3 | 0.07 | 0.38 | 0.24 |
| 3 | 116973408 | rs938170 | 0.825 | -0.287 | -1.701 | -1.222 | 0 | -0.002 | -0.002 | 0 | 0.09 | 0.08 | 0.13 | 0.1 | 0.07 | 0.09 |
| 3 | 116974370 | rs1218626 | -0.559 | -0.29 | -1.67 | -1.256 | 0.004 | -0.014 | -0.014 | 0 | 0.76 | 0.67 | 0.77 | 0.8 | 0.72 | 0.73 |
| 3 | 116982057 | rs1513179 | -0.035 | -0.488 | -1.691 | -1.263 | 0.035 | 0.028 | 0.028 | 0 | 0.7 | 0.86 | 0.63 | 0.27 | 0.78 | 0.69 |
| 3 | 116983850 | rs4855899 | -0.723 | -0.482 | -1.687 | -1.026 | 0.001 | -0.004 | -0.004 | 0.992 | 0.85 | 0.88 | 0.85 | 0.8 | 0.88 | 0.88 |
| 3 | 116990925 | rs6807085 | -0.528 | -0.174 | -1.468 | -1.557 | 0.026 | 0.027 | 0.027 | 0 | 0.69 | 0.8 | 0.62 | 0.26 | 0.78 | 0.66 |
| 3 | 117000128 | rs3849471 | 0.057 | 0.168 | -1.418 | -1.477 | 0 | -0.073 | -0.073 | 0 | 0.47 | 0.47 | 0.41 | 0.07 | 0.51 | 0.4 |
| 3 | 117010780 | rs1513171 | 0.153 | 0.214 | -1.568 | -1.357 | 0.004 | -0.125 | -0.125 | 0 | 0.41 | 0.49 | 0.36 | 0.07 | 0.43 | 0.39 |
| 3 | 117014065 | rs4855941 | 0.145 | 0.043 | -2.341 | -0.553 | -0.058 | 0.07 | 0.07 | 0 | 0.41 | 0.08 | 0.36 | 0.27 | 0.52 | 0.41 |
| 3 | 117015399 | rs11716697 | -0.255 | -0.469 | -1.923 | -0.2 | -0.013 | 0.16 | 0.16 | 0 | 0.77 | 0.57 | 0.56 | 0.49 | 0.8 | 0.71 |
| 3 | 117017073 | rs16826629 | 1.984 | -0.611 | -1.953 | -0.332 | 0 | -0.01 | -0.01 | 0 | 0.24 | 0.27 | 0.13 | 0.1 | 0.2 | 0.16 |
| 3 | 117017795 | rs9815346 | -0.701 | -0.025 | -1.417 | 0.331 | -0.025 | -0.034 | -0.034 | 0 | 0.32 | 0.2 | 0.24 | 0.4 | 0.38 | 0.31 |
| 3 | 117023557 | rs1398634 | -0.415 | -0.726 | -0.516 | 1.251 | -0.005 | -0.041 | -0.041 | 0.185 | 0.16 | 0.06 | 0.37 | 0.43 | 0.15 | 0.23 |
| 3 | 117025312 | rs3860542 | 0.753 | -0.692 | -0.519 | 0.391 | -0.008 | 0.098 | 0.098 | 0.01 | 0.77 | 0.57 | 0.54 | 0.42 | 0.78 | 0.66 |
| 3 | 117036955 | rs1398630 | 0.665 | -1.088 | -0.279 | 0.928 | -0.028 | 0.227 | 0.227 | 0 | 0.86 | 0.62 | 0.72 | 0.42 | 0.89 | 0.72 |
| 3 | 117045006 | rs6800555 | 0.45 | -1.036 | -0.257 | 0.82 | -0.009 | 0.14 | 0.14 | 0 | 0.82 | 0.62 | 0.68 | 0.41 | 0.85 | 0.72 |
| 3 | 117049817 | rs6438389 | 0.93 | -0.821 | -0.733 | 0.444 | -0.035 | -0.005 | -0.005 | 0 | 0.7 | 0.47 | 0.63 | 0.32 | 0.79 | 0.65 |
| 3 | 117077219 | rs1398641 | 0.64 | -0.788 | -1.405 | 0.676 | 0.073 | -0.098 | -0.098 | 0 | 0.59 | 0.1 | 0.52 | 0.51 | 0.69 | 0.68 |
| 3 | 117078853 | rs9878146 | -0.318 | -1.019 | -0.688 | 0.603 | 0.004 | -0.02 | -0.02 | 0 | 0.25 | 0.22 | 0.41 | 0.42 | 0.27 | 0.28 |
| 3 | 117087224 | rs7609678 | -0.423 | -1.037 | 0 | 0.585 | -0.01 | 0.051 | 0.051 | 0 | 0.26 | 0.35 | 0.42 | 0.42 | 0.27 | 0.28 |
| 3 | 117088533 | rs12497924 | -0.895 | -0.957 | -0.256 | 0.747 | -0.002 | -0.028 | -0.028 | 0 | 0.15 | 0.08 | 0.3 | 0.57 | 0.17 | 0.3 |
| 3 | 117093435 | rs7623774 | 0.065 | -1.189 | 0.612 | 0.52 | -0.036 | -0.067 | -0.067 | 0 | 0.12 | 0.18 | 0.04 | 0 | 0.05 | 0.04 |
| 3 | 117094777 | rs7615693 | 0.719 | -1.233 | 0.975 | 0.477 | -0.004 | -0.044 | -0.044 | 0.125 | 0.68 | 0.39 | 0.79 | 0.95 | 0.76 | 0.83 |
| 3 | 117094804 | rs13094218 | 0.432 | -1.181 | 0.927 | 0.55 | -0.011 | -0.029 | -0.029 | 0 | 0.09 | 0.02 | 0.05 | 0.21 | 0.1 | 0.16 |
| 3 | 117095341 | rs2048610 | -0.331 | -1.232 | 0.957 | 0.28 | -0.011 | -0.097 | -0.097 | 0 | 0.8 | 0.57 | 0.83 | 0.95 | 0.81 | 0.87 |
| 3 | 117102416 | rs1849474 | 2.124 | -0.391 | 0.413 | 0.09 | -0.067 | 0.13 | 0.13 | 0.25 | 0.39 | 0.02 | 0.34 | 0.21 | 0.36 | 0.38 |
| 3 | 117111635 | rs1398629 | 1.376 | -1.251 | 1.188 | 0.626 | 0.058 | -0.115 | -0.115 | 0 | 0.6 | 0.08 | 0.52 | 0.56 | 0.68 | 0.68 |
| 3 | 117116725 | rs9846338 | -0.028 | -1.056 | 0.892 | 0.759 | 0 | 0.002 | 0.002 | 0 | 0.05 | 0.05 | 0.05 | 0.03 | 0.05 | 0.09 |
| 3 | 117117949 | rs12638372 | -0.197 | -1.037 | 0.944 | 0.868 | 0 | 0.078 | 0.078 | 0 | 0.07 | 0.06 | 0.25 | 0.4 | 0.08 | 0.16 |
| 3 | 117118513 | rs2869501 | 1.165 | -1.192 | 1.219 | 0.757 | -0.037 | -0.068 | -0.068 | 0 | 0.8 | 0.34 | 0.87 | 0.97 | 0.9 | 0.86 |
| 3 | 117126566 | rs4855946 | 0.649 | -1.171 | 1.271 | 0.768 | -0.062 | -0.104 | -0.104 | 0.674 | 0.85 | 0.46 | 0.94 | 1 | 0.95 | 0.95 |
| 3 | 117132295 | rs12631527 | -0.182 | -1.104 | 1.055 | 0.911 | -0.004 | 0.146 | 0.146 | 0 | 0.07 | 0.11 | 0.25 | 0.4 | 0.08 | 0.16 |
| 3 | 117132512 | rs1355713 | 1.715 | -1.294 | 0.929 | 0.776 | 0.062 | 0.058 | 0.058 | 0 | 0.51 | 0.13 | 0.48 | 0.36 | 0.58 | 0.53 |
| 3 | 117133546 | rs1398625 | -0.121 | -1.332 | 1.13 | 0.754 | -0.041 | -0.077 | -0.077 | 0 | 0.88 | 0.71 | 0.95 | 1 | 0.95 | 0.96 |
| 3 | 117137202 | rs6777348 | 0.232 | -1.312 | 1.133 | 0.753 | 0 | -0.008 | -0.008 | 0 | 0.22 | 0.26 | 0.16 | 0.2 | 0.24 | 0.19 |
| 3 | 117137446 | rs7635558 | -0.117 | -1.316 | 1.126 | 0.754 | -0.038 | -0.071 | -0.071 | 0 | 0.88 | 0.75 | 0.96 | 1 | 0.95 | 0.96 |
| 3 | 117141277 | rs6797385 | 1.313 | -1.313 | 0.932 | 0.747 | 0.051 | 0.136 | 0.136 | 0 | 0.58 | 0.16 | 0.72 | 0.77 | 0.67 | 0.68 |
| 3 | 117141815 | rs9847858 | 1.568 | -1.216 | 0.599 | 0.504 | 0.007 | 0.016 | 0.016 | 0 | 0.5 | 0.03 | 0.46 | 0.36 | 0.58 | 0.52 |
| 3 | 117146882 | rs4073041 | -0.349 | -0.852 | 0.835 | 0.683 | -0.004 | 0.021 | 0.021 | 0 | 0.3 | 0.39 | 0.41 | 0.61 | 0.32 | 0.36 |
| 3 | 117152802 | rs16826914 | -0.911 | -0.713 | 1.148 | 0.942 | -0.041 | -0.078 | -0.078 | 0.174 | 0.12 | 0.28 | 0.05 | 0 | 0.05 | 0.04 |
| 3 | 117156240 | rs6806321 | 1.036 | -0.544 | 0.91 | 1.65 | -0.017 | 0.248 | 0.248 | 0.107 | 0.84 | 0.62 | 0.66 | 0.53 | 0.86 | 0.76 |
| 3 | 117156904 | rs4536834 | -0.856 | -0.034 | 0.115 | 0.8 | -0.036 | 0.089 | 0.089 | 0.102 | 0.2 | 0.04 | 0.25 | 0 | 0.21 | 0.16 |
| 3 | 117158450 | rs9881096 | 1.616 | -0.097 | 0.66 | 1.223 | -0.003 | 0.005 | 0.005 | 0 | 0.62 | 0.36 | 0.37 | 0.31 | 0.57 | 0.56 |

| | | | | | | | | | | | | | | | | |
|---|---|---|---|---|---|---|---|---|---|---|---|---|---|---|---|---|
| 3 | 117162784 | rs4075039 | -1.406 | -0.197 | 1.099 | 1.652 | -0.001 | 0.001 | 0.001 | 0.13 | 0.15 | 0.12 | 0.33 | 0.54 | 0.2 | 0.26 |
| 3 | 117166593 | rs1870916 | -1.936 | -0.228 | 1.654 | 2.114 | 0.049 | 0.43 | 0.43 | 0 | 0.07 | 0.3 | 0.24 | 0.41 | 0.07 | 0.05 |
| 3 | 117172164 | rs6776358 | 1.218 | -0.222 | 1.646 | 2.089 | 0.019 | 0.075 | 0.075 | 0.208 | 0.72 | 0.46 | 0.54 | 0.45 | 0.69 | 0.8 |
| 3 | 117172333 | rs10511371 | -0.959 | -0.488 | 1.623 | 2.074 | -0.01 | 0.04 | 0.04 | 0.913 | 0.07 | 0.31 | 0.07 | 0.06 | 0.03 | 0.1 |
| 3 | 117172973 | rs17662573 | -0.373 | -0.466 | 1.26 | 1.989 | -0.031 | 0.039 | 0.039 | 0.229 | 0.19 | 0.08 | 0.26 | 0 | 0.24 | 0.11 |
| 3 | 117174568 | rs11917701 | 2.247 | -0.456 | 1.241 | 1.993 | -0.007 | -0.022 | -0.022 | 0 | 0.57 | 0.12 | 0.41 | 0.39 | 0.6 | 0.63 |
| 3 | 117177909 | rs11715287 | -0.126 | -0.555 | 0.81 | 1.595 | -0.02 | 0.019 | 0.019 | 0.065 | 0.16 | 0.07 | 0.24 | 0 | 0.21 | 0.09 |
| 3 | 117191259 | rs2937685 | 0.565 | -0.586 | 0.795 | 1.793 | 0.018 | -0.005 | -0.005 | 0.264 | 0.77 | 0.44 | 0.59 | 0.76 | 0.76 | 0.74 |
| 3 | 117194656 | rs2927278 | 0.786 | -0.592 | 0.786 | 1.777 | 0.015 | -0.014 | -0.014 | 0 | 0.75 | 0.42 | 0.52 | 0.76 | 0.73 | 0.73 |
| 3 | 117200255 | rs7632761 | -2.023 | -0.87 | 0.579 | 1.894 | -0.047 | 0.093 | 0.093 | 0.206 | 0.22 | 0.38 | 0.22 | 0.53 | 0.15 | 0.25 |
| 3 | 117200934 | rs2927279 | 0.305 | -0.839 | 0.588 | 1.895 | 0.022 | 0 | 0 | 0.069 | 0.78 | 0.47 | 0.61 | 0.77 | 0.77 | 0.73 |
| 3 | 117205693 | rs2927280 | 2.435 | -0.932 | 0.341 | 1.478 | -0.002 | 0.205 | 0.205 | 1.48 | 0.5 | 0.05 | 0.3 | 0.22 | 0.56 | 0.49 |
| 3 | 117208691 | rs16827045 | -0.668 | -0.891 | 0.69 | 1.926 | -0.014 | 0.022 | 0.022 | 0 | 0.08 | 0.09 | 0.04 | 0.15 | 0.03 | 0.12 |
| 3 | 117212184 | rs2937678 | 0.269 | -0.788 | 0.65 | 1.97 | 0.03 | -0.069 | -0.069 | 0 | 0.79 | 0.47 | 0.63 | 0.92 | 0.78 | 0.78 |
| 3 | 117217583 | rs9849429 | -0.842 | -0.566 | -0.323 | 2.169 | 0.002 | -0.003 | -0.003 | 0 | 0.06 | 0.06 | 0.04 | 0.11 | 0.04 | 0.05 |
| 3 | 117224118 | rs9830523 | -0.877 | -0.578 | -0.324 | 2.168 | 0.005 | -0.006 | -0.006 | 0 | 0.06 | 0.06 | 0.03 | 0.11 | 0.04 | 0.08 |
| 3 | 117234839 | rs1676232 | 1.153 | -0.532 | -0.484 | 0.688 | 0.036 | 0.082 | 0.082 | 1.601 | 0.57 | 0.15 | 0.38 | 0.29 | 0.64 | 0.5 |
| 3 | 117237561 | rs2937670 | 1.441 | -0.349 | 0.49 | 0.487 | -0.008 | -0.016 | -0.016 | 0 | 0.85 | 0.4 | 0.73 | 0.91 | 0.87 | 0.81 |
| 3 | 117237625 | rs2927287 | 1.024 | -0.586 | 0.396 | 0.17 | -0.039 | -0.063 | -0.063 | 0 | 0.8 | 0.65 | 0.89 | 0.9 | 0.87 | 0.86 |
| 3 | 117238369 | rs6792959 | -1.172 | -0.009 | 0.025 | 0.434 | 0.001 | 0.001 | 0.001 | 0 | 0.11 | 0.12 | 0.09 | 0.15 | 0.15 | 0.08 |
| 3 | 117239197 | rs1979867 | -0.532 | -0.305 | -0.115 | 0.485 | 0.002 | -0.002 | -0.002 | 0 | 0.05 | 0.05 | 0.01 | 0.08 | 0.01 | 0.04 |
| 3 | 117239341 | rs1979868 | 0.603 | -0.376 | -1.051 | -0.358 | -0.052 | 0.034 | 0.034 | 0 | 0.39 | 0.19 | 0.48 | 0.7 | 0.43 | 0.52 |
| 3 | 117239883 | rs4616651 | -0.267 | -0.382 | -0.588 | -0.599 | -0.01 | 0.099 | 0.099 | 0.021 | 0.53 | 0.28 | 0.47 | 0.22 | 0.56 | 0.43 |
| 3 | 117244009 | rs17720543 | 0.113 | -0.674 | -1.035 | -0.735 | -0.03 | 0.034 | 0.034 | 0 | 0.18 | 0.02 | 0.08 | 0.1 | 0.2 | 0.14 |
| 3 | 117248700 | rs2927271 | 0.408 | -0.589 | -1.01 | -1.008 | -0.023 | 0.153 | 0.153 | 0 | 0.45 | 0.16 | 0.41 | 0.8 | 0.55 | 0.57 |
| 3 | 117250719 | rs1910044 | 0.167 | -1.042 | -0.111 | -1.184 | -0.016 | 0.076 | 0.076 | 0 | 0.38 | 0.19 | 0.51 | 0.16 | 0.4 | 0.35 |
| 3 | 117254425 | rs13314246 | -1.171 | -1.472 | 0.476 | -1.418 | 0.013 | 0.057 | 0.057 | 0 | 0.45 | 0.64 | 0.57 | 0.2 | 0.4 | 0.35 |
| 3 | 117259093 | rs4456868 | -0.98 | -1.5 | 0.893 | -1.43 | 0.019 | 0.09 | 0.09 | 0 | 0.45 | 0.69 | 0.58 | 0.2 | 0.4 | 0.34 |
| 3 | 117270806 | rs1466416 | 0.461 | -1.06 | 0.573 | -1.241 | 0.024 | 0.004 | 0.004 | 0 | 0.06 | 0.01 | 0.01 | 0.04 | 0.01 | 0.04 |
| 3 | 117280848 | rs7644537 | 0.256 | -1.257 | 0.683 | -0.499 | -0.023 | -0.019 | -0.019 | 0 | 0.4 | 0.12 | 0.53 | 0.41 | 0.41 | 0.57 |
| 3 | 117284829 | rs7620877 | -0.067 | -1.28 | 0.756 | -0.404 | -0.001 | -0.018 | -0.018 | 0.022 | 0.42 | 0.19 | 0.58 | 0.47 | 0.43 | 0.6 |
| 3 | 117298203 | rs1393190 | -0.904 | -1.371 | 0.891 | -0.323 | 0.016 | 0.003 | 0.003 | 0.277 | 0.53 | 0.67 | 0.63 | 0.54 | 0.44 | 0.65 |
| 3 | 117301697 | rs1501870 | -1.016 | -1.374 | 0.966 | -0.323 | 0.013 | -0.004 | -0.004 | 0.178 | 0.54 | 0.76 | 0.64 | 0.55 | 0.44 | 0.65 |
| 3 | 117302160 | rs9849920 | -0.916 | -1.349 | 0.765 | -0.312 | 0.007 | 0.002 | 0.002 | 0.247 | 0.53 | 0.57 | 0.62 | 0.55 | 0.44 | 0.65 |
| 3 | 117304479 | rs1501868 | -0.209 | -0.836 | 0.195 | -0.088 | 0.022 | -0.015 | -0.015 | 0.789 | 0.47 | 0.21 | 0.59 | 0.51 | 0.43 | 0.6 |
| 3 | 117307567 | rs1875517 | -0.177 | -0.515 | -0.081 | 0.164 | 0.016 | -0.04 | -0.04 | 0 | 0.46 | 0.15 | 0.51 | 0.5 | 0.41 | 0.62 |
| 3 | 117308818 | rs2869787 | -0.316 | -0.53 | -0.002 | 0.157 | 0.027 | -0.019 | -0.019 | 0 | 0.48 | 0.2 | 0.51 | 0.5 | 0.41 | 0.62 |
| 3 | 117311967 | rs4516643 | -0.349 | 0.19 | -0.087 | -0.002 | 0.01 | -0.033 | -0.033 | 0 | 0.62 | 0.75 | 0.62 | 0.53 | 0.65 | 0.51 |
| 3 | 117315806 | rs11719399 | -0.281 | 0.278 | 0.027 | 0.031 | 0.016 | -0.021 | -0.021 | 0 | 0.59 | 0.72 | 0.61 | 0.53 | 0.64 | 0.5 |
| 3 | 117317414 | rs9867393 | -0.643 | 0.208 | -0.106 | -0.002 | -0.023 | -0.11 | -0.11 | 0 | 0.65 | 0.89 | 0.63 | 0.53 | 0.66 | 0.53 |
| 3 | 117325977 | rs10511370 | -0.688 | -0.047 | 0.367 | 0.023 | 0.031 | -0.054 | -0.054 | 0.318 | 0.16 | 0.43 | 0.16 | 0.06 | 0.15 | 0.11 |
| 3 | 117329656 | rs11721241 | 0.889 | 0.009 | -0.447 | 0.405 | -0.102 | -0.123 | -0.123 | 0 | 0.38 | 0.03 | 0.42 | 0.44 | 0.46 | 0.37 |
| 3 | 117330716 | rs1501871 | 0.502 | 0.189 | 0.084 | 0.356 | -0.001 | 0 | 0 | 0 | 0.46 | 0.46 | 0.48 | 0.52 | 0.5 | 0.43 |
| 3 | 117330858 | rs1847023 | 0.037 | 0.167 | -0.011 | 0.36 | 0.008 | 0.004 | 0.004 | 0 | 0.51 | 0.59 | 0.5 | 0.52 | 0.51 | 0.46 |
| 3 | 117332915 | rs6809736 | -0.009 | 0.755 | -0.554 | 0.8 | -0.028 | -0.021 | -0.021 | 0 | 0.95 | 0.99 | 0.94 | 0.79 | 0.89 | 0.94 |
| 3 | 117338327 | rs13059762 | 0.107 | 0.432 | -0.346 | 0.611 | 0 | 0 | 0 | 0.406 | 0.48 | 0.41 | 0.48 | 0.53 | 0.5 | 0.43 |
| 3 | 117346618 | rs2904196 | 0.296 | 0.154 | -0.175 | 1.168 | -0.007 | 0.019 | 0.019 | 0 | 0.73 | 0.64 | 0.79 | 0.58 | 0.73 | 0.81 |
| 3 | 117354726 | rs6773314 | 0.976 | -0.738 | -0.624 | 1.44 | 0.115 | 0.153 | 0.153 | 0.792 | 0.51 | 0.09 | 0.54 | 0.21 | 0.63 | 0.55 |
| 3 | 117356239 | rs6804100 | 0.774 | -1.033 | 1.25 | 1.526 | -0.034 | 0.023 | 0.023 | 0 | 0.73 | 0.48 | 0.67 | 0.32 | 0.77 | 0.65 |
| 3 | 117357839 | rs6772369 | 0.34 | -1.123 | 1.232 | 1.85 | 0.003 | -0.015 | -0.015 | 2.252 | 0.81 | 0.87 | 0.72 | 0.44 | 0.81 | 0.68 |
| 3 | 117366248 | rs9816414 | 1.444 | -0.506 | 1.023 | 1.027 | -0.006 | 0.122 | 0.122 | 0 | 0.59 | 0.22 | 0.53 | 0.17 | 0.59 | 0.56 |
| 3 | 117366881 | rs6799173 | 1.074 | -0.464 | 1.035 | 1.059 | 0.003 | -0.041 | -0.041 | 0 | 0.63 | 0.59 | 0.56 | 0.18 | 0.59 | 0.58 |
| 3 | 117379540 | rs9877923 | 0.978 | -0.396 | 1.295 | 1.25 | 0.007 | 0.008 | 0.008 | 0 | 0.19 | 0.16 | 0.12 | 0.11 | 0.15 | 0.07 |
| 3 | 117379876 | rs9882233 | 0.683 | -0.4 | 1.741 | 1.236 | -0.028 | -0.052 | -0.052 | 0 | 0.21 | 0.33 | 0.19 | 0.12 | 0.16 | 0.07 |
| 3 | 117380644 | rs4440150 | 0.129 | -0.561 | 1.672 | 1.46 | -0.018 | 0.006 | 0.006 | 0 | 0.7 | 0.61 | 0.78 | 0.6 | 0.74 | 0.77 |
| 3 | 117383588 | rs12631137 | 0.066 | -0.53 | 1.486 | 1.572 | -0.001 | 0.039 | 0.039 | 0 | 0.09 | 0.08 | 0.11 | 0.29 | 0.06 | 0.19 |
| 3 | 117393911 | rs6764568 | 0.955 | -0.674 | 1.37 | 1.462 | 0.018 | -0.003 | -0.003 | 0 | 0.61 | 0.36 | 0.74 | 0.51 | 0.68 | 0.73 |
| 3 | 117401316 | rs13097890 | 0.993 | -0.718 | 1.513 | 1.356 | 0.01 | 0.013 | 0.013 | 0.207 | 0.2 | 0.11 | 0.12 | 0.11 | 0.15 | 0.08 |
| 3 | 117407287 | rs3814059 | -0.2 | -0.574 | 1.324 | 1.398 | -0.008 | -0.018 | -0.018 | 0.366 | 0.17 | 0.13 | 0.22 | 0.5 | 0.21 | 0.23 |
| 3 | 117410498 | rs3923050 | 1.477 | -0.637 | 1.065 | 1.399 | 0.108 | 0.106 | 0.106 | 0.497 | 0.51 | 0.23 | 0.63 | 0.27 | 0.64 | 0.56 |
| 3 | 117413844 | rs4234649 | 1.3 | -0.559 | 1.062 | 1.42 | -0.025 | 0.069 | 0.069 | 0 | 0.8 | 0.41 | 0.85 | 0.65 | 0.83 | 0.81 |
| 3 | 117420673 | rs4301042 | -0.014 | -0.509 | 0.684 | 1.427 | -0.018 | -0.045 | -0.045 | 0.342 | 0.91 | 0.98 | 0.91 | 0.73 | 0.87 | 0.84 |
| 3 | 117425366 | rs9289086 | -0.142 | -0.614 | 0.959 | 1.327 | -0.02 | 0.012 | 0.012 | 0.006 | 0.31 | 0.36 | 0.24 | 0.39 | 0.21 | 0.27 |
| 3 | 117427214 | rs13325341 | 1.903 | -0.512 | 0.338 | 0.837 | -0.097 | 0.249 | 0.249 | 1.063 | 0.42 | 0.09 | 0.49 | 0.05 | 0.44 | 0.49 |
| 3 | 117427801 | rs4345097 | 0.004 | -0.256 | 0.396 | 1.076 | -0.017 | -0.03 | -0.03 | 0 | 0.9 | 0.94 | 0.89 | 0.73 | 0.83 | 0.86 |
| 3 | 117429566 | rs6438403 | 0.059 | 0.006 | 0.388 | 0.937 | -0.024 | 0.011 | 0.011 | 0 | 0.9 | 0.94 | 0.91 | 0.88 | 0.82 | 0.85 |
| 3 | 117431660 | rs7646668 | 1.218 | 0.15 | 0.566 | 1.141 | -0.004 | 0.081 | 0.081 | 0.145 | 0.51 | 0.29 | 0.59 | 0.25 | 0.46 | 0.66 |
| 3 | 117435160 | rs12629100 | 0.102 | 0.024 | 0.664 | 1.235 | -0.001 | 0.018 | 0.018 | 0 | 0.07 | 0.1 | 0.09 | 0.22 | 0.06 | 0.19 |
| 3 | 117436989 | rs4635741 | 1.348 | -0.054 | 0.699 | 1.246 | 0.039 | 0.035 | 0.035 | 0.071 | 0.52 | 0.25 | 0.63 | 0.31 | 0.57 | 0.54 |

| | | | | | | | | | | | | | | | | |
|---|---|---|---|---|---|---|---|---|---|---|---|---|---|---|---|---|
| 3 | 117438965 | rs9289087 | -0.995 | 0.406 | 0.932 | 1.338 | 0.002 | 0.128 | 0.128 | 0 | 0.1 | 0.28 | 0.11 | 0.24 | 0.06 | 0.2 |
| 3 | 117441231 | rs4568174 | 1.141 | 0.472 | 0.528 | 1.122 | 0.019 | 0 | 0 | 0 | 0.87 | 0.74 | 0.89 | 0.82 | 0.82 | 0.83 |
| 3 | 117448084 | rs13325314 | 0.282 | 0.432 | 0.351 | 0.982 | 0.01 | -0.002 | -0.002 | 0 | 0.06 | 0.09 | 0.03 | 0.04 | 0.07 | 0.02 |
| 3 | 117449309 | rs9876009 | 0.282 | 0.469 | 0.34 | 0.982 | 0.006 | -0.002 | -0.002 | 0 | 0.06 | 0.06 | 0.03 | 0.04 | 0.07 | 0.02 |
| 3 | 117454822 | rs6809420 | 0.557 | 0.47 | 0.278 | 0.853 | 0.016 | -0.033 | -0.033 | 0.078 | 0.89 | 0.81 | 0.91 | 0.95 | 0.82 | 0.83 |
| 3 | 117456698 | rs7647501 | 0.63 | 0.069 | 0.119 | 0.813 | -0.001 | 0.006 | 0.006 | 0 | 0.62 | 0.65 | 0.62 | 0.28 | 0.56 | 0.67 |
| 3 | 117461709 | rs7646416 | 0.923 | 0.131 | 0.025 | 0.952 | 0.014 | 0.024 | 0.024 | 0 | 0.8 | 0.74 | 0.78 | 0.42 | 0.66 | 0.62 |
| 3 | 117464639 | rs9833119 | 1.852 | -0.02 | -0.103 | 0.879 | -0.073 | 0.311 | 0.311 | 1.409 | 0.45 | 0.08 | 0.51 | 0.03 | 0.44 | 0.48 |
| 3 | 117465114 | rs13059541 | 0.557 | 0.064 | -0.125 | 1.114 | 0.062 | -0.08 | -0.08 | 0.001 | 0.23 | 0.03 | 0.19 | 0.36 | 0.14 | 0.11 |
| 3 | 117473977 | rs4687996 | 0.934 | 0.241 | 0.072 | 0.714 | 0 | 0.017 | 0.017 | 0 | 0.55 | 0.6 | 0.57 | 0.06 | 0.51 | 0.5 |
| 3 | 117479214 | rs4687856 | 0.712 | -0.136 | 0.082 | 0.675 | 0.01 | -0.001 | -0.001 | 0 | 0.58 | 0.62 | 0.66 | 0.36 | 0.62 | 0.57 |
| 3 | 117484067 | rs6808588 | -0.179 | 0.023 | -0.147 | 0.371 | 0.004 | 0 | 0 | 0.099 | 0.07 | 0.09 | 0.03 | 0 | 0.06 | 0.08 |
| 3 | 117488858 | rs4687866 | 0.878 | -0.786 | -1.008 | -0.565 | 0.07 | 0.092 | 0.092 | 6.798 | 0.28 | 0.08 | 0.32 | 0 | 0.17 | 0.1 |
| 3 | 117489779 | rs4320082 | -0.554 | -0.292 | -0.341 | -0.24 | -0.002 | 0.007 | 0.007 | 0 | 0.67 | 0.59 | 0.75 | 0.57 | 0.66 | 0.73 |
| 3 | 117490114 | rs4499636 | 0.605 | -0.257 | -0.59 | -0.357 | 0.041 | 0.01 | 0.01 | 2.9 | 0.34 | 0.21 | 0.43 | 0.3 | 0.22 | 0.21 |
| 3 | 117491325 | rs4610260 | 1.143 | 0.705 | -0.238 | 0.131 | -0.012 | 0.01 | 0.01 | 0.063 | 0.22 | 0.18 | 0.18 | 0.02 | 0.26 | 0.39 |
| 3 | 117505310 | rs7635561 | 0.202 | 0.584 | -0.255 | 0.634 | -0.011 | -0.013 | -0.013 | 0 | 0.1 | 0.08 | 0.14 | 0.3 | 0.11 | 0.11 |
| 3 | 117521471 | rs9864015 | 0.355 | -0.085 | -0.034 | -0.41 | 0.004 | -0.013 | -0.013 | 0 | 0.33 | 0.33 | 0.36 | 0.06 | 0.2 | 0.11 |
| 3 | 117522951 | rs12489939 | 0.745 | -0.122 | -0.03 | -0.271 | 0.046 | -0.029 | -0.029 | 1.094 | 0.32 | 0.1 | 0.26 | 0.41 | 0.27 | 0.2 |
| 3 | 117524434 | rs9842336 | 0.849 | -0.167 | 0.035 | -0.313 | 0.02 | 0.022 | 0.022 | 1.987 | 0.32 | 0.26 | 0.36 | 0.06 | 0.2 | 0.11 |
| 3 | 117526264 | rs12491422 | 0.334 | 0.345 | 0.046 | -0.032 | 0.006 | -0.074 | -0.074 | 1.377 | 0.37 | 0.12 | 0.34 | 0.54 | 0.4 | 0.31 |
| 3 | 117532231 | rs9856456 | -0.432 | 0.328 | 0.972 | -0.038 | 0.026 | 0.339 | 0.339 | 0.296 | 0.94 | 0.75 | 0.93 | 0.64 | 0.97 | 0.88 |
| 3 | 117537439 | rs4687889 | 0.252 | 0.877 | 1.182 | -0.098 | -0.001 | -0.011 | -0.011 | 0 | 0.55 | 0.6 | 0.54 | 0.09 | 0.44 | 0.5 |
| 3 | 117539255 | rs7431379 | -0.187 | 0.759 | 1.318 | -0.124 | 0.004 | -0.018 | -0.018 | 0 | 0.57 | 0.23 | 0.5 | 0.57 | 0.64 | 0.67 |
| 3 | 117542954 | rs9831027 | 0.852 | 0.635 | 1.357 | -0.213 | 0.043 | 0.07 | 0.07 | 2.26 | 0.34 | 0.22 | 0.35 | 0.06 | 0.21 | 0.11 |
| 3 | 117544977 | rs6438414 | -0.601 | 0.675 | 1.669 | -0.156 | 0.069 | 0.044 | 0.044 | 1.141 | 0.71 | 0.45 | 0.71 | 0.61 | 0.59 | 0.44 |
| 3 | 117547273 | rs6438416 | -0.872 | 0.701 | 1.615 | -0.084 | 0.032 | 0.309 | 0.309 | 0.169 | 0.95 | 0.76 | 0.93 | 0.67 | 0.97 | 0.88 |
| 3 | 117547988 | rs6438418 | 0.389 | 0.718 | 1.645 | -0.14 | -0.005 | -0.078 | -0.078 | 0 | 0.39 | 0.47 | 0.42 | 0.07 | 0.34 | 0.25 |
| 3 | 117549152 | rs9834986 | 0.747 | 0.56 | 1.788 | -0.138 | 0.006 | -0.012 | -0.012 | 0 | 0.36 | 0.35 | 0.37 | 0.06 | 0.2 | 0.13 |
| 3 | 117549667 | rs13066603 | 0.348 | 0.589 | 1.755 | -0.092 | 0 | 0.027 | 0.027 | 0.017 | 0.51 | 0.28 | 0.44 | 0.46 | 0.51 | 0.53 |
| 3 | 117550431 | rs6438419 | 1.019 | 0.585 | 1.748 | -0.1 | 0.041 | -0.031 | -0.031 | 0 | 0.31 | 0.11 | 0.27 | 0.41 | 0.27 | 0.2 |
| 3 | 117551526 | rs4688064 | 2.022 | 0.493 | 1.736 | -0.139 | -0.031 | 0.048 | 0.048 | 0.018 | 0.19 | 0.1 | 0.16 | 0.05 | 0.24 | 0.32 |
| 3 | 117552111 | rs10934420 | 0.078 | 0.504 | 1.69 | -0.117 | -0.026 | -0.023 | -0.023 | 0 | 0.39 | 0.23 | 0.39 | 0.56 | 0.52 | 0.42 |
| 3 | 117554999 | rs9838478 | 0.751 | 0.471 | 1.639 | -0.152 | 0.018 | 0.009 | 0.009 | 1.607 | 0.35 | 0.31 | 0.36 | 0.06 | 0.2 | 0.14 |
| 3 | 117562436 | rs11715566 | 0.232 | 0.534 | 1.539 | -0.11 | -0.042 | -0.042 | -0.042 | 0.422 | 0.38 | 0.18 | 0.39 | 0.55 | 0.52 | 0.42 |
| 3 | 117567340 | rs7641471 | -0.343 | 0.314 | 1.624 | -0.121 | 0.002 | -0.002 | -0.002 | 0 | 0.61 | 0.38 | 0.56 | 0.61 | 0.77 | 0.75 |
| 3 | 117584724 | rs4688067 | -0.565 | 0.375 | 1.752 | -0.05 | 0.042 | 0.369 | 0.369 | 0 | 0.95 | 0.75 | 0.92 | 0.66 | 0.97 | 0.88 |
| 3 | 117589477 | rs6788088 | -0.523 | 0.307 | 1.714 | -0.081 | 0.013 | 0.261 | 0.261 | 0 | 0.94 | 0.8 | 0.94 | 0.66 | 0.97 | 0.88 |
| 3 | 117589610 | rs6788211 | -0.516 | 0.289 | 1.751 | -0.147 | -0.034 | 0.013 | 0.013 | 0.525 | 0.62 | 0.46 | 0.57 | 0.6 | 0.77 | 0.75 |
| 3 | 117593588 | rs12054269 | 0.711 | 0.309 | 1.492 | -0.191 | 0.05 | -0.014 | -0.014 | 0 | 0.34 | 0.14 | 0.29 | 0.4 | 0.27 | 0.21 |
| 3 | 117594641 | rs9812067 | 1.843 | 0.139 | 1.263 | -0.364 | 0.044 | 0.081 | 0.081 | 1.863 | 0.24 | 0.11 | 0.3 | 0 | 0.16 | 0.09 |
| 3 | 117598159 | rs9860111 | 1.692 | 0.155 | 1.291 | -0.356 | 0.031 | 0.029 | 0.029 | 1.551 | 0.25 | 0.19 | 0.32 | 0 | 0.16 | 0.09 |
| 3 | 117599116 | rs9809760 | -0.702 | 0.25 | 1.459 | -0.295 | 0.015 | 0.016 | 0.016 | 2.24 | 0.68 | 0.57 | 0.66 | 0.46 | 0.48 | 0.35 |
| 3 | 117599362 | rs9289092 | -0.033 | 0.301 | 1.22 | -0.268 | 0.003 | -0.017 | -0.017 | 0 | 0.41 | 0.44 | 0.48 | 0.2 | 0.45 | 0.34 |
| 3 | 117618668 | rs6791432 | -1.099 | 0.623 | 1.191 | 0.019 | 0.011 | 0.028 | 0.028 | 0.425 | 0.76 | 0.66 | 0.75 | 0.61 | 0.73 | 0.53 |
| 3 | 117638221 | rs2045307 | 0.875 | 0.836 | 1.563 | 0.067 | 0.101 | 0.202 | 0.202 | 0 | 0.84 | 0.63 | 0.83 | 0.62 | 0.8 | 0.75 |
| 3 | 117640199 | rs6805565 | 1.717 | 0.355 | 1 | -0.246 | 0.012 | -0.081 | -0.081 | 0 | 0.21 | 0.05 | 0.21 | 0.38 | 0.19 | 0.12 |
| 3 | 117642675 | rs6795060 | -0.219 | 0.512 | 1.095 | -0.149 | 0.034 | -0.016 | -0.016 | 0 | 0.11 | 0.13 | 0.08 | 0.06 | 0.17 | 0.12 |
| 3 | 117652206 | rs16827737 | 0.75 | 0.163 | 0.839 | -0.467 | -0.009 | -0.011 | -0.011 | 0.027 | 0.16 | 0.07 | 0.12 | 0.19 | 0.21 | 0.19 |
| 3 | 117653416 | rs1378835 | 0.017 | 0.22 | 1.224 | -0.511 | -0.001 | -0.001 | -0.001 | 0 | 0.63 | 0.39 | 0.66 | 0.58 | 0.56 | 0.4 |
| 3 | 117656521 | rs1563438 | 1.886 | 0.328 | 0.804 | -0.625 | -0.053 | 0.082 | 0.082 | 0 | 0.2 | 0.08 | 0.15 | 0.05 | 0.24 | 0.36 |
| 3 | 117671762 | rs17722838 | 0.472 | 0.289 | 1.063 | -0.609 | 0.001 | -0.054 | -0.054 | 0 | 0.3 | 0.33 | 0.19 | 0.07 | 0.28 | 0.4 |
| 3 | 117690978 | rs9849010 | 0.441 | 0.07 | 0.588 | -0.61 | 0.007 | -0.05 | -0.05 | 0 | 0.37 | 0.12 | 0.33 | 0.56 | 0.39 | 0.31 |
| 3 | 117698800 | rs2605532 | 1.575 | -0.132 | 0.52 | -0.819 | 0.034 | 0.043 | 0.043 | 0.622 | 0.26 | 0.16 | 0.31 | 0 | 0.17 | 0.09 |
| 3 | 117709714 | rs11709110 | 0.651 | 0.038 | 0.81 | -0.744 | -0.006 | 0.014 | 0.014 | 0 | 0.29 | 0.25 | 0.19 | 0.07 | 0.28 | 0.4 |
| 3 | 117721983 | rs10934427 | 0.076 | 0.132 | 1.129 | -0.591 | 0.003 | 0.021 | 0.021 | 0 | 0.35 | 0.46 | 0.28 | 0.43 | 0.31 | 0.52 |
| 3 | 117731061 | rs10511373 | 2.111 | -0.075 | 0.821 | -0.624 | -0.006 | -0.081 | -0.081 | 0 | 0.18 | 0.05 | 0.18 | 0.38 | 0.17 | 0.11 |
| 3 | 117738666 | rs6798512 | -0.235 | 0.364 | 1.047 | -0.592 | -0.002 | -0.056 | -0.056 | 0 | 0.1 | 0.26 | 0.04 | 0.02 | 0.04 | 0.04 |
| 3 | 117750008 | rs817510 | 1.514 | 0.286 | 0.87 | -0.275 | 0.041 | 0.066 | 0.066 | 0 | 0.43 | 0.3 | 0.51 | 0.23 | 0.33 | 0.18 |
| 3 | 117755084 | rs11711199 | -0.225 | 0.577 | 0.672 | -0.169 | -0.011 | 0.058 | 0.058 | 0.044 | 0.1 | 0 | 0.03 | 0.02 | 0.05 | 0.04 |
| 3 | 117755434 | rs7618215 | 1.212 | 0.712 | 0.947 | -0.075 | -0.005 | 0 | 0 | 0.289 | 0.19 | 0.21 | 0.16 | 0.06 | 0.23 | 0.34 |
| 3 | 117773472 | rs1499981 | -0.482 | 0.732 | 1 | 0.051 | -0.009 | -0.01 | -0.01 | 0 | 0.75 | 0.76 | 0.73 | 0.58 | 0.62 | 0.46 |
| 3 | 117773652 | rs12695358 | -0.356 | 0.715 | 1.181 | 0.042 | 0.027 | 0.03 | 0.03 | 1.712 | 0.74 | 0.64 | 0.72 | 0.58 | 0.62 | 0.46 |
| 3 | 117774395 | rs11915336 | 0.353 | 0.614 | 1.098 | 0.017 | 0.017 | 0.01 | 0.01 | 0 | 0.64 | 0.57 | 0.68 | 0.56 | 0.57 | 0.42 |
| 3 | 117794284 | rs11925566 | 0.38 | 0.804 | 0.739 | 0.148 | -0.001 | 0.098 | 0.098 | 0 | 0.05 | 0.06 | 0.09 | 0.35 | 0.03 | 0.11 |
| 3 | 117800545 | rs1392265 | 0.368 | 0.746 | 0.827 | 0.052 | 0.023 | 0.015 | 0.015 | 0 | 0.65 | 0.54 | 0.68 | 0.57 | 0.57 | 0.43 |
| 3 | 117801330 | rs1499976 | 0.212 | 0.943 | 0.22 | -0.018 | -0.018 | 0.013 | 0.013 | 0 | 0.91 | 0.96 | 0.93 | 0.98 | 0.84 | 0.88 |
| 3 | 117822657 | rs10049077 | 0.481 | 0.587 | -0.053 | 0.389 | -0.014 | -0.029 | -0.029 | 0 | 0.23 | 0.18 | 0.24 | 0.37 | 0.26 | 0.42 |
| 3 | 117822894 | rs1032164 | -0.231 | 0.149 | -0.123 | 0.197 | 0.013 | -0.027 | -0.027 | 0 | 0.19 | 0.15 | 0.14 | 0.31 | 0.18 | 0.26 |

| | | | | | | | | | | | | | | | | |
|---|---|---|---|---|---|---|---|---|---|---|---|---|---|---|---|---|
| 3 | 117825077 | rs6770134 | -0.248 | -0.139 | -0.157 | -0.139 | -0.007 | 0.003 | 0.003 | 0 | 0.39 | 0.46 | 0.42 | 0.57 | 0.31 | 0.39 |
| 3 | 117825677 | rs10433401 | 0.823 | -0.134 | -0.253 | -0.129 | -0.014 | -0.032 | -0.032 | 0 | 0.07 | 0.01 | 0.07 | 0.29 | 0.1 | 0.08 |
| 3 | 117825865 | rs4688076 | -0.286 | -0.102 | -0.031 | -0.091 | -0.013 | 0.006 | 0.006 | 0 | 0.39 | 0.48 | 0.47 | 0.58 | 0.31 | 0.4 |
| 3 | 117827857 | rs9289095 | -0.236 | -0.287 | 0.416 | 0.335 | 0.003 | 0.001 | 0.001 | 0 | 0.6 | 0.43 | 0.67 | 0.59 | 0.64 | 0.65 |
| 3 | 117831077 | rs9846164 | 0.287 | -0.119 | 0.192 | -0.075 | -0.008 | -0.019 | -0.019 | 0 | 0.19 | 0.15 | 0.2 | 0.33 | 0.26 | 0.25 |
| 3 | 117831093 | rs4688077 | -1.078 | -0.22 | 0.161 | -0.075 | -0.001 | -0.059 | -0.059 | 0.145 | 0.69 | 0.66 | 0.77 | 0.93 | 0.67 | 0.81 |
| 3 | 117832779 | rs16827959 | 1.177 | -0.02 | -0.35 | 0.079 | 0.003 | 0.004 | 0.004 | 0 | 0.06 | 0.02 | 0.05 | 0.29 | 0.07 | 0.06 |
| 3 | 117833398 | rs16827961 | 1.13 | -0.021 | -0.35 | 0.079 | 0.003 | 0.002 | 0.002 | 0 | 0.06 | 0.02 | 0.05 | 0.29 | 0.07 | 0.06 |
| 3 | 117838063 | rs1499984 | -0.505 | -0.129 | 0.106 | -0.557 | -0.013 | -0.07 | -0.07 | 0.719 | 0.63 | 0.55 | 0.72 | 0.94 | 0.67 | 0.72 |
| 3 | 117840295 | rs2129229 | 0.486 | -0.221 | 0.044 | -0.266 | -0.004 | 0.107 | 0.107 | 0 | 0.07 | 0.09 | 0.12 | 0.39 | 0.04 | 0.17 |
| 3 | 117850135 | rs4688079 | -0.536 | -0.286 | 0.206 | -0.425 | -0.015 | -0.028 | -0.028 | 1.093 | 0.29 | 0.36 | 0.23 | 0.13 | 0.25 | 0.31 |
| 3 | 117852018 | rs11917814 | 1.351 | -0.082 | 0.168 | -0.319 | -0.024 | -0.028 | -0.028 | 0 | 0.32 | 0.22 | 0.41 | 0.46 | 0.38 | 0.33 |
| 3 | 117891733 | rs6438441 | -1.215 | -0.747 | 0.211 | 0.107 | -0.068 | 0.087 | 0.087 | 1.083 | 0.23 | 0.35 | 0.22 | 0.5 | 0.08 | 0.45 |
| 3 | 117904003 | rs1486336 | -0.189 | 0.267 | 0.417 | -0.248 | 0.104 | -0.073 | -0.073 | 0 | 0.82 | 0.59 | 0.79 | 0.92 | 0.68 | 0.75 |
| 3 | 117913539 | rs6795313 | 0.588 | -0.255 | 0.215 | -0.617 | -0.003 | 0.005 | 0.005 | 0 | 0.46 | 0.38 | 0.62 | 0.88 | 0.46 | 0.53 |
| 3 | 117915454 | rs4688081 | 1.065 | -0.322 | 0.159 | -0.724 | 0.011 | -0.037 | -0.037 | 0 | 0.67 | 0.37 | 0.72 | 0.86 | 0.62 | 0.6 |
| 3 | 117919392 | rs9846497 | -0.117 | -0.309 | 0.127 | -0.734 | 0.085 | -0.12 | -0.12 | 0 | 0.8 | 0.6 | 0.76 | 0.98 | 0.64 | 0.88 |
| 3 | 117919948 | rs1906041 | 1.558 | -0.306 | 0.21 | -0.555 | -0.053 | -0.074 | -0.074 | 1.91 | 0.34 | 0.1 | 0.44 | 0.44 | 0.37 | 0.38 |
| 3 | 117928767 | rs9849225 | 1.649 | -0.379 | 0.236 | -0.61 | 0.02 | 0.183 | 0.183 | 0.159 | 0.26 | 0.02 | 0.13 | 0 | 0.19 | 0.08 |
| 3 | 117939731 | rs9821669 | 0.134 | -0.246 | 0.452 | -0.544 | 0.052 | -0.075 | -0.075 | 1.227 | 0.76 | 0.56 | 0.77 | 0.94 | 0.66 | 0.77 |
| 3 | 117948652 | rs4428171 | 0.883 | -0.13 | 0.456 | -0.56 | 0.042 | -0.059 | -0.059 | 0.297 | 0.71 | 0.43 | 0.75 | 0.87 | 0.6 | 0.61 |
| 3 | 117952509 | rs4475034 | 1.538 | -0.189 | 0.365 | -0.524 | -0.049 | -0.066 | -0.066 | 0 | 0.34 | 0.12 | 0.45 | 0.45 | 0.36 | 0.38 |
| 3 | 117958399 | rs9828714 | 1.81 | -0.231 | 0.559 | -0.42 | -0.032 | -0.043 | -0.043 | 1.065 | 0.6 | 0.2 | 0.57 | 0.52 | 0.53 | 0.6 |
| 3 | 117958778 | rs6804074 | 0.978 | -0.072 | 0.663 | -0.407 | 0.031 | -0.068 | -0.068 | 0 | 0.79 | 0.5 | 0.86 | 0.93 | 0.8 | 0.76 |
| 3 | 117959502 | rs9853921 | 0.175 | -0.704 | 0.745 | -0.329 | 0.092 | -0.118 | -0.118 | 1.373 | 0.18 | 0.41 | 0.22 | 0.01 | 0.33 | 0.12 |
| 3 | 117972228 | rs4688091 | 0.894 | -0.642 | 0.67 | -0.301 | 0.077 | -0.08 | -0.08 | 0 | 0.15 | 0.27 | 0.2 | 0.01 | 0.33 | 0.11 |
| 3 | 117973455 | rs4447753 | 1.471 | -0.644 | 0.272 | -0.25 | -0.029 | -0.036 | -0.036 | 0.31 | 0.62 | 0.23 | 0.58 | 0.52 | 0.54 | 0.61 |
| 3 | 117974999 | rs12636586 | 0.185 | -0.27 | 0.269 | 0.402 | 0.005 | 0.257 | 0.257 | 0 | 0.05 | 0.16 | 0.12 | 0.38 | 0.03 | 0.14 |
| 3 | 117975155 | rs12636589 | -0.499 | -0.258 | 0.267 | 0.396 | -0.013 | 0.081 | 0.081 | 1.964 | 0.14 | 0.2 | 0.15 | 0.43 | 0.08 | 0.23 |
| 3 | 117992898 | rs9874084 | -1.263 | -0.26 | 0.246 | 0.397 | -0.04 | 0.158 | 0.158 | 0.809 | 0.2 | 0.4 | 0.18 | 0.47 | 0.11 | 0.25 |
| 3 | 117995184 | rs17281325 | 0.072 | 0.111 | -0.357 | 0.447 | 0.005 | 0.002 | 0.002 | 1.126 | 0.12 | 0.1 | 0.06 | 0.05 | 0.08 | 0.13 |
| 3 | 117998391 | rs12486432 | -0.113 | 0.661 | -0.492 | 0.823 | -0.006 | -0.006 | -0.006 | 0.968 | 0.43 | 0.37 | 0.51 | 0.51 | 0.45 | 0.4 |
| 3 | 118000812 | rs6802983 | 0.416 | 0.831 | -0.547 | 1.129 | -0.009 | -0.002 | -0.002 | 0 | 0.6 | 0.62 | 0.52 | 0.41 | 0.56 | 0.53 |
| 3 | 118001529 | rs9831693 | -0.625 | 0.6 | -0.575 | 0.86 | 0 | -0.013 | -0.013 | 1.705 | 0.07 | 0.06 | 0.04 | 0.01 | 0.05 | 0.06 |
| 3 | 118016233 | rs6779441 | -1.706 | 0.118 | 1.247 | 0.939 | -0.025 | 0.033 | 0.033 | 0.712 | 0.12 | 0.52 | 0.13 | 0.15 | 0.05 | 0.19 |
| 3 | 118020932 | rs705260 | -1.116 | 0.737 | 1.129 | 1.147 | -0.003 | -0.015 | -0.015 | 0 | 0.34 | 0.47 | 0.35 | 0.34 | 0.34 | 0.49 |
| 3 | 118021980 | rs4688095 | 1.711 | 0.638 | 1.131 | 1.207 | 0.024 | 0.118 | 0.118 | 0.053 | 0.49 | 0.18 | 0.39 | 0.28 | 0.47 | 0.36 |
| 3 | 118023515 | rs779206 | 0.985 | 0.736 | 1.565 | 1.448 | 0.037 | 0.075 | 0.075 | 0 | 0.8 | 0.66 | 0.68 | 0.65 | 0.77 | 0.81 |
| 3 | 118026811 | rs705257 | -0.431 | 0.687 | 1.826 | 1.41 | 0 | -0.003 | -0.003 | 0 | 0.29 | 0.28 | 0.27 | 0.38 | 0.29 | 0.45 |
| 3 | 118031202 | rs779596 | 0.256 | 0.691 | 1.826 | 1.418 | -0.004 | 0.012 | 0.012 | 0 | 0.23 | 0.17 | 0.22 | 0.19 | 0.26 | 0.28 |
| 3 | 118045905 | rs9865837 | -1.496 | 0.679 | 1.944 | 1.43 | 0.007 | 0.054 | 0.054 | 0.436 | 0.11 | 0.32 | 0.13 | 0.15 | 0.07 | 0.23 |
| 3 | 118053399 | rs779607 | -0.446 | 0.682 | 2.148 | 1.432 | -0.004 | 0.001 | 0.001 | 0 | 0.3 | 0.38 | 0.29 | 0.37 | 0.29 | 0.43 |
| 3 | 118062305 | rs705266 | -0.46 | 0.654 | 2.224 | 1.656 | 0 | -0.001 | -0.001 | 0.524 | 0.3 | 0.29 | 0.29 | 0.44 | 0.29 | 0.49 |
| 3 | 118062786 | rs705265 | -0.287 | 0.737 | 1.94 | 1.704 | 0.011 | 0.039 | 0.039 | 0 | 0.9 | 0.88 | 0.77 | 0.74 | 0.83 | 0.94 |
| 3 | 118081025 | rs2166753 | 0.722 | 0.679 | 1.919 | 1.609 | 0.024 | 0.018 | 0.018 | 0 | 0.64 | 0.54 | 0.5 | 0.48 | 0.55 | 0.58 |
| 3 | 118091164 | rs1447594 | -1.593 | 0.623 | 1.911 | 1.635 | -0.023 | 0.026 | 0.026 | 0 | 0.14 | 0.31 | 0.11 | 0.19 | 0.08 | 0.21 |
| 3 | 118095784 | rs9856360 | 2.12 | 0.096 | 0.836 | 0.756 | 0.021 | -0.019 | -0.019 | 0 | 0.29 | 0.1 | 0.29 | 0.28 | 0.27 | 0.25 |
| 3 | 118098264 | rs1374760 | 0.985 | 0.066 | 0.977 | 0.66 | -0.01 | 0.005 | 0.005 | 0 | 0.39 | 0.55 | 0.4 | 0.48 | 0.32 | 0.44 |
| 3 | 118100550 | rs536720 | -0.145 | 0.229 | 0.814 | 0.675 | -0.008 | -0.006 | -0.006 | 0 | 0.31 | 0.31 | 0.35 | 0.32 | 0.37 | 0.22 |
| 3 | 118106643 | rs1587453 | 1.276 | 0.437 | 0.551 | 0.508 | 0.007 | 0.001 | 0.001 | 1.428 | 0.14 | 0.11 | 0.05 | 0 | 0.1 | 0.02 |
| 3 | 118115160 | rs4687917 | 2.009 | 0.246 | 0.344 | 0.694 | 0.031 | -0.017 | -0.017 | 0.581 | 0.32 | 0.1 | 0.3 | 0.29 | 0.3 | 0.32 |
| 3 | 118115968 | rs1696485 | -0.566 | 0.502 | 0.72 | 0.839 | 0.045 | 0.037 | 0.037 | 0.334 | 0.3 | 0.51 | 0.39 | 0.34 | 0.34 | 0.15 |
| 3 | 118119682 | rs524438 | -1.034 | 0.301 | 0.937 | 0.991 | 0.007 | 0.014 | 0.014 | 0.84 | 0.37 | 0.56 | 0.35 | 0.52 | 0.35 | 0.3 |
| 3 | 118121335 | rs488628 | -0.093 | 0.289 | 1.368 | 1.28 | 0.004 | 0.007 | 0.007 | 0.914 | 0.25 | 0.3 | 0.26 | 0.27 | 0.24 | 0.08 |
| 3 | 118122816 | rs492668 | 0.761 | 0.357 | 1.433 | 1.287 | 0.011 | 0.031 | 0.031 | 0 | 0.1 | 0.11 | 0.21 | 0.26 | 0.15 | 0.06 |
| 3 | 118125046 | rs480491 | -0.55 | 0.2 | 1.391 | 1.11 | -0.041 | 0.021 | 0.021 | 0 | 0.75 | 0.84 | 0.71 | 0.79 | 0.64 | 0.67 |
| 3 | 118142356 | rs2090713 | -0.212 | 0.079 | 1.557 | 1.051 | -0.009 | -0.043 | -0.043 | 0.786 | 0.08 | 0.14 | 0.05 | 0.01 | 0.05 | 0.07 |
| 3 | 118146818 | rs16828422 | 0.288 | 0.05 | 1.525 | 1.118 | 0.003 | 0.005 | 0.005 | 0 | 0.06 | 0.04 | 0.01 | 0.2 | 0.03 | 0.14 |
| 3 | 118147502 | rs513437 | 0.61 | 0.105 | 1.729 | 1.056 | 0.054 | 0.101 | 0.101 | 0 | 0.11 | 0.23 | 0.23 | 0.27 | 0.16 | 0.06 |
| 3 | 118154526 | rs4688104 | 1.998 | 0.029 | 1.662 | 1.005 | 0.027 | 0.002 | 0.002 | 3.689 | 0.35 | 0.25 | 0.36 | 0.31 | 0.31 | 0.37 |
| 3 | 118154937 | rs553070 | -0.623 | 0.14 | 1.662 | 1.032 | 0.035 | 0.053 | 0.053 | 0.27 | 0.18 | 0.31 | 0.27 | 0.27 | 0.2 | 0.13 |
| 3 | 118157897 | rs577196 | 0.316 | 0.144 | 1.469 | 1.006 | -0.01 | -0.016 | -0.016 | 3.898 | 0.63 | 0.69 | 0.48 | 0.43 | 0.48 | 0.49 |
| 3 | 118162623 | rs10511378 | -0.022 | -0.017 | 1.644 | 0.722 | -0.005 | -0.045 | -0.045 | 1.005 | 0.23 | 0.21 | 0.09 | 0.03 | 0.15 | 0.09 |
| 3 | 118163149 | rs499789 | 2.01 | -0.059 | 1.575 | 0.757 | -0.004 | -0.001 | -0.001 | 0 | 0.34 | 0.39 | 0.38 | 0.29 | 0.31 | 0.37 |
| 3 | 118165434 | rs9834139 | 2.304 | -0.022 | 1.192 | 0.746 | 0.018 | -0.013 | -0.013 | 0 | 0.32 | 0.07 | 0.33 | 0.25 | 0.3 | 0.32 |
| 3 | 118166055 | rs553769 | -0.017 | 0.047 | 1.325 | 0.827 | -0.027 | -0.019 | -0.019 | 0 | 0.69 | 0.77 | 0.7 | 0.58 | 0.61 | 0.52 |
| 3 | 118183783 | rs1949542 | 0.803 | 0.655 | 1.392 | 0.302 | 0.064 | 0.024 | 0.024 | 0.106 | 0.73 | 0.38 | 0.55 | 0.64 | 0.6 | 0.74 |
| 3 | 118184249 | rs7632701 | 0.652 | 0.872 | 1.381 | 0.309 | 0.041 | 0.064 | 0.064 | 0 | 0.1 | 0.17 | 0.23 | 0.27 | 0.16 | 0.05 |
| 3 | 118186591 | rs1401955 | 1.751 | 0.531 | 1.042 | 0.067 | 0.044 | 0.055 | 0.055 | 0.011 | 0.5 | 0.08 | 0.39 | 0.62 | 0.42 | 0.53 |

| | | | | | | | | | | | | | | | | |
|---|---|---|---|---|---|---|---|---|---|---|---|---|---|---|---|---|
| 3 | 118187821 | rs2970148 | -0.355 | 0.567 | 0.991 | -0.183 | -0.001 | 0 | 0 | 0 | 0.85 | 0.85 | 0.8 | 0.92 | 0.77 | 0.8 |
| 3 | 118193612 | rs17676122 | -0.434 | 0.372 | 0.832 | -0.274 | -0.002 | -0.033 | -0.033 | 0 | 0.23 | 0.19 | 0.16 | 0.03 | 0.18 | 0.2 |
| 3 | 118196219 | rs1401957 | 1.043 | 0.454 | 0.729 | -0.252 | 0.007 | 0.091 | 0.091 | 0 | 0.6 | 0.26 | 0.63 | 0.89 | 0.6 | 0.6 |
| 3 | 118198045 | rs9849379 | 2.398 | 0.304 | 0.336 | 0.005 | -0.034 | 0.097 | 0.097 | 3.148 | 0.31 | 0.04 | 0.32 | 0.14 | 0.31 | 0.32 |
| 3 | 118201229 | rs1464311 | -0.184 | 0.624 | 0.638 | 0.247 | 0.005 | 0.008 | 0.008 | 0.01 | 0.12 | 0.08 | 0.1 | 0.03 | 0.15 | 0.08 |
| 3 | 118202086 | rs7618451 | 1.528 | 0.244 | 0.336 | 0.375 | -0.009 | 0.014 | 0.014 | 0 | 0.59 | 0.31 | 0.46 | 0.63 | 0.48 | 0.68 |
| 3 | 118204601 | rs2139547 | 0.703 | 0.146 | 0.105 | 0.439 | 0.032 | -0.117 | -0.117 | 0 | 0.21 | 0.05 | 0.07 | 0.48 | 0.12 | 0.23 |
| 3 | 118206941 | rs2970120 | -0.816 | 0.639 | 0.704 | 0.284 | 0.04 | -0.098 | -0.098 | 1.918 | 0.17 | 0.39 | 0.15 | 0.03 | 0.23 | 0.09 |
| 3 | 118217387 | rs2946818 | 0.224 | 0.746 | 0.08 | 0.718 | -0.012 | -0.012 | -0.012 | 0 | 0.77 | 0.8 | 0.63 | 0.65 | 0.71 | 0.77 |
| 3 | 118218911 | rs2946819 | -0.255 | 0.517 | 0.463 | 0.534 | -0.003 | -0.016 | -0.016 | 5.129 | 0.88 | 0.83 | 0.94 | 0.93 | 0.91 | 0.88 |
| 3 | 118221011 | rs11710875 | -0.202 | 0.775 | 0.793 | 0.582 | -0.001 | 0.046 | 0.046 | 6.381 | 0.27 | 0.29 | 0.24 | 0.5 | 0.24 | 0.27 |
| 3 | 118226026 | rs1401951 | -1.405 | 0.733 | 0.828 | 0.493 | 0.001 | 0.004 | 0.004 | 6.135 | 0.4 | 0.55 | 0.43 | 0.58 | 0.4 | 0.4 |
| 3 | 118229370 | rs939316 | -0.382 | 0.449 | 0.335 | 0.522 | 0.001 | -0.001 | -0.001 | 0 | 0.1 | 0.09 | 0.14 | 0.1 | 0.11 | 0.13 |
| 3 | 118232666 | rs16828610 | -0.366 | 0.457 | 0.292 | 0.455 | -0.005 | -0.024 | -0.024 | 0 | 0.1 | 0.17 | 0.1 | 0.03 | 0.1 | 0.11 |
| 3 | 118252847 | rs4630940 | 1.56 | 0.578 | 0.133 | 0.419 | 0.016 | -0.024 | -0.024 | 0 | 0.56 | 0.27 | 0.59 | 0.46 | 0.61 | 0.62 |
| 3 | 118253469 | rs6807699 | 1.064 | 0.558 | 0.151 | 0.421 | 0.01 | -0.029 | -0.029 | 0.016 | 0.6 | 0.3 | 0.63 | 0.47 | 0.64 | 0.7 |
| 3 | 118257029 | rs16828633 | -0.478 | 0.536 | 0.438 | 0.393 | -0.01 | -0.031 | -0.031 | 0 | 0.1 | 0.14 | 0.1 | 0.01 | 0.1 | 0.12 |
| 3 | 118263121 | rs4074439 | 0.482 | 0.524 | 0.43 | 0.39 | -0.003 | 0.023 | 0.023 | 5.07 | 0.67 | 0.41 | 0.69 | 0.47 | 0.72 | 0.74 |
| 3 | 118273774 | rs9875662 | 0.884 | 0.571 | 0.38 | 0.197 | -0.03 | 0.014 | 0.014 | 0 | 0.11 | 0.01 | 0.05 | 0.06 | 0.11 | 0.12 |
| 3 | 118274165 | rs4295164 | 1.227 | 0.399 | 0.929 | 0.392 | 0.08 | 0.03 | 0.03 | 2.046 | 0.9 | 0.49 | 0.91 | 0.92 | 0.9 | 0.97 |
| 3 | 118281297 | rs9814010 | 1.804 | 0.4 | 0.614 | 0.423 | -0.006 | -0.013 | -0.013 | 0 | 0.79 | 0.34 | 0.77 | 0.84 | 0.79 | 0.83 |
| 3 | 118293560 | rs6775374 | 1.105 | 0.45 | 0.257 | 0.423 | 0.02 | -0.023 | -0.023 | 0.46 | 0.62 | 0.3 | 0.72 | 0.44 | 0.7 | 0.64 |
| 3 | 118297492 | rs6801870 | 0.791 | 0.255 | 0.247 | 0.471 | -0.011 | 0.031 | 0.031 | 0 | 0.06 | 0.01 | 0.05 | 0 | 0.08 | 0.05 |
| 3 | 118310117 | rs6798298 | 1.394 | 0.318 | 0.284 | 0.421 | 0.006 | -0.011 | -0.011 | 1.774 | 0.58 | 0.37 | 0.61 | 0.47 | 0.62 | 0.61 |
| 3 | 118322916 | rs9822639 | 0.87 | 0.214 | 0.349 | 0.492 | -0.006 | 0.017 | 0.017 | 2.516 | 0.66 | 0.45 | 0.66 | 0.52 | 0.71 | 0.66 |
| 3 | 118329449 | rs1846641 | 0.184 | 0.195 | 0.767 | 0.517 | -0.012 | 0.068 | 0.068 | 3.971 | 0.74 | 0.58 | 0.75 | 0.53 | 0.81 | 0.76 |
| 3 | 118335040 | rs9830308 | 0.467 | 0.174 | 0.85 | 0.499 | -0.012 | 0.036 | 0.036 | 3.801 | 0.7 | 0.47 | 0.71 | 0.52 | 0.78 | 0.7 |
| 3 | 118336471 | rs9836543 | 1.479 | 0.157 | 0.87 | 0.52 | 0.018 | 0.012 | 0.012 | 4.669 | 0.57 | 0.26 | 0.55 | 0.39 | 0.62 | 0.6 |
| 3 | 118345006 | rs1500084 | 0.437 | 0.192 | 0.815 | 0.607 | -0.006 | 0.07 | 0.07 | 3.889 | 0.71 | 0.57 | 0.74 | 0.48 | 0.77 | 0.72 |
| 3 | 118356071 | rs13083186 | 1.195 | 0.165 | 0.815 | 0.668 | 0.008 | -0.021 | -0.021 | 0 | 0.59 | 0.33 | 0.61 | 0.49 | 0.62 | 0.63 |
| 3 | 118359465 | rs1909616 | -1.514 | 0.036 | 1.13 | 0.611 | 0.005 | -0.007 | -0.007 | 0 | 0.43 | 0.58 | 0.4 | 0.51 | 0.38 | 0.38 |
| 3 | 118381709 | rs9832838 | 1.262 | 0.049 | 1.004 | 0.583 | 0.017 | -0.002 | -0.002 | 0.827 | 0.57 | 0.24 | 0.56 | 0.42 | 0.62 | 0.62 |
| 3 | 118382257 | rs9817827 | -0.986 | 0.114 | 1.061 | 0.615 | 0.034 | 0.05 | 0.05 | 0.862 | 0.06 | 0.31 | 0.04 | 0.05 | 0.04 | 0.04 |
| 3 | 118395847 | rs9831518 | -0.932 | 0.218 | 0.583 | 0.505 | -0.018 | -0.005 | -0.005 | 0.144 | 0.21 | 0.53 | 0.1 | 0.2 | 0.15 | 0.19 |
| 3 | 118401982 | rs6769257 | -0.211 | 0.128 | 0.436 | 0.338 | 0.003 | 0.01 | 0.01 | 0 | 0.1 | 0.07 | 0.1 | 0.01 | 0.11 | 0.1 |
| 3 | 118402091 | rs17202866 | 1.702 | -0.127 | 0.084 | 0.388 | -0.157 | 0.035 | 0.035 | 2.343 | 0.43 | 0.04 | 0.49 | 0.26 | 0.51 | 0.45 |
| 3 | 118402543 | rs6438471 | -0.313 | -0.219 | 0.925 | 0.518 | -0.009 | -0.038 | -0.038 | 0 | 0.16 | 0.32 | 0.07 | 0.1 | 0.11 | 0.15 |
| 3 | 118402562 | rs6438472 | -1.141 | -0.223 | 0.562 | 0.601 | -0.014 | 0.025 | 0.025 | 0.086 | 0.23 | 0.61 | 0.18 | 0.29 | 0.19 | 0.21 |
| 3 | 118408200 | rs6765745 | -1.573 | -0.293 | 0.258 | 0.384 | -0.013 | -0.021 | -0.021 | 0 | 0.08 | 0.07 | 0.08 | 0.04 | 0.05 | 0.1 |
| 3 | 118411464 | rs6782741 | 0.948 | -0.155 | -0.467 | 0.424 | -0.04 | 0.029 | 0.029 | 1.061 | 0.34 | 0.13 | 0.4 | 0.27 | 0.36 | 0.22 |
| 3 | 118413478 | rs4687935 | -0.348 | -0.328 | -0.214 | 0.508 | 0 | 0 | 0 | 0.102 | 0.42 | 0.4 | 0.26 | 0.44 | 0.37 | 0.57 |
| 3 | 118418924 | rs16828784 | 0.885 | -0.383 | -0.195 | 0.579 | -0.005 | -0.008 | -0.008 | 0.14 | 0.2 | 0.16 | 0.21 | 0.17 | 0.25 | 0.17 |
| 3 | 118418948 | rs13091270 | 0.257 | -0.374 | -0.312 | 0.481 | 0.009 | -0.013 | -0.013 | 0.643 | 0.33 | 0.17 | 0.15 | 0.41 | 0.26 | 0.46 |
| 3 | 118422245 | rs4687937 | 0.199 | -0.302 | -0.035 | 0.533 | -0.001 | 0.01 | 0.01 | 0 | 0.68 | 0.39 | 0.57 | 0.68 | 0.63 | 0.69 |
| 3 | 118423392 | rs9859270 | -0.017 | -0.352 | -0.307 | 0.608 | 0.01 | -0.02 | -0.02 | 0 | 0.33 | 0.22 | 0.16 | 0.45 | 0.26 | 0.46 |
| 3 | 118440360 | rs2684311 | -0.582 | -0.331 | -0.173 | 0.549 | -0.002 | -0.001 | -0.001 | 0.022 | 0.47 | 0.51 | 0.42 | 0.42 | 0.45 | 0.46 |
| 3 | 118442496 | rs982040 | 0.692 | -0.41 | -0.16 | 0.549 | -0.001 | -0.002 | -0.002 | 0.092 | 0.19 | 0.21 | 0.08 | 0.15 | 0.15 | 0.2 |
| 3 | 118449242 | rs6809627 | 0.466 | -0.435 | -0.162 | 0.56 | -0.022 | -0.006 | -0.006 | 0 | 0.14 | 0.23 | 0.16 | 0.11 | 0.12 | 0.14 |
| 3 | 118451648 | rs705245 | -1.71 | -0.458 | 0.04 | 0.6 | -0.008 | 0.055 | 0.055 | 0 | 0.87 | 0.74 | 0.91 | 0.72 | 0.9 | 0.73 |
| 3 | 118456547 | rs798401 | 0.085 | -0.455 | -0.364 | 0.578 | -0.001 | -0.003 | -0.003 | 0 | 0.33 | 0.25 | 0.25 | 0.31 | 0.33 | 0.32 |
| 3 | 118461040 | rs705236 | -0.663 | -0.47 | -0.35 | 0.557 | -0.001 | 0 | 0 | 0 | 0.47 | 0.49 | 0.4 | 0.42 | 0.45 | 0.46 |
| 3 | 118463829 | rs697378 | -1.607 | -0.433 | -0.284 | 0.552 | 0.007 | 0.075 | 0.075 | 0 | 0.86 | 0.72 | 0.9 | 0.72 | 0.89 | 0.73 |
| 3 | 118464116 | rs808887 | -1.022 | -0.537 | -0.545 | 0.548 | -0.006 | 0.08 | 0.08 | 0.29 | 0.82 | 0.58 | 0.83 | 0.69 | 0.86 | 0.67 |
| 3 | 118467574 | rs798256 | 0.604 | -0.642 | -0.586 | 0.56 | -0.003 | -0.007 | -0.007 | 0.004 | 0.2 | 0.23 | 0.08 | 0.15 | 0.15 | 0.2 |
| 3 | 118468626 | rs798257 | 1.56 | -0.778 | -0.661 | 0.254 | -0.037 | 0.004 | 0.004 | 0.044 | 0.14 | 0.01 | 0.06 | 0.11 | 0.14 | 0.14 |
| 3 | 118468712 | rs4687827 | 1.254 | -0.684 | -0.613 | 0.27 | -0.085 | 0.043 | 0.043 | 0.388 | 0.39 | 0.13 | 0.49 | 0.3 | 0.46 | 0.27 |
| 3 | 118474092 | rs2650158 | 0.117 | -0.586 | -0.892 | 0.345 | -0.001 | -0.002 | -0.002 | 0 | 0.33 | 0.25 | 0.24 | 0.31 | 0.33 | 0.33 |
| 3 | 118482997 | rs798593 | -0.079 | -0.628 | -0.881 | 0.35 | 0 | -0.022 | -0.022 | 0 | 0.35 | 0.27 | 0.33 | 0.42 | 0.35 | 0.37 |
| 3 | 118488359 | rs2037430 | 2.247 | -0.25 | -1.082 | 0.252 | -0.074 | 0.099 | 0.099 | 0.01 | 0.26 | 0.02 | 0.15 | 0.06 | 0.26 | 0.15 |
| 3 | 118499493 | rs705225 | 1.761 | -0.574 | -0.329 | 0.729 | -0.039 | -0.055 | -0.055 | 0 | 0.13 | 0.02 | 0.16 | 0.17 | 0.18 | 0.12 |
| 3 | 118501868 | rs798577 | -1.183 | -0.745 | 0.047 | 0.641 | -0.04 | 0.046 | 0.046 | 0.603 | 0.59 | 0.77 | 0.49 | 0.7 | 0.53 | 0.72 |
| 3 | 118515016 | rs6770279 | -1.322 | -0.719 | 0.468 | 0.798 | -0.036 | -0.019 | -0.019 | 0 | 0.17 | 0.3 | 0.18 | 0.11 | 0.11 | 0.15 |
| 3 | 118519824 | rs1392230 | -1.411 | -0.337 | 0.371 | 0.466 | -0.041 | 0.019 | 0.019 | 0.017 | 0.24 | 0.48 | 0.15 | 0.28 | 0.19 | 0.35 |
| 3 | 118525556 | rs7610551 | -0.803 | -0.056 | -0.332 | -0.092 | -0.023 | -0.057 | -0.057 | 0.523 | 0.11 | 0.21 | 0.07 | 0 | 0.09 | 0.1 |
| 3 | 118530692 | rs881604 | -0.045 | 0.045 | -0.575 | -0.256 | -0.011 | -0.046 | -0.046 | 0 | 0.37 | 0.42 | 0.24 | 0.15 | 0.3 | 0.36 |
| 3 | 118537079 | rs17465252 | -0.198 | -0.195 | -0.849 | -0.275 | 0.006 | 0.006 | 0.006 | 0 | 0.09 | 0.04 | 0.1 | 0.07 | 0.08 | 0.09 |
| 3 | 118539108 | rs614673 | 0.501 | 0.267 | -1.681 | -0.369 | -0.06 | 0.025 | 0.025 | 0 | 0.24 | 0.04 | 0.2 | 0.18 | 0.31 | 0.33 |
| 3 | 118543021 | rs12631973 | 1.374 | 1.051 | -1.526 | 0.558 | -0.039 | -0.04 | -0.04 | 0 | 0.09 | 0.02 | 0.06 | 0.29 | 0.14 | 0.1 |
| 3 | 118546044 | rs1081891 | -0.03 | 0.74 | -1.136 | 0.568 | -0.011 | -0.029 | -0.029 | 0 | 0.28 | 0.2 | 0.47 | 0.35 | 0.31 | 0.21 |

| | | | | | | | | | | | | | | | | |
|---|---|---|---|---|---|---|---|---|---|---|---|---|---|---|---|---|
| 3 | 118549344 | rs1081896 | -0.554 | 0.721 | -0.993 | 0.554 | -0.041 | 0.013 | 0.013 | 0 | 0.37 | 0.23 | 0.53 | 0.64 | 0.45 | 0.31 |
| 3 | 118550230 | rs7610864 | -0.137 | 0.757 | -1.021 | 0.577 | -0.007 | -0.012 | -0.012 | 0.002 | 0.18 | 0.14 | 0.17 | 0.36 | 0.22 | 0.2 |
| 3 | 118551721 | rs1081900 | 1.01 | 0.48 | -1.145 | 0.41 | -0.063 | -0.094 | -0.094 | 0 | 0.19 | 0.05 | 0.37 | 0.28 | 0.24 | 0.12 |
| 3 | 118553190 | rs7625631 | 0.844 | 0.384 | -1.118 | 0.406 | -0.002 | -0.003 | -0.003 | 0 | 0.09 | 0.01 | 0.1 | 0.07 | 0.07 | 0.09 |
| 3 | 118557806 | rs13062633 | 1.113 | 0.559 | -1.118 | 0.482 | -0.049 | -0.083 | -0.083 | 0.013 | 0.18 | 0.06 | 0.36 | 0.28 | 0.22 | 0.13 |
| 3 | 118559855 | rs1521290 | 0.181 | 0.376 | -0.4 | 0.412 | -0.04 | -0.027 | -0.027 | 0.082 | 0.36 | 0.25 | 0.3 | 0.46 | 0.47 | 0.44 |
| 3 | 118561751 | rs2649882 | -1.436 | 0.319 | -0.088 | 0.369 | -0.034 | -0.059 | -0.059 | 0 | 0.64 | 0.54 | 0.77 | 0.82 | 0.76 | 0.66 |
| 3 | 118561959 | rs2649881 | 0.116 | 0.293 | -0.272 | 0.415 | -0.04 | -0.028 | -0.028 | 0 | 0.36 | 0.24 | 0.3 | 0.46 | 0.47 | 0.44 |
| 3 | 118578547 | rs9829595 | 0.62 | 0.028 | -1.034 | 0.077 | -0.096 | -0.047 | -0.047 | 0 | 0.27 | 0.06 | 0.42 | 0.27 | 0.37 | 0.2 |
| 3 | 118578656 | rs1589441 | -0.953 | -0.037 | -0.577 | 0.374 | 0.061 | 0.053 | 0.053 | 0.319 | 0.87 | 0.66 | 0.9 | 0.8 | 0.84 | 0.89 |
| 3 | 118581688 | rs1088694 | -1.001 | -1.015 | -0.989 | -0.1 | 0.018 | -0.006 | -0.006 | 0 | 0.75 | 0.23 | 0.89 | 0.99 | 0.94 | 0.77 |
| 3 | 118586272 | rs950336 | 0.327 | -0.79 | -0.961 | 0.004 | -0.008 | -0.051 | -0.051 | 0.02 | 0.39 | 0.13 | 0.41 | 0.52 | 0.41 | 0.46 |
| 3 | 118587348 | rs6769280 | 2.043 | -0.844 | -0.836 | -0.036 | -0.049 | -0.094 | -0.094 | 0.042 | 0.18 | 0.29 | 0.07 | 0.01 | 0.06 | 0.23 |
| 3 | 118588401 | rs12486625 | 0.434 | -0.489 | -1.532 | 0.186 | 0.046 | 0.097 | 0.097 | 1.512 | 0.47 | 0.81 | 0.51 | 0.27 | 0.37 | 0.38 |
| 3 | 118594133 | rs1402578 | 0.453 | -0.34 | -1.375 | 0.382 | -0.007 | -0.057 | -0.057 | 0 | 0.32 | 0.13 | 0.31 | 0.44 | 0.34 | 0.37 |
| 3 | 118610916 | rs9818140 | 0.727 | 0.203 | -1.228 | 0.54 | -0.016 | 0.012 | 0.012 | 0 | 0.22 | 0.13 | 0.17 | 0.16 | 0.26 | 0.16 |
| 3 | 118615463 | rs1879553 | 0.797 | -0.039 | -1.194 | 0.657 | 0.001 | -0.003 | -0.003 | 0 | 0.08 | 0.05 | 0.09 | 0.13 | 0.08 | 0.07 |
| 3 | 118618170 | rs1521293 | -0.366 | -0.218 | -1.276 | 0.537 | 0.022 | 0.009 | 0.009 | 0 | 0.53 | 0.36 | 0.53 | 0.58 | 0.64 | 0.46 |
| 3 | 118618642 | rs10934467 | 0.253 | -0.261 | -1.293 | 0.518 | -0.021 | -0.019 | -0.019 | 0 | 0.44 | 0.16 | 0.44 | 0.45 | 0.56 | 0.39 |
| 3 | 118618860 | rs6438476 | -1.597 | -0.516 | -1.451 | 0.493 | -0.055 | -0.039 | -0.039 | 0 | 0.88 | 0.96 | 0.93 | 0.8 | 0.78 | 0.85 |
| 3 | 118623248 | rs11927719 | 0.67 | -1.148 | -1.244 | 0.404 | -0.038 | -0.028 | -0.028 | 0 | 0.33 | 0.57 | 0.34 | 0.13 | 0.13 | 0.35 |
| 3 | 118624631 | rs6782002 | 0.231 | -1.125 | -1.195 | 0.388 | -0.009 | -0.003 | -0.003 | 0 | 0.37 | 0.61 | 0.41 | 0.21 | 0.19 | 0.42 |
| 3 | 118641879 | rs2160052 | 0.42 | -1.151 | -1.249 | 0.404 | 0.044 | 0.029 | 0.029 | 0 | 0.35 | 0.72 | 0.39 | 0.21 | 0.14 | 0.39 |
| 3 | 118643250 | rs11709968 | 0.659 | -0.991 | -1.188 | 0.48 | -0.003 | 0.005 | 0.005 | 0 | 0.09 | 0.09 | 0.08 | 0.13 | 0.08 | 0.06 |
| 3 | 118649060 | rs2903250 | -1.696 | -0.943 | -1.423 | 0.508 | -0.068 | -0.051 | -0.051 | 0.033 | 0.81 | 0.98 | 0.87 | 0.8 | 0.76 | 0.86 |
| 3 | 118655664 | rs9837571 | -0.062 | -1.081 | -1 | 0.472 | 0.031 | 0.073 | 0.073 | 0 | 0.08 | 0.31 | 0.29 | 0.13 | 0.08 | 0.06 |
| 3 | 118659679 | rs9878706 | -0.234 | -1.266 | -1.272 | 0.391 | 0.141 | 0.034 | 0.034 | 0 | 0.46 | 0.83 | 0.5 | 0.34 | 0.25 | 0.46 |
| 3 | 118667405 | rs7651619 | 1.12 | -1.3 | -1.119 | 0.338 | -0.096 | -0.083 | -0.083 | 5.454 | 0.28 | 0.38 | 0.12 | 0.08 | 0.08 | 0.34 |
| 3 | 118676492 | rs4687833 | -1.233 | -0.713 | -0.991 | 0.606 | -0.012 | -0.001 | -0.001 | 0 | 0.8 | 0.88 | 0.86 | 0.82 | 0.76 | 0.84 |
| 3 | 118706164 | rs1016284 | 0.162 | -0.38 | -0.782 | 0.764 | -0.002 | 0.003 | 0.003 | 0 | 0.09 | 0.11 | 0.1 | 0.13 | 0.08 | 0.06 |
| 3 | 118719480 | rs4687835 | -0.806 | -1.064 | -0.842 | 0.783 | 0.058 | -0.002 | -0.002 | 0 | 0.36 | 0.79 | 0.48 | 0.33 | 0.25 | 0.45 |
| 3 | 118730933 | rs4687836 | -0.826 | -0.856 | -0.45 | 0.851 | -0.018 | 0.05 | 0.05 | 0.305 | 0.16 | 0.41 | 0.18 | 0.2 | 0.14 | 0.14 |
| 3 | 118742848 | rs39688 | -0.337 | -0.014 | -0.659 | 0.815 | -0.003 | -0.001 | -0.001 | 1.306 | 0.54 | 0.52 | 0.64 | 0.58 | 0.57 | 0.67 |
| 3 | 118747001 | rs9840095 | -0.142 | -0.013 | -0.619 | 0.833 | -0.002 | 0.047 | 0.047 | 0 | 0.14 | 0.28 | 0.19 | 0.2 | 0.16 | 0.14 |
| 3 | 118747851 | rs859018 | -1.719 | 0.028 | -0.886 | 0.537 | 0.001 | 0.02 | 0.02 | 2.281 | 0.69 | 0.92 | 0.84 | 0.79 | 0.73 | 0.81 |
| 3 | 118751053 | rs35859 | -1.437 | 0.064 | -0.881 | 0.548 | 0.004 | 0.026 | 0.026 | 2.243 | 0.69 | 0.92 | 0.84 | 0.78 | 0.73 | 0.8 |
| 3 | 118755319 | rs35862 | -1.015 | 0.231 | -0.767 | 0.641 | -0.005 | 0.005 | 0.005 | 0.468 | 0.51 | 0.67 | 0.6 | 0.47 | 0.48 | 0.56 |
| 3 | 118757870 | rs251448 | -0.985 | 0.137 | -0.376 | 0.633 | 0.003 | 0.024 | 0.024 | 1.059 | 0.61 | 0.68 | 0.7 | 0.76 | 0.58 | 0.76 |
| 3 | 118760722 | rs12636639 | 0.443 | -0.254 | -0.066 | 0.848 | -0.037 | 0.018 | 0.018 | 0 | 0.19 | 0.31 | 0.29 | 0.2 | 0.11 | 0.33 |
| 3 | 118779507 | rs1369583 | 0.444 | -0.254 | -0.069 | 0.856 | -0.036 | 0.019 | 0.019 | 0 | 0.19 | 0.3 | 0.29 | 0.21 | 0.11 | 0.33 |
| 3 | 118781005 | rs7638463 | 0.105 | -0.305 | 0.025 | 0.833 | -0.055 | 0.026 | 0.026 | 0 | 0.2 | 0.43 | 0.3 | 0.21 | 0.11 | 0.33 |
| 3 | 118786845 | rs1347449 | 1.445 | -0.011 | -0.337 | 0.664 | -0.008 | 0.024 | 0.024 | 1.08 | 0.34 | 0.17 | 0.27 | 0.25 | 0.38 | 0.19 |
| 3 | 118793832 | rs9840510 | 2.656 | 0.244 | -0.6 | 0.764 | -0.08 | -0.019 | -0.019 | 0 | 0.17 | 0.01 | 0.15 | 0.14 | 0.21 | 0.18 |
| 3 | 118799592 | rs4435634 | 1.584 | 0.208 | -0.359 | 0.748 | 0 | 0.011 | 0.011 | 0.636 | 0.34 | 0.3 | 0.28 | 0.25 | 0.38 | 0.19 |
| 3 | 118799682 | rs4449310 | 0.745 | 0.074 | -0.148 | 0.623 | -0.001 | 0.01 | 0.01 | 0.583 | 0.42 | 0.5 | 0.55 | 0.48 | 0.45 | 0.28 |
| 3 | 118801854 | rs16829334 | 1.304 | -0.058 | -0.037 | 0.599 | -0.058 | 0.003 | 0.003 | 0 | 0.16 | 0.29 | 0.09 | 0.14 | 0.05 | 0.33 |
| 3 | 118803278 | rs1369574 | -0.482 | 0.168 | -0.374 | 0.612 | -0.044 | 0.003 | 0.003 | 1.006 | 0.76 | 0.93 | 0.8 | 0.76 | 0.71 | 0.79 |
| 3 | 118812027 | rs2917080 | -0.047 | 0.158 | -0.434 | 0.603 | -0.022 | 0.031 | 0.031 | 1.352 | 0.43 | 0.21 | 0.36 | 0.38 | 0.5 | 0.39 |
| 3 | 118823141 | rs1347448 | 1.747 | 0.188 | -0.099 | 0.754 | -0.031 | -0.014 | -0.014 | 0.16 | 0.22 | 0.09 | 0.19 | 0.2 | 0.23 | 0.28 |
| 3 | 118824418 | rs6779584 | 0.432 | 0.375 | -0.043 | 0.504 | -0.022 | -0.066 | -0.066 | 0 | 0.66 | 0.54 | 0.56 | 0.84 | 0.72 | 0.6 |
| 3 | 118864700 | rs9289121 | 0.027 | 0.443 | 0.093 | 0.528 | 0.012 | 0.016 | 0.016 | 0 | 0.39 | 0.48 | 0.47 | 0.51 | 0.42 | 0.43 |
| 3 | 118864912 | rs10049288 | 1.658 | 0.426 | 0.168 | 0.53 | -0.004 | 0.003 | 0.003 | 0 | 0.24 | 0.2 | 0.19 | 0.2 | 0.23 | 0.28 |
| 3 | 118865332 | rs11550908 | 2.078 | 0.425 | 0.158 | 0.53 | 0.005 | 0.012 | 0.012 | 0.086 | 0.11 | 0.13 | 0.07 | 0.24 | 0.15 | 0.11 |
| 3 | 118865609 | rs10934483 | 2.078 | 0.425 | 0.158 | 0.53 | 0.005 | 0.012 | 0.012 | 0.086 | 0.11 | 0.13 | 0.07 | 0.24 | 0.15 | 0.11 |
| 3 | 118865801 | rs4077931 | 1.575 | 0.338 | -0.347 | 0.392 | -0.004 | 0.003 | 0.003 | 0 | 0.24 | 0.21 | 0.2 | 0.2 | 0.23 | 0.28 |
| 3 | 118865970 | rs4077930 | 0.19 | -0.168 | -0.318 | 0.433 | -0.008 | -0.027 | -0.027 | 0.009 | 0.26 | 0.26 | 0.19 | 0.08 | 0.17 | 0.35 |
| 3 | 118870356 | rs4133080 | 1.815 | 0.289 | -0.759 | 0.956 | -0.083 | -0.014 | -0.014 | 0.492 | 0.25 | 0.02 | 0.19 | 0.18 | 0.28 | 0.12 |
| 3 | 118876771 | rs6783216 | 0.61 | 0.182 | -0.51 | 1.064 | -0.018 | 0.021 | 0.021 | 0 | 0.06 | 0.14 | 0.03 | 0.06 | 0.03 | 0.1 |
| 3 | 118891246 | rs2903301 | -0.798 | 0.606 | -0.815 | 1.1 | 0.001 | -0.01 | -0.01 | 2.001 | 0.65 | 0.72 | 0.67 | 0.6 | 0.62 | 0.77 |
| 3 | 118901013 | rs9814507 | 0.335 | 0.668 | -0.903 | 1.07 | 0.002 | -0.019 | -0.019 | 0.099 | 0.31 | 0.29 | 0.28 | 0.41 | 0.27 | 0.33 |
| 3 | 118902844 | rs4455314 | 0.329 | 0.988 | -0.356 | 1.183 | -0.002 | -0.006 | -0.006 | 0.001 | 0.77 | 0.79 | 0.82 | 0.66 | 0.76 | 0.64 |
| 3 | 118903100 | rs12107009 | -1.1 | 0.37 | 0.129 | 0.872 | -0.01 | -0.031 | -0.031 | 0 | 0.09 | 0.16 | 0.14 | 0.05 | 0.08 | 0.13 |
| 3 | 118906442 | rs10934486 | -1.26 | 0.675 | 1.191 | 1.414 | 0.044 | 0.089 | 0.089 | 0.013 | 0.12 | 0.34 | 0.31 | 0.23 | 0.1 | 0.22 |
| 3 | 118908376 | rs6772525 | -0.63 | 0.357 | 0.774 | 1.465 | -0.005 | -0.02 | -0.02 | 0 | 0.41 | 0.29 | 0.32 | 0.47 | 0.6 | 0.47 |
| 3 | 118909159 | rs9840317 | -0.386 | 0.089 | 0.856 | 1.445 | -0.015 | 0 | 0 | 0 | 0.93 | 0.89 | 0.92 | 0.93 | 0.97 | 0.87 |
| 3 | 118909538 | rs9878996 | 1.446 | 0.281 | 1.038 | 1.706 | -0.017 | 0.05 | 0.05 | 0 | 0.76 | 0.66 | 0.62 | 0.52 | 0.82 | 0.7 |
| 3 | 118909987 | rs3796357 | 0.073 | 0.286 | 0.973 | 1.684 | -0.039 | 0.007 | 0.007 | 0 | 0.91 | 0.78 | 0.91 | 0.93 | 0.97 | 0.87 |
| 3 | 118910184 | rs3796358 | 0.47 | 0.344 | 0.696 | 1.68 | -0.016 | -0.048 | -0.048 | 0 | 0.1 | 0.02 | 0.07 | 0.25 | 0.11 | 0.1 |
| 3 | 118913337 | rs7628485 | 0.092 | 0.362 | 0.745 | 1.527 | -0.06 | 0.014 | 0.014 | 0.013 | 0.3 | 0.17 | 0.24 | 0.22 | 0.5 | 0.38 |

| | | | | | | | | | | | | | | | | |
|---|---|---|---|---|---|---|---|---|---|---|---|---|---|---|---|---|
| 3 | 118916627 | rs12632242 | 2.01 | 0.306 | 0.648 | 1.417 | 0.04 | 0.091 | 0.091 | 1.603 | 0.4 | 0.08 | 0.31 | 0.22 | 0.3 | 0.17 |
| 3 | 118917422 | rs13315549 | 0.221 | 0.46 | 1.183 | 1.338 | 0.034 | 0.046 | 0.046 | 0.011 | 0.06 | 0.19 | 0.07 | 0.08 | 0.06 | 0.15 |
| 3 | 118926750 | rs11927939 | 0.118 | 0.415 | 1.087 | 1.358 | -0.029 | -0.001 | -0.001 | 0 | 0.08 | 0.12 | 0.06 | 0.06 | 0.03 | 0.15 |
| 3 | 118930884 | rs6774 | -0.739 | 0.491 | 0.979 | 1.338 | 0.033 | 0.015 | 0.015 | 0 | 0.09 | 0.26 | 0.28 | 0.07 | 0.09 | 0.18 |
| 3 | 118940383 | rs6766503 | 0.517 | 0.501 | 1.02 | 1.348 | 0.016 | 0.023 | 0.023 | 0 | 0.05 | 0.11 | 0.06 | 0.07 | 0.06 | 0.14 |
| 3 | 118940400 | rs3936077 | -1.243 | 0.516 | 0.99 | 1.453 | 0.014 | 0.044 | 0.044 | 0.437 | 0.19 | 0.3 | 0.35 | 0.33 | 0.2 | 0.31 |
| 3 | 118941029 | rs10511382 | -0.143 | 0.433 | 1.079 | 1.425 | -0.029 | 0.013 | 0.013 | 0.265 | 0.09 | 0.32 | 0.08 | 0.06 | 0.03 | 0.13 |
| 3 | 118941098 | rs4314161 | -2.177 | 0.4 | 0.436 | 1.443 | 0.045 | 0.071 | 0.071 | 1.813 | 0.61 | 0.9 | 0.68 | 0.76 | 0.7 | 0.85 |
| 3 | 118941441 | rs4528932 | -2.273 | 0.402 | 0.426 | 1.444 | 0.052 | 0.074 | 0.074 | 1.503 | 0.6 | 0.86 | 0.68 | 0.76 | 0.69 | 0.84 |
| 3 | 118945614 | rs3752376 | 0.458 | 0.527 | 0.021 | 1.401 | -0.003 | -0.018 | -0.018 | 0.31 | 0.1 | 0.05 | 0.07 | 0.26 | 0.11 | 0.12 |
| 3 | 118948256 | rs4568126 | -0.064 | 0.555 | -0.033 | 1.429 | -0.032 | -0.002 | -0.002 | 0 | 0.32 | 0.24 | 0.26 | 0.35 | 0.47 | 0.37 |
| 3 | 118953569 | rs4687841 | -1.713 | 0.631 | -0.241 | 1.379 | 0.01 | 0.007 | 0.007 | 0.244 | 0.56 | 0.9 | 0.64 | 0.68 | 0.64 | 0.71 |
| 3 | 118955888 | rs3806698 | 0.17 | 1.068 | 0.495 | 1.254 | 0.018 | -0.011 | -0.011 | 0.861 | 0.2 | 0.26 | 0.2 | 0.14 | 0.32 | 0.18 |
| 3 | 118962626 | rs9869645 | -0.91 | 0.733 | 0.473 | 1.3 | 0 | 0.007 | 0.007 | 0 | 0.14 | 0.18 | 0.29 | 0.26 | 0.14 | 0.17 |
| 3 | 118963980 | rs2594731 | 0.516 | 0.763 | 0.136 | 0.727 | 0.011 | 0.022 | 0.022 | 0 | 0.95 | 0.9 | 0.94 | 0.91 | 0.94 | 0.86 |
| 3 | 118979065 | rs2717265 | 0.687 | 0.586 | 0.149 | 0.477 | -0.001 | -0.006 | -0.006 | 0 | 0.84 | 0.81 | 0.9 | 0.9 | 0.85 | 0.85 |
| 3 | 118980188 | rs2971544 | 1.7 | 0.339 | 0.803 | 0.305 | -0.019 | -0.038 | -0.038 | 0 | 0.69 | 0.38 | 0.61 | 0.78 | 0.76 | 0.66 |
| 3 | 118982422 | rs2717263 | 1.882 | 0.298 | 1.134 | 0.513 | 0.016 | 0.011 | 0.011 | 0.639 | 0.68 | 0.24 | 0.61 | 0.71 | 0.77 | 0.6 |
| 3 | 118985011 | rs6781116 | -1.894 | 0.359 | 1.137 | 0.523 | -0.031 | -0.046 | -0.046 | 0.05 | 0.29 | 0.59 | 0.37 | 0.21 | 0.2 | 0.31 |
| 3 | 118993976 | rs16829641 | -1.049 | 0.362 | 1.279 | 0.525 | 0.036 | 0.092 | 0.092 | 0 | 0.07 | 0.23 | 0.23 | 0.14 | 0.07 | 0.12 |
| 3 | 118994546 | rs10511385 | -0.243 | 0.28 | 1.067 | 0.508 | -0.014 | -0.007 | -0.007 | 0 | 0.11 | 0.01 | 0.06 | 0.1 | 0.11 | 0.1 |
| 3 | 118996122 | rs6438515 | -1.672 | 0.249 | 1.235 | 0.481 | -0.013 | 0.04 | 0.04 | 0.008 | 0.14 | 0.55 | 0.09 | 0.18 | 0.08 | 0.25 |
| 3 | 119000376 | rs9827972 | -2.028 | 0.189 | 0.996 | 0.564 | -0.005 | -0.001 | -0.001 | 0 | 0.27 | 0.68 | 0.16 | 0.28 | 0.19 | 0.35 |
| 3 | 119023971 | rs7622994 | -2.249 | 0.215 | 0.875 | 0.431 | 0.007 | 0.008 | 0.008 | 0.097 | 0.3 | 0.72 | 0.4 | 0.22 | 0.21 | 0.35 |
| 3 | 119024937 | rs11918598 | -1.14 | 0.117 | 1.149 | 0.441 | -0.002 | -0.016 | -0.016 | 0 | 0.16 | 0.31 | 0.31 | 0.12 | 0.16 | 0.21 |
| 3 | 119027344 | rs7610546 | -0.307 | -0.069 | 0.351 | 0.177 | 0 | -0.005 | -0.005 | 0 | 0.4 | 0.47 | 0.28 | 0.35 | 0.41 | 0.52 |
| 3 | 119033157 | rs9834750 | -0.049 | -0.075 | 0.406 | 0.175 | 0.007 | -0.002 | -0.002 | 0 | 0.37 | 0.37 | 0.25 | 0.32 | 0.41 | 0.43 |
| 3 | 119034756 | rs7650193 | -1.887 | 0.203 | 0.249 | 0.184 | -0.067 | -0.145 | -0.145 | 0.867 | 0.63 | 0.96 | 0.65 | 0.48 | 0.64 | 0.76 |
| 3 | 119049071 | rs9845965 | -0.336 | -0.218 | 0.975 | 0.173 | 0.041 | -0.014 | -0.014 | 0.692 | 0.07 | 0.31 | 0.06 | 0.01 | 0.07 | 0.09 |
| 3 | 119050732 | rs2044621 | -2.172 | -0.312 | 0.763 | 0.295 | -0.014 | -0.013 | -0.013 | 0.001 | 0.29 | 0.66 | 0.37 | 0.24 | 0.23 | 0.38 |
| 3 | 119054711 | rs4687995 | -2.372 | -0.335 | 0.713 | 0.348 | -0.002 | -0.001 | -0.001 | 0 | 0.33 | 0.73 | 0.39 | 0.34 | 0.27 | 0.44 |
| 3 | 119055547 | rs9289128 | -0.478 | -0.353 | 0.788 | 0.328 | 0.008 | -0.048 | -0.048 | 1.717 | 0.1 | 0.41 | 0.06 | 0.02 | 0.07 | 0.18 |
| 3 | 119056040 | rs9835104 | -1.6 | -0.355 | 0.447 | 0.351 | -0.028 | -0.02 | -0.02 | 0 | 0.24 | 0.63 | 0.15 | 0.18 | 0.19 | 0.3 |
| 3 | 119060080 | rs12493383 | -1.763 | -0.34 | 0.311 | 0.373 | -0.007 | -0.002 | -0.002 | 0 | 0.31 | 0.71 | 0.38 | 0.3 | 0.25 | 0.4 |
| 3 | 119066079 | rs9860092 | -0.058 | -0.298 | -0.217 | 0.348 | 0.005 | 0.001 | 0.001 | 0 | 0.11 | 0.13 | 0.06 | 0.1 | 0.1 | 0.09 |
| 3 | 119066610 | rs7612016 | -1.851 | -0.555 | -0.732 | 0.027 | 0.057 | 0.006 | 0.006 | 0.79 | 0.49 | 0.83 | 0.45 | 0.42 | 0.38 | 0.65 |
| 3 | 119067735 | rs4687848 | -0.329 | -0.492 | -0.713 | 0.061 | -0.062 | -0.035 | -0.035 | 0 | 0.19 | 0.41 | 0.09 | 0.11 | 0.1 | 0.31 |
| 3 | 119089770 | rs10934492 | -0.363 | -0.106 | -1.069 | 0.192 | 0.004 | 0.011 | 0.011 | 0 | 0.2 | 0.2 | 0.28 | 0.11 | 0.14 | 0.29 |
| 3 | 119090325 | rs4687998 | -0.27 | 0.313 | -1.324 | 0.415 | -0.026 | -0.03 | -0.03 | 0 | 0.35 | 0.5 | 0.38 | 0.2 | 0.23 | 0.48 |
| 3 | 119095812 | rs17203069 | -0.322 | 1.22 | -1.858 | 0.624 | -0.07 | 0.068 | 0.068 | 0 | 0.21 | 0.07 | 0.28 | 0.06 | 0.24 | 0.19 |
| 3 | 119099049 | rs9837853 | 0.209 | 0.839 | -1.173 | 1.419 | 0.002 | 0.038 | 0.038 | 0 | 0.81 | 0.85 | 0.68 | 0.43 | 0.75 | 0.76 |
| 3 | 119107277 | rs13078855 | -0.552 | 0.286 | -1.634 | 1.434 | 0.018 | -0.048 | -0.048 | 0 | 0.19 | 0.11 | 0.09 | 0.33 | 0.16 | 0.19 |
| 3 | 119110006 | rs2241994 | -0.001 | -0.013 | -2.508 | 1.12 | 0 | -0.049 | -0.049 | 0 | 0.24 | 0.02 | 0.08 | 0.23 | 0.14 | 0.21 |
| 3 | 119110957 | rs2873688 | -0.81 | 0.872 | -2.582 | 0.378 | -0.016 | 0.026 | 0.026 | 0 | 0.8 | 0.87 | 0.69 | 0.9 | 0.76 | 0.85 |
| 3 | 119111762 | rs9831023 | -0.08 | 0.676 | -1.869 | 0.904 | -0.013 | 0.043 | 0.043 | 0.187 | 0.28 | 0.61 | 0.19 | 0.47 | 0.24 | 0.33 |
| 3 | 119111783 | rs2121994 | 0.996 | 1.326 | -2.147 | 0.758 | -0.036 | -0.03 | -0.03 | 0 | 0.06 | 0.01 | 0.2 | 0.11 | 0.11 | 0.07 |
| 3 | 119112559 | rs935616 | -0.548 | 0.928 | -2.22 | 0.466 | -0.02 | 0.058 | 0.058 | 0.073 | 0.6 | 0.72 | 0.47 | 0.82 | 0.51 | 0.64 |
| 3 | 119112898 | rs2029034 | 0.136 | 0.943 | -1.39 | 0.976 | 0 | 0.012 | 0.012 | 0 | 0.39 | 0.5 | 0.36 | 0.47 | 0.35 | 0.4 |
| 3 | 119113762 | rs1000199 | 1.142 | 0.967 | -1.589 | 0.994 | -0.036 | -0.033 | -0.033 | 0 | 0.06 | 0.01 | 0.2 | 0.12 | 0.1 | 0.07 |
| 3 | 119113820 | rs1000198 | 0.396 | 1.354 | -1.54 | 0.672 | -0.02 | -0.01 | -0.01 | 0 | 0.46 | 0.23 | 0.51 | 0.45 | 0.49 | 0.48 |
| 3 | 119118497 | rs11715698 | 0.933 | 1.298 | -1.647 | 0.556 | -0.098 | -0.076 | -0.076 | 0 | 0.33 | 0.05 | 0.39 | 0.35 | 0.39 | 0.36 |
| 3 | 119118796 | rs11712165 | 0.993 | 1.268 | -1.665 | 0.553 | -0.133 | -0.105 | -0.105 | 0 | 0.32 | 0.03 | 0.39 | 0.34 | 0.39 | 0.3 |
| 3 | 119122820 | rs12494314 | 1.269 | 0.956 | -1.54 | 0.589 | 0.019 | -0.044 | -0.044 | 0 | 0.2 | 0.09 | 0.1 | 0.38 | 0.16 | 0.23 |
| 3 | 119123344 | rs6772534 | 0.975 | 0.838 | -0.936 | 0.518 | -0.023 | -0.038 | -0.038 | 0 | 0.17 | 0.3 | 0.15 | 0.1 | 0.15 | 0.22 |
| 3 | 119123814 | rs6773050 | 0.259 | 0.851 | -0.936 | 0.52 | 0 | 0.015 | 0.015 | 0 | 0.49 | 0.33 | 0.54 | 0.44 | 0.53 | 0.52 |
| 3 | 119123999 | rs17281612 | 1.666 | 0.806 | -0.836 | 0.508 | 0.003 | 0.007 | 0.007 | 0 | 0.14 | 0.11 | 0.13 | 0.09 | 0.15 | 0.16 |
| 3 | 119128398 | rs2305249 | 1.246 | 0.797 | -0.836 | 0.541 | 0.02 | -0.042 | -0.042 | 0 | 0.2 | 0.09 | 0.09 | 0.37 | 0.16 | 0.23 |
| 3 | 119129803 | rs11714363 | 0.784 | 0.835 | -0.832 | 0.542 | 0.015 | 0.012 | 0.012 | 0 | 0.41 | 0.32 | 0.31 | 0.32 | 0.37 | 0.4 |
| 3 | 119132554 | rs4687855 | -1.832 | 0.967 | -0.676 | 0.488 | -0.027 | 0.014 | 0.014 | 0 | 0.8 | 0.84 | 0.75 | 0.85 | 0.78 | 0.86 |
| 3 | 119133183 | rs3732413 | -1.929 | 0.945 | -0.7 | 0.478 | -0.055 | 0.009 | 0.009 | 0 | 0.81 | 0.93 | 0.76 | 0.85 | 0.79 | 0.86 |
| 3 | 119137466 | rs12631504 | 1.582 | 0.829 | -0.689 | 0.477 | 0.003 | 0.005 | 0.005 | 0 | 0.14 | 0.13 | 0.13 | 0.09 | 0.15 | 0.16 |
| 3 | 119141369 | rs11706042 | 2.211 | 0.785 | -0.852 | 0.473 | -0.041 | -0.069 | -0.069 | 0 | 0.23 | 0.01 | 0.16 | 0.22 | 0.22 | 0.13 |
| 3 | 119141643 | rs10511391 | 1.394 | 0.896 | -0.896 | 0.486 | -0.054 | -0.036 | -0.036 | 3.356 | 0.09 | 0.01 | 0.23 | 0.13 | 0.17 | 0.13 |
| 3 | 119145512 | rs6763561 | 1.064 | 0.834 | -0.84 | 0.471 | 0.036 | 0.013 | 0.013 | 1.83 | 0.44 | 0.18 | 0.26 | 0.59 | 0.37 | 0.37 |
| 3 | 119150089 | rs3732421 | 1.434 | 0.826 | -0.766 | 0.474 | 0.02 | -0.046 | -0.046 | 0 | 0.2 | 0.08 | 0.09 | 0.36 | 0.16 | 0.24 |
| 3 | 119150836 | rs1132200 | 1.841 | 0.723 | -0.93 | 0.566 | -0.006 | 0.018 | 0.018 | 0.037 | 0.16 | 0.01 | 0.07 | 0.07 | 0.14 | 0.17 |
| 3 | 119157615 | rs3195852 | 0.803 | 0.776 | -0.879 | 0.406 | 0.026 | 0.031 | 0.031 | 0.862 | 0.48 | 0.59 | 0.31 | 0.62 | 0.4 | 0.45 |
| 3 | 119161982 | rs7430156 | -0.004 | 0.784 | -0.857 | 0.402 | -0.009 | 0.062 | 0.062 | 0 | 0.57 | 0.64 | 0.55 | 0.76 | 0.56 | 0.64 |
| 3 | 119165650 | rs16829853 | 1.485 | 0.792 | -0.85 | 0.413 | -0.054 | -0.036 | -0.036 | 0.189 | 0.09 | 0.01 | 0.23 | 0.13 | 0.17 | 0.13 |

| | | | | | | | | | | | | | | | | |
|---|---|---|---|---|---|---|---|---|---|---|---|---|---|---|---|---|
| 3 | 119182259 | rs2282175 | 1.793 | 0.8 | -0.909 | 0.405 | 0.004 | 0.009 | 0.009 | 0 | 0.14 | 0.12 | 0.13 | 0.09 | 0.15 | 0.16 |
| 3 | 119190976 | rs12493965 | 1.253 | 0.84 | -0.925 | 0.414 | -0.012 | -0.006 | -0.006 | 0.035 | 0.09 | 0.1 | 0.24 | 0.12 | 0.17 | 0.13 |
| 3 | 119196166 | rs3732420 | 1.428 | 0.84 | -0.927 | 0.414 | -0.054 | -0.036 | -0.036 | 0.133 | 0.09 | 0.01 | 0.23 | 0.12 | 0.17 | 0.13 |
| 3 | 119196334 | rs3732419 | 0.019 | 0.829 | -0.935 | 0.399 | -0.037 | 0.036 | 0.036 | 0.021 | 0.62 | 0.73 | 0.45 | 0.72 | 0.54 | 0.67 |
| 3 | 119197285 | rs4688005 | -1.582 | 0.828 | -0.961 | 0.376 | -0.008 | 0.008 | 0.008 | 0 | 0.79 | 0.79 | 0.74 | 0.85 | 0.78 | 0.86 |
| 3 | 119198873 | rs4688007 | -1.721 | 0.832 | -1.022 | 0.378 | -0.024 | 0.02 | 0.02 | 0 | 0.79 | 0.88 | 0.75 | 0.85 | 0.78 | 0.86 |
| 3 | 119201144 | rs6780684 | 0.44 | 0.792 | -0.975 | 0.387 | 0.041 | 0.051 | 0.051 | 0 | 0.5 | 0.73 | 0.42 | 0.36 | 0.46 | 0.49 |
| 3 | 119211294 | rs1997341 | 1.344 | 0.893 | -1.02 | 0.431 | -0.054 | -0.036 | -0.036 | 0 | 0.09 | 0.01 | 0.23 | 0.12 | 0.17 | 0.13 |
| 3 | 119214549 | rs3755579 | 1.421 | 0.84 | -0.56 | 0.335 | -0.015 | -0.005 | -0.005 | 0 | 0.32 | 0.05 | 0.23 | 0.23 | 0.28 | 0.24 |
| 3 | 119216554 | rs2049500 | 0.83 | 0.82 | -0.471 | 0.328 | -0.001 | -0.001 | -0.001 | 0 | 0.37 | 0.61 | 0.29 | 0.26 | 0.31 | 0.34 |
| 3 | 119217312 | rs2280670 | 2.291 | 0.81 | -0.482 | 0.318 | -0.031 | -0.063 | -0.063 | 0 | 0.24 | 0.02 | 0.16 | 0.23 | 0.21 | 0.17 |
| 3 | 119219573 | rs11539377 | 0.656 | 0.812 | -0.481 | 0.305 | 0.06 | 0.056 | 0.056 | 0.908 | 0.06 | 0.56 | 0.06 | 0.03 | 0.02 | 0.1 |
| 3 | 119233558 | rs9869760 | -0.71 | 1.313 | -1.084 | 0.179 | 0.008 | 0.018 | 0.018 | 0 | 0.49 | 0.37 | 0.5 | 0.6 | 0.56 | 0.52 |
| 3 | 119235102 | rs12695388 | -0.319 | 1.312 | -1.077 | 0.161 | -0.014 | -0.003 | -0.003 | 0 | 0.42 | 0.23 | 0.46 | 0.59 | 0.48 | 0.51 |
| 3 | 119242930 | rs11916272 | 0.771 | 1.297 | -1.062 | 0.231 | 0.06 | 0.115 | 0.115 | 0.908 | 0.06 | 0.56 | 0.06 | 0.06 | 0.02 | 0.1 |
| 3 | 119243152 | rs11916344 | 0.771 | 1.297 | -1.062 | 0.222 | 0.06 | 0.056 | 0.056 | 0.908 | 0.06 | 0.56 | 0.06 | 0.03 | 0.02 | 0.1 |
| 3 | 119243549 | rs17281703 | 2.136 | 1.29 | -1.162 | 0.249 | -0.002 | -0.002 | -0.002 | 0 | 0.13 | 0.04 | 0.13 | 0.11 | 0.14 | 0.16 |
| 3 | 119243934 | rs1599796 | 2.384 | 1.294 | -1.081 | 0.176 | -0.036 | -0.064 | -0.064 | 0 | 0.24 | 0.01 | 0.16 | 0.23 | 0.21 | 0.17 |
| 3 | 119244421 | rs7628626 | -1.178 | 1.488 | -0.748 | 0.09 | -0.007 | 0.01 | 0.01 | 0 | 0.8 | 0.84 | 0.75 | 0.85 | 0.81 | 0.85 |
| 3 | 119245957 | rs10934493 | 0.38 | 1.488 | -0.741 | 0.093 | 0.008 | -0.026 | -0.026 | 0 | 0.06 | 0.15 | 0.04 | 0.01 | 0.06 | 0.01 |
| 3 | 119246827 | rs6778945 | -2.167 | 1.472 | -0.852 | 0.119 | -0.029 | -0.014 | -0.014 | 0.001 | 0.86 | 0.97 | 0.79 | 0.86 | 0.86 | 0.86 |
| 3 | 119250890 | rs4688014 | 0.401 | 1.459 | -0.812 | 0.088 | -0.003 | 0.044 | 0.044 | 0.313 | 0.66 | 0.24 | 0.61 | 0.82 | 0.69 | 0.67 |
| 3 | 119251785 | rs610902 | -0.086 | 1.382 | -0.997 | 0.164 | -0.034 | 0.023 | 0.023 | 0 | 0.68 | 0.8 | 0.51 | 0.74 | 0.63 | 0.71 |
| 3 | 119257079 | rs624035 | -0.416 | 0.823 | -1.1 | 0.213 | -0.001 | -0.003 | -0.003 | 0 | 0.67 | 0.72 | 0.66 | 0.78 | 0.71 | 0.63 |
| 3 | 119257188 | rs491407 | -1.751 | 0.845 | -1.132 | 0.37 | 0 | -0.004 | -0.004 | 0 | 0.71 | 0.83 | 0.8 | 0.73 | 0.74 | 0.79 |
| 3 | 119260944 | rs6804441 | 0.829 | 0.94 | -1.108 | 0.379 | 0.014 | -0.046 | -0.046 | 3.509 | 0.18 | 0.11 | 0.09 | 0.35 | 0.15 | 0.24 |
| 3 | 119261375 | rs7648642 | -0.794 | 0.765 | -1.189 | 0.344 | 0.01 | 0.021 | 0.021 | 0 | 0.39 | 0.55 | 0.45 | 0.51 | 0.45 | 0.42 |
| 3 | 119263680 | rs2228017 | 1.749 | 0.726 | -1.165 | 0.4 | -0.001 | -0.004 | -0.004 | 0 | 0.29 | 0.17 | 0.21 | 0.27 | 0.26 | 0.21 |
| 3 | 119267302 | rs16829957 | -0.43 | 0.768 | -1.097 | 0.485 | -0.001 | 0.068 | 0.068 | 0 | 0.32 | 0.53 | 0.36 | 0.5 | 0.3 | 0.37 |
| 3 | 119269112 | rs527004 | -0.665 | 0.782 | -1.665 | 0.087 | 0.01 | 0.035 | 0.035 | 0 | 0.71 | 0.86 | 0.69 | 0.85 | 0.73 | 0.7 |
| 3 | 119272883 | rs626364 | 0.252 | 0.386 | -1.572 | 0.466 | -0.006 | 0.049 | 0.049 | 0 | 0.56 | 0.75 | 0.38 | 0.63 | 0.45 | 0.52 |
| 3 | 119274749 | rs1852211 | -0.722 | 0.114 | -1.179 | 0.425 | -0.012 | 0.016 | 0.016 | 0.154 | 0.15 | 0.13 | 0.27 | 0.06 | 0.21 | 0.18 |
| 3 | 119274841 | rs6807532 | -0.076 | -0.958 | -0.854 | 0.625 | -0.002 | -0.023 | -0.023 | 0.507 | 0.06 | 0.12 | 0.05 | 0 | 0.09 | 0.02 |
| 3 | 119275098 | rs6810204 | -0.413 | -1.195 | -0.82 | 1.161 | -0.034 | 0.084 | 0.084 | 0 | 0.17 | 0.29 | 0.09 | 0.33 | 0.1 | 0.26 |
| 3 | 119277846 | rs6438537 | -0.307 | -0.998 | -1.167 | 0.829 | -0.028 | 0.024 | 0.024 | 0 | 0.21 | 0.28 | 0.1 | 0.29 | 0.12 | 0.31 |
| 3 | 119278848 | rs1880661 | -1.106 | -1.041 | -1.369 | 0.76 | -0.076 | 0.093 | 0.093 | 0 | 0.63 | 0.88 | 0.39 | 0.78 | 0.42 | 0.67 |
| 3 | 119279281 | rs3915166 | -0.181 | -1.751 | -1.279 | 0.812 | -0.029 | 0.022 | 0.022 | 0 | 0.22 | 0.29 | 0.1 | 0.29 | 0.12 | 0.31 |
| 3 | 119288848 | rs4330287 | -0.1 | -1.696 | -1.378 | 0.836 | -0.024 | 0.021 | 0.021 | 0 | 0.21 | 0.26 | 0.1 | 0.29 | 0.12 | 0.3 |
| 3 | 119291419 | rs12487334 | -0.093 | -1.688 | -1.751 | 0.769 | -0.012 | 0.007 | 0.007 | 0 | 0.21 | 0.21 | 0.1 | 0.28 | 0.11 | 0.3 |
| 3 | 119295128 | rs529650 | -0.588 | -0.651 | -2.34 | 0.832 | -0.035 | 0.092 | 0.092 | 0 | 0.63 | 0.87 | 0.38 | 0.78 | 0.36 | 0.67 |
| 3 | 119300655 | rs1485331 | -0.455 | -0.076 | -1.212 | 1.488 | -0.004 | 0.015 | 0.015 | 0 | 0.11 | 0.13 | 0.05 | 0.26 | 0.06 | 0.22 |
| 3 | 119305075 | rs2242494 | -0.015 | 0.061 | -1.497 | 1.163 | -0.003 | -0.051 | -0.051 | 0.798 | 0.17 | 0.02 | 0.13 | 0.23 | 0.11 | 0.19 |
| 3 | 119309260 | rs538136 | -0.611 | -0.505 | -0.347 | 1.406 | -0.03 | 0.143 | 0.143 | 0 | 0.22 | 0.51 | 0.19 | 0.44 | 0.18 | 0.32 |
| 3 | 119312613 | rs2171025 | -0.536 | -0.702 | 0.283 | 1.299 | -0.005 | 0.271 | 0.271 | 0 | 0.15 | 0.36 | 0.14 | 0.44 | 0.16 | 0.27 |
| 3 | 119321563 | rs2688638 | -1.489 | -0.511 | 0.409 | 1.069 | -0.027 | 0.096 | 0.096 | 0 | 0.22 | 0.6 | 0.2 | 0.44 | 0.18 | 0.31 |
| 3 | 119325907 | rs2251566 | -0.752 | -0.453 | 0.478 | 1.137 | -0.003 | 0.312 | 0.312 | 0 | 0.16 | 0.42 | 0.15 | 0.45 | 0.16 | 0.27 |
| 3 | 119334330 | rs579809 | -1.312 | -0.567 | 0.698 | 1.573 | 0.027 | 0.234 | 0.234 | 0.878 | 0.08 | 0.47 | 0.07 | 0.15 | 0.03 | 0.17 |
| 3 | 119334986 | rs2247660 | -1.009 | -0.507 | -0.522 | 0.604 | -0.046 | 0.155 | 0.155 | 0 | 0.57 | 0.83 | 0.6 | 0.9 | 0.49 | 0.62 |
| 3 | 119344693 | rs483349 | -1.066 | -0.573 | -0.542 | 0.559 | -0.059 | 0.173 | 0.173 | 0 | 0.58 | 0.86 | 0.6 | 0.9 | 0.49 | 0.62 |
| 3 | 119347400 | rs2280580 | -0.496 | -0.869 | 0.059 | 0.965 | 0.004 | 0.13 | 0.13 | 0 | 0.11 | 0.26 | 0.12 | 0.34 | 0.13 | 0.09 |
| 3 | 119348586 | rs12023 | -0.108 | -1.519 | -0.67 | 0.86 | 0.05 | -0.045 | -0.045 | 0 | 0.33 | 0.14 | 0.15 | 0.41 | 0.22 | 0.43 |
| 3 | 119353844 | rs2692618 | -1.61 | -1.68 | 0.426 | 0.614 | -0.054 | -0.003 | -0.003 | 0 | 0.69 | 0.83 | 0.59 | 0.67 | 0.56 | 0.69 |
| 3 | 119353893 | rs16830097 | 0.359 | -1.644 | 0.611 | -0.017 | 0.008 | -0.033 | -0.033 | 0 | 0.08 | 0.13 | 0.09 | 0 | 0.11 | 0.03 |
| 3 | 119355411 | rs17203235 | -0.316 | -1.857 | 0.529 | -0.124 | -0.027 | -0.018 | -0.018 | 0 | 0.28 | 0.15 | 0.39 | 0.25 | 0.34 | 0.21 |
| 3 | 119359045 | rs571352 | -1.918 | -1.934 | 0.514 | -0.131 | -0.047 | -0.034 | -0.034 | 0 | 0.69 | 0.82 | 0.59 | 0.34 | 0.57 | 0.64 |
| 3 | 119360867 | rs6792223 | 0.952 | -2.019 | 0.538 | -0.176 | 0.063 | 0.132 | 0.132 | 0 | 0.3 | 0.07 | 0.12 | 0.09 | 0.19 | 0.38 |
| 3 | 119361250 | rs8007 | -0.953 | -2.004 | 0.993 | -0.169 | 0.007 | 0 | 0 | 0.993 | 0.6 | 0.51 | 0.48 | 0.34 | 0.46 | 0.61 |
| 3 | 119363566 | rs4688021 | -0.142 | -1.885 | 1.079 | -0.093 | -0.013 | -0.025 | -0.025 | 0 | 0.26 | 0.15 | 0.32 | 0.25 | 0.27 | 0.23 |
| 3 | 119369652 | rs2670292 | 0.283 | -1.889 | 0.874 | -0.091 | 0 | 0.005 | 0.005 | 0 | 0.73 | 0.65 | 0.66 | 0.75 | 0.73 | 0.77 |
| 3 | 119373315 | rs2688629 | 1.433 | -1.754 | 0.784 | -0.01 | -0.002 | 0 | 0 | 0 | 0.66 | 0.35 | 0.54 | 0.75 | 0.63 | 0.74 |
| 3 | 119377511 | rs11914676 | -0.094 | -1.751 | 0.766 | 0.003 | -0.01 | -0.017 | -0.017 | 0.473 | 0.18 | 0.1 | 0.32 | 0.24 | 0.24 | 0.17 |
| 3 | 119378872 | rs1880040 | -0.459 | -1.893 | 0.762 | -0.079 | 0.027 | 0.033 | 0.033 | 0.17 | 0.49 | 0.29 | 0.46 | 0.33 | 0.43 | 0.55 |
| 3 | 119378977 | rs2873788 | 1.573 | -2.121 | 0.625 | -0.211 | 0.032 | 0.143 | 0.143 | 0 | 0.22 | 0.02 | 0.1 | 0 | 0.16 | 0.17 |
| 3 | 119379186 | rs4688023 | -0.408 | -1.807 | 0.807 | -0.017 | 0.027 | 0.032 | 0.032 | 0.17 | 0.49 | 0.29 | 0.46 | 0.33 | 0.43 | 0.55 |
| 3 | 119379352 | rs4688024 | 0.452 | -1.777 | 0.748 | 0.005 | -0.053 | -0.072 | -0.072 | 0 | 0.16 | 0.03 | 0.31 | 0.24 | 0.24 | 0.17 |
| 3 | 119383278 | rs16830286 | 0.954 | -1.647 | 0.691 | 0.109 | -0.017 | -0.026 | -0.026 | 0 | 0.06 | 0 | 0.07 | 0.18 | 0.08 | 0.13 |
| 3 | 119388666 | rs3772134 | 1.685 | -1.712 | 0.788 | 0.098 | -0.049 | -0.03 | -0.03 | 0 | 0.35 | 0.19 | 0.41 | 0.66 | 0.43 | 0.37 |
| 3 | 119390418 | rs6807079 | 1.667 | -1.718 | 0.76 | 0.102 | -0.039 | -0.024 | -0.024 | 1.442 | 0.35 | 0.22 | 0.41 | 0.66 | 0.43 | 0.37 |
| 3 | 119391423 | rs6803814 | -0.843 | -1.715 | 0.416 | 0.16 | -0.016 | -0.011 | -0.011 | 2.061 | 0.25 | 0.17 | 0.4 | 0.24 | 0.35 | 0.2 |

| | | | | | | | | | | | | | | | | |
|---|---|---|---|---|---|---|---|---|---|---|---|---|---|---|---|---|
| 3 | 119392101 | rs13323756 | 0.723 | -1.723 | 0.437 | 0.154 | 0.044 | 0.061 | 0.061 | 0 | 0.32 | 0.19 | 0.14 | 0.09 | 0.19 | 0.38 |
| 3 | 119395799 | rs2280578 | -0.948 | -1.46 | 0.204 | 0.319 | -0.015 | -0.011 | -0.011 | 0 | 0.25 | 0.18 | 0.4 | 0.24 | 0.35 | 0.15 |
| 3 | 119403145 | rs6809361 | 0.69 | -1.549 | 0.465 | 0.366 | 0 | 0 | 0 | 0 | 0.46 | 0.51 | 0.68 | 0.69 | 0.57 | 0.41 |
| 3 | 119404233 | rs34775086 | 0.624 | -1.673 | -0.224 | 0.014 | 0.074 | 0.124 | 0.124 | 0.012 | 0.32 | 0.07 | 0.12 | 0.1 | 0.2 | 0.37 |
| 3 | 119407403 | rs2137619 | -0.871 | -1.591 | -0.184 | 0.156 | -0.046 | -0.017 | -0.017 | 0 | 0.27 | 0.17 | 0.39 | 0.23 | 0.36 | 0.24 |
| 3 | 119411243 | rs11720605 | 1.551 | -1.707 | -0.007 | 0.084 | -0.052 | -0.028 | -0.028 | 1.659 | 0.36 | 0.15 | 0.42 | 0.66 | 0.44 | 0.38 |
| 3 | 119413445 | rs4585237 | -0.784 | -1.012 | 0.423 | 0.238 | -0.039 | -0.026 | -0.026 | 0 | 0.22 | 0.15 | 0.38 | 0.23 | 0.33 | 0.2 |
| 3 | 119413823 | rs6766563 | 1.303 | -1.006 | 0.67 | 0.262 | 0.019 | -0.002 | -0.002 | 0 | 0.7 | 0.43 | 0.57 | 0.76 | 0.64 | 0.76 |
| 3 | 119414604 | rs7626940 | -1.068 | -0.839 | -0.364 | 0.345 | -0.037 | -0.016 | -0.016 | 0.667 | 0.27 | 0.19 | 0.39 | 0.24 | 0.36 | 0.24 |
| 3 | 119418531 | rs7617287 | -0.558 | -0.963 | -0.058 | 0.389 | 0.002 | 0.026 | 0.026 | 0 | 0.14 | 0.17 | 0.24 | 0.18 | 0.15 | 0.2 |
| 3 | 119420914 | rs13082557 | -0.537 | -0.962 | -0.058 | 0.389 | 0.003 | 0.027 | 0.027 | 0 | 0.14 | 0.17 | 0.24 | 0.18 | 0.15 | 0.2 |
| 3 | 119422106 | rs7612954 | -0.581 | -0.976 | -0.065 | 0.382 | 0.002 | 0.025 | 0.025 | 0 | 0.14 | 0.17 | 0.23 | 0.18 | 0.15 | 0.2 |
| 3 | 119424822 | rs4688028 | -0.178 | -0.984 | -0.074 | 0.378 | -0.007 | -0.019 | -0.019 | 0 | 0.12 | 0.06 | 0.22 | 0.18 | 0.15 | 0.2 |
| 3 | 119426307 | rs9848716 | 1.893 | -1.488 | -0.041 | -0.057 | 0.003 | 0.014 | 0.014 | 0 | 0.7 | 0.18 | 0.64 | 0.81 | 0.73 | 0.72 |
| 3 | 119430289 | rs7638068 | -0.136 | -1.513 | 0.85 | -0.086 | 0.016 | -0.055 | -0.055 | 0 | 0.1 | 0.26 | 0.1 | 0 | 0.11 | 0.03 |
| 3 | 119433300 | rs10934496 | 1.048 | -1.506 | 0.88 | -0.058 | -0.01 | 0.025 | 0.025 | 0 | 0.8 | 0.46 | 0.73 | 0.81 | 0.84 | 0.75 |
| 3 | 119437131 | rs6763266 | -0.213 | -1.925 | 0.136 | 0.042 | 0.019 | 0.006 | 0.006 | 0 | 0.42 | 0.35 | 0.29 | 0.41 | 0.39 | 0.39 |
| 3 | 119438608 | rs6778570 | 0.761 | -2.086 | 0.547 | 0.083 | -0.011 | 0.024 | 0.024 | 0 | 0.79 | 0.45 | 0.73 | 0.81 | 0.84 | 0.75 |
| 3 | 119442572 | rs9830785 | 0.209 | -1.941 | 0.731 | 0.094 | 0.03 | -0.008 | -0.008 | 0.102 | 0.36 | 0.26 | 0.24 | 0.4 | 0.28 | 0.32 |
| 3 | 119445735 | rs11711386 | 2.095 | -2.42 | 0.838 | 0.052 | -0.192 | 0.041 | 0.041 | 0.084 | 0.33 | 0.03 | 0.43 | 0.19 | 0.48 | 0.26 |
| 3 | 119446441 | rs9841230 | 0.5 | -2.454 | 0.987 | 0.061 | 0.031 | -0.008 | -0.008 | 0.321 | 0.36 | 0.26 | 0.23 | 0.4 | 0.28 | 0.32 |
| 3 | 119458851 | rs4688033 | 0.194 | -2.453 | 0.863 | 0.091 | 0.024 | 0.028 | 0.028 | 0.028 | 0.17 | 0.28 | 0.29 | 0.2 | 0.21 | 0.24 |
| 3 | 119469439 | rs2472662 | 2.173 | -2.468 | 0.824 | 0.085 | -0.193 | 0.039 | 0.039 | 0.102 | 0.33 | 0.03 | 0.43 | 0.19 | 0.48 | 0.26 |
| 3 | 119471473 | rs1464599 | 0.126 | -2.405 | 0.877 | 0.017 | 0.022 | 0.026 | 0.026 | 0.019 | 0.17 | 0.26 | 0.29 | 0.2 | 0.21 | 0.24 |
| 3 | 119473719 | rs9289134 | -1.47 | -2.421 | 0.876 | 0.01 | -0.001 | 0 | 0 | 0 | 0.27 | 0.76 | 0.34 | 0.2 | 0.23 | 0.31 |
| 3 | 119478763 | rs9815093 | -1.47 | -2.421 | 0.876 | 0.01 | -0.001 | 0 | 0 | 0 | 0.27 | 0.76 | 0.34 | 0.2 | 0.23 | 0.31 |
| 3 | 119479987 | rs4687882 | -0.889 | -2.393 | 0.803 | 0.023 | 0.033 | 0.027 | 0.027 | 0 | 0.23 | 0.51 | 0.32 | 0.2 | 0.23 | 0.31 |
| 3 | 119483231 | rs9682652 | 0.113 | -2.345 | 0.647 | 0.048 | 0.023 | 0.027 | 0.027 | 0.019 | 0.17 | 0.28 | 0.3 | 0.2 | 0.22 | 0.24 |
| 3 | 119483267 | rs11712308 | 1.486 | -2.378 | 0.598 | 0.042 | -0.189 | -0.066 | -0.066 | 0.455 | 0.38 | 0.03 | 0.44 | 0.4 | 0.49 | 0.37 |
| 3 | 119484601 | rs10511394 | -0.175 | -1.971 | 1.162 | 0.046 | 0.023 | 0.027 | 0.027 | 0.019 | 0.17 | 0.27 | 0.29 | 0.2 | 0.21 | 0.24 |
| 3 | 119495499 | rs1581451 | 0.77 | -2.655 | 0.739 | -0.384 | 0.017 | 0.031 | 0.031 | 0 | 0.58 | 0.32 | 0.57 | 0.78 | 0.63 | 0.6 |
| 3 | 119497045 | rs7643038 | 0.763 | -2.657 | 0.738 | -0.385 | 0.016 | 0.03 | 0.03 | 0 | 0.58 | 0.32 | 0.57 | 0.78 | 0.63 | 0.6 |
| 3 | 119499507 | rs1523130 | 1.212 | -2.813 | 0.052 | -0.48 | 0.041 | 0.128 | 0.128 | 0 | 0.52 | 0.07 | 0.55 | 0.78 | 0.62 | 0.55 |
| 3 | 119500035 | rs3814055 | -0.357 | -2.707 | 0.111 | -0.409 | 0.009 | 0.026 | 0.026 | 0 | 0.38 | 0.31 | 0.4 | 0.22 | 0.37 | 0.35 |
| 3 | 119501039 | rs1523127 | 0.888 | -2.639 | 0.099 | -0.402 | 0.049 | 0.101 | 0.101 | 0.129 | 0.55 | 0.12 | 0.55 | 0.78 | 0.63 | 0.6 |
| 3 | 119503005 | rs2472670 | -0.985 | -2.253 | -0.366 | -0.539 | -0.053 | -0.119 | -0.119 | 0.934 | 0.77 | 0.38 | 0.86 | 1 | 0.86 | 0.85 |
| 3 | 119503045 | rs2472671 | -0.985 | -2.253 | -0.366 | -0.539 | -0.053 | -0.119 | -0.119 | 0.934 | 0.77 | 0.38 | 0.86 | 1 | 0.86 | 0.85 |
| 3 | 119503971 | rs13071341 | -0.976 | -2.249 | -0.39 | -0.537 | -0.053 | -0.118 | -0.118 | 0.04 | 0.77 | 0.38 | 0.86 | 1 | 0.86 | 0.85 |
| 3 | 119504140 | rs11712211 | -0.976 | -2.192 | -0.39 | -0.537 | -0.051 | -0.118 | -0.118 | 0 | 0.77 | 0.38 | 0.86 | 1 | 0.86 | 0.85 |
| 3 | 119504381 | rs10934498 | 1.151 | -2.252 | -0.402 | -0.526 | 0.072 | 0.146 | 0.146 | 0 | 0.5 | 0.13 | 0.55 | 0.78 | 0.61 | 0.55 |
| 3 | 119506462 | rs2056530 | -0.363 | -2.058 | 0.19 | -0.575 | -0.053 | -0.118 | -0.118 | 0 | 0.77 | 0.38 | 0.86 | 1 | 0.86 | 0.85 |
| 3 | 119506744 | rs2873951 | -0.363 | -2.054 | 0.19 | -0.575 | -0.051 | -0.118 | -0.118 | 0.149 | 0.77 | 0.38 | 0.86 | 1 | 0.86 | 0.85 |
| 3 | 119506880 | rs1403527 | -0.363 | -2.051 | 0.19 | -0.575 | -0.051 | -0.118 | -0.118 | 0 | 0.77 | 0.38 | 0.86 | 1 | 0.86 | 0.85 |
| 3 | 119509381 | rs1403526 | -0.886 | -1.116 | -0.085 | 0.155 | -0.01 | 0.002 | 0.002 | 0 | 0.36 | 0.45 | 0.45 | 0.38 | 0.26 | 0.36 |
| 3 | 119512832 | rs6773295 | -0.117 | -1.12 | -0.006 | 0.073 | -0.021 | -0.036 | -0.036 | 0 | 0.08 | 0.15 | 0.02 | 0.01 | 0.02 | 0.04 |
| 3 | 119514556 | rs1357459 | -0.204 | -1.316 | -0.159 | -0.019 | -0.057 | -0.1 | -0.1 | 0.224 | 0.26 | 0.04 | 0.42 | 0.36 | 0.26 | 0.34 |
| 3 | 119514628 | rs4687885 | -0.815 | -1.256 | 0.144 | 0.063 | -0.037 | -0.054 | -0.054 | 0.763 | 0.1 | 0.33 | 0.03 | 0.01 | 0.02 | 0.04 |
| 3 | 119514789 | rs4687886 | -0.872 | -1.257 | 0.143 | 0.055 | -0.039 | -0.061 | -0.061 | 0.289 | 0.1 | 0.33 | 0.03 | 0.01 | 0.02 | 0.04 |
| 3 | 119515585 | rs4687887 | -0.765 | -1.296 | 0.14 | 0.058 | -0.029 | -0.051 | -0.051 | 0 | 0.09 | 0.32 | 0.02 | 0.01 | 0.02 | 0.04 |
| 3 | 119518898 | rs4440154 | 0.768 | -1.31 | 0.314 | -0.175 | -0.001 | 0 | 0 | 0 | 0.65 | 0.66 | 0.51 | 0.64 | 0.68 | 0.62 |
| 3 | 119519464 | rs6772976 | 0.076 | -1.37 | 0.148 | -0.188 | -0.048 | -0.102 | -0.102 | 2.292 | 0.24 | 0.04 | 0.4 | 0.36 | 0.24 | 0.33 |
| 3 | 119520125 | rs2461823 | 0.803 | -1.384 | 0.231 | -0.138 | -0.005 | -0.004 | -0.004 | 0 | 0.64 | 0.51 | 0.5 | 0.64 | 0.68 | 0.62 |
| 3 | 119520165 | rs2461822 | 0.809 | -1.38 | 0.233 | -0.136 | -0.005 | 0.001 | 0.001 | 0 | 0.64 | 0.51 | 0.5 | 0.64 | 0.68 | 0.62 |
| 3 | 119520558 | rs2461821 | 0.805 | -1.378 | 0.234 | -0.183 | -0.005 | -0.004 | -0.004 | 0 | 0.64 | 0.51 | 0.5 | 0.64 | 0.68 | 0.62 |
| 3 | 119521055 | rs13059232 | 0.795 | -1.39 | 0.353 | -0.188 | -0.004 | -0.004 | -0.004 | 1.256 | 0.64 | 0.51 | 0.5 | 0.64 | 0.68 | 0.62 |
| 3 | 119523746 | rs6438549 | 0.444 | -1.365 | 0.124 | -0.202 | 0.019 | -0.077 | -0.077 | 0 | 0.88 | 0.54 | 0.9 | 1 | 0.92 | 0.95 |
| 3 | 119523872 | rs2461818 | 0.679 | -1.407 | 0.103 | -0.222 | 0.007 | -0.088 | -0.088 | 0.726 | 0.87 | 0.54 | 0.9 | 1 | 0.92 | 0.95 |
| 3 | 119524166 | rs6771638 | 0.064 | -1.318 | 0.052 | -0.133 | -0.008 | -0.042 | -0.042 | 0 | 0.25 | 0.14 | 0.41 | 0.37 | 0.24 | 0.33 |
| 3 | 119524651 | rs2461817 | 0.942 | -1.161 | 0.044 | 0.001 | 0.007 | 0.009 | 0.009 | 0 | 0.57 | 0.39 | 0.49 | 0.63 | 0.67 | 0.58 |
| 3 | 119527776 | rs2472680 | 0.214 | -0.955 | 0.594 | 0.427 | -0.001 | 0.032 | 0.032 | 0.077 | 0.93 | 0.8 | 0.96 | 0.88 | 0.96 | 0.9 |
| 3 | 119530858 | rs3732357 | 1.086 | -1.736 | -0.308 | 0.64 | 0.041 | -0.005 | -0.005 | 0 | 0.63 | 0.26 | 0.72 | 0.62 | 0.73 | 0.58 |
| 3 | 119532652 | rs2472682 | 1.44 | -1.333 | -0.592 | 0.768 | 0.067 | -0.023 | -0.023 | 0 | 0.57 | 0.15 | 0.65 | 0.38 | 0.66 | 0.55 |
| 3 | 119533346 | rs11917714 | -0.606 | -1.619 | 0.183 | 0.648 | -0.04 | 0.051 | 0.051 | 0 | 0.25 | 0.35 | 0.18 | 0.47 | 0.17 | 0.29 |
| 3 | 119533733 | rs6785049 | 1.895 | -1.689 | -0.102 | 0.536 | 0.035 | -0.11 | -0.11 | 0.563 | 0.52 | 0.04 | 0.62 | 0.4 | 0.62 | 0.51 |
| 3 | 119536429 | rs3732359 | 1.073 | -1.799 | 0.602 | 0.52 | 0.032 | -0.026 | -0.026 | 0 | 0.67 | 0.31 | 0.77 | 0.41 | 0.8 | 0.57 |
| 3 | 119536559 | rs10511395 | -0.393 | -1.788 | 0.585 | 0.495 | -0.004 | -0.016 | -0.016 | 2.514 | 0.13 | 0.15 | 0.14 | 0 | 0.17 | 0.05 |
| 3 | 119536581 | rs3732360 | 1.304 | -1.769 | 0.575 | 0.487 | 0.043 | -0.03 | -0.03 | 0 | 0.65 | 0.3 | 0.75 | 0.41 | 0.77 | 0.55 |
| 3 | 119536718 | rs1054190 | 0.177 | -1.74 | 0.597 | 0.481 | -0.033 | 0.05 | 0.05 | 0.579 | 0.11 | 0.04 | 0.1 | 0 | 0.14 | 0.04 |
| 3 | 119536817 | rs6438550 | -0.894 | -1.761 | 0.919 | 0.484 | -0.004 | 0.022 | 0.022 | 0 | 0.94 | 0.87 | 0.96 | 0.88 | 0.97 | 0.85 |

| | | | | | | | | | | | | | | | | |
|---|---|---|---|---|---|---|---|---|---|---|---|---|---|---|---|---|
| 3 | 119536897 | rs1054191 | -0.59 | -1.753 | 0.887 | 0.481 | -0.004 | -0.016 | -0.016 | 0 | 0.13 | 0.15 | 0.14 | 0 | 0.17 | 0.05 |
| 3 | 119537254 | rs3814057 | -0.596 | -1.78 | 0.994 | 0.513 | -0.086 | 0.107 | 0.107 | 0 | 0.26 | 0.52 | 0.19 | 0.47 | 0.17 | 0.29 |
| 3 | 119542297 | rs3732361 | 1.881 | -1.75 | 0.962 | 0.546 | 0.075 | -0.026 | -0.026 | 0 | 0.54 | 0.15 | 0.64 | 0.41 | 0.63 | 0.51 |
| 3 | 119544615 | rs2037547 | 0.943 | -1.738 | 1.042 | 0.547 | 0.015 | -0.058 | -0.058 | 0.069 | 0.81 | 0.68 | 0.82 | 0.88 | 0.79 | 0.82 |
| 3 | 119546197 | rs4261913 | 2.006 | -1.778 | 0.938 | 0.542 | 0.055 | -0.072 | -0.072 | 0 | 0.52 | 0.05 | 0.62 | 0.4 | 0.62 | 0.51 |
| 3 | 119547062 | rs2873950 | -0.497 | -1.789 | 1.049 | 0.561 | -0.049 | 0.084 | 0.084 | 0 | 0.24 | 0.39 | 0.18 | 0.49 | 0.17 | 0.29 |
| 3 | 119578880 | rs1719888 | -0.904 | -1.775 | 0.882 | 0.521 | -0.002 | 0.001 | 0.001 | 0.05 | 0.92 | 0.87 | 0.94 | 0.88 | 0.94 | 0.84 |
| 3 | 119594488 | rs3108749 | -0.432 | -1.79 | 0.888 | 0.527 | -0.002 | 0.006 | 0.006 | 0.006 | 0.91 | 0.82 | 0.93 | 0.88 | 0.94 | 0.84 |
| 3 | 119606078 | rs16830594 | 0.493 | -1.957 | 0.879 | 0.707 | 0.034 | -0.067 | -0.067 | 0 | 0.15 | 0.04 | 0.07 | 0.34 | 0.11 | 0.14 |
| 3 | 119610793 | rs6782799 | -2.171 | -1.812 | 0.788 | 0.566 | 0.056 | -0.089 | -0.089 | 0.357 | 0.47 | 0.95 | 0.38 | 0.59 | 0.38 | 0.49 |
| 3 | 119622014 | rs4340737 | -0.544 | -1.812 | 0.779 | 0.572 | -0.043 | 0.066 | 0.066 | 0 | 0.24 | 0.36 | 0.18 | 0.47 | 0.17 | 0.3 |
| 3 | 119631098 | rs13314595 | -1.633 | -1.813 | 0.718 | 0.564 | -0.005 | -0.074 | -0.074 | 0 | 0.23 | 0.56 | 0.2 | 0.13 | 0.21 | 0.2 |
| 3 | 119631814 | rs6438552 | -2.036 | -1.766 | 0.559 | 0.579 | 0.056 | -0.08 | -0.08 | 0 | 0.47 | 0.94 | 0.37 | 0.6 | 0.38 | 0.49 |
| 3 | 119675051 | rs17810235 | 1.634 | -0.884 | -0.116 | 0.116 | -0.117 | 0.107 | 0.107 | 0.008 | 0.26 | 0.01 | 0.29 | 0.06 | 0.32 | 0.29 |
| 3 | 119685477 | rs17810302 | 0.459 | -0.754 | -0.063 | 0.142 | -0.031 | 0.033 | 0.033 | 0 | 0.09 | 0.04 | 0.09 | 0 | 0.14 | 0.03 |
| 3 | 119693611 | rs6771023 | -0.648 | -0.791 | 0.049 | 0.14 | 0.024 | -0.052 | -0.052 | 0 | 0.17 | 0.31 | 0.17 | 0.12 | 0.2 | 0.19 |
| 3 | 119776880 | rs17204878 | -1.725 | -0.754 | -1.152 | 0.252 | 0.032 | -0.101 | -0.101 | 0 | 0.45 | 0.95 | 0.37 | 0.59 | 0.37 | 0.49 |
| 3 | 119778489 | rs2199503 | -2.098 | -0.625 | -1.094 | 0.173 | -0.107 | -0.138 | -0.138 | 0 | 0.73 | 0.98 | 0.82 | 0.65 | 0.69 | 0.79 |
| 3 | 119790194 | rs182839 | 0.598 | -0.451 | -0.364 | 0.291 | -0.003 | 0 | 0 | 0.055 | 0.08 | 0.13 | 0.06 | 0.12 | 0.06 | 0.16 |
| 3 | 119792474 | rs17205095 | -0.224 | -0.236 | -0.886 | 0.274 | 0.006 | -0.025 | -0.025 | 0 | 0.09 | 0.13 | 0.09 | 0 | 0.14 | 0.03 |
| 3 | 119813282 | rs334558 | -1.399 | -0.804 | -1.295 | 0.435 | 0.078 | -0.037 | -0.037 | 0 | 0.46 | 0.94 | 0.34 | 0.65 | 0.33 | 0.55 |
| 3 | 119822335 | rs11925868 | -1.122 | -0.418 | -0.768 | 0.257 | -0.004 | 0.02 | 0.02 | 0 | 0.13 | 0.13 | 0.15 | 0.08 | 0.16 | 0.19 |
| 3 | 119824037 | rs11927974 | -0.142 | -0.345 | -0.793 | 0.255 | -0.014 | 0.016 | 0.016 | 0 | 0.08 | 0.05 | 0.06 | 0 | 0.1 | 0.04 |
| 3 | 119824166 | rs16830720 | -0.252 | -0.561 | -0.71 | 0.479 | 0.004 | -0.006 | -0.006 | 0 | 0.05 | 0.06 | 0.08 | 0.13 | 0.06 | 0.17 |
| 3 | 119828512 | rs16830730 | -0.736 | -0.91 | -0.24 | 0.474 | -0.027 | 0.043 | 0.043 | 0 | 0.2 | 0.29 | 0.1 | 0.57 | 0.15 | 0.3 |
| 3 | 119832621 | rs334543 | -1.447 | -0.64 | -0.949 | 0.119 | -0.053 | -0.045 | -0.045 | 0 | 0.72 | 0.9 | 0.59 | 0.71 | 0.69 | 0.8 |
| 3 | 119839546 | rs9861883 | -0.06 | -0.586 | -0.575 | -0.03 | 0.001 | 0.004 | 0.004 | 0 | 0.18 | 0.17 | 0.09 | 0.53 | 0.15 | 0.22 |
| 3 | 119839869 | rs334547 | -0.36 | -0.391 | -0.294 | -0.377 | 0.005 | -0.013 | -0.013 | 1.719 | 0.14 | 0.3 | 0.11 | 0.12 | 0.11 | 0.17 |
| 3 | 119842143 | rs6792814 | 1.206 | -0.068 | -0.721 | -0.596 | -0.075 | 0.143 | 0.143 | 0 | 0.28 | 0.09 | 0.34 | 0.01 | 0.35 | 0.24 |
| 3 | 119842377 | rs3112282 | 0.315 | 0.133 | -0.346 | -0.155 | -0.006 | 0.002 | 0.002 | 0.02 | 0.09 | 0.16 | 0.06 | 0.11 | 0.07 | 0.15 |
| 3 | 119844557 | rs13086387 | 1.171 | 0.098 | -0.57 | -0.326 | -0.052 | 0.065 | 0.065 | 0 | 0.31 | 0.11 | 0.21 | 0.22 | 0.36 | 0.26 |
| 3 | 119855410 | rs923380 | -0.286 | 0.019 | -0.209 | -0.447 | -0.01 | -0.021 | -0.021 | 0 | 0.9 | 0.84 | 0.93 | 0.96 | 0.93 | 0.86 |
| 3 | 119867803 | rs6794478 | -0.296 | 0.062 | -0.046 | -0.436 | -0.006 | -0.012 | -0.012 | 0 | 0.9 | 0.88 | 0.93 | 0.94 | 0.93 | 0.86 |
| 3 | 119869525 | rs4563433 | 0.765 | -0.15 | 0.402 | -0.716 | 0.005 | -0.052 | -0.052 | 0.041 | 0.34 | 0.47 | 0.39 | 0.16 | 0.39 | 0.39 |
| 3 | 119876676 | rs12489527 | 0.047 | -0.688 | 0.171 | -0.572 | 0.026 | -0.061 | -0.061 | 0 | 0.19 | 0.08 | 0.28 | 0.59 | 0.14 | 0.23 |
| 3 | 119877871 | rs11712811 | 1.148 | -0.4 | 0.333 | -0.646 | -0.013 | 0.053 | 0.053 | 0.007 | 0.36 | 0.26 | 0.24 | 0.18 | 0.38 | 0.25 |
| 3 | 119884407 | rs13092071 | 1.762 | -0.436 | 0.254 | -0.695 | -0.017 | 0.046 | 0.046 | 0.007 | 0.29 | 0.16 | 0.2 | 0.17 | 0.33 | 0.22 |
| 3 | 119884663 | rs1488763 | 1.053 | -0.677 | 0.457 | -0.718 | 0.007 | 0.006 | 0.006 | 1.038 | 0.63 | 0.39 | 0.58 | 0.19 | 0.73 | 0.47 |
| 3 | 119893257 | rs1488762 | -0.788 | -0.614 | 0.445 | -0.673 | -0.006 | -0.006 | -0.006 | 0.728 | 0.9 | 0.88 | 0.93 | 0.93 | 0.93 | 0.86 |
| 3 | 119893904 | rs2319540 | 0.388 | -0.673 | 0.929 | -0.65 | -0.002 | 0.125 | 0.125 | 0 | 0.07 | 0.27 | 0.05 | 0.15 | 0.05 | 0.15 |
| 3 | 119895964 | rs7634938 | 1.683 | -0.651 | 0.773 | -0.761 | -0.072 | 0.117 | 0.117 | 0 | 0.28 | 0.11 | 0.34 | 0.02 | 0.35 | 0.23 |
| 3 | 119896657 | rs7635647 | 0.144 | -0.698 | 0.774 | -0.728 | 0.023 | -0.049 | -0.049 | 0 | 0.19 | 0.09 | 0.28 | 0.59 | 0.14 | 0.23 |
| 3 | 119899392 | rs979683 | 0.645 | -0.68 | 0.832 | -0.686 | -0.007 | 0.01 | 0.01 | 0 | 0.06 | 0.05 | 0.03 | 0 | 0.05 | 0.02 |
| 3 | 119900785 | rs11710115 | 0.906 | -0.913 | 1.002 | -0.529 | -0.003 | -0.015 | -0.015 | 0.044 | 0.36 | 0.5 | 0.26 | 0.32 | 0.39 | 0.38 |
| 3 | 119915437 | rs2035669 | 1.623 | -0.813 | 0.604 | -0.501 | -0.015 | 0.037 | 0.037 | 0 | 0.29 | 0.16 | 0.2 | 0.17 | 0.33 | 0.2 |
| 3 | 119930433 | rs9878040 | -0.016 | -1.5 | 0.379 | -0.349 | 0.012 | -0.023 | -0.023 | 0 | 0.18 | 0.12 | 0.28 | 0.58 | 0.14 | 0.23 |
| 3 | 119943453 | rs11929146 | 0.647 | -1.674 | 0.591 | -0.58 | -0.032 | 0.036 | 0.036 | 0.061 | 0.77 | 0.7 | 0.69 | 0.36 | 0.88 | 0.65 |
| 3 | 119946098 | rs7648922 | 1.819 | -1.165 | 0.055 | -0.69 | 0.014 | 0.028 | 0.028 | 0.172 | 0.28 | 0.1 | 0.18 | 0.19 | 0.29 | 0.17 |
| 3 | 119947503 | rs6785180 | 0.047 | -0.861 | -0.133 | -0.546 | -0.001 | -0.023 | -0.023 | 0 | 0.08 | 0.12 | 0.06 | 0.01 | 0.1 | 0.09 |
| 3 | 119975699 | rs9826773 | -0.059 | -0.763 | -0.316 | -0.398 | -0.024 | 0.083 | 0.083 | 0 | 0.13 | 0.2 | 0.24 | 0.58 | 0.05 | 0.21 |
| 3 | 119984085 | rs1873652 | -0.2 | -0.769 | -0.322 | -0.333 | -0.03 | 0.1 | 0.1 | 0 | 0.14 | 0.21 | 0.24 | 0.57 | 0.05 | 0.21 |
| 3 | 120001235 | rs6795868 | -0.126 | -0.806 | -0.409 | -0.515 | 0.025 | 0.018 | 0.018 | 0 | 0.48 | 0.38 | 0.65 | 0.58 | 0.49 | 0.42 |
| 3 | 120004825 | rs9849695 | -0.299 | -0.787 | -0.3 | -0.2 | -0.029 | 0.1 | 0.1 | 0 | 0.14 | 0.22 | 0.24 | 0.56 | 0.05 | 0.2 |
| 3 | 120020619 | rs2971211 | -1.212 | -0.764 | -0.266 | -0.974 | -0.005 | -0.053 | -0.053 | 0 | 0.19 | 0.39 | 0.15 | 0.08 | 0.19 | 0.24 |
| 3 | 120028644 | rs6803846 | 1.358 | -0.771 | -0.891 | -1.388 | -0.164 | 0.224 | 0.224 | 0.018 | 0.32 | 0.04 | 0.36 | 0.02 | 0.43 | 0.24 |
| 3 | 120032474 | rs787193 | 0.061 | -0.838 | -0.862 | -1.351 | -0.007 | 0.017 | 0.017 | 0 | 0.09 | 0.09 | 0.08 | 0.01 | 0.13 | 0.1 |
| 3 | 120040449 | rs787209 | -1.596 | -1.131 | -0.559 | -1.414 | -0.048 | -0.081 | -0.081 | 0.296 | 0.3 | 0.56 | 0.39 | 0.11 | 0.21 | 0.27 |
| 3 | 120043023 | rs16831033 | -0.84 | -1.081 | -0.424 | -1.426 | -0.058 | -0.035 | -0.035 | 0 | 0.18 | 0.39 | 0.3 | 0.1 | 0.08 | 0.17 |
| 3 | 120043836 | rs10511400 | -0.84 | -1.081 | -0.424 | -1.426 | -0.058 | -0.035 | -0.035 | 0.01 | 0.18 | 0.39 | 0.3 | 0.1 | 0.08 | 0.16 |
| 3 | 120049117 | rs6438575 | 1.385 | -1.017 | -0.772 | -1.512 | -0.176 | 0.246 | 0.246 | 0.023 | 0.33 | 0.04 | 0.36 | 0.02 | 0.44 | 0.25 |
| 3 | 120049292 | rs2638597 | -0.093 | -0.873 | -0.479 | -1.241 | -0.001 | -0.032 | -0.032 | 0 | 0.11 | 0.17 | 0.09 | 0 | 0.13 | 0.1 |
| 3 | 120071567 | rs7622363 | -0.836 | -0.917 | -0.957 | -1.457 | 0.006 | 0.133 | 0.133 | 0.136 | 0.56 | 0.8 | 0.59 | 0.79 | 0.55 | 0.69 |
| 3 | 120074964 | rs13075278 | 0.603 | -1.118 | -0.803 | -1.435 | 0.013 | -0.007 | -0.007 | 0 | 0.38 | 0.4 | 0.29 | 0.68 | 0.47 | 0.53 |
| 3 | 120079894 | rs12495760 | 0.209 | -1.109 | -0.835 | -1.461 | -0.002 | 0 | 0 | 0 | 0.13 | 0.13 | 0.26 | 0.09 | 0.07 | 0.12 |
| 3 | 120083854 | rs1147683 | -1.846 | -1.023 | -1.034 | -1.518 | -0.036 | 0.036 | 0.036 | 0.621 | 0.67 | 0.98 | 0.67 | 0.81 | 0.68 | 0.8 |
| 3 | 120099799 | rs11708297 | -0.384 | -1.102 | -0.936 | -1.584 | 0.037 | 0.139 | 0.139 | 0.32 | 0.49 | 0.75 | 0.38 | 0.8 | 0.52 | 0.67 |
| 3 | 120100054 | rs1733354 | -1.869 | -1.014 | -1.061 | -1.533 | -0.036 | 0.036 | 0.036 | 0.621 | 0.67 | 0.98 | 0.67 | 0.81 | 0.68 | 0.8 |
| 3 | 120106539 | rs4676776 | 1.413 | -0.839 | -0.874 | -1.617 | -0.107 | 0.134 | 0.134 | 0 | 0.28 | 0.09 | 0.2 | 0.02 | 0.37 | 0.24 |
| 3 | 120113286 | rs2488 | -0.357 | -0.731 | -0.345 | -1.582 | 0.007 | 0.042 | 0.042 | 0 | 0.48 | 0.43 | 0.51 | 0.04 | 0.52 | 0.41 |

| | | | | | | | | | | | | | | | | |
|---|---|---|---|---|---|---|---|---|---|---|---|---|---|---|---|---|
| 3 | 120115385 | rs11708686 | 0.88 | -0.734 | -0.326 | -1.094 | 0.017 | 0.319 | 0.319 | 0.493 | 0.12 | 0.37 | 0.09 | 0.6 | 0.1 | 0.3 |
| 3 | 120119479 | rs1147696 | -0.132 | -1.002 | -0.958 | -1.657 | -0.005 | 0.104 | 0.104 | 0 | 0.45 | 0.32 | 0.49 | 0.04 | 0.51 | 0.35 |
| 3 | 120122624 | rs9880550 | 0.058 | -1 | -0.976 | -1.656 | -0.005 | 0.104 | 0.104 | 0 | 0.45 | 0.31 | 0.49 | 0.04 | 0.51 | 0.35 |
| 3 | 120127340 | rs6799005 | 0.144 | -0.722 | -1.039 | -1.559 | -0.001 | -0.038 | -0.038 | 1.631 | 0.12 | 0.19 | 0.09 | 0.01 | 0.13 | 0.08 |
| 3 | 120131762 | rs13097755 | 0.464 | -0.558 | -1.417 | -1.45 | -0.026 | 0.064 | 0.064 | 0 | 0.37 | 0.29 | 0.42 | 0.03 | 0.39 | 0.33 |
| 3 | 120141209 | rs1259297 | 0.799 | -1.107 | -1.73 | -1.884 | -0.056 | 0.051 | 0.051 | 0.293 | 0.34 | 0.03 | 0.34 | 0.21 | 0.34 | 0.2 |
| 3 | 120144064 | rs916512 | 0.788 | -1.079 | -1.741 | -1.845 | -0.048 | 0.052 | 0.052 | 0.365 | 0.35 | 0.03 | 0.34 | 0.22 | 0.34 | 0.2 |
| 3 | 120147864 | rs1259299 | 0.598 | -1.246 | -1.62 | -1.894 | 0.027 | 0.019 | 0.019 | 0 | 0.38 | 0.16 | 0.36 | 0.31 | 0.33 | 0.31 |
| 3 | 120152222 | rs1624195 | 0.224 | -1.882 | -0.269 | -2.044 | -0.035 | 0.02 | 0.02 | 0 | 0.91 | 0.75 | 0.74 | 0.92 | 0.98 | 0.94 |
| 3 | 120154181 | rs1733302 | 0.738 | -1.837 | -0.321 | -2.133 | -0.026 | -0.051 | -0.051 | 0 | 0.83 | 0.73 | 0.68 | 0.92 | 0.89 | 0.91 |
| 3 | 120165097 | rs1267675 | 2.318 | -1.886 | -0.331 | -2.303 | 0.018 | 0.049 | 0.049 | 0.303 | 0.64 | 0.28 | 0.52 | 0.81 | 0.71 | 0.73 |
| 3 | 120166480 | rs13326852 | -1.14 | -1.896 | -0.293 | -2.345 | 0.018 | -0.133 | -0.133 | 0 | 0.21 | 0.47 | 0.2 | 0.02 | 0.23 | 0.13 |
| 3 | 120167006 | rs1259339 | 0.681 | -1.662 | -0.252 | -2.168 | -0.043 | 0.033 | 0.033 | 0 | 0.85 | 0.75 | 0.72 | 0.83 | 0.94 | 0.85 |
| 3 | 120179752 | rs1259309 | -2.05 | -1.566 | -0.56 | -2.269 | 0.019 | 0.148 | 0.148 | 0.002 | 0.47 | 0.86 | 0.51 | 0.25 | 0.35 | 0.44 |
| 3 | 120179828 | rs1259308 | -0.38 | -1.926 | -0.303 | -2.225 | -0.049 | 0.002 | 0.002 | 0 | 0.17 | 0.29 | 0.27 | 0.15 | 0.07 | 0.12 |
| 3 | 120194175 | rs4676787 | -0.516 | -1.913 | -0.164 | -2.251 | 0 | -0.013 | -0.013 | 1.971 | 0.17 | 0.19 | 0.16 | 0.06 | 0.2 | 0.19 |
| 3 | 120194615 | rs1259342 | 0.06 | -1.951 | -0.37 | -2.256 | -0.057 | 0.032 | 0.032 | 0 | 0.13 | 0.27 | 0.25 | 0.15 | 0.05 | 0.12 |
| 3 | 120199413 | rs1262395 | -2.326 | -1.955 | -0.395 | -2.271 | 0.035 | 0.106 | 0.106 | 0 | 0.43 | 0.81 | 0.49 | 0.25 | 0.31 | 0.42 |
| 3 | 120199467 | rs1267693 | -1.259 | -1.953 | -0.387 | -2.28 | -0.026 | -0.016 | -0.016 | 0.019 | 0.28 | 0.32 | 0.35 | 0.22 | 0.16 | 0.31 |
| 3 | 120204169 | rs13099637 | -1.475 | -1.937 | -0.43 | -2.34 | 0.017 | -0.102 | -0.102 | 0.832 | 0.23 | 0.48 | 0.22 | 0.06 | 0.23 | 0.19 |
| 3 | 120206372 | rs1147720 | -0.281 | -1.902 | -1.647 | -2.323 | -0.003 | -0.001 | -0.001 | 0 | 0.17 | 0.13 | 0.26 | 0.18 | 0.08 | 0.22 |
| 3 | 120206415 | rs893460 | 0.738 | -1.869 | -1.8 | -2.297 | -0.032 | 0.009 | 0.009 | 0 | 0.09 | 0.01 | 0.07 | 0.04 | 0.08 | 0.09 |
| 3 | 120206843 | rs988924 | -2.251 | -1.867 | -1.79 | -2.299 | 0.031 | 0.153 | 0.153 | 0.175 | 0.47 | 0.89 | 0.51 | 0.24 | 0.33 | 0.43 |
| 3 | 120210509 | rs4676791 | 0.017 | -1.774 | -1.694 | -2.229 | -0.002 | -0.005 | -0.005 | 0 | 0.08 | 0.08 | 0.09 | 0.02 | 0.12 | 0.1 |
| 3 | 120214132 | rs1259313 | 0.48 | -1.778 | -1.654 | -2.257 | -0.019 | -0.033 | -0.033 | 0 | 0.11 | 0.12 | 0.2 | 0.01 | 0.05 | 0.13 |
| 3 | 120214270 | rs1147721 | -1.195 | -1.682 | -1.586 | -2.185 | -0.007 | -0.012 | -0.012 | 0.785 | 0.28 | 0.32 | 0.38 | 0.2 | 0.22 | 0.33 |
| 3 | 120216892 | rs1272257 | -1.817 | -1.68 | -1.707 | -2.129 | 0.04 | 0.108 | 0.108 | 0.025 | 0.44 | 0.84 | 0.49 | 0.24 | 0.31 | 0.42 |
| 3 | 120225355 | rs1259340 | 0.769 | -1.37 | -1.118 | -1.939 | 0.007 | 0 | 0 | 0 | 0.1 | 0.07 | 0.06 | 0.01 | 0.08 | 0.15 |
| 3 | 120232831 | rs3901452 | -0.869 | -1.291 | -1.155 | -1.992 | 0.003 | -0.003 | -0.003 | 0 | 0.14 | 0.15 | 0.31 | 0.11 | 0.18 | 0.13 |
| 3 | 120234219 | rs1259475 | -2.071 | -1.307 | -1.286 | -2.094 | -0.011 | 0.226 | 0.226 | 0 | 0.48 | 0.85 | 0.53 | 0.16 | 0.46 | 0.39 |
| 3 | 120234541 | rs1259474 | -0.594 | -1.28 | -1.247 | -2.011 | 0.007 | 0.018 | 0.018 | 0 | 0.27 | 0.22 | 0.17 | 0.05 | 0.25 | 0.25 |
| 3 | 120236329 | rs9289146 | 0.201 | -1.216 | -1.204 | -1.959 | 0 | -0.009 | -0.009 | 0.06 | 0.17 | 0.18 | 0.12 | 0.04 | 0.17 | 0.1 |
| 3 | 120237730 | rs1259495 | 0.758 | -1.124 | -0.833 | -1.546 | 0.006 | 0.002 | 0.002 | 0.145 | 0.11 | 0.07 | 0.07 | 0.01 | 0.09 | 0.14 |
| 3 | 120238431 | rs1259494 | -0.167 | -1.16 | -0.9 | -1.598 | 0.01 | 0.027 | 0.027 | 0 | 0.26 | 0.18 | 0.19 | 0.05 | 0.24 | 0.23 |
| 3 | 120239317 | rs2319822 | 0.752 | -0.878 | -0.708 | -1.465 | 0.078 | 0.068 | 0.068 | 2.723 | 0.05 | 0.47 | 0.03 | 0 | 0 | 0 |
| 3 | 120244109 | rs16831397 | -1.564 | -0.902 | -0.838 | -1.476 | -0.003 | -0.039 | -0.039 | 0 | 0.23 | 0.63 | 0.19 | 0.11 | 0.19 | 0.14 |
| 3 | 120250343 | rs4286452 | 1.009 | -0.859 | -1.092 | -1.459 | -0.005 | 0.052 | 0.052 | 0.348 | 0.14 | 0.05 | 0.12 | 0.04 | 0.15 | 0.09 |
| 3 | 120250723 | rs1595481 | -2.037 | -0.249 | -0.73 | -0.986 | -0.009 | -0.024 | -0.024 | 0 | 0.24 | 0.58 | 0.23 | 0.15 | 0.2 | 0.19 |
| 3 | 120269643 | rs4676799 | -0.964 | -0.21 | -0.78 | -0.801 | 0.005 | 0.131 | 0.131 | 0 | 0.54 | 0.82 | 0.43 | 0.2 | 0.45 | 0.42 |
| 3 | 120272358 | rs9853121 | -1.487 | -0.217 | -0.778 | -0.802 | 0.001 | -0.024 | -0.024 | 0 | 0.14 | 0.52 | 0.12 | 0.05 | 0.07 | 0.07 |
| 3 | 120277583 | rs701989 | 0.492 | 0.399 | -0.871 | -0.367 | 0.003 | -0.016 | -0.016 | 0.215 | 0.38 | 0.33 | 0.26 | 0.07 | 0.34 | 0.3 |
| 3 | 120280776 | rs805009 | 0.012 | 0.673 | -0.664 | -0.001 | 0.009 | 0.034 | 0.034 | 0 | 0.49 | 0.41 | 0.39 | 0.2 | 0.45 | 0.42 |
| 3 | 120316203 | rs16831523 | -0.647 | 0.799 | -1.172 | 1.058 | -0.008 | -0.063 | -0.063 | 0 | 0.23 | 0.11 | 0.3 | 0.57 | 0.22 | 0.36 |
| 3 | 120421729 | rs9850589 | 0.22 | 0.369 | -0.582 | 0.459 | -0.011 | -0.024 | -0.024 | 0 | 0.93 | 0.84 | 0.96 | 0.99 | 0.95 | 0.98 |
| 3 | 120425331 | rs13077101 | 0.335 | 0.152 | -1.593 | -1.139 | -0.005 | 0.176 | 0.176 | 0.132 | 0.37 | 0.1 | 0.2 | 0.03 | 0.33 | 0.34 |
| 3 | 120428621 | rs11720353 | 0.666 | 0.614 | -1.201 | -1.352 | -0.004 | 0.003 | 0.003 | 0 | 0.23 | 0.18 | 0.22 | 0.13 | 0.34 | 0.15 |
| 3 | 120430134 | rs12489104 | 0.361 | 0.113 | -1.222 | -1.521 | -0.064 | 0.022 | 0.022 | 0.048 | 0.24 | 0.05 | 0.49 | 0.83 | 0.27 | 0.37 |
| 3 | 120436898 | rs11708525 | 0.821 | 0.107 | -1.214 | -1.525 | -0.004 | 0.003 | 0.003 | 0 | 0.23 | 0.18 | 0.22 | 0.13 | 0.34 | 0.15 |
| 3 | 120437310 | rs6787364 | -0.425 | 0.373 | -1.138 | -0.093 | -0.01 | -0.015 | -0.015 | 0 | 0.45 | 0.25 | 0.38 | 0.49 | 0.57 | 0.44 |
| 3 | 120446626 | rs9985346 | 0.531 | 0.664 | -1.267 | -0.212 | -0.016 | -0.083 | -0.083 | 0.001 | 0.21 | 0.07 | 0.15 | 0.37 | 0.23 | 0.29 |
| 3 | 120457838 | rs9854081 | 0.62 | 0.668 | -1.277 | -0.21 | -0.028 | -0.101 | -0.101 | 0 | 0.2 | 0.04 | 0.15 | 0.36 | 0.23 | 0.29 |
| 3 | 120461576 | rs7647895 | 0.808 | 0.634 | 0.144 | -0.224 | -0.004 | 0.003 | 0.003 | 0 | 0.23 | 0.18 | 0.22 | 0.13 | 0.34 | 0.15 |
| 3 | 120465622 | rs1466960 | -1.528 | 0.086 | -0.599 | -1.024 | -0.054 | 0.229 | 0.229 | 0.167 | 0.68 | 0.97 | 0.82 | 0.99 | 0.68 | 0.79 |
| 3 | 120473570 | rs17243218 | 1.16 | 0.342 | -0.303 | -0.804 | -0.044 | 0.018 | 0.018 | 0.173 | 0.19 | 0.07 | 0.19 | 0.13 | 0.3 | 0.13 |
| 3 | 120485709 | rs378759 | -0.022 | 0.267 | -0.257 | -0.774 | 0.012 | 0.269 | 0.269 | 0 | 0.13 | 0.39 | 0.39 | 0.47 | 0.09 | 0.1 |
| 3 | 120492428 | rs6769985 | 0.522 | 0.285 | -0.181 | -0.772 | -0.035 | 0.036 | 0.036 | 0 | 0.23 | 0.11 | 0.22 | 0.13 | 0.34 | 0.15 |
| 3 | 120507751 | rs9858561 | 0.923 | 0.574 | -0.693 | -0.362 | -0.012 | -0.042 | -0.042 | 0.038 | 0.11 | 0.02 | 0.05 | 0.17 | 0.1 | 0.21 |
| 3 | 120511490 | rs373188 | -0.323 | 0.752 | -0.368 | -0.153 | 0.017 | 0.012 | 0.012 | 0 | 0.36 | 0.52 | 0.61 | 0.6 | 0.42 | 0.23 |
| 3 | 120522716 | rs12629908 | -0.784 | 0.689 | -0.354 | 0.775 | -0.001 | 0.086 | 0.086 | 0 | 0.09 | 0.1 | 0.37 | 0.4 | 0.05 | 0.09 |
| 3 | 120524205 | rs434559 | -0.056 | 0.569 | -0.055 | -0.008 | -0.005 | -0.016 | -0.016 | 0 | 0.13 | 0.27 | 0.28 | 0.08 | 0.12 | 0.12 |
| 3 | 120525029 | rs322459 | -0.997 | 0.692 | -0.701 | -0.524 | 0.024 | 0.049 | 0.049 | 0 | 0.59 | 0.79 | 0.71 | 0.35 | 0.61 | 0.58 |
| 3 | 120526289 | rs186522 | -0.646 | 1.129 | -0.085 | -0.887 | 0.019 | -0.072 | -0.072 | 1.245 | 0.33 | 0.52 | 0.46 | 0.09 | 0.38 | 0.29 |
| 3 | 120529229 | rs413163 | 0.714 | 1.102 | 0.08 | -0.845 | 0 | -0.019 | -0.019 | 0.234 | 0.14 | 0.39 | 0.29 | 0.08 | 0.13 | 0.12 |
| 3 | 120531127 | rs399491 | -1.044 | 0.864 | -0.195 | -1 | -0.012 | 0.061 | 0.061 | 0 | 0.37 | 0.69 | 0.48 | 0.11 | 0.39 | 0.3 |
| 3 | 120533211 | rs559461 | -0.045 | 0.863 | -0.027 | -0.968 | -0.02 | -0.042 | -0.042 | 0 | 0.19 | 0.54 | 0.34 | 0.1 | 0.18 | 0.13 |
| 3 | 120535520 | rs2016491 | -1.328 | 0.92 | -0.56 | -1.158 | -0.007 | 0.173 | 0.173 | 0 | 0.44 | 0.79 | 0.51 | 0.12 | 0.43 | 0.35 |
| 3 | 120536346 | rs444107 | 0.426 | 0.693 | -0.049 | -1.176 | -0.011 | -0.033 | -0.033 | 0.166 | 0.2 | 0.34 | 0.31 | 0.11 | 0.19 | 0.17 |
| 3 | 120542461 | rs561375 | -1.754 | 0.779 | -0.963 | -1.224 | 0.001 | 0.033 | 0.033 | 0 | 0.62 | 0.9 | 0.73 | 0.34 | 0.65 | 0.64 |
| 3 | 120551100 | rs646244 | 0.06 | 0.607 | -0.569 | -1.109 | -0.01 | -0.018 | -0.018 | 2.098 | 0.25 | 0.7 | 0.35 | 0.11 | 0.19 | 0.18 |

| | | | | | | | | | | | | | | | | |
|---|---|---|---|---|---|---|---|---|---|---|---|---|---|---|---|---|
| 3 | 120558109 | rs7619694 | 0.121 | 1.356 | -0.972 | -0.679 | -0.012 | -0.002 | -0.002 | 0 | 0.19 | 0.12 | 0.18 | 0.01 | 0.27 | 0.17 |
| 3 | 120561365 | rs7612901 | 0.094 | 1.359 | -0.987 | -0.677 | -0.003 | -0.019 | -0.019 | 0 | 0.19 | 0.16 | 0.18 | 0.01 | 0.27 | 0.17 |
| 3 | 120572921 | rs371768 | -0.594 | 0.789 | -1.651 | -0.79 | 0.039 | 0.103 | 0.103 | 0.01 | 0.57 | 0.85 | 0.73 | 0.35 | 0.68 | 0.57 |
| 3 | 120574135 | rs2171167 | 1.227 | 0.674 | -1.951 | -1.229 | -0.065 | 0.082 | 0.082 | 0 | 0.14 | 0.01 | 0.1 | 0 | 0.21 | 0.05 |
| 3 | 120575246 | rs9852152 | -0.299 | 0.371 | -1.148 | -1.231 | 0.006 | 0.02 | 0.02 | 2.045 | 0.05 | 0.21 | 0.26 | 0.07 | 0.04 | 0.05 |
| 3 | 120577143 | rs697011 | -0.185 | -0.18 | -1.645 | -1.665 | 0.003 | 0.122 | 0.122 | 0.243 | 0.52 | 0.7 | 0.67 | 0.24 | 0.46 | 0.35 |
| 3 | 120585315 | rs322458 | 0.357 | -0.455 | -1.752 | -1.861 | 0.001 | -0.035 | -0.035 | 0.013 | 0.38 | 0.46 | 0.34 | 0.17 | 0.36 | 0.26 |
| 3 | 120589727 | rs322457 | 0.472 | -0.478 | -1.48 | -1.86 | 0.02 | 0.022 | 0.022 | 0 | 0.29 | 0.32 | 0.32 | 0.53 | 0.36 | 0.38 |
| 3 | 120595986 | rs420621 | -0.505 | -0.647 | -1.553 | -2.128 | -0.005 | 0.095 | 0.095 | 0.336 | 0.53 | 0.7 | 0.69 | 0.25 | 0.47 | 0.35 |
| 3 | 120597030 | rs9856996 | 0.1 | -0.853 | -1.419 | -2.137 | -0.041 | -0.051 | -0.051 | 0 | 0.3 | 0.44 | 0.4 | 0.18 | 0.2 | 0.2 |
| 3 | 120601829 | rs16831912 | 2.665 | -0.718 | -1.486 | -2.038 | -0.035 | 0.026 | 0.026 | 5.295 | 0.06 | 0.01 | 0.05 | 0.47 | 0.1 | 0.24 |
| 3 | 120604574 | rs11927953 | 1.846 | -0.83 | -1.31 | -2.112 | 0.006 | 0.007 | 0.007 | 0 | 0.14 | 0.08 | 0.07 | 0.11 | 0.11 | 0.1 |
| 3 | 120606438 | rs10451894 | 1.771 | -0.835 | -1.163 | -2.114 | -0.011 | -0.013 | -0.013 | 0 | 0.15 | 0.23 | 0.09 | 0.11 | 0.11 | 0.1 |
| 3 | 120626429 | rs470931 | -0.304 | -0.937 | -0.867 | -2.234 | 0.001 | 0.061 | 0.061 | 0 | 0.49 | 0.58 | 0.66 | 0.25 | 0.47 | 0.33 |
| 3 | 120674527 | rs662467 | -1.328 | -1.38 | -1.124 | -2.416 | -0.023 | -0.012 | -0.012 | 0 | 0.85 | 0.8 | 0.71 | 0.87 | 0.91 | 0.84 |
| 3 | 120726942 | rs1403719 | 2.701 | -1.49 | -1.434 | -2.548 | -0.046 | -0.043 | -0.043 | 1.219 | 0.15 | 0.01 | 0.11 | 0.22 | 0.17 | 0.16 |
| 3 | 120728401 | rs605537 | -2.538 | -1.526 | -1.333 | -2.537 | 0.019 | 0.011 | 0.011 | 0 | 0.91 | 0.93 | 0.95 | 0.94 | 0.95 | 0.92 |
| 3 | 120761771 | rs9831480 | 1.323 | -1.528 | -1.294 | -2.539 | -0.007 | 0.006 | 0.006 | 1.298 | 0.06 | 0.2 | 0.25 | 0.07 | 0.04 | 0.08 |
| 3 | 120782246 | rs6438601 | -0.351 | -1.519 | -1.229 | -2.565 | 0.002 | 0.021 | 0.021 | 0.01 | 0.64 | 0.68 | 0.75 | 0.26 | 0.67 | 0.38 |
| 3 | 120792926 | rs7615991 | 1.985 | -1.519 | -1.244 | -2.597 | 0.028 | -0.049 | -0.049 | 0 | 0.2 | 0.28 | 0.27 | 0.07 | 0.26 | 0.12 |
| 3 | 120806790 | rs9848328 | 0.867 | -1.541 | -1.275 | -2.608 | 0.008 | -0.03 | -0.03 | 2.174 | 0.35 | 0.37 | 0.34 | 0.18 | 0.37 | 0.23 |
| 3 | 120817040 | rs2331522 | -1.258 | -1.557 | -1.245 | -2.599 | -0.004 | 0.024 | 0.024 | 0.411 | 0.81 | 0.74 | 0.87 | 0.48 | 0.85 | 0.67 |
| 3 | 120817063 | rs10212452 | -0.022 | -1.521 | -1.221 | -2.63 | 0.007 | 0.059 | 0.059 | 0 | 0.49 | 0.57 | 0.65 | 0.25 | 0.47 | 0.33 |
| 3 | 120844696 | rs17677959 | 2.976 | -1.511 | -1.263 | -2.63 | -0.038 | 0.06 | 0.06 | 0 | 0.14 | 0.03 | 0.1 | 0 | 0.2 | 0.05 |
| 3 | 120845812 | rs3856576 | 2.372 | -1.518 | -1.244 | -2.612 | 0.006 | 0.005 | 0.005 | 0 | 0.14 | 0.08 | 0.07 | 0.12 | 0.1 | 0.1 |
| 3 | 120847203 | rs9289163 | 1.606 | -1.501 | -1.226 | -2.585 | -0.023 | -0.03 | -0.03 | 0 | 0.14 | 0.2 | 0.31 | 0.07 | 0.1 | 0.11 |
| 3 | 120851373 | rs3905999 | -2.666 | -1.392 | -1.33 | -2.514 | 0.013 | 0 | 0 | 0.16 | 0.91 | 1 | 0.96 | 0.94 | 0.95 | 0.92 |
| 3 | 120871957 | rs2331641 | -0.516 | -1.249 | -1.126 | -2.495 | 0.002 | 0.021 | 0.021 | 0 | 0.64 | 0.68 | 0.75 | 0.26 | 0.67 | 0.38 |
| 3 | 120977383 | rs9875986 | 2.968 | -0.525 | -0.882 | -2.116 | -0.05 | 0.06 | 0.06 | 0.051 | 0.1 | 0.01 | 0.1 | 0 | 0.15 | 0.06 |
| 3 | 120987767 | rs11921531 | 1.901 | -0.581 | -0.844 | -2.104 | 0.006 | 0.004 | 0.004 | 0 | 0.14 | 0.09 | 0.07 | 0.12 | 0.1 | 0.1 |
| 3 | 121003444 | rs338986 | -1.615 | -0.602 | -0.712 | -2.187 | 0.029 | -0.049 | -0.049 | 0.057 | 0.8 | 0.72 | 0.73 | 0.93 | 0.73 | 0.88 |
| 3 | 121009772 | rs338997 | -0.403 | -0.602 | -0.72 | -2.187 | 0.008 | -0.029 | -0.029 | 0.057 | 0.65 | 0.63 | 0.66 | 0.82 | 0.63 | 0.78 |
| 3 | 121015824 | rs13324292 | 1.046 | -0.653 | -0.667 | -2.208 | -0.006 | 0.007 | 0.007 | 1.156 | 0.06 | 0.2 | 0.25 | 0.07 | 0.05 | 0.08 |
| 3 | 121039333 | rs6787064 | 0.576 | -0.725 | -0.683 | -2.258 | 0.012 | 0.088 | 0.088 | 0 | 0.5 | 0.33 | 0.35 | 0.75 | 0.53 | 0.67 |
| 3 | 121080249 | rs7649908 | -1.611 | -0.429 | -0.479 | -2.092 | 0.036 | -0.052 | -0.052 | 1.193 | 0.8 | 0.68 | 0.74 | 0.93 | 0.74 | 0.88 |
| 3 | 121084087 | rs12489855 | 0.742 | -0.473 | -0.576 | -1.893 | -0.012 | 0.029 | 0.029 | 0.643 | 0.07 | 0.2 | 0.25 | 0.09 | 0.05 | 0.08 |
| 3 | 121100283 | rs17740066 | 2.722 | -0.335 | -0.884 | -2.087 | -0.028 | 0.026 | 0.026 | 0.059 | 0.07 | 0.01 | 0.15 | 0.07 | 0.1 | 0.06 |
| 3 | 121103502 | rs3914164 | 1.398 | -1.001 | -0.722 | -1.839 | 0.019 | 0.017 | 0.017 | 0 | 0.19 | 0.05 | 0.08 | 0.14 | 0.13 | 0.15 |
| 3 | 121119101 | rs6766105 | -0.303 | -0.277 | 0.003 | -1.238 | -0.004 | 0.002 | 0.002 | 0 | 0.26 | 0.23 | 0.42 | 0.73 | 0.3 | 0.64 |
| 3 | 121121958 | rs9878946 | 1.391 | -0.115 | 0.068 | -1.216 | 0.029 | -0.032 | -0.032 | 0 | 0.07 | 0.21 | 0.05 | 0 | 0.08 | 0.01 |
| 3 | 121137976 | rs6782025 | -1.629 | -0.104 | 0.018 | -1.249 | 0.039 | -0.052 | -0.052 | 1.372 | 0.8 | 0.64 | 0.74 | 0.88 | 0.75 | 0.88 |
| 3 | 121138015 | rs6782033 | -1.629 | -0.104 | 0.018 | -1.249 | 0.039 | -0.052 | -0.052 | 0 | 0.8 | 0.64 | 0.74 | 0.88 | 0.75 | 0.88 |
| 3 | 121141096 | rs17678695 | 3.028 | -0.074 | -0.076 | -1.292 | -0.008 | 0.036 | 0.036 | 0 | 0.07 | 0.01 | 0.06 | 0 | 0.06 | 0.01 |
| 3 | 121141718 | rs7632907 | -1.918 | -0.246 | 0.048 | -1.229 | 0.012 | -0.019 | -0.019 | 0 | 0.81 | 0.8 | 0.75 | 0.88 | 0.75 | 0.89 |
| 3 | 121146061 | rs2169302 | 1.849 | -0.222 | -0.071 | -1.102 | -0.019 | -0.02 | -0.02 | 0 | 0.12 | 0.04 | 0.11 | 0.18 | 0.14 | 0.25 |
| 3 | 121149009 | rs9862879 | 0.712 | -0.319 | -0.096 | -1.178 | -0.015 | -0.088 | -0.088 | 2.218 | 0.16 | 0.3 | 0.12 | 0 | 0.14 | 0.04 |
| 3 | 121154193 | rs6800901 | 1.72 | -0.282 | -0.176 | -1.188 | 0.02 | 0.137 | 0.137 | 0 | 0.38 | 0.11 | 0.21 | 0.15 | 0.32 | 0.21 |
| 3 | 121154974 | rs1381057 | 0.038 | -0.129 | 0.677 | -0.456 | 0.033 | -0.07 | -0.07 | 0 | 0.74 | 0.63 | 0.69 | 0.88 | 0.67 | 0.88 |
| 3 | 121155198 | rs2030531 | 0.778 | -0.126 | 0.681 | -0.454 | 0.012 | -0.095 | -0.095 | 0.279 | 0.65 | 0.54 | 0.63 | 0.88 | 0.61 | 0.85 |
| 3 | 121167632 | rs11709726 | 0.873 | -0.078 | 0.721 | -0.425 | -0.003 | -0.001 | -0.001 | 0 | 0.64 | 0.37 | 0.62 | 0.87 | 0.61 | 0.85 |
| 3 | 121186422 | rs532411 | 0.73 | -0.018 | 0.471 | -0.48 | 0.028 | -0.034 | -0.034 | 0 | 0.07 | 0.22 | 0.06 | 0 | 0.08 | 0.01 |
| 3 | 121186543 | rs1381058 | 0.826 | -0.023 | 0.473 | -0.488 | 0.007 | -0.069 | -0.069 | 0.721 | 0.64 | 0.48 | 0.62 | 0.88 | 0.61 | 0.85 |
| 3 | 121214952 | rs11719937 | 0.638 | 0.859 | 0.32 | 0.039 | 0.002 | 0.089 | 0.089 | 5.856 | 0.1 | 0.1 | 0.07 | 0.48 | 0.11 | 0.31 |
| 3 | 121228960 | rs487848 | -0.068 | 1.19 | -0.049 | -0.638 | 0.032 | -0.034 | -0.034 | 0 | 0.07 | 0.22 | 0.05 | 0 | 0.08 | 0.01 |
| 3 | 121239027 | rs702019 | -0.423 | 1.175 | -1.018 | -0.44 | -0.002 | 0 | 0 | 0 | 0.81 | 0.84 | 0.75 | 0.88 | 0.75 | 0.89 |
| 3 | 121280139 | rs4676681 | 0.807 | 1.295 | -1.554 | -1.369 | 0 | 0.081 | 0.081 | 0 | 0.53 | 0.33 | 0.3 | 0.3 | 0.48 | 0.54 |
| 3 | 121280791 | rs616966 | -0.308 | 0.908 | -1.217 | -1.078 | -0.005 | 0.001 | 0.001 | 0.188 | 0.29 | 0.6 | 0.25 | 0.61 | 0.24 | 0.35 |
| 3 | 121285031 | rs3101345 | 0.015 | 1.998 | -1.486 | -1.152 | -0.053 | 0.064 | 0.064 | 0 | 0.82 | 0.93 | 0.53 | 0.9 | 0.72 | 0.89 |
| 3 | 121288890 | rs9810670 | -0.193 | 1.678 | -1.261 | -1.389 | 0.004 | 0.031 | 0.031 | 0.001 | 0.51 | 0.38 | 0.68 | 0.65 | 0.55 | 0.47 |
| 3 | 121289499 | rs12485719 | -0.1 | 1.949 | -1.341 | -1.645 | -0.02 | 0.031 | 0.031 | 0 | 0.17 | 0.12 | 0.45 | 0.06 | 0.26 | 0.1 |
| 3 | 121294785 | rs4437078 | -1.046 | 2.347 | -0.949 | -1.844 | 0.011 | 0.016 | 0.016 | 0.059 | 0.6 | 0.61 | 0.76 | 0.75 | 0.7 | 0.61 |
| 3 | 121302410 | rs13065635 | 0.686 | 2.43 | -0.995 | -1.802 | -0.021 | -0.015 | -0.015 | 0 | 0.07 | 0.03 | 0.07 | 0.1 | 0.15 | 0.14 |
| 3 | 121302677 | rs11714812 | 0.95 | 2.377 | -0.978 | -1.834 | 0.002 | -0.001 | -0.001 | 0 | 0.33 | 0.21 | 0.23 | 0.58 | 0.29 | 0.38 |
| 3 | 121315813 | rs7631876 | 0.475 | 2.619 | -0.938 | -2.125 | -0.002 | 0.012 | 0.012 | 0.172 | 0.38 | 0.24 | 0.29 | 0.62 | 0.4 | 0.41 |
| 3 | 121319824 | rs6775899 | 1.531 | 2.41 | -0.972 | -2.431 | 0.009 | 0.01 | 0.01 | 0.011 | 0.29 | 0.32 | 0.15 | 0.19 | 0.18 | 0.36 |
| 3 | 121320486 | rs4676736 | -0.31 | 2.565 | -1.152 | -2.431 | 0.004 | 0.014 | 0.014 | 0.017 | 0.53 | 0.49 | 0.7 | 0.68 | 0.59 | 0.49 |
| 3 | 121323465 | rs6808258 | 0.092 | 2.575 | -1.161 | -2.426 | 0.01 | -0.037 | -0.037 | 0.589 | 0.15 | 0.21 | 0.42 | 0.05 | 0.19 | 0.08 |
| 3 | 121324643 | rs1464099 | -1.189 | 2.558 | -1.379 | -2.327 | 0.003 | 0.014 | 0.014 | 0 | 0.89 | 0.95 | 0.91 | 0.96 | 0.88 | 0.96 |
| 3 | 121324957 | rs17691111 | 1.403 | 2.09 | -1.575 | -2.323 | -0.052 | 0.12 | 0.12 | 0.287 | 0.25 | 0.01 | 0.16 | 0.05 | 0.26 | 0.15 |

| | | | | | | | | | | | | | | | | |
|---|---|---|---|---|---|---|---|---|---|---|---|---|---|---|---|---|
| 3 | 121327184 | rs6773252 | 0.819 | 1.294 | -1.155 | -2.098 | -0.033 | -0.068 | -0.068 | 0.004 | 0.33 | 0.55 | 0.18 | 0.19 | 0.21 | 0.39 |
| 3 | 121329372 | rs12491882 | 0.025 | 1.416 | -1.353 | -1.58 | -0.015 | 0.021 | 0.021 | 0 | 0.4 | 0.21 | 0.3 | 0.64 | 0.46 | 0.39 |
| 3 | 121332245 | rs3845862 | -0.552 | 1.012 | -1.307 | -1.611 | 0.002 | -0.058 | -0.058 | 0 | 0.43 | 0.54 | 0.63 | 0.1 | 0.48 | 0.21 |
| 3 | 121335146 | rs938391 | -0.325 | 1.137 | -1.155 | -1.47 | -0.016 | 0.12 | 0.12 | 0 | 0.65 | 0.86 | 0.77 | 0.93 | 0.61 | 0.81 |
| 3 | 121335461 | rs1567990 | -1.008 | 1.022 | -0.319 | -0.209 | 0.013 | -0.039 | -0.039 | 0 | 0.15 | 0.41 | 0.43 | 0.08 | 0.16 | 0.06 |
| 3 | 121339354 | rs7640859 | 0.317 | 0.779 | -0.913 | -0.092 | 0.004 | 0.018 | 0.018 | 0 | 0.9 | 0.98 | 0.96 | 0.95 | 0.92 | 0.86 |
| 3 | 121343500 | rs1492178 | 1.082 | 0.63 | -1.055 | 0.429 | 0.004 | -0.015 | -0.015 | 0 | 0.76 | 0.81 | 0.82 | 0.72 | 0.78 | 0.69 |
| 3 | 121343534 | rs1492177 | 0.126 | 0.811 | -1.039 | -0.038 | -0.001 | 0.004 | 0.004 | 0 | 0.24 | 0.17 | 0.5 | 0.14 | 0.25 | 0.41 |
| 3 | 121344140 | rs3772130 | 1.052 | 1.035 | -1.116 | -0.314 | -0.015 | 0.011 | 0.011 | 0.045 | 0.19 | 0.13 | 0.14 | 0.09 | 0.28 | 0.09 |
| 3 | 121346656 | rs2141488 | -0.2 | 0.874 | 0.063 | -0.385 | 0.004 | -0.016 | -0.016 | 0 | 0.76 | 0.81 | 0.82 | 0.72 | 0.78 | 0.69 |
| 3 | 121346930 | rs9810952 | 0.721 | 0.873 | 0.063 | -0.355 | 0.002 | 0.007 | 0.007 | 0 | 0.5 | 0.43 | 0.31 | 0.57 | 0.53 | 0.28 |
| 3 | 121348499 | rs1873645 | 0.675 | 0.832 | 0.164 | -0.373 | 0 | -0.057 | -0.057 | 2.623 | 0.26 | 0.43 | 0.52 | 0.15 | 0.25 | 0.41 |
| 3 | 121350593 | rs1128158 | 0.071 | 0.831 | 0.162 | -0.374 | 0.004 | -0.016 | -0.016 | 1.7 | 0.24 | 0.18 | 0.18 | 0.28 | 0.22 | 0.3 |
| 3 | 121350616 | rs1128154 | 0.057 | 0.831 | 0.162 | -0.374 | 0.004 | -0.016 | -0.016 | 0 | 0.24 | 0.19 | 0.18 | 0.28 | 0.22 | 0.3 |
| 3 | 121351338 | rs2070180 | 1.119 | 0.842 | 0.147 | -0.425 | -0.015 | 0.011 | 0.011 | 0.045 | 0.19 | 0.13 | 0.14 | 0.09 | 0.28 | 0.1 |
| 3 | 121353254 | rs2070179 | -0.645 | 0.729 | 0.176 | -0.457 | -0.004 | -0.056 | -0.056 | 1.06 | 0.73 | 0.57 | 0.48 | 0.85 | 0.75 | 0.58 |
| 3 | 121387784 | rs9819530 | 0.136 | 0.677 | 0.108 | -0.377 | -0.03 | -0.014 | -0.014 | 0 | 0.3 | 0.2 | 0.26 | 0.3 | 0.41 | 0.24 |
| 3 | 121394749 | rs6766737 | 0.451 | 0.54 | 0.276 | -0.399 | -0.006 | -0.018 | -0.018 | 0 | 0.14 | 0.19 | 0.06 | 0.07 | 0.09 | 0.16 |
| 3 | 121412892 | rs11915822 | 0.451 | 0.54 | 0.346 | -0.399 | -0.008 | -0.023 | -0.023 | 0.181 | 0.14 | 0.19 | 0.06 | 0.07 | 0.09 | 0.16 |
| 3 | 121414061 | rs1127412 | 1.785 | 0.537 | 0.41 | -0.426 | 0.01 | -0.039 | -0.039 | 1.782 | 0.29 | 0.16 | 0.16 | 0.48 | 0.25 | 0.19 |
| 3 | 121415610 | rs33988592 | 1.213 | 0.588 | 0.39 | -0.463 | -0.014 | 0.01 | 0.01 | 0.003 | 0.19 | 0.13 | 0.14 | 0.09 | 0.28 | 0.09 |
| 3 | 121415720 | rs3732410 | 1.885 | 0.529 | 0.356 | -0.476 | 0 | -0.128 | -0.128 | 0 | 0.28 | 0.05 | 0.16 | 0.48 | 0.25 | 0.19 |
| 3 | 121459589 | rs12498138 | 1.679 | 0.701 | 0.084 | -0.713 | -0.011 | -0.023 | -0.023 | 0 | 0.05 | 0.01 | 0.36 | 0.08 | 0.07 | 0.04 |
| 3 | 121465220 | rs9849087 | 1.207 | 0.687 | 0.144 | -0.68 | -0.004 | -0.037 | -0.037 | 0.249 | 0.11 | 0.07 | 0.12 | 0.21 | 0.13 | 0.14 |
| 3 | 121469417 | rs1919552 | -0.543 | 0.718 | 0.423 | -1.078 | -0.001 | -0.043 | -0.043 | 0 | 0.73 | 0.59 | 0.48 | 0.85 | 0.75 | 0.58 |
| 3 | 121480038 | rs10934559 | 0.613 | 1.36 | 0.39 | -1.19 | 0.104 | -0.004 | -0.004 | 0 | 0.07 | 0.23 | 0.06 | 0 | 0.13 | 0.02 |
| 3 | 121481801 | rs4676744 | -1.294 | 1.139 | 0.732 | -1.007 | 0.023 | 0.028 | 0.028 | 0 | 0.59 | 0.76 | 0.7 | 0.33 | 0.62 | 0.68 |
| 3 | 121488353 | rs9878086 | 0.708 | 1.076 | 0.937 | -1.06 | 0.038 | -0.031 | -0.031 | 0 | 0.21 | 0.27 | 0.14 | 0.09 | 0.28 | 0.1 |
| 3 | 121499506 | rs12492861 | 1.121 | 1.076 | 0.99 | -1.066 | 0.003 | 0.009 | 0.009 | 0 | 0.12 | 0.1 | 0.05 | 0.07 | 0.09 | 0.15 |
| 3 | 121509881 | rs6808500 | -1.336 | 1.076 | 0.99 | -1.066 | 0.023 | 0.029 | 0.029 | 0 | 0.59 | 0.76 | 0.7 | 0.33 | 0.62 | 0.68 |
| 3 | 121513185 | rs12489511 | 0.593 | 1.128 | 0.987 | -1.069 | -0.004 | -0.023 | -0.023 | 0.049 | 0.27 | 0.39 | 0.51 | 0.16 | 0.25 | 0.44 |
| 3 | 121514161 | rs4311252 | 0.85 | 1.101 | 0.993 | -1.066 | 0.043 | -0.029 | -0.029 | 0.27 | 0.2 | 0.27 | 0.14 | 0.09 | 0.28 | 0.09 |
| 3 | 121526107 | rs4280673 | 0.795 | 1.072 | 0.97 | -1.079 | 0.041 | -0.03 | -0.03 | 0 | 0.21 | 0.27 | 0.14 | 0.09 | 0.28 | 0.09 |
| 3 | 121526849 | rs6770719 | 0.702 | 1.087 | 1.009 | -1.083 | 0 | -0.005 | -0.005 | 0 | 0.26 | 0.32 | 0.51 | 0.16 | 0.25 | 0.44 |
| 3 | 121536497 | rs4482697 | -0.369 | 1.087 | 0.996 | -1.083 | -0.003 | -0.017 | -0.017 | 0.172 | 0.38 | 0.49 | 0.56 | 0.23 | 0.34 | 0.59 |
| 3 | 121554329 | rs3805125 | 0.809 | 1.154 | 0.971 | -0.991 | -0.001 | -0.005 | -0.005 | 0 | 0.13 | 0.16 | 0.06 | 0.07 | 0.09 | 0.15 |
| 3 | 121556219 | rs9826473 | 0.732 | 1.187 | 0.874 | -0.908 | 0.044 | -0.023 | -0.023 | 3.304 | 0.2 | 0.27 | 0.14 | 0.11 | 0.28 | 0.09 |
| 3 | 121563355 | rs9884018 | 0.614 | 0.928 | 0.69 | -0.72 | 0.001 | 0.012 | 0.012 | 0 | 0.14 | 0.13 | 0.15 | 0.62 | 0.14 | 0.2 |
| 3 | 121564989 | rs6767409 | -0.321 | 0.886 | 0.781 | -0.851 | 0.025 | -0.047 | -0.047 | 0 | 0.34 | 0.44 | 0.2 | 0.18 | 0.37 | 0.25 |
| 3 | 121568595 | rs10511407 | 1.019 | 0.955 | 0.685 | -1.165 | 0.003 | 0.016 | 0.016 | 0 | 0.12 | 0.1 | 0.05 | 0.07 | 0.09 | 0.15 |
| 3 | 121573850 | rs7637495 | -1.999 | 0.912 | 0.563 | -1.257 | 0.007 | 0.117 | 0.117 | 0 | 0.72 | 0.91 | 0.63 | 0.95 | 0.74 | 0.88 |
| 3 | 121574215 | rs7340674 | 0.029 | 0.884 | 0.882 | -1.227 | -0.016 | 0.009 | 0.009 | 0 | 0.33 | 0.24 | 0.28 | 0.72 | 0.41 | 0.29 |
| 3 | 121584999 | rs2700413 | 0.789 | 0.825 | 0.906 | -1.225 | -0.002 | -0.022 | -0.022 | 0.117 | 0.13 | 0.25 | 0.06 | 0.08 | 0.09 | 0.15 |
| 3 | 121593769 | rs1920299 | 1.927 | 0.82 | 0.816 | -1.242 | 0.008 | 0.116 | 0.116 | 0 | 0.28 | 0.1 | 0.37 | 0.05 | 0.26 | 0.12 |
| 3 | 121597022 | rs7620562 | -1.137 | 0.833 | 0.844 | -1.235 | 0.002 | 0.008 | 0.008 | 0 | 0.88 | 0.88 | 0.95 | 0.92 | 0.91 | 0.85 |
| 3 | 121597888 | rs1920298 | 1.92 | 0.805 | 0.799 | -1.246 | 0.009 | 0.112 | 0.112 | 0 | 0.28 | 0.11 | 0.37 | 0.05 | 0.26 | 0.12 |
| 3 | 121610395 | rs1920307 | -0.525 | 0.777 | 0.93 | -1.124 | -0.007 | -0.03 | -0.03 | 0 | 0.86 | 0.68 | 0.93 | 0.92 | 0.91 | 0.85 |
| 3 | 121613054 | rs10470412 | -1.147 | 0.787 | 0.843 | -1.119 | 0.001 | 0.012 | 0.012 | 0 | 0.88 | 0.91 | 0.95 | 0.92 | 0.91 | 0.85 |
| 3 | 121614175 | rs1881998 | -0.444 | 0.826 | 0.931 | -1.126 | -0.006 | -0.032 | -0.032 | 0 | 0.86 | 0.68 | 0.93 | 0.92 | 0.91 | 0.85 |
| 3 | 121618056 | rs3805134 | -0.397 | 0.906 | 0.825 | -1.359 | 0.009 | -0.019 | -0.019 | 0 | 0.38 | 0.49 | 0.3 | 0.69 | 0.42 | 0.3 |
| 3 | 121618371 | rs9830721 | -0.397 | 0.906 | 0.825 | -1.359 | 0.009 | -0.019 | -0.019 | 0 | 0.38 | 0.49 | 0.3 | 0.69 | 0.42 | 0.3 |
| 3 | 121620787 | rs2141615 | -0.397 | 0.906 | 0.825 | -1.359 | 0.009 | -0.019 | -0.019 | 1.422 | 0.38 | 0.49 | 0.3 | 0.69 | 0.42 | 0.3 |
| 3 | 121630194 | rs6803757 | -0.397 | 0.906 | 0.827 | -1.359 | 0.009 | -0.02 | -0.02 | 0 | 0.38 | 0.49 | 0.29 | 0.69 | 0.42 | 0.3 |
| 3 | 121630328 | rs6438687 | -0.397 | 0.906 | 0.827 | -1.359 | 0.009 | -0.02 | -0.02 | 0 | 0.38 | 0.49 | 0.29 | 0.69 | 0.42 | 0.3 |
| 3 | 121632432 | rs866927 | -0.397 | 0.906 | 0.827 | -1.359 | 0.009 | -0.02 | -0.02 | 0.017 | 0.38 | 0.49 | 0.29 | 0.69 | 0.42 | 0.3 |
| 3 | 121632718 | rs953730 | -0.942 | 0.631 | 0.905 | -1.61 | 0.014 | -0.062 | -0.062 | 0.688 | 0.8 | 0.69 | 0.72 | 0.93 | 0.77 | 0.61 |
| 3 | 121633503 | rs9862977 | -0.283 | 0.6 | 0.927 | -1.58 | 0.009 | -0.021 | -0.021 | 0 | 0.38 | 0.48 | 0.29 | 0.7 | 0.42 | 0.3 |
| 3 | 121633904 | rs2049330 | -0.283 | 0.6 | 0.927 | -1.58 | 0.009 | -0.021 | -0.021 | 0 | 0.38 | 0.48 | 0.29 | 0.7 | 0.42 | 0.3 |
| 3 | 121633964 | rs2049331 | -0.283 | 0.6 | 0.927 | -1.57 | 0.009 | -0.02 | -0.02 | 0.593 | 0.38 | 0.48 | 0.29 | 0.7 | 0.42 | 0.3 |
| 3 | 121634887 | rs1357531 | -0.283 | 0.6 | 0.927 | -1.587 | 0.009 | -0.021 | -0.021 | 1.417 | 0.38 | 0.48 | 0.29 | 0.7 | 0.42 | 0.3 |
| 3 | 121635230 | rs2700420 | -0.283 | 0.6 | 0.927 | -1.587 | 0.009 | -0.021 | -0.021 | 0 | 0.38 | 0.48 | 0.29 | 0.7 | 0.42 | 0.3 |
| 3 | 121636223 | rs11914993 | -0.283 | 0.596 | 1.058 | -1.589 | 0.009 | -0.021 | -0.021 | 0 | 0.38 | 0.48 | 0.29 | 0.7 | 0.42 | 0.3 |
| 3 | 121641352 | rs11920521 | -0.274 | 0.594 | 1.057 | -1.59 | 0.009 | -0.021 | -0.021 | 0.231 | 0.38 | 0.48 | 0.29 | 0.7 | 0.42 | 0.3 |
| 3 | 121641528 | rs7637569 | -0.283 | 0.596 | 1.079 | -1.589 | 0.009 | -0.021 | -0.021 | 0 | 0.38 | 0.48 | 0.29 | 0.7 | 0.42 | 0.3 |
| 3 | 121643699 | rs2257132 | -0.283 | 0.596 | 1.065 | -1.589 | 0.009 | -0.021 | -0.021 | 0 | 0.38 | 0.48 | 0.29 | 0.7 | 0.42 | 0.3 |
| 3 | 121643804 | rs2257212 | -0.283 | 0.596 | 1.065 | -1.589 | 0.009 | -0.021 | -0.021 | 0 | 0.38 | 0.48 | 0.29 | 0.7 | 0.42 | 0.3 |
| 3 | 121643893 | rs2257214 | -0.283 | 0.596 | 1.065 | -1.589 | 0.009 | -0.021 | -0.021 | 0.984 | 0.38 | 0.48 | 0.29 | 0.7 | 0.42 | 0.3 |
| 3 | 121644417 | rs1920319 | -0.283 | 0.596 | 1.065 | -1.589 | 0.009 | -0.021 | -0.021 | 0.284 | 0.38 | 0.48 | 0.29 | 0.7 | 0.42 | 0.3 |
| 3 | 121646641 | rs1143670 | -0.243 | 0.596 | 1.141 | -1.589 | 0.01 | -0.022 | -0.022 | 0 | 0.37 | 0.43 | 0.29 | 0.7 | 0.42 | 0.3 |

| | | | | | | | | | | | | | | | | |
|---|---|---|---|---|---|---|---|---|---|---|---|---|---|---|---|---|
| 3 | 121646759 | rs3762819 | -0.284 | 0.596 | 1.104 | -1.589 | 0.009 | -0.021 | -0.021 | 0 | 0.38 | 0.48 | 0.29 | 0.7 | 0.42 | 0.3 |
| 3 | 121646886 | rs3762821 | 0.032 | 0.586 | 1.174 | -1.593 | -0.018 | 0.019 | 0.019 | 0 | 0.35 | 0.27 | 0.28 | 0.7 | 0.42 | 0.3 |
| 3 | 121647286 | rs1143671 | -0.291 | 0.587 | 1.164 | -1.593 | 0.009 | -0.021 | -0.021 | 0.225 | 0.38 | 0.48 | 0.29 | 0.7 | 0.42 | 0.3 |
| 3 | 121648168 | rs1143672 | -0.291 | 0.587 | 1.164 | -1.593 | 0.009 | -0.021 | -0.021 | 0 | 0.38 | 0.48 | 0.29 | 0.7 | 0.42 | 0.3 |
| 3 | 121657967 | rs874742 | -0.187 | 0.592 | 1.165 | -1.59 | 0.013 | -0.024 | -0.024 | 0 | 0.37 | 0.47 | 0.29 | 0.7 | 0.43 | 0.3 |
| 3 | 121658082 | rs874741 | -0.162 | 0.592 | 1.165 | -1.59 | 0.013 | -0.024 | -0.024 | 0 | 0.37 | 0.47 | 0.29 | 0.7 | 0.43 | 0.3 |
| 3 | 121659197 | rs3817599 | -0.553 | 0.655 | 1.135 | -1.615 | 0 | -0.027 | -0.027 | 0.357 | 0.76 | 0.7 | 0.7 | 0.87 | 0.76 | 0.6 |
| 3 | 121662400 | rs1920310 | 1.316 | 0.478 | 1.093 | -1.585 | 0.005 | 0.004 | 0.004 | 0 | 0.11 | 0.09 | 0.03 | 0.1 | 0.06 | 0.16 |
| 3 | 121663962 | rs9829181 | -0.192 | 0.455 | 1.135 | -1.6 | -0.008 | -0.084 | -0.084 | 4.786 | 0.73 | 0.55 | 0.69 | 0.87 | 0.76 | 0.6 |
| 3 | 121664112 | rs9867460 | 0.194 | 0.474 | 1.107 | -1.604 | -0.018 | 0.018 | 0.018 | 0 | 0.35 | 0.27 | 0.28 | 0.7 | 0.43 | 0.3 |
| 3 | 121665702 | rs6438689 | -0.512 | 0.519 | 1.069 | -1.57 | 0 | -0.027 | -0.027 | 0 | 0.76 | 0.7 | 0.7 | 0.87 | 0.76 | 0.6 |
| 3 | 121665950 | rs7615571 | 2.099 | 0.519 | 0.953 | -1.571 | 0.005 | 0.114 | 0.114 | 0 | 0.28 | 0.12 | 0.38 | 0.07 | 0.27 | 0.14 |
| 3 | 121670405 | rs9871743 | -0.405 | 0.526 | 0.963 | -1.562 | -0.002 | 0.006 | 0.006 | 0 | 0.46 | 0.44 | 0.31 | 0.8 | 0.49 | 0.46 |
| 3 | 121706021 | rs12491991 | 1.949 | 0.614 | 0.437 | -1.465 | 0.002 | 0.12 | 0.12 | 0 | 0.28 | 0.11 | 0.38 | 0.07 | 0.27 | 0.14 |
| 3 | 121712051 | rs2877561 | -1.739 | 0.604 | 0.524 | -1.498 | 0.005 | 0.04 | 0.04 | 0 | 0.71 | 0.83 | 0.6 | 0.92 | 0.71 | 0.8 |
| 3 | 121715319 | rs9868257 | 0.089 | 0.839 | 0.163 | -1.138 | -0.031 | 0.013 | 0.013 | 0 | 0.31 | 0.24 | 0.25 | 0.71 | 0.39 | 0.31 |
| 3 | 121717577 | rs9824120 | -0.846 | 0.952 | 0.189 | -1.175 | -0.001 | 0.042 | 0.042 | 0 | 0.47 | 0.49 | 0.33 | 0.79 | 0.5 | 0.4 |
| 3 | 121723700 | rs12489988 | 0.087 | 0.779 | 0.016 | -1.032 | -0.003 | -0.01 | -0.01 | 0.014 | 0.15 | 0.23 | 0.08 | 0.11 | 0.1 | 0.11 |
| 3 | 121728846 | rs2001212 | 0.06 | 1.049 | 0.037 | -1.103 | -0.001 | 0 | 0 | 0 | 0.67 | 0.64 | 0.74 | 0.28 | 0.6 | 0.73 |
| 3 | 121729899 | rs11929195 | -0.082 | 1.264 | -0.365 | -1.079 | -0.009 | 0.012 | 0.012 | 0 | 0.69 | 0.75 | 0.74 | 0.3 | 0.67 | 0.72 |
| 3 | 121740065 | rs7627252 | -0.024 | 1.418 | -0.993 | -1.539 | -0.005 | 0.002 | 0.002 | 0 | 0.21 | 0.16 | 0.15 | 0.05 | 0.29 | 0.28 |
| 3 | 121740304 | rs9289186 | -0.667 | 1.124 | -0.474 | -1.682 | 0.021 | -0.041 | -0.041 | 0 | 0.89 | 0.83 | 0.93 | 0.99 | 0.91 | 0.97 |
| 3 | 121744849 | rs4350947 | 1.454 | 0.78 | -1.098 | -1.994 | -0.006 | 0.056 | 0.056 | 0 | 0.32 | 0.14 | 0.38 | 0.23 | 0.28 | 0.33 |
| 3 | 121751531 | rs1962532 | -1.357 | 0.138 | -0.96 | -1.699 | 0.013 | 0.143 | 0.143 | 0.795 | 0.49 | 0.75 | 0.31 | 0.28 | 0.5 | 0.63 |
| 3 | 121767054 | rs12106790 | 1.169 | 0.087 | -0.81 | -1.539 | 0.004 | 0 | 0 | 0.018 | 0.19 | 0.15 | 0.12 | 0.04 | 0.2 | 0.25 |
| 3 | 121770858 | rs2715267 | 0.542 | 0.175 | -0.51 | -1.462 | 0.021 | 0.047 | 0.047 | 0.932 | 0.6 | 0.37 | 0.76 | 0.73 | 0.63 | 0.5 |
| 3 | 121775134 | rs13082681 | 0.449 | 0.166 | 0.189 | -0.78 | -0.001 | -0.001 | -0.001 | 0 | 0.07 | 0.09 | 0.06 | 0.1 | 0.08 | 0.09 |
| 3 | 121778970 | rs2681404 | 0.684 | 0.221 | 0.511 | -0.853 | 0.022 | -0.042 | -0.042 | 1.043 | 0.11 | 0.23 | 0.08 | 0.01 | 0.12 | 0.04 |
| 3 | 121781386 | rs2715276 | 0.213 | 0.206 | 0.036 | -0.835 | -0.007 | 0.022 | 0.022 | 0 | 0.15 | 0.11 | 0.11 | 0.01 | 0.17 | 0.17 |
| 3 | 121781426 | rs17203397 | 2.044 | -0.029 | -0.096 | -0.62 | 0.026 | 0.081 | 0.081 | 0.331 | 0.55 | 0.19 | 0.64 | 0.33 | 0.62 | 0.53 |
| 3 | 121781723 | rs2715274 | 0.203 | 0.08 | -0.183 | -0.467 | 0.001 | -0.036 | -0.036 | 0.251 | 0.16 | 0.2 | 0.12 | 0.01 | 0.17 | 0.17 |
| 3 | 121784718 | rs13075281 | 0.101 | 0.08 | -0.206 | -0.423 | -0.003 | 0.018 | 0.018 | 0 | 0.09 | 0.18 | 0.07 | 0.1 | 0.08 | 0.1 |
| 3 | 121785631 | rs2681411 | 1.703 | 0.065 | -0.038 | -0.452 | 0.013 | 0.023 | 0.023 | 0 | 0.58 | 0.31 | 0.68 | 0.33 | 0.64 | 0.55 |
| 3 | 121786069 | rs13074706 | 0.574 | 0.034 | -0.126 | -0.409 | -0.001 | -0.001 | -0.001 | 0 | 0.08 | 0.08 | 0.06 | 0.1 | 0.08 | 0.1 |
| 3 | 121796768 | rs9282641 | 1.473 | -0.002 | -0.167 | -0.37 | 0.005 | 0.052 | 0.052 | 0.711 | 0.12 | 0.02 | 0.08 | 0 | 0.09 | 0.09 |
| 3 | 121799554 | rs2681422 | -0.923 | 0.056 | -0.114 | -0.288 | -0.028 | 0.183 | 0.183 | 0.005 | 0.83 | 0.63 | 0.85 | 0.45 | 0.89 | 0.83 |
| 3 | 121801230 | rs2681426 | 0.785 | 0 | -0.131 | -0.299 | -0.002 | -0.001 | -0.001 | 0.263 | 0.11 | 0.11 | 0.08 | 0.01 | 0.12 | 0.05 |
| 3 | 121806825 | rs11717893 | -1.106 | -0.073 | -0.189 | -0.295 | 0 | -0.013 | -0.013 | 0.009 | 0.22 | 0.31 | 0.17 | 0.12 | 0.22 | 0.21 |
| 3 | 121811811 | rs9836399 | 1.248 | 0.081 | -0.612 | -0.065 | -0.005 | -0.024 | -0.024 | 0 | 0.59 | 0.47 | 0.66 | 0.33 | 0.6 | 0.58 |
| 3 | 121815615 | rs2681415 | -0.291 | -0.205 | -0.242 | 0.451 | -0.038 | 0.178 | 0.178 | 0.021 | 0.82 | 0.61 | 0.84 | 0.47 | 0.87 | 0.81 |
| 3 | 121819586 | rs3792285 | -0.316 | 0.201 | -0.494 | -0.036 | 0.002 | -0.008 | -0.008 | 0 | 0.1 | 0.09 | 0.08 | 0.11 | 0.1 | 0.11 |
| 3 | 121825197 | rs2681417 | 0.445 | 0.14 | 0.019 | -0.398 | -0.001 | -0.055 | -0.055 | 0 | 0.9 | 0.66 | 0.94 | 1 | 0.94 | 0.88 |
| 3 | 121828837 | rs2254911 | 0.054 | 0.126 | 0.178 | -0.405 | 0.006 | -0.039 | -0.039 | 0 | 0.92 | 0.8 | 0.95 | 1 | 0.94 | 0.9 |
| 3 | 121831497 | rs9848900 | 0.756 | -0.215 | -0.389 | -0.162 | -0.033 | 0.019 | 0.019 | 0 | 0.64 | 0.48 | 0.76 | 0.39 | 0.73 | 0.41 |
| 3 | 121834057 | rs2681420 | -0.063 | -0.023 | -1.09 | -0.047 | 0.024 | -0.063 | -0.063 | 0.308 | 0.26 | 0.18 | 0.19 | 0.6 | 0.21 | 0.47 |
| 3 | 121838319 | rs1129055 | -0.182 | 0.309 | -1.213 | -0.104 | -0.009 | -0.076 | -0.076 | 0 | 0.27 | 0.16 | 0.21 | 0.6 | 0.28 | 0.48 |
| 3 | 121838791 | rs1915087 | -0.608 | 0.355 | -1.179 | -0.098 | -0.004 | -0.061 | -0.061 | 0 | 0.34 | 0.17 | 0.23 | 0.6 | 0.33 | 0.6 |
| 3 | 121843127 | rs2681401 | -0.5 | 0.36 | -1.237 | -1.105 | -0.019 | 0.062 | 0.062 | 0.012 | 0.59 | 0.73 | 0.42 | 0.1 | 0.56 | 0.37 |
| 3 | 121850297 | rs6438697 | 0.555 | 0.098 | -1.044 | -0.652 | 0.011 | -0.014 | -0.014 | 0 | 0.7 | 0.76 | 0.84 | 0.36 | 0.73 | 0.52 |
| 3 | 121850554 | rs9818482 | 1.045 | -0.063 | -0.635 | -1.133 | -0.004 | -0.004 | -0.004 | 0 | 0.56 | 0.47 | 0.74 | 0.34 | 0.59 | 0.35 |
| 3 | 121854657 | rs9839875 | -0.368 | -0.212 | -0.727 | -1.125 | 0.014 | -0.04 | -0.04 | 1.849 | 0.78 | 0.89 | 0.87 | 0.37 | 0.78 | 0.56 |
| 3 | 121865014 | rs937629 | -0.029 | -0.241 | -0.641 | -1.137 | -0.012 | 0.022 | 0.022 | 0 | 0.75 | 0.73 | 0.86 | 0.37 | 0.78 | 0.57 |
| 3 | 121873433 | rs11721321 | 1.392 | -0.242 | -0.969 | -1.705 | -0.041 | -0.019 | -0.019 | 0 | 0.34 | 0.2 | 0.6 | 0.34 | 0.43 | 0.23 |
| 3 | 121889088 | rs7652589 | 0.158 | -0.327 | -0.581 | -1.609 | -0.003 | 0.011 | 0.011 | 0.823 | 0.66 | 0.59 | 0.79 | 0.36 | 0.64 | 0.45 |
| 3 | 121899848 | rs7648041 | 0.691 | 0.188 | -0.817 | -1.941 | -0.017 | 0.007 | 0.007 | 0 | 0.22 | 0.16 | 0.19 | 0.01 | 0.31 | 0.17 |
| 3 | 121900462 | rs4678013 | -0.734 | 0.18 | -0.817 | -1.693 | -0.01 | 0.025 | 0.025 | 0 | 0.79 | 0.8 | 0.89 | 0.39 | 0.82 | 0.58 |
| 3 | 121902021 | rs34028592 | 1.451 | 0.147 | -0.76 | -1.671 | 0.013 | -0.042 | -0.042 | 0.022 | 0.13 | 0.22 | 0.1 | 0.02 | 0.15 | 0.13 |
| 3 | 121902907 | rs1048213 | 1.953 | 0.129 | -0.867 | -1.704 | -0.064 | -0.133 | -0.133 | 1.407 | 0.23 | 0.03 | 0.45 | 0.35 | 0.23 | 0.16 |
| 3 | 121905657 | rs6762782 | 0.079 | 0.116 | -0.851 | -1.71 | -0.009 | 0.018 | 0.018 | 1.26 | 0.67 | 0.57 | 0.79 | 0.36 | 0.67 | 0.45 |
| 3 | 121908434 | rs4678029 | 1.68 | 0.239 | -0.834 | -1.394 | -0.064 | -0.133 | -0.133 | 0 | 0.23 | 0.03 | 0.45 | 0.35 | 0.23 | 0.16 |
| 3 | 121913370 | rs937627 | 1.575 | 0.307 | -1.18 | -1.381 | -0.041 | -0.11 | -0.11 | 4.42 | 0.24 | 0.06 | 0.45 | 0.35 | 0.23 | 0.16 |
| 3 | 121916109 | rs6438706 | 1.134 | 0.312 | -1.107 | -1.427 | 0.013 | -0.045 | -0.045 | 0 | 0.13 | 0.23 | 0.1 | 0.02 | 0.15 | 0.13 |
| 3 | 121916325 | rs6438707 | -1.413 | 0.066 | -1.179 | -1.311 | 0.003 | 0.03 | 0.03 | 0 | 0.88 | 0.93 | 0.94 | 0.45 | 0.92 | 0.62 |
| 3 | 121918491 | rs1814740 | -0.135 | 0.359 | -1.08 | -1.66 | 0.001 | 0.078 | 0.078 | 0 | 0.52 | 0.64 | 0.39 | 0.08 | 0.54 | 0.33 |
| 3 | 121934161 | rs4678126 | 0.786 | 0.59 | -0.763 | -1.175 | 0.007 | -0.023 | -0.023 | 0 | 0.13 | 0.17 | 0.1 | 0.04 | 0.15 | 0.14 |
| 3 | 121937943 | rs35274320 | 0.94 | 0.614 | -0.778 | -1.19 | 0.003 | -0.019 | -0.019 | 0.009 | 0.12 | 0.16 | 0.09 | 0.04 | 0.14 | 0.08 |
| 3 | 121954077 | rs1354162 | 1.435 | 0.303 | -1.228 | -1.54 | -0.002 | 0.108 | 0.108 | 0.132 | 0.18 | 0.02 | 0.11 | 0 | 0.12 | 0.1 |
| 3 | 121963877 | rs6438715 | -0.24 | 0.391 | -0.568 | -0.946 | -0.011 | 0.048 | 0.048 | 0 | 0.82 | 0.79 | 0.9 | 0.49 | 0.87 | 0.69 |
| 3 | 121965745 | rs7632399 | -1.247 | 0.383 | -0.576 | -0.908 | -0.018 | 0.113 | 0.113 | 0 | 0.9 | 0.9 | 0.95 | 0.5 | 0.92 | 0.78 |

| | | | | | | | | | | | | | | | | |
|---|---|---|---|---|---|---|---|---|---|---|---|---|---|---|---|---|
| 3 | 121966952 | rs7637874 | 0.82 | 0.583 | -0.353 | -1.146 | 0.002 | -0.035 | -0.035 | 1.233 | 0.16 | 0.21 | 0.18 | 0 | 0.25 | 0.1 |
| 3 | 121968267 | rs34345120 | 1.254 | 0.231 | -0.159 | -0.558 | -0.004 | 0.02 | 0.02 | 0 | 0.06 | 0.03 | 0.03 | 0.01 | 0.05 | 0.09 |
| 3 | 121969937 | rs1463890 | -0.105 | 0.228 | -0.154 | -0.547 | -0.004 | 0.034 | 0.034 | 0 | 0.83 | 0.86 | 0.91 | 0.49 | 0.87 | 0.69 |
| 3 | 121970020 | rs7647446 | 0.634 | 0.255 | -0.093 | -0.588 | 0.011 | -0.025 | -0.025 | 0 | 0.13 | 0.18 | 0.1 | 0.04 | 0.16 | 0.14 |
| 3 | 121970301 | rs1965358 | 0.65 | 0.238 | -0.195 | -0.515 | -0.005 | 0.033 | 0.033 | 0.773 | 0.7 | 0.62 | 0.8 | 0.45 | 0.71 | 0.53 |
| 3 | 121971512 | rs937625 | -0.684 | 0.048 | -0.377 | 0.153 | 0.003 | 0.046 | 0.046 | 0 | 0.9 | 0.95 | 0.95 | 0.5 | 0.92 | 0.78 |
| 3 | 121976926 | rs10222633 | 0.115 | 0.875 | 0.234 | -0.519 | 0.006 | 0.003 | 0.003 | 0 | 0.38 | 0.4 | 0.34 | 0.4 | 0.51 | 0.25 |
| 3 | 121977125 | rs10222469 | -0.559 | 0.907 | 0.043 | -0.913 | 0.079 | -0.097 | -0.097 | 4.084 | 0.85 | 0.48 | 0.7 | 0.95 | 0.83 | 0.8 |
| 3 | 121977282 | rs10934578 | 0.412 | 0.862 | -0.039 | -0.907 | 0.084 | -0.051 | -0.051 | 0 | 0.46 | 0.07 | 0.36 | 0.56 | 0.32 | 0.51 |
| 3 | 121979229 | rs3804592 | 0.369 | 0.893 | 0.095 | -0.876 | 0.033 | -0.052 | -0.052 | 0.011 | 0.11 | 0.3 | 0.27 | 0.03 | 0.14 | 0.16 |
| 3 | 121984021 | rs3804593 | 2.143 | 0.723 | -0.062 | -1.405 | 0.051 | 0.242 | 0.242 | 2.944 | 0.1 | 0.24 | 0.11 | 0.39 | 0.14 | 0.06 |
| 3 | 121987932 | rs1393198 | 2.065 | 0.646 | -0.176 | -1.462 | -0.004 | 0.095 | 0.095 | 0.108 | 0.15 | 0.02 | 0.06 | 0 | 0.1 | 0.07 |
| 3 | 121993247 | rs17251221 | 1.47 | 0.41 | -0.358 | -1.605 | 0.047 | 0.161 | 0.161 | 0 | 0.25 | 0.01 | 0.11 | 0.04 | 0.14 | 0.21 |
| 3 | 122000241 | rs7647405 | 1.963 | 0.561 | -0.072 | -1.413 | 0 | -0.011 | -0.011 | 0 | 0.06 | 0.07 | 0.21 | 0.02 | 0.04 | 0.14 |
| 3 | 122003757 | rs1801725 | 1.473 | 0.558 | -0.055 | -1.446 | 0.043 | 0.161 | 0.161 | 0 | 0.25 | 0.01 | 0.11 | 0.04 | 0.15 | 0.21 |
| 3 | 122003769 | rs1042636 | 1.043 | 0.899 | 0.176 | -1.172 | -0.003 | 0.088 | 0.088 | 0 | 0.06 | 0.03 | 0.2 | 0.52 | 0.07 | 0.26 |
| 3 | 122005053 | rs9740 | 1.43 | 1.157 | 0.015 | -1.195 | -0.028 | 0.074 | 0.074 | 0.462 | 0.28 | 0.15 | 0.23 | 0 | 0.36 | 0.19 |
| 3 | 122005131 | rs1802757 | 1.952 | 1.237 | -0.06 | -0.917 | 0.052 | 0.245 | 0.245 | 2.944 | 0.1 | 0.24 | 0.11 | 0.39 | 0.15 | 0.06 |
| 3 | 122014566 | rs13083990 | 0.225 | 0.451 | -0.906 | -0.912 | 0.109 | -0.017 | -0.017 | 0 | 0.49 | 0.08 | 0.4 | 0.56 | 0.35 | 0.56 |
| 3 | 122017186 | rs6808422 | 0.116 | 0.683 | -0.283 | -1.121 | 0.011 | -0.091 | -0.091 | 1.541 | 0.41 | 0.57 | 0.51 | 0.11 | 0.51 | 0.38 |
| 3 | 122017232 | rs6438720 | -0.143 | 0.627 | 0.011 | -0.79 | 0.022 | -0.002 | -0.002 | 0 | 0.47 | 0.64 | 0.57 | 0.5 | 0.62 | 0.42 |
| 3 | 122020194 | rs9851884 | -0.333 | 0.317 | -0.056 | -0.856 | -0.025 | 0.015 | 0.015 | 0 | 0.62 | 0.58 | 0.63 | 0.51 | 0.71 | 0.48 |
| 3 | 122031859 | rs4247197 | 1.344 | 0.301 | -0.173 | -0.791 | -0.019 | 0.048 | 0.048 | 0 | 0.05 | 0 | 0.21 | 0.43 | 0.08 | 0.27 |
| 3 | 122032436 | rs11718426 | 1.139 | 0.501 | 0.153 | -1.049 | -0.002 | 0.009 | 0.009 | 0.36 | 0.25 | 0.25 | 0.33 | 0.09 | 0.23 | 0.21 |
| 3 | 122034854 | rs4678179 | -0.35 | 0.43 | 0.245 | -1.091 | -0.001 | -0.038 | -0.038 | 0.045 | 0.68 | 0.33 | 0.61 | 0.89 | 0.73 | 0.75 |
| 3 | 122040062 | rs9864290 | 0.02 | 0.177 | 0.327 | -0.694 | -0.007 | 0.002 | 0.002 | 0 | 0.6 | 0.48 | 0.63 | 0.51 | 0.71 | 0.48 |
| 3 | 122040283 | rs9864835 | 2.201 | 0.281 | 0.286 | -0.71 | -0.019 | 0.008 | 0.008 | 3.174 | 0.1 | 0.06 | 0.08 | 0.43 | 0.15 | 0.09 |
| 3 | 122053715 | rs6438722 | 0.276 | 0.405 | -0.038 | -0.589 | 0.017 | 0.031 | 0.031 | 0.784 | 0.51 | 0.43 | 0.55 | 0.09 | 0.56 | 0.39 |
| 3 | 122056432 | rs17589 | 0.292 | 0.411 | -0.149 | -0.586 | 0.023 | 0.062 | 0.062 | 0.744 | 0.5 | 0.41 | 0.55 | 0.09 | 0.56 | 0.39 |
| 3 | 122057873 | rs12491981 | 0.605 | 0.638 | -0.505 | 0.095 | -0.017 | 0.034 | 0.034 | 0.008 | 0.06 | 0 | 0.2 | 0.43 | 0.08 | 0.27 |
| 3 | 122063811 | rs9856248 | 0.163 | 0.15 | -0.099 | -0.229 | -0.014 | 0.018 | 0.018 | 0 | 0.36 | 0.44 | 0.24 | 0.46 | 0.3 | 0.38 |
| 3 | 122066340 | rs10934586 | 0.239 | 0.531 | -0.39 | -0.162 | 0.025 | 0.06 | 0.06 | 0 | 0.49 | 0.28 | 0.33 | 0.28 | 0.46 | 0.43 |
| 3 | 122074704 | rs6798997 | 0.506 | 0.612 | -0.421 | 0.292 | -0.061 | -0.054 | -0.054 | 0.252 | 0.3 | 0.14 | 0.29 | 0.56 | 0.41 | 0.34 |
| 3 | 122078323 | rs4678189 | -0.427 | 0.509 | 0.185 | 0.234 | -0.002 | 0.01 | 0.01 | 0 | 0.59 | 0.59 | 0.46 | 0.69 | 0.59 | 0.63 |
| 3 | 122079181 | rs13081171 | 1.169 | 0.438 | -0.055 | 0.023 | -0.058 | 0.123 | 0.123 | 0.186 | 0.29 | 0.08 | 0.37 | 0.11 | 0.3 | 0.25 |
| 3 | 122080747 | rs4491840 | 1.357 | 0.487 | 0.18 | 0.078 | -0.015 | -0.05 | -0.05 | 0 | 0.27 | 0.26 | 0.16 | 0.05 | 0.17 | 0.18 |
| 3 | 122082241 | rs7628690 | 1.368 | 1.188 | -0.039 | 0.026 | -0.006 | 0.027 | 0.027 | 0 | 0.06 | 0.04 | 0.13 | 0.24 | 0.14 | 0.14 |
| 3 | 122089359 | rs9834317 | 1.275 | 1.11 | -0.07 | -0.041 | -0.055 | -0.115 | -0.115 | 1.951 | 0.28 | 0.36 | 0.16 | 0.05 | 0.17 | 0.18 |
| 3 | 122101721 | rs6771201 | 0.109 | 1.192 | -0.187 | 0.042 | -0.061 | -0.055 | -0.055 | 1.599 | 0.3 | 0.15 | 0.29 | 0.56 | 0.41 | 0.33 |
| 3 | 122102710 | rs3828361 | 1.189 | 1.134 | -0.268 | 0.008 | -0.035 | -0.015 | -0.015 | 0 | 0.06 | 0 | 0.12 | 0.24 | 0.14 | 0.17 |
| 3 | 122104122 | rs1961957 | -0.622 | 0.842 | -0.134 | -0.004 | -0.003 | 0.011 | 0.011 | 0.305 | 0.61 | 0.61 | 0.47 | 0.69 | 0.61 | 0.63 |
| 3 | 122110921 | rs6438726 | 0.817 | 1.109 | -0.412 | 0.121 | -0.029 | -0.035 | -0.035 | 0.722 | 0.22 | 0.08 | 0.2 | 0.24 | 0.26 | 0.22 |
| 3 | 122121238 | rs4678191 | -0.342 | 1.059 | -0.306 | -0.041 | -0.017 | 0.004 | 0.004 | 0 | 0.35 | 0.31 | 0.31 | 0.66 | 0.44 | 0.46 |
| 3 | 122127592 | rs4678192 | 0.234 | 0.846 | -0.612 | 0.059 | 0.001 | -0.005 | -0.005 | 0 | 0.4 | 0.3 | 0.37 | 0.44 | 0.39 | 0.28 |
| 3 | 122127789 | rs10511409 | 0.301 | 0.146 | -0.666 | 0.104 | -0.006 | -0.009 | -0.009 | 0 | 0.38 | 0.35 | 0.27 | 0.3 | 0.31 | 0.34 |
| 3 | 122127822 | rs13320117 | -0.386 | 0.193 | 0.056 | 0.482 | 0.001 | -0.006 | -0.006 | 0 | 0.43 | 0.38 | 0.32 | 0.5 | 0.42 | 0.25 |
| 3 | 122127926 | rs17266628 | 0.59 | 0.131 | 0.041 | 0.319 | 0.045 | 0.006 | 0.006 | 0 | 0.33 | 0.14 | 0.19 | 0.27 | 0.24 | 0.21 |
| 3 | 122128394 | rs1127343 | 0.332 | 0.002 | 0.425 | 0.285 | -0.004 | -0.06 | -0.06 | 0.014 | 0.59 | 0.57 | 0.72 | 0.79 | 0.59 | 0.67 |
| 3 | 122133830 | rs3749213 | 0.992 | 0.301 | 0.47 | 0.857 | -0.018 | -0.011 | -0.011 | 0 | 0.22 | 0.11 | 0.21 | 0.22 | 0.26 | 0.22 |
| 3 | 122135242 | rs10934592 | 0.95 | 0.295 | 0.471 | 0.854 | -0.017 | -0.01 | -0.01 | 0.723 | 0.22 | 0.11 | 0.21 | 0.22 | 0.26 | 0.22 |
| 3 | 122136480 | rs9879508 | 2.21 | 0.426 | 0.521 | 0.946 | -0.009 | 0.027 | 0.027 | 0 | 0.08 | 0.06 | 0.08 | 0.32 | 0.15 | 0.1 |
| 3 | 122141135 | rs11932 | 1.746 | 0.425 | 0.64 | 0.941 | 0.051 | 0.146 | 0.146 | 0 | 0.09 | 0.18 | 0.09 | 0.32 | 0.15 | 0.1 |
| 3 | 122145324 | rs3762637 | 2.078 | 0.455 | 0.612 | 0.976 | 0.037 | 0.121 | 0.121 | 0 | 0.08 | 0.13 | 0.08 | 0.32 | 0.15 | 0.1 |
| 3 | 122145470 | rs17201246 | 1.245 | 0.242 | 0.68 | 0.711 | 0.056 | 0.08 | 0.08 | 0.529 | 0.27 | 0.09 | 0.12 | 0.11 | 0.15 | 0.18 |
| 3 | 122178602 | rs12487598 | 0.246 | 0.255 | 0.689 | 0.719 | -0.008 | -0.009 | -0.009 | 0.021 | 0.3 | 0.28 | 0.3 | 0.54 | 0.41 | 0.31 |
| 3 | 122195922 | rs10934599 | -0.416 | 0.399 | 0.885 | 0.651 | -0.004 | 0.011 | 0.011 | 0 | 0.57 | 0.37 | 0.42 | 0.65 | 0.56 | 0.49 |
| 3 | 122208293 | rs10934601 | -0.412 | 0.397 | 0.966 | 0.65 | -0.004 | 0.011 | 0.011 | 0 | 0.57 | 0.37 | 0.42 | 0.65 | 0.56 | 0.49 |
| 3 | 122227471 | rs13316042 | -0.727 | 0.506 | 0.585 | 0.701 | -0.011 | 0.037 | 0.037 | 0.053 | 0.61 | 0.68 | 0.47 | 0.77 | 0.58 | 0.62 |
| 3 | 122228252 | rs4470444 | -0.427 | 0.528 | 0.273 | 0.699 | -0.004 | 0.011 | 0.011 | 0 | 0.57 | 0.37 | 0.42 | 0.65 | 0.56 | 0.49 |
| 3 | 122259606 | rs9851180 | -0.434 | 0.581 | 0.193 | 0.696 | 0.001 | -0.006 | -0.006 | 0 | 0.57 | 0.42 | 0.41 | 0.65 | 0.56 | 0.49 |
| 3 | 122264619 | rs6780181 | -0.419 | 0.538 | -0.198 | 0.707 | -0.002 | 0.013 | 0.013 | 0.003 | 0.57 | 0.36 | 0.41 | 0.65 | 0.56 | 0.49 |
| 3 | 122277034 | rs9289197 | -0.612 | 0.546 | -0.385 | 0.638 | 0 | 0.026 | 0.026 | 0.198 | 0.59 | 0.36 | 0.42 | 0.79 | 0.58 | 0.63 |
| 3 | 122285005 | rs12485294 | -0.467 | 0.67 | -0.269 | 0.762 | 0.001 | -0.011 | -0.011 | 0.105 | 0.57 | 0.47 | 0.59 | 0.66 | 0.56 | 0.49 |
| 3 | 122287289 | rs9839782 | -0.009 | 0.661 | -0.367 | 0.863 | 0.001 | 0.02 | 0.02 | 0 | 0.27 | 0.34 | 0.4 | 0.36 | 0.29 | 0.36 |
| 3 | 122288210 | rs2332285 | 0.014 | 0.49 | -0.456 | 0.475 | 0.001 | -0.011 | -0.011 | 0 | 0.57 | 0.46 | 0.59 | 0.66 | 0.56 | 0.49 |
| 3 | 122290579 | rs2036342 | 0.037 | 0.33 | -0.452 | 0.666 | 0.009 | -0.006 | -0.006 | 0 | 0.37 | 0.27 | 0.26 | 0.44 | 0.32 | 0.42 |
| 3 | 122291353 | rs7616403 | -1.383 | 0.462 | 0.609 | 1.419 | -0.001 | 0.004 | 0.004 | 0 | 0.14 | 0.13 | 0.1 | 0.23 | 0.14 | 0.23 |
| 3 | 122293209 | rs1045089 | -1.515 | 0.554 | 0.469 | 1.513 | -0.018 | -0.048 | -0.048 | 1.645 | 0.26 | 0.19 | 0.4 | 0.48 | 0.34 | 0.38 |
| 3 | 122295235 | rs6796228 | -0.152 | 0.501 | 1.097 | 1.485 | 0.054 | 0.093 | 0.093 | 0 | 0.14 | 0.35 | 0.34 | 0.25 | 0.21 | 0.15 |

| | | | | | | | | | | | | | | | | |
|---|---|---|---|---|---|---|---|---|---|---|---|---|---|---|---|---|
| 3 | 122295780 | rs7651414 | -1.936 | 0.452 | 1.177 | 1.475 | 0.015 | 0.081 | 0.081 | 0 | 0.3 | 0.53 | 0.45 | 0.49 | 0.33 | 0.38 |
| 3 | 122296647 | rs7632426 | -1.126 | 0.395 | 1.164 | 1.475 | -0.002 | 0.009 | 0.009 | 0 | 0.14 | 0.15 | 0.09 | 0.23 | 0.12 | 0.23 |
| 3 | 122296834 | rs2332240 | -2.055 | 0.625 | 1.169 | 1.444 | 0.028 | 0.026 | 0.026 | 0 | 0.34 | 0.57 | 0.58 | 0.54 | 0.43 | 0.44 |
| 3 | 122303929 | rs7624614 | -1.019 | 0.725 | 1.098 | 1.43 | 0.089 | 0.081 | 0.081 | 0 | 0.2 | 0.47 | 0.48 | 0.32 | 0.31 | 0.2 |
| 3 | 122310203 | rs4677964 | 1.905 | -0.745 | 1.294 | 0.958 | -0.001 | 0.004 | 0.004 | 0 | 0.69 | 0.64 | 0.73 | 0.64 | 0.76 | 0.65 |
| 3 | 122310266 | rs4678200 | 1.248 | -0.588 | 1.668 | 0.179 | 0 | -0.009 | -0.009 | 0 | 0.83 | 0.73 | 0.91 | 0.91 | 0.87 | 0.84 |
| 3 | 122313712 | rs1553247 | 1.351 | -0.659 | 1.547 | 0.133 | -0.022 | -0.032 | -0.032 | 0 | 0.75 | 0.71 | 0.92 | 0.93 | 0.9 | 0.82 |
| 3 | 122314874 | rs6762418 | -0.014 | -0.667 | 1.466 | 0.143 | -0.049 | -0.039 | -0.039 | 0 | 0.85 | 0.8 | 0.95 | 0.98 | 0.98 | 0.89 |
| 3 | 122315893 | rs9882797 | 1.779 | -0.592 | 1.016 | 0.173 | 0.009 | 0.015 | 0.015 | 1.507 | 0.67 | 0.31 | 0.85 | 0.9 | 0.82 | 0.76 |
| 3 | 122325467 | rs1909808 | -2.186 | 0.111 | 1.406 | 0.733 | -0.017 | 0.005 | 0.005 | 1.114 | 0.15 | 0.74 | 0.09 | 0.15 | 0.07 | 0.12 |
| 3 | 122325735 | rs16833242 | -1.001 | 0.133 | 1.486 | 0.746 | 0.052 | 0.202 | 0.202 | 6.529 | 0.06 | 0.48 | 0.04 | 0.1 | 0.01 | 0.06 |
| 3 | 122325769 | rs1909807 | -1.448 | 0.14 | 1.355 | 0.741 | 0 | -0.007 | -0.007 | 0.377 | 0.09 | 0.15 | 0.04 | 0.05 | 0.07 | 0.07 |
| 3 | 122328753 | rs2173764 | -1.14 | -0.103 | 1.209 | 0.83 | 0.02 | 0.152 | 0.152 | 8.221 | 0.06 | 0.57 | 0.05 | 0.1 | 0.01 | 0.06 |
| 3 | 122340626 | rs12636277 | -1.045 | 0.075 | -0.121 | 0.523 | -0.002 | -0.003 | -0.003 | 1.115 | 0.11 | 0.14 | 0.06 | 0.07 | 0.08 | 0.07 |
| 3 | 122341401 | rs9827225 | -1.318 | 0.261 | -0.51 | 0.5 | -0.001 | -0.005 | -0.005 | 0 | 0.22 | 0.73 | 0.2 | 0.2 | 0.19 | 0.2 |
| 3 | 122343439 | rs4677965 | 0.086 | -0.071 | -1.529 | 0.246 | 0.046 | 0.094 | 0.094 | 0 | 0.54 | 0.2 | 0.43 | 0.69 | 0.65 | 0.51 |
| 3 | 122345870 | rs17208928 | 0.167 | -0.494 | -1.979 | -0.364 | -0.049 | 0.141 | 0.141 | 2.426 | 0.23 | 0.01 | 0.09 | 0.01 | 0.22 | 0.12 |
| 3 | 122347389 | rs13314756 | -0.483 | -1.349 | -2.302 | 0.163 | 0.038 | -0.074 | -0.074 | 0.075 | 0.21 | 0.1 | 0.25 | 0.49 | 0.13 | 0.19 |
| 3 | 122348423 | rs12632799 | -0.624 | -1.389 | -1.864 | 0.172 | 0.005 | -0.006 | -0.006 | 0 | 0.21 | 0.2 | 0.24 | 0.48 | 0.13 | 0.12 |
| 3 | 122349815 | rs9880413 | -0.467 | -1.388 | -1.92 | 0.166 | 0.039 | -0.072 | -0.072 | 0 | 0.21 | 0.1 | 0.25 | 0.49 | 0.13 | 0.19 |
| 3 | 122351791 | rs7612991 | -1.205 | -1.61 | -0.913 | 0.014 | 0.054 | 0.004 | 0.004 | 1.957 | 0.49 | 0.71 | 0.42 | 0.5 | 0.38 | 0.44 |
| 3 | 122354037 | rs1106346 | -0.843 | -1.554 | -0.566 | 0.036 | 0.044 | -0.01 | -0.01 | 0 | 0.46 | 0.26 | 0.39 | 0.5 | 0.38 | 0.31 |
| 3 | 122360083 | rs17209173 | 0.564 | -1.458 | -0.782 | -0.431 | -0.032 | 0.12 | 0.12 | 1.894 | 0.24 | 0.06 | 0.13 | 0.01 | 0.25 | 0.12 |
| 3 | 122362079 | rs9844986 | -0.477 | -1.638 | -0.487 | -0.269 | 0.016 | 0.016 | 0.016 | 0.22 | 0.11 | 0.11 | 0.07 | 0.03 | 0.05 | 0.14 |
| 3 | 122368402 | rs12485657 | -0.562 | -1.903 | -0.206 | -0.146 | -0.013 | 0.021 | 0.021 | 0 | 0.23 | 0.31 | 0.51 | 0.64 | 0.24 | 0.29 |
| 3 | 122377674 | rs1256208 | -1.077 | -2.295 | 0.223 | -0.011 | -0.029 | -0.003 | -0.003 | 0 | 0.18 | 0.44 | 0.22 | 0.17 | 0.15 | 0.12 |
| 3 | 122377999 | rs7652006 | 0.018 | -2.314 | 0.209 | 0.021 | -0.013 | 0.1 | 0.1 | 0 | 0.16 | 0.24 | 0.36 | 0.49 | 0.15 | 0.23 |
| 3 | 122379262 | rs17269087 | 1.072 | -2.261 | 0.162 | -0.225 | 0.001 | 0.071 | 0.071 | 2.521 | 0.26 | 0.11 | 0.1 | 0 | 0.22 | 0.13 |
| 3 | 122380614 | rs790118 | 0.004 | -2.232 | 0.595 | -0.203 | 0.007 | -0.016 | -0.016 | 0 | 0.36 | 0.27 | 0.15 | 0.02 | 0.28 | 0.19 |
| 3 | 122380722 | rs6438753 | 0.285 | -2.228 | 0.649 | -0.178 | 0.019 | 0.088 | 0.088 | 0 | 0.07 | 0.28 | 0.17 | 0.15 | 0.09 | 0.06 |
| 3 | 122385402 | rs796892 | -0.034 | -2.214 | 0.654 | -0.192 | 0.008 | -0.014 | -0.014 | 0.021 | 0.37 | 0.27 | 0.15 | 0.02 | 0.28 | 0.19 |
| 3 | 122388312 | rs1877377 | 0.129 | -2.292 | 0.587 | -0.165 | -0.005 | 0.233 | 0.233 | 0.134 | 0.1 | 0.23 | 0.32 | 0.48 | 0.12 | 0.22 |
| 3 | 122388464 | rs790125 | -0.807 | -2.375 | 0.252 | -0.169 | 0.036 | 0.021 | 0.021 | 0 | 0.49 | 0.61 | 0.49 | 0.53 | 0.41 | 0.53 |
| 3 | 122394381 | rs2971372 | -0.516 | -2.555 | 0.226 | -0.09 | -0.025 | 0.249 | 0.249 | 0.034 | 0.15 | 0.32 | 0.33 | 0.47 | 0.14 | 0.24 |
| 3 | 122396818 | rs2650961 | -1.571 | -2.3 | -0.052 | -0.126 | -0.097 | 0.012 | 0.012 | 0 | 0.6 | 0.91 | 0.66 | 0.67 | 0.5 | 0.58 |
| 3 | 122397064 | rs1355562 | -0.885 | -2.258 | 0.092 | -0.13 | 0.01 | 0.006 | 0.006 | 0 | 0.51 | 0.52 | 0.49 | 0.53 | 0.41 | 0.54 |
| 3 | 122397717 | rs2280070 | -0.399 | -1.931 | -0.199 | 0.061 | 0.002 | 0.004 | 0.004 | 0 | 0.11 | 0.1 | 0.05 | 0.06 | 0.07 | 0.18 |
| 3 | 122403480 | rs4678207 | -1.118 | -2.015 | -0.1 | -0.011 | -0.021 | 0.028 | 0.028 | 0 | 0.33 | 0.44 | 0.45 | 0.52 | 0.24 | 0.45 |
| 3 | 122403544 | rs1256197 | -1.431 | -2.087 | -0.945 | -0.346 | -0.021 | 0.02 | 0.02 | 0 | 0.55 | 0.96 | 0.63 | 0.67 | 0.42 | 0.55 |
| 3 | 122406077 | rs790131 | -0.643 | -2.555 | -0.635 | -0.367 | 0.079 | -0.001 | -0.001 | 0 | 0.41 | 0.69 | 0.42 | 0.56 | 0.23 | 0.49 |
| 3 | 122406265 | rs11711456 | 0.311 | -2.544 | 0.425 | -0.312 | -0.03 | 0.162 | 0.162 | 0 | 0.13 | 0.23 | 0.32 | 0.49 | 0.09 | 0.24 |
| 3 | 122407374 | rs17270460 | 1.553 | -2.479 | 0.201 | -0.345 | -0.001 | -0.01 | -0.01 | 5.208 | 0.07 | 0.05 | 0.06 | 0 | 0.07 | 0.01 |
| 3 | 122410208 | rs10934611 | -0.013 | -2.485 | 0.051 | -0.422 | 0.003 | -0.007 | -0.007 | 0 | 0.1 | 0.04 | 0.14 | 0.13 | 0.14 | 0.07 |
| 3 | 122412901 | rs1256200 | -0.483 | -2.44 | 0.111 | -0.444 | 0.075 | -0.02 | -0.02 | 0.736 | 0.38 | 0.74 | 0.42 | 0.55 | 0.23 | 0.45 |
| 3 | 122416425 | rs812046 | 1.749 | -2.176 | 0.614 | -0.536 | -0.004 | -0.049 | -0.049 | 0.011 | 0.16 | 0.27 | 0.08 | 0.05 | 0.12 | 0.14 |
| 3 | 122418750 | rs11719086 | -0.592 | -2.239 | 0.459 | -0.579 | 0.004 | 0.012 | 0.012 | 0 | 0.18 | 0.18 | 0.22 | 0.13 | 0.21 | 0.09 |
| 3 | 122419083 | rs13093808 | 0.842 | -2.232 | 0.542 | -0.565 | 0.002 | -0.006 | -0.006 | 0 | 0.07 | 0.04 | 0.11 | 0.13 | 0.09 | 0.05 |
| 3 | 122423256 | rs2288618 | 0.842 | -2.204 | 0.617 | -0.53 | 0.002 | -0.006 | -0.006 | 0 | 0.07 | 0.04 | 0.11 | 0.13 | 0.09 | 0.05 |
| 3 | 122433930 | rs10433340 | 2.874 | -1.898 | 0.143 | -0.582 | -0.135 | -0.059 | -0.059 | 0 | 0.28 | 0.02 | 0.29 | 0.29 | 0.39 | 0.26 |
| 3 | 122437321 | rs7645033 | -2.019 | -1.76 | 1.137 | -0.067 | -0.002 | -0.023 | -0.023 | 0 | 0.85 | 0.86 | 0.94 | 0.98 | 0.88 | 0.96 |
| 3 | 122437834 | rs3821688 | 1.838 | -1.759 | 1.372 | -0.09 | 0.013 | 0.056 | 0.056 | 0 | 0.46 | 0.32 | 0.38 | 0.32 | 0.54 | 0.56 |
| 3 | 122440043 | rs16833457 | 0.769 | -1.835 | 1.36 | -0.081 | 0.002 | -0.003 | -0.003 | 0.072 | 0.07 | 0.04 | 0.11 | 0.13 | 0.09 | 0.05 |
| 3 | 122447574 | rs3732832 | 1.839 | -1.834 | 1.418 | -0.088 | 0.015 | 0.056 | 0.056 | 0 | 0.46 | 0.32 | 0.38 | 0.32 | 0.54 | 0.56 |
| 3 | 122447911 | rs11557048 | -0.264 | -1.834 | 1.4 | -0.088 | -0.005 | -0.002 | -0.002 | 0.375 | 0.11 | 0.16 | 0.16 | 0.13 | 0.14 | 0.08 |
| 3 | 122448203 | rs10934615 | 0.71 | -1.873 | 1.387 | 0.295 | 0.013 | -0.018 | -0.018 | 6.493 | 0.11 | 0.06 | 0.15 | 0.35 | 0.08 | 0.14 |
| 3 | 122448914 | rs6774933 | 1.939 | -1.837 | 1.375 | 0.238 | 0.013 | 0.063 | 0.063 | 0 | 0.45 | 0.28 | 0.37 | 0.32 | 0.54 | 0.56 |
| 3 | 122449657 | rs6726 | 1.924 | -2.024 | 1.382 | 0.245 | 0.014 | 0.064 | 0.064 | 0 | 0.45 | 0.28 | 0.37 | 0.32 | 0.54 | 0.57 |
| 3 | 122458887 | rs3115 | -0.361 | -1.953 | 1.416 | 0.272 | -0.009 | -0.001 | -0.001 | 0 | 0.11 | 0.23 | 0.17 | 0.13 | 0.14 | 0.08 |
| 3 | 122473354 | rs12496363 | 0.112 | -2.025 | 1.589 | 0.259 | -0.028 | 0.005 | 0.005 | 0 | 0.65 | 0.55 | 0.61 | 0.44 | 0.75 | 0.66 |
| 3 | 122477795 | rs2137210 | 1.161 | -2.035 | 1.622 | 0.235 | 0.031 | 0.027 | 0.027 | 0 | 0.53 | 0.37 | 0.49 | 0.45 | 0.63 | 0.62 |
| 3 | 122479371 | rs16833512 | 0.068 | -2.068 | 1.761 | 0.245 | -0.021 | 0.152 | 0.152 | 0 | 0.13 | 0.25 | 0.31 | 0.51 | 0.1 | 0.23 |
| 3 | 122504041 | rs11712252 | 0.169 | -2.076 | 1.717 | 0.237 | -0.028 | 0.004 | 0.004 | 0 | 0.64 | 0.57 | 0.61 | 0.45 | 0.75 | 0.66 |
| 3 | 122504474 | rs13095775 | 0.62 | -2.037 | 1.75 | 0.255 | -0.003 | 0.003 | 0.003 | 0 | 0.07 | 0.09 | 0.11 | 0.13 | 0.09 | 0.05 |
| 3 | 122509518 | rs9812646 | 1.911 | -2.047 | 1.756 | 0.251 | 0.011 | 0.084 | 0.084 | 0 | 0.45 | 0.23 | 0.37 | 0.32 | 0.54 | 0.57 |
| 3 | 122510763 | rs9855669 | 2.051 | -2.034 | 1.79 | 0.255 | 0.002 | 0.011 | 0.011 | 1.596 | 0.14 | 0.08 | 0.06 | 0.02 | 0.12 | 0.04 |
| 3 | 122512631 | rs3806643 | 0.173 | -2.024 | 1.659 | 0.258 | -0.028 | 0.005 | 0.005 | 0 | 0.64 | 0.57 | 0.61 | 0.45 | 0.75 | 0.66 |
| 3 | 122524265 | rs16833535 | 0.587 | -2.007 | 1.574 | 0.311 | 0.01 | -0.007 | -0.007 | 0.019 | 0.11 | 0.09 | 0.15 | 0.32 | 0.08 | 0.13 |
| 3 | 122531377 | rs9872909 | 0.879 | -1.997 | 1.608 | 0.28 | 0 | 0 | 0 | 0 | 0.57 | 0.48 | 0.5 | 0.32 | 0.66 | 0.61 |
| 3 | 122545494 | rs9853452 | 1.825 | -1.953 | 1.394 | 0.299 | 0.005 | 0.084 | 0.084 | 0 | 0.45 | 0.19 | 0.37 | 0.32 | 0.54 | 0.57 |

| | | | | | | | | | | | | | | | | |
|---|---|---|---|---|---|---|---|---|---|---|---|---|---|---|---|---|
| 3 | 122550989 | rs9847351 | 1.975 | -1.64 | 1.39 | 0.438 | -0.002 | 0.043 | 0.043 | 2.169 | 0.14 | 0.03 | 0.05 | 0.02 | 0.12 | 0.04 |
| 3 | 122567173 | rs10934619 | 0.153 | -1.616 | 1.465 | 0.47 | -0.009 | 0.121 | 0.121 | 0.164 | 0.11 | 0.27 | 0.16 | 0.33 | 0.09 | 0.13 |
| 3 | 122571435 | rs7650313 | -0.802 | -1.557 | 1.283 | 0.27 | 0 | -0.009 | -0.009 | 0 | 0.78 | 0.81 | 0.92 | 0.96 | 0.85 | 0.89 |
| 3 | 122595777 | rs6780615 | 2.279 | -1.296 | 1.994 | 0.943 | 0.028 | 0.025 | 0.025 | 1.026 | 0.61 | 0.24 | 0.38 | 0.29 | 0.65 | 0.61 |
| 3 | 122598291 | rs2288677 | -1.861 | -1.303 | 1.992 | 0.965 | -0.029 | 0.017 | 0.017 | 3.097 | 0.31 | 0.57 | 0.56 | 0.69 | 0.29 | 0.38 |
| 3 | 122600983 | rs9859230 | 1.998 | -1.272 | 1.852 | 0.87 | 0.018 | 0.017 | 0.017 | 1.152 | 0.62 | 0.29 | 0.39 | 0.29 | 0.65 | 0.61 |
| 3 | 122605544 | rs9864868 | 1.998 | -1.272 | 1.852 | 0.87 | 0.018 | 0.017 | 0.017 | 1.152 | 0.62 | 0.29 | 0.39 | 0.29 | 0.65 | 0.61 |
| 3 | 122612470 | rs9681491 | -0.295 | -1.032 | 1.877 | 1.168 | 0.001 | 0.023 | 0.023 | 0 | 0.11 | 0.11 | 0.21 | 0.42 | 0.11 | 0.14 |
| 3 | 122617445 | rs13076885 | -1.184 | -0.847 | 1.969 | 1.187 | -0.007 | 0.063 | 0.063 | 0.343 | 0.17 | 0.24 | 0.26 | 0.48 | 0.19 | 0.18 |
| 3 | 122639941 | rs13088040 | -0.312 | -0.771 | 1.873 | 1.242 | 0 | 0.114 | 0.114 | 0.001 | 0.1 | 0.14 | 0.16 | 0.42 | 0.1 | 0.14 |
| 3 | 122640541 | rs3755750 | 2.067 | -0.929 | 1.443 | 0.778 | 0.014 | 0.031 | 0.031 | 0.362 | 0.62 | 0.24 | 0.44 | 0.29 | 0.65 | 0.59 |
| 3 | 122642850 | rs9846967 | 1.228 | -0.983 | 1.348 | 0.85 | -0.007 | 0.076 | 0.076 | 0 | 0.78 | 0.43 | 0.58 | 0.41 | 0.8 | 0.74 |
| 3 | 122643492 | rs11710442 | 0.36 | -0.832 | 1.118 | 0.795 | 0 | -0.003 | -0.003 | 0.248 | 0.06 | 0.06 | 0.04 | 0.05 | 0.08 | 0.04 |
| 3 | 122645277 | rs2276777 | -0.157 | -0.904 | 1.073 | 0.799 | 0.009 | 0.012 | 0.012 | 0.645 | 0.11 | 0.07 | 0.1 | 0.06 | 0.07 | 0.11 |
| 3 | 122645304 | rs2276776 | 0.493 | -0.898 | 1.057 | 0.793 | 0.007 | 0.004 | 0.004 | 0.041 | 0.08 | 0.06 | 0.06 | 0.06 | 0.03 | 0.08 |
| 3 | 122646734 | rs2276775 | -0.346 | -0.899 | 1.065 | 0.773 | 0.005 | 0.033 | 0.033 | 0.118 | 0.15 | 0.07 | 0.12 | 0.06 | 0.1 | 0.09 |
| 3 | 122646828 | rs2276774 | -0.908 | -0.913 | 1.275 | 0.77 | 0 | -0.005 | -0.005 | 0 | 0.18 | 0.18 | 0.17 | 0.06 | 0.14 | 0.12 |
| 3 | 122648199 | rs9867255 | -1.638 | -0.919 | 1.253 | 0.77 | 0 | -0.005 | -0.005 | 0 | 0.25 | 0.23 | 0.21 | 0.11 | 0.22 | 0.16 |
| 3 | 122648823 | rs17281161 | 2.211 | -0.989 | 0.747 | 0.466 | -0.074 | 0.025 | 0.025 | 1.77 | 0.39 | 0.16 | 0.47 | 0.3 | 0.49 | 0.39 |
| 3 | 122650890 | rs13099843 | -1.514 | -0.896 | 0.909 | 0.392 | -0.002 | -0.016 | -0.016 | 0 | 0.24 | 0.23 | 0.21 | 0.11 | 0.22 | 0.16 |
| 3 | 122652533 | rs11718661 | 0.062 | -0.716 | 0.744 | 0.483 | 0.012 | -0.01 | -0.01 | 0 | 0.06 | 0.18 | 0.06 | 0.05 | 0.08 | 0.05 |
| 3 | 122656016 | rs4677976 | -0.238 | -0.753 | 0.529 | 0.768 | -0.004 | -0.037 | -0.037 | 0 | 0.28 | 0.11 | 0.17 | 0.33 | 0.26 | 0.33 |
| 3 | 122657204 | rs2060090 | 2.101 | -0.861 | 0.596 | 0.838 | -0.047 | 0.051 | 0.051 | 1.267 | 0.42 | 0.18 | 0.49 | 0.29 | 0.5 | 0.43 |
| 3 | 122658417 | rs3732827 | -0.114 | -0.799 | 1.023 | 0.892 | 0.005 | 0.09 | 0.09 | 0.052 | 0.11 | 0.2 | 0.17 | 0.33 | 0.1 | 0.12 |
| 3 | 122659829 | rs10934622 | -0.114 | -0.799 | 1.023 | 0.892 | 0.005 | 0.09 | 0.09 | 0.052 | 0.11 | 0.2 | 0.17 | 0.33 | 0.1 | 0.12 |
| 3 | 122660687 | rs6803981 | -0.175 | -0.836 | 0.65 | 0.903 | 0.002 | -0.005 | -0.005 | 0 | 0.29 | 0.25 | 0.18 | 0.33 | 0.27 | 0.33 |
| 3 | 122661637 | rs2332410 | 0.361 | -0.704 | 0.806 | 0.938 | -0.009 | -0.008 | -0.008 | 0 | 0.09 | 0.17 | 0.07 | 0.06 | 0.03 | 0.08 |
| 3 | 122661863 | rs11719594 | 1.121 | -0.671 | 0.556 | 0.962 | -0.001 | 0.002 | 0.002 | 0.001 | 0.74 | 0.32 | 0.7 | 0.61 | 0.8 | 0.8 |
| 3 | 122664852 | rs4677868 | 1.721 | -0.677 | 0.265 | 1.081 | -0.008 | 0.024 | 0.024 | 0.481 | 0.41 | 0.32 | 0.52 | 0.27 | 0.5 | 0.4 |
| 3 | 122666048 | rs9831044 | 1.168 | -0.614 | 0.385 | 1.063 | 0.003 | 0.01 | 0.01 | 1.862 | 0.5 | 0.42 | 0.69 | 0.61 | 0.6 | 0.51 |
| 3 | 122667277 | rs2288678 | -0.157 | -0.535 | 0.139 | 0.58 | 0.012 | 0.009 | 0.009 | 0 | 0.15 | 0.08 | 0.08 | 0.13 | 0.09 | 0.19 |
| 3 | 122676523 | rs10934626 | -0.082 | -0.59 | 0.3 | 0.62 | -0.005 | 0.005 | 0.005 | 0 | 0.16 | 0.22 | 0.08 | 0.29 | 0.09 | 0.25 |
| 3 | 122676671 | rs9857254 | -1.409 | -0.637 | -0.267 | 0.565 | 0 | -0.001 | -0.001 | 0 | 0.51 | 0.52 | 0.28 | 0.29 | 0.38 | 0.48 |
| 3 | 122676705 | rs9839593 | -0.758 | -0.639 | -0.311 | 0.488 | -0.009 | -0.02 | -0.02 | 0.018 | 0.32 | 0.38 | 0.16 | 0.29 | 0.22 | 0.39 |
| 3 | 122678507 | rs4478030 | -1.611 | -0.329 | -0.333 | 0.828 | -0.022 | 0.021 | 0.021 | 1.095 | 0.59 | 0.7 | 0.47 | 0.71 | 0.49 | 0.51 |
| 3 | 122680141 | rs13094003 | -0.641 | 0.027 | -0.534 | 0.936 | 0 | -0.003 | -0.003 | 0 | 0.23 | 0.22 | 0.27 | 0.35 | 0.24 | 0.21 |
| 3 | 122683207 | rs12635724 | -0.632 | -0.259 | -0.559 | 0.362 | -0.007 | -0.018 | -0.018 | 1.212 | 0.33 | 0.61 | 0.21 | 0.18 | 0.23 | 0.28 |
| 3 | 122686113 | rs6764971 | 0.193 | -0.451 | -0.987 | 0.522 | 0.021 | 0.003 | 0.003 | 0 | 0.3 | 0.2 | 0.22 | 0.27 | 0.25 | 0.22 |
| 3 | 122688170 | rs9816759 | -0.698 | -0.698 | -1.357 | 0.788 | 0.086 | 0.129 | 0.129 | 0.764 | 0.49 | 0.83 | 0.41 | 0.72 | 0.42 | 0.53 |
| 3 | 122689398 | rs9880238 | -0.465 | -1.079 | -0.354 | 1.14 | -0.044 | 0.176 | 0.176 | 0.237 | 0.12 | 0.37 | 0.05 | 0.32 | 0.04 | 0.22 |
| 3 | 122689412 | rs6800093 | 0.606 | -0.739 | -0.957 | 0.909 | 0.023 | -0.013 | -0.013 | 0 | 0.27 | 0.13 | 0.21 | 0.27 | 0.24 | 0.19 |
| 3 | 122693800 | rs4234209 | 0.495 | -0.26 | -0.863 | 0.74 | 0.003 | -0.026 | -0.026 | 0.217 | 0.65 | 0.25 | 0.47 | 0.54 | 0.71 | 0.62 |
| 3 | 122694400 | rs895074 | 0.087 | -0.1 | -1.208 | 0.205 | -0.06 | -0.03 | -0.03 | 0 | 0.15 | 0.03 | 0.36 | 0.13 | 0.18 | 0.22 |
| 3 | 122694527 | rs895075 | -0.536 | -0.433 | -0.749 | 0.043 | 0.011 | 0.03 | 0.03 | 0 | 0.13 | 0.08 | 0.11 | 0.03 | 0.11 | 0.16 |
| 3 | 122694930 | rs9968193 | -0.662 | -0.589 | -0.016 | -0.041 | -0.017 | -0.016 | -0.016 | 0 | 0.13 | 0.46 | 0.07 | 0.11 | 0.07 | 0.12 |
| 3 | 122698761 | rs1470399 | -1.028 | -0.393 | 0.19 | -0.04 | -0.021 | -0.111 | -0.111 | 0 | 0.19 | 0.43 | 0.16 | 0.04 | 0.14 | 0.18 |
| 3 | 122707433 | rs998945 | 1.08 | -0.045 | -1.142 | -0.058 | -0.082 | 0.057 | 0.057 | 0.003 | 0.43 | 0.13 | 0.58 | 0.69 | 0.44 | 0.51 |
| 3 | 122709853 | rs6769548 | -0.428 | -0.402 | -0.2 | -0.12 | -0.025 | 0.025 | 0.025 | 0.017 | 0.17 | 0.49 | 0.1 | 0.15 | 0.1 | 0.14 |
| 3 | 122711344 | rs1444820 | -1.444 | -0.541 | -0.869 | -0.089 | 0.056 | -0.041 | -0.041 | 0.153 | 0.28 | 0.86 | 0.22 | 0.31 | 0.16 | 0.31 |
| 3 | 122713511 | rs6786625 | -0.477 | -0.4 | -0.97 | -0.715 | -0.099 | 0.107 | 0.107 | 0 | 0.72 | 0.98 | 0.78 | 0.93 | 0.64 | 0.84 |
| 3 | 122714241 | rs16833750 | 0.203 | -0.336 | -0.63 | -0.567 | 0.005 | 0.003 | 0.003 | 0 | 0.09 | 0.07 | 0.04 | 0.09 | 0.06 | 0.17 |
| 3 | 122715812 | rs2120806 | -0.126 | -0.135 | -0.295 | -0.548 | 0.043 | -0.043 | -0.043 | 1.569 | 0.12 | 0.44 | 0.1 | 0.03 | 0.08 | 0.06 |
| 3 | 122719728 | rs9834538 | 0.261 | -0.074 | -0.952 | -0.518 | 0.026 | -0.036 | -0.036 | 0.028 | 0.1 | 0.32 | 0.08 | 0.03 | 0.08 | 0.06 |
| 3 | 122723191 | rs11715232 | 0.754 | 0.009 | -1.379 | -0.602 | -0.039 | 0.067 | 0.067 | 0 | 0.11 | 0.01 | 0.06 | 0 | 0.13 | 0.05 |
| 3 | 122729859 | rs9863672 | 0.141 | 0.052 | -1.19 | -0.57 | 0.035 | -0.042 | -0.042 | 0 | 0.11 | 0.41 | 0.09 | 0.03 | 0.08 | 0.06 |
| 3 | 122750703 | rs9839656 | -0.051 | 0.132 | -2.629 | -0.645 | -0.076 | 0.167 | 0.167 | 0 | 0.31 | 0.04 | 0.21 | 0.06 | 0.36 | 0.18 |
| 3 | 122754753 | rs13074864 | 0.1 | 0.12 | -2.389 | 0.024 | -0.006 | -0.003 | -0.003 | 0 | 0.06 | 0.04 | 0.34 | 0.23 | 0.07 | 0.14 |
| 3 | 122755775 | rs12107644 | -0.723 | -0.155 | -2.261 | -0.661 | 0.05 | 0.06 | 0.06 | 3.029 | 0.25 | 0.87 | 0.18 | 0.16 | 0.14 | 0.33 |
| 3 | 122759622 | rs6799522 | -1.303 | -0.171 | -2.291 | -0.691 | 0.013 | 0.137 | 0.137 | 1.068 | 0.35 | 0.89 | 0.23 | 0.15 | 0.27 | 0.38 |
| 3 | 122760119 | rs2877681 | -0.946 | -0.425 | -1.881 | -0.429 | 0.049 | 0.065 | 0.065 | 3.084 | 0.25 | 0.87 | 0.18 | 0.13 | 0.14 | 0.32 |
| 3 | 122775487 | rs12485921 | 0.144 | 0.411 | -1.784 | -0.613 | -0.045 | 0.134 | 0.134 | 0.37 | 0.41 | 0.19 | 0.43 | 0.19 | 0.41 | 0.39 |
| 3 | 122780164 | rs6809947 | 0.513 | 0.446 | -1.16 | -0.207 | 0.012 | 0.003 | 0.003 | 0 | 0.47 | 0.39 | 0.38 | 0.57 | 0.38 | 0.4 |
| 3 | 122780413 | rs10934637 | 0.165 | 0.692 | -0.922 | -0.374 | 0.031 | -0.01 | -0.01 | 0 | 0.05 | 0.08 | 0.27 | 0.02 | 0.11 | 0.05 |
| 3 | 122780470 | rs6786428 | 0.433 | 0.603 | -0.495 | -0.44 | 0.001 | 0.021 | 0.021 | 0 | 0.58 | 0.4 | 0.53 | 0.9 | 0.6 | 0.77 |
| 3 | 122782715 | rs9850019 | 0.491 | 0.549 | -0.754 | -0.34 | -0.003 | -0.001 | -0.001 | 0 | 0.14 | 0.12 | 0.11 | 0.07 | 0.17 | 0.12 |
| 3 | 122783953 | rs6781240 | 0.202 | 0.402 | -0.452 | -0.446 | -0.007 | -0.025 | -0.025 | 0 | 0.65 | 0.64 | 0.8 | 0.92 | 0.7 | 0.81 |
| 3 | 122788359 | rs9873177 | 0.443 | 0.633 | -0.854 | -0.084 | -0.015 | -0.003 | -0.003 | 0 | 0.16 | 0.06 | 0.1 | 0.07 | 0.18 | 0.15 |
| 3 | 122792686 | rs13434289 | 0.751 | 0.797 | -0.746 | 0.053 | -0.022 | -0.023 | -0.023 | 0.036 | 0.13 | 0.04 | 0.09 | 0.08 | 0.18 | 0.14 |
| 3 | 122796461 | rs9812472 | -0.093 | 1.004 | -0.658 | 0.143 | -0.027 | -0.011 | -0.011 | 0.086 | 0.18 | 0.15 | 0.35 | 0.12 | 0.29 | 0.18 |

| | | | | | | | | | | | | | | | | |
|---|---|---|---|---|---|---|---|---|---|---|---|---|---|---|---|---|
| 3 | 122801981 | rs13075358 | 1.086 | 0.234 | -0.981 | 0.414 | -0.092 | -0.064 | -0.064 | 0 | 0.37 | 0.02 | 0.26 | 0.38 | 0.38 | 0.28 |
| 3 | 122809400 | rs11709496 | 0.099 | 0.337 | 0.506 | 0.768 | -0.009 | -0.025 | -0.025 | 0.469 | 0.06 | 0.17 | 0.03 | 0 | 0.02 | 0.04 |
| 3 | 122819587 | rs2589597 | 0.417 | 0.318 | 0.679 | 0.982 | 0.001 | 0.027 | 0.027 | 0 | 0.81 | 0.75 | 0.74 | 0.69 | 0.77 | 0.67 |
| 3 | 122821671 | rs2241963 | 1.064 | 0.28 | 0.581 | 0.973 | -0.02 | 0.099 | 0.099 | 0 | 0.88 | 0.65 | 0.81 | 0.76 | 0.94 | 0.76 |
| 3 | 122824513 | rs1808988 | 0.288 | 0.633 | -0.007 | 1.01 | -0.001 | -0.006 | -0.006 | 0.054 | 0.81 | 0.81 | 0.75 | 0.68 | 0.77 | 0.67 |
| 3 | 122825040 | rs1078982 | 0.308 | 0.635 | -0.024 | 1.011 | -0.001 | -0.001 | -0.001 | 0 | 0.81 | 0.79 | 0.75 | 0.68 | 0.77 | 0.65 |
| 3 | 122828749 | rs2589596 | 0.332 | 0.711 | 0.187 | 1.02 | 0.005 | 0.084 | 0.084 | 0 | 0.8 | 0.6 | 0.51 | 0.63 | 0.77 | 0.6 |
| 3 | 122834810 | rs836838 | 0.183 | 0.358 | -0.196 | 0.423 | 0.014 | 0.028 | 0.028 | 0 | 0.27 | 0.2 | 0.14 | 0.09 | 0.19 | 0.15 |
| 3 | 122835232 | rs2278668 | 0.793 | 0.592 | -0.258 | 0.722 | 0.008 | -0.013 | -0.013 | 0 | 0.49 | 0.15 | 0.57 | 0.43 | 0.59 | 0.47 |
| 3 | 122841099 | rs836852 | 0.094 | 0.588 | 0.096 | 0.752 | 0.007 | 0.101 | 0.101 | 0 | 0.76 | 0.47 | 0.72 | 0.52 | 0.78 | 0.62 |
| 3 | 122841794 | rs836853 | -0.138 | 0.531 | 0.591 | 0.726 | 0.015 | 0.104 | 0.104 | 0 | 0.78 | 0.6 | 0.73 | 0.52 | 0.78 | 0.6 |
| 3 | 122843110 | rs861375 | 0.2 | 0.532 | 0.449 | 0.741 | 0.006 | 0.102 | 0.102 | 0 | 0.76 | 0.47 | 0.72 | 0.51 | 0.78 | 0.6 |
| 3 | 122845165 | rs3792379 | 1.018 | 0.875 | 0.321 | 0.713 | -0.044 | -0.01 | -0.01 | 0 | 0.46 | 0.12 | 0.53 | 0.4 | 0.51 | 0.44 |
| 3 | 122846881 | rs9820435 | 0.786 | 0.643 | 0.372 | 0.688 | 0.015 | 0.008 | 0.008 | 0 | 0.49 | 0.32 | 0.58 | 0.42 | 0.59 | 0.44 |
| 3 | 122848468 | rs702030 | 0.717 | 0.66 | 0.23 | 0.684 | 0.022 | 0.075 | 0.075 | 0 | 0.26 | 0.1 | 0.14 | 0.09 | 0.19 | 0.15 |
| 3 | 122850576 | rs2589601 | 0.017 | 0.739 | 0.14 | 0.737 | 0.016 | 0.131 | 0.131 | 0 | 0.77 | 0.5 | 0.72 | 0.52 | 0.78 | 0.6 |
| 3 | 122850938 | rs3792382 | 0.613 | 0.868 | -0.335 | 0.792 | 0.023 | 0.008 | 0.008 | 0 | 0.49 | 0.26 | 0.57 | 0.43 | 0.59 | 0.45 |
| 3 | 122853805 | rs836832 | 0.087 | 1.056 | -0.348 | 1.102 | 0.013 | 0.07 | 0.07 | 0 | 0.79 | 0.67 | 0.74 | 0.52 | 0.79 | 0.61 |
| 3 | 122857849 | rs3804755 | 0.308 | 1.034 | -1.201 | 0.78 | -0.004 | 0.051 | 0.051 | 0 | 0.49 | 0.12 | 0.45 | 0.59 | 0.5 | 0.54 |
| 3 | 122858671 | rs920899 | 0.871 | 1.966 | -0.084 | 0.555 | 0.131 | -0.028 | -0.028 | 0 | 0.91 | 0.74 | 0.92 | 0.96 | 0.85 | 0.9 |
| 3 | 122860081 | rs3792390 | -0.433 | 1.504 | -1.085 | 0.311 | -0.014 | 0.005 | 0.005 | 0 | 0.21 | 0.05 | 0.24 | 0.17 | 0.23 | 0.28 |
| 3 | 122860765 | rs836871 | 0.468 | 1.674 | -0.884 | 0.055 | 0 | 0 | 0 | 0 | 0.3 | 0.34 | 0.27 | 0.63 | 0.26 | 0.4 |
| 3 | 122863027 | rs1969262 | -0.862 | 1.276 | -1.219 | 0.256 | -0.035 | 0.129 | 0.129 | 0 | 0.62 | 0.87 | 0.6 | 0.81 | 0.58 | 0.71 |
| 3 | 122870152 | rs2717220 | 0.082 | 0.505 | -1.366 | 0.233 | -0.021 | 0.025 | 0.025 | 0 | 0.25 | 0.03 | 0.21 | 0.16 | 0.25 | 0.33 |
| 3 | 122870545 | rs13062598 | 1.049 | 0.298 | -1.872 | 0.322 | -0.023 | -0.044 | -0.044 | 1.57 | 0.16 | 0.03 | 0.13 | 0.17 | 0.22 | 0.17 |
| 3 | 122871926 | rs2717224 | -0.901 | -0.124 | -1.628 | 0.553 | 0.045 | 0.024 | 0.024 | 0.114 | 0.51 | 0.79 | 0.48 | 0.52 | 0.51 | 0.57 |
| 3 | 122872354 | rs2673341 | -0.413 | -0.128 | -0.746 | 0.6 | 0.028 | -0.004 | -0.004 | 0 | 0.45 | 0.35 | 0.39 | 0.5 | 0.47 | 0.56 |
| 3 | 122877390 | rs872554 | 1.34 | -0.164 | -0.814 | 0.551 | -0.023 | -0.044 | -0.044 | 0 | 0.16 | 0.03 | 0.13 | 0.17 | 0.21 | 0.15 |
| 3 | 122878019 | rs870429 | -0.687 | -0.164 | -0.163 | 0.541 | -0.004 | -0.002 | -0.002 | 0 | 0.48 | 0.5 | 0.45 | 0.52 | 0.51 | 0.58 |
| 3 | 122880118 | rs3828393 | 0.261 | -0.142 | -0.536 | 0.455 | 0.004 | -0.007 | -0.007 | 0 | 0.31 | 0.29 | 0.3 | 0.35 | 0.29 | 0.43 |
| 3 | 122885247 | rs2673338 | -0.316 | -0.476 | -0.565 | 0.216 | 0.049 | -0.01 | -0.01 | 0 | 0.33 | 0.8 | 0.34 | 0.32 | 0.26 | 0.41 |
| 3 | 122886174 | rs2717228 | -0.302 | -0.489 | -0.646 | 0.094 | 0.01 | 0.01 | 0.01 | 0 | 0.41 | 0.36 | 0.4 | 0.32 | 0.46 | 0.48 |
| 3 | 122888637 | rs2667469 | -0.343 | -0.446 | -1.295 | 0.046 | 0.058 | -0.023 | -0.023 | 0 | 0.34 | 0.81 | 0.34 | 0.33 | 0.27 | 0.41 |
| 3 | 122894253 | rs1515372 | -0.047 | -0.336 | -1.522 | -0.169 | -0.042 | 0.087 | 0.087 | 0 | 0.39 | 0.05 | 0.3 | 0.22 | 0.44 | 0.28 |
| 3 | 122897812 | rs1511580 | 0.418 | -1.646 | -0.724 | 0.03 | 0.043 | 0.057 | 0.057 | 0 | 0.27 | 0.12 | 0.21 | 0.13 | 0.22 | 0.24 |
| 3 | 122898729 | rs6438779 | -1.049 | -2.2 | -1.516 | -0.374 | 0.023 | -0.156 | -0.156 | 0 | 0.78 | 0.97 | 0.8 | 0.5 | 0.83 | 0.78 |
| 3 | 122902918 | rs7614713 | 0.133 | -2.194 | -0.542 | -0.649 | -0.036 | -0.065 | -0.065 | 0.309 | 0.33 | 0.53 | 0.27 | 0.13 | 0.26 | 0.25 |
| 3 | 122912292 | rs9821373 | 0.583 | -2.205 | -0.381 | -0.592 | -0.004 | -0.006 | -0.006 | 0 | 0.29 | 0.3 | 0.22 | 0.13 | 0.21 | 0.24 |
| 3 | 122914169 | rs13064484 | 1.449 | -1.591 | -1.015 | -0.711 | -0.156 | 0.009 | 0.009 | 0 | 0.25 | 0.02 | 0.21 | 0.17 | 0.38 | 0.22 |
| 3 | 122915756 | rs6767379 | 1.239 | -1.81 | -0.789 | -0.772 | 0.021 | 0.028 | 0.028 | 0 | 0.25 | 0.2 | 0.19 | 0.12 | 0.21 | 0.22 |
| 3 | 122920007 | rs28413179 | 1.239 | -1.81 | -0.763 | -0.775 | 0.02 | 0.026 | 0.026 | 0 | 0.25 | 0.2 | 0.19 | 0.11 | 0.21 | 0.22 |
| 3 | 122923429 | rs6800592 | 1.101 | -1.802 | -0.773 | -0.774 | 0.021 | 0.028 | 0.028 | 0 | 0.25 | 0.2 | 0.19 | 0.12 | 0.21 | 0.22 |
| 3 | 122929056 | rs2332472 | -1.225 | -2.069 | -1.08 | -0.989 | 0.007 | 0.106 | 0.106 | 0 | 0.77 | 0.98 | 0.88 | 0.95 | 0.84 | 0.89 |
| 3 | 122942402 | rs1546604 | 1.223 | -1.946 | -0.931 | -1.018 | 0.021 | 0.028 | 0.028 | 0 | 0.25 | 0.2 | 0.19 | 0.11 | 0.21 | 0.22 |
| 3 | 122947572 | rs7642344 | 1.06 | -1.96 | -0.486 | -1.05 | 0.04 | -0.05 | -0.05 | 0 | 0.31 | 0.48 | 0.27 | 0.17 | 0.41 | 0.22 |
| 3 | 122956591 | rs9839589 | 0.877 | -1.958 | -0.3 | -1.069 | -0.022 | -0.042 | -0.042 | 4.262 | 0.29 | 0.43 | 0.24 | 0.12 | 0.24 | 0.22 |
| 3 | 122964428 | rs11720939 | 1.643 | -2.016 | -0.423 | -1.102 | -0.079 | 0.089 | 0.089 | 0.164 | 0.16 | 0.02 | 0.27 | 0.01 | 0.17 | 0.21 |
| 3 | 122966184 | rs13101217 | 0.004 | -1.994 | -0.022 | -0.939 | 0.005 | 0.006 | 0.006 | 0 | 0.56 | 0.71 | 0.49 | 0.45 | 0.63 | 0.54 |
| 3 | 122982226 | rs6796833 | 0.97 | -1.951 | -0.375 | -0.924 | -0.014 | -0.022 | -0.022 | 0 | 0.27 | 0.36 | 0.2 | 0.12 | 0.21 | 0.22 |
| 3 | 122990702 | rs1137117 | 1.321 | -1.888 | -0.483 | -0.897 | 0.019 | 0.026 | 0.026 | 0 | 0.25 | 0.2 | 0.19 | 0.12 | 0.21 | 0.22 |
| 3 | 122991161 | rs1137140 | 1.321 | -1.888 | -0.483 | -0.897 | 0.019 | 0.026 | 0.026 | 0 | 0.25 | 0.2 | 0.19 | 0.12 | 0.21 | 0.22 |
| 3 | 122991235 | rs1204 | 1.321 | -1.888 | -0.483 | -0.897 | 0.019 | 0.026 | 0.026 | 0 | 0.25 | 0.2 | 0.19 | 0.12 | 0.21 | 0.22 |
| 3 | 123001494 | rs6784930 | 1.447 | -1.575 | -0.483 | -0.813 | -0.079 | 0.089 | 0.089 | 0.161 | 0.16 | 0.02 | 0.26 | 0.01 | 0.17 | 0.21 |
| 3 | 123006453 | rs6777397 | -0.348 | -1.685 | -0.19 | -0.838 | 0.024 | 0.048 | 0.048 | 0 | 0.54 | 0.38 | 0.46 | 0.82 | 0.43 | 0.55 |
| 3 | 123007207 | rs4678001 | -1.182 | -1.363 | -0.216 | -0.73 | 0.013 | -0.021 | -0.021 | 0.122 | 0.71 | 0.64 | 0.76 | 0.84 | 0.64 | 0.78 |
| 3 | 123013637 | rs4678003 | 0.62 | -1.814 | -0.402 | -0.585 | 0.03 | -0.049 | -0.049 | 0 | 0.24 | 0.1 | 0.25 | 0.69 | 0.18 | 0.32 |
| 3 | 123018963 | rs4482616 | -1.376 | -1.798 | -0.127 | -0.63 | 0 | -0.005 | -0.005 | 0 | 0.75 | 0.79 | 0.79 | 0.88 | 0.79 | 0.78 |
| 3 | 123021298 | rs4678005 | -0.443 | -1.691 | -0.152 | -0.631 | -0.01 | -0.018 | -0.018 | 0.013 | 0.41 | 0.49 | 0.36 | 0.72 | 0.34 | 0.51 |
| 3 | 123025165 | rs4528887 | 1.069 | -2.135 | -0.707 | -0.387 | 0.025 | 0.058 | 0.058 | 0 | 0.24 | 0.04 | 0.18 | 0.13 | 0.21 | 0.22 |
| 3 | 123033693 | rs4234214 | 0.465 | -2.085 | -0.512 | 0.112 | 0.028 | -0.094 | -0.094 | 0 | 0.21 | 0.09 | 0.21 | 0.56 | 0.19 | 0.22 |
| 3 | 123039496 | rs3935566 | -0.423 | -2.215 | -0.004 | 0.084 | 0.012 | 0.174 | 0.174 | 5.311 | 0.08 | 0.49 | 0.08 | 0.14 | 0.01 | 0.21 |
| 3 | 123042911 | rs6806851 | 0.42 | -2.109 | -0.31 | -0.132 | -0.017 | -0.002 | -0.002 | 1.806 | 0.69 | 0.36 | 0.69 | 0.28 | 0.76 | 0.56 |
| 3 | 123045055 | rs9826728 | 0.434 | -2.018 | -0.625 | -0.22 | 0.034 | 0.087 | 0.087 | 0 | 0.43 | 0.22 | 0.5 | 0.25 | 0.55 | 0.35 |
| 3 | 123049938 | rs6806529 | -0.617 | -1.712 | -0.545 | -0.508 | -0.001 | 0.011 | 0.011 | 0 | 0.57 | 0.69 | 0.49 | 0.96 | 0.61 | 0.65 |
| 3 | 123055019 | rs4073374 | 0.898 | -1.85 | -0.994 | 0.134 | 0.004 | -0.095 | -0.095 | 0 | 0.16 | 0.04 | 0.16 | 0.47 | 0.16 | 0.26 |
| 3 | 123056153 | rs4678010 | -0.173 | -1.799 | -0.933 | 0.119 | 0.017 | -0.102 | -0.102 | 0 | 0.25 | 0.11 | 0.24 | 0.49 | 0.26 | 0.31 |
| 3 | 123064831 | rs9289218 | -0.016 | -1.64 | -0.454 | 0.191 | 0.019 | 0.118 | 0.118 | 0 | 0.49 | 0.24 | 0.41 | 0.69 | 0.57 | 0.43 |
| 3 | 123065778 | rs11708067 | 1.328 | -1.718 | -0.523 | -0.142 | -0.007 | -0.003 | -0.003 | 0 | 0.16 | 0.13 | 0.31 | 0 | 0.17 | 0.19 |
| 3 | 123066010 | rs11708903 | 0.374 | -1.766 | -0.42 | -0.151 | -0.009 | 0.129 | 0.129 | 0.079 | 0.43 | 0.2 | 0.53 | 0.03 | 0.42 | 0.36 |

| | | | | | | | | | | | | | | | | |
|---|---|---|---|---|---|---|---|---|---|---|---|---|---|---|---|---|
| 3 | 123068359 | rs4383453 | 1.82 | -1.763 | -0.711 | -0.152 | 0.001 | 0.074 | 0.074 | 0 | 0.23 | 0.07 | 0.18 | 0.03 | 0.21 | 0.12 |
| 3 | 123068743 | rs4630884 | 0.067 | -1.311 | -0.484 | 0.287 | 0.027 | 0.098 | 0.098 | 0 | 0.51 | 0.74 | 0.59 | 0.32 | 0.43 | 0.57 |
| 3 | 123082416 | rs9881942 | -0.012 | -1.312 | -0.628 | 0.281 | -0.014 | 0.094 | 0.094 | 0 | 0.47 | 0.15 | 0.44 | 0.68 | 0.57 | 0.44 |
| 3 | 123094451 | rs2877716 | -0.775 | -1.348 | -0.557 | 0.236 | 0.012 | -0.064 | -0.064 | 0 | 0.82 | 0.75 | 0.67 | 1 | 0.81 | 0.81 |
| 3 | 123095312 | rs6798189 | 1.022 | -1.214 | -0.528 | 0.297 | -0.023 | 0.031 | 0.031 | 0.54 | 0.16 | 0.08 | 0.31 | 0 | 0.19 | 0.19 |
| 3 | 123098965 | rs931574 | 0.239 | -1.301 | -0.655 | 0.715 | 0.018 | -0.052 | -0.052 | 0.398 | 0.2 | 0.11 | 0.2 | 0.36 | 0.18 | 0.26 |
| 3 | 123100223 | rs4677887 | -0.158 | -0.953 | -0.817 | 0.284 | 0.014 | 0.02 | 0.02 | 0 | 0.56 | 0.65 | 0.44 | 0.68 | 0.44 | 0.64 |
| 3 | 123105721 | rs17295246 | 0.702 | -0.664 | -1.305 | -0.168 | 0.034 | 0.133 | 0.133 | 0.079 | 0.31 | 0.07 | 0.19 | 0.07 | 0.25 | 0.21 |
| 3 | 123121286 | rs12496583 | 0.528 | -0.362 | -1.073 | 0.269 | 0.051 | 0.042 | 0.042 | 0.018 | 0.37 | 0.1 | 0.23 | 0.29 | 0.29 | 0.32 |
| 3 | 123124137 | rs7635110 | 0.677 | -0.301 | -1.346 | -0.081 | 0.017 | 0.18 | 0.18 | 0.344 | 0.34 | 0.03 | 0.18 | 0.08 | 0.29 | 0.23 |
| 3 | 123128413 | rs17295401 | 0.66 | -0.304 | -1.336 | -0.067 | 0.015 | 0.179 | 0.179 | 0.676 | 0.35 | 0.03 | 0.18 | 0.08 | 0.29 | 0.23 |
| 3 | 123131254 | rs17361324 | -0.457 | 0.363 | -0.164 | 0.776 | -0.02 | 0.028 | 0.028 | 0 | 0.17 | 0.1 | 0.31 | 0.01 | 0.2 | 0.19 |
| 3 | 123134810 | rs7616545 | -1.51 | -0.13 | 0.639 | 1.643 | 0.003 | 0.359 | 0.359 | 0.77 | 0.07 | 0.38 | 0.1 | 0.35 | 0.01 | 0.23 |
| 3 | 123143272 | rs4677889 | 0.666 | -0.051 | 0.344 | 1.176 | -0.013 | 0.019 | 0.019 | 0 | 0.73 | 0.46 | 0.64 | 0.72 | 0.78 | 0.8 |
| 3 | 123152942 | rs9847700 | 0.243 | 0.481 | 0.29 | 0.821 | -0.046 | 0.047 | 0.047 | 0 | 0.08 | 0.01 | 0.07 | 0.03 | 0.16 | 0.05 |
| 3 | 123157477 | rs9841477 | 0.968 | 0.048 | 0.343 | 1.343 | 0.009 | 0.062 | 0.062 | 0.069 | 0.56 | 0.25 | 0.42 | 0.35 | 0.6 | 0.5 |
| 3 | 123173119 | rs13074111 | -0.794 | -0.24 | 0.61 | 1.425 | -0.002 | -0.009 | -0.009 | 0 | 0.39 | 0.45 | 0.4 | 0.27 | 0.35 | 0.33 |
| 3 | 123176747 | rs4678030 | 0.841 | -0.268 | 0.708 | 1.416 | 0.009 | 0.016 | 0.016 | 0.415 | 0.58 | 0.33 | 0.58 | 0.69 | 0.66 | 0.65 |
| 3 | 123179907 | rs11714453 | -0.219 | -0.366 | 0.619 | 1.223 | -0.015 | 0.135 | 0.135 | 1.17 | 0.3 | 0.1 | 0.32 | 0.05 | 0.29 | 0.29 |
| 3 | 123183651 | rs6798524 | -1.402 | -0.281 | 0.746 | 1.568 | 0.022 | 0.473 | 0.473 | 0 | 0.06 | 0.33 | 0.18 | 0.37 | 0.01 | 0.15 |
| 3 | 123189192 | rs9851728 | 1.079 | -0.233 | 0.611 | 1.589 | 0.052 | 0.564 | 0.564 | 0.285 | 0.95 | 0.62 | 0.82 | 0.63 | 0.99 | 0.89 |
| 3 | 123195946 | rs4677896 | 0.817 | -0.027 | -0.303 | 0.794 | 0.022 | 0.073 | 0.073 | 0.434 | 0.53 | 0.18 | 0.4 | 0.36 | 0.62 | 0.5 |
| 3 | 123196692 | rs9861395 | 0.78 | -0.99 | 0.1 | 1.02 | -0.047 | 0.036 | 0.036 | 0 | 0.74 | 0.43 | 0.73 | 0.39 | 0.86 | 0.68 |
| 3 | 123209551 | rs1039043 | 0.55 | -1.324 | 0.421 | 0.894 | -0.055 | 0.068 | 0.068 | 0 | 0.8 | 0.48 | 0.85 | 0.7 | 0.9 | 0.86 |
| 3 | 123210664 | rs9881425 | 0.865 | -1.412 | 0.611 | 0.847 | -0.001 | -0.007 | -0.007 | 0 | 0.05 | 0.04 | 0.02 | 0.01 | 0.03 | 0.01 |
| 3 | 123211064 | rs10668 | 1.071 | -1.495 | -0.58 | -0.242 | -0.191 | 0.207 | 0.207 | 1.088 | 0.32 | 0.03 | 0.33 | 0.05 | 0.49 | 0.32 |
| 3 | 123212563 | rs1055137 | 0.636 | -1.893 | -0.684 | 0.184 | -0.031 | -0.093 | -0.093 | 0.303 | 0.19 | 0.07 | 0.31 | 0.36 | 0.2 | 0.2 |
| 3 | 123212832 | rs9816876 | 1.782 | -1.885 | -0.682 | 0.183 | -0.001 | -0.007 | -0.007 | 0 | 0.05 | 0.04 | 0.02 | 0.01 | 0.03 | 0.01 |
| 3 | 123217769 | rs16834365 | 0.672 | -1.938 | -0.043 | 0.217 | -0.022 | -0.032 | -0.032 | 0 | 0.11 | 0.21 | 0.07 | 0.02 | 0.06 | 0.02 |
| 3 | 123229780 | rs12495900 | 0.249 | -2.049 | 0.366 | 0.168 | 0 | 0.028 | 0.028 | 0 | 0.22 | 0.28 | 0.32 | 0.37 | 0.21 | 0.2 |
| 3 | 123232693 | rs9811572 | 1.864 | -2.05 | 0.314 | 0.156 | 0.002 | 0.001 | 0.001 | 0.642 | 0.05 | 0.04 | 0.02 | 0.01 | 0.03 | 0.01 |
| 3 | 123243426 | rs3100788 | 1.309 | -2.083 | 0.372 | 0.167 | 0.048 | 0.062 | 0.062 | 1.265 | 0.58 | 0.13 | 0.52 | 0.34 | 0.69 | 0.65 |
| 3 | 123251793 | rs11927656 | 0.953 | -2.173 | 0.424 | 0.125 | -0.011 | 0.007 | 0.007 | 0 | 0.17 | 0.12 | 0.11 | 0.01 | 0.2 | 0.11 |
| 3 | 123253000 | rs16834404 | 0.031 | -2.158 | 0.658 | 0.158 | -0.035 | -0.063 | -0.063 | 0.013 | 0.16 | 0.3 | 0.1 | 0.03 | 0.09 | 0.03 |
| 3 | 123261968 | rs4678041 | -0.548 | -2.16 | 0.657 | 0.173 | -0.043 | 0.097 | 0.097 | 0 | 0.19 | 0.3 | 0.3 | 0.39 | 0.1 | 0.12 |
| 3 | 123280101 | rs6780507 | -1.12 | -2.089 | 0.122 | 0.117 | 0.025 | 0.027 | 0.027 | 0.168 | 0.38 | 0.81 | 0.28 | 0.3 | 0.31 | 0.25 |
| 3 | 123287366 | rs2049713 | -0.868 | -2.153 | 0.107 | 0.123 | -0.004 | -0.004 | -0.004 | 0 | 0.36 | 0.58 | 0.26 | 0.3 | 0.3 | 0.25 |
| 3 | 123289679 | rs2697520 | -1.695 | -2.836 | -0.234 | -0.006 | 0.158 | -0.017 | -0.017 | 0 | 0.34 | 0.91 | 0.41 | 0.65 | 0.15 | 0.31 |
| 3 | 123298615 | rs3109650 | -1.133 | -2.327 | 0.327 | 0.281 | -0.058 | 0.026 | 0.026 | 3.281 | 0.28 | 0.48 | 0.39 | 0.61 | 0.18 | 0.32 |
| 3 | 123318520 | rs861890 | -1.213 | -2.337 | 0.233 | 0.25 | 0 | -0.001 | -0.001 | 0 | 0.31 | 0.69 | 0.41 | 0.63 | 0.18 | 0.33 |
| 3 | 123329932 | rs2626028 | -1.597 | -2.385 | 0.128 | 0.229 | 0.093 | -0.005 | -0.005 | 0 | 0.34 | 0.84 | 0.42 | 0.63 | 0.18 | 0.34 |
| 3 | 123332340 | rs9422 | -1.449 | -2.37 | 0.135 | 0.228 | 0.08 | -0.005 | -0.005 | 0 | 0.33 | 0.8 | 0.41 | 0.63 | 0.18 | 0.33 |
| 3 | 123339834 | rs3845915 | -0.928 | -2.292 | 0.136 | 0.358 | -0.026 | 0.026 | 0.026 | 0.255 | 0.25 | 0.65 | 0.35 | 0.38 | 0.14 | 0.22 |
| 3 | 123351308 | rs16834493 | 0.367 | -1.933 | 0.014 | 0.824 | -0.021 | 0.308 | 0.308 | 0.144 | 0.06 | 0.21 | 0.23 | 0.37 | 0.01 | 0.13 |
| 3 | 123357037 | rs820463 | -0.625 | -1.99 | -0.257 | 0.77 | -0.11 | 0.073 | 0.073 | 3.728 | 0.22 | 0.47 | 0.33 | 0.64 | 0.08 | 0.3 |
| 3 | 123380781 | rs11709947 | 0.472 | -1.01 | -2.07 | -0.745 | -0.115 | 0.302 | 0.302 | 0.734 | 0.41 | 0.03 | 0.38 | 0.04 | 0.55 | 0.36 |
| 3 | 123389855 | rs17299206 | 0.197 | -0.988 | -0.703 | 0.211 | 0.002 | -0.009 | -0.009 | 0.118 | 0.11 | 0.07 | 0.06 | 0.01 | 0.09 | 0.08 |
| 3 | 123393133 | rs820360 | -0.38 | -0.412 | -0.857 | 0.382 | -0.109 | 0.179 | 0.179 | 0 | 0.35 | 0.08 | 0.32 | 0.08 | 0.37 | 0.24 |
| 3 | 123426984 | rs7641248 | -0.72 | -1.52 | 1.592 | 1.475 | -0.001 | 0.02 | 0.02 | 0.455 | 0.17 | 0.14 | 0.38 | 0.48 | 0.13 | 0.1 |
| 3 | 123430265 | rs820339 | 1.288 | -1.079 | 2.462 | 0.496 | -0.047 | -0.008 | -0.008 | 0.014 | 0.9 | 0.38 | 0.94 | 0.95 | 0.97 | 0.87 |
| 3 | 123463708 | rs11718105 | -0.459 | 0.079 | 0.791 | 0.033 | -0.076 | 0.08 | 0.08 | 0.031 | 0.17 | 0.05 | 0.21 | 0.01 | 0.35 | 0.14 |
| 3 | 123473110 | rs9837215 | -1.568 | -1.118 | 1.296 | 0.379 | 0 | -0.006 | -0.006 | 0 | 0.08 | 0.08 | 0.07 | 0.06 | 0.06 | 0.07 |
| 3 | 123493190 | rs9867910 | -1.744 | -1.443 | 2.221 | 0.386 | -0.028 | -0.026 | -0.026 | 4.048 | 0.18 | 0.73 | 0.14 | 0.1 | 0.1 | 0.23 |
| 3 | 123513478 | rs9846863 | -1.746 | -1.517 | 1.63 | 0.421 | -0.065 | -0.129 | -0.129 | 0.131 | 0.18 | 0.44 | 0.1 | 0.06 | 0.09 | 0.23 |
| 3 | 123516240 | rs4678060 | 1.43 | -1.475 | 0.782 | 0.443 | -0.011 | -0.033 | -0.033 | 0 | 0.81 | 0.29 | 0.87 | 0.94 | 0.91 | 0.74 |
| 3 | 123526911 | rs2124508 | -1.091 | -1.573 | 0.4 | 0.67 | 0.01 | -0.012 | -0.012 | 0.313 | 0.36 | 0.33 | 0.51 | 0.59 | 0.33 | 0.38 |
| 3 | 123532046 | rs10511418 | -0.746 | -1.921 | 0.256 | 0.721 | 0.005 | 0.008 | 0.008 | 0 | 0.16 | 0.12 | 0.11 | 0.09 | 0.06 | 0.12 |
| 3 | 123534283 | rs17373884 | -0.563 | -1.941 | 0.191 | 0.773 | -0.076 | 0.076 | 0.076 | 0 | 0.15 | 0.03 | 0.11 | 0.03 | 0.25 | 0.17 |
| 3 | 123535336 | rs10511417 | 0.486 | -2.139 | 0.64 | 0.912 | -0.012 | 0.356 | 0.356 | 0.007 | 0.05 | 0.2 | 0.3 | 0.46 | 0.01 | 0.08 |
| 3 | 123538843 | rs3911406 | -1.487 | -2.224 | 0.847 | 0.848 | -0.083 | 0.104 | 0.104 | 0 | 0.21 | 0.36 | 0.4 | 0.55 | 0.08 | 0.2 |
| 3 | 123544576 | rs2682215 | -0.861 | -1.941 | 0.753 | 0.629 | -0.013 | 0.025 | 0.025 | 0 | 0.16 | 0.1 | 0.12 | 0.06 | 0.21 | 0.25 |
| 3 | 123550463 | rs2700352 | -0.861 | -1.967 | 1.134 | 0.617 | -0.013 | 0.025 | 0.025 | 0 | 0.16 | 0.1 | 0.12 | 0.06 | 0.21 | 0.25 |
| 3 | 123584231 | rs4118366 | 0.71 | -1.99 | 1.371 | 0.598 | -0.001 | -0.01 | -0.01 | 0 | 0.84 | 0.8 | 0.88 | 0.96 | 0.79 | 0.76 |
| 3 | 123584488 | rs16834826 | 2.005 | -2.378 | 1.412 | 0.337 | 0.002 | 0.001 | 0.001 | 0.234 | 0.54 | 0.43 | 0.45 | 0.39 | 0.64 | 0.51 |
| 3 | 123587569 | rs3863980 | -0.215 | -2.622 | 1.886 | 0.107 | -0.009 | 0.169 | 0.169 | 0.616 | 0.07 | 0.19 | 0.08 | 0.36 | 0.06 | 0.18 |
| 3 | 123593561 | rs4118370 | 0.505 | -2.724 | 2.032 | 0.152 | -0.028 | 0.288 | 0.288 | 0 | 0.06 | 0.19 | 0.3 | 0.46 | 0.01 | 0.09 |
| 3 | 123595432 | rs2700408 | 0.948 | -2.775 | 1.89 | 0.11 | 0 | 0.014 | 0.014 | 0 | 0.77 | 0.71 | 0.8 | 0.59 | 0.72 | 0.56 |
| 3 | 123624252 | rs12496579 | 2.678 | -2.798 | 1.921 | -0.055 | 0.005 | 0.264 | 0.264 | 0.049 | 0.45 | 0.12 | 0.39 | 0.1 | 0.56 | 0.36 |
| 3 | 123634046 | rs2700373 | 0.243 | -2.626 | 2.288 | 0.324 | -0.056 | 0.029 | 0.029 | 0 | 0.84 | 0.95 | 0.87 | 0.89 | 0.71 | 0.76 |

| | | | | | | | | | | | | | | | | |
|---|---|---|---|---|---|---|---|---|---|---|---|---|---|---|---|---|
| 3 | 123645834 | rs2091662 | -0.409 | -2.675 | 2.758 | 0.455 | -0.025 | 0.364 | 0.364 | 0.1 | 0.08 | 0.27 | 0.31 | 0.41 | 0.01 | 0.1 |
| 3 | 123671866 | rs2700361 | -0.64 | -2.688 | 2.927 | 0.452 | -0.01 | 0.356 | 0.356 | 0.559 | 0.07 | 0.35 | 0.08 | 0.35 | 0.01 | 0.16 |
| 3 | 123672008 | rs2682198 | -0.827 | -2.728 | 2.909 | 0.434 | -0.019 | 0.333 | 0.333 | 1.001 | 0.07 | 0.36 | 0.08 | 0.35 | 0.01 | 0.16 |
| 3 | 123672378 | rs2700357 | 0.234 | -2.732 | 2.895 | 0.422 | -0.058 | 0.029 | 0.029 | 0 | 0.84 | 0.95 | 0.87 | 0.89 | 0.72 | 0.75 |
| 3 | 123672874 | rs2700351 | 0.234 | -2.732 | 2.895 | 0.422 | -0.058 | 0.029 | 0.029 | 0 | 0.84 | 0.95 | 0.87 | 0.89 | 0.72 | 0.75 |
| 3 | 123673213 | rs4677909 | 0.675 | -2.767 | 2.929 | 0.428 | -0.025 | 0.284 | 0.284 | 0 | 0.06 | 0.16 | 0.3 | 0.41 | 0.01 | 0.1 |
| 3 | 123674627 | rs2700409 | -0.833 | -2.738 | 2.769 | 0.433 | -0.019 | 0.333 | 0.333 | 1.001 | 0.07 | 0.36 | 0.08 | 0.35 | 0.01 | 0.16 |
| 3 | 123683440 | rs7633013 | 0.42 | -2.649 | 2.414 | 0.346 | -0.06 | 0.034 | 0.034 | 0 | 0.83 | 0.95 | 0.86 | 0.89 | 0.7 | 0.76 |
| 3 | 123686157 | rs12695434 | 0.332 | -2.615 | 2.428 | 0.358 | -0.06 | 0.03 | 0.03 | 0 | 0.84 | 0.95 | 0.86 | 0.89 | 0.7 | 0.76 |
| 3 | 123699823 | rs17310770 | -0.619 | -3.032 | 2.47 | 0.15 | 0.003 | 0.047 | 0.047 | 0 | 0.21 | 0.02 | 0.09 | 0.12 | 0.16 | 0.21 |
| 3 | 123699904 | rs16834952 | -1.298 | -3.033 | 2.62 | 0.219 | 0.017 | 0.492 | 0.492 | 0 | 0.05 | 0.25 | 0.26 | 0.42 | 0.01 | 0.08 |
| 3 | 123702716 | rs7621877 | -0.296 | -2.993 | 2.48 | 0.137 | -0.005 | -0.006 | -0.006 | 1.096 | 0.89 | 0.93 | 0.92 | 1 | 0.84 | 0.89 |
| 3 | 123702960 | rs9881117 | 2.249 | -3.06 | 2.388 | 0.133 | 0.092 | 0.178 | 0.178 | 0.821 | 0.52 | 0.25 | 0.43 | 0.1 | 0.65 | 0.47 |
| 3 | 123704039 | rs6770339 | -1.2 | -2.742 | 2.589 | 0.513 | -0.009 | 0.374 | 0.374 | 1.132 | 0.07 | 0.37 | 0.14 | 0.35 | 0.01 | 0.12 |
| 3 | 123708048 | rs2332717 | -0.59 | -2.729 | 1.726 | 0.564 | 0.01 | -0.029 | -0.029 | 0 | 0.13 | 0.06 | 0.17 | 0.38 | 0.11 | 0.2 |
| 3 | 123712966 | rs2332719 | -2.434 | -3.063 | 1.683 | 0.323 | 0.077 | -0.064 | -0.064 | 0 | 0.4 | 0.82 | 0.36 | 0.5 | 0.31 | 0.36 |
| 3 | 123719219 | rs9289229 | -0.875 | -3.084 | 2.352 | 0.316 | 0.031 | -0.014 | -0.014 | 0 | 0.06 | 0.29 | 0.07 | 0.01 | 0.08 | 0.02 |
| 3 | 123728685 | rs9866970 | -0.601 | -3.129 | 2.509 | 0.305 | -0.038 | -0.1 | -0.1 | 0 | 0.17 | 0.34 | 0.09 | 0.01 | 0.1 | 0.08 |
| 3 | 123734690 | rs9823891 | -2.602 | -3.155 | 2.501 | 0.315 | 0.063 | -0.045 | -0.045 | 0.111 | 0.39 | 0.78 | 0.35 | 0.5 | 0.31 | 0.4 |
| 3 | 123738355 | rs4485668 | -2.664 | -3.163 | 2.395 | 0.308 | 0.086 | -0.083 | -0.083 | 0.09 | 0.4 | 0.87 | 0.36 | 0.5 | 0.31 | 0.4 |
| 3 | 123744470 | rs12634530 | -1.987 | -2.999 | 2.368 | 0.412 | -0.031 | 0.133 | 0.133 | 0 | 0.22 | 0.44 | 0.26 | 0.48 | 0.21 | 0.31 |
| 3 | 123764705 | rs2332740 | 1.339 | -2.906 | 2.436 | 0.34 | 0.041 | -0.005 | -0.005 | 0 | 0.93 | 0.61 | 0.92 | 0.99 | 0.93 | 0.99 |
| 3 | 123767922 | rs2332690 | 1.769 | -2.977 | 2.489 | 0.41 | -0.048 | 0.163 | 0.163 | 0 | 0.77 | 0.51 | 0.73 | 0.5 | 0.81 | 0.73 |
| 3 | 123774078 | rs9289231 | -0.344 | -3.192 | 2.572 | 0.283 | -0.028 | -0.087 | -0.087 | 0 | 0.15 | 0.3 | 0.08 | 0.01 | 0.09 | 0.09 |
| 3 | 123774706 | rs13083500 | 2.213 | -3.171 | 2.528 | 0.293 | 0.067 | -0.053 | -0.053 | 0 | 0.62 | 0.22 | 0.65 | 0.49 | 0.71 | 0.65 |
| 3 | 123776606 | rs16835038 | -0.674 | -3.07 | 2.441 | 0.373 | 0.002 | -0.01 | -0.01 | 0 | 0.07 | 0.01 | 0.24 | 0.39 | 0.07 | 0.16 |
| 3 | 123805421 | rs6773078 | -1.805 | -2.928 | 2.79 | 0.384 | -0.055 | 0.074 | 0.074 | 0 | 0.3 | 0.46 | 0.28 | 0.47 | 0.22 | 0.34 |
| 3 | 123807012 | rs11721323 | -0.684 | -2.935 | 2.778 | 0.382 | 0.01 | -0.016 | -0.016 | 0 | 0.14 | 0.09 | 0.19 | 0.45 | 0.13 | 0.26 |
| 3 | 123810460 | rs4678082 | 0.909 | -2.936 | 2.792 | 0.375 | 0.039 | -0.004 | -0.004 | 0 | 0.93 | 0.67 | 0.92 | 1 | 0.93 | 0.99 |
| 3 | 123812836 | rs10934657 | 2.36 | -2.936 | 2.583 | 0.378 | 0.073 | -0.058 | -0.058 | 0 | 0.63 | 0.2 | 0.65 | 0.51 | 0.72 | 0.65 |
| 3 | 123824333 | rs16835100 | -0.885 | -2.92 | 2.492 | 0.427 | -0.01 | 0.069 | 0.069 | 0 | 0.15 | 0.17 | 0.2 | 0.46 | 0.13 | 0.26 |
| 3 | 123831367 | rs4678085 | -1.679 | -2.804 | 2.595 | 0.438 | -0.034 | 0.037 | 0.037 | 0 | 0.31 | 0.54 | 0.28 | 0.47 | 0.22 | 0.34 |
| 3 | 123834347 | rs1320182 | 1.775 | -2.187 | 2.126 | 0.267 | 0.069 | -0.072 | -0.072 | 0 | 0.59 | 0.18 | 0.61 | 0.51 | 0.66 | 0.61 |
| 3 | 123858749 | rs10512620 | -1.615 | -2.201 | 2.476 | 0.257 | -0.02 | 0.023 | 0.023 | 0 | 0.37 | 0.5 | 0.34 | 0.48 | 0.34 | 0.39 |
| 3 | 123878673 | rs6786840 | -1.343 | -2.208 | 2.256 | 0.253 | 0.02 | -0.084 | -0.084 | 0 | 0.34 | 0.14 | 0.31 | 0.47 | 0.34 | 0.38 |
| 3 | 123880441 | rs4678090 | -1.333 | -2.204 | 2.208 | 0.253 | 0.024 | -0.095 | -0.095 | 0.356 | 0.34 | 0.12 | 0.31 | 0.47 | 0.33 | 0.38 |
| 3 | 123895464 | rs1920609 | -1.395 | -2.199 | 2.247 | 0.228 | 0.022 | -0.081 | -0.081 | 0.041 | 0.35 | 0.13 | 0.31 | 0.47 | 0.34 | 0.38 |
| 3 | 123897358 | rs12488684 | 1.73 | -2.181 | 2.007 | 0.236 | 0.064 | -0.06 | -0.06 | 0 | 0.58 | 0.2 | 0.6 | 0.52 | 0.66 | 0.61 |
| 3 | 123916071 | rs2141665 | -1.454 | -2.009 | 1.916 | 0.259 | 0.001 | -0.003 | -0.003 | 0 | 0.37 | 0.37 | 0.34 | 0.55 | 0.34 | 0.39 |
| 3 | 123923922 | rs1444768 | 1.52 | -2.048 | 0.256 | 0.057 | 0.101 | 0 | 0 | 0 | 0.5 | 0.16 | 0.59 | 0.44 | 0.62 | 0.53 |
| 3 | 123925295 | rs1822791 | -0.136 | -2.117 | 0.134 | 0.185 | -0.018 | -0.1 | -0.1 | 0 | 0.13 | 0.02 | 0.18 | 0.42 | 0.11 | 0.24 |
| 3 | 123927948 | rs2084017 | 0.532 | -2.118 | 0.473 | 0.185 | -0.023 | 0.169 | 0.169 | 0 | 0.83 | 0.6 | 0.77 | 0.61 | 0.88 | 0.76 |
| 3 | 123930542 | rs1444757 | 1.373 | -1.903 | -0.469 | 0.212 | 0.094 | 0 | 0 | 0 | 0.5 | 0.14 | 0.57 | 0.47 | 0.62 | 0.53 |
| 3 | 123939607 | rs1444755 | -0.924 | -1.535 | 0.238 | 0.698 | -0.008 | 0.104 | 0.104 | 0 | 0.15 | 0.22 | 0.2 | 0.46 | 0.12 | 0.23 |
| 3 | 123940692 | rs9843963 | -1.83 | -2.147 | 0.37 | 0.897 | 0.001 | -0.005 | -0.005 | 0 | 0.17 | 0.19 | 0.29 | 0.75 | 0.13 | 0.33 |
| 3 | 123941100 | rs1920623 | -2.075 | -2.168 | 0.508 | 0.902 | -0.063 | 0.066 | 0.066 | 0 | 0.18 | 0.36 | 0.31 | 0.72 | 0.13 | 0.31 |
| 3 | 123942353 | rs6789224 | -1.411 | -2.221 | 0.716 | 1.023 | -0.017 | 0.098 | 0.098 | 0 | 0.09 | 0.15 | 0.19 | 0.56 | 0.08 | 0.19 |
| 3 | 123943740 | rs9859929 | -1.719 | -2.237 | 0.78 | 1.059 | -0.044 | 0.152 | 0.152 | 0 | 0.11 | 0.23 | 0.24 | 0.63 | 0.06 | 0.25 |
| 3 | 123944955 | rs9858797 | -2.467 | -2.239 | 0.778 | 1.054 | -0.066 | 0.074 | 0.074 | 0 | 0.18 | 0.37 | 0.31 | 0.72 | 0.13 | 0.31 |
| 3 | 123952217 | rs17286604 | 0.76 | -1.604 | -0.289 | 0.09 | -0.118 | 0.248 | 0.248 | 0.387 | 0.36 | 0.07 | 0.19 | 0.02 | 0.38 | 0.32 |
| 3 | 123952276 | rs3772728 | 0.262 | -1.462 | -0.163 | 0.273 | 0.021 | -0.025 | -0.025 | 2.224 | 0.07 | 0.02 | 0.11 | 0.16 | 0.05 | 0.12 |
| 3 | 123954939 | rs9846518 | 0.109 | -1.229 | -0.028 | 0.336 | -0.015 | 0.011 | 0.011 | 0 | 0.07 | 0.14 | 0.06 | 0.09 | 0.07 | 0.06 |
| 3 | 123956596 | rs4678095 | 0.772 | -1.228 | 0.069 | 0.298 | -0.078 | 0.077 | 0.077 | 0 | 0.41 | 0.13 | 0.44 | 0.23 | 0.49 | 0.35 |
| 3 | 123956804 | rs4321476 | 0.779 | -1.141 | 1.094 | 1.015 | 0.021 | 0.504 | 0.504 | 0 | 0.95 | 0.7 | 0.83 | 0.53 | 0.98 | 0.87 |
| 3 | 123959585 | rs7616435 | -1.002 | -1.344 | 1.433 | 0.95 | -0.032 | 0.022 | 0.022 | 0 | 0.11 | 0.29 | 0.09 | 0.09 | 0.07 | 0.08 |
| 3 | 123960730 | rs4678097 | -1.749 | -1.565 | 0.933 | 0.639 | -0.034 | 0.025 | 0.025 | 0 | 0.27 | 0.58 | 0.3 | 0.59 | 0.2 | 0.39 |
| 3 | 123966566 | rs2332768 | 1.191 | -1.186 | -0.036 | 0.348 | -0.055 | 0 | 0 | 0.915 | 0.3 | 0.07 | 0.39 | 0.23 | 0.36 | 0.23 |
| 3 | 123967952 | rs6781700 | -0.068 | -1.065 | 0.125 | 0.23 | -0.002 | -0.001 | -0.001 | 0.076 | 0.13 | 0.13 | 0.15 | 0.19 | 0.09 | 0.24 |
| 3 | 123970685 | rs6438837 | -0.348 | -1.086 | 0.275 | 0.215 | -0.009 | -0.002 | -0.002 | 0.02 | 0.14 | 0.18 | 0.15 | 0.18 | 0.09 | 0.24 |
| 3 | 123970864 | rs13321072 | 0.467 | -0.993 | 0.24 | 0.27 | 0.001 | -0.003 | -0.003 | 0.454 | 0.07 | 0.07 | 0.07 | 0.09 | 0.08 | 0.07 |
| 3 | 123974407 | rs13320544 | 0.427 | -1.001 | 0.216 | 0.266 | 0.003 | -0.005 | -0.005 | 1.215 | 0.07 | 0.06 | 0.06 | 0.09 | 0.08 | 0.06 |
| 3 | 123974576 | rs2120789 | 1.503 | -0.955 | 0.475 | 0.201 | 0.02 | 0.029 | 0.029 | 0.027 | 0.65 | 0.24 | 0.68 | 0.71 | 0.72 | 0.6 |
| 3 | 123976794 | rs7636752 | 0.167 | -0.86 | 0.674 | 0.512 | -0.006 | 0.2 | 0.2 | 0.315 | 0.84 | 0.38 | 0.8 | 0.51 | 0.88 | 0.78 |
| 3 | 123980608 | rs1997422 | 0.328 | -0.89 | 0.577 | 0.295 | 0.001 | -0.011 | -0.011 | 0 | 0.43 | 0.45 | 0.47 | 0.26 | 0.46 | 0.33 |
| 3 | 123983604 | rs2276739 | -0.365 | -0.793 | 0.586 | 0.379 | -0.001 | 0.009 | 0.009 | 0 | 0.23 | 0.18 | 0.21 | 0.18 | 0.2 | 0.34 |
| 3 | 123985016 | rs7636024 | 0.516 | -0.756 | 0.63 | 0.328 | -0.005 | -0.004 | -0.004 | 0 | 0.4 | 0.47 | 0.46 | 0.33 | 0.42 | 0.31 |
| 3 | 123988039 | rs2272486 | -1.107 | -0.767 | 0.562 | 0.354 | -0.003 | -0.008 | -0.008 | 0 | 0.63 | 0.65 | 0.66 | 0.53 | 0.62 | 0.65 |
| 3 | 124003228 | rs2332769 | -1.535 | -0.871 | 0.935 | 0.328 | 0.002 | -0.021 | -0.021 | 0.184 | 0.7 | 0.52 | 0.59 | 0.79 | 0.66 | 0.8 |
| 3 | 124013947 | rs1373606 | -0.49 | -0.811 | 0.661 | 0.38 | 0.005 | 0.007 | 0.007 | 0 | 0.58 | 0.36 | 0.52 | 0.66 | 0.57 | 0.68 |

| | | | | | | | | | | | | | | | | |
|---|---|---|---|---|---|---|---|---|---|---|---|---|---|---|---|---|
| 3 | 124017996 | rs12695436 | -0.38 | -0.653 | 0.68 | 0.491 | -0.002 | 0.008 | 0.008 | 0 | 0.22 | 0.19 | 0.21 | 0.18 | 0.19 | 0.32 |
| 3 | 124039542 | rs13067494 | 0.017 | -0.53 | 0.599 | 0.653 | 0.009 | -0.039 | -0.039 | 0.001 | 0.56 | 0.21 | 0.45 | 0.56 | 0.55 | 0.67 |
| 3 | 124044949 | rs2289778 | 1.194 | -0.418 | 1.301 | 0.801 | -0.021 | 0.085 | 0.085 | 0 | 0.86 | 0.5 | 0.81 | 0.78 | 0.89 | 0.88 |
| 3 | 124048764 | rs2276740 | -0.734 | -0.433 | 1.278 | 0.673 | -0.009 | 0.01 | 0.01 | 0 | 0.11 | 0.16 | 0.1 | 0.15 | 0.08 | 0.1 |
| 3 | 124053501 | rs6438839 | -1.144 | -0.459 | 1.675 | 0.637 | -0.006 | -0.056 | -0.056 | 0 | 0.12 | 0.47 | 0.11 | 0.01 | 0.07 | 0.09 |
| 3 | 124055024 | rs3821525 | 0.386 | 0.181 | 1.153 | 0.842 | -0.044 | -0.065 | -0.065 | 0.058 | 0.81 | 0.89 | 0.71 | 0.48 | 0.72 | 0.61 |
| 3 | 124058705 | rs3772731 | 0.592 | -0.234 | 1.066 | 0.397 | -0.008 | -0.011 | -0.011 | 0 | 0.39 | 0.55 | 0.33 | 0.23 | 0.31 | 0.23 |
| 3 | 124061586 | rs11708466 | 0.985 | -0.254 | 0.592 | 0.425 | 0.032 | 0.032 | 0.032 | 0.03 | 0.35 | 0.13 | 0.27 | 0.23 | 0.29 | 0.22 |
| 3 | 124066969 | rs1444750 | -0.177 | 0.079 | 0.476 | 0.508 | -0.01 | 0.003 | 0.003 | 0 | 0.87 | 0.89 | 0.76 | 0.47 | 0.81 | 0.71 |
| 3 | 124070129 | rs10934667 | 1.098 | -0.211 | 0.324 | 0.363 | 0.05 | 0.017 | 0.017 | 0.728 | 0.32 | 0.13 | 0.18 | 0.22 | 0.22 | 0.2 |
| 3 | 124078675 | rs1444756 | -0.698 | 0.006 | 0.946 | -0.024 | 0.001 | 0.021 | 0.021 | 0 | 0.56 | 0.59 | 0.63 | 0.89 | 0.59 | 0.63 |
| 3 | 124078870 | rs10511423 | 0.986 | -0.098 | 0.921 | -0.084 | -0.039 | 0.191 | 0.191 | 0.337 | 0.4 | 0.14 | 0.27 | 0 | 0.4 | 0.29 |
| 3 | 124079272 | rs9878723 | -0.888 | -0.097 | 0.967 | -0.076 | -0.009 | 0.13 | 0.13 | 0 | 0.58 | 0.75 | 0.63 | 0.9 | 0.59 | 0.63 |
| 3 | 124092145 | rs659450 | 0.031 | -0.094 | 0.965 | -0.025 | 0.026 | 0.096 | 0.096 | 0 | 0.44 | 0.19 | 0.41 | 0.75 | 0.4 | 0.52 |
| 3 | 124095793 | rs498536 | -0.022 | -0.083 | 1.215 | -0.022 | -0.005 | -0.018 | -0.018 | 0 | 0.47 | 0.47 | 0.43 | 0.76 | 0.4 | 0.52 |
| 3 | 124100519 | rs895286 | -0.005 | 0.65 | 1.448 | 0.708 | 0.038 | 0.019 | 0.019 | 0 | 0.1 | 0.19 | 0.19 | 0.14 | 0.19 | 0.11 |
| 3 | 124102388 | rs9869539 | -0.127 | 0.755 | 0.756 | 0.688 | -0.027 | 0.001 | 0.001 | 0 | 0.1 | 0.08 | 0.23 | 0.12 | 0.19 | 0.18 |
| 3 | 124121911 | rs13068267 | -0.372 | 0.701 | 0.224 | 1.027 | -0.023 | -0.034 | -0.034 | 0 | 0.13 | 0.05 | 0.24 | 0.63 | 0.19 | 0.33 |
| 3 | 124125522 | rs4677925 | 0.924 | 0.563 | 0.289 | 0.362 | 0.015 | -0.018 | -0.018 | 0 | 0.75 | 0.74 | 0.72 | 0.86 | 0.67 | 0.74 |
| 3 | 124127961 | rs661798 | 0.282 | -0.306 | -0.068 | -0.032 | 0.002 | 0.039 | 0.039 | 0 | 0.51 | 0.64 | 0.57 | 0.75 | 0.5 | 0.55 |
| 3 | 124131844 | rs3887233 | 0.407 | 0.163 | 0.202 | 0.193 | 0.002 | -0.001 | -0.001 | 0 | 0.45 | 0.53 | 0.48 | 0.53 | 0.45 | 0.46 |
| 3 | 124132568 | rs6762346 | 0.344 | 0.106 | 0.356 | 0.168 | -0.006 | 0.057 | 0.057 | 0 | 0.47 | 0.67 | 0.6 | 0.71 | 0.47 | 0.51 |
| 3 | 124136699 | rs7632785 | 0.596 | 0.093 | 0.752 | 0.16 | -0.007 | 0.015 | 0.015 | 0 | 0.43 | 0.36 | 0.46 | 0.65 | 0.44 | 0.49 |
| 3 | 124141503 | rs1373928 | 0.899 | 0.064 | 0.831 | 0.461 | -0.019 | -0.03 | -0.03 | 0 | 0.41 | 0.25 | 0.43 | 0.46 | 0.42 | 0.47 |
| 3 | 124145425 | rs3772753 | 0.474 | 0.367 | 0.396 | 0.124 | -0.004 | 0.03 | 0.03 | 0 | 0.07 | 0.09 | 0.07 | 0.19 | 0.07 | 0.04 |
| 3 | 124149174 | rs630760 | 0.317 | 0.579 | 0.125 | -0.127 | 0.059 | 0.128 | 0.128 | 0 | 0.43 | 0.17 | 0.24 | 0.22 | 0.35 | 0.39 |
| 3 | 124154852 | rs717955 | 0.201 | 0.473 | 0.26 | 0.018 | -0.008 | 0.062 | 0.062 | 0 | 0.46 | 0.68 | 0.61 | 0.76 | 0.46 | 0.52 |
| 3 | 124163162 | rs6798951 | 0.441 | 0.727 | 0.294 | 0.732 | 0.02 | 0.184 | 0.184 | 0 | 0.06 | 0.13 | 0.26 | 0.5 | 0.13 | 0.19 |
| 3 | 124165217 | rs624450 | -0.264 | 0.222 | 0.3 | 0.019 | -0.014 | -0.014 | -0.014 | 0 | 0.43 | 0.5 | 0.31 | 0.31 | 0.37 | 0.41 |
| 3 | 124178163 | rs572209 | 0.448 | 0.063 | 0.156 | -0.475 | 0 | -0.002 | -0.002 | 0 | 0.15 | 0.15 | 0.08 | 0.08 | 0.12 | 0.18 |
| 3 | 124178276 | rs573186 | -0.063 | -0.101 | 0.303 | -0.761 | -0.007 | -0.054 | -0.054 | 0 | 0.63 | 0.5 | 0.41 | 0.12 | 0.6 | 0.56 |
| 3 | 124181261 | rs535121 | 0.6 | -0.11 | 0.34 | -0.736 | -0.006 | -0.022 | -0.022 | 1.052 | 0.16 | 0.24 | 0.1 | 0.08 | 0.12 | 0.18 |
| 3 | 124181454 | rs2289839 | 1.625 | -0.178 | -0.03 | -0.697 | 0.007 | -0.027 | -0.027 | 0 | 0.06 | 0.01 | 0.04 | 0.13 | 0.05 | 0.05 |
| 3 | 124181739 | rs2289840 | 1.034 | -0.181 | 0.08 | -0.667 | 0 | -0.031 | -0.031 | 0 | 0.26 | 0.2 | 0.42 | 0.61 | 0.32 | 0.37 |
| 3 | 124182136 | rs9817308 | 0.987 | 0.037 | -0.127 | -0.643 | -0.001 | 0.157 | 0.157 | 0 | 0.47 | 0.25 | 0.31 | 0.05 | 0.48 | 0.37 |
| 3 | 124189656 | rs6770075 | 0.46 | 0.586 | -0.254 | -0.646 | -0.005 | 0.089 | 0.089 | 0 | 0.52 | 0.33 | 0.35 | 0.18 | 0.49 | 0.46 |
| 3 | 124194149 | rs7623685 | 0.516 | 0.591 | -0.355 | -0.642 | -0.002 | 0.128 | 0.128 | 0 | 0.51 | 0.27 | 0.34 | 0.18 | 0.49 | 0.46 |
| 3 | 124197610 | rs3755674 | 1.005 | 0.968 | -0.427 | -0.108 | -0.024 | 0.061 | 0.061 | 0 | 0.06 | 0.02 | 0.25 | 0.5 | 0.16 | 0.17 |
| 3 | 124209438 | rs2289841 | -0.687 | 0.436 | -0.437 | -0.549 | 0.03 | 0.01 | 0.01 | 0 | 0.63 | 0.75 | 0.72 | 0.72 | 0.69 | 0.58 |
| 3 | 124210544 | rs2076716 | -0.456 | 0.791 | -0.164 | -0.39 | 0.01 | 0.02 | 0.02 | 0 | 0.49 | 0.61 | 0.59 | 0.69 | 0.5 | 0.54 |
| 3 | 124211050 | rs3755678 | -0.36 | 1.026 | 0.178 | -0.15 | 0.028 | 0.153 | 0.153 | 0 | 0.13 | 0.21 | 0.25 | 0.49 | 0.15 | 0.22 |
| 3 | 124219712 | rs1445637 | -0.784 | 0.689 | 0.105 | -0.539 | 0.012 | 0.015 | 0.015 | 0 | 0.52 | 0.59 | 0.43 | 0.38 | 0.48 | 0.56 |
| 3 | 124222935 | rs9859074 | 0.667 | 0.948 | -0.122 | -0.709 | 0.01 | 0.007 | 0.007 | 0 | 0.32 | 0.29 | 0.16 | 0.28 | 0.24 | 0.37 |
| 3 | 124223298 | rs522554 | 0.055 | 0.886 | 0.193 | -0.677 | 0.017 | 0.018 | 0.018 | 1.13 | 0.89 | 0.54 | 0.84 | 0.92 | 0.95 | 0.91 |
| 3 | 124224993 | rs485887 | 0.258 | 0.788 | 0.142 | -0.68 | 0.038 | -0.01 | -0.01 | 0 | 0.87 | 0.52 | 0.82 | 0.92 | 0.92 | 0.91 |
| 3 | 124229502 | rs556031 | 0.99 | 0.845 | -0.384 | -0.685 | 0.018 | 0.036 | 0.036 | 0 | 0.46 | 0.22 | 0.54 | 0.61 | 0.52 | 0.44 |
| 3 | 124229890 | rs6805335 | 1.816 | 0.779 | -0.422 | -0.834 | 0.014 | 0.018 | 0.018 | 0 | 0.2 | 0.12 | 0.08 | 0 | 0.14 | 0.18 |
| 3 | 124236829 | rs1867647 | -1.215 | 0.951 | -0.739 | -0.723 | 0.022 | 0.026 | 0.026 | 0 | 0.54 | 0.77 | 0.47 | 0.39 | 0.49 | 0.56 |
| 3 | 124236917 | rs33981978 | 0.242 | 1.015 | -0.667 | -0.753 | -0.032 | -0.029 | -0.029 | 0 | 0.19 | 0.12 | 0.19 | 0.31 | 0.26 | 0.28 |
| 3 | 124237591 | rs6765634 | 0.432 | 0.965 | -0.578 | -0.733 | -0.013 | -0.087 | -0.087 | 1.569 | 0.32 | 0.53 | 0.26 | 0.08 | 0.22 | 0.28 |
| 3 | 124238130 | rs7613442 | 0.43 | 0.977 | -0.573 | -0.729 | -0.012 | -0.087 | -0.087 | 0.107 | 0.32 | 0.53 | 0.26 | 0.08 | 0.22 | 0.28 |
| 3 | 124244479 | rs16835545 | 0.977 | 1.077 | -0.8 | -0.631 | 0.019 | -0.039 | -0.039 | 0 | 0.08 | 0.1 | 0.05 | 0 | 0.06 | 0.08 |
| 3 | 124247149 | rs332414 | -1.831 | 0.316 | -1.76 | -0.747 | -0.021 | -0.091 | -0.091 | 0 | 0.79 | 0.94 | 0.59 | 0.4 | 0.69 | 0.65 |
| 3 | 124258080 | rs332408 | -1.132 | 0.657 | -0.954 | -0.374 | 0.022 | 0.027 | 0.027 | 0 | 0.53 | 0.73 | 0.38 | 0.4 | 0.48 | 0.55 |
| 3 | 124263881 | rs6800965 | -0.215 | 1.056 | -0.932 | -0.41 | 0.097 | 0.033 | 0.033 | 0 | 0.27 | 0.48 | 0.27 | 0.29 | 0.33 | 0.35 |
| 3 | 124272171 | rs2713651 | 0.035 | 0.221 | -0.454 | -0.325 | -0.019 | -0.09 | -0.09 | 0 | 0.23 | 0.34 | 0.11 | 0 | 0.15 | 0.15 |
| 3 | 124281604 | rs2279784 | -0.407 | 0.3 | -0.891 | 0.542 | 0.027 | -0.043 | -0.043 | 0.035 | 0.19 | 0.1 | 0.31 | 0.65 | 0.12 | 0.21 |
| 3 | 124281980 | rs2256831 | -0.875 | 0.843 | -1.354 | -0.196 | -0.044 | 0.257 | 0.257 | 2.791 | 0.52 | 0.85 | 0.57 | 0.86 | 0.5 | 0.42 |
| 3 | 124283993 | rs986911 | -1.002 | 1.067 | -0.368 | 0.424 | 0.023 | 0.086 | 0.086 | 0.151 | 0.29 | 0.49 | 0.32 | 0.51 | 0.3 | 0.27 |
| 3 | 124285457 | rs7633543 | -0.53 | 1.162 | -0.187 | 0.424 | 0.014 | 0.167 | 0.167 | 0.16 | 0.23 | 0.49 | 0.28 | 0.51 | 0.19 | 0.14 |
| 3 | 124288962 | rs16835606 | -0.919 | 1.03 | 0.003 | 0.453 | 0.015 | 0.061 | 0.061 | 0.314 | 0.3 | 0.6 | 0.35 | 0.51 | 0.3 | 0.26 |
| 3 | 124290306 | rs1532668 | -0.875 | 1.06 | 0.036 | 0.476 | 0.022 | 0.085 | 0.085 | 0.043 | 0.29 | 0.52 | 0.33 | 0.51 | 0.3 | 0.26 |
| 3 | 124292219 | rs2713649 | 0.236 | 1.208 | 0.213 | 0.545 | -0.001 | 0.037 | 0.037 | 0.439 | 0.28 | 0.4 | 0.33 | 0.39 | 0.3 | 0.22 |
| 3 | 124297645 | rs6438847 | -0.072 | 1.216 | 0.246 | 0.584 | 0 | -0.003 | -0.003 | 0.023 | 0.31 | 0.6 | 0.46 | 0.8 | 0.35 | 0.25 |
| 3 | 124298174 | rs6766077 | 0.455 | 1.212 | -0.057 | 0.612 | -0.028 | -0.05 | -0.05 | 0 | 0.25 | 0.07 | 0.28 | 0.34 | 0.29 | 0.2 |
| 3 | 124299968 | rs6772598 | 0.455 | 1.212 | -0.057 | 0.612 | -0.028 | -0.05 | -0.05 | 0 | 0.25 | 0.07 | 0.28 | 0.34 | 0.29 | 0.2 |
| 3 | 124305845 | rs4677935 | -0.304 | 1.167 | -0.27 | 0.468 | 0 | 0.101 | 0.101 | 0 | 0.39 | 0.1 | 0.3 | 0.21 | 0.36 | 0.28 |
| 3 | 124309338 | rs2651620 | -1.1 | 1.361 | 0.284 | 0.44 | -0.001 | -0.001 | -0.001 | 2.897 | 0.3 | 0.68 | 0.34 | 0.51 | 0.31 | 0.26 |
| 3 | 124314560 | rs7639225 | -0.636 | 1.29 | 0.431 | 0.471 | -0.005 | 0.105 | 0.105 | 0 | 0.25 | 0.44 | 0.28 | 0.52 | 0.24 | 0.24 |

| | | | | | | | | | | | | | | | | |
|---|---|---|---|---|---|---|---|---|---|---|---|---|---|---|---|---|
| 3 | 124318900 | rs6797559 | 0.162 | 1.278 | 0.299 | -0.128 | -0.007 | 0.064 | 0.064 | 0 | 0.19 | 0.11 | 0.12 | 0 | 0.19 | 0.17 |
| 3 | 124323191 | rs4678130 | -1.302 | 1.28 | 0.309 | -0.18 | 0.004 | 0.002 | 0.002 | 0 | 0.54 | 0.58 | 0.58 | 0.85 | 0.59 | 0.47 |
| 3 | 124326801 | rs333338 | -1.082 | 1.298 | -0.081 | -0.138 | -0.03 | 0.233 | 0.233 | 0 | 0.52 | 0.85 | 0.58 | 0.85 | 0.53 | 0.46 |
| 3 | 124327305 | rs333335 | -1.275 | 0.914 | -0.504 | -0.534 | -0.052 | 0.051 | 0.051 | 0 | 0.64 | 0.78 | 0.59 | 0.76 | 0.56 | 0.46 |
| 3 | 124328494 | rs333332 | 0.015 | 0.768 | -0.529 | -0.311 | 0.031 | 0.015 | 0.015 | 0 | 0.42 | 0.67 | 0.44 | 0.64 | 0.34 | 0.24 |
| 3 | 124328506 | rs1561578 | 0.055 | 0.788 | -0.477 | -0.285 | 0.001 | 0.018 | 0.018 | 0 | 0.23 | 0.19 | 0.17 | 0.12 | 0.22 | 0.23 |
| 3 | 124330744 | rs333327 | -1.1 | 0.859 | -0.521 | -0.353 | -0.036 | 0.081 | 0.081 | 0.175 | 0.6 | 0.78 | 0.53 | 0.76 | 0.46 | 0.33 |
| 3 | 124331600 | rs333321 | -1.792 | 0.78 | -0.516 | -0.362 | -0.04 | 0.112 | 0.112 | 0 | 0.7 | 0.91 | 0.7 | 0.89 | 0.68 | 0.57 |
| 3 | 124334599 | rs333318 | -0.302 | 0.755 | -0.461 | -0.315 | 0.024 | 0.045 | 0.045 | 0 | 0.47 | 0.74 | 0.53 | 0.64 | 0.44 | 0.36 |
| 3 | 124341535 | rs582288 | 1.092 | 0.583 | -0.198 | -0.167 | -0.032 | 0.098 | 0.098 | 0.003 | 0.23 | 0.51 | 0.35 | 0.61 | 0.19 | 0.12 |
| 3 | 124341806 | rs10512628 | -0.911 | 0.68 | -0.337 | -0.227 | -0.005 | 0.044 | 0.044 | 0 | 0.47 | 0.35 | 0.35 | 0.28 | 0.49 | 0.45 |
| 3 | 124345166 | rs2119185 | -1.578 | 0.62 | -0.401 | -0.202 | -0.014 | 0.053 | 0.053 | 0 | 0.71 | 0.81 | 0.69 | 0.9 | 0.68 | 0.6 |
| 3 | 124345877 | rs619460 | -1.528 | 0.629 | -0.394 | -0.12 | -0.011 | 0.035 | 0.035 | 0 | 0.68 | 0.78 | 0.64 | 0.81 | 0.66 | 0.56 |
| 3 | 124345910 | rs589663 | -0.456 | 0.62 | -0.318 | -0.059 | -0.003 | 0 | 0 | 2.52 | 0.27 | 0.63 | 0.23 | 0.22 | 0.27 | 0.31 |
| 3 | 124345983 | rs13071295 | 1.043 | 0.899 | -0.268 | 0.11 | 0.032 | -0.033 | -0.033 | 0 | 0.07 | 0.11 | 0.11 | 0 | 0.13 | 0.16 |
| 3 | 124348999 | rs16835749 | -0.286 | 0.232 | -0.542 | 0.25 | -0.007 | 0.037 | 0.037 | 0.143 | 0.44 | 0.17 | 0.46 | 0.68 | 0.41 | 0.28 |
| 3 | 124349120 | rs16835753 | -0.295 | 0.232 | -0.542 | 0.25 | -0.006 | 0.037 | 0.037 | 0.131 | 0.44 | 0.18 | 0.46 | 0.68 | 0.41 | 0.28 |
| 3 | 124350438 | rs6764025 | -0.104 | 0.229 | -0.485 | 0.112 | 0.013 | 0.018 | 0.018 | 0 | 0.21 | 0.16 | 0.15 | 0.11 | 0.16 | 0.13 |
| 3 | 124351316 | rs1708303 | -1.272 | 0.338 | -0.37 | -0.185 | -0.021 | 0.121 | 0.121 | 0 | 0.6 | 0.8 | 0.69 | 0.92 | 0.6 | 0.48 |
| 3 | 124352135 | rs13087377 | 0.483 | 0.216 | 0.538 | 0.501 | 0.001 | 0.006 | 0.006 | 0 | 0.33 | 0.39 | 0.39 | 0.43 | 0.38 | 0.29 |
| 3 | 124352728 | rs652863 | 1.132 | 0.022 | 0.71 | 0.331 | -0.034 | -0.055 | -0.055 | 0 | 0.1 | 0.2 | 0.04 | 0 | 0.05 | 0.02 |
| 3 | 124356462 | rs667883 | -0.804 | 0.269 | 0.975 | 0.277 | 0.003 | 0.03 | 0.03 | 0.104 | 0.53 | 0.63 | 0.58 | 0.92 | 0.5 | 0.38 |
| 3 | 124358726 | rs644007 | -0.487 | 0.212 | 1.01 | 0.529 | -0.015 | 0.032 | 0.032 | 0 | 0.26 | 0.3 | 0.29 | 0.49 | 0.22 | 0.19 |
| 3 | 124366662 | rs1392912 | -0.974 | 0.268 | 1.071 | 0.5 | 0.001 | 0.037 | 0.037 | 0 | 0.32 | 0.41 | 0.4 | 0.49 | 0.32 | 0.28 |
| 3 | 124371035 | rs333306 | -0.724 | 0.266 | 0.937 | 0.578 | 0.004 | 0.012 | 0.012 | 0.117 | 0.52 | 0.59 | 0.44 | 0.57 | 0.48 | 0.35 |
| 3 | 124372075 | rs333304 | -1.382 | 0.276 | 0.832 | 0.538 | -0.008 | 0.078 | 0.078 | 0 | 0.59 | 0.73 | 0.68 | 0.92 | 0.61 | 0.48 |
| 3 | 124372449 | rs3772777 | 1.129 | 0.365 | 0.749 | 0.604 | 0.001 | -0.039 | -0.039 | 0.039 | 0.26 | 0.24 | 0.27 | 0.43 | 0.28 | 0.2 |
| 3 | 124374215 | rs11711670 | 1.152 | 0.362 | 0.757 | 0.656 | 0 | -0.035 | -0.035 | 0 | 0.26 | 0.21 | 0.26 | 0.4 | 0.28 | 0.2 |
| 3 | 124377632 | rs333300 | -0.658 | 0.307 | 0.808 | 0.812 | 0.007 | 0.008 | 0.008 | 0.798 | 0.52 | 0.63 | 0.42 | 0.5 | 0.47 | 0.32 |
| 3 | 124377732 | rs16835783 | 1.178 | 0.345 | 0.831 | 0.795 | 0.066 | 0.034 | 0.034 | 0 | 0.06 | 0.17 | 0.11 | 0.08 | 0.1 | 0.1 |
| 3 | 124379817 | rs333289 | -1.266 | 0.331 | 0.781 | 0.796 | -0.015 | 0.028 | 0.028 | 0.019 | 0.58 | 0.8 | 0.53 | 0.59 | 0.57 | 0.42 |
| 3 | 124380793 | rs665927 | -0.631 | 0.307 | 0.993 | 1.036 | -0.024 | -0.002 | -0.002 | 0 | 0.27 | 0.53 | 0.18 | 0.19 | 0.24 | 0.16 |
| 3 | 124381439 | rs333287 | -1.398 | 0.286 | 0.75 | 1.025 | -0.001 | -0.003 | -0.003 | 0 | 0.35 | 0.77 | 0.31 | 0.27 | 0.34 | 0.26 |
| 3 | 124393490 | rs9845054 | 1.145 | 0.062 | 1.155 | 0.941 | -0.03 | -0.049 | -0.049 | 0 | 0.09 | 0.24 | 0.05 | 0 | 0.03 | 0.02 |
| 3 | 124393610 | rs333354 | -1.358 | 0.349 | 1.069 | 0.998 | -0.002 | 0.015 | 0.015 | 0 | 0.58 | 0.81 | 0.55 | 0.58 | 0.6 | 0.44 |
| 3 | 124393940 | rs333352 | -1.424 | 0.343 | 1.035 | 0.995 | -0.013 | 0.007 | 0.007 | 0.016 | 0.59 | 0.85 | 0.56 | 0.58 | 0.6 | 0.44 |
| 3 | 124394667 | rs3772785 | 0.228 | 0.408 | 1.11 | 1.036 | 0.003 | 0.006 | 0.006 | 0 | 0.18 | 0.26 | 0.14 | 0.18 | 0.23 | 0.16 |
| 3 | 124395138 | rs189151 | 0.887 | 0.447 | 1.139 | 1.049 | 0.095 | 0.059 | 0.059 | 0 | 0.07 | 0.23 | 0.12 | 0.08 | 0.1 | 0.1 |
| 3 | 124395387 | rs333350 | -2.265 | 0.005 | 0.744 | 0.805 | -0.118 | 0.143 | 0.143 | 0.018 | 0.69 | 0.99 | 0.72 | 0.93 | 0.64 | 0.54 |
| 3 | 124396338 | rs333349 | -0.154 | 0.015 | 0.775 | 0.842 | 0.002 | 0.001 | 0.001 | 0 | 0.47 | 0.55 | 0.45 | 0.5 | 0.54 | 0.39 |
| 3 | 124401225 | rs3772790 | -0.39 | 0.036 | 0.783 | 0.879 | -0.056 | 0.053 | 0.053 | 0.002 | 0.22 | 0.42 | 0.27 | 0.42 | 0.1 | 0.16 |
| 3 | 124401692 | rs586395 | -1.335 | 0.05 | 0.703 | 0.872 | 0.007 | -0.011 | -0.011 | 0 | 0.3 | 0.71 | 0.39 | 0.5 | 0.2 | 0.25 |
| 3 | 124402365 | rs680822 | -1.252 | 0.057 | 0.745 | 0.862 | -0.007 | 0.005 | 0.005 | 0 | 0.38 | 0.6 | 0.37 | 0.52 | 0.31 | 0.28 |
| 3 | 124403275 | rs1708320 | -2.043 | 0.099 | 0.676 | 0.772 | -0.017 | 0.064 | 0.064 | 0 | 0.68 | 0.81 | 0.72 | 0.92 | 0.64 | 0.54 |
| 3 | 124403643 | rs1501187 | -1.352 | 0.101 | 0.778 | 0.862 | -0.001 | -0.002 | -0.002 | 1.506 | 0.34 | 0.62 | 0.36 | 0.35 | 0.38 | 0.31 |
| 3 | 124404514 | rs2713637 | -1.332 | 0.112 | 0.811 | 0.868 | 0 | 0 | 0 | 0 | 0.34 | 0.6 | 0.35 | 0.35 | 0.38 | 0.31 |
| 3 | 124404632 | rs3755699 | 1.049 | 0.384 | 0.67 | 1.014 | 0.067 | 0.038 | 0.038 | 0 | 0.06 | 0.18 | 0.12 | 0.08 | 0.1 | 0.1 |
| 3 | 124411690 | rs333285 | -0.935 | 0.327 | 0.548 | 1.011 | 0.004 | -0.011 | -0.011 | 0 | 0.3 | 0.25 | 0.35 | 0.6 | 0.29 | 0.29 |
| 3 | 124412153 | rs333284 | -0.672 | 0.302 | 0.51 | 0.952 | 0.003 | 0.121 | 0.121 | 0 | 0.51 | 0.31 | 0.54 | 0.84 | 0.5 | 0.42 |
| 3 | 124412524 | rs2289428 | 1.748 | 0.3 | 0.341 | 0.959 | -0.016 | -0.028 | -0.028 | 0 | 0.22 | 0.07 | 0.19 | 0.24 | 0.21 | 0.13 |
| 3 | 124414454 | rs3755702 | -1.096 | 0.333 | 0.528 | 0.947 | 0.001 | 0.002 | 0.002 | 0 | 0.33 | 0.36 | 0.42 | 0.61 | 0.37 | 0.35 |
| 3 | 124416186 | rs16845650 | 0.524 | -0.177 | 0.691 | 1.215 | -0.012 | 0.021 | 0.021 | 0 | 0.11 | 0.15 | 0.17 | 0.34 | 0.05 | 0.1 |
| 3 | 124422005 | rs7621976 | 0.242 | -0.172 | 0.771 | 1.17 | 0.064 | 0.033 | 0.033 | 0 | 0.07 | 0.22 | 0.12 | 0.08 | 0.1 | 0.12 |
| 3 | 124428896 | rs184559 | -0.506 | -0.297 | 0.585 | 1.015 | -0.003 | -0.008 | -0.008 | 0 | 0.42 | 0.54 | 0.54 | 0.74 | 0.39 | 0.36 |
| 3 | 124433512 | rs3863064 | 0.269 | -0.32 | 0.191 | 0.869 | 0.004 | 0.001 | 0.001 | 0 | 0.1 | 0.12 | 0.17 | 0.34 | 0.05 | 0.1 |
| 3 | 124438329 | rs4234221 | -0.214 | -0.245 | 0.493 | 0.543 | -0.016 | 0.003 | 0.003 | 0 | 0.16 | 0.31 | 0.2 | 0.14 | 0.11 | 0.21 |
| 3 | 124438439 | rs332512 | -0.486 | -0.33 | 0.19 | 0.233 | -0.018 | -0.016 | -0.016 | 0 | 0.34 | 0.54 | 0.37 | 0.27 | 0.29 | 0.46 |
| 3 | 124439685 | rs1162 | -0.486 | -0.33 | 0.19 | 0.233 | -0.018 | -0.016 | -0.016 | 0.432 | 0.34 | 0.54 | 0.37 | 0.27 | 0.29 | 0.46 |
| 3 | 124441237 | rs162740 | 1.095 | -0.054 | -0.375 | 0.107 | 0 | -0.015 | -0.015 | 0.398 | 0.34 | 0.26 | 0.28 | 0.37 | 0.28 | 0.2 |
| 3 | 124443522 | rs3736388 | 0.733 | -0.253 | -0.68 | -0.009 | -0.027 | -0.041 | -0.041 | 0 | 0.44 | 0.15 | 0.37 | 0.42 | 0.42 | 0.39 |
| 3 | 124445904 | rs16835912 | -0.358 | -0.307 | -0.326 | -0.001 | -0.006 | 0.004 | 0.004 | 2.649 | 0.63 | 0.34 | 0.67 | 0.64 | 0.57 | 0.66 |
| 3 | 124446077 | rs17282057 | 2.133 | -0.287 | -0.395 | -0.012 | -0.011 | 0.011 | 0.011 | 0 | 0.12 | 0.01 | 0.09 | 0.05 | 0.13 | 0.18 |
| 3 | 124454174 | rs3772804 | 1.49 | -0.225 | -0.145 | 0.071 | -0.059 | -0.092 | -0.092 | 0 | 0.32 | 0.07 | 0.26 | 0.37 | 0.29 | 0.2 |
| 3 | 124462808 | rs13146 | 0.849 | -0.235 | -0.104 | 0.08 | 0.001 | 0.002 | 0.002 | 0 | 0.18 | 0.15 | 0.26 | 0.17 | 0.15 | 0.24 |
| 3 | 124478734 | rs13082230 | 0.057 | -0.272 | -0.091 | -0.168 | -0.007 | 0.029 | 0.029 | 0.251 | 0.51 | 0.34 | 0.6 | 0.59 | 0.45 | 0.48 |
| 3 | 124479212 | rs13087417 | 1.451 | -0.267 | -0.102 | -0.164 | -0.019 | -0.058 | -0.058 | 0 | 0.32 | 0.17 | 0.29 | 0.4 | 0.29 | 0.2 |
| 3 | 124481052 | rs14386 | 0.627 | -0.607 | 0.185 | 0.141 | 0.004 | 0.006 | 0.006 | 0 | 0.18 | 0.12 | 0.26 | 0.16 | 0.15 | 0.23 |
| 3 | 124481987 | rs2676 | 0.439 | -0.614 | 0.234 | 0.138 | -0.002 | -0.003 | -0.003 | 0 | 0.19 | 0.2 | 0.27 | 0.16 | 0.15 | 0.23 |
| 3 | 124482869 | rs585021 | 1.015 | -0.852 | 0.34 | -0.006 | -0.014 | 0.047 | 0.047 | 2.34 | 0.21 | 0.34 | 0.25 | 0.37 | 0.14 | 0.18 |

| | | | | | | | | | | | | | | | | |
|---|---|---|---|---|---|---|---|---|---|---|---|---|---|---|---|---|
| 3 | 124485235 | rs2291081 | 0.768 | -0.51 | 0.908 | 0.394 | 0.001 | 0 | 0 | 0 | 0.16 | 0.13 | 0.26 | 0.16 | 0.14 | 0.18 |
| 3 | 124486993 | rs614664 | -1.533 | -0.575 | 1.268 | 0.336 | 0.033 | 0.028 | 0.028 | 0 | 0.42 | 0.71 | 0.52 | 0.37 | 0.35 | 0.52 |
| 3 | 124494030 | rs621209 | -0.549 | -0.485 | 1.347 | 0.348 | -0.035 | -0.036 | -0.036 | 0.097 | 0.27 | 0.59 | 0.27 | 0.21 | 0.22 | 0.35 |
| 3 | 124498173 | rs9879289 | 1.155 | -0.423 | 1.241 | 0.027 | -0.016 | 0.051 | 0.051 | 0 | 0.14 | 0.01 | 0.14 | 0.03 | 0.12 | 0.13 |
| 3 | 124499685 | rs3772827 | 0.352 | -0.418 | 1.435 | 0.037 | 0.013 | 0.074 | 0.074 | 0.466 | 0.06 | 0.27 | 0.05 | 0.12 | 0.03 | 0.04 |
| 3 | 124501222 | rs2619328 | -1.584 | -0.403 | 1.412 | 0.028 | 0.032 | 0.03 | 0.03 | 0 | 0.42 | 0.72 | 0.51 | 0.37 | 0.36 | 0.51 |
| 3 | 124515308 | rs1803825 | -0.823 | -0.387 | 1.182 | 0.212 | -0.03 | 0.012 | 0.012 | 0 | 0.24 | 0.48 | 0.35 | 0.32 | 0.19 | 0.31 |
| 3 | 124515314 | rs2291085 | 0.621 | -0.388 | 1.211 | 0.219 | -0.013 | 0.051 | 0.051 | 0.247 | 0.05 | 0.22 | 0.03 | 0.13 | 0.01 | 0.04 |
| 3 | 124515509 | rs2291088 | 0.907 | -0.359 | 1.07 | 0.227 | 0.002 | 0 | 0 | 0.189 | 0.16 | 0.12 | 0.26 | 0.17 | 0.15 | 0.17 |
| 3 | 124521035 | rs12487905 | -0.622 | -0.355 | 1.084 | 0.217 | -0.012 | 0.009 | 0.009 | 0 | 0.24 | 0.37 | 0.3 | 0.33 | 0.2 | 0.26 |
| 3 | 124521263 | rs17282085 | 1.523 | -0.433 | 1.009 | 0.167 | 0.01 | 0.026 | 0.026 | 0 | 0.14 | 0.01 | 0.13 | 0.05 | 0.12 | 0.19 |
| 3 | 124521357 | rs12695446 | -0.622 | -0.346 | 1.135 | 0.236 | -0.011 | 0.003 | 0.003 | 0 | 0.24 | 0.37 | 0.3 | 0.29 | 0.2 | 0.26 |
| 3 | 124525071 | rs4678164 | 0.805 | -0.317 | 1.04 | 0.222 | 0 | -0.001 | -0.001 | 0 | 0.16 | 0.13 | 0.26 | 0.19 | 0.17 | 0.18 |
| 3 | 124533315 | rs7650893 | -0.474 | -0.317 | 1.209 | 0.229 | -0.005 | 0.087 | 0.087 | 0.585 | 0.07 | 0.36 | 0.1 | 0.16 | 0.02 | 0.08 |
| 3 | 124535635 | rs3772836 | -1.409 | -0.349 | 1.447 | 0.202 | 0.002 | 0.072 | 0.072 | 1.186 | 0.1 | 0.53 | 0.12 | 0.15 | 0.04 | 0.13 |
| 3 | 124539303 | rs3772839 | 0.693 | -0.264 | 1.252 | 0.236 | -0.014 | 0.045 | 0.045 | 0.027 | 0.05 | 0.19 | 0.03 | 0.13 | 0.01 | 0.03 |
| 3 | 124540556 | rs4678166 | 1.441 | -0.256 | 0.933 | 0.143 | 0.004 | 0.038 | 0.038 | 0 | 0.14 | 0.04 | 0.13 | 0.06 | 0.13 | 0.2 |
| 3 | 124540832 | rs4678167 | 0.851 | -0.289 | 1.127 | 0.125 | -0.012 | -0.042 | -0.042 | 0 | 0.17 | 0.28 | 0.16 | 0.06 | 0.14 | 0.25 |
| 3 | 124544266 | rs7624392 | -1.012 | -0.259 | 0.891 | 0.165 | 0.026 | 0.018 | 0.018 | 0 | 0.41 | 0.72 | 0.52 | 0.39 | 0.33 | 0.51 |
| 3 | 124545750 | rs6771896 | 0.165 | 0.081 | 0.761 | 0.218 | 0 | 0 | 0 | 3.999 | 0.18 | 0.2 | 0.32 | 0.21 | 0.18 | 0.23 |
| 3 | 124546047 | rs4677946 | -0.344 | 0.068 | 0.971 | 0.202 | -0.009 | -0.022 | -0.022 | 0 | 0.22 | 0.43 | 0.35 | 0.19 | 0.19 | 0.27 |
| 3 | 124547871 | rs2076722 | 0.691 | 0.575 | 0.551 | 0.317 | 0.004 | 0.023 | 0.023 | 2.848 | 0.18 | 0.25 | 0.25 | 0.37 | 0.25 | 0.19 |
| 3 | 124548684 | rs16836080 | -0.037 | 0.593 | 0.795 | -0.127 | 0.031 | -0.067 | -0.067 | 0 | 0.23 | 0.53 | 0.29 | 0.17 | 0.29 | 0.22 |
| 3 | 124550364 | rs3772840 | 0.841 | 0.264 | 0.788 | -0.271 | -0.001 | 0.002 | 0.002 | 0.055 | 0.48 | 0.49 | 0.39 | 0.73 | 0.43 | 0.39 |
| 3 | 124554130 | rs3772841 | -1.716 | 0.27 | 0.601 | -0.304 | 0.001 | 0.002 | 0.002 | 0 | 0.83 | 0.84 | 0.83 | 0.93 | 0.82 | 0.79 |
| 3 | 124560890 | rs3772845 | -1.808 | 0.395 | 0.563 | -0.265 | -0.012 | 0.028 | 0.028 | 0 | 0.82 | 0.91 | 0.82 | 0.91 | 0.79 | 0.77 |
| 3 | 124562515 | rs3772847 | 0.186 | 0.358 | 0.581 | -0.248 | -0.003 | -0.019 | -0.019 | 1.005 | 0.19 | 0.21 | 0.23 | 0.06 | 0.16 | 0.23 |
| 3 | 124577555 | rs11928651 | 0.97 | 0.383 | 0.687 | -0.257 | -0.001 | 0.017 | 0.017 | 0 | 0.46 | 0.4 | 0.38 | 0.71 | 0.42 | 0.39 |
| 3 | 124577826 | rs6778643 | 0.703 | 0.388 | 0.501 | -0.254 | 0.005 | 0.101 | 0.101 | 0.095 | 0.49 | 0.66 | 0.42 | 0.72 | 0.42 | 0.39 |
| 3 | 124588929 | rs13091878 | 0.363 | 0.436 | 0.46 | -0.263 | 0.003 | 0.047 | 0.047 | 0 | 0.18 | 0.1 | 0.22 | 0.06 | 0.16 | 0.23 |
| 3 | 124592700 | rs1265646 | 0.175 | 0.41 | 0.478 | -0.301 | -0.001 | 0.087 | 0.087 | 0 | 0.63 | 0.75 | 0.59 | 0.85 | 0.63 | 0.55 |
| 3 | 124594803 | rs2047574 | -1.468 | 0.414 | 0.401 | -0.304 | -0.003 | 0.003 | 0.003 | 2.411 | 0.85 | 0.86 | 0.81 | 0.86 | 0.79 | 0.85 |
| 3 | 124598499 | rs10804563 | 1.633 | 0.687 | 0.07 | -0.207 | -0.037 | -0.083 | -0.083 | 0 | 0.34 | 0.1 | 0.29 | 0.39 | 0.33 | 0.28 |
| 3 | 124601972 | rs9968182 | 1.061 | 0.579 | 0.48 | -1.14 | 0 | 0.001 | 0.001 | 0 | 0.47 | 0.5 | 0.38 | 0.71 | 0.42 | 0.39 |
| 3 | 124609945 | rs9847614 | 0.815 | 0.558 | 0.053 | -1.102 | -0.002 | 0.044 | 0.044 | 0 | 0.14 | 0.03 | 0.18 | 0.06 | 0.12 | 0.18 |
| 3 | 124610238 | rs7373878 | 0.476 | 0.566 | 0.306 | -1.08 | -0.002 | -0.017 | -0.017 | 0 | 0.64 | 0.41 | 0.54 | 0.79 | 0.62 | 0.62 |
| 3 | 124624568 | rs12732 | 0.964 | 0.705 | -0.554 | -1.102 | -0.035 | 0.032 | 0.032 | 0 | 0.18 | 0.06 | 0.18 | 0.09 | 0.21 | 0.23 |
| 3 | 124635512 | rs4679166 | 0.085 | 0.677 | -0.619 | -1.131 | 0.007 | -0.052 | -0.052 | 0 | 0.73 | 0.61 | 0.74 | 0.89 | 0.7 | 0.67 |
| 3 | 124645034 | rs2877822 | -0.854 | 0.605 | -0.342 | -0.742 | -0.002 | -0.039 | -0.039 | 0.209 | 0.23 | 0.24 | 0.25 | 0.05 | 0.21 | 0.3 |
| 3 | 124646594 | rs4679392 | -1.068 | 0.602 | -0.517 | -0.661 | 0.001 | -0.028 | -0.028 | 0 | 0.38 | 0.43 | 0.45 | 0.22 | 0.43 | 0.45 |
| 3 | 124646837 | rs4679394 | -0.46 | 0.404 | -0.998 | 0.158 | -0.033 | 0.09 | 0.09 | 0 | 0.16 | 0.02 | 0.2 | 0.03 | 0.15 | 0.19 |
| 3 | 124653868 | rs10934702 | 0.59 | 0.528 | -0.358 | 0.296 | -0.002 | 0.01 | 0.01 | 0 | 0.87 | 0.72 | 0.83 | 0.86 | 0.9 | 0.81 |
| 3 | 124656695 | rs7612467 | 0.489 | 0.691 | -1.063 | 0.039 | -0.012 | 0.026 | 0.026 | 3.183 | 0.23 | 0.13 | 0.27 | 0.15 | 0.28 | 0.29 |
| 3 | 124659978 | rs7644047 | -0.114 | 0.246 | -0.488 | 0.257 | -0.018 | -0.042 | -0.042 | 0 | 0.65 | 0.42 | 0.57 | 0.79 | 0.69 | 0.67 |
| 3 | 124664036 | rs9880945 | -0.664 | 0.23 | 0.058 | 1.439 | 0.008 | 0.095 | 0.095 | 0 | 0.21 | 0.25 | 0.2 | 0.5 | 0.19 | 0.18 |
| 3 | 124665328 | rs1969056 | -0.609 | 0.332 | 0.119 | 1.489 | 0.012 | 0.114 | 0.114 | 0 | 0.2 | 0.25 | 0.2 | 0.5 | 0.19 | 0.17 |
| 3 | 124672358 | rs6793951 | 0.667 | 0.348 | -0.123 | 1.129 | -0.014 | 0.057 | 0.057 | 1.016 | 0.92 | 0.72 | 0.85 | 0.87 | 0.96 | 0.88 |
| 3 | 124686733 | rs3184121 | -1.004 | 0.797 | -0.493 | 1.322 | 0 | 0.001 | 0.001 | 0.613 | 0.2 | 0.19 | 0.27 | 0.65 | 0.24 | 0.31 |
| 3 | 124687432 | rs17237607 | -0.798 | 0.659 | -0.642 | 1.286 | -0.024 | 0.003 | 0.003 | 0 | 0.07 | 0.01 | 0.09 | 0.05 | 0.09 | 0.12 |
| 3 | 124687767 | rs1909586 | -0.955 | 0.58 | -1.423 | 0.393 | -0.031 | 0.062 | 0.062 | 0 | 0.62 | 0.85 | 0.61 | 0.75 | 0.62 | 0.57 |
| 3 | 124688269 | rs16836316 | -0.367 | 0.293 | -0.674 | 1.045 | -0.001 | 0 | 0 | 0 | 0.29 | 0.25 | 0.29 | 0.39 | 0.24 | 0.25 |
| 3 | 124688416 | rs16836319 | -0.248 | 0.199 | -0.741 | 1.378 | 0.007 | -0.01 | -0.01 | 3.996 | 0.06 | 0.03 | 0.06 | 0.1 | 0.05 | 0.07 |
| 3 | 124691470 | rs3732610 | -1.868 | 0.112 | -0.749 | 1.3 | -0.048 | 0.073 | 0.073 | 0 | 0.62 | 0.88 | 0.61 | 0.78 | 0.61 | 0.58 |
| 3 | 124692194 | rs6438864 | 0.092 | -0.158 | -0.139 | 1.29 | 0.011 | 0.026 | 0.026 | 0.336 | 0.25 | 0.36 | 0.23 | 0.3 | 0.33 | 0.25 |
| 3 | 124693518 | rs9790108 | -0.165 | -0.161 | -0.077 | 1.252 | -0.003 | 0.001 | 0.001 | 0.053 | 0.27 | 0.25 | 0.28 | 0.37 | 0.23 | 0.24 |
| 3 | 124694303 | rs3821541 | -0.989 | -0.211 | 0.103 | 1.23 | -0.01 | 0.005 | 0.005 | 0.461 | 0.37 | 0.55 | 0.36 | 0.49 | 0.28 | 0.32 |
| 3 | 124702008 | rs6763492 | -0.899 | -0.343 | 0.245 | 1.193 | -0.021 | 0.014 | 0.014 | 0.243 | 0.36 | 0.45 | 0.35 | 0.49 | 0.28 | 0.32 |
| 3 | 124705111 | rs6438865 | -0.339 | -0.362 | 0.215 | 1.175 | 0.002 | 0.006 | 0.006 | 1.972 | 0.75 | 0.72 | 0.78 | 0.71 | 0.67 | 0.75 |
| 3 | 124715526 | rs10934705 | -0.499 | -0.481 | 0.396 | 1.143 | -0.02 | 0.006 | 0.006 | 1.381 | 0.3 | 0.39 | 0.29 | 0.37 | 0.23 | 0.25 |
| 3 | 124719687 | rs4286466 | -1.198 | -0.55 | 0.34 | 1.126 | 0.019 | -0.01 | -0.01 | 0 | 0.38 | 0.67 | 0.37 | 0.49 | 0.27 | 0.32 |
| 3 | 124721791 | rs6796048 | -0.439 | -0.521 | 0.305 | 1.051 | -0.018 | 0.004 | 0.004 | 2.337 | 0.3 | 0.51 | 0.31 | 0.37 | 0.22 | 0.23 |
| 3 | 124722021 | rs4287948 | 2 | -0.63 | 0.058 | 0.991 | -0.115 | 0.046 | 0.046 | 0.072 | 0.36 | 0.03 | 0.38 | 0.23 | 0.4 | 0.43 |
| 3 | 124725659 | rs4618262 | 0.866 | -0.364 | 0.521 | 1.121 | 0.002 | -0.007 | -0.007 | 0 | 0.63 | 0.29 | 0.65 | 0.6 | 0.62 | 0.66 |
| 3 | 124728626 | rs6438869 | 0.485 | -0.274 | 0.517 | 1.129 | 0 | 0.044 | 0.044 | 0 | 0.69 | 0.51 | 0.72 | 0.6 | 0.65 | 0.73 |
| 3 | 124729193 | rs6784454 | 0.868 | -0.281 | 0.391 | 1.345 | -0.001 | -0.001 | -0.001 | 0 | 0.66 | 0.32 | 0.63 | 0.37 | 0.66 | 0.72 |
| 3 | 124732618 | rs6790837 | 0.997 | 0.013 | 0.463 | 1.32 | -0.022 | -0.024 | -0.024 | 0.745 | 0.53 | 0.23 | 0.44 | 0.47 | 0.54 | 0.61 |
| 3 | 124732842 | rs6438871 | 0.231 | -0.032 | 1.095 | 1.488 | 0.004 | -0.034 | -0.034 | 0.36 | 0.65 | 0.63 | 0.73 | 0.93 | 0.62 | 0.81 |
| 3 | 124739892 | rs6438874 | 1.118 | -0.049 | 1.372 | 1.495 | -0.003 | -0.003 | -0.003 | 0.385 | 0.54 | 0.46 | 0.48 | 0.48 | 0.54 | 0.6 |
| 3 | 124741456 | rs2948791 | -1.223 | -0.055 | 1.44 | 1.581 | -0.008 | 0.122 | 0.122 | 0 | 0.11 | 0.21 | 0.22 | 0.41 | 0.08 | 0.2 |

| | | | | | | | | | | | | | | | | |
|---|---|---|---|---|---|---|---|---|---|---|---|---|---|---|---|---|
| 3 | 124742610 | rs2981548 | 1.142 | -0.052 | 1.298 | 1.579 | -0.007 | -0.006 | -0.006 | 0.391 | 0.54 | 0.4 | 0.47 | 0.48 | 0.54 | 0.6 |
| 3 | 124746049 | rs2981546 | 1.103 | -0.053 | 1.199 | 1.579 | -0.012 | -0.011 | -0.011 | 0.056 | 0.54 | 0.37 | 0.48 | 0.48 | 0.55 | 0.61 |
| 3 | 124746347 | rs4404487 | 0.098 | -0.06 | 1.254 | 1.574 | -0.013 | 0.026 | 0.026 | 0 | 0.66 | 0.79 | 0.72 | 0.89 | 0.62 | 0.8 |
| 3 | 124748226 | rs2860440 | 1.033 | 0.006 | 0.758 | 1.529 | -0.011 | -0.01 | -0.01 | 0 | 0.54 | 0.38 | 0.47 | 0.48 | 0.55 | 0.6 |
| 3 | 124756771 | rs4679258 | -1.265 | -0.07 | 0.39 | 1.335 | -0.011 | 0.03 | 0.03 | 0 | 0.28 | 0.61 | 0.42 | 0.53 | 0.23 | 0.36 |
| 3 | 124757318 | rs1574733 | -0.717 | 0.094 | 0.242 | 1.202 | -0.002 | 0 | 0 | 0.001 | 0.83 | 0.82 | 0.9 | 0.82 | 0.86 | 0.88 |
| 3 | 124757777 | rs7626291 | -1.07 | 0.021 | 0.544 | 1.303 | 0.075 | 0.198 | 0.198 | 3.081 | 0.06 | 0.46 | 0.09 | 0.12 | 0.01 | 0.06 |
| 3 | 124757888 | rs7429439 | 1.057 | 0.255 | -0.106 | 1.025 | -0.037 | 0.061 | 0.061 | 0 | 0.58 | 0.15 | 0.62 | 0.7 | 0.59 | 0.72 |
| 3 | 124759846 | rs13067154 | 0.332 | 0.853 | 0.178 | 0.982 | -0.049 | 0.045 | 0.045 | 0 | 0.18 | 0.06 | 0.17 | 0 | 0.26 | 0.1 |
| 3 | 124760086 | rs2333033 | -1.112 | 0.676 | 0.39 | 1.146 | -0.025 | 0.023 | 0.023 | 0 | 0.39 | 0.31 | 0.51 | 0.35 | 0.46 | 0.37 |
| 3 | 124764110 | rs2981534 | -1.496 | 1.301 | 0.617 | 1.305 | 0.1 | 0.115 | 0.115 | 1.941 | 0.12 | 0.37 | 0.35 | 0.24 | 0.2 | 0.21 |
| 3 | 124768366 | rs2333038 | 1.151 | 0.882 | -1.106 | 0.682 | -0.13 | -0.043 | -0.043 | 0 | 0.55 | 0.03 | 0.5 | 0.41 | 0.5 | 0.51 |
| 3 | 124772082 | rs2001425 | 1.194 | 1.283 | -0.454 | 0.295 | 0.023 | 0 | 0 | 0 | 0.45 | 0.36 | 0.42 | 0.57 | 0.37 | 0.49 |
| 3 | 124776845 | rs2877839 | 0.68 | 2.059 | 0.7 | 0.996 | 0.016 | 0.052 | 0.052 | 0 | 0.71 | 0.53 | 0.65 | 0.6 | 0.71 | 0.69 |
| 3 | 124785277 | rs11718242 | 1.272 | 1.729 | 1.098 | 1.082 | 0.013 | 0.129 | 0.129 | 0.008 | 0.8 | 0.49 | 0.68 | 0.61 | 0.77 | 0.7 |
| 3 | 124788013 | rs2981496 | 0.806 | 1.66 | 0.915 | 1.153 | -0.015 | 0.108 | 0.108 | 0 | 0.85 | 0.71 | 0.79 | 0.62 | 0.87 | 0.81 |
| 3 | 124788161 | rs2981495 | 0.877 | 1.966 | 0.951 | 1.237 | 0.005 | 0.114 | 0.114 | 0 | 0.87 | 0.76 | 0.8 | 0.62 | 0.85 | 0.82 |
| 3 | 124788634 | rs7629128 | 0.943 | 1.837 | 1.105 | 1.239 | 0 | 0.189 | 0.189 | 0 | 0.86 | 0.66 | 0.79 | 0.62 | 0.87 | 0.82 |
| 3 | 124794877 | rs2333034 | 1.396 | 1.926 | 0.88 | 1.25 | 0.005 | 0.061 | 0.061 | 0 | 0.78 | 0.68 | 0.69 | 0.55 | 0.74 | 0.64 |
| 3 | 124799357 | rs2948819 | -0.711 | 1.944 | 0.776 | 1.085 | 0.012 | 0.016 | 0.016 | 0 | 0.06 | 0.11 | 0.08 | 0.13 | 0.09 | 0.16 |
| 3 | 124799400 | rs2333035 | -0.447 | 2.106 | 1 | 0.74 | 0.006 | -0.058 | -0.058 | 0 | 0.24 | 0.33 | 0.19 | 0.04 | 0.24 | 0.13 |
| 3 | 124801799 | rs2981486 | -1.284 | 1.902 | 0.66 | 0.842 | 0 | 0.013 | 0.013 | 0 | 0.09 | 0.11 | 0.08 | 0.21 | 0.1 | 0.22 |
| 3 | 124801903 | rs2981485 | -2.013 | 2.394 | 0.666 | 0.917 | -0.004 | 0.024 | 0.024 | 0 | 0.47 | 0.84 | 0.6 | 0.59 | 0.54 | 0.53 |
| 3 | 124801998 | rs2948818 | -1.096 | 2.4 | 0.641 | 0.797 | 0 | 0.013 | 0.013 | 0 | 0.09 | 0.11 | 0.08 | 0.21 | 0.1 | 0.22 |
| 3 | 124802231 | rs13975 | -1.922 | 2.399 | 0.567 | 0.73 | 0.004 | 0.015 | 0.015 | 0 | 0.47 | 0.8 | 0.6 | 0.58 | 0.54 | 0.53 |
| 3 | 124802677 | rs1574340 | -1.922 | 2.399 | 0.57 | 0.73 | 0.004 | 0.016 | 0.016 | 0 | 0.47 | 0.8 | 0.6 | 0.58 | 0.54 | 0.53 |
| 3 | 124802888 | rs2981482 | -1.905 | 2.398 | 0.57 | 0.729 | 0.003 | 0.015 | 0.015 | 0 | 0.47 | 0.8 | 0.6 | 0.58 | 0.54 | 0.52 |
| 3 | 124803791 | rs17239028 | -0.112 | 2.509 | 0.711 | 0.696 | -0.002 | -0.006 | -0.006 | 0 | 0.23 | 0.22 | 0.23 | 0.04 | 0.31 | 0.13 |
| 3 | 124804204 | rs2948814 | -1.49 | 2.296 | 0.559 | 0.938 | 0.015 | 0.208 | 0.208 | 0.57 | 0.2 | 0.53 | 0.36 | 0.49 | 0.2 | 0.34 |
| 3 | 124811314 | rs651630 | 1.301 | 2.473 | -0.717 | 0.427 | 0.008 | 0.013 | 0.013 | 0.512 | 0.52 | 0.18 | 0.38 | 0.45 | 0.47 | 0.56 |
| 3 | 124814347 | rs535575 | 1.073 | 1.869 | -0.957 | 0.457 | 0.021 | 0.004 | 0.004 | 0 | 0.52 | 0.27 | 0.43 | 0.46 | 0.45 | 0.57 |
| 3 | 124814690 | rs532106 | 0.757 | 1.923 | -0.859 | 0.46 | -0.008 | -0.009 | -0.009 | 0.089 | 0.58 | 0.35 | 0.46 | 0.54 | 0.51 | 0.67 |
| 3 | 124815896 | rs2993639 | -0.199 | 1.936 | -1.059 | 0.535 | -0.101 | 0.056 | 0.056 | 0 | 0.3 | 0.11 | 0.28 | 0.14 | 0.42 | 0.18 |
| 3 | 124818338 | rs2670170 | 0.167 | 1.433 | -1.085 | 0.391 | -0.014 | -0.014 | -0.014 | 0 | 0.6 | 0.35 | 0.51 | 0.54 | 0.52 | 0.71 |
| 3 | 124820293 | rs2788462 | 0.612 | 1.213 | -1.228 | 0.5 | 0.016 | -0.05 | -0.05 | 0 | 0.37 | 0.14 | 0.37 | 0.45 | 0.35 | 0.46 |
| 3 | 124820470 | rs13325275 | -1.191 | 1.483 | -0.28 | 0.355 | 0.007 | -0.017 | -0.017 | 0.044 | 0.11 | 0.31 | 0.12 | 0.11 | 0.1 | 0.2 |
| 3 | 124820822 | rs2948798 | -1.08 | 2.05 | -0.828 | 0.428 | -0.029 | 0.011 | 0.011 | 0 | 0.13 | 0.07 | 0.17 | 0.06 | 0.22 | 0.11 |
| 3 | 124831048 | rs12487908 | -0.523 | 2.01 | -0.535 | 1.152 | -0.052 | -0.059 | -0.059 | 1.032 | 0.09 | 0 | 0.08 | 0.23 | 0.15 | 0.11 |
| 3 | 124831327 | rs2993638 | -0.12 | 1.673 | -0.973 | 0.548 | 0 | -0.001 | -0.001 | 0.232 | 0.35 | 0.37 | 0.44 | 0.32 | 0.42 | 0.34 |
| 3 | 124831414 | rs2993637 | -0.061 | 1.879 | -1.127 | 0.555 | -0.034 | -0.01 | -0.01 | 0 | 0.09 | 0.03 | 0.21 | 0.1 | 0.2 | 0.13 |
| 3 | 124832747 | rs11714323 | 0.695 | 1.741 | -1.136 | 0.509 | 0.024 | 0.004 | 0.004 | 0.957 | 0.21 | 0.04 | 0.18 | 0.16 | 0.19 | 0.07 |
| 3 | 124833968 | rs2948797 | -0.412 | 1.602 | -1.017 | 0.576 | -0.002 | 0 | 0 | 0.788 | 0.35 | 0.35 | 0.43 | 0.25 | 0.42 | 0.22 |
| 3 | 124835779 | rs591803 | -0.292 | 1.152 | -0.973 | 0.593 | 0 | -0.002 | -0.002 | 0.125 | 0.31 | 0.27 | 0.36 | 0.68 | 0.27 | 0.47 |
| 3 | 124838938 | rs2942819 | -0.96 | 1.743 | -0.585 | 1.114 | 0.025 | 0.015 | 0.015 | 0.4 | 0.26 | 0.29 | 0.32 | 0.39 | 0.43 | 0.38 |
| 3 | 124840018 | rs6790154 | -0.249 | 1.112 | -0.2 | 1.127 | -0.004 | 0.017 | 0.017 | 0.201 | 0.33 | 0.38 | 0.27 | 0.47 | 0.32 | 0.19 |
| 3 | 124841661 | rs1877581 | 1.177 | 0.933 | -0.467 | 1.19 | 0.008 | 0 | 0 | 0 | 0.17 | 0.04 | 0.15 | 0.16 | 0.16 | 0.06 |
| 3 | 124842747 | rs12486816 | 0.537 | 0.898 | -0.407 | 1.216 | -0.031 | -0.057 | -0.057 | 0 | 0.12 | 0.03 | 0.11 | 0.33 | 0.16 | 0.12 |
| 3 | 124843489 | rs2945118 | -0.255 | 1.164 | -0.532 | 0.986 | -0.024 | 0.013 | 0.013 | 0 | 0.08 | 0.04 | 0.18 | 0.04 | 0.17 | 0.1 |
| 3 | 124847902 | rs516355 | 1.08 | 0.842 | 0.047 | 1.021 | -0.008 | -0.022 | -0.022 | 0 | 0.55 | 0.27 | 0.49 | 0.46 | 0.48 | 0.69 |
| 3 | 124848755 | rs2228677 | 0.667 | 0.298 | -0.562 | 0.158 | -0.011 | 0.028 | 0.028 | 0 | 0.42 | 0.3 | 0.23 | 0.34 | 0.46 | 0.27 |
| 3 | 124849770 | rs702045 | 1.051 | 0.592 | -0.149 | 0.194 | 0.083 | 0.051 | 0.051 | 0.213 | 0.89 | 0.54 | 0.82 | 0.93 | 0.89 | 0.84 |
| 3 | 124850245 | rs9813946 | 1.578 | 0.557 | -0.166 | 0.165 | 0.004 | -0.015 | -0.015 | 0 | 0.31 | 0.28 | 0.15 | 0.01 | 0.29 | 0.17 |
| 3 | 124851233 | rs652746 | 0.208 | 0.571 | 0.081 | 0.217 | 0.053 | 0.109 | 0.109 | 0.253 | 0.94 | 0.62 | 0.95 | 0.97 | 0.99 | 0.91 |
| 3 | 124853253 | rs1068926 | 0.743 | 0.579 | -0.145 | 0.234 | 0.026 | -0.059 | -0.059 | 0 | 0.24 | 0.12 | 0.34 | 0.41 | 0.18 | 0.46 |
| 3 | 124854158 | rs16836516 | 1.463 | 0.531 | -0.142 | 0.165 | 0.004 | 0.014 | 0.014 | 0 | 0.29 | 0.24 | 0.14 | 0.01 | 0.26 | 0.15 |
| 3 | 124859482 | rs673693 | -0.498 | 0.387 | 0.141 | 0.817 | 0.001 | 0.013 | 0.013 | 2.091 | 0.42 | 0.58 | 0.51 | 0.65 | 0.4 | 0.62 |
| 3 | 124860138 | rs816151 | -0.126 | 0.231 | -0.301 | 0.669 | 0.002 | 0.084 | 0.084 | 0 | 0.14 | 0.22 | 0.29 | 0.28 | 0.12 | 0.33 |
| 3 | 124862868 | rs684030 | 0.16 | 0.568 | -0.929 | 0.605 | 0.009 | 0.086 | 0.086 | 0 | 0.51 | 0.77 | 0.49 | 0.29 | 0.44 | 0.56 |
| 3 | 124867634 | rs938226 | 0.833 | 0.642 | -1.059 | 0.553 | -0.002 | 0.023 | 0.023 | 0 | 0.2 | 0.1 | 0.18 | 0.16 | 0.21 | 0.1 |
| 3 | 124868722 | rs13067346 | 0.566 | 0.631 | -1.506 | 0.867 | -0.057 | -0.09 | -0.09 | 0 | 0.27 | 0.03 | 0.2 | 0.37 | 0.31 | 0.24 |
| 3 | 124868891 | rs6786220 | -0.432 | 0.809 | -0.311 | 0.582 | 0.074 | 0.085 | 0.085 | 0 | 0.1 | 0.38 | 0.22 | 0.18 | 0.13 | 0.14 |
| 3 | 124869928 | rs679990 | -1.509 | 0.423 | -0.242 | 0.37 | 0.039 | -0.014 | -0.014 | 0 | 0.94 | 0.89 | 0.88 | 0.97 | 0.93 | 0.94 |
| 3 | 124871467 | rs581557 | -0.128 | 0.03 | -0.022 | 0.423 | 0.087 | 0.092 | 0.092 | 0 | 0.73 | 0.54 | 0.67 | 0.51 | 0.62 | 0.7 |
| 3 | 124872058 | rs564809 | 0.521 | -0.388 | -0.428 | 0.305 | 0.041 | -0.07 | -0.07 | 0.363 | 0.24 | 0.04 | 0.3 | 0.36 | 0.17 | 0.44 |
| 3 | 124872379 | rs630439 | 0.521 | -0.388 | -0.428 | 0.305 | 0.041 | -0.07 | -0.07 | 0 | 0.24 | 0.04 | 0.3 | 0.36 | 0.17 | 0.44 |
| 3 | 124873295 | rs1464172 | 0.553 | -0.415 | 0.009 | -0.034 | 0.008 | 0.017 | 0.017 | 0 | 0.49 | 0.47 | 0.37 | 0.15 | 0.45 | 0.24 |
| 3 | 124874835 | rs633055 | -1.152 | -0.609 | -0.143 | 0.55 | 0 | -0.001 | -0.001 | 0.171 | 0.82 | 0.86 | 0.81 | 0.69 | 0.83 | 0.72 |
| 3 | 124875043 | rs6788986 | 0.511 | -0.683 | 0.342 | 0.017 | 0.004 | 0.01 | 0.01 | 0.019 | 0.08 | 0.3 | 0.04 | 0.07 | 0.04 | 0.04 |
| 3 | 124876757 | rs16836571 | 0.173 | -0.732 | 0.273 | 0.008 | 0.009 | 0.02 | 0.02 | 0.194 | 0.09 | 0.39 | 0.05 | 0.07 | 0.04 | 0.04 |

| | | | | | | | | | | | | | | | | |
|---|---|---|---|---|---|---|---|---|---|---|---|---|---|---|---|---|
| 3 | 124879996 | rs632682 | -1.205 | -0.696 | 0.137 | 0.058 | -0.003 | -0.003 | -0.003 | 0 | 0.82 | 0.84 | 0.81 | 0.82 | 0.83 | 0.72 |
| 3 | 124881156 | rs484728 | -1.205 | -0.696 | 0.078 | 0.058 | 0 | 0 | 0 | 0 | 0.82 | 0.86 | 0.81 | 0.82 | 0.83 | 0.72 |
| 3 | 124883952 | rs9852163 | -0.014 | -0.755 | 0.141 | 0.039 | 0 | 0.011 | 0.011 | 0.716 | 0.09 | 0.41 | 0.05 | 0.07 | 0.04 | 0.04 |
| 3 | 124884508 | rs2076726 | 1.584 | -1.042 | -0.653 | -0.139 | 0.009 | -0.004 | -0.004 | 0 | 0.22 | 0.04 | 0.22 | 0.18 | 0.16 | 0.19 |
| 3 | 124885287 | rs9829910 | -0.1 | -0.728 | 0.104 | -0.027 | -0.011 | 0 | 0 | 0 | 0.09 | 0.35 | 0.04 | 0.07 | 0.04 | 0.04 |
| 3 | 124887233 | rs6763338 | 0.949 | -0.743 | -0.602 | -0.358 | 0.018 | 0.117 | 0.117 | 0.012 | 0.22 | 0.03 | 0.09 | 0 | 0.18 | 0.15 |
| 3 | 124888091 | rs682870 | -0.081 | -0.915 | 0.006 | -0.168 | -0.033 | 0.051 | 0.051 | 0.712 | 0.89 | 0.76 | 0.96 | 0.8 | 0.95 | 0.74 |
| 3 | 124891424 | rs13316796 | 0.209 | -0.95 | 0.026 | -0.47 | 0.014 | 0.006 | 0.006 | 0 | 0.38 | 0.33 | 0.29 | 0.05 | 0.35 | 0.21 |
| 3 | 124892546 | rs503875 | -0.047 | -0.856 | 0.012 | -0.142 | -0.032 | 0.057 | 0.057 | 0.712 | 0.89 | 0.75 | 0.96 | 0.8 | 0.95 | 0.74 |
| 3 | 124894481 | rs13074637 | 0.276 | -0.816 | 0.095 | -0.395 | -0.018 | -0.069 | -0.069 | 3.716 | 0.31 | 0.3 | 0.15 | 0.07 | 0.24 | 0.2 |
| 3 | 124897700 | rs602770 | 0.768 | -0.635 | -0.188 | 0.775 | 0.006 | 0.031 | 0.031 | 0.123 | 0.07 | 0.06 | 0.1 | 0.35 | 0.02 | 0.24 |
| 3 | 124903161 | rs6777718 | 0.186 | -0.117 | 0.092 | 0.706 | -0.016 | 0.033 | 0.033 | 0.03 | 0.68 | 0.46 | 0.53 | 0.4 | 0.75 | 0.53 |
| 3 | 124905511 | rs9816529 | 0.747 | -1.071 | -0.992 | 0.256 | -0.112 | -0.069 | -0.069 | 0.223 | 0.27 | 0.1 | 0.24 | 0.32 | 0.43 | 0.19 |
| 3 | 124909178 | rs4679159 | 0.703 | -0.736 | -0.487 | 0.638 | 0.001 | 0.005 | 0.005 | 0 | 0.08 | 0.13 | 0.13 | 0.12 | 0.08 | 0.02 |
| 3 | 124909866 | rs867805 | 0.62 | -1.396 | -0.428 | 0.447 | 0.002 | -0.001 | -0.001 | 0 | 0.59 | 0.41 | 0.49 | 0.39 | 0.58 | 0.45 |
| 3 | 124910439 | rs1464065 | 0.912 | -1.383 | -0.458 | 0.414 | 0.007 | 0.002 | 0.002 | 1.286 | 0.56 | 0.4 | 0.44 | 0.38 | 0.56 | 0.39 |
| 3 | 124911288 | rs702043 | 0.104 | -1.294 | -0.175 | 0.584 | 0.013 | -0.019 | -0.019 | 0 | 0.29 | 0.2 | 0.27 | 0.44 | 0.3 | 0.49 |
| 3 | 124911841 | rs1980080 | 0.336 | -1.267 | -0.132 | 0.64 | -0.016 | 0.023 | 0.023 | 0.849 | 0.63 | 0.55 | 0.56 | 0.5 | 0.64 | 0.4 |
| 3 | 124915818 | rs816341 | -0.121 | -1.452 | -0.142 | 1.003 | -0.017 | 0.114 | 0.114 | 0.074 | 0.11 | 0.17 | 0.15 | 0.44 | 0.07 | 0.31 |
| 3 | 124918754 | rs6801331 | -0.437 | -1.179 | -0.181 | 0.44 | -0.004 | 0.054 | 0.054 | 0 | 0.95 | 0.83 | 0.93 | 0.9 | 0.98 | 0.92 |
| 3 | 124918879 | rs6801517 | -0.364 | -1.334 | -0.447 | 0.297 | -0.014 | 0.016 | 0.016 | 1.523 | 0.65 | 0.52 | 0.56 | 0.39 | 0.66 | 0.5 |
| 3 | 124919384 | rs6766250 | -0.452 | -1.159 | -0.529 | 0.409 | -0.015 | 0.016 | 0.016 | 0.969 | 0.66 | 0.44 | 0.61 | 0.4 | 0.67 | 0.51 |
| 3 | 124921457 | rs9873519 | -0.254 | -0.464 | -0.463 | 0.769 | -0.011 | 0.018 | 0.018 | 0 | 0.4 | 0.27 | 0.47 | 0.32 | 0.54 | 0.24 |
| 3 | 124922985 | rs13100784 | -0.41 | -0.095 | -0.677 | 0.562 | -0.012 | 0.032 | 0.032 | 0 | 0.05 | 0.16 | 0.03 | 0.09 | 0.02 | 0.08 |
| 3 | 124926054 | rs9833435 | 0.273 | -0.096 | -1.182 | 0.515 | 0.018 | 0.017 | 0.017 | 0 | 0.53 | 0.66 | 0.57 | 0.57 | 0.42 | 0.63 |
| 3 | 124929748 | rs9832863 | 0.354 | -0.093 | -1.254 | 0.363 | -0.013 | -0.002 | -0.002 | 0 | 0.86 | 1 | 0.86 | 0.87 | 0.89 | 0.9 |
| 3 | 124938103 | rs6438884 | -0.927 | -1.007 | -0.64 | -0.173 | -0.048 | -0.083 | -0.083 | 0 | 0.26 | 0.6 | 0.16 | 0.05 | 0.11 | 0.13 |
| 3 | 124948552 | rs4679372 | 0.028 | -0.042 | -0.886 | 0.696 | -0.014 | -0.013 | -0.013 | 0 | 0.83 | 0.9 | 0.63 | 0.53 | 0.76 | 0.66 |
| 3 | 124951821 | rs1053738 | 0.109 | -0.054 | -0.777 | 0.691 | -0.006 | 0.002 | 0.002 | 0 | 0.83 | 0.88 | 0.63 | 0.53 | 0.75 | 0.66 |
| 3 | 124959834 | rs2011442 | 0.537 | 0.768 | -0.777 | 0.865 | 0.001 | 0.008 | 0.008 | 0 | 0.8 | 0.83 | 0.57 | 0.46 | 0.66 | 0.57 |
| 3 | 124963914 | rs9858110 | -0.491 | 0.69 | -0.805 | 1.294 | -0.065 | -0.065 | -0.065 | 0.81 | 0.88 | 0.98 | 0.73 | 0.71 | 0.77 | 0.69 |
| 3 | 124967313 | rs11709450 | 0.033 | 0.754 | -0.774 | 0.983 | -0.006 | 0 | 0 | 0 | 0.82 | 0.88 | 0.63 | 0.53 | 0.75 | 0.66 |
| 3 | 124979785 | rs17264366 | 0.938 | 0.93 | -1.305 | 0.796 | -0.031 | -0.019 | -0.019 | 0 | 0.13 | 0.02 | 0.15 | 0.13 | 0.18 | 0.14 |
| 3 | 124989394 | rs7638240 | 1.966 | 0.191 | -0.996 | 0.132 | 0.016 | 0.01 | 0.01 | 0 | 0.36 | 0.29 | 0.22 | 0.01 | 0.32 | 0.23 |
| 3 | 125030381 | rs1975760 | -0.533 | 0.495 | -1.235 | 0.439 | -0.006 | 0 | 0 | 0 | 0.82 | 0.87 | 0.63 | 0.53 | 0.75 | 0.66 |
| 3 | 125054369 | rs10512647 | 0.328 | 0.2 | -1.353 | 0.425 | -0.014 | -0.008 | -0.008 | 0.046 | 0.16 | 0.02 | 0.11 | 0.14 | 0.12 | 0.08 |
| 3 | 125070745 | rs4508729 | -0.033 | 0.394 | -1.119 | 0.488 | -0.029 | -0.04 | -0.04 | 3.555 | 0.8 | 0.9 | 0.62 | 0.55 | 0.73 | 0.66 |
| 3 | 125075013 | rs7609684 | 1.041 | 0.29 | -0.78 | 0.427 | 0.003 | 0.01 | 0.01 | 0.118 | 0.67 | 0.37 | 0.5 | 0.48 | 0.62 | 0.52 |
| 3 | 125079521 | rs13074986 | -0.18 | 0.917 | -1.053 | 0.737 | -0.071 | -0.093 | -0.093 | 0.039 | 0.81 | 0.95 | 0.63 | 0.54 | 0.73 | 0.65 |
| 3 | 125118082 | rs7640441 | 0.081 | 0.972 | -0.47 | 0.222 | 0.024 | 0.016 | 0.016 | 0.178 | 0.15 | 0.17 | 0.21 | 0.2 | 0.26 | 0.2 |
| 3 | 125128677 | rs10934713 | 1.008 | 0.518 | -0.803 | -0.312 | 0.015 | 0.071 | 0.071 | 0 | 0.38 | 0.28 | 0.23 | 0.08 | 0.35 | 0.3 |
| 3 | 125131761 | rs2333226 | 0.406 | 0.704 | -0.372 | 0.327 | -0.002 | -0.006 | -0.006 | 0 | 0.12 | 0.11 | 0.16 | 0.15 | 0.15 | 0.07 |
| 3 | 125148476 | rs7620749 | -0.817 | 0.096 | -0.157 | 0.074 | 0 | 0 | 0 | 4.311 | 0.22 | 0.25 | 0.19 | 0.15 | 0.17 | 0.14 |
| 3 | 125153891 | rs6798446 | -1.273 | -0.904 | -0.69 | -0.131 | 0.116 | 0.048 | 0.048 | 3.3 | 0.47 | 0.76 | 0.33 | 0.36 | 0.26 | 0.37 |
| 3 | 125154388 | rs4679393 | -1.109 | -0.67 | -0.106 | -0.177 | 0.001 | 0.003 | 0.003 | 0 | 0.39 | 0.64 | 0.42 | 0.28 | 0.31 | 0.4 |
| 3 | 125161022 | rs9838753 | 0.035 | -0.724 | -0.049 | -0.156 | 0.008 | 0.008 | 0.008 | 0 | 0.17 | 0.22 | 0.29 | 0.22 | 0.2 | 0.23 |
| 3 | 125161649 | rs7618106 | -1.362 | -0.578 | -0.153 | -0.395 | 0.015 | 0.051 | 0.051 | 0 | 0.43 | 0.74 | 0.45 | 0.29 | 0.35 | 0.43 |
| 3 | 125173964 | rs4679170 | -0.287 | -0.688 | 0.23 | -0.409 | 0.023 | 0.044 | 0.044 | 0 | 0.25 | 0.16 | 0.17 | 0.13 | 0.17 | 0.19 |
| 3 | 125181941 | rs7613952 | -1.735 | -0.738 | 0.357 | -0.371 | 0.001 | 0.019 | 0.019 | 0 | 0.52 | 0.8 | 0.58 | 0.42 | 0.43 | 0.51 |
| 3 | 125182246 | rs7636435 | -1.638 | -0.744 | 0.688 | -0.375 | 0.007 | 0.017 | 0.017 | 0 | 0.51 | 0.65 | 0.56 | 0.42 | 0.43 | 0.51 |
| 3 | 125193134 | rs7610077 | -1.377 | -0.88 | 0.614 | -0.53 | 0.017 | 0.02 | 0.02 | 0 | 0.46 | 0.64 | 0.49 | 0.37 | 0.36 | 0.48 |
| 3 | 125194108 | rs4679397 | -0.237 | -0.81 | 0.609 | -0.496 | 0.019 | -0.01 | -0.01 | 0 | 0.31 | 0.18 | 0.35 | 0.32 | 0.24 | 0.29 |
| 3 | 125197444 | rs3772214 | -1.515 | -0.813 | 0.994 | -0.494 | 0 | 0.002 | 0.002 | 0 | 0.49 | 0.5 | 0.55 | 0.41 | 0.42 | 0.46 |
| 3 | 125201519 | rs9864550 | -2.33 | -0.807 | 0.945 | -0.487 | -0.03 | 0.004 | 0.004 | 0.011 | 0.57 | 0.83 | 0.59 | 0.42 | 0.44 | 0.53 |
| 3 | 125204577 | rs17331632 | 0.085 | -0.796 | 0.89 | -0.411 | 0.023 | 0.027 | 0.027 | 0 | 0.27 | 0.16 | 0.23 | 0.21 | 0.22 | 0.24 |
| 3 | 125205106 | rs9848399 | 0.017 | -0.642 | 0.967 | -0.407 | 0.019 | 0.022 | 0.022 | 0.096 | 0.06 | 0.08 | 0.19 | 0.15 | 0.12 | 0.15 |
| 3 | 125238099 | rs4630895 | -1.443 | -0.621 | 0.951 | -0.506 | 0.004 | 0.016 | 0.016 | 0 | 0.5 | 0.57 | 0.53 | 0.3 | 0.43 | 0.43 |
| 3 | 125239869 | rs28384489 | -1.314 | -0.793 | 0.993 | -0.474 | 0.038 | 0.038 | 0.038 | 0 | 0.47 | 0.72 | 0.51 | 0.38 | 0.34 | 0.45 |
| 3 | 125243742 | rs1839063 | -0.016 | -0.866 | 0.761 | -0.651 | 0 | 0.001 | 0.001 | 0 | 0.25 | 0.26 | 0.21 | 0.18 | 0.21 | 0.25 |
| 3 | 125246363 | rs9841525 | -0.029 | -0.857 | 0.765 | -0.647 | 0 | 0 | 0 | 0 | 0.24 | 0.26 | 0.21 | 0.18 | 0.21 | 0.25 |
| 3 | 125247295 | rs6804431 | -2.419 | -0.809 | 0.691 | -0.617 | -0.06 | 0.036 | 0.036 | 0.457 | 0.57 | 0.92 | 0.54 | 0.36 | 0.43 | 0.51 |
| 3 | 125249340 | rs2294053 | 1.166 | -0.688 | 0.688 | -0.534 | -0.014 | 0.038 | 0.038 | 0 | 0.11 | 0 | 0.09 | 0.04 | 0.07 | 0.03 |
| 3 | 125250956 | rs9867731 | -0.284 | -0.711 | 1.05 | -0.541 | -0.038 | -0.065 | -0.065 | 0 | 0.19 | 0.37 | 0.17 | 0.11 | 0.1 | 0.14 |
| 3 | 125255110 | rs7644750 | -0.992 | -0.707 | 1.022 | -0.539 | -0.03 | -0.043 | -0.043 | 0.131 | 0.23 | 0.5 | 0.27 | 0.18 | 0.13 | 0.19 |
| 3 | 125272159 | rs9848230 | 0.29 | -0.6 | 0.939 | -0.529 | 0.008 | -0.045 | -0.045 | 0 | 0.1 | 0.19 | 0.08 | 0 | 0.12 | 0.07 |
| 3 | 125278095 | rs2922188 | -2.418 | -0.747 | 0.908 | -0.543 | -0.044 | 0.024 | 0.024 | 0.016 | 0.59 | 0.83 | 0.55 | 0.36 | 0.47 | 0.51 |
| 3 | 125280473 | rs2922180 | 0.545 | -0.493 | 1.493 | -0.254 | 0.016 | 0.037 | 0.037 | 0 | 0.06 | 0.27 | 0.07 | 0.07 | 0.03 | 0.09 |
| 3 | 125303459 | rs9853814 | 0.26 | -0.256 | 1.026 | -0.3 | 0.007 | -0.046 | -0.046 | 0 | 0.1 | 0.19 | 0.07 | 0 | 0.13 | 0.08 |
| 3 | 125312130 | rs2979306 | 0.292 | -0.246 | 0.991 | -0.556 | -0.008 | -0.045 | -0.045 | 0.769 | 0.17 | 0.33 | 0.17 | 0.09 | 0.18 | 0.18 |

| | | | | | | | | | | | | | | | | |
|---|---|---|---|---|---|---|---|---|---|---|---|---|---|---|---|---|
| 3 | 125313904 | rs7625936 | -0.559 | -0.059 | 0.854 | -0.508 | 0.001 | -0.012 | -0.012 | 0.006 | 0.28 | 0.37 | 0.25 | 0.16 | 0.3 | 0.25 |
| 3 | 125319251 | rs9855951 | -0.538 | -0.075 | 0.781 | -0.491 | 0.002 | -0.02 | -0.02 | 0 | 0.28 | 0.38 | 0.29 | 0.16 | 0.3 | 0.25 |
| 3 | 125326749 | rs2922195 | -1.873 | -0.177 | 0.571 | -0.52 | -0.035 | 0.091 | 0.091 | 0 | 0.47 | 0.85 | 0.54 | 0.32 | 0.49 | 0.46 |
| 3 | 125332263 | rs2922194 | 0.534 | -0.393 | 0.716 | -0.4 | -0.004 | -0.027 | -0.027 | 0 | 0.16 | 0.25 | 0.14 | 0.1 | 0.16 | 0.16 |
| 3 | 125351276 | rs17273179 | 1.233 | -0.395 | 0.277 | -0.407 | -0.018 | -0.013 | -0.013 | 0 | 0.14 | 0 | 0.16 | 0.1 | 0.13 | 0.1 |
| 3 | 125352873 | rs4679177 | -0.207 | -0.327 | 0.672 | -0.453 | 0.01 | -0.036 | -0.036 | 0 | 0.21 | 0.46 | 0.26 | 0.16 | 0.2 | 0.21 |
| 3 | 125355558 | rs4679428 | -0.246 | -0.332 | 0.675 | -0.428 | 0.008 | -0.034 | -0.034 | 0 | 0.21 | 0.46 | 0.27 | 0.16 | 0.2 | 0.21 |
| 3 | 125379972 | rs13323469 | -0.818 | -0.538 | 0.595 | -0.587 | -0.009 | -0.021 | -0.021 | 0 | 0.3 | 0.39 | 0.28 | 0.16 | 0.24 | 0.25 |
| 3 | 125384035 | rs2922167 | 0.736 | -0.575 | 0.676 | -0.58 | -0.004 | -0.026 | -0.026 | 0.743 | 0.16 | 0.21 | 0.14 | 0.09 | 0.17 | 0.16 |
| 3 | 125391858 | rs13094641 | 2.053 | -0.572 | 0.613 | -0.58 | 0.008 | 0.1 | 0.1 | 0 | 0.49 | 0.13 | 0.46 | 0.68 | 0.56 | 0.54 |
| 3 | 125392975 | rs6784346 | 1.45 | -0.092 | 0.263 | -0.439 | -0.024 | -0.004 | -0.004 | 0 | 0.2 | 0.11 | 0.18 | 0.22 | 0.29 | 0.12 |
| 3 | 125393494 | rs16836952 | -0.027 | -0.518 | 1.079 | -0.394 | 0.036 | 0.051 | 0.051 | 0.424 | 0.06 | 0.25 | 0.12 | 0.07 | 0.03 | 0.05 |
| 3 | 125403248 | rs2001665 | 0.44 | -0.505 | 0.778 | -0.208 | 0.004 | -0.014 | -0.014 | 0.203 | 0.18 | 0.14 | 0.22 | 0.22 | 0.16 | 0.17 |
| 3 | 125410843 | rs2979311 | -0.955 | -0.432 | 1.181 | -0.384 | -0.007 | -0.031 | -0.031 | 0.068 | 0.22 | 0.47 | 0.26 | 0.16 | 0.2 | 0.21 |
| 3 | 125435432 | rs9289256 | 1.061 | -0.511 | 0.85 | -0.32 | -0.01 | -0.037 | -0.037 | 0 | 0.73 | 0.61 | 0.73 | 0.84 | 0.76 | 0.71 |
| 3 | 125452841 | rs2947630 | -0.892 | -0.426 | 0.89 | -0.406 | 0.015 | 0.117 | 0.117 | 0.273 | 0.11 | 0.47 | 0.12 | 0.18 | 0.08 | 0.24 |
| 3 | 125477762 | rs2735247 | -0.167 | -0.467 | 0.383 | -0.523 | 0.001 | 0.004 | 0.004 | 0 | 0.9 | 0.93 | 0.95 | 1 | 0.92 | 0.93 |
| 3 | 125479062 | rs2735246 | 0.732 | -0.407 | 0.549 | -0.558 | 0 | -0.023 | -0.023 | 0 | 0.1 | 0.11 | 0.1 | 0.01 | 0.11 | 0.05 |
| 3 | 125481779 | rs9827398 | -0.762 | -0.475 | 1.146 | -0.5 | 0.022 | 0.131 | 0.131 | 0.105 | 0.1 | 0.44 | 0.1 | 0.17 | 0.08 | 0.25 |
| 3 | 125486820 | rs2735232 | 0.859 | -0.614 | 0.633 | -0.654 | -0.01 | -0.013 | -0.013 | 0.482 | 0.68 | 0.42 | 0.64 | 0.71 | 0.71 | 0.6 |
| 3 | 125504275 | rs2735226 | -0.561 | -0.429 | 0.852 | -0.666 | -0.007 | -0.054 | -0.054 | 4.656 | 0.12 | 0.18 | 0.05 | 0 | 0.08 | 0.08 |
| 3 | 125540308 | rs17334039 | -1.155 | -0.234 | 0.971 | -0.518 | -0.005 | -0.005 | -0.005 | 0.086 | 0.2 | 0.28 | 0.22 | 0.21 | 0.2 | 0.33 |
| 3 | 125541332 | rs2979929 | 0.457 | -0.239 | 1.229 | -0.852 | -0.007 | -0.029 | -0.029 | 0 | 0.86 | 0.84 | 0.94 | 0.98 | 0.91 | 0.91 |
| 3 | 125552742 | rs2979948 | 0.862 | 0.209 | 1.369 | -0.811 | 0.007 | -0.047 | -0.047 | 0 | 0.82 | 0.43 | 0.83 | 0.92 | 0.84 | 0.89 |
| 3 | 125620850 | rs9821905 | -0.07 | 0.437 | 1.296 | -0.599 | -0.001 | -0.026 | -0.026 | 1.692 | 0.12 | 0.19 | 0.11 | 0.08 | 0.14 | 0.1 |
| 3 | 125669719 | rs2017409 | -1.35 | 0.436 | 1.398 | -0.679 | -0.012 | -0.116 | -0.116 | 0 | 0.24 | 0.58 | 0.18 | 0.08 | 0.21 | 0.13 |
| 3 | 125677514 | rs7611945 | -1.302 | 0.442 | 1.377 | -0.676 | -0.01 | -0.139 | -0.139 | 0 | 0.23 | 0.49 | 0.17 | 0.08 | 0.21 | 0.13 |
| 3 | 125681401 | rs4340655 | -1.029 | 0.772 | 0.477 | -0.604 | -0.099 | -0.066 | -0.066 | 0.027 | 0.64 | 0.92 | 0.52 | 0.51 | 0.56 | 0.53 |
| 3 | 125682330 | rs7630575 | -0.631 | 0.745 | 0.785 | -0.669 | -0.007 | -0.092 | -0.092 | 0 | 0.2 | 0.5 | 0.19 | 0.13 | 0.19 | 0.18 |
| 3 | 125683874 | rs1827492 | -1.004 | 0.875 | 0.17 | -0.533 | -0.116 | -0.081 | -0.081 | 0 | 0.64 | 0.94 | 0.52 | 0.51 | 0.56 | 0.54 |
| 3 | 125707977 | rs7652883 | 0.151 | 0.617 | 0.957 | -0.739 | 0.004 | 0.024 | 0.024 | 0 | 0.21 | 0.17 | 0.14 | 0.15 | 0.22 | 0.24 |
| 3 | 125725652 | rs1044176 | 0.756 | 0.894 | 1.03 | -0.774 | 0 | -0.014 | -0.014 | 1.67 | 0.07 | 0.04 | 0.03 | 0 | 0.05 | 0.03 |
| 3 | 125733081 | rs883207 | 0.677 | 0.754 | 1.118 | -0.834 | -0.012 | -0.032 | -0.032 | 0 | 0.38 | 0.54 | 0.29 | 0.18 | 0.28 | 0.26 |
| 3 | 125737621 | rs3804587 | 1.037 | 0.707 | 1.247 | -0.855 | 0.026 | 0.074 | 0.074 | 0.154 | 0.34 | 0.19 | 0.23 | 0.18 | 0.28 | 0.26 |
| 3 | 125740393 | rs11925965 | 0.926 | 0.749 | 1.088 | -0.836 | 0.001 | -0.004 | -0.004 | 0 | 0.35 | 0.36 | 0.26 | 0.18 | 0.28 | 0.26 |
| 3 | 125748213 | rs7623362 | -1.305 | 1.148 | 0.966 | -0.724 | -0.102 | -0.048 | -0.048 | 0.113 | 0.72 | 0.93 | 0.42 | 0.37 | 0.5 | 0.46 |
| 3 | 125758232 | rs1112001 | -0.716 | 0.824 | 0.876 | -0.735 | 0.001 | 0.005 | 0.005 | 0 | 0.34 | 0.36 | 0.26 | 0.3 | 0.34 | 0.33 |
| 3 | 125760349 | rs12633421 | 0.793 | 0.683 | 0.692 | -0.729 | -0.007 | -0.022 | -0.022 | 0.351 | 0.36 | 0.38 | 0.26 | 0.18 | 0.28 | 0.26 |
| 3 | 125769757 | rs4077563 | 0.962 | 1.021 | 0.706 | -0.672 | 0.003 | 0.005 | 0.005 | 0 | 0.3 | 0.25 | 0.45 | 0.51 | 0.33 | 0.4 |
| 3 | 125771698 | rs12495947 | 0.457 | 0.929 | 0.701 | -0.761 | 0.004 | 0.038 | 0.038 | 0 | 0.21 | 0.13 | 0.14 | 0.12 | 0.22 | 0.24 |
| 3 | 125773630 | rs6798580 | -0.1 | 0.907 | 0.895 | -0.79 | 0.002 | -0.042 | -0.042 | 0 | 0.13 | 0.19 | 0.1 | 0.03 | 0.12 | 0.06 |
| 3 | 125776438 | rs4502539 | -0.871 | 0.9 | 0.892 | -0.79 | 0.002 | 0.031 | 0.031 | 0 | 0.34 | 0.32 | 0.23 | 0.15 | 0.34 | 0.3 |
| 3 | 125783086 | rs7632497 | 0.49 | 0.925 | 0.774 | -0.764 | 0.004 | 0.037 | 0.037 | 0 | 0.21 | 0.13 | 0.14 | 0.13 | 0.22 | 0.24 |
| 3 | 125784995 | rs3936031 | -0.436 | 0.82 | 0.945 | -0.814 | 0.035 | 0.169 | 0.169 | 1.714 | 0.05 | 0.27 | 0.09 | 0.15 | 0.06 | 0.05 |
| 3 | 125786760 | rs4308226 | -0.917 | 0.822 | 0.924 | -0.832 | -0.01 | -0.007 | -0.007 | 0 | 0.63 | 0.57 | 0.68 | 0.66 | 0.66 | 0.7 |
| 3 | 125787632 | rs4314124 | 1.425 | 0.845 | 0.58 | -0.842 | 0.015 | 0.076 | 0.076 | 0.474 | 0.31 | 0.13 | 0.23 | 0.18 | 0.29 | 0.25 |
| 3 | 125797913 | rs6796610 | 1.425 | 0.845 | 0.58 | -0.842 | 0.015 | 0.076 | 0.076 | 0.474 | 0.31 | 0.13 | 0.23 | 0.18 | 0.29 | 0.25 |
| 3 | 125800761 | rs11717532 | 1.114 | 0.923 | 0.524 | -0.787 | -0.001 | -0.01 | -0.01 | 0.073 | 0.07 | 0.04 | 0.03 | 0 | 0.06 | 0.03 |
| 3 | 125809971 | rs1806651 | -1.453 | 0.881 | 0.438 | -0.727 | -0.087 | -0.111 | -0.111 | 0.081 | 0.73 | 0.94 | 0.58 | 0.5 | 0.68 | 0.6 |
| 3 | 125817218 | rs2365011 | 1.471 | 0.829 | 0.454 | -0.76 | 0.008 | 0.073 | 0.073 | 0.659 | 0.3 | 0.09 | 0.23 | 0.19 | 0.29 | 0.25 |
| 3 | 125818091 | rs9840118 | -0.636 | 1.009 | 0.859 | -0.533 | -0.011 | -0.011 | -0.011 | 0 | 0.66 | 0.74 | 0.5 | 0.47 | 0.61 | 0.57 |
| 3 | 125822003 | rs2003334 | 0.56 | 1.012 | 1.017 | -0.515 | 0.009 | 0.021 | 0.021 | 0 | 0.43 | 0.36 | 0.29 | 0.18 | 0.36 | 0.29 |
| 3 | 125822272 | rs4646761 | 0.494 | 1.024 | 0.898 | -0.507 | 0.01 | 0.019 | 0.019 | 0.093 | 0.45 | 0.62 | 0.37 | 0.34 | 0.4 | 0.33 |
| 3 | 125824899 | rs4646754 | 0.52 | 0.973 | 0.715 | -0.653 | 0.002 | -0.006 | -0.006 | 0.186 | 0.43 | 0.42 | 0.29 | 0.18 | 0.36 | 0.3 |
| 3 | 125826003 | rs4646750 | 0.676 | 0.955 | 0.472 | -0.456 | -0.003 | -0.016 | -0.016 | 0.015 | 0.06 | 0.11 | 0.07 | 0.03 | 0.06 | 0.03 |
| 3 | 125826059 | rs1127717 | -0.012 | 0.998 | 0.204 | -0.443 | -0.002 | -0.01 | -0.01 | 0.061 | 0.21 | 0.23 | 0.16 | 0.13 | 0.22 | 0.24 |
| 3 | 125826243 | rs2276727 | 0.111 | 0.997 | -0.054 | -0.43 | 0.004 | 0.032 | 0.032 | 0 | 0.2 | 0.12 | 0.13 | 0.13 | 0.21 | 0.24 |
| 3 | 125826914 | rs3772431 | -0.799 | 1.549 | 0.049 | -0.17 | 0.004 | 0.014 | 0.014 | 0.603 | 0.73 | 0.75 | 0.58 | 0.5 | 0.7 | 0.54 |
| 3 | 125827399 | rs4646745 | -0.418 | 1.122 | -0.019 | -0.544 | 0.002 | -0.058 | -0.058 | 1.875 | 0.18 | 0.2 | 0.17 | 0.03 | 0.22 | 0.12 |
| 3 | 125829505 | rs3821466 | 0.51 | 0.929 | -0.45 | -0.281 | 0.04 | 0.028 | 0.028 | 1.262 | 0.4 | 0.11 | 0.19 | 0.36 | 0.3 | 0.27 |
| 3 | 125829541 | rs3772430 | -0.962 | 1.553 | -0.458 | -0.228 | -0.074 | -0.117 | -0.117 | 0.03 | 0.66 | 0.91 | 0.55 | 0.56 | 0.56 | 0.45 |
| 3 | 125831017 | rs3772424 | -0.125 | 1.777 | 0.043 | -0.314 | 0.053 | 0.035 | 0.035 | 0 | 0.11 | 0.21 | 0.24 | 0.21 | 0.22 | 0.15 |
| 3 | 125833260 | rs2290053 | -0.197 | 1.626 | 0.476 | -0.386 | -0.004 | 0.013 | 0.013 | 0.832 | 0.53 | 0.49 | 0.4 | 0.37 | 0.48 | 0.38 |
| 3 | 125836673 | rs6792028 | -1.258 | 1.552 | 0.584 | -0.471 | -0.016 | -0.004 | -0.004 | 0 | 0.36 | 0.72 | 0.4 | 0.42 | 0.39 | 0.38 |
| 3 | 125839224 | rs11924478 | 0.376 | 1.588 | 0.51 | -0.449 | 0.003 | 0 | 0 | 1.76 | 0.14 | 0.16 | 0.21 | 0.24 | 0.26 | 0.28 |
| 3 | 125842439 | rs6804272 | -0.71 | 1.563 | 0.921 | -0.352 | -0.007 | -0.016 | -0.016 | 0.039 | 0.63 | 0.69 | 0.52 | 0.54 | 0.56 | 0.46 |
| 3 | 125844718 | rs2278756 | 0.821 | 1.47 | 0.477 | -0.396 | -0.037 | 0.03 | 0.03 | 1.489 | 0.41 | 0.16 | 0.32 | 0.36 | 0.43 | 0.36 |
| 3 | 125846325 | rs3772419 | -0.991 | 1.565 | 0.528 | -0.175 | -0.001 | -0.006 | -0.006 | 0 | 0.67 | 0.65 | 0.57 | 0.6 | 0.64 | 0.48 |
| 3 | 125847121 | rs2365004 | -0.991 | 1.603 | 0.526 | -0.175 | -0.001 | -0.007 | -0.007 | 2.366 | 0.67 | 0.66 | 0.58 | 0.6 | 0.64 | 0.48 |

| | | | | | | | | | | | | | | | | |
|---|---|---|---|---|---|---|---|---|---|---|---|---|---|---|---|---|
| 3 | 125847468 | rs10934746 | -0.991 | 1.603 | 0.526 | -0.175 | -0.001 | -0.007 | -0.007 | 0.018 | 0.67 | 0.66 | 0.58 | 0.6 | 0.64 | 0.48 |
| 3 | 125848067 | rs6763254 | -0.991 | 1.603 | 0.526 | -0.175 | -0.001 | -0.007 | -0.007 | 0 | 0.67 | 0.66 | 0.58 | 0.6 | 0.64 | 0.48 |
| 3 | 125848710 | rs1868126 | -1.131 | 1.6 | 0.533 | -0.176 | -0.003 | -0.016 | -0.016 | 0 | 0.68 | 0.74 | 0.58 | 0.6 | 0.64 | 0.48 |
| 3 | 125849031 | rs873696 | -0.986 | 1.601 | 0.53 | -0.176 | -0.001 | -0.007 | -0.007 | 0 | 0.67 | 0.66 | 0.58 | 0.6 | 0.64 | 0.48 |
| 3 | 125849714 | rs1562475 | -0.986 | 1.601 | 0.53 | -0.176 | -0.001 | -0.007 | -0.007 | 0.084 | 0.67 | 0.66 | 0.58 | 0.6 | 0.64 | 0.48 |
| 3 | 125849928 | rs3821453 | -0.986 | 1.601 | 0.53 | -0.176 | -0.001 | -0.007 | -0.007 | 0 | 0.67 | 0.66 | 0.58 | 0.6 | 0.64 | 0.48 |
| 3 | 125850381 | rs2305228 | -1.335 | 1.598 | 0.536 | -0.177 | -0.031 | -0.08 | -0.08 | 0 | 0.7 | 0.9 | 0.61 | 0.6 | 0.65 | 0.48 |
| 3 | 125850702 | rs4646731 | -0.977 | 1.601 | 0.518 | -0.151 | -0.001 | -0.007 | -0.007 | 0 | 0.67 | 0.66 | 0.62 | 0.6 | 0.64 | 0.48 |
| 3 | 125853019 | rs4646728 | -1.185 | 1.599 | 0.554 | -0.153 | 0.003 | 0.019 | 0.019 | 2.983 | 0.68 | 0.73 | 0.58 | 0.59 | 0.65 | 0.47 |
| 3 | 125854409 | rs2276724 | 0.639 | 0.71 | 0.563 | -0.189 | 0 | 0.004 | 0.004 | 0.005 | 0.12 | 0.13 | 0.15 | 0.24 | 0.17 | 0.08 |
| 3 | 125855160 | rs3772413 | 0.18 | 0.735 | 0.529 | -0.189 | 0 | 0.01 | 0.01 | 0.579 | 0.15 | 0.21 | 0.16 | 0.23 | 0.17 | 0.1 |
| 3 | 125856695 | rs2305230 | -0.071 | 0.735 | 0.523 | -0.19 | 0.014 | 0.065 | 0.065 | 0.795 | 0.16 | 0.44 | 0.19 | 0.23 | 0.17 | 0.11 |
| 3 | 125857342 | rs6800400 | -0.071 | 0.735 | 0.523 | -0.19 | 0.014 | 0.065 | 0.065 | 0.795 | 0.16 | 0.44 | 0.19 | 0.23 | 0.17 | 0.11 |
| 3 | 125859367 | rs11923466 | 0.122 | 0.808 | 0.479 | -0.162 | 0.042 | 0.083 | 0.083 | 0.248 | 0.51 | 0.29 | 0.38 | 0.38 | 0.38 | 0.37 |
| 3 | 125865117 | rs11715574 | -0.114 | 0.816 | 0.488 | -0.158 | 0.012 | 0.061 | 0.061 | 0.883 | 0.16 | 0.44 | 0.19 | 0.23 | 0.17 | 0.11 |
| 3 | 125865766 | rs2886059 | 0.008 | 0.82 | 0.435 | -0.156 | 0.019 | 0.071 | 0.071 | 0.69 | 0.16 | 0.42 | 0.19 | 0.23 | 0.17 | 0.11 |
| 3 | 125866026 | rs1965848 | -1.347 | 0.854 | -0.251 | -0.153 | -0.125 | -0.123 | -0.123 | 0 | 0.69 | 0.96 | 0.64 | 0.62 | 0.55 | 0.49 |
| 3 | 125868513 | rs10934751 | -0.15 | 0.95 | -0.282 | -0.13 | 0.021 | -0.005 | -0.005 | 0 | 0.52 | 0.29 | 0.39 | 0.6 | 0.46 | 0.38 |
| 3 | 125870798 | rs4646717 | -0.004 | 1.208 | -0.306 | -0.022 | 0 | 0 | 0 | 0 | 0.49 | 0.48 | 0.33 | 0.36 | 0.34 | 0.3 |
| 3 | 125872293 | rs16837178 | -0.173 | 1.473 | -0.445 | 0.207 | -0.001 | -0.021 | -0.021 | 0.695 | 0.1 | 0.13 | 0.06 | 0 | 0.04 | 0.02 |
| 3 | 125875579 | rs4234274 | 0.814 | 1.584 | -0.732 | 0.71 | -0.003 | -0.003 | -0.003 | 0 | 0.85 | 0.85 | 0.78 | 0.75 | 0.81 | 0.88 |
| 3 | 125876862 | rs4646703 | -0.98 | 1.389 | -0.991 | 0.492 | -0.015 | 0.016 | 0.016 | 0 | 0.18 | 0.1 | 0.11 | 0.1 | 0.16 | 0.15 |
| 3 | 125879456 | rs4646701 | -0.378 | 1.29 | -0.669 | 0.902 | -0.007 | -0.012 | -0.012 | 0.087 | 0.63 | 0.38 | 0.46 | 0.48 | 0.59 | 0.54 |
| 3 | 125880208 | rs16837183 | 0.587 | 0.878 | -0.88 | 0.463 | -0.001 | -0.02 | -0.02 | 0.141 | 0.1 | 0.1 | 0.04 | 0 | 0.05 | 0.03 |
| 3 | 125881122 | rs4679208 | -0.956 | 1.001 | 0.317 | 1.022 | 0.004 | 0.003 | 0.003 | 0.005 | 0.67 | 0.64 | 0.64 | 0.62 | 0.62 | 0.58 |
| 3 | 125883511 | rs4521172 | -0.523 | 1.079 | 0.228 | 1.074 | -0.01 | -0.009 | -0.009 | 0 | 0.63 | 0.37 | 0.41 | 0.43 | 0.58 | 0.53 |
| 3 | 125890744 | rs7617733 | -0.618 | 0.312 | 1.235 | 0.869 | 0.045 | 0.069 | 0.069 | 0 | 0.14 | 0.43 | 0.27 | 0.23 | 0.16 | 0.13 |
| 3 | 125890830 | rs7639712 | 0.166 | 0.303 | 1.185 | 0.856 | -0.006 | -0.057 | -0.057 | 1.689 | 0.11 | 0.25 | 0.09 | 0.04 | 0.12 | 0.08 |
| 3 | 125896572 | rs9842910 | -0.772 | 0.271 | 1.239 | 0.854 | 0.016 | 0.035 | 0.035 | 0 | 0.15 | 0.53 | 0.28 | 0.23 | 0.16 | 0.13 |
| 3 | 125897538 | rs13060596 | 0.404 | 0.192 | 1.145 | 0.745 | 0.035 | -0.023 | -0.023 | 1.649 | 0.4 | 0.2 | 0.28 | 0.39 | 0.36 | 0.38 |
| 3 | 125902417 | rs9874508 | -0.529 | -0.074 | 1.076 | 0.859 | 0.041 | 0.005 | 0.005 | 1.142 | 0.5 | 0.73 | 0.54 | 0.49 | 0.48 | 0.36 |
| 3 | 125905389 | rs2077523 | -0.67 | -0.087 | 1.546 | 0.831 | -0.002 | 0.001 | 0.001 | 2.778 | 0.55 | 0.43 | 0.39 | 0.49 | 0.49 | 0.37 |
| 3 | 125908195 | rs6795005 | -1.092 | 0.365 | 1.655 | 0.87 | 0.043 | 0.063 | 0.063 | 0 | 0.21 | 0.44 | 0.29 | 0.3 | 0.22 | 0.24 |
| 3 | 125908310 | rs9851577 | -0.002 | 0.345 | 1.726 | 0.86 | 0.04 | 0.062 | 0.062 | 1.961 | 0.44 | 0.19 | 0.31 | 0.31 | 0.43 | 0.3 |
| 3 | 125908841 | rs4591460 | 0.181 | 0.041 | 1.476 | 0.796 | -0.002 | -0.012 | -0.012 | 0 | 0.16 | 0.18 | 0.13 | 0.08 | 0.19 | 0.2 |
| 3 | 125908975 | rs13069815 | -0.558 | 0.054 | 1.508 | 0.802 | 0.017 | 0.03 | 0.03 | 0.012 | 0.09 | 0.23 | 0.2 | 0.12 | 0.08 | 0.17 |
| 3 | 125909415 | rs13070856 | 0.4 | 0.043 | 1.676 | 0.791 | 0.027 | 0.004 | 0.004 | 4.668 | 0.34 | 0.09 | 0.23 | 0.26 | 0.32 | 0.23 |
| 3 | 125909525 | rs4679103 | -1.041 | 0.063 | 1.662 | 0.795 | -0.003 | -0.02 | -0.02 | 0 | 0.26 | 0.41 | 0.33 | 0.21 | 0.27 | 0.37 |
| 3 | 125910955 | rs13082246 | -1.862 | 0.084 | 1.626 | 0.811 | 0 | 0 | 0 | 0.238 | 0.64 | 0.63 | 0.6 | 0.61 | 0.65 | 0.54 |
| 3 | 125913438 | rs4679213 | -0.571 | 0.105 | 1.547 | 0.818 | 0.022 | 0.037 | 0.037 | 0.001 | 0.09 | 0.23 | 0.2 | 0.12 | 0.08 | 0.17 |
| 3 | 125919451 | rs6765882 | -1.791 | 0.116 | 1.559 | 0.881 | 0.003 | 0 | 0 | 0.318 | 0.64 | 0.61 | 0.6 | 0.63 | 0.65 | 0.55 |
| 3 | 125919839 | rs6793040 | -0.818 | 0.583 | 1.887 | 1.242 | 0.022 | 0.047 | 0.047 | 0 | 0.09 | 0.23 | 0.21 | 0.14 | 0.08 | 0.17 |
| 3 | 125920064 | rs4374483 | -0.75 | 0.567 | 1.907 | 1.221 | -0.001 | 0.004 | 0.004 | 3.362 | 0.46 | 0.39 | 0.47 | 0.54 | 0.46 | 0.35 |
| 3 | 125920117 | rs1447659 | 0.695 | 0.604 | 1.777 | 1.182 | 0.014 | 0.016 | 0.016 | 0 | 0.11 | 0.15 | 0.08 | 0.18 | 0.15 | 0.07 |
| 3 | 125927761 | rs9822040 | -1.769 | 0.47 | 1.522 | 1.153 | -0.048 | 0.003 | 0.003 | 0.352 | 0.58 | 0.8 | 0.56 | 0.62 | 0.54 | 0.51 |
| 3 | 125929182 | rs1447656 | -0.085 | 0.552 | 1.687 | 0.782 | 0.023 | -0.013 | -0.013 | 0.643 | 0.1 | 0.32 | 0.24 | 0.08 | 0.09 | 0.18 |
| 3 | 125929471 | rs1447657 | -0.359 | 0.499 | 1.644 | 1.084 | 0.007 | -0.003 | -0.003 | 1.27 | 0.38 | 0.33 | 0.46 | 0.41 | 0.34 | 0.3 |
| 3 | 125931750 | rs4679219 | -0.491 | 0.421 | 1.836 | 1.088 | 0.004 | -0.005 | -0.005 | 0 | 0.39 | 0.33 | 0.44 | 0.52 | 0.34 | 0.18 |
| 3 | 125934193 | rs16837285 | -0.107 | 0.309 | 1.735 | 1.003 | 0.022 | -0.041 | -0.041 | 2.082 | 0.3 | 0.16 | 0.26 | 0.41 | 0.3 | 0.23 |
| 3 | 125934221 | rs11924509 | -0.644 | 0.337 | 1.828 | 0.997 | 0.044 | 0.005 | 0.005 | 0.413 | 0.19 | 0.45 | 0.27 | 0.18 | 0.19 | 0.09 |
| 3 | 125943511 | rs2365020 | -1.3 | 0.381 | 1.514 | 0.938 | 0.01 | 0.023 | 0.023 | 0.02 | 0.48 | 0.6 | 0.53 | 0.59 | 0.49 | 0.33 |
| 3 | 125954785 | rs6767551 | -0.807 | 0.378 | 1.53 | 0.938 | -0.013 | 0 | 0 | 0.006 | 0.23 | 0.43 | 0.29 | 0.25 | 0.17 | 0.23 |
| 3 | 125956224 | rs1839029 | 0.393 | 0.334 | 1.272 | 0.976 | -0.001 | -0.021 | -0.021 | 0 | 0.25 | 0.17 | 0.24 | 0.34 | 0.32 | 0.1 |
| 3 | 125970734 | rs1816949 | -0.721 | 0.39 | 1.054 | 0.332 | 0.005 | 0.002 | 0.002 | 0.212 | 0.46 | 0.51 | 0.34 | 0.57 | 0.39 | 0.31 |
| 3 | 125973585 | rs10934761 | 0.045 | 0.376 | 0.865 | 0.263 | -0.023 | -0.077 | -0.077 | 0.72 | 0.29 | 0.1 | 0.27 | 0.56 | 0.37 | 0.28 |
| 3 | 125973658 | rs2121745 | -1.468 | 0.368 | 0.937 | 0.188 | -0.006 | 0.158 | 0.158 | 0.262 | 0.51 | 0.79 | 0.55 | 0.76 | 0.5 | 0.48 |
| 3 | 125978418 | rs11708255 | -1.024 | 0.419 | 1.253 | 0.33 | -0.015 | -0.008 | -0.008 | 0.016 | 0.22 | 0.5 | 0.26 | 0.17 | 0.15 | 0.18 |
| 3 | 125983084 | rs10470318 | -1.329 | 0.52 | 1.191 | 0.309 | 0.014 | 0.014 | 0.014 | 0 | 0.24 | 0.5 | 0.31 | 0.26 | 0.22 | 0.22 |
| 3 | 125983428 | rs10470321 | -1.532 | 0.468 | 0.781 | 0.286 | 0.001 | 0.002 | 0.002 | 0.358 | 0.27 | 0.8 | 0.34 | 0.26 | 0.22 | 0.22 |
| 3 | 125987924 | rs9825772 | -0.917 | 0.506 | 0.508 | 0.249 | 0.001 | 0.012 | 0.012 | 0.032 | 0.46 | 0.79 | 0.42 | 0.58 | 0.46 | 0.34 |
| 3 | 125989724 | rs2044579 | 0.578 | 0.547 | 0.572 | 0.206 | -0.002 | -0.07 | -0.07 | 0 | 0.11 | 0.27 | 0.07 | 0 | 0.06 | 0.02 |
| 3 | 125990017 | rs2044577 | -1.1 | 0.626 | 0.112 | 0.371 | -0.009 | 0.182 | 0.182 | 0.18 | 0.52 | 0.81 | 0.57 | 0.78 | 0.54 | 0.49 |
| 3 | 125990463 | rs6809287 | -1.376 | 0.506 | 0.931 | 0.88 | -0.007 | 0.133 | 0.133 | 0.582 | 0.17 | 0.49 | 0.25 | 0.27 | 0.1 | 0.19 |
| 3 | 125992478 | rs731221 | -1.856 | 0.836 | 0.696 | 0.703 | 0.163 | 0.06 | 0.06 | 0.413 | 0.08 | 0.47 | 0.26 | 0.05 | 0.12 | 0.12 |
| 3 | 125993159 | rs747037 | -0.786 | 0.578 | -0.212 | 0.937 | -0.018 | -0.071 | -0.071 | 0 | 0.11 | 0 | 0.08 | 0.2 | 0.08 | 0.13 |
| 3 | 125994799 | rs6807689 | 0.377 | 0.382 | -0.654 | 0.565 | 0.02 | -0.003 | -0.003 | 0 | 0.75 | 0.9 | 0.72 | 0.78 | 0.82 | 0.8 |
| 3 | 125997198 | rs4679115 | 0.797 | 0.287 | -1.021 | -0.327 | -0.023 | 0.05 | 0.05 | 0.195 | 0.45 | 0.29 | 0.44 | 0.7 | 0.49 | 0.58 |
| 3 | 125999565 | rs10934762 | 0.018 | 0.329 | -0.43 | -0.68 | 0.001 | -0.088 | -0.088 | 0 | 0.32 | 0.53 | 0.27 | 0.09 | 0.33 | 0.27 |
| 3 | 126006518 | rs7612364 | 0.149 | 0.412 | -0.971 | -1.242 | 0.001 | 0.099 | 0.099 | 0.326 | 0.52 | 0.67 | 0.38 | 0.09 | 0.5 | 0.52 |

| | | | | | | | | | | | | | | | | |
|---|---|---|---|---|---|---|---|---|---|---|---|---|---|---|---|---|
| 3 | 126007124 | rs9844302 | 0.752 | 0.17 | -1.082 | -1.074 | 0.004 | 0.006 | 0.006 | 0.093 | 0.15 | 0.09 | 0.2 | 0.1 | 0.12 | 0.15 |
| 3 | 126009008 | rs6797108 | 0.535 | 0.504 | -0.579 | -1.522 | 0.005 | -0.07 | -0.07 | 0.332 | 0.29 | 0.46 | 0.24 | 0.08 | 0.28 | 0.42 |
| 3 | 126012852 | rs6804946 | 1.811 | 0.518 | -0.427 | -1.673 | -0.024 | 0.03 | 0.03 | 0 | 0.15 | 0.08 | 0.15 | 0 | 0.19 | 0.15 |
| 3 | 126019929 | rs6767688 | 0.647 | 0.359 | -0.382 | -1.506 | -0.021 | 0.036 | 0.036 | 0 | 0.09 | 0 | 0.07 | 0.01 | 0.07 | 0.19 |
| 3 | 126020366 | rs10934765 | 1.231 | 0.213 | -0.691 | -1.067 | -0.027 | 0.064 | 0.064 | 0 | 0.35 | 0.15 | 0.3 | 0.27 | 0.42 | 0.29 |
| 3 | 126023648 | rs11918049 | 0.979 | 0.159 | -0.212 | -0.991 | 0.003 | -0.035 | -0.035 | 0.802 | 0.38 | 0.48 | 0.32 | 0.27 | 0.43 | 0.3 |
| 3 | 126028764 | rs7626340 | -0.277 | 0.282 | 0.276 | -0.705 | 0.006 | -0.004 | -0.004 | 1.333 | 0.59 | 0.7 | 0.61 | 0.45 | 0.61 | 0.69 |
| 3 | 126033522 | rs9851868 | -0.269 | 0.29 | 0.286 | -0.692 | 0.009 | -0.003 | -0.003 | 0 | 0.59 | 0.71 | 0.61 | 0.45 | 0.61 | 0.69 |
| 3 | 126035952 | rs12633128 | 0.829 | 0.361 | 0.319 | -0.654 | -0.004 | -0.008 | -0.008 | 0 | 0.09 | 0.12 | 0.07 | 0.04 | 0.05 | 0.15 |
| 3 | 126036904 | rs11712964 | 0.709 | 0.517 | 0.26 | -0.607 | 0.022 | 0.03 | 0.03 | 1.149 | 0.12 | 0.24 | 0.22 | 0.15 | 0.13 | 0.2 |
| 3 | 126039033 | rs9854816 | -0.186 | 0.291 | 0.061 | -0.219 | -0.003 | -0.049 | -0.049 | 0.862 | 0.61 | 0.84 | 0.62 | 0.47 | 0.62 | 0.7 |
| 3 | 126040602 | rs6785183 | 1.051 | 0.295 | 0.506 | -0.275 | -0.001 | 0.003 | 0.003 | 0 | 0.42 | 0.42 | 0.35 | 0.31 | 0.46 | 0.39 |
| 3 | 126041824 | rs3856649 | -0.195 | 0.379 | 0.528 | 0.052 | -0.008 | 0.006 | 0.006 | 0.109 | 0.61 | 0.56 | 0.62 | 0.56 | 0.63 | 0.7 |
| 3 | 126049305 | rs9880406 | -0.08 | 0.65 | 0.369 | 0.015 | 0 | -0.001 | -0.001 | 0 | 0.2 | 0.21 | 0.3 | 0.25 | 0.2 | 0.32 |
| 3 | 126057797 | rs7623133 | -0.24 | 0.362 | 0.168 | 0.141 | 0.018 | -0.03 | -0.03 | 0.02 | 0.62 | 0.83 | 0.63 | 0.58 | 0.63 | 0.71 |
| 3 | 126059958 | rs6783805 | 1.331 | 0.347 | 0.136 | 0.127 | -0.008 | 0.038 | 0.038 | 0 | 0.4 | 0.32 | 0.34 | 0.27 | 0.46 | 0.32 |
| 3 | 126060617 | rs7615776 | -0.645 | 0.33 | -0.054 | 0.024 | 0 | 0.003 | 0.003 | 0.28 | 0.32 | 0.37 | 0.37 | 0.3 | 0.29 | 0.43 |
| 3 | 126066220 | rs9838915 | -0.251 | 0.173 | -0.041 | 0.157 | -0.001 | -0.004 | -0.004 | 0 | 0.2 | 0.18 | 0.28 | 0.25 | 0.15 | 0.32 |
| 3 | 126068779 | rs9850626 | -0.489 | 0.201 | 0.007 | 0.3 | -0.005 | 0.032 | 0.032 | 0 | 0.73 | 0.67 | 0.7 | 0.59 | 0.73 | 0.74 |
| 3 | 126078890 | rs1358087 | 0.38 | 0.07 | -0.055 | 0.186 | 0 | 0 | 0 | 0 | 0.6 | 0.65 | 0.56 | 0.48 | 0.62 | 0.6 |
| 3 | 126083378 | rs9846116 | -1.473 | 0.189 | -0.32 | 0.842 | -0.014 | -0.032 | -0.032 | 0 | 0.85 | 0.95 | 0.83 | 0.82 | 0.84 | 0.88 |
| 3 | 126088338 | rs6777173 | 1.045 | 0.351 | 0.134 | 0.155 | -0.033 | 0.089 | 0.089 | 0.341 | 0.41 | 0.24 | 0.31 | 0.31 | 0.46 | 0.4 |
| 3 | 126089186 | rs938385 | 0.794 | 0.425 | 0.006 | 0.187 | -0.003 | 0.012 | 0.012 | 0.156 | 0.44 | 0.63 | 0.34 | 0.31 | 0.46 | 0.4 |
| 3 | 126092351 | rs9865702 | -0.43 | 0.75 | -0.107 | 0.314 | -0.003 | 0.001 | 0.001 | 0.106 | 0.31 | 0.35 | 0.39 | 0.28 | 0.27 | 0.36 |
| 3 | 126096047 | rs4299436 | -0.26 | 0.787 | -0.389 | 0.273 | 0.003 | 0.001 | 0.001 | 0 | 0.28 | 0.3 | 0.39 | 0.27 | 0.32 | 0.4 |
| 3 | 126097849 | rs1877966 | 0.209 | 0.85 | -0.943 | 0.236 | -0.043 | 0.039 | 0.039 | 0 | 0.27 | 0.01 | 0.28 | 0.12 | 0.22 | 0.31 |
| 3 | 126099612 | rs11707482 | 0.189 | 1.414 | 0.11 | 1.028 | -0.007 | -0.076 | -0.076 | 0 | 0.73 | 0.87 | 0.7 | 0.52 | 0.7 | 0.68 |
| 3 | 126099847 | rs1849900 | 0.909 | 1.548 | 0.33 | 0.968 | -0.008 | -0.054 | -0.054 | 0 | 0.65 | 0.12 | 0.57 | 0.5 | 0.62 | 0.66 |
| 3 | 126101047 | rs9864977 | 0.314 | 0.868 | 0.124 | 1.18 | -0.002 | -0.049 | -0.049 | 0 | 0.73 | 0.83 | 0.69 | 0.51 | 0.7 | 0.68 |
| 3 | 126105301 | rs6795344 | -1.677 | 1.078 | 0.863 | 0.613 | -0.03 | -0.075 | -0.075 | 0 | 0.19 | 0.48 | 0.18 | 0.08 | 0.16 | 0.09 |
| 3 | 126109485 | rs6798944 | 1.221 | 1.3 | 0.834 | 0.854 | 0.023 | 0.091 | 0.091 | 0.119 | 0.81 | 0.34 | 0.75 | 0.59 | 0.79 | 0.81 |
| 3 | 126111104 | rs12695485 | -0.195 | 1.278 | 0.734 | 0.876 | 0.007 | 0.06 | 0.06 | 0 | 0.91 | 0.9 | 0.86 | 0.65 | 0.89 | 0.83 |
| 3 | 126112299 | rs6801075 | 1.526 | 1.208 | 0.235 | 0.797 | 0 | -0.003 | -0.003 | 0.905 | 0.71 | 0.25 | 0.68 | 0.57 | 0.73 | 0.74 |
| 3 | 126113127 | rs2280677 | 0.707 | 1.954 | 0.257 | 1.283 | 0.062 | 0.195 | 0.195 | 0 | 0.82 | 0.5 | 0.77 | 0.59 | 0.79 | 0.8 |
| 3 | 126116410 | rs883052 | -1.107 | 2.144 | -0.177 | 0.66 | 0.063 | 0.009 | 0.009 | 0 | 0.05 | 0.1 | 0.07 | 0.06 | 0.14 | 0.05 |
| 3 | 126118561 | rs11919855 | -0.006 | 1.851 | -0.6 | 0.499 | -0.027 | 0.027 | 0.027 | 0 | 0.05 | 0.01 | 0.05 | 0 | 0.08 | 0.02 |
| 3 | 126129646 | rs9828868 | 0.377 | 1.237 | -0.664 | 0.393 | 0.03 | 0.045 | 0.045 | 0 | 0.5 | 0.7 | 0.52 | 0.42 | 0.42 | 0.34 |
| 3 | 126132031 | rs9863762 | 0.213 | 0.77 | -1.162 | -0.05 | 0.001 | -0.058 | -0.058 | 1.183 | 0.65 | 0.87 | 0.73 | 0.58 | 0.7 | 0.57 |
| 3 | 126134636 | rs6794488 | 0.471 | 0.704 | -1.585 | -0.432 | -0.036 | -0.027 | -0.027 | 0 | 0.55 | 0.86 | 0.68 | 0.58 | 0.52 | 0.5 |
| 3 | 126135774 | rs4679241 | -0.013 | 0.531 | -0.87 | -0.771 | 0 | -0.022 | -0.022 | 0 | 0.23 | 0.36 | 0.27 | 0.13 | 0.19 | 0.27 |
| 3 | 126138778 | rs2271630 | -0.827 | -0.047 | -0.713 | -0.955 | -0.02 | 0.088 | 0.088 | 0 | 0.88 | 0.89 | 0.86 | 0.59 | 0.93 | 0.79 |
| 3 | 126139620 | rs4410406 | -0.021 | 0.074 | 0.075 | -1.03 | -0.004 | -0.059 | -0.059 | 0.28 | 0.24 | 0.5 | 0.28 | 0.13 | 0.19 | 0.26 |
| 3 | 126140654 | rs9859308 | -0.613 | -0.255 | 0.064 | -1.339 | 0.007 | -0.042 | -0.042 | 0 | 0.32 | 0.45 | 0.43 | 0.2 | 0.32 | 0.33 |
| 3 | 126141173 | rs13100907 | 1.439 | -0.216 | 0.326 | -1.019 | 0.025 | -0.026 | -0.026 | 0 | 0.06 | 0.11 | 0.06 | 0 | 0.09 | 0.01 |
| 3 | 126141857 | rs13083581 | 0.288 | -0.391 | 0.322 | -1.03 | -0.005 | 0.001 | 0.001 | 0 | 0.37 | 0.27 | 0.38 | 0.4 | 0.4 | 0.28 |
| 3 | 126141893 | rs9868566 | -0.156 | -0.545 | 0.354 | -1.11 | -0.009 | -0.032 | -0.032 | 0.588 | 0.07 | 0.17 | 0.1 | 0.01 | 0.06 | 0.06 |
| 3 | 126142871 | rs6791445 | 0.971 | -0.311 | 0.422 | -0.994 | 0.003 | 0.001 | 0.001 | 0 | 0.29 | 0.28 | 0.31 | 0.32 | 0.34 | 0.19 |
| 3 | 126143297 | rs11711483 | 1.729 | -0.275 | 0.418 | -1.004 | -0.007 | -0.016 | -0.016 | 0.015 | 0.22 | 0.14 | 0.25 | 0.32 | 0.26 | 0.17 |
| 3 | 126143727 | rs9875507 | -0.587 | -0.397 | 0.553 | -1.108 | -0.003 | 0.008 | 0.008 | 0 | 0.49 | 0.79 | 0.55 | 0.52 | 0.5 | 0.47 |
| 3 | 126149705 | rs13316625 | -0.116 | -0.179 | 0.659 | -0.969 | -0.008 | 0.006 | 0.006 | 0 | 0.05 | 0.18 | 0.11 | 0.07 | 0.05 | 0.17 |
| 3 | 126151680 | rs7647378 | 1.611 | -0.093 | 0.529 | -0.956 | -0.007 | -0.017 | -0.017 | 0 | 0.23 | 0.14 | 0.25 | 0.32 | 0.26 | 0.17 |
| 3 | 126152517 | rs11916460 | 1.244 | -0.152 | 0.542 | -0.967 | 0.037 | -0.026 | -0.026 | 0 | 0.06 | 0.14 | 0.06 | 0 | 0.09 | 0.02 |
| 3 | 126153361 | rs13099144 | 0.716 | -0.139 | 0.593 | -0.97 | 0.01 | 0.013 | 0.013 | 0 | 0.31 | 0.39 | 0.35 | 0.4 | 0.36 | 0.22 |
| 3 | 126154201 | rs12631841 | 1.684 | -0.098 | 0.53 | -0.965 | -0.013 | -0.025 | -0.025 | 1.52 | 0.21 | 0.1 | 0.17 | 0.31 | 0.25 | 0.14 |
| 3 | 126155325 | rs1055451 | 1.517 | 0.101 | 0.75 | -0.943 | 0.038 | -0.022 | -0.022 | 0 | 0.05 | 0.12 | 0.06 | 0 | 0.08 | 0.01 |
| 3 | 126160601 | rs4679121 | -0.626 | -0.523 | 0.632 | -0.782 | 0.011 | 0.007 | 0.007 | 0 | 0.07 | 0.21 | 0.23 | 0.09 | 0.09 | 0.07 |
| 3 | 126168529 | rs6795359 | 1.23 | -0.653 | 0.644 | -0.855 | 0.012 | -0.028 | -0.028 | 0 | 0.24 | 0.17 | 0.18 | 0.35 | 0.22 | 0.17 |
| 3 | 126175401 | rs812368 | -0.625 | -0.602 | 0.695 | -0.878 | -0.013 | 0.015 | 0.015 | 0 | 0.67 | 0.43 | 0.59 | 0.57 | 0.67 | 0.74 |
| 3 | 126180088 | rs3230 | 0.799 | -0.683 | 0.262 | -1.196 | -0.044 | -0.023 | -0.023 | 0.268 | 0.41 | 0.14 | 0.42 | 0.51 | 0.48 | 0.4 |
| 3 | 126180121 | rs3231 | 0.235 | -0.438 | 0.447 | -0.949 | 0.038 | -0.004 | -0.004 | 0 | 0.56 | 0.19 | 0.52 | 0.51 | 0.62 | 0.54 |
| 3 | 126182920 | rs1687462 | 0.235 | -0.431 | 0.451 | -0.946 | 0.038 | -0.004 | -0.004 | 0 | 0.56 | 0.19 | 0.52 | 0.51 | 0.62 | 0.54 |
| 3 | 126184450 | rs4679245 | -0.601 | -0.439 | 0.529 | -0.95 | 0.005 | -0.002 | -0.002 | 0 | 0.65 | 0.27 | 0.57 | 0.57 | 0.67 | 0.74 |
| 3 | 126200291 | rs800950 | 0.61 | -0.354 | -0.553 | -0.856 | 0.019 | 0.012 | 0.012 | 0 | 0.48 | 0.19 | 0.47 | 0.51 | 0.53 | 0.46 |
| 3 | 126200403 | rs1799398 | 0.088 | -0.35 | -0.54 | -0.846 | 0.036 | -0.004 | -0.004 | 0 | 0.56 | 0.19 | 0.52 | 0.51 | 0.62 | 0.54 |
| 3 | 126202257 | rs1687477 | 0.512 | -0.284 | -0.433 | -0.681 | 0.016 | -0.003 | -0.003 | 0 | 0.47 | 0.19 | 0.47 | 0.49 | 0.52 | 0.47 |
| 3 | 126207222 | rs777483 | -0.855 | -0.225 | -0.066 | 0.008 | -0.016 | 0.02 | 0.02 | 0 | 0.71 | 0.56 | 0.62 | 0.59 | 0.74 | 0.73 |
| 3 | 126207595 | rs777481 | 0.779 | -0.345 | -0.134 | -0.078 | 0.001 | -0.003 | -0.003 | 1.872 | 0.46 | 0.18 | 0.43 | 0.51 | 0.51 | 0.44 |
| 3 | 126207792 | rs748465 | -0.806 | -0.112 | -0.05 | -0.142 | -0.016 | 0.018 | 0.018 | 0 | 0.71 | 0.56 | 0.62 | 0.59 | 0.75 | 0.74 |
| 3 | 126211946 | rs708907 | -0.029 | 0.402 | -0.784 | 0.34 | 0.021 | 0.026 | 0.026 | 0 | 0.49 | 0.22 | 0.45 | 0.53 | 0.52 | 0.58 |

| | | | | | | | | | | | | | | | | |
|---|---|---|---|---|---|---|---|---|---|---|---|---|---|---|---|---|
| 3 | 126215694 | rs1687482 | 1.057 | 0.656 | 0.13 | 0.599 | 0.044 | 0.13 | 0.13 | 1.327 | 0.86 | 0.48 | 0.76 | 0.69 | 0.83 | 0.74 |
| 3 | 126216273 | rs17742502 | 0.37 | 0.854 | -0.531 | -0.639 | -0.008 | 0.019 | 0.019 | 0 | 0.29 | 0.21 | 0.2 | 0.01 | 0.31 | 0.29 |
| 3 | 126216386 | rs4383463 | -0.045 | 0.584 | -0.775 | -0.655 | -0.001 | 0 | 0 | 0 | 0.57 | 0.5 | 0.45 | 0.52 | 0.57 | 0.66 |
| 3 | 126216566 | rs4322954 | 1.17 | 0.856 | -0.772 | -0.644 | 0.003 | -0.011 | -0.011 | 0 | 0.27 | 0.26 | 0.28 | 0.52 | 0.24 | 0.34 |
| 3 | 126220539 | rs9844797 | 0.262 | 0.578 | -0.495 | -0.421 | 0.003 | 0.004 | 0.004 | 0 | 0.61 | 0.51 | 0.51 | 0.52 | 0.58 | 0.65 |
| 3 | 126222084 | rs4679251 | 0.248 | 0.514 | -0.446 | -0.454 | 0 | -0.001 | -0.001 | 0 | 0.59 | 0.44 | 0.5 | 0.51 | 0.57 | 0.65 |
| 3 | 126222357 | rs1799390 | 0.114 | 0.497 | -0.172 | -0.607 | -0.021 | -0.009 | -0.009 | 0 | 0.27 | 0.18 | 0.32 | 0.29 | 0.3 | 0.18 |
| 3 | 126225147 | rs1799388 | 0.445 | 0.082 | 0.621 | -0.532 | 0.008 | -0.017 | -0.017 | 0 | 0.92 | 0.74 | 0.94 | 0.95 | 0.94 | 0.92 |
| 3 | 126227185 | rs16837531 | 1.365 | -0.214 | 0.367 | -0.864 | 0.01 | 0.034 | 0.034 | 2.031 | 0.6 | 0.22 | 0.59 | 0.68 | 0.62 | 0.8 |
| 3 | 126228953 | rs1873388 | -0.81 | -0.289 | 0.49 | -0.855 | 0.016 | 0.033 | 0.033 | 0 | 0.28 | 0.39 | 0.42 | 0.4 | 0.35 | 0.31 |
| 3 | 126230104 | rs777490 | -0.529 | -0.238 | 0.537 | -0.808 | 0.007 | 0.029 | 0.029 | 0 | 0.26 | 0.32 | 0.4 | 0.41 | 0.31 | 0.3 |
| 3 | 126230343 | rs16837548 | 1.966 | -0.351 | 0.485 | -0.869 | -0.032 | -0.049 | -0.049 | 0 | 0.56 | 0.1 | 0.48 | 0.54 | 0.57 | 0.61 |
| 3 | 126230388 | rs777488 | -0.862 | -0.212 | 0.537 | -0.759 | 0.015 | 0.034 | 0.034 | 0 | 0.28 | 0.37 | 0.42 | 0.41 | 0.35 | 0.31 |
| 3 | 126230886 | rs777487 | -0.621 | -0.215 | 0.464 | -0.75 | 0.009 | 0.037 | 0.037 | 0 | 0.26 | 0.35 | 0.41 | 0.41 | 0.31 | 0.3 |
| 3 | 126233910 | rs777486 | 0.003 | -0.177 | 0.361 | -0.736 | -0.004 | -0.007 | -0.007 | 0 | 0.22 | 0.2 | 0.29 | 0.26 | 0.26 | 0.12 |
| 3 | 126236080 | rs1687481 | -0.363 | -0.192 | 0.457 | -0.724 | 0.013 | 0.034 | 0.034 | 0.261 | 0.24 | 0.29 | 0.39 | 0.41 | 0.31 | 0.24 |
| 3 | 126236761 | rs12493410 | -1.002 | -0.23 | 0.447 | -0.723 | 0.005 | 0.022 | 0.022 | 0 | 0.31 | 0.36 | 0.42 | 0.41 | 0.34 | 0.31 |
| 3 | 126239083 | rs9813564 | -2.068 | -0.23 | 0.139 | -1.059 | 0.006 | -0.012 | -0.012 | 0 | 0.45 | 0.84 | 0.52 | 0.46 | 0.43 | 0.39 |
| 3 | 126241795 | rs6786437 | 1.635 | -0.287 | 0.013 | -1.171 | -0.058 | -0.057 | -0.057 | 0.004 | 0.09 | 0 | 0.16 | 0.26 | 0.19 | 0.06 |
| 3 | 126243419 | rs7613347 | -2.323 | -0.394 | 0.098 | -1.169 | 0.008 | -0.007 | -0.007 | 2.422 | 0.45 | 0.83 | 0.52 | 0.46 | 0.42 | 0.4 |
| 3 | 126244591 | rs6796854 | -0.444 | -0.409 | 0.099 | -1.133 | -0.068 | -0.067 | -0.067 | 0 | 0.22 | 0.04 | 0.3 | 0.26 | 0.28 | 0.12 |
| 3 | 126246635 | rs3845960 | -1.714 | -0.535 | 0.826 | -1.161 | 0.002 | 0.006 | 0.006 | 0 | 0.37 | 0.58 | 0.48 | 0.45 | 0.41 | 0.42 |
| 3 | 126252506 | rs4679256 | 0.422 | -0.409 | 0.552 | -1.215 | -0.064 | -0.066 | -0.066 | 0 | 0.16 | 0.03 | 0.27 | 0.26 | 0.25 | 0.14 |
| 3 | 126256669 | rs6783962 | -2.148 | -0.569 | 0.722 | -1.29 | -0.002 | 0.018 | 0.018 | 2.297 | 0.38 | 0.79 | 0.41 | 0.32 | 0.35 | 0.26 |
| 3 | 126257236 | rs1873400 | -1.229 | -0.557 | 1.13 | -1.286 | 0.04 | 0.031 | 0.031 | 0 | 0.24 | 0.46 | 0.34 | 0.26 | 0.3 | 0.21 |
| 3 | 126257356 | rs1873399 | -0.56 | -0.548 | 1.126 | -1.276 | 0.022 | 0 | 0 | 0.901 | 0.08 | 0.32 | 0.07 | 0.06 | 0.05 | 0.06 |
| 3 | 126260163 | rs6781798 | -0.067 | -0.528 | 1.259 | -1.267 | 0.022 | 0.024 | 0.024 | 0 | 0.18 | 0.27 | 0.3 | 0.26 | 0.25 | 0.14 |
| 3 | 126261345 | rs1056522 | -1.406 | -0.544 | 1.245 | -1.275 | 0.019 | 0.01 | 0.01 | 0 | 0.28 | 0.46 | 0.34 | 0.26 | 0.3 | 0.21 |
| 3 | 126264060 | rs1552641 | 0.517 | -0.518 | 0.985 | -1.254 | -0.064 | -0.066 | -0.066 | 0 | 0.16 | 0.03 | 0.27 | 0.26 | 0.25 | 0.13 |
| 3 | 126265325 | rs12492352 | 1.968 | -0.661 | 0.347 | -1.715 | -0.046 | -0.144 | -0.144 | 0.164 | 0.22 | 0.01 | 0.27 | 0.58 | 0.21 | 0.29 |
| 3 | 126265706 | rs1123373 | 2.088 | -0.579 | 1.427 | -1.044 | -0.041 | -0.044 | -0.044 | 0 | 0.78 | 0.42 | 0.8 | 0.84 | 0.84 | 0.72 |
| 3 | 126267231 | rs12490724 | 1.79 | -0.471 | 1.485 | -0.987 | -0.03 | -0.027 | -0.027 | 0 | 0.82 | 0.5 | 0.81 | 0.84 | 0.85 | 0.74 |
| 3 | 126269232 | rs1873393 | 1.151 | -0.417 | 1.258 | -0.944 | -0.012 | 0.008 | 0.008 | 0 | 0.89 | 0.7 | 0.87 | 0.9 | 0.91 | 0.78 |
| 3 | 126269316 | rs1873392 | 1.134 | -0.418 | 1.258 | -0.944 | -0.012 | 0.009 | 0.009 | 0 | 0.9 | 0.7 | 0.87 | 0.9 | 0.91 | 0.79 |
| 3 | 126270484 | rs6782739 | 1.156 | -0.417 | 1.169 | -0.944 | -0.015 | 0.005 | 0.005 | 0 | 0.89 | 0.71 | 0.87 | 0.89 | 0.91 | 0.79 |
| 3 | 126276710 | rs12636448 | 1.734 | -0.337 | 0.6 | -0.973 | 0.003 | 0.019 | 0.019 | 2.269 | 0.62 | 0.25 | 0.59 | 0.69 | 0.65 | 0.74 |
| 3 | 126283079 | rs7617257 | 2.164 | -0.458 | 0.588 | -0.86 | 0.009 | -0.006 | -0.006 | 0 | 0.55 | 0.19 | 0.46 | 0.56 | 0.57 | 0.59 |
| 3 | 126287018 | rs3888972 | -1.059 | -0.354 | 0.73 | -0.959 | -0.002 | -0.049 | -0.049 | 0.057 | 0.27 | 0.6 | 0.33 | 0.1 | 0.26 | 0.12 |
| 3 | 126287440 | rs2363947 | 0.364 | -0.179 | 0.474 | -0.891 | 0.009 | -0.007 | -0.007 | 0.304 | 0.17 | 0.23 | 0.25 | 0.1 | 0.2 | 0.09 |
| 3 | 126288409 | rs17743103 | -0.744 | -0.254 | -0.078 | -1.025 | -0.043 | 0.077 | 0.077 | 0.262 | 0.26 | 0.06 | 0.31 | 0.1 | 0.29 | 0.12 |
| 3 | 126289091 | rs2363948 | 2.025 | -0.893 | 0.146 | -1.38 | 0.041 | 0.105 | 0.105 | 0 | 0.56 | 0.27 | 0.56 | 0.85 | 0.65 | 0.56 |
| 3 | 126289362 | rs1873389 | 1.703 | -0.799 | 0.113 | -1.328 | 0.02 | 0.06 | 0.06 | 0 | 0.6 | 0.25 | 0.55 | 0.78 | 0.65 | 0.56 |
| 3 | 126292703 | rs4679269 | 2.147 | -0.654 | -0.122 | -1.156 | 0.054 | 0.065 | 0.065 | 0 | 0.5 | 0.21 | 0.52 | 0.59 | 0.59 | 0.5 |
| 3 | 126299279 | rs796573 | 0.336 | -0.618 | -0.101 | -1.348 | -0.011 | 0.078 | 0.078 | 0 | 0.22 | 0.1 | 0.3 | 0.05 | 0.21 | 0.12 |
| 3 | 126299757 | rs777216 | 0.82 | -0.719 | 0.531 | -1.396 | -0.033 | -0.044 | -0.044 | 0 | 0.81 | 0.58 | 0.89 | 0.84 | 0.87 | 0.71 |
| 3 | 126304038 | rs777230 | 0.306 | -0.641 | 0.404 | -1.334 | -0.005 | 0.068 | 0.068 | 0 | 0.23 | 0.13 | 0.31 | 0.05 | 0.21 | 0.12 |
| 3 | 126323541 | rs11921028 | 1.08 | -0.724 | 0.786 | -1.333 | -0.01 | -0.009 | -0.009 | 0 | 0.6 | 0.47 | 0.54 | 0.6 | 0.65 | 0.56 |
| 3 | 126329910 | rs777241 | -1.011 | -0.632 | 0.822 | -1.178 | -0.011 | 0.017 | 0.017 | 0 | 0.95 | 0.93 | 0.97 | 0.85 | 0.95 | 0.77 |
| 3 | 126365316 | rs777237 | 0.033 | -0.656 | 0.691 | -1.181 | -0.004 | 0.072 | 0.072 | 0 | 0.92 | 0.64 | 0.95 | 0.85 | 0.95 | 0.77 |
| 3 | 126367640 | rs777238 | 1.484 | -0.726 | 0.473 | -1.223 | -0.046 | 0.042 | 0.042 | 0 | 0.79 | 0.44 | 0.84 | 0.64 | 0.85 | 0.67 |
| 3 | 126370166 | rs9637365 | 1.73 | -0.623 | 0.051 | -1.288 | 0.015 | 0.058 | 0.058 | 0 | 0.47 | 0.22 | 0.47 | 0.6 | 0.57 | 0.51 |
| 3 | 126372919 | rs11718498 | 2.512 | -0.337 | -0.399 | -1.462 | -0.05 | -0.009 | -0.009 | 0 | 0.37 | 0.17 | 0.37 | 0.58 | 0.42 | 0.37 |
| 3 | 126374855 | rs9815125 | -1.398 | -0.567 | -0.065 | -1.164 | -0.048 | 0.05 | 0.05 | 0 | 0.24 | 0.46 | 0.16 | 0.37 | 0.17 | 0.32 |
| 3 | 126377499 | rs2010479 | -1.097 | -0.572 | -0.289 | -0.799 | -0.045 | 0.122 | 0.122 | 0 | 0.14 | 0.35 | 0.1 | 0.37 | 0.08 | 0.3 |
| 3 | 126380804 | rs4679128 | -0.24 | -0.557 | -0.406 | -0.821 | -0.011 | 0.097 | 0.097 | 0 | 0.78 | 0.92 | 0.79 | 0.95 | 0.8 | 0.87 |
| 3 | 126384464 | rs9863992 | -0.619 | -0.31 | 0.039 | -0.32 | -0.023 | 0.051 | 0.051 | 0 | 0.08 | 0.25 | 0.04 | 0.18 | 0.05 | 0.23 |
| 3 | 126387875 | rs2139747 | -1.365 | -0.287 | -0.299 | -0.403 | -0.026 | 0.009 | 0.009 | 0.231 | 0.92 | 1 | 0.87 | 0.96 | 0.9 | 0.95 |
| 3 | 126394025 | rs9850754 | -1.078 | -0.316 | 0.126 | -0.18 | -0.047 | 0.131 | 0.131 | 0 | 0.15 | 0.36 | 0.1 | 0.37 | 0.08 | 0.3 |
| 3 | 126411836 | rs9860180 | -1.111 | 0.157 | -0.12 | -0.53 | 0.003 | -0.005 | -0.005 | 0.291 | 0.91 | 0.9 | 0.87 | 0.95 | 0.9 | 0.96 |
| 3 | 126422362 | rs9814583 | -0.877 | -0.136 | 0.114 | -0.42 | 0.004 | -0.006 | -0.006 | 1.471 | 0.12 | 0.11 | 0.18 | 0.21 | 0.15 | 0.33 |
| 3 | 126425661 | rs951739 | -1.327 | -0.129 | 0.31 | -0.374 | -0.002 | 0.013 | 0.013 | 0.037 | 0.15 | 0.16 | 0.24 | 0.35 | 0.16 | 0.4 |
| 3 | 126430248 | rs16837708 | -2.083 | -0.583 | 1.063 | -0.618 | -0.045 | 0.029 | 0.029 | 0 | 0.25 | 0.53 | 0.29 | 0.35 | 0.19 | 0.4 |
| 3 | 126432392 | rs9839501 | -1.392 | -0.571 | 1.273 | -0.611 | -0.013 | 0.048 | 0.048 | 0 | 0.16 | 0.25 | 0.26 | 0.35 | 0.16 | 0.4 |
| 3 | 126435975 | rs6806049 | -0.695 | -0.677 | 0.506 | -1.474 | 0.065 | 0.097 | 0.097 | 0 | 0.48 | 0.74 | 0.41 | 0.36 | 0.37 | 0.45 |
| 3 | 126439532 | rs13061453 | 1.986 | -0.624 | 0.075 | -1.636 | -0.083 | -0.099 | -0.099 | 0 | 0.21 | 0.04 | 0.29 | 0.59 | 0.31 | 0.31 |
| 3 | 126445381 | rs9852440 | -1.838 | -0.404 | 0.555 | -1.398 | -0.02 | -0.001 | -0.001 | 0 | 0.3 | 0.59 | 0.32 | 0.35 | 0.23 | 0.42 |
| 3 | 126451937 | rs2272487 | -0.847 | -0.568 | 0.635 | -1.385 | -0.013 | 0.048 | 0.048 | 0 | 0.16 | 0.25 | 0.26 | 0.35 | 0.17 | 0.4 |
| 3 | 126457276 | rs1550991 | -2.203 | -0.558 | 0.625 | -1.406 | -0.003 | 0.003 | 0.003 | 0.953 | 0.91 | 0.95 | 0.87 | 0.95 | 0.9 | 0.96 |
| 3 | 126479286 | rs6808646 | 1.714 | -0.558 | 0.668 | -1.427 | -0.006 | -0.019 | -0.019 | 0.644 | 0.05 | 0.04 | 0.01 | 0 | 0.02 | 0 |

| | | | | | | | | | | | | | | | | |
|---|---|---|---|---|---|---|---|---|---|---|---|---|---|---|---|---|
| 3 | 126483156 | rs6779670 | -0.923 | -0.627 | 0.775 | -1.5 | 0.046 | 0.067 | 0.067 | 0 | 0.48 | 0.67 | 0.39 | 0.35 | 0.37 | 0.45 |
| 3 | 126495306 | rs2053820 | -0.914 | -0.598 | 0.788 | -1.487 | 0.045 | 0.066 | 0.066 | 0 | 0.48 | 0.67 | 0.4 | 0.35 | 0.37 | 0.45 |
| 3 | 126504141 | rs4679293 | -0.653 | -0.606 | 1.039 | -1.474 | -0.028 | 0.021 | 0.021 | 0.006 | 0.15 | 0.33 | 0.21 | 0.2 | 0.15 | 0.32 |
| 3 | 126529804 | rs9858948 | -1.904 | -0.632 | 0.977 | -1.493 | -0.001 | 0 | 0 | 0.618 | 0.31 | 0.66 | 0.32 | 0.34 | 0.23 | 0.41 |
| 3 | 126565847 | rs1530691 | 1.897 | -0.842 | 0.846 | -1.566 | -0.057 | 0.108 | 0.108 | 0 | 0.17 | 0.01 | 0.14 | 0.01 | 0.21 | 0.14 |
| 3 | 126573995 | rs11714878 | 1.43 | -1.231 | 1.895 | -0.617 | 0.001 | 0.034 | 0.034 | 0.029 | 0.17 | 0.08 | 0.08 | 0 | 0.15 | 0.03 |
| 3 | 126584800 | rs9860191 | -1.762 | -1.232 | 1.932 | -0.606 | -0.045 | 0 | 0 | 1.21 | 0.19 | 0.35 | 0.2 | 0.21 | 0.14 | 0.31 |
| 3 | 126585852 | rs17750671 | 1.528 | -1.19 | 1.782 | -0.614 | -0.032 | 0.108 | 0.108 | 0.018 | 0.17 | 0.01 | 0.07 | 0 | 0.14 | 0.03 |
| 3 | 126620149 | rs6785036 | -0.249 | -1.175 | 1.8 | -0.53 | -0.003 | 0.007 | 0.007 | 0 | 0.08 | 0.06 | 0.09 | 0.15 | 0.04 | 0.08 |
| 3 | 126623316 | rs9872353 | -1.131 | -1.216 | 1.852 | -0.604 | -0.003 | -0.012 | -0.012 | 1.309 | 0.4 | 0.6 | 0.32 | 0.2 | 0.32 | 0.38 |
| 3 | 126627781 | rs4679302 | 1.595 | -1.25 | 1.924 | -0.568 | -0.055 | -0.025 | -0.025 | 0 | 0.78 | 0.41 | 0.76 | 0.8 | 0.83 | 0.66 |
| 3 | 126664392 | rs11721085 | 1.492 | -1.191 | 1.794 | -0.555 | 0.072 | 0.096 | 0.096 | 0 | 0.52 | 0.24 | 0.57 | 0.66 | 0.66 | 0.54 |
| 3 | 126669726 | rs4679311 | 0.967 | -1.161 | 1.899 | -0.485 | 0.017 | 0.016 | 0.016 | 0 | 0.6 | 0.35 | 0.61 | 0.66 | 0.69 | 0.54 |
| 3 | 126674224 | rs746139 | -1.228 | -1.378 | 2.157 | -0.295 | -0.004 | -0.054 | -0.054 | 0 | 0.81 | 0.77 | 0.91 | 0.99 | 0.85 | 0.97 |
| 3 | 126675154 | rs4624521 | -1.525 | -1.279 | 1.986 | -0.233 | -0.038 | 0.018 | 0.018 | 0.034 | 0.15 | 0.33 | 0.2 | 0.19 | 0.12 | 0.33 |
| 3 | 126680355 | rs1007390 | 1.141 | -1.237 | 1.6 | -0.228 | 0.053 | 0.058 | 0.058 | 0 | 0.54 | 0.29 | 0.58 | 0.66 | 0.67 | 0.54 |
| 3 | 126685000 | rs4679316 | 0.812 | -1.124 | 1.656 | 0.006 | 0.003 | 0.002 | 0.002 | 3.191 | 0.61 | 0.34 | 0.73 | 0.61 | 0.73 | 0.72 |
| 3 | 126688559 | rs9851250 | -1.576 | -0.862 | 1.533 | 0.369 | 0.003 | 0.023 | 0.023 | 0.319 | 0.12 | 0.12 | 0.18 | 0.4 | 0.13 | 0.39 |
| 3 | 126695927 | rs7646841 | -0.496 | -0.742 | 1.792 | 0.417 | -0.005 | 0.112 | 0.112 | 0 | 0.08 | 0.14 | 0.17 | 0.22 | 0.07 | 0.15 |
| 3 | 126709202 | rs6764158 | -1.457 | -0.668 | 1.798 | 0.373 | -0.008 | 0.005 | 0.005 | 0 | 0.39 | 0.61 | 0.34 | 0.42 | 0.34 | 0.43 |
| 3 | 126712758 | rs732737 | -1.379 | -0.655 | 1.634 | 0.372 | -0.021 | 0.031 | 0.031 | 0 | 0.09 | 0.24 | 0.07 | 0.2 | 0.08 | 0.25 |
| 3 | 126720564 | rs747967 | -1.162 | -0.657 | 1.227 | 0.398 | 0.005 | -0.005 | -0.005 | 0 | 0.11 | 0.1 | 0.05 | 0.21 | 0.12 | 0.26 |
| 3 | 126732047 | rs1347003 | -1.449 | -0.652 | 1.145 | 0.389 | 0.021 | -0.016 | -0.016 | 0 | 0.4 | 0.72 | 0.35 | 0.43 | 0.34 | 0.43 |
| 3 | 126741011 | rs9289290 | -0.149 | -1.048 | 1.111 | 0.493 | -0.013 | 0.019 | 0.019 | 0 | 0.05 | 0.06 | 0.03 | 0.15 | 0.06 | 0.17 |
| 3 | 126741478 | rs4679325 | 0.941 | -1.037 | 1.028 | 0.454 | -0.005 | 0.007 | 0.007 | 1.327 | 0.09 | 0.03 | 0.14 | 0.05 | 0.09 | 0.04 |
| 3 | 126746690 | rs9851451 | -1.289 | -1.076 | 1.36 | 0.651 | -0.014 | 0.001 | 0.001 | 0.33 | 0.34 | 0.64 | 0.2 | 0.28 | 0.26 | 0.23 |
| 3 | 126746731 | rs9833827 | -0.818 | -1.03 | 1.38 | 0.672 | -0.025 | 0.013 | 0.013 | 0 | 0.3 | 0.64 | 0.19 | 0.28 | 0.21 | 0.22 |
| 3 | 126755134 | rs3749395 | -1.137 | -0.952 | 0.919 | 0.602 | -0.02 | 0.074 | 0.074 | 0 | 0.15 | 0.2 | 0.08 | 0.33 | 0.15 | 0.26 |
| 3 | 126756304 | rs11711837 | 0.051 | -0.496 | 0.568 | -0.187 | 0.005 | 0.016 | 0.016 | 1.184 | 0.18 | 0.09 | 0.08 | 0 | 0.14 | 0.03 |
| 3 | 126757949 | rs9848753 | -1.235 | -0.183 | 1.026 | 0.673 | -0.004 | 0.006 | 0.006 | 0.078 | 0.06 | 0.05 | 0.04 | 0.13 | 0.08 | 0.18 |
| 3 | 126778239 | rs4234280 | -1.304 | -0.192 | 0.493 | 0.66 | -0.011 | 0.086 | 0.086 | 0.522 | 0.1 | 0.59 | 0.1 | 0.2 | 0.1 | 0.26 |
| 3 | 126783989 | rs900429 | 0.683 | 0.109 | 0.154 | 1.127 | 0.005 | 0.074 | 0.074 | 0.017 | 0.87 | 0.76 | 0.72 | 0.7 | 0.87 | 0.78 |
| 3 | 126784415 | rs4679328 | 0.02 | 0.014 | -0.2 | 0.612 | -0.012 | 0.06 | 0.06 | 0 | 0.28 | 0.52 | 0.34 | 0.51 | 0.25 | 0.34 |
| 3 | 126784752 | rs4679329 | -0.583 | -0.51 | -0.964 | 0.177 | -0.041 | -0.111 | -0.111 | 0 | 0.17 | 0.01 | 0.31 | 0.38 | 0.19 | 0.23 |
| 3 | 126786864 | rs6789402 | -0.98 | -0.296 | -1.59 | -0.19 | 0.004 | 0.044 | 0.044 | 0 | 0.69 | 0.8 | 0.83 | 0.91 | 0.7 | 0.82 |
| 3 | 126787159 | rs11921451 | 1.041 | 0.122 | -1.545 | -0.268 | -0.092 | 0.07 | 0.07 | 0 | 0.24 | 0.02 | 0.15 | 0.11 | 0.26 | 0.26 |
| 3 | 126790130 | rs9859428 | 0.228 | 0.208 | -1.106 | 0.043 | -0.009 | 0.042 | 0.042 | 0 | 0.54 | 0.77 | 0.52 | 0.62 | 0.51 | 0.61 |
| 3 | 126798493 | rs6439033 | -0.167 | 0.262 | -1.123 | 0.038 | 0.014 | 0.122 | 0.122 | 0.009 | 0.62 | 0.81 | 0.78 | 0.91 | 0.64 | 0.76 |
| 3 | 126800773 | rs9810372 | -0.023 | 0.305 | -1.217 | 0.194 | -0.013 | 0.038 | 0.038 | 0 | 0.55 | 0.78 | 0.52 | 0.61 | 0.52 | 0.61 |
| 3 | 126805061 | rs10934803 | -0.77 | 0.815 | -0.464 | 0.153 | 0.019 | -0.035 | -0.035 | 0 | 0.22 | 0.33 | 0.28 | 0.14 | 0.28 | 0.29 |
| 3 | 126807142 | rs11914882 | 0.265 | 1.281 | -0.577 | -0.175 | 0.012 | 0.116 | 0.116 | 0.246 | 0.52 | 0.27 | 0.47 | 0.18 | 0.48 | 0.28 |
| 3 | 126812577 | rs1471659 | 0.643 | 1.663 | 0.343 | 1.051 | 0.068 | 0.275 | 0.275 | 0.586 | 0.88 | 0.44 | 0.82 | 0.74 | 0.89 | 0.87 |
| 3 | 126815345 | rs16837988 | 0.073 | 2.218 | 0.271 | 1.514 | -0.001 | 0.005 | 0.005 | 0 | 0.26 | 0.23 | 0.21 | 0.4 | 0.26 | 0.42 |
| 3 | 126816310 | rs16837992 | 0.923 | 2.267 | 0.012 | 1.634 | -0.015 | -0.032 | -0.032 | 0 | 0.11 | 0 | 0.07 | 0.11 | 0.09 | 0.21 |
| 3 | 126817583 | rs4679336 | 1.59 | 2.366 | 0.093 | 1.506 | -0.018 | 0.014 | 0.014 | 0 | 0.6 | 0.3 | 0.59 | 0.35 | 0.58 | 0.45 |
| 3 | 126820688 | rs13066695 | -1.566 | 2.518 | 0.214 | 1.571 | 0.013 | 0.032 | 0.032 | 0 | 0.35 | 0.51 | 0.31 | 0.54 | 0.37 | 0.53 |
| 3 | 126821906 | rs4679337 | -1.469 | 2.644 | 0.195 | 1.635 | 0.028 | 0.029 | 0.029 | 0 | 0.35 | 0.46 | 0.38 | 0.57 | 0.41 | 0.54 |
| 3 | 126822685 | rs9844620 | -0.666 | 2.297 | 0.177 | 1.611 | -0.003 | -0.004 | -0.004 | 0 | 0.26 | 0.19 | 0.3 | 0.43 | 0.31 | 0.43 |
| 3 | 126824804 | rs7627218 | -1.288 | 2.327 | 0.85 | 1.583 | -0.002 | 0.022 | 0.022 | 0 | 0.32 | 0.41 | 0.3 | 0.41 | 0.29 | 0.49 |
| 3 | 126831861 | rs9868010 | -0.669 | 2.231 | 0.878 | 1.597 | -0.001 | 0.001 | 0.001 | 0 | 0.26 | 0.17 | 0.26 | 0.4 | 0.26 | 0.42 |
| 3 | 126857090 | rs9879157 | 0.601 | 1.978 | -0.013 | 1.744 | 0.002 | -0.049 | -0.049 | 0 | 0.11 | 0.01 | 0.07 | 0.18 | 0.07 | 0.2 |
| 3 | 126857142 | rs4599251 | -1.629 | 2.133 | 0.045 | 1.885 | -0.006 | 0.069 | 0.069 | 0 | 0.07 | 0.12 | 0.15 | 0.37 | 0.11 | 0.13 |
| 3 | 126860724 | rs6807545 | -1.318 | 2.133 | -0.071 | 1.053 | 0 | 0.002 | 0.002 | 0 | 0.45 | 0.54 | 0.41 | 0.42 | 0.46 | 0.49 |
| 3 | 126862321 | rs4679343 | 0.685 | 2.231 | 0.024 | 1.055 | 0.024 | 0.022 | 0.022 | 0 | 0.16 | 0.19 | 0.19 | 0.25 | 0.2 | 0.21 |
| 3 | 126863808 | rs12487078 | -1.466 | 2.412 | -0.146 | 0.956 | 0.001 | -0.003 | -0.003 | 1.115 | 0.43 | 0.49 | 0.39 | 0.43 | 0.46 | 0.48 |
| 3 | 126864404 | rs12490870 | -0.697 | 2.219 | -0.054 | 1.048 | 0.021 | 0.034 | 0.034 | 0 | 0.3 | 0.44 | 0.32 | 0.42 | 0.31 | 0.44 |
| 3 | 126877020 | rs729154 | 0.271 | 2.225 | -0.33 | 1.015 | -0.014 | -0.031 | -0.031 | 0 | 0.1 | 0.04 | 0.11 | 0.19 | 0.12 | 0.2 |
| 3 | 126877085 | rs729153 | -0.856 | 1.712 | -1.246 | 0.484 | -0.014 | -0.024 | -0.024 | 0 | 0.66 | 0.72 | 0.5 | 0.51 | 0.61 | 0.68 |
| 3 | 126885815 | rs6784566 | 1.52 | 0.713 | -1.241 | -0.014 | -0.027 | -0.07 | -0.07 | 0 | 0.23 | 0.28 | 0.12 | 0.02 | 0.18 | 0.15 |
| 3 | 126887252 | rs1482921 | -2.216 | 0.725 | -1.521 | -0.049 | -0.025 | 0.04 | 0.04 | 0 | 0.75 | 0.83 | 0.62 | 0.92 | 0.67 | 0.88 |
| 3 | 126887617 | rs881839 | -0.159 | 0.647 | -1.42 | 0.001 | -0.017 | -0.008 | -0.008 | 0 | 0.41 | 0.25 | 0.36 | 0.5 | 0.42 | 0.52 |
| 3 | 126888869 | rs1351666 | 2.415 | 0.841 | -1.446 | -0.056 | -0.055 | 0.079 | 0.079 | 0 | 0.22 | 0.05 | 0.33 | 0.08 | 0.27 | 0.1 |
| 3 | 126890358 | rs1384567 | 0.697 | 0.84 | -1.384 | -0.04 | -0.005 | 0.003 | 0.003 | 0 | 0.3 | 0.22 | 0.25 | 0.3 | 0.3 | 0.32 |
| 3 | 126897431 | rs11717754 | 2.342 | 0.814 | -1.423 | -0.006 | -0.054 | 0.038 | 0.038 | 0 | 0.23 | 0.05 | 0.33 | 0.15 | 0.27 | 0.1 |
| 3 | 126898751 | rs12638109 | 1.621 | 0.704 | -1.359 | 0.533 | 0.02 | -0.033 | -0.033 | 0 | 0.08 | 0.05 | 0.04 | 0.16 | 0.04 | 0.13 |
| 3 | 126901114 | rs11710590 | -0.769 | 1.115 | -1.161 | 0.399 | -0.006 | -0.006 | -0.006 | 0 | 0.68 | 0.39 | 0.69 | 0.74 | 0.68 | 0.73 |
| 3 | 126904293 | rs6793657 | 0.715 | 1.334 | -1.106 | 0.344 | -0.001 | 0.001 | 0.001 | 0 | 0.28 | 0.21 | 0.21 | 0.23 | 0.24 | 0.31 |
| 3 | 126912432 | rs6802930 | -0.179 | 0.931 | -0.968 | 0.724 | -0.012 | -0.001 | -0.001 | 4.654 | 0.59 | 0.27 | 0.42 | 0.45 | 0.54 | 0.67 |
| 3 | 126917584 | rs4679349 | -0.335 | 1.137 | -0.625 | 1.157 | 0.016 | 0.098 | 0.098 | 0 | 0.93 | 0.85 | 0.86 | 0.75 | 0.91 | 0.91 |

| | | | | | | | | | | | | | | | | |
|---|---|---|---|---|---|---|---|---|---|---|---|---|---|---|---|---|
| 3 | 126920448 | rs1482923 | -0.245 | 1.127 | -0.634 | 1.152 | 0.01 | 0.082 | 0.082 | 0 | 0.93 | 0.86 | 0.86 | 0.75 | 0.91 | 0.91 |
| 3 | 126923982 | rs11923216 | 1.097 | 0.909 | -0.981 | 0.846 | -0.004 | -0.003 | -0.003 | 0 | 0.62 | 0.3 | 0.45 | 0.46 | 0.57 | 0.69 |
| 3 | 126926181 | rs873094 | 1.67 | 0.209 | -1.125 | 0.622 | 0.062 | 0.148 | 0.148 | 0 | 0.37 | 0.1 | 0.2 | 0.1 | 0.32 | 0.24 |
| 3 | 126926547 | rs900419 | 0.287 | 0.715 | -0.016 | 1.198 | 0.042 | 0.174 | 0.174 | 0 | 0.92 | 0.71 | 0.85 | 0.75 | 0.91 | 0.91 |
| 3 | 126927416 | rs11920916 | 0.083 | 0.502 | -0.177 | 0.588 | -0.047 | 0.037 | 0.037 | 0 | 0.13 | 0.25 | 0.08 | 0.16 | 0.05 | 0.12 |
| 3 | 126935726 | rs4234283 | 0.352 | 1.197 | -0.187 | 0.655 | 0.035 | 0.036 | 0.036 | 0.098 | 0.83 | 0.42 | 0.86 | 0.84 | 0.85 | 0.84 |
| 3 | 126940784 | rs10934807 | 0.375 | 1.025 | -0.827 | 0.356 | 0.014 | 0.039 | 0.039 | 0 | 0.43 | 0.34 | 0.51 | 0.34 | 0.44 | 0.26 |
| 3 | 126942965 | rs13095826 | 0.447 | 1.09 | -0.702 | -0.162 | -0.006 | 0.047 | 0.047 | 0.054 | 0.36 | 0.27 | 0.29 | 0.24 | 0.39 | 0.29 |
| 3 | 126945750 | rs1994128 | -0.722 | 1.367 | -0.15 | 0.182 | -0.002 | 0.036 | 0.036 | 0 | 0.77 | 0.69 | 0.56 | 0.57 | 0.75 | 0.74 |
| 3 | 126947703 | rs4679354 | 0.551 | 1.21 | -0.29 | 0.074 | -0.01 | 0.097 | 0.097 | 0 | 0.39 | 0.18 | 0.29 | 0.24 | 0.41 | 0.32 |
| 3 | 126950126 | rs4234284 | 0.551 | 1.21 | -0.224 | 0.074 | -0.004 | 0.078 | 0.078 | 0 | 0.39 | 0.2 | 0.29 | 0.24 | 0.4 | 0.32 |
| 3 | 126956890 | rs9289293 | 0.499 | 1.18 | -0.052 | 0.037 | -0.007 | 0 | 0 | 0 | 0.35 | 0.21 | 0.25 | 0.33 | 0.34 | 0.43 |
| 3 | 126966294 | rs2358766 | 0.596 | 1.352 | -0.055 | 0.066 | -0.009 | 0.091 | 0.091 | 0 | 0.37 | 0.23 | 0.28 | 0.06 | 0.41 | 0.21 |
| 3 | 126977097 | rs16838209 | 0.276 | 1.359 | 0.047 | 0.191 | -0.001 | -0.01 | -0.01 | 0 | 0.4 | 0.44 | 0.3 | 0.27 | 0.4 | 0.32 |
| 3 | 126980437 | rs7627541 | 1.256 | 1.545 | -0.315 | -0.715 | 0.023 | 0.098 | 0.098 | 0.128 | 0.29 | 0.11 | 0.19 | 0.05 | 0.27 | 0.17 |
| 3 | 126986209 | rs13087651 | -0.175 | 1.384 | -0.069 | -0.596 | 0.013 | 0.025 | 0.025 | 0 | 0.33 | 0.45 | 0.5 | 0.61 | 0.37 | 0.39 |
| 3 | 126990806 | rs6439045 | -0.967 | 1.492 | -0.292 | -0.842 | -0.006 | -0.046 | -0.046 | 0 | 0.8 | 0.85 | 0.82 | 0.71 | 0.8 | 0.71 |
| 3 | 126992473 | rs12495245 | 0.782 | 1.475 | -0.275 | -0.855 | -0.003 | -0.046 | -0.046 | 0.132 | 0.47 | 0.4 | 0.32 | 0.11 | 0.43 | 0.33 |
| 3 | 126993099 | rs9871760 | 0.781 | 1.486 | -0.242 | -0.79 | -0.005 | 0.024 | 0.024 | 0 | 0.25 | 0.29 | 0.42 | 0.6 | 0.24 | 0.36 |
| 3 | 126996775 | rs9860652 | 0.781 | 1.486 | -0.242 | -0.79 | -0.005 | 0.024 | 0.024 | 0 | 0.25 | 0.29 | 0.42 | 0.6 | 0.24 | 0.36 |
| 3 | 127001159 | rs9681626 | -0.833 | 1.729 | -0.217 | -0.719 | -0.012 | -0.047 | -0.047 | 0 | 0.81 | 0.9 | 0.83 | 0.73 | 0.8 | 0.72 |
| 3 | 127003972 | rs9826609 | 0.012 | 1.755 | -0.183 | -0.814 | -0.034 | -0.02 | -0.02 | 0 | 0.73 | 0.79 | 0.75 | 0.72 | 0.65 | 0.69 |
| 3 | 127011798 | rs6414306 | -0.142 | 1.693 | -0.178 | -1.26 | 0 | 0.033 | 0.033 | 0 | 0.43 | 0.64 | 0.56 | 0.77 | 0.4 | 0.57 |
| 3 | 127024201 | rs2001774 | 1.561 | 1.634 | -0.168 | -1.245 | 0.054 | 0.082 | 0.082 | 2.52 | 0.3 | 0.11 | 0.19 | 0.01 | 0.24 | 0.12 |
| 3 | 127024461 | rs6439055 | -1.336 | 1.689 | -0.176 | -1.244 | -0.012 | -0.022 | -0.022 | 0 | 0.83 | 0.87 | 0.83 | 0.77 | 0.8 | 0.75 |
| 3 | 127028929 | rs11922489 | 1.576 | 1.501 | -0.187 | -1.341 | 0.05 | 0.072 | 0.072 | 2.733 | 0.3 | 0.12 | 0.19 | 0.01 | 0.24 | 0.12 |
| 3 | 127036142 | rs7610764 | 0.87 | 1.736 | 0.044 | -1.218 | -0.01 | 0.02 | 0.02 | 0 | 0.26 | 0.37 | 0.36 | 0.67 | 0.21 | 0.34 |
| 3 | 127037677 | rs6439060 | -1.298 | 1.968 | -0.385 | -1.133 | -0.055 | -0.072 | -0.072 | 0.341 | 0.83 | 0.96 | 0.83 | 0.76 | 0.8 | 0.75 |
| 3 | 127046788 | rs2358559 | 0.622 | 1.404 | -0.222 | -1.221 | 0.005 | 0.01 | 0.01 | 1.067 | 0.39 | 0.26 | 0.26 | 0.01 | 0.4 | 0.16 |
| 3 | 127055016 | rs2884486 | 1.433 | 1.185 | -0.25 | -1.187 | 0.033 | 0.027 | 0.027 | 2.282 | 0.31 | 0.18 | 0.18 | 0.01 | 0.25 | 0.13 |
| 3 | 127066341 | rs12106895 | 0.175 | 1.248 | -0.037 | -0.912 | -0.022 | 0.005 | 0.005 | 0 | 0.31 | 0.61 | 0.45 | 0.72 | 0.27 | 0.43 |
| 3 | 127067194 | rs13077884 | 0.257 | 1.415 | -0.194 | -0.838 | -0.013 | 0.01 | 0.01 | 0 | 0.29 | 0.48 | 0.44 | 0.71 | 0.26 | 0.42 |
| 3 | 127068225 | rs7627270 | 0.3 | 2.62 | -0.369 | -0.011 | -0.015 | 0.002 | 0.002 | 0.395 | 0.69 | 0.8 | 0.71 | 0.73 | 0.66 | 0.61 |
| 3 | 127068660 | rs9990353 | -0.648 | 2.318 | 0.007 | -0.279 | 0.075 | -0.1 | -0.1 | 0.197 | 0.28 | 0.48 | 0.3 | 0.09 | 0.41 | 0.28 |
| 3 | 127071136 | rs2358545 | -1.02 | 1.84 | 0.234 | 0.471 | 0.098 | 0.142 | 0.142 | 0 | 0.08 | 0.33 | 0.18 | 0.14 | 0.11 | 0.16 |
| 3 | 127077801 | rs2358544 | 0.56 | 1.671 | -0.764 | 0.298 | -0.005 | 0.008 | 0.008 | 0.016 | 0.56 | 0.46 | 0.35 | 0.62 | 0.53 | 0.68 |
| 3 | 127079760 | rs6439064 | -0.035 | 1.418 | -1.174 | 0.635 | -0.015 | -0.052 | -0.052 | 0 | 0.91 | 0.99 | 0.7 | 0.78 | 0.93 | 0.93 |
| 3 | 127081915 | rs6439065 | 0.007 | 1.338 | -1.057 | 0.732 | 0.006 | -0.029 | -0.029 | 0 | 0.9 | 0.95 | 0.71 | 0.76 | 0.93 | 0.92 |
| 3 | 127082168 | rs6768941 | 0.329 | 1.068 | -1.962 | -0.256 | -0.024 | 0.153 | 0.153 | 0.541 | 0.34 | 0.13 | 0.2 | 0.04 | 0.36 | 0.2 |
| 3 | 127084030 | rs12634857 | 0.97 | 0.229 | -1.524 | -0.004 | 0.033 | 0.022 | 0.022 | 0.001 | 0.28 | 0.2 | 0.1 | 0.23 | 0.18 | 0.22 |
| 3 | 127097277 | rs6802625 | -0.96 | 0.146 | -1.626 | 0.022 | -0.004 | -0.064 | -0.064 | 0.807 | 0.89 | 0.99 | 0.71 | 0.76 | 0.92 | 0.89 |
| 3 | 127103555 | rs4247199 | -0.463 | 0.39 | -1.349 | 0.246 | 0.05 | 0.02 | 0.02 | 0 | 0.63 | 0.79 | 0.59 | 0.65 | 0.72 | 0.71 |
| 3 | 127104054 | rs11709061 | 0.247 | 0.278 | -0.916 | 0.173 | -0.071 | -0.089 | -0.089 | 0 | 0.24 | 0.12 | 0.35 | 0.45 | 0.37 | 0.48 |
| 3 | 127112049 | rs4632497 | -0.603 | 0.136 | -0.847 | 0.26 | 0.055 | 0.008 | 0.008 | 0 | 0.64 | 0.81 | 0.6 | 0.63 | 0.72 | 0.72 |
| 3 | 127112273 | rs1806680 | 0.959 | 0.318 | -0.034 | 0.256 | -0.001 | 0.022 | 0.022 | 0.325 | 0.35 | 0.28 | 0.21 | 0.19 | 0.37 | 0.24 |
| 3 | 127113121 | rs9818768 | 0.105 | 0.532 | 0.172 | 0.408 | -0.035 | -0.046 | -0.046 | 0 | 0.24 | 0.18 | 0.31 | 0.42 | 0.35 | 0.47 |
| 3 | 127116269 | rs4679364 | -0.23 | 1.099 | 0.742 | 0.743 | 0.061 | 0.049 | 0.049 | 0 | 0.09 | 0.33 | 0.21 | 0.18 | 0.12 | 0.24 |
| 3 | 127124699 | rs2044614 | -1.283 | 1.029 | 0.585 | 1.375 | -0.024 | -0.059 | -0.059 | 0 | 0.91 | 0.99 | 0.72 | 0.68 | 0.91 | 0.92 |
| 3 | 127124826 | rs11706462 | 1.512 | 0.705 | 0.134 | 0.847 | -0.012 | 0.082 | 0.082 | 0.105 | 0.2 | 0.03 | 0.1 | 0.03 | 0.18 | 0.16 |
| 3 | 127125439 | rs2594228 | 0.38 | 0.589 | 0.589 | 0.809 | 0.001 | -0.01 | -0.01 | 0.11 | 0.58 | 0.31 | 0.4 | 0.46 | 0.61 | 0.51 |
| 3 | 127125589 | rs10804580 | 0.421 | 0.626 | 0.541 | 0.774 | 0.001 | -0.007 | -0.007 | 0 | 0.28 | 0.33 | 0.3 | 0.22 | 0.3 | 0.4 |
| 3 | 127129934 | rs2594203 | 0.082 | 0.544 | 0.649 | 0.794 | 0.018 | 0.175 | 0.175 | 0 | 0.8 | 0.51 | 0.51 | 0.5 | 0.82 | 0.68 |
| 3 | 127138277 | rs11710009 | -0.209 | 0.557 | 0.828 | 0.799 | 0.012 | 0.103 | 0.103 | 0 | 0.82 | 0.64 | 0.53 | 0.5 | 0.82 | 0.68 |
| 3 | 127139117 | rs10934813 | 0.311 | 0.399 | 0.359 | 0.494 | -0.002 | 0.02 | 0.02 | 0.374 | 0.35 | 0.26 | 0.22 | 0.22 | 0.37 | 0.25 |
| 3 | 127142913 | rs6767818 | 1.115 | 0.406 | 0.326 | 0.162 | -0.009 | -0.059 | -0.059 | 0 | 0.22 | 0.26 | 0.11 | 0.03 | 0.18 | 0.16 |
| 3 | 127145694 | rs12487492 | 1.484 | 0.449 | -0.042 | 0.245 | -0.055 | 0.078 | 0.078 | 0 | 0.13 | 0.01 | 0.1 | 0 | 0.18 | 0.06 |
| 3 | 127150025 | rs4637240 | 1.21 | 0.104 | 0.267 | 0.406 | -0.008 | -0.015 | -0.015 | 0.312 | 0.29 | 0.27 | 0.12 | 0.11 | 0.19 | 0.2 |
| 3 | 127158718 | rs9647334 | 0.045 | 0.262 | 0.401 | 0.658 | -0.001 | 0.005 | 0.005 | 0.197 | 0.59 | 0.65 | 0.41 | 0.34 | 0.46 | 0.61 |
| 3 | 127159199 | rs6767638 | 1.14 | 0.177 | 0.576 | 0.63 | 0.001 | 0.002 | 0.002 | 0.212 | 0.23 | 0.24 | 0.16 | 0.11 | 0.19 | 0.22 |
| 3 | 127159701 | rs10934815 | -1.493 | 0.334 | 0.788 | 1.11 | 0 | 0.068 | 0.068 | 0 | 0.9 | 0.88 | 0.7 | 0.67 | 0.91 | 0.92 |
| 3 | 127161090 | rs2884483 | -1.12 | 0.31 | 0.847 | 1.111 | 0.004 | 0.201 | 0.201 | 0 | 0.88 | 0.69 | 0.69 | 0.6 | 0.91 | 0.88 |
| 3 | 127164816 | rs1374924 | 0.191 | 0.19 | 0.779 | 0.996 | 0.03 | 0.013 | 0.013 | 0 | 0.32 | 0.36 | 0.32 | 0.32 | 0.47 | 0.31 |
| 3 | 127166771 | rs6776442 | -1.703 | 0.067 | 0.685 | 1.234 | 0.001 | -0.003 | -0.003 | 0.556 | 0.91 | 0.94 | 0.74 | 0.74 | 0.93 | 0.92 |
| 3 | 127166819 | rs13070949 | 0.677 | 0.073 | 0.742 | 1.186 | 0.001 | 0.001 | 0.001 | 0 | 0.07 | 0.08 | 0.11 | 0.12 | 0.08 | 0.2 |
| 3 | 127171638 | rs6799846 | -0.78 | -0.08 | 0.794 | 1.098 | 0.011 | 0.001 | 0.001 | 0 | 0.63 | 0.65 | 0.61 | 0.62 | 0.74 | 0.76 |
| 3 | 127183990 | rs7620937 | 1.194 | 0.303 | 0.595 | 1.28 | 0.003 | 0.004 | 0.004 | 0 | 0.14 | 0.09 | 0.25 | 0.21 | 0.25 | 0.19 |
| 3 | 127186461 | rs11919024 | 1.818 | 0.25 | 0.306 | 0.997 | -0.008 | 0.057 | 0.057 | 0.533 | 0.2 | 0.03 | 0.16 | 0.09 | 0.16 | 0.15 |
| 3 | 127188015 | rs10804581 | 0.258 | 0.298 | 0.617 | 1.045 | -0.002 | 0.015 | 0.015 | 0 | 0.41 | 0.35 | 0.31 | 0.28 | 0.41 | 0.34 |
| 3 | 127189935 | rs2594215 | -1.808 | 0.617 | 0.534 | 1.253 | 0.001 | -0.019 | -0.019 | 1.121 | 0.92 | 0.96 | 0.82 | 0.82 | 0.94 | 0.93 |

| | | | | | | | | | | | | | | | | |
|---|---|---|---|---|---|---|---|---|---|---|---|---|---|---|---|---|
| 3 | 127190795 | rs2594216 | -1.828 | 0.598 | 0.546 | 1.257 | 0.003 | -0.013 | -0.013 | 0 | 0.92 | 0.95 | 0.82 | 0.82 | 0.95 | 0.93 |
| 3 | 127191375 | rs9858161 | -0.184 | 0.505 | 0.739 | 1.07 | 0.077 | 0.073 | 0.073 | 0 | 0.22 | 0.34 | 0.38 | 0.34 | 0.34 | 0.42 |
| 3 | 127193427 | rs2594220 | 1.047 | -0.585 | 0.488 | 0.45 | -0.005 | -0.005 | -0.005 | 0.044 | 0.17 | 0.22 | 0.11 | 0.13 | 0.16 | 0.13 |
| 3 | 127202167 | rs2594227 | 0.383 | -0.227 | 0.59 | 0.37 | 0.015 | -0.003 | -0.003 | 0 | 0.87 | 0.62 | 0.84 | 0.88 | 0.9 | 0.72 |
| 3 | 127206375 | rs1562531 | 0.39 | -0.36 | 0.309 | 0.374 | 0.005 | -0.023 | -0.023 | 0 | 0.66 | 0.62 | 0.56 | 0.77 | 0.64 | 0.55 |
| 3 | 127206836 | rs1562532 | 0.474 | -0.357 | 0.31 | 0.376 | 0.006 | -0.037 | -0.037 | 0.272 | 0.65 | 0.57 | 0.57 | 0.77 | 0.64 | 0.55 |
| 3 | 127209361 | rs1868240 | -0.21 | -0.666 | 0.437 | 0.995 | -0.013 | 0.207 | 0.207 | 0.012 | 0.17 | 0.42 | 0.19 | 0.49 | 0.17 | 0.19 |
| 3 | 127220987 | rs2720251 | 0.369 | -0.639 | -0.281 | 0.697 | -0.012 | -0.011 | -0.011 | 0 | 0.77 | 0.65 | 0.67 | 0.77 | 0.84 | 0.68 |
| 3 | 127236915 | rs11917019 | 1.717 | -0.128 | -1.159 | 0.312 | -0.042 | 0.06 | 0.06 | 1.661 | 0.24 | 0.03 | 0.18 | 0.11 | 0.24 | 0.16 |
| 3 | 127247374 | rs2720240 | 0.266 | -0.243 | -0.434 | 0.636 | -0.011 | -0.008 | -0.008 | 0 | 0.78 | 0.71 | 0.69 | 0.77 | 0.84 | 0.68 |
| 3 | 127251527 | rs9831691 | 0.8 | -0.093 | -0.166 | 0.929 | -0.008 | 0.03 | 0.03 | 0 | 0.72 | 0.57 | 0.54 | 0.29 | 0.73 | 0.56 |
| 3 | 127253553 | rs6773160 | 0.736 | -0.04 | -0.222 | 1.002 | -0.002 | 0.012 | 0.012 | 0 | 0.72 | 0.64 | 0.54 | 0.36 | 0.73 | 0.63 |
| 3 | 127254061 | rs2594205 | -0.1 | -0.257 | -0.324 | 1.518 | -0.002 | 0.037 | 0.037 | 0 | 0.87 | 0.84 | 0.73 | 0.51 | 0.9 | 0.83 |
| 3 | 127262757 | rs2594210 | 0.916 | 0.246 | -0.692 | 0.958 | -0.007 | 0.007 | 0.007 | 0.76 | 0.54 | 0.58 | 0.42 | 0.33 | 0.49 | 0.42 |
| 3 | 127264909 | rs768854 | 0.403 | 0.23 | -0.915 | 0.949 | -0.006 | -0.01 | -0.01 | 0.362 | 0.22 | 0.18 | 0.13 | 0.28 | 0.26 | 0.21 |
| 3 | 127281890 | rs12489933 | 0.752 | 0.066 | -0.901 | 1.052 | 0.002 | -0.016 | -0.016 | 1.673 | 0.1 | 0.08 | 0.27 | 0.25 | 0.09 | 0.19 |
| 3 | 127281914 | rs12489937 | 0.657 | 0.006 | -0.825 | 1.023 | 0.002 | -0.007 | -0.007 | 0 | 0.1 | 0.11 | 0.28 | 0.25 | 0.09 | 0.19 |
| 3 | 127292788 | rs4974445 | 1.971 | -0.018 | -1.499 | 0.47 | -0.072 | 0.106 | 0.106 | 1.489 | 0.27 | 0.04 | 0.25 | 0.1 | 0.29 | 0.16 |
| 3 | 127295084 | rs920233 | 0.324 | -0.126 | -1.927 | 0.137 | -0.037 | -0.032 | -0.032 | 0.793 | 0.23 | 0.02 | 0.33 | 0.26 | 0.2 | 0.32 |
| 3 | 127304957 | rs920230 | 0.324 | -0.129 | -1.929 | 0.181 | -0.037 | -0.039 | -0.039 | 0 | 0.23 | 0.02 | 0.33 | 0.27 | 0.2 | 0.32 |
| 3 | 127351492 | rs13082666 | 1.209 | -0.291 | -0.861 | 0.724 | -0.054 | 0.095 | 0.095 | 5.641 | 0.4 | 0.11 | 0.32 | 0.19 | 0.42 | 0.27 |
| 3 | 127373586 | rs11714815 | 0.909 | -0.161 | -0.435 | 0.696 | 0 | 0.001 | 0.001 | 0.282 | 0.13 | 0.18 | 0.17 | 0.18 | 0.14 | 0.18 |
| 3 | 127379270 | rs920232 | 0.909 | -0.161 | -0.435 | 0.696 | 0 | 0.001 | 0.001 | 0.282 | 0.13 | 0.18 | 0.17 | 0.18 | 0.14 | 0.18 |
| 3 | 127385752 | rs9861025 | -0.879 | -0.174 | -0.448 | 0.674 | 0.056 | 0.058 | 0.058 | 2.391 | 0.05 | 0.36 | 0.03 | 0.01 | 0.01 | 0.06 |
| 3 | 127388150 | rs2304551 | 1.432 | -0.04 | -0.87 | 0.741 | -0.038 | -0.034 | -0.034 | 0 | 0.06 | 0 | 0.06 | 0.09 | 0.11 | 0.1 |
| 3 | 127403849 | rs782446 | -0.352 | -0.431 | -0.479 | 0.537 | 0.005 | -0.006 | -0.006 | 0.009 | 0.75 | 0.78 | 0.65 | 0.64 | 0.78 | 0.61 |
| 3 | 127404314 | rs2336408 | 0.199 | -0.345 | -0.481 | 0.244 | -0.006 | -0.111 | -0.111 | 0 | 0.21 | 0.35 | 0.15 | 0.02 | 0.24 | 0.12 |
| 3 | 127407753 | rs6801421 | 0.924 | -0.359 | -0.624 | 0.238 | -0.043 | 0.099 | 0.099 | 0 | 0.17 | 0.03 | 0.12 | 0.01 | 0.24 | 0.09 |
| 3 | 127408909 | rs6439081 | -0.495 | -0.322 | -0.591 | 0.341 | 0.01 | -0.01 | -0.01 | 0 | 0.75 | 0.84 | 0.65 | 0.67 | 0.79 | 0.61 |
| 3 | 127409941 | rs782444 | 0.765 | -0.324 | -0.556 | 0.336 | 0.004 | 0.01 | 0.01 | 0 | 0.53 | 0.45 | 0.5 | 0.65 | 0.54 | 0.46 |
| 3 | 127410959 | rs814130 | -0.463 | -0.321 | -0.556 | 0.339 | 0.01 | -0.007 | -0.007 | 0 | 0.75 | 0.82 | 0.65 | 0.67 | 0.79 | 0.62 |
| 3 | 127411436 | rs812156 | 0.367 | -0.127 | -0.563 | 0.626 | 0.005 | 0.01 | 0.01 | 0 | 0.53 | 0.45 | 0.5 | 0.65 | 0.54 | 0.46 |
| 3 | 127412573 | rs782441 | -0.225 | -0.39 | -0.328 | 0.79 | 0.001 | 0.056 | 0.056 | 0.234 | 0.19 | 0.45 | 0.26 | 0.27 | 0.2 | 0.26 |
| 3 | 127413881 | rs4881 | 1.197 | -0.535 | -0.058 | 0.681 | 0.046 | -0.02 | -0.02 | 0 | 0.06 | 0.2 | 0.08 | 0.01 | 0.09 | 0.08 |
| 3 | 127415047 | rs13086586 | 1.123 | -0.876 | -0.522 | 0.433 | -0.024 | 0.107 | 0.107 | 0 | 0.35 | 0.12 | 0.25 | 0.13 | 0.32 | 0.26 |
| 3 | 127416473 | rs6765071 | 2.306 | -0.861 | -0.589 | 0.354 | 0.001 | 0.026 | 0.026 | 0 | 0.23 | 0.12 | 0.19 | 0.13 | 0.19 | 0.17 |
| 3 | 127417202 | rs782437 | -0.499 | -1.032 | -0.525 | 0.764 | -0.108 | 0.081 | 0.081 | 0.996 | 0.56 | 0.93 | 0.43 | 0.61 | 0.46 | 0.56 |
| 3 | 127419578 | rs7633484 | 1.117 | -0.971 | -0.319 | 1.065 | 0.052 | 0.262 | 0.262 | 0.817 | 0.08 | 0.29 | 0.08 | 0.28 | 0.1 | 0.11 |
| 3 | 127421486 | rs9828443 | 1.27 | -1.004 | -0.363 | 1.008 | -0.015 | 0.1 | 0.1 | 0 | 0.35 | 0.17 | 0.25 | 0.13 | 0.32 | 0.25 |
| 3 | 127426929 | rs1071912 | -0.409 | -1.119 | -0.245 | 0.985 | 0.001 | -0.006 | -0.006 | 0 | 0.54 | 0.54 | 0.39 | 0.52 | 0.46 | 0.48 |
| 3 | 127426999 | rs2078637 | 0.567 | -1.08 | -0.243 | 0.999 | -0.001 | 0 | 0 | 0 | 0.43 | 0.46 | 0.34 | 0.41 | 0.42 | 0.36 |
| 3 | 127427592 | rs1437553 | 0.567 | -1.08 | -0.228 | 0.999 | 0 | 0.001 | 0.001 | 0 | 0.43 | 0.48 | 0.34 | 0.41 | 0.42 | 0.36 |
| 3 | 127430886 | rs3773133 | 0.776 | -1.12 | -0.416 | 0.911 | 0.009 | -0.008 | -0.008 | 0.032 | 0.09 | 0.08 | 0.05 | 0.11 | 0.04 | 0.11 |
| 3 | 127432559 | rs597956 | -1.958 | -1.111 | -0.607 | 0.911 | -0.007 | -0.029 | -0.029 | 0 | 0.92 | 0.98 | 0.83 | 0.8 | 0.91 | 0.8 |
| 3 | 127433542 | rs534018 | -0.018 | -1.128 | -0.241 | 0.873 | -0.004 | 0.022 | 0.022 | 0 | 0.47 | 0.37 | 0.46 | 0.38 | 0.5 | 0.44 |
| 3 | 127435100 | rs479504 | -0.539 | -1.114 | -0.243 | 0.889 | 0.009 | 0.149 | 0.149 | 0.811 | 0.84 | 0.69 | 0.74 | 0.51 | 0.81 | 0.69 |
| 3 | 127435191 | rs3773146 | 0.999 | -1.075 | -0.722 | 0.537 | -0.137 | 0.101 | 0.101 | 1.181 | 0.33 | 0.05 | 0.38 | 0.19 | 0.45 | 0.23 |
| 3 | 127435806 | rs782449 | -0.85 | -1.056 | -0.585 | 0.47 | -0.004 | 0.067 | 0.067 | 0 | 0.56 | 0.78 | 0.55 | 0.66 | 0.59 | 0.55 |
| 3 | 127438528 | rs3773155 | -0.236 | -1.073 | -0.201 | 0.519 | -0.041 | -0.002 | -0.002 | 0 | 0.19 | 0.39 | 0.13 | 0.14 | 0.13 | 0.19 |
| 3 | 127438887 | rs9811688 | 1.234 | -1.074 | -0.443 | 0.495 | -0.003 | 0.068 | 0.068 | 0 | 0.36 | 0.23 | 0.29 | 0.13 | 0.32 | 0.26 |
| 3 | 127438938 | rs3773159 | 2.299 | -0.831 | -0.906 | 0.183 | 0.001 | 0.022 | 0.022 | 0 | 0.23 | 0.12 | 0.23 | 0.13 | 0.19 | 0.17 |
| 3 | 127445938 | rs16826716 | 1.307 | -0.117 | -0.639 | 0.071 | 0.036 | -0.013 | -0.013 | 0 | 0.07 | 0.24 | 0.08 | 0.02 | 0.09 | 0.1 |
| 3 | 127447884 | rs11713989 | 0.2 | 0.05 | -1.532 | 0.198 | -0.059 | 0.076 | 0.076 | 0.288 | 0.34 | 0.13 | 0.37 | 0.21 | 0.42 | 0.22 |
| 3 | 127448986 | rs594323 | 0.413 | 0.06 | -1.824 | 0.233 | -0.033 | 0.035 | 0.035 | 0.005 | 0.58 | 0.77 | 0.46 | 0.58 | 0.49 | 0.56 |
| 3 | 127449008 | rs497423 | -0.039 | 0.034 | -1.049 | 0.647 | -0.006 | 0.018 | 0.018 | 0 | 0.91 | 0.89 | 0.83 | 0.79 | 0.91 | 0.78 |
| 3 | 127463608 | rs6787155 | 0.375 | -0.012 | -1.778 | 0.431 | 0.005 | 0.017 | 0.017 | 0 | 0.77 | 0.86 | 0.77 | 0.76 | 0.79 | 0.71 |
| 3 | 127469895 | rs605188 | 0.459 | 0.13 | -1.682 | 0.948 | 0.019 | 0.004 | 0.004 | 1.512 | 0.55 | 0.39 | 0.64 | 0.41 | 0.61 | 0.58 |
| 3 | 127470166 | rs7651814 | -0.301 | 0.015 | -0.863 | 1.176 | -0.03 | 0.017 | 0.017 | 2.231 | 0.22 | 0.59 | 0.14 | 0.26 | 0.15 | 0.15 |
| 3 | 127473488 | rs6778770 | 0.759 | 0.472 | -1.483 | 0.96 | -0.007 | 0.019 | 0.019 | 0 | 0.31 | 0.05 | 0.17 | 0.25 | 0.25 | 0.22 |
| 3 | 127474030 | rs664910 | 0.317 | 0.808 | -0.264 | 1.393 | -0.009 | -0.012 | -0.012 | 0.074 | 0.71 | 0.47 | 0.79 | 0.31 | 0.72 | 0.6 |
| 3 | 127474541 | rs17282181 | -0.17 | 0.514 | -0.729 | 1.234 | -0.063 | -0.018 | -0.018 | 0 | 0.18 | 0.03 | 0.33 | 0.17 | 0.25 | 0.29 |
| 3 | 127475626 | rs7653663 | -0.435 | 0.194 | 0.38 | 1.81 | -0.028 | 0.112 | 0.112 | 0 | 0.23 | 0.44 | 0.14 | 0.39 | 0.17 | 0.31 |
| 3 | 127480473 | rs7652615 | 0.673 | 0.6 | 1.315 | 2.819 | -0.006 | 0.063 | 0.063 | 0 | 0.8 | 0.65 | 0.87 | 0.68 | 0.84 | 0.68 |
| 3 | 127484757 | rs608318 | 0.49 | 0.858 | 0.517 | 2.702 | -0.022 | 0.052 | 0.052 | 0 | 0.81 | 0.66 | 0.8 | 0.71 | 0.86 | 0.83 |
| 3 | 127491331 | rs13066225 | -0.894 | 1.748 | 0.033 | 2.359 | -0.019 | -0.016 | -0.016 | 0 | 0.13 | 0.06 | 0.23 | 0.23 | 0.18 | 0.24 |
| 3 | 127493864 | rs567384 | 1.316 | 1.642 | 0.618 | 2.91 | 0.015 | 0.135 | 0.135 | 0 | 0.93 | 0.9 | 0.79 | 0.57 | 0.88 | 0.74 |
| 3 | 127495915 | rs587779 | 1.372 | 1.192 | 1.433 | 2.427 | -0.006 | 0.093 | 0.093 | 0 | 0.83 | 0.67 | 0.84 | 0.67 | 0.85 | 0.69 |
| 3 | 127496076 | rs541855 | 1.939 | 1.199 | 2.21 | 2.419 | 0.016 | 0.095 | 0.095 | 0 | 0.8 | 0.44 | 0.78 | 0.66 | 0.8 | 0.53 |
| 3 | 127500291 | rs3773161 | 1.279 | 1.194 | 2.464 | 2.397 | -0.002 | 0.16 | 0.16 | 0 | 0.95 | 0.86 | 0.94 | 0.73 | 0.98 | 0.93 |

| | | | | | | | | | | | | | | | | |
|---|---|---|---|---|---|---|---|---|---|---|---|---|---|---|---|---|
| 3 | 127502549 | rs9846048 | -2.531 | 1.026 | 2.317 | 2.324 | 0.002 | 0.06 | 0.06 | 0 | 0.21 | 0.58 | 0.22 | 0.35 | 0.2 | 0.47 |
| 3 | 127502920 | rs9289316 | -0.725 | 1.031 | 1.716 | 1.81 | -0.018 | 0.036 | 0.036 | 0 | 0.1 | 0.02 | 0.09 | 0.04 | 0.12 | 0.16 |
| 3 | 127515984 | rs11715363 | -1.891 | 0.878 | 1.684 | 1.601 | -0.01 | 0 | 0 | 1.798 | 0.25 | 0.34 | 0.16 | 0.24 | 0.18 | 0.29 |
| 3 | 127518962 | rs1474332 | -1.351 | 0.954 | 1.529 | 1.648 | 0 | 0.014 | 0.014 | 0 | 0.16 | 0.24 | 0.14 | 0.25 | 0.15 | 0.28 |
| 3 | 127521678 | rs9825729 | -1.254 | 0.961 | 1.375 | 1.651 | 0 | 0.001 | 0.001 | 0 | 0.16 | 0.17 | 0.14 | 0.24 | 0.15 | 0.28 |
| 3 | 127538412 | rs874546 | -1.806 | 0.875 | 1.423 | 1.568 | -0.001 | 0 | 0 | 3.862 | 0.27 | 0.7 | 0.19 | 0.24 | 0.18 | 0.33 |
| 3 | 127551630 | rs11716997 | -1.229 | 1.005 | 0.508 | 0.965 | 0.002 | -0.017 | -0.017 | 0 | 0.59 | 0.74 | 0.5 | 0.51 | 0.61 | 0.75 |
| 3 | 127568933 | rs10439996 | 0.144 | 0.944 | 0.575 | 1.015 | 0.001 | -0.006 | -0.006 | 0.134 | 0.38 | 0.23 | 0.34 | 0.39 | 0.41 | 0.49 |
| 3 | 127577789 | rs4974416 | -1.021 | 0.715 | 0.42 | 0.987 | -0.034 | -0.018 | -0.018 | 0 | 0.56 | 0.82 | 0.47 | 0.49 | 0.49 | 0.7 |
| 3 | 127590236 | rs6439086 | -1.461 | 0.684 | 1.148 | 0.86 | 0.021 | -0.054 | -0.054 | 0 | 0.18 | 0.48 | 0.14 | 0.1 | 0.14 | 0.23 |
| 3 | 127594748 | rs11926127 | -1.115 | 0.956 | 0.345 | 0.661 | 0.01 | -0.057 | -0.057 | 0 | 0.15 | 0.22 | 0.11 | 0.05 | 0.14 | 0.21 |
| 3 | 127611229 | rs4577421 | -1.359 | 0.783 | -0.114 | 1.169 | 0.004 | 0.004 | 0.004 | 0 | 0.07 | 0.1 | 0.04 | 0.16 | 0.09 | 0.06 |
| 3 | 127617605 | rs7634887 | 0.994 | -0.162 | 0.104 | 1.263 | -0.029 | 0.012 | 0.012 | 0 | 0.72 | 0.38 | 0.85 | 0.72 | 0.77 | 0.68 |
| 3 | 127618032 | rs6439093 | 0.757 | -0.342 | 0.797 | 1.448 | -0.019 | 0.101 | 0.101 | 0 | 0.92 | 0.79 | 0.95 | 0.85 | 0.95 | 0.91 |
| 3 | 127622736 | rs891885 | -0.798 | -0.276 | -0.025 | 1.239 | -0.036 | -0.013 | -0.013 | 0.024 | 0.68 | 0.78 | 0.48 | 0.67 | 0.57 | 0.77 |
| 3 | 127627327 | rs9865083 | 0.595 | -0.95 | 0.171 | 0.954 | -0.002 | 0.01 | 0.01 | 0 | 0.35 | 0.34 | 0.34 | 0.18 | 0.34 | 0.31 |
| 3 | 127631561 | rs6439099 | -0.2 | -1.164 | 0.677 | 1.156 | -0.006 | 0.103 | 0.103 | 0 | 0.1 | 0.18 | 0.05 | 0.25 | 0.1 | 0.13 |
| 3 | 127633543 | rs13073655 | 0.726 | -1.338 | 0.996 | 1.129 | -0.008 | 0.031 | 0.031 | 0 | 0.34 | 0.26 | 0.32 | 0.19 | 0.34 | 0.31 |
| 3 | 127639072 | rs16838982 | -0.549 | -1.44 | 1.199 | 1.2 | -0.02 | 0.019 | 0.019 | 0 | 0.24 | 0.26 | 0.11 | 0.39 | 0.17 | 0.4 |
| 3 | 127641368 | rs16839044 | 0.427 | -1.408 | 1.069 | 1.215 | -0.007 | 0.011 | 0.011 | 0 | 0.07 | 0.1 | 0.03 | 0.1 | 0.06 | 0.06 |
| 3 | 127641968 | rs4141499 | -1.798 | -1.415 | 1.083 | 1.214 | -0.083 | -0.039 | -0.039 | 0 | 0.67 | 0.85 | 0.49 | 0.67 | 0.57 | 0.77 |
| 3 | 127651629 | rs2217628 | -1.073 | -1.503 | 1.175 | 0.971 | -0.026 | -0.003 | -0.003 | 0 | 0.29 | 0.3 | 0.11 | 0.32 | 0.16 | 0.39 |
| 3 | 127655873 | rs2878603 | 0.694 | -1.534 | 1.346 | 0.939 | -0.002 | 0.007 | 0.007 | 0 | 0.35 | 0.32 | 0.34 | 0.19 | 0.34 | 0.31 |
| 3 | 127662082 | rs6766308 | 1.368 | -1.575 | 0.99 | 0.784 | -0.087 | 0.031 | 0.031 | 0 | 0.25 | 0.02 | 0.19 | 0.11 | 0.26 | 0.19 |
| 3 | 127663013 | rs4857882 | -1.874 | -1.549 | 1.058 | 0.799 | -0.043 | -0.043 | -0.043 | 0 | 0.65 | 0.78 | 0.47 | 0.51 | 0.53 | 0.7 |
| 3 | 127669950 | rs9830149 | 0.593 | -1.508 | 1.107 | 0.816 | -0.006 | 0.009 | 0.009 | 0 | 0.07 | 0.09 | 0.03 | 0.1 | 0.06 | 0.06 |
| 3 | 127684748 | rs17203687 | 1.662 | -1.45 | 0.969 | 0.819 | -0.023 | -0.01 | -0.01 | 0 | 0.05 | 0 | 0.1 | 0.07 | 0.08 | 0.06 |
| 3 | 127685088 | rs13066952 | 0.28 | -1.527 | 1.075 | 0.804 | 0.002 | -0.027 | -0.027 | 0 | 0.35 | 0.42 | 0.34 | 0.19 | 0.34 | 0.3 |
| 3 | 127689918 | rs7633735 | -1.202 | -1.456 | 1.072 | 0.946 | -0.014 | 0.011 | 0.011 | 0 | 0.58 | 0.68 | 0.44 | 0.42 | 0.46 | 0.64 |
| 3 | 127693417 | rs16839633 | -1.372 | -1.682 | 1.268 | 1.292 | -0.08 | 0.069 | 0.069 | 0.02 | 0.22 | 0.53 | 0.11 | 0.33 | 0.11 | 0.33 |
| 3 | 127694318 | rs16839641 | 0.04 | -1.526 | 0.822 | 1.201 | -0.028 | 0 | 0 | 0 | 0.13 | 0.2 | 0.05 | 0.2 | 0.05 | 0.24 |
| 3 | 127694583 | rs16839675 | -0.015 | -0.797 | 0.379 | 1.341 | -0.006 | 0.02 | 0.02 | 0 | 0.07 | 0.09 | 0.03 | 0.12 | 0.07 | 0.04 |
| 3 | 127701413 | rs6798286 | -0.392 | -0.584 | 0.035 | 0.976 | -0.002 | 0.007 | 0.007 | 0 | 0.31 | 0.27 | 0.26 | 0.18 | 0.31 | 0.38 |
| 3 | 127703085 | rs13100951 | 0.007 | -0.132 | -0.356 | 1.397 | -0.023 | -0.03 | -0.03 | 0.02 | 0.19 | 0.12 | 0.3 | 0.29 | 0.26 | 0.18 |
| 3 | 127709698 | rs2878602 | 0.728 | -0.383 | -0.415 | 0.908 | -0.004 | 0.003 | 0.003 | 0.476 | 0.7 | 0.42 | 0.57 | 0.31 | 0.75 | 0.72 |
| 3 | 127710292 | rs4857926 | -0.626 | 0.14 | -0.755 | 0.637 | -0.004 | -0.018 | -0.018 | 0 | 0.65 | 0.7 | 0.63 | 0.6 | 0.63 | 0.65 |
| 3 | 127714611 | rs1549876 | -0.577 | -0.341 | -1.083 | 0.232 | 0.053 | -0.012 | -0.012 | 0 | 0.5 | 0.79 | 0.44 | 0.56 | 0.42 | 0.51 |
| 3 | 127720740 | rs1469659 | 0.31 | -1.276 | -0.542 | 0.214 | -0.053 | 0.06 | 0.06 | 0 | 0.25 | 0.54 | 0.25 | 0.47 | 0.16 | 0.24 |
| 3 | 127727752 | rs11717102 | 0.82 | -1.026 | -0.841 | 0.01 | 0.018 | 0.04 | 0.04 | 0 | 0.55 | 0.66 | 0.63 | 0.81 | 0.61 | 0.54 |
| 3 | 127747841 | rs6764517 | -1.039 | -1.026 | -0.905 | 0.036 | 0.028 | 0.025 | 0.025 | 0 | 0.8 | 0.93 | 0.82 | 0.9 | 0.85 | 0.8 |
| 3 | 127760006 | rs2053014 | 0.041 | -1.07 | -0.849 | -0.032 | -0.002 | -0.02 | -0.02 | 0 | 0.25 | 0.26 | 0.19 | 0.09 | 0.24 | 0.26 |
| 3 | 127773676 | rs1030655 | 0.857 | -1.091 | -0.895 | 0.029 | 0.027 | 0.06 | 0.06 | 0 | 0.56 | 0.73 | 0.65 | 0.8 | 0.61 | 0.54 |
| 3 | 127779633 | rs9849210 | 0.029 | -1.079 | -0.811 | 0.033 | -0.001 | -0.009 | -0.009 | 0 | 0.24 | 0.23 | 0.18 | 0.09 | 0.24 | 0.26 |
| 3 | 127780082 | rs759945 | -1.344 | -1.114 | -0.828 | 0.018 | 0.021 | 0.009 | 0.009 | 0 | 0.81 | 0.96 | 0.82 | 0.88 | 0.85 | 0.79 |
| 3 | 127788955 | rs10934847 | -0.036 | -0.975 | -0.726 | 0.056 | 0 | -0.006 | -0.006 | 0 | 0.24 | 0.22 | 0.18 | 0.09 | 0.24 | 0.26 |
| 3 | 127789695 | rs1042912 | -0.03 | -0.98 | -0.728 | 0.054 | -0.001 | -0.009 | -0.009 | 0 | 0.24 | 0.23 | 0.18 | 0.09 | 0.24 | 0.26 |
| 3 | 127799155 | rs9860614 | 1.092 | -1.052 | -0.835 | -0.043 | -0.042 | -0.027 | -0.027 | 1.277 | 0.1 | 0.01 | 0.05 | 0.08 | 0.12 | 0.09 |
| 3 | 127799983 | rs1057220 | -0.324 | -1.038 | -0.812 | -0.027 | 0 | -0.006 | -0.006 | 0 | 0.24 | 0.21 | 0.18 | 0.09 | 0.24 | 0.26 |
| 3 | 127801896 | rs1007232 | -0.346 | -1.033 | -0.811 | -0.025 | -0.001 | -0.009 | -0.009 | 0 | 0.24 | 0.23 | 0.18 | 0.09 | 0.24 | 0.26 |
| 3 | 127803922 | rs11706281 | -0.346 | -1.033 | -0.814 | -0.025 | -0.001 | -0.008 | -0.008 | 0.353 | 0.24 | 0.22 | 0.18 | 0.09 | 0.24 | 0.26 |
| 3 | 127804368 | rs11714256 | -0.346 | -1.033 | -0.814 | -0.025 | 0 | -0.006 | -0.006 | 0 | 0.24 | 0.22 | 0.18 | 0.09 | 0.24 | 0.26 |
| 3 | 127826618 | rs11924142 | -0.846 | -1.206 | -0.592 | 0.246 | 0.036 | -0.077 | -0.077 | 0 | 0.79 | 0.91 | 0.73 | 0.53 | 0.85 | 0.75 |
| 3 | 127834003 | rs7650365 | 1.949 | -1.372 | -1.061 | 0.04 | -0.18 | -0.12 | -0.12 | 0 | 0.32 | 0.02 | 0.41 | 0.37 | 0.48 | 0.26 |
| 3 | 127872175 | rs11920225 | 1.062 | -1.252 | -0.665 | -0.073 | 0.022 | 0.016 | 0.016 | 0 | 0.2 | 0.11 | 0.19 | 0.1 | 0.15 | 0.2 |
| 3 | 127886957 | rs10934850 | 0.851 | -1.315 | -0.658 | -0.119 | -0.012 | -0.06 | -0.06 | 0 | 0.13 | 0.21 | 0.13 | 0.01 | 0.11 | 0.16 |
| 3 | 127888694 | rs2687731 | 1.214 | -1.328 | -0.721 | -0.124 | 0.024 | 0.015 | 0.015 | 0 | 0.19 | 0.05 | 0.18 | 0.1 | 0.15 | 0.21 |
| 3 | 127889375 | rs3122174 | 1.11 | -1.323 | -0.723 | -0.122 | 0.024 | 0.018 | 0.018 | 0 | 0.2 | 0.1 | 0.19 | 0.1 | 0.15 | 0.21 |
| 3 | 127892037 | rs2999052 | -0.05 | -1.207 | -0.665 | -0.049 | -0.014 | -0.028 | -0.028 | 0 | 0.33 | 0.35 | 0.32 | 0.11 | 0.26 | 0.37 |
| 3 | 127911077 | rs17401543 | 1.782 | -1.052 | -0.918 | -0.058 | -0.039 | 0.046 | 0.046 | 0 | 0.11 | 0.01 | 0.07 | 0.02 | 0.12 | 0.1 |
| 3 | 127929120 | rs4293718 | -0.85 | -1.395 | -0.898 | 0.075 | -0.003 | 0.005 | 0.005 | 0.249 | 0.34 | 0.65 | 0.28 | 0.53 | 0.26 | 0.37 |
| 3 | 127931340 | rs2999090 | -1.33 | -1.57 | -0.726 | -0.135 | -0.003 | -0.035 | -0.035 | 0.325 | 0.89 | 0.88 | 0.88 | 0.99 | 0.89 | 0.83 |
| 3 | 127937326 | rs2254379 | 1.562 | -1.668 | -0.955 | -0.182 | 0.019 | 0.01 | 0.01 | 0 | 0.19 | 0.04 | 0.18 | 0.1 | 0.15 | 0.21 |
| 3 | 127965570 | rs10512698 | 2.085 | -1.604 | -0.923 | -0.186 | -0.039 | 0.046 | 0.046 | 0 | 0.11 | 0.01 | 0.07 | 0.02 | 0.12 | 0.1 |
| 3 | 127977978 | rs2999068 | -0.465 | -1.654 | -0.343 | -0.183 | -0.018 | -0.086 | -0.086 | 0 | 0.85 | 0.7 | 0.86 | 0.99 | 0.88 | 0.83 |
| 3 | 127983495 | rs2811544 | -0.484 | -1.648 | -0.34 | -0.181 | -0.017 | -0.085 | -0.085 | 0 | 0.86 | 0.7 | 0.86 | 0.99 | 0.88 | 0.83 |
| 3 | 127991527 | rs2811415 | 0.099 | -1.7 | -0.411 | -0.26 | -0.019 | -0.104 | -0.104 | 1.158 | 0.8 | 0.69 | 0.84 | 0.99 | 0.84 | 0.82 |
| 3 | 128015236 | rs2955103 | 0.971 | -1.733 | -0.413 | -0.096 | 0.059 | 0.004 | 0.004 | 0.013 | 0.49 | 0.1 | 0.48 | 0.49 | 0.61 | 0.49 |
| 3 | 128038373 | rs10934853 | -0.652 | -1.627 | -0.173 | -0.097 | 0.062 | -0.026 | -0.026 | 0 | 0.38 | 0.78 | 0.45 | 0.49 | 0.28 | 0.47 |
| 3 | 128038834 | rs12486156 | 1.591 | -1.756 | -0.215 | -0.128 | -0.002 | -0.017 | -0.017 | 0 | 0.19 | 0.03 | 0.19 | 0.1 | 0.16 | 0.27 |

| | | | | | | | | | | | | | | | | |
|---|---|---|---|---|---|---|---|---|---|---|---|---|---|---|---|---|
| 3 | 128046643 | rs4857841 | -0.757 | -1.768 | -0.173 | -0.125 | 0.062 | -0.026 | -0.026 | 0.026 | 0.38 | 0.78 | 0.45 | 0.49 | 0.28 | 0.47 |
| 3 | 128053487 | rs16844002 | 0.357 | -1.78 | 0.004 | -0.139 | -0.016 | -0.092 | -0.092 | 0 | 0.14 | 0.36 | 0.15 | 0.01 | 0.11 | 0.16 |
| 3 | 128059610 | rs1735558 | 1.535 | -1.751 | -0.018 | -0.124 | 0.003 | -0.016 | -0.016 | 3.271 | 0.19 | 0.03 | 0.19 | 0.1 | 0.16 | 0.27 |
| 3 | 128064118 | rs4857879 | 0.605 | -1.727 | 0.07 | -0.134 | -0.044 | 0.088 | 0.088 | 0 | 0.24 | 0.45 | 0.31 | 0.48 | 0.17 | 0.33 |
| 3 | 128066610 | rs1735557 | -2.24 | -1.634 | -0.221 | -0.14 | -0.036 | 0.046 | 0.046 | 0 | 0.89 | 0.99 | 0.93 | 0.98 | 0.88 | 0.9 |
| 3 | 128075398 | rs1735546 | -0.631 | -1.733 | -0.21 | -0.137 | 0.088 | -0.04 | -0.04 | 0 | 0.38 | 0.81 | 0.44 | 0.49 | 0.25 | 0.47 |
| 3 | 128085090 | rs1108313 | 1.906 | -1.531 | -0.401 | -0.133 | -0.18 | -0.128 | -0.128 | 0 | 0.35 | 0.02 | 0.4 | 0.37 | 0.47 | 0.3 |
| 3 | 128092102 | rs1735538 | -0.464 | -1.399 | -0.153 | -0.212 | 0.077 | -0.038 | -0.038 | 0.16 | 0.38 | 0.8 | 0.45 | 0.49 | 0.25 | 0.42 |
| 3 | 128095652 | rs3021461 | 0.201 | -1.525 | 0.127 | -0.276 | -0.026 | 0.03 | 0.03 | 0 | 0.29 | 0.4 | 0.38 | 0.49 | 0.22 | 0.37 |
| 3 | 128097496 | rs2293947 | 0.963 | -1.484 | 0.088 | -0.226 | -0.009 | -0.015 | -0.015 | 0 | 0.06 | 0.02 | 0.16 | 0.38 | 0.07 | 0.17 |
| 3 | 128104562 | rs6806480 | 0.583 | -1.511 | 0.178 | -0.332 | 0.024 | -0.015 | -0.015 | 1.587 | 0.08 | 0.39 | 0.06 | 0 | 0.04 | 0.02 |
| 3 | 128106283 | rs1010786 | -2.192 | -1.429 | -0.046 | -0.339 | -0.037 | 0.048 | 0.048 | 0 | 0.89 | 0.99 | 0.92 | 0.98 | 0.86 | 0.9 |
| 3 | 128111201 | rs1702134 | 0.994 | -1.57 | 0.036 | -0.315 | -0.003 | -0.001 | -0.001 | 1.926 | 0.21 | 0.25 | 0.21 | 0.1 | 0.13 | 0.22 |
| 3 | 128112307 | rs1620440 | 1.08 | -1.566 | -0.067 | -0.313 | 0.012 | 0.004 | 0.004 | 0 | 0.2 | 0.2 | 0.2 | 0.1 | 0.13 | 0.22 |
| 3 | 128117121 | rs1735562 | -1.528 | -1.49 | -0.36 | -0.334 | -0.014 | 0.048 | 0.048 | 0 | 0.84 | 0.91 | 0.91 | 0.98 | 0.84 | 0.89 |
| 3 | 128119874 | rs11705709 | 1.197 | -1.022 | -0.147 | -0.151 | -0.02 | -0.009 | -0.009 | 0 | 0.06 | 0.03 | 0.16 | 0.38 | 0.09 | 0.13 |
| 3 | 128120355 | rs11705891 | 0.679 | -1.024 | -0.144 | -0.152 | 0.029 | 0.138 | 0.138 | 0 | 0.07 | 0.16 | 0.17 | 0.38 | 0.09 | 0.16 |
| 3 | 128122820 | rs1735537 | 0.008 | -1.091 | -0.423 | -0.253 | -0.024 | 0.023 | 0.023 | 0 | 0.31 | 0.57 | 0.4 | 0.48 | 0.23 | 0.39 |
| 3 | 128123786 | rs2977564 | 2.21 | -1.393 | -1.517 | -0.154 | -0.165 | 0.003 | 0.003 | 0.036 | 0.24 | 0.02 | 0.35 | 0.17 | 0.41 | 0.14 |
| 3 | 128127643 | rs2939820 | 0.072 | -1.619 | -1.491 | -0.248 | -0.037 | -0.109 | -0.109 | 0.055 | 0.28 | 0.1 | 0.27 | 0.46 | 0.25 | 0.42 |
| 3 | 128128254 | rs3828417 | -0.982 | -1.353 | -1.118 | -0.615 | 0.005 | -0.007 | -0.007 | 0 | 0.2 | 0.76 | 0.25 | 0.4 | 0.08 | 0.14 |
| 3 | 128133108 | rs2811366 | -1.491 | -0.849 | -1.462 | -0.525 | -0.191 | 0.236 | 0.236 | 0 | 0.68 | 0.98 | 0.73 | 1 | 0.5 | 0.87 |
| 3 | 128138198 | rs4521245 | -0.637 | -0.616 | -0.167 | 0.413 | -0.016 | 0.167 | 0.167 | 0 | 0.09 | 0.27 | 0.17 | 0.4 | 0.04 | 0.15 |
| 3 | 128148525 | rs7644598 | -1.056 | -0.797 | -0.217 | 0.155 | -0.02 | 0.032 | 0.032 | 0 | 0.27 | 0.4 | 0.28 | 0.51 | 0.18 | 0.35 |
| 3 | 128149590 | rs2981030 | 0.317 | -1.344 | -0.466 | -0.351 | 0 | -0.001 | -0.001 | 0 | 0.32 | 0.32 | 0.43 | 0.47 | 0.28 | 0.45 |
| 3 | 128152714 | rs1962408 | -0.253 | -1.725 | -0.422 | -0.702 | -0.006 | 0.013 | 0.013 | 0 | 0.37 | 0.28 | 0.49 | 0.59 | 0.35 | 0.54 |
| 3 | 128159359 | rs2713585 | -0.491 | -2.041 | -0.121 | -0.86 | -0.008 | -0.071 | -0.071 | 0 | 0.41 | 0.48 | 0.59 | 0.87 | 0.36 | 0.55 |
| 3 | 128162235 | rs12491486 | 0.435 | -1.945 | -0.523 | -0.775 | -0.016 | -0.017 | -0.017 | 0 | 0.31 | 0.2 | 0.44 | 0.52 | 0.31 | 0.41 |
| 3 | 128163890 | rs9855048 | 0.562 | -1.678 | -0.225 | -0.719 | -0.004 | -0.002 | -0.002 | 0 | 0.08 | 0.07 | 0.04 | 0.08 | 0.04 | 0.17 |
| 3 | 128168267 | rs13061811 | 0.633 | -1.975 | -0.739 | -0.668 | 0.005 | -0.021 | -0.021 | 0.416 | 0.4 | 0.12 | 0.43 | 0.53 | 0.34 | 0.45 |
| 3 | 128168298 | rs6781634 | 0.948 | -2.109 | -0.814 | -0.994 | -0.076 | 0.11 | 0.11 | 0 | 0.34 | 0.16 | 0.29 | 0.1 | 0.51 | 0.33 |
| 3 | 128169862 | rs2659703 | 0.084 | -2.14 | -0.273 | -1.006 | 0.002 | -0.028 | -0.028 | 2.338 | 0.51 | 0.52 | 0.55 | 0.89 | 0.39 | 0.52 |
| 3 | 128170341 | rs2953130 | -0.141 | -2.228 | 0.076 | -0.775 | -0.018 | -0.029 | -0.029 | 0 | 0.88 | 0.72 | 0.84 | 0.97 | 0.9 | 0.85 |
| 3 | 128172058 | rs4857895 | 1.382 | -2.379 | 0.082 | -0.74 | 0.012 | -0.008 | -0.008 | 0 | 0.67 | 0.37 | 0.7 | 0.53 | 0.82 | 0.61 |
| 3 | 128172844 | rs2001309 | 0.394 | -2.18 | -0.195 | -1.206 | 0 | 0 | 0 | 0 | 0.79 | 0.8 | 0.8 | 0.89 | 0.85 | 0.7 |
| 3 | 128174018 | rs9880311 | -0.154 | -1.789 | -0.169 | -0.934 | 0.002 | 0.005 | 0.005 | 0 | 0.48 | 0.54 | 0.4 | 0.32 | 0.56 | 0.42 |
| 3 | 128174742 | rs11713004 | 1.497 | -1.673 | -0.369 | -1.047 | -0.104 | 0.121 | 0.121 | 0 | 0.27 | 0.11 | 0.25 | 0.02 | 0.47 | 0.18 |
| 3 | 128177123 | rs9850678 | -0.222 | -1.557 | -0.191 | -0.872 | -0.056 | -0.011 | -0.011 | 0 | 0.21 | 0.44 | 0.1 | 0.24 | 0.09 | 0.23 |
| 3 | 128177991 | rs2659694 | 0.056 | -1.446 | -0.738 | -0.729 | 0.005 | 0.111 | 0.111 | 0 | 0.47 | 0.22 | 0.56 | 0.66 | 0.44 | 0.57 |
| 3 | 128182074 | rs2981026 | 0.081 | -1.043 | -0.846 | 0.155 | -0.009 | -0.08 | -0.08 | 0 | 0.3 | 0.13 | 0.36 | 0.49 | 0.28 | 0.36 |
| 3 | 128182378 | rs13071640 | 0.337 | 0.328 | -0.13 | 0.065 | 0.005 | -0.001 | -0.001 | 0 | 0.15 | 0.27 | 0.16 | 0.22 | 0.16 | 0.17 |
| 3 | 128185399 | rs11713343 | -0.704 | 0.312 | -0.056 | -0.059 | -0.006 | -0.038 | -0.038 | 0 | 0.23 | 0.64 | 0.2 | 0.07 | 0.19 | 0.18 |
| 3 | 128185532 | rs12634139 | 0.645 | 0.094 | 0.317 | 0.084 | -0.01 | -0.006 | -0.006 | 4.862 | 0.75 | 0.32 | 0.74 | 0.71 | 0.81 | 0.7 |
| 3 | 128188343 | rs2981013 | -0.591 | 0.573 | -0.081 | 0 | 0.006 | -0.023 | -0.023 | 0 | 0.78 | 0.6 | 0.8 | 0.78 | 0.84 | 0.84 |
| 3 | 128189572 | rs2953128 | -0.381 | 0.565 | -1.281 | 0.234 | 0.005 | -0.005 | -0.005 | 0 | 0.53 | 0.72 | 0.44 | 0.54 | 0.44 | 0.58 |
| 3 | 128190941 | rs7431368 | -0.733 | 0.452 | -0.913 | 0.511 | -0.002 | 0.004 | 0.004 | 0 | 0.8 | 0.81 | 0.83 | 0.78 | 0.84 | 0.85 |
| 3 | 128192653 | rs10934854 | -0.226 | 0.469 | -1.322 | 0.486 | 0.002 | 0.004 | 0.004 | 0 | 0.73 | 0.39 | 0.72 | 0.64 | 0.78 | 0.79 |
| 3 | 128192862 | rs10934856 | -0.028 | 0.193 | -1.771 | 0.595 | 0.01 | -0.085 | -0.085 | 0 | 0.29 | 0.06 | 0.18 | 0.35 | 0.22 | 0.34 |
| 3 | 128195814 | rs11718692 | -0.021 | 0.777 | -1.18 | 0.191 | -0.004 | 0.004 | 0.004 | 0 | 0.16 | 0.16 | 0.13 | 0.06 | 0.19 | 0.06 |
| 3 | 128196500 | rs2713594 | 0.3 | 0.845 | -1.111 | 0.334 | 0.009 | -0.001 | -0.001 | 0 | 0.53 | 0.56 | 0.51 | 0.53 | 0.45 | 0.53 |
| 3 | 128197282 | rs1573949 | -0.979 | 1.092 | -1.145 | 0.416 | 0.001 | 0.001 | 0.001 | 0.423 | 0.75 | 0.77 | 0.72 | 0.67 | 0.71 | 0.61 |
| 3 | 128199380 | rs3803 | 0.702 | 1.111 | -1.131 | 0.395 | -0.005 | 0.007 | 0.007 | 0 | 0.16 | 0.16 | 0.13 | 0.06 | 0.2 | 0.06 |
| 3 | 128200459 | rs2713604 | -0.713 | 0.969 | -1.225 | 0.364 | -0.003 | -0.003 | -0.003 | 0 | 0.73 | 0.78 | 0.69 | 0.68 | 0.7 | 0.57 |
| 3 | 128204951 | rs2335052 | 0.648 | 0.028 | -0.838 | 0.468 | -0.004 | 0.017 | 0.017 | 0 | 0.2 | 0.22 | 0.24 | 0.38 | 0.18 | 0.15 |
| 3 | 128209667 | rs2860228 | 0.074 | 0.047 | -0.543 | 0.292 | 0.019 | 0.041 | 0.041 | 0.052 | 0.5 | 0.45 | 0.4 | 0.24 | 0.43 | 0.44 |
| 3 | 128212526 | rs9851497 | -1.537 | 0.066 | -0.608 | 0.235 | -0.007 | -0.004 | -0.004 | 0 | 0.75 | 0.82 | 0.71 | 0.7 | 0.67 | 0.62 |
| 3 | 128218410 | rs7629791 | -0.486 | 0.068 | -0.639 | 0.073 | 0 | 0.002 | 0.002 | 0.104 | 0.55 | 0.62 | 0.47 | 0.31 | 0.49 | 0.47 |
| 3 | 128222777 | rs7628169 | 0.662 | -0.559 | -0.216 | 0.025 | 0.005 | 0.017 | 0.017 | 0 | 0.13 | 0.07 | 0.07 | 0 | 0.12 | 0.02 |
| 3 | 128229571 | rs2659695 | -1.197 | -0.711 | 0.042 | 0.236 | 0.009 | -0.002 | -0.002 | 0 | 0.44 | 0.61 | 0.42 | 0.49 | 0.4 | 0.19 |
| 3 | 128240954 | rs7433900 | -0.452 | -1.063 | 0.443 | 0.102 | -0.009 | 0.012 | 0.012 | 0.597 | 0.27 | 0.28 | 0.32 | 0.42 | 0.23 | 0.17 |
| 3 | 128241281 | rs4613470 | -1.34 | -0.988 | 0.239 | 0.085 | 0.009 | 0.002 | 0.002 | 1.056 | 0.43 | 0.62 | 0.42 | 0.42 | 0.37 | 0.19 |
| 3 | 128243631 | rs6769565 | -1.254 | -1.241 | 0.243 | 0.113 | 0.006 | 0 | 0 | 1.151 | 0.42 | 0.62 | 0.41 | 0.42 | 0.34 | 0.19 |
| 3 | 128245696 | rs9819395 | -0.145 | -1.237 | 0.373 | 0.286 | -0.015 | 0.065 | 0.065 | 0.771 | 0.21 | 0.33 | 0.29 | 0.4 | 0.19 | 0.16 |
| 3 | 128248399 | rs7629705 | -0.809 | -1.373 | 0.424 | 0.228 | -0.013 | 0.01 | 0.01 | 0 | 0.33 | 0.45 | 0.36 | 0.45 | 0.28 | 0.29 |
| 3 | 128257214 | rs9854612 | -1.253 | -1.503 | 0.621 | 0.159 | 0.01 | -0.002 | -0.002 | 0 | 0.41 | 0.68 | 0.4 | 0.45 | 0.32 | 0.29 |
| 3 | 128260550 | rs4857855 | 0.153 | -1.499 | 0.351 | 0.329 | 0.004 | -0.035 | -0.035 | 0.07 | 0.19 | 0.13 | 0.24 | 0.33 | 0.19 | 0.23 |
| 3 | 128272481 | rs7635061 | -1.449 | -1.651 | 0.882 | 0.21 | -0.012 | 0.001 | 0.001 | 0.464 | 0.4 | 0.61 | 0.36 | 0.44 | 0.31 | 0.27 |
| 3 | 128276332 | rs9824329 | -0.104 | -1.77 | 0.883 | 0.15 | -0.051 | -0.109 | -0.109 | 0.004 | 0.17 | 0.48 | 0.09 | 0.02 | 0.1 | 0.07 |
| 3 | 128278803 | rs2335050 | -0.012 | -1.442 | 0.467 | 0.413 | 0.008 | -0.07 | -0.07 | 0 | 0.19 | 0.11 | 0.25 | 0.43 | 0.19 | 0.23 |

| | | | | | | | | | | | | | | | | |
|---|---|---|---|---|---|---|---|---|---|---|---|---|---|---|---|---|
| 3 | 128281563 | rs12495912 | -0.665 | -1.508 | 0.589 | 0.378 | -0.038 | 0.006 | 0.006 | 0.885 | 0.09 | 0.27 | 0.06 | 0.08 | 0.04 | 0.02 |
| 3 | 128282697 | rs6806253 | -0.153 | -1.253 | 0.558 | 0.357 | 0.009 | -0.014 | -0.014 | 0 | 0.2 | 0.16 | 0.22 | 0.32 | 0.18 | 0.23 |
| 3 | 128287743 | rs2712421 | -0.463 | -0.846 | 0.371 | -0.189 | 0.012 | -0.054 | -0.054 | 0 | 0.75 | 0.75 | 0.79 | 0.97 | 0.69 | 0.79 |
| 3 | 128290208 | rs2713589 | -0.227 | -0.978 | 0.091 | 0.196 | -0.023 | 0.029 | 0.029 | 0.31 | 0.2 | 0.37 | 0.21 | 0.32 | 0.16 | 0.23 |
| 3 | 128292213 | rs2734046 | -0.53 | -0.821 | -0.484 | 0.269 | -0.011 | -0.087 | -0.087 | 0.071 | 0.83 | 0.99 | 0.81 | 0.68 | 0.84 | 0.77 |
| 3 | 128294355 | rs2713575 | -0.653 | -0.912 | -0.168 | 0.034 | 0.001 | -0.003 | -0.003 | 0.275 | 0.45 | 0.55 | 0.43 | 0.44 | 0.45 | 0.53 |
| 3 | 128294897 | rs2465354 | -0.063 | -0.727 | 0.045 | 0.23 | -0.023 | 0.063 | 0.063 | 0.002 | 0.16 | 0.26 | 0.18 | 0.35 | 0.09 | 0.23 |
| 3 | 128303373 | rs6798431 | 0.901 | -1.002 | -0.215 | 0.379 | 0.006 | 0.017 | 0.017 | 1.163 | 0.39 | 0.32 | 0.23 | 0.33 | 0.38 | 0.31 |
| 3 | 128306418 | rs6772849 | 0.908 | -1.012 | -0.219 | 0.375 | 0.005 | 0.017 | 0.017 | 1.172 | 0.38 | 0.33 | 0.23 | 0.33 | 0.38 | 0.3 |
| 3 | 128311781 | rs1806722 | 0.377 | -0.986 | -0.293 | 0.19 | 0.008 | 0.035 | 0.035 | 0 | 0.24 | 0.13 | 0.2 | 0.1 | 0.23 | 0.19 |
| 3 | 128313880 | rs4857857 | 0.568 | -0.434 | 0.279 | 0.528 | 0.043 | 0.008 | 0.008 | 0 | 0.2 | 0.29 | 0.36 | 0.2 | 0.26 | 0.29 |
| 3 | 128316939 | rs9864772 | 0.179 | -0.449 | 0.196 | 0.219 | 0.03 | 0.028 | 0.028 | 0 | 0.66 | 0.52 | 0.48 | 0.44 | 0.64 | 0.49 |
| 3 | 128324161 | rs1962040 | 0.955 | -0.913 | -0.131 | -0.088 | -0.001 | 0.003 | 0.003 | 0.376 | 0.36 | 0.3 | 0.25 | 0.31 | 0.39 | 0.34 |
| 3 | 128336023 | rs13325747 | 0.615 | -1.034 | -0.039 | -0.138 | 0 | 0.003 | 0.003 | 0.084 | 0.4 | 0.38 | 0.37 | 0.34 | 0.42 | 0.34 |
| 3 | 128338748 | rs6782407 | 1.137 | -0.723 | 0.217 | -0.102 | 0.012 | -0.011 | -0.011 | 0 | 0.19 | 0.21 | 0.24 | 0.1 | 0.26 | 0.2 |
| 3 | 128344786 | rs1126828 | 0.809 | -0.909 | 0.067 | 0.109 | 0.006 | -0.088 | -0.088 | 0 | 0.25 | 0.14 | 0.2 | 0.41 | 0.21 | 0.24 |
| 3 | 128351054 | rs9868579 | 1.548 | -0.929 | 0.186 | 0.096 | -0.003 | 0.007 | 0.007 | 0.27 | 0.36 | 0.26 | 0.34 | 0.34 | 0.39 | 0.34 |
| 3 | 128354016 | rs9289331 | 0.827 | -0.807 | -0.09 | 0.161 | 0.004 | -0.088 | -0.088 | 0.478 | 0.24 | 0.14 | 0.2 | 0.41 | 0.21 | 0.24 |
| 3 | 128356852 | rs1697 | 0.292 | -0.802 | 0.111 | 0.132 | 0.012 | -0.021 | -0.021 | 0 | 0.66 | 0.62 | 0.59 | 0.75 | 0.63 | 0.57 |
| 3 | 128391789 | rs6806687 | -0.051 | -0.67 | 0.072 | 0.201 | 0.021 | -0.022 | -0.022 | 0 | 0.69 | 0.58 | 0.6 | 0.75 | 0.66 | 0.58 |
| 3 | 128404269 | rs11712115 | 0.934 | -0.652 | 0.03 | 0.163 | 0.024 | -0.022 | -0.022 | 1.585 | 0.2 | 0.29 | 0.28 | 0.1 | 0.28 | 0.2 |
| 3 | 128409680 | rs10048945 | 0.067 | -0.656 | -0.365 | 0.199 | 0.023 | -0.027 | -0.027 | 0 | 0.67 | 0.57 | 0.57 | 0.75 | 0.63 | 0.58 |
| 3 | 128412024 | rs6785206 | -1.088 | -0.511 | -1.258 | 0.304 | 0.004 | -0.007 | -0.007 | 0 | 0.89 | 0.94 | 0.85 | 0.84 | 0.92 | 0.78 |
| 3 | 128417293 | rs7624610 | 0.583 | -0.397 | -1.403 | 0.373 | 0.001 | -0.08 | -0.08 | 0 | 0.27 | 0.16 | 0.2 | 0.39 | 0.25 | 0.24 |
| 3 | 128421166 | rs4521246 | 0.99 | -0.322 | -1.319 | 0.386 | -0.01 | -0.005 | -0.005 | 0 | 0.34 | 0.1 | 0.31 | 0.33 | 0.35 | 0.33 |
| 3 | 128424456 | rs4634140 | 0.254 | -0.111 | -0.921 | 0.302 | -0.015 | 0.023 | 0.023 | 0.099 | 0.65 | 0.25 | 0.54 | 0.73 | 0.63 | 0.57 |
| 3 | 128427739 | rs10934861 | 0.023 | 0.295 | -0.776 | 0.224 | 0.003 | -0.014 | -0.014 | 0 | 0.44 | 0.29 | 0.54 | 0.5 | 0.54 | 0.58 |
| 3 | 128430941 | rs2811502 | 0.164 | 0.514 | -0.169 | 0.095 | -0.001 | 0 | 0 | 0 | 0.42 | 0.59 | 0.5 | 0.46 | 0.44 | 0.57 |
| 3 | 128434420 | rs4443195 | -0.119 | 0.497 | -0.117 | 0.084 | -0.003 | 0.003 | 0.003 | 0 | 0.45 | 0.6 | 0.5 | 0.45 | 0.5 | 0.57 |
| 3 | 128446412 | rs4384970 | -0.077 | 0.263 | 0.347 | 0.184 | -0.025 | 0.034 | 0.034 | 0 | 0.16 | 0.49 | 0.24 | 0.2 | 0.12 | 0.31 |
| 3 | 128499195 | rs7372263 | 0.959 | -0.699 | 0.247 | -0.174 | 0.098 | 0.08 | 0.08 | 0 | 0.5 | 0.15 | 0.54 | 0.33 | 0.61 | 0.44 |
| 3 | 128503532 | rs6762861 | -1.037 | -0.418 | 0.95 | -0.204 | -0.016 | -0.01 | -0.01 | 0 | 0.23 | 0.67 | 0.3 | 0.22 | 0.2 | 0.33 |
| 3 | 128505829 | rs12054283 | 0.324 | -0.34 | 0.542 | -0.17 | -0.015 | 0.014 | 0.014 | 0 | 0.14 | 0.27 | 0.21 | 0.2 | 0.11 | 0.31 |
| 3 | 128516142 | rs3772158 | 0.324 | -0.34 | 0.542 | -0.17 | -0.015 | 0.014 | 0.014 | 0 | 0.14 | 0.27 | 0.21 | 0.2 | 0.11 | 0.31 |
| 3 | 128541260 | rs12106864 | -0.132 | -0.365 | 0.387 | -0.125 | -0.03 | 0.012 | 0.012 | 0.01 | 0.17 | 0.43 | 0.23 | 0.2 | 0.11 | 0.32 |
| 3 | 128542465 | rs6775459 | -0.163 | -0.265 | 0.097 | -0.196 | -0.002 | -0.044 | -0.044 | 0 | 0.17 | 0.33 | 0.17 | 0.07 | 0.15 | 0.15 |
| 3 | 128563551 | rs9843124 | -1.118 | -0.134 | 0.366 | -0.058 | 0.033 | 0.084 | 0.084 | 0 | 0.12 | 0.48 | 0.17 | 0.15 | 0.11 | 0.19 |
| 3 | 128566324 | rs9830429 | 0.937 | -0.414 | -0.741 | -0.508 | 0.007 | 0.069 | 0.069 | 0 | 0.48 | 0.05 | 0.49 | 0.31 | 0.58 | 0.45 |
| 3 | 128566942 | rs13088146 | 0.754 | -0.407 | -0.446 | -0.625 | 0 | 0.004 | 0.004 | 0.054 | 0.13 | 0.11 | 0.14 | 0.06 | 0.15 | 0.14 |
| 3 | 128568202 | rs9840633 | 0.043 | -0.335 | 0.133 | -0.499 | 0.058 | -0.021 | -0.021 | 0 | 0.08 | 0.38 | 0.08 | 0 | 0.09 | 0.02 |
| 3 | 128585707 | rs789242 | -0.562 | -0.443 | -0.465 | -0.704 | 0.041 | -0.002 | -0.002 | 0 | 0.39 | 0.85 | 0.43 | 0.63 | 0.29 | 0.38 |
| 3 | 128593040 | rs796862 | 0.627 | -0.531 | -0.177 | -0.646 | -0.011 | 0.025 | 0.025 | 0 | 0.28 | 0.36 | 0.27 | 0.56 | 0.19 | 0.24 |
| 3 | 128593201 | rs789217 | 0.742 | -0.533 | -0.172 | -0.646 | -0.004 | 0.011 | 0.011 | 0 | 0.27 | 0.29 | 0.26 | 0.56 | 0.19 | 0.24 |
| 3 | 128598231 | rs789224 | -0.616 | -0.499 | -0.182 | -0.653 | 0.04 | -0.014 | -0.014 | 0 | 0.38 | 0.85 | 0.37 | 0.62 | 0.29 | 0.38 |
| 3 | 128608341 | rs2929859 | 0.598 | -0.499 | -0.197 | -0.644 | -0.01 | 0.024 | 0.024 | 0 | 0.28 | 0.35 | 0.27 | 0.56 | 0.19 | 0.24 |
| 3 | 128614185 | rs1680778 | -0.614 | -0.495 | -0.309 | -0.657 | 0.042 | -0.001 | -0.001 | 0.225 | 0.39 | 0.85 | 0.43 | 0.62 | 0.29 | 0.38 |
| 3 | 128627021 | rs1683787 | -0.562 | -0.534 | -0.219 | -0.692 | 0.038 | -0.006 | -0.006 | 0.387 | 0.38 | 0.79 | 0.37 | 0.62 | 0.3 | 0.38 |
| 3 | 128627933 | rs876755 | 0.682 | -0.451 | -0.113 | -0.697 | -0.004 | 0.008 | 0.008 | 0 | 0.28 | 0.29 | 0.27 | 0.57 | 0.19 | 0.24 |
| 3 | 128636961 | rs789237 | 0.731 | -0.511 | 0.305 | -0.747 | 0.003 | -0.007 | -0.007 | 0 | 0.27 | 0.24 | 0.26 | 0.57 | 0.19 | 0.24 |
| 3 | 128648117 | rs6767104 | -0.315 | -0.509 | 0.267 | -0.747 | 0.013 | -0.016 | -0.016 | 0 | 0.36 | 0.64 | 0.35 | 0.57 | 0.28 | 0.25 |
| 3 | 128652553 | rs9810890 | 0.582 | -0.411 | 0.291 | -0.686 | 0.062 | -0.015 | -0.015 | 0 | 0.08 | 0.32 | 0.08 | 0 | 0.09 | 0.02 |
| 3 | 128655765 | rs2879784 | 1.055 | -0.514 | -0.333 | -0.794 | 0.001 | -0.001 | -0.001 | 0 | 0.16 | 0.16 | 0.14 | 0.06 | 0.15 | 0.14 |
| 3 | 128663159 | rs789236 | 0.915 | -0.629 | -0.447 | -0.879 | -0.003 | -0.012 | -0.012 | 0 | 0.18 | 0.11 | 0.22 | 0.57 | 0.14 | 0.2 |
| 3 | 128666029 | rs1683813 | -1.768 | -0.71 | -0.513 | -0.796 | -0.088 | 0.107 | 0.107 | 0 | 0.55 | 0.95 | 0.53 | 0.76 | 0.46 | 0.52 |
| 3 | 128666129 | rs1680777 | 0.038 | -0.692 | -0.112 | -0.865 | 0 | 0.017 | 0.017 | 0 | 0.23 | 0.27 | 0.27 | 0.62 | 0.2 | 0.33 |
| 3 | 128669652 | rs4596167 | 1.306 | -0.677 | -0.361 | -1.021 | 0.001 | 0.049 | 0.049 | 0 | 0.17 | 0.07 | 0.14 | 0.05 | 0.16 | 0.13 |
| 3 | 128670446 | rs789222 | -1.768 | -0.514 | -0.546 | -1.112 | -0.141 | -0.009 | -0.009 | 8.543 | 0.57 | 0.95 | 0.51 | 0.67 | 0.47 | 0.53 |
| 3 | 128677098 | rs2630262 | -1.699 | -0.603 | -0.357 | -1.149 | -0.078 | -0.023 | -0.023 | 0.003 | 0.64 | 0.93 | 0.58 | 0.67 | 0.59 | 0.5 |
| 3 | 128686912 | rs789212 | -0.398 | -0.629 | -0.064 | -1.093 | 0.001 | 0.011 | 0.011 | 0 | 0.32 | 0.28 | 0.31 | 0.57 | 0.3 | 0.28 |
| 3 | 128688499 | rs718613 | 1.216 | -0.7 | 0.044 | -0.882 | 0.004 | 0.087 | 0.087 | 0 | 0.25 | 0.09 | 0.21 | 0.05 | 0.27 | 0.15 |
| 3 | 128691941 | rs2929860 | -1.326 | 0.332 | -0.034 | -0.41 | -0.072 | 0.033 | 0.033 | 0 | 0.87 | 0.99 | 0.8 | 0.95 | 0.82 | 0.84 |
| 3 | 128704891 | rs13072370 | 0.109 | -0.455 | 0.224 | 0.134 | 0.003 | -0.018 | -0.018 | 0 | 0.17 | 0.1 | 0.2 | 0.57 | 0.11 | 0.22 |
| 3 | 128712378 | rs11916893 | 0.375 | -1.015 | 0.012 | -0.143 | -0.009 | -0.008 | -0.008 | 0 | 0.15 | 0.03 | 0.1 | 0.13 | 0.11 | 0.18 |
| 3 | 128741642 | rs419463 | 0.326 | -1.129 | -0.256 | -0.429 | 0.019 | 0.039 | 0.039 | 0 | 0.35 | 0.17 | 0.17 | 0.03 | 0.32 | 0.22 |
| 3 | 128743450 | rs693476 | 0.362 | -1.138 | -0.256 | -0.433 | 0.012 | -0.001 | -0.001 | 0 | 0.35 | 0.21 | 0.17 | 0.03 | 0.32 | 0.22 |
| 3 | 128744213 | rs413183 | -0.667 | -0.976 | -0.566 | 0.153 | -0.021 | 0.058 | 0.058 | 0 | 0.59 | 0.85 | 0.4 | 0.25 | 0.48 | 0.48 |
| 3 | 128744735 | rs395020 | -0.654 | -0.956 | -0.548 | 0.162 | -0.019 | 0.059 | 0.059 | 0 | 0.59 | 0.84 | 0.4 | 0.25 | 0.48 | 0.48 |
| 3 | 128748148 | rs1531807 | -0.629 | -0.93 | -0.533 | 0.17 | -0.011 | 0.055 | 0.055 | 0 | 0.58 | 0.81 | 0.4 | 0.25 | 0.48 | 0.48 |
| 3 | 128755953 | rs3732430 | 0.343 | -0.729 | -0.747 | 0.138 | 0 | 0.042 | 0.042 | 0 | 0.41 | 0.35 | 0.28 | 0.21 | 0.44 | 0.31 |

| | | | | | | | | | | | | | | | | |
|---|---|---|---|---|---|---|---|---|---|---|---|---|---|---|---|---|
| 3 | 128762833 | rs6439154 | -0.482 | -0.131 | -1.369 | 0.136 | -0.063 | -0.01 | -0.01 | 0 | 0.59 | 0.91 | 0.54 | 0.64 | 0.48 | 0.49 |
| 3 | 128778267 | rs4927926 | -0.345 | -0.467 | -0.103 | -0.551 | -0.032 | -0.026 | -0.026 | 0.012 | 0.92 | 0.82 | 0.92 | 0.99 | 0.99 | 0.94 |
| 3 | 128792751 | rs4927903 | -0.081 | -0.538 | 0.218 | -0.576 | -0.04 | -0.03 | -0.03 | 2.167 | 0.91 | 0.71 | 0.92 | 0.99 | 0.99 | 0.96 |
| 3 | 128796544 | rs9826969 | -0.359 | -0.57 | -0.058 | -0.59 | -0.031 | -0.023 | -0.023 | 0.658 | 0.92 | 0.81 | 0.92 | 0.99 | 0.99 | 0.96 |
| 3 | 128803534 | rs9826460 | -0.523 | -0.579 | 0.344 | -0.594 | -0.034 | -0.025 | -0.025 | 0 | 0.92 | 0.84 | 0.92 | 0.99 | 0.99 | 0.96 |
| 3 | 128818712 | rs9861780 | 1.087 | -0.874 | 1.001 | -0.466 | -0.066 | 0.012 | 0.012 | 0.127 | 0.86 | 0.34 | 0.86 | 0.89 | 0.98 | 0.85 |
| 3 | 128880416 | rs4927905 | -0.786 | -0.865 | 1.319 | -0.491 | -0.028 | -0.02 | -0.02 | 0 | 0.93 | 0.91 | 0.93 | 0.99 | 0.99 | 0.96 |
| 3 | 128886264 | rs9875392 | 0.396 | -0.857 | 1.219 | -0.475 | -0.053 | 0.071 | 0.071 | 0 | 0.89 | 0.65 | 0.9 | 0.89 | 0.99 | 0.86 |
| 3 | 128887527 | rs12633752 | -0.105 | -0.856 | 1.161 | -0.483 | -0.037 | 0.066 | 0.066 | 0 | 0.91 | 0.78 | 0.91 | 0.89 | 0.99 | 0.86 |
| 3 | 128890350 | rs4303883 | 0.826 | -0.849 | 0.975 | -0.48 | -0.068 | 0.02 | 0.02 | 0.278 | 0.86 | 0.36 | 0.87 | 0.89 | 0.98 | 0.86 |
| 3 | 128915591 | rs6805004 | 0.41 | -0.336 | -0.003 | -0.86 | -0.013 | 0.036 | 0.036 | 0 | 0.46 | 0.11 | 0.54 | 0.59 | 0.54 | 0.57 |
| 3 | 128937637 | rs7628487 | 0.92 | -0.964 | 0.313 | -0.559 | 0.049 | -0.016 | -0.016 | 0 | 0.67 | 0.13 | 0.66 | 0.62 | 0.8 | 0.76 |
| 3 | 128959313 | rs6439158 | 0.317 | -1.063 | 0.425 | -0.511 | -0.057 | -0.022 | -0.022 | 0.111 | 0.8 | 0.4 | 0.83 | 0.9 | 0.91 | 0.84 |
| 3 | 128976396 | rs4927910 | -0.53 | -1.099 | 0.414 | -0.509 | -0.058 | -0.019 | -0.019 | 0.029 | 0.2 | 0.61 | 0.17 | 0.11 | 0.08 | 0.17 |
| 3 | 128979008 | rs4927943 | 0.59 | -0.298 | 0.138 | -0.248 | -0.05 | -0.023 | -0.023 | 0 | 0.08 | 0.01 | 0.3 | 0.56 | 0.15 | 0.25 |
| 3 | 128979108 | rs4927906 | -1.278 | -0.591 | 0.571 | -0.234 | 0.041 | 0.186 | 0.186 | 1.071 | 0.06 | 0.4 | 0.05 | 0.1 | 0.01 | 0.12 |
| 3 | 128987730 | rs10934875 | -0.495 | -0.93 | 0.413 | -0.44 | -0.03 | -0.014 | -0.014 | 0 | 0.25 | 0.66 | 0.16 | 0.11 | 0.1 | 0.18 |
| 3 | 128988045 | rs9784321 | -0.613 | -0.937 | 0.627 | -0.444 | -0.075 | -0.041 | -0.041 | 0 | 0.25 | 0.6 | 0.16 | 0.11 | 0.1 | 0.18 |
| 3 | 128988225 | rs6767119 | 0.159 | -0.96 | 0.767 | -0.454 | -0.01 | -0.032 | -0.032 | 3.913 | 0.2 | 0.21 | 0.11 | 0.01 | 0.09 | 0.07 |
| 3 | 128996677 | rs7637014 | -0.786 | -1.011 | 0.753 | -0.475 | -0.014 | -0.004 | -0.004 | 0.187 | 0.27 | 0.68 | 0.17 | 0.11 | 0.1 | 0.19 |
| 3 | 129002120 | rs7434264 | 1.31 | -0.728 | 0.226 | -0.957 | -0.118 | 0.145 | 0.145 | 0 | 0.25 | 0.02 | 0.18 | 0.03 | 0.42 | 0.21 |
| 3 | 129008303 | rs4075182 | -0.482 | -1.436 | 0.526 | -0.804 | -0.021 | -0.008 | -0.008 | 0 | 0.26 | 0.67 | 0.17 | 0.11 | 0.1 | 0.19 |
| 3 | 129012744 | rs10934878 | -0.53 | -1.437 | 0.53 | -0.804 | -0.016 | -0.005 | -0.005 | 0 | 0.26 | 0.68 | 0.17 | 0.11 | 0.1 | 0.19 |
| 3 | 129020778 | rs6765930 | -1.108 | -1.571 | 0.242 | -1.353 | -0.019 | -0.012 | -0.012 | 0 | 0.78 | 0.9 | 0.77 | 0.7 | 0.81 | 0.82 |
| 3 | 129027158 | rs6775488 | -0.319 | -1.719 | 0.296 | -1.382 | -0.04 | -0.069 | -0.069 | 0.741 | 0.25 | 0.65 | 0.17 | 0.01 | 0.1 | 0.08 |
| 3 | 129032876 | rs7623849 | 0.288 | -1.705 | 0.264 | -1.376 | -0.022 | -0.048 | -0.048 | 1.134 | 0.21 | 0.25 | 0.12 | 0.01 | 0.09 | 0.08 |
| 3 | 129041882 | rs6790635 | -1.711 | -1.647 | -0.236 | -1.379 | -0.071 | 0.048 | 0.048 | 3.272 | 0.83 | 0.99 | 0.81 | 0.93 | 0.84 | 0.87 |
| 3 | 129043582 | rs10934881 | -1.362 | -1.464 | -0.07 | -1.241 | -0.02 | 0.066 | 0.066 | 2.924 | 0.78 | 0.91 | 0.79 | 0.92 | 0.81 | 0.85 |
| 3 | 129044478 | rs6787676 | -0.141 | -1.488 | 0.061 | -1.203 | -0.09 | -0.01 | -0.01 | 0 | 0.23 | 0.5 | 0.16 | 0.25 | 0.1 | 0.11 |
| 3 | 129045906 | rs7636293 | 0.341 | -1.468 | -0.007 | -1.174 | 0.044 | 0.059 | 0.059 | 0.11 | 0.54 | 0.32 | 0.62 | 0.66 | 0.71 | 0.74 |
| 3 | 129050943 | rs6439168 | -0.975 | -1.423 | -0.055 | -1.693 | -0.002 | 0.039 | 0.039 | 1.868 | 0.76 | 0.86 | 0.78 | 0.91 | 0.81 | 0.84 |
| 3 | 129054338 | rs7612685 | -0.358 | -1.035 | 0.617 | -1.132 | -0.084 | -0.127 | -0.127 | 0 | 0.22 | 0.53 | 0.15 | 0.03 | 0.1 | 0.08 |
| 3 | 129060180 | rs6439170 | -0.329 | -0.09 | 0.335 | -0.613 | 0.046 | -0.126 | -0.126 | 0.096 | 0.3 | 0.58 | 0.28 | 0.03 | 0.44 | 0.22 |
| 3 | 129064902 | rs9853279 | -0.414 | -0.047 | -0.424 | -0.594 | -0.013 | -0.006 | -0.006 | 0.069 | 0.31 | 0.74 | 0.29 | 0.03 | 0.44 | 0.23 |
| 3 | 129068655 | rs7430161 | 0.067 | 0.353 | -1.286 | -0.333 | -0.13 | 0.141 | 0.141 | 0 | 0.25 | 0.03 | 0.21 | 0.02 | 0.43 | 0.25 |
| 3 | 129083281 | rs6810203 | -0.838 | -1.136 | -0.078 | 0.796 | -0.077 | 0.068 | 0.068 | 0 | 0.21 | 0.53 | 0.14 | 0.39 | 0.07 | 0.25 |
| 3 | 129109507 | rs3821918 | -1.099 | -1.319 | -0.045 | 1.158 | 0 | -0.003 | -0.003 | 0 | 0.32 | 0.7 | 0.28 | 0.58 | 0.17 | 0.39 |
| 3 | 129118583 | rs16859134 | -0.426 | -1.527 | 0.537 | 1.043 | -0.027 | -0.042 | -0.042 | 0 | 0.13 | 0.24 | 0.05 | 0.02 | 0.05 | 0.06 |
| 3 | 129122292 | rs7615711 | -1.745 | -1.402 | 1.196 | 1.228 | -0.012 | 0.007 | 0.007 | 0 | 0.32 | 0.67 | 0.28 | 0.63 | 0.2 | 0.39 |
| 3 | 129125231 | rs7647218 | -0.478 | -1.462 | 1.357 | 1.339 | -0.004 | -0.021 | -0.021 | 1.76 | 0.13 | 0.22 | 0.1 | 0.03 | 0.11 | 0.15 |
| 3 | 129149021 | rs3138373 | -0.728 | -1.508 | 2.016 | 1.649 | -0.006 | -0.009 | -0.009 | 0 | 0.15 | 0.23 | 0.09 | 0.09 | 0.11 | 0.27 |
| 3 | 129151667 | rs2005618 | -1.433 | -1.553 | 2.472 | 1.813 | -0.035 | 0.185 | 0.185 | 0.094 | 0.18 | 0.55 | 0.24 | 0.51 | 0.11 | 0.3 |
| 3 | 129152144 | rs140695 | -1.217 | -2.235 | 1.169 | 0.711 | 0.008 | 0.227 | 0.227 | 0 | 0.66 | 0.98 | 0.81 | 0.98 | 0.69 | 0.76 |
| 3 | 129170305 | rs4688850 | 0.921 | -2.016 | 0.821 | 0.731 | -0.034 | -0.051 | -0.051 | 0 | 0.09 | 0.01 | 0.18 | 0.29 | 0.11 | 0.17 |
| 3 | 129186270 | rs10934884 | 0.766 | -3.1 | 0.647 | 0.442 | 0.009 | 0.228 | 0.228 | 0 | 0.34 | 0.02 | 0.19 | 0.02 | 0.32 | 0.24 |
| 3 | 129188325 | rs7612530 | -1.496 | -2.87 | 1.644 | 1.155 | 0.042 | -0.024 | -0.024 | 0 | 0.32 | 0.8 | 0.29 | 0.64 | 0.16 | 0.39 |
| 3 | 129190001 | rs6439183 | -0.1 | -2.87 | 1.645 | 1.143 | -0.006 | -0.009 | -0.009 | 0 | 0.15 | 0.24 | 0.09 | 0.09 | 0.11 | 0.27 |
| 3 | 129195663 | rs2301570 | 0.179 | -2.852 | 1.625 | 1.143 | -0.037 | 0.018 | 0.018 | 0 | 0.13 | 0.24 | 0.05 | 0.12 | 0.05 | 0.09 |
| 3 | 129201692 | rs7628918 | -0.677 | -2.848 | 1.634 | 1.152 | -0.034 | 0.203 | 0.203 | 0.187 | 0.18 | 0.56 | 0.24 | 0.52 | 0.11 | 0.3 |
| 3 | 129210867 | rs2285353 | 0.162 | -2.814 | 1.515 | 1.12 | -0.037 | 0.018 | 0.018 | 0 | 0.13 | 0.24 | 0.05 | 0.12 | 0.05 | 0.09 |
| 3 | 129218944 | rs6799876 | -0.125 | -2.658 | 1.208 | 1.173 | -0.006 | -0.009 | -0.009 | 0 | 0.15 | 0.24 | 0.09 | 0.09 | 0.11 | 0.27 |
| 3 | 129263140 | rs1872111 | 0.415 | -2.494 | 0.601 | 1.037 | 0 | -0.006 | -0.006 | 0 | 0.12 | 0.17 | 0.09 | 0.06 | 0.12 | 0.11 |
| 3 | 129265270 | rs2625956 | -1.517 | -2.866 | 0.477 | 1.179 | 0.089 | -0.082 | -0.082 | 0 | 0.33 | 0.91 | 0.29 | 0.63 | 0.17 | 0.39 |
| 3 | 129274363 | rs1052910 | -0.441 | -2.284 | 1.032 | 1.276 | 0.003 | 0.053 | 0.053 | 0.051 | 0.14 | 0.26 | 0.2 | 0.21 | 0.14 | 0.24 |
| 3 | 129274456 | rs2824 | -0.005 | -2.288 | 1.062 | 1.29 | 0.008 | -0.064 | -0.064 | 0 | 0.13 | 0.21 | 0.1 | 0 | 0.14 | 0.09 |
| 3 | 129274655 | rs1052883 | -0.549 | -2.289 | 1.128 | 1.268 | 0.006 | 0.067 | 0.067 | 0.033 | 0.15 | 0.31 | 0.2 | 0.21 | 0.14 | 0.24 |
| 3 | 129275206 | rs11920311 | -0.484 | -2.293 | 1.193 | 1.266 | 0.006 | 0.075 | 0.075 | 0.079 | 0.15 | 0.33 | 0.2 | 0.21 | 0.14 | 0.24 |
| 3 | 129284818 | rs2625973 | -0.158 | -2.524 | 0.79 | 0.639 | 0 | -0.023 | -0.023 | 0 | 0.29 | 0.22 | 0.19 | 0 | 0.31 | 0.16 |
| 3 | 129285175 | rs11925462 | -0.63 | -3.015 | 1.01 | 0.608 | -0.05 | 0.029 | 0.029 | 1.841 | 0.13 | 0.3 | 0.05 | 0.11 | 0.04 | 0.1 |
| 3 | 129286635 | rs2245278 | -0.103 | -2.897 | 0.58 | 1.202 | 0 | -0.001 | -0.001 | 0 | 0.29 | 0.23 | 0.19 | 0.39 | 0.31 | 0.2 |
| 3 | 129292065 | rs2625976 | -0.069 | -2.892 | 0.569 | 1.209 | 0 | -0.006 | -0.006 | 0 | 0.29 | 0.21 | 0.23 | 0.41 | 0.31 | 0.2 |
| 3 | 129293256 | rs2255703 | -1.435 | -3.115 | 0.277 | 1.074 | 0.052 | -0.009 | -0.009 | 0 | 0.53 | 0.72 | 0.36 | 0.51 | 0.41 | 0.44 |
| 3 | 129298994 | rs10934885 | 0.339 | -2.813 | 0.47 | 1.229 | 0.018 | -0.005 | -0.005 | 0 | 0.71 | 0.78 | 0.8 | 0.59 | 0.77 | 0.81 |
| 3 | 129300291 | rs7610060 | 1.791 | -3.01 | 0.254 | 1.427 | 0.11 | 0.167 | 0.167 | 0.019 | 0.5 | 0.14 | 0.58 | 0.25 | 0.64 | 0.55 |
| 3 | 129302159 | rs2285369 | -0.47 | -2.703 | 0.955 | 2.135 | -0.002 | 0.001 | 0.001 | 0.007 | 0.09 | 0.09 | 0.03 | 0.11 | 0.04 | 0.1 |
| 3 | 129303072 | rs2285371 | -0.441 | -2.703 | 0.966 | 2.135 | -0.004 | 0.004 | 0.004 | 0 | 0.09 | 0.08 | 0.03 | 0.11 | 0.04 | 0.1 |
| 3 | 129304306 | rs10934887 | -0.472 | -2.681 | 1.012 | 2.12 | -0.007 | -0.017 | -0.017 | 0 | 0.1 | 0.06 | 0.13 | 0.24 | 0.1 | 0.17 |
| 3 | 129305460 | rs2301572 | -0.657 | -2.672 | 1.174 | 2.035 | -0.021 | 0.024 | 0.024 | 0.064 | 0.09 | 0.13 | 0.03 | 0.11 | 0.04 | 0.1 |
| 3 | 129312989 | rs2811450 | -1.787 | -2.878 | 1.244 | 1.959 | 0.013 | 0.011 | 0.011 | 0 | 0.37 | 0.32 | 0.31 | 0.75 | 0.24 | 0.39 |

| | | | | | | | | | | | | | | | | |
|---|---|---|---|---|---|---|---|---|---|---|---|---|---|---|---|---|
| 3 | 129317585 | rs9847824 | 0.113 | -2.731 | 0.935 | 2.123 | 0.018 | -0.021 | -0.021 | 0 | 0.17 | 0.14 | 0.14 | 0.4 | 0.1 | 0.12 |
| 3 | 129322824 | rs1872113 | -1.595 | -2.699 | 1.142 | 1.911 | -0.006 | -0.002 | -0.002 | 0 | 0.35 | 0.67 | 0.34 | 0.74 | 0.23 | 0.37 |
| 3 | 129327291 | rs2625978 | -0.466 | -2.564 | 0.697 | 2.277 | -0.01 | 0.105 | 0.105 | 0.428 | 0.07 | 0.08 | 0.09 | 0.47 | 0.03 | 0.12 |
| 3 | 129334233 | rs10804591 | 1.426 | -2.724 | 0.906 | 1.847 | 0.055 | 0.027 | 0.027 | 0.32 | 0.63 | 0.3 | 0.65 | 0.26 | 0.78 | 0.63 |
| 3 | 129339615 | rs2811434 | -1.646 | -2.836 | 0.897 | 1.834 | 0.064 | 0.037 | 0.037 | 0.286 | 0.37 | 0.78 | 0.37 | 0.75 | 0.22 | 0.38 |
| 3 | 129345155 | rs2625989 | 1.456 | -2.687 | 0.814 | 2.46 | 0.088 | -0.154 | -0.154 | 0.236 | 0.7 | 0.07 | 0.67 | 0.43 | 0.81 | 0.77 |
| 3 | 129350751 | rs1316964 | -0.143 | -2.819 | 1.341 | 2.321 | -0.047 | -0.071 | -0.071 | 0 | 0.14 | 0.25 | 0.06 | 0 | 0.08 | 0.05 |
| 3 | 129355240 | rs12636474 | -0.067 | -2.844 | 1.477 | 2.649 | -0.005 | -0.015 | -0.015 | 0 | 0.09 | 0.05 | 0.13 | 0.19 | 0.09 | 0.11 |
| 3 | 129366185 | rs2625946 | 1.943 | -2.109 | 1.542 | 2.71 | 0.082 | -0.099 | -0.099 | 0.261 | 0.7 | 0.11 | 0.67 | 0.41 | 0.81 | 0.77 |
| 3 | 129366963 | rs1136553 | 0.05 | -2.109 | 1.546 | 2.711 | -0.01 | -0.024 | -0.024 | 0 | 0.09 | 0.03 | 0.13 | 0.19 | 0.09 | 0.11 |
| 3 | 129374592 | rs2811425 | 1.958 | -2.108 | 1.835 | 2.737 | 0.085 | -0.105 | -0.105 | 1.281 | 0.7 | 0.11 | 0.67 | 0.41 | 0.81 | 0.77 |
| 3 | 129401505 | rs6780938 | 1.951 | -2.134 | 2.258 | 2.728 | 0.087 | -0.147 | -0.147 | 0.067 | 0.7 | 0.07 | 0.67 | 0.43 | 0.81 | 0.77 |
| 3 | 129432347 | rs4688772 | -0.882 | -2.112 | 2.304 | 2.738 | 0 | 0.039 | 0.039 | 0 | 0.12 | 0.13 | 0.21 | 0.59 | 0.11 | 0.18 |
| 3 | 129436362 | rs7617899 | -0.072 | -2.112 | 2.316 | 2.714 | -0.014 | -0.032 | -0.032 | 0.184 | 0.14 | 0.18 | 0.06 | 0 | 0.08 | 0.05 |
| 3 | 129463505 | rs4075143 | -0.721 | -1.954 | 2.018 | 2.605 | 0 | -0.1 | -0.1 | 0.479 | 0.13 | 0.01 | 0.16 | 0.35 | 0.09 | 0.3 |
| 3 | 129490835 | rs4688813 | -0.921 | -1.524 | 2.469 | 2.97 | -0.001 | 0.061 | 0.061 | 0 | 0.12 | 0.17 | 0.21 | 0.59 | 0.11 | 0.18 |
| 3 | 129494907 | rs12485251 | 0.072 | -1.556 | 2.401 | 2.95 | -0.01 | -0.023 | -0.023 | 0 | 0.09 | 0.03 | 0.13 | 0.18 | 0.09 | 0.11 |
| 3 | 129501662 | rs10934893 | -0.063 | -1.556 | 2.409 | 2.942 | -0.064 | -0.087 | -0.087 | 0 | 0.85 | 0.67 | 0.93 | 1 | 0.92 | 0.95 |
| 3 | 129517529 | rs16863175 | 0.07 | -1.547 | 2.202 | 3.078 | -0.01 | -0.029 | -0.029 | 0 | 0.09 | 0.03 | 0.14 | 0.21 | 0.09 | 0.12 |
| 3 | 129546197 | rs784688 | -0.082 | -1.625 | 2.082 | 2.93 | -0.064 | -0.088 | -0.088 | 0 | 0.85 | 0.67 | 0.93 | 1 | 0.92 | 0.95 |
| 3 | 129604169 | rs2669833 | -1.671 | -1.458 | 1.147 | 2.798 | 0.088 | -0.088 | -0.088 | 0 | 0.3 | 0.88 | 0.33 | 0.59 | 0.18 | 0.23 |
| 3 | 129646689 | rs9818312 | -1.446 | -0.331 | 1.372 | 2.724 | 0.003 | -0.011 | -0.011 | 0 | 0.26 | 0.09 | 0.22 | 0.25 | 0.25 | 0.27 |
| 3 | 129647645 | rs10934895 | 2.465 | -0.379 | 1.349 | 2.88 | 0.045 | -0.021 | -0.021 | 0 | 0.61 | 0.17 | 0.6 | 0.34 | 0.69 | 0.55 |
| 3 | 129651554 | rs13316898 | -1.12 | -0.376 | 1.334 | 2.865 | 0.012 | -0.026 | -0.026 | 0 | 0.23 | 0.07 | 0.21 | 0.24 | 0.22 | 0.28 |
| 3 | 129658498 | rs2137574 | 1.173 | -0.337 | 2.164 | 2.922 | -0.026 | 0.056 | 0.056 | 0 | 0.75 | 0.51 | 0.77 | 0.58 | 0.77 | 0.73 |
| 3 | 129658706 | rs938196 | -0.466 | -0.321 | 2.194 | 2.983 | -0.015 | 0.022 | 0.022 | 0 | 0.19 | 0.28 | 0.18 | 0.34 | 0.18 | 0.16 |
| 3 | 129662273 | rs2896579 | 0.311 | -0.321 | 2.083 | 2.983 | -0.027 | 0.049 | 0.049 | 0 | 0.81 | 0.6 | 0.81 | 0.66 | 0.81 | 0.84 |
| 3 | 129663810 | rs9880871 | 0.295 | -0.321 | 2.066 | 2.977 | -0.024 | 0.042 | 0.042 | 0 | 0.81 | 0.64 | 0.81 | 0.66 | 0.81 | 0.81 |
| 3 | 129664970 | rs4688822 | 0.332 | -0.316 | 2.007 | 2.985 | -0.027 | 0.017 | 0.017 | 0 | 0.8 | 0.6 | 0.81 | 0.71 | 0.81 | 0.81 |
| 3 | 129666819 | rs4688828 | -0.401 | -0.26 | 1.054 | 3.064 | 0 | 0.041 | 0.041 | 0 | 0.07 | 0.05 | 0.13 | 0.28 | 0.05 | 0.18 |
| 3 | 129668516 | rs10934897 | 1.444 | -0.241 | 1.094 | 2.795 | 0.008 | -0.001 | -0.001 | 0 | 0.72 | 0.25 | 0.75 | 0.63 | 0.76 | 0.73 |
| 3 | 129672976 | rs1708884 | -1.481 | -0.308 | 1.173 | 2.712 | -0.012 | 0.023 | 0.023 | 0 | 0.27 | 0.64 | 0.33 | 0.58 | 0.23 | 0.38 |
| 3 | 129674580 | rs884697 | -1.521 | -0.284 | 1.143 | 2.873 | -0.002 | 0.004 | 0.004 | 0 | 0.27 | 0.7 | 0.37 | 0.63 | 0.26 | 0.3 |
| 3 | 129676089 | rs1708950 | -2.283 | 0.177 | 1.623 | 2.737 | -0.001 | 0.222 | 0.222 | 0 | 0.19 | 0.51 | 0.37 | 0.41 | 0.16 | 0.27 |
| 3 | 129676824 | rs1708227 | -0.347 | 0.186 | 1.529 | 2.745 | 0.004 | 0.006 | 0.006 | 0 | 0.07 | 0.1 | 0.16 | 0.22 | 0.03 | 0.08 |
| 3 | 129687018 | rs784718 | 0.449 | 0.308 | 2.273 | 2.637 | 0.054 | 0.265 | 0.265 | 0 | 0.94 | 0.67 | 0.85 | 0.77 | 0.95 | 0.89 |
| 3 | 129687948 | rs784717 | 0.758 | 0.385 | 1.659 | 2.384 | -0.003 | -0.009 | -0.009 | 0.492 | 0.68 | 0.73 | 0.73 | 0.57 | 0.61 | 0.66 |
| 3 | 129693337 | rs784707 | -1.49 | 0.052 | 1.858 | 2.56 | 0.007 | 0.088 | 0.088 | 0 | 0.17 | 0.36 | 0.37 | 0.37 | 0.18 | 0.18 |
| 3 | 129695373 | rs7645772 | 2.277 | 0.285 | 1.467 | 2.374 | -0.067 | 0.095 | 0.095 | 0.664 | 0.45 | 0.07 | 0.35 | 0.2 | 0.46 | 0.37 |
| 3 | 129698175 | rs2670898 | 1.01 | 0.29 | 1.259 | 2.464 | 0.06 | 0.146 | 0.146 | 0.166 | 0.92 | 0.59 | 0.91 | 0.82 | 0.95 | 0.94 |
| 3 | 129699722 | rs2670897 | -1.218 | 0.357 | 0.841 | 2.528 | 0.001 | 0.02 | 0.02 | 0 | 0.16 | 0.24 | 0.35 | 0.37 | 0.18 | 0.17 |
| 3 | 129706392 | rs6791923 | 1.239 | 0.556 | 0.93 | 2.49 | 0.021 | 0.003 | 0.003 | 0.44 | 0.54 | 0.29 | 0.56 | 0.46 | 0.59 | 0.49 |
| 3 | 129706641 | rs13059321 | 0.96 | 0.244 | 0.962 | 2.03 | 0.001 | -0.001 | -0.001 | 0 | 0.55 | 0.42 | 0.43 | 0.4 | 0.5 | 0.46 |
| 3 | 129709647 | rs925613 | 1.368 | 0.501 | -0.11 | 1.827 | -0.042 | 0.081 | 0.081 | 0.474 | 0.46 | 0.11 | 0.36 | 0.22 | 0.45 | 0.37 |
| 3 | 129823705 | rs4299520 | -1.428 | 0.533 | 0.624 | 1.811 | 0.017 | 0.105 | 0.105 | 0 | 0.06 | 0.16 | 0.05 | 0.16 | 0.06 | 0.15 |
| 3 | 129844461 | rs9757129 | 0.881 | 0.408 | 0.045 | 1.355 | 0.01 | 0.016 | 0.016 | 0.064 | 0.5 | 0.36 | 0.41 | 0.41 | 0.48 | 0.5 |
| 3 | 129920258 | rs2587988 | -1.376 | -0.662 | -0.238 | 1.664 | -0.042 | -0.113 | -0.113 | 0.375 | 0.29 | 0.1 | 0.4 | 0.51 | 0.29 | 0.39 |
| 3 | 129933246 | rs9855392 | -1.116 | -0.796 | 0.143 | 1.215 | 0 | -0.001 | -0.001 | 1.744 | 0.2 | 0.25 | 0.19 | 0.2 | 0.16 | 0.21 |
| 3 | 129933648 | rs9855777 | 2.143 | -1.104 | 0.685 | 1.325 | 0.019 | 0.038 | 0.038 | 1.555 | 0.61 | 0.17 | 0.49 | 0.27 | 0.71 | 0.54 |
| 3 | 129939787 | rs6802606 | -1.483 | -1.102 | 1.941 | 1.282 | -0.028 | 0.193 | 0.193 | 0 | 0.2 | 0.53 | 0.33 | 0.52 | 0.13 | 0.2 |
| 3 | 129944129 | rs9289367 | 1.785 | -1.074 | 2.085 | 1.303 | -0.007 | -0.013 | -0.013 | 0 | 0.64 | 0.43 | 0.51 | 0.28 | 0.71 | 0.54 |
| 3 | 129946531 | rs4336159 | 0.24 | -1.261 | 2.248 | 1.292 | -0.005 | 0.092 | 0.092 | 0 | 0.89 | 0.85 | 0.74 | 0.48 | 0.92 | 0.78 |
| 3 | 129969817 | rs6777871 | 0.648 | -1.742 | 2.486 | 0.917 | 0.007 | -0.01 | -0.01 | 0 | 0.78 | 0.83 | 0.71 | 0.48 | 0.87 | 0.75 |
| 3 | 129970462 | rs9864582 | -1.263 | -1.657 | 2.486 | 0.872 | -0.002 | -0.016 | -0.016 | 0.207 | 0.25 | 0.39 | 0.22 | 0.18 | 0.2 | 0.26 |
| 3 | 129976048 | rs6804874 | -1.284 | -1.664 | 2.526 | 0.871 | 0 | 0.013 | 0.013 | 0.187 | 0.25 | 0.29 | 0.21 | 0.18 | 0.2 | 0.26 |
| 3 | 129977006 | rs7619208 | 1.401 | -1.662 | 2.394 | 0.872 | -0.006 | -0.028 | -0.028 | 0 | 0.72 | 0.34 | 0.76 | 0.82 | 0.8 | 0.68 |
| 3 | 129985198 | rs9849611 | -0.917 | -1.732 | 2.391 | 1.007 | -0.019 | 0.041 | 0.041 | 0.045 | 0.22 | 0.25 | 0.3 | 0.52 | 0.13 | 0.25 |
| 3 | 129990413 | rs4688871 | 1.385 | -1.835 | 2.051 | 1.03 | 0 | -0.003 | -0.003 | 0 | 0.72 | 0.39 | 0.65 | 0.35 | 0.83 | 0.6 |
| 3 | 130001692 | rs1453240 | -1.767 | -1.617 | 1.84 | 0.962 | -0.012 | -0.008 | -0.008 | 0 | 0.36 | 0.55 | 0.46 | 0.69 | 0.28 | 0.46 |
| 3 | 130005239 | rs10512760 | -0.2 | -1.637 | 1.377 | 1.007 | -0.008 | 0.103 | 0.103 | 0 | 0.1 | 0.14 | 0.26 | 0.52 | 0.06 | 0.2 |
| 3 | 130016770 | rs9829426 | -0.4 | -1.547 | -0.156 | 1.011 | -0.008 | 0.014 | 0.014 | 0 | 0.18 | 0.11 | 0.16 | 0.17 | 0.16 | 0.23 |
| 3 | 130018358 | rs9821802 | 0.178 | -1.3 | -0.402 | 0.87 | 0.04 | -0.019 | -0.019 | 0 | 0.13 | 0.02 | 0.05 | 0.13 | 0.08 | 0.14 |
| 3 | 130019843 | rs13080253 | 1.236 | -0.869 | -0.102 | 0.632 | 0.032 | 0.188 | 0.188 | 0.019 | 0.48 | 0.08 | 0.44 | 0.17 | 0.64 | 0.34 |
| 3 | 130024270 | rs9853619 | -1.464 | -0.348 | 1.203 | 1.253 | 0.117 | 0.127 | 0.127 | 2.029 | 0.06 | 0.26 | 0.13 | 0.15 | 0.08 | 0.12 |
| 3 | 130026739 | rs2403381 | -1.311 | -0.775 | 1.233 | 0.752 | 0.01 | 0.031 | 0.031 | 0.581 | 0.45 | 0.6 | 0.47 | 0.87 | 0.36 | 0.52 |
| 3 | 130028717 | rs7649424 | -1.648 | -0.957 | 1.23 | 0.62 | 0.043 | 0.217 | 0.217 | 0.095 | 0.49 | 0.77 | 0.5 | 0.87 | 0.37 | 0.6 |
| 3 | 130036205 | rs4688874 | 1.616 | -0.965 | 1.206 | 0.616 | 0.041 | 0.259 | 0.259 | 0.135 | 0.5 | 0.18 | 0.49 | 0.13 | 0.63 | 0.4 |
| 3 | 130041509 | rs9868118 | 1.286 | -1.071 | 1.918 | 1.232 | 0.006 | 0.007 | 0.007 | 0 | 0.58 | 0.47 | 0.75 | 0.77 | 0.69 | 0.53 |
| 3 | 130045057 | rs9880602 | 1.568 | -1.094 | 1.877 | 1.154 | 0.028 | -0.061 | -0.061 | 0.061 | 0.9 | 0.76 | 0.9 | 1 | 0.93 | 0.9 |

| | | | | | | | | | | | | | | | | |
|---|---|---|---|---|---|---|---|---|---|---|---|---|---|---|---|---|
| 3 | 130047316 | rs1453245 | 1.651 | -1.094 | 1.861 | 1.154 | 0.032 | -0.062 | -0.062 | 0 | 0.9 | 0.72 | 0.89 | 1 | 0.93 | 0.9 |
| 3 | 130048207 | rs7609940 | -0.645 | -1.069 | 1.411 | 1.204 | 0.041 | -0.012 | -0.012 | 1.622 | 0.31 | 0.06 | 0.13 | 0.22 | 0.24 | 0.38 |
| 3 | 130061040 | rs9879726 | 1.655 | -0.693 | 2.356 | 1.683 | 0.07 | 0.056 | 0.056 | 0 | 0.93 | 0.72 | 0.91 | 0.93 | 0.94 | 0.92 |
| 3 | 130064743 | rs1869319 | 2.482 | -0.736 | 2.18 | 1.661 | 0.014 | -0.008 | -0.008 | 0 | 0.87 | 0.54 | 0.88 | 0.92 | 0.93 | 0.84 |
| 3 | 130087842 | rs10804598 | -1.994 | -0.484 | 1.556 | 1.918 | -0.013 | -0.008 | -0.008 | 0.033 | 0.07 | 0.05 | 0.18 | 0.18 | 0.11 | 0.1 |
| 3 | 130096569 | rs9841662 | -2.291 | -0.622 | 2.005 | 2.643 | 0.006 | 0.123 | 0.123 | 0 | 0.12 | 0.28 | 0.29 | 0.34 | 0.1 | 0.14 |
| 3 | 130096881 | rs16826935 | -2.786 | -0.782 | 2.014 | 2.487 | 0.015 | 0.125 | 0.125 | 0 | 0.21 | 0.43 | 0.46 | 0.52 | 0.2 | 0.31 |
| 3 | 130104206 | rs10212372 | -2.786 | -0.838 | 2.303 | 2.515 | 0.022 | 0.162 | 0.162 | 0.006 | 0.22 | 0.49 | 0.46 | 0.52 | 0.2 | 0.33 |
| 3 | 130105223 | rs4688794 | 0.697 | -1.886 | 2.235 | 2.429 | -0.024 | 0.067 | 0.067 | 0 | 0.84 | 0.78 | 0.89 | 0.71 | 0.89 | 0.77 |
| 3 | 130106394 | rs1391062 | -2.43 | -2.029 | 2.34 | 2.444 | -0.014 | 0.183 | 0.183 | 0 | 0.2 | 0.48 | 0.42 | 0.54 | 0.15 | 0.32 |
| 3 | 130109668 | rs924193 | 2.78 | -2.671 | 2.217 | 2.21 | 0.046 | 0.07 | 0.07 | 0 | 0.59 | 0.29 | 0.45 | 0.17 | 0.71 | 0.44 |
| 3 | 130110550 | rs11917356 | -2.482 | -2.572 | 2.198 | 2.413 | 0.004 | 0.113 | 0.113 | 0 | 0.18 | 0.4 | 0.43 | 0.63 | 0.15 | 0.28 |
| 3 | 130113635 | rs10934938 | -2.808 | -2.579 | 2.191 | 2.41 | -0.021 | 0.046 | 0.046 | 0.054 | 0.25 | 0.42 | 0.45 | 0.63 | 0.19 | 0.36 |
| 3 | 130122125 | rs10934939 | -2.173 | -2.547 | 2.273 | 2.602 | -0.045 | 0.066 | 0.066 | 0 | 0.18 | 0.42 | 0.22 | 0.29 | 0.11 | 0.29 |
| 3 | 130130573 | rs1566139 | -1.464 | -2.723 | 2.254 | 2.568 | 0.001 | 0 | 0 | 0 | 0.13 | 0.09 | 0.1 | 0.11 | 0.11 | 0.13 |
| 3 | 130131647 | rs16827439 | -1.228 | -2.663 | 2.286 | 2.601 | 0.011 | 0.018 | 0.018 | 0 | 0.07 | 0.09 | 0.09 | 0.1 | 0.07 | 0.05 |
| 3 | 130135456 | rs6439233 | 1.041 | -2.826 | 2.821 | 2.705 | -0.059 | 0.262 | 0.262 | 0.581 | 0.83 | 0.57 | 0.73 | 0.46 | 0.9 | 0.74 |
| 3 | 130137471 | rs1542829 | -1.199 | -2.798 | 2.546 | 2.639 | 0.003 | -0.011 | -0.011 | 0.026 | 0.09 | 0.1 | 0.15 | 0.17 | 0.06 | 0.16 |
| 3 | 130138117 | rs6769205 | 0.593 | -2.78 | 2.557 | 2.574 | -0.031 | 0.013 | 0.013 | 0.227 | 0.84 | 0.77 | 0.93 | 0.8 | 0.92 | 0.8 |
| 3 | 130139884 | rs12636289 | -1.548 | -2.783 | 2.434 | 2.513 | -0.001 | -0.016 | -0.016 | 0.015 | 0.18 | 0.11 | 0.14 | 0.27 | 0.15 | 0.24 |
| 3 | 130141306 | rs976323 | -2.192 | -2.85 | 2.514 | 2.498 | 0.013 | -0.005 | -0.005 | 0 | 0.25 | 0.15 | 0.17 | 0.29 | 0.19 | 0.33 |
| 3 | 130144198 | rs16827628 | -0.003 | -2.846 | 2.532 | 2.475 | -0.003 | -0.012 | -0.012 | 0 | 0.08 | 0.09 | 0.03 | 0.02 | 0.04 | 0.09 |
| 3 | 130145541 | rs9821748 | -1.432 | -2.843 | 2.486 | 2.502 | -0.002 | -0.02 | -0.02 | 0 | 0.17 | 0.09 | 0.14 | 0.27 | 0.15 | 0.24 |
| 3 | 130145718 | rs981548 | 2.242 | -2.922 | 2.95 | 2.437 | -0.051 | 0.057 | 0.057 | 0 | 0.78 | 0.39 | 0.68 | 0.57 | 0.88 | 0.73 |
| 3 | 130147332 | rs16827675 | -2.133 | -2.859 | 3.245 | 2.495 | 0.023 | 0.419 | 0.419 | 0 | 0.07 | 0.35 | 0.22 | 0.36 | 0.03 | 0.07 |
| 3 | 130147409 | rs6770893 | -2.675 | -2.969 | 3.249 | 2.463 | -0.078 | 0.104 | 0.104 | 0 | 0.23 | 0.53 | 0.32 | 0.42 | 0.12 | 0.26 |
| 3 | 130147813 | rs6785181 | -1.855 | -2.876 | 3.111 | 2.549 | 0.022 | 0.393 | 0.393 | 0 | 0.06 | 0.31 | 0.2 | 0.36 | 0.03 | 0.07 |
| 3 | 130148510 | rs9875499 | -2.094 | -3.275 | 2.502 | 2.398 | 0.036 | -0.066 | -0.066 | 0.061 | 0.28 | 0.05 | 0.22 | 0.34 | 0.21 | 0.33 |
| 3 | 130149333 | rs16827728 | -1.542 | -3.282 | 2.44 | 2.396 | -0.016 | -0.114 | -0.114 | 0 | 0.21 | 0.02 | 0.2 | 0.32 | 0.18 | 0.25 |
| 3 | 130149845 | rs1391050 | 2.888 | -3.324 | 3.152 | 2.446 | -0.006 | 0.001 | 0.001 | 0 | 0.58 | 0.48 | 0.64 | 0.51 | 0.72 | 0.49 |
| 3 | 130162395 | rs9883988 | 1.645 | -3.315 | 3.321 | 2.317 | -0.006 | 0.075 | 0.075 | 0 | 0.91 | 0.74 | 0.9 | 0.85 | 0.95 | 0.88 |
| 3 | 130172754 | rs864701 | 2.085 | -3.297 | 3.55 | 2.278 | -0.01 | 0.063 | 0.063 | 0 | 0.87 | 0.53 | 0.86 | 0.85 | 0.94 | 0.85 |
| 3 | 130172792 | rs819086 | 1.423 | -3.294 | 3.317 | 2.28 | -0.013 | 0.043 | 0.043 | 0 | 0.89 | 0.74 | 0.89 | 0.85 | 0.94 | 0.86 |
| 3 | 130174334 | rs819085 | 0.821 | -3.291 | 3.305 | 2.244 | -0.007 | -0.01 | -0.01 | 0 | 0.91 | 0.83 | 0.9 | 0.96 | 0.94 | 0.91 |
| 3 | 130177740 | rs2047366 | 3.248 | -3.415 | 3.154 | 2.225 | 0.073 | 0.111 | 0.111 | 0 | 0.56 | 0.26 | 0.42 | 0.18 | 0.7 | 0.43 |
| 3 | 130181587 | rs9682426 | 2.595 | -3.773 | 3.038 | 2.285 | -0.025 | 0.02 | 0.02 | 1.141 | 0.66 | 0.56 | 0.62 | 0.6 | 0.76 | 0.66 |
| 3 | 130199738 | rs2971565 | 2.998 | -3.731 | 3.166 | 2.314 | 0.04 | -0.023 | -0.023 | 6.853 | 0.64 | 0.27 | 0.6 | 0.41 | 0.79 | 0.6 |
| 3 | 130203742 | rs7624712 | 2.365 | -3.742 | 3.264 | 2.334 | -0.036 | 0.137 | 0.137 | 0.345 | 0.79 | 0.45 | 0.69 | 0.48 | 0.87 | 0.82 |
| 3 | 130203904 | rs10934943 | -0.885 | -3.669 | 3.131 | 2.319 | -0.008 | -0.009 | -0.009 | 0 | 0.09 | 0.18 | 0.08 | 0.04 | 0.05 | 0.1 |
| 3 | 130207777 | rs1391054 | -1.221 | -3.673 | 2.757 | 2.217 | -0.005 | -0.007 | -0.007 | 4.493 | 0.15 | 0.18 | 0.1 | 0.06 | 0.09 | 0.21 |
| 3 | 130216566 | rs9883486 | -2.403 | -3.711 | 2.567 | 2.213 | -0.015 | 0.084 | 0.084 | 0 | 0.21 | 0.58 | 0.33 | 0.46 | 0.16 | 0.22 |
| 3 | 130220079 | rs1497317 | 3.458 | -3.831 | 2.323 | 2.102 | 0.116 | 0.167 | 0.167 | 0.529 | 0.54 | 0.19 | 0.42 | 0.17 | 0.72 | 0.42 |
| 3 | 130224642 | rs2645950 | 2.216 | -3.751 | 2.385 | 1.742 | -0.021 | 0.044 | 0.044 | 0.301 | 0.72 | 0.5 | 0.63 | 0.54 | 0.78 | 0.74 |
| 3 | 130227216 | rs2661936 | 2.056 | -3.725 | 2.396 | 1.755 | -0.058 | 0.061 | 0.061 | 0 | 0.73 | 0.5 | 0.66 | 0.52 | 0.83 | 0.64 |
| 3 | 130229942 | rs2645961 | 0.481 | -3.682 | 2.575 | 1.841 | -0.022 | 0.218 | 0.218 | 0 | 0.89 | 0.73 | 0.77 | 0.58 | 0.92 | 0.86 |
| 3 | 130231320 | rs2645959 | 2.379 | -3.565 | 2.297 | 1.884 | 0.097 | -0.018 | -0.018 | 0.64 | 0.55 | 0.28 | 0.56 | 0.41 | 0.73 | 0.51 |
| 3 | 130231509 | rs16828872 | -1.204 | -3.202 | 2.132 | 0.976 | 0 | -0.015 | -0.015 | 1.57 | 0.1 | 0.21 | 0.1 | 0.03 | 0.07 | 0.1 |
| 3 | 130239724 | rs6799357 | -1.475 | -3.452 | 1.949 | 0.783 | -0.017 | -0.03 | -0.03 | 0.201 | 0.36 | 0.6 | 0.2 | 0.17 | 0.24 | 0.22 |
| 3 | 130243249 | rs4682614 | -0.925 | -3.069 | 1.611 | 0.986 | -0.035 | -0.031 | -0.031 | 0 | 0.26 | 0.44 | 0.22 | 0.17 | 0.2 | 0.25 |
| 3 | 130244120 | rs6795367 | -1.325 | -2.771 | 1.351 | 1.143 | 0.11 | 0.179 | 0.179 | 0.543 | 0.48 | 0.79 | 0.32 | 0.21 | 0.33 | 0.37 |
| 3 | 130253804 | rs3993281 | 2.46 | -2.581 | 2.396 | 2.175 | 0.018 | -0.018 | -0.018 | 0 | 0.7 | 0.24 | 0.75 | 0.41 | 0.83 | 0.64 |
| 3 | 130255483 | rs937907 | 1.635 | -2.617 | 3.195 | 2.318 | -0.017 | 0.128 | 0.128 | 0 | 0.81 | 0.72 | 0.8 | 0.5 | 0.84 | 0.71 |
| 3 | 130255698 | rs1396330 | 1.468 | -2.622 | 3.395 | 2.316 | 0.007 | -0.032 | -0.032 | 0 | 0.82 | 0.9 | 0.82 | 0.5 | 0.84 | 0.71 |
| 3 | 130255839 | rs1396329 | 1.785 | -2.623 | 3.597 | 2.314 | -0.029 | 0.148 | 0.148 | 0 | 0.79 | 0.58 | 0.79 | 0.44 | 0.84 | 0.69 |
| 3 | 130277221 | rs2174944 | 2.776 | -2.627 | 3.428 | 2.299 | 0.056 | -0.062 | -0.062 | 0.101 | 0.7 | 0.14 | 0.74 | 0.41 | 0.82 | 0.64 |
| 3 | 130284284 | rs9830253 | 2.696 | -2.644 | 3.292 | 2.289 | 0.059 | -0.084 | -0.084 | 0 | 0.7 | 0.14 | 0.75 | 0.44 | 0.82 | 0.64 |
| 3 | 130284840 | rs4682599 | 1.607 | -2.581 | 3.482 | 2.298 | 0.01 | -0.021 | -0.021 | 0 | 0.8 | 0.88 | 0.83 | 0.52 | 0.85 | 0.69 |
| 3 | 130289617 | rs10934949 | 1.511 | -2.411 | 3.736 | 2.34 | 0.048 | 0.068 | 0.068 | 0.147 | 0.92 | 0.46 | 0.95 | 0.92 | 0.98 | 0.96 |
| 3 | 130291211 | rs4682600 | 2.09 | -2.488 | 2.969 | 2.388 | 0.051 | -0.088 | -0.088 | 0.837 | 0.71 | 0.15 | 0.75 | 0.46 | 0.8 | 0.64 |
| 3 | 130297259 | rs16829695 | -2.504 | -1.503 | 3.296 | 2.829 | -0.003 | 0.259 | 0.259 | 0 | 0.14 | 0.34 | 0.18 | 0.48 | 0.12 | 0.3 |
| 3 | 130322553 | rs6781724 | -1.607 | -1.28 | 2.679 | 2.156 | 0.109 | 0.128 | 0.128 | 0.769 | 0.07 | 0.36 | 0.13 | 0.09 | 0.07 | 0.18 |
| 3 | 130328911 | rs7636350 | -2.757 | -1.551 | 2.178 | 2.325 | 0.017 | -0.028 | -0.028 | 0 | 0.27 | 0.79 | 0.53 | 0.34 | 0.23 | 0.37 |
| 3 | 130330135 | rs1508518 | -2.169 | -1.554 | 2.376 | 2.341 | -0.036 | 0.096 | 0.096 | 0 | 0.21 | 0.47 | 0.38 | 0.28 | 0.18 | 0.22 |
| 3 | 130333768 | rs6439251 | -2.187 | -1.343 | 2.141 | 2.188 | -0.012 | 0.064 | 0.064 | 0.435 | 0.22 | 0.6 | 0.51 | 0.3 | 0.21 | 0.33 |
| 3 | 130336698 | rs16830219 | -0.787 | -1.359 | 2.15 | 2.216 | -0.02 | 0.031 | 0.031 | 0 | 0.09 | 0.15 | 0.2 | 0.15 | 0.05 | 0.07 |
| 3 | 130357328 | rs1484937 | -1.931 | -1.275 | 0.103 | 2.075 | 0.003 | -0.007 | -0.007 | 0.804 | 0.25 | 0.73 | 0.53 | 0.3 | 0.24 | 0.35 |
| 3 | 130360904 | rs965122 | -1.005 | -1.445 | -0.638 | 1.867 | 0.04 | -0.092 | -0.092 | 0.014 | 0.26 | 0.06 | 0.34 | 0.27 | 0.21 | 0.38 |
| 3 | 130361856 | rs16830494 | -0.499 | -1.478 | -0.567 | 2.177 | -0.002 | -0.019 | -0.019 | 0 | 0.09 | 0.05 | 0.16 | 0.19 | 0.09 | 0.16 |
| 3 | 130363293 | rs6798745 | -1.638 | -1.69 | 0.688 | 2.27 | -0.012 | 0.137 | 0.137 | 0 | 0.14 | 0.51 | 0.2 | 0.25 | 0.1 | 0.19 |

| | | | | | | | | | | | | | | | | |
|---|---|---|---|---|---|---|---|---|---|---|---|---|---|---|---|---|
| 3 | 130368069 | rs7614116 | -2.076 | -1.902 | 0.886 | 2.287 | 0.067 | 0.02 | 0.02 | 0 | 0.41 | 0.81 | 0.57 | 0.53 | 0.31 | 0.57 |
| 3 | 130368301 | rs7636656 | -1.309 | -1.806 | 1.239 | 2.288 | -0.014 | 0.074 | 0.074 | 0 | 0.13 | 0.35 | 0.19 | 0.24 | 0.1 | 0.18 |
| 3 | 130370748 | rs17748304 | 0.015 | -1.76 | 1.108 | 2.209 | -0.002 | 0.01 | 0.01 | 0.016 | 0.09 | 0.01 | 0.04 | 0.04 | 0.07 | 0.24 |
| 3 | 130377034 | rs1396331 | -2.214 | -1.887 | 1.298 | 2.286 | 0.014 | -0.014 | -0.014 | 0 | 0.33 | 0.73 | 0.41 | 0.37 | 0.22 | 0.41 |
| 3 | 130386977 | rs1872378 | -1.586 | -1.777 | 1.666 | 2.583 | 0.018 | 0.002 | 0.002 | 0 | 0.24 | 0.14 | 0.34 | 0.24 | 0.21 | 0.38 |
| 3 | 130394728 | rs17748751 | -1.599 | -1.733 | 1.823 | 2.583 | 0 | 0 | 0 | 0 | 0.24 | 0.19 | 0.35 | 0.24 | 0.21 | 0.38 |
| 3 | 130428146 | rs10934954 | -1.587 | -1.757 | 1.884 | 2.479 | 0.017 | 0.002 | 0.002 | 0 | 0.24 | 0.14 | 0.34 | 0.24 | 0.21 | 0.38 |
| 3 | 130449222 | rs2170990 | -1.6 | -1.749 | 1.9 | 2.424 | 0.022 | 0.011 | 0.011 | 0 | 0.24 | 0.12 | 0.31 | 0.21 | 0.21 | 0.37 |
| 3 | 130465474 | rs2293179 | -2.154 | -1.719 | 1.968 | 2.426 | -0.001 | -0.002 | -0.002 | 0 | 0.31 | 0.72 | 0.41 | 0.37 | 0.21 | 0.4 |
| 3 | 130477566 | rs16845869 | -0.183 | -1.765 | 1.822 | 2.397 | 0.016 | -0.011 | -0.011 | 1.916 | 0.1 | 0.02 | 0.24 | 0.11 | 0.07 | 0.09 |
| 3 | 130481527 | rs2090994 | -0.869 | -1.698 | 1.849 | 2.385 | 0.015 | 0.008 | 0.008 | 0 | 0.2 | 0.09 | 0.29 | 0.15 | 0.15 | 0.33 |
| 3 | 130491805 | rs4682640 | -1.715 | -1.647 | 1.757 | 2.34 | -0.055 | 0.07 | 0.07 | 0 | 0.29 | 0.51 | 0.38 | 0.37 | 0.21 | 0.4 |
| 3 | 130497531 | rs10512789 | -1.751 | -1.635 | 1.057 | 2.326 | 0.01 | -0.012 | -0.012 | 1.484 | 0.32 | 0.72 | 0.43 | 0.36 | 0.23 | 0.39 |
| 3 | 130499302 | rs16831670 | -1.373 | -1.218 | 0.438 | 2.148 | 0.009 | -0.016 | -0.016 | 0 | 0.28 | 0.24 | 0.35 | 0.47 | 0.26 | 0.41 |
| 3 | 130511038 | rs11708087 | -0.478 | -1.2 | 0.691 | 1.597 | -0.019 | -0.033 | -0.033 | 0 | 0.17 | 0.28 | 0.17 | 0.09 | 0.16 | 0.15 |
| 3 | 130513372 | rs11710411 | 1.287 | -0.716 | 0.24 | 1.424 | 0.049 | 0.069 | 0.069 | 0 | 0.43 | 0.27 | 0.29 | 0.29 | 0.31 | 0.29 |
| 3 | 130513617 | rs13064499 | 0.242 | -0.704 | 0.299 | 1.405 | -0.02 | -0.035 | -0.035 | 1.643 | 0.17 | 0.31 | 0.17 | 0.09 | 0.16 | 0.15 |
| 3 | 130514590 | rs12493491 | -0.006 | -0.693 | 0.036 | 2.038 | -0.111 | -0.05 | -0.05 | 0 | 0.64 | 0.9 | 0.52 | 0.71 | 0.48 | 0.48 |
| 3 | 130521374 | rs16831979 | 0.597 | -0.76 | 0.174 | 1.978 | -0.01 | -0.019 | -0.019 | 0 | 0.16 | 0.19 | 0.16 | 0.08 | 0.16 | 0.15 |
| 3 | 130522321 | rs13063749 | 1.289 | -0.811 | 0.184 | 2.076 | 0.042 | 0.018 | 0.018 | 0 | 0.43 | 0.31 | 0.31 | 0.47 | 0.32 | 0.3 |
| 3 | 130529906 | rs4682644 | 0.508 | -0.792 | -0.091 | 1.804 | -0.018 | -0.029 | -0.029 | 0 | 0.16 | 0.22 | 0.16 | 0.08 | 0.16 | 0.15 |
| 3 | 130530685 | rs6439260 | -0.568 | 0.373 | -0.799 | 1.832 | -0.139 | 0.025 | 0.025 | 0.478 | 0.76 | 0.98 | 0.63 | 0.9 | 0.64 | 0.81 |
| 3 | 130539871 | rs9814702 | 1.211 | -0.351 | -0.393 | 1.755 | 0.049 | 0.068 | 0.068 | 0 | 0.42 | 0.16 | 0.27 | 0.3 | 0.31 | 0.29 |
| 3 | 130546582 | rs10934962 | -0.418 | -0.058 | -0.814 | 1.499 | 0.02 | 0.042 | 0.042 | 0.421 | 0.76 | 0.89 | 0.82 | 0.92 | 0.78 | 0.8 |
| 3 | 130547086 | rs4487219 | -0.394 | -0.04 | -0.878 | 1.507 | 0.002 | -0.004 | -0.004 | 0 | 0.76 | 0.8 | 0.81 | 0.92 | 0.78 | 0.8 |
| 3 | 130547366 | rs9824635 | -0.025 | 0.129 | -1.507 | 1.699 | -0.156 | -0.078 | -0.078 | 0 | 0.27 | 0.08 | 0.45 | 0.3 | 0.46 | 0.47 |
| 3 | 130555052 | rs12493708 | 1.119 | -0.362 | -1.641 | 0.726 | 0.084 | 0.02 | 0.02 | 0.838 | 0.31 | 0.01 | 0.14 | 0.21 | 0.16 | 0.22 |
| 3 | 130559316 | rs6789666 | -0.391 | 0.819 | -0.57 | 1.199 | 0.001 | -0.001 | -0.001 | 0 | 0.31 | 0.31 | 0.53 | 0.6 | 0.47 | 0.51 |
| 3 | 130572374 | rs2669864 | -1.016 | 0.815 | -0.359 | 1.411 | -0.008 | -0.019 | -0.019 | 0.294 | 0.29 | 0.21 | 0.34 | 0.37 | 0.33 | 0.37 |
| 3 | 130573787 | rs852211 | 1.348 | 0.446 | 0.397 | 1.61 | 0.001 | 0.003 | 0.003 | 0 | 0.63 | 0.43 | 0.5 | 0.4 | 0.62 | 0.51 |
| 3 | 130588475 | rs1606490 | 0.395 | 0.497 | 0.602 | 1.725 | -0.002 | 0.067 | 0.067 | 0 | 0.94 | 0.83 | 0.85 | 0.79 | 0.94 | 0.9 |
| 3 | 130606850 | rs2760254 | -0.585 | 0.508 | 1.294 | 1.794 | -0.001 | -0.028 | -0.028 | 0 | 0.2 | 0.21 | 0.15 | 0.11 | 0.2 | 0.22 |
| 3 | 130612466 | rs2760252 | -0.491 | 0.474 | 1.249 | 1.78 | -0.001 | 0.01 | 0.01 | 2.253 | 0.19 | 0.12 | 0.13 | 0.04 | 0.19 | 0.17 |
| 3 | 130624071 | rs2669865 | -0.601 | 0.222 | 1.155 | 2.118 | 0 | -0.06 | -0.06 | 0 | 0.19 | 0.17 | 0.13 | 0 | 0.2 | 0.2 |
| 3 | 130657525 | rs1745663 | 0.488 | 0.193 | 1.65 | 2.11 | -0.006 | 0.056 | 0.056 | 0 | 0.93 | 0.8 | 0.85 | 0.8 | 0.93 | 0.89 |
| 3 | 130678263 | rs2760271 | -0.975 | 0.026 | 2.011 | 1.816 | -0.008 | -0.075 | -0.075 | 0.165 | 0.22 | 0.31 | 0.14 | 0.04 | 0.2 | 0.21 |
| 3 | 130693691 | rs218489 | 2.255 | 0.075 | 1.97 | 2.046 | 0.016 | -0.025 | -0.025 | 0 | 0.66 | 0.22 | 0.61 | 0.45 | 0.7 | 0.65 |
| 3 | 130709466 | rs218482 | 0.641 | 0.205 | 1.746 | 2.143 | -0.002 | 0.212 | 0.212 | 0 | 0.93 | 0.78 | 0.8 | 0.58 | 0.93 | 0.89 |
| 3 | 130715748 | rs3773814 | 0.089 | 0.289 | 1.41 | 1.911 | -0.033 | -0.033 | -0.033 | 0 | 0.06 | 0 | 0.14 | 0.1 | 0.14 | 0.12 |
| 3 | 130734061 | rs7641006 | -0.583 | 0.148 | 1.559 | 1.805 | -0.003 | 0.034 | 0.034 | 1.599 | 0.21 | 0.08 | 0.13 | 0.13 | 0.22 | 0.24 |
| 3 | 130737632 | rs218503 | 1.808 | 0.09 | 1.86 | 1.949 | -0.01 | 0.03 | 0.03 | 1.438 | 0.69 | 0.37 | 0.63 | 0.45 | 0.71 | 0.65 |
| 3 | 130738613 | rs3773816 | -0.672 | 0.12 | 1.771 | 1.959 | 0.001 | 0.008 | 0.008 | 2.507 | 0.21 | 0.13 | 0.14 | 0.13 | 0.22 | 0.24 |
| 3 | 130742029 | rs13073173 | -0.672 | 0.12 | 1.771 | 1.959 | 0.001 | 0.008 | 0.008 | 2.507 | 0.21 | 0.13 | 0.14 | 0.13 | 0.22 | 0.24 |
| 3 | 130742557 | rs13074752 | 0.252 | 0.117 | 1.904 | 1.964 | -0.001 | 0.207 | 0.207 | 0 | 0.93 | 0.8 | 0.8 | 0.58 | 0.93 | 0.89 |
| 3 | 130744384 | rs13066072 | -0.683 | 0.141 | 1.77 | 1.857 | 0.001 | 0.008 | 0.008 | 2.507 | 0.21 | 0.13 | 0.14 | 0.13 | 0.22 | 0.24 |
| 3 | 130764130 | rs11709564 | 1.42 | 0.054 | 1.219 | 1.533 | -0.017 | 0.038 | 0.038 | 0.35 | 0.08 | 0 | 0.04 | 0 | 0.08 | 0.01 |
| 3 | 130768474 | rs10428246 | -1.006 | 0.052 | 1.631 | 1.671 | -0.025 | -0.027 | -0.027 | 0 | 0.07 | 0.24 | 0.03 | 0 | 0.01 | 0.08 |
| 3 | 130786277 | rs2220775 | -0.831 | 0.231 | 1.677 | 1.788 | 0.024 | 0.023 | 0.023 | 0 | 0.08 | 0.16 | 0.16 | 0.15 | 0.15 | 0.18 |
| 3 | 130804076 | rs2029684 | -0.33 | 0.28 | 1.499 | 1.821 | -0.002 | 0.001 | 0.001 | 1.864 | 0.18 | 0.14 | 0.14 | 0.14 | 0.21 | 0.16 |
| 3 | 130830371 | rs16835887 | -0.388 | 0.273 | 1.514 | 1.855 | -0.001 | 0 | 0 | 0.077 | 0.18 | 0.13 | 0.14 | 0.15 | 0.21 | 0.16 |
| 3 | 130841101 | rs1033221 | 0.131 | 0.394 | 1.432 | 2.062 | -0.001 | 0.137 | 0.137 | 0 | 0.94 | 0.89 | 0.8 | 0.58 | 0.94 | 0.91 |
| 3 | 130884169 | rs959072 | -1.816 | 0.365 | 1.488 | 1.948 | 0.027 | 0.026 | 0.026 | 0 | 0.13 | 0.33 | 0.18 | 0.15 | 0.15 | 0.22 |
| 3 | 130893844 | rs7639697 | -1.816 | 0.365 | 1.488 | 1.948 | 0.027 | 0.026 | 0.026 | 0 | 0.13 | 0.33 | 0.18 | 0.15 | 0.15 | 0.22 |
| 3 | 130923834 | rs885076 | 0.607 | 0.476 | 1.001 | 1.945 | -0.007 | 0.015 | 0.015 | 0 | 0.89 | 0.89 | 0.76 | 0.54 | 0.86 | 0.9 |
| 3 | 130936691 | rs16836134 | -0.408 | 0.178 | 1.474 | 1.868 | 0.017 | 0.194 | 0.194 | 0 | 0.05 | 0.25 | 0.16 | 0.18 | 0.03 | 0.02 |
| 3 | 130947273 | rs3737999 | -2.441 | -0.147 | 1.791 | 1.994 | -0.035 | 0.206 | 0.206 | 1.771 | 0.1 | 0.6 | 0.22 | 0.3 | 0.05 | 0.07 |
| 3 | 130954093 | rs3965152 | -2.348 | -0.149 | 1.911 | 2.001 | -0.023 | 0.201 | 0.201 | 1.406 | 0.1 | 0.53 | 0.19 | 0.19 | 0.04 | 0.04 |
| 3 | 130957814 | rs10512803 | -1.457 | -0.133 | 2.263 | 2.008 | -0.005 | 0.21 | 0.21 | 0.108 | 0.08 | 0.32 | 0.16 | 0.19 | 0.04 | 0.04 |
| 3 | 130968017 | rs1991443 | -0.374 | 0.208 | 2.072 | 2.241 | -0.042 | -0.041 | -0.041 | 2.657 | 0.07 | 0.01 | 0.23 | 0.17 | 0.15 | 0.19 |
| 3 | 130972087 | rs1549600 | 1.413 | -0.047 | 2.141 | 2.27 | 0.001 | 0.006 | 0.006 | 0 | 0.82 | 0.78 | 0.81 | 0.7 | 0.75 | 0.79 |
| 3 | 130992385 | rs16836266 | -1.457 | 0.004 | 2.357 | 2.323 | 0.006 | 0.307 | 0.307 | 0.016 | 0.08 | 0.31 | 0.18 | 0.28 | 0.05 | 0.07 |
| 3 | 131009133 | rs6439279 | 1.332 | 0.027 | 2.399 | 2.315 | 0.007 | 0.022 | 0.022 | 0.047 | 0.83 | 0.76 | 0.8 | 0.7 | 0.76 | 0.79 |
| 3 | 131035264 | rs17334851 | -0.12 | -0.053 | 2.53 | 1.958 | 0.004 | -0.018 | -0.018 | 0 | 0.06 | 0.09 | 0.08 | 0 | 0.1 | 0.12 |
| 3 | 131041928 | rs748155 | -1.702 | -0.249 | 2.545 | 2.135 | 0.006 | 0.309 | 0.309 | 0.012 | 0.07 | 0.29 | 0.18 | 0.29 | 0.05 | 0.07 |
| 3 | 131047910 | rs17275526 | 0.452 | 0.067 | 2.113 | 1.791 | -0.004 | -0.001 | -0.001 | 0 | 0.13 | 0.12 | 0.07 | 0.03 | 0.14 | 0.22 |
| 3 | 131057916 | rs755568 | -1.591 | -0.002 | 2.196 | 2.1 | -0.007 | 0.254 | 0.254 | 0.241 | 0.07 | 0.25 | 0.17 | 0.29 | 0.05 | 0.08 |
| 3 | 131082104 | rs16836493 | -2.491 | -0.011 | 2.359 | 2.064 | -0.003 | 0.297 | 0.297 | 0 | 0.09 | 0.56 | 0.22 | 0.28 | 0.05 | 0.07 |
| 3 | 131082489 | rs3749392 | -2.861 | 0.065 | 2.311 | 1.968 | 0.067 | 0.245 | 0.245 | 0.279 | 0.05 | 0.38 | 0.07 | 0.12 | 0.02 | 0.05 |
| 3 | 131104016 | rs12631539 | -0.804 | 0.088 | 2.046 | 2.095 | 0.006 | 0.249 | 0.249 | 0 | 0.05 | 0.21 | 0.15 | 0.23 | 0.03 | 0.04 |

| | | | | | | | | | | | | | | | | |
|---|---|---|---|---|---|---|---|---|---|---|---|---|---|---|---|---|
| 3 | 131136797 | rs1352108 | 1.564 | 0.344 | 2.571 | 2.697 | 0.009 | 0.246 | 0.246 | 1.818 | 0.95 | 0.83 | 0.93 | 0.71 | 0.96 | 0.91 |
| 3 | 131141501 | rs4854560 | 3.015 | 0.409 | 2.457 | 2.711 | -0.01 | 0.278 | 0.278 | 0.372 | 0.91 | 0.39 | 0.86 | 0.63 | 0.95 | 0.9 |
| 3 | 131150266 | rs11710141 | 0.798 | 0.296 | 2.15 | 2.333 | -0.029 | 0.045 | 0.045 | 0 | 0.78 | 0.37 | 0.7 | 0.45 | 0.86 | 0.83 |
| 3 | 131183784 | rs7622001 | 0.172 | 0.761 | 1.711 | 2.034 | -0.042 | -0.099 | -0.099 | 0 | 0.75 | 0.96 | 0.54 | 0.39 | 0.73 | 0.58 |
| 3 | 131188528 | rs2291382 | 0.127 | 0.971 | 1.543 | 1.854 | -0.042 | 0.032 | 0.032 | 0 | 0.13 | 0.02 | 0.07 | 0.06 | 0.16 | 0.24 |
| 3 | 131190769 | rs11923992 | -2.808 | 0.858 | 2.506 | 2.334 | 0.082 | 0.124 | 0.124 | 1.488 | 0.05 | 0.32 | 0.05 | 0.09 | 0.04 | 0.05 |
| 3 | 131195753 | rs1486230 | 0.403 | 0.921 | 2.221 | 2.381 | -0.036 | -0.096 | -0.096 | 0.015 | 0.75 | 0.94 | 0.54 | 0.39 | 0.72 | 0.58 |
| 3 | 131224541 | rs11706454 | -0.523 | 1.278 | 2.072 | 2.074 | 0 | -0.008 | -0.008 | 0 | 0.06 | 0.09 | 0.03 | 0 | 0.07 | 0.02 |
| 3 | 131237247 | rs1843604 | -0.029 | 1.309 | 2.068 | 2.381 | -0.023 | -0.108 | -0.108 | 2.045 | 0.21 | 0.04 | 0.39 | 0.36 | 0.2 | 0.23 |
| 3 | 131239070 | rs9289391 | 1.562 | 1.292 | 2.28 | 2.386 | 0.065 | 0.423 | 0.423 | 0 | 0.95 | 0.68 | 0.84 | 0.67 | 0.95 | 0.81 |
| 3 | 131239666 | rs1054473 | -0.006 | 1.399 | 1.319 | 2.086 | -0.003 | -0.061 | -0.061 | 0 | 0.21 | 0.1 | 0.38 | 0.37 | 0.22 | 0.2 |
| 3 | 131240642 | rs7618892 | -1.091 | 1.454 | 1.358 | 1.79 | 0.016 | -0.027 | -0.027 | 0 | 0.06 | 0.18 | 0.07 | 0.01 | 0.05 | 0.11 |
| 3 | 131242502 | rs1385969 | -0.15 | 1.189 | 0.819 | 2.063 | 0.001 | -0.022 | -0.022 | 0.154 | 0.21 | 0.19 | 0.34 | 0.39 | 0.22 | 0.2 |
| 3 | 131249129 | rs4274722 | 1.069 | 1.264 | 1.189 | 1.908 | 0.028 | 0.026 | 0.026 | 0 | 0.8 | 0.4 | 0.76 | 0.71 | 0.74 | 0.63 |
| 3 | 131251079 | rs9942077 | -0.56 | 1.037 | 1.423 | 1.999 | -0.026 | -0.031 | -0.031 | 0 | 0.06 | 0.16 | 0.04 | 0 | 0.01 | 0.05 |
| 3 | 131254432 | rs4241351 | -0.579 | 1.62 | 1.323 | 2.398 | 0.013 | -0.119 | -0.119 | 2.394 | 0.23 | 0.07 | 0.43 | 0.45 | 0.19 | 0.26 |
| 3 | 131273951 | rs12106976 | -2.939 | 1.557 | 2.937 | 2.836 | 0.031 | -0.001 | -0.001 | 0.788 | 0.17 | 0.67 | 0.2 | 0.16 | 0.18 | 0.26 |
| 3 | 131278974 | rs7622111 | -2.792 | 1.497 | 3.025 | 2.752 | 0.052 | -0.043 | -0.043 | 0.676 | 0.16 | 0.54 | 0.18 | 0.1 | 0.17 | 0.17 |
| 3 | 131280659 | rs7611550 | -2.405 | 0.918 | 2.986 | 2.411 | -0.006 | -0.023 | -0.023 | 0 | 0.08 | 0.37 | 0.08 | 0.02 | 0.04 | 0.02 |
| 3 | 131282084 | rs7637123 | -2.358 | 0.918 | 2.986 | 2.412 | -0.005 | -0.022 | -0.022 | 0 | 0.08 | 0.37 | 0.08 | 0.02 | 0.04 | 0.02 |
| 3 | 131290201 | rs6771189 | -2.236 | 0.917 | 2.985 | 2.408 | -0.016 | -0.035 | -0.035 | 0 | 0.08 | 0.31 | 0.07 | 0.02 | 0.04 | 0.02 |
| 3 | 131290291 | rs6808429 | -3.521 | 0.936 | 3.623 | 2.715 | -0.009 | 0.024 | 0.024 | 0 | 0.23 | 0.63 | 0.25 | 0.3 | 0.16 | 0.33 |
| 3 | 131290333 | rs6808442 | -3.113 | 1.178 | 3.773 | 2.273 | -0.021 | -0.06 | -0.06 | 0 | 0.11 | 0.43 | 0.1 | 0.02 | 0.05 | 0.03 |
| 3 | 131290631 | rs16837140 | -1.951 | 1.131 | 3.68 | 2.346 | 0.075 | 0.035 | 0.035 | 0 | 0.07 | 0.23 | 0.06 | 0.06 | 0.07 | 0.12 |
| 3 | 131307353 | rs16837158 | -1.314 | 1.166 | 3.128 | 2.358 | 0.113 | 0.124 | 0.124 | 0 | 0.06 | 0.25 | 0.06 | 0.1 | 0.07 | 0.12 |
| 3 | 131319165 | rs6799539 | -1.691 | 1.175 | 3.09 | 2.364 | 0.122 | 0.134 | 0.134 | 0 | 0.07 | 0.31 | 0.07 | 0.12 | 0.07 | 0.18 |
| 3 | 131321322 | rs12632185 | -3.552 | 1.131 | 3.194 | 2.338 | 0.003 | 0.007 | 0.007 | 0 | 0.22 | 0.8 | 0.18 | 0.19 | 0.15 | 0.25 |
| 3 | 131322261 | rs6767491 | -1.714 | 1.145 | 3.217 | 2.343 | 0.115 | 0.121 | 0.121 | 0 | 0.07 | 0.31 | 0.07 | 0.12 | 0.07 | 0.18 |
| 3 | 131325437 | rs1502665 | -3.302 | 1.157 | 3.22 | 2.36 | 0 | 0 | 0 | 0 | 0.2 | 0.77 | 0.18 | 0.19 | 0.15 | 0.23 |
| 3 | 131330902 | rs10512820 | -2.482 | 1.264 | 3.252 | 2.405 | 0.053 | 0.044 | 0.044 | 0 | 0.11 | 0.32 | 0.1 | 0.13 | 0.11 | 0.17 |
| 3 | 131342265 | rs7621659 | -2.776 | 1.509 | 2.787 | 2.576 | 0.003 | -0.002 | -0.002 | 2.323 | 0.17 | 0.73 | 0.15 | 0.18 | 0.13 | 0.21 |
| 3 | 131346138 | rs7612052 | -2.776 | 1.131 | 2.844 | 1.71 | 0.003 | -0.003 | -0.003 | 0.38 | 0.17 | 0.73 | 0.15 | 0.18 | 0.13 | 0.21 |
| 3 | 131349427 | rs6439308 | -2.778 | 1.199 | 2.691 | 1.738 | 0.008 | -0.002 | -0.002 | 0 | 0.17 | 0.73 | 0.16 | 0.17 | 0.13 | 0.21 |
| 3 | 131352286 | rs6780136 | -2.752 | 1.21 | 2.696 | 1.743 | 0.008 | -0.001 | -0.001 | 0 | 0.17 | 0.72 | 0.16 | 0.17 | 0.12 | 0.21 |
| 3 | 131354026 | rs12634351 | -2.702 | 1.237 | 2.688 | 1.835 | 0.009 | 0.026 | 0.026 | 0 | 0.17 | 0.72 | 0.23 | 0.24 | 0.14 | 0.22 |
| 3 | 131363382 | rs3891970 | -1.68 | 1.294 | 2.182 | 1.806 | 0.083 | 0.096 | 0.096 | 0 | 0.08 | 0.29 | 0.15 | 0.15 | 0.09 | 0.14 |
| 3 | 131364117 | rs989765 | -1.736 | 1.297 | 2.164 | 1.707 | 0.09 | 0.153 | 0.153 | 0 | 0.09 | 0.32 | 0.16 | 0.19 | 0.09 | 0.14 |
| 3 | 131372608 | rs6766639 | -1.487 | 1.242 | 1.21 | 1.543 | 0.081 | 0.078 | 0.078 | 0 | 0.14 | 0.5 | 0.21 | 0.18 | 0.16 | 0.2 |
| 3 | 131374290 | rs9822908 | -0.898 | 0.889 | 0.636 | 1.544 | 0.016 | 0.079 | 0.079 | 0 | 0.2 | 0.31 | 0.22 | 0.37 | 0.24 | 0.22 |
| 3 | 131375337 | rs16837371 | -1.132 | 0.126 | 0.975 | 0.911 | -0.036 | -0.064 | -0.064 | 2.989 | 0.11 | 0.28 | 0.05 | 0.02 | 0.03 | 0.08 |
| 3 | 131377277 | rs2221308 | 0.874 | 0.305 | 1.077 | 1.836 | 0 | -0.013 | -0.013 | 0 | 0.92 | 0.88 | 0.9 | 0.96 | 0.86 | 0.89 |
| 3 | 131382627 | rs11924321 | -0.234 | 0.343 | 0.634 | 1.078 | -0.003 | -0.036 | -0.036 | 0 | 0.38 | 0.46 | 0.28 | 0.17 | 0.31 | 0.26 |
| 3 | 131383862 | rs17293396 | 1.017 | -0.34 | 0.569 | 0.867 | -0.099 | 0.246 | 0.246 | 0.61 | 0.42 | 0.07 | 0.32 | 0.06 | 0.5 | 0.41 |
| 3 | 131385562 | rs4854775 | 0.38 | -0.378 | 0.807 | 0.841 | -0.001 | 0.046 | 0.046 | 0 | 0.36 | 0.26 | 0.25 | 0.16 | 0.3 | 0.26 |
| 3 | 131387073 | rs4854782 | -0.451 | -0.385 | 0.792 | 0.839 | -0.003 | 0.006 | 0.006 | 0 | 0.93 | 0.93 | 0.9 | 0.96 | 0.86 | 0.89 |
| 3 | 131387344 | rs10512815 | 0.228 | -0.386 | 0.86 | 0.839 | -0.001 | -0.021 | -0.021 | 0 | 0.37 | 0.4 | 0.26 | 0.16 | 0.3 | 0.26 |
| 3 | 131399961 | rs9876228 | 0.202 | -0.409 | 1.582 | 0.959 | 0 | -0.01 | -0.01 | 0 | 0.54 | 0.44 | 0.62 | 0.79 | 0.55 | 0.62 |
| 3 | 131405652 | rs4854827 | 0.395 | -0.314 | 1.786 | 0.976 | 0 | 0.025 | 0.025 | 0 | 0.35 | 0.25 | 0.24 | 0.09 | 0.29 | 0.22 |
| 3 | 131408143 | rs1393560 | 0.354 | -0.265 | 1.657 | 1.086 | 0.014 | 0.136 | 0.136 | 0 | 0.9 | 0.81 | 0.73 | 0.6 | 0.83 | 0.8 |
| 3 | 131409530 | rs7619962 | 0.637 | -0.263 | 1.634 | 1.088 | 0.01 | 0.122 | 0.122 | 0 | 0.89 | 0.77 | 0.73 | 0.6 | 0.82 | 0.8 |
| 3 | 131412454 | rs6799202 | 0.825 | -0.294 | 1.582 | 1.296 | 0.038 | 0.306 | 0.306 | 0.143 | 0.89 | 0.51 | 0.75 | 0.68 | 0.85 | 0.83 |
| 3 | 131414045 | rs6769500 | -0.996 | -0.199 | 0.705 | -0.008 | -0.002 | 0.002 | 0.002 | 0 | 0.29 | 0.28 | 0.22 | 0.19 | 0.34 | 0.27 |
| 3 | 131414382 | rs1566695 | -1.324 | -0.225 | 0.691 | -0.001 | 0.001 | 0.003 | 0.003 | 0 | 0.64 | 0.67 | 0.48 | 0.31 | 0.64 | 0.53 |
| 3 | 131414469 | rs7433923 | 0.63 | -0.186 | 0.495 | -0.039 | -0.033 | 0.072 | 0.072 | 6.085 | 0.15 | 0.04 | 0.1 | 0.02 | 0.19 | 0.11 |
| 3 | 131415307 | rs11707675 | 0.16 | -0.052 | 0.754 | 0.221 | 0.014 | -0.032 | -0.032 | 1.942 | 0.08 | 0.16 | 0.09 | 0.02 | 0.12 | 0.11 |
| 3 | 131418966 | rs3883881 | 0.522 | -0.113 | 0.608 | 0.119 | 0 | -0.008 | -0.008 | 0 | 0.32 | 0.33 | 0.24 | 0.16 | 0.3 | 0.24 |
| 3 | 131420495 | rs6762188 | -0.651 | -0.042 | 0.237 | -0.148 | 0.001 | -0.033 | -0.033 | 0 | 0.45 | 0.52 | 0.36 | 0.22 | 0.44 | 0.4 |
| 3 | 131429611 | rs7637944 | -0.911 | 0.169 | 0.107 | 0.392 | 0.001 | -0.019 | -0.019 | 0 | 0.13 | 0.23 | 0.12 | 0.12 | 0.15 | 0.16 |
| 3 | 131433077 | rs995540 | 0.007 | -0.035 | -0.03 | 0.485 | 0.012 | 0.025 | 0.025 | 0 | 0.5 | 0.48 | 0.34 | 0.32 | 0.42 | 0.54 |
| 3 | 131433362 | rs7653478 | -0.416 | -0.283 | -0.232 | 0.247 | 0.004 | -0.046 | -0.046 | 0 | 0.14 | 0.43 | 0.08 | 0.05 | 0.11 | 0.15 |
| 3 | 131433979 | rs7611999 | -0.359 | -0.577 | 0.171 | 0.785 | -0.017 | 0.106 | 0.106 | 0.108 | 0.89 | 0.79 | 0.78 | 0.68 | 0.92 | 0.88 |
| 3 | 131434259 | rs6802186 | 1.006 | -0.544 | -0.108 | 0.536 | -0.03 | -0.021 | -0.021 | 0 | 0.32 | 0.17 | 0.34 | 0.35 | 0.37 | 0.23 |
| 3 | 131439834 | rs2061871 | 0.412 | -0.516 | 0.424 | 0.656 | -0.005 | 0.047 | 0.047 | 0.32 | 0.79 | 0.67 | 0.69 | 0.65 | 0.79 | 0.68 |
| 3 | 131443898 | rs7429462 | 0.955 | -0.7 | -0.335 | 0.364 | 0.003 | 0.026 | 0.026 | 0.121 | 0.32 | 0.14 | 0.26 | 0.23 | 0.31 | 0.34 |
| 3 | 131445951 | rs17294121 | -1.202 | -0.5 | 0.251 | 0.316 | 0.02 | -0.007 | -0.007 | 0.358 | 0.06 | 0.17 | 0.06 | 0.03 | 0.08 | 0.03 |
| 3 | 131450887 | rs7610027 | 0.474 | -0.803 | -0.45 | 0.008 | 0.018 | 0.032 | 0.032 | 0 | 0.21 | 0.1 | 0.12 | 0.11 | 0.16 | 0.16 |
| 3 | 131455600 | rs17294289 | 1.069 | -0.688 | -0.365 | 0.03 | -0.001 | 0.013 | 0.013 | 0 | 0.48 | 0.48 | 0.52 | 0.64 | 0.48 | 0.49 |
| 3 | 131456327 | rs4854621 | -0.325 | -0.853 | -0.391 | -0.024 | -0.023 | -0.04 | -0.04 | 0 | 0.68 | 0.51 | 0.79 | 0.82 | 0.76 | 0.71 |
| 3 | 131460188 | rs6806286 | 0.752 | -0.964 | -0.425 | 0.24 | 0.012 | 0.009 | 0.009 | 0.196 | 0.52 | 0.36 | 0.61 | 0.56 | 0.61 | 0.52 |

| | | | | | | | | | | | | | | | | |
|---|---|---|---|---|---|---|---|---|---|---|---|---|---|---|---|---|
| 3 | 131460681 | rs9850812 | 1.18 | -0.68 | -0.666 | 0.274 | -0.053 | -0.023 | -0.023 | 0.895 | 0.19 | 0.03 | 0.27 | 0.18 | 0.28 | 0.21 |
| 3 | 131464049 | rs6801905 | 0.148 | -0.433 | -0.087 | 0.552 | -0.001 | 0 | 0 | 0.016 | 0.34 | 0.31 | 0.27 | 0.41 | 0.32 | 0.44 |
| 3 | 131470173 | rs1913288 | -2.39 | -0.521 | 0.221 | 0.498 | -0.031 | -0.017 | -0.017 | 0 | 0.94 | 0.87 | 0.97 | 0.96 | 0.99 | 0.97 |
| 3 | 131470703 | rs9875838 | 1.458 | -0.472 | -0.087 | 0.472 | -0.063 | -0.001 | -0.001 | 0 | 0.19 | 0.02 | 0.28 | 0.15 | 0.27 | 0.19 |
| 3 | 131480369 | rs1513375 | 0.334 | -0.688 | 0.428 | 0.538 | 0.002 | 0.013 | 0.013 | 0.42 | 0.74 | 0.43 | 0.66 | 0.68 | 0.76 | 0.7 |
| 3 | 131481625 | rs1398766 | 0.814 | -0.628 | 0.064 | 0.578 | -0.088 | -0.053 | -0.053 | 0 | 0.25 | 0.04 | 0.4 | 0.3 | 0.35 | 0.26 |
| 3 | 131483386 | rs2036998 | 2.883 | -0.845 | 0.076 | 0.469 | -0.04 | -0.056 | -0.056 | 0 | 0.27 | 0.06 | 0.14 | 0.32 | 0.27 | 0.31 |
| 3 | 131483721 | rs1513380 | 0.663 | -0.675 | 0.367 | 0.494 | 0.005 | 0 | 0 | 0 | 0.58 | 0.51 | 0.4 | 0.57 | 0.52 | 0.58 |
| 3 | 131487158 | rs1877589 | 1.145 | -0.766 | 0.329 | 0.477 | -0.001 | 0 | 0 | 0 | 0.54 | 0.53 | 0.34 | 0.56 | 0.42 | 0.56 |
| 3 | 131487666 | rs1355782 | 1.896 | -0.769 | 0.196 | 0.484 | 0.029 | -0.01 | -0.01 | 0.29 | 0.44 | 0.21 | 0.23 | 0.42 | 0.36 | 0.43 |
| 3 | 131491981 | rs1913285 | -2.173 | -0.684 | 0.481 | 0.229 | -0.023 | -0.025 | -0.025 | 0 | 0.93 | 0.74 | 0.97 | 1 | 0.99 | 1 |
| 3 | 131495940 | rs9853154 | 1.961 | -0.601 | 0.153 | 0.24 | -0.06 | -0.006 | -0.006 | 0 | 0.18 | 0.02 | 0.27 | 0.14 | 0.26 | 0.19 |
| 3 | 131502605 | rs9848181 | -1.683 | -0.751 | 0.328 | 0.256 | -0.004 | 0.004 | 0.004 | 0 | 0.64 | 0.44 | 0.75 | 0.53 | 0.71 | 0.58 |
| 3 | 131503343 | rs1355784 | 0.835 | -0.565 | -0.077 | 0.322 | -0.018 | -0.061 | -0.061 | 0 | 0.32 | 0.1 | 0.36 | 0.38 | 0.31 | 0.33 |
| 3 | 131503502 | rs9835665 | -2.56 | -0.447 | 0.003 | 0.232 | -0.029 | -0.031 | -0.031 | 0 | 0.94 | 0.83 | 0.98 | 1 | 0.99 | 1 |
| 3 | 131505266 | rs6767389 | 2.469 | -0.219 | -0.517 | 0.357 | -0.012 | -0.017 | -0.017 | 0 | 0.26 | 0.13 | 0.14 | 0.28 | 0.27 | 0.3 |
| 3 | 131514365 | rs938241 | 0.777 | -0.052 | -0.561 | 0.434 | -0.037 | -0.045 | -0.045 | 3.255 | 0.09 | 0.01 | 0.09 | 0.17 | 0.14 | 0.07 |
| 3 | 131522552 | rs1112706 | -0.981 | -0.169 | -0.241 | 0.367 | 0.004 | -0.002 | -0.002 | 0 | 0.45 | 0.68 | 0.56 | 0.54 | 0.43 | 0.48 |
| 3 | 131522925 | rs16837851 | 0.914 | -0.33 | -0.278 | 0.356 | 0.006 | 0.032 | 0.032 | 0 | 0.15 | 0.21 | 0.33 | 0.35 | 0.19 | 0.26 |
| 3 | 131527944 | rs1913292 | -1.551 | -0.431 | -0.6 | 0.19 | 0.01 | 0.012 | 0.012 | 10.914 | 0.91 | 0.95 | 0.97 | 1 | 0.95 | 0.98 |
| 3 | 131532115 | rs885250 | 2.547 | -0.412 | -0.707 | 0.252 | -0.021 | -0.042 | -0.042 | 0 | 0.28 | 0.09 | 0.15 | 0.32 | 0.26 | 0.32 |
| 3 | 131533805 | rs9826180 | 1.616 | -0.273 | -0.677 | 0.144 | -0.068 | 0.007 | 0.007 | 0.857 | 0.18 | 0.02 | 0.26 | 0.14 | 0.26 | 0.19 |
| 3 | 131537007 | rs2137606 | -0.079 | -0.468 | -0.135 | 0.114 | 0.002 | 0 | 0 | 0 | 0.61 | 0.45 | 0.61 | 0.8 | 0.63 | 0.61 |
| 3 | 131541210 | rs16837878 | 0.749 | -0.217 | -0.383 | 0.381 | -0.037 | -0.045 | -0.045 | 3.109 | 0.18 | 0.12 | 0.37 | 0.29 | 0.25 | 0.24 |
| 3 | 131559574 | rs938243 | -2.145 | -0.376 | -0.011 | 0.11 | -0.027 | -0.074 | -0.074 | 0 | 0.82 | 0.78 | 0.9 | 1 | 0.9 | 0.92 |
| 3 | 131561519 | rs1513385 | 0.116 | -0.221 | -0.194 | 0.2 | 0.012 | 0.019 | 0.019 | 0 | 0.54 | 0.42 | 0.6 | 0.85 | 0.62 | 0.72 |
| 3 | 131563646 | rs13077768 | 2.239 | -0.01 | -0.586 | 0.665 | -0.013 | -0.055 | -0.055 | 1.559 | 0.28 | 0.08 | 0.14 | 0.31 | 0.26 | 0.33 |
| 3 | 131574307 | rs16837915 | -0.023 | -0.176 | -0.054 | 0.635 | -0.016 | -0.021 | -0.021 | 0 | 0.37 | 0.47 | 0.43 | 0.27 | 0.28 | 0.38 |
| 3 | 131577637 | rs10512830 | -0.023 | -0.176 | -0.054 | 0.635 | -0.016 | -0.021 | -0.021 | 0 | 0.37 | 0.47 | 0.43 | 0.27 | 0.28 | 0.38 |
| 3 | 131578097 | rs9848661 | 1.537 | -0.067 | -0.161 | 0.611 | -0.059 | 0.027 | 0.027 | 1.022 | 0.18 | 0.03 | 0.26 | 0.13 | 0.26 | 0.2 |
| 3 | 131578561 | rs1513378 | -0.503 | -0.038 | -0.141 | 0.733 | -0.004 | -0.001 | -0.001 | 0 | 0.43 | 0.48 | 0.49 | 0.45 | 0.38 | 0.41 |
| 3 | 131581274 | rs12629059 | 1.437 | 0.188 | -0.391 | 0.865 | -0.024 | -0.029 | -0.029 | 1.188 | 0.06 | 0.01 | 0.07 | 0.18 | 0.1 | 0.04 |
| 3 | 131585192 | rs7621543 | -1.675 | 0.035 | -0.185 | 0.817 | -0.003 | -0.02 | -0.02 | 0 | 0.64 | 0.75 | 0.79 | 0.56 | 0.64 | 0.54 |
| 3 | 131589527 | rs4854845 | 0.558 | -0.253 | 0.08 | 0.684 | -0.002 | -0.005 | -0.005 | 0 | 0.29 | 0.33 | 0.42 | 0.21 | 0.26 | 0.25 |
| 3 | 131600425 | rs6808555 | 1.174 | -0.035 | -0.198 | 0.834 | -0.002 | -0.001 | -0.001 | 0 | 0.07 | 0.06 | 0.06 | 0.1 | 0.09 | 0.09 |
| 3 | 131602528 | rs9864995 | 0.191 | -0.184 | -0.255 | 0.795 | -0.071 | -0.029 | -0.029 | 0.501 | 0.24 | 0.08 | 0.34 | 0.3 | 0.36 | 0.21 |
| 3 | 131607539 | rs2199351 | -0.168 | -0.322 | -0.049 | 0.685 | -0.012 | -0.017 | -0.017 | 0 | 0.39 | 0.58 | 0.44 | 0.27 | 0.28 | 0.31 |
| 3 | 131619401 | rs1461143 | -2.415 | -0.261 | 0.021 | 0.94 | 0 | -0.012 | -0.012 | 1.467 | 0.72 | 0.84 | 0.85 | 0.71 | 0.73 | 0.62 |
| 3 | 131621309 | rs1381073 | -0.78 | -0.284 | 0.035 | 0.968 | 0.009 | -0.002 | -0.002 | 0 | 0.46 | 0.63 | 0.51 | 0.49 | 0.39 | 0.35 |
| 3 | 131627205 | rs1225003 | -2.366 | -0.231 | 0.018 | 1.051 | 0.001 | 0.004 | 0.004 | 0 | 0.72 | 0.76 | 0.87 | 0.79 | 0.75 | 0.61 |
| 3 | 131627288 | rs883370 | -1.352 | -0.237 | 0.026 | 1.011 | -0.002 | 0.023 | 0.023 | 0.093 | 0.54 | 0.64 | 0.59 | 0.67 | 0.48 | 0.43 |
| 3 | 131627680 | rs1966947 | -0.392 | -0.413 | 0.378 | 0.959 | -0.024 | -0.004 | -0.004 | 0.275 | 0.17 | 0.31 | 0.09 | 0.15 | 0.11 | 0.14 |
| 3 | 131635559 | rs1225062 | -2.378 | -0.325 | 0.484 | 0.937 | 0 | 0.006 | 0.006 | 0 | 0.71 | 0.76 | 0.87 | 0.79 | 0.74 | 0.61 |
| 3 | 131638046 | rs876423 | -1.119 | -0.282 | 0.455 | 1.013 | 0.001 | -0.024 | -0.024 | 0 | 0.37 | 0.53 | 0.37 | 0.28 | 0.38 | 0.33 |
| 3 | 131639140 | rs2035095 | -0.148 | -0.367 | 0.617 | 1.008 | -0.006 | -0.001 | -0.001 | 0 | 0.16 | 0.17 | 0.07 | 0.15 | 0.11 | 0.14 |
| 3 | 131646464 | rs1225047 | -0.855 | -0.277 | 0.525 | 1.102 | 0.007 | 0.005 | 0.005 | 0 | 0.38 | 0.42 | 0.42 | 0.46 | 0.47 | 0.36 |
| 3 | 131647122 | rs1225041 | -2.255 | -0.247 | 0.566 | 1.146 | -0.003 | 0.016 | 0.016 | 0 | 0.72 | 0.87 | 0.87 | 0.78 | 0.74 | 0.61 |
| 3 | 131647348 | rs2369231 | -0.719 | -0.287 | 0.685 | 1.16 | -0.009 | 0.019 | 0.019 | 0.966 | 0.21 | 0.38 | 0.15 | 0.27 | 0.19 | 0.18 |
| 3 | 131647784 | rs1225037 | -0.055 | -0.469 | 0.353 | 1.022 | 0.001 | 0.015 | 0.015 | 0 | 0.43 | 0.36 | 0.4 | 0.34 | 0.43 | 0.36 |
| 3 | 131649935 | rs7610136 | -1.059 | -0.901 | 0.331 | 1.112 | 0.021 | 0.076 | 0.076 | 0.389 | 0.29 | 0.48 | 0.49 | 0.44 | 0.32 | 0.29 |
| 3 | 131650094 | rs16838123 | 0.479 | -0.798 | 0.31 | 1.129 | 0.001 | 0.014 | 0.014 | 0.833 | 0.15 | 0.18 | 0.39 | 0.3 | 0.19 | 0.16 |
| 3 | 131654264 | rs12633498 | 0.664 | -0.795 | 0.26 | 1.131 | -0.011 | -0.016 | -0.016 | 0.91 | 0.14 | 0.12 | 0.38 | 0.3 | 0.19 | 0.16 |
| 3 | 131659109 | rs7610352 | 0.678 | -0.794 | 0.261 | 1.168 | -0.011 | -0.013 | -0.013 | 0 | 0.14 | 0.12 | 0.38 | 0.33 | 0.19 | 0.17 |
| 3 | 131659745 | rs1225028 | -0.053 | -1.03 | 0.086 | 0.866 | -0.065 | -0.032 | -0.032 | 0 | 0.44 | 0.16 | 0.69 | 0.54 | 0.5 | 0.41 |
| 3 | 131663289 | rs6806898 | -0.715 | -0.919 | 0.325 | 0.843 | 0.005 | 0.004 | 0.004 | 0 | 0.15 | 0.37 | 0.11 | 0.13 | 0.14 | 0.12 |
| 3 | 131663987 | rs6785853 | -0.76 | -0.911 | 0.36 | 0.832 | -0.001 | -0.002 | -0.002 | 0 | 0.57 | 0.48 | 0.77 | 0.65 | 0.6 | 0.52 |
| 3 | 131664194 | rs1381077 | -0.724 | -1.316 | 0.364 | 0.867 | 0.008 | 0.015 | 0.015 | 0 | 0.15 | 0.31 | 0.1 | 0.14 | 0.14 | 0.13 |
| 3 | 131668941 | rs1225020 | -2.457 | -1.274 | 0.386 | 0.703 | 0.001 | -0.016 | -0.016 | 0 | 0.72 | 0.64 | 0.86 | 0.78 | 0.74 | 0.65 |
| 3 | 131673153 | rs6798778 | -0.643 | -1.31 | 0.28 | 0.648 | 0.006 | 0 | 0 | 0 | 0.15 | 0.27 | 0.1 | 0.13 | 0.15 | 0.13 |
| 3 | 131674049 | rs2613981 | -2.689 | -1.315 | 0.204 | 0.644 | -0.001 | 0.001 | 0.001 | 0 | 0.73 | 0.78 | 0.87 | 0.78 | 0.74 | 0.65 |
| 3 | 131674718 | rs1870712 | -0.768 | -1.299 | 0.031 | 0.621 | -0.004 | 0.016 | 0.016 | 0 | 0.57 | 0.34 | 0.76 | 0.65 | 0.6 | 0.52 |
| 3 | 131676584 | rs1919987 | -0.118 | -1.345 | -0.214 | 0.615 | -0.019 | -0.023 | -0.023 | 0 | 0.47 | 0.25 | 0.56 | 0.51 | 0.5 | 0.46 |
| 3 | 131680868 | rs9882252 | 0.983 | -1.355 | -0.492 | 0.406 | -0.057 | 0.027 | 0.027 | 0 | 0.29 | 0.01 | 0.17 | 0.2 | 0.3 | 0.24 |
| 3 | 131682092 | rs13082797 | -0.171 | -1.369 | -0.308 | 0.401 | 0.004 | 0.12 | 0.12 | 0.016 | 0.43 | 0.18 | 0.27 | 0.22 | 0.44 | 0.4 |
| 3 | 131689596 | rs1381083 | -1.854 | -1.238 | 0.233 | 0.437 | 0.054 | 0.044 | 0.044 | 2.07 | 0.06 | 0.5 | 0.03 | 0 | 0 | 0 |
| 3 | 131693006 | rs748731 | -0.902 | -1.063 | -0.573 | 0.54 | -0.011 | 0.013 | 0.013 | 0 | 0.18 | 0.1 | 0.26 | 0.15 | 0.21 | 0.22 |
| 3 | 131693200 | rs2077315 | -1.756 | -1.173 | -0.5 | 0.664 | -0.003 | 0.003 | 0.003 | 0 | 0.74 | 0.82 | 0.89 | 0.79 | 0.75 | 0.63 |
| 3 | 131713566 | rs1320900 | -0.593 | -1 | 0.366 | 1.195 | -0.029 | -0.033 | -0.033 | 0 | 0.31 | 0.13 | 0.45 | 0.35 | 0.33 | 0.36 |
| 3 | 131716105 | rs9834560 | -1.254 | -1.162 | 0.66 | 0.857 | -0.001 | 0 | 0 | 0 | 0.43 | 0.49 | 0.52 | 0.49 | 0.41 | 0.4 |

| | | | | | | | | | | | | | | | | |
|---|---|---|---|---|---|---|---|---|---|---|---|---|---|---|---|---|
| 3 | 131718729 | rs6806453 | -0.538 | -1.193 | 0.712 | 0.194 | -0.033 | -0.058 | -0.058 | 0.086 | 0.25 | 0.51 | 0.15 | 0.15 | 0.2 | 0.22 |
| 3 | 131723626 | rs4854866 | 0.051 | -1.151 | 0.771 | 0.475 | -0.004 | 0.193 | 0.193 | 0.11 | 0.84 | 0.6 | 0.7 | 0.64 | 0.86 | 0.9 |
| 3 | 131725225 | rs2001306 | -0.105 | -1.16 | 0.838 | 0.504 | 0.037 | 0.183 | 0.183 | 1.904 | 0.9 | 0.65 | 0.75 | 0.76 | 0.92 | 0.93 |
| 3 | 131725318 | rs11715147 | 0.997 | -1.11 | 0.62 | 0.221 | -0.068 | 0.07 | 0.07 | 0.003 | 0.14 | 0.01 | 0.13 | 0.01 | 0.2 | 0.09 |
| 3 | 131726722 | rs7647029 | 0.865 | -1.249 | 0.77 | 0.176 | -0.001 | -0.025 | -0.025 | 5.592 | 0.62 | 0.61 | 0.34 | 0.25 | 0.62 | 0.71 |
| 3 | 131730242 | rs10512839 | 0.274 | -1.251 | 0.792 | 0.205 | 0.015 | 0.036 | 0.036 | 1.134 | 0.72 | 0.63 | 0.38 | 0.38 | 0.68 | 0.75 |
| 3 | 131733945 | rs16838297 | 0.777 | -1.294 | 0.957 | 0.169 | -0.001 | -0.004 | -0.004 | 0 | 0.16 | 0.21 | 0.11 | 0.13 | 0.19 | 0.15 |
| 3 | 131734203 | rs9827101 | 1.194 | -1.281 | 0.926 | 0.141 | 0 | -0.019 | -0.019 | 0 | 0.53 | 0.54 | 0.28 | 0.25 | 0.51 | 0.6 |
| 3 | 131734429 | rs9864905 | 0.596 | -1.239 | 1.09 | 0.092 | 0.006 | -0.051 | -0.051 | 0 | 0.1 | 0.16 | 0.07 | 0 | 0.11 | 0.11 |
| 3 | 131743254 | rs2369364 | 0.804 | -1.259 | 1.075 | 0.124 | 0.009 | 0.008 | 0.008 | 0.159 | 0.62 | 0.57 | 0.33 | 0.39 | 0.57 | 0.64 |
| 3 | 131743312 | rs9847058 | 1.175 | -1.258 | 1.04 | 0.121 | -0.01 | 0.019 | 0.019 | 0 | 0.08 | 0.03 | 0.05 | 0 | 0.11 | 0.11 |
| 3 | 131743694 | rs2887465 | 0.204 | -1.26 | 1.048 | 0.138 | 0.002 | 0.007 | 0.007 | 1.299 | 0.71 | 0.69 | 0.4 | 0.4 | 0.68 | 0.75 |
| 3 | 131743988 | rs6439335 | -0.94 | -1.245 | 0.992 | 0.189 | 0 | -0.011 | -0.011 | 0 | 0.92 | 0.96 | 0.76 | 0.74 | 0.92 | 0.93 |
| 3 | 131751649 | rs10934987 | 0.776 | -1.284 | 1.06 | 0.162 | 0 | 0.008 | 0.008 | 0 | 0.67 | 0.62 | 0.58 | 0.6 | 0.63 | 0.67 |
| 3 | 131751775 | rs13076253 | 1.346 | -1.286 | 1.046 | 0.147 | -0.003 | 0.001 | 0.001 | 0.073 | 0.13 | 0.19 | 0.09 | 0.12 | 0.15 | 0.12 |
| 3 | 131753688 | rs7649058 | -0.034 | -1.312 | 1.023 | 0.215 | 0.004 | -0.031 | -0.031 | 0 | 0.25 | 0.34 | 0.18 | 0.13 | 0.3 | 0.26 |
| 3 | 131754721 | rs1569283 | 0.588 | -1.323 | 0.978 | 0.215 | 0.008 | 0.022 | 0.022 | 0 | 0.21 | 0.26 | 0.37 | 0.35 | 0.25 | 0.18 |
| 3 | 131756012 | rs16838396 | 1.68 | -1.35 | 0.931 | 0.199 | -0.001 | -0.05 | -0.05 | 1.761 | 0.08 | 0 | 0.04 | 0.13 | 0.06 | 0.03 |
| 3 | 131756349 | rs1569282 | 1.451 | -1.341 | 1.012 | 0.205 | 0.01 | 0.034 | 0.034 | 0 | 0.46 | 0.37 | 0.22 | 0.26 | 0.38 | 0.49 |
| 3 | 131758077 | rs1320903 | 0.043 | -1.35 | 1.018 | 0.201 | 0.003 | -0.03 | -0.03 | 0 | 0.25 | 0.33 | 0.18 | 0.14 | 0.3 | 0.26 |
| 3 | 131761368 | rs2951406 | -1.119 | -1.289 | 0.827 | 0.271 | -0.01 | -0.036 | -0.036 | 0 | 0.93 | 0.99 | 0.77 | 0.74 | 0.93 | 0.93 |
| 3 | 131766671 | rs1881913 | -0.328 | -1.348 | 0.989 | 0.226 | -0.002 | 0.017 | 0.017 | 0 | 0.33 | 0.35 | 0.43 | 0.5 | 0.31 | 0.28 |
| 3 | 131776192 | rs3009505 | 0.44 | -1.497 | 1.009 | 0.5 | 0.001 | -0.006 | -0.006 | 0 | 0.08 | 0.05 | 0.24 | 0.26 | 0.08 | 0.08 |
| 3 | 131783593 | rs6792706 | 0.426 | -1.498 | 0.965 | 0.404 | 0 | -0.003 | -0.003 | 0 | 0.18 | 0.22 | 0.13 | 0.14 | 0.22 | 0.17 |
| 3 | 131785594 | rs3009501 | 0.304 | -1.549 | 0.997 | 0.353 | 0 | -0.002 | -0.002 | 0 | 0.59 | 0.43 | 0.54 | 0.54 | 0.62 | 0.63 |
| 3 | 131786302 | rs11924777 | 0.832 | -1.515 | 1.091 | 0.362 | -0.017 | -0.02 | -0.02 | 0 | 0.25 | 0.17 | 0.37 | 0.34 | 0.31 | 0.28 |
| 3 | 131786554 | rs3009500 | 1.089 | -1.138 | 0.921 | 0.404 | -0.012 | 0.024 | 0.024 | 0 | 0.41 | 0.21 | 0.43 | 0.37 | 0.4 | 0.46 |
| 3 | 131789220 | rs13091881 | 1.746 | -1.092 | 0.512 | 0.098 | -0.017 | 0.081 | 0.081 | 0 | 0.33 | 0.17 | 0.19 | 0.11 | 0.33 | 0.38 |
| 3 | 131789864 | rs9842509 | 0.027 | -0.721 | 0.172 | 0.329 | 0 | -0.002 | -0.002 | 0 | 0.19 | 0.22 | 0.13 | 0.17 | 0.23 | 0.18 |
| 3 | 131797114 | rs3009491 | -1.801 | -0.601 | -0.186 | 0.218 | -0.006 | 0.018 | 0.018 | 0 | 0.69 | 0.8 | 0.83 | 0.89 | 0.68 | 0.64 |
| 3 | 131797500 | rs6439340 | -0.168 | 0.256 | 0.402 | 0.719 | 0.003 | -0.017 | -0.017 | 0 | 0.19 | 0.51 | 0.16 | 0.14 | 0.2 | 0.15 |
| 3 | 131797662 | rs3009490 | -0.683 | 0.23 | 0.373 | 0.894 | 0.005 | -0.012 | -0.012 | 0.101 | 0.1 | 0.09 | 0.21 | 0.24 | 0.08 | 0.08 |
| 3 | 131800249 | rs9289416 | -1.042 | 0.233 | 0.355 | 0.898 | -0.018 | 0.049 | 0.049 | 0.004 | 0.28 | 0.51 | 0.35 | 0.39 | 0.27 | 0.28 |
| 3 | 131801556 | rs6765085 | 0.422 | 0.39 | -0.382 | 0.98 | -0.002 | 0.002 | 0.002 | 0 | 0.16 | 0.13 | 0.13 | 0.14 | 0.2 | 0.15 |
| 3 | 131807866 | rs9818159 | 0.314 | 0.395 | -0.688 | 0.902 | -0.079 | -0.082 | -0.082 | 0.082 | 0.23 | 0.05 | 0.35 | 0.35 | 0.31 | 0.26 |
| 3 | 131808995 | rs2091650 | 0.03 | 0.094 | -0.277 | 0.619 | -0.037 | -0.033 | -0.033 | 0.051 | 0.39 | 0.23 | 0.48 | 0.49 | 0.52 | 0.41 |
| 3 | 131832227 | rs4513497 | -0.193 | 0.418 | -0.565 | -0.071 | 0.024 | 0.08 | 0.08 | 0 | 0.51 | 0.34 | 0.72 | 0.75 | 0.6 | 0.53 |
| 3 | 131835569 | rs13074860 | 0.103 | 0.604 | -1.038 | 0.029 | 0.009 | 0.094 | 0.094 | 0.358 | 0.49 | 0.29 | 0.28 | 0.25 | 0.42 | 0.48 |
| 3 | 131837290 | rs9813389 | -1.133 | 0.867 | -0.992 | -0.138 | -0.007 | 0.007 | 0.007 | 0 | 0.08 | 0.08 | 0.04 | 0 | 0.08 | 0.08 |
| 3 | 131838077 | rs1356802 | -0.758 | 0.275 | -0.961 | -0.844 | 0.017 | 0.04 | 0.04 | 0.021 | 0.42 | 0.63 | 0.24 | 0.21 | 0.31 | 0.56 |
| 3 | 131842612 | rs2369372 | -0.452 | 0.668 | -1.009 | -0.75 | 0.012 | -0.026 | -0.026 | 0 | 0.62 | 0.72 | 0.62 | 0.5 | 0.65 | 0.66 |
| 3 | 131842788 | rs2369373 | 1.351 | 1.356 | -0.674 | -0.812 | -0.006 | -0.032 | -0.032 | 0 | 0.25 | 0.14 | 0.3 | 0.45 | 0.29 | 0.12 |
| 3 | 131843104 | rs16838754 | 0.068 | 1.046 | -0.146 | -1.029 | -0.009 | -0.091 | -0.091 | 1.235 | 0.15 | 0.32 | 0.1 | 0 | 0.12 | 0.16 |
| 3 | 131843203 | rs6785388 | -0.57 | 1.05 | -0.099 | -0.994 | -0.031 | -0.019 | -0.019 | 0.243 | 0.22 | 0.5 | 0.14 | 0.16 | 0.16 | 0.19 |
| 3 | 131849410 | rs11714053 | -0.576 | 1.03 | 0.01 | -1.003 | -0.03 | -0.019 | -0.019 | 5.614 | 0.22 | 0.47 | 0.14 | 0.16 | 0.16 | 0.19 |
| 3 | 131852707 | rs3851997 | 0.346 | 1.045 | -0.022 | -1.017 | 0 | -0.001 | -0.001 | 0 | 0.46 | 0.41 | 0.52 | 0.49 | 0.53 | 0.5 |
| 3 | 131853424 | rs9681094 | -1.249 | 1.088 | -0.183 | -0.99 | -0.016 | -0.011 | -0.011 | 1.397 | 0.68 | 0.91 | 0.66 | 0.66 | 0.69 | 0.7 |
| 3 | 131854840 | rs1521286 | -1.009 | 0.959 | 0.32 | -0.969 | -0.006 | 0.004 | 0.004 | 0 | 0.67 | 0.75 | 0.53 | 0.65 | 0.63 | 0.64 |
| 3 | 131865787 | rs4083132 | -0.016 | 1.001 | 0.234 | -1.076 | -0.003 | 0.047 | 0.047 | 0 | 0.47 | 0.39 | 0.45 | 0.62 | 0.49 | 0.33 |
| 3 | 131867527 | rs3914906 | 0.998 | 0.955 | 0.125 | -1.044 | -0.001 | 0.015 | 0.015 | 0 | 0.19 | 0.1 | 0.1 | 0.05 | 0.16 | 0.33 |
| 3 | 131874596 | rs12486037 | -0.75 | 1.234 | 0.453 | -0.808 | 0.001 | -0.001 | -0.001 | 0 | 0.67 | 0.7 | 0.64 | 0.38 | 0.65 | 0.76 |
| 3 | 131875067 | rs3914305 | 1.297 | 1.011 | 0.46 | -0.759 | 0.01 | -0.066 | -0.066 | 0 | 0.26 | 0.11 | 0.31 | 0.45 | 0.27 | 0.2 |
| 3 | 131879029 | rs3844275 | 0.87 | 1.207 | 0.962 | -0.957 | -0.006 | -0.009 | -0.009 | 0 | 0.35 | 0.41 | 0.38 | 0.34 | 0.31 | 0.31 |
| 3 | 131879796 | rs3900591 | 0.731 | 1.176 | 1.091 | -0.971 | -0.002 | -0.002 | -0.002 | 0 | 0.36 | 0.53 | 0.39 | 0.34 | 0.31 | 0.31 |
| 3 | 131884654 | rs3914303 | 0.208 | 1.276 | 1.144 | -1.003 | -0.003 | 0.054 | 0.054 | 0.028 | 0.63 | 0.69 | 0.75 | 0.79 | 0.66 | 0.47 |
| 3 | 131884794 | rs9876304 | 1.21 | 1.262 | 1.192 | -0.984 | -0.001 | -0.001 | -0.001 | 0 | 0.32 | 0.29 | 0.35 | 0.34 | 0.3 | 0.24 |
| 3 | 131895292 | rs11720059 | 0.398 | 1.286 | 1.095 | -0.975 | -0.027 | -0.041 | -0.041 | 0.879 | 0.07 | 0 | 0.07 | 0.17 | 0.1 | 0.11 |
| 3 | 131901570 | rs1505807 | -2.531 | 1.217 | 1.191 | -1.016 | 0.028 | -0.051 | -0.051 | 0 | 0.09 | 0.62 | 0.15 | 0.01 | 0.12 | 0.11 |
| 3 | 131906594 | rs1395006 | 0.857 | 1.186 | 1.194 | -1.068 | 0.003 | -0.032 | -0.032 | 0 | 0.56 | 0.43 | 0.63 | 0.78 | 0.54 | 0.35 |
| 3 | 131908305 | rs7630827 | 0.845 | 1.046 | 0.885 | -1.149 | -0.002 | 0.008 | 0.008 | 0.431 | 0.56 | 0.58 | 0.65 | 0.79 | 0.54 | 0.37 |
| 3 | 131916052 | rs9831157 | 1.312 | 1.002 | 1.339 | -1.147 | -0.009 | -0.008 | -0.008 | 0.622 | 0.29 | 0.21 | 0.33 | 0.34 | 0.29 | 0.25 |
| 3 | 131919483 | rs9289423 | 1.433 | 1.001 | 1.233 | -1.147 | -0.033 | -0.031 | -0.031 | 0 | 0.28 | 0.12 | 0.33 | 0.34 | 0.29 | 0.25 |
| 3 | 131932053 | rs884431 | -0.197 | 1.029 | 1.317 | -1.154 | -0.022 | -0.015 | -0.015 | 0 | 0.44 | 0.68 | 0.52 | 0.52 | 0.5 | 0.38 |
| 3 | 131937682 | rs11710548 | 1.915 | 1.089 | 0.951 | -1.131 | -0.045 | -0.18 | -0.18 | 0 | 0.23 | 0.02 | 0.27 | 0.45 | 0.25 | 0.12 |
| 3 | 131938708 | rs10934991 | -0.911 | 1.116 | 1.09 | -1.1 | 0.011 | 0.008 | 0.008 | 0 | 0.12 | 0.19 | 0.16 | 0.18 | 0.21 | 0.13 |
| 3 | 131944637 | rs11714139 | -0.582 | 1.059 | 0.984 | -1.099 | 0.001 | -0.014 | -0.014 | 0 | 0.11 | 0.06 | 0.1 | 0.17 | 0.12 | 0.18 |
| 3 | 131947937 | rs9878957 | 1.332 | 1.046 | 1.054 | -1.12 | 0.028 | 0.014 | 0.014 | 0.433 | 0.83 | 0.56 | 0.83 | 0.82 | 0.79 | 0.8 |
| 3 | 131952108 | rs12494585 | 1.369 | 1.002 | 0.48 | -1.076 | -0.048 | -0.045 | -0.045 | 0 | 0.28 | 0.04 | 0.37 | 0.33 | 0.3 | 0.24 |
| 3 | 131962828 | rs10512846 | 0.408 | 0.999 | 0.566 | -1.186 | -0.011 | 0.119 | 0.119 | 0.164 | 0.29 | 0.13 | 0.16 | 0.04 | 0.24 | 0.44 |

| | | | | | | | | | | | | | | | | |
|---|---|---|---|---|---|---|---|---|---|---|---|---|---|---|---|---|
| 3 | 131964467 | rs7653315 | -0.841 | 1.005 | 0.727 | -1.147 | 0.009 | 0.049 | 0.049 | 1.186 | 0.43 | 0.67 | 0.3 | 0.21 | 0.36 | 0.62 |
| 3 | 131980800 | rs4854895 | -0.152 | 0.899 | 0.555 | -1.068 | -0.001 | 0 | 0 | 0 | 0.59 | 0.44 | 0.45 | 0.57 | 0.6 | 0.58 |
| 3 | 131981255 | rs9858118 | 1.272 | 0.999 | -0.029 | -1.075 | -0.017 | 0.008 | 0.008 | 0 | 0.27 | 0.12 | 0.36 | 0.26 | 0.27 | 0.22 |
| 3 | 131995246 | rs2310228 | -0.497 | 1.015 | 0.192 | -1.2 | -0.002 | 0.005 | 0.005 | 0.138 | 0.38 | 0.67 | 0.28 | 0.06 | 0.36 | 0.54 |
| 3 | 132000576 | rs3931393 | 0.048 | 0.882 | -0.055 | -1.004 | -0.001 | -0.003 | -0.003 | 0 | 0.9 | 0.92 | 0.86 | 0.85 | 0.89 | 0.88 |
| 3 | 132002082 | rs11927890 | 0.163 | 0.839 | 0.027 | -1.217 | 0.001 | 0.077 | 0.077 | 0.007 | 0.3 | 0.19 | 0.19 | 0.06 | 0.27 | 0.46 |
| 3 | 132003117 | rs2684303 | 0.941 | 0.639 | -0.408 | -1.226 | 0.026 | -0.038 | -0.038 | 0 | 0.36 | 0.05 | 0.31 | 0.59 | 0.28 | 0.4 |
| 3 | 132006118 | rs1505829 | -0.457 | 0.5 | -0.101 | -1.269 | -0.002 | 0.005 | 0.005 | 0.069 | 0.44 | 0.59 | 0.58 | 0.41 | 0.55 | 0.38 |
| 3 | 132010325 | rs1910955 | 0.935 | 0.5 | 0.219 | -1.36 | -0.008 | -0.002 | -0.002 | 0 | 0.28 | 0.25 | 0.43 | 0.3 | 0.38 | 0.24 |
| 3 | 132012079 | rs2670104 | 1.423 | 0.092 | 0.25 | -1.572 | -0.002 | -0.016 | -0.016 | 0 | 0.38 | 0.2 | 0.34 | 0.57 | 0.37 | 0.45 |
| 3 | 132021906 | rs1100396 | -1.343 | 0.28 | 0.464 | -1.765 | -0.016 | -0.062 | -0.062 | 0.189 | 0.34 | 0.55 | 0.22 | 0.13 | 0.25 | 0.31 |
| 3 | 132031058 | rs11709247 | -0.067 | 0.488 | 0.49 | -1.413 | 0.03 | -0.028 | -0.028 | 0 | 0.16 | 0.34 | 0.14 | 0.11 | 0.18 | 0.15 |
| 3 | 132034199 | rs3844501 | 0.452 | 0.245 | 0.403 | -1.371 | 0.003 | -0.007 | -0.007 | 0 | 0.13 | 0.21 | 0.1 | 0.11 | 0.14 | 0.14 |
| 3 | 132054049 | rs310001 | -1.149 | 0.424 | -0.338 | -1.156 | -0.026 | 0.078 | 0.078 | 0.009 | 0.65 | 0.78 | 0.72 | 0.94 | 0.61 | 0.69 |
| 3 | 132059239 | rs2310230 | -0.592 | -0.467 | -0.38 | -0.167 | 0.006 | -0.002 | -0.002 | 0 | 0.33 | 0.26 | 0.35 | 0.63 | 0.23 | 0.45 |
| 3 | 132059280 | rs2310229 | -0.969 | -0.712 | -0.463 | -0.224 | 0.069 | 0.032 | 0.032 | 0 | 0.42 | 0.2 | 0.37 | 0.63 | 0.22 | 0.46 |
| 3 | 132063643 | rs972775 | 0.998 | 0.467 | 0.579 | 0.151 | -0.001 | -0.005 | -0.005 | 0 | 0.61 | 0.39 | 0.49 | 0.16 | 0.55 | 0.54 |
| 3 | 132065564 | rs16839122 | 0.684 | 0.396 | 0.707 | 0.425 | 0.002 | 0.004 | 0.004 | 0 | 0.63 | 0.45 | 0.57 | 0.6 | 0.57 | 0.67 |
| 3 | 132066160 | rs17241868 | 1.979 | 0.515 | 0.313 | 0.008 | -0.002 | 0.105 | 0.105 | 1.298 | 0.43 | 0.25 | 0.38 | 0.08 | 0.38 | 0.38 |
| 3 | 132075417 | rs3853148 | 1.055 | 0.522 | 0.353 | 0.066 | -0.002 | 0.031 | 0.031 | 0 | 0.58 | 0.36 | 0.48 | 0.12 | 0.54 | 0.52 |
| 3 | 132075746 | rs1042330 | 1.055 | 0.522 | 0.353 | 0.066 | -0.002 | 0.031 | 0.031 | 0 | 0.58 | 0.36 | 0.48 | 0.12 | 0.54 | 0.52 |
| 3 | 132076551 | rs14192 | 1.055 | 0.582 | 0.086 | 0.059 | -0.002 | 0.031 | 0.031 | 0 | 0.58 | 0.36 | 0.48 | 0.12 | 0.54 | 0.52 |
| 3 | 132080342 | rs16839139 | -0.649 | 1.016 | 0.311 | 0.787 | -0.014 | 0.294 | 0.294 | 0 | 0.05 | 0.17 | 0.09 | 0.49 | 0.02 | 0.14 |
| 3 | 132081831 | rs3905051 | -0.469 | 1.022 | 0.284 | 0.744 | 0.001 | -0.002 | -0.002 | 2.268 | 0.2 | 0.2 | 0.2 | 0.53 | 0.18 | 0.27 |
| 3 | 132085885 | rs9873881 | -0.047 | 1.111 | 0.376 | 0.746 | -0.032 | -0.045 | -0.045 | 0.134 | 0.22 | 0.12 | 0.29 | 0.29 | 0.27 | 0.28 |
| 3 | 132102933 | rs6776932 | -1.545 | 1 | 0.373 | 0.724 | -0.001 | -0.022 | -0.022 | 0 | 0.46 | 0.62 | 0.56 | 0.86 | 0.54 | 0.58 |
| 3 | 132111213 | rs2310232 | -1.017 | 0.821 | -0.011 | 1.457 | 0 | 0.008 | 0.008 | 1.979 | 0.21 | 0.23 | 0.21 | 0.53 | 0.18 | 0.27 |
| 3 | 132126527 | rs7620445 | -0.959 | 1.155 | -0.001 | 1.452 | -0.004 | 0.004 | 0.004 | 0 | 0.11 | 0.06 | 0.26 | 0.3 | 0.16 | 0.28 |
| 3 | 132129504 | rs17244458 | -0.856 | 0.923 | 0.694 | 1.098 | -0.017 | -0.012 | -0.012 | 0 | 0.22 | 0.35 | 0.19 | 0.17 | 0.16 | 0.17 |
| 3 | 132132699 | rs11706084 | -0.816 | 0.907 | 1.052 | 1.091 | -0.013 | -0.009 | -0.009 | 0 | 0.21 | 0.35 | 0.19 | 0.17 | 0.16 | 0.17 |
| 3 | 132144164 | rs9882900 | -0.555 | 0.807 | 1.572 | 1.038 | -0.003 | 0.005 | 0.005 | 0 | 0.07 | 0.16 | 0.05 | 0.05 | 0.06 | 0.07 |
| 3 | 132149551 | rs6792616 | -0.677 | 0.803 | 1.503 | 1.036 | -0.013 | -0.009 | -0.009 | 0.086 | 0.21 | 0.35 | 0.19 | 0.18 | 0.16 | 0.17 |
| 3 | 132150183 | rs3860499 | -2.141 | 0.666 | 1.468 | 1.047 | -0.006 | -0.082 | -0.082 | 0.586 | 0.41 | 0.9 | 0.52 | 0.47 | 0.37 | 0.52 |
| 3 | 132157429 | rs3849409 | -2.628 | 0.613 | 1.115 | 0.81 | -0.052 | 0.307 | 0.307 | 0 | 0.52 | 0.92 | 0.52 | 0.86 | 0.49 | 0.69 |
| 3 | 132160713 | rs10490862 | -0.076 | 0.516 | 0.948 | 0.838 | -0.008 | -0.034 | -0.034 | 0 | 0.17 | 0.01 | 0.14 | 0.15 | 0.13 | 0.17 |
| 3 | 132163522 | rs9835254 | -2.008 | 0.537 | 1.288 | 0.775 | 0.004 | 0.087 | 0.087 | 0 | 0.46 | 0.7 | 0.45 | 0.81 | 0.43 | 0.62 |
| 3 | 132172395 | rs7613098 | -0.13 | 0.65 | 1.218 | 0.931 | -0.011 | -0.014 | -0.014 | 0.171 | 0.1 | 0.05 | 0.19 | 0.18 | 0.15 | 0.28 |
| 3 | 132176651 | rs11717545 | -0.97 | 0.571 | 1.352 | 0.932 | 0.006 | 0.206 | 0.206 | 0.027 | 0.17 | 0.4 | 0.1 | 0.49 | 0.15 | 0.18 |
| 3 | 132211993 | rs9833058 | -1.01 | 0.777 | 1.417 | 0.881 | 0.013 | 0.025 | 0.025 | 0 | 0.17 | 0.21 | 0.25 | 0.23 | 0.23 | 0.34 |
| 3 | 132218883 | rs16839317 | -0.428 | 0.375 | 1.422 | 0.814 | -0.018 | -0.009 | -0.009 | 4.364 | 0.18 | 0.3 | 0.17 | 0.15 | 0.13 | 0.15 |
| 3 | 132219554 | rs11925382 | -0.249 | 0.369 | 0.853 | 0.818 | 0.022 | 0.005 | 0.005 | 0 | 0.17 | 0.1 | 0.14 | 0.15 | 0.11 | 0.14 |
| 3 | 132240339 | rs7622472 | -2.444 | 0.441 | 0.747 | 0.671 | -0.066 | 0.319 | 0.319 | 0 | 0.52 | 0.91 | 0.5 | 0.85 | 0.48 | 0.68 |
| 3 | 132242642 | rs9833627 | -1.157 | 0.613 | 0.952 | 0.922 | 0.009 | 0.026 | 0.026 | 0 | 0.17 | 0.22 | 0.25 | 0.23 | 0.22 | 0.34 |
| 3 | 132244313 | rs9843883 | -0.318 | 0.612 | 0.621 | 0.936 | -0.041 | -0.064 | -0.064 | 0 | 0.11 | 0.01 | 0.18 | 0.19 | 0.14 | 0.29 |
| 3 | 132255721 | rs2133692 | -2.223 | 0.453 | 0.524 | 0.668 | -0.047 | 0.32 | 0.32 | 0 | 0.51 | 0.9 | 0.49 | 0.85 | 0.48 | 0.67 |
| 3 | 132257747 | rs6671 | -1.118 | 0.442 | 1.397 | 0.713 | 0.031 | 0.081 | 0.081 | 0 | 0.19 | 0.44 | 0.26 | 0.23 | 0.22 | 0.34 |
| 3 | 132268469 | rs10935018 | -0.916 | 0.007 | 2.08 | 0.993 | -0.028 | -0.034 | -0.034 | 0.003 | 0.2 | 0.33 | 0.18 | 0.11 | 0.14 | 0.14 |
| 3 | 132269435 | rs7619350 | -1.007 | -0.015 | 2.279 | 1.047 | -0.008 | 0.174 | 0.174 | 0.172 | 0.15 | 0.38 | 0.08 | 0.54 | 0.12 | 0.19 |
| 3 | 132269761 | rs6439360 | -2.051 | -0.018 | 2.072 | 0.987 | 0.029 | -0.008 | -0.008 | 0 | 0.35 | 0.76 | 0.25 | 0.65 | 0.26 | 0.34 |
| 3 | 132274513 | rs12639254 | -1.482 | -0.049 | 2.187 | 1.014 | 0.006 | -0.055 | -0.055 | 1.662 | 0.24 | 0.14 | 0.12 | 0.51 | 0.2 | 0.27 |
| 3 | 132276356 | rs12488259 | -0.096 | -0.056 | 2.094 | 1.007 | -0.024 | -0.059 | -0.059 | 0 | 0.11 | 0.02 | 0.05 | 0.51 | 0.12 | 0.19 |
| 3 | 132277866 | rs2270801 | -0.827 | -0.082 | 2.078 | 1.002 | 0.001 | -0.002 | -0.002 | 0.42 | 0.17 | 0.01 | 0.13 | 0.1 | 0.11 | 0.14 |
| 3 | 132280930 | rs6777072 | -2.503 | -0.109 | 2.264 | 1.052 | 0.026 | 0.012 | 0.012 | 0 | 0.37 | 0.73 | 0.25 | 0.69 | 0.27 | 0.43 |
| 3 | 132283715 | rs10935019 | -0.105 | -0.091 | 2.067 | 1.086 | -0.023 | -0.053 | -0.053 | 0.871 | 0.11 | 0.02 | 0.05 | 0.51 | 0.12 | 0.19 |
| 3 | 132300293 | rs12487584 | -2.405 | -0.13 | 2.24 | 1.089 | 0.02 | 0.007 | 0.007 | 0 | 0.36 | 0.73 | 0.24 | 0.68 | 0.25 | 0.43 |
| 3 | 132306191 | rs11927882 | -2.463 | -0.122 | 2.241 | 1.093 | 0.016 | 0.004 | 0.004 | 0 | 0.35 | 0.73 | 0.24 | 0.69 | 0.25 | 0.43 |
| 3 | 132310651 | rs10804610 | -1.409 | -0.123 | 2.395 | 1.101 | -0.013 | 0.184 | 0.184 | 0.202 | 0.16 | 0.62 | 0.09 | 0.5 | 0.12 | 0.21 |
| 3 | 132322262 | rs2305623 | -0.608 | -0.091 | 2.346 | 1.104 | 0 | -0.02 | -0.02 | 0.454 | 0.13 | 0.09 | 0.05 | 0.5 | 0.12 | 0.21 |
| 3 | 132364731 | rs17348055 | -0.776 | -0.165 | 2.467 | 1.235 | -0.001 | -0.022 | -0.022 | 0 | 0.17 | 0.01 | 0.13 | 0.12 | 0.11 | 0.14 |
| 3 | 132366498 | rs6776500 | -2.371 | -0.168 | 2.514 | 1.242 | 0.02 | 0.017 | 0.017 | 0 | 0.37 | 0.74 | 0.26 | 0.7 | 0.29 | 0.44 |
| 3 | 132369134 | rs16839479 | -0.328 | -0.146 | 2.504 | 1.253 | -0.003 | -0.01 | -0.01 | 0 | 0.12 | 0.07 | 0.05 | 0.51 | 0.12 | 0.21 |
| 3 | 132371778 | rs1001122 | -2.36 | -0.144 | 2.501 | 1.249 | 0.027 | 0.023 | 0.023 | 0 | 0.36 | 0.76 | 0.26 | 0.69 | 0.29 | 0.44 |
| 3 | 132379065 | rs2168435 | -1.142 | -0.096 | 2.609 | 1.143 | 0.003 | 0.268 | 0.268 | 0.022 | 0.15 | 0.38 | 0.08 | 0.57 | 0.12 | 0.21 |
| 3 | 132379314 | rs3749272 | 0.239 | -0.112 | 2.125 | 1.143 | 0.012 | -0.033 | -0.033 | 0 | 0.05 | 0 | 0.07 | 0.12 | 0.04 | 0.06 |
| 3 | 132390336 | rs12495862 | -0.254 | -0.024 | 2.234 | 1.194 | 0.001 | 0.077 | 0.077 | 0 | 0.11 | 0.14 | 0.06 | 0.57 | 0.12 | 0.21 |
| 3 | 132394556 | rs2305629 | -2.039 | -0.087 | 1.894 | 1.172 | 0.032 | 0.034 | 0.034 | 0 | 0.37 | 0.76 | 0.27 | 0.76 | 0.3 | 0.43 |
| 3 | 132401600 | rs6794496 | -0.419 | 0.006 | 1.195 | 1.164 | -0.005 | 0.012 | 0.012 | 0 | 0.12 | 0.09 | 0.07 | 0.57 | 0.16 | 0.22 |
| 3 | 132413996 | rs10935028 | -0.686 | 0.048 | 1.154 | 1.156 | -0.014 | 0.131 | 0.131 | 0.096 | 0.12 | 0.19 | 0.06 | 0.58 | 0.1 | 0.2 |
| 3 | 132414454 | rs12636390 | -1.551 | 0.073 | 0.468 | 0.697 | 0.025 | 0.03 | 0.03 | 0 | 0.37 | 0.74 | 0.26 | 0.76 | 0.29 | 0.42 |

| | | | | | | | | | | | | | | | | |
|---|---|---|---|---|---|---|---|---|---|---|---|---|---|---|---|---|
| 3 | 132424780 | rs3860501 | -0.306 | 0.203 | -0.13 | 0.895 | -0.02 | -0.036 | -0.036 | 0 | 0.08 | 0.01 | 0.2 | 0.15 | 0.08 | 0.18 |
| 3 | 132429522 | rs12636883 | 0.224 | 0.3 | 0.069 | 0.898 | -0.006 | -0.012 | -0.012 | 0 | 0.15 | 0.16 | 0.15 | 0.64 | 0.19 | 0.29 |
| 3 | 132430804 | rs12486323 | -0.354 | 0.178 | 0 | 0.83 | 0 | -0.046 | -0.046 | 0 | 0.21 | 0.16 | 0.15 | 0.64 | 0.21 | 0.3 |
| 3 | 132434156 | rs17414558 | -0.306 | 0.147 | -0.275 | 0.65 | -0.007 | 0.006 | 0.006 | 0 | 0.18 | 0.01 | 0.14 | 0.11 | 0.12 | 0.13 |
| 3 | 132435823 | rs11718115 | -1.724 | 0.083 | 0.223 | 0.607 | -0.011 | -0.024 | -0.024 | 0.278 | 0.43 | 0.51 | 0.33 | 0.75 | 0.37 | 0.43 |
| 3 | 132438301 | rs3860503 | -1.076 | 0.428 | -0.153 | 1.002 | 0.006 | -0.025 | -0.025 | 0 | 0.27 | 0.24 | 0.16 | 0.57 | 0.23 | 0.35 |
| 3 | 132443661 | rs747328 | -0.683 | 0.758 | -0.067 | 1.176 | 0 | 0.016 | 0.016 | 0.031 | 0.22 | 0.25 | 0.16 | 0.5 | 0.24 | 0.17 |
| 3 | 132447104 | rs10935030 | -0.399 | 0.723 | -0.64 | 1.228 | -0.055 | -0.146 | -0.146 | 0 | 0.2 | 0.01 | 0.13 | 0.48 | 0.2 | 0.16 |
| 3 | 132451038 | rs9811369 | -1.038 | 0.795 | -0.631 | 0.437 | -0.015 | 0.05 | 0.05 | 0 | 0.58 | 0.73 | 0.5 | 0.77 | 0.5 | 0.61 |
| 3 | 132457395 | rs899432 | 0.249 | 0.64 | -1.156 | 0.824 | 0.03 | -0.168 | -0.168 | 1.234 | 0.22 | 0.01 | 0.06 | 0.5 | 0.12 | 0.19 |
| 3 | 132476678 | rs7624974 | 0.134 | 1.103 | -0.827 | 0.767 | 0.02 | 0.018 | 0.018 | 0 | 0.08 | 0.11 | 0.2 | 0.19 | 0.17 | 0.13 |
| 3 | 132488754 | rs9879204 | -0.715 | 0.851 | -0.063 | 0.672 | 0.047 | -0.009 | -0.009 | 1.579 | 0.45 | 0.31 | 0.27 | 0.56 | 0.33 | 0.41 |
| 3 | 132489686 | rs13317201 | -0.186 | 0.889 | 0.018 | 0.572 | 0.014 | -0.049 | -0.049 | 0 | 0.28 | 0.17 | 0.12 | 0.5 | 0.21 | 0.31 |
| 3 | 132491096 | rs3849410 | -0.841 | 0.966 | 0.253 | 0.555 | 0.042 | -0.018 | -0.018 | 0 | 0.44 | 0.25 | 0.25 | 0.57 | 0.33 | 0.42 |
| 3 | 132491910 | rs6807073 | -1.995 | 1.144 | 0.505 | 0.373 | -0.011 | 0.049 | 0.049 | 0 | 0.59 | 0.75 | 0.5 | 0.77 | 0.52 | 0.6 |
| 3 | 132494523 | rs6779839 | 0.013 | 1.182 | 0.261 | 0.394 | 0.06 | -0.028 | -0.028 | 1.757 | 0.06 | 0.19 | 0.06 | 0.01 | 0.09 | 0.13 |
| 3 | 132502340 | rs4082181 | -0.998 | 1.116 | 0.501 | 0.417 | 0.054 | 0.028 | 0.028 | 0 | 0.48 | 0.31 | 0.27 | 0.61 | 0.35 | 0.48 |
| 3 | 132503440 | rs6810216 | -1.221 | 0.967 | 0.678 | 0.361 | 0.017 | -0.028 | -0.028 | 0.182 | 0.36 | 0.52 | 0.4 | 0.25 | 0.38 | 0.4 |
| 3 | 132509832 | rs2053954 | -0.518 | 0.978 | 0.557 | 0.418 | 0.083 | -0.019 | -0.019 | 0 | 0.42 | 0.21 | 0.21 | 0.59 | 0.26 | 0.3 |
| 3 | 132512459 | rs1398722 | -1.679 | 1.093 | 0.758 | 0.392 | 0.002 | -0.031 | -0.031 | 0 | 0.57 | 0.52 | 0.47 | 0.77 | 0.52 | 0.6 |
| 3 | 132522262 | rs11706847 | -1.076 | 1.168 | 0.766 | 0.428 | 0.098 | 0.021 | 0.021 | 0 | 0.14 | 0.31 | 0.26 | 0.17 | 0.25 | 0.27 |
| 3 | 132525101 | rs4343624 | -0.584 | 1.018 | 0.701 | 0.423 | 0.084 | -0.01 | -0.01 | 0 | 0.43 | 0.23 | 0.22 | 0.59 | 0.26 | 0.3 |
| 3 | 132533091 | rs1349606 | -1.27 | 1.174 | 0.836 | 0.392 | 0.002 | 0.023 | 0.023 | 0 | 0.52 | 0.46 | 0.43 | 0.75 | 0.44 | 0.44 |
| 3 | 132542963 | rs6764659 | -0.985 | 1.075 | 0.736 | 0.383 | 0.043 | 0.021 | 0.021 | 0 | 0.49 | 0.34 | 0.27 | 0.6 | 0.35 | 0.43 |
| 3 | 132548645 | rs16839679 | 0.356 | 0.86 | 0.432 | 0.376 | 0.041 | -0.108 | -0.108 | 0 | 0.23 | 0.08 | 0.08 | 0.53 | 0.14 | 0.19 |
| 3 | 132548805 | rs7652595 | -1.152 | 0.601 | 0.628 | 0.224 | 0.115 | 0.014 | 0.014 | 0 | 0.16 | 0.41 | 0.27 | 0.16 | 0.26 | 0.28 |
| 3 | 132554134 | rs9844515 | -1.996 | 0.644 | 1.098 | 0.234 | -0.031 | 0.057 | 0.057 | 0 | 0.59 | 0.78 | 0.51 | 0.76 | 0.51 | 0.6 |
| 3 | 132558241 | rs9818010 | -0.81 | 0.636 | 1.051 | 0.252 | 0.085 | 0.003 | 0.003 | 0 | 0.44 | 0.23 | 0.25 | 0.6 | 0.29 | 0.33 |
| 3 | 132560225 | rs17351032 | 0.504 | 0.644 | 0.78 | 0.25 | 0.002 | 0.06 | 0.06 | 0.007 | 0.2 | 0.01 | 0.15 | 0.07 | 0.15 | 0.11 |
| 3 | 132562866 | rs9881437 | -1.598 | 0.887 | 1.037 | 0.326 | -0.01 | 0.022 | 0.022 | 0 | 0.58 | 0.67 | 0.49 | 0.76 | 0.51 | 0.6 |
| 3 | 132574744 | rs1452152 | -0.718 | 0.844 | 1.061 | 0.298 | 0.021 | 0.006 | 0.006 | 0 | 0.48 | 0.43 | 0.29 | 0.59 | 0.35 | 0.43 |
| 3 | 132584189 | rs1452146 | -1.525 | 0.901 | 0.818 | 0.234 | -0.011 | 0.022 | 0.022 | 0 | 0.58 | 0.68 | 0.49 | 0.75 | 0.51 | 0.6 |
| 3 | 132609896 | rs11714577 | -0.061 | 1.014 | 0.363 | 0.407 | -0.014 | -0.037 | -0.037 | 0 | 0.08 | 0.02 | 0.16 | 0.15 | 0.09 | 0.14 |
| 3 | 132609907 | rs9851851 | 0.709 | 0.957 | 0.432 | 0.025 | 0.003 | 0.003 | 0.003 | 0 | 0.48 | 0.55 | 0.68 | 0.4 | 0.56 | 0.54 |
| 3 | 132610596 | rs9852993 | 0.256 | 0.788 | 0.108 | -0.333 | -0.014 | 0.089 | 0.089 | 0 | 0.25 | 0.01 | 0.18 | 0.07 | 0.21 | 0.16 |
| 3 | 132610752 | rs6439371 | 1.124 | 0.773 | 0.237 | -0.352 | -0.001 | -0.004 | -0.004 | 0 | 0.59 | 0.59 | 0.6 | 0.76 | 0.64 | 0.55 |
| 3 | 132614374 | rs1869151 | 1.555 | 1.33 | 0.467 | -0.689 | -0.022 | 0.063 | 0.063 | 0 | 0.37 | 0.25 | 0.48 | 0.07 | 0.44 | 0.34 |
| 3 | 132614582 | rs2600961 | 0.72 | 1.191 | 0.679 | -0.491 | 0.041 | -0.014 | -0.014 | 0.927 | 0.93 | 0.8 | 0.95 | 0.98 | 0.93 | 0.8 |
| 3 | 132622722 | rs9813321 | 0.332 | 1.285 | 0.383 | -0.301 | -0.009 | -0.04 | -0.04 | 0 | 0.07 | 0.01 | 0.15 | 0.16 | 0.07 | 0.11 |
| 3 | 132624113 | rs2600956 | 1.075 | 1.278 | 0.833 | -0.352 | 0.072 | 0.004 | 0.004 | 0.129 | 0.92 | 0.65 | 0.93 | 0.97 | 0.93 | 0.81 |
| 3 | 132624322 | rs10512870 | 1.032 | 1.212 | 0.212 | -0.399 | 0.042 | -0.108 | -0.108 | 0 | 0.22 | 0.05 | 0.08 | 0.57 | 0.14 | 0.22 |
| 3 | 132624354 | rs10512869 | 0.383 | 1.688 | -0.016 | -0.599 | -0.058 | 0.082 | 0.082 | 0 | 0.23 | 0.01 | 0.2 | 0.07 | 0.22 | 0.13 |
| 3 | 132638242 | rs9878846 | 1.196 | 1.751 | 0.726 | -0.612 | 0.006 | -0.012 | -0.012 | 2.399 | 0.6 | 0.47 | 0.53 | 0.73 | 0.55 | 0.53 |
| 3 | 132639561 | rs2600968 | 1.441 | 1.78 | 0.774 | -0.598 | 0.081 | 0.033 | 0.033 | 0 | 0.82 | 0.54 | 0.73 | 0.8 | 0.77 | 0.66 |
| 3 | 132640698 | rs11927414 | -1.264 | 1.777 | 0.855 | -0.737 | 0.012 | -0.036 | -0.036 | 1.304 | 0.6 | 0.55 | 0.5 | 0.82 | 0.5 | 0.66 |
| 3 | 132640879 | rs10512871 | -0.393 | 1.783 | 0.824 | -0.757 | 0.005 | -0.09 | -0.09 | 0.161 | 0.3 | 0.53 | 0.27 | 0.09 | 0.29 | 0.32 |
| 3 | 132641236 | rs7648145 | 0.72 | 1.684 | 0.543 | -0.741 | -0.006 | 0 | 0 | 0.023 | 0.66 | 0.34 | 0.68 | 0.89 | 0.62 | 0.65 |
| 3 | 132642627 | rs7355965 | 1.484 | 1.707 | 0.631 | -0.688 | -0.004 | 0.019 | 0.019 | 0 | 0.36 | 0.32 | 0.45 | 0.16 | 0.41 | 0.31 |
| 3 | 132643409 | rs9868404 | 0.143 | 1.268 | 0.113 | -0.582 | 0.034 | -0.037 | -0.037 | 0.039 | 0.32 | 0.02 | 0.24 | 0.73 | 0.21 | 0.38 |
| 3 | 132646256 | rs4479596 | 0.358 | 1.91 | 0.802 | -0.547 | -0.001 | 0.04 | 0.04 | 0.124 | 0.58 | 0.23 | 0.42 | 0.68 | 0.45 | 0.47 |
| 3 | 132646367 | rs4538372 | -0.233 | 2.034 | 1.051 | -0.608 | -0.025 | 0.008 | 0.008 | 0 | 0.67 | 0.74 | 0.52 | 0.71 | 0.55 | 0.64 |
| 3 | 132653045 | rs6767367 | -0.037 | 1.63 | 1.135 | -0.347 | 0.019 | 0.008 | 0.008 | 0 | 0.06 | 0.13 | 0.07 | 0.13 | 0.1 | 0.16 |
| 3 | 132653345 | rs10804615 | 0.742 | 1.773 | 0.914 | -0.324 | 0.004 | -0.018 | -0.018 | 0.068 | 0.56 | 0.2 | 0.37 | 0.51 | 0.45 | 0.42 |
| 3 | 132672981 | rs17353336 | -0.232 | 1.799 | 0.79 | -0.266 | -0.003 | 0 | 0 | 0 | 0.67 | 0.7 | 0.54 | 0.71 | 0.58 | 0.64 |
| 3 | 132682335 | rs6764114 | 0.605 | 2.246 | 0.512 | -0.466 | 0.006 | 0.067 | 0.067 | 0.1 | 0.56 | 0.24 | 0.52 | 0.69 | 0.58 | 0.52 |
| 3 | 132685171 | rs7653397 | -1.744 | 2.262 | 1.036 | -0.328 | 0.083 | 0.133 | 0.133 | 0.47 | 0.17 | 0.36 | 0.29 | 0.4 | 0.24 | 0.36 |
| 3 | 132690957 | rs6763720 | 0.053 | 2.184 | 0.758 | -0.504 | -0.011 | -0.106 | -0.106 | 0.767 | 0.38 | 0.49 | 0.37 | 0.08 | 0.34 | 0.27 |
| 3 | 132694401 | rs7620792 | -0.443 | 2.31 | 0.595 | -0.425 | -0.005 | -0.005 | -0.005 | 0 | 0.09 | 0.08 | 0.07 | 0.16 | 0.1 | 0.14 |
| 3 | 132701065 | rs7632365 | -1.391 | 1.832 | 0.489 | -0.36 | 0.034 | -0.013 | -0.013 | 0 | 0.56 | 0.77 | 0.66 | 0.48 | 0.6 | 0.64 |
| 3 | 132702211 | rs7625328 | 0.608 | 1.939 | 0.926 | -0.387 | 0 | -0.002 | -0.002 | 0 | 0.58 | 0.44 | 0.34 | 0.73 | 0.53 | 0.54 |
| 3 | 132704649 | rs6789188 | -1.424 | 1.687 | 0.608 | -0.322 | 0.003 | 0.055 | 0.055 | 0 | 0.49 | 0.77 | 0.57 | 0.41 | 0.49 | 0.5 |
| 3 | 132728006 | rs6770608 | -0.961 | 1.695 | 0.742 | -0.218 | 0.038 | 0.057 | 0.057 | 1.154 | 0.11 | 0.38 | 0.09 | 0.18 | 0.1 | 0.14 |
| 3 | 132743344 | rs9866375 | -0.397 | 1.692 | 0.792 | -0.338 | -0.016 | 0.044 | 0.044 | 0 | 0.37 | 0.2 | 0.47 | 0.24 | 0.41 | 0.37 |
| 3 | 132751699 | rs1464403 | -0.664 | 1.679 | 0.876 | -0.345 | 0.003 | -0.025 | -0.025 | 0.102 | 0.39 | 0.43 | 0.49 | 0.24 | 0.41 | 0.37 |
| 3 | 132756182 | rs7633217 | -1.708 | 1.677 | 0.439 | -0.309 | 0.011 | 0.019 | 0.019 | 0 | 0.52 | 0.82 | 0.63 | 0.48 | 0.53 | 0.57 |
| 3 | 132761684 | rs2030444 | -0.649 | 1.535 | 0.656 | -0.333 | 0 | -0.021 | -0.021 | 0.176 | 0.4 | 0.54 | 0.5 | 0.25 | 0.41 | 0.37 |
| 3 | 132765826 | rs1402457 | 1.175 | 1.491 | 0.145 | -0.467 | 0.012 | 0.026 | 0.026 | 0.008 | 0.11 | 0.04 | 0.07 | 0.01 | 0.08 | 0.04 |
| 3 | 132766222 | rs13061327 | 0.163 | 1.544 | 0.192 | -0.437 | -0.007 | -0.007 | -0.007 | 0 | 0.07 | 0.07 | 0.06 | 0.19 | 0.09 | 0.13 |
| 3 | 132766843 | rs12330756 | -0.48 | 1.787 | 0.063 | -0.475 | -0.009 | 0.164 | 0.164 | 0 | 0.32 | 0.06 | 0.33 | 0.06 | 0.3 | 0.22 |

| | | | | | | | | | | | | | | | | |
|---|---|---|---|---|---|---|---|---|---|---|---|---|---|---|---|---|
| 3 | 132768296 | rs6787057 | -0.525 | 1.786 | 0.129 | -0.458 | 0.013 | 0.031 | 0.031 | 0.37 | 0.09 | 0.21 | 0.07 | 0.19 | 0.09 | 0.13 |
| 3 | 132768387 | rs9839659 | 0.616 | 1.761 | -0.368 | -0.462 | -0.05 | 0.086 | 0.086 | 0 | 0.21 | 0.01 | 0.25 | 0.05 | 0.23 | 0.19 |
| 3 | 132777314 | rs9879280 | -1.026 | 1.877 | -0.08 | -0.329 | -0.022 | 0.042 | 0.042 | 0 | 0.4 | 0.23 | 0.54 | 0.3 | 0.44 | 0.43 |
| 3 | 132778027 | rs6802908 | 0.317 | 1.912 | -0.2 | -0.083 | 0.02 | 0.023 | 0.023 | 0 | 0.06 | 0.09 | 0.14 | 0.17 | 0.12 | 0.12 |
| 3 | 132799209 | rs1357019 | -1.113 | 1.613 | -0.12 | -0.306 | 0.009 | 0.042 | 0.042 | 0 | 0.46 | 0.63 | 0.49 | 0.3 | 0.42 | 0.38 |
| 3 | 132800435 | rs10512887 | 0.266 | 1.989 | -0.086 | -0.173 | 0.038 | 0.046 | 0.046 | 0 | 0.07 | 0.15 | 0.17 | 0.19 | 0.15 | 0.17 |
| 3 | 132806802 | rs6782366 | -0.204 | 1.873 | -0.225 | -0.299 | -0.016 | -0.016 | -0.016 | 0 | 0.21 | 0.17 | 0.26 | 0.34 | 0.26 | 0.28 |
| 3 | 132808392 | rs1965461 | 0.833 | 1.929 | -0.224 | -0.308 | 0.005 | 0.003 | 0.003 | 0 | 0.14 | 0.16 | 0.17 | 0.19 | 0.16 | 0.19 |
| 3 | 132810967 | rs6802205 | -1.32 | 1.725 | -0.311 | -0.459 | 0.007 | 0.033 | 0.033 | 0 | 0.46 | 0.23 | 0.55 | 0.43 | 0.55 | 0.49 |
| 3 | 132815704 | rs6807522 | -1.32 | 1.725 | -0.311 | -0.459 | 0.007 | 0.033 | 0.033 | 0 | 0.46 | 0.23 | 0.55 | 0.43 | 0.55 | 0.49 |
| 3 | 132825695 | rs1450045 | -2.315 | 1.762 | -0.335 | -0.475 | 0.02 | 0.003 | 0.003 | 0 | 0.55 | 0.79 | 0.67 | 0.48 | 0.57 | 0.55 |
| 3 | 132834411 | rs12497903 | -0.113 | 1.938 | -0.545 | -0.422 | -0.031 | -0.038 | -0.038 | 0.853 | 0.08 | 0.03 | 0.19 | 0.18 | 0.11 | 0.16 |
| 3 | 132837567 | rs3811649 | 0.166 | 1.937 | -0.553 | -0.422 | -0.034 | -0.034 | -0.034 | 0.068 | 0.07 | 0.01 | 0.18 | 0.15 | 0.11 | 0.13 |
| 3 | 132837961 | rs7631163 | -0.936 | 1.941 | -0.001 | -0.446 | 0.028 | 0.026 | 0.026 | 0 | 0.27 | 0.41 | 0.43 | 0.39 | 0.29 | 0.36 |
| 3 | 132844398 | rs9870361 | 0.032 | 1.963 | -0.701 | -0.446 | -0.032 | 0.09 | 0.09 | 0 | 0.24 | 0.05 | 0.2 | 0.09 | 0.28 | 0.19 |
| 3 | 132846319 | rs17361457 | 1.009 | 1.844 | -0.619 | -0.427 | -0.015 | -0.007 | -0.007 | 0.252 | 0.14 | 0.11 | 0.16 | 0.17 | 0.15 | 0.14 |
| 3 | 132846881 | rs1464401 | -0.785 | 1.681 | -0.646 | -0.492 | -0.069 | 0.033 | 0.033 | 0 | 0.33 | 0.07 | 0.39 | 0.26 | 0.39 | 0.37 |
| 3 | 132856691 | rs3897016 | -0.697 | 1.493 | -0.349 | -0.672 | -0.009 | 0.045 | 0.045 | 0 | 0.4 | 0.71 | 0.6 | 0.23 | 0.45 | 0.35 |
| 3 | 132860651 | rs9289439 | -0.169 | 1.759 | 0.066 | -0.752 | 0.048 | -0.015 | -0.015 | 0 | 0.12 | 0.25 | 0.36 | 0.13 | 0.18 | 0.15 |
| 3 | 132862516 | rs9854387 | 0.293 | 1.367 | -0.109 | -0.691 | -0.003 | 0.059 | 0.059 | 0 | 0.25 | 0.12 | 0.18 | 0.09 | 0.26 | 0.17 |
| 3 | 132879712 | rs6784028 | -0.793 | 0.907 | -0.681 | -0.386 | 0.012 | -0.001 | -0.001 | 0.074 | 0.61 | 0.79 | 0.68 | 0.42 | 0.63 | 0.68 |
| 3 | 132884215 | rs1450053 | -0.046 | 0.979 | -0.346 | -0.078 | -0.003 | 0.023 | 0.023 | 0 | 0.67 | 0.42 | 0.7 | 0.51 | 0.69 | 0.71 |
| 3 | 132888446 | rs6794137 | 0.969 | 0.939 | -0.89 | -0.79 | -0.004 | 0.025 | 0.025 | 0 | 0.21 | 0.12 | 0.13 | 0.04 | 0.2 | 0.19 |
| 3 | 132890677 | rs4854697 | 0.582 | 1.431 | -0.75 | -1.217 | -0.007 | 0.029 | 0.029 | 0 | 0.55 | 0.34 | 0.66 | 0.85 | 0.53 | 0.5 |
| 3 | 132891280 | rs1450055 | -0.173 | 1.412 | -0.681 | -1.197 | -0.002 | -0.006 | -0.006 | 0.574 | 0.36 | 0.34 | 0.27 | 0.15 | 0.37 | 0.42 |
| 3 | 132892428 | rs1450054 | 1.037 | 1.445 | -0.276 | -1.129 | -0.002 | -0.021 | -0.021 | 0.158 | 0.31 | 0.24 | 0.22 | 0.04 | 0.32 | 0.22 |
| 3 | 132899962 | rs6767920 | 0.084 | 1.394 | -0.221 | -1.138 | -0.003 | -0.001 | -0.001 | 1.039 | 0.46 | 0.39 | 0.34 | 0.14 | 0.46 | 0.44 |
| 3 | 132902053 | rs12493846 | 1.478 | 1.404 | -0.583 | -1.03 | -0.039 | -0.157 | -0.157 | 0 | 0.22 | 0.02 | 0.37 | 0.39 | 0.23 | 0.29 |
| 3 | 132905128 | rs7642901 | 1.337 | 1.306 | -0.114 | -1.008 | -0.006 | 0.03 | 0.03 | 0 | 0.23 | 0.24 | 0.22 | 0.47 | 0.2 | 0.2 |
| 3 | 132905331 | rs9858099 | 0.841 | 1.583 | -0.181 | -0.852 | -0.021 | 0.099 | 0.099 | 0 | 0.3 | 0.13 | 0.19 | 0.04 | 0.31 | 0.22 |
| 3 | 132908793 | rs12490128 | 0.413 | 1.551 | -0.144 | -0.789 | -0.021 | 0.035 | 0.035 | 0 | 0.14 | 0.05 | 0.12 | 0.1 | 0.16 | 0.23 |
| 3 | 132913122 | rs1868602 | 0.705 | 1.774 | 0.228 | -0.677 | -0.004 | 0.015 | 0.015 | 0 | 0.29 | 0.19 | 0.21 | 0.04 | 0.29 | 0.21 |
| 3 | 132915738 | rs7638408 | -1.339 | 1.709 | 0.581 | -0.563 | 0.004 | 0.028 | 0.028 | 0 | 0.35 | 0.62 | 0.42 | 0.48 | 0.35 | 0.54 |
| 3 | 132916612 | rs6439389 | 1.225 | 1.67 | 0.592 | -0.595 | 0.021 | -0.023 | -0.023 | 0.339 | 0.39 | 0.14 | 0.38 | 0.46 | 0.38 | 0.28 |
| 3 | 132934607 | rs12639218 | 0.758 | 1.748 | 0.593 | -0.613 | -0.003 | 0.013 | 0.013 | 0 | 0.1 | 0.05 | 0.26 | 0.36 | 0.12 | 0.15 |
| 3 | 132938130 | rs4854701 | -1.444 | 1.758 | 0.637 | -0.626 | -0.013 | 0.041 | 0.041 | 0 | 0.74 | 0.87 | 0.81 | 0.95 | 0.74 | 0.82 |
| 3 | 132941928 | rs1901039 | -1.371 | 1.758 | 0.656 | -0.626 | -0.007 | 0.015 | 0.015 | 0 | 0.74 | 0.81 | 0.81 | 0.95 | 0.74 | 0.82 |
| 3 | 132948929 | rs4854576 | 0.786 | 1.683 | 0.614 | -0.493 | -0.004 | 0.018 | 0.018 | 0 | 0.1 | 0.05 | 0.26 | 0.37 | 0.12 | 0.15 |
| 3 | 132978969 | rs9866337 | -0.984 | 1.682 | 1.04 | -0.498 | -0.003 | 0 | 0 | 0.091 | 0.57 | 0.6 | 0.6 | 0.53 | 0.62 | 0.68 |
| 3 | 132984117 | rs7624858 | -1.04 | 1.679 | 1.061 | -0.5 | 0.009 | -0.002 | -0.002 | 1.943 | 0.57 | 0.66 | 0.61 | 0.53 | 0.62 | 0.69 |
| 3 | 132993565 | rs7639076 | -0.992 | 1.679 | 1.06 | -0.499 | 0.002 | -0.001 | -0.001 | 0 | 0.57 | 0.62 | 0.6 | 0.53 | 0.62 | 0.68 |
| 3 | 132995726 | rs16840161 | 0.463 | 1.686 | 1.143 | -0.535 | -0.003 | 0.005 | 0.005 | 0 | 0.14 | 0.13 | 0.1 | 0.1 | 0.15 | 0.25 |
| 3 | 133000230 | rs6439400 | 0.158 | 1.68 | 1.194 | -0.537 | -0.008 | -0.008 | -0.008 | 0 | 0.54 | 0.3 | 0.48 | 0.57 | 0.53 | 0.56 |
| 3 | 133004714 | rs11706654 | 1.286 | 1.669 | 1.149 | -0.528 | 0.021 | -0.025 | -0.025 | 0.018 | 0.38 | 0.14 | 0.37 | 0.46 | 0.36 | 0.28 |
| 3 | 133005613 | rs6776275 | -0.182 | 1.693 | 1.21 | -0.58 | 0 | -0.033 | -0.033 | 0 | 0.44 | 0.47 | 0.32 | 0.16 | 0.43 | 0.48 |
| 3 | 133009814 | rs2370409 | 0.22 | 1.706 | 1.018 | -0.575 | -0.001 | 0.151 | 0.151 | 0 | 0.52 | 0.21 | 0.65 | 0.84 | 0.56 | 0.51 |
| 3 | 133014479 | rs9867089 | 1.028 | 1.713 | 0.979 | -0.579 | -0.002 | -0.06 | -0.06 | 0.206 | 0.3 | 0.36 | 0.22 | 0.06 | 0.28 | 0.23 |
| 3 | 133023568 | rs1708368 | 0.843 | 1.776 | 0.856 | -0.526 | -0.007 | 0.006 | 0.006 | 0 | 0.1 | 0.04 | 0.25 | 0.37 | 0.12 | 0.15 |
| 3 | 133032892 | rs2699869 | -0.883 | 1.801 | 0.885 | -0.516 | -0.001 | -0.001 | -0.001 | 0 | 0.54 | 0.57 | 0.58 | 0.53 | 0.55 | 0.63 |
| 3 | 133035774 | rs3849406 | 1.071 | 1.814 | 0.626 | -0.501 | 0.02 | -0.015 | -0.015 | 0 | 0.41 | 0.2 | 0.38 | 0.47 | 0.38 | 0.32 |
| 3 | 133051572 | rs9876399 | 1.392 | 2.048 | 0.044 | -0.752 | -0.022 | 0.089 | 0.089 | 0 | 0.26 | 0.07 | 0.2 | 0.05 | 0.28 | 0.17 |
| 3 | 133071425 | rs1920021 | 0.332 | 2.212 | 0.03 | -0.75 | 0.002 | -0.013 | -0.013 | 0 | 0.14 | 0.16 | 0.1 | 0.1 | 0.15 | 0.25 |
| 3 | 133074428 | rs9289443 | 1.267 | 2.234 | -0.031 | -0.648 | -0.018 | 0.079 | 0.079 | 0 | 0.27 | 0.09 | 0.2 | 0.05 | 0.28 | 0.17 |
| 3 | 133088362 | rs938229 | 0.366 | 2.251 | 0.024 | -0.187 | -0.015 | -0.018 | -0.018 | 0.44 | 0.37 | 0.27 | 0.48 | 0.39 | 0.41 | 0.35 |
| 3 | 133088541 | rs938230 | -0.288 | 2.307 | 0.089 | -0.202 | 0.004 | -0.001 | -0.001 | 0 | 0.46 | 0.65 | 0.54 | 0.4 | 0.52 | 0.42 |
| 3 | 133090973 | rs4479595 | 2.157 | 1.798 | -0.361 | -0.209 | -0.007 | -0.016 | -0.016 | 0 | 0.16 | 0.08 | 0.18 | 0.29 | 0.18 | 0.17 |
| 3 | 133093153 | rs13318906 | 0.906 | 2.743 | 0.359 | 0.002 | -0.013 | 0.11 | 0.11 | 0 | 0.26 | 0.06 | 0.18 | 0.05 | 0.25 | 0.16 |
| 3 | 133094559 | rs1114630 | 0.016 | 2.76 | 0.466 | 0.026 | 0.001 | -0.037 | -0.037 | 1.923 | 0.34 | 0.25 | 0.43 | 0.39 | 0.34 | 0.32 |
| 3 | 133097119 | rs4854719 | -0.4 | 2.533 | 0.504 | 0.021 | -0.006 | -0.013 | -0.013 | 0 | 0.39 | 0.38 | 0.46 | 0.39 | 0.4 | 0.38 |
| 3 | 133099245 | rs1197314 | -0.499 | 1.913 | 0.01 | 0.147 | -0.038 | -0.026 | -0.026 | 0 | 0.4 | 0.26 | 0.49 | 0.39 | 0.45 | 0.39 |
| 3 | 133100452 | rs1197313 | -0.293 | 1.935 | 0.021 | 0.139 | -0.048 | -0.039 | -0.039 | 0.283 | 0.38 | 0.22 | 0.46 | 0.39 | 0.43 | 0.38 |
| 3 | 133101068 | rs1197309 | 0.071 | 1.97 | 0.007 | 0.138 | -0.041 | -0.063 | -0.063 | 4.194 | 0.34 | 0.15 | 0.44 | 0.39 | 0.38 | 0.33 |
| 3 | 133101390 | rs6807356 | 1.662 | 1.9 | -0.072 | 0.101 | 0.024 | -0.019 | -0.019 | 0 | 0.41 | 0.25 | 0.4 | 0.47 | 0.38 | 0.34 |
| 3 | 133105513 | rs1795107 | -0.128 | 1.957 | -0.342 | 0.11 | -0.024 | -0.025 | -0.025 | 0 | 0.37 | 0.28 | 0.47 | 0.39 | 0.42 | 0.34 |
| 3 | 133105729 | rs1197298 | -0.405 | 1.933 | -0.457 | 0.119 | -0.036 | -0.037 | -0.037 | 0 | 0.4 | 0.19 | 0.47 | 0.39 | 0.48 | 0.4 |
| 3 | 133109469 | rs3732572 | 0.866 | 2.024 | -0.441 | 0.166 | 0.019 | 0.001 | 0.001 | 0 | 0.5 | 0.39 | 0.48 | 0.58 | 0.48 | 0.56 |
| 3 | 133109606 | rs16840309 | 0.099 | 2.424 | -1.664 | 0.316 | -0.006 | 0.011 | 0.011 | 0 | 0.11 | 0.07 | 0.08 | 0.13 | 0.12 | 0.21 |
| 3 | 133110636 | rs2670179 | -0.306 | 2.509 | -1.521 | 0.722 | -0.087 | -0.105 | -0.105 | 0 | 0.36 | 0.07 | 0.44 | 0.39 | 0.4 | 0.32 |
| 3 | 133110681 | rs2699882 | -0.359 | 2.607 | -1.468 | 0.766 | -0.054 | -0.071 | -0.071 | 0 | 0.37 | 0.12 | 0.44 | 0.39 | 0.4 | 0.32 |

| | | | | | | | | | | | | | | | | |
|---|---|---|---|---|---|---|---|---|---|---|---|---|---|---|---|---|
| 3 | 133112197 | rs1567788 | 0.921 | 2.585 | -1.354 | 1.01 | -0.022 | 0.03 | 0.03 | 0 | 0.05 | 0.01 | 0.22 | 0.3 | 0.09 | 0.07 |
| 3 | 133115340 | rs2699880 | -0.674 | 2.083 | -1.165 | 0.708 | 0 | -0.002 | -0.002 | 0 | 0.45 | 0.45 | 0.5 | 0.41 | 0.51 | 0.38 |
| 3 | 133118054 | rs517255 | -0.73 | 2.083 | -1.593 | 0.58 | -0.003 | -0.005 | -0.005 | 0 | 0.46 | 0.36 | 0.5 | 0.5 | 0.49 | 0.48 |
| 3 | 133125597 | rs16840382 | -0.176 | 1.9 | -2.17 | 0.298 | 0.003 | -0.012 | -0.012 | 0 | 0.14 | 0.16 | 0.08 | 0.09 | 0.15 | 0.2 |
| 3 | 133125713 | rs1197312 | -0.367 | 0.846 | -1.846 | 0.6 | 0.036 | 0.011 | 0.011 | 0 | 0.49 | 0.28 | 0.51 | 0.49 | 0.55 | 0.51 |
| 3 | 133130469 | rs17365742 | -0.969 | 1.174 | -1.641 | 0.547 | -0.045 | -0.011 | -0.011 | 0 | 0.13 | 0.01 | 0.2 | 0.1 | 0.18 | 0.1 |
| 3 | 133137284 | rs4854585 | 0.488 | 0.785 | -1.47 | -0.374 | -0.002 | -0.002 | -0.002 | 0 | 0.62 | 0.37 | 0.73 | 0.92 | 0.59 | 0.7 |
| 3 | 133137434 | rs10935060 | 0.768 | 0.784 | -0.816 | -0.219 | -0.005 | -0.024 | -0.024 | 0 | 0.68 | 0.37 | 0.65 | 0.82 | 0.67 | 0.68 |
| 3 | 133143318 | rs13073286 | 1.169 | 1.396 | -0.952 | 0.559 | -0.002 | 0.003 | 0.003 | 0 | 0.92 | 0.93 | 0.8 | 0.82 | 0.8 | 0.83 |
| 3 | 133156692 | rs6762405 | -0.096 | 0.559 | -1.71 | 0.441 | 0.089 | 0.043 | 0.043 | 0.132 | 0.47 | 0.71 | 0.23 | 0.36 | 0.26 | 0.3 |
| 3 | 133167363 | rs2276737 | 0.341 | 0.474 | -2.231 | 0.097 | 0.014 | 0.049 | 0.049 | 2.318 | 0.5 | 0.1 | 0.38 | 0.59 | 0.57 | 0.64 |
| 3 | 133172767 | rs10935064 | -0.059 | 0.252 | -2.26 | 0.003 | 0.005 | -0.039 | -0.039 | 0 | 0.63 | 0.13 | 0.53 | 0.59 | 0.68 | 0.72 |
| 3 | 133179580 | rs931099 | 0.78 | 1.193 | -2.433 | -0.551 | -0.028 | -0.007 | -0.007 | 0 | 0.43 | 0.12 | 0.4 | 0.57 | 0.38 | 0.61 |
| 3 | 133180449 | rs12632048 | 0.626 | 0.699 | -1.431 | -0.787 | -0.007 | 0.051 | 0.051 | 0 | 0.15 | 0.32 | 0.2 | 0.25 | 0.11 | 0.14 |
| 3 | 133184552 | rs1479179 | -0.925 | 0.477 | -0.118 | -0.993 | -0.014 | -0.011 | -0.011 | 0 | 0.85 | 0.82 | 0.84 | 0.91 | 0.9 | 0.89 |
| 3 | 133189192 | rs7628262 | -1.182 | 0.699 | 0.886 | -1.044 | 0.013 | -0.012 | -0.012 | 0 | 0.39 | 0.46 | 0.43 | 0.33 | 0.52 | 0.28 |
| 3 | 133189316 | rs9289448 | -0.051 | 0.945 | 0.643 | -1.025 | 0.004 | 0.007 | 0.007 | 0 | 0.17 | 0.18 | 0.24 | 0.29 | 0.23 | 0.15 |
| 3 | 133191093 | rs899458 | -2.1 | 1.244 | 0.427 | -1.21 | -0.026 | 0.059 | 0.059 | 0.621 | 0.43 | 0.73 | 0.42 | 0.32 | 0.51 | 0.3 |
| 3 | 133191453 | rs7628499 | 1.218 | 1.266 | 0.715 | -0.984 | -0.005 | 0.001 | 0.001 | 0.812 | 0.68 | 0.36 | 0.59 | 0.68 | 0.73 | 0.8 |
| 3 | 133197271 | rs7625229 | 0.226 | 1.815 | 0.433 | -0.957 | -0.016 | 0.006 | 0.006 | 0 | 0.1 | 0.06 | 0.14 | 0 | 0.26 | 0.05 |
| 3 | 133208595 | rs6439411 | 1.736 | 1.413 | 0.525 | -0.936 | -0.001 | 0.003 | 0.003 | 0 | 0.57 | 0.3 | 0.43 | 0.6 | 0.44 | 0.72 |
| 3 | 133209377 | rs7652889 | 1.054 | 1.587 | 0.087 | -0.863 | -0.016 | -0.009 | -0.009 | 0 | 0.62 | 0.72 | 0.48 | 0.6 | 0.46 | 0.75 |
| 3 | 133221952 | rs13073410 | 0.483 | 1.644 | 0.131 | -0.992 | -0.013 | -0.024 | -0.024 | 0 | 0.76 | 0.84 | 0.63 | 0.66 | 0.74 | 0.81 |
| 3 | 133222283 | rs10935066 | -0.679 | 1.761 | 0.384 | -1.145 | 0.05 | -0.08 | -0.08 | 0 | 0.15 | 0.27 | 0.18 | 0 | 0.3 | 0.08 |
| 3 | 133226149 | rs10433342 | 1.439 | 1.272 | 0.288 | -1.104 | 0.012 | 0.022 | 0.022 | 0 | 0.58 | 0.3 | 0.41 | 0.65 | 0.38 | 0.69 |
| 3 | 133227463 | rs6762168 | -1.49 | 0.607 | 0.297 | -2.056 | -0.016 | 0.16 | 0.16 | 0.017 | 0.45 | 0.78 | 0.31 | 0.09 | 0.5 | 0.27 |
| 3 | 133227514 | rs9876777 | -1.633 | 0.368 | -0.147 | -2.052 | -0.015 | 0.207 | 0.207 | 1.274 | 0.53 | 0.84 | 0.36 | 0.17 | 0.48 | 0.35 |
| 3 | 133230966 | rs6439419 | -0.925 | 0.776 | 0.319 | -2.085 | 0 | -0.096 | -0.096 | 0.12 | 0.36 | 0.5 | 0.22 | 0.07 | 0.35 | 0.29 |
| 3 | 133234857 | rs9814681 | -0.679 | 0.627 | 0.431 | -1.834 | -0.002 | 0.061 | 0.061 | 0 | 0.24 | 0.15 | 0.16 | 0.06 | 0.25 | 0.13 |
| 3 | 133235272 | rs13065952 | -0.193 | 0.804 | 0.579 | -1.911 | 0.008 | -0.086 | -0.086 | 0.007 | 0.18 | 0.29 | 0.14 | 0.02 | 0.19 | 0.03 |
| 3 | 133236066 | rs9820205 | -1.352 | 0.571 | 0.124 | -1.886 | 0.007 | 0.13 | 0.13 | 0.851 | 0.47 | 0.72 | 0.33 | 0.07 | 0.46 | 0.21 |
| 3 | 133236819 | rs13095648 | 0.051 | 0.436 | 0.205 | -1.112 | -0.014 | -0.051 | -0.051 | 0 | 0.13 | 0.22 | 0.06 | 0.02 | 0.08 | 0.02 |
| 3 | 133241106 | rs13059279 | -0.875 | 0.65 | 0.189 | -0.909 | 0.003 | -0.053 | -0.053 | 0 | 0.22 | 0.41 | 0.14 | 0.08 | 0.22 | 0.14 |
| 3 | 133243896 | rs9882521 | -1.045 | 1.119 | -0.37 | -0.61 | 0 | -0.001 | -0.001 | 0 | 0.43 | 0.44 | 0.41 | 0.38 | 0.53 | 0.34 |
| 3 | 133244619 | rs9825996 | -1.035 | 0.375 | -0.39 | -0.751 | 0.005 | -0.107 | -0.107 | 0.075 | 0.16 | 0.43 | 0.13 | 0.01 | 0.15 | 0.14 |
| 3 | 133245961 | rs7647249 | 0.039 | 1.205 | -1.341 | -0.243 | 0.005 | -0.032 | -0.032 | 0.345 | 0.21 | 0.08 | 0.19 | 0.31 | 0.21 | 0.12 |
| 3 | 133250747 | rs11706487 | -0.644 | 1.454 | -0.85 | -0.353 | -0.002 | -0.003 | -0.003 | 0.024 | 0.07 | 0.05 | 0.09 | 0 | 0.14 | 0.04 |
| 3 | 133252369 | rs12630919 | -0.449 | 0.943 | -1.013 | -0.349 | 0 | 0.007 | 0.007 | 0 | 0.24 | 0.2 | 0.2 | 0.13 | 0.24 | 0.25 |
| 3 | 133254258 | rs4854723 | 0.405 | 1.371 | -1.031 | -0.668 | -0.014 | 0.02 | 0.02 | 0 | 0.71 | 0.25 | 0.66 | 0.87 | 0.64 | 0.81 |
| 3 | 133257885 | rs4854727 | 0.153 | 1.557 | 0.189 | -0.755 | 0.037 | -0.048 | -0.048 | 1.789 | 0.76 | 0.64 | 0.73 | 0.87 | 0.68 | 0.83 |
| 3 | 133258597 | rs4533654 | 0.953 | 0.722 | -0.33 | -0.771 | 0.005 | 0.046 | 0.046 | 0 | 0.63 | 0.25 | 0.63 | 0.77 | 0.63 | 0.67 |
| 3 | 133264251 | rs7614812 | 0.558 | 0.665 | -0.229 | -0.894 | 0.009 | -0.045 | -0.045 | 0.394 | 0.68 | 0.65 | 0.69 | 0.87 | 0.63 | 0.83 |
| 3 | 133265532 | rs872134 | 0.784 | 0.637 | 0.212 | -0.906 | 0.01 | -0.068 | -0.068 | 0 | 0.66 | 0.47 | 0.66 | 0.87 | 0.63 | 0.81 |
| 3 | 133268364 | rs16840524 | 0.086 | 0.27 | -0.623 | -0.846 | -0.013 | 0.005 | 0.005 | 0 | 0.29 | 0.56 | 0.26 | 0.34 | 0.25 | 0.37 |
| 3 | 133277735 | rs11714566 | 0.814 | 0.469 | -1.027 | -0.901 | -0.041 | -0.055 | -0.055 | 0.058 | 0.46 | 0.03 | 0.45 | 0.53 | 0.42 | 0.47 |
| 3 | 133287065 | rs4854733 | -0.16 | 0.559 | -0.452 | -0.844 | 0.042 | -0.056 | -0.056 | 0.797 | 0.75 | 0.44 | 0.71 | 0.87 | 0.67 | 0.84 |
| 3 | 133291621 | rs10049041 | 0.199 | -0.16 | -0.431 | -0.877 | -0.012 | -0.037 | -0.037 | 0 | 0.14 | 0.2 | 0.08 | 0.01 | 0.08 | 0.03 |
| 3 | 133293618 | rs3821508 | 0.943 | -0.003 | -1.124 | -1.03 | 0.076 | 0.03 | 0.03 | 1.339 | 0.55 | 0.07 | 0.52 | 0.62 | 0.65 | 0.53 |
| 3 | 133294203 | rs10049438 | -0.541 | -0.134 | -0.317 | -1.1 | -0.007 | 0 | 0 | 0 | 0.31 | 0.71 | 0.31 | 0.33 | 0.19 | 0.4 |
| 3 | 133301508 | rs9847556 | -0.174 | 0.322 | -0.082 | -1.08 | 0.015 | -0.028 | -0.028 | 0 | 0.86 | 0.77 | 0.83 | 0.95 | 0.84 | 0.93 |
| 3 | 133307491 | rs6792846 | -1.055 | 0.343 | -0.008 | -1.052 | 0.017 | -0.005 | -0.005 | 0 | 0.94 | 0.91 | 0.87 | 0.98 | 0.9 | 0.94 |
| 3 | 133316003 | rs13098257 | -0.713 | 0.909 | 0.071 | -1.493 | 0 | -0.038 | -0.038 | 4.711 | 0.29 | 0.28 | 0.2 | 0.04 | 0.31 | 0.09 |
| 3 | 133327457 | rs1051772 | 0.009 | 0.851 | 0.083 | -1.496 | -0.002 | -0.007 | -0.007 | 0 | 0.14 | 0.17 | 0.07 | 0.05 | 0.1 | 0.04 |
| 3 | 133329943 | rs1444601 | -0.227 | 0.751 | 0.432 | -1.573 | -0.001 | -0.004 | -0.004 | 0 | 0.75 | 0.78 | 0.63 | 0.63 | 0.75 | 0.57 |
| 3 | 133338862 | rs4854588 | -1.108 | 0.76 | 0.438 | -1.594 | 0.002 | 0.001 | 0.001 | 0 | 0.92 | 0.93 | 0.83 | 0.98 | 0.85 | 0.91 |
| 3 | 133339340 | rs2887747 | -1.357 | 0.723 | 0.441 | -1.591 | 0.001 | 0 | 0 | 0 | 0.93 | 0.93 | 0.84 | 0.98 | 0.87 | 0.93 |
| 3 | 133341988 | rs10935070 | 2.381 | 0.628 | 0.036 | -1.754 | -0.051 | 0.253 | 0.253 | 0.241 | 0.36 | 0.01 | 0.3 | 0.03 | 0.36 | 0.34 |
| 3 | 133347034 | rs1550742 | -0.538 | 0.199 | 0.296 | -1.615 | 0 | 0.001 | 0.001 | 0 | 0.78 | 0.79 | 0.64 | 0.63 | 0.77 | 0.6 |
| 3 | 133356790 | rs17301766 | 1.192 | 0.22 | 0.318 | -1.622 | -0.02 | 0.013 | 0.013 | 0 | 0.14 | 0.07 | 0.12 | 0 | 0.18 | 0.05 |
| 3 | 133357145 | rs10512907 | 1.192 | 0.22 | 0.318 | -1.622 | -0.02 | 0.013 | 0.013 | 0 | 0.14 | 0.07 | 0.12 | 0 | 0.18 | 0.05 |
| 3 | 133366227 | rs1822744 | 1.574 | 0.185 | 0.83 | -1.625 | -0.003 | 0.014 | 0.014 | 1.435 | 0.46 | 0.25 | 0.42 | 0.55 | 0.47 | 0.52 |
| 3 | 133367858 | rs12634591 | 0.972 | 0.216 | 0.706 | -1.561 | 0.003 | -0.005 | -0.005 | 0.305 | 0.17 | 0.13 | 0.2 | 0.35 | 0.1 | 0.34 |
| 3 | 133368362 | rs3192149 | -0.576 | 0.332 | 0.745 | -1.498 | 0 | 0.001 | 0.001 | 0 | 0.78 | 0.79 | 0.64 | 0.63 | 0.77 | 0.6 |
| 3 | 133392191 | rs11713248 | 1.093 | 0.39 | 0.749 | -1.634 | 0.005 | -0.028 | -0.028 | 0 | 0.14 | 0.1 | 0.12 | 0 | 0.18 | 0.05 |
| 3 | 133394478 | rs11924101 | 1.681 | 0.173 | 0.517 | -1.258 | -0.034 | -0.053 | -0.053 | 0.093 | 0.43 | 0.15 | 0.42 | 0.48 | 0.47 | 0.52 |
| 3 | 133399702 | rs2718812 | -1.02 | 0.164 | 0.618 | -1.238 | -0.011 | -0.006 | -0.006 | 0.175 | 0.43 | 0.73 | 0.51 | 0.46 | 0.42 | 0.44 |
| 3 | 133406341 | rs7630957 | 0.906 | 0.381 | 0.634 | -1.254 | 0.038 | -0.082 | -0.082 | 0.233 | 0.14 | 0.27 | 0.13 | 0 | 0.17 | 0.05 |
| 3 | 133411478 | rs1006097 | -0.604 | 0.306 | 0.043 | -1.128 | -0.009 | -0.01 | -0.01 | 0 | 0.4 | 0.64 | 0.5 | 0.47 | 0.41 | 0.45 |
| 3 | 133411605 | rs1006098 | -1.465 | 0.385 | -0.131 | -0.954 | -0.028 | -0.037 | -0.037 | 0 | 0.56 | 0.78 | 0.57 | 0.53 | 0.51 | 0.48 |

| | | | | | | | | | | | | | | | | |
|---|---|---|---|---|---|---|---|---|---|---|---|---|---|---|---|---|
| 3 | 133412094 | rs1444598 | 0.604 | -0.07 | -0.217 | -1.44 | -0.017 | 0.067 | 0.067 | 0 | 0.56 | 0.35 | 0.61 | 0.93 | 0.58 | 0.81 |
| 3 | 133412277 | rs13061966 | 0.504 | 0.274 | -0.023 | -0.834 | -0.008 | 0.001 | 0.001 | 4.464 | 0.41 | 0.31 | 0.57 | 0.55 | 0.57 | 0.54 |
| 3 | 133413516 | rs1005933 | -0.753 | 0.129 | -0.265 | -0.925 | 0.049 | -0.055 | -0.055 | 0 | 0.73 | 0.87 | 0.78 | 0.57 | 0.81 | 0.66 |
| 3 | 133415371 | rs4854742 | 0.632 | 0.177 | -0.36 | -0.973 | -0.021 | -0.005 | -0.005 | 0 | 0.4 | 0.28 | 0.61 | 0.52 | 0.55 | 0.53 |
| 3 | 133419018 | rs13071381 | 0.443 | 0.16 | 0.11 | -0.998 | 0 | -0.052 | -0.052 | 7.759 | 0.32 | 0.35 | 0.16 | 0.05 | 0.26 | 0.13 |
| 3 | 133423473 | rs4419374 | -0.605 | 0.09 | -0.403 | -0.913 | -0.035 | 0.023 | 0.023 | 3.941 | 0.72 | 0.61 | 0.76 | 0.57 | 0.81 | 0.66 |
| 3 | 133427224 | rs4854593 | 0.439 | 0.186 | -0.405 | -0.931 | 0.004 | -0.004 | -0.004 | 0 | 0.42 | 0.48 | 0.62 | 0.56 | 0.57 | 0.54 |
| 3 | 133429086 | rs13080154 | 0.256 | 0.173 | -0.637 | -0.386 | 0.032 | -0.016 | -0.016 | 0 | 0.57 | 0.19 | 0.55 | 0.48 | 0.63 | 0.53 |
| 3 | 133436136 | rs4525863 | 0.797 | 1.172 | -0.317 | 0.423 | 0.037 | 0.053 | 0.053 | 0.985 | 0.21 | 0.29 | 0.39 | 0.42 | 0.32 | 0.43 |
| 3 | 133439450 | rs4441652 | -0.654 | 0.381 | -1.297 | -0.364 | 0.001 | 0.158 | 0.158 | 1.139 | 0.62 | 0.84 | 0.81 | 0.94 | 0.66 | 0.84 |
| 3 | 133452444 | rs4241356 | 0.076 | -0.18 | -1.278 | -0.118 | 0.003 | -0.009 | -0.009 | 0 | 0.21 | 0.15 | 0.27 | 0.27 | 0.19 | 0.23 |
| 3 | 133452671 | rs4854755 | -0.054 | -0.689 | -1.684 | -0.481 | 0.033 | 0.012 | 0.012 | 0 | 0.32 | 0.15 | 0.28 | 0.27 | 0.22 | 0.28 |
| 3 | 133464102 | rs8177313 | 0.415 | -0.705 | -1.544 | -0.31 | 0.035 | 0.017 | 0.017 | 0 | 0.11 | 0.02 | 0.05 | 0.05 | 0.06 | 0.13 |
| 3 | 133464618 | rs8177185 | 1.279 | -0.71 | -1.511 | -0.256 | -0.017 | 0.058 | 0.058 | 0 | 0.08 | 0.12 | 0.12 | 0.25 | 0.07 | 0.15 |
| 3 | 133466222 | rs6796795 | 0.183 | -0.791 | -1.135 | -0.326 | -0.018 | -0.011 | -0.011 | 0 | 0.12 | 0.16 | 0.06 | 0.05 | 0.06 | 0.13 |
| 3 | 133466374 | rs4428180 | 0.167 | -0.268 | -1.328 | -0.157 | 0.003 | 0.031 | 0.031 | 0 | 0.12 | 0.14 | 0.23 | 0.26 | 0.14 | 0.19 |
| 3 | 133467696 | rs8177190 | 0.649 | 0.324 | -1.864 | -0.397 | -0.039 | 0.017 | 0.017 | 0.013 | 0.08 | 0.01 | 0.12 | 0.03 | 0.1 | 0.08 |
| 3 | 133468139 | rs8177191 | 0.968 | 0.096 | -1.624 | -0.272 | 0.003 | -0.003 | -0.003 | 0.071 | 0.17 | 0.07 | 0.12 | 0.2 | 0.15 | 0.09 |
| 3 | 133474328 | rs12769 | 1.438 | 0.607 | -1.502 | 0.037 | -0.058 | -0.078 | -0.078 | 0 | 0.19 | 0.09 | 0.38 | 0.45 | 0.32 | 0.42 |
| 3 | 133476852 | rs3811658 | 1.052 | 0.083 | -1.397 | -0.069 | -0.051 | -0.057 | -0.057 | 0.01 | 0.32 | 0.12 | 0.39 | 0.43 | 0.35 | 0.41 |
| 3 | 133480588 | rs8177261 | 1.008 | 0.106 | -1.24 | -0.077 | -0.02 | -0.026 | -0.026 | 0 | 0.32 | 0.2 | 0.39 | 0.43 | 0.35 | 0.41 |
| 3 | 133480817 | rs6762719 | 0.976 | 0.069 | -1.169 | -0.093 | -0.015 | -0.021 | -0.021 | 0.604 | 0.33 | 0.21 | 0.39 | 0.43 | 0.35 | 0.41 |
| 3 | 133482230 | rs8177271 | 1.105 | 0.169 | -1.175 | -0.052 | -0.054 | -0.051 | -0.051 | 0 | 0.31 | 0.12 | 0.39 | 0.43 | 0.35 | 0.41 |
| 3 | 133483696 | rs1880669 | 0.527 | 0.182 | -1.19 | -0.219 | 0.052 | 0.001 | 0.001 | 0 | 0.5 | 0.35 | 0.56 | 0.5 | 0.61 | 0.56 |
| 3 | 133484029 | rs3811647 | 1.286 | 0.154 | -1.286 | -0.219 | -0.018 | -0.018 | -0.018 | 0 | 0.33 | 0.19 | 0.39 | 0.43 | 0.35 | 0.41 |
| 3 | 133484188 | rs1358024 | 1.93 | 0.433 | -1.581 | -0.182 | -0.016 | -0.109 | -0.109 | 2.928 | 0.25 | 0.01 | 0.17 | 0.39 | 0.19 | 0.27 |
| 3 | 133485830 | rs2715632 | 1.432 | 0.366 | -0.646 | -0.219 | 0.055 | -0.019 | -0.019 | 0 | 0.18 | 0.34 | 0.18 | 0.07 | 0.26 | 0.14 |
| 3 | 133490033 | rs1525889 | 1.077 | 0.384 | -1.247 | -0.193 | -0.034 | -0.029 | -0.029 | 0.236 | 0.33 | 0.14 | 0.39 | 0.42 | 0.35 | 0.41 |
| 3 | 133492471 | rs9824452 | 0.837 | 0.601 | -1.235 | -0.452 | -0.021 | 0.019 | 0.019 | 0 | 0.09 | 0.04 | 0.13 | 0.04 | 0.12 | 0.11 |
| 3 | 133495673 | rs2715628 | 1.711 | 0.553 | -1.006 | -0.384 | -0.019 | 0.005 | 0.005 | 0 | 0.16 | 0.12 | 0.16 | 0.07 | 0.26 | 0.15 |
| 3 | 133518198 | rs9881405 | 0.423 | 0.286 | -1.225 | -0.483 | -0.017 | -0.008 | -0.008 | 0 | 0.24 | 0.08 | 0.22 | 0.23 | 0.2 | 0.16 |
| 3 | 133519135 | rs13061203 | 2.197 | 0.523 | -1.225 | -0.49 | -0.015 | 0.014 | 0.014 | 0 | 0.06 | 0.04 | 0.22 | 0.03 | 0.16 | 0.14 |
| 3 | 133519981 | rs9853615 | -0.395 | 0.13 | -0.842 | -0.622 | 0.003 | 0.036 | 0.036 | 0 | 0.37 | 0.31 | 0.3 | 0.24 | 0.34 | 0.26 |
| 3 | 133526403 | rs3817524 | -0.47 | 0.065 | -0.731 | -0.639 | 0.002 | 0.017 | 0.017 | 0 | 0.38 | 0.39 | 0.33 | 0.24 | 0.35 | 0.26 |
| 3 | 133527169 | rs7650925 | -0.662 | -0.125 | -0.429 | -0.439 | -0.029 | -0.008 | -0.008 | 1.825 | 0.71 | 0.78 | 0.54 | 0.72 | 0.52 | 0.63 |
| 3 | 133529594 | rs6439441 | -0.702 | -0.076 | -0.551 | -0.437 | -0.091 | -0.035 | -0.035 | 0.136 | 0.71 | 0.87 | 0.56 | 0.72 | 0.53 | 0.64 |
| 3 | 133532054 | rs6806081 | 0.68 | -0.41 | -0.134 | -0.469 | 0 | 0.041 | 0.041 | 0 | 0.06 | 0.24 | 0.03 | 0.05 | 0.01 | 0.13 |
| 3 | 133537067 | rs1534166 | -0.16 | -0.37 | -0.568 | -0.562 | 0.024 | 0.041 | 0.041 | 0 | 0.34 | 0.29 | 0.22 | 0.22 | 0.28 | 0.22 |
| 3 | 133538015 | rs11707556 | 0.696 | -0.12 | -1.161 | -0.635 | -0.071 | 0.092 | 0.092 | 0 | 0.25 | 0.07 | 0.2 | 0.05 | 0.36 | 0.14 |
| 3 | 133539574 | rs10512913 | 1.881 | 0.171 | -1.081 | -0.489 | -0.089 | 0.003 | 0.003 | 0 | 0.15 | 0.01 | 0.15 | 0.04 | 0.27 | 0.07 |
| 3 | 133540359 | rs6439442 | -1.094 | 0.487 | -0.749 | -0.204 | -0.053 | -0.009 | -0.009 | 0 | 0.66 | 0.86 | 0.66 | 0.72 | 0.56 | 0.63 |
| 3 | 133541701 | rs6786236 | 0.267 | 0.183 | -0.357 | -0.213 | -0.002 | 0.002 | 0.002 | 0 | 0.38 | 0.39 | 0.44 | 0.45 | 0.32 | 0.33 |
| 3 | 133543917 | rs7626433 | 1.359 | -0.014 | -0.273 | 0 | -0.016 | 0.013 | 0.013 | 0 | 0.26 | 0.29 | 0.3 | 0.39 | 0.18 | 0.25 |
| 3 | 133544998 | rs1043147 | 0.778 | -0.005 | -0.391 | 0.002 | 0.006 | -0.003 | -0.003 | 0 | 0.31 | 0.25 | 0.34 | 0.39 | 0.25 | 0.3 |
| 3 | 133546297 | rs12637730 | 1.516 | 0.064 | -1.029 | 0.049 | 0.016 | -0.103 | -0.103 | 5.758 | 0.24 | 0.02 | 0.27 | 0.39 | 0.17 | 0.26 |
| 3 | 133547216 | rs2293375 | 0.879 | 0.041 | -1.009 | 0.052 | 0.034 | -0.078 | -0.078 | 0 | 0.3 | 0.11 | 0.32 | 0.43 | 0.24 | 0.3 |
| 3 | 133547245 | rs2293374 | 1.645 | 0.057 | -1.163 | 0.014 | 0.006 | -0.113 | -0.113 | 0 | 0.24 | 0.01 | 0.27 | 0.39 | 0.17 | 0.26 |
| 3 | 133547692 | rs2293373 | 1.659 | 0.058 | -1.163 | 0.015 | 0.005 | -0.113 | -0.113 | 0.512 | 0.24 | 0.01 | 0.27 | 0.39 | 0.17 | 0.26 |
| 3 | 133549938 | rs2715609 | -1.116 | 0.201 | -1.043 | 0.066 | 0.004 | -0.093 | -0.093 | 0 | 0.71 | 0.97 | 0.64 | 0.6 | 0.76 | 0.7 |
| 3 | 133552883 | rs1880666 | -0.636 | 0.11 | -0.545 | -0.126 | -0.012 | -0.011 | -0.011 | 2.691 | 0.2 | 0.66 | 0.23 | 0.17 | 0.14 | 0.25 |
| 3 | 133553272 | rs6793673 | 0.938 | 0.295 | -0.702 | -0.234 | 0.002 | -0.029 | -0.029 | 0 | 0.14 | 0.19 | 0.1 | 0.04 | 0.12 | 0.08 |
| 3 | 133553527 | rs2280672 | -0.259 | 0.381 | -0.648 | -0.091 | -0.007 | 0.002 | 0.002 | 0 | 0.56 | 0.53 | 0.47 | 0.4 | 0.63 | 0.59 |
| 3 | 133555500 | rs2692672 | -0.413 | 1.005 | -0.943 | -0.204 | 0.001 | -0.003 | -0.003 | 0 | 0.23 | 0.28 | 0.25 | 0.16 | 0.21 | 0.17 |
| 3 | 133556345 | rs9840108 | 0.248 | 1.075 | -1.194 | 0.123 | -0.027 | 0.001 | 0.001 | 0 | 0.35 | 0.27 | 0.32 | 0.28 | 0.5 | 0.36 |
| 3 | 133558698 | rs6765093 | -0.313 | 0.594 | -2.147 | 0.016 | 0.044 | -0.02 | -0.02 | 0 | 0.47 | 0.84 | 0.47 | 0.39 | 0.59 | 0.44 |
| 3 | 133561095 | rs1105877 | 0.151 | 0.532 | -2.049 | -0.06 | -0.069 | -0.04 | -0.04 | 0.015 | 0.3 | 0.04 | 0.19 | 0.27 | 0.31 | 0.23 |
| 3 | 133576458 | rs9815265 | -0.018 | 1.313 | -1.69 | -0.716 | -0.072 | 0.079 | 0.079 | 0.172 | 0.24 | 0.11 | 0.25 | 0.07 | 0.38 | 0.26 |
| 3 | 133577269 | rs1534167 | -0.611 | 0.401 | -1.883 | -1.474 | 0 | 0.043 | 0.043 | 0.157 | 0.62 | 0.68 | 0.54 | 0.13 | 0.66 | 0.56 |
| 3 | 133579775 | rs17376789 | 0.562 | -0.115 | -2.637 | -1.609 | 0.002 | 0.17 | 0.17 | 0 | 0.27 | 0.01 | 0.22 | 0.03 | 0.23 | 0.21 |
| 3 | 133581770 | rs1113479 | -0.507 | -0.208 | -2.526 | -0.721 | -0.036 | 0.016 | 0.016 | 0 | 0.49 | 0.88 | 0.47 | 0.53 | 0.49 | 0.59 |
| 3 | 133585781 | rs9835459 | 0.325 | 0.599 | -2.315 | -0.884 | -0.063 | 0.049 | 0.049 | 0 | 0.27 | 0.11 | 0.28 | 0.17 | 0.41 | 0.33 |
| 3 | 133589324 | rs9835973 | 0.041 | 0.036 | -2.029 | -0.52 | -0.012 | 0.009 | 0.009 | 0 | 0.21 | 0.01 | 0.25 | 0.13 | 0.18 | 0.19 |
| 3 | 133594691 | rs2692682 | -0.304 | -0.377 | -1.055 | -0.65 | -0.046 | -0.052 | -0.052 | 0 | 0.23 | 0.37 | 0.11 | 0.12 | 0.1 | 0.24 |
| 3 | 133595618 | rs2692681 | -1.156 | -0.112 | -1.343 | -1.347 | -0.019 | 0.033 | 0.033 | 0 | 0.6 | 0.76 | 0.51 | 0.33 | 0.58 | 0.57 |
| 3 | 133595969 | rs9844944 | 0.871 | 0.004 | -1.44 | -1.459 | -0.048 | -0.022 | -0.022 | 0 | 0.13 | 0.03 | 0.2 | 0.14 | 0.22 | 0.15 |
| 3 | 133600082 | rs1917779 | 0.327 | -0.113 | -1.431 | -1.486 | -0.032 | -0.054 | -0.054 | 0 | 0.25 | 0.37 | 0.13 | 0.12 | 0.15 | 0.26 |
| 3 | 133602195 | rs1949881 | -0.371 | -0.297 | -0.862 | -1.63 | -0.01 | -0.021 | -0.021 | 0 | 0.37 | 0.47 | 0.31 | 0.28 | 0.35 | 0.44 |
| 3 | 133603480 | rs2715638 | -0.966 | -0.198 | -0.987 | -1.67 | 0.046 | 0.19 | 0.19 | 0 | 0.51 | 0.8 | 0.35 | 0.2 | 0.41 | 0.44 |
| 3 | 133604743 | rs9813363 | 0.995 | -0.109 | -0.712 | -1.558 | 0.022 | 0.014 | 0.014 | 0 | 0.12 | 0.2 | 0.2 | 0.14 | 0.17 | 0.13 |

| | | | | | | | | | | | | | | | | |
|---|---|---|---|---|---|---|---|---|---|---|---|---|---|---|---|---|
| 3 | 133604787 | rs17311500 | 1.471 | -0.084 | -0.75 | -1.618 | -0.033 | 0.05 | 0.05 | 0 | 0.16 | 0.05 | 0.15 | 0.05 | 0.2 | 0.12 |
| 3 | 133611938 | rs7636726 | 0.825 | -0.123 | -0.511 | -1.381 | 0.023 | 0.016 | 0.016 | 0 | 0.12 | 0.23 | 0.21 | 0.14 | 0.17 | 0.13 |
| 3 | 133616371 | rs940898 | -0.938 | -0.189 | -0.625 | -1.504 | 0.001 | 0.001 | 0.001 | 0 | 0.43 | 0.43 | 0.34 | 0.27 | 0.36 | 0.44 |
| 3 | 133618581 | rs9824788 | 0.385 | -0.283 | -1.006 | -1.544 | -0.029 | 0.01 | 0.01 | 0.246 | 0.2 | 0.06 | 0.22 | 0.14 | 0.26 | 0.21 |
| 3 | 133629112 | rs2692686 | 0.018 | -0.347 | -0.544 | -1.412 | -0.032 | -0.034 | -0.034 | 0 | 0.25 | 0.62 | 0.15 | 0.14 | 0.13 | 0.23 |
| 3 | 133636566 | rs6439445 | -0.16 | -0.315 | -0.354 | -1.249 | -0.001 | 0.023 | 0.023 | 0 | 0.5 | 0.53 | 0.48 | 0.31 | 0.52 | 0.55 |
| 3 | 133638387 | rs6809337 | -1.104 | 0.026 | -0.703 | -0.941 | -0.038 | 0.003 | 0.003 | 0 | 0.81 | 0.9 | 0.76 | 0.82 | 0.74 | 0.84 |
| 3 | 133639221 | rs9790196 | 0.215 | 0.002 | -0.631 | -0.952 | -0.001 | 0.04 | 0.04 | 0 | 0.5 | 0.59 | 0.49 | 0.3 | 0.52 | 0.55 |
| 3 | 133644169 | rs6763132 | -1.073 | 0.103 | -0.775 | -0.911 | -0.012 | 0.003 | 0.003 | 0 | 0.81 | 0.82 | 0.75 | 0.82 | 0.74 | 0.83 |
| 3 | 133644893 | rs7623757 | 0.873 | 0.2 | -0.808 | -0.559 | -0.001 | 0.002 | 0.002 | 0 | 0.3 | 0.27 | 0.26 | 0.52 | 0.23 | 0.29 |
| 3 | 133646650 | rs6808547 | 1.044 | 0.804 | -0.957 | -0.571 | -0.029 | 0.039 | 0.039 | 0 | 0.14 | 0.08 | 0.14 | 0.04 | 0.17 | 0.13 |
| 3 | 133649908 | rs11915399 | 0.964 | 0.806 | -0.99 | -0.563 | 0.012 | -0.041 | -0.041 | 0 | 0.16 | 0.25 | 0.15 | 0.04 | 0.17 | 0.13 |
| 3 | 133650996 | rs9819520 | 0.607 | 0.1 | -1.486 | -0.328 | -0.035 | -0.023 | -0.023 | 0 | 0.14 | 0.02 | 0.18 | 0.14 | 0.18 | 0.12 |
| 3 | 133652322 | rs1131598 | -0.293 | -1.242 | -1.218 | 0.417 | -0.005 | 0.007 | 0.007 | 0 | 0.27 | 0.25 | 0.28 | 0.48 | 0.2 | 0.45 |
| 3 | 133654846 | rs7616492 | 1.035 | -1.51 | -1.521 | 0.391 | -0.137 | 0.03 | 0.03 | 0 | 0.36 | 0.04 | 0.32 | 0.25 | 0.39 | 0.23 |
| 3 | 133655001 | rs4502579 | 0.481 | -1.592 | -0.952 | 0.578 | -0.022 | 0.18 | 0.18 | 0 | 0.17 | 0.4 | 0.23 | 0.44 | 0.11 | 0.28 |
| 3 | 133656433 | rs6439448 | -1.041 | -1.177 | -0.765 | 0.357 | 0.029 | 0.016 | 0.016 | 0.732 | 0.83 | 0.63 | 0.77 | 0.83 | 0.8 | 0.86 |
| 3 | 133658566 | rs11710021 | 0.916 | -1.103 | -1.408 | 0.141 | -0.009 | 0.055 | 0.055 | 0 | 0.14 | 0 | 0.09 | 0.05 | 0.1 | 0.13 |
| 3 | 133663681 | rs9818609 | 0.953 | -1.048 | -1.28 | 0.253 | -0.042 | 0.13 | 0.13 | 0.546 | 0.42 | 0.13 | 0.36 | 0.18 | 0.41 | 0.22 |
| 3 | 133664543 | rs11720811 | 1.637 | -0.952 | -1.366 | 0.253 | -0.011 | 0.027 | 0.027 | 0.054 | 0.11 | 0.06 | 0.07 | 0 | 0.18 | 0.13 |
| 3 | 133670073 | rs6767412 | 0.58 | -1.174 | -1.023 | 0.471 | -0.004 | 0.009 | 0.009 | 0 | 0.21 | 0.32 | 0.24 | 0.28 | 0.19 | 0.21 |
| 3 | 133670391 | rs4854769 | 0.654 | -1.175 | -1.041 | 0.471 | -0.003 | 0.007 | 0.007 | 0 | 0.2 | 0.3 | 0.24 | 0.28 | 0.19 | 0.21 |
| 3 | 133670663 | rs3924031 | 1.375 | -1.219 | -1.143 | 0.426 | -0.104 | 0.095 | 0.095 | 0.007 | 0.37 | 0.05 | 0.35 | 0.18 | 0.41 | 0.21 |
| 3 | 133674895 | rs7340717 | -0.359 | -1.286 | -0.505 | 0.975 | -0.001 | 0.001 | 0.001 | 0 | 0.3 | 0.32 | 0.31 | 0.29 | 0.37 | 0.33 |
| 3 | 133674985 | rs7340718 | -0.229 | -1.288 | -0.511 | 1.128 | -0.002 | -0.006 | -0.006 | 0 | 0.3 | 0.31 | 0.31 | 0.35 | 0.37 | 0.34 |
| 3 | 133681395 | rs6804798 | 0.587 | -1.179 | -0.454 | 0.961 | -0.001 | -0.039 | -0.039 | 0 | 0.14 | 0.23 | 0.08 | 0.01 | 0.18 | 0.13 |
| 3 | 133681762 | rs9820625 | -0.805 | -1.38 | -0.492 | 1.049 | 0.006 | 0.01 | 0.01 | 0 | 0.52 | 0.47 | 0.46 | 0.43 | 0.53 | 0.55 |
| 3 | 133685254 | rs9917636 | -0.797 | -1.382 | -0.493 | 1.048 | 0.006 | 0.01 | 0.01 | 0 | 0.52 | 0.47 | 0.46 | 0.43 | 0.53 | 0.55 |
| 3 | 133687864 | rs4854776 | -0.789 | -1.489 | -1.201 | 1.031 | 0.03 | 0.082 | 0.082 | 0 | 0.53 | 0.28 | 0.53 | 0.79 | 0.44 | 0.76 |
| 3 | 133690038 | rs13319548 | -0.237 | -1.558 | -0.993 | 1.115 | -0.016 | 0.028 | 0.028 | 0 | 0.28 | 0.52 | 0.26 | 0.39 | 0.28 | 0.47 |
| 3 | 133690376 | rs13093879 | 1.878 | -1.66 | -1.057 | 1.003 | 0.033 | 0.036 | 0.036 | 0 | 0.16 | 0.06 | 0.07 | 0.06 | 0.12 | 0.12 |
| 3 | 133691147 | rs4854777 | -1.631 | -1.419 | -0.945 | 0.999 | -0.005 | 0.003 | 0.003 | 0.076 | 0.66 | 0.68 | 0.62 | 0.81 | 0.62 | 0.89 |
| 3 | 133691165 | rs9882333 | 0.769 | -1.301 | -0.727 | 0.969 | 0.037 | -0.069 | -0.069 | 0 | 0.13 | 0.42 | 0.1 | 0.02 | 0.18 | 0.13 |
| 3 | 133692743 | rs4241362 | -1.238 | -1.197 | -1.08 | 1.014 | -0.007 | 0.013 | 0.013 | 0.117 | 0.88 | 0.92 | 0.92 | 0.98 | 0.82 | 0.87 |
| 3 | 133693859 | rs6795214 | 0.516 | -1.03 | -1.009 | 1.176 | 0.004 | -0.003 | -0.003 | 0 | 0.15 | 0.1 | 0.16 | 0.38 | 0.1 | 0.35 |
| 3 | 133696285 | rs4241363 | -0.382 | -0.975 | -1.023 | 0.804 | 0.027 | 0.021 | 0.021 | 0 | 0.47 | 0.63 | 0.47 | 0.69 | 0.41 | 0.64 |
| 3 | 133698349 | rs10935090 | -0.02 | -0.259 | -1.102 | 1.063 | 0.002 | 0.003 | 0.003 | 0 | 0.16 | 0.12 | 0.19 | 0.54 | 0.13 | 0.32 |
| 3 | 133699345 | rs3924290 | -0.17 | -0.36 | -1.3 | 0.86 | -0.017 | 0.025 | 0.025 | 0 | 0.82 | 0.75 | 0.85 | 0.57 | 0.89 | 0.65 |
| 3 | 133699575 | rs7646392 | 0.017 | 0.58 | -1.769 | 0.344 | -0.034 | 0.004 | 0.004 | 0.253 | 0.35 | 0.22 | 0.41 | 0.26 | 0.39 | 0.21 |
| 3 | 133700071 | rs5013525 | -0.036 | 0.634 | -1.745 | 0.345 | -0.034 | 0.005 | 0.005 | 0.345 | 0.35 | 0.21 | 0.42 | 0.25 | 0.39 | 0.19 |
| 3 | 133701960 | rs4327389 | 0 | 0.685 | -1.89 | 0.317 | -0.044 | 0.007 | 0.007 | 0.441 | 0.34 | 0.18 | 0.41 | 0.25 | 0.39 | 0.19 |
| 3 | 133709523 | rs4331673 | -0.479 | 0.521 | -1.521 | 0.203 | -0.02 | 0.092 | 0.092 | 0 | 0.15 | 0.03 | 0.08 | 0 | 0.18 | 0.09 |
| 3 | 133717948 | rs4854785 | -0.804 | 0.768 | -1.113 | 0.986 | -0.002 | -0.007 | -0.007 | 0 | 0.24 | 0.2 | 0.35 | 0.46 | 0.23 | 0.36 |
| 3 | 133719806 | rs13320795 | -0.465 | 0.307 | -1.538 | 0.246 | 0.018 | 0.091 | 0.091 | 0 | 0.15 | 0.02 | 0.08 | 0.01 | 0.09 | 0.2 |
| 3 | 133720809 | rs4854786 | 0.654 | -0.23 | -0.585 | 0.046 | -0.033 | -0.057 | -0.057 | 0 | 0.77 | 0.76 | 0.88 | 0.99 | 0.91 | 0.67 |
| 3 | 133723028 | rs9790150 | 1.272 | 0.793 | 0.079 | -0.255 | -0.009 | -0.095 | -0.095 | 0 | 0.69 | 0.35 | 0.73 | 0.98 | 0.72 | 0.65 |
| 3 | 133725592 | rs7428024 | 0.891 | 0.647 | 0.623 | 0.921 | -0.059 | 0.06 | 0.06 | 0 | 0.74 | 0.59 | 0.68 | 0.68 | 0.83 | 0.55 |
| 3 | 133727866 | rs7625035 | 1.08 | 0.718 | 0.576 | 0.857 | -0.018 | 0.011 | 0.011 | 0 | 0.71 | 0.73 | 0.65 | 0.66 | 0.79 | 0.54 |
| 3 | 133730608 | rs4854787 | -1.339 | 1.225 | 0.975 | 1.621 | -0.003 | 0.028 | 0.028 | 0 | 0.13 | 0.13 | 0.14 | 0.37 | 0.07 | 0.14 |
| 3 | 133730932 | rs7629416 | 2.226 | 1.816 | 1.492 | 1.467 | 0.045 | 0.052 | 0.052 | 0 | 0.78 | 0.43 | 0.7 | 0.62 | 0.66 | 0.79 |
| 3 | 133731842 | rs4854789 | -1.177 | 1.83 | 1.239 | 1.487 | -0.003 | 0.033 | 0.033 | 0.103 | 0.12 | 0.13 | 0.11 | 0.36 | 0.08 | 0.15 |
| 3 | 133735861 | rs9841380 | 2.02 | 1.588 | 0.978 | 1.421 | -0.007 | -0.013 | -0.013 | 0.315 | 0.66 | 0.26 | 0.57 | 0.56 | 0.6 | 0.68 |
| 3 | 133738545 | rs4241365 | 0.846 | 1.509 | 1.279 | 1.416 | 0.162 | -0.025 | -0.025 | 0 | 0.93 | 0.75 | 0.83 | 0.99 | 0.76 | 0.94 |
| 3 | 133747404 | rs9834727 | 1.545 | 1.379 | 0.504 | 1.495 | -0.005 | -0.013 | -0.013 | 0 | 0.67 | 0.27 | 0.57 | 0.56 | 0.61 | 0.67 |
| 3 | 133756102 | rs9816473 | 1.293 | 1.226 | 0.333 | 1.554 | -0.003 | -0.007 | -0.007 | 0 | 0.68 | 0.31 | 0.6 | 0.35 | 0.65 | 0.77 |
| 3 | 133759848 | rs4508778 | 1.541 | 2.356 | 0.953 | 1.69 | 0.041 | 0.038 | 0.038 | 0 | 0.72 | 0.41 | 0.45 | 0.38 | 0.64 | 0.77 |
| 3 | 133760537 | rs4528941 | -1.759 | 2.432 | 1.664 | 2.414 | 0.04 | 0.054 | 0.054 | 0 | 0.27 | 0.36 | 0.44 | 0.54 | 0.38 | 0.23 |
| 3 | 133760908 | rs4301004 | -1.292 | 2.942 | 1.569 | 2.551 | 0.015 | 0.015 | 0.015 | 0 | 0.17 | 0.2 | 0.22 | 0.31 | 0.22 | 0.14 |
| 3 | 133767223 | rs7431223 | -1.612 | 2.747 | 1.825 | 2.558 | 0.087 | 0.242 | 0.242 | 0 | 0.1 | 0.33 | 0.33 | 0.29 | 0.17 | 0.11 |
| 3 | 133769778 | rs9858660 | -0.042 | 1.949 | 1.136 | 1.683 | -0.005 | -0.005 | -0.005 | 0 | 0.83 | 0.8 | 0.89 | 0.91 | 0.83 | 0.59 |
| 3 | 133778591 | rs6765810 | 1.416 | 2.314 | 1.172 | 1.909 | 0.073 | 0.079 | 0.079 | 0 | 0.73 | 0.49 | 0.46 | 0.42 | 0.63 | 0.77 |
| 3 | 133780301 | rs4854605 | -1.319 | 2.124 | 1.3 | 1.742 | -0.004 | 0.013 | 0.013 | 0 | 0.25 | 0.3 | 0.4 | 0.6 | 0.25 | 0.55 |
| 3 | 133784988 | rs7615471 | 0.417 | 2.464 | 1.41 | 1.796 | 0.046 | 0.033 | 0.033 | 0 | 0.93 | 0.83 | 0.82 | 0.82 | 0.84 | 0.93 |
| 3 | 133793121 | rs4241367 | -1.731 | 2.056 | 0.941 | 0.962 | 0 | 0.124 | 0.124 | 0 | 0.47 | 0.71 | 0.59 | 0.79 | 0.44 | 0.72 |
| 3 | 133794199 | rs4241370 | -1.704 | 2.05 | 1.373 | 0.951 | 0.001 | 0.054 | 0.054 | 0 | 0.46 | 0.65 | 0.58 | 0.79 | 0.44 | 0.72 |
| 3 | 133796681 | rs7431119 | -1.539 | 2.04 | 1.294 | 1.325 | -0.001 | 0.01 | 0.01 | 0 | 0.4 | 0.53 | 0.46 | 0.43 | 0.37 | 0.61 |
| 3 | 133798207 | rs12635144 | 0.687 | 1.91 | 1.175 | 1.382 | 0.025 | -0.003 | -0.003 | 0.407 | 0.06 | 0 | 0.19 | 0.26 | 0.02 | 0.29 |
| 3 | 133798818 | rs6439455 | -0.545 | 1.825 | 1.21 | 1.276 | 0.06 | -0.024 | -0.024 | 0 | 0.94 | 0.86 | 0.86 | 0.99 | 0.85 | 0.99 |
| 3 | 133800471 | rs12631386 | -0.465 | 1.72 | 1.366 | 1.288 | -0.005 | 0.082 | 0.082 | 0 | 0.18 | 0.27 | 0.29 | 0.29 | 0.18 | 0.43 |

| | | | | | | | | | | | | | | | | |
|---|---|---|---|---|---|---|---|---|---|---|---|---|---|---|---|---|
| 3 | 133800995 | rs6770237 | 0.157 | 1.721 | 0.939 | 1.062 | -0.006 | 0.001 | 0.001 | 0 | 0.15 | 0.12 | 0.08 | 0 | 0.13 | 0.12 |
| 3 | 133804070 | rs6414331 | -0.838 | 1.655 | 1.043 | 1.102 | 0 | -0.023 | -0.023 | 0 | 0.22 | 0.32 | 0.18 | 0.14 | 0.18 | 0.16 |
| 3 | 133804728 | rs6783575 | -0.069 | 1.923 | 0.982 | 0.913 | 0.034 | -0.015 | -0.015 | 0 | 0.92 | 0.91 | 0.8 | 0.99 | 0.7 | 0.96 |
| 3 | 133805349 | rs4854610 | -0.637 | 1.514 | 1.011 | 1.23 | -0.001 | 0 | 0 | 0 | 0.21 | 0.19 | 0.2 | 0.35 | 0.19 | 0.21 |
| 3 | 133807007 | rs4426678 | -0.623 | 1.515 | 0.943 | 1.23 | -0.012 | -0.031 | -0.031 | 0 | 0.21 | 0.15 | 0.19 | 0.35 | 0.19 | 0.21 |
| 3 | 133813666 | rs4854811 | -0.684 | 1.414 | 1.235 | 1.278 | 0.026 | 0.238 | 0.238 | 0 | 0.07 | 0.22 | 0.12 | 0.37 | 0.07 | 0.1 |
| 3 | 133830753 | rs4854818 | -0.532 | 1.416 | 1.201 | 1.279 | 0.003 | 0.123 | 0.123 | 0 | 0.07 | 0.12 | 0.12 | 0.37 | 0.07 | 0.1 |
| 3 | 133853335 | rs7630560 | 0.805 | 1.632 | 1.119 | 1.131 | 0.034 | -0.017 | -0.017 | 1.576 | 0.83 | 0.75 | 0.7 | 0.88 | 0.67 | 0.89 |
| 3 | 133866728 | rs9878348 | -0.233 | 1.606 | 1.154 | 1.202 | 0.018 | 0.201 | 0.201 | 0 | 0.06 | 0.16 | 0.12 | 0.37 | 0.07 | 0.1 |
| 3 | 133868797 | rs4072181 | 0.122 | 1.587 | 1.168 | 1.143 | -0.001 | 0 | 0 | 0 | 0.06 | 0.04 | 0.08 | 0.11 | 0.05 | 0.06 |
| 3 | 133874566 | rs9283588 | -0.568 | 1.425 | 1.276 | 1.163 | 0.078 | 0.352 | 0.352 | 0 | 0.07 | 0.22 | 0.2 | 0.48 | 0.12 | 0.16 |
| 3 | 133877570 | rs4854617 | 0.186 | 1.491 | 0.971 | 1.121 | 0.056 | -0.053 | -0.053 | 0 | 0.88 | 0.76 | 0.78 | 1 | 0.72 | 0.95 |
| 3 | 133882151 | rs4470517 | 0.04 | 1.561 | 0.957 | 1.124 | 0.048 | -0.041 | -0.041 | 0 | 0.89 | 0.78 | 0.78 | 1 | 0.7 | 0.95 |
| 3 | 133896121 | rs6762690 | -1.147 | 1.371 | 0.85 | 1.107 | 0 | 0.03 | 0.03 | 0 | 0.14 | 0.12 | 0.2 | 0.48 | 0.12 | 0.16 |
| 3 | 133949713 | rs4339087 | 0.366 | 1.265 | 0.6 | 0.943 | 0 | -0.029 | -0.029 | 0 | 0.15 | 0.1 | 0.09 | 0 | 0.13 | 0.12 |
| 3 | 133980309 | rs6439470 | 0.946 | 1.074 | 0.315 | 1.094 | 0.025 | 0.047 | 0.047 | 0 | 0.76 | 0.59 | 0.59 | 0.67 | 0.68 | 0.57 |
| 3 | 133989068 | rs6780295 | -1.309 | 1.633 | 0.466 | 1.082 | -0.003 | 0.016 | 0.016 | 0 | 0.3 | 0.58 | 0.32 | 0.48 | 0.25 | 0.27 |
| 3 | 133990426 | rs9289462 | -0.537 | 2.01 | 0.299 | 1.505 | 0 | 0.063 | 0.063 | 0 | 0.07 | 0.11 | 0.06 | 0.33 | 0.07 | 0.12 |
| 3 | 133992299 | rs7619206 | 0.546 | 2.096 | 0.1 | 0.789 | 0.084 | -0.062 | -0.062 | 0 | 0.88 | 0.77 | 0.8 | 1 | 0.7 | 0.95 |
| 3 | 133992984 | rs10935106 | 0.507 | 2.106 | -0.154 | 0.793 | 0.029 | -0.033 | -0.033 | 0.594 | 0.88 | 0.82 | 0.8 | 1 | 0.7 | 0.96 |
| 3 | 133995154 | rs12185973 | -0.513 | 1.79 | -0.367 | 1.044 | -0.002 | -0.039 | -0.039 | 1.135 | 0.07 | 0.01 | 0.15 | 0.13 | 0.05 | 0.06 |
| 3 | 133995452 | rs9870797 | 0.507 | 2.128 | 0.027 | 0.529 | 0.08 | -0.062 | -0.062 | 0 | 0.88 | 0.79 | 0.79 | 1 | 0.71 | 0.95 |
| 3 | 133997360 | rs4634087 | 0.646 | 2.088 | -0.056 | 0.983 | 0.047 | 0.017 | 0.017 | 0 | 0.87 | 0.82 | 0.64 | 0.83 | 0.71 | 0.92 |
| 3 | 134010124 | rs6763231 | 0.947 | 0.849 | -0.344 | -0.043 | -0.002 | -0.06 | -0.06 | 0.299 | 0.42 | 0.43 | 0.28 | 0.15 | 0.43 | 0.28 |
| 3 | 134013144 | rs4535199 | 0.662 | 1.068 | -0.292 | 0.28 | -0.006 | -0.006 | -0.006 | 0.169 | 0.74 | 0.71 | 0.57 | 0.47 | 0.61 | 0.69 |
| 3 | 134020548 | rs4504146 | 0.388 | 0.624 | -0.54 | -1.305 | 0.001 | 0.005 | 0.005 | 0 | 0.34 | 0.28 | 0.25 | 0.08 | 0.38 | 0.21 |
| 3 | 134029172 | rs13080059 | 0.149 | 0.661 | -0.055 | -1.783 | -0.006 | -0.05 | -0.05 | 3.129 | 0.43 | 0.45 | 0.34 | 0.15 | 0.34 | 0.39 |
| 3 | 134033585 | rs4533618 | -0.726 | 0.856 | 0.039 | -2.04 | 0.001 | -0.021 | -0.021 | 0 | 0.61 | 0.41 | 0.68 | 0.93 | 0.59 | 0.79 |
| 3 | 134035007 | rs4682662 | 0.325 | 0.816 | 0.036 | -1.794 | -0.008 | 0.002 | 0.002 | 0 | 0.2 | 0.18 | 0.37 | 0.81 | 0.26 | 0.4 |
| 3 | 134043422 | rs7636249 | 0.369 | 0.457 | 0.065 | -2.234 | -0.003 | -0.156 | -0.156 | 0.047 | 0.25 | 0.46 | 0.14 | 0.01 | 0.22 | 0.09 |
| 3 | 134043922 | rs11928672 | 0.369 | 0.457 | 0.065 | -2.234 | -0.003 | -0.156 | -0.156 | 0 | 0.25 | 0.46 | 0.14 | 0.01 | 0.22 | 0.09 |
| 3 | 134043988 | rs11917363 | 0.369 | 0.457 | 0.065 | -2.234 | -0.003 | -0.156 | -0.156 | 0 | 0.25 | 0.46 | 0.14 | 0.01 | 0.22 | 0.09 |
| 3 | 134053382 | rs4974489 | -0.004 | 0.653 | -0.35 | -2.278 | -0.004 | 0.064 | 0.064 | 0 | 0.35 | 0.25 | 0.52 | 0.92 | 0.34 | 0.71 |
| 3 | 134063742 | rs4974496 | -0.558 | 0.49 | -0.412 | -2.154 | 0.003 | 0.037 | 0.037 | 0 | 0.63 | 0.73 | 0.69 | 0.93 | 0.61 | 0.81 |
| 3 | 134075522 | rs4974498 | -0.642 | 0.544 | -1.449 | -2.109 | -0.07 | 0.188 | 0.188 | 0 | 0.64 | 0.96 | 0.72 | 0.93 | 0.61 | 0.79 |
| 3 | 134076154 | rs3732852 | -0.653 | 0.55 | -1.445 | -2.106 | -0.07 | 0.187 | 0.187 | 0 | 0.64 | 0.96 | 0.72 | 0.92 | 0.61 | 0.79 |
| 3 | 134083642 | rs3773820 | 0.664 | 0.693 | -1.24 | -2.089 | -0.055 | 0.107 | 0.107 | 0 | 0.29 | 0.11 | 0.26 | 0.08 | 0.37 | 0.19 |
| 3 | 134090703 | rs11919447 | 0.61 | 0.723 | -1.228 | -2.303 | 0.002 | 0.051 | 0.051 | 0 | 0.28 | 0.15 | 0.16 | 0.01 | 0.25 | 0.09 |
| 3 | 134090900 | rs10755085 | 0.03 | 1.01 | -0.426 | -1.78 | -0.002 | -0.015 | -0.015 | 0.19 | 0.62 | 0.66 | 0.67 | 0.92 | 0.56 | 0.78 |
| 3 | 134106494 | rs2400273 | 0.217 | 1.335 | -0.866 | -1.618 | -0.055 | 0.097 | 0.097 | 0 | 0.29 | 0.14 | 0.27 | 0.08 | 0.39 | 0.2 |
| 3 | 134108539 | rs9682157 | -0.079 | 1.009 | -0.655 | -0.902 | 0.007 | -0.011 | -0.011 | 0 | 0.45 | 0.39 | 0.45 | 0.49 | 0.42 | 0.66 |
| 3 | 134109630 | rs4974500 | -1.089 | 1.107 | -0.262 | -0.983 | -0.005 | -0.006 | -0.006 | 0 | 0.54 | 0.7 | 0.53 | 0.5 | 0.52 | 0.66 |
| 3 | 134109861 | rs4974501 | -0.076 | 1.134 | -0.597 | -0.796 | -0.042 | -0.036 | -0.036 | 0 | 0.35 | 0.21 | 0.44 | 0.39 | 0.42 | 0.46 |
| 3 | 134120357 | rs3846051 | 0.452 | 0.816 | -0.636 | -0.675 | -0.011 | 0.062 | 0.062 | 0 | 0.28 | 0.12 | 0.18 | 0.1 | 0.3 | 0.32 |
| 3 | 134134236 | rs10433423 | -0.434 | 0.71 | -0.472 | -0.449 | 0.023 | 0.01 | 0.01 | 0 | 0.49 | 0.65 | 0.38 | 0.43 | 0.4 | 0.66 |
| 3 | 134138009 | rs3846052 | -0.526 | 0.539 | -0.499 | -1.847 | 0.003 | -0.04 | -0.04 | 0 | 0.69 | 0.76 | 0.82 | 0.99 | 0.76 | 0.86 |
| 3 | 134140357 | rs16841633 | -0.437 | -0.124 | 0.305 | -0.923 | -0.012 | 0.045 | 0.045 | 1.388 | 0.19 | 0.35 | 0.08 | 0.32 | 0.11 | 0.48 |
| 3 | 134142093 | rs4602357 | -1.042 | -0.162 | 0.134 | -1.146 | -0.001 | 0.014 | 0.014 | 0.371 | 0.49 | 0.8 | 0.35 | 0.46 | 0.44 | 0.66 |
| 3 | 134143907 | rs9839295 | 0.592 | -0.35 | 0.651 | -1.211 | -0.014 | 0.034 | 0.034 | 0 | 0.26 | 0.18 | 0.24 | 0.1 | 0.33 | 0.15 |
| 3 | 134143919 | rs16841638 | -0.123 | -0.404 | 1.094 | -1.173 | -0.012 | 0.048 | 0.048 | 2.543 | 0.2 | 0.31 | 0.07 | 0.36 | 0.11 | 0.5 |
| 3 | 134144721 | rs6796705 | 0.394 | -0.237 | 1.06 | -1.473 | -0.002 | -0.005 | -0.005 | 0 | 0.27 | 0.27 | 0.24 | 0.1 | 0.33 | 0.15 |
| 3 | 134146709 | rs13097818 | 0.107 | -0.236 | 1.41 | -1.46 | 0.015 | -0.073 | -0.073 | 0 | 0.28 | 0.45 | 0.27 | 0.12 | 0.34 | 0.16 |
| 3 | 134146915 | rs6779053 | -0.115 | -0.44 | 1.572 | -1.395 | -0.033 | 0.02 | 0.02 | 0.869 | 0.18 | 0.45 | 0.08 | 0.18 | 0.08 | 0.29 |
| 3 | 134151959 | rs9790169 | -1.081 | -0.34 | 1.299 | -1.423 | -0.004 | 0.049 | 0.049 | 0.026 | 0.45 | 0.71 | 0.34 | 0.28 | 0.44 | 0.42 |
| 3 | 134152631 | rs4312629 | 0.328 | -0.423 | 1.02 | -1.413 | -0.028 | 0.051 | 0.051 | 0 | 0.26 | 0.17 | 0.24 | 0.1 | 0.36 | 0.16 |
| 3 | 134152990 | rs11916983 | 0.234 | -0.401 | 1.033 | -1.403 | 0 | -0.013 | -0.013 | 2.415 | 0.27 | 0.3 | 0.25 | 0.1 | 0.36 | 0.15 |
| 3 | 134160593 | rs9851770 | 0.286 | -0.499 | 0.924 | -1.489 | 0.01 | -0.036 | -0.036 | 0 | 0.26 | 0.33 | 0.25 | 0.1 | 0.35 | 0.11 |
| 3 | 134166224 | rs9835166 | 0.237 | -0.522 | 0.975 | -1.499 | 0.015 | -0.049 | -0.049 | 0 | 0.27 | 0.34 | 0.25 | 0.09 | 0.35 | 0.11 |
| 3 | 134170971 | rs7650509 | 0.865 | -0.76 | 0.837 | -1.594 | -0.002 | 0.026 | 0.026 | 0 | 0.57 | 0.33 | 0.67 | 0.74 | 0.59 | 0.59 |
| 3 | 134172488 | rs1498719 | -1.269 | -0.706 | 0.681 | -1.507 | 0 | -0.011 | -0.011 | 0.053 | 0.91 | 0.99 | 0.97 | 0.92 | 0.95 | 0.85 |
| 3 | 134191938 | rs9841212 | 0.457 | -0.642 | 0.686 | -1.563 | -0.002 | -0.016 | -0.016 | 0 | 0.65 | 0.46 | 0.73 | 0.74 | 0.69 | 0.6 |
| 3 | 134192338 | rs9842113 | 1.478 | -0.763 | 0.392 | -1.708 | -0.113 | -0.024 | -0.024 | 0.783 | 0.5 | 0.03 | 0.63 | 0.57 | 0.52 | 0.39 |
| 3 | 134192469 | rs9842140 | 2.179 | -1.29 | -0.003 | -2.104 | -0.105 | -0.149 | -0.149 | 0 | 0.29 | 0.02 | 0.53 | 0.56 | 0.38 | 0.27 |
| 3 | 134195885 | rs934093 | -0.157 | -1.101 | 0.239 | -2.225 | 0.002 | -0.036 | -0.036 | 0.86 | 0.68 | 0.58 | 0.79 | 0.83 | 0.69 | 0.7 |
| 3 | 134200205 | rs1863913 | 0.105 | -1.1 | 0.231 | -2.225 | -0.003 | -0.039 | -0.039 | 0 | 0.66 | 0.47 | 0.77 | 0.82 | 0.69 | 0.6 |
| 3 | 134205071 | rs1874883 | -0.248 | -1.017 | 0.155 | -2.187 | 0.004 | -0.037 | -0.037 | 0 | 0.68 | 0.58 | 0.79 | 0.82 | 0.69 | 0.7 |
| 3 | 134205920 | rs1498718 | 0.012 | -1.018 | 0.057 | -2.188 | -0.001 | -0.047 | -0.047 | 0.756 | 0.66 | 0.49 | 0.77 | 0.82 | 0.69 | 0.6 |
| 3 | 134206069 | rs3932818 | 0.012 | -1.018 | 0.057 | -2.188 | -0.001 | -0.047 | -0.047 | 0.02 | 0.66 | 0.49 | 0.77 | 0.82 | 0.69 | 0.6 |
| 3 | 134217904 | rs10935116 | 0.903 | -1.091 | -0.048 | -2.271 | -0.001 | 0.149 | 0.149 | 0 | 0.23 | 0.01 | 0.1 | 0.01 | 0.15 | 0.12 |

| | | | | | | | | | | | | | | | | |
|---|---|---|---|---|---|---|---|---|---|---|---|---|---|---|---|---|
| 3 | 134218557 | rs9814567 | -0.431 | -0.846 | 0.295 | -2.001 | 0.001 | -0.013 | -0.013 | 0 | 0.69 | 0.65 | 0.79 | 0.83 | 0.69 | 0.7 |
| 3 | 134225969 | rs9827878 | 0.473 | -0.818 | 0.462 | -1.981 | 0.003 | 0.004 | 0.004 | 0 | 0.13 | 0.14 | 0.12 | 0.22 | 0.17 | 0.21 |
| 3 | 134247601 | rs12636917 | 1.918 | -0.625 | -0.095 | -1.914 | -0.003 | -0.026 | -0.026 | 0 | 0.09 | 0.01 | 0.02 | 0.09 | 0.05 | 0.17 |
| 3 | 134261654 | rs7427492 | -0.326 | -0.528 | -0.054 | -1.855 | 0.003 | -0.036 | -0.036 | 0 | 0.68 | 0.58 | 0.79 | 0.83 | 0.69 | 0.7 |
| 3 | 134270886 | rs9821505 | 1.558 | -0.516 | -0.06 | -1.852 | -0.04 | -0.047 | -0.047 | 0 | 0.08 | 0.01 | 0.1 | 0.21 | 0.14 | 0.15 |
| 3 | 134280948 | rs9809619 | 0.584 | -1.059 | 0.436 | -2.536 | -0.001 | -0.035 | -0.035 | 1.869 | 0.66 | 0.58 | 0.78 | 0.82 | 0.69 | 0.6 |
| 3 | 134317337 | rs4955526 | 0.797 | -0.812 | 0.294 | -2.696 | 0 | -0.067 | -0.067 | 0 | 0.63 | 0.54 | 0.76 | 0.88 | 0.67 | 0.66 |
| 3 | 134322742 | rs2293293 | 0.779 | -0.806 | 0.315 | -2.677 | 0 | -0.069 | -0.069 | 2.802 | 0.63 | 0.53 | 0.75 | 0.88 | 0.67 | 0.65 |
| 3 | 134325536 | rs4074521 | -0.819 | -0.805 | 0.294 | -2.677 | 0.001 | -0.043 | -0.043 | 0.229 | 0.36 | 0.41 | 0.23 | 0.11 | 0.33 | 0.34 |
| 3 | 134325626 | rs4074522 | -0.749 | -0.806 | 0.276 | -2.677 | -0.002 | 0.001 | 0.001 | 0.239 | 0.36 | 0.33 | 0.22 | 0.11 | 0.33 | 0.34 |
| 3 | 134331048 | rs7633868 | 0.832 | -0.741 | 0.161 | -2.653 | 0.002 | -0.055 | -0.055 | 0 | 0.62 | 0.55 | 0.74 | 0.88 | 0.65 | 0.65 |
| 3 | 134342758 | rs9809344 | 0.771 | -0.595 | 0.031 | -2.575 | 0.007 | -0.073 | -0.073 | 2.762 | 0.63 | 0.5 | 0.74 | 0.88 | 0.65 | 0.64 |
| 3 | 134348796 | rs13083717 | 0.449 | -0.614 | -0.035 | -2.673 | -0.089 | 0.106 | 0.106 | 0 | 0.2 | 0.02 | 0.16 | 0.02 | 0.26 | 0.1 |
| 3 | 134351724 | rs4543053 | 0.637 | -0.514 | 0.077 | -2.561 | 0.008 | -0.065 | -0.065 | 1.302 | 0.63 | 0.5 | 0.74 | 0.86 | 0.65 | 0.65 |
| 3 | 134358947 | rs6775611 | -0.556 | -0.459 | 0.007 | -2.378 | 0.002 | -0.017 | -0.017 | 0 | 0.33 | 0.36 | 0.23 | 0.13 | 0.32 | 0.3 |
| 3 | 134363340 | rs6771583 | -0.16 | -0.252 | -0.248 | -1.997 | 0.023 | 0.019 | 0.019 | 0 | 0.2 | 0.15 | 0.09 | 0.01 | 0.12 | 0.07 |
| 3 | 134365064 | rs766492 | 1.019 | -0.232 | -0.203 | -2.024 | -0.001 | -0.033 | -0.033 | 0.047 | 0.58 | 0.5 | 0.71 | 0.9 | 0.61 | 0.8 |
| 3 | 134369716 | rs13060869 | -0.068 | 0.134 | -0.11 | -1.611 | -0.067 | 0.095 | 0.095 | 0 | 0.22 | 0.11 | 0.18 | 0.02 | 0.29 | 0.14 |
| 3 | 134370556 | rs6439506 | -0.114 | 0.122 | -0.281 | -1.615 | -0.02 | -0.049 | -0.049 | 0.066 | 0.82 | 0.78 | 0.9 | 0.99 | 0.9 | 0.95 |
| 3 | 134372486 | rs9883955 | 0.379 | 0.633 | -0.595 | -1.208 | 0.001 | -0.003 | -0.003 | 0 | 0.57 | 0.36 | 0.69 | 0.89 | 0.61 | 0.78 |
| 3 | 134393397 | rs2895694 | 0.425 | 0.559 | -0.521 | -1.271 | 0.001 | -0.065 | -0.065 | 0 | 0.67 | 0.55 | 0.71 | 0.93 | 0.66 | 0.82 |
| 3 | 134395599 | rs6781259 | 0.116 | 0.499 | -0.454 | -1.428 | -0.002 | -0.024 | -0.024 | 0.268 | 0.92 | 0.86 | 0.93 | 1 | 0.92 | 0.91 |
| 3 | 134397858 | rs6763799 | -0.462 | 0.338 | 0.23 | -1.531 | 0.01 | -0.102 | -0.102 | 1.167 | 0.27 | 0.43 | 0.23 | 0.07 | 0.27 | 0.1 |
| 3 | 134400448 | rs11923418 | -0.789 | 0.435 | 0.017 | -1.188 | -0.002 | 0.007 | 0.007 | 0 | 0.46 | 0.57 | 0.43 | 0.31 | 0.44 | 0.3 |
| 3 | 134401030 | rs7632935 | 1.661 | 0.085 | -0.56 | -1.486 | -0.002 | -0.002 | -0.002 | 0 | 0.34 | 0.29 | 0.44 | 0.64 | 0.42 | 0.33 |
| 3 | 134402587 | rs10935125 | 1.646 | 0.579 | -0.843 | -1.713 | -0.025 | 0.024 | 0.024 | 0 | 0.06 | 0.01 | 0.13 | 0 | 0.23 | 0.07 |
| 3 | 134402835 | rs10212493 | 0.294 | -0.232 | -0.396 | -1.696 | -0.031 | -0.12 | -0.12 | 0.016 | 0.84 | 0.54 | 0.88 | 1 | 0.88 | 0.89 |
| 3 | 134403049 | rs17694541 | 1.124 | -0.185 | -0.4 | -1.651 | -0.008 | 0.007 | 0.007 | 0.395 | 0.12 | 0.01 | 0.06 | 0.06 | 0.06 | 0.3 |
| 3 | 134405873 | rs1562489 | 0.372 | -0.279 | -0.272 | -1.787 | 0.006 | -0.068 | -0.068 | 0.743 | 0.69 | 0.44 | 0.71 | 0.91 | 0.67 | 0.84 |
| 3 | 134407891 | rs1562487 | 0.66 | -0.289 | -0.289 | -1.777 | 0 | -0.018 | -0.018 | 0 | 0.22 | 0.15 | 0.21 | 0.23 | 0.2 | 0.21 |
| 3 | 134409348 | rs4245900 | -0.076 | -0.285 | -0.278 | -1.778 | 0.004 | -0.024 | -0.024 | 0 | 0.26 | 0.28 | 0.25 | 0.09 | 0.3 | 0.15 |
| 3 | 134419118 | rs1447690 | 1.189 | -0.238 | -0.305 | -1.699 | -0.005 | -0.002 | -0.002 | 0 | 0.42 | 0.29 | 0.5 | 0.65 | 0.5 | 0.41 |
| 3 | 134428788 | rs11915953 | 0.891 | -0.857 | 0.828 | -1.632 | 0.031 | 0 | 0 | 0 | 0.93 | 0.61 | 0.96 | 1 | 0.96 | 0.98 |
| 3 | 134431256 | rs4955476 | 0.603 | -0.927 | 1.012 | -1.678 | -0.049 | -0.12 | -0.12 | 0 | 0.79 | 0.49 | 0.89 | 0.97 | 0.9 | 0.67 |
| 3 | 134437773 | rs7644265 | -0.063 | -0.904 | 0.754 | -1.62 | 0.003 | 0.002 | 0.002 | 0 | 0.91 | 0.93 | 0.93 | 1 | 0.92 | 0.94 |
| 3 | 134437955 | rs9289483 | 1.578 | -0.925 | 0.686 | -1.578 | 0.025 | 0.129 | 0.129 | 0 | 0.48 | 0.12 | 0.59 | 0.75 | 0.62 | 0.41 |
| 3 | 134439849 | rs9851083 | 1.45 | -0.924 | 0.716 | -1.577 | 0.044 | 0.1 | 0.1 | 0 | 0.5 | 0.27 | 0.6 | 0.75 | 0.62 | 0.41 |
| 3 | 134446125 | rs7619098 | -0.778 | -1.109 | 0.731 | -1.424 | 0.008 | 0.017 | 0.017 | 0.317 | 0.37 | 0.34 | 0.31 | 0.24 | 0.28 | 0.54 |
| 3 | 134460268 | rs1447699 | -0.126 | -1.021 | 0.612 | -1.391 | -0.008 | -0.005 | -0.005 | 0 | 0.26 | 0.45 | 0.25 | 0.22 | 0.23 | 0.22 |
| 3 | 134460368 | rs1447698 | -0.049 | -1.018 | 0.561 | -1.389 | -0.004 | -0.002 | -0.002 | 0 | 0.26 | 0.37 | 0.24 | 0.22 | 0.23 | 0.22 |
| 3 | 134467688 | rs11711304 | -0.235 | -0.994 | 0.216 | -1.445 | 0.008 | 0.061 | 0.061 | 0.407 | 0.19 | 0.01 | 0.12 | 0.04 | 0.12 | 0.36 |
| 3 | 134471654 | rs9819372 | 0.842 | -0.511 | -1.585 | -1.158 | 0.059 | 0.178 | 0.178 | 0.036 | 0.52 | 0.08 | 0.6 | 0.8 | 0.67 | 0.42 |
| 3 | 134477303 | rs13097181 | -0.836 | -0.84 | -1.195 | -1.01 | 0.031 | 0.036 | 0.036 | 0 | 0.2 | 0.01 | 0.09 | 0.07 | 0.1 | 0.3 |
| 3 | 134482381 | rs6439517 | 1.515 | -0.93 | 0.577 | -1.391 | -0.005 | -0.018 | -0.018 | 2.723 | 0.64 | 0.43 | 0.69 | 0.85 | 0.75 | 0.52 |
| 3 | 134489905 | rs9825801 | 1.569 | -1.039 | 1.26 | -1.428 | 0.007 | 0.005 | 0.005 | 0 | 0.64 | 0.39 | 0.69 | 0.85 | 0.75 | 0.52 |
| 3 | 134491843 | rs1554112 | 0.712 | -0.88 | 1.355 | -1.353 | 0.02 | -0.017 | -0.017 | 1.025 | 0.89 | 0.81 | 0.82 | 0.91 | 0.86 | 0.89 |
| 3 | 134492589 | rs16842042 | -0.689 | -0.992 | 1.333 | -1.397 | -0.061 | -0.043 | -0.043 | 0 | 0.16 | 0.42 | 0.09 | 0.06 | 0.05 | 0.34 |
| 3 | 134497786 | rs10154906 | 1.607 | -0.991 | 1.273 | -1.396 | -0.015 | -0.035 | -0.035 | 0 | 0.73 | 0.42 | 0.74 | 0.86 | 0.81 | 0.55 |
| 3 | 134508366 | rs5008643 | -0.869 | -0.849 | 1.174 | -1.321 | 0.01 | -0.01 | -0.01 | 0 | 0.1 | 0.12 | 0.17 | 0.1 | 0.14 | 0.09 |
| 3 | 134513356 | rs11918092 | 1.66 | -1.258 | 1.655 | -1.398 | -0.019 | -0.009 | -0.009 | 0.459 | 0.75 | 0.34 | 0.81 | 0.89 | 0.89 | 0.58 |
| 3 | 134518101 | rs11921721 | 0.59 | -1.116 | 1.58 | -1.394 | -0.012 | -0.041 | -0.041 | 0 | 0.91 | 0.88 | 0.93 | 0.99 | 0.94 | 0.97 |
| 3 | 134519445 | rs1004551 | 2.823 | -1.028 | 1.323 | -1.473 | 0.036 | 0.081 | 0.081 | 0.148 | 0.65 | 0.2 | 0.7 | 0.86 | 0.76 | 0.56 |
| 3 | 134523608 | rs7648359 | -1.667 | -0.917 | 1.34 | -1.173 | -0.042 | -0.001 | -0.001 | 1.719 | 0.2 | 0.68 | 0.14 | 0.19 | 0.06 | 0.42 |
| 3 | 134529708 | rs9818551 | 0.999 | -0.897 | 1.268 | -1.211 | 0.013 | -0.03 | -0.03 | 0.035 | 0.91 | 0.86 | 0.88 | 0.96 | 0.87 | 0.96 |
| 3 | 134530843 | rs6439522 | 2.116 | -0.946 | 1.152 | -1.236 | -0.003 | -0.085 | -0.085 | 0 | 0.76 | 0.42 | 0.77 | 0.85 | 0.79 | 0.8 |
| 3 | 134532637 | rs4955455 | 2.545 | -0.947 | 1.192 | -1.22 | 0.007 | 0.01 | 0.01 | 0 | 0.65 | 0.3 | 0.7 | 0.75 | 0.75 | 0.5 |
| 3 | 134532784 | rs4955502 | 2.336 | -0.892 | 1.084 | -1.222 | -0.01 | -0.032 | -0.032 | 2.175 | 0.67 | 0.38 | 0.73 | 0.79 | 0.75 | 0.58 |
| 3 | 134536972 | rs1913685 | 1.582 | -0.894 | 0.519 | -1.131 | -0.015 | -0.008 | -0.008 | 0 | 0.82 | 0.28 | 0.85 | 0.85 | 0.91 | 0.75 |
| 3 | 134538797 | rs4955504 | 1.839 | -0.954 | 0.502 | -1.158 | -0.014 | -0.008 | -0.008 | 0.072 | 0.81 | 0.28 | 0.85 | 0.85 | 0.91 | 0.74 |
| 3 | 134544299 | rs6802089 | 1.536 | -1.078 | -0.141 | -1.238 | 0.071 | 0.105 | 0.105 | 0.104 | 0.59 | 0.18 | 0.7 | 0.78 | 0.7 | 0.31 |
| 3 | 134551528 | rs885594 | -1.239 | -1.147 | 0.874 | -1.293 | -0.025 | -0.03 | -0.03 | 0 | 0.33 | 0.53 | 0.22 | 0.18 | 0.25 | 0.53 |
| 3 | 134551713 | rs1949305 | -1.322 | -1.208 | 1.545 | -1.338 | 0.048 | 0.044 | 0.044 | 0 | 0.08 | 0.34 | 0.11 | 0.08 | 0.1 | 0.1 |
| 3 | 134554163 | rs2138213 | -0.78 | -1.204 | 1.41 | -1.333 | 0.009 | 0.01 | 0.01 | 0 | 0.25 | 0.16 | 0.11 | 0.1 | 0.15 | 0.42 |
| 3 | 134560109 | rs6777246 | -1.273 | -1.21 | 1.421 | -1.336 | 0.019 | 0.028 | 0.028 | 0 | 0.3 | 0.18 | 0.17 | 0.13 | 0.2 | 0.46 |
| 3 | 134563277 | rs6793813 | -1.477 | -1.269 | 1.988 | -1.346 | -0.057 | -0.07 | -0.07 | 0 | 0.26 | 0.44 | 0.17 | 0.12 | 0.18 | 0.37 |
| 3 | 134565434 | rs13073658 | -1.382 | -1.219 | 2.05 | -1.341 | 0.039 | 0.04 | 0.04 | 0 | 0.08 | 0.3 | 0.12 | 0.08 | 0.1 | 0.1 |
| 3 | 134571470 | rs10935135 | 1.197 | -1.202 | 2.337 | -1.34 | 0.051 | 0.062 | 0.062 | 0.792 | 0.92 | 0.61 | 0.94 | 0.94 | 0.94 | 0.86 |
| 3 | 134573223 | rs13078906 | -2.279 | -1.208 | 2.004 | -1.379 | -0.01 | -0.027 | -0.027 | 0 | 0.39 | 0.45 | 0.3 | 0.2 | 0.33 | 0.58 |
| 3 | 134574719 | rs12489360 | 2.276 | -1.205 | 1.801 | -1.377 | 0.044 | 0.105 | 0.105 | 0 | 0.59 | 0.19 | 0.67 | 0.8 | 0.67 | 0.42 |

| | | | | | | | | | | | | | | | | |
|---|---|---|---|---|---|---|---|---|---|---|---|---|---|---|---|---|
| 3 | 134579223 | rs868130 | 2.346 | -1.209 | 1.763 | -1.379 | 0.05 | 0.116 | 0.116 | 0 | 0.58 | 0.18 | 0.67 | 0.8 | 0.67 | 0.43 |
| 3 | 134591138 | rs6762261 | -0.86 | -1.227 | 2.007 | -1.384 | -0.001 | -0.002 | -0.002 | 0 | 0.23 | 0.24 | 0.16 | 0.11 | 0.17 | 0.32 |
| 3 | 134592046 | rs6439532 | -2.81 | -1.256 | 1.936 | -1.401 | 0.056 | 0.124 | 0.124 | 0 | 0.43 | 0.8 | 0.33 | 0.21 | 0.33 | 0.59 |
| 3 | 134592440 | rs1515367 | -1.842 | -1.243 | 2.026 | -1.397 | 0.002 | -0.004 | -0.004 | 0.205 | 0.33 | 0.34 | 0.25 | 0.14 | 0.27 | 0.44 |
| 3 | 134599236 | rs7646638 | -1.762 | -1.235 | 2.127 | -1.333 | 0.028 | 0.039 | 0.039 | 1.786 | 0.1 | 0.47 | 0.07 | 0.07 | 0.06 | 0.16 |
| 3 | 134607243 | rs9868348 | -1.954 | 0.208 | 1.733 | -1.627 | 0.003 | -0.039 | -0.039 | 0 | 0.4 | 0.46 | 0.38 | 0.17 | 0.46 | 0.48 |
| 3 | 134608500 | rs9874454 | -0.324 | 0.228 | 1.61 | -1.627 | -0.06 | 0.067 | 0.067 | 0 | 0.16 | 0.05 | 0.21 | 0.06 | 0.29 | 0.17 |
| 3 | 134611198 | rs1515364 | -2.263 | 0.114 | 1.539 | -1.478 | 0.003 | -0.027 | -0.027 | 0 | 0.45 | 0.52 | 0.4 | 0.26 | 0.5 | 0.59 |
| 3 | 134618819 | rs1996242 | 2.021 | 0.316 | 0.388 | -1.188 | -0.103 | 0.087 | 0.087 | 0 | 0.55 | 0.06 | 0.61 | 0.7 | 0.51 | 0.35 |
| 3 | 134618953 | rs16842235 | -0.917 | -0.098 | 1.117 | -0.904 | -0.023 | -0.009 | -0.009 | 0.008 | 0.15 | 0.33 | 0.09 | 0.12 | 0.1 | 0.34 |
| 3 | 134619724 | rs16842240 | -0.592 | -0.081 | 0.977 | -0.895 | -0.015 | -0.003 | -0.003 | 0.084 | 0.13 | 0.28 | 0.1 | 0.12 | 0.1 | 0.24 |
| 3 | 134622297 | rs1515360 | 0.87 | 0.167 | 0.943 | -0.755 | -0.011 | -0.014 | -0.014 | 0 | 0.89 | 0.72 | 0.93 | 0.93 | 0.94 | 0.74 |
| 3 | 134624640 | rs36221 | 1.224 | 0.664 | 0.812 | -0.502 | 0.014 | 0.017 | 0.017 | 0.56 | 0.8 | 0.38 | 0.84 | 0.77 | 0.76 | 0.7 |
| 3 | 134624806 | rs882340 | -1.248 | -0.07 | 0.527 | -0.928 | -0.001 | -0.006 | -0.006 | 0 | 0.18 | 0.18 | 0.1 | 0.12 | 0.13 | 0.18 |
| 3 | 134624858 | rs6782258 | 0.878 | -0.14 | -0.715 | -1.094 | -0.049 | 0.054 | 0.054 | 0 | 0.24 | 0.06 | 0.14 | 0.11 | 0.26 | 0.17 |
| 3 | 134625255 | rs10512935 | 0.631 | -0.211 | -0.62 | -0.968 | -0.036 | 0.02 | 0.02 | 0 | 0.19 | 0.05 | 0.12 | 0.19 | 0.21 | 0.12 |
| 3 | 134625555 | rs7630597 | 0.442 | -0.032 | 0.027 | -0.612 | 0.022 | 0.023 | 0.023 | 0.033 | 0.16 | 0.2 | 0.29 | 0.21 | 0.19 | 0.2 |
| 3 | 134625614 | rs9870698 | 0.234 | -0.927 | -0.242 | 0.003 | -0.033 | 0.009 | 0.009 | 0.125 | 0.28 | 0.36 | 0.16 | 0.32 | 0.1 | 0.34 |
| 3 | 134626339 | rs36218 | -0.378 | -0.488 | -0.59 | 0.059 | 0.04 | -0.015 | -0.015 | 0.312 | 0.4 | 0.79 | 0.46 | 0.53 | 0.27 | 0.47 |
| 3 | 134629174 | rs11707204 | 0.527 | -0.346 | -0.969 | -0.088 | -0.036 | 0.066 | 0.066 | 4.97 | 0.26 | 0.04 | 0.13 | 0.2 | 0.22 | 0.16 |
| 3 | 134629492 | rs36214 | -0.195 | -0.775 | -0.76 | -0.006 | 0.028 | -0.021 | -0.021 | 0.123 | 0.33 | 0.75 | 0.27 | 0.41 | 0.2 | 0.45 |
| 3 | 134630329 | rs36211 | -0.655 | -0.743 | -0.419 | -0.073 | 0.001 | 0 | 0 | 0.023 | 0.42 | 0.6 | 0.52 | 0.56 | 0.28 | 0.61 |
| 3 | 134632521 | rs9682984 | 0.613 | -0.526 | -0.433 | 0.046 | 0.005 | 0.025 | 0.025 | 0.115 | 0.13 | 0.17 | 0.35 | 0.19 | 0.14 | 0.18 |
| 3 | 134636524 | rs11709198 | -0.085 | -0.371 | -0.867 | -0.308 | 0.007 | 0.032 | 0.032 | 1.454 | 0.33 | 0.22 | 0.2 | 0.22 | 0.25 | 0.25 |
| 3 | 134636796 | rs36198 | -1.588 | -0.151 | -1.177 | -0.194 | -0.163 | -0.056 | -0.056 | 0.037 | 0.72 | 0.97 | 0.68 | 0.76 | 0.51 | 0.78 |
| 3 | 134637946 | rs11719058 | 0.495 | -0.226 | -0.699 | -0.231 | -0.009 | 0.01 | 0.01 | 0.287 | 0.24 | 0.19 | 0.16 | 0.2 | 0.26 | 0.21 |
| 3 | 134638836 | rs36193 | -0.488 | -0.736 | -0.423 | 0.048 | 0.024 | 0.003 | 0.003 | 0.802 | 0.38 | 0.7 | 0.49 | 0.62 | 0.24 | 0.54 |
| 3 | 134642667 | rs11920780 | -0.256 | -1.099 | -0.422 | 0.109 | -0.014 | 0 | 0 | 0.434 | 0.36 | 0.61 | 0.44 | 0.62 | 0.18 | 0.52 |
| 3 | 134643468 | rs4459937 | 0.481 | -0.717 | -1.008 | 0.684 | 0.007 | -0.01 | -0.01 | 2.096 | 0.06 | 0.02 | 0.27 | 0.2 | 0.06 | 0.09 |
| 3 | 134643602 | rs40439 | -1.259 | -0.267 | -1.508 | 0.327 | -0.157 | -0.049 | -0.049 | 0 | 0.71 | 0.97 | 0.67 | 0.75 | 0.51 | 0.78 |
| 3 | 134645401 | rs36183 | -0.362 | -0.486 | -1.405 | 0.215 | -0.055 | -0.055 | -0.055 | 1.679 | 0.59 | 0.83 | 0.45 | 0.55 | 0.47 | 0.64 |
| 3 | 134647169 | rs36178 | -0.721 | -0.199 | -1.338 | 0.303 | -0.103 | -0.116 | -0.116 | 0 | 0.65 | 0.91 | 0.41 | 0.52 | 0.47 | 0.69 |
| 3 | 134647849 | rs4955518 | 0.471 | -0.255 | -1.275 | 0.239 | 0.008 | -0.032 | -0.032 | 0.101 | 0.07 | 0.02 | 0.28 | 0.19 | 0.06 | 0.1 |
| 3 | 134650932 | rs40437 | -0.745 | -0.246 | -1.062 | 0.179 | -0.06 | -0.072 | -0.072 | 0 | 0.64 | 0.86 | 0.42 | 0.58 | 0.48 | 0.67 |
| 3 | 134652757 | rs36167 | -0.74 | -0.316 | -0.952 | -1.203 | -0.047 | 0.104 | 0.104 | 0.401 | 0.64 | 0.82 | 0.41 | 0.2 | 0.48 | 0.58 |
| 3 | 134657852 | rs984709 | 0.789 | -0.944 | -0.227 | -1.648 | -0.056 | -0.104 | -0.104 | 0 | 0.21 | 0.3 | 0.11 | 0.04 | 0.09 | 0.26 |
| 3 | 134659551 | rs9874260 | 0.193 | -0.942 | -0.303 | -1.836 | -0.006 | -0.045 | -0.045 | 0 | 0.44 | 0.39 | 0.46 | 0.17 | 0.32 | 0.34 |
| 3 | 134661430 | rs983401 | 0.798 | -0.82 | -0.097 | -1.84 | 0 | -0.047 | -0.047 | 2.011 | 0.12 | 0.14 | 0.05 | 0 | 0.1 | 0.11 |
| 3 | 134666452 | rs16842308 | 0.879 | -0.831 | -0.099 | -1.843 | 0 | -0.032 | -0.032 | 0 | 0.11 | 0.1 | 0.05 | 0 | 0.1 | 0.11 |
| 3 | 134666475 | rs975149 | -1.302 | -0.942 | -0.174 | -1.928 | -0.071 | 0.112 | 0.112 | 0.345 | 0.64 | 0.85 | 0.6 | 0.22 | 0.48 | 0.58 |
| 3 | 134667951 | rs11719020 | -1.016 | -0.988 | -0.17 | -1.923 | -0.02 | 0.058 | 0.058 | 0.615 | 0.6 | 0.65 | 0.57 | 0.21 | 0.44 | 0.53 |
| 3 | 134669241 | rs4955460 | 0.249 | -1.004 | -0.145 | -1.934 | -0.007 | -0.04 | -0.04 | 0 | 0.44 | 0.4 | 0.47 | 0.18 | 0.32 | 0.35 |
| 3 | 134670524 | rs7644369 | -0.976 | -0.967 | -0.408 | -1.905 | -0.018 | 0.054 | 0.054 | 0.518 | 0.6 | 0.64 | 0.57 | 0.21 | 0.44 | 0.53 |
| 3 | 134676076 | rs36132 | -0.043 | -0.627 | -0.335 | -1.486 | 0.012 | 0.014 | 0.014 | 0 | 0.13 | 0.3 | 0.15 | 0.15 | 0.14 | 0.15 |
| 3 | 134682799 | rs3940695 | -0.36 | -0.674 | -0.32 | -1.845 | -0.014 | -0.059 | -0.059 | 0.014 | 0.45 | 0.43 | 0.46 | 0.18 | 0.32 | 0.34 |
| 3 | 134684918 | rs6782004 | -0.35 | -0.677 | -0.328 | -1.846 | -0.013 | -0.056 | -0.056 | 0 | 0.45 | 0.44 | 0.46 | 0.18 | 0.32 | 0.34 |
| 3 | 134687294 | rs6789372 | 0.613 | -0.278 | -0.7 | -1.646 | -0.012 | 0.042 | 0.042 | 2.825 | 0.26 | 0.14 | 0.38 | 0.13 | 0.27 | 0.15 |
| 3 | 134693052 | rs7636504 | -0.282 | -0.184 | -0.396 | -1.672 | 0.012 | 0.049 | 0.049 | 3.172 | 0.5 | 0.59 | 0.51 | 0.21 | 0.39 | 0.42 |
| 3 | 134694660 | rs16842415 | -1.049 | 0.011 | 0.156 | -2.499 | 0.004 | -0.096 | -0.096 | 0 | 0.26 | 0.47 | 0.2 | 0.13 | 0.23 | 0.27 |
| 3 | 134694729 | rs36124 | -1.516 | -0.123 | 0.355 | -2.535 | -0.001 | -0.027 | -0.027 | 0 | 0.28 | 0.65 | 0.23 | 0.13 | 0.23 | 0.28 |
| 3 | 134695446 | rs574719 | -0.648 | 0.295 | -0.17 | -3.079 | -0.056 | 0.119 | 0.119 | 1.86 | 0.68 | 0.87 | 0.53 | 0.17 | 0.6 | 0.48 |
| 3 | 134698869 | rs6776069 | -0.777 | 0.294 | 0.072 | -3.128 | 0.008 | -0.105 | -0.105 | 0 | 0.27 | 0.41 | 0.2 | 0.12 | 0.25 | 0.28 |
| 3 | 134699750 | rs36112 | -0.654 | 0.389 | -0.068 | -3.047 | -0.051 | 0.116 | 0.116 | 2.086 | 0.68 | 0.87 | 0.53 | 0.17 | 0.6 | 0.48 |
| 3 | 134707191 | rs9866987 | 0.173 | 0.335 | 0.106 | -3.026 | 0.003 | -0.008 | -0.008 | 0.046 | 0.11 | 0.15 | 0.1 | 0.12 | 0.09 | 0.17 |
| 3 | 134709951 | rs7613213 | 1.386 | 0.306 | -0.184 | -3.088 | 0.015 | 0.074 | 0.074 | 4.914 | 0.4 | 0.25 | 0.3 | 0.05 | 0.37 | 0.2 |
| 3 | 134716279 | rs16842460 | 0.048 | 0.089 | 0.286 | -3.036 | -0.001 | -0.03 | -0.03 | 0 | 0.14 | 0.15 | 0.09 | 0.01 | 0.14 | 0.11 |
| 3 | 134719138 | rs40524 | -0.49 | 0.143 | 0.356 | -3.004 | 0.016 | -0.021 | -0.021 | 3.964 | 0.13 | 0.52 | 0.13 | 0.12 | 0.09 | 0.16 |
| 3 | 134719415 | rs36074 | -0.122 | 0.156 | 0.351 | -2.998 | 0.02 | -0.012 | -0.012 | 0 | 0.12 | 0.34 | 0.11 | 0.12 | 0.09 | 0.16 |
| 3 | 134727833 | rs36064 | -1.006 | 0.144 | -0.346 | -2.954 | -0.11 | 0.013 | 0.013 | 0.794 | 0.75 | 0.96 | 0.58 | 0.27 | 0.65 | 0.64 |
| 3 | 134732732 | rs36059 | -1.023 | -0.628 | 0.124 | -2.843 | -0.002 | -0.027 | -0.027 | 0 | 0.27 | 0.67 | 0.23 | 0.12 | 0.22 | 0.27 |
| 3 | 134733516 | rs40347 | -0.269 | -0.377 | -0.073 | -2.553 | 0.003 | -0.002 | -0.002 | 0 | 0.92 | 0.98 | 0.94 | 0.9 | 0.93 | 0.81 |
| 3 | 134736671 | rs36057 | 0.843 | -0.057 | 0.047 | -2.759 | 0 | -0.025 | -0.025 | 0 | 0.44 | 0.4 | 0.36 | 0.05 | 0.43 | 0.19 |
| 3 | 134740452 | rs36054 | 0.366 | 0.509 | 0.242 | -2.146 | 0.005 | -0.013 | -0.013 | 0 | 0.25 | 0.41 | 0.35 | 0.68 | 0.3 | 0.34 |
| 3 | 134753629 | rs36037 | 0.13 | 0.657 | 0.318 | -1.897 | 0.068 | -0.019 | -0.019 | 0 | 0.93 | 0.73 | 0.95 | 1 | 0.95 | 0.92 |
| 3 | 134755591 | rs36039 | 0.436 | 0.414 | -0.812 | -1.964 | -0.04 | 0.183 | 0.183 | 0 | 0.4 | 0.12 | 0.44 | 0.21 | 0.4 | 0.32 |
| 3 | 134763460 | rs6791224 | -1.852 | 0.112 | 0.671 | -1.301 | -0.02 | -0.011 | -0.011 | 0.548 | 0.14 | 0.46 | 0.17 | 0.16 | 0.11 | 0.16 |
| 3 | 134763517 | rs6793611 | -1.379 | 0.214 | 0.733 | -1.267 | 0.004 | -0.05 | -0.05 | 0 | 0.09 | 0.22 | 0.08 | 0 | 0.09 | 0.06 |
| 3 | 134770520 | rs12639432 | -0.224 | 0.646 | -0.015 | -1.139 | -0.024 | -0.001 | -0.001 | 0 | 0.26 | 0.09 | 0.4 | 0.7 | 0.3 | 0.39 |
| 3 | 134778492 | rs6778165 | 1.292 | 0.771 | 0.128 | -1.311 | 0.001 | -0.051 | -0.051 | 0.022 | 0.56 | 0.44 | 0.4 | 0.15 | 0.51 | 0.47 |

| | | | | | | | | | | | | | | | | |
|---|---|---|---|---|---|---|---|---|---|---|---|---|---|---|---|---|
| 3 | 134779021 | rs9826733 | 0.325 | 0.158 | 0.394 | -0.977 | -0.008 | -0.048 | -0.048 | 0 | 0.91 | 0.8 | 0.94 | 1 | 0.94 | 0.97 |
| 3 | 134792844 | rs6439552 | 0.908 | 0.171 | 0.257 | -0.953 | -0.023 | -0.002 | -0.002 | 0 | 0.87 | 0.55 | 0.89 | 0.85 | 0.92 | 0.87 |
| 3 | 134805581 | rs6439553 | 0.977 | 0.095 | -0.131 | -0.806 | 0.005 | 0.004 | 0.004 | 0.532 | 0.62 | 0.34 | 0.53 | 0.18 | 0.68 | 0.51 |
| 3 | 134805772 | rs959562 | -1.146 | 0.5 | -0.353 | -0.249 | -0.021 | -0.003 | -0.003 | 0 | 0.11 | 0.24 | 0.07 | 0.13 | 0.06 | 0.14 |
| 3 | 134816062 | rs7372166 | 0.608 | 1.327 | -1.161 | -0.697 | -0.062 | 0.097 | 0.097 | 0 | 0.73 | 0.87 | 0.8 | 0.95 | 0.66 | 0.86 |
| 3 | 134816417 | rs17765097 | 0.807 | 0.844 | -1.561 | -1.115 | 0.014 | 0.086 | 0.086 | 0.129 | 0.4 | 0.23 | 0.35 | 0.2 | 0.39 | 0.43 |
| 3 | 134818733 | rs9821105 | 0.829 | 0.942 | -1.664 | -1.154 | 0.009 | 0.08 | 0.08 | 0.124 | 0.39 | 0.23 | 0.36 | 0.2 | 0.38 | 0.42 |
| 3 | 134820091 | rs7373984 | -0.795 | 0.996 | -0.576 | -1.221 | -0.003 | -0.039 | -0.039 | 0.035 | 0.56 | 0.51 | 0.58 | 0.79 | 0.56 | 0.57 |
| 3 | 134821129 | rs11711351 | 0.058 | 0.859 | -0.048 | -1.437 | -0.009 | -0.017 | -0.017 | 0 | 0.28 | 0.19 | 0.18 | 0.01 | 0.33 | 0.15 |
| 3 | 134825988 | rs7374961 | 0.336 | 0.546 | -0.127 | -1.537 | -0.023 | -0.009 | -0.009 | 3.055 | 0.25 | 0.03 | 0.37 | 0.77 | 0.22 | 0.41 |
| 3 | 134826917 | rs1554675 | -0.572 | 0.495 | 0.018 | -1.561 | 0.048 | 0.009 | 0.009 | 0 | 0.94 | 0.75 | 0.95 | 0.98 | 0.92 | 0.99 |
| 3 | 134829538 | rs7374193 | 0.876 | 0.343 | -0.287 | -0.168 | -0.024 | -0.088 | -0.088 | 0.143 | 0.14 | 0.05 | 0.3 | 0.4 | 0.18 | 0.24 |
| 3 | 134829562 | rs9813334 | 0.868 | 0.331 | -0.243 | -0.14 | 0.007 | -0.008 | -0.008 | 0.019 | 0.1 | 0.02 | 0.05 | 0.12 | 0.05 | 0.16 |
| 3 | 134830532 | rs10935154 | -0.644 | 0.362 | 0.065 | -0.143 | -0.018 | 0.027 | 0.027 | 0.005 | 0.42 | 0.29 | 0.51 | 0.67 | 0.51 | 0.4 |
| 3 | 134837339 | rs7647433 | 0.174 | 0.584 | 0.596 | -0.271 | 0.097 | 0.021 | 0.021 | 0.068 | 0.92 | 0.54 | 0.91 | 0.98 | 0.93 | 0.99 |
| 3 | 134837615 | rs16842632 | -0.222 | 0.579 | 0.56 | 0 | 0.023 | 0.078 | 0.078 | 0 | 0.17 | 0.29 | 0.4 | 0.61 | 0.24 | 0.26 |
| 3 | 134841452 | rs11706256 | -0.947 | 0.837 | 0.411 | -0.268 | -0.068 | 0.141 | 0.141 | 0.053 | 0.44 | 0.15 | 0.53 | 0.77 | 0.49 | 0.58 |
| 3 | 134842423 | rs940639 | 0.94 | 0.803 | 0.357 | -0.232 | -0.016 | -0.045 | -0.045 | 0 | 0.12 | 0.08 | 0.33 | 0.6 | 0.17 | 0.25 |
| 3 | 134843286 | rs2140766 | 1.546 | 0.741 | 0.607 | -0.323 | 0.035 | 0.143 | 0.143 | 0.14 | 0.43 | 0.14 | 0.35 | 0.2 | 0.38 | 0.38 |
| 3 | 134845428 | rs13096641 | 1.503 | -0.146 | -0.412 | -1.056 | -0.005 | 0.015 | 0.015 | 0 | 0.22 | 0.07 | 0.22 | 0.19 | 0.22 | 0.27 |
| 3 | 134845693 | rs7625747 | 1.977 | -0.22 | -0.342 | -1.048 | 0 | -0.032 | -0.032 | 0 | 0.2 | 0.2 | 0.12 | 0.01 | 0.14 | 0.1 |
| 3 | 134846285 | rs7626314 | 0.953 | -0.137 | -0.346 | -0.848 | 0.008 | -0.014 | -0.014 | 0.662 | 0.32 | 0.26 | 0.46 | 0.59 | 0.33 | 0.34 |
| 3 | 134850771 | rs940638 | 0.991 | -0.206 | -0.516 | -1.027 | -0.024 | 0.018 | 0.018 | 0 | 0.26 | 0.08 | 0.25 | 0.23 | 0.26 | 0.28 |
| 3 | 134852689 | rs6772933 | 0.229 | -0.482 | 0.591 | -1.193 | 0.022 | -0.025 | -0.025 | 0 | 0.91 | 0.73 | 0.93 | 0.98 | 0.92 | 0.97 |
| 3 | 134853762 | rs11713704 | 1.228 | -0.001 | 0.212 | -1.362 | -0.077 | 0.15 | 0.15 | 0 | 0.22 | 0.04 | 0.15 | 0 | 0.24 | 0.13 |
| 3 | 134854943 | rs12488683 | -0.007 | -0.005 | 0.467 | -1.329 | -0.005 | 0.045 | 0.045 | 0 | 0.34 | 0.26 | 0.24 | 0.17 | 0.31 | 0.38 |
| 3 | 134857076 | rs6762543 | 0.495 | -0.032 | 0.534 | -1.393 | -0.024 | 0.064 | 0.064 | 0 | 0.38 | 0.24 | 0.58 | 0.81 | 0.45 | 0.49 |
| 3 | 134857783 | rs9840863 | 1.503 | -0.045 | 0.518 | -1.311 | -0.007 | -0.039 | -0.039 | 0 | 0.07 | 0.02 | 0.05 | 0.16 | 0.04 | 0.19 |
| 3 | 134859834 | rs868909 | -1.022 | -0.064 | 0.565 | -1.339 | -0.026 | -0.124 | -0.124 | 0.019 | 0.71 | 0.48 | 0.81 | 0.97 | 0.76 | 0.86 |
| 3 | 134864529 | rs3772647 | -1.169 | 0.304 | 0.166 | -1.345 | -0.055 | -0.08 | -0.08 | 0 | 0.83 | 0.94 | 0.63 | 0.42 | 0.72 | 0.73 |
| 3 | 134867952 | rs3772646 | 0.61 | 0.014 | 0.115 | -1.334 | 0.006 | 0.011 | 0.011 | 0 | 0.11 | 0.18 | 0.11 | 0.18 | 0.1 | 0.2 |
| 3 | 134870058 | rs6764768 | 0.954 | -0.289 | 0.128 | -1.389 | -0.004 | -0.015 | -0.015 | 0 | 0.32 | 0.35 | 0.23 | 0.19 | 0.26 | 0.35 |
| 3 | 134874705 | rs1112766 | 0.968 | 0.398 | -0.177 | -1.275 | -0.061 | 0.152 | 0.152 | 0 | 0.22 | 0.04 | 0.13 | 0 | 0.25 | 0.15 |
| 3 | 134877635 | rs7642038 | 0.701 | 0.538 | -0.106 | -1.14 | -0.068 | -0.084 | -0.084 | 0 | 0.18 | 0.05 | 0.39 | 0.58 | 0.3 | 0.28 |
| 3 | 134879621 | rs3772643 | 0.339 | 0.51 | 0.142 | -1.14 | -0.004 | 0.016 | 0.016 | 0 | 0.11 | 0.19 | 0.08 | 0.17 | 0.06 | 0.17 |
| 3 | 134881518 | rs1392918 | 0.184 | 0.651 | 0.397 | -1.046 | 0.005 | 0.014 | 0.014 | 0.859 | 0.21 | 0.27 | 0.4 | 0.57 | 0.3 | 0.33 |
| 3 | 134882233 | rs905047 | -0.638 | 0.357 | 0.581 | -1.245 | 0 | -0.112 | -0.112 | 1.561 | 0.77 | 0.67 | 0.85 | 1 | 0.76 | 0.86 |
| 3 | 134886890 | rs4894248 | -0.799 | 0.401 | 0.413 | -1.232 | -0.001 | -0.047 | -0.047 | 0 | 0.78 | 0.8 | 0.85 | 1 | 0.77 | 0.86 |
| 3 | 134891032 | rs1501219 | 0.726 | 0.138 | 0.591 | -1.067 | 0.001 | -0.017 | -0.017 | 0.324 | 0.92 | 0.65 | 0.96 | 1 | 0.97 | 0.95 |
| 3 | 134892486 | rs6801620 | -0.227 | 0.203 | -0.21 | -1.399 | -0.014 | 0.061 | 0.061 | 0 | 0.44 | 0.32 | 0.64 | 0.86 | 0.53 | 0.56 |
| 3 | 134892683 | rs4894252 | -0.105 | 0.388 | -0.328 | -1.013 | 0.011 | -0.032 | -0.032 | 0 | 0.39 | 0.49 | 0.6 | 0.78 | 0.46 | 0.48 |
| 3 | 134898344 | rs10512944 | -0.207 | 0.097 | -0.349 | -0.561 | 0.025 | 0.165 | 0.165 | 0.001 | 0.08 | 0.25 | 0.15 | 0.25 | 0.05 | 0.21 |
| 3 | 134900383 | rs16842755 | 1.124 | 0.131 | -0.712 | -0.887 | -0.015 | -0.029 | -0.029 | 0 | 0.2 | 0.02 | 0.2 | 0.24 | 0.2 | 0.19 |
| 3 | 134904385 | rs875368 | -0.449 | 0.503 | 0.161 | -0.306 | 0.023 | -0.02 | -0.02 | 0 | 0.33 | 0.5 | 0.49 | 0.71 | 0.39 | 0.55 |
| 3 | 134911917 | rs6776570 | -0.603 | 0.55 | -0.347 | -0.127 | -0.004 | 0.001 | 0.001 | 0.018 | 0.36 | 0.64 | 0.51 | 0.65 | 0.39 | 0.56 |
| 3 | 134912567 | rs3821502 | -0.243 | 0.505 | -0.025 | -0.155 | -0.027 | -0.007 | -0.007 | 0 | 0.33 | 0.26 | 0.48 | 0.65 | 0.38 | 0.56 |
| 3 | 134916043 | rs6766459 | -1.383 | 0.396 | -0.742 | -0.504 | -0.008 | 0.005 | 0.005 | 0.053 | 0.62 | 0.87 | 0.68 | 0.65 | 0.63 | 0.73 |
| 3 | 134918522 | rs7622312 | 0.074 | 0.612 | -0.288 | -0.543 | 0.003 | -0.147 | -0.147 | 4.759 | 0.21 | 0.37 | 0.16 | 0 | 0.22 | 0.11 |
| 3 | 134924970 | rs2036367 | -1.201 | 0.894 | -0.571 | -0.145 | -0.004 | -0.014 | -0.014 | 0.099 | 0.53 | 0.58 | 0.67 | 0.7 | 0.58 | 0.68 |
| 3 | 134928717 | rs6439563 | 0.641 | 1.131 | -0.489 | 0.097 | 0.001 | -0.001 | -0.001 | 0 | 0.53 | 0.44 | 0.45 | 0.51 | 0.49 | 0.47 |
| 3 | 134933616 | rs4309752 | -0.475 | 1.562 | -1.328 | 0.117 | -0.018 | 0.01 | 0.01 | 0 | 0.28 | 0.25 | 0.29 | 0.23 | 0.33 | 0.25 |
| 3 | 134936583 | rs9858763 | 0.885 | 1.37 | -1.329 | 0.054 | 0.008 | -0.01 | -0.01 | 0 | 0.53 | 0.29 | 0.45 | 0.52 | 0.46 | 0.46 |
| 3 | 134937333 | rs6777929 | -0.502 | 1.44 | -1.316 | 0.033 | -0.002 | 0.082 | 0.082 | 0 | 0.77 | 0.43 | 0.61 | 0.52 | 0.76 | 0.68 |
| 3 | 134944521 | rs16842829 | 0.398 | 1.04 | -1.562 | 0.244 | -0.001 | -0.008 | -0.008 | 0 | 0.09 | 0.12 | 0.04 | 0 | 0.08 | 0.11 |
| 3 | 134945866 | rs7613488 | 0.261 | 1.057 | -1.329 | 0.572 | -0.035 | 0.063 | 0.063 | 0 | 0.59 | 0.28 | 0.36 | 0.29 | 0.5 | 0.53 |
| 3 | 134955715 | rs9865620 | -0.301 | 0.204 | -0.892 | 0.471 | -0.008 | 0.003 | 0.003 | 0 | 0.62 | 0.48 | 0.66 | 0.53 | 0.71 | 0.76 |
| 3 | 134955912 | rs6806284 | -0.306 | 1.051 | -0.863 | -0.482 | 0.002 | -0.057 | -0.057 | 0.02 | 0.26 | 0.23 | 0.2 | 0 | 0.32 | 0.18 |
| 3 | 134957661 | rs11719703 | -0.504 | 1.069 | -1.238 | -0.204 | -0.018 | 0.024 | 0.024 | 0 | 0.42 | 0.22 | 0.31 | 0.34 | 0.42 | 0.6 |
| 3 | 134960470 | rs6793828 | -0.477 | 1.1 | -0.786 | -0.271 | -0.004 | -0.001 | -0.001 | 1.035 | 0.34 | 0.34 | 0.3 | 0.34 | 0.37 | 0.49 |
| 3 | 134961213 | rs1501202 | 0.704 | 0.416 | -1.043 | -0.239 | 0.019 | 0.011 | 0.011 | 0 | 0.27 | 0.17 | 0.21 | 0.25 | 0.19 | 0.25 |
| 3 | 134962423 | rs17712024 | 0.883 | 0.485 | -1.13 | -0.359 | -0.004 | 0.006 | 0.006 | 0 | 0.13 | 0.11 | 0.07 | 0 | 0.11 | 0.13 |
| 3 | 134967368 | rs11717042 | 0.517 | 0.579 | -1.11 | -0.297 | 0.002 | 0.016 | 0.016 | 0.025 | 0.2 | 0.26 | 0.22 | 0.32 | 0.23 | 0.36 |
| 3 | 134978236 | rs3182239 | 0.962 | 0.464 | -0.439 | -0.361 | -0.034 | 0.081 | 0.081 | 0 | 0.13 | 0.02 | 0.06 | 0 | 0.11 | 0.14 |
| 3 | 134979169 | rs10608 | 0.719 | 0.45 | -0.305 | -0.309 | 0 | -0.028 | -0.028 | 0 | 0.15 | 0.19 | 0.07 | 0 | 0.11 | 0.14 |
| 3 | 134987090 | rs10935158 | 0.426 | 0.404 | -0.212 | -0.286 | 0.005 | -0.005 | -0.005 | 0 | 0.65 | 0.44 | 0.7 | 0.69 | 0.65 | 0.51 |
| 3 | 134996284 | rs2120927 | 0.281 | 0.604 | -0.238 | -0.419 | 0.011 | -0.087 | -0.087 | 0.43 | 0.75 | 0.59 | 0.79 | 0.98 | 0.75 | 0.85 |
| 3 | 134996955 | rs1021752 | 1.333 | 0.669 | -0.36 | -0.277 | 0.002 | 0.042 | 0.042 | 0 | 0.13 | 0.18 | 0.15 | 0.29 | 0.15 | 0.34 |
| 3 | 134999888 | rs1596707 | 1.26 | 0.435 | -0.512 | -0.357 | -0.097 | -0.099 | -0.099 | 0.538 | 0.36 | 0.07 | 0.43 | 0.45 | 0.41 | 0.27 |
| 3 | 135018895 | rs10935162 | -0.387 | 0.518 | -0.439 | -0.422 | -0.006 | 0.003 | 0.003 | 0.653 | 0.25 | 0.23 | 0.19 | 0.02 | 0.25 | 0.14 |

| | | | | | | | | | | | | | | | | |
|---|---|---|---|---|---|---|---|---|---|---|---|---|---|---|---|---|
| 3 | 135023570 | rs1553283 | 0.049 | 0.498 | -0.643 | -0.355 | -0.008 | -0.02 | -0.02 | 0 | 0.27 | 0.16 | 0.2 | 0.3 | 0.26 | 0.47 |
| 3 | 135025212 | rs4894277 | 1.209 | 0.316 | -0.548 | 0.109 | -0.032 | 0.026 | 0.026 | 0 | 0.36 | 0.2 | 0.43 | 0.27 | 0.41 | 0.24 |
| 3 | 135028331 | rs12494371 | 1.151 | 0.262 | -0.364 | 0.129 | 0.025 | -0.024 | -0.024 | 1.462 | 0.26 | 0.17 | 0.21 | 0.41 | 0.19 | 0.27 |
| 3 | 135032852 | rs1445216 | -0.579 | 0.229 | -0.216 | 0.042 | -0.02 | -0.004 | -0.004 | 0 | 0.34 | 0.24 | 0.32 | 0.31 | 0.39 | 0.49 |
| 3 | 135043714 | rs864800 | -0.991 | 0.22 | -0.456 | -0.258 | 0.006 | 0.015 | 0.015 | 0.699 | 0.67 | 0.83 | 0.71 | 0.67 | 0.66 | 0.65 |
| 3 | 135044357 | rs844024 | 1.298 | 0.563 | -0.349 | -0.096 | -0.035 | 0.016 | 0.016 | 0.037 | 0.08 | 0.02 | 0.12 | 0.02 | 0.13 | 0.02 |
| 3 | 135045376 | rs10935163 | 0.913 | 0.552 | -0.273 | 0.169 | 0 | 0.052 | 0.052 | 0 | 0.14 | 0.18 | 0.16 | 0.49 | 0.16 | 0.31 |
| 3 | 135045517 | rs11928139 | 0.647 | 0.366 | -0.191 | 0.003 | 0.001 | 0.114 | 0.114 | 0 | 0.46 | 0.24 | 0.43 | 0.17 | 0.42 | 0.33 |
| 3 | 135046428 | rs7612961 | -1.077 | 0.623 | -0.548 | 0.122 | 0 | -0.004 | -0.004 | 0 | 0.73 | 0.78 | 0.72 | 0.66 | 0.71 | 0.67 |
| 3 | 135046768 | rs9826781 | -0.193 | 0.656 | -0.491 | 0.531 | -0.004 | -0.003 | -0.003 | 0 | 0.22 | 0.23 | 0.27 | 0.48 | 0.29 | 0.32 |
| 3 | 135049624 | rs1445221 | 0.575 | 0.734 | -0.512 | 0.161 | -0.01 | -0.013 | -0.013 | 0 | 0.33 | 0.34 | 0.34 | 0.25 | 0.25 | 0.2 |
| 3 | 135052959 | rs823956 | 0.405 | 1.247 | -0.515 | -0.17 | 0.004 | -0.005 | -0.005 | 0.217 | 0.11 | 0.13 | 0.16 | 0.02 | 0.2 | 0.03 |
| 3 | 135067701 | rs823966 | 0.489 | 1.185 | -0.515 | -0.198 | 0.005 | -0.008 | -0.008 | 0.645 | 0.11 | 0.12 | 0.16 | 0.02 | 0.2 | 0.03 |
| 3 | 135075532 | rs16842953 | 1.257 | 1.113 | -0.692 | 0.189 | -0.018 | -0.043 | -0.043 | 0 | 0.08 | 0 | 0.11 | 0.25 | 0.09 | 0.16 |
| 3 | 135077155 | rs711933 | -0.632 | 0.726 | -0.376 | -0.904 | -0.033 | 0.115 | 0.115 | 3.098 | 0.55 | 0.81 | 0.56 | 0.2 | 0.49 | 0.24 |
| 3 | 135083504 | rs823970 | -0.7 | 0.781 | -0.162 | -0.911 | 0.001 | -0.019 | -0.019 | 0 | 0.55 | 0.52 | 0.54 | 0.2 | 0.53 | 0.25 |
| 3 | 135086592 | rs711936 | -0.264 | 0.736 | -0.019 | -0.876 | 0.003 | 0.092 | 0.092 | 0.021 | 0.47 | 0.64 | 0.51 | 0.18 | 0.45 | 0.22 |
| 3 | 135098822 | rs824070 | 0.099 | 0.733 | 0.071 | -0.998 | 0.003 | 0.099 | 0.099 | 0.423 | 0.53 | 0.37 | 0.4 | 0.84 | 0.54 | 0.78 |
| 3 | 135103031 | rs998591 | 0.517 | 0.755 | 0.239 | -1.109 | -0.017 | -0.051 | -0.051 | 0.931 | 0.35 | 0.37 | 0.35 | 0.14 | 0.26 | 0.19 |
| 3 | 135110411 | rs711943 | 0.34 | 0.724 | 0.091 | -1.088 | 0.003 | 0.165 | 0.165 | 0.328 | 0.52 | 0.22 | 0.39 | 0.84 | 0.54 | 0.78 |
| 3 | 135112615 | rs711948 | -0.591 | 1.025 | 0.35 | -0.752 | -0.002 | 0.002 | 0.002 | 0 | 0.9 | 0.86 | 0.77 | 0.98 | 0.8 | 0.98 |
| 3 | 135123375 | rs698266 | 0.207 | 0.727 | 0.631 | -1.041 | 0 | -0.016 | -0.016 | 0.253 | 0.62 | 0.43 | 0.45 | 0.84 | 0.63 | 0.78 |
| 3 | 135130301 | rs1579819 | 0.757 | 0.664 | 0.876 | -1.049 | -0.01 | -0.016 | -0.016 | 1.474 | 0.26 | 0.23 | 0.29 | 0.13 | 0.16 | 0.19 |
| 3 | 135136722 | rs711959 | -0.818 | 0.711 | 0.756 | -1.017 | 0.045 | -0.026 | -0.026 | 0.712 | 0.9 | 0.76 | 0.76 | 0.98 | 0.8 | 0.98 |
| 3 | 135140925 | rs1526944 | 0.666 | 0.578 | 0.579 | -1.015 | -0.04 | -0.056 | -0.056 | 0.318 | 0.27 | 0.32 | 0.29 | 0.13 | 0.16 | 0.19 |
| 3 | 135145304 | rs1094787 | 0.294 | 0.672 | -0.066 | -1.073 | 0.009 | 0.066 | 0.066 | 0 | 0.61 | 0.27 | 0.45 | 0.85 | 0.63 | 0.79 |
| 3 | 135153284 | rs824087 | 0.287 | 0.706 | -0.074 | -1.078 | 0.014 | 0.067 | 0.067 | 0 | 0.61 | 0.27 | 0.45 | 0.85 | 0.64 | 0.79 |
| 3 | 135167677 | rs12494120 | 0.213 | 1.32 | 0.081 | -1.275 | 0.099 | -0.073 | -0.073 | 0 | 0.87 | 0.6 | 0.74 | 0.99 | 0.79 | 0.98 |
| 3 | 135176294 | rs824097 | 0.281 | 1.587 | -0.745 | -1.662 | -0.097 | 0.118 | 0.118 | 0 | 0.75 | 0.92 | 0.7 | 1 | 0.61 | 0.91 |
| 3 | 135176959 | rs16843013 | 0.995 | 0.978 | -0.642 | -1.935 | -0.012 | -0.05 | -0.05 | 0 | 0.2 | 0.25 | 0.13 | 0.07 | 0.15 | 0.14 |
| 3 | 135186269 | rs711962 | 0.561 | 1.06 | -0.772 | -1.827 | -0.049 | 0.077 | 0.077 | 0.431 | 0.14 | 0.02 | 0.22 | 0 | 0.2 | 0.04 |
| 3 | 135186481 | rs7628597 | 0.448 | 1.193 | -0.622 | -1.897 | 0.007 | -0.069 | -0.069 | 1.138 | 0.3 | 0.34 | 0.21 | 0.07 | 0.33 | 0.19 |
| 3 | 135189819 | rs711963 | -0.723 | 1.185 | -0.35 | -1.927 | 0.006 | 0.14 | 0.14 | 0.536 | 0.47 | 0.73 | 0.47 | 0.07 | 0.55 | 0.23 |
| 3 | 135193301 | rs11706224 | 0.381 | 1.042 | -0.259 | -1.87 | 0.008 | -0.067 | -0.067 | 0 | 0.31 | 0.58 | 0.23 | 0.07 | 0.34 | 0.2 |
| 3 | 135199973 | rs7653465 | 1.666 | 0.93 | -0.776 | -1.377 | -0.046 | -0.118 | -0.118 | 0 | 0.17 | 0.02 | 0.29 | 0.5 | 0.2 | 0.41 |
| 3 | 135213840 | rs11915808 | 0.246 | 0.948 | -0.495 | -1.487 | 0.016 | -0.093 | -0.093 | 3.694 | 0.17 | 0.33 | 0.25 | 0 | 0.21 | 0.04 |
| 3 | 135221504 | rs6789221 | 0.047 | 0.717 | -0.698 | -1.466 | 0.002 | -0.01 | -0.01 | 0 | 0.46 | 0.38 | 0.43 | 0.07 | 0.55 | 0.23 |
| 3 | 135223002 | rs2062735 | 0.18 | 0.894 | -1.57 | -0.872 | -0.022 | -0.1 | -0.1 | 0 | 0.3 | 0.09 | 0.35 | 0.51 | 0.3 | 0.45 |
| 3 | 135225249 | rs11712655 | 0.09 | 0.852 | -1.564 | -0.84 | -0.015 | -0.101 | -0.101 | 0 | 0.3 | 0.09 | 0.35 | 0.51 | 0.29 | 0.45 |
| 3 | 135238918 | rs9868429 | 0.029 | -0.085 | -1.139 | -1.087 | 0.008 | -0.003 | -0.003 | 0 | 0.36 | 0.3 | 0.27 | 0.32 | 0.22 | 0.38 |
| 3 | 135238987 | rs4894292 | 0.319 | 0.491 | -1.605 | -1.158 | 0.012 | 0.042 | 0.042 | 0 | 0.51 | 0.18 | 0.61 | 0.67 | 0.58 | 0.54 |
| 3 | 135244629 | rs1502186 | 0.688 | 0.462 | -1.739 | -1.289 | -0.067 | 0.054 | 0.054 | 0 | 0.46 | 0.11 | 0.49 | 0.69 | 0.52 | 0.5 |
| 3 | 135245818 | rs4894293 | 0.205 | -0.147 | -1.47 | -1.248 | 0.055 | 0.079 | 0.079 | 0 | 0.53 | 0.24 | 0.62 | 0.71 | 0.63 | 0.57 |
| 3 | 135249401 | rs6786506 | 0.838 | -0.549 | -1.703 | -1.312 | 0.011 | 0.053 | 0.053 | 0.266 | 0.2 | 0.04 | 0.13 | 0.08 | 0.18 | 0.1 |
| 3 | 135251491 | rs1393318 | -0.313 | -0.565 | -1.702 | -1.374 | 0.017 | 0.077 | 0.077 | 0 | 0.75 | 0.89 | 0.83 | 0.92 | 0.77 | 0.88 |
| 3 | 135254047 | rs10512967 | 0.651 | -1.236 | -1.302 | -1.514 | -0.061 | -0.006 | -0.006 | 0 | 0.22 | 0.36 | 0.18 | 0.17 | 0.11 | 0.14 |
| 3 | 135255401 | rs7622460 | 0.392 | -1.094 | -1.278 | -1.659 | 0.003 | -0.089 | -0.089 | 0 | 0.63 | 0.54 | 0.73 | 0.92 | 0.56 | 0.8 |
| 3 | 135257140 | rs6782791 | 0.634 | -1.014 | -1.191 | -1.602 | 0.096 | -0.079 | -0.079 | 0 | 0.12 | 0.35 | 0.11 | 0 | 0.21 | 0.08 |
| 3 | 135259530 | rs13320368 | 1.241 | -1.01 | -1.363 | -1.507 | -0.006 | -0.031 | -0.031 | 0 | 0.07 | 0.01 | 0.05 | 0.11 | 0.06 | 0.21 |
| 3 | 135259765 | rs6792890 | 0.633 | -1.1 | -1.136 | -1.635 | -0.059 | -0.005 | -0.005 | 0.244 | 0.22 | 0.35 | 0.18 | 0.17 | 0.1 | 0.14 |
| 3 | 135260047 | rs11926571 | -0.108 | -0.514 | -0.5 | -1.405 | 0.053 | -0.119 | -0.119 | 0 | 0.83 | 0.62 | 0.86 | 1 | 0.75 | 0.9 |
| 3 | 135267861 | rs7645988 | 0.467 | -0.261 | -1.228 | -0.945 | -0.029 | 0.05 | 0.05 | 0 | 0.41 | 0.23 | 0.55 | 0.74 | 0.46 | 0.65 |
| 3 | 135268608 | rs9826153 | 0.709 | -0.894 | -0.707 | -0.745 | -0.026 | -0.011 | -0.011 | 0.061 | 0.75 | 0.37 | 0.88 | 0.87 | 0.85 | 0.73 |
| 3 | 135269091 | rs1502174 | -0.366 | -0.448 | -0.672 | -0.895 | -0.001 | -0.005 | -0.005 | 0.242 | 0.43 | 0.49 | 0.5 | 0.36 | 0.46 | 0.45 |
| 3 | 135270611 | rs9289494 | 0.566 | 0.073 | -0.93 | -1.557 | -0.008 | -0.019 | -0.019 | 0 | 0.35 | 0.28 | 0.39 | 0.55 | 0.37 | 0.58 |
| 3 | 135274088 | rs10512957 | -0.315 | 0.009 | -0.721 | -1.726 | -0.002 | 0.001 | 0.001 | 0 | 0.17 | 0.11 | 0.1 | 0.06 | 0.22 | 0.26 |
| 3 | 135274692 | rs9840594 | 0.641 | -0.115 | -0.32 | -1.776 | 0.006 | 0.015 | 0.015 | 0 | 0.37 | 0.41 | 0.4 | 0.53 | 0.37 | 0.57 |
| 3 | 135274861 | rs6771993 | 0.154 | -0.287 | -0.24 | -1.843 | 0 | -0.001 | -0.001 | 0 | 0.35 | 0.32 | 0.43 | 0.4 | 0.37 | 0.42 |
| 3 | 135275666 | rs10512956 | 0.58 | -0.237 | -0.297 | -1.852 | 0.001 | -0.022 | -0.022 | 0 | 0.13 | 0.11 | 0.09 | 0.03 | 0.2 | 0.23 |
| 3 | 135275693 | rs10935172 | 0.929 | -0.454 | -0.475 | -1.985 | -0.044 | 0.06 | 0.06 | 0 | 0.22 | 0.05 | 0.15 | 0.11 | 0.26 | 0.14 |
| 3 | 135276133 | rs9289495 | 0.274 | -0.453 | -0.33 | -1.951 | 0.006 | 0.01 | 0.01 | 0.173 | 0.12 | 0.09 | 0.05 | 0.02 | 0.08 | 0.04 |
| 3 | 135276559 | rs9878270 | -0.572 | -0.453 | -0.269 | -1.96 | -0.002 | -0.042 | -0.042 | 0 | 0.26 | 0.28 | 0.13 | 0.05 | 0.28 | 0.27 |
| 3 | 135276789 | rs1540734 | 1.482 | -0.419 | -0.242 | -1.873 | -0.001 | -0.002 | -0.002 | 0 | 0.29 | 0.27 | 0.36 | 0.52 | 0.3 | 0.41 |
| 3 | 135278405 | rs6770007 | 1.482 | -0.392 | -0.32 | -1.87 | 0.003 | 0.024 | 0.024 | 0 | 0.29 | 0.36 | 0.36 | 0.52 | 0.29 | 0.41 |
| 3 | 135280770 | rs6439578 | -1.605 | -0.482 | -0.278 | -1.967 | -0.025 | 0.051 | 0.051 | 0 | 0.77 | 0.61 | 0.64 | 0.68 | 0.84 | 0.81 |
| 3 | 135281668 | rs7612415 | -0.746 | -0.445 | -0.349 | -1.973 | -0.017 | 0.157 | 0.157 | 0.395 | 0.47 | 0.19 | 0.28 | 0.15 | 0.53 | 0.41 |
| 3 | 135282024 | rs9833932 | -0.278 | -0.519 | -0.637 | -1.884 | 0.001 | 0.021 | 0.021 | 0 | 0.36 | 0.29 | 0.21 | 0.12 | 0.34 | 0.17 |
| 3 | 135283689 | rs13094401 | -0.424 | -0.226 | -0.724 | -1.741 | -0.003 | -0.003 | -0.003 | 0 | 0.56 | 0.43 | 0.55 | 0.44 | 0.5 | 0.36 |
| 3 | 135283812 | rs13095064 | -0.424 | -0.226 | -0.724 | -1.741 | -0.003 | -0.003 | -0.003 | 0 | 0.56 | 0.43 | 0.55 | 0.44 | 0.5 | 0.36 |

| | | | | | | | | | | | | | | | | |
|---|---|---|---|---|---|---|---|---|---|---|---|---|---|---|---|---|
| 3 | 135283979 | rs16843144 | 0.169 | -0.752 | -0.474 | -1.632 | -0.006 | 0.001 | 0.001 | 0 | 0.35 | 0.38 | 0.4 | 0.34 | 0.26 | 0.22 |
| 3 | 135285923 | rs16846019 | -0.122 | -0.067 | -0.307 | -1.436 | 0.038 | 0.013 | 0.013 | 0 | 0.51 | 0.25 | 0.45 | 0.6 | 0.56 | 0.54 |
| 3 | 135287325 | rs6806781 | 0.105 | 0.032 | -0.407 | -1.287 | -0.019 | 0.079 | 0.079 | 0 | 0.26 | 0.07 | 0.17 | 0.14 | 0.26 | 0.16 |
| 3 | 135288025 | rs6785912 | 0.185 | 0.314 | 0.07 | -1.129 | 0.011 | 0.01 | 0.01 | 0 | 0.4 | 0.58 | 0.49 | 0.48 | 0.43 | 0.45 |
| 3 | 135289506 | rs17721287 | 0.244 | 0.718 | 0.028 | -1.449 | -0.01 | 0.027 | 0.027 | 0 | 0.18 | 0.03 | 0.1 | 0.14 | 0.17 | 0.13 |
| 3 | 135291733 | rs6439579 | -1.054 | 0.742 | 0.558 | -1.618 | 0 | 0.022 | 0.022 | 0 | 0.5 | 0.56 | 0.38 | 0.2 | 0.5 | 0.39 |
| 3 | 135310068 | rs6808137 | -1.411 | 0.618 | 1.547 | -1.273 | -0.009 | -0.132 | -0.132 | 0 | 0.33 | 0.51 | 0.3 | 0.04 | 0.29 | 0.29 |
| 3 | 135313029 | rs974709 | -1.125 | 0.636 | 1.469 | -1.266 | -0.004 | -0.072 | -0.072 | 0 | 0.31 | 0.38 | 0.26 | 0.03 | 0.26 | 0.26 |
| 3 | 135337113 | rs6762033 | -0.366 | 0.674 | 1.306 | -1.292 | 0.007 | 0.049 | 0.049 | 0 | 0.21 | 0.1 | 0.09 | 0 | 0.14 | 0.18 |
| 3 | 135345493 | rs1447598 | -2.282 | 1.034 | 1.59 | -0.681 | 0.044 | -0.038 | -0.038 | 0 | 0.13 | 0.61 | 0.22 | 0.03 | 0.16 | 0.11 |
| 3 | 135351730 | rs3850168 | -2.297 | 0.998 | 1.59 | -0.405 | 0.038 | 0.038 | 0.038 | 1.59 | 0.09 | 0.48 | 0.08 | 0.04 | 0.06 | 0.01 |
| 3 | 135361053 | rs9814548 | 1.898 | 1.335 | 0.951 | -0.415 | 0.059 | 0.02 | 0.02 | 1.301 | 0.93 | 0.65 | 0.93 | 0.98 | 0.94 | 0.99 |
| 3 | 135377187 | rs2218625 | 0.995 | 1.396 | 0.765 | -0.396 | 0.053 | 0.01 | 0.01 | 0.79 | 0.95 | 0.75 | 0.94 | 0.99 | 0.94 | 0.99 |
| 3 | 135434791 | rs17248276 | -0.803 | 1.58 | 0.111 | -0.356 | -0.037 | 0.016 | 0.016 | 0 | 0.06 | 0 | 0.13 | 0.01 | 0.15 | 0.03 |
| 3 | 135447944 | rs11922097 | -0.811 | 1.212 | 0.263 | -0.501 | 0.024 | -0.105 | -0.105 | 2.146 | 0.28 | 0.46 | 0.26 | 0.05 | 0.28 | 0.33 |
| 3 | 135457984 | rs1447600 | -0.924 | 0.962 | -0.746 | 0.999 | -0.015 | -0.056 | -0.056 | 0 | 0.16 | 0.09 | 0.24 | 0.62 | 0.23 | 0.39 |
| 3 | 135459917 | rs7613386 | 0.041 | 0.12 | -0.927 | -0.663 | 0.003 | -0.048 | -0.048 | 1.689 | 0.62 | 0.64 | 0.74 | 0.98 | 0.68 | 0.76 |
| 3 | 135461655 | rs9851816 | 0.127 | 0.051 | -1.723 | -0.076 | -0.015 | -0.144 | -0.144 | 0.152 | 0.23 | 0.05 | 0.36 | 0.53 | 0.24 | 0.34 |
| 3 | 135464712 | rs11717874 | 0.45 | -0.585 | -1.84 | -0.92 | 0.002 | 0.031 | 0.031 | 0 | 0.22 | 0.15 | 0.2 | 0.01 | 0.24 | 0.12 |
| 3 | 135475923 | rs4894319 | 0.27 | -1.15 | -1.801 | -0.647 | -0.008 | -0.169 | -0.169 | 0.621 | 0.25 | 0.02 | 0.27 | 0.57 | 0.19 | 0.37 |
| 3 | 135483586 | rs7427644 | -0.715 | -1.402 | -1.407 | -1.737 | -0.016 | -0.101 | -0.101 | 0 | 0.33 | 0.42 | 0.21 | 0.02 | 0.27 | 0.24 |
| 3 | 135502373 | rs1401547 | 0.706 | -1.18 | -1.378 | -1.531 | 0.001 | 0.014 | 0.014 | 0.177 | 0.09 | 0.02 | 0.03 | 0 | 0.07 | 0.11 |
| 3 | 135514304 | rs6766026 | 0.11 | -1.19 | -1.141 | -1.534 | -0.01 | -0.037 | -0.037 | 0.267 | 0.11 | 0.24 | 0.04 | 0.02 | 0.08 | 0.12 |
| 3 | 135548749 | rs16843405 | -0.393 | -1.375 | -0.534 | -1.83 | -0.012 | -0.139 | -0.139 | 0 | 0.22 | 0.54 | 0.21 | 0.01 | 0.21 | 0.13 |
| 3 | 135561715 | rs7646013 | -0.406 | -1.354 | -0.525 | -1.833 | -0.01 | -0.133 | -0.133 | 0 | 0.22 | 0.54 | 0.21 | 0.01 | 0.21 | 0.13 |
| 3 | 135575079 | rs7648592 | -0.057 | -1.384 | -0.371 | -1.843 | -0.002 | -0.074 | -0.074 | 0.688 | 0.19 | 0.2 | 0.14 | 0.01 | 0.21 | 0.13 |
| 3 | 135580066 | rs1519260 | -0.942 | -0.639 | -0.926 | -2.244 | -0.126 | 0.128 | 0.128 | 0.165 | 0.64 | 0.97 | 0.45 | 0.16 | 0.57 | 0.54 |
| 3 | 135582293 | rs6762218 | 0.001 | -0.547 | -0.547 | -2.101 | -0.004 | -0.131 | -0.131 | 1.295 | 0.2 | 0.35 | 0.15 | 0.01 | 0.21 | 0.13 |
| 3 | 135587811 | rs1401543 | -0.061 | -0.548 | -0.436 | -2.101 | -0.005 | -0.149 | -0.149 | 0 | 0.21 | 0.41 | 0.15 | 0.01 | 0.21 | 0.13 |
| 3 | 135591867 | rs6791082 | -0.26 | -0.493 | -0.506 | -2.107 | -0.011 | -0.138 | -0.138 | 0 | 0.22 | 0.56 | 0.16 | 0.01 | 0.21 | 0.13 |
| 3 | 135594236 | rs1607504 | 0.038 | -0.467 | -0.582 | -2.088 | -0.003 | -0.102 | -0.102 | 0.062 | 0.2 | 0.28 | 0.14 | 0.01 | 0.21 | 0.12 |
| 3 | 135604118 | rs10512975 | -0.662 | -0.427 | -0.882 | -2.178 | -0.052 | 0.228 | 0.228 | 0 | 0.54 | 0.89 | 0.41 | 0.14 | 0.5 | 0.39 |
| 3 | 135604830 | rs17196014 | 0.197 | -0.511 | -0.852 | -2.201 | -0.004 | -0.069 | -0.069 | 0 | 0.19 | 0.22 | 0.14 | 0.01 | 0.2 | 0.13 |
| 3 | 135607701 | rs6772366 | 0.563 | -0.5 | -0.962 | -2.203 | -0.008 | -0.084 | -0.084 | 0 | 0.34 | 0.48 | 0.25 | 0.13 | 0.3 | 0.27 |
| 3 | 135607768 | rs6772454 | -0.591 | -0.395 | -1.178 | -2.06 | -0.077 | 0.243 | 0.243 | 0 | 0.55 | 0.93 | 0.4 | 0.14 | 0.5 | 0.39 |
| 3 | 135609010 | rs13074773 | -0.38 | -0.382 | -1.021 | -1.901 | -0.001 | -0.005 | -0.005 | 0 | 0.43 | 0.56 | 0.37 | 0.21 | 0.48 | 0.38 |
| 3 | 135609461 | rs6776731 | 0.435 | -0.259 | -1.432 | -1.879 | -0.001 | 0.039 | 0.039 | 0 | 0.08 | 0.04 | 0.05 | 0.01 | 0.07 | 0.05 |
| 3 | 135609551 | rs13075467 | 0.821 | 0.412 | -1.402 | -1.933 | -0.079 | 0.111 | 0.111 | 0 | 0.18 | 0.01 | 0.15 | 0.01 | 0.24 | 0.16 |
| 3 | 135623469 | rs6803679 | -1.101 | -0.016 | -1.662 | -2.131 | -0.132 | 0.074 | 0.074 | 2.742 | 0.61 | 0.97 | 0.47 | 0.23 | 0.57 | 0.55 |
| 3 | 135624570 | rs9819988 | -0.334 | 0.04 | -0.599 | -2.589 | -0.019 | -0.15 | -0.15 | 0 | 0.34 | 0.53 | 0.19 | 0.03 | 0.29 | 0.27 |
| 3 | 135625601 | rs13065836 | 1.524 | 0.03 | -0.682 | -2.64 | -0.035 | 0.047 | 0.047 | 0 | 0.09 | 0.01 | 0.07 | 0 | 0.12 | 0.02 |
| 3 | 135641904 | rs9875948 | 0.181 | -0.101 | -0.673 | -2.594 | -0.017 | 0.177 | 0.177 | 1.657 | 0.31 | 0.06 | 0.15 | 0.03 | 0.29 | 0.25 |
| 3 | 135643303 | rs13434196 | -0.134 | -0.094 | -0.669 | -2.591 | 0.002 | -0.034 | -0.034 | 0.21 | 0.33 | 0.3 | 0.18 | 0.03 | 0.29 | 0.27 |
| 3 | 135648685 | rs1568350 | 0.076 | -0.047 | -0.728 | -2.576 | 0.006 | 0.102 | 0.102 | 1.55 | 0.32 | 0.13 | 0.15 | 0.03 | 0.29 | 0.25 |
| 3 | 135669685 | rs16843560 | -0.156 | 0.209 | 0.161 | -2.41 | 0 | -0.019 | -0.019 | 0 | 0.25 | 0.37 | 0.26 | 0.19 | 0.27 | 0.3 |
| 3 | 135686355 | rs1523599 | -0.416 | 0.054 | -0.058 | -2.481 | -0.012 | -0.031 | -0.031 | 0 | 0.31 | 0.56 | 0.28 | 0.19 | 0.28 | 0.34 |
| 3 | 135702704 | rs6802942 | 1.183 | 0.476 | -0.362 | -2.433 | 0.006 | -0.024 | -0.024 | 0 | 0.07 | 0.09 | 0.05 | 0 | 0.11 | 0.09 |
| 3 | 135714105 | rs6774289 | 0.13 | 0.433 | -0.764 | -2.588 | -0.009 | 0.064 | 0.064 | 2.923 | 0.28 | 0.15 | 0.14 | 0.03 | 0.28 | 0.26 |
| 3 | 135720673 | rs9872542 | 0.063 | 0.419 | -0.695 | -2.594 | -0.002 | 0.009 | 0.009 | 2.3 | 0.29 | 0.2 | 0.15 | 0.03 | 0.28 | 0.26 |
| 3 | 135722264 | rs17197552 | 0.123 | 0.427 | -0.723 | -2.591 | -0.008 | 0.065 | 0.065 | 2.962 | 0.28 | 0.15 | 0.14 | 0.03 | 0.28 | 0.26 |
| 3 | 135731314 | rs6771997 | 0.168 | 0.515 | -0.737 | -2.593 | -0.016 | 0.099 | 0.099 | 3.075 | 0.28 | 0.1 | 0.14 | 0.03 | 0.27 | 0.27 |
| 3 | 135738428 | rs13066914 | 1.519 | 0.679 | -0.81 | -2.596 | -0.04 | 0.042 | 0.042 | 0 | 0.08 | 0 | 0.05 | 0 | 0.11 | 0.04 |
| 3 | 135741565 | rs2177407 | -0.433 | 0.515 | -0.653 | -2.574 | -0.016 | -0.041 | -0.041 | 0.063 | 0.31 | 0.49 | 0.28 | 0.2 | 0.29 | 0.33 |
| 3 | 135769006 | rs13072433 | -0.461 | 0.622 | -1.267 | -2.566 | 0.004 | 0.007 | 0.007 | 0 | 0.31 | 0.65 | 0.29 | 0.2 | 0.28 | 0.33 |
| 3 | 135785491 | rs10048942 | 0.586 | 1.069 | -1.801 | -2.679 | 0.003 | 0.019 | 0.019 | 0.666 | 0.24 | 0.17 | 0.11 | 0.02 | 0.21 | 0.2 |
| 3 | 135787529 | rs6439623 | -1.037 | 1.091 | -1.547 | -2.191 | 0.005 | 0.097 | 0.097 | 0 | 0.37 | 0.81 | 0.33 | 0.2 | 0.35 | 0.39 |
| 3 | 135823077 | rs10512987 | -0.321 | 0.711 | -0.365 | -2.639 | -0.016 | -0.124 | -0.124 | 3.096 | 0.32 | 0.46 | 0.18 | 0.06 | 0.29 | 0.29 |
| 3 | 135824130 | rs7634492 | 0.884 | 0.777 | -0.174 | -2.453 | 0.028 | -0.009 | -0.009 | 0 | 0.05 | 0.16 | 0.03 | 0.01 | 0.07 | 0.07 |
| 3 | 135889651 | rs13065565 | 2.081 | 0.882 | -0.681 | -2.683 | -0.021 | 0.036 | 0.036 | 0 | 0.07 | 0.01 | 0.07 | 0 | 0.1 | 0.01 |
| 3 | 135893295 | rs10935178 | 0.628 | 0.797 | -0.104 | -2.652 | 0.059 | 0.017 | 0.017 | 0.104 | 0.06 | 0.27 | 0.04 | 0.01 | 0.06 | 0.06 |
| 3 | 135914476 | rs9845457 | 0.947 | 0.609 | -1.206 | -3.108 | -0.009 | 0.206 | 0.206 | 0.472 | 0.55 | 0.05 | 0.68 | 0.79 | 0.61 | 0.52 |
| 3 | 135917197 | rs35087366 | 1.08 | 0.483 | -1.22 | -3.136 | -0.022 | 0.077 | 0.077 | 0 | 0.13 | 0.01 | 0.1 | 0.01 | 0.13 | 0.16 |
| 3 | 135930406 | rs9873767 | 2.644 | 0.994 | -1.777 | -2.362 | -0.056 | -0.156 | -0.156 | 0 | 0.3 | 0.02 | 0.32 | 0.53 | 0.27 | 0.33 |
| 3 | 135932359 | rs687339 | -0.065 | 0.773 | -1.229 | -2.475 | -0.002 | -0.027 | -0.027 | 0 | 0.76 | 0.71 | 0.81 | 0.85 | 0.77 | 0.74 |
| 3 | 135933264 | rs1279831 | 1.543 | 1.007 | -1.397 | -2.368 | -0.115 | 0.17 | 0.17 | 0.971 | 0.41 | 0.06 | 0.61 | 0.79 | 0.47 | 0.45 |
| 3 | 135953476 | rs661739 | 0.719 | 1.126 | -1.221 | -2.367 | 0.021 | 0.062 | 0.062 | 0.502 | 0.5 | 0.37 | 0.7 | 0.8 | 0.57 | 0.59 |
| 3 | 135960485 | rs561307 | 1.565 | 1.317 | -1.432 | -2.273 | -0.056 | -0.063 | -0.063 | 0 | 0.11 | 0.01 | 0.3 | 0.25 | 0.2 | 0.11 |
| 3 | 135977305 | rs7650071 | 2.686 | 1.124 | -1.73 | -2.511 | -0.025 | 0.042 | 0.042 | 0.008 | 0.08 | 0.01 | 0.07 | 0 | 0.11 | 0.01 |
| 3 | 135980673 | rs2290131 | 3.032 | 0.853 | -1.921 | -2.495 | -0.058 | -0.155 | -0.155 | 0 | 0.28 | 0.01 | 0.3 | 0.54 | 0.25 | 0.39 |

| | | | | | | | | | | | | | | | | |
|---|---|---|---|---|---|---|---|---|---|---|---|---|---|---|---|---|
| 3 | 135983089 | rs9826146 | -0.104 | 0.104 | -0.982 | -2.891 | -0.008 | -0.076 | -0.076 | 2.978 | 0.34 | 0.57 | 0.2 | 0.06 | 0.3 | 0.3 |
| 3 | 136002809 | rs3821445 | 0.588 | 0.029 | -0.802 | -2.94 | -0.02 | 0.051 | 0.051 | 0 | 0.19 | 0.07 | 0.14 | 0.01 | 0.2 | 0.15 |
| 3 | 136003194 | rs16843879 | 1.068 | 0.07 | -0.606 | -2.923 | 0.131 | 0.019 | 0.019 | 0 | 0.08 | 0.39 | 0.07 | 0.01 | 0.1 | 0.07 |
| 3 | 136004925 | rs1153877 | 2.019 | 0.054 | -0.717 | -2.915 | -0.116 | 0.142 | 0.142 | 0.808 | 0.4 | 0.06 | 0.6 | 0.78 | 0.45 | 0.5 |
| 3 | 136006576 | rs1279840 | 0.178 | 0.099 | -0.534 | -2.819 | -0.002 | -0.017 | -0.017 | 3.024 | 0.74 | 0.69 | 0.81 | 0.85 | 0.75 | 0.8 |
| 3 | 136020541 | rs1711171 | 0.018 | 0.098 | -0.429 | -2.815 | -0.001 | -0.003 | -0.003 | 0.811 | 0.76 | 0.77 | 0.82 | 0.85 | 0.76 | 0.8 |
| 3 | 136047189 | rs696081 | 1.803 | 0.241 | -0.572 | -2.84 | -0.075 | 0.142 | 0.142 | 0.993 | 0.4 | 0.11 | 0.6 | 0.78 | 0.43 | 0.51 |
| 3 | 136054982 | rs10513002 | 2.089 | 0.267 | -0.469 | -2.803 | 0.037 | -0.013 | -0.013 | 0 | 0.05 | 0.07 | 0.04 | 0.01 | 0.1 | 0.06 |
| 3 | 136056861 | rs696520 | 1.853 | 0.233 | -0.762 | -2.864 | -0.112 | 0.148 | 0.148 | 0.553 | 0.4 | 0.05 | 0.61 | 0.79 | 0.45 | 0.51 |
| 3 | 136064547 | rs695983 | 0.441 | 0.248 | -0.786 | -2.951 | -0.002 | 0.006 | 0.006 | 0 | 0.72 | 0.74 | 0.79 | 0.84 | 0.7 | 0.8 |
| 3 | 136067232 | rs17200216 | 0.885 | 0.134 | -0.699 | -2.952 | 0.04 | 0.081 | 0.081 | 0.254 | 0.27 | 0.14 | 0.13 | 0 | 0.2 | 0.22 |
| 3 | 136067655 | rs696517 | -0.806 | 0.165 | -0.753 | -2.899 | 0.006 | -0.067 | -0.067 | 0 | 0.81 | 0.73 | 0.85 | 0.99 | 0.8 | 0.85 |
| 3 | 136069472 | rs667920 | -0.176 | 0.155 | -0.754 | -2.89 | -0.001 | -0.009 | -0.009 | 0 | 0.77 | 0.74 | 0.82 | 0.85 | 0.77 | 0.81 |
| 3 | 136088038 | rs9860801 | -0.028 | 0.13 | -0.731 | -2.905 | -0.008 | -0.071 | -0.071 | 0 | 0.35 | 0.55 | 0.2 | 0.06 | 0.3 | 0.29 |
| 3 | 136097576 | rs655836 | 1.512 | 0.182 | -0.872 | -2.91 | -0.12 | 0.16 | 0.16 | 0.384 | 0.42 | 0.06 | 0.61 | 0.79 | 0.46 | 0.52 |
| 3 | 136111917 | rs1145106 | 0.151 | 0.18 | -0.785 | -2.868 | 0.001 | -0.009 | -0.009 | 0 | 0.76 | 0.71 | 0.81 | 0.85 | 0.72 | 0.8 |
| 3 | 136132705 | rs7620314 | -0.751 | 0.039 | -0.793 | -2.913 | 0.02 | 0.038 | 0.038 | 0.126 | 0.43 | 0.6 | 0.26 | 0.2 | 0.36 | 0.34 |
| 3 | 136149139 | rs10935183 | 2.081 | 0.085 | -0.81 | -2.886 | -0.081 | 0.115 | 0.115 | 0.804 | 0.38 | 0.09 | 0.59 | 0.78 | 0.41 | 0.51 |
| 3 | 136175525 | rs35375192 | 0.015 | 0.056 | -0.728 | -2.889 | -0.007 | -0.089 | -0.089 | 0 | 0.34 | 0.54 | 0.2 | 0.01 | 0.3 | 0.29 |
| 3 | 136205007 | rs6768743 | 0.88 | -0.008 | -0.794 | -2.923 | 0.038 | 0.098 | 0.098 | 0.145 | 0.26 | 0.1 | 0.12 | 0 | 0.2 | 0.23 |
| 3 | 136208788 | rs9883916 | -0.647 | 0.002 | -0.877 | -2.969 | -0.012 | 0.005 | 0.005 | 0 | 0.79 | 0.86 | 0.84 | 0.98 | 0.76 | 0.85 |
| 3 | 136235908 | rs711975 | 2.313 | -0.11 | -1.078 | -3.056 | -0.122 | 0.123 | 0.123 | 0.91 | 0.38 | 0.04 | 0.59 | 0.79 | 0.44 | 0.51 |
| 3 | 136238860 | rs9812319 | -0.225 | -0.153 | -0.908 | -3.067 | 0 | -0.039 | -0.039 | 0 | 0.35 | 0.61 | 0.2 | 0.06 | 0.3 | 0.29 |
| 3 | 136255169 | rs833752 | 0.146 | -0.169 | -0.793 | -3.055 | -0.001 | -0.018 | -0.018 | 0 | 0.73 | 0.66 | 0.8 | 0.86 | 0.74 | 0.8 |
| 3 | 136260722 | rs9813691 | 0.829 | -0.19 | -0.888 | -3.068 | 0.027 | 0.154 | 0.154 | 0.191 | 0.25 | 0.04 | 0.1 | 0 | 0.19 | 0.22 |
| 3 | 136268818 | rs34967300 | 0.833 | -1.031 | -1.352 | -3.061 | -0.018 | 0.024 | 0.024 | 0 | 0.21 | 0.1 | 0.16 | 0.01 | 0.24 | 0.15 |
| 3 | 136279390 | rs9857966 | 0.982 | -1.005 | -1.43 | -3.032 | 0.027 | 0.146 | 0.146 | 0.122 | 0.25 | 0.04 | 0.1 | 0 | 0.19 | 0.22 |
| 3 | 136309710 | rs940174 | 1.938 | -0.979 | -1.485 | -3.035 | -0.122 | 0.118 | 0.118 | 0.885 | 0.38 | 0.04 | 0.59 | 0.79 | 0.44 | 0.51 |
| 3 | 136335795 | rs6782151 | -0.046 | -1.001 | -1.364 | -3.053 | -0.004 | -0.055 | -0.055 | 0.008 | 0.35 | 0.57 | 0.2 | 0.06 | 0.3 | 0.29 |
| 3 | 136343945 | rs10513016 | 2.714 | -1.003 | -1.298 | -3.052 | 0.044 | -0.012 | -0.012 | 2.711 | 0.05 | 0.07 | 0.04 | 0.01 | 0.1 | 0.08 |
| 3 | 136346206 | rs10935186 | 1.681 | -0.805 | -1.585 | -2.974 | -0.122 | 0.124 | 0.124 | 0.877 | 0.38 | 0.04 | 0.59 | 0.79 | 0.43 | 0.51 |
| 3 | 136356464 | rs9845762 | 0.261 | -0.853 | -1.59 | -2.987 | 0.003 | -0.029 | -0.029 | 0 | 0.72 | 0.62 | 0.79 | 0.85 | 0.69 | 0.8 |
| 3 | 136377377 | rs6791142 | -1.821 | -0.836 | -1.612 | -2.972 | -0.113 | 0.11 | 0.11 | 0.873 | 0.62 | 0.96 | 0.41 | 0.21 | 0.59 | 0.49 |
| 3 | 136425600 | rs13072691 | 2.991 | -0.796 | -1.649 | -3.007 | -0.036 | 0.049 | 0.049 | 0.078 | 0.09 | 0.01 | 0.07 | 0 | 0.14 | 0.01 |
| 3 | 136444589 | rs7616204 | -1.652 | -0.866 | -1.423 | -2.863 | -0.112 | 0.111 | 0.111 | 0.918 | 0.62 | 0.96 | 0.41 | 0.21 | 0.59 | 0.49 |
| 3 | 136450593 | rs6771713 | -0.217 | -0.706 | -0.967 | -2.408 | -0.007 | -0.061 | -0.061 | 0 | 0.34 | 0.58 | 0.19 | 0.06 | 0.28 | 0.29 |
| 3 | 136451746 | rs9289513 | 0.78 | -0.714 | -0.922 | -2.409 | 0.046 | 0.102 | 0.102 | 0.448 | 0.25 | 0.11 | 0.12 | 0 | 0.18 | 0.2 |
| 3 | 136458866 | rs6799699 | -0.206 | -0.7 | -1.018 | -2.402 | -0.007 | -0.061 | -0.061 | 0 | 0.34 | 0.58 | 0.19 | 0.06 | 0.28 | 0.29 |
| 3 | 136468637 | rs34992220 | -0.556 | -0.707 | -1.009 | -2.398 | -0.002 | -0.013 | -0.013 | 0 | 0.38 | 0.56 | 0.23 | 0.15 | 0.33 | 0.33 |
| 3 | 136472573 | rs9845862 | 0.648 | -0.551 | -1.077 | -2.325 | 0.043 | 0.121 | 0.121 | 0.399 | 0.25 | 0.07 | 0.11 | 0 | 0.18 | 0.2 |
| 3 | 136476862 | rs6789329 | -0.438 | -0.547 | -1.038 | -2.292 | 0.007 | 0.01 | 0.01 | 0.16 | 0.37 | 0.61 | 0.24 | 0.2 | 0.31 | 0.33 |
| 3 | 136513282 | rs4420814 | 0.718 | -0.114 | -0.997 | -2.109 | 0 | -0.016 | -0.016 | 0 | 0.42 | 0.52 | 0.66 | 0.85 | 0.46 | 0.53 |
| 3 | 136537044 | rs6439663 | -0.326 | -0.167 | -1.1 | -2.112 | 0.008 | -0.035 | -0.035 | 0 | 0.68 | 0.57 | 0.76 | 0.85 | 0.66 | 0.75 |
| 3 | 136539689 | rs9682783 | -0.76 | -0.14 | -1.079 | -2.141 | 0 | -0.027 | -0.027 | 0 | 0.73 | 0.64 | 0.81 | 0.92 | 0.76 | 0.83 |
| 3 | 136551620 | rs9842577 | -0.262 | -0.128 | -1.066 | -2.137 | 0.007 | -0.03 | -0.03 | 0 | 0.68 | 0.57 | 0.77 | 0.85 | 0.66 | 0.74 |
| 3 | 136553923 | rs1406539 | -0.754 | -0.127 | -1.068 | -2.137 | 0 | -0.023 | -0.023 | 0 | 0.73 | 0.64 | 0.81 | 0.92 | 0.76 | 0.83 |
| 3 | 136558759 | rs1569228 | -0.286 | -0.136 | -1.068 | -2.132 | 0.005 | -0.025 | -0.025 | 0.671 | 0.68 | 0.6 | 0.77 | 0.85 | 0.66 | 0.74 |
| 3 | 136573440 | rs9827648 | -0.263 | -0.136 | -1.068 | -2.132 | 0.007 | -0.03 | -0.03 | 0 | 0.68 | 0.57 | 0.77 | 0.85 | 0.66 | 0.74 |
| 3 | 136580524 | rs9836807 | -0.206 | -0.051 | -0.961 | -2.025 | 0.007 | -0.031 | -0.031 | 0 | 0.68 | 0.57 | 0.77 | 0.85 | 0.66 | 0.74 |
| 3 | 136587117 | rs9845460 | -0.742 | -0.03 | -0.951 | -2.025 | 0 | -0.025 | -0.025 | 0.295 | 0.74 | 0.64 | 0.82 | 0.92 | 0.76 | 0.83 |
| 3 | 136604697 | rs9815771 | -0.199 | -0.033 | -0.96 | -2.024 | 0.007 | -0.031 | -0.031 | 0 | 0.68 | 0.57 | 0.77 | 0.85 | 0.66 | 0.75 |
| 3 | 136606271 | rs9846691 | -0.145 | -0.044 | -0.889 | -2.029 | 0.013 | -0.052 | -0.052 | 0 | 0.68 | 0.52 | 0.77 | 0.85 | 0.66 | 0.75 |
| 3 | 136612149 | rs9878367 | -0.204 | -0.025 | -0.99 | -2.021 | 0.007 | -0.031 | -0.031 | 0 | 0.68 | 0.57 | 0.77 | 0.85 | 0.66 | 0.74 |
| 3 | 136618214 | rs7645773 | -0.194 | -0.029 | -0.991 | -2.023 | 0.007 | -0.031 | -0.031 | 0 | 0.68 | 0.57 | 0.77 | 0.85 | 0.66 | 0.74 |
| 3 | 136626697 | rs1347209 | 0.171 | -0.102 | -1.043 | -2.085 | 0.001 | -0.015 | -0.015 | 0 | 0.63 | 0.57 | 0.73 | 0.85 | 0.59 | 0.71 |
| 3 | 136635171 | rs16844241 | 0.57 | 0.135 | -0.71 | -1.831 | -0.006 | 0.038 | 0.038 | 0 | 0.25 | 0.19 | 0.18 | 0.08 | 0.24 | 0.17 |
| 3 | 136639237 | rs4678274 | -0.012 | 0.136 | -0.786 | -1.838 | 0.007 | -0.035 | -0.035 | 0 | 0.68 | 0.56 | 0.77 | 0.85 | 0.66 | 0.74 |
| 3 | 136647246 | rs3772388 | -0.064 | 0.229 | -0.78 | -1.815 | 0.008 | -0.031 | -0.031 | 0 | 0.68 | 0.57 | 0.76 | 0.85 | 0.66 | 0.74 |
| 3 | 136662529 | rs17301365 | 0.535 | 0.909 | -0.412 | -1.001 | 0.007 | -0.001 | -0.001 | 0 | 0.06 | 0.04 | 0.04 | 0.07 | 0.1 | 0.09 |
| 3 | 136674500 | rs835636 | -0.451 | 0.599 | -0.316 | -1.166 | -0.005 | 0.036 | 0.036 | 0 | 0.47 | 0.56 | 0.69 | 0.91 | 0.5 | 0.59 |
| 3 | 136676303 | rs835634 | -0.302 | 1.066 | 0.238 | 0.215 | 0.015 | -0.001 | -0.001 | 1.337 | 0.38 | 0.5 | 0.55 | 0.68 | 0.43 | 0.41 |
| 3 | 136693459 | rs10935213 | -0.309 | 1.045 | 0.227 | 0.216 | 0.014 | -0.001 | -0.001 | 0 | 0.38 | 0.51 | 0.56 | 0.69 | 0.43 | 0.42 |
| 3 | 136700455 | rs75976 | -0.265 | 1.051 | 0.228 | 0.219 | 0.014 | 0 | 0 | 0.028 | 0.37 | 0.43 | 0.56 | 0.69 | 0.43 | 0.41 |
| 3 | 136715468 | rs9845076 | 1.048 | 0.966 | -0.026 | -0.078 | 0.026 | 0.145 | 0.145 | 0 | 0.44 | 0.18 | 0.27 | 0.1 | 0.38 | 0.38 |
| 3 | 136746022 | rs835631 | -1.05 | 1.182 | 0.059 | 0.006 | 0.005 | -0.015 | -0.015 | 0 | 0.52 | 0.51 | 0.7 | 0.91 | 0.62 | 0.6 |
| 3 | 136751674 | rs900949 | 0.784 | 1.314 | 0.184 | 0.315 | 0.006 | 0.006 | 0.006 | 0 | 0.5 | 0.49 | 0.29 | 0.24 | 0.4 | 0.44 |
| 3 | 136755982 | rs1470142 | 1.025 | 1.384 | 0.069 | 0.225 | 0.015 | 0.043 | 0.043 | 0 | 0.47 | 0.37 | 0.28 | 0.09 | 0.39 | 0.4 |
| 3 | 136757858 | rs6780453 | -1.212 | 1.62 | 0.274 | 0.702 | 0.038 | 0.165 | 0.165 | 0.106 | 0.18 | 0.43 | 0.33 | 0.38 | 0.23 | 0.25 |
| 3 | 136760945 | rs6772642 | 0.066 | 1.351 | 0.122 | 0.875 | 0.015 | -0.001 | -0.001 | 1.029 | 0.62 | 0.47 | 0.56 | 0.61 | 0.58 | 0.66 |

| | | | | | | | | | | | | | | | | |
|---|---|---|---|---|---|---|---|---|---|---|---|---|---|---|---|---|
| 3 | 136761075 | rs9845921 | 0.537 | 1.353 | 0.033 | 0.593 | 0.001 | -0.009 | -0.009 | 0.047 | 0.54 | 0.48 | 0.33 | 0.3 | 0.51 | 0.47 |
| 3 | 136762937 | rs12489946 | -0.437 | 1.133 | 0.135 | 0.7 | -0.006 | 0.071 | 0.071 | 0.065 | 0.15 | 0.32 | 0.28 | 0.26 | 0.16 | 0.21 |
| 3 | 136763653 | rs6439680 | -0.113 | 1.418 | -0.563 | 0.708 | -0.046 | -0.041 | -0.041 | 0.264 | 0.65 | 0.84 | 0.58 | 0.55 | 0.59 | 0.64 |
| 3 | 136765859 | rs894004 | -0.795 | 1.617 | -0.694 | 0.795 | -0.027 | -0.099 | -0.099 | 0 | 0.78 | 0.93 | 0.73 | 0.51 | 0.78 | 0.83 |
| 3 | 136769944 | rs4678222 | 0.745 | 1.6 | -0.46 | 0.693 | -0.002 | 0.006 | 0.006 | 0 | 0.63 | 0.64 | 0.46 | 0.26 | 0.6 | 0.62 |
| 3 | 136770481 | rs6775490 | -0.784 | 1.603 | -0.847 | 0.825 | -0.038 | -0.118 | -0.118 | 0 | 0.78 | 0.96 | 0.73 | 0.51 | 0.78 | 0.82 |
| 3 | 136771382 | rs7622879 | -0.77 | 1.603 | -0.809 | 0.825 | -0.012 | -0.054 | -0.054 | 0 | 0.78 | 0.91 | 0.73 | 0.51 | 0.78 | 0.82 |
| 3 | 136781083 | rs4678285 | -1.441 | 1.669 | -0.777 | 0.893 | -0.009 | -0.039 | -0.039 | 0 | 0.82 | 0.88 | 0.76 | 0.65 | 0.83 | 0.85 |
| 3 | 136792237 | rs752923 | -0.329 | 1.577 | -0.582 | 0.713 | 0.029 | 0.086 | 0.086 | 0 | 0.05 | 0.23 | 0.04 | 0.15 | 0.05 | 0.03 |
| 3 | 136795007 | rs9818918 | 0.969 | 1.641 | -0.67 | 0.497 | 0.018 | 0.032 | 0.032 | 0 | 0.51 | 0.56 | 0.31 | 0.3 | 0.4 | 0.49 |
| 3 | 136795393 | rs9819250 | 1.37 | 1.634 | -0.668 | 0.455 | 0.005 | 0.001 | 0.001 | 0 | 0.47 | 0.53 | 0.28 | 0.16 | 0.36 | 0.45 |
| 3 | 136796050 | rs17303046 | 0.646 | 1.842 | -0.869 | 0.464 | -0.032 | 0.008 | 0.008 | 0 | 0.12 | 0.08 | 0.13 | 0.1 | 0.25 | 0.16 |
| 3 | 136804213 | rs6439683 | 0.915 | 1.357 | -0.828 | 0.447 | 0.019 | 0.032 | 0.032 | 0 | 0.5 | 0.55 | 0.29 | 0.31 | 0.4 | 0.49 |
| 3 | 136804986 | rs1484794 | 0.82 | 1.357 | -0.822 | 0.45 | 0.032 | 0.057 | 0.057 | 0 | 0.51 | 0.63 | 0.3 | 0.32 | 0.4 | 0.49 |
| 3 | 136811406 | rs6439686 | 0.732 | 1.407 | -0.922 | 0.453 | 0.01 | 0.016 | 0.016 | 0 | 0.52 | 0.52 | 0.3 | 0.31 | 0.4 | 0.52 |
| 3 | 136828838 | rs7628560 | -1.178 | 1.673 | -1.46 | 0.664 | -0.096 | -0.115 | -0.115 | 0 | 0.78 | 0.98 | 0.74 | 0.66 | 0.75 | 0.81 |
| 3 | 136835237 | rs9855204 | 1.293 | 1.531 | -1.259 | 0.298 | -0.006 | -0.016 | -0.016 | 0 | 0.45 | 0.47 | 0.33 | 0.17 | 0.33 | 0.46 |
| 3 | 136837555 | rs1837603 | -0.693 | 1.445 | -1.273 | 0.475 | -0.025 | -0.048 | -0.048 | 0 | 0.74 | 0.8 | 0.5 | 0.42 | 0.72 | 0.72 |
| 3 | 136841449 | rs6788460 | 0.612 | 1.676 | -1.26 | 0.429 | -0.029 | 0.043 | 0.043 | 0 | 0.6 | 0.76 | 0.43 | 0.27 | 0.55 | 0.57 |
| 3 | 136841553 | rs10935222 | -0.213 | 1.539 | -1.398 | 0.302 | -0.016 | 0.017 | 0.017 | 0.011 | 0.16 | 0.03 | 0.09 | 0.15 | 0.18 | 0.15 |
| 3 | 136841721 | rs10935223 | -0.302 | 1.587 | -1.307 | 0.373 | -0.001 | 0.004 | 0.004 | 0 | 0.15 | 0.12 | 0.08 | 0.15 | 0.17 | 0.15 |
| 3 | 136844600 | rs2053439 | -0.846 | 1.696 | -1.456 | 0.452 | -0.031 | -0.054 | -0.054 | 0 | 0.75 | 0.88 | 0.58 | 0.42 | 0.72 | 0.72 |
| 3 | 136849972 | rs9831130 | 0.203 | 1.703 | -1.53 | 0.319 | -0.056 | 0.036 | 0.036 | 0 | 0.63 | 0.78 | 0.5 | 0.26 | 0.55 | 0.6 |
| 3 | 136854978 | rs1440699 | -0.188 | 1.65 | -1.487 | 0.613 | -0.014 | -0.056 | -0.056 | 0.151 | 0.15 | 0.07 | 0.22 | 0.4 | 0.2 | 0.2 |
| 3 | 136863079 | rs7624691 | 0.265 | 1.107 | -1.764 | -0.06 | 0.02 | 0.019 | 0.019 | 0 | 0.51 | 0.59 | 0.53 | 0.58 | 0.47 | 0.54 |
| 3 | 136867313 | rs7647718 | 0.571 | 1.353 | -1.949 | -0.081 | 0.051 | -0.027 | -0.027 | 0 | 0.14 | 0.3 | 0.17 | 0.09 | 0.23 | 0.12 |
| 3 | 136875691 | rs1440697 | -0.876 | 1.097 | -2.016 | 0.03 | -0.044 | -0.047 | -0.047 | 0.018 | 0.66 | 0.96 | 0.69 | 0.67 | 0.69 | 0.69 |
| 3 | 136879530 | rs7626849 | -0.661 | 1.174 | -1.791 | -0.089 | 0.006 | -0.009 | -0.009 | 0 | 0.28 | 0.36 | 0.25 | 0.19 | 0.41 | 0.24 |
| 3 | 136882119 | rs9289530 | -1.232 | 0.996 | -1.995 | -0.126 | -0.061 | -0.102 | -0.102 | 0.107 | 0.67 | 0.96 | 0.7 | 0.66 | 0.7 | 0.76 |
| 3 | 136885813 | rs9830729 | -0.383 | 1.057 | -1.569 | -0.113 | -0.022 | -0.014 | -0.014 | 0 | 0.54 | 0.74 | 0.61 | 0.52 | 0.53 | 0.64 |
| 3 | 136893356 | rs10935230 | -0.162 | 1.245 | -1.35 | -0.019 | 0.063 | 0.1 | 0.1 | 0 | 0.13 | 0.25 | 0.25 | 0.35 | 0.2 | 0.2 |
| 3 | 136902177 | rs2722546 | -0.167 | 1.197 | -1.81 | -0.182 | -0.003 | -0.01 | -0.01 | 0.039 | 0.49 | 0.75 | 0.58 | 0.52 | 0.48 | 0.59 |
| 3 | 136905916 | rs10513028 | 0.007 | 1.27 | -1.677 | -0.339 | -0.032 | 0.034 | 0.034 | 0 | 0.14 | 0.11 | 0.11 | 0.11 | 0.21 | 0.14 |
| 3 | 136907111 | rs4678297 | 0.569 | 0.953 | -1.309 | -0.384 | -0.004 | -0.007 | -0.007 | 0.002 | 0.38 | 0.51 | 0.36 | 0.17 | 0.33 | 0.43 |
| 3 | 136909995 | rs428788 | -0.754 | 0.981 | -1.054 | -0.244 | 0.037 | 0.039 | 0.039 | 0 | 0.28 | 0.46 | 0.25 | 0.45 | 0.35 | 0.35 |
| 3 | 136923798 | rs979604 | 1.649 | 0.866 | -1.467 | -0.275 | -0.071 | -0.158 | -0.158 | 1.575 | 0.28 | 0.01 | 0.35 | 0.38 | 0.25 | 0.17 |
| 3 | 136938031 | rs9869035 | 0.237 | 0.819 | -1.083 | -0.443 | -0.004 | -0.021 | -0.021 | 0.014 | 0.46 | 0.53 | 0.39 | 0.17 | 0.4 | 0.49 |
| 3 | 136947138 | rs1381251 | 0.443 | 0.374 | -0.893 | -0.415 | -0.032 | -0.035 | -0.035 | 0 | 0.27 | 0.36 | 0.24 | 0.17 | 0.18 | 0.45 |
| 3 | 136949826 | rs2880858 | 0.905 | 0.626 | -1.409 | -0.217 | 0.026 | 0.096 | 0.096 | 0 | 0.1 | 0.14 | 0.07 | 0.3 | 0.14 | 0.16 |
| 3 | 136957752 | rs1478312 | 0.695 | 0.558 | -1.555 | -0.352 | 0.026 | 0.103 | 0.103 | 1.642 | 0.54 | 0.21 | 0.49 | 0.69 | 0.62 | 0.36 |
| 3 | 136963967 | rs9838168 | 1.447 | 0.532 | -1.66 | -0.329 | -0.1 | -0.089 | -0.089 | 0.037 | 0.44 | 0.03 | 0.41 | 0.38 | 0.47 | 0.2 |
| 3 | 136978661 | rs1382270 | 0.706 | 0.609 | -1.488 | -0.431 | 0.019 | 0.087 | 0.087 | 1.88 | 0.55 | 0.22 | 0.49 | 0.68 | 0.61 | 0.35 |
| 3 | 136984847 | rs17311950 | 0.364 | 0.566 | -1.745 | -0.467 | -0.069 | 0.022 | 0.022 | 0 | 0.14 | 0.04 | 0.22 | 0.14 | 0.23 | 0.2 |
| 3 | 136985329 | rs1039617 | 0.523 | 0.45 | -1.239 | -0.495 | -0.007 | 0.038 | 0.038 | 1.784 | 0.59 | 0.59 | 0.54 | 0.68 | 0.59 | 0.35 |
| 3 | 136992840 | rs12107468 | -0.014 | 0.411 | -1.134 | -0.072 | -0.011 | 0.104 | 0.104 | 0 | 0.17 | 0.54 | 0.14 | 0.33 | 0.14 | 0.19 |
| 3 | 137002382 | rs9853224 | -0.146 | 0.401 | -1.218 | -0.075 | -0.015 | 0.091 | 0.091 | 0 | 0.83 | 0.41 | 0.86 | 0.66 | 0.86 | 0.81 |
| 3 | 137003783 | rs11916892 | 1.798 | 0.584 | -1.661 | -0.398 | -0.098 | 0.13 | 0.13 | 0 | 0.3 | 0.06 | 0.3 | 0.15 | 0.36 | 0.2 |
| 3 | 137012388 | rs9851969 | 1.129 | 0.473 | -0.714 | 0.548 | 0.033 | -0.092 | -0.092 | 4.049 | 0.61 | 0.15 | 0.64 | 0.49 | 0.7 | 0.39 |
| 3 | 137015487 | rs6439699 | 0.188 | 0.46 | -0.641 | 0.536 | -0.006 | -0.005 | -0.005 | 0 | 0.21 | 0.2 | 0.21 | 0.17 | 0.16 | 0.42 |
| 3 | 137031237 | rs7616632 | 1.677 | 0.191 | -0.538 | 0.611 | -0.019 | 0.055 | 0.055 | 4.11 | 0.48 | 0.23 | 0.54 | 0.35 | 0.49 | 0.23 |
| 3 | 137032092 | rs1564186 | -0.034 | 0.178 | -0.369 | 0.729 | 0.008 | 0.028 | 0.028 | 0 | 0.85 | 0.59 | 0.89 | 0.86 | 0.87 | 0.85 |
| 3 | 137051116 | rs3912121 | 0.734 | 0.17 | -0.449 | 0.742 | -0.015 | 0.073 | 0.073 | 0 | 0.82 | 0.31 | 0.85 | 0.66 | 0.87 | 0.82 |
| 3 | 137052096 | rs3845928 | -0.014 | 0.177 | -0.437 | 0.746 | 0.013 | 0.029 | 0.029 | 0 | 0.85 | 0.59 | 0.89 | 0.86 | 0.87 | 0.86 |
| 3 | 137055154 | rs6439704 | -0.824 | 0.186 | -0.512 | 0.725 | 0.001 | 0.008 | 0.008 | 1.077 | 0.38 | 0.48 | 0.36 | 0.5 | 0.39 | 0.63 |
| 3 | 137065862 | rs1542535 | 0.793 | 0.327 | -0.485 | 0.817 | -0.016 | 0.082 | 0.082 | 0 | 0.82 | 0.34 | 0.85 | 0.66 | 0.87 | 0.84 |
| 3 | 137071042 | rs11921899 | -1.329 | 0.147 | -0.73 | 0.467 | -0.017 | -0.081 | -0.081 | 0 | 0.45 | 0.88 | 0.42 | 0.51 | 0.39 | 0.64 |
| 3 | 137100966 | rs13100602 | -0.858 | 0.221 | -0.723 | 0.5 | -0.001 | 0 | 0 | 0 | 0.41 | 0.4 | 0.37 | 0.51 | 0.39 | 0.62 |
| 3 | 137107585 | rs6792394 | -1.15 | 0.401 | -1.574 | 0.594 | 0.001 | -0.052 | -0.052 | 0 | 0.44 | 0.75 | 0.41 | 0.51 | 0.4 | 0.62 |
| 3 | 137130750 | rs9834972 | 1.378 | 0.026 | -1.57 | 0.721 | -0.02 | 0.046 | 0.046 | 0.387 | 0.49 | 0.1 | 0.56 | 0.35 | 0.56 | 0.31 |
| 3 | 137159572 | rs9809649 | 1.413 | -0.053 | -1.164 | 0.697 | 0.035 | 0.074 | 0.074 | 0.329 | 0.5 | 0.21 | 0.56 | 0.35 | 0.57 | 0.31 |
| 3 | 137185210 | rs10804650 | -0.06 | 0.013 | -1.369 | 0.766 | -0.025 | -0.031 | -0.031 | 0.566 | 0.14 | 0.07 | 0.17 | 0.36 | 0.24 | 0.18 |
| 3 | 137190765 | rs4678242 | -0.22 | -0.101 | -1.504 | 0.862 | -0.025 | -0.046 | -0.046 | 0.081 | 0.88 | 0.96 | 0.85 | 0.81 | 0.87 | 0.83 |
| 3 | 137196162 | rs12490164 | -0.018 | -0.285 | -1.073 | 0.755 | -0.045 | -0.025 | -0.025 | 0 | 0.13 | 0.23 | 0.06 | 0.08 | 0.04 | 0.29 |
| 3 | 137198193 | rs13324482 | 1.906 | -0.139 | -1.474 | 0.804 | -0.026 | -0.021 | -0.021 | 0 | 0.5 | 0.06 | 0.55 | 0.37 | 0.57 | 0.31 |
| 3 | 137199714 | rs4478034 | -1.233 | -0.119 | -0.771 | 0.605 | -0.001 | 0.046 | 0.046 | 0.501 | 0.28 | 0.42 | 0.29 | 0.45 | 0.27 | 0.48 |
| 3 | 137208725 | rs9289539 | 1.615 | -0.27 | -1.097 | 0.499 | 0.094 | -0.012 | -0.012 | 0 | 0.59 | 0.13 | 0.72 | 0.6 | 0.76 | 0.42 |
| 3 | 137211259 | rs12633755 | 0.878 | -0.4 | -0.86 | 0.475 | -0.017 | -0.018 | -0.018 | 0 | 0.76 | 0.68 | 0.85 | 0.81 | 0.83 | 0.53 |
| 3 | 137214174 | rs3922442 | -0.309 | -0.409 | -0.811 | 0.473 | -0.071 | 0.048 | 0.048 | 0 | 0.16 | 0.5 | 0.12 | 0.21 | 0.07 | 0.11 |
| 3 | 137214696 | rs6439732 | 0.927 | -0.41 | -0.738 | 0.472 | -0.022 | -0.023 | -0.023 | 0 | 0.75 | 0.59 | 0.84 | 0.81 | 0.83 | 0.53 |

| | | | | | | | | | | | | | | | | |
|---|---|---|---|---|---|---|---|---|---|---|---|---|---|---|---|---|
| 3 | 137215756 | rs4314115 | 1.797 | -0.38 | -0.864 | 0.476 | 0.091 | -0.007 | -0.007 | 0 | 0.59 | 0.15 | 0.73 | 0.6 | 0.76 | 0.42 |
| 3 | 137232860 | rs9682216 | -0.095 | 0.375 | -0.375 | 0.461 | -0.029 | 0 | 0 | 0.02 | 0.16 | 0.14 | 0.14 | 0.18 | 0.24 | 0.19 |
| 3 | 137235880 | rs9815948 | 1.831 | 0.316 | -0.412 | 0.444 | 0.003 | -0.015 | -0.015 | 0 | 0.59 | 0.5 | 0.71 | 0.63 | 0.58 | 0.36 |
| 3 | 137239115 | rs1372737 | -0.198 | 0.753 | -0.749 | 0.626 | -0.012 | -0.009 | -0.009 | 0 | 0.9 | 0.94 | 0.93 | 0.89 | 0.89 | 0.88 |
| 3 | 137246402 | rs1442629 | 0.764 | 0.637 | -0.746 | 0.613 | -0.065 | -0.018 | -0.018 | 0.003 | 0.74 | 0.88 | 0.77 | 0.71 | 0.63 | 0.69 |
| 3 | 137247183 | rs1442593 | 0.746 | 0.678 | -0.774 | 0.614 | -0.072 | -0.021 | -0.021 | 0.194 | 0.74 | 0.9 | 0.77 | 0.71 | 0.63 | 0.69 |
| 3 | 137248353 | rs12492700 | 0.159 | 0.514 | 0.012 | 0.563 | 0.04 | 0.095 | 0.095 | 0 | 0.11 | 0.27 | 0.12 | 0.24 | 0.09 | 0.18 |
| 3 | 137249069 | rs16847101 | -0.453 | 0.477 | 0.101 | 0.573 | -0.006 | 0 | 0 | 0 | 0.19 | 0.45 | 0.16 | 0.17 | 0.15 | 0.24 |
| 3 | 137253713 | rs6797178 | -1.536 | 0.235 | 0.021 | 0.343 | 0.048 | 0.002 | 0.002 | 0 | 0.46 | 0.82 | 0.35 | 0.47 | 0.34 | 0.58 |
| 3 | 137254071 | rs10935254 | -0.774 | 0.428 | -0.011 | 0.395 | -0.018 | 0.001 | 0.001 | 0 | 0.29 | 0.55 | 0.22 | 0.31 | 0.21 | 0.34 |
| 3 | 137261477 | rs11707619 | 0.065 | 0.554 | -0.145 | 0.595 | 0.001 | 0 | 0 | 2.062 | 0.15 | 0.13 | 0.12 | 0.16 | 0.13 | 0.24 |
| 3 | 137282985 | rs12630965 | -0.218 | 0.551 | -0.012 | 0.606 | -0.004 | 0.014 | 0.014 | 0 | 0.22 | 0.23 | 0.16 | 0.31 | 0.16 | 0.34 |
| 3 | 137290019 | rs6439749 | -0.336 | 0.781 | -0.122 | 0.447 | -0.002 | -0.001 | -0.001 | 0 | 0.16 | 0.17 | 0.13 | 0.17 | 0.13 | 0.24 |
| 3 | 137292798 | rs959177 | 0.491 | 1.294 | -0.458 | 0.252 | -0.013 | 0.037 | 0.037 | 0 | 0.77 | 0.88 | 0.78 | 0.92 | 0.67 | 0.88 |
| 3 | 137301511 | rs10804653 | 0.943 | 1.334 | -0.025 | 0.78 | -0.006 | -0.016 | -0.016 | 0 | 0.58 | 0.25 | 0.61 | 0.51 | 0.56 | 0.56 |
| 3 | 137301734 | rs1610111 | 0.092 | 1.427 | 0.569 | 0.848 | -0.001 | 0.003 | 0.003 | 0 | 0.22 | 0.21 | 0.11 | 0.34 | 0.2 | 0.32 |
| 3 | 137306262 | rs6795373 | 1.357 | 1.55 | 0.505 | 0.878 | -0.113 | -0.065 | -0.065 | 0.148 | 0.53 | 0.04 | 0.56 | 0.39 | 0.52 | 0.47 |
| 3 | 137311307 | rs6782909 | 1.382 | 1.039 | 1.166 | 0.573 | 0.095 | 0.06 | 0.06 | 4.661 | 0.94 | 0.48 | 0.95 | 0.99 | 0.98 | 0.98 |
| 3 | 137312571 | rs10513037 | -0.858 | 1.036 | 1.279 | 0.587 | -0.014 | -0.026 | -0.026 | 0.829 | 0.36 | 0.19 | 0.35 | 0.46 | 0.42 | 0.44 |
| 3 | 137321590 | rs984491 | -0.894 | 0.924 | 1.635 | 0.65 | 0.002 | 0.016 | 0.016 | 0 | 0.29 | 0.59 | 0.16 | 0.35 | 0.27 | 0.37 |
| 3 | 137324323 | rs2165694 | 0.761 | 0.923 | 1.554 | 0.65 | 0 | -0.001 | -0.001 | 0 | 0.7 | 0.26 | 0.82 | 0.64 | 0.72 | 0.62 |
| 3 | 137330345 | rs7637509 | -0.507 | 1.032 | 1.233 | 0.673 | -0.003 | 0.005 | 0.005 | 0 | 0.28 | 0.36 | 0.14 | 0.35 | 0.26 | 0.36 |
| 3 | 137336749 | rs10935257 | 0.649 | 1.091 | 0.949 | 0.699 | -0.004 | 0.002 | 0.002 | 0 | 0.69 | 0.34 | 0.83 | 0.65 | 0.72 | 0.63 |
| 3 | 137345119 | rs1899450 | 0.516 | 1.235 | 0.126 | 0.792 | -0.005 | 0.011 | 0.011 | 0 | 0.72 | 0.61 | 0.86 | 0.65 | 0.74 | 0.64 |
| 3 | 137350423 | rs1540518 | 0.516 | 1.235 | 0.113 | 0.792 | -0.005 | 0.012 | 0.012 | 0 | 0.72 | 0.6 | 0.85 | 0.65 | 0.74 | 0.63 |
| 3 | 137363976 | rs9851498 | -0.411 | 1.379 | -0.321 | 0.892 | 0 | 0.002 | 0.002 | 0.003 | 0.26 | 0.23 | 0.14 | 0.35 | 0.26 | 0.37 |
| 3 | 137367809 | rs6777144 | 0.377 | 1.315 | -0.847 | 0.419 | -0.028 | 0.051 | 0.051 | 0 | 0.56 | 0.69 | 0.58 | 0.71 | 0.54 | 0.64 |
| 3 | 137372132 | rs17203979 | -0.882 | 1.468 | -0.463 | 0.069 | -0.001 | 0.016 | 0.016 | 0 | 0.25 | 0.18 | 0.15 | 0.09 | 0.19 | 0.19 |
| 3 | 137372151 | rs4678365 | -1.037 | 1.666 | 0.166 | 0.488 | 0.06 | 0.116 | 0.116 | 0.04 | 0.05 | 0.23 | 0.21 | 0.13 | 0.1 | 0.1 |
| 3 | 137389062 | rs1372733 | -1.253 | 1.367 | 0.685 | 0.315 | -0.01 | -0.027 | -0.027 | 0.697 | 0.31 | 0.56 | 0.37 | 0.21 | 0.24 | 0.3 |
| 3 | 137393428 | rs1074338 | -0.743 | 1.345 | 0.96 | 0.29 | -0.001 | -0.005 | -0.005 | 0 | 0.24 | 0.21 | 0.16 | 0.08 | 0.19 | 0.2 |
| 3 | 137406508 | rs6772470 | -1.671 | 2.101 | 0.702 | 0.599 | 0.017 | 0 | 0 | 0 | 0.38 | 0.51 | 0.48 | 0.36 | 0.42 | 0.38 |
| 3 | 137408592 | rs12636521 | 0.069 | 2.154 | 0.693 | 0.488 | -0.003 | 0.036 | 0.036 | 0.657 | 0.16 | 0.08 | 0.1 | 0.06 | 0.12 | 0.08 |
| 3 | 137409366 | rs12637203 | 0.107 | 2.164 | 0.698 | 0.492 | -0.004 | 0.035 | 0.035 | 0.846 | 0.16 | 0.08 | 0.1 | 0.06 | 0.12 | 0.08 |
| 3 | 137410101 | rs17203993 | 0.013 | 2.161 | 0.708 | 0.531 | 0.003 | -0.007 | -0.007 | 0.323 | 0.08 | 0.08 | 0.06 | 0.06 | 0.07 | 0.12 |
| 3 | 137411496 | rs917934 | -2.108 | 2.144 | 0.696 | 0.543 | 0.043 | 0.059 | 0.059 | 1.027 | 0.22 | 0.43 | 0.38 | 0.3 | 0.3 | 0.29 |
| 3 | 137413421 | rs1837955 | -1.662 | 2.121 | 0.721 | 0.536 | 0.019 | 0.057 | 0.057 | 0.136 | 0.14 | 0.26 | 0.31 | 0.28 | 0.23 | 0.17 |
| 3 | 137413940 | rs17468050 | -0.227 | 1.538 | 0.895 | 0.411 | 0.002 | -0.026 | -0.026 | 0 | 0.08 | 0.09 | 0.06 | 0.01 | 0.07 | 0.12 |
| 3 | 137415813 | rs10935263 | -1.179 | 1.534 | 0.979 | 0.443 | 0.002 | 0.109 | 0.109 | 0.002 | 0.05 | 0.16 | 0.19 | 0.14 | 0.05 | 0.1 |
| 3 | 137416355 | rs2017141 | 0.01 | 1.536 | 0.956 | 0.435 | -0.008 | 0.038 | 0.038 | 0.844 | 0.15 | 0.07 | 0.1 | 0.06 | 0.12 | 0.08 |
| 3 | 137419421 | rs3845931 | -1.938 | 1.412 | 1.041 | 0.448 | 0.008 | 0.053 | 0.053 | 0.695 | 0.14 | 0.42 | 0.26 | 0.15 | 0.12 | 0.22 |
| 3 | 137419702 | rs3856631 | -0.44 | 1.41 | 0.911 | 0.287 | 0.006 | -0.039 | -0.039 | 0 | 0.08 | 0.14 | 0.07 | 0.01 | 0.07 | 0.12 |
| 3 | 137422031 | rs17546951 | -0.466 | 1.403 | 0.898 | 0.284 | 0.004 | -0.035 | -0.035 | 0 | 0.08 | 0.14 | 0.06 | 0.01 | 0.07 | 0.12 |
| 3 | 137429433 | rs16847385 | -1.344 | 1.304 | 1.439 | 0.444 | 0.015 | 0.053 | 0.053 | 0.201 | 0.13 | 0.38 | 0.24 | 0.16 | 0.1 | 0.17 |
| 3 | 137434886 | rs10935266 | -1.73 | 1.243 | 1.174 | 0.412 | -0.005 | 0.006 | 0.006 | 0.497 | 0.15 | 0.63 | 0.29 | 0.15 | 0.11 | 0.17 |
| 3 | 137441140 | rs6768778 | -1.557 | 1.28 | 1.111 | 0.348 | -0.011 | -0.028 | -0.028 | 0 | 0.28 | 0.64 | 0.35 | 0.16 | 0.2 | 0.19 |
| 3 | 137447485 | rs10935268 | -0.845 | 0.558 | 1.209 | 0.081 | -0.034 | -0.046 | -0.046 | 2.388 | 0.22 | 0.36 | 0.12 | 0.06 | 0.11 | 0.07 |
| 3 | 137453126 | rs201689 | 1.798 | 0.398 | 1.556 | -0.05 | 0.012 | 0.019 | 0.019 | 0 | 0.59 | 0.32 | 0.72 | 0.76 | 0.67 | 0.71 |
| 3 | 137457771 | rs10490795 | -0.796 | 0.366 | 1.684 | -0.062 | -0.025 | -0.023 | -0.023 | 0 | 0.14 | 0.15 | 0.06 | 0.07 | 0.08 | 0.15 |
| 3 | 137462071 | rs6804350 | -1.954 | 0.359 | 1.696 | -0.115 | -0.013 | -0.034 | -0.034 | 0 | 0.41 | 0.54 | 0.27 | 0.2 | 0.33 | 0.29 |
| 3 | 137466335 | rs16847421 | -0.505 | 0.396 | 1.792 | -0.175 | -0.017 | 0.013 | 0.013 | 0.137 | 0.06 | 0.25 | 0.04 | 0.04 | 0.02 | 0.06 |
| 3 | 137481342 | rs16847433 | -0.331 | 0.392 | 1.794 | -0.175 | -0.016 | -0.002 | -0.002 | 0 | 0.06 | 0.1 | 0.03 | 0.04 | 0.02 | 0.06 |
| 3 | 137481510 | rs11717337 | 1.173 | 0.065 | 1.098 | -0.342 | -0.021 | -0.023 | -0.023 | 0.432 | 0.11 | 0.01 | 0.09 | 0.08 | 0.14 | 0.08 |
| 3 | 137482363 | rs149863 | 0.755 | 0.118 | 1.282 | -0.435 | 0.001 | -0.08 | -0.08 | 0 | 0.73 | 0.43 | 0.79 | 0.87 | 0.74 | 0.86 |
| 3 | 137483362 | rs201718 | 1.976 | 0.091 | 1.184 | -0.421 | 0.033 | 0.114 | 0.114 | 0 | 0.49 | 0.24 | 0.62 | 0.78 | 0.53 | 0.63 |
| 3 | 137487504 | rs157280 | 0.064 | 0.446 | 1.353 | -0.384 | 0.009 | -0.037 | -0.037 | 0.117 | 0.79 | 0.63 | 0.82 | 0.91 | 0.76 | 0.91 |
| 3 | 137498659 | rs201733 | 0.792 | 0.084 | 1.234 | -0.335 | -0.072 | -0.106 | -0.106 | 0 | 0.84 | 0.48 | 0.91 | 0.97 | 0.93 | 0.86 |
| 3 | 137502980 | rs149869 | 0.785 | 0.111 | 1.133 | -0.331 | -0.064 | -0.1 | -0.1 | 0 | 0.84 | 0.43 | 0.9 | 0.97 | 0.93 | 0.86 |
| 3 | 137504684 | rs6764959 | -0.642 | 0.652 | 0.451 | -0.293 | -0.018 | 0.056 | 0.056 | 0.017 | 0.3 | 0.2 | 0.27 | 0.14 | 0.37 | 0.16 |
| 3 | 137508790 | rs201739 | 0.343 | 0.223 | 0.574 | -0.033 | -0.014 | -0.04 | -0.04 | 0 | 0.91 | 0.86 | 0.94 | 0.99 | 0.94 | 0.87 |
| 3 | 137511135 | rs1442583 | -1.375 | 0.537 | 0.737 | 0.832 | 0.009 | 0.103 | 0.103 | 0.009 | 0.07 | 0.16 | 0.18 | 0.27 | 0.07 | 0.11 |
| 3 | 137512736 | rs6798439 | -0.567 | 0.489 | 0.633 | 0.323 | 0.009 | 0.009 | 0.009 | 0 | 0.25 | 0.21 | 0.19 | 0.05 | 0.27 | 0.13 |
| 3 | 137515661 | rs13083385 | -0.487 | 0.496 | 0.629 | 0.315 | 0.013 | 0.02 | 0.02 | 0 | 0.24 | 0.17 | 0.18 | 0.05 | 0.27 | 0.13 |
| 3 | 137515989 | rs1372729 | -2.048 | 0.412 | 0.94 | 0.211 | 0.038 | 0.043 | 0.043 | 0 | 0.45 | 0.74 | 0.53 | 0.39 | 0.43 | 0.45 |
| 3 | 137519670 | rs1442585 | -2.006 | 0.863 | 0.67 | -0.046 | 0.014 | 0.07 | 0.07 | 0 | 0.49 | 0.74 | 0.55 | 0.4 | 0.5 | 0.49 |
| 3 | 137521204 | rs17548687 | -0.209 | 0.888 | 0.859 | -0.011 | -0.001 | -0.011 | -0.011 | 0 | 0.12 | 0.17 | 0.16 | 0.07 | 0.11 | 0.21 |
| 3 | 137522609 | rs2024369 | -2.027 | 0.864 | 0.95 | 0.713 | 0.036 | 0.063 | 0.063 | 0 | 0.51 | 0.72 | 0.55 | 0.65 | 0.47 | 0.48 |
| 3 | 137524754 | rs2024370 | -0.063 | 0.833 | 1.04 | 0.732 | 0 | -0.003 | -0.003 | 0 | 0.1 | 0.09 | 0.11 | 0.05 | 0.18 | 0.1 |
| 3 | 137526940 | rs12488404 | -1.5 | 0.801 | 1.19 | 0.791 | -0.005 | 0.091 | 0.091 | 0 | 0.24 | 0.44 | 0.29 | 0.39 | 0.29 | 0.34 |

| | | | | | | | | | | | | | | | | |
|---|---|---|---|---|---|---|---|---|---|---|---|---|---|---|---|---|
| 3 | 137534327 | rs157275 | 0.337 | 0.876 | 0.931 | 0.65 | 0.003 | 0.009 | 0.009 | 0 | 0.86 | 0.75 | 0.8 | 0.8 | 0.89 | 0.84 |
| 3 | 137535110 | rs157276 | 2.521 | 0.941 | 0.647 | 0.419 | -0.03 | 0.093 | 0.093 | 0 | 0.45 | 0.22 | 0.41 | 0.28 | 0.46 | 0.42 |
| 3 | 137537074 | rs201679 | 1.461 | 0.796 | 1.088 | -0.134 | 0.003 | 0.013 | 0.013 | 0.015 | 0.6 | 0.37 | 0.62 | 0.77 | 0.6 | 0.72 |
| 3 | 137538158 | rs201681 | 0.721 | 0.72 | 1.138 | -0.138 | -0.034 | -0.055 | -0.055 | 1.088 | 0.71 | 0.5 | 0.72 | 0.8 | 0.8 | 0.83 |
| 3 | 137540972 | rs9873940 | 0.653 | 0.515 | 1.357 | 0.088 | -0.032 | -0.075 | -0.075 | 0 | 0.14 | 0.22 | 0.08 | 0 | 0.09 | 0.01 |
| 3 | 137543708 | rs4678382 | -0.268 | 0.3 | 0.73 | 0.076 | -0.011 | -0.002 | -0.002 | 0 | 0.11 | 0.13 | 0.07 | 0.1 | 0.05 | 0.11 |
| 3 | 137543903 | rs11706822 | -0.153 | 0.327 | 0.822 | 0.044 | 0.011 | -0.032 | -0.032 | 0 | 0.11 | 0.14 | 0.11 | 0.03 | 0.19 | 0.12 |
| 3 | 137544378 | rs149999 | 2.113 | 0.375 | 0.839 | 0.03 | 0.077 | 0.119 | 0.119 | 0.806 | 0.53 | 0.18 | 0.55 | 0.67 | 0.59 | 0.65 |
| 3 | 137546102 | rs9872573 | 1.208 | 0.781 | 0.501 | -0.301 | -0.059 | 0.032 | 0.032 | 0 | 0.1 | 0.01 | 0.09 | 0.05 | 0.16 | 0.07 |
| 3 | 137552109 | rs10935269 | -1.546 | 0.105 | 1.079 | 0.113 | -0.009 | -0.029 | -0.029 | 0 | 0.22 | 0.28 | 0.18 | 0.13 | 0.25 | 0.22 |
| 3 | 137568499 | rs6791229 | 0.536 | 0.182 | 1.334 | 0.301 | -0.028 | -0.097 | -0.097 | 0.081 | 0.14 | 0.33 | 0.08 | 0 | 0.09 | 0.01 |
| 3 | 137568920 | rs16847557 | -1.143 | 0.385 | 0.889 | 0.646 | 0.019 | -0.024 | -0.024 | 0 | 0.19 | 0.1 | 0.23 | 0.33 | 0.12 | 0.24 |
| 3 | 137573307 | rs1902576 | 1.884 | 0.661 | 1.047 | 0.216 | 0.006 | 0.009 | 0.009 | 0 | 0.61 | 0.4 | 0.61 | 0.7 | 0.67 | 0.66 |
| 3 | 137574640 | rs2124865 | -0.666 | 0.655 | 1.051 | 0.186 | 0.023 | -0.032 | -0.032 | 0 | 0.08 | 0.13 | 0.06 | 0.02 | 0.11 | 0.11 |
| 3 | 137584058 | rs1350287 | 1.959 | 0.646 | 0.81 | 0.194 | 0.034 | 0.083 | 0.083 | 0.151 | 0.6 | 0.16 | 0.6 | 0.7 | 0.67 | 0.66 |
| 3 | 137600246 | rs4678385 | 0.512 | 0.582 | 0.813 | 0.318 | 0.002 | -0.002 | -0.002 | 0.018 | 0.83 | 0.84 | 0.76 | 0.74 | 0.87 | 0.79 |
| 3 | 137623229 | rs9832826 | 0.3 | 0.557 | 0.96 | 0.249 | -0.029 | -0.05 | -0.05 | 0.002 | 0.14 | 0.26 | 0.08 | 0.03 | 0.08 | 0.02 |
| 3 | 137624652 | rs1978983 | 1.275 | 0.642 | 0.992 | 0.257 | -0.029 | -0.022 | -0.022 | 1.169 | 0.69 | 0.57 | 0.67 | 0.71 | 0.77 | 0.77 |
| 3 | 137637478 | rs988140 | 0.479 | 0.625 | 0.862 | 0.294 | 0.002 | -0.002 | -0.002 | 0 | 0.83 | 0.84 | 0.76 | 0.73 | 0.87 | 0.79 |
| 3 | 137644546 | rs6439768 | 2.007 | 0.848 | 0.908 | 0.167 | 0.01 | 0.026 | 0.026 | 0 | 0.59 | 0.34 | 0.59 | 0.7 | 0.64 | 0.64 |
| 3 | 137644967 | rs1429775 | 2.007 | 0.848 | 0.908 | 0.167 | 0.01 | 0.026 | 0.026 | 0 | 0.59 | 0.34 | 0.59 | 0.7 | 0.64 | 0.64 |
| 3 | 137646826 | rs16847604 | -0.777 | 0.871 | 1.028 | 0.157 | 0.038 | -0.045 | -0.045 | 5.405 | 0.09 | 0.22 | 0.07 | 0.01 | 0.12 | 0.11 |
| 3 | 137650736 | rs16847609 | 0.376 | 0.852 | 1.082 | 0.146 | -0.029 | -0.047 | -0.047 | 0 | 0.14 | 0.26 | 0.09 | 0.04 | 0.08 | 0.02 |
| 3 | 137653568 | rs17282551 | 0.348 | 0.833 | 1.048 | 0.148 | -0.022 | 0.001 | 0.001 | 0 | 0.08 | 0 | 0.08 | 0.05 | 0.08 | 0.04 |
| 3 | 137655693 | rs16847638 | 0.661 | 0.83 | 1.048 | 0.15 | -0.007 | -0.04 | -0.04 | 0 | 0.08 | 0 | 0.16 | 0.2 | 0.06 | 0.18 |
| 3 | 137657512 | rs13078457 | -0.916 | 0.844 | 1.141 | 0.144 | -0.011 | -0.111 | -0.111 | 0 | 0.23 | 0.38 | 0.15 | 0.05 | 0.2 | 0.13 |
| 3 | 137659603 | rs1429760 | 2.029 | 0.881 | 0.97 | 0.164 | 0.01 | 0.026 | 0.026 | 0 | 0.59 | 0.34 | 0.59 | 0.7 | 0.64 | 0.64 |
| 3 | 137664132 | rs1348564 | 1.964 | 0.833 | 0.98 | 0.17 | 0 | -0.001 | -0.001 | 0 | 0.6 | 0.43 | 0.59 | 0.7 | 0.64 | 0.64 |
| 3 | 137666147 | rs1444576 | 1.146 | 0.694 | 1.011 | 0.188 | -0.025 | -0.017 | -0.017 | 0 | 0.69 | 0.58 | 0.66 | 0.7 | 0.76 | 0.76 |
| 3 | 137669856 | rs1429764 | 1.999 | 0.734 | 0.883 | 0.184 | 0.011 | 0.022 | 0.022 | 0 | 0.6 | 0.34 | 0.59 | 0.7 | 0.65 | 0.65 |
| 3 | 137678882 | rs1530442 | -0.979 | 0.732 | 0.692 | 0.203 | 0.027 | -0.047 | -0.047 | 0 | 0.09 | 0.2 | 0.06 | 0.01 | 0.12 | 0.11 |
| 3 | 137691793 | rs11717942 | -0.61 | 0.831 | 0.63 | 0.21 | 0.024 | -0.037 | -0.037 | 0 | 0.08 | 0.1 | 0.06 | 0.01 | 0.12 | 0.11 |
| 3 | 137708155 | rs12635524 | 1.832 | 0.785 | 0.551 | 0.228 | 0.017 | 0.09 | 0.09 | 0.248 | 0.6 | 0.15 | 0.58 | 0.7 | 0.64 | 0.67 |
| 3 | 137721875 | rs4678398 | -1.751 | 0.941 | 0.084 | 0.467 | -0.001 | -0.002 | -0.002 | 0.03 | 0.25 | 0.24 | 0.3 | 0.26 | 0.27 | 0.33 |
| 3 | 137726704 | rs3755753 | 0.582 | 1.14 | -0.394 | 0.933 | 0.002 | -0.02 | -0.02 | 0 | 0.79 | 0.82 | 0.63 | 0.57 | 0.77 | 0.7 |
| 3 | 137736469 | rs7637370 | 1.746 | 1.282 | -1.222 | 0.393 | -0.029 | 0.128 | 0.128 | 0.244 | 0.52 | 0.16 | 0.3 | 0.29 | 0.47 | 0.44 |
| 3 | 137744531 | rs6766201 | -0.035 | 1.361 | -1.15 | 0.419 | 0.002 | -0.003 | -0.003 | 0 | 0.07 | 0.07 | 0.1 | 0.03 | 0.13 | 0.11 |
| 3 | 137746121 | rs12495548 | -0.207 | 1.37 | -1.2 | 0.423 | 0.027 | -0.028 | -0.028 | 0 | 0.08 | 0.13 | 0.11 | 0.03 | 0.13 | 0.11 |
| 3 | 137748979 | rs6780412 | 1.235 | 1.362 | -1.223 | 0.419 | 0.013 | 0.042 | 0.042 | 0.022 | 0.6 | 0.24 | 0.4 | 0.33 | 0.6 | 0.55 |
| 3 | 137750672 | rs11707222 | -0.285 | 1.575 | -1.177 | 0.426 | 0.027 | -0.029 | -0.029 | 0.019 | 0.09 | 0.14 | 0.11 | 0.03 | 0.13 | 0.11 |
| 3 | 137760156 | rs329129 | 1.232 | 1.233 | -1.801 | 0.637 | -0.019 | 0.056 | 0.056 | 0.128 | 0.54 | 0.27 | 0.31 | 0.34 | 0.48 | 0.49 |
| 3 | 137764127 | rs11925123 | 0.212 | 1.246 | -1.88 | 0.615 | -0.036 | 0.024 | 0.024 | 0.073 | 0.17 | 0.01 | 0.13 | 0.07 | 0.16 | 0.06 |
| 3 | 137767073 | rs329138 | 0.583 | 0.928 | -1.469 | 0.683 | -0.019 | 0.061 | 0.061 | 0.208 | 0.54 | 0.27 | 0.31 | 0.33 | 0.48 | 0.49 |
| 3 | 137771152 | rs922688 | 0.552 | 1.003 | -1.888 | 0.65 | -0.104 | 0.076 | 0.076 | 0.891 | 0.57 | 0.07 | 0.48 | 0.77 | 0.51 | 0.68 |
| 3 | 137776539 | rs436145 | 0.925 | -0.076 | -0.958 | 0.975 | -0.047 | 0.04 | 0.04 | 0.033 | 0.8 | 0.33 | 0.78 | 0.63 | 0.86 | 0.72 |
| 3 | 137779492 | rs2724713 | 0.923 | -0.084 | -0.962 | 0.971 | -0.047 | 0.041 | 0.041 | 0.058 | 0.8 | 0.33 | 0.78 | 0.64 | 0.86 | 0.72 |
| 3 | 137781923 | rs329388 | 1.023 | -0.139 | -0.736 | 0.948 | -0.018 | 0.009 | 0.009 | 0.119 | 0.79 | 0.24 | 0.77 | 0.63 | 0.86 | 0.72 |
| 3 | 137783700 | rs179282 | 1.778 | -0.264 | -1.344 | 0.33 | 0.041 | 0.009 | 0.009 | 0 | 0.66 | 0.19 | 0.68 | 0.62 | 0.69 | 0.65 |
| 3 | 137786093 | rs365504 | 0.435 | -0.363 | -0.723 | 0.341 | 0.007 | -0.009 | -0.009 | 1.204 | 0.85 | 0.82 | 0.84 | 0.63 | 0.87 | 0.72 |
| 3 | 137787192 | rs446644 | 0.962 | -0.39 | 0.739 | 0.329 | -0.042 | 0.052 | 0.052 | 0.107 | 0.81 | 0.31 | 0.78 | 0.63 | 0.86 | 0.72 |
| 3 | 137787282 | rs419075 | 0.774 | -0.39 | 1.112 | 0.329 | -0.042 | 0.073 | 0.073 | 0.042 | 0.82 | 0.55 | 0.8 | 0.63 | 0.86 | 0.72 |
| 3 | 137789903 | rs2622704 | 1.749 | -0.418 | 1.057 | 0.302 | -0.006 | 0 | 0 | 0 | 0.69 | 0.29 | 0.69 | 0.61 | 0.69 | 0.65 |
| 3 | 137798155 | rs2724697 | 1.764 | -0.571 | 1.038 | 0.237 | -0.035 | 0 | 0 | 0 | 0.69 | 0.39 | 0.7 | 0.61 | 0.69 | 0.67 |
| 3 | 137799498 | rs405602 | 1.693 | -0.571 | 0.999 | 0.237 | -0.029 | 0.001 | 0.001 | 0 | 0.69 | 0.46 | 0.71 | 0.61 | 0.69 | 0.67 |
| 3 | 137807259 | rs2724693 | 2.019 | -0.612 | 0.953 | 0.255 | 0.036 | 0.001 | 0.001 | 0 | 0.67 | 0.19 | 0.69 | 0.61 | 0.69 | 0.67 |
| 3 | 137814703 | rs405265 | 0.382 | -0.615 | 0.903 | 0.253 | 0.009 | -0.012 | -0.012 | 0 | 0.86 | 0.83 | 0.84 | 0.63 | 0.87 | 0.74 |
| 3 | 137817829 | rs2622723 | 1.859 | -0.656 | -0.552 | -0.44 | -0.168 | -0.167 | -0.167 | 0 | 0.36 | 0.02 | 0.49 | 0.51 | 0.41 | 0.47 |
| 3 | 137818099 | rs11928535 | -0.082 | -0.672 | -0.286 | -0.651 | -0.009 | -0.078 | -0.078 | 0 | 0.16 | 0.32 | 0.12 | 0.01 | 0.18 | 0.07 |
| 3 | 137835746 | rs9857908 | 0.755 | -0.972 | -0.603 | -1.032 | 0.056 | 0.074 | 0.074 | 0 | 0.56 | 0.13 | 0.64 | 0.63 | 0.6 | 0.58 |
| 3 | 137843106 | rs2170309 | 0.161 | -0.926 | -0.315 | -0.999 | -0.001 | -0.001 | -0.001 | 0 | 0.67 | 0.64 | 0.72 | 0.64 | 0.67 | 0.62 |
| 3 | 137844645 | rs2622695 | 0.31 | -0.948 | -0.046 | -0.766 | -0.016 | -0.001 | -0.001 | 0 | 0.66 | 0.46 | 0.71 | 0.59 | 0.68 | 0.62 |
| 3 | 137850003 | rs2724691 | 0.18 | -0.761 | -0.181 | -0.666 | -0.006 | 0.006 | 0.006 | 0 | 0.65 | 0.59 | 0.72 | 0.55 | 0.66 | 0.58 |
| 3 | 137854258 | rs1158066 | 0.921 | -0.348 | -0.334 | -0.696 | -0.043 | 0.142 | 0.142 | 0 | 0.32 | 0.06 | 0.22 | 0.05 | 0.32 | 0.19 |
| 3 | 137867667 | rs2622708 | 0.109 | -0.433 | 0.113 | -0.43 | 0.003 | -0.039 | -0.039 | 0 | 0.69 | 0.81 | 0.75 | 0.55 | 0.68 | 0.6 |
| 3 | 137880743 | rs2622736 | 0.306 | -0.441 | 0.181 | -0.502 | -0.011 | 0.017 | 0.017 | 0 | 0.68 | 0.46 | 0.7 | 0.55 | 0.68 | 0.6 |
| 3 | 137882949 | rs2724712 | 0.296 | -0.431 | 0.155 | -0.505 | -0.013 | 0.02 | 0.02 | 0 | 0.68 | 0.48 | 0.71 | 0.55 | 0.68 | 0.6 |
| 3 | 137897905 | rs774807 | 0.297 | -0.405 | -0.105 | -0.482 | -0.008 | 0.016 | 0.016 | 0 | 0.68 | 0.56 | 0.71 | 0.54 | 0.68 | 0.6 |
| 3 | 137899898 | rs774813 | 0.225 | -0.383 | -0.29 | -0.471 | -0.001 | 0.001 | 0.001 | 0 | 0.68 | 0.63 | 0.72 | 0.54 | 0.68 | 0.59 |
| 3 | 137921391 | rs17204145 | 0.026 | -0.08 | -1.049 | -1.152 | -0.068 | 0 | 0 | 0 | 0.22 | 0.02 | 0.16 | 0.16 | 0.29 | 0.15 |

| | | | | | | | | | | | | | | | | |
|---|---|---|---|---|---|---|---|---|---|---|---|---|---|---|---|---|
| 3 | 137925562 | rs13314286 | 1.82 | -0.007 | -1.254 | -1.22 | -0.116 | -0.078 | -0.078 | 0 | 0.26 | 0.01 | 0.45 | 0.72 | 0.34 | 0.43 |
| 3 | 137940324 | rs9862439 | 0.397 | -0.38 | -0.744 | -1.837 | -0.001 | 0.066 | 0.066 | 0 | 0.49 | 0.63 | 0.7 | 0.78 | 0.52 | 0.55 |
| 3 | 137970907 | rs7631734 | 0.634 | -0.43 | -0.592 | -1.874 | -0.03 | 0.186 | 0.186 | 0 | 0.46 | 0.14 | 0.65 | 0.83 | 0.53 | 0.56 |
| 3 | 137979007 | rs7612622 | 0.398 | -0.288 | -0.789 | -1.718 | -0.007 | 0.111 | 0.111 | 0 | 0.48 | 0.7 | 0.68 | 0.78 | 0.5 | 0.55 |
| 3 | 138008140 | rs3732840 | 0.21 | -0.854 | -0.249 | -1.181 | 0.018 | 0.049 | 0.049 | 0 | 0.63 | 0.73 | 0.83 | 0.85 | 0.7 | 0.74 |
| 3 | 138025136 | rs3856636 | 0.423 | -1.026 | -0.17 | -1.24 | 0.04 | 0.118 | 0.118 | 0 | 0.62 | 0.8 | 0.84 | 0.85 | 0.71 | 0.73 |
| 3 | 138025396 | rs9846480 | 1.119 | -0.947 | -0.127 | -1.249 | 0.041 | 0.203 | 0.203 | 0 | 0.47 | 0.19 | 0.71 | 0.82 | 0.64 | 0.56 |
| 3 | 138052754 | rs6807945 | -0.378 | 0.047 | 0.606 | -0.635 | -0.01 | -0.01 | -0.01 | 0.169 | 0.87 | 0.89 | 0.93 | 0.76 | 0.85 | 0.91 |
| 3 | 138053187 | rs4678408 | 1.571 | -0.367 | 0.239 | -1.022 | -0.003 | 0.149 | 0.149 | 0.007 | 0.56 | 0.16 | 0.72 | 0.77 | 0.64 | 0.71 |
| 3 | 138057719 | rs4678409 | 1.305 | 0.04 | 1.172 | -0.793 | 0.007 | -0.033 | -0.033 | 0 | 0.66 | 0.59 | 0.78 | 0.73 | 0.67 | 0.71 |
| 3 | 138076206 | rs1201763 | 0.185 | 0.057 | 1.071 | -0.847 | -0.035 | 0.101 | 0.101 | 0 | 0.77 | 0.95 | 0.85 | 0.97 | 0.77 | 0.8 |
| 3 | 138078198 | rs730158 | 1.23 | 0.039 | 1.222 | -0.844 | -0.011 | 0.026 | 0.026 | 0 | 0.67 | 0.77 | 0.8 | 0.73 | 0.67 | 0.71 |
| 3 | 138091140 | rs1199334 | -0.078 | 0.087 | 1.338 | -0.816 | -0.001 | -0.051 | -0.051 | 0.444 | 0.8 | 0.72 | 0.83 | 0.97 | 0.82 | 0.81 |
| 3 | 138091701 | rs1199333 | -0.295 | 0.086 | 1.35 | -0.816 | -0.007 | 0.052 | 0.052 | 0 | 0.81 | 0.94 | 0.86 | 0.97 | 0.82 | 0.81 |
| 3 | 138094605 | rs940233 | 1.422 | 0.101 | 1.447 | -0.795 | -0.002 | 0.011 | 0.011 | 0 | 0.65 | 0.38 | 0.75 | 0.72 | 0.67 | 0.71 |
| 3 | 138096097 | rs12695685 | -0.048 | 0.16 | 1.285 | -0.789 | 0.002 | -0.028 | -0.028 | 0.044 | 0.12 | 0.14 | 0.07 | 0.03 | 0.15 | 0.09 |
| 3 | 138096512 | rs4678417 | 1.505 | 0.149 | 1.263 | -0.792 | 0.005 | -0.02 | -0.02 | 1.961 | 0.66 | 0.51 | 0.76 | 0.72 | 0.66 | 0.71 |
| 3 | 138097313 | rs253662 | -0.307 | 0.145 | 1.199 | -0.789 | -0.009 | 0.052 | 0.052 | 0 | 0.81 | 0.94 | 0.86 | 0.97 | 0.82 | 0.81 |
| 3 | 138101529 | rs1199330 | -0.785 | 0.176 | 1.211 | -0.776 | 0.052 | 0.114 | 0.114 | 0.21 | 0.11 | 0.37 | 0.08 | 0.24 | 0.11 | 0.09 |
| 3 | 138106698 | rs1209710 | 0.22 | 0.191 | 0.627 | -0.814 | -0.037 | 0.113 | 0.113 | 0 | 0.77 | 0.95 | 0.85 | 0.98 | 0.77 | 0.8 |
| 3 | 138119952 | rs2306374 | -0.03 | 0.211 | 0.697 | -0.747 | -0.027 | 0.04 | 0.04 | 0 | 0.12 | 0.07 | 0.06 | 0.03 | 0.15 | 0.09 |
| 3 | 138122122 | rs9818870 | -0.03 | 0.163 | 0.707 | -0.747 | -0.019 | 0.027 | 0.027 | 0 | 0.12 | 0.1 | 0.07 | 0.03 | 0.14 | 0.09 |
| 3 | 138124114 | rs2293251 | -0.025 | 0.181 | 0.508 | -0.841 | -0.025 | 0.04 | 0.04 | 0 | 0.12 | 0.07 | 0.07 | 0.03 | 0.14 | 0.09 |
| 3 | 138126481 | rs1720827 | 0.255 | 0.059 | 0.529 | -0.73 | -0.032 | 0.099 | 0.099 | 0 | 0.78 | 0.95 | 0.85 | 0.98 | 0.78 | 0.79 |
| 3 | 138130484 | rs7626349 | 0.63 | 0.044 | 0.789 | -0.44 | 0.003 | 0.015 | 0.015 | 0.005 | 0.71 | 0.56 | 0.75 | 0.63 | 0.69 | 0.77 |
| 3 | 138133412 | rs1564406 | 0.532 | 0.039 | 0.696 | -0.413 | -0.002 | 0.009 | 0.009 | 0.087 | 0.7 | 0.51 | 0.74 | 0.63 | 0.68 | 0.77 |
| 3 | 138151971 | rs1679153 | -1.539 | -0.129 | 0.966 | 0.018 | -0.002 | 0.202 | 0.202 | 0 | 0.14 | 0.36 | 0.07 | 0.39 | 0.09 | 0.14 |
| 3 | 138152940 | rs1679181 | -1.269 | -0.085 | 1.062 | 0.137 | 0.028 | 0.352 | 0.352 | 0.003 | 0.09 | 0.38 | 0.06 | 0.4 | 0.05 | 0.13 |
| 3 | 138177119 | rs808925 | 0.026 | -0.195 | 0.456 | -0.679 | 0 | -0.026 | -0.026 | 0.01 | 0.71 | 0.78 | 0.82 | 0.94 | 0.74 | 0.74 |
| 3 | 138184885 | rs11709290 | 0.7 | -0.548 | 0.921 | -0.52 | -0.012 | 0 | 0 | 0.171 | 0.62 | 0.48 | 0.78 | 0.55 | 0.72 | 0.64 |
| 3 | 138188485 | rs13316499 | 1.149 | -0.089 | 0.742 | -0.866 | -0.007 | -0.008 | -0.008 | 4.528 | 0.11 | 0.07 | 0.1 | 0 | 0.2 | 0.03 |
| 3 | 138189031 | rs2347469 | -0.415 | -0.129 | 0.656 | -0.883 | -0.006 | -0.058 | -0.058 | 0.146 | 0.7 | 0.72 | 0.82 | 0.94 | 0.75 | 0.75 |
| 3 | 138189658 | rs3813264 | 0.757 | -0.222 | 0.978 | -0.909 | 0.051 | -0.002 | -0.002 | 0 | 0.6 | 0.31 | 0.76 | 0.55 | 0.72 | 0.64 |
| 3 | 138191232 | rs10935282 | 2.088 | -0.867 | 0.122 | -1.241 | -0.145 | -0.095 | -0.095 | 0.004 | 0.36 | 0.05 | 0.54 | 0.53 | 0.44 | 0.48 |
| 3 | 138193977 | rs12630060 | -0.037 | -0.725 | 0.334 | -1.043 | -0.022 | 0.121 | 0.121 | 0 | 0.07 | 0.13 | 0.03 | 0.4 | 0.03 | 0.16 |
| 3 | 138194435 | rs1618357 | 0.74 | -0.45 | -0.236 | -1.737 | 0.013 | 0.148 | 0.148 | 0 | 0.52 | 0.34 | 0.71 | 0.91 | 0.64 | 0.6 |
| 3 | 138218709 | rs7653815 | -0.512 | -1.775 | 0.134 | -0.94 | 0.002 | -0.078 | -0.078 | 0.021 | 0.43 | 0.4 | 0.3 | 0.06 | 0.47 | 0.29 |
| 3 | 138223014 | rs7627685 | -0.497 | -1.777 | 0.133 | -0.867 | 0.002 | -0.065 | -0.065 | 0 | 0.43 | 0.41 | 0.29 | 0.09 | 0.47 | 0.29 |
| 3 | 138244400 | rs9878436 | -0.488 | -1.766 | 0.139 | -0.927 | 0.002 | -0.094 | -0.094 | 0.704 | 0.42 | 0.42 | 0.29 | 0.06 | 0.45 | 0.29 |
| 3 | 138254278 | rs1672927 | -1.426 | -1.978 | 0.224 | -0.731 | -0.044 | 0.042 | 0.042 | 2.3 | 0.52 | 0.81 | 0.36 | 0.45 | 0.5 | 0.4 |
| 3 | 138266596 | rs12632793 | -0.492 | -1.981 | 0.442 | -0.744 | -0.023 | 0.127 | 0.127 | 0 | 0.07 | 0.14 | 0.03 | 0.39 | 0.03 | 0.11 |
| 3 | 138266635 | rs12632794 | -1.107 | -1.975 | 0.308 | -0.741 | -0.024 | 0.251 | 0.251 | 0 | 0.09 | 0.32 | 0.06 | 0.39 | 0.03 | 0.11 |
| 3 | 138289163 | rs1673608 | 1.23 | -1.971 | 0.362 | -0.748 | -0.018 | 0.056 | 0.056 | 2.446 | 0.48 | 0.24 | 0.64 | 0.55 | 0.51 | 0.61 |
| 3 | 138289221 | rs1673607 | 0.272 | -1.948 | 0.213 | -0.774 | 0.002 | -0.082 | -0.082 | 0.367 | 0.57 | 0.59 | 0.71 | 0.94 | 0.53 | 0.71 |
| 3 | 138300769 | rs11916525 | -1.234 | -1.864 | 0.24 | -0.647 | -0.02 | 0.284 | 0.284 | 0 | 0.09 | 0.36 | 0.06 | 0.4 | 0.03 | 0.11 |
| 3 | 138329862 | rs811322 | 0.212 | -1.982 | 0.107 | -0.793 | 0.004 | -0.14 | -0.14 | 0 | 0.58 | 0.55 | 0.73 | 0.98 | 0.56 | 0.78 |
| 3 | 138334927 | rs1672928 | 0.997 | -1.993 | 0.059 | -0.747 | -0.043 | 0.057 | 0.057 | 3.189 | 0.49 | 0.16 | 0.65 | 0.58 | 0.53 | 0.67 |
| 3 | 138339907 | rs3773748 | -1.476 | -1.667 | 0.611 | -0.155 | -0.016 | 0.312 | 0.312 | 0.125 | 0.09 | 0.36 | 0.06 | 0.39 | 0.03 | 0.11 |
| 3 | 138347957 | rs641320 | -2.416 | -1.751 | 0.576 | -0.785 | -0.023 | -0.039 | -0.039 | 0 | 0.14 | 0.75 | 0.1 | 0.02 | 0.06 | 0.05 |
| 3 | 138347988 | rs13043 | -2.691 | -1.796 | 1.378 | -0.729 | 0.012 | -0.001 | -0.001 | 0 | 0.07 | 0.63 | 0.07 | 0 | 0.01 | 0 |
| 3 | 138348081 | rs470678 | -0.539 | -2.16 | 0.658 | -0.356 | -0.008 | 0.01 | 0.01 | 0 | 0.26 | 0.7 | 0.33 | 0.46 | 0.2 | 0.29 |
| 3 | 138348310 | rs17282752 | -0.717 | -1.856 | 0.625 | -1.006 | 0.036 | 0.092 | 0.092 | 0.358 | 0.37 | 0.76 | 0.26 | 0.01 | 0.28 | 0.22 |
| 3 | 138349332 | rs601032 | -0.793 | -1.752 | 0.787 | -0.731 | -0.019 | 0.182 | 0.182 | 0 | 0.73 | 0.99 | 0.86 | 0.98 | 0.78 | 0.73 |
| 3 | 138372984 | rs17548850 | 1.185 | -1.459 | 1.136 | -0.577 | -0.066 | 0.061 | 0.061 | 0 | 0.11 | 0.02 | 0.09 | 0 | 0.21 | 0.03 |
| 3 | 138380069 | rs559160 | 0.687 | -1.449 | 1.251 | -0.57 | -0.02 | 0.234 | 0.234 | 1.511 | 0.48 | 0.2 | 0.64 | 0.98 | 0.51 | 0.69 |
| 3 | 138384073 | rs2305268 | 0.303 | -1.555 | 1.202 | -0.289 | 0.018 | 0.011 | 0.011 | 0.093 | 0.09 | 0.05 | 0.02 | 0.45 | 0.02 | 0.11 |
| 3 | 138400332 | rs12493155 | 1.852 | -1.513 | 1.159 | -0.503 | -0.168 | -0.174 | -0.174 | 0 | 0.33 | 0.02 | 0.57 | 0.44 | 0.43 | 0.47 |
| 3 | 138405597 | rs524164 | 0.943 | -0.793 | 1.696 | -1.411 | -0.001 | 0.077 | 0.077 | 1.057 | 0.49 | 0.35 | 0.66 | 0.98 | 0.52 | 0.69 |
| 3 | 138435158 | rs385254 | 0.918 | -0.78 | 1.852 | -1.405 | 0 | -0.023 | -0.023 | 13.761 | 0.49 | 0.41 | 0.67 | 0.98 | 0.52 | 0.69 |
| 3 | 138443067 | rs361059 | 0.773 | -0.753 | 1.358 | -1.392 | 0 | 0.026 | 0.026 | 0.834 | 0.51 | 0.64 | 0.7 | 0.98 | 0.52 | 0.68 |
| 3 | 138451352 | rs10513055 | 0.029 | -0.605 | 1.324 | -1.33 | -0.024 | 0.035 | 0.035 | 1.595 | 0.18 | 0.11 | 0.12 | 0 | 0.25 | 0.04 |
| 3 | 138460203 | rs500687 | 0.271 | -0.179 | 1.665 | -0.862 | 0.002 | 0.003 | 0.003 | 1.472 | 0.5 | 0.6 | 0.69 | 0.98 | 0.52 | 0.68 |
| 3 | 138568591 | rs7613579 | 0.318 | -0.046 | 0.675 | -0.797 | 0.002 | -0.046 | -0.046 | 0 | 0.5 | 0.54 | 0.68 | 0.98 | 0.52 | 0.69 |
| 3 | 138599529 | rs9882957 | -0.348 | -0.021 | 0.094 | -0.603 | -0.001 | -0.09 | -0.09 | 1.21 | 0.48 | 0.46 | 0.31 | 0.02 | 0.47 | 0.35 |
| 3 | 138638081 | rs17485052 | 0.455 | 0.529 | -1.161 | -0.324 | -0.057 | 0.066 | 0.066 | 0 | 0.12 | 0.02 | 0.09 | 0 | 0.21 | 0.07 |
| 3 | 138640704 | rs11706377 | 0.032 | 0.303 | -0.763 | 0.019 | -0.062 | 0.209 | 0.209 | 1.043 | 0.43 | 0.13 | 0.6 | 0.91 | 0.45 | 0.57 |
| 3 | 138658195 | rs750321 | -1.35 | 0.145 | 0.411 | 1.744 | -0.022 | 0.244 | 0.244 | 0.013 | 0.05 | 0.12 | 0.03 | 0.56 | 0.01 | 0.03 |
| 3 | 138661115 | rs1482976 | -2.211 | 0.021 | 1.12 | 1.719 | -0.002 | 0.474 | 0.474 | 2.207 | 0.08 | 0.38 | 0.27 | 0.6 | 0.01 | 0.06 |
| 3 | 138666922 | rs13064974 | -2.203 | 0.188 | 1.572 | 1.012 | 0.045 | -0.024 | -0.024 | 0 | 0.09 | 0.42 | 0.05 | 0.05 | 0.04 | 0.07 |

| | | | | | | | | | | | | | | | | |
|---|---|---|---|---|---|---|---|---|---|---|---|---|---|---|---|---|
| 3 | 138667669 | rs931250 | -2.183 | -0.062 | 1.596 | 1.013 | -0.012 | -0.022 | -0.022 | 0 | 0.09 | 0.42 | 0.05 | 0.05 | 0.02 | 0.07 |
| 3 | 138673922 | rs2018794 | -2.642 | -0.124 | 1.601 | 1.084 | -0.046 | -0.012 | -0.012 | 0 | 0.11 | 0.45 | 0.29 | 0.08 | 0.02 | 0.2 |
| 3 | 138691912 | rs360711 | 1.872 | 0.754 | 1.416 | 0.836 | 0.011 | -0.056 | -0.056 | 0 | 0.84 | 0.67 | 0.63 | 0.92 | 0.85 | 0.76 |
| 3 | 138693471 | rs360710 | 1.763 | 0.657 | 1.366 | 0.779 | -0.011 | -0.072 | -0.072 | 0 | 0.85 | 0.71 | 0.89 | 0.95 | 0.89 | 0.88 |
| 3 | 138705061 | rs6799036 | -1.514 | 0.63 | 1.236 | 1.447 | 0.003 | 0.043 | 0.043 | 0 | 0.09 | 0.13 | 0.28 | 0.27 | 0.12 | 0.14 |
| 3 | 138707522 | rs1039944 | 1.499 | 0.344 | 2.121 | 1.354 | 0.003 | -0.033 | -0.033 | 0.054 | 0.85 | 0.6 | 0.89 | 0.9 | 0.88 | 0.88 |
| 3 | 138730737 | rs9860483 | 1.359 | 0.322 | 2.258 | 1.25 | -0.009 | -0.068 | -0.068 | 1.826 | 0.86 | 0.77 | 0.91 | 0.96 | 0.89 | 0.9 |
| 3 | 138734214 | rs7640145 | 1.8 | 0.308 | 2.349 | 1.243 | -0.015 | -0.103 | -0.103 | 0.889 | 0.81 | 0.7 | 0.86 | 0.96 | 0.82 | 0.89 |
| 3 | 138736641 | rs9884065 | 1.241 | 0.323 | 2.275 | 1.257 | -0.006 | -0.052 | -0.052 | 0 | 0.87 | 0.83 | 0.92 | 0.96 | 0.89 | 0.9 |
| 3 | 138739617 | rs1720230 | 0.981 | 0.33 | 2.224 | 1.252 | 0.003 | 0.001 | 0.001 | 0 | 0.88 | 0.91 | 0.93 | 0.96 | 0.89 | 0.9 |
| 3 | 138757635 | rs619202 | -1.529 | 0.27 | 2.201 | 1.851 | -0.007 | -0.018 | -0.018 | 0 | 0.14 | 0.11 | 0.12 | 0.35 | 0.16 | 0.13 |
| 3 | 138764229 | rs608841 | -2.477 | 0.202 | 2.246 | 1.83 | 0 | -0.004 | -0.004 | 0 | 0.22 | 0.22 | 0.39 | 0.64 | 0.25 | 0.27 |
| 3 | 138775377 | rs696638 | -1.571 | 0.558 | 2.198 | 1.776 | -0.014 | -0.022 | -0.022 | 0 | 0.15 | 0.11 | 0.13 | 0.37 | 0.18 | 0.13 |
| 3 | 138809935 | rs683094 | 1.626 | -0.586 | 2.442 | 1.306 | -0.067 | 0.08 | 0.08 | 0 | 0.82 | 0.66 | 0.7 | 0.68 | 0.89 | 0.85 |
| 3 | 138814520 | rs610949 | 1.714 | -0.455 | 2.504 | 1.347 | -0.028 | 0.211 | 0.211 | 0 | 0.89 | 0.67 | 0.72 | 0.68 | 0.91 | 0.87 |
| 3 | 138823398 | rs2667303 | 2.356 | -0.479 | 2.394 | 1.417 | -0.02 | 0.005 | 0.005 | 0 | 0.74 | 0.41 | 0.61 | 0.27 | 0.79 | 0.73 |
| 3 | 138831086 | rs584059 | 2.06 | -0.435 | 2.242 | 1.191 | -0.042 | 0.205 | 0.205 | 0.071 | 0.88 | 0.5 | 0.7 | 0.67 | 0.91 | 0.86 |
| 3 | 138832331 | rs676102 | 2.015 | -0.365 | 2.197 | 1.195 | -0.032 | 0.246 | 0.246 | 0.085 | 0.88 | 0.54 | 0.7 | 0.67 | 0.91 | 0.86 |
| 3 | 138840886 | rs940189 | 1.732 | -0.428 | 2.013 | 0.683 | -0.019 | -0.106 | -0.106 | 0.68 | 0.72 | 0.45 | 0.8 | 0.96 | 0.78 | 0.87 |
| 3 | 138841476 | rs939646 | 1.82 | -0.462 | 1.865 | 0.669 | -0.002 | -0.05 | -0.05 | 0 | 0.71 | 0.37 | 0.79 | 0.96 | 0.78 | 0.87 |
| 3 | 138842704 | rs646315 | -1.235 | -0.349 | 1.783 | 0.719 | -0.017 | -0.092 | -0.092 | 0 | 0.21 | 0.31 | 0.13 | 0.04 | 0.14 | 0.12 |
| 3 | 138843703 | rs661149 | -0.98 | -0.469 | 1.007 | 0.621 | 0.005 | 0.036 | 0.036 | 1.933 | 0.22 | 0.16 | 0.15 | 0.03 | 0.21 | 0.11 |
| 3 | 138843870 | rs10935299 | 0.39 | -0.074 | -0.245 | 0.13 | -0.024 | -0.032 | -0.032 | 0.445 | 0.29 | 0.12 | 0.38 | 0.31 | 0.29 | 0.26 |
| 3 | 138847358 | rs685220 | -0.946 | -0.352 | 0.349 | 0.195 | -0.015 | 0.021 | 0.021 | 0.42 | 0.28 | 0.64 | 0.23 | 0.4 | 0.22 | 0.15 |
| 3 | 138848435 | rs17487338 | -0.08 | 0.377 | -0.005 | -0.099 | -0.017 | 0.008 | 0.008 | 0 | 0.09 | 0.06 | 0.09 | 0 | 0.15 | 0.06 |
| 3 | 138848899 | rs16848785 | -0.487 | 0.22 | 0.166 | -0.164 | 0.02 | -0.021 | -0.021 | 0 | 0.2 | 0.3 | 0.41 | 0.11 | 0.29 | 0.22 |
| 3 | 138850756 | rs4894381 | 0.557 | 0.515 | -0.195 | -0.597 | -0.055 | 0.083 | 0.083 | 0 | 0.27 | 0.16 | 0.18 | 0.04 | 0.29 | 0.22 |
| 3 | 138855190 | rs12632107 | 0.538 | 0.195 | -0.206 | -0.301 | -0.08 | -0.085 | -0.085 | 2.608 | 0.31 | 0.07 | 0.2 | 0.4 | 0.31 | 0.28 |
| 3 | 138876040 | rs7615670 | 0.573 | 0.233 | -0.083 | -0.287 | -0.051 | -0.06 | -0.06 | 1.433 | 0.31 | 0.13 | 0.2 | 0.41 | 0.31 | 0.28 |
| 3 | 138877924 | rs6775443 | -0.14 | 0.168 | 0.171 | -0.385 | 0.012 | 0.007 | 0.007 | 0 | 0.38 | 0.49 | 0.28 | 0.41 | 0.38 | 0.29 |
| 3 | 138884724 | rs7626828 | -0.483 | 0.21 | -0.634 | -0.418 | 0.094 | -0.002 | -0.002 | 0.157 | 0.49 | 0.21 | 0.58 | 0.53 | 0.57 | 0.49 |
| 3 | 138885236 | rs12489068 | 0.272 | 0.218 | -0.64 | -0.414 | -0.081 | 0.01 | 0.01 | 0 | 0.18 | 0.05 | 0.38 | 0.12 | 0.26 | 0.21 |
| 3 | 138893426 | rs11719269 | 0.674 | 0.344 | -0.616 | -0.344 | -0.04 | -0.048 | -0.048 | 0 | 0.31 | 0.16 | 0.21 | 0.41 | 0.31 | 0.28 |
| 3 | 138903294 | rs4894396 | -1.214 | 0.153 | -0.786 | -0.418 | 0.055 | -0.053 | -0.053 | 0.421 | 0.61 | 0.85 | 0.69 | 0.54 | 0.66 | 0.57 |
| 3 | 138903842 | rs11925186 | 0.428 | 0.457 | -0.758 | -0.118 | -0.039 | -0.048 | -0.048 | 0 | 0.3 | 0.16 | 0.21 | 0.41 | 0.31 | 0.28 |
| 3 | 138903985 | rs7638110 | -0.325 | 0.3 | -0.171 | 0.28 | -0.003 | -0.037 | -0.037 | 0 | 0.07 | 0.15 | 0.05 | 0 | 0.07 | 0.01 |
| 3 | 138905786 | rs16848873 | -0.213 | 0.327 | -0.197 | 0.29 | -0.001 | -0.035 | -0.035 | 0.066 | 0.07 | 0.15 | 0.05 | 0 | 0.08 | 0.01 |
| 3 | 138916393 | rs11706018 | -0.899 | 0.252 | -0.004 | 0.661 | 0.004 | 0.276 | 0.276 | 0 | 0.08 | 0.29 | 0.09 | 0.37 | 0.08 | 0.03 |
| 3 | 138922545 | rs10804664 | 0.406 | 0.491 | -0.537 | 0.205 | -0.026 | -0.061 | -0.061 | 0 | 0.3 | 0.13 | 0.2 | 0.53 | 0.29 | 0.29 |
| 3 | 138928207 | rs1521558 | -0.427 | 0.317 | -0.602 | -0.115 | 0.017 | 0.035 | 0.035 | 0 | 0.23 | 0.32 | 0.43 | 0.64 | 0.28 | 0.31 |
| 3 | 138934384 | rs13074105 | 0.754 | 0.241 | -0.723 | -0.088 | -0.026 | -0.061 | -0.061 | 0 | 0.3 | 0.13 | 0.2 | 0.53 | 0.29 | 0.29 |
| 3 | 138940053 | rs9825837 | -1.288 | 0.074 | -0.648 | -0.145 | 0.028 | 0.007 | 0.007 | 0.098 | 0.62 | 0.86 | 0.68 | 0.67 | 0.65 | 0.58 |
| 3 | 138954012 | rs7623058 | -0.168 | 0.039 | -0.922 | -0.439 | 0.04 | 0.186 | 0.186 | 0.839 | 0.44 | 0.21 | 0.32 | 0.04 | 0.53 | 0.32 |
| 3 | 138954773 | rs4298067 | -1.718 | -0.583 | -1.807 | -0.715 | -0.024 | -0.107 | -0.107 | 0 | 0.74 | 0.99 | 0.73 | 0.67 | 0.72 | 0.77 |
| 3 | 138959151 | rs4455342 | 1.002 | -0.391 | -1.314 | -0.538 | 0.037 | -0.009 | -0.009 | 0 | 0.1 | 0.04 | 0.27 | 0.11 | 0.09 | 0.16 |
| 3 | 138973582 | rs11712639 | 1.043 | -0.224 | -1.178 | -0.661 | -0.029 | -0.071 | -0.071 | 0 | 0.31 | 0.17 | 0.39 | 0.54 | 0.28 | 0.28 |
| 3 | 138980265 | rs6439841 | -1.106 | -0.552 | -0.686 | -0.738 | -0.003 | 0.049 | 0.049 | 0 | 0.28 | 0.39 | 0.22 | 0.54 | 0.28 | 0.26 |
| 3 | 138981318 | rs3892677 | 0.182 | -0.239 | -0.517 | -0.582 | -0.004 | -0.011 | -0.011 | 0 | 0.12 | 0.11 | 0.11 | 0 | 0.18 | 0.07 |
| 3 | 138985300 | rs1602631 | -0.545 | -0.169 | -1.406 | -0.393 | 0.021 | -0.065 | -0.065 | 0.003 | 0.64 | 0.88 | 0.48 | 0.55 | 0.66 | 0.51 |
| 3 | 138997688 | rs9866054 | 0.219 | -0.246 | -1.346 | -0.261 | 0.008 | -0.136 | -0.136 | 0.66 | 0.3 | 0.04 | 0.52 | 0.46 | 0.26 | 0.45 |
| 3 | 138998584 | rs11919329 | 0.623 | -0.099 | -1.103 | -0.052 | 0.004 | 0 | 0 | 0 | 0.09 | 0.1 | 0.1 | 0.09 | 0.1 | 0.05 |
| 3 | 139004920 | rs10212419 | 0.377 | -0.129 | -0.861 | -0.014 | 0.022 | -0.094 | -0.094 | 1.762 | 0.28 | 0.09 | 0.5 | 0.44 | 0.23 | 0.42 |
| 3 | 139009532 | rs2046965 | 1.138 | 0.072 | -0.999 | -0.149 | -0.039 | -0.014 | -0.014 | 0 | 0.4 | 0.2 | 0.22 | 0.43 | 0.45 | 0.35 |
| 3 | 139021139 | rs4894405 | 0.067 | 0.153 | -0.481 | 0.041 | 0 | 0 | 0 | 0 | 0.72 | 0.74 | 0.44 | 0.56 | 0.71 | 0.58 |
| 3 | 139026525 | rs10935317 | -0.32 | 0.227 | -0.431 | -0.32 | 0.005 | 0.011 | 0.011 | 0 | 0.35 | 0.39 | 0.63 | 0.54 | 0.4 | 0.47 |
| 3 | 139027710 | rs6764916 | 0.852 | 0.247 | -0.132 | 0.414 | -0.011 | 0.196 | 0.196 | 0 | 0.94 | 0.87 | 0.95 | 0.59 | 0.95 | 0.89 |
| 3 | 139028722 | rs1602629 | 1.123 | 0.581 | -0.025 | 0.388 | 0.004 | 0.1 | 0.1 | 0 | 0.83 | 0.66 | 0.86 | 0.58 | 0.81 | 0.82 |
| 3 | 139030783 | rs6781671 | -0.003 | 0.555 | -0.302 | 0.263 | -0.003 | -0.01 | -0.01 | 0.019 | 0.1 | 0.12 | 0.08 | 0.01 | 0.13 | 0.06 |
| 3 | 139031115 | rs11716220 | 0.703 | 0.579 | -0.312 | 0.363 | 0.01 | 0.087 | 0.087 | 0 | 0.83 | 0.66 | 0.85 | 0.59 | 0.79 | 0.82 |
| 3 | 139034049 | rs4894345 | 0.52 | -0.165 | -1.009 | -0.752 | 0.033 | 0.071 | 0.071 | 0 | 0.7 | 0.83 | 0.8 | 0.89 | 0.75 | 0.8 |
| 3 | 139035269 | rs11707053 | 0.536 | -0.359 | -0.882 | -1.563 | 0.017 | 0.049 | 0.049 | 0 | 0.48 | 0.62 | 0.33 | 0.08 | 0.56 | 0.38 |
| 3 | 139040591 | rs6810170 | 0.01 | -0.297 | -0.451 | -1.951 | -0.018 | -0.065 | -0.065 | 0 | 0.26 | 0.4 | 0.15 | 0.08 | 0.19 | 0.17 |
| 3 | 139041241 | rs6798217 | 0.346 | -0.559 | -0.631 | -2.067 | -0.014 | 0.004 | 0.004 | 0 | 0.2 | 0.16 | 0.5 | 0.8 | 0.23 | 0.35 |
| 3 | 139041889 | rs9814754 | 0.444 | -0.258 | -0.411 | -2.25 | -0.005 | -0.015 | -0.015 | 0 | 0.4 | 0.37 | 0.27 | 0.09 | 0.51 | 0.33 |
| 3 | 139042283 | rs6439849 | 0.944 | -0.27 | -0.322 | -2.226 | -0.003 | -0.033 | -0.033 | 0 | 0.13 | 0.19 | 0.09 | 0.04 | 0.1 | 0.07 |
| 3 | 139043477 | rs9820915 | 1.463 | -0.262 | -0.303 | -2.209 | -0.047 | 0.062 | 0.062 | 0 | 0.26 | 0.13 | 0.18 | 0.05 | 0.4 | 0.26 |
| 3 | 139056959 | rs12107249 | 0.52 | -0.322 | 0.138 | -1.54 | -0.014 | -0.041 | -0.041 | 2.071 | 0.13 | 0.18 | 0.07 | 0.03 | 0.09 | 0.07 |
| 3 | 139059066 | rs9848910 | 0.843 | -0.854 | 0.233 | -1.886 | 0.019 | 0.021 | 0.021 | 0 | 0.66 | 0.28 | 0.81 | 0.91 | 0.79 | 0.73 |
| 3 | 139061844 | rs11924964 | 0.465 | -0.726 | 0.394 | -1.876 | -0.021 | -0.053 | -0.053 | 0 | 0.14 | 0.25 | 0.07 | 0.03 | 0.09 | 0.07 |

| | | | | | | | | | | | | | | | | |
|---|---|---|---|---|---|---|---|---|---|---|---|---|---|---|---|---|
| 3 | 139064374 | rs1602628 | 1.841 | -0.765 | 0.218 | -1.898 | 0.002 | 0.009 | 0.009 | 0 | 0.1 | 0.06 | 0.04 | 0.03 | 0.08 | 0.08 |
| 3 | 139067890 | rs13083195 | 0.436 | -0.487 | -0.086 | -2.158 | -0.047 | 0.023 | 0.023 | 0 | 0.2 | 0.08 | 0.5 | 0.87 | 0.23 | 0.33 |
| 3 | 139075938 | rs7833 | 1.679 | -0.593 | -0.09 | -2.224 | -0.087 | 0.312 | 0.312 | 0 | 0.45 | 0.06 | 0.29 | 0.04 | 0.55 | 0.4 |
| 3 | 139083770 | rs1602626 | 2.186 | -0.602 | -0.058 | -2.218 | 0.001 | 0.006 | 0.006 | 0 | 0.1 | 0.07 | 0.04 | 0.03 | 0.08 | 0.08 |
| 3 | 139090736 | rs7373116 | 1.686 | -0.605 | -0.076 | -2.23 | -0.087 | 0.313 | 0.313 | 0 | 0.45 | 0.06 | 0.29 | 0.04 | 0.55 | 0.4 |
| 3 | 139101777 | rs13060697 | 0.365 | -0.565 | 0.049 | -2.196 | -0.01 | 0.01 | 0.01 | 0 | 0.2 | 0.16 | 0.51 | 0.87 | 0.23 | 0.34 |
| 3 | 139110135 | rs10935323 | 0.4 | -0.554 | 0.017 | -2.191 | -0.053 | 0.026 | 0.026 | 0 | 0.2 | 0.08 | 0.5 | 0.87 | 0.23 | 0.33 |
| 3 | 139116581 | rs1599744 | -0.69 | -0.601 | 0.318 | -2.183 | -0.013 | -0.019 | -0.019 | 0 | 0.94 | 0.85 | 0.96 | 0.98 | 0.96 | 0.96 |
| 3 | 139152480 | rs13092507 | 0.412 | -0.581 | -0.053 | -2.189 | -0.045 | 0.026 | 0.026 | 0 | 0.2 | 0.08 | 0.5 | 0.87 | 0.23 | 0.33 |
| 3 | 139163707 | rs7635513 | 0.951 | -0.662 | 0.122 | -2.208 | -0.018 | -0.053 | -0.053 | 0 | 0.1 | 0.22 | 0.04 | 0 | 0.06 | 0.02 |
| 3 | 139164266 | rs2289348 | 2.033 | -0.694 | -0.146 | -2.243 | 0.001 | 0.007 | 0.007 | 0 | 0.11 | 0.07 | 0.05 | 0.03 | 0.09 | 0.1 |
| 3 | 139165108 | rs12495890 | -1.985 | -0.672 | -0.174 | -2.251 | -0.098 | 0.309 | 0.309 | 0 | 0.56 | 0.94 | 0.72 | 0.96 | 0.46 | 0.6 |
| 3 | 139169369 | rs11919679 | -1.066 | -0.648 | -0.127 | -2.242 | -0.036 | -0.073 | -0.073 | 0 | 0.23 | 0.66 | 0.14 | 0.06 | 0.13 | 0.17 |
| 3 | 139176535 | rs9882014 | 0.329 | -0.644 | -0.173 | -2.252 | -0.01 | 0.012 | 0.012 | 0 | 0.21 | 0.15 | 0.51 | 0.87 | 0.23 | 0.34 |
| 3 | 139212730 | rs444264 | -1.913 | -0.694 | -0.257 | -2.057 | -0.089 | 0.278 | 0.278 | 0 | 0.56 | 0.93 | 0.71 | 0.9 | 0.47 | 0.61 |
| 3 | 139217951 | rs2882080 | 1.663 | -0.694 | -0.055 | -1.911 | 0.001 | 0.001 | 0.001 | 0 | 0.12 | 0.11 | 0.06 | 0.03 | 0.09 | 0.09 |
| 3 | 139227224 | rs176986 | -0.278 | -0.679 | 0.089 | -1.856 | -0.004 | -0.027 | -0.027 | 1.762 | 0.09 | 0.36 | 0.06 | 0.02 | 0.04 | 0.11 |
| 3 | 139234514 | rs211584 | 0.936 | -0.759 | 0.057 | -1.918 | -0.041 | -0.072 | -0.072 | 0.788 | 0.77 | 0.33 | 0.86 | 0.95 | 0.87 | 0.83 |
| 3 | 139234533 | rs167186 | 0.141 | -0.687 | 0.089 | -1.869 | -0.003 | -0.027 | -0.027 | 0 | 0.91 | 0.64 | 0.94 | 0.98 | 0.96 | 0.89 |
| 3 | 139238368 | rs184156 | 0.713 | -0.625 | -0.139 | -1.945 | 0.035 | 0.052 | 0.052 | 0 | 0.68 | 0.21 | 0.82 | 0.95 | 0.81 | 0.78 |
| 3 | 139239271 | rs397940 | -0.174 | -0.583 | -0.107 | -1.913 | -0.033 | -0.098 | -0.098 | 0 | 0.79 | 0.53 | 0.87 | 0.95 | 0.86 | 0.8 |
| 3 | 139239323 | rs295488 | 1.307 | -0.391 | -0.576 | -1.94 | -0.058 | 0.326 | 0.326 | 0 | 0.47 | 0.06 | 0.3 | 0.08 | 0.57 | 0.45 |
| 3 | 139246382 | rs295493 | 1.008 | -0.423 | -0.432 | -1.448 | 0.033 | 0.088 | 0.088 | 0 | 0.6 | 0.18 | 0.57 | 0.83 | 0.7 | 0.69 |
| 3 | 139248983 | rs4683502 | 0.562 | -0.041 | -0.396 | -0.169 | -0.015 | 0.025 | 0.025 | 0.02 | 0.09 | 0.05 | 0.22 | 0.46 | 0.13 | 0.13 |
| 3 | 139249598 | rs9821204 | 0.626 | -0.201 | -0.51 | -0.294 | -0.052 | 0.101 | 0.101 | 0 | 0.22 | 0.03 | 0.13 | 0.03 | 0.21 | 0.14 |
| 3 | 139250649 | rs893703 | 0.421 | -0.201 | -0.481 | -0.284 | 0.026 | 0.151 | 0.151 | 0.018 | 0.1 | 0.14 | 0.23 | 0.46 | 0.13 | 0.13 |
| 3 | 139252215 | rs893704 | -1.318 | -0.298 | -0.317 | -0.338 | 0 | -0.002 | -0.002 | 0 | 0.42 | 0.59 | 0.42 | 0.52 | 0.38 | 0.32 |
| 3 | 139253871 | rs12485273 | -0.996 | -0.27 | -0.256 | -0.134 | 0.007 | -0.06 | -0.06 | 0 | 0.19 | 0.47 | 0.19 | 0.12 | 0.16 | 0.13 |
| 3 | 139257603 | rs2071387 | -1.098 | -0.252 | -0.264 | -0.083 | -0.006 | -0.005 | -0.005 | 1.757 | 0.22 | 0.47 | 0.37 | 0.22 | 0.19 | 0.18 |
| 3 | 139259787 | rs176990 | 1.362 | -0.398 | -0.537 | -0.005 | 0.003 | 0.082 | 0.082 | 0 | 0.51 | 0.1 | 0.36 | 0.38 | 0.58 | 0.56 |
| 3 | 139271591 | rs9826668 | -1.24 | -0.25 | -0.543 | -0.374 | -0.001 | -0.025 | -0.025 | 0.175 | 0.44 | 0.54 | 0.5 | 0.25 | 0.4 | 0.32 |
| 3 | 139274323 | rs9822458 | -0.247 | 0.354 | -1.172 | -0.401 | -0.046 | 0.09 | 0.09 | 0 | 0.3 | 0.14 | 0.25 | 0.11 | 0.31 | 0.2 |
| 3 | 139280685 | rs169825 | 0.606 | 0.863 | -1.656 | 0.062 | -0.045 | 0.053 | 0.053 | 0.65 | 0.54 | 0.07 | 0.33 | 0.39 | 0.45 | 0.63 |
| 3 | 139281153 | rs211587 | -0.421 | 1.472 | -0.932 | -0.148 | 0.021 | -0.064 | -0.064 | 0 | 0.84 | 0.72 | 0.84 | 0.99 | 0.8 | 0.97 |
| 3 | 139283462 | rs1497266 | -0.353 | 1.418 | -0.637 | -0.156 | 0.011 | 0.008 | 0.008 | 0 | 0.3 | 0.59 | 0.51 | 0.6 | 0.36 | 0.38 |
| 3 | 139294516 | rs10935336 | -0.134 | 1.702 | -0.047 | -0.16 | 0.031 | 0.082 | 0.082 | 0 | 0.21 | 0.43 | 0.45 | 0.58 | 0.25 | 0.28 |
| 3 | 139295138 | rs167189 | 0.422 | 1.761 | -0.059 | -0.537 | 0.036 | -0.118 | -0.118 | 0.009 | 0.76 | 0.5 | 0.78 | 0.97 | 0.7 | 0.9 |
| 3 | 139298567 | rs12485554 | -0.043 | 1.509 | 0.395 | 0.27 | -0.001 | -0.001 | -0.001 | 0.06 | 0.19 | 0.2 | 0.4 | 0.2 | 0.23 | 0.18 |
| 3 | 139306828 | rs295465 | -0.313 | 1.223 | 0.343 | 0.198 | 0.023 | -0.028 | -0.028 | 0 | 0.88 | 0.6 | 0.91 | 0.97 | 0.89 | 0.92 |
| 3 | 139326484 | rs7639753 | -0.5 | 1.316 | -0.358 | 0.525 | 0.022 | 0.019 | 0.019 | 0 | 0.33 | 0.42 | 0.51 | 0.37 | 0.44 | 0.31 |
| 3 | 139327261 | rs10804666 | -0.439 | 0.972 | -0.754 | 0.539 | -0.01 | 0.056 | 0.056 | 0.011 | 0.81 | 0.46 | 0.8 | 0.73 | 0.84 | 0.85 |
| 3 | 139328475 | rs11920622 | 1.91 | 0.859 | -0.412 | 0.101 | 0.018 | -0.023 | -0.023 | 0.016 | 0.05 | 0.17 | 0.04 | 0 | 0.03 | 0.03 |
| 3 | 139331061 | rs12635533 | 1.009 | 0.754 | -0.218 | 0.081 | 0.012 | -0.011 | -0.011 | 0 | 0.49 | 0.08 | 0.36 | 0.63 | 0.42 | 0.67 |
| 3 | 139336381 | rs9873488 | -0.608 | 0.771 | -0.195 | 0.09 | 0 | -0.001 | -0.001 | 0 | 0.46 | 0.46 | 0.6 | 0.38 | 0.57 | 0.33 |
| 3 | 139348518 | rs16849220 | 0.005 | 0.926 | 0.27 | 0.046 | 0.003 | -0.023 | -0.023 | 1.294 | 0.75 | 0.44 | 0.8 | 0.84 | 0.75 | 0.88 |
| 3 | 139351191 | rs6786606 | 0.946 | 0.902 | 0.214 | -0.086 | 0 | -0.037 | -0.037 | 1.647 | 0.16 | 0.14 | 0.12 | 0 | 0.17 | 0.02 |
| 3 | 139351199 | rs16849222 | 0.378 | 0.822 | 0.067 | -0.244 | 0.002 | 0.008 | 0.008 | 0 | 0.32 | 0.4 | 0.23 | 0.33 | 0.31 | 0.34 |
| 3 | 139353664 | rs2292412 | -0.046 | 0.648 | 0.177 | 0.014 | 0.003 | 0.006 | 0.006 | 0 | 0.5 | 0.53 | 0.59 | 0.71 | 0.47 | 0.61 |
| 3 | 139356796 | rs2291803 | 1.337 | 0.581 | -0.952 | 0.247 | -0.025 | -0.133 | -0.133 | 0.604 | 0.3 | 0.03 | 0.38 | 0.47 | 0.28 | 0.39 |
| 3 | 139359410 | rs4683613 | -0.867 | 0.64 | -0.81 | 0.125 | -0.045 | 0.096 | 0.096 | 0 | 0.6 | 0.79 | 0.65 | 0.82 | 0.58 | 0.71 |
| 3 | 139377406 | rs7625738 | -0.871 | 0.6 | -0.729 | 0.296 | -0.007 | 0.053 | 0.053 | 0 | 0.54 | 0.7 | 0.69 | 0.73 | 0.51 | 0.65 |
| 3 | 139379385 | rs11707887 | 0.636 | 1.077 | -1.549 | 0.079 | -0.028 | -0.132 | -0.132 | 0.048 | 0.29 | 0.03 | 0.37 | 0.48 | 0.28 | 0.37 |
| 3 | 139379940 | rs13075367 | 0.743 | 1.231 | -0.65 | -0.382 | 0 | -0.028 | -0.028 | 0 | 0.13 | 0.09 | 0.1 | 0 | 0.14 | 0.02 |
| 3 | 139387857 | rs7631422 | 0.794 | 0.605 | -0.402 | -0.109 | -0.012 | -0.046 | -0.046 | 0.133 | 0.08 | 0.24 | 0.05 | 0 | 0.06 | 0.06 |
| 3 | 139392368 | rs7639511 | -0.721 | 0.731 | -0.694 | -0.708 | 0.002 | -0.047 | -0.047 | 0 | 0.68 | 0.63 | 0.8 | 0.83 | 0.74 | 0.75 |
| 3 | 139393833 | rs16849307 | -0.225 | 0.924 | -1.236 | -0.551 | 0.002 | -0.003 | -0.003 | 0 | 0.34 | 0.39 | 0.35 | 0.24 | 0.45 | 0.35 |
| 3 | 139404169 | rs4518082 | 0.722 | 0.981 | -1.245 | -0.52 | -0.011 | -0.006 | -0.006 | 0 | 0.26 | 0.2 | 0.26 | 0.24 | 0.37 | 0.28 |
| 3 | 139404858 | rs9845929 | 2.085 | 1.26 | -1.408 | -0.702 | -0.01 | 0.029 | 0.029 | 0 | 0.12 | 0.04 | 0.11 | 0 | 0.17 | 0.02 |
| 3 | 139404982 | rs12634170 | 0.47 | 1.259 | -1.394 | -0.699 | -0.024 | -0.014 | -0.014 | 0.631 | 0.26 | 0.15 | 0.25 | 0.24 | 0.37 | 0.28 |
| 3 | 139405342 | rs10935342 | -0.745 | 1.119 | -1.5 | -0.827 | 0.004 | 0.037 | 0.037 | 2.535 | 0.61 | 0.34 | 0.75 | 0.84 | 0.69 | 0.71 |
| 3 | 139408355 | rs10154999 | 2.21 | 1.02 | -1.467 | -0.738 | -0.007 | 0.017 | 0.017 | 0 | 0.12 | 0.05 | 0.11 | 0 | 0.17 | 0.02 |
| 3 | 139409436 | rs9790066 | 1.438 | 0.714 | -1.854 | -0.462 | -0.048 | -0.081 | -0.081 | 0.09 | 0.13 | 0.02 | 0.29 | 0.33 | 0.19 | 0.13 |
| 3 | 139411031 | rs11720063 | 0.345 | 0.662 | -1.817 | -0.505 | 0.014 | -0.038 | -0.038 | 0 | 0.35 | 0.21 | 0.51 | 0.6 | 0.32 | 0.43 |
| 3 | 139423824 | rs4504138 | -0.082 | 0.679 | -1.765 | -0.685 | -0.016 | 0.185 | 0.185 | 0.605 | 0.46 | 0.21 | 0.61 | 0.83 | 0.48 | 0.61 |
| 3 | 139429937 | rs7648532 | 1.332 | 0.839 | -1.617 | -0.784 | 0.004 | -0.05 | -0.05 | 0 | 0.33 | 0.4 | 0.23 | 0.17 | 0.29 | 0.31 |
| 3 | 139435050 | rs11713380 | -0.055 | 0.834 | -2.258 | -0.692 | 0.011 | 0.189 | 0.189 | 0.468 | 0.46 | 0.2 | 0.6 | 0.83 | 0.44 | 0.61 |
| 3 | 139437905 | rs4389442 | 1.24 | 0.874 | -2.025 | -0.333 | -0.004 | -0.08 | -0.08 | 0.104 | 0.13 | 0.02 | 0.27 | 0.34 | 0.14 | 0.13 |
| 3 | 139447500 | rs10935348 | -0.079 | 0.948 | -1.747 | -0.892 | -0.019 | 0.168 | 0.168 | 3.342 | 0.55 | 0.21 | 0.7 | 0.83 | 0.6 | 0.63 |
| 3 | 139454159 | rs4683686 | 0.02 | 0.857 | -1.287 | -1.121 | 0.007 | -0.07 | -0.07 | 0.758 | 0.78 | 0.34 | 0.81 | 0.98 | 0.76 | 0.76 |

| | | | | | | | | | | | | | | | | |
|---|---|---|---|---|---|---|---|---|---|---|---|---|---|---|---|---|
| 3 | 139455326 | rs13072543 | 0.451 | 0.604 | -1.993 | -1.298 | 0.044 | 0.003 | 0.003 | 0 | 0.44 | 0.21 | 0.6 | 0.61 | 0.41 | 0.42 |
| 3 | 139459271 | rs4073897 | 1.018 | 0.634 | -2.058 | -1.294 | -0.08 | -0.071 | -0.071 | 0 | 0.34 | 0.08 | 0.21 | 0.37 | 0.36 | 0.35 |
| 3 | 139460879 | rs12635210 | 1.43 | 0.371 | -2.266 | -1.505 | 0.022 | -0.093 | -0.093 | 0 | 0.27 | 0.06 | 0.48 | 0.61 | 0.22 | 0.36 |
| 3 | 139464203 | rs9813586 | 1.137 | 0.391 | -1.477 | -1.543 | 0.044 | -0.078 | -0.078 | 2.569 | 0.14 | 0.32 | 0.14 | 0.02 | 0.17 | 0.2 |
| 3 | 139464938 | rs4267617 | -2.421 | 0.372 | -1.43 | -1.526 | 0.018 | -0.035 | -0.035 | 0 | 0.93 | 0.75 | 0.96 | 1 | 0.93 | 0.94 |
| 3 | 139469121 | rs4683428 | 1.229 | 0.328 | -1.546 | -1.548 | -0.083 | -0.071 | -0.071 | 1.755 | 0.34 | 0.07 | 0.21 | 0.37 | 0.36 | 0.34 |
| 3 | 139472488 | rs4568080 | 1.469 | 0.326 | -1.563 | -1.562 | 0.015 | -0.093 | -0.093 | 0 | 0.27 | 0.06 | 0.48 | 0.61 | 0.23 | 0.36 |
| 3 | 139475253 | rs12695704 | 1.167 | 0.275 | -1.021 | -1.306 | 0.023 | -0.165 | -0.165 | 2.224 | 0.27 | 0.48 | 0.24 | 0.02 | 0.3 | 0.22 |
| 3 | 139475978 | rs12487317 | -0.563 | 0.245 | -1.042 | -1.325 | -0.003 | 0.018 | 0.018 | 0 | 0.66 | 0.28 | 0.72 | 0.98 | 0.63 | 0.72 |
| 3 | 139476592 | rs13093713 | 0.407 | -0.193 | -0.632 | -0.763 | -0.011 | 0.001 | 0.001 | 0 | 0.6 | 0.55 | 0.45 | 0.39 | 0.66 | 0.57 |
| 3 | 139477769 | rs12488425 | 0.218 | -0.113 | -1.031 | -0.671 | 0.029 | -0.064 | -0.064 | 0 | 0.32 | 0.1 | 0.5 | 0.61 | 0.27 | 0.37 |
| 3 | 139480123 | rs1374828 | 1.011 | 0.093 | -0.713 | -0.569 | -0.009 | 0.073 | 0.073 | 0 | 0.46 | 0.32 | 0.37 | 0.24 | 0.49 | 0.39 |
| 3 | 139481170 | rs9847705 | 0.55 | 0.793 | -0.251 | -0.069 | 0.029 | 0.037 | 0.037 | 0 | 0.77 | 0.62 | 0.83 | 0.77 | 0.74 | 0.73 |
| 3 | 139483668 | rs9876670 | -0.918 | 0.821 | -0.067 | -0.211 | 0.025 | -0.031 | -0.031 | 0 | 0.93 | 0.74 | 0.95 | 1 | 0.93 | 0.94 |
| 3 | 139489458 | rs11921303 | -0.448 | 0.996 | 0.162 | 0.031 | 0.035 | 0.03 | 0.03 | 0 | 0.19 | 0.32 | 0.14 | 0.22 | 0.23 | 0.24 |
| 3 | 139496471 | rs6772801 | -0.463 | 1.04 | -0.032 | 0.023 | -0.015 | -0.009 | -0.009 | 0 | 0.21 | 0.13 | 0.15 | 0.22 | 0.29 | 0.28 |
| 3 | 139499055 | rs1900627 | -0.647 | 0.972 | 0.03 | 0.205 | 0 | 0 | 0 | 0 | 0.44 | 0.43 | 0.34 | 0.48 | 0.5 | 0.46 |
| 3 | 139500341 | rs1374831 | -1.434 | 0.766 | -0.224 | 0.157 | -0.002 | 0.032 | 0.032 | 0 | 0.84 | 0.97 | 0.92 | 0.87 | 0.88 | 0.79 |
| 3 | 139502118 | rs9289585 | 2.084 | 0.645 | 0.039 | 0.163 | 0.004 | -0.002 | -0.002 | 0 | 0.31 | 0.32 | 0.52 | 0.28 | 0.29 | 0.27 |
| 3 | 139505788 | rs13098819 | 1.082 | 0.337 | 0.544 | 0.52 | 0 | -0.002 | -0.002 | 1.957 | 0.22 | 0.17 | 0.18 | 0.26 | 0.21 | 0.18 |
| 3 | 139506968 | rs2028994 | 0.54 | 0.378 | 0.4 | 0.532 | 0 | 0 | 0 | 0 | 0.53 | 0.53 | 0.68 | 0.5 | 0.58 | 0.57 |
| 3 | 139511746 | rs6771183 | -0.945 | 0.336 | 0.359 | 0.515 | -0.013 | -0.009 | -0.009 | 0.109 | 0.83 | 0.96 | 0.91 | 0.76 | 0.84 | 0.76 |
| 3 | 139513569 | rs9849159 | 1.72 | 0.383 | 0.176 | 0.499 | 0.002 | 0.004 | 0.004 | 0 | 0.37 | 0.32 | 0.57 | 0.28 | 0.4 | 0.34 |
| 3 | 139516804 | rs6777158 | 1.091 | 0.537 | -0.217 | 0.561 | -0.01 | -0.011 | -0.011 | 0.126 | 0.61 | 0.71 | 0.75 | 0.54 | 0.61 | 0.55 |
| 3 | 139521148 | rs2882296 | -1.251 | 0.642 | -0.446 | 0.531 | -0.025 | 0.039 | 0.039 | 0 | 0.85 | 0.99 | 0.94 | 0.91 | 0.89 | 0.79 |
| 3 | 139530894 | rs6789124 | -0.011 | 0.747 | 0.009 | 0.641 | 0.004 | 0.01 | 0.01 | 0 | 0.14 | 0.14 | 0.12 | 0.23 | 0.18 | 0.2 |
| 3 | 139533909 | rs10935353 | -0.656 | 0.744 | -0.174 | 0.645 | -0.003 | 0.003 | 0.003 | 0.112 | 0.46 | 0.59 | 0.37 | 0.59 | 0.47 | 0.42 |
| 3 | 139547504 | rs11716660 | -0.496 | 0.813 | -0.539 | 0.469 | -0.009 | -0.012 | -0.012 | 0 | 0.14 | 0.12 | 0.12 | 0.23 | 0.18 | 0.2 |
| 3 | 139549939 | rs11707795 | -0.515 | 0.793 | -0.66 | 0.225 | -0.005 | 0.034 | 0.034 | 0 | 0.84 | 0.95 | 0.93 | 0.88 | 0.88 | 0.79 |
| 3 | 139550666 | rs1447726 | 0.977 | 0.749 | -0.096 | 0.35 | 0.002 | 0.003 | 0.003 | 0 | 0.66 | 0.58 | 0.75 | 0.6 | 0.67 | 0.55 |
| 3 | 139554677 | rs9823958 | -0.342 | 0.636 | -0.321 | 0.035 | -0.001 | -0.003 | -0.003 | 0 | 0.92 | 0.95 | 0.95 | 1 | 0.91 | 0.91 |
| 3 | 139555222 | rs1868170 | -0.717 | 0.529 | -0.086 | -0.105 | 0.007 | 0.013 | 0.013 | 0 | 0.54 | 0.56 | 0.56 | 0.68 | 0.54 | 0.47 |
| 3 | 139558777 | rs9872548 | 0.172 | 0.469 | 0.056 | -0.346 | 0.005 | 0.008 | 0.008 | 0 | 0.19 | 0.15 | 0.06 | 0.04 | 0.14 | 0.15 |
| 3 | 139560314 | rs9844671 | 1.31 | 0.497 | -0.065 | -0.321 | -0.019 | 0.05 | 0.05 | 0 | 0.12 | 0.01 | 0.03 | 0.04 | 0.08 | 0.14 |
| 3 | 139565057 | rs1374825 | 1.457 | 0.462 | 0.113 | -0.286 | 0.003 | 0.005 | 0.005 | 0 | 0.36 | 0.36 | 0.36 | 0.27 | 0.36 | 0.34 |
| 3 | 139567313 | rs13315295 | -0.223 | 0.278 | 0.085 | -0.314 | 0.007 | 0.016 | 0.016 | 0 | 0.21 | 0.16 | 0.08 | 0.1 | 0.17 | 0.19 |
| 3 | 139567666 | rs6800179 | 0.542 | 0.286 | 0.18 | -0.345 | -0.012 | -0.038 | -0.038 | 0 | 0.87 | 0.79 | 0.92 | 0.94 | 0.88 | 0.88 |
| 3 | 139568279 | rs1374832 | -1.358 | 0.365 | 0.105 | -0.368 | -0.023 | 0.038 | 0.038 | 0 | 0.61 | 0.87 | 0.62 | 0.72 | 0.56 | 0.61 |
| 3 | 139570504 | rs1597079 | -0.028 | 0.652 | 0.689 | -0.139 | 0.01 | 0.004 | 0.004 | 0 | 0.37 | 0.54 | 0.51 | 0.57 | 0.37 | 0.33 |
| 3 | 139573880 | rs12635255 | -0.028 | 0.652 | 0.691 | -0.142 | 0.012 | 0.007 | 0.007 | 0 | 0.37 | 0.52 | 0.51 | 0.57 | 0.37 | 0.33 |
| 3 | 139581879 | rs10935358 | 0.031 | 0.599 | 0.843 | -0.149 | 0.005 | 0.003 | 0.003 | 0 | 0.37 | 0.4 | 0.5 | 0.58 | 0.37 | 0.35 |
| 3 | 139585439 | rs16849528 | 1.261 | 0.827 | 0.448 | 0.151 | -0.034 | -0.087 | -0.087 | 0.818 | 0.21 | 0.04 | 0.36 | 0.34 | 0.21 | 0.16 |
| 3 | 139585818 | rs16846176 | 0.013 | 0.784 | 0.437 | 0.026 | -0.025 | -0.027 | -0.027 | 0.145 | 0.17 | 0.32 | 0.09 | 0.15 | 0.11 | 0.24 |
| 3 | 139589227 | rs9838863 | 0.333 | 0.815 | 0.087 | 0.04 | 0.008 | 0.008 | 0.008 | 0.059 | 0.16 | 0.11 | 0.07 | 0.15 | 0.11 | 0.24 |
| 3 | 139590722 | rs1562494 | -1.832 | 0.913 | -0.025 | 0.073 | -0.019 | 0.026 | 0.026 | 0 | 0.95 | 0.99 | 0.95 | 1 | 0.92 | 0.95 |
| 3 | 139595530 | rs9881047 | -0.412 | 0.871 | -0.023 | 0.024 | -0.001 | 0.018 | 0.018 | 0 | 0.22 | 0.16 | 0.11 | 0.17 | 0.2 | 0.28 |
| 3 | 139606794 | rs2554152 | -0.155 | 0.787 | 0.074 | 0.042 | -0.014 | 0.018 | 0.018 | 1.819 | 0.18 | 0.62 | 0.15 | 0.22 | 0.13 | 0.18 |
| 3 | 139610720 | rs787267 | -1.611 | 0.839 | -0.468 | 0.046 | -0.097 | 0.013 | 0.013 | 0.006 | 0.59 | 0.97 | 0.61 | 0.73 | 0.53 | 0.62 |
| 3 | 139611392 | rs17336118 | 0.914 | 1.326 | 0.117 | 0.368 | -0.003 | 0.005 | 0.005 | 0.047 | 0.06 | 0.04 | 0.02 | 0.05 | 0.05 | 0.12 |
| 3 | 139612162 | rs16849559 | 0.97 | 1.507 | 0.096 | 0.508 | -0.039 | -0.086 | -0.086 | 2.372 | 0.18 | 0.04 | 0.36 | 0.33 | 0.2 | 0.2 |
| 3 | 139614254 | rs13065203 | 1.218 | 1.529 | 0.102 | 0.509 | -0.069 | -0.113 | -0.113 | 0 | 0.17 | 0.02 | 0.35 | 0.32 | 0.2 | 0.16 |
| 3 | 139614923 | rs2349987 | 0.135 | 1.554 | 0.662 | 0.453 | 0.067 | 0.014 | 0.014 | 0 | 0.95 | 0.72 | 0.95 | 1 | 0.95 | 1 |
| 3 | 139615610 | rs983323 | 0.68 | 1.546 | 0.27 | 0.429 | -0.008 | -0.017 | -0.017 | 0 | 0.76 | 0.38 | 0.81 | 0.78 | 0.79 | 0.79 |
| 3 | 139619069 | rs1426053 | 1.353 | 1.383 | 0.052 | 0.349 | -0.065 | 0.086 | 0.086 | 0.006 | 0.5 | 0.05 | 0.39 | 0.29 | 0.53 | 0.44 |
| 3 | 139620397 | rs6439885 | 1.117 | 1.419 | 1.033 | 0.161 | -0.014 | -0.009 | -0.009 | 0 | 0.76 | 0.42 | 0.83 | 0.75 | 0.81 | 0.84 |
| 3 | 139628356 | rs16849578 | -2.293 | 1.502 | 1.075 | 0.222 | 0.006 | -0.007 | -0.007 | 0 | 0.17 | 0.5 | 0.12 | 0.16 | 0.11 | 0.24 |
| 3 | 139632175 | rs11714886 | -1.72 | 1.514 | 1.096 | 0.121 | -0.003 | -0.003 | -0.003 | 0 | 0.28 | 0.41 | 0.2 | 0.29 | 0.23 | 0.38 |
| 3 | 139632361 | rs11707263 | -1.061 | 1.393 | 1.151 | 0.126 | -0.002 | -0.002 | -0.002 | 0 | 0.23 | 0.25 | 0.15 | 0.22 | 0.18 | 0.25 |
| 3 | 139635017 | rs970929 | -1.834 | 1.609 | 1.108 | 0.045 | 0.008 | -0.007 | -0.007 | 0.252 | 0.35 | 0.58 | 0.28 | 0.3 | 0.37 | 0.4 |
| 3 | 139636462 | rs4613415 | -1.294 | 1.661 | 0.688 | 0.065 | -0.009 | 0.007 | 0.007 | 0 | 0.28 | 0.24 | 0.2 | 0.23 | 0.32 | 0.29 |
| 3 | 139641010 | rs9855416 | -1.766 | 1.695 | 0.678 | 0.082 | 0.02 | -0.008 | -0.008 | 0 | 0.33 | 0.38 | 0.25 | 0.3 | 0.35 | 0.41 |
| 3 | 139648874 | rs9869519 | -1.242 | 1.729 | 0.247 | 0.096 | -0.032 | 0.021 | 0.021 | 0 | 0.27 | 0.18 | 0.19 | 0.23 | 0.3 | 0.28 |
| 3 | 139661938 | rs9835329 | 0.786 | 1.446 | 0.125 | 0.066 | -0.016 | 0.02 | 0.02 | 0 | 0.06 | 0.02 | 0.03 | 0 | 0.07 | 0.05 |
| 3 | 139667733 | rs7642046 | 1.099 | 1.265 | 0.523 | -0.054 | 0.006 | -0.024 | -0.024 | 0 | 0.9 | 0.46 | 0.91 | 0.95 | 0.93 | 0.96 |
| 3 | 139667945 | rs7642260 | 2.202 | 1.5 | 0.407 | 0.03 | -0.035 | 0.02 | 0.02 | 0.558 | 0.75 | 0.12 | 0.76 | 0.79 | 0.72 | 0.77 |
| 3 | 139669929 | rs769307 | -0.364 | 1.388 | 0.333 | 0.013 | 0.007 | 0.019 | 0.019 | 0 | 0.19 | 0.09 | 0.11 | 0.13 | 0.13 | 0.21 |
| 3 | 139677838 | rs9860322 | 0.623 | 1.469 | 0.224 | 0.02 | -0.008 | 0.002 | 0.002 | 0 | 0.07 | 0.05 | 0.04 | 0 | 0.08 | 0.05 |
| 3 | 139677993 | rs2007640 | 2.255 | 1.446 | 0.207 | 0.009 | -0.071 | 0.06 | 0.06 | 1.035 | 0.56 | 0.14 | 0.66 | 0.66 | 0.6 | 0.57 |
| 3 | 139678136 | rs2007634 | 2.334 | 1.457 | 0.209 | 0.014 | -0.041 | 0.066 | 0.066 | 1.049 | 0.56 | 0.21 | 0.67 | 0.66 | 0.59 | 0.57 |

| | | | | | | | | | | | | | | | | |
|---|---|---|---|---|---|---|---|---|---|---|---|---|---|---|---|---|
| 3 | 139679469 | rs9870376 | 0.361 | 2.032 | 0.559 | 0.286 | -0.013 | 0.009 | 0.009 | 0 | 0.07 | 0.05 | 0.04 | 0 | 0.08 | 0.05 |
| 3 | 139679506 | rs9870392 | -0.024 | 2.033 | 0.563 | 0.287 | 0.014 | -0.033 | -0.033 | 0 | 0.07 | 0.14 | 0.04 | 0 | 0.09 | 0.05 |
| 3 | 139682048 | rs966866 | -2.088 | 2.092 | 0.641 | 0.369 | 0.038 | 0.013 | 0.013 | 0 | 0.19 | 0.28 | 0.15 | 0.28 | 0.26 | 0.26 |
| 3 | 139682495 | rs6439886 | -0.681 | 2.088 | 0.658 | 0.366 | 0.02 | -0.014 | -0.014 | 0 | 0.1 | 0.19 | 0.05 | 0.07 | 0.12 | 0.1 |
| 3 | 139684873 | rs17337414 | 2.471 | 2.082 | 0.898 | 0.346 | 0.013 | -0.001 | -0.001 | 1.637 | 0.75 | 0.26 | 0.77 | 0.72 | 0.68 | 0.73 |
| 3 | 139686400 | rs9821561 | -0.221 | 2.171 | 0.236 | 0.482 | -0.002 | 0.002 | 0.002 | 0 | 0.08 | 0.08 | 0.04 | 0.08 | 0.12 | 0.09 |
| 3 | 139688415 | rs6806737 | -0.623 | 2.174 | 0.205 | 0.483 | 0.017 | -0.012 | -0.012 | 0 | 0.1 | 0.18 | 0.05 | 0.07 | 0.12 | 0.09 |
| 3 | 139688912 | rs6439887 | -2.06 | 2.211 | 0.158 | 0.492 | 0.043 | 0.014 | 0.014 | 0 | 0.19 | 0.32 | 0.16 | 0.28 | 0.26 | 0.26 |
| 3 | 139691884 | rs1426044 | -0.503 | 2.158 | -0.458 | 0.626 | -0.036 | -0.034 | -0.034 | 0 | 0.07 | 0.03 | 0.09 | 0.16 | 0.14 | 0.14 |
| 3 | 139695953 | rs1559795 | 1.085 | 1.556 | -0.411 | 0.335 | -0.001 | 0.032 | 0.032 | 0.869 | 0.66 | 0.27 | 0.7 | 0.74 | 0.7 | 0.65 |
| 3 | 139696209 | rs6791621 | 0.945 | 1.446 | -0.088 | 0.748 | -0.001 | 0.004 | 0.004 | 0.879 | 0.7 | 0.33 | 0.67 | 0.61 | 0.72 | 0.61 |
| 3 | 139696765 | rs7628696 | 1.413 | 1.324 | -0.714 | 0.61 | -0.015 | -0.018 | -0.018 | 0 | 0.61 | 0.21 | 0.6 | 0.52 | 0.62 | 0.57 |
| 3 | 139697073 | rs1426041 | 0.474 | 1.22 | -0.12 | 0.557 | 0.005 | -0.019 | -0.019 | 0 | 0.82 | 0.36 | 0.82 | 0.88 | 0.86 | 0.8 |
| 3 | 139699811 | rs9289594 | 1.244 | 1.279 | -0.103 | 1.367 | -0.004 | 0.009 | 0.009 | 0.289 | 0.66 | 0.4 | 0.67 | 0.47 | 0.75 | 0.58 |
| 3 | 139700967 | rs4683773 | 0.103 | 1.716 | -0.787 | 0.917 | 0.023 | 0.052 | 0.052 | 0 | 0.2 | 0.27 | 0.38 | 0.4 | 0.26 | 0.18 |
| 3 | 139701495 | rs4683465 | -0.047 | 1.833 | -0.81 | 0.887 | 0.037 | 0.061 | 0.061 | 0 | 0.21 | 0.33 | 0.41 | 0.4 | 0.31 | 0.19 |
| 3 | 139704174 | rs4683774 | 0.141 | 1.78 | -0.886 | 0.863 | 0.008 | 0.013 | 0.013 | 0 | 0.08 | 0.14 | 0.07 | 0.14 | 0.11 | 0.08 |
| 3 | 139707965 | rs1426034 | -0.074 | 1.762 | -0.83 | 0.909 | 0.011 | 0.023 | 0.023 | 0 | 0.91 | 0.84 | 0.93 | 0.85 | 0.89 | 0.92 |
| 3 | 139710090 | rs1426033 | 1.186 | 1.627 | -0.935 | 0.782 | -0.028 | 0.003 | 0.003 | 0.023 | 0.63 | 0.19 | 0.62 | 0.33 | 0.63 | 0.57 |
| 3 | 139713959 | rs9812091 | 0.881 | 1.312 | -1.243 | 0.489 | -0.004 | 0.018 | 0.018 | 0 | 0.6 | 0.26 | 0.63 | 0.33 | 0.64 | 0.57 |
| 3 | 139723808 | rs987193 | 0.676 | 1.282 | -1.223 | 0.528 | -0.007 | -0.017 | -0.017 | 1.877 | 0.18 | 0.15 | 0.36 | 0.26 | 0.19 | 0.17 |
| 3 | 139724111 | rs9875299 | 1.853 | 1.339 | -1.304 | 0.389 | -0.034 | 0.266 | 0.266 | 0 | 0.44 | 0.09 | 0.28 | 0.07 | 0.48 | 0.39 |
| 3 | 139726964 | rs986561 | -1.286 | 1.283 | -1.277 | 0.425 | -0.014 | 0.145 | 0.145 | 0 | 0.49 | 0.76 | 0.67 | 0.79 | 0.47 | 0.54 |
| 3 | 139727401 | rs9863642 | 2.129 | 1.212 | -1.625 | 0.093 | -0.049 | 0.263 | 0.263 | 0 | 0.37 | 0.02 | 0.22 | 0.03 | 0.36 | 0.34 |
| 3 | 139727563 | rs1426028 | -0.329 | 1.23 | -1.219 | 0.478 | 0.014 | 0.022 | 0.022 | 0 | 0.91 | 0.82 | 0.93 | 0.86 | 0.89 | 0.93 |
| 3 | 139728371 | rs10513093 | 1.01 | 1.006 | -1.243 | 0.999 | -0.055 | -0.061 | -0.061 | 0 | 0.13 | 0.01 | 0.11 | 0.19 | 0.15 | 0.2 |
| 3 | 139729281 | rs12638188 | -0.259 | 1.044 | -1.091 | 1.024 | -0.009 | 0.036 | 0.036 | 0 | 0.29 | 0.57 | 0.27 | 0.51 | 0.22 | 0.34 |
| 3 | 139731000 | rs2194938 | 0.391 | 1.159 | -0.609 | 1.134 | 0.01 | 0.114 | 0.114 | 0.379 | 0.83 | 0.65 | 0.59 | 0.47 | 0.84 | 0.68 |
| 3 | 139731266 | rs1365152 | 1.286 | 1.486 | -0.712 | 0.789 | -0.02 | 0.008 | 0.008 | 0 | 0.06 | 0.02 | 0.09 | 0.03 | 0.09 | 0.05 |
| 3 | 139742354 | rs10935364 | 1.784 | 1.552 | -0.332 | 0.878 | -0.004 | -0.007 | -0.007 | 0 | 0.66 | 0.28 | 0.48 | 0.57 | 0.66 | 0.73 |
| 3 | 139744460 | rs11714306 | 1.23 | 1.429 | -0.77 | 0.622 | -0.038 | -0.032 | -0.032 | 0.162 | 0.4 | 0.04 | 0.29 | 0.38 | 0.38 | 0.51 |
| 3 | 139745438 | rs4271881 | 0.35 | 1.451 | -0.329 | 0.613 | 0.01 | 0.012 | 0.012 | 0 | 0.13 | 0.16 | 0.31 | 0.2 | 0.15 | 0.12 |
| 3 | 139750888 | rs9820731 | -0.058 | 1.517 | -0.224 | 0.522 | 0.021 | -0.025 | -0.025 | 1.098 | 0.1 | 0.22 | 0.14 | 0.03 | 0.16 | 0.08 |
| 3 | 139753307 | rs9832111 | 0.392 | 1.169 | -0.507 | 0.137 | -0.018 | 0.017 | 0.017 | 0 | 0.38 | 0.37 | 0.37 | 0.23 | 0.52 | 0.28 |
| 3 | 139753703 | rs768337 | 1.49 | 1.55 | -0.042 | 0.733 | -0.001 | -0.001 | -0.001 | 0 | 0.68 | 0.29 | 0.46 | 0.58 | 0.6 | 0.75 |
| 3 | 139754061 | rs4467410 | 0.561 | 1.578 | -0.043 | 0.704 | 0.073 | -0.002 | -0.002 | 0 | 0.77 | 0.49 | 0.53 | 0.78 | 0.68 | 0.81 |
| 3 | 139761163 | rs4683474 | 0.977 | 1.483 | -0.251 | 0.764 | 0.009 | 0.038 | 0.038 | 0 | 0.06 | 0.1 | 0.06 | 0.15 | 0.08 | 0.04 |
| 3 | 139766782 | rs6769310 | 1.113 | 1.269 | -0.242 | 0.555 | -0.003 | -0.009 | -0.009 | 1.481 | 0.27 | 0.25 | 0.17 | 0.17 | 0.23 | 0.25 |
| 3 | 139769880 | rs1157820 | 1.047 | 1.589 | 0.089 | 0.892 | 0.016 | 0.035 | 0.035 | 0 | 0.74 | 0.35 | 0.55 | 0.59 | 0.69 | 0.82 |
| 3 | 139779073 | rs16849766 | -0.148 | 1.418 | 0.257 | 1.165 | -0.001 | 0.008 | 0.008 | 0 | 0.12 | 0.15 | 0.27 | 0.19 | 0.11 | 0.13 |
| 3 | 139779359 | rs2114167 | 0.735 | 1.463 | 0.211 | 0.876 | 0.028 | -0.017 | -0.017 | 0 | 0.07 | 0.17 | 0.1 | 0.03 | 0.08 | 0.04 |
| 3 | 139782290 | rs4683795 | -0.095 | 0.757 | -0.005 | 1.003 | 0.004 | 0.01 | 0.01 | 1.352 | 0.52 | 0.28 | 0.44 | 0.47 | 0.54 | 0.65 |
| 3 | 139783002 | rs7633131 | -0.784 | 0.159 | 0.638 | 1.46 | -0.004 | 0.009 | 0.009 | 0 | 0.13 | 0.11 | 0.36 | 0.45 | 0.16 | 0.14 |
| 3 | 139784874 | rs2350488 | -1.889 | -0.053 | 0.825 | 1.223 | 0.042 | 0.029 | 0.029 | 0.055 | 0.48 | 0.81 | 0.65 | 0.61 | 0.46 | 0.46 |
| 3 | 139785389 | rs2350487 | -1.926 | -0.057 | 1.066 | 1.31 | 0.004 | -0.013 | -0.013 | 0 | 0.31 | 0.73 | 0.48 | 0.54 | 0.31 | 0.21 |
| 3 | 139785564 | rs4600787 | -0.105 | -0.044 | 1.22 | 1.319 | -0.024 | -0.004 | -0.004 | 0 | 0.12 | 0.29 | 0.05 | 0.09 | 0.06 | 0.07 |
| 3 | 139787303 | rs10935368 | 0.591 | -0.004 | 1.071 | 1.369 | -0.027 | -0.044 | -0.044 | 0 | 0.08 | 0.02 | 0.3 | 0.3 | 0.1 | 0.09 |
| 3 | 139788146 | rs16849801 | -1.143 | -0.008 | 1.154 | 1.365 | -0.006 | -0.014 | -0.014 | 0.268 | 0.18 | 0.16 | 0.4 | 0.45 | 0.24 | 0.14 |
| 3 | 139793375 | rs1317468 | 0.316 | -0.147 | 1.103 | 1.368 | -0.002 | 0.022 | 0.022 | 0 | 0.08 | 0.13 | 0.08 | 0.15 | 0.07 | 0.05 |
| 3 | 139797844 | rs16849828 | -1.836 | -0.175 | 1.206 | 1.35 | -0.025 | 0.076 | 0.076 | 0.845 | 0.29 | 0.51 | 0.43 | 0.54 | 0.23 | 0.21 |
| 3 | 139813731 | rs4683797 | -1.86 | -0.278 | 0.796 | 1.357 | -0.012 | 0.02 | 0.02 | 0.942 | 0.3 | 0.61 | 0.41 | 0.56 | 0.22 | 0.21 |
| 3 | 139814577 | rs9790156 | 0.6 | -0.258 | 0.627 | 1.268 | -0.016 | -0.047 | -0.047 | 0.653 | 0.08 | 0.02 | 0.25 | 0.28 | 0.09 | 0.09 |
| 3 | 139817289 | rs7652623 | -1.957 | -0.258 | 0.611 | 1.14 | 0.043 | 0.009 | 0.009 | 0.272 | 0.47 | 0.81 | 0.62 | 0.62 | 0.44 | 0.46 |
| 3 | 139818305 | rs4683798 | -0.082 | 0.141 | 0.897 | 1.888 | 0.006 | 0.053 | 0.053 | 0 | 0.08 | 0.16 | 0.08 | 0.16 | 0.06 | 0.05 |
| 3 | 139826735 | rs1878979 | -1.561 | 0.198 | 0.822 | 1.392 | -0.005 | 0.033 | 0.033 | 0.058 | 0.27 | 0.35 | 0.39 | 0.51 | 0.22 | 0.2 |
| 3 | 139826751 | rs4462908 | -0.144 | 0.145 | 0.387 | 1.371 | 0.004 | 0.005 | 0.005 | 0 | 0.1 | 0.08 | 0.03 | 0.06 | 0.06 | 0.06 |
| 3 | 139832777 | rs4368461 | -1.65 | 0.131 | 0.128 | 1.405 | -0.02 | 0.049 | 0.049 | 0.782 | 0.29 | 0.56 | 0.4 | 0.55 | 0.22 | 0.21 |
| 3 | 139832841 | rs4568079 | -0.761 | 0.907 | -0.756 | 1.684 | -0.001 | 0.004 | 0.004 | 0 | 0.15 | 0.13 | 0.34 | 0.45 | 0.15 | 0.14 |
| 3 | 139835865 | rs2882429 | -0.376 | 0.895 | -0.753 | 1.115 | 0.001 | -0.001 | -0.001 | 11.549 | 0.37 | 0.37 | 0.35 | 0.39 | 0.32 | 0.26 |
| 3 | 139835921 | rs2350498 | -0.48 | 1.179 | -0.439 | 0.641 | 0.007 | -0.021 | -0.021 | 0 | 0.44 | 0.46 | 0.4 | 0.28 | 0.51 | 0.41 |
| 3 | 139839000 | rs2350501 | 0.37 | 1.88 | -0.077 | 0.455 | -0.006 | 0.007 | 0.007 | 0 | 0.74 | 0.72 | 0.8 | 0.79 | 0.71 | 0.72 |
| 3 | 139840886 | rs4605505 | 0.122 | 1.412 | 0.418 | 0.912 | 0.022 | 0.162 | 0.162 | 0 | 0.15 | 0.35 | 0.14 | 0.36 | 0.13 | 0.25 |
| 3 | 139844136 | rs4470469 | 0.026 | 1.531 | 0.551 | 0.83 | 0 | -0.007 | -0.007 | 0.24 | 0.75 | 0.71 | 0.84 | 0.8 | 0.74 | 0.72 |
| 3 | 139844670 | rs4683806 | 1.271 | 1.58 | 0.352 | 0.759 | -0.008 | 0.029 | 0.029 | 0 | 0.52 | 0.21 | 0.6 | 0.35 | 0.53 | 0.4 |
| 3 | 139845375 | rs10460847 | -0.095 | 1.129 | 0.765 | 1.048 | 0.02 | 0.12 | 0.12 | 0 | 0.14 | 0.27 | 0.14 | 0.35 | 0.13 | 0.25 |
| 3 | 139846802 | rs2350279 | -0.832 | 1.088 | 0.357 | 0.767 | 0.002 | 0.056 | 0.056 | 0 | 0.25 | 0.6 | 0.26 | 0.45 | 0.23 | 0.33 |
| 3 | 139850795 | rs4683808 | 0.652 | 0.672 | 0.81 | -0.248 | -0.013 | -0.12 | -0.12 | 0.82 | 0.79 | 0.44 | 0.85 | 0.93 | 0.85 | 0.88 |
| 3 | 139854998 | rs1320260 | 0.24 | 0.637 | 1.164 | -0.263 | 0.002 | -0.043 | -0.043 | 0 | 0.84 | 0.77 | 0.89 | 0.93 | 0.85 | 0.88 |
| 3 | 139861050 | rs4638924 | -0.407 | 0.777 | 0.545 | -0.129 | -0.05 | -0.138 | -0.138 | 0 | 0.27 | 0.04 | 0.18 | 0.59 | 0.25 | 0.37 |

| | | | | | | | | | | | | | | | | |
|---|---|---|---|---|---|---|---|---|---|---|---|---|---|---|---|---|
| 3 | 139863001 | rs4317058 | 0.825 | 0.449 | 1.093 | -1.387 | 0.01 | 0.044 | 0.044 | 0.456 | 0.79 | 0.18 | 0.83 | 0.92 | 0.85 | 0.89 |
| 3 | 139882137 | rs9862768 | 0.846 | 0.427 | 0.825 | -1.386 | 0.007 | 0.012 | 0.012 | 0.394 | 0.79 | 0.21 | 0.82 | 0.92 | 0.84 | 0.88 |
| 3 | 139887425 | rs6776629 | -1.003 | 0.43 | 1.033 | -1.046 | -0.002 | -0.016 | -0.016 | 0 | 0.05 | 0.2 | 0.02 | 0 | 0.01 | 0.02 |
| 3 | 139887968 | rs12637089 | -0.634 | 0.508 | 0.376 | -0.945 | -0.038 | -0.019 | -0.019 | 0 | 0.11 | 0.01 | 0.04 | 0.12 | 0.11 | 0.09 |
| 3 | 139890882 | rs9289600 | 1.403 | 0.254 | 1.037 | -1.251 | -0.001 | -0.033 | -0.033 | 0 | 0.78 | 0.52 | 0.88 | 0.86 | 0.85 | 0.88 |
| 3 | 139893732 | rs9818712 | -0.208 | 0.172 | 1.07 | -1.837 | 0.002 | -0.053 | -0.053 | 0 | 0.28 | 0.35 | 0.14 | 0.09 | 0.25 | 0.19 |
| 3 | 139894697 | rs4305391 | -0.637 | -0.051 | 0.963 | -2.059 | -0.004 | -0.092 | -0.092 | 0 | 0.37 | 0.49 | 0.2 | 0.1 | 0.33 | 0.22 |
| 3 | 139900975 | rs4361242 | -0.069 | 0.142 | 0.268 | -1.956 | -0.013 | 0.067 | 0.067 | 0 | 0.29 | 0.2 | 0.14 | 0.09 | 0.26 | 0.19 |
| 3 | 139915713 | rs9825100 | 0.307 | 0.266 | 0.302 | -1.849 | -0.024 | 0.058 | 0.058 | 0 | 0.25 | 0.17 | 0.12 | 0.09 | 0.25 | 0.18 |
| 3 | 139922531 | rs4594581 | -0.074 | 0.21 | 0.002 | -1.772 | -0.069 | 0.188 | 0.188 | 0 | 0.37 | 0.09 | 0.2 | 0.09 | 0.39 | 0.25 |
| 3 | 139922915 | rs4532091 | -0.196 | 0.162 | 0.34 | -1.748 | -0.014 | 0.095 | 0.095 | 0 | 0.39 | 0.23 | 0.22 | 0.09 | 0.41 | 0.29 |
| 3 | 139931034 | rs6808021 | 0.089 | 0.129 | 0.712 | -1.743 | 0 | -0.051 | -0.051 | 0 | 0.59 | 0.59 | 0.78 | 0.91 | 0.6 | 0.73 |
| 3 | 139938425 | rs9815072 | -0.08 | 0.245 | -0.555 | -1.649 | -0.04 | 0.165 | 0.165 | 0 | 0.38 | 0.13 | 0.21 | 0.09 | 0.4 | 0.27 |
| 3 | 139940965 | rs7610536 | 0.049 | 0.762 | -0.428 | -1.122 | -0.021 | 0.02 | 0.02 | 0 | 0.36 | 0.22 | 0.6 | 0.7 | 0.44 | 0.39 |
| 3 | 139948261 | rs2350512 | 0.132 | 0.539 | 0.154 | -1.14 | 0 | -0.031 | -0.031 | 0 | 0.55 | 0.47 | 0.7 | 0.88 | 0.47 | 0.65 |
| 3 | 139950502 | rs2350513 | 0.033 | 0.482 | 0.142 | -1.146 | -0.067 | 0.175 | 0.175 | 0 | 0.41 | 0.1 | 0.23 | 0.11 | 0.51 | 0.33 |
| 3 | 139952343 | rs1878982 | 0.099 | 0.558 | 0.028 | -1.076 | 0.002 | -0.053 | -0.053 | 0.097 | 0.55 | 0.52 | 0.72 | 0.88 | 0.48 | 0.67 |
| 3 | 139979100 | rs9844835 | -0.198 | 0.6 | -0.264 | -1.183 | -0.007 | 0.045 | 0.045 | 0 | 0.68 | 0.25 | 0.78 | 0.98 | 0.67 | 0.62 |
| 3 | 139992571 | rs9811609 | -0.592 | 0.246 | -0.192 | -1.576 | 0.012 | 0.173 | 0.173 | 0 | 0.5 | 0.3 | 0.57 | 0.97 | 0.52 | 0.55 |
| 3 | 139993059 | rs9853327 | 0.229 | 0.413 | -0.626 | -0.951 | -0.032 | -0.055 | -0.055 | 0 | 0.11 | 0.01 | 0.04 | 0.2 | 0.11 | 0.08 |
| 3 | 140000677 | rs4583599 | 0.59 | 0.235 | 0.572 | -1.144 | 0.017 | 0.014 | 0.014 | 0.042 | 0.93 | 0.74 | 0.97 | 0.97 | 0.97 | 0.91 |
| 3 | 140007588 | rs967861 | -1.269 | 0.023 | 0.786 | -1.381 | 0.003 | -0.033 | -0.033 | 0.146 | 0.56 | 0.44 | 0.74 | 0.97 | 0.59 | 0.57 |
| 3 | 140007919 | rs922566 | -0.448 | 0.424 | 0.828 | -0.208 | 0.003 | 0.025 | 0.025 | 0 | 0.19 | 0.19 | 0.25 | 0.58 | 0.23 | 0.24 |
| 3 | 140010660 | rs1001518 | 1.381 | 0.275 | 0.511 | -0.077 | 0.004 | -0.008 | -0.008 | 0.596 | 0.07 | 0.05 | 0.11 | 0.13 | 0.05 | 0.1 |
| 3 | 140011602 | rs2061929 | 1.858 | 0.205 | 0.627 | -0.304 | -0.001 | -0.08 | -0.08 | 0 | 0.46 | 0.47 | 0.27 | 0.03 | 0.42 | 0.42 |
| 3 | 140014763 | rs1351313 | 1.836 | 0.225 | 0.813 | -0.332 | -0.006 | 0.1 | 0.1 | 0.019 | 0.47 | 0.29 | 0.27 | 0 | 0.48 | 0.34 |
| 3 | 140018489 | rs9858663 | -0.748 | 0.225 | 0.713 | 0.504 | -0.004 | -0.081 | -0.081 | 0.308 | 0.22 | 0.12 | 0.21 | 0.49 | 0.21 | 0.21 |
| 3 | 140019495 | rs4327349 | 2.361 | 0.219 | 0.682 | 0.085 | -0.02 | 0.162 | 0.162 | 0 | 0.41 | 0.16 | 0.22 | 0 | 0.41 | 0.32 |
| 3 | 140020694 | rs6439916 | 0.366 | 0.256 | 0.639 | 0.216 | 0.007 | 0.014 | 0.014 | 0 | 0.67 | 0.59 | 0.48 | 0.38 | 0.64 | 0.62 |
| 3 | 140022990 | rs1011757 | -0.872 | 0.035 | -0.172 | 0.496 | -0.017 | -0.094 | -0.094 | 0.006 | 0.22 | 0.07 | 0.21 | 0.38 | 0.21 | 0.21 |
| 3 | 140032477 | rs1479873 | 0.823 | 0.027 | 0.129 | 0.548 | 0.028 | -0.023 | -0.023 | 0.498 | 0.64 | 0.24 | 0.49 | 0.37 | 0.69 | 0.52 |
| 3 | 140036034 | rs7632227 | -0.184 | 0.08 | 0.178 | 0.622 | 0.008 | -0.014 | -0.014 | 1.791 | 0.09 | 0.1 | 0.13 | 0.18 | 0.08 | 0.21 |
| 3 | 140043640 | rs1479875 | 0.999 | -0.031 | 0.874 | 0.434 | 0 | 0.018 | 0.018 | 0 | 0.6 | 0.4 | 0.42 | 0.23 | 0.61 | 0.55 |
| 3 | 140047836 | rs7630996 | 2.128 | 0.125 | 0.985 | 0.242 | -0.005 | 0.007 | 0.007 | 0 | 0.44 | 0.36 | 0.27 | 0.03 | 0.46 | 0.33 |
| 3 | 140049319 | rs987486 | 1.915 | 0.138 | 0.986 | 0.238 | -0.002 | 0.005 | 0.005 | 0 | 0.47 | 0.46 | 0.29 | 0.09 | 0.49 | 0.42 |
| 3 | 140052260 | rs879716 | 1.023 | 0.201 | 1.015 | 0.285 | 0 | 0.001 | 0.001 | 0 | 0.6 | 0.42 | 0.43 | 0.21 | 0.6 | 0.55 |
| 3 | 140052671 | rs985439 | 1.03 | 0.188 | 1.05 | 0.278 | -0.002 | -0.021 | -0.021 | 0 | 0.61 | 0.54 | 0.45 | 0.21 | 0.6 | 0.55 |
| 3 | 140053766 | rs1006877 | 2.44 | 0.068 | 0.926 | 0.125 | -0.068 | 0.22 | 0.22 | 0 | 0.41 | 0.11 | 0.25 | 0.03 | 0.46 | 0.34 |
| 3 | 140054568 | rs1154766 | 2.375 | 0.214 | 0.981 | 0.244 | -0.035 | 0.169 | 0.169 | 0 | 0.42 | 0.2 | 0.26 | 0.04 | 0.46 | 0.34 |
| 3 | 140055784 | rs999819 | 1.084 | 0.253 | 1.479 | 0.405 | 0 | -0.023 | -0.023 | 0 | 0.61 | 0.53 | 0.45 | 0.19 | 0.6 | 0.55 |
| 3 | 140058622 | rs1479878 | 0.361 | 0.294 | 1.466 | 0.639 | 0 | 0.018 | 0.018 | 0 | 0.7 | 0.65 | 0.54 | 0.36 | 0.67 | 0.6 |
| 3 | 140060165 | rs16850177 | 0.965 | 0.328 | 1.205 | 0.536 | -0.024 | -0.045 | -0.045 | 0 | 0.08 | 0.01 | 0.08 | 0.19 | 0.07 | 0.07 |
| 3 | 140063820 | rs17345089 | -0.319 | 0.316 | 1.278 | 0.429 | 0.002 | 0.005 | 0.005 | 0 | 0.14 | 0.12 | 0.15 | 0.08 | 0.12 | 0.11 |
| 3 | 140064908 | rs9871764 | 2.036 | 0.32 | 1.31 | 0.428 | -0.005 | 0.029 | 0.029 | 0 | 0.47 | 0.37 | 0.29 | 0.09 | 0.48 | 0.42 |
| 3 | 140066293 | rs7610675 | 1.908 | 0.314 | 1.298 | 0.416 | 0 | -0.039 | -0.039 | 0.046 | 0.48 | 0.49 | 0.31 | 0.08 | 0.48 | 0.43 |
| 3 | 140071249 | rs6810326 | 2.351 | 0.472 | 1.204 | 0.372 | -0.007 | 0.077 | 0.077 | 0 | 0.43 | 0.28 | 0.26 | 0.03 | 0.44 | 0.34 |
| 3 | 140074782 | rs1542567 | 1.62 | 0.353 | 1.162 | 0.394 | -0.001 | 0.036 | 0.036 | 0 | 0.56 | 0.39 | 0.41 | 0.12 | 0.57 | 0.45 |
| 3 | 140075657 | rs11716346 | 1.674 | 0.248 | 1.137 | 0.398 | 0.002 | -0.001 | -0.001 | 0 | 0.52 | 0.47 | 0.34 | 0.09 | 0.57 | 0.42 |
| 3 | 140079839 | rs1479851 | 1.631 | 0.256 | 1.051 | 0.375 | 0.007 | 0.051 | 0.051 | 0 | 0.53 | 0.39 | 0.42 | 0.14 | 0.57 | 0.44 |
| 3 | 140080015 | rs9810564 | -0.519 | 0.201 | 1.169 | 0.016 | 0.008 | -0.033 | -0.033 | 4.543 | 0.16 | 0.05 | 0.13 | 0.22 | 0.12 | 0.11 |
| 3 | 140081011 | rs6790930 | 2.077 | 0.352 | 1.281 | -0.115 | -0.035 | 0.139 | 0.139 | 0 | 0.42 | 0.23 | 0.27 | 0.04 | 0.47 | 0.33 |
| 3 | 140081692 | rs9874598 | 2.097 | 0.35 | 1.262 | -0.098 | -0.075 | 0.211 | 0.211 | 0 | 0.41 | 0.12 | 0.26 | 0.05 | 0.48 | 0.33 |
| 3 | 140086874 | rs11719890 | 2.079 | 0.304 | 1.208 | -0.125 | -0.013 | 0.04 | 0.04 | 0 | 0.41 | 0.33 | 0.28 | 0.04 | 0.48 | 0.34 |
| 3 | 140088658 | rs1026779 | 2.051 | 0.332 | 1.177 | -0.129 | -0.006 | -0.001 | -0.001 | 0 | 0.42 | 0.38 | 0.3 | 0.04 | 0.47 | 0.34 |
| 3 | 140089029 | rs11915085 | 0.156 | 0.532 | 1.19 | -0.081 | 0.035 | -0.032 | -0.032 | 0.916 | 0.06 | 0.12 | 0.05 | 0 | 0.08 | 0.02 |
| 3 | 140089611 | rs899622 | -1.795 | 0.457 | 1.294 | -0.199 | 0.008 | 0.065 | 0.065 | 0 | 0.5 | 0.38 | 0.67 | 0.91 | 0.48 | 0.62 |
| 3 | 140090936 | rs1383016 | 2.495 | 0.349 | 1.291 | -0.311 | -0.059 | 0.186 | 0.186 | 0 | 0.4 | 0.14 | 0.27 | 0.03 | 0.47 | 0.3 |
| 3 | 140094984 | rs7618673 | 1.885 | 0.534 | 1.457 | -0.216 | 0.002 | 0.073 | 0.073 | 0 | 0.47 | 0.38 | 0.3 | 0.1 | 0.52 | 0.38 |
| 3 | 140100521 | rs10454214 | 0.161 | 0.192 | 0.021 | 0.164 | -0.015 | -0.063 | -0.063 | 0 | 0.15 | 0.01 | 0.15 | 0.24 | 0.14 | 0.15 |
| 3 | 140101406 | rs6793803 | -0.609 | 0.243 | 0.213 | 0.21 | -0.003 | 0.01 | 0.01 | 0 | 0.24 | 0.29 | 0.46 | 0.67 | 0.24 | 0.43 |
| 3 | 140101504 | rs6771702 | -1.75 | 0.311 | 0.054 | 0.072 | 0.044 | 0.127 | 0.127 | 0 | 0.45 | 0.3 | 0.62 | 0.92 | 0.4 | 0.68 |
| 3 | 140101652 | rs7640776 | -0.942 | 0.405 | 0.235 | 0.551 | 0.024 | -0.005 | -0.005 | 0.608 | 0.2 | 0.1 | 0.15 | 0.23 | 0.15 | 0.16 |
| 3 | 140101953 | rs1871493 | 1.356 | 0.926 | 0.58 | 0.5 | 0.002 | -0.003 | -0.003 | 0 | 0.72 | 0.68 | 0.78 | 0.74 | 0.72 | 0.77 |
| 3 | 140102984 | rs4683491 | -1.561 | 0.954 | 0.802 | 0.684 | 0.018 | -0.003 | -0.003 | 0 | 0.31 | 0.46 | 0.42 | 0.36 | 0.33 | 0.28 |
| 3 | 140103136 | rs4683829 | -1.558 | 0.984 | 1.22 | 0.67 | 0.012 | -0.01 | -0.01 | 0 | 0.27 | 0.47 | 0.37 | 0.27 | 0.26 | 0.25 |
| 3 | 140112407 | rs1479863 | 0.712 | 1.223 | 1.162 | 1.368 | 0.003 | 0.017 | 0.017 | 0 | 0.77 | 0.75 | 0.8 | 0.59 | 0.75 | 0.77 |
| 3 | 140114551 | rs2170059 | 2.383 | 1.31 | 0.736 | 1.145 | -0.08 | 0.065 | 0.065 | 0 | 0.51 | 0.06 | 0.54 | 0.35 | 0.51 | 0.59 |
| 3 | 140116226 | rs4683832 | 1.317 | 1.38 | 0.979 | 1.129 | 0.005 | 0.003 | 0.003 | 0 | 0.69 | 0.65 | 0.66 | 0.37 | 0.65 | 0.68 |
| 3 | 140119555 | rs13096759 | 0.743 | 1.371 | 1.441 | 1.039 | 0.075 | -0.036 | -0.036 | 0.102 | 0.07 | 0.21 | 0.06 | 0 | 0.09 | 0.06 |

| | | | | | | | | | | | | | | | | |
|---|---|---|---|---|---|---|---|---|---|---|---|---|---|---|---|---|
| 3 | 140120742 | rs993691 | 1.38 | 1.335 | 1.701 | 1.067 | 0.022 | 0.005 | 0.005 | 0 | 0.69 | 0.44 | 0.65 | 0.37 | 0.65 | 0.76 |
| 3 | 140124454 | rs13064082 | 2.139 | 1.307 | 1.408 | 1.069 | 0.001 | 0.006 | 0.006 | 0 | 0.59 | 0.35 | 0.61 | 0.3 | 0.6 | 0.69 |
| 3 | 140125714 | rs993483 | 0.353 | 1.428 | 0.66 | 1.236 | -0.003 | -0.001 | -0.001 | 0 | 0.78 | 0.81 | 0.81 | 0.49 | 0.75 | 0.77 |
| 3 | 140126067 | rs993484 | 1.927 | 1.323 | 0.715 | 1.08 | -0.047 | 0.027 | 0.027 | 0 | 0.61 | 0.21 | 0.58 | 0.35 | 0.55 | 0.69 |
| 3 | 140127332 | rs7619971 | 1.185 | 1.305 | 0.082 | 1.188 | -0.001 | -0.001 | -0.001 | 0 | 0.71 | 0.68 | 0.68 | 0.35 | 0.65 | 0.75 |
| 3 | 140137723 | rs13099443 | -1.157 | 1.401 | 0.004 | 1.428 | 0.039 | -0.027 | -0.027 | 0.03 | 0.35 | 0.24 | 0.29 | 0.43 | 0.3 | 0.34 |
| 3 | 140140442 | rs13073891 | -2.212 | 1.495 | -0.066 | 1.36 | -0.094 | 0.05 | 0.05 | 0 | 0.49 | 0.94 | 0.46 | 0.65 | 0.49 | 0.4 |
| 3 | 140141925 | rs347320 | -1.69 | 1.507 | -0.019 | 1.363 | -0.011 | 0.038 | 0.038 | 0 | 0.42 | 0.26 | 0.38 | 0.67 | 0.44 | 0.4 |
| 3 | 140142144 | rs347322 | -2.163 | 1.49 | -0.163 | 1.381 | -0.094 | 0.05 | 0.05 | 0 | 0.49 | 0.94 | 0.46 | 0.65 | 0.5 | 0.4 |
| 3 | 140147286 | rs347328 | -1.431 | 1.213 | -0.121 | 1.274 | 0.031 | 0.031 | 0.031 | 0 | 0.41 | 0.24 | 0.54 | 0.71 | 0.4 | 0.59 |
| 3 | 140149805 | rs347332 | -1.323 | 1.393 | 0.614 | 1.359 | 0.001 | 0 | 0 | 0 | 0.34 | 0.64 | 0.36 | 0.65 | 0.33 | 0.28 |
| 3 | 140150969 | rs347335 | -1.016 | 1.531 | 0.16 | 1.433 | -0.001 | -0.002 | -0.002 | 0 | 0.31 | 0.25 | 0.33 | 0.66 | 0.33 | 0.29 |
| 3 | 140160906 | rs9877176 | -1.068 | 1.522 | -0.054 | 1.488 | -0.018 | -0.036 | -0.036 | 0 | 0.13 | 0.07 | 0.29 | 0.27 | 0.18 | 0.36 |
| 3 | 140164183 | rs2114860 | 0.751 | 1.41 | 0.358 | 1.808 | -0.017 | -0.041 | -0.041 | 0 | 0.7 | 0.81 | 0.72 | 0.55 | 0.69 | 0.76 |
| 3 | 140167131 | rs6439923 | -1.993 | 1.764 | 0.316 | 2.042 | -0.009 | -0.045 | -0.045 | 0 | 0.38 | 0.81 | 0.54 | 0.59 | 0.38 | 0.41 |
| 3 | 140167536 | rs2081803 | 0.158 | 1.775 | 0.291 | 2.049 | -0.053 | -0.052 | -0.052 | 0 | 0.91 | 0.99 | 0.86 | 0.65 | 0.86 | 0.83 |
| 3 | 140169657 | rs6439924 | -0.859 | 1.695 | 1.302 | 1.862 | -0.022 | -0.065 | -0.065 | 0.242 | 0.23 | 0.37 | 0.16 | 0.1 | 0.17 | 0.08 |
| 3 | 140171892 | rs6439925 | -1.916 | 1.488 | 0.935 | 1.351 | 0.003 | 0.002 | 0.002 | 0 | 0.34 | 0.79 | 0.47 | 0.35 | 0.31 | 0.37 |
| 3 | 140173355 | rs1427781 | -1.277 | 1.599 | 0.742 | 1.478 | -0.023 | -0.027 | -0.027 | 0.628 | 0.08 | 0.04 | 0.28 | 0.26 | 0.14 | 0.29 |
| 3 | 140176199 | rs6791077 | -1.044 | 1.742 | 0.807 | 1.476 | -0.013 | -0.024 | -0.024 | 0.404 | 0.25 | 0.12 | 0.4 | 0.33 | 0.25 | 0.33 |
| 3 | 140180563 | rs6439928 | -1.348 | 1.506 | 1.171 | 1.021 | 0.001 | -0.052 | -0.052 | 1.001 | 0.34 | 0.5 | 0.45 | 0.16 | 0.31 | 0.38 |
| 3 | 140182046 | rs10935384 | -1.583 | 1.394 | 1.065 | 1.124 | -0.013 | 0.045 | 0.045 | 0 | 0.41 | 0.67 | 0.52 | 0.24 | 0.39 | 0.43 |
| 3 | 140185248 | rs534395 | 1.902 | 1.174 | 1.944 | 1.88 | 0.024 | 0.058 | 0.058 | 0 | 0.71 | 0.61 | 0.56 | 0.54 | 0.69 | 0.69 |
| 3 | 140185294 | rs533657 | -1.474 | 1.279 | 2.049 | 2.039 | 0.034 | 0.037 | 0.037 | 0 | 0.22 | 0.32 | 0.38 | 0.33 | 0.25 | 0.28 |
| 3 | 140189148 | rs524554 | -2.485 | 1.369 | 2.165 | 2.058 | -0.012 | -0.005 | -0.005 | 0 | 0.36 | 0.67 | 0.57 | 0.58 | 0.36 | 0.37 |
| 3 | 140190071 | rs12632435 | -0.608 | 1.413 | 1.99 | 2.052 | -0.024 | -0.042 | -0.042 | 0.13 | 0.1 | 0.01 | 0.09 | 0.14 | 0.1 | 0.03 |
| 3 | 140196275 | rs531617 | -2.58 | 1.343 | 2.271 | 2.094 | 0.007 | -0.017 | -0.017 | 0 | 0.34 | 0.54 | 0.31 | 0.27 | 0.33 | 0.25 |
| 3 | 140198492 | rs546655 | -1.842 | 1.341 | 2.31 | 2.069 | 0.025 | -0.069 | -0.069 | 0 | 0.23 | 0.52 | 0.22 | 0.13 | 0.23 | 0.21 |
| 3 | 140218207 | rs349507 | -2.765 | 0.908 | 2.147 | 1.955 | -0.013 | 0.017 | 0.017 | 0 | 0.43 | 0.74 | 0.6 | 0.58 | 0.4 | 0.39 |
| 3 | 140223794 | rs9862730 | -1.252 | 0.958 | 2.426 | 1.869 | 0.139 | 0.065 | 0.065 | 0 | 0.06 | 0.33 | 0.08 | 0.06 | 0.09 | 0.02 |
| 3 | 140226452 | rs6793607 | -1.132 | 0.937 | 2.409 | 1.93 | 0.007 | 0.091 | 0.091 | 0 | 0.16 | 0.26 | 0.36 | 0.41 | 0.16 | 0.18 |
| 3 | 140229445 | rs349504 | -2.215 | 1.004 | 2.184 | 1.851 | -0.005 | -0.029 | -0.029 | 1.278 | 0.28 | 0.72 | 0.53 | 0.55 | 0.27 | 0.25 |
| 3 | 140241720 | rs180971 | -1.165 | 1.268 | 1.932 | 1.817 | 0.051 | 0.194 | 0.194 | 0 | 0.05 | 0.12 | 0.3 | 0.36 | 0.12 | 0.14 |
| 3 | 140252017 | rs349538 | -1.599 | 1.286 | 1.832 | 1.431 | 0.193 | 0.361 | 0.361 | 0 | 0.06 | 0.34 | 0.31 | 0.29 | 0.13 | 0.15 |
| 3 | 140257037 | rs12638211 | -1.611 | 0.841 | 1.016 | 0.731 | -0.004 | 0.057 | 0.057 | 0 | 0.31 | 0.16 | 0.17 | 0.12 | 0.26 | 0.2 |
| 3 | 140260276 | rs349554 | -1.173 | 1 | 0.979 | 0.977 | 0.026 | 0.01 | 0.01 | 0 | 0.2 | 0.39 | 0.17 | 0.18 | 0.2 | 0.07 |
| 3 | 140262088 | rs6808158 | -1.207 | 0.8 | -0.06 | 0.842 | -0.027 | -0.007 | -0.007 | 0 | 0.15 | 0.03 | 0.08 | 0.14 | 0.15 | 0.2 |
| 3 | 140262974 | rs10513121 | -1.541 | 0.706 | 0.635 | 0.478 | 0.008 | -0.031 | -0.031 | 0 | 0.16 | 0.24 | 0.11 | 0.06 | 0.17 | 0.14 |
| 3 | 140266593 | rs16850488 | -0.818 | 0.356 | 1.004 | 0.238 | -0.025 | -0.065 | -0.065 | 0.005 | 0.22 | 0.49 | 0.18 | 0.11 | 0.18 | 0.08 |
| 3 | 140272445 | rs9682918 | -0.164 | 0.42 | 0.587 | 0.231 | -0.006 | 0 | 0 | 0 | 0.06 | 0.05 | 0.03 | 0 | 0.08 | 0.03 |
| 3 | 140274442 | rs17350052 | -2.059 | 0.343 | 0.588 | 0.196 | 0.016 | 0.048 | 0.048 | 0 | 0.38 | 0.73 | 0.28 | 0.17 | 0.33 | 0.23 |
| 3 | 140274759 | rs6773067 | -1.32 | 0.452 | 0.742 | 0.155 | -0.001 | -0.015 | -0.015 | 0 | 0.08 | 0.07 | 0.05 | 0 | 0.11 | 0.05 |
| 3 | 140276484 | rs907489 | -1.398 | 0.103 | 0.738 | 0.28 | -0.002 | -0.009 | -0.009 | 0 | 0.27 | 0.73 | 0.22 | 0.11 | 0.2 | 0.11 |
| 3 | 140277903 | rs7629162 | -2.272 | 0.464 | 1.59 | 0.697 | 0.055 | 0.05 | 0.05 | 0 | 0.08 | 0.44 | 0.11 | 0.07 | 0.08 | 0.04 |
| 3 | 140288049 | rs16846209 | -1.352 | 0.496 | 1.312 | 0.527 | 0.003 | -0.032 | -0.032 | 0.765 | 0.07 | 0.07 | 0.05 | 0 | 0.1 | 0.04 |
| 3 | 140298808 | rs1834869 | -0.914 | 1.029 | 2.329 | 0.221 | -0.004 | -0.01 | -0.01 | 0 | 0.09 | 0.09 | 0.09 | 0 | 0.12 | 0.14 |
| 3 | 140299573 | rs347957 | -1.77 | 0.766 | 2.4 | 0.901 | -0.001 | 0.011 | 0.011 | 0 | 0.18 | 0.29 | 0.33 | 0.72 | 0.19 | 0.39 |
| 3 | 140299603 | rs1845476 | -0.904 | 0.892 | 2.336 | 0.794 | -0.012 | 0.001 | 0.001 | 0 | 0.11 | 0.07 | 0.1 | 0 | 0.14 | 0.17 |
| 3 | 140311901 | rs347967 | -0.278 | 0.462 | 1.748 | 1.115 | 0.021 | -0.041 | -0.041 | 0 | 0.64 | 0.72 | 0.53 | 0.57 | 0.67 | 0.67 |
| 3 | 140315715 | rs16850633 | 0.42 | 0.419 | 1.746 | 1.118 | -0.006 | -0.122 | -0.122 | 0.203 | 0.33 | 0.11 | 0.45 | 0.46 | 0.33 | 0.33 |
| 3 | 140316851 | rs724761 | 0.673 | 0.8 | 1.785 | 1.4 | -0.059 | 0.152 | 0.152 | 0.004 | 0.5 | 0.08 | 0.35 | 0.27 | 0.52 | 0.38 |
| 3 | 140336271 | rs6808213 | -0.505 | 0.842 | 2.278 | 1.387 | 0.033 | -0.043 | -0.043 | 0 | 0.08 | 0.16 | 0.11 | 0 | 0.12 | 0.1 |
| 3 | 140350567 | rs1494524 | 0.059 | 0.932 | 1.976 | 1.589 | -0.007 | -0.002 | -0.002 | 0 | 0.38 | 0.49 | 0.49 | 0.59 | 0.36 | 0.34 |
| 3 | 140351826 | rs1696200 | 0.079 | 0.95 | 1.975 | 1.588 | -0.009 | -0.002 | -0.002 | 0 | 0.38 | 0.5 | 0.49 | 0.59 | 0.33 | 0.34 |
| 3 | 140355911 | rs1690863 | 0.226 | 0.825 | 1.945 | 1.587 | 0.009 | 0.001 | 0.001 | 0 | 0.37 | 0.33 | 0.48 | 0.6 | 0.35 | 0.34 |
| 3 | 140366189 | rs698661 | 0.226 | 0.825 | 1.945 | 1.587 | 0.01 | 0.001 | 0.001 | 0 | 0.37 | 0.33 | 0.48 | 0.6 | 0.35 | 0.34 |
| 3 | 140366948 | rs698664 | 0.321 | 0.829 | 1.895 | 1.628 | 0.021 | -0.044 | -0.044 | 0 | 0.36 | 0.27 | 0.46 | 0.48 | 0.33 | 0.33 |
| 3 | 140375145 | rs837662 | 1.001 | 0.833 | 1.92 | 1.715 | 0.135 | 0.096 | 0.096 | 0 | 0.9 | 0.53 | 0.85 | 0.82 | 0.87 | 0.72 |
| 3 | 140396022 | rs7651388 | 0.433 | 0.883 | 1.732 | 1.719 | -0.016 | 0.126 | 0.126 | 0 | 0.52 | 0.25 | 0.37 | 0.21 | 0.51 | 0.38 |
| 3 | 140399199 | rs2047563 | 0.371 | 0.883 | 1.731 | 1.724 | -0.007 | 0.046 | 0.046 | 0 | 0.53 | 0.26 | 0.39 | 0.34 | 0.53 | 0.39 |
| 3 | 140404389 | rs4683376 | 0.523 | 0.868 | 1.395 | 1.893 | -0.028 | 0.143 | 0.143 | 0 | 0.52 | 0.19 | 0.49 | 0.27 | 0.51 | 0.38 |
| 3 | 140407260 | rs9876490 | 0.425 | 0.866 | 1.492 | 1.877 | -0.004 | 0.029 | 0.029 | 0 | 0.52 | 0.27 | 0.51 | 0.39 | 0.53 | 0.38 |
| 3 | 140408467 | rs698677 | 0.304 | 0.866 | 1.491 | 1.87 | 0.02 | -0.028 | -0.028 | 0 | 0.37 | 0.27 | 0.34 | 0.42 | 0.33 | 0.33 |
| 3 | 140414730 | rs961554 | 0.533 | 0.952 | 0.647 | 1.517 | -0.072 | 0.166 | 0.166 | 0 | 0.5 | 0.06 | 0.48 | 0.28 | 0.51 | 0.38 |
| 3 | 140419715 | rs1540506 | -0.234 | 0.949 | 0.693 | 1.522 | 0.013 | 0.002 | 0.002 | 0 | 0.59 | 0.36 | 0.62 | 0.39 | 0.66 | 0.49 |
| 3 | 140422914 | rs12493620 | 0.505 | 1.03 | 0.234 | 1.527 | -0.039 | 0.165 | 0.165 | 0 | 0.49 | 0.09 | 0.48 | 0.28 | 0.51 | 0.38 |
| 3 | 140429298 | rs7645465 | -0.045 | 1.052 | 0.282 | 1.497 | -0.009 | 0.004 | 0.004 | 0 | 0.06 | 0.05 | 0.1 | 0 | 0.12 | 0.1 |
| 3 | 140435469 | rs7627814 | -0.084 | 1.017 | 0.284 | 1.499 | -0.01 | 0.004 | 0.004 | 0 | 0.06 | 0.05 | 0.1 | 0 | 0.11 | 0.1 |
| 3 | 140448219 | rs869394 | 0.214 | 0.764 | 0.192 | 1.595 | 0.02 | 0.06 | 0.06 | 0 | 0.52 | 0.19 | 0.63 | 0.38 | 0.58 | 0.41 |

| | | | | | | | | | | | | | | | |
|---|---|---|---|---|---|---|---|---|---|---|---|---|---|---|---|
| 3 | 140454836 | rs2044078 | 0.029 | 0.977 | -0.19 | 1.244 | -0.1 | 0.157 | 0.157 | 0 | 0.38 | 0.05 | 0.46 | 0.13 | 0.42 | 0.35 |
| 3 | 140455487 | rs1561764 | -0.635 | 0.839 | 0.557 | 1.623 | -0.002 | 0.011 | 0.011 | 0 | 0.48 | 0.64 | 0.56 | 0.37 | 0.49 | 0.42 |
| 3 | 140458433 | rs9856981 | -0.082 | 0.847 | 0.633 | 1.654 | -0.014 | 0.044 | 0.044 | 0 | 0.4 | 0.23 | 0.47 | 0.23 | 0.42 | 0.35 |
| 3 | 140459247 | rs9861826 | -0.048 | 0.848 | 0.566 | 1.626 | -0.039 | 0.143 | 0.143 | 0 | 0.39 | 0.14 | 0.46 | 0.13 | 0.42 | 0.34 |
| 3 | 140466634 | rs4571221 | -0.622 | 0.815 | 0.663 | 1.644 | -0.004 | 0.007 | 0.007 | 0 | 0.48 | 0.62 | 0.56 | 0.37 | 0.48 | 0.42 |
| 3 | 140468370 | rs7624520 | -0.054 | 0.718 | 0.579 | 1.485 | -0.059 | 0.149 | 0.149 | 0 | 0.38 | 0.1 | 0.45 | 0.13 | 0.41 | 0.3 |
| 3 | 140476375 | rs7643274 | -0.303 | 0.768 | 0.505 | 1.344 | 0 | -0.06 | -0.06 | 0 | 0.42 | 0.49 | 0.49 | 0.13 | 0.41 | 0.3 |
| 3 | 140484942 | rs7632490 | -0.711 | 0.638 | 0.791 | 1.858 | 0 | -0.001 | -0.001 | 0 | 0.38 | 0.41 | 0.54 | 0.49 | 0.36 | 0.39 |
| 3 | 140485038 | rs7644202 | 0.612 | 0.602 | 0.591 | 1.742 | -0.033 | 0.069 | 0.069 | 0 | 0.19 | 0.08 | 0.34 | 0.05 | 0.22 | 0.21 |
| 3 | 140488941 | rs9861350 | -0.368 | 0.595 | 0.586 | 1.755 | -0.022 | 0.082 | 0.082 | 0 | 0.31 | 0.14 | 0.42 | 0.15 | 0.33 | 0.26 |
| 3 | 140492007 | rs4465920 | -0.928 | 0.597 | 0.635 | 1.776 | 0.007 | -0.01 | -0.01 | 0 | 0.37 | 0.33 | 0.53 | 0.42 | 0.36 | 0.4 |
| 3 | 140492239 | rs4428155 | -1.143 | 0.591 | 0.646 | 1.732 | -0.012 | 0.014 | 0.014 | 0 | 0.39 | 0.55 | 0.56 | 0.49 | 0.36 | 0.4 |
| 3 | 140511157 | rs11707850 | 0.518 | 0.325 | -0.202 | 1.7 | -0.041 | 0.072 | 0.072 | 0 | 0.19 | 0.08 | 0.34 | 0.05 | 0.22 | 0.21 |
| 3 | 140526480 | rs2139266 | -0.942 | 0.697 | -0.183 | 1.497 | 0.002 | 0.001 | 0.001 | 0 | 0.46 | 0.47 | 0.61 | 0.51 | 0.44 | 0.48 |
| 3 | 140533823 | rs11712892 | -0.447 | 0.398 | -0.564 | 1.24 | 0.013 | 0.019 | 0.019 | 0 | 0.33 | 0.25 | 0.42 | 0.21 | 0.3 | 0.26 |
| 3 | 140540718 | rs908821 | -0.655 | 0.846 | -0.235 | 1.77 | 0 | 0.011 | 0.011 | 0 | 0.26 | 0.27 | 0.39 | 0.47 | 0.26 | 0.35 |
| 3 | 140542380 | rs6808549 | 0.982 | 0.799 | -0.046 | 1.895 | 0 | 0.008 | 0.008 | 0 | 0.66 | 0.72 | 0.66 | 0.33 | 0.68 | 0.58 |
| 3 | 140561465 | rs2139269 | 0.474 | 0.854 | 0.599 | 2.164 | 0.004 | 0.049 | 0.049 | 0 | 0.81 | 0.69 | 0.76 | 0.68 | 0.83 | 0.76 |
| 3 | 140570721 | rs9814403 | -0.034 | 0.847 | 0.665 | 2.169 | -0.006 | 0.01 | 0.01 | 0.016 | 0.85 | 0.87 | 0.85 | 0.94 | 0.84 | 0.79 |
| 3 | 140590632 | rs9864312 | -0.031 | 0.869 | 0.633 | 2.097 | 0.002 | -0.006 | -0.006 | 0 | 0.84 | 0.79 | 0.83 | 0.94 | 0.81 | 0.78 |
| 3 | 140603142 | rs6793954 | 0.016 | 0.87 | 0.599 | 2.098 | -0.032 | 0.031 | 0.031 | 0.255 | 0.85 | 0.91 | 0.84 | 0.94 | 0.81 | 0.78 |
| 3 | 140618640 | rs7433538 | 0.562 | 0.861 | 0.648 | 2.125 | 0.008 | 0.035 | 0.035 | 0 | 0.81 | 0.71 | 0.74 | 0.64 | 0.8 | 0.75 |
| 3 | 140632237 | rs4683387 | 1.484 | 0.554 | 0.691 | 1.475 | 0.003 | 0 | 0 | 0 | 0.69 | 0.17 | 0.71 | 0.68 | 0.71 | 0.78 |
| 3 | 140636790 | rs7650615 | -2.386 | 0.653 | 1.357 | 1.49 | 0.07 | 0.175 | 0.175 | 0 | 0.07 | 0.29 | 0.11 | 0.22 | 0.12 | 0.07 |
| 3 | 140636886 | rs7628638 | -2.499 | 0.652 | 1.398 | 1.49 | 0.091 | 0.211 | 0.211 | 0 | 0.07 | 0.37 | 0.11 | 0.22 | 0.12 | 0.07 |
| 3 | 140639942 | rs1401509 | 1.548 | 0.507 | 1.364 | 1.38 | 0.006 | 0.001 | 0.001 | 0 | 0.7 | 0.23 | 0.74 | 0.69 | 0.76 | 0.81 |
| 3 | 140659454 | rs7620524 | -1.551 | 0.285 | 2.496 | 1.458 | -0.023 | -0.078 | -0.078 | 0.084 | 0.12 | 0.48 | 0.08 | 0 | 0.07 | 0.01 |
| 3 | 140669440 | rs13099216 | -0.658 | 0.241 | 2.975 | 1.464 | -0.009 | -0.033 | -0.033 | 0.06 | 0.09 | 0.09 | 0.05 | 0 | 0.07 | 0.01 |
| 3 | 140669825 | rs9845648 | 0.057 | 0.356 | 1.97 | 0.897 | -0.086 | 0.144 | 0.144 | 0.772 | 0.22 | 0.02 | 0.17 | 0.02 | 0.31 | 0.08 |
| 3 | 140676957 | rs7643434 | -2.756 | -0.093 | 2.573 | 1.421 | -0.014 | 0.002 | 0.002 | 0.887 | 0.17 | 0.77 | 0.16 | 0.21 | 0.13 | 0.05 |
| 3 | 140684003 | rs7616174 | -0.985 | -0.091 | 2.547 | 1.42 | -0.01 | -0.061 | -0.061 | 0.014 | 0.1 | 0.33 | 0.07 | 0 | 0.07 | 0.01 |
| 3 | 140692486 | rs2555707 | 2.007 | -0.09 | 2.538 | 1.423 | -0.031 | 0.013 | 0.013 | 0.614 | 0.83 | 0.31 | 0.85 | 0.79 | 0.87 | 0.95 |
| 3 | 140726843 | rs4583620 | -2.02 | -0.033 | 2.317 | 1.285 | -0.021 | 0.025 | 0.025 | 0.135 | 0.15 | 0.58 | 0.13 | 0.17 | 0.12 | 0.05 |
| 3 | 140732654 | rs6767867 | -1.946 | -0.013 | 2.128 | 1.282 | -0.012 | 0.052 | 0.052 | 0.056 | 0.14 | 0.49 | 0.12 | 0.18 | 0.12 | 0.05 |
| 3 | 140746321 | rs11923192 | -1.525 | -0.02 | 2.065 | 1.298 | -0.011 | 0.044 | 0.044 | 0 | 0.13 | 0.36 | 0.11 | 0.18 | 0.12 | 0.05 |
| 3 | 140755988 | rs9876562 | 1.512 | -0.365 | 1.463 | 1.201 | -0.032 | -0.004 | -0.004 | 0 | 0.84 | 0.27 | 0.86 | 0.82 | 0.92 | 0.95 |
| 3 | 140761280 | rs6810025 | 1.454 | -0.46 | 1.366 | 1.459 | -0.058 | 0.073 | 0.073 | 0.903 | 0.83 | 0.37 | 0.71 | 0.69 | 0.91 | 0.94 |
| 3 | 140766208 | rs9826671 | 1.366 | -0.339 | 1.242 | 1.341 | -0.024 | 0.003 | 0.003 | 4.602 | 0.72 | 0.36 | 0.65 | 0.68 | 0.78 | 0.91 |
| 3 | 140778763 | rs890528 | -1.904 | -0.257 | 1.982 | 1.617 | -0.002 | 0 | 0 | 0 | 0.22 | 0.75 | 0.35 | 0.34 | 0.14 | 0.08 |
| 3 | 140780992 | rs6793789 | -1.69 | -0.248 | 2.615 | 1.602 | -0.046 | 0.071 | 0.071 | 2.131 | 0.2 | 0.5 | 0.3 | 0.31 | 0.14 | 0.09 |
| 3 | 140784435 | rs6804474 | -1.948 | -0.248 | 2.579 | 1.538 | -0.024 | 0.018 | 0.018 | 2.69 | 0.22 | 0.69 | 0.33 | 0.32 | 0.14 | 0.09 |
| 3 | 140785834 | rs10935417 | -1.798 | -0.257 | 3.089 | 1.52 | -0.033 | 0.039 | 0.039 | 2.196 | 0.2 | 0.35 | 0.28 | 0.29 | 0.14 | 0.09 |
| 3 | 140792365 | rs16851040 | -3.073 | -0.313 | 2.998 | 1.234 | 0.08 | 0.07 | 0.07 | 1.623 | 0.05 | 0.53 | 0.05 | 0 | 0.01 | 0 |
| 3 | 140799213 | rs16851055 | -2.101 | -0.466 | 3.056 | 1.468 | -0.046 | -0.01 | -0.01 | 0.903 | 0.22 | 0.45 | 0.17 | 0.19 | 0.18 | 0.13 |
| 3 | 140804298 | rs4683550 | -0.936 | -0.42 | 2.801 | 1.504 | -0.009 | -0.004 | -0.004 | 0 | 0.13 | 0.14 | 0.1 | 0.11 | 0.12 | 0.05 |
| 3 | 140804947 | rs7620622 | -2.862 | -0.428 | 2.612 | 1.842 | 0.009 | -0.011 | -0.011 | 0 | 0.25 | 0.77 | 0.34 | 0.34 | 0.19 | 0.13 |
| 3 | 140810146 | rs6791826 | -2.568 | 0.045 | 2.103 | 1.506 | 0.002 | -0.011 | -0.011 | 0 | 0.39 | 0.67 | 0.46 | 0.44 | 0.41 | 0.31 |
| 3 | 140819229 | rs11918400 | -2.119 | -0.624 | 1.783 | 2.026 | -0.024 | 0.12 | 0.12 | 0 | 0.19 | 0.51 | 0.41 | 0.29 | 0.16 | 0.07 |
| 3 | 140828801 | rs899348 | -2.64 | -0.637 | 1.628 | 1.809 | -0.01 | 0.014 | 0.014 | 0 | 0.28 | 0.6 | 0.47 | 0.37 | 0.26 | 0.23 |
| 3 | 140845584 | rs2086180 | -0.347 | -0.578 | 1.478 | 1.822 | 0.002 | -0.012 | -0.012 | 0 | 0.11 | 0.04 | 0.09 | 0.14 | 0.12 | 0.05 |
| 3 | 140848571 | rs7647349 | -0.842 | -0.543 | 1.316 | 1.765 | -0.001 | 0.007 | 0.007 | 0 | 0.11 | 0.08 | 0.07 | 0.11 | 0.1 | 0.17 |
| 3 | 140849754 | rs4683395 | -1.602 | -0.519 | 1.496 | 1.855 | -0.011 | 0.111 | 0.111 | 0 | 0.18 | 0.51 | 0.4 | 0.27 | 0.17 | 0.07 |
| 3 | 140863405 | rs7614592 | -1.254 | -0.888 | 0.845 | 1.235 | -0.048 | -0.023 | -0.023 | 0 | 0.29 | 0.48 | 0.15 | 0.24 | 0.18 | 0.29 |
| 3 | 140866631 | rs2291043 | -1.284 | -0.971 | 1.442 | 1.37 | -0.027 | -0.006 | -0.006 | 0.175 | 0.15 | 0.32 | 0.23 | 0.15 | 0.13 | 0.05 |
| 3 | 140866728 | rs9647343 | -0.797 | -0.891 | 1.766 | 1.262 | -0.003 | -0.046 | -0.046 | 0 | 0.11 | 0.23 | 0.11 | 0.02 | 0.11 | 0.01 |
| 3 | 140867094 | rs2291042 | -2.06 | -1.218 | 2.025 | 1.651 | -0.029 | 0.047 | 0.047 | 0 | 0.13 | 0.32 | 0.1 | 0.22 | 0.06 | 0.09 |
| 3 | 140867938 | rs875870 | -1.521 | -1.612 | 2.046 | 1.463 | -0.047 | 0.006 | 0.006 | 0 | 0.21 | 0.37 | 0.23 | 0.16 | 0.1 | 0.1 |
| 3 | 140868715 | rs1108693 | 2.71 | -1 | 1.473 | 1.905 | -0.006 | -0.113 | -0.113 | 0 | 0.59 | 0.03 | 0.46 | 0.6 | 0.63 | 0.82 |
| 3 | 140875814 | rs13080353 | 2.73 | -0.64 | 1.661 | 2.1 | 0.002 | -0.122 | -0.122 | 0.231 | 0.63 | 0.03 | 0.47 | 0.62 | 0.66 | 0.83 |
| 3 | 140890184 | rs2871782 | 1.055 | -0.715 | 2.635 | 2.112 | -0.001 | 0.044 | 0.044 | 0 | 0.84 | 0.65 | 0.72 | 0.79 | 0.81 | 0.91 |
| 3 | 140896329 | rs12494491 | 2.734 | -0.655 | 2.683 | 2.088 | 0.01 | 0.022 | 0.022 | 0.65 | 0.68 | 0.22 | 0.62 | 0.77 | 0.68 | 0.85 |
| 3 | 140902668 | rs7613134 | 0.77 | -0.509 | 2.762 | 2.111 | -0.003 | 0.007 | 0.007 | 0.046 | 0.87 | 0.89 | 0.81 | 0.95 | 0.83 | 0.93 |
| 3 | 140905177 | rs1564257 | 3.036 | -0.599 | 2.73 | 2.178 | 0.026 | -0.057 | -0.057 | 1.918 | 0.63 | 0.15 | 0.48 | 0.61 | 0.66 | 0.84 |
| 3 | 140906678 | rs1533128 | -2.822 | -0.595 | 3.305 | 2.209 | -0.019 | 0.307 | 0.307 | 3.709 | 0.1 | 0.49 | 0.25 | 0.31 | 0.04 | 0.04 |
| 3 | 140907232 | rs1511385 | 0.703 | -0.59 | 3.266 | 2.193 | -0.019 | 0.009 | 0.009 | 0 | 0.87 | 0.96 | 0.82 | 0.95 | 0.83 | 0.93 |
| 3 | 140907784 | rs1533130 | -3.438 | -0.641 | 3.319 | 2.211 | 0.066 | 0.236 | 0.236 | 5.088 | 0.06 | 0.44 | 0.12 | 0.16 | 0.02 | 0.03 |
| 3 | 140908639 | rs2048456 | 2.745 | -0.581 | 3.307 | 2.107 | -0.002 | -0.002 | -0.002 | 0.029 | 0.67 | 0.33 | 0.56 | 0.77 | 0.66 | 0.85 |
| 3 | 140914251 | rs2086182 | 2.471 | -0.532 | 3.109 | 2.113 | -0.011 | -0.013 | -0.013 | 0 | 0.69 | 0.49 | 0.59 | 0.78 | 0.68 | 0.88 |
| 3 | 140920166 | rs7650287 | -1.043 | -0.682 | 2.642 | 1.972 | -0.02 | -0.028 | -0.028 | 0 | 0.07 | 0.11 | 0.03 | 0 | 0.03 | 0.02 |

| | | | | | | | | | | | | | | | | |
|---|---|---|---|---|---|---|---|---|---|---|---|---|---|---|---|---|
| 3 | 140938546 | rs1877222 | 2.118 | -0.972 | 2.632 | 2.209 | 0.006 | 0.006 | 0.006 | 0 | 0.66 | 0.46 | 0.62 | 0.63 | 0.63 | 0.72 |
| 3 | 140940921 | rs1511370 | 1.1 | -0.956 | 2.554 | 2.239 | -0.039 | 0.039 | 0.039 | 0 | 0.77 | 0.97 | 0.8 | 0.89 | 0.76 | 0.78 |
| 3 | 140941407 | rs1511372 | 2.123 | -0.932 | 2.702 | 2.233 | 0.01 | 0.012 | 0.012 | 0 | 0.66 | 0.53 | 0.63 | 0.62 | 0.63 | 0.72 |
| 3 | 140942963 | rs4683586 | 2.551 | -0.921 | 2.249 | 2.251 | 0.018 | -0.017 | -0.017 | 0 | 0.55 | 0.27 | 0.47 | 0.54 | 0.58 | 0.63 |
| 3 | 140956014 | rs6439985 | 0.504 | -0.714 | 2.013 | 2.287 | -0.033 | 0.006 | 0.006 | 0 | 0.87 | 0.97 | 0.83 | 0.92 | 0.79 | 0.93 |
| 3 | 140956600 | rs13069904 | 2.691 | -0.781 | 2.056 | 2.323 | 0.014 | -0.049 | -0.049 | 0 | 0.54 | 0.09 | 0.45 | 0.55 | 0.57 | 0.64 |
| 3 | 140958061 | rs6779870 | 2.435 | -0.72 | 2.234 | 2.153 | 0.001 | 0.003 | 0.003 | 0 | 0.6 | 0.3 | 0.59 | 0.62 | 0.59 | 0.7 |
| 3 | 140960116 | rs9819541 | -1.589 | -0.685 | 2.153 | 2.031 | 0.014 | -0.001 | -0.001 | 0.288 | 0.17 | 0.26 | 0.21 | 0.15 | 0.22 | 0.06 |
| 3 | 140964265 | rs7638594 | -1.195 | -0.772 | 2.171 | 2.149 | -0.023 | 0.009 | 0.009 | 0.11 | 0.07 | 0.15 | 0.14 | 0.1 | 0.02 | 0.08 |
| 3 | 140965583 | rs1877219 | 0.389 | -0.67 | 2.05 | 2.485 | -0.054 | -0.007 | -0.007 | 0 | 0.87 | 0.99 | 0.83 | 0.89 | 0.78 | 0.93 |
| 3 | 140970556 | rs11713226 | -1.313 | -0.601 | 2.306 | 2.428 | 0.051 | -0.029 | -0.029 | 0 | 0.16 | 0.47 | 0.13 | 0.09 | 0.17 | 0.07 |
| 3 | 140978098 | rs9682606 | -1.201 | -0.605 | 2.302 | 2.451 | 0.037 | -0.018 | -0.018 | 0 | 0.15 | 0.36 | 0.11 | 0.09 | 0.16 | 0.05 |
| 3 | 140983891 | rs1026716 | 2.821 | -0.766 | 1.687 | 2.548 | -0.011 | -0.049 | -0.049 | 1.383 | 0.52 | 0.08 | 0.43 | 0.43 | 0.56 | 0.59 |
| 3 | 140984705 | rs1318221 | 2.051 | -0.776 | 1.808 | 2.461 | -0.02 | 0.023 | 0.023 | 0.106 | 0.68 | 0.59 | 0.56 | 0.52 | 0.72 | 0.66 |
| 3 | 140988528 | rs6769909 | -2.397 | -0.747 | 0.786 | 2.497 | 0.032 | -0.032 | -0.032 | 0 | 0.35 | 0.81 | 0.38 | 0.47 | 0.23 | 0.34 |
| 3 | 140990456 | rs9819731 | 2.404 | -0.616 | 0.585 | 2.11 | -0.01 | -0.013 | -0.013 | 0 | 0.58 | 0.2 | 0.57 | 0.53 | 0.57 | 0.67 |
| 3 | 140992753 | rs4683592 | 0.347 | -0.726 | 0.587 | 2.11 | -0.054 | -0.039 | -0.039 | 0 | 0.87 | 0.99 | 0.83 | 0.8 | 0.79 | 0.92 |
| 3 | 140995898 | rs11921252 | -1.811 | -0.778 | 0.673 | 2.41 | -0.013 | 0.027 | 0.027 | 0.027 | 0.28 | 0.63 | 0.25 | 0.38 | 0.18 | 0.25 |
| 3 | 140996410 | rs10935425 | 2.787 | -0.778 | 0.616 | 2.399 | -0.01 | 0.015 | 0.015 | 1.576 | 0.52 | 0.09 | 0.54 | 0.38 | 0.57 | 0.58 |
| 3 | 140998214 | rs13087169 | 2.004 | -0.215 | -0.209 | 2.399 | 0.009 | 0.022 | 0.022 | 0.78 | 0.67 | 0.4 | 0.66 | 0.54 | 0.64 | 0.69 |
| 3 | 140999116 | rs6786620 | -0.44 | -0.191 | -0.396 | 2.012 | 0.01 | 0 | 0 | 0 | 0.1 | 0.06 | 0.05 | 0.07 | 0.03 | 0.13 |
| 3 | 141002564 | rs10935427 | 2.217 | -0.121 | -0.552 | 2.136 | 0.004 | 0.011 | 0.011 | 0.184 | 0.58 | 0.23 | 0.56 | 0.41 | 0.59 | 0.66 |
| 3 | 141003094 | rs9880195 | 0.622 | 0.043 | -0.86 | 1.624 | -0.053 | -0.055 | -0.055 | 0 | 0.9 | 0.99 | 0.83 | 0.79 | 0.81 | 0.92 |
| 3 | 141007003 | rs6779015 | -1.443 | -0.487 | -0.351 | 1.105 | -0.025 | -0.011 | -0.011 | 3.162 | 0.13 | 0.18 | 0.06 | 0.08 | 0.07 | 0.13 |
| 3 | 141011630 | rs3210458 | -1.286 | -0.039 | -0.821 | 1.177 | 0.007 | 0 | 0 | 0 | 0.15 | 0.1 | 0.07 | 0.12 | 0.11 | 0.1 |
| 3 | 141022270 | rs4683401 | -1.66 | 0.035 | -1.158 | 1.558 | 0.015 | -0.052 | -0.052 | 0.234 | 0.22 | 0.07 | 0.1 | 0.28 | 0.18 | 0.14 |
| 3 | 141022487 | rs11917422 | -0.846 | 0.227 | -1.951 | 1.458 | -0.035 | 0.006 | 0.006 | 0.116 | 0.23 | 0.86 | 0.24 | 0.2 | 0.24 | 0.23 |
| 3 | 141023325 | rs9809435 | -1.681 | -0.603 | -2.072 | 1.421 | 0.089 | -0.014 | -0.014 | 0 | 0.43 | 0.85 | 0.29 | 0.39 | 0.25 | 0.3 |
| 3 | 141026385 | rs11927391 | -1.629 | -0.655 | -0.561 | 1.548 | -0.001 | 0.057 | 0.057 | 0 | 0.14 | 0.36 | 0.08 | 0.16 | 0.09 | 0.16 |
| 3 | 141027614 | rs9968231 | -1.358 | -0.634 | -0.269 | 1.328 | 0.002 | -0.013 | -0.013 | 0 | 0.12 | 0.22 | 0.06 | 0.06 | 0.09 | 0.15 |
| 3 | 141030543 | rs12630390 | -1.065 | -0.746 | -0.335 | 1.357 | 0.008 | 0 | 0 | 0 | 0.09 | 0.05 | 0.04 | 0.05 | 0.04 | 0.14 |
| 3 | 141042222 | rs6777886 | -0.849 | -0.275 | 0.081 | 0.43 | -0.004 | 0.025 | 0.025 | 0 | 0.34 | 0.27 | 0.21 | 0 | 0.37 | 0.2 |
| 3 | 141046859 | rs4683600 | -1.891 | -0.517 | 0.209 | 0.721 | 0.002 | 0.083 | 0.083 | 0 | 0.51 | 0.87 | 0.48 | 0.39 | 0.46 | 0.38 |
| 3 | 141049791 | rs1983234 | 0.631 | -0.408 | -0.053 | 0.659 | -0.023 | 0.054 | 0.054 | 0 | 0.1 | 0 | 0.05 | 0.01 | 0.08 | 0.05 |
| 3 | 141055901 | rs6807533 | -1.933 | -0.345 | 0.241 | 0.872 | -0.012 | 0.038 | 0.038 | 0.053 | 0.16 | 0.7 | 0.26 | 0.39 | 0.09 | 0.19 |
| 3 | 141058687 | rs4683602 | -1.471 | -0.348 | 0.604 | 0.799 | 0.075 | 0.182 | 0.182 | 0.402 | 0.08 | 0.39 | 0.21 | 0.15 | 0.05 | 0.03 |
| 3 | 141061438 | rs9823785 | -0.552 | -0.638 | -0.473 | 0.564 | -0.001 | 0.029 | 0.029 | 0 | 0.26 | 0.24 | 0.18 | 0.06 | 0.3 | 0.15 |
| 3 | 141063953 | rs9838625 | -2.057 | -0.642 | -0.65 | 0.666 | -0.047 | 0.011 | 0.011 | 0 | 0.44 | 0.97 | 0.44 | 0.34 | 0.42 | 0.34 |
| 3 | 141072289 | rs1346408 | -2.007 | -0.345 | -0.688 | 0.257 | -0.067 | 0.105 | 0.105 | 0.031 | 0.42 | 0.97 | 0.43 | 0.23 | 0.42 | 0.33 |
| 3 | 141073675 | rs6784404 | -1.556 | -0.35 | -0.645 | 0.259 | -0.055 | 0.154 | 0.154 | 0.057 | 0.39 | 0.94 | 0.41 | 0.14 | 0.42 | 0.27 |
| 3 | 141074962 | rs13095453 | -0.487 | -0.342 | -0.575 | 0.168 | -0.001 | 0.003 | 0.003 | 0 | 0.26 | 0.22 | 0.17 | 0 | 0.31 | 0.16 |
| 3 | 141079309 | rs7613516 | -1.858 | -0.234 | -0.986 | 0.671 | -0.058 | -0.023 | -0.023 | 0 | 0.43 | 0.97 | 0.44 | 0.34 | 0.42 | 0.33 |
| 3 | 141093285 | rs7624084 | -1.755 | 0.28 | -0.881 | 0.689 | -0.046 | 0.026 | 0.026 | 0 | 0.42 | 0.88 | 0.45 | 0.33 | 0.43 | 0.36 |
| 3 | 141094209 | rs6440003 | -1.474 | 0.272 | -0.689 | 0.671 | -0.057 | -0.013 | -0.013 | 0 | 0.36 | 0.9 | 0.43 | 0.32 | 0.43 | 0.31 |
| 3 | 141096736 | rs9822195 | -0.939 | 0.305 | 0.224 | 0.599 | 0.002 | -0.098 | -0.098 | 0 | 0.29 | 0.38 | 0.18 | 0 | 0.32 | 0.15 |
| 3 | 141100280 | rs6764769 | -1.848 | 0.312 | 0.142 | 0.614 | -0.056 | -0.012 | -0.012 | 0 | 0.36 | 0.9 | 0.43 | 0.33 | 0.42 | 0.29 |
| 3 | 141102833 | rs6763931 | -1.856 | 0.313 | 0.142 | 0.614 | -0.056 | -0.011 | -0.011 | 0.209 | 0.36 | 0.89 | 0.43 | 0.33 | 0.43 | 0.29 |
| 3 | 141105570 | rs724016 | -1.883 | 0.316 | 0.191 | 0.563 | -0.04 | 0.003 | 0.003 | 0 | 0.35 | 0.82 | 0.42 | 0.32 | 0.42 | 0.27 |
| 3 | 141107612 | rs10513136 | 0.738 | 0.708 | -0.074 | 0.318 | -0.024 | 0.04 | 0.04 | 0 | 0.07 | 0 | 0.03 | 0 | 0.06 | 0.05 |
| 3 | 141121814 | rs2871960 | -1.603 | 0.647 | -0.273 | 0.347 | -0.069 | -0.01 | -0.01 | 0.012 | 0.36 | 0.9 | 0.43 | 0.33 | 0.43 | 0.27 |
| 3 | 141128804 | rs13099193 | -0.451 | 0.673 | -0.117 | 0.317 | -0.035 | 0.067 | 0.067 | 0 | 0.24 | 0.12 | 0.17 | 0 | 0.31 | 0.12 |
| 3 | 141133960 | rs13091182 | -0.391 | 0.619 | -0.495 | 0.293 | -0.048 | 0.093 | 0.093 | 0 | 0.23 | 0.09 | 0.17 | 0 | 0.31 | 0.12 |
| 3 | 141135126 | rs11919595 | -2.091 | 0.642 | -0.205 | 0.38 | 0.085 | 0.081 | 0.081 | 1.865 | 0.06 | 0.6 | 0.16 | 0.09 | 0.07 | 0.05 |
| 3 | 141137035 | rs9825379 | -2.161 | 0.62 | 0.287 | 0.444 | 0.135 | 0.296 | 0.296 | 0.632 | 0.05 | 0.35 | 0.09 | 0.18 | 0.07 | 0.06 |
| 3 | 141140968 | rs9846396 | -1.741 | 0.585 | 0.363 | 0.415 | -0.02 | 0.033 | 0.033 | 0 | 0.33 | 0.74 | 0.41 | 0.23 | 0.42 | 0.28 |
| 3 | 141145315 | rs6762826 | -2.911 | 0.624 | 0.659 | 0.632 | 0.137 | 0.394 | 0.394 | 0.558 | 0.07 | 0.45 | 0.14 | 0.28 | 0.09 | 0.1 |
| 3 | 141158212 | rs7612543 | -1.985 | 0.773 | 0.421 | 0.466 | -0.008 | 0.022 | 0.022 | 0 | 0.38 | 0.69 | 0.44 | 0.2 | 0.42 | 0.29 |
| 3 | 141158614 | rs7636914 | -0.461 | 0.862 | 0.094 | 0.386 | -0.048 | 0.07 | 0.07 | 0 | 0.22 | 0.12 | 0.18 | 0 | 0.31 | 0.14 |
| 3 | 141160882 | rs7626871 | 0.363 | 0.737 | 0.23 | 0.625 | -0.03 | -0.055 | -0.055 | 0 | 0.09 | 0.01 | 0.2 | 0.22 | 0.09 | 0.1 |
| 3 | 141162185 | rs16851435 | -0.486 | 0.734 | 0.261 | 0.625 | -0.001 | -0.012 | -0.012 | 0 | 0.12 | 0.09 | 0.22 | 0.31 | 0.11 | 0.16 |
| 3 | 141205185 | rs3821712 | -1.98 | 0.351 | 0.264 | 0.174 | 0 | 0.008 | 0.008 | 0 | 0.39 | 0.67 | 0.46 | 0.29 | 0.39 | 0.32 |
| 3 | 141208684 | rs11717842 | -0.186 | 0.397 | -0.074 | -0.178 | -0.091 | 0.141 | 0.141 | 0 | 0.21 | 0.02 | 0.16 | 0 | 0.29 | 0.13 |
| 3 | 141218057 | rs10513140 | 2.275 | -0.099 | -0.661 | 0.03 | -0.079 | -0.012 | -0.012 | 0 | 0.34 | 0.01 | 0.3 | 0.32 | 0.32 | 0.41 |
| 3 | 141272216 | rs9850416 | -0.762 | -0.169 | 0.117 | -0.087 | -0.002 | 0.001 | 0.001 | 0 | 0.26 | 0.25 | 0.2 | 0.08 | 0.32 | 0.22 |
| 3 | 141275436 | rs16851483 | 0.993 | -0.153 | 0.078 | -0.06 | -0.025 | -0.055 | -0.055 | 0 | 0.09 | 0 | 0.23 | 0.21 | 0.06 | 0.08 |
| 3 | 141301451 | rs3821710 | -1.624 | -0.155 | 0.108 | -0.055 | 0 | -0.001 | -0.001 | 0 | 0.38 | 0.57 | 0.45 | 0.29 | 0.38 | 0.28 |
| 3 | 141306013 | rs2035935 | -0.715 | -0.051 | 0.03 | 0.368 | -0.001 | 0.112 | 0.112 | 0 | 0.13 | 0.3 | 0.26 | 0.21 | 0.07 | 0.09 |
| 3 | 141326602 | rs295322 | 0.054 | 0.11 | -0.15 | 0.44 | 0.007 | 0.111 | 0.111 | 0 | 0.11 | 0.25 | 0.25 | 0.21 | 0.07 | 0.08 |
| 3 | 141338641 | rs295301 | -0.579 | -0.098 | -0.84 | 0.11 | -0.002 | -0.001 | -0.001 | 0.019 | 0.41 | 0.45 | 0.43 | 0.34 | 0.34 | 0.29 |

| | | | | | | | | | | | | | | | | |
|---|---|---|---|---|---|---|---|---|---|---|---|---|---|---|---|---|
| 3 | 141338997 | rs295302 | 0.464 | 0.387 | -1.3 | -0.119 | -0.015 | 0.043 | 0.043 | 0 | 0.78 | 0.93 | 0.8 | 0.88 | 0.78 | 0.81 |
| 3 | 141341000 | rs295303 | -0.118 | 0.737 | -0.38 | -0.184 | 0.026 | -0.041 | -0.041 | 0 | 0.83 | 0.61 | 0.85 | 0.94 | 0.8 | 0.85 |
| 3 | 141344198 | rs13072855 | 0.876 | 0.765 | -0.275 | 0.389 | 0.001 | -0.002 | -0.002 | 0 | 0.09 | 0.11 | 0.11 | 0.19 | 0.08 | 0.07 |
| 3 | 141345102 | rs2971542 | 0.43 | 0.583 | -0.829 | -0.951 | 0 | -0.003 | -0.003 | 0 | 0.44 | 0.48 | 0.59 | 0.76 | 0.44 | 0.59 |
| 3 | 141345678 | rs11711597 | 0.441 | 0.773 | -1.376 | -1.459 | 0.005 | -0.003 | -0.003 | 0 | 0.25 | 0.16 | 0.43 | 0.74 | 0.23 | 0.48 |
| 3 | 141346188 | rs2131495 | -0.308 | 0.69 | -1.594 | -1.81 | 0.007 | -0.016 | -0.016 | 0 | 0.62 | 0.46 | 0.65 | 0.77 | 0.53 | 0.64 |
| 3 | 141347053 | rs2640006 | 0.111 | 0.533 | -1.701 | -1.92 | -0.001 | 0.009 | 0.009 | 0 | 0.43 | 0.42 | 0.57 | 0.77 | 0.41 | 0.58 |
| 3 | 141349381 | rs2677420 | -0.308 | 0.283 | -1.506 | -2.232 | -0.008 | 0.026 | 0.026 | 0 | 0.57 | 0.59 | 0.62 | 0.78 | 0.48 | 0.69 |
| 3 | 141361654 | rs295313 | -0.799 | 0.384 | -1.571 | -2.375 | -0.003 | -0.042 | -0.042 | 0 | 0.66 | 0.64 | 0.73 | 0.98 | 0.59 | 0.78 |
| 3 | 141362225 | rs295315 | 0.688 | 0.356 | -1.608 | -2.436 | -0.004 | -0.008 | -0.008 | 0 | 0.17 | 0.13 | 0.4 | 0.76 | 0.17 | 0.46 |
| 3 | 141365883 | rs7618625 | 0.327 | 0.053 | -1.666 | -2.351 | 0.006 | 0.017 | 0.017 | 0.235 | 0.49 | 0.51 | 0.33 | 0.22 | 0.43 | 0.32 |
| 3 | 141367440 | rs953265 | 2.365 | 0.009 | -1.905 | -2.394 | 0.03 | 0.098 | 0.098 | 0.196 | 0.19 | 0.03 | 0.07 | 0 | 0.12 | 0.07 |
| 3 | 141368245 | rs1994846 | 1.469 | 0.035 | -1.858 | -2.38 | -0.003 | -0.125 | -0.125 | 0 | 0.21 | 0.42 | 0.15 | 0.03 | 0.19 | 0.17 |
| 3 | 141371399 | rs2131494 | 1.541 | 0.048 | -1.908 | -2.374 | -0.002 | -0.11 | -0.11 | 0.369 | 0.2 | 0.39 | 0.14 | 0.03 | 0.19 | 0.17 |
| 3 | 141371896 | rs13060724 | 2.082 | -0.213 | -2.615 | -1.878 | -0.039 | -0.049 | -0.049 | 0.148 | 0.08 | 0.01 | 0.1 | 0.19 | 0.12 | 0.09 |
| 3 | 141377091 | rs17195414 | 1.65 | -0.403 | -2.376 | -1.694 | 0.032 | 0.094 | 0.094 | 0.09 | 0.19 | 0.03 | 0.07 | 0 | 0.12 | 0.07 |
| 3 | 141377330 | rs868767 | 0.226 | 0.109 | -2.004 | -1.132 | 0.009 | -0.055 | -0.055 | 0 | 0.13 | 0.04 | 0.35 | 0.74 | 0.1 | 0.42 |
| 3 | 141382051 | rs2030184 | 0.35 | 0.123 | -2.077 | -1.184 | -0.008 | 0.046 | 0.046 | 0.128 | 0.54 | 0.24 | 0.46 | 0.39 | 0.53 | 0.42 |
| 3 | 141385690 | rs2312500 | 1.171 | 0.44 | -2.589 | -2.176 | -0.003 | -0.002 | -0.002 | 2.368 | 0.32 | 0.22 | 0.5 | 0.7 | 0.29 | 0.43 |
| 3 | 141387271 | rs4683618 | 0.505 | 0.486 | -2.666 | -2.269 | -0.001 | 0.059 | 0.059 | 0 | 0.43 | 0.24 | 0.57 | 0.73 | 0.4 | 0.53 |
| 3 | 141395810 | rs12485838 | 0.28 | 0.229 | -2.401 | -1.997 | -0.002 | 0.004 | 0.004 | 0.056 | 0.67 | 0.35 | 0.69 | 0.73 | 0.65 | 0.65 |
| 3 | 141400107 | rs1122929 | 1.062 | 0.201 | -2.694 | -1.603 | -0.005 | 0.037 | 0.037 | 0.781 | 0.11 | 0.03 | 0.07 | 0.02 | 0.12 | 0.1 |
| 3 | 141402645 | rs4683619 | 0.2 | 0.365 | -2.157 | -1.165 | 0.002 | 0.004 | 0.004 | 0 | 0.77 | 0.68 | 0.76 | 0.75 | 0.73 | 0.7 |
| 3 | 141404210 | rs925861 | 0.361 | 0.212 | -2.69 | -0.716 | -0.002 | -0.011 | -0.011 | 0 | 0.11 | 0.1 | 0.12 | 0.22 | 0.13 | 0.1 |
| 3 | 141405708 | rs12485563 | 0.074 | 0.212 | -2.763 | -0.574 | -0.014 | -0.018 | -0.018 | 0.056 | 0.29 | 0.16 | 0.5 | 0.69 | 0.27 | 0.4 |
| 3 | 141408287 | rs6791307 | -0.631 | -0.534 | -2.586 | -1.009 | 0.05 | 0.095 | 0.095 | 0 | 0.24 | 0.09 | 0.22 | 0 | 0.16 | 0.12 |
| 3 | 141408691 | rs6440031 | 0.16 | -0.011 | -2.309 | -0.649 | 0.008 | 0.023 | 0.023 | 0 | 0.89 | 0.98 | 0.93 | 0.96 | 0.92 | 0.88 |
| 3 | 141409701 | rs17195428 | 0.287 | -0.146 | -2.746 | -0.753 | -0.036 | 0.003 | 0.003 | 0.118 | 0.37 | 0.08 | 0.25 | 0.28 | 0.35 | 0.3 |
| 3 | 141410864 | rs4683620 | 0.459 | -0.211 | -1.681 | -0.618 | 0.004 | -0.015 | -0.015 | 0 | 0.83 | 0.9 | 0.93 | 0.99 | 0.91 | 0.81 |
| 3 | 141417021 | rs4683623 | -0.025 | 0.145 | -1.411 | -0.561 | 0.004 | 0.021 | 0.021 | 0 | 0.53 | 0.59 | 0.38 | 0.29 | 0.46 | 0.5 |
| 3 | 141417155 | rs11721133 | 0.213 | -0.452 | -1.206 | -1.228 | -0.008 | -0.015 | -0.015 | 0.119 | 0.46 | 0.48 | 0.31 | 0.27 | 0.34 | 0.37 |
| 3 | 141418843 | rs6772861 | -0.58 | -0.372 | -1.634 | -1.543 | -0.032 | 0.066 | 0.066 | 0 | 0.85 | 0.97 | 0.89 | 0.97 | 0.85 | 0.89 |
| 3 | 141418907 | rs9855944 | -0.217 | -0.869 | -0.972 | -2.037 | -0.009 | -0.062 | -0.062 | 1.767 | 0.41 | 0.48 | 0.29 | 0.14 | 0.36 | 0.35 |
| 3 | 141419203 | rs9876068 | 0.777 | -0.793 | -0.893 | -1.929 | 0.002 | -0.007 | -0.007 | 0.289 | 0.08 | 0.11 | 0.06 | 0 | 0.05 | 0.02 |
| 3 | 141420562 | rs9882031 | -0.369 | -0.96 | -0.453 | -1.69 | -0.029 | -0.005 | -0.005 | 0 | 0.19 | 0.33 | 0.12 | 0.13 | 0.12 | 0.14 |
| 3 | 141420575 | rs9824350 | 0.922 | -0.875 | -0.612 | -1.564 | -0.005 | -0.026 | -0.026 | 0.541 | 0.18 | 0.26 | 0.14 | 0.09 | 0.15 | 0.15 |
| 3 | 141425418 | rs6440032 | 0.013 | -1.043 | -0.77 | -1.768 | -0.001 | -0.017 | -0.017 | 0 | 0.32 | 0.31 | 0.2 | 0.13 | 0.26 | 0.36 |
| 3 | 141426135 | rs16851667 | -0.584 | -1.221 | -0.91 | -1.674 | -0.008 | -0.013 | -0.013 | 0 | 0.43 | 0.45 | 0.27 | 0.26 | 0.34 | 0.46 |
| 3 | 141426649 | rs17787760 | 0.582 | -0.9 | -1.319 | -1.44 | -0.001 | 0.065 | 0.065 | 0.322 | 0.15 | 0.02 | 0.06 | 0.04 | 0.11 | 0.18 |
| 3 | 141428703 | rs9821235 | -0.662 | -0.082 | -1.199 | -0.67 | -0.006 | -0.013 | -0.013 | 0.079 | 0.65 | 0.68 | 0.48 | 0.52 | 0.6 | 0.55 |
| 3 | 141428857 | rs9842908 | 0.03 | -0.407 | -1.177 | -0.617 | 0.012 | 0.016 | 0.016 | 0 | 0.21 | 0.14 | 0.12 | 0.2 | 0.16 | 0.18 |
| 3 | 141440311 | rs7630533 | 0.239 | -0.043 | -0.565 | -0.529 | 0.003 | -0.096 | -0.096 | 0 | 0.32 | 0.51 | 0.22 | 0.13 | 0.3 | 0.28 |
| 3 | 141440993 | rs6782796 | 0.874 | 0.067 | -0.222 | -0.629 | 0.003 | -0.128 | -0.128 | 0.197 | 0.23 | 0.37 | 0.14 | 0.01 | 0.2 | 0.13 |
| 3 | 141442131 | rs9847061 | 0.97 | -0.183 | -0.955 | -0.157 | 0.01 | -0.043 | -0.043 | 0 | 0.21 | 0.11 | 0.34 | 0.38 | 0.17 | 0.38 |
| 3 | 141444530 | rs10433346 | -0.198 | -0.055 | -0.193 | 0.151 | -0.001 | -0.009 | -0.009 | 0 | 0.34 | 0.62 | 0.25 | 0.21 | 0.31 | 0.29 |
| 3 | 141448766 | rs1105082 | 0.068 | -0.057 | -0.303 | -0.18 | -0.009 | -0.096 | -0.096 | 0 | 0.32 | 0.54 | 0.22 | 0.1 | 0.27 | 0.27 |
| 3 | 141457998 | rs1980190 | -0.332 | -0.088 | -0.649 | -0.185 | 0.03 | 0.095 | 0.095 | 0 | 0.37 | 0.78 | 0.26 | 0.17 | 0.27 | 0.27 |
| 3 | 141463298 | rs16851720 | 0.573 | -0.251 | 0.555 | -0.285 | -0.002 | -0.048 | -0.048 | 0.79 | 0.25 | 0.3 | 0.16 | 0.08 | 0.2 | 0.14 |
| 3 | 141463803 | rs17195449 | 1.682 | -0.257 | 0.434 | -0.052 | -0.019 | -0.028 | -0.028 | 0 | 0.05 | 0 | 0.06 | 0.11 | 0.06 | 0.13 |
| 3 | 141466147 | rs1055690 | -1.242 | -0.187 | 0.803 | -0.122 | -0.003 | 0.001 | 0.001 | 0 | 0.14 | 0.38 | 0.13 | 0.15 | 0.12 | 0.16 |
| 3 | 141469078 | rs16851736 | 1.091 | -0.283 | 0.636 | -0.137 | -0.005 | -0.066 | -0.066 | 0.025 | 0.21 | 0.27 | 0.12 | 0.02 | 0.15 | 0.11 |
| 3 | 141469631 | rs16851740 | 0.491 | -0.659 | 0.363 | 0.005 | -0.004 | 0.001 | 0.001 | 0 | 0.07 | 0.09 | 0.06 | 0.09 | 0.06 | 0.13 |
| 3 | 141484860 | rs2677421 | 0.624 | -0.222 | -0.184 | 0.286 | 0.011 | 0.009 | 0.009 | 0 | 0.5 | 0.3 | 0.5 | 0.5 | 0.45 | 0.5 |
| 3 | 141492170 | rs7616846 | -0.139 | -0.238 | 0.204 | 0.645 | -0.006 | -0.047 | -0.047 | 0.04 | 0.8 | 0.92 | 0.73 | 0.72 | 0.76 | 0.73 |
| 3 | 141500037 | rs9835566 | 1.446 | -0.358 | 0.137 | 0.367 | 0.032 | 0.053 | 0.053 | 0 | 0.54 | 0.22 | 0.55 | 0.59 | 0.59 | 0.52 |
| 3 | 141501812 | rs2681699 | 0.477 | -0.461 | 0.321 | 0.244 | 0.002 | 0.008 | 0.008 | 0 | 0.13 | 0.09 | 0.07 | 0.02 | 0.11 | 0.05 |
| 3 | 141512513 | rs11921607 | -1.408 | -0.496 | 0.229 | 0.256 | 0.03 | 0.042 | 0.042 | 0.111 | 0.29 | 0.77 | 0.18 | 0.11 | 0.17 | 0.21 |
| 3 | 141515136 | rs6806847 | -0.351 | -0.332 | 0.149 | 0.329 | -0.038 | -0.096 | -0.096 | 0 | 0.8 | 0.98 | 0.74 | 0.69 | 0.77 | 0.72 |
| 3 | 141526995 | rs13317804 | -0.131 | -0.373 | 0.495 | 0.4 | 0.005 | -0.02 | -0.02 | 0.01 | 0.78 | 0.86 | 0.71 | 0.67 | 0.76 | 0.76 |
| 3 | 141528376 | rs9835447 | 1.562 | -0.383 | 0.068 | 0.278 | 0.052 | 0.013 | 0.013 | 0 | 0.53 | 0.2 | 0.52 | 0.55 | 0.59 | 0.58 |
| 3 | 141540962 | rs7615804 | -0.27 | -0.005 | -0.315 | 0.375 | -0.022 | -0.112 | -0.112 | 0 | 0.8 | 0.97 | 0.71 | 0.64 | 0.76 | 0.77 |
| 3 | 141541105 | rs13071985 | 0.314 | 0.004 | -0.251 | 0.145 | 0 | 0.013 | 0.013 | 0 | 0.13 | 0.1 | 0.11 | 0.1 | 0.13 | 0.08 |
| 3 | 141546876 | rs2861381 | 1.033 | 0.067 | 0.088 | 0.197 | 0.008 | -0.004 | -0.004 | 0 | 0.58 | 0.65 | 0.58 | 0.55 | 0.59 | 0.59 |
| 3 | 141551574 | rs4586785 | 0.017 | 0.277 | 0.277 | -0.322 | 0 | -0.009 | -0.009 | 0 | 0.07 | 0.07 | 0.03 | 0.02 | 0.06 | 0.06 |
| 3 | 141551881 | rs6440041 | -1.361 | 0.133 | 0.772 | -0.458 | -0.004 | -0.027 | -0.027 | 0.292 | 0.32 | 0.62 | 0.22 | 0.15 | 0.27 | 0.16 |
| 3 | 141552043 | rs13083686 | -1.039 | 0.068 | 0.285 | -0.314 | -0.016 | -0.068 | -0.068 | 0.046 | 0.24 | 0.34 | 0.11 | 0.08 | 0.17 | 0.11 |
| 3 | 141555626 | rs6801834 | 0.204 | 0.306 | -0.396 | -0.136 | 0.002 | -0.006 | -0.006 | 0 | 0.11 | 0.08 | 0.05 | 0 | 0.1 | 0.04 |
| 3 | 141563276 | rs4683630 | -0.502 | 0.518 | -0.268 | -0.121 | 0 | -0.016 | -0.016 | 0 | 0.45 | 0.56 | 0.35 | 0.19 | 0.48 | 0.3 |
| 3 | 141564786 | rs12107859 | 0.301 | 0.268 | -0.202 | 0.177 | -0.007 | -0.011 | -0.011 | 0 | 0.14 | 0.22 | 0.09 | 0.09 | 0.11 | 0.15 |

| | | | | | | | | | | | | | | | | |
|---|---|---|---|---|---|---|---|---|---|---|---|---|---|---|---|---|
| 3 | 141566263 | rs16846283 | -0.313 | -0.303 | -0.312 | 1.103 | -0.029 | 0.099 | 0.099 | 0 | 0.09 | 0.17 | 0.18 | 0.31 | 0.04 | 0.25 |
| 3 | 141568628 | rs6792646 | 0.934 | -0.055 | -0.559 | 0.586 | -0.002 | -0.008 | -0.008 | 0 | 0.35 | 0.27 | 0.37 | 0.41 | 0.36 | 0.31 |
| 3 | 141569773 | rs12485634 | -0.601 | -0.165 | -0.638 | 0.794 | 0.006 | -0.02 | -0.02 | 0 | 0.25 | 0.22 | 0.32 | 0.38 | 0.23 | 0.32 |
| 3 | 141570294 | rs11710724 | 0.304 | 0.031 | -1.016 | 0.597 | -0.019 | -0.016 | -0.016 | 0 | 0.06 | 0.01 | 0.07 | 0.11 | 0.06 | 0.09 |
| 3 | 141572495 | rs13078635 | -1.162 | -0.426 | -0.951 | 0.484 | 0.026 | -0.015 | -0.015 | 0.002 | 0.35 | 0.72 | 0.41 | 0.51 | 0.27 | 0.34 |
| 3 | 141573761 | rs9832234 | 0.667 | -0.179 | -1.102 | 0.027 | -0.019 | 0.066 | 0.066 | 0.002 | 0.15 | 0.06 | 0.17 | 0.03 | 0.16 | 0.05 |
| 3 | 141580483 | rs4683634 | -1.067 | 0.592 | -0.717 | 0.59 | -0.005 | -0.026 | -0.026 | 0 | 0.23 | 0.19 | 0.23 | 0.52 | 0.24 | 0.31 |
| 3 | 141589505 | rs6440043 | -1.56 | 0.201 | -0.652 | 0.197 | 0.004 | 0.01 | 0.01 | 0 | 0.48 | 0.83 | 0.48 | 0.59 | 0.45 | 0.44 |
| 3 | 141596848 | rs1560413 | -1.511 | 0.195 | -0.035 | 0.216 | 0.009 | 0.015 | 0.015 | 0 | 0.48 | 0.66 | 0.45 | 0.59 | 0.45 | 0.44 |
| 3 | 141602770 | rs11721024 | 0.149 | 0.132 | -0.185 | 0.396 | -0.029 | -0.025 | -0.025 | 0 | 0.11 | 0.01 | 0.13 | 0.14 | 0.12 | 0.08 |
| 3 | 141605961 | rs1897139 | -0.005 | 0.112 | 0.226 | 0.395 | -0.035 | -0.047 | -0.047 | 0 | 0.1 | 0.02 | 0.1 | 0.39 | 0.13 | 0.23 |
| 3 | 141606244 | rs9810380 | -0.308 | 0.113 | 0.331 | 0.373 | -0.004 | 0.008 | 0.008 | 0 | 0.12 | 0.1 | 0.1 | 0.4 | 0.13 | 0.23 |
| 3 | 141610807 | rs4683645 | -1.628 | 0.139 | 0.215 | 0.316 | -0.007 | 0.007 | 0.007 | 0 | 0.48 | 0.85 | 0.48 | 0.59 | 0.45 | 0.44 |
| 3 | 141622055 | rs6440048 | 0.004 | -0.093 | 0.514 | -0.506 | -0.003 | -0.042 | -0.042 | 0 | 0.23 | 0.24 | 0.2 | 0.04 | 0.2 | 0.13 |
| 3 | 141630680 | rs16851955 | 1.137 | 0.081 | 0.163 | -0.897 | -0.001 | -0.019 | -0.019 | 0 | 0.09 | 0.07 | 0.09 | 0 | 0.09 | 0.01 |
| 3 | 141630722 | rs6799078 | 1.633 | -0.026 | -0.054 | -1.215 | -0.005 | 0.001 | 0.001 | 1.332 | 0.35 | 0.35 | 0.39 | 0.65 | 0.38 | 0.58 |
| 3 | 141635598 | rs11712817 | 1.698 | -0.033 | 0.007 | -1.332 | -0.004 | 0.001 | 0.001 | 0.702 | 0.34 | 0.33 | 0.38 | 0.67 | 0.38 | 0.57 |
| 3 | 141644808 | rs1050190 | 0.215 | -0.228 | 0.067 | -1.573 | -0.01 | -0.074 | -0.074 | 0 | 0.3 | 0.34 | 0.24 | 0.04 | 0.25 | 0.22 |
| 3 | 141646962 | rs16852004 | 1.497 | -0.138 | 0.226 | -1.417 | 0.003 | -0.008 | -0.008 | 0 | 0.36 | 0.46 | 0.39 | 0.68 | 0.38 | 0.57 |
| 3 | 141647298 | rs13319758 | 0.899 | -0.186 | 0.292 | -1.443 | 0.004 | -0.032 | -0.032 | 0 | 0.44 | 0.52 | 0.49 | 0.83 | 0.47 | 0.66 |
| 3 | 141647359 | rs12695746 | -0.376 | -0.03 | 0.548 | -0.911 | -0.049 | -0.027 | -0.027 | 0 | 0.84 | 0.98 | 0.84 | 0.87 | 0.8 | 0.88 |
| 3 | 141654200 | rs10935445 | 0.549 | -0.143 | 0.609 | -0.937 | -0.018 | -0.033 | -0.033 | 0 | 0.06 | 0.01 | 0.08 | 0.15 | 0.06 | 0.03 |
| 3 | 141658875 | rs11711748 | 0.549 | -0.2 | 0.609 | -0.937 | -0.02 | -0.033 | -0.033 | 0.363 | 0.06 | 0.01 | 0.09 | 0.15 | 0.08 | 0.03 |
| 3 | 141659241 | rs6440052 | 1.112 | -0.234 | 1.043 | -1 | 0 | 0.001 | 0.001 | 0 | 0.74 | 0.68 | 0.72 | 0.75 | 0.74 | 0.83 |
| 3 | 141663752 | rs9942 | -0.25 | -0.244 | 0.565 | -1 | -0.009 | 0.003 | 0.003 | 0 | 0.83 | 0.94 | 0.82 | 0.9 | 0.83 | 0.87 |
| 3 | 141668453 | rs2163294 | 1.489 | -0.08 | 0.378 | -1.243 | 0.001 | 0.084 | 0.084 | 0 | 0.58 | 0.21 | 0.59 | 0.8 | 0.6 | 0.71 |
| 3 | 141669685 | rs11569291 | 0.29 | 0.531 | 0.793 | -0.51 | -0.022 | -0.013 | -0.013 | 0 | 0.06 | 0.01 | 0.06 | 0.05 | 0.08 | 0.04 |
| 3 | 141683097 | rs4683653 | -2.194 | 0.492 | 1.031 | -0.585 | -0.001 | -0.001 | -0.001 | 0.362 | 0.32 | 0.62 | 0.32 | 0.26 | 0.31 | 0.28 |
| 3 | 141691407 | rs3804771 | -1.351 | 0.664 | 1.263 | -0.485 | 0.018 | 0.034 | 0.034 | 1.549 | 0.17 | 0.55 | 0.2 | 0.18 | 0.18 | 0.14 |
| 3 | 141699787 | rs11705778 | 0.268 | 0.505 | 0.703 | -0.528 | -0.023 | -0.011 | -0.011 | 1.974 | 0.06 | 0.01 | 0.06 | 0.05 | 0.08 | 0.04 |
| 3 | 141733434 | rs6794224 | -0.602 | 0.453 | 1.377 | -0.682 | 0.024 | -0.021 | -0.021 | 0.244 | 0.06 | 0.19 | 0.03 | 0 | 0.05 | 0.07 |
| 3 | 141743951 | rs7644383 | 1.134 | 0.469 | 0.996 | -0.717 | -0.035 | -0.035 | -0.035 | 0 | 0.79 | 0.96 | 0.83 | 0.77 | 0.73 | 0.84 |
| 3 | 141747640 | rs10935453 | 2.477 | 0.383 | 1.282 | -0.751 | 0 | 0.002 | 0.002 | 0 | 0.69 | 0.35 | 0.76 | 0.76 | 0.68 | 0.75 |
| 3 | 141750539 | rs963396 | 0.521 | 0.487 | 0.729 | -0.648 | -0.048 | -0.071 | -0.071 | 0 | 0.84 | 0.99 | 0.88 | 0.8 | 0.81 | 0.88 |
| 3 | 141754558 | rs6795301 | 0.424 | 0.501 | 0.735 | -0.642 | -0.055 | -0.072 | -0.072 | 0 | 0.85 | 0.99 | 0.87 | 0.8 | 0.79 | 0.88 |
| 3 | 141756541 | rs1543064 | 0.462 | 0.501 | 0.735 | -0.642 | -0.055 | -0.071 | -0.071 | 1.031 | 0.85 | 0.99 | 0.87 | 0.8 | 0.78 | 0.88 |
| 3 | 141763540 | rs9289641 | 0.462 | 0.501 | 0.735 | -0.595 | -0.055 | -0.079 | -0.079 | 0.15 | 0.85 | 0.99 | 0.87 | 0.78 | 0.79 | 0.88 |
| 3 | 141767853 | rs1355438 | 1.498 | 0.536 | 1.012 | -0.626 | 0.005 | 0.007 | 0.007 | 0 | 0.77 | 0.76 | 0.81 | 0.75 | 0.72 | 0.83 |
| 3 | 141770001 | rs2862124 | 0.487 | 0.558 | 0.788 | -0.613 | -0.057 | -0.079 | -0.079 | 0.035 | 0.84 | 0.99 | 0.87 | 0.78 | 0.78 | 0.88 |
| 3 | 141775278 | rs1704990 | 0.174 | 0.61 | 0.814 | -0.569 | -0.039 | -0.068 | -0.068 | 0.562 | 0.9 | 0.99 | 0.88 | 0.75 | 0.83 | 0.89 |
| 3 | 141777113 | rs4683663 | -0.006 | 0.6 | 0.805 | -0.572 | -0.038 | 0.03 | 0.03 | 0.035 | 0.1 | 0 | 0.06 | 0.03 | 0.11 | 0.04 |
| 3 | 141786073 | rs1705619 | 0.17 | 0.6 | 0.806 | -0.564 | -0.039 | -0.068 | -0.068 | 0 | 0.9 | 0.99 | 0.88 | 0.75 | 0.83 | 0.89 |
| 3 | 141791408 | rs4683419 | -0.008 | 0.597 | 0.808 | -0.569 | -0.038 | 0.03 | 0.03 | 0.035 | 0.1 | 0 | 0.06 | 0.03 | 0.11 | 0.04 |
| 3 | 141795573 | rs7635444 | -1.733 | 0.625 | 0.802 | -0.941 | 0.01 | 0.013 | 0.013 | 2.431 | 0.37 | 0.76 | 0.34 | 0.28 | 0.37 | 0.15 |
| 3 | 141806655 | rs347684 | 0.35 | 0.659 | 0.487 | -0.865 | -0.04 | -0.069 | -0.069 | 0 | 0.89 | 0.99 | 0.88 | 0.77 | 0.83 | 0.89 |
| 3 | 141809984 | rs1705612 | 0.283 | 0.655 | 0.487 | -0.863 | -0.038 | -0.069 | -0.069 | 0 | 0.9 | 0.99 | 0.88 | 0.75 | 0.83 | 0.89 |
| 3 | 141818850 | rs11705932 | -0.889 | 0.63 | 0.721 | -0.853 | -0.007 | 0.057 | 0.057 | 0.217 | 0.18 | 0.26 | 0.16 | 0.29 | 0.18 | 0.09 |
| 3 | 141835715 | rs6789998 | -1.645 | 0.67 | 0.16 | -0.845 | 0 | 0.046 | 0.046 | 0.84 | 0.43 | 0.86 | 0.36 | 0.28 | 0.42 | 0.21 |
| 3 | 141845000 | rs5028811 | 1.493 | 0.608 | 0.467 | -0.825 | -0.008 | 0.012 | 0.012 | 0 | 0.56 | 0.38 | 0.61 | 0.7 | 0.48 | 0.73 |
| 3 | 141867638 | rs6771905 | -0.227 | 0.235 | 0.719 | -0.69 | 0.007 | -0.099 | -0.099 | 0 | 0.18 | 0.43 | 0.16 | 0.02 | 0.19 | 0.06 |
| 3 | 141870908 | rs9784370 | -1.106 | 0.193 | 0.539 | -0.744 | 0.003 | -0.101 | -0.101 | 0 | 0.33 | 0.56 | 0.25 | 0.04 | 0.34 | 0.16 |
| 3 | 141871000 | rs9289642 | -1.273 | 0.003 | -0.377 | -0.61 | 0.003 | 0.028 | 0.028 | 0 | 0.48 | 0.78 | 0.36 | 0.34 | 0.47 | 0.37 |
| 3 | 141881351 | rs6769556 | -1.273 | -0.139 | -0.371 | -0.929 | 0.013 | 0.069 | 0.069 | 0 | 0.42 | 0.69 | 0.33 | 0.18 | 0.4 | 0.25 |
| 3 | 141882459 | rs6808709 | -1.312 | -0.347 | 0.02 | -0.74 | -0.015 | -0.029 | -0.029 | 0 | 0.21 | 0.44 | 0.16 | 0.12 | 0.2 | 0.16 |
| 3 | 141889009 | rs13067957 | -0.766 | -0.397 | -0.751 | -0.564 | 0.006 | -0.008 | -0.008 | 0 | 0.11 | 0.07 | 0.07 | 0.12 | 0.1 | 0.09 |
| 3 | 141893901 | rs7628622 | -0.877 | -0.782 | -1.008 | -0.981 | 0.078 | 0.103 | 0.103 | 1.002 | 0.28 | 0.86 | 0.2 | 0.04 | 0.18 | 0.28 |
| 3 | 141898980 | rs6764361 | -0.003 | -0.331 | -0.81 | -1.11 | -0.013 | -0.042 | -0.042 | 0 | 0.38 | 0.55 | 0.33 | 0.17 | 0.27 | 0.31 |
| 3 | 141903905 | rs4577487 | -0.614 | -0.101 | -1.754 | -1.06 | 0.054 | 0.287 | 0.287 | 0 | 0.5 | 0.89 | 0.42 | 0.17 | 0.39 | 0.4 |
| 3 | 141909481 | rs7623942 | -0.608 | -0.101 | -1.731 | -1.06 | 0.058 | 0.276 | 0.276 | 0 | 0.5 | 0.87 | 0.42 | 0.17 | 0.39 | 0.4 |
| 3 | 141920434 | rs9814627 | -0.019 | 0.499 | -1.723 | -0.015 | -0.051 | -0.058 | -0.058 | 0.174 | 0.63 | 0.8 | 0.6 | 0.5 | 0.56 | 0.43 |
| 3 | 141925039 | rs4485736 | -0.301 | 0.807 | -1.857 | 0.421 | -0.022 | -0.056 | -0.056 | 0.856 | 0.67 | 0.85 | 0.66 | 0.53 | 0.66 | 0.45 |
| 3 | 141932356 | rs6440072 | -0.297 | 0.801 | -1.684 | 0.403 | -0.023 | -0.054 | -0.054 | 0 | 0.66 | 0.86 | 0.66 | 0.53 | 0.66 | 0.45 |
| 3 | 141933649 | rs6781598 | -0.047 | 0.266 | -0.937 | 0.312 | -0.029 | -0.01 | -0.01 | 0 | 0.25 | 0.42 | 0.2 | 0.23 | 0.21 | 0.19 |
| 3 | 141943537 | rs9842771 | -0.335 | 0.259 | -0.611 | 0.395 | -0.002 | 0.003 | 0.003 | 0.174 | 0.63 | 0.59 | 0.64 | 0.38 | 0.67 | 0.37 |
| 3 | 141952543 | rs6774637 | -0.439 | 0.247 | -0.647 | 0.39 | 0 | -0.004 | -0.004 | 1.894 | 0.64 | 0.69 | 0.65 | 0.38 | 0.67 | 0.37 |
| 3 | 141962168 | rs13325146 | 1.168 | 0.241 | -0.18 | 0.166 | -0.006 | 0.046 | 0.046 | 0.363 | 0.17 | 0.41 | 0.22 | 0.24 | 0.14 | 0.14 |
| 3 | 141963015 | rs13067734 | -0.136 | 0.201 | -0.068 | 0.141 | -0.001 | 0.015 | 0.015 | 0 | 0.49 | 0.52 | 0.45 | 0.31 | 0.53 | 0.3 |
| 3 | 141965461 | rs10935457 | -0.173 | 0.146 | -0.085 | 0.117 | -0.001 | 0.019 | 0.019 | 0 | 0.5 | 0.53 | 0.45 | 0.31 | 0.53 | 0.3 |
| 3 | 141979191 | rs11721308 | 1.041 | 0.133 | -0.098 | -0.036 | -0.033 | -0.031 | -0.031 | 0.166 | 0.27 | 0.12 | 0.26 | 0.25 | 0.32 | 0.18 |

| | | | | | | | | | | | | | | | | |
|---|---|---|---|---|---|---|---|---|---|---|---|---|---|---|---|---|
| 3 | 141989460 | rs9871394 | 2.072 | 0.075 | -0.189 | -0.033 | -0.03 | 0.034 | 0.034 | 0.321 | 0.13 | 0.02 | 0.18 | 0.07 | 0.13 | 0.05 |
| 3 | 141992053 | rs7621485 | -0.032 | -0.031 | -0.104 | -0.135 | -0.002 | 0.01 | 0.01 | 0 | 0.47 | 0.51 | 0.45 | 0.31 | 0.54 | 0.3 |
| 3 | 142000804 | rs10428215 | -0.114 | -0.015 | 0.009 | -0.119 | -0.001 | 0.01 | 0.01 | 0 | 0.48 | 0.52 | 0.45 | 0.31 | 0.54 | 0.3 |
| 3 | 142015590 | rs1552340 | 0.164 | 0.049 | 0.412 | -0.261 | 0.002 | -0.006 | -0.006 | 0.392 | 0.41 | 0.38 | 0.39 | 0.38 | 0.41 | 0.34 |
| 3 | 142027811 | rs3762799 | 0.118 | 0.039 | 0.414 | -0.247 | 0.002 | -0.005 | -0.005 | 0 | 0.41 | 0.38 | 0.38 | 0.38 | 0.41 | 0.34 |
| 3 | 142070013 | rs11708870 | 0.907 | 0.124 | 0.541 | -0.144 | -0.003 | 0 | 0 | 0 | 0.1 | 0.11 | 0.09 | 0.1 | 0.15 | 0.08 |
| 3 | 142076059 | rs3816805 | -1.032 | 0.09 | 0.596 | -0.159 | -0.071 | -0.032 | -0.032 | 0.175 | 0.55 | 0.91 | 0.51 | 0.48 | 0.56 | 0.42 |
| 3 | 142110818 | rs7642634 | 0.803 | 0.099 | 0.648 | -0.153 | 0.003 | -0.003 | -0.003 | 0.534 | 0.11 | 0.15 | 0.09 | 0.1 | 0.15 | 0.08 |
| 3 | 142114406 | rs6440082 | -0.95 | 0.13 | 0.628 | -0.165 | -0.064 | -0.02 | -0.02 | 0.105 | 0.54 | 0.86 | 0.5 | 0.48 | 0.55 | 0.42 |
| 3 | 142120786 | rs13061823 | -0.889 | 0.101 | 0.636 | -0.151 | -0.04 | -0.006 | -0.006 | 0 | 0.53 | 0.84 | 0.49 | 0.48 | 0.56 | 0.42 |
| 3 | 142123384 | rs1351965 | -1.476 | 0.18 | 0.275 | -0.062 | -0.097 | -0.078 | -0.078 | 0.994 | 0.69 | 0.97 | 0.76 | 0.64 | 0.67 | 0.52 |
| 3 | 142123841 | rs2306700 | 0.685 | 0.191 | 0.224 | -0.076 | 0.002 | -0.004 | -0.004 | 0 | 0.11 | 0.16 | 0.1 | 0.1 | 0.15 | 0.08 |
| 3 | 142124169 | rs7349528 | 1.19 | 0.017 | -0.077 | -0.146 | -0.025 | 0.096 | 0.096 | 0 | 0.2 | 0.02 | 0.15 | 0.06 | 0.21 | 0.1 |
| 3 | 142146261 | rs9826786 | -0.966 | 0 | -0.265 | -0.088 | -0.077 | -0.035 | -0.035 | 0 | 0.54 | 0.91 | 0.51 | 0.48 | 0.56 | 0.42 |
| 3 | 142164735 | rs12635931 | -2.317 | 0.111 | -0.057 | -0.053 | -0.028 | 0.047 | 0.047 | 0.972 | 0.86 | 0.99 | 0.81 | 0.93 | 0.87 | 0.95 |
| 3 | 142168331 | rs1802904 | -2.347 | 0.1 | -0.065 | -0.048 | -0.035 | 0.041 | 0.041 | 0.635 | 0.86 | 0.99 | 0.81 | 0.93 | 0.87 | 0.94 |
| 3 | 142178144 | rs2229032 | 1.034 | 0.346 | 0.116 | 0.045 | -0.021 | -0.002 | -0.002 | 0 | 0.1 | 0.07 | 0.1 | 0.1 | 0.17 | 0.08 |
| 3 | 142216916 | rs6805118 | 1.959 | 0.904 | -0.101 | -0.142 | -0.041 | 0.035 | 0.035 | 0 | 0.07 | 0 | 0.07 | 0 | 0.12 | 0.1 |
| 3 | 142219542 | rs4256152 | -0.226 | -0.073 | 0.466 | 0.1 | -0.003 | 0.002 | 0.002 | 0.112 | 0.41 | 0.56 | 0.37 | 0.37 | 0.39 | 0.34 |
| 3 | 142234534 | rs10804682 | 1.115 | -0.162 | 0.557 | 0.043 | -0.019 | 0.098 | 0.098 | 0 | 0.21 | 0.03 | 0.15 | 0.07 | 0.2 | 0.11 |
| 3 | 142281612 | rs2227928 | -1.206 | -0.137 | 0.309 | 0.054 | -0.038 | -0.034 | -0.034 | 0 | 0.56 | 0.85 | 0.51 | 0.47 | 0.59 | 0.46 |
| 3 | 142287627 | rs11707731 | 2.077 | -0.105 | 0.109 | 0.101 | -0.021 | -0.052 | -0.052 | 0.026 | 0.16 | 0.05 | 0.19 | 0.15 | 0.18 | 0.14 |
| 3 | 142290490 | rs11923612 | -0.001 | -0.082 | 0.204 | -0.001 | -0.01 | -0.049 | -0.049 | 0.169 | 0.39 | 0.17 | 0.35 | 0.36 | 0.39 | 0.3 |
| 3 | 142322055 | rs7647212 | 0.147 | 0.061 | 0.371 | -0.42 | 0.059 | 0.002 | 0.002 | 0 | 0.14 | 0.44 | 0.13 | 0.12 | 0.18 | 0.15 |
| 3 | 142331360 | rs6440096 | 0.03 | 0.045 | 0.239 | -0.428 | 0.036 | -0.011 | -0.011 | 0 | 0.15 | 0.58 | 0.14 | 0.12 | 0.18 | 0.15 |
| 3 | 142337430 | rs2049325 | -1.221 | -0.103 | 0.095 | -0.365 | -0.061 | -0.039 | -0.039 | 0 | 0.58 | 0.92 | 0.46 | 0.41 | 0.59 | 0.47 |
| 3 | 142342112 | rs34392719 | -0.251 | -0.103 | 0.113 | -0.497 | 0.007 | 0.02 | 0.02 | 0 | 0.43 | 0.34 | 0.32 | 0.29 | 0.42 | 0.33 |
| 3 | 142344984 | rs16852464 | 2.101 | -0.09 | -0.115 | -0.448 | -0.025 | -0.037 | -0.037 | 0 | 0.16 | 0.05 | 0.13 | 0.14 | 0.18 | 0.15 |
| 3 | 142345817 | rs6800573 | -0.041 | 0.051 | 0.176 | -0.448 | 0.044 | -0.027 | -0.027 | 0.069 | 0.15 | 0.53 | 0.14 | 0.1 | 0.18 | 0.15 |
| 3 | 142347757 | rs1983434 | -1.085 | 0.026 | 0.159 | -0.47 | -0.016 | 0.007 | 0.007 | 0.008 | 0.57 | 0.8 | 0.44 | 0.4 | 0.59 | 0.47 |
| 3 | 142352978 | rs13074172 | 0.074 | 0.031 | 0.083 | -0.432 | 0.006 | 0.04 | 0.04 | 0 | 0.42 | 0.21 | 0.31 | 0.3 | 0.42 | 0.33 |
| 3 | 142362179 | rs17552506 | 2.31 | -0.078 | -0.125 | -0.53 | -0.036 | 0.07 | 0.07 | 0.407 | 0.12 | 0.01 | 0.09 | 0 | 0.13 | 0.01 |
| 3 | 142363432 | rs16852486 | -0.634 | 0.042 | 0.558 | -0.28 | 0.003 | 0.002 | 0.002 | 0 | 0.56 | 0.63 | 0.43 | 0.4 | 0.65 | 0.48 |
| 3 | 142365769 | rs6808205 | -0.841 | 0.054 | 0.623 | -0.28 | 0.011 | 0.007 | 0.007 | 0 | 0.57 | 0.7 | 0.44 | 0.4 | 0.66 | 0.46 |
| 3 | 142375817 | rs11715610 | -0.642 | 0.121 | -0.163 | -0.193 | -0.002 | 0.019 | 0.019 | 0 | 0.55 | 0.79 | 0.44 | 0.39 | 0.61 | 0.41 |
| 3 | 142387809 | rs4637328 | -1.539 | 0.069 | -0.934 | -0.177 | -0.088 | -0.17 | -0.17 | 0 | 0.73 | 0.99 | 0.63 | 0.5 | 0.73 | 0.52 |
| 3 | 142391847 | rs2140437 | -0.058 | 0.577 | -0.341 | 0.035 | -0.018 | 0.054 | 0.054 | 0 | 0.43 | 0.21 | 0.31 | 0.29 | 0.47 | 0.31 |
| 3 | 142405348 | rs11714087 | 1.179 | 0.632 | -0.367 | 0.08 | 0.005 | -0.002 | -0.002 | 0 | 0.08 | 0.09 | 0.1 | 0.09 | 0.14 | 0.07 |
| 3 | 142409228 | rs4683427 | -0.181 | 0.528 | -0.578 | 0.049 | -0.009 | 0.046 | 0.046 | 0 | 0.43 | 0.24 | 0.31 | 0.29 | 0.47 | 0.31 |
| 3 | 142411200 | rs11929107 | 0.254 | 0.504 | -1.05 | -0.063 | -0.036 | 0.052 | 0.052 | 0 | 0.26 | 0.06 | 0.17 | 0.14 | 0.29 | 0.16 |
| 3 | 142411287 | rs9823492 | 0.017 | 0.968 | -0.845 | 0.001 | 0.002 | 0 | 0 | 0 | 0.13 | 0.14 | 0.14 | 0.19 | 0.2 | 0.17 |
| 3 | 142412224 | rs1916638 | 0.751 | 0.557 | -0.625 | 0.148 | -0.008 | 0.047 | 0.047 | 0 | 0.22 | 0.11 | 0.16 | 0 | 0.24 | 0.11 |
| 3 | 142420683 | rs6440104 | -0.852 | 0.499 | -0.73 | 0.375 | -0.039 | 0.071 | 0.071 | 0 | 0.4 | 0.79 | 0.34 | 0.31 | 0.46 | 0.3 |
| 3 | 142428926 | rs7623762 | -0.82 | 0.506 | -0.57 | 0.378 | -0.025 | 0.055 | 0.055 | 0 | 0.39 | 0.74 | 0.34 | 0.31 | 0.46 | 0.3 |
| 3 | 142444625 | rs13094259 | 1.052 | 0.562 | -0.673 | 0.419 | -0.028 | 0.107 | 0.107 | 0 | 0.2 | 0.02 | 0.14 | 0 | 0.21 | 0.11 |
| 3 | 142446205 | rs953239 | -0.632 | 0.625 | -0.534 | 0.457 | -0.005 | 0.006 | 0.006 | 0 | 0.36 | 0.63 | 0.32 | 0.31 | 0.43 | 0.29 |
| 3 | 142503195 | rs13086677 | 0.811 | 1.094 | -0.621 | 0.73 | -0.022 | -0.008 | -0.008 | 0.167 | 0.17 | 0.06 | 0.14 | 0.13 | 0.19 | 0.15 |
| 3 | 142523349 | rs3821647 | 0.045 | 1.109 | -0.53 | 0.718 | 0.032 | 0.027 | 0.027 | 0 | 0.12 | 0.22 | 0.14 | 0.24 | 0.2 | 0.15 |
| 3 | 142526594 | rs4627 | 0.651 | 0.987 | -1.068 | 0.551 | -0.021 | -0.03 | -0.03 | 0 | 0.18 | 0.06 | 0.14 | 0.18 | 0.19 | 0.15 |
| 3 | 142541687 | rs9841007 | -0.841 | 1.106 | -0.032 | 0.502 | 0.021 | 0.012 | 0.012 | 0.127 | 0.18 | 0.66 | 0.19 | 0.29 | 0.26 | 0.16 |
| 3 | 142545647 | rs2608069 | 0.841 | 0.402 | -0.886 | 0.38 | 0.009 | -0.051 | -0.051 | 0 | 0.13 | 0.01 | 0.2 | 0.18 | 0.08 | 0.06 |
| 3 | 142547417 | rs9681405 | -0.57 | 0.799 | -0.194 | -0.172 | 0.001 | 0.012 | 0.012 | 0 | 0.51 | 0.58 | 0.45 | 0.33 | 0.55 | 0.43 |
| 3 | 142551850 | rs9680986 | 0.144 | 0.981 | 0.331 | -0.689 | 0.013 | -0.116 | -0.116 | 0.519 | 0.2 | 0.34 | 0.16 | 0.01 | 0.2 | 0.08 |
| 3 | 142555228 | rs13092560 | -0.632 | 0.937 | 0.041 | -0.795 | -0.027 | 0.065 | 0.065 | 0.03 | 0.33 | 0.23 | 0.27 | 0.11 | 0.42 | 0.26 |
| 3 | 142555846 | rs11720678 | -0.097 | 0.625 | 0.254 | -0.214 | -0.031 | 0.011 | 0.011 | 0 | 0.16 | 0.06 | 0.12 | 0.1 | 0.22 | 0.18 |
| 3 | 142561195 | rs16852706 | -1.106 | 0.679 | 0.699 | -0.214 | 0.013 | -0.037 | -0.037 | 0 | 0.24 | 0.49 | 0.2 | 0.16 | 0.29 | 0.21 |
| 3 | 142564853 | rs7624783 | -1.596 | 0.308 | 0.809 | -0.142 | -0.003 | 0.018 | 0.018 | 0 | 0.15 | 0.26 | 0.18 | 0.17 | 0.16 | 0.09 |
| 3 | 142567030 | rs3816690 | -0.257 | 0.062 | 0.554 | -0.074 | 0.001 | 0 | 0 | 0.023 | 0.09 | 0.05 | 0.04 | 0.05 | 0.06 | 0.12 |
| 3 | 142567327 | rs893883 | -0.981 | -0.017 | 0.543 | -0.342 | -0.001 | -0.001 | -0.001 | 0.385 | 0.59 | 0.62 | 0.49 | 0.33 | 0.7 | 0.46 |
| 3 | 142573789 | rs7647899 | 0.13 | -0.001 | 0.33 | -0.342 | 0.018 | 0.064 | 0.064 | 0.279 | 0.46 | 0.25 | 0.42 | 0.28 | 0.64 | 0.34 |
| 3 | 142574456 | rs4324442 | -0.697 | -0.211 | 0.175 | -0.24 | -0.001 | -0.001 | -0.001 | 0.133 | 0.57 | 0.43 | 0.48 | 0.33 | 0.69 | 0.45 |
| 3 | 142576455 | rs6440116 | -0.888 | -0.145 | -0.192 | -0.211 | -0.006 | -0.005 | -0.005 | 1.523 | 0.59 | 0.57 | 0.49 | 0.33 | 0.69 | 0.46 |
| 3 | 142585802 | rs7624656 | -1.574 | -0.018 | 0.489 | 0.129 | -0.047 | -0.052 | -0.052 | 0 | 0.17 | 0.5 | 0.1 | 0.06 | 0.09 | 0.14 |
| 3 | 142599285 | rs7637508 | -1.205 | 0.444 | 0.483 | 0.137 | 0.065 | -0.038 | -0.038 | 0 | 0.1 | 0.43 | 0.09 | 0.02 | 0.08 | 0.12 |
| 3 | 142600087 | rs2125125 | -0.31 | 0.066 | -0.649 | 0.241 | -0.041 | -0.034 | -0.034 | 0 | 0.79 | 0.97 | 0.71 | 0.78 | 0.82 | 0.65 |
| 3 | 142603110 | rs7639314 | -0.323 | -0.236 | -1.11 | 0.363 | 0.006 | -0.01 | -0.01 | 0.771 | 0.22 | 0.08 | 0.16 | 0.21 | 0.2 | 0.27 |
| 3 | 142603915 | rs11718990 | -0.404 | -0.266 | -1.457 | 0.506 | 0.008 | 0.017 | 0.017 | 0 | 0.47 | 0.23 | 0.41 | 0.36 | 0.58 | 0.42 |
| 3 | 142604476 | rs2707982 | 0.053 | -0.312 | -1.219 | 0.727 | 0.018 | -0.011 | -0.011 | 0 | 0.62 | 0.73 | 0.6 | 0.56 | 0.68 | 0.47 |
| 3 | 142610610 | rs6778966 | 1.345 | -0.296 | -0.966 | 0.673 | -0.027 | 0.015 | 0.015 | 0 | 0.32 | 0.52 | 0.31 | 0.36 | 0.23 | 0.24 |

| | | | | | | | | | | | | | | | | |
|---|---|---|---|---|---|---|---|---|---|---|---|---|---|---|---|---|
| 3 | 142611657 | rs2707975 | -0.966 | -0.255 | -1.015 | 0.68 | 0.004 | -0.006 | -0.006 | 0 | 0.78 | 0.81 | 0.73 | 0.72 | 0.81 | 0.66 |
| 3 | 142617138 | rs4683702 | 1.08 | -0.29 | -1.023 | 0.653 | -0.08 | 0.075 | 0.075 | 0 | 0.23 | 0.08 | 0.25 | 0.06 | 0.39 | 0.13 |
| 3 | 142625778 | rs1470121 | 1.048 | -0.372 | -1.334 | 0.587 | -0.084 | 0.087 | 0.087 | 0 | 0.23 | 0.08 | 0.23 | 0.04 | 0.39 | 0.13 |
| 3 | 142628616 | rs2581618 | -0.109 | -0.413 | -1.255 | 0.69 | 0.003 | 0.001 | 0.001 | 0 | 0.71 | 0.76 | 0.65 | 0.66 | 0.75 | 0.58 |
| 3 | 142629862 | rs7645967 | 0.218 | -0.395 | -1.342 | 0.689 | -0.001 | 0.007 | 0.007 | 0 | 0.07 | 0.05 | 0.06 | 0.06 | 0.06 | 0.08 |
| 3 | 142630330 | rs2581620 | -1.165 | -0.4 | -1.298 | 0.697 | 0.002 | -0.013 | -0.013 | 0.762 | 0.79 | 0.88 | 0.71 | 0.72 | 0.82 | 0.66 |
| 3 | 142630633 | rs1479136 | -1.194 | -0.397 | -1.365 | 0.698 | -0.015 | -0.04 | -0.04 | 0 | 0.79 | 0.93 | 0.72 | 0.72 | 0.82 | 0.66 |
| 3 | 142632418 | rs2581623 | -0.305 | -0.38 | -1.332 | 0.695 | 0.006 | 0.042 | 0.042 | 0 | 0.55 | 0.67 | 0.48 | 0.68 | 0.43 | 0.52 |
| 3 | 142632679 | rs17554998 | 0.169 | -0.609 | -1.43 | 0.682 | -0.001 | 0.005 | 0.005 | 0 | 0.07 | 0.04 | 0.06 | 0.06 | 0.06 | 0.08 |
| 3 | 142633319 | rs16852815 | 0.938 | -0.813 | -1.496 | 0.789 | -0.018 | -0.068 | -0.068 | 0.097 | 0.15 | 0.02 | 0.09 | 0.18 | 0.14 | 0.18 |
| 3 | 142633960 | rs2581625 | 0.347 | -0.519 | -1.358 | 0.415 | 0.033 | 0.028 | 0.028 | 0 | 0.49 | 0.62 | 0.43 | 0.6 | 0.39 | 0.45 |
| 3 | 142636762 | rs2581628 | -0.875 | -0.252 | -1.102 | 0.653 | -0.013 | -0.031 | -0.031 | 0 | 0.81 | 0.88 | 0.73 | 0.73 | 0.84 | 0.67 |
| 3 | 142645905 | rs2581631 | 0.362 | -0.113 | -0.658 | 0.331 | 0.002 | 0.002 | 0.002 | 0 | 0.53 | 0.47 | 0.54 | 0.23 | 0.65 | 0.34 |
| 3 | 142647219 | rs16852846 | -0.888 | 0.614 | -0.516 | 0.549 | -0.001 | 0.016 | 0.016 | 0 | 0.07 | 0.13 | 0.06 | 0.14 | 0.05 | 0.07 |
| 3 | 142652559 | rs4683438 | -0.086 | 0.909 | -0.415 | -0.075 | 0.045 | -0.116 | -0.116 | 0.55 | 0.27 | 0.43 | 0.25 | 0.02 | 0.36 | 0.1 |
| 3 | 142661350 | rs3262 | -0.251 | 1.085 | -1.46 | 0.487 | -0.038 | -0.058 | -0.058 | 1.545 | 0.09 | 0.01 | 0.21 | 0.29 | 0.15 | 0.16 |
| 3 | 142664512 | rs7612265 | -0.725 | 0.41 | -1.351 | 0.273 | -0.017 | 0.004 | 0.004 | 0 | 0.45 | 0.79 | 0.55 | 0.37 | 0.54 | 0.29 |
| 3 | 142666745 | rs9812607 | -0.761 | 0.447 | -1.215 | 0.353 | -0.046 | 0.007 | 0.007 | 0.196 | 0.42 | 0.81 | 0.5 | 0.34 | 0.5 | 0.3 |
| 3 | 142670056 | rs6769643 | -0.891 | 0.402 | -1.293 | 0.281 | -0.05 | 0.023 | 0.023 | 0.083 | 0.45 | 0.82 | 0.48 | 0.35 | 0.53 | 0.35 |
| 3 | 142670616 | rs2270768 | -0.642 | 0.906 | -1.251 | 0.443 | -0.037 | -0.063 | -0.063 | 0.132 | 0.15 | 0.06 | 0.25 | 0.26 | 0.23 | 0.13 |
| 3 | 142672769 | rs9844683 | -0.936 | 0.548 | -0.816 | 0.594 | -0.013 | 0.1 | 0.1 | 0 | 0.06 | 0.16 | 0.15 | 0.21 | 0.04 | 0.15 |
| 3 | 142677280 | rs36294 | -0.287 | 0.432 | -0.883 | -0.183 | 0.003 | -0.066 | -0.066 | 0 | 0.38 | 0.4 | 0.29 | 0.06 | 0.39 | 0.17 |
| 3 | 142688622 | rs36287 | 0.118 | 0.788 | -1.529 | -0.675 | -0.095 | 0.176 | 0.176 | 0 | 0.31 | 0.07 | 0.23 | 0.02 | 0.37 | 0.09 |
| 3 | 142691972 | rs11711259 | -0.113 | 0.075 | -1.125 | -0.497 | -0.017 | -0.016 | -0.016 | 0.317 | 0.35 | 0.44 | 0.28 | 0.33 | 0.27 | 0.43 |
| 3 | 142693563 | rs1355760 | -0.426 | -0.143 | -0.906 | -1.174 | 0.021 | -0.126 | -0.126 | 0.432 | 0.25 | 0.47 | 0.22 | 0.05 | 0.29 | 0.21 |
| 3 | 142695801 | rs40160 | -1.329 | -0.197 | -1.245 | -0.623 | -0.022 | -0.016 | -0.016 | 0 | 0.76 | 0.82 | 0.7 | 0.68 | 0.67 | 0.72 |
| 3 | 142696524 | rs2354165 | -0.378 | -0.846 | -1.357 | -0.903 | -0.003 | -0.114 | -0.114 | 0 | 0.28 | 0.5 | 0.25 | 0.06 | 0.29 | 0.26 |
| 3 | 142699541 | rs251489 | -0.826 | -0.753 | -1.596 | -0.638 | -0.057 | -0.013 | -0.013 | 0 | 0.68 | 0.84 | 0.65 | 0.67 | 0.57 | 0.65 |
| 3 | 142700273 | rs1353887 | 0.919 | -1.245 | -1.644 | -0.34 | 0.023 | -0.047 | -0.047 | 0 | 0.22 | 0.13 | 0.23 | 0.36 | 0.15 | 0.14 |
| 3 | 142703764 | rs251484 | -0.912 | -0.958 | -1.658 | -0.349 | -0.078 | -0.014 | -0.014 | 2.52 | 0.67 | 0.84 | 0.64 | 0.68 | 0.55 | 0.61 |
| 3 | 142710064 | rs4683441 | -0.207 | -0.964 | -0.744 | -0.191 | 0.002 | 0.007 | 0.007 | 0 | 0.47 | 0.61 | 0.46 | 0.42 | 0.47 | 0.48 |
| 3 | 142714537 | rs12492904 | -1.4 | -1.018 | -0.67 | -0.13 | -0.081 | -0.028 | -0.028 | 0.04 | 0.72 | 0.88 | 0.69 | 0.72 | 0.62 | 0.6 |
| 3 | 142716921 | rs1814565 | -0.892 | -1.231 | -0.144 | -0.225 | 0.02 | -0.002 | -0.002 | 0.04 | 0.66 | 0.5 | 0.66 | 0.69 | 0.61 | 0.6 |
| 3 | 142720348 | rs766360 | -0.849 | -1.213 | -0.137 | -0.218 | 0.017 | -0.004 | -0.004 | 0 | 0.65 | 0.5 | 0.66 | 0.68 | 0.61 | 0.6 |
| 3 | 142739103 | rs6440128 | -0.033 | -1.317 | -0.134 | -0.458 | 0.026 | -0.007 | -0.007 | 0 | 0.36 | 0.72 | 0.34 | 0.4 | 0.25 | 0.27 |
| 3 | 142743889 | rs7611721 | -1.6 | -1.306 | -0.182 | -0.467 | -0.05 | -0.01 | -0.01 | 0 | 0.71 | 0.8 | 0.69 | 0.72 | 0.62 | 0.6 |
| 3 | 142746876 | rs1449868 | -1.724 | -1.318 | -0.186 | -0.467 | -0.097 | -0.037 | -0.037 | 0 | 0.72 | 0.89 | 0.7 | 0.72 | 0.62 | 0.62 |
| 3 | 142746988 | rs1839406 | -1.618 | -1.316 | -0.193 | -0.467 | -0.05 | -0.011 | -0.011 | 0 | 0.71 | 0.8 | 0.69 | 0.72 | 0.62 | 0.62 |
| 3 | 142748023 | rs16852909 | 1.667 | -1.358 | -0.213 | -0.492 | 0.022 | -0.071 | -0.071 | 0 | 0.2 | 0.04 | 0.22 | 0.33 | 0.15 | 0.1 |
| 3 | 142756907 | rs11929440 | 1.301 | -1.319 | -0.263 | -0.501 | -0.055 | 0.097 | 0.097 | 0 | 0.15 | 0.01 | 0.14 | 0 | 0.22 | 0.23 |
| 3 | 142770314 | rs1053950 | -1.296 | -1.005 | 0.244 | -0.099 | -0.05 | -0.01 | -0.01 | 0 | 0.71 | 0.8 | 0.69 | 0.72 | 0.62 | 0.6 |
| 3 | 142776151 | rs1043898 | -1.296 | -1.005 | 0.244 | -0.099 | -0.05 | -0.01 | -0.01 | 0 | 0.71 | 0.8 | 0.69 | 0.72 | 0.62 | 0.6 |
| 3 | 142778355 | rs17633378 | 0.556 | -0.976 | 0.253 | -0.096 | 0.009 | 0.012 | 0.012 | 0 | 0.36 | 0.18 | 0.36 | 0.33 | 0.37 | 0.33 |
| 3 | 142784321 | rs13094293 | -0.985 | -0.784 | 0.252 | -0.044 | -0.022 | 0.003 | 0.003 | 0 | 0.67 | 0.72 | 0.68 | 0.72 | 0.62 | 0.6 |
| 3 | 142795437 | rs11708528 | 0.17 | -0.875 | 0.22 | 0.097 | -0.033 | 0.007 | 0.007 | 0 | 0.34 | 0.48 | 0.32 | 0.36 | 0.24 | 0.29 |
| 3 | 142796422 | rs11925953 | 0.591 | -0.767 | -0.144 | 0.127 | 0.007 | 0.01 | 0.01 | 0 | 0.36 | 0.24 | 0.37 | 0.32 | 0.37 | 0.31 |
| 3 | 142800051 | rs750870 | 0.75 | -0.77 | -0.242 | 0.153 | 0.005 | -0.042 | -0.042 | 0 | 0.27 | 0.13 | 0.29 | 0.36 | 0.23 | 0.29 |
| 3 | 142819277 | rs936198 | 0.434 | -0.631 | -0.206 | 0.129 | 0.01 | 0.014 | 0.014 | 0 | 0.37 | 0.22 | 0.37 | 0.33 | 0.38 | 0.3 |
| 3 | 142834022 | rs7640077 | -0.286 | -0.828 | 0.319 | 0.006 | 0.023 | 0.002 | 0.002 | 0 | 0.39 | 0.7 | 0.35 | 0.41 | 0.27 | 0.32 |
| 3 | 142836966 | rs3755740 | 0.076 | -0.837 | 0.384 | -0.006 | 0.014 | -0.001 | -0.001 | 0.483 | 0.36 | 0.3 | 0.31 | 0.38 | 0.26 | 0.31 |
| 3 | 142837272 | rs3755739 | -0.298 | -0.79 | -0.133 | 0.01 | 0.023 | 0.003 | 0.003 | 0 | 0.39 | 0.71 | 0.35 | 0.41 | 0.27 | 0.33 |
| 3 | 142841740 | rs6664 | -0.254 | -0.686 | -0.014 | -0.045 | 0.007 | 0.004 | 0.004 | 0 | 0.39 | 0.66 | 0.31 | 0.34 | 0.29 | 0.42 |
| 3 | 142842195 | rs4149496 | -1.686 | -0.615 | -0.488 | 0.081 | -0.072 | -0.029 | -0.029 | 3.899 | 0.76 | 0.93 | 0.75 | 0.72 | 0.64 | 0.66 |
| 3 | 142842764 | rs2292509 | 0.887 | -0.686 | -0.504 | 0.053 | 0.001 | -0.065 | -0.065 | 5.742 | 0.28 | 0.05 | 0.24 | 0.32 | 0.21 | 0.35 |
| 3 | 142844327 | rs11718140 | 0.429 | -0.588 | -0.159 | 0.254 | -0.009 | -0.004 | -0.004 | 0 | 0.36 | 0.16 | 0.33 | 0.32 | 0.36 | 0.45 |
| 3 | 142844453 | rs7611217 | -1.652 | -0.574 | -0.21 | 0.265 | -0.078 | -0.036 | -0.036 | 0 | 0.76 | 0.93 | 0.74 | 0.72 | 0.64 | 0.65 |
| 3 | 142845585 | rs4683448 | 0.156 | -0.579 | 0.35 | 0.181 | 0.003 | 0.005 | 0.005 | 0 | 0.39 | 0.34 | 0.36 | 0.31 | 0.36 | 0.45 |
| 3 | 142846285 | rs6770489 | 0.156 | -0.579 | 0.368 | 0.181 | 0.001 | 0.002 | 0.002 | 0 | 0.39 | 0.34 | 0.36 | 0.31 | 0.36 | 0.45 |
| 3 | 142846370 | rs6794836 | 0.116 | -0.583 | 0.366 | 0.179 | 0.002 | 0.002 | 0.002 | 0 | 0.39 | 0.34 | 0.36 | 0.31 | 0.36 | 0.45 |
| 3 | 142846718 | rs11919563 | 0.513 | -0.544 | 0.442 | 0.215 | 0.023 | -0.005 | -0.005 | 0 | 0.33 | 0.15 | 0.35 | 0.34 | 0.27 | 0.18 |
| 3 | 142846800 | rs4683449 | -0.296 | -0.918 | 0.934 | -0.199 | 0.003 | 0.004 | 0.004 | 0 | 0.39 | 0.35 | 0.36 | 0.31 | 0.36 | 0.45 |
| 3 | 142849107 | rs9829651 | -1.017 | -0.829 | 0.212 | -0.748 | -0.038 | 0.067 | 0.067 | 0 | 0.56 | 0.86 | 0.46 | 0.31 | 0.47 | 0.53 |
| 3 | 142850990 | rs1056360 | -0.199 | -0.447 | 0.726 | -0.217 | -0.002 | 0.039 | 0.039 | 0.426 | 0.36 | 0.19 | 0.34 | 0.26 | 0.34 | 0.36 |
| 3 | 142852015 | rs6440136 | -0.621 | -0.35 | 0.337 | -0.208 | 0.016 | 0.042 | 0.042 | 0 | 0.41 | 0.71 | 0.38 | 0.29 | 0.35 | 0.4 |
| 3 | 142857148 | rs16853045 | -0.665 | -0.292 | 0.069 | -0.282 | 0.005 | 0.01 | 0.01 | 0 | 0.42 | 0.61 | 0.37 | 0.25 | 0.35 | 0.37 |
| 3 | 142860107 | rs726737 | -0.551 | 0.018 | -0.428 | -0.147 | -0.005 | -0.019 | -0.019 | 0.028 | 0.39 | 0.46 | 0.36 | 0.25 | 0.35 | 0.37 |
| 3 | 142864702 | rs9790318 | -0.831 | 0.377 | -0.56 | 0.063 | -0.002 | -0.004 | -0.004 | 0 | 0.22 | 0.23 | 0.21 | 0.17 | 0.2 | 0.3 |
| 3 | 142865022 | rs4683456 | -0.764 | 0.453 | -0.682 | 0.018 | 0.003 | 0.008 | 0.008 | 0 | 0.22 | 0.15 | 0.18 | 0.17 | 0.22 | 0.33 |
| 3 | 142865637 | rs4683458 | -0.213 | 0.601 | 0.01 | -0.313 | 0.032 | -0.019 | -0.019 | 0.092 | 0.13 | 0.38 | 0.1 | 0.09 | 0.12 | 0.08 |

| | | | | | | | | | | | | | | | | |
|---|---|---|---|---|---|---|---|---|---|---|---|---|---|---|---|---|
| 3 | 142866513 | rs6805378 | -0.514 | 0.887 | -0.604 | -0.356 | 0 | -0.004 | -0.004 | 0 | 0.47 | 0.48 | 0.58 | 0.63 | 0.45 | 0.46 |
| 3 | 142867543 | rs4683749 | -0.659 | 0.912 | -0.557 | -0.384 | 0 | 0.003 | 0.003 | 1.532 | 0.49 | 0.55 | 0.6 | 0.64 | 0.47 | 0.47 |
| 3 | 142867790 | rs10804686 | -0.248 | 0.824 | -0.666 | -0.496 | -0.009 | 0.01 | 0.01 | 1.32 | 0.4 | 0.28 | 0.52 | 0.63 | 0.37 | 0.4 |
| 3 | 142869470 | rs4683751 | -0.377 | 0.973 | -0.196 | -0.303 | -0.016 | -0.036 | -0.036 | 0.062 | 0.25 | 0.55 | 0.24 | 0.17 | 0.22 | 0.35 |
| 3 | 142871062 | rs6801461 | -0.771 | 0.897 | -0.265 | -0.34 | -0.019 | -0.032 | -0.032 | 0 | 0.28 | 0.61 | 0.26 | 0.18 | 0.22 | 0.38 |
| 3 | 142872107 | rs2352734 | -1.536 | 0.863 | -0.625 | -0.521 | -0.053 | 0.044 | 0.044 | 0.346 | 0.6 | 0.89 | 0.68 | 0.71 | 0.56 | 0.54 |
| 3 | 142877227 | rs10804687 | 0.107 | 0.946 | -0.517 | -0.339 | -0.001 | -0.001 | -0.001 | 0.757 | 0.32 | 0.29 | 0.43 | 0.53 | 0.34 | 0.16 |
| 3 | 142888043 | rs17572584 | 0.265 | 0.799 | -0.573 | -0.336 | 0.004 | -0.008 | -0.008 | 0.943 | 0.07 | 0.11 | 0.08 | 0.04 | 0.11 | 0.05 |
| 3 | 142888606 | rs1530266 | -0.674 | 0.912 | -0.656 | -0.426 | 0.001 | 0.002 | 0.002 | 0.438 | 0.28 | 0.65 | 0.27 | 0.24 | 0.26 | 0.39 |
| 3 | 142889135 | rs9874259 | -1.477 | 1.13 | -1.178 | -0.44 | -0.057 | 0.001 | 0.001 | 1.052 | 0.62 | 0.95 | 0.72 | 0.71 | 0.61 | 0.56 |
| 3 | 142890392 | rs10935483 | 0.398 | 1.039 | -0.553 | -0.122 | -0.004 | 0 | 0 | 0 | 0.18 | 0.22 | 0.17 | 0.2 | 0.14 | 0.32 |
| 3 | 142891007 | rs11717285 | 0.273 | 1.015 | -1.087 | 0.11 | -0.015 | -0.06 | -0.06 | 0 | 0.23 | 0.12 | 0.38 | 0.45 | 0.25 | 0.1 |
| 3 | 142896286 | rs7646486 | -0.704 | 0.521 | -1.092 | -0.465 | 0.038 | 0.032 | 0.032 | 0 | 0.65 | 0.79 | 0.74 | 0.78 | 0.7 | 0.58 |
| 3 | 142901537 | rs1372288 | 0.008 | 1.746 | -0.744 | -0.543 | 0.005 | -0.016 | -0.016 | 0.106 | 0.1 | 0.14 | 0.11 | 0.02 | 0.2 | 0.07 |
| 3 | 142902953 | rs4371486 | -1.259 | 1.23 | -1.666 | -1.03 | 0.015 | 0.025 | 0.025 | 0 | 0.66 | 0.96 | 0.76 | 0.78 | 0.7 | 0.58 |
| 3 | 142903366 | rs4683461 | 0.249 | 1.273 | -1.103 | -1.192 | -0.001 | 0.028 | 0.028 | 0 | 0.37 | 0.23 | 0.53 | 0.74 | 0.37 | 0.38 |
| 3 | 142905461 | rs294470 | -0.749 | 1.188 | -0.604 | -1.204 | 0.029 | 0.143 | 0.143 | 0 | 0.5 | 0.78 | 0.67 | 0.76 | 0.57 | 0.45 |
| 3 | 142905661 | rs6776634 | -1.553 | 1.078 | -0.578 | -1.167 | 0.048 | 0.054 | 0.054 | 0 | 0.65 | 0.87 | 0.73 | 0.77 | 0.7 | 0.56 |
| 3 | 142906885 | rs294468 | -1.74 | 1.131 | -0.577 | -0.946 | 0.009 | -0.018 | -0.018 | 0.103 | 0.66 | 0.97 | 0.69 | 0.75 | 0.7 | 0.58 |
| 3 | 142907815 | rs294466 | -0.998 | 1.087 | -0.585 | -0.772 | -0.041 | 0.078 | 0.078 | 0.03 | 0.4 | 0.79 | 0.32 | 0.25 | 0.45 | 0.41 |
| 3 | 142915227 | rs4839656 | -1.76 | 0.826 | -0.529 | -0.726 | 0.044 | 0.049 | 0.049 | 0 | 0.65 | 0.86 | 0.75 | 0.77 | 0.71 | 0.56 |
| 3 | 142916200 | rs9812350 | 0.46 | 1.015 | 0.208 | -0.819 | 0.038 | -0.089 | -0.089 | 2.742 | 0.25 | 0.4 | 0.19 | 0.03 | 0.33 | 0.17 |
| 3 | 142921964 | rs4839620 | 0.944 | 1.019 | 0.431 | -0.741 | -0.009 | -0.021 | -0.021 | 0 | 0.27 | 0.19 | 0.44 | 0.52 | 0.3 | 0.16 |
| 3 | 142923599 | rs6777882 | -1.509 | 0.531 | 0.503 | -0.619 | 0.038 | 0.042 | 0.042 | 0 | 0.66 | 0.85 | 0.74 | 0.77 | 0.71 | 0.56 |
| 3 | 142926765 | rs990739 | 0.689 | 0.661 | 0.464 | -0.636 | 0.057 | -0.064 | -0.064 | 0 | 0.11 | 0.3 | 0.13 | 0.02 | 0.2 | 0.06 |
| 3 | 142927446 | rs4839629 | -1.258 | 0.105 | 0.324 | -0.481 | 0.024 | 0.025 | 0.025 | 0 | 0.65 | 0.77 | 0.74 | 0.77 | 0.71 | 0.56 |
| 3 | 142928972 | rs6795908 | -1.293 | 0.097 | 0.235 | -0.471 | 0.025 | 0.027 | 0.027 | 0 | 0.65 | 0.8 | 0.75 | 0.77 | 0.71 | 0.56 |
| 3 | 142929441 | rs12635345 | 0.203 | 0.125 | 0.013 | -0.249 | -0.012 | 0.008 | 0.008 | 0 | 0.11 | 0.15 | 0.11 | 0.21 | 0.07 | 0.22 |
| 3 | 142947872 | rs11708440 | -1.414 | 0.308 | -0.559 | -0.469 | 0.009 | 0.032 | 0.032 | 0 | 0.67 | 0.86 | 0.72 | 0.77 | 0.67 | 0.56 |
| 3 | 142949459 | rs11709751 | 1.471 | 0.695 | -0.778 | -0.755 | -0.047 | 0.057 | 0.057 | 0 | 0.18 | 0.09 | 0.13 | 0.01 | 0.22 | 0.17 |
| 3 | 142951172 | rs12633118 | 1.335 | 0.467 | -0.688 | -0.36 | 0.009 | -0.057 | -0.057 | 0 | 0.21 | 0.07 | 0.23 | 0.43 | 0.19 | 0.12 |
| 3 | 142951838 | rs9839394 | -0.396 | 0.344 | -0.24 | -0.333 | 0.007 | -0.024 | -0.024 | 0 | 0.39 | 0.45 | 0.43 | 0.25 | 0.45 | 0.38 |
| 3 | 142954177 | rs7624504 | 1.679 | 0.234 | -0.207 | -0.204 | 0.006 | -0.047 | -0.047 | 0.04 | 0.16 | 0.05 | 0.2 | 0.42 | 0.14 | 0.11 |
| 3 | 142954515 | rs7648972 | 1.621 | 0.23 | -0.209 | -0.23 | 0.007 | -0.049 | -0.049 | 0 | 0.16 | 0.05 | 0.2 | 0.43 | 0.14 | 0.11 |
| 3 | 142955098 | rs9831427 | -0.774 | 0.156 | -0.144 | -0.195 | -0.02 | -0.004 | -0.004 | 0 | 0.61 | 0.5 | 0.66 | 0.67 | 0.65 | 0.52 |
| 3 | 142957007 | rs17470719 | -1.677 | 0.348 | -1.224 | -0.206 | -0.018 | 0.013 | 0.013 | 0 | 0.68 | 0.94 | 0.74 | 0.77 | 0.67 | 0.58 |
| 3 | 142960273 | rs4839604 | 0.268 | 0.889 | -0.914 | -0.093 | -0.001 | 0.001 | 0.001 | 0 | 0.12 | 0.15 | 0.21 | 0.11 | 0.18 | 0.1 |
| 3 | 142961223 | rs13317029 | 0.247 | 0.886 | -0.914 | -0.178 | -0.001 | 0.001 | 0.001 | 0 | 0.12 | 0.15 | 0.1 | 0.09 | 0.18 | 0.09 |
| 3 | 142962075 | rs961413 | 1.029 | 0.597 | -1.234 | -0.285 | -0.059 | 0.094 | 0.094 | 0 | 0.24 | 0.1 | 0.18 | 0.02 | 0.29 | 0.16 |
| 3 | 142963642 | rs9875090 | -0.806 | -0.176 | -0.658 | 0.531 | 0.007 | -0.058 | -0.058 | 0 | 0.86 | 0.96 | 0.7 | 0.56 | 0.88 | 0.89 |
| 3 | 142964700 | rs11712677 | -0.242 | -0.248 | -0.37 | 0.243 | -0.021 | -0.005 | -0.005 | 0 | 0.09 | 0.12 | 0.08 | 0.13 | 0.05 | 0.15 |
| 3 | 142965442 | rs6440155 | -0.196 | -0.256 | -0.838 | 0.586 | 0.026 | -0.099 | -0.099 | 0 | 0.79 | 0.97 | 0.7 | 0.62 | 0.86 | 0.79 |
| 3 | 142966143 | rs9862139 | -1.027 | 0.223 | -0.051 | 0.275 | 0.072 | 0.078 | 0.078 | 0.216 | 0.13 | 0.37 | 0.14 | 0.23 | 0.15 | 0.11 |
| 3 | 142971279 | rs9289659 | -0.025 | 0.291 | -0.254 | 0.293 | 0.007 | 0.004 | 0.004 | 0 | 0.08 | 0.05 | 0.09 | 0.21 | 0.13 | 0.07 |
| 3 | 142972148 | rs9856144 | -0.507 | 0.108 | -0.416 | 0.187 | -0.024 | -0.017 | -0.017 | 0 | 0.11 | 0.04 | 0.1 | 0.21 | 0.17 | 0.09 |
| 3 | 142976429 | rs6769212 | 1.259 | -0.146 | -0.18 | 0.133 | -0.002 | 0.002 | 0.002 | 0 | 0.62 | 0.47 | 0.49 | 0.38 | 0.64 | 0.62 |
| 3 | 142976751 | rs6414353 | -0.198 | -0.029 | -0.673 | 0.194 | 0.024 | -0.067 | -0.067 | 0 | 0.78 | 0.87 | 0.65 | 0.58 | 0.84 | 0.75 |
| 3 | 142979264 | rs7625382 | -0.503 | 0.346 | -0.629 | 0.162 | -0.026 | -0.011 | -0.011 | 0 | 0.11 | 0.05 | 0.11 | 0.17 | 0.19 | 0.1 |
| 3 | 142980880 | rs9857832 | -0.123 | -1.115 | -0.554 | -0.28 | -0.019 | 0.018 | 0.018 | 0 | 0.66 | 0.43 | 0.75 | 0.6 | 0.73 | 0.72 |
| 3 | 142982422 | rs1992426 | -0.073 | -1.323 | -0.925 | -0.396 | 0.003 | -0.008 | -0.008 | 0.043 | 0.23 | 0.16 | 0.39 | 0.44 | 0.22 | 0.29 |
| 3 | 142982837 | rs7621065 | 0.645 | -1.414 | -0.506 | -0.256 | -0.01 | -0.001 | -0.001 | 0 | 0.07 | 0.12 | 0.03 | 0.09 | 0.02 | 0.05 |
| 3 | 142982899 | rs9810857 | 0.81 | -1.457 | -0.727 | -0.557 | -0.055 | 0.195 | 0.195 | 0.167 | 0.43 | 0.13 | 0.36 | 0.13 | 0.5 | 0.37 |
| 3 | 142982930 | rs9757063 | 0.201 | -0.759 | -0.195 | -0.161 | -0.004 | 0.008 | 0.008 | 0 | 0.65 | 0.42 | 0.71 | 0.66 | 0.62 | 0.65 |
| 3 | 142984522 | rs3796229 | -1.428 | -0.947 | 0.616 | -0.56 | 0.028 | -0.03 | -0.03 | 0 | 0.1 | 0.37 | 0.04 | 0.03 | 0.06 | 0.07 |
| 3 | 142984711 | rs3796228 | -0.451 | -1.349 | -0.282 | -0.705 | 0.005 | 0.038 | 0.038 | 0 | 0.16 | 0.08 | 0.09 | 0.06 | 0.14 | 0.07 |
| 3 | 142985100 | rs1046706 | 0.62 | -1.626 | -0.086 | -0.901 | 0.006 | -0.006 | -0.006 | 0 | 0.54 | 0.57 | 0.37 | 0.1 | 0.58 | 0.44 |
| 3 | 142989332 | rs920572 | -1.718 | -1.887 | 0.176 | -0.007 | 0.007 | 0 | 0 | 0.029 | 0.29 | 0.68 | 0.19 | 0.29 | 0.16 | 0.35 |
| 3 | 142991226 | rs7641350 | -2.038 | -1.987 | 0.085 | -0.099 | 0.121 | 0.022 | 0.022 | 0.631 | 0.36 | 0.88 | 0.24 | 0.29 | 0.2 | 0.36 |
| 3 | 142991989 | rs9813097 | -1.951 | -1.709 | 0.386 | -0.469 | 0.016 | 0.016 | 0.016 | 1.053 | 0.28 | 0.74 | 0.16 | 0.1 | 0.12 | 0.21 |
| 3 | 142994194 | rs7639109 | -1.288 | -1.529 | 0.487 | -0.39 | -0.059 | -0.127 | -0.127 | 0 | 0.2 | 0.52 | 0.09 | 0.01 | 0.06 | 0.1 |
| 3 | 142994822 | rs1501626 | -1.265 | -1.518 | 0.546 | -0.381 | -0.068 | -0.069 | -0.069 | 0 | 0.2 | 0.47 | 0.11 | 0.09 | 0.07 | 0.11 |
| 3 | 143009413 | rs6786757 | -1.025 | -1.225 | -0.74 | -0.084 | -0.02 | -0.012 | -0.012 | 0 | 0.24 | 0.66 | 0.13 | 0.09 | 0.09 | 0.16 |
| 3 | 143016478 | rs1135438 | 0.508 | 0.03 | -1.213 | 0.558 | -0.021 | -0.044 | -0.044 | 3.035 | 0.77 | 0.93 | 0.73 | 0.35 | 0.73 | 0.58 |
| 3 | 143017187 | rs986376 | 0.991 | 0.023 | -0.372 | 0.374 | 0.002 | 0.008 | 0.008 | 0.551 | 0.72 | 0.57 | 0.68 | 0.34 | 0.69 | 0.57 |
| 3 | 143020269 | rs6440162 | 0.504 | 0.149 | -1.448 | -0.308 | -0.008 | 0.163 | 0.163 | 0.313 | 0.55 | 0.85 | 0.41 | 0.19 | 0.58 | 0.36 |
| 3 | 143020870 | rs1393072 | -0.571 | 0.087 | -1.497 | -0.874 | 0.002 | 0.007 | 0.007 | 0 | 0.76 | 0.81 | 0.84 | 0.97 | 0.76 | 0.79 |
| 3 | 143021578 | rs16853385 | -0.184 | -0.37 | -1.264 | 0.083 | 0.004 | 0.001 | 0.001 | 0 | 0.11 | 0.08 | 0.11 | 0.22 | 0.09 | 0.2 |
| 3 | 143023597 | rs7428843 | -0.465 | -0.25 | -0.794 | -0.241 | 0.009 | 0.013 | 0.013 | 0.064 | 0.67 | 0.69 | 0.8 | 0.93 | 0.73 | 0.76 |
| 3 | 143025766 | rs7641069 | -0.706 | -0.036 | -0.178 | 0.148 | -0.016 | 0.04 | 0.04 | 0 | 0.15 | 0.22 | 0.13 | 0.28 | 0.11 | 0.21 |

| | | | | | | | | | | | | | | | | |
|---|---|---|---|---|---|---|---|---|---|---|---|---|---|---|---|---|
| 3 | 143029022 | rs1995356 | 0.93 | 0.561 | -0.169 | -0.541 | -0.002 | -0.089 | -0.089 | 0 | 0.28 | 0.3 | 0.15 | 0 | 0.25 | 0.07 |
| 3 | 143029090 | rs7431684 | -1.055 | 0.706 | -0.8 | 0.098 | -0.084 | -0.081 | -0.081 | 0.125 | 0.64 | 0.97 | 0.67 | 0.56 | 0.64 | 0.49 |
| 3 | 143031250 | rs6796444 | -1.001 | 0.771 | -0.253 | 0.313 | -0.008 | -0.014 | -0.014 | 0.723 | 0.31 | 0.77 | 0.48 | 0.38 | 0.36 | 0.39 |
| 3 | 143044440 | rs12488603 | 0.83 | 1.04 | 0.349 | 0.48 | 0.021 | 0.003 | 0.003 | 0.154 | 0.1 | 0.16 | 0.28 | 0.12 | 0.18 | 0.09 |
| 3 | 143045530 | rs1812200 | 1.191 | 0.768 | 0.018 | 0.031 | -0.003 | 0.031 | 0.031 | 0 | 0.27 | 0.17 | 0.14 | 0 | 0.24 | 0.08 |
| 3 | 143049769 | rs7431637 | -1.049 | 0.763 | -0.158 | 0.066 | -0.034 | -0.048 | -0.048 | 0 | 0.6 | 0.96 | 0.66 | 0.4 | 0.62 | 0.47 |
| 3 | 143050502 | rs9332458 | -0.303 | 0.787 | 0.222 | 0.019 | 0.061 | 0.023 | 0.023 | 0.216 | 0.18 | 0.54 | 0.41 | 0.23 | 0.25 | 0.18 |
| 3 | 143052348 | rs6440164 | -0.458 | 0.751 | 0.169 | -0.009 | 0.014 | 0.196 | 0.196 | 0 | 0.49 | 0.85 | 0.57 | 0.23 | 0.51 | 0.3 |
| 3 | 143056467 | rs7632299 | 0.683 | 0.788 | 0.316 | 0.03 | 0.115 | 0.099 | 0.099 | 0.031 | 0.13 | 0.37 | 0.37 | 0.23 | 0.21 | 0.15 |
| 3 | 143058731 | rs6769555 | -0.402 | 0.758 | 0.193 | 0.002 | 0.016 | 0.153 | 0.153 | 0 | 0.49 | 0.82 | 0.57 | 0.23 | 0.51 | 0.3 |
| 3 | 143059497 | rs16853457 | 0.475 | 0.695 | 0.137 | 0.053 | 0.004 | -0.017 | -0.017 | 0 | 0.1 | 0.05 | 0.07 | 0.16 | 0.11 | 0.16 |
| 3 | 143060387 | rs7642361 | 1.205 | 0.677 | 0.159 | 0.013 | 0 | -0.011 | -0.011 | 0 | 0.28 | 0.24 | 0.14 | 0.01 | 0.24 | 0.08 |
| 3 | 143066827 | rs6779058 | -0.474 | 0.62 | -0.045 | 0.212 | -0.005 | 0.001 | 0.001 | 0 | 0.15 | 0.19 | 0.11 | 0.17 | 0.13 | 0.21 |
| 3 | 143068250 | rs10446322 | -0.649 | 0.717 | -0.434 | 0.202 | 0.01 | 0.028 | 0.028 | 0 | 0.56 | 0.86 | 0.63 | 0.39 | 0.59 | 0.43 |
| 3 | 143070312 | rs6765636 | -1.289 | 0.78 | -0.051 | 0.575 | 0.013 | 0.033 | 0.033 | 0.074 | 0.31 | 0.58 | 0.51 | 0.5 | 0.36 | 0.38 |
| 3 | 143074208 | rs4839623 | 0.163 | 0.659 | -0.144 | 0.753 | 0.006 | -0.007 | -0.007 | 2.819 | 0.11 | 0.08 | 0.07 | 0.16 | 0.12 | 0.18 |
| 3 | 143082012 | rs7628692 | -0.417 | 0.639 | 0.14 | 0.674 | 0.037 | 0.034 | 0.034 | 1.298 | 0.23 | 0.36 | 0.44 | 0.34 | 0.31 | 0.34 |
| 3 | 143085345 | rs868702 | -1.117 | 0.654 | -0.005 | 0.662 | 0.004 | 0 | 0 | 1.384 | 0.29 | 0.67 | 0.47 | 0.33 | 0.34 | 0.37 |
| 3 | 143086325 | rs4839628 | -1.006 | 0.452 | -0.566 | 0.237 | -0.004 | -0.066 | -0.066 | 0 | 0.6 | 0.94 | 0.69 | 0.52 | 0.64 | 0.54 |
| 3 | 143090003 | rs7619769 | -0.526 | 0.483 | -0.301 | 0.231 | 0.016 | 0.004 | 0.004 | 0 | 0.52 | 0.63 | 0.63 | 0.49 | 0.59 | 0.51 |
| 3 | 143090698 | rs11715693 | -0.542 | 0.491 | -0.355 | 0.233 | 0.021 | 0.005 | 0.005 | 0 | 0.52 | 0.66 | 0.62 | 0.48 | 0.59 | 0.51 |
| 3 | 143094640 | rs4839631 | 0.272 | 0.835 | -0.226 | 0.25 | 0.005 | 0.015 | 0.015 | 0 | 0.81 | 0.84 | 0.8 | 0.61 | 0.77 | 0.78 |
| 3 | 143097448 | rs6440165 | 0.425 | 0.765 | -0.192 | 0.119 | 0.003 | 0.015 | 0.015 | 0 | 0.8 | 0.82 | 0.79 | 0.56 | 0.78 | 0.78 |
| 3 | 143097665 | rs9828435 | 0.975 | 0.834 | -0.208 | -0.015 | 0.005 | -0.007 | -0.007 | 0.749 | 0.42 | 0.35 | 0.5 | 0.41 | 0.41 | 0.46 |
| 3 | 143100626 | rs13099124 | 1.38 | 0.695 | -0.295 | -0.075 | -0.022 | 0.097 | 0.097 | 0 | 0.25 | 0.12 | 0.16 | 0 | 0.25 | 0.08 |
| 3 | 143101313 | rs6782465 | 0.89 | 0.692 | -0.262 | -0.006 | -0.001 | -0.006 | -0.006 | 0 | 0.09 | 0.05 | 0.1 | 0.15 | 0.1 | 0.18 |
| 3 | 143103734 | rs2293328 | 1.048 | 0.564 | -0.171 | -0.071 | 0.002 | -0.01 | -0.01 | 0.304 | 0.42 | 0.29 | 0.49 | 0.4 | 0.41 | 0.46 |
| 3 | 143106228 | rs6440168 | 0.879 | 0.798 | -0.567 | 0.029 | 0.011 | -0.001 | -0.001 | 0 | 0.46 | 0.6 | 0.52 | 0.41 | 0.42 | 0.48 |
| 3 | 143109003 | rs11707857 | 0.313 | 0.822 | -0.801 | -0.027 | -0.004 | 0.04 | 0.04 | 0 | 0.34 | 0.25 | 0.26 | 0.16 | 0.35 | 0.27 |
| 3 | 143124710 | rs11915421 | 1.19 | 0.739 | -0.907 | -0.096 | -0.036 | 0.125 | 0.125 | 0 | 0.24 | 0.07 | 0.15 | 0 | 0.25 | 0.09 |
| 3 | 143127587 | rs4839633 | -0.602 | 0.867 | -0.881 | 0.121 | -0.021 | -0.03 | -0.03 | 0.239 | 0.87 | 0.95 | 0.85 | 0.61 | 0.83 | 0.82 |
| 3 | 143129061 | rs9849797 | -0.497 | 0.895 | -0.735 | 0.125 | -0.008 | -0.005 | -0.005 | 1.38 | 0.87 | 0.88 | 0.85 | 0.6 | 0.83 | 0.8 |
| 3 | 143131978 | rs17757146 | 0.146 | 0.708 | -0.761 | -0.001 | 0.001 | 0.033 | 0.033 | 0 | 0.35 | 0.28 | 0.27 | 0.16 | 0.36 | 0.31 |
| 3 | 143132725 | rs1875459 | 0.393 | 0.971 | -1.406 | 0.121 | -0.024 | 0.093 | 0.093 | 0 | 0.32 | 0.11 | 0.25 | 0.16 | 0.34 | 0.25 |
| 3 | 143133504 | rs11718728 | 1.146 | 1.016 | -1.394 | 0.108 | -0.034 | 0.115 | 0.115 | 0 | 0.24 | 0.07 | 0.16 | 0 | 0.25 | 0.09 |
| 3 | 143136819 | rs4839634 | -0.631 | 0.887 | -1.091 | 0.532 | -0.005 | 0.01 | 0.01 | 0 | 0.13 | 0.16 | 0.15 | 0.2 | 0.14 | 0.19 |
| 3 | 143137224 | rs2037676 | -0.528 | 0.768 | -1.604 | 0.297 | -0.075 | -0.051 | -0.051 | 0 | 0.68 | 0.93 | 0.66 | 0.62 | 0.6 | 0.75 |
| 3 | 143137747 | rs6805667 | 0.054 | 0.724 | -1.526 | 0.279 | -0.004 | -0.001 | -0.001 | 0.488 | 0.06 | 0.11 | 0.06 | 0.04 | 0.05 | 0.02 |
| 3 | 143142681 | rs6440172 | -0.046 | 0.88 | -1.508 | 0.292 | -0.01 | -0.003 | -0.003 | 0 | 0.61 | 0.71 | 0.61 | 0.56 | 0.56 | 0.69 |
| 3 | 143144568 | rs1554672 | -0.174 | 1.017 | -1.679 | 0.363 | -0.013 | -0.008 | -0.008 | 0 | 0.62 | 0.75 | 0.58 | 0.56 | 0.58 | 0.67 |
| 3 | 143145568 | rs3860228 | 0.285 | 0.942 | -1.678 | 0.685 | -0.018 | -0.035 | -0.035 | 0 | 0.07 | 0.01 | 0.09 | 0.14 | 0.1 | 0.19 |
| 3 | 143155026 | rs905142 | -0.354 | 0.878 | -2.12 | 0.475 | -0.001 | -0.075 | -0.075 | 0 | 0.62 | 0.84 | 0.65 | 0.56 | 0.64 | 0.5 |
| 3 | 143157753 | rs12636237 | -0.306 | 1.343 | -1.947 | 0.845 | -0.003 | -0.005 | -0.005 | 0 | 0.1 | 0.07 | 0.13 | 0.17 | 0.1 | 0.11 |
| 3 | 143160538 | rs11923950 | 0.815 | 1.516 | -2.322 | 0.33 | -0.047 | -0.069 | -0.069 | 0.821 | 0.21 | 0.05 | 0.27 | 0.39 | 0.25 | 0.23 |
| 3 | 143161737 | rs9861304 | 1.035 | 1.678 | -2.27 | -0.037 | -0.041 | 0.042 | 0.042 | 0 | 0.09 | 0.02 | 0.12 | 0 | 0.18 | 0.09 |
| 3 | 143162091 | rs10222419 | -0.755 | 1.19 | -2.09 | -0.269 | -0.001 | 0.001 | 0.001 | 0 | 0.24 | 0.23 | 0.2 | 0.15 | 0.28 | 0.17 |
| 3 | 143163169 | rs4839595 | -0.775 | 0.536 | -1.957 | -0.253 | 0.014 | 0.006 | 0.006 | 0.012 | 0.5 | 0.71 | 0.53 | 0.57 | 0.44 | 0.55 |
| 3 | 143163381 | rs9871107 | -0.462 | 0.714 | -1.231 | -0.517 | 0.004 | -0.015 | -0.015 | 0 | 0.33 | 0.34 | 0.32 | 0.15 | 0.41 | 0.21 |
| 3 | 143163868 | rs12496742 | 0.033 | 0.212 | -1.203 | -0.092 | 0.003 | -0.002 | -0.002 | 0 | 0.16 | 0.13 | 0.15 | 0.15 | 0.15 | 0.11 |
| 3 | 143165502 | rs1875463 | -0.903 | -0.403 | -1.303 | 0.359 | -0.018 | -0.092 | -0.092 | 0.842 | 0.74 | 0.92 | 0.74 | 0.6 | 0.75 | 0.69 |
| 3 | 143178526 | rs838610 | -0.477 | -1.018 | -0.91 | 0.082 | -0.01 | -0.057 | -0.057 | 0.305 | 0.63 | 0.85 | 0.61 | 0.5 | 0.66 | 0.56 |
| 3 | 143179552 | rs6804097 | 0.739 | -1.307 | -0.169 | -0.149 | -0.007 | 0.023 | 0.023 | 0 | 0.28 | 0.32 | 0.33 | 0.5 | 0.23 | 0.32 |
| 3 | 143181685 | rs11919836 | 0.293 | -1.443 | 0.001 | -0.329 | -0.035 | -0.117 | -0.117 | 0 | 0.16 | 0.44 | 0.08 | 0 | 0.1 | 0.09 |
| 3 | 143192433 | rs838626 | -0.25 | -1.202 | 0.135 | -0.062 | 0.011 | 0.009 | 0.009 | 0 | 0.49 | 0.39 | 0.51 | 0.5 | 0.56 | 0.48 |
| 3 | 143195455 | rs838631 | -1.136 | -1.256 | -0.154 | -0.056 | -0.007 | 0.005 | 0.005 | 0 | 0.69 | 0.81 | 0.71 | 0.65 | 0.68 | 0.76 |
| 3 | 143199328 | rs838638 | -1.17 | -1.264 | -0.141 | -0.071 | -0.007 | 0.004 | 0.004 | 0 | 0.69 | 0.79 | 0.71 | 0.65 | 0.68 | 0.76 |
| 3 | 143200129 | rs10513205 | -0.42 | -1.375 | -0.208 | 0.104 | 0 | -0.001 | -0.001 | 0 | 0.49 | 0.54 | 0.51 | 0.64 | 0.35 | 0.61 |
| 3 | 143201591 | rs12488945 | 0.252 | -0.968 | -0.314 | 0.63 | -0.03 | 0.024 | 0.024 | 0 | 0.31 | 0.54 | 0.33 | 0.37 | 0.22 | 0.3 |
| 3 | 143209571 | rs838606 | 0.139 | -0.747 | -1 | 0.565 | -0.002 | -0.003 | -0.003 | 0 | 0.45 | 0.22 | 0.51 | 0.49 | 0.54 | 0.47 |
| 3 | 143210189 | rs1404568 | 0.562 | -0.436 | -1.042 | 0.799 | 0.009 | -0.037 | -0.037 | 0 | 0.15 | 0.02 | 0.15 | 0.15 | 0.11 | 0.26 |
| 3 | 143214114 | rs838598 | -0.428 | -0.294 | -0.427 | 1.057 | -0.001 | -0.001 | -0.001 | 0 | 0.36 | 0.37 | 0.43 | 0.4 | 0.42 | 0.44 |
| 3 | 143214462 | rs10935496 | 0.586 | -0.286 | -0.642 | 1.081 | -0.006 | 0.007 | 0.007 | 0 | 0.16 | 0.16 | 0.13 | 0.26 | 0.09 | 0.25 |
| 3 | 143215072 | rs838599 | 1.013 | 0.005 | -0.805 | 0.735 | -0.104 | 0.108 | 0.108 | 0 | 0.18 | 0.02 | 0.17 | 0 | 0.32 | 0.16 |
| 3 | 143220697 | rs838602 | 0.751 | 0.067 | -0.793 | 0.875 | -0.095 | -0.031 | -0.031 | 0 | 0.19 | 0.04 | 0.27 | 0.15 | 0.33 | 0.18 |
| 3 | 143222874 | rs6771449 | 0.519 | -0.042 | -0.766 | 0.328 | -0.07 | 0.062 | 0.062 | 0.559 | 0.19 | 0.06 | 0.19 | 0.01 | 0.36 | 0.22 |
| 3 | 143223146 | rs9844522 | 0.954 | -0.038 | -0.778 | 0.326 | -0.112 | 0.111 | 0.111 | 0 | 0.17 | 0.02 | 0.18 | 0 | 0.32 | 0.19 |
| 3 | 143223366 | rs9844871 | 0.311 | -0.111 | -0.656 | 0.318 | -0.029 | 0.006 | 0.006 | 0.315 | 0.21 | 0.15 | 0.2 | 0.01 | 0.36 | 0.22 |
| 3 | 143227959 | rs6440177 | -0.346 | -0.633 | -0.155 | 0.891 | -0.06 | 0.084 | 0.084 | 0 | 0.82 | 0.6 | 0.83 | 0.7 | 0.92 | 0.77 |
| 3 | 143237419 | rs2049504 | 0.253 | -0.883 | -0.479 | 0.618 | -0.062 | 0.034 | 0.034 | 0 | 0.74 | 0.52 | 0.79 | 0.7 | 0.86 | 0.73 |

| | | | | | | | | | | | | | | | | |
|---|---|---|---|---|---|---|---|---|---|---|---|---|---|---|---|---|
| 3 | 143237653 | rs2119380 | 0.688 | -0.674 | -0.531 | 0.593 | 0.002 | -0.006 | -0.006 | 0 | 0.56 | 0.52 | 0.61 | 0.69 | 0.53 | 0.53 |
| 3 | 143238874 | rs9872790 | 0.743 | -0.594 | -0.36 | 0.403 | -0.008 | -0.007 | -0.007 | 0 | 0.58 | 0.37 | 0.52 | 0.55 | 0.51 | 0.47 |
| 3 | 143241871 | rs12489418 | 0.569 | -0.93 | -0.89 | 0.419 | 0.02 | -0.008 | -0.008 | 0 | 0.22 | 0.03 | 0.13 | 0.17 | 0.15 | 0.34 |
| 3 | 143245008 | rs9821051 | 0.823 | -0.033 | 0.006 | 0.661 | 0.017 | 0.017 | 0.017 | 0 | 0.63 | 0.44 | 0.54 | 0.56 | 0.54 | 0.55 |
| 3 | 143247158 | rs1405019 | 0.36 | -0.845 | 0.159 | 0.062 | -0.038 | -0.089 | -0.089 | 0.102 | 0.84 | 0.72 | 0.9 | 0.98 | 0.91 | 0.8 |
| 3 | 143255258 | rs11922063 | -0.674 | -0.771 | 0.248 | 0.539 | -0.031 | 0.065 | 0.065 | 0 | 0.09 | 0.25 | 0.12 | 0.18 | 0.03 | 0.17 |
| 3 | 143257069 | rs7642430 | 0.325 | -0.742 | -0.391 | 0.495 | 0.005 | 0.002 | 0.002 | 0 | 0.16 | 0.06 | 0.15 | 0.15 | 0.15 | 0.13 |
| 3 | 143267083 | rs17579952 | 0.855 | -0.535 | -0.659 | 0.182 | -0.006 | 0.011 | 0.011 | 0.159 | 0.19 | 0.16 | 0.2 | 0.1 | 0.2 | 0.16 |
| 3 | 143269658 | rs10513209 | 0.989 | -0.412 | -0.626 | -0.109 | -0.002 | 0.006 | 0.006 | 0 | 0.12 | 0.13 | 0.09 | 0.04 | 0.14 | 0.09 |
| 3 | 143271365 | rs1525861 | 1.202 | -0.991 | -0.225 | -0.559 | -0.033 | 0.009 | 0.009 | 0 | 0.16 | 0.28 | 0.17 | 0.17 | 0.09 | 0.19 |
| 3 | 143274521 | rs17641385 | 1.462 | -0.725 | -0.437 | -0.445 | 0.001 | -0.009 | -0.009 | 0 | 0.19 | 0.24 | 0.21 | 0.1 | 0.2 | 0.17 |
| 3 | 143281041 | rs17641479 | 1.398 | -0.74 | -0.557 | -0.452 | -0.002 | 0.006 | 0.006 | 0.052 | 0.2 | 0.19 | 0.21 | 0.1 | 0.21 | 0.18 |
| 3 | 143283221 | rs13098869 | 1.66 | -0.883 | -0.787 | -0.642 | 0.01 | 0.03 | 0.03 | 0 | 0.06 | 0 | 0.01 | 0 | 0.04 | 0.01 |
| 3 | 143286467 | rs13085041 | -1.008 | -0.821 | -0.624 | -0.565 | -0.055 | -0.072 | -0.072 | 0.129 | 0.7 | 0.85 | 0.64 | 0.47 | 0.6 | 0.67 |
| 3 | 143288932 | rs10935498 | -0.589 | -0.985 | -0.449 | -0.671 | 0.052 | 0.101 | 0.101 | 0 | 0.51 | 0.66 | 0.44 | 0.26 | 0.39 | 0.5 |
| 3 | 143293012 | rs6763202 | -0.214 | -1.037 | 0.168 | -0.663 | -0.025 | -0.027 | -0.027 | 0 | 0.39 | 0.44 | 0.33 | 0.25 | 0.26 | 0.38 |
| 3 | 143293348 | rs6763515 | 0.274 | -1.157 | 0.167 | -0.748 | -0.075 | -0.1 | -0.1 | 0.935 | 0.24 | 0.45 | 0.2 | 0.08 | 0.11 | 0.24 |
| 3 | 143294547 | rs5003886 | 1.432 | -1.233 | 0.248 | -0.791 | -0.005 | -0.034 | -0.034 | 1.22 | 0.16 | 0.19 | 0.09 | 0 | 0.08 | 0.13 |
| 3 | 143294669 | rs727114 | -0.918 | -1.232 | -0.026 | -0.768 | 0.048 | 0.099 | 0.099 | 0 | 0.52 | 0.71 | 0.43 | 0.29 | 0.39 | 0.49 |
| 3 | 143296624 | rs6776465 | 1.105 | -1.199 | -0.05 | -0.607 | -0.024 | 0.001 | 0.001 | 0 | 0.07 | 0.12 | 0.1 | 0.08 | 0.02 | 0.11 |
| 3 | 143297591 | rs13076949 | 1.589 | -1.213 | -0.588 | -0.644 | -0.025 | -0.036 | -0.036 | 0.349 | 0.15 | 0 | 0.13 | 0.16 | 0.15 | 0.14 |
| 3 | 143305613 | rs959821 | -1.387 | -0.582 | -0.791 | -0.103 | -0.002 | -0.071 | -0.071 | 0 | 0.79 | 0.93 | 0.77 | 0.68 | 0.79 | 0.71 |
| 3 | 143307072 | rs13074737 | 1.684 | -0.62 | -0.686 | -0.117 | 0.002 | -0.002 | -0.002 | 0 | 0.14 | 0.13 | 0.13 | 0.17 | 0.13 | 0.12 |
| 3 | 143310347 | rs16853885 | 1.755 | -0.675 | -0.278 | -0.081 | 0.013 | 0.07 | 0.07 | 0 | 0.07 | 0.25 | 0.11 | 0.09 | 0.02 | 0.11 |
| 3 | 143310941 | rs6771556 | -0.414 | -0.649 | -0.268 | -0.224 | 0.025 | 0.032 | 0.032 | 0 | 0.48 | 0.56 | 0.41 | 0.3 | 0.38 | 0.46 |
| 3 | 143311260 | rs4839603 | 0.713 | -0.692 | -0.402 | -0.268 | 0.005 | 0.006 | 0.006 | 0 | 0.3 | 0.27 | 0.22 | 0.17 | 0.21 | 0.26 |
| 3 | 143311453 | rs17582102 | 1.424 | -0.545 | -0.382 | -0.194 | -0.005 | 0.016 | 0.016 | 0 | 0.13 | 0.07 | 0.08 | 0.04 | 0.14 | 0.1 |
| 3 | 143313963 | rs6770565 | -0.942 | 0.079 | -0.655 | -0.141 | -0.069 | -0.097 | -0.097 | 0 | 0.74 | 0.88 | 0.64 | 0.46 | 0.64 | 0.65 |
| 3 | 143319444 | rs16853934 | -0.158 | 0.12 | -0.1 | 0.108 | -0.021 | -0.043 | -0.043 | 0.454 | 0.22 | 0.44 | 0.2 | 0.13 | 0.17 | 0.21 |
| 3 | 143319916 | rs9846132 | 0.214 | 0.221 | -0.367 | 0.161 | -0.059 | 0.034 | 0.034 | 0 | 0.33 | 0.17 | 0.29 | 0.26 | 0.38 | 0.29 |
| 3 | 143330927 | rs6779888 | -0.977 | 0.591 | -0.631 | 0.317 | 0.01 | -0.015 | -0.015 | 0 | 0.58 | 0.73 | 0.56 | 0.48 | 0.58 | 0.51 |
| 3 | 143333030 | rs16853938 | 0.882 | 0.539 | -0.591 | 0.304 | 0.018 | -0.007 | -0.007 | 0.021 | 0.28 | 0.41 | 0.32 | 0.29 | 0.27 | 0.29 |
| 3 | 143334681 | rs13089318 | -1.06 | 0.615 | -0.78 | 0.325 | 0.012 | -0.035 | -0.035 | 0 | 0.59 | 0.72 | 0.56 | 0.52 | 0.6 | 0.51 |
| 3 | 143335479 | rs11714312 | 0.916 | 0.624 | -0.753 | 0.237 | 0.014 | -0.005 | -0.005 | 0 | 0.28 | 0.37 | 0.31 | 0.29 | 0.27 | 0.29 |
| 3 | 143347484 | rs9811274 | 0.44 | 0.382 | -1.638 | 0.445 | -0.02 | 0.019 | 0.019 | 0 | 0.43 | 0.72 | 0.44 | 0.37 | 0.41 | 0.39 |
| 3 | 143351588 | rs4425252 | -0.314 | -0.028 | -1.488 | 0.486 | 0.009 | -0.037 | -0.037 | 0 | 0.59 | 0.72 | 0.57 | 0.51 | 0.59 | 0.51 |
| 3 | 143354691 | rs13061355 | 0.345 | 0.014 | -1.38 | 0.301 | -0.007 | -0.049 | -0.049 | 0.099 | 0.15 | 0.01 | 0.13 | 0.15 | 0.16 | 0.12 |
| 3 | 143355137 | rs9862274 | 0.122 | -0.827 | -1.305 | -0.326 | 0.035 | 0.041 | 0.041 | 0 | 0.64 | 0.74 | 0.68 | 0.81 | 0.72 | 0.76 |
| 3 | 143355892 | rs9829661 | -0.567 | -0.45 | -0.142 | -0.288 | 0.05 | 0.014 | 0.014 | 0 | 0.07 | 0.34 | 0.07 | 0.06 | 0.07 | 0.03 |
| 3 | 143357546 | rs9873621 | -0.121 | -0.924 | -0.321 | -0.432 | 0.01 | -0.004 | -0.004 | 0 | 0.68 | 0.7 | 0.65 | 0.63 | 0.72 | 0.54 |
| 3 | 143358785 | rs7613679 | -0.558 | -0.954 | -0.365 | -0.484 | 0 | -0.008 | -0.008 | 1.718 | 0.76 | 0.72 | 0.7 | 0.63 | 0.76 | 0.58 |
| 3 | 143360568 | rs12635131 | 0.465 | -1.27 | -0.203 | -0.688 | 0.005 | 0.007 | 0.007 | 0 | 0.42 | 0.5 | 0.47 | 0.47 | 0.5 | 0.39 |
| 3 | 143363920 | rs16853964 | 0.299 | -1.242 | -0.233 | -0.689 | -0.023 | -0.049 | -0.049 | 0 | 0.25 | 0.57 | 0.23 | 0.12 | 0.21 | 0.24 |
| 3 | 143364025 | rs11916000 | 1.44 | -1.408 | 0.212 | -0.751 | -0.041 | -0.009 | -0.009 | 0 | 0.14 | 0.2 | 0.14 | 0.12 | 0.06 | 0.15 |
| 3 | 143365765 | rs1992866 | 1.395 | -1.468 | 0.045 | -0.851 | -0.057 | -0.054 | -0.054 | 0 | 0.23 | 0.12 | 0.27 | 0.36 | 0.34 | 0.19 |
| 3 | 143366830 | rs1242075 | -1.586 | -1.674 | -0.046 | -0.948 | -0.002 | -0.018 | -0.018 | 0 | 0.71 | 0.82 | 0.65 | 0.64 | 0.71 | 0.55 |
| 3 | 143367794 | rs12152544 | 0.726 | -1.708 | 0.006 | -0.931 | -0.003 | -0.007 | -0.007 | 0 | 0.09 | 0.08 | 0.06 | 0 | 0.13 | 0.09 |
| 3 | 143372127 | rs1242065 | -1.239 | -1.695 | -0.004 | -0.811 | -0.003 | -0.013 | -0.013 | 0 | 0.66 | 0.74 | 0.61 | 0.59 | 0.65 | 0.53 |
| 3 | 143372266 | rs1242064 | -0.438 | -1.664 | 0 | -0.807 | -0.002 | 0 | 0 | 0 | 0.54 | 0.52 | 0.53 | 0.49 | 0.58 | 0.45 |
| 3 | 143380236 | rs6440181 | 0.326 | -1.909 | 0.295 | -0.757 | -0.024 | 0.025 | 0.025 | 0 | 0.13 | 0.32 | 0.09 | 0.11 | 0.07 | 0.08 |
| 3 | 143387743 | rs1447738 | 0.488 | -1.762 | 0.089 | -0.688 | 0.008 | 0.008 | 0.008 | 0.096 | 0.61 | 0.42 | 0.69 | 0.72 | 0.78 | 0.73 |
| 3 | 143389999 | rs1530650 | 0.176 | -1.69 | 0.124 | -0.645 | -0.03 | -0.011 | -0.011 | 0.075 | 0.65 | 0.48 | 0.72 | 0.77 | 0.82 | 0.75 |
| 3 | 143397460 | rs7644727 | 0.587 | -1.264 | -0.028 | -0.505 | 0.047 | 0.051 | 0.051 | 0 | 0.52 | 0.39 | 0.62 | 0.71 | 0.63 | 0.63 |
| 3 | 143398613 | rs7650371 | -0.196 | -0.497 | 0.559 | -0.293 | -0.061 | -0.017 | -0.017 | 0.054 | 0.83 | 0.66 | 0.85 | 0.89 | 0.92 | 0.94 |
| 3 | 143400556 | rs10935500 | -0.196 | -0.497 | 0.559 | -0.293 | -0.061 | -0.017 | -0.017 | 0 | 0.83 | 0.67 | 0.85 | 0.89 | 0.92 | 0.94 |
| 3 | 143402200 | rs2121768 | -0.252 | -0.209 | 0.523 | -0.213 | -0.003 | -0.003 | -0.003 | 0.017 | 0.35 | 0.39 | 0.35 | 0.38 | 0.49 | 0.29 |
| 3 | 143403277 | rs7631319 | 0.842 | -0.063 | 0.294 | -0.115 | 0.032 | 0.007 | 0.007 | 0 | 0.49 | 0.3 | 0.51 | 0.51 | 0.43 | 0.65 |
| 3 | 143405250 | rs1868174 | 0.126 | -0.05 | 0.348 | -0.148 | -0.048 | -0.061 | -0.061 | 0 | 0.75 | 0.51 | 0.83 | 0.89 | 0.84 | 0.83 |
| 3 | 143405663 | rs2360867 | 0.896 | -0.057 | 0.346 | -0.152 | 0.018 | 0.004 | 0.004 | 0 | 0.49 | 0.34 | 0.52 | 0.51 | 0.43 | 0.65 |
| 3 | 143405698 | rs10513217 | 0.952 | -0.066 | 0.264 | -0.156 | 0.025 | 0.005 | 0.005 | 0 | 0.48 | 0.31 | 0.52 | 0.51 | 0.43 | 0.65 |
| 3 | 143409272 | rs6782243 | 0.413 | 0.078 | -0.468 | -0.025 | -0.062 | -0.051 | -0.051 | 0 | 0.26 | 0.2 | 0.3 | 0.38 | 0.4 | 0.18 |
| 3 | 143418390 | rs12489656 | 0.578 | 0.099 | -0.424 | -0.09 | 0.013 | 0.002 | 0.002 | 0 | 0.51 | 0.55 | 0.56 | 0.51 | 0.44 | 0.67 |
| 3 | 143419181 | rs6774032 | 0.471 | -0.3 | -0.26 | -0.713 | -0.031 | -0.066 | -0.066 | 0 | 0.76 | 0.62 | 0.83 | 0.93 | 0.85 | 0.85 |
| 3 | 143421221 | rs1900645 | 0.047 | 0.247 | -0.735 | -0.97 | 0.003 | -0.021 | -0.021 | 0 | 0.44 | 0.47 | 0.38 | 0.28 | 0.47 | 0.31 |
| 3 | 143421571 | rs12637424 | 0.052 | 0.071 | -1.067 | -1.036 | -0.006 | -0.007 | -0.007 | 0.55 | 0.42 | 0.47 | 0.53 | 0.66 | 0.36 | 0.64 |
| 3 | 143422497 | rs9836173 | 0.236 | 0.102 | -1.091 | -1.052 | 0.006 | 0.016 | 0.016 | 0.075 | 0.5 | 0.53 | 0.45 | 0.34 | 0.56 | 0.36 |
| 3 | 143422775 | rs11927652 | 0.254 | 0.082 | -1.1 | -1.061 | 0.007 | 0.017 | 0.017 | 0.043 | 0.5 | 0.53 | 0.45 | 0.34 | 0.56 | 0.36 |
| 3 | 143424822 | rs2800 | -0.123 | -0.206 | -1.1 | -1.19 | -0.002 | -0.004 | -0.004 | 0 | 0.66 | 0.71 | 0.64 | 0.4 | 0.66 | 0.59 |
| 3 | 143427040 | rs12233450 | 1.16 | -0.262 | -1.292 | -1.112 | -0.011 | 0.003 | 0.003 | 0.029 | 0.08 | 0.02 | 0.21 | 0.54 | 0.1 | 0.28 |

| | | | | | | | | | | | | | | | | |
|---|---|---|---|---|---|---|---|---|---|---|---|---|---|---|---|---|
| 3 | 143427194 | rs1124328 | 1.088 | -0.36 | -1.146 | -1.17 | -0.006 | -0.031 | -0.031 | 0.275 | 0.19 | 0.28 | 0.12 | 0.05 | 0.16 | 0.13 |
| 3 | 143427928 | rs7609691 | 0.641 | -0.513 | -1.047 | -1.186 | -0.012 | -0.018 | -0.018 | 0 | 0.13 | 0.19 | 0.19 | 0.07 | 0.1 | 0.21 |
| 3 | 143427951 | rs4839655 | 0.287 | 0.199 | -1.198 | -1.287 | -0.003 | 0.002 | 0.002 | 0.359 | 0.6 | 0.61 | 0.58 | 0.4 | 0.61 | 0.53 |
| 3 | 143428240 | rs765723 | -0.269 | -0.158 | -1.387 | -1.582 | 0.013 | 0.018 | 0.018 | 0 | 0.33 | 0.68 | 0.3 | 0.13 | 0.24 | 0.32 |
| 3 | 143428737 | rs1900649 | -0.852 | 0.549 | -1.124 | -1.124 | 0.001 | 0.015 | 0.015 | 2.232 | 0.91 | 0.97 | 0.97 | 1 | 0.92 | 1 |
| 3 | 143438124 | rs13058809 | 0.244 | 0.621 | -1.101 | -1.298 | -0.002 | -0.019 | -0.019 | 0 | 0.15 | 0.2 | 0.08 | 0 | 0.15 | 0.07 |
| 3 | 143438609 | rs7649414 | 0.915 | 0.484 | -1.441 | -1.383 | -0.027 | 0.12 | 0.12 | 0 | 0.49 | 0.17 | 0.41 | 0.35 | 0.45 | 0.55 |
| 3 | 143440367 | rs9819844 | 0.214 | 0.782 | -0.454 | -1.523 | 0.015 | -0.145 | -0.145 | 0.215 | 0.72 | 0.58 | 0.84 | 1 | 0.73 | 0.88 |
| 3 | 143440698 | rs6807319 | 0.406 | 0.664 | -0.543 | -1.637 | 0.009 | -0.119 | -0.119 | 0 | 0.7 | 0.6 | 0.81 | 1 | 0.71 | 0.86 |
| 3 | 143441481 | rs11712783 | 1.027 | 0.78 | -0.812 | -1.579 | 0.024 | 0.065 | 0.065 | 0 | 0.49 | 0.3 | 0.43 | 0.37 | 0.45 | 0.43 |
| 3 | 143442254 | rs12629217 | -0.134 | 0.889 | -0.197 | -1.157 | 0.006 | 0.08 | 0.08 | 1.434 | 0.2 | 0.32 | 0.4 | 0.63 | 0.24 | 0.42 |
| 3 | 143442774 | rs7645701 | -0.271 | 0.82 | -0.244 | -1.216 | -0.002 | 0.058 | 0.058 | 1.23 | 0.22 | 0.32 | 0.42 | 0.63 | 0.27 | 0.42 |
| 3 | 143443103 | rs7645841 | -0.753 | 1.208 | -0.184 | -1.217 | 0.026 | -0.049 | -0.049 | 0 | 0.09 | 0.27 | 0.08 | 0 | 0.07 | 0.07 |
| 3 | 143443933 | rs6780721 | -0.375 | 1.516 | -0.584 | -1.06 | 0.004 | 0.015 | 0.015 | 0 | 0.2 | 0.27 | 0.41 | 0.34 | 0.26 | 0.24 |
| 3 | 143445815 | rs6763423 | -0.67 | 1.456 | -0.841 | -1.042 | 0.003 | -0.025 | -0.025 | 0 | 0.91 | 0.92 | 0.96 | 1 | 0.88 | 0.95 |
| 3 | 143447722 | rs9829503 | -0.247 | 1.067 | -1.373 | -1.168 | -0.04 | 0.052 | 0.052 | 0 | 0.41 | 0.15 | 0.56 | 0.33 | 0.55 | 0.35 |
| 3 | 143450856 | rs12492227 | 1.335 | 0.787 | -1.208 | -0.881 | -0.038 | -0.025 | -0.025 | 0 | 0.34 | 0.02 | 0.21 | 0.22 | 0.27 | 0.43 |
| 3 | 143453195 | rs7616384 | -0.566 | 1.077 | 0.246 | -0.12 | -0.008 | -0.071 | -0.071 | 0.038 | 0.12 | 0.26 | 0.09 | 0 | 0.07 | 0.01 |
| 3 | 143464437 | rs17591646 | 0.511 | 1.259 | -0.32 | -0.053 | 0 | -0.011 | -0.011 | 0 | 0.08 | 0.06 | 0.06 | 0 | 0.07 | 0.01 |
| 3 | 143473442 | rs11714850 | -1.008 | 1.278 | 0.059 | 0.293 | -0.016 | 0.009 | 0.009 | 0.823 | 0.35 | 0.25 | 0.6 | 0.78 | 0.51 | 0.5 |
| 3 | 143478454 | rs4839657 | -1.043 | 1.398 | 0.214 | 0.363 | 0.001 | -0.001 | -0.001 | 0.057 | 0.27 | 0.7 | 0.38 | 0.29 | 0.32 | 0.2 |
| 3 | 143481878 | rs7613141 | -0.293 | 1.536 | 0.113 | 0.574 | 0.016 | 0.011 | 0.011 | 0 | 0.21 | 0.3 | 0.34 | 0.29 | 0.29 | 0.15 |
| 3 | 143482873 | rs1349001 | -0.451 | 1.228 | -0.274 | 0.599 | 0.047 | -0.003 | -0.003 | 0 | 0.26 | 0.56 | 0.4 | 0.3 | 0.38 | 0.16 |
| 3 | 143488471 | rs5025389 | -0.608 | 1.311 | -1.432 | 0.654 | -0.015 | 0.003 | 0.003 | 0.051 | 0.29 | 0.78 | 0.41 | 0.31 | 0.38 | 0.16 |
| 3 | 143493643 | rs16854267 | -0.824 | 0.583 | -1.465 | 1.47 | 0 | -0.041 | -0.041 | 0 | 0.14 | 0.06 | 0.26 | 0.46 | 0.18 | 0.33 |
| 3 | 143498566 | rs6805483 | -0.049 | 0.273 | -2.064 | 0.738 | -0.037 | 0.086 | 0.086 | 0.757 | 0.59 | 0.9 | 0.45 | 0.72 | 0.4 | 0.7 |
| 3 | 143500636 | rs16854291 | 0.116 | -0.937 | -1.053 | 1.15 | -0.045 | 0.035 | 0.035 | 0.206 | 0.26 | 0.42 | 0.21 | 0.29 | 0.14 | 0.27 |
| 3 | 143501570 | rs7626610 | -0.706 | -0.923 | -1.184 | 1.025 | -0.013 | 0.09 | 0.09 | 0.572 | 0.59 | 0.85 | 0.45 | 0.72 | 0.4 | 0.7 |
| 3 | 143502226 | rs12491888 | -0.78 | -0.905 | -1.21 | 1.033 | -0.042 | 0.079 | 0.079 | 1.005 | 0.59 | 0.9 | 0.45 | 0.72 | 0.4 | 0.69 |
| 3 | 143506434 | rs6808041 | 0.536 | -0.788 | -0.486 | 1.375 | -0.005 | -0.002 | -0.002 | 0 | 0.32 | 0.35 | 0.21 | 0.28 | 0.26 | 0.4 |
| 3 | 143510594 | rs2007192 | 0.251 | -0.323 | -0.17 | 1.296 | -0.012 | -0.014 | -0.014 | 0 | 0.36 | 0.44 | 0.3 | 0.29 | 0.3 | 0.42 |
| 3 | 143511130 | rs12233446 | 0.339 | -0.963 | -0.659 | 1.247 | 0.012 | -0.046 | -0.046 | 0.198 | 0.23 | 0.04 | 0.18 | 0.25 | 0.14 | 0.28 |
| 3 | 143526250 | rs4839610 | -0.036 | -1.352 | -0.171 | 1.043 | 0.051 | -0.027 | -0.027 | 0 | 0.41 | 0.73 | 0.28 | 0.54 | 0.27 | 0.39 |
| 3 | 143526787 | rs1868182 | -0.841 | -1.111 | 0.484 | 0.559 | 0.029 | 0.008 | 0.008 | 0 | 0.07 | 0.37 | 0.07 | 0 | 0.02 | 0.01 |
| 3 | 143533737 | rs13070587 | 0.454 | -0.822 | -0.235 | 0.567 | 0.013 | 0.049 | 0.049 | 0 | 0.35 | 0.14 | 0.19 | 0.24 | 0.35 | 0.39 |
| 3 | 143534874 | rs10513219 | -0.038 | -0.795 | 0.674 | 0.951 | 0.005 | -0.021 | -0.021 | 0 | 0.23 | 0.14 | 0.28 | 0.26 | 0.21 | 0.37 |
| 3 | 143535544 | rs6769847 | 0.829 | -0.796 | 1.456 | 0.926 | 0.043 | 0.076 | 0.076 | 0 | 0.93 | 0.66 | 0.95 | 0.91 | 0.95 | 0.95 |
| 3 | 143535626 | rs6794755 | -0.165 | -0.848 | 1.671 | 0.932 | -0.004 | 0.008 | 0.008 | 0.003 | 0.25 | 0.27 | 0.31 | 0.27 | 0.21 | 0.37 |
| 3 | 143537964 | rs9817066 | 0.926 | -0.916 | 1.658 | 0.91 | 0.034 | -0.045 | -0.045 | 0 | 0.64 | 0.16 | 0.62 | 0.62 | 0.73 | 0.56 |
| 3 | 143538156 | rs2361199 | 1.377 | -0.893 | 1.778 | 0.901 | 0.037 | 0.054 | 0.054 | 0 | 0.91 | 0.54 | 0.93 | 0.91 | 0.94 | 0.94 |
| 3 | 143542008 | rs7641634 | -0.596 | -0.879 | 1.962 | 0.94 | -0.02 | 0.011 | 0.011 | 0.563 | 0.32 | 0.47 | 0.34 | 0.3 | 0.25 | 0.43 |
| 3 | 143545699 | rs9847107 | 1.79 | -0.608 | 2.415 | 0.659 | -0.003 | 0.072 | 0.072 | 3.061 | 0.92 | 0.41 | 0.92 | 0.88 | 0.98 | 0.97 |
| 3 | 143547182 | rs4839670 | 2.044 | -0.63 | 2.258 | 0.652 | -0.036 | 0.005 | 0.005 | 4.101 | 0.89 | 0.27 | 0.9 | 0.88 | 0.97 | 0.96 |
| 3 | 143570871 | rs9832471 | 2.043 | -0.639 | 2.427 | 0.661 | -0.029 | 0 | 0 | 0 | 0.89 | 0.22 | 0.89 | 0.88 | 0.96 | 0.96 |
| 3 | 143574386 | rs6780027 | 1.3 | -0.518 | 2.153 | 0.667 | 0.04 | 0.036 | 0.036 | 0.707 | 0.94 | 0.62 | 0.94 | 0.97 | 0.97 | 0.99 |
| 3 | 143575195 | rs9289669 | 1.432 | -0.543 | 2.186 | 0.621 | 0.003 | -0.013 | -0.013 | 0.17 | 0.94 | 0.72 | 0.95 | 0.99 | 0.97 | 0.99 |
| 3 | 143576347 | rs4839614 | 2.135 | -0.582 | 2.373 | 0.623 | 0.006 | 0.056 | 0.056 | 1.581 | 0.9 | 0.49 | 0.91 | 0.91 | 0.96 | 0.95 |
| 3 | 143582463 | rs10513226 | -2.25 | -0.577 | 2.317 | 0.696 | 0.026 | 0.101 | 0.101 | 1.817 | 0.07 | 0.44 | 0.04 | 0.09 | 0.02 | 0.04 |
| 3 | 143585240 | rs6788160 | -1.658 | -0.562 | 2.292 | 0.663 | 0.063 | 0.042 | 0.042 | 0.648 | 0.06 | 0.43 | 0.05 | 0.01 | 0.02 | 0.02 |
| 3 | 143600997 | rs16854607 | -1.946 | -0.582 | 2.313 | 0.683 | 0.038 | 0.113 | 0.113 | 2.093 | 0.07 | 0.52 | 0.05 | 0.08 | 0.02 | 0.03 |
| 3 | 143605831 | rs9289670 | -0.049 | -0.112 | 0.982 | 0.383 | -0.054 | -0.011 | -0.011 | 0 | 0.26 | 0.07 | 0.21 | 0.28 | 0.33 | 0.25 |
| 3 | 143617422 | rs6440204 | -0.873 | -0.137 | -0.27 | 0.377 | -0.036 | -0.021 | -0.021 | 0 | 0.58 | 0.89 | 0.61 | 0.57 | 0.59 | 0.56 |
| 3 | 143620809 | rs6803662 | -1.083 | -0.566 | 0.35 | 0.211 | -0.037 | -0.056 | -0.056 | 4.036 | 0.11 | 0.6 | 0.08 | 0.01 | 0.03 | 0.06 |
| 3 | 143637187 | rs1027726 | -0.936 | -0.18 | -0.923 | 0.223 | -0.119 | -0.115 | -0.115 | 2.093 | 0.55 | 0.97 | 0.46 | 0.52 | 0.49 | 0.51 |
| 3 | 143656905 | rs1910691 | -0.257 | -0.397 | -0.079 | 0.212 | -0.001 | -0.025 | -0.025 | 0 | 0.36 | 0.56 | 0.32 | 0.26 | 0.39 | 0.26 |
| 3 | 143689222 | rs1992093 | -0.634 | -0.451 | 0.173 | 0.155 | -0.004 | 0.044 | 0.044 | 1.512 | 0.41 | 0.75 | 0.35 | 0.26 | 0.42 | 0.3 |
| 3 | 143690077 | rs4431106 | -1.38 | -0.559 | 0.202 | 0.217 | -0.032 | -0.038 | -0.038 | 0 | 0.54 | 0.85 | 0.45 | 0.52 | 0.48 | 0.51 |
| 3 | 143690416 | rs7631704 | -1.333 | -0.588 | 0.572 | 0.205 | -0.018 | -0.024 | -0.024 | 0 | 0.53 | 0.76 | 0.45 | 0.52 | 0.48 | 0.51 |
| 3 | 143697002 | rs6767811 | -1.388 | -0.605 | 0.483 | 0.217 | -0.063 | -0.069 | -0.069 | 0 | 0.54 | 0.92 | 0.45 | 0.52 | 0.48 | 0.52 |
| 3 | 143698664 | rs10513235 | 0.313 | -0.692 | 0.807 | 0.114 | -0.029 | 0.013 | 0.013 | 0.04 | 0.29 | 0.13 | 0.25 | 0.25 | 0.36 | 0.22 |
| 3 | 143703750 | rs7616037 | -1.134 | -0.655 | 1.202 | 0.131 | 0.111 | 0.027 | 0.027 | 0 | 0.06 | 0.41 | 0.06 | 0 | 0.05 | 0.04 |
| 3 | 143704516 | rs9847681 | 1.132 | -0.68 | 1.165 | 0.215 | 0.027 | -0.073 | -0.073 | 1.114 | 0.13 | 0 | 0.09 | 0.2 | 0.06 | 0.2 |
| 3 | 143706302 | rs9879081 | 0.978 | -0.676 | 1.194 | 0.207 | -0.007 | 0.011 | 0.011 | 0.367 | 0.13 | 0.11 | 0.1 | 0.26 | 0.06 | 0.22 |
| 3 | 143709368 | rs17667256 | -0.782 | -0.679 | 1.199 | -0.039 | -0.01 | 0.066 | 0.066 | 1.539 | 0.41 | 0.8 | 0.35 | 0.26 | 0.42 | 0.3 |
| 3 | 143711298 | rs7610378 | -0.329 | -0.825 | 1.135 | -0.016 | 0 | -0.015 | -0.015 | 0.101 | 0.34 | 0.39 | 0.29 | 0.26 | 0.36 | 0.26 |
| 3 | 143735572 | rs4561830 | -0.696 | -0.861 | 0.891 | 0.036 | -0.002 | 0.076 | 0.076 | 1.258 | 0.38 | 0.8 | 0.3 | 0.2 | 0.39 | 0.29 |
| 3 | 143762420 | rs11706025 | 1.024 | -1.095 | 0.768 | 0.216 | 0.007 | -0.008 | -0.008 | 0.934 | 0.13 | 0.08 | 0.15 | 0.22 | 0.04 | 0.2 |
| 3 | 143772247 | rs13068298 | 0.057 | -0.962 | 1.096 | 0.077 | -0.008 | -0.038 | -0.038 | 0 | 0.3 | 0.54 | 0.22 | 0.18 | 0.3 | 0.21 |
| 3 | 143774531 | rs16854874 | 1.113 | -0.96 | 1.107 | 0.078 | 0.018 | -0.02 | -0.02 | 0 | 0.13 | 0.07 | 0.14 | 0.22 | 0.04 | 0.2 |

| | | | | | | | | | | | | | | | | |
|---|---|---|---|---|---|---|---|---|---|---|---|---|---|---|---|---|
| 3 | 143774592 | rs2053211 | -1.462 | -0.978 | 1.115 | 0.046 | 0 | -0.05 | -0.05 | 0 | 0.51 | 0.96 | 0.43 | 0.42 | 0.4 | 0.49 |
| 3 | 143776096 | rs1898265 | -0.891 | -0.961 | 1.126 | 0.039 | 0.007 | 0.117 | 0.117 | 0.936 | 0.39 | 0.89 | 0.28 | 0.19 | 0.35 | 0.26 |
| 3 | 143778708 | rs16854884 | 0.374 | -1.086 | 1.283 | 0.039 | 0.001 | -0.024 | -0.024 | 0 | 0.28 | 0.4 | 0.2 | 0.18 | 0.3 | 0.21 |
| 3 | 143794370 | rs17668478 | 0.592 | -1.11 | 1.116 | 0.004 | -0.032 | 0.037 | 0.037 | 0.181 | 0.26 | 0.06 | 0.17 | 0.16 | 0.3 | 0.23 |
| 3 | 143796305 | rs990320 | -1.395 | -1.154 | 0.961 | -0.007 | 0.026 | -0.021 | -0.021 | 0 | 0.51 | 0.96 | 0.43 | 0.4 | 0.38 | 0.49 |
| 3 | 143801696 | rs13083224 | -0.48 | -1.064 | 1.253 | -0.03 | -0.01 | -0.036 | -0.036 | 1.564 | 0.34 | 0.53 | 0.24 | 0.25 | 0.33 | 0.27 |
| 3 | 143804040 | rs6804060 | -0.548 | -1.013 | 1.226 | -0.025 | -0.001 | -0.005 | -0.005 | 0 | 0.35 | 0.64 | 0.26 | 0.24 | 0.32 | 0.27 |
| 3 | 143817528 | rs6784938 | -0.495 | -1.143 | 1.162 | -0.008 | -0.002 | -0.003 | -0.003 | 0 | 0.36 | 0.6 | 0.3 | 0.27 | 0.33 | 0.3 |
| 3 | 143819506 | rs1530477 | -0.688 | -1.141 | 0.734 | -0.007 | 0.026 | 0.055 | 0.055 | 1.781 | 0.38 | 0.81 | 0.32 | 0.27 | 0.33 | 0.3 |
| 3 | 143825439 | rs9809103 | 1.245 | -1.133 | 0.63 | 0.056 | 0.014 | -0.011 | -0.011 | 0.196 | 0.12 | 0.07 | 0.1 | 0.14 | 0.04 | 0.16 |
| 3 | 143828642 | rs9825324 | 0.75 | -1.135 | 0.758 | 0.055 | -0.032 | 0.012 | 0.012 | 2.486 | 0.13 | 0.19 | 0.11 | 0.14 | 0.04 | 0.16 |
| 3 | 143831886 | rs13320658 | -1.273 | -1.101 | 0.171 | -0.019 | 0.029 | -0.036 | -0.036 | 0 | 0.51 | 0.94 | 0.43 | 0.41 | 0.39 | 0.51 |
| 3 | 143853149 | rs13078077 | -1.064 | -1.238 | 0.548 | 0.26 | 0.066 | -0.027 | -0.027 | 0 | 0.45 | 0.8 | 0.39 | 0.5 | 0.33 | 0.4 |
| 3 | 143853187 | rs11717074 | -0.205 | -1.736 | 1.245 | 0.317 | -0.085 | 0.018 | 0.018 | 4.352 | 0.21 | 0.55 | 0.19 | 0.26 | 0.05 | 0.18 |
| 3 | 143858564 | rs9862907 | 0.63 | -1.629 | 0.888 | 0.29 | 0.004 | -0.001 | -0.001 | 0 | 0.16 | 0.12 | 0.12 | 0.18 | 0.05 | 0.15 |
| 3 | 143888701 | rs7648286 | -1.502 | -0.956 | 0.926 | 0.297 | -0.012 | 0.01 | 0.01 | 2.498 | 0.13 | 0.69 | 0.14 | 0.21 | 0.05 | 0.17 |
| 3 | 143892876 | rs6800503 | -0.09 | -0.941 | 0.459 | 0.06 | 0.018 | 0.089 | 0.089 | 0.159 | 0.43 | 0.15 | 0.32 | 0.25 | 0.42 | 0.27 |
| 3 | 143905694 | rs13086045 | -0.09 | -0.941 | 0.459 | 0.115 | 0.018 | 0.107 | 0.107 | 0.159 | 0.43 | 0.15 | 0.32 | 0.22 | 0.42 | 0.27 |
| 3 | 143916142 | rs9824917 | -1.294 | -0.864 | 0.513 | 0.126 | -0.014 | 0.016 | 0.016 | 2.498 | 0.13 | 0.67 | 0.14 | 0.21 | 0.05 | 0.17 |
| 3 | 143922257 | rs4431107 | -1.302 | -0.831 | 0.238 | 0.098 | -0.088 | -0.053 | -0.053 | 0.042 | 0.58 | 0.89 | 0.47 | 0.46 | 0.48 | 0.44 |
| 3 | 143949314 | rs9841242 | 0.002 | -0.823 | 0.253 | 0.108 | -0.001 | 0.004 | 0.004 | 0 | 0.07 | 0.09 | 0.1 | 0.15 | 0.05 | 0.15 |
| 3 | 143952189 | rs1503335 | -0.426 | -0.706 | 0.354 | 0.14 | 0.028 | 0.076 | 0.076 | 0.665 | 0.5 | 0.33 | 0.4 | 0.33 | 0.45 | 0.42 |
| 3 | 143958863 | rs9289673 | -0.975 | -0.661 | 0.017 | 0.175 | -0.108 | -0.069 | -0.069 | 0 | 0.59 | 0.92 | 0.48 | 0.46 | 0.49 | 0.45 |
| 3 | 143970983 | rs7356114 | -0.354 | -0.237 | -0.395 | 0.699 | 0.005 | 0.051 | 0.051 | 0 | 0.05 | 0.11 | 0.04 | 0.22 | 0.05 | 0.14 |
| 3 | 143985479 | rs1354316 | 0.241 | -0.664 | -1.855 | 0.402 | -0.067 | -0.054 | -0.054 | 0.314 | 0.42 | 0.06 | 0.59 | 0.64 | 0.58 | 0.42 |
| 3 | 143992630 | rs6763060 | 0.652 | -1.186 | -1.224 | -0.043 | -0.003 | -0.016 | -0.016 | 0 | 0.2 | 0.32 | 0.14 | 0.12 | 0.19 | 0.22 |
| 3 | 144000439 | rs4681527 | -0.875 | -1.782 | -0.838 | -0.214 | 0.003 | -0.003 | -0.003 | 0 | 0.78 | 0.83 | 0.81 | 0.77 | 0.81 | 0.71 |
| 3 | 144007968 | rs1568979 | -0.06 | -1.563 | -0.572 | -0.728 | -0.001 | -0.014 | -0.014 | 0 | 0.54 | 0.49 | 0.7 | 0.84 | 0.55 | 0.59 |
| 3 | 144009300 | rs6770845 | 1.379 | -1.883 | -0.433 | -0.216 | -0.062 | -0.007 | -0.007 | 0 | 0.32 | 0.14 | 0.52 | 0.34 | 0.43 | 0.31 |
| 3 | 144019763 | rs13094155 | 1.104 | -1.677 | -0.241 | -0.706 | -0.001 | -0.017 | -0.017 | 0 | 0.22 | 0.29 | 0.18 | 0.07 | 0.22 | 0.2 |
| 3 | 144027934 | rs931773 | -1.231 | -1.668 | -0.258 | -0.72 | 0.001 | -0.007 | -0.007 | 0.079 | 0.77 | 0.79 | 0.88 | 0.97 | 0.79 | 0.81 |
| 3 | 144028498 | rs1358171 | 0.638 | -1.749 | -0.154 | -0.734 | 0.007 | 0.049 | 0.049 | 0 | 0.46 | 0.34 | 0.68 | 0.85 | 0.54 | 0.59 |
| 3 | 144039116 | rs197848 | -0.852 | -1.765 | 0.236 | -0.469 | 0.006 | 0.025 | 0.025 | 0 | 0.75 | 0.8 | 0.88 | 0.97 | 0.78 | 0.81 |
| 3 | 144039257 | rs197849 | 0.255 | -1.903 | 0.256 | -0.507 | -0.008 | -0.031 | -0.031 | 0 | 0.63 | 0.36 | 0.83 | 0.92 | 0.76 | 0.78 |
| 3 | 144050897 | rs197825 | -0.318 | -1.892 | 0.314 | -0.511 | -0.011 | -0.091 | -0.091 | 0 | 0.71 | 0.51 | 0.84 | 0.97 | 0.77 | 0.8 |
| 3 | 144058395 | rs197831 | -0.182 | -1.836 | 0.136 | -0.483 | -0.012 | -0.046 | -0.046 | 0 | 0.66 | 0.62 | 0.85 | 0.92 | 0.74 | 0.77 |
| 3 | 144067955 | rs197840 | 0.035 | -1.212 | 0.024 | -0.427 | -0.022 | -0.033 | -0.033 | 0.002 | 0.14 | 0.36 | 0.08 | 0.06 | 0.07 | 0.08 |
| 3 | 144072672 | rs197858 | -0.744 | -1.522 | -0.098 | -0.683 | 0.006 | -0.03 | -0.03 | 0 | 0.71 | 0.72 | 0.86 | 1 | 0.78 | 0.81 |
| 3 | 144080620 | rs1019950 | -0.423 | -1.571 | -0.099 | -0.7 | 0.016 | -0.023 | -0.023 | 0 | 0.64 | 0.68 | 0.85 | 1 | 0.76 | 0.8 |
| 3 | 144082862 | rs16855296 | 0.993 | -1.752 | -0.311 | -0.345 | -0.001 | -0.065 | -0.065 | 0 | 0.1 | 0.01 | 0.07 | 0.24 | 0.07 | 0.23 |
| 3 | 144084535 | rs856797 | 0.672 | -1.59 | -0.019 | -0.806 | -0.003 | -0.064 | -0.064 | 0 | 0.3 | 0.34 | 0.16 | 0.02 | 0.26 | 0.24 |
| 3 | 144087856 | rs979000 | 0.218 | -1.525 | -0.089 | -0.764 | 0.028 | 0.034 | 0.034 | 1.808 | 0.57 | 0.31 | 0.79 | 0.94 | 0.71 | 0.73 |
| 3 | 144100011 | rs728053 | -0.391 | -1.275 | -1.119 | 0.111 | 0.032 | -0.061 | -0.061 | 0.549 | 0.79 | 0.9 | 0.85 | 0.5 | 0.87 | 0.68 |
| 3 | 144106346 | rs892263 | -0.722 | -1.279 | -1.108 | 0.118 | 0.02 | -0.036 | -0.036 | 0 | 0.82 | 0.89 | 0.87 | 0.5 | 0.89 | 0.71 |
| 3 | 144118360 | rs9852498 | 1.522 | -0.142 | -1.521 | -0.048 | -0.112 | -0.015 | -0.015 | 0 | 0.16 | 0.01 | 0.28 | 0.14 | 0.29 | 0.12 |
| 3 | 144126216 | rs16855401 | 0.517 | -0.953 | -0.937 | -0.108 | 0.001 | 0.001 | 0.001 | 0.158 | 0.51 | 0.56 | 0.57 | 0.4 | 0.59 | 0.37 |
| 3 | 144130645 | rs4681292 | 1.336 | -0.856 | -1.204 | -0.338 | 0 | 0.023 | 0.023 | 0.591 | 0.23 | 0.11 | 0.24 | 0.08 | 0.23 | 0.22 |
| 3 | 144134473 | rs6440251 | -0.659 | -0.855 | -1.158 | -0.235 | 0.023 | -0.037 | -0.037 | 0 | 0.8 | 0.88 | 0.79 | 0.46 | 0.86 | 0.68 |
| 3 | 144146926 | rs2569948 | -0.061 | -0.249 | -0.797 | -0.081 | 0.007 | 0.01 | 0.01 | 0.233 | 0.52 | 0.62 | 0.52 | 0.36 | 0.58 | 0.37 |
| 3 | 144146945 | rs2569949 | -0.296 | 0.007 | -0.953 | 0.03 | 0.026 | 0.016 | 0.016 | 0.086 | 0.57 | 0.78 | 0.55 | 0.37 | 0.63 | 0.46 |
| 3 | 144167396 | rs2717399 | -0.18 | 0.164 | -1.181 | 0.205 | 0.029 | 0.055 | 0.055 | 0 | 0.5 | 0.72 | 0.51 | 0.36 | 0.56 | 0.36 |
| 3 | 144170066 | rs13068539 | 0.135 | 0.351 | -1.278 | -0.101 | -0.019 | -0.003 | -0.003 | 0 | 0.68 | 0.69 | 0.54 | 0.3 | 0.62 | 0.69 |
| 3 | 144171765 | rs13075539 | 0.449 | -0.48 | -0.844 | -0.797 | -0.046 | -0.138 | -0.138 | 0 | 0.21 | 0.35 | 0.11 | 0 | 0.15 | 0.2 |
| 3 | 144173107 | rs10513242 | 0.592 | -0.025 | -1.07 | -0.495 | -0.013 | 0.044 | 0.044 | 0 | 0.08 | 0.01 | 0.05 | 0 | 0.09 | 0.03 |
| 3 | 144203082 | rs9833354 | 0.122 | 0.015 | -1.134 | -0.384 | -0.073 | -0.052 | -0.052 | 0 | 0.7 | 0.9 | 0.63 | 0.31 | 0.6 | 0.66 |
| 3 | 144213414 | rs7611785 | 0.088 | -0.418 | -0.947 | -0.681 | -0.02 | 0.224 | 0.224 | 0 | 0.13 | 0.41 | 0.17 | 0.29 | 0.09 | 0.3 |
| 3 | 144215405 | rs9852425 | 1.068 | -0.481 | -0.938 | -0.725 | 0.001 | -0.102 | -0.102 | 0 | 0.56 | 0.46 | 0.46 | 0.02 | 0.52 | 0.36 |
| 3 | 144216555 | rs9840395 | -0.018 | -0.471 | -1.022 | -0.711 | -0.059 | -0.034 | -0.034 | 0 | 0.69 | 0.87 | 0.63 | 0.31 | 0.6 | 0.67 |
| 3 | 144216886 | rs9844730 | 1.202 | -0.789 | -1.101 | -0.866 | -0.026 | 0.11 | 0.11 | 0.07 | 0.51 | 0.13 | 0.47 | 0.27 | 0.52 | 0.61 |
| 3 | 144230196 | rs1527237 | 0.664 | -0.725 | -1.08 | -0.894 | -0.043 | 0.136 | 0.136 | 0 | 0.58 | 0.81 | 0.54 | 0.05 | 0.53 | 0.38 |
| 3 | 144241161 | rs1178897 | 0.536 | -0.722 | -1.1 | -0.826 | -0.071 | 0.183 | 0.183 | 0 | 0.59 | 0.87 | 0.54 | 0.09 | 0.54 | 0.45 |
| 3 | 144242847 | rs1657010 | 1.372 | -0.823 | -1.168 | -0.827 | -0.083 | 0.054 | 0.054 | 0.16 | 0.5 | 0.05 | 0.46 | 0.31 | 0.51 | 0.53 |
| 3 | 144244265 | rs1657012 | 0.593 | -0.693 | -1.072 | -0.769 | -0.072 | 0.189 | 0.189 | 0 | 0.59 | 0.88 | 0.54 | 0.09 | 0.54 | 0.45 |
| 3 | 144249504 | rs1726536 | -0.172 | -0.437 | -1.24 | -0.685 | -0.083 | -0.065 | -0.065 | 0.114 | 0.69 | 0.9 | 0.63 | 0.34 | 0.6 | 0.66 |
| 3 | 144252161 | rs1726526 | 0.758 | -0.429 | -1.204 | -0.786 | -0.049 | 0.156 | 0.156 | 0 | 0.58 | 0.8 | 0.54 | 0.09 | 0.53 | 0.38 |
| 3 | 144256185 | rs1880901 | -0.482 | -0.421 | -1.177 | -0.251 | -0.103 | -0.066 | -0.066 | 0 | 0.76 | 0.98 | 0.67 | 0.68 | 0.64 | 0.75 |
| 3 | 144256399 | rs1880902 | 1.698 | -0.969 | -1.096 | -0.438 | -0.048 | 0.322 | 0.322 | 0 | 0.45 | 0.12 | 0.38 | 0.05 | 0.46 | 0.32 |
| 3 | 144259158 | rs2717381 | 1.823 | -0.968 | -1.09 | -0.438 | -0.007 | 0.226 | 0.226 | 0 | 0.45 | 0.18 | 0.39 | 0.05 | 0.46 | 0.32 |
| 3 | 144262510 | rs10490826 | 0.977 | -0.962 | -1.013 | -0.269 | 0.014 | -0.038 | -0.038 | 0.019 | 0.1 | 0.03 | 0.09 | 0.26 | 0.06 | 0.22 |

| | | | | | | | | | | | | | | | | |
|---|---|---|---|---|---|---|---|---|---|---|---|---|---|---|---|---|
| 3 | 144265840 | rs12637065 | -0.873 | -0.98 | -0.904 | -0.292 | -0.014 | 0.006 | 0.006 | 0 | 0.26 | 0.63 | 0.2 | 0.26 | 0.15 | 0.34 |
| 3 | 144268855 | rs12631899 | 0.49 | -0.949 | -0.693 | -0.292 | -0.009 | 0.177 | 0.177 | 0 | 0.12 | 0.34 | 0.13 | 0.25 | 0.09 | 0.23 |
| 3 | 144271654 | rs2010067 | -0.026 | -0.825 | -0.625 | -0.263 | -0.024 | -0.098 | -0.098 | 0 | 0.15 | 0.3 | 0.09 | 0 | 0.08 | 0.11 |
| 3 | 144277333 | rs2717389 | 1.088 | -0.822 | -0.7 | -0.281 | 0.004 | -0.004 | -0.004 | 0 | 0.56 | 0.55 | 0.55 | 0.35 | 0.54 | 0.56 |
| 3 | 144286112 | rs1989624 | 0.003 | -1.108 | -0.545 | -0.226 | -0.025 | -0.096 | -0.096 | 0.983 | 0.15 | 0.3 | 0.09 | 0 | 0.07 | 0.11 |
| 3 | 144287355 | rs4681363 | -0.099 | -1.101 | -0.623 | -0.235 | -0.094 | -0.023 | -0.023 | 0.476 | 0.33 | 0.11 | 0.38 | 0.36 | 0.44 | 0.31 |
| 3 | 144291254 | rs12637885 | 0.001 | -1.134 | -0.632 | -0.225 | -0.119 | -0.043 | -0.043 | 0 | 0.32 | 0.07 | 0.38 | 0.36 | 0.44 | 0.31 |
| 3 | 144298314 | rs12631326 | 1.135 | -1.191 | -0.636 | -0.223 | 0.005 | 0.001 | 0.001 | 0 | 0.1 | 0.07 | 0.09 | 0.26 | 0.08 | 0.17 |
| 3 | 144301659 | rs16855659 | 1.054 | -1.192 | -0.637 | -0.223 | 0.006 | -0.006 | -0.006 | 0 | 0.1 | 0.07 | 0.09 | 0.26 | 0.08 | 0.17 |
| 3 | 144307096 | rs9289678 | -0.075 | -1.121 | -0.379 | -0.077 | -0.119 | -0.039 | -0.039 | 2.142 | 0.32 | 0.07 | 0.38 | 0.36 | 0.44 | 0.3 |
| 3 | 144308606 | rs7613909 | -0.714 | -1.129 | -0.362 | -0.099 | -0.003 | 0 | 0 | 0 | 0.38 | 0.39 | 0.46 | 0.69 | 0.45 | 0.32 |
| 3 | 144311843 | rs1007564 | -0.283 | -1.109 | -0.368 | -0.112 | -0.077 | -0.037 | -0.037 | 0 | 0.34 | 0.16 | 0.43 | 0.4 | 0.45 | 0.32 |
| 3 | 144314964 | rs6793949 | 0.097 | -1.046 | -0.357 | -0.144 | -0.018 | -0.095 | -0.095 | 0 | 0.15 | 0.27 | 0.08 | 0 | 0.08 | 0.11 |
| 3 | 144323993 | rs16855734 | -0.001 | -1.139 | -0.379 | -0.145 | -0.027 | -0.12 | -0.12 | 0.055 | 0.16 | 0.47 | 0.1 | 0 | 0.08 | 0.19 |
| 3 | 144331004 | rs11706820 | 1.341 | -0.755 | -0.657 | 0.028 | -0.046 | -0.041 | -0.041 | 0 | 0.09 | 0.01 | 0.16 | 0.13 | 0.11 | 0.13 |
| 3 | 144339980 | rs800081 | -1.434 | -1.563 | -0.439 | -0.081 | 0.007 | 0.045 | 0.045 | 0 | 0.53 | 0.69 | 0.55 | 0.69 | 0.52 | 0.43 |
| 3 | 144340205 | rs800082 | -0.104 | -1.373 | -0.476 | -0.063 | -0.037 | -0.001 | -0.001 | 0 | 0.34 | 0.2 | 0.4 | 0.36 | 0.44 | 0.3 |
| 3 | 144342672 | rs800086 | -1.373 | -1.336 | -0.115 | 0.199 | 0.004 | 0.008 | 0.008 | 0.248 | 0.51 | 0.5 | 0.52 | 0.58 | 0.51 | 0.56 |
| 3 | 144347471 | rs800072 | -0.952 | -1.466 | 0.179 | 0.104 | 0.003 | -0.062 | -0.062 | 0 | 0.42 | 0.6 | 0.51 | 0.9 | 0.45 | 0.46 |
| 3 | 144347744 | rs800075 | -1.982 | -1.452 | 0.134 | 0.121 | -0.033 | 0.192 | 0.192 | 0.022 | 0.58 | 0.84 | 0.59 | 0.9 | 0.51 | 0.58 |
| 3 | 144352296 | rs1151556 | -0.535 | -1.352 | 0.236 | 0.253 | 0.014 | 0.002 | 0.002 | 0 | 0.33 | 0.56 | 0.46 | 0.37 | 0.39 | 0.31 |
| 3 | 144354430 | rs1092103 | -0.976 | -1.256 | 0.273 | 0.319 | -0.009 | 0.003 | 0.003 | 0 | 0.41 | 0.58 | 0.48 | 0.65 | 0.43 | 0.43 |
| 3 | 144375349 | rs941214 | 0.33 | -1.049 | -0.121 | 0.477 | 0.01 | -0.01 | -0.01 | 0.446 | 0.19 | 0.18 | 0.12 | 0.25 | 0.11 | 0.16 |
| 3 | 144382589 | rs1178938 | -0.615 | -0.801 | -0.644 | 0.542 | 0.019 | 0 | 0 | 0 | 0.35 | 0.36 | 0.44 | 0.37 | 0.4 | 0.35 |
| 3 | 144388175 | rs800065 | -0.532 | -0.679 | -1.147 | 0.425 | -0.006 | 0.001 | 0.001 | 0 | 0.34 | 0.24 | 0.39 | 0.33 | 0.4 | 0.34 |
| 3 | 144391772 | rs1178943 | 0.515 | -0.098 | -2.2 | 0.779 | -0.141 | -0.101 | -0.101 | 1.307 | 0.21 | 0.01 | 0.32 | 0.33 | 0.34 | 0.22 |
| 3 | 144407872 | rs1151564 | -0.464 | 0.295 | -1.539 | -0.601 | -0.034 | 0.212 | 0.212 | 0.02 | 0.57 | 0.84 | 0.59 | 0.9 | 0.51 | 0.57 |
| 3 | 144413287 | rs16855880 | -1.068 | -0.44 | -0.128 | 0.028 | -0.02 | -0.013 | -0.013 | 0 | 0.13 | 0.69 | 0.09 | 0.09 | 0.09 | 0.15 |
| 3 | 144416508 | rs2117999 | -1.088 | -0.447 | -0.137 | 1.086 | -0.016 | 0.052 | 0.052 | 0.133 | 0.13 | 0.73 | 0.11 | 0.31 | 0.09 | 0.16 |
| 3 | 144428012 | rs7639659 | -0.701 | -0.58 | 0.546 | 1.094 | 0 | 0.049 | 0.049 | 0 | 0.08 | 0.23 | 0.07 | 0.12 | 0.07 | 0.2 |
| 3 | 144441451 | rs1512775 | 0.041 | -1.628 | -0.782 | 0.628 | 0.056 | 0.057 | 0.057 | 0 | 0.25 | 0.05 | 0.13 | 0.11 | 0.19 | 0.21 |
| 3 | 144456570 | rs7647656 | 0.004 | -1.374 | -1.049 | 0.549 | 0.043 | -0.126 | -0.126 | 0.231 | 0.72 | 0.94 | 0.74 | 0.52 | 0.81 | 0.77 |
| 3 | 144466929 | rs1512779 | -0.527 | -0.551 | -1.12 | 0.275 | -0.014 | -0.061 | -0.061 | 0 | 0.53 | 0.93 | 0.61 | 0.5 | 0.6 | 0.59 |
| 3 | 144473663 | rs12634654 | 0.906 | -0.787 | -1.084 | 0.424 | -0.013 | -0.097 | -0.097 | 0.551 | 0.14 | 0.01 | 0.22 | 0.28 | 0.11 | 0.18 |
| 3 | 144479727 | rs7630559 | -1.008 | -1.036 | -0.873 | -0.289 | 0.051 | 0.038 | 0.038 | 1.459 | 0.2 | 0.88 | 0.16 | 0.11 | 0.11 | 0.23 |
| 3 | 144486315 | rs7611272 | -1.139 | -1.026 | -0.887 | -0.285 | 0.054 | 0.039 | 0.039 | 1.598 | 0.2 | 0.9 | 0.16 | 0.11 | 0.11 | 0.21 |
| 3 | 144492101 | rs6790749 | -0.554 | -1.034 | -0.196 | -0.287 | -0.012 | 0.035 | 0.035 | 0.305 | 0.11 | 0.41 | 0.1 | 0.11 | 0.08 | 0.2 |
| 3 | 144496334 | rs7615606 | 0.672 | -0.652 | -0.875 | -0.385 | -0.13 | -0.115 | -0.115 | 0.599 | 0.26 | 0.03 | 0.44 | 0.4 | 0.46 | 0.26 |
| 3 | 144500151 | rs1355661 | -0.993 | -0.789 | 0.68 | -0.554 | -0.036 | -0.026 | -0.026 | 0 | 0.17 | 0.7 | 0.14 | 0.11 | 0.08 | 0.21 |
| 3 | 144511555 | rs6805089 | -1.486 | -0.782 | 0.316 | -0.658 | 0.021 | 0.028 | 0.028 | 0.949 | 0.27 | 0.77 | 0.15 | 0.12 | 0.14 | 0.3 |
| 3 | 144538795 | rs2889113 | -0.106 | -0.176 | 0.356 | -0.285 | -0.001 | -0.007 | -0.007 | 2.911 | 0.24 | 0.19 | 0.25 | 0.27 | 0.23 | 0.28 |
| 3 | 144539001 | rs6789186 | -1.6 | -0.317 | 0.137 | -0.48 | 0.019 | -0.006 | -0.006 | 0 | 0.45 | 0.86 | 0.39 | 0.39 | 0.37 | 0.5 |
| 3 | 144556283 | rs10935515 | -0.889 | -0.32 | 0.614 | -0.489 | -0.002 | 0.002 | 0.002 | 0.46 | 0.33 | 0.52 | 0.35 | 0.3 | 0.33 | 0.37 |
| 3 | 144577620 | rs2203774 | 0.068 | -0.363 | 0.495 | -0.532 | -0.012 | 0.007 | 0.007 | 0 | 0.1 | 0.41 | 0.06 | 0.09 | 0.04 | 0.12 |
| 3 | 144586275 | rs7632306 | -0.179 | -0.336 | 0.051 | -0.573 | 0.007 | -0.029 | -0.029 | 0 | 0.09 | 0.22 | 0.08 | 0.03 | 0.1 | 0.09 |
| 3 | 144589580 | rs7644213 | 0.032 | -0.331 | 0.06 | -0.571 | 0 | -0.023 | -0.023 | 0 | 0.09 | 0.16 | 0.07 | 0.03 | 0.1 | 0.09 |
| 3 | 144590550 | rs4401402 | -0.936 | -0.357 | 0.045 | -0.571 | -0.004 | 0.002 | 0.002 | 0.387 | 0.34 | 0.48 | 0.34 | 0.3 | 0.33 | 0.37 |
| 3 | 144613189 | rs11708824 | -2.052 | -0.462 | 1.079 | -0.291 | -0.043 | -0.025 | -0.025 | 0 | 0.21 | 0.54 | 0.14 | 0.16 | 0.14 | 0.22 |
| 3 | 144617422 | rs953859 | -1.955 | -0.371 | -0.018 | -0.271 | 0.012 | 0.013 | 0.013 | 0.45 | 0.27 | 0.79 | 0.17 | 0.16 | 0.17 | 0.23 |
| 3 | 144630219 | rs7632741 | -0.778 | 0.101 | -0.009 | -0.067 | -0.004 | -0.056 | -0.056 | 1.057 | 0.1 | 0.21 | 0.05 | 0 | 0.1 | 0.06 |
| 3 | 144642293 | rs6440278 | -1.222 | 0.052 | -0.544 | 0.02 | -0.005 | -0.005 | -0.005 | 0 | 0.21 | 0.78 | 0.17 | 0.16 | 0.16 | 0.19 |
| 3 | 144643319 | rs6783105 | -0.19 | 0.026 | -1.558 | -0.763 | -0.003 | 0.139 | 0.139 | 0 | 0.59 | 0.85 | 0.69 | 0.85 | 0.47 | 0.68 |
| 3 | 144646012 | rs13061557 | 0.341 | -0.146 | -1.293 | -1.668 | 0.006 | 0.033 | 0.033 | 0 | 0.59 | 0.36 | 0.59 | 0.85 | 0.66 | 0.69 |
| 3 | 144658558 | rs4308318 | 0.634 | -0.393 | -1.255 | -1.705 | -0.032 | 0.022 | 0.022 | 0 | 0.32 | 0.19 | 0.4 | 0.23 | 0.37 | 0.18 |
| 3 | 144661211 | rs4611857 | 0.25 | -0.803 | -1.108 | -1.649 | -0.041 | 0.068 | 0.068 | 0.017 | 0.45 | 0.22 | 0.44 | 0.7 | 0.52 | 0.7 |
| 3 | 144663231 | rs10454765 | 0.391 | -0.812 | -1.02 | -1.652 | -0.028 | 0.074 | 0.074 | 0 | 0.45 | 0.25 | 0.48 | 0.71 | 0.52 | 0.71 |
| 3 | 144664456 | rs6794760 | -1.071 | -0.583 | -0.852 | -1.411 | 0.008 | 0.027 | 0.027 | 2.27 | 0.89 | 0.95 | 0.95 | 1 | 0.92 | 0.96 |
| 3 | 144668698 | rs9835486 | 0.134 | -0.585 | -0.801 | -0.999 | 0.002 | -0.005 | -0.005 | 0 | 0.25 | 0.21 | 0.22 | 0.31 | 0.24 | 0.29 |
| 3 | 144669044 | rs4681640 | 1.491 | -0.629 | -1.135 | -1.079 | -0.097 | 0.049 | 0.049 | 0 | 0.37 | 0.07 | 0.38 | 0.25 | 0.43 | 0.15 |
| 3 | 144672307 | rs6779715 | 0.369 | -0.342 | -0.023 | -0.703 | -0.006 | -0.038 | -0.038 | 0 | 0.67 | 0.81 | 0.69 | 0.57 | 0.67 | 0.46 |
| 3 | 144677749 | rs4681641 | 0.5 | -0.383 | -0.062 | -0.706 | -0.011 | -0.04 | -0.04 | 0 | 0.21 | 0.13 | 0.26 | 0.43 | 0.25 | 0.42 |
| 3 | 144681790 | rs3924267 | 0.645 | -0.355 | 0.225 | -0.69 | -0.005 | 0.011 | 0.011 | 0 | 0.64 | 0.54 | 0.67 | 0.57 | 0.67 | 0.47 |
| 3 | 144689230 | rs13065774 | -0.311 | -0.322 | 0.163 | -0.699 | -0.004 | -0.015 | -0.015 | 0 | 0.06 | 0.27 | 0.04 | 0 | 0.01 | 0 |
| 3 | 144698368 | rs9834819 | 0.602 | -0.249 | -0.234 | -0.661 | -0.008 | 0.022 | 0.022 | 0 | 0.64 | 0.47 | 0.63 | 0.56 | 0.67 | 0.51 |
| 3 | 144701909 | rs6440283 | -0.909 | -0.249 | -0.679 | -0.691 | -0.015 | 0.058 | 0.058 | 1.165 | 0.9 | 1 | 0.96 | 1 | 0.92 | 0.98 |
| 3 | 144719412 | rs4608734 | -0.217 | -0.194 | -0.411 | -0.499 | -0.001 | 0.001 | 0.001 | 0 | 0.23 | 0.22 | 0.2 | 0.25 | 0.22 | 0.27 |
| 3 | 144720051 | rs11706681 | 1.382 | -0.529 | -0.301 | -0.524 | 0.003 | 0.113 | 0.113 | 0 | 0.46 | 0.23 | 0.47 | 0.22 | 0.56 | 0.25 |
| 3 | 144728277 | rs9883600 | 1.424 | -0.557 | -0.274 | -0.515 | -0.014 | 0.148 | 0.148 | 0 | 0.46 | 0.13 | 0.47 | 0.22 | 0.56 | 0.25 |
| 3 | 144734068 | rs9859874 | -0.415 | -0.654 | 0.306 | -0.567 | -0.013 | -0.044 | -0.044 | 0.007 | 0.87 | 0.78 | 0.92 | 0.99 | 0.94 | 0.89 |

| | | | | | | | | | | | | | | | | |
|---|---|---|---|---|---|---|---|---|---|---|---|---|---|---|---|---|
| 3 | 144744693 | rs6414361 | -0.693 | -0.637 | 0.462 | -0.573 | -0.012 | -0.063 | -0.063 | 0.217 | 0.88 | 0.75 | 0.94 | 1 | 0.94 | 0.9 |
| 3 | 144750754 | rs6440299 | -0.125 | -0.547 | 0.404 | -0.383 | -0.007 | 0.005 | 0.005 | 0 | 0.24 | 0.33 | 0.22 | 0.26 | 0.22 | 0.27 |
| 3 | 144764442 | rs7427602 | 0.013 | -0.577 | 0.231 | -0.391 | -0.002 | 0 | 0 | 0 | 0.23 | 0.22 | 0.2 | 0.25 | 0.22 | 0.27 |
| 3 | 144782354 | rs6440306 | 0.231 | -0.877 | 0.649 | -0.162 | -0.026 | 0.006 | 0.006 | 0 | 0.1 | 0.25 | 0.08 | 0.06 | 0.05 | 0.04 |
| 3 | 144798660 | rs7373412 | 1.16 | -1.049 | 0.717 | -0.2 | 0.029 | -0.004 | -0.004 | 0 | 0.6 | 0.28 | 0.63 | 0.44 | 0.72 | 0.5 |
| 3 | 144806160 | rs7632966 | -0.145 | -0.959 | 0.748 | -0.231 | -0.035 | 0.002 | 0.002 | 0 | 0.12 | 0.36 | 0.09 | 0.06 | 0.05 | 0.04 |
| 3 | 144812596 | rs774 | 1.278 | -0.7 | -0.094 | -0.352 | -0.053 | -0.014 | -0.014 | 0.039 | 0.17 | 0.02 | 0.17 | 0.17 | 0.2 | 0.14 |
| 3 | 144815506 | rs7433098 | 0.992 | -0.997 | 1.165 | 0.18 | -0.042 | -0.057 | -0.057 | 0.152 | 0.76 | 0.42 | 0.85 | 0.94 | 0.89 | 0.87 |
| 3 | 144843921 | rs9863274 | 0.993 | -0.953 | 1.119 | 0.202 | -0.023 | -0.053 | -0.053 | 0.124 | 0.76 | 0.38 | 0.84 | 0.94 | 0.87 | 0.84 |
| 3 | 144850047 | rs7433333 | -0.23 | -0.913 | 1.089 | 0.218 | -0.013 | -0.064 | -0.064 | 0 | 0.88 | 0.77 | 0.94 | 1 | 0.94 | 0.9 |
| 3 | 144864268 | rs11928035 | 0.491 | -1.002 | 0.919 | 0.298 | 0.007 | -0.001 | -0.001 | 0.23 | 0.15 | 0.11 | 0.22 | 0.51 | 0.14 | 0.34 |
| 3 | 144868289 | rs9868905 | -1.102 | -0.954 | 1.076 | 0.266 | -0.012 | 0.013 | 0.013 | 0 | 0.28 | 0.61 | 0.33 | 0.56 | 0.21 | 0.39 |
| 3 | 144869013 | rs7432214 | 0.761 | -0.875 | 0.983 | 0.309 | -0.02 | -0.055 | -0.055 | 0 | 0.76 | 0.42 | 0.81 | 0.93 | 0.85 | 0.84 |
| 3 | 144880588 | rs9867653 | -1.067 | -0.65 | 0.643 | 0.395 | 0.012 | -0.001 | -0.001 | 0 | 0.39 | 0.64 | 0.4 | 0.57 | 0.29 | 0.51 |
| 3 | 144881442 | rs6803080 | -1.244 | -0.601 | 0.335 | 0.273 | 0.053 | 0.019 | 0.019 | 0.586 | 0.42 | 0.77 | 0.42 | 0.6 | 0.29 | 0.54 |
| 3 | 144883424 | rs13100466 | -0.126 | -0.208 | 0.36 | 0.275 | 0.001 | 0.009 | 0.009 | 0 | 0.1 | 0.27 | 0.09 | 0.06 | 0.07 | 0.06 |
| 3 | 144884332 | rs7622285 | -0.827 | -0.177 | -0.454 | 0.141 | 0.029 | 0.016 | 0.016 | 0.064 | 0.4 | 0.7 | 0.41 | 0.63 | 0.29 | 0.54 |
| 3 | 144895451 | rs12489832 | -0.174 | 0.812 | -0.223 | 0.013 | 0.023 | -0.011 | -0.011 | 0.105 | 0.27 | 0.28 | 0.37 | 0.16 | 0.44 | 0.26 |
| 3 | 144896643 | rs12493808 | -0.267 | 0.765 | 0.007 | -0.002 | 0.049 | -0.024 | -0.024 | 0 | 0.28 | 0.34 | 0.37 | 0.16 | 0.44 | 0.32 |
| 3 | 144898708 | rs2120986 | 0.291 | 0.227 | -0.201 | 0.193 | -0.027 | -0.045 | -0.045 | 0 | 0.8 | 0.98 | 0.83 | 0.82 | 0.82 | 0.83 |
| 3 | 144927723 | rs1867602 | 1.177 | 0.474 | 0.298 | -0.115 | -0.009 | -0.061 | -0.061 | 0.046 | 0.7 | 0.55 | 0.78 | 0.88 | 0.77 | 0.75 |
| 3 | 144928287 | rs1822906 | 0.601 | 0.285 | 0.321 | -0.095 | 0.006 | 0.023 | 0.023 | 0.076 | 0.75 | 0.87 | 0.81 | 0.88 | 0.8 | 0.87 |
| 3 | 144933520 | rs1348656 | 0.825 | 0.341 | 0.088 | -0.09 | -0.02 | -0.053 | -0.053 | 0 | 0.73 | 0.6 | 0.79 | 0.88 | 0.8 | 0.83 |
| 3 | 144949453 | rs6800639 | 0.581 | 0.925 | -0.265 | 0.061 | 0.016 | -0.049 | -0.049 | 0 | 0.77 | 0.65 | 0.8 | 0.91 | 0.75 | 0.8 |
| 3 | 144955389 | rs6778538 | -0.182 | 0.573 | -1.149 | 0.218 | 0.015 | 0.033 | 0.033 | 0 | 0.55 | 0.76 | 0.48 | 0.41 | 0.54 | 0.37 |
| 3 | 144955867 | rs9868910 | 0.619 | 1.095 | -1.553 | -0.202 | -0.012 | 0.021 | 0.021 | 3.174 | 0.23 | 0.06 | 0.19 | 0.23 | 0.21 | 0.15 |
| 3 | 144959753 | rs1822907 | -0.541 | 1.576 | -0.576 | -0.283 | -0.001 | -0.008 | -0.008 | 0 | 0.49 | 0.51 | 0.42 | 0.35 | 0.47 | 0.29 |
| 3 | 144962452 | rs4681216 | 0.848 | 1.351 | -0.696 | -0.409 | -0.089 | 0.016 | 0.016 | 0 | 0.19 | 0.02 | 0.19 | 0.1 | 0.26 | 0.14 |
| 3 | 144966507 | rs6800765 | 0.661 | 1.032 | -1 | 0.165 | -0.002 | 0.001 | 0.001 | 1.224 | 0.18 | 0.21 | 0.2 | 0.4 | 0.2 | 0.3 |
| 3 | 144968864 | rs9834421 | 0.167 | 0.976 | -0.472 | 0.087 | -0.006 | 0.072 | 0.072 | 0.089 | 0.47 | 0.3 | 0.4 | 0.33 | 0.47 | 0.29 |
| 3 | 144968890 | rs12637929 | 0.497 | 1.059 | -0.592 | 0.009 | -0.04 | 0.089 | 0.089 | 0.021 | 0.41 | 0.13 | 0.28 | 0.27 | 0.38 | 0.22 |
| 3 | 144976679 | rs9845170 | 1.592 | 0.811 | -0.659 | 0.059 | 0.011 | 0.012 | 0.012 | 2.551 | 0.19 | 0.07 | 0.09 | 0.19 | 0.12 | 0.08 |
| 3 | 144980846 | rs7611509 | 1.622 | 0.391 | -1.169 | -0.094 | -0.011 | -0.06 | -0.06 | 0 | 0.22 | 0.02 | 0.29 | 0.24 | 0.23 | 0.28 |
| 3 | 144993940 | rs1445367 | 1.613 | 0.66 | -0.877 | -0.135 | -0.092 | 0.036 | 0.036 | 0 | 0.21 | 0.02 | 0.18 | 0.1 | 0.26 | 0.14 |
| 3 | 144993976 | rs9856022 | 1.505 | 0.42 | -0.689 | -0.131 | 0.003 | 0.003 | 0.003 | 0.932 | 0.2 | 0.16 | 0.09 | 0.19 | 0.12 | 0.08 |
| 3 | 145007309 | rs7644653 | -0.71 | 0.403 | -0.668 | -0.138 | -0.008 | -0.002 | -0.002 | 0 | 0.66 | 0.65 | 0.61 | 0.72 | 0.68 | 0.6 |
| 3 | 145010745 | rs16856619 | 1.805 | 0.418 | -0.727 | -0.128 | 0.009 | 0.007 | 0.007 | 1.072 | 0.18 | 0.06 | 0.09 | 0.19 | 0.12 | 0.08 |
| 3 | 145012642 | rs7646194 | 1.28 | 0.408 | -0.687 | -0.132 | -0.019 | -0.017 | -0.017 | 0.905 | 0.21 | 0.31 | 0.11 | 0.19 | 0.12 | 0.08 |
| 3 | 145023840 | rs1584153 | 0.488 | 0.459 | -0.745 | -0.135 | -0.08 | -0.085 | -0.085 | 0.033 | 0.39 | 0.16 | 0.38 | 0.48 | 0.47 | 0.44 |
| 3 | 145024118 | rs1822905 | 1.597 | 0.432 | -0.679 | -0.139 | -0.008 | -0.008 | -0.008 | 0.054 | 0.19 | 0.21 | 0.1 | 0.19 | 0.12 | 0.08 |
| 3 | 145024569 | rs1373855 | -0.561 | 0.282 | -0.783 | -0.224 | -0.025 | -0.006 | -0.006 | 0 | 0.66 | 0.55 | 0.61 | 0.72 | 0.68 | 0.62 |
| 3 | 145025613 | rs9883003 | 0.639 | 0.567 | -0.59 | -0.146 | -0.004 | -0.011 | -0.011 | 0 | 0.26 | 0.32 | 0.23 | 0.24 | 0.21 | 0.18 |
| 3 | 145028127 | rs9878196 | 1.737 | 0.464 | -0.652 | -0.133 | 0.01 | 0.003 | 0.003 | 1.415 | 0.18 | 0.09 | 0.09 | 0.17 | 0.11 | 0.09 |
| 3 | 145030556 | rs10935547 | 0.455 | 0.197 | -0.77 | -0.263 | -0.006 | 0.004 | 0.004 | 0 | 0.29 | 0.32 | 0.38 | 0.31 | 0.32 | 0.38 |
| 3 | 145031681 | rs6806923 | 0.744 | 0.178 | -0.913 | -0.266 | -0.07 | -0.066 | -0.066 | 0 | 0.41 | 0.14 | 0.38 | 0.47 | 0.47 | 0.44 |
| 3 | 145037522 | rs4637241 | -1.099 | 0.199 | -0.887 | -0.247 | -0.012 | 0.008 | 0.008 | 0 | 0.94 | 0.97 | 0.89 | 0.95 | 0.9 | 0.9 |
| 3 | 145049826 | rs9859461 | -1.138 | 0.213 | -0.855 | -0.241 | 0.002 | -0.003 | -0.003 | 0 | 0.94 | 0.94 | 0.88 | 0.95 | 0.9 | 0.9 |
| 3 | 145066106 | rs1320914 | -0.211 | 0.146 | -0.793 | -0.249 | 0.041 | 0.001 | 0.001 | 0.424 | 0.49 | 0.67 | 0.5 | 0.48 | 0.43 | 0.46 |
| 3 | 145066128 | rs12490099 | 0.509 | 0.153 | -0.669 | -0.242 | -0.008 | 0.006 | 0.006 | 0 | 0.29 | 0.32 | 0.39 | 0.31 | 0.31 | 0.38 |
| 3 | 145067341 | rs1373859 | 0.44 | 0.165 | -0.608 | -0.192 | 0.03 | -0.001 | -0.001 | 0 | 0.51 | 0.3 | 0.43 | 0.46 | 0.54 | 0.47 |
| 3 | 145072970 | rs11927695 | 1.512 | 0.451 | -0.808 | -0.312 | -0.099 | 0.118 | 0.118 | 0.829 | 0.31 | 0.02 | 0.21 | 0.09 | 0.33 | 0.15 |
| 3 | 145079739 | rs13072906 | 0.808 | 0.431 | -0.689 | -0.274 | -0.009 | 0.045 | 0.045 | 0 | 0.21 | 0.31 | 0.34 | 0.32 | 0.24 | 0.33 |
| 3 | 145092747 | rs6810154 | 0.465 | 0.501 | -0.969 | -0.164 | 0.016 | -0.021 | -0.021 | 0 | 0.5 | 0.16 | 0.42 | 0.49 | 0.54 | 0.48 |
| 3 | 145102498 | rs345056 | -0.049 | 0.517 | -0.907 | -0.16 | -0.002 | 0.004 | 0.004 | 0.583 | 0.7 | 0.41 | 0.53 | 0.64 | 0.66 | 0.57 |
| 3 | 145106442 | rs2166819 | 0.749 | 0.47 | -0.868 | -0.126 | -0.012 | 0.068 | 0.068 | 0 | 0.22 | 0.39 | 0.34 | 0.32 | 0.24 | 0.3 |
| 3 | 145107340 | rs345062 | 1.824 | 0.479 | -0.957 | -0.043 | 0.005 | 0.003 | 0.003 | 1.204 | 0.18 | 0.07 | 0.09 | 0.15 | 0.12 | 0.09 |
| 3 | 145114811 | rs11716171 | 0.909 | 0.523 | -0.955 | -0.022 | -0.006 | -0.028 | -0.028 | 0 | 0.18 | 0.14 | 0.22 | 0.4 | 0.21 | 0.33 |
| 3 | 145147420 | rs1621465 | 1.525 | 0.527 | -0.825 | 0.022 | 0.004 | 0.004 | 0.004 | 1.323 | 0.19 | 0.17 | 0.1 | 0.15 | 0.12 | 0.09 |
| 3 | 145157591 | rs1117544 | 1.01 | 0.635 | -0.842 | 0.074 | -0.005 | -0.023 | -0.023 | 0.397 | 0.17 | 0.13 | 0.21 | 0.4 | 0.19 | 0.33 |
| 3 | 145171975 | rs10935557 | 0.621 | 0.768 | -0.882 | 0.381 | -0.02 | 0.07 | 0.07 | 2.645 | 0.48 | 0.22 | 0.33 | 0.33 | 0.49 | 0.44 |
| 3 | 145181910 | rs686865 | 0.141 | 1.27 | -0.424 | 0.543 | 0.029 | 0.078 | 0.078 | 0.239 | 0.7 | 0.49 | 0.42 | 0.46 | 0.63 | 0.52 |
| 3 | 145185102 | rs345015 | 1.361 | 1.399 | -0.751 | 0.406 | 0.012 | 0.037 | 0.037 | 0.003 | 0.24 | 0.05 | 0.12 | 0.14 | 0.15 | 0.14 |
| 3 | 145186466 | rs12233570 | 0.665 | 1.385 | -0.667 | 0.345 | -0.024 | 0.036 | 0.036 | 0.986 | 0.43 | 0.18 | 0.27 | 0.32 | 0.48 | 0.38 |
| 3 | 145187073 | rs11710718 | 0.66 | 1.376 | -0.668 | 0.341 | -0.024 | 0.036 | 0.036 | 0.991 | 0.43 | 0.18 | 0.27 | 0.32 | 0.48 | 0.38 |
| 3 | 145189041 | rs345020 | 0.033 | 1.345 | -0.288 | 0.503 | 0.03 | 0.043 | 0.043 | 0.044 | 0.89 | 0.75 | 0.77 | 0.8 | 0.84 | 0.83 |
| 3 | 145214966 | rs1026826 | 0.084 | 1.36 | -0.481 | 0.517 | -0.011 | 0.013 | 0.013 | 0.243 | 0.26 | 0.52 | 0.37 | 0.32 | 0.22 | 0.31 |
| 3 | 145215363 | rs1476335 | 1.662 | 1.485 | -0.723 | 0.658 | -0.017 | 0.109 | 0.109 | 1.697 | 0.29 | 0.08 | 0.16 | 0.11 | 0.27 | 0.16 |
| 3 | 145225110 | rs2077807 | 0.698 | 1.456 | -0.718 | 0.724 | 0.009 | 0.02 | 0.02 | 0 | 0.11 | 0.14 | 0.12 | 0.21 | 0.18 | 0.25 |
| 3 | 145232366 | rs624226 | -0.694 | 1.581 | -0.743 | 0.72 | -0.003 | -0.003 | -0.003 | 1.735 | 0.92 | 0.92 | 0.78 | 0.8 | 0.83 | 0.85 |

| | | | | | | | | | | | | | | | | |
|---|---|---|---|---|---|---|---|---|---|---|---|---|---|---|---|---|
| 3 | 145238164 | rs1481074 | 0.212 | 1.524 | -0.713 | 0.687 | -0.007 | 0.004 | 0.004 | 0 | 0.56 | 0.59 | 0.53 | 0.43 | 0.49 | 0.47 |
| 3 | 145247365 | rs12497842 | 0.221 | 1.41 | -0.562 | 0.676 | -0.006 | 0.015 | 0.015 | 0.056 | 0.26 | 0.4 | 0.37 | 0.32 | 0.22 | 0.31 |
| 3 | 145250034 | rs17289140 | -0.853 | 1.471 | -0.765 | 0.679 | -0.044 | -0.069 | -0.069 | 1.005 | 0.69 | 0.91 | 0.66 | 0.64 | 0.67 | 0.72 |
| 3 | 145254142 | rs17825849 | 1.676 | 1.46 | -0.713 | 0.688 | 0.01 | 0.059 | 0.059 | 1.217 | 0.3 | 0.19 | 0.16 | 0.11 | 0.27 | 0.16 |
| 3 | 145256049 | rs548453 | -0.735 | 1.074 | -0.734 | 0.739 | -0.002 | -0.002 | -0.002 | 0 | 0.92 | 0.92 | 0.78 | 0.8 | 0.83 | 0.86 |
| 3 | 145258296 | rs1481070 | 0.175 | 1.046 | -0.727 | 0.69 | -0.016 | 0.009 | 0.009 | 0 | 0.56 | 0.65 | 0.54 | 0.43 | 0.49 | 0.47 |
| 3 | 145275185 | rs2687748 | 0.376 | 1.128 | -0.946 | 0.663 | 0.011 | 0.031 | 0.031 | 0.295 | 0.66 | 0.5 | 0.43 | 0.5 | 0.63 | 0.56 |
| 3 | 145286224 | rs6440359 | 0.108 | 1.333 | -0.824 | 0.778 | 0 | 0.009 | 0.009 | 0 | 0.14 | 0.19 | 0.12 | 0.23 | 0.17 | 0.24 |
| 3 | 145287372 | rs2695775 | 1.486 | 1.323 | -0.867 | 0.742 | 0.013 | 0.008 | 0.008 | 0 | 0.22 | 0.13 | 0.13 | 0.17 | 0.18 | 0.14 |
| 3 | 145291677 | rs1881036 | -0.518 | 1.215 | -0.859 | 0.797 | -0.009 | -0.011 | -0.011 | 0.096 | 0.69 | 0.72 | 0.62 | 0.62 | 0.63 | 0.7 |
| 3 | 145295342 | rs1731413 | -0.399 | 1.244 | -0.919 | 0.82 | 0.02 | 0.031 | 0.031 | 0 | 0.91 | 0.88 | 0.77 | 0.79 | 0.83 | 0.84 |
| 3 | 145295363 | rs11721325 | 0.233 | 1.164 | -0.902 | 0.8 | -0.001 | 0.006 | 0.006 | 0 | 0.13 | 0.15 | 0.1 | 0.23 | 0.15 | 0.24 |
| 3 | 145300490 | rs13088893 | 0.479 | 1.163 | -0.925 | 0.776 | -0.005 | 0.007 | 0.007 | 0 | 0.56 | 0.55 | 0.51 | 0.38 | 0.48 | 0.47 |
| 3 | 145312260 | rs13099610 | 0.841 | 1.102 | -0.955 | 0.711 | 0.004 | 0.014 | 0.014 | 0.922 | 0.47 | 0.53 | 0.46 | 0.35 | 0.41 | 0.41 |
| 3 | 145313719 | rs11922887 | 0.493 | 1.149 | -0.888 | 0.785 | -0.005 | 0.007 | 0.007 | 0 | 0.56 | 0.56 | 0.51 | 0.38 | 0.48 | 0.47 |
| 3 | 145316935 | rs7633943 | -0.043 | 1.214 | -0.85 | 0.801 | -0.004 | 0.027 | 0.027 | 0 | 0.14 | 0.24 | 0.11 | 0.24 | 0.16 | 0.24 |
| 3 | 145322930 | rs13070115 | 0.457 | 1.257 | -0.825 | 0.765 | -0.021 | 0.001 | 0.001 | 0.676 | 0.25 | 0.52 | 0.36 | 0.28 | 0.19 | 0.3 |
| 3 | 145324049 | rs1608311 | 1.285 | 1.328 | -1.439 | 0.75 | -0.016 | 0.138 | 0.138 | 0.945 | 0.32 | 0.13 | 0.17 | 0.09 | 0.29 | 0.16 |
| 3 | 145328543 | rs6440364 | -0.845 | 0.867 | -1.291 | 0.865 | -0.029 | 0.043 | 0.043 | 0.167 | 0.58 | 0.75 | 0.59 | 0.69 | 0.53 | 0.68 |
| 3 | 145335395 | rs13327693 | 1.314 | 0.975 | -1.304 | 0.801 | -0.022 | -0.035 | -0.035 | 0 | 0.2 | 0.03 | 0.13 | 0.18 | 0.17 | 0.15 |
| 3 | 145341579 | rs1403495 | -0.076 | 0.818 | -0.324 | 0.921 | -0.003 | 0.005 | 0.005 | 0 | 0.23 | 0.34 | 0.35 | 0.28 | 0.21 | 0.29 |
| 3 | 145347449 | rs9863210 | 0.607 | 0.954 | -0.181 | 1.089 | 0.015 | 0.072 | 0.072 | 0 | 0.8 | 0.64 | 0.77 | 0.69 | 0.76 | 0.66 |
| 3 | 145357593 | rs9868350 | -0.018 | 0.554 | -0.478 | 0.663 | 0.011 | 0.005 | 0.005 | 0 | 0.45 | 0.62 | 0.59 | 0.58 | 0.42 | 0.4 |
| 3 | 145359343 | rs17825933 | 0.816 | 1.068 | -1.111 | 0.353 | -0.058 | 0.046 | 0.046 | 0 | 0.23 | 0.04 | 0.13 | 0.1 | 0.23 | 0.09 |
| 3 | 145362754 | rs1405858 | -0.533 | 1.041 | -0.897 | 0.408 | -0.028 | -0.011 | -0.011 | 0 | 0.73 | 0.86 | 0.75 | 0.68 | 0.66 | 0.5 |
| 3 | 145370402 | rs9834043 | -0.426 | 1.065 | -0.75 | 0.418 | -0.008 | -0.002 | -0.002 | 3.127 | 0.72 | 0.78 | 0.75 | 0.68 | 0.66 | 0.5 |
| 3 | 145372322 | rs9289702 | 0.92 | 0.779 | -0.372 | 0.584 | 0.007 | 0.003 | 0.003 | 0 | 0.37 | 0.3 | 0.39 | 0.38 | 0.26 | 0.24 |
| 3 | 145372738 | rs1523076 | 0.546 | 0.855 | -0.363 | 0.644 | 0.006 | 0.03 | 0.03 | 0 | 0.07 | 0.11 | 0.19 | 0.17 | 0.15 | 0.14 |
| 3 | 145377618 | rs10935579 | 0.607 | 0.331 | -0.148 | 0.537 | -0.047 | -0.027 | -0.027 | 0 | 0.31 | 0.15 | 0.32 | 0.28 | 0.39 | 0.25 |
| 3 | 145387917 | rs1609195 | -0.607 | 0.654 | -0.271 | 0.656 | -0.082 | -0.05 | -0.05 | 0 | 0.73 | 0.96 | 0.76 | 0.68 | 0.66 | 0.5 |
| 3 | 145409452 | rs7647052 | -0.693 | 0.73 | -0.681 | 0.682 | -0.067 | -0.032 | -0.032 | 0 | 0.75 | 0.94 | 0.77 | 0.7 | 0.67 | 0.52 |
| 3 | 145409857 | rs9869039 | -0.404 | 0.762 | -0.178 | 0.303 | 0.002 | -0.001 | -0.001 | 0.572 | 0.4 | 0.54 | 0.36 | 0.3 | 0.4 | 0.26 |
| 3 | 145420382 | rs3844271 | -0.448 | 0.809 | -0.726 | 0.278 | -0.016 | 0.013 | 0.013 | 0 | 0.42 | 0.71 | 0.4 | 0.29 | 0.43 | 0.27 |
| 3 | 145422134 | rs2461807 | -0.089 | 0.071 | -0.596 | -0.164 | -0.036 | 0.02 | 0.02 | 0 | 0.16 | 0.26 | 0.08 | 0.23 | 0.07 | 0.12 |
| 3 | 145423008 | rs2687864 | -0.648 | 0.748 | -1.228 | -0.839 | 0.013 | 0.05 | 0.05 | 0 | 0.36 | 0.23 | 0.26 | 0.22 | 0.35 | 0.2 |
| 3 | 145426265 | rs2459133 | -0.049 | 0.74 | -1.507 | -0.777 | -0.001 | 0.003 | 0.003 | 0.188 | 0.25 | 0.02 | 0.12 | 0.19 | 0.2 | 0.14 |
| 3 | 145436914 | rs2673900 | -1.14 | 0.835 | -2.319 | -1.281 | -0.089 | -0.065 | -0.065 | 0 | 0.7 | 0.97 | 0.55 | 0.38 | 0.66 | 0.57 |
| 3 | 145443276 | rs17236193 | 0.093 | 0.793 | -1.867 | -1.066 | 0.004 | 0.02 | 0.02 | 0 | 0.27 | 0.09 | 0.16 | 0.26 | 0.25 | 0.26 |
| 3 | 145449231 | rs2459139 | 0.25 | 0.525 | -0.771 | -1.274 | -0.022 | -0.037 | -0.037 | 0.557 | 0.23 | 0.4 | 0.21 | 0.15 | 0.18 | 0.19 |
| 3 | 145450531 | rs2687863 | -0.722 | 1.023 | -0.625 | -1.758 | -0.006 | 0.013 | 0.013 | 0 | 0.57 | 0.6 | 0.43 | 0.29 | 0.5 | 0.38 |
| 3 | 145458948 | rs2459126 | -1.246 | 1.329 | -0.693 | -1.884 | -0.036 | 0.014 | 0.014 | 0 | 0.66 | 0.86 | 0.52 | 0.34 | 0.64 | 0.49 |
| 3 | 145460835 | rs7643465 | 1.534 | 1.281 | -0.738 | -1.996 | 0.008 | 0.042 | 0.042 | 0.61 | 0.22 | 0.08 | 0.13 | 0.12 | 0.21 | 0.14 |
| 3 | 145464401 | rs2687868 | 0.437 | 1.12 | -0.243 | -1.961 | 0.017 | 0.027 | 0.027 | 0 | 0.27 | 0.19 | 0.21 | 0.15 | 0.2 | 0.2 |
| 3 | 145464550 | rs6764100 | -0.042 | 1.199 | -0.165 | -1.825 | 0.08 | -0.003 | -0.003 | 0 | 0.09 | 0.27 | 0.09 | 0.05 | 0.15 | 0.11 |
| 3 | 145468854 | rs2687870 | -1.436 | 1.23 | -0.549 | -2.261 | -0.045 | 0.005 | 0.005 | 0 | 0.7 | 0.96 | 0.54 | 0.31 | 0.69 | 0.54 |
| 3 | 145471762 | rs6770324 | 1.147 | 1.169 | -0.423 | -2.383 | -0.005 | 0.038 | 0.038 | 1.037 | 0.27 | 0.14 | 0.2 | 0.14 | 0.29 | 0.18 |
| 3 | 145471923 | rs2461810 | 0.532 | 1.215 | -0.367 | -2.345 | -0.015 | 0.049 | 0.049 | 0 | 0.29 | 0.18 | 0.23 | 0.16 | 0.28 | 0.34 |
| 3 | 145475756 | rs2459129 | 0.53 | 1.209 | -0.369 | -2.27 | -0.015 | 0.037 | 0.037 | 0 | 0.29 | 0.18 | 0.23 | 0.19 | 0.28 | 0.34 |
| 3 | 145477425 | rs16857193 | 0.252 | 1.178 | -0.261 | -2.386 | 0.088 | 0.017 | 0.017 | 0.778 | 0.05 | 0.29 | 0.06 | 0 | 0.07 | 0.01 |
| 3 | 145484110 | rs3863994 | 0.468 | 1.206 | -0.529 | -2.435 | -0.009 | 0.04 | 0.04 | 0 | 0.29 | 0.21 | 0.23 | 0.16 | 0.29 | 0.34 |
| 3 | 145484296 | rs12108087 | 0.105 | 1.336 | -0.458 | -2.24 | 0.112 | 0.035 | 0.035 | 1.138 | 0.05 | 0.34 | 0.06 | 0 | 0.07 | 0.01 |
| 3 | 145487982 | rs6807108 | -1.235 | 1.346 | -1.401 | -2.223 | -0.044 | 0.025 | 0.025 | 0 | 0.69 | 0.92 | 0.53 | 0.3 | 0.67 | 0.54 |
| 3 | 145492901 | rs6806150 | 0.245 | 1.054 | -0.704 | -1.733 | -0.001 | -0.022 | -0.022 | 0.896 | 0.28 | 0.26 | 0.21 | 0.14 | 0.29 | 0.18 |
| 3 | 145499439 | rs9846710 | 0.463 | 1.066 | -1.1 | -1.495 | -0.021 | 0.122 | 0.122 | 0 | 0.44 | 0.06 | 0.49 | 0.74 | 0.4 | 0.54 |
| 3 | 145504317 | rs7652768 | -1.024 | 1.34 | -0.356 | -1.603 | 0.053 | -0.062 | -0.062 | 0.085 | 0.13 | 0.36 | 0.13 | 0.03 | 0.18 | 0.07 |
| 3 | 145504887 | rs9839546 | -0.331 | 1.314 | -0.437 | -1.649 | 0.002 | 0.041 | 0.041 | 0 | 0.57 | 0.37 | 0.62 | 0.76 | 0.58 | 0.61 |
| 3 | 145505755 | rs9878603 | 0.675 | 1.347 | -0.445 | -1.636 | -0.005 | 0.115 | 0.115 | 0 | 0.43 | 0.08 | 0.49 | 0.74 | 0.4 | 0.48 |
| 3 | 145506173 | rs7614629 | -0.207 | 1.233 | -0.326 | -1.496 | 0.007 | 0.06 | 0.06 | 0 | 0.54 | 0.35 | 0.62 | 0.77 | 0.58 | 0.55 |
| 3 | 145506580 | rs9866001 | 0.206 | 1.265 | -0.331 | -1.499 | 0.002 | 0.087 | 0.087 | 0 | 0.5 | 0.35 | 0.57 | 0.74 | 0.5 | 0.5 |
| 3 | 145508776 | rs6440373 | -0.074 | 1.115 | -0.554 | -1.36 | -0.002 | 0.019 | 0.019 | 0.009 | 0.54 | 0.6 | 0.61 | 0.74 | 0.54 | 0.53 |
| 3 | 145509794 | rs4681260 | -0.248 | 0.793 | -0.419 | -1.273 | 0 | -0.004 | -0.004 | 0 | 0.59 | 0.6 | 0.66 | 0.77 | 0.61 | 0.58 |
| 3 | 145517239 | rs9875815 | -0.008 | 0.739 | -0.72 | -0.768 | -0.009 | 0.023 | 0.023 | 0.106 | 0.44 | 0.32 | 0.52 | 0.6 | 0.44 | 0.3 |
| 3 | 145518144 | rs11709200 | -0.341 | 0.187 | -0.298 | -1.234 | -0.025 | -0.056 | -0.056 | 1.057 | 0.3 | 0.48 | 0.19 | 0.22 | 0.23 | 0.33 |
| 3 | 145519571 | rs9831692 | 0.574 | 0.152 | -0.682 | -1.478 | -0.037 | 0.076 | 0.076 | 0 | 0.42 | 0.17 | 0.55 | 0.73 | 0.48 | 0.51 |
| 3 | 145524923 | rs9814478 | -0.153 | -0.092 | -0.188 | -1.54 | -0.003 | -0.046 | -0.046 | 0 | 0.61 | 0.41 | 0.66 | 0.89 | 0.64 | 0.74 |
| 3 | 145525989 | rs7643638 | -0.017 | 0.367 | -1.014 | -1.128 | -0.009 | 0.049 | 0.049 | 0.06 | 0.44 | 0.61 | 0.61 | 0.81 | 0.53 | 0.54 |
| 3 | 145530567 | rs4681269 | 0.045 | -0.027 | -1.446 | -1.16 | 0.047 | 0.115 | 0.115 | 0 | 0.37 | 0.12 | 0.18 | 0.16 | 0.29 | 0.4 |
| 3 | 145530708 | rs4681270 | 0.364 | 0.025 | -1.556 | -1.249 | 0.03 | 0.088 | 0.088 | 0 | 0.32 | 0.11 | 0.17 | 0.14 | 0.28 | 0.34 |
| 3 | 145542558 | rs7630694 | -0.638 | -0.074 | -0.518 | -1.438 | 0.005 | -0.031 | -0.031 | 0.017 | 0.1 | 0.22 | 0.05 | 0.02 | 0.07 | 0.18 |

| | | | | | | | | | | | | | | | | |
|---|---|---|---|---|---|---|---|---|---|---|---|---|---|---|---|---|
| 3 | 145547356 | rs1398775 | 0.654 | 0.101 | -0.199 | -1.099 | -0.003 | 0.094 | 0.094 | 0 | 0.47 | 0.26 | 0.57 | 0.7 | 0.55 | 0.44 |
| 3 | 145563316 | rs13071352 | 0.839 | 0.229 | -0.3 | -1.116 | -0.081 | -0.073 | -0.073 | 0.051 | 0.23 | 0.08 | 0.35 | 0.63 | 0.38 | 0.33 |
| 3 | 145572381 | rs10513264 | -1.495 | -0.161 | -0.321 | -1.139 | 0.003 | 0.004 | 0.004 | 0 | 0.7 | 0.74 | 0.82 | 0.81 | 0.71 | 0.67 |
| 3 | 145581255 | rs1398777 | 0.748 | 0.807 | -0.03 | -0.855 | 0 | 0.02 | 0.02 | 0 | 0.18 | 0.21 | 0.24 | 0.55 | 0.25 | 0.3 |
| 3 | 145585459 | rs7652584 | -0.37 | 0.476 | 0.005 | -1.126 | -0.001 | 0.026 | 0.026 | 0 | 0.45 | 0.63 | 0.52 | 0.19 | 0.45 | 0.35 |
| 3 | 145598921 | rs5020216 | -0.164 | 0.535 | 0.091 | -1.174 | 0 | -0.02 | -0.02 | 0 | 0.39 | 0.43 | 0.36 | 0.11 | 0.38 | 0.32 |
| 3 | 145600492 | rs13061647 | 0.911 | 0.341 | -0.096 | -1.261 | -0.002 | -0.018 | -0.018 | 0.23 | 0.21 | 0.16 | 0.29 | 0.62 | 0.29 | 0.33 |
| 3 | 145603219 | rs16857489 | 3.095 | 0.299 | -0.092 | -1.265 | -0.016 | -0.017 | -0.017 | 0 | 0.05 | 0.06 | 0.02 | 0 | 0.01 | 0 |
| 3 | 145607628 | rs9849753 | 1.495 | 0.351 | -0.031 | -1.254 | 0.034 | 0.016 | 0.016 | 0 | 0.06 | 0.14 | 0.17 | 0.09 | 0.07 | 0.03 |
| 3 | 145608233 | rs7641056 | -1.961 | 0.322 | -0.16 | -1.273 | 0.01 | 0.021 | 0.021 | 0 | 0.72 | 0.8 | 0.83 | 0.81 | 0.75 | 0.69 |
| 3 | 145609432 | rs1355789 | 0.434 | 0.388 | -0.299 | -1.286 | -0.003 | -0.027 | -0.027 | 0 | 0.25 | 0.16 | 0.31 | 0.62 | 0.32 | 0.44 |
| 3 | 145612756 | rs7639600 | -2.027 | 0.37 | -0.272 | -1.286 | 0.008 | 0.018 | 0.018 | 0 | 0.72 | 0.8 | 0.84 | 0.81 | 0.75 | 0.69 |
| 3 | 145614120 | rs9810981 | 1.935 | 0.422 | -0.311 | -1.268 | -0.029 | -0.019 | -0.019 | 0 | 0.05 | 0 | 0.16 | 0.09 | 0.07 | 0.04 |
| 3 | 145616506 | rs7652032 | 0.916 | 0.392 | -0.207 | -1.338 | -0.015 | 0.022 | 0.022 | 0 | 0.28 | 0.27 | 0.3 | 0.09 | 0.31 | 0.16 |
| 3 | 145616862 | rs4318523 | 0.359 | 0.017 | -0.348 | -1.302 | -0.001 | -0.012 | -0.012 | 0 | 0.25 | 0.22 | 0.32 | 0.62 | 0.32 | 0.44 |
| 3 | 145618127 | rs13316664 | 1.944 | 0.358 | -0.31 | -1.4 | -0.015 | 0.043 | 0.043 | 0 | 0.19 | 0.09 | 0.2 | 0.08 | 0.19 | 0.11 |
| 3 | 145623801 | rs6440401 | -0.248 | 0.344 | -0.253 | -1.328 | -0.003 | 0.001 | 0.001 | 0 | 0.42 | 0.62 | 0.5 | 0.18 | 0.42 | 0.24 |
| 3 | 145624626 | rs10935590 | 1.085 | 0.356 | -0.39 | -0.436 | -0.002 | 0.005 | 0.005 | 0 | 0.17 | 0.16 | 0.19 | 0.33 | 0.16 | 0.21 |
| 3 | 145629083 | rs9289707 | -0.046 | 0.389 | -0.322 | -0.434 | 0.011 | -0.046 | -0.046 | 0 | 0.41 | 0.51 | 0.49 | 0.19 | 0.42 | 0.24 |
| 3 | 145632742 | rs12629663 | 2.158 | 0.421 | -0.646 | -0.364 | -0.024 | -0.067 | -0.067 | 0.162 | 0.1 | 0.01 | 0.17 | 0.32 | 0.16 | 0.21 |
| 3 | 145635676 | rs1813256 | -1.387 | 0.402 | -0.628 | -0.394 | 0.007 | -0.045 | -0.045 | 0.007 | 0.63 | 0.8 | 0.71 | 0.51 | 0.62 | 0.58 |
| 3 | 145643853 | rs6762748 | -1.399 | 0.414 | -0.622 | -0.4 | 0.006 | -0.046 | -0.046 | 0 | 0.63 | 0.8 | 0.71 | 0.51 | 0.62 | 0.58 |
| 3 | 145653627 | rs1398776 | 0.999 | 0.343 | -0.6 | -0.295 | 0.001 | 0 | 0 | 0 | 0.16 | 0.15 | 0.19 | 0.32 | 0.16 | 0.2 |
| 3 | 145654103 | rs6766386 | 0.115 | 0.262 | -0.683 | -0.741 | -0.001 | -0.024 | -0.024 | 0 | 0.34 | 0.38 | 0.25 | 0.09 | 0.27 | 0.29 |
| 3 | 145655364 | rs578641 | 0.914 | 0.608 | -0.897 | -0.567 | 0.008 | 0.107 | 0.107 | 0 | 0.3 | 0.15 | 0.21 | 0.09 | 0.23 | 0.17 |
| 3 | 145656927 | rs6440405 | 0.883 | 0.593 | -0.582 | -0.312 | -0.014 | 0.173 | 0.173 | 0 | 0.15 | 0.47 | 0.22 | 0.31 | 0.16 | 0.19 |
| 3 | 145660362 | rs1553977 | 0.977 | 1.247 | -1.579 | -0.115 | 0.044 | -0.01 | -0.01 | 0 | 0.13 | 0.19 | 0.22 | 0.1 | 0.18 | 0.07 |
| 3 | 145662227 | rs13062906 | 1.055 | 0.728 | -2.231 | 0.129 | 0.001 | -0.067 | -0.067 | 0.486 | 0.17 | 0.06 | 0.2 | 0.32 | 0.19 | 0.22 |
| 3 | 145662242 | rs13314579 | 0.478 | 0.913 | -1.701 | -0.122 | 0.002 | 0.066 | 0.066 | 0 | 0.24 | 0.14 | 0.19 | 0.08 | 0.2 | 0.13 |
| 3 | 145662790 | rs16857644 | 0.277 | 0.762 | -1.358 | 0.34 | -0.033 | -0.06 | -0.06 | 2.093 | 0.09 | 0 | 0.12 | 0.33 | 0.13 | 0.09 |
| 3 | 145664091 | rs17401096 | -0.578 | 0.484 | -1.021 | -0.292 | -0.002 | -0.02 | -0.02 | 0 | 0.06 | 0.13 | 0.04 | 0.01 | 0.04 | 0.13 |
| 3 | 145664135 | rs12494193 | 0.671 | 0.386 | -1.466 | -0.368 | -0.025 | 0.044 | 0.044 | 0 | 0.2 | 0.01 | 0.17 | 0.09 | 0.17 | 0.13 |
| 3 | 145664959 | rs16857652 | -0.766 | 0.364 | -0.729 | -0.349 | -0.005 | -0.008 | -0.008 | 0 | 0.3 | 0.62 | 0.22 | 0.21 | 0.24 | 0.22 |
| 3 | 145665253 | rs13092361 | -0.72 | 0.39 | 0.172 | -0.174 | 0.006 | 0.03 | 0.03 | 0 | 0.16 | 0.27 | 0.15 | 0.23 | 0.18 | 0.13 |
| 3 | 145671953 | rs6791851 | 0.103 | 0.818 | -0.477 | -0.422 | 0.001 | 0.025 | 0.025 | 0 | 0.6 | 0.64 | 0.66 | 0.81 | 0.62 | 0.57 |
| 3 | 145675927 | rs10935594 | 1.292 | 0.814 | -1.176 | -0.281 | -0.033 | -0.016 | -0.016 | 0 | 0.4 | 0.07 | 0.46 | 0.55 | 0.39 | 0.43 |
| 3 | 145678873 | rs1513420 | -0.194 | 0.816 | -1.151 | -0.395 | -0.05 | 0.131 | 0.131 | 0 | 0.65 | 0.92 | 0.7 | 0.81 | 0.65 | 0.67 |
| 3 | 145687305 | rs9289708 | -0.873 | 0.743 | 0.295 | -0.24 | 0.003 | 0.043 | 0.043 | 0.129 | 0.2 | 0.51 | 0.19 | 0.25 | 0.18 | 0.21 |
| 3 | 145687497 | rs502645 | 1.405 | 0.656 | -0.036 | -0.359 | 0.002 | -0.007 | -0.007 | 0 | 0.4 | 0.15 | 0.45 | 0.53 | 0.39 | 0.43 |
| 3 | 145697193 | rs1355792 | -0.12 | 0.714 | -0.122 | -0.365 | -0.069 | 0.13 | 0.13 | 0 | 0.63 | 0.88 | 0.67 | 0.79 | 0.6 | 0.66 |
| 3 | 145699202 | rs7611414 | -0.27 | 0.559 | 0.705 | -0.381 | 0.006 | 0.024 | 0.024 | 0 | 0.17 | 0.24 | 0.17 | 0.25 | 0.18 | 0.22 |
| 3 | 145701146 | rs5029231 | -2.234 | 0.54 | 0.832 | -0.377 | 0.02 | -0.009 | -0.009 | 0 | 0.07 | 0.6 | 0.05 | 0 | 0.02 | 0.03 |
| 3 | 145709314 | rs6763296 | -2.119 | 0.55 | 1.065 | -0.364 | 0.058 | 0.03 | 0.03 | 3.235 | 0.07 | 0.52 | 0.05 | 0 | 0.02 | 0.01 |
| 3 | 145709551 | rs4681280 | 0.467 | 0.58 | 0.817 | -0.589 | -0.001 | 0.024 | 0.024 | 0 | 0.64 | 0.8 | 0.72 | 0.73 | 0.65 | 0.67 |
| 3 | 145731553 | rs9844407 | 0.491 | 0.592 | 0.785 | -0.658 | -0.014 | 0.077 | 0.077 | 0 | 0.58 | 0.77 | 0.67 | 0.79 | 0.59 | 0.64 |
| 3 | 145732933 | rs16857741 | 1.818 | 0.678 | 0.661 | -0.29 | -0.009 | -0.063 | -0.063 | 0.514 | 0.39 | 0.1 | 0.49 | 0.5 | 0.39 | 0.44 |
| 3 | 145735049 | rs6808907 | 0.432 | 0.712 | 0.773 | -0.401 | -0.015 | 0.102 | 0.102 | 0 | 0.59 | 0.79 | 0.69 | 0.79 | 0.6 | 0.64 |
| 3 | 145737357 | rs1449452 | 0.276 | 0.707 | 1.059 | -0.309 | -0.008 | -0.027 | -0.027 | 0 | 0.64 | 0.64 | 0.79 | 0.77 | 0.69 | 0.69 |
| 3 | 145744022 | rs6797906 | 1.387 | 0.782 | 1.131 | -0.251 | 0.002 | 0.009 | 0.009 | 0 | 0.43 | 0.37 | 0.53 | 0.56 | 0.43 | 0.46 |
| 3 | 145746162 | rs988501 | 0.482 | 0.462 | 1.396 | -0.317 | 0.001 | 0.004 | 0.004 | 0 | 0.66 | 0.71 | 0.79 | 0.77 | 0.69 | 0.67 |
| 3 | 145758307 | rs993521 | 0.646 | 0.498 | 1.091 | -0.286 | -0.013 | -0.039 | -0.039 | 0.36 | 0.63 | 0.53 | 0.78 | 0.77 | 0.69 | 0.68 |
| 3 | 145763446 | rs4681290 | -0.241 | 0.975 | 0.868 | 0.125 | -0.02 | 0.001 | 0.001 | 0.579 | 0.27 | 0.18 | 0.4 | 0.26 | 0.38 | 0.44 |
| 3 | 145767684 | rs9867807 | 0.859 | 1.015 | 0.835 | 0.46 | 0.002 | -0.011 | -0.011 | 0.05 | 0.57 | 0.34 | 0.41 | 0.5 | 0.48 | 0.43 |
| 3 | 145776673 | rs7651139 | 0.664 | 1.067 | 1.066 | 0.562 | 0.008 | 0.01 | 0.01 | 0 | 0.59 | 0.53 | 0.48 | 0.52 | 0.5 | 0.44 |
| 3 | 145787657 | rs3792347 | 1.104 | 1.183 | 0.807 | 0.276 | -0.01 | -0.022 | -0.022 | 0 | 0.11 | 0.07 | 0.14 | 0.21 | 0.1 | 0.11 |
| 3 | 145790803 | rs1449444 | 0.862 | 1.056 | 0.732 | 0.253 | -0.003 | -0.005 | -0.005 | 0 | 0.58 | 0.49 | 0.46 | 0.57 | 0.47 | 0.4 |
| 3 | 145818406 | rs9820945 | -0.906 | 1.213 | 0.503 | 0.237 | 0 | -0.001 | -0.001 | 0 | 0.82 | 0.82 | 0.73 | 0.84 | 0.77 | 0.73 |
| 3 | 145824025 | rs9289713 | 1.06 | 1.182 | 0.518 | 0.22 | 0.02 | 0.023 | 0.023 | 0 | 0.55 | 0.25 | 0.42 | 0.57 | 0.46 | 0.4 |
| 3 | 145836188 | rs6787634 | -0.37 | 1.207 | 0.078 | 0.263 | 0.008 | 0.002 | 0.002 | 0 | 0.22 | 0.22 | 0.25 | 0.26 | 0.27 | 0.28 |
| 3 | 145838895 | rs4681297 | -0.884 | 1.226 | 0.088 | 0.266 | 0 | -0.001 | -0.001 | 0 | 0.82 | 0.82 | 0.73 | 0.84 | 0.77 | 0.73 |
| 3 | 145844373 | rs4681298 | 0.921 | 1.233 | 0.042 | 0.267 | -0.03 | 0.024 | 0.024 | 0 | 0.09 | 0.01 | 0.08 | 0.03 | 0.13 | 0.18 |
| 3 | 145849908 | rs12632001 | -0.483 | 1.218 | 0.066 | 0.262 | -0.005 | -0.001 | -0.001 | 0 | 0.23 | 0.18 | 0.26 | 0.26 | 0.28 | 0.29 |
| 3 | 145861305 | rs1707469 | 0.186 | 1.034 | -0.015 | 0.222 | -0.007 | 0.006 | 0.006 | 0.316 | 0.72 | 0.77 | 0.65 | 0.8 | 0.64 | 0.55 |
| 3 | 145868148 | rs10935606 | -0.242 | 1.048 | -0.243 | 0.235 | -0.016 | 0.014 | 0.014 | 0.243 | 0.27 | 0.19 | 0.35 | 0.2 | 0.36 | 0.44 |
| 3 | 145885467 | rs1913186 | -0.103 | 0.823 | -0.23 | 0.289 | 0.037 | -0.03 | -0.03 | 0 | 0.29 | 0.4 | 0.37 | 0.18 | 0.4 | 0.47 |
| 3 | 145886725 | rs11923433 | -1.511 | 0.412 | -0.691 | 0.227 | -0.036 | -0.165 | -0.165 | 0.149 | 0.62 | 0.97 | 0.64 | 0.53 | 0.62 | 0.71 |
| 3 | 145888457 | rs1913185 | 0.731 | 0.54 | -0.445 | 0.353 | -0.005 | -0.006 | -0.006 | 2.21 | 0.15 | 0.12 | 0.18 | 0.26 | 0.15 | 0.13 |
| 3 | 145888493 | rs4342076 | -0.341 | 0.574 | -0.653 | 0.369 | -0.004 | 0.003 | 0.003 | 0.393 | 0.34 | 0.61 | 0.4 | 0.25 | 0.41 | 0.5 |
| 3 | 145890816 | rs4681303 | -2.067 | 0.633 | -1.365 | 0.525 | -0.016 | -0.03 | -0.03 | 0 | 0.73 | 0.99 | 0.85 | 0.79 | 0.78 | 0.8 |

| | | | | | | | | | | | | | | | | |
|---|---|---|---|---|---|---|---|---|---|---|---|---|---|---|---|---|
| 3 | 145892711 | rs1355685 | -1.494 | 1.608 | -0.399 | 0.393 | 0.076 | -0.002 | -0.002 | 0 | 0.51 | 0.85 | 0.58 | 0.48 | 0.56 | 0.61 |
| 3 | 145894415 | rs6793582 | -0.16 | 2.043 | 0.017 | 0.354 | 0.053 | -0.001 | -0.001 | 0 | 0.21 | 0.38 | 0.32 | 0.19 | 0.29 | 0.32 |
| 3 | 145903426 | rs11928424 | -0.516 | 1.46 | -0.486 | 0.448 | 0.002 | -0.009 | -0.009 | 0 | 0.1 | 0.11 | 0.08 | 0.03 | 0.12 | 0.17 |
| 3 | 145905301 | rs16857999 | -0.537 | 1.354 | -0.146 | 0.789 | 0.011 | 0.047 | 0.047 | 0 | 0.16 | 0.29 | 0.18 | 0.26 | 0.14 | 0.12 |
| 3 | 145905391 | rs7628805 | 0.223 | 1.996 | -0.666 | 0.745 | -0.041 | -0.016 | -0.016 | 0 | 0.13 | 0.04 | 0.24 | 0.14 | 0.22 | 0.22 |
| 3 | 145910430 | rs1059836 | -0.354 | 1.823 | -0.141 | 0.149 | 0.007 | -0.042 | -0.042 | 0.669 | 0.17 | 0.21 | 0.14 | 0.06 | 0.19 | 0.11 |
| 3 | 145911164 | rs3749281 | -0.3 | 1.826 | -0.135 | 0.089 | 0.012 | -0.02 | -0.02 | 0 | 0.08 | 0.1 | 0.09 | 0.03 | 0.13 | 0.04 |
| 3 | 145911733 | rs9985294 | 0.98 | 1.822 | -0.014 | -0.139 | 0 | -0.001 | -0.001 | 6.88 | 0.64 | 0.4 | 0.61 | 0.71 | 0.62 | 0.62 |
| 3 | 145912177 | rs9815240 | -0.312 | 1.833 | -0.279 | -0.052 | 0.011 | -0.039 | -0.039 | 0 | 0.13 | 0.26 | 0.09 | 0.06 | 0.14 | 0.09 |
| 3 | 145912299 | rs1512902 | 0.304 | 1.611 | -0.785 | -0.231 | -0.048 | 0.032 | 0.032 | 0 | 0.18 | 0.06 | 0.15 | 0.06 | 0.27 | 0.21 |
| 3 | 145912548 | rs1512903 | 0.677 | 0.88 | -0.558 | -0.338 | 0 | -0.003 | -0.003 | 10.42 | 0.6 | 0.64 | 0.58 | 0.69 | 0.58 | 0.64 |
| 3 | 145915013 | rs1828652 | 0.302 | 0.235 | -0.608 | -0.135 | 0.007 | 0.011 | 0.011 | 0 | 0.45 | 0.55 | 0.3 | 0.32 | 0.4 | 0.32 |
| 3 | 145917761 | rs1061409 | 0.153 | -0.104 | -0.934 | -0.073 | -0.066 | -0.013 | -0.013 | 0 | 0.25 | 0.1 | 0.38 | 0.19 | 0.34 | 0.27 |
| 3 | 145921145 | rs3920230 | 0.988 | -1.222 | -1.136 | -0.245 | -0.095 | -0.137 | -0.137 | 0 | 0.33 | 0.05 | 0.29 | 0.46 | 0.36 | 0.46 |
| 3 | 145921954 | rs9839874 | -0.58 | -1.348 | -0.431 | -0.263 | -0.008 | -0.001 | -0.001 | 2.544 | 0.7 | 0.69 | 0.75 | 0.66 | 0.76 | 0.83 |
| 3 | 145925572 | rs17349248 | 0.984 | -1.515 | -0.588 | -0.331 | 0.014 | 0.032 | 0.032 | 0.144 | 0.2 | 0.12 | 0.1 | 0.09 | 0.14 | 0.08 |
| 3 | 145935896 | rs3804654 | -0.433 | -1.13 | -0.546 | -0.215 | -0.027 | -0.008 | -0.008 | 0 | 0.67 | 0.61 | 0.81 | 0.66 | 0.76 | 0.83 |
| 3 | 145938619 | rs3762685 | 0.603 | -0.573 | -0.343 | -0.09 | 0.018 | -0.002 | -0.002 | 0.589 | 0.26 | 0.35 | 0.42 | 0.19 | 0.34 | 0.27 |
| 3 | 145948249 | rs9836419 | 0.729 | -1.453 | -0.456 | -0.291 | 0.103 | 0.086 | 0.086 | 0 | 0.55 | 0.22 | 0.75 | 0.65 | 0.7 | 0.71 |
| 3 | 145960310 | rs10737947 | -0.089 | -1.894 | 0.242 | -0.23 | -0.089 | 0.014 | 0.014 | 0 | 0.74 | 0.53 | 0.85 | 0.66 | 0.86 | 0.84 |
| 3 | 145967173 | rs9863647 | -0.087 | -1.863 | 0.259 | -0.245 | -0.08 | 0.011 | 0.011 | 0 | 0.74 | 0.54 | 0.85 | 0.67 | 0.85 | 0.84 |
| 3 | 145968098 | rs4475004 | 0.69 | -1.983 | 0.365 | -0.317 | 0.011 | 0.006 | 0.006 | 0 | 0.65 | 0.35 | 0.82 | 0.65 | 0.8 | 0.75 |
| 3 | 145975420 | rs13072399 | -0.355 | -1.874 | 0.296 | -0.359 | -0.021 | -0.056 | -0.056 | 0.72 | 0.12 | 0.2 | 0.05 | 0.01 | 0.06 | 0.11 |
| 3 | 145979875 | rs1828560 | -0.101 | -2.023 | 0.51 | -0.305 | -0.033 | 0.008 | 0.008 | 0 | 0.79 | 0.67 | 0.84 | 0.75 | 0.86 | 0.87 |
| 3 | 145982150 | rs4681310 | 0.746 | -1.825 | 0.233 | -0.224 | 0.038 | 0.022 | 0.022 | 0 | 0.62 | 0.29 | 0.73 | 0.67 | 0.78 | 0.72 |
| 3 | 145988231 | rs7645100 | -0.189 | -1.737 | 0.944 | -0.21 | -0.053 | -0.003 | -0.003 | 0 | 0.75 | 0.63 | 0.85 | 0.7 | 0.84 | 0.84 |
| 3 | 145996473 | rs13092543 | -0.475 | -1.678 | 0.912 | -0.278 | -0.022 | -0.076 | -0.076 | 1.434 | 0.12 | 0.34 | 0.05 | 0.01 | 0.07 | 0.12 |
| 3 | 146014404 | rs12497460 | 0.926 | -1.097 | -0.035 | -0.118 | -0.12 | -0.087 | -0.087 | 0 | 0.22 | 0.02 | 0.41 | 0.21 | 0.32 | 0.22 |
| 3 | 146026957 | rs13067886 | 0.637 | -1.326 | 0.266 | -0.002 | 0.003 | 0.008 | 0.008 | 0 | 0.07 | 0.04 | 0.09 | 0.02 | 0.07 | 0.08 |
| 3 | 146031241 | rs1849260 | 0.576 | -1.617 | 0.516 | 0.105 | -0.033 | -0.006 | -0.006 | 0 | 0.65 | 0.51 | 0.76 | 0.66 | 0.74 | 0.74 |
| 3 | 146032938 | rs1160640 | -0.533 | -1.643 | 0.581 | 0.094 | -0.022 | 0.008 | 0.008 | 0.125 | 0.75 | 0.73 | 0.87 | 0.68 | 0.83 | 0.85 |
| 3 | 146035144 | rs2203129 | 0.133 | -1.641 | 0.594 | 0.095 | -0.027 | 0.001 | 0.001 | 0 | 0.69 | 0.64 | 0.77 | 0.66 | 0.75 | 0.77 |
| 3 | 146043900 | rs13075285 | 0.687 | -1.607 | 0.488 | -0.088 | -0.003 | -0.007 | -0.007 | 0 | 0.07 | 0.09 | 0.1 | 0.02 | 0.07 | 0.08 |
| 3 | 146047142 | rs1512068 | 0.399 | -1.613 | 0.537 | 0.243 | -0.014 | 0.001 | 0.001 | 0.012 | 0.65 | 0.44 | 0.75 | 0.62 | 0.73 | 0.72 |
| 3 | 146049601 | rs1912892 | 0.485 | -1.627 | 0.802 | 0.237 | -0.03 | 0.003 | 0.003 | 0 | 0.65 | 0.48 | 0.75 | 0.63 | 0.73 | 0.72 |
| 3 | 146055242 | rs9881632 | 0.458 | -1.645 | 0.321 | 0.23 | -0.001 | -0.001 | -0.001 | 0 | 0.63 | 0.4 | 0.74 | 0.63 | 0.73 | 0.72 |
| 3 | 146062738 | rs7642687 | -0.215 | -1.635 | 0.426 | 0.228 | -0.053 | -0.013 | -0.013 | 0.191 | 0.7 | 0.6 | 0.85 | 0.68 | 0.8 | 0.81 |
| 3 | 146067740 | rs9842818 | 0.332 | -1.665 | 0.41 | 0.161 | 0.002 | 0.002 | 0.002 | 0 | 0.66 | 0.39 | 0.74 | 0.66 | 0.76 | 0.77 |
| 3 | 146091891 | rs907491 | -0.283 | -0.919 | 0.471 | -0.047 | 0 | 0 | 0 | 0 | 0.7 | 0.76 | 0.77 | 0.68 | 0.71 | 0.81 |
| 3 | 146093034 | rs17417377 | 0.543 | -0.975 | -0.181 | -0.061 | -0.054 | 0.005 | 0.005 | 0 | 0.4 | 0.22 | 0.34 | 0.39 | 0.45 | 0.46 |
| 3 | 146106142 | rs2867865 | -0.118 | -0.536 | -0.081 | -0.199 | -0.001 | -0.007 | -0.007 | 0 | 0.58 | 0.51 | 0.62 | 0.65 | 0.58 | 0.76 |
| 3 | 146109133 | rs6770000 | -0.295 | -0.358 | -0.392 | -0.272 | 0.001 | -0.011 | -0.011 | 0.521 | 0.58 | 0.53 | 0.61 | 0.63 | 0.56 | 0.76 |
| 3 | 146109354 | rs13064298 | -0.13 | -0.386 | -0.589 | -0.284 | -0.002 | 0.028 | 0.028 | 0 | 0.56 | 0.37 | 0.59 | 0.63 | 0.56 | 0.74 |
| 3 | 146115594 | rs11715366 | 0.675 | -0.369 | -0.977 | -0.285 | -0.06 | -0.039 | -0.039 | 0.279 | 0.28 | 0.1 | 0.22 | 0.31 | 0.31 | 0.47 |
| 3 | 146117041 | rs6803636 | -2.061 | -0.567 | -1.32 | -0.363 | -0.041 | -0.065 | -0.065 | 1.113 | 0.87 | 0.99 | 0.88 | 0.73 | 0.82 | 0.92 |
| 3 | 146117770 | rs7650361 | -0.772 | -0.569 | -0.665 | -0.039 | -0.001 | 0.007 | 0.007 | 0 | 0.56 | 0.45 | 0.59 | 0.6 | 0.56 | 0.74 |
| 3 | 146118568 | rs4476473 | -0.721 | -0.569 | -0.672 | -0.039 | -0.002 | 0.041 | 0.041 | 0 | 0.55 | 0.36 | 0.59 | 0.6 | 0.56 | 0.72 |
| 3 | 146119295 | rs13092702 | -0.721 | -0.569 | -0.672 | -0.039 | -0.002 | 0.041 | 0.041 | 0 | 0.55 | 0.36 | 0.59 | 0.6 | 0.56 | 0.72 |
| 3 | 146151175 | rs1054736 | 0.333 | -0.469 | -0.734 | -0.014 | -0.044 | -0.024 | -0.024 | 0 | 0.28 | 0.12 | 0.22 | 0.32 | 0.31 | 0.47 |
| 3 | 146152479 | rs13068178 | -1.233 | -0.527 | -0.692 | -0.043 | 0 | -0.016 | -0.016 | 0 | 0.6 | 0.55 | 0.69 | 0.63 | 0.6 | 0.79 |
| 3 | 146159847 | rs4444684 | -0.8 | -0.524 | -0.698 | -0.046 | -0.001 | 0.066 | 0.066 | 0 | 0.55 | 0.28 | 0.59 | 0.6 | 0.57 | 0.73 |
| 3 | 146162424 | rs1589674 | 0.349 | -0.558 | -0.684 | -0.034 | -0.044 | -0.024 | -0.024 | 0 | 0.28 | 0.12 | 0.22 | 0.32 | 0.31 | 0.47 |
| 3 | 146179536 | rs6440440 | -1.253 | -0.244 | -0.22 | 0.317 | 0.036 | 0.024 | 0.024 | 0 | 0.05 | 0.24 | 0.1 | 0.02 | 0.04 | 0.06 |
| 3 | 146183649 | rs12633694 | -0.534 | -0.189 | -0.405 | 0.233 | -0.02 | 0.079 | 0.079 | 0 | 0.47 | 0.28 | 0.56 | 0.6 | 0.49 | 0.64 |
| 3 | 146191013 | rs506791 | -1.137 | 0.277 | -0.273 | 0.496 | 0.002 | 0.021 | 0.021 | 0 | 0.5 | 0.56 | 0.67 | 0.63 | 0.51 | 0.66 |
| 3 | 146214126 | rs575834 | -1.058 | 0.747 | 0.59 | 0.535 | 0.012 | 0.01 | 0.01 | 0 | 0.28 | 0.38 | 0.31 | 0.33 | 0.28 | 0.44 |
| 3 | 146216915 | rs491900 | -0.41 | 0.703 | 0.295 | 0.558 | -0.013 | -0.012 | -0.012 | 0.07 | 0.23 | 0.17 | 0.21 | 0.32 | 0.24 | 0.39 |
| 3 | 146219740 | rs4681327 | -1.375 | 1.02 | 0.461 | 0.495 | 0.02 | 0.069 | 0.069 | 0 | 0.28 | 0.5 | 0.37 | 0.38 | 0.27 | 0.4 |
| 3 | 146221640 | rs7612740 | -1.587 | 0.497 | 0.424 | -0.509 | 0 | -0.007 | -0.007 | 0 | 0.06 | 0.27 | 0.03 | 0 | 0.01 | 0.01 |
| 3 | 146231988 | rs1870424 | -0.438 | 1.207 | -0.352 | -0.536 | -0.021 | 0.024 | 0.024 | 0 | 0.16 | 0.16 | 0.1 | 0.08 | 0.19 | 0.18 |
| 3 | 146232469 | rs454645 | -0.667 | 1.176 | -0.144 | -0.649 | -0.002 | -0.055 | -0.055 | 0 | 0.1 | 0.19 | 0.07 | 0 | 0.07 | 0.03 |
| 3 | 146235394 | rs343319 | -1.444 | 0.89 | 0.01 | -0.855 | 0 | -0.028 | -0.028 | 0 | 0.41 | 0.59 | 0.32 | 0.15 | 0.35 | 0.26 |
| 3 | 146236161 | rs343317 | -0.896 | 1.086 | -0.125 | -0.361 | 0 | -0.058 | -0.058 | 0 | 0.1 | 0.2 | 0.07 | 0 | 0.07 | 0.03 |
| 3 | 146242961 | rs2587028 | -0.83 | 1.615 | -0.536 | -0.08 | -0.026 | -0.006 | -0.006 | 0.493 | 0.1 | 0.04 | 0.14 | 0.15 | 0.14 | 0.15 |
| 3 | 146243369 | rs2688693 | -1.54 | 1.351 | -0.578 | 0 | -0.015 | -0.011 | -0.011 | 0.153 | 0.17 | 0.07 | 0.2 | 0.2 | 0.17 | 0.16 |
| 3 | 146245506 | rs2688692 | -1.954 | 1.888 | -0.074 | -0.517 | 0.023 | -0.033 | -0.033 | 0 | 0.34 | 0.45 | 0.37 | 0.23 | 0.39 | 0.33 |
| 3 | 146246892 | rs2738908 | -1.235 | 1.681 | -0.111 | -0.239 | -0.01 | -0.008 | -0.008 | 0 | 0.16 | 0.09 | 0.21 | 0.2 | 0.17 | 0.16 |
| 3 | 146258699 | rs2587011 | -1.801 | 2.208 | 0.814 | -0.494 | 0.05 | -0.03 | -0.03 | 0 | 0.2 | 0.35 | 0.22 | 0.17 | 0.26 | 0.22 |
| 3 | 146261279 | rs1997602 | -0.025 | 2.168 | 0.936 | -0.501 | -0.032 | 0.017 | 0.017 | 0.957 | 0.07 | 0.02 | 0.12 | 0.05 | 0.1 | 0.11 |

| | | | | | | | | | | | | | | | | |
|---|---|---|---|---|---|---|---|---|---|---|---|---|---|---|---|---|
| 3 | 146262344 | rs1061307 | -0.868 | 2.085 | 1.042 | -0.467 | -0.012 | 0.043 | 0.043 | 0 | 0.1 | 0.34 | 0.06 | 0.13 | 0.08 | 0.11 |
| 3 | 146262781 | rs1079369 | -2.236 | 1.954 | 1.215 | -0.538 | 0.039 | -0.045 | -0.045 | 0 | 0.22 | 0.47 | 0.23 | 0.17 | 0.25 | 0.23 |
| 3 | 146265311 | rs7625713 | -1.151 | 1.858 | 1.106 | -0.488 | 0.003 | -0.004 | -0.004 | 0 | 0.12 | 0.16 | 0.09 | 0.13 | 0.15 | 0.12 |
| 3 | 146270331 | rs16858488 | -0.535 | 1.649 | 1.072 | -0.517 | -0.024 | -0.049 | -0.049 | 0 | 0.09 | 0.28 | 0.05 | 0 | 0.05 | 0.08 |
| 3 | 146271303 | rs343340 | 0.017 | 1.758 | 0.101 | -0.123 | 0.011 | 0.041 | 0.041 | 0 | 0.24 | 0.35 | 0.27 | 0.44 | 0.29 | 0.27 |
| 3 | 146276607 | rs977759 | -1.077 | 1.048 | 0.048 | -0.486 | -0.001 | -0.017 | -0.017 | 0 | 0.37 | 0.56 | 0.37 | 0.21 | 0.37 | 0.36 |
| 3 | 146288645 | rs342920 | -0.83 | 1.319 | 0.26 | -0.598 | 0 | 0.001 | 0.001 | 0 | 0.56 | 0.55 | 0.57 | 0.59 | 0.65 | 0.55 |
| 3 | 146288792 | rs342921 | -0.151 | 1.519 | 0.167 | -0.923 | 0.007 | 0.008 | 0.008 | 0 | 0.33 | 0.38 | 0.42 | 0.59 | 0.42 | 0.41 |
| 3 | 146288803 | rs9819872 | 0.383 | 1.548 | 0.129 | -0.942 | 0.002 | -0.004 | -0.004 | 0.033 | 0.2 | 0.14 | 0.1 | 0 | 0.18 | 0.12 |
| 3 | 146289450 | rs17431978 | 0.797 | 1.538 | 0.076 | -0.94 | -0.02 | 0.11 | 0.11 | 0 | 0.17 | 0.01 | 0.06 | 0 | 0.14 | 0.1 |
| 3 | 146294737 | rs17432334 | -0.63 | 1.363 | 0.202 | -0.946 | 0.004 | 0.015 | 0.015 | 0.078 | 0.3 | 0.29 | 0.17 | 0.16 | 0.27 | 0.24 |
| 3 | 146296485 | rs343292 | 0.03 | 1.375 | 0.224 | -0.96 | 0.001 | 0.002 | 0.002 | 0 | 0.33 | 0.34 | 0.42 | 0.58 | 0.42 | 0.41 |
| 3 | 146297818 | rs17354140 | 0.396 | 1.36 | 0.225 | -0.959 | -0.002 | 0.073 | 0.073 | 0 | 0.2 | 0.12 | 0.12 | 0.01 | 0.19 | 0.13 |
| 3 | 146297951 | rs17354168 | -0.618 | 1.367 | 0.191 | -0.956 | 0.004 | 0.015 | 0.015 | 0.086 | 0.3 | 0.29 | 0.17 | 0.16 | 0.27 | 0.24 |
| 3 | 146302296 | rs342903 | -2.23 | 1.329 | 0.233 | -0.979 | 0.006 | 0.019 | 0.019 | 0 | 0.66 | 0.76 | 0.63 | 0.75 | 0.73 | 0.67 |
| 3 | 146302786 | rs10513292 | 0.458 | 1.284 | 0.243 | -0.941 | -0.002 | -0.015 | -0.015 | 0.045 | 0.08 | 0.05 | 0.05 | 0.16 | 0.08 | 0.12 |
| 3 | 146303165 | rs10513294 | -0.555 | 1.281 | 0.253 | -0.93 | 0.003 | 0.013 | 0.013 | 0.103 | 0.29 | 0.29 | 0.17 | 0.16 | 0.27 | 0.24 |
| 3 | 146310721 | rs343281 | -0.058 | 1.312 | 0.197 | -0.916 | 0.004 | 0.005 | 0.005 | 0 | 0.34 | 0.35 | 0.42 | 0.58 | 0.42 | 0.41 |
| 3 | 146312941 | rs6787813 | -0.636 | 1.251 | 0.269 | -0.911 | -0.005 | -0.019 | -0.019 | 0 | 0.3 | 0.38 | 0.19 | 0.16 | 0.27 | 0.24 |
| 3 | 146315380 | rs6807700 | 0.346 | 1.237 | 0.252 | -0.939 | 0.002 | 0.03 | 0.03 | 0 | 0.2 | 0.14 | 0.12 | 0.01 | 0.19 | 0.13 |
| 3 | 146319946 | rs423266 | -1.485 | 1.293 | 0.254 | -0.961 | -0.003 | -0.03 | -0.03 | 0 | 0.58 | 0.53 | 0.58 | 0.73 | 0.65 | 0.58 |
| 3 | 146323885 | rs12491246 | 0.412 | 1.303 | 0.38 | -0.985 | 0.002 | 0.003 | 0.003 | 0 | 0.2 | 0.16 | 0.13 | 0.01 | 0.19 | 0.13 |
| 3 | 146325770 | rs9822212 | 0.004 | 1.345 | 0.495 | -1.026 | 0.038 | -0.033 | -0.033 | 0.152 | 0.08 | 0.24 | 0.06 | 0.01 | 0.08 | 0.08 |
| 3 | 146327284 | rs342887 | -2.221 | 1.25 | 0.505 | -1.068 | 0.009 | 0.026 | 0.026 | 0 | 0.65 | 0.76 | 0.64 | 0.75 | 0.73 | 0.66 |
| 3 | 146328944 | rs342885 | 0.44 | 1.55 | 0.675 | -1.149 | -0.013 | -0.027 | -0.027 | 0 | 0.26 | 0.19 | 0.35 | 0.58 | 0.34 | 0.29 |
| 3 | 146330953 | rs342897 | -0.608 | 1.504 | 0.688 | -1.164 | 0.004 | -0.012 | -0.012 | 0.151 | 0.37 | 0.42 | 0.45 | 0.73 | 0.46 | 0.46 |
| 3 | 146332304 | rs342896 | -0.868 | 1.541 | 0.516 | -1.153 | 0.002 | -0.001 | -0.001 | 0 | 0.4 | 0.43 | 0.45 | 0.6 | 0.48 | 0.49 |
| 3 | 146334631 | rs342892 | 0.563 | 1.634 | 0.308 | 0.193 | -0.027 | -0.066 | -0.066 | 0 | 0.1 | 0.01 | 0.13 | 0.25 | 0.1 | 0.18 |
| 3 | 146338040 | rs163731 | -1.682 | 1.669 | 0.467 | 0.102 | 0.002 | 0.007 | 0.007 | 0 | 0.49 | 0.58 | 0.51 | 0.66 | 0.54 | 0.48 |
| 3 | 146338648 | rs342913 | -2.505 | 1.614 | 0.404 | 0.084 | -0.004 | 0.043 | 0.043 | 0.189 | 0.58 | 0.91 | 0.6 | 0.68 | 0.62 | 0.6 |
| 3 | 146341576 | rs343278 | -1.783 | 1.933 | 0.352 | 0.232 | 0.022 | 0.002 | 0.002 | 0 | 0.5 | 0.63 | 0.54 | 0.52 | 0.58 | 0.52 |
| 3 | 146343322 | rs342907 | -0.755 | 1.908 | 0.16 | 0.324 | -0.025 | -0.037 | -0.037 | 0.004 | 0.35 | 0.25 | 0.38 | 0.49 | 0.39 | 0.32 |
| 3 | 146350346 | rs342938 | 0.3 | 2.034 | 0.334 | 0.448 | 0.017 | 0.121 | 0.121 | 0 | 0.11 | 0.19 | 0.24 | 0.48 | 0.16 | 0.17 |
| 3 | 146354136 | rs3852001 | 0.141 | 1.974 | 0.048 | 0.074 | -0.058 | 0.144 | 0.144 | 1.323 | 0.23 | 0.03 | 0.1 | 0.01 | 0.22 | 0.17 |
| 3 | 146368187 | rs9842222 | -1.323 | 1.73 | 0.619 | 0.339 | 0.035 | 0.026 | 0.026 | 0 | 0.19 | 0.4 | 0.18 | 0.17 | 0.23 | 0.24 |
| 3 | 146370996 | rs11916605 | -1.362 | 1.63 | 0.733 | 0.299 | 0.035 | -0.007 | -0.007 | 0.029 | 0.24 | 0.42 | 0.19 | 0.19 | 0.3 | 0.32 |
| 3 | 146371033 | rs4681329 | 1.655 | 1.505 | 0.742 | 0.457 | 0.032 | 0.036 | 0.036 | 0 | 0.46 | 0.29 | 0.5 | 0.33 | 0.38 | 0.42 |
| 3 | 146371578 | rs11924592 | -1.278 | 1.48 | 0.741 | 0.445 | 0.032 | -0.009 | -0.009 | 0 | 0.25 | 0.42 | 0.19 | 0.19 | 0.3 | 0.32 |
| 3 | 146375713 | rs11915439 | 0.024 | 1.445 | 0.046 | 0.53 | -0.003 | -0.008 | -0.008 | 0.021 | 0.71 | 0.71 | 0.69 | 0.51 | 0.67 | 0.73 |
| 3 | 146376541 | rs1858326 | -0.079 | 1.481 | -0.414 | 0.496 | -0.025 | -0.041 | -0.041 | 0.059 | 0.14 | 0.02 | 0.07 | 0.17 | 0.15 | 0.16 |
| 3 | 146381657 | rs7644114 | 1.03 | 1.435 | -0.094 | 0.495 | 0.006 | -0.032 | -0.032 | 0.038 | 0.07 | 0.11 | 0.07 | 0.01 | 0.08 | 0.13 |
| 3 | 146385081 | rs1511559 | 0.117 | 1.404 | -0.458 | 0.639 | -0.023 | -0.057 | -0.057 | 0 | 0.72 | 0.85 | 0.7 | 0.5 | 0.68 | 0.73 |
| 3 | 146390895 | rs7623328 | 0.117 | 1.404 | -0.449 | 0.639 | -0.022 | -0.055 | -0.055 | 0 | 0.72 | 0.85 | 0.7 | 0.5 | 0.67 | 0.73 |
| 3 | 146391374 | rs2868452 | -0.569 | 1.494 | -0.479 | 0.627 | 0.002 | 0.056 | 0.056 | 0 | 0.89 | 0.85 | 0.79 | 0.5 | 0.85 | 0.82 |
| 3 | 146394094 | rs10935638 | 1.574 | 0.89 | -1.008 | 0.243 | -0.011 | -0.02 | -0.02 | 0.02 | 0.36 | 0.07 | 0.34 | 0.3 | 0.31 | 0.33 |
| 3 | 146396006 | rs11718101 | -0.549 | 0.942 | -0.629 | 0.39 | -0.001 | 0.008 | 0.008 | 0 | 0.79 | 0.75 | 0.62 | 0.49 | 0.72 | 0.74 |
| 3 | 146397951 | rs16858754 | -0.287 | 0.617 | -0.287 | 0.38 | -0.002 | -0.013 | -0.013 | 0 | 0.15 | 0.12 | 0.09 | 0.19 | 0.15 | 0.15 |
| 3 | 146401145 | rs17370295 | 0.061 | 0.732 | -0.083 | 0.468 | -0.038 | -0.058 | -0.058 | 0 | 0.12 | 0.01 | 0.08 | 0.19 | 0.11 | 0.11 |
| 3 | 146401686 | rs986173 | -0.736 | 0.774 | 0.533 | 0.529 | 0.065 | 0.171 | 0.171 | 0 | 0.15 | 0.46 | 0.15 | 0.24 | 0.19 | 0.29 |
| 3 | 146402108 | rs986171 | 0.48 | 0.599 | 0.263 | 0.475 | 0.005 | 0.007 | 0.007 | 0 | 0.5 | 0.53 | 0.51 | 0.48 | 0.55 | 0.52 |
| 3 | 146402171 | rs1912785 | 0.505 | 0.599 | 0.255 | 0.475 | 0.009 | 0.013 | 0.013 | 0 | 0.49 | 0.57 | 0.52 | 0.48 | 0.55 | 0.52 |
| 3 | 146402194 | rs986170 | 0.839 | 0.582 | 0.356 | 0.522 | -0.023 | 0.06 | 0.06 | 0 | 0.46 | 0.22 | 0.45 | 0.27 | 0.51 | 0.48 |
| 3 | 146405051 | rs1877259 | -0.168 | 0.609 | 0.155 | 0.472 | 0.007 | -0.036 | -0.036 | 0.053 | 0.68 | 0.77 | 0.61 | 0.48 | 0.69 | 0.61 |
| 3 | 146414212 | rs9872768 | -0.036 | 0.657 | 0.265 | 0.411 | -0.003 | 0.003 | 0.003 | 0 | 0.66 | 0.58 | 0.65 | 0.56 | 0.69 | 0.61 |
| 3 | 146419523 | rs9871995 | 0.253 | 0.698 | 0.463 | -0.108 | 0.001 | -0.013 | -0.013 | 0 | 0.62 | 0.53 | 0.52 | 0.25 | 0.6 | 0.44 |
| 3 | 146424531 | rs12330695 | 1.421 | 0.706 | 0.374 | -0.177 | -0.003 | 0.008 | 0.008 | 0 | 0.16 | 0.1 | 0.08 | 0 | 0.14 | 0.09 |
| 3 | 146429560 | rs9859125 | 1.248 | 0.681 | 0.362 | -0.161 | -0.001 | 0.023 | 0.023 | 0 | 0.17 | 0.1 | 0.08 | 0 | 0.14 | 0.08 |
| 3 | 146429761 | rs2174783 | 1.893 | 0.552 | 0.38 | -0.367 | -0.02 | 0.037 | 0.037 | 0 | 0.31 | 0.14 | 0.35 | 0.22 | 0.35 | 0.22 |
| 3 | 146432172 | rs9836760 | -0.143 | 0.804 | 0.807 | -0.167 | -0.001 | -0.001 | -0.001 | 0 | 0.64 | 0.59 | 0.55 | 0.25 | 0.65 | 0.56 |
| 3 | 146432343 | rs9857093 | 1.416 | 0.764 | 0.654 | -0.151 | -0.002 | 0.014 | 0.014 | 0 | 0.16 | 0.1 | 0.08 | 0 | 0.14 | 0.08 |
| 3 | 146437116 | rs1355476 | 1.298 | 0.808 | 0.333 | -0.135 | 0 | 0 | 0 | 0 | 0.08 | 0.08 | 0.06 | 0.03 | 0.08 | 0.12 |
| 3 | 146437590 | rs12695826 | 1.027 | 0.8 | 0.369 | -0.089 | -0.013 | 0.064 | 0.064 | 0 | 0.43 | 0.22 | 0.42 | 0.25 | 0.45 | 0.35 |
| 3 | 146444219 | rs7613056 | 0.949 | 0.861 | 0.153 | -0.019 | -0.001 | -0.052 | -0.052 | 0 | 0.17 | 0.19 | 0.09 | 0 | 0.14 | 0.08 |
| 3 | 146468430 | rs12497166 | 0.455 | 0.976 | 0.134 | 0.01 | 0.004 | 0.024 | 0.024 | 0 | 0.54 | 0.36 | 0.46 | 0.23 | 0.55 | 0.43 |
| 3 | 146477052 | rs6440456 | -0.091 | 0.918 | 0.59 | 0.374 | 0.003 | -0.01 | -0.01 | 1.419 | 0.64 | 0.67 | 0.64 | 0.53 | 0.66 | 0.56 |
| 3 | 146479144 | rs9870986 | -0.375 | 0.9 | 0.543 | 0.451 | 0.007 | -0.014 | -0.014 | 0 | 0.68 | 0.75 | 0.68 | 0.57 | 0.7 | 0.6 |
| 3 | 146484999 | rs9836412 | 0.57 | 0.898 | 0.257 | 0.206 | 0.005 | 0.029 | 0.029 | 0 | 0.53 | 0.35 | 0.46 | 0.23 | 0.55 | 0.42 |
| 3 | 146487468 | rs7629060 | 1.749 | 0.988 | 0.077 | 0.037 | -0.008 | 0.034 | 0.034 | 0 | 0.33 | 0.18 | 0.34 | 0.22 | 0.36 | 0.21 |
| 3 | 146488309 | rs9289731 | 1.409 | 0.988 | 0.064 | 0.036 | -0.007 | 0.019 | 0.019 | 0 | 0.36 | 0.26 | 0.37 | 0.23 | 0.41 | 0.33 |

| | | | | | | | | | | | | | | | | |
|---|---|---|---|---|---|---|---|---|---|---|---|---|---|---|---|---|
| 3 | 146495703 | rs1019897 | 0.481 | 0.758 | 0.079 | 0.071 | 0.01 | 0.078 | 0.078 | 0 | 0.53 | 0.31 | 0.45 | 0.23 | 0.55 | 0.42 |
| 3 | 146496302 | rs16858887 | 1.005 | 0.764 | -0.239 | 0.144 | -0.008 | 0.005 | 0.005 | 0 | 0.09 | 0 | 0.03 | 0.07 | 0.05 | 0.07 |
| 3 | 146504254 | rs9834616 | 0.184 | 0.785 | 0.148 | 0.18 | 0.007 | 0.067 | 0.067 | 0 | 0.54 | 0.33 | 0.46 | 0.23 | 0.55 | 0.42 |
| 3 | 146507727 | rs12152233 | 0.761 | 0.88 | 0.191 | 0.36 | 0 | 0.035 | 0.035 | 0 | 0.09 | 0.09 | 0.14 | 0.3 | 0.09 | 0.14 |
| 3 | 146524417 | rs6781142 | 0.84 | 1.114 | 0.417 | 0.556 | -0.011 | 0.019 | 0.019 | 0 | 0.36 | 0.27 | 0.39 | 0.27 | 0.42 | 0.32 |
| 3 | 146539373 | rs10049183 | 1.163 | 0.947 | 0.38 | 0.554 | -0.007 | 0.02 | 0.02 | 0.071 | 0.32 | 0.22 | 0.36 | 0.23 | 0.37 | 0.21 |
| 3 | 146541654 | rs1431969 | -1.218 | 1.033 | -0.23 | 0.537 | -0.025 | 0.107 | 0.107 | 0.117 | 0.83 | 0.99 | 0.92 | 1 | 0.86 | 0.9 |
| 3 | 146556091 | rs1608002 | -0.117 | 1.083 | -0.027 | 0.554 | -0.002 | 0.006 | 0.006 | 0 | 0.48 | 0.5 | 0.54 | 0.77 | 0.49 | 0.69 |
| 3 | 146562424 | rs2013690 | 0.07 | 0.827 | -0.11 | 0.587 | -0.001 | -0.014 | -0.014 | 0 | 0.46 | 0.52 | 0.52 | 0.77 | 0.45 | 0.58 |
| 3 | 146568369 | rs4408842 | -0.902 | 0.63 | -0.578 | 0.22 | 0.008 | 0.015 | 0.015 | 0.151 | 0.82 | 0.89 | 0.91 | 1 | 0.85 | 0.9 |
| 3 | 146570867 | rs1978683 | -0.168 | 0.76 | -0.556 | 0.542 | 0 | -0.004 | -0.004 | 0 | 0.43 | 0.39 | 0.43 | 0.44 | 0.42 | 0.52 |
| 3 | 146579888 | rs17438440 | 0.395 | 0.793 | -1.137 | 0.218 | -0.011 | 0.09 | 0.09 | 0.353 | 0.15 | 0.01 | 0.06 | 0 | 0.12 | 0.1 |
| 3 | 146584464 | rs9877252 | -1.098 | 0.91 | -1.419 | -0.197 | -0.001 | 0.038 | 0.038 | 0.283 | 0.74 | 0.83 | 0.72 | 0.89 | 0.74 | 0.9 |
| 3 | 146594746 | rs4318522 | 0.514 | 1.422 | -1.566 | 0.728 | -0.029 | -0.04 | -0.04 | 0 | 0.11 | 0.04 | 0.07 | 0.18 | 0.13 | 0.17 |
| 3 | 146598479 | rs13091393 | 0.408 | 1.255 | -0.854 | 0.771 | -0.002 | 0.002 | 0.002 | 0.745 | 0.4 | 0.39 | 0.49 | 0.51 | 0.41 | 0.5 |
| 3 | 146601295 | rs1464446 | -0.76 | 1.308 | -0.714 | 1.495 | -0.028 | -0.039 | -0.039 | 0 | 0.14 | 0.08 | 0.11 | 0.38 | 0.18 | 0.24 |
| 3 | 146617655 | rs11924113 | -0.911 | 1.056 | 0.193 | 1.242 | 0.075 | 0.128 | 0.128 | 0 | 0.08 | 0.28 | 0.09 | 0.21 | 0.1 | 0.17 |
| 3 | 146622509 | rs7632942 | -0.377 | 0.853 | -0.193 | 1.295 | 0.002 | -0.003 | -0.003 | 0.169 | 0.16 | 0.14 | 0.27 | 0.25 | 0.13 | 0.14 |
| 3 | 146622901 | rs6440468 | -1.928 | 0.633 | -0.081 | 1.096 | -0.001 | -0.047 | -0.047 | 0 | 0.37 | 0.78 | 0.45 | 0.5 | 0.32 | 0.37 |
| 3 | 146628382 | rs1357130 | -0.105 | 0.514 | 0.25 | 1.087 | -0.007 | -0.023 | -0.023 | 1.886 | 0.12 | 0.21 | 0.07 | 0.04 | 0.1 | 0.05 |
| 3 | 146633580 | rs11708123 | -1.127 | 0.525 | 0.174 | 1.103 | -0.01 | 0 | 0 | 0 | 0.29 | 0.36 | 0.35 | 0.29 | 0.22 | 0.19 |
| 3 | 146634035 | rs11928953 | -1.129 | 0.499 | -0.323 | 1.105 | 0.007 | -0.041 | -0.041 | 0.045 | 0.3 | 0.77 | 0.42 | 0.49 | 0.24 | 0.34 |
| 3 | 146647304 | rs4643673 | -0.778 | 0.423 | -1.386 | 0.686 | -0.078 | -0.126 | -0.126 | 0 | 0.74 | 0.99 | 0.67 | 0.64 | 0.7 | 0.71 |
| 3 | 146666094 | rs11721056 | -0.963 | 0.253 | -0.71 | 0.36 | 0.002 | -0.03 | -0.03 | 0.721 | 0.59 | 0.75 | 0.53 | 0.49 | 0.57 | 0.62 |
| 3 | 146683672 | rs7624556 | -0.133 | -0.256 | -0.316 | 0.135 | -0.036 | 0.083 | 0.083 | 0 | 0.15 | 0.23 | 0.24 | 0.42 | 0.07 | 0.29 |
| 3 | 146687502 | rs13098281 | -0.27 | -0.204 | -0.32 | 0.082 | 0 | -0.084 | -0.084 | 0 | 0.43 | 0.54 | 0.3 | 0.07 | 0.5 | 0.31 |
| 3 | 146690215 | rs7611207 | -0.029 | -0.176 | -0.207 | -0.028 | -0.002 | -0.032 | -0.032 | 3.125 | 0.38 | 0.34 | 0.27 | 0.05 | 0.47 | 0.23 |
| 3 | 146695770 | rs13072250 | 0.566 | 0.302 | -0.297 | -0.164 | 0.022 | -0.041 | -0.041 | 0.104 | 0.1 | 0.2 | 0.09 | 0.01 | 0.14 | 0.13 |
| 3 | 146700377 | rs4131188 | 0.714 | 0.182 | -0.559 | 0.127 | 0.03 | -0.029 | -0.029 | 0 | 0.1 | 0.04 | 0.21 | 0.18 | 0.04 | 0.1 |
| 3 | 146709009 | rs1354197 | 0.158 | 0.557 | -0.372 | 0.163 | 0.026 | -0.059 | -0.059 | 0 | 0.14 | 0.32 | 0.11 | 0.03 | 0.17 | 0.22 |
| 3 | 146713828 | rs6774215 | -0.012 | 0.558 | -1.218 | 0.199 | 0.028 | 0.067 | 0.067 | 0.357 | 0.2 | 0.55 | 0.15 | 0.27 | 0.2 | 0.43 |
| 3 | 146719217 | rs1995392 | 0.407 | 0.572 | -1.324 | 0.215 | 0.001 | -0.005 | -0.005 | 0 | 0.23 | 0.11 | 0.31 | 0.22 | 0.2 | 0.14 |
| 3 | 146722198 | rs1502122 | -0.414 | 0.431 | -1.192 | 0.231 | 0.006 | -0.01 | -0.01 | 0.065 | 0.58 | 0.69 | 0.53 | 0.52 | 0.57 | 0.62 |
| 3 | 146746263 | rs7615695 | -1.405 | 0.744 | -0.807 | 0.108 | 0 | 0 | 0 | 0 | 0.35 | 0.69 | 0.23 | 0.34 | 0.28 | 0.33 |
| 3 | 146758405 | rs4274714 | -1.45 | 0.955 | -1.751 | -0.249 | -0.002 | -0.064 | -0.064 | 1.58 | 0.57 | 0.82 | 0.5 | 0.52 | 0.6 | 0.49 |
| 3 | 146763391 | rs9873630 | 0.703 | 0.972 | -1.33 | -0.228 | -0.014 | 0.022 | 0.022 | 0 | 0.23 | 0.11 | 0.28 | 0.17 | 0.32 | 0.16 |
| 3 | 146809521 | rs9289736 | -1.624 | 1.136 | -0.755 | -0.712 | 0.008 | -0.014 | -0.014 | 0 | 0.69 | 0.72 | 0.55 | 0.56 | 0.73 | 0.58 |
| 3 | 146810327 | rs9881485 | -0.038 | 1.298 | -0.285 | -0.716 | 0.003 | 0.004 | 0.004 | 0.433 | 0.41 | 0.42 | 0.38 | 0.5 | 0.5 | 0.4 |
| 3 | 146823099 | rs2318315 | -0.59 | 1.196 | 0.064 | -0.758 | 0.013 | 0.004 | 0.004 | 0.445 | 0.51 | 0.44 | 0.45 | 0.56 | 0.61 | 0.5 |
| 3 | 146827987 | rs7635583 | -0.183 | 1.202 | 0.073 | -0.742 | 0 | 0.002 | 0.002 | 0.006 | 0.43 | 0.43 | 0.39 | 0.5 | 0.53 | 0.4 |
| 3 | 146830477 | rs10513303 | 2.291 | 1.101 | -0.232 | -0.717 | -0.049 | 0.033 | 0.033 | 0 | 0.11 | 0.01 | 0.05 | 0.03 | 0.14 | 0.09 |
| 3 | 146835182 | rs9289738 | 1.599 | 1.254 | -0.256 | -0.814 | -0.001 | -0.001 | -0.001 | 0 | 0.08 | 0.08 | 0.2 | 0.17 | 0.12 | 0.09 |
| 3 | 146837929 | rs9814423 | -0.595 | 1.136 | 0.02 | -0.704 | 0 | 0 | 0 | 0 | 0.5 | 0.47 | 0.44 | 0.53 | 0.56 | 0.41 |
| 3 | 146848626 | rs10513306 | 1.47 | 1.098 | -0.187 | -0.815 | -0.048 | 0.085 | 0.085 | 0 | 0.15 | 0.02 | 0.09 | 0 | 0.15 | 0.04 |
| 3 | 146867800 | rs9865951 | -1.186 | 1.009 | 0.048 | -0.78 | -0.008 | 0.019 | 0.019 | 0 | 0.65 | 0.56 | 0.53 | 0.53 | 0.7 | 0.46 |
| 3 | 146871560 | rs4309688 | -1.144 | 0.965 | -0.095 | -1.166 | -0.02 | 0.011 | 0.011 | 0 | 0.58 | 0.72 | 0.4 | 0.39 | 0.57 | 0.42 |
| 3 | 146875745 | rs17564438 | 0.718 | 0.855 | 0.019 | -1.33 | 0.005 | 0.045 | 0.045 | 0 | 0.26 | 0.16 | 0.22 | 0.04 | 0.3 | 0.24 |
| 3 | 146880458 | rs1502111 | -0.628 | 0.843 | 0.458 | -1.325 | 0 | -0.029 | -0.029 | 0 | 0.5 | 0.44 | 0.37 | 0.1 | 0.54 | 0.34 |
| 3 | 146881054 | rs11719030 | 0.684 | 0.9 | 0.538 | -1.286 | 0.006 | 0.027 | 0.027 | 0 | 0.21 | 0.3 | 0.1 | 0.3 | 0.2 | 0.24 |
| 3 | 146884607 | rs7625438 | 1.382 | 0.879 | 0.436 | -1.314 | -0.001 | -0.052 | -0.052 | 0 | 0.17 | 0.22 | 0.08 | 0.02 | 0.16 | 0.11 |
| 3 | 146905640 | rs6773881 | -1.897 | 0.917 | -0.434 | -1.283 | 0.006 | -0.023 | -0.023 | 0.687 | 0.71 | 0.78 | 0.48 | 0.39 | 0.74 | 0.58 |
| 3 | 146912522 | rs9815177 | -0.971 | 0.964 | -0.559 | -1.413 | 0 | 0.041 | 0.041 | 0 | 0.55 | 0.75 | 0.4 | 0.35 | 0.59 | 0.47 |
| 3 | 146914749 | rs11717071 | 2.361 | 0.966 | -0.558 | -1.425 | -0.016 | 0.03 | 0.03 | 0 | 0.12 | 0.04 | 0.06 | 0 | 0.14 | 0.09 |
| 3 | 146916848 | rs17565826 | 1.521 | 0.89 | -0.584 | -1.407 | -0.021 | 0.013 | 0.013 | 0 | 0.08 | 0.01 | 0.05 | 0.04 | 0.08 | 0.06 |
| 3 | 146920104 | rs6780114 | -0.259 | 1.007 | -0.497 | -1.341 | 0.002 | -0.013 | -0.013 | 0 | 0.4 | 0.59 | 0.31 | 0.1 | 0.4 | 0.28 |
| 3 | 146922987 | rs2202054 | -0.642 | 1 | -0.748 | -1.084 | -0.016 | 0.015 | 0.015 | 0 | 0.57 | 0.76 | 0.39 | 0.38 | 0.59 | 0.5 |
| 3 | 146943112 | rs9839538 | 0.849 | 0.859 | -1.016 | -1.206 | -0.001 | -0.02 | -0.02 | 2.321 | 0.2 | 0.17 | 0.19 | 0.05 | 0.24 | 0.16 |
| 3 | 146947340 | rs9833001 | -0.791 | 0.942 | -1.409 | -0.788 | 0.003 | -0.018 | -0.018 | 0 | 0.64 | 0.81 | 0.45 | 0.39 | 0.67 | 0.52 |
| 3 | 146964828 | rs12496794 | 0.837 | 1.1 | -0.882 | -0.386 | -0.024 | 0.03 | 0.03 | 0 | 0.07 | 0.01 | 0.06 | 0.01 | 0.08 | 0.03 |
| 3 | 146969974 | rs9878692 | -0.426 | 0.779 | -0.875 | -0.252 | 0 | 0.014 | 0.014 | 0 | 0.45 | 0.63 | 0.4 | 0.61 | 0.46 | 0.52 |
| 3 | 146973383 | rs9857512 | -0.527 | 0.782 | -0.97 | -0.963 | -0.001 | -0.036 | -0.036 | 0.051 | 0.37 | 0.61 | 0.34 | 0.07 | 0.47 | 0.19 |
| 3 | 146983089 | rs17566714 | 0.377 | 0.899 | -0.844 | -0.768 | -0.01 | 0.005 | 0.005 | 0 | 0.22 | 0.17 | 0.21 | 0.04 | 0.31 | 0.13 |
| 3 | 146997009 | rs7645827 | -1.672 | 0.679 | -1.21 | -0.344 | -0.012 | -0.038 | -0.038 | 1.616 | 0.53 | 0.92 | 0.47 | 0.56 | 0.59 | 0.55 |
| 3 | 147000266 | rs1814982 | -1.265 | 0.588 | -0.833 | -0.192 | -0.035 | 0.064 | 0.064 | 0 | 0.42 | 0.76 | 0.37 | 0.25 | 0.51 | 0.3 |
| 3 | 147035587 | rs6440477 | -1.687 | 0.225 | -1.347 | -0.172 | -0.018 | -0.192 | -0.192 | 0 | 0.74 | 0.99 | 0.54 | 0.56 | 0.78 | 0.77 |
| 3 | 147038181 | rs9811539 | 0.818 | 0.336 | -0.748 | -0.268 | 0.026 | -0.033 | -0.033 | 0.117 | 0.08 | 0.27 | 0.08 | 0.02 | 0.11 | 0.05 |
| 3 | 147039856 | rs16846545 | 0.607 | 0.381 | -0.682 | -0.214 | 0.018 | 0.124 | 0.124 | 0 | 0.06 | 0.16 | 0.03 | 0.18 | 0.05 | 0.1 |
| 3 | 147048365 | rs9289746 | -0.058 | 0.36 | -0.752 | -0.5 | 0.028 | -0.106 | -0.106 | 0 | 0.3 | 0.44 | 0.3 | 0.06 | 0.41 | 0.17 |
| 3 | 147050225 | rs17567518 | 1.113 | 0.55 | -0.74 | -0.362 | 0.002 | 0.019 | 0.019 | 0 | 0.32 | 0.21 | 0.15 | 0.3 | 0.28 | 0.49 |

| | | | | | | | | | | | | | | | | |
|---|---|---|---|---|---|---|---|---|---|---|---|---|---|---|---|---|
| 3 | 147058367 | rs1354054 | -0.933 | 0.338 | -0.678 | -0.381 | -0.003 | 0 | 0 | 0 | 0.6 | 0.59 | 0.44 | 0.51 | 0.67 | 0.59 |
| 3 | 147061844 | rs2651315 | -0.719 | 0.354 | -0.757 | -0.389 | -0.007 | -0.027 | -0.027 | 0.173 | 0.65 | 0.79 | 0.82 | 0.53 | 0.67 | 0.55 |
| 3 | 147063796 | rs2651314 | 0.827 | 0.336 | -0.743 | -0.357 | -0.005 | 0.001 | 0.001 | 0 | 0.4 | 0.41 | 0.55 | 0.51 | 0.32 | 0.41 |
| 3 | 147066342 | rs7642125 | 0.502 | 0.347 | -0.762 | -0.339 | 0.016 | 0.005 | 0.005 | 0 | 0.1 | 0.3 | 0.09 | 0.09 | 0.11 | 0.17 |
| 3 | 147076615 | rs11720889 | 0.18 | 0.386 | -1.31 | -0.337 | -0.022 | -0.005 | -0.005 | 0.043 | 0.24 | 0.2 | 0.24 | 0.24 | 0.35 | 0.22 |
| 3 | 147076791 | rs884370 | -1.388 | 0.38 | -1.468 | -0.399 | -0.001 | 0.031 | 0.031 | 0 | 0.69 | 0.8 | 0.84 | 0.75 | 0.72 | 0.64 |
| 3 | 147081271 | rs11718368 | 0.713 | 0.426 | -1.662 | -0.34 | -0.031 | -0.051 | -0.051 | 0 | 0.43 | 0.16 | 0.64 | 0.45 | 0.45 | 0.31 |
| 3 | 147082063 | rs1503732 | 0.607 | 0.396 | -1.9 | -0.213 | -0.021 | -0.042 | -0.042 | 0 | 0.1 | 0.01 | 0.08 | 0.32 | 0.13 | 0.26 |
| 3 | 147082427 | rs1503733 | -1.017 | 0.387 | -1.891 | -0.245 | -0.048 | 0.017 | 0.017 | 0.087 | 0.4 | 0.79 | 0.39 | 0.36 | 0.5 | 0.4 |
| 3 | 147082740 | rs4681356 | -0.869 | 0.388 | -1.842 | -0.231 | -0.007 | 0.075 | 0.075 | 0 | 0.64 | 0.82 | 0.82 | 0.77 | 0.65 | 0.59 |
| 3 | 147085744 | rs1503737 | -0.232 | 0.691 | -1.341 | -0.08 | 0.015 | 0.154 | 0.154 | 0.286 | 0.16 | 0.51 | 0.13 | 0.32 | 0.15 | 0.28 |
| 3 | 147086268 | rs1910466 | -0.793 | 0.715 | -2.406 | 0.07 | -0.056 | 0.012 | 0.012 | 0.035 | 0.41 | 0.79 | 0.39 | 0.36 | 0.51 | 0.41 |
| 3 | 147090343 | rs1532769 | 0.514 | 0.858 | -2.266 | 0.139 | 0 | -0.008 | -0.008 | 0 | 0.47 | 0.66 | 0.6 | 0.46 | 0.47 | 0.44 |
| 3 | 147093600 | rs1503738 | -0.379 | 0.864 | -2.523 | 0.006 | 0.009 | 0.054 | 0.054 | 0 | 0.63 | 0.76 | 0.79 | 0.72 | 0.64 | 0.64 |
| 3 | 147096847 | rs1394041 | 0.742 | 0.892 | -2.634 | 0.044 | -0.016 | -0.021 | -0.021 | 0 | 0.1 | 0.02 | 0.08 | 0.15 | 0.11 | 0.12 |
| 3 | 147099798 | rs16859387 | 0.087 | 0.84 | -2.711 | 0.064 | 0 | 0.005 | 0.005 | 0 | 0.1 | 0.05 | 0.11 | 0.25 | 0.08 | 0.19 |
| 3 | 147101640 | rs2279832 | -1.77 | 0.688 | -3.093 | -0.211 | 0.005 | -0.012 | -0.012 | 0 | 0.35 | 0.3 | 0.29 | 0.53 | 0.32 | 0.47 |
| 3 | 147105790 | rs2279830 | -2.148 | 0.821 | -3.595 | -0.387 | -0.076 | -0.155 | -0.155 | 0 | 0.58 | 0.98 | 0.51 | 0.58 | 0.62 | 0.6 |
| 3 | 147106319 | rs2279829 | -0.658 | 0.678 | -2.545 | 0.06 | -0.001 | 0 | 0 | 0.073 | 0.2 | 0.22 | 0.14 | 0.28 | 0.21 | 0.27 |
| 3 | 147134032 | rs9833875 | -2.203 | 0.7 | -1.986 | 0.661 | 0.004 | -0.013 | -0.013 | 0.453 | 0.36 | 0.3 | 0.3 | 0.52 | 0.36 | 0.53 |
| 3 | 147137976 | rs9289748 | -2.321 | 0.718 | -1.96 | 0.668 | 0.004 | -0.009 | -0.009 | 0 | 0.37 | 0.32 | 0.3 | 0.52 | 0.36 | 0.53 |
| 3 | 147141344 | rs16859444 | -0.196 | 0.716 | -1.962 | 0.688 | -0.01 | 0.007 | 0.007 | 0 | 0.15 | 0.23 | 0.11 | 0.24 | 0.13 | 0.22 |
| 3 | 147142513 | rs9289749 | -2.425 | 0.725 | -1.978 | 0.702 | 0.004 | -0.009 | -0.009 | 0 | 0.37 | 0.31 | 0.31 | 0.51 | 0.36 | 0.53 |
| 3 | 147161075 | rs16859457 | -1.415 | 0.745 | -1.962 | 0.711 | -0.009 | 0.027 | 0.027 | 0 | 0.26 | 0.3 | 0.23 | 0.47 | 0.22 | 0.41 |
| 3 | 147174864 | rs9818183 | -2.361 | 0.762 | -2.087 | 0.709 | 0.009 | -0.023 | -0.023 | 0 | 0.37 | 0.27 | 0.3 | 0.51 | 0.36 | 0.53 |
| 3 | 147175324 | rs12630408 | -0.276 | 0.857 | -1.968 | 0.8 | -0.009 | 0.007 | 0.007 | 0 | 0.15 | 0.2 | 0.11 | 0.25 | 0.13 | 0.22 |
| 3 | 147209179 | rs12629668 | -1.681 | 0.867 | -1.961 | 0.81 | 0.008 | -0.022 | -0.022 | 0 | 0.31 | 0.23 | 0.29 | 0.47 | 0.29 | 0.47 |
| 3 | 147214792 | rs1027696 | -0.374 | 1.059 | -1.807 | 1.024 | -0.012 | 0.015 | 0.015 | 0 | 0.15 | 0.19 | 0.11 | 0.24 | 0.13 | 0.23 |
| 3 | 147215564 | rs1027695 | -1.514 | 1.056 | -1.812 | 1.022 | 0.001 | -0.002 | -0.002 | 0 | 0.25 | 0.2 | 0.23 | 0.46 | 0.22 | 0.45 |
| 3 | 147217797 | rs10513315 | -0.449 | 1.083 | -1.896 | 0.982 | -0.017 | -0.048 | -0.048 | 0 | 0.1 | 0.01 | 0.12 | 0.22 | 0.09 | 0.22 |
| 3 | 147237708 | rs9821069 | -0.742 | 1.228 | -1.907 | 0.872 | -0.055 | -0.073 | -0.073 | 0.312 | 0.13 | 0.01 | 0.13 | 0.22 | 0.18 | 0.29 |
| 3 | 147237920 | rs7613624 | -2.053 | 1.131 | -1.87 | 0.875 | -0.014 | -0.03 | -0.03 | 0.787 | 0.35 | 0.22 | 0.31 | 0.48 | 0.42 | 0.53 |
| 3 | 147246362 | rs11928555 | -0.42 | 0.964 | -1.842 | 0.883 | -0.006 | 0.006 | 0.006 | 0 | 0.17 | 0.17 | 0.11 | 0.26 | 0.14 | 0.23 |
| 3 | 147250714 | rs4561800 | -0.063 | 0.928 | -1.805 | 0.901 | -0.031 | -0.059 | -0.059 | 0 | 0.1 | 0.01 | 0.11 | 0.21 | 0.11 | 0.22 |
| 3 | 147268583 | rs11917877 | -1.546 | 0.96 | -1.559 | 0.814 | -0.008 | -0.017 | -0.017 | 0 | 0.34 | 0.28 | 0.31 | 0.48 | 0.42 | 0.52 |
| 3 | 147270026 | rs9873611 | -1.767 | 1.111 | -1.322 | 0.575 | -0.002 | -0.003 | -0.003 | 0 | 0.44 | 0.4 | 0.46 | 0.5 | 0.55 | 0.62 |
| 3 | 147275430 | rs11919670 | 0.195 | 1.38 | -1.405 | 0.618 | -0.049 | 0.054 | 0.054 | 0 | 0.11 | 0.02 | 0.14 | 0 | 0.2 | 0.05 |
| 3 | 147275754 | rs9817905 | 0.136 | 1.41 | -1.492 | 0.713 | -0.044 | -0.057 | -0.057 | 1.965 | 0.11 | 0.01 | 0.1 | 0.17 | 0.13 | 0.23 |
| 3 | 147276761 | rs9878930 | -1.304 | 1.262 | -1.342 | 0.673 | -0.002 | 0.028 | 0.028 | 0 | 0.27 | 0.32 | 0.27 | 0.48 | 0.25 | 0.47 |
| 3 | 147285912 | rs7624879 | -0.062 | 1.216 | -1.289 | 0.68 | -0.005 | 0.036 | 0.036 | 0 | 0.13 | 0.15 | 0.16 | 0.32 | 0.11 | 0.24 |
| 3 | 147292393 | rs4596113 | -1.593 | 1.244 | -1.303 | 0.592 | -0.003 | -0.003 | -0.003 | 0 | 0.37 | 0.37 | 0.42 | 0.49 | 0.44 | 0.52 |
| 3 | 147299274 | rs13325052 | -0.906 | 1.065 | -1.302 | 0.571 | 0 | 0.01 | 0.01 | 0 | 0.25 | 0.29 | 0.27 | 0.33 | 0.25 | 0.35 |
| 3 | 147305595 | rs13058918 | -0.593 | 1.265 | -1.514 | 0.306 | -0.027 | 0.043 | 0.043 | 0 | 0.91 | 0.98 | 0.9 | 1 | 0.87 | 0.96 |
| 3 | 147305791 | rs6440487 | 0.88 | 1.081 | -1.581 | 0.086 | 0.007 | 0.016 | 0.016 | 0.265 | 0.1 | 0.09 | 0.03 | 0.05 | 0.06 | 0.12 |
| 3 | 147310700 | rs11719156 | 1.503 | 0.626 | -1.626 | -0.081 | 0.015 | 0.066 | 0.066 | 0.003 | 0.5 | 0.18 | 0.61 | 0.62 | 0.58 | 0.37 |
| 3 | 147317832 | rs9882625 | 0.86 | 0.572 | -1.329 | -0.115 | -0.002 | -0.009 | -0.009 | 0 | 0.68 | 0.33 | 0.76 | 0.83 | 0.75 | 0.7 |
| 3 | 147320598 | rs13074490 | 1.644 | 0.585 | -1.313 | -0.104 | 0.004 | 0.064 | 0.064 | 0.032 | 0.5 | 0.12 | 0.61 | 0.62 | 0.58 | 0.38 |
| 3 | 147323069 | rs11715857 | 0.274 | 0.479 | -1.092 | 0.099 | -0.004 | -0.038 | -0.038 | 0.118 | 0.78 | 0.44 | 0.79 | 0.88 | 0.81 | 0.81 |
| 3 | 147339776 | rs13063052 | 1.365 | 0.571 | -1.022 | 0.051 | 0.014 | 0.039 | 0.039 | 0 | 0.5 | 0.31 | 0.62 | 0.62 | 0.55 | 0.38 |
| 3 | 147340091 | rs4681364 | -1.029 | 0.552 | -0.999 | 0.107 | 0.011 | 0.005 | 0.005 | 0 | 0.41 | 0.68 | 0.28 | 0.38 | 0.32 | 0.58 |
| 3 | 147341672 | rs6796711 | 0.518 | 0.379 | -0.922 | 0.124 | 0.003 | 0.05 | 0.05 | 1.121 | 0.11 | 0.39 | 0.09 | 0.12 | 0.07 | 0.14 |
| 3 | 147345411 | rs7651055 | -0.307 | 1.056 | -0.671 | 0.201 | 0 | -0.034 | -0.034 | 0 | 0.31 | 0.21 | 0.36 | 0.47 | 0.33 | 0.42 |
| 3 | 147353269 | rs9838382 | 1.082 | 0.611 | -0.961 | 0.367 | 0 | -0.003 | -0.003 | 0 | 0.46 | 0.42 | 0.48 | 0.46 | 0.43 | 0.37 |
| 3 | 147359950 | rs9847153 | 1.064 | 0.465 | -1.363 | 0.422 | -0.002 | 0 | 0 | 0 | 0.46 | 0.49 | 0.48 | 0.46 | 0.41 | 0.36 |
| 3 | 147360072 | rs16859608 | -0.037 | 0.149 | -1.634 | 0.372 | 0.006 | 0.01 | 0.01 | 1.107 | 0.09 | 0.08 | 0.03 | 0.05 | 0.06 | 0.1 |
| 3 | 147361337 | rs10513322 | -0.6 | 0.148 | -1.8 | 0.596 | 0.001 | -0.054 | -0.054 | 0 | 0.18 | 0.01 | 0.13 | 0.2 | 0.14 | 0.25 |
| 3 | 147362266 | rs2319336 | 0.095 | 0.55 | -1.375 | 0.274 | 0.007 | -0.031 | -0.031 | 0 | 0.64 | 0.5 | 0.62 | 0.79 | 0.65 | 0.74 |
| 3 | 147364266 | rs2873949 | 0.077 | 0.836 | -1.7 | -0.341 | -0.013 | 0.018 | 0.018 | 0.704 | 0.17 | 0.08 | 0.13 | 0 | 0.21 | 0.08 |
| 3 | 147364330 | rs2319335 | -0.398 | 0.53 | -2.029 | -0.094 | 0.027 | 0.009 | 0.009 | 0 | 0.63 | 0.8 | 0.67 | 0.66 | 0.73 | 0.68 |
| 3 | 147368416 | rs9862216 | -0.073 | 0.647 | -1.533 | 0.116 | -0.001 | 0 | 0 | 0 | 0.47 | 0.51 | 0.48 | 0.49 | 0.52 | 0.45 |
| 3 | 147377697 | rs2319333 | 0.072 | -0.252 | -1.79 | 0.242 | 0.04 | -0.012 | -0.012 | 0.26 | 0.36 | 0.27 | 0.41 | 0.61 | 0.23 | 0.43 |
| 3 | 147388753 | rs6788109 | 0.686 | -0.6 | -1.954 | 0.594 | -0.002 | 0 | 0 | 0 | 0.24 | 0.26 | 0.23 | 0.3 | 0.16 | 0.25 |
| 3 | 147389227 | rs13075781 | 1.452 | 0.077 | -2.653 | 0.321 | -0.059 | 0.089 | 0.089 | 0.253 | 0.14 | 0.01 | 0.1 | 0 | 0.19 | 0.07 |
| 3 | 147390505 | rs7622746 | -1.394 | 0.248 | -2.767 | 0.359 | -0.029 | 0.012 | 0.012 | 1.366 | 0.83 | 0.98 | 0.88 | 0.89 | 0.82 | 0.91 |
| 3 | 147390532 | rs11914463 | -0.48 | 0.308 | -2.465 | 0.457 | -0.005 | 0.006 | 0.006 | 0.031 | 0.73 | 0.83 | 0.73 | 0.78 | 0.71 | 0.77 |
| 3 | 147394548 | rs6802980 | 0.835 | 0.463 | -1.827 | 0.438 | -0.018 | 0.003 | 0.003 | 0 | 0.28 | 0.56 | 0.28 | 0.31 | 0.23 | 0.24 |
| 3 | 147394647 | rs16859648 | -0.436 | 0.837 | -2.161 | 0.986 | -0.017 | -0.122 | -0.122 | 0 | 0.27 | 0.03 | 0.17 | 0.35 | 0.21 | 0.26 |
| 3 | 147403067 | rs2873943 | -0.465 | 1.038 | -1.799 | 0.484 | -0.023 | -0.061 | -0.061 | 0 | 0.29 | 0.14 | 0.33 | 0.57 | 0.3 | 0.47 |
| 3 | 147409986 | rs12629902 | -0.306 | 0.811 | -1.252 | 0.008 | 0 | -0.013 | -0.013 | 0 | 0.33 | 0.4 | 0.42 | 0.71 | 0.33 | 0.39 |

| | | | | | | | | | | | | | | | | |
|---|---|---|---|---|---|---|---|---|---|---|---|---|---|---|---|---|
| 3 | 147413101 | rs7643549 | 0.449 | -0.003 | -1.697 | -0.252 | 0.016 | 0.011 | 0.011 | 0 | 0.26 | 0.16 | 0.3 | 0.23 | 0.2 | 0.33 |
| 3 | 147414845 | rs9855367 | 1.573 | 0.35 | -2.315 | -0.942 | -0.118 | 0.128 | 0.128 | 0 | 0.22 | 0.03 | 0.16 | 0.01 | 0.36 | 0.1 |
| 3 | 147417621 | rs12639104 | -0.141 | -0.154 | -1.977 | -0.974 | -0.041 | 0.057 | 0.057 | 0.059 | 0.23 | 0.54 | 0.27 | 0.61 | 0.14 | 0.3 |
| 3 | 147418936 | rs4681369 | -0.937 | 0.799 | -2.17 | -1.233 | 0.014 | 0.02 | 0.02 | 0 | 0.77 | 0.84 | 0.81 | 0.88 | 0.8 | 0.77 |
| 3 | 147421382 | rs7652411 | -0.28 | 0.718 | -2.05 | -1.112 | -0.004 | 0.039 | 0.039 | 0 | 0.66 | 0.76 | 0.68 | 0.88 | 0.62 | 0.73 |
| 3 | 147422935 | rs11711379 | 1.052 | 1.363 | -2.183 | -0.857 | -0.119 | 0.122 | 0.122 | 0 | 0.21 | 0.03 | 0.15 | 0.01 | 0.34 | 0.09 |
| 3 | 147423509 | rs2319306 | -0.28 | -0.103 | -2.231 | -0.451 | 0.042 | 0.066 | 0.066 | 0 | 0.52 | 0.77 | 0.46 | 0.43 | 0.56 | 0.53 |
| 3 | 147424585 | rs17511826 | 0.992 | 0.222 | -2.019 | -0.986 | -0.141 | 0.192 | 0.192 | 0 | 0.29 | 0.03 | 0.18 | 0.01 | 0.39 | 0.14 |
| 3 | 147431755 | rs9653947 | 0.262 | -0.168 | -1.672 | -0.498 | 0 | 0.044 | 0.044 | 0 | 0.13 | 0.16 | 0.24 | 0.56 | 0.17 | 0.33 |
| 3 | 147432019 | rs4681371 | 0.014 | -0.314 | -1.32 | -0.519 | 0.053 | 0.017 | 0.017 | 0 | 0.49 | 0.32 | 0.48 | 0.57 | 0.62 | 0.47 |
| 3 | 147441796 | rs2319319 | -0.063 | -0.642 | -0.338 | -0.682 | 0 | -0.04 | -0.04 | 0 | 0.91 | 0.41 | 0.94 | 1 | 0.94 | 0.99 |
| 3 | 147443667 | rs6777677 | -0.313 | -0.543 | -0.647 | -1.004 | -0.03 | 0.047 | 0.047 | 0 | 0.6 | 0.32 | 0.75 | 0.97 | 0.56 | 0.84 |
| 3 | 147461796 | rs4279060 | -0.622 | -0.697 | -0.225 | -0.948 | -0.001 | -0.059 | -0.059 | 0.098 | 0.62 | 0.62 | 0.75 | 0.97 | 0.53 | 0.84 |
| 3 | 147463055 | rs9861579 | 0.068 | -1.29 | -0.337 | -0.933 | 0.018 | 0.021 | 0.021 | 0 | 0.35 | 0.25 | 0.34 | 0.7 | 0.33 | 0.5 |
| 3 | 147469466 | rs16859725 | 1.203 | -1.299 | -0.32 | -0.945 | 0.039 | -0.01 | -0.01 | 0 | 0.07 | 0.36 | 0.05 | 0 | 0.06 | 0.01 |
| 3 | 147470053 | rs6770996 | 0.723 | -1.274 | -0.325 | -0.899 | 0 | 0 | 0 | 0.264 | 0.26 | 0.23 | 0.4 | 0.27 | 0.2 | 0.35 |
| 3 | 147472174 | rs2319327 | -0.956 | -1.217 | -0.432 | -0.905 | 0.001 | -0.069 | -0.069 | 0.35 | 0.62 | 0.57 | 0.75 | 0.97 | 0.54 | 0.84 |
| 3 | 147472375 | rs10513320 | 1.472 | -1.407 | -0.855 | -0.639 | -0.044 | -0.09 | -0.09 | 0 | 0.13 | 0.03 | 0.2 | 0.45 | 0.23 | 0.31 |
| 3 | 147473912 | rs9835933 | -0.987 | -1.239 | -0.567 | -1.063 | -0.011 | -0.031 | -0.031 | 0 | 0.63 | 0.63 | 0.75 | 0.97 | 0.54 | 0.84 |
| 3 | 147475370 | rs2319323 | -0.934 | -0.994 | -0.232 | 0.015 | -0.01 | 0.005 | 0.005 | 0 | 0.24 | 0.23 | 0.18 | 0.25 | 0.16 | 0.21 |
| 3 | 147481380 | rs6808403 | -0.643 | -0.795 | -0.641 | -0.189 | 0.009 | -0.003 | -0.003 | 0.012 | 0.21 | 0.15 | 0.13 | 0.21 | 0.15 | 0.2 |
| 3 | 147484611 | rs937780 | 0.252 | -0.44 | -0.588 | -0.793 | 0.006 | 0.023 | 0.023 | 0 | 0.79 | 0.88 | 0.88 | 0.94 | 0.83 | 0.85 |
| 3 | 147492616 | rs12695850 | -0.303 | -0.354 | -0.043 | -0.247 | -0.012 | -0.074 | -0.074 | 1.894 | 0.12 | 0.35 | 0.08 | 0 | 0.1 | 0.05 |
| 3 | 147492839 | rs11928247 | -0.421 | -0.137 | -0.757 | -0.081 | -0.001 | -0.005 | -0.005 | 0 | 0.36 | 0.65 | 0.46 | 0.79 | 0.35 | 0.37 |
| 3 | 147494512 | rs9829326 | -0.072 | 0.277 | -0.963 | 0.369 | -0.013 | 0.014 | 0.014 | 0 | 0.57 | 0.64 | 0.52 | 0.58 | 0.58 | 0.55 |
| 3 | 147494530 | rs2873925 | -1.422 | 0.185 | -0.665 | 0.804 | -0.001 | 0.006 | 0.006 | 2.229 | 0.1 | 0.09 | 0.11 | 0.25 | 0.06 | 0.35 |
| 3 | 147495559 | rs9682099 | -0.916 | -0.216 | -0.174 | 0.702 | -0.019 | 0.024 | 0.024 | 0.864 | 0.22 | 0.56 | 0.19 | 0.23 | 0.16 | 0.15 |
| 3 | 147496499 | rs4681375 | 1.13 | -0.33 | -0.505 | 0.744 | 0.013 | -0.024 | -0.024 | 0 | 0.64 | 0.3 | 0.59 | 0.47 | 0.67 | 0.69 |
| 3 | 147502818 | rs7627412 | -0.856 | -0.599 | 0.056 | 0.588 | -0.002 | 0.013 | 0.013 | 0 | 0.21 | 0.18 | 0.32 | 0.41 | 0.21 | 0.19 |
| 3 | 147505136 | rs1391796 | 1.288 | -0.694 | 0.48 | 0.448 | -0.017 | 0.018 | 0.018 | 0 | 0.69 | 0.54 | 0.65 | 0.63 | 0.71 | 0.76 |
| 3 | 147508791 | rs1391795 | 0.699 | -1.724 | -0.092 | -0.262 | 0.035 | 0.107 | 0.107 | 0 | 0.8 | 0.95 | 0.97 | 0.99 | 0.89 | 0.87 |
| 3 | 147513235 | rs1909357 | -2 | -1.697 | -0.151 | -0.326 | 0.077 | -0.012 | -0.012 | 0 | 0.51 | 0.89 | 0.47 | 0.51 | 0.42 | 0.39 |
| 3 | 147513921 | rs4681376 | -0.46 | -1.889 | 0.564 | -0.024 | -0.008 | 0.104 | 0.104 | 0 | 0.21 | 0.31 | 0.35 | 0.5 | 0.24 | 0.24 |
| 3 | 147533843 | rs6785065 | -1.03 | -1.921 | 1.107 | -0.031 | -0.014 | 0.043 | 0.043 | 0.347 | 0.27 | 0.62 | 0.39 | 0.49 | 0.27 | 0.25 |
| 3 | 147536474 | rs11922072 | -1.321 | -2.033 | 0.92 | -0.221 | -0.027 | -0.064 | -0.064 | 0.14 | 0.15 | 0.25 | 0.04 | 0.01 | 0.06 | 0.1 |
| 3 | 147546076 | rs4681379 | -0.593 | -1.296 | 1.345 | 0.237 | -0.005 | 0.045 | 0.045 | 0 | 0.22 | 0.24 | 0.34 | 0.51 | 0.23 | 0.26 |
| 3 | 147554766 | rs994371 | -1.245 | -1.318 | 1.427 | 0.226 | -0.02 | 0.056 | 0.056 | 0.667 | 0.28 | 0.62 | 0.38 | 0.51 | 0.26 | 0.28 |
| 3 | 147557097 | rs4681137 | -1.149 | -1.315 | 1.253 | 0.227 | -0.024 | 0.081 | 0.081 | 0.244 | 0.27 | 0.48 | 0.38 | 0.51 | 0.26 | 0.28 |
| 3 | 147564316 | rs1498899 | -1.353 | -1.32 | 1.051 | 0.225 | 0 | -0.001 | -0.001 | 0 | 0.28 | 0.72 | 0.39 | 0.51 | 0.26 | 0.28 |
| 3 | 147596280 | rs1498891 | -1.18 | -1.078 | 0.397 | 0.364 | -0.003 | 0.005 | 0.005 | 0 | 0.32 | 0.37 | 0.37 | 0.51 | 0.29 | 0.29 |
| 3 | 147599246 | rs1116439 | -1.059 | -1.139 | 0.369 | 0.386 | -0.002 | 0.002 | 0.002 | 0 | 0.31 | 0.35 | 0.36 | 0.51 | 0.27 | 0.28 |
| 3 | 147610313 | rs4681386 | -0.923 | -0.87 | 0.254 | 0.461 | -0.001 | 0.003 | 0.003 | 0 | 0.28 | 0.26 | 0.34 | 0.49 | 0.26 | 0.29 |
| 3 | 147613651 | rs6774903 | -0.889 | -0.673 | 0.123 | 0.439 | 0 | -0.022 | -0.022 | 0.638 | 0.31 | 0.21 | 0.38 | 0.48 | 0.35 | 0.29 |
| 3 | 147623914 | rs4681390 | -0.653 | -0.32 | 0.343 | 0.399 | 0.012 | 0.055 | 0.055 | 0.146 | 0.21 | 0.39 | 0.2 | 0.27 | 0.21 | 0.15 |
| 3 | 147624267 | rs4681141 | -1.314 | -0.19 | 0.504 | 0.289 | 0 | 0 | 0 | 1.061 | 0.26 | 0.64 | 0.27 | 0.27 | 0.26 | 0.15 |
| 3 | 147625933 | rs6798441 | -1.036 | 0.314 | -0.135 | -0.006 | 0.054 | -0.001 | -0.001 | 0.968 | 0.05 | 0.26 | 0.07 | 0.01 | 0.05 | 0 |
| 3 | 147631763 | rs6775904 | -0.915 | 0.124 | -0.951 | -0.126 | -0.143 | 0.115 | 0.115 | 0.41 | 0.61 | 0.97 | 0.74 | 0.79 | 0.52 | 0.65 |
| 3 | 147653591 | rs7613691 | -1.766 | 0.588 | 0.451 | 0.255 | 0.098 | 0.01 | 0.01 | 0.592 | 0.07 | 0.34 | 0.07 | 0.02 | 0.05 | 0.01 |
| 3 | 147677770 | rs11711809 | -1.558 | 0.418 | -0.478 | 0.42 | 0 | 0 | 0 | 1.287 | 0.14 | 0.14 | 0.1 | 0.09 | 0.13 | 0.07 |
| 3 | 147679289 | rs11713814 | -0.939 | 0.504 | -0.633 | 0.486 | -0.007 | -0.009 | -0.009 | 2.175 | 0.11 | 0.08 | 0.09 | 0.1 | 0.12 | 0.07 |
| 3 | 147681840 | rs11708371 | -0.414 | 0.367 | -0.897 | 0.394 | 0.022 | -0.033 | -0.033 | 3.237 | 0.18 | 0.11 | 0.09 | 0.27 | 0.13 | 0.28 |
| 3 | 147682334 | rs1462654 | -1.263 | 0.893 | -0.452 | 0.401 | 0 | -0.009 | -0.009 | 0 | 0.12 | 0.16 | 0.12 | 0.03 | 0.15 | 0.04 |
| 3 | 147703442 | rs17529042 | -0.741 | 0.911 | -0.544 | 0.406 | -0.028 | 0.03 | 0.03 | 0.939 | 0.1 | 0.03 | 0.09 | 0.02 | 0.12 | 0.04 |
| 3 | 147704388 | rs6771140 | -1.599 | 0.72 | -0.102 | 0.761 | 0.013 | 0.133 | 0.133 | 1.895 | 0.17 | 0.32 | 0.36 | 0.45 | 0.17 | 0.22 |
| 3 | 147727960 | rs16860017 | -1.192 | 1.378 | 0.635 | -0.216 | 0.116 | 0.016 | 0.016 | 0 | 0.1 | 0.49 | 0.13 | 0.06 | 0.16 | 0.06 |
| 3 | 147730134 | rs16860023 | -0.907 | 1.379 | 0.552 | -0.214 | 0.106 | 0.05 | 0.05 | 0 | 0.07 | 0.36 | 0.1 | 0.04 | 0.12 | 0.05 |
| 3 | 147746622 | rs16860030 | -1.784 | 1.31 | 0.263 | -0.218 | 0.057 | -0.009 | -0.009 | 0 | 0.11 | 0.6 | 0.14 | 0.06 | 0.17 | 0.06 |
| 3 | 147750355 | rs734873 | 0.148 | 2.012 | -0.272 | 0.019 | -0.044 | -0.007 | -0.007 | 0.029 | 0.08 | 0.01 | 0.29 | 0.43 | 0.11 | 0.26 |
| 3 | 147751494 | rs9289763 | 0.472 | 1.408 | -0.549 | -1.11 | -0.051 | 0.05 | 0.05 | 0 | 0.25 | 0.01 | 0.12 | 0.11 | 0.26 | 0.21 |
| 3 | 147756686 | rs1841770 | 0.515 | 1.163 | 0.523 | -0.909 | -0.064 | 0.086 | 0.086 | 0.406 | 0.68 | 0.86 | 0.65 | 0.82 | 0.55 | 0.8 |
| 3 | 147774712 | rs9848851 | 1.082 | 1.335 | 1.116 | -0.282 | 0.006 | -0.011 | -0.011 | 0.103 | 0.75 | 0.31 | 0.7 | 0.8 | 0.65 | 0.8 |
| 3 | 147787204 | rs6764882 | 0.751 | 1.449 | 0.442 | -0.184 | 0.042 | -0.036 | -0.036 | 0 | 0.78 | 0.63 | 0.72 | 0.82 | 0.67 | 0.8 |
| 3 | 147787591 | rs1870000 | 0.462 | 1.649 | -0.76 | -0.096 | -0.042 | 0.024 | 0.024 | 0.006 | 0.8 | 0.88 | 0.74 | 0.82 | 0.67 | 0.8 |
| 3 | 147794002 | rs10513327 | 0.722 | 1.291 | -0.323 | -0.2 | 0.008 | -0.039 | -0.039 | 0.168 | 0.76 | 0.66 | 0.79 | 0.85 | 0.74 | 0.82 |
| 3 | 147803350 | rs1869998 | -0.385 | 1.082 | -0.198 | 0.121 | -0.001 | -0.001 | -0.001 | 0.119 | 0.31 | 0.3 | 0.41 | 0.31 | 0.33 | 0.35 |
| 3 | 147805662 | rs15606 | -0.931 | 0.884 | 0.311 | 0.019 | 0.002 | -0.009 | -0.009 | 0 | 0.37 | 0.73 | 0.45 | 0.33 | 0.37 | 0.38 |
| 3 | 147818813 | rs7635164 | -1.171 | 0.729 | 0.485 | 0.059 | -0.002 | 0.01 | 0.01 | 0 | 0.19 | 0.21 | 0.23 | 0.25 | 0.18 | 0.25 |
| 3 | 147829201 | rs4591472 | -1.252 | 0.684 | 0.643 | 0.113 | -0.009 | 0.017 | 0.017 | 0 | 0.33 | 0.44 | 0.43 | 0.31 | 0.33 | 0.34 |
| 3 | 147845656 | rs1456682 | 0.467 | 0.715 | 0.597 | 0.119 | 0.006 | -0.029 | -0.029 | 0 | 0.17 | 0.24 | 0.2 | 0.08 | 0.18 | 0.13 |

| | | | | | | | | | | | | | | | | |
|---|---|---|---|---|---|---|---|---|---|---|---|---|---|---|---|---|
| 3 | 147850902 | rs4269055 | 0.081 | 0.638 | 0.452 | 0.093 | -0.028 | -0.051 | -0.051 | 0 | 0.09 | 0 | 0.19 | 0.22 | 0.12 | 0.16 |
| 3 | 147859661 | rs1456679 | -1.26 | 0.609 | 0.633 | 0.174 | -0.009 | 0.017 | 0.017 | 0 | 0.33 | 0.56 | 0.42 | 0.3 | 0.31 | 0.34 |
| 3 | 147866816 | rs12497438 | 0.557 | 0.465 | 0.139 | 0.16 | -0.031 | 0.017 | 0.017 | 0 | 0.16 | 0.02 | 0.14 | 0.15 | 0.16 | 0.16 |
| 3 | 147867286 | rs7614394 | -1.442 | 0.497 | 0.466 | 0.388 | 0.004 | -0.014 | -0.014 | 0.461 | 0.35 | 0.74 | 0.45 | 0.36 | 0.35 | 0.37 |
| 3 | 147867628 | rs7628508 | 0.03 | 0.815 | 0.139 | 0.33 | 0.016 | -0.043 | -0.043 | 0.003 | 0.17 | 0.36 | 0.21 | 0.08 | 0.18 | 0.14 |
| 3 | 147869237 | rs980933 | -0.273 | 0.803 | 0.113 | 0.329 | 0.025 | -0.031 | -0.031 | 0 | 0.19 | 0.38 | 0.23 | 0.14 | 0.22 | 0.17 |
| 3 | 147884168 | rs12495960 | 0.178 | 0.855 | -0.526 | 0.426 | -0.01 | 0.01 | 0.01 | 0 | 0.15 | 0.09 | 0.19 | 0.1 | 0.17 | 0.12 |
| 3 | 147888040 | rs4681394 | -0.675 | 0.744 | -0.356 | 0.622 | -0.003 | 0.065 | 0.065 | 0 | 0.13 | 0.27 | 0.22 | 0.22 | 0.13 | 0.2 |
| 3 | 147895516 | rs16860163 | -1.353 | 0.832 | -0.287 | 0.619 | 0.005 | 0.03 | 0.03 | 0.026 | 0.31 | 0.45 | 0.44 | 0.36 | 0.33 | 0.37 |
| 3 | 147901443 | rs6770020 | -1.186 | 0.799 | -0.691 | 0.616 | 0 | 0.006 | 0.006 | 0 | 0.28 | 0.34 | 0.42 | 0.31 | 0.3 | 0.33 |
| 3 | 147911005 | rs6810197 | 0.265 | 0.836 | -1.005 | 0.628 | -0.016 | 0.014 | 0.014 | 0 | 0.14 | 0.07 | 0.19 | 0.09 | 0.17 | 0.13 |
| 3 | 147918738 | rs9838737 | -1.45 | 0.788 | -1.762 | 0.434 | -0.081 | -0.047 | -0.047 | 0 | 0.65 | 0.98 | 0.7 | 0.67 | 0.65 | 0.68 |
| 3 | 147927668 | rs7645488 | -1.22 | 0.903 | -0.791 | 0.675 | -0.002 | -0.013 | -0.013 | 0 | 0.27 | 0.26 | 0.41 | 0.31 | 0.3 | 0.33 |
| 3 | 147935700 | rs897666 | -0.504 | 0.958 | -0.984 | 0.701 | 0.002 | -0.014 | -0.014 | 0 | 0.12 | 0.1 | 0.21 | 0.21 | 0.13 | 0.2 |
| 3 | 147939350 | rs1379078 | -0.093 | 1.027 | -0.734 | 0.18 | -0.008 | 0.008 | 0.008 | 0 | 0.14 | 0.11 | 0.19 | 0.09 | 0.17 | 0.12 |
| 3 | 147943603 | rs6771516 | -0.093 | 1.049 | -0.739 | 0.18 | -0.002 | 0.003 | 0.003 | 0 | 0.15 | 0.16 | 0.19 | 0.09 | 0.17 | 0.12 |
| 3 | 147951658 | rs7638279 | -0.098 | 0.94 | -0.751 | 0.01 | -0.005 | 0.02 | 0.02 | 0 | 0.45 | 0.58 | 0.34 | 0.19 | 0.46 | 0.24 |
| 3 | 147954392 | rs1495331 | 0.15 | 1.001 | -0.63 | 0.008 | -0.001 | 0 | 0 | 0 | 0.89 | 0.91 | 0.9 | 0.96 | 0.86 | 0.94 |
| 3 | 147955974 | rs1874295 | 1.287 | 0.153 | -1.063 | 0.354 | 0.021 | -0.054 | -0.054 | 0.63 | 0.6 | 0.14 | 0.54 | 0.46 | 0.66 | 0.61 |
| 3 | 147957721 | rs4355248 | 0.368 | -0.164 | 0.311 | 0.472 | 0 | 0.007 | 0.007 | 0 | 0.77 | 0.78 | 0.68 | 0.69 | 0.77 | 0.78 |
| 3 | 147959124 | rs1495328 | 0.496 | -0.221 | 1.077 | 0.411 | 0.018 | -0.037 | -0.037 | 0 | 0.07 | 0.16 | 0.04 | 0 | 0.08 | 0.03 |
| 3 | 147960665 | rs6769088 | -0.619 | -0.184 | 1.146 | 0.465 | 0.008 | -0.004 | -0.004 | 0 | 0.88 | 0.87 | 0.83 | 0.89 | 0.86 | 0.94 |
| 3 | 147963098 | rs9851377 | 1.293 | -0.222 | 1.437 | 0.475 | -0.018 | 0.06 | 0.06 | 0 | 0.7 | 0.49 | 0.61 | 0.47 | 0.73 | 0.65 |
| 3 | 147978935 | rs7611797 | 0.772 | -0.186 | 1.263 | 0.357 | 0.008 | -0.017 | -0.017 | 0 | 0.11 | 0.06 | 0.15 | 0.2 | 0.09 | 0.16 |
| 3 | 147987077 | rs4578991 | -0.803 | -0.116 | 1.182 | 0.333 | -0.03 | 0.015 | 0.015 | 0 | 0.9 | 0.99 | 0.9 | 0.95 | 0.86 | 0.94 |
| 3 | 147989624 | rs10804724 | -1.377 | -0.304 | 1.452 | 0.347 | 0.004 | 0.001 | 0.001 | 0.91 | 0.24 | 0.72 | 0.31 | 0.26 | 0.18 | 0.19 |
| 3 | 147995347 | rs16860281 | -0.404 | -0.163 | 1.688 | 0.284 | 0.017 | -0.053 | -0.053 | 1.25 | 0.09 | 0.24 | 0.06 | 0 | 0.08 | 0.03 |
| 3 | 147997618 | rs7615701 | -1.916 | -0.459 | 1.828 | 0.504 | -0.038 | 0.074 | 0.074 | 1.969 | 0.2 | 0.52 | 0.2 | 0.32 | 0.14 | 0.2 |
| 3 | 148001131 | rs10935692 | -1.201 | -0.41 | 1.643 | 0.59 | -0.03 | 0.094 | 0.094 | 0 | 0.11 | 0.27 | 0.14 | 0.32 | 0.06 | 0.17 |
| 3 | 148004138 | rs11712977 | -1.201 | -0.41 | 1.643 | 0.59 | -0.03 | 0.094 | 0.094 | 0 | 0.11 | 0.27 | 0.14 | 0.32 | 0.06 | 0.17 |
| 3 | 148008513 | rs10433444 | -1.574 | -0.428 | 1.724 | 0.582 | -0.038 | 0.174 | 0.174 | 0 | 0.12 | 0.44 | 0.15 | 0.32 | 0.06 | 0.16 |
| 3 | 148020990 | rs2029807 | -0.064 | -0.335 | 1.368 | 0.604 | -0.003 | 0.041 | 0.041 | 0 | 0.06 | 0.14 | 0.06 | 0.27 | 0.05 | 0.16 |
| 3 | 148037315 | rs7612531 | -1.044 | 0.196 | 1.173 | 0.679 | -0.027 | 0.093 | 0.093 | 0 | 0.22 | 0.41 | 0.26 | 0.57 | 0.17 | 0.33 |
| 3 | 148079188 | rs6440537 | -0.516 | 0.156 | 1.417 | 0.687 | 0.002 | 0.168 | 0.168 | 0 | 0.08 | 0.35 | 0.08 | 0.26 | 0.05 | 0.16 |
| 3 | 148085522 | rs4528908 | 2.121 | 0.21 | 1.127 | 0.503 | 0.01 | -0.031 | -0.031 | 0.674 | 0.54 | 0.07 | 0.48 | 0.36 | 0.6 | 0.59 |
| 3 | 148088465 | rs4643674 | 0.34 | 0.017 | 1.267 | 0.639 | 0.014 | -0.066 | -0.066 | 0.079 | 0.13 | 0.02 | 0.19 | 0.29 | 0.11 | 0.17 |
| 3 | 148103369 | rs7610876 | -1.033 | 0.006 | 1.653 | 0.689 | 0.009 | 0.232 | 0.232 | 0 | 0.09 | 0.43 | 0.09 | 0.27 | 0.06 | 0.16 |
| 3 | 148114124 | rs10935703 | 0.143 | 0.004 | 1.607 | 0.689 | 0.018 | -0.023 | -0.023 | 0 | 0.14 | 0.08 | 0.2 | 0.28 | 0.11 | 0.17 |
| 3 | 148115945 | rs4374498 | -0.432 | 0.028 | 1.587 | 0.838 | -0.008 | 0.123 | 0.123 | 0 | 0.07 | 0.26 | 0.07 | 0.26 | 0.05 | 0.16 |
| 3 | 148120514 | rs4681408 | -1.345 | 0.066 | 1.565 | 0.785 | -0.027 | 0.095 | 0.095 | 0 | 0.11 | 0.29 | 0.15 | 0.31 | 0.06 | 0.16 |
| 3 | 148124129 | rs6440539 | -2.316 | -0.052 | 1.602 | 0.692 | -0.011 | 0.016 | 0.016 | 1.406 | 0.29 | 0.71 | 0.38 | 0.6 | 0.23 | 0.35 |
| 3 | 148150193 | rs12634011 | -0.827 | 0.078 | 1.509 | 0.706 | -0.005 | 0.157 | 0.157 | 0 | 0.07 | 0.27 | 0.07 | 0.28 | 0.05 | 0.16 |
| 3 | 148188630 | rs3851993 | -0.568 | 0.307 | 1.026 | 0.404 | -0.003 | -0.004 | -0.004 | 0.81 | 0.95 | 0.95 | 0.95 | 1 | 0.88 | 0.98 |
| 3 | 148189939 | rs1916224 | 0.878 | -0.148 | 0.151 | 0.023 | -0.048 | 0.002 | 0.002 | 0.064 | 0.46 | 0.15 | 0.51 | 0.34 | 0.52 | 0.42 |
| 3 | 148195726 | rs13084464 | 1.007 | -0.409 | -0.274 | -0.1 | -0.071 | 0.005 | 0.005 | 0 | 0.4 | 0.13 | 0.39 | 0.29 | 0.46 | 0.32 |
| 3 | 148202707 | rs3915004 | 0.546 | -0.586 | -0.006 | -0.096 | -0.004 | 0.005 | 0.005 | 0 | 0.33 | 0.43 | 0.25 | 0.38 | 0.29 | 0.45 |
| 3 | 148210054 | rs3863998 | 0.639 | -0.461 | -0.161 | -0.015 | -0.008 | -0.049 | -0.049 | 0 | 0.11 | 0.03 | 0.18 | 0.23 | 0.1 | 0.17 |
| 3 | 148219044 | rs7631173 | 1.002 | -0.671 | -0.208 | -0.074 | -0.036 | -0.012 | -0.012 | 0 | 0.47 | 0.15 | 0.51 | 0.37 | 0.51 | 0.39 |
| 3 | 148220105 | rs987558 | -0.185 | -0.646 | -0.069 | -0.078 | 0.03 | -0.049 | -0.049 | 0 | 0.91 | 0.77 | 0.93 | 1 | 0.87 | 0.97 |
| 3 | 148221464 | rs35942196 | 0.626 | -0.668 | 0.021 | -0.081 | -0.009 | 0.018 | 0.018 | 0 | 0.3 | 0.4 | 0.22 | 0.38 | 0.25 | 0.38 |
| 3 | 148228410 | rs6440552 | -0.049 | -0.594 | -0.054 | -0.074 | 0.035 | -0.051 | -0.051 | 0 | 0.91 | 0.73 | 0.93 | 1 | 0.87 | 0.97 |
| 3 | 148228808 | rs4349458 | -0.116 | -0.599 | -0.068 | -0.076 | 0.029 | -0.05 | -0.05 | 0.52 | 0.91 | 0.77 | 0.93 | 1 | 0.87 | 0.97 |
| 3 | 148233254 | rs1403430 | 0.833 | -0.6 | -0.076 | -0.069 | 0 | -0.049 | -0.049 | 0.444 | 0.1 | 0.02 | 0.17 | 0.25 | 0.09 | 0.17 |
| 3 | 148234314 | rs1159882 | -0.343 | -0.57 | -0.03 | -0.094 | 0.012 | -0.036 | -0.036 | 0 | 0.92 | 0.87 | 0.94 | 1 | 0.88 | 0.97 |
| 3 | 148236575 | rs7631739 | -0.178 | -0.571 | -0.039 | -0.095 | 0.032 | -0.048 | -0.048 | 0 | 0.92 | 0.77 | 0.93 | 1 | 0.87 | 0.97 |
| 3 | 148240207 | rs1589634 | -0.048 | -0.567 | -0.064 | -0.093 | 0.034 | -0.051 | -0.051 | 1.45 | 0.91 | 0.74 | 0.93 | 1 | 0.87 | 0.97 |
| 3 | 148251079 | rs9848403 | -0.031 | -0.622 | -0.377 | -0.027 | 0.018 | -0.052 | -0.052 | 0 | 0.15 | 0.05 | 0.19 | 0.26 | 0.11 | 0.2 |
| 3 | 148254350 | rs7630439 | 0.194 | -0.585 | -0.286 | -0.062 | -0.015 | 0.004 | 0.004 | 0 | 0.34 | 0.41 | 0.25 | 0.39 | 0.26 | 0.41 |
| 3 | 148258098 | rs1522951 | 0.373 | -0.639 | -0.496 | -0.017 | 0.006 | 0.046 | 0.046 | 0.023 | 0.55 | 0.2 | 0.65 | 0.61 | 0.59 | 0.55 |
| 3 | 148263249 | rs13058910 | 0.548 | -0.223 | -0.052 | 0.038 | 0.02 | -0.008 | -0.008 | 0 | 0.76 | 0.61 | 0.73 | 0.76 | 0.71 | 0.8 |
| 3 | 148266924 | rs4681425 | 0.548 | -0.223 | -0.052 | 0.017 | 0.02 | -0.003 | -0.003 | 0 | 0.76 | 0.62 | 0.73 | 0.75 | 0.7 | 0.79 |
| 3 | 148272024 | rs4681426 | 0.568 | -0.479 | -0.719 | 0.089 | 0.01 | -0.028 | -0.028 | 0 | 0.33 | 0.25 | 0.32 | 0.48 | 0.24 | 0.44 |
| 3 | 148285788 | rs3920359 | -0.69 | 0.028 | -1.246 | 0.364 | 0.046 | -0.061 | -0.061 | 0.136 | 0.89 | 0.85 | 0.92 | 1 | 0.82 | 0.95 |
| 3 | 148301245 | rs3103760 | 0.389 | -0.007 | -1.265 | 0.389 | 0.007 | 0.006 | 0.006 | 0.095 | 0.79 | 0.79 | 0.75 | 0.74 | 0.73 | 0.79 |
| 3 | 148303859 | rs10513333 | 1.065 | 0.088 | -1.341 | 0.338 | -0.055 | -0.001 | -0.001 | 0 | 0.36 | 0.1 | 0.35 | 0.27 | 0.43 | 0.31 |
| 3 | 148308873 | rs2639369 | 1.04 | 0.301 | -0.457 | 0.481 | 0.007 | 0.007 | 0.007 | 0.1 | 0.79 | 0.79 | 0.75 | 0.74 | 0.73 | 0.79 |
| 3 | 148315687 | rs1522940 | -0.187 | -0.211 | -1.075 | 0.937 | 0.011 | -0.027 | -0.027 | 0 | 0.1 | 0.06 | 0.17 | 0.26 | 0.09 | 0.16 |
| 3 | 148321254 | rs2172347 | 0.08 | -0.246 | -0.892 | 0.581 | -0.016 | -0.001 | -0.001 | 0 | 0.4 | 0.48 | 0.42 | 0.4 | 0.32 | 0.33 |
| 3 | 148328711 | rs6800835 | 0.669 | -0.318 | -1.219 | 0.162 | 0.009 | 0.005 | 0.005 | 0 | 0.38 | 0.42 | 0.45 | 0.56 | 0.49 | 0.54 |

| | | | | | | | | | | | | | | | | |
|---|---|---|---|---|---|---|---|---|---|---|---|---|---|---|---|---|
| 3 | 148328996 | rs397650 | -0.576 | -0.423 | -1.316 | 0.068 | 0.004 | -0.006 | -0.006 | 0.833 | 0.9 | 0.91 | 0.94 | 1 | 0.9 | 0.92 |
| 3 | 148335030 | rs6803324 | 0.293 | -0.491 | -0.94 | 0.435 | -0.018 | 0.078 | 0.078 | 0 | 0.05 | 0.14 | 0.2 | 0.17 | 0.02 | 0.05 |
| 3 | 148336197 | rs188019 | -0.481 | -0.443 | -0.993 | 0.047 | 0.011 | 0.031 | 0.031 | 0 | 0.56 | 0.69 | 0.72 | 0.71 | 0.61 | 0.64 |
| 3 | 148337463 | rs4306816 | 0.123 | -0.079 | -0.922 | 0.304 | 0.002 | -0.01 | -0.01 | 0 | 0.43 | 0.53 | 0.65 | 0.73 | 0.51 | 0.59 |
| 3 | 148343517 | rs1879976 | 0.149 | -0.069 | -0.926 | 0.224 | 0.002 | -0.007 | -0.007 | 0 | 0.43 | 0.53 | 0.64 | 0.68 | 0.51 | 0.59 |
| 3 | 148345526 | rs275690 | -0.709 | -0.274 | -0.949 | 0.195 | 0.002 | 0.006 | 0.006 | 0 | 0.1 | 0.09 | 0.06 | 0.06 | 0.08 | 0.03 |
| 3 | 148346933 | rs9825391 | 0.638 | -0.237 | -1.073 | 0.229 | 0 | -0.005 | -0.005 | 3.522 | 0.46 | 0.4 | 0.61 | 0.6 | 0.54 | 0.63 |
| 3 | 148347742 | rs275685 | -0.287 | -0.445 | -0.617 | -0.598 | -0.006 | -0.044 | -0.044 | 0.277 | 0.29 | 0.33 | 0.14 | 0.07 | 0.23 | 0.12 |
| 3 | 148353740 | rs275697 | 0.539 | -0.999 | -0.436 | -0.428 | 0.002 | 0.017 | 0.017 | 0 | 0.94 | 0.81 | 0.9 | 0.95 | 0.94 | 0.94 |
| 3 | 148356615 | rs9846257 | 0.31 | -1.617 | -1.08 | -1.137 | -0.004 | 0.066 | 0.066 | 0 | 0.41 | 0.31 | 0.75 | 0.8 | 0.63 | 0.55 |
| 3 | 148358582 | rs6440561 | -0.453 | -2.218 | -1.222 | -1.552 | 0 | 0.043 | 0.043 | 0 | 0.74 | 0.85 | 0.88 | 0.92 | 0.77 | 0.92 |
| 3 | 148358840 | rs13070340 | 0.481 | -2.296 | -0.841 | -1.5 | -0.025 | -0.015 | -0.015 | 0 | 0.08 | 0.16 | 0.02 | 0.04 | 0.02 | 0.06 |
| 3 | 148361474 | rs1873901 | 0.827 | -2.298 | -0.842 | -1.525 | -0.024 | 0.069 | 0.069 | 0 | 0.39 | 0.23 | 0.7 | 0.79 | 0.6 | 0.53 |
| 3 | 148362977 | rs388488 | -0.077 | -2.551 | -0.984 | -1.77 | -0.017 | -0.09 | -0.09 | 1.046 | 0.71 | 0.58 | 0.84 | 0.92 | 0.79 | 0.7 |
| 3 | 148367386 | rs275680 | -0.119 | -2.453 | -1.743 | -1.585 | 0.022 | 0.032 | 0.032 | 0 | 0.34 | 0.25 | 0.13 | 0.13 | 0.2 | 0.18 |
| 3 | 148375111 | rs7617587 | 0.61 | -2.361 | -1.726 | -1.41 | -0.029 | -0.005 | -0.005 | 0 | 0.06 | 0.11 | 0.02 | 0.05 | 0 | 0.03 |
| 3 | 148378780 | rs10804730 | 0.942 | -2.588 | -2.119 | -1.597 | -0.027 | 0.128 | 0.128 | 0 | 0.41 | 0.16 | 0.71 | 0.8 | 0.63 | 0.53 |
| 3 | 148379707 | rs9836959 | 0.868 | -2.62 | -2.113 | -1.575 | -0.016 | 0.136 | 0.136 | 0 | 0.42 | 0.13 | 0.72 | 0.8 | 0.65 | 0.55 |
| 3 | 148383072 | rs17762633 | 0.547 | -2.822 | -2.207 | -1.683 | -0.007 | 0.033 | 0.033 | 0 | 0.12 | 0.01 | 0.07 | 0.07 | 0.1 | 0.2 |
| 3 | 148383381 | rs275701 | 0.485 | -2.832 | -2.128 | -1.696 | 0.087 | 0.141 | 0.141 | 0.582 | 0.6 | 0.24 | 0.8 | 0.89 | 0.79 | 0.63 |
| 3 | 148386360 | rs389571 | 0.074 | -2.807 | -1.988 | -1.688 | -0.006 | -0.047 | -0.047 | 0 | 0.66 | 0.46 | 0.83 | 0.93 | 0.8 | 0.67 |
| 3 | 148387876 | rs275657 | 1.071 | -2.702 | -2.358 | -1.631 | 0.004 | 0.028 | 0.028 | 0 | 0.21 | 0.12 | 0.09 | 0.09 | 0.16 | 0.09 |
| 3 | 148392710 | rs275667 | 0.555 | -2.518 | -2.213 | -1.566 | 0.062 | 0.087 | 0.087 | 0.462 | 0.6 | 0.3 | 0.82 | 0.89 | 0.8 | 0.63 |
| 3 | 148424143 | rs2131127 | -0.514 | -1.442 | -2.417 | -0.692 | 0.03 | -0.014 | -0.014 | 0 | 0.41 | 0.72 | 0.36 | 0.36 | 0.33 | 0.42 |
| 3 | 148430767 | rs931490 | -0.951 | -1.864 | -2.087 | -1.63 | -0.015 | 0.034 | 0.034 | 0 | 0.79 | 0.97 | 0.92 | 0.92 | 0.83 | 0.84 |
| 3 | 148432469 | rs4681443 | -0.863 | -1.506 | -1.102 | -1.133 | 0.024 | 0.031 | 0.031 | 0.343 | 0.3 | 0.68 | 0.18 | 0.06 | 0.18 | 0.24 |
| 3 | 148432574 | rs4681444 | -0.89 | -1.489 | -1.103 | -1.134 | 0.025 | 0.033 | 0.033 | 0.244 | 0.3 | 0.68 | 0.18 | 0.06 | 0.18 | 0.24 |
| 3 | 148438191 | rs3772616 | -1.149 | -0.946 | -1.059 | -0.749 | 0.04 | 0.065 | 0.065 | 0.032 | 0.32 | 0.8 | 0.28 | 0.2 | 0.18 | 0.33 |
| 3 | 148439788 | rs4524238 | -0.867 | -0.623 | -0.733 | -0.711 | 0 | -0.008 | -0.008 | 0.833 | 0.26 | 0.69 | 0.27 | 0.13 | 0.18 | 0.28 |
| 3 | 148448418 | rs12695902 | -0.723 | -0.017 | -0.746 | -0.479 | -0.001 | -0.013 | -0.013 | 0 | 0.11 | 0.08 | 0.07 | 0 | 0.1 | 0.04 |
| 3 | 148455537 | rs6801836 | -0.883 | -0.395 | -0.51 | -0.779 | -0.007 | -0.03 | -0.03 | 0.086 | 0.33 | 0.4 | 0.17 | 0.1 | 0.22 | 0.14 |
| 3 | 148457642 | rs1800766 | -0.461 | -0.541 | -0.721 | -0.775 | 0.026 | 0.008 | 0.008 | 0 | 0.25 | 0.08 | 0.09 | 0.16 | 0.14 | 0.13 |
| 3 | 148459884 | rs5183 | -0.966 | -0.727 | 0.205 | -1.189 | 0.008 | -0.036 | -0.036 | 0 | 0.08 | 0.32 | 0.03 | 0 | 0.05 | 0.03 |
| 3 | 148459988 | rs5186 | 0.288 | 0.453 | -0.486 | -1.737 | -0.082 | 0.048 | 0.048 | 0 | 0.15 | 0.02 | 0.23 | 0.06 | 0.27 | 0.07 |
| 3 | 148464454 | rs275645 | 1.261 | -0.49 | -0.61 | -2.173 | -0.014 | 0.049 | 0.049 | 0.127 | 0.4 | 0.26 | 0.57 | 0.75 | 0.47 | 0.55 |
| 3 | 148465540 | rs275644 | -0.633 | -0.753 | -0.585 | -2.147 | 0.013 | -0.008 | -0.008 | 0 | 0.9 | 0.94 | 0.97 | 0.83 | 0.95 | 0.77 |
| 3 | 148466363 | rs427832 | 1.319 | -0.781 | -0.376 | -2.263 | -0.005 | 0.027 | 0.027 | 0 | 0.45 | 0.34 | 0.63 | 0.76 | 0.55 | 0.57 |
| 3 | 148468746 | rs13097326 | 0.806 | -0.593 | -0.019 | -2.12 | -0.079 | 0.047 | 0.047 | 0 | 0.15 | 0.02 | 0.23 | 0.06 | 0.26 | 0.06 |
| 3 | 148469590 | rs10513338 | -0.062 | -0.694 | 0.119 | -2.066 | 0.037 | -0.012 | -0.012 | 0 | 0.07 | 0.39 | 0.04 | 0 | 0.05 | 0.03 |
| 3 | 148469627 | rs2675513 | -0.613 | -0.679 | -0.246 | -2.045 | 0.012 | -0.002 | -0.002 | 0.742 | 0.9 | 0.92 | 0.97 | 0.85 | 0.95 | 0.77 |
| 3 | 148476957 | rs1492081 | 0.847 | -0.773 | 0.199 | -2.091 | -0.061 | -0.031 | -0.031 | 0 | 0.81 | 0.31 | 0.92 | 0.85 | 0.92 | 0.74 |
| 3 | 148477713 | rs17713681 | 1.131 | -0.668 | 0.129 | -2.148 | -0.001 | 0.002 | 0.002 | 0.082 | 0.66 | 0.3 | 0.68 | 0.78 | 0.64 | 0.67 |
| 3 | 148481248 | rs6772419 | -0.876 | -0.267 | 0.116 | -2.206 | 0.047 | -0.073 | -0.073 | 0 | 0.12 | 0.45 | 0.09 | 0 | 0.13 | 0.09 |
| 3 | 148483203 | rs16860871 | 0.308 | -0.33 | 0.226 | -2.178 | 0.022 | -0.018 | -0.018 | 0.056 | 0.06 | 0.29 | 0.04 | 0 | 0.05 | 0.03 |
| 3 | 148485532 | rs2934084 | -0.741 | -0.572 | 0.098 | -2.258 | 0.012 | -0.006 | -0.006 | 0 | 0.9 | 0.94 | 0.98 | 0.84 | 0.97 | 0.77 |
| 3 | 148490358 | rs2934086 | 0.661 | -0.711 | -0.137 | -2.487 | -0.027 | -0.048 | -0.048 | 0.007 | 0.58 | 0.47 | 0.83 | 0.81 | 0.76 | 0.59 |
| 3 | 148496554 | rs11718189 | 0.534 | -0.494 | -0.364 | -2.405 | -0.008 | 0.081 | 0.081 | 0 | 0.25 | 0.13 | 0.13 | 0.04 | 0.21 | 0.15 |
| 3 | 148504871 | rs9822689 | -0.074 | 0.098 | 0.127 | -1.97 | -0.04 | 0.048 | 0.048 | 0 | 0.21 | 0.14 | 0.26 | 0.06 | 0.31 | 0.1 |
| 3 | 148506993 | rs2934128 | 1.574 | -0.345 | 0.339 | -2.18 | -0.016 | 0.102 | 0.102 | 0 | 0.57 | 0.22 | 0.63 | 0.78 | 0.56 | 0.6 |
| 3 | 148509827 | rs13099960 | 0.431 | -0.385 | 0.3 | -2.187 | 0.014 | -0.004 | -0.004 | 0 | 0.11 | 0.06 | 0.02 | 0.15 | 0.03 | 0.23 |
| 3 | 148510503 | rs1492087 | -0.859 | -0.366 | 0.452 | -2.227 | 0.002 | -0.006 | -0.006 | 1.526 | 0.83 | 0.85 | 0.76 | 0.94 | 0.73 | 0.93 |
| 3 | 148510797 | rs6803439 | -1.065 | -0.448 | 0.711 | -2 | -0.011 | -0.01 | -0.01 | 0 | 0.18 | 0.21 | 0.08 | 0.16 | 0.13 | 0.28 |
| 3 | 148513931 | rs11710606 | -1.959 | -0.406 | 0.438 | -2.078 | -0.007 | -0.016 | -0.016 | 0 | 0.28 | 0.68 | 0.14 | 0.16 | 0.19 | 0.33 |
| 3 | 148515005 | rs10935726 | -1.457 | -0.215 | 0.433 | -1.93 | -0.036 | -0.046 | -0.046 | 0 | 0.21 | 0.51 | 0.11 | 0.16 | 0.14 | 0.29 |
| 3 | 148531358 | rs11918693 | -0.878 | 0.255 | -0.097 | -1.899 | -0.003 | -0.007 | -0.007 | 1.342 | 0.09 | 0.09 | 0.07 | 0 | 0.11 | 0.06 |
| 3 | 148538324 | rs11711897 | -0.12 | 0.168 | 0.114 | -1.829 | 0.01 | -0.027 | -0.027 | 0 | 0.06 | 0.15 | 0.04 | 0 | 0.06 | 0.04 |
| 3 | 148539284 | rs4681158 | 0.438 | -0.002 | -0.23 | -1.454 | 0 | -0.001 | -0.001 | 0.184 | 0.57 | 0.37 | 0.43 | 0.66 | 0.34 | 0.65 |
| 3 | 148540154 | rs9849625 | 1.472 | 0.582 | -0.635 | -0.634 | -0.07 | -0.018 | -0.018 | 0.162 | 0.58 | 0.09 | 0.56 | 0.66 | 0.55 | 0.6 |
| 3 | 148543411 | rs1352897 | -1.444 | 1.214 | -0.379 | -0.455 | -0.064 | 0.029 | 0.029 | 0.365 | 0.89 | 0.98 | 0.77 | 0.96 | 0.76 | 0.9 |
| 3 | 148546001 | rs1492094 | 0.78 | 1.2 | -0.188 | -0.554 | 0.073 | -0.094 | -0.094 | 0 | 0.75 | 0.37 | 0.65 | 0.96 | 0.6 | 0.83 |
| 3 | 148547435 | rs9858818 | -0.675 | 1.249 | -0.091 | -0.541 | 0.113 | -0.062 | -0.062 | 0.716 | 0.85 | 0.65 | 0.71 | 0.96 | 0.69 | 0.88 |
| 3 | 148547780 | rs9821558 | 0.569 | 1.156 | -0.756 | -0.402 | 0.051 | 0.047 | 0.047 | 0.243 | 0.73 | 0.35 | 0.61 | 0.67 | 0.63 | 0.64 |
| 3 | 148548024 | rs11717064 | -0.34 | 1.378 | -1.629 | -0.238 | 0.039 | -0.048 | -0.048 | 1.847 | 0.1 | 0.38 | 0.06 | 0 | 0.1 | 0.05 |
| 3 | 148550522 | rs7617340 | 0.636 | 0.803 | -2.707 | 0.266 | -0.06 | -0.007 | -0.007 | 0 | 0.26 | 0.11 | 0.28 | 0.33 | 0.36 | 0.25 |
| 3 | 148550827 | rs11711637 | 0.832 | 0.685 | -2.803 | 0.278 | 0.094 | 0.014 | 0.014 | 0.831 | 0.42 | 0.08 | 0.31 | 0.33 | 0.28 | 0.41 |
| 3 | 148551733 | rs9842791 | -0.104 | 1.327 | -2.053 | 0.32 | -0.073 | -0.014 | -0.014 | 0 | 0.27 | 0.09 | 0.28 | 0.33 | 0.36 | 0.27 |
| 3 | 148553480 | rs13066553 | 0.308 | 0.753 | -2.152 | 0.081 | 0.015 | 0.1 | 0.1 | 0 | 0.4 | 0.14 | 0.28 | 0.2 | 0.36 | 0.29 |
| 3 | 148553560 | rs1318579 | -0.239 | 0.347 | -1.536 | 0.586 | -0.012 | -0.008 | -0.008 | 0.317 | 0.41 | 0.3 | 0.5 | 0.6 | 0.51 | 0.56 |
| 3 | 148554668 | rs3772602 | 0.975 | 0.794 | -1.787 | 1.159 | 0.004 | 0.152 | 0.152 | 0.166 | 0.47 | 0.14 | 0.43 | 0.23 | 0.45 | 0.34 |

| | | | | | | | | | | | | | | | | |
|---|---|---|---|---|---|---|---|---|---|---|---|---|---|---|---|---|
| 3 | 148554759 | rs3772601 | 0.401 | 1.437 | -1.938 | 0.644 | -0.009 | 0.008 | 0.008 | 0 | 0.06 | 0.03 | 0.03 | 0 | 0.08 | 0.08 |
| 3 | 148558051 | rs7639856 | 0.493 | 1.324 | -1.526 | 0.5 | 0.007 | -0.052 | -0.052 | 0.05 | 0.1 | 0.18 | 0.03 | 0 | 0.09 | 0.02 |
| 3 | 148558447 | rs2291671 | -0.228 | 1.306 | -0.837 | 1.108 | 0.037 | 0.05 | 0.05 | 0 | 0.16 | 0.35 | 0.14 | 0.28 | 0.2 | 0.28 |
| 3 | 148560974 | rs1492104 | -1.137 | 1.267 | -0.866 | 1.127 | 0.001 | 0.009 | 0.009 | 0 | 0.32 | 0.36 | 0.32 | 0.55 | 0.35 | 0.42 |
| 3 | 148562310 | rs1059502 | -0.454 | 1.303 | -0.447 | 0.734 | 0.037 | 0.057 | 0.057 | 0 | 0.16 | 0.35 | 0.14 | 0.3 | 0.2 | 0.3 |
| 3 | 148571953 | rs1850288 | -0.153 | 0.96 | -0.447 | 0.409 | -0.021 | -0.011 | -0.011 | 0 | 0.79 | 0.83 | 0.68 | 0.74 | 0.66 | 0.55 |
| 3 | 148575156 | rs2331407 | 1.007 | 0.465 | -0.053 | 0.403 | -0.011 | -0.035 | -0.035 | 0 | 0.64 | 0.21 | 0.56 | 0.52 | 0.59 | 0.51 |
| 3 | 148576599 | rs4521178 | 0.434 | 0.466 | 0.271 | 0.408 | 0.008 | 0.017 | 0.017 | 0 | 0.72 | 0.68 | 0.65 | 0.57 | 0.64 | 0.53 |
| 3 | 148578446 | rs2001988 | -0.297 | 0.474 | 0.25 | 0.408 | 0.004 | 0.01 | 0.01 | 0 | 0.26 | 0.28 | 0.34 | 0.43 | 0.34 | 0.45 |
| 3 | 148578728 | rs9826662 | 1.394 | -0.07 | -0.145 | -0.173 | -0.01 | -0.02 | -0.02 | 0.135 | 0.44 | 0.23 | 0.49 | 0.52 | 0.46 | 0.39 |
| 3 | 148579686 | rs1356258 | -0.713 | -0.129 | 0.089 | 0.085 | -0.004 | -0.006 | -0.006 | 0.733 | 0.34 | 0.31 | 0.36 | 0.43 | 0.38 | 0.54 |
| 3 | 148586327 | rs7633618 | 1.444 | -0.74 | -0.165 | 0.285 | 0.053 | -0.039 | -0.039 | 0 | 0.49 | 0.1 | 0.52 | 0.51 | 0.59 | 0.41 |
| 3 | 148587489 | rs6807270 | 0.447 | -0.71 | 0.149 | 0.311 | -0.001 | -0.001 | -0.001 | 0 | 0.64 | 0.33 | 0.61 | 0.66 | 0.64 | 0.46 |
| 3 | 148590428 | rs7646258 | 0.143 | -0.515 | 0.595 | 0.293 | -0.011 | -0.014 | -0.014 | 0 | 0.69 | 0.71 | 0.59 | 0.61 | 0.63 | 0.44 |
| 3 | 148597482 | rs1032465 | 0.99 | -0.531 | 0.61 | 0.275 | 0 | -0.001 | -0.001 | 6.787 | 0.58 | 0.47 | 0.55 | 0.61 | 0.61 | 0.42 |
| 3 | 148602499 | rs3772577 | 0.981 | -0.518 | 0.614 | 0.296 | 0 | -0.001 | -0.001 | 0 | 0.58 | 0.47 | 0.53 | 0.59 | 0.6 | 0.42 |
| 3 | 148606254 | rs3772575 | 0.738 | -0.48 | 0.339 | 0.307 | -0.001 | -0.001 | -0.001 | 0 | 0.21 | 0.24 | 0.3 | 0.27 | 0.28 | 0.43 |
| 3 | 148609045 | rs4681161 | 1.247 | -0.413 | 0.094 | 0.251 | -0.009 | -0.019 | -0.019 | 0 | 0.46 | 0.13 | 0.46 | 0.44 | 0.56 | 0.35 |
| 3 | 148615287 | rs3819340 | -0.359 | -0.097 | 0.277 | 0.251 | -0.008 | 0.032 | 0.032 | 0 | 0.11 | 0.4 | 0.12 | 0.09 | 0.05 | 0.1 |
| 3 | 148615695 | rs1440640 | 0.489 | 0.083 | -0.724 | 0.32 | -0.008 | -0.001 | -0.001 | 0 | 0.56 | 0.2 | 0.5 | 0.59 | 0.6 | 0.4 |
| 3 | 148624027 | rs7647772 | 0.324 | 0.06 | -0.631 | 0.327 | 0.001 | -0.006 | -0.006 | 0 | 0.58 | 0.32 | 0.53 | 0.58 | 0.61 | 0.4 |
| 3 | 148630929 | rs10804733 | -0.655 | 0.709 | -0.883 | 0.091 | -0.015 | 0.038 | 0.038 | 0 | 0.54 | 0.74 | 0.63 | 0.66 | 0.52 | 0.65 |
| 3 | 148631606 | rs7623208 | 0.383 | 0.873 | -0.683 | -0.049 | -0.009 | 0.007 | 0.007 | 0 | 0.92 | 0.96 | 0.95 | 1 | 0.88 | 0.96 |
| 3 | 148633264 | rs10935735 | -0.919 | 0.471 | -0.52 | 0.321 | -0.005 | -0.009 | -0.009 | 0 | 0.14 | 0.2 | 0.14 | 0.1 | 0.1 | 0.09 |
| 3 | 148636416 | rs9289775 | 0.647 | 0.666 | -0.524 | 0.267 | 0.006 | -0.017 | -0.017 | 0 | 0.74 | 0.66 | 0.82 | 0.87 | 0.74 | 0.82 |
| 3 | 148637771 | rs6785178 | 0.35 | 0.374 | -1.596 | 0.204 | -0.003 | 0.029 | 0.029 | 0.081 | 0.34 | 0.09 | 0.32 | 0.27 | 0.28 | 0.16 |
| 3 | 148648193 | rs9653951 | 0.098 | 0.381 | -1.687 | 0.352 | -0.044 | -0.043 | -0.043 | 0 | 0.31 | 0.22 | 0.43 | 0.46 | 0.5 | 0.58 |
| 3 | 148656318 | rs16861176 | 0.077 | 0.602 | -0.98 | 0.247 | 0.025 | -0.032 | -0.032 | 3.382 | 0.18 | 0.32 | 0.27 | 0.11 | 0.26 | 0.21 |
| 3 | 148657780 | rs9883066 | 0.478 | 0.975 | -1.061 | 0.708 | -0.037 | -0.05 | -0.05 | 1.829 | 0.15 | 0.08 | 0.18 | 0.35 | 0.25 | 0.33 |
| 3 | 148661164 | rs4681163 | 0.512 | 0.433 | -0.506 | 0.158 | 0.007 | -0.004 | -0.004 | 0.511 | 0.8 | 0.67 | 0.68 | 0.76 | 0.78 | 0.71 |
| 3 | 148661836 | rs7611664 | -0.59 | 0.132 | -0.619 | -0.057 | 0.006 | 0.008 | 0.008 | 0 | 0.27 | 0.26 | 0.16 | 0.14 | 0.21 | 0.26 |
| 3 | 148662845 | rs11710943 | 0.119 | 0.536 | -0.444 | 0.067 | 0.004 | -0.019 | -0.019 | 0 | 0.14 | 0.18 | 0.2 | 0.03 | 0.2 | 0.25 |
| 3 | 148666886 | rs16861192 | 0.291 | 0.341 | -0.366 | 0.332 | 0.007 | 0.009 | 0.009 | 0 | 0.09 | 0.1 | 0.13 | 0.18 | 0.14 | 0.12 |
| 3 | 148672002 | rs1440642 | -0.43 | 0.131 | -0.399 | 0.176 | -0.002 | -0.017 | -0.017 | 0 | 0.92 | 0.93 | 0.97 | 1 | 0.95 | 0.9 |
| 3 | 148679329 | rs6780596 | 1.291 | 0.075 | -0.393 | 0.76 | -0.001 | 0.002 | 0.002 | 0.005 | 0.56 | 0.55 | 0.4 | 0.37 | 0.45 | 0.45 |
| 3 | 148681019 | rs6806078 | 0.339 | 0.097 | -0.57 | 0.713 | 0.006 | 0.008 | 0.008 | 0 | 0.11 | 0.06 | 0.07 | 0.03 | 0.11 | 0.06 |
| 3 | 148683013 | rs2165342 | 1.82 | -0.236 | -0.184 | 0.684 | 0.016 | 0.049 | 0.049 | 0 | 0.5 | 0.31 | 0.38 | 0.36 | 0.45 | 0.44 |
| 3 | 148683203 | rs2029756 | -0.963 | -0.079 | -0.261 | 0.928 | -0.063 | -0.088 | -0.088 | 0 | 0.85 | 0.97 | 0.74 | 0.58 | 0.75 | 0.75 |
| 3 | 148686636 | rs6764973 | -1.393 | -0.369 | 0.21 | 0.911 | 0 | 0.027 | 0.027 | 0 | 0.23 | 0.56 | 0.37 | 0.29 | 0.19 | 0.25 |
| 3 | 148686894 | rs6765244 | 0.351 | -0.843 | 0.113 | 0.895 | -0.008 | -0.001 | -0.001 | 0 | 0.13 | 0.13 | 0.05 | 0.07 | 0.06 | 0.1 |
| 3 | 148693400 | rs7615371 | 1.958 | -0.487 | 0.24 | 0.659 | 0.038 | 0.052 | 0.052 | 0.008 | 0.48 | 0.22 | 0.37 | 0.36 | 0.39 | 0.26 |
| 3 | 148693509 | rs7617449 | -0.571 | -0.078 | 0.626 | 0.467 | 0.008 | -0.079 | -0.079 | 4.115 | 0.12 | 0.42 | 0.16 | 0 | 0.07 | 0.1 |
| 3 | 148694336 | rs4681473 | -1.082 | -0.007 | 0.429 | 0.543 | -0.045 | -0.053 | -0.053 | 4.649 | 0.89 | 0.97 | 0.85 | 0.7 | 0.78 | 0.71 |
| 3 | 148695855 | rs10935737 | 0.813 | -0.057 | 0.557 | 0.552 | 0.045 | 0.044 | 0.044 | 0 | 0.76 | 0.48 | 0.68 | 0.72 | 0.72 | 0.6 |
| 3 | 148698898 | rs13075612 | -1.191 | -0.197 | 0.512 | 0.405 | -0.005 | -0.009 | -0.009 | 0 | 0.33 | 0.39 | 0.32 | 0.27 | 0.33 | 0.27 |
| 3 | 148703787 | rs3732559 | -1.434 | -0.52 | 0.125 | 0.356 | 0.017 | 0.007 | 0.007 | 0.025 | 0.39 | 0.68 | 0.34 | 0.36 | 0.35 | 0.24 |
| 3 | 148708411 | rs6784845 | -1.337 | -0.368 | -0.096 | 0.418 | -0.062 | -0.066 | -0.066 | 0.055 | 0.9 | 0.99 | 0.85 | 0.7 | 0.81 | 0.71 |
| 3 | 148710801 | rs3772583 | 0.327 | -0.496 | 0.083 | 0.272 | -0.022 | -0.023 | -0.023 | 0 | 0.14 | 0.07 | 0.24 | 0.24 | 0.25 | 0.13 |
| 3 | 148728842 | rs3772573 | 3.3 | -0.55 | -0.189 | 0.224 | -0.044 | -0.079 | -0.079 | 0.518 | 0.34 | 0.07 | 0.42 | 0.4 | 0.34 | 0.18 |
| 3 | 148734744 | rs3772572 | 0.358 | -0.569 | -0.091 | -0.082 | -0.024 | -0.033 | -0.033 | 0 | 0.81 | 0.86 | 0.81 | 0.71 | 0.74 | 0.55 |
| 3 | 148736576 | rs3772569 | -2.089 | -0.72 | -0.071 | -0.156 | 0.034 | 0.046 | 0.046 | 0 | 0.43 | 0.73 | 0.37 | 0.34 | 0.38 | 0.32 |
| 3 | 148747119 | rs13068009 | -0.784 | -0.63 | 0.488 | -0.319 | 0 | 0.004 | 0.004 | 0 | 0.25 | 0.28 | 0.29 | 0.34 | 0.3 | 0.21 |
| 3 | 148748380 | rs3182285 | -1.401 | -0.634 | 0.505 | -0.304 | 0 | 0 | 0 | 0 | 0.32 | 0.28 | 0.31 | 0.34 | 0.34 | 0.27 |
| 3 | 148758135 | rs1457604 | -1.447 | -0.613 | 0.381 | -0.287 | -0.064 | -0.069 | -0.069 | 1.571 | 0.9 | 0.99 | 0.85 | 0.72 | 0.8 | 0.71 |
| 3 | 148759325 | rs2290725 | -1.413 | -0.616 | 0.438 | -0.267 | -0.001 | -0.002 | -0.002 | 0.103 | 0.32 | 0.32 | 0.31 | 0.34 | 0.34 | 0.27 |
| 3 | 148761815 | rs9864549 | 3.59 | -0.63 | 0.399 | -0.281 | -0.043 | -0.046 | -0.046 | 0 | 0.34 | 0.06 | 0.4 | 0.38 | 0.34 | 0.17 |
| 3 | 148772455 | rs7637603 | -1.574 | -0.533 | 0.557 | -0.187 | -0.007 | -0.01 | -0.01 | 0 | 0.33 | 0.43 | 0.31 | 0.34 | 0.34 | 0.27 |
| 3 | 148776611 | rs9850529 | 2.843 | -0.516 | 0.24 | -0.264 | -0.002 | 0.008 | 0.008 | 0.41 | 0.4 | 0.12 | 0.44 | 0.38 | 0.37 | 0.23 |
| 3 | 148786085 | rs2305868 | 1.477 | -0.424 | 0.312 | -0.463 | 0.028 | 0.029 | 0.029 | 0 | 0.07 | 0 | 0.03 | 0.03 | 0.03 | 0.02 |
| 3 | 148820658 | rs7647629 | 0.926 | -0.168 | 0.291 | -0.396 | 0.004 | -0.026 | -0.026 | 0 | 0.07 | 0.1 | 0.05 | 0 | 0.06 | 0.14 |
| 3 | 148835753 | rs6788799 | -0.858 | -0.368 | 0.37 | -0.408 | -0.001 | -0.002 | -0.002 | 0 | 0.3 | 0.28 | 0.26 | 0.25 | 0.28 | 0.26 |
| 3 | 148836573 | rs6792168 | -1.613 | -0.307 | 0.29 | -0.505 | 0.024 | 0.032 | 0.032 | 0.401 | 0.4 | 0.71 | 0.3 | 0.27 | 0.32 | 0.34 |
| 3 | 148837122 | rs772805 | 2.497 | -0.217 | 0.105 | -0.43 | 0.002 | -0.017 | -0.017 | 0.011 | 0.41 | 0.11 | 0.45 | 0.44 | 0.38 | 0.24 |
| 3 | 148845710 | rs10935741 | -0.526 | -0.216 | 0.213 | -0.555 | -0.018 | -0.033 | -0.033 | 0 | 0.19 | 0.26 | 0.1 | 0.1 | 0.14 | 0.13 |
| 3 | 148846090 | rs6805170 | 0.355 | -0.19 | 0.256 | -0.507 | -0.009 | -0.005 | -0.005 | 0.604 | 0.13 | 0.26 | 0.09 | 0.1 | 0.1 | 0.07 |
| 3 | 148852455 | rs4681169 | -1.411 | -0.199 | 0.246 | -0.514 | -0.012 | -0.066 | -0.066 | 1.252 | 0.33 | 0.54 | 0.32 | 0.17 | 0.31 | 0.41 |
| 3 | 148853477 | rs4681487 | -1.988 | -0.176 | 0.001 | -0.458 | -0.007 | -0.005 | -0.005 | 0.008 | 0.46 | 0.68 | 0.48 | 0.46 | 0.51 | 0.69 |
| 3 | 148854505 | rs17786967 | 1.864 | -0.687 | 0.158 | -0.542 | -0.016 | -0.026 | -0.026 | 0 | 0.06 | 0.09 | 0.03 | 0 | 0.02 | 0.05 |
| 3 | 148863151 | rs11718908 | -0.5 | -0.6 | 0.234 | -0.504 | -0.036 | -0.043 | -0.043 | 0 | 0.25 | 0.45 | 0.24 | 0.17 | 0.2 | 0.26 |

| | | | | | | | | | | | | | | | | |
|---|---|---|---|---|---|---|---|---|---|---|---|---|---|---|---|---|
| 3 | 148863395 | rs11715291 | -0.5 | -0.6 | 0.234 | -0.504 | -0.036 | -0.043 | -0.043 | 0 | 0.25 | 0.45 | 0.24 | 0.17 | 0.2 | 0.26 |
| 3 | 148865528 | rs2689224 | -1.519 | -0.56 | 0.172 | -0.454 | 0 | 0.01 | 0.01 | 0.831 | 0.38 | 0.59 | 0.4 | 0.45 | 0.4 | 0.58 |
| 3 | 148866675 | rs12487869 | -0.432 | -0.565 | 0.197 | -0.455 | -0.033 | -0.039 | -0.039 | 0 | 0.25 | 0.45 | 0.24 | 0.17 | 0.2 | 0.26 |
| 3 | 148866708 | rs2689225 | -1.971 | -0.571 | 0.097 | -0.474 | -0.003 | -0.004 | -0.004 | 0.26 | 0.44 | 0.67 | 0.44 | 0.45 | 0.46 | 0.71 |
| 3 | 148867126 | rs17787392 | 0.349 | -0.537 | 0.061 | -0.326 | -0.008 | -0.007 | -0.007 | 0 | 0.19 | 0.27 | 0.2 | 0.17 | 0.17 | 0.22 |
| 3 | 148867141 | rs2689227 | -0.164 | -0.465 | -0.24 | -0.291 | -0.006 | -0.002 | -0.002 | 0 | 0.13 | 0.14 | 0.16 | 0.27 | 0.2 | 0.32 |
| 3 | 148868247 | rs2254913 | -1.892 | -0.344 | -0.566 | -0.215 | -0.059 | -0.073 | -0.073 | 0 | 0.61 | 0.9 | 0.53 | 0.54 | 0.59 | 0.81 |
| 3 | 148872987 | rs6440589 | -1.564 | -0.03 | 0.308 | -0.097 | -0.01 | 0.006 | 0.006 | 0.596 | 0.44 | 0.7 | 0.35 | 0.36 | 0.43 | 0.5 |
| 3 | 148874252 | rs6785780 | -1.277 | 0.17 | -0.189 | -0.008 | 0.006 | 0 | 0 | 0.161 | 0.41 | 0.49 | 0.33 | 0.37 | 0.42 | 0.5 |
| 3 | 148881673 | rs3732557 | -0.407 | 0.479 | -0.55 | 0.797 | -0.047 | -0.036 | -0.036 | 0 | 0.08 | 0.01 | 0.14 | 0.27 | 0.18 | 0.21 |
| 3 | 148896820 | rs16861588 | -0.565 | 0.18 | -0.522 | 0.795 | -0.046 | -0.034 | -0.034 | 0 | 0.08 | 0.01 | 0.13 | 0.28 | 0.18 | 0.21 |
| 3 | 148900046 | rs16861598 | -1.306 | -0.627 | 0.485 | 0.011 | 0.057 | 0.029 | 0.029 | 7.133 | 0.06 | 0.51 | 0.05 | 0 | 0.02 | 0.01 |
| 3 | 148902714 | rs701755 | -0.378 | -0.903 | -0.085 | -0.608 | -0.011 | 0.021 | 0.021 | 0 | 0.17 | 0.11 | 0.17 | 0 | 0.17 | 0.19 |
| 3 | 148903923 | rs4974389 | 1.62 | -1.967 | -0.95 | -0.672 | -0.04 | -0.065 | -0.065 | 0.109 | 0.4 | 0.1 | 0.53 | 0.58 | 0.56 | 0.39 |
| 3 | 148906027 | rs6799507 | -0.54 | -1.749 | 0.148 | -0.898 | 0.072 | 0.044 | 0.044 | 5.8 | 0.06 | 0.45 | 0.04 | 0 | 0.02 | 0.01 |
| 3 | 148912996 | rs773050 | -0.505 | -2.172 | -0.042 | -1.096 | -0.053 | -0.071 | -0.071 | 2.506 | 0.85 | 0.79 | 0.97 | 1 | 0.96 | 0.88 |
| 3 | 148913126 | rs13072552 | -0.79 | -2.08 | 0.133 | -1.066 | -0.03 | -0.083 | -0.083 | 0 | 0.13 | 0.48 | 0.09 | 0 | 0.07 | 0.03 |
| 3 | 148923656 | rs772908 | 0.015 | -2.196 | -0.453 | -1.203 | -0.002 | -0.022 | -0.022 | 0 | 0.23 | 0.2 | 0.21 | 0.01 | 0.25 | 0.27 |
| 3 | 148931980 | rs11709714 | 1.861 | -1.799 | -0.755 | -0.874 | -0.072 | -0.055 | -0.055 | 0 | 0.4 | 0.08 | 0.57 | 0.54 | 0.51 | 0.4 |
| 3 | 148938950 | rs7652826 | 1.485 | -1.788 | -0.708 | -1.11 | -0.024 | 0.03 | 0.03 | 0.054 | 0.47 | 0.07 | 0.59 | 0.62 | 0.56 | 0.41 |
| 3 | 148939861 | rs17838831 | -0.42 | -1.779 | -0.384 | -0.89 | 0.002 | 0.059 | 0.059 | 0 | 0.12 | 0.16 | 0.2 | 0.31 | 0.15 | 0.21 |
| 3 | 148941958 | rs7623663 | 0.134 | -1.802 | -0.404 | -0.963 | 0.004 | 0.064 | 0.064 | 0 | 0.09 | 0.11 | 0.17 | 0.27 | 0.11 | 0.19 |
| 3 | 148947188 | rs16861645 | 0.123 | -2.084 | 0.098 | -1.722 | -0.066 | -0.089 | -0.089 | 0.132 | 0.16 | 0.25 | 0.04 | 0 | 0.04 | 0.13 |
| 3 | 148950780 | rs16861650 | 0.225 | -2.064 | -0.003 | -1.713 | -0.046 | -0.066 | -0.066 | 0 | 0.15 | 0.19 | 0.03 | 0 | 0.04 | 0.14 |
| 3 | 148957233 | rs701741 | -1.066 | -2.132 | 0.056 | -1.845 | -0.017 | -0.097 | -0.097 | 0.358 | 0.37 | 0.38 | 0.16 | 0.07 | 0.26 | 0.38 |
| 3 | 148958367 | rs9844628 | -0.422 | -1.565 | -0.52 | -1.563 | -0.003 | -0.067 | -0.067 | 0 | 0.23 | 0.25 | 0.14 | 0.07 | 0.22 | 0.24 |
| 3 | 148962041 | rs9827666 | -0.25 | -1.503 | -0.966 | -1.536 | 0 | -0.014 | -0.014 | 0 | 0.22 | 0.17 | 0.12 | 0.07 | 0.21 | 0.24 |
| 3 | 148964692 | rs7618020 | -0.325 | -1.509 | -0.968 | -1.536 | 0.001 | -0.013 | -0.013 | 0 | 0.22 | 0.18 | 0.13 | 0.07 | 0.21 | 0.24 |
| 3 | 148966357 | rs35969781 | 0.189 | -1.162 | -1.401 | -1.355 | -0.003 | 0.039 | 0.039 | 0 | 0.07 | 0 | 0.03 | 0 | 0.04 | 0.02 |
| 3 | 148966687 | rs6777996 | -1.127 | -1.677 | -1.683 | -1.684 | 0.085 | 0.397 | 0.397 | 0 | 0.51 | 0.93 | 0.24 | 0.08 | 0.33 | 0.41 |
| 3 | 148967335 | rs7624706 | -1.066 | -0.804 | -0.265 | -1.065 | 0.01 | -0.105 | -0.105 | 0.447 | 0.18 | 0.41 | 0.13 | 0.05 | 0.16 | 0.16 |
| 3 | 148973465 | rs13072463 | -0.257 | -0.836 | -0.81 | -1.458 | 0.015 | 0.02 | 0.02 | 0 | 0.4 | 0.69 | 0.25 | 0.06 | 0.35 | 0.37 |
| 3 | 148974301 | rs951776 | -0.366 | -0.755 | -1.6 | -0.367 | -0.014 | -0.094 | -0.094 | 0.001 | 0.87 | 0.99 | 0.89 | 0.62 | 0.88 | 0.73 |
| 3 | 148977314 | rs7617219 | -0.488 | -0.267 | -1.43 | -0.316 | 0.038 | 0 | 0 | 0.504 | 0.49 | 0.9 | 0.57 | 0.45 | 0.42 | 0.47 |
| 3 | 148988153 | rs7613093 | -0.934 | 0.058 | 0.056 | 0.167 | 0.035 | 0.031 | 0.031 | 0.167 | 0.07 | 0.42 | 0.07 | 0.03 | 0.04 | 0.06 |
| 3 | 148993079 | rs7633905 | -0.73 | 0.414 | -1.469 | -0.027 | -0.043 | -0.011 | -0.011 | 0 | 0.51 | 0.94 | 0.66 | 0.46 | 0.47 | 0.51 |
| 3 | 148993158 | rs11919380 | -0.789 | 0.265 | -0.735 | 0.146 | 0.003 | -0.005 | -0.005 | 0 | 0.31 | 0.74 | 0.53 | 0.37 | 0.32 | 0.26 |
| 3 | 148993230 | rs7634024 | -0.249 | 0.28 | -0.162 | 0.215 | 0.03 | 0.114 | 0.114 | 0.884 | 0.05 | 0.15 | 0.19 | 0.15 | 0.08 | 0.09 |
| 3 | 148998726 | rs12152496 | -0.965 | 0.289 | -0.551 | 0.054 | 0.009 | 0.013 | 0.013 | 0.219 | 0.29 | 0.73 | 0.4 | 0.19 | 0.26 | 0.18 |
| 3 | 149000201 | rs6788873 | -1.426 | 0.791 | -0.038 | 0.72 | 0.04 | 0.15 | 0.15 | 0.854 | 0.13 | 0.37 | 0.45 | 0.37 | 0.22 | 0.19 |
| 3 | 149009839 | rs6787602 | 0.064 | 0.889 | -0.848 | -0.311 | 0.006 | -0.072 | -0.072 | 0 | 0.32 | 0.44 | 0.38 | 0.14 | 0.4 | 0.26 |
| 3 | 149014143 | rs6783464 | -0.162 | 0.327 | -0.768 | -0.513 | 0.002 | -0.003 | -0.003 | 0 | 0.15 | 0.19 | 0.09 | 0.03 | 0.1 | 0.17 |
| 3 | 149014360 | rs2030612 | -0.073 | 0.735 | -1.099 | 0.003 | -0.005 | -0.011 | -0.011 | 0.332 | 0.75 | 0.73 | 0.79 | 0.8 | 0.78 | 0.92 |
| 3 | 149019086 | rs1526787 | 0.085 | 1.442 | -1.681 | 0.335 | -0.061 | -0.069 | -0.069 | 0.477 | 0.72 | 0.92 | 0.62 | 0.42 | 0.65 | 0.46 |
| 3 | 149022622 | rs12487547 | 0.936 | 1.2 | -1.318 | -0.671 | -0.004 | -0.055 | -0.055 | 0 | 0.33 | 0.39 | 0.35 | 0.03 | 0.29 | 0.15 |
| 3 | 149022765 | rs1908052 | -0.243 | 1.324 | -1.238 | -0.623 | -0.029 | 0.075 | 0.075 | 0 | 0.65 | 0.83 | 0.55 | 0.16 | 0.56 | 0.26 |
| 3 | 149025861 | rs13084366 | 0.688 | 1.578 | -1.579 | -0.802 | -0.051 | -0.008 | -0.008 | 0 | 0.12 | 0.01 | 0.08 | 0.11 | 0.14 | 0.08 |
| 3 | 149026545 | rs9917642 | 0.325 | 1.306 | -1.086 | -0.955 | -0.02 | -0.111 | -0.111 | 0 | 0.19 | 0.4 | 0.1 | 0.03 | 0.11 | 0.04 |
| 3 | 149033096 | rs9831865 | 0.423 | 1.735 | -0.756 | -0.711 | 0.063 | 0.128 | 0.128 | 0 | 0.1 | 0.27 | 0.15 | 0.28 | 0.18 | 0.22 |
| 3 | 149036714 | rs13353411 | -1.437 | 1.441 | -1.113 | -0.934 | 0.01 | 0.032 | 0.032 | 0.298 | 0.71 | 0.77 | 0.67 | 0.98 | 0.72 | 0.87 |
| 3 | 149040646 | rs16861768 | 0.222 | 1.221 | -0.763 | -0.69 | -0.003 | 0.05 | 0.05 | 0 | 0.15 | 0.06 | 0.07 | 0.03 | 0.11 | 0.04 |
| 3 | 149042363 | rs2291603 | -0.05 | 1.196 | -0.539 | -0.568 | 0 | -0.02 | -0.02 | 0 | 0.84 | 0.77 | 0.9 | 0.89 | 0.86 | 0.92 |
| 3 | 149043131 | rs17787898 | -0.336 | 1.62 | -1.054 | -0.508 | -0.051 | 0.081 | 0.081 | 0 | 0.18 | 0.05 | 0.13 | 0.04 | 0.21 | 0.07 |
| 3 | 149045840 | rs7634903 | -0.111 | 0.678 | -0.764 | 0.338 | -0.004 | 0.017 | 0.017 | 0 | 0.17 | 0.21 | 0.14 | 0.31 | 0.15 | 0.18 |
| 3 | 149050925 | rs1526786 | 0.903 | 0.542 | -0.598 | 0.068 | -0.001 | 0 | 0 | 0 | 0.57 | 0.43 | 0.71 | 0.49 | 0.64 | 0.65 |
| 3 | 149057323 | rs17195589 | 1.638 | 0.533 | -1.074 | -0.561 | -0.049 | 0.094 | 0.094 | 0 | 0.28 | 0.13 | 0.45 | 0.04 | 0.37 | 0.21 |
| 3 | 149058025 | rs4681171 | 1.358 | 0.317 | -1.419 | -0.155 | -0.026 | -0.073 | -0.073 | 0.849 | 0.11 | 0.01 | 0.15 | 0.4 | 0.09 | 0.36 |
| 3 | 149058404 | rs7623948 | 0.917 | 0.198 | -0.95 | -0.369 | -0.001 | -0.027 | -0.027 | 1.411 | 0.4 | 0.56 | 0.54 | 0.08 | 0.47 | 0.24 |
| 3 | 149060020 | rs7618084 | 0.933 | -0.029 | -1.116 | -0.289 | 0.011 | -0.013 | -0.013 | 0.048 | 0.16 | 0.14 | 0.13 | 0.34 | 0.13 | 0.2 |
| 3 | 149061297 | rs11916903 | 1.323 | -0.043 | -0.468 | -0.326 | 0.045 | -0.005 | -0.005 | 0 | 0.06 | 0.16 | 0.05 | 0.03 | 0.07 | 0.02 |
| 3 | 149062515 | rs7622141 | -0.256 | -0.166 | -0.343 | -0.325 | -0.035 | 0.077 | 0.077 | 0 | 0.24 | 0.38 | 0.16 | 0.34 | 0.16 | 0.21 |
| 3 | 149064800 | rs9875422 | -0.549 | -0.175 | -0.619 | -0.445 | -0.02 | -0.028 | -0.028 | 0 | 0.39 | 0.55 | 0.36 | 0.74 | 0.31 | 0.62 |
| 3 | 149066126 | rs7632041 | 0.369 | -0.211 | -0.652 | -0.537 | -0.001 | -0.002 | -0.002 | 0 | 0.59 | 0.65 | 0.66 | 0.42 | 0.6 | 0.43 |
| 3 | 149066998 | rs7623431 | 0.838 | -0.19 | -0.819 | -0.466 | -0.001 | 0.018 | 0.018 | 0 | 0.17 | 0.21 | 0.13 | 0.34 | 0.13 | 0.22 |
| 3 | 149070226 | rs977333 | -0.762 | -0.069 | -0.981 | -0.538 | -0.068 | -0.001 | -0.001 | 0 | 0.8 | 0.98 | 0.89 | 0.82 | 0.77 | 0.83 |
| 3 | 149072123 | rs6440601 | -0.872 | -0.301 | -0.753 | -0.482 | 0.015 | 0.024 | 0.024 | 0 | 0.43 | 0.61 | 0.39 | 0.74 | 0.34 | 0.64 |
| 3 | 149074545 | rs6786289 | 0.797 | -0.268 | -1.082 | -0.232 | 0.017 | -0.037 | -0.037 | 0 | 0.16 | 0.11 | 0.12 | 0.37 | 0.13 | 0.21 |
| 3 | 149084352 | rs13314474 | 0.69 | -0.631 | -0.941 | -0.41 | -0.003 | 0.015 | 0.015 | 0 | 0.51 | 0.4 | 0.59 | 0.42 | 0.53 | 0.4 |
| 3 | 149087404 | rs8479 | -0.253 | -0.899 | -1.015 | -0.381 | 0.004 | 0.01 | 0.01 | 0 | 0.8 | 0.82 | 0.91 | 0.86 | 0.83 | 0.87 |

| | | | | | | | | | | | | | | | |
|---|---|---|---|---|---|---|---|---|---|---|---|---|---|---|---|
| 3 | 149088190 | rs9842403 | 0.766 | -0.951 | -1.129 | -0.333 | 0.029 | -0.06 | -0.06 | 0.63 | 0.17 | 0.14 | 0.11 | 0.38 | 0.13 | 0.22 |
| 3 | 149090724 | rs6440603 | 0.087 | -0.995 | -0.921 | -0.395 | 0.009 | -0.015 | -0.015 | 0 | 0.22 | 0.26 | 0.14 | 0.38 | 0.15 | 0.24 |
| 3 | 149090907 | rs6440605 | -0.186 | -0.66 | -1.212 | -0.932 | 0.002 | 0.005 | 0.005 | 0 | 0.8 | 0.84 | 0.91 | 0.86 | 0.83 | 0.87 |
| 3 | 149097084 | rs4681502 | -0.719 | -0.543 | -1.48 | -1.144 | 0.01 | 0.112 | 0.112 | 0 | 0.48 | 0.7 | 0.43 | 0.89 | 0.45 | 0.71 |
| 3 | 149099575 | rs10935749 | -0.133 | -0.741 | -2.147 | -0.758 | -0.045 | -0.032 | -0.032 | 0 | 0.34 | 0.08 | 0.36 | 0.69 | 0.35 | 0.59 |
| 3 | 149101811 | rs2004107 | -0.719 | -0.743 | -2.375 | -1.133 | -0.035 | 0.051 | 0.051 | 0 | 0.82 | 0.99 | 0.92 | 0.94 | 0.83 | 0.89 |
| 3 | 149104849 | rs2059991 | 2.036 | -0.944 | -2.065 | -1.201 | -0.016 | 0.026 | 0.026 | 0 | 0.28 | 0.21 | 0.41 | 0.06 | 0.36 | 0.17 |
| 3 | 149112810 | rs10935751 | 0.991 | -1.565 | -2.333 | -0.929 | 0.1 | -0.03 | -0.03 | 0 | 0.53 | 0.15 | 0.75 | 0.62 | 0.68 | 0.62 |
| 3 | 149114614 | rs976052 | -1.047 | -1.485 | -2.358 | -0.899 | -0.012 | 0.029 | 0.029 | 0.041 | 0.85 | 0.99 | 0.93 | 0.94 | 0.88 | 0.9 |
| 3 | 149121095 | rs4317046 | 0.602 | -1.136 | -1.808 | -0.849 | 0.06 | 0.002 | 0.002 | 0.38 | 0.53 | 0.3 | 0.76 | 0.63 | 0.68 | 0.61 |
| 3 | 149121931 | rs1304026 | 0.514 | -0.511 | -1.789 | -0.409 | -0.002 | -0.028 | -0.028 | 0.357 | 0.12 | 0.07 | 0.05 | 0.18 | 0.12 | 0.14 |
| 3 | 149122161 | rs12638370 | 0.115 | -0.236 | -1.671 | -0.46 | -0.053 | -0.131 | -0.131 | 0 | 0.16 | 0.02 | 0.29 | 0.57 | 0.18 | 0.41 |
| 3 | 149134372 | rs6766558 | -0.546 | -0.136 | -1.08 | -1.769 | -0.003 | -0.021 | -0.021 | 0 | 0.69 | 0.7 | 0.53 | 0.92 | 0.55 | 0.71 |
| 3 | 149141886 | rs9869237 | -0.226 | -1.055 | -1.064 | -2.102 | -0.016 | 0.074 | 0.074 | 0.22 | 0.58 | 0.25 | 0.46 | 0.85 | 0.47 | 0.69 |
| 3 | 149144558 | rs1574462 | -0.008 | -0.98 | -0.614 | -1.902 | 0.036 | 0.084 | 0.084 | 0 | 0.07 | 0.32 | 0.05 | 0.1 | 0.06 | 0.04 |
| 3 | 149145414 | rs881950 | 0.707 | -1.01 | -1.495 | -2.098 | 0.008 | 0.042 | 0.042 | 0 | 0.33 | 0.15 | 0.37 | 0.8 | 0.31 | 0.56 |
| 3 | 149148302 | rs6808859 | -0.542 | -0.391 | -1.701 | -2.02 | -0.093 | 0.17 | 0.17 | 0 | 0.65 | 0.89 | 0.62 | 0.97 | 0.53 | 0.69 |
| 3 | 149151285 | rs941084 | -0.229 | -0.428 | -1.6 | -1.801 | -0.032 | 0.225 | 0.225 | 0 | 0.57 | 0.84 | 0.54 | 0.94 | 0.47 | 0.74 |
| 3 | 149151571 | rs9832360 | 0.401 | -0.553 | -1.224 | -1.623 | 0.004 | 0.038 | 0.038 | 0 | 0.38 | 0.27 | 0.42 | 0.8 | 0.37 | 0.64 |
| 3 | 149151611 | rs7636153 | -0.287 | -1.017 | -0.463 | -1.117 | -0.027 | -0.03 | -0.03 | 0 | 0.3 | 0.54 | 0.22 | 0.18 | 0.21 | 0.13 |
| 3 | 149152329 | rs6440611 | -0.4 | -0.82 | -0.788 | -0.677 | 0.002 | 0.002 | 0.002 | 1.033 | 0.45 | 0.55 | 0.25 | 0.27 | 0.29 | 0.2 |
| 3 | 149159718 | rs9829432 | 0.556 | -0.129 | -0.804 | -0.722 | 0.003 | 0.016 | 0.016 | 0.383 | 0.53 | 0.61 | 0.6 | 0.15 | 0.57 | 0.33 |
| 3 | 149160028 | rs9833550 | 0.598 | -0.165 | -0.704 | -0.738 | 0.002 | -0.005 | -0.005 | 0 | 0.53 | 0.57 | 0.6 | 0.15 | 0.57 | 0.33 |
| 3 | 149172012 | rs7621657 | 0.151 | -0.59 | -0.778 | -0.82 | -0.013 | -0.057 | -0.057 | 0 | 0.2 | 0.23 | 0.11 | 0.02 | 0.14 | 0.03 |
| 3 | 149174034 | rs9822199 | -0.195 | -0.638 | -0.748 | -0.702 | -0.028 | -0.074 | -0.074 | 0.003 | 0.23 | 0.33 | 0.12 | 0.07 | 0.17 | 0.08 |
| 3 | 149177380 | rs4681173 | 0.003 | -0.523 | -0.896 | 0.275 | -0.017 | -0.037 | -0.037 | 0 | 0.1 | 0.02 | 0.18 | 0.52 | 0.1 | 0.43 |
| 3 | 149178661 | rs4405858 | -0.123 | -0.517 | -0.804 | 0.025 | -0.003 | -0.019 | -0.019 | 0 | 0.93 | 0.98 | 0.98 | 0.77 | 0.93 | 0.89 |
| 3 | 149180484 | rs6440614 | 0.751 | -0.553 | -0.718 | -0.019 | -0.019 | -0.003 | -0.003 | 0 | 0.7 | 0.65 | 0.86 | 0.71 | 0.76 | 0.8 |
| 3 | 149180923 | rs9868841 | -0.216 | -0.029 | -0.695 | -0.389 | 0 | -0.046 | -0.046 | 0 | 0.27 | 0.26 | 0.19 | 0.02 | 0.25 | 0.12 |
| 3 | 149181661 | rs4456805 | 0.273 | -0.812 | -0.184 | 0.103 | -0.028 | 0.075 | 0.075 | 0.015 | 0.86 | 0.56 | 0.85 | 0.77 | 0.91 | 0.7 |
| 3 | 149182555 | rs6794608 | 0.553 | -0.59 | -0.48 | -0.38 | -0.01 | 0.001 | 0.001 | 1.124 | 0.25 | 0.14 | 0.21 | 0 | 0.26 | 0.07 |
| 3 | 149182882 | rs6794817 | -0.771 | -0.692 | -0.413 | -0.004 | 0.009 | 0.176 | 0.176 | 0.188 | 0.13 | 0.32 | 0.13 | 0.6 | 0.14 | 0.37 |
| 3 | 149183359 | rs9880421 | 2.148 | -0.895 | -1.458 | -0.56 | -0.115 | 0.225 | 0.225 | 0.002 | 0.34 | 0.06 | 0.36 | 0.02 | 0.4 | 0.2 |
| 3 | 149190879 | rs13069729 | 0.619 | -0.806 | -1.212 | -0.537 | -0.032 | 0.073 | 0.073 | 0.01 | 0.24 | 0.08 | 0.21 | 0 | 0.29 | 0.07 |
| 3 | 149193684 | rs1802837 | -0.539 | -0.868 | -1.192 | -0.552 | -0.003 | 0.062 | 0.062 | 0 | 0.37 | 0.23 | 0.3 | 0.07 | 0.4 | 0.17 |
| 3 | 149201983 | rs13062142 | -0.337 | -0.88 | -0.948 | -0.316 | -0.001 | -0.003 | -0.003 | 0 | 0.35 | 0.27 | 0.28 | 0.14 | 0.4 | 0.13 |
| 3 | 149203000 | rs4681177 | 0.285 | -0.787 | -1.044 | -0.249 | -0.007 | 0.012 | 0.012 | 0 | 0.28 | 0.17 | 0.25 | 0.14 | 0.35 | 0.1 |
| 3 | 149207934 | rs9289788 | 0.697 | -0.804 | -1.563 | -0.096 | 0.003 | -0.038 | -0.038 | 0 | 0.52 | 0.15 | 0.59 | 0.52 | 0.57 | 0.64 |
| 3 | 149211387 | rs9843304 | 1.07 | -0.805 | -1.648 | -0.103 | -0.055 | -0.076 | -0.076 | 0 | 0.47 | 0.08 | 0.55 | 0.51 | 0.53 | 0.61 |
| 3 | 149213337 | rs4681179 | 0.387 | -0.527 | -1.374 | 0.187 | -0.008 | 0.056 | 0.056 | 0 | 0.06 | 0.02 | 0.15 | 0.42 | 0.07 | 0.35 |
| 3 | 149219070 | rs10935763 | 0.302 | -0.743 | -1.038 | -0.018 | -0.003 | 0.003 | 0.003 | 0 | 0.29 | 0.23 | 0.26 | 0.14 | 0.33 | 0.09 |
| 3 | 149220517 | rs12636725 | -1.167 | -1.023 | -0.51 | 0.745 | 0 | 0.005 | 0.005 | 0 | 0.11 | 0.08 | 0.16 | 0.42 | 0.1 | 0.4 |
| 3 | 149220940 | rs12631874 | 0.037 | -1.009 | -0.564 | 0.792 | -0.021 | 0.039 | 0.039 | 0.053 | 0.06 | 0 | 0.14 | 0.42 | 0.07 | 0.36 |
| 3 | 149225903 | rs6779678 | 1.273 | -1.274 | -0.293 | 0.323 | -0.026 | 0.01 | 0.01 | 0 | 0.75 | 0.32 | 0.82 | 0.65 | 0.88 | 0.72 |
| 3 | 149230353 | rs1435621 | 0.755 | -0.809 | -0.378 | 0.095 | 0.001 | -0.008 | -0.008 | 0 | 0.92 | 0.93 | 0.97 | 1 | 0.95 | 0.94 |
| 3 | 149232489 | rs7652887 | -1.006 | -0.988 | 0.608 | 0.414 | 0.021 | 0.264 | 0.264 | 1.925 | 0.08 | 0.42 | 0.12 | 0.21 | 0.02 | 0.1 |
| 3 | 149235707 | rs7640098 | 0.893 | -1.14 | 0.176 | -0.161 | 0.007 | 0.016 | 0.016 | 0 | 0.76 | 0.8 | 0.84 | 0.89 | 0.8 | 0.77 |
| 3 | 149242016 | rs17195624 | -0.775 | -1.172 | 0.365 | -0.205 | 0.004 | 0.001 | 0.001 | 0 | 0.2 | 0.19 | 0.11 | 0.11 | 0.15 | 0.19 |
| 3 | 149242596 | rs17195631 | 0.24 | -1.13 | 0.31 | -0.201 | -0.003 | 0.001 | 0.001 | 0 | 0.07 | 0.05 | 0.07 | 0.06 | 0.1 | 0.05 |
| 3 | 149242957 | rs9289790 | 1.712 | -0.893 | 0.124 | -0.47 | -0.005 | -0.018 | -0.018 | 0.112 | 0.57 | 0.52 | 0.61 | 0.25 | 0.61 | 0.36 |
| 3 | 149244786 | rs10935766 | 0.655 | -0.571 | 0.281 | 0.178 | 0.011 | 0.013 | 0.013 | 0 | 0.82 | 0.74 | 0.83 | 0.81 | 0.8 | 0.89 |
| 3 | 149246213 | rs10513353 | -0.664 | -0.503 | 0.232 | -0.032 | -0.004 | 0.099 | 0.099 | 0 | 0.14 | 0.43 | 0.19 | 0.22 | 0.09 | 0.12 |
| 3 | 149247639 | rs7616772 | 0.764 | -0.501 | 0.15 | -0.055 | 0.005 | 0 | 0 | 0 | 0.74 | 0.73 | 0.77 | 0.74 | 0.7 | 0.83 |
| 3 | 149250601 | rs4681524 | 1.536 | -0.437 | -0.058 | -0.099 | 0.002 | -0.027 | -0.027 | 0 | 0.58 | 0.28 | 0.57 | 0.52 | 0.62 | 0.71 |
| 3 | 149252801 | rs10446281 | 0.191 | -0.476 | -0.019 | -0.165 | 0.001 | 0.007 | 0.007 | 0 | 0.81 | 0.84 | 0.85 | 0.94 | 0.81 | 0.87 |
| 3 | 149254700 | rs16861954 | 0.046 | -0.218 | -0.245 | -0.149 | 0.002 | -0.011 | -0.011 | 4.597 | 0.08 | 0.12 | 0.09 | 0.05 | 0.07 | 0.03 |
| 3 | 149255315 | rs4681525 | 0.762 | -0.223 | -0.402 | 0.091 | -0.003 | -0.003 | -0.003 | 0 | 0.65 | 0.39 | 0.65 | 0.7 | 0.72 | 0.77 |
| 3 | 149257404 | rs13325694 | 1.896 | 0.045 | -0.899 | -0.523 | -0.009 | 0.155 | 0.155 | 0 | 0.37 | 0.13 | 0.35 | 0.04 | 0.4 | 0.2 |
| 3 | 149259169 | rs1865604 | 0.759 | 0.178 | -0.57 | 0.087 | 0 | -0.002 | -0.002 | 0 | 0.52 | 0.38 | 0.54 | 0.52 | 0.52 | 0.68 |
| 3 | 149260786 | rs2278476 | 0.773 | 0.201 | -0.84 | 0.098 | 0.001 | -0.003 | -0.003 | 0 | 0.52 | 0.31 | 0.54 | 0.52 | 0.52 | 0.68 |
| 3 | 149266671 | rs9836653 | 0.499 | -0.089 | -1.247 | -0.285 | 0.024 | 0.173 | 0.173 | 0.03 | 0.51 | 0.18 | 0.49 | 0.19 | 0.62 | 0.35 |
| 3 | 149267986 | rs7624541 | -1.109 | -0.334 | -1.183 | -0.688 | -0.08 | 0.204 | 0.204 | 0.112 | 0.6 | 0.85 | 0.58 | 0.96 | 0.52 | 0.74 |
| 3 | 149271087 | rs11716804 | -0.679 | -0.771 | -0.297 | -0.075 | -0.005 | -0.002 | -0.002 | 0 | 0.38 | 0.49 | 0.23 | 0.35 | 0.36 | 0.56 |
| 3 | 149274090 | rs2005066 | 0.677 | -0.681 | 0.388 | 0.019 | 0.009 | 0.107 | 0.107 | 0 | 0.12 | 0.2 | 0.09 | 0.28 | 0.11 | 0.37 |
| 3 | 149277773 | rs713372 | -0.015 | -0.763 | 0.553 | 0.103 | -0.002 | -0.049 | -0.049 | 0.234 | 0.24 | 0.29 | 0.13 | 0.02 | 0.23 | 0.16 |
| 3 | 149284432 | rs4681526 | -1.451 | -0.578 | 0.334 | 0.213 | -0.024 | 0.108 | 0.108 | 0 | 0.56 | 0.73 | 0.53 | 0.79 | 0.51 | 0.63 |
| 3 | 149289808 | rs16862004 | -0.326 | 0.134 | 0.775 | 0.876 | 0.018 | 0.136 | 0.136 | 0 | 0.13 | 0.2 | 0.26 | 0.48 | 0.15 | 0.14 |
| 3 | 149292509 | rs7652697 | -0.71 | 0.06 | 0.755 | 0.238 | -0.008 | 0.006 | 0.006 | 0 | 0.3 | 0.37 | 0.22 | 0.29 | 0.27 | 0.45 |
| 3 | 149293316 | rs16862007 | 0.398 | -0.032 | 0.725 | 0.566 | -0.005 | 0.07 | 0.07 | 0 | 0.12 | 0.17 | 0.08 | 0.23 | 0.07 | 0.33 |

| | | | | | | | | | | | | | | | | |
|---|---|---|---|---|---|---|---|---|---|---|---|---|---|---|---|---|
| 3 | 149301095 | rs2864418 | -0.324 | 0.259 | 0.347 | 0.703 | -0.003 | -0.011 | -0.011 | 0 | 0.4 | 0.52 | 0.29 | 0.32 | 0.35 | 0.51 |
| 3 | 149301605 | rs13098021 | -0.772 | 0.278 | -0.322 | 0.737 | -0.002 | 0.047 | 0.047 | 0 | 0.51 | 0.7 | 0.4 | 0.34 | 0.48 | 0.57 |
| 3 | 149302530 | rs1344817 | -0.778 | 0.286 | -0.318 | 0.743 | -0.002 | 0.046 | 0.046 | 0 | 0.51 | 0.7 | 0.4 | 0.34 | 0.48 | 0.57 |
| 3 | 149308504 | rs7626565 | -0.703 | 0.385 | 0.093 | 1.001 | -0.003 | -0.016 | -0.016 | 0 | 0.11 | 0.18 | 0.1 | 0.06 | 0.12 | 0.07 |
| 3 | 149340813 | rs1344815 | 0.364 | 0.589 | 0.616 | 1.244 | -0.001 | 0 | 0 | 0.741 | 0.65 | 0.35 | 0.48 | 0.33 | 0.65 | 0.54 |
| 3 | 149358539 | rs9820339 | 0.607 | 0.701 | -0.012 | 1.15 | 0 | 0.051 | 0.051 | 0 | 0.53 | 0.14 | 0.3 | 0.37 | 0.52 | 0.46 |
| 3 | 149362405 | rs9821781 | 0.366 | 0.784 | 0.858 | 1.453 | 0.007 | -0.011 | -0.011 | 0.143 | 0.59 | 0.64 | 0.37 | 0.57 | 0.56 | 0.52 |
| 3 | 149363508 | rs2117754 | -0.036 | 0.642 | 0.511 | 1.486 | -0.007 | 0.026 | 0.026 | 0 | 0.6 | 0.69 | 0.6 | 0.64 | 0.54 | 0.6 |
| 3 | 149364473 | rs6779455 | 0.707 | 0.616 | 0.977 | 1.419 | -0.003 | 0.01 | 0.01 | 0 | 0.46 | 0.33 | 0.31 | 0.38 | 0.44 | 0.39 |
| 3 | 149374873 | rs1055153 | 0.534 | 0.77 | 0.453 | 1.286 | 0.004 | 0.008 | 0.008 | 0 | 0.07 | 0.06 | 0.04 | 0.15 | 0.07 | 0.05 |
| 3 | 149375289 | rs6783790 | 0.662 | 0.985 | 0.782 | 1.482 | 0.009 | -0.003 | -0.003 | 0.575 | 0.42 | 0.28 | 0.46 | 0.46 | 0.38 | 0.42 |
| 3 | 149382718 | rs4681535 | 0.438 | 0.797 | 1.11 | 1.603 | 0.001 | 0.002 | 0.002 | 0 | 0.48 | 0.4 | 0.35 | 0.44 | 0.42 | 0.44 |
| 3 | 149387303 | rs11710532 | 0.627 | 0.759 | 1.174 | 1.598 | 0.006 | 0.043 | 0.043 | 0 | 0.46 | 0.31 | 0.32 | 0.29 | 0.41 | 0.42 |
| 3 | 149387681 | rs3811715 | 0.325 | 0.891 | 0.879 | 1.48 | 0.01 | 0.062 | 0.062 | 0 | 0.11 | 0.19 | 0.34 | 0.22 | 0.17 | 0.13 |
| 3 | 149389695 | rs17195721 | 0.158 | 0.874 | 1.042 | 1.458 | 0 | 0.002 | 0.002 | 0 | 0.56 | 0.45 | 0.65 | 0.51 | 0.58 | 0.56 |
| 3 | 149390031 | rs9810250 | 0.158 | 0.874 | 1.042 | 1.458 | 0 | 0.002 | 0.002 | 0 | 0.56 | 0.45 | 0.65 | 0.51 | 0.58 | 0.56 |
| 3 | 149393280 | rs4681542 | 0.922 | 0.899 | 0.96 | 1.424 | 0.007 | 0.057 | 0.057 | 0 | 0.45 | 0.24 | 0.32 | 0.29 | 0.41 | 0.42 |
| 3 | 149395320 | rs9809185 | -0.009 | 0.921 | 0.881 | 1.308 | 0 | -0.005 | -0.005 | 0 | 0.58 | 0.53 | 0.67 | 0.66 | 0.59 | 0.57 |
| 3 | 149399908 | rs12632606 | -1.245 | 0.876 | 1.016 | 1.338 | 0.007 | 0.045 | 0.045 | 0 | 0.21 | 0.36 | 0.42 | 0.28 | 0.25 | 0.27 |
| 3 | 149401665 | rs6804325 | -0.643 | 0.863 | 1.041 | 1.318 | 0.023 | -0.018 | -0.018 | 0 | 0.1 | 0.28 | 0.08 | 0.05 | 0.09 | 0.04 |
| 3 | 149403563 | rs13064993 | -0.017 | 0.857 | 0.971 | 1.283 | 0.002 | -0.043 | -0.043 | 0 | 0.08 | 0.17 | 0.05 | 0 | 0.07 | 0.01 |
| 3 | 149421197 | rs6785251 | 0.239 | 0.92 | 0.938 | 1.318 | 0 | -0.004 | -0.004 | 2.495 | 0.35 | 0.4 | 0.28 | 0.34 | 0.33 | 0.42 |
| 3 | 149422599 | rs9870573 | 0.027 | 1.039 | 0.863 | 1.318 | -0.001 | -0.004 | -0.004 | 0 | 0.38 | 0.43 | 0.33 | 0.38 | 0.35 | 0.54 |
| 3 | 149425773 | rs4681549 | 1.371 | 1.038 | 0.818 | 1.376 | -0.01 | 0.05 | 0.05 | 0 | 0.41 | 0.14 | 0.26 | 0.25 | 0.39 | 0.29 |
| 3 | 149434570 | rs13321700 | 0.01 | 1.024 | 1.009 | 1.423 | 0 | -0.001 | -0.001 | 0 | 0.38 | 0.58 | 0.32 | 0.47 | 0.36 | 0.43 |
| 3 | 149449946 | rs1002896 | 0.072 | 0.79 | 1.105 | 1.421 | -0.012 | 0.002 | 0.002 | 0 | 0.55 | 0.65 | 0.38 | 0.46 | 0.47 | 0.44 |
| 3 | 149456154 | rs12330718 | 0.308 | 0.745 | 1.166 | 1.417 | -0.002 | 0.032 | 0.032 | 0 | 0.51 | 0.39 | 0.35 | 0.3 | 0.47 | 0.43 |
| 3 | 149456926 | rs10935774 | 0.283 | 0.738 | 1.162 | 1.413 | -0.002 | 0.02 | 0.02 | 0 | 0.51 | 0.41 | 0.35 | 0.31 | 0.47 | 0.43 |
| 3 | 149459341 | rs7633238 | -0.775 | 0.693 | 0.567 | 1.211 | -0.073 | -0.039 | -0.039 | 0 | 0.72 | 0.97 | 0.6 | 0.82 | 0.67 | 0.72 |
| 3 | 149468530 | rs9843784 | 1.186 | 0.54 | 0.354 | 1.173 | -0.095 | 0.03 | 0.03 | 0 | 0.42 | 0.04 | 0.27 | 0.25 | 0.41 | 0.3 |
| 3 | 149470818 | rs7626954 | -1.252 | 0.53 | 0.427 | 1.19 | -0.007 | 0.059 | 0.059 | 0.839 | 0.13 | 0.61 | 0.14 | 0.26 | 0.08 | 0.17 |
| 3 | 149476779 | rs13319447 | 0.392 | 0.64 | 0.439 | 1.194 | -0.029 | -0.064 | -0.064 | 0.042 | 0.11 | 0.01 | 0.11 | 0.31 | 0.12 | 0.22 |
| 3 | 149479301 | rs954714 | -0.04 | 0.565 | 0.182 | 0.792 | -0.011 | -0.016 | -0.016 | 0 | 0.56 | 0.65 | 0.4 | 0.51 | 0.49 | 0.46 |
| 3 | 149485293 | rs7610425 | 0.651 | 0.701 | -0.546 | 0.485 | -0.011 | 0.05 | 0.05 | 0 | 0.4 | 0.17 | 0.26 | 0.25 | 0.39 | 0.28 |
| 3 | 149488309 | rs7630033 | -0.32 | 0.442 | -0.504 | -0.381 | 0.001 | -0.074 | -0.074 | 0.026 | 0.32 | 0.65 | 0.36 | 0.02 | 0.29 | 0.23 |
| 3 | 149493646 | rs2130319 | -1.024 | -0.073 | -0.946 | 0.104 | 0.011 | 0.091 | 0.091 | 0.247 | 0.46 | 0.85 | 0.49 | 0.3 | 0.4 | 0.4 |
| 3 | 149495855 | rs1403028 | 0.389 | -0.337 | -0.689 | 0.058 | -0.015 | -0.045 | -0.045 | 0 | 0.74 | 0.55 | 0.81 | 0.89 | 0.8 | 0.79 |
| 3 | 149505805 | rs9843236 | 0.75 | -0.48 | -0.382 | -0.144 | -0.013 | -0.047 | -0.047 | 0 | 0.68 | 0.44 | 0.77 | 0.89 | 0.72 | 0.75 |
| 3 | 149512110 | rs1522244 | 0.583 | -0.606 | -0.397 | -0.202 | -0.006 | -0.019 | -0.019 | 0 | 0.71 | 0.65 | 0.8 | 0.89 | 0.75 | 0.77 |
| 3 | 149512284 | rs2222400 | 0.171 | -0.639 | -0.387 | -0.223 | -0.012 | -0.03 | -0.03 | 0 | 0.74 | 0.63 | 0.81 | 0.89 | 0.79 | 0.78 |
| 3 | 149514788 | rs988987 | 1.854 | -0.486 | -0.775 | 0.72 | -0.101 | -0.062 | -0.062 | 0 | 0.33 | 0.03 | 0.28 | 0.29 | 0.4 | 0.35 |
| 3 | 149519777 | rs9814585 | -1.24 | -0.524 | -0.649 | 0.692 | -0.002 | -0.004 | -0.004 | 0 | 0.85 | 0.93 | 0.88 | 0.89 | 0.87 | 0.85 |
| 3 | 149519843 | rs1552596 | -0.172 | -0.485 | -0.56 | 0.743 | -0.001 | -0.011 | -0.011 | 0 | 0.06 | 0.09 | 0.03 | 0.01 | 0.07 | 0.03 |
| 3 | 149532826 | rs9875202 | 1.503 | -0.619 | -0.279 | 0.735 | -0.015 | 0.011 | 0.011 | 0.126 | 0.67 | 0.51 | 0.7 | 0.65 | 0.71 | 0.65 |
| 3 | 149540465 | rs7614059 | 0.653 | -0.605 | -0.049 | 0.728 | -0.008 | -0.041 | -0.041 | 0.031 | 0.09 | 0.17 | 0.05 | 0 | 0.07 | 0.04 |
| 3 | 149544358 | rs3772200 | -0.6 | -0.613 | -0.053 | 0.742 | -0.014 | 0.012 | 0.012 | 0 | 0.13 | 0.24 | 0.15 | 0.23 | 0.1 | 0.15 |
| 3 | 149579776 | rs6762307 | -0.853 | -0.609 | -0.023 | 0.525 | -0.02 | -0.081 | -0.081 | 0 | 0.15 | 0.38 | 0.09 | 0.01 | 0.11 | 0.09 |
| 3 | 149599826 | rs2035741 | -0.821 | -0.667 | 0.776 | 0.506 | -0.006 | -0.002 | -0.002 | 0.242 | 0.85 | 0.93 | 0.87 | 0.89 | 0.84 | 0.84 |
| 3 | 149604799 | rs9815364 | -1.647 | -0.609 | 0.759 | 0.511 | -0.011 | -0.013 | -0.013 | 0 | 0.89 | 0.95 | 0.89 | 0.89 | 0.88 | 0.85 |
| 3 | 149624839 | rs16862294 | 1.997 | -0.701 | 0.373 | 0.43 | 0.001 | -0.011 | -0.011 | 0 | 0.06 | 0.02 | 0.02 | 0.21 | 0.04 | 0.2 |
| 3 | 149660475 | rs9879862 | 0.904 | -0.751 | 0.746 | -0.158 | -0.03 | 0.088 | 0.088 | 0.222 | 0.29 | 0.13 | 0.32 | 0.06 | 0.29 | 0.22 |
| 3 | 149668016 | rs2100633 | 0.81 | -0.744 | 1.208 | -0.04 | -0.017 | -0.068 | -0.068 | 0 | 0.69 | 0.45 | 0.78 | 0.91 | 0.73 | 0.75 |
| 3 | 149672139 | rs3911489 | -2.05 | -0.627 | 1.655 | 0.031 | 0.015 | 0.091 | 0.091 | 0 | 0.11 | 0.4 | 0.11 | 0.15 | 0.09 | 0.09 |
| 3 | 149678534 | rs6768054 | 1.038 | -0.652 | 1.175 | -0.138 | -0.01 | -0.037 | -0.037 | 0.052 | 0.67 | 0.45 | 0.72 | 0.87 | 0.71 | 0.69 |
| 3 | 149681694 | rs9860302 | 0.61 | -0.757 | 0.989 | -0.098 | -0.001 | 0.006 | 0.006 | 0.119 | 0.32 | 0.27 | 0.36 | 0.27 | 0.29 | 0.26 |
| 3 | 149709004 | rs3915852 | 0.445 | -0.679 | 1.022 | -0.058 | 0.001 | -0.035 | -0.035 | 0 | 0.33 | 0.42 | 0.38 | 0.24 | 0.29 | 0.28 |
| 3 | 149710906 | rs9815028 | -0.304 | -0.706 | 0.981 | -0.038 | -0.002 | -0.004 | -0.004 | 0 | 0.78 | 0.75 | 0.78 | 0.85 | 0.8 | 0.73 |
| 3 | 149723269 | rs6440644 | -0.439 | -0.698 | 0.874 | -0.048 | -0.001 | -0.016 | -0.016 | 0.71 | 0.05 | 0.19 | 0.03 | 0 | 0.03 | 0.01 |
| 3 | 149732017 | rs13063435 | 1.68 | -0.704 | 0.697 | 0.052 | -0.077 | -0.117 | -0.117 | 0 | 0.35 | 0.07 | 0.35 | 0.56 | 0.42 | 0.4 |
| 3 | 149732367 | rs16862366 | 0.298 | -0.663 | 0.939 | -0.067 | -0.011 | -0.029 | -0.029 | 0 | 0.05 | 0.12 | 0.04 | 0 | 0.05 | 0.02 |
| 3 | 149737819 | rs9812618 | 0.332 | -0.65 | 0.983 | -0.03 | 0.001 | -0.006 | -0.006 | 0.324 | 0.33 | 0.35 | 0.34 | 0.26 | 0.31 | 0.3 |
| 3 | 149752251 | rs1542568 | -0.723 | -0.649 | 0.62 | 0.012 | 0.004 | 0.007 | 0.007 | 0 | 0.38 | 0.62 | 0.39 | 0.32 | 0.33 | 0.39 |
| 3 | 149752398 | rs12695946 | 0.73 | -0.398 | 0.431 | -0.588 | -0.002 | -0.001 | -0.001 | 0 | 0.19 | 0.21 | 0.24 | 0.23 | 0.17 | 0.26 |
| 3 | 149754422 | rs1545782 | 0.465 | -0.37 | 0.321 | -0.461 | -0.012 | -0.025 | -0.025 | 0.035 | 0.06 | 0.11 | 0.03 | 0 | 0.05 | 0.02 |
| 3 | 149765785 | rs6789518 | -0.123 | -0.398 | 0.449 | -0.477 | -0.005 | -0.03 | -0.03 | 0.011 | 0.07 | 0.24 | 0.05 | 0 | 0.05 | 0.02 |
| 3 | 149775072 | rs11925509 | -0.173 | -0.344 | -0.028 | -0.475 | -0.008 | -0.03 | -0.03 | 0 | 0.35 | 0.52 | 0.38 | 0.23 | 0.29 | 0.33 |
| 3 | 149777272 | rs4681564 | -0.673 | -0.231 | -0.783 | -0.315 | 0.012 | 0.156 | 0.156 | 0 | 0.51 | 0.77 | 0.5 | 0.29 | 0.43 | 0.57 |
| 3 | 149777669 | rs9837142 | -0.193 | -0.202 | -0.994 | -0.3 | -0.003 | -0.015 | -0.015 | 0 | 0.43 | 0.51 | 0.47 | 0.29 | 0.39 | 0.5 |
| 3 | 149779984 | rs1979425 | 0.653 | 0.315 | -2.215 | -0.033 | 0.001 | 0 | 0 | 0 | 0.1 | 0.09 | 0.15 | 0.13 | 0.06 | 0.07 |

| | | | | | | | | | | | | | | | | |
|---|---|---|---|---|---|---|---|---|---|---|---|---|---|---|---|---|
| 3 | 149780516 | rs1994374 | 0.734 | 0.853 | -2.28 | 0.349 | -0.136 | -0.098 | -0.098 | 0.11 | 0.35 | 0.03 | 0.38 | 0.61 | 0.45 | 0.3 |
| 3 | 149782159 | rs962179 | -0.319 | 1.082 | -1.13 | 0.714 | -0.007 | -0.005 | -0.005 | 0 | 0.88 | 0.95 | 0.9 | 0.91 | 0.88 | 0.88 |
| 3 | 149787516 | rs11921628 | -0.224 | 1.13 | -1.243 | 0.435 | -0.027 | 0.034 | 0.034 | 0 | 0.09 | 0.02 | 0.07 | 0 | 0.15 | 0.05 |
| 3 | 149788430 | rs4681194 | -0.563 | 0.745 | -0.768 | 0.857 | -0.007 | 0.006 | 0.006 | 0.03 | 0.26 | 0.55 | 0.23 | 0.68 | 0.19 | 0.38 |
| 3 | 149808060 | rs4484141 | 0.25 | 1.446 | -1.582 | 1.122 | -0.053 | -0.006 | -0.006 | 0.101 | 0.67 | 0.84 | 0.6 | 0.27 | 0.49 | 0.61 |
| 3 | 149809746 | rs10513359 | -0.517 | 0.887 | -2.175 | 0.49 | 0.009 | 0.063 | 0.063 | 0 | 0.32 | 0.14 | 0.32 | 0.19 | 0.25 | 0.24 |
| 3 | 149809872 | rs12639497 | 0.122 | 1.188 | -2.407 | 0.691 | -0.063 | 0.054 | 0.054 | 0.001 | 0.29 | 0.08 | 0.31 | 0.2 | 0.42 | 0.2 |
| 3 | 149812073 | rs1531640 | 0.22 | 0.858 | -1.715 | 1.18 | 0.005 | 0.009 | 0.009 | 2.182 | 0.15 | 0.18 | 0.25 | 0.24 | 0.16 | 0.21 |
| 3 | 149812267 | rs11927779 | 0.231 | 0.659 | -1.998 | 0.801 | -0.024 | 0.141 | 0.141 | 0.009 | 0.4 | 0.15 | 0.36 | 0.16 | 0.47 | 0.24 |
| 3 | 149816159 | rs17195840 | 2.121 | 0.629 | -2.359 | 0.859 | -0.035 | -0.04 | -0.04 | 0.974 | 0.08 | 0.01 | 0.04 | 0.07 | 0.1 | 0.06 |
| 3 | 149823546 | rs1552037 | -1.097 | 0.023 | -1.631 | 0.825 | 0.002 | -0.003 | -0.003 | 0 | 0.71 | 0.68 | 0.72 | 0.6 | 0.8 | 0.66 |
| 3 | 149824124 | rs1564996 | 0.662 | 0.031 | -1.754 | 0.57 | -0.041 | 0.151 | 0.151 | 0.431 | 0.39 | 0.1 | 0.35 | 0.16 | 0.47 | 0.22 |
| 3 | 149831459 | rs9844449 | 0.873 | -0.532 | -1.158 | 1.121 | 0.023 | 0.036 | 0.036 | 0.244 | 0.25 | 0.18 | 0.18 | 0.12 | 0.16 | 0.19 |
| 3 | 149832485 | rs2046542 | 0.689 | 0.146 | -0.71 | 1.725 | 0.001 | 0.086 | 0.086 | 0 | 0.94 | 0.91 | 0.89 | 0.66 | 0.94 | 0.86 |
| 3 | 149834544 | rs10513360 | -0.299 | -0.025 | -0.486 | 1.164 | -0.007 | 0.086 | 0.086 | 0.081 | 0.06 | 0.16 | 0.04 | 0.15 | 0.04 | 0.14 |
| 3 | 149835069 | rs6785346 | 0.066 | -0.272 | -1.055 | 0.607 | 0.021 | 0.083 | 0.083 | 0 | 0.52 | 0.16 | 0.38 | 0.36 | 0.41 | 0.47 |
| 3 | 149835786 | rs763438 | 0.71 | -0.484 | -1.199 | 0.456 | 0.011 | 0.005 | 0.005 | 0 | 0.19 | 0.13 | 0.12 | 0.16 | 0.15 | 0.19 |
| 3 | 149838890 | rs9857853 | -0.26 | -0.449 | -0.818 | 0.558 | -0.001 | 0.001 | 0.001 | 0 | 0.56 | 0.42 | 0.4 | 0.43 | 0.44 | 0.59 |
| 3 | 149839156 | rs4681581 | -1.136 | -0.229 | -1.063 | 0.682 | -0.048 | -0.089 | -0.089 | 0 | 0.84 | 0.96 | 0.84 | 0.67 | 0.77 | 0.77 |
| 3 | 149844021 | rs1386287 | -0.877 | -0.24 | -0.876 | 0.677 | 0.003 | 0.022 | 0.022 | 0 | 0.83 | 0.83 | 0.83 | 0.67 | 0.77 | 0.77 |
| 3 | 149844435 | rs12485454 | 0.957 | -0.286 | -1.028 | 0.616 | -0.052 | 0.073 | 0.073 | 0 | 0.36 | 0.04 | 0.29 | 0.25 | 0.31 | 0.31 |
| 3 | 149844734 | rs9848025 | -0.159 | -0.37 | -0.944 | 0.585 | -0.001 | 0.008 | 0.008 | 0 | 0.55 | 0.31 | 0.4 | 0.43 | 0.44 | 0.59 |
| 3 | 149845539 | rs9862910 | 0.99 | -0.179 | -1.441 | 0.616 | -0.043 | 0.085 | 0.085 | 0 | 0.34 | 0.04 | 0.27 | 0.21 | 0.29 | 0.3 |
| 3 | 149847927 | rs13326757 | 0.995 | -0.18 | -1.405 | 0.615 | -0.037 | 0.088 | 0.088 | 0 | 0.34 | 0.05 | 0.27 | 0.21 | 0.29 | 0.3 |
| 3 | 149850328 | rs6440651 | 0.558 | -0.227 | -0.986 | 0.659 | -0.034 | 0.014 | 0.014 | 0.196 | 0.22 | 0.38 | 0.13 | 0.21 | 0.15 | 0.28 |
| 3 | 149855012 | rs2171266 | 1.255 | -0.174 | -1.039 | 0.547 | -0.038 | 0.032 | 0.032 | 0.711 | 0.24 | 0.12 | 0.37 | 0.2 | 0.31 | 0.16 |
| 3 | 149858232 | rs12634425 | 0.762 | -0.167 | -0.96 | 0.538 | -0.028 | -0.01 | -0.01 | 0 | 0.2 | 0.32 | 0.11 | 0.13 | 0.13 | 0.17 |
| 3 | 149871159 | rs2867840 | 1.037 | 0.142 | -1.356 | 0.495 | 0.015 | 0.067 | 0.067 | 0 | 0.28 | 0.12 | 0.2 | 0.15 | 0.22 | 0.27 |
| 3 | 149878428 | rs2903684 | 0.479 | 0.528 | -0.996 | 0.973 | -0.038 | -0.049 | -0.049 | 0.004 | 0.75 | 0.88 | 0.7 | 0.59 | 0.64 | 0.71 |
| 3 | 149882962 | rs10935794 | -0.201 | 0.526 | -1.035 | 1.089 | -0.034 | -0.057 | -0.057 | 0 | 0.82 | 0.94 | 0.79 | 0.67 | 0.72 | 0.74 |
| 3 | 149889584 | rs9853551 | 0.733 | 0.012 | -0.404 | 0.811 | 0.001 | -0.004 | -0.004 | 5.699 | 0.59 | 0.52 | 0.62 | 0.43 | 0.58 | 0.62 |
| 3 | 149893873 | rs17210844 | 1.346 | -0.26 | -0.729 | 0.501 | -0.075 | 0.062 | 0.062 | 0.341 | 0.3 | 0.07 | 0.4 | 0.19 | 0.4 | 0.22 |
| 3 | 149894219 | rs1386291 | 1.281 | -0.208 | -0.631 | 0.487 | -0.012 | -0.023 | -0.023 | 1.419 | 0.26 | 0.33 | 0.19 | 0.19 | 0.18 | 0.2 |
| 3 | 149894929 | rs4681586 | 1.037 | -0.128 | -0.65 | 0.565 | -0.032 | 0.034 | 0.034 | 0.053 | 0.33 | 0.2 | 0.42 | 0.24 | 0.4 | 0.42 |
| 3 | 149895807 | rs10513366 | 0.832 | -0.033 | -0.78 | 0.973 | -0.018 | -0.03 | -0.03 | 0 | 0.08 | 0.01 | 0.05 | 0.07 | 0.1 | 0.12 |
| 3 | 149896384 | rs6764451 | 0.694 | -0.243 | -0.815 | 0.686 | -0.002 | 0.002 | 0.002 | 0 | 0.29 | 0.29 | 0.28 | 0.28 | 0.25 | 0.2 |
| 3 | 149897772 | rs9289805 | -0.497 | -0.138 | -0.792 | 0.968 | -0.012 | -0.015 | -0.015 | 0 | 0.89 | 0.95 | 0.82 | 0.72 | 0.8 | 0.84 |
| 3 | 149897783 | rs9289806 | 0.809 | -0.359 | -0.335 | 1.016 | 0.017 | 0.02 | 0.02 | 0 | 0.23 | 0.17 | 0.12 | 0.09 | 0.16 | 0.18 |
| 3 | 149900326 | rs9790045 | 1.171 | -0.099 | -0.259 | 1.148 | -0.031 | -0.015 | -0.015 | 0 | 0.33 | 0.2 | 0.42 | 0.3 | 0.4 | 0.45 |
| 3 | 149913817 | rs6772257 | 0.993 | -0.101 | -0.15 | 1.147 | -0.002 | -0.001 | -0.001 | 0.28 | 0.35 | 0.33 | 0.44 | 0.3 | 0.4 | 0.45 |
| 3 | 149914790 | rs2903688 | 0.929 | 0.536 | 0.042 | 1.219 | 0.012 | -0.001 | -0.001 | 0 | 0.07 | 0.12 | 0.1 | 0.09 | 0.08 | 0.03 |
| 3 | 149917021 | rs4681593 | -0.053 | 0.442 | 0.083 | 1.166 | 0.01 | 0.039 | 0.039 | 0 | 0.87 | 0.81 | 0.78 | 0.62 | 0.82 | 0.87 |
| 3 | 149919818 | rs901462 | 0.438 | 0.242 | 0.042 | 1.21 | 0.034 | 0.082 | 0.082 | 0 | 0.77 | 0.58 | 0.71 | 0.56 | 0.72 | 0.79 |
| 3 | 149926130 | rs7629819 | -0.077 | -0.035 | -0.745 | 0.805 | -0.06 | -0.049 | -0.049 | 0 | 0.4 | 0.14 | 0.37 | 0.38 | 0.46 | 0.39 |
| 3 | 149935215 | rs12233598 | 0.058 | -0.585 | 0.03 | 0.762 | -0.032 | 0.088 | 0.088 | 0 | 0.43 | 0.24 | 0.35 | 0.3 | 0.45 | 0.48 |
| 3 | 149936122 | rs12638737 | -0.363 | -0.471 | 0.34 | 0.837 | 0 | 0 | 0 | 0 | 0.55 | 0.42 | 0.41 | 0.45 | 0.52 | 0.61 |
| 3 | 149936213 | rs13323544 | -0.645 | -0.398 | 0.074 | 0.82 | -0.004 | -0.008 | -0.008 | 0 | 0.59 | 0.64 | 0.56 | 0.53 | 0.55 | 0.66 |
| 3 | 149939444 | rs6440654 | -0.001 | 0.034 | -0.119 | 0.412 | -0.002 | 0.001 | 0.001 | 0 | 0.39 | 0.37 | 0.38 | 0.38 | 0.45 | 0.46 |
| 3 | 149944462 | rs6772814 | -0.344 | -0.122 | -0.09 | 0.215 | 0.002 | 0.019 | 0.019 | 0 | 0.52 | 0.44 | 0.39 | 0.4 | 0.56 | 0.39 |
| 3 | 149948279 | rs6786486 | -0.689 | -0.29 | -0.161 | 0.281 | -0.02 | 0.003 | 0.003 | 0 | 0.57 | 0.78 | 0.58 | 0.49 | 0.59 | 0.51 |
| 3 | 149954300 | rs7623697 | 0.263 | 0.057 | -0.556 | 0.331 | -0.051 | -0.003 | -0.003 | 0.625 | 0.36 | 0.17 | 0.32 | 0.44 | 0.46 | 0.36 |
| 3 | 149960105 | rs6809105 | 0.051 | 0.08 | -0.572 | -0.033 | -0.046 | 0.05 | 0.05 | 1.139 | 0.39 | 0.23 | 0.43 | 0.35 | 0.52 | 0.38 |
| 3 | 149965315 | rs9835316 | 0.203 | 0.003 | -0.343 | -0.068 | -0.095 | 0.058 | 0.058 | 1.055 | 0.37 | 0.12 | 0.39 | 0.36 | 0.51 | 0.33 |
| 3 | 149968000 | rs9813089 | 0.035 | 0.023 | -0.381 | -0.105 | -0.057 | 0.072 | 0.072 | 0.919 | 0.39 | 0.12 | 0.4 | 0.36 | 0.55 | 0.33 |
| 3 | 149972433 | rs7430967 | -0.557 | -0.169 | -0.266 | -0.069 | -0.001 | -0.001 | -0.001 | 0 | 0.49 | 0.53 | 0.48 | 0.45 | 0.62 | 0.47 |
| 3 | 149979591 | rs6440662 | -0.213 | 0.006 | -0.547 | -0.069 | -0.028 | 0.022 | 0.022 | 1.955 | 0.11 | 0.57 | 0.09 | 0.1 | 0.06 | 0.13 |
| 3 | 149980548 | rs7432940 | 0.497 | 0.805 | -1.384 | -0.254 | -0.125 | 0.046 | 0.046 | 0.014 | 0.21 | 0.01 | 0.29 | 0.12 | 0.36 | 0.15 |
| 3 | 149986210 | rs7430029 | 0.014 | -0.004 | -0.985 | 0.252 | -0.022 | -0.026 | -0.026 | 0 | 0.19 | 0.01 | 0.13 | 0.13 | 0.15 | 0.21 |
| 3 | 149994882 | rs9834503 | 0.086 | 0.648 | -1.563 | 0.174 | -0.1 | 0.077 | 0.077 | 0.815 | 0.39 | 0.09 | 0.4 | 0.31 | 0.52 | 0.32 |
| 3 | 149995170 | rs11718906 | -0.44 | 0.331 | -1.027 | 0.607 | -0.004 | 0.019 | 0.019 | 0 | 0.66 | 0.59 | 0.63 | 0.58 | 0.68 | 0.59 |
| 3 | 150006338 | rs7430433 | 0.352 | 1.329 | -1.603 | 0.114 | -0.059 | 0.053 | 0.053 | 0 | 0.82 | 0.94 | 0.72 | 0.91 | 0.76 | 0.88 |
| 3 | 150012931 | rs9838013 | 0.236 | 1.05 | -1.739 | 0.153 | -0.002 | 0.01 | 0.01 | 0 | 0.49 | 0.34 | 0.46 | 0.59 | 0.54 | 0.34 |
| 3 | 150013987 | rs9881412 | 0.847 | 0.972 | -1.739 | 0.156 | 0.003 | -0.01 | -0.01 | 0 | 0.41 | 0.3 | 0.37 | 0.58 | 0.45 | 0.31 |
| 3 | 150021485 | rs7648426 | 0.39 | 0.416 | -1.231 | 0.16 | -0.048 | 0.009 | 0.009 | 0 | 0.17 | 0.4 | 0.2 | 0.17 | 0.09 | 0.18 |
| 3 | 150022818 | rs7428155 | 1.603 | 0.533 | -1.681 | 0.521 | -0.03 | -0.062 | -0.062 | 0 | 0.1 | 0 | 0.1 | 0.3 | 0.08 | 0.13 |
| 3 | 150023454 | rs7652403 | 0.839 | 0.512 | -1.564 | 0.487 | -0.002 | 0.002 | 0.002 | 0 | 0.24 | 0.21 | 0.15 | 0.14 | 0.22 | 0.38 |
| 3 | 150024518 | rs6440670 | -0.933 | 0.639 | -1.823 | 0.284 | -0.067 | 0.027 | 0.027 | 0.067 | 0.87 | 0.98 | 0.78 | 0.92 | 0.82 | 0.91 |
| 3 | 150024765 | rs7427160 | 0.598 | 0.73 | -1.623 | 0.439 | 0.003 | 0.007 | 0.007 | 0 | 0.48 | 0.3 | 0.44 | 0.59 | 0.51 | 0.33 |
| 3 | 150028555 | rs7647971 | -0.117 | 0.317 | -1.302 | 0.738 | -0.004 | 0.005 | 0.005 | 0 | 0.31 | 0.38 | 0.28 | 0.32 | 0.24 | 0.55 |

| | | | | | | | | | | | | | | | | |
|---|---|---|---|---|---|---|---|---|---|---|---|---|---|---|---|---|
| 3 | 150028979 | rs11710003 | 0.6 | 0.779 | -1.383 | 0.897 | -0.03 | -0.061 | -0.061 | 0 | 0.1 | 0 | 0.1 | 0.3 | 0.08 | 0.13 |
| 3 | 150031832 | rs4681607 | -0.204 | 0.629 | -0.889 | 0.317 | 0.002 | -0.006 | -0.006 | 0 | 0.22 | 0.34 | 0.21 | 0.16 | 0.2 | 0.39 |
| 3 | 150034556 | rs1463230 | 0.253 | 0.583 | -1.688 | 0.08 | -0.082 | 0.012 | 0.012 | 0 | 0.57 | 0.89 | 0.39 | 0.24 | 0.49 | 0.39 |
| 3 | 150035821 | rs13087661 | 0.828 | -0.088 | -2.101 | -0.39 | 0.002 | 0.001 | 0.001 | 0 | 0.39 | 0.62 | 0.31 | 0.24 | 0.33 | 0.28 |
| 3 | 150042618 | rs4301033 | 0.794 | -0.192 | -1.669 | -0.285 | -0.022 | -0.067 | -0.067 | 0 | 0.13 | 0.22 | 0.05 | 0 | 0.1 | 0.06 |
| 3 | 150047765 | rs1901099 | 0.882 | -0.58 | -1.489 | -0.464 | -0.003 | -0.004 | -0.004 | 0 | 0.19 | 0.17 | 0.05 | 0.07 | 0.11 | 0.1 |
| 3 | 150051590 | rs9917701 | 1.741 | -0.326 | -0.933 | -0.176 | -0.005 | 0.129 | 0.129 | 0.564 | 0.06 | 0.15 | 0.13 | 0.17 | 0.07 | 0.08 |
| 3 | 150052807 | rs16862610 | 0.673 | -0.456 | -0.405 | -0.426 | -0.032 | -0.094 | -0.094 | 0 | 0.15 | 0.39 | 0.06 | 0 | 0.1 | 0.11 |
| 3 | 150056041 | rs9813497 | 0.204 | 0.175 | -1.212 | -0.255 | 0.005 | -0.016 | -0.016 | 0 | 0.24 | 0.19 | 0.29 | 0.26 | 0.2 | 0.41 |
| 3 | 150062028 | rs9845738 | 0.181 | 0.605 | -1.238 | -0.441 | -0.002 | -0.016 | -0.016 | 0.464 | 0.27 | 0.23 | 0.34 | 0.28 | 0.29 | 0.39 |
| 3 | 150064115 | rs4681610 | 1.465 | 1.09 | -1.435 | -0.536 | -0.025 | 0.024 | 0.024 | 0 | 0.18 | 0.14 | 0.14 | 0.08 | 0.23 | 0.16 |
| 3 | 150065494 | rs9871507 | -0.539 | 1.063 | -1.566 | -0.469 | -0.031 | -0.039 | -0.039 | 0 | 0.7 | 0.75 | 0.58 | 0.41 | 0.63 | 0.67 |
| 3 | 150066331 | rs1562789 | -1.313 | 0.937 | -1.315 | -0.344 | -0.004 | 0.013 | 0.013 | 0 | 0.88 | 0.89 | 0.72 | 0.48 | 0.86 | 0.82 |
| 3 | 150069785 | rs4681202 | -1.079 | 1.104 | -1.182 | -0.335 | 0.015 | 0.051 | 0.051 | 0.01 | 0.87 | 0.82 | 0.71 | 0.48 | 0.8 | 0.82 |
| 3 | 150072153 | rs7633948 | 0.523 | 0.07 | -1.024 | -0.868 | -0.033 | -0.054 | -0.054 | 0 | 0.34 | 0.38 | 0.2 | 0.09 | 0.22 | 0.19 |
| 3 | 150073637 | rs11719816 | 1.195 | 0.493 | -1.385 | -0.984 | -0.004 | -0.023 | -0.023 | 0 | 0.23 | 0.24 | 0.15 | 0.07 | 0.21 | 0.22 |
| 3 | 150077967 | rs9283603 | 0.781 | 1.025 | -1.587 | -0.641 | -0.01 | 0.018 | 0.018 | 0 | 0.22 | 0.14 | 0.22 | 0.13 | 0.23 | 0.32 |
| 3 | 150078836 | rs12638422 | -0.171 | 0.505 | -1.498 | -0.566 | -0.052 | -0.052 | -0.052 | 1.067 | 0.62 | 0.81 | 0.47 | 0.34 | 0.52 | 0.48 |
| 3 | 150079669 | rs13097358 | 0.757 | -0.075 | -1.251 | -1.058 | 0.003 | -0.023 | -0.023 | 0.331 | 0.26 | 0.2 | 0.14 | 0.03 | 0.2 | 0.22 |
| 3 | 150087008 | rs1012583 | 0.552 | 0.688 | -1.539 | -0.703 | -0.041 | 0.058 | 0.058 | 0 | 0.22 | 0.1 | 0.22 | 0.08 | 0.25 | 0.38 |
| 3 | 150094926 | rs10804747 | -1.209 | 0.819 | -1.68 | -0.389 | -0.037 | -0.075 | -0.075 | 0 | 0.83 | 0.95 | 0.7 | 0.48 | 0.78 | 0.85 |
| 3 | 150114504 | rs12639090 | 0.806 | 0.334 | -0.712 | -0.754 | 0.003 | -0.033 | -0.033 | 0 | 0.31 | 0.19 | 0.33 | 0.36 | 0.3 | 0.27 |
| 3 | 150115046 | rs7620193 | 1.407 | 0.104 | -0.303 | -0.931 | -0.008 | -0.051 | -0.051 | 0 | 0.15 | 0.4 | 0.09 | 0.03 | 0.15 | 0.12 |
| 3 | 150116149 | rs9862778 | 1.99 | 0.005 | -0.285 | -0.941 | 0.005 | 0.01 | 0.01 | 0 | 0.12 | 0.07 | 0.05 | 0 | 0.12 | 0.08 |
| 3 | 150122121 | rs10513373 | 1.836 | 0.049 | -0.429 | -0.921 | -0.046 | -0.076 | -0.076 | 0.229 | 0.11 | 0.02 | 0.21 | 0.31 | 0.15 | 0.12 |
| 3 | 150146026 | rs4681618 | 0.222 | -0.014 | -0.315 | -0.886 | 0.008 | -0.013 | -0.013 | 0 | 0.43 | 0.26 | 0.38 | 0.36 | 0.43 | 0.35 |
| 3 | 150153887 | rs10513372 | 1.972 | 0.302 | -0.231 | -0.997 | 0.03 | -0.004 | -0.004 | 0 | 0.07 | 0.08 | 0.1 | 0.03 | 0.17 | 0.1 |
| 3 | 150157229 | rs10513370 | 1.003 | 0.28 | -0.112 | -1.005 | 0.107 | -0.009 | -0.009 | 0 | 0.09 | 0.3 | 0.11 | 0.03 | 0.17 | 0.1 |
| 3 | 150159767 | rs9784335 | 0.191 | 0.174 | -0.002 | -1.039 | 0 | -0.001 | -0.001 | 0 | 0.44 | 0.41 | 0.4 | 0.37 | 0.43 | 0.38 |
| 3 | 150161522 | rs4323025 | -1.428 | 0.119 | -0.073 | -0.997 | -0.003 | -0.007 | -0.007 | 0 | 0.76 | 0.82 | 0.67 | 0.5 | 0.74 | 0.84 |
| 3 | 150166817 | rs1376027 | 2.115 | 0.16 | -0.187 | -1.003 | 0.028 | -0.004 | -0.004 | 0 | 0.07 | 0.08 | 0.1 | 0.03 | 0.17 | 0.1 |
| 3 | 150167144 | rs2903834 | -0.494 | 0.067 | -0.271 | -0.994 | 0.045 | 0.021 | 0.021 | 0 | 0.67 | 0.53 | 0.56 | 0.47 | 0.57 | 0.73 |
| 3 | 150171628 | rs12485291 | 1.831 | 0.248 | -0.575 | -0.817 | 0.024 | 0.011 | 0.011 | 0 | 0.16 | 0.04 | 0.14 | 0.1 | 0.1 | 0.33 |
| 3 | 150173434 | rs6767155 | 1.975 | 0.214 | -0.568 | -0.896 | 0.002 | -0.012 | -0.012 | 0 | 0.13 | 0.1 | 0.05 | 0 | 0.12 | 0.08 |
| 3 | 150178939 | rs16862676 | 1.772 | 0.473 | -0.858 | -0.614 | -0.006 | -0.07 | -0.07 | 0 | 0.11 | 0.02 | 0.19 | 0.31 | 0.11 | 0.11 |
| 3 | 150179597 | rs10513368 | 1.618 | 0.78 | -0.613 | -0.471 | 0.006 | 0.002 | 0.002 | 0 | 0.08 | 0.09 | 0.1 | 0.03 | 0.17 | 0.11 |
| 3 | 150181413 | rs4681620 | 0.101 | 0.749 | -0.556 | -0.525 | 0.02 | 0.009 | 0.009 | 0.121 | 0.61 | 0.51 | 0.54 | 0.46 | 0.53 | 0.67 |
| 3 | 150187314 | rs1868673 | -0.196 | 0.367 | -0.702 | -0.472 | 0.013 | -0.004 | -0.004 | 5.206 | 0.54 | 0.68 | 0.5 | 0.4 | 0.6 | 0.49 |
| 3 | 150203023 | rs4459923 | -0.25 | 0.135 | -0.808 | -0.769 | 0.023 | 0.06 | 0.06 | 0.094 | 0.45 | 0.43 | 0.64 | 0.86 | 0.59 | 0.44 |
| 3 | 150205466 | rs4681623 | -0.837 | -0.385 | -0.806 | -0.957 | -0.048 | -0.126 | -0.126 | 0 | 0.79 | 0.53 | 0.9 | 1 | 0.86 | 0.87 |
| 3 | 150209313 | rs4277711 | -0.041 | 0.719 | -0.594 | -0.496 | -0.026 | 0.03 | 0.03 | 0 | 0.38 | 0.25 | 0.55 | 0.79 | 0.48 | 0.35 |
| 3 | 150212698 | rs7634891 | -0.041 | 0.703 | -0.594 | -0.463 | -0.027 | 0.028 | 0.028 | 0 | 0.38 | 0.26 | 0.55 | 0.79 | 0.48 | 0.35 |
| 3 | 150218614 | rs4447788 | -0.893 | 0.141 | -0.513 | -0.492 | -0.014 | -0.052 | -0.052 | 0.896 | 0.64 | 0.66 | 0.79 | 0.93 | 0.7 | 0.79 |
| 3 | 150226136 | rs9810770 | -0.229 | 0.473 | -0.678 | -0.117 | 0.01 | 0.01 | 0.01 | 0 | 0.49 | 0.47 | 0.56 | 0.53 | 0.54 | 0.59 |
| 3 | 150229958 | rs9289815 | -0.788 | 0.318 | -0.23 | 0.055 | -0.024 | -0.089 | -0.089 | 0 | 0.86 | 0.7 | 0.92 | 1 | 0.89 | 0.93 |
| 3 | 150237962 | rs9865803 | -0.454 | 0.303 | -0.289 | 0.73 | -0.021 | 0.131 | 0.131 | 0.246 | 0.83 | 0.7 | 0.77 | 0.47 | 0.85 | 0.73 |
| 3 | 150238321 | rs9846474 | 0.635 | 0.611 | -0.393 | -0.032 | -0.011 | -0.021 | -0.021 | 1.528 | 0.23 | 0.11 | 0.29 | 0.42 | 0.27 | 0.15 |
| 3 | 150238549 | rs6806768 | -0.136 | 0.818 | -0.654 | 0.136 | -0.006 | -0.015 | -0.015 | 0 | 0.63 | 0.6 | 0.63 | 0.58 | 0.63 | 0.78 |
| 3 | 150239275 | rs4072809 | 0.221 | 1.321 | -0.882 | 0.177 | -0.012 | -0.005 | -0.005 | 3.447 | 0.16 | 0.14 | 0.3 | 0.45 | 0.27 | 0.38 |
| 3 | 150239458 | rs4234379 | 0.339 | 0.864 | -0.592 | -0.154 | 0.018 | 0.019 | 0.019 | 0 | 0.58 | 0.37 | 0.62 | 0.58 | 0.61 | 0.75 |
| 3 | 150243878 | rs7638277 | 0.305 | 0.878 | -0.726 | -0.067 | -0.003 | -0.005 | -0.005 | 0 | 0.58 | 0.47 | 0.61 | 0.58 | 0.59 | 0.74 |
| 3 | 150257082 | rs10935807 | -0.327 | 0.99 | -1.144 | -0.321 | -0.015 | -0.023 | -0.023 | 0 | 0.31 | 0.2 | 0.35 | 0.4 | 0.38 | 0.34 |
| 3 | 150259668 | rs11922157 | -0.809 | 1.556 | -0.671 | -0.605 | 0.007 | 0.001 | 0.001 | 0 | 0.41 | 0.51 | 0.5 | 0.42 | 0.52 | 0.44 |
| 3 | 150259942 | rs8052 | -0.886 | 1.259 | -0.615 | -0.668 | -0.003 | 0 | 0 | 0.924 | 0.86 | 0.67 | 0.86 | 0.91 | 0.88 | 0.91 |
| 3 | 150260853 | rs10272 | 1.332 | 1.249 | -0.622 | -0.671 | 0.049 | -0.033 | -0.033 | 0 | 0.45 | 0.16 | 0.36 | 0.49 | 0.36 | 0.47 |
| 3 | 150264428 | rs6790726 | -0.805 | 1.254 | -0.52 | -0.675 | -0.002 | 0.002 | 0.002 | 0.303 | 0.86 | 0.63 | 0.86 | 0.91 | 0.88 | 0.91 |
| 3 | 150264779 | rs17280260 | 1.473 | 0.883 | -0.816 | -0.818 | 0.007 | 0.004 | 0.004 | 0 | 0.11 | 0.1 | 0.03 | 0.09 | 0.07 | 0.12 |
| 3 | 150272305 | rs4681636 | 1.647 | 1.303 | -0.937 | -0.944 | -0.044 | 0.044 | 0.044 | 0 | 0.08 | 0.01 | 0.13 | 0 | 0.15 | 0.03 |
| 3 | 150285520 | rs2293151 | 1.71 | 1.32 | -0.803 | -0.942 | 0.046 | -0.025 | -0.025 | 1.462 | 0.08 | 0.16 | 0.14 | 0.02 | 0.16 | 0.11 |
| 3 | 150285664 | rs2293150 | 1.77 | 1.318 | -0.777 | -0.943 | 0.031 | -0.019 | -0.019 | 0 | 0.08 | 0.12 | 0.13 | 0.02 | 0.16 | 0.11 |
| 3 | 150288309 | rs9840992 | 1.273 | 1.284 | -0.797 | -0.933 | 0.051 | -0.007 | -0.007 | 0 | 0.47 | 0.27 | 0.37 | 0.49 | 0.36 | 0.47 |
| 3 | 150298317 | rs9883613 | 0.839 | 1.214 | -0.771 | -0.847 | -0.002 | -0.004 | -0.004 | 0 | 0.55 | 0.43 | 0.51 | 0.51 | 0.52 | 0.57 |
| 3 | 150302160 | rs7638500 | -0.003 | 1.275 | -0.796 | -0.88 | 0.04 | -0.058 | -0.058 | 0.004 | 0.78 | 0.6 | 0.73 | 0.91 | 0.73 | 0.88 |
| 3 | 150307263 | rs2090916 | -0.395 | 0.961 | -0.941 | -0.977 | -0.013 | -0.037 | -0.037 | 0 | 0.8 | 0.67 | 0.85 | 0.91 | 0.86 | 0.86 |
| 3 | 150316866 | rs1915944 | 0.586 | 0.878 | -1.012 | -0.873 | -0.003 | 0.008 | 0.008 | 0 | 0.23 | 0.27 | 0.23 | 0.4 | 0.21 | 0.3 |
| 3 | 150317636 | rs9809820 | -0.275 | 1.09 | -1.023 | -0.996 | 0.05 | -0.072 | -0.072 | 0.612 | 0.77 | 0.57 | 0.75 | 0.92 | 0.73 | 0.87 |
| 3 | 150320322 | rs2204113 | 0.247 | 1.175 | -1.082 | -1.061 | 0.039 | -0.05 | -0.05 | 0 | 0.69 | 0.45 | 0.61 | 0.9 | 0.56 | 0.77 |
| 3 | 150332011 | rs1568585 | 2.648 | 0.971 | -1.232 | -1.472 | 0.011 | -0.086 | -0.086 | 0.976 | 0.29 | 0.06 | 0.32 | 0.4 | 0.26 | 0.3 |
| 3 | 150340114 | rs2177148 | 0.546 | 1.143 | -1.07 | -1.417 | -0.006 | -0.013 | -0.013 | 0 | 0.54 | 0.33 | 0.53 | 0.52 | 0.52 | 0.57 |

| | | | | | | | | | | | | | | | | |
|---|---|---|---|---|---|---|---|---|---|---|---|---|---|---|---|---|
| 3 | 150342895 | rs1464517 | 0.18 | 1.163 | -1.114 | -1.45 | 0.036 | -0.166 | -0.166 | 0 | 0.73 | 0.58 | 0.78 | 1 | 0.7 | 0.85 |
| 3 | 150345664 | rs11548666 | 1.923 | 1.218 | -1.156 | -1.248 | 0.009 | -0.004 | -0.004 | 0 | 0.08 | 0.07 | 0.14 | 0.02 | 0.17 | 0.11 |
| 3 | 150346654 | rs6777911 | 0.888 | 0.503 | -1.192 | -1.142 | 0.063 | -0.039 | -0.039 | 3.442 | 0.46 | 0.16 | 0.39 | 0.5 | 0.36 | 0.47 |
| 3 | 150353177 | rs6440688 | 0.189 | 0.821 | -0.593 | -1.005 | -0.007 | -0.09 | -0.09 | 0 | 0.14 | 0.28 | 0.08 | 0 | 0.1 | 0.08 |
| 3 | 150357070 | rs13064363 | 2.076 | 0.621 | -1.348 | -1.07 | 0.054 | -0.067 | -0.067 | 0 | 0.32 | 0.06 | 0.3 | 0.39 | 0.23 | 0.29 |
| 3 | 150360152 | rs9849937 | -0.115 | 0.314 | -0.902 | -0.86 | 0.009 | 0.03 | 0.03 | 0 | 0.74 | 0.77 | 0.87 | 0.9 | 0.76 | 0.8 |
| 3 | 150365815 | rs6783018 | -0.609 | 0.414 | -0.662 | -0.826 | -0.016 | 0.115 | 0.115 | 0 | 0.26 | 0.48 | 0.3 | 0.49 | 0.22 | 0.37 |
| 3 | 150367481 | rs12695959 | 0.838 | 0.571 | -0.718 | -0.727 | 0.035 | 0.026 | 0.026 | 0 | 0.53 | 0.26 | 0.57 | 0.41 | 0.58 | 0.48 |
| 3 | 150369151 | rs1403086 | 0.446 | 1.281 | -0.123 | -0.24 | -0.011 | 0.002 | 0.002 | 0 | 0.87 | 0.89 | 0.87 | 0.99 | 0.82 | 0.89 |
| 3 | 150378858 | rs6773558 | -1.991 | 1.28 | 0.123 | -0.24 | 0.003 | 0.012 | 0.012 | 0 | 0.35 | 0.64 | 0.4 | 0.52 | 0.34 | 0.4 |
| 3 | 150379145 | rs6809409 | 0.099 | 0.477 | 0.474 | -0.73 | -0.007 | -0.037 | -0.037 | 0 | 0.07 | 0.14 | 0.02 | 0 | 0.03 | 0.05 |
| 3 | 150380030 | rs16862792 | -1.523 | 0.76 | 0.434 | -0.405 | -0.001 | 0.003 | 0.003 | 0.502 | 0.13 | 0.12 | 0.13 | 0.06 | 0.15 | 0.1 |
| 3 | 150384307 | rs6808787 | -1.45 | 0.722 | 0.464 | -0.25 | 0.027 | -0.012 | -0.012 | 0 | 0.13 | 0.51 | 0.14 | 0.06 | 0.13 | 0.09 |
| 3 | 150384807 | rs6440691 | 0.452 | 0.685 | -0.318 | -0.084 | -0.014 | 0.023 | 0.023 | 0 | 0.92 | 0.97 | 0.96 | 1 | 0.91 | 0.96 |
| 3 | 150402481 | rs4680090 | -0.57 | 0.329 | 0.103 | 0.407 | 0.013 | 0.05 | 0.05 | 0 | 0.26 | 0.35 | 0.4 | 0.47 | 0.29 | 0.23 |
| 3 | 150403749 | rs4680107 | -0.57 | 0.329 | 0.119 | 0.407 | 0.014 | 0.052 | 0.052 | 0 | 0.26 | 0.36 | 0.4 | 0.47 | 0.29 | 0.23 |
| 3 | 150404970 | rs2116682 | 0.907 | -0.63 | -0.741 | -0.198 | -0.123 | -0.153 | -0.153 | 1.537 | 0.43 | 0.06 | 0.48 | 0.51 | 0.53 | 0.49 |
| 3 | 150406250 | rs9832682 | -0.55 | -1.01 | 0.125 | -0.294 | -0.034 | -0.086 | -0.086 | 6.356 | 0.71 | 0.66 | 0.9 | 0.97 | 0.82 | 0.77 |
| 3 | 150415498 | rs6440693 | 0.189 | -0.746 | 0.302 | -0.03 | 0.015 | 0.01 | 0.01 | 0 | 0.52 | 0.43 | 0.69 | 0.64 | 0.67 | 0.6 |
| 3 | 150417238 | rs6766256 | 0.608 | -0.557 | 0.697 | -0.027 | -0.004 | -0.004 | -0.004 | 0.038 | 0.45 | 0.41 | 0.42 | 0.49 | 0.46 | 0.54 |
| 3 | 150423391 | rs7648424 | 0.959 | -0.732 | 0.664 | -0.2 | -0.014 | -0.032 | -0.032 | 0 | 0.32 | 0.24 | 0.42 | 0.6 | 0.35 | 0.51 |
| 3 | 150427570 | rs4680240 | -0.708 | -0.808 | 0.699 | -0.245 | 0.001 | -0.012 | -0.012 | 1.437 | 0.73 | 0.72 | 0.9 | 0.96 | 0.82 | 0.77 |
| 3 | 150427673 | rs13086215 | 0.203 | -0.671 | 0.523 | -0.2 | -0.087 | 0.153 | 0.153 | 0 | 0.37 | 0.12 | 0.32 | 0 | 0.46 | 0.13 |
| 3 | 150429131 | rs13074670 | 0.029 | -0.591 | 0.501 | -0.146 | -0.036 | 0.003 | 0.003 | 0 | 0.39 | 0.23 | 0.46 | 0.36 | 0.46 | 0.19 |
| 3 | 150441617 | rs9859732 | -0.797 | -0.996 | 0.537 | -0.254 | 0.035 | 0.052 | 0.052 | 0 | 0.77 | 0.84 | 0.92 | 0.96 | 0.84 | 0.77 |
| 3 | 150449883 | rs476663 | -0.608 | 0.052 | 0.923 | 0.449 | -0.006 | 0.001 | 0.001 | 0 | 0.41 | 0.34 | 0.44 | 0.45 | 0.42 | 0.24 |
| 3 | 150452545 | rs1520818 | -0.782 | -0.563 | 1.815 | 0.84 | -0.027 | -0.109 | -0.109 | 0 | 0.21 | 0.4 | 0.11 | 0.05 | 0.13 | 0.17 |
| 3 | 150455755 | rs502941 | 1.881 | -0.468 | 2.287 | 1.49 | -0.024 | 0.115 | 0.115 | 0 | 0.85 | 0.61 | 0.82 | 0.61 | 0.88 | 0.74 |
| 3 | 150458543 | rs481236 | 2.121 | -0.461 | 2.44 | 1.474 | -0.033 | 0.149 | 0.149 | 0 | 0.83 | 0.46 | 0.81 | 0.61 | 0.88 | 0.74 |
| 3 | 150467808 | rs6788895 | -2.652 | -0.339 | 2.491 | 1.73 | 0.006 | 0.368 | 0.368 | 1.639 | 0.09 | 0.54 | 0.16 | 0.4 | 0.03 | 0.19 |
| 3 | 150487945 | rs2014279 | -2.42 | -0.199 | 2.28 | 2.006 | 0.001 | 0.357 | 0.357 | 1.536 | 0.08 | 0.44 | 0.14 | 0.45 | 0.03 | 0.18 |
| 3 | 150510292 | rs1220472 | 0.191 | 0.137 | 1.136 | 2.096 | -0.002 | 0.026 | 0.026 | 0 | 0.95 | 0.97 | 0.88 | 0.82 | 0.92 | 0.87 |
| 3 | 150527120 | rs6768306 | -1.794 | 0.051 | 0.853 | 1.667 | 0.031 | 0.37 | 0.37 | 0 | 0.06 | 0.36 | 0.13 | 0.41 | 0.04 | 0.17 |
| 3 | 150530740 | rs4679980 | -0.716 | -0.293 | 0.235 | 0.322 | -0.026 | -0.079 | -0.079 | 0 | 0.36 | 0.49 | 0.19 | 0.12 | 0.25 | 0.38 |
| 3 | 150530994 | rs4679683 | -0.955 | -0.41 | 0.7 | 0.162 | -0.005 | -0.047 | -0.047 | 2.979 | 0.25 | 0.41 | 0.15 | 0.11 | 0.2 | 0.32 |
| 3 | 150531601 | rs12495611 | -0.099 | 0.732 | 0.664 | 1.144 | 0.029 | 0.162 | 0.162 | 0 | 0.1 | 0.16 | 0.12 | 0.44 | 0.14 | 0.1 |
| 3 | 150541363 | rs10935819 | -2.049 | 0.674 | 0.256 | 0.804 | 0.003 | 0.05 | 0.05 | 0 | 0.28 | 0.56 | 0.25 | 0.46 | 0.23 | 0.4 |
| 3 | 150545285 | rs2904193 | -0.995 | 0.683 | -0.557 | 0.263 | 0.003 | 0.026 | 0.026 | 0.15 | 0.15 | 0.09 | 0.11 | 0.02 | 0.1 | 0.3 |
| 3 | 150580747 | rs4679690 | -0.818 | 0.986 | -0.865 | -0.178 | 0.013 | 0.076 | 0.076 | 0.323 | 0.34 | 0.22 | 0.23 | 0.02 | 0.28 | 0.45 |
| 3 | 150582992 | rs2037409 | 0.543 | 1.301 | -1.496 | 0.348 | -0.024 | 0.033 | 0.033 | 0 | 0.08 | 0.04 | 0.08 | 0.44 | 0.13 | 0.11 |
| 3 | 150585344 | rs4680005 | -1.461 | 0.819 | -1.481 | 0.373 | -0.021 | 0.019 | 0.019 | 0.981 | 0.53 | 0.93 | 0.52 | 0.59 | 0.52 | 0.65 |
| 3 | 150590168 | rs1444200 | -0.916 | 0.571 | -1.006 | 0.343 | -0.008 | 0.001 | 0.001 | 0.152 | 0.46 | 0.72 | 0.34 | 0.48 | 0.42 | 0.56 |
| 3 | 150594033 | rs1220544 | 0.575 | 0.401 | -1.663 | 0.393 | -0.048 | -0.104 | -0.104 | 3.302 | 0.58 | 0.09 | 0.68 | 0.56 | 0.65 | 0.54 |
| 3 | 150594685 | rs574445 | 0.606 | 0.417 | -1.374 | 0.252 | -0.032 | 0.085 | 0.085 | 1.14 | 0.5 | 0.13 | 0.44 | 0.39 | 0.56 | 0.5 |
| 3 | 150600047 | rs639693 | 0.557 | -0.019 | -0.136 | 0.575 | -0.002 | 0.01 | 0.01 | 0 | 0.74 | 0.75 | 0.71 | 0.44 | 0.81 | 0.78 |
| 3 | 150601083 | rs6440703 | 0.662 | 0.003 | -0.441 | 0.343 | -0.007 | -0.037 | -0.037 | 0 | 0.62 | 0.8 | 0.58 | 0.42 | 0.65 | 0.72 |
| 3 | 150602373 | rs16862966 | 0.382 | 0.106 | -0.651 | 0.281 | -0.004 | -0.018 | -0.018 | 0 | 0.06 | 0.1 | 0.03 | 0 | 0.03 | 0.06 |
| 3 | 150603913 | rs520976 | 1.174 | 0.057 | -0.168 | 0.254 | 0.002 | -0.001 | -0.001 | 0 | 0.55 | 0.37 | 0.54 | 0.39 | 0.62 | 0.63 |
| 3 | 150606046 | rs569220 | -0.006 | 0.519 | -0.643 | 0.484 | -0.003 | -0.021 | -0.021 | 1.029 | 0.07 | 0.12 | 0.04 | 0.01 | 0.04 | 0.08 |
| 3 | 150606263 | rs4680029 | -0.048 | 0.517 | -0.625 | 0.483 | -0.005 | -0.026 | -0.026 | 0.595 | 0.07 | 0.13 | 0.04 | 0.01 | 0.04 | 0.08 |
| 3 | 150609866 | rs6768408 | 0.764 | 0.602 | -0.803 | 0.557 | 0.009 | 0.012 | 0.012 | 0 | 0.5 | 0.65 | 0.42 | 0.41 | 0.5 | 0.57 |
| 3 | 150609925 | rs628383 | 0.22 | 0.816 | -0.924 | 1.409 | -0.083 | -0.122 | -0.122 | 0 | 0.83 | 0.98 | 0.66 | 0.61 | 0.77 | 0.82 |
| 3 | 150614723 | rs6767398 | -1.325 | 0.072 | 0.241 | 1.267 | -0.006 | 0.086 | 0.086 | 0 | 0.11 | 0.39 | 0.17 | 0.17 | 0.07 | 0.14 |
| 3 | 150617682 | rs17219034 | -0.768 | -0.363 | 0.116 | 0.629 | 0.012 | 0.002 | 0.002 | 0 | 0.13 | 0.06 | 0.04 | 0 | 0.08 | 0.05 |
| 3 | 150619023 | rs3732753 | -1.806 | -0.345 | 0.336 | 1.267 | -0.032 | 0.037 | 0.037 | 0 | 0.32 | 0.56 | 0.44 | 0.4 | 0.22 | 0.27 |
| 3 | 150622735 | rs11717908 | -1.351 | 0.264 | 0.499 | 1.198 | 0.017 | -0.035 | -0.035 | 0 | 0.07 | 0.15 | 0.05 | 0 | 0.08 | 0.1 |
| 3 | 150625775 | rs1000079 | -1.701 | -0.123 | 0.95 | 1.477 | -0.018 | 0.064 | 0.064 | 0.618 | 0.18 | 0.66 | 0.31 | 0.36 | 0.12 | 0.19 |
| 3 | 150632774 | rs2061201 | -0.607 | -0.053 | 0.14 | 1.232 | 0.032 | -0.012 | -0.012 | 0 | 0.45 | 0.68 | 0.57 | 0.5 | 0.39 | 0.37 |
| 3 | 150636224 | rs936189 | -0.544 | -0.425 | 0.433 | 1.125 | -0.007 | -0.013 | -0.013 | 0 | 0.4 | 0.44 | 0.38 | 0.32 | 0.32 | 0.33 |
| 3 | 150636345 | rs732289 | -0.714 | -1.081 | -0.138 | 0.849 | 0.061 | 0.084 | 0.084 | 0.342 | 0.45 | 0.15 | 0.34 | 0.34 | 0.31 | 0.36 |
| 3 | 150641304 | rs1456139 | -1.108 | -1.22 | -0.06 | 1.425 | 0.069 | 0.028 | 0.028 | 0.018 | 0.52 | 0.82 | 0.46 | 0.52 | 0.38 | 0.41 |
| 3 | 150643373 | rs10513380 | 0.758 | 0.157 | 0.036 | 1.349 | -0.001 | 0.028 | 0.028 | 0 | 0.52 | 0.36 | 0.57 | 0.78 | 0.55 | 0.61 |
| 3 | 150644839 | rs12635299 | -1.674 | -0.089 | -0.421 | 2.155 | -0.008 | 0.165 | 0.165 | 0 | 0.05 | 0.18 | 0.12 | 0.35 | 0.05 | 0.05 |
| 3 | 150645351 | rs4680058 | -0.029 | 0.056 | -0.841 | 1.466 | -0.056 | 0.032 | 0.032 | 0 | 0.66 | 0.95 | 0.72 | 0.79 | 0.7 | 0.65 |
| 3 | 150647216 | rs9289826 | 0.687 | -0.129 | -0.317 | 1.709 | 0.045 | 0.021 | 0.021 | 0 | 0.55 | 0.29 | 0.64 | 0.58 | 0.69 | 0.62 |
| 3 | 150648913 | rs1456136 | 0.617 | 0.026 | -0.218 | 1.457 | 0.027 | 0.037 | 0.037 | 0.006 | 0.56 | 0.31 | 0.65 | 0.68 | 0.68 | 0.62 |
| 3 | 150654154 | rs12635944 | -0.402 | 0.096 | -0.416 | 1.264 | -0.004 | 0.032 | 0.032 | 0 | 0.36 | 0.27 | 0.36 | 0.31 | 0.35 | 0.38 |
| 3 | 150656671 | rs11720116 | -0.184 | 0.542 | -0.792 | 1.48 | -0.012 | -0.075 | -0.075 | 0.54 | 0.33 | 0.07 | 0.32 | 0.44 | 0.29 | 0.43 |
| 3 | 150661842 | rs11914988 | -0.063 | 0.291 | -0.767 | 1.497 | 0.016 | -0.078 | -0.078 | 4.168 | 0.31 | 0.08 | 0.27 | 0.43 | 0.25 | 0.39 |

| | | | | | | | | | | | | | | | | |
|---|---|---|---|---|---|---|---|---|---|---|---|---|---|---|---|---|
| 3 | 150662914 | rs13092829 | 0.261 | 0.432 | -0.168 | 1.433 | 0.016 | -0.055 | -0.055 | 0.165 | 0.9 | 0.72 | 0.94 | 1 | 0.92 | 0.97 |
| 3 | 150663334 | rs10935822 | -0.063 | 0.389 | -0.533 | 1.463 | 0.014 | -0.077 | -0.077 | 0.229 | 0.31 | 0.08 | 0.27 | 0.43 | 0.26 | 0.39 |
| 3 | 150664390 | rs13099797 | 1.075 | 0.373 | -0.644 | 1.579 | -0.064 | 0.072 | 0.072 | 0.238 | 0.49 | 0.04 | 0.48 | 0.3 | 0.56 | 0.5 |
| 3 | 150673377 | rs1350154 | -0.084 | -1.022 | -0.259 | 1.399 | -0.045 | -0.005 | -0.005 | 0 | 0.37 | 0.49 | 0.21 | 0.33 | 0.19 | 0.42 |
| 3 | 150674627 | rs11714757 | -1.033 | -0.705 | -0.487 | 1.145 | 0.042 | 0.049 | 0.049 | 0.133 | 0.55 | 0.68 | 0.34 | 0.75 | 0.33 | 0.6 |
| 3 | 150675887 | rs9814253 | 0.936 | -1.011 | -0.813 | 0.89 | -0.066 | 0.021 | 0.021 | 0 | 0.39 | 0.14 | 0.49 | 0.28 | 0.48 | 0.41 |
| 3 | 150675977 | rs7617623 | 0.615 | -1.243 | -0.641 | 1.114 | -0.002 | 0.026 | 0.026 | 0 | 0.09 | 0.1 | 0.14 | 0.24 | 0.14 | 0.08 |
| 3 | 150678233 | rs9825721 | 0.259 | -1.321 | -0.614 | 1.026 | 0.021 | 0.005 | 0.005 | 0.006 | 0.49 | 0.46 | 0.66 | 0.52 | 0.63 | 0.49 |
| 3 | 150685372 | rs6796933 | 0.015 | -1.337 | -0.587 | 1.062 | 0.003 | 0.001 | 0.001 | 0.024 | 0.53 | 0.56 | 0.59 | 0.43 | 0.53 | 0.55 |
| 3 | 150690018 | rs12497559 | 0.75 | -1.281 | -0.72 | 1.104 | -0.022 | -0.055 | -0.055 | 0 | 0.09 | 0.01 | 0.13 | 0.24 | 0.14 | 0.08 |
| 3 | 150690566 | rs3796240 | 0.323 | -1.328 | -0.717 | 1.06 | 0.038 | 0.022 | 0.022 | 0 | 0.5 | 0.33 | 0.58 | 0.44 | 0.53 | 0.55 |
| 3 | 150691112 | rs6786256 | 0.789 | -1.111 | -0.846 | 0.833 | -0.005 | 0.036 | 0.036 | 0 | 0.43 | 0.36 | 0.51 | 0.29 | 0.5 | 0.46 |
| 3 | 150692298 | rs7612632 | -0.328 | -0.599 | -1.043 | 1.068 | -0.012 | -0.005 | -0.005 | 0 | 0.62 | 0.65 | 0.74 | 0.67 | 0.67 | 0.63 |
| 3 | 150697208 | rs16863118 | -0.911 | -0.441 | -1.097 | 1.471 | -0.006 | 0.029 | 0.029 | 0 | 0.16 | 0.23 | 0.21 | 0.38 | 0.16 | 0.14 |
| 3 | 150707804 | rs11710513 | -0.698 | -0.838 | -1.689 | 0.962 | 0.047 | -0.03 | -0.03 | 0 | 0.2 | 0.02 | 0.05 | 0.13 | 0.1 | 0.32 |
| 3 | 150707822 | rs7652387 | -0.958 | -1.033 | -0.954 | 0.836 | -0.02 | -0.003 | -0.003 | 0.783 | 0.2 | 0.23 | 0.06 | 0.12 | 0.09 | 0.29 |
| 3 | 150709018 | rs1014691 | -0.53 | -1.171 | -1.499 | 0.606 | 0.054 | 0.042 | 0.042 | 2.725 | 0.51 | 0.71 | 0.37 | 0.4 | 0.32 | 0.44 |
| 3 | 150711491 | rs7625740 | -0.114 | -0.712 | -2.015 | 0.515 | 0.027 | 0.011 | 0.011 | 0 | 0.49 | 0.27 | 0.49 | 0.41 | 0.58 | 0.56 |
| 3 | 150714425 | rs6763580 | -0.003 | -0.396 | -2.133 | 0.544 | -0.01 | 0.03 | 0.03 | 0 | 0.48 | 0.21 | 0.41 | 0.35 | 0.53 | 0.54 |
| 3 | 150716919 | rs957231 | 0.755 | -0.694 | -1.422 | 0.635 | -0.035 | 0.061 | 0.061 | 0.65 | 0.25 | 0.43 | 0.26 | 0.38 | 0.18 | 0.25 |
| 3 | 150721400 | rs9840169 | 0.58 | 0.186 | -1.74 | 0.782 | -0.09 | -0.088 | -0.088 | 0.163 | 0.27 | 0.07 | 0.4 | 0.36 | 0.46 | 0.36 |
| 3 | 150730241 | rs6800123 | 0.015 | 0.14 | -0.983 | 0.994 | 0 | 0 | 0 | 3.595 | 0.75 | 0.74 | 0.72 | 0.77 | 0.75 | 0.74 |
| 3 | 150732589 | rs3105873 | -0.097 | -0.036 | -1.028 | 0.92 | 0 | 0 | 0 | 0 | 0.76 | 0.77 | 0.74 | 0.81 | 0.78 | 0.76 |
| 3 | 150732819 | rs3100792 | 0.025 | -0.247 | -1.303 | 1.167 | 0.004 | -0.035 | -0.035 | 0 | 0.25 | 0.17 | 0.26 | 0.42 | 0.22 | 0.16 |
| 3 | 150745041 | rs6766361 | 0.014 | -0.827 | -0.882 | 0.76 | 0.028 | 0.026 | 0.026 | 1.983 | 0.55 | 0.55 | 0.31 | 0.23 | 0.33 | 0.49 |
| 3 | 150750799 | rs625164 | 1.117 | -0.802 | 0.269 | 1.577 | 0.041 | 0.187 | 0.187 | 0 | 0.92 | 0.67 | 0.79 | 0.79 | 0.93 | 0.87 |
| 3 | 150763869 | rs1456130 | 0.363 | -1.155 | -0.495 | 1.652 | 0.023 | -0.088 | -0.088 | 0 | 0.81 | 0.95 | 0.73 | 0.62 | 0.84 | 0.78 |
| 3 | 150776673 | rs16863147 | -0.586 | -1.297 | 0.375 | 1.969 | 0.018 | 0.321 | 0.321 | 0.926 | 0.1 | 0.47 | 0.19 | 0.42 | 0.06 | 0.21 |
| 3 | 150778251 | rs7634663 | -0.961 | -1.199 | 0.33 | 1.963 | 0.028 | 0.309 | 0.309 | 0.138 | 0.12 | 0.47 | 0.2 | 0.44 | 0.08 | 0.25 |
| 3 | 150782182 | rs358967 | 1.286 | -1.484 | 0.304 | 1.811 | -0.02 | -0.002 | -0.002 | 0.6 | 0.68 | 0.34 | 0.56 | 0.31 | 0.77 | 0.55 |
| 3 | 150792444 | rs6440717 | 0.026 | -1.646 | 0.023 | 1.168 | 0 | -0.003 | -0.003 | 0 | 0.19 | 0.18 | 0.13 | 0.11 | 0.19 | 0.26 |
| 3 | 150801068 | rs7629773 | 0.221 | -0.961 | -1.089 | 0.3 | -0.048 | 0.211 | 0.211 | 0.474 | 0.56 | 0.91 | 0.6 | 0.79 | 0.46 | 0.49 |
| 3 | 150806332 | rs7645665 | 0.775 | -0.611 | -0.744 | 0.91 | 0.01 | 0.024 | 0.024 | 0 | 0.13 | 0.2 | 0.2 | 0.27 | 0.18 | 0.18 |
| 3 | 150818075 | rs2116290 | -1.191 | -0.625 | -1.073 | 0.756 | -0.133 | 0.13 | 0.13 | 0 | 0.75 | 0.98 | 0.77 | 0.92 | 0.65 | 0.77 |
| 3 | 150819083 | rs12634353 | 0.053 | -1.809 | -1.254 | 0.972 | -0.021 | -0.027 | -0.027 | 0.049 | 0.4 | 0.78 | 0.43 | 0.52 | 0.42 | 0.37 |
| 3 | 150822068 | rs970235 | 0.137 | -2.106 | -1.378 | 0.884 | -0.025 | -0.058 | -0.058 | 0 | 0.3 | 0.77 | 0.36 | 0.47 | 0.36 | 0.3 |
| 3 | 150824794 | rs1346943 | 0.153 | -2.17 | -1.366 | 0.917 | -0.026 | -0.06 | -0.06 | 0.359 | 0.3 | 0.77 | 0.36 | 0.47 | 0.36 | 0.3 |
| 3 | 150834015 | rs4680158 | -0.868 | -1.829 | -1.433 | 0.993 | 0.031 | 0.001 | 0.001 | 0.193 | 0.72 | 0.87 | 0.8 | 0.76 | 0.78 | 0.85 |
| 3 | 150848022 | rs12636173 | 1.64 | -1.69 | -1.287 | 1.109 | 0.035 | -0.008 | -0.008 | 0 | 0.11 | 0.06 | 0.06 | 0.08 | 0.05 | 0.09 |
| 3 | 150854520 | rs3915228 | 0.995 | -1.668 | -0.334 | 1.112 | 0.064 | 0.129 | 0.129 | 0 | 0.15 | 0.39 | 0.22 | 0.27 | 0.18 | 0.18 |
| 3 | 150857406 | rs1920392 | 0.814 | -1.674 | -0.098 | 1.096 | 0.063 | 0.125 | 0.125 | 0 | 0.16 | 0.5 | 0.23 | 0.27 | 0.19 | 0.24 |
| 3 | 150867063 | rs6780053 | -0.606 | -1.659 | -0.148 | 1.067 | 0.005 | -0.016 | -0.016 | 0 | 0.3 | 0.68 | 0.33 | 0.54 | 0.27 | 0.35 |
| 3 | 150878729 | rs9823077 | 1.01 | -1.472 | -0.242 | 1.123 | 0.037 | 0.08 | 0.08 | 0 | 0.14 | 0.3 | 0.21 | 0.26 | 0.18 | 0.17 |
| 3 | 150880908 | rs3116549 | 2.094 | -1.307 | -1.052 | 0.808 | -0.056 | 0.015 | 0.015 | 0.008 | 0.19 | 0.01 | 0.24 | 0.14 | 0.21 | 0.27 |
| 3 | 150883600 | rs2567322 | 1.166 | -1.511 | -0.836 | 1.018 | -0.079 | 0.183 | 0.183 | 0.463 | 0.43 | 0.05 | 0.46 | 0.22 | 0.54 | 0.51 |
| 3 | 150886786 | rs1005726 | 1.414 | -1.431 | -0.397 | 1.159 | 0.02 | 0.044 | 0.044 | 0 | 0.13 | 0.26 | 0.2 | 0.27 | 0.18 | 0.17 |
| 3 | 150898634 | rs1017348 | -0.656 | -1.58 | -1.132 | 1.322 | -0.012 | 0.017 | 0.017 | 0.746 | 0.72 | 0.66 | 0.77 | 0.61 | 0.79 | 0.77 |
| 3 | 150911683 | rs1221002 | -1.095 | -1.557 | -1.404 | 1.23 | 0.017 | -0.023 | -0.023 | 0 | 0.75 | 0.93 | 0.81 | 0.76 | 0.81 | 0.88 |
| 3 | 150917000 | rs3773614 | 0.866 | -1.135 | -1.417 | 0.283 | -0.031 | -0.006 | -0.006 | 0.637 | 0.13 | 0.22 | 0.09 | 0.07 | 0.07 | 0.11 |
| 3 | 150928122 | rs12497969 | 0.452 | -0.618 | -1.197 | 0.19 | -0.05 | 0.184 | 0.184 | 0.402 | 0.43 | 0.1 | 0.46 | 0.23 | 0.53 | 0.51 |
| 3 | 150929865 | rs1220996 | 0.719 | -0.472 | -1.197 | 0.279 | -0.089 | 0.162 | 0.162 | 0.212 | 0.41 | 0.06 | 0.45 | 0.23 | 0.51 | 0.49 |
| 3 | 150939858 | rs9881116 | 1.972 | 0.243 | -0.921 | 0.516 | -0.05 | 0.044 | 0.044 | 0 | 0.08 | 0.01 | 0.1 | 0 | 0.18 | 0.04 |
| 3 | 150942578 | rs9817216 | 1.632 | 0.206 | -0.568 | 0.496 | 0.017 | -0.027 | -0.027 | 0 | 0.09 | 0.1 | 0.11 | 0 | 0.18 | 0.04 |
| 3 | 150943539 | rs9289834 | -0.504 | -0.618 | -1.229 | 0.639 | 0.121 | 0 | 0 | 0 | 0.51 | 0.75 | 0.59 | 0.45 | 0.7 | 0.5 |
| 3 | 150944534 | rs16863264 | 0.708 | -0.148 | -0.989 | 0.983 | -0.006 | -0.005 | -0.005 | 0 | 0.24 | 0.23 | 0.25 | 0.25 | 0.33 | 0.3 |
| 3 | 150945243 | rs2870518 | -0.431 | 0.012 | -1.722 | 0.996 | 0.005 | -0.108 | -0.108 | 0 | 0.69 | 0.88 | 0.81 | 0.49 | 0.75 | 0.7 |
| 3 | 150946047 | rs4680254 | 0.202 | -0.505 | -1.236 | 0.924 | -0.031 | -0.019 | -0.019 | 0 | 0.61 | 0.3 | 0.57 | 0.62 | 0.51 | 0.6 |
| 3 | 150946450 | rs13317873 | 0.208 | -0.515 | -1.073 | 0.901 | -0.02 | -0.01 | -0.01 | 0 | 0.61 | 0.36 | 0.58 | 0.62 | 0.51 | 0.6 |
| 3 | 150967269 | rs1882013 | 0.729 | -0.316 | -0.286 | 0.914 | 0.006 | -0.004 | -0.004 | 0 | 0.26 | 0.25 | 0.26 | 0.22 | 0.31 | 0.29 |
| 3 | 150968277 | rs6782715 | 0.367 | -0.368 | -0.108 | 0.889 | -0.007 | -0.004 | -0.004 | 0.111 | 0.61 | 0.37 | 0.57 | 0.62 | 0.51 | 0.59 |
| 3 | 150970104 | rs2141631 | -1.578 | -0.291 | -0.246 | 0.951 | -0.041 | -0.025 | -0.025 | 0 | 0.89 | 0.98 | 0.9 | 0.9 | 0.84 | 0.9 |
| 3 | 150970686 | rs1920395 | 0.519 | -0.564 | -0.115 | 0.582 | 0.048 | 0.01 | 0.01 | 0 | 0.53 | 0.32 | 0.46 | 0.62 | 0.33 | 0.55 |
| 3 | 150977411 | rs9815520 | 0.23 | -0.362 | 0.191 | 0.717 | 0.005 | 0.002 | 0.002 | 0 | 0.62 | 0.44 | 0.59 | 0.63 | 0.5 | 0.58 |
| 3 | 150989323 | rs9870740 | 0.624 | -0.429 | 0.136 | 0.98 | -0.006 | 0.004 | 0.004 | 0.104 | 0.58 | 0.38 | 0.56 | 0.51 | 0.48 | 0.5 |
| 3 | 150990042 | rs7653603 | 0.543 | -0.429 | 0.194 | 0.943 | 0.006 | -0.003 | -0.003 | 0 | 0.59 | 0.47 | 0.57 | 0.51 | 0.48 | 0.5 |
| 3 | 150990510 | rs6440732 | -1.89 | -0.135 | 0.112 | 1.061 | -0.023 | -0.021 | -0.021 | 2.638 | 0.95 | 0.99 | 0.92 | 0.85 | 0.83 | 0.9 |
| 3 | 150996434 | rs3773621 | 0.417 | -0.227 | 0.16 | 0.975 | 0.01 | -0.006 | -0.006 | 0 | 0.27 | 0.35 | 0.3 | 0.21 | 0.32 | 0.31 |
| 3 | 150998157 | rs10935835 | 0.609 | -0.234 | 0.177 | 0.971 | -0.027 | 0.003 | 0.003 | 0 | 0.25 | 0.15 | 0.28 | 0.21 | 0.32 | 0.3 |
| 3 | 151004814 | rs7617677 | 0.772 | -0.267 | 0.044 | 0.985 | -0.005 | 0.004 | 0.004 | 0.102 | 0.57 | 0.35 | 0.56 | 0.52 | 0.48 | 0.5 |

| | | | | | | | | | | | | | | | | |
|---|---|---|---|---|---|---|---|---|---|---|---|---|---|---|---|---|
| 3 | 151007310 | rs9863983 | 0.396 | -0.285 | -0.037 | 0.964 | 0.055 | 0.158 | 0.158 | 1.138 | 0.82 | 0.58 | 0.83 | 0.72 | 0.78 | 0.79 |
| 3 | 151011944 | rs12492699 | 1.405 | -0.875 | -0.449 | 0.775 | -0.052 | 0.069 | 0.069 | 0.042 | 0.25 | 0.01 | 0.22 | 0.16 | 0.23 | 0.35 |
| 3 | 151011969 | rs1048408 | 0.567 | -0.819 | -0.381 | 0.814 | -0.001 | 0 | 0 | 0 | 0.57 | 0.46 | 0.57 | 0.51 | 0.46 | 0.51 |
| 3 | 151012497 | rs2242353 | -0.175 | -0.804 | -0.361 | 0.825 | 0 | 0.01 | 0.01 | 2.276 | 0.83 | 0.87 | 0.86 | 0.72 | 0.78 | 0.79 |
| 3 | 151021983 | rs11719936 | 0.559 | -0.849 | -0.382 | 0.767 | 0 | -0.002 | -0.002 | 0 | 0.57 | 0.48 | 0.57 | 0.51 | 0.47 | 0.53 |
| 3 | 151027359 | rs1125541 | 1.518 | -0.855 | -0.388 | 0.733 | 0 | 0.05 | 0.05 | 0.011 | 0.25 | 0.09 | 0.22 | 0.16 | 0.23 | 0.35 |
| 3 | 151027834 | rs1565573 | 1.518 | -0.855 | -0.388 | 0.733 | 0 | 0.05 | 0.05 | 0.011 | 0.25 | 0.08 | 0.22 | 0.15 | 0.23 | 0.35 |
| 3 | 151034429 | rs3821668 | 0.853 | -0.541 | -0.229 | 1.024 | 0.046 | -0.049 | -0.049 | 0 | 0.11 | 0.01 | 0.03 | 0.13 | 0.05 | 0.1 |
| 3 | 151034602 | rs1472122 | 0.747 | -0.534 | -0.139 | 1.02 | -0.001 | 0 | 0 | 0.745 | 0.57 | 0.54 | 0.58 | 0.51 | 0.46 | 0.53 |
| 3 | 151041513 | rs3971191 | -0.503 | -0.402 | -0.425 | 1.079 | 0.089 | 0.09 | 0.09 | 0 | 0.93 | 0.82 | 0.89 | 0.85 | 0.83 | 0.87 |
| 3 | 151044732 | rs6440735 | 0.034 | -0.398 | -0.573 | 1.115 | -0.059 | -0.106 | -0.106 | 2.967 | 0.83 | 0.97 | 0.87 | 0.72 | 0.78 | 0.8 |
| 3 | 151044903 | rs12497019 | 0.65 | -0.41 | -0.585 | 1.1 | -0.023 | 0.005 | 0.005 | 0 | 0.25 | 0.17 | 0.28 | 0.21 | 0.32 | 0.28 |
| 3 | 151046308 | rs1466684 | -1.691 | -0.363 | -0.656 | 1.169 | -0.011 | -0.008 | -0.008 | 1.734 | 0.95 | 0.97 | 0.91 | 0.85 | 0.83 | 0.9 |
| 3 | 151046604 | rs3732757 | 0.76 | -0.382 | -0.658 | 1.165 | 0.046 | -0.049 | -0.049 | 0 | 0.11 | 0.01 | 0.03 | 0.13 | 0.05 | 0.09 |
| 3 | 151048544 | rs4146770 | -0.212 | -0.378 | -0.828 | 0.961 | -0.052 | -0.071 | -0.071 | 0.003 | 0.84 | 0.96 | 0.88 | 0.79 | 0.8 | 0.83 |
| 3 | 151051278 | rs10935837 | 1.974 | -0.193 | -0.521 | 0.823 | 0.003 | -0.002 | -0.002 | 0 | 0.06 | 0.06 | 0.06 | 0.07 | 0.05 | 0.04 |
| 3 | 151053898 | rs7644001 | 0.054 | -0.029 | -0.305 | 0.695 | -0.008 | 0.006 | 0.006 | 0.096 | 0.38 | 0.5 | 0.34 | 0.34 | 0.36 | 0.37 |
| 3 | 151058441 | rs3821667 | -0.42 | 0.339 | 0.292 | 1.425 | 0.097 | 0.095 | 0.095 | 0 | 0.94 | 0.87 | 0.9 | 0.85 | 0.83 | 0.9 |
| 3 | 151079436 | rs10755105 | -0.078 | -0.101 | -0.04 | 1.329 | -0.025 | -0.047 | -0.047 | 0 | 0.64 | 0.76 | 0.6 | 0.5 | 0.6 | 0.73 |
| 3 | 151082923 | rs1463725 | 0.737 | -1.236 | 0.338 | 1.116 | -0.025 | -0.001 | -0.001 | 0.541 | 0.22 | 0.22 | 0.2 | 0.35 | 0.04 | 0.13 |
| 3 | 151088006 | rs7646193 | 1.171 | -1.042 | -0.337 | 0.96 | 0.017 | 0.008 | 0.008 | 0.229 | 0.18 | 0.09 | 0.04 | 0.17 | 0.03 | 0.13 |
| 3 | 151089226 | rs7637803 | -0.035 | -1.053 | -0.002 | 0.573 | -0.029 | -0.041 | -0.041 | 2.191 | 0.34 | 0.38 | 0.19 | 0.17 | 0.2 | 0.17 |
| 3 | 151090424 | rs3732765 | 1.333 | -0.584 | -0.615 | 0.16 | -0.104 | 0.125 | 0.125 | 0 | 0.27 | 0.03 | 0.31 | 0.14 | 0.38 | 0.25 |
| 3 | 151090963 | rs9859538 | 0.425 | -0.591 | -0.566 | 0.17 | -0.083 | 0.245 | 0.245 | 0 | 0.41 | 0.1 | 0.43 | 0.14 | 0.56 | 0.33 |
| 3 | 151097391 | rs17283010 | 1.018 | -0.576 | -0.462 | 0.183 | 0.002 | -0.033 | -0.033 | 0.283 | 0.3 | 0.33 | 0.34 | 0.14 | 0.38 | 0.25 |
| 3 | 151100956 | rs11713504 | 1.192 | -0.565 | -0.448 | 0.188 | -0.062 | 0.114 | 0.114 | 0 | 0.28 | 0.12 | 0.31 | 0.14 | 0.38 | 0.26 |
| 3 | 151101083 | rs10935840 | 1.204 | -0.564 | -0.448 | 0.188 | -0.062 | 0.113 | 0.113 | 0 | 0.28 | 0.13 | 0.31 | 0.14 | 0.38 | 0.26 |
| 3 | 151101167 | rs7429509 | 0.989 | -0.303 | -0.162 | 0.946 | -0.009 | 0.012 | 0.012 | 0 | 0.24 | 0.33 | 0.22 | 0.28 | 0.2 | 0.37 |
| 3 | 151104838 | rs1907637 | -1.364 | -0.202 | -0.185 | 1.254 | -0.014 | -0.011 | -0.011 | 0 | 0.95 | 0.98 | 0.92 | 0.83 | 0.86 | 0.91 |
| 3 | 151111790 | rs9653953 | 0.175 | -0.263 | -0.123 | 1.388 | 0.011 | -0.006 | -0.006 | 0.11 | 0.63 | 0.52 | 0.65 | 0.45 | 0.59 | 0.47 |
| 3 | 151112568 | rs3732768 | 0.931 | -0.556 | -0.42 | 1.343 | -0.017 | 0.024 | 0.024 | 0 | 0.24 | 0.37 | 0.22 | 0.28 | 0.2 | 0.38 |
| 3 | 151115991 | rs765281 | 0.534 | -0.775 | -0.588 | 1.158 | 0.033 | 0.12 | 0.12 | 0 | 0.46 | 0.23 | 0.36 | 0.31 | 0.41 | 0.42 |
| 3 | 151128895 | rs10935844 | -0.273 | -0.662 | -0.546 | 1.079 | 0.001 | 0.006 | 0.006 | 0 | 0.56 | 0.51 | 0.58 | 0.55 | 0.48 | 0.49 |
| 3 | 151132206 | rs16863374 | -0.368 | -0.843 | 0.026 | 1.469 | 0.002 | 0.115 | 0.115 | 0 | 0.09 | 0.26 | 0.07 | 0.13 | 0.06 | 0.07 |
| 3 | 151133180 | rs16863376 | 0.973 | -0.92 | -0.597 | 1.225 | -0.018 | -0.003 | -0.003 | 0 | 0.18 | 0.13 | 0.19 | 0.3 | 0.03 | 0.13 |
| 3 | 151133816 | rs3821665 | -0.292 | 0.649 | -0.454 | 1.224 | -0.021 | 0.01 | 0.01 | 0 | 0.41 | 0.3 | 0.53 | 0.43 | 0.52 | 0.52 |
| 3 | 151135126 | rs11714493 | 1.183 | 1.486 | 0.171 | 1.337 | -0.021 | 0.005 | 0.005 | 0 | 0.62 | 0.28 | 0.41 | 0.43 | 0.54 | 0.53 |
| 3 | 151140556 | rs9840738 | 0.273 | 1.64 | 0.345 | 1.851 | 0.039 | 0.005 | 0.005 | 0 | 0.93 | 0.86 | 0.83 | 0.9 | 0.85 | 0.81 |
| 3 | 151147968 | rs6772196 | 0.166 | 1.647 | 0.295 | 1.973 | 0.007 | 0.224 | 0.224 | 3.338 | 0.08 | 0.19 | 0.21 | 0.41 | 0.09 | 0.17 |
| 3 | 151151220 | rs7611956 | 0.354 | 1.694 | -0.247 | 1.077 | 0.01 | 0 | 0 | 0.052 | 0.93 | 0.91 | 0.83 | 0.9 | 0.85 | 0.81 |
| 3 | 151161112 | rs12487205 | -0.66 | 1.267 | -0.428 | 0.995 | 0.021 | 0.04 | 0.04 | 0 | 0.08 | 0.01 | 0.05 | 0.02 | 0.04 | 0.03 |
| 3 | 151165678 | rs9871952 | 0.888 | 1.85 | -0.398 | 1.169 | 0.05 | 0.031 | 0.031 | 0 | 0.91 | 0.82 | 0.77 | 0.76 | 0.77 | 0.79 |
| 3 | 151166058 | rs17204557 | -0.399 | 1.804 | -0.431 | 1.148 | 0.015 | 0.036 | 0.036 | 0 | 0.08 | 0 | 0.05 | 0.02 | 0.03 | 0.13 |
| 3 | 151166422 | rs6798252 | 1.767 | 1.791 | -0.339 | 1.143 | 0.024 | 0.008 | 0.008 | 0 | 0.82 | 0.71 | 0.71 | 0.74 | 0.74 | 0.66 |
| 3 | 151168919 | rs4680442 | 1.145 | 2.182 | -0.375 | 1.42 | 0.054 | 0.036 | 0.036 | 0 | 0.91 | 0.81 | 0.78 | 0.76 | 0.77 | 0.81 |
| 3 | 151171542 | rs7621591 | 0.707 | 2.271 | -0.604 | 1.444 | 0.017 | 0.005 | 0.005 | 0 | 0.92 | 0.89 | 0.78 | 0.76 | 0.77 | 0.81 |
| 3 | 151184862 | rs6790903 | 0.626 | 1.108 | -0.613 | 1.337 | 0.002 | 0.017 | 0.017 | 0.734 | 0.62 | 0.46 | 0.57 | 0.45 | 0.57 | 0.61 |
| 3 | 151187290 | rs4564957 | -0.531 | 1.411 | -1.066 | 1.238 | -0.002 | -0.003 | -0.003 | 0.025 | 0.37 | 0.34 | 0.38 | 0.35 | 0.43 | 0.26 |
| 3 | 151194880 | rs9289840 | 1.284 | 0.543 | -1.607 | 1.087 | 0.001 | -0.004 | -0.004 | 0.599 | 0.4 | 0.6 | 0.45 | 0.44 | 0.35 | 0.46 |
| 3 | 151194924 | rs4312631 | 0.311 | 0.841 | -1.37 | 1.155 | -0.022 | -0.003 | -0.003 | 0 | 0.25 | 0.17 | 0.29 | 0.27 | 0.38 | 0.16 |
| 3 | 151196321 | rs6773513 | -0.737 | 0.754 | -1.404 | 1.147 | -0.049 | -0.016 | -0.016 | 0 | 0.9 | 0.97 | 0.87 | 0.9 | 0.84 | 0.83 |
| 3 | 151199905 | rs9877416 | 0.593 | 0.758 | -1.26 | 1.158 | 0.001 | 0 | 0 | 0 | 0.78 | 0.69 | 0.77 | 0.82 | 0.8 | 0.72 |
| 3 | 151210190 | rs9876367 | -0.598 | 0.848 | -1.272 | 1.328 | -0.046 | -0.049 | -0.049 | 0.094 | 0.9 | 0.94 | 0.85 | 0.75 | 0.84 | 0.82 |
| 3 | 151217514 | rs7617486 | 1.03 | 0.808 | -1.18 | 1.288 | -0.004 | 0.024 | 0.024 | 0.452 | 0.55 | 0.68 | 0.55 | 0.61 | 0.43 | 0.61 |
| 3 | 151219429 | rs11920961 | -0.139 | 0.777 | -1.271 | 1.227 | -0.032 | 0.01 | 0.01 | 0 | 0.3 | 0.16 | 0.29 | 0.27 | 0.4 | 0.18 |
| 3 | 151220186 | rs4679848 | -0.139 | 0.76 | -1.24 | 1.239 | 0.01 | 0.011 | 0.011 | 0 | 0.84 | 0.73 | 0.79 | 0.83 | 0.83 | 0.76 |
| 3 | 151220624 | rs7622119 | -0.116 | 0.799 | -1.229 | 1.071 | -0.033 | 0.079 | 0.079 | 0 | 0.3 | 0.16 | 0.28 | 0.12 | 0.4 | 0.18 |
| 3 | 151227212 | rs7629867 | 0.68 | 0.254 | -1.094 | 1.023 | 0.005 | 0.001 | 0.001 | 0.39 | 0.51 | 0.52 | 0.49 | 0.55 | 0.41 | 0.54 |
| 3 | 151245039 | rs9836445 | 0.686 | 0.274 | -1.3 | 0.988 | 0.016 | 0.006 | 0.006 | 0.262 | 0.51 | 0.57 | 0.48 | 0.56 | 0.39 | 0.56 |
| 3 | 151262987 | rs7631760 | 0.081 | 0.615 | -1.372 | 0.973 | -0.058 | -0.013 | -0.013 | 0 | 0.65 | 0.81 | 0.62 | 0.66 | 0.5 | 0.67 |
| 3 | 151272986 | rs4383491 | -0.189 | 0.46 | -1.543 | 0.448 | -0.005 | 0.028 | 0.028 | 0 | 0.22 | 0.16 | 0.11 | 0.13 | 0.17 | 0.11 |
| 3 | 151274346 | rs7649833 | -0.476 | 0.753 | -1.284 | 0.867 | 0.004 | 0.008 | 0.008 | 0.25 | 0.75 | 0.68 | 0.69 | 0.68 | 0.65 | 0.7 |
| 3 | 151283646 | rs7631281 | -1.009 | 0.729 | -1.262 | 0.904 | -0.006 | -0.001 | -0.001 | 0.018 | 0.78 | 0.77 | 0.76 | 0.8 | 0.7 | 0.78 |
| 3 | 151283691 | rs9842591 | 0.743 | 0.619 | -0.738 | 0.862 | -0.002 | 0 | 0 | 0.198 | 0.52 | 0.47 | 0.57 | 0.54 | 0.47 | 0.59 |
| 3 | 151287356 | rs7629779 | -0.851 | 0.848 | -0.735 | 0.833 | -0.015 | -0.001 | -0.001 | 4.778 | 0.78 | 0.82 | 0.77 | 0.79 | 0.7 | 0.78 |
| 3 | 151314855 | rs6783939 | 0.301 | 0.877 | -0.782 | 0.452 | -0.008 | 0 | 0 | 0 | 0.11 | 0.07 | 0.07 | 0 | 0.14 | 0.06 |
| 3 | 151319088 | rs7620281 | 0.248 | -0.161 | -0.866 | 0.434 | -0.021 | -0.008 | -0.008 | 0 | 0.27 | 0.38 | 0.34 | 0.26 | 0.26 | 0.13 |
| 3 | 151320010 | rs9289851 | -0.046 | 0.035 | -0.926 | 0.432 | -0.006 | -0.005 | -0.005 | 0.884 | 0.58 | 0.57 | 0.6 | 0.58 | 0.52 | 0.64 |
| 3 | 151323327 | rs7616331 | -0.197 | -0.159 | -0.453 | 0.149 | 0.016 | 0.064 | 0.064 | 0 | 0.22 | 0.55 | 0.29 | 0.28 | 0.28 | 0.14 |

| | | | | | | | | | | | | | | | | |
|---|---|---|---|---|---|---|---|---|---|---|---|---|---|---|---|---|
| 3 | 151325890 | rs7637081 | 0.43 | -0.32 | -0.027 | -0.151 | -0.02 | -0.023 | -0.023 | 0.185 | 0.13 | 0.29 | 0.18 | 0.05 | 0.1 | 0.05 |
| 3 | 151328453 | rs1521592 | -0.815 | 0.172 | -0.741 | -0.337 | -0.017 | 0.084 | 0.084 | 0 | 0.57 | 0.78 | 0.59 | 0.69 | 0.56 | 0.71 |
| 3 | 151333454 | rs16863517 | 0.146 | 0.753 | -0.217 | 0.013 | 0.032 | -0.048 | -0.048 | 0.041 | 0.09 | 0.25 | 0.09 | 0 | 0.1 | 0.02 |
| 3 | 151335734 | rs4680567 | 0.025 | 0.522 | -0.998 | 0.205 | -0.016 | -0.009 | -0.009 | 0 | 0.3 | 0.22 | 0.3 | 0.63 | 0.34 | 0.56 |
| 3 | 151342245 | rs1915717 | -0.601 | 0.766 | -1.253 | 0.112 | -0.001 | -0.006 | -0.006 | 0 | 0.62 | 0.66 | 0.6 | 0.77 | 0.64 | 0.64 |
| 3 | 151345293 | rs4680575 | -0.333 | 0.885 | -1.848 | 0.117 | -0.02 | 0.078 | 0.078 | 0.505 | 0.35 | 0.14 | 0.31 | 0.23 | 0.34 | 0.22 |
| 3 | 151359752 | rs9882483 | -0.673 | 0.926 | -1.315 | 0.582 | -0.003 | -0.008 | -0.008 | 0.022 | 0.15 | 0.14 | 0.13 | 0.14 | 0.17 | 0.06 |
| 3 | 151360593 | rs17283150 | 0.519 | 1.135 | -1.484 | 0.267 | -0.01 | 0.014 | 0.014 | 0 | 0.07 | 0.02 | 0.04 | 0 | 0.07 | 0.06 |
| 3 | 151361249 | rs4680584 | 0.491 | 1.76 | -1.072 | 0.468 | 0.026 | 0.063 | 0.063 | 0 | 0.19 | 0.27 | 0.33 | 0.57 | 0.26 | 0.3 |
| 3 | 151370083 | rs1402740 | -0.193 | 1.056 | -0.801 | -0.285 | 0.006 | -0.012 | -0.012 | 0.752 | 0.42 | 0.49 | 0.55 | 0.74 | 0.45 | 0.62 |
| 3 | 151371789 | rs12330894 | -0.506 | 0.851 | -0.273 | -0.497 | 0.005 | -0.007 | -0.007 | 0 | 0.23 | 0.28 | 0.21 | 0.14 | 0.28 | 0.14 |
| 3 | 151371825 | rs12491104 | 1.359 | 0.906 | -0.543 | -0.361 | -0.063 | -0.09 | -0.09 | 0 | 0.18 | 0.07 | 0.31 | 0.55 | 0.27 | 0.3 |
| 3 | 151374176 | rs12630593 | 0.988 | 0.767 | -0.409 | -0.455 | 0.012 | 0.016 | 0.016 | 0 | 0.22 | 0.15 | 0.22 | 0.2 | 0.18 | 0.31 |
| 3 | 151375352 | rs17204620 | 0.957 | 0.78 | -0.459 | -0.469 | -0.017 | 0.03 | 0.03 | 0 | 0.09 | 0.02 | 0.08 | 0 | 0.11 | 0.09 |
| 3 | 151375575 | rs4679898 | -0.871 | 0.679 | -0.369 | -0.46 | 0.002 | 0.018 | 0.018 | 0 | 0.43 | 0.65 | 0.55 | 0.68 | 0.55 | 0.45 |
| 3 | 151377883 | rs7628408 | -1.48 | 0.739 | -0.319 | -0.423 | 0.015 | 0.014 | 0.014 | 0.011 | 0.66 | 0.75 | 0.77 | 0.88 | 0.75 | 0.79 |
| 3 | 151393037 | rs10935868 | 0.413 | 0.987 | -0.576 | -0.227 | -0.087 | -0.11 | -0.11 | 0.795 | 0.25 | 0.08 | 0.33 | 0.55 | 0.34 | 0.33 |
| 3 | 151393494 | rs13080679 | 0.048 | 0.74 | -0.553 | -0.253 | -0.026 | 0.112 | 0.112 | 0 | 0.36 | 0.15 | 0.35 | 0.18 | 0.39 | 0.42 |
| 3 | 151394601 | rs11718642 | 0.879 | 0.782 | -0.624 | -0.258 | -0.005 | 0.049 | 0.049 | 0 | 0.27 | 0.12 | 0.27 | 0.19 | 0.28 | 0.34 |
| 3 | 151396537 | rs1521590 | -1.469 | 0.78 | -0.455 | -0.229 | -0.043 | -0.011 | -0.011 | 0 | 0.65 | 0.56 | 0.71 | 0.74 | 0.75 | 0.78 |
| 3 | 151398073 | rs10222585 | -1.459 | 0.78 | -0.455 | -0.229 | -0.043 | -0.011 | -0.011 | 0 | 0.65 | 0.56 | 0.71 | 0.74 | 0.75 | 0.78 |
| 3 | 151399344 | rs12495578 | 0.197 | 0.963 | -0.685 | -0.15 | -0.042 | -0.047 | -0.047 | 0 | 0.27 | 0.19 | 0.35 | 0.56 | 0.36 | 0.36 |
| 3 | 151412554 | rs7644103 | 0.18 | 0.786 | -0.356 | 0.407 | -0.006 | -0.004 | -0.004 | 0 | 0.29 | 0.3 | 0.3 | 0.42 | 0.4 | 0.41 |
| 3 | 151414310 | rs7648113 | -1.322 | 0.197 | -0.863 | -0.056 | 0.062 | 0.097 | 0.097 | 0.142 | 0.67 | 0.89 | 0.77 | 0.84 | 0.75 | 0.79 |
| 3 | 151414394 | rs7648055 | -1.216 | 0.23 | -0.804 | -0.041 | 0.049 | 0.069 | 0.069 | 0.015 | 0.66 | 0.82 | 0.77 | 0.84 | 0.75 | 0.79 |
| 3 | 151425097 | rs991866 | -0.503 | 0.759 | -0.437 | 0.31 | -0.01 | 0.027 | 0.027 | 0 | 0.4 | 0.7 | 0.51 | 0.65 | 0.48 | 0.46 |
| 3 | 151430857 | rs3988161 | -0.073 | 0.422 | -0.389 | 0.362 | 0.002 | 0.003 | 0.003 | 0 | 0.76 | 0.71 | 0.6 | 0.75 | 0.68 | 0.76 |
| 3 | 151436104 | rs1474251 | -0.154 | 0.407 | -0.268 | 0.363 | -0.007 | -0.012 | -0.012 | 0 | 0.76 | 0.81 | 0.61 | 0.75 | 0.68 | 0.76 |
| 3 | 151438247 | rs16846748 | 1.062 | -0.371 | -0.485 | -0.477 | 0.025 | 0.028 | 0.028 | 0.053 | 0.23 | 0.16 | 0.1 | 0.02 | 0.11 | 0.14 |
| 3 | 151441983 | rs16863583 | 1.713 | -0.859 | -0.056 | -0.209 | -0.041 | 0.035 | 0.035 | 0 | 0.09 | 0.18 | 0.04 | 0.15 | 0.01 | 0.03 |
| 3 | 151445322 | rs1850945 | 1.575 | -0.667 | 0.007 | -0.134 | -0.01 | -0.011 | -0.011 | 0.313 | 0.19 | 0.14 | 0.17 | 0.4 | 0.27 | 0.23 |
| 3 | 151452440 | rs1520132 | 1.412 | -0.624 | -0.07 | -0.201 | -0.006 | 0.019 | 0.019 | 0 | 0.23 | 0.13 | 0.27 | 0.17 | 0.26 | 0.32 |
| 3 | 151460532 | rs1401962 | 0.644 | -0.379 | 0.041 | -0.018 | 0.032 | 0.094 | 0.094 | 0 | 0.46 | 0.3 | 0.37 | 0.2 | 0.37 | 0.46 |
| 3 | 151461258 | rs9815308 | -0.667 | -0.309 | -0.081 | -0.077 | -0.018 | -0.01 | -0.01 | 0 | 0.74 | 0.81 | 0.53 | 0.75 | 0.64 | 0.73 |
| 3 | 151461385 | rs10513418 | 0.389 | -0.253 | -0.091 | 0.018 | 0.024 | 0.056 | 0.056 | 0 | 0.29 | 0.41 | 0.22 | 0.55 | 0.31 | 0.3 |
| 3 | 151462665 | rs1401961 | -0.378 | -0.098 | 0.036 | 0.224 | 0.004 | 0.011 | 0.011 | 0 | 0.75 | 0.67 | 0.59 | 0.74 | 0.68 | 0.76 |
| 3 | 151463421 | rs1972977 | -0.467 | -0.13 | -0.109 | 0.645 | -0.008 | -0.026 | -0.026 | 0 | 0.76 | 0.82 | 0.65 | 0.62 | 0.67 | 0.72 |
| 3 | 151464131 | rs1120606 | 0.253 | -0.354 | -0.066 | 0.407 | -0.008 | -0.007 | -0.007 | 0 | 0.52 | 0.45 | 0.33 | 0.35 | 0.35 | 0.47 |
| 3 | 151465100 | rs9846265 | 1.288 | -0.063 | -0.16 | 0.255 | 0.018 | 0.018 | 0.018 | 0.336 | 0.27 | 0.16 | 0.12 | 0.18 | 0.18 | 0.11 |
| 3 | 151471137 | rs977547 | 0.888 | -0.114 | -0.053 | 0.253 | -0.035 | -0.012 | -0.012 | 0.045 | 0.29 | 0.35 | 0.16 | 0.25 | 0.18 | 0.13 |
| 3 | 151476514 | rs9839054 | 1.321 | -0.09 | -0.186 | 0.265 | 0.021 | 0.006 | 0.006 | 0.284 | 0.27 | 0.16 | 0.13 | 0.25 | 0.18 | 0.13 |
| 3 | 151486525 | rs9826691 | 1.321 | -0.09 | -0.166 | 0.265 | 0.028 | 0.008 | 0.008 | 0.284 | 0.27 | 0.15 | 0.13 | 0.25 | 0.18 | 0.13 |
| 3 | 151488333 | rs6809359 | 0.776 | 0.074 | 0.078 | 0.263 | 0.04 | 0.015 | 0.015 | 0 | 0.2 | 0.39 | 0.24 | 0.26 | 0.19 | 0.23 |
| 3 | 151493391 | rs12152279 | 1.147 | 0.088 | -0.016 | 0.271 | 0.001 | 0 | 0 | 0 | 0.18 | 0.23 | 0.22 | 0.26 | 0.19 | 0.23 |
| 3 | 151499361 | rs4679938 | 0.095 | 0.047 | -0.04 | 0.16 | 0.045 | -0.02 | -0.02 | 0 | 0.29 | 0.5 | 0.3 | 0.26 | 0.28 | 0.3 |
| 3 | 151500579 | rs7644262 | -0.215 | 0.022 | -0.074 | 0.168 | -0.004 | -0.003 | -0.003 | 0.007 | 0.55 | 0.66 | 0.43 | 0.51 | 0.46 | 0.42 |
| 3 | 151502077 | rs1401967 | 0.087 | 0.071 | -0.028 | 0.174 | 0.045 | -0.02 | -0.02 | 0 | 0.29 | 0.5 | 0.3 | 0.26 | 0.28 | 0.3 |
| 3 | 151503678 | rs16863663 | 1.47 | -0.019 | -0.373 | 0.287 | 0.027 | 0.008 | 0.008 | 0.347 | 0.26 | 0.15 | 0.13 | 0.25 | 0.18 | 0.13 |
| 3 | 151505556 | rs6440785 | 0.961 | 0.224 | -0.565 | 0.156 | -0.001 | -0.011 | -0.011 | 1.156 | 0.12 | 0.13 | 0.15 | 0.02 | 0.12 | 0.05 |
| 3 | 151507750 | rs1401970 | 1.331 | 0.55 | -0.594 | 0.395 | -0.027 | -0.028 | -0.028 | 0 | 0.18 | 0.11 | 0.22 | 0.28 | 0.17 | 0.2 |
| 3 | 151508246 | rs1520149 | -0.266 | 0.459 | -0.302 | 0.502 | 0.002 | 0.02 | 0.02 | 0 | 0.65 | 0.6 | 0.58 | 0.54 | 0.56 | 0.47 |
| 3 | 151510257 | rs6440786 | -1.262 | 0.586 | -0.858 | -0.112 | -0.041 | 0.035 | 0.035 | 0 | 0.88 | 0.98 | 0.84 | 0.95 | 0.79 | 0.82 |
| 3 | 151514590 | rs17204697 | 1.105 | 0.349 | -0.584 | -0.092 | 0.001 | -0.036 | -0.036 | 0.061 | 0.09 | 0.13 | 0.07 | 0 | 0.09 | 0.06 |
| 3 | 151514705 | rs9874357 | 0.994 | 0.275 | -0.665 | -0.17 | 0.002 | 0.006 | 0.006 | 0 | 0.2 | 0.15 | 0.17 | 0.1 | 0.22 | 0.28 |
| 3 | 151517293 | rs954710 | -0.097 | 0.837 | -0.664 | 0.01 | -0.01 | 0.009 | 0.009 | 0 | 0.4 | 0.31 | 0.44 | 0.33 | 0.38 | 0.31 |
| 3 | 151518154 | rs13081526 | 0.888 | 1.003 | -0.447 | 0.189 | -0.004 | 0.008 | 0.008 | 0 | 0.16 | 0.25 | 0.15 | 0.15 | 0.15 | 0.21 |
| 3 | 151521001 | rs9862443 | 1.32 | 0.93 | -0.508 | 0.024 | -0.001 | -0.002 | -0.002 | 0.238 | 0.18 | 0.18 | 0.17 | 0.1 | 0.22 | 0.24 |
| 3 | 151525324 | rs12107596 | 0.679 | 0.863 | -0.576 | -0.041 | 0.002 | 0.003 | 0.003 | 0 | 0.31 | 0.31 | 0.36 | 0.33 | 0.29 | 0.26 |
| 3 | 151530121 | rs2120982 | -1.099 | 0.96 | -0.662 | -0.075 | 0.004 | -0.012 | -0.012 | 0.184 | 0.66 | 0.79 | 0.69 | 0.58 | 0.68 | 0.73 |
| 3 | 151531894 | rs2293004 | 0.68 | 0.895 | -0.676 | -0.051 | 0.002 | 0.003 | 0.003 | 0 | 0.31 | 0.31 | 0.36 | 0.33 | 0.3 | 0.26 |
| 3 | 151533341 | rs1520137 | -0.007 | 0.846 | -0.701 | 0.026 | 0 | 0.001 | 0.001 | 0.772 | 0.49 | 0.47 | 0.52 | 0.42 | 0.52 | 0.51 |
| 3 | 151535422 | rs2271942 | -2.087 | 0.849 | -0.629 | 0.038 | 0.002 | -0.008 | -0.008 | 0 | 0.76 | 0.79 | 0.76 | 0.58 | 0.77 | 0.8 |
| 3 | 151537717 | rs8193027 | 1.29 | 0.847 | -0.614 | 0.016 | -0.005 | 0.004 | 0.004 | 0 | 0.15 | 0.22 | 0.14 | 0.15 | 0.14 | 0.21 |
| 3 | 151538039 | rs2271940 | -0.549 | 0.854 | -0.616 | 0.015 | -0.003 | 0.002 | 0.002 | 0.742 | 0.6 | 0.55 | 0.61 | 0.43 | 0.63 | 0.59 |
| 3 | 151545601 | rs1803155 | -2.002 | 0.867 | -0.629 | 0.125 | 0.002 | -0.007 | -0.007 | 1.181 | 0.76 | 0.77 | 0.75 | 0.58 | 0.77 | 0.8 |
| 3 | 151546041 | rs1042201 | -0.117 | 0.905 | -0.607 | 0.11 | -0.001 | 0 | 0 | 0.806 | 0.49 | 0.47 | 0.52 | 0.43 | 0.51 | 0.5 |
| 3 | 151554749 | rs6440788 | -0.81 | 1.433 | -0.436 | 0.395 | 0.001 | -0.002 | -0.002 | 0.045 | 0.57 | 0.6 | 0.58 | 0.51 | 0.54 | 0.55 |
| 3 | 151561598 | rs1881988 | -0.882 | 1.989 | -0.31 | 0.433 | -0.032 | -0.012 | -0.012 | 0 | 0.38 | 0.71 | 0.49 | 0.36 | 0.38 | 0.31 |
| 3 | 151574286 | rs10804755 | -1.681 | 2.046 | -0.564 | 0.368 | 0.053 | -0.013 | -0.013 | 0 | 0.5 | 0.84 | 0.61 | 0.41 | 0.56 | 0.47 |

| | | | | | | | | | | | | | | | | |
|---|---|---|---|---|---|---|---|---|---|---|---|---|---|---|---|---|
| 3 | 151575600 | rs12152407 | 0.839 | 2.458 | -0.44 | 0.395 | -0.034 | 0.001 | 0.001 | 0.405 | 0.19 | 0.07 | 0.2 | 0.19 | 0.2 | 0.18 |
| 3 | 151582195 | rs2410833 | 0.416 | 2.856 | 0.214 | 0.607 | 0 | -0.001 | -0.001 | 0 | 0.07 | 0.06 | 0.05 | 0.02 | 0.08 | 0.05 |
| 3 | 151584657 | rs9823831 | -1.233 | 3.105 | 0.205 | 0.488 | -0.004 | 0 | 0 | 0.013 | 0.23 | 0.21 | 0.3 | 0.08 | 0.35 | 0.28 |
| 3 | 151585619 | rs6800968 | -0.418 | 3.476 | 0.447 | 0.971 | -0.018 | -0.004 | -0.004 | 0 | 0.25 | 0.17 | 0.37 | 0.24 | 0.3 | 0.24 |
| 3 | 151587637 | rs6783742 | -0.476 | 3.605 | 0.323 | 0.92 | -0.05 | -0.023 | -0.023 | 0 | 0.81 | 0.94 | 0.71 | 0.78 | 0.77 | 0.82 |
| 3 | 151589418 | rs6763405 | -0.984 | 3.468 | 0.31 | 0.936 | -0.033 | 0.006 | 0.006 | 0.066 | 0.93 | 1 | 0.87 | 0.97 | 0.92 | 0.94 |
| 3 | 151589783 | rs1402012 | 0.029 | 3.256 | 0.633 | 1.064 | 0.004 | -0.009 | -0.009 | 0.535 | 0.1 | 0.11 | 0.15 | 0.02 | 0.14 | 0.08 |
| 3 | 151596588 | rs9811297 | -0.029 | 3.103 | 0.806 | 0.711 | 0.003 | -0.008 | -0.008 | 0 | 0.1 | 0.11 | 0.15 | 0.02 | 0.14 | 0.08 |
| 3 | 151597310 | rs13315275 | -1.684 | 3.383 | 1.006 | 0.661 | 0.025 | -0.009 | -0.009 | 0.02 | 0.33 | 0.34 | 0.44 | 0.25 | 0.39 | 0.31 |
| 3 | 151599393 | rs13079080 | -1.487 | 3.375 | 2.02 | 0.87 | 0.001 | 0 | 0 | 0 | 0.35 | 0.57 | 0.44 | 0.3 | 0.38 | 0.35 |
| 3 | 151602690 | rs2120978 | 1.444 | 3.472 | 1.987 | 0.85 | 0.007 | -0.023 | -0.023 | 0 | 0.06 | 0.1 | 0.05 | 0 | 0.07 | 0.04 |
| 3 | 151607481 | rs6786530 | -0.06 | 3.411 | 1.839 | 0.869 | 0.003 | -0.006 | -0.006 | 0 | 0.1 | 0.1 | 0.15 | 0.03 | 0.13 | 0.09 |
| 3 | 151617573 | rs17204725 | -0.666 | 3.398 | 1.454 | 0.863 | -0.008 | -0.002 | -0.002 | 0 | 0.26 | 0.19 | 0.32 | 0.23 | 0.29 | 0.24 |
| 3 | 151618335 | rs9865108 | 0.155 | 3.494 | 1.582 | 0.831 | 0.119 | 0.042 | 0.042 | 0 | 0.08 | 0.33 | 0.11 | 0.07 | 0.1 | 0.11 |
| 3 | 151618571 | rs17283255 | 1.059 | 3.268 | 1.555 | 0.889 | -0.027 | -0.04 | -0.04 | 0 | 0.16 | 0.05 | 0.17 | 0.2 | 0.17 | 0.16 |
| 3 | 151619375 | rs4679951 | -1.304 | 3.329 | 1.626 | 0.871 | 0.074 | -0.034 | -0.034 | 0 | 0.18 | 0.46 | 0.26 | 0.09 | 0.21 | 0.19 |
| 3 | 151619732 | rs12495929 | 0.925 | 3.255 | 1.457 | 0.864 | -0.026 | -0.04 | -0.04 | 0.038 | 0.16 | 0.05 | 0.17 | 0.2 | 0.17 | 0.16 |
| 3 | 151624629 | rs3107329 | -0.469 | 3.429 | 1.498 | 0.637 | 0.012 | -0.015 | -0.015 | 0 | 0.92 | 0.84 | 0.86 | 0.95 | 0.81 | 0.86 |
| 3 | 151627988 | rs6806340 | -0.089 | 3.161 | 1.529 | 1.11 | 0.033 | 0.115 | 0.115 | 0 | 0.19 | 0.39 | 0.27 | 0.34 | 0.19 | 0.23 |
| 3 | 151631698 | rs12489140 | -1.422 | 2.989 | 1.226 | 1.093 | 0.001 | 0 | 0 | 0 | 0.34 | 0.56 | 0.46 | 0.37 | 0.38 | 0.35 |
| 3 | 151639765 | rs7648585 | -1.189 | 2.847 | 1.315 | 1.406 | -0.001 | 0 | 0 | 0 | 0.33 | 0.56 | 0.45 | 0.48 | 0.36 | 0.34 |
| 3 | 151653029 | rs4679954 | -0.25 | 2.536 | 1.114 | 1.219 | -0.012 | 0.011 | 0.011 | 0 | 0.08 | 0.05 | 0.14 | 0.02 | 0.11 | 0.09 |
| 3 | 151653472 | rs17283269 | 0.067 | 2.746 | 1.02 | 1.244 | -0.069 | -0.061 | -0.061 | 0.692 | 0.21 | 0.04 | 0.25 | 0.27 | 0.27 | 0.17 |
| 3 | 151655279 | rs10935880 | -0.169 | 2.777 | 1.1 | 1.199 | 0.03 | 0.057 | 0.057 | 0 | 0.22 | 0.28 | 0.3 | 0.37 | 0.27 | 0.21 |
| 3 | 151660625 | rs12497640 | -0.844 | 2.62 | 0.939 | 1.098 | 0.016 | -0.023 | -0.023 | 0 | 0.09 | 0.19 | 0.15 | 0.02 | 0.11 | 0.09 |
| 3 | 151670949 | rs17204732 | -0.943 | 2.587 | 0.799 | 1.205 | 0.037 | 0.03 | 0.03 | 0 | 0.31 | 0.48 | 0.46 | 0.38 | 0.38 | 0.31 |
| 3 | 151673589 | rs7622233 | -0.906 | 2.454 | 0.926 | 1.052 | 0.066 | 0.012 | 0.012 | 0 | 0.15 | 0.4 | 0.26 | 0.11 | 0.17 | 0.16 |
| 3 | 151674477 | rs16863765 | -0.595 | 2.494 | 1 | 1.276 | 0.026 | 0.059 | 0.059 | 0.693 | 0.25 | 0.51 | 0.38 | 0.37 | 0.28 | 0.25 |
| 3 | 151676322 | rs2723362 | -0.122 | 2.175 | 1.051 | 0.902 | 0.061 | 0.003 | 0.003 | 0 | 0.84 | 0.63 | 0.74 | 0.86 | 0.76 | 0.82 |
| 3 | 151676355 | rs2590702 | 1.304 | 2.031 | 0.898 | 0.851 | 0.056 | 0.022 | 0.022 | 0 | 0.51 | 0.31 | 0.4 | 0.59 | 0.38 | 0.54 |
| 3 | 151678985 | rs3109798 | -0.809 | 2.079 | 0.944 | 1.063 | -0.02 | 0.02 | 0.02 | 0 | 0.87 | 0.92 | 0.8 | 0.94 | 0.76 | 0.86 |
| 3 | 151680669 | rs10428197 | 0.329 | 2.047 | 0.965 | 1.036 | -0.063 | -0.015 | -0.015 | 0 | 0.24 | 0.1 | 0.2 | 0.25 | 0.33 | 0.21 |
| 3 | 151681018 | rs989769 | 0.744 | 2.003 | 1.045 | 1.042 | -0.004 | -0.001 | -0.001 | 0.044 | 0.6 | 0.53 | 0.53 | 0.61 | 0.42 | 0.61 |
| 3 | 151697625 | rs12486319 | -0.378 | 2.033 | 1.009 | 0.925 | -0.064 | 0.014 | 0.014 | 0 | 0.32 | 0.14 | 0.31 | 0.24 | 0.37 | 0.28 |
| 3 | 151698181 | rs2046034 | 0.351 | 2.04 | 0.974 | 0.928 | -0.08 | -0.014 | -0.014 | 0 | 0.24 | 0.07 | 0.2 | 0.22 | 0.33 | 0.21 |
| 3 | 151708414 | rs13433939 | 1.704 | 1.23 | 0.311 | 0.945 | 0.012 | -0.012 | -0.012 | 0 | 0.3 | 0.26 | 0.26 | 0.46 | 0.18 | 0.24 |
| 3 | 151710325 | rs6804426 | -1.311 | 1.411 | 0.028 | 0.923 | -0.018 | 0.016 | 0.016 | 0 | 0.88 | 0.93 | 0.8 | 0.93 | 0.76 | 0.85 |
| 3 | 151722893 | rs13064648 | -0.139 | 1.378 | 0.116 | 0.953 | -0.004 | 0.002 | 0.002 | 0 | 0.33 | 0.34 | 0.32 | 0.23 | 0.38 | 0.28 |
| 3 | 151724954 | rs17283297 | 0.055 | 1.377 | -0.086 | 0.966 | -0.062 | 0.017 | 0.017 | 0 | 0.32 | 0.16 | 0.32 | 0.23 | 0.37 | 0.28 |
| 3 | 151731717 | rs1987989 | 0.057 | 1.055 | -0.06 | 0.856 | 0 | 0.001 | 0.001 | 0 | 0.59 | 0.65 | 0.59 | 0.69 | 0.55 | 0.52 |
| 3 | 151732641 | rs7647733 | 0.511 | 1.018 | -0.056 | 0.754 | -0.008 | -0.001 | -0.001 | 0.399 | 0.29 | 0.24 | 0.2 | 0.24 | 0.21 | 0.34 |
| 3 | 151739498 | rs1600350 | -0.301 | 1.459 | 0.304 | 1.074 | 0.011 | 0.028 | 0.028 | 0 | 0.81 | 0.69 | 0.72 | 0.8 | 0.75 | 0.81 |
| 3 | 151740395 | rs13317725 | 0.814 | 1.414 | 0.217 | 1.027 | 0.033 | 0.045 | 0.045 | 0 | 0.31 | 0.16 | 0.27 | 0.14 | 0.26 | 0.35 |
| 3 | 151743406 | rs2724675 | -0.099 | 1.411 | 0.227 | 1.032 | -0.002 | 0.032 | 0.032 | 0 | 0.57 | 0.79 | 0.53 | 0.73 | 0.5 | 0.5 |
| 3 | 151744900 | rs13078935 | 0.795 | 1.584 | 0.228 | 0.985 | -0.051 | 0.031 | 0.031 | 0 | 0.23 | 0.09 | 0.18 | 0.16 | 0.32 | 0.2 |
| 3 | 151761488 | rs2898579 | -0.498 | 1.564 | 0.33 | 1.218 | 0.009 | -0.008 | -0.008 | 1.832 | 0.54 | 0.37 | 0.5 | 0.46 | 0.58 | 0.56 |
| 3 | 151763504 | rs1678483 | 1.817 | 1.536 | 0.384 | 1.209 | 0.01 | 0.004 | 0.004 | 0 | 0.26 | 0.23 | 0.21 | 0.32 | 0.17 | 0.24 |
| 3 | 151769549 | rs842500 | -0.763 | 1.541 | 0.381 | 1.213 | 0.006 | 0.019 | 0.019 | 0 | 0.82 | 0.78 | 0.73 | 0.77 | 0.75 | 0.81 |
| 3 | 151769599 | rs842501 | 1.807 | 1.564 | 0.251 | 1.223 | 0.007 | 0.003 | 0.003 | 0 | 0.26 | 0.25 | 0.22 | 0.32 | 0.17 | 0.24 |
| 3 | 151771309 | rs6765457 | 0.691 | 1.578 | 0.151 | 1.209 | -0.019 | -0.012 | -0.012 | 0.779 | 0.23 | 0.17 | 0.22 | 0.31 | 0.32 | 0.22 |
| 3 | 151775979 | rs12489232 | 0.54 | 1.649 | -0.028 | 1.012 | 0.037 | 0.052 | 0.052 | 0 | 0.3 | 0.13 | 0.27 | 0.14 | 0.26 | 0.35 |
| 3 | 151781244 | rs1715648 | -0.44 | 1.933 | 0.453 | 1.306 | 0.012 | 0.039 | 0.039 | 0 | 0.2 | 0.38 | 0.3 | 0.23 | 0.25 | 0.19 |
| 3 | 151786022 | rs842497 | -0.568 | 2.199 | -0.103 | 0.88 | 0.007 | 0.026 | 0.026 | 0.602 | 0.51 | 0.41 | 0.58 | 0.31 | 0.58 | 0.57 |
| 3 | 151790057 | rs10935885 | 1.072 | 2.274 | 0.043 | 0.979 | -0.006 | 0.037 | 0.037 | 0 | 0.22 | 0.28 | 0.19 | 0.38 | 0.17 | 0.22 |
| 3 | 151795212 | rs502775 | -1.357 | 2.282 | 0.059 | 0.983 | -0.001 | 0 | 0 | 0 | 0.73 | 0.71 | 0.77 | 0.68 | 0.75 | 0.79 |
| 3 | 151795988 | rs551508 | -0.111 | 2.646 | -0.128 | 0.904 | 0.01 | 0.008 | 0.008 | 0.026 | 0.43 | 0.39 | 0.46 | 0.3 | 0.37 | 0.5 |
| 3 | 151799506 | rs1494016 | 0.555 | 2.6 | -0.243 | 0.858 | -0.003 | -0.015 | -0.015 | 0 | 0.29 | 0.32 | 0.28 | 0.1 | 0.21 | 0.39 |
| 3 | 151803306 | rs556925 | 1.038 | 2.623 | -0.274 | 0.834 | 0.001 | -0.035 | -0.035 | 0 | 0.14 | 0.05 | 0.18 | 0.2 | 0.16 | 0.11 |
| 3 | 151818217 | rs1993937 | 1.233 | 2.227 | -0.21 | 0.754 | -0.004 | 0.034 | 0.034 | 0 | 0.22 | 0.26 | 0.18 | 0.36 | 0.17 | 0.22 |
| 3 | 151828999 | rs6807750 | 0.647 | 2.198 | -0.352 | 0.668 | 0.023 | 0.053 | 0.053 | 0.576 | 0.28 | 0.19 | 0.25 | 0.1 | 0.2 | 0.38 |
| 3 | 151846308 | rs325738 | -0.291 | 2.26 | -0.057 | 1.107 | -0.035 | -0.039 | -0.039 | 0.847 | 0.23 | 0.16 | 0.32 | 0.31 | 0.37 | 0.22 |
| 3 | 151850421 | rs325731 | -0.263 | 2.26 | -0.076 | 1.084 | -0.043 | -0.04 | -0.04 | 0 | 0.23 | 0.14 | 0.32 | 0.29 | 0.37 | 0.22 |
| 3 | 151850483 | rs226750 | -1.152 | 2.226 | -0.024 | 1.081 | -0.002 | 0.005 | 0.005 | 0.597 | 0.73 | 0.67 | 0.77 | 0.66 | 0.75 | 0.79 |
| 3 | 151880172 | rs325786 | 0.867 | 2.066 | -0.11 | 1.086 | -0.025 | -0.077 | -0.077 | 0 | 0.14 | 0.01 | 0.19 | 0.2 | 0.16 | 0.1 |
| 3 | 151883510 | rs1494014 | 0.72 | 2.115 | -0.054 | 1.082 | 0.02 | -0.001 | -0.001 | 0.002 | 0.09 | 0.12 | 0.13 | 0.09 | 0.21 | 0.11 |
| 3 | 151883726 | rs325781 | 0.862 | 2.118 | -0.068 | 1.092 | -0.014 | -0.064 | -0.064 | 0 | 0.14 | 0.02 | 0.19 | 0.2 | 0.16 | 0.1 |
| 3 | 151887185 | rs325772 | 0.377 | 1.718 | 0.107 | 1.151 | -0.017 | -0.016 | -0.016 | 0 | 0.24 | 0.21 | 0.32 | 0.29 | 0.37 | 0.22 |
| 3 | 151888934 | rs2098930 | 0.921 | 1.752 | -0.012 | 1.101 | 0.003 | -0.003 | -0.003 | 2.264 | 0.28 | 0.21 | 0.22 | 0.28 | 0.26 | 0.2 |
| 3 | 151892847 | rs9854676 | 0.845 | 1.512 | 0.059 | 0.63 | -0.002 | 0.026 | 0.026 | 0 | 0.29 | 0.34 | 0.25 | 0.36 | 0.28 | 0.21 |

| | | | | | | | | | | | | | | | | |
|---|---|---|---|---|---|---|---|---|---|---|---|---|---|---|---|---|
| 3 | 151894165 | rs325761 | 0.616 | 1.457 | -0.08 | 0.663 | -0.016 | -0.014 | -0.014 | 0 | 0.23 | 0.2 | 0.34 | 0.32 | 0.37 | 0.22 |
| 3 | 151899661 | rs16863988 | 0.834 | 1.502 | -0.541 | 0.544 | 0.022 | 0.031 | 0.031 | 0.441 | 0.22 | 0.14 | 0.2 | 0.09 | 0.11 | 0.36 |
| 3 | 151899724 | rs323614 | 0.513 | 1.899 | -0.446 | 0.961 | 0 | 0 | 0 | 0 | 0.25 | 0.27 | 0.22 | 0.26 | 0.24 | 0.2 |
| 3 | 151900862 | rs9831800 | -0.48 | 2.083 | -0.484 | 1.498 | 0 | -0.018 | -0.018 | 0 | 0.29 | 0.21 | 0.22 | 0.37 | 0.28 | 0.22 |
| 3 | 151911863 | rs323613 | 0.877 | 1.909 | -0.081 | 1.451 | 0.036 | 0.06 | 0.06 | 0 | 0.85 | 0.77 | 0.8 | 0.67 | 0.76 | 0.86 |
| 3 | 151920995 | rs1402008 | -0.452 | 0.831 | 0.469 | 1.382 | 0.027 | -0.009 | -0.009 | 0 | 0.12 | 0.04 | 0.19 | 0.11 | 0.09 | 0.15 |
| 3 | 151921122 | rs545144 | 0.905 | 0.537 | 0.45 | 1.194 | -0.001 | 0.003 | 0.003 | 0 | 0.86 | 0.88 | 0.81 | 0.68 | 0.78 | 0.88 |
| 3 | 151921374 | rs325798 | 2.06 | 0.177 | 0.48 | 1.139 | 0.005 | -0.102 | -0.102 | 0 | 0.62 | 0.11 | 0.51 | 0.5 | 0.62 | 0.64 |
| 3 | 151929158 | rs10513419 | -1.036 | 0.27 | 0.651 | 0.887 | 0.017 | -0.01 | -0.01 | 0 | 0.19 | 0.07 | 0.26 | 0.17 | 0.2 | 0.23 |
| 3 | 151960936 | rs323618 | 1.398 | 0.336 | 0.97 | 0.698 | 0.004 | -0.007 | -0.007 | 0.015 | 0.66 | 0.74 | 0.63 | 0.56 | 0.63 | 0.63 |
| 3 | 151976553 | rs370612 | 1.217 | 0.076 | 1.854 | 0.846 | -0.001 | 0 | 0 | 0.427 | 0.68 | 0.34 | 0.58 | 0.48 | 0.67 | 0.65 |
| 3 | 151978186 | rs323623 | 1.168 | 0.038 | 1.845 | 0.83 | 0 | 0.005 | 0.005 | 0.093 | 0.69 | 0.34 | 0.58 | 0.48 | 0.67 | 0.65 |
| 3 | 151983948 | rs842049 | 1.904 | -0.042 | 1.75 | 0.79 | -0.008 | -0.017 | -0.017 | 0.204 | 0.56 | 0.18 | 0.49 | 0.47 | 0.55 | 0.59 |
| 3 | 151998053 | rs1426385 | -0.048 | 0.022 | 1.169 | 0.584 | 0.009 | 0.015 | 0.015 | 0 | 0.64 | 0.61 | 0.49 | 0.55 | 0.57 | 0.58 |
| 3 | 152016969 | rs1426383 | 0.543 | -0.05 | 0.646 | -0.225 | 0.011 | -0.012 | -0.012 | 0 | 0.36 | 0.35 | 0.31 | 0.53 | 0.28 | 0.46 |
| 3 | 152025423 | rs11705943 | 0.562 | 0.057 | 0.51 | -0.056 | -0.016 | -0.017 | -0.017 | 0 | 0.12 | 0.09 | 0.24 | 0.27 | 0.19 | 0.1 |
| 3 | 152029167 | rs13063739 | -0.297 | 0.072 | 0.518 | -0.072 | 0.01 | 0.016 | 0.016 | 0 | 0.64 | 0.61 | 0.49 | 0.56 | 0.57 | 0.58 |
| 3 | 152039322 | rs2681437 | 1.668 | -0.146 | 0.343 | -0.188 | -0.016 | -0.027 | -0.027 | 0 | 0.06 | 0 | 0.03 | 0.07 | 0.06 | 0.1 |
| 3 | 152042244 | rs3913371 | -2.14 | -0.209 | 0.349 | -0.244 | -0.053 | 0.003 | 0.003 | 0 | 0.83 | 0.99 | 0.78 | 0.89 | 0.81 | 0.78 |
| 3 | 152053250 | rs4472028 | -0.329 | -0.334 | 0.642 | 0.396 | 0.01 | 0.015 | 0.015 | 0 | 0.64 | 0.61 | 0.49 | 0.57 | 0.57 | 0.58 |
| 3 | 152058449 | rs13077912 | 0.52 | 0.504 | -0.344 | 0.238 | -0.025 | 0.051 | 0.051 | 0 | 0.13 | 0.05 | 0.09 | 0.01 | 0.16 | 0.05 |
| 3 | 152062632 | rs17435143 | 0.469 | 0.484 | -0.336 | 0.224 | -0.014 | 0.026 | 0.026 | 0 | 0.27 | 0.16 | 0.17 | 0.01 | 0.28 | 0.12 |
| 3 | 152066793 | rs2036291 | -0.02 | 0.589 | -0.366 | 0.235 | 0.022 | -0.001 | -0.001 | 1.588 | 0.49 | 0.41 | 0.4 | 0.57 | 0.45 | 0.53 |
| 3 | 152070404 | rs6440800 | 0.582 | 0.606 | -0.429 | 0.241 | -0.027 | 0.05 | 0.05 | 0 | 0.13 | 0.05 | 0.09 | 0.01 | 0.16 | 0.05 |
| 3 | 152073055 | rs3920075 | 0.894 | 0.49 | -0.268 | 0.367 | 0.023 | 0.018 | 0.018 | 0 | 0.33 | 0.21 | 0.3 | 0.22 | 0.29 | 0.31 |
| 3 | 152084243 | rs7633673 | 0.281 | 0.559 | -0.321 | 0.349 | 0.01 | 0.002 | 0.002 | 0 | 0.5 | 0.47 | 0.33 | 0.51 | 0.41 | 0.53 |
| 3 | 152095318 | rs7651564 | 1.687 | 0.679 | -0.015 | 0.094 | 0.001 | -0.088 | -0.088 | 0 | 0.16 | 0.27 | 0.1 | 0 | 0.11 | 0.07 |
| 3 | 152095576 | rs12632591 | 0.776 | 0.25 | -0.197 | -0.29 | 0.018 | -0.005 | -0.005 | 0 | 0.28 | 0.12 | 0.25 | 0.23 | 0.25 | 0.29 |
| 3 | 152103745 | rs13094115 | 0.145 | 1.209 | -0.228 | -0.286 | 0.003 | 0.005 | 0.005 | 0 | 0.44 | 0.39 | 0.34 | 0.23 | 0.36 | 0.36 |
| 3 | 152104237 | rs13099394 | 0.726 | 1.321 | -0.317 | -0.487 | -0.016 | 0.031 | 0.031 | 0 | 0.12 | 0.05 | 0.09 | 0.01 | 0.16 | 0.05 |
| 3 | 152116652 | rs13079208 | -0.393 | 1.192 | -0.076 | -0.611 | -0.001 | 0 | 0 | 0 | 0.57 | 0.54 | 0.36 | 0.2 | 0.52 | 0.41 |
| 3 | 152127354 | rs7642123 | 1.919 | 1.123 | -0.022 | -0.607 | -0.002 | -0.088 | -0.088 | 0.014 | 0.16 | 0.26 | 0.1 | 0 | 0.11 | 0.07 |
| 3 | 152129990 | rs7629065 | 0.291 | 1.175 | -0.163 | -0.61 | 0.003 | 0.006 | 0.006 | 0 | 0.44 | 0.38 | 0.34 | 0.23 | 0.36 | 0.36 |
| 3 | 152135108 | rs3773908 | -1.351 | 1.029 | -0.236 | -0.492 | -0.011 | 0.037 | 0.037 | 0 | 0.72 | 0.92 | 0.7 | 0.84 | 0.72 | 0.66 |
| 3 | 152137036 | rs6440804 | -0.331 | 1.124 | -0.384 | -0.51 | -0.001 | -0.001 | -0.001 | 0 | 0.57 | 0.55 | 0.43 | 0.24 | 0.53 | 0.41 |
| 3 | 152177708 | rs2019689 | 1.044 | 0.869 | -0.789 | -0.431 | 0.02 | 0.004 | 0.004 | 0 | 0.27 | 0.12 | 0.17 | 0.19 | 0.24 | 0.27 |
| 3 | 152180329 | rs17437560 | 2.288 | 1.023 | -1.12 | -0.619 | 0 | 0.016 | 0.016 | 0 | 0.14 | 0.08 | 0.06 | 0 | 0.11 | 0.05 |
| 3 | 152181040 | rs1801769 | 0.986 | 1.015 | -1.178 | -0.476 | 0.02 | -0.007 | -0.007 | 0 | 0.27 | 0.12 | 0.25 | 0.23 | 0.24 | 0.28 |
| 3 | 152183512 | rs7221 | 1.018 | 1.018 | -1.238 | -0.475 | 0.017 | -0.016 | -0.016 | 0 | 0.27 | 0.08 | 0.25 | 0.23 | 0.24 | 0.28 |
| 3 | 152187431 | rs13069559 | 0.54 | 1.412 | -1.081 | -0.61 | -0.02 | 0.034 | 0.034 | 0 | 0.13 | 0.05 | 0.09 | 0.01 | 0.17 | 0.07 |
| 3 | 152188507 | rs7615064 | -1.253 | 0.976 | -1.115 | -0.46 | -0.07 | -0.012 | -0.012 | 0.43 | 0.76 | 0.99 | 0.75 | 0.85 | 0.74 | 0.68 |
| 3 | 152190321 | rs16864316 | -0.807 | 0.857 | -0.349 | -0.662 | -0.005 | 0.005 | 0.005 | 1.023 | 0.58 | 0.53 | 0.59 | 0.48 | 0.58 | 0.41 |
| 3 | 152190918 | rs1434794 | -0.587 | 0.855 | -0.416 | -0.803 | 0.03 | -0.018 | -0.018 | 0 | 0.32 | 0.47 | 0.4 | 0.25 | 0.41 | 0.17 |
| 3 | 152193063 | rs1946671 | -1.444 | 0.864 | -0.553 | -1.004 | 0.005 | -0.017 | -0.017 | 0 | 0.6 | 0.71 | 0.61 | 0.49 | 0.59 | 0.43 |
| 3 | 152193819 | rs17372345 | 1.245 | 1.103 | -0.704 | -1.071 | -0.012 | 0.015 | 0.015 | 0 | 0.08 | 0.03 | 0.1 | 0.02 | 0.1 | 0.01 |
| 3 | 152203657 | rs17372441 | -0.844 | 0.577 | -0.686 | -1.094 | 0.015 | 0.006 | 0.006 | 0.414 | 0.54 | 0.65 | 0.57 | 0.48 | 0.53 | 0.37 |
| 3 | 152206062 | rs9990017 | 2.219 | 1.117 | -0.41 | -1.749 | -0.005 | 0.012 | 0.012 | 0 | 0.14 | 0.08 | 0.08 | 0 | 0.12 | 0.06 |
| 3 | 152206633 | rs13326290 | -0.123 | 1.144 | -0.287 | -1.616 | -0.056 | 0.074 | 0.074 | 0.361 | 0.36 | 0.18 | 0.35 | 0.14 | 0.43 | 0.11 |
| 3 | 152217569 | rs723661 | -0.921 | 1.131 | -0.196 | -1.613 | -0.005 | 0.02 | 0.02 | 0.452 | 0.44 | 0.56 | 0.41 | 0.14 | 0.49 | 0.19 |
| 3 | 152222143 | rs1915948 | -0.955 | 1.703 | 0.717 | -0.853 | 0.054 | -0.015 | -0.015 | 0 | 0.22 | 0.36 | 0.29 | 0.13 | 0.3 | 0.05 |
| 3 | 152222960 | rs7650390 | -0.315 | 1.632 | 0.83 | -0.863 | 0.033 | -0.005 | -0.005 | 0 | 0.08 | 0.41 | 0.07 | 0.01 | 0.06 | 0.1 |
| 3 | 152227812 | rs17372844 | -0.78 | 1.798 | 0.125 | -0.787 | -0.052 | 0.002 | 0.002 | 0 | 0.2 | 0.08 | 0.27 | 0.13 | 0.3 | 0.05 |
| 3 | 152234694 | rs1278095 | 1.57 | 2.068 | 0.232 | -0.728 | 0.014 | -0.031 | -0.031 | 1.561 | 0.63 | 0.47 | 0.54 | 0.83 | 0.48 | 0.75 |
| 3 | 152237084 | rs7611727 | -1.293 | 1.962 | 0.687 | -0.147 | 0.071 | 0.009 | 0.009 | 0 | 0.18 | 0.57 | 0.32 | 0.15 | 0.29 | 0.06 |
| 3 | 152266836 | rs7633109 | -1.545 | 1.2 | 0.36 | 0.056 | 0.009 | -0.004 | -0.004 | 0 | 0.29 | 0.52 | 0.29 | 0.21 | 0.32 | 0.26 |
| 3 | 152271261 | rs13100355 | -0.813 | 1.566 | 1.018 | 0.556 | 0.042 | 0.067 | 0.067 | 0 | 0.09 | 0.17 | 0.14 | 0.17 | 0.15 | 0.04 |
| 3 | 152285415 | rs11925597 | -1.112 | 1.592 | 0.826 | 0.559 | 0.141 | 0.197 | 0.197 | 0.105 | 0.11 | 0.5 | 0.18 | 0.18 | 0.16 | 0.04 |
| 3 | 152304617 | rs939871 | -0.885 | 1.019 | 0.749 | 0.253 | -0.013 | -0.031 | -0.031 | 0.34 | 0.19 | 0.39 | 0.11 | 0.09 | 0.14 | 0.15 |
| 3 | 152310614 | rs4635681 | -0.774 | 1.116 | 0.719 | 0.193 | 0.001 | 0.01 | 0.01 | 0.876 | 0.18 | 0.17 | 0.1 | 0.06 | 0.16 | 0.15 |
| 3 | 152311489 | rs13076788 | -1.406 | 1.302 | 0.663 | -0.252 | 0.004 | 0.057 | 0.057 | 0 | 0.5 | 0.66 | 0.39 | 0.27 | 0.44 | 0.41 |
| 3 | 152321858 | rs11923583 | -1.009 | 1.619 | 0.768 | -0.433 | 0.001 | -0.002 | -0.002 | 0.373 | 0.34 | 0.32 | 0.25 | 0.1 | 0.27 | 0.32 |
| 3 | 152324182 | rs2178405 | -1.125 | 1.604 | 0.801 | -0.446 | -0.001 | -0.018 | -0.018 | 0.056 | 0.35 | 0.36 | 0.26 | 0.1 | 0.27 | 0.37 |
| 3 | 152324451 | rs13080295 | 0.829 | 1.598 | 0.632 | -0.179 | -0.032 | -0.016 | -0.016 | 0 | 0.13 | 0.01 | 0.08 | 0.1 | 0.13 | 0.04 |
| 3 | 152328890 | rs7646238 | -1.788 | 1.936 | 0.771 | -0.128 | 0.003 | 0.033 | 0.033 | 0 | 0.49 | 0.67 | 0.38 | 0.35 | 0.44 | 0.42 |
| 3 | 152329006 | rs7646341 | -0.061 | 1.798 | 0.802 | -0.202 | 0.001 | 0.002 | 0.002 | 0.077 | 0.2 | 0.2 | 0.09 | 0.06 | 0.15 | 0.21 |
| 3 | 152339970 | rs1878434 | -1.895 | 2.073 | 0.837 | 0.108 | 0.008 | 0.016 | 0.016 | 0 | 0.5 | 0.67 | 0.49 | 0.41 | 0.51 | 0.43 |
| 3 | 152345234 | rs6772600 | -0.409 | 2.069 | 0.878 | 0.093 | 0.001 | 0.01 | 0.01 | 0.114 | 0.81 | 0.83 | 0.91 | 0.94 | 0.86 | 0.85 |
| 3 | 152347482 | rs2203686 | 0.95 | 2.077 | 0.962 | 0.086 | -0.009 | 0.036 | 0.036 | 0 | 0.66 | 0.71 | 0.72 | 0.9 | 0.65 | 0.68 |
| 3 | 152350457 | rs9844440 | 1.88 | 1.127 | 0.398 | 0.279 | -0.03 | -0.005 | -0.005 | 0 | 0.27 | 0.1 | 0.32 | 0.22 | 0.32 | 0.36 |

| | | | | | | | | | | | | | | | | |
|---|---|---|---|---|---|---|---|---|---|---|---|---|---|---|---|---|
| 3 | 152350499 | rs1400612 | 0.273 | 1.36 | 0.758 | 0.231 | -0.055 | 0.068 | 0.068 | 0 | 0.14 | 0.02 | 0.18 | 0.04 | 0.21 | 0.17 |
| 3 | 152350527 | rs1517250 | 0.248 | 1.354 | 0.858 | 0.267 | 0.004 | 0.087 | 0.087 | 0 | 0.16 | 0.31 | 0.2 | 0.31 | 0.16 | 0.05 |
| 3 | 152351739 | rs4679687 | 1.459 | 1.346 | 0.697 | 0.328 | 0.007 | 0.019 | 0.019 | 0 | 0.5 | 0.33 | 0.51 | 0.59 | 0.49 | 0.62 |
| 3 | 152352602 | rs9289868 | 1.459 | 1.346 | 0.697 | 0.328 | 0.007 | 0.019 | 0.019 | 0 | 0.5 | 0.33 | 0.51 | 0.59 | 0.49 | 0.62 |
| 3 | 152375596 | rs1983338 | 0.591 | 1.256 | 0.501 | 0.526 | -0.001 | 0.024 | 0.024 | 0 | 0.68 | 0.76 | 0.75 | 0.77 | 0.72 | 0.81 |
| 3 | 152382636 | rs1517240 | 0.567 | 1.26 | 0.482 | 0.528 | -0.003 | 0.025 | 0.025 | 0 | 0.68 | 0.79 | 0.75 | 0.77 | 0.72 | 0.81 |
| 3 | 152385404 | rs13096682 | 0.673 | 1.282 | 0.568 | 0.556 | -0.022 | -0.038 | -0.038 | 0 | 0.13 | 0.03 | 0.16 | 0.18 | 0.14 | 0.04 |
| 3 | 152385842 | rs12233485 | 1.097 | 1.489 | 1.058 | 0.509 | 0.004 | 0.002 | 0.002 | 0.078 | 0.63 | 0.38 | 0.67 | 0.61 | 0.64 | 0.71 |
| 3 | 152387723 | rs9873341 | 0.607 | 1.232 | 0.983 | 0 | -0.002 | 0.023 | 0.023 | 0 | 0.68 | 0.78 | 0.75 | 0.77 | 0.72 | 0.81 |
| 3 | 152404607 | rs4679689 | 1.073 | 1.226 | 1.058 | -0.023 | 0.003 | -0.004 | -0.004 | 0 | 0.63 | 0.37 | 0.68 | 0.63 | 0.65 | 0.72 |
| 3 | 152405102 | rs9865555 | 1.09 | 1.221 | 1.107 | -0.025 | 0 | -0.001 | -0.001 | 0 | 0.62 | 0.35 | 0.68 | 0.63 | 0.65 | 0.72 |
| 3 | 152424154 | rs2063948 | 1.084 | 1.228 | 1.114 | -0.045 | 0 | -0.001 | -0.001 | 0 | 0.63 | 0.34 | 0.68 | 0.63 | 0.65 | 0.72 |
| 3 | 152439040 | rs13061814 | 0.397 | 1.185 | 1.325 | -0.026 | 0.002 | 0.044 | 0.044 | 0 | 0.14 | 0.24 | 0.19 | 0.3 | 0.15 | 0.05 |
| 3 | 152439912 | rs4680002 | -0.255 | 1.141 | 1.433 | -0.082 | 0 | -0.003 | -0.003 | 0 | 0.81 | 0.8 | 0.9 | 0.95 | 0.86 | 0.85 |
| 3 | 152441612 | rs9289869 | 1.273 | 1.147 | 1.53 | -0.087 | -0.005 | 0.004 | 0.004 | 0.272 | 0.62 | 0.28 | 0.67 | 0.63 | 0.65 | 0.72 |
| 3 | 152444315 | rs9875868 | 0.39 | 1.151 | 1.533 | -0.088 | 0.022 | -0.084 | -0.084 | 0.005 | 0.77 | 0.56 | 0.87 | 0.94 | 0.8 | 0.77 |
| 3 | 152450333 | rs9842398 | -0.231 | 0.927 | 1.541 | -0.082 | 0 | -0.001 | -0.001 | 0 | 0.81 | 0.81 | 0.9 | 0.94 | 0.86 | 0.85 |
| 3 | 152454727 | rs12488256 | 0.379 | 0.927 | 1.557 | -0.079 | 0.001 | 0.04 | 0.04 | 0 | 0.15 | 0.23 | 0.19 | 0.31 | 0.15 | 0.05 |
| 3 | 152455261 | rs13094696 | 0.538 | 0.943 | 1.086 | -0.52 | -0.053 | 0.057 | 0.057 | 0 | 0.12 | 0.01 | 0.15 | 0.03 | 0.2 | 0.11 |
| 3 | 152460701 | rs1461169 | 0.095 | 0.619 | 1.444 | -0.266 | -0.002 | -0.001 | -0.001 | 0 | 0.8 | 0.79 | 0.88 | 0.94 | 0.82 | 0.83 |
| 3 | 152463662 | rs1461167 | 0.366 | 0.627 | 1.456 | -0.262 | 0.012 | -0.084 | -0.084 | 0 | 0.78 | 0.56 | 0.87 | 0.94 | 0.82 | 0.83 |
| 3 | 152471575 | rs4234292 | -0.377 | 0.461 | 1.062 | -0.299 | -0.007 | 0.013 | 0.013 | 0.034 | 0.86 | 0.88 | 0.92 | 0.94 | 0.86 | 0.85 |
| 3 | 152471753 | rs721661 | 0.298 | 0.472 | 1.016 | -0.267 | 0 | 0.002 | 0.002 | 0.82 | 0.81 | 0.81 | 0.9 | 0.93 | 0.86 | 0.85 |
| 3 | 152481845 | rs12491993 | 0.627 | 0.559 | 0.91 | -0.233 | -0.024 | -0.05 | -0.05 | 0.753 | 0.13 | 0.02 | 0.17 | 0.19 | 0.15 | 0.04 |
| 3 | 152490171 | rs6804565 | 0.353 | 0.5 | 1.093 | -0.326 | -0.002 | -0.023 | -0.023 | 0.549 | 0.81 | 0.75 | 0.9 | 0.94 | 0.86 | 0.86 |
| 3 | 152492432 | rs1350794 | 0.066 | 0.475 | 0.86 | -0.36 | -0.004 | 0.01 | 0.01 | 0 | 0.82 | 0.86 | 0.88 | 0.96 | 0.81 | 0.86 |
| 3 | 152496566 | rs6440822 | -0.594 | 0.358 | 1.108 | -0.339 | 0.005 | 0.001 | 0.001 | 1.747 | 0.07 | 0.38 | 0.04 | 0 | 0.01 | 0 |
| 3 | 152505305 | rs9825291 | 1.397 | 0.462 | 0.765 | -0.271 | -0.002 | 0.021 | 0.021 | 0.177 | 0.62 | 0.26 | 0.67 | 0.76 | 0.66 | 0.75 |
| 3 | 152515862 | rs6773552 | -0.772 | 0.193 | 1.211 | 0.07 | 0.015 | 0.085 | 0.085 | 0.301 | 0.17 | 0.55 | 0.21 | 0.22 | 0.14 | 0.18 |
| 3 | 152532750 | rs2028282 | 0.555 | 0.22 | 0.787 | 0.075 | -0.001 | 0.01 | 0.01 | 0 | 0.11 | 0.12 | 0.14 | 0.19 | 0.1 | 0.09 |
| 3 | 152535365 | rs10513432 | -1.374 | 0.14 | 1.12 | -0.094 | 0.097 | 0.058 | 0.058 | 0.06 | 0.06 | 0.38 | 0.06 | 0.03 | 0.05 | 0.11 |
| 3 | 152544329 | rs16864594 | -0.252 | -0.051 | 1.177 | -0.143 | 0.018 | 0.016 | 0.016 | 1.759 | 0.05 | 0.3 | 0.03 | 0 | 0 | 0 |
| 3 | 152552190 | rs1371097 | -1.614 | -0.045 | 1.012 | -0.097 | 0.008 | -0.006 | -0.006 | 0 | 0.24 | 0.78 | 0.26 | 0.25 | 0.16 | 0.22 |
| 3 | 152554357 | rs701265 | -1.616 | -0.063 | 1.018 | -0.105 | 0.009 | -0.007 | -0.007 | 0.025 | 0.24 | 0.79 | 0.27 | 0.25 | 0.16 | 0.2 |
| 3 | 152567908 | rs6763988 | -0.615 | -0.171 | 0.432 | -0.631 | 0.014 | 0.011 | 0.011 | 2.758 | 0.06 | 0.33 | 0.03 | 0 | 0.01 | 0 |
| 3 | 152577715 | rs1473285 | -0.277 | -0.457 | -0.476 | -0.801 | 0.006 | 0.01 | 0.01 | 0 | 0.86 | 0.99 | 0.93 | 0.88 | 0.88 | 0.89 |
| 3 | 152586415 | rs9880812 | -0.112 | -0.142 | -0.985 | -1.243 | -0.01 | 0.008 | 0.008 | 0 | 0.39 | 0.34 | 0.32 | 0.11 | 0.44 | 0.2 |
| 3 | 152589597 | rs7356076 | 0.534 | -0.301 | -1.006 | -1.175 | 0.021 | 0.041 | 0.041 | 0 | 0.45 | 0.65 | 0.47 | 0.72 | 0.34 | 0.54 |
| 3 | 152589723 | rs7356077 | 1.964 | -0.585 | -1.277 | -1.492 | 0.037 | 0.043 | 0.043 | 0.013 | 0.23 | 0.11 | 0.1 | 0 | 0.13 | 0.11 |
| 3 | 152597534 | rs9835953 | 0.234 | -0.367 | -1.281 | -1.362 | 0.038 | 0.069 | 0.069 | 0 | 0.47 | 0.7 | 0.48 | 0.72 | 0.35 | 0.58 |
| 3 | 152611324 | rs9855770 | 1.647 | -0.572 | -1.241 | -1.501 | 0.05 | 0.043 | 0.043 | 0.145 | 0.26 | 0.15 | 0.1 | 0.01 | 0.12 | 0.13 |
| 3 | 152611465 | rs7649494 | 0.347 | 0.321 | -1.25 | -0.963 | -0.081 | 0.068 | 0.068 | 0 | 0.31 | 0.12 | 0.34 | 0.17 | 0.42 | 0.25 |
| 3 | 152627736 | rs1371101 | -0.11 | 0.039 | -0.848 | -0.278 | -0.013 | -0.039 | -0.039 | 2.86 | 0.79 | 0.69 | 0.85 | 0.92 | 0.82 | 0.79 |
| 3 | 152630713 | rs10935907 | 0.002 | 0.55 | -1.496 | 0.08 | -0.064 | -0.033 | -0.033 | 0 | 0.18 | 0.02 | 0.23 | 0.16 | 0.24 | 0.19 |
| 3 | 152631729 | rs11718068 | -0.164 | 0.155 | -0.91 | 0.507 | -0.011 | 0.004 | 0.004 | 0 | 0.58 | 0.43 | 0.56 | 0.63 | 0.61 | 0.48 |
| 3 | 152632751 | rs12632803 | 0.65 | 0.114 | -0.834 | 0.246 | -0.037 | 0.001 | 0.001 | 0 | 0.39 | 0.16 | 0.31 | 0.46 | 0.43 | 0.32 |
| 3 | 152635958 | rs7625688 | -0.795 | -0.201 | -0.42 | 0.032 | -0.008 | -0.04 | -0.04 | 0 | 0.69 | 0.58 | 0.8 | 0.91 | 0.72 | 0.73 |
| 3 | 152643911 | rs16864714 | 1.262 | -0.323 | -0.911 | 0.386 | 0.023 | -0.055 | -0.055 | 0 | 0.24 | 0.07 | 0.25 | 0.28 | 0.18 | 0.24 |
| 3 | 152652737 | rs17452880 | 0.513 | -0.45 | -0.928 | 0.318 | 0.016 | 0.015 | 0.015 | 0 | 0.19 | 0.09 | 0.21 | 0.14 | 0.15 | 0.24 |
| 3 | 152655130 | rs2043853 | -1.069 | 0.245 | -0.704 | 0.502 | -0.078 | -0.068 | -0.068 | 0 | 0.81 | 0.97 | 0.78 | 0.67 | 0.71 | 0.8 |
| 3 | 152655157 | rs9817893 | -0.799 | -0.46 | 0.249 | 0.455 | 0.003 | 0.002 | 0.002 | 0 | 0.53 | 0.65 | 0.63 | 0.56 | 0.46 | 0.55 |
| 3 | 152658125 | rs10513437 | 0.289 | -0.172 | 0.296 | 0.284 | 0.027 | 0.003 | 0.003 | 0 | 0.07 | 0.1 | 0.22 | 0.16 | 0.2 | 0.12 |
| 3 | 152663631 | rs2043854 | -1.025 | -0.241 | 0.388 | 0.271 | 0.005 | -0.01 | -0.01 | 2.121 | 0.3 | 0.38 | 0.48 | 0.3 | 0.35 | 0.36 |
| 3 | 152665425 | rs9823698 | 0.072 | 0.089 | 0.433 | 0.539 | 0.016 | 0 | 0 | 0 | 0.07 | 0.09 | 0.23 | 0.16 | 0.2 | 0.11 |
| 3 | 152674921 | rs1403501 | 1.004 | -0.08 | 0.385 | 0.554 | -0.009 | -0.015 | -0.015 | 0.04 | 0.34 | 0.29 | 0.34 | 0.49 | 0.41 | 0.43 |
| 3 | 152676283 | rs6771615 | 1.83 | -0.743 | -0.229 | 0.431 | 0.003 | -0.001 | -0.001 | 0.275 | 0.19 | 0.11 | 0.12 | 0.19 | 0.11 | 0.12 |
| 3 | 152679420 | rs6440836 | -0.044 | -0.684 | -0.389 | 0.273 | 0.023 | 0.106 | 0.106 | 0 | 0.68 | 0.93 | 0.72 | 0.86 | 0.72 | 0.71 |
| 3 | 152687097 | rs4679695 | 0.359 | -0.504 | -0.264 | 0.529 | 0.028 | 0.022 | 0.022 | 0.09 | 0.48 | 0.77 | 0.6 | 0.6 | 0.6 | 0.6 |
| 3 | 152691847 | rs9872160 | 1.645 | -0.34 | -0.253 | 0.589 | -0.005 | 0.005 | 0.005 | 0 | 0.2 | 0.15 | 0.12 | 0.26 | 0.11 | 0.1 |
| 3 | 152708841 | rs1533913 | 0.256 | 0.44 | 0.008 | 0.94 | -0.017 | -0.023 | -0.023 | 0 | 0.39 | 0.33 | 0.46 | 0.53 | 0.53 | 0.51 |
| 3 | 152716014 | rs6795065 | 0.633 | 0.455 | -0.416 | 0.996 | 0.039 | 0.015 | 0.015 | 0 | 0.39 | 0.17 | 0.21 | 0.34 | 0.26 | 0.26 |
| 3 | 152721975 | rs9883654 | 0.004 | 0.81 | -0.145 | 0.985 | 0.01 | 0.005 | 0.005 | 0 | 0.51 | 0.64 | 0.46 | 0.47 | 0.42 | 0.39 |
| 3 | 152724407 | rs13070843 | 0.113 | 0.538 | -0.358 | 0.902 | 0.011 | -0.009 | -0.009 | 1.29 | 0.8 | 0.63 | 0.76 | 0.82 | 0.77 | 0.8 |
| 3 | 152726146 | rs9839183 | 0.309 | 0.405 | -0.745 | 0.907 | 0.011 | 0.007 | 0.007 | 0 | 0.4 | 0.3 | 0.22 | 0.31 | 0.24 | 0.26 |
| 3 | 152737318 | rs1464593 | 0.361 | 0.854 | -0.844 | 1.504 | -0.002 | -0.021 | -0.021 | 0.052 | 0.85 | 0.9 | 0.78 | 0.72 | 0.85 | 0.83 |
| 3 | 152743124 | rs12631439 | 0.02 | 1.121 | -1.017 | 1.74 | -0.017 | -0.067 | -0.067 | 0 | 0.17 | 0.06 | 0.24 | 0.37 | 0.21 | 0.2 |
| 3 | 152746223 | rs9814144 | -0.375 | 0.878 | -1.048 | 1.322 | 0.007 | -0.043 | -0.043 | 0 | 0.32 | 0.22 | 0.32 | 0.49 | 0.35 | 0.29 |
| 3 | 152760119 | rs13097238 | 0.389 | 0.782 | -0.817 | 0.528 | -0.005 | 0.082 | 0.082 | 0 | 0.46 | 0.74 | 0.48 | 0.25 | 0.46 | 0.33 |
| 3 | 152761774 | rs9829175 | 0.803 | 0.847 | -0.055 | 0.375 | 0.004 | 0.023 | 0.023 | 0 | 0.38 | 0.32 | 0.33 | 0.14 | 0.34 | 0.24 |

| | | | | | | | | | | | | | | | | |
|---|---|---|---|---|---|---|---|---|---|---|---|---|---|---|---|---|
| 3 | 152762462 | rs9854001 | 0.864 | 0.677 | -0.289 | 0.137 | 0.012 | 0.028 | 0.028 | 0 | 0.18 | 0.11 | 0.05 | 0.04 | 0.13 | 0.26 |
| 3 | 152769540 | rs559751 | -0.336 | 0.729 | -0.184 | 0.139 | 0 | -0.009 | -0.009 | 0.059 | 0.44 | 0.51 | 0.61 | 0.82 | 0.52 | 0.5 |
| 3 | 152770183 | rs843790 | 0.051 | 0.715 | 0.008 | 0.135 | -0.036 | 0.054 | 0.054 | 0 | 0.4 | 0.21 | 0.58 | 0.78 | 0.52 | 0.49 |
| 3 | 152778561 | rs579638 | -1.047 | 0.682 | -0.314 | 0.177 | 0.006 | 0.024 | 0.024 | 0 | 0.82 | 0.89 | 0.95 | 0.96 | 0.86 | 0.74 |
| 3 | 152785719 | rs506400 | -0.38 | 0.722 | -0.242 | 0.228 | 0.004 | 0.022 | 0.022 | 0 | 0.47 | 0.53 | 0.63 | 0.85 | 0.58 | 0.54 |
| 3 | 152790138 | rs1717662 | -1.459 | 0.716 | -0.265 | 0.205 | 0.008 | 0.06 | 0.06 | 0.008 | 0.86 | 0.96 | 0.97 | 1 | 0.92 | 0.77 |
| 3 | 152791742 | rs511669 | -0.352 | 0.689 | -0.287 | 0.31 | 0.003 | 0.023 | 0.023 | 0 | 0.47 | 0.55 | 0.63 | 0.8 | 0.58 | 0.52 |
| 3 | 152792177 | rs13086307 | 1.057 | 0.718 | -0.354 | 0.367 | 0.005 | 0.016 | 0.016 | 0 | 0.38 | 0.34 | 0.33 | 0.2 | 0.34 | 0.26 |
| 3 | 152805309 | rs528581 | 0.081 | 0.686 | -0.393 | 0.449 | -0.001 | 0.001 | 0.001 | 0 | 0.38 | 0.31 | 0.48 | 0.67 | 0.41 | 0.36 |
| 3 | 152816505 | rs815035 | -1.056 | 0.553 | -1.05 | 0.374 | -0.002 | 0.075 | 0.075 | 0.017 | 0.85 | 0.97 | 0.96 | 0.99 | 0.91 | 0.77 |
| 3 | 152823845 | rs568925 | 0.733 | 0.115 | -1.422 | 0.934 | 0.013 | -0.11 | -0.11 | 0 | 0.63 | 0.07 | 0.58 | 0.47 | 0.72 | 0.48 |
| 3 | 152831259 | rs17468698 | 0.442 | 0.323 | -0.825 | 0.923 | 0.02 | 0.001 | 0.001 | 0 | 0.05 | 0.09 | 0.13 | 0.07 | 0.11 | 0.07 |
| 3 | 152831552 | rs9827303 | 0.015 | 0.219 | -0.061 | 0.929 | 0.019 | 0.002 | 0.002 | 0 | 0.06 | 0.17 | 0.04 | 0.03 | 0.07 | 0.07 |
| 3 | 152833338 | rs9837387 | -0.219 | 0.388 | 0.095 | 1.094 | 0.001 | 0.001 | 0.001 | 0 | 0.53 | 0.52 | 0.53 | 0.41 | 0.59 | 0.48 |
| 3 | 152833680 | rs10935915 | 1.085 | 0.301 | -0.307 | 0.843 | -0.083 | 0.134 | 0.134 | 0.29 | 0.38 | 0.06 | 0.32 | 0.13 | 0.41 | 0.26 |
| 3 | 152861999 | rs6776457 | -1.007 | 0.269 | 0.587 | 1.501 | 0.032 | 0.436 | 0.436 | 7.619 | 0.06 | 0.48 | 0.04 | 0.27 | 0.01 | 0.07 |
| 3 | 152864622 | rs9843465 | -1.823 | 0.108 | -0.166 | 0.867 | -0.019 | 0.151 | 0.151 | 0 | 0.78 | 0.99 | 0.95 | 0.99 | 0.88 | 0.8 |
| 3 | 152867288 | rs9830448 | 0.81 | 0.39 | -0.154 | 0.902 | -0.083 | 0.061 | 0.061 | 0.287 | 0.3 | 0.02 | 0.29 | 0.12 | 0.35 | 0.23 |
| 3 | 152868246 | rs9855757 | -0.034 | 0.331 | 0.114 | 0.701 | 0.007 | -0.025 | -0.025 | 5.439 | 0.34 | 0.71 | 0.34 | 0.44 | 0.33 | 0.27 |
| 3 | 152878719 | rs894802 | -0.822 | 0.485 | 0.468 | 0.842 | 0.087 | 0.071 | 0.071 | 0.08 | 0.1 | 0.44 | 0.13 | 0.11 | 0.12 | 0.08 |
| 3 | 152878838 | rs894801 | 1.152 | 0.477 | 0.373 | 0.818 | -0.009 | 0.007 | 0.007 | 0 | 0.05 | 0.02 | 0.09 | 0.01 | 0.11 | 0.06 |
| 3 | 152885818 | rs6809490 | -0.641 | 0.465 | 0.465 | 0.837 | 0.082 | 0.102 | 0.102 | 0.446 | 0.1 | 0.51 | 0.14 | 0.14 | 0.11 | 0.08 |
| 3 | 152890607 | rs524562 | 1.091 | 0.401 | -0.774 | 0.895 | -0.107 | -0.091 | -0.091 | 0.057 | 0.26 | 0.02 | 0.28 | 0.36 | 0.33 | 0.29 |
| 3 | 152894141 | rs6770653 | 0.783 | 0.214 | -0.157 | 1.164 | -0.008 | 0.023 | 0.023 | 0.183 | 0.23 | 0.28 | 0.25 | 0.26 | 0.23 | 0.23 |
| 3 | 152915925 | rs7637356 | -0.664 | 0.385 | -1.139 | 0.713 | 0.005 | 0.027 | 0.027 | 0.638 | 0.44 | 0.93 | 0.56 | 0.59 | 0.43 | 0.47 |
| 3 | 152927952 | rs16822423 | 0.81 | 0.839 | -0.881 | 0.629 | -0.11 | -0.084 | -0.084 | 0 | 0.26 | 0.02 | 0.26 | 0.35 | 0.32 | 0.33 |
| 3 | 152929955 | rs9856931 | -0.735 | 0.779 | -1.152 | 0.305 | -0.09 | 0.084 | 0.084 | 0.281 | 0.5 | 0.97 | 0.68 | 0.62 | 0.54 | 0.52 |
| 3 | 152930576 | rs1899695 | -0.375 | 0.691 | -0.342 | 0.859 | -0.002 | 0.233 | 0.233 | 0 | 0.1 | 0.29 | 0.23 | 0.33 | 0.07 | 0.15 |
| 3 | 152933001 | rs1025917 | 1.089 | 0.659 | -0.104 | 0.725 | -0.002 | -0.001 | -0.001 | 0 | 0.23 | 0.23 | 0.27 | 0.22 | 0.31 | 0.17 |
| 3 | 152935128 | rs16822433 | -0.166 | 0.601 | 0.174 | 0.721 | -0.005 | 0.147 | 0.147 | 0 | 0.09 | 0.17 | 0.22 | 0.33 | 0.07 | 0.15 |
| 3 | 152935967 | rs6800715 | -0.222 | 0.713 | 0.179 | 0.597 | 0.038 | -0.063 | -0.063 | 0.071 | 0.15 | 0.31 | 0.17 | 0.04 | 0.21 | 0.21 |
| 3 | 152937218 | rs6440853 | 0.536 | 0.674 | 0.108 | 0.542 | 0.009 | -0.058 | -0.058 | 0.537 | 0.1 | 0.26 | 0.08 | 0 | 0.08 | 0.06 |
| 3 | 152953343 | rs9811019 | -1.264 | 0.939 | -0.507 | 0.66 | -0.047 | 0.093 | 0.093 | 0.02 | 0.51 | 0.94 | 0.68 | 0.63 | 0.55 | 0.53 |
| 3 | 152960022 | rs7620292 | 0.886 | 0.883 | -0.296 | 0.581 | 0 | 0.001 | 0.001 | 0.128 | 0.23 | 0.33 | 0.28 | 0.23 | 0.25 | 0.17 |
| 3 | 152976450 | rs6793048 | -1.153 | 1.092 | -0.983 | 0.689 | -0.106 | 0.006 | 0.006 | 0.063 | 0.51 | 0.98 | 0.69 | 0.6 | 0.55 | 0.54 |
| 3 | 152987718 | rs16822470 | -0.015 | 0.978 | -0.71 | 0.831 | 0 | 0.039 | 0.039 | 0 | 0.08 | 0.1 | 0.21 | 0.19 | 0.07 | 0.08 |
| 3 | 152992604 | rs6778334 | 1.198 | 1.067 | -1.072 | 0.988 | -0.075 | -0.072 | -0.072 | 0 | 0.25 | 0.02 | 0.25 | 0.22 | 0.3 | 0.26 |
| 3 | 152994772 | rs7355869 | -0.275 | 1.079 | -0.941 | 0.916 | -0.022 | -0.03 | -0.03 | 0.805 | 0.41 | 0.76 | 0.47 | 0.59 | 0.5 | 0.5 |
| 3 | 152996074 | rs9822986 | -0.373 | 1.105 | -1.492 | 0.91 | -0.053 | -0.068 | -0.068 | 0 | 0.42 | 0.88 | 0.48 | 0.59 | 0.49 | 0.5 |
| 3 | 153014098 | rs12053979 | 0.925 | 1.18 | -1.348 | 0.883 | -0.079 | -0.075 | -0.075 | 0 | 0.25 | 0.02 | 0.25 | 0.26 | 0.31 | 0.27 |
| 3 | 153017762 | rs971385 | -0.826 | 1.381 | 0.11 | 0.557 | 0.079 | 0.026 | 0.026 | 0.649 | 0.08 | 0.49 | 0.09 | 0.02 | 0.06 | 0.03 |
| 3 | 153017793 | rs9815437 | 0.199 | 1.539 | -0.697 | 0.751 | -0.011 | -0.009 | -0.009 | 0.132 | 0.22 | 0.18 | 0.28 | 0.27 | 0.3 | 0.16 |
| 3 | 153022780 | rs9824588 | 0.086 | 1.044 | -1.157 | 0.181 | -0.056 | 0.091 | 0.091 | 0.002 | 0.77 | 0.97 | 0.84 | 0.89 | 0.78 | 0.77 |
| 3 | 153029536 | rs12629639 | -0.825 | 1.009 | -0.591 | 0.769 | 0.009 | 0.293 | 0.293 | 0 | 0.07 | 0.21 | 0.29 | 0.43 | 0.08 | 0.16 |
| 3 | 153035536 | rs11923708 | 1.3 | 1.108 | -0.325 | 0.413 | 0.007 | -0.029 | -0.029 | 0 | 0.15 | 0.27 | 0.12 | 0.08 | 0.17 | 0.11 |
| 3 | 153037911 | rs16822529 | -0.336 | 1.386 | -0.498 | 0.497 | 0.019 | 0.124 | 0.124 | 0 | 0.12 | 0.18 | 0.34 | 0.43 | 0.16 | 0.2 |
| 3 | 153038982 | rs2174748 | -0.763 | 1.3 | -0.717 | 0.147 | -0.034 | 0.07 | 0.07 | 0.262 | 0.81 | 0.97 | 0.85 | 0.9 | 0.79 | 0.81 |
| 3 | 153040189 | rs6764891 | 0.294 | 1.318 | -0.525 | 0.351 | -0.009 | -0.007 | -0.007 | 0 | 0.7 | 0.78 | 0.57 | 0.61 | 0.67 | 0.69 |
| 3 | 153040808 | rs9289901 | 1.261 | 1.11 | -0.699 | 0.128 | 0.004 | -0.004 | -0.004 | 0 | 0.3 | 0.23 | 0.14 | 0.03 | 0.29 | 0.2 |
| 3 | 153041225 | rs9848440 | 1.299 | 1.123 | -1.028 | 0.115 | 0.004 | 0.04 | 0.04 | 0 | 0.29 | 0.13 | 0.13 | 0.01 | 0.29 | 0.19 |
| 3 | 153048676 | rs2221582 | -0.15 | 0.673 | -1.566 | 0.43 | -0.008 | -0.084 | -0.084 | 0 | 0.71 | 0.87 | 0.71 | 0.52 | 0.73 | 0.57 |
| 3 | 153048865 | rs9862609 | 0.382 | 0.799 | -1.403 | 0.309 | -0.001 | -0.012 | -0.012 | 0 | 0.29 | 0.39 | 0.29 | 0.21 | 0.31 | 0.25 |
| 3 | 153054244 | rs7622837 | 0.236 | 1.195 | -1.914 | 0.35 | -0.017 | -0.007 | -0.007 | 0 | 0.52 | 0.82 | 0.35 | 0.5 | 0.44 | 0.44 |
| 3 | 153060454 | rs1507157 | -0.007 | 1.149 | -1.608 | -0.46 | -0.013 | 0.048 | 0.048 | 0 | 0.27 | 0.13 | 0.14 | 0.02 | 0.3 | 0.19 |
| 3 | 153066164 | rs1507163 | 0.596 | 0.907 | -1.952 | -0.225 | 0 | 0.007 | 0.007 | 0 | 0.69 | 0.18 | 0.74 | 0.74 | 0.72 | 0.61 |
| 3 | 153066493 | rs6778259 | 0.074 | 0.747 | -1.43 | -0.361 | -0.005 | -0.033 | -0.033 | 1.945 | 0.13 | 0.12 | 0.04 | 0 | 0.08 | 0.09 |
| 3 | 153068547 | rs9852831 | -0.014 | 0.767 | 0.25 | -0.218 | -0.012 | -0.004 | -0.004 | 0 | 0.87 | 0.9 | 0.84 | 0.86 | 0.81 | 0.78 |
| 3 | 153071132 | rs9810406 | 0.142 | 0.903 | -0.449 | -0.56 | -0.004 | 0.002 | 0.002 | 0 | 0.14 | 0.11 | 0.13 | 0.08 | 0.22 | 0.11 |
| 3 | 153096783 | rs9822378 | 0.344 | 0.359 | -1.047 | -0.116 | 0.039 | 0.001 | 0.001 | 0 | 0.57 | 0.2 | 0.67 | 0.41 | 0.64 | 0.39 |
| 3 | 153099036 | rs955013 | 0.876 | 0.503 | -1 | -0.198 | 0.004 | -0.005 | -0.005 | 0 | 0.4 | 0.12 | 0.22 | 0.41 | 0.36 | 0.43 |
| 3 | 153099453 | rs955012 | -0.484 | 0.217 | -1.193 | -0.404 | -0.049 | -0.012 | -0.012 | 0.002 | 0.83 | 0.99 | 0.84 | 0.88 | 0.81 | 0.8 |
| 3 | 153100477 | rs9843491 | 0.209 | 0.307 | -0.998 | -0.36 | -0.028 | -0.022 | -0.022 | 2.018 | 0.36 | 0.19 | 0.55 | 0.43 | 0.43 | 0.27 |
| 3 | 153116888 | rs6794116 | -0.29 | 0.308 | -1.289 | -0.281 | -0.036 | -0.065 | -0.065 | 0 | 0.43 | 0.87 | 0.62 | 0.49 | 0.46 | 0.37 |
| 3 | 153120549 | rs2862071 | 0.996 | 0.132 | -1.297 | -0.031 | -0.003 | -0.037 | -0.037 | 0 | 0.24 | 0.1 | 0.33 | 0.3 | 0.22 | 0.17 |
| 3 | 153121917 | rs6440877 | 0.436 | 0.123 | -1.306 | -0.053 | -0.041 | 0.01 | 0.01 | 0 | 0.7 | 0.86 | 0.61 | 0.75 | 0.6 | 0.68 |
| 3 | 153124222 | rs16822612 | 0.987 | 0.181 | -1.361 | -0.042 | -0.016 | -0.057 | -0.057 | 3.503 | 0.24 | 0.06 | 0.32 | 0.29 | 0.21 | 0.16 |
| 3 | 153128470 | rs2862827 | -0.975 | 0.339 | -1.314 | -0.114 | 0.021 | -0.005 | -0.005 | 0 | 0.85 | 0.81 | 0.74 | 0.84 | 0.74 | 0.89 |
| 3 | 153130204 | rs10433456 | 1.234 | 0.287 | -1.314 | 0.052 | 0.013 | 0.041 | 0.041 | 0 | 0.41 | 0.11 | 0.22 | 0.36 | 0.35 | 0.43 |
| 3 | 153132672 | rs7622713 | 1.015 | 0.651 | -1.612 | 0.035 | 0.033 | 0.025 | 0.025 | 0 | 0.46 | 0.74 | 0.28 | 0.41 | 0.37 | 0.52 |

| | | | | | | | | | | | | | | | | |
|---|---|---|---|---|---|---|---|---|---|---|---|---|---|---|---|---|
| 3 | 153132716 | rs4680062 | 0.002 | 0.873 | -1.582 | 0.051 | -0.024 | -0.039 | -0.039 | 0.056 | 0.15 | 0.01 | 0.14 | 0.13 | 0.15 | 0.2 |
| 3 | 153139795 | rs7644682 | -0.452 | 1.069 | -1.063 | 0.084 | 0.008 | -0.002 | -0.002 | 0 | 0.85 | 0.85 | 0.74 | 0.84 | 0.74 | 0.89 |
| 3 | 153157523 | rs7611860 | 0.266 | 0.603 | -1.203 | 0.281 | -0.023 | -0.04 | -0.04 | 0 | 0.24 | 0.07 | 0.32 | 0.29 | 0.23 | 0.2 |
| 3 | 153175186 | rs11920638 | -0.637 | 0.173 | 0.383 | 0.44 | -0.003 | 0.139 | 0.139 | 0 | 0.13 | 0.47 | 0.21 | 0.2 | 0.12 | 0.18 |
| 3 | 153175351 | rs9839852 | -0.068 | 1.029 | 0.189 | 0.807 | -0.026 | -0.013 | -0.013 | 0.101 | 0.12 | 0.1 | 0.25 | 0.15 | 0.25 | 0.12 |
| 3 | 153177445 | rs1881950 | 0.496 | 0.674 | 0.646 | 0.591 | 0.001 | -0.012 | -0.012 | 0 | 0.57 | 0.47 | 0.51 | 0.67 | 0.49 | 0.66 |
| 3 | 153177649 | rs4643686 | 0.768 | 0.735 | 0.516 | 0.558 | -0.024 | 0.016 | 0.016 | 0 | 0.1 | 0.06 | 0.13 | 0 | 0.22 | 0.05 |
| 3 | 153177762 | rs4611796 | 0.397 | 0.487 | 0.66 | 0.65 | -0.001 | 0.009 | 0.009 | 0 | 0.57 | 0.59 | 0.45 | 0.64 | 0.47 | 0.62 |
| 3 | 153185593 | rs17792167 | 0.355 | 0.474 | 1.048 | 0.613 | -0.009 | 0.022 | 0.022 | 0.166 | 0.08 | 0.11 | 0.09 | 0.09 | 0.06 | 0.14 |
| 3 | 153186265 | rs17735715 | 1.149 | 0.416 | 0.617 | 0.578 | 0.035 | 0.032 | 0.032 | 0 | 0.47 | 0.14 | 0.29 | 0.43 | 0.4 | 0.44 |
| 3 | 153186281 | rs6782397 | -0.204 | 0.582 | 0.558 | 0.423 | -0.06 | 0.078 | 0.078 | 0 | 0.37 | 0.17 | 0.35 | 0.19 | 0.46 | 0.2 |
| 3 | 153188510 | rs6767634 | -0.204 | 0.581 | 0.499 | 0.448 | -0.039 | 0.057 | 0.057 | 0 | 0.37 | 0.25 | 0.36 | 0.19 | 0.46 | 0.2 |
| 3 | 153212332 | rs2862078 | 0.82 | 0.399 | 0.49 | 0.434 | 0 | -0.002 | -0.002 | 0 | 0.57 | 0.5 | 0.43 | 0.63 | 0.48 | 0.58 |
| 3 | 153220021 | rs6778736 | -0.641 | 0.213 | 0.513 | 0.741 | 0.024 | 0.371 | 0.371 | 0.262 | 0.09 | 0.43 | 0.19 | 0.29 | 0.06 | 0.28 |
| 3 | 153221215 | rs1881955 | 0.612 | 0.424 | -0.279 | 0.198 | 0.003 | 0.001 | 0.001 | 0 | 0.19 | 0.17 | 0.19 | 0.2 | 0.18 | 0.11 |
| 3 | 153223622 | rs2862073 | -0.074 | 0.208 | -0.581 | 0.094 | 0.007 | 0.005 | 0.005 | 0.147 | 0.86 | 0.88 | 0.89 | 0.98 | 0.91 | 0.86 |
| 3 | 153244664 | rs16822688 | -1.018 | 0.193 | -0.639 | 0.114 | 0.023 | 0.027 | 0.027 | 0 | 0.2 | 0.13 | 0.14 | 0.09 | 0.14 | 0.29 |
| 3 | 153245938 | rs958322 | 1.043 | 0.435 | -0.358 | -0.08 | 0.017 | -0.042 | -0.042 | 0 | 0.71 | 0.45 | 0.63 | 0.8 | 0.66 | 0.69 |
| 3 | 153252942 | rs6440892 | 0.672 | 1.284 | -0.3 | 0.335 | 0.102 | 0.01 | 0.01 | 0.521 | 0.84 | 0.64 | 0.67 | 0.8 | 0.67 | 0.81 |
| 3 | 153256247 | rs10049379 | 0.147 | 0.738 | -1.686 | 0.23 | -0.056 | 0.04 | 0.04 | 0.011 | 0.65 | 0.88 | 0.77 | 0.76 | 0.53 | 0.64 |
| 3 | 153258272 | rs6764190 | -0.382 | -0.13 | -2.085 | 0.409 | 0.06 | -0.054 | -0.054 | 0.093 | 0.44 | 0.91 | 0.55 | 0.4 | 0.29 | 0.49 |
| 3 | 153259376 | rs6767565 | -0.257 | -0.617 | -1.162 | 0.271 | -0.041 | -0.02 | -0.02 | 0 | 0.43 | 0.51 | 0.39 | 0.21 | 0.18 | 0.32 |
| 3 | 153271806 | rs1388008 | -0.941 | -0.366 | -0.983 | 0.345 | 0.048 | -0.009 | -0.009 | 0.123 | 0.55 | 0.82 | 0.57 | 0.42 | 0.35 | 0.57 |
| 3 | 153280557 | rs7632990 | 0.366 | -0.387 | -0.813 | 0.372 | -0.003 | 0.004 | 0.004 | 0 | 0.09 | 0.11 | 0.05 | 0.12 | 0.04 | 0.09 |
| 3 | 153286204 | rs4680074 | -0.053 | -0.548 | -0.341 | -0.131 | -0.014 | -0.037 | -0.037 | 0 | 0.41 | 0.52 | 0.37 | 0.15 | 0.26 | 0.33 |
| 3 | 153289393 | rs718894 | -0.351 | -0.345 | -0.186 | 0.009 | 0.036 | 0.043 | 0.043 | 0.097 | 0.47 | 0.42 | 0.4 | 0.22 | 0.3 | 0.4 |
| 3 | 153294641 | rs6781989 | -0.663 | 0.501 | -0.132 | -0.067 | -0.007 | -0.006 | -0.006 | 0 | 0.71 | 0.76 | 0.72 | 0.42 | 0.59 | 0.69 |
| 3 | 153297048 | rs6776300 | -0.652 | 0.429 | 0.108 | -0.107 | 0.029 | 0.024 | 0.024 | 0 | 0.69 | 0.61 | 0.7 | 0.42 | 0.58 | 0.69 |
| 3 | 153297924 | rs902432 | 0.185 | 0.151 | 0.024 | -0.276 | -0.047 | 0.008 | 0.008 | 0 | 0.35 | 0.18 | 0.4 | 0.34 | 0.4 | 0.45 |
| 3 | 153300061 | rs11920558 | 0.058 | 0.138 | 0.053 | -0.281 | -0.011 | 0.011 | 0.011 | 0 | 0.37 | 0.32 | 0.41 | 0.34 | 0.4 | 0.44 |
| 3 | 153312759 | rs989181 | 0.226 | 0.14 | -0.047 | -0.281 | -0.03 | 0.013 | 0.013 | 0 | 0.36 | 0.21 | 0.4 | 0.34 | 0.4 | 0.45 |
| 3 | 153320659 | rs9863960 | 1.118 | 0.099 | -0.269 | 0.302 | -0.017 | -0.017 | -0.017 | 0.261 | 0.06 | 0 | 0.09 | 0.24 | 0.11 | 0.12 |
| 3 | 153328203 | rs1873354 | 1.143 | -0.11 | -0.215 | 0.107 | -0.061 | -0.074 | -0.074 | 0 | 0.31 | 0.4 | 0.3 | 0.07 | 0.15 | 0.19 |
| 3 | 153329877 | rs6769589 | 0.617 | -0.071 | -0.214 | 0.118 | -0.059 | -0.088 | -0.088 | 0 | 0.35 | 0.49 | 0.31 | 0.08 | 0.18 | 0.24 |
| 3 | 153335441 | rs7616085 | 0.31 | 0.145 | -0.436 | 0.242 | -0.054 | 0.013 | 0.013 | 0 | 0.35 | 0.15 | 0.39 | 0.34 | 0.4 | 0.46 |
| 3 | 153349772 | rs13323208 | 0.685 | 0.131 | 0.144 | 0.275 | 0.015 | 0.054 | 0.054 | 0 | 0.07 | 0.23 | 0.17 | 0.08 | 0.07 | 0.04 |
| 3 | 153352722 | rs6806502 | -0.656 | 0.814 | 0.213 | 0.181 | 0.109 | 0.008 | 0.008 | 1.305 | 0.95 | 0.79 | 0.95 | 0.98 | 0.88 | 0.96 |
| 3 | 153354747 | rs13321012 | 0.898 | 0.781 | 0.347 | 0.191 | 0.012 | 0.035 | 0.035 | 0 | 0.07 | 0.2 | 0.17 | 0.07 | 0.07 | 0.05 |
| 3 | 153355197 | rs10935939 | 0.774 | 0.689 | -0.405 | 0.107 | -0.07 | -0.134 | -0.134 | 0 | 0.24 | 0.03 | 0.23 | 0.56 | 0.3 | 0.27 |
| 3 | 153356213 | rs6780167 | 0.347 | 0.389 | -0.571 | -0.118 | -0.024 | 0.014 | 0.014 | 0 | 0.35 | 0.4 | 0.39 | 0.46 | 0.25 | 0.42 |
| 3 | 153367793 | rs9822824 | 0.123 | 0.773 | -0.509 | -0.097 | 0.062 | 0.048 | 0.048 | 0 | 0.83 | 0.63 | 0.7 | 0.84 | 0.76 | 0.76 |
| 3 | 153368311 | rs13079953 | 1.873 | 0.857 | -0.607 | -0.09 | -0.05 | -0.089 | -0.089 | 0 | 0.18 | 0.02 | 0.15 | 0.37 | 0.2 | 0.12 |
| 3 | 153370666 | rs11711889 | 0.972 | 0.803 | -0.651 | -0.118 | -0.042 | -0.072 | -0.072 | 0.074 | 0.11 | 0 | 0.13 | 0.25 | 0.16 | 0.14 |
| 3 | 153374978 | rs1006977 | 0.782 | 0.663 | -0.593 | -0.302 | -0.021 | 0.033 | 0.033 | 0 | 0.38 | 0.12 | 0.31 | 0.29 | 0.44 | 0.29 |
| 3 | 153375167 | rs1554560 | -0.043 | 0.667 | -0.687 | -0.168 | 0.033 | 0.021 | 0.021 | 0 | 0.64 | 0.51 | 0.55 | 0.47 | 0.56 | 0.64 |
| 3 | 153387494 | rs7621258 | 0.757 | 0.776 | -1.072 | -0.096 | -0.016 | -0.034 | -0.034 | 0 | 0.11 | 0.08 | 0.13 | 0.26 | 0.17 | 0.14 |
| 3 | 153388774 | rs6778630 | 0.549 | 0.607 | -0.953 | -0.348 | 0 | 0.004 | 0.004 | 0 | 0.21 | 0.3 | 0.24 | 0.33 | 0.25 | 0.37 |
| 3 | 153390148 | rs17744453 | 1.425 | 0.424 | -0.798 | -0.696 | 0.034 | 0.021 | 0.021 | 0.331 | 0.19 | 0.12 | 0.09 | 0 | 0.07 | 0.14 |
| 3 | 153390848 | rs9817352 | 0.332 | 1.807 | -0.481 | 0.087 | 0.072 | 0.026 | 0.026 | 0 | 0.89 | 0.75 | 0.79 | 0.91 | 0.81 | 0.91 |
| 3 | 153392603 | rs3846078 | -0.277 | 1.591 | -0.328 | 0.206 | 0.142 | 0.12 | 0.12 | 0 | 0.94 | 0.75 | 0.82 | 0.91 | 0.89 | 0.93 |
| 3 | 153396096 | rs1490543 | 0.431 | 0.74 | -0.8 | -0.182 | 0.004 | -0.011 | -0.011 | 2.996 | 0.51 | 0.44 | 0.4 | 0.2 | 0.38 | 0.42 |
| 3 | 153404708 | rs9879759 | -0.119 | 1.146 | -1.162 | 0.441 | -0.029 | -0.054 | -0.054 | 0 | 0.24 | 0.11 | 0.18 | 0.37 | 0.27 | 0.18 |
| 3 | 153408662 | rs952122 | -0.356 | 0.612 | -0.794 | -0.722 | -0.001 | -0.041 | -0.041 | 0 | 0.46 | 0.44 | 0.43 | 0.1 | 0.65 | 0.32 |
| 3 | 153409421 | rs2037556 | -0.482 | 0.664 | -0.597 | -0.823 | 0.032 | 0.088 | 0.088 | 1.233 | 0.44 | 0.3 | 0.41 | 0.08 | 0.63 | 0.31 |
| 3 | 153418139 | rs7638828 | 0.143 | 0.869 | -0.041 | -1.046 | 0.03 | -0.092 | -0.092 | 0 | 0.13 | 0.43 | 0.09 | 0.01 | 0.09 | 0.03 |
| 3 | 153418189 | rs12696007 | -0.024 | 0.842 | -0.216 | -1.093 | -0.051 | 0.09 | 0.09 | 0.225 | 0.32 | 0.15 | 0.35 | 0.07 | 0.54 | 0.29 |
| 3 | 153420934 | rs10935945 | -0.015 | 0.849 | -0.213 | -1.09 | -0.051 | 0.09 | 0.09 | 0.185 | 0.32 | 0.15 | 0.35 | 0.07 | 0.54 | 0.3 |
| 3 | 153422774 | rs17744868 | 0.771 | 0.901 | -0.411 | -1.031 | -0.004 | 0.021 | 0.021 | 0 | 0.11 | 0.06 | 0.06 | 0.01 | 0.09 | 0.03 |
| 3 | 153423761 | rs9823065 | -0.355 | 1.74 | -0.519 | -1.072 | -0.003 | -0.063 | -0.063 | 0.178 | 0.36 | 0.55 | 0.25 | 0.01 | 0.44 | 0.2 |
| 3 | 153430648 | rs7645891 | -1.039 | 0.3 | -0.621 | -0.928 | 0.09 | 0.146 | 0.146 | 0.248 | 0.52 | 0.69 | 0.47 | 0.09 | 0.7 | 0.38 |
| 3 | 153437261 | rs9857982 | -0.804 | 0.228 | -0.567 | -0.033 | 0.006 | 0.016 | 0.016 | 0.134 | 0.54 | 0.64 | 0.71 | 0.6 | 0.55 | 0.45 |
| 3 | 153440291 | rs11715526 | -0.941 | 0.075 | -0.612 | -0.056 | 0.007 | 0.009 | 0.009 | 0 | 0.57 | 0.63 | 0.75 | 0.62 | 0.63 | 0.54 |
| 3 | 153443023 | rs6801094 | 0.403 | -0.135 | -0.812 | -0.141 | -0.096 | 0.087 | 0.087 | 0 | 0.27 | 0.07 | 0.31 | 0.1 | 0.4 | 0.26 |
| 3 | 153447119 | rs9865106 | -0.146 | 0.039 | -0.688 | 0.511 | 0.001 | -0.029 | -0.029 | 3.821 | 0.19 | 0.13 | 0.22 | 0.26 | 0.21 | 0.23 |
| 3 | 153449080 | rs579695 | 0.899 | -0.046 | -0.688 | 0.574 | 0.007 | 0.009 | 0.009 | 0 | 0.43 | 0.35 | 0.25 | 0.4 | 0.36 | 0.46 |
| 3 | 153453684 | rs1032844 | -0.019 | -0.008 | -0.884 | 0.864 | 0.011 | -0.047 | -0.047 | 0 | 0.14 | 0.06 | 0.19 | 0.26 | 0.13 | 0.17 |
| 3 | 153459400 | rs532478 | -0.654 | -0.001 | -0.41 | 0.67 | -0.009 | 0.018 | 0.018 | 0.065 | 0.36 | 0.5 | 0.27 | 0.36 | 0.34 | 0.33 |
| 3 | 153460899 | rs1532795 | -0.809 | -0.228 | -0.578 | 0.563 | -0.015 | 0.01 | 0.01 | 0.342 | 0.4 | 0.52 | 0.28 | 0.37 | 0.35 | 0.34 |
| 3 | 153464278 | rs579468 | -0.798 | -0.341 | -0.797 | 0.14 | 0.09 | -0.086 | -0.086 | 0 | 0.76 | 0.6 | 0.66 | 0.94 | 0.63 | 0.82 |

| | | | | | | | | | | | | | | | | |
|---|---|---|---|---|---|---|---|---|---|---|---|---|---|---|---|---|
| 3 | 153467805 | rs12490796 | 0.792 | -0.282 | -1.024 | 0.292 | 0.063 | -0.059 | -0.059 | 0 | 0.38 | 0.17 | 0.38 | 0.59 | 0.27 | 0.49 |
| 3 | 153470222 | rs514105 | -0.451 | -0.068 | -0.811 | 0.442 | -0.002 | 0.004 | 0.004 | 0 | 0.28 | 0.24 | 0.24 | 0.19 | 0.34 | 0.27 |
| 3 | 153474220 | rs510173 | -0.18 | -0.333 | -0.695 | 0.601 | -0.002 | -0.001 | -0.001 | 0 | 0.22 | 0.22 | 0.18 | 0.19 | 0.19 | 0.21 |
| 3 | 153476041 | rs9877084 | 0.351 | -0.646 | -0.602 | 0.872 | -0.015 | 0.049 | 0.049 | 0.352 | 0.15 | 0.23 | 0.25 | 0.37 | 0.12 | 0.14 |
| 3 | 153477976 | rs9812437 | 0.324 | -0.821 | -1.111 | 0.268 | 0.034 | 0.056 | 0.056 | 0 | 0.35 | 0.22 | 0.16 | 0.13 | 0.25 | 0.4 |
| 3 | 153481697 | rs663256 | -0.023 | -0.631 | -0.987 | 0.566 | -0.003 | 0 | 0 | 0 | 0.4 | 0.38 | 0.38 | 0.47 | 0.33 | 0.53 |
| 3 | 153482959 | rs482953 | 0.41 | -0.415 | -0.942 | 0.708 | 0.013 | 0.108 | 0.108 | 0 | 0.05 | 0.1 | 0.21 | 0.34 | 0.09 | 0.14 |
| 3 | 153496242 | rs12486336 | 0.018 | 0.724 | -1.621 | 0.164 | -0.072 | -0.063 | -0.063 | 0.91 | 0.23 | 0.13 | 0.48 | 0.4 | 0.47 | 0.34 |
| 3 | 153497897 | rs1524543 | -0.679 | 0.519 | -1.148 | -0.448 | -0.012 | -0.06 | -0.06 | 0.462 | 0.74 | 0.71 | 0.83 | 0.94 | 0.84 | 0.9 |
| 3 | 153504560 | rs4680092 | 0.882 | -0.036 | -1.614 | -0.952 | 0.001 | 0.013 | 0.013 | 0.042 | 0.21 | 0.18 | 0.12 | 0.06 | 0.22 | 0.3 |
| 3 | 153504639 | rs16823172 | 1.808 | -0.035 | -1.589 | -0.453 | -0.001 | -0.014 | -0.014 | 0 | 0.1 | 0.07 | 0.14 | 0.28 | 0.07 | 0.07 |
| 3 | 153505042 | rs1524541 | 1.536 | -0.286 | -1.7 | -0.643 | 0.005 | 0.003 | 0.003 | 0.025 | 0.19 | 0.18 | 0.08 | 0.05 | 0.12 | 0.27 |
| 3 | 153505779 | rs16823176 | 2.224 | -0.25 | -1.881 | -0.459 | -0.008 | -0.043 | -0.043 | 0 | 0.11 | 0.03 | 0.06 | 0.2 | 0.1 | 0.13 |
| 3 | 153507426 | rs4679721 | 0.14 | 0.056 | -1.34 | -0.488 | -0.003 | -0.003 | -0.003 | 0 | 0.29 | 0.28 | 0.62 | 0.59 | 0.46 | 0.37 |
| 3 | 153513139 | rs7622338 | 0.838 | 0.282 | -1.095 | -0.495 | 0.032 | 0.022 | 0.022 | 0.693 | 0.2 | 0.26 | 0.49 | 0.33 | 0.39 | 0.3 |
| 3 | 153521473 | rs13071232 | 0.154 | 0.238 | -1.043 | -0.541 | 0.05 | 0.018 | 0.018 | 0.612 | 0.26 | 0.37 | 0.53 | 0.34 | 0.49 | 0.33 |
| 3 | 153524432 | rs10935951 | -0.201 | -0.007 | -1.081 | -0.742 | -0.001 | -0.002 | -0.002 | 0 | 0.45 | 0.56 | 0.64 | 0.39 | 0.61 | 0.6 |
| 3 | 153525308 | rs10935952 | 1.23 | 0.284 | -1.083 | -0.55 | -0.013 | -0.012 | -0.012 | 0 | 0.18 | 0.13 | 0.49 | 0.33 | 0.39 | 0.33 |
| 3 | 153526188 | rs12490250 | 0.194 | 0.235 | -1.062 | -0.553 | 0.001 | -0.001 | -0.001 | 0 | 0.25 | 0.24 | 0.53 | 0.34 | 0.48 | 0.36 |
| 3 | 153536704 | rs13098484 | 1.184 | 0.034 | -1.077 | -0.459 | -0.012 | -0.011 | -0.011 | 0.464 | 0.18 | 0.16 | 0.46 | 0.34 | 0.36 | 0.29 |
| 3 | 153538262 | rs12486542 | -0.426 | -0.452 | -0.87 | -0.802 | 0.003 | 0.003 | 0.003 | 0.861 | 0.5 | 0.55 | 0.65 | 0.42 | 0.64 | 0.62 |
| 3 | 153538769 | rs2174299 | -0.998 | -0.41 | -0.851 | -0.684 | 0.006 | 0.002 | 0.002 | 0 | 0.58 | 0.65 | 0.78 | 0.67 | 0.69 | 0.67 |
| 3 | 153538939 | rs1524538 | 0.217 | 0.022 | -0.656 | -0.906 | -0.004 | -0.001 | -0.001 | 0 | 0.36 | 0.37 | 0.57 | 0.42 | 0.51 | 0.5 |
| 3 | 153548571 | rs17803063 | -1.183 | 0.266 | -0.28 | -0.401 | -0.039 | -0.041 | -0.041 | 0.062 | 0.81 | 0.9 | 0.55 | 0.64 | 0.63 | 0.73 |
| 3 | 153550577 | rs4680104 | 0.631 | 0.221 | 0.297 | -0.55 | 0.001 | 0 | 0 | 0 | 0.65 | 0.36 | 0.45 | 0.64 | 0.51 | 0.65 |
| 3 | 153553501 | rs6794969 | -0.332 | 0.131 | 0.456 | -0.544 | -0.008 | -0.08 | -0.08 | 0.016 | 0.14 | 0.31 | 0.09 | 0 | 0.11 | 0.05 |
| 3 | 153555839 | rs7645911 | 0.164 | 0.274 | -0.005 | -0.355 | 0.022 | -0.003 | -0.003 | 0 | 0.64 | 0.46 | 0.46 | 0.63 | 0.49 | 0.64 |
| 3 | 153567441 | rs10513451 | 0.787 | -0.677 | -0.316 | -1.176 | 0.003 | -0.012 | -0.012 | 0 | 0.33 | 0.3 | 0.2 | 0.08 | 0.19 | 0.28 |
| 3 | 153572537 | rs6788015 | -0.124 | -0.667 | 0.009 | -1.386 | 0.014 | 0.004 | 0.004 | 0.033 | 0.45 | 0.56 | 0.28 | 0.08 | 0.33 | 0.35 |
| 3 | 153573828 | rs6803060 | 0.953 | -0.566 | -0.332 | -1.342 | 0.002 | -0.015 | -0.015 | 0.054 | 0.33 | 0.3 | 0.2 | 0.08 | 0.19 | 0.28 |
| 3 | 153578076 | rs7637630 | 0.112 | -0.507 | -0.238 | -1.235 | 0.016 | 0.031 | 0.031 | 0.018 | 0.42 | 0.31 | 0.26 | 0.08 | 0.33 | 0.31 |
| 3 | 153578641 | rs9824066 | 1.116 | -0.497 | -0.676 | -1.187 | 0.005 | 0.057 | 0.057 | 0.066 | 0.13 | 0.04 | 0.05 | 0.01 | 0.11 | 0.12 |
| 3 | 153598526 | rs11915835 | -0.101 | -0.62 | -0.207 | -1.122 | 0.001 | -0.034 | -0.034 | 0 | 0.43 | 0.5 | 0.28 | 0.08 | 0.34 | 0.35 |
| 3 | 153603294 | rs16823270 | 1.559 | -0.64 | -0.692 | -0.698 | -0.01 | -0.075 | -0.075 | 0 | 0.15 | 0.04 | 0.09 | 0.25 | 0.12 | 0.15 |
| 3 | 153610263 | rs3112291 | -1.065 | -0.745 | -0.276 | -1 | 0.001 | -0.016 | -0.016 | 0 | 0.68 | 0.71 | 0.8 | 0.92 | 0.8 | 0.72 |
| 3 | 153610297 | rs11925488 | 0.989 | -0.678 | -0.299 | -0.949 | -0.028 | 0.045 | 0.045 | 0 | 0.1 | 0.01 | 0.06 | 0 | 0.14 | 0.04 |
| 3 | 153616295 | rs787964 | -1.012 | -0.766 | -0.204 | -0.988 | -0.009 | -0.04 | -0.04 | 0 | 0.67 | 0.66 | 0.8 | 0.92 | 0.8 | 0.72 |
| 3 | 153620022 | rs17815430 | 0.932 | -0.655 | -0.55 | -0.959 | -0.012 | 0.017 | 0.017 | 0 | 0.1 | 0.04 | 0.06 | 0 | 0.14 | 0.04 |
| 3 | 153627929 | rs701263 | -1.266 | -0.769 | -0.395 | -0.988 | 0.037 | 0.094 | 0.094 | 0.505 | 0.7 | 0.87 | 0.81 | 0.92 | 0.8 | 0.72 |
| 3 | 153640591 | rs10513453 | 0.143 | -0.324 | 0.233 | -0.805 | 0.026 | -0.078 | -0.078 | 2.006 | 0.12 | 0.29 | 0.08 | 0 | 0.15 | 0.07 |
| 3 | 153656818 | rs1879557 | -1.333 | -0.654 | -0.415 | -0.882 | 0.014 | 0.033 | 0.033 | 0 | 0.75 | 0.85 | 0.82 | 0.93 | 0.84 | 0.76 |
| 3 | 153663422 | rs1357087 | 1.37 | -0.501 | -0.328 | -0.322 | -0.002 | 0.003 | 0.003 | 0 | 0.15 | 0.13 | 0.16 | 0.23 | 0.09 | 0.13 |
| 3 | 153671400 | rs1533750 | -0.547 | -0.668 | -0.234 | -0.569 | 0.003 | 0.08 | 0.08 | 0 | 0.63 | 0.78 | 0.74 | 0.92 | 0.65 | 0.67 |
| 3 | 153672174 | rs1467266 | -0.476 | -0.676 | 0.057 | -0.57 | 0 | -0.027 | -0.027 | 0 | 0.63 | 0.64 | 0.73 | 0.92 | 0.65 | 0.67 |
| 3 | 153673498 | rs16823342 | 1.088 | -0.635 | 0 | -0.578 | 0.007 | -0.042 | -0.042 | 0.045 | 0.08 | 0.15 | 0.05 | 0 | 0.08 | 0.03 |
| 3 | 153673681 | rs1568489 | 0.377 | -0.651 | -0.138 | -0.567 | -0.001 | -0.003 | -0.003 | 3.428 | 0.47 | 0.51 | 0.58 | 0.69 | 0.56 | 0.54 |
| 3 | 153677863 | rs6440914 | -1.047 | -0.438 | -0.608 | -0.555 | 0.003 | 0.084 | 0.084 | 0 | 0.71 | 0.87 | 0.79 | 0.92 | 0.74 | 0.7 |
| 3 | 153680401 | rs2900641 | -0.662 | -0.491 | -0.542 | -0.505 | -0.008 | 0.131 | 0.131 | 0 | 0.64 | 0.85 | 0.75 | 0.92 | 0.65 | 0.67 |
| 3 | 153684680 | rs16823359 | 1.677 | -0.656 | -0.616 | -0.417 | 0.017 | -0.041 | -0.041 | 0 | 0.15 | 0.03 | 0.15 | 0.19 | 0.1 | 0.12 |
| 3 | 153685843 | rs6440917 | -0.729 | -0.631 | -0.532 | -0.406 | 0 | 0.106 | 0.106 | 0 | 0.64 | 0.81 | 0.74 | 0.92 | 0.65 | 0.67 |
| 3 | 153688870 | rs1521303 | 1.159 | -0.525 | -0.049 | -0.074 | -0.023 | 0.032 | 0.032 | 0 | 0.17 | 0.23 | 0.16 | 0.23 | 0.1 | 0.15 |
| 3 | 153690297 | rs12633006 | 1.001 | 0.277 | -0.305 | 0.27 | -0.01 | -0.015 | -0.015 | 0 | 0.1 | 0.07 | 0.08 | 0.26 | 0.09 | 0.14 |
| 3 | 153704356 | rs6440920 | -0.337 | -0.099 | -0.699 | 0.074 | -0.008 | 0.118 | 0.118 | 0.016 | 0.55 | 0.91 | 0.69 | 0.73 | 0.59 | 0.55 |
| 3 | 153712602 | rs883228 | -0.444 | -0.091 | -0.531 | 0.079 | 0.017 | 0.111 | 0.111 | 0 | 0.55 | 0.84 | 0.68 | 0.72 | 0.6 | 0.55 |
| 3 | 153713874 | rs9844300 | 1.296 | -0.177 | -0.406 | -0.09 | -0.004 | -0.016 | -0.016 | 0.041 | 0.09 | 0.12 | 0.02 | 0.02 | 0.05 | 0.08 |
| 3 | 153715341 | rs7611756 | -0.323 | -0.067 | -0.556 | 0.11 | -0.012 | 0.108 | 0.108 | 0.04 | 0.56 | 0.91 | 0.69 | 0.72 | 0.6 | 0.55 |
| 3 | 153725622 | rs6794747 | -0.612 | -0.211 | -0.138 | 0.27 | 0.031 | 0.002 | 0.002 | 0 | 0.62 | 0.71 | 0.73 | 0.6 | 0.74 | 0.58 |
| 3 | 153730237 | rs4613417 | -0.733 | -0.185 | -0.546 | 0.109 | 0.058 | 0.041 | 0.041 | 0.011 | 0.63 | 0.93 | 0.75 | 0.72 | 0.74 | 0.58 |
| 3 | 153732845 | rs4234301 | -0.834 | -0.339 | -0.487 | 0.175 | 0.072 | 0.021 | 0.021 | 0.016 | 0.64 | 0.9 | 0.76 | 0.7 | 0.75 | 0.58 |
| 3 | 153754091 | rs11923161 | 1.234 | -0.311 | -0.012 | 0.671 | -0.014 | 0.067 | 0.067 | 0 | 0.17 | 0.2 | 0.28 | 0.3 | 0.1 | 0.21 |
| 3 | 153754124 | rs6440932 | -0.993 | -0.435 | 0.131 | 0.555 | 0.064 | 0.03 | 0.03 | 0 | 0.64 | 0.84 | 0.76 | 0.69 | 0.74 | 0.58 |
| 3 | 153755820 | rs4256090 | 0.221 | -0.244 | 0.42 | 0.532 | 0.007 | 0.103 | 0.103 | 0 | 0.26 | 0.42 | 0.36 | 0.42 | 0.24 | 0.24 |
| 3 | 153761914 | rs13080778 | -0.348 | -0.206 | 0.393 | 0.41 | 0.022 | 0.028 | 0.028 | 0 | 0.54 | 0.68 | 0.69 | 0.58 | 0.6 | 0.55 |
| 3 | 153766961 | rs7646922 | -0.056 | -0.181 | 0.504 | 0.386 | 0.017 | 0.029 | 0.029 | 0 | 0.45 | 0.66 | 0.48 | 0.38 | 0.63 | 0.36 |
| 3 | 153774462 | rs9882680 | 0.514 | -0.254 | 0.482 | 0.387 | 0.024 | -0.038 | -0.038 | 0 | 0.37 | 0.46 | 0.41 | 0.26 | 0.5 | 0.34 |
| 3 | 153776149 | rs13079099 | 1.179 | -0.228 | 0.618 | 0.74 | 0.007 | -0.011 | -0.011 | 0 | 0.16 | 0.1 | 0.27 | 0.33 | 0.1 | 0.21 |
| 3 | 153779868 | rs17817774 | 0.519 | -0.036 | 0.513 | 0.62 | 0.02 | 0.004 | 0.004 | 0 | 0.08 | 0.2 | 0.08 | 0.11 | 0.13 | 0.02 |
| 3 | 153809316 | rs1713795 | 0.069 | -0.281 | 0.835 | 0.639 | -0.007 | 0.1 | 0.1 | 0 | 0.73 | 0.49 | 0.6 | 0.67 | 0.77 | 0.76 |
| 3 | 153818962 | rs1617481 | 0.119 | -0.324 | 1.091 | 0.638 | -0.007 | 0.073 | 0.073 | 0.008 | 0.73 | 0.48 | 0.6 | 0.55 | 0.77 | 0.77 |

| | | | | | | | | | | | | | | | | |
|---|---|---|---|---|---|---|---|---|---|---|---|---|---|---|---|---|
| 3 | 153831289 | rs1470666 | 0.677 | -0.313 | 1.121 | 0.967 | -0.038 | 0.146 | 0.146 | 0.08 | 0.2 | 0.46 | 0.27 | 0.31 | 0.11 | 0.2 |
| 3 | 153843667 | rs1727889 | 0.254 | -0.368 | 1.212 | 0.375 | -0.008 | 0.055 | 0.055 | 0 | 0.72 | 0.43 | 0.59 | 0.55 | 0.76 | 0.77 |
| 3 | 153846481 | rs10935975 | 0.671 | -0.361 | 1.231 | 0.382 | -0.039 | 0.207 | 0.207 | 0.063 | 0.2 | 0.49 | 0.34 | 0.45 | 0.11 | 0.21 |
| 3 | 153846700 | rs1463217 | 0.24 | -0.365 | 1.25 | 0.383 | -0.006 | 0.038 | 0.038 | 0.036 | 0.72 | 0.4 | 0.59 | 0.55 | 0.76 | 0.77 |
| 3 | 153857679 | rs7622562 | 1.121 | -0.197 | 0.63 | -0.021 | -0.123 | 0.021 | 0.021 | 0 | 0.29 | 0.05 | 0.36 | 0.24 | 0.45 | 0.26 |
| 3 | 153860779 | rs12496189 | 0.625 | -0.317 | 1.125 | 0.31 | -0.038 | 0.179 | 0.179 | 0.063 | 0.19 | 0.4 | 0.34 | 0.45 | 0.11 | 0.21 |
| 3 | 153870252 | rs2044757 | 1.056 | -0.315 | 1.293 | 0.32 | -0.003 | 0.023 | 0.023 | 0 | 0.17 | 0.15 | 0.31 | 0.34 | 0.11 | 0.19 |
| 3 | 153873574 | rs1962674 | 0.843 | -0.318 | 1.314 | 0.318 | -0.022 | 0.092 | 0.092 | 0 | 0.18 | 0.27 | 0.33 | 0.34 | 0.11 | 0.19 |
| 3 | 153875713 | rs1727949 | -0.377 | -0.282 | 1.143 | 0.135 | 0.036 | 0.015 | 0.015 | 0 | 0.91 | 0.76 | 0.92 | 0.88 | 0.87 | 0.97 |
| 3 | 153891686 | rs357478 | 0.231 | -0.572 | 0.748 | 0.006 | 0 | 0.001 | 0.001 | 0.008 | 0.46 | 0.49 | 0.44 | 0.48 | 0.54 | 0.43 |
| 3 | 153894996 | rs7639613 | 1.093 | -0.645 | 0.588 | 0.035 | -0.02 | 0.075 | 0.075 | 0 | 0.18 | 0.21 | 0.32 | 0.38 | 0.11 | 0.18 |
| 3 | 153898669 | rs6796038 | 1.093 | -0.649 | 0.698 | -0.075 | -0.023 | 0.1 | 0.1 | 0 | 0.18 | 0.23 | 0.32 | 0.42 | 0.11 | 0.17 |
| 3 | 153902784 | rs9289938 | 0.846 | -0.704 | 0.518 | -0.248 | 0.009 | 0.013 | 0.013 | 0 | 0.31 | 0.25 | 0.19 | 0.14 | 0.23 | 0.38 |
| 3 | 153907658 | rs380401 | 1.177 | -0.487 | 0.059 | -0.441 | -0.044 | 0.023 | 0.023 | 0.484 | 0.27 | 0.18 | 0.36 | 0.21 | 0.44 | 0.3 |
| 3 | 153912641 | rs12330779 | 0.998 | -0.48 | 0.002 | -0.443 | 0.02 | 0.084 | 0.084 | 0 | 0.28 | 0.08 | 0.16 | 0.14 | 0.22 | 0.38 |
| 3 | 153914902 | rs386893 | 0.397 | -0.388 | 0.003 | -0.389 | 0 | 0.001 | 0.001 | 0 | 0.46 | 0.48 | 0.45 | 0.36 | 0.54 | 0.43 |
| 3 | 153927792 | rs399118 | 0.087 | -0.249 | 0.126 | -0.695 | -0.007 | -0.048 | -0.048 | 0 | 0.61 | 0.51 | 0.74 | 0.85 | 0.64 | 0.62 |
| 3 | 153930808 | rs2122365 | -0.802 | 0.026 | -0.641 | -0.516 | 0.008 | -0.026 | -0.026 | 0 | 0.93 | 0.88 | 0.93 | 1 | 0.87 | 0.98 |
| 3 | 153949975 | rs9844663 | 0.541 | -0.18 | -0.869 | -0.594 | 0.013 | 0.076 | 0.076 | 0 | 0.29 | 0.12 | 0.17 | 0.14 | 0.24 | 0.38 |
| 3 | 153964496 | rs355748 | 0.461 | -0.464 | -0.355 | -0.486 | 0 | 0 | 0 | 0.093 | 0.42 | 0.45 | 0.41 | 0.49 | 0.4 | 0.38 |
| 3 | 153967490 | rs12491012 | 1.387 | -0.586 | -0.193 | -0.557 | 0 | 0.022 | 0.022 | 0.017 | 0.15 | 0.12 | 0.3 | 0.37 | 0.1 | 0.17 |
| 3 | 153973408 | rs3172332 | 0.764 | -0.493 | -0.24 | -0.635 | -0.042 | 0.1 | 0.1 | 0 | 0.33 | 0.1 | 0.23 | 0.15 | 0.37 | 0.42 |
| 3 | 153974029 | rs13074879 | 1.036 | -0.4 | 0.36 | -0.552 | -0.032 | 0.148 | 0.148 | 0.356 | 0.17 | 0.3 | 0.31 | 0.37 | 0.1 | 0.17 |
| 3 | 153977253 | rs355779 | -0.07 | -0.398 | 0.36 | -0.564 | -0.001 | 0.106 | 0.106 | 0.127 | 0.75 | 0.57 | 0.63 | 0.63 | 0.77 | 0.79 |
| 3 | 153992431 | rs6771155 | 1.409 | -0.387 | 0.325 | -0.585 | -0.005 | 0.04 | 0.04 | 0.007 | 0.15 | 0.14 | 0.3 | 0.37 | 0.1 | 0.17 |
| 3 | 154010991 | rs355761 | 2.343 | -0.461 | -0.189 | -0.703 | -0.076 | -0.049 | -0.049 | 0 | 0.15 | 0.01 | 0.28 | 0.19 | 0.24 | 0.18 |
| 3 | 154012614 | rs355762 | 0.248 | 0.048 | 0.563 | -0.319 | -0.006 | 0.108 | 0.108 | 0.277 | 0.74 | 0.52 | 0.63 | 0.63 | 0.77 | 0.79 |
| 3 | 154018690 | rs3732652 | 0.545 | 0.084 | 0.4 | -0.345 | -0.036 | 0.079 | 0.079 | 0 | 0.31 | 0.12 | 0.22 | 0.15 | 0.36 | 0.4 |
| 3 | 154045001 | rs355768 | -0.973 | 0.314 | 0.47 | -0.137 | -0.043 | 0.041 | 0.041 | 0 | 0.92 | 1 | 0.95 | 1 | 0.87 | 0.96 |
| 3 | 154052774 | rs696856 | 0.917 | 0.246 | 0.547 | -0.199 | 0.01 | 0.02 | 0.02 | 0.079 | 0.5 | 0.39 | 0.44 | 0.52 | 0.48 | 0.44 |
| 3 | 154054799 | rs701145 | -1.144 | 0.575 | 0.112 | -0.194 | -0.041 | 0.036 | 0.036 | 0 | 0.93 | 1 | 0.95 | 1 | 0.87 | 0.95 |
| 3 | 154064740 | rs789296 | 0.044 | 0.445 | 0.261 | -0.275 | -0.004 | -0.004 | -0.004 | 0.894 | 0.68 | 0.7 | 0.75 | 0.89 | 0.58 | 0.62 |
| 3 | 154070312 | rs701133 | -1.173 | 0.568 | -0.155 | -0.242 | -0.041 | 0.037 | 0.037 | 0 | 0.93 | 1 | 0.95 | 1 | 0.87 | 0.96 |
| 3 | 154076157 | rs2037580 | 0.787 | 0.323 | 0.187 | 0.095 | -0.01 | 0.057 | 0.057 | 0.025 | 0.16 | 0.17 | 0.3 | 0.46 | 0.1 | 0.17 |
| 3 | 154082012 | rs13315200 | 0.439 | 0.488 | -0.203 | -0.094 | -0.016 | 0.025 | 0.025 | 0 | 0.23 | 0.16 | 0.19 | 0.11 | 0.29 | 0.33 |
| 3 | 154091066 | rs10935984 | 0.551 | 0.201 | 0.142 | 0.453 | -0.025 | 0.142 | 0.142 | 0 | 0.13 | 0.23 | 0.21 | 0.44 | 0.08 | 0.09 |
| 3 | 154095084 | rs2689326 | 0.471 | 0.236 | 0.334 | 0.246 | -0.014 | 0.027 | 0.027 | 0 | 0.74 | 0.58 | 0.58 | 0.34 | 0.78 | 0.79 |
| 3 | 154095778 | rs12488867 | -0.517 | 0.251 | 0.293 | 0.207 | -0.016 | 0.011 | 0.011 | 0 | 0.25 | 0.33 | 0.4 | 0.66 | 0.17 | 0.21 |
| 3 | 154100758 | rs701121 | -0.295 | 0.527 | 0.475 | 0.289 | 0.005 | 0.014 | 0.014 | 0 | 0.87 | 0.8 | 0.79 | 0.77 | 0.86 | 0.88 |
| 3 | 154102981 | rs696853 | -0.25 | 0.526 | 0.474 | 0.289 | 0.004 | 0.013 | 0.013 | 0 | 0.87 | 0.8 | 0.79 | 0.77 | 0.86 | 0.88 |
| 3 | 154106880 | rs7643022 | 1.739 | 0.401 | -0.005 | -0.183 | -0.037 | 0.045 | 0.045 | 0.112 | 0.27 | 0.09 | 0.27 | 0.17 | 0.29 | 0.33 |
| 3 | 154111414 | rs2136837 | 1.157 | 1.159 | 1.289 | -0.244 | -0.012 | 0.001 | 0.001 | 0 | 0.61 | 0.35 | 0.58 | 0.56 | 0.57 | 0.56 |
| 3 | 154115627 | rs6762664 | 0.446 | 1.258 | 1.321 | -0.634 | 0.041 | -0.04 | -0.04 | 0.003 | 0.72 | 0.57 | 0.78 | 0.78 | 0.65 | 0.65 |
| 3 | 154133729 | rs13080773 | 0.02 | 0.99 | 1.374 | -0.063 | 0.002 | -0.008 | -0.008 | 0 | 0.17 | 0.11 | 0.21 | 0.27 | 0.14 | 0.17 |
| 3 | 154134232 | rs10513457 | -0.352 | 1.04 | 1.448 | -0.258 | 0.01 | -0.004 | -0.004 | 0 | 0.21 | 0.21 | 0.18 | 0.22 | 0.29 | 0.27 |
| 3 | 154134251 | rs17369867 | -0.169 | 1.037 | 1.193 | -0.261 | -0.068 | 0.032 | 0.032 | 0 | 0.2 | 0.05 | 0.17 | 0.13 | 0.29 | 0.27 |
| 3 | 154144067 | rs7614655 | -1.555 | 0.955 | 1.441 | -0.184 | -0.009 | 0.001 | 0.001 | 0 | 0.4 | 0.62 | 0.41 | 0.48 | 0.43 | 0.44 |
| 3 | 154149215 | rs16824038 | -0.948 | 0.129 | 1.224 | 0.191 | -0.005 | 0.04 | 0.04 | 0.057 | 0.2 | 0.39 | 0.24 | 0.36 | 0.16 | 0.19 |
| 3 | 154149405 | rs1996256 | -0.1 | 0.165 | 0.896 | 0.208 | -0.011 | -0.035 | -0.035 | 0 | 0.16 | 0.04 | 0.2 | 0.24 | 0.14 | 0.14 |
| 3 | 154157941 | rs4680132 | -0.864 | 0.175 | 0.935 | 0.204 | -0.005 | 0.029 | 0.029 | 0.03 | 0.19 | 0.29 | 0.23 | 0.36 | 0.16 | 0.2 |
| 3 | 154172230 | rs4679735 | -0.198 | 0.231 | -0.097 | -0.039 | -0.012 | -0.036 | -0.036 | 0 | 0.16 | 0.04 | 0.2 | 0.24 | 0.17 | 0.14 |
| 3 | 154186021 | rs2314039 | -1.075 | 0.112 | 0.925 | -0.431 | -0.023 | -0.036 | -0.036 | 0 | 0.13 | 0.29 | 0.11 | 0.08 | 0.08 | 0.15 |
| 3 | 154186823 | rs7638340 | -1.347 | 0.066 | 1.444 | -0.451 | -0.023 | -0.041 | -0.041 | 0 | 0.14 | 0.38 | 0.11 | 0.08 | 0.08 | 0.15 |
| 3 | 154194897 | rs16824085 | -0.792 | 0.039 | 1.632 | -0.462 | -0.023 | -0.024 | -0.024 | 0 | 0.11 | 0.12 | 0.06 | 0.06 | 0.07 | 0.1 |
| 3 | 154200833 | rs4292196 | -1.807 | -0.088 | 2.414 | -0.499 | -0.042 | -0.049 | -0.049 | 0 | 0.15 | 0.56 | 0.13 | 0.09 | 0.09 | 0.15 |
| 3 | 154240489 | rs1519568 | -1.791 | 0.141 | 2.713 | -0.542 | 0.002 | -0.054 | -0.054 | 0 | 0.15 | 0.44 | 0.13 | 0.08 | 0.12 | 0.17 |
| 3 | 154252288 | rs11915244 | -1.918 | 0.305 | 2.615 | -0.702 | -0.019 | -0.055 | -0.055 | 0 | 0.16 | 0.57 | 0.15 | 0.1 | 0.12 | 0.17 |
| 3 | 154258918 | rs1160335 | -0.876 | 0.311 | 2.585 | -0.699 | -0.001 | -0.007 | -0.007 | 0 | 0.12 | 0.1 | 0.07 | 0.06 | 0.11 | 0.11 |
| 3 | 154267617 | rs7644760 | -2.055 | 0.325 | 2.845 | -0.69 | -0.019 | -0.042 | -0.042 | 0.581 | 0.17 | 0.65 | 0.15 | 0.1 | 0.12 | 0.16 |
| 3 | 154268934 | rs9809000 | -0.095 | 0.263 | 2.977 | -0.911 | 0.002 | -0.03 | -0.03 | 0 | 0.13 | 0.19 | 0.1 | 0.04 | 0.12 | 0.14 |
| 3 | 154277932 | rs2314058 | -1.104 | 0.251 | 3.168 | -0.707 | -0.012 | -0.04 | -0.04 | 0 | 0.13 | 0.29 | 0.09 | 0.06 | 0.11 | 0.11 |
| 3 | 154279750 | rs2313950 | -1.081 | 0.252 | 3.156 | -0.707 | -0.012 | -0.039 | -0.039 | 0 | 0.13 | 0.28 | 0.09 | 0.06 | 0.11 | 0.11 |
| 3 | 154281430 | rs10513461 | -1.081 | 0.252 | 3.156 | -0.707 | -0.012 | -0.039 | -0.039 | 0 | 0.13 | 0.28 | 0.09 | 0.06 | 0.11 | 0.11 |
| 3 | 154287700 | rs7630004 | -1.957 | 0.265 | 3.217 | -0.723 | -0.011 | -0.056 | -0.056 | 0 | 0.17 | 0.57 | 0.14 | 0.09 | 0.12 | 0.16 |
| 3 | 154291478 | rs7614327 | -1.034 | 0.281 | 2.873 | -0.71 | -0.009 | -0.024 | -0.024 | 0 | 0.13 | 0.17 | 0.08 | 0.06 | 0.11 | 0.11 |
| 3 | 154292518 | rs6440947 | -0.691 | 0.315 | 1.923 | -0.952 | -0.01 | 0.07 | 0.07 | 0 | 0.58 | 0.88 | 0.55 | 0.33 | 0.57 | 0.56 |
| 3 | 154309977 | rs17372655 | -0.313 | 0.208 | 1.922 | -1.054 | 0.001 | 0.001 | 0.001 | 0 | 0.11 | 0.09 | 0.05 | 0.06 | 0.09 | 0.1 |
| 3 | 154316007 | rs6809316 | -1.166 | 0.208 | 2.458 | -1.065 | 0.015 | -0.041 | -0.041 | 0 | 0.14 | 0.43 | 0.11 | 0.07 | 0.11 | 0.16 |
| 3 | 154317529 | rs7646874 | -0.713 | 0.233 | 2.497 | -1.064 | -0.046 | 0.085 | 0.085 | 0 | 0.58 | 0.92 | 0.53 | 0.31 | 0.57 | 0.55 |

| | | | | | | | | | | | | | | | | |
|---|---|---|---|---|---|---|---|---|---|---|---|---|---|---|---|---|
| 3 | 154325046 | rs7620094 | -0.686 | -0.024 | 2.668 | -0.999 | -0.014 | -0.042 | -0.042 | 0 | 0.12 | 0.29 | 0.07 | 0.06 | 0.09 | 0.1 |
| 3 | 154326754 | rs7614620 | -1.333 | -0.023 | 2.666 | -0.999 | 0.003 | -0.051 | -0.051 | 0 | 0.15 | 0.43 | 0.11 | 0.07 | 0.11 | 0.16 |
| 3 | 154330618 | rs12107040 | -1.331 | -0.028 | 2.664 | -1.001 | 0.003 | -0.051 | -0.051 | 0 | 0.15 | 0.43 | 0.11 | 0.07 | 0.11 | 0.16 |
| 3 | 154353017 | rs11915900 | -0.696 | -0.012 | 2.554 | -0.985 | -0.014 | -0.042 | -0.042 | 0 | 0.12 | 0.29 | 0.07 | 0.06 | 0.1 | 0.1 |
| 3 | 154381756 | rs4603925 | -0.528 | 0.173 | 2.106 | -0.906 | -0.033 | 0.085 | 0.085 | 0 | 0.56 | 0.88 | 0.51 | 0.34 | 0.53 | 0.55 |
| 3 | 154405414 | rs6776293 | -2.568 | 0.015 | 2.413 | -1.038 | 0.079 | 0.076 | 0.076 | 5.037 | 0.05 | 0.59 | 0.06 | 0.06 | 0.02 | 0.05 |
| 3 | 154412877 | rs4423728 | -0.614 | -0.123 | 1.839 | -1.158 | -0.043 | 0.094 | 0.094 | 0 | 0.58 | 0.85 | 0.37 | 0.29 | 0.51 | 0.56 |
| 3 | 154425941 | rs11928410 | -1.041 | -0.092 | 1.889 | -0.866 | -0.013 | -0.021 | -0.021 | 0 | 0.13 | 0.36 | 0.06 | 0.09 | 0.07 | 0.19 |
| 3 | 154432075 | rs11927311 | -1.041 | -0.092 | 1.889 | -0.866 | -0.013 | -0.021 | -0.021 | 2.169 | 0.13 | 0.36 | 0.06 | 0.09 | 0.07 | 0.19 |
| 3 | 154443871 | rs7611483 | -0.634 | 0.057 | 1.653 | -0.767 | -0.008 | -0.012 | -0.012 | 0.144 | 0.1 | 0.2 | 0.04 | 0.1 | 0.06 | 0.19 |
| 3 | 154449684 | rs7629959 | -0.521 | 0.005 | 0.981 | -0.867 | 0.014 | 0.064 | 0.064 | 0 | 0.54 | 0.71 | 0.29 | 0.3 | 0.44 | 0.55 |
| 3 | 154454529 | rs7651148 | -1.616 | 0.319 | 1.589 | -0.068 | 0.007 | 0.002 | 0.002 | 0.833 | 0.12 | 0.48 | 0.06 | 0.1 | 0.06 | 0.19 |
| 3 | 154488411 | rs7616989 | -1.254 | 0.487 | 1.499 | 0.022 | 0.092 | 0.118 | 0.118 | 1.745 | 0.06 | 0.4 | 0.03 | 0.13 | 0.04 | 0.19 |
| 3 | 154492161 | rs6789924 | -1.123 | 0.872 | 1.404 | -0.582 | -0.001 | -0.049 | -0.049 | 1.198 | 0.25 | 0.62 | 0.17 | 0.15 | 0.24 | 0.29 |
| 3 | 154493381 | rs10935990 | -0.495 | 0.861 | 1.503 | -0.59 | -0.005 | -0.001 | -0.001 | 0 | 0.19 | 0.17 | 0.14 | 0.02 | 0.21 | 0.13 |
| 3 | 154494952 | rs6440957 | -1.638 | 1.36 | 0.937 | -0.028 | -0.092 | -0.045 | -0.045 | 0 | 0.43 | 0.92 | 0.48 | 0.42 | 0.44 | 0.5 |
| 3 | 154496091 | rs6803413 | -0.446 | 1.331 | 0.682 | -0.527 | -0.032 | -0.064 | -0.064 | 0 | 0.88 | 0.97 | 0.69 | 0.56 | 0.83 | 0.82 |
| 3 | 154508800 | rs16824293 | -0.987 | 0.75 | 1.375 | -1.379 | -0.001 | -0.049 | -0.049 | 0 | 0.25 | 0.62 | 0.17 | 0.15 | 0.24 | 0.29 |
| 3 | 154511264 | rs997183 | -0.872 | 0.77 | 1.198 | -1.366 | 0.007 | -0.08 | -0.08 | 1.197 | 0.23 | 0.36 | 0.15 | 0.14 | 0.22 | 0.26 |
| 3 | 154518276 | rs7645606 | -1.047 | 0.736 | 1.086 | -1.365 | -0.002 | -0.042 | -0.042 | 0 | 0.26 | 0.64 | 0.18 | 0.15 | 0.24 | 0.29 |
| 3 | 154548410 | rs16824330 | -1.064 | 0.666 | 1.941 | -0.848 | 0.077 | 0.116 | 0.116 | 2.209 | 0.06 | 0.41 | 0.03 | 0.13 | 0.03 | 0.16 |
| 3 | 154551924 | rs11921486 | -0.842 | 0.801 | 1.14 | -0.948 | 0.003 | -0.052 | -0.052 | 0 | 0.19 | 0.24 | 0.14 | 0.02 | 0.21 | 0.13 |
| 3 | 154559907 | rs11924507 | -0.842 | 0.801 | 1.141 | -0.948 | 0.003 | -0.049 | -0.049 | 0.086 | 0.19 | 0.24 | 0.14 | 0.02 | 0.21 | 0.13 |
| 3 | 154578366 | rs6801607 | -0.781 | 0.86 | 0.597 | -0.909 | 0.002 | -0.041 | -0.041 | 0 | 0.19 | 0.21 | 0.14 | 0.02 | 0.21 | 0.13 |
| 3 | 154587654 | rs6440961 | -1.353 | 0.781 | 1.159 | -0.608 | 0.063 | -0.016 | -0.016 | 0.093 | 0.1 | 0.44 | 0.11 | 0.04 | 0.11 | 0.14 |
| 3 | 154672849 | rs16824435 | -1.633 | 0.731 | 1.507 | -0.648 | 0.08 | -0.009 | -0.009 | 0 | 0.1 | 0.46 | 0.1 | 0.03 | 0.1 | 0.1 |
| 3 | 154676122 | rs1608001 | -0.893 | 0.488 | 0.201 | -1.367 | 0.008 | 0.133 | 0.133 | 0.078 | 0.43 | 0.75 | 0.29 | 0.06 | 0.38 | 0.26 |
| 3 | 154681770 | rs1525865 | -0.342 | 0.342 | 0.258 | -1.346 | -0.004 | -0.07 | -0.07 | 0 | 0.35 | 0.56 | 0.21 | 0.04 | 0.27 | 0.19 |
| 3 | 154715596 | rs7635887 | -0.674 | 0.107 | -0.627 | -0.6 | -0.061 | -0.058 | -0.058 | 0 | 0.58 | 0.83 | 0.47 | 0.57 | 0.44 | 0.63 |
| 3 | 154716357 | rs2314612 | 0.531 | -0.742 | -0.889 | -0.426 | -0.053 | -0.053 | -0.053 | 2.064 | 0.43 | 0.06 | 0.53 | 0.52 | 0.6 | 0.36 |
| 3 | 154719349 | rs7623854 | 0.52 | -0.64 | -0.586 | -0.464 | 0.02 | -0.005 | -0.005 | 0 | 0.46 | 0.37 | 0.56 | 0.48 | 0.6 | 0.38 |
| 3 | 154737184 | rs16824513 | 0.094 | -0.709 | 0.174 | -0.613 | -0.022 | -0.029 | -0.029 | 0 | 0.08 | 0.14 | 0.03 | 0 | 0.03 | 0.08 |
| 3 | 154739536 | rs13326335 | -0.598 | -0.025 | -0.042 | -0.262 | 0.033 | 0.048 | 0.048 | 0 | 0.13 | 0.01 | 0.03 | 0.03 | 0.05 | 0.04 |
| 3 | 154800995 | rs9829757 | 0.76 | 0.803 | -0.052 | -0.191 | -0.007 | -0.082 | -0.082 | 3.215 | 0.78 | 0.55 | 0.86 | 0.87 | 0.79 | 0.87 |
| 3 | 154801365 | rs989692 | 0.657 | 1.416 | -0.155 | -0.624 | -0.001 | -0.003 | -0.003 | 0 | 0.61 | 0.44 | 0.55 | 0.8 | 0.53 | 0.74 |
| 3 | 154806647 | rs1370031 | -0.792 | 2.992 | -0.669 | -0.732 | -0.074 | 0.007 | 0.007 | 0 | 0.11 | 0.01 | 0.11 | 0.04 | 0.24 | 0.05 |
| 3 | 154809102 | rs1836915 | 0.182 | 2.677 | -0.978 | -0.928 | 0.001 | 0.013 | 0.013 | 1.752 | 0.15 | 0.08 | 0.08 | 0.11 | 0.15 | 0.11 |
| 3 | 154812696 | rs13065680 | 0.207 | 3.094 | -1.175 | -0.83 | 0 | 0.026 | 0.026 | 2.025 | 0.15 | 0.05 | 0.08 | 0.11 | 0.14 | 0.11 |
| 3 | 154813986 | rs17443031 | -1.217 | 3.03 | -0.864 | -0.861 | -0.029 | 0.03 | 0.03 | 0.887 | 0.27 | 0.16 | 0.19 | 0.15 | 0.38 | 0.15 |
| 3 | 154816911 | rs968628 | -1.49 | 2.165 | -1.223 | -1.462 | -0.021 | 0.069 | 0.069 | 0 | 0.59 | 0.79 | 0.38 | 0.33 | 0.56 | 0.42 |
| 3 | 154819781 | rs1436634 | 1.638 | 1.395 | -1.661 | -0.828 | 0.023 | -0.029 | -0.029 | 0 | 0.1 | 0.03 | 0.02 | 0.14 | 0.07 | 0.16 |
| 3 | 154821348 | rs9834487 | 0.982 | 0.859 | -1.833 | -1.293 | 0.003 | 0.042 | 0.042 | 0 | 0.14 | 0.05 | 0.08 | 0.08 | 0.12 | 0.19 |
| 3 | 154824652 | rs1436631 | 1.285 | 0.752 | -0.532 | -1.211 | 0.055 | 0.085 | 0.085 | 0 | 0.06 | 0.33 | 0.05 | 0.07 | 0.02 | 0.05 |
| 3 | 154825213 | rs9827313 | 0.447 | 0.788 | -0.896 | -1.271 | 0.004 | 0.011 | 0.011 | 0.993 | 0.18 | 0.11 | 0.1 | 0.11 | 0.16 | 0.12 |
| 3 | 154825385 | rs9827586 | -0.522 | 0.794 | -0.937 | -1.277 | -0.02 | 0.031 | 0.031 | 0.965 | 0.3 | 0.18 | 0.2 | 0.15 | 0.39 | 0.18 |
| 3 | 154827008 | rs3773895 | -0.032 | 0.137 | -1.236 | -0.972 | 0.018 | 0.006 | 0.006 | 0 | 0.16 | 0.13 | 0.08 | 0.12 | 0.16 | 0.17 |
| 3 | 154827787 | rs1025192 | 0.484 | -0.494 | -1.055 | -0.313 | 0.006 | -0.003 | -0.003 | 0 | 0.47 | 0.46 | 0.59 | 0.5 | 0.52 | 0.64 |
| 3 | 154829907 | rs9853221 | 0.641 | -0.657 | -1.267 | -0.401 | -0.009 | -0.019 | -0.019 | 0 | 0.18 | 0.24 | 0.12 | 0.12 | 0.16 | 0.11 |
| 3 | 154829999 | rs1436630 | -0.247 | -0.656 | -1.521 | -0.423 | -0.014 | 0.007 | 0.007 | 2.685 | 0.65 | 0.62 | 0.7 | 0.62 | 0.69 | 0.74 |
| 3 | 154830469 | rs10935998 | 1.324 | -0.196 | -2.248 | 0.08 | 0.037 | -0.039 | -0.039 | 0 | 0.11 | 0.03 | 0.03 | 0.16 | 0.08 | 0.16 |
| 3 | 154835722 | rs10513469 | 0.043 | 0.985 | -1.826 | 0.137 | -0.023 | -0.038 | -0.038 | 2.155 | 0.42 | 0.25 | 0.5 | 0.47 | 0.39 | 0.56 |
| 3 | 154847174 | rs10513470 | 0.458 | 1.036 | -1.727 | 0.233 | -0.031 | -0.01 | -0.01 | 0 | 0.05 | 0.01 | 0.07 | 0.03 | 0.14 | 0.08 |
| 3 | 154857146 | rs11719420 | 0.689 | 0.555 | -1.737 | 0.112 | -0.101 | -0.123 | -0.123 | 1.243 | 0.28 | 0.1 | 0.45 | 0.44 | 0.34 | 0.39 |
| 3 | 154858689 | rs2016846 | 0.208 | 1.193 | -1.39 | 0.422 | -0.016 | -0.038 | -0.038 | 0.165 | 0.42 | 0.25 | 0.49 | 0.47 | 0.38 | 0.56 |
| 3 | 154863088 | rs3821719 | -0.716 | 1.771 | -0.754 | 0.517 | 0.014 | -0.001 | -0.001 | 0.82 | 0.07 | 0.08 | 0.08 | 0.03 | 0.15 | 0.09 |
| 3 | 154866453 | rs1436633 | -0.312 | 1.162 | -0.334 | 0.39 | 0.019 | -0.051 | -0.051 | 0.015 | 0.32 | 0.46 | 0.25 | 0.21 | 0.36 | 0.32 |
| 3 | 154871679 | rs9829347 | -0.613 | 1.011 | -0.126 | 0.768 | -0.004 | 0.002 | 0.002 | 0.021 | 0.34 | 0.65 | 0.28 | 0.35 | 0.38 | 0.42 |
| 3 | 154882287 | rs3773876 | 0.122 | 1.154 | -0.005 | 0.666 | 0.008 | -0.075 | -0.075 | 0 | 0.27 | 0.33 | 0.18 | 0.08 | 0.29 | 0.17 |
| 3 | 154882950 | rs1385480 | -1.169 | 0.569 | -0.833 | 0.796 | -0.083 | -0.094 | -0.094 | 0 | 0.5 | 0.97 | 0.36 | 0.52 | 0.46 | 0.57 |
| 3 | 154894805 | rs3773874 | -0.377 | 1.22 | -0.098 | 1 | 0.021 | 0.075 | 0.075 | 0.007 | 0.05 | 0.13 | 0.07 | 0.15 | 0.07 | 0.16 |
| 3 | 154898407 | rs701109 | -0.555 | 1.099 | -0.358 | 0.945 | 0.005 | 0.002 | 0.002 | 0.018 | 0.34 | 0.62 | 0.28 | 0.38 | 0.38 | 0.41 |
| 3 | 154900690 | rs12765 | -0.164 | 0.849 | 0.321 | 0.971 | 0.001 | 0.001 | 0.001 | 0 | 0.68 | 0.71 | 0.7 | 0.73 | 0.69 | 0.61 |
| 3 | 154901205 | rs6665 | -0.12 | 0.87 | 0.446 | 1 | -0.006 | -0.009 | -0.009 | 0 | 0.67 | 0.65 | 0.7 | 0.72 | 0.69 | 0.61 |
| 3 | 154920664 | rs17386464 | -0.147 | 1.351 | 0.854 | 1.443 | 0.018 | 0.016 | 0.016 | 0 | 0.67 | 0.55 | 0.59 | 0.63 | 0.64 | 0.68 |
| 3 | 154925064 | rs8180028 | -0.515 | 1.713 | 0.673 | 1.549 | -0.071 | -0.046 | -0.046 | 0 | 0.73 | 0.98 | 0.65 | 0.75 | 0.73 | 0.74 |
| 3 | 154933041 | rs16824714 | 1.164 | 1.548 | 0.412 | 1.422 | -0.002 | 0.057 | 0.057 | 0 | 0.4 | 0.13 | 0.38 | 0.35 | 0.4 | 0.55 |
| 3 | 154934868 | rs16824723 | 0.214 | 1.53 | 0.683 | 1.332 | -0.001 | -0.03 | -0.03 | 0.321 | 0.23 | 0.27 | 0.16 | 0.12 | 0.24 | 0.1 |
| 3 | 154934897 | rs9869815 | -0.652 | 1.579 | 0.832 | 1.353 | -0.008 | 0.005 | 0.005 | 0 | 0.71 | 0.75 | 0.61 | 0.75 | 0.69 | 0.73 |
| 3 | 154949850 | rs7624578 | 0.553 | 1.559 | 0.838 | 1.367 | 0 | -0.001 | -0.001 | 0.675 | 0.48 | 0.47 | 0.44 | 0.63 | 0.45 | 0.63 |

| | | | | | | | | | | | | | | | | |
|---|---|---|---|---|---|---|---|---|---|---|---|---|---|---|---|---|
| 3 | 154977461 | rs987771 | 1.178 | 1.475 | 0.885 | 1.305 | -0.009 | 0.038 | 0.038 | 0 | 0.38 | 0.14 | 0.28 | 0.36 | 0.4 | 0.54 |
| 3 | 154980236 | rs1385482 | -0.274 | 1.534 | 0.885 | 1.325 | 0.091 | 0.17 | 0.17 | 1.411 | 0.06 | 0.34 | 0.15 | 0.12 | 0.07 | 0.06 |
| 3 | 154982968 | rs1385481 | 0.894 | 1.386 | 0.645 | 1.123 | -0.001 | 0.026 | 0.026 | 0 | 0.15 | 0.12 | 0.07 | 0.02 | 0.16 | 0.05 |
| 3 | 154984971 | rs1021213 | -0.451 | 1.382 | 0.401 | 1.279 | -0.003 | -0.024 | -0.024 | 0 | 0.67 | 0.75 | 0.58 | 0.6 | 0.69 | 0.7 |
| 3 | 154986954 | rs9855353 | 1.049 | 1.104 | 0.752 | 1.111 | 0.003 | 0.004 | 0.004 | 0 | 0.39 | 0.34 | 0.27 | 0.36 | 0.35 | 0.55 |
| 3 | 155003301 | rs9289952 | -0.601 | 1.392 | 0.067 | 1.113 | -0.032 | 0.015 | 0.015 | 0 | 0.71 | 0.82 | 0.6 | 0.77 | 0.65 | 0.73 |
| 3 | 155012313 | rs10513474 | 0.478 | 1.275 | -0.022 | 1.149 | 0.011 | -0.012 | -0.012 | 0 | 0.08 | 0.15 | 0.07 | 0.08 | 0.09 | 0.04 |
| 3 | 155015174 | rs1601383 | 1.644 | 1.234 | -0.279 | 1.162 | -0.042 | -0.022 | -0.022 | 0.065 | 0.34 | 0.02 | 0.23 | 0.31 | 0.33 | 0.29 |
| 3 | 155019933 | rs2200392 | 0.882 | 1.319 | 0.085 | 1.127 | 0.009 | 0.029 | 0.029 | 0 | 0.46 | 0.25 | 0.33 | 0.45 | 0.45 | 0.61 |
| 3 | 155021779 | rs3796271 | 0.674 | 1.265 | 0.089 | 1.052 | 0.009 | 0.008 | 0.008 | 0 | 0.47 | 0.28 | 0.44 | 0.59 | 0.47 | 0.63 |
| 3 | 155021827 | rs3796270 | 0.62 | 1.291 | 0.052 | 1.063 | 0.009 | 0.009 | 0.009 | 0 | 0.48 | 0.33 | 0.44 | 0.59 | 0.47 | 0.63 |
| 3 | 155031155 | rs6803323 | 0.492 | 0.949 | -0.213 | 0.629 | -0.006 | -0.007 | -0.007 | 0 | 0.56 | 0.64 | 0.4 | 0.53 | 0.51 | 0.67 |
| 3 | 155034166 | rs6440971 | 0.31 | 1.054 | -0.495 | 0.751 | -0.034 | -0.033 | -0.033 | 0 | 0.31 | 0.19 | 0.51 | 0.36 | 0.37 | 0.28 |
| 3 | 155037006 | rs7643746 | -0.524 | 0.29 | -0.306 | 0.784 | 0.002 | -0.001 | -0.001 | 0 | 0.52 | 0.57 | 0.64 | 0.48 | 0.6 | 0.36 |
| 3 | 155042640 | rs13314317 | 0.45 | 0.087 | -0.903 | 0.441 | 0.065 | -0.02 | -0.02 | 0 | 0.44 | 0.3 | 0.54 | 0.49 | 0.35 | 0.63 |
| 3 | 155045430 | rs6440972 | -0.261 | 0.404 | -0.28 | -0.05 | 0.011 | -0.043 | -0.043 | 0.016 | 0.15 | 0.36 | 0.07 | 0.1 | 0.13 | 0.08 |
| 3 | 155046798 | rs9824811 | 0.302 | -0.064 | -0.817 | -0.248 | -0.006 | -0.005 | -0.005 | 1.118 | 0.75 | 0.26 | 0.71 | 0.83 | 0.85 | 0.86 |
| 3 | 155048352 | rs6783837 | 1.041 | -0.268 | -1.702 | -0.196 | 0.048 | -0.069 | -0.069 | 0 | 0.36 | 0.02 | 0.38 | 0.63 | 0.28 | 0.58 |
| 3 | 155050565 | rs9845749 | 0.541 | 0.296 | -1.495 | -0.309 | 0.033 | 0.086 | 0.086 | 0 | 0.45 | 0.2 | 0.43 | 0.71 | 0.41 | 0.62 |
| 3 | 155055548 | rs13320391 | 0.435 | 0.518 | -1.212 | -0.213 | -0.009 | 0.006 | 0.006 | 0 | 0.11 | 0.07 | 0.06 | 0.08 | 0.14 | 0.07 |
| 3 | 155055610 | rs9844189 | 0.028 | 0.665 | -0.896 | -0.591 | -0.006 | -0.034 | -0.034 | 0 | 0.16 | 0.25 | 0.08 | 0.11 | 0.1 | 0.11 |
| 3 | 155061124 | rs9817455 | 0.028 | 0.665 | -0.896 | -0.591 | -0.006 | -0.034 | -0.034 | 0 | 0.16 | 0.25 | 0.08 | 0.11 | 0.1 | 0.11 |
| 3 | 155064239 | rs9833591 | 0.028 | 0.665 | -0.896 | -0.591 | -0.006 | -0.034 | -0.034 | 0 | 0.16 | 0.25 | 0.08 | 0.11 | 0.1 | 0.11 |
| 3 | 155070591 | rs6440975 | 1.263 | 0.879 | -0.15 | -0.779 | -0.001 | 0.024 | 0.024 | 0.028 | 0.58 | 0.34 | 0.64 | 0.72 | 0.63 | 0.74 |
| 3 | 155075068 | rs7619671 | -0.085 | 0.614 | -0.099 | -1.014 | 0.004 | -0.014 | -0.014 | 0 | 0.3 | 0.37 | 0.29 | 0.12 | 0.38 | 0.22 |
| 3 | 155077398 | rs13320892 | -0.381 | 0.672 | -0.217 | -0.899 | -0.003 | 0.008 | 0.008 | 0.775 | 0.26 | 0.23 | 0.14 | 0.19 | 0.25 | 0.18 |
| 3 | 155078062 | rs4455360 | -0.833 | 0.91 | -0.179 | -0.535 | -0.03 | 0.002 | 0.002 | 0 | 0.93 | 0.99 | 0.81 | 0.95 | 0.88 | 0.94 |
| 3 | 155081940 | rs4600855 | 1.574 | 1.142 | 0.126 | -0.279 | -0.006 | -0.001 | -0.001 | 0 | 0.46 | 0.56 | 0.45 | 0.52 | 0.41 | 0.51 |
| 3 | 155082806 | rs6769372 | -0.253 | 1.056 | -0.15 | -0.352 | -0.041 | 0.012 | 0.012 | 0 | 0.84 | 0.96 | 0.75 | 0.84 | 0.78 | 0.82 |
| 3 | 155087563 | rs6786301 | -0.421 | 1.501 | 0.306 | -0.249 | 0.025 | 0.026 | 0.026 | 0 | 0.24 | 0.42 | 0.29 | 0.27 | 0.25 | 0.22 |
| 3 | 155088059 | rs13092986 | -0.158 | 1.516 | 0.151 | -0.253 | -0.029 | -0.024 | -0.024 | 0 | 0.23 | 0.1 | 0.28 | 0.27 | 0.27 | 0.22 |
| 3 | 155088878 | rs6790153 | 2.043 | 1.451 | 0.001 | -0.333 | 0.005 | -0.063 | -0.063 | 0.264 | 0.42 | 0.13 | 0.37 | 0.5 | 0.43 | 0.49 |
| 3 | 155089985 | rs9849682 | -2.099 | 1.418 | -0.137 | 0.431 | 0.021 | 0.059 | 0.059 | 0 | 0.57 | 0.71 | 0.71 | 0.75 | 0.58 | 0.74 |
| 3 | 155095455 | rs12696030 | -0.652 | 1.375 | -0.153 | 0.029 | -0.001 | -0.001 | -0.001 | 0.107 | 0.28 | 0.27 | 0.28 | 0.29 | 0.28 | 0.3 |
| 3 | 155101831 | rs4234306 | -1.883 | 1.248 | -0.433 | -0.061 | -0.066 | -0.082 | -0.082 | 0 | 0.56 | 0.94 | 0.65 | 0.5 | 0.53 | 0.53 |
| 3 | 155110950 | rs6794214 | -1.667 | 1.322 | 0.306 | 0.529 | -0.02 | -0.002 | -0.002 | 0.788 | 0.39 | 0.83 | 0.44 | 0.41 | 0.34 | 0.39 |
| 3 | 155122855 | rs4680174 | -1.659 | 1.488 | 0.64 | 0.578 | -0.01 | -0.005 | -0.005 | 0 | 0.39 | 0.69 | 0.45 | 0.43 | 0.36 | 0.43 |
| 3 | 155128968 | rs6802130 | -0.601 | 1.407 | 1.063 | 0.531 | -0.001 | -0.017 | -0.017 | 0 | 0.07 | 0.29 | 0.03 | 0 | 0.01 | 0.04 |
| 3 | 155139253 | rs13083852 | -1.093 | 1.418 | 0.498 | 0.519 | -0.083 | -0.081 | -0.081 | 0 | 0.3 | 0.09 | 0.36 | 0.42 | 0.34 | 0.37 |
| 3 | 155142297 | rs13081656 | -1.123 | 1.42 | 0.533 | 0.519 | -0.073 | -0.072 | -0.072 | 0 | 0.3 | 0.09 | 0.37 | 0.42 | 0.33 | 0.37 |
| 3 | 155152505 | rs7615026 | -1.417 | 1.427 | 1.055 | 0.579 | 0 | 0.002 | 0.002 | 0 | 0.33 | 0.6 | 0.39 | 0.4 | 0.29 | 0.33 |
| 3 | 155171784 | rs12631218 | -2.055 | 1.275 | 1.358 | 0.959 | 0.004 | -0.036 | -0.036 | 0 | 0.27 | 0.84 | 0.27 | 0.4 | 0.25 | 0.33 |
| 3 | 155177333 | rs11709568 | -1.275 | 1.314 | 1.801 | 0.981 | 0.017 | 0.099 | 0.099 | 0 | 0.22 | 0.49 | 0.21 | 0.39 | 0.22 | 0.26 |
| 3 | 155181831 | rs9829532 | -1.194 | 1.314 | 1.917 | 0.979 | 0.009 | 0.068 | 0.068 | 0 | 0.21 | 0.4 | 0.19 | 0.39 | 0.22 | 0.26 |
| 3 | 155183770 | rs35259851 | -1.235 | 1.296 | 1.91 | 0.971 | 0.007 | 0.065 | 0.065 | 0 | 0.21 | 0.4 | 0.19 | 0.39 | 0.22 | 0.26 |
| 3 | 155184048 | rs7618495 | -1.328 | 1.294 | 1.887 | 0.97 | 0.014 | 0.09 | 0.09 | 0 | 0.22 | 0.48 | 0.21 | 0.39 | 0.22 | 0.26 |
| 3 | 155204939 | rs1125539 | 0.606 | 1.701 | 1.404 | 0.613 | -0.048 | -0.005 | -0.005 | 0 | 0.88 | 0.98 | 0.8 | 0.94 | 0.8 | 0.85 |
| 3 | 155234715 | rs6790399 | -0.316 | 1.65 | 1.463 | 0.622 | 0 | -0.017 | -0.017 | 3.07 | 0.08 | 0.12 | 0.06 | 0.01 | 0.11 | 0.07 |
| 3 | 155240638 | rs7637342 | 2.396 | 1.603 | 1.632 | 0.591 | 0.013 | -0.034 | -0.034 | 0.5 | 0.72 | 0.37 | 0.66 | 0.9 | 0.65 | 0.73 |
| 3 | 155241740 | rs9840234 | 1.895 | 1.555 | 1.308 | 0.602 | 0.026 | -0.048 | -0.048 | 0.393 | 0.78 | 0.69 | 0.78 | 0.93 | 0.7 | 0.79 |
| 3 | 155244835 | rs6770715 | -2.249 | 1.391 | 1.192 | 0.992 | -0.015 | 0.005 | 0.005 | 0 | 0.27 | 0.66 | 0.25 | 0.35 | 0.21 | 0.35 |
| 3 | 155267180 | rs16825084 | -0.022 | 1.472 | 0.789 | 0.852 | -0.02 | 0.043 | 0.043 | 0 | 0.08 | 0 | 0.04 | 0.01 | 0.08 | 0.07 |
| 3 | 155267440 | rs3851357 | -0.788 | 1.551 | 0.838 | 0.89 | -0.042 | 0.012 | 0.012 | 0 | 0.11 | 0.02 | 0.13 | 0.04 | 0.18 | 0.14 |
| 3 | 155306781 | rs13085233 | 0.456 | 0.939 | 0.706 | 0.818 | 0.007 | 0.001 | 0.001 | 0 | 0.06 | 0.01 | 0.07 | 0.03 | 0.05 | 0.02 |
| 3 | 155347040 | rs9823723 | -0.985 | 0.639 | 1.101 | 0.846 | -0.04 | 0.036 | 0.036 | 0.529 | 0.17 | 0.41 | 0.14 | 0.25 | 0.1 | 0.23 |
| 3 | 155350269 | rs932207 | -0.985 | 0.639 | 1.101 | 0.846 | -0.04 | 0.036 | 0.036 | 0.529 | 0.17 | 0.41 | 0.14 | 0.25 | 0.1 | 0.23 |
| 3 | 155355295 | rs1519534 | -1.756 | 0.856 | 0.945 | 1.375 | 0.02 | 0.087 | 0.087 | 0 | 0.21 | 0.37 | 0.37 | 0.44 | 0.24 | 0.44 |
| 3 | 155367330 | rs16825206 | -2.244 | 0.592 | 0.831 | 1.217 | 0.022 | 0.015 | 0.015 | 0 | 0.37 | 0.68 | 0.31 | 0.33 | 0.29 | 0.4 |
| 3 | 155370090 | rs6806485 | -0.071 | 0.615 | 0.942 | 1.238 | 0 | 0.006 | 0.006 | 0 | 0.08 | 0.04 | 0.09 | 0.03 | 0.08 | 0.09 |
| 3 | 155371659 | rs9838305 | -2.003 | 0.574 | 0.922 | 1.237 | -0.003 | -0.001 | -0.001 | 0 | 0.27 | 0.67 | 0.25 | 0.31 | 0.2 | 0.36 |
| 3 | 155382134 | rs7610374 | -1.112 | 0.63 | 0.982 | 1.186 | -0.001 | -0.016 | -0.016 | 0 | 0.16 | 0.13 | 0.14 | 0.03 | 0.16 | 0.12 |
| 3 | 155383148 | rs13089059 | -0.506 | 0.856 | 0.806 | 1.448 | -0.033 | -0.032 | -0.032 | 0.037 | 0.1 | 0.07 | 0.29 | 0.44 | 0.15 | 0.35 |
| 3 | 155386745 | rs16825229 | -0.119 | 0.726 | 1.377 | 1.448 | -0.009 | 0.041 | 0.041 | 0 | 0.1 | 0.17 | 0.11 | 0.2 | 0.08 | 0.23 |
| 3 | 155408234 | rs16825274 | -1.893 | 0.696 | 1.72 | 1.391 | -0.019 | -0.061 | -0.061 | 1.617 | 0.24 | 0.38 | 0.17 | 0.1 | 0.19 | 0.16 |
| 3 | 155412668 | rs393081 | -2.49 | 0.793 | 1.824 | 1.524 | 0.002 | 0.025 | 0.025 | 0 | 0.35 | 0.57 | 0.47 | 0.51 | 0.33 | 0.5 |
| 3 | 155420079 | rs183088 | -2.886 | 0.768 | 1.645 | 1.451 | 0.013 | 0.105 | 0.105 | 0 | 0.47 | 0.78 | 0.6 | 0.74 | 0.43 | 0.74 |
| 3 | 155423302 | rs1835212 | -1.988 | 0.743 | 1.507 | 1.471 | -0.001 | -0.036 | -0.036 | 0 | 0.26 | 0.26 | 0.39 | 0.5 | 0.25 | 0.46 |
| 3 | 155425831 | rs12053896 | 2.298 | 0.805 | 1.561 | 1.355 | 0 | 0 | 0 | 0 | 0.62 | 0.6 | 0.5 | 0.31 | 0.65 | 0.32 |
| 3 | 155425866 | rs193313 | -2.587 | 0.679 | 1.728 | 1.391 | 0.007 | 0.025 | 0.025 | 0 | 0.45 | 0.63 | 0.56 | 0.69 | 0.42 | 0.71 |

| | | | | | | | | | | | | | | | | |
|---|---|---|---|---|---|---|---|---|---|---|---|---|---|---|---|---|
| 3 | 155426459 | rs117442 | -2.153 | 0.942 | 1.917 | 1.769 | 0.001 | 0.066 | 0.066 | 0 | 0.31 | 0.45 | 0.46 | 0.57 | 0.31 | 0.59 |
| 3 | 155433793 | rs4680214 | -0.693 | 0.832 | 1.903 | 1.566 | -0.009 | 0.038 | 0.038 | 0 | 0.1 | 0.22 | 0.1 | 0.15 | 0.06 | 0.18 |
| 3 | 155438022 | rs10428198 | -1.536 | 0.885 | 1.834 | 1.509 | -0.002 | -0.031 | -0.031 | 0 | 0.21 | 0.24 | 0.17 | 0.15 | 0.2 | 0.27 |
| 3 | 155470585 | rs6765861 | -0.179 | 0.989 | 1.464 | 1.437 | -0.034 | 0.057 | 0.057 | 0 | 0.11 | 0.02 | 0.07 | 0 | 0.14 | 0.09 |
| 3 | 155483319 | rs358734 | -1.609 | 0.96 | 1.765 | 1.833 | 0 | 0.121 | 0.121 | 0 | 0.14 | 0.22 | 0.36 | 0.51 | 0.14 | 0.42 |
| 3 | 155485302 | rs358733 | -0.532 | 1.39 | 1.396 | 1.6 | 0.002 | -0.005 | -0.005 | 0 | 0.83 | 0.83 | 0.85 | 0.91 | 0.77 | 0.84 |
| 3 | 155487290 | rs358732 | -0.322 | 1.376 | 1.338 | 1.722 | -0.011 | 0.038 | 0.038 | 0.006 | 0.05 | 0.05 | 0.25 | 0.39 | 0.08 | 0.25 |
| 3 | 155490066 | rs17349490 | -0.634 | 1.376 | 1.36 | 1.161 | -0.005 | -0.004 | -0.004 | 0 | 0.15 | 0.12 | 0.09 | 0.03 | 0.18 | 0.14 |
| 3 | 155492177 | rs358730 | -0.5 | 1.333 | 1.166 | 1.36 | -0.03 | 0.033 | 0.033 | 0 | 0.83 | 0.92 | 0.86 | 0.92 | 0.78 | 0.84 |
| 3 | 155505597 | rs11712034 | -0.372 | 1.333 | 1.205 | 1.36 | 0.001 | -0.005 | -0.005 | 0 | 0.83 | 0.84 | 0.85 | 0.92 | 0.78 | 0.84 |
| 3 | 155518043 | rs416063 | -0.052 | 1.136 | 1.571 | 1.581 | 0.02 | -0.001 | -0.001 | 0 | 0.81 | 0.68 | 0.78 | 0.8 | 0.79 | 0.71 |
| 3 | 155520717 | rs404570 | -1.133 | 1.192 | 1.628 | 1.793 | -0.012 | -0.029 | -0.029 | 0.008 | 0.19 | 0.18 | 0.34 | 0.38 | 0.23 | 0.32 |
| 3 | 155520927 | rs389893 | -0.366 | 1.171 | 1.712 | 1.09 | -0.007 | 0.004 | 0.004 | 0 | 0.82 | 0.87 | 0.81 | 0.84 | 0.79 | 0.71 |
| 3 | 155525798 | rs358715 | 1.595 | 1.101 | 1.741 | 1.018 | -0.04 | 0.074 | 0.074 | 0.369 | 0.62 | 0.18 | 0.67 | 0.74 | 0.61 | 0.53 |
| 3 | 155529965 | rs168587 | -0.314 | 1.06 | 1.781 | 1.044 | 0.001 | -0.002 | -0.002 | 0 | 0.82 | 0.81 | 0.8 | 0.83 | 0.79 | 0.71 |
| 3 | 155539266 | rs11921267 | -1.221 | 0.798 | 1.435 | 1.149 | -0.026 | 0.023 | 0.023 | 0.266 | 0.26 | 0.06 | 0.2 | 0.2 | 0.26 | 0.39 |
| 3 | 155542475 | rs358702 | -2.119 | 0.759 | 1.489 | 1.229 | 0.007 | 0.009 | 0.009 | 0.693 | 0.33 | 0.82 | 0.44 | 0.32 | 0.29 | 0.45 |
| 3 | 155548882 | rs1025341 | -1.98 | 0.725 | 1.325 | 1.27 | 0.031 | 0.094 | 0.094 | 3.258 | 0.06 | 0.71 | 0.23 | 0.11 | 0.02 | 0.06 |
| 3 | 155551990 | rs6809229 | 1.722 | 0.737 | 0.436 | 1.224 | -0.048 | -0.034 | -0.034 | 0 | 0.49 | 0.07 | 0.37 | 0.51 | 0.5 | 0.28 |
| 3 | 155572129 | rs1131925 | 1.166 | 0.882 | 1.536 | 1.385 | 0.008 | 0.021 | 0.021 | 0 | 0.63 | 0.53 | 0.47 | 0.46 | 0.6 | 0.36 |
| 3 | 155588148 | rs11706359 | 1.703 | 0.804 | 1.479 | 1.238 | -0.029 | 0.037 | 0.037 | 0 | 0.54 | 0.2 | 0.46 | 0.58 | 0.57 | 0.42 |
| 3 | 155622422 | rs9882341 | -0.234 | 0.804 | 2.121 | 1.228 | 0.007 | -0.074 | -0.074 | 2.547 | 0.12 | 0.24 | 0.07 | 0 | 0.1 | 0.12 |
| 3 | 155624605 | rs6441028 | 1.26 | 0.858 | 2.094 | 1.23 | 0.002 | 0.006 | 0.006 | 0 | 0.63 | 0.64 | 0.48 | 0.43 | 0.6 | 0.35 |
| 3 | 155628877 | rs10936021 | -0.501 | 0.846 | 2.212 | 1.265 | 0.037 | 0.035 | 0.035 | 0 | 0.21 | 0.25 | 0.26 | 0.42 | 0.3 | 0.47 |
| 3 | 155630845 | rs2067877 | 0.375 | 0.847 | 2.214 | 1.26 | 0.044 | 0.017 | 0.017 | 0 | 0.17 | 0.22 | 0.19 | 0.26 | 0.23 | 0.29 |
| 3 | 155654017 | rs10804774 | 1.625 | 0.847 | 2.195 | 1.28 | 0.002 | 0.006 | 0.006 | 0 | 0.63 | 0.64 | 0.48 | 0.42 | 0.6 | 0.36 |
| 3 | 155658252 | rs4679758 | 0.458 | 0.755 | 2.338 | 1.29 | 0.004 | 0.001 | 0.001 | 0 | 0.16 | 0.15 | 0.18 | 0.26 | 0.23 | 0.29 |
| 3 | 155682269 | rs701112 | 2.154 | 0.518 | 2.304 | 1.139 | -0.021 | -0.012 | -0.012 | 1.614 | 0.64 | 0.51 | 0.58 | 0.68 | 0.73 | 0.49 |
| 3 | 155682770 | rs819834 | 1.255 | 0.802 | 2.544 | 1.285 | 0.005 | -0.007 | -0.007 | 0.516 | 0.79 | 0.79 | 0.66 | 0.69 | 0.84 | 0.65 |
| 3 | 155685093 | rs1516500 | 1.475 | 0.802 | 2.539 | 1.292 | -0.024 | 0.037 | 0.037 | 0 | 0.79 | 0.67 | 0.64 | 0.69 | 0.83 | 0.65 |
| 3 | 155687546 | rs819855 | 0.28 | 0.795 | 2.377 | 1.208 | 0.025 | -0.009 | -0.009 | 0 | 0.84 | 0.9 | 0.73 | 0.84 | 0.9 | 0.83 |
| 3 | 155700066 | rs1711084 | -0.999 | 0.753 | 2.551 | 1.135 | 0.035 | -0.03 | -0.03 | 0 | 0.29 | 0.47 | 0.26 | 0.27 | 0.33 | 0.4 |
| 3 | 155704794 | rs7629523 | 2.554 | 0.817 | 2.259 | 1.236 | -0.083 | -0.08 | -0.08 | 0.233 | 0.51 | 0.08 | 0.39 | 0.41 | 0.53 | 0.25 |
| 3 | 155706888 | rs819851 | 0.113 | 0.782 | 2.355 | 1.254 | -0.015 | 0.025 | 0.025 | 0 | 0.17 | 0.09 | 0.11 | 0.07 | 0.21 | 0.16 |
| 3 | 155707094 | rs9808909 | -2.176 | 0.825 | 2.259 | 1.315 | 0.013 | 0.139 | 0.139 | 0 | 0.08 | 0.25 | 0.27 | 0.28 | 0.08 | 0.19 |
| 3 | 155723064 | rs819867 | 0.915 | 0.72 | 2.06 | 1.242 | -0.011 | 0.01 | 0.01 | 0.911 | 0.87 | 0.81 | 0.72 | 0.84 | 0.89 | 0.79 |
| 3 | 155725030 | rs2200707 | 2.55 | 0.733 | 1.61 | 1.273 | -0.07 | -0.046 | -0.046 | 0.37 | 0.5 | 0.09 | 0.4 | 0.4 | 0.52 | 0.23 |
| 3 | 155726800 | rs13067477 | 2.55 | 0.729 | 1.594 | 1.273 | -0.072 | -0.052 | -0.052 | 0 | 0.5 | 0.08 | 0.39 | 0.4 | 0.52 | 0.23 |
| 3 | 155730790 | rs1093955 | -1.498 | 0.834 | 1.816 | 1.174 | 0.019 | 0.001 | 0.001 | 0.421 | 0.34 | 0.53 | 0.3 | 0.44 | 0.36 | 0.54 |
| 3 | 155734056 | rs1685748 | -0.636 | 0.901 | 1.763 | 1.346 | 0.012 | 0.03 | 0.03 | 0 | 0.21 | 0.41 | 0.22 | 0.35 | 0.22 | 0.25 |
| 3 | 155736404 | rs819849 | 0.907 | 0.706 | 1.476 | 0.969 | -0.002 | 0.006 | 0.006 | 0.769 | 0.86 | 0.9 | 0.86 | 0.91 | 0.84 | 0.77 |
| 3 | 155738576 | rs12330401 | -0.645 | 0.261 | 1.982 | 1.095 | -0.012 | 0.007 | 0.007 | 0 | 0.11 | 0.24 | 0.06 | 0.13 | 0.08 | 0.29 |
| 3 | 155743407 | rs10936023 | 1.325 | 0.199 | 1.857 | 0.823 | 0.006 | -0.002 | -0.002 | 0 | 0.63 | 0.52 | 0.58 | 0.64 | 0.6 | 0.62 |
| 3 | 155748694 | rs4680230 | 2.339 | 0.153 | 1.62 | 0.764 | -0.085 | -0.073 | -0.073 | 0.351 | 0.49 | 0.07 | 0.49 | 0.5 | 0.5 | 0.29 |
| 3 | 155751484 | rs6781558 | -0.328 | 0.075 | 1.708 | 0.469 | -0.008 | -0.059 | -0.059 | 0 | 0.1 | 0.21 | 0.04 | 0 | 0.08 | 0.08 |
| 3 | 155755327 | rs9848466 | -0.9 | 0.139 | 1.767 | 0.497 | -0.003 | 0.003 | 0.003 | 0 | 0.12 | 0.3 | 0.08 | 0.13 | 0.1 | 0.31 |
| 3 | 155769380 | rs901862 | 1.469 | 0.1 | 1.767 | 0.522 | 0 | 0.001 | 0.001 | 0 | 0.62 | 0.41 | 0.41 | 0.54 | 0.6 | 0.62 |
| 3 | 155811323 | rs2013676 | -0.907 | -0.301 | 1.984 | -0.615 | 0.009 | -0.072 | -0.072 | 0 | 0.12 | 0.27 | 0.05 | 0 | 0.1 | 0.12 |
| 3 | 155822097 | rs12493813 | 1.25 | -0.328 | 2.301 | -0.7 | 0.001 | -0.001 | -0.001 | 0 | 0.73 | 0.72 | 0.79 | 0.76 | 0.67 | 0.59 |
| 3 | 155830631 | rs9873706 | 2.174 | -0.335 | 2.073 | -0.707 | -0.046 | 0.054 | 0.054 | 0 | 0.59 | 0.11 | 0.71 | 0.71 | 0.57 | 0.48 |
| 3 | 155836231 | rs13093154 | -1.321 | -0.313 | 2.227 | -0.703 | -0.009 | 0.007 | 0.007 | 0 | 0.22 | 0.26 | 0.11 | 0.28 | 0.18 | 0.39 |
| 3 | 155838900 | rs1488425 | 2.196 | -0.27 | 1.916 | -0.702 | -0.014 | 0.06 | 0.06 | 0 | 0.59 | 0.24 | 0.7 | 0.72 | 0.57 | 0.45 |
| 3 | 155843745 | rs13073556 | 2.162 | -0.234 | 1.874 | -0.673 | -0.019 | 0.054 | 0.054 | 0 | 0.59 | 0.2 | 0.7 | 0.71 | 0.57 | 0.45 |
| 3 | 155849212 | rs9862554 | -0.028 | -0.119 | 1.997 | -0.655 | 0.045 | -0.066 | -0.066 | 0 | 0.87 | 0.78 | 0.85 | 0.99 | 0.78 | 0.91 |
| 3 | 155862593 | rs4359774 | 2.152 | -0.264 | 1.885 | -0.653 | -0.011 | 0.044 | 0.044 | 0 | 0.59 | 0.2 | 0.7 | 0.69 | 0.56 | 0.37 |
| 3 | 155873269 | rs4680238 | 0.498 | -0.303 | 2.023 | -1.161 | 0.012 | -0.014 | -0.014 | 0.458 | 0.84 | 0.78 | 0.83 | 0.91 | 0.74 | 0.6 |
| 3 | 155894892 | rs4680239 | 1.82 | -0.189 | 1.935 | -1.246 | 0.057 | -0.106 | -0.106 | 0.083 | 0.72 | 0.51 | 0.76 | 0.91 | 0.6 | 0.45 |
| 3 | 155899665 | rs4577423 | 2.14 | -0.405 | 1.434 | -1.341 | 0.006 | -0.049 | -0.049 | 0.55 | 0.63 | 0.46 | 0.71 | 0.88 | 0.57 | 0.41 |
| 3 | 155901863 | rs13085540 | -1.25 | -0.379 | 1.429 | -1.333 | -0.002 | -0.046 | -0.046 | 0 | 0.19 | 0.17 | 0.09 | 0 | 0.15 | 0.17 |
| 3 | 155902491 | rs13086539 | -0.745 | -0.371 | 1.326 | -1.329 | 0.003 | 0.004 | 0.004 | 0 | 0.17 | 0.08 | 0.07 | 0 | 0.14 | 0.14 |
| 3 | 155907740 | rs1353812 | 1.122 | -0.371 | 1.345 | -1.323 | 0.05 | -0.055 | -0.055 | 0 | 0.79 | 0.54 | 0.78 | 0.88 | 0.7 | 0.54 |
| 3 | 155929451 | rs4505650 | 1.746 | -0.263 | 1.05 | -1.208 | 0 | -0.022 | -0.022 | 0 | 0.61 | 0.44 | 0.71 | 0.88 | 0.56 | 0.41 |
| 3 | 155955485 | rs9864967 | 1.732 | -0.266 | 0.99 | -1.257 | 0.004 | -0.05 | -0.05 | 0.09 | 0.61 | 0.49 | 0.72 | 0.89 | 0.56 | 0.41 |
| 3 | 155965750 | rs9870819 | 1.195 | -0.213 | 0.394 | -1.223 | 0.005 | 0.011 | 0.011 | 0 | 0.58 | 0.42 | 0.69 | 0.77 | 0.61 | 0.52 |
| 3 | 155984606 | rs11706532 | -0.389 | 0.086 | -0.275 | -1.308 | 0.016 | -0.061 | -0.061 | 4.08 | 0.13 | 0.24 | 0.06 | 0.02 | 0.15 | 0.17 |
| 3 | 155991348 | rs1160221 | 0.165 | 0.259 | -0.643 | -1.21 | -0.032 | 0.102 | 0.102 | 0 | 0.75 | 0.92 | 0.79 | 0.98 | 0.7 | 0.88 |
| 3 | 156004954 | rs13082914 | -1.091 | -0.084 | 0.201 | -0.45 | -0.012 | -0.066 | -0.066 | 0 | 0.26 | 0.46 | 0.13 | 0.17 | 0.22 | 0.46 |
| 3 | 156012679 | rs882089 | -0.842 | -0.97 | -0.848 | -0.688 | 0.081 | 0.098 | 0.098 | 0.493 | 0.4 | 0.72 | 0.25 | 0.19 | 0.23 | 0.32 |
| 3 | 156019961 | rs9844584 | -0.969 | -1.183 | -0.087 | -0.743 | 0.03 | 0.016 | 0.016 | 1.153 | 0.07 | 0.53 | 0.05 | 0.03 | 0.02 | 0.12 |

| | | | | | | | | | | | | | | | | |
|---|---|---|---|---|---|---|---|---|---|---|---|---|---|---|---|---|
| 3 | 156032178 | rs6770663 | -1.55 | -1.953 | -0.118 | -1.057 | -0.006 | -0.002 | -0.002 | 0 | 0.21 | 0.71 | 0.16 | 0.16 | 0.11 | 0.22 |
| 3 | 156034647 | rs1874952 | -1.703 | -2.099 | -0.174 | -1.071 | 0.115 | 0.12 | 0.12 | 0.731 | 0.42 | 0.78 | 0.27 | 0.24 | 0.24 | 0.42 |
| 3 | 156034741 | rs498033 | -2.185 | -1.504 | -0.926 | -1.3 | -0.158 | -0.016 | -0.016 | 0 | 0.61 | 0.96 | 0.42 | 0.34 | 0.51 | 0.64 |
| 3 | 156037384 | rs4680259 | -1.016 | -1.619 | -0.48 | -1.133 | 0.077 | 0.086 | 0.086 | 1.265 | 0.35 | 0.11 | 0.22 | 0.22 | 0.23 | 0.3 |
| 3 | 156045286 | rs3821415 | -0.891 | -1.619 | -0.48 | -1.094 | 0.081 | 0.079 | 0.079 | 1.074 | 0.35 | 0.12 | 0.22 | 0.22 | 0.23 | 0.33 |
| 3 | 156050641 | rs3821416 | -1.522 | -1.617 | -0.351 | -1.088 | 0.103 | 0.108 | 0.108 | 0.51 | 0.42 | 0.75 | 0.27 | 0.24 | 0.25 | 0.44 |
| 3 | 156061497 | rs3772238 | -0.19 | -1.777 | -0.314 | -1.15 | 0.058 | 0.072 | 0.072 | 6.874 | 0.23 | 0.07 | 0.08 | 0.07 | 0.12 | 0.21 |
| 3 | 156067894 | rs6441061 | -0.97 | -1.771 | 0.063 | -1.176 | -0.018 | -0.035 | -0.035 | 0 | 0.3 | 0.62 | 0.13 | 0.09 | 0.15 | 0.33 |
| 3 | 156068886 | rs3772242 | -0.118 | -1.683 | -0.288 | -1.112 | 0.066 | 0.098 | 0.098 | 3.968 | 0.23 | 0.04 | 0.07 | 0.07 | 0.12 | 0.21 |
| 3 | 156078627 | rs6786247 | -0.334 | -1.581 | 0.095 | -0.885 | -0.01 | 0.004 | 0.004 | 0 | 0.13 | 0.11 | 0.09 | 0.2 | 0.08 | 0.27 |
| 3 | 156079516 | rs9859443 | -1.805 | -1.667 | 0.02 | -1.111 | 0.104 | 0.129 | 0.129 | 0.165 | 0.44 | 0.77 | 0.25 | 0.25 | 0.28 | 0.47 |
| 3 | 156098920 | rs9850840 | -0.7 | -1.437 | 0.665 | -1.11 | -0.042 | -0.106 | -0.106 | 0.014 | 0.26 | 0.55 | 0.14 | 0.05 | 0.18 | 0.15 |
| 3 | 156105173 | rs11926141 | -1.014 | -1.448 | 0.366 | -1.114 | -0.005 | -0.035 | -0.035 | 0 | 0.3 | 0.66 | 0.16 | 0.05 | 0.2 | 0.18 |
| 3 | 156112914 | rs16826083 | -1.175 | -1.44 | 0.01 | -1.088 | 0.047 | 0.074 | 0.074 | 2.031 | 0.32 | 0.76 | 0.17 | 0.09 | 0.2 | 0.3 |
| 3 | 156129973 | rs13324876 | -1.404 | -1.307 | 0.11 | -1.043 | 0.012 | 0.016 | 0.016 | 1.349 | 0.27 | 0.72 | 0.15 | 0.09 | 0.18 | 0.3 |
| 3 | 156148662 | rs16826108 | -0.786 | -1.191 | 0.563 | -0.987 | -0.026 | -0.101 | -0.101 | 0 | 0.21 | 0.38 | 0.12 | 0.05 | 0.17 | 0.14 |
| 3 | 156150082 | rs997876 | -1.492 | -0.959 | -0.144 | -1.181 | 0.009 | 0.053 | 0.053 | 0.031 | 0.34 | 0.75 | 0.24 | 0.13 | 0.3 | 0.38 |
| 3 | 156150661 | rs2122418 | 0.607 | -0.411 | -0.862 | -0.687 | -0.081 | -0.005 | -0.005 | 0 | 0.65 | 0.96 | 0.77 | 0.69 | 0.63 | 0.72 |
| 3 | 156156285 | rs4680266 | 0.855 | -0.037 | -0.146 | -0.466 | -0.016 | 0.033 | 0.033 | 0 | 0.65 | 0.76 | 0.76 | 0.77 | 0.62 | 0.67 |
| 3 | 156157738 | rs1357432 | 1.438 | -0.021 | 0.559 | 0.504 | 0 | 0.001 | 0.001 | 0 | 0.55 | 0.43 | 0.62 | 0.51 | 0.56 | 0.42 |
| 3 | 156158932 | rs6777084 | -1.142 | -0.255 | 1.267 | 1.097 | -0.002 | 0.017 | 0.017 | 0 | 0.06 | 0.16 | 0.11 | 0.07 | 0.04 | 0.01 |
| 3 | 156162373 | rs11917301 | -2.331 | -0.064 | 1.498 | 1.227 | 0.015 | 0.116 | 0.116 | 0.12 | 0.13 | 0.52 | 0.17 | 0.2 | 0.1 | 0.14 |
| 3 | 156162567 | rs6772280 | -2.342 | -0.228 | 1.503 | 1.026 | -0.004 | 0.047 | 0.047 | 0.037 | 0.13 | 0.55 | 0.09 | 0.14 | 0.09 | 0.14 |
| 3 | 156170443 | rs16826199 | -1.937 | -0.248 | 1.281 | 1.135 | 0.006 | 0.12 | 0.12 | 0 | 0.1 | 0.4 | 0.08 | 0.19 | 0.07 | 0.15 |
| 3 | 156175584 | rs2293198 | -1.436 | -0.432 | 1.1 | 1.494 | 0.001 | 0.001 | 0.001 | 0 | 0.5 | 0.54 | 0.36 | 0.54 | 0.48 | 0.57 |
| 3 | 156176734 | rs3772265 | -0.476 | -0.581 | 0.768 | 1.413 | -0.059 | -0.012 | -0.012 | 0.02 | 0.27 | 0.04 | 0.16 | 0.22 | 0.28 | 0.39 |
| 3 | 156176979 | rs3772266 | -0.886 | -0.619 | 0.942 | 1.387 | -0.028 | -0.068 | -0.068 | 0 | 0.3 | 0.12 | 0.19 | 0.37 | 0.33 | 0.42 |
| 3 | 156178206 | rs2280561 | -2.25 | -0.639 | 1.411 | 1.382 | -0.018 | -0.013 | -0.013 | 0 | 0.54 | 0.78 | 0.37 | 0.44 | 0.5 | 0.63 |
| 3 | 156182208 | rs13315136 | -1.172 | -0.633 | 1.921 | 1.359 | 0.047 | 0.022 | 0.022 | 0 | 0.09 | 0.44 | 0.06 | 0.06 | 0.07 | 0.08 |
| 3 | 156182956 | rs7648656 | -1.038 | -0.608 | 1.524 | 1.308 | 0.06 | -0.012 | -0.012 | 0 | 0.09 | 0.44 | 0.06 | 0.01 | 0.07 | 0.02 |
| 3 | 156185892 | rs1546750 | -1.563 | -0.472 | 0.985 | 1.412 | 0.015 | -0.004 | -0.004 | 0.304 | 0.43 | 0.31 | 0.29 | 0.39 | 0.41 | 0.51 |
| 3 | 156204177 | rs7644005 | -1.941 | -0.572 | 0.924 | 1.895 | 0.019 | -0.014 | -0.014 | 1.808 | 0.31 | 0.78 | 0.24 | 0.33 | 0.24 | 0.33 |
| 3 | 156205797 | rs11918294 | -1.178 | -0.286 | 1.433 | 1.812 | 0.028 | 0.093 | 0.093 | 0 | 0.08 | 0.27 | 0.04 | 0.15 | 0.09 | 0.1 |
| 3 | 156218007 | rs16826325 | -1.426 | -0.394 | 1.604 | 1.692 | -0.001 | -0.001 | -0.001 | 0 | 0.24 | 0.29 | 0.09 | 0.21 | 0.19 | 0.19 |
| 3 | 156220466 | rs12490937 | -1.666 | -0.301 | 1.327 | 1.893 | -0.006 | -0.039 | -0.039 | 0 | 0.24 | 0.14 | 0.23 | 0.36 | 0.28 | 0.38 |
| 3 | 156225491 | rs4257505 | 0.628 | -0.265 | 1.129 | 1.952 | -0.016 | -0.021 | -0.021 | 0 | 0.05 | 0 | 0.08 | 0.03 | 0.08 | 0.06 |
| 3 | 156227002 | rs12485831 | -2.236 | -0.338 | 1.429 | 2.016 | -0.005 | 0.024 | 0.024 | 0.049 | 0.27 | 0.64 | 0.25 | 0.46 | 0.28 | 0.48 |
| 3 | 156227643 | rs6772323 | -2.071 | -0.27 | 1.663 | 2.057 | 0.006 | -0.002 | -0.002 | 0 | 0.42 | 0.56 | 0.29 | 0.43 | 0.37 | 0.47 |
| 3 | 156231801 | rs9942113 | -2.041 | -0.24 | 1.704 | 2.045 | -0.004 | 0 | 0 | 0 | 0.42 | 0.55 | 0.28 | 0.56 | 0.37 | 0.47 |
| 3 | 156234268 | rs4295133 | -2.493 | -0.214 | 1.708 | 2.079 | 0.007 | 0.001 | 0.001 | 0 | 0.49 | 0.63 | 0.33 | 0.51 | 0.48 | 0.56 |
| 3 | 156234773 | rs17239426 | -0.661 | -0.291 | 1.444 | 1.422 | -0.001 | -0.005 | -0.005 | 0 | 0.08 | 0.09 | 0.05 | 0.01 | 0.12 | 0.09 |
| 3 | 156237352 | rs13092077 | -1.832 | -0.342 | 1.74 | 1.466 | -0.003 | -0.008 | -0.008 | 0 | 0.36 | 0.47 | 0.23 | 0.24 | 0.34 | 0.31 |
| 3 | 156238535 | rs4254605 | -1.218 | -0.326 | 1.595 | 1.396 | -0.014 | 0.002 | 0.002 | 0 | 0.27 | 0.38 | 0.17 | 0.23 | 0.22 | 0.16 |
| 3 | 156239255 | rs9848416 | 1.52 | 0.011 | 1.461 | 0.661 | -0.005 | -0.011 | -0.011 | 1.058 | 0.65 | 0.66 | 0.72 | 0.66 | 0.64 | 0.5 |
| 3 | 156239308 | rs9830768 | 1.175 | 0.206 | 1.206 | 0.894 | -0.001 | 0.016 | 0.016 | 0 | 0.92 | 0.93 | 0.92 | 0.72 | 0.88 | 0.78 |
| 3 | 156241306 | rs6441075 | -1.228 | 0.317 | 1.147 | 0.428 | -0.003 | -0.041 | -0.041 | 0 | 0.42 | 0.44 | 0.29 | 0.06 | 0.39 | 0.29 |
| 3 | 156241965 | rs2280299 | 2.129 | 0.094 | 1.028 | 0.505 | -0.012 | -0.004 | -0.004 | 0.006 | 0.42 | 0.33 | 0.61 | 0.58 | 0.46 | 0.36 |
| 3 | 156245781 | rs4680277 | 0.99 | 0.103 | 0.726 | 0.647 | 0.002 | 0.035 | 0.035 | 0 | 0.9 | 0.88 | 0.91 | 0.65 | 0.86 | 0.73 |
| 3 | 156247836 | rs875582 | 1.209 | 0.275 | 0.932 | 0.358 | -0.001 | -0.023 | -0.023 | 0.307 | 0.56 | 0.52 | 0.68 | 0.81 | 0.57 | 0.57 |
| 3 | 156248127 | rs875581 | 1.005 | 0.288 | 0.744 | 0.241 | 0.001 | 0.005 | 0.005 | 0 | 0.67 | 0.74 | 0.73 | 0.77 | 0.69 | 0.6 |
| 3 | 156248263 | rs13083246 | -0.009 | 0.495 | 0.78 | 0.328 | -0.008 | -0.007 | -0.007 | 0 | 0.07 | 0.08 | 0.12 | 0.04 | 0.13 | 0.06 |
| 3 | 156248426 | rs9830714 | -1.598 | 0.486 | 0.841 | 0.325 | 0.007 | 0.017 | 0.017 | 0 | 0.16 | 0.21 | 0.27 | 0.26 | 0.24 | 0.26 |
| 3 | 156249214 | rs2280031 | -1.77 | 0.464 | 0.78 | 0.479 | 0.001 | 0.008 | 0.008 | 0 | 0.17 | 0.22 | 0.27 | 0.32 | 0.24 | 0.27 |
| 3 | 156254213 | rs908807 | 0.35 | 0.409 | 0.767 | 0.152 | 0.003 | 0.003 | 0.003 | 0 | 0.71 | 0.78 | 0.87 | 0.87 | 0.79 | 0.74 |
| 3 | 156255087 | rs907060 | -0.015 | 0.397 | 0.702 | 0.077 | 0.011 | 0.139 | 0.139 | 0 | 0.74 | 0.95 | 0.88 | 0.98 | 0.81 | 0.81 |
| 3 | 156255689 | rs1027915 | 0.243 | 0.409 | 0.718 | 0.083 | 0.018 | 0.031 | 0.031 | 0 | 0.72 | 0.85 | 0.87 | 0.98 | 0.79 | 0.8 |
| 3 | 156256407 | rs10936027 | 0.203 | 0.427 | 0.719 | 0.092 | -0.008 | -0.006 | -0.006 | 4.308 | 0.07 | 0.08 | 0.12 | 0.04 | 0.13 | 0.06 |
| 3 | 156257121 | rs1510278 | 0.239 | 0.416 | 0.718 | 0.085 | 0.018 | 0.031 | 0.031 | 0 | 0.72 | 0.85 | 0.87 | 0.98 | 0.79 | 0.8 |
| 3 | 156259178 | rs6764992 | 0.186 | 0.444 | 0.707 | 0.116 | -0.008 | -0.006 | -0.006 | 0 | 0.07 | 0.08 | 0.12 | 0.04 | 0.13 | 0.06 |
| 3 | 156259485 | rs8665 | 0.868 | 0.414 | 0.738 | 0.093 | 0.007 | 0.053 | 0.053 | 0 | 0.66 | 0.77 | 0.75 | 0.94 | 0.66 | 0.75 |
| 3 | 156260245 | rs1510277 | 0.127 | 0.443 | 0.599 | 0.066 | 0.017 | 0.135 | 0.135 | 0 | 0.73 | 0.91 | 0.89 | 0.98 | 0.81 | 0.81 |
| 3 | 156262460 | rs6795731 | -1.513 | 0.447 | 0.714 | 0.261 | 0.024 | 0.092 | 0.092 | 0.134 | 0.12 | 0.19 | 0.16 | 0.34 | 0.15 | 0.26 |
| 3 | 156266880 | rs3773716 | 2.287 | 0.302 | 0.812 | 0.105 | -0.062 | -0.016 | -0.016 | 0 | 0.41 | 0.2 | 0.54 | 0.6 | 0.46 | 0.34 |
| 3 | 156268070 | rs12631297 | -1.658 | 0.698 | 0.949 | 0.399 | -0.001 | -0.004 | -0.004 | 0 | 0.46 | 0.43 | 0.4 | 0.38 | 0.46 | 0.44 |
| 3 | 156271608 | rs2292338 | -1.955 | 0.985 | 1.221 | 0.894 | 0.016 | 0.031 | 0.031 | 0.035 | 0.11 | 0.15 | 0.15 | 0.23 | 0.15 | 0.19 |
| 3 | 156281184 | rs16826539 | -1.613 | 0.934 | 0.9 | 0.563 | -0.023 | -0.017 | -0.017 | 0.01 | 0.09 | 0.04 | 0.15 | 0.17 | 0.12 | 0.2 |
| 3 | 156287509 | rs13063568 | -1.138 | 1.435 | 1.125 | 0.492 | 0.03 | 0.002 | 0.002 | 0 | 0.09 | 0.23 | 0.1 | 0.06 | 0.13 | 0.09 |
| 3 | 156292361 | rs907059 | -2.628 | 1.424 | 1.611 | 0.568 | 0.044 | 0.059 | 0.059 | 0 | 0.2 | 0.45 | 0.27 | 0.33 | 0.24 | 0.36 |
| 3 | 156295341 | rs9826775 | -1.496 | 1.411 | 1.601 | 0.585 | 0.039 | 0.038 | 0.038 | 0 | 0.1 | 0.26 | 0.11 | 0.17 | 0.13 | 0.17 |

| | | | | | | | | | | | | | | | | |
|---|---|---|---|---|---|---|---|---|---|---|---|---|---|---|---|---|
| 3 | 156296255 | rs9870053 | -1.091 | 1.412 | 1.583 | 0.583 | 0.03 | -0.002 | -0.002 | 0 | 0.09 | 0.23 | 0.1 | 0.05 | 0.13 | 0.08 |
| 3 | 156300724 | rs1510272 | -2.426 | 1.467 | 1.51 | 0.579 | 0.024 | 0.021 | 0.021 | 0 | 0.19 | 0.19 | 0.26 | 0.34 | 0.26 | 0.36 |
| 3 | 156320230 | rs10513491 | -2.448 | 1.413 | 1.708 | 0.601 | 0.043 | 0.059 | 0.059 | 0 | 0.21 | 0.52 | 0.26 | 0.35 | 0.24 | 0.35 |
| 3 | 156332037 | rs11927843 | -2.036 | 1.286 | 0.842 | 0.606 | -0.007 | -0.007 | -0.007 | 0 | 0.1 | 0.08 | 0.15 | 0.18 | 0.14 | 0.18 |
| 3 | 156334430 | rs12492052 | -1.721 | 1.233 | 0.825 | 0.608 | -0.007 | -0.01 | -0.01 | 0.094 | 0.09 | 0.06 | 0.14 | 0.18 | 0.11 | 0.18 |
| 3 | 156337285 | rs11919879 | -1.809 | 1.187 | 0.946 | 0.793 | -0.003 | 0.068 | 0.068 | 0 | 0.07 | 0.1 | 0.05 | 0.29 | 0.07 | 0.22 |
| 3 | 156337595 | rs10446282 | -2.026 | 0.821 | 1.393 | 0.345 | -0.006 | 0.03 | 0.03 | 0.248 | 0.17 | 0.5 | 0.11 | 0.19 | 0.14 | 0.22 |
| 3 | 156339632 | rs7638936 | 0.43 | 0.406 | 0.648 | -0.746 | 0.008 | 0.041 | 0.041 | 0 | 0.52 | 0.64 | 0.65 | 0.89 | 0.59 | 0.63 |
| 3 | 156342849 | rs2102506 | 1.473 | 1.499 | 0.589 | -0.636 | 0.007 | -0.011 | -0.011 | 0.532 | 0.7 | 0.65 | 0.67 | 0.81 | 0.64 | 0.67 |
| 3 | 156351846 | rs9811767 | -0.965 | 1.485 | 0.366 | -0.49 | -0.003 | -0.001 | -0.001 | 0 | 0.19 | 0.18 | 0.2 | 0.16 | 0.2 | 0.15 |
| 3 | 156355404 | rs13092225 | 0.902 | 1.086 | 0.62 | -0.881 | 0.002 | 0.007 | 0.007 | 0 | 0.62 | 0.36 | 0.77 | 0.85 | 0.67 | 0.45 |
| 3 | 156356839 | rs4680294 | -0.547 | 1.079 | 1.044 | -0.904 | -0.014 | -0.051 | -0.051 | 0 | 0.32 | 0.41 | 0.18 | 0.11 | 0.23 | 0.42 |
| 3 | 156357328 | rs11926561 | 1.239 | 1.088 | 0.962 | -0.874 | 0.049 | 0.015 | 0.015 | 0 | 0.94 | 0.79 | 0.95 | 0.96 | 0.91 | 0.87 |
| 3 | 156370877 | rs10428096 | 2.058 | 1.488 | 0.814 | -0.849 | -0.01 | 0.042 | 0.042 | 0 | 0.41 | 0.17 | 0.6 | 0.73 | 0.43 | 0.36 |
| 3 | 156372177 | rs6807894 | 0.809 | 1.381 | 0.909 | -0.895 | -0.007 | -0.052 | -0.052 | 0 | 0.63 | 0.5 | 0.78 | 0.85 | 0.68 | 0.45 |
| 3 | 156374086 | rs6790457 | 0.587 | 1.247 | 1.003 | -0.909 | -0.016 | -0.043 | -0.043 | 0.195 | 0.67 | 0.57 | 0.82 | 0.86 | 0.74 | 0.55 |
| 3 | 156376555 | rs1397190 | -0.815 | 1.163 | 0.895 | -1.18 | -0.016 | -0.058 | -0.058 | 0 | 0.36 | 0.57 | 0.2 | 0.12 | 0.24 | 0.44 |
| 3 | 156389947 | rs9289971 | -0.121 | 1.509 | 0.706 | -1.123 | -0.046 | -0.006 | -0.006 | 1.415 | 0.14 | 0.01 | 0.17 | 0.09 | 0.15 | 0.07 |
| 3 | 156395329 | rs345986 | -1.547 | 1.385 | 1.011 | -1.252 | -0.004 | 0.122 | 0.122 | 0 | 0.44 | 0.25 | 0.33 | 0.1 | 0.38 | 0.41 |
| 3 | 156418756 | rs1367311 | -0.008 | 1.473 | 0.89 | -1.221 | -0.041 | -0.003 | -0.003 | 0 | 0.14 | 0.01 | 0.17 | 0.09 | 0.14 | 0.07 |
| 3 | 156427827 | rs1469536 | 1.718 | 1.479 | 1.051 | -1.213 | 0.003 | 0.067 | 0.067 | 0 | 0.49 | 0.33 | 0.6 | 0.76 | 0.53 | 0.4 |
| 3 | 156444383 | rs237250 | -1.312 | 1.611 | 0.832 | -1.403 | -0.016 | 0.127 | 0.127 | 0 | 0.46 | 0.29 | 0.36 | 0.11 | 0.43 | 0.41 |
| 3 | 156470955 | rs13068223 | 1.775 | 1.576 | 0.859 | -1.386 | 0.01 | 0.06 | 0.06 | 0 | 0.47 | 0.31 | 0.57 | 0.75 | 0.49 | 0.4 |
| 3 | 156480263 | rs391833 | 0.112 | 1.771 | 0.634 | -1.381 | 0 | 0.043 | 0.043 | 0 | 0.13 | 0.02 | 0.06 | 0 | 0.11 | 0.06 |
| 3 | 156481051 | rs344045 | 1.249 | 1.724 | 1.063 | -1.355 | 0.082 | 0.046 | 0.046 | 0.337 | 0.87 | 0.5 | 0.86 | 0.83 | 0.82 | 0.78 |
| 3 | 156489509 | rs1430408 | 2.086 | 1.693 | 0.955 | -1.451 | -0.019 | 0.16 | 0.16 | 0 | 0.57 | 0.17 | 0.7 | 0.82 | 0.6 | 0.45 |
| 3 | 156489778 | rs344019 | -1.04 | 1.7 | 1.039 | -1.455 | 0.001 | -0.026 | -0.026 | 1.375 | 0.29 | 0.27 | 0.16 | 0.01 | 0.22 | 0.33 |
| 3 | 156491160 | rs9867857 | 2.385 | 1.591 | 0.756 | -1.437 | 0.004 | 0.126 | 0.126 | 0 | 0.46 | 0.16 | 0.54 | 0.72 | 0.48 | 0.41 |
| 3 | 156499981 | rs2054880 | -1.564 | 1.805 | 0.758 | -1.387 | -0.006 | 0.01 | 0.01 | 0 | 0.42 | 0.37 | 0.36 | 0.22 | 0.41 | 0.46 |
| 3 | 156524220 | rs9818684 | 1.738 | 2.429 | 0.862 | -0.995 | -0.059 | 0.067 | 0.067 | 0.373 | 0.6 | 0.19 | 0.63 | 0.73 | 0.56 | 0.6 |
| 3 | 156525562 | rs1430410 | 1.534 | 2.461 | 1.06 | -1.037 | -0.011 | 0.029 | 0.029 | 0.133 | 0.63 | 0.33 | 0.67 | 0.84 | 0.62 | 0.7 |
| 3 | 156555984 | rs344081 | -1.102 | 2.387 | 1.274 | -1.217 | -0.02 | -0.086 | -0.086 | 0 | 0.2 | 0.58 | 0.1 | 0.03 | 0.12 | 0.24 |
| 3 | 156558120 | rs344085 | 0.042 | 2.351 | 1.505 | -1.235 | -0.002 | 0.01 | 0.01 | 0 | 0.12 | 0.07 | 0.06 | 0.03 | 0.11 | 0.22 |
| 3 | 156574076 | rs491423 | -1.476 | 2.612 | 1.342 | -1.222 | 0 | -0.02 | -0.02 | 0.804 | 0.26 | 0.7 | 0.17 | 0.03 | 0.25 | 0.25 |
| 3 | 156574252 | rs453219 | 0.029 | 2.676 | 1.317 | -0.917 | -0.032 | -0.03 | -0.03 | 0 | 0.87 | 0.93 | 0.82 | 0.78 | 0.82 | 0.87 |
| 3 | 156577669 | rs13070271 | 0.146 | 2.675 | 1.317 | -0.917 | -0.041 | -0.031 | -0.031 | 0 | 0.87 | 0.93 | 0.8 | 0.78 | 0.77 | 0.87 |
| 3 | 156580766 | rs16826869 | 0.271 | 2.658 | 1.314 | -0.929 | 0.001 | -0.03 | -0.03 | 0.033 | 0.06 | 0.11 | 0.06 | 0 | 0.08 | 0.01 |
| 3 | 156581034 | rs7616633 | 1.297 | 2.654 | 1.524 | -0.847 | 0.028 | -0.075 | -0.075 | 1.059 | 0.71 | 0.61 | 0.76 | 0.88 | 0.71 | 0.71 |
| 3 | 156601655 | rs2321291 | -0.248 | 2.661 | 1.306 | -1.007 | -0.064 | 0.033 | 0.033 | 0 | 0.88 | 0.99 | 0.83 | 0.94 | 0.83 | 0.95 |
| 3 | 156642634 | rs7650127 | -1.397 | 2.528 | 1.349 | -1.019 | 0.001 | -0.028 | -0.028 | 0 | 0.26 | 0.68 | 0.16 | 0.03 | 0.19 | 0.24 |
| 3 | 156677025 | rs10936040 | 1.965 | 2.487 | 1.253 | -1.004 | -0.045 | 0.059 | 0.059 | 0.216 | 0.61 | 0.24 | 0.64 | 0.74 | 0.58 | 0.62 |
| 3 | 156681774 | rs16826948 | -0.484 | 2.502 | 1.189 | -0.995 | -0.014 | -0.077 | -0.077 | 1.954 | 0.18 | 0.26 | 0.07 | 0.03 | 0.12 | 0.23 |
| 3 | 156682397 | rs2874495 | 0.997 | 2.293 | 0.505 | -0.889 | -0.001 | 0 | 0 | 0 | 0.67 | 0.36 | 0.7 | 0.75 | 0.65 | 0.63 |
| 3 | 156696506 | rs10936043 | 1.676 | 2.177 | 0.1 | -0.878 | -0.019 | 0.132 | 0.132 | 0.111 | 0.53 | 0.11 | 0.59 | 0.74 | 0.52 | 0.5 |
| 3 | 156697097 | rs6780427 | 1.265 | 2.279 | 0.17 | -0.803 | -0.046 | 0.06 | 0.06 | 0.216 | 0.61 | 0.23 | 0.64 | 0.74 | 0.57 | 0.62 |
| 3 | 156699828 | rs9878410 | -1.932 | 2.441 | 0.788 | -0.138 | 0.058 | 0.077 | 0.077 | 0.029 | 0.16 | 0.59 | 0.23 | 0.25 | 0.18 | 0.14 |
| 3 | 156700416 | rs7635876 | -0.53 | 2.494 | 1.234 | -0.131 | 0.123 | 0.065 | 0.065 | 0 | 0.07 | 0.27 | 0.13 | 0.06 | 0.11 | 0.07 |
| 3 | 156700986 | rs1871967 | -0.146 | 2.302 | 1.311 | -0.123 | 0.014 | 0.012 | 0.012 | 0 | 0.07 | 0.1 | 0.11 | 0.06 | 0.06 | 0.06 |
| 3 | 156704760 | rs10513501 | -0.478 | 2.29 | 1.394 | -0.136 | 0.001 | -0.017 | -0.017 | 0.184 | 0.05 | 0.08 | 0.04 | 0 | 0.06 | 0.01 |
| 3 | 156707330 | rs1032044 | 1.429 | 2.575 | 0.987 | -0.473 | -0.086 | 0.012 | 0.012 | 0.878 | 0.64 | 0.16 | 0.6 | 0.7 | 0.57 | 0.8 |
| 3 | 156709318 | rs11920497 | -0.644 | 2.561 | 1.225 | -0.403 | 0.041 | 0.148 | 0.148 | 0 | 0.08 | 0.2 | 0.16 | 0.24 | 0.08 | 0.13 |
| 3 | 156715450 | rs4680326 | -0.292 | 2.313 | 1.047 | -0.558 | 0.013 | -0.04 | -0.04 | 3.625 | 0.79 | 0.76 | 0.81 | 0.95 | 0.71 | 0.94 |
| 3 | 156728057 | rs1842840 | 0.836 | 2.266 | 0.523 | -0.536 | 0.032 | -0.021 | -0.021 | 0 | 0.71 | 0.42 | 0.67 | 0.79 | 0.64 | 0.81 |
| 3 | 156728610 | rs7616559 | -0.251 | 2.279 | 0.474 | -0.542 | 0.056 | -0.093 | -0.093 | 0 | 0.79 | 0.6 | 0.81 | 0.94 | 0.71 | 0.94 |
| 3 | 156733218 | rs2321458 | 0.162 | 2.264 | 0.463 | -0.516 | 0.012 | 0.012 | 0.012 | 0 | 0.07 | 0.1 | 0.11 | 0.06 | 0.06 | 0.06 |
| 3 | 156739425 | rs9839037 | -0.258 | 2.266 | 0.45 | -0.551 | 0.057 | -0.093 | -0.093 | 0.032 | 0.79 | 0.6 | 0.81 | 0.94 | 0.71 | 0.94 |
| 3 | 156739808 | rs10936051 | 0.166 | 2.319 | 0.133 | -0.538 | 0.013 | 0.01 | 0.01 | 0 | 0.07 | 0.1 | 0.11 | 0.06 | 0.07 | 0.06 |
| 3 | 156745105 | rs1032041 | 1.351 | 2.316 | -0.54 | -0.583 | -0.082 | 0.079 | 0.079 | 1.073 | 0.63 | 0.15 | 0.62 | 0.78 | 0.59 | 0.8 |
| 3 | 156753029 | rs4680332 | 1.401 | 2.13 | -0.751 | -0.571 | -0.092 | 0.072 | 0.072 | 0.207 | 0.62 | 0.09 | 0.66 | 0.76 | 0.56 | 0.73 |
| 3 | 156754441 | rs1482856 | 0.273 | 1.786 | -0.562 | -0.468 | -0.022 | 0.01 | 0.01 | 0.609 | 0.75 | 0.19 | 0.76 | 0.78 | 0.69 | 0.79 |
| 3 | 156759307 | rs9851046 | 0.386 | 1.202 | -1.383 | -0.39 | -0.115 | -0.053 | -0.053 | 0.061 | 0.27 | 0.02 | 0.26 | 0.21 | 0.33 | 0.38 |
| 3 | 156777225 | rs9990369 | -1.393 | 1.386 | -0.167 | -0.343 | -0.025 | 0.019 | 0.019 | 0.146 | 0.09 | 0.35 | 0.05 | 0.1 | 0.01 | 0.12 |
| 3 | 156781395 | rs931245 | 0.824 | 1.676 | -1.483 | -0.282 | -0.056 | 0.058 | 0.058 | 0.928 | 0.66 | 0.11 | 0.71 | 0.78 | 0.64 | 0.77 |
| 3 | 156784535 | rs10936058 | 0.75 | 1.544 | -1.472 | -0.331 | -0.035 | 0.094 | 0.094 | 1.021 | 0.67 | 0.11 | 0.73 | 0.85 | 0.66 | 0.84 |
| 3 | 156789192 | rs2128535 | 0.555 | 1.313 | -1.165 | -0.04 | -0.008 | 0.026 | 0.026 | 1.66 | 0.67 | 0.25 | 0.74 | 0.77 | 0.66 | 0.76 |
| 3 | 156797609 | rs17451107 | 0.099 | 1.013 | -1.024 | 0.133 | 0.004 | -0.003 | -0.003 | 0 | 0.43 | 0.34 | 0.45 | 0.48 | 0.38 | 0.24 |
| 3 | 156800360 | rs11720805 | 0.039 | 0.757 | -1.552 | -0.218 | 0.031 | -0.06 | -0.06 | 0 | 0.59 | 0.08 | 0.56 | 0.49 | 0.68 | 0.4 |
| 3 | 156800384 | rs11712937 | -0.042 | 0.784 | -1.548 | -0.219 | 0.021 | -0.014 | -0.014 | 0 | 0.61 | 0.24 | 0.58 | 0.49 | 0.68 | 0.39 |
| 3 | 156802256 | rs11719286 | 0.073 | 0.695 | -1.885 | -0.087 | 0.046 | 0.02 | 0.02 | 0 | 0.59 | 0.08 | 0.63 | 0.57 | 0.68 | 0.37 |

| | | | | | | | | | | | | | | | | |
|---|---|---|---|---|---|---|---|---|---|---|---|---|---|---|---|---|
| 3 | 156803565 | rs9822326 | 0.652 | 1.019 | -1.615 | 0.078 | -0.006 | -0.01 | -0.01 | 0.832 | 0.4 | 0.18 | 0.49 | 0.52 | 0.4 | 0.28 |
| 3 | 156812581 | rs35649103 | -0.542 | 0.75 | -1.35 | 0.358 | 0 | -0.008 | -0.008 | 0.115 | 0.12 | 0.11 | 0.15 | 0.22 | 0.1 | 0.43 |
| 3 | 156815221 | rs9860743 | -0.6 | 1.218 | -1.023 | 0.049 | 0.029 | 0.033 | 0.033 | 0 | 0.76 | 0.57 | 0.73 | 0.67 | 0.71 | 0.87 |
| 3 | 156817662 | rs2306531 | -0.258 | 0.997 | -1.371 | -0.081 | -0.024 | 0.069 | 0.069 | 0.293 | 0.29 | 0.15 | 0.19 | 0.09 | 0.32 | 0.3 |
| 3 | 156820489 | rs9867835 | -0.505 | 0.532 | -1.024 | -0.13 | -0.016 | 0.003 | 0.003 | 0 | 0.69 | 0.74 | 0.73 | 0.66 | 0.65 | 0.88 |
| 3 | 156821808 | rs6441111 | -0.627 | 0.387 | -1.123 | 0.162 | -0.011 | 0.015 | 0.015 | 0 | 0.55 | 0.67 | 0.47 | 0.63 | 0.49 | 0.65 |
| 3 | 156827918 | rs13098118 | -0.598 | -0.251 | -0.871 | 0.432 | 0.019 | -0.017 | -0.017 | 0 | 0.13 | 0.03 | 0.12 | 0.15 | 0.09 | 0.18 |
| 3 | 156828823 | rs729257 | 0.006 | -0.716 | -0.576 | 0.116 | 0.013 | 0.007 | 0.007 | 0 | 0.45 | 0.59 | 0.56 | 0.56 | 0.41 | 0.57 |
| 3 | 156832894 | rs7623382 | -0.738 | -0.935 | -0.502 | 0.018 | 0.009 | 0.056 | 0.056 | 0 | 0.56 | 0.82 | 0.6 | 0.64 | 0.44 | 0.69 |
| 3 | 156833616 | rs16827244 | -0.017 | -1.047 | -0.433 | 0.036 | 0.001 | 0 | 0 | 0 | 0.12 | 0.09 | 0.07 | 0.13 | 0.09 | 0.35 |
| 3 | 156836517 | rs2086667 | 0.918 | -0.862 | -0.159 | 0.204 | -0.028 | 0.038 | 0.038 | 0 | 0.29 | 0.5 | 0.39 | 0.36 | 0.23 | 0.12 |
| 3 | 156848637 | rs7644860 | -1.381 | -0.931 | -0.167 | 0.137 | -0.008 | 0.002 | 0.002 | 0 | 0.27 | 0.67 | 0.19 | 0.28 | 0.15 | 0.49 |
| 3 | 156849811 | rs6804404 | -0.708 | -0.715 | 0.056 | 0.148 | -0.028 | 0.02 | 0.02 | 0 | 0.19 | 0.23 | 0.08 | 0.22 | 0.09 | 0.41 |
| 3 | 156852205 | rs12487629 | -0.134 | -0.581 | -0.04 | 0.155 | 0.003 | 0.001 | 0.001 | 0 | 0.41 | 0.32 | 0.54 | 0.58 | 0.3 | 0.55 |
| 3 | 156861460 | rs4680340 | 0.689 | -0.622 | 0.464 | 0.313 | -0.002 | 0.005 | 0.005 | 0 | 0.27 | 0.31 | 0.49 | 0.33 | 0.26 | 0.19 |
| 3 | 156862145 | rs6809394 | -0.431 | -0.862 | 0.367 | 0.326 | -0.008 | -0.003 | -0.003 | 0 | 0.41 | 0.49 | 0.63 | 0.55 | 0.37 | 0.38 |
| 3 | 156880883 | rs10936063 | 1.223 | -0.849 | 0.307 | 0.308 | -0.002 | -0.062 | -0.062 | 0 | 0.24 | 0.08 | 0.48 | 0.33 | 0.26 | 0.19 |
| 3 | 156901525 | rs9867618 | -0.959 | -0.595 | 0.634 | 0.207 | 0.063 | 0.102 | 0.102 | 0 | 0.49 | 0.72 | 0.66 | 0.67 | 0.39 | 0.62 |
| 3 | 156919138 | rs16827308 | 0.112 | -0.591 | 0.605 | 0.242 | -0.003 | 0.003 | 0.003 | 0 | 0.38 | 0.42 | 0.53 | 0.46 | 0.31 | 0.3 |
| 3 | 156946257 | rs12491378 | 1.489 | -0.143 | 0.172 | -0.028 | -0.071 | 0.063 | 0.063 | 0.124 | 0.12 | 0.02 | 0.13 | 0.03 | 0.25 | 0.15 |
| 3 | 156952986 | rs17387112 | 1.197 | -0.269 | 0.208 | 0.046 | 0.013 | -0.023 | -0.023 | 0 | 0.09 | 0.02 | 0.03 | 0.09 | 0.04 | 0.09 |
| 3 | 156964032 | rs7613866 | -0.066 | -0.322 | 0.366 | 0.05 | -0.005 | 0.047 | 0.047 | 0 | 0.06 | 0.26 | 0.1 | 0.09 | 0.05 | 0.11 |
| 3 | 156966235 | rs7642179 | -0.051 | -0.36 | 0.382 | 0.04 | -0.003 | -0.001 | -0.001 | 0 | 0.4 | 0.46 | 0.62 | 0.54 | 0.36 | 0.39 |
| 3 | 156967195 | rs4680348 | -0.79 | -0.387 | 0.424 | 0.038 | 0.063 | 0.099 | 0.099 | 0 | 0.49 | 0.72 | 0.66 | 0.67 | 0.39 | 0.62 |
| 3 | 156973544 | rs4680349 | 1.654 | -0.379 | 0.275 | 0.052 | -0.013 | -0.089 | -0.089 | 0 | 0.24 | 0.04 | 0.47 | 0.33 | 0.26 | 0.18 |
| 3 | 156978173 | rs1046938 | -0.321 | -0.386 | 0.427 | 0.044 | -0.01 | -0.002 | -0.002 | 1.142 | 0.42 | 0.55 | 0.63 | 0.54 | 0.35 | 0.39 |
| 3 | 156983414 | rs7652298 | 1.606 | -0.279 | 0.132 | 0.058 | -0.008 | -0.081 | -0.081 | 0.086 | 0.24 | 0.05 | 0.47 | 0.33 | 0.26 | 0.18 |
| 3 | 156986803 | rs1403104 | 0.376 | -0.313 | 0.163 | 0.043 | 0.02 | -0.027 | -0.027 | 0 | 0.37 | 0.3 | 0.51 | 0.42 | 0.31 | 0.28 |
| 3 | 156987482 | rs1915952 | -0.899 | -0.375 | 0.329 | -0.069 | -0.007 | 0.007 | 0.007 | 0 | 0.31 | 0.25 | 0.19 | 0.25 | 0.33 | 0.49 |
| 3 | 156987632 | rs13326287 | -0.448 | -0.374 | 0.401 | -0.067 | 0.008 | 0.072 | 0.072 | 0.01 | 0.07 | 0.38 | 0.11 | 0.09 | 0.05 | 0.11 |
| 3 | 156988661 | rs7636413 | -1.427 | -0.334 | 0.328 | -0.081 | -0.001 | 0.003 | 0.003 | 0 | 0.59 | 0.64 | 0.69 | 0.62 | 0.59 | 0.67 |
| 3 | 156994234 | rs12490031 | 0.823 | -0.194 | -0.124 | -0.073 | 0.027 | -0.054 | -0.054 | 0 | 0.16 | 0.22 | 0.15 | 0.04 | 0.25 | 0.16 |
| 3 | 156998998 | rs12637932 | -0.417 | -0.314 | -0.076 | 0.013 | 0.009 | 0.008 | 0.008 | 0 | 0.49 | 0.3 | 0.65 | 0.5 | 0.55 | 0.43 |
| 3 | 157004323 | rs4680353 | 0.315 | -0.156 | -0.101 | 0.055 | -0.033 | -0.013 | -0.013 | 0 | 0.39 | 0.23 | 0.62 | 0.41 | 0.51 | 0.33 |
| 3 | 157028417 | rs1522393 | -0.063 | -0.187 | -0.26 | 0.045 | -0.035 | -0.026 | -0.026 | 0 | 0.43 | 0.15 | 0.63 | 0.58 | 0.55 | 0.55 |
| 3 | 157029522 | rs6769530 | -0.194 | -0.439 | 0.188 | -0.03 | -0.061 | 0.041 | 0.041 | 0.027 | 0.17 | 0.43 | 0.15 | 0.17 | 0.09 | 0.21 |
| 3 | 157044184 | rs1500913 | 0.142 | -0.183 | -0.168 | 0.039 | -0.024 | 0.002 | 0.002 | 0 | 0.18 | 0.11 | 0.15 | 0.18 | 0.28 | 0.39 |
| 3 | 157045959 | rs12494423 | -0.002 | -0.279 | -0.133 | 0.035 | -0.029 | -0.037 | -0.037 | 3.156 | 0.42 | 0.17 | 0.63 | 0.53 | 0.55 | 0.55 |
| 3 | 157054271 | rs988587 | -1.394 | -0.448 | 0.244 | 0.037 | -0.005 | 0.02 | 0.02 | 0 | 0.63 | 0.73 | 0.81 | 0.75 | 0.64 | 0.79 |
| 3 | 157057390 | rs13325944 | 0.274 | -0.576 | 0.053 | 0.191 | 0.003 | -0.004 | -0.004 | 0.189 | 0.13 | 0.12 | 0.12 | 0.17 | 0.09 | 0.21 |
| 3 | 157076854 | rs10513512 | 1.276 | -0.485 | -0.063 | 0.191 | 0.009 | -0.077 | -0.077 | 0.001 | 0.28 | 0.1 | 0.5 | 0.39 | 0.27 | 0.17 |
| 3 | 157078151 | rs16827528 | -0.617 | -0.518 | 0.075 | 0.167 | -0.014 | 0.019 | 0.019 | 0 | 0.2 | 0.21 | 0.14 | 0.31 | 0.13 | 0.45 |
| 3 | 157080331 | rs9883822 | -0.507 | -0.517 | 0.073 | 0.14 | 0.046 | 0.074 | 0.074 | 0 | 0.48 | 0.32 | 0.65 | 0.71 | 0.4 | 0.64 |
| 3 | 157081324 | rs11918974 | 1.302 | -0.482 | 0.023 | 0.169 | 0.009 | -0.061 | -0.061 | 0 | 0.28 | 0.09 | 0.5 | 0.35 | 0.27 | 0.17 |
| 3 | 157088982 | rs9883103 | -0.444 | -0.387 | 0.096 | 0.07 | 0.049 | 0.074 | 0.074 | 0 | 0.48 | 0.31 | 0.65 | 0.7 | 0.39 | 0.67 |
| 3 | 157094479 | rs4680360 | -1.47 | -0.566 | 0.167 | 0.242 | -0.05 | 0.062 | 0.062 | 0 | 0.66 | 0.9 | 0.8 | 0.75 | 0.62 | 0.77 |
| 3 | 157113276 | rs12486427 | -1.213 | -0.582 | 0.527 | 0.252 | -0.006 | 0.031 | 0.031 | 0 | 0.64 | 0.74 | 0.73 | 0.75 | 0.62 | 0.77 |
| 3 | 157123357 | rs1869857 | -0.369 | -0.668 | 0.689 | 0.297 | 0.003 | 0.008 | 0.008 | 0 | 0.53 | 0.56 | 0.68 | 0.61 | 0.58 | 0.68 |
| 3 | 157125584 | rs6803628 | -1.431 | -0.675 | 0.631 | 0.276 | -0.014 | 0.041 | 0.041 | 0 | 0.65 | 0.77 | 0.73 | 0.74 | 0.63 | 0.78 |
| 3 | 157126631 | rs1456104 | 0.538 | -0.735 | 0.717 | 0.374 | -0.001 | 0 | 0 | 0 | 0.41 | 0.39 | 0.62 | 0.36 | 0.5 | 0.26 |
| 3 | 157131788 | rs1378796 | -0.2 | -0.693 | 0.567 | 0.411 | -0.018 | 0.018 | 0.018 | 0 | 0.15 | 0.22 | 0.08 | 0.23 | 0.1 | 0.3 |
| 3 | 157132577 | rs1074864 | 0.564 | -0.722 | 0.827 | 0.372 | -0.039 | 0.006 | 0.006 | 0 | 0.4 | 0.23 | 0.61 | 0.37 | 0.49 | 0.23 |
| 3 | 157135595 | rs7639767 | 0.299 | -0.858 | 0.649 | 0.602 | -0.008 | 0.021 | 0.021 | 0.153 | 0.08 | 0.13 | 0.03 | 0.14 | 0.04 | 0.25 |
| 3 | 157138405 | rs16827620 | 0.299 | -0.858 | 0.649 | 0.602 | -0.008 | 0.021 | 0.021 | 0.153 | 0.08 | 0.13 | 0.03 | 0.14 | 0.04 | 0.25 |
| 3 | 157139541 | rs11919151 | -1.418 | -0.823 | 0.536 | 0.582 | -0.031 | 0.035 | 0.035 | 0.053 | 0.23 | 0.35 | 0.1 | 0.33 | 0.14 | 0.55 |
| 3 | 157143996 | rs9681270 | -0.536 | -0.302 | 0.425 | 0.363 | 0.008 | 0.045 | 0.045 | 0 | 0.52 | 0.66 | 0.69 | 0.66 | 0.57 | 0.67 |
| 3 | 157145903 | rs7632702 | 0.19 | -0.206 | 0.356 | 0.486 | -0.02 | -0.032 | -0.032 | 0 | 0.42 | 0.24 | 0.62 | 0.48 | 0.51 | 0.37 |
| 3 | 157159694 | rs3845978 | -0.431 | -0.018 | 0.681 | 0.622 | 0.014 | 0.086 | 0.086 | 0 | 0.08 | 0.24 | 0.05 | 0.15 | 0.05 | 0.26 |
| 3 | 157163635 | rs16827657 | -0.946 | -0.059 | 0.145 | 0.604 | 0.032 | 0.109 | 0.109 | 0.198 | 0.11 | 0.53 | 0.07 | 0.16 | 0.06 | 0.3 |
| 3 | 157166123 | rs4680367 | -0.656 | 0.031 | -0.554 | 0.13 | -0.005 | 0.064 | 0.064 | 0 | 0.59 | 0.74 | 0.72 | 0.76 | 0.6 | 0.74 |
| 3 | 157168586 | rs16827662 | -0.579 | -0.266 | -0.037 | 0.193 | -0.021 | -0.047 | -0.047 | 0 | 0.21 | 0.44 | 0.11 | 0.12 | 0.12 | 0.09 |
| 3 | 157169652 | rs1378797 | -0.795 | -0.775 | -0.631 | 0.241 | 0.013 | 0.065 | 0.065 | 0 | 0.56 | 0.8 | 0.7 | 0.7 | 0.58 | 0.6 |
| 3 | 157170524 | rs1456103 | -1.613 | -0.807 | -0.907 | 0.109 | -0.045 | 0.164 | 0.164 | 0 | 0.68 | 0.99 | 0.84 | 0.91 | 0.7 | 0.81 |
| 3 | 157172112 | rs12634412 | 1.926 | -0.181 | -0.897 | 0.709 | -0.05 | -0.102 | -0.102 | 5.896 | 0.18 | 0.03 | 0.31 | 0.36 | 0.21 | 0.11 |
| 3 | 157182981 | rs9877620 | 0.837 | -0.214 | -0.776 | 0.674 | -0.018 | -0.015 | -0.015 | 0.416 | 0.17 | 0.1 | 0.27 | 0.2 | 0.24 | 0.37 |
| 3 | 157186678 | rs4679796 | -0.003 | -0.285 | -0.489 | 0.632 | -0.001 | 0.005 | 0.005 | 0 | 0.37 | 0.42 | 0.42 | 0.49 | 0.34 | 0.2 |
| 3 | 157188045 | rs1456113 | -1.145 | -0.209 | -0.562 | 0.666 | 0.01 | 0.056 | 0.056 | 0 | 0.57 | 0.81 | 0.7 | 0.67 | 0.57 | 0.6 |
| 3 | 157188378 | rs1011060 | -0.163 | -0.258 | -0.562 | 0.676 | 0.002 | -0.006 | -0.006 | 0 | 0.39 | 0.69 | 0.43 | 0.39 | 0.33 | 0.22 |
| 3 | 157189235 | rs12631456 | 0.546 | -0.25 | -0.563 | 0.679 | -0.017 | -0.026 | -0.026 | 0 | 0.18 | 0.12 | 0.27 | 0.28 | 0.24 | 0.38 |

| | | | | | | | | | | | | | | | | |
|---|---|---|---|---|---|---|---|---|---|---|---|---|---|---|---|---|
| 3 | 157195091 | rs12485456 | -0.949 | -0.174 | -0.578 | 0.702 | 0.015 | 0.035 | 0.035 | 0 | 0.55 | 0.81 | 0.7 | 0.59 | 0.56 | 0.6 |
| 3 | 157209768 | rs4680372 | 1.528 | -0.15 | -0.526 | 0.688 | 0.003 | 0.026 | 0.026 | 0 | 0.21 | 0.26 | 0.32 | 0.35 | 0.21 | 0.12 |
| 3 | 157216017 | rs1456095 | -0.971 | -0.142 | -0.637 | 0.553 | 0.012 | 0.046 | 0.046 | 0 | 0.56 | 0.74 | 0.69 | 0.67 | 0.57 | 0.57 |
| 3 | 157219233 | rs10513515 | 0.285 | -0.039 | -0.314 | 0.702 | 0.001 | -0.041 | -0.041 | 0.622 | 0.15 | 0.27 | 0.08 | 0.04 | 0.11 | 0.08 |
| 3 | 157221966 | rs16827747 | 1.38 | 0.002 | -0.953 | 0.872 | -0.048 | -0.106 | -0.106 | 0.04 | 0.2 | 0.03 | 0.31 | 0.35 | 0.21 | 0.1 |
| 3 | 157226796 | rs1838141 | -0.362 | -0.17 | -0.583 | 0.691 | 0.003 | 0.002 | 0.002 | 0 | 0.29 | 0.37 | 0.23 | 0.31 | 0.29 | 0.28 |
| 3 | 157233698 | rs10049206 | -1.442 | -0.065 | -0.528 | 0.444 | -0.01 | -0.119 | -0.119 | 0 | 0.68 | 0.52 | 0.8 | 0.91 | 0.68 | 0.78 |
| 3 | 157240416 | rs9855702 | 1.168 | 0.034 | -0.602 | 0.924 | -0.009 | -0.048 | -0.048 | 0 | 0.13 | 0.02 | 0.21 | 0.21 | 0.15 | 0.19 |
| 3 | 157246226 | rs16827796 | 0.908 | 0.006 | -0.579 | 0.941 | 0.009 | 0.082 | 0.082 | 0 | 0.16 | 0.32 | 0.17 | 0.28 | 0.18 | 0.24 |
| 3 | 157247946 | rs6793453 | -0.358 | -0.005 | -0.64 | 0.98 | -0.003 | -0.002 | -0.002 | 0.642 | 0.31 | 0.42 | 0.43 | 0.67 | 0.34 | 0.61 |
| 3 | 157248336 | rs11714198 | 0.803 | 0.046 | -1.424 | 1.031 | -0.007 | -0.069 | -0.069 | 2.833 | 0.14 | 0.03 | 0.26 | 0.31 | 0.16 | 0.37 |
| 3 | 157250987 | rs4680375 | -0.684 | -0.006 | -1.361 | 0.906 | 0 | 0.033 | 0.033 | 0 | 0.57 | 0.63 | 0.77 | 0.81 | 0.58 | 0.74 |
| 3 | 157251358 | rs1840675 | 0.196 | 0.189 | -1.405 | 1.032 | -0.005 | 0.002 | 0.002 | 0.014 | 0.42 | 0.62 | 0.51 | 0.56 | 0.43 | 0.38 |
| 3 | 157251387 | rs1840674 | -0.629 | 0.159 | -1.382 | 0.991 | 0.012 | 0.035 | 0.035 | 1.285 | 0.54 | 0.76 | 0.56 | 0.6 | 0.52 | 0.41 |
| 3 | 157257016 | rs6441134 | 0.231 | 0.176 | -1.374 | 1.01 | -0.004 | -0.008 | -0.008 | 0 | 0.42 | 0.62 | 0.51 | 0.49 | 0.42 | 0.38 |
| 3 | 157263695 | rs749422 | 0.005 | 0.272 | -1.407 | 0.745 | -0.001 | -0.001 | -0.001 | 0 | 0.45 | 0.57 | 0.55 | 0.59 | 0.43 | 0.56 |
| 3 | 157265066 | rs9884066 | -0.54 | 0.346 | -2.153 | 0.787 | 0.005 | 0.094 | 0.094 | 0.51 | 0.56 | 0.83 | 0.61 | 0.69 | 0.53 | 0.59 |
| 3 | 157278882 | rs1579771 | -1.606 | 0.332 | -2.367 | 0.795 | -0.024 | 0.043 | 0.043 | 0 | 0.7 | 0.9 | 0.8 | 0.8 | 0.67 | 0.8 |
| 3 | 157283248 | rs11928144 | 0.222 | 0.449 | -1.556 | 1.101 | 0.005 | -0.023 | -0.023 | 0 | 0.14 | 0.06 | 0.21 | 0.22 | 0.15 | 0.22 |
| 3 | 157288885 | rs16827846 | -0.13 | 0.703 | -1.163 | 0.843 | 0.01 | -0.031 | -0.031 | 0 | 0.1 | 0.13 | 0.05 | 0.03 | 0.1 | 0.04 |
| 3 | 157290908 | rs9834772 | -0.418 | 0.701 | -1.144 | 0.861 | 0.004 | -0.023 | -0.023 | 0 | 0.18 | 0.11 | 0.15 | 0.28 | 0.18 | 0.24 |
| 3 | 157292403 | rs7649021 | -0.641 | 0.8 | -0.928 | 0.466 | 0.01 | -0.031 | -0.031 | 0 | 0.1 | 0.13 | 0.05 | 0.03 | 0.1 | 0.04 |
| 3 | 157312876 | rs6414386 | -0.079 | 0.386 | -1.622 | 0.81 | -0.01 | -0.077 | -0.077 | 0 | 0.79 | 0.98 | 0.74 | 0.72 | 0.8 | 0.61 |
| 3 | 157313589 | rs9812718 | 0.41 | 0.156 | -1.003 | 0.137 | -0.005 | -0.009 | -0.009 | 0 | 0.56 | 0.55 | 0.57 | 0.7 | 0.63 | 0.41 |
| 3 | 157318743 | rs4472012 | 0.127 | 0.216 | -0.855 | 0.067 | 0.025 | 0.042 | 0.042 | 0.018 | 0.6 | 0.78 | 0.6 | 0.7 | 0.65 | 0.4 |
| 3 | 157324261 | rs4680382 | 0.248 | 0.212 | -0.841 | 0.074 | 0.023 | 0.049 | 0.049 | 0.017 | 0.6 | 0.81 | 0.6 | 0.7 | 0.65 | 0.4 |
| 3 | 157332411 | rs10936094 | 0.901 | 0.845 | -0.795 | 0.413 | -0.002 | -0.007 | -0.007 | 0 | 0.33 | 0.28 | 0.42 | 0.35 | 0.35 | 0.29 |
| 3 | 157340576 | rs9854256 | 1.299 | 0.651 | -1.058 | 0.581 | -0.072 | 0.058 | 0.058 | 0 | 0.27 | 0.01 | 0.21 | 0.15 | 0.32 | 0.19 |
| 3 | 157343492 | rs2649728 | -0.426 | 0.389 | -0.591 | 0.581 | 0.006 | 0.01 | 0.01 | 0.001 | 0.56 | 0.65 | 0.63 | 0.71 | 0.63 | 0.43 |
| 3 | 157359662 | rs1015389 | 1.045 | 0.407 | -0.405 | 0.554 | -0.002 | 0.021 | 0.021 | 0 | 0.31 | 0.25 | 0.28 | 0.16 | 0.33 | 0.25 |
| 3 | 157365321 | rs1392187 | -0.536 | 0.186 | -0.275 | 0.512 | 0.029 | -0.021 | -0.021 | 0 | 0.57 | 0.7 | 0.51 | 0.53 | 0.63 | 0.4 |
| 3 | 157366084 | rs1392188 | -0.145 | 0.232 | -0.495 | 0.541 | 0.028 | 0 | 0 | 0 | 0.52 | 0.37 | 0.47 | 0.53 | 0.61 | 0.39 |
| 3 | 157375233 | rs792746 | 0.598 | 0.444 | -0.761 | 0.716 | -0.054 | -0.057 | -0.057 | 0 | 0.19 | 0.07 | 0.18 | 0.36 | 0.28 | 0.14 |
| 3 | 157386185 | rs11925333 | -1.056 | 0.586 | -0.902 | 0.666 | -0.007 | -0.155 | -0.155 | 0 | 0.31 | 0.95 | 0.37 | 0.52 | 0.25 | 0.17 |
| 3 | 157393770 | rs963354 | -1.35 | 0.407 | -1.086 | 0.489 | -0.094 | -0.078 | -0.078 | 0 | 0.75 | 0.97 | 0.7 | 0.69 | 0.7 | 0.52 |
| 3 | 157419690 | rs9289990 | -0.543 | 0.124 | -0.258 | 0.596 | 0.005 | -0.021 | -0.021 | 0 | 0.38 | 0.43 | 0.56 | 0.67 | 0.42 | 0.34 |
| 3 | 157421151 | rs6783747 | 0.632 | 0.154 | -0.265 | 0.749 | -0.011 | -0.038 | -0.038 | 0 | 0.17 | 0.1 | 0.13 | 0.37 | 0.19 | 0.12 |
| 3 | 157421613 | rs1008893 | -1.175 | 0.083 | -0.001 | 0.599 | 0.007 | 0.108 | 0.108 | 0.017 | 0.46 | 0.77 | 0.6 | 0.72 | 0.45 | 0.38 |
| 3 | 157425014 | rs1001102 | -0.228 | 0.092 | -0.018 | 0.614 | -0.006 | -0.013 | -0.013 | 0 | 0.08 | 0.34 | 0.05 | 0.02 | 0.03 | 0.04 |
| 3 | 157425796 | rs1604027 | -0.228 | 0.092 | -0.018 | 0.614 | -0.006 | -0.013 | -0.013 | 0 | 0.08 | 0.34 | 0.05 | 0.02 | 0.03 | 0.04 |
| 3 | 157428862 | rs9838511 | -0.228 | 0.092 | -0.018 | 0.614 | -0.006 | -0.013 | -0.013 | 0 | 0.08 | 0.34 | 0.05 | 0.02 | 0.03 | 0.04 |
| 3 | 157434826 | rs9829465 | -0.228 | 0.092 | -0.018 | 0.614 | -0.006 | -0.013 | -0.013 | 0 | 0.08 | 0.34 | 0.05 | 0.02 | 0.03 | 0.04 |
| 3 | 157435503 | rs9289992 | -0.228 | 0.092 | -0.018 | 0.614 | -0.006 | -0.013 | -0.013 | 0 | 0.08 | 0.34 | 0.05 | 0.02 | 0.03 | 0.04 |
| 3 | 157441860 | rs4680395 | -1.741 | 0.118 | -0.279 | 0.526 | -0.034 | 0.077 | 0.077 | 0.825 | 0.6 | 0.81 | 0.66 | 0.73 | 0.59 | 0.49 |
| 3 | 157451200 | rs10154880 | 0.991 | 0.053 | -0.085 | 0.761 | -0.005 | 0.002 | 0.002 | 0 | 0.17 | 0.11 | 0.27 | 0.15 | 0.2 | 0.18 |
| 3 | 157466626 | rs718144 | 0.937 | 0.017 | -0.252 | 0.941 | -0.002 | -0.002 | -0.002 | 0 | 0.1 | 0.09 | 0.08 | 0.17 | 0.13 | 0.06 |
| 3 | 157471663 | rs9837835 | -0.834 | -0.078 | -0.215 | 0.781 | 0 | -0.013 | -0.013 | 0 | 0.39 | 0.52 | 0.55 | 0.7 | 0.42 | 0.34 |
| 3 | 157474123 | rs1696007 | 0.235 | -0.46 | 0.054 | 0.942 | -0.025 | -0.012 | -0.012 | 2.765 | 0.73 | 0.59 | 0.84 | 0.74 | 0.8 | 0.82 |
| 3 | 157475101 | rs792742 | 0.972 | -0.451 | 0.425 | 0.997 | -0.015 | -0.003 | -0.003 | 0 | 0.57 | 0.45 | 0.58 | 0.58 | 0.62 | 0.67 |
| 3 | 157479716 | rs6797803 | 0.786 | -0.25 | 0.175 | 1.165 | 0.005 | 0.013 | 0.013 | 0 | 0.08 | 0.12 | 0.08 | 0.17 | 0.1 | 0.09 |
| 3 | 157493146 | rs703174 | -1.061 | -0.353 | 0.227 | 1.169 | 0.001 | -0.002 | -0.002 | 0.804 | 0.82 | 0.83 | 0.78 | 0.85 | 0.83 | 0.87 |
| 3 | 157493952 | rs4585149 | -0.931 | -0.378 | -0.07 | 1.072 | -0.008 | 0.001 | 0.001 | 0 | 0.84 | 0.87 | 0.87 | 0.83 | 0.81 | 0.9 |
| 3 | 157494573 | rs792739 | 1.291 | -0.292 | 0.126 | 0.951 | 0.001 | 0.011 | 0.011 | 0.057 | 0.52 | 0.47 | 0.39 | 0.26 | 0.51 | 0.55 |
| 3 | 157505518 | rs1994480 | 1.315 | -0.512 | 0.138 | 0.958 | 0.002 | 0.009 | 0.009 | 0.062 | 0.51 | 0.47 | 0.4 | 0.27 | 0.52 | 0.55 |
| 3 | 157514666 | rs16828327 | 0.02 | -0.856 | 0.227 | 1.156 | -0.002 | 0.003 | 0.003 | 0.009 | 0.18 | 0.23 | 0.26 | 0.19 | 0.19 | 0.18 |
| 3 | 157531347 | rs10513518 | 0.057 | -0.824 | 0.062 | 1.214 | -0.037 | 0.083 | 0.083 | 0 | 0.08 | 0.19 | 0.05 | 0.23 | 0.01 | 0.05 |
| 3 | 157536183 | rs16828352 | 0.244 | -0.642 | -0.223 | 1.114 | -0.001 | -0.003 | -0.003 | 0 | 0.16 | 0.16 | 0.26 | 0.17 | 0.19 | 0.22 |
| 3 | 157537988 | rs1877593 | 1.19 | -1.114 | -0.328 | 0.935 | 0.036 | -0.012 | -0.012 | 0 | 0.57 | 0.29 | 0.63 | 0.49 | 0.66 | 0.67 |
| 3 | 157538373 | rs2361516 | 1.524 | -1.362 | -0.263 | 1.229 | 0.055 | 0.123 | 0.123 | 0 | 0.48 | 0.21 | 0.4 | 0.27 | 0.58 | 0.53 |
| 3 | 157542303 | rs6794184 | 0.719 | -1.591 | -0.176 | 1.423 | 0.01 | -0.013 | -0.013 | 0.981 | 0.63 | 0.34 | 0.67 | 0.53 | 0.74 | 0.69 |
| 3 | 157545506 | rs6806806 | -1.188 | -1.661 | -0.224 | 1.416 | -0.057 | 0.108 | 0.108 | 0 | 0.23 | 0.63 | 0.26 | 0.46 | 0.12 | 0.22 |
| 3 | 157547513 | rs2137611 | 0.116 | -1.523 | -0.204 | 1.635 | -0.019 | 0.057 | 0.057 | 0 | 0.1 | 0.27 | 0.06 | 0.17 | 0.06 | 0.11 |
| 3 | 157553448 | rs1515473 | 0.995 | -1.139 | -0.922 | 1.51 | -0.006 | -0.073 | -0.073 | 0 | 0.11 | 0.01 | 0.23 | 0.27 | 0.13 | 0.15 |
| 3 | 157559512 | rs1399759 | -0.003 | -0.697 | 0.006 | 1.502 | -0.031 | 0.07 | 0.07 | 0.458 | 0.07 | 0.18 | 0.04 | 0.16 | 0.01 | 0.03 |
| 3 | 157567963 | rs953427 | -0.105 | -1.025 | -0.549 | 1.452 | 0.004 | -0.01 | -0.01 | 0.56 | 0.65 | 0.68 | 0.49 | 0.55 | 0.64 | 0.68 |
| 3 | 157570503 | rs16828412 | -0.281 | -1.334 | -0.451 | 1.385 | -0.061 | 0.032 | 0.032 | 0 | 0.25 | 0.53 | 0.15 | 0.31 | 0.14 | 0.16 |
| 3 | 157570753 | rs10936110 | 0.23 | -1.459 | -0.164 | 1.413 | -0.056 | -0.003 | -0.003 | 0.075 | 0.19 | 0.37 | 0.1 | 0.16 | 0.08 | 0.13 |
| 3 | 157570795 | rs2056123 | -0.701 | -1.369 | -0.533 | 1.355 | 0.007 | 0.008 | 0.008 | 0 | 0.66 | 0.75 | 0.69 | 0.67 | 0.65 | 0.71 |
| 3 | 157572952 | rs11713033 | 0.971 | -1.534 | -1.27 | 1.233 | 0 | -0.013 | -0.013 | 0 | 0.14 | 0.13 | 0.19 | 0.31 | 0.18 | 0.15 |

| | | | | | | | | | | | | | | | | |
|---|---|---|---|---|---|---|---|---|---|---|---|---|---|---|---|---|
| 3 | 157574195 | rs7622880 | 1.511 | -1.507 | -1.302 | 1.05 | -0.028 | 0.066 | 0.066 | 0 | 0.4 | 0.2 | 0.34 | 0.24 | 0.49 | 0.52 |
| 3 | 157578673 | rs4680401 | 0.259 | -1.517 | -1.194 | 1.227 | 0 | -0.028 | -0.028 | 0.388 | 0.2 | 0.18 | 0.24 | 0.45 | 0.25 | 0.19 |
| 3 | 157579469 | rs16828444 | 0.311 | -1.743 | -1.444 | 0.886 | 0.054 | -0.019 | -0.019 | 2.21 | 0.2 | 0.03 | 0.06 | 0.17 | 0.1 | 0.13 |
| 3 | 157579641 | rs7640829 | 1.304 | -0.944 | -1.313 | 0.739 | -0.049 | 0.026 | 0.026 | 0 | 0.36 | 0.19 | 0.34 | 0.29 | 0.47 | 0.48 |
| 3 | 157581467 | rs9841945 | -0.317 | -0.743 | -1.545 | 0.657 | -0.001 | 0.067 | 0.067 | 0 | 0.66 | 0.82 | 0.72 | 0.77 | 0.65 | 0.67 |
| 3 | 157582078 | rs9864196 | 0.228 | -1.503 | -1.97 | 0.731 | 0.092 | -0.095 | -0.095 | 0.079 | 0.24 | 0.04 | 0.23 | 0.32 | 0.12 | 0.19 |
| 3 | 157586847 | rs10936111 | 0.685 | -0.474 | -1.811 | 0.228 | 0.007 | 0.01 | 0.01 | 0 | 0.31 | 0.39 | 0.39 | 0.58 | 0.42 | 0.45 |
| 3 | 157591106 | rs17618255 | 0.538 | -0.327 | -1.479 | 0.099 | 0.008 | 0.001 | 0.001 | 0 | 0.07 | 0.08 | 0.07 | 0.13 | 0.12 | 0.02 |
| 3 | 157593238 | rs9290004 | -0.097 | -0.551 | -1.632 | 0.024 | 0.006 | 0.017 | 0.017 | 0 | 0.48 | 0.49 | 0.37 | 0.28 | 0.39 | 0.36 |
| 3 | 157594704 | rs13078073 | 1.618 | 0.065 | -1.41 | 0.392 | 0 | 0.006 | 0.006 | 0 | 0.14 | 0.15 | 0.19 | 0.35 | 0.19 | 0.17 |
| 3 | 157608860 | rs9859533 | 1.263 | 0.562 | -1.14 | 0.395 | 0.03 | 0.082 | 0.082 | 0 | 0.16 | 0.32 | 0.32 | 0.36 | 0.21 | 0.3 |
| 3 | 157609802 | rs1515464 | 0.763 | 0.556 | -1.069 | -0.301 | 0.046 | -0.077 | -0.077 | 0 | 0.26 | 0.44 | 0.25 | 0.15 | 0.3 | 0.14 |
| 3 | 157643014 | rs2673549 | -2.248 | 0.183 | -1.579 | -0.3 | 0.011 | 0.059 | 0.059 | 0 | 0.84 | 0.95 | 0.92 | 1 | 0.86 | 0.86 |
| 3 | 157653791 | rs10936114 | -0.526 | -0.052 | -1.514 | -0.477 | 0.007 | -0.009 | -0.009 | 0.169 | 0.63 | 0.6 | 0.55 | 0.47 | 0.62 | 0.65 |
| 3 | 157672947 | rs9825122 | -1.466 | 0.105 | -1.502 | -0.463 | 0 | -0.003 | -0.003 | 0 | 0.8 | 0.79 | 0.63 | 0.47 | 0.75 | 0.8 |
| 3 | 157675069 | rs9290009 | 1.68 | -0.335 | -1.911 | -0.947 | 0.033 | -0.016 | -0.016 | 0.552 | 0.19 | 0.01 | 0.15 | 0.14 | 0.11 | 0.17 |
| 3 | 157697874 | rs4387925 | 0.123 | -0.028 | -1.37 | -1.029 | -0.028 | 0 | 0 | 0 | 0.4 | 0.27 | 0.37 | 0.41 | 0.43 | 0.51 |
| 3 | 157705221 | rs7642284 | 1.726 | 0.081 | -1.195 | -1.017 | 0.022 | -0.132 | -0.132 | 0.01 | 0.18 | 0.38 | 0.16 | 0.02 | 0.18 | 0.14 |
| 3 | 157723257 | rs9866741 | 1.069 | 0.218 | -1.607 | -0.939 | -0.009 | -0.012 | -0.012 | 0 | 0.23 | 0.07 | 0.2 | 0.24 | 0.22 | 0.24 |
| 3 | 157746923 | rs17681602 | 1.507 | -0.146 | -1.322 | -0.795 | 0.004 | 0.133 | 0.133 | 0.288 | 0.2 | 0.01 | 0.08 | 0 | 0.15 | 0.14 |
| 3 | 157794275 | rs11924696 | 1.667 | -0.071 | -0.84 | -0.84 | 0.014 | -0.091 | -0.091 | 0 | 0.16 | 0.25 | 0.13 | 0.02 | 0.15 | 0.14 |
| 3 | 157804409 | rs4679819 | -0.048 | 0.03 | -1.684 | -0.8 | -0.002 | -0.005 | -0.005 | 0.834 | 0.38 | 0.69 | 0.52 | 0.59 | 0.38 | 0.35 |
| 3 | 157813401 | rs1123115 | 0.85 | 0.029 | -1.863 | -0.133 | -0.017 | -0.085 | -0.085 | 1.82 | 0.19 | 0.08 | 0.35 | 0.57 | 0.24 | 0.2 |
| 3 | 157815088 | rs6441173 | -1.075 | -0.033 | -1.44 | -0.485 | 0.009 | 0.133 | 0.133 | 0.515 | 0.8 | 0.98 | 0.92 | 1 | 0.85 | 0.86 |
| 3 | 157815380 | rs1608114 | -1.062 | -0.032 | -1.434 | -0.484 | 0.025 | 0.102 | 0.102 | 0.523 | 0.8 | 0.94 | 0.91 | 0.99 | 0.85 | 0.84 |
| 3 | 157822669 | rs1915940 | -1.022 | -0.013 | -1.437 | -0.475 | 0.028 | 0.094 | 0.094 | 0.257 | 0.79 | 0.93 | 0.91 | 0.99 | 0.85 | 0.84 |
| 3 | 157828322 | rs11713363 | 0.194 | 0.524 | -0.433 | 0.307 | 0.003 | 0.104 | 0.104 | 0 | 0.22 | 0.42 | 0.36 | 0.57 | 0.24 | 0.22 |
| 3 | 157832974 | rs12696067 | -0.415 | 0.454 | -0.233 | -0.517 | 0 | -0.004 | -0.004 | 0 | 0.62 | 0.62 | 0.7 | 0.82 | 0.67 | 0.58 |
| 3 | 157845417 | rs4542974 | 1.642 | 0.686 | -0.823 | -0.829 | -0.04 | 0.047 | 0.047 | 0 | 0.19 | 0.02 | 0.2 | 0.09 | 0.18 | 0.2 |
| 3 | 157846181 | rs12485440 | -1.123 | 0.51 | -0.567 | -0.859 | -0.007 | 0.08 | 0.08 | 0 | 0.65 | 0.88 | 0.72 | 0.82 | 0.67 | 0.59 |
| 3 | 157851223 | rs7624303 | 1.522 | 0.237 | -0.776 | -1.077 | -0.032 | 0.018 | 0.018 | 0 | 0.24 | 0.06 | 0.23 | 0.18 | 0.24 | 0.26 |
| 3 | 157877607 | rs13086348 | -0.02 | 0.205 | -0.32 | -1.19 | 0.005 | 0.019 | 0.019 | 0 | 0.57 | 0.62 | 0.66 | 0.69 | 0.61 | 0.46 |
| 3 | 157887272 | rs9857883 | 0.885 | 0.216 | 0.192 | -1.264 | -0.015 | 0.007 | 0.007 | 0 | 0.36 | 0.22 | 0.43 | 0.63 | 0.44 | 0.26 |
| 3 | 157924373 | rs852110 | 0.075 | 0.212 | -0.011 | -1.254 | -0.008 | -0.03 | -0.03 | 0.199 | 0.57 | 0.5 | 0.64 | 0.71 | 0.62 | 0.47 |
| 3 | 157934971 | rs4680421 | 1.608 | 0.205 | -0.377 | -1.263 | -0.008 | 0.023 | 0.023 | 0 | 0.25 | 0.15 | 0.23 | 0.17 | 0.24 | 0.27 |
| 3 | 157935626 | rs10513524 | 1.011 | 0.206 | -0.338 | -1.226 | -0.005 | 0.006 | 0.006 | 0 | 0.06 | 0.1 | 0.05 | 0.12 | 0.06 | 0.13 |
| 3 | 157944984 | rs2362248 | 1.265 | 0.213 | -0.335 | -1.224 | -0.003 | 0.007 | 0.007 | 0 | 0.06 | 0.1 | 0.05 | 0.12 | 0.06 | 0.13 |
| 3 | 157989979 | rs1213048 | 0.524 | 0.2 | -0.343 | -1.284 | -0.001 | 0 | 0 | 0 | 0.39 | 0.6 | 0.47 | 0.62 | 0.44 | 0.28 |
| 3 | 157992363 | rs1851062 | 1.718 | 0.178 | -0.363 | -1.316 | -0.023 | 0.031 | 0.031 | 0 | 0.24 | 0.09 | 0.23 | 0.17 | 0.24 | 0.26 |
| 3 | 157992814 | rs7643792 | 0.481 | 0.27 | -0.321 | -1.225 | -0.001 | 0 | 0 | 0.979 | 0.39 | 0.6 | 0.47 | 0.62 | 0.44 | 0.28 |
| 3 | 157993838 | rs2682405 | 0.481 | 0.27 | -0.321 | -1.225 | -0.001 | 0 | 0 | 0 | 0.39 | 0.6 | 0.47 | 0.62 | 0.44 | 0.28 |
| 3 | 158001555 | rs2693540 | -0.919 | 0.298 | -0.417 | -1.165 | 0.004 | 0.062 | 0.062 | 0 | 0.64 | 0.78 | 0.72 | 0.83 | 0.67 | 0.61 |
| 3 | 158007807 | rs1210359 | 1.002 | 0.575 | -0.646 | -1.01 | -0.035 | -0.044 | -0.044 | 1.294 | 0.24 | 0.08 | 0.32 | 0.62 | 0.32 | 0.24 |
| 3 | 158029931 | rs10513525 | 0.8 | 0.35 | -0.432 | -0.733 | -0.003 | 0.008 | 0.008 | 0 | 0.06 | 0.11 | 0.06 | 0.12 | 0.06 | 0.13 |
| 3 | 158036690 | rs17683861 | 0.822 | 0.305 | -0.535 | -0.865 | 0.003 | 0.018 | 0.018 | 0 | 0.25 | 0.18 | 0.26 | 0.22 | 0.23 | 0.33 |
| 3 | 158054568 | rs827183 | -2.294 | 0.217 | -0.594 | -0.944 | 0.009 | 0.016 | 0.016 | 0.403 | 0.9 | 0.95 | 0.95 | 1 | 0.91 | 0.88 |
| 3 | 158062256 | rs4411841 | 0.986 | 0.175 | -0.574 | -0.835 | -0.003 | 0.007 | 0.007 | 0 | 0.06 | 0.08 | 0.06 | 0.13 | 0.06 | 0.13 |
| 3 | 158070858 | rs10936131 | 1.563 | 0.19 | -0.758 | -1.01 | -0.034 | 0.039 | 0.039 | 0 | 0.24 | 0.06 | 0.23 | 0.15 | 0.24 | 0.27 |
| 3 | 158092801 | rs6774461 | -1.174 | 0.035 | -0.279 | -1.112 | -0.017 | 0.001 | 0.001 | 0 | 0.64 | 0.78 | 0.57 | 0.39 | 0.56 | 0.72 |
| 3 | 158104706 | rs7648196 | -0.487 | -0.056 | -0.085 | -1.096 | -0.002 | 0.003 | 0.003 | 0 | 0.55 | 0.61 | 0.52 | 0.38 | 0.47 | 0.6 |
| 3 | 158110709 | rs7640075 | 1.61 | 0.073 | -0.154 | -1.045 | -0.034 | 0.044 | 0.044 | 0 | 0.24 | 0.06 | 0.23 | 0.15 | 0.24 | 0.27 |
| 3 | 158115253 | rs939114 | 0.005 | 0.061 | -0.105 | -1.073 | -0.012 | 0.11 | 0.11 | 0 | 0.49 | 0.21 | 0.48 | 0.34 | 0.47 | 0.6 |
| 3 | 158172133 | rs12637243 | 1.841 | -0.183 | -0.122 | -0.968 | -0.006 | 0.05 | 0.05 | 0 | 0.19 | 0.08 | 0.19 | 0.07 | 0.16 | 0.19 |
| 3 | 158172405 | rs10936142 | 0.618 | 0.155 | -0.163 | -1.06 | -0.01 | 0.13 | 0.13 | 0 | 0.43 | 0.13 | 0.41 | 0.21 | 0.37 | 0.46 |
| 3 | 158178245 | rs6806967 | 1.628 | 0.216 | -0.249 | -0.998 | -0.007 | 0.061 | 0.061 | 0 | 0.23 | 0.06 | 0.22 | 0.11 | 0.21 | 0.26 |
| 3 | 158190313 | rs7649463 | -0.412 | 0.255 | -0.087 | -0.938 | -0.002 | 0.028 | 0.028 | 0 | 0.53 | 0.66 | 0.5 | 0.36 | 0.45 | 0.61 |
| 3 | 158198722 | rs6769314 | 1.988 | 0.138 | 0.244 | -1.027 | -0.001 | -0.012 | -0.012 | 0.967 | 0.19 | 0.22 | 0.21 | 0.1 | 0.17 | 0.19 |
| 3 | 158203468 | rs10936146 | -0.508 | 0.154 | 0.099 | -1.019 | -0.002 | 0.027 | 0.027 | 0 | 0.53 | 0.66 | 0.5 | 0.36 | 0.45 | 0.61 |
| 3 | 158206114 | rs11710813 | 1.713 | 0.111 | 0.005 | -1.059 | -0.015 | 0.058 | 0.058 | 0 | 0.23 | 0.06 | 0.22 | 0.11 | 0.23 | 0.25 |
| 3 | 158236860 | rs2117764 | 0.481 | 0.173 | 0.079 | -0.669 | 0.001 | 0.015 | 0.015 | 0 | 0.32 | 0.21 | 0.28 | 0.31 | 0.29 | 0.41 |
| 3 | 158239143 | rs9849688 | 1.653 | 0.214 | 0.184 | -0.615 | -0.014 | -0.055 | -0.055 | 0 | 0.2 | 0.43 | 0.21 | 0.1 | 0.16 | 0.2 |
| 3 | 158247867 | rs16829064 | 0.58 | 0.314 | 0.016 | -0.495 | -0.002 | 0 | 0 | 0 | 0.08 | 0.08 | 0.05 | 0.14 | 0.06 | 0.13 |
| 3 | 158270299 | rs1630524 | 0.129 | 0.15 | 0.167 | -0.662 | 0 | 0.013 | 0.013 | 0.019 | 0.52 | 0.49 | 0.47 | 0.27 | 0.48 | 0.55 |
| 3 | 158273096 | rs1714524 | -0.683 | 0.195 | 0.079 | -0.626 | -0.015 | 0.004 | 0.004 | 0.273 | 0.62 | 0.77 | 0.54 | 0.41 | 0.54 | 0.67 |
| 3 | 158284861 | rs1714521 | 0.072 | 0.07 | 0.061 | -0.411 | 0.015 | 0.025 | 0.025 | 0 | 0.4 | 0.31 | 0.33 | 0.36 | 0.38 | 0.46 |
| 3 | 158285175 | rs1714520 | 0.271 | 0.154 | -0.132 | -0.692 | 0.005 | 0.034 | 0.034 | 0 | 0.21 | 0.17 | 0.16 | 0.1 | 0.21 | 0.25 |
| 3 | 158285280 | rs1656377 | -0.811 | 0.504 | -0.044 | -0.845 | -0.007 | 0.003 | 0.003 | 0.274 | 0.62 | 0.74 | 0.55 | 0.38 | 0.58 | 0.69 |
| 3 | 158286742 | rs7638518 | 1.869 | 0.463 | 0.112 | -0.784 | -0.005 | -0.05 | -0.05 | 0.068 | 0.21 | 0.32 | 0.21 | 0.06 | 0.17 | 0.23 |
| 3 | 158301682 | rs10513527 | 2.174 | 0.47 | 0.113 | -0.778 | 0 | -0.016 | -0.016 | 0 | 0.19 | 0.22 | 0.18 | 0.06 | 0.16 | 0.23 |

| | | | | | | | | | | | | | | | | |
|---|---|---|---|---|---|---|---|---|---|---|---|---|---|---|---|---|
| 3 | 158312667 | rs16829122 | 1.783 | 0.444 | -0.095 | -0.854 | -0.014 | -0.039 | -0.039 | 0 | 0.16 | 0.04 | 0.19 | 0.2 | 0.16 | 0.21 |
| 3 | 158313003 | rs1435643 | 0.2 | 0.573 | -0.359 | -1.096 | -0.009 | 0.011 | 0.011 | 0 | 0.35 | 0.26 | 0.33 | 0.33 | 0.35 | 0.36 |
| 3 | 158314188 | rs6804230 | 1.068 | 0.592 | -0.387 | -1.065 | -0.024 | 0.036 | 0.036 | 0 | 0.18 | 0.07 | 0.13 | 0.13 | 0.2 | 0.15 |
| 3 | 158317068 | rs12107742 | 2.009 | 0.402 | -0.215 | -1.014 | -0.002 | -0.032 | -0.032 | 0.608 | 0.21 | 0.29 | 0.19 | 0.06 | 0.16 | 0.23 |
| 3 | 158320597 | rs4875 | -0.582 | 0.586 | -0.193 | -1.091 | -0.001 | 0.015 | 0.015 | 0 | 0.58 | 0.76 | 0.53 | 0.38 | 0.54 | 0.6 |
| 3 | 158328274 | rs13315529 | 1.812 | 0.586 | -0.102 | -1.026 | -0.015 | -0.111 | -0.111 | 0.045 | 0.22 | 0.5 | 0.19 | 0.06 | 0.16 | 0.23 |
| 3 | 158342928 | rs2566332 | 0.972 | 0.638 | -0.267 | -0.976 | -0.001 | -0.001 | -0.001 | 0.275 | 0.32 | 0.25 | 0.39 | 0.62 | 0.33 | 0.29 |
| 3 | 158362038 | rs3806642 | 2.331 | 0.606 | -0.209 | -0.977 | 0 | -0.018 | -0.018 | 0 | 0.19 | 0.22 | 0.18 | 0.06 | 0.16 | 0.23 |
| 3 | 158364309 | rs2303910 | 1.802 | 0.638 | -0.982 | -0.313 | -0.035 | -0.109 | -0.109 | 0.239 | 0.14 | 0 | 0.23 | 0.38 | 0.16 | 0.16 |
| 3 | 158366900 | rs2303909 | -0.467 | 0.588 | -0.731 | -0.563 | -0.002 | 0.017 | 0.017 | 0 | 0.58 | 0.77 | 0.53 | 0.38 | 0.54 | 0.61 |
| 3 | 158368524 | rs12639052 | -0.483 | 0.592 | -0.73 | -0.111 | -0.002 | 0.01 | 0.01 | 0 | 0.58 | 0.77 | 0.66 | 0.58 | 0.55 | 0.62 |
| 3 | 158379649 | rs9873667 | 0.44 | 0.613 | -0.694 | -1.54 | -0.001 | -0.012 | -0.012 | 0 | 0.41 | 0.64 | 0.34 | 0.19 | 0.39 | 0.4 |
| 3 | 158388780 | rs8455 | -0.324 | 0.552 | -0.727 | -1.58 | 0 | 0.02 | 0.02 | 0 | 0.57 | 0.75 | 0.53 | 0.38 | 0.54 | 0.61 |
| 3 | 158404267 | rs10936158 | -0.324 | 0.552 | -0.727 | -1.58 | 0 | 0.02 | 0.02 | 0 | 0.57 | 0.75 | 0.53 | 0.38 | 0.54 | 0.61 |
| 3 | 158415344 | rs1051109 | 1.679 | 0.738 | -0.994 | -1.315 | 0.001 | -0.011 | -0.011 | 0 | 0.17 | 0.13 | 0.19 | 0.2 | 0.16 | 0.21 |
| 3 | 158415840 | rs3867391 | 1.006 | 0.766 | -0.984 | -1.371 | -0.022 | 0.033 | 0.033 | 0 | 0.19 | 0.11 | 0.14 | 0.13 | 0.22 | 0.15 |
| 3 | 158421824 | rs12488447 | 1.659 | 0.72 | -0.875 | -1.141 | -0.001 | -0.011 | -0.011 | 0 | 0.16 | 0.11 | 0.19 | 0.2 | 0.16 | 0.21 |
| 3 | 158424331 | rs13074694 | -1.032 | 0.994 | -0.5 | -0.966 | -0.004 | -0.032 | -0.032 | 0 | 0.79 | 0.9 | 0.71 | 0.38 | 0.78 | 0.74 |
| 3 | 158427059 | rs17642880 | 2.277 | 1.206 | -0.589 | -1.332 | -0.008 | 0.016 | 0.016 | 1.951 | 0.09 | 0.02 | 0.06 | 0 | 0.08 | 0.02 |
| 3 | 158428642 | rs2307064 | 0.541 | 1.214 | -0.503 | -1.433 | -0.007 | 0.015 | 0.015 | 0 | 0.2 | 0.12 | 0.14 | 0.15 | 0.24 | 0.16 |
| 3 | 158429009 | rs9840571 | -0.143 | 1.146 | -0.222 | -1.541 | 0 | -0.007 | -0.007 | 0 | 0.58 | 0.5 | 0.54 | 0.4 | 0.54 | 0.63 |
| 3 | 158430706 | rs9864589 | 2.47 | 0.917 | -0.347 | -1.592 | 0.007 | 0.037 | 0.037 | 0 | 0.21 | 0.16 | 0.2 | 0.05 | 0.16 | 0.25 |
| 3 | 158434137 | rs9847482 | 1.081 | 1.016 | -0.33 | -1.54 | 0.004 | 0.016 | 0.016 | 0 | 0.37 | 0.34 | 0.39 | 0.25 | 0.31 | 0.47 |
| 3 | 158435374 | rs9825140 | 1.91 | 1.101 | -0.394 | -1.513 | -0.01 | -0.025 | -0.025 | 0.16 | 0.14 | 0.05 | 0.18 | 0.2 | 0.15 | 0.21 |
| 3 | 158453279 | rs7646881 | 1.816 | 0.924 | -0.113 | -1.48 | -0.003 | -0.056 | -0.056 | 4.699 | 0.24 | 0.3 | 0.13 | 0.02 | 0.19 | 0.24 |
| 3 | 158459976 | rs7635993 | -0.452 | 0.826 | -0.16 | -1.391 | -0.005 | -0.001 | -0.001 | 0 | 0.65 | 0.78 | 0.48 | 0.37 | 0.6 | 0.65 |
| 3 | 158463046 | rs9861938 | -0.13 | 0.757 | 0.071 | -1.42 | -0.003 | -0.007 | -0.007 | 0.003 | 0.62 | 0.51 | 0.45 | 0.37 | 0.6 | 0.63 |
| 3 | 158463101 | rs7617304 | 1.629 | 0.771 | 0.026 | -1.576 | -0.002 | -0.049 | -0.049 | 0 | 0.31 | 0.32 | 0.14 | 0.02 | 0.22 | 0.24 |
| 3 | 158470321 | rs6441225 | -0.109 | 0.787 | -0.368 | -1.477 | 0.002 | 0 | 0 | 0.443 | 0.33 | 0.37 | 0.32 | 0.35 | 0.36 | 0.4 |
| 3 | 158477796 | rs9861289 | 1.409 | 0.875 | -0.552 | -1.505 | -0.019 | -0.017 | -0.017 | 1.91 | 0.19 | 0.07 | 0.2 | 0.19 | 0.21 | 0.24 |
| 3 | 158484933 | rs17700475 | 0.969 | 0.846 | -0.465 | -1.359 | 0.013 | -0.002 | -0.002 | 1.147 | 0.12 | 0.16 | 0.11 | 0.15 | 0.15 | 0.15 |
| 3 | 158486700 | rs3863076 | 1.021 | 0.861 | -0.555 | -1.353 | 0.003 | -0.001 | -0.001 | 0 | 0.12 | 0.12 | 0.11 | 0.15 | 0.16 | 0.15 |
| 3 | 158489641 | rs1560418 | 0.647 | 0.833 | -0.646 | -1.44 | -0.005 | 0.006 | 0.006 | 0.018 | 0.17 | 0.12 | 0.15 | 0.15 | 0.22 | 0.15 |
| 3 | 158490420 | rs1346834 | -0.731 | 0.828 | -0.585 | -1.433 | -0.001 | -0.001 | -0.001 | 0.236 | 0.7 | 0.73 | 0.5 | 0.38 | 0.66 | 0.66 |
| 3 | 158503232 | rs7616425 | 0.28 | 0.712 | -0.559 | -1.466 | -0.001 | 0.017 | 0.017 | 0 | 0.53 | 0.63 | 0.37 | 0.23 | 0.45 | 0.5 |
| 3 | 158515499 | rs2713734 | 1.619 | -0.045 | -0.542 | -2.223 | -0.003 | -0.032 | -0.032 | 0 | 0.11 | 0.1 | 0.07 | 0 | 0.11 | 0.08 |
| 3 | 158518963 | rs1381462 | 0.89 | -0.209 | -0.512 | -2.013 | -0.001 | -0.004 | -0.004 | 0.161 | 0.19 | 0.17 | 0.2 | 0.08 | 0.18 | 0.17 |
| 3 | 158523819 | rs2061617 | 1.174 | -0.156 | -0.703 | -2.129 | 0.013 | 0.03 | 0.03 | 0 | 0.31 | 0.26 | 0.16 | 0.05 | 0.23 | 0.25 |
| 3 | 158524569 | rs10513528 | 0.938 | 0.088 | -0.752 | -1.83 | 0.007 | 0.01 | 0.01 | 0 | 0.11 | 0.12 | 0.13 | 0.19 | 0.16 | 0.11 |
| 3 | 158525574 | rs3816637 | 0.455 | -0.133 | -0.892 | -1.887 | -0.026 | 0.102 | 0.102 | 0 | 0.27 | 0.04 | 0.23 | 0.08 | 0.25 | 0.19 |
| 3 | 158526223 | rs6441226 | 1.051 | -0.095 | -0.81 | -1.863 | 0.007 | 0.023 | 0.023 | 0 | 0.11 | 0.18 | 0.11 | 0.19 | 0.12 | 0.11 |
| 3 | 158527448 | rs12497958 | 0.709 | -0.089 | -0.831 | -1.858 | 0.007 | 0.05 | 0.05 | 0 | 0.12 | 0.26 | 0.12 | 0.21 | 0.12 | 0.13 |
| 3 | 158529117 | rs12493280 | -0.404 | -0.09 | -0.796 | -1.945 | -0.001 | 0.015 | 0.015 | 0 | 0.38 | 0.3 | 0.35 | 0.29 | 0.37 | 0.31 |
| 3 | 158537443 | rs3765083 | -0.636 | -0.013 | -0.559 | -1.647 | -0.002 | -0.003 | -0.003 | 0 | 0.7 | 0.72 | 0.53 | 0.37 | 0.61 | 0.5 |
| 3 | 158539294 | rs1548092 | 0.28 | -0.077 | -0.508 | -1.768 | 0.012 | 0.082 | 0.082 | 0 | 0.48 | 0.36 | 0.35 | 0.15 | 0.4 | 0.4 |
| 3 | 158545195 | rs6792827 | 1.416 | -0.162 | -0.509 | -1.725 | -0.004 | -0.04 | -0.04 | 0 | 0.26 | 0.3 | 0.14 | 0.05 | 0.17 | 0.23 |
| 3 | 158550968 | rs2364910 | -0.226 | -0.077 | -0.496 | -1.669 | -0.001 | 0.004 | 0.004 | 0 | 0.46 | 0.42 | 0.41 | 0.34 | 0.44 | 0.31 |
| 3 | 158554978 | rs1870829 | -0.494 | 0.289 | -0.457 | -1.24 | -0.004 | -0.006 | -0.006 | 0 | 0.7 | 0.75 | 0.54 | 0.38 | 0.61 | 0.55 |
| 3 | 158562133 | rs6778714 | -0.542 | 0.665 | -0.438 | -0.675 | -0.002 | -0.001 | -0.001 | 0.24 | 0.34 | 0.33 | 0.36 | 0.33 | 0.37 | 0.29 |
| 3 | 158567174 | rs1503100 | 0.074 | 0.727 | -0.504 | -0.619 | -0.001 | 0 | 0 | 0 | 0.19 | 0.19 | 0.17 | 0.19 | 0.21 | 0.12 |
| 3 | 158568590 | rs340313 | -0.347 | 0.367 | -0.022 | -1.181 | 0.006 | -0.033 | -0.033 | 0.87 | 0.3 | 0.34 | 0.31 | 0.14 | 0.36 | 0.31 |
| 3 | 158570108 | rs340309 | 0.537 | 0.422 | 0.102 | -1.09 | 0.027 | -0.028 | -0.028 | 0 | 0.13 | 0.22 | 0.14 | 0.06 | 0.18 | 0.11 |
| 3 | 158578044 | rs187345 | 0.378 | 0.329 | 0.24 | -1.136 | 0.017 | -0.037 | -0.037 | 0 | 0.15 | 0.26 | 0.14 | 0.06 | 0.18 | 0.11 |
| 3 | 158578714 | rs17645548 | 0.569 | 0.263 | -0.267 | -1.327 | -0.015 | -0.017 | -0.017 | 0.257 | 0.2 | 0.15 | 0.37 | 0.76 | 0.25 | 0.44 |
| 3 | 158583584 | rs340284 | -0.964 | 0.026 | 0.016 | -1.348 | 0.001 | -0.014 | -0.014 | 0 | 0.52 | 0.48 | 0.64 | 0.88 | 0.59 | 0.67 |
| 3 | 158585531 | rs340280 | -1.514 | -0.127 | 0.097 | -1.446 | -0.013 | 0.038 | 0.038 | 0 | 0.65 | 0.77 | 0.76 | 0.99 | 0.64 | 0.72 |
| 3 | 158592935 | rs340271 | -0.501 | -0.054 | 0.115 | -1.29 | 0 | -0.002 | -0.002 | 0 | 0.43 | 0.44 | 0.49 | 0.73 | 0.44 | 0.52 |
| 3 | 158593799 | rs4679840 | 1.116 | -0.09 | -0.259 | -1.309 | -0.009 | -0.02 | -0.02 | 1.96 | 0.15 | 0.05 | 0.18 | 0.15 | 0.16 | 0.15 |
| 3 | 158594051 | rs340270 | -1.38 | -0.094 | -0.226 | -1.302 | -0.004 | 0.038 | 0.038 | 0 | 0.61 | 0.71 | 0.7 | 0.92 | 0.6 | 0.7 |
| 3 | 158594656 | rs340268 | -1.361 | -0.061 | -0.212 | -1.325 | -0.002 | 0.023 | 0.023 | 0 | 0.61 | 0.67 | 0.69 | 0.89 | 0.6 | 0.7 |
| 3 | 158595198 | rs10936163 | -0.461 | -0.073 | -0.273 | -1.342 | -0.005 | -0.029 | -0.029 | 0 | 0.35 | 0.51 | 0.34 | 0.21 | 0.34 | 0.28 |
| 3 | 158596415 | rs12494894 | -0.474 | 0.367 | -1.181 | -1.158 | -0.011 | 0.085 | 0.085 | 0 | 0.41 | 0.25 | 0.54 | 0.83 | 0.41 | 0.53 |
| 3 | 158599864 | rs340251 | 0.315 | 0.557 | -1.142 | -0.65 | 0.002 | -0.027 | -0.027 | 0.037 | 0.16 | 0.18 | 0.13 | 0.06 | 0.18 | 0.13 |
| 3 | 158601397 | rs7630650 | -0.419 | 0.53 | -1.472 | 0.102 | -0.018 | -0.066 | -0.066 | 0 | 0.25 | 0.1 | 0.34 | 0.64 | 0.25 | 0.37 |
| 3 | 158613419 | rs1393757 | -0.439 | 0.784 | -1.509 | 0.162 | -0.044 | -0.079 | -0.079 | 0 | 0.21 | 0.05 | 0.35 | 0.66 | 0.23 | 0.39 |
| 3 | 158614804 | rs10513532 | 0.07 | 0.572 | -1.521 | -0.079 | 0.005 | -0.012 | -0.012 | 0.556 | 0.13 | 0.08 | 0.09 | 0.15 | 0.1 | 0.14 |
| 3 | 158616233 | rs6769725 | -0.31 | 0.377 | -0.742 | -0.045 | -0.014 | 0.037 | 0.037 | 0 | 0.14 | 0.26 | 0.13 | 0.21 | 0.1 | 0.15 |
| 3 | 158624462 | rs6794838 | -0.432 | 0.326 | -0.416 | -0.013 | -0.016 | 0.06 | 0.06 | 0 | 0.15 | 0.34 | 0.13 | 0.22 | 0.1 | 0.15 |
| 3 | 158624804 | rs414357 | -0.302 | 0.585 | -0.377 | -0.941 | -0.012 | 0.008 | 0.008 | 0 | 0.38 | 0.27 | 0.62 | 0.7 | 0.53 | 0.48 |

| | | | | | | | | | | | | | | | | |
|---|---|---|---|---|---|---|---|---|---|---|---|---|---|---|---|---|
| 3 | 158625596 | rs9833196 | 1.249 | 0.861 | -0.317 | -1.13 | -0.001 | -0.068 | -0.068 | 0 | 0.13 | 0.17 | 0.08 | 0 | 0.13 | 0.02 |
| 3 | 158626899 | rs340233 | -1.612 | 0.226 | -0.623 | -1.669 | 0.008 | 0.012 | 0.012 | 0 | 0.66 | 0.72 | 0.83 | 0.94 | 0.72 | 0.71 |
| 3 | 158626971 | rs340232 | -1.457 | 0.187 | -0.599 | -1.68 | -0.001 | -0.034 | -0.034 | 0 | 0.64 | 0.61 | 0.82 | 0.94 | 0.72 | 0.71 |
| 3 | 158628264 | rs340228 | 1.338 | 0.419 | -0.845 | -1.66 | -0.029 | 0.011 | 0.011 | 0 | 0.1 | 0.05 | 0.23 | 0.06 | 0.2 | 0.12 |
| 3 | 158633726 | rs12496761 | 0.332 | 0.173 | -0.673 | -1.69 | -0.004 | -0.009 | -0.009 | 0 | 0.26 | 0.25 | 0.17 | 0.17 | 0.23 | 0.16 |
| 3 | 158634689 | rs340260 | 0.356 | 0.297 | -0.599 | -1.596 | -0.031 | -0.018 | -0.018 | 0 | 0.25 | 0.13 | 0.54 | 0.69 | 0.4 | 0.46 |
| 3 | 158636057 | rs340261 | -1.133 | 0.181 | -0.561 | -1.495 | -0.008 | -0.063 | -0.063 | 0 | 0.64 | 0.57 | 0.81 | 0.94 | 0.72 | 0.71 |
| 3 | 158650962 | rs6768619 | 0.972 | 0.292 | -0.824 | -1.203 | 0.007 | -0.018 | -0.018 | 0.127 | 0.1 | 0.05 | 0.07 | 0.15 | 0.07 | 0.11 |
| 3 | 158651538 | rs340295 | -0.134 | 1.003 | -1.363 | -0.818 | -0.049 | -0.054 | -0.054 | 0 | 0.76 | 0.85 | 0.61 | 0.36 | 0.68 | 0.55 |
| 3 | 158653634 | rs1604643 | 1.581 | 0.634 | -0.929 | -1.481 | 0 | -0.024 | -0.024 | 0 | 0.13 | 0.11 | 0.08 | 0 | 0.12 | 0.02 |
| 3 | 158661559 | rs340241 | -1.044 | 0.64 | -0.442 | -1.655 | -0.012 | 0.034 | 0.034 | 0 | 0.56 | 0.65 | 0.5 | 0.21 | 0.54 | 0.34 |
| 3 | 158664527 | rs10513530 | 0.108 | 0.526 | -0.466 | -1.78 | -0.024 | -0.002 | -0.002 | 0 | 0.24 | 0.13 | 0.29 | 0.2 | 0.3 | 0.23 |
| 3 | 158664659 | rs11712844 | 0.369 | 0.515 | -0.692 | -1.782 | -0.089 | -0.028 | -0.028 | 0 | 0.23 | 0.02 | 0.28 | 0.16 | 0.3 | 0.22 |
| 3 | 158665225 | rs11707952 | -0.721 | 0.468 | -0.651 | -1.804 | -0.001 | -0.019 | -0.019 | 0 | 0.4 | 0.43 | 0.4 | 0.16 | 0.42 | 0.31 |
| 3 | 158669688 | rs12696090 | 0.006 | 0.04 | -0.42 | -1.727 | -0.041 | -0.147 | -0.147 | 0 | 0.31 | 0.5 | 0.21 | 0.04 | 0.25 | 0.1 |
| 3 | 158671184 | rs1503114 | 1.033 | 0.089 | -0.809 | -1.445 | 0.01 | 0.015 | 0.015 | 0 | 0.14 | 0.1 | 0.1 | 0 | 0.12 | 0.07 |
| 3 | 158674592 | rs9830992 | -0.162 | 0.36 | -0.869 | -1.221 | -0.02 | -0.005 | -0.005 | 0 | 0.35 | 0.52 | 0.25 | 0.28 | 0.27 | 0.22 |
| 3 | 158681384 | rs1875689 | -0.492 | 0.734 | -1.499 | -1.116 | -0.017 | 0.103 | 0.103 | 0 | 0.71 | 0.92 | 0.82 | 0.91 | 0.73 | 0.72 |
| 3 | 158684240 | rs992089 | 0.453 | 1.008 | -1.167 | -0.332 | 0.005 | 0.001 | 0.001 | 0 | 0.38 | 0.26 | 0.58 | 0.62 | 0.33 | 0.55 |
| 3 | 158687391 | rs13077666 | 0.963 | 1.067 | -0.601 | -0.518 | 0.001 | -0.001 | -0.001 | 0.46 | 0.11 | 0.08 | 0.09 | 0 | 0.12 | 0.04 |
| 3 | 158714381 | rs1846897 | 0.749 | 1.346 | -0.781 | -0.616 | -0.003 | 0.07 | 0.07 | 0 | 0.55 | 0.29 | 0.64 | 0.77 | 0.53 | 0.69 |
| 3 | 158715865 | rs9811068 | 0.103 | 1.319 | -0.235 | -0.574 | 0.011 | -0.039 | -0.039 | 0.189 | 0.67 | 0.46 | 0.73 | 0.8 | 0.64 | 0.74 |
| 3 | 158721128 | rs7614679 | 0.305 | 1.393 | -0.813 | -0.541 | -0.001 | -0.002 | -0.002 | 0 | 0.66 | 0.36 | 0.73 | 0.8 | 0.64 | 0.75 |
| 3 | 158722357 | rs16829746 | 0.376 | 1.184 | -0.474 | -0.225 | -0.004 | -0.024 | -0.024 | 0.216 | 0.12 | 0.16 | 0.1 | 0.03 | 0.12 | 0.06 |
| 3 | 158724532 | rs2047675 | 0.724 | 1.415 | -0.697 | -0.069 | -0.013 | 0.023 | 0.023 | 0.344 | 0.59 | 0.28 | 0.65 | 0.72 | 0.54 | 0.72 |
| 3 | 158725874 | rs16829759 | 0.476 | 1.349 | -0.028 | -0.295 | 0.002 | -0.058 | -0.058 | 0 | 0.67 | 0.39 | 0.76 | 0.88 | 0.65 | 0.74 |
| 3 | 158726484 | rs9836856 | 0.499 | 1.492 | -0.39 | -0.488 | -0.019 | 0.022 | 0.022 | 0 | 0.77 | 0.79 | 0.81 | 0.92 | 0.68 | 0.79 |
| 3 | 158726789 | rs12632916 | 1.074 | 0.617 | -0.309 | -0.514 | 0.009 | 0.125 | 0.125 | 1.899 | 0.61 | 0.2 | 0.74 | 0.86 | 0.63 | 0.68 |
| 3 | 158726922 | rs9841375 | 0.649 | 0.261 | 0.28 | -0.415 | 0.003 | -0.056 | -0.056 | 0 | 0.11 | 0.23 | 0.09 | 0 | 0.14 | 0.06 |
| 3 | 158730892 | rs16829770 | -0.097 | -0.14 | -0.623 | -0.356 | -0.037 | -0.008 | -0.008 | 0 | 0.33 | 0.05 | 0.51 | 0.72 | 0.32 | 0.57 |
| 3 | 158731993 | rs7610202 | -0.036 | -0.272 | 0.143 | -0.501 | -0.007 | -0.037 | -0.037 | 0 | 0.72 | 0.63 | 0.83 | 0.88 | 0.73 | 0.75 |
| 3 | 158736714 | rs7647847 | 0.297 | -0.277 | 0.436 | -0.502 | 0.012 | 0.024 | 0.024 | 1.765 | 0.66 | 0.28 | 0.77 | 0.87 | 0.72 | 0.75 |
| 3 | 158743012 | rs1112924 | 0.33 | -0.277 | 0.291 | -0.501 | 0.022 | 0.048 | 0.048 | 2.14 | 0.66 | 0.23 | 0.77 | 0.87 | 0.72 | 0.75 |
| 3 | 158755620 | rs12489127 | 1.313 | 0.285 | -0.247 | -0.441 | -0.103 | 0.103 | 0.103 | 0.035 | 0.32 | 0.07 | 0.25 | 0.15 | 0.41 | 0.18 |
| 3 | 158759012 | rs7639579 | -0.104 | 0.121 | 0.025 | -0.009 | -0.066 | 0.001 | 0.001 | 0 | 0.13 | 0.01 | 0.21 | 0.09 | 0.18 | 0.15 |
| 3 | 158768420 | rs1566550 | 0.097 | -0.735 | 0.586 | 0.692 | -0.012 | 0.021 | 0.021 | 0 | 0.3 | 0.38 | 0.29 | 0.39 | 0.23 | 0.44 |
| 3 | 158776705 | rs2364931 | 0.546 | -0.795 | 0.435 | 0.717 | -0.008 | -0.044 | -0.044 | 0.296 | 0.72 | 0.86 | 0.61 | 0.59 | 0.69 | 0.67 |
| 3 | 158778092 | rs7643629 | 1.408 | -0.778 | 0.585 | 0.713 | -0.012 | 0.025 | 0.025 | 0.034 | 0.4 | 0.35 | 0.31 | 0.19 | 0.46 | 0.23 |
| 3 | 158778215 | rs1501281 | -1.117 | -0.605 | -0.167 | 0.525 | -0.002 | -0.01 | -0.01 | 0 | 0.89 | 0.9 | 0.94 | 1 | 0.86 | 0.93 |
| 3 | 158781278 | rs16829848 | 0.345 | -0.606 | -0.142 | 0.751 | 0.016 | -0.043 | -0.043 | 0.479 | 0.28 | 0.17 | 0.28 | 0.39 | 0.23 | 0.44 |
| 3 | 158790222 | rs6798734 | -0.141 | -0.468 | -0.193 | 0.686 | 0.003 | -0.035 | -0.035 | 1.092 | 0.75 | 0.67 | 0.8 | 0.86 | 0.73 | 0.76 |
| 3 | 158791891 | rs974155 | 0.585 | -0.349 | -1.089 | 0.74 | -0.031 | -0.091 | -0.091 | 0 | 0.25 | 0.02 | 0.33 | 0.31 | 0.22 | 0.43 |
| 3 | 158798149 | rs10513534 | 0.178 | -0.433 | -1.694 | 0.828 | 0.02 | -0.108 | -0.108 | 0.519 | 0.64 | 0.16 | 0.62 | 0.49 | 0.68 | 0.65 |
| 3 | 158819701 | rs2364930 | 0.691 | -0.026 | -1.65 | 0.431 | -0.141 | 0.106 | 0.106 | 0.376 | 0.38 | 0.08 | 0.29 | 0.18 | 0.46 | 0.22 |
| 3 | 158830704 | rs9876582 | 0.247 | -0.342 | -0.918 | 1.067 | 0.008 | -0.017 | -0.017 | 0.15 | 0.71 | 0.79 | 0.74 | 0.68 | 0.71 | 0.65 |
| 3 | 158831323 | rs12497650 | 0.199 | -0.412 | -0.442 | 1.048 | 0.002 | -0.003 | -0.003 | 0 | 0.7 | 0.69 | 0.73 | 0.67 | 0.7 | 0.65 |
| 3 | 158831718 | rs6785766 | 0.194 | -0.411 | -0.442 | 1.049 | 0.002 | -0.003 | -0.003 | 0.053 | 0.7 | 0.69 | 0.73 | 0.67 | 0.7 | 0.65 |
| 3 | 158833189 | rs9816422 | 0.283 | -0.441 | 0.507 | 1.383 | -0.011 | 0.044 | 0.044 | 0 | 0.69 | 0.56 | 0.63 | 0.48 | 0.7 | 0.62 |
| 3 | 158839647 | rs1566548 | 0.279 | -0.484 | 0.678 | 1.634 | -0.002 | 0.001 | 0.001 | 0 | 0.71 | 0.65 | 0.63 | 0.72 | 0.7 | 0.62 |
| 3 | 158840694 | rs17782680 | 0.892 | -0.504 | 0.736 | 1.136 | -0.021 | 0.046 | 0.046 | 0 | 0.36 | 0.27 | 0.27 | 0.17 | 0.39 | 0.16 |
| 3 | 158846028 | rs10513535 | 1.058 | -0.582 | 0.665 | 1.101 | -0.098 | 0.093 | 0.093 | 0 | 0.34 | 0.09 | 0.26 | 0.17 | 0.39 | 0.16 |
| 3 | 158863882 | rs6792979 | 1.091 | -0.602 | 0.693 | 1.092 | -0.073 | 0.085 | 0.085 | 0 | 0.34 | 0.11 | 0.26 | 0.17 | 0.39 | 0.16 |
| 3 | 158864418 | rs1392949 | 0.692 | -0.809 | 0.759 | 1.105 | -0.055 | -0.037 | -0.037 | 0.128 | 0.31 | 0.02 | 0.33 | 0.3 | 0.3 | 0.45 |
| 3 | 158867887 | rs13318193 | -0.144 | -0.823 | 1.095 | 0.99 | -0.015 | 0.012 | 0.012 | 0 | 0.38 | 0.56 | 0.37 | 0.54 | 0.3 | 0.46 |
| 3 | 158870237 | rs6765003 | 0.013 | -1.069 | 0.748 | 0.453 | 0.044 | -0.006 | -0.006 | 0 | 0.46 | 0.18 | 0.58 | 0.47 | 0.41 | 0.63 |
| 3 | 158870253 | rs6789328 | -0.164 | -1.066 | 1.2 | 0.442 | 0.014 | 0.003 | 0.003 | 0 | 0.47 | 0.38 | 0.6 | 0.48 | 0.41 | 0.62 |
| 3 | 158873992 | rs13078847 | 0.728 | -0.891 | 0.771 | 0.421 | -0.038 | -0.004 | -0.004 | 0 | 0.16 | 0.06 | 0.12 | 0.12 | 0.19 | 0.18 |
| 3 | 158892264 | rs4501157 | -1.279 | -0.99 | 1.095 | 0.387 | -0.039 | -0.027 | -0.027 | 0 | 0.66 | 0.84 | 0.74 | 0.61 | 0.6 | 0.8 |
| 3 | 158892812 | rs4269119 | 0.839 | -1.046 | 0.95 | 0.38 | 0.049 | -0.005 | -0.005 | 0.455 | 0.32 | 0.14 | 0.33 | 0.37 | 0.25 | 0.43 |
| 3 | 158901791 | rs1962071 | 0.681 | -1.056 | 1.282 | 0.37 | 0.018 | 0.001 | 0.001 | 0 | 0.33 | 0.23 | 0.34 | 0.38 | 0.25 | 0.43 |
| 3 | 158903157 | rs10470473 | 0.55 | -0.68 | 1.696 | 0.611 | -0.008 | -0.002 | -0.002 | 0 | 0.13 | 0.11 | 0.25 | 0.11 | 0.15 | 0.16 |
| 3 | 158915760 | rs13075120 | 0.474 | -0.616 | 0.598 | 0.644 | -0.039 | -0.009 | -0.009 | 0 | 0.16 | 0.06 | 0.12 | 0.12 | 0.2 | 0.19 |
| 3 | 158923515 | rs1449017 | -0.716 | -0.645 | 0.801 | 0.627 | -0.02 | 0.005 | 0.005 | 0 | 0.29 | 0.23 | 0.38 | 0.23 | 0.35 | 0.35 |
| 3 | 158935342 | rs2621277 | 1.264 | -1.308 | 0.016 | 0.046 | -0.081 | -0.034 | -0.034 | 0 | 0.42 | 0.11 | 0.31 | 0.4 | 0.42 | 0.2 |
| 3 | 158937870 | rs6781001 | 0.798 | -1.171 | -0.017 | 0.068 | -0.03 | 0.004 | 0.004 | 0 | 0.17 | 0.06 | 0.12 | 0.12 | 0.2 | 0.2 |
| 3 | 158943008 | rs10936176 | 0.93 | -1.063 | 0.106 | 0.088 | -0.024 | 0.002 | 0.002 | 0.08 | 0.13 | 0.07 | 0.14 | 0.09 | 0.15 | 0.16 |
| 3 | 158962830 | rs2595233 | -0.444 | -1.604 | -0.062 | -0.117 | 0.062 | -0.158 | -0.158 | 3.32 | 0.69 | 0.94 | 0.57 | 0.5 | 0.74 | 0.44 |
| 3 | 158970212 | rs2621293 | 0.979 | -1.599 | -0.229 | -0.345 | 0.078 | 0.002 | 0.002 | 0.2 | 0.49 | 0.18 | 0.44 | 0.49 | 0.55 | 0.36 |
| 3 | 158974294 | rs9862769 | 1.151 | -1.535 | -0.169 | -0.159 | -0.027 | -0.007 | -0.007 | 0 | 0.13 | 0.07 | 0.14 | 0.11 | 0.16 | 0.16 |

| | | | | | | | | | | | | | | | | |
|---|---|---|---|---|---|---|---|---|---|---|---|---|---|---|---|---|
| 3 | 158975626 | rs7624604 | 0.097 | -1.627 | 0.491 | -0.183 | -0.037 | 0.018 | 0.018 | 0 | 0.66 | 0.48 | 0.54 | 0.49 | 0.74 | 0.44 |
| 3 | 158977434 | rs2621306 | 0.72 | -1.559 | 0.44 | -0.146 | 0.04 | 0.019 | 0.019 | 0.121 | 0.52 | 0.38 | 0.39 | 0.39 | 0.58 | 0.27 |
| 3 | 158985073 | rs9827048 | 0.056 | -1.54 | 0.627 | -0.181 | 0.02 | -0.103 | -0.103 | 0 | 0.16 | 0.39 | 0.1 | 0 | 0.17 | 0.07 |
| 3 | 158990095 | rs2595241 | -0.52 | -1.783 | 0.393 | -0.296 | 0.061 | -0.157 | -0.157 | 0 | 0.72 | 0.96 | 0.58 | 0.5 | 0.76 | 0.45 |
| 3 | 158993239 | rs1448996 | 1.168 | -1.748 | 0.564 | -0.293 | 0.013 | -0.05 | -0.05 | 0 | 0.12 | 0.18 | 0.09 | 0 | 0.16 | 0.06 |
| 3 | 158994949 | rs2595243 | 1.324 | -1.764 | 0.535 | -0.311 | -0.006 | 0.001 | 0.001 | 0 | 0.43 | 0.42 | 0.33 | 0.39 | 0.42 | 0.2 |
| 3 | 158998151 | rs13083244 | 1.437 | -1.68 | 0.35 | -0.304 | -0.008 | -0.006 | -0.006 | 0 | 0.11 | 0.09 | 0.07 | 0 | 0.16 | 0.06 |
| 3 | 159017010 | rs2595247 | 1.566 | -1.507 | 0.265 | -0.184 | -0.077 | -0.037 | -0.037 | 0 | 0.39 | 0.12 | 0.31 | 0.4 | 0.42 | 0.21 |
| 3 | 159018527 | rs2595248 | 1.072 | -1.438 | 0.499 | -0.167 | 0.004 | -0.004 | -0.004 | 0.785 | 0.45 | 0.51 | 0.35 | 0.4 | 0.43 | 0.22 |
| 3 | 159019946 | rs2595249 | -0.165 | -1.478 | 0.1 | 0.028 | 0.035 | -0.174 | -0.174 | 0 | 0.72 | 0.97 | 0.59 | 0.51 | 0.77 | 0.46 |
| 3 | 159047257 | rs2365820 | 0.581 | -0.991 | 0.837 | -0.169 | -0.004 | -0.014 | -0.014 | 0 | 0.22 | 0.37 | 0.26 | 0.17 | 0.17 | 0.21 |
| 3 | 159056759 | rs9290036 | 0.995 | -0.994 | 0.795 | -0.172 | -0.003 | 0.006 | 0.006 | 0 | 0.19 | 0.14 | 0.24 | 0.17 | 0.17 | 0.21 |
| 3 | 159072815 | rs7646959 | 1.581 | -1.1 | 0.698 | -0.134 | -0.057 | -0.061 | -0.061 | 0 | 0.4 | 0.15 | 0.37 | 0.45 | 0.47 | 0.24 |
| 3 | 159080907 | rs895877 | -0.383 | -1.165 | 0.321 | -0.168 | 0.053 | -0.05 | -0.05 | 0.845 | 0.74 | 0.89 | 0.7 | 0.62 | 0.79 | 0.53 |
| 3 | 159081357 | rs1449004 | 0.932 | -1.141 | 0.473 | -0.186 | 0.004 | 0.004 | 0.004 | 0.145 | 0.61 | 0.41 | 0.62 | 0.62 | 0.63 | 0.44 |
| 3 | 159085425 | rs11917829 | -1.043 | -1.304 | 0.37 | -0.18 | 0.025 | -0.07 | -0.07 | 0 | 0.19 | 0.44 | 0.09 | 0.13 | 0.19 | 0.14 |
| 3 | 159092308 | rs4680500 | 1.877 | -1.251 | 0.131 | -0.189 | -0.081 | -0.098 | -0.098 | 3.075 | 0.4 | 0.1 | 0.37 | 0.45 | 0.46 | 0.24 |
| 3 | 159096538 | rs2365835 | -1.17 | -1.236 | 0.187 | -0.103 | 0.023 | -0.03 | -0.03 | 0 | 0.81 | 0.92 | 0.72 | 0.75 | 0.82 | 0.58 |
| 3 | 159098080 | rs4440163 | -0.297 | -1.214 | 0.296 | -0.175 | -0.014 | -0.009 | -0.009 | 0.006 | 0.29 | 0.42 | 0.27 | 0.29 | 0.19 | 0.29 |
| 3 | 159098147 | rs1109313 | -0.933 | -1.184 | 0.367 | -0.234 | -0.003 | 0.029 | 0.029 | 0 | 0.38 | 0.68 | 0.34 | 0.27 | 0.35 | 0.35 |
| 3 | 159098996 | rs9871574 | 1.007 | -1.201 | 0.334 | -0.235 | -0.012 | 0.014 | 0.014 | 0 | 0.2 | 0.06 | 0.23 | 0.15 | 0.16 | 0.21 |
| 3 | 159099226 | rs878382 | -1.154 | -1.189 | 0.306 | -0.146 | 0.016 | -0.014 | -0.014 | 0 | 0.81 | 0.87 | 0.71 | 0.75 | 0.82 | 0.59 |
| 3 | 159104487 | rs7649056 | 1.631 | -1.188 | 0.23 | -0.135 | -0.067 | -0.051 | -0.051 | 1.481 | 0.42 | 0.13 | 0.37 | 0.48 | 0.47 | 0.24 |
| 3 | 159105475 | rs7640445 | -0.317 | -1.127 | 0.324 | -0.237 | -0.001 | -0.001 | -0.001 | 0 | 0.3 | 0.37 | 0.27 | 0.27 | 0.2 | 0.32 |
| 3 | 159133317 | rs7631740 | 0.264 | -0.945 | 0.107 | -0.177 | 0.071 | -0.046 | -0.046 | 0 | 0.84 | 0.58 | 0.91 | 0.88 | 0.78 | 0.85 |
| 3 | 159137792 | rs12487721 | 0.805 | -1.167 | -0.054 | 0.053 | 0.014 | -0.014 | -0.014 | 0.38 | 0.21 | 0.16 | 0.3 | 0.29 | 0.15 | 0.42 |
| 3 | 159140615 | rs9290042 | 0.89 | -1.066 | -0.283 | 0.044 | -0.002 | 0.012 | 0.012 | 0 | 0.2 | 0.17 | 0.22 | 0.14 | 0.17 | 0.2 |
| 3 | 159140638 | rs4680505 | 1.224 | -0.971 | -0.679 | 0.162 | -0.034 | -0.029 | -0.029 | 0 | 0.43 | 0.25 | 0.39 | 0.44 | 0.48 | 0.24 |
| 3 | 159151726 | rs1489919 | 0.23 | -0.934 | -0.885 | 0.154 | -0.009 | 0.029 | 0.029 | 0 | 0.63 | 0.33 | 0.68 | 0.73 | 0.63 | 0.69 |
| 3 | 159152259 | rs4680509 | 0.3 | -0.989 | -0.786 | 0.391 | 0.036 | -0.034 | -0.034 | 0 | 0.21 | 0.11 | 0.29 | 0.29 | 0.15 | 0.42 |
| 3 | 159155377 | rs16830368 | 0.077 | -0.068 | -0.681 | -0.179 | 0.001 | -0.012 | -0.012 | 0.736 | 0.41 | 0.51 | 0.48 | 0.21 | 0.5 | 0.19 |
| 3 | 159155504 | rs2171846 | 0.851 | -0.076 | -0.154 | 0.024 | 0.046 | -0.008 | -0.008 | 0 | 0.82 | 0.75 | 0.87 | 0.8 | 0.74 | 0.9 |
| 3 | 159158217 | rs12636257 | 0.588 | -0.416 | -0.687 | 0.489 | -0.016 | -0.038 | -0.038 | 0 | 0.16 | 0.15 | 0.2 | 0.32 | 0.19 | 0.39 |
| 3 | 159168091 | rs6803496 | -0.197 | -0.853 | -0.628 | 0.333 | -0.027 | -0.029 | -0.029 | 0.645 | 0.25 | 0.53 | 0.16 | 0.19 | 0.19 | 0.17 |
| 3 | 159173074 | rs4142769 | -0.937 | -0.738 | -1.117 | 0.364 | 0.021 | -0.033 | -0.033 | 0 | 0.41 | 0.77 | 0.36 | 0.51 | 0.38 | 0.54 |
| 3 | 159180395 | rs7635938 | 0.833 | -0.755 | -1.405 | 0.222 | -0.034 | -0.078 | -0.078 | 0 | 0.24 | 0.08 | 0.19 | 0.35 | 0.23 | 0.26 |
| 3 | 159180640 | rs1489920 | -1.793 | -0.813 | -1.601 | 0.013 | -0.081 | 0.068 | 0.068 | 0 | 0.66 | 0.9 | 0.58 | 0.8 | 0.59 | 0.78 |
| 3 | 159184569 | rs7634185 | 0.198 | -0.41 | -1.032 | 0.282 | -0.006 | -0.011 | -0.011 | 0.051 | 0.2 | 0.19 | 0.09 | 0.14 | 0.17 | 0.11 |
| 3 | 159200834 | rs17798579 | 0.59 | -0.454 | -0.984 | 0.227 | -0.02 | -0.054 | -0.054 | 0 | 0.19 | 0.11 | 0.17 | 0.28 | 0.21 | 0.4 |
| 3 | 159216270 | rs6787578 | 0.976 | -0.529 | -0.933 | 0.038 | 0.002 | 0.002 | 0.002 | 0.128 | 0.15 | 0.14 | 0.14 | 0.09 | 0.14 | 0.11 |
| 3 | 159217153 | rs1565352 | -1.643 | -0.535 | -1.263 | 0.002 | -0.015 | 0.037 | 0.037 | 0 | 0.68 | 0.8 | 0.58 | 0.8 | 0.63 | 0.79 |
| 3 | 159221461 | rs10470336 | -1.651 | -0.531 | -1.541 | 0.004 | -0.025 | 0.045 | 0.045 | 0 | 0.68 | 0.84 | 0.59 | 0.8 | 0.63 | 0.79 |
| 3 | 159232929 | rs1826118 | -0.251 | -0.541 | -1.753 | 0.001 | -0.011 | 0.13 | 0.13 | 0 | 0.48 | 0.26 | 0.42 | 0.7 | 0.49 | 0.64 |
| 3 | 159234376 | rs9870894 | -1.489 | -0.402 | -1.833 | 0.085 | -0.007 | 0.013 | 0.013 | 0 | 0.68 | 0.74 | 0.58 | 0.8 | 0.63 | 0.79 |
| 3 | 159239866 | rs9835769 | -0.165 | -0.192 | -1.857 | 0.502 | -0.024 | -0.078 | -0.078 | 1.995 | 0.29 | 0.14 | 0.25 | 0.43 | 0.28 | 0.24 |
| 3 | 159245263 | rs1387696 | 0.592 | -0.013 | -1.94 | 0.184 | 0 | 0.038 | 0.038 | 2.157 | 0.13 | 0.01 | 0.06 | 0.07 | 0.13 | 0.11 |
| 3 | 159252752 | rs2196619 | -0.515 | 0.329 | -1.353 | 0.648 | -0.036 | -0.091 | -0.091 | 0 | 0.28 | 0.11 | 0.25 | 0.41 | 0.28 | 0.24 |
| 3 | 159269142 | rs11710258 | 0.312 | -0.123 | -0.849 | 0.424 | 0.014 | 0.044 | 0.044 | 1.148 | 0.58 | 0.29 | 0.67 | 0.68 | 0.63 | 0.76 |
| 3 | 159276026 | rs13326718 | -1.369 | -0.599 | 0.325 | 0.532 | 0.004 | -0.008 | -0.008 | 0 | 0.09 | 0.24 | 0.1 | 0.08 | 0.11 | 0.05 |
| 3 | 159276226 | rs9839862 | -0.815 | -1.029 | 0.617 | 0.826 | -0.005 | 0.01 | 0.01 | 0.129 | 0.17 | 0.2 | 0.16 | 0.21 | 0.1 | 0.23 |
| 3 | 159277496 | rs6782081 | -1.999 | -1.158 | 1.212 | 1.061 | -0.019 | 0.038 | 0.038 | 0.144 | 0.29 | 0.65 | 0.29 | 0.47 | 0.22 | 0.3 |
| 3 | 159298036 | rs12108170 | -1.573 | -1.187 | 1.606 | 1.116 | -0.016 | 0.04 | 0.04 | 2.08 | 0.23 | 0.7 | 0.2 | 0.39 | 0.13 | 0.25 |
| 3 | 159302556 | rs7629269 | -2.028 | -1.226 | 1.885 | 0.773 | -0.034 | 0.026 | 0.026 | 2.43 | 0.1 | 0.73 | 0.06 | 0.18 | 0.03 | 0.05 |
| 3 | 159316263 | rs1581342 | -2.561 | -1.326 | 1.841 | 0.729 | -0.04 | 0.013 | 0.013 | 2.566 | 0.12 | 0.73 | 0.06 | 0.18 | 0.03 | 0.05 |
| 3 | 159321775 | rs9866095 | -2.102 | -1.232 | 1.883 | 0.771 | -0.036 | 0.023 | 0.023 | 3.095 | 0.11 | 0.73 | 0.06 | 0.18 | 0.03 | 0.05 |
| 3 | 159327356 | rs6805934 | -1.396 | -1.669 | 1.337 | 0.351 | 0.098 | 0.071 | 0.071 | 1.423 | 0.33 | 0.85 | 0.2 | 0.25 | 0.19 | 0.21 |
| 3 | 159332305 | rs12107648 | -2.251 | -1.005 | 1.986 | 0.892 | -0.043 | 0.07 | 0.07 | 2.668 | 0.1 | 0.62 | 0.05 | 0.17 | 0.01 | 0.04 |
| 3 | 159336334 | rs16830532 | -0.978 | -0.824 | 1.547 | 0.942 | 0 | -0.002 | -0.002 | 0.077 | 0.05 | 0.25 | 0.02 | 0 | 0.01 | 0 |
| 3 | 159351238 | rs9827448 | -0.275 | -0.693 | 0.596 | 1.144 | -0.029 | -0.071 | -0.071 | 0 | 0.11 | 0.01 | 0.14 | 0.17 | 0.09 | 0.21 |
| 3 | 159359371 | rs9842465 | -1.809 | -0.858 | 1.613 | 0.809 | 0.009 | 0.119 | 0.119 | 0.954 | 0.08 | 0.45 | 0.03 | 0.16 | 0.01 | 0.01 |
| 3 | 159380949 | rs7630289 | -1.268 | -0.851 | 1.455 | 0.62 | -0.004 | -0.002 | -0.002 | 0 | 0.06 | 0.31 | 0.06 | 0.01 | 0.01 | 0.02 |
| 3 | 159386571 | rs9826416 | -2.14 | -0.794 | 1.361 | 0.612 | -0.003 | 0.009 | 0.009 | 2.323 | 0.1 | 0.55 | 0.08 | 0.07 | 0.03 | 0.03 |
| 3 | 159399494 | rs7628548 | -0.909 | -1.734 | 0.607 | 0.307 | 0.009 | 0.011 | 0.011 | 0.161 | 0.29 | 0.69 | 0.32 | 0.21 | 0.14 | 0.22 |
| 3 | 159402180 | rs9845025 | -0.302 | -1.804 | 1.338 | 0.294 | -0.015 | -0.015 | -0.015 | 0 | 0.22 | 0.21 | 0.25 | 0.15 | 0.13 | 0.17 |
| 3 | 159413737 | rs9812406 | -1.034 | -1.793 | 1.309 | 0.296 | 0.001 | 0.006 | 0.006 | 0.161 | 0.29 | 0.68 | 0.32 | 0.21 | 0.14 | 0.22 |
| 3 | 159452704 | rs9851867 | 0.284 | -0.718 | 0.433 | 0.091 | 0.002 | -0.003 | -0.003 | 1.488 | 0.43 | 0.65 | 0.39 | 0.44 | 0.38 | 0.43 |
| 3 | 159452813 | rs9869514 | 0.097 | -0.629 | 0.529 | 0.231 | -0.003 | 0 | 0 | 0 | 0.24 | 0.21 | 0.25 | 0.2 | 0.13 | 0.18 |
| 3 | 159454827 | rs4602438 | -0.843 | -0.361 | 0.379 | 0.217 | -0.008 | 0.037 | 0.037 | 1.502 | 0.56 | 0.67 | 0.61 | 0.72 | 0.53 | 0.57 |
| 3 | 159455394 | rs12487057 | -0.71 | 0.144 | 0.523 | 0.861 | 0.01 | 0.063 | 0.063 | 0.262 | 0.84 | 0.83 | 0.74 | 0.55 | 0.8 | 0.66 |

| | | | | | | | | | | | | | | | | |
|---|---|---|---|---|---|---|---|---|---|---|---|---|---|---|---|---|
| 3 | 159459059 | rs7632574 | -0.483 | 0.104 | 0.354 | 0.872 | -0.013 | -0.034 | -0.034 | 0 | 0.82 | 0.87 | 0.76 | 0.59 | 0.81 | 0.67 |
| 3 | 159459178 | rs9840100 | 0.011 | 0.301 | 0.334 | 0.759 | -0.007 | -0.005 | -0.005 | 0.113 | 0.28 | 0.4 | 0.36 | 0.26 | 0.26 | 0.36 |
| 3 | 159461456 | rs4680514 | -0.323 | 0.284 | 0.351 | 0.749 | 0.001 | 0.018 | 0.018 | 0 | 0.79 | 0.72 | 0.72 | 0.57 | 0.8 | 0.64 |
| 3 | 159461558 | rs11915303 | 0.001 | 0.44 | -0.165 | 0.73 | -0.006 | -0.012 | -0.012 | 0 | 0.28 | 0.34 | 0.33 | 0.24 | 0.26 | 0.33 |
| 3 | 159462903 | rs1461100 | 0 | 0.112 | -0.004 | 0.889 | 0.017 | 0.112 | 0.112 | 0 | 0.79 | 0.57 | 0.74 | 0.59 | 0.81 | 0.67 |
| 3 | 159477493 | rs719070 | -0.26 | -0.814 | -0.209 | 0.342 | -0.046 | -0.023 | -0.023 | 0 | 0.26 | 0.41 | 0.27 | 0.18 | 0.18 | 0.28 |
| 3 | 159486818 | rs6780187 | 0.122 | -0.947 | -0.641 | 0.517 | 0.028 | -0.042 | -0.042 | 0 | 0.31 | 0.2 | 0.39 | 0.48 | 0.24 | 0.35 |
| 3 | 159489875 | rs6803317 | -0.442 | -1.808 | -0.509 | -0.041 | 0.02 | 0.008 | 0.008 | 0 | 0.56 | 0.58 | 0.53 | 0.62 | 0.43 | 0.49 |
| 3 | 159491807 | rs16830748 | 0.704 | -1.874 | -0.274 | -0.018 | 0.026 | 0.026 | 0.026 | 0 | 0.21 | 0.06 | 0.11 | 0.14 | 0.17 | 0.14 |
| 3 | 159509380 | rs2169305 | -0.544 | -1.395 | -0.337 | -0.487 | 0.012 | -0.019 | -0.019 | 0.986 | 0.39 | 0.3 | 0.25 | 0.48 | 0.37 | 0.41 |
| 3 | 159512199 | rs16830780 | 0.505 | -1.368 | -0.505 | -0.458 | 0.021 | -0.012 | -0.012 | 4.548 | 0.26 | 0.13 | 0.15 | 0.27 | 0.22 | 0.35 |
| 3 | 159523497 | rs6792870 | 0.095 | -1.421 | -0.272 | -0.52 | -0.024 | -0.01 | -0.01 | 5.325 | 0.3 | 0.3 | 0.18 | 0.23 | 0.22 | 0.35 |
| 3 | 159532411 | rs13066560 | 1.071 | -1.455 | -0.382 | -0.536 | 0.027 | 0.039 | 0.039 | 0 | 0.21 | 0.05 | 0.1 | 0.1 | 0.17 | 0.12 |
| 3 | 159551651 | rs10513546 | -0.277 | -1.426 | -0.169 | -0.599 | 0.012 | 0.006 | 0.006 | 0 | 0.35 | 0.27 | 0.2 | 0.27 | 0.33 | 0.2 |
| 3 | 159554958 | rs13088856 | -0.548 | -1.394 | -0.414 | -0.586 | 0.002 | 0.002 | 0.002 | 0 | 0.38 | 0.58 | 0.24 | 0.27 | 0.34 | 0.21 |
| 3 | 159557538 | rs11925758 | 0.357 | -1.404 | -0.44 | -0.584 | 0.014 | 0.074 | 0.074 | 0.343 | 0.16 | 0.48 | 0.13 | 0.2 | 0.16 | 0.08 |
| 3 | 159557953 | rs13064339 | 0.929 | -1.272 | -0.585 | -0.788 | 0.032 | 0.065 | 0.065 | 0.113 | 0.22 | 0.03 | 0.1 | 0.07 | 0.17 | 0.13 |
| 3 | 159567428 | rs7640429 | 0.026 | -0.923 | -0.692 | -0.942 | 0.016 | 0.094 | 0.094 | 0.058 | 0.52 | 0.64 | 0.55 | 0.78 | 0.48 | 0.48 |
| 3 | 159580653 | rs2366405 | 0.502 | -1.1 | -1.152 | -0.974 | 0.036 | 0.019 | 0.019 | 0 | 0.27 | 0.12 | 0.16 | 0.19 | 0.22 | 0.16 |
| 3 | 159582382 | rs1675497 | -1.682 | -0.763 | -1.184 | -0.939 | -0.095 | 0.092 | 0.092 | 0.55 | 0.81 | 0.96 | 0.71 | 0.99 | 0.7 | 0.84 |
| 3 | 159583518 | rs1675496 | 1.114 | -1.077 | -0.707 | -0.627 | 0.019 | -0.014 | -0.014 | 0 | 0.33 | 0.24 | 0.4 | 0.61 | 0.25 | 0.35 |
| 3 | 159599190 | rs33051 | 0.452 | -1.296 | -1.131 | -1.345 | 0.074 | 0.188 | 0.188 | 0 | 0.44 | 0.15 | 0.45 | 0.9 | 0.35 | 0.64 |
| 3 | 159608812 | rs33058 | 0.702 | -1.29 | -1.045 | -1.392 | 0.006 | 0.007 | 0.007 | 0 | 0.4 | 0.35 | 0.48 | 0.74 | 0.31 | 0.6 |
| 3 | 159610911 | rs3773786 | 0.132 | -0.465 | -1.292 | -1.841 | 0.004 | -0.03 | -0.03 | 0 | 0.21 | 0.26 | 0.14 | 0.05 | 0.26 | 0.11 |
| 3 | 159611291 | rs26298 | -0.539 | -0.098 | -1.569 | -1.637 | -0.159 | 0.167 | 0.167 | 0 | 0.74 | 0.97 | 0.66 | 1 | 0.61 | 0.78 |
| 3 | 159613455 | rs6785153 | -0.4 | -0.123 | -1.085 | -1.306 | 0.006 | -0.03 | -0.03 | 0.221 | 0.13 | 0.17 | 0.1 | 0.02 | 0.17 | 0.05 |
| 3 | 159625545 | rs9872563 | 0.5 | 0.278 | -1.264 | -1.543 | 0.001 | -0.007 | -0.007 | 0 | 0.15 | 0.17 | 0.07 | 0 | 0.13 | 0.08 |
| 3 | 159629241 | rs6808758 | 1.129 | -0.109 | -1.222 | -1.535 | -0.004 | -0.001 | -0.001 | 0 | 0.07 | 0.07 | 0.02 | 0.06 | 0.04 | 0.08 |
| 3 | 159631796 | rs7644697 | 0.95 | 0.568 | -1.674 | -1.677 | -0.013 | 0.002 | 0.002 | 0 | 0.12 | 0.11 | 0.17 | 0.07 | 0.15 | 0.06 |
| 3 | 159634474 | rs7631387 | -0.906 | 0.334 | -1.336 | -2.126 | -0.007 | 0.021 | 0.021 | 0 | 0.6 | 0.65 | 0.36 | 0.19 | 0.45 | 0.57 |
| 3 | 159635428 | rs9877650 | -1.756 | 0.205 | -0.97 | -2.154 | 0.003 | -0.004 | -0.004 | 0 | 0.65 | 0.7 | 0.52 | 0.26 | 0.57 | 0.57 |
| 3 | 159640152 | rs6792701 | -0.561 | -0.174 | -0.875 | -2.452 | -0.002 | -0.032 | -0.032 | 0.712 | 0.43 | 0.58 | 0.41 | 0.2 | 0.39 | 0.46 |
| 3 | 159644297 | rs6799788 | 0.313 | -0.333 | -0.932 | -2.555 | -0.018 | 0.073 | 0.073 | 0 | 0.23 | 0.08 | 0.19 | 0.08 | 0.21 | 0.38 |
| 3 | 159645671 | rs6805758 | -0.768 | -0.391 | -0.704 | -2.591 | -0.005 | -0.019 | -0.019 | 0 | 0.4 | 0.53 | 0.3 | 0.2 | 0.33 | 0.46 |
| 3 | 159655667 | rs9814871 | 1.97 | -0.161 | -0.793 | -2.482 | -0.017 | 0.045 | 0.045 | 0 | 0.09 | 0.01 | 0.12 | 0 | 0.09 | 0.06 |
| 3 | 159663317 | rs550662 | -0.808 | -0.249 | -0.566 | -2.574 | 0 | 0.086 | 0.086 | 0 | 0.46 | 0.33 | 0.36 | 0.15 | 0.44 | 0.44 |
| 3 | 159669994 | rs11918254 | 1.413 | -0.434 | -0.634 | -2.511 | -0.011 | 0.012 | 0.012 | 0 | 0.14 | 0.14 | 0.19 | 0.07 | 0.18 | 0.08 |
| 3 | 159672597 | rs10513547 | 1.458 | -0.465 | -0.631 | -2.53 | 0.01 | 0.043 | 0.043 | 0 | 0.17 | 0.06 | 0.09 | 0 | 0.15 | 0.04 |
| 3 | 159675391 | rs10513548 | 0.81 | -0.397 | -0.365 | -2.411 | -0.002 | -0.038 | -0.038 | 0 | 0.13 | 0.37 | 0.15 | 0.05 | 0.1 | 0.07 |
| 3 | 159678641 | rs7636840 | 0.596 | -0.4 | -0.315 | -2.014 | 0.01 | 0.037 | 0.037 | 0.045 | 0.18 | 0.5 | 0.22 | 0.18 | 0.18 | 0.08 |
| 3 | 159678736 | rs12494630 | 1.336 | -0.331 | -0.514 | -1.995 | -0.011 | -0.011 | -0.011 | 0 | 0.14 | 0.14 | 0.19 | 0.13 | 0.18 | 0.08 |
| 3 | 159686068 | rs2561285 | 1.078 | -0.342 | -0.645 | -2.044 | -0.005 | 0.022 | 0.022 | 0 | 0.14 | 0.07 | 0.08 | 0.08 | 0.11 | 0.32 |
| 3 | 159686345 | rs7610160 | -0.942 | -0.356 | -0.554 | -2.141 | -0.003 | -0.025 | -0.025 | 0.531 | 0.44 | 0.55 | 0.42 | 0.18 | 0.42 | 0.19 |
| 3 | 159688029 | rs17811014 | 1.222 | -0.278 | -0.54 | -2.078 | 0.004 | 0.061 | 0.061 | 0 | 0.18 | 0.04 | 0.09 | 0 | 0.16 | 0.04 |
| 3 | 159688294 | rs681588 | -1.283 | -0.303 | -0.482 | -1.998 | -0.001 | 0.01 | 0.01 | 0 | 0.51 | 0.61 | 0.44 | 0.26 | 0.46 | 0.47 |
| 3 | 159691049 | rs479952 | -1.708 | -0.209 | -0.494 | -1.678 | 0.001 | 0.001 | 0.001 | 0 | 0.62 | 0.69 | 0.45 | 0.35 | 0.55 | 0.55 |
| 3 | 159691112 | rs1014486 | -0.737 | -0.264 | -0.607 | -1.728 | 0.017 | 0.087 | 0.087 | 0 | 0.48 | 0.33 | 0.36 | 0.23 | 0.46 | 0.23 |
| 3 | 159692776 | rs609907 | -0.028 | -0.28 | -0.973 | -1.934 | -0.005 | 0.078 | 0.078 | 0 | 0.3 | 0.13 | 0.14 | 0.13 | 0.25 | 0.37 |
| 3 | 159692986 | rs540722 | 0.423 | -0.281 | -1.229 | -1.869 | -0.041 | 0.089 | 0.089 | 0 | 0.22 | 0.03 | 0.21 | 0.07 | 0.2 | 0.18 |
| 3 | 159695933 | rs2886666 | 1.15 | -0.232 | -1.222 | -1.81 | -0.051 | -0.058 | -0.058 | 0 | 0.15 | 0.02 | 0.2 | 0.15 | 0.19 | 0.07 |
| 3 | 159696099 | rs2366408 | -0.25 | -0.324 | -0.554 | -1.821 | -0.002 | -0.011 | -0.011 | 0.007 | 0.37 | 0.47 | 0.33 | 0.15 | 0.35 | 0.12 |
| 3 | 159696998 | rs9811792 | -1.713 | -0.116 | -0.871 | -1.61 | -0.01 | -0.003 | -0.003 | 0 | 0.64 | 0.74 | 0.48 | 0.33 | 0.56 | 0.55 |
| 3 | 159699077 | rs6441282 | -1.005 | 0.122 | -0.924 | -1.506 | 0.004 | 0.054 | 0.054 | 0 | 0.5 | 0.68 | 0.4 | 0.24 | 0.45 | 0.43 |
| 3 | 159699573 | rs692890 | -0.875 | -0.155 | -0.592 | -1.291 | 0.007 | 0.016 | 0.016 | 0 | 0.41 | 0.67 | 0.32 | 0.16 | 0.29 | 0.4 |
| 3 | 159700325 | rs755004 | 0.692 | 0.223 | -0.98 | -1.03 | -0.024 | -0.015 | -0.015 | 0 | 0.08 | 0.01 | 0.13 | 0.08 | 0.12 | 0.09 |
| 3 | 159701231 | rs662959 | 0.044 | -0.178 | -0.619 | -1.247 | -0.001 | -0.017 | -0.017 | 0 | 0.15 | 0.21 | 0.07 | 0.12 | 0.12 | 0.37 |
| 3 | 159706280 | rs2243115 | 0.939 | -0.571 | -0.96 | -1.449 | 0.023 | 0.034 | 0.034 | 0 | 0.18 | 0.08 | 0.09 | 0.09 | 0.13 | 0.12 |
| 3 | 159709651 | rs2243123 | 0.214 | -0.315 | -1.029 | -1.498 | -0.001 | 0 | 0 | 0 | 0.17 | 0.14 | 0.31 | 0.09 | 0.25 | 0.08 |
| 3 | 159712273 | rs2243133 | 1.761 | -0.468 | -1.502 | -1.32 | 0.015 | 0.01 | 0.01 | 0 | 0.08 | 0.03 | 0.01 | 0 | 0.03 | 0.02 |
| 3 | 159713467 | rs568408 | 0.343 | -0.48 | -1.485 | -1.313 | -0.003 | -0.016 | -0.016 | 0 | 0.16 | 0.21 | 0.07 | 0.12 | 0.12 | 0.37 |
| 3 | 159714802 | rs2243143 | -0.328 | -0.377 | -1.28 | -1.272 | 0 | -0.011 | -0.011 | 0.481 | 0.38 | 0.56 | 0.42 | 0.18 | 0.38 | 0.2 |
| 3 | 159715551 | rs668998 | -1.679 | -0.37 | -1.623 | -1.233 | -0.055 | 0.052 | 0.052 | 0 | 0.61 | 0.86 | 0.5 | 0.3 | 0.53 | 0.57 |
| 3 | 159718268 | rs6441284 | -0.538 | -0.163 | -1.349 | -1.081 | -0.003 | 0.018 | 0.018 | 0 | 0.39 | 0.61 | 0.42 | 0.18 | 0.38 | 0.2 |
| 3 | 159719132 | rs485497 | -0.935 | -0.009 | -1.593 | -0.877 | -0.043 | 0.16 | 0.16 | 0 | 0.53 | 0.84 | 0.37 | 0.24 | 0.47 | 0.5 |
| 3 | 159720271 | rs4680536 | -0.499 | 0.731 | -1.075 | -0.425 | -0.013 | 0.011 | 0.011 | 0 | 0.26 | 0.26 | 0.24 | 0.14 | 0.38 | 0.19 |
| 3 | 159720817 | rs598638 | -1.108 | 0.485 | -1.609 | -0.143 | -0.013 | -0.023 | -0.023 | 0.044 | 0.7 | 0.99 | 0.88 | 0.81 | 0.8 | 0.84 |
| 3 | 159728878 | rs6441286 | 0.953 | 0.735 | -0.825 | 0.179 | -0.013 | -0.049 | -0.049 | 0 | 0.35 | 0.21 | 0.46 | 0.46 | 0.42 | 0.42 |
| 3 | 159728987 | rs564799 | -0.188 | 0.753 | -0.726 | 0.156 | -0.047 | 0.037 | 0.037 | 0 | 0.25 | 0.18 | 0.23 | 0.14 | 0.35 | 0.17 |
| 3 | 159732983 | rs574808 | -0.361 | 0.756 | -0.694 | 0.159 | 0.035 | -0.021 | -0.021 | 0.467 | 0.27 | 0.45 | 0.36 | 0.22 | 0.36 | 0.18 |

| | | | | | | | | | | | | | | | |
|---|---|---|---|---|---|---|---|---|---|---|---|---|---|---|---|
| 3 | 159743231 | rs2647926 | -0.307 | 0.091 | -0.172 | 0.331 | -0.012 | -0.004 | -0.004 | 0 | 0.62 | 0.66 | 0.82 | 0.66 | 0.78 | 0.6 |
| 3 | 159743883 | rs2936298 | -0.51 | 0.224 | -0.902 | 0.343 | 0.031 | 0.014 | 0.014 | 3.008 | 0.63 | 0.33 | 0.7 | 0.71 | 0.79 | 0.84 |
| 3 | 159744022 | rs2914117 | -0.216 | 0.383 | -1.021 | 0.316 | 0.07 | 0.008 | 0.008 | 1.111 | 0.58 | 0.29 | 0.68 | 0.59 | 0.77 | 0.6 |
| 3 | 159751822 | rs6783320 | -1.24 | 1.449 | -0.526 | 0.426 | 0.015 | 0.044 | 0.044 | 0 | 0.15 | 0.31 | 0.21 | 0.3 | 0.19 | 0.25 |
| 3 | 159754507 | rs10513549 | -0.548 | 1.725 | -0.804 | 0.081 | -0.031 | 0.028 | 0.028 | 0 | 0.16 | 0.11 | 0.18 | 0 | 0.27 | 0.08 |
| 3 | 159756451 | rs1075498 | -1.599 | 1.132 | -0.69 | 0.13 | -0.002 | -0.002 | -0.002 | 0 | 0.26 | 0.37 | 0.31 | 0.25 | 0.29 | 0.31 |
| 3 | 159759401 | rs7618691 | -1.648 | 1.014 | 0.037 | 0.479 | -0.006 | 0.097 | 0.097 | 0.175 | 0.07 | 0.31 | 0.13 | 0.16 | 0.03 | 0.08 |
| 3 | 159767453 | rs16831052 | -0.615 | 1.019 | 0.05 | 0.491 | -0.003 | 0.063 | 0.063 | 0 | 0.06 | 0.19 | 0.14 | 0.13 | 0.07 | 0.07 |
| 3 | 159772812 | rs1818023 | -0.569 | 1.02 | 0.056 | 0.489 | -0.002 | 0.065 | 0.065 | 0 | 0.06 | 0.19 | 0.14 | 0.13 | 0.07 | 0.07 |
| 3 | 159780373 | rs12492688 | -0.525 | 1.02 | 0.056 | 0.489 | -0.001 | 0.067 | 0.067 | 0 | 0.06 | 0.19 | 0.14 | 0.13 | 0.07 | 0.07 |
| 3 | 159791628 | rs12491474 | -0.525 | 1.02 | 0.056 | 0.489 | -0.001 | 0.067 | 0.067 | 0 | 0.06 | 0.19 | 0.14 | 0.13 | 0.07 | 0.07 |
| 3 | 159792242 | rs13062665 | 0.646 | 1.318 | -0.482 | 0.384 | -0.068 | 0.019 | 0.019 | 0.002 | 0.19 | 0.02 | 0.12 | 0.15 | 0.24 | 0.28 |
| 3 | 159793827 | rs9818492 | 1.965 | 1.281 | 0.407 | 0.692 | 0.043 | 0.031 | 0.031 | 0.238 | 0.74 | 0.44 | 0.72 | 0.67 | 0.7 | 0.64 |
| 3 | 159796991 | rs9847609 | 1.752 | 1.333 | 0.293 | 0.72 | 0.029 | -0.013 | -0.013 | 0 | 0.77 | 0.64 | 0.83 | 0.79 | 0.71 | 0.7 |
| 3 | 159803697 | rs9880666 | 0.567 | 1.358 | 0.281 | 0.688 | 0.007 | 0 | 0 | 0 | 0.86 | 0.88 | 0.88 | 0.85 | 0.79 | 0.74 |
| 3 | 159804528 | rs12634498 | 1.967 | 1.358 | 0.224 | 0.696 | 0.045 | 0.032 | 0.032 | 0.107 | 0.75 | 0.51 | 0.72 | 0.66 | 0.7 | 0.66 |
| 3 | 159804903 | rs12496326 | -0.353 | 1.345 | 0.019 | 0.688 | -0.007 | 0.029 | 0.029 | 0 | 0.06 | 0.15 | 0.13 | 0.1 | 0.07 | 0.07 |
| 3 | 159814546 | rs9866988 | 1.249 | 1.461 | -0.211 | 0.546 | 0.052 | 0.014 | 0.014 | 0.087 | 0.79 | 0.57 | 0.73 | 0.75 | 0.72 | 0.67 |
| 3 | 159819567 | rs11921898 | -1.216 | 0.186 | -0.13 | 0.746 | -0.009 | -0.002 | -0.002 | 0 | 0.1 | 0.22 | 0.05 | 0.08 | 0.08 | 0.05 |
| 3 | 159820753 | rs11922988 | -1.721 | 0.114 | -0.046 | 0.836 | -0.038 | 0.082 | 0.082 | 0 | 0.17 | 0.49 | 0.21 | 0.29 | 0.12 | 0.26 |
| 3 | 159822667 | rs9861636 | -0.382 | -0.686 | -1.211 | 0.151 | 0.101 | 0.21 | 0.21 | 4.551 | 0.56 | 0.95 | 0.52 | 0.81 | 0.28 | 0.47 |
| 3 | 159834050 | rs11916869 | -1.664 | 0.08 | 0.478 | 0.262 | 0.02 | 0.003 | 0.003 | 0 | 0.17 | 0.62 | 0.15 | 0.17 | 0.17 | 0.2 |
| 3 | 159843433 | rs4541448 | -1.107 | 0.047 | 0.731 | 0.268 | -0.005 | 0.065 | 0.065 | 0 | 0.12 | 0.55 | 0.09 | 0.17 | 0.06 | 0.16 |
| 3 | 159845146 | rs7622728 | -0.592 | 0.081 | 0.649 | 0.268 | 0.006 | 0 | 0 | 0.06 | 0.07 | 0.13 | 0.1 | 0.08 | 0.14 | 0.06 |
| 3 | 159846158 | rs11709882 | -1.819 | 0.061 | 0.833 | 0.262 | 0.05 | 0.021 | 0.021 | 0 | 0.16 | 0.53 | 0.16 | 0.17 | 0.17 | 0.19 |
| 3 | 159861147 | rs9880646 | -0.631 | -0.323 | 0.281 | 0.747 | -0.03 | -0.073 | -0.073 | 2.929 | 0.62 | 0.89 | 0.51 | 0.57 | 0.39 | 0.49 |
| 3 | 159876691 | rs4130284 | -0.997 | -0.876 | 0.245 | 0.666 | -0.05 | 0.016 | 0.016 | 0.162 | 0.18 | 0.64 | 0.16 | 0.2 | 0.07 | 0.15 |
| 3 | 159877251 | rs9853972 | 0.585 | -1.057 | 0.321 | 0.579 | 0.055 | 0.026 | 0.026 | 0.118 | 0.42 | 0.14 | 0.34 | 0.38 | 0.32 | 0.34 |
| 3 | 159880446 | rs7638698 | -1.246 | -1.214 | 0.467 | 0.236 | -0.051 | 0.013 | 0.013 | 0.125 | 0.19 | 0.65 | 0.16 | 0.2 | 0.07 | 0.15 |
| 3 | 159888261 | rs12631488 | -0.948 | -0.911 | 0.61 | 0.266 | -0.017 | 0.041 | 0.041 | 0 | 0.12 | 0.27 | 0.11 | 0.14 | 0.06 | 0.13 |
| 3 | 159892169 | rs6441301 | -0.553 | -0.72 | 0.056 | 0.062 | 0.036 | 0.052 | 0.052 | 1.449 | 0.58 | 0.76 | 0.46 | 0.33 | 0.35 | 0.41 |
| 3 | 159899731 | rs9840016 | 0.409 | -1.138 | -0.394 | -0.07 | 0.049 | 0.035 | 0.035 | 0.115 | 0.37 | 0.24 | 0.36 | 0.27 | 0.21 | 0.25 |
| 3 | 159911288 | rs868916 | -1.425 | -0.624 | 0.083 | 0.155 | -0.016 | -0.007 | -0.007 | 0 | 0.27 | 0.66 | 0.21 | 0.2 | 0.15 | 0.23 |
| 3 | 159912709 | rs2276750 | -0.481 | -0.606 | 0.016 | 0.179 | 0.01 | -0.015 | -0.015 | 0.485 | 0.15 | 0.09 | 0.15 | 0.18 | 0.09 | 0.18 |
| 3 | 159931416 | rs11928737 | -0.263 | -0.623 | 0.045 | 0.171 | 0.008 | -0.014 | -0.014 | 0 | 0.13 | 0.07 | 0.15 | 0.14 | 0.08 | 0.05 |
| 3 | 159932610 | rs6775953 | -1.774 | 0.602 | -0.135 | -0.074 | -0.013 | 0.017 | 0.017 | 0 | 0.35 | 0.7 | 0.34 | 0.2 | 0.39 | 0.3 |
| 3 | 159942445 | rs2152426 | -1.557 | 0.63 | 0.367 | -0.034 | 0.031 | -0.02 | -0.02 | 0 | 0.29 | 0.59 | 0.33 | 0.2 | 0.36 | 0.29 |
| 3 | 159947951 | rs11711568 | -1.536 | 0.623 | 0.856 | -0.076 | 0.049 | -0.022 | -0.022 | 0 | 0.23 | 0.43 | 0.2 | 0.14 | 0.33 | 0.24 |
| 3 | 159950498 | rs4679876 | -1.797 | 0.156 | 0.017 | -0.701 | -0.004 | 0.121 | 0.121 | 0 | 0.49 | 0.74 | 0.33 | 0.16 | 0.49 | 0.33 |
| 3 | 159952941 | rs10513552 | -1.609 | -0.273 | 0.305 | -0.691 | -0.006 | -0.027 | -0.027 | 0 | 0.29 | 0.66 | 0.16 | 0.05 | 0.18 | 0.1 |
| 3 | 159956559 | rs6797926 | -1.86 | 0.066 | 0.262 | -0.633 | 0.01 | 0.025 | 0.025 | 0.39 | 0.35 | 0.71 | 0.23 | 0.13 | 0.31 | 0.19 |
| 3 | 159973102 | rs4532176 | -1.413 | -0.365 | 0.734 | -0.746 | -0.034 | -0.046 | -0.046 | 0 | 0.22 | 0.32 | 0.1 | 0.04 | 0.14 | 0.08 |
| 3 | 159976271 | rs16831149 | -1.13 | -0.447 | 1.245 | -0.828 | -0.04 | -0.054 | -0.054 | 0 | 0.12 | 0.53 | 0.08 | 0.03 | 0.05 | 0.02 |
| 3 | 159993490 | rs4679880 | -1.514 | -0.127 | 1.519 | -0.896 | 0.004 | -0.08 | -0.08 | 0 | 0.26 | 0.56 | 0.18 | 0.07 | 0.23 | 0.14 |
| 3 | 159994945 | rs10936199 | -1.502 | -0.127 | 1.52 | -0.909 | 0.004 | -0.083 | -0.083 | 0.137 | 0.25 | 0.56 | 0.19 | 0.07 | 0.24 | 0.14 |
| 3 | 160065568 | rs6809313 | -2.429 | -0.336 | 1.225 | -0.837 | -0.044 | 0.028 | 0.028 | 0 | 0.57 | 0.87 | 0.5 | 0.31 | 0.58 | 0.43 |
| 3 | 160073391 | rs7614701 | -0.973 | -0.449 | 1.331 | -0.909 | -0.02 | 0.003 | 0.003 | 0 | 0.09 | 0.32 | 0.06 | 0.07 | 0.08 | 0.05 |
| 3 | 160083918 | rs34182424 | 1.766 | -0.468 | 1.371 | -0.92 | 0.001 | -0.006 | -0.006 | 0 | 0.06 | 0.05 | 0.03 | 0 | 0.02 | 0 |
| 3 | 160089896 | rs12233589 | -0.973 | -0.456 | 1.371 | -0.911 | -0.02 | 0.003 | 0.003 | 0 | 0.09 | 0.32 | 0.06 | 0.07 | 0.08 | 0.05 |
| 3 | 160116174 | rs9843737 | 1.381 | -0.473 | 1.434 | -0.92 | -0.027 | -0.032 | -0.032 | 0 | 0.07 | 0.12 | 0.03 | 0 | 0.02 | 0.01 |
| 3 | 160121839 | rs10936201 | 0.678 | -0.484 | 1.45 | -0.917 | -0.012 | 0.014 | 0.014 | 0.133 | 0.18 | 0.12 | 0.25 | 0.13 | 0.24 | 0.2 |
| 3 | 160130110 | rs2305407 | -1.758 | -0.435 | 1.484 | -0.92 | -0.001 | -0.004 | -0.004 | 0 | 0.47 | 0.57 | 0.45 | 0.24 | 0.52 | 0.36 |
| 3 | 160135286 | rs4680582 | -0.18 | -0.211 | 1.547 | -0.805 | -0.031 | 0.054 | 0.054 | 0.877 | 0.19 | 0.09 | 0.13 | 0.04 | 0.2 | 0.12 |
| 3 | 160135488 | rs9813697 | 1.321 | -0.234 | 1.589 | -0.815 | -0.031 | -0.037 | -0.037 | 0 | 0.07 | 0.14 | 0.03 | 0 | 0.02 | 0.02 |
| 3 | 160152076 | rs3773371 | -0.056 | -0.137 | 1.493 | -0.737 | -0.01 | -0.005 | -0.005 | 0 | 0.07 | 0.17 | 0.04 | 0.04 | 0.07 | 0.05 |
| 3 | 160152696 | rs10923 | 0.867 | -0.174 | 1.464 | -0.777 | -0.013 | 0.008 | 0.008 | 0.925 | 0.16 | 0.12 | 0.25 | 0.13 | 0.23 | 0.2 |
| 3 | 160154258 | rs1047210 | -0.921 | -0.178 | 1.472 | -0.762 | -0.034 | 0.047 | 0.047 | 0 | 0.35 | 0.24 | 0.38 | 0.17 | 0.43 | 0.31 |
| 3 | 160154747 | rs4679885 | -1.689 | -0.179 | 1.472 | -0.761 | 0.001 | -0.011 | -0.011 | 1.881 | 0.45 | 0.57 | 0.44 | 0.24 | 0.51 | 0.36 |
| 3 | 160159921 | rs11706810 | -0.953 | -0.167 | 1.401 | -0.759 | -0.017 | 0.023 | 0.023 | 0 | 0.36 | 0.26 | 0.38 | 0.17 | 0.43 | 0.31 |
| 3 | 160168292 | rs2279457 | -1.625 | -0.189 | 1.321 | -0.731 | 0.003 | -0.017 | -0.017 | 0 | 0.45 | 0.56 | 0.44 | 0.25 | 0.51 | 0.36 |
| 3 | 160171926 | rs17236487 | 0.279 | -0.135 | 1.215 | -0.694 | -0.008 | 0.007 | 0.007 | 0.067 | 0.15 | 0.15 | 0.12 | 0.07 | 0.19 | 0.12 |
| 3 | 160191374 | rs3851366 | -1.373 | -0.084 | 1.235 | -0.664 | -0.033 | 0.07 | 0.07 | 0 | 0.42 | 0.25 | 0.41 | 0.21 | 0.51 | 0.36 |
| 3 | 160192518 | rs4680588 | -1.524 | -0.081 | 1.296 | -0.659 | -0.003 | 0.011 | 0.011 | 0 | 0.45 | 0.61 | 0.45 | 0.25 | 0.51 | 0.36 |
| 3 | 160208092 | rs10936203 | 0.753 | 0.081 | 1.128 | -0.68 | -0.013 | 0.006 | 0.006 | 0.968 | 0.17 | 0.12 | 0.25 | 0.14 | 0.25 | 0.2 |
| 3 | 160213102 | rs1056669 | -2.044 | -0.196 | 1.479 | -0.541 | 0 | 0.038 | 0.038 | 0 | 0.52 | 0.66 | 0.47 | 0.25 | 0.55 | 0.38 |
| 3 | 160217483 | rs7627756 | -2.07 | -0.195 | 1.411 | -0.54 | -0.001 | 0.067 | 0.067 | 0 | 0.52 | 0.71 | 0.47 | 0.25 | 0.55 | 0.38 |
| 3 | 160227606 | rs2305401 | 0.444 | -0.055 | 0.94 | -0.47 | 0.001 | 0.002 | 0.002 | 0 | 0.06 | 0.05 | 0.03 | 0.03 | 0.07 | 0.04 |
| 3 | 160232432 | rs3773372 | -1.65 | -0.08 | 1.059 | -0.493 | 0.001 | -0.012 | -0.012 | 0 | 0.45 | 0.5 | 0.43 | 0.25 | 0.52 | 0.37 |
| 3 | 160245980 | rs4679890 | -1.501 | -0.082 | 0.935 | -0.485 | -0.017 | 0.062 | 0.062 | 0 | 0.44 | 0.32 | 0.41 | 0.25 | 0.52 | 0.37 |

| | | | | | | | | | | | | | | | | |
|---|---|---|---|---|---|---|---|---|---|---|---|---|---|---|---|---|
| 3 | 160283815 | rs3806629 | 0.431 | -0.186 | 0.673 | -0.48 | 0.001 | 0.002 | 0.002 | 0 | 0.06 | 0.05 | 0.03 | 0.03 | 0.07 | 0.04 |
| 3 | 160294561 | rs17570467 | -1.196 | -0.144 | 0.665 | -0.432 | -0.01 | 0.025 | 0.025 | 0 | 0.42 | 0.4 | 0.42 | 0.21 | 0.52 | 0.37 |
| 3 | 160324015 | rs7629423 | 1.834 | -0.996 | 0.536 | -0.11 | 0.008 | 0.069 | 0.069 | 0 | 0.49 | 0.14 | 0.57 | 0.7 | 0.51 | 0.61 |
| 3 | 160324463 | rs16831289 | 0.699 | -1.073 | 1.049 | -0.159 | -0.036 | -0.041 | -0.041 | 0 | 0.07 | 0.24 | 0.04 | 0 | 0.02 | 0.01 |
| 3 | 160325458 | rs4320089 | 1.135 | -1.18 | 1.123 | -0.215 | 0.002 | 0.041 | 0.041 | 0 | 0.63 | 0.26 | 0.65 | 0.82 | 0.62 | 0.69 |
| 3 | 160327589 | rs6797357 | 0.706 | -1.075 | 1.284 | -0.197 | -0.033 | -0.06 | -0.06 | 0.008 | 0.79 | 0.34 | 0.88 | 0.97 | 0.88 | 0.85 |
| 3 | 160328337 | rs7641978 | 0.58 | -1.073 | 1.491 | -0.196 | -0.054 | -0.1 | -0.1 | 0.062 | 0.8 | 0.46 | 0.89 | 0.97 | 0.88 | 0.85 |
| 3 | 160334529 | rs6778401 | 1.244 | -1.025 | 1.263 | -0.163 | 0.001 | -0.003 | -0.003 | 0.462 | 0.06 | 0.08 | 0.03 | 0 | 0.02 | 0.01 |
| 3 | 160336211 | rs17505069 | -0.809 | -1.016 | 1.227 | -0.147 | 0.003 | 0 | 0 | 0 | 0.13 | 0.12 | 0.08 | 0.09 | 0.12 | 0.08 |
| 3 | 160340896 | rs4679904 | 0.755 | -0.998 | 0.998 | -0.182 | -0.035 | -0.003 | -0.003 | 0 | 0.16 | 0.06 | 0.24 | 0.15 | 0.26 | 0.15 |
| 3 | 160351021 | rs7631510 | 0.025 | -1.063 | 1.297 | -0.17 | -0.04 | -0.049 | -0.049 | 0 | 0.1 | 0.39 | 0.05 | 0 | 0.02 | 0.01 |
| 3 | 160352568 | rs1920660 | 1.834 | -1.008 | 0.616 | -0.086 | 0.014 | 0.074 | 0.074 | 0 | 0.5 | 0.14 | 0.57 | 0.7 | 0.51 | 0.62 |
| 3 | 160358220 | rs1920657 | 1.472 | -1.001 | 0.497 | -0.073 | -0.008 | 0.043 | 0.043 | 0 | 0.56 | 0.27 | 0.61 | 0.74 | 0.52 | 0.64 |
| 3 | 160358415 | rs2141671 | 1.534 | -1.196 | 0.81 | -0.12 | -0.055 | -0.066 | -0.066 | 0 | 0.71 | 0.45 | 0.84 | 0.9 | 0.81 | 0.82 |
| 3 | 160358462 | rs2141670 | 0.5 | -1.127 | 0.234 | -0.086 | -0.004 | -0.015 | -0.015 | 0.238 | 0.82 | 0.95 | 0.9 | 0.88 | 0.81 | 0.8 |
| 3 | 160359711 | rs4635753 | -0.199 | -1.084 | 0.211 | -0.147 | 0.019 | 0.027 | 0.027 | 0 | 0.83 | 0.91 | 0.91 | 0.93 | 0.89 | 0.86 |
| 3 | 160367991 | rs1920656 | -0.108 | -1.06 | 0.967 | 0.176 | 0.07 | -0.015 | -0.015 | 0 | 0.19 | 0.44 | 0.27 | 0.14 | 0.27 | 0.17 |
| 3 | 160376851 | rs4679909 | -0.994 | -1.082 | 0.857 | 0.605 | 0.004 | 0.004 | 0.004 | 1.202 | 0.26 | 0.61 | 0.31 | 0.28 | 0.34 | 0.21 |
| 3 | 160377195 | rs17236586 | -1.425 | -1.088 | 0.844 | 0.597 | 0 | -0.001 | -0.001 | 0.549 | 0.3 | 0.62 | 0.33 | 0.28 | 0.35 | 0.21 |
| 3 | 160385610 | rs6765579 | -1.18 | -1.215 | 0.916 | 0.665 | -0.038 | 0.021 | 0.021 | 0 | 0.11 | 0.25 | 0.07 | 0.13 | 0.05 | 0.07 |
| 3 | 160394315 | rs35498840 | 0.282 | -1.137 | 0.988 | 0.655 | 0.049 | 0 | 0 | 0.146 | 0.16 | 0.32 | 0.21 | 0.14 | 0.24 | 0.15 |
| 3 | 160398885 | rs2367209 | -1.185 | -1.255 | 0.916 | 0.598 | 0.001 | 0.031 | 0.031 | 0.122 | 0.27 | 0.54 | 0.3 | 0.32 | 0.26 | 0.22 |
| 3 | 160400751 | rs6763892 | -1.684 | -1.186 | 0.646 | 0.588 | 0.014 | -0.003 | -0.003 | 1.061 | 0.39 | 0.71 | 0.36 | 0.36 | 0.35 | 0.34 |
| 3 | 160401027 | rs13072556 | -0.234 | -0.147 | 0.862 | 0.913 | 0.046 | 0.025 | 0.025 | 0.097 | 0.16 | 0.32 | 0.2 | 0.19 | 0.24 | 0.16 |
| 3 | 160412175 | rs7651855 | -0.825 | -0.794 | 0.353 | 1.013 | 0.003 | 0.04 | 0.04 | 0.004 | 0.34 | 0.44 | 0.37 | 0.55 | 0.36 | 0.34 |
| 3 | 160416023 | rs11707740 | -0.752 | -0.805 | 0.668 | 1.04 | 0.006 | 0.024 | 0.024 | 0 | 0.34 | 0.37 | 0.38 | 0.5 | 0.38 | 0.31 |
| 3 | 160421402 | rs720320 | -0.705 | -1.16 | 0.705 | 1.12 | -0.009 | -0.003 | -0.003 | 0 | 0.06 | 0.19 | 0.16 | 0.04 | 0.03 | 0.1 |
| 3 | 160427758 | rs13062999 | -0.71 | -1.273 | 0.831 | 1.247 | -0.002 | 0.023 | 0.023 | 0.046 | 0.36 | 0.48 | 0.36 | 0.55 | 0.36 | 0.33 |
| 3 | 160429869 | rs7624766 | -1.13 | -1.308 | 0.511 | 1.137 | 0.011 | -0.002 | -0.002 | 0.142 | 0.41 | 0.76 | 0.41 | 0.61 | 0.38 | 0.36 |
| 3 | 160461351 | rs7633816 | -1.457 | -1.247 | 0.131 | 0.976 | -0.025 | 0.054 | 0.054 | 0.954 | 0.47 | 0.95 | 0.57 | 0.66 | 0.4 | 0.44 |
| 3 | 160462363 | rs9837945 | -1.228 | -1.184 | 0.494 | 0.976 | -0.009 | -0.002 | -0.002 | 0.186 | 0.1 | 0.19 | 0.05 | 0.07 | 0.05 | 0.08 |
| 3 | 160466753 | rs9869775 | -1.01 | -1.291 | 0.503 | 0.863 | -0.034 | -0.058 | -0.058 | 0.288 | 0.21 | 0.3 | 0.09 | 0.06 | 0.1 | 0.11 |
| 3 | 160469056 | rs959960 | -1.722 | -1.304 | -0.005 | 0.845 | 0.016 | 0.039 | 0.039 | 1.12 | 0.45 | 0.93 | 0.57 | 0.66 | 0.38 | 0.43 |
| 3 | 160499192 | rs16831457 | -1.852 | -1.34 | 0.427 | 0.932 | 0.008 | 0.037 | 0.037 | 0.149 | 0.09 | 0.37 | 0.06 | 0.07 | 0.04 | 0.07 |
| 3 | 160519718 | rs17826376 | -0.309 | -1.331 | 0.371 | 0.937 | 0.003 | -0.006 | -0.006 | 0 | 0.06 | 0.04 | 0.04 | 0.07 | 0.04 | 0.07 |
| 3 | 160520986 | rs7618082 | -1.412 | -1.331 | 0.398 | 0.937 | 0 | 0.025 | 0.025 | 0 | 0.08 | 0.28 | 0.05 | 0.07 | 0.04 | 0.07 |
| 3 | 160531628 | rs4390979 | -1.126 | -1.321 | 0.373 | 0.935 | -0.006 | 0.001 | 0.001 | 0.225 | 0.08 | 0.23 | 0.05 | 0.06 | 0.04 | 0.07 |
| 3 | 160535721 | rs12487861 | -0.73 | -1.307 | 0.253 | 0.913 | 0.001 | 0.05 | 0.05 | 0 | 0.27 | 0.55 | 0.47 | 0.59 | 0.28 | 0.35 |
| 3 | 160557063 | rs13080397 | -0.733 | -1.385 | 0.009 | 0.852 | -0.005 | 0.026 | 0.026 | 0 | 0.27 | 0.55 | 0.49 | 0.6 | 0.28 | 0.35 |
| 3 | 160562018 | rs6766368 | -0.524 | -1.116 | 0.283 | 1.009 | -0.006 | -0.015 | -0.015 | 0.079 | 0.06 | 0.22 | 0.03 | 0.02 | 0.03 | 0.03 |
| 3 | 160570195 | rs9869116 | -0.833 | -1.083 | 0.021 | 0.928 | 0.004 | -0.001 | -0.001 | 0.769 | 0.4 | 0.58 | 0.55 | 0.59 | 0.36 | 0.4 |
| 3 | 160578472 | rs1839019 | -1.33 | -1.263 | -0.16 | 0.956 | 0.039 | 0.002 | 0.002 | 0.758 | 0.47 | 0.95 | 0.58 | 0.62 | 0.37 | 0.44 |
| 3 | 160579248 | rs9831287 | -0.997 | -1.196 | 0.07 | 0.966 | 0.005 | 0 | 0 | 0 | 0.42 | 0.56 | 0.46 | 0.58 | 0.36 | 0.42 |
| 3 | 160580103 | rs1965186 | -0.127 | -1.315 | 0.666 | 1.303 | -0.018 | 0.041 | 0.041 | 0 | 0.28 | 0.35 | 0.26 | 0.46 | 0.23 | 0.24 |
| 3 | 160589852 | rs11708813 | 1.548 | -1.245 | -0.036 | 0.57 | -0.106 | 0.267 | 0.267 | 0 | 0.41 | 0.04 | 0.3 | 0.07 | 0.54 | 0.21 |
| 3 | 160594496 | rs10513557 | 0.361 | -1.359 | 0.402 | 0.524 | -0.015 | -0.016 | -0.016 | 0 | 0.1 | 0.14 | 0.14 | 0.07 | 0.06 | 0.03 |
| 3 | 160595284 | rs6801330 | 1.235 | -1.449 | 0.735 | 0.757 | -0.003 | 0.027 | 0.027 | 0 | 0.77 | 0.67 | 0.63 | 0.6 | 0.79 | 0.57 |
| 3 | 160601327 | rs6785844 | 0.213 | -1.458 | 0.954 | 0.863 | 0 | 0.012 | 0.012 | 0 | 0.9 | 0.89 | 0.79 | 0.7 | 0.87 | 0.65 |
| 3 | 160616852 | rs1447622 | 0.023 | -1.542 | 1.127 | 0.819 | -0.032 | 0.026 | 0.026 | 0.373 | 0.33 | 0.44 | 0.31 | 0.5 | 0.25 | 0.33 |
| 3 | 160620006 | rs11718966 | -0.051 | -1.543 | 1.156 | 0.819 | -0.034 | 0.024 | 0.024 | 0.478 | 0.33 | 0.46 | 0.31 | 0.5 | 0.25 | 0.33 |
| 3 | 160626929 | rs9881807 | -0.833 | -1.545 | 1.065 | 0.809 | 0.027 | 0.012 | 0.012 | 0 | 0.43 | 0.59 | 0.45 | 0.57 | 0.3 | 0.36 |
| 3 | 160630267 | rs6776120 | 0.744 | -1.533 | 1.095 | 0.805 | 0.002 | 0.018 | 0.018 | 0 | 0.87 | 0.8 | 0.77 | 0.68 | 0.85 | 0.62 |
| 3 | 160630477 | rs6809177 | 1.711 | -1.522 | 0.952 | 0.739 | -0.031 | 0.203 | 0.203 | 0 | 0.42 | 0.15 | 0.31 | 0.11 | 0.54 | 0.26 |
| 3 | 160631965 | rs13086250 | 1.641 | -1.491 | 0.958 | 0.75 | -0.013 | 0.147 | 0.147 | 0 | 0.43 | 0.22 | 0.31 | 0.11 | 0.54 | 0.26 |
| 3 | 160635849 | rs9845943 | 0.362 | -1.382 | 1.423 | 0.846 | 0.017 | -0.04 | -0.04 | 1.1 | 0.28 | 0.17 | 0.29 | 0.5 | 0.24 | 0.32 |
| 3 | 160641930 | rs12696110 | 0.99 | -1.339 | 1.594 | 0.834 | 0.007 | 0.046 | 0.046 | 0 | 0.87 | 0.8 | 0.78 | 0.65 | 0.85 | 0.63 |
| 3 | 160643787 | rs11705811 | 1.002 | -1.395 | 1.803 | 0.681 | 0.01 | 0.016 | 0.016 | 0.55 | 0.54 | 0.36 | 0.47 | 0.15 | 0.61 | 0.3 |
| 3 | 160647912 | rs12690576 | -0.495 | -1.472 | 1.772 | 0.974 | 0.009 | -0.007 | -0.007 | 0 | 0.39 | 0.28 | 0.34 | 0.47 | 0.32 | 0.38 |
| 3 | 160664954 | rs2044571 | 0.43 | -1.426 | 1.76 | 0.65 | 0.011 | -0.012 | -0.012 | 1.498 | 0.29 | 0.2 | 0.3 | 0.6 | 0.25 | 0.32 |
| 3 | 160668220 | rs1374795 | 1.488 | -1.376 | 1.878 | 0.675 | 0.001 | 0.078 | 0.078 | 0.111 | 0.82 | 0.42 | 0.64 | 0.68 | 0.85 | 0.61 |
| 3 | 160681614 | rs1562464 | 0.366 | -1.383 | 1.742 | 0.687 | 0.011 | -0.011 | -0.011 | 0 | 0.29 | 0.21 | 0.31 | 0.6 | 0.25 | 0.32 |
| 3 | 160682500 | rs1868109 | -0.482 | -1.375 | 1.814 | 0.653 | 0.003 | 0.003 | 0.003 | 0 | 0.41 | 0.36 | 0.36 | 0.66 | 0.33 | 0.39 |
| 3 | 160682599 | rs11709802 | 0.103 | -1.429 | 1.926 | 0.655 | 0.01 | 0.03 | 0.03 | 0 | 0.92 | 0.88 | 0.79 | 0.77 | 0.88 | 0.69 |
| 3 | 160695770 | rs7624185 | 0.397 | -1.421 | 1.92 | 0.698 | 0.011 | -0.022 | -0.022 | 0 | 0.29 | 0.21 | 0.31 | 0.55 | 0.25 | 0.32 |
| 3 | 160696867 | rs6441345 | -1.175 | -1.417 | 1.822 | 0.657 | 0.071 | 0.09 | 0.09 | 0 | 0.49 | 0.68 | 0.49 | 0.7 | 0.36 | 0.47 |
| 3 | 160700144 | rs1374787 | -0.421 | -1.417 | 1.837 | 0.658 | 0.003 | 0.001 | 0.001 | 0 | 0.41 | 0.36 | 0.36 | 0.61 | 0.33 | 0.39 |
| 3 | 160700958 | rs1562466 | 0.144 | -1.387 | 1.48 | 0.668 | 0.01 | 0.054 | 0.054 | 0 | 0.92 | 0.88 | 0.79 | 0.72 | 0.88 | 0.69 |
| 3 | 160701442 | rs1562465 | -1.149 | -1.391 | 1.456 | 0.669 | 0.07 | 0.09 | 0.09 | 0 | 0.49 | 0.68 | 0.49 | 0.7 | 0.36 | 0.47 |
| 3 | 160701866 | rs12487156 | -0.349 | -1.372 | 1.499 | 0.66 | -0.013 | 0.015 | 0.015 | 0 | 0.07 | 0.16 | 0.12 | 0.09 | 0.03 | 0.07 |

| | | | | | | | | | | | | | | | | |
|---|---|---|---|---|---|---|---|---|---|---|---|---|---|---|---|---|
| 3 | 160706041 | rs778643 | -1.921 | -1.367 | 1.453 | 0.566 | -0.063 | 0.264 | 0.264 | 0.077 | 0.58 | 0.91 | 0.71 | 0.98 | 0.48 | 0.78 |
| 3 | 160710141 | rs778650 | -0.517 | -1.332 | 1.498 | 0.63 | 0.025 | 0.1 | 0.1 | 0 | 0.09 | 0.22 | 0.22 | 0.28 | 0.12 | 0.3 |
| 3 | 160716041 | rs778653 | -1.658 | -1.276 | 1.424 | 0.596 | -0.006 | 0.15 | 0.15 | 0 | 0.55 | 0.75 | 0.6 | 0.92 | 0.48 | 0.76 |
| 3 | 160735255 | rs16831756 | -1.197 | -1.299 | 1.533 | 0.599 | -0.004 | 0.021 | 0.021 | 0 | 0.2 | 0.29 | 0.27 | 0.28 | 0.2 | 0.34 |
| 3 | 160744571 | rs898677 | -0.206 | -1.33 | 1.485 | 0.644 | -0.018 | -0.006 | -0.006 | 0.104 | 0.35 | 0.43 | 0.34 | 0.64 | 0.28 | 0.42 |
| 3 | 160767278 | rs1580254 | -0.713 | -1.27 | 1.214 | 0.613 | 0.033 | 0.036 | 0.036 | 0 | 0.45 | 0.64 | 0.49 | 0.69 | 0.34 | 0.4 |
| 3 | 160768149 | rs12496196 | -0.287 | -1.165 | 1.164 | 0.607 | 0.018 | 0.067 | 0.067 | 0 | 0.08 | 0.13 | 0.21 | 0.26 | 0.13 | 0.31 |
| 3 | 160768962 | rs9860182 | -1.146 | -1.201 | 1.181 | 0.657 | 0.075 | 0.112 | 0.112 | 0.011 | 0.49 | 0.74 | 0.51 | 0.71 | 0.36 | 0.45 |
| 3 | 160774244 | rs16831830 | -1.009 | -0.979 | 1.497 | 0.818 | -0.042 | -0.009 | -0.009 | 1.704 | 0.17 | 0.39 | 0.18 | 0.14 | 0.1 | 0.14 |
| 3 | 160783445 | rs6763838 | -1.237 | -0.812 | 1.473 | 0.911 | -0.003 | 0.028 | 0.028 | 0 | 0.22 | 0.33 | 0.33 | 0.3 | 0.21 | 0.26 |
| 3 | 160787502 | rs2306061 | -1.79 | -1.094 | 1.492 | 0.573 | -0.022 | 0.075 | 0.075 | 0 | 0.59 | 0.75 | 0.71 | 0.9 | 0.51 | 0.73 |
| 3 | 160789129 | rs2279106 | -1.358 | -1.182 | 1.704 | 0.242 | -0.016 | 0.04 | 0.04 | 0 | 0.24 | 0.45 | 0.34 | 0.31 | 0.21 | 0.26 |
| 3 | 160796505 | rs13256 | 0.47 | -1.302 | 1.699 | 0.221 | 0.015 | -0.059 | -0.059 | 0 | 0.3 | 0.15 | 0.3 | 0.54 | 0.27 | 0.33 |
| 3 | 160797282 | rs9829882 | -1.046 | -0.781 | 1.777 | 0.247 | 0.002 | 0.018 | 0.018 | 0 | 0.22 | 0.28 | 0.33 | 0.31 | 0.22 | 0.27 |
| 3 | 160800002 | rs1599384 | -0.822 | -0.793 | 1.774 | 0.28 | 0.006 | 0.014 | 0.014 | 0 | 0.19 | 0.22 | 0.22 | 0.26 | 0.22 | 0.25 |
| 3 | 160802353 | rs35599916 | -0.493 | -0.782 | 1.774 | 0.273 | 0.016 | 0.03 | 0.03 | 0 | 0.17 | 0.22 | 0.21 | 0.26 | 0.22 | 0.26 |
| 3 | 160803242 | rs9861850 | -1.185 | -0.821 | 1.963 | 0.249 | 0.041 | 0.115 | 0.115 | 0 | 0.2 | 0.5 | 0.34 | 0.32 | 0.22 | 0.27 |
| 3 | 160805352 | rs2231252 | -0.486 | -0.814 | 1.929 | 0.249 | 0.016 | 0.031 | 0.031 | 0 | 0.17 | 0.22 | 0.21 | 0.27 | 0.22 | 0.26 |
| 3 | 160810269 | rs36011360 | 0.519 | -0.786 | 1.961 | 0.239 | -0.01 | -0.014 | -0.014 | 0 | 0.06 | 0.11 | 0.07 | 0.04 | 0.03 | 0.14 |
| 3 | 160810355 | rs11717272 | 0.277 | -0.84 | 1.895 | 0.192 | 0.033 | -0.066 | -0.066 | 0 | 0.33 | 0.15 | 0.3 | 0.54 | 0.27 | 0.32 |
| 3 | 160812752 | rs4521287 | 0.489 | -0.678 | 1.941 | 0.149 | 0.031 | -0.074 | -0.074 | 3.254 | 0.32 | 0.13 | 0.3 | 0.54 | 0.27 | 0.32 |
| 3 | 160813597 | rs7630674 | -1.552 | -0.608 | 1.847 | 0.078 | -0.025 | 0.092 | 0.092 | 0 | 0.58 | 0.77 | 0.72 | 0.88 | 0.49 | 0.69 |
| 3 | 160821929 | rs7651087 | -0.889 | -0.565 | 1.781 | 0.129 | 0.003 | 0.039 | 0.039 | 0 | 0.5 | 0.41 | 0.52 | 0.8 | 0.49 | 0.58 |
| 3 | 160826404 | rs6441354 | -1.163 | -0.571 | 1.449 | 0.102 | -0.004 | 0.076 | 0.076 | 0 | 0.53 | 0.68 | 0.65 | 0.86 | 0.49 | 0.61 |
| 3 | 160853401 | rs9810737 | -0.401 | -0.404 | 1.173 | 0.286 | -0.025 | -0.037 | -0.037 | 0 | 0.14 | 0.05 | 0.2 | 0.25 | 0.19 | 0.2 |
| 3 | 160878126 | rs9878365 | -0.507 | -0.527 | 1.409 | 0.186 | 0.012 | 0.034 | 0.034 | 0 | 0.16 | 0.22 | 0.22 | 0.26 | 0.18 | 0.2 |
| 3 | 160883062 | rs12631850 | -1.209 | -0.64 | 1.528 | 0.04 | 0 | -0.025 | -0.025 | 0 | 0.54 | 0.55 | 0.55 | 0.79 | 0.48 | 0.61 |
| 3 | 160887654 | rs1026572 | -1.302 | -0.649 | 1.601 | 0.031 | -0.006 | 0.062 | 0.062 | 0 | 0.55 | 0.69 | 0.66 | 0.85 | 0.48 | 0.62 |
| 3 | 160897701 | rs6795586 | -0.209 | -0.69 | 1.757 | 0.111 | -0.009 | -0.002 | -0.002 | 0 | 0.39 | 0.6 | 0.46 | 0.6 | 0.3 | 0.42 |
| 3 | 160897780 | rs16831981 | 0.329 | -0.541 | 1.667 | 0.117 | 0.023 | -0.039 | -0.039 | 1.138 | 0.34 | 0.2 | 0.32 | 0.52 | 0.29 | 0.33 |
| 3 | 160907163 | rs12634256 | -1.6 | -0.709 | 1.663 | 0.107 | -0.046 | -0.029 | -0.029 | 0 | 0.24 | 0.51 | 0.27 | 0.2 | 0.15 | 0.31 |
| 3 | 160936364 | rs6799805 | -0.434 | -0.614 | 1.501 | -0.006 | 0.052 | 0.059 | 0.059 | 0 | 0.45 | 0.75 | 0.52 | 0.71 | 0.33 | 0.58 |
| 3 | 160941318 | rs6774276 | -1.236 | -0.378 | 1.487 | 0.015 | -0.068 | 0.185 | 0.185 | 0 | 0.6 | 0.88 | 0.74 | 0.94 | 0.49 | 0.78 |
| 3 | 160946769 | rs6794601 | -1.346 | -0.381 | 1.516 | 0.012 | -0.026 | 0.061 | 0.061 | 0 | 0.6 | 0.73 | 0.63 | 0.91 | 0.51 | 0.78 |
| 3 | 160959896 | rs4487256 | -0.542 | -0.421 | 1.552 | 0.029 | -0.002 | 0.009 | 0.009 | 0 | 0.1 | 0.08 | 0.19 | 0.24 | 0.15 | 0.19 |
| 3 | 160987158 | rs9876553 | -0.44 | -0.424 | 1.566 | 0.036 | -0.018 | -0.025 | -0.025 | 0 | 0.1 | 0.04 | 0.18 | 0.24 | 0.15 | 0.19 |
| 3 | 161003124 | rs7633186 | -1.295 | -0.425 | 1.539 | 0.016 | -0.07 | 0.19 | 0.19 | 0 | 0.6 | 0.88 | 0.74 | 0.94 | 0.49 | 0.78 |
| 3 | 161008125 | rs13083229 | -0.508 | -0.527 | 1.839 | 0.055 | 0.001 | -0.008 | -0.008 | 0 | 0.49 | 0.52 | 0.55 | 0.76 | 0.46 | 0.54 |
| 3 | 161014276 | rs1478568 | -0.681 | -0.526 | 1.816 | 0.054 | 0.01 | 0.115 | 0.115 | 0 | 0.51 | 0.7 | 0.67 | 0.82 | 0.46 | 0.58 |
| 3 | 161015153 | rs1382429 | 0.472 | -0.554 | 1.908 | 0.051 | 0.024 | -0.073 | -0.073 | 0 | 0.37 | 0.13 | 0.33 | 0.52 | 0.3 | 0.35 |
| 3 | 161017192 | rs9850455 | -1.444 | -0.536 | 1.923 | 0.033 | -0.026 | 0.061 | 0.061 | 0 | 0.6 | 0.73 | 0.63 | 0.91 | 0.51 | 0.76 |
| 3 | 161036373 | rs978933 | -0.656 | -0.516 | 2.056 | 0.087 | 0.055 | 0.127 | 0.127 | 0 | 0.11 | 0.24 | 0.2 | 0.23 | 0.15 | 0.19 |
| 3 | 161043869 | rs10936227 | -0.546 | -0.558 | 2.045 | 0.061 | 0.011 | 0.013 | 0.013 | 0 | 0.5 | 0.53 | 0.43 | 0.67 | 0.36 | 0.6 |
| 3 | 161047467 | rs1993891 | -1.485 | -0.543 | 2.03 | 0.063 | -0.107 | 0.226 | 0.226 | 0 | 0.62 | 0.91 | 0.75 | 0.96 | 0.51 | 0.8 |
| 3 | 161047720 | rs4856703 | 0.373 | -0.564 | 1.978 | 0.07 | 0.029 | -0.036 | -0.036 | 0.146 | 0.38 | 0.24 | 0.34 | 0.53 | 0.31 | 0.35 |
| 3 | 161049534 | rs12636434 | -0.439 | -0.557 | 1.996 | 0.075 | -0.025 | 0.006 | 0.006 | 0 | 0.09 | 0.16 | 0.06 | 0.12 | 0.03 | 0.2 |
| 3 | 161058726 | rs6780536 | 0.328 | -0.569 | 1.965 | 0.039 | 0.026 | -0.024 | -0.024 | 0.035 | 0.38 | 0.32 | 0.35 | 0.54 | 0.31 | 0.35 |
| 3 | 161060507 | rs336545 | -0.547 | -0.523 | 1.904 | 0.079 | -0.001 | 0.009 | 0.009 | 0 | 0.11 | 0.08 | 0.19 | 0.23 | 0.15 | 0.22 |
| 3 | 161062331 | rs7652447 | -0.727 | -0.549 | 1.928 | 0.054 | -0.037 | 0.003 | 0.003 | 0 | 0.1 | 0.28 | 0.08 | 0.11 | 0.03 | 0.16 |
| 3 | 161062724 | rs163327 | -1.011 | -0.483 | 1.94 | 0.001 | -0.005 | 0.008 | 0.008 | 0 | 0.54 | 0.71 | 0.68 | 0.96 | 0.46 | 0.78 |
| 3 | 161071947 | rs6800515 | -0.453 | -0.19 | 1.863 | 0.184 | -0.027 | 0.005 | 0.005 | 0 | 0.09 | 0.27 | 0.18 | 0.1 | 0.03 | 0.15 |
| 3 | 161077630 | rs1450522 | 0.374 | -0.458 | 1.663 | 0.14 | 0.034 | -0.059 | -0.059 | 0 | 0.39 | 0.13 | 0.35 | 0.54 | 0.31 | 0.34 |
| 3 | 161087857 | rs1376122 | -0.347 | -0.469 | 1.393 | 0.098 | -0.002 | -0.001 | -0.001 | 0.51 | 0.48 | 0.52 | 0.53 | 0.64 | 0.34 | 0.5 |
| 3 | 161090616 | rs2054710 | 0.278 | -0.356 | 0.912 | 0.098 | 0.024 | -0.019 | -0.019 | 0.029 | 0.39 | 0.23 | 0.35 | 0.54 | 0.31 | 0.34 |
| 3 | 161097268 | rs456611 | -0.984 | -0.303 | 0.681 | -0.202 | -0.042 | 0.102 | 0.102 | 0 | 0.61 | 0.77 | 0.73 | 0.97 | 0.51 | 0.78 |
| 3 | 161100429 | rs465539 | -1.319 | -0.142 | 1.416 | 0.535 | 0.01 | 0.19 | 0.19 | 0.541 | 0.13 | 0.49 | 0.37 | 0.3 | 0.1 | 0.2 |
| 3 | 161102097 | rs393076 | -1.306 | -0.117 | 1.419 | 0.538 | 0.02 | 0.192 | 0.192 | 0.486 | 0.13 | 0.5 | 0.38 | 0.31 | 0.11 | 0.21 |
| 3 | 161102127 | rs420081 | -0.645 | -0.144 | 1.179 | 0.535 | 0 | 0.012 | 0.012 | 0 | 0.1 | 0.14 | 0.23 | 0.2 | 0.1 | 0.19 |
| 3 | 161102861 | rs459679 | -0.645 | -0.144 | 1.179 | 0.535 | 0 | 0.012 | 0.012 | 0 | 0.1 | 0.14 | 0.23 | 0.2 | 0.1 | 0.18 |
| 3 | 161112867 | rs336550 | -1.432 | -0.151 | 1.504 | 0.569 | 0.032 | 0.177 | 0.177 | 1.388 | 0.14 | 0.44 | 0.26 | 0.3 | 0.12 | 0.23 |
| 3 | 161130900 | rs1811483 | 0.144 | -0.337 | 0.623 | 0.341 | 0.02 | -0.062 | -0.062 | 0.166 | 0.39 | 0.13 | 0.27 | 0.55 | 0.35 | 0.39 |
| 3 | 161148190 | rs7618391 | 0.373 | 0.373 | 0.498 | -0.382 | -0.001 | 0.013 | 0.013 | 0 | 0.4 | 0.37 | 0.28 | 0.06 | 0.42 | 0.38 |
| 3 | 161150771 | rs12163614 | 0.788 | 0.28 | 0.534 | -0.478 | -0.001 | -0.001 | -0.001 | 0 | 0.09 | 0.08 | 0.07 | 0.1 | 0.11 | 0.21 |
| 3 | 161151441 | rs1450525 | 0.343 | 0.243 | 0.947 | -0.368 | -0.006 | 0.073 | 0.073 | 0 | 0.4 | 0.31 | 0.32 | 0.14 | 0.4 | 0.3 |
| 3 | 161153006 | rs971515 | -0.082 | 0.224 | 1.239 | -0.239 | 0.004 | 0.018 | 0.018 | 0 | 0.48 | 0.42 | 0.5 | 0.35 | 0.47 | 0.44 |
| 3 | 161158310 | rs7611369 | 0.142 | 0.231 | 1.21 | -0.225 | -0.009 | 0.05 | 0.05 | 0 | 0.08 | 0.11 | 0.19 | 0.21 | 0.06 | 0.14 |
| 3 | 161160475 | rs7625210 | -0.624 | 0.202 | 1.359 | -0.245 | 0 | 0.002 | 0.002 | 0 | 0.53 | 0.61 | 0.56 | 0.42 | 0.48 | 0.63 |
| 3 | 161162664 | rs6762110 | 0.223 | 0.142 | 1.241 | -0.266 | -0.024 | 0.022 | 0.022 | 0.154 | 0.43 | 0.13 | 0.48 | 0.33 | 0.43 | 0.39 |
| 3 | 161164355 | rs16832376 | -2.131 | 0.113 | 1.519 | -0.278 | 0.033 | 0.11 | 0.11 | 0.974 | 0.07 | 0.36 | 0.07 | 0.07 | 0.02 | 0.21 |

| | | | | | | | | | | | | | | | | |
|---|---|---|---|---|---|---|---|---|---|---|---|---|---|---|---|---|
| 3 | 161164453 | rs12632716 | 0.193 | 0.121 | 1.477 | -0.278 | 0 | 0.033 | 0.033 | 0.134 | 0.43 | 0.17 | 0.48 | 0.33 | 0.43 | 0.39 |
| 3 | 161170611 | rs12633334 | -0.62 | 0.142 | 1.474 | -0.238 | 0.001 | 0 | 0 | 0 | 0.53 | 0.6 | 0.56 | 0.44 | 0.48 | 0.64 |
| 3 | 161176756 | rs9843804 | 0.913 | 0.162 | 1.232 | -0.241 | -0.085 | 0.093 | 0.093 | 0 | 0.35 | 0.03 | 0.29 | 0.14 | 0.36 | 0.25 |
| 3 | 161193325 | rs16832441 | 0.326 | 0.25 | 1.146 | -0.138 | 0.004 | -0.003 | -0.003 | 0 | 0.07 | 0.06 | 0.18 | 0.22 | 0.06 | 0.14 |
| 3 | 161195729 | rs6441378 | 0.107 | 0.158 | 1.008 | -0.411 | 0.006 | 0.021 | 0.021 | 0 | 0.46 | 0.59 | 0.4 | 0.23 | 0.43 | 0.58 |
| 3 | 161203534 | rs17478663 | -0.51 | 0.161 | 1.013 | -0.415 | 0 | -0.001 | -0.001 | 0 | 0.11 | 0.08 | 0.1 | 0.07 | 0.1 | 0.31 |
| 3 | 161209750 | rs17478991 | 0.934 | 0.157 | 0.994 | -0.413 | -0.003 | 0.004 | 0.004 | 0.316 | 0.06 | 0.01 | 0.08 | 0.03 | 0.06 | 0.26 |
| 3 | 161213636 | rs6441380 | 1.051 | 0.158 | 1.114 | -0.416 | 0 | -0.004 | -0.004 | 0 | 0.34 | 0.35 | 0.29 | 0.15 | 0.33 | 0.26 |
| 3 | 161214536 | rs12630808 | -0.329 | 0.291 | 0.986 | -0.331 | -0.003 | 0.01 | 0.01 | 0 | 0.12 | 0.12 | 0.2 | 0.21 | 0.1 | 0.14 |
| 3 | 161221705 | rs3921595 | -0.271 | 0.2 | 0.764 | -0.458 | 0.01 | 0.074 | 0.074 | 0 | 0.51 | 0.65 | 0.42 | 0.27 | 0.47 | 0.59 |
| 3 | 161226185 | rs4287919 | 0.179 | 0.197 | 0.76 | -0.37 | 0.018 | 0.034 | 0.034 | 0.247 | 0.45 | 0.24 | 0.37 | 0.32 | 0.42 | 0.41 |
| 3 | 161237020 | rs9821185 | 0.835 | 0.208 | 0.747 | -0.625 | -0.003 | 0.105 | 0.105 | 0 | 0.37 | 0.17 | 0.2 | 0.11 | 0.37 | 0.24 |
| 3 | 161268611 | rs6781716 | 0.201 | 0.192 | 0.76 | -0.571 | 0.016 | 0.058 | 0.058 | 0.342 | 0.44 | 0.24 | 0.37 | 0.27 | 0.42 | 0.39 |
| 3 | 161268974 | rs10513567 | 0.429 | 0.17 | 0.73 | -0.467 | 0.004 | -0.002 | -0.002 | 0 | 0.08 | 0.07 | 0.17 | 0.16 | 0.05 | 0.16 |
| 3 | 161274019 | rs6780945 | -0.935 | 0.218 | 1.049 | -0.506 | 0.014 | 0.009 | 0.009 | 0.63 | 0.13 | 0.35 | 0.12 | 0.12 | 0.1 | 0.3 |
| 3 | 161275848 | rs17421738 | -0.364 | 0.24 | 0.788 | -0.509 | -0.002 | -0.001 | -0.001 | 0 | 0.1 | 0.06 | 0.1 | 0.11 | 0.1 | 0.3 |
| 3 | 161328281 | rs2674222 | 0.772 | 0.234 | 0.794 | -0.552 | -0.084 | -0.036 | -0.036 | 0 | 0.4 | 0.1 | 0.47 | 0.61 | 0.47 | 0.3 |
| 3 | 161328660 | rs1515735 | 1.354 | 0.714 | 0.474 | -0.659 | -0.034 | 0.033 | 0.033 | 0 | 0.07 | 0.02 | 0.1 | 0 | 0.13 | 0.01 |
| 3 | 161339062 | rs6441392 | -0.882 | 0.27 | 0.921 | -0.464 | -0.026 | -0.001 | -0.001 | 0.013 | 0.58 | 0.73 | 0.51 | 0.39 | 0.53 | 0.7 |
| 3 | 161340855 | rs9840911 | 1.18 | 0.252 | 0.979 | -0.509 | -0.002 | 0.105 | 0.105 | 0.897 | 0.33 | 0.11 | 0.19 | 0.1 | 0.34 | 0.24 |
| 3 | 161348584 | rs9851175 | 0.31 | 0.268 | 1.04 | -0.492 | 0.003 | 0.005 | 0.005 | 1.765 | 0.42 | 0.41 | 0.38 | 0.28 | 0.39 | 0.36 |
| 3 | 161359375 | rs1399916 | -0.244 | 0.221 | 0.823 | -0.522 | 0.026 | 0.061 | 0.061 | 0.188 | 0.5 | 0.68 | 0.44 | 0.29 | 0.44 | 0.44 |
| 3 | 161362119 | rs4269094 | 0.797 | 0.414 | 0.701 | -0.506 | 0 | -0.009 | -0.009 | 0 | 0.35 | 0.36 | 0.21 | 0.1 | 0.34 | 0.25 |
| 3 | 161380386 | rs9881042 | 0.078 | 0.339 | 0.689 | -0.425 | -0.001 | 0.002 | 0.002 | 0 | 0.47 | 0.45 | 0.33 | 0.14 | 0.47 | 0.52 |
| 3 | 161381008 | rs16833090 | 0.765 | 0.433 | 0.43 | -0.238 | -0.011 | -0.017 | -0.017 | 0 | 0.06 | 0 | 0.08 | 0.04 | 0.06 | 0.22 |
| 3 | 161385281 | rs9879283 | -0.484 | 0.366 | 0.672 | -0.154 | -0.012 | 0.016 | 0.016 | 0 | 0.55 | 0.69 | 0.55 | 0.34 | 0.53 | 0.69 |
| 3 | 161393049 | rs9863808 | -0.389 | 0.337 | 0.478 | 0.141 | -0.005 | 0.012 | 0.012 | 0 | 0.11 | 0.15 | 0.23 | 0.2 | 0.09 | 0.15 |
| 3 | 161393266 | rs2404572 | -0.65 | 0.292 | 0.729 | -0.126 | -0.002 | -0.024 | -0.024 | 0.971 | 0.13 | 0.27 | 0.11 | 0.07 | 0.1 | 0.27 |
| 3 | 161393745 | rs9827593 | -1.042 | 0.355 | 0.235 | -0.07 | -0.026 | 0.012 | 0.012 | 0 | 0.55 | 0.84 | 0.53 | 0.39 | 0.55 | 0.72 |
| 3 | 161394052 | rs4635724 | -1.159 | 0.56 | 0.306 | -0.223 | -0.012 | -0.03 | -0.03 | 0 | 0.54 | 0.86 | 0.56 | 0.43 | 0.53 | 0.72 |
| 3 | 161396297 | rs4298056 | -0.629 | 0.58 | 1.079 | -0.318 | -0.004 | 0.05 | 0.05 | 0 | 0.16 | 0.34 | 0.16 | 0.18 | 0.12 | 0.2 |
| 3 | 161396942 | rs6773928 | -0.188 | 0.574 | 0.82 | -0.321 | -0.001 | -0.026 | -0.026 | 0 | 0.13 | 0.1 | 0.15 | 0.18 | 0.11 | 0.16 |
| 3 | 161405094 | rs9845108 | -1.325 | 0.575 | 1.168 | -0.346 | 0.006 | 0.054 | 0.054 | 0 | 0.19 | 0.55 | 0.17 | 0.21 | 0.13 | 0.24 |
| 3 | 161417866 | rs9848511 | -1.478 | 0.505 | 1.153 | -0.467 | 0.009 | 0.016 | 0.016 | 0 | 0.22 | 0.34 | 0.19 | 0.23 | 0.19 | 0.33 |
| 3 | 161421122 | rs10084648 | 0.646 | 0.41 | 0.786 | -0.79 | -0.012 | 0.092 | 0.092 | 0 | 0.35 | 0.25 | 0.38 | 0.11 | 0.39 | 0.4 |
| 3 | 161445796 | rs4856718 | -1.162 | 0.421 | 0.91 | -0.692 | 0.001 | 0.002 | 0.002 | 0 | 0.56 | 0.72 | 0.57 | 0.39 | 0.55 | 0.72 |
| 3 | 161447835 | rs16833265 | 0.031 | 0.36 | 0.832 | -0.656 | -0.004 | 0.019 | 0.019 | 0.021 | 0.06 | 0.19 | 0.04 | 0.04 | 0.03 | 0.2 |
| 3 | 161455195 | rs9819763 | -0.275 | 0.369 | 0.375 | -0.682 | -0.065 | 0.092 | 0.092 | 0 | 0.44 | 0.13 | 0.48 | 0.27 | 0.46 | 0.5 |
| 3 | 161460016 | rs16833312 | -1.929 | 0.337 | 0.657 | -0.642 | 0.081 | 0.198 | 0.198 | 1.913 | 0.05 | 0.49 | 0.04 | 0.08 | 0.01 | 0.05 |
| 3 | 161474909 | rs12489261 | 0.268 | 0.457 | 0.018 | -0.619 | -0.01 | -0.056 | -0.056 | 0.035 | 0.13 | 0.01 | 0.12 | 0.15 | 0.09 | 0.14 |
| 3 | 161475537 | rs9832859 | -0.234 | 0.45 | 0.046 | -0.657 | -0.009 | 0.07 | 0.07 | 0 | 0.45 | 0.29 | 0.49 | 0.27 | 0.46 | 0.5 |
| 3 | 161475954 | rs17424987 | 0.292 | 0.425 | 0.04 | -0.65 | -0.01 | -0.056 | -0.056 | 0 | 0.13 | 0.01 | 0.12 | 0.15 | 0.09 | 0.14 |
| 3 | 161482645 | rs308690 | -0.082 | 0.556 | 0.176 | -0.552 | -0.007 | 0.07 | 0.07 | 0 | 0.45 | 0.29 | 0.49 | 0.27 | 0.45 | 0.5 |
| 3 | 161490079 | rs1515730 | -0.834 | 0.771 | 0.77 | -0.562 | 0.055 | 0.068 | 0.068 | 0 | 0.09 | 0.25 | 0.06 | 0.15 | 0.1 | 0.19 |
| 3 | 161498506 | rs2404822 | -0.023 | 0.583 | 0.482 | -1.489 | -0.001 | -0.055 | -0.055 | 0 | 0.55 | 0.58 | 0.54 | 0.91 | 0.49 | 0.58 |
| 3 | 161504007 | rs1996264 | 0.469 | 0.62 | 0.288 | -1.483 | 0.025 | 0.097 | 0.097 | 0.256 | 0.5 | 0.34 | 0.48 | 0.84 | 0.46 | 0.4 |
| 3 | 161514758 | rs6441399 | -0.194 | 0.698 | 0.228 | -1.595 | 0.027 | -0.08 | -0.08 | 0.635 | 0.86 | 0.69 | 0.88 | 1 | 0.87 | 0.92 |
| 3 | 161515001 | rs1370087 | 0.784 | 0.678 | 0.045 | -1.626 | -0.017 | 0.04 | 0.04 | 0 | 0.32 | 0.19 | 0.34 | 0.08 | 0.38 | 0.33 |
| 3 | 161515014 | rs1370086 | 1.087 | 0.645 | -0.653 | -1.575 | 0.017 | 0.183 | 0.183 | 0.176 | 0.45 | 0.13 | 0.44 | 0.81 | 0.4 | 0.37 |
| 3 | 161517861 | rs12639469 | 0.684 | 0.61 | -0.73 | -1.569 | 0.025 | 0.111 | 0.111 | 0.058 | 0.5 | 0.32 | 0.48 | 0.85 | 0.46 | 0.41 |
| 3 | 161520740 | rs1436742 | 1.077 | 0.632 | -0.966 | -1.6 | -0.133 | 0.111 | 0.111 | 0 | 0.26 | 0.02 | 0.32 | 0.09 | 0.36 | 0.27 |
| 3 | 161522742 | rs1436740 | 0.247 | 0.314 | -0.681 | -1.375 | -0.001 | 0 | 0 | 0 | 0.09 | 0.1 | 0.07 | 0.05 | 0.06 | 0.23 |
| 3 | 161523244 | rs2118097 | 0.367 | 0.247 | -0.79 | -1.471 | 0.028 | 0.124 | 0.124 | 0.214 | 0.5 | 0.31 | 0.49 | 0.87 | 0.45 | 0.41 |
| 3 | 161523859 | rs9859426 | -0.333 | 0.007 | -0.888 | -0.9 | 0 | 0 | 0 | 0 | 0.09 | 0.08 | 0.06 | 0.05 | 0.06 | 0.23 |
| 3 | 161525058 | rs2118096 | -0.003 | 0.452 | -1.445 | -0.979 | -0.061 | 0.029 | 0.029 | 0 | 0.36 | 0.12 | 0.32 | 0.26 | 0.41 | 0.26 |
| 3 | 161526024 | rs2404825 | 1.077 | 0.165 | -1.897 | -1.385 | -0.007 | 0.1 | 0.1 | 0.084 | 0.29 | 0.07 | 0.17 | 0.15 | 0.29 | 0.49 |
| 3 | 161535407 | rs2033686 | 0.656 | 0.181 | -1.781 | -1.18 | -0.015 | -0.013 | -0.013 | 0 | 0.15 | 0.09 | 0.21 | 0.23 | 0.24 | 0.16 |
| 3 | 161537899 | rs17427131 | 0.721 | 0.169 | -1.638 | -1.195 | 0.007 | 0.002 | 0.002 | 0 | 0.15 | 0.14 | 0.21 | 0.23 | 0.24 | 0.16 |
| 3 | 161540640 | rs6779918 | -0.898 | -0.076 | -0.989 | -1.394 | -0.001 | -0.026 | -0.026 | 0 | 0.55 | 0.56 | 0.73 | 0.84 | 0.59 | 0.49 |
| 3 | 161540839 | rs11926881 | 1.022 | -0.779 | -0.824 | -0.882 | -0.001 | -0.003 | -0.003 | 0.002 | 0.15 | 0.12 | 0.39 | 0.6 | 0.17 | 0.22 |
| 3 | 161544997 | rs13061797 | 0.062 | -0.718 | -0.837 | -1.088 | 0.014 | 0.002 | 0.002 | 0.746 | 0.21 | 0.15 | 0.1 | 0.01 | 0.16 | 0.1 |
| 3 | 161556214 | rs1583563 | -0.208 | -0.752 | -0.699 | -1.077 | -0.015 | -0.072 | -0.072 | 0.479 | 0.2 | 0.22 | 0.11 | 0.01 | 0.18 | 0.13 |
| 3 | 161563784 | rs2404829 | -0.221 | -0.45 | -0.789 | -0.581 | -0.007 | -0.021 | -0.021 | 0 | 0.22 | 0.18 | 0.16 | 0.35 | 0.23 | 0.37 |
| 3 | 161564087 | rs1115517 | 0.478 | -0.937 | -0.693 | -1.171 | 0.008 | 0.014 | 0.014 | 0 | 0.56 | 0.63 | 0.71 | 0.65 | 0.61 | 0.61 |
| 3 | 161564168 | rs2019291 | 0.616 | -0.977 | -0.479 | -1.189 | -0.004 | -0.006 | -0.006 | 0 | 0.55 | 0.51 | 0.7 | 0.64 | 0.61 | 0.6 |
| 3 | 161568993 | rs7433024 | -1.314 | -0.995 | -0.498 | -1.198 | 0.004 | -0.028 | -0.028 | 0.035 | 0.8 | 0.84 | 0.89 | 1 | 0.85 | 0.98 |
| 3 | 161570645 | rs1972828 | 0.152 | -0.842 | -0.476 | -1.061 | -0.017 | -0.04 | -0.04 | 0 | 0.22 | 0.12 | 0.15 | 0.35 | 0.23 | 0.37 |
| 3 | 161574584 | rs1388699 | 0.436 | -0.966 | -0.538 | -0.931 | -0.003 | -0.002 | -0.002 | 0 | 0.49 | 0.47 | 0.69 | 0.59 | 0.61 | 0.61 |
| 3 | 161581684 | rs1492146 | 0.157 | -1.031 | -0.796 | -0.974 | -0.031 | -0.096 | -0.096 | 1.1 | 0.26 | 0.1 | 0.17 | 0.4 | 0.25 | 0.38 |

| | | | | | | | | | | | | | | | | |
|---|---|---|---|---|---|---|---|---|---|---|---|---|---|---|---|---|
| 3 | 161582755 | rs4856730 | -1.358 | -0.986 | -0.692 | -1.01 | 0.022 | 0.001 | 0.001 | 1.926 | 0.77 | 0.88 | 0.88 | 1 | 0.85 | 0.98 |
| 3 | 161587817 | rs1994742 | 0.506 | -0.408 | -0.44 | -0.791 | 0.004 | 0 | 0 | 0 | 0.44 | 0.42 | 0.62 | 0.56 | 0.4 | 0.33 |
| 3 | 161587855 | rs17501198 | 0.96 | -0.323 | -0.498 | -0.835 | 0.012 | -0.047 | -0.047 | 0 | 0.27 | 0.26 | 0.19 | 0.05 | 0.35 | 0.29 |
| 3 | 161593739 | rs4093549 | 0.575 | -0.274 | -0.569 | -0.77 | 0.002 | 0 | 0 | 0.03 | 0.44 | 0.4 | 0.62 | 0.56 | 0.4 | 0.33 |
| 3 | 161604866 | rs9840287 | 0.545 | -0.046 | -0.482 | -0.04 | 0 | -0.005 | -0.005 | 0 | 0.72 | 0.68 | 0.82 | 0.6 | 0.76 | 0.62 |
| 3 | 161616585 | rs11707244 | 0.545 | -0.009 | -0.529 | -0.001 | -0.002 | 0.008 | 0.008 | 0 | 0.72 | 0.68 | 0.82 | 0.6 | 0.76 | 0.61 |
| 3 | 161620429 | rs722711 | -0.748 | 0.223 | -0.9 | 0.385 | 0.001 | -0.021 | -0.021 | 0 | 0.22 | 0.18 | 0.39 | 0.71 | 0.22 | 0.34 |
| 3 | 161625084 | rs2405046 | 0.559 | 0.318 | -0.178 | -0.01 | 0.014 | 0.012 | 0.012 | 0 | 0.94 | 0.8 | 0.95 | 0.94 | 0.95 | 0.84 |
| 3 | 161633104 | rs7626487 | 0.117 | 0.514 | -0.087 | -0.279 | -0.002 | -0.003 | -0.003 | 0 | 0.54 | 0.51 | 0.38 | 0.43 | 0.58 | 0.4 |
| 3 | 161652269 | rs6441419 | 0.667 | 1.16 | -0.287 | -0.643 | 0.017 | 0.006 | 0.006 | 0 | 0.43 | 0.22 | 0.27 | 0.43 | 0.39 | 0.31 |
| 3 | 161653188 | rs6788766 | 0.436 | 1.251 | -0.124 | -0.67 | -0.001 | 0.014 | 0.014 | 0.02 | 0.32 | 0.35 | 0.5 | 0.53 | 0.31 | 0.33 |
| 3 | 161653669 | rs2101653 | -0.298 | 1.392 | -0.487 | -0.792 | -0.062 | -0.031 | -0.031 | 0.003 | 0.46 | 0.86 | 0.64 | 0.53 | 0.48 | 0.41 |
| 3 | 161657384 | rs6778365 | 0.708 | 2.107 | 0.157 | -0.568 | 0.031 | -0.043 | -0.043 | 0 | 0.1 | 0.15 | 0.1 | 0.01 | 0.19 | 0.09 |
| 3 | 161662483 | rs11927460 | 0.193 | 1.489 | -0.185 | -0.472 | 0.001 | 0.005 | 0.005 | 1.616 | 0.08 | 0.18 | 0.16 | 0.06 | 0.05 | 0.05 |
| 3 | 161665740 | rs9878666 | 0.368 | 1.632 | -0.311 | -0.468 | -0.018 | -0.008 | -0.008 | 0 | 0.11 | 0.06 | 0.09 | 0.41 | 0.15 | 0.19 |
| 3 | 161669809 | rs6441421 | -0.558 | 1.424 | -0.391 | -0.54 | 0.002 | 0.008 | 0.008 | 0 | 0.26 | 0.24 | 0.28 | 0.48 | 0.31 | 0.3 |
| 3 | 161678110 | rs1352913 | 0.17 | 1.034 | -0.451 | -0.289 | -0.007 | -0.007 | -0.007 | 0 | 0.9 | 0.86 | 0.92 | 0.93 | 0.94 | 0.76 |
| 3 | 161695760 | rs4575917 | -0.321 | 1.315 | -0.612 | -0.134 | -0.001 | -0.041 | -0.041 | 0 | 0.65 | 0.84 | 0.58 | 0.56 | 0.68 | 0.65 |
| 3 | 161700441 | rs9867137 | -0.338 | 1.305 | -0.369 | -0.064 | 0 | 0.016 | 0.016 | 0 | 0.16 | 0.2 | 0.14 | 0.42 | 0.21 | 0.24 |
| 3 | 161700564 | rs4299495 | 0.822 | 1.271 | -0.198 | -0.434 | 0.003 | -0.006 | -0.006 | 0 | 0.42 | 0.54 | 0.4 | 0.11 | 0.35 | 0.32 |
| 3 | 161702108 | rs9821852 | -0.76 | 1.685 | -0.243 | -0.653 | -0.037 | -0.028 | -0.028 | 0 | 0.3 | 0.27 | 0.25 | 0.44 | 0.48 | 0.4 |
| 3 | 161708249 | rs9855315 | 1.237 | 1.702 | -0.271 | -0.673 | -0.09 | 0.078 | 0.078 | 0.094 | 0.14 | 0.01 | 0.11 | 0.01 | 0.27 | 0.16 |
| 3 | 161708628 | rs4856741 | 1.968 | 1.506 | -0.057 | -0.721 | 0.037 | 0.081 | 0.081 | 0 | 0.29 | 0.14 | 0.18 | 0.1 | 0.22 | 0.24 |
| 3 | 161709943 | rs4425278 | -0.241 | 1.605 | -0.109 | -0.66 | -0.017 | -0.023 | -0.023 | 0 | 0.58 | 0.81 | 0.54 | 0.53 | 0.56 | 0.56 |
| 3 | 161710141 | rs4256145 | 0.82 | 1.566 | -0.087 | -0.65 | 0.009 | 0.016 | 0.016 | 0 | 0.42 | 0.58 | 0.4 | 0.14 | 0.35 | 0.32 |
| 3 | 161717623 | rs9875486 | 0.372 | 1.573 | -0.345 | -0.619 | -0.003 | 0.007 | 0.007 | 0 | 0.15 | 0.13 | 0.12 | 0.39 | 0.16 | 0.21 |
| 3 | 161747143 | rs4856666 | 0.932 | 1.421 | -0.319 | -0.599 | 0 | -0.046 | -0.046 | 0 | 0.41 | 0.19 | 0.29 | 0.5 | 0.38 | 0.45 |
| 3 | 161770119 | rs4856746 | -0.571 | 1.451 | 0.078 | -0.603 | -0.02 | 0.016 | 0.016 | 0.808 | 0.67 | 0.63 | 0.6 | 0.56 | 0.76 | 0.69 |
| 3 | 161771382 | rs9871102 | -0.448 | 1.569 | -0.105 | -0.681 | 0.046 | -0.05 | -0.05 | 0 | 0.15 | 0.48 | 0.23 | 0.04 | 0.18 | 0.11 |
| 3 | 161778666 | rs9825150 | -0.482 | 1.346 | -0.322 | -0.627 | 0.033 | -0.067 | -0.067 | 0 | 0.29 | 0.62 | 0.35 | 0.06 | 0.45 | 0.27 |
| 3 | 161778975 | rs1397222 | 0.648 | 1.286 | -0.444 | -0.182 | -0.001 | 0.029 | 0.029 | 0 | 0.11 | 0.11 | 0.09 | 0.36 | 0.11 | 0.06 |
| 3 | 161791399 | rs12696134 | -0.734 | 0.839 | -0.496 | -0.347 | 0.048 | 0.09 | 0.09 | 0.618 | 0.76 | 0.93 | 0.91 | 0.94 | 0.85 | 0.8 |
| 3 | 161801528 | rs6441433 | -0.092 | 1.038 | -0.229 | -0.119 | 0.024 | -0.027 | -0.027 | 0.114 | 0.67 | 0.78 | 0.61 | 0.53 | 0.76 | 0.56 |
| 3 | 161807012 | rs6804961 | 0.05 | 1.136 | -0.088 | -0.117 | 0.003 | -0.004 | -0.004 | 0.541 | 0.63 | 0.68 | 0.57 | 0.53 | 0.7 | 0.53 |
| 3 | 161812412 | rs4465957 | 0.363 | 1.113 | 0.141 | -0.135 | -0.011 | 0.003 | 0.003 | 1.597 | 0.57 | 0.51 | 0.54 | 0.53 | 0.65 | 0.49 |
| 3 | 161817702 | rs4856761 | 0.916 | 1.393 | 0.085 | -0.286 | 0.015 | -0.014 | -0.014 | 0 | 0.43 | 0.25 | 0.42 | 0.51 | 0.38 | 0.33 |
| 3 | 161823150 | rs7609974 | 1.05 | 1.264 | 0.138 | -0.231 | 0.005 | 0.011 | 0.011 | 0.036 | 0.05 | 0.1 | 0.11 | 0.04 | 0.06 | 0.05 |
| 3 | 161824408 | rs13088846 | 1.165 | 1.285 | -0.276 | -0.244 | 0.014 | -0.049 | -0.049 | 0 | 0.38 | 0.19 | 0.34 | 0.48 | 0.34 | 0.31 |
| 3 | 161825807 | rs13063743 | 0.381 | 1.515 | -0.29 | -0.334 | 0.018 | 0.082 | 0.082 | 1.227 | 0.55 | 0.33 | 0.7 | 0.87 | 0.59 | 0.54 |
| 3 | 161826690 | rs4539953 | 1.312 | 1.564 | -0.833 | -0.362 | -0.076 | 0.071 | 0.071 | 0 | 0.12 | 0.01 | 0.09 | 0.01 | 0.23 | 0.14 |
| 3 | 161831242 | rs6771785 | -0.375 | 1.429 | -0.73 | -0.31 | 0.005 | 0.039 | 0.039 | 0 | 0.66 | 0.76 | 0.83 | 0.91 | 0.68 | 0.63 |
| 3 | 161832815 | rs11922869 | 0.841 | 1.182 | -0.854 | 0.033 | 0.012 | -0.064 | -0.064 | 0 | 0.37 | 0.15 | 0.33 | 0.47 | 0.34 | 0.31 |
| 3 | 161845898 | rs9829131 | -0.252 | 0.905 | -0.514 | -0.159 | 0.002 | -0.033 | -0.033 | 1.948 | 0.89 | 0.81 | 0.91 | 0.97 | 0.88 | 0.94 |
| 3 | 161846859 | rs1510329 | 0.419 | 0.726 | -1.385 | -0.258 | 0.034 | 0.023 | 0.023 | 1.465 | 0.27 | 0.11 | 0.15 | 0.12 | 0.2 | 0.23 |
| 3 | 161847016 | rs1510328 | -0.231 | 1.669 | -1.601 | 0.183 | -0.05 | -0.048 | -0.048 | 0.021 | 0.36 | 0.18 | 0.32 | 0.41 | 0.42 | 0.49 |
| 3 | 161850511 | rs17749916 | 2.081 | 1.423 | -2.231 | 0.133 | -0.018 | 0.064 | 0.064 | 0.14 | 0.29 | 0.04 | 0.2 | 0.14 | 0.25 | 0.18 |
| 3 | 161853873 | rs10513578 | 1.156 | 1.144 | -1.981 | 0.29 | 0.004 | 0.101 | 0.101 | 0.353 | 0.22 | 0.03 | 0.07 | 0.04 | 0.16 | 0.19 |
| 3 | 161854329 | rs1397216 | 0.258 | 1.519 | -1.372 | 0.848 | -0.013 | -0.014 | -0.014 | 0.623 | 0.1 | 0.06 | 0.16 | 0.29 | 0.22 | 0.26 |
| 3 | 161866740 | rs7648209 | 0.929 | 0.145 | -0.979 | 0.76 | 0.032 | -0.042 | -0.042 | 0 | 0.74 | 0.26 | 0.59 | 0.55 | 0.8 | 0.76 |
| 3 | 161883401 | rs9867116 | 0.78 | 0.226 | -0.82 | 0.838 | 0.007 | -0.01 | -0.01 | 1.146 | 0.76 | 0.32 | 0.59 | 0.55 | 0.82 | 0.79 |
| 3 | 161885828 | rs1567455 | 0.667 | 0.669 | -0.418 | 0.568 | -0.001 | 0.017 | 0.017 | 0 | 0.81 | 0.68 | 0.81 | 0.85 | 0.8 | 0.85 |
| 3 | 161888626 | rs1876994 | -0.914 | 0.376 | -1.23 | 0.027 | -0.028 | 0.094 | 0.094 | 0.137 | 0.8 | 0.96 | 0.92 | 0.97 | 0.8 | 0.84 |
| 3 | 161893088 | rs12496424 | 1.125 | 0.755 | -1.442 | 0.127 | 0.001 | -0.002 | -0.002 | 0 | 0.47 | 0.37 | 0.63 | 0.51 | 0.47 | 0.35 |
| 3 | 161895309 | rs6798308 | 0.311 | 0.443 | -1.794 | -0.118 | 0 | -0.016 | -0.016 | 0 | 0.59 | 0.72 | 0.56 | 0.52 | 0.58 | 0.49 |
| 3 | 161895538 | rs16850784 | -0.086 | 0.261 | -1.745 | -0.351 | 0.005 | -0.009 | -0.009 | 0 | 0.64 | 0.68 | 0.54 | 0.49 | 0.7 | 0.53 |
| 3 | 161911123 | rs985703 | 0.008 | 0.268 | -1.101 | -0.332 | 0.007 | 0.096 | 0.096 | 0 | 0.12 | 0.26 | 0.13 | 0.23 | 0.1 | 0.18 |
| 3 | 161913102 | rs13094424 | 2.31 | 0.313 | -1.575 | -0.592 | -0.119 | 0.225 | 0.225 | 0 | 0.31 | 0.02 | 0.25 | 0 | 0.36 | 0.06 |
| 3 | 161913373 | rs9823100 | 0.912 | 0.321 | -1.331 | -0.624 | 0.002 | 0.107 | 0.107 | 0 | 0.08 | 0.21 | 0.07 | 0.26 | 0.06 | 0.2 |
| 3 | 161913772 | rs13095680 | -0.352 | 0.246 | -1.11 | -0.693 | 0.015 | -0.026 | -0.026 | 0 | 0.64 | 0.73 | 0.54 | 0.49 | 0.69 | 0.53 |
| 3 | 161914718 | rs12485327 | 1.015 | 0.104 | -1.507 | -0.56 | -0.008 | -0.062 | -0.062 | 0 | 0.1 | 0.01 | 0.33 | 0.42 | 0.08 | 0.17 |
| 3 | 161914770 | rs11914348 | 2.286 | 0.361 | -1.262 | -0.868 | -0.075 | 0.167 | 0.167 | 0 | 0.31 | 0.06 | 0.25 | 0 | 0.36 | 0.06 |
| 3 | 161915857 | rs9827542 | 1.937 | 0.414 | -1.249 | -0.71 | -0.042 | 0.066 | 0.066 | 1.163 | 0.11 | 0.01 | 0.09 | 0 | 0.16 | 0.11 |
| 3 | 161916714 | rs13081918 | 0.479 | 0.064 | -1.03 | -0.531 | -0.006 | -0.013 | -0.013 | 0.061 | 0.56 | 0.55 | 0.48 | 0.23 | 0.63 | 0.34 |
| 3 | 161921368 | rs7624830 | -0.49 | 0.419 | -0.923 | -0.972 | 0.007 | -0.093 | -0.093 | 0.162 | 0.62 | 0.63 | 0.7 | 0.99 | 0.52 | 0.75 |
| 3 | 161922679 | rs9843284 | 0.767 | 0.657 | -0.72 | -0.173 | -0.005 | 0.023 | 0.023 | 0 | 0.63 | 0.56 | 0.73 | 0.57 | 0.67 | 0.55 |
| 3 | 161927159 | rs16854366 | 0.412 | 0.633 | -0.426 | 0.428 | -0.007 | 0.018 | 0.018 | 1.264 | 0.1 | 0.1 | 0.12 | 0.14 | 0.08 | 0.14 |
| 3 | 161932536 | rs7340714 | -0.126 | 1.373 | -0.373 | 0.404 | 0.016 | 0.02 | 0.02 | 0 | 0.18 | 0.26 | 0.15 | 0.25 | 0.23 | 0.2 |
| 3 | 161935875 | rs6787636 | 0.009 | 1.102 | -0.353 | 0.482 | 0.003 | 0.016 | 0.016 | 1.414 | 0.13 | 0.13 | 0.14 | 0.14 | 0.13 | 0.19 |
| 3 | 161937414 | rs9846433 | -0.346 | 1.07 | -0.08 | 0.361 | -0.014 | 0.025 | 0.025 | 0 | 0.2 | 0.25 | 0.21 | 0.22 | 0.2 | 0.38 |

| | | | | | | | | | | | | | | | | |
|---|---|---|---|---|---|---|---|---|---|---|---|---|---|---|---|---|
| 3 | 161937965 | rs1387010 | -1.866 | 1.178 | -0.561 | 0.444 | -0.008 | 0 | 0 | 0 | 0.91 | 0.91 | 0.84 | 0.85 | 0.88 | 0.91 |
| 3 | 161939246 | rs10513581 | 0.855 | 1.171 | -0.555 | 0.438 | 0 | -0.004 | -0.004 | 0 | 0.71 | 0.75 | 0.63 | 0.63 | 0.68 | 0.52 |
| 3 | 161944949 | rs9880055 | 0.457 | 1.168 | -0.492 | 0.458 | -0.009 | 0.025 | 0.025 | 0 | 0.14 | 0.21 | 0.07 | 0.25 | 0.1 | 0.19 |
| 3 | 161949284 | rs1488583 | -0.405 | 1.442 | -0.772 | 0.248 | -0.001 | 0.002 | 0.002 | 0 | 0.2 | 0.16 | 0.21 | 0.22 | 0.2 | 0.38 |
| 3 | 161951039 | rs6802741 | 0.762 | 1.468 | -0.744 | 0.235 | -0.002 | -0.002 | -0.002 | 2.179 | 0.68 | 0.71 | 0.6 | 0.63 | 0.62 | 0.47 |
| 3 | 161972497 | rs11927519 | 1.675 | 1.501 | -1.373 | -0.087 | 0.034 | -0.008 | -0.008 | 3.637 | 0.39 | 0.2 | 0.39 | 0.61 | 0.31 | 0.33 |
| 3 | 161984543 | rs13086855 | 0.944 | 1.684 | -1.15 | -0.122 | -0.014 | 0.009 | 0.009 | 2.424 | 0.13 | 0.06 | 0.09 | 0 | 0.17 | 0.04 |
| 3 | 161993708 | rs12635341 | 1.948 | 1.596 | -1.217 | -0.135 | 0.034 | 0 | 0 | 0 | 0.42 | 0.2 | 0.4 | 0.61 | 0.34 | 0.33 |
| 3 | 161994639 | rs13072299 | 1.384 | 1.878 | -1.015 | 0.099 | 0.005 | 0.04 | 0.04 | 1.322 | 0.1 | 0.01 | 0.02 | 0 | 0.06 | 0.02 |
| 3 | 162003134 | rs2405563 | 1.027 | 1.796 | -0.453 | 0.408 | -0.007 | -0.069 | -0.069 | 0.175 | 0.12 | 0.25 | 0.06 | 0 | 0.1 | 0.06 |
| 3 | 162004273 | rs9883161 | -0.101 | 2.328 | -0.264 | 0.901 | 0.004 | 0.102 | 0.102 | 2.113 | 0.82 | 0.63 | 0.7 | 0.63 | 0.83 | 0.64 |
| 3 | 162049171 | rs7611892 | 1.149 | 2.17 | -0.276 | 0.955 | 0.014 | 0.012 | 0.012 | 0 | 0.67 | 0.56 | 0.6 | 0.63 | 0.65 | 0.57 |
| 3 | 162050329 | rs13060652 | 0.436 | 2.149 | -0.324 | 0.954 | -0.028 | 0.038 | 0.038 | 2.633 | 0.15 | 0.03 | 0.1 | 0 | 0.18 | 0.05 |
| 3 | 162053213 | rs13100767 | 1.341 | 2.109 | -0.411 | 1.226 | 0.002 | 0.002 | 0.002 | 0.944 | 0.66 | 0.59 | 0.52 | 0.42 | 0.57 | 0.34 |
| 3 | 162055990 | rs9833336 | -0.542 | 1.867 | -0.774 | 0.994 | -0.04 | -0.096 | -0.096 | 0 | 0.86 | 0.98 | 0.75 | 0.69 | 0.86 | 0.77 |
| 3 | 162058440 | rs11917604 | 0.992 | 1.502 | -0.584 | 1.065 | -0.002 | -0.001 | -0.001 | 0 | 0.55 | 0.47 | 0.47 | 0.49 | 0.44 | 0.43 |
| 3 | 162063995 | rs4413326 | 0.714 | 0.945 | -0.682 | 1.524 | 0 | -0.018 | -0.018 | 0 | 0.06 | 0.01 | 0.08 | 0.16 | 0.03 | 0.11 |
| 3 | 162074782 | rs6441449 | 1.157 | 1.093 | -0.492 | 0.867 | 0 | 0.001 | 0.001 | 0 | 0.5 | 0.45 | 0.39 | 0.23 | 0.5 | 0.43 |
| 3 | 162083007 | rs13100275 | 0.323 | 1.162 | -0.656 | 0.996 | -0.011 | 0.017 | 0.017 | 0 | 0.17 | 0.02 | 0.19 | 0.15 | 0.15 | 0.18 |
| 3 | 162084947 | rs2405749 | 0.267 | 1.238 | -0.576 | 1.699 | -0.024 | 0.031 | 0.031 | 0 | 0.05 | 0.01 | 0.17 | 0.34 | 0.05 | 0.05 |
| 3 | 162085820 | rs12488754 | -1.106 | 1.387 | -0.267 | 1.622 | 0.02 | 0.139 | 0.139 | 0 | 0.09 | 0.17 | 0.26 | 0.38 | 0.11 | 0.2 |
| 3 | 162089122 | rs2897221 | -0.854 | 1.36 | -0.254 | 1.636 | 0.035 | 0.131 | 0.131 | 0 | 0.05 | 0.16 | 0.09 | 0.15 | 0.07 | 0.18 |
| 3 | 162094430 | rs2897222 | -0.281 | 1.42 | -0.335 | 1.582 | 0.001 | -0.019 | -0.019 | 0 | 0.17 | 0.18 | 0.16 | 0.37 | 0.15 | 0.25 |
| 3 | 162112120 | rs6800943 | -0.017 | 1.721 | -0.472 | 1.62 | -0.079 | -0.109 | -0.109 | 0.302 | 0.85 | 0.98 | 0.75 | 0.64 | 0.8 | 0.84 |
| 3 | 162113116 | rs4393923 | 0.463 | 1.69 | -0.453 | 1.608 | -0.005 | 0 | 0 | 0 | 0.81 | 0.82 | 0.66 | 0.6 | 0.74 | 0.69 |
| 3 | 162116546 | rs16848682 | 0.847 | 1.634 | -0.613 | 1.849 | -0.007 | -0.028 | -0.028 | 0 | 0.06 | 0 | 0.01 | 0.18 | 0.04 | 0.12 |
| 3 | 162117467 | rs2405755 | 0.684 | 1.593 | -0.319 | 1.359 | -0.012 | -0.005 | -0.005 | 0 | 0.69 | 0.81 | 0.64 | 0.68 | 0.62 | 0.53 |
| 3 | 162131408 | rs12636148 | -0.546 | 1.888 | -0.026 | 1.49 | -0.039 | -0.031 | -0.031 | 0.39 | 0.07 | 0.02 | 0.27 | 0.26 | 0.13 | 0.16 |
| 3 | 162142475 | rs6803545 | 0.786 | 1.673 | -0.058 | 1.406 | 0.002 | 0.018 | 0.018 | 0 | 0.1 | 0.25 | 0.18 | 0.15 | 0.11 | 0.09 |
| 3 | 162155076 | rs11929273 | -0.512 | 1.623 | -0.12 | 1.318 | 0.003 | -0.028 | -0.028 | 0.193 | 0.18 | 0.18 | 0.16 | 0.39 | 0.15 | 0.25 |
| 3 | 162156517 | rs2318630 | 1.639 | 1.633 | -0.064 | 1.173 | -0.009 | 0.047 | 0.047 | 0 | 0.59 | 0.65 | 0.57 | 0.27 | 0.59 | 0.51 |
| 3 | 162161616 | rs11924705 | 2.34 | 1.447 | 0.022 | 1.011 | 0.012 | 0.15 | 0.15 | 0 | 0.45 | 0.21 | 0.31 | 0.08 | 0.43 | 0.28 |
| 3 | 162186802 | rs1478161 | -1.015 | 1.408 | 0.05 | 1.117 | -0.006 | 0.043 | 0.043 | 0.065 | 0.28 | 0.42 | 0.34 | 0.54 | 0.25 | 0.34 |
| 3 | 162188734 | rs16849242 | -1.098 | 1.184 | -0.395 | 0.995 | 0.016 | -0.026 | -0.026 | 0 | 0.27 | 0.25 | 0.17 | 0.5 | 0.2 | 0.36 |
| 3 | 162198135 | rs7349544 | 1.014 | 1.314 | -0.507 | 0.616 | -0.048 | 0.047 | 0.047 | 0 | 0.09 | 0.01 | 0.08 | 0.01 | 0.15 | 0.11 |
| 3 | 162216666 | rs1351091 | -1.378 | 1.071 | -0.431 | 0.924 | -0.01 | 0.028 | 0.028 | 0.757 | 0.44 | 0.66 | 0.65 | 0.78 | 0.47 | 0.6 |
| 3 | 162237495 | rs1031837 | -1.539 | 0.995 | -0.571 | 0.624 | 0.017 | 0.117 | 0.117 | 0 | 0.5 | 0.73 | 0.68 | 0.86 | 0.53 | 0.69 |
| 3 | 162242541 | rs9868015 | 1.163 | 0.907 | -0.709 | 0.772 | 0.004 | -0.004 | -0.004 | 0 | 0.07 | 0.03 | 0.02 | 0 | 0.04 | 0.08 |
| 3 | 162243751 | rs1478146 | 1.149 | 1.185 | -0.739 | 1.3 | -0.007 | 0.011 | 0.011 | 0 | 0.06 | 0.07 | 0.19 | 0.22 | 0.07 | 0.03 |
| 3 | 162259706 | rs35780213 | -0.646 | 1.099 | -0.727 | 1.53 | 0.001 | -0.044 | -0.044 | 0 | 0.16 | 0.11 | 0.18 | 0.45 | 0.15 | 0.36 |
| 3 | 162264998 | rs12488833 | 1.13 | 1.094 | -0.676 | 1.518 | -0.003 | 0.015 | 0.015 | 0 | 0.06 | 0.06 | 0.19 | 0.23 | 0.08 | 0.04 |
| 3 | 162275610 | rs16850590 | -0.588 | 0.88 | -0.647 | 1.456 | 0.002 | 0.045 | 0.045 | 0 | 0.1 | 0.1 | 0.1 | 0.42 | 0.06 | 0.16 |
| 3 | 162276129 | rs6800542 | -1.9 | 1.098 | -0.371 | 1.057 | -0.009 | 0.012 | 0.012 | 0 | 0.33 | 0.48 | 0.4 | 0.62 | 0.31 | 0.5 |
| 3 | 162276175 | rs6764816 | -1.449 | 1.08 | -0.283 | 1.026 | -0.012 | 0.044 | 0.044 | 0 | 0.25 | 0.44 | 0.37 | 0.62 | 0.24 | 0.42 |
| 3 | 162282328 | rs7647075 | -2.29 | 1.138 | -0.448 | 0.832 | 0 | -0.005 | -0.005 | 0 | 0.43 | 0.5 | 0.48 | 0.64 | 0.44 | 0.6 |
| 3 | 162283234 | rs6806429 | 0.955 | 1.105 | -0.113 | 0.802 | 0.01 | -0.039 | -0.039 | 0 | 0.11 | 0.18 | 0.09 | 0.02 | 0.14 | 0.1 |
| 3 | 162283321 | rs6770712 | -1.077 | 1.1 | -0.12 | 0.801 | 0.014 | -0.048 | -0.048 | 0.395 | 0.29 | 0.5 | 0.32 | 0.21 | 0.31 | 0.3 |
| 3 | 162286357 | rs13083177 | 1.32 | 1.055 | -0.189 | 1.076 | 0 | -0.001 | -0.001 | 0 | 0.08 | 0.04 | 0.02 | 0 | 0.07 | 0.08 |
| 3 | 162286692 | rs7617688 | -2.438 | 1.059 | -0.184 | 1.093 | 0 | 0.01 | 0.01 | 0 | 0.45 | 0.66 | 0.49 | 0.64 | 0.44 | 0.6 |
| 3 | 162287573 | rs9816189 | -0.326 | 1.048 | -0.328 | 0.999 | 0.002 | -0.047 | -0.047 | 0.022 | 0.2 | 0.22 | 0.11 | 0.02 | 0.21 | 0.18 |
| 3 | 162305516 | rs16850831 | -1.324 | 0.764 | -0.451 | 1.067 | 0.006 | -0.032 | -0.032 | 0.836 | 0.15 | 0.11 | 0.16 | 0.43 | 0.14 | 0.3 |
| 3 | 162314038 | rs9647350 | -0.299 | 0.853 | -0.521 | 0.751 | -0.016 | 0.029 | 0.029 | 0 | 0.06 | 0.01 | 0.06 | 0.01 | 0.08 | 0.14 |
| 3 | 162322748 | rs1121055 | 1.254 | 0.881 | -0.551 | 0.55 | -0.045 | 0.038 | 0.038 | 0 | 0.09 | 0.01 | 0.08 | 0.02 | 0.14 | 0.08 |
| 3 | 162330008 | rs6804879 | 0.187 | 0.278 | -1.369 | -0.335 | -0.01 | 0.088 | 0.088 | 0.146 | 0.47 | 0.33 | 0.48 | 0.25 | 0.49 | 0.44 |
| 3 | 162370984 | rs13096777 | 0.988 | 0.129 | -1.062 | -0.206 | 0.002 | 0.001 | 0.001 | 0.53 | 0.36 | 0.35 | 0.41 | 0.33 | 0.44 | 0.34 |
| 3 | 162373246 | rs1496737 | 1.131 | 0.086 | -1.263 | -0.62 | 0.01 | 0.026 | 0.026 | 0.249 | 0.34 | 0.27 | 0.35 | 0.23 | 0.27 | 0.2 |
| 3 | 162376870 | rs903943 | 0.556 | 0.131 | -1.276 | -0.584 | 0.006 | 0.039 | 0.039 | 0 | 0.4 | 0.32 | 0.41 | 0.24 | 0.35 | 0.36 |
| 3 | 162389205 | rs12491408 | -0.232 | 0.238 | -1.143 | -0.599 | -0.037 | 0.132 | 0.132 | 0.487 | 0.18 | 0.48 | 0.22 | 0.34 | 0.12 | 0.25 |
| 3 | 162391801 | rs12489920 | -1.54 | 0.212 | -1.291 | -0.622 | -0.019 | 0.048 | 0.048 | 0 | 0.32 | 0.52 | 0.38 | 0.44 | 0.26 | 0.53 |
| 3 | 162396246 | rs12492947 | 1 | 0.309 | -1.235 | -0.658 | 0.006 | -0.025 | -0.025 | 0 | 0.12 | 0.29 | 0.13 | 0.05 | 0.1 | 0.13 |
| 3 | 162397319 | rs10936299 | -0.231 | 0.589 | -1.687 | -0.627 | -0.037 | 0.053 | 0.053 | 0.007 | 0.8 | 0.89 | 0.77 | 0.95 | 0.74 | 0.66 |
| 3 | 162399269 | rs12634787 | -0.626 | 0.567 | -1.678 | -0.605 | -0.001 | 0.046 | 0.046 | 0 | 0.09 | 0.13 | 0.1 | 0.36 | 0.05 | 0.16 |
| 3 | 162400620 | rs9290132 | 1.052 | 0.572 | -1.678 | -0.612 | -0.012 | -0.026 | -0.026 | 0 | 0.71 | 0.73 | 0.67 | 0.6 | 0.68 | 0.5 |
| 3 | 162410741 | rs6441541 | -1.574 | 0.31 | -1.49 | -0.419 | -0.027 | 0.029 | 0.029 | 0 | 0.91 | 0.96 | 0.91 | 0.98 | 0.85 | 0.91 |
| 3 | 162423035 | rs7433221 | 0.165 | -0.038 | -1.805 | -0.883 | 0.013 | 0.013 | 0.013 | 0 | 0.55 | 0.64 | 0.51 | 0.6 | 0.39 | 0.37 |
| 3 | 162424485 | rs4858909 | 0.411 | 0.09 | -1.644 | -0.63 | 0.002 | 0.012 | 0.012 | 0 | 0.81 | 0.75 | 0.81 | 0.63 | 0.79 | 0.74 |
| 3 | 162424689 | rs12631404 | 0.208 | 0.126 | -1.536 | -0.629 | -0.017 | 0.126 | 0.126 | 0.012 | 0.05 | 0.15 | 0.09 | 0.29 | 0.02 | 0.12 |
| 3 | 162429810 | rs9844124 | -0.284 | 0.158 | -1.548 | -0.62 | 0.006 | 0.029 | 0.029 | 0 | 0.85 | 0.76 | 0.83 | 0.69 | 0.83 | 0.78 |
| 3 | 162449000 | rs9835079 | -0.987 | -0.279 | -1.717 | -0.728 | -0.012 | -0.005 | -0.005 | 0.499 | 0.71 | 0.78 | 0.75 | 0.68 | 0.7 | 0.74 |

| | | | | | | | | | | | | | | | | |
|---|---|---|---|---|---|---|---|---|---|---|---|---|---|---|---|---|
| 3 | 162453428 | rs12490517 | 0.698 | -0.35 | -2.031 | -1.089 | -0.058 | 0.133 | 0.133 | 0 | 0.29 | 0.1 | 0.3 | 0.03 | 0.4 | 0.37 |
| 3 | 162460891 | rs2201659 | -0.798 | -0.405 | -2.097 | -1.152 | 0.051 | 0.324 | 0.324 | 0 | 0.51 | 0.86 | 0.55 | 0.95 | 0.36 | 0.53 |
| 3 | 162461144 | rs1908780 | -2.342 | -0.343 | -2.071 | -1.107 | -0.061 | 0.114 | 0.114 | 0 | 0.79 | 0.97 | 0.85 | 0.98 | 0.76 | 0.89 |
| 3 | 162482732 | rs9290133 | -0.329 | -0.681 | -1.62 | -1.106 | 0.001 | -0.013 | -0.013 | 0 | 0.39 | 0.61 | 0.44 | 0.89 | 0.29 | 0.4 |
| 3 | 162486664 | rs9682116 | 0.988 | -0.728 | -1.558 | -1.006 | -0.018 | -0.02 | -0.02 | 0 | 0.11 | 0.26 | 0.11 | 0.06 | 0.07 | 0.12 |
| 3 | 162491025 | rs206906 | 0.362 | -0.748 | -1.472 | -0.882 | -0.002 | 0.004 | 0.004 | 0 | 0.3 | 0.3 | 0.33 | 0.61 | 0.24 | 0.26 |
| 3 | 162495102 | rs3853175 | 0.858 | -0.664 | -1.677 | -1.037 | -0.075 | 0.148 | 0.148 | 0 | 0.28 | 0.08 | 0.3 | 0.03 | 0.4 | 0.36 |
| 3 | 162496237 | rs9851876 | 0.026 | -0.687 | -1.495 | -1.014 | -0.004 | 0.008 | 0.008 | 0 | 0.41 | 0.35 | 0.41 | 0.09 | 0.48 | 0.52 |
| 3 | 162498022 | rs4140866 | 1.998 | -0.717 | -1.718 | -0.669 | -0.039 | -0.066 | -0.066 | 0 | 0.1 | 0.01 | 0.22 | 0.41 | 0.12 | 0.08 |
| 3 | 162507286 | rs206285 | -2.47 | -0.655 | -1.73 | -0.799 | -0.054 | 0.118 | 0.118 | 0 | 0.79 | 0.97 | 0.85 | 0.98 | 0.76 | 0.89 |
| 3 | 162511139 | rs206280 | 0.602 | -0.82 | -1.472 | -0.637 | -0.005 | 0.028 | 0.028 | 0 | 0.25 | 0.26 | 0.31 | 0.54 | 0.2 | 0.2 |
| 3 | 162626585 | rs206276 | -0.522 | -0.81 | -1.439 | -0.746 | -0.002 | -0.022 | -0.022 | 0.22 | 0.37 | 0.6 | 0.45 | 0.89 | 0.28 | 0.35 |
| 3 | 162632001 | rs206868 | -0.897 | -0.789 | -1.67 | -0.906 | 0.001 | 0.008 | 0.008 | 0 | 0.61 | 0.62 | 0.66 | 0.71 | 0.6 | 0.69 |
| 3 | 162636681 | rs206265 | -0.905 | -0.791 | -1.671 | -0.907 | 0.001 | 0.008 | 0.008 | 0 | 0.61 | 0.62 | 0.66 | 0.71 | 0.6 | 0.69 |
| 3 | 162638937 | rs16847382 | 1.246 | -0.822 | -1.675 | -0.936 | 0.014 | 0.02 | 0.02 | 0 | 0.12 | 0.04 | 0.09 | 0.06 | 0.07 | 0.13 |
| 3 | 162640469 | rs206262 | 2.216 | -0.891 | -1.783 | -0.907 | 0.002 | -0.034 | -0.034 | 0 | 0.12 | 0.01 | 0.06 | 0.14 | 0.07 | 0.06 |
| 3 | 162649358 | rs9290153 | -0.109 | -0.756 | -1.664 | -0.948 | -0.002 | -0.001 | -0.001 | 0 | 0.35 | 0.35 | 0.35 | 0.15 | 0.4 | 0.48 |
| 3 | 162650452 | rs9836765 | -0.1 | -0.756 | -1.664 | -0.948 | -0.002 | -0.002 | -0.002 | 0 | 0.35 | 0.35 | 0.35 | 0.15 | 0.4 | 0.48 |
| 3 | 162654424 | rs3912608 | 1.651 | -0.74 | -1.683 | -0.96 | -0.082 | 0.076 | 0.076 | 0.348 | 0.17 | 0.04 | 0.22 | 0.03 | 0.3 | 0.27 |
| 3 | 162656280 | rs6777496 | 0.549 | -0.862 | -1.461 | -0.885 | -0.006 | 0.195 | 0.195 | 0 | 0.06 | 0.22 | 0.11 | 0.27 | 0.04 | 0.09 |
| 3 | 162666172 | rs9865194 | 0.96 | -0.767 | -1.447 | -0.886 | -0.021 | -0.03 | -0.03 | 0 | 0.13 | 0.28 | 0.11 | 0.06 | 0.07 | 0.16 |
| 3 | 162671443 | rs13067901 | 1.715 | -0.567 | -1.375 | -0.784 | -0.001 | 0.037 | 0.037 | 1.643 | 0.11 | 0.02 | 0.08 | 0 | 0.12 | 0.08 |
| 3 | 162672041 | rs1386866 | -0.227 | -0.612 | -1.405 | -0.821 | -0.012 | 0.012 | 0.012 | 1.395 | 0.38 | 0.42 | 0.4 | 0.56 | 0.32 | 0.29 |
| 3 | 162675279 | rs11918594 | 1.881 | -0.641 | -1.283 | -0.835 | 0.007 | 0.118 | 0.118 | 0.388 | 0.13 | 0.2 | 0.24 | 0.42 | 0.13 | 0.08 |
| 3 | 162675520 | rs13060008 | -1.609 | -0.453 | -1.298 | -1.009 | -0.002 | -0.005 | -0.005 | 0 | 0.73 | 0.76 | 0.74 | 0.7 | 0.72 | 0.77 |
| 3 | 162678220 | rs11921809 | -1.609 | -0.453 | -1.298 | -1.009 | -0.002 | -0.005 | -0.005 | 0 | 0.73 | 0.76 | 0.74 | 0.7 | 0.72 | 0.77 |
| 3 | 162682265 | rs9847141 | 0.663 | -0.421 | -1.288 | -1.011 | 0.003 | -0.024 | -0.024 | 0 | 0.3 | 0.32 | 0.32 | 0.09 | 0.36 | 0.43 |
| 3 | 162687835 | rs13080866 | 0.111 | -0.378 | -1.372 | -0.974 | 0.004 | -0.008 | -0.008 | 0.017 | 0.36 | 0.28 | 0.39 | 0.55 | 0.32 | 0.29 |
| 3 | 162689869 | rs1386856 | -0.002 | -0.327 | -1.427 | -0.968 | 0.001 | 0.002 | 0.002 | 1.507 | 0.39 | 0.43 | 0.45 | 0.43 | 0.44 | 0.57 |
| 3 | 162695483 | rs9860284 | -0.002 | -0.34 | -1.244 | -0.841 | 0.001 | 0.002 | 0.002 | 1.465 | 0.39 | 0.43 | 0.45 | 0.43 | 0.44 | 0.57 |
| 3 | 162698133 | rs1955090 | 0.006 | -0.336 | -1.242 | -0.839 | 0.001 | 0.002 | 0.002 | 1.485 | 0.39 | 0.43 | 0.45 | 0.43 | 0.44 | 0.57 |
| 3 | 162704020 | rs10513598 | 1.394 | -0.335 | -1.235 | -0.838 | 0.002 | -0.009 | -0.009 | 0 | 0.11 | 0.08 | 0.09 | 0 | 0.12 | 0.08 |
| 3 | 162707978 | rs1011306 | -0.09 | -0.199 | -1.234 | -0.702 | 0.001 | 0.002 | 0.002 | 1.485 | 0.39 | 0.43 | 0.45 | 0.43 | 0.44 | 0.57 |
| 3 | 162724542 | rs1352369 | 0.113 | -0.224 | -1.243 | -0.736 | -0.011 | 0.014 | 0.014 | 1.631 | 0.38 | 0.41 | 0.39 | 0.55 | 0.32 | 0.29 |
| 3 | 162729163 | rs6802688 | -1.768 | -0.124 | -1.235 | -0.707 | -0.008 | 0.026 | 0.026 | 0 | 0.77 | 0.83 | 0.85 | 0.98 | 0.75 | 0.86 |
| 3 | 162733793 | rs9838708 | -0.023 | -0.308 | -1.212 | -0.666 | 0.001 | 0.004 | 0.004 | 1.721 | 0.38 | 0.41 | 0.45 | 0.43 | 0.41 | 0.56 |
| 3 | 162735677 | rs1586501 | 0.158 | -0.311 | -1.224 | -0.697 | -0.01 | 0.013 | 0.013 | 2.425 | 0.38 | 0.41 | 0.39 | 0.55 | 0.32 | 0.3 |
| 3 | 162741452 | rs2046696 | -1.737 | -0.168 | -1.389 | -0.816 | -0.013 | 0.021 | 0.021 | 0 | 0.77 | 0.82 | 0.84 | 0.98 | 0.74 | 0.87 |
| 3 | 162749836 | rs7641571 | 0.926 | -0.008 | -1.18 | -0.453 | -0.002 | -0.003 | -0.003 | 3.742 | 0.08 | 0.09 | 0.08 | 0.06 | 0.05 | 0.13 |
| 3 | 162756477 | rs13092443 | 0.607 | 0.058 | -1.633 | -0.586 | -0.094 | 0.065 | 0.065 | 0.66 | 0.25 | 0.03 | 0.24 | 0.09 | 0.32 | 0.34 |
| 3 | 162773033 | rs6548375 | -0.373 | -0.065 | -1.213 | -0.422 | 0.002 | 0.006 | 0.006 | 1.661 | 0.39 | 0.46 | 0.45 | 0.42 | 0.41 | 0.56 |
| 3 | 162781486 | rs6796209 | 1.146 | -0.289 | -1.194 | -0.438 | 0.003 | 0.007 | 0.007 | 1.401 | 0.11 | 0.05 | 0.08 | 0 | 0.12 | 0.08 |
| 3 | 162789598 | rs13090583 | 0.107 | -0.395 | -1.098 | -0.434 | -0.006 | 0.007 | 0.007 | 2.545 | 0.37 | 0.36 | 0.39 | 0.56 | 0.32 | 0.3 |
| 3 | 162802753 | rs7427962 | 0.165 | -0.389 | -1.103 | -0.529 | 0 | 0.001 | 0.001 | 0.08 | 0.37 | 0.37 | 0.43 | 0.59 | 0.4 | 0.31 |
| 3 | 162809107 | rs6548383 | -0.457 | -0.37 | -0.875 | -0.599 | -0.027 | 0.082 | 0.082 | 0 | 0.2 | 0.51 | 0.22 | 0.32 | 0.14 | 0.3 |
| 3 | 162836008 | rs7641999 | 1.089 | -0.349 | -0.782 | -0.584 | 0.049 | 0.03 | 0.03 | 0 | 0.08 | 0.31 | 0.09 | 0.05 | 0.05 | 0.1 |
| 3 | 162856377 | rs4387975 | -0.043 | -0.447 | -1.025 | -0.817 | -0.003 | 0.021 | 0.021 | 0 | 0.57 | 0.32 | 0.61 | 0.67 | 0.59 | 0.56 |
| 3 | 162857478 | rs4589940 | 1.223 | -0.434 | -1.06 | -0.801 | 0.048 | 0.029 | 0.029 | 0 | 0.08 | 0.31 | 0.09 | 0.05 | 0.05 | 0.1 |
| 3 | 162859171 | rs6794545 | 0.041 | -0.471 | -1.044 | -0.788 | -0.013 | 0.028 | 0.028 | 0.283 | 0.13 | 0.22 | 0.14 | 0.26 | 0.09 | 0.2 |
| 3 | 162860260 | rs12330861 | 1.506 | -0.461 | -1.341 | -0.93 | -0.022 | 0.051 | 0.051 | 0.002 | 0.19 | 0.01 | 0.08 | 0.07 | 0.17 | 0.18 |
| 3 | 162861639 | rs6767542 | -1.206 | -0.391 | -1.218 | -0.848 | 0.017 | -0.069 | -0.069 | 0.021 | 0.72 | 0.67 | 0.76 | 0.93 | 0.68 | 0.76 |
| 3 | 162871454 | rs11915237 | -1.149 | -0.392 | -1.218 | -0.848 | 0.016 | -0.068 | -0.068 | 12.082 | 0.71 | 0.66 | 0.76 | 0.93 | 0.68 | 0.76 |
| 3 | 162876062 | rs6774828 | 0.127 | -0.398 | -1.222 | -0.709 | -0.004 | 0.002 | 0.002 | 0.244 | 0.13 | 0.2 | 0.13 | 0.2 | 0.09 | 0.19 |
| 3 | 162878626 | rs12636292 | 0.183 | -0.397 | -1.221 | -0.708 | -0.004 | 0.003 | 0.003 | 0.33 | 0.13 | 0.2 | 0.13 | 0.2 | 0.09 | 0.19 |
| 3 | 162879193 | rs4855272 | 1.526 | -0.54 | -1.279 | -0.574 | 0.009 | 0.038 | 0.038 | 0.337 | 0.2 | 0.21 | 0.32 | 0.45 | 0.28 | 0.09 |
| 3 | 162879873 | rs6788517 | 0.167 | -0.491 | -1.234 | -0.547 | -0.003 | 0.002 | 0.002 | 0.355 | 0.13 | 0.2 | 0.14 | 0.2 | 0.09 | 0.19 |
| 3 | 162893823 | rs9874856 | -0.331 | -0.499 | -1.266 | -0.56 | 0.002 | -0.01 | -0.01 | 0 | 0.58 | 0.46 | 0.6 | 0.67 | 0.59 | 0.58 |
| 3 | 162899930 | rs16845642 | 1.132 | -0.655 | -1.279 | -0.479 | 0.017 | -0.02 | -0.02 | 1.685 | 0.09 | 0.05 | 0.1 | 0.2 | 0.04 | 0.12 |
| 3 | 162918387 | rs4021819 | -1.27 | -0.529 | -1.253 | -0.577 | 0.016 | -0.042 | -0.042 | 0 | 0.72 | 0.65 | 0.73 | 0.87 | 0.69 | 0.76 |
| 3 | 162919329 | rs1492171 | -1.237 | -0.532 | -1.254 | -0.578 | 0.015 | -0.042 | -0.042 | 0.152 | 0.71 | 0.65 | 0.73 | 0.87 | 0.69 | 0.76 |
| 3 | 162921554 | rs1492172 | -1.237 | -0.532 | -1.254 | -0.578 | 0.015 | -0.042 | -0.042 | 0 | 0.71 | 0.65 | 0.73 | 0.87 | 0.69 | 0.76 |
| 3 | 162925738 | rs4324469 | -0.168 | -0.575 | -1.192 | -0.61 | 0 | 0.021 | 0.021 | 0 | 0.57 | 0.33 | 0.6 | 0.68 | 0.6 | 0.58 |
| 3 | 162932624 | rs4855204 | -1.243 | -0.523 | -1.061 | -0.595 | 0.015 | -0.042 | -0.042 | 0 | 0.71 | 0.65 | 0.73 | 0.87 | 0.69 | 0.76 |
| 3 | 162935109 | rs6809992 | 1.54 | -0.581 | -1.164 | -0.622 | 0.011 | 0.038 | 0.038 | 0.282 | 0.2 | 0.21 | 0.32 | 0.45 | 0.29 | 0.09 |
| 3 | 162935921 | rs2087740 | -1.282 | -0.496 | -1.105 | -0.65 | 0.015 | -0.042 | -0.042 | 1.455 | 0.71 | 0.65 | 0.73 | 0.87 | 0.69 | 0.77 |
| 3 | 162937939 | rs4021822 | -1.141 | -0.497 | -1.092 | -0.65 | 0.02 | -0.063 | -0.063 | 0 | 0.7 | 0.52 | 0.73 | 0.87 | 0.69 | 0.76 |
| 3 | 162941903 | rs1994750 | -1.277 | -0.49 | -1.116 | -0.647 | 0.015 | -0.042 | -0.042 | 0 | 0.71 | 0.65 | 0.73 | 0.87 | 0.69 | 0.76 |
| 3 | 162950711 | rs12696185 | 0.365 | -0.432 | -1.05 | -0.575 | 0.054 | 0.048 | 0.048 | 0 | 0.11 | 0.42 | 0.15 | 0.11 | 0.1 | 0.17 |
| 3 | 162976133 | rs843057 | 1.253 | -0.582 | -1.25 | -0.64 | 0.028 | -0.013 | -0.013 | 0 | 0.14 | 0.07 | 0.06 | 0.15 | 0.07 | 0.07 |

| | | | | | | | | | | | | | | | | |
|---|---|---|---|---|---|---|---|---|---|---|---|---|---|---|---|---|
| 3 | 162997262 | rs9809962 | 0.954 | -0.268 | -1.22 | -0.686 | -0.036 | 0.089 | 0.089 | 0.322 | 0.23 | 0.05 | 0.22 | 0.08 | 0.25 | 0.32 |
| 3 | 163022631 | rs1099110 | -2.038 | -0.209 | -1.25 | -0.735 | -0.093 | 0.131 | 0.131 | 0 | 0.78 | 0.99 | 0.82 | 0.98 | 0.73 | 0.85 |
| 3 | 163033205 | rs1186529 | 0.738 | -0.299 | -1.151 | -0.64 | 0 | -0.005 | -0.005 | 0 | 0.34 | 0.29 | 0.36 | 0.6 | 0.35 | 0.25 |
| 3 | 163048842 | rs7632852 | 1.158 | -0.292 | -1.136 | -0.639 | 0.01 | -0.04 | -0.04 | 0.331 | 0.3 | 0.34 | 0.27 | 0.13 | 0.31 | 0.25 |
| 3 | 163058603 | rs1834723 | -0.56 | -0.209 | -1.185 | -0.555 | -0.002 | -0.003 | -0.003 | 0 | 0.63 | 0.69 | 0.62 | 0.4 | 0.6 | 0.68 |
| 3 | 163075070 | rs1365702 | -0.032 | -0.153 | -0.846 | -0.365 | -0.028 | 0.132 | 0.132 | 0 | 0.16 | 0.4 | 0.21 | 0.31 | 0.08 | 0.2 |
| 3 | 163096848 | rs6796340 | -0.157 | -0.125 | -1.017 | -0.495 | 0 | 0.015 | 0.015 | 0 | 0.43 | 0.63 | 0.42 | 0.32 | 0.39 | 0.43 |
| 3 | 163139188 | rs1380781 | 1.472 | -0.027 | -1.026 | -0.474 | -0.017 | 0.035 | 0.035 | 0 | 0.23 | 0.12 | 0.17 | 0.02 | 0.26 | 0.15 |
| 3 | 163159314 | rs13060071 | 0.638 | -0.045 | -0.94 | -0.328 | 0.006 | -0.017 | -0.017 | 0.009 | 0.11 | 0.06 | 0.15 | 0.19 | 0.09 | 0.19 |
| 3 | 163178778 | rs2711774 | -1.439 | -0.022 | -0.988 | -0.183 | 0.007 | 0.026 | 0.026 | 0 | 0.78 | 0.77 | 0.68 | 0.58 | 0.71 | 0.89 |
| 3 | 163214096 | rs9283631 | 0.348 | 0.244 | -0.539 | -0.42 | 0.055 | 0.144 | 0.144 | 0 | 0.08 | 0.44 | 0.1 | 0.11 | 0.04 | 0.09 |
| 3 | 163223847 | rs1599271 | 0.077 | 0.322 | -0.582 | -0.445 | 0.062 | 0.086 | 0.086 | 0 | 0.11 | 0.45 | 0.21 | 0.14 | 0.13 | 0.24 |
| 3 | 163224607 | rs996784 | -1.418 | 0.36 | -1.132 | -0.604 | -0.093 | 0.125 | 0.125 | 0 | 0.78 | 0.99 | 0.83 | 0.98 | 0.74 | 0.85 |
| 3 | 163233510 | rs6765213 | -0.027 | 0.251 | -0.795 | -0.18 | -0.008 | 0.026 | 0.026 | 0 | 0.53 | 0.32 | 0.47 | 0.63 | 0.51 | 0.44 |
| 3 | 163240717 | rs6810129 | 0.35 | 0.375 | -1.228 | 0.015 | 0 | 0.01 | 0.01 | 0 | 0.38 | 0.49 | 0.43 | 0.56 | 0.41 | 0.2 |
| 3 | 163242030 | rs9811637 | -0.017 | 0.805 | -1.287 | -0.038 | -0.012 | 0.033 | 0.033 | 0 | 0.66 | 0.73 | 0.68 | 0.78 | 0.63 | 0.66 |
| 3 | 163257974 | rs1460434 | 0.049 | 0.988 | -1.044 | -0.097 | 0.006 | -0.02 | -0.02 | 0.413 | 0.6 | 0.49 | 0.58 | 0.75 | 0.58 | 0.5 |
| 3 | 163263886 | rs12632458 | -0.005 | 0.944 | -0.808 | -0.1 | 0.014 | -0.029 | -0.029 | 0.268 | 0.61 | 0.58 | 0.58 | 0.74 | 0.57 | 0.51 |
| 3 | 163281470 | rs12492596 | 0.308 | 0.447 | -0.916 | 0.681 | -0.015 | 0.111 | 0.111 | 0 | 0.13 | 0.21 | 0.25 | 0.45 | 0.07 | 0.17 |
| 3 | 163281927 | rs1471271 | 0.469 | 0.604 | -1.517 | 0.249 | -0.003 | 0.022 | 0.022 | 0 | 0.37 | 0.34 | 0.3 | 0.17 | 0.45 | 0.22 |
| 3 | 163313230 | rs1460421 | 0.873 | 0.246 | -1.731 | -0.132 | -0.026 | 0.019 | 0.019 | 0 | 0.24 | 0.07 | 0.17 | 0.18 | 0.27 | 0.11 |
| 3 | 163323028 | rs4855274 | -0.644 | 0.108 | -1.529 | -0.194 | -0.003 | -0.044 | -0.044 | 0 | 0.9 | 0.83 | 0.91 | 1 | 0.88 | 0.95 |
| 3 | 163338289 | rs16847040 | 1.139 | 0.039 | -1.539 | -0.044 | 0.006 | 0.013 | 0.013 | 0 | 0.13 | 0.09 | 0.25 | 0.45 | 0.07 | 0.2 |
| 3 | 163342357 | rs6767112 | 0.057 | 0.06 | -1.475 | -0.098 | 0.004 | -0.006 | -0.006 | 0.615 | 0.36 | 0.17 | 0.42 | 0.63 | 0.34 | 0.31 |
| 3 | 163347630 | rs2364424 | -0.799 | 0.046 | -1.599 | -0.187 | 0.001 | -0.004 | -0.004 | 0 | 0.89 | 0.91 | 0.92 | 0.88 | 0.88 | 0.94 |
| 3 | 163350517 | rs7622802 | 1.201 | 0.076 | -1.564 | -0.262 | 0.002 | -0.013 | -0.013 | 0 | 0.44 | 0.42 | 0.43 | 0.25 | 0.46 | 0.58 |
| 3 | 163366652 | rs6764070 | 0.477 | 0.385 | -1.748 | -0.748 | -0.035 | 0.046 | 0.046 | 0 | 0.22 | 0.12 | 0.35 | 0.09 | 0.24 | 0.3 |
| 3 | 163370769 | rs11708018 | 0.853 | 0.164 | -1.678 | -0.26 | -0.003 | -0.007 | -0.007 | 0 | 0.58 | 0.44 | 0.71 | 0.65 | 0.58 | 0.74 |
| 3 | 163398158 | rs9825896 | -0.921 | 0.147 | -1.681 | -0.211 | 0 | -0.002 | -0.002 | 0.643 | 0.89 | 0.88 | 0.91 | 0.85 | 0.88 | 0.94 |
| 3 | 163427533 | rs13085094 | 0.789 | 0.224 | -1.717 | -0.227 | -0.024 | 0.012 | 0.012 | 0 | 0.23 | 0.11 | 0.17 | 0.19 | 0.28 | 0.13 |
| 3 | 163461849 | rs2365142 | -0.964 | 0.323 | -1.688 | -0.095 | -0.002 | -0.018 | -0.018 | 0 | 0.9 | 0.95 | 0.91 | 0.85 | 0.88 | 0.94 |
| 3 | 163494059 | rs16847256 | 1.386 | 0.405 | -1.652 | 0.062 | -0.025 | -0.068 | -0.068 | 0 | 0.11 | 0.01 | 0.25 | 0.42 | 0.09 | 0.16 |
| 3 | 163525723 | rs1881969 | 0.52 | 0.43 | -1.459 | -0.492 | 0 | -0.008 | -0.008 | 0 | 0.45 | 0.43 | 0.46 | 0.23 | 0.46 | 0.55 |
| 3 | 163535387 | rs7634506 | -0.162 | 0.423 | -1.361 | -0.208 | -0.007 | -0.031 | -0.031 | 0 | 0.06 | 0.15 | 0.03 | 0 | 0.03 | 0.05 |
| 3 | 163543844 | rs13081339 | 0.162 | 0.445 | -1.36 | -0.437 | -0.001 | -0.017 | -0.017 | 0 | 0.4 | 0.56 | 0.32 | 0.12 | 0.33 | 0.36 |
| 3 | 163546423 | rs1907860 | 1.038 | 0.679 | -1.58 | -0.189 | -0.024 | -0.039 | -0.039 | 0 | 0.07 | 0.01 | 0.04 | 0.14 | 0.07 | 0.03 |
| 3 | 163549965 | rs1907858 | 1.08 | 0.763 | -1.847 | 0.224 | 0.004 | -0.009 | -0.009 | 0 | 0.26 | 0.24 | 0.41 | 0.48 | 0.19 | 0.27 |
| 3 | 163554331 | rs1492791 | 0.191 | 0.789 | -1.86 | 0.241 | -0.063 | -0.028 | -0.028 | 0 | 0.66 | 0.82 | 0.73 | 0.61 | 0.53 | 0.65 |
| 3 | 163563267 | rs1492782 | 0.591 | 0.59 | -1.521 | 0.082 | -0.003 | 0.006 | 0.006 | 0 | 0.56 | 0.66 | 0.66 | 0.57 | 0.45 | 0.59 |
| 3 | 163567068 | rs13059483 | 1.556 | 0.566 | -1.481 | 0.087 | 0 | 0.003 | 0.003 | 0 | 0.08 | 0.07 | 0.04 | 0.14 | 0.07 | 0.03 |
| 3 | 163571753 | rs1492789 | 0.608 | 0.586 | -1.497 | 0.073 | -0.002 | 0.006 | 0.006 | 0 | 0.56 | 0.66 | 0.66 | 0.57 | 0.45 | 0.59 |
| 3 | 163572006 | rs1881970 | -0.613 | 0.712 | -1.482 | -0.091 | -0.07 | -0.009 | -0.009 | 1.788 | 0.74 | 0.87 | 0.77 | 0.74 | 0.6 | 0.68 |
| 3 | 163572618 | rs1948418 | 1.262 | 0.684 | -1.413 | -0.067 | -0.016 | -0.066 | -0.066 | 0.863 | 0.3 | 0.42 | 0.25 | 0.11 | 0.25 | 0.32 |
| 3 | 163572907 | rs6774925 | 0.439 | 0.664 | -1.447 | 0.127 | 0.002 | -0.004 | -0.004 | 0.331 | 0.34 | 0.29 | 0.45 | 0.61 | 0.26 | 0.3 |
| 3 | 163578711 | rs4643712 | -0.253 | 1.309 | -1.536 | 0.104 | -0.039 | -0.01 | -0.01 | 0.592 | 0.76 | 0.83 | 0.7 | 0.73 | 0.57 | 0.67 |
| 3 | 163582602 | rs10936368 | 1.406 | 0.862 | -1.775 | 0.201 | 0.044 | -0.101 | -0.101 | 0 | 0.29 | 0.07 | 0.31 | 0.47 | 0.19 | 0.31 |
| 3 | 163585643 | rs1389014 | 0.388 | 0.842 | -1.628 | 0.15 | 0.009 | 0.013 | 0.013 | 0 | 0.39 | 0.64 | 0.26 | 0.13 | 0.31 | 0.34 |
| 3 | 163585938 | rs1389017 | 1.569 | 0.771 | -1.691 | 0.067 | 0.003 | 0.025 | 0.025 | 0 | 0.1 | 0.06 | 0.06 | 0.02 | 0.09 | 0.05 |
| 3 | 163599603 | rs1389021 | -0.214 | 0.778 | -1.732 | 0.065 | 0.051 | 0.091 | 0.091 | 0.052 | 0.47 | 0.76 | 0.39 | 0.26 | 0.38 | 0.36 |
| 3 | 163602176 | rs6772132 | 1.414 | 0.688 | -1.354 | 0.023 | -0.026 | -0.064 | -0.064 | 1.083 | 0.27 | 0.41 | 0.18 | 0.11 | 0.22 | 0.28 |
| 3 | 163606337 | rs4855202 | -0.257 | 0.709 | -1.664 | 0.031 | 0.047 | 0.09 | 0.09 | 0.083 | 0.47 | 0.76 | 0.39 | 0.26 | 0.38 | 0.36 |
| 3 | 163607266 | rs16847343 | 1.354 | 0.586 | -1.592 | 0.009 | -0.031 | -0.075 | -0.075 | 2.284 | 0.27 | 0.44 | 0.19 | 0.11 | 0.22 | 0.28 |
| 3 | 163607772 | rs7626120 | 1.482 | 0.378 | -1.707 | 0.044 | -0.006 | 0.03 | 0.03 | 0 | 0.1 | 0.04 | 0.06 | 0.02 | 0.09 | 0.05 |
| 3 | 163609772 | rs7611011 | 1.56 | 0.392 | -1.635 | 0.141 | 0.043 | -0.107 | -0.107 | 1.18 | 0.29 | 0.07 | 0.31 | 0.47 | 0.19 | 0.31 |
| 3 | 163616880 | rs2262960 | 1.04 | 1.41 | -1.64 | -0.004 | -0.002 | -0.006 | -0.006 | 0 | 0.15 | 0.13 | 0.13 | 0.04 | 0.22 | 0.09 |
| 3 | 163617092 | rs2720014 | 0.616 | 1.246 | -1.758 | 0.089 | 0.003 | -0.019 | -0.019 | 0 | 0.45 | 0.2 | 0.53 | 0.5 | 0.41 | 0.39 |
| 3 | 163633168 | rs3926031 | 0.251 | 1.616 | -1.828 | -0.222 | -0.023 | -0.015 | -0.015 | 0.212 | 0.69 | 0.79 | 0.53 | 0.61 | 0.53 | 0.48 |
| 3 | 163636679 | rs9878887 | 1.606 | 1.544 | -1.876 | -0.333 | 0.042 | -0.113 | -0.113 | 0.097 | 0.34 | 0.09 | 0.31 | 0.57 | 0.22 | 0.32 |
| 3 | 163641971 | rs9834331 | 2.034 | 1.546 | -1.875 | -0.301 | 0 | 0.003 | 0.003 | 0 | 0.05 | 0.07 | 0.02 | 0.11 | 0.04 | 0.01 |
| 3 | 163643042 | rs6774001 | 0.181 | 1.533 | -1.881 | -0.307 | -0.024 | -0.015 | -0.015 | 0 | 0.69 | 0.79 | 0.53 | 0.61 | 0.52 | 0.48 |
| 3 | 163645437 | rs9822851 | 2.035 | 1.529 | -1.875 | -0.311 | 0 | 0.003 | 0.003 | 0 | 0.05 | 0.07 | 0.02 | 0.11 | 0.04 | 0.01 |
| 3 | 163653210 | rs1150433 | 1.272 | 1.476 | -1.907 | -0.341 | 0.007 | -0.042 | -0.042 | 0.001 | 0.16 | 0.17 | 0.13 | 0.05 | 0.2 | 0.07 |
| 3 | 163658357 | rs16847384 | 1.748 | 1.389 | -1.786 | -0.356 | -0.025 | -0.141 | -0.141 | 0 | 0.22 | 0.46 | 0.13 | 0.02 | 0.18 | 0.1 |
| 3 | 163661532 | rs7613590 | 1.257 | 1.378 | -1.863 | -0.356 | -0.006 | -0.08 | -0.08 | 0 | 0.24 | 0.66 | 0.16 | 0.02 | 0.21 | 0.12 |
| 3 | 163662721 | rs7619158 | 1.244 | 1.378 | -1.863 | -0.356 | -0.006 | -0.08 | -0.08 | 0 | 0.24 | 0.66 | 0.16 | 0.02 | 0.21 | 0.14 |
| 3 | 163664285 | rs9857844 | 0.784 | 1.379 | -1.868 | -0.349 | 0.055 | -0.027 | -0.027 | 0 | 0.44 | 0.13 | 0.38 | 0.59 | 0.32 | 0.35 |
| 3 | 163671071 | rs9829001 | 0.782 | 1.384 | -1.892 | -0.348 | 0.054 | -0.029 | -0.029 | 0.37 | 0.44 | 0.13 | 0.38 | 0.59 | 0.32 | 0.35 |
| 3 | 163672702 | rs12489788 | 2.202 | 1.378 | -1.902 | -0.348 | 0.022 | -0.147 | -0.147 | 0.236 | 0.28 | 0.02 | 0.29 | 0.46 | 0.18 | 0.3 |
| 3 | 163678177 | rs3903745 | 0.838 | 1.443 | -1.725 | -0.507 | 0.06 | -0.016 | -0.016 | 0 | 0.44 | 0.15 | 0.38 | 0.59 | 0.32 | 0.35 |

| | | | | | | | | | | | | | | | | |
|---|---|---|---|---|---|---|---|---|---|---|---|---|---|---|---|---|
| 3 | 163681978 | rs3860532 | 0.159 | 1.449 | -1.721 | -0.507 | -0.023 | -0.015 | -0.015 | 0.131 | 0.69 | 0.79 | 0.53 | 0.61 | 0.52 | 0.48 |
| 3 | 163687895 | rs1581740 | 0.17 | 1.45 | -1.721 | -0.507 | -0.024 | -0.015 | -0.015 | 0.119 | 0.69 | 0.79 | 0.53 | 0.61 | 0.52 | 0.47 |
| 3 | 163693304 | rs1407537 | 0.833 | 1.435 | -1.729 | -0.503 | 0.06 | -0.016 | -0.016 | 0 | 0.44 | 0.15 | 0.38 | 0.59 | 0.32 | 0.35 |
| 3 | 163714630 | rs9864398 | 0.839 | 1.429 | -1.732 | -0.509 | 0.059 | -0.003 | -0.003 | 0.117 | 0.44 | 0.14 | 0.38 | 0.61 | 0.32 | 0.35 |
| 3 | 163719325 | rs36074517 | 1.098 | 1.485 | -1.58 | -0.472 | -0.008 | -0.096 | -0.096 | 0 | 0.24 | 0.64 | 0.15 | 0.02 | 0.21 | 0.12 |
| 3 | 163723996 | rs7614697 | 1.105 | 1.5 | -1.606 | -0.455 | -0.006 | -0.102 | -0.102 | 2.086 | 0.24 | 0.62 | 0.16 | 0.02 | 0.23 | 0.12 |
| 3 | 163725853 | rs7626982 | 1.549 | 1.454 | -1.716 | -0.435 | 0.048 | -0.068 | -0.068 | 0 | 0.34 | 0.11 | 0.32 | 0.58 | 0.22 | 0.31 |
| 3 | 163729593 | rs13083232 | 1.559 | 1.454 | -1.725 | -0.434 | 0.048 | -0.073 | -0.073 | 0.319 | 0.34 | 0.11 | 0.32 | 0.58 | 0.22 | 0.3 |
| 3 | 163730330 | rs6772713 | 1.335 | 1.504 | -1.481 | -0.251 | -0.019 | -0.13 | -0.13 | 0.069 | 0.22 | 0.44 | 0.14 | 0.02 | 0.2 | 0.1 |
| 3 | 163735214 | rs9815702 | -0.331 | 1.73 | -1.688 | -0.027 | -0.091 | -0.107 | -0.107 | 0 | 0.84 | 0.98 | 0.66 | 0.66 | 0.72 | 0.54 |
| 3 | 163761825 | rs13095640 | 0.409 | 1.342 | -1.41 | -0.144 | -0.012 | 0.028 | 0.028 | 0.505 | 0.55 | 0.66 | 0.46 | 0.67 | 0.45 | 0.48 |
| 3 | 163761964 | rs7427021 | 0.409 | 1.342 | -1.41 | -0.144 | -0.012 | 0.028 | 0.028 | 0.505 | 0.55 | 0.66 | 0.46 | 0.67 | 0.45 | 0.48 |
| 3 | 163769857 | rs12490884 | 1.115 | 1.275 | -1.448 | -0.205 | -0.007 | -0.046 | -0.046 | 1.653 | 0.35 | 0.17 | 0.31 | 0.56 | 0.27 | 0.37 |
| 3 | 163777235 | rs13084099 | 0.297 | 1.271 | -1.443 | -0.226 | -0.012 | 0.029 | 0.029 | 0.505 | 0.55 | 0.66 | 0.46 | 0.67 | 0.45 | 0.48 |
| 3 | 163781290 | rs6808152 | 1.212 | 1.247 | -1.355 | -0.218 | -0.023 | -0.046 | -0.046 | 0 | 0.2 | 0.41 | 0.14 | 0.11 | 0.18 | 0.12 |
| 3 | 163785690 | rs10936400 | 1.025 | 1.299 | -1.419 | -0.126 | -0.002 | -0.03 | -0.03 | 0 | 0.35 | 0.21 | 0.32 | 0.56 | 0.27 | 0.36 |
| 3 | 163795233 | rs12489384 | 1.025 | 1.299 | -1.436 | -0.126 | -0.003 | -0.032 | -0.032 | 0 | 0.35 | 0.21 | 0.32 | 0.55 | 0.27 | 0.36 |
| 3 | 163795964 | rs2198280 | 1.792 | 1.109 | -1.409 | 0.068 | 0.021 | -0.108 | -0.108 | 1.7 | 0.27 | 0.03 | 0.27 | 0.43 | 0.18 | 0.32 |
| 3 | 163810778 | rs10936401 | 1.462 | 1.188 | -1.348 | 0.032 | 0.004 | -0.014 | -0.014 | 0.91 | 0.29 | 0.25 | 0.3 | 0.55 | 0.21 | 0.34 |
| 3 | 163817702 | rs1839943 | 1.459 | 1.198 | -1.364 | 0.036 | 0.006 | -0.019 | -0.019 | 0 | 0.29 | 0.22 | 0.3 | 0.55 | 0.21 | 0.34 |
| 3 | 163827733 | rs1449938 | 1.459 | 1.198 | -1.368 | 0.036 | 0.005 | -0.016 | -0.016 | 1.409 | 0.29 | 0.23 | 0.3 | 0.55 | 0.21 | 0.34 |
| 3 | 163832988 | rs12629383 | 1.459 | 1.198 | -1.368 | 0.036 | 0.005 | -0.016 | -0.016 | 2.041 | 0.29 | 0.23 | 0.3 | 0.55 | 0.21 | 0.34 |
| 3 | 163838015 | rs9290226 | 1.566 | 1.116 | -1.549 | -0.127 | 0.01 | 0.052 | 0.052 | 0 | 0.13 | 0.04 | 0.1 | 0.02 | 0.1 | 0.04 |
| 3 | 163849580 | rs1992987 | 1.607 | 1.134 | -1.521 | -0.107 | 0.005 | -0.016 | -0.016 | 4.099 | 0.29 | 0.23 | 0.3 | 0.55 | 0.21 | 0.34 |
| 3 | 163857437 | rs1839412 | 1.607 | 1.134 | -1.52 | -0.107 | 0.005 | -0.018 | -0.018 | 1.871 | 0.29 | 0.22 | 0.3 | 0.55 | 0.21 | 0.34 |
| 3 | 163861651 | rs1947243 | -0.11 | 1.531 | -1.531 | -0.213 | 0.004 | 0.023 | 0.023 | 0 | 0.63 | 0.73 | 0.65 | 0.79 | 0.61 | 0.69 |
| 3 | 163864862 | rs11718123 | 1.656 | 1.509 | -1.443 | -0.107 | 0.005 | -0.016 | -0.016 | 0 | 0.29 | 0.23 | 0.3 | 0.55 | 0.21 | 0.34 |
| 3 | 163870667 | rs10936408 | 1.656 | 1.509 | -1.443 | -0.114 | 0.005 | -0.016 | -0.016 | 0 | 0.29 | 0.22 | 0.3 | 0.55 | 0.21 | 0.34 |
| 3 | 163885473 | rs2218935 | 0.531 | 1.411 | -1.515 | -0.189 | 0.004 | 0.049 | 0.049 | 0.113 | 0.52 | 0.64 | 0.45 | 0.67 | 0.42 | 0.47 |
| 3 | 163895047 | rs12495540 | 2.218 | 1.355 | -1.548 | -0.111 | 0.013 | -0.109 | -0.109 | 0 | 0.26 | 0.03 | 0.27 | 0.43 | 0.17 | 0.31 |
| 3 | 163898428 | rs710855 | -0.596 | 1.439 | -1.486 | -0.544 | 0.002 | -0.023 | -0.023 | 0 | 0.46 | 0.54 | 0.45 | 0.26 | 0.49 | 0.39 |
| 3 | 163902187 | rs6548335 | 0.929 | 1.458 | -1.505 | -0.57 | 0.011 | -0.058 | -0.058 | 0 | 0.23 | 0.42 | 0.15 | 0.12 | 0.23 | 0.13 |
| 3 | 163904370 | rs788895 | -0.628 | 1.444 | -1.477 | -0.553 | 0.001 | -0.016 | -0.016 | 0 | 0.46 | 0.55 | 0.45 | 0.26 | 0.49 | 0.39 |
| 3 | 163907728 | rs572190 | -0.775 | 1.443 | -1.487 | -0.55 | 0.001 | -0.003 | -0.003 | 0.187 | 0.75 | 0.76 | 0.74 | 0.81 | 0.68 | 0.74 |
| 3 | 163910768 | rs558949 | -0.775 | 1.443 | -1.478 | -0.55 | 0.001 | -0.003 | -0.003 | 0 | 0.75 | 0.76 | 0.74 | 0.81 | 0.68 | 0.74 |
| 3 | 163914791 | rs622209 | -0.597 | 1.445 | -1.478 | -0.549 | 0.002 | -0.023 | -0.023 | 0 | 0.46 | 0.54 | 0.45 | 0.26 | 0.49 | 0.39 |
| 3 | 163917329 | rs7629729 | -0.722 | 1.44 | -1.456 | -0.552 | -0.004 | 0.037 | 0.037 | 0 | 0.47 | 0.65 | 0.45 | 0.26 | 0.49 | 0.4 |
| 3 | 163922738 | rs1321548 | -0.605 | 1.47 | -1.501 | -0.538 | 0.002 | -0.023 | -0.023 | 0.404 | 0.46 | 0.54 | 0.45 | 0.26 | 0.49 | 0.39 |
| 3 | 163930675 | rs1667822 | -0.583 | 1.52 | -1.547 | -0.418 | 0.001 | -0.003 | -0.003 | 0 | 0.75 | 0.76 | 0.74 | 0.81 | 0.68 | 0.74 |
| 3 | 163934363 | rs1086092 | -0.496 | 1.415 | -1.455 | -0.388 | -0.011 | 0.041 | 0.041 | 2.7 | 0.23 | 0.16 | 0.29 | 0.14 | 0.26 | 0.26 |
| 3 | 163938800 | rs686697 | -0.481 | 1.415 | -1.455 | -0.388 | -0.011 | 0.041 | 0.041 | 2.705 | 0.23 | 0.16 | 0.29 | 0.14 | 0.26 | 0.26 |
| 3 | 163941553 | rs483558 | -0.488 | 1.418 | -1.453 | -0.387 | -0.011 | 0.041 | 0.041 | 2.7 | 0.23 | 0.16 | 0.29 | 0.14 | 0.26 | 0.26 |
| 3 | 163957118 | rs663534 | -0.054 | 1.495 | -1.731 | -0.365 | -0.05 | -0.021 | -0.021 | 0.315 | 0.09 | 0.01 | 0.2 | 0.12 | 0.16 | 0.22 |
| 3 | 163961014 | rs9818705 | 0.99 | 1.299 | -1.639 | -0.346 | 0.01 | 0.056 | 0.056 | 0 | 0.13 | 0.04 | 0.1 | 0.02 | 0.1 | 0.04 |
| 3 | 163970655 | rs789324 | -0.16 | 1.61 | -1.573 | -0.22 | -0.05 | -0.021 | -0.021 | 0 | 0.09 | 0.01 | 0.19 | 0.12 | 0.16 | 0.22 |
| 3 | 163983338 | rs13073971 | -0.927 | 1.585 | -1.407 | -0.529 | -0.001 | 0.001 | 0.001 | 0 | 0.47 | 0.54 | 0.45 | 0.25 | 0.5 | 0.4 |
| 3 | 163994508 | rs1575160 | -0.228 | 1.685 | -1.581 | -0.624 | -0.068 | 0.051 | 0.051 | 1.395 | 0.24 | 0.05 | 0.31 | 0.14 | 0.3 | 0.25 |
| 3 | 164001278 | rs13098760 | -1.184 | 1.676 | -1.549 | -0.671 | 0 | 0.011 | 0.011 | 0 | 0.48 | 0.54 | 0.46 | 0.25 | 0.54 | 0.4 |
| 3 | 164004812 | rs4855216 | -0.028 | 1.74 | -1.594 | -0.639 | -0.053 | -0.02 | -0.02 | 0 | 0.09 | 0.01 | 0.2 | 0.12 | 0.16 | 0.2 |
| 3 | 164016016 | rs6785566 | 0.779 | 1.575 | -1.349 | -0.654 | -0.013 | -0.04 | -0.04 | 0 | 0.19 | 0.4 | 0.13 | 0.12 | 0.18 | 0.14 |
| 3 | 164037502 | rs7611861 | 0.318 | 1.507 | -1.304 | -0.604 | 0.011 | -0.06 | -0.06 | 0 | 0.23 | 0.43 | 0.16 | 0.11 | 0.24 | 0.15 |
| 3 | 164073788 | rs6796508 | 0.398 | 1.738 | -0.816 | -0.52 | -0.021 | 0.019 | 0.019 | 0 | 0.18 | 0.12 | 0.13 | 0.11 | 0.23 | 0.15 |
| 3 | 164076198 | rs6548344 | -1.115 | 1.791 | -0.894 | -0.564 | -0.001 | -0.045 | -0.045 | 0 | 0.4 | 0.52 | 0.28 | 0.19 | 0.41 | 0.34 |
| 3 | 164083080 | rs13071561 | 0.942 | 1.463 | -0.993 | -0.444 | 0.006 | 0.071 | 0.071 | 0 | 0.15 | 0.03 | 0.11 | 0.03 | 0.14 | 0.05 |
| 3 | 164087419 | rs2141853 | -1.768 | 1.584 | -1.097 | -0.206 | 0.023 | -0.074 | -0.074 | 0 | 0.61 | 0.84 | 0.73 | 0.5 | 0.7 | 0.52 |
| 3 | 164088664 | rs13070763 | 0.509 | 1.84 | -0.961 | -0.147 | -0.008 | 0.012 | 0.012 | 0 | 0.18 | 0.16 | 0.14 | 0.12 | 0.23 | 0.13 |
| 3 | 164101193 | rs721976 | -0.983 | 1.802 | -0.95 | -0.149 | 0.006 | -0.023 | -0.023 | 2.321 | 0.46 | 0.8 | 0.62 | 0.48 | 0.56 | 0.47 |
| 3 | 164101429 | rs10513611 | 0.337 | 1.849 | -0.697 | -0.182 | 0.002 | -0.009 | -0.009 | 0 | 0.19 | 0.23 | 0.14 | 0.12 | 0.23 | 0.13 |
| 3 | 164102274 | rs1821575 | -0.325 | 1.639 | -0.139 | -0.099 | 0.026 | 0.028 | 0.028 | 0 | 0.26 | 0.4 | 0.46 | 0.36 | 0.33 | 0.33 |
| 3 | 164114823 | rs9857988 | -0.59 | 1.631 | -0.129 | -0.101 | 0.003 | 0.004 | 0.004 | 0.279 | 0.28 | 0.64 | 0.48 | 0.36 | 0.33 | 0.33 |
| 3 | 164123367 | rs16848141 | -0.427 | 1.689 | -0.237 | -0.076 | 0.028 | 0.031 | 0.031 | 0.016 | 0.27 | 0.57 | 0.48 | 0.36 | 0.33 | 0.33 |
| 3 | 164156963 | rs9849665 | -1.063 | 1.603 | -0.315 | -0.16 | -0.001 | 0 | 0 | 0.101 | 0.42 | 0.44 | 0.57 | 0.38 | 0.47 | 0.38 |
| 3 | 164170151 | rs892765 | -0.215 | 1.834 | -0.312 | -0.067 | 0.007 | 0.002 | 0.002 | 0.217 | 0.23 | 0.31 | 0.37 | 0.24 | 0.27 | 0.13 |
| 3 | 164172830 | rs810692 | 0.251 | 1.811 | -0.372 | -0.058 | 0.003 | -0.013 | -0.013 | 0.742 | 0.2 | 0.31 | 0.21 | 0.15 | 0.2 | 0.25 |
| 3 | 164182926 | rs1708504 | -0.984 | 1.794 | -0.753 | -0.007 | -0.004 | 0.002 | 0.002 | 0.158 | 0.43 | 0.61 | 0.58 | 0.39 | 0.47 | 0.37 |
| 3 | 164200220 | rs4855182 | -0.515 | 1.789 | -0.759 | 0.04 | -0.005 | -0.001 | -0.001 | 0 | 0.06 | 0.06 | 0.23 | 0.09 | 0.12 | 0.08 |
| 3 | 164209389 | rs298750 | -0.074 | 1.335 | -0.27 | -0.133 | -0.006 | -0.051 | -0.051 | 0 | 0.18 | 0.31 | 0.13 | 0.1 | 0.15 | 0.27 |
| 3 | 164209573 | rs1865482 | -0.653 | 1.363 | -0.647 | -0.156 | 0.018 | 0.014 | 0.014 | 0 | 0.34 | 0.49 | 0.43 | 0.38 | 0.44 | 0.34 |
| 3 | 164210971 | rs1478001 | -0.982 | 1.218 | -0.691 | -0.281 | -0.001 | -0.001 | -0.001 | 0.195 | 0.41 | 0.45 | 0.46 | 0.44 | 0.49 | 0.5 |

| | | | | | | | | | | | | | | | | |
|---|---|---|---|---|---|---|---|---|---|---|---|---|---|---|---|---|
| 3 | 164210992 | rs13070146 | -0.279 | 1.326 | -0.785 | -0.316 | -0.055 | 0.035 | 0.035 | 0 | 0.28 | 0.08 | 0.27 | 0.19 | 0.34 | 0.31 |
| 3 | 164211352 | rs298748 | -2.778 | 1.439 | -0.896 | -0.424 | 0.009 | -0.113 | -0.113 | 1.077 | 0.67 | 0.93 | 0.74 | 0.56 | 0.73 | 0.73 |
| 3 | 164212378 | rs1351067 | -0.978 | 1.398 | -0.605 | -0.554 | -0.035 | 0.085 | 0.085 | 0 | 0.39 | 0.27 | 0.43 | 0.17 | 0.47 | 0.44 |
| 3 | 164217297 | rs13074881 | 0.129 | 1.387 | -0.573 | -0.539 | -0.027 | 0.048 | 0.048 | 0.068 | 0.25 | 0.19 | 0.27 | 0.13 | 0.33 | 0.31 |
| 3 | 164238956 | rs892761 | -1.843 | 1.4 | -0.552 | -0.443 | 0.013 | 0.018 | 0.018 | 0 | 0.52 | 0.88 | 0.55 | 0.43 | 0.58 | 0.62 |
| 3 | 164244419 | rs967422 | 0.651 | 1.238 | -0.489 | -0.499 | -0.005 | 0.019 | 0.019 | 0 | 0.14 | 0.07 | 0.17 | 0.04 | 0.14 | 0.12 |
| 3 | 164248306 | rs9828418 | 0.809 | 1.194 | -0.367 | -0.342 | 0.03 | -0.007 | -0.007 | 0 | 0.1 | 0.31 | 0.09 | 0.05 | 0.1 | 0.11 |
| 3 | 164249319 | rs9853726 | 0.368 | 1.197 | -0.59 | -0.322 | 0.046 | 0.185 | 0.185 | 0.221 | 0.12 | 0.46 | 0.12 | 0.25 | 0.11 | 0.18 |
| 3 | 164267629 | rs6764920 | -2.499 | 1.262 | -1.052 | -0.319 | -0.044 | -0.175 | -0.175 | 0 | 0.7 | 0.98 | 0.75 | 0.51 | 0.75 | 0.72 |
| 3 | 164269269 | rs17503925 | -0.111 | 1.469 | -0.655 | -0.271 | -0.025 | 0.03 | 0.03 | 0 | 0.22 | 0.19 | 0.26 | 0.12 | 0.32 | 0.14 |
| 3 | 164272414 | rs12633984 | -0.863 | 1.489 | -0.61 | -0.261 | 0.02 | 0.012 | 0.012 | 0.215 | 0.3 | 0.5 | 0.31 | 0.34 | 0.37 | 0.39 |
| 3 | 164280365 | rs6775887 | -0.015 | 1.578 | -1.007 | -0.217 | -0.052 | -0.013 | -0.013 | 1.001 | 0.13 | 0.03 | 0.16 | 0.12 | 0.17 | 0.11 |
| 3 | 164294002 | rs2035199 | -0.656 | 1.444 | -1.069 | -0.596 | 0.016 | 0.103 | 0.103 | 0 | 0.48 | 0.67 | 0.42 | 0.3 | 0.48 | 0.38 |
| 3 | 164297552 | rs9290240 | 0.41 | 1.393 | -1.109 | -0.821 | -0.015 | -0.066 | -0.066 | 0 | 0.3 | 0.53 | 0.21 | 0.05 | 0.26 | 0.17 |
| 3 | 164305622 | rs973273 | -2.32 | 1.521 | -1.604 | -0.297 | -0.018 | -0.109 | -0.109 | 0 | 0.7 | 0.93 | 0.72 | 0.58 | 0.72 | 0.71 |
| 3 | 164347792 | rs13073673 | 0.367 | 0.881 | -1.083 | -0.587 | -0.052 | 0.049 | 0.049 | 0 | 0.14 | 0.02 | 0.17 | 0.04 | 0.19 | 0.12 |
| 3 | 164359700 | rs9858061 | 0.156 | 0.62 | -0.544 | -0.716 | 0 | -0.024 | -0.024 | 0.992 | 0.29 | 0.26 | 0.19 | 0.05 | 0.26 | 0.17 |
| 3 | 164359759 | rs17445697 | 0.367 | 0.712 | -0.588 | -0.737 | -0.024 | 0.043 | 0.043 | 0 | 0.14 | 0.05 | 0.17 | 0.04 | 0.19 | 0.12 |
| 3 | 164360829 | rs9863328 | -0.857 | 0.822 | -0.307 | -0.623 | 0.001 | 0.019 | 0.019 | 0 | 0.45 | 0.36 | 0.41 | 0.27 | 0.48 | 0.36 |
| 3 | 164361106 | rs9867481 | -0.914 | 0.796 | -0.283 | -0.708 | -0.001 | 0.023 | 0.023 | 0 | 0.43 | 0.31 | 0.37 | 0.07 | 0.47 | 0.29 |
| 3 | 164361328 | rs9829997 | -0.005 | 0.834 | -0.279 | -0.788 | 0.002 | -0.005 | -0.005 | 3.103 | 0.3 | 0.24 | 0.21 | 0.04 | 0.3 | 0.18 |
| 3 | 164361859 | rs7616852 | -0.784 | 0.537 | -0.007 | -0.884 | -0.001 | -0.007 | -0.007 | 0 | 0.22 | 0.28 | 0.23 | 0.1 | 0.26 | 0.28 |
| 3 | 164363372 | rs9878865 | -2.419 | 0.621 | -0.215 | -0.788 | -0.059 | -0.06 | -0.06 | 0 | 0.65 | 0.92 | 0.54 | 0.41 | 0.66 | 0.62 |
| 3 | 164363695 | rs6776744 | -1.524 | 0.668 | -0.064 | -0.775 | -0.004 | 0.022 | 0.022 | 0.183 | 0.53 | 0.6 | 0.47 | 0.31 | 0.57 | 0.53 |
| 3 | 164369406 | rs1390044 | 0.296 | 0.636 | -0.071 | -0.732 | 0.002 | -0.031 | -0.031 | 0.685 | 0.3 | 0.22 | 0.21 | 0.01 | 0.29 | 0.19 |
| 3 | 164370044 | rs1546148 | 0.53 | 0.805 | 0.428 | -0.153 | -0.006 | -0.005 | -0.005 | 0 | 0.1 | 0.05 | 0.06 | 0.1 | 0.1 | 0.09 |
| 3 | 164375880 | rs16848471 | -0.878 | 0.781 | 0.732 | -0.056 | 0.015 | 0.169 | 0.169 | 0 | 0.11 | 0.45 | 0.1 | 0.24 | 0.1 | 0.23 |
| 3 | 164380029 | rs9856794 | -1.27 | 0.702 | 1.112 | -0.094 | -0.002 | -0.019 | -0.019 | 0.267 | 0.24 | 0.23 | 0.15 | 0.13 | 0.26 | 0.27 |
| 3 | 164382627 | rs1495063 | -0.266 | 0.723 | 1.033 | -0.051 | 0.031 | 0.146 | 0.146 | 0 | 0.18 | 0.49 | 0.32 | 0.37 | 0.21 | 0.23 |
| 3 | 164384338 | rs2722633 | -2.161 | 0.707 | 1.022 | -0.055 | -0.012 | -0.046 | -0.046 | 0 | 0.43 | 0.81 | 0.48 | 0.5 | 0.47 | 0.5 |
| 3 | 164399225 | rs9881897 | -1.267 | 0.62 | 1.003 | 0.048 | 0 | -0.007 | -0.007 | 0 | 0.27 | 0.21 | 0.2 | 0.13 | 0.31 | 0.28 |
| 3 | 164411535 | rs1495045 | -1.081 | 0.618 | 0.845 | 0.049 | -0.011 | 0.031 | 0.031 | 0 | 0.26 | 0.13 | 0.2 | 0.14 | 0.31 | 0.28 |
| 3 | 164412364 | rs16847459 | -1.415 | 0.606 | 0.69 | 0.084 | 0.001 | 0 | 0 | 0.019 | 0.31 | 0.56 | 0.26 | 0.33 | 0.33 | 0.35 |
| 3 | 164415896 | rs9290242 | -1.047 | 0.448 | 0.821 | 0.019 | -0.001 | 0.001 | 0.001 | 0.016 | 0.21 | 0.66 | 0.2 | 0.24 | 0.22 | 0.27 |
| 3 | 164420232 | rs2036034 | 0.203 | 0.352 | 0.149 | -0.242 | -0.043 | 0.04 | 0.04 | 0 | 0.28 | 0.05 | 0.35 | 0.16 | 0.31 | 0.28 |
| 3 | 164430644 | rs1112188 | -1.445 | 0.318 | 0.204 | -0.164 | 0.006 | 0.033 | 0.033 | 0 | 0.49 | 0.76 | 0.55 | 0.44 | 0.53 | 0.55 |
| 3 | 164432677 | rs2643190 | 0.225 | 0.374 | -0.013 | -0.221 | -0.06 | 0.019 | 0.019 | 0 | 0.27 | 0.03 | 0.33 | 0.18 | 0.3 | 0.28 |
| 3 | 164435533 | rs6548369 | -0.903 | 0.101 | 0.499 | -0.498 | -0.008 | 0.018 | 0.018 | 0.195 | 0.19 | 0.64 | 0.24 | 0.26 | 0.19 | 0.26 |
| 3 | 164438520 | rs6763526 | 1.187 | 0.221 | 0.336 | -0.511 | -0.002 | 0.071 | 0.071 | 0.015 | 0.53 | 0.24 | 0.51 | 0.65 | 0.49 | 0.44 |
| 3 | 164449337 | rs9850374 | -0.209 | 0.254 | 0.701 | -1.025 | 0.006 | -0.025 | -0.025 | 0.031 | 0.15 | 0.23 | 0.18 | 0.07 | 0.18 | 0.17 |
| 3 | 164468993 | rs1390040 | -0.165 | 0.244 | 0.467 | -1.03 | -0.005 | 0.01 | 0.01 | 0 | 0.14 | 0.11 | 0.17 | 0.07 | 0.18 | 0.16 |
| 3 | 164474373 | rs11718253 | 1.293 | 0.231 | 0.336 | -1.11 | -0.003 | 0.097 | 0.097 | 0.163 | 0.51 | 0.27 | 0.51 | 0.74 | 0.44 | 0.41 |
| 3 | 164515528 | rs4387973 | -0.092 | 0.335 | 0.091 | -0.985 | -0.049 | 0.122 | 0.122 | 0 | 0.72 | 0.94 | 0.75 | 0.92 | 0.68 | 0.67 |
| 3 | 164537423 | rs4560300 | 0.742 | 0.197 | 0.844 | -1.1 | 0 | 0.05 | 0.05 | 1.88 | 0.59 | 0.38 | 0.76 | 0.84 | 0.6 | 0.64 |
| 3 | 164541172 | rs9812705 | 1.034 | 0.362 | 0.844 | -1.116 | -0.042 | 0.138 | 0.138 | 3.001 | 0.55 | 0.21 | 0.69 | 0.76 | 0.53 | 0.52 |
| 3 | 164558595 | rs9847462 | 0.506 | 0.355 | 1.127 | -1.145 | 0.037 | 0.018 | 0.018 | 0.024 | 0.11 | 0.25 | 0.08 | 0.1 | 0.09 | 0.13 |
| 3 | 164559726 | rs7652863 | -0.319 | 0.348 | 0.888 | -1.153 | -0.026 | 0.099 | 0.099 | 0.021 | 0.73 | 0.92 | 0.85 | 0.93 | 0.7 | 0.78 |
| 3 | 164582258 | rs9290246 | 0.544 | 0.307 | 0.749 | -1.157 | 0.001 | -0.05 | -0.05 | 0 | 0.65 | 0.44 | 0.77 | 0.85 | 0.63 | 0.65 |
| 3 | 164586522 | rs4618222 | -1.437 | 0.436 | 0.317 | -0.87 | -0.006 | 0.015 | 0.015 | 5.122 | 0.2 | 0.87 | 0.17 | 0.18 | 0.16 | 0.25 |
| 3 | 164588702 | rs9810313 | -0.66 | 0.507 | 0.391 | -0.841 | -0.004 | -0.017 | -0.017 | 0 | 0.14 | 0.62 | 0.1 | 0.09 | 0.1 | 0.13 |
| 3 | 164589184 | rs7636613 | -0.103 | 0.527 | 0.182 | -0.563 | 0.019 | 0.027 | 0.027 | 0 | 0.05 | 0.15 | 0.06 | 0.1 | 0.07 | 0.12 |
| 3 | 164600590 | rs1993522 | 1.562 | 0.216 | -0.782 | -0.951 | -0.117 | -0.064 | -0.064 | 1.036 | 0.3 | 0.03 | 0.43 | 0.7 | 0.33 | 0.34 |
| 3 | 164613717 | rs6789934 | -0.057 | -0.112 | -0.036 | -1.318 | -0.012 | -0.003 | -0.003 | 0.022 | 0.14 | 0.57 | 0.1 | 0.11 | 0.1 | 0.15 |
| 3 | 164631530 | rs6781516 | 0.63 | -0.14 | 0.694 | -1.328 | -0.001 | -0.002 | -0.002 | 1.24 | 0.11 | 0.18 | 0.07 | 0.08 | 0.1 | 0.13 |
| 3 | 164654374 | rs7619352 | -0.683 | -0.253 | 0.313 | -1.678 | 0.001 | -0.03 | -0.03 | 0 | 0.29 | 0.56 | 0.18 | 0.14 | 0.29 | 0.22 |
| 3 | 164658624 | rs6548380 | -0.099 | -0.158 | 0.552 | -1.532 | -0.013 | -0.01 | -0.01 | 0 | 0.14 | 0.63 | 0.1 | 0.11 | 0.1 | 0.14 |
| 3 | 164688034 | rs7630103 | -0.137 | -0.229 | 0.802 | -1.589 | -0.008 | -0.041 | -0.041 | 0 | 0.15 | 0.56 | 0.1 | 0.07 | 0.11 | 0.15 |
| 3 | 164694254 | rs6774322 | 0.176 | -0.234 | 1.114 | -1.591 | 0.019 | -0.025 | -0.025 | 0 | 0.14 | 0.44 | 0.09 | 0.07 | 0.11 | 0.15 |
| 3 | 164695827 | rs6806082 | 0.721 | -0.234 | 1.283 | -1.591 | -0.001 | -0.009 | -0.009 | 0 | 0.12 | 0.14 | 0.07 | 0.06 | 0.11 | 0.15 |
| 3 | 164696120 | rs13072967 | 0.268 | -0.424 | 1.022 | -1.791 | 0.002 | 0.067 | 0.067 | 0 | 0.31 | 0.15 | 0.17 | 0.07 | 0.32 | 0.16 |
| 3 | 164706642 | rs6788137 | -0.09 | -0.333 | 0.609 | -1.754 | -0.016 | -0.043 | -0.043 | 0 | 0.16 | 0.63 | 0.11 | 0.07 | 0.11 | 0.15 |
| 3 | 164711387 | rs16848834 | -0.128 | -0.335 | 0.608 | -1.755 | -0.018 | -0.045 | -0.045 | 0 | 0.16 | 0.63 | 0.11 | 0.07 | 0.11 | 0.17 |
| 3 | 164724567 | rs7618643 | 0.745 | -0.15 | 0.129 | -1.644 | -0.001 | -0.006 | -0.006 | 0.956 | 0.12 | 0.13 | 0.07 | 0.06 | 0.11 | 0.17 |
| 3 | 164747930 | rs4557160 | 1.156 | -0.248 | 0.07 | -1.719 | -0.012 | 0.106 | 0.106 | 0 | 0.47 | 0.31 | 0.56 | 0.78 | 0.5 | 0.45 |
| 3 | 164748156 | rs13069479 | 0.186 | -0.269 | 0.388 | -1.54 | -0.005 | 0.087 | 0.087 | 0 | 0.3 | 0.12 | 0.17 | 0.08 | 0.31 | 0.16 |
| 3 | 164759707 | rs13062602 | 0.445 | 0.055 | 0.465 | -1.466 | 0.001 | -0.036 | -0.036 | 0 | 0.63 | 0.47 | 0.62 | 0.83 | 0.63 | 0.54 |
| 3 | 164764719 | rs9838509 | 0.454 | 0.054 | 0.465 | -1.465 | 0 | -0.034 | -0.034 | 0.544 | 0.63 | 0.47 | 0.62 | 0.83 | 0.63 | 0.54 |
| 3 | 164773057 | rs9290257 | 0.304 | 0.16 | 0.501 | -1.431 | 0.003 | -0.042 | -0.042 | 1.442 | 0.65 | 0.47 | 0.62 | 0.83 | 0.63 | 0.58 |
| 3 | 164783165 | rs9283633 | 0.312 | 0.176 | 0.508 | -1.437 | 0.003 | -0.039 | -0.039 | 0 | 0.65 | 0.46 | 0.62 | 0.83 | 0.63 | 0.58 |

| | | | | | | | | | | | | | | | | |
|---|---|---|---|---|---|---|---|---|---|---|---|---|---|---|---|---|
| 3 | 164793758 | rs9290264 | 0.518 | 0.222 | 0.438 | -1.46 | -0.003 | 0.015 | 0.015 | 0 | 0.25 | 0.21 | 0.18 | 0.11 | 0.28 | 0.21 |
| 3 | 164796640 | rs7641586 | -0.004 | 0.24 | 0.177 | -1.481 | -0.007 | 0.072 | 0.072 | 0 | 0.29 | 0.13 | 0.17 | 0.09 | 0.31 | 0.15 |
| 3 | 164799079 | rs9756306 | -0.067 | 0.38 | -0.037 | -1.229 | -0.002 | -0.059 | -0.059 | 0.119 | 0.3 | 0.35 | 0.21 | 0.12 | 0.3 | 0.32 |
| 3 | 164807679 | rs9825346 | 0.446 | 0.374 | -0.173 | -1.205 | 0.008 | -0.069 | -0.069 | 0 | 0.65 | 0.53 | 0.63 | 0.83 | 0.63 | 0.58 |
| 3 | 164871580 | rs12696221 | 0.237 | 0.233 | -0.834 | -1.101 | -0.011 | 0.087 | 0.087 | 0 | 0.36 | 0.22 | 0.25 | 0.12 | 0.38 | 0.28 |
| 3 | 164876267 | rs4858792 | 1.069 | -0.185 | -0.783 | -0.821 | -0.025 | 0.034 | 0.034 | 0 | 0.33 | 0.07 | 0.34 | 0.75 | 0.31 | 0.39 |
| 3 | 164891218 | rs920417 | 0.188 | -0.211 | -0.697 | -0.866 | 0.015 | 0.143 | 0.143 | 0.003 | 0.48 | 0.25 | 0.42 | 0.17 | 0.56 | 0.35 |
| 3 | 164895453 | rs2053371 | -0.251 | 0.016 | -1.144 | -0.675 | 0 | 0.003 | 0.003 | 0 | 0.08 | 0.08 | 0.16 | 0.05 | 0.08 | 0.14 |
| 3 | 164903829 | rs9856989 | 0.956 | -0.025 | -1.251 | -0.718 | -0.031 | 0.026 | 0.026 | 0 | 0.33 | 0.07 | 0.34 | 0.75 | 0.3 | 0.39 |
| 3 | 164904433 | rs4858782 | -0.358 | 0.283 | -0.741 | -0.326 | -0.001 | 0.002 | 0.002 | 0 | 0.08 | 0.06 | 0.16 | 0.05 | 0.08 | 0.14 |
| 3 | 164905570 | rs11922760 | -0.628 | 0.282 | -0.737 | -0.365 | 0 | 0.004 | 0.004 | 0 | 0.08 | 0.11 | 0.16 | 0.05 | 0.08 | 0.14 |
| 3 | 164905688 | rs17449404 | -0.558 | 0.288 | -0.75 | -0.362 | -0.001 | 0.004 | 0.004 | 0 | 0.08 | 0.09 | 0.16 | 0.05 | 0.08 | 0.14 |
| 3 | 164910800 | rs11928265 | -0.789 | 0.32 | -0.844 | -0.167 | 0.003 | 0.021 | 0.021 | 0 | 0.13 | 0.1 | 0.18 | 0.06 | 0.13 | 0.15 |
| 3 | 164926605 | rs13082820 | -0.399 | 0.239 | -0.715 | -0.138 | 0.029 | 0.106 | 0.106 | 0.02 | 0.53 | 0.25 | 0.57 | 0.23 | 0.6 | 0.46 |
| 3 | 164941049 | rs2033857 | -0.738 | 0.237 | 0.024 | 0.032 | -0.002 | -0.004 | -0.004 | 0 | 0.6 | 0.51 | 0.6 | 0.34 | 0.63 | 0.57 |
| 3 | 164963899 | rs9871479 | -0.666 | 0.485 | 0.416 | 0.298 | 0.004 | 0.021 | 0.021 | 0 | 0.51 | 0.64 | 0.52 | 0.4 | 0.43 | 0.52 |
| 3 | 164967758 | rs12494164 | -0.348 | 0.682 | 0.549 | 0.189 | 0.015 | -0.002 | -0.002 | 0 | 0.14 | 0.24 | 0.26 | 0.16 | 0.17 | 0.12 |
| 3 | 164979912 | rs16848988 | 0.094 | -0.069 | 0.779 | 0.165 | 0.018 | 0.012 | 0.012 | 0 | 0.31 | 0.13 | 0.24 | 0.22 | 0.25 | 0.37 |
| 3 | 164992076 | rs1371567 | -0.482 | -0.029 | 0.876 | 0.101 | 0.012 | 0.016 | 0.016 | 0 | 0.38 | 0.64 | 0.28 | 0.21 | 0.27 | 0.41 |
| 3 | 165002300 | rs16849016 | -0.427 | -0.021 | 0.976 | 0.093 | -0.003 | -0.005 | -0.005 | 0 | 0.37 | 0.57 | 0.28 | 0.21 | 0.27 | 0.41 |
| 3 | 165020180 | rs1525383 | -0.516 | 0.098 | 0.889 | 0.09 | 0.009 | 0.015 | 0.015 | 0 | 0.39 | 0.62 | 0.29 | 0.21 | 0.28 | 0.39 |
| 3 | 165026679 | rs10936438 | -0.472 | 0.1 | 0.899 | 0.091 | -0.006 | -0.014 | -0.014 | 0 | 0.39 | 0.58 | 0.29 | 0.21 | 0.28 | 0.39 |
| 3 | 165041116 | rs9841819 | 0.105 | 0.159 | 0.527 | 0.254 | 0.005 | 0.006 | 0.006 | 0 | 0.32 | 0.21 | 0.25 | 0.24 | 0.26 | 0.37 |
| 3 | 165041791 | rs9862939 | -0.413 | 0.108 | 0.645 | 0.232 | -0.012 | -0.013 | -0.013 | 0 | 0.37 | 0.51 | 0.27 | 0.26 | 0.27 | 0.4 |
| 3 | 165043405 | rs9815183 | 0.075 | 0.163 | 0.448 | 0.244 | 0.008 | 0.01 | 0.01 | 0 | 0.32 | 0.18 | 0.24 | 0.24 | 0.26 | 0.38 |
| 3 | 165054691 | rs7619601 | -0.244 | 0.304 | 0.081 | 0.308 | -0.013 | -0.014 | -0.014 | 0.006 | 0.37 | 0.51 | 0.27 | 0.26 | 0.27 | 0.41 |
| 3 | 165056353 | rs822715 | 0.157 | 0.508 | -0.301 | 0.042 | -0.006 | 0.009 | 0.009 | 0 | 0.83 | 0.84 | 0.73 | 0.89 | 0.78 | 0.87 |
| 3 | 165068119 | rs1723543 | 0.303 | 0.3 | -0.179 | -0.072 | 0.007 | -0.007 | -0.007 | 0.007 | 0.85 | 0.76 | 0.74 | 0.88 | 0.78 | 0.9 |
| 3 | 165073006 | rs822718 | 1.087 | 0.141 | -1.339 | 0.135 | -0.157 | -0.168 | -0.168 | 0 | 0.35 | 0.02 | 0.39 | 0.51 | 0.49 | 0.41 |
| 3 | 165110392 | rs2312457 | -0.495 | 0.111 | -0.511 | -0.01 | 0.003 | -0.007 | -0.007 | 0.028 | 0.19 | 0.26 | 0.28 | 0.13 | 0.24 | 0.1 |
| 3 | 165116509 | rs9839242 | -0.752 | -0.287 | -0.539 | 0.009 | -0.022 | 0.039 | 0.039 | 0 | 0.52 | 0.85 | 0.56 | 0.33 | 0.44 | 0.32 |
| 3 | 165118985 | rs1404782 | -0.746 | -0.282 | -0.539 | 0.012 | -0.023 | 0.04 | 0.04 | 0 | 0.52 | 0.85 | 0.56 | 0.33 | 0.44 | 0.32 |
| 3 | 165125159 | rs9878522 | 0.23 | -0.572 | 0.189 | -0.076 | -0.009 | -0.004 | -0.004 | 0.043 | 0.34 | 0.34 | 0.24 | 0.23 | 0.21 | 0.25 |
| 3 | 165131390 | rs11923404 | 0.636 | -0.566 | -0.003 | -0.04 | -0.017 | -0.005 | -0.005 | 0.598 | 0.29 | 0.37 | 0.23 | 0.21 | 0.19 | 0.21 |
| 3 | 165132797 | rs2312406 | 0.745 | -0.444 | 0.027 | -0.046 | -0.008 | 0.004 | 0.004 | 0 | 0.27 | 0.37 | 0.24 | 0.22 | 0.2 | 0.21 |
| 3 | 165134957 | rs6444811 | 0.259 | -0.424 | 0.006 | -0.023 | -0.013 | 0.001 | 0.001 | 0.299 | 0.32 | 0.53 | 0.26 | 0.26 | 0.22 | 0.25 |
| 3 | 165136797 | rs4626117 | 0.192 | -0.217 | -0.156 | -0.021 | 0.001 | -0.003 | -0.003 | 0 | 0.82 | 0.73 | 0.71 | 0.87 | 0.78 | 0.9 |
| 3 | 165160646 | rs10513615 | 0.234 | -0.104 | -0.996 | -0.317 | -0.037 | -0.001 | -0.001 | 0 | 0.44 | 0.14 | 0.38 | 0.65 | 0.44 | 0.68 |
| 3 | 165168292 | rs13076475 | 0.803 | 0.022 | -1.179 | -0.392 | -0.081 | 0.074 | 0.074 | 3.104 | 0.22 | 0.02 | 0.18 | 0.06 | 0.31 | 0.1 |
| 3 | 165177057 | rs6803772 | -0.181 | -0.141 | -0.186 | -0.088 | 0.002 | 0.007 | 0.007 | 0 | 0.48 | 0.53 | 0.41 | 0.66 | 0.44 | 0.68 |
| 3 | 165182446 | rs4593053 | 0.47 | -0.185 | -0.724 | 0.188 | -0.06 | -0.111 | -0.111 | 0.964 | 0.34 | 0.08 | 0.33 | 0.53 | 0.38 | 0.57 |
| 3 | 165211894 | rs12152542 | 0.54 | -0.223 | -0.866 | 0.214 | -0.078 | -0.138 | -0.138 | 1.05 | 0.32 | 0.08 | 0.32 | 0.51 | 0.38 | 0.43 |
| 3 | 165219144 | rs7643628 | 0.165 | -0.312 | -0.825 | 0.194 | -0.001 | -0.049 | -0.049 | 0 | 0.31 | 0.17 | 0.28 | 0.56 | 0.31 | 0.4 |
| 3 | 165219707 | rs4452350 | -0.278 | -0.043 | -0.445 | 0.24 | 0.007 | 0.04 | 0.04 | 0 | 0.05 | 0.08 | 0.05 | 0.18 | 0.06 | 0.1 |
| 3 | 165245914 | rs1464623 | 0.592 | -0.215 | -0.372 | -0.103 | -0.016 | 0.038 | 0.038 | 3.968 | 0.25 | 0.43 | 0.38 | 0.33 | 0.22 | 0.2 |
| 3 | 165253400 | rs6444873 | -0.962 | -0.333 | -1.178 | -0.509 | 0 | -0.067 | -0.067 | 0.009 | 0.73 | 0.82 | 0.68 | 0.46 | 0.7 | 0.63 |
| 3 | 165255278 | rs1403617 | 0.167 | -0.114 | -0.644 | -0.232 | -0.004 | 0.013 | 0.013 | 3.049 | 0.3 | 0.6 | 0.42 | 0.44 | 0.27 | 0.32 |
| 3 | 165265189 | rs4282082 | -1.191 | 0.07 | -1.047 | -0.158 | -0.012 | -0.053 | -0.053 | 0 | 0.78 | 0.81 | 0.7 | 0.46 | 0.74 | 0.63 |
| 3 | 165267123 | rs6794521 | -0.135 | -0.335 | -1.664 | -0.907 | -0.018 | 0.063 | 0.063 | 0 | 0.56 | 0.76 | 0.53 | 0.31 | 0.52 | 0.42 |
| 3 | 165267688 | rs1568701 | 1.782 | -0.514 | -1.356 | -1.164 | -0.001 | 0 | 0 | 0 | 0.19 | 0.21 | 0.32 | 0.18 | 0.18 | 0.16 |
| 3 | 165270444 | rs4634149 | -1.898 | -0.506 | -1.555 | -1.219 | -0.024 | -0.066 | -0.066 | 0.313 | 0.75 | 0.81 | 0.67 | 0.45 | 0.7 | 0.56 |
| 3 | 165277899 | rs9847083 | -1.835 | -0.791 | -1.44 | -1.289 | -0.035 | -0.073 | -0.073 | 0.064 | 0.71 | 0.8 | 0.62 | 0.44 | 0.63 | 0.52 |
| 3 | 165289424 | rs2222464 | 0.077 | -0.294 | -0.647 | -1.261 | -0.009 | -0.021 | -0.021 | 1.537 | 0.33 | 0.55 | 0.37 | 0.18 | 0.27 | 0.24 |
| 3 | 165303141 | rs7622979 | 0.432 | -0.524 | -0.661 | -1.348 | -0.021 | 0.009 | 0.009 | 0 | 0.09 | 0.14 | 0.04 | 0.12 | 0.04 | 0.08 |
| 3 | 165304697 | rs6786218 | 1.365 | -0.481 | -0.562 | -1.345 | 0.005 | -0.013 | -0.013 | 0 | 0.13 | 0.08 | 0.05 | 0 | 0.08 | 0.07 |
| 3 | 165310746 | rs11923311 | -1.316 | -0.374 | -0.601 | -1.331 | -0.021 | 0.063 | 0.063 | 0 | 0.56 | 0.76 | 0.54 | 0.3 | 0.52 | 0.41 |
| 3 | 165311604 | rs6444909 | 1.277 | -0.909 | -0.034 | -1.184 | -0.001 | -0.003 | -0.003 | 0.511 | 0.19 | 0.22 | 0.32 | 0.15 | 0.18 | 0.11 |
| 3 | 165314375 | rs7432504 | 0.299 | -0.463 | 0.072 | -1.024 | 0.07 | 0.074 | 0.074 | 0.226 | 0.58 | 0.25 | 0.58 | 0.72 | 0.72 | 0.73 |
| 3 | 165339415 | rs7620921 | -0.448 | 0.359 | 1.316 | -0.593 | 0.078 | 0.131 | 0.131 | 0 | 0.05 | 0.34 | 0.18 | 0.12 | 0.05 | 0.09 |
| 3 | 165342529 | rs606353 | 0.475 | 0.263 | 0.877 | -0.713 | 0.014 | 0.039 | 0.039 | 0 | 0.12 | 0.03 | 0.04 | 0 | 0.08 | 0.11 |
| 3 | 165351210 | rs553149 | -0.463 | 0.255 | 0.963 | -0.714 | -0.022 | -0.048 | -0.048 | 0 | 0.19 | 0.43 | 0.22 | 0.12 | 0.12 | 0.19 |
| 3 | 165374980 | rs17706250 | -0.301 | 0.265 | 0.853 | -0.606 | 0.017 | -0.022 | -0.022 | 0 | 0.14 | 0.08 | 0.07 | 0.16 | 0.11 | 0.12 |
| 3 | 165385580 | rs10513618 | 0.43 | 0.655 | 0.721 | -0.732 | -0.075 | 0.071 | 0.071 | 0.08 | 0.13 | 0.01 | 0.14 | 0.01 | 0.21 | 0.09 |
| 3 | 165388035 | rs3107699 | -0.771 | 0.579 | 0.916 | -0.657 | 0.01 | -0.048 | -0.048 | 0 | 0.32 | 0.4 | 0.36 | 0.12 | 0.34 | 0.23 |
| 3 | 165392256 | rs17706693 | -0.27 | 0.507 | 0.781 | -0.622 | 0.015 | -0.027 | -0.027 | 3.159 | 0.14 | 0.08 | 0.08 | 0.16 | 0.11 | 0.12 |
| 3 | 165403192 | rs694955 | -1.765 | 0.566 | 0.981 | -0.633 | -0.014 | 0.058 | 0.058 | 0 | 0.47 | 0.76 | 0.46 | 0.28 | 0.47 | 0.35 |
| 3 | 165405491 | rs632101 | -1.019 | 0.823 | 1.12 | -0.742 | 0 | -0.015 | -0.015 | 0 | 0.33 | 0.68 | 0.38 | 0.12 | 0.36 | 0.23 |
| 3 | 165435642 | rs697355 | -1.602 | 0.828 | -0.157 | -0.35 | 0.005 | 0.054 | 0.054 | 0 | 0.52 | 0.73 | 0.4 | 0.33 | 0.52 | 0.41 |
| 3 | 165435695 | rs11928838 | 0.197 | 0.91 | 0.03 | -0.651 | -0.019 | 0.014 | 0.014 | 0 | 0.12 | 0.1 | 0.15 | 0 | 0.21 | 0.08 |

| | | | | | | | | | | | | | | | | |
|---|---|---|---|---|---|---|---|---|---|---|---|---|---|---|---|---|
| 3 | 165436169 | rs6771420 | -1.767 | 0.18 | 0.739 | -0.458 | 0.004 | 0.008 | 0.008 | 0 | 0.35 | 0.63 | 0.26 | 0.13 | 0.33 | 0.21 |
| 3 | 165438286 | rs1707706 | -1.188 | 0.356 | 1.019 | -0.396 | -0.002 | -0.026 | -0.026 | 0 | 0.16 | 0.25 | 0.15 | 0.08 | 0.18 | 0.13 |
| 3 | 165439372 | rs17459269 | -0.187 | 0.224 | 1.627 | -0.455 | -0.007 | 0.018 | 0.018 | 0 | 0.11 | 0.19 | 0.06 | 0.13 | 0.1 | 0.09 |
| 3 | 165441595 | rs1707713 | -2.886 | 0.171 | 1.525 | -0.307 | -0.029 | -0.013 | -0.013 | 0 | 0.45 | 0.93 | 0.46 | 0.36 | 0.42 | 0.35 |
| 3 | 165442455 | rs1707714 | -2.543 | 0.08 | 2.414 | -0.329 | -0.005 | -0.01 | -0.01 | 0.011 | 0.34 | 0.63 | 0.26 | 0.21 | 0.33 | 0.26 |
| 3 | 165442777 | rs1706444 | 0.426 | 0.086 | 2.305 | -0.317 | -0.004 | 0.004 | 0.004 | 0 | 0.06 | 0.11 | 0.03 | 0.07 | 0.03 | 0.08 |
| 3 | 165442806 | rs1398616 | -1.936 | 0.086 | 2.267 | -0.318 | -0.01 | -0.008 | -0.008 | 0 | 0.27 | 0.44 | 0.21 | 0.21 | 0.28 | 0.22 |
| 3 | 165445557 | rs17519507 | -0.165 | 0.095 | 2.046 | -0.296 | -0.007 | 0.02 | 0.02 | 0 | 0.11 | 0.19 | 0.06 | 0.13 | 0.11 | 0.09 |
| 3 | 165447997 | rs13098460 | -0.062 | 0.129 | 1.828 | -0.325 | -0.027 | 0.058 | 0.058 | 1.702 | 0.1 | 0 | 0.1 | 0 | 0.15 | 0.04 |
| 3 | 165468445 | rs1398089 | -1.284 | 0.113 | 2.231 | -0.324 | 0.003 | -0.028 | -0.028 | 0 | 0.14 | 0.52 | 0.1 | 0.07 | 0.08 | 0.12 |
| 3 | 165468999 | rs1403015 | -2.588 | 0.081 | 2.308 | -0.341 | 0.008 | 0.015 | 0.015 | 0.845 | 0.37 | 0.67 | 0.27 | 0.21 | 0.34 | 0.29 |
| 3 | 165476026 | rs4113819 | -0.345 | 0.103 | 2.42 | -0.313 | 0.016 | -0.033 | -0.033 | 0.102 | 0.07 | 0.2 | 0.05 | 0 | 0.05 | 0.04 |
| 3 | 165486145 | rs11720167 | -1.37 | 0.105 | 2.447 | -0.309 | 0.001 | 0 | 0 | 0 | 0.19 | 0.18 | 0.12 | 0.13 | 0.19 | 0.16 |
| 3 | 165489497 | rs10049010 | -0.333 | 0.111 | 2.324 | -0.325 | 0.014 | -0.035 | -0.035 | 1.012 | 0.07 | 0.2 | 0.05 | 0 | 0.05 | 0.04 |
| 3 | 165494567 | rs2668209 | 2.295 | 0.101 | 2.33 | -0.327 | -0.002 | -0.003 | -0.003 | 0 | 0.67 | 0.35 | 0.79 | 0.79 | 0.74 | 0.72 |
| 3 | 165500160 | rs16849623 | -0.203 | 0.132 | 2.442 | -0.31 | -0.001 | -0.037 | -0.037 | 0.146 | 0.07 | 0.14 | 0.05 | 0 | 0.05 | 0.04 |
| 3 | 165500685 | rs3863086 | -0.103 | 0.131 | 2.461 | -0.311 | -0.003 | -0.029 | -0.029 | 0 | 0.07 | 0.11 | 0.05 | 0 | 0.05 | 0.04 |
| 3 | 165502724 | rs829508 | 2.086 | 0.115 | 2.512 | -0.308 | 0 | 0.001 | 0.001 | 0.192 | 0.74 | 0.31 | 0.67 | 0.77 | 0.76 | 0.78 |
| 3 | 165505177 | rs1355538 | 2.996 | 0.152 | 1.991 | -0.352 | -0.056 | -0.008 | -0.008 | 0 | 0.52 | 0.05 | 0.53 | 0.64 | 0.54 | 0.6 |
| 3 | 165538468 | rs1511982 | 1.074 | 0.269 | 2.466 | -0.112 | -0.009 | -0.032 | -0.032 | 0 | 0.86 | 0.63 | 0.92 | 0.93 | 0.91 | 0.88 |
| 3 | 165539122 | rs9878990 | 2.733 | 0.254 | 2.22 | -0.124 | -0.021 | -0.04 | -0.04 | 0.079 | 0.61 | 0.1 | 0.6 | 0.65 | 0.65 | 0.66 |
| 3 | 165553646 | rs4680608 | 1.46 | 0.247 | 2.069 | -0.092 | -0.008 | 0.003 | 0.003 | 0 | 0.78 | 0.71 | 0.86 | 0.8 | 0.82 | 0.8 |
| 3 | 165555125 | rs1126680 | 0.651 | 0.292 | 1.527 | -0.086 | -0.017 | 0.025 | 0.025 | 0 | 0.05 | 0.02 | 0.05 | 0 | 0.08 | 0.04 |
| 3 | 165586274 | rs4602440 | 0.954 | 0.275 | 1.595 | -0.009 | 0.001 | -0.004 | -0.004 | 0 | 0.85 | 0.86 | 0.91 | 0.8 | 0.87 | 0.84 |
| 3 | 165606177 | rs9879188 | -2.15 | 0.717 | 1.7 | -0.071 | 0.002 | -0.002 | -0.002 | 0 | 0.33 | 0.53 | 0.37 | 0.28 | 0.35 | 0.26 |
| 3 | 165612248 | rs12489886 | -0.739 | 0.953 | 1.354 | 0.095 | 0.031 | 0.031 | 0.031 | 0 | 0.07 | 0.13 | 0.2 | 0.15 | 0.12 | 0.07 |
| 3 | 165637529 | rs4365667 | -1.678 | 0.762 | 1.601 | -0.082 | 0.015 | -0.031 | -0.031 | 0.026 | 0.21 | 0.59 | 0.28 | 0.15 | 0.29 | 0.15 |
| 3 | 165654797 | rs830202 | 0.367 | 0.879 | 1.485 | -0.069 | -0.002 | -0.001 | -0.001 | 0 | 0.06 | 0.03 | 0.03 | 0.08 | 0.05 | 0.08 |
| 3 | 165655877 | rs16849898 | -0.215 | 0.901 | 1.734 | -0.081 | 0 | -0.001 | -0.001 | 0 | 0.06 | 0.05 | 0.15 | 0.15 | 0.12 | 0.07 |
| 3 | 165666554 | rs7627395 | -0.139 | 0.813 | 1.567 | -0.072 | -0.001 | -0.021 | -0.021 | 0 | 0.12 | 0.03 | 0.06 | 0.13 | 0.12 | 0.09 |
| 3 | 165669181 | rs9819444 | -0.441 | 0.826 | 2.419 | -0.22 | -0.006 | -0.071 | -0.071 | 0 | 0.12 | 0.29 | 0.1 | 0 | 0.14 | 0.04 |
| 3 | 165683741 | rs9829437 | -2.326 | 0.74 | 2.193 | -0.233 | -0.009 | 0.035 | 0.035 | 0.382 | 0.36 | 0.77 | 0.36 | 0.25 | 0.41 | 0.27 |
| 3 | 165706855 | rs12491503 | -2.428 | 0.695 | 1.638 | -0.226 | -0.075 | -0.052 | -0.052 | 0.897 | 0.34 | 0.93 | 0.32 | 0.33 | 0.32 | 0.31 |
| 3 | 165711873 | rs1900912 | -1.962 | 0.651 | 2.132 | -0.136 | 0.001 | 0.004 | 0.004 | 0.119 | 0.32 | 0.69 | 0.23 | 0.2 | 0.33 | 0.25 |
| 3 | 165751527 | rs9854178 | -1.865 | 0.62 | 2.157 | -0.139 | 0.003 | -0.017 | -0.017 | 0 | 0.34 | 0.53 | 0.34 | 0.25 | 0.38 | 0.26 |
| 3 | 165756573 | rs2872065 | 0.066 | 0.785 | 2.063 | 0.376 | -0.001 | 0.005 | 0.005 | 0 | 0.06 | 0.11 | 0.03 | 0.14 | 0.04 | 0.08 |
| 3 | 165761918 | rs1868531 | -2.418 | 0.758 | 2.547 | 0.119 | -0.004 | 0 | 0 | 0 | 0.39 | 0.64 | 0.48 | 0.4 | 0.44 | 0.33 |
| 3 | 165766451 | rs6777981 | -2.05 | 0.796 | 2.692 | 0.127 | -0.006 | -0.013 | -0.013 | 0 | 0.32 | 0.58 | 0.24 | 0.26 | 0.32 | 0.23 |
| 3 | 165767741 | rs16849971 | -1.925 | 0.786 | 2.955 | 0.122 | 0 | 0 | 0 | 0.365 | 0.29 | 0.33 | 0.2 | 0.25 | 0.3 | 0.19 |
| 3 | 165769683 | rs13069682 | -2.425 | 0.742 | 2.697 | 0.093 | -0.004 | 0 | 0 | 0.938 | 0.39 | 0.64 | 0.48 | 0.4 | 0.44 | 0.33 |
| 3 | 165778002 | rs11920160 | -0.879 | 0.675 | 2.399 | 0.061 | 0.006 | -0.003 | -0.003 | 0 | 0.16 | 0.12 | 0.11 | 0 | 0.18 | 0.05 |
| 3 | 165778330 | rs7632669 | -2.063 | 0.362 | 1.94 | -0.018 | -0.035 | 0.001 | 0.001 | 0 | 0.47 | 0.87 | 0.55 | 0.45 | 0.51 | 0.36 |
| 3 | 165785410 | rs7613317 | -2.059 | 0.312 | 1.2 | 0.04 | -0.01 | 0.009 | 0.009 | 0.41 | 0.5 | 0.79 | 0.39 | 0.47 | 0.53 | 0.3 |
| 3 | 165786026 | rs1551041 | -1.633 | 0.268 | 1.804 | 0.015 | 0.04 | 0.031 | 0.031 | 0 | 0.25 | 0.39 | 0.32 | 0.26 | 0.35 | 0.28 |
| 3 | 165790002 | rs2084525 | -2.601 | 0.347 | 1.671 | 0.016 | -0.022 | -0.012 | -0.012 | 0 | 0.51 | 0.84 | 0.47 | 0.49 | 0.56 | 0.35 |
| 3 | 165791692 | rs1900911 | -0.135 | 0.479 | 1.786 | 0.027 | 0 | -0.026 | -0.026 | 0.769 | 0.09 | 0.08 | 0.08 | 0 | 0.12 | 0.03 |
| 3 | 165799452 | rs6767888 | -0.041 | 0.526 | 1.907 | 0.105 | 0.003 | 0.024 | 0.024 | 0 | 0.08 | 0.1 | 0.12 | 0.12 | 0.11 | 0.1 |
| 3 | 165805972 | rs7431160 | -0.25 | 0.553 | 1.827 | 0.122 | 0.042 | 0.044 | 0.044 | 0 | 0.07 | 0.14 | 0.11 | 0.14 | 0.12 | 0.12 |
| 3 | 165824337 | rs11922912 | -1.867 | 0.482 | 2.038 | 0.054 | 0.021 | 0.002 | 0.002 | 0 | 0.33 | 0.48 | 0.39 | 0.3 | 0.44 | 0.31 |
| 3 | 165827721 | rs4632505 | -0.522 | 0.375 | 2.2 | 0.025 | -0.009 | 0.009 | 0.009 | 0 | 0.17 | 0.34 | 0.13 | 0.14 | 0.17 | 0.12 |
| 3 | 165836041 | rs13091389 | -1.135 | 0.404 | 2.125 | 0.035 | 0 | -0.001 | -0.001 | 0 | 0.15 | 0.11 | 0.18 | 0.14 | 0.24 | 0.15 |
| 3 | 165843402 | rs13091795 | -1.077 | 0.405 | 1.959 | 0.035 | -0.014 | 0 | 0 | 0 | 0.15 | 0.08 | 0.18 | 0.14 | 0.24 | 0.15 |
| 3 | 165852207 | rs7653493 | -0.53 | 0.388 | 1.919 | 0.042 | -0.009 | 0.005 | 0.005 | 0 | 0.17 | 0.3 | 0.13 | 0.14 | 0.17 | 0.12 |
| 3 | 165859107 | rs13061243 | -1.945 | 0.544 | 1.732 | 0.041 | -0.005 | 0 | 0 | 0 | 0.34 | 0.63 | 0.39 | 0.3 | 0.43 | 0.31 |
| 3 | 165867004 | rs4680618 | -0.016 | 0.378 | 1.608 | 0.029 | 0 | -0.003 | -0.003 | 0 | 0.86 | 0.9 | 0.88 | 0.78 | 0.85 | 0.88 |
| 3 | 165868939 | rs17662011 | 0.025 | 0.643 | 1.205 | -0.133 | -0.011 | 0.01 | 0.01 | 0 | 0.09 | 0.08 | 0.07 | 0.04 | 0.12 | 0.04 |
| 3 | 165879515 | rs9879491 | -0.754 | -0.444 | 1.477 | -0.157 | -0.043 | -0.024 | -0.024 | 0.873 | 0.26 | 0.56 | 0.21 | 0.16 | 0.18 | 0.18 |
| 3 | 165894868 | rs1899169 | 2.036 | -0.503 | 1.16 | -0.116 | 0.056 | 0.026 | 0.026 | 4.147 | 0.51 | 0.21 | 0.59 | 0.61 | 0.56 | 0.65 |
| 3 | 165909425 | rs12638042 | 2.097 | -0.517 | 1.141 | -0.083 | 0.052 | 0.008 | 0.008 | 3.976 | 0.51 | 0.15 | 0.59 | 0.6 | 0.56 | 0.65 |
| 3 | 165914538 | rs1440650 | -0.365 | -0.514 | 1.343 | -0.07 | 0.017 | 0.007 | 0.007 | 0.081 | 0.22 | 0.16 | 0.12 | 0.16 | 0.16 | 0.13 |
| 3 | 165917057 | rs6776482 | -0.662 | -0.445 | 0.721 | -0.042 | -0.015 | -0.007 | -0.007 | 0 | 0.27 | 0.68 | 0.22 | 0.17 | 0.18 | 0.18 |
| 3 | 165925527 | rs10936474 | 0.879 | -0.257 | 0.43 | -0.03 | 0.042 | -0.014 | -0.014 | 0 | 0.6 | 0.27 | 0.66 | 0.6 | 0.67 | 0.7 |
| 3 | 165931222 | rs11924406 | -0.826 | 0.096 | 0.694 | -0.013 | -0.001 | -0.009 | -0.009 | 0 | 0.09 | 0.07 | 0.07 | 0 | 0.12 | 0.05 |
| 3 | 165936570 | rs9857044 | 0.736 | -0.448 | 0.735 | 0.398 | -0.032 | 0.014 | 0.014 | 0 | 0.76 | 0.56 | 0.77 | 0.75 | 0.79 | 0.81 |
| 3 | 165940003 | rs9877342 | -0.62 | 0.237 | 0.754 | 0.006 | 0.006 | -0.023 | -0.023 | 0.042 | 0.19 | 0.28 | 0.14 | 0.11 | 0.17 | 0.13 |
| 3 | 165942272 | rs4413279 | 0.103 | 0.365 | 0.479 | -0.001 | -0.014 | -0.005 | -0.005 | 0 | 0.68 | 0.77 | 0.67 | 0.7 | 0.67 | 0.8 |
| 3 | 165952297 | rs9855808 | 1.295 | 0.378 | 0.765 | 0.017 | 0.002 | 0.001 | 0.001 | 2.925 | 0.51 | 0.46 | 0.52 | 0.59 | 0.51 | 0.67 |
| 3 | 165961111 | rs1371901 | 0.097 | 0.398 | 0.789 | 0.028 | 0.001 | 0 | 0 | 0 | 0.69 | 0.66 | 0.66 | 0.7 | 0.68 | 0.8 |
| 3 | 165980721 | rs4634069 | -1.059 | 0.209 | 0.888 | -0.111 | -0.006 | -0.021 | -0.021 | 0 | 0.27 | 0.67 | 0.21 | 0.13 | 0.25 | 0.17 |

| | | | | | | | | | | | | | | | | |
|---|---|---|---|---|---|---|---|---|---|---|---|---|---|---|---|---|
| 3 | 165986370 | rs638307 | 0.447 | 0.327 | 1.2 | -0.154 | 0.001 | -0.038 | -0.038 | 1.457 | 0.14 | 0.2 | 0.13 | 0.03 | 0.12 | 0.09 |
| 3 | 165988394 | rs9821978 | 0.941 | 0.319 | 1.026 | -0.172 | 0 | -0.009 | -0.009 | 0 | 0.53 | 0.52 | 0.53 | 0.68 | 0.55 | 0.68 |
| 3 | 165992588 | rs1542072 | 0.941 | 0.319 | 1.026 | -0.172 | 0 | -0.009 | -0.009 | 0.15 | 0.53 | 0.52 | 0.53 | 0.68 | 0.55 | 0.68 |
| 3 | 166003323 | rs9873795 | 0.281 | 0.289 | 0.986 | -0.181 | -0.029 | -0.034 | -0.034 | 0 | 0.08 | 0.19 | 0.05 | 0.01 | 0.08 | 0.03 |
| 3 | 166003976 | rs478024 | 0.523 | 0.481 | 0.839 | -0.116 | -0.019 | 0.051 | 0.051 | 0 | 0.13 | 0.02 | 0.11 | 0.03 | 0.12 | 0.09 |
| 3 | 166012938 | rs636225 | -0.324 | 0.488 | 1.156 | -0.13 | -0.024 | -0.09 | -0.09 | 0 | 0.8 | 0.6 | 0.88 | 0.97 | 0.81 | 0.87 |
| 3 | 166022534 | rs649748 | 0.217 | 0.482 | 1.168 | -0.126 | -0.022 | -0.068 | -0.068 | 0.043 | 0.19 | 0.33 | 0.11 | 0.03 | 0.17 | 0.12 |
| 3 | 166032924 | rs9283636 | 1.42 | 0.354 | 1.085 | -0.178 | -0.007 | 0.023 | 0.023 | 4.053 | 0.44 | 0.32 | 0.48 | 0.66 | 0.46 | 0.63 |
| 3 | 166040108 | rs4336070 | 0.539 | 0.377 | 1.078 | -0.169 | -0.007 | 0.004 | 0.004 | 0.304 | 0.64 | 0.69 | 0.59 | 0.7 | 0.63 | 0.75 |
| 3 | 166046267 | rs4494879 | 0.938 | 0.428 | 1.171 | -0.117 | 0.005 | -0.001 | -0.001 | 0 | 0.6 | 0.57 | 0.57 | 0.61 | 0.58 | 0.68 |
| 3 | 166047504 | rs1574560 | 1.474 | 0.44 | 0.994 | -0.103 | -0.041 | 0.031 | 0.031 | 3.965 | 0.43 | 0.16 | 0.47 | 0.66 | 0.46 | 0.63 |
| 3 | 166065099 | rs2863381 | 2.109 | 0.225 | 0.357 | -0.183 | -0.075 | -0.158 | -0.158 | 0.011 | 0.33 | 0.03 | 0.3 | 0.51 | 0.34 | 0.52 |
| 3 | 166076333 | rs4680690 | -1.022 | 0.347 | 0.601 | -0.207 | 0 | -0.015 | -0.015 | 0 | 0.86 | 0.86 | 0.88 | 0.96 | 0.89 | 0.91 |
| 3 | 166085645 | rs2863384 | 0.907 | 0.403 | 0.263 | -0.162 | -0.056 | -0.055 | -0.055 | 0 | 0.09 | 0.01 | 0.23 | 0.17 | 0.17 | 0.1 |
| 3 | 166103246 | rs6801797 | 0.163 | 0.322 | 0.693 | -0.134 | 0.101 | 0.138 | 0.138 | 0 | 0.05 | 0.31 | 0.04 | 0.09 | 0.06 | 0.06 |
| 3 | 166124934 | rs1996483 | 1.009 | 0.227 | 0.338 | -0.195 | -0.03 | 0.009 | 0.009 | 0.53 | 0.58 | 0.27 | 0.65 | 0.63 | 0.57 | 0.68 |
| 3 | 166148892 | rs2863389 | 1.056 | 0.15 | 0.315 | -0.228 | -0.002 | -0.008 | -0.008 | 0.013 | 0.19 | 0.2 | 0.2 | 0.06 | 0.17 | 0.12 |
| 3 | 166161902 | rs9876783 | -0.963 | 0.203 | 0.517 | -0.228 | -0.001 | 0.002 | 0.002 | 1.986 | 0.45 | 0.6 | 0.41 | 0.38 | 0.47 | 0.34 |
| 3 | 166166063 | rs9832900 | 0.138 | 0.202 | 0.748 | -0.221 | 0.07 | 0.08 | 0.08 | 0 | 0.05 | 0.21 | 0.04 | 0.09 | 0.07 | 0.08 |
| 3 | 166170696 | rs4396843 | 2.726 | 0.01 | -0.112 | 0.003 | -0.072 | -0.015 | -0.015 | 0 | 0.21 | 0.01 | 0.14 | 0.22 | 0.23 | 0.3 |
| 3 | 166173120 | rs13085064 | 0.273 | -0.019 | 0.478 | -0.441 | -0.012 | -0.01 | -0.01 | 0.172 | 0.22 | 0.19 | 0.18 | 0.24 | 0.23 | 0.15 |
| 3 | 166178844 | rs9873387 | 1.16 | 0.007 | 0.566 | -0.379 | -0.023 | -0.043 | -0.043 | 0 | 0.19 | 0.29 | 0.2 | 0.05 | 0.17 | 0.12 |
| 3 | 166179549 | rs9819099 | 0.439 | 0.02 | 0.428 | -0.369 | -0.017 | -0.036 | -0.036 | 0 | 0.23 | 0.41 | 0.23 | 0.14 | 0.24 | 0.2 |
| 3 | 166193793 | rs16850660 | 1.979 | 0.018 | 0.066 | -0.297 | -0.075 | -0.16 | -0.16 | 0 | 0.33 | 0.03 | 0.3 | 0.52 | 0.34 | 0.51 |
| 3 | 166202007 | rs10936492 | 1.506 | 0.321 | 0.571 | -0.435 | -0.005 | -0.018 | -0.018 | 0 | 0.41 | 0.18 | 0.46 | 0.58 | 0.41 | 0.56 |
| 3 | 166217046 | rs6781717 | -0.1 | 0.219 | 0.742 | -0.547 | 0.015 | -0.054 | -0.054 | 0 | 0.73 | 0.6 | 0.73 | 0.89 | 0.69 | 0.77 |
| 3 | 166217236 | rs6806752 | -0.402 | 0.218 | 0.778 | -0.499 | 0.002 | -0.011 | -0.011 | 0 | 0.32 | 0.25 | 0.26 | 0.31 | 0.28 | 0.23 |
| 3 | 166221093 | rs6766252 | -0.807 | 0.231 | 0.647 | -0.494 | -0.003 | 0.026 | 0.026 | 0 | 0.78 | 0.81 | 0.78 | 0.95 | 0.76 | 0.85 |
| 3 | 166249372 | rs12638016 | 1.642 | 0.211 | 0.571 | -0.472 | 0.002 | -0.025 | -0.025 | 0 | 0.4 | 0.22 | 0.42 | 0.57 | 0.39 | 0.53 |
| 3 | 166267241 | rs9836632 | -0.554 | 0.195 | 0.687 | -0.35 | 0.004 | -0.008 | -0.008 | 0 | 0.32 | 0.39 | 0.29 | 0.25 | 0.31 | 0.22 |
| 3 | 166298091 | rs13095617 | -0.56 | 0.145 | 0.706 | -0.371 | -0.011 | 0.011 | 0.011 | 0.644 | 0.32 | 0.2 | 0.29 | 0.25 | 0.31 | 0.23 |
| 3 | 166298493 | rs10513630 | 1.158 | 0.354 | 0.325 | -0.424 | 0.003 | 0.01 | 0.01 | 0.113 | 0.1 | 0.07 | 0.11 | 0.03 | 0.08 | 0.09 |
| 3 | 166298513 | rs12696252 | 0.722 | 0.289 | 0.297 | -0.378 | -0.053 | -0.048 | -0.048 | 0 | 0.08 | 0 | 0.12 | 0.13 | 0.16 | 0.09 |
| 3 | 166304564 | rs1403779 | -0.019 | 0.148 | 0.474 | -0.479 | -0.003 | -0.025 | -0.025 | 0.474 | 0.72 | 0.49 | 0.7 | 0.82 | 0.7 | 0.76 |
| 3 | 166309393 | rs537548 | 0.833 | 0.186 | -0.02 | -0.588 | 0.001 | -0.003 | -0.003 | 0 | 0.2 | 0.26 | 0.21 | 0.11 | 0.19 | 0.14 |
| 3 | 166315090 | rs10936499 | 0.356 | 0.093 | -0.16 | -0.66 | -0.058 | -0.022 | -0.022 | 0 | 0.15 | 0.04 | 0.15 | 0.13 | 0.21 | 0.1 |
| 3 | 166319387 | rs600723 | 0.818 | 0.108 | 0.113 | -0.567 | -0.001 | 0.001 | 0.001 | 0 | 0.2 | 0.22 | 0.21 | 0.11 | 0.19 | 0.14 |
| 3 | 166322997 | rs9836306 | -0.77 | 0.186 | 0.052 | -0.575 | 0.017 | 0.018 | 0.018 | 1.183 | 0.25 | 0.21 | 0.16 | 0.12 | 0.18 | 0.14 |
| 3 | 166341617 | rs11916399 | 0.515 | 0.506 | -0.023 | -0.478 | 0.004 | 0.007 | 0.007 | 0 | 0.11 | 0.07 | 0.11 | 0.03 | 0.13 | 0.1 |
| 3 | 166388061 | rs7648668 | -1.538 | 0.777 | -0.407 | -0.693 | -0.107 | 0.047 | 0.047 | 0.029 | 0.38 | 0.88 | 0.39 | 0.26 | 0.5 | 0.35 |
| 3 | 166389520 | rs9820644 | -1.351 | 0.853 | -0.217 | -0.701 | -0.054 | 0.038 | 0.038 | 0 | 0.37 | 0.82 | 0.34 | 0.26 | 0.45 | 0.31 |
| 3 | 166399946 | rs4680626 | -1.535 | 0.899 | -0.217 | -0.697 | -0.073 | 0.047 | 0.047 | 0 | 0.38 | 0.85 | 0.39 | 0.26 | 0.5 | 0.39 |
| 3 | 166403296 | rs13081649 | 0.244 | 0.961 | 0.11 | -0.717 | 0.011 | -0.03 | -0.03 | 0 | 0.1 | 0.19 | 0.14 | 0.03 | 0.13 | 0.1 |
| 3 | 166416014 | rs9872777 | -1.156 | 1.053 | 0.373 | -0.705 | 0.007 | -0.035 | -0.035 | 0.472 | 0.31 | 0.52 | 0.3 | 0.19 | 0.37 | 0.3 |
| 3 | 166420044 | rs9874501 | -0.721 | 1.092 | 0.007 | -0.704 | 0.036 | -0.028 | -0.028 | 2.734 | 0.2 | 0.3 | 0.26 | 0.11 | 0.31 | 0.28 |
| 3 | 166431551 | rs9867633 | 0.314 | 1.022 | -0.926 | -0.625 | -0.005 | 0 | 0 | 0 | 0.1 | 0.07 | 0.12 | 0.08 | 0.17 | 0.18 |
| 3 | 166431713 | rs12488985 | 0.897 | 0.558 | -1.083 | -0.533 | -0.001 | -0.006 | -0.006 | 0 | 0.11 | 0.13 | 0.07 | 0.08 | 0.11 | 0.06 |
| 3 | 166444013 | rs1517057 | -0.708 | 0.346 | -1.282 | -0.555 | -0.035 | 0.011 | 0.011 | 0 | 0.31 | 0.75 | 0.34 | 0.25 | 0.4 | 0.4 |
| 3 | 166444743 | rs2863837 | 0.707 | -0.235 | -0.674 | -0.386 | -0.006 | 0.002 | 0.002 | 0.534 | 0.1 | 0.16 | 0.03 | 0.15 | 0.05 | 0.02 |
| 3 | 166450664 | rs501102 | 0.292 | -0.155 | -0.934 | -0.58 | 0.062 | 0.026 | 0.026 | 0.889 | 0.23 | 0.08 | 0.18 | 0.13 | 0.12 | 0.14 |
| 3 | 166459187 | rs10513629 | -1.183 | 0.452 | 0.313 | -0.481 | -0.002 | 0.062 | 0.062 | 0.292 | 0.1 | 0.32 | 0.08 | 0.18 | 0.07 | 0.09 |
| 3 | 166459814 | rs16850869 | 0.137 | 0.447 | 0.152 | -0.497 | 0.01 | -0.011 | -0.011 | 0 | 0.05 | 0 | 0.01 | 0.08 | 0.02 | 0.07 |
| 3 | 166460951 | rs2863836 | -1.436 | 0.959 | 0.625 | -0.611 | 0.02 | -0.003 | -0.003 | 0 | 0.14 | 0.59 | 0.13 | 0.18 | 0.13 | 0.14 |
| 3 | 166462794 | rs17679734 | -0.382 | 1.429 | 0.345 | -0.656 | 0.005 | 0 | 0 | 0 | 0.07 | 0.09 | 0.1 | 0.07 | 0.11 | 0.14 |
| 3 | 166480559 | rs9874962 | 0.644 | 1.442 | -0.436 | -1.247 | 0.003 | 0.185 | 0.185 | 0.326 | 0.37 | 0.08 | 0.35 | 0.08 | 0.34 | 0.32 |
| 3 | 166509376 | rs7430710 | -1.003 | 1.532 | -0.445 | -1.132 | -0.064 | 0.033 | 0.033 | 0.14 | 0.6 | 0.95 | 0.61 | 0.33 | 0.59 | 0.6 |
| 3 | 166517229 | rs6778565 | 0.156 | 1.576 | -0.312 | -1.152 | 0.017 | 0.001 | 0.001 | 0 | 0.07 | 0.11 | 0.09 | 0.07 | 0.1 | 0.13 |
| 3 | 166517250 | rs6767590 | -0.98 | 1.575 | -0.397 | -1.143 | -0.03 | 0.056 | 0.056 | 0.078 | 0.59 | 0.87 | 0.6 | 0.32 | 0.59 | 0.6 |
| 3 | 166522847 | rs13062144 | -0.245 | 1.339 | -0.236 | -1.078 | 0.002 | 0.035 | 0.035 | 0.016 | 0.51 | 0.57 | 0.5 | 0.26 | 0.49 | 0.47 |
| 3 | 166528151 | rs7615817 | -1.048 | 1.855 | -0.34 | -1.037 | 0.062 | -0.025 | -0.025 | 0 | 0.27 | 0.41 | 0.27 | 0.25 | 0.39 | 0.36 |
| 3 | 166538324 | rs6787835 | -1.04 | 1.968 | -0.247 | -1.224 | 0.033 | -0.029 | -0.029 | 0.253 | 0.3 | 0.39 | 0.31 | 0.25 | 0.46 | 0.38 |
| 3 | 166555665 | rs13433888 | -0.837 | 2.056 | -0.033 | -1.167 | 0.066 | 0.044 | 0.044 | 0 | 0.12 | 0.23 | 0.13 | 0.24 | 0.18 | 0.21 |
| 3 | 166572963 | rs9290300 | -0.91 | 1.324 | -0.727 | -1.798 | 0.081 | 0.083 | 0.083 | 0 | 0.48 | 0.3 | 0.49 | 0.28 | 0.69 | 0.63 |
| 3 | 166576022 | rs13062023 | 0.344 | 1.451 | -1.247 | -1.756 | -0.187 | 0.203 | 0.203 | 0 | 0.32 | 0.03 | 0.31 | 0.03 | 0.46 | 0.36 |
| 3 | 166576103 | rs9812298 | -1.089 | 0.231 | -0.996 | -1.641 | 0.045 | -0.024 | -0.024 | 0 | 0.71 | 0.87 | 0.73 | 0.34 | 0.81 | 0.75 |
| 3 | 166580001 | rs1916286 | -0.406 | 0.174 | -0.893 | -1.782 | 0.129 | 0.198 | 0.198 | 0 | 0.46 | 0.22 | 0.46 | 0.14 | 0.69 | 0.61 |
| 3 | 166585179 | rs9831825 | -0.647 | 1.634 | -0.213 | -1.642 | -0.042 | -0.01 | -0.01 | 3.668 | 0.12 | 0.06 | 0.11 | 0.16 | 0.23 | 0.19 |
| 3 | 166589013 | rs7649772 | -0.912 | 1.051 | -0.418 | -1.589 | 0.01 | -0.093 | -0.093 | 0 | 0.73 | 0.97 | 0.73 | 0.39 | 0.81 | 0.76 |
| 3 | 166613118 | rs10212208 | 0.042 | 1.045 | 0.131 | -1.69 | 0.055 | 0.068 | 0.068 | 0 | 0.62 | 0.21 | 0.6 | 0.24 | 0.73 | 0.64 |

| | | | | | | | | | | | | | | | | |
|---|---|---|---|---|---|---|---|---|---|---|---|---|---|---|---|---|
| 3 | 166666870 | rs2037546 | -0.689 | 1.27 | 0.562 | -1.586 | 0.007 | -0.019 | -0.019 | 1.523 | 0.68 | 0.79 | 0.71 | 0.4 | 0.77 | 0.73 |
| 3 | 166679062 | rs17471644 | 1.033 | 1.546 | 0.509 | -1.782 | -0.002 | 0.301 | 0.301 | 0.018 | 0.48 | 0.07 | 0.47 | 0.09 | 0.48 | 0.44 |
| 3 | 166679453 | rs9290306 | -0.677 | 1.703 | 0.757 | -1.776 | -0.004 | -0.005 | -0.005 | 0 | 0.67 | 0.71 | 0.69 | 0.39 | 0.75 | 0.71 |
| 3 | 166681458 | rs9857493 | -0.413 | 1.664 | 0.484 | -1.754 | -0.026 | -0.003 | -0.003 | 0 | 0.09 | 0.07 | 0.11 | 0.08 | 0.19 | 0.11 |
| 3 | 166687065 | rs9840614 | -0.413 | 1.664 | 0.484 | -1.754 | -0.026 | -0.003 | -0.003 | 0 | 0.09 | 0.07 | 0.11 | 0.08 | 0.19 | 0.11 |
| 3 | 166689010 | rs16850977 | 1.103 | 1.356 | 0.702 | -1.722 | 0.034 | 0.128 | 0.128 | 3.563 | 0.05 | 0.2 | 0.05 | 0.13 | 0.03 | 0.05 |
| 3 | 166695625 | rs1608323 | 0.856 | 1.448 | -0.071 | -1.785 | 0.002 | 0.301 | 0.301 | 0.018 | 0.48 | 0.07 | 0.46 | 0.09 | 0.48 | 0.44 |
| 3 | 166715641 | rs1607650 | -1.756 | 1.552 | 1.195 | -1.228 | 0.026 | 0.06 | 0.06 | 0 | 0.22 | 0.5 | 0.21 | 0.32 | 0.29 | 0.27 |
| 3 | 166726765 | rs6781033 | -1.029 | 1.661 | 1.267 | -1.219 | 0.068 | 0.038 | 0.038 | 0 | 0.13 | 0.36 | 0.13 | 0.16 | 0.21 | 0.2 |
| 3 | 166739585 | rs2203919 | 0.672 | 1.27 | 0.439 | -1.668 | -0.049 | 0.144 | 0.144 | 0 | 0.36 | 0.12 | 0.48 | 0.11 | 0.41 | 0.41 |
| 3 | 166745838 | rs12631237 | 0.878 | 1.502 | 0.775 | -1.616 | 0.011 | 0.004 | 0.004 | 0 | 0.42 | 0.27 | 0.32 | 0.58 | 0.34 | 0.33 |
| 3 | 166746608 | rs1356768 | 0.081 | 1.509 | 0.794 | -1.626 | 0.002 | 0.018 | 0.018 | 0.271 | 0.45 | 0.49 | 0.55 | 0.26 | 0.45 | 0.48 |
| 3 | 166747878 | rs1356769 | 0.347 | 1.342 | 0.762 | -1.611 | 0.007 | -0.008 | -0.008 | 0.055 | 0.79 | 0.72 | 0.8 | 0.83 | 0.73 | 0.79 |
| 3 | 166749739 | rs10460872 | 1.168 | 1.829 | 0.914 | -1.239 | 0.013 | 0.006 | 0.006 | 0.017 | 0.42 | 0.25 | 0.32 | 0.58 | 0.34 | 0.35 |
| 3 | 166769968 | rs1519943 | 0.861 | 1.953 | 0.934 | -1.272 | -0.029 | 0.014 | 0.014 | 0 | 0.85 | 0.9 | 0.87 | 0.93 | 0.76 | 0.87 |
| 3 | 166770445 | rs1401866 | 1.397 | 1.922 | 0.849 | -1.243 | 0.03 | -0.015 | -0.015 | 0 | 0.39 | 0.24 | 0.31 | 0.58 | 0.31 | 0.34 |
| 3 | 166782814 | rs2030352 | 1.563 | 2.386 | 1.099 | -1.057 | 0.046 | -0.006 | -0.006 | 0 | 0.82 | 0.64 | 0.83 | 0.84 | 0.73 | 0.81 |
| 3 | 166819223 | rs1356771 | 0.569 | 2.34 | 0.558 | -0.779 | -0.057 | -0.041 | -0.041 | 1.423 | 0.88 | 0.97 | 0.83 | 0.85 | 0.76 | 0.84 |
| 3 | 166827141 | rs13091161 | 0.327 | 2.156 | 0.38 | -0.906 | -0.094 | 0.167 | 0.167 | 0 | 0.37 | 0.04 | 0.31 | 0.08 | 0.39 | 0.37 |
| 3 | 166855373 | rs6797201 | -1.147 | 2.212 | 0.503 | -0.847 | -0.067 | -0.017 | -0.017 | 0 | 0.15 | 0.04 | 0.19 | 0.11 | 0.28 | 0.18 |
| 3 | 166861678 | rs4522766 | 0.183 | 2.208 | 1.149 | -0.88 | 0.006 | -0.045 | -0.045 | 0 | 0.57 | 0.54 | 0.62 | 0.82 | 0.56 | 0.62 |
| 3 | 166872713 | rs4541410 | -1.211 | 2.165 | 1.181 | -0.786 | -0.034 | -0.002 | -0.002 | 0.047 | 0.16 | 0.09 | 0.2 | 0.11 | 0.28 | 0.18 |
| 3 | 166889466 | rs9848597 | -0.759 | 2.179 | 1.248 | -0.775 | -0.009 | -0.004 | -0.004 | 0 | 0.13 | 0.11 | 0.19 | 0.1 | 0.26 | 0.17 |
| 3 | 166903382 | rs9857004 | -0.759 | 2.179 | 1.233 | -0.775 | -0.042 | -0.016 | -0.016 | 0 | 0.13 | 0.05 | 0.19 | 0.1 | 0.27 | 0.17 |
| 3 | 166941144 | rs6776832 | 0.367 | 2.177 | 1.373 | -0.725 | -0.041 | -0.033 | -0.033 | 0 | 0.07 | 0.01 | 0.13 | 0.07 | 0.19 | 0.08 |
| 3 | 166944651 | rs11925091 | 0.281 | 1.9 | 1.392 | -0.744 | -0.095 | 0.146 | 0.146 | 0 | 0.36 | 0.03 | 0.3 | 0.11 | 0.38 | 0.36 |
| 3 | 166948241 | rs13067249 | 0.321 | 1.89 | 1.58 | -0.661 | 0.069 | 0.044 | 0.044 | 0 | 0.86 | 0.65 | 0.77 | 0.83 | 0.74 | 0.75 |
| 3 | 166956806 | rs4245910 | -0.139 | 1.848 | 1.333 | -0.659 | -0.045 | -0.017 | -0.017 | 0 | 0.87 | 0.95 | 0.81 | 0.9 | 0.74 | 0.83 |
| 3 | 166958824 | rs9683261 | 0.274 | 1.812 | 1.421 | -0.647 | 0.026 | 0.017 | 0.017 | 0 | 0.86 | 0.79 | 0.79 | 0.83 | 0.74 | 0.75 |
| 3 | 166964218 | rs6770878 | 1.452 | 1.826 | 1.356 | -0.68 | 0.048 | 0.005 | 0.005 | 0 | 0.41 | 0.27 | 0.42 | 0.64 | 0.3 | 0.4 |
| 3 | 166965063 | rs13085286 | -0.231 | 1.846 | 1.277 | -0.656 | -0.05 | -0.022 | -0.022 | 0.07 | 0.87 | 0.96 | 0.81 | 0.9 | 0.74 | 0.83 |
| 3 | 166979675 | rs10936523 | -0.135 | 1.842 | 1.413 | -0.658 | 0.003 | 0 | 0 | 0 | 0.87 | 0.86 | 0.8 | 0.9 | 0.74 | 0.83 |
| 3 | 167000720 | rs7641073 | -0.592 | 1.81 | 1.349 | -0.684 | -0.013 | 0.088 | 0.088 | 0 | 0.46 | 0.69 | 0.4 | 0.26 | 0.44 | 0.43 |
| 3 | 167013777 | rs13068321 | 0.248 | 2.195 | 0.985 | -0.638 | -0.05 | 0.088 | 0.088 | 0 | 0.34 | 0.07 | 0.2 | 0.18 | 0.31 | 0.31 |
| 3 | 167023279 | rs7643215 | -0.234 | 2.203 | 1.005 | -0.621 | -0.05 | -0.02 | -0.02 | 2.937 | 0.87 | 0.96 | 0.81 | 0.9 | 0.74 | 0.83 |
| 3 | 167034814 | rs13068132 | -0.334 | 2.302 | 1.274 | -0.594 | -0.029 | 0.079 | 0.079 | 0 | 0.39 | 0.21 | 0.34 | 0.19 | 0.41 | 0.37 |
| 3 | 167037939 | rs10451920 | -0.219 | 2.295 | 1.142 | -0.611 | -0.014 | -0.001 | -0.001 | 0 | 0.86 | 0.88 | 0.81 | 0.9 | 0.74 | 0.83 |
| 3 | 167050814 | rs6791786 | 1.624 | 1.945 | 1.248 | -0.617 | 0.03 | 0.003 | 0.003 | 0 | 0.4 | 0.25 | 0.42 | 0.64 | 0.34 | 0.39 |
| 3 | 167051726 | rs10936530 | 1.436 | 2.027 | 0.77 | -0.584 | 0 | 0 | 0 | 0 | 0.42 | 0.46 | 0.44 | 0.66 | 0.34 | 0.39 |
| 3 | 167051769 | rs11923054 | -0.164 | 2.029 | 0.649 | -0.585 | -0.098 | 0.126 | 0.126 | 0 | 0.38 | 0.07 | 0.33 | 0.18 | 0.41 | 0.37 |
| 3 | 167057794 | rs4368502 | 0.66 | 2.089 | 0.501 | -0.586 | -0.069 | 0.09 | 0.09 | 0 | 0.29 | 0.02 | 0.18 | 0.11 | 0.27 | 0.29 |
| 3 | 167062663 | rs7647253 | -0.085 | 1.883 | 0.763 | -0.412 | 0.015 | -0.011 | -0.011 | 0 | 0.86 | 0.8 | 0.81 | 0.93 | 0.77 | 0.83 |
| 3 | 167062902 | rs13433693 | 1.022 | 1.898 | 0.553 | -0.413 | -0.003 | -0.002 | -0.002 | 1.335 | 0.42 | 0.46 | 0.44 | 0.66 | 0.34 | 0.39 |
| 3 | 167074305 | rs6762521 | -0.054 | 1.8 | 0.216 | -0.708 | -0.006 | 0.002 | 0.002 | 0 | 0.85 | 0.87 | 0.82 | 0.91 | 0.77 | 0.83 |
| 3 | 167074873 | rs7634221 | 0.574 | 1.015 | 0.106 | -0.388 | 0.009 | 0.011 | 0.011 | 0 | 0.42 | 0.4 | 0.45 | 0.73 | 0.34 | 0.45 |
| 3 | 167088475 | rs2900891 | -0.355 | 1.083 | 0.442 | -0.03 | -0.006 | -0.049 | -0.049 | 4.316 | 0.31 | 0.53 | 0.25 | 0.22 | 0.28 | 0.3 |
| 3 | 167097421 | rs13073265 | -0.859 | 1.199 | 0.444 | -0.1 | 0.013 | -0.025 | -0.025 | 0.071 | 0.06 | 0.11 | 0.14 | 0.01 | 0.1 | 0.04 |
| 3 | 167102124 | rs1554568 | -0.275 | 1.175 | 0.457 | -0.124 | -0.004 | -0.029 | -0.029 | 0 | 0.31 | 0.39 | 0.25 | 0.22 | 0.29 | 0.31 |
| 3 | 167105744 | rs13060971 | -0.841 | 1.275 | -0.114 | -0.094 | 0 | -0.019 | -0.019 | 0 | 0.37 | 0.62 | 0.39 | 0.22 | 0.38 | 0.35 |
| 3 | 167113205 | rs1464634 | -1.333 | 1.559 | -0.428 | 0.326 | -0.032 | 0.035 | 0.035 | 0 | 0.07 | 0.01 | 0.15 | 0 | 0.15 | 0.06 |
| 3 | 167117774 | rs2419135 | 0.869 | 1.68 | -0.538 | 0.97 | -0.032 | -0.054 | -0.054 | 0 | 0.86 | 0.96 | 0.81 | 0.77 | 0.78 | 0.82 |
| 3 | 167121000 | rs1403646 | -1.363 | 1.439 | 0.265 | 1.25 | 0.004 | 0.126 | 0.126 | 0.566 | 0.1 | 0.17 | 0.22 | 0.3 | 0.13 | 0.09 |
| 3 | 167132319 | rs10936537 | -1.691 | 1.333 | 1.153 | 0.858 | 0.037 | -0.091 | -0.091 | 0.64 | 0.13 | 0.45 | 0.12 | 0 | 0.16 | 0.03 |
| 3 | 167138546 | rs13078395 | -0.933 | 1.324 | 1.305 | 0.853 | 0.001 | -0.015 | -0.015 | 0 | 0.06 | 0.07 | 0.06 | 0 | 0.11 | 0.03 |
| 3 | 167139670 | rs9869341 | -1.824 | 1.154 | 0.927 | 0.509 | -0.023 | 0 | 0 | 0 | 0.47 | 0.84 | 0.47 | 0.42 | 0.52 | 0.44 |
| 3 | 167156003 | rs9846882 | -2.228 | 1.042 | 1.708 | 0.552 | -0.017 | 0.113 | 0.113 | 0 | 0.2 | 0.49 | 0.29 | 0.29 | 0.2 | 0.11 |
| 3 | 167156429 | rs11918193 | -1.067 | 1.036 | 1.861 | 0.575 | 0.008 | -0.072 | -0.072 | 0.25 | 0.34 | 0.46 | 0.25 | 0.13 | 0.43 | 0.36 |
| 3 | 167156958 | rs11919181 | -1.067 | 1.036 | 1.861 | 0.59 | 0.008 | -0.082 | -0.082 | 0.528 | 0.34 | 0.46 | 0.25 | 0.1 | 0.43 | 0.36 |
| 3 | 167159831 | rs9864094 | -1.435 | 0.978 | 1.952 | 0.653 | -0.006 | 0.246 | 0.246 | 0.487 | 0.14 | 0.43 | 0.23 | 0.29 | 0.09 | 0.08 |
| 3 | 167159834 | rs9844202 | -1.493 | 0.972 | 1.955 | 0.651 | -0.009 | 0.242 | 0.242 | 0.501 | 0.14 | 0.43 | 0.23 | 0.29 | 0.09 | 0.08 |
| 3 | 167160291 | rs9848915 | -1.431 | 0.994 | 1.964 | 0.66 | -0.006 | 0.246 | 0.246 | 0.535 | 0.14 | 0.43 | 0.23 | 0.29 | 0.09 | 0.08 |
| 3 | 167160385 | rs9811119 | -2.617 | 0.953 | 2.048 | 0.639 | 0.02 | -0.046 | -0.046 | 0 | 0.22 | 0.82 | 0.31 | 0.33 | 0.2 | 0.11 |
| 3 | 167164187 | rs2272138 | -1.436 | 0.971 | 1.996 | 0.66 | -0.006 | 0.246 | 0.246 | 0.535 | 0.14 | 0.43 | 0.23 | 0.29 | 0.09 | 0.08 |
| 3 | 167168148 | rs9863185 | -0.874 | 1.005 | 2.003 | 0.696 | 0.007 | 0.172 | 0.172 | 0.601 | 0.1 | 0.21 | 0.21 | 0.28 | 0.09 | 0.07 |
| 3 | 167180161 | rs13074993 | -1.186 | 0.907 | 1.953 | 0.544 | -0.003 | -0.026 | -0.026 | 0 | 0.09 | 0.11 | 0.06 | 0 | 0.11 | 0.03 |
| 3 | 167184878 | rs9841174 | -1.286 | 0.931 | 2.111 | 0.946 | 0 | 0.007 | 0.007 | 0 | 0.37 | 0.37 | 0.38 | 0.51 | 0.4 | 0.4 |
| 3 | 167189607 | rs17246389 | 0.036 | 0.95 | 1.909 | 0.94 | -0.044 | 0.089 | 0.089 | 0.407 | 0.22 | 0.06 | 0.13 | 0.1 | 0.26 | 0.32 |
| 3 | 167189890 | rs6444350 | -1.88 | 0.924 | 2.011 | 0.936 | -0.004 | 0.004 | 0.004 | 0 | 0.46 | 0.76 | 0.46 | 0.54 | 0.51 | 0.44 |
| 3 | 167189923 | rs11922309 | -0.58 | 0.91 | 1.944 | 0.749 | 0 | -0.017 | -0.017 | 7.862 | 0.06 | 0.07 | 0.05 | 0 | 0.11 | 0.03 |

| | | | | | | | | | | | | | | | | |
|---|---|---|---|---|---|---|---|---|---|---|---|---|---|---|---|---|
| 3 | 167207837 | rs12493588 | -1.579 | 0.88 | 2.163 | 0.74 | 0.004 | -0.038 | -0.038 | 0 | 0.1 | 0.29 | 0.07 | 0.03 | 0.11 | 0.03 |
| 3 | 167208012 | rs9881165 | -0.806 | 0.87 | 2.035 | 0.845 | -0.008 | 0.174 | 0.174 | 0.371 | 0.1 | 0.18 | 0.22 | 0.4 | 0.09 | 0.07 |
| 3 | 167212340 | rs11918205 | -1.621 | 0.887 | 2.057 | 0.833 | 0.005 | -0.014 | -0.014 | 2.447 | 0.42 | 0.62 | 0.42 | 0.52 | 0.46 | 0.43 |
| 3 | 167212764 | rs9290315 | -1.894 | 0.893 | 1.891 | 0.833 | 0.018 | -0.011 | -0.011 | 0 | 0.44 | 0.71 | 0.43 | 0.53 | 0.46 | 0.43 |
| 3 | 167218543 | rs2140364 | -1.121 | 0.94 | 1.674 | 0.77 | -0.005 | -0.029 | -0.029 | 0.401 | 0.34 | 0.28 | 0.37 | 0.48 | 0.4 | 0.39 |
| 3 | 167225951 | rs11709117 | -0.666 | 0.877 | 1.669 | 0.01 | -0.002 | -0.065 | -0.065 | 0 | 0.28 | 0.38 | 0.16 | 0.13 | 0.29 | 0.33 |
| 3 | 167240938 | rs4305424 | -1.694 | 0.891 | 1.702 | 0.094 | 0.003 | 0.011 | 0.011 | 0.014 | 0.19 | 0.49 | 0.22 | 0.19 | 0.21 | 0.1 |
| 3 | 167243869 | rs9855001 | -1.887 | 0.898 | 1.226 | 0.104 | -0.026 | 0.104 | 0.104 | 0 | 0.47 | 0.89 | 0.39 | 0.31 | 0.5 | 0.43 |
| 3 | 167253580 | rs6791262 | -1.257 | 0.862 | 1.328 | 0.204 | -0.01 | 0.011 | 0.011 | 0.308 | 0.16 | 0.27 | 0.19 | 0.19 | 0.16 | 0.09 |
| 3 | 167269394 | rs7650514 | -1.055 | 0.987 | 1.317 | 0.008 | 0.001 | 0 | 0 | 0 | 0.3 | 0.27 | 0.16 | 0.13 | 0.31 | 0.35 |
| 3 | 167275119 | rs6778707 | -0.827 | 0.968 | 1.264 | -0.022 | 0.001 | 0.008 | 0.008 | 0.018 | 0.28 | 0.22 | 0.16 | 0.1 | 0.31 | 0.35 |
| 3 | 167279650 | rs6774632 | -1.424 | 0.119 | 1.453 | -0.226 | -0.02 | -0.026 | -0.026 | 0 | 0.36 | 0.4 | 0.25 | 0.2 | 0.31 | 0.37 |
| 3 | 167283904 | rs6788731 | -1.788 | 0.521 | 1.857 | -0.11 | -0.004 | -0.009 | -0.009 | 0 | 0.35 | 0.61 | 0.37 | 0.25 | 0.36 | 0.37 |
| 3 | 167284846 | rs10936541 | -0.806 | 0.23 | 1.828 | -0.15 | 0.004 | 0.003 | 0.003 | 0.042 | 0.22 | 0.15 | 0.13 | 0.05 | 0.23 | 0.29 |
| 3 | 167287678 | rs4552347 | -1.983 | 0.284 | 1.901 | -0.155 | -0.007 | -0.009 | -0.009 | 0 | 0.34 | 0.6 | 0.32 | 0.25 | 0.3 | 0.37 |
| 3 | 167290798 | rs6780019 | -0.929 | 0.366 | 1.073 | -0.391 | 0.003 | 0 | 0 | 0 | 0.23 | 0.15 | 0.13 | 0.05 | 0.23 | 0.29 |
| 3 | 167300830 | rs13318752 | -1.967 | 1.418 | 1.445 | -0.257 | 0.064 | -0.026 | -0.026 | 0.882 | 0.18 | 0.62 | 0.25 | 0.12 | 0.28 | 0.1 |
| 3 | 167303485 | rs4274741 | 1.995 | 1.493 | 1.456 | 0.162 | 0.018 | 0.02 | 0.02 | 1.385 | 0.78 | 0.28 | 0.77 | 0.71 | 0.68 | 0.86 |
| 3 | 167336554 | rs7614964 | -2.019 | 1.235 | 2.247 | 0.179 | 0.053 | -0.068 | -0.068 | 0.011 | 0.17 | 0.37 | 0.22 | 0.04 | 0.26 | 0.11 |
| 3 | 167348838 | rs7627289 | -0.847 | 1.004 | 2.001 | -0.057 | 0.003 | 0.015 | 0.015 | 0.601 | 0.2 | 0.14 | 0.14 | 0.07 | 0.19 | 0.24 |
| 3 | 167349441 | rs6807003 | -1.812 | 1.119 | 1.93 | 0.024 | 0.059 | 0.071 | 0.071 | 0 | 0.17 | 0.4 | 0.22 | 0.25 | 0.27 | 0.1 |
| 3 | 167364471 | rs9829009 | -2.109 | 1.119 | 2.04 | 0.057 | 0.064 | -0.013 | -0.013 | 0 | 0.18 | 0.46 | 0.24 | 0.15 | 0.26 | 0.13 |
| 3 | 167375967 | rs7626799 | -2.539 | 1.121 | 2.021 | 0.037 | -0.068 | -0.044 | -0.044 | 0 | 0.4 | 0.95 | 0.41 | 0.42 | 0.46 | 0.38 |
| 3 | 167400290 | rs9870066 | -1.86 | 1.16 | 2.183 | -0.015 | 0.16 | -0.049 | -0.049 | 0.005 | 0.12 | 0.44 | 0.12 | 0.02 | 0.26 | 0.08 |
| 3 | 167402444 | rs7640968 | -1.549 | 1.158 | 2.357 | -0.016 | 0.092 | -0.043 | -0.043 | 0 | 0.11 | 0.27 | 0.12 | 0.02 | 0.26 | 0.08 |
| 3 | 167403273 | rs9848373 | -2.489 | 1.156 | 2.453 | -0.01 | 0.094 | 0.147 | 0.147 | 0.947 | 0.14 | 0.65 | 0.16 | 0.34 | 0.26 | 0.12 |
| 3 | 167407896 | rs16851407 | -1.86 | 1.147 | 2.428 | -0.026 | 0.158 | -0.048 | -0.048 | 0 | 0.12 | 0.43 | 0.12 | 0.02 | 0.26 | 0.08 |
| 3 | 167425194 | rs6777835 | -1.674 | 1.153 | 2.435 | -0.024 | 0.153 | -0.043 | -0.043 | 0 | 0.11 | 0.38 | 0.12 | 0.02 | 0.26 | 0.08 |
| 3 | 167435193 | rs7615915 | -1.648 | 1.151 | 2.49 | -0.025 | 0.132 | -0.043 | -0.043 | 0 | 0.11 | 0.32 | 0.12 | 0.02 | 0.26 | 0.08 |
| 3 | 167439397 | rs6444571 | -2.087 | 1.152 | 2.295 | -0.021 | 0.147 | 0.017 | 0.017 | 0.146 | 0.13 | 0.45 | 0.14 | 0.13 | 0.26 | 0.12 |
| 3 | 167441991 | rs9841137 | -2.422 | 1.177 | 2.292 | -0.016 | 0.094 | 0.147 | 0.147 | 0.947 | 0.14 | 0.65 | 0.16 | 0.34 | 0.26 | 0.12 |
| 3 | 167452504 | rs3792293 | -2.104 | 1.2 | 2.142 | -0.016 | 0.158 | 0.215 | 0.215 | 0.298 | 0.13 | 0.45 | 0.15 | 0.34 | 0.27 | 0.12 |
| 3 | 167460383 | rs11921535 | -2.052 | 1.094 | 1.274 | -0.037 | -0.061 | -0.029 | -0.029 | 0 | 0.36 | 0.92 | 0.31 | 0.4 | 0.46 | 0.37 |
| 3 | 167466514 | rs6444589 | -2.028 | 1.129 | 1.312 | -0.08 | -0.06 | -0.021 | -0.021 | 0.202 | 0.35 | 0.92 | 0.31 | 0.34 | 0.46 | 0.36 |
| 3 | 167474916 | rs9828581 | -1.788 | 1.318 | 1.857 | 0.099 | 0.089 | 0.092 | 0.092 | 0.06 | 0.1 | 0.27 | 0.15 | 0.23 | 0.21 | 0.1 |
| 3 | 167478327 | rs9883327 | -2.768 | 1.222 | 1.618 | 0.133 | 0.033 | 0.054 | 0.054 | 0.133 | 0.2 | 0.63 | 0.3 | 0.28 | 0.27 | 0.13 |
| 3 | 167488623 | rs11927026 | -1.299 | 1.173 | 1.652 | 0.127 | 0.003 | -0.008 | -0.008 | 0 | 0.09 | 0.21 | 0.16 | 0.05 | 0.1 | 0.04 |
| 3 | 167497444 | rs9815034 | -2.341 | 1.301 | 0.863 | 0.013 | 0.005 | 0.004 | 0.004 | 0.093 | 0.31 | 0.64 | 0.36 | 0.34 | 0.38 | 0.27 |
| 3 | 167504672 | rs9852595 | -1.478 | 1.43 | -0.118 | -0.555 | -0.072 | 0.055 | 0.055 | 0.193 | 0.18 | 0.04 | 0.18 | 0.07 | 0.27 | 0.21 |
| 3 | 167518303 | rs2131114 | -0.677 | 0.861 | 0.011 | -1.187 | -0.022 | -0.105 | -0.105 | 0 | 0.32 | 0.51 | 0.2 | 0.05 | 0.32 | 0.19 |
| 3 | 167521341 | rs9858935 | -1.464 | 1.186 | -0.191 | -1.654 | -0.001 | 0.051 | 0.051 | 0 | 0.48 | 0.66 | 0.4 | 0.08 | 0.53 | 0.25 |
| 3 | 167526429 | rs13093535 | -0.439 | 1.154 | -0.17 | -1.684 | -0.021 | -0.116 | -0.116 | 0.778 | 0.33 | 0.58 | 0.2 | 0.02 | 0.33 | 0.2 |
| 3 | 167527371 | rs3804617 | -0.671 | 1.313 | -0.344 | -1.333 | -0.018 | 0.023 | 0.023 | 0 | 0.16 | 0.07 | 0.2 | 0.03 | 0.21 | 0.05 |
| 3 | 167527846 | rs11706323 | -0.323 | 1.065 | -0.244 | -1.326 | 0.004 | 0.016 | 0.016 | 0 | 0.12 | 0.06 | 0.05 | 0.02 | 0.13 | 0.15 |
| 3 | 167530216 | rs16851498 | -1.11 | 1.057 | 0.538 | -1.648 | 0.012 | -0.131 | -0.131 | 1.172 | 0.2 | 0.51 | 0.2 | 0.01 | 0.25 | 0.04 |
| 3 | 167530444 | rs2055027 | 0.622 | 0.794 | 0.251 | -1.867 | 0.003 | -0.004 | -0.004 | 0 | 0.8 | 0.73 | 0.92 | 0.97 | 0.83 | 0.86 |
| 3 | 167531607 | rs7641394 | -1.293 | 1.281 | 0.945 | -1.614 | 0.023 | -0.017 | -0.017 | 0 | 0.14 | 0.55 | 0.22 | 0.11 | 0.18 | 0.04 |
| 3 | 167534712 | rs16851502 | -1.243 | 1.347 | 0.906 | -1.684 | 0.015 | -0.029 | -0.029 | 0 | 0.17 | 0.27 | 0.2 | 0.08 | 0.23 | 0.06 |
| 3 | 167534993 | rs7651327 | -1.42 | 1.266 | 0.886 | -1.751 | -0.018 | -0.058 | -0.058 | 0 | 0.22 | 0.62 | 0.15 | 0.07 | 0.22 | 0.12 |
| 3 | 167537664 | rs6797197 | -1.358 | 1.273 | 1.087 | -1.752 | -0.023 | -0.096 | -0.096 | 0 | 0.22 | 0.53 | 0.14 | 0.04 | 0.22 | 0.12 |
| 3 | 167552548 | rs11914302 | -0.019 | 1.249 | 1.691 | -1.749 | -0.007 | -0.041 | -0.041 | 0 | 0.09 | 0.24 | 0.08 | 0 | 0.1 | 0.02 |
| 3 | 167552712 | rs16851520 | -0.239 | 1.321 | 1.583 | -1.748 | 0.003 | -0.024 | -0.024 | 0 | 0.11 | 0.12 | 0.1 | 0.01 | 0.15 | 0.03 |
| 3 | 167558356 | rs1492024 | -1.614 | 1.564 | 1.584 | -1.772 | 0.013 | -0.061 | -0.061 | 0 | 0.26 | 0.58 | 0.26 | 0.11 | 0.29 | 0.15 |
| 3 | 167568177 | rs1392326 | -2.186 | 1.559 | 1.326 | -1.77 | -0.012 | 0.115 | 0.115 | 0 | 0.36 | 0.87 | 0.34 | 0.14 | 0.39 | 0.16 |
| 3 | 167578227 | rs1027242 | -2.183 | 1.678 | 0.696 | -1.479 | 0.007 | 0.024 | 0.024 | 0 | 0.49 | 0.66 | 0.68 | 0.36 | 0.59 | 0.28 |
| 3 | 167589870 | rs2131112 | -0.002 | 1.689 | 0.15 | -1.775 | 0.002 | -0.014 | -0.014 | 0 | 0.08 | 0.09 | 0.09 | 0.01 | 0.14 | 0.02 |
| 3 | 167594951 | rs11928048 | 0.118 | 1.467 | -0.236 | -1.705 | -0.011 | 0.021 | 0.021 | 0 | 0.11 | 0.03 | 0.04 | 0.07 | 0.12 | 0.1 |
| 3 | 167604119 | rs6763376 | -1.411 | 1.666 | -0.591 | -1.779 | -0.045 | 0.129 | 0.129 | 0 | 0.36 | 0.9 | 0.41 | 0.16 | 0.42 | 0.17 |
| 3 | 167607869 | rs882751 | 0.055 | 1.848 | -0.745 | -1.818 | -0.022 | 0.02 | 0.02 | 0 | 0.88 | 0.96 | 0.84 | 0.91 | 0.86 | 0.9 |
| 3 | 167610853 | rs6794207 | -1.094 | 1.772 | -0.099 | -1.877 | 0.008 | -0.072 | -0.072 | 0.071 | 0.29 | 0.53 | 0.3 | 0.04 | 0.38 | 0.13 |
| 3 | 167617797 | rs4955779 | 1.809 | 1.832 | -0.842 | -1.784 | -0.05 | 0.119 | 0.119 | 0.191 | 0.47 | 0.04 | 0.41 | 0.73 | 0.39 | 0.69 |
| 3 | 167626220 | rs17417881 | 0.287 | 1.814 | -0.754 | -1.744 | -0.042 | 0.046 | 0.046 | 0 | 0.09 | 0.01 | 0.08 | 0.02 | 0.17 | 0.04 |
| 3 | 167631414 | rs6444727 | -0.746 | 1.48 | 0.059 | -1.821 | -0.009 | -0.073 | -0.073 | 0 | 0.24 | 0.43 | 0.28 | 0.07 | 0.24 | 0.16 |
| 3 | 167633424 | rs6800444 | -2.026 | 1.512 | -0.621 | -2.312 | 0.013 | 0.106 | 0.106 | 0.044 | 0.57 | 0.89 | 0.65 | 0.26 | 0.68 | 0.34 |
| 3 | 167635818 | rs17418357 | 0.311 | 1.452 | -0.969 | -2.404 | -0.053 | 0.07 | 0.07 | 0 | 0.2 | 0.04 | 0.25 | 0.07 | 0.25 | 0.14 |
| 3 | 167635899 | rs9824464 | -2.171 | 1.378 | -1.076 | -2.455 | -0.024 | 0.146 | 0.146 | 0.017 | 0.58 | 0.97 | 0.6 | 0.15 | 0.69 | 0.34 |
| 3 | 167635958 | rs4955597 | -0.483 | 1.426 | -0.862 | -2.432 | 0.017 | -0.069 | -0.069 | 0 | 0.35 | 0.44 | 0.31 | 0.08 | 0.44 | 0.19 |
| 3 | 167640021 | rs2055362 | -1.81 | 1.422 | -0.857 | -2.427 | 0.004 | -0.002 | -0.002 | 0 | 0.55 | 0.49 | 0.55 | 0.15 | 0.69 | 0.34 |
| 3 | 167646960 | rs972730 | -0.757 | 1.316 | -0.553 | -2.339 | 0.021 | -0.081 | -0.081 | 0.911 | 0.35 | 0.43 | 0.31 | 0.07 | 0.44 | 0.19 |

| | | | | | | | | | | | | | | | | |
|---|---|---|---|---|---|---|---|---|---|---|---|---|---|---|---|---|
| 3 | 167661136 | rs12639288 | -0.211 | 1.396 | -0.719 | -2.373 | -0.026 | 0.118 | 0.118 | 0 | 0.29 | 0.09 | 0.15 | 0.06 | 0.33 | 0.17 |
| 3 | 167666657 | rs7428224 | 0.055 | 1.016 | -0.604 | -2.129 | 0.002 | -0.033 | -0.033 | 0.011 | 0.45 | 0.45 | 0.33 | 0.19 | 0.46 | 0.41 |
| 3 | 167669335 | rs7632605 | -0.437 | 1.298 | 0.292 | -1.757 | 0.013 | -0.052 | -0.052 | 0 | 0.29 | 0.54 | 0.27 | 0.2 | 0.33 | 0.14 |
| 3 | 167680850 | rs13069589 | 0.02 | 1.299 | -0.241 | -1.782 | -0.037 | 0.097 | 0.097 | 0 | 0.27 | 0.09 | 0.17 | 0.06 | 0.34 | 0.14 |
| 3 | 167692392 | rs12492208 | -0.1 | 0.935 | 0.097 | -1.555 | -0.024 | 0.048 | 0.048 | 0 | 0.31 | 0.22 | 0.22 | 0.04 | 0.39 | 0.18 |
| 3 | 167696174 | rs4431164 | -1.125 | 0.96 | 0.36 | -1.614 | 0.004 | 0.002 | 0.002 | 1.232 | 0.12 | 0.7 | 0.21 | 0.18 | 0.1 | 0.08 |
| 3 | 167696683 | rs4356862 | 0.03 | 0.937 | 0.317 | -1.661 | -0.034 | 0.069 | 0.069 | 0 | 0.31 | 0.2 | 0.21 | 0.04 | 0.4 | 0.19 |
| 3 | 167696838 | rs6444773 | -0.678 | 0.714 | 1.095 | -1.382 | 0.062 | -0.037 | -0.037 | 0 | 0.09 | 0.37 | 0.07 | 0.04 | 0.09 | 0.08 |
| 3 | 167699702 | rs7631039 | -1.654 | 0.747 | 0.635 | -1.358 | 0 | 0 | 0 | 0.005 | 0.13 | 0.8 | 0.22 | 0.18 | 0.1 | 0.08 |
| 3 | 167702376 | rs7626620 | -1.753 | 0.71 | 0.583 | -1.352 | 0.002 | 0.002 | 0.002 | 4.156 | 0.13 | 0.85 | 0.22 | 0.17 | 0.1 | 0.08 |
| 3 | 167721648 | rs10936548 | -1.427 | 0.783 | 0.666 | -1.312 | 0.005 | 0.005 | 0.005 | 3.658 | 0.11 | 0.72 | 0.21 | 0.17 | 0.1 | 0.07 |
| 3 | 167722697 | rs10936549 | -0.345 | 0.789 | 1.355 | -1.311 | 0.083 | 0.162 | 0.162 | 2.629 | 0.07 | 0.43 | 0.18 | 0.13 | 0.04 | 0.04 |
| 3 | 167725349 | rs7632500 | -0.493 | 0.769 | 1.279 | -1.332 | 0.066 | 0.133 | 0.133 | 1.537 | 0.07 | 0.48 | 0.18 | 0.13 | 0.04 | 0.04 |
| 3 | 167725408 | rs7634939 | -0.493 | 0.769 | 1.279 | -1.332 | 0.066 | 0.133 | 0.133 | 1.537 | 0.07 | 0.48 | 0.18 | 0.13 | 0.04 | 0.04 |
| 3 | 167726096 | rs9873288 | -1.113 | 0.893 | 0.503 | -1.662 | -0.056 | 0.175 | 0.175 | 0 | 0.43 | 0.88 | 0.44 | 0.2 | 0.51 | 0.28 |
| 3 | 167727436 | rs4254706 | -1.079 | 0.887 | 0.988 | -1.245 | 0.011 | 0.063 | 0.063 | 3.317 | 0.09 | 0.59 | 0.19 | 0.14 | 0.04 | 0.04 |
| 3 | 167737813 | rs11914431 | -1.628 | 0.913 | 0.503 | -1.241 | 0 | -0.002 | -0.002 | 0.421 | 0.12 | 0.79 | 0.22 | 0.17 | 0.1 | 0.07 |
| 3 | 167749515 | rs9845075 | -0.712 | 0.784 | 0.206 | -1.787 | -0.012 | 0.153 | 0.153 | 0 | 0.52 | 0.71 | 0.33 | 0.08 | 0.48 | 0.3 |
| 3 | 167749543 | rs9845202 | 0.386 | 0.855 | 0.105 | -1.97 | -0.035 | 0.092 | 0.092 | 0 | 0.25 | 0.07 | 0.15 | 0.03 | 0.26 | 0.15 |
| 3 | 167766491 | rs7639014 | -0.893 | 0.712 | 0.54 | -1.978 | 0.002 | 0.098 | 0.098 | 0 | 0.51 | 0.65 | 0.34 | 0.08 | 0.5 | 0.3 |
| 3 | 167770115 | rs9824277 | -1.225 | 0.816 | 0.272 | -1.979 | 0 | 0.218 | 0.218 | 0 | 0.54 | 0.81 | 0.36 | 0.09 | 0.55 | 0.33 |
| 3 | 167775964 | rs1522379 | 0.16 | 0.903 | 0.446 | -1.809 | 0.024 | -0.116 | -0.116 | 0.616 | 0.28 | 0.44 | 0.19 | 0.04 | 0.3 | 0.15 |
| 3 | 167783178 | rs1522380 | -0.369 | 0.612 | 0.46 | -1.005 | 0.002 | -0.015 | -0.015 | 0 | 0.23 | 0.21 | 0.15 | 0.04 | 0.2 | 0.15 |
| 3 | 167799902 | rs877439 | -0.496 | 0.903 | 0.448 | 0.318 | 0.006 | 0.016 | 0.016 | 0 | 0.49 | 0.72 | 0.5 | 0.61 | 0.5 | 0.54 |
| 3 | 167805891 | rs10513641 | -1.065 | 0.928 | 1.288 | 0.045 | 0.075 | 0.001 | 0.001 | 0.492 | 0.09 | 0.45 | 0.06 | 0.02 | 0.09 | 0.06 |
| 3 | 167823611 | rs7651464 | -1.459 | 0.889 | 1.227 | 0.028 | 0.03 | -0.032 | -0.032 | 0 | 0.1 | 0.54 | 0.07 | 0.02 | 0.09 | 0.06 |
| 3 | 167837748 | rs4345115 | 0.125 | 0.883 | 0.67 | 0.346 | -0.01 | -0.06 | -0.06 | 0 | 0.39 | 0.15 | 0.46 | 0.56 | 0.41 | 0.46 |
| 3 | 167852482 | rs10513642 | -0.224 | 0.794 | 1.07 | 0.371 | -0.001 | -0.002 | -0.002 | 0.187 | 0.05 | 0.06 | 0.03 | 0.01 | 0.09 | 0.06 |
| 3 | 167860563 | rs11916217 | -0.622 | 0.795 | 1.361 | 0.367 | 0.047 | -0.022 | -0.022 | 0 | 0.06 | 0.22 | 0.04 | 0.01 | 0.09 | 0.06 |
| 3 | 167893774 | rs6807050 | -0.919 | 0.783 | 1.268 | 0.362 | 0.059 | -0.02 | -0.02 | 4.481 | 0.07 | 0.29 | 0.04 | 0.01 | 0.09 | 0.06 |
| 3 | 167896868 | rs2593420 | 0.496 | 0.751 | 0.632 | 0.7 | 0.003 | -0.018 | -0.018 | 0 | 0.29 | 0.21 | 0.4 | 0.65 | 0.27 | 0.5 |
| 3 | 167899270 | rs2593423 | -0.012 | 0.658 | 0.029 | 0.579 | 0.001 | 0.003 | 0.003 | 0 | 0.37 | 0.63 | 0.45 | 0.7 | 0.37 | 0.56 |
| 3 | 167904845 | rs6444798 | 0.615 | 0.973 | 0.157 | 0.365 | -0.002 | 0 | 0 | 0 | 0.65 | 0.69 | 0.73 | 0.73 | 0.56 | 0.79 |
| 3 | 167907755 | rs6767716 | -1.387 | 0.292 | 0.279 | -0.128 | 0.014 | 0.041 | 0.041 | 8.688 | 0.2 | 0.82 | 0.15 | 0.01 | 0.16 | 0.17 |
| 3 | 167913564 | rs7643342 | -0.943 | 0.628 | 0.343 | 0.126 | 0.017 | -0.048 | -0.048 | 0 | 0.16 | 0.41 | 0.13 | 0.06 | 0.15 | 0.16 |
| 3 | 167918046 | rs2871744 | -0.23 | -0.032 | -0.886 | -0.113 | 0.031 | 0.054 | 0.054 | 0.202 | 0.36 | 0.13 | 0.47 | 0.26 | 0.28 | 0.36 |
| 3 | 167925722 | rs13100658 | -0.268 | 0.014 | 0.274 | -0.269 | 0.024 | 0.049 | 0.049 | 0.11 | 0.36 | 0.19 | 0.46 | 0.25 | 0.3 | 0.35 |
| 3 | 167928439 | rs6800871 | -1.15 | -0.242 | 1.405 | -0.407 | -0.024 | -0.041 | -0.041 | 1.203 | 0.09 | 0.73 | 0.08 | 0.02 | 0.04 | 0.13 |
| 3 | 167937888 | rs7624807 | -0.756 | -0.629 | 1.234 | -0.696 | -0.002 | -0.034 | -0.034 | 0 | 0.24 | 0.73 | 0.16 | 0.03 | 0.18 | 0.29 |
| 3 | 167940314 | rs2067678 | -0.45 | -0.621 | 1.443 | -0.693 | -0.009 | -0.109 | -0.109 | 0.005 | 0.22 | 0.57 | 0.15 | 0.03 | 0.18 | 0.29 |
| 3 | 167940789 | rs3914953 | -0.265 | -0.623 | 1.717 | -0.694 | -0.002 | -0.066 | -0.066 | 0.116 | 0.2 | 0.27 | 0.12 | 0.03 | 0.18 | 0.27 |
| 3 | 167947516 | rs16852036 | -0.312 | -0.623 | 1.891 | -0.694 | -0.001 | -0.095 | -0.095 | 0 | 0.2 | 0.38 | 0.13 | 0.03 | 0.18 | 0.28 |
| 3 | 167974061 | rs1605389 | -1.467 | -0.19 | 0.552 | -0.669 | -0.039 | -0.111 | -0.111 | 1.032 | 0.66 | 0.96 | 0.67 | 0.51 | 0.63 | 0.67 |
| 3 | 167976376 | rs727800 | -1.367 | -0.437 | 1.024 | -0.495 | 0.063 | 0.11 | 0.11 | 0.038 | 0.44 | 0.88 | 0.52 | 0.29 | 0.32 | 0.48 |
| 3 | 168044074 | rs16852171 | 0.885 | -0.563 | 1.331 | -0.286 | 0.033 | -0.041 | -0.041 | 0 | 0.14 | 0.06 | 0.34 | 0.25 | 0.09 | 0.1 |
| 3 | 168064070 | rs11915706 | 0.057 | -0.45 | 1.233 | -0.596 | 0.001 | -0.075 | -0.075 | 0 | 0.16 | 0.3 | 0.11 | 0.03 | 0.15 | 0.24 |
| 3 | 168066580 | rs3908793 | 1.102 | -0.554 | 0.973 | -0.492 | 0.016 | -0.049 | -0.049 | 6.923 | 0.14 | 0.01 | 0.33 | 0.21 | 0.1 | 0.07 |
| 3 | 168093655 | rs11706167 | 1.181 | -0.503 | 0.646 | 0.008 | -0.004 | -0.01 | -0.01 | 0 | 0.1 | 0.04 | 0.18 | 0.31 | 0.11 | 0.14 |
| 3 | 168125439 | rs9879247 | -1.486 | -0.611 | 0.979 | -0.534 | 0.058 | 0.132 | 0.132 | 0.007 | 0.41 | 0.9 | 0.5 | 0.23 | 0.29 | 0.41 |
| 3 | 168132678 | rs9879751 | -0.72 | -0.63 | 1.152 | -0.535 | -0.053 | -0.028 | -0.028 | 0 | 0.25 | 0.57 | 0.39 | 0.21 | 0.14 | 0.17 |
| 3 | 168136690 | rs7614661 | -1.243 | -0.488 | 1.116 | -0.459 | 0.039 | 0.083 | 0.083 | 0.144 | 0.35 | 0.86 | 0.48 | 0.23 | 0.26 | 0.32 |
| 3 | 168161589 | rs6798963 | -0.099 | -0.484 | 1.304 | -0.448 | 0.019 | -0.098 | -0.098 | 0 | 0.16 | 0.4 | 0.11 | 0.02 | 0.15 | 0.24 |
| 3 | 168167082 | rs6780394 | 0.137 | -0.412 | 1.293 | -0.351 | -0.045 | 0.007 | 0.007 | 0.08 | 0.18 | 0.4 | 0.36 | 0.21 | 0.11 | 0.08 |
| 3 | 168175261 | rs863492 | -0.343 | -0.251 | 1.418 | -0.543 | 0.004 | -0.076 | -0.076 | 0.136 | 0.17 | 0.28 | 0.1 | 0.02 | 0.17 | 0.23 |
| 3 | 168181535 | rs11926516 | -0.47 | -0.331 | 1.802 | -0.609 | 0.044 | -0.003 | -0.003 | 0 | 0.07 | 0.29 | 0.05 | 0 | 0.06 | 0.08 |
| 3 | 168183506 | rs329072 | -1.523 | -0.294 | 1.725 | -0.582 | 0.025 | 0.077 | 0.077 | 0 | 0.38 | 0.75 | 0.46 | 0.19 | 0.31 | 0.38 |
| 3 | 168183879 | rs97529 | -0.862 | -0.315 | 1.667 | -0.583 | -0.009 | -0.022 | -0.022 | 0 | 0.31 | 0.39 | 0.4 | 0.19 | 0.24 | 0.29 |
| 3 | 168194499 | rs16852344 | 0.779 | -0.258 | 0.984 | -0.524 | 0.022 | -0.011 | -0.011 | 0 | 0.14 | 0.08 | 0.29 | 0.17 | 0.07 | 0.07 |
| 3 | 168196092 | rs184502 | -1.543 | -0.327 | 0.787 | -0.553 | 0.007 | 0.135 | 0.135 | 0.126 | 0.41 | 0.96 | 0.47 | 0.19 | 0.3 | 0.38 |
| 3 | 168201575 | rs2595924 | -1.511 | -0.105 | 0.818 | -0.552 | 0.02 | 0.148 | 0.148 | 0 | 0.41 | 0.9 | 0.48 | 0.19 | 0.32 | 0.39 |
| 3 | 168202823 | rs865772 | -1.464 | -0.041 | 0.908 | -0.526 | 0.023 | 0.085 | 0.085 | 0 | 0.4 | 0.76 | 0.47 | 0.19 | 0.32 | 0.39 |
| 3 | 168205232 | rs3906954 | 0.703 | -0.005 | 0.781 | -0.393 | 0.015 | -0.036 | -0.036 | 0 | 0.14 | 0.02 | 0.3 | 0.17 | 0.09 | 0.08 |
| 3 | 168212344 | rs1025217 | -1.292 | 0.379 | 0.727 | -0.428 | -0.039 | 0.091 | 0.091 | 0 | 0.53 | 0.81 | 0.59 | 0.3 | 0.5 | 0.42 |
| 3 | 168219437 | rs13089730 | -0.385 | 0.443 | 0.944 | -0.582 | 0.012 | -0.118 | -0.118 | 0 | 0.16 | 0.43 | 0.11 | 0.01 | 0.15 | 0.09 |
| 3 | 168220116 | rs329082 | -0.09 | 0.379 | 0.745 | -0.596 | 0.009 | -0.038 | -0.038 | 2.323 | 0.08 | 0.2 | 0.05 | 0.02 | 0.07 | 0.19 |
| 3 | 168223132 | rs7642311 | -0.192 | 0.39 | 0.855 | -0.582 | 0 | -0.083 | -0.083 | 0 | 0.15 | 0.28 | 0.1 | 0.01 | 0.13 | 0.09 |
| 3 | 168231158 | rs11718699 | 0.496 | 0.352 | 0.487 | -0.369 | -0.063 | -0.06 | -0.06 | 0.29 | 0.26 | 0.02 | 0.42 | 0.26 | 0.28 | 0.13 |
| 3 | 168233747 | rs329079 | -0.255 | 0.461 | 0.652 | -0.324 | 0.012 | -0.038 | -0.038 | 0 | 0.08 | 0.2 | 0.05 | 0.02 | 0.07 | 0.18 |
| 3 | 168234218 | rs884828 | 0.048 | 0.429 | 0.263 | -0.255 | -0.002 | -0.004 | -0.004 | 1.042 | 0.3 | 0.34 | 0.43 | 0.26 | 0.3 | 0.13 |

| | | | | | | | | | | | | | | | | |
|---|---|---|---|---|---|---|---|---|---|---|---|---|---|---|---|---|
| 3 | 168245830 | rs2555499 | -0.361 | 0.19 | 0.021 | -0.055 | -0.002 | 0 | 0 | 0.118 | 0.41 | 0.58 | 0.5 | 0.45 | 0.36 | 0.28 |
| 3 | 168254003 | rs2053021 | -0.145 | 0.231 | 0.005 | -0.045 | 0.021 | -0.023 | -0.023 | 0.15 | 0.07 | 0.29 | 0.05 | 0.01 | 0.03 | 0.13 |
| 3 | 168266180 | rs3853728 | -0.153 | 0.27 | -0.632 | 0.141 | 0 | -0.001 | -0.001 | 0.446 | 0.41 | 0.61 | 0.5 | 0.45 | 0.36 | 0.28 |
| 3 | 168267259 | rs13091735 | -0.615 | 0.52 | -0.753 | 0.114 | -0.026 | 0.027 | 0.027 | 0 | 0.12 | 0.05 | 0.09 | 0.01 | 0.18 | 0.07 |
| 3 | 168273670 | rs4091213 | 0.793 | -0.556 | -0.613 | 0.651 | 0.042 | 0.011 | 0.011 | 1.42 | 0.58 | 0.24 | 0.61 | 0.43 | 0.66 | 0.37 |
| 3 | 168281038 | rs3911093 | 0.394 | -0.871 | -0.603 | 1.112 | 0.015 | -0.1 | -0.1 | 0 | 0.67 | 0.9 | 0.67 | 0.53 | 0.68 | 0.49 |
| 3 | 168283582 | rs3853730 | 0.99 | -0.878 | 0.123 | 1.156 | -0.001 | 0 | 0 | 0 | 0.6 | 0.6 | 0.61 | 0.39 | 0.66 | 0.37 |
| 3 | 168302190 | rs3853733 | 0.249 | -0.96 | -0.54 | 1.144 | -0.021 | -0.116 | -0.116 | 0 | 0.17 | 0.01 | 0.16 | 0.35 | 0.16 | 0.24 |
| 3 | 168306182 | rs3914114 | 0.196 | -0.792 | -0.359 | 1.209 | -0.001 | -0.001 | -0.001 | 0.463 | 0.05 | 0.07 | 0.04 | 0.09 | 0.08 | 0.16 |
| 3 | 168311682 | rs8179961 | 1.478 | -0.883 | -0.409 | 1.157 | 0.056 | 0.02 | 0.02 | 2.271 | 0.57 | 0.19 | 0.61 | 0.42 | 0.65 | 0.36 |
| 3 | 168314500 | rs7630157 | -1.347 | -0.159 | 0.244 | 1.084 | 0.028 | 0.268 | 0.268 | 0.988 | 0.07 | 0.5 | 0.06 | 0.13 | 0.01 | 0.13 |
| 3 | 168326057 | rs16852499 | -1.285 | 0.352 | -0.933 | 0.629 | -0.014 | 0.008 | 0.008 | 0 | 0.1 | 0.04 | 0.06 | 0.09 | 0.12 | 0.17 |
| 3 | 168335051 | rs3863110 | -0.965 | 0.531 | -1.14 | 0.848 | -0.011 | -0.026 | -0.026 | 0.021 | 0.05 | 0.01 | 0.25 | 0.11 | 0.05 | 0.17 |
| 3 | 168335438 | rs9290340 | -0.761 | 0.285 | -0.67 | 0.95 | -0.016 | 0.037 | 0.037 | 0 | 0.25 | 0.36 | 0.23 | 0.3 | 0.23 | 0.25 |
| 3 | 168337826 | rs3914117 | -1.157 | 0.006 | -0.237 | 0.779 | 0.011 | -0.007 | -0.007 | 0.525 | 0.29 | 0.72 | 0.29 | 0.3 | 0.23 | 0.29 |
| 3 | 168338128 | rs6766202 | -1.001 | 0.185 | -0.612 | 0.479 | -0.006 | -0.059 | -0.059 | 1.108 | 0.2 | 0.4 | 0.16 | 0.11 | 0.15 | 0.3 |
| 3 | 168341985 | rs17736747 | -0.535 | -0.077 | -1.305 | 0.662 | -0.024 | -0.059 | -0.059 | 0 | 0.18 | 0.02 | 0.11 | 0.21 | 0.18 | 0.16 |
| 3 | 168344334 | rs13069078 | 0.465 | 0.105 | -2.173 | -0.642 | -0.112 | 0.241 | 0.241 | 2.092 | 0.35 | 0.03 | 0.19 | 0.03 | 0.39 | 0.21 |
| 3 | 168350231 | rs16847695 | -0.176 | -0.575 | -1.526 | -0.333 | -0.051 | 0.011 | 0.011 | 0 | 0.31 | 0.5 | 0.42 | 0.64 | 0.22 | 0.33 |
| 3 | 168351860 | rs12629405 | 0.223 | -0.611 | -1.239 | -0.015 | 0 | -0.002 | -0.002 | 0 | 0.55 | 0.55 | 0.4 | 0.56 | 0.5 | 0.5 |
| 3 | 168363205 | rs16852525 | 0.205 | -0.649 | -1.125 | 0.045 | -0.019 | 0.047 | 0.047 | 0 | 0.19 | 0.47 | 0.37 | 0.23 | 0.17 | 0.15 |
| 3 | 168366063 | rs16852530 | 0.71 | -0.664 | -1.102 | 0.058 | 0.002 | 0.009 | 0.009 | 0 | 0.16 | 0.15 | 0.14 | 0.06 | 0.16 | 0.1 |
| 3 | 168369375 | rs10804826 | -0.781 | -0.696 | -1.139 | -0.012 | -0.046 | 0.043 | 0.043 | 0 | 0.72 | 0.89 | 0.76 | 0.79 | 0.66 | 0.61 |
| 3 | 168381002 | rs16852539 | 0.586 | -0.768 | -1.266 | 0.106 | 0.028 | -0.073 | -0.073 | 0 | 0.12 | 0.02 | 0.25 | 0.54 | 0.07 | 0.22 |
| 3 | 168383190 | rs4955617 | 1.694 | -0.506 | -0.924 | -0.303 | -0.031 | 0.13 | 0.13 | 1.726 | 0.35 | 0.12 | 0.19 | 0.03 | 0.37 | 0.18 |
| 3 | 168383288 | rs6807896 | 0.77 | -0.379 | -0.869 | -0.208 | 0.002 | 0.005 | 0.005 | 0 | 0.16 | 0.15 | 0.14 | 0.06 | 0.15 | 0.09 |
| 3 | 168389435 | rs6444819 | -0.371 | -0.18 | -0.63 | -0.014 | -0.003 | 0 | 0 | 0.014 | 0.71 | 0.75 | 0.53 | 0.64 | 0.65 | 0.55 |
| 3 | 168391645 | rs988230 | -0.607 | -0.075 | -0.888 | -0.144 | -0.089 | 0.036 | 0.036 | 0.047 | 0.72 | 0.94 | 0.75 | 0.79 | 0.65 | 0.58 |
| 3 | 168413945 | rs10049401 | 1.381 | -0.313 | -0.759 | -0.494 | -0.003 | -0.018 | -0.018 | 0.82 | 0.26 | 0.2 | 0.15 | 0.04 | 0.28 | 0.14 |
| 3 | 168417194 | rs9841841 | -0.872 | -0.218 | -0.859 | -0.472 | -0.038 | 0.022 | 0.022 | 0 | 0.72 | 0.89 | 0.7 | 0.81 | 0.66 | 0.7 |
| 3 | 168419832 | rs16852581 | -0.437 | -0.199 | -0.525 | -0.209 | -0.003 | 0 | 0 | 0 | 0.44 | 0.48 | 0.32 | 0.52 | 0.38 | 0.45 |
| 3 | 168424384 | rs9821618 | -0.996 | -0.174 | -0.898 | -0.822 | -0.07 | 0.091 | 0.091 | 0.039 | 0.73 | 0.96 | 0.79 | 0.88 | 0.66 | 0.71 |
| 3 | 168430783 | rs925363 | -0.242 | -0.34 | -0.999 | -0.848 | -0.04 | 0.006 | 0.006 | 0 | 0.62 | 0.94 | 0.52 | 0.32 | 0.6 | 0.52 |
| 3 | 168435479 | rs9862098 | -1.063 | -0.338 | -0.999 | -0.849 | -0.07 | 0.091 | 0.091 | 0.039 | 0.73 | 0.96 | 0.79 | 0.88 | 0.66 | 0.71 |
| 3 | 168437510 | rs1522982 | -0.072 | -0.346 | -0.824 | -0.851 | 0.006 | 0.086 | 0.086 | 0 | 0.6 | 0.78 | 0.4 | 0.14 | 0.6 | 0.41 |
| 3 | 168438555 | rs13074219 | 1.63 | -0.186 | -0.944 | -1.093 | -0.072 | 0.02 | 0.02 | 0 | 0.19 | 0.01 | 0.11 | 0.12 | 0.24 | 0.24 |
| 3 | 168455787 | rs2421468 | -0.837 | -0.204 | -0.729 | -0.927 | -0.004 | 0.001 | 0.001 | 0.398 | 0.67 | 0.73 | 0.67 | 0.72 | 0.63 | 0.6 |
| 3 | 168458567 | rs17800311 | 1.143 | -0.23 | -1.026 | -0.843 | -0.017 | 0.078 | 0.078 | 0 | 0.23 | 0.1 | 0.14 | 0.05 | 0.23 | 0.16 |
| 3 | 168463089 | rs6763367 | 1.076 | -0.173 | -0.98 | -0.811 | -0.006 | 0.047 | 0.047 | 0 | 0.24 | 0.17 | 0.15 | 0.05 | 0.22 | 0.16 |
| 3 | 168465419 | rs6770546 | 0.836 | 0.277 | -1.554 | -0.691 | 0.03 | -0.054 | -0.054 | 0 | 0.11 | 0.01 | 0.28 | 0.58 | 0.06 | 0.2 |
| 3 | 168469608 | rs2687707 | 1.009 | 0.299 | -1.559 | -0.758 | -0.075 | 0.092 | 0.092 | 0.05 | 0.27 | 0.03 | 0.21 | 0.11 | 0.34 | 0.29 |
| 3 | 168493689 | rs516215 | 0.064 | 0.6 | -1.398 | -0.614 | 0.01 | 0.092 | 0.092 | 0 | 0.4 | 0.17 | 0.52 | 0.8 | 0.4 | 0.6 |
| 3 | 168496750 | rs535028 | -1.315 | 0.582 | -1.384 | -0.638 | -0.005 | -0.001 | -0.001 | 0 | 0.74 | 0.8 | 0.85 | 0.96 | 0.72 | 0.87 |
| 3 | 168520843 | rs496741 | 0.064 | 0.662 | -1.647 | -0.596 | 0.011 | 0.091 | 0.091 | 0 | 0.4 | 0.17 | 0.52 | 0.8 | 0.4 | 0.6 |
| 3 | 168538993 | rs603638 | -1.344 | 0.701 | -1.67 | -0.706 | -0.01 | 0.029 | 0.029 | 0 | 0.75 | 0.86 | 0.85 | 0.96 | 0.72 | 0.87 |
| 3 | 168546914 | rs678690 | -0.666 | 0.083 | -1.423 | 0.366 | 0.012 | 0.08 | 0.08 | 0 | 0.41 | 0.23 | 0.61 | 0.88 | 0.4 | 0.61 |
| 3 | 168564812 | rs1513431 | -0.488 | 0.045 | -0.475 | 1.554 | -0.025 | 0.087 | 0.087 | 0 | 0.1 | 0.15 | 0.3 | 0.68 | 0.05 | 0.26 |
| 3 | 168581491 | rs7633280 | 0.153 | 0.455 | -0.877 | 0.496 | 0.042 | -0.011 | -0.011 | 0 | 0.47 | 0.68 | 0.37 | 0.51 | 0.38 | 0.5 |
| 3 | 168583606 | rs9851042 | -0.519 | 1.252 | -1.364 | 0.182 | -0.008 | -0.019 | -0.019 | 0 | 0.79 | 0.84 | 0.66 | 0.6 | 0.75 | 0.72 |
| 3 | 168585686 | rs11718651 | 0.392 | 1.442 | -1.508 | 0.14 | -0.05 | 0.121 | 0.121 | 1.141 | 0.31 | 0.11 | 0.29 | 0.09 | 0.34 | 0.22 |
| 3 | 168586930 | rs9825517 | 0.172 | 1.469 | -0.89 | 0.029 | 0.035 | 0.06 | 0.06 | 0 | 0.68 | 0.55 | 0.5 | 0.54 | 0.63 | 0.63 |
| 3 | 168595470 | rs986738 | -0.04 | 0.885 | -1.534 | -0.093 | -0.058 | 0.077 | 0.077 | 0 | 0.3 | 0.05 | 0.24 | 0.2 | 0.33 | 0.31 |
| 3 | 168610857 | rs1588880 | 0.584 | 0.534 | -0.935 | 0.136 | -0.019 | -0.029 | -0.029 | 0 | 0.57 | 0.72 | 0.46 | 0.56 | 0.44 | 0.49 |
| 3 | 168615125 | rs4565022 | 0.329 | 0.214 | 0.011 | 0.151 | 0.007 | 0.282 | 0.282 | 0 | 0.07 | 0.29 | 0.11 | 0.29 | 0.03 | 0.06 |
| 3 | 168621773 | rs1877267 | 0.423 | 0.22 | 0.16 | 0.142 | -0.012 | -0.019 | -0.019 | 0 | 0.57 | 0.66 | 0.46 | 0.56 | 0.44 | 0.49 |
| 3 | 168627721 | rs11921316 | -0.035 | 0.334 | 0.047 | -0.077 | -0.001 | -0.022 | -0.022 | 0 | 0.22 | 0.25 | 0.17 | 0.03 | 0.29 | 0.2 |
| 3 | 168640814 | rs1511598 | -0.189 | 0.411 | 0.08 | -0.036 | -0.022 | -0.051 | -0.051 | 0 | 0.78 | 0.84 | 0.63 | 0.59 | 0.75 | 0.69 |
| 3 | 168645035 | rs1498444 | 1.709 | 0.17 | 0.19 | -0.293 | -0.084 | 0.224 | 0.224 | 0.159 | 0.46 | 0.07 | 0.33 | 0.25 | 0.44 | 0.34 |
| 3 | 168645151 | rs2280063 | -0.026 | 0.279 | 0.297 | -0.279 | 0.001 | -0.003 | -0.003 | 0 | 0.22 | 0.23 | 0.17 | 0.03 | 0.28 | 0.19 |
| 3 | 168646471 | rs1511603 | -0.474 | 0.264 | 0.311 | -0.142 | -0.04 | -0.077 | -0.077 | 0 | 0.8 | 0.91 | 0.63 | 0.59 | 0.75 | 0.69 |
| 3 | 168656997 | rs10513652 | -0.074 | -0.342 | 0.771 | 0.049 | -0.018 | -0.06 | -0.06 | 1.635 | 0.12 | 0.26 | 0.07 | 0.01 | 0.11 | 0.14 |
| 3 | 168669946 | rs9845424 | -0.759 | -0.213 | 1.096 | 0.262 | -0.001 | -0.024 | -0.024 | 0 | 0.25 | 0.28 | 0.16 | 0.03 | 0.3 | 0.2 |
| 3 | 168670179 | rs9290348 | -0.617 | -0.181 | 1.034 | 0.23 | 0.024 | 0.225 | 0.225 | 0 | 0.12 | 0.28 | 0.1 | 0.39 | 0.11 | 0.28 |
| 3 | 168684844 | rs11918821 | -1.349 | -0.358 | 1.002 | 0.117 | -0.009 | 0.004 | 0.004 | 0 | 0.34 | 0.61 | 0.48 | 0.35 | 0.34 | 0.36 |
| 3 | 168692457 | rs9290352 | -1.388 | -0.342 | 1.2 | 0.377 | 0.012 | 0.222 | 0.222 | 0.863 | 0.14 | 0.33 | 0.12 | 0.41 | 0.13 | 0.43 |
| 3 | 168692989 | rs9290353 | -0.455 | -0.552 | 1.279 | 0.403 | -0.011 | 0.126 | 0.126 | 0 | 0.09 | 0.13 | 0.09 | 0.38 | 0.07 | 0.3 |
| 3 | 168693089 | rs9835962 | -1.294 | -0.552 | 1.099 | 0.39 | -0.026 | 0.271 | 0.271 | 0.362 | 0.13 | 0.33 | 0.1 | 0.41 | 0.08 | 0.4 |
| 3 | 168698189 | rs6802894 | 1.503 | -1.071 | 0.936 | 0.472 | 0.006 | 0.025 | 0.025 | 0 | 0.54 | 0.42 | 0.41 | 0.28 | 0.57 | 0.35 |
| 3 | 168698220 | rs9834862 | -0.645 | -1.109 | 0.935 | 0.648 | -0.037 | 0.098 | 0.098 | 0 | 0.19 | 0.25 | 0.12 | 0.4 | 0.12 | 0.44 |

| | | | | | | | | | | | | | | | | |
|---|---|---|---|---|---|---|---|---|---|---|---|---|---|---|---|---|
| 3 | 168715048 | rs16852963 | 0.128 | -1.131 | 0.482 | 1.212 | 0.051 | -0.068 | -0.068 | 1.521 | 0.15 | 0.02 | 0.07 | 0.22 | 0.08 | 0.16 |
| 3 | 168726724 | rs13089423 | 0.858 | -2.104 | 0.864 | 0.894 | -0.041 | -0.022 | -0.022 | 0 | 0.74 | 0.48 | 0.84 | 0.82 | 0.81 | 0.67 |
| 3 | 168741644 | rs1397899 | 1.95 | -2.261 | 0.925 | 1.006 | 0.077 | -0.006 | -0.006 | 0 | 0.54 | 0.29 | 0.75 | 0.6 | 0.71 | 0.44 |
| 3 | 168742461 | rs1420469 | 0.813 | -2.323 | 0.627 | 0.961 | 0.036 | 0.029 | 0.029 | 0 | 0.1 | 0.05 | 0.04 | 0.05 | 0.03 | 0.06 |
| 3 | 168743995 | rs9809610 | -1.28 | -2.224 | 0.627 | 0.946 | 0.001 | -0.035 | -0.035 | 0.124 | 0.36 | 0.23 | 0.26 | 0.38 | 0.36 | 0.44 |
| 3 | 168746694 | rs974942 | -1.221 | -2.19 | 0.719 | 0.922 | -0.003 | -0.041 | -0.041 | 0 | 0.92 | 0.8 | 0.95 | 1 | 0.92 | 0.96 |
| 3 | 168746967 | rs974944 | 1.235 | -2.155 | 0.463 | 0.953 | -0.05 | 0 | 0 | 0 | 0.12 | 0.04 | 0.12 | 0.07 | 0.19 | 0.07 |
| 3 | 168753276 | rs7647582 | -0.825 | -2.109 | 0.541 | 0.978 | 0.002 | -0.004 | -0.004 | 0 | 0.24 | 0.21 | 0.14 | 0.31 | 0.17 | 0.37 |
| 3 | 168754246 | rs12635194 | 0.742 | -2.104 | 0.362 | 0.974 | 0.056 | -0.055 | -0.055 | 0 | 0.15 | 0.02 | 0.07 | 0.18 | 0.08 | 0.15 |
| 3 | 168757650 | rs3851376 | 2.627 | -2.049 | 0.376 | 0.923 | -0.126 | 0.105 | 0.105 | 0 | 0.38 | 0.05 | 0.31 | 0.24 | 0.49 | 0.24 |
| 3 | 168778197 | rs7631699 | -1.581 | -2.362 | 0.597 | 0.935 | 0.009 | -0.002 | -0.002 | 0 | 0.36 | 0.36 | 0.19 | 0.34 | 0.2 | 0.44 |
| 3 | 168781277 | rs6444837 | 0.67 | -2.234 | 0.948 | 1.312 | -0.002 | 0.005 | 0.005 | 0 | 0.7 | 0.64 | 0.54 | 0.62 | 0.73 | 0.77 |
| 3 | 168784088 | rs721128 | 2.615 | -2.328 | 0.564 | 1.263 | -0.099 | 0.083 | 0.083 | 0 | 0.39 | 0.03 | 0.36 | 0.24 | 0.56 | 0.25 |
| 3 | 168785079 | rs12637241 | 1.855 | -2.302 | 0.629 | 1.35 | 0.015 | -0.005 | -0.005 | 1.887 | 0.55 | 0.08 | 0.43 | 0.4 | 0.64 | 0.4 |
| 3 | 168789705 | rs6806825 | 1.391 | -2.253 | 0.781 | 1.334 | 0.035 | -0.015 | -0.015 | 0.145 | 0.56 | 0.72 | 0.48 | 0.47 | 0.64 | 0.62 |
| 3 | 168789847 | rs10513658 | -1.454 | -2.17 | 1.057 | 1.177 | -0.042 | 0.09 | 0.09 | 0 | 0.15 | 0.53 | 0.12 | 0.21 | 0.09 | 0.38 |
| 3 | 168795475 | rs9837244 | -1.126 | -2.141 | 0.46 | 0.998 | -0.005 | -0.012 | -0.012 | 0.239 | 0.92 | 0.92 | 0.98 | 0.99 | 0.96 | 0.96 |
| 3 | 168810874 | rs17466625 | 0.916 | -2.125 | 0.388 | 1.014 | -0.067 | 0.007 | 0.007 | 0 | 0.16 | 0.02 | 0.1 | 0.07 | 0.2 | 0.09 |
| 3 | 168811043 | rs12634370 | -0.972 | -2.169 | 0.482 | 1.037 | 0.016 | 0.017 | 0.017 | 0 | 0.33 | 0.17 | 0.19 | 0.23 | 0.29 | 0.26 |
| 3 | 168814909 | rs13094430 | -0.512 | -2.155 | 0.552 | 1.05 | 0.008 | 0.009 | 0.009 | 0.046 | 0.3 | 0.17 | 0.19 | 0.23 | 0.28 | 0.24 |
| 3 | 168815099 | rs4955637 | 1.642 | -2.176 | 0.635 | 1.158 | 0.032 | 0.003 | 0.003 | 0 | 0.56 | 0.66 | 0.73 | 0.63 | 0.62 | 0.48 |
| 3 | 168820427 | rs2111082 | -0.447 | -2.192 | 0.592 | 1.157 | 0.008 | -0.027 | -0.027 | 0 | 0.09 | 0.04 | 0.06 | 0.18 | 0.08 | 0.27 |
| 3 | 168822441 | rs1362521 | -0.935 | -1.806 | 0.925 | 1.289 | 0.017 | 0.017 | 0.017 | 0 | 0.32 | 0.18 | 0.19 | 0.23 | 0.29 | 0.27 |
| 3 | 168828505 | rs1564356 | -0.96 | -1.701 | 0.925 | 1.316 | -0.012 | -0.022 | -0.022 | 0 | 0.91 | 0.86 | 0.98 | 0.99 | 0.96 | 0.96 |
| 3 | 168832107 | rs9847631 | 1.675 | -1.719 | 0.928 | 1.336 | -0.002 | -0.001 | -0.001 | 0 | 0.54 | 0.52 | 0.72 | 0.63 | 0.62 | 0.48 |
| 3 | 168832835 | rs16853173 | -0.052 | -1.295 | 0.83 | 1.903 | 0.041 | -0.032 | -0.032 | 0 | 0.14 | 0.03 | 0.07 | 0.16 | 0.08 | 0.15 |
| 3 | 168837949 | rs3851378 | 0.54 | -1.293 | 0.842 | 1.904 | -0.006 | 0.003 | 0.003 | 0.286 | 0.88 | 0.84 | 0.93 | 0.85 | 0.91 | 0.74 |
| 3 | 168840179 | rs16853198 | -0.172 | -1.119 | 0.787 | 1.99 | 0.041 | -0.032 | -0.032 | 0 | 0.14 | 0.03 | 0.07 | 0.16 | 0.08 | 0.15 |
| 3 | 168864643 | rs7633965 | 0.433 | -0.4 | 0.885 | 1.561 | 0.009 | -0.013 | -0.013 | 0 | 0.86 | 0.92 | 0.67 | 0.73 | 0.92 | 0.88 |
| 3 | 168867120 | rs6796432 | -1.684 | 0.309 | 1.715 | 1.467 | 0.021 | 0.018 | 0.018 | 0 | 0.05 | 0.22 | 0.02 | 0.01 | 0.01 | 0.08 |
| 3 | 168874071 | rs13324022 | -1.206 | 0.209 | 1.784 | 1.403 | 0.02 | 0.006 | 0.006 | 0 | 0.08 | 0.39 | 0.05 | 0.02 | 0.02 | 0.12 |
| 3 | 168885028 | rs9809961 | -1.859 | 1.176 | 1.589 | 1.64 | 0.071 | -0.05 | -0.05 | 0 | 0.14 | 0.5 | 0.15 | 0.05 | 0.18 | 0.18 |
| 3 | 168885707 | rs7426660 | -0.036 | 1.011 | 1.372 | 2.158 | 0.011 | -0.008 | -0.008 | 0.252 | 0.09 | 0.08 | 0.06 | 0.3 | 0.06 | 0.12 |
| 3 | 168889604 | rs12629466 | -1.524 | 0.971 | 1.438 | 2.129 | 0.004 | -0.006 | -0.006 | 0 | 0.16 | 0.15 | 0.12 | 0.41 | 0.14 | 0.35 |
| 3 | 168903973 | rs7632657 | -2.139 | 1.06 | 1.439 | 2.127 | 0.05 | -0.002 | -0.002 | 0 | 0.21 | 0.52 | 0.21 | 0.18 | 0.27 | 0.38 |
| 3 | 168904516 | rs6444847 | -2.194 | 1.034 | 1.041 | 1.97 | -0.001 | -0.003 | -0.003 | 0 | 0.3 | 0.72 | 0.29 | 0.5 | 0.33 | 0.52 |
| 3 | 168913273 | rs7652391 | -1.299 | 1.677 | 0.95 | 1.998 | 0.065 | 0.087 | 0.087 | 0 | 0.16 | 0.46 | 0.32 | 0.24 | 0.2 | 0.16 |
| 3 | 168915658 | rs2220393 | -1.131 | 1.848 | 0.675 | 2.075 | -0.012 | -0.036 | -0.036 | 0.179 | 0.17 | 0.1 | 0.22 | 0.32 | 0.22 | 0.17 |
| 3 | 168916240 | rs9864370 | -0.411 | 1.894 | 0.78 | 2.16 | 0.008 | 0.02 | 0.02 | 0 | 0.05 | 0.07 | 0.06 | 0.11 | 0.07 | 0.23 |
| 3 | 168917632 | rs1825886 | 1.734 | 1.867 | 0.464 | 1.744 | -0.003 | -0.001 | -0.001 | 0 | 0.7 | 0.69 | 0.7 | 0.68 | 0.66 | 0.76 |
| 3 | 168924023 | rs3863100 | -0.314 | 1.826 | 0.495 | 1.713 | 0 | -0.015 | -0.015 | 0.176 | 0.89 | 0.87 | 0.93 | 1 | 0.9 | 0.94 |
| 3 | 168932039 | rs2220392 | -0.18 | 1.913 | 0.464 | 1.743 | -0.007 | -0.002 | -0.002 | 0 | 0.09 | 0.06 | 0.09 | 0 | 0.13 | 0.05 |
| 3 | 168932906 | rs16853318 | 0.353 | 1.912 | 0.373 | 1.897 | 0.004 | -0.011 | -0.011 | 2.361 | 0.09 | 0.07 | 0.13 | 0.28 | 0.09 | 0.13 |
| 3 | 168933691 | rs13085351 | 2.38 | 1.913 | 0.385 | 1.514 | 0.036 | 0.07 | 0.07 | 0 | 0.51 | 0.16 | 0.31 | 0.34 | 0.4 | 0.37 |
| 3 | 168942339 | rs6797224 | -1.39 | 0.981 | 0.621 | 1.676 | 0.042 | 0.03 | 0.03 | 0 | 0.17 | 0.48 | 0.33 | 0.23 | 0.21 | 0.14 |
| 3 | 168950999 | rs784280 | -0.257 | 0.47 | 0.667 | 1.592 | -0.017 | -0.069 | -0.069 | 0 | 0.12 | 0.29 | 0.07 | 0 | 0.08 | 0.06 |
| 3 | 168954774 | rs11706512 | 2.192 | 0.58 | 0.457 | 1.6 | -0.012 | 0.057 | 0.057 | 0 | 0.51 | 0.12 | 0.38 | 0.32 | 0.51 | 0.38 |
| 3 | 168959330 | rs1386827 | 2.221 | 0.589 | 0.436 | 1.6 | -0.043 | 0.033 | 0.033 | 0 | 0.5 | 0.06 | 0.37 | 0.33 | 0.5 | 0.38 |
| 3 | 168962600 | rs784281 | 0.114 | 0.525 | 0.579 | 1.616 | -0.01 | -0.047 | -0.047 | 0.492 | 0.11 | 0.17 | 0.07 | 0 | 0.08 | 0.06 |
| 3 | 168963285 | rs1352340 | -0.514 | 0.539 | 0.14 | 1.65 | -0.024 | 0.028 | 0.028 | 0 | 0.11 | 0.28 | 0.25 | 0.14 | 0.06 | 0.08 |
| 3 | 168967606 | rs784285 | 0.891 | 0.776 | -0.456 | 1.397 | 0.001 | 0.001 | 0.001 | 0 | 0.63 | 0.45 | 0.47 | 0.33 | 0.59 | 0.45 |
| 3 | 168972802 | rs998749 | 1.018 | 0.894 | -1.595 | 1.598 | -0.022 | 0.082 | 0.082 | 0 | 0.51 | 0.11 | 0.39 | 0.3 | 0.51 | 0.39 |
| 3 | 168976870 | rs10513664 | -1.466 | 0.912 | -0.902 | 1.756 | -0.027 | -0.026 | -0.026 | 0.065 | 0.06 | 0.01 | 0.05 | 0.11 | 0.08 | 0.08 |
| 3 | 168982572 | rs6444848 | -1.12 | 0.57 | -0.832 | 2.134 | -0.003 | -0.038 | -0.038 | 0.015 | 0.33 | 0.78 | 0.33 | 0.61 | 0.29 | 0.48 |
| 3 | 168983547 | rs4955642 | -1.735 | 0.241 | -0.404 | 2.769 | 0.001 | 0.031 | 0.031 | 0 | 0.1 | 0.09 | 0.11 | 0.31 | 0.12 | 0.23 |
| 3 | 168983985 | rs987254 | -1.578 | -0.135 | -0.504 | 2.612 | 0.014 | -0.047 | -0.047 | 0 | 0.23 | 0.16 | 0.19 | 0.58 | 0.17 | 0.31 |
| 3 | 169001968 | rs3851384 | -1.07 | -0.871 | -0.253 | 2.301 | 0.055 | -0.053 | -0.053 | 0 | 0.16 | 0.01 | 0.14 | 0.22 | 0.07 | 0.2 |
| 3 | 169004819 | rs7634654 | -1.083 | -0.788 | 0.767 | 2.737 | 0.028 | 0.266 | 0.266 | 1.023 | 0.06 | 0.33 | 0.06 | 0.23 | 0.02 | 0.02 |
| 3 | 169008194 | rs9848313 | -2.21 | -1.087 | 1.096 | 2.822 | -0.046 | 0.156 | 0.156 | 5.241 | 0.13 | 0.71 | 0.1 | 0.38 | 0.04 | 0.1 |
| 3 | 169008292 | rs9865969 | -0.828 | -0.963 | 1.16 | 2.737 | 0.016 | 0.061 | 0.061 | 0.266 | 0.06 | 0.25 | 0.05 | 0.08 | 0.02 | 0.01 |
| 3 | 169013588 | rs17251130 | 0.203 | -1.075 | 0.847 | 2.605 | -0.001 | 0 | 0 | 0 | 0.07 | 0.05 | 0.03 | 0.05 | 0.06 | 0.02 |
| 3 | 169017471 | rs7612163 | -1.542 | -1.114 | 0.381 | 2.275 | 0.057 | 0.291 | 0.291 | 0 | 0.5 | 0.85 | 0.64 | 0.87 | 0.42 | 0.47 |
| 3 | 169024814 | rs6806711 | -0.686 | -1.129 | 0.488 | 2.346 | -0.017 | 0.023 | 0.023 | 0.136 | 0.12 | 0.22 | 0.08 | 0.13 | 0.08 | 0.02 |
| 3 | 169031754 | rs16853481 | -1.656 | -1.298 | 0.178 | 2.525 | -0.06 | 0.251 | 0.251 | 0.377 | 0.18 | 0.46 | 0.1 | 0.47 | 0.09 | 0.11 |
| 3 | 169032006 | rs16853486 | -0.276 | -1.018 | 0.021 | 2.59 | -0.001 | 0.043 | 0.043 | 0.525 | 0.09 | 0.09 | 0.04 | 0.34 | 0.06 | 0.09 |
| 3 | 169033482 | rs6780761 | -0.364 | -0.946 | 0.214 | 2.28 | 0 | -0.001 | -0.001 | 1.889 | 0.33 | 0.31 | 0.54 | 0.39 | 0.33 | 0.36 |
| 3 | 169035053 | rs12632175 | -0.334 | -0.579 | 0.013 | 2.057 | -0.002 | 0.007 | 0.007 | 1.873 | 0.08 | 0.07 | 0.04 | 0.19 | 0.06 | 0.08 |
| 3 | 169040917 | rs17679846 | -0.924 | -0.304 | 0.431 | 1.997 | 0.002 | 0.006 | 0.006 | 0 | 0.4 | 0.6 | 0.62 | 0.78 | 0.41 | 0.44 |
| 3 | 169042352 | rs9849779 | -0.195 | -0.24 | 0.46 | 2.575 | -0.003 | 0.014 | 0.014 | 0 | 0.15 | 0.24 | 0.4 | 0.29 | 0.16 | 0.21 |

| | | | | | | | | | | | | | | | | |
|---|---|---|---|---|---|---|---|---|---|---|---|---|---|---|---|---|
| 3 | 169042446 | rs9869781 | -1.323 | -0.191 | 0.566 | 2.844 | -0.001 | 0.081 | 0.081 | 0.137 | 0.22 | 0.46 | 0.51 | 0.6 | 0.25 | 0.28 |
| 3 | 169042979 | rs9854404 | -0.208 | -0.182 | 0.39 | 2.792 | 0.003 | -0.025 | -0.025 | 0 | 0.15 | 0.1 | 0.39 | 0.3 | 0.16 | 0.21 |
| 3 | 169063121 | rs1879796 | -1.504 | -0.284 | 0.383 | 2.728 | -0.011 | 0.081 | 0.081 | 1.399 | 0.27 | 0.61 | 0.53 | 0.63 | 0.27 | 0.28 |
| 3 | 169071370 | rs13324208 | -1.194 | -0.39 | 0.312 | 2.856 | -0.003 | 0.002 | 0.002 | 0 | 0.35 | 0.46 | 0.57 | 0.43 | 0.36 | 0.35 |
| 3 | 169079193 | rs1430435 | -1.925 | 0.42 | 0.296 | 2.786 | 0.002 | -0.006 | -0.006 | 0 | 0.43 | 0.64 | 0.67 | 0.71 | 0.5 | 0.43 |
| 3 | 169080185 | rs9883650 | -1.119 | 0.422 | 0.306 | 2.819 | 0.032 | 0.114 | 0.114 | 0 | 0.05 | 0.14 | 0.09 | 0.23 | 0.09 | 0.03 |
| 3 | 169081584 | rs7617754 | -0.999 | 0.419 | 0.286 | 2.822 | 0.009 | 0.09 | 0.09 | 0.682 | 0.23 | 0.48 | 0.53 | 0.49 | 0.26 | 0.24 |
| 3 | 169082175 | rs6763243 | -1.247 | 0.457 | 0.327 | 2.811 | 0.001 | -0.002 | -0.002 | 0 | 0.35 | 0.34 | 0.56 | 0.43 | 0.36 | 0.35 |
| 3 | 169082633 | rs6774494 | -1.667 | 0.405 | 0.313 | 2.8 | 0.004 | 0.055 | 0.055 | 1.11 | 0.3 | 0.63 | 0.56 | 0.6 | 0.32 | 0.28 |
| 3 | 169083095 | rs4955568 | -0.741 | 0.318 | 0.055 | 2.598 | -0.014 | 0.008 | 0.008 | 0.32 | 0.22 | 0.33 | 0.44 | 0.32 | 0.18 | 0.21 |
| 3 | 169084827 | rs756644 | -1.076 | 0.427 | 0.047 | 2.374 | 0.004 | 0.01 | 0.01 | 0 | 0.37 | 0.49 | 0.64 | 0.59 | 0.44 | 0.38 |
| 3 | 169091584 | rs1290791 | -0.582 | 0.621 | -0.99 | 1.814 | -0.094 | -0.073 | -0.073 | 0.283 | 0.77 | 0.97 | 0.85 | 0.78 | 0.76 | 0.71 |
| 3 | 169098791 | rs988397 | 1.001 | 0.557 | -0.404 | 1.761 | 0.012 | 0.042 | 0.042 | 0 | 0.52 | 0.55 | 0.36 | 0.15 | 0.45 | 0.4 |
| 3 | 169098992 | rs7622799 | -0.708 | 0.655 | -0.041 | 2.205 | 0.061 | 0.067 | 0.067 | 0 | 0.05 | 0.14 | 0.33 | 0.16 | 0.12 | 0.08 |
| 3 | 169100117 | rs16853620 | -1.248 | 0.653 | -0.056 | 2.261 | 0.016 | 0.086 | 0.086 | 0 | 0.23 | 0.4 | 0.48 | 0.58 | 0.3 | 0.29 |
| 3 | 169103185 | rs1835374 | -0.955 | 0.641 | -0.035 | 2.359 | 0.021 | 0.071 | 0.071 | 0 | 0.21 | 0.32 | 0.48 | 0.51 | 0.3 | 0.29 |
| 3 | 169118660 | rs9814713 | -0.567 | 0.187 | 0.112 | 2.148 | -0.032 | 0.042 | 0.042 | 0.265 | 0.08 | 0.18 | 0.03 | 0.15 | 0.01 | 0.02 |
| 3 | 169128274 | rs1918966 | -1.443 | 0.278 | -0.02 | 1.941 | -0.001 | -0.039 | -0.039 | 0.011 | 0.48 | 0.58 | 0.65 | 0.89 | 0.53 | 0.55 |
| 3 | 169139890 | rs1918969 | -1.094 | 0.466 | -0.038 | 2.063 | -0.017 | 0.023 | 0.023 | 0 | 0.38 | 0.36 | 0.58 | 0.74 | 0.45 | 0.44 |
| 3 | 169146497 | rs2014590 | -1.427 | 0.504 | -0.069 | 2.05 | 0.01 | -0.041 | -0.041 | 0 | 0.42 | 0.55 | 0.61 | 0.82 | 0.48 | 0.46 |
| 3 | 169148435 | rs9637451 | 0.829 | 0.378 | -0.563 | 2.197 | -0.077 | -0.07 | -0.07 | 0.903 | 0.14 | 0.04 | 0.15 | 0.22 | 0.25 | 0.18 |
| 3 | 169156443 | rs16853741 | -0.557 | 0.249 | -0.529 | 2.283 | 0.019 | -0.078 | -0.078 | 0 | 0.16 | 0.08 | 0.37 | 0.48 | 0.14 | 0.19 |
| 3 | 169171699 | rs4955660 | 0.352 | -0.053 | -0.382 | 1.855 | 0.007 | 0.012 | 0.012 | 0 | 0.07 | 0.07 | 0.29 | 0.27 | 0.02 | 0.02 |
| 3 | 169172788 | rs4245909 | -1.588 | -0.01 | -0.363 | 1.736 | -0.003 | 0.008 | 0.008 | 0 | 0.45 | 0.49 | 0.63 | 0.81 | 0.51 | 0.53 |
| 3 | 169177924 | rs9850919 | -0.014 | -0.014 | -1.055 | 1.724 | -0.087 | -0.055 | -0.055 | 0 | 0.7 | 0.9 | 0.76 | 0.63 | 0.6 | 0.6 |
| 3 | 169179876 | rs1918975 | -0.028 | 0.031 | -1.244 | 1.724 | -0.133 | -0.096 | -0.096 | 0 | 0.7 | 0.96 | 0.77 | 0.62 | 0.6 | 0.6 |
| 3 | 169183443 | rs17559919 | 0.207 | 0.132 | -0.9 | 1.33 | 0 | -0.005 | -0.005 | 0 | 0.08 | 0.05 | 0.05 | 0.09 | 0.06 | 0.09 |
| 3 | 169188577 | rs3851386 | -0.632 | -0.002 | -0.433 | 1.098 | -0.029 | 0.149 | 0.149 | 0.152 | 0.14 | 0.45 | 0.38 | 0.29 | 0.09 | 0.09 |
| 3 | 169192040 | rs6809660 | 0.518 | 0.029 | -0.941 | 0.119 | 0.007 | 0.043 | 0.043 | 0 | 0.48 | 0.38 | 0.34 | 0.21 | 0.46 | 0.45 |
| 3 | 169195349 | rs2178175 | 0.027 | -0.8 | -1.457 | -0.197 | 0.049 | 0.081 | 0.081 | 4.465 | 0.83 | 0.97 | 0.87 | 0.95 | 0.89 | 0.94 |
| 3 | 169196978 | rs9820408 | -0.14 | -0.994 | -1.102 | -0.674 | -0.011 | -0.058 | -0.058 | 0 | 0.14 | 0.19 | 0.05 | 0 | 0.11 | 0.11 |
| 3 | 169198024 | rs7652987 | 0.988 | -1.273 | -1.025 | -0.416 | -0.003 | -0.015 | -0.015 | 0 | 0.4 | 0.31 | 0.53 | 0.48 | 0.51 | 0.56 |
| 3 | 169209276 | rs9867896 | -0.239 | -1.471 | -1.358 | -0.348 | 0.024 | 0.065 | 0.065 | 1.553 | 0.61 | 0.91 | 0.65 | 0.74 | 0.66 | 0.74 |
| 3 | 169209897 | rs6783972 | -0.015 | -0.997 | -0.569 | -0.256 | -0.021 | -0.079 | -0.079 | 0 | 0.16 | 0.26 | 0.05 | 0 | 0.11 | 0.11 |
| 3 | 169211767 | rs1918964 | -0.212 | -0.302 | -0.718 | -0.226 | -0.031 | 0.052 | 0.052 | 0 | 0.34 | 0.22 | 0.25 | 0.23 | 0.42 | 0.23 |
| 3 | 169212367 | rs7638569 | -0.335 | -0.707 | 0.067 | -0.155 | -0.02 | -0.015 | -0.015 | 0 | 0.88 | 0.5 | 0.85 | 0.91 | 0.91 | 0.94 |
| 3 | 169213004 | rs12634968 | -0.241 | -0.99 | -0.048 | -0.468 | -0.022 | 0.074 | 0.074 | 0 | 0.36 | 0.3 | 0.3 | 0.21 | 0.44 | 0.23 |
| 3 | 169213559 | rs6797464 | -0.1 | -1.233 | 0.413 | -0.399 | 0.018 | -0.012 | -0.012 | 0.118 | 0.92 | 0.67 | 0.96 | 0.98 | 0.94 | 0.94 |
| 3 | 169216068 | rs5004899 | 0.653 | -1.187 | 0.206 | 0.161 | 0.004 | 0.034 | 0.034 | 0 | 0.06 | 0.06 | 0.09 | 0.33 | 0.03 | 0.02 |
| 3 | 169221557 | rs1881305 | -0.564 | -1.474 | 0.444 | -0.062 | 0.018 | -0.018 | -0.018 | 0 | 0.2 | 0.22 | 0.14 | 0.35 | 0.15 | 0.17 |
| 3 | 169227560 | rs9815003 | -0.492 | -1.317 | 0.566 | -0.183 | -0.011 | 0.049 | 0.049 | 0 | 0.38 | 0.7 | 0.31 | 0.21 | 0.43 | 0.27 |
| 3 | 169233179 | rs7613029 | -0.118 | -1.294 | 0.671 | -0.17 | 0.001 | 0.006 | 0.006 | 0 | 0.3 | 0.62 | 0.36 | 0.52 | 0.34 | 0.13 |
| 3 | 169238015 | rs1997280 | -1.068 | -1.507 | 0.154 | -0.154 | -0.006 | 0.099 | 0.099 | 0 | 0.56 | 0.84 | 0.62 | 0.75 | 0.6 | 0.47 |
| 3 | 169242142 | rs1343034 | -0.649 | -1.415 | 0.729 | -0.62 | 0 | 0.061 | 0.061 | 0 | 0.44 | 0.68 | 0.34 | 0.19 | 0.46 | 0.28 |
| 3 | 169244024 | rs9876326 | -0.273 | -1.397 | 0.745 | -0.607 | -0.003 | 0.01 | 0.01 | 0 | 0.39 | 0.63 | 0.31 | 0.17 | 0.43 | 0.28 |
| 3 | 169249745 | rs3909247 | 0.604 | -1.051 | 0.482 | -0.464 | -0.01 | 0.013 | 0.013 | 0 | 0.26 | 0.23 | 0.27 | 0.16 | 0.33 | 0.12 |
| 3 | 169253993 | rs6767836 | 0.507 | -0.91 | 0.349 | -0.357 | 0.025 | -0.04 | -0.04 | 0.168 | 0.27 | 0.4 | 0.29 | 0.16 | 0.33 | 0.13 |
| 3 | 169257233 | rs1362772 | 0.053 | -1.128 | -0.484 | -0.214 | 0.003 | -0.025 | -0.025 | 0 | 0.24 | 0.09 | 0.29 | 0.68 | 0.21 | 0.3 |
| 3 | 169259913 | rs16854053 | 1.766 | -1.146 | -0.516 | 0.183 | 0.005 | -0.038 | -0.038 | 0 | 0.08 | 0 | 0.04 | 0.33 | 0.05 | 0.12 |
| 3 | 169262998 | rs12631412 | 1.254 | -1.205 | -0.217 | -0.026 | 0.019 | -0.016 | -0.016 | 0 | 0.22 | 0.26 | 0.32 | 0.17 | 0.28 | 0.15 |
| 3 | 169263506 | rs7626930 | 0.997 | -0.9 | -0.823 | 0.256 | -0.006 | -0.036 | -0.036 | 0 | 0.15 | 0.08 | 0.25 | 0.34 | 0.15 | 0.18 |
| 3 | 169267796 | rs1421012 | 0.49 | -0.959 | -0.714 | 0.214 | 0.002 | -0.027 | -0.027 | 0 | 0.18 | 0.12 | 0.26 | 0.36 | 0.16 | 0.22 |
| 3 | 169272121 | rs11713381 | 0.05 | -0.856 | -0.643 | -0.328 | 0.015 | -0.006 | -0.006 | 0 | 0.26 | 0.12 | 0.3 | 0.69 | 0.21 | 0.34 |
| 3 | 169283495 | rs3863105 | -0.621 | -0.686 | -0.889 | -0.314 | -0.073 | 0.087 | 0.087 | 0 | 0.73 | 0.96 | 0.79 | 0.88 | 0.67 | 0.6 |
| 3 | 169302430 | rs6774008 | -0.627 | -0.565 | -0.459 | -0.36 | 0.003 | -0.027 | -0.027 | 0 | 0.8 | 0.76 | 0.84 | 0.94 | 0.75 | 0.88 |
| 3 | 169304895 | rs7631725 | -0.084 | -0.55 | -0.048 | -0.153 | 0.03 | 0.011 | 0.011 | 0 | 0.09 | 0.32 | 0.13 | 0.1 | 0.08 | 0.05 |
| 3 | 169307123 | rs6791154 | 0.695 | -0.766 | 0.235 | -0.419 | 0.002 | 0.051 | 0.051 | 0 | 0.51 | 0.34 | 0.44 | 0.2 | 0.5 | 0.44 |
| 3 | 169308593 | rs10513677 | 0.994 | -1.138 | 0.091 | -0.669 | -0.024 | -0.061 | -0.061 | 0.211 | 0.16 | 0.18 | 0.09 | 0.03 | 0.1 | 0.14 |
| 3 | 169309227 | rs7635759 | 1.193 | -1.064 | 0.068 | -0.621 | 0.03 | 0.013 | 0.013 | 0 | 0.08 | 0.24 | 0.13 | 0.1 | 0.08 | 0.05 |
| 3 | 169312608 | rs9647377 | 1.778 | -1.123 | -0.205 | 0.063 | -0.02 | -0.056 | -0.056 | 0.721 | 0.12 | 0.03 | 0.18 | 0.25 | 0.12 | 0.23 |
| 3 | 169316020 | rs6803541 | -0.433 | -1.298 | -0.129 | 0.137 | -0.004 | 0.002 | 0.002 | 0 | 0.34 | 0.28 | 0.34 | 0.57 | 0.29 | 0.49 |
| 3 | 169320058 | rs11716015 | -0.504 | -1.305 | 0.02 | -0.044 | 0.013 | -0.035 | -0.035 | 1.344 | 0.69 | 0.54 | 0.69 | 0.76 | 0.65 | 0.74 |
| 3 | 169322958 | rs6763377 | 1.298 | -1.212 | -0.067 | 0.093 | -0.005 | 0.049 | 0.049 | 0 | 0.35 | 0.2 | 0.35 | 0.19 | 0.36 | 0.24 |
| 3 | 169325234 | rs2032701 | 1.398 | -1.009 | -0.155 | 0.3 | -0.018 | -0.066 | -0.066 | 0 | 0.13 | 0.03 | 0.18 | 0.26 | 0.13 | 0.25 |
| 3 | 169327132 | rs1895030 | 0.685 | -1.073 | -0.027 | 0.196 | -0.004 | 0.013 | 0.013 | 0 | 0.45 | 0.46 | 0.5 | 0.3 | 0.49 | 0.37 |
| 3 | 169327436 | rs6762435 | 0.588 | -1.078 | 0.046 | 0.215 | -0.009 | 0.041 | 0.041 | 0 | 0.19 | 0.24 | 0.22 | 0.27 | 0.16 | 0.25 |
| 3 | 169330644 | rs7626686 | 0.306 | -0.738 | -0.08 | 0.374 | -0.004 | -0.004 | -0.004 | 1.11 | 0.46 | 0.46 | 0.53 | 0.57 | 0.49 | 0.41 |
| 3 | 169343223 | rs6804374 | 0.382 | -0.5 | -0.426 | 0.506 | 0.001 | 0.002 | 0.002 | 0 | 0.41 | 0.58 | 0.4 | 0.44 | 0.42 | 0.34 |
| 3 | 169344717 | rs933605 | -0.549 | -0.402 | -0.838 | 0.576 | -0.006 | 0.018 | 0.018 | 0 | 0.68 | 0.74 | 0.67 | 0.74 | 0.65 | 0.72 |

| | | | | | | | | | | | | | | | | |
|---|---|---|---|---|---|---|---|---|---|---|---|---|---|---|---|---|
| 3 | 169347832 | rs7625357 | 0.676 | 0.069 | -1.01 | 0.735 | -0.007 | 0.022 | 0.022 | 0 | 0.3 | 0.25 | 0.28 | 0.18 | 0.3 | 0.23 |
| 3 | 169348816 | rs16854236 | -1.026 | 0.139 | -0.796 | 1.473 | -0.001 | 0.058 | 0.058 | 0 | 0.05 | 0.07 | 0.08 | 0.25 | 0.07 | 0.1 |
| 3 | 169355082 | rs4955666 | -0.814 | 0.048 | -1.196 | 0.827 | 0.003 | 0.031 | 0.031 | 0.056 | 0.42 | 0.83 | 0.54 | 0.59 | 0.4 | 0.58 |
| 3 | 169360455 | rs16854266 | -0.211 | -0.262 | -1.088 | 1.013 | -0.046 | -0.112 | -0.112 | 0.07 | 0.27 | 0.07 | 0.38 | 0.44 | 0.34 | 0.27 |
| 3 | 169366536 | rs2421748 | -1.57 | -0.566 | -0.748 | 0.814 | 0.017 | 0.021 | 0.021 | 0.255 | 0.45 | 0.82 | 0.5 | 0.52 | 0.42 | 0.58 |
| 3 | 169372156 | rs7644900 | -1.273 | -0.684 | 0.29 | 0.611 | -0.047 | -0.1 | -0.1 | 0.425 | 0.16 | 0.62 | 0.08 | 0 | 0.07 | 0.28 |
| 3 | 169379797 | rs9874200 | -1.376 | -0.725 | 0.566 | 0.561 | -0.05 | -0.017 | -0.017 | 0 | 0.16 | 0.62 | 0.11 | 0.08 | 0.07 | 0.15 |
| 3 | 169396070 | rs4955669 | -1.602 | -0.277 | 0.272 | 0.784 | 0 | -0.006 | -0.006 | 0.189 | 0.45 | 0.67 | 0.49 | 0.49 | 0.43 | 0.57 |
| 3 | 169406484 | rs10936590 | -0.375 | -0.372 | 0.251 | 0.68 | -0.038 | -0.056 | -0.056 | 0 | 0.3 | 0.1 | 0.38 | 0.37 | 0.34 | 0.35 |
| 3 | 169411792 | rs1421018 | -1.479 | -0.371 | 0.866 | 0.68 | -0.001 | -0.004 | -0.004 | 0.678 | 0.43 | 0.5 | 0.44 | 0.38 | 0.42 | 0.54 |
| 3 | 169416569 | rs16854319 | -0.249 | -0.457 | 0.435 | 0.894 | -0.028 | -0.081 | -0.081 | 0 | 0.27 | 0.05 | 0.34 | 0.37 | 0.3 | 0.22 |
| 3 | 169420117 | rs1997397 | -1.127 | -0.409 | 0.61 | 0.87 | -0.001 | 0.001 | 0.001 | 0.023 | 0.38 | 0.62 | 0.4 | 0.45 | 0.33 | 0.42 |
| 3 | 169442363 | rs9880534 | -1.176 | -0.385 | 0.479 | 0.877 | 0.002 | -0.012 | -0.012 | 0.019 | 0.39 | 0.65 | 0.41 | 0.45 | 0.34 | 0.54 |
| 3 | 169461571 | rs16854335 | -0.506 | -0.461 | 0.375 | 0.924 | -0.007 | -0.026 | -0.026 | 0 | 0.31 | 0.18 | 0.31 | 0.4 | 0.32 | 0.26 |
| 3 | 169477506 | rs12638862 | -0.832 | -0.89 | 0.462 | 0.75 | 0.019 | -0.045 | -0.045 | 0 | 0.33 | 0.13 | 0.37 | 0.59 | 0.25 | 0.26 |
| 3 | 169480204 | rs12630450 | -1.273 | -1.452 | 0.986 | 0.533 | 0.004 | 0.004 | 0.004 | 0 | 0.41 | 0.41 | 0.43 | 0.68 | 0.26 | 0.37 |
| 3 | 169486144 | rs9822885 | -1.277 | -1.458 | 0.965 | 0.534 | 0.009 | 0.009 | 0.009 | 0 | 0.41 | 0.39 | 0.43 | 0.68 | 0.26 | 0.39 |
| 3 | 169489937 | rs35100673 | 0.668 | -1.537 | 0.979 | 0.497 | 0 | 0.026 | 0.026 | 0 | 0.08 | 0.01 | 0.03 | 0.03 | 0.05 | 0.03 |
| 3 | 169489946 | rs3821383 | -1.243 | -1.453 | 1.078 | 0.497 | 0.008 | 0.008 | 0.008 | 0 | 0.41 | 0.39 | 0.43 | 0.68 | 0.27 | 0.36 |
| 3 | 169492101 | rs10936599 | -0.221 | -1.451 | 0.906 | 0.574 | -0.039 | -0.13 | -0.13 | 0 | 0.28 | 0.04 | 0.35 | 0.58 | 0.24 | 0.24 |
| 3 | 169497585 | rs1317082 | -0.221 | -1.472 | 0.934 | 0.575 | -0.039 | -0.127 | -0.127 | 0 | 0.28 | 0.04 | 0.35 | 0.58 | 0.24 | 0.22 |
| 3 | 169508272 | rs13069553 | -0.194 | -1.471 | 0.935 | 0.571 | -0.04 | -0.13 | -0.13 | 5.295 | 0.28 | 0.04 | 0.35 | 0.58 | 0.24 | 0.22 |
| 3 | 169509244 | rs7633750 | -1.454 | -1.551 | 1.257 | 0.536 | 0.003 | 0.003 | 0.003 | 0 | 0.42 | 0.58 | 0.44 | 0.68 | 0.27 | 0.34 |
| 3 | 169512145 | rs7621631 | -0.168 | -1.483 | 1.104 | 0.54 | -0.044 | -0.13 | -0.13 | 1.305 | 0.27 | 0.04 | 0.35 | 0.58 | 0.24 | 0.22 |
| 3 | 169514585 | rs10936600 | -0.219 | -1.533 | 1.131 | 0.543 | -0.02 | -0.104 | -0.104 | 0 | 0.28 | 0.06 | 0.35 | 0.58 | 0.24 | 0.22 |
| 3 | 169528449 | rs10936601 | -1.246 | -1.409 | 1.164 | 0.191 | -0.007 | -0.006 | -0.006 | 0.065 | 0.42 | 0.57 | 0.43 | 0.68 | 0.27 | 0.34 |
| 3 | 169529958 | rs9878797 | -1.269 | -1.366 | 0.909 | -0.406 | 0.009 | 0.067 | 0.067 | 0 | 0.56 | 0.71 | 0.59 | 0.79 | 0.43 | 0.55 |
| 3 | 169536637 | rs10936602 | 0.083 | -1.957 | 1.262 | -0.245 | 0.016 | -0.003 | -0.003 | 0 | 0.33 | 0.23 | 0.38 | 0.66 | 0.25 | 0.22 |
| 3 | 169540397 | rs1920119 | -1.343 | -2.053 | 1.381 | -0.32 | 0.023 | 0.058 | 0.058 | 0 | 0.54 | 0.67 | 0.61 | 0.76 | 0.4 | 0.52 |
| 3 | 169545652 | rs10936603 | -0.013 | -1.815 | 1.372 | -0.144 | 0.017 | -0.002 | -0.002 | 0 | 0.33 | 0.23 | 0.38 | 0.66 | 0.26 | 0.22 |
| 3 | 169553498 | rs9833035 | -0.117 | -1.713 | 0.968 | -0.114 | 0.002 | 0 | 0 | 0 | 0.36 | 0.35 | 0.44 | 0.65 | 0.26 | 0.4 |
| 3 | 169558025 | rs3732451 | -1.111 | -1.779 | 0.974 | -0.157 | -0.002 | -0.01 | -0.01 | 0.256 | 0.51 | 0.55 | 0.6 | 0.75 | 0.39 | 0.51 |
| 3 | 169558821 | rs12489230 | -0.068 | -1.679 | 0.844 | -0.124 | 0.024 | -0.004 | -0.004 | 0.032 | 0.34 | 0.25 | 0.38 | 0.64 | 0.26 | 0.27 |
| 3 | 169565571 | rs13074500 | 0.045 | -1.624 | 0.275 | -0.154 | 0.027 | -0.023 | -0.023 | 0 | 0.33 | 0.14 | 0.37 | 0.64 | 0.26 | 0.25 |
| 3 | 169570241 | rs11709840 | -0.004 | -1.53 | 0.383 | -0.11 | 0.028 | 0.006 | 0.006 | 0 | 0.37 | 0.22 | 0.4 | 0.64 | 0.29 | 0.33 |
| 3 | 169571145 | rs16854469 | 0.927 | -1.536 | 0.353 | -0.124 | 0.001 | 0.014 | 0.014 | 0 | 0.09 | 0.04 | 0.1 | 0.05 | 0.08 | 0.06 |
| 3 | 169574465 | rs11919269 | 0.046 | -1.456 | -0.032 | -0.085 | 0.024 | 0.002 | 0.002 | 0 | 0.37 | 0.2 | 0.4 | 0.64 | 0.29 | 0.31 |
| 3 | 169582223 | rs7647589 | -0.904 | -1.254 | -0.014 | -0.498 | 0.009 | 0.037 | 0.037 | 0 | 0.55 | 0.66 | 0.62 | 0.78 | 0.43 | 0.58 |
| 3 | 169589918 | rs9841073 | -1.333 | -0.643 | -0.328 | -0.455 | -0.002 | -0.011 | -0.011 | 0.027 | 0.29 | 0.69 | 0.3 | 0.13 | 0.24 | 0.29 |
| 3 | 169595485 | rs7614694 | -0.63 | -0.545 | -0.314 | 0.231 | 0.008 | 0.015 | 0.015 | 0 | 0.48 | 0.58 | 0.41 | 0.68 | 0.41 | 0.53 |
| 3 | 169597186 | rs7628292 | -0.98 | -0.267 | -0.022 | 0.513 | -0.026 | 0.012 | 0.012 | 0 | 0.08 | 0.18 | 0.09 | 0.1 | 0.07 | 0.15 |
| 3 | 169606936 | rs7635067 | -0.073 | 0.035 | -0.73 | 0.378 | -0.011 | 0.019 | 0.019 | 0 | 0.43 | 0.31 | 0.39 | 0.67 | 0.46 | 0.32 |
| 3 | 169608873 | rs16854516 | -0.656 | -0.237 | -0.026 | 0.317 | -0.019 | -0.005 | -0.005 | 0 | 0.1 | 0.23 | 0.15 | 0.05 | 0.07 | 0.16 |
| 3 | 169611035 | rs6444879 | 0.275 | 0.027 | -0.175 | 0.021 | -0.005 | -0.007 | -0.007 | 0 | 0.9 | 0.92 | 0.94 | 1 | 0.86 | 0.83 |
| 3 | 169616462 | rs4955686 | 0.274 | 0.045 | 0.179 | 0.39 | 0.017 | -0.003 | -0.003 | 0 | 0.9 | 0.86 | 0.91 | 0.88 | 0.84 | 0.8 |
| 3 | 169617342 | rs9874716 | 0.76 | -0.131 | 0.7 | 0.106 | 0.003 | 0.005 | 0.005 | 0.054 | 0.35 | 0.24 | 0.35 | 0.25 | 0.32 | 0.29 |
| 3 | 169617405 | rs16854522 | 0.595 | -0.296 | 0.905 | 0.263 | -0.022 | -0.021 | -0.021 | 5.634 | 0.44 | 0.28 | 0.42 | 0.55 | 0.48 | 0.38 |
| 3 | 169617842 | rs7633907 | -0.133 | -0.302 | 0.966 | 0.266 | 0.022 | 0.003 | 0.003 | 0 | 0.9 | 0.83 | 0.91 | 0.87 | 0.83 | 0.8 |
| 3 | 169624528 | rs9283638 | 0.17 | -0.359 | 1.183 | 0.223 | 0.004 | 0.004 | 0.004 | 0.812 | 0.43 | 0.35 | 0.45 | 0.3 | 0.35 | 0.4 |
| 3 | 169630589 | rs7613633 | 0.397 | -0.333 | 1.347 | 0.228 | 0.035 | 0.013 | 0.013 | 0.143 | 0.88 | 0.65 | 0.88 | 0.85 | 0.83 | 0.79 |
| 3 | 169633705 | rs9876612 | 0.837 | -0.396 | 1.351 | 0.447 | 0.005 | 0.008 | 0.008 | 0.18 | 0.36 | 0.25 | 0.36 | 0.25 | 0.33 | 0.29 |
| 3 | 169633958 | rs4955695 | 0.664 | -0.39 | 1.207 | 0.448 | -0.057 | -0.047 | -0.047 | 0.013 | 0.43 | 0.19 | 0.42 | 0.55 | 0.48 | 0.39 |
| 3 | 169634639 | rs7614795 | 0.653 | -0.514 | 1.524 | 0.499 | 0.035 | 0.013 | 0.013 | 0.143 | 0.88 | 0.65 | 0.88 | 0.85 | 0.83 | 0.79 |
| 3 | 169643955 | rs1290670 | -1.561 | -0.825 | 1.016 | 0.604 | -0.006 | -0.008 | -0.008 | 0 | 0.3 | 0.36 | 0.22 | 0.17 | 0.25 | 0.31 |
| 3 | 169644710 | rs10513680 | -0.807 | -0.841 | 0.833 | 0.669 | 0.01 | -0.003 | -0.003 | 0 | 0.19 | 0.14 | 0.17 | 0.17 | 0.12 | 0.14 |
| 3 | 169651974 | rs6765033 | -0.808 | -0.82 | 0.915 | 0.706 | -0.009 | 0.001 | 0.001 | 0.841 | 0.15 | 0.15 | 0.11 | 0.16 | 0.08 | 0.09 |
| 3 | 169652186 | rs6790443 | -0.808 | -0.82 | 0.915 | 0.731 | -0.009 | 0.003 | 0.003 | 0.841 | 0.15 | 0.15 | 0.11 | 0.19 | 0.08 | 0.09 |
| 3 | 169654216 | rs9836170 | -0.894 | -0.887 | 0.956 | 0.761 | -0.023 | 0.008 | 0.008 | 1.139 | 0.13 | 0.15 | 0.11 | 0.16 | 0.04 | 0.09 |
| 3 | 169655805 | rs9851636 | -2.129 | -0.994 | 1.767 | 0.961 | -0.037 | 0.028 | 0.028 | 0 | 0.25 | 0.57 | 0.2 | 0.33 | 0.19 | 0.3 |
| 3 | 169659270 | rs960353 | -2.025 | -0.929 | 1.758 | 0.964 | -0.032 | 0.028 | 0.028 | 0 | 0.26 | 0.58 | 0.22 | 0.34 | 0.22 | 0.32 |
| 3 | 169662411 | rs6773121 | -1.157 | -0.96 | 1.806 | 0.964 | -0.027 | 0.117 | 0.117 | 0.773 | 0.11 | 0.45 | 0.13 | 0.16 | 0.04 | 0.06 |
| 3 | 169676977 | rs12107990 | -1.185 | -0.924 | 1.7 | 0.926 | -0.005 | 0.001 | 0.001 | 0.035 | 0.13 | 0.17 | 0.12 | 0.21 | 0.09 | 0.13 |
| 3 | 169679950 | rs6795822 | -2.231 | -0.832 | 1.863 | 0.909 | -0.026 | 0.031 | 0.031 | 0 | 0.25 | 0.59 | 0.22 | 0.34 | 0.24 | 0.34 |
| 3 | 169680664 | rs13315158 | -0.498 | -0.774 | 1.948 | 0.889 | 0.04 | -0.066 | -0.066 | 0 | 0.11 | 0.28 | 0.06 | 0 | 0.15 | 0.17 |
| 3 | 169682202 | rs3755608 | -0.226 | -0.78 | 1.925 | 0.894 | 0.009 | -0.01 | -0.01 | 0 | 0.09 | 0.09 | 0.1 | 0.16 | 0.05 | 0.08 |
| 3 | 169683036 | rs6808317 | -0.697 | -0.724 | 1.945 | 0.86 | 0.062 | -0.07 | -0.07 | 0 | 0.11 | 0.38 | 0.07 | 0 | 0.15 | 0.17 |
| 3 | 169697376 | rs9811291 | -1.357 | -0.766 | 1.842 | 1.087 | -0.006 | 0.048 | 0.048 | 0 | 0.19 | 0.32 | 0.16 | 0.3 | 0.2 | 0.29 |
| 3 | 169698932 | rs3772179 | -1.485 | -0.779 | 1.953 | 1.08 | -0.01 | 0.068 | 0.068 | 0 | 0.21 | 0.54 | 0.18 | 0.3 | 0.2 | 0.29 |
| 3 | 169713483 | rs1132237 | -1.485 | -0.719 | 1.781 | 1.111 | -0.01 | 0.068 | 0.068 | 0 | 0.21 | 0.54 | 0.18 | 0.3 | 0.2 | 0.28 |

| | | | | | | | | | | | | | | | | |
|---|---|---|---|---|---|---|---|---|---|---|---|---|---|---|---|---|
| 3 | 169713732 | rs9416 | -1.404 | -0.719 | 1.787 | 1.11 | -0.006 | 0.083 | 0.083 | 0 | 0.2 | 0.51 | 0.18 | 0.3 | 0.2 | 0.29 |
| 3 | 169714142 | rs3348 | -1.378 | -0.727 | 1.725 | 1.142 | 0 | 0.054 | 0.054 | 0 | 0.19 | 0.54 | 0.13 | 0.26 | 0.2 | 0.28 |
| 3 | 169716639 | rs16854711 | -0.563 | -0.649 | 1.261 | 1.044 | -0.02 | -0.037 | -0.037 | 0 | 0.16 | 0.04 | 0.15 | 0.17 | 0.2 | 0.25 |
| 3 | 169718125 | rs718205 | -0.618 | -0.649 | 1.278 | 1.044 | -0.003 | -0.012 | -0.012 | 0 | 0.17 | 0.12 | 0.15 | 0.17 | 0.2 | 0.25 |
| 3 | 169722804 | rs6804761 | -0.654 | -0.287 | 0.732 | 0.618 | -0.007 | 0.012 | 0.012 | 0.404 | 0.64 | 0.7 | 0.72 | 0.79 | 0.6 | 0.67 |
| 3 | 169723927 | rs2901585 | 0.455 | -0.512 | 0.652 | 0.964 | 0.009 | 0.009 | 0.009 | 0 | 0.47 | 0.41 | 0.39 | 0.39 | 0.37 | 0.35 |
| 3 | 169731027 | rs12636720 | -0.483 | -0.847 | 0.423 | 0.401 | -0.004 | -0.014 | -0.014 | 0 | 0.39 | 0.65 | 0.55 | 0.74 | 0.37 | 0.47 |
| 3 | 169737301 | rs4955705 | 1.071 | -0.788 | 0.078 | 0.501 | 0.012 | 0.02 | 0.02 | 0.078 | 0.08 | 0.19 | 0.15 | 0.13 | 0.08 | 0.05 |
| 3 | 169738278 | rs7651441 | -0.584 | -0.875 | 0.256 | 0.426 | -0.001 | -0.005 | -0.005 | 0.195 | 0.37 | 0.4 | 0.54 | 0.74 | 0.36 | 0.47 |
| 3 | 169741250 | rs9868869 | -0.013 | -1.064 | -0.298 | 0.557 | 0.026 | -0.089 | -0.089 | 0.389 | 0.32 | 0.13 | 0.38 | 0.54 | 0.28 | 0.41 |
| 3 | 169741291 | rs9868633 | -1.266 | -1.336 | -0.076 | 0.345 | 0.011 | 0.019 | 0.019 | 1.622 | 0.48 | 0.64 | 0.58 | 0.75 | 0.38 | 0.47 |
| 3 | 169741925 | rs9818038 | 0.067 | -1.277 | -0.577 | 0.364 | 0.027 | -0.086 | -0.086 | 0.021 | 0.32 | 0.14 | 0.38 | 0.54 | 0.28 | 0.41 |
| 3 | 169756846 | rs2287483 | 1.083 | -1.095 | -0.148 | 0.546 | -0.001 | -0.001 | -0.001 | 0 | 0.17 | 0.16 | 0.18 | 0.16 | 0.1 | 0.04 |
| 3 | 169772965 | rs4955585 | 0.096 | -1.029 | -0.294 | 0.394 | -0.013 | -0.134 | -0.134 | 0.009 | 0.34 | 0.09 | 0.39 | 0.51 | 0.34 | 0.44 |
| 3 | 169774383 | rs16854812 | 1.469 | -1.022 | -0.132 | 0.335 | -0.006 | 0.001 | 0.001 | 1.694 | 0.16 | 0.21 | 0.19 | 0.16 | 0.1 | 0.04 |
| 3 | 169774821 | rs6444891 | -1.286 | -0.984 | -0.674 | -0.098 | -0.003 | 0.083 | 0.083 | 0 | 0.85 | 0.98 | 0.89 | 0.98 | 0.83 | 0.91 |
| 3 | 169777735 | rs6787395 | 0.097 | -0.891 | -0.458 | 0.201 | 0.017 | -0.017 | -0.017 | 0 | 0.37 | 0.3 | 0.4 | 0.6 | 0.34 | 0.45 |
| 3 | 169782204 | rs6804193 | -0.464 | -0.937 | -0.366 | 0.227 | 0.009 | 0.02 | 0.02 | 1.696 | 0.51 | 0.46 | 0.55 | 0.7 | 0.43 | 0.46 |
| 3 | 169784616 | rs6798945 | -0.203 | -0.926 | -0.529 | 0.219 | 0.011 | 0.076 | 0.076 | 1.371 | 0.48 | 0.19 | 0.53 | 0.7 | 0.43 | 0.46 |
| 3 | 169785660 | rs2287481 | 0.414 | -0.922 | -0.61 | 0.265 | -0.013 | -0.125 | -0.125 | 0.076 | 0.34 | 0.09 | 0.35 | 0.5 | 0.32 | 0.42 |
| 3 | 169801658 | rs6444895 | -0.344 | -0.916 | -0.399 | 0.226 | 0.015 | 0.032 | 0.032 | 0.383 | 0.49 | 0.44 | 0.57 | 0.7 | 0.43 | 0.45 |
| 3 | 169802930 | rs1861935 | 0.343 | -0.916 | -0.391 | 0.208 | 0.009 | -0.046 | -0.046 | 0.021 | 0.35 | 0.24 | 0.38 | 0.49 | 0.32 | 0.4 |
| 3 | 169806170 | rs7615039 | 2.219 | -0.896 | -0.367 | 0.223 | 0.001 | 0.006 | 0.006 | 3.047 | 0.13 | 0.09 | 0.18 | 0.1 | 0.12 | 0.04 |
| 3 | 169807839 | rs6804888 | -0.447 | -0.924 | 0.099 | 0.259 | 0.008 | 0.015 | 0.015 | 0.906 | 0.5 | 0.59 | 0.58 | 0.7 | 0.43 | 0.45 |
| 3 | 169808114 | rs10936611 | 2.26 | -0.62 | -0.332 | -0.069 | -0.114 | -0.036 | -0.036 | 5.084 | 0.24 | 0.03 | 0.22 | 0.22 | 0.31 | 0.35 |
| 3 | 169812115 | rs879161 | 2.151 | -0.517 | -0.141 | 0.098 | -0.004 | -0.008 | -0.008 | 0 | 0.14 | 0.21 | 0.19 | 0.1 | 0.12 | 0.04 |
| 3 | 169813340 | rs7643249 | -0.034 | -0.444 | -0.086 | 0.043 | -0.002 | -0.004 | -0.004 | 0.064 | 0.49 | 0.51 | 0.5 | 0.68 | 0.44 | 0.45 |
| 3 | 169832116 | rs6444898 | 0.565 | -0.655 | -0.055 | 0.117 | 0.012 | -0.048 | -0.048 | 0.233 | 0.34 | 0.24 | 0.38 | 0.49 | 0.31 | 0.4 |
| 3 | 169838000 | rs7638591 | 0.146 | -0.668 | -0.013 | 0.103 | 0.016 | 0.017 | 0.017 | 0.423 | 0.47 | 0.28 | 0.55 | 0.6 | 0.43 | 0.44 |
| 3 | 169857007 | rs10804838 | -0.729 | -0.692 | 0.364 | 0.049 | 0.012 | 0.022 | 0.022 | 0 | 0.94 | 0.91 | 0.95 | 0.93 | 0.91 | 0.96 |
| 3 | 169861358 | rs6775783 | -0.231 | -0.809 | 0.354 | -0.051 | -0.002 | 0.006 | 0.006 | 0 | 0.59 | 0.59 | 0.57 | 0.51 | 0.62 | 0.57 |
| 3 | 169887114 | rs1566426 | 0.493 | -1.193 | -0.208 | 0.115 | 0.011 | -0.031 | -0.031 | 0 | 0.35 | 0.28 | 0.38 | 0.5 | 0.32 | 0.4 |
| 3 | 169902079 | rs17200223 | 0.197 | -1.1 | -0.598 | 0.154 | 0.001 | 0.003 | 0.003 | 0.577 | 0.46 | 0.17 | 0.54 | 0.6 | 0.43 | 0.44 |
| 3 | 169908121 | rs12492192 | 0.103 | -1.053 | -0.571 | 0.154 | 0.01 | 0.057 | 0.057 | 0.102 | 0.47 | 0.28 | 0.56 | 0.72 | 0.44 | 0.45 |
| 3 | 169918311 | rs1625949 | 0.522 | -0.807 | -0.989 | 0.262 | -0.035 | -0.137 | -0.137 | 0 | 0.32 | 0.08 | 0.38 | 0.51 | 0.33 | 0.39 |
| 3 | 169925664 | rs1705589 | 0.262 | -0.353 | -0.224 | 0.613 | -0.054 | 0.093 | 0.093 | 0.001 | 0.82 | 0.96 | 0.83 | 0.98 | 0.78 | 0.92 |
| 3 | 169933134 | rs1705603 | 0.29 | -0.817 | 0.266 | 1.049 | 0 | 0.008 | 0.008 | 0.445 | 0.54 | 0.63 | 0.61 | 0.76 | 0.48 | 0.51 |
| 3 | 169938385 | rs1082977 | 0.863 | -0.686 | -0.195 | 1.111 | -0.039 | -0.027 | -0.027 | 0 | 0.47 | 0.11 | 0.55 | 0.59 | 0.46 | 0.46 |
| 3 | 169938890 | rs1082976 | 0.973 | -0.704 | -0.212 | 1.103 | -0.067 | -0.054 | -0.054 | 0.1 | 0.46 | 0.07 | 0.55 | 0.59 | 0.46 | 0.46 |
| 3 | 169944100 | rs546950 | 0.82 | -0.672 | -0.187 | 1.104 | -0.033 | -0.046 | -0.046 | 0.109 | 0.47 | 0.12 | 0.48 | 0.56 | 0.46 | 0.46 |
| 3 | 169947906 | rs1082969 | 0.912 | -0.103 | 0.578 | 1.3 | 0.001 | -0.005 | -0.005 | 0 | 0.82 | 0.68 | 0.83 | 0.88 | 0.82 | 0.89 |
| 3 | 169958552 | rs1684885 | 0.525 | -0.113 | 0.603 | 1.295 | 0 | -0.001 | -0.001 | 0 | 0.84 | 0.84 | 0.84 | 0.89 | 0.81 | 0.89 |
| 3 | 169966100 | rs516741 | -2.137 | 0.002 | 0.843 | 1.875 | 0.021 | 0.084 | 0.084 | 0 | 0.24 | 0.54 | 0.38 | 0.36 | 0.26 | 0.3 |
| 3 | 169972081 | rs2650229 | -1.414 | -0.727 | 0.955 | 1.708 | 0.013 | 0.041 | 0.041 | 0 | 0.18 | 0.43 | 0.2 | 0.19 | 0.2 | 0.12 |
| 3 | 169990410 | rs2140825 | 0.906 | -0.819 | 0.732 | 1.594 | -0.126 | -0.124 | -0.124 | 0 | 0.29 | 0.06 | 0.44 | 0.45 | 0.4 | 0.44 |
| 3 | 170010234 | rs1039929 | -0.89 | -0.397 | 0.562 | 1.324 | 0.001 | -0.001 | -0.001 | 1.755 | 0.16 | 0.18 | 0.18 | 0.11 | 0.19 | 0.11 |
| 3 | 170022495 | rs2650220 | -0.182 | -0.217 | 0.194 | 1.247 | -0.053 | 0.041 | 0.041 | 0 | 0.1 | 0.01 | 0.16 | 0.02 | 0.19 | 0.08 |
| 3 | 170032677 | rs1997061 | 0.347 | -0.174 | 0.54 | 1.54 | 0.005 | -0.001 | -0.001 | 0 | 0.36 | 0.33 | 0.5 | 0.39 | 0.42 | 0.47 |
| 3 | 170061756 | rs6444914 | -0.756 | -0.862 | 0.767 | 1.518 | 0.043 | -0.014 | -0.014 | 0 | 0.53 | 0.85 | 0.59 | 0.48 | 0.62 | 0.75 |
| 3 | 170062936 | rs17826519 | 0.673 | -0.834 | 1.007 | 1.14 | -0.06 | 0.068 | 0.068 | 5.027 | 0.35 | 0.19 | 0.43 | 0.25 | 0.43 | 0.4 |
| 3 | 170064776 | rs9875109 | -1.46 | -0.914 | 1.284 | 1.188 | -0.034 | 0.013 | 0.013 | 1.273 | 0.18 | 0.6 | 0.15 | 0.15 | 0.15 | 0.3 |
| 3 | 170072864 | rs4955588 | 0.807 | -0.889 | 1.522 | 1.193 | -0.017 | -0.046 | -0.046 | 0 | 0.84 | 0.57 | 0.88 | 0.94 | 0.86 | 0.71 |
| 3 | 170077713 | rs3772172 | 1.284 | -1.135 | 1.613 | 1.175 | -0.034 | -0.002 | -0.002 | 0.188 | 0.79 | 0.55 | 0.83 | 0.84 | 0.82 | 0.67 |
| 3 | 170078232 | rs3772173 | 1.37 | -1.137 | 1.624 | 1.174 | -0.04 | -0.004 | -0.004 | 0.08 | 0.78 | 0.43 | 0.82 | 0.84 | 0.82 | 0.67 |
| 3 | 170091902 | rs9682041 | 0.681 | -1.111 | 1.502 | 0.957 | -0.017 | -0.022 | -0.022 | 0 | 0.86 | 0.71 | 0.89 | 0.94 | 0.88 | 0.84 |
| 3 | 170105378 | rs6786542 | 1.148 | -1.097 | 1.335 | 0.461 | -0.035 | -0.064 | -0.064 | 2.159 | 0.79 | 0.56 | 0.85 | 0.93 | 0.83 | 0.68 |
| 3 | 170128750 | rs13096001 | 0.537 | -1.034 | 0.641 | 0.481 | 0.043 | 0.05 | 0.05 | 0.043 | 0.46 | 0.36 | 0.45 | 0.67 | 0.35 | 0.3 |
| 3 | 170133639 | rs6782252 | -0.032 | -0.91 | 0.556 | 0.474 | -0.004 | -0.056 | -0.056 | 0 | 0.89 | 0.85 | 0.93 | 0.99 | 0.88 | 0.89 |
| 3 | 170136084 | rs11711130 | 0.865 | -0.981 | -0.141 | 0.725 | 0.022 | -0.007 | -0.007 | 0 | 0.05 | 0 | 0.07 | 0.22 | 0.02 | 0.01 |
| 3 | 170140792 | rs1477211 | 0.855 | -1.014 | 0.046 | 0.427 | -0.027 | -0.14 | -0.14 | 0 | 0.81 | 0.53 | 0.81 | 0.99 | 0.82 | 0.71 |
| 3 | 170147991 | rs6794467 | 0.49 | -0.422 | -0.475 | 0.895 | -0.001 | -0.001 | -0.001 | 0 | 0.36 | 0.31 | 0.36 | 0.32 | 0.49 | 0.45 |
| 3 | 170150076 | rs6444929 | 0.096 | -0.125 | -0.223 | 0.531 | 0 | -0.01 | -0.01 | 0.07 | 0.9 | 0.87 | 0.92 | 0.99 | 0.88 | 0.87 |
| 3 | 170158128 | rs10936633 | 0.123 | 0.079 | -0.363 | 0.786 | -0.003 | -0.007 | -0.007 | 0 | 0.41 | 0.32 | 0.42 | 0.52 | 0.5 | 0.47 |
| 3 | 170160493 | rs1035298 | 0.053 | -0.151 | -0.248 | 0.875 | -0.004 | 0 | 0 | 0 | 0.45 | 0.42 | 0.4 | 0.41 | 0.38 | 0.4 |
| 3 | 170162684 | rs6444930 | 1.116 | 0.278 | 0.152 | 1.442 | 0.007 | 0.03 | 0.03 | 0 | 0.88 | 0.82 | 0.91 | 0.77 | 0.87 | 0.94 |
| 3 | 170166091 | rs6764438 | 0.933 | 0.22 | 0.293 | 1.41 | 0.011 | 0.04 | 0.04 | 0.401 | 0.88 | 0.82 | 0.88 | 0.74 | 0.86 | 0.87 |
| 3 | 170166138 | rs6789670 | 1.115 | 0.175 | 0.367 | 1.443 | 0.019 | 0.122 | 0.122 | 2.494 | 0.82 | 0.5 | 0.82 | 0.7 | 0.86 | 0.84 |
| 3 | 170171835 | rs7647823 | -0.785 | -0.246 | -0.192 | 1.037 | -0.012 | -0.006 | -0.006 | 0 | 0.46 | 0.61 | 0.47 | 0.62 | 0.34 | 0.53 |
| 3 | 170172774 | rs6769887 | 0.295 | -0.222 | 0.78 | 1.164 | 0.024 | 0.014 | 0.014 | 0.057 | 0.82 | 0.56 | 0.72 | 0.8 | 0.8 | 0.71 |

| | | | | | | | | | | | | | | | | |
|---|---|---|---|---|---|---|---|---|---|---|---|---|---|---|---|---|
| 3 | 170176173 | rs6414536 | 0.462 | -0.149 | 0.832 | 1.042 | 0 | -0.001 | -0.001 | 0 | 0.8 | 0.76 | 0.68 | 0.8 | 0.75 | 0.73 |
| 3 | 170178057 | rs2287489 | 0.393 | -0.164 | 1.017 | 1.224 | -0.001 | 0 | 0 | 0 | 0.81 | 0.78 | 0.69 | 0.8 | 0.75 | 0.73 |
| 3 | 170178305 | rs2287490 | 1.271 | -0.449 | 0.783 | 1.618 | 0 | -0.012 | -0.012 | 0.329 | 0.71 | 0.33 | 0.47 | 0.56 | 0.67 | 0.56 |
| 3 | 170179621 | rs4955730 | -0.011 | -0.462 | 0.226 | 1.639 | -0.029 | 0.014 | 0.014 | 0.032 | 0.82 | 0.94 | 0.69 | 0.83 | 0.75 | 0.75 |
| 3 | 170179772 | rs12633223 | 1.913 | -0.761 | -0.264 | 1.569 | -0.059 | 0.059 | 0.059 | 0 | 0.46 | 0.12 | 0.31 | 0.31 | 0.48 | 0.32 |
| 3 | 170180811 | rs16855088 | 0.21 | -0.848 | -0.318 | 1.494 | 0.007 | -0.071 | -0.071 | 0 | 0.18 | 0.03 | 0.14 | 0.29 | 0.14 | 0.19 |
| 3 | 170184132 | rs10936636 | 1.743 | -0.695 | -0.179 | 1.527 | -0.064 | 0.062 | 0.062 | 0 | 0.46 | 0.12 | 0.3 | 0.28 | 0.48 | 0.31 |
| 3 | 170185361 | rs9290392 | -0.323 | -0.362 | 0.659 | 1.659 | 0.012 | 0.126 | 0.126 | 0 | 0.08 | 0.25 | 0.2 | 0.23 | 0.08 | 0.17 |
| 3 | 170187805 | rs13089480 | 1.105 | -0.116 | 0.042 | 1.354 | -0.002 | -0.015 | -0.015 | 0.007 | 0.65 | 0.38 | 0.51 | 0.55 | 0.64 | 0.56 |
| 3 | 170194623 | rs759712 | 0.679 | 0.326 | 0.09 | 1.193 | 0.033 | 0.018 | 0.018 | 1.051 | 0.9 | 0.77 | 0.75 | 0.83 | 0.83 | 0.81 |
| 3 | 170198898 | rs3732449 | -0.682 | 0.181 | 0.17 | 1.336 | 0.009 | 0.162 | 0.162 | 0 | 0.19 | 0.52 | 0.26 | 0.38 | 0.2 | 0.24 |
| 3 | 170203820 | rs3732448 | -1.317 | 0.173 | -0.117 | 0.789 | 0.003 | 0.012 | 0.012 | 0 | 0.43 | 0.66 | 0.5 | 0.37 | 0.44 | 0.47 |
| 3 | 170204909 | rs9828037 | 0.235 | 0.15 | 0.134 | 1.067 | 0.021 | 0.171 | 0.171 | 0 | 0.08 | 0.21 | 0.12 | 0.17 | 0.06 | 0.1 |
| 3 | 170209234 | rs10490803 | -0.054 | 0.167 | 0.03 | 0.595 | -0.026 | -0.028 | -0.028 | 0 | 0.07 | 0.03 | 0.22 | 0.18 | 0.14 | 0.14 |
| 3 | 170217126 | rs11929657 | -0.96 | -0.054 | 0.11 | 0.271 | -0.03 | -0.14 | -0.14 | 0 | 0.2 | 0.53 | 0.11 | 0 | 0.13 | 0.11 |
| 3 | 170217252 | rs13078511 | 0.218 | -0.112 | 0.696 | 0.235 | 0.004 | -0.035 | -0.035 | 0 | 0.07 | 0.26 | 0.05 | 0 | 0.06 | 0.01 |
| 3 | 170217912 | rs11915263 | -2.133 | -0.171 | 0.62 | 0.264 | -0.003 | -0.017 | -0.017 | 0 | 0.28 | 0.73 | 0.33 | 0.18 | 0.27 | 0.25 |
| 3 | 170220967 | rs6767520 | 0.044 | -0.118 | 0.576 | 0.249 | 0.003 | -0.004 | -0.004 | 0 | 0.11 | 0.07 | 0.04 | 0 | 0.07 | 0.09 |
| 3 | 170225104 | rs12490693 | -1.105 | -0.102 | 0.618 | 0.294 | 0.021 | 0.009 | 0.009 | 0.456 | 0.15 | 0.42 | 0.28 | 0.18 | 0.2 | 0.15 |
| 3 | 170231016 | rs1861940 | 0.898 | 0.133 | -0.075 | -0.019 | -0.011 | 0.011 | 0.011 | 0.231 | 0.62 | 0.28 | 0.46 | 0.36 | 0.51 | 0.58 |
| 3 | 170234819 | rs6799974 | -0.644 | 0.171 | -0.282 | 0.077 | 0.012 | 0.081 | 0.081 | 0 | 0.78 | 0.75 | 0.76 | 0.54 | 0.71 | 0.75 |
| 3 | 170241620 | rs6802900 | 0.262 | 0.204 | -0.272 | -0.007 | 0.011 | 0.004 | 0.004 | 0 | 0.71 | 0.72 | 0.55 | 0.35 | 0.57 | 0.6 |
| 3 | 170244995 | rs11915593 | 0.099 | 0.168 | -0.184 | 0.092 | 0 | -0.022 | -0.022 | 0 | 0.11 | 0.11 | 0.04 | 0 | 0.07 | 0.09 |
| 3 | 170246133 | rs11924505 | 0.123 | 0.177 | -0.189 | 0.096 | 0.001 | -0.011 | -0.011 | 0 | 0.11 | 0.08 | 0.04 | 0 | 0.07 | 0.09 |
| 3 | 170250400 | rs9880460 | -0.311 | -0.087 | -0.189 | 0.087 | 0.009 | 0.004 | 0.004 | 0.104 | 0.36 | 0.49 | 0.47 | 0.62 | 0.47 | 0.4 |
| 3 | 170251603 | rs10513683 | -0.929 | 0.07 | -0.482 | -0.142 | 0 | -0.006 | -0.006 | 0 | 0.33 | 0.34 | 0.23 | 0.23 | 0.29 | 0.29 |
| 3 | 170255506 | rs1023599 | 0.751 | 0.155 | -0.614 | -0.027 | 0.004 | 0.077 | 0.077 | 0 | 0.27 | 0.33 | 0.26 | 0.46 | 0.33 | 0.24 |
| 3 | 170263320 | rs6790988 | -2.289 | 0.417 | -0.619 | -0.127 | -0.051 | 0.058 | 0.058 | 0.021 | 0.7 | 0.98 | 0.71 | 0.84 | 0.76 | 0.69 |
| 3 | 170274083 | rs7611581 | -0.889 | 0.063 | -0.344 | 0.204 | -0.014 | 0.029 | 0.029 | 0.727 | 0.49 | 0.74 | 0.54 | 0.59 | 0.54 | 0.48 |
| 3 | 170275098 | rs1468977 | -0.288 | 0.159 | -0.281 | 0.246 | -0.001 | 0.007 | 0.007 | 0 | 0.39 | 0.41 | 0.31 | 0.4 | 0.4 | 0.33 |
| 3 | 170275580 | rs2422040 | 0.364 | 0.269 | -0.118 | 0.467 | -0.002 | 0.039 | 0.039 | 0 | 0.29 | 0.35 | 0.26 | 0.4 | 0.33 | 0.24 |
| 3 | 170293574 | rs2052398 | -2.023 | 0.092 | 0.405 | 0.822 | -0.012 | -0.042 | -0.042 | 0 | 0.37 | 0.24 | 0.38 | 0.48 | 0.39 | 0.5 |
| 3 | 170293715 | rs4955590 | -0.459 | 0.127 | 0.25 | 0.881 | -0.029 | -0.093 | -0.093 | 0.005 | 0.19 | 0 | 0.1 | 0.26 | 0.18 | 0.2 |
| 3 | 170306455 | rs12152317 | -1.369 | 0.116 | 0.635 | 0.851 | 0.07 | 0.086 | 0.086 | 0.007 | 0.1 | 0.39 | 0.25 | 0.21 | 0.15 | 0.21 |
| 3 | 170315805 | rs4955737 | -2.097 | 0.013 | 0.599 | 0.971 | -0.001 | 0.019 | 0.019 | 0 | 0.36 | 0.51 | 0.36 | 0.47 | 0.36 | 0.47 |
| 3 | 170323055 | rs874590 | -1.525 | 0.059 | -0.362 | 0.965 | -0.002 | -0.023 | -0.023 | 0.466 | 0.29 | 0.07 | 0.14 | 0.26 | 0.25 | 0.29 |
| 3 | 170332956 | rs17289598 | -1.138 | 0.794 | 0.212 | 1.468 | -0.008 | -0.006 | -0.006 | 0 | 0.06 | 0.04 | 0.24 | 0.22 | 0.12 | 0.2 |
| 3 | 170341683 | rs17826561 | -1.577 | 0.763 | 0.127 | 1.474 | -0.004 | -0.019 | -0.019 | 0.203 | 0.18 | 0.15 | 0.12 | 0.27 | 0.18 | 0.27 |
| 3 | 170347683 | rs4955593 | -1.147 | 0.519 | 0.102 | 1.278 | -0.032 | -0.031 | -0.031 | 0 | 0.05 | 0.01 | 0.23 | 0.22 | 0.12 | 0.19 |
| 3 | 170362619 | rs1011295 | -1.67 | -0.155 | 0.462 | 1.229 | -0.022 | -0.013 | -0.013 | 0 | 0.17 | 0.65 | 0.26 | 0.19 | 0.12 | 0.24 |
| 3 | 170377367 | rs9862393 | -1.585 | 0.159 | 0.619 | 1.718 | 0.016 | 0.277 | 0.277 | 0.806 | 0.14 | 0.5 | 0.24 | 0.42 | 0.13 | 0.3 |
| 3 | 170396745 | rs17842406 | -0.907 | -0.073 | 0.267 | 1.369 | 0 | 0 | 0 | 0.853 | 0.15 | 0.17 | 0.13 | 0.13 | 0.12 | 0.14 |
| 3 | 170408158 | rs499061 | -1.538 | 0.546 | 0.515 | 1.907 | 0.03 | 0.098 | 0.098 | 0 | 0.05 | 0.1 | 0.04 | 0.2 | 0.1 | 0.16 |
| 3 | 170408254 | rs9841319 | -1.174 | 0.216 | 1.314 | 1.835 | -0.019 | -0.002 | -0.002 | 0 | 0.17 | 0.31 | 0.14 | 0.16 | 0.12 | 0.14 |
| 3 | 170410059 | rs558514 | -2.112 | 0.419 | 1.225 | 1.764 | 0.027 | 0.097 | 0.097 | 0 | 0.23 | 0.41 | 0.19 | 0.37 | 0.24 | 0.3 |
| 3 | 170415627 | rs9870546 | -1.208 | 0.408 | 1.253 | 1.725 | -0.007 | -0.009 | -0.009 | 0 | 0.18 | 0.25 | 0.15 | 0.16 | 0.14 | 0.15 |
| 3 | 170417673 | rs546601 | -0.644 | -0.307 | 0.636 | 1.41 | -0.016 | -0.059 | -0.059 | 4.88 | 0.12 | 0.02 | 0.05 | 0.21 | 0.14 | 0.17 |
| 3 | 170438978 | rs2088672 | -0.978 | -0.475 | 1.086 | 1.436 | -0.012 | -0.002 | -0.002 | 0.85 | 0.17 | 0.25 | 0.17 | 0.19 | 0.11 | 0.17 |
| 3 | 170440005 | rs479645 | -2.08 | -0.538 | 1.383 | 1.476 | -0.016 | 0.02 | 0.02 | 0 | 0.32 | 0.47 | 0.27 | 0.4 | 0.27 | 0.34 |
| 3 | 170440352 | rs1532591 | -0.322 | -0.53 | 1.057 | 1.45 | 0.002 | -0.007 | -0.007 | 0 | 0.13 | 0.08 | 0.16 | 0.19 | 0.11 | 0.16 |
| 3 | 170442722 | rs502233 | -1.392 | -0.448 | 1.073 | 1.526 | 0.006 | -0.009 | -0.009 | 0 | 0.17 | 0.18 | 0.09 | 0.17 | 0.14 | 0.18 |
| 3 | 170451587 | rs559311 | -2.51 | -0.515 | 1.199 | 2.432 | 0.016 | 0.154 | 0.154 | 0 | 0.09 | 0.22 | 0.11 | 0.4 | 0.11 | 0.28 |
| 3 | 170462589 | rs474588 | -2.373 | -1.164 | 1.288 | 2.265 | -0.012 | 0.033 | 0.033 | 0 | 0.27 | 0.43 | 0.26 | 0.59 | 0.21 | 0.4 |
| 3 | 170463622 | rs17236893 | 0.074 | -1.152 | 1.158 | 2.256 | -0.015 | -0.042 | -0.042 | 0 | 0.12 | 0 | 0.12 | 0.19 | 0.1 | 0.1 |
| 3 | 170469336 | rs472617 | -1.684 | -1.146 | 1.203 | 2.213 | 0.005 | -0.02 | -0.02 | 0 | 0.23 | 0.24 | 0.23 | 0.58 | 0.19 | 0.4 |
| 3 | 170479343 | rs9871548 | 1.926 | -1.164 | 1.439 | 2.247 | -0.025 | 0.081 | 0.081 | 0 | 0.73 | 0.49 | 0.74 | 0.42 | 0.79 | 0.6 |
| 3 | 170494409 | rs4955755 | 2.2 | -1.42 | 1.796 | 1.886 | 0.03 | -0.021 | -0.021 | 0.064 | 0.62 | 0.2 | 0.59 | 0.41 | 0.7 | 0.54 |
| 3 | 170510715 | rs4499602 | -1.771 | -1.349 | 1.991 | 1.947 | -0.023 | 0.06 | 0.06 | 0 | 0.27 | 0.61 | 0.22 | 0.45 | 0.18 | 0.26 |
| 3 | 170519746 | rs9851496 | -1.606 | -1.437 | 1.971 | 1.922 | -0.024 | 0.067 | 0.067 | 0 | 0.26 | 0.45 | 0.21 | 0.45 | 0.18 | 0.27 |
| 3 | 170522227 | rs12485248 | 0.843 | -1.287 | 0.544 | 1.355 | -0.169 | -0.023 | -0.023 | 0 | 0.34 | 0.03 | 0.45 | 0.25 | 0.46 | 0.39 |
| 3 | 170527300 | rs9845497 | -1.768 | -1.501 | 1.237 | 1.692 | -0.01 | 0.113 | 0.113 | 0 | 0.13 | 0.34 | 0.09 | 0.26 | 0.09 | 0.16 |
| 3 | 170530581 | rs6779973 | -1.779 | -1.809 | 1.086 | 1.631 | -0.021 | 0.002 | 0.002 | 0 | 0.3 | 0.67 | 0.29 | 0.67 | 0.2 | 0.33 |
| 3 | 170533587 | rs6790486 | -1.1 | -1.15 | 0.915 | 1.407 | -0.024 | 0.048 | 0.048 | 0.474 | 0.31 | 0.48 | 0.25 | 0.42 | 0.29 | 0.26 |
| 3 | 170561866 | rs4955756 | -0.738 | -1.131 | 0.68 | 0.979 | -0.001 | -0.001 | -0.001 | 4.533 | 0.57 | 0.39 | 0.57 | 0.76 | 0.51 | 0.62 |
| 3 | 170561907 | rs12495896 | -0.136 | -1.317 | 0.711 | 0.95 | 0.021 | 0.031 | 0.031 | 0.004 | 0.24 | 0.13 | 0.16 | 0.18 | 0.21 | 0.17 |
| 3 | 170568001 | rs10936656 | 0.951 | -1.313 | 0.734 | 0.988 | 0.016 | -0.017 | -0.017 | 0.08 | 0.3 | 0.17 | 0.37 | 0.38 | 0.23 | 0.38 |
| 3 | 170568341 | rs10936658 | -0.713 | -1.298 | 0.757 | 0.987 | 0.005 | 0 | 0 | 4.167 | 0.54 | 0.37 | 0.53 | 0.56 | 0.45 | 0.55 |
| 3 | 170572112 | rs10936661 | -1.153 | -0.969 | 0.784 | 0.827 | 0.026 | -0.065 | -0.065 | 0 | 0.15 | 0.5 | 0.17 | 0.05 | 0.14 | 0.14 |
| 3 | 170572896 | rs12488260 | -1.754 | -0.971 | 0.82 | 0.828 | -0.062 | 0.049 | 0.049 | 0.029 | 0.66 | 0.82 | 0.71 | 0.8 | 0.62 | 0.71 |

| | | | | | | | | | | | | | | | | |
|---|---|---|---|---|---|---|---|---|---|---|---|---|---|---|---|---|
| 3 | 170574165 | rs11919048 | -1.761 | -0.301 | 1.817 | 1.236 | 0.027 | -0.064 | -0.064 | 0.176 | 0.15 | 0.5 | 0.17 | 0.05 | 0.14 | 0.14 |
| 3 | 170582888 | rs13063927 | -0.417 | -0.305 | 1.468 | 1.235 | -0.03 | -0.09 | -0.09 | 0 | 0.2 | 0.04 | 0.18 | 0.38 | 0.23 | 0.18 |
| 3 | 170589304 | rs12486780 | -0.878 | -0.315 | 1.502 | 1.225 | 0.011 | 0.018 | 0.018 | 0.052 | 0.24 | 0.17 | 0.21 | 0.18 | 0.22 | 0.17 |
| 3 | 170610316 | rs6792607 | -2.339 | -0.301 | 1.563 | 1.217 | 0.005 | 0 | 0 | 1.68 | 0.38 | 0.78 | 0.4 | 0.42 | 0.39 | 0.32 |
| 3 | 170611490 | rs13065440 | -1.024 | -0.387 | 1.359 | 1.179 | 0.008 | 0.007 | 0.007 | 0.012 | 0.25 | 0.23 | 0.23 | 0.22 | 0.21 | 0.19 |
| 3 | 170618763 | rs1356012 | -1.053 | -0.37 | 1.366 | 1.179 | 0.005 | 0.005 | 0.005 | 0 | 0.25 | 0.24 | 0.23 | 0.22 | 0.22 | 0.19 |
| 3 | 170619974 | rs35280911 | -0.365 | -0.432 | 1.274 | 1.194 | 0.002 | 0 | 0 | 0.105 | 0.19 | 0.17 | 0.17 | 0.22 | 0.17 | 0.13 |
| 3 | 170622714 | rs1879441 | -0.354 | -0.428 | 1.259 | 1.196 | 0.003 | 0 | 0 | 0.091 | 0.19 | 0.17 | 0.17 | 0.22 | 0.17 | 0.13 |
| 3 | 170625806 | rs2271998 | -2.238 | -0.393 | 1.348 | 1.238 | 0.006 | 0.001 | 0.001 | 1.03 | 0.39 | 0.77 | 0.39 | 0.42 | 0.39 | 0.32 |
| 3 | 170628132 | rs6769079 | -2.337 | 0.45 | 1.476 | 1.582 | 0.002 | 0.003 | 0.003 | 0.419 | 0.33 | 0.66 | 0.36 | 0.45 | 0.36 | 0.32 |
| 3 | 170628770 | rs6444952 | -1.686 | 0.393 | 1.596 | 1.581 | 0.007 | 0.071 | 0.071 | 0.081 | 0.24 | 0.45 | 0.26 | 0.45 | 0.26 | 0.26 |
| 3 | 170659019 | rs10936665 | -0.951 | -0.057 | 1.523 | 1.178 | 0.005 | 0.063 | 0.063 | 0 | 0.11 | 0.2 | 0.18 | 0.22 | 0.13 | 0.14 |
| 3 | 170670279 | rs16855567 | -0.483 | -0.058 | 1.6 | -0.095 | 0.012 | -0.045 | -0.045 | 0.151 | 0.09 | 0.17 | 0.09 | 0 | 0.1 | 0.05 |
| 3 | 170671309 | rs7642480 | 0.01 | -0.142 | 1.627 | -0.098 | -0.001 | -0.002 | -0.002 | 0.775 | 0.14 | 0.11 | 0.13 | 0.2 | 0.15 | 0.1 |
| 3 | 170671429 | rs7628878 | -1.585 | -0.145 | 1.627 | -0.095 | 0.003 | -0.006 | -0.006 | 0 | 0.23 | 0.31 | 0.25 | 0.25 | 0.25 | 0.18 |
| 3 | 170693708 | rs16855591 | -0.246 | -0.054 | 1.549 | -0.143 | 0.001 | -0.046 | -0.046 | 0 | 0.11 | 0.17 | 0.11 | 0 | 0.11 | 0.09 |
| 3 | 170694497 | rs11709140 | -1.321 | -0.05 | 1.423 | -0.13 | -0.003 | -0.072 | -0.072 | 0 | 0.17 | 0.4 | 0.17 | 0.05 | 0.14 | 0.19 |
| 3 | 170715830 | rs5398 | -2 | -0.077 | 1.463 | -0.146 | -0.005 | -0.02 | -0.02 | 0.57 | 0.32 | 0.64 | 0.31 | 0.24 | 0.29 | 0.28 |
| 3 | 170717521 | rs11920090 | -1.057 | 0.002 | 1.497 | -0.139 | 0.004 | -0.081 | -0.081 | 0.196 | 0.15 | 0.37 | 0.17 | 0.02 | 0.14 | 0.16 |
| 3 | 170717996 | rs11924648 | -1.296 | 0 | 1.443 | -0.14 | 0.006 | -0.102 | -0.102 | 0.042 | 0.16 | 0.45 | 0.17 | 0.02 | 0.14 | 0.16 |
| 3 | 170722448 | rs1996220 | -1.151 | 0.001 | 1.176 | -0.14 | 0.01 | -0.094 | -0.094 | 0.219 | 0.16 | 0.43 | 0.17 | 0.02 | 0.14 | 0.16 |
| 3 | 170724729 | rs1499821 | -0.04 | 0.001 | 0.933 | -0.134 | -0.002 | -0.003 | -0.003 | 0.02 | 0.15 | 0.12 | 0.13 | 0.19 | 0.15 | 0.1 |
| 3 | 170732300 | rs5400 | -1.158 | 0.104 | 0.995 | -0.093 | 0.015 | -0.099 | -0.099 | 0 | 0.16 | 0.49 | 0.17 | 0.02 | 0.14 | 0.16 |
| 3 | 170735099 | rs11924032 | -1.277 | 0.013 | 0.913 | -0.058 | 0.003 | -0.028 | -0.028 | 0 | 0.27 | 0.41 | 0.26 | 0.21 | 0.26 | 0.23 |
| 3 | 170747810 | rs6800180 | -1.487 | -0.132 | 0.581 | -0.125 | -0.013 | -0.082 | -0.082 | 0 | 0.17 | 0.57 | 0.15 | 0.04 | 0.12 | 0.18 |
| 3 | 170767302 | rs9681556 | -0.694 | -0.028 | -0.127 | -0.02 | -0.003 | -0.06 | -0.06 | 0 | 0.12 | 0.38 | 0.09 | 0.02 | 0.08 | 0.11 |
| 3 | 170769171 | rs4133045 | -0.251 | -0.434 | -1.529 | -0.013 | 0.02 | 0.008 | 0.008 | 0 | 0.69 | 0.97 | 0.79 | 0.78 | 0.78 | 0.74 |
| 3 | 170777684 | rs16855718 | -0.737 | -0.57 | -0.985 | -0.275 | -0.008 | -0.036 | -0.036 | 0 | 0.1 | 0.13 | 0.07 | 0 | 0.09 | 0.07 |
| 3 | 170811667 | rs16855789 | -0.968 | -0.809 | -0.564 | -0.437 | -0.01 | -0.034 | -0.034 | 0 | 0.12 | 0.15 | 0.07 | 0.01 | 0.09 | 0.1 |
| 3 | 170814299 | rs9827202 | -1.541 | -1.247 | 0.489 | -0.559 | -0.063 | -0.1 | -0.1 | 0 | 0.17 | 0.52 | 0.17 | 0.05 | 0.1 | 0.13 |
| 3 | 170823923 | rs9789980 | -0.341 | -0.751 | -0.447 | -0.346 | 0 | 0 | 0 | 0.001 | 0.55 | 0.46 | 0.54 | 0.68 | 0.58 | 0.54 |
| 3 | 170825920 | rs2291900 | 0.882 | -0.995 | -0.514 | -0.464 | -0.046 | -0.019 | -0.019 | 0 | 0.26 | 0.08 | 0.24 | 0.19 | 0.29 | 0.24 |
| 3 | 170832832 | rs999979 | -0.048 | -0.879 | -0.512 | -0.121 | 0 | 0.003 | 0.003 | 0 | 0.53 | 0.42 | 0.53 | 0.54 | 0.56 | 0.53 |
| 3 | 170839690 | rs6767021 | 0.963 | -0.251 | -1.224 | -0.175 | -0.033 | -0.108 | -0.108 | 0.786 | 0.21 | 0.1 | 0.22 | 0.5 | 0.23 | 0.24 |
| 3 | 170849546 | rs4955768 | 0.233 | -0.284 | -0.588 | -0.472 | -0.029 | -0.067 | -0.067 | 0.368 | 0.14 | 0.28 | 0.08 | 0.03 | 0.1 | 0.1 |
| 3 | 170849988 | rs12488990 | 1.441 | 0.059 | -0.892 | -0.521 | -0.07 | 0.013 | 0.013 | 0 | 0.12 | 0.01 | 0.18 | 0.05 | 0.2 | 0.12 |
| 3 | 170859893 | rs13315469 | 0.773 | -0.215 | -0.713 | -0.643 | 0.002 | -0.052 | -0.052 | 0 | 0.4 | 0.21 | 0.43 | 0.45 | 0.37 | 0.43 |
| 3 | 170861559 | rs6791187 | -0.737 | 0.539 | 0.056 | -0.684 | 0.004 | -0.044 | -0.044 | 1.391 | 0.28 | 0.32 | 0.29 | 0.12 | 0.33 | 0.27 |
| 3 | 170865850 | rs7372276 | -0.396 | 0.278 | 0.193 | -0.604 | -0.043 | -0.088 | -0.088 | 0 | 0.16 | 0.31 | 0.09 | 0.04 | 0.1 | 0.14 |
| 3 | 170874398 | rs6444967 | -0.536 | 0.994 | 0.376 | -0.641 | 0.032 | -0.098 | -0.098 | 0.005 | 0.17 | 0.35 | 0.17 | 0 | 0.22 | 0.12 |
| 3 | 170876501 | rs6788393 | 0.47 | 0.034 | 0.005 | -0.625 | -0.017 | 0.051 | 0.051 | 0 | 0.72 | 0.82 | 0.69 | 0.88 | 0.65 | 0.76 |
| 3 | 170886171 | rs6789167 | -1.008 | 0.159 | -0.365 | -0.562 | -0.06 | 0.008 | 0.008 | 0 | 0.85 | 0.98 | 0.78 | 0.89 | 0.75 | 0.85 |
| 3 | 170893070 | rs12637875 | 0.784 | -0.174 | -0.096 | -0.692 | 0.003 | 0.073 | 0.073 | 0 | 0.62 | 0.27 | 0.6 | 0.87 | 0.58 | 0.63 |
| 3 | 170908762 | rs12495729 | -0.152 | 0.106 | 0.139 | 0.092 | 0 | 0.002 | 0.002 | 0 | 0.33 | 0.32 | 0.39 | 0.5 | 0.32 | 0.43 |
| 3 | 170916577 | rs2422193 | 0.091 | 0.146 | 0.089 | 0.186 | -0.002 | -0.012 | -0.012 | 4.25 | 0.31 | 0.25 | 0.33 | 0.47 | 0.29 | 0.36 |
| 3 | 170921178 | rs6783543 | 1.287 | 0.119 | 0.202 | 0.062 | 0.003 | 0.007 | 0.007 | 0 | 0.39 | 0.36 | 0.29 | 0.38 | 0.31 | 0.36 |
| 3 | 170922054 | rs11716192 | -0.256 | 0.116 | 0.118 | 0.003 | -0.014 | -0.038 | -0.038 | 0 | 0.39 | 0.12 | 0.38 | 0.48 | 0.38 | 0.41 |
| 3 | 170928797 | rs6444972 | -1.361 | 0.275 | 0.359 | -0.034 | 0.045 | -0.015 | -0.015 | 0.189 | 0.86 | 0.78 | 0.74 | 0.89 | 0.72 | 0.85 |
| 3 | 170929190 | rs2292004 | -0.002 | 0.202 | 0.339 | 0.004 | -0.012 | -0.002 | -0.002 | 0 | 0.34 | 0.21 | 0.4 | 0.58 | 0.34 | 0.43 |
| 3 | 170931099 | rs10936674 | 0.982 | -0.935 | 0.724 | 0.058 | 0.006 | 0.013 | 0.013 | 0 | 0.38 | 0.32 | 0.26 | 0.31 | 0.28 | 0.33 |
| 3 | 170934496 | rs13091938 | -0.204 | -0.939 | 0.62 | 0.399 | 0.013 | -0.017 | -0.017 | 0 | 0.18 | 0.1 | 0.15 | 0.24 | 0.11 | 0.09 |
| 3 | 170935941 | rs1124184 | -1.168 | -0.923 | 0.499 | 0.409 | -0.142 | -0.094 | -0.094 | 0.723 | 0.67 | 0.91 | 0.53 | 0.62 | 0.49 | 0.55 |
| 3 | 170941083 | rs7374644 | 1.045 | -1.347 | 0.479 | 0.471 | 0.026 | 0.05 | 0.05 | 0 | 0.38 | 0.16 | 0.25 | 0.34 | 0.28 | 0.34 |
| 3 | 170943009 | rs905128 | -1.435 | -1.219 | 0.284 | 0.387 | -0.049 | 0.029 | 0.029 | 0 | 0.27 | 0.56 | 0.26 | 0.3 | 0.17 | 0.19 |
| 3 | 170943712 | rs871468 | 0.538 | -0.747 | -0.643 | 0.597 | -0.011 | 0.143 | 0.143 | 0 | 0.08 | 0.14 | 0.1 | 0.25 | 0.03 | 0.05 |
| 3 | 170948750 | rs6787205 | -0.611 | -0.302 | -0.911 | -0.376 | 0 | -0.002 | -0.002 | 0.222 | 0.56 | 0.55 | 0.43 | 0.38 | 0.45 | 0.51 |
| 3 | 170949276 | rs7355905 | -1.397 | -0.184 | -1.076 | 0.007 | 0.017 | 0.024 | 0.024 | 0 | 0.16 | 0.07 | 0.12 | 0.07 | 0.13 | 0.14 |
| 3 | 170963758 | rs13086608 | -0.746 | -0.012 | -1.186 | -0.047 | 0.019 | 0.063 | 0.063 | 0 | 0.12 | 0.01 | 0.07 | 0 | 0.08 | 0.07 |
| 3 | 170964909 | rs16856044 | -0.821 | -0.015 | -1.117 | -0.049 | 0.019 | 0.018 | 0.018 | 0 | 0.12 | 0.09 | 0.07 | 0 | 0.08 | 0.07 |
| 3 | 170965875 | rs9819421 | -0.752 | -0.062 | -1.062 | 0.049 | 0.001 | 0 | 0 | 0 | 0.24 | 0.23 | 0.23 | 0.22 | 0.21 | 0.19 |
| 3 | 170970433 | rs13075838 | -0.423 | 0.191 | -1.705 | -0.725 | -0.017 | 0.106 | 0.106 | 0 | 0.2 | 0.02 | 0.11 | 0 | 0.18 | 0.08 |
| 3 | 170974549 | rs16856054 | -0.643 | 0.115 | -0.871 | -1.099 | 0.002 | -0.003 | -0.003 | 0 | 0.09 | 0.08 | 0.06 | 0.01 | 0.06 | 0.05 |
| 3 | 170977841 | rs6772621 | -1.099 | 0.226 | -0.849 | -1.137 | -0.005 | 0.037 | 0.037 | 0 | 0.57 | 0.69 | 0.43 | 0.31 | 0.52 | 0.43 |
| 3 | 170981057 | rs4894643 | 0.83 | 0.187 | -0.44 | -1.302 | -0.016 | -0.013 | -0.013 | 0 | 0.22 | 0.17 | 0.2 | 0.3 | 0.25 | 0.23 |
| 3 | 170984600 | rs11927009 | -1.381 | 0.234 | -0.303 | -1.397 | -0.013 | 0.101 | 0.101 | 0 | 0.54 | 0.69 | 0.43 | 0.12 | 0.49 | 0.41 |
| 3 | 170987337 | rs6806583 | 1.027 | 0.559 | -0.456 | -1.515 | 0.027 | -0.024 | -0.024 | 0 | 0.06 | 0.14 | 0.09 | 0.01 | 0.1 | 0.07 |
| 3 | 170991647 | rs6785520 | 0.769 | 0.441 | -0.079 | -1.551 | 0.004 | 0.017 | 0.017 | 0 | 0.21 | 0.15 | 0.12 | 0 | 0.18 | 0.08 |
| 3 | 170993127 | rs11915443 | -1.148 | 0.426 | 0.07 | -1.489 | -0.002 | -0.074 | -0.074 | 0 | 0.33 | 0.52 | 0.31 | 0.12 | 0.31 | 0.33 |
| 3 | 170995335 | rs13065441 | -0.131 | 0.407 | -0.033 | -1.497 | 0.004 | -0.001 | -0.001 | 0 | 0.13 | 0.08 | 0.1 | 0.01 | 0.09 | 0.06 |

| | | | | | | | | | | | | | | | | |
|---|---|---|---|---|---|---|---|---|---|---|---|---|---|---|---|---|
| 3 | 170998771 | rs7646338 | 1.111 | 0.241 | -0.458 | -1.101 | -0.014 | -0.048 | -0.048 | 0.603 | 0.12 | 0.01 | 0.15 | 0.2 | 0.13 | 0.28 |
| 3 | 171000533 | rs10513694 | 0.11 | 0.307 | -0.146 | -1.149 | -0.008 | 0.014 | 0.014 | 0 | 0.19 | 0.16 | 0.19 | 0.11 | 0.22 | 0.24 |
| 3 | 171021756 | rs2172394 | 0.858 | -0.131 | -0.385 | -1.236 | 0.014 | 0.01 | 0.01 | 0 | 0.16 | 0 | 0.11 | 0.1 | 0.11 | 0.16 |
| 3 | 171024522 | rs4455343 | -0.288 | 0.513 | -0.258 | -1.257 | 0 | 0.116 | 0.116 | 0.114 | 0.43 | 0.23 | 0.33 | 0.13 | 0.38 | 0.27 |
| 3 | 171027615 | rs4894521 | -0.378 | 0.507 | -0.123 | -1.258 | 0.002 | 0.014 | 0.014 | 0.035 | 0.44 | 0.34 | 0.32 | 0.13 | 0.38 | 0.27 |
| 3 | 171027683 | rs4894793 | -0.77 | 0.77 | -0.191 | -1.024 | -0.016 | 0.099 | 0.099 | 0 | 0.57 | 0.71 | 0.44 | 0.13 | 0.5 | 0.4 |
| 3 | 171038437 | rs9829436 | -0.141 | 0.084 | -0.213 | -0.243 | 0.008 | 0.005 | 0.005 | 0 | 0.44 | 0.35 | 0.31 | 0.37 | 0.29 | 0.39 |
| 3 | 171039539 | rs6798267 | -0.066 | 0.153 | -0.363 | -0.246 | 0.046 | 0.034 | 0.034 | 0 | 0.43 | 0.29 | 0.31 | 0.35 | 0.29 | 0.39 |
| 3 | 171040813 | rs9823183 | -0.791 | 0.849 | 0.378 | -0.306 | 0.062 | -0.099 | -0.099 | 0 | 0.15 | 0.45 | 0.14 | 0.02 | 0.16 | 0.11 |
| 3 | 171041055 | rs10513695 | -0.873 | 0.674 | 0.174 | 0.236 | 0.022 | 0.02 | 0.02 | 0 | 0.21 | 0.39 | 0.2 | 0.22 | 0.25 | 0.31 |
| 3 | 171041464 | rs7615965 | -0.88 | 0.957 | 0.141 | 0.225 | 0.038 | 0.011 | 0.011 | 0 | 0.21 | 0.38 | 0.22 | 0.19 | 0.28 | 0.33 |
| 3 | 171048165 | rs6444980 | -0.902 | 1.001 | -0.02 | 0.511 | 0.004 | 0.002 | 0.002 | 0 | 0.23 | 0.26 | 0.21 | 0.25 | 0.33 | 0.31 |
| 3 | 171049177 | rs2172390 | -0.996 | 1.051 | -0.616 | 0.596 | 0.019 | 0.084 | 0.084 | 0.003 | 0.47 | 0.74 | 0.54 | 0.82 | 0.6 | 0.59 |
| 3 | 171056156 | rs10936678 | 0.248 | 0.507 | -0.543 | 0.906 | 0.001 | 0.021 | 0.021 | 0 | 0.19 | 0.18 | 0.27 | 0.51 | 0.27 | 0.2 |
| 3 | 171057320 | rs2010071 | -0.11 | 0.6 | -0.52 | 0.687 | -0.003 | -0.011 | -0.011 | 0 | 0.23 | 0.2 | 0.28 | 0.47 | 0.28 | 0.28 |
| 3 | 171061886 | rs9824791 | -0.291 | 0.346 | -1.013 | 0.196 | -0.024 | 0.06 | 0.06 | 0 | 0.53 | 0.78 | 0.62 | 0.59 | 0.53 | 0.55 |
| 3 | 171071204 | rs6768644 | 0.26 | 0.315 | -0.637 | 0.331 | -0.003 | -0.008 | -0.008 | 0 | 0.26 | 0.21 | 0.27 | 0.42 | 0.3 | 0.45 |
| 3 | 171072227 | rs6780532 | 0.982 | -0.13 | -0.428 | -0.149 | -0.01 | -0.085 | -0.085 | 0 | 0.31 | 0.14 | 0.39 | 0.53 | 0.35 | 0.27 |
| 3 | 171073436 | rs931497 | -0.742 | -0.134 | -0.101 | -0.269 | 0.006 | 0.064 | 0.064 | 0 | 0.6 | 0.74 | 0.72 | 0.97 | 0.65 | 0.74 |
| 3 | 171073616 | rs931498 | 0.315 | 0.217 | -0.305 | -0.023 | -0.004 | -0.008 | -0.008 | 0 | 0.26 | 0.22 | 0.27 | 0.42 | 0.3 | 0.45 |
| 3 | 171076593 | rs1038883 | 0.044 | -0.87 | -0.326 | -0.984 | 0.009 | 0.017 | 0.017 | 0 | 0.52 | 0.59 | 0.64 | 0.71 | 0.63 | 0.68 |
| 3 | 171080678 | rs1881961 | -0.251 | -1.363 | -0.311 | -0.694 | -0.062 | 0.006 | 0.006 | 0.821 | 0.8 | 0.62 | 0.85 | 0.76 | 0.91 | 0.9 |
| 3 | 171081370 | rs12495787 | 0.721 | -1.349 | -1.225 | -0.785 | -0.01 | 0.046 | 0.046 | 0.392 | 0.11 | 0.01 | 0.11 | 0.02 | 0.08 | 0.16 |
| 3 | 171083708 | rs9290409 | 0.548 | -1.314 | -1.315 | -0.616 | 0 | -0.049 | -0.049 | 0.117 | 0.12 | 0.05 | 0.08 | 0.31 | 0.1 | 0.18 |
| 3 | 171087469 | rs3796295 | 0.794 | -1.217 | -1.391 | -0.667 | 0.079 | -0.075 | -0.075 | 0.173 | 0.55 | 0.08 | 0.63 | 0.43 | 0.73 | 0.56 |
| 3 | 171088709 | rs902952 | 0.01 | -0.415 | -0.175 | -1.038 | -0.047 | 0.008 | 0.008 | 0.008 | 0.82 | 0.7 | 0.86 | 0.76 | 0.92 | 0.9 |
| 3 | 171092504 | rs4894572 | 0.167 | 0.593 | -0.769 | -1.797 | -0.036 | 0.099 | 0.099 | 0 | 0.35 | 0.22 | 0.64 | 0.93 | 0.48 | 0.56 |
| 3 | 171093738 | rs7636254 | 0.746 | 0.406 | -0.624 | -2.178 | -0.002 | -0.009 | -0.009 | 0.226 | 0.32 | 0.3 | 0.23 | 0.02 | 0.35 | 0.19 |
| 3 | 171098402 | rs9811628 | -0.924 | 0.75 | -1.117 | -2.197 | 0 | -0.041 | -0.041 | 0 | 0.84 | 0.85 | 0.9 | 1 | 0.87 | 0.79 |
| 3 | 171098694 | rs902954 | -0.024 | 0.734 | -1.321 | -2.08 | -0.003 | -0.048 | -0.048 | 0.018 | 0.65 | 0.72 | 0.84 | 0.94 | 0.73 | 0.72 |
| 3 | 171098888 | rs2901705 | 0.139 | 1.049 | -1.955 | -1.157 | -0.038 | 0.045 | 0.045 | 0 | 0.3 | 0.14 | 0.59 | 0.84 | 0.38 | 0.52 |
| 3 | 171100675 | rs882172 | 0.01 | 0.684 | -1.842 | -1.302 | -0.028 | 0.215 | 0.215 | 0 | 0.55 | 0.82 | 0.31 | 0.07 | 0.48 | 0.27 |
| 3 | 171101556 | rs9883206 | 0.227 | -0.324 | -1.374 | -1.168 | -0.017 | -0.016 | -0.016 | 0 | 0.65 | 0.38 | 0.75 | 0.85 | 0.77 | 0.83 |
| 3 | 171103807 | rs902956 | -0.283 | 0.273 | -2.272 | -1.676 | -0.134 | 0.164 | 0.164 | 0 | 0.7 | 0.98 | 0.76 | 0.9 | 0.59 | 0.72 |
| 3 | 171109146 | rs4894432 | -0.034 | -0.219 | -1.606 | -1.823 | 0.009 | 0.011 | 0.011 | 0 | 0.36 | 0.28 | 0.19 | 0.14 | 0.25 | 0.38 |
| 3 | 171112827 | rs6807555 | 0.23 | -0.394 | -1.534 | -1.91 | -0.014 | 0.058 | 0.058 | 0.903 | 0.61 | 0.72 | 0.37 | 0.16 | 0.56 | 0.4 |
| 3 | 171116588 | rs6787776 | -1.257 | -0.419 | -1.625 | -1.912 | -0.027 | -0.018 | -0.018 | 0 | 0.83 | 0.88 | 0.5 | 0.25 | 0.72 | 0.74 |
| 3 | 171118709 | rs6785069 | 0.661 | -0.37 | -0.997 | -2.239 | 0.006 | -0.095 | -0.095 | 0.068 | 0.43 | 0.48 | 0.3 | 0.11 | 0.47 | 0.32 |
| 3 | 171119213 | rs4894435 | -0.095 | -0.336 | -0.75 | -1.669 | -0.002 | -0.011 | -0.011 | 0 | 0.67 | 0.68 | 0.41 | 0.11 | 0.64 | 0.65 |
| 3 | 171130793 | rs10936684 | 0.102 | -0.245 | -0.805 | -1.629 | -0.001 | -0.037 | -0.037 | 2.618 | 0.63 | 0.66 | 0.41 | 0.11 | 0.64 | 0.65 |
| 3 | 171135807 | rs10936685 | 0.377 | 0.045 | -0.899 | -0.185 | -0.002 | -0.002 | -0.002 | 0 | 0.78 | 0.81 | 0.79 | 0.67 | 0.72 | 0.62 |
| 3 | 171139467 | rs7650598 | -0.174 | 0.146 | -1.066 | -0.118 | -0.006 | -0.009 | -0.009 | 0 | 0.83 | 0.86 | 0.8 | 0.69 | 0.74 | 0.67 |
| 3 | 171145711 | rs9875030 | -0.11 | -0.118 | -1.32 | -0.966 | -0.045 | 0.055 | 0.055 | 0 | 0.71 | 0.91 | 0.55 | 0.17 | 0.64 | 0.51 |
| 3 | 171146626 | rs6444987 | 0.308 | 0.014 | -0.91 | -1.051 | 0 | -0.066 | -0.066 | 0 | 0.45 | 0.47 | 0.39 | 0.05 | 0.47 | 0.42 |
| 3 | 171160434 | rs721912 | -0.874 | -0.419 | -1.02 | -0.103 | 0.03 | 0.019 | 0.019 | 0.31 | 0.28 | 0.77 | 0.3 | 0.21 | 0.18 | 0.3 |
| 3 | 171165273 | rs13314339 | -0.586 | -0.458 | -0.465 | -0.346 | -0.062 | -0.117 | -0.117 | 0 | 0.2 | 0.53 | 0.1 | 0.01 | 0.11 | 0.18 |
| 3 | 171173555 | rs10936688 | -0.104 | 0.945 | -1.645 | -0.292 | -0.125 | -0.028 | -0.028 | 0.406 | 0.77 | 0.95 | 0.7 | 0.76 | 0.64 | 0.78 |
| 3 | 171173627 | rs1491664 | 0.337 | 0.527 | -1.644 | -0.823 | -0.001 | -0.013 | -0.013 | 0.378 | 0.22 | 0.27 | 0.16 | 0.14 | 0.19 | 0.14 |
| 3 | 171174213 | rs1038828 | -0.39 | 0.752 | -1.56 | -0.626 | -0.123 | -0.025 | -0.025 | 0.162 | 0.76 | 0.93 | 0.7 | 0.76 | 0.64 | 0.78 |
| 3 | 171180685 | rs12634193 | -0.026 | 0.501 | -1.504 | -0.467 | -0.004 | -0.003 | -0.003 | 0 | 0.75 | 0.86 | 0.73 | 0.76 | 0.75 | 0.71 |
| 3 | 171183791 | rs7645984 | 0.324 | 0.275 | -0.485 | -0.582 | 0.002 | -0.008 | -0.008 | 0 | 0.72 | 0.62 | 0.69 | 0.74 | 0.74 | 0.71 |
| 3 | 171188929 | rs9844740 | 0.146 | -0.207 | -0.133 | -0.749 | -0.045 | -0.039 | -0.039 | 0 | 0.25 | 0.53 | 0.19 | 0.13 | 0.2 | 0.15 |
| 3 | 171192425 | rs10936689 | 0.906 | -0.23 | -0.168 | -0.738 | -0.009 | 0.009 | 0.009 | 1.349 | 0.47 | 0.15 | 0.49 | 0.6 | 0.54 | 0.57 |
| 3 | 171196599 | rs11916004 | -0.357 | -0.343 | -0.082 | -0.843 | 0.008 | 0.008 | 0.008 | 1.483 | 0.31 | 0.66 | 0.22 | 0.13 | 0.22 | 0.17 |
| 3 | 171199203 | rs6777074 | -0.564 | 0.166 | 0.095 | -0.93 | -0.005 | -0.006 | -0.006 | 0 | 0.35 | 0.58 | 0.49 | 0.65 | 0.32 | 0.39 |
| 3 | 171199970 | rs13091534 | 1.496 | 0.147 | -0.202 | -1.124 | -0.047 | 0.213 | 0.213 | 0.603 | 0.37 | 0.1 | 0.2 | 0.09 | 0.39 | 0.32 |
| 3 | 171201400 | rs9825902 | -1.514 | 0.481 | 0.081 | -0.855 | -0.028 | -0.033 | -0.033 | 0.748 | 0.82 | 0.93 | 0.75 | 0.75 | 0.79 | 0.77 |
| 3 | 171203303 | rs6805933 | -0.335 | 0.458 | 0.85 | -0.855 | 0.01 | 0 | 0 | 0 | 0.75 | 0.5 | 0.71 | 0.74 | 0.74 | 0.71 |
| 3 | 171212317 | rs918310 | -0.876 | 0.72 | 0.392 | -0.756 | -0.004 | -0.003 | -0.003 | 3.962 | 0.78 | 0.79 | 0.73 | 0.74 | 0.77 | 0.74 |
| 3 | 171220439 | rs6788198 | 1.093 | 0.563 | 0.18 | -0.874 | -0.003 | -0.023 | -0.023 | 0.808 | 0.16 | 0.02 | 0.15 | 0.13 | 0.17 | 0.09 |
| 3 | 171226045 | rs7651813 | -0.129 | 0.424 | 0.613 | -0.444 | 0 | 0.002 | 0.002 | 0 | 0.72 | 0.64 | 0.58 | 0.69 | 0.76 | 0.72 |
| 3 | 171226979 | rs9283639 | 1.083 | 0.508 | 0.895 | -0.685 | 0 | -0.066 | -0.066 | 0 | 0.46 | 0.56 | 0.25 | 0.1 | 0.43 | 0.37 |
| 3 | 171227112 | rs2075360 | 0.483 | 0.685 | 0.748 | -0.584 | -0.036 | -0.077 | -0.077 | 0 | 0.21 | 0.04 | 0.34 | 0.62 | 0.25 | 0.39 |
| 3 | 171227756 | rs2075361 | -0.106 | 0.628 | 0.913 | -0.539 | 0.006 | -0.007 | -0.007 | 0.34 | 0.7 | 0.55 | 0.6 | 0.72 | 0.71 | 0.76 |
| 3 | 171231125 | rs1388464 | -0.708 | 0.663 | 0.908 | -0.513 | 0.025 | 0.024 | 0.024 | 0 | 0.77 | 0.66 | 0.64 | 0.72 | 0.76 | 0.8 |
| 3 | 171232768 | rs9861012 | -0.189 | 0.839 | 0.794 | -0.544 | -0.033 | 0.012 | 0.012 | 0 | 0.72 | 0.8 | 0.45 | 0.26 | 0.62 | 0.48 |
| 3 | 171238322 | rs6804896 | 0.475 | 0.746 | 1.085 | -0.916 | -0.001 | -0.049 | -0.049 | 0 | 0.55 | 0.45 | 0.27 | 0.1 | 0.48 | 0.4 |
| 3 | 171241012 | rs6445008 | -0.26 | 0.741 | 1.125 | -0.905 | 0.004 | -0.067 | -0.067 | 0 | 0.11 | 0.34 | 0.07 | 0 | 0.08 | 0.05 |
| 3 | 171241762 | rs400721 | -0.569 | 0.727 | 0.672 | -0.782 | -0.002 | 0.009 | 0.009 | 0 | 0.34 | 0.72 | 0.42 | 0.26 | 0.31 | 0.24 |

| | | | | | | | | | | | | | | | | |
|---|---|---|---|---|---|---|---|---|---|---|---|---|---|---|---|---|
| 3 | 171245526 | rs260774 | 1.59 | 0.729 | 0.648 | -0.771 | -0.03 | -0.033 | -0.033 | 0 | 0.18 | 0.08 | 0.35 | 0.25 | 0.21 | 0.18 |
| 3 | 171253756 | rs7645103 | 1.289 | 0.718 | 0.394 | -0.932 | 0.015 | 0.111 | 0.111 | 0 | 0.4 | 0.21 | 0.21 | 0.13 | 0.36 | 0.33 |
| 3 | 171255914 | rs7641259 | 0.173 | 0.637 | 0.418 | -0.737 | -0.022 | -0.053 | -0.053 | 0.147 | 0.1 | 0.24 | 0.06 | 0 | 0.06 | 0.04 |
| 3 | 171264618 | rs16856450 | -1.066 | 0.735 | 0.025 | -0.658 | 0.023 | -0.026 | -0.026 | 0 | 0.06 | 0.25 | 0.04 | 0.01 | 0.05 | 0.02 |
| 3 | 171269484 | rs2061065 | 1.147 | 0.777 | -0.295 | -0.839 | 0.008 | 0.052 | 0.052 | 0 | 0.41 | 0.26 | 0.22 | 0.13 | 0.39 | 0.32 |
| 3 | 171270126 | rs360458 | 0.042 | 0.646 | 0.167 | -0.228 | 0.001 | 0.006 | 0.006 | 1.326 | 0.73 | 0.7 | 0.66 | 0.37 | 0.68 | 0.56 |
| 3 | 171272488 | rs2160992 | 0.056 | 0.83 | -0.627 | -0.165 | 0.007 | -0.024 | -0.024 | 6.041 | 0.06 | 0.13 | 0.03 | 0.01 | 0.05 | 0.01 |
| 3 | 171277982 | rs360430 | -0.05 | 0.755 | -0.898 | -0.158 | -0.025 | -0.044 | -0.044 | 0.02 | 0.74 | 0.9 | 0.67 | 0.37 | 0.69 | 0.55 |
| 3 | 171284593 | rs682455 | -0.177 | 0.761 | -1.044 | -0.014 | -0.058 | -0.059 | -0.059 | 0 | 0.75 | 0.96 | 0.66 | 0.34 | 0.7 | 0.54 |
| 3 | 171286713 | rs360414 | 1.084 | 0.864 | -0.735 | -0.003 | -0.002 | -0.024 | -0.024 | 0 | 0.56 | 0.45 | 0.47 | 0.3 | 0.56 | 0.41 |
| 3 | 171309620 | rs360402 | -0.068 | 0.917 | -1.935 | 0.37 | -0.046 | -0.062 | -0.062 | 0 | 0.69 | 0.94 | 0.65 | 0.38 | 0.69 | 0.42 |
| 3 | 171311729 | rs83615 | -0.544 | 0.859 | -1.941 | 0.66 | 0.021 | -0.042 | -0.042 | 0.027 | 0.76 | 0.97 | 0.82 | 0.78 | 0.83 | 0.75 |
| 3 | 171311936 | rs83616 | 0.392 | 0.883 | -1.947 | 0.021 | -0.047 | 0.124 | 0.124 | 0 | 0.54 | 0.1 | 0.44 | 0.31 | 0.58 | 0.38 |
| 3 | 171312880 | rs6780561 | -0.299 | 0.964 | -1.481 | 0.227 | -0.007 | 0.061 | 0.061 | 0.389 | 0.45 | 0.21 | 0.52 | 0.33 | 0.55 | 0.4 |
| 3 | 171314225 | rs10936694 | -0.033 | 2.068 | -0.042 | 0.784 | 0.106 | 0.124 | 0.124 | 0 | 0.09 | 0.22 | 0.33 | 0.22 | 0.19 | 0.07 |
| 3 | 171317259 | rs13327509 | -1.179 | 1.741 | 0.148 | 0.521 | -0.021 | -0.006 | -0.006 | 0 | 0.33 | 0.72 | 0.46 | 0.35 | 0.4 | 0.33 |
| 3 | 171362977 | rs360399 | -0.518 | 1.999 | 0.261 | 1.003 | -0.04 | -0.101 | -0.101 | 0.009 | 0.83 | 0.97 | 0.84 | 0.58 | 0.83 | 0.61 |
| 3 | 171369776 | rs4894707 | -1.144 | 1.961 | 0.879 | 0.722 | -0.038 | 0.001 | 0.001 | 0 | 0.31 | 0.76 | 0.47 | 0.32 | 0.39 | 0.33 |
| 3 | 171380525 | rs3821745 | -0.107 | 2.006 | 0.932 | 0.734 | 0.178 | 0.163 | 0.163 | 0 | 0.11 | 0.36 | 0.34 | 0.19 | 0.19 | 0.07 |
| 3 | 171386231 | rs1473377 | -1.147 | 1.908 | 1.114 | 0.76 | 0.015 | 0.004 | 0.004 | 0 | 0.29 | 0.58 | 0.44 | 0.32 | 0.39 | 0.31 |
| 3 | 171395468 | rs2124147 | -1.285 | 1.918 | 1.066 | 0.763 | -0.024 | -0.002 | -0.002 | 0.138 | 0.31 | 0.73 | 0.46 | 0.32 | 0.39 | 0.33 |
| 3 | 171404478 | rs2290480 | 0.183 | 1.948 | 1.048 | 0.765 | -0.006 | 0.021 | 0.021 | 0 | 0.17 | 0.14 | 0.11 | 0.13 | 0.2 | 0.25 |
| 3 | 171409603 | rs7649974 | -0.893 | 1.944 | 1.119 | 0.766 | 0.018 | 0.007 | 0.007 | 0 | 0.26 | 0.34 | 0.43 | 0.32 | 0.38 | 0.32 |
| 3 | 171419209 | rs12632647 | 0.132 | 1.984 | 1.156 | 0.812 | 0.084 | 0.085 | 0.085 | 0 | 0.1 | 0.2 | 0.33 | 0.19 | 0.18 | 0.07 |
| 3 | 171436724 | rs16856586 | -0.273 | 1.443 | 0.594 | 0.807 | 0.218 | 0.127 | 0.127 | 0 | 0.13 | 0.4 | 0.36 | 0.19 | 0.25 | 0.09 |
| 3 | 171437153 | rs9290432 | -1.232 | 1.336 | -0.112 | 0.775 | -0.035 | -0.01 | -0.01 | 0 | 0.29 | 0.76 | 0.45 | 0.32 | 0.36 | 0.32 |
| 3 | 171452645 | rs9869368 | 0.222 | 1.327 | -0.091 | 0.748 | 0.02 | 0.018 | 0.018 | 0 | 0.16 | 0.14 | 0.09 | 0.13 | 0.14 | 0.24 |
| 3 | 171463272 | rs6801138 | -0.637 | 1.414 | -0.063 | 0.807 | 0.05 | 0.021 | 0.021 | 0 | 0.15 | 0.61 | 0.38 | 0.2 | 0.26 | 0.09 |
| 3 | 171483668 | rs11715665 | -0.67 | 1.299 | -0.132 | 0.852 | -0.002 | -0.001 | -0.001 | 0.137 | 0.31 | 0.61 | 0.45 | 0.36 | 0.39 | 0.34 |
| 3 | 171498922 | rs7631361 | 0.031 | 1.605 | -0.262 | 1.365 | -0.009 | -0.001 | -0.001 | 0.286 | 0.88 | 0.9 | 0.85 | 0.59 | 0.86 | 0.68 |
| 3 | 171500490 | rs9968111 | 0.559 | 1.547 | 0.172 | 1.33 | 0.019 | 0.105 | 0.105 | 0 | 0.86 | 0.76 | 0.84 | 0.59 | 0.86 | 0.68 |
| 3 | 171510238 | rs3934620 | -0.108 | 1.917 | 0.516 | 1.276 | 0.034 | 0.019 | 0.019 | 0 | 0.16 | 0.29 | 0.39 | 0.23 | 0.27 | 0.18 |
| 3 | 171514103 | rs11714531 | 0.596 | 1.707 | 0.012 | 1.048 | -0.024 | -0.048 | -0.048 | 0 | 0.14 | 0.03 | 0.2 | 0.21 | 0.17 | 0.09 |
| 3 | 171518678 | rs4894771 | -0.623 | 1.881 | 0.195 | 1.167 | -0.023 | -0.031 | -0.031 | 0 | 0.89 | 0.94 | 0.89 | 0.63 | 0.86 | 0.69 |
| 3 | 171519793 | rs13321742 | 0.179 | 1.919 | 0.254 | 1.193 | 0.026 | 0.024 | 0.024 | 0 | 0.16 | 0.27 | 0.39 | 0.29 | 0.27 | 0.19 |
| 3 | 171520989 | rs6787821 | 0.075 | 1.824 | 0.46 | 1.165 | 0.02 | 0.074 | 0.074 | 0 | 0.86 | 0.76 | 0.87 | 0.63 | 0.85 | 0.69 |
| 3 | 171522415 | rs6445029 | 0.846 | 0.922 | -0.065 | 0.015 | 0.039 | 0.037 | 0.037 | 0 | 0.53 | 0.21 | 0.36 | 0.33 | 0.42 | 0.39 |
| 3 | 171523099 | rs4128634 | 1.142 | 0.873 | -0.172 | -0.147 | -0.006 | -0.047 | -0.047 | 3.355 | 0.15 | 0.18 | 0.11 | 0 | 0.14 | 0.1 |
| 3 | 171531746 | rs10936701 | 0.321 | 1.299 | -0.235 | 0.523 | 0.006 | 0.13 | 0.13 | 0.003 | 0.07 | 0.14 | 0.16 | 0.45 | 0.08 | 0.31 |
| 3 | 171533283 | rs3843979 | 0.54 | 1.86 | -0.463 | 0.428 | -0.05 | -0.048 | -0.048 | 0 | 0.11 | 0.05 | 0.34 | 0.26 | 0.25 | 0.11 |
| 3 | 171536255 | rs4515081 | 1.245 | 0.909 | -0.551 | -0.386 | 0.052 | 0.059 | 0.059 | 0 | 0.34 | 0.19 | 0.13 | 0.13 | 0.19 | 0.27 |
| 3 | 171537110 | rs12486202 | 1.451 | 0.724 | -0.65 | -0.574 | 0.009 | -0.003 | -0.003 | 0.269 | 0.19 | 0.14 | 0.11 | 0.01 | 0.18 | 0.11 |
| 3 | 171537598 | rs7652788 | 1.132 | 0.661 | -0.337 | -0.391 | 0.019 | 0.019 | 0.019 | 0 | 0.35 | 0.27 | 0.13 | 0.13 | 0.18 | 0.27 |
| 3 | 171545219 | rs9848505 | -0.032 | 0.953 | -0.189 | -0.697 | -0.001 | -0.061 | -0.061 | 1.45 | 0.43 | 0.47 | 0.35 | 0.14 | 0.46 | 0.33 |
| 3 | 171546804 | rs9835008 | 0.682 | 1.042 | -0.211 | -0.695 | 0.007 | -0.024 | -0.024 | 0 | 0.18 | 0.31 | 0.19 | 0.14 | 0.18 | 0.17 |
| 3 | 171550202 | rs9817267 | -0.088 | 1.143 | -0.412 | -0.749 | 0 | -0.034 | -0.034 | 0.306 | 0.45 | 0.54 | 0.37 | 0.14 | 0.49 | 0.34 |
| 3 | 171551241 | rs6788037 | 0.846 | 0.043 | -0.802 | -0.806 | 0.004 | 0.034 | 0.034 | 0 | 0.26 | 0.17 | 0.17 | 0.01 | 0.27 | 0.16 |
| 3 | 171558472 | rs4894514 | -0.089 | 1.263 | -0.663 | 0.165 | 0.003 | 0 | 0 | 0 | 0.27 | 0.28 | 0.52 | 0.4 | 0.45 | 0.28 |
| 3 | 171559827 | rs3850188 | -0.153 | 0.926 | -0.495 | 0.409 | 0.015 | 0.183 | 0.183 | 0 | 0.89 | 0.8 | 0.82 | 0.47 | 0.89 | 0.72 |
| 3 | 171562199 | rs3929518 | -0.36 | 0.909 | -1.179 | 0.417 | -0.02 | -0.01 | -0.01 | 0 | 0.31 | 0.27 | 0.53 | 0.4 | 0.47 | 0.3 |
| 3 | 171567390 | rs17479963 | -0.057 | -0.406 | -1.536 | -0.227 | 0.048 | 0.084 | 0.084 | 0.883 | 0.34 | 0.14 | 0.22 | 0.12 | 0.23 | 0.21 |
| 3 | 171580943 | rs7632751 | -0.718 | -0.364 | -1.995 | -0.005 | -0.089 | -0.023 | -0.023 | 0 | 0.76 | 0.84 | 0.59 | 0.7 | 0.52 | 0.77 |
| 3 | 171582012 | rs17421040 | 0.531 | -1.052 | -1.829 | -0.532 | 0.017 | 0.094 | 0.094 | 0 | 0.27 | 0.08 | 0.18 | 0.05 | 0.22 | 0.24 |
| 3 | 171593340 | rs3850193 | -0.188 | -0.849 | -1.081 | -0.598 | 0.006 | -0.018 | -0.018 | 0 | 0.6 | 0.48 | 0.39 | 0.19 | 0.52 | 0.42 |
| 3 | 171599454 | rs9825237 | 1.019 | -0.761 | -1.258 | -0.689 | 0.014 | 0.05 | 0.05 | 0.339 | 0.29 | 0.18 | 0.19 | 0.14 | 0.26 | 0.16 |
| 3 | 171606858 | rs11919089 | 0.476 | -0.691 | -0.794 | -0.723 | 0 | -0.04 | -0.04 | 6.066 | 0.21 | 0.22 | 0.13 | 0.01 | 0.2 | 0.14 |
| 3 | 171611313 | rs4618247 | 0.73 | -0.781 | -0.929 | -0.693 | 0.008 | 0.042 | 0.042 | 0 | 0.19 | 0.09 | 0.11 | 0.01 | 0.17 | 0.12 |
| 3 | 171622499 | rs9826915 | 0.154 | -1.259 | -0.385 | -0.097 | -0.004 | 0 | 0 | 0 | 0.46 | 0.49 | 0.38 | 0.5 | 0.38 | 0.55 |
| 3 | 171629269 | rs9873456 | 0.126 | -1.313 | -0.289 | -0.12 | -0.004 | -0.001 | -0.001 | 0.733 | 0.47 | 0.49 | 0.37 | 0.48 | 0.37 | 0.54 |
| 3 | 171645174 | rs4894786 | -1.084 | -1.243 | -0.519 | -0.174 | -0.086 | -0.083 | -0.083 | 0 | 0.73 | 0.86 | 0.5 | 0.56 | 0.53 | 0.73 |
| 3 | 171646042 | rs9832169 | 1.142 | -1.775 | -0.228 | -0.032 | 0.01 | 0.075 | 0.075 | 0.464 | 0.26 | 0.13 | 0.17 | 0.12 | 0.25 | 0.14 |
| 3 | 171653903 | rs3913363 | -0.434 | -1.498 | -0.101 | -0.099 | 0.114 | -0.016 | -0.016 | 0.197 | 0.5 | 0.8 | 0.4 | 0.53 | 0.37 | 0.57 |
| 3 | 171655674 | rs6792542 | 0.203 | -0.914 | 0.066 | 0.137 | 0.008 | -0.008 | -0.008 | 1.811 | 0.33 | 0.33 | 0.27 | 0.43 | 0.29 | 0.38 |
| 3 | 171680513 | rs7619106 | 0.31 | -0.161 | -0.693 | -0.604 | -0.014 | -0.04 | -0.04 | 0 | 0.42 | 0.41 | 0.23 | 0.16 | 0.29 | 0.29 |
| 3 | 171684515 | rs2901755 | -0.609 | 0.321 | -1.239 | -0.265 | -0.039 | -0.067 | -0.067 | 0 | 0.75 | 0.88 | 0.63 | 0.66 | 0.61 | 0.78 |
| 3 | 171684613 | rs4243421 | -0.358 | 0.319 | -0.62 | -0.103 | -0.012 | 0.024 | 0.024 | 0 | 0.28 | 0.31 | 0.24 | 0.46 | 0.16 | 0.35 |
| 3 | 171690432 | rs6782709 | 0.103 | 0.389 | -0.798 | -1.229 | 0.003 | -0.003 | -0.003 | 0 | 0.56 | 0.43 | 0.75 | 0.89 | 0.61 | 0.62 |
| 3 | 171691679 | rs3913364 | 0.129 | 0.699 | -0.858 | -1.326 | 0.002 | 0.023 | 0.023 | 0 | 0.53 | 0.42 | 0.69 | 0.89 | 0.54 | 0.6 |
| 3 | 171696155 | rs6794800 | 0.407 | 0.197 | -1.189 | -1.927 | 0.018 | 0.001 | 0.001 | 0 | 0.33 | 0.22 | 0.15 | 0 | 0.24 | 0.15 |

| | | | | | | | | | | | | | | | | |
|---|---|---|---|---|---|---|---|---|---|---|---|---|---|---|---|---|
| 3 | 171705816 | rs1499814 | -0.169 | 0.575 | -0.184 | -1.212 | 0.004 | -0.069 | -0.069 | 0.492 | 0.8 | 0.7 | 0.88 | 0.94 | 0.78 | 0.81 |
| 3 | 171726910 | rs476651 | -0.096 | 0.438 | -0.402 | -1.147 | -0.01 | -0.057 | -0.057 | 0 | 0.19 | 0.25 | 0.12 | 0.05 | 0.17 | 0.2 |
| 3 | 171731061 | rs817797 | 0.1 | 0.819 | -0.56 | -0.754 | -0.001 | -0.002 | -0.002 | 0 | 0.62 | 0.64 | 0.35 | 0.18 | 0.52 | 0.47 |
| 3 | 171733155 | rs555654 | -0.214 | 1.216 | -0.848 | 0.118 | -0.044 | -0.031 | -0.031 | 0 | 0.64 | 0.81 | 0.56 | 0.56 | 0.62 | 0.73 |
| 3 | 171736443 | rs12497673 | 0.494 | 0.467 | -0.384 | -0.518 | -0.019 | -0.064 | -0.064 | 0.744 | 0.19 | 0.33 | 0.1 | 0.05 | 0.16 | 0.1 |
| 3 | 171742181 | rs579806 | 0.164 | 0.85 | -0.408 | -0.023 | 0.002 | 0.036 | 0.036 | 0 | 0.31 | 0.45 | 0.4 | 0.47 | 0.34 | 0.3 |
| 3 | 171747356 | rs6781921 | 0.553 | 1.117 | -0.802 | 0.143 | -0.013 | 0.058 | 0.058 | 0 | 0.36 | 0.24 | 0.25 | 0.1 | 0.38 | 0.38 |
| 3 | 171748643 | rs705028 | 0.391 | 0.974 | -0.695 | 0.268 | -0.003 | -0.02 | -0.02 | 0 | 0.3 | 0.32 | 0.39 | 0.47 | 0.34 | 0.31 |
| 3 | 171750699 | rs9852803 | 0.367 | 1.114 | -0.915 | 0.027 | -0.003 | 0.023 | 0.023 | 0 | 0.33 | 0.26 | 0.23 | 0.1 | 0.33 | 0.36 |
| 3 | 171755243 | rs559235 | -0.223 | 1.022 | -0.884 | 0.809 | -0.009 | -0.044 | -0.044 | 0.001 | 0.3 | 0.24 | 0.39 | 0.46 | 0.35 | 0.31 |
| 3 | 171779873 | rs6773362 | -0.59 | 0.67 | -0.307 | 0.64 | 0.01 | 0.012 | 0.012 | 0 | 0.53 | 0.58 | 0.75 | 0.8 | 0.63 | 0.51 |
| 3 | 171780763 | rs4894796 | -0.214 | 0.682 | -0.301 | 1.233 | 0.021 | -0.004 | -0.004 | 1.504 | 0.47 | 0.38 | 0.6 | 0.55 | 0.59 | 0.44 |
| 3 | 171791275 | rs6808166 | -0.176 | 0.634 | -0.796 | 1.189 | 0.015 | -0.057 | -0.057 | 0 | 0.44 | 0.15 | 0.58 | 0.55 | 0.58 | 0.42 |
| 3 | 171800256 | rs4894803 | 0.054 | 1.422 | -1.096 | 1.623 | -0.092 | -0.026 | -0.026 | 2.553 | 0.28 | 0.08 | 0.37 | 0.28 | 0.41 | 0.23 |
| 3 | 171803226 | rs9862371 | -0.245 | 0.867 | -0.646 | 1.501 | 0.029 | -0.005 | -0.005 | 1.872 | 0.44 | 0.21 | 0.57 | 0.59 | 0.58 | 0.4 |
| 3 | 171808897 | rs7620170 | -0.158 | 0.876 | -0.428 | 1.434 | 0.011 | -0.006 | -0.006 | 0 | 0.46 | 0.41 | 0.63 | 0.56 | 0.58 | 0.41 |
| 3 | 171821513 | rs7651931 | -0.642 | 0.757 | -0.815 | 0.867 | 0.034 | 0.07 | 0.07 | 0 | 0.54 | 0.7 | 0.74 | 0.79 | 0.63 | 0.59 |
| 3 | 171842567 | rs6787115 | -0.448 | 0.402 | -1.787 | 0.866 | 0.091 | 0.115 | 0.115 | 0.154 | 0.54 | 0.24 | 0.68 | 0.8 | 0.7 | 0.62 |
| 3 | 171847196 | rs6763884 | -0.15 | 0.602 | -1.72 | 1.042 | 0.077 | -0.003 | -0.003 | 0 | 0.47 | 0.24 | 0.49 | 0.46 | 0.61 | 0.46 |
| 3 | 171854170 | rs7616583 | -0.228 | 1.247 | -1.779 | 1.065 | -0.059 | -0.085 | -0.085 | 0 | 0.18 | 0.01 | 0.09 | 0.3 | 0.21 | 0.2 |
| 3 | 171858078 | rs6765198 | 0.19 | 1.224 | -1.672 | 0.748 | 0.003 | 0.013 | 0.013 | 0 | 0.48 | 0.63 | 0.46 | 0.45 | 0.49 | 0.49 |
| 3 | 171861100 | rs4894813 | -0.37 | 1.342 | -1.569 | 0.3 | -0.017 | -0.003 | -0.003 | 0 | 0.44 | 0.29 | 0.47 | 0.59 | 0.54 | 0.45 |
| 3 | 171863080 | rs3980306 | -0.399 | 1.368 | -1.315 | 0.291 | -0.01 | 0 | 0 | 0 | 0.45 | 0.39 | 0.48 | 0.59 | 0.54 | 0.45 |
| 3 | 171863425 | rs11714903 | -0.373 | 1.32 | -1.314 | 0.292 | -0.009 | 0 | 0 | 0 | 0.44 | 0.39 | 0.48 | 0.59 | 0.54 | 0.45 |
| 3 | 171870344 | rs9825202 | -0.076 | 1.203 | -1.168 | -1.014 | 0.017 | 0.054 | 0.054 | 0 | 0.49 | 0.61 | 0.32 | 0.06 | 0.59 | 0.4 |
| 3 | 171872996 | rs6790174 | -0.349 | 0.679 | -1.243 | -1.069 | -0.016 | -0.016 | -0.016 | 0 | 0.68 | 0.64 | 0.64 | 0.2 | 0.78 | 0.55 |
| 3 | 171876079 | rs10936713 | 0.05 | 1.127 | -0.528 | -1.227 | 0.013 | -0.03 | -0.03 | 0 | 0.25 | 0.41 | 0.34 | 0.14 | 0.27 | 0.17 |
| 3 | 171878103 | rs9839226 | 0.785 | 1.609 | -1.224 | -1.658 | -0.04 | 0.096 | 0.096 | 1.381 | 0.37 | 0.19 | 0.28 | 0.04 | 0.45 | 0.31 |
| 3 | 171880571 | rs4435670 | -0.462 | 0.982 | -1.221 | -1.315 | 0.009 | 0.053 | 0.053 | 0 | 0.73 | 0.88 | 0.59 | 0.13 | 0.77 | 0.49 |
| 3 | 171886438 | rs13089740 | -0.44 | 0.442 | -0.531 | -1.367 | 0.003 | -0.033 | -0.033 | 0 | 0.39 | 0.34 | 0.29 | 0.06 | 0.25 | 0.28 |
| 3 | 171889467 | rs4450850 | -0.169 | 1.196 | -0.347 | -0.092 | 0.03 | 0.105 | 0.105 | 0 | 0.84 | 0.72 | 0.69 | 0.57 | 0.79 | 0.76 |
| 3 | 171893935 | rs6768553 | -0.289 | 1.157 | -0.567 | -0.084 | 0.003 | 0.034 | 0.034 | 0 | 0.86 | 0.85 | 0.69 | 0.57 | 0.79 | 0.76 |
| 3 | 171898476 | rs4894819 | 0.77 | 1.255 | -0.687 | -0.344 | -0.03 | -0.035 | -0.035 | 0 | 0.75 | 0.81 | 0.56 | 0.61 | 0.65 | 0.72 |
| 3 | 171904227 | rs7620401 | -0.622 | 0.697 | -0.679 | -0.729 | 0.052 | 0.119 | 0.119 | 0 | 0.5 | 0.73 | 0.44 | 0.27 | 0.38 | 0.38 |
| 3 | 171909101 | rs9837802 | -0.802 | 0.069 | 0.258 | -0.483 | -0.035 | -0.022 | -0.022 | 0 | 0.23 | 0.59 | 0.13 | 0.19 | 0.15 | 0.14 |
| 3 | 171915214 | rs7640569 | -0.155 | 0.259 | 0.416 | -0.052 | 0.002 | 0.086 | 0.086 | 0 | 0.89 | 0.8 | 0.76 | 0.65 | 0.9 | 0.87 |
| 3 | 171915776 | rs7643363 | 0.917 | 0.12 | 0.957 | -0.126 | 0.008 | 0.161 | 0.161 | 0.442 | 0.83 | 0.56 | 0.72 | 0.65 | 0.86 | 0.86 |
| 3 | 171917715 | rs9853317 | -1.287 | 0.235 | 0.541 | -0.037 | -0.013 | 0.005 | 0.005 | 0 | 0.93 | 0.96 | 0.78 | 0.65 | 0.91 | 0.87 |
| 3 | 171919214 | rs4639001 | 1.464 | 0.375 | 0.406 | -0.311 | 0.013 | 0.015 | 0.015 | 0.074 | 0.49 | 0.18 | 0.43 | 0.39 | 0.47 | 0.46 |
| 3 | 171937014 | rs6445047 | 0.254 | 0.386 | 0.056 | -0.47 | 0.001 | -0.047 | -0.047 | 1.427 | 0.71 | 0.81 | 0.68 | 0.53 | 0.75 | 0.65 |
| 3 | 171956421 | rs7614432 | -0.004 | 0.356 | 0.165 | -0.557 | -0.002 | -0.005 | -0.005 | 1.075 | 0.22 | 0.26 | 0.11 | 0.14 | 0.14 | 0.2 |
| 3 | 171956536 | rs7622134 | 0.154 | 0.354 | 0.165 | -0.574 | 0.01 | 0.015 | 0.015 | 0 | 0.21 | 0.15 | 0.1 | 0.14 | 0.14 | 0.2 |
| 3 | 171964223 | rs12635823 | -0.831 | 0.329 | 0.168 | -0.552 | 0.007 | -0.026 | -0.026 | 0.482 | 0.33 | 0.26 | 0.34 | 0.47 | 0.26 | 0.36 |
| 3 | 171965790 | rs11719016 | 0.005 | 0.366 | 0.543 | -0.725 | 0.052 | -0.069 | -0.069 | 0 | 0.17 | 0.38 | 0.2 | 0.08 | 0.24 | 0.19 |
| 3 | 171967050 | rs13099331 | 1.297 | 0.452 | 0.453 | -0.622 | -0.03 | -0.049 | -0.049 | 0.546 | 0.1 | 0.02 | 0.24 | 0.33 | 0.12 | 0.15 |
| 3 | 171973601 | rs13094089 | -1.258 | 0.438 | 0.448 | -0.665 | -0.031 | -0.033 | -0.033 | 0.022 | 0.41 | 0.31 | 0.49 | 0.55 | 0.46 | 0.54 |
| 3 | 171973943 | rs13098737 | 1.045 | 0.302 | -0.096 | -0.796 | -0.067 | 0.021 | 0.021 | 0 | 0.11 | 0.01 | 0.17 | 0.08 | 0.21 | 0.19 |
| 3 | 171978832 | rs9876039 | -0.698 | -0.505 | 0.447 | -0.467 | -0.033 | -0.073 | -0.073 | 0 | 0.23 | 0.48 | 0.14 | 0.14 | 0.17 | 0.2 |
| 3 | 171995605 | rs4894535 | 0.291 | -0.34 | -0.297 | -0.188 | -0.003 | 0.054 | 0.054 | 0.022 | 0.17 | 0.23 | 0.26 | 0.34 | 0.15 | 0.2 |
| 3 | 171997463 | rs4894413 | -1.586 | -0.34 | -0.542 | -0.337 | 0.03 | -0.006 | -0.006 | 0.279 | 0.48 | 0.77 | 0.46 | 0.57 | 0.4 | 0.56 |
| 3 | 171998350 | rs9812681 | -0.855 | -0.84 | -0.601 | -0.899 | -0.013 | -0.035 | -0.035 | 0 | 0.25 | 0.57 | 0.16 | 0.23 | 0.22 | 0.35 |
| 3 | 172002955 | rs9838734 | -0.744 | -0.82 | -1.595 | -0.883 | 0.054 | 0.106 | 0.106 | 3.581 | 0.38 | 0.77 | 0.19 | 0.15 | 0.23 | 0.26 |
| 3 | 172006714 | rs7630681 | -0.788 | -0.892 | -1.187 | -0.75 | 0.023 | 0.065 | 0.065 | 1.238 | 0.34 | 0.74 | 0.21 | 0.22 | 0.27 | 0.39 |
| 3 | 172011774 | rs12494991 | -0.77 | -1.203 | -0.467 | -0.197 | -0.003 | 0.001 | 0.001 | 0 | 0.3 | 0.29 | 0.35 | 0.34 | 0.25 | 0.23 |
| 3 | 172013848 | rs8180024 | -0.807 | -1.217 | -0.266 | -0.243 | -0.004 | 0 | 0 | 0 | 0.32 | 0.36 | 0.36 | 0.34 | 0.28 | 0.23 |
| 3 | 172016725 | rs16845307 | 0.004 | -1.687 | -0.331 | -0.405 | 0.007 | 0.02 | 0.02 | 0 | 0.32 | 0.28 | 0.12 | 0.18 | 0.24 | 0.45 |
| 3 | 172020156 | rs16845310 | -0.221 | -1.653 | -0.564 | -0.322 | 0.016 | -0.009 | -0.009 | 0 | 0.28 | 0.18 | 0.34 | 0.32 | 0.24 | 0.24 |
| 3 | 172022010 | rs17461979 | -1.378 | -1.606 | -0.67 | -0.082 | -0.107 | -0.101 | -0.101 | 1.993 | 0.64 | 0.88 | 0.47 | 0.54 | 0.46 | 0.73 |
| 3 | 172024832 | rs3733027 | 0.511 | -1.193 | -0.459 | -0.063 | 0.009 | -0.048 | -0.048 | 1.142 | 0.12 | 0.04 | 0.22 | 0.3 | 0.1 | 0.14 |
| 3 | 172026710 | rs7613643 | -0.006 | -1.303 | -0.06 | -0.399 | -0.005 | -0.072 | -0.072 | 0 | 0.25 | 0.33 | 0.12 | 0.07 | 0.2 | 0.32 |
| 3 | 172031363 | rs6773689 | -0.7 | -1.464 | -0.357 | -0.352 | -0.013 | -0.052 | -0.052 | 0.447 | 0.4 | 0.44 | 0.18 | 0.16 | 0.3 | 0.41 |
| 3 | 172034271 | rs9837821 | -0.74 | -1.431 | -1.194 | -0.327 | 0.022 | 0.055 | 0.055 | 0 | 0.42 | 0.64 | 0.2 | 0.21 | 0.32 | 0.49 |
| 3 | 172046113 | rs7642511 | -0.582 | -1.192 | -0.487 | 0.255 | 0.02 | -0.012 | -0.012 | 0.091 | 0.38 | 0.64 | 0.36 | 0.44 | 0.26 | 0.35 |
| 3 | 172048596 | rs6774744 | -1.581 | -0.733 | 0.249 | -0.062 | 0.006 | -0.033 | -0.033 | 0 | 0.08 | 0.3 | 0.06 | 0 | 0.06 | 0.03 |
| 3 | 172048734 | rs2172839 | -0.617 | -1.64 | 0.043 | -0.605 | -0.025 | -0.042 | -0.042 | 0 | 0.4 | 0.45 | 0.19 | 0.15 | 0.28 | 0.31 |
| 3 | 172053664 | rs669552 | -0.737 | -1.659 | 0.125 | -1.121 | -0.014 | -0.095 | -0.095 | 0 | 0.2 | 0.33 | 0.09 | 0.04 | 0.14 | 0.15 |
| 3 | 172059121 | rs570549 | 0.913 | -2.141 | -0.014 | -1.321 | 0.023 | 0.031 | 0.031 | 0 | 0.49 | 0.44 | 0.75 | 0.8 | 0.67 | 0.65 |
| 3 | 172060700 | rs606797 | 1.559 | -2.548 | -0.587 | -0.592 | -0.067 | -0.002 | -0.002 | 0.045 | 0.35 | 0.13 | 0.4 | 0.29 | 0.54 | 0.35 |
| 3 | 172061303 | rs593739 | -0.095 | -2.73 | -0.485 | -0.817 | 0.029 | 0.036 | 0.036 | 0 | 0.63 | 0.72 | 0.81 | 0.84 | 0.76 | 0.69 |

| | | | | | | | | | | | | | | | | |
|---|---|---|---|---|---|---|---|---|---|---|---|---|---|---|---|---|
| 3 | 172065559 | rs10513701 | 0.319 | -2.865 | -0.854 | -0.834 | 0.004 | -0.008 | -0.008 | 0.297 | 0.14 | 0.14 | 0.06 | 0.04 | 0.09 | 0.04 |
| 3 | 172069865 | rs9821137 | 0.888 | -2.929 | -1.029 | -0.877 | 0.014 | 0.019 | 0.019 | 0.396 | 0.23 | 0.05 | 0.12 | 0.15 | 0.2 | 0.19 |
| 3 | 172081022 | rs7639108 | -0.573 | -2.67 | -0.685 | -0.687 | 0.01 | -0.024 | -0.024 | 0 | 0.19 | 0.16 | 0.35 | 0.57 | 0.16 | 0.36 |
| 3 | 172087898 | rs660239 | 1.697 | -2.866 | -0.74 | -0.732 | -0.05 | 0.011 | 0.011 | 0.026 | 0.34 | 0.16 | 0.42 | 0.27 | 0.51 | 0.31 |
| 3 | 172091717 | rs4894544 | 0.083 | -2.4 | -0.319 | -0.836 | 0.059 | -0.042 | -0.042 | 0.612 | 0.9 | 0.63 | 0.92 | 1 | 0.9 | 0.93 |
| 3 | 172097617 | rs953883 | 0.517 | -2.586 | -0.415 | -0.887 | -0.008 | -0.036 | -0.036 | 0.69 | 0.14 | 0.23 | 0.08 | 0.04 | 0.09 | 0.04 |
| 3 | 172101711 | rs609424 | 0.889 | -2.823 | -0.517 | -0.659 | -0.034 | 0.079 | 0.079 | 0 | 0.76 | 0.52 | 0.59 | 0.41 | 0.79 | 0.6 |
| 3 | 172102729 | rs9870218 | -0.166 | -2.494 | -0.875 | -0.446 | 0.004 | 0.009 | 0.009 | 0 | 0.14 | 0.12 | 0.33 | 0.59 | 0.12 | 0.34 |
| 3 | 172108555 | rs9810131 | 0.242 | -2.042 | -0.941 | -0.628 | -0.005 | -0.025 | -0.025 | 7.346 | 0.14 | 0.22 | 0.07 | 0.04 | 0.08 | 0.04 |
| 3 | 172113324 | rs1285082 | 0.688 | -1.543 | -1.52 | -0.485 | -0.044 | 0.086 | 0.086 | 0.049 | 0.39 | 0.16 | 0.39 | 0.21 | 0.5 | 0.37 |
| 3 | 172115902 | rs12897 | 0.418 | -1.151 | -1.662 | -0.133 | 0.032 | 0.148 | 0.148 | 0.351 | 0.48 | 0.12 | 0.52 | 0.26 | 0.58 | 0.39 |
| 3 | 172120847 | rs9812159 | -0.903 | -0.396 | -1.258 | 0.725 | 0 | 0.019 | 0.019 | 0 | 0.14 | 0.14 | 0.27 | 0.63 | 0.14 | 0.27 |
| 3 | 172122521 | rs9856558 | 0.23 | 0.031 | -1.141 | 0.484 | -0.012 | -0.001 | -0.001 | 4.287 | 0.58 | 0.31 | 0.45 | 0.47 | 0.51 | 0.54 |
| 3 | 172131456 | rs493789 | 0.154 | -0.081 | -0.424 | 0.069 | -0.037 | -0.136 | -0.136 | 0.292 | 0.82 | 0.54 | 0.9 | 1 | 0.88 | 0.85 |
| 3 | 172132236 | rs522101 | -0.014 | -0.124 | 0.148 | 0.029 | -0.015 | -0.074 | -0.074 | 0 | 0.87 | 0.7 | 0.93 | 1 | 0.91 | 0.88 |
| 3 | 172133060 | rs551228 | 0.326 | 0.11 | -0.113 | 0.254 | -0.002 | 0.003 | 0.003 | 0.462 | 0.21 | 0.24 | 0.41 | 0.45 | 0.34 | 0.23 |
| 3 | 172133487 | rs2593858 | 0.257 | 0.102 | -0.013 | 0.245 | 0.013 | 0.025 | 0.025 | 0.575 | 0.21 | 0.28 | 0.41 | 0.45 | 0.33 | 0.23 |
| 3 | 172138843 | rs520418 | 1.059 | -0.083 | 0.241 | 0.284 | -0.013 | 0.01 | 0.01 | 0.077 | 0.73 | 0.39 | 0.7 | 0.61 | 0.77 | 0.43 |
| 3 | 172145026 | rs16845548 | 0.138 | 0.101 | 0.12 | 0.428 | 0.001 | 0.096 | 0.096 | 0 | 0.09 | 0.12 | 0.19 | 0.38 | 0.11 | 0.4 |
| 3 | 172157110 | rs558572 | 0.765 | -0.23 | 0.008 | -0.113 | -0.045 | -0.168 | -0.168 | 0 | 0.73 | 0.56 | 0.85 | 0.99 | 0.81 | 0.8 |
| 3 | 172162117 | rs565105 | 0.559 | -0.214 | -0.04 | 0.035 | -0.009 | 0.044 | 0.044 | 0 | 0.74 | 0.53 | 0.68 | 0.62 | 0.73 | 0.42 |
| 3 | 172163449 | rs509035 | 0.987 | 0.071 | -0.445 | 0.066 | -0.073 | -0.104 | -0.104 | 0 | 0.17 | 0.06 | 0.29 | 0.42 | 0.28 | 0.22 |
| 3 | 172165727 | rs572169 | 0.972 | 0.061 | -0.45 | 0.08 | -0.073 | -0.103 | -0.103 | 0 | 0.17 | 0.06 | 0.29 | 0.42 | 0.28 | 0.22 |
| 3 | 172166033 | rs495225 | 1.241 | -0.166 | 0.151 | 0.103 | -0.001 | -0.001 | -0.001 | 0 | 0.64 | 0.4 | 0.66 | 0.62 | 0.7 | 0.39 |
| 3 | 172170104 | rs9819506 | -0.746 | -0.006 | 0.256 | 0.037 | -0.006 | 0.021 | 0.021 | 0 | 0.29 | 0.52 | 0.35 | 0.39 | 0.31 | 0.51 |
| 3 | 172185376 | rs9881097 | -1.318 | 0.024 | 0.108 | -0.272 | -0.006 | 0.01 | 0.01 | 0.131 | 0.55 | 0.66 | 0.6 | 0.41 | 0.56 | 0.64 |
| 3 | 172193632 | rs6445063 | 0.081 | -0.049 | 0.025 | -0.614 | 0 | -0.028 | -0.028 | 0 | 0.15 | 0.16 | 0.1 | 0 | 0.15 | 0.15 |
| 3 | 172202876 | rs2100143 | -1.243 | -0.1 | 0.107 | -0.301 | 0.004 | 0.002 | 0.002 | 0 | 0.51 | 0.57 | 0.49 | 0.54 | 0.5 | 0.6 |
| 3 | 172202999 | rs2084373 | -0.022 | 0.539 | 0.113 | -0.16 | 0.073 | 0.128 | 0.128 | 0 | 0.82 | 0.66 | 0.61 | 0.64 | 0.71 | 0.76 |
| 3 | 172215575 | rs2041693 | -0.886 | 0.102 | 0.175 | -0.154 | -0.002 | 0.002 | 0.002 | 0 | 0.41 | 0.53 | 0.41 | 0.53 | 0.43 | 0.48 |
| 3 | 172215637 | rs2041692 | 0.415 | 0.153 | -0.032 | -0.271 | 0.001 | 0.054 | 0.054 | 0 | 0.18 | 0.18 | 0.23 | 0.51 | 0.21 | 0.39 |
| 3 | 172220976 | rs1992868 | -0.147 | 0.466 | -0.21 | -0.403 | -0.016 | -0.019 | -0.019 | 0 | 0.76 | 0.78 | 0.56 | 0.65 | 0.69 | 0.64 |
| 3 | 172223376 | rs11720451 | -0.117 | 0.461 | -0.211 | -0.405 | -0.015 | -0.017 | -0.017 | 0 | 0.76 | 0.77 | 0.56 | 0.65 | 0.69 | 0.64 |
| 3 | 172224075 | rs1131535 | -1.134 | 0.374 | -0.195 | -0.455 | 0.009 | -0.029 | -0.029 | 0 | 0.42 | 0.67 | 0.42 | 0.54 | 0.43 | 0.47 |
| 3 | 172226240 | rs3136601 | -1.169 | 0.343 | -0.166 | -0.362 | 0.012 | -0.016 | -0.016 | 0 | 0.42 | 0.66 | 0.41 | 0.57 | 0.43 | 0.47 |
| 3 | 172227199 | rs3815496 | 0.05 | 1.175 | 0.049 | -0.065 | -0.015 | -0.014 | -0.014 | 0.039 | 0.76 | 0.77 | 0.56 | 0.69 | 0.69 | 0.64 |
| 3 | 172228544 | rs3136597 | -0.517 | 1.106 | 0.12 | -0.055 | -0.004 | -0.105 | -0.105 | 0 | 0.22 | 0.47 | 0.21 | 0.06 | 0.23 | 0.08 |
| 3 | 172228827 | rs1126161 | 0.196 | 1.231 | -0.213 | 0.188 | -0.014 | -0.01 | -0.01 | 0 | 0.76 | 0.77 | 0.59 | 0.74 | 0.69 | 0.64 |
| 3 | 172231986 | rs3774315 | 1.896 | 1.141 | -0.487 | -0.036 | 0.027 | 0.127 | 0.127 | 0 | 0.37 | 0.09 | 0.2 | 0.16 | 0.32 | 0.17 |
| 3 | 172233369 | rs4894559 | -0.22 | 1.374 | -0.236 | 0.383 | -0.05 | -0.059 | -0.059 | 0 | 0.84 | 0.92 | 0.65 | 0.74 | 0.76 | 0.76 |
| 3 | 172234844 | rs231986 | -0.154 | 1.181 | 0.218 | 0.105 | -0.008 | -0.069 | -0.069 | 0 | 0.84 | 0.72 | 0.89 | 1 | 0.86 | 0.85 |
| 3 | 172240999 | rs2270418 | 0.365 | 1.328 | -0.004 | 0.351 | -0.006 | -0.038 | -0.038 | 0.059 | 0.16 | 0.09 | 0.2 | 0.55 | 0.15 | 0.39 |
| 3 | 172241866 | rs365238 | -0.082 | 1.641 | 0.688 | 0.575 | 0.001 | 0.024 | 0.024 | 0 | 0.86 | 0.9 | 0.91 | 1 | 0.86 | 0.84 |
| 3 | 172242747 | rs388515 | -1.386 | 1.838 | 1.308 | 0.964 | 0.025 | 0.084 | 0.084 | 0 | 0.19 | 0.58 | 0.37 | 0.29 | 0.19 | 0.23 |
| 3 | 172245403 | rs17601515 | -0.81 | 0.965 | 1.175 | 0.586 | -0.014 | -0.004 | -0.004 | 0 | 0.09 | 0.08 | 0.13 | 0.17 | 0.18 | 0.06 |
| 3 | 172245463 | rs9854603 | -2.233 | 0.75 | 1.882 | 0.699 | 0.024 | 0.004 | 0.004 | 0 | 0.14 | 0.46 | 0.12 | 0.17 | 0.16 | 0.05 |
| 3 | 172247790 | rs11925997 | -0.915 | 1.125 | 1.423 | 0.845 | 0.033 | 0.046 | 0.046 | 0.043 | 0.07 | 0.24 | 0.08 | 0.16 | 0.1 | 0.04 |
| 3 | 172258131 | rs17601879 | -1.327 | 1.072 | 1.386 | 0.81 | 0.005 | 0.018 | 0.018 | 0 | 0.08 | 0.25 | 0.08 | 0.16 | 0.09 | 0.04 |
| 3 | 172262053 | rs13316580 | -1.449 | 1.155 | 1.37 | 0.839 | 0.065 | 0.045 | 0.045 | 0 | 0.1 | 0.43 | 0.11 | 0.16 | 0.12 | 0.05 |
| 3 | 172270248 | rs3843973 | 0.44 | 1.259 | 0.874 | 1.223 | 0.022 | 0.074 | 0.074 | 0 | 0.76 | 0.65 | 0.6 | 0.55 | 0.68 | 0.54 |
| 3 | 172270437 | rs3900917 | -0.267 | 0.972 | 0.489 | 0.718 | -0.001 | -0.013 | -0.013 | 0.783 | 0.22 | 0.19 | 0.27 | 0.31 | 0.21 | 0.29 |
| 3 | 172273512 | rs231997 | -0.634 | 1.103 | 0.342 | 1.086 | 0.002 | 0.085 | 0.085 | 0 | 0.13 | 0.21 | 0.29 | 0.41 | 0.16 | 0.24 |
| 3 | 172276754 | rs231989 | -0.175 | 1.149 | 0.483 | 0.961 | 0.01 | 0 | 0 | 0 | 0.8 | 0.77 | 0.85 | 0.82 | 0.76 | 0.77 |
| 3 | 172281430 | rs9868605 | 0.159 | 0.95 | 0.143 | 0.765 | -0.004 | -0.016 | -0.016 | 0 | 0.18 | 0.12 | 0.25 | 0.34 | 0.18 | 0.27 |
| 3 | 172291604 | rs234063 | -0.118 | 1.04 | -0.208 | 0.687 | 0.016 | -0.021 | -0.021 | 0 | 0.77 | 0.69 | 0.84 | 0.82 | 0.73 | 0.76 |
| 3 | 172292484 | rs2286983 | -0.771 | 0.769 | -0.176 | 0.819 | -0.011 | 0.049 | 0.049 | 0 | 0.25 | 0.44 | 0.29 | 0.33 | 0.2 | 0.29 |
| 3 | 172294880 | rs234061 | 0.529 | 0.944 | -0.563 | 0.646 | -0.002 | 0.009 | 0.009 | 0 | 0.72 | 0.32 | 0.81 | 0.84 | 0.74 | 0.75 |
| 3 | 172294944 | rs6805167 | 0.757 | 0.815 | -0.986 | 0.766 | -0.001 | -0.058 | -0.058 | 0.589 | 0.61 | 0.17 | 0.55 | 0.45 | 0.65 | 0.57 |
| 3 | 172301933 | rs11918360 | -0.123 | 0.651 | -0.897 | -0.089 | 0.002 | 0.018 | 0.018 | 0 | 0.45 | 0.43 | 0.38 | 0.28 | 0.4 | 0.4 |
| 3 | 172302780 | rs17608660 | 0.835 | 0.582 | -1.444 | -0.419 | -0.053 | -0.036 | -0.036 | 0 | 0.22 | 0.03 | 0.23 | 0.25 | 0.24 | 0.19 |
| 3 | 172304658 | rs16845868 | -0.102 | 0.49 | -1.092 | -0.469 | -0.003 | -0.048 | -0.048 | 0.003 | 0.16 | 0.17 | 0.07 | 0.02 | 0.12 | 0.17 |
| 3 | 172310820 | rs234055 | -0.41 | 0.407 | -0.759 | -0.659 | 0.002 | 0.017 | 0.017 | 0 | 0.69 | 0.78 | 0.73 | 0.76 | 0.71 | 0.72 |
| 3 | 172311712 | rs234051 | -0.404 | 0.379 | -0.586 | -0.602 | 0.002 | 0.018 | 0.018 | 0 | 0.69 | 0.79 | 0.74 | 0.75 | 0.72 | 0.72 |
| 3 | 172313268 | rs234044 | 0.981 | 0.354 | -1.067 | -0.441 | -0.029 | -0.004 | -0.004 | 0 | 0.24 | 0.2 | 0.23 | 0.25 | 0.37 | 0.17 |
| 3 | 172313367 | rs234043 | -0.757 | 0.416 | -1.231 | -0.248 | 0.001 | 0.008 | 0.008 | 0 | 0.7 | 0.8 | 0.74 | 0.74 | 0.72 | 0.72 |
| 3 | 172314227 | rs234041 | 0.833 | 0.6 | -1.291 | -0.237 | -0.072 | -0.03 | -0.03 | 0 | 0.22 | 0.08 | 0.19 | 0.25 | 0.36 | 0.15 |
| 3 | 172314679 | rs16845879 | -0.342 | -0.139 | -0.622 | -0.184 | -0.03 | -0.046 | -0.046 | 0 | 0.12 | 0.19 | 0.03 | 0.03 | 0.03 | 0.15 |
| 3 | 172318864 | rs4894568 | 0.11 | 0.449 | -0.827 | 0.15 | 0.005 | -0.004 | -0.004 | 0 | 0.37 | 0.32 | 0.42 | 0.39 | 0.36 | 0.31 |
| 3 | 172319115 | rs6784737 | 0.804 | 0.259 | -1.102 | 0.059 | -0.001 | 0.054 | 0.054 | 0.011 | 0.27 | 0.03 | 0.19 | 0.16 | 0.22 | 0.16 |

| | | | | | | | | | | | | | | | | |
|---|---|---|---|---|---|---|---|---|---|---|---|---|---|---|---|---|
| 3 | 172321159 | rs6779133 | 0.823 | 0.252 | -1.104 | 0.054 | -0.002 | 0.061 | 0.061 | 0.061 | 0.27 | 0.03 | 0.19 | 0.15 | 0.22 | 0.16 |
| 3 | 172321434 | rs16845891 | 1.316 | -0.098 | -1.063 | 0.129 | 0.009 | -0.021 | -0.021 | 0 | 0.18 | 0.14 | 0.3 | 0.38 | 0.18 | 0.34 |
| 3 | 172321698 | rs9848146 | 0.902 | 0.133 | -0.828 | -0.432 | 0 | -0.024 | -0.024 | 0 | 0.07 | 0.07 | 0.03 | 0 | 0.05 | 0.06 |
| 3 | 172321768 | rs17524400 | 1.372 | 0.174 | -0.857 | -0.416 | -0.002 | 0.062 | 0.062 | 0.069 | 0.27 | 0.03 | 0.19 | 0.15 | 0.22 | 0.16 |
| 3 | 172321976 | rs9872004 | 0.248 | 0.327 | -0.192 | -0.375 | -0.006 | -0.056 | -0.056 | 0 | 0.1 | 0.16 | 0.04 | 0 | 0.06 | 0.07 |
| 3 | 172331985 | rs374777 | -0.387 | 1.2 | -0.06 | -0.177 | 0.005 | -0.008 | -0.008 | 0 | 0.71 | 0.68 | 0.76 | 0.82 | 0.67 | 0.77 |
| 3 | 172339752 | rs6771424 | 0.529 | 0.72 | 0.023 | -0.562 | -0.009 | -0.083 | -0.083 | 0 | 0.32 | 0.37 | 0.25 | 0.13 | 0.27 | 0.15 |
| 3 | 172340607 | rs234025 | -0.293 | 1.024 | -0.152 | -0.342 | 0.029 | 0.052 | 0.052 | 0 | 0.53 | 0.71 | 0.53 | 0.41 | 0.61 | 0.35 |
| 3 | 172347134 | rs234016 | -0.08 | 0.962 | -0.09 | -0.322 | 0.007 | 0.038 | 0.038 | 0 | 0.49 | 0.62 | 0.48 | 0.4 | 0.56 | 0.33 |
| 3 | 172349216 | rs3205144 | 0.822 | 1.036 | -0.447 | -0.25 | -0.004 | -0.013 | -0.013 | 0.651 | 0.28 | 0.29 | 0.22 | 0.19 | 0.23 | 0.13 |
| 3 | 172351006 | rs3733023 | 0.279 | 1.144 | -0.751 | -0.326 | 0.001 | 0.003 | 0.003 | 0 | 0.32 | 0.3 | 0.25 | 0.18 | 0.32 | 0.15 |
| 3 | 172359864 | rs234003 | -0.502 | 1.452 | -0.299 | 0.063 | 0.004 | -0.002 | -0.002 | 0.784 | 0.32 | 0.39 | 0.31 | 0.3 | 0.42 | 0.25 |
| 3 | 172362707 | rs98106 | 0.18 | 1.695 | -0.911 | 0.414 | -0.031 | -0.029 | -0.029 | 0 | 0.23 | 0.18 | 0.26 | 0.3 | 0.32 | 0.22 |
| 3 | 172365619 | rs9865941 | 0.288 | 1.497 | -0.784 | 0.299 | -0.007 | -0.008 | -0.008 | 0 | 0.67 | 0.66 | 0.62 | 0.54 | 0.55 | 0.57 |
| 3 | 172366481 | rs170883 | -0.324 | 0.979 | -1.485 | -0.68 | -0.002 | 0.156 | 0.156 | 0.098 | 0.67 | 0.9 | 0.82 | 0.94 | 0.69 | 0.69 |
| 3 | 172368701 | rs6778941 | 0.053 | 1.093 | -0.965 | -0.389 | 0.02 | 0.012 | 0.012 | 0 | 0.33 | 0.37 | 0.42 | 0.44 | 0.45 | 0.27 |
| 3 | 172369828 | rs2901811 | -0.493 | 0.995 | -0.929 | -0.655 | 0.001 | -0.033 | -0.033 | 0 | 0.58 | 0.55 | 0.76 | 0.94 | 0.64 | 0.69 |
| 3 | 172371929 | rs17756228 | -0.011 | 1.397 | -1.316 | -0.636 | 0 | 0.055 | 0.055 | 0 | 0.49 | 0.34 | 0.6 | 0.74 | 0.46 | 0.56 |
| 3 | 172372521 | rs10513704 | 0.565 | 1.803 | -1.572 | -0.072 | -0.003 | -0.003 | -0.003 | 0 | 0.09 | 0.08 | 0.16 | 0.21 | 0.17 | 0.13 |
| 3 | 172374119 | rs6796724 | 0.528 | 1.522 | -1.538 | -0.148 | -0.031 | -0.015 | -0.015 | 0 | 0.2 | 0.1 | 0.26 | 0.22 | 0.28 | 0.15 |
| 3 | 172374658 | rs669376 | -0.043 | 0.788 | -1.472 | -0.726 | 0.006 | 0.093 | 0.093 | 0 | 0.33 | 0.2 | 0.18 | 0.06 | 0.3 | 0.25 |
| 3 | 172377148 | rs3902414 | -1.251 | 0.859 | -1.118 | -0.038 | -0.019 | -0.009 | -0.009 | 0 | 0.39 | 0.82 | 0.34 | 0.44 | 0.39 | 0.4 |
| 3 | 172384590 | rs9883329 | -1.191 | 0.965 | -0.845 | -0.205 | -0.005 | 0.013 | 0.013 | 0 | 0.38 | 0.67 | 0.33 | 0.36 | 0.39 | 0.39 |
| 3 | 172385693 | rs11712263 | -0.449 | 1.295 | -1.514 | -0.062 | -0.053 | 0.018 | 0.018 | 0 | 0.23 | 0.08 | 0.14 | 0.15 | 0.28 | 0.24 |
| 3 | 172389700 | rs9881902 | -0.512 | 0.799 | -0.728 | -0.079 | -0.001 | 0.016 | 0.016 | 0 | 0.1 | 0.2 | 0.15 | 0.2 | 0.1 | 0.15 |
| 3 | 172394890 | rs492748 | 0.172 | 1.359 | -1.262 | -0.651 | -0.041 | 0.061 | 0.061 | 0.047 | 0.78 | 0.92 | 0.86 | 1 | 0.71 | 0.88 |
| 3 | 172395351 | rs497316 | 0.821 | 0.825 | -0.997 | -0.371 | 0.001 | 0.039 | 0.039 | 0.974 | 0.55 | 0.38 | 0.64 | 0.67 | 0.53 | 0.58 |
| 3 | 172397868 | rs483772 | 0.489 | 1.379 | -0.794 | -0.576 | -0.007 | 0.01 | 0.01 | 0 | 0.62 | 0.38 | 0.6 | 0.77 | 0.5 | 0.68 |
| 3 | 172399955 | rs4894576 | -0.649 | 1.471 | -0.414 | -0.805 | 0.025 | -0.014 | -0.014 | 0 | 0.27 | 0.4 | 0.24 | 0.25 | 0.33 | 0.41 |
| 3 | 172400899 | rs9821237 | 0.238 | 1.454 | -0.445 | -0.822 | -0.007 | -0.007 | -0.007 | 0 | 0.17 | 0.27 | 0.12 | 0.17 | 0.15 | 0.28 |
| 3 | 172404542 | rs494694 | -1.26 | 1.607 | -0.867 | -1.165 | -0.049 | 0.174 | 0.174 | 0.221 | 0.48 | 0.9 | 0.39 | 0.25 | 0.54 | 0.51 |
| 3 | 172412887 | rs651165 | 0.463 | 1.248 | -0.668 | -1.163 | -0.024 | 0.052 | 0.052 | 0 | 0.17 | 0.08 | 0.12 | 0 | 0.21 | 0.12 |
| 3 | 172421663 | rs487255 | -0.975 | 1.356 | 0.052 | -1.265 | 0.02 | -0.063 | -0.063 | 0 | 0.3 | 0.56 | 0.29 | 0.14 | 0.39 | 0.32 |
| 3 | 172421968 | rs1105152 | 0.058 | 1.335 | -0.021 | -1.249 | -0.004 | -0.014 | -0.014 | 0 | 0.2 | 0.19 | 0.12 | 0 | 0.22 | 0.13 |
| 3 | 172428782 | rs2276805 | 0.953 | 1.23 | -0.059 | -1.3 | 0.023 | 0.047 | 0.047 | 0 | 0.64 | 0.18 | 0.75 | 0.72 | 0.69 | 0.56 |
| 3 | 172449817 | rs4894588 | 0.829 | 1.2 | 0.157 | -1.26 | -0.014 | 0.061 | 0.061 | 0 | 0.11 | 0.31 | 0.08 | 0.15 | 0.06 | 0.16 |
| 3 | 172455843 | rs9863121 | 0.374 | 1.229 | 0.181 | -1.295 | 0.002 | -0.066 | -0.066 | 0 | 0.75 | 0.56 | 0.83 | 0.86 | 0.75 | 0.72 |
| 3 | 172457287 | rs7612516 | -0.349 | 1.159 | 0.138 | -1.222 | 0.014 | 0.002 | 0.002 | 0 | 0.17 | 0.32 | 0.19 | 0.16 | 0.21 | 0.24 |
| 3 | 172462291 | rs11719562 | 1.532 | 1.293 | 0.006 | -1.266 | -0.053 | 0.087 | 0.087 | 0.192 | 0.57 | 0.05 | 0.63 | 0.71 | 0.56 | 0.5 |
| 3 | 172471316 | rs10936737 | 1.5 | 1.153 | 0.027 | -1.232 | -0.011 | 0.086 | 0.086 | 0.142 | 0.57 | 0.07 | 0.65 | 0.7 | 0.58 | 0.51 |
| 3 | 172476419 | rs7646507 | -0.723 | 1.157 | 0.086 | -1.265 | -0.042 | 0.032 | 0.032 | 0 | 0.87 | 0.71 | 0.92 | 0.85 | 0.94 | 0.82 |
| 3 | 172498216 | rs4459929 | 1.536 | 1.163 | 0.11 | -1.256 | 0.008 | 0.087 | 0.087 | 0.019 | 0.58 | 0.12 | 0.65 | 0.7 | 0.58 | 0.51 |
| 3 | 172514294 | rs10804849 | 0.116 | 1.174 | 0.106 | -1.229 | -0.003 | -0.009 | -0.009 | 0 | 0.81 | 0.7 | 0.82 | 0.85 | 0.79 | 0.74 |
| 3 | 172527977 | rs11714381 | 0.101 | 1.182 | 0.11 | -1.225 | -0.003 | -0.009 | -0.009 | 0 | 0.81 | 0.71 | 0.83 | 0.85 | 0.79 | 0.74 |
| 3 | 172533725 | rs4491935 | 0.646 | 1.275 | 0.133 | -1.207 | 0 | -0.032 | -0.032 | 0 | 0.16 | 0.19 | 0.11 | 0 | 0.17 | 0.07 |
| 3 | 172558312 | rs9813760 | -0.91 | 1.118 | -0.506 | -1.142 | -0.02 | -0.003 | -0.003 | 0 | 0.27 | 0.54 | 0.21 | 0.24 | 0.24 | 0.43 |
| 3 | 172558337 | rs9833643 | -0.93 | 0.973 | -1.411 | -1.036 | -0.007 | 0.122 | 0.122 | 0.071 | 0.46 | 0.94 | 0.36 | 0.25 | 0.42 | 0.57 |
| 3 | 172564677 | rs7433573 | -0.193 | 1.32 | -1.493 | -1.024 | 0.003 | -0.016 | -0.016 | 0.516 | 0.29 | 0.35 | 0.29 | 0.21 | 0.29 | 0.3 |
| 3 | 172564788 | rs9290453 | -0.678 | 1.158 | -2.11 | -0.199 | -0.04 | -0.054 | -0.054 | 0 | 0.59 | 0.95 | 0.64 | 0.44 | 0.64 | 0.65 |
| 3 | 172576619 | rs13066810 | -0.885 | 1.422 | -1.19 | -0.081 | -0.002 | 0.007 | 0.007 | 0 | 0.15 | 0.06 | 0.11 | 0.17 | 0.14 | 0.14 |
| 3 | 172582602 | rs1499626 | 0.486 | 1.855 | -0.991 | 0.545 | 0.005 | 0.012 | 0.012 | 0 | 0.92 | 0.93 | 0.89 | 0.84 | 0.85 | 0.93 |
| 3 | 172584312 | rs6787188 | -0.327 | 1.381 | -1.016 | 0.079 | 0.011 | 0.024 | 0.024 | 0.771 | 0.25 | 0.18 | 0.19 | 0.08 | 0.2 | 0.08 |
| 3 | 172587874 | rs869258 | 0.537 | 1.605 | -0.834 | 0.508 | 0.002 | 0.001 | 0.001 | 0 | 0.92 | 0.93 | 0.89 | 0.87 | 0.85 | 0.93 |
| 3 | 172591359 | rs6799845 | 0.784 | 1.23 | -0.764 | 0.388 | 0.001 | 0.003 | 0.003 | 2.125 | 0.69 | 0.8 | 0.73 | 0.73 | 0.71 | 0.85 |
| 3 | 172592311 | rs7639187 | 1.636 | 1.045 | -0.533 | 0.047 | 0.003 | 0.062 | 0.062 | 0.109 | 0.51 | 0.18 | 0.41 | 0.64 | 0.49 | 0.58 |
| 3 | 172594253 | rs4299505 | 1.552 | 0.851 | -0.214 | 0.026 | 0.02 | 0.048 | 0.048 | 0.202 | 0.53 | 0.26 | 0.45 | 0.67 | 0.55 | 0.66 |
| 3 | 172601502 | rs12638320 | -0.133 | 0.857 | 0.282 | -0.12 | -0.013 | -0.009 | -0.009 | 0 | 0.09 | 0.14 | 0.11 | 0.05 | 0.07 | 0.11 |
| 3 | 172603682 | rs1156615 | 1.795 | 0.71 | 0.341 | -0.161 | 0.011 | 0.104 | 0.104 | 0.925 | 0.52 | 0.1 | 0.44 | 0.69 | 0.55 | 0.67 |
| 3 | 172604675 | rs12637859 | -1.162 | -0.038 | 1.004 | -0.199 | -0.034 | -0.041 | -0.041 | 0.012 | 0.13 | 0.34 | 0.15 | 0.05 | 0.1 | 0.12 |
| 3 | 172623001 | rs12629617 | -0.138 | 0.001 | 0.941 | -0.174 | -0.014 | -0.01 | -0.01 | 0 | 0.09 | 0.15 | 0.11 | 0.05 | 0.07 | 0.11 |
| 3 | 172624328 | rs13070200 | 1.62 | 0.15 | 0.745 | -0.175 | 0.038 | 0.05 | 0.05 | 0 | 0.56 | 0.23 | 0.48 | 0.69 | 0.6 | 0.71 |
| 3 | 172632849 | rs1499630 | 1.847 | 0.129 | 0.686 | -0.185 | 0.057 | 0.099 | 0.099 | 1.228 | 0.52 | 0.11 | 0.45 | 0.69 | 0.58 | 0.7 |
| 3 | 172648167 | rs13099104 | 1.718 | 0.115 | 0.671 | 0.009 | 0.054 | 0.053 | 0.053 | 0 | 0.57 | 0.12 | 0.5 | 0.69 | 0.64 | 0.68 |
| 3 | 172650339 | rs1499632 | 0.857 | 0.238 | 1.063 | 0.021 | -0.01 | -0.025 | -0.025 | 0.213 | 0.66 | 0.51 | 0.61 | 0.76 | 0.68 | 0.82 |
| 3 | 172659200 | rs1587420 | -0.301 | 0.294 | 0.909 | 0.256 | 0.054 | 0.021 | 0.021 | 1.53 | 0.29 | 0.13 | 0.27 | 0.26 | 0.2 | 0.22 |
| 3 | 172661239 | rs1353868 | -1.594 | 0.266 | 1.257 | 0.388 | 0.001 | 0 | 0 | 0 | 0.4 | 0.42 | 0.43 | 0.38 | 0.38 | 0.37 |
| 3 | 172672695 | rs12636168 | 0.165 | 0.668 | 1.32 | 0.362 | -0.019 | -0.023 | -0.023 | 0 | 0.05 | 0.02 | 0.09 | 0.15 | 0.11 | 0.1 |
| 3 | 172672792 | rs6781018 | -1.248 | 0.923 | 1.236 | 0.386 | 0.001 | 0.001 | 0.001 | 0 | 0.45 | 0.6 | 0.4 | 0.33 | 0.38 | 0.38 |
| 3 | 172673482 | rs9864296 | -1.77 | 1.065 | 1.355 | 0.34 | -0.003 | 0.001 | 0.001 | 0 | 0.53 | 0.59 | 0.54 | 0.47 | 0.56 | 0.52 |

| | | | | | | | | | | | | | | | | |
|---|---|---|---|---|---|---|---|---|---|---|---|---|---|---|---|---|
| 3 | 172681942 | rs12497957 | -0.567 | 1.077 | 1.033 | 0.322 | 0.04 | 0.025 | 0.025 | 0.065 | 0.39 | 0.12 | 0.35 | 0.32 | 0.35 | 0.34 |
| 3 | 172695355 | rs6445091 | -1.552 | 0.685 | 1.073 | 0.496 | -0.006 | -0.005 | -0.005 | 0 | 0.55 | 0.69 | 0.55 | 0.47 | 0.56 | 0.53 |
| 3 | 172718955 | rs12629188 | -0.771 | 0.167 | 1.263 | 1.189 | -0.008 | 0.011 | 0.011 | 0 | 0.19 | 0.26 | 0.22 | 0.34 | 0.14 | 0.23 |
| 3 | 172721786 | rs1515438 | -1.146 | 0.161 | 1.271 | 1.048 | -0.013 | 0.107 | 0.107 | 0.044 | 0.25 | 0.65 | 0.36 | 0.48 | 0.24 | 0.35 |
| 3 | 172728171 | rs791832 | -0.922 | 0.116 | 1.249 | 0.688 | 0.005 | -0.004 | -0.004 | 0 | 0.18 | 0.12 | 0.12 | 0 | 0.19 | 0.07 |
| 3 | 172735766 | rs575511 | -1.274 | 0.28 | 2.073 | 1.6 | 0.009 | 0.087 | 0.087 | 0.358 | 0.11 | 0.26 | 0.16 | 0.14 | 0.13 | 0.16 |
| 3 | 172751927 | rs16846434 | -0.749 | 0.482 | 1.533 | 1.46 | -0.021 | -0.006 | -0.006 | 0 | 0.09 | 0.02 | 0.06 | 0.08 | 0.11 | 0.09 |
| 3 | 172766822 | rs508508 | -1.091 | 0.222 | 2.271 | 1.623 | 0.002 | 0.049 | 0.049 | 0.034 | 0.1 | 0.22 | 0.18 | 0.14 | 0.15 | 0.15 |
| 3 | 172772994 | rs10936741 | -0.771 | 0.168 | 2.271 | 1.214 | 0.007 | -0.036 | -0.036 | 0.668 | 0.07 | 0.17 | 0.04 | 0.01 | 0.05 | 0.06 |
| 3 | 172775311 | rs1398455 | -0.91 | 0.208 | 2.361 | 1.589 | -0.005 | 0.021 | 0.021 | 0 | 0.11 | 0.12 | 0.14 | 0.22 | 0.09 | 0.09 |
| 3 | 172780554 | rs7620311 | -0.553 | 0.186 | 2.296 | 1.64 | -0.002 | -0.002 | -0.002 | 0 | 0.12 | 0.16 | 0.19 | 0.16 | 0.19 | 0.26 |
| 3 | 172782231 | rs7645546 | -2.084 | 0.104 | 2.713 | 1.608 | 0.001 | -0.007 | -0.007 | 0 | 0.31 | 0.72 | 0.39 | 0.39 | 0.32 | 0.42 |
| 3 | 172782527 | rs7638139 | -1.482 | 0.209 | 2.692 | 1.58 | -0.022 | 0.041 | 0.041 | 0 | 0.19 | 0.57 | 0.2 | 0.23 | 0.13 | 0.15 |
| 3 | 172786148 | rs6766715 | -0.89 | 0.357 | 2.14 | 1.626 | -0.008 | 0.061 | 0.061 | 0 | 0.16 | 0.35 | 0.19 | 0.24 | 0.13 | 0.13 |
| 3 | 172795631 | rs12630529 | -0.729 | 0.27 | 1.399 | 1.416 | -0.004 | -0.011 | -0.011 | 0 | 0.15 | 0.08 | 0.18 | 0.16 | 0.18 | 0.32 |
| 3 | 172798800 | rs1878007 | -0.731 | 0.286 | 1.189 | 1.423 | -0.027 | -0.041 | -0.041 | 0 | 0.14 | 0.01 | 0.17 | 0.16 | 0.18 | 0.32 |
| 3 | 172823151 | rs6778909 | -1.498 | -0.004 | 2.263 | 1.178 | -0.031 | 0.049 | 0.049 | 0 | 0.2 | 0.42 | 0.2 | 0.24 | 0.1 | 0.18 |
| 3 | 172828084 | rs2861071 | -2.103 | 0.169 | 2.702 | 1.103 | -0.006 | 0.003 | 0.003 | 0 | 0.31 | 0.66 | 0.31 | 0.3 | 0.24 | 0.31 |
| 3 | 172830063 | rs13070480 | -2.187 | 0.154 | 2.889 | 1.103 | -0.003 | 0.012 | 0.012 | 0 | 0.3 | 0.39 | 0.29 | 0.36 | 0.25 | 0.33 |
| 3 | 172832917 | rs1355653 | -1.314 | 0.128 | 3.011 | 1.115 | 0 | -0.002 | -0.002 | 0 | 0.19 | 0.19 | 0.21 | 0.31 | 0.12 | 0.24 |
| 3 | 172835082 | rs16846616 | -0.332 | 0.113 | 2.955 | 1.105 | -0.016 | -0.074 | -0.074 | 0 | 0.13 | 0.02 | 0.17 | 0.31 | 0.09 | 0.14 |
| 3 | 172835125 | rs1515442 | -2.447 | 0.125 | 3.021 | 1.124 | 0.002 | -0.005 | -0.005 | 0 | 0.32 | 0.72 | 0.31 | 0.37 | 0.25 | 0.36 |
| 3 | 172835290 | rs1515441 | -1.135 | 0.18 | 2.911 | 1.13 | 0.014 | -0.048 | -0.048 | 0 | 0.17 | 0.1 | 0.18 | 0.31 | 0.1 | 0.19 |
| 3 | 172843082 | rs11708501 | -1.838 | 0.164 | 2.946 | 1.107 | 0.001 | -0.07 | -0.07 | 0 | 0.18 | 0.45 | 0.13 | 0.07 | 0.17 | 0.2 |
| 3 | 172845803 | rs11929145 | -0.665 | 0.208 | 2.629 | 1.13 | 0.005 | -0.016 | -0.016 | 0 | 0.12 | 0.21 | 0.09 | 0.06 | 0.14 | 0.12 |
| 3 | 172851177 | rs1512747 | -2.109 | 0.19 | 2.632 | 1.142 | 0.005 | -0.01 | -0.01 | 0 | 0.34 | 0.75 | 0.32 | 0.37 | 0.27 | 0.34 |
| 3 | 172855271 | rs16846652 | -1.317 | -0.247 | 2.251 | 0.645 | -0.001 | -0.008 | -0.008 | 0 | 0.23 | 0.33 | 0.16 | 0.18 | 0.18 | 0.23 |
| 3 | 172858094 | rs10460879 | -2.209 | -0.216 | 2.441 | 0.768 | 0.001 | -0.005 | -0.005 | 0 | 0.35 | 0.66 | 0.28 | 0.35 | 0.32 | 0.36 |
| 3 | 172859705 | rs7652619 | -1.32 | -0.208 | 2.431 | 0.621 | -0.001 | 0.002 | 0.002 | 0 | 0.23 | 0.22 | 0.16 | 0.18 | 0.18 | 0.23 |
| 3 | 172863513 | rs1606845 | -1.356 | -0.203 | 2.432 | 0.584 | -0.001 | 0.001 | 0.001 | 0 | 0.23 | 0.27 | 0.16 | 0.23 | 0.18 | 0.23 |
| 3 | 172881415 | rs1512745 | -0.845 | 0.393 | 2.673 | 0.484 | 0.005 | 0.001 | 0.001 | 0.221 | 0.16 | 0.21 | 0.12 | 0.17 | 0.18 | 0.17 |
| 3 | 172881577 | rs1512744 | -0.845 | 0.393 | 2.673 | 0.484 | 0.005 | 0.001 | 0.001 | 0.221 | 0.16 | 0.21 | 0.12 | 0.17 | 0.18 | 0.17 |
| 3 | 172882261 | rs16826562 | -1.393 | 0.331 | 2.836 | 0.442 | 0.005 | 0.003 | 0.003 | 0.042 | 0.07 | 0.24 | 0.05 | 0.03 | 0.07 | 0.04 |
| 3 | 172883801 | rs7650308 | -1.868 | 0.33 | 2.155 | 0.441 | 0.007 | -0.007 | -0.007 | 0 | 0.25 | 0.59 | 0.18 | 0.2 | 0.24 | 0.21 |
| 3 | 172884734 | rs1512743 | -1.749 | 0.335 | 1.667 | 0.443 | 0.001 | -0.004 | -0.004 | 0 | 0.25 | 0.66 | 0.18 | 0.2 | 0.24 | 0.21 |
| 3 | 172888973 | rs7611388 | -1.285 | 0.241 | 1.226 | 0.429 | 0.017 | 0.012 | 0.012 | 2.868 | 0.09 | 0.51 | 0.07 | 0.03 | 0.06 | 0.03 |
| 3 | 172917785 | rs1607765 | -0.831 | 0.298 | 0.496 | 0.475 | 0.001 | 0.008 | 0.008 | 4.509 | 0.09 | 0.52 | 0.07 | 0.03 | 0.06 | 0.02 |
| 3 | 172950553 | rs2861203 | -0.824 | 0.519 | -0.832 | 0.516 | 0.006 | 0.024 | 0.024 | 0.431 | 0.34 | 0.79 | 0.26 | 0.24 | 0.32 | 0.22 |
| 3 | 172973714 | rs792127 | -0.824 | 0.536 | -0.344 | 0.668 | -0.007 | -0.001 | -0.001 | 0 | 0.26 | 0.31 | 0.14 | 0.24 | 0.22 | 0.21 |
| 3 | 172973907 | rs792128 | -1.058 | 0.93 | -0.265 | 0.908 | 0 | -0.001 | -0.001 | 0.012 | 0.38 | 0.62 | 0.26 | 0.36 | 0.4 | 0.32 |
| 3 | 172974531 | rs10936746 | -1.198 | 2.222 | -0.018 | 0.775 | 0.036 | 0.001 | 0.001 | 1.828 | 0.09 | 0.08 | 0.2 | 0.09 | 0.27 | 0.12 |
| 3 | 172982314 | rs9840514 | -1.311 | 1.705 | 0.593 | 0.686 | -0.016 | 0.02 | 0.02 | 1.348 | 0.41 | 0.62 | 0.58 | 0.66 | 0.48 | 0.51 |
| 3 | 172986183 | rs9290468 | -1.416 | 1.605 | 0.867 | 0.609 | -0.031 | 0.041 | 0.041 | 1.213 | 0.42 | 0.75 | 0.59 | 0.65 | 0.49 | 0.51 |
| 3 | 172993091 | rs9853065 | -0.994 | 2.179 | 0.922 | 0.647 | -0.002 | 0.002 | 0.002 | 0.464 | 0.36 | 0.62 | 0.44 | 0.56 | 0.31 | 0.4 |
| 3 | 173001387 | rs10936748 | -0.565 | 2.183 | 1.01 | 0.554 | 0.039 | -0.026 | -0.026 | 0 | 0.1 | 0.16 | 0.2 | 0.03 | 0.24 | 0.14 |
| 3 | 173004184 | rs6772821 | -1.597 | 2.161 | 1.03 | 0.568 | -0.001 | 0.036 | 0.036 | 1.241 | 0.45 | 0.79 | 0.61 | 0.6 | 0.55 | 0.54 |
| 3 | 173005973 | rs2030398 | -0.595 | 1.879 | 0.965 | 0.559 | 0.034 | -0.023 | -0.023 | 0 | 0.1 | 0.16 | 0.18 | 0.03 | 0.24 | 0.14 |
| 3 | 173007330 | rs9825385 | -1.066 | 0.756 | 1.206 | 0.872 | 0.015 | -0.038 | -0.038 | 0.308 | 0.16 | 0.48 | 0.15 | 0.09 | 0.11 | 0.08 |
| 3 | 173008491 | rs12489095 | -1.401 | 0.865 | 0.97 | 0.874 | -0.004 | 0.018 | 0.018 | 0.147 | 0.32 | 0.33 | 0.4 | 0.54 | 0.28 | 0.39 |
| 3 | 173010075 | rs9840933 | -0.554 | 1.014 | 0.685 | 0.929 | 0.015 | -0.028 | -0.028 | 0 | 0.13 | 0.23 | 0.13 | 0.09 | 0.11 | 0.09 |
| 3 | 173012476 | rs4894444 | -1.113 | 1.026 | 0.454 | 0.931 | -0.012 | 0.023 | 0.023 | 0.495 | 0.34 | 0.5 | 0.42 | 0.54 | 0.28 | 0.39 |
| 3 | 173018314 | rs705041 | 0.748 | 1.013 | 0.336 | 0.792 | -0.007 | -0.004 | -0.004 | 0 | 0.62 | 0.52 | 0.61 | 0.68 | 0.67 | 0.71 |
| 3 | 173026064 | rs16863019 | -2.13 | 0.703 | 1.351 | 0.645 | 0.041 | 0.032 | 0.032 | 0.025 | 0.05 | 0.38 | 0.03 | 0 | 0.01 | 0.01 |
| 3 | 173036814 | rs13071945 | -0.766 | 1.449 | 0.973 | 1.363 | -0.018 | -0.003 | -0.003 | 0 | 0.11 | 0.08 | 0.25 | 0.48 | 0.17 | 0.35 |
| 3 | 173052338 | rs1402216 | -1.965 | 1.386 | 1.27 | 1.267 | 0.004 | -0.02 | -0.02 | 0 | 0.07 | 0.28 | 0.03 | 0.01 | 0.02 | 0.09 |
| 3 | 173055512 | rs7629349 | -1.549 | 1.42 | 1.419 | 1.329 | 0.034 | 0.152 | 0.152 | 0 | 0.14 | 0.31 | 0.27 | 0.46 | 0.16 | 0.31 |
| 3 | 173065435 | rs11915539 | -1.301 | 1.353 | 1.43 | 1.26 | 0.008 | -0.019 | -0.019 | 0.044 | 0.07 | 0.35 | 0.03 | 0.01 | 0.02 | 0.09 |
| 3 | 173079012 | rs10513713 | -1.982 | 1.318 | 1.474 | 1.288 | 0.036 | 0.193 | 0.193 | 0 | 0.19 | 0.48 | 0.28 | 0.48 | 0.18 | 0.37 |
| 3 | 173089355 | rs7632429 | -0.656 | 1.384 | 0.801 | 1.333 | -0.032 | -0.028 | -0.028 | 0 | 0.11 | 0.04 | 0.23 | 0.47 | 0.15 | 0.3 |
| 3 | 173107443 | rs247975 | -1.219 | 0.585 | -0.665 | 0.135 | -0.009 | 0.018 | 0.018 | 0 | 0.55 | 0.74 | 0.57 | 0.62 | 0.53 | 0.65 |
| 3 | 173124923 | rs497383 | -0.192 | 0.339 | -1.302 | -0.562 | -0.013 | 0.058 | 0.058 | 0.301 | 0.28 | 0.11 | 0.23 | 0.12 | 0.27 | 0.18 |
| 3 | 173130026 | rs532021 | 0.375 | 0.439 | -1.322 | -0.833 | -0.033 | 0.084 | 0.084 | 0 | 0.15 | 0.02 | 0.16 | 0 | 0.15 | 0.09 |
| 3 | 173130573 | rs621369 | -0.95 | 0.143 | -1.002 | -1.314 | 0.003 | 0.008 | 0.008 | 0 | 0.53 | 0.57 | 0.69 | 0.78 | 0.56 | 0.67 |
| 3 | 173131044 | rs619092 | 0.105 | -0.461 | -0.919 | -1.452 | 0.018 | 0.051 | 0.051 | 1.025 | 0.58 | 0.28 | 0.73 | 0.79 | 0.65 | 0.78 |
| 3 | 173131865 | rs605211 | -0.987 | -1.732 | -0.575 | -1.863 | 0.025 | 0.074 | 0.074 | 2.158 | 0.55 | 0.7 | 0.44 | 0.19 | 0.38 | 0.4 |
| 3 | 173133190 | rs686535 | -0.79 | -1.33 | -0.232 | -1.387 | -0.021 | -0.045 | -0.045 | 0 | 0.4 | 0.54 | 0.34 | 0.18 | 0.27 | 0.29 |
| 3 | 173133322 | rs548757 | -0.768 | -1.218 | -0.856 | -1.338 | -0.003 | -0.014 | -0.014 | 0 | 0.39 | 0.38 | 0.33 | 0.18 | 0.26 | 0.29 |
| 3 | 173136240 | rs569221 | -0.068 | -0.847 | -1.982 | -1.056 | 0.042 | 0.084 | 0.084 | 4.422 | 0.28 | 0.09 | 0.28 | 0.15 | 0.19 | 0.2 |
| 3 | 173152759 | rs247979 | -0.325 | -0.577 | -1.788 | -0.945 | 0.074 | 0.127 | 0.127 | 2.346 | 0.46 | 0.8 | 0.41 | 0.25 | 0.3 | 0.38 |

| | | | | | | | | | | | | | | | | |
|---|---|---|---|---|---|---|---|---|---|---|---|---|---|---|---|---|
| 3 | 173153693 | rs584633 | 0.413 | -1.194 | -1.259 | -1.963 | -0.022 | -0.127 | -0.127 | 0 | 0.57 | 0.53 | 0.82 | 0.97 | 0.73 | 0.78 |
| 3 | 173154109 | rs586444 | 0.154 | -1.705 | -1.084 | -2.167 | -0.001 | -0.024 | -0.024 | 0.064 | 0.87 | 0.86 | 0.94 | 0.96 | 0.88 | 0.9 |
| 3 | 173155275 | rs569009 | -0.469 | -1.727 | -0.844 | -2.256 | 0.007 | -0.034 | -0.034 | 0.049 | 0.13 | 0.28 | 0.19 | 0.06 | 0.11 | 0.1 |
| 3 | 173158613 | rs658132 | -0.922 | -1.556 | -0.931 | -2.38 | 0.029 | 0.065 | 0.065 | 0 | 0.37 | 0.72 | 0.3 | 0.07 | 0.24 | 0.19 |
| 3 | 173160531 | rs626337 | 0.88 | -1.541 | -0.955 | -2.827 | 0.051 | 0.145 | 0.145 | 0 | 0.57 | 0.27 | 0.77 | 0.91 | 0.65 | 0.68 |
| 3 | 173161912 | rs583173 | 0.801 | -1.586 | 0.143 | -2.861 | -0.003 | -0.053 | -0.053 | 0.181 | 0.58 | 0.42 | 0.79 | 0.91 | 0.66 | 0.69 |
| 3 | 173162477 | rs12488070 | -0.179 | -1.937 | 0.547 | -3.041 | -0.029 | -0.13 | -0.13 | 0.134 | 0.79 | 0.56 | 0.87 | 0.97 | 0.84 | 0.91 |
| 3 | 173171411 | rs247984 | -0.299 | -1.955 | 0.884 | -3.053 | -0.024 | -0.111 | -0.111 | 0.923 | 0.8 | 0.64 | 0.87 | 0.97 | 0.84 | 0.91 |
| 3 | 173176376 | rs247968 | 1.513 | -1.955 | 1.039 | -2.99 | 0.069 | 0.197 | 0.197 | 0 | 0.56 | 0.22 | 0.77 | 0.91 | 0.65 | 0.68 |
| 3 | 173179298 | rs17318925 | -0.9 | -1.999 | 1.149 | -3.006 | -0.009 | -0.033 | -0.033 | 0.363 | 0.22 | 0.21 | 0.09 | 0.06 | 0.18 | 0.23 |
| 3 | 173180633 | rs11709498 | 1.985 | -2.113 | 1.115 | -3.064 | 0.054 | 0.256 | 0.256 | 0 | 0.47 | 0.18 | 0.63 | 0.91 | 0.61 | 0.69 |
| 3 | 173180781 | rs665913 | 0.21 | -2.117 | 1.016 | -3.056 | -0.008 | 0.025 | 0.025 | 0 | 0.12 | 0.05 | 0.17 | 0.06 | 0.12 | 0.11 |
| 3 | 173185376 | rs6771377 | 2.118 | -2.142 | 0.979 | -3.037 | -0.023 | 0.205 | 0.205 | 0 | 0.44 | 0.16 | 0.61 | 0.85 | 0.56 | 0.58 |
| 3 | 173209781 | rs548436 | -0.138 | -1.828 | 1.266 | -2.852 | -0.02 | -0.121 | -0.121 | 0 | 0.79 | 0.58 | 0.87 | 0.97 | 0.83 | 0.91 |
| 3 | 173211632 | rs7646820 | 0.908 | -1.767 | 0.901 | -2.764 | 0.005 | -0.036 | -0.036 | 0 | 0.63 | 0.47 | 0.56 | 0.79 | 0.57 | 0.54 |
| 3 | 173223419 | rs484341 | 0.23 | -1.518 | 0.228 | -2.911 | 0.036 | 0.137 | 0.137 | 0 | 0.7 | 0.89 | 0.86 | 0.97 | 0.78 | 0.89 |
| 3 | 173224367 | rs6767159 | 0.887 | -1.142 | 0.434 | -3.158 | 0.002 | 0.062 | 0.062 | 0 | 0.51 | 0.6 | 0.51 | 0.88 | 0.47 | 0.7 |
| 3 | 173233354 | rs6779841 | -0.505 | -1.204 | 0.846 | -3.103 | 0.038 | 0.051 | 0.051 | 0.21 | 0.07 | 0.34 | 0.04 | 0.05 | 0.05 | 0.09 |
| 3 | 173239453 | rs2901898 | 0.578 | -1.294 | 0.949 | -3.219 | 0 | -0.06 | -0.06 | 0 | 0.56 | 0.49 | 0.51 | 0.88 | 0.53 | 0.73 |
| 3 | 173246813 | rs7652570 | 0.172 | -1.498 | 0.832 | -3.318 | 0.029 | 0.023 | 0.023 | 0 | 0.14 | 0.06 | 0.03 | 0 | 0.07 | 0.1 |
| 3 | 173248181 | rs16827533 | 0.106 | -1.49 | 0.836 | -3.313 | 0.031 | 0.031 | 0.031 | 0 | 0.14 | 0.04 | 0.02 | 0.05 | 0.07 | 0.08 |
| 3 | 173256811 | rs4289370 | 0.228 | -1.292 | 0.716 | -3.315 | -0.019 | 0.175 | 0.175 | 0 | 0.55 | 0.8 | 0.52 | 0.86 | 0.5 | 0.71 |
| 3 | 173259356 | rs10936759 | 0.543 | -1.321 | 0.699 | -3.33 | -0.013 | 0.207 | 0.207 | 0 | 0.52 | 0.78 | 0.52 | 0.86 | 0.48 | 0.71 |
| 3 | 173259658 | rs11713253 | 2.208 | -1.299 | 0.641 | -3.315 | -0.095 | 0.107 | 0.107 | 0 | 0.33 | 0.1 | 0.43 | 0.81 | 0.38 | 0.53 |
| 3 | 173264141 | rs9833723 | 0.33 | -1.28 | 0.738 | -3.29 | -0.003 | 0.09 | 0.09 | 0 | 0.54 | 0.7 | 0.51 | 0.81 | 0.5 | 0.65 |
| 3 | 173267784 | rs766004 | 0.561 | -1.284 | 0.537 | -3.291 | 0 | -0.002 | -0.002 | 0 | 0.52 | 0.52 | 0.49 | 0.81 | 0.5 | 0.64 |
| 3 | 173272363 | rs6445111 | 0.302 | -1.25 | 0.328 | -3.281 | -0.006 | 0.119 | 0.119 | 0 | 0.54 | 0.72 | 0.51 | 0.81 | 0.5 | 0.65 |
| 3 | 173276758 | rs9842389 | 1.644 | -1.058 | 0.095 | -2.862 | 0.025 | -0.045 | -0.045 | 0 | 0.35 | 0.43 | 0.43 | 0.74 | 0.37 | 0.52 |
| 3 | 173281266 | rs1080310 | 1.974 | -1.112 | -0.427 | -2.763 | -0.083 | 0.048 | 0.048 | 0 | 0.32 | 0.12 | 0.42 | 0.73 | 0.38 | 0.52 |
| 3 | 173281958 | rs6808060 | 0.55 | -1.188 | -0.319 | -2.811 | 0 | -0.046 | -0.046 | 0 | 0.49 | 0.44 | 0.48 | 0.86 | 0.46 | 0.68 |
| 3 | 173289281 | rs9844612 | 0.619 | -1.161 | -0.405 | -2.744 | 0 | -0.015 | -0.015 | 0.676 | 0.48 | 0.43 | 0.47 | 0.73 | 0.46 | 0.62 |
| 3 | 173299666 | rs6803916 | 1.496 | -0.726 | -1.165 | -2.478 | -0.07 | 0.048 | 0.048 | 0 | 0.33 | 0.15 | 0.44 | 0.73 | 0.4 | 0.54 |
| 3 | 173301037 | rs3774047 | -0.482 | -0.05 | -0.969 | -2.023 | -0.025 | 0 | 0 | 0 | 0.08 | 0.03 | 0.23 | 0.09 | 0.12 | 0.1 |
| 3 | 173309839 | rs1983060 | 0.453 | 0.083 | -0.824 | -2.209 | 0.006 | 0.012 | 0.012 | 0 | 0.54 | 0.47 | 0.51 | 0.85 | 0.51 | 0.74 |
| 3 | 173310353 | rs17178976 | 0.496 | 0.778 | -1.202 | -1.177 | 0.001 | 0.013 | 0.013 | 0 | 0.51 | 0.43 | 0.57 | 0.74 | 0.5 | 0.6 |
| 3 | 173310458 | rs1980296 | -0.795 | 0.966 | -1.599 | -0.196 | -0.085 | -0.097 | -0.097 | 0.345 | 0.19 | 0.02 | 0.24 | 0.28 | 0.28 | 0.22 |
| 3 | 173314411 | rs6801149 | 0.742 | 0.286 | -1.171 | 0.027 | -0.001 | 0.003 | 0.003 | 0 | 0.89 | 0.87 | 0.84 | 0.94 | 0.89 | 0.94 |
| 3 | 173317852 | rs16828127 | 0.503 | -0.613 | -1.414 | 0.066 | 0.051 | 0.006 | 0.006 | 0 | 0.2 | 0.06 | 0.11 | 0.15 | 0.1 | 0.22 |
| 3 | 173322248 | rs3844523 | 0.558 | -0.807 | -0.775 | -0.03 | 0.011 | 0.007 | 0.007 | 0 | 0.69 | 0.69 | 0.73 | 0.79 | 0.8 | 0.74 |
| 3 | 173322277 | rs11718812 | 0.767 | -0.801 | -0.919 | -0.051 | 0.059 | 0.035 | 0.035 | 0 | 0.18 | 0.01 | 0.04 | 0.08 | 0.08 | 0.19 |
| 3 | 173323726 | rs10513715 | 1.073 | -1.365 | -1.353 | -0.701 | 0.027 | 0.073 | 0.073 | 0.418 | 0.19 | 0.08 | 0.09 | 0.04 | 0.13 | 0.1 |
| 3 | 173328600 | rs9820667 | 0.924 | -1.828 | -0.574 | -0.539 | 0.009 | 0.007 | 0.007 | 1.02 | 0.46 | 0.38 | 0.62 | 0.72 | 0.68 | 0.66 |
| 3 | 173332780 | rs11712636 | 0.841 | -1.959 | -0.436 | -0.563 | 0.023 | 0.002 | 0.002 | 0 | 0.22 | 0.15 | 0.06 | 0.16 | 0.09 | 0.21 |
| 3 | 173336649 | rs10936764 | 0.849 | -1.938 | -0.37 | -0.589 | -0.008 | -0.005 | -0.005 | 0.006 | 0.45 | 0.41 | 0.62 | 0.72 | 0.67 | 0.63 |
| 3 | 173336718 | rs9852917 | -0.242 | -1.989 | -0.291 | -0.616 | 0.02 | 0.01 | 0.01 | 0 | 0.32 | 0.3 | 0.25 | 0.2 | 0.14 | 0.25 |
| 3 | 173339777 | rs9868927 | 1.361 | -1.514 | -0.058 | -0.543 | -0.014 | 0.001 | 0.001 | 0 | 0.33 | 0.24 | 0.4 | 0.26 | 0.53 | 0.31 |
| 3 | 173340221 | rs4894620 | -0.208 | -1.526 | 0.457 | -1.307 | -0.031 | -0.043 | -0.043 | 2.44 | 0.74 | 0.57 | 0.7 | 0.87 | 0.81 | 0.84 |
| 3 | 173342387 | rs2901925 | -1.212 | -1.303 | 0.88 | -1.469 | 0.03 | 0.048 | 0.048 | 0.336 | 0.53 | 0.67 | 0.37 | 0.24 | 0.34 | 0.37 |
| 3 | 173344536 | rs6801163 | -0.557 | -1.175 | 0.847 | -1.36 | -0.02 | -0.016 | -0.016 | 0.73 | 0.33 | 0.42 | 0.27 | 0.2 | 0.21 | 0.27 |
| 3 | 173353739 | rs16828799 | 0.959 | -1.089 | 0.349 | -1.338 | -0.005 | -0.019 | -0.019 | 0 | 0.2 | 0.02 | 0.06 | 0.15 | 0.15 | 0.22 |
| 3 | 173362365 | rs16828950 | 0.451 | -0.969 | 0.582 | -1.278 | -0.02 | -0.016 | -0.016 | 0 | 0.25 | 0.35 | 0.09 | 0.15 | 0.16 | 0.22 |
| 3 | 173379185 | rs9290476 | -0.145 | -1.047 | 0.342 | -1.397 | 0.018 | 0.03 | 0.03 | 0 | 0.28 | 0.27 | 0.27 | 0.08 | 0.19 | 0.14 |
| 3 | 173379289 | rs9290477 | -1.052 | -1.01 | 0.386 | -1.369 | 0.009 | 0.016 | 0.016 | 0.198 | 0.53 | 0.63 | 0.37 | 0.24 | 0.36 | 0.37 |
| 3 | 173386910 | rs1553121 | -1.038 | -1.003 | 0.385 | -1.369 | 0.007 | 0.012 | 0.012 | 0.125 | 0.53 | 0.61 | 0.36 | 0.24 | 0.36 | 0.37 |
| 3 | 173397582 | rs11708138 | -0.277 | -1.037 | 0.46 | -1.387 | -0.026 | -0.056 | -0.056 | 0 | 0.33 | 0.45 | 0.13 | 0.13 | 0.24 | 0.23 |
| 3 | 173401090 | rs1502465 | 1.092 | -0.962 | -0.098 | -1.385 | 0.024 | 0.062 | 0.062 | 0.065 | 0.18 | 0.07 | 0.09 | 0.04 | 0.12 | 0.1 |
| 3 | 173402170 | rs1393450 | 0.055 | -0.868 | 0.015 | -1.329 | -0.014 | -0.026 | -0.026 | 0 | 0.31 | 0.36 | 0.12 | 0.15 | 0.22 | 0.23 |
| 3 | 173406816 | rs1502463 | -0.199 | -0.531 | -0.172 | -1.287 | 0.002 | 0 | 0 | 0 | 0.23 | 0.28 | 0.25 | 0.08 | 0.15 | 0.14 |
| 3 | 173410323 | rs1502461 | -0.46 | -0.59 | -0.292 | -1.291 | 0.024 | 0.116 | 0.116 | 0 | 0.26 | 0.09 | 0.12 | 0.04 | 0.19 | 0.11 |
| 3 | 173413519 | rs7638716 | -1.089 | 0.313 | 0.336 | -0.661 | -0.007 | -0.006 | -0.006 | 2.3 | 0.08 | 0.19 | 0.18 | 0.04 | 0.05 | 0.04 |
| 3 | 173418466 | rs987741 | 0.687 | 0.663 | 0.317 | -0.983 | 0 | 0.029 | 0.029 | 0 | 0.62 | 0.32 | 0.7 | 0.79 | 0.61 | 0.71 |
| 3 | 173420348 | rs2901924 | 0.05 | 0.75 | 0.388 | -1.112 | -0.026 | 0.024 | 0.024 | 0 | 0.16 | 0.09 | 0.22 | 0.08 | 0.24 | 0.16 |
| 3 | 173422417 | rs959148 | 1.502 | 0.708 | 0.63 | -1.14 | -0.005 | 0.047 | 0.047 | 0 | 0.57 | 0.33 | 0.6 | 0.74 | 0.58 | 0.65 |
| 3 | 173424373 | rs4894626 | 0.663 | 0.703 | 0.713 | -1.134 | -0.017 | -0.028 | -0.028 | 0 | 0.74 | 0.41 | 0.83 | 0.82 | 0.82 | 0.82 |
| 3 | 173431705 | rs9815508 | 0.988 | 0.636 | 1.137 | -1.337 | 0.001 | -0.055 | -0.055 | 0 | 0.65 | 0.54 | 0.73 | 0.78 | 0.66 | 0.75 |
| 3 | 173432639 | rs16829911 | 0.305 | 0.673 | 0.893 | -1.316 | -0.017 | 0.028 | 0.028 | 0 | 0.15 | 0.06 | 0.13 | 0.07 | 0.18 | 0.15 |
| 3 | 173438357 | rs3980098 | 1.514 | 0.749 | 0.939 | -1.301 | -0.012 | 0.081 | 0.081 | 0.007 | 0.57 | 0.26 | 0.6 | 0.75 | 0.58 | 0.66 |
| 3 | 173440437 | rs13087558 | -0.158 | 1.219 | 0.67 | -1.2 | -0.036 | 0.034 | 0.034 | 0 | 0.18 | 0.11 | 0.24 | 0.08 | 0.27 | 0.16 |
| 3 | 173440913 | rs3850180 | -0.186 | 0.711 | 1.174 | -0.971 | -0.002 | -0.068 | -0.068 | 0 | 0.84 | 0.76 | 0.9 | 1 | 0.86 | 0.93 |

| | | | | | | | | | | | | | | | | |
|---|---|---|---|---|---|---|---|---|---|---|---|---|---|---|---|---|
| 3 | 173441676 | rs9877840 | -0.063 | 0.616 | 0.939 | -0.893 | -0.028 | -0.033 | -0.033 | 0 | 0.83 | 0.55 | 0.87 | 0.86 | 0.88 | 0.91 |
| 3 | 173453868 | rs1488564 | 0.153 | 0.701 | 0.142 | -0.739 | 0.007 | 0.018 | 0.018 | 0 | 0.54 | 0.58 | 0.71 | 0.75 | 0.62 | 0.68 |
| 3 | 173462988 | rs6779753 | 0.105 | 0.571 | -0.496 | -0.767 | -0.005 | 0.068 | 0.068 | 0 | 0.54 | 0.66 | 0.62 | 0.77 | 0.57 | 0.67 |
| 3 | 173466849 | rs722472 | -0.604 | -0.183 | -0.218 | -0.806 | -0.014 | -0.056 | -0.056 | 0.008 | 0.23 | 0.31 | 0.13 | 0.1 | 0.17 | 0.31 |
| 3 | 173468188 | rs1038644 | -0.202 | -0.236 | 0.001 | -0.826 | -0.008 | -0.034 | -0.034 | 0.864 | 0.16 | 0.24 | 0.1 | 0.1 | 0.11 | 0.3 |
| 3 | 173469241 | rs1387004 | 0.667 | -0.634 | -1.042 | -1.014 | 0.021 | 0.147 | 0.147 | 0 | 0.49 | 0.1 | 0.62 | 0.64 | 0.6 | 0.47 |
| 3 | 173472327 | rs2046718 | 0.12 | -1.047 | -0.498 | -1.288 | -0.011 | -0.085 | -0.085 | 0 | 0.68 | 0.42 | 0.8 | 0.93 | 0.77 | 0.62 |
| 3 | 173475674 | rs991932 | 0.641 | -0.798 | -0.67 | -1.108 | 0.051 | 0.138 | 0.138 | 0 | 0.5 | 0.22 | 0.62 | 0.66 | 0.6 | 0.46 |
| 3 | 173477704 | rs9863894 | 0.763 | -1.186 | -0.063 | -1.233 | 0.015 | 0.005 | 0.005 | 0 | 0.57 | 0.62 | 0.65 | 0.05 | 0.67 | 0.46 |
| 3 | 173485371 | rs976683 | 0.965 | -1.571 | 0.492 | -0.378 | 0.042 | 0.021 | 0.021 | 0.941 | 0.53 | 0.38 | 0.67 | 0.34 | 0.7 | 0.48 |
| 3 | 173487044 | rs901884 | -0.058 | -1.9 | 0.771 | -0.112 | 0.039 | -0.037 | -0.037 | 0.255 | 0.68 | 0.8 | 0.74 | 0.4 | 0.82 | 0.58 |
| 3 | 173489454 | rs9879266 | 1.287 | -1.626 | 0.876 | -0.262 | 0.003 | 0.003 | 0.003 | 0.338 | 0.52 | 0.48 | 0.64 | 0.34 | 0.65 | 0.48 |
| 3 | 173491396 | rs1948161 | 0.681 | -1.593 | 1.109 | -0.167 | -0.008 | -0.001 | -0.001 | 1.66 | 0.6 | 0.59 | 0.7 | 0.37 | 0.76 | 0.51 |
| 3 | 173494302 | rs9830510 | 0.356 | -1.774 | 1.349 | 0.091 | 0.003 | -0.003 | -0.003 | 0.019 | 0.68 | 0.7 | 0.72 | 0.36 | 0.82 | 0.52 |
| 3 | 173497299 | rs9818020 | 1.256 | -1.825 | 1.514 | 0.052 | 0.016 | -0.004 | -0.004 | 0 | 0.58 | 0.37 | 0.66 | 0.38 | 0.71 | 0.51 |
| 3 | 173507395 | rs4499633 | 0.658 | -1.694 | 1.331 | 0.152 | -0.003 | 0.002 | 0.002 | 0 | 0.65 | 0.67 | 0.69 | 0.42 | 0.74 | 0.56 |
| 3 | 173510129 | rs10513719 | 0.853 | -1.721 | 1.476 | 0.182 | -0.024 | 0.016 | 0.016 | 0 | 0.62 | 0.57 | 0.68 | 0.39 | 0.72 | 0.51 |
| 3 | 173514219 | rs7640492 | 0.132 | -1.75 | 1.153 | 0.195 | 0.026 | -0.032 | -0.032 | 0 | 0.72 | 0.8 | 0.75 | 0.42 | 0.85 | 0.57 |
| 3 | 173514581 | rs13059427 | -0.748 | -1.518 | 1.083 | 0.216 | 0.02 | -0.033 | -0.033 | 0.032 | 0.06 | 0.11 | 0.05 | 0 | 0.11 | 0.02 |
| 3 | 173533557 | rs6790407 | 0.221 | -1.772 | 0.807 | 0.373 | 0.061 | 0.032 | 0.032 | 0 | 0.69 | 0.8 | 0.73 | 0.23 | 0.82 | 0.57 |
| 3 | 173534651 | rs6782728 | 0.2 | -1.768 | 0.655 | 0.378 | 0.081 | 0.03 | 0.03 | 0 | 0.69 | 0.86 | 0.73 | 0.23 | 0.83 | 0.57 |
| 3 | 173538547 | rs7612948 | -1.073 | -1.601 | 0.703 | 0.369 | 0.002 | -0.007 | -0.007 | 3.865 | 0.08 | 0.06 | 0.05 | 0.03 | 0.13 | 0.06 |
| 3 | 173553008 | rs1948160 | 1.006 | -1.629 | 0.843 | 0.342 | 0.01 | 0.012 | 0.012 | 0 | 0.6 | 0.68 | 0.67 | 0.19 | 0.71 | 0.5 |
| 3 | 173553783 | rs9829309 | 0.935 | -1.717 | 0.703 | 0.354 | 0.034 | -0.023 | -0.023 | 0 | 0.61 | 0.81 | 0.67 | 0.39 | 0.7 | 0.49 |
| 3 | 173556777 | rs7619910 | 0.275 | -2.058 | 0.619 | 0.497 | 0.035 | 0.015 | 0.015 | 0 | 0.69 | 0.78 | 0.73 | 0.23 | 0.84 | 0.57 |
| 3 | 173557837 | rs1948162 | 0.104 | -2.011 | 0.367 | 0.453 | 0.076 | -0.102 | -0.102 | 0 | 0.72 | 0.91 | 0.75 | 0.43 | 0.85 | 0.57 |
| 3 | 173562636 | rs1906569 | -0.027 | -1.94 | 0.318 | 1.051 | 0.059 | -0.072 | -0.072 | 0 | 0.73 | 0.89 | 0.75 | 0.54 | 0.85 | 0.63 |
| 3 | 173563692 | rs13067039 | -1.258 | -1.947 | 0.57 | 1.149 | 0.008 | 0.139 | 0.139 | 0 | 0.12 | 0.25 | 0.07 | 0.31 | 0.14 | 0.07 |
| 3 | 173563936 | rs6779415 | -0.111 | -1.96 | 0.505 | 1.159 | 0.057 | -0.076 | -0.076 | 0 | 0.73 | 0.89 | 0.75 | 0.54 | 0.84 | 0.63 |
| 3 | 173564727 | rs4355309 | -1.794 | -1.829 | 0.547 | 1.074 | 0.002 | 0.104 | 0.104 | 0 | 0.14 | 0.26 | 0.08 | 0.31 | 0.15 | 0.07 |
| 3 | 173565119 | rs2861593 | 1.624 | -2.15 | 0.42 | 0.75 | 0.094 | 0.158 | 0.158 | 0 | 0.5 | 0.29 | 0.61 | 0.19 | 0.65 | 0.51 |
| 3 | 173573365 | rs1545991 | 0.196 | -1.559 | 0.974 | 1.002 | 0.019 | -0.03 | -0.03 | 0.463 | 0.8 | 0.84 | 0.66 | 0.59 | 0.87 | 0.71 |
| 3 | 173573842 | rs9883293 | -0.281 | -1.584 | 0.915 | 0.992 | 0.018 | -0.052 | -0.052 | 1.045 | 0.83 | 0.91 | 0.68 | 0.58 | 0.88 | 0.7 |
| 3 | 173573991 | rs13100306 | 2.51 | -1.838 | 0.965 | 0.733 | 0.051 | 0.085 | 0.085 | 0 | 0.57 | 0.32 | 0.53 | 0.22 | 0.68 | 0.54 |
| 3 | 173588268 | rs4621354 | -1.425 | -1.649 | 1.354 | 0.892 | 0.021 | 0.181 | 0.181 | 0 | 0.14 | 0.33 | 0.09 | 0.34 | 0.18 | 0.14 |
| 3 | 173589827 | rs1021303 | -0.423 | -1.557 | 1.121 | 0.926 | 0.022 | -0.05 | -0.05 | 0.042 | 0.84 | 0.92 | 0.68 | 0.59 | 0.89 | 0.7 |
| 3 | 173620010 | rs16831765 | -1.308 | -1.315 | 1.228 | 0.922 | 0.007 | 0.067 | 0.067 | 0 | 0.12 | 0.16 | 0.09 | 0.35 | 0.18 | 0.1 |
| 3 | 173623133 | rs4280663 | -1.945 | -1.387 | 1.204 | 0.744 | 0.022 | 0.033 | 0.033 | 0 | 0.39 | 0.66 | 0.46 | 0.78 | 0.32 | 0.44 |
| 3 | 173626250 | rs4513478 | -1.266 | -0.967 | 1.338 | 1.075 | -0.017 | 0.028 | 0.028 | 0 | 0.29 | 0.37 | 0.37 | 0.59 | 0.28 | 0.42 |
| 3 | 173632989 | rs9851272 | -1.869 | -0.22 | 1.59 | 1.178 | 0.01 | 0.078 | 0.078 | 0 | 0.19 | 0.29 | 0.13 | 0.36 | 0.23 | 0.19 |
| 3 | 173641361 | rs1873586 | -2.109 | -0.351 | 1.947 | 0.986 | -0.007 | 0.076 | 0.076 | 0.208 | 0.21 | 0.55 | 0.15 | 0.35 | 0.23 | 0.25 |
| 3 | 173644485 | rs9882442 | -1.227 | -0.344 | 2.157 | 0.916 | -0.009 | -0.003 | -0.003 | 0 | 0.08 | 0.25 | 0.05 | 0.03 | 0.03 | 0.06 |
| 3 | 173648400 | rs9875618 | -0.697 | -0.342 | 2.031 | 0.892 | -0.014 | -0.03 | -0.03 | 0 | 0.07 | 0.1 | 0.02 | 0 | 0.02 | 0.06 |
| 3 | 173649154 | rs13097226 | -1.451 | 0.209 | 2.107 | 0.992 | 0.097 | 0.191 | 0.191 | 0 | 0.15 | 0.37 | 0.15 | 0.35 | 0.26 | 0.17 |
| 3 | 173659090 | rs11926346 | -1.485 | 0.135 | 2.092 | 0.966 | 0.036 | 0.19 | 0.19 | 0 | 0.15 | 0.36 | 0.12 | 0.35 | 0.21 | 0.18 |
| 3 | 173661286 | rs12490818 | -2.201 | 0.121 | 1.837 | 0.96 | 0.016 | 0.068 | 0.068 | 0.313 | 0.22 | 0.55 | 0.18 | 0.35 | 0.28 | 0.25 |
| 3 | 173665677 | rs13067869 | -1.022 | 0.229 | 1.569 | 1.039 | -0.026 | -0.029 | -0.029 | 0 | 0.05 | 0.01 | 0.04 | 0.15 | 0.08 | 0.12 |
| 3 | 173666966 | rs9844274 | -2.105 | 0.069 | 1.727 | 1.06 | 0.015 | 0.058 | 0.058 | 0 | 0.21 | 0.34 | 0.16 | 0.35 | 0.28 | 0.25 |
| 3 | 173681906 | rs963350 | -0.184 | 0.135 | 1.466 | 1.243 | -0.002 | 0.035 | 0.035 | 0 | 0.14 | 0.13 | 0.3 | 0.37 | 0.15 | 0.26 |
| 3 | 173685923 | rs1038832 | -1.193 | -0.185 | 1.404 | 0.711 | 0.014 | 0.059 | 0.059 | 0 | 0.28 | 0.5 | 0.21 | 0.4 | 0.33 | 0.3 |
| 3 | 173689779 | rs9290488 | -1.074 | -0.184 | 1.471 | 0.703 | -0.014 | 0.07 | 0.07 | 0 | 0.27 | 0.52 | 0.18 | 0.4 | 0.28 | 0.29 |
| 3 | 173697708 | rs6792822 | 1.825 | -0.117 | 1.598 | 0.55 | 0.018 | 0.112 | 0.112 | 0.179 | 0.54 | 0.28 | 0.5 | 0.22 | 0.55 | 0.44 |
| 3 | 173699986 | rs12494045 | -0.544 | -0.103 | 1.599 | 0.634 | 0.028 | 0.11 | 0.11 | 0 | 0.11 | 0.23 | 0.11 | 0.33 | 0.19 | 0.17 |
| 3 | 173712563 | rs6789981 | 1.367 | -0.002 | 1.524 | 0.613 | -0.022 | 0.059 | 0.059 | 0 | 0.11 | 0.01 | 0.04 | 0 | 0.08 | 0.06 |
| 3 | 173712693 | rs1388478 | 0.355 | 0.125 | 1.664 | 0.825 | 0 | 0.003 | 0.003 | 0 | 0.08 | 0.09 | 0.06 | 0.18 | 0.12 | 0.08 |
| 3 | 173713299 | rs12496665 | -0.313 | 0.118 | 1.678 | 0.806 | 0.018 | 0.079 | 0.079 | 0 | 0.11 | 0.19 | 0.11 | 0.34 | 0.19 | 0.17 |
| 3 | 173716814 | rs13078193 | -0.558 | 0.126 | 1.661 | 0.816 | 0.033 | 0.078 | 0.078 | 0 | 0.12 | 0.22 | 0.14 | 0.34 | 0.24 | 0.19 |
| 3 | 173724146 | rs291933 | 0.019 | 0.07 | 1.568 | 0.829 | 0.001 | -0.024 | -0.024 | 0 | 0.27 | 0.16 | 0.39 | 0.46 | 0.24 | 0.33 |
| 3 | 173727983 | rs291928 | -0.17 | -0.342 | 1.561 | 0.69 | 0 | -0.007 | -0.007 | 0 | 0.94 | 0.92 | 0.96 | 0.98 | 0.94 | 0.94 |
| 3 | 173733659 | rs9812924 | -1.421 | -0.544 | 1.739 | 0.791 | 0.039 | 0.221 | 0.221 | 0 | 0.15 | 0.5 | 0.14 | 0.39 | 0.2 | 0.24 |
| 3 | 173737594 | rs4894458 | 1.819 | -0.491 | 1.616 | 0.79 | 0.003 | 0.102 | 0.102 | 0.076 | 0.46 | 0.28 | 0.42 | 0.22 | 0.44 | 0.37 |
| 3 | 173741221 | rs2241302 | -1.986 | -0.994 | 2.153 | 1.425 | 0.032 | 0.17 | 0.17 | 0 | 0.15 | 0.43 | 0.13 | 0.37 | 0.2 | 0.24 |
| 3 | 173752898 | rs1997060 | -0.932 | -0.741 | 1.9 | 1.44 | 0.018 | 0.077 | 0.077 | 0 | 0.11 | 0.15 | 0.1 | 0.35 | 0.2 | 0.17 |
| 3 | 173758062 | rs6767848 | -1.943 | -1.224 | 2.105 | 1.016 | -0.021 | 0.052 | 0.052 | 0 | 0.16 | 0.45 | 0.12 | 0.23 | 0.14 | 0.17 |
| 3 | 173758553 | rs11921475 | -1.229 | -1.246 | 2.327 | 1.001 | 0.002 | -0.001 | -0.001 | 1.825 | 0.07 | 0.24 | 0.07 | 0.05 | 0.05 | 0.02 |
| 3 | 173769511 | rs9854235 | -0.895 | -1.158 | 2.153 | 1.047 | 0.002 | 0.058 | 0.058 | 0 | 0.1 | 0.28 | 0.06 | 0.18 | 0.1 | 0.15 |
| 3 | 173785874 | rs16833012 | -1.816 | -1.161 | 1.811 | 1.038 | -0.017 | 0.056 | 0.056 | 0 | 0.16 | 0.49 | 0.12 | 0.23 | 0.14 | 0.16 |
| 3 | 173789058 | rs1917638 | -0.457 | -1.095 | 1.294 | 1.173 | 0.002 | -0.024 | -0.024 | 0 | 0.26 | 0.16 | 0.24 | 0.48 | 0.24 | 0.36 |
| 3 | 173810957 | rs16833234 | -0.19 | -0.943 | 1.222 | 1.006 | 0.002 | -0.021 | -0.021 | 0 | 0.22 | 0.13 | 0.21 | 0.38 | 0.21 | 0.24 |

| | | | | | | | | | | | | | | | | |
|---|---|---|---|---|---|---|---|---|---|---|---|---|---|---|---|---|
| 3 | 173818395 | rs9872442 | -1.57 | -0.663 | 1.769 | 0.696 | 0.046 | 0.189 | 0.189 | 0 | 0.1 | 0.44 | 0.1 | 0.21 | 0.11 | 0.1 |
| 3 | 173844235 | rs4140699 | -2.234 | -0.776 | 1.662 | 0.598 | -0.042 | -0.025 | -0.025 | 0 | 0.23 | 0.59 | 0.14 | 0.18 | 0.18 | 0.19 |
| 3 | 173847402 | rs7626528 | -1.416 | -0.745 | 1.73 | 0.563 | -0.045 | -0.101 | -0.101 | 0.323 | 0.15 | 0.35 | 0.07 | 0 | 0.07 | 0.1 |
| 3 | 173852415 | rs13315608 | -0.207 | -0.725 | 1.682 | 0.574 | 0.008 | -0.003 | -0.003 | 0 | 0.1 | 0.1 | 0.02 | 0 | 0.04 | 0.09 |
| 3 | 173853784 | rs6445141 | -1.569 | -0.727 | 1.699 | 0.588 | -0.02 | 0.009 | 0.009 | 0 | 0.18 | 0.33 | 0.1 | 0.22 | 0.16 | 0.19 |
| 3 | 173855250 | rs9843125 | -1.675 | -0.723 | 1.67 | 0.59 | -0.03 | 0.013 | 0.013 | 0 | 0.18 | 0.39 | 0.1 | 0.22 | 0.16 | 0.19 |
| 3 | 173855560 | rs12491232 | -0.856 | -0.693 | 1.565 | 0.603 | 0.007 | 0.081 | 0.081 | 0 | 0.08 | 0.24 | 0.07 | 0.21 | 0.1 | 0.1 |
| 3 | 173858380 | rs9821957 | -1.081 | -0.728 | 1.522 | 0.827 | -0.053 | 0.067 | 0.067 | 1.958 | 0.14 | 0.42 | 0.06 | 0.22 | 0.05 | 0.1 |
| 3 | 173859622 | rs9290494 | -1.318 | -0.689 | 0.786 | 0.846 | -0.057 | 0.036 | 0.036 | 6.036 | 0.18 | 0.6 | 0.1 | 0.22 | 0.08 | 0.11 |
| 3 | 173866424 | rs11919911 | -1.656 | -0.298 | 0.959 | 1.103 | 0.045 | 0.251 | 0.251 | 0.565 | 0.08 | 0.42 | 0.06 | 0.24 | 0.05 | 0.02 |
| 3 | 173868469 | rs6798464 | -1.195 | -0.844 | 0.412 | 0.938 | 0.018 | -0.009 | -0.009 | 0 | 0.4 | 0.64 | 0.31 | 0.56 | 0.28 | 0.27 |
| 3 | 173878934 | rs6768717 | -0.979 | -1.45 | 0.608 | -0.037 | -0.043 | -0.112 | -0.112 | 0.181 | 0.29 | 0.54 | 0.2 | 0.07 | 0.15 | 0.2 |
| 3 | 173880681 | rs9883269 | -0.573 | -1.549 | 0.625 | -0.083 | -0.043 | -0.042 | -0.042 | 0.072 | 0.13 | 0.52 | 0.07 | 0.07 | 0.05 | 0.11 |
| 3 | 173884293 | rs13340122 | -0.705 | -1.49 | 0.982 | -0.09 | -0.041 | -0.045 | -0.045 | 0 | 0.13 | 0.44 | 0.07 | 0.07 | 0.05 | 0.12 |
| 3 | 173889865 | rs7625721 | -0.418 | -1.406 | 1.001 | -0.09 | -0.029 | -0.038 | -0.038 | 0.367 | 0.13 | 0.45 | 0.06 | 0.07 | 0.05 | 0.11 |
| 3 | 173890458 | rs6771925 | 1.488 | -1.47 | -0.149 | -0.591 | -0.123 | 0.023 | 0.023 | 0 | 0.26 | 0.07 | 0.39 | 0.13 | 0.43 | 0.29 |
| 3 | 173894275 | rs16857409 | 0.861 | -1.785 | -0.008 | -0.468 | -0.002 | 0.007 | 0.007 | 0 | 0.21 | 0.23 | 0.2 | 0.63 | 0.18 | 0.15 |
| 3 | 173899303 | rs2160977 | -1.141 | -1.908 | -0.053 | -0.508 | 0.017 | 0.039 | 0.039 | 0.811 | 0.74 | 0.87 | 0.77 | 0.79 | 0.76 | 0.64 |
| 3 | 173900788 | rs16857557 | -0.307 | -2.084 | 0.676 | -0.606 | -0.029 | -0.09 | -0.09 | 0.02 | 0.27 | 0.39 | 0.19 | 0.03 | 0.15 | 0.19 |
| 3 | 173901722 | rs4894645 | 1.068 | -2.093 | 0.878 | -0.583 | 0.001 | -0.008 | -0.008 | 0 | 0.21 | 0.2 | 0.21 | 0.59 | 0.18 | 0.15 |
| 3 | 173902385 | rs4894650 | -0.67 | -2.085 | 1.214 | -0.682 | -0.023 | -0.036 | -0.036 | 0 | 0.71 | 0.65 | 0.78 | 0.76 | 0.75 | 0.65 |
| 3 | 173903820 | rs1345537 | -1.014 | -2.078 | 1.201 | -0.679 | -0.002 | -0.003 | -0.003 | 0 | 0.73 | 0.75 | 0.79 | 0.76 | 0.75 | 0.65 |
| 3 | 173904977 | rs2193744 | 0.618 | -2.209 | 1.347 | -0.761 | 0.027 | 0.058 | 0.058 | 0 | 0.48 | 0.37 | 0.6 | 0.79 | 0.61 | 0.46 |
| 3 | 173905478 | rs1421432 | -1.245 | -2.248 | 1.296 | -0.771 | 0.013 | 0.034 | 0.034 | 0.144 | 0.75 | 0.86 | 0.8 | 0.8 | 0.75 | 0.62 |
| 3 | 173908675 | rs9883446 | -0.054 | -2.069 | 1.382 | -0.82 | -0.01 | -0.076 | -0.076 | 0 | 0.24 | 0.32 | 0.19 | 0 | 0.16 | 0.2 |
| 3 | 173909881 | rs6445149 | -0.999 | -1.882 | 1.408 | -0.794 | -0.001 | -0.013 | -0.013 | 0 | 0.73 | 0.78 | 0.79 | 0.83 | 0.74 | 0.61 |
| 3 | 173917669 | rs1549100 | -0.676 | -1.86 | 1.544 | -0.797 | -0.015 | -0.046 | -0.046 | 0 | 0.71 | 0.7 | 0.78 | 0.83 | 0.74 | 0.61 |
| 3 | 173920993 | rs4494938 | -0.207 | -1.851 | 1.55 | -0.837 | -0.022 | -0.088 | -0.088 | 0 | 0.25 | 0.38 | 0.2 | 0.03 | 0.16 | 0.2 |
| 3 | 173925625 | rs1421420 | -0.813 | -1.886 | 1.52 | -0.781 | -0.012 | -0.032 | -0.032 | 0 | 0.71 | 0.74 | 0.78 | 0.8 | 0.74 | 0.6 |
| 3 | 173934318 | rs13064752 | 0.099 | -1.917 | 1.5 | -0.798 | 0.023 | 0.025 | 0.025 | 0 | 0.6 | 0.33 | 0.73 | 0.8 | 0.71 | 0.5 |
| 3 | 173939218 | rs9290496 | 1.027 | -1.867 | 1.492 | -0.769 | -0.038 | -0.054 | -0.054 | 0.11 | 0.09 | 0.34 | 0.03 | 0.01 | 0.03 | 0.11 |
| 3 | 173952876 | rs1345534 | 0.008 | -1.642 | 1.152 | -0.522 | -0.021 | -0.029 | -0.029 | 0 | 0.37 | 0.16 | 0.37 | 0.51 | 0.37 | 0.21 |
| 3 | 173957297 | rs4894467 | -0.662 | -1.942 | 1.459 | -0.587 | -0.001 | -0.003 | -0.003 | 0.16 | 0.48 | 0.48 | 0.42 | 0.57 | 0.41 | 0.48 |
| 3 | 173958362 | rs16858344 | 0.736 | -1.916 | 1.496 | -0.582 | -0.041 | -0.04 | -0.04 | 0 | 0.11 | 0.3 | 0.04 | 0.06 | 0.04 | 0.27 |
| 3 | 173960436 | rs961879 | -0.268 | -2.019 | 1.304 | -0.646 | -0.019 | 0.003 | 0.003 | 0.052 | 0.36 | 0.52 | 0.34 | 0.58 | 0.27 | 0.41 |
| 3 | 173961129 | rs1362957 | 1.501 | -1.888 | 1.224 | -0.737 | 0.021 | 0.057 | 0.057 | 0.733 | 0.22 | 0.1 | 0.19 | 0.14 | 0.2 | 0.22 |
| 3 | 173965464 | rs1421425 | -1.15 | -1.836 | 1.36 | -0.574 | 0.018 | 0.037 | 0.037 | 0 | 0.52 | 0.71 | 0.43 | 0.58 | 0.42 | 0.49 |
| 3 | 173966738 | rs13088246 | 0.772 | -1.794 | 1.309 | -0.514 | 0.001 | -0.021 | -0.021 | 0.653 | 0.24 | 0.16 | 0.28 | 0.5 | 0.22 | 0.14 |
| 3 | 173967286 | rs16858417 | 0.453 | -1.751 | 1.219 | -0.504 | -0.042 | -0.012 | -0.012 | 0 | 0.12 | 0.33 | 0.05 | 0.08 | 0.05 | 0.26 |
| 3 | 173971715 | rs4894661 | 0.478 | -1.648 | 0.83 | -0.415 | 0.002 | -0.056 | -0.056 | 0 | 0.26 | 0.15 | 0.29 | 0.49 | 0.23 | 0.14 |
| 3 | 173976450 | rs808079 | 0.854 | -1.499 | 0.823 | -0.586 | 0.007 | -0.077 | -0.077 | 0.621 | 0.14 | 0.17 | 0.09 | 0 | 0.14 | 0.08 |
| 3 | 173985441 | rs13095909 | -0.021 | -1.465 | 0.291 | -0.542 | 0.003 | -0.012 | -0.012 | 0.449 | 0.08 | 0.07 | 0.05 | 0.05 | 0.11 | 0.13 |
| 3 | 173990588 | rs4504175 | 0.395 | -1.599 | -0.028 | -0.104 | 0.017 | -0.058 | -0.058 | 0 | 0.15 | 0.07 | 0.15 | 0.54 | 0.11 | 0.2 |
| 3 | 173997153 | rs7646919 | 1.243 | -1.744 | -0.132 | -0.523 | -0.006 | -0.028 | -0.028 | 0 | 0.27 | 0.24 | 0.14 | 0.09 | 0.2 | 0.24 |
| 3 | 174000154 | rs9855544 | 0.742 | -1.391 | -0.425 | -0.543 | 0.017 | -0.058 | -0.058 | 0.902 | 0.15 | 0.07 | 0.15 | 0.54 | 0.11 | 0.21 |
| 3 | 174010051 | rs811197 | -0.826 | -1.46 | -0.157 | -1.019 | -0.028 | 0.028 | 0.028 | 0 | 0.62 | 0.73 | 0.44 | 0.69 | 0.48 | 0.54 |
| 3 | 174022055 | rs11714567 | 0.549 | -0.496 | -0.457 | -1.314 | 0.001 | -0.053 | -0.053 | 0.578 | 0.2 | 0.21 | 0.15 | 0.01 | 0.24 | 0.13 |
| 3 | 174024904 | rs9821632 | -1.045 | -0.562 | -0.269 | -1.206 | -0.01 | -0.055 | -0.055 | 0 | 0.61 | 0.63 | 0.76 | 0.85 | 0.67 | 0.63 |
| 3 | 174025863 | rs6445164 | 1.821 | -0.591 | -0.247 | -1.208 | 0.011 | 0.018 | 0.018 | 0.061 | 0.27 | 0.18 | 0.15 | 0.11 | 0.2 | 0.24 |
| 3 | 174036130 | rs10084713 | 0.513 | -0.627 | -0.364 | -1.174 | 0.022 | 0.016 | 0.016 | 0.207 | 0.43 | 0.34 | 0.3 | 0.65 | 0.31 | 0.44 |
| 3 | 174038312 | rs13098004 | -0.077 | -0.555 | -0.319 | -1.155 | -0.004 | 0.02 | 0.02 | 0 | 0.43 | 0.43 | 0.57 | 0.31 | 0.53 | 0.42 |
| 3 | 174048651 | rs11926733 | -0.03 | -0.415 | -0.244 | -0.95 | -0.006 | 0.002 | 0.002 | 0 | 0.71 | 0.71 | 0.76 | 0.38 | 0.77 | 0.67 |
| 3 | 174052468 | rs1014345 | -1.775 | 0.161 | 0.009 | -0.784 | 0.013 | -0.061 | -0.061 | 0 | 0.88 | 0.8 | 0.9 | 0.99 | 0.87 | 0.91 |
| 3 | 174058551 | rs7614469 | -0.17 | 0.046 | 0.056 | -0.356 | 0.002 | 0.083 | 0.083 | 0 | 0.76 | 0.49 | 0.77 | 0.46 | 0.78 | 0.71 |
| 3 | 174062083 | rs11716098 | -0.547 | 0.187 | -0.785 | -0.296 | 0.002 | 0.057 | 0.057 | 0 | 0.78 | 0.75 | 0.8 | 0.5 | 0.79 | 0.71 |
| 3 | 174062834 | rs5018958 | 2.183 | 0.327 | -1.038 | -0.415 | -0.051 | -0.016 | -0.016 | 0.806 | 0.18 | 0.1 | 0.41 | 0.22 | 0.29 | 0.3 |
| 3 | 174077631 | rs9863756 | 1.317 | 0.271 | -0.717 | -0.027 | 0.001 | 0.037 | 0.037 | 0 | 0.1 | 0.09 | 0.11 | 0.51 | 0.09 | 0.2 |
| 3 | 174077949 | rs4894469 | -1.799 | 0.62 | -0.734 | -0.399 | 0.012 | -0.056 | -0.056 | 0 | 0.88 | 0.84 | 0.91 | 0.99 | 0.87 | 0.91 |
| 3 | 174084394 | rs939320 | 0.728 | 0.645 | -1.525 | -0.362 | -0.028 | 0.023 | 0.023 | 0 | 0.31 | 0.24 | 0.5 | 0.23 | 0.43 | 0.34 |
| 3 | 174085014 | rs9821469 | 1.261 | 0.475 | -1.533 | -0.323 | -0.012 | 0.021 | 0.021 | 0 | 0.23 | 0.1 | 0.18 | 0.18 | 0.23 | 0.27 |
| 3 | 174088909 | rs7610632 | 0.429 | 0.691 | -1.085 | -0.146 | 0.015 | -0.037 | -0.037 | 0 | 0.07 | 0.13 | 0.08 | 0.01 | 0.12 | 0.03 |
| 3 | 174092252 | rs7620258 | 0.202 | -0.021 | -1.534 | 0.099 | 0.004 | -0.039 | -0.039 | 0 | 0.63 | 0.25 | 0.6 | 0.54 | 0.66 | 0.65 |
| 3 | 174092341 | rs11712043 | 1.262 | -0.421 | -2.055 | -0.252 | -0.047 | 0.006 | 0.006 | 0 | 0.26 | 0.11 | 0.32 | 0.24 | 0.33 | 0.33 |
| 3 | 174092928 | rs9844366 | -0.843 | -0.91 | -2.4 | -0.612 | -0.052 | 0.085 | 0.085 | 0.009 | 0.59 | 0.89 | 0.66 | 0.77 | 0.58 | 0.66 |
| 3 | 174093339 | rs9882210 | 0.651 | -1.039 | -1.924 | -0.393 | 0.02 | -0.046 | -0.046 | 0.073 | 0.22 | 0.15 | 0.26 | 0.5 | 0.19 | 0.26 |
| 3 | 174095188 | rs13079516 | 1.81 | -1.036 | -1.961 | -0.479 | -0.003 | 0 | 0 | 0 | 0.19 | 0.14 | 0.2 | 0.18 | 0.18 | 0.17 |
| 3 | 174104641 | rs1522249 | 0.004 | -1.493 | -1.427 | -0.25 | 0.027 | -0.088 | -0.088 | 0.032 | 0.24 | 0.12 | 0.28 | 0.5 | 0.22 | 0.3 |
| 3 | 174105112 | rs1520168 | -0.902 | -1.56 | -0.404 | -0.359 | 0.001 | -0.006 | -0.006 | 0 | 0.58 | 0.59 | 0.65 | 0.74 | 0.58 | 0.71 |
| 3 | 174108632 | rs1356816 | 0.792 | -1.552 | -0.271 | -0.39 | 0.005 | -0.012 | -0.012 | 0 | 0.33 | 0.38 | 0.37 | 0.24 | 0.38 | 0.4 |

| | | | | | | | | | | | | | | | | |
|---|---|---|---|---|---|---|---|---|---|---|---|---|---|---|---|---|
| 3 | 174109138 | rs11713639 | 1.134 | -1.583 | 0.002 | -0.418 | -0.015 | 0.016 | 0.016 | 0 | 0.3 | 0.2 | 0.36 | 0.24 | 0.35 | 0.4 |
| 3 | 174112212 | rs1915102 | -0.952 | -1.585 | -0.029 | -0.386 | -0.043 | 0.075 | 0.075 | 0 | 0.6 | 0.84 | 0.67 | 0.77 | 0.59 | 0.71 |
| 3 | 174112354 | rs1915103 | 0.344 | -0.929 | 0.247 | 0.069 | 0.004 | -0.012 | -0.012 | 0 | 0.34 | 0.39 | 0.37 | 0.24 | 0.38 | 0.4 |
| 3 | 174124164 | rs6783143 | -1.053 | -0.676 | 0.249 | -0.171 | -0.009 | 0.052 | 0.052 | 0 | 0.56 | 0.73 | 0.63 | 0.77 | 0.53 | 0.76 |
| 3 | 174127710 | rs12629482 | -1.074 | -0.591 | 0.42 | -0.09 | -0.01 | 0.087 | 0.087 | 0.146 | 0.53 | 0.73 | 0.62 | 0.77 | 0.52 | 0.76 |
| 3 | 174136298 | rs6650901 | -0.436 | -1.072 | 1.143 | 0.439 | -0.004 | 0.01 | 0.01 | 0 | 0.36 | 0.43 | 0.36 | 0.49 | 0.32 | 0.46 |
| 3 | 174142441 | rs9863476 | 0.427 | -1.083 | 1.365 | 0.505 | -0.01 | 0.171 | 0.171 | 0 | 0.06 | 0.16 | 0.17 | 0.31 | 0.03 | 0.16 |
| 3 | 174143382 | rs11706616 | 0.994 | -1.123 | 1.262 | 0.44 | -0.039 | -0.063 | -0.063 | 1.142 | 0.21 | 0.05 | 0.26 | 0.27 | 0.22 | 0.31 |
| 3 | 174146956 | rs7620744 | -1.035 | -1.115 | 1.339 | 0.374 | 0.017 | 0.03 | 0.03 | 0 | 0.42 | 0.67 | 0.53 | 0.79 | 0.35 | 0.61 |
| 3 | 174152030 | rs9812312 | -0.924 | -0.873 | 1.13 | 0.473 | -0.058 | 0.116 | 0.116 | 0 | 0.2 | 0.4 | 0.24 | 0.45 | 0.12 | 0.31 |
| 3 | 174164942 | rs7632482 | 0.02 | -0.58 | 1.129 | 0.201 | -0.007 | 0.236 | 0.236 | 0.013 | 0.06 | 0.18 | 0.26 | 0.43 | 0.04 | 0.11 |
| 3 | 174166224 | rs9830993 | -1.098 | -0.642 | 0.987 | -0.094 | 0.018 | 0.137 | 0.137 | 0.022 | 0.52 | 0.74 | 0.68 | 0.94 | 0.46 | 0.69 |
| 3 | 174177349 | rs12631086 | 0.199 | -0.581 | 0.769 | 0.225 | 0.005 | -0.055 | -0.055 | 0 | 0.3 | 0.15 | 0.33 | 0.43 | 0.3 | 0.46 |
| 3 | 174190024 | rs4894675 | -0.431 | -0.567 | 0.677 | 0.213 | -0.009 | -0.043 | -0.043 | 0 | 0.37 | 0.45 | 0.61 | 0.9 | 0.34 | 0.57 |
| 3 | 174194243 | rs10936797 | 0.236 | -0.592 | 0.652 | 0.247 | 0.001 | -0.077 | -0.077 | 0 | 0.29 | 0.14 | 0.33 | 0.43 | 0.29 | 0.46 |
| 3 | 174198101 | rs9290501 | -0.758 | -0.332 | 0.954 | 0.746 | 0.005 | 0.375 | 0.375 | 0.623 | 0.08 | 0.33 | 0.28 | 0.47 | 0.04 | 0.11 |
| 3 | 174213188 | rs938441 | -0.307 | -0.313 | 0.398 | -0.046 | 0.008 | 0.038 | 0.038 | 0 | 0.39 | 0.32 | 0.62 | 0.91 | 0.39 | 0.57 |
| 3 | 174225345 | rs12632844 | -0.928 | -0.381 | 0.461 | -0.311 | 0.01 | 0.009 | 0.009 | 0 | 0.6 | 0.56 | 0.49 | 0.51 | 0.59 | 0.65 |
| 3 | 174231077 | rs7619222 | -1.378 | -0.366 | 0.404 | -0.452 | -0.003 | -0.005 | -0.005 | 0.059 | 0.65 | 0.7 | 0.76 | 0.9 | 0.62 | 0.76 |
| 3 | 174241461 | rs1515592 | -0.217 | -0.335 | 0.467 | -0.33 | -0.005 | -0.021 | -0.021 | 6.568 | 0.27 | 0.34 | 0.13 | 0.12 | 0.26 | 0.29 |
| 3 | 174243684 | rs12637269 | -1.004 | -0.008 | 0.586 | 0.013 | -0.008 | 0.012 | 0.012 | 0 | 0.64 | 0.7 | 0.76 | 0.9 | 0.63 | 0.71 |
| 3 | 174245721 | rs6780558 | -1.282 | 0.35 | 0.93 | 0.816 | -0.004 | 0.13 | 0.13 | 0 | 0.22 | 0.47 | 0.38 | 0.5 | 0.24 | 0.27 |
| 3 | 174245949 | rs6772350 | -0.482 | 0.4 | 0.307 | 0.456 | -0.001 | 0.003 | 0.003 | 0 | 0.15 | 0.18 | 0.11 | 0.12 | 0.2 | 0.17 |
| 3 | 174252463 | rs2861835 | 0.958 | 0.646 | 0.19 | -0.128 | 0.004 | 0.051 | 0.051 | 0 | 0.58 | 0.31 | 0.43 | 0.11 | 0.6 | 0.51 |
| 3 | 174256126 | rs1515604 | -0.538 | 0.175 | 0.663 | 0.248 | -0.05 | 0.211 | 0.211 | 0 | 0.16 | 0.37 | 0.33 | 0.53 | 0.11 | 0.26 |
| 3 | 174256743 | rs6776623 | -1.015 | 1.131 | 0.84 | 0.067 | 0.016 | 0.004 | 0.004 | 0 | 0.3 | 0.46 | 0.25 | 0.35 | 0.34 | 0.28 |
| 3 | 174260303 | rs6793059 | -0.201 | 1.254 | 0.313 | 0.155 | -0.004 | -0.01 | -0.01 | 0 | 0.19 | 0.17 | 0.17 | 0.35 | 0.2 | 0.22 |
| 3 | 174273043 | rs1913733 | -0.097 | 1.21 | 0.048 | 0.079 | 0.027 | -0.044 | -0.044 | 0.916 | 0.08 | 0.17 | 0.07 | 0 | 0.12 | 0.04 |
| 3 | 174273874 | rs9290503 | -0.551 | 1.11 | -0.454 | 0.251 | 0 | -0.001 | -0.001 | 0 | 0.35 | 0.66 | 0.33 | 0.39 | 0.39 | 0.29 |
| 3 | 174274011 | rs9871610 | -0.776 | 0.946 | -0.564 | 0.295 | -0.015 | 0.005 | 0.005 | 0 | 0.4 | 0.7 | 0.38 | 0.42 | 0.47 | 0.29 |
| 3 | 174276798 | rs1515585 | -0.525 | 0.625 | -0.883 | 0.213 | 0.018 | 0.043 | 0.043 | 0.018 | 0.42 | 0.75 | 0.61 | 0.76 | 0.36 | 0.51 |
| 3 | 174277472 | rs9822073 | -0.602 | 0.801 | -1.062 | 0.005 | 0.016 | 0.098 | 0.098 | 0.03 | 0.47 | 0.78 | 0.66 | 0.79 | 0.44 | 0.61 |
| 3 | 174303113 | rs9290506 | -0.784 | 0.707 | -0.067 | -0.022 | -0.005 | 0 | 0 | 0 | 0.24 | 0.34 | 0.48 | 0.72 | 0.18 | 0.43 |
| 3 | 174308012 | rs685963 | -0.762 | 0.349 | 0.365 | -0.323 | -0.002 | -0.076 | -0.076 | 0 | 0.17 | 0.33 | 0.14 | 0.08 | 0.15 | 0.16 |
| 3 | 174316915 | rs536223 | -0.614 | 0.279 | 0.792 | -0.225 | -0.004 | -0.048 | -0.048 | 0.134 | 0.12 | 0.25 | 0.12 | 0.08 | 0.11 | 0.16 |
| 3 | 174348263 | rs556852 | 0.116 | 0.045 | 0.911 | 0.67 | -0.015 | 0.049 | 0.049 | 0.534 | 0.83 | 0.7 | 0.84 | 0.7 | 0.88 | 0.7 |
| 3 | 174351626 | rs545495 | 1.659 | 0.197 | 1.116 | 0.946 | 0 | -0.002 | -0.002 | 0 | 0.66 | 0.34 | 0.68 | 0.55 | 0.71 | 0.39 |
| 3 | 174354964 | rs10936812 | -0.333 | 0.312 | 1.117 | 1.008 | -0.007 | -0.008 | -0.008 | 0.877 | 0.13 | 0.08 | 0.14 | 0.15 | 0.16 | 0.27 |
| 3 | 174357197 | rs4365637 | 1.552 | 1.054 | 1.033 | 0.9 | 0.02 | 0.042 | 0.042 | 0.022 | 0.68 | 0.52 | 0.37 | 0.4 | 0.56 | 0.55 |
| 3 | 174360371 | rs588400 | -0.709 | 0.652 | 0.973 | 0.559 | 0.008 | 0.01 | 0.01 | 0 | 0.14 | 0.32 | 0.17 | 0.15 | 0.17 | 0.28 |
| 3 | 174364200 | rs614699 | -0.296 | 0.936 | 0.304 | 0.353 | -0.022 | 0.025 | 0.025 | 0 | 0.17 | 0.07 | 0.13 | 0.11 | 0.23 | 0.26 |
| 3 | 174367708 | rs693471 | -1.412 | 0.365 | 0.787 | 0.512 | -0.002 | 0 | 0 | 0 | 0.34 | 0.63 | 0.27 | 0.34 | 0.29 | 0.57 |
| 3 | 174383294 | rs488776 | 1.133 | 0.08 | 0.958 | 0.381 | -0.018 | -0.002 | -0.002 | 0 | 0.64 | 0.48 | 0.74 | 0.64 | 0.7 | 0.43 |
| 3 | 174385622 | rs7629875 | -0.129 | -0.01 | 1.011 | 0.371 | -0.027 | 0.032 | 0.032 | 0 | 0.14 | 0.2 | 0.11 | 0.26 | 0.06 | 0.3 |
| 3 | 174389212 | rs4611834 | 1.403 | -0.114 | 0.995 | 0.507 | -0.015 | -0.004 | -0.004 | 1.492 | 0.63 | 0.47 | 0.72 | 0.65 | 0.7 | 0.41 |
| 3 | 174399432 | rs560409 | 0.44 | 0.318 | -0.085 | 0.187 | -0.032 | 0.015 | 0.015 | 0.311 | 0.19 | 0.07 | 0.27 | 0.13 | 0.25 | 0.19 |
| 3 | 174399571 | rs559369 | 0.175 | 0.019 | 0.3 | 0.328 | 0 | -0.037 | -0.037 | 0 | 0.77 | 0.67 | 0.83 | 0.89 | 0.76 | 0.71 |
| 3 | 174400892 | rs12486421 | 0.371 | 0.19 | -0.293 | 0.415 | -0.004 | 0.005 | 0.005 | 0 | 0.13 | 0.1 | 0.1 | 0.26 | 0.06 | 0.31 |
| 3 | 174405392 | rs4414863 | -0.257 | 0.432 | -0.315 | 0.208 | -0.014 | 0.035 | 0.035 | 0 | 0.81 | 0.89 | 0.85 | 0.91 | 0.77 | 0.74 |
| 3 | 174411312 | rs8180018 | -0.672 | -0.095 | 0.271 | 0.795 | -0.041 | 0.16 | 0.16 | 0 | 0.13 | 0.33 | 0.17 | 0.37 | 0.04 | 0.27 |
| 3 | 174413204 | rs537932 | 0.632 | 0.041 | -0.732 | 0.481 | -0.061 | -0.015 | -0.015 | 0 | 0.37 | 0.12 | 0.34 | 0.33 | 0.43 | 0.21 |
| 3 | 174413915 | rs565334 | -0.028 | 0.059 | -0.7 | 0.419 | -0.005 | 0.088 | 0.088 | 0 | 0.47 | 0.25 | 0.49 | 0.71 | 0.46 | 0.44 |
| 3 | 174415345 | rs552008 | -0.399 | 0.021 | -0.981 | 0.168 | 0.001 | -0.031 | -0.031 | 0 | 0.56 | 0.54 | 0.63 | 0.82 | 0.54 | 0.54 |
| 3 | 174417912 | rs12494312 | -0.318 | 0.058 | -0.771 | 0.625 | -0.042 | 0.201 | 0.201 | 0.088 | 0.19 | 0.46 | 0.3 | 0.48 | 0.11 | 0.33 |
| 3 | 174418262 | rs4367080 | 0.725 | 0.822 | -1.652 | 0.579 | -0.06 | -0.073 | -0.073 | 0.256 | 0.24 | 0.05 | 0.21 | 0.32 | 0.28 | 0.17 |
| 3 | 174418538 | rs7639982 | 0.18 | 0.803 | -1.112 | 0.323 | -0.015 | -0.004 | -0.004 | 0 | 0.68 | 0.7 | 0.58 | 0.63 | 0.6 | 0.51 |
| 3 | 174419856 | rs6774160 | -0.776 | 0.276 | -0.833 | 0.265 | 0.001 | 0 | 0 | 0 | 0.45 | 0.51 | 0.39 | 0.46 | 0.41 | 0.56 |
| 3 | 174420162 | rs9834476 | -0.314 | 0.054 | -0.844 | 0.041 | -0.016 | 0.025 | 0.025 | 0 | 0.33 | 0.46 | 0.32 | 0.46 | 0.29 | 0.49 |
| 3 | 174422011 | rs654146 | 1.896 | 0.206 | -0.988 | -0.09 | -0.035 | 0.077 | 0.077 | 0 | 0.36 | 0.16 | 0.3 | 0.17 | 0.38 | 0.24 |
| 3 | 174425811 | rs6783729 | 0.792 | 0.13 | -0.802 | -0.128 | 0.003 | -0.051 | -0.051 | 0 | 0.12 | 0.25 | 0.08 | 0.01 | 0.13 | 0.08 |
| 3 | 174426336 | rs4410453 | -1.387 | 0.147 | -0.877 | -0.115 | -0.007 | 0.001 | 0.001 | 0 | 0.88 | 0.89 | 0.72 | 0.63 | 0.83 | 0.84 |
| 3 | 174426402 | rs6784378 | 1.942 | -0.123 | -0.769 | -0.201 | 0.021 | -0.027 | -0.027 | 0 | 0.11 | 0.07 | 0.06 | 0.34 | 0.07 | 0.22 |
| 3 | 174426709 | rs11719844 | -0.121 | -0.084 | -0.463 | -0.258 | -0.011 | 0.053 | 0.053 | 0 | 0.26 | 0.38 | 0.25 | 0.44 | 0.24 | 0.44 |
| 3 | 174434433 | rs630690 | 1.399 | -0.029 | -0.524 | -0.25 | -0.002 | 0.059 | 0.059 | 0 | 0.48 | 0.39 | 0.39 | 0.2 | 0.5 | 0.32 |
| 3 | 174443661 | rs1447782 | 1.011 | 0.025 | -0.432 | -0.34 | -0.001 | -0.047 | -0.047 | 0 | 0.12 | 0.24 | 0.08 | 0.01 | 0.12 | 0.07 |
| 3 | 174444171 | rs13325005 | -0.861 | -0.017 | 0.244 | -0.398 | -0.009 | -0.157 | -0.157 | 0 | 0.24 | 0.43 | 0.2 | 0.01 | 0.23 | 0.17 |
| 3 | 174452796 | rs6809168 | 0.535 | 0.035 | 0.281 | -0.264 | -0.003 | 0.033 | 0.033 | 0 | 0.2 | 0.3 | 0.19 | 0.39 | 0.18 | 0.39 |
| 3 | 174461025 | rs9846937 | -1.665 | 0.056 | 0.104 | -0.297 | 0.016 | 0.06 | 0.06 | 0 | 0.5 | 0.76 | 0.41 | 0.41 | 0.47 | 0.59 |
| 3 | 174463389 | rs490042 | 2.164 | 0.046 | -0.08 | -0.27 | -0.029 | 0.071 | 0.071 | 0 | 0.36 | 0.14 | 0.31 | 0.21 | 0.36 | 0.25 |

| | | | | | | | | | | | | | | | | |
|---|---|---|---|---|---|---|---|---|---|---|---|---|---|---|---|---|
| 3 | 174471345 | rs1184856 | 0.637 | -0.216 | 0.174 | -0.18 | -0.016 | 0.022 | 0.022 | 0 | 0.7 | 0.68 | 0.6 | 0.62 | 0.76 | 0.72 |
| 3 | 174474613 | rs545012 | -0.91 | -0.17 | 0.115 | -0.044 | -0.012 | -0.027 | -0.027 | 0.026 | 0.84 | 0.87 | 0.7 | 0.62 | 0.78 | 0.8 |
| 3 | 174476314 | rs6778906 | 1.296 | -0.349 | 0.268 | -0.048 | -0.032 | 0.065 | 0.065 | 0 | 0.13 | 0.27 | 0.08 | 0.29 | 0.07 | 0.24 |
| 3 | 174486282 | rs16864198 | 0.947 | 0.074 | 0.089 | -0.374 | -0.01 | 0.013 | 0.013 | 0 | 0.1 | 0.05 | 0.09 | 0.01 | 0.15 | 0.06 |
| 3 | 174494183 | rs538340 | 0.171 | 0.069 | 0.239 | -0.401 | 0.006 | -0.033 | -0.033 | 0.624 | 0.62 | 0.46 | 0.48 | 0.22 | 0.58 | 0.43 |
| 3 | 174495959 | rs9290515 | -0.554 | 0.051 | 0.192 | -0.427 | -0.004 | -0.074 | -0.074 | 0.09 | 0.24 | 0.24 | 0.18 | 0.01 | 0.26 | 0.14 |
| 3 | 174504462 | rs564378 | 0.867 | 0.08 | 0.118 | -0.151 | -0.002 | -0.01 | -0.01 | 0.082 | 0.59 | 0.38 | 0.51 | 0.62 | 0.57 | 0.69 |
| 3 | 174506772 | rs7648550 | 0.443 | 0.039 | -0.026 | -0.26 | -0.001 | -0.003 | -0.003 | 0.049 | 0.26 | 0.32 | 0.22 | 0.46 | 0.28 | 0.42 |
| 3 | 174511549 | rs2140285 | -2.07 | -0.156 | -0.061 | -0.283 | -0.043 | 0.087 | 0.087 | 0 | 0.65 | 0.9 | 0.72 | 0.82 | 0.68 | 0.73 |
| 3 | 174519856 | rs7609643 | -0.391 | -0.239 | 0.442 | -0.357 | 0.008 | 0.032 | 0.032 | 0 | 0.29 | 0.33 | 0.39 | 0.36 | 0.34 | 0.28 |
| 3 | 174522942 | rs7619062 | -1.079 | -0.221 | 0.372 | -0.309 | 0 | -0.003 | -0.003 | 0 | 0.42 | 0.54 | 0.48 | 0.37 | 0.43 | 0.33 |
| 3 | 174523783 | rs9290518 | -1.846 | -0.216 | 0.197 | -0.32 | -0.025 | 0.1 | 0.1 | 0 | 0.63 | 0.87 | 0.69 | 0.82 | 0.67 | 0.72 |
| 3 | 174528506 | rs6774585 | -0.921 | -0.218 | 0.521 | -0.306 | 0 | 0.001 | 0.001 | 0 | 0.41 | 0.37 | 0.48 | 0.37 | 0.43 | 0.33 |
| 3 | 174532637 | rs9867956 | 0.397 | -0.244 | 0.604 | -0.302 | 0 | 0.045 | 0.045 | 0 | 0.24 | 0.38 | 0.24 | 0.45 | 0.25 | 0.4 |
| 3 | 174533443 | rs6445183 | -1.191 | -0.308 | 0.135 | -0.247 | 0.003 | 0.029 | 0.029 | 0 | 0.49 | 0.8 | 0.43 | 0.48 | 0.5 | 0.52 |
| 3 | 174536522 | rs940058 | 0.086 | -0.798 | 0.5 | -0.265 | -0.015 | -0.045 | -0.045 | 1.154 | 0.12 | 0.15 | 0.08 | 0.01 | 0.09 | 0.05 |
| 3 | 174540260 | rs6797551 | 0.532 | -0.713 | 0.505 | -0.24 | 0 | -0.017 | -0.017 | 0 | 0.11 | 0.11 | 0.07 | 0 | 0.12 | 0.05 |
| 3 | 174563022 | rs9872466 | -0.697 | -0.961 | 0.682 | -0.019 | -0.018 | 0.041 | 0.041 | 0 | 0.32 | 0.4 | 0.39 | 0.51 | 0.31 | 0.26 |
| 3 | 174563280 | rs16865032 | 0.49 | -1.013 | 0.579 | 0.084 | 0.009 | 0.012 | 0.012 | 0.598 | 0.11 | 0.06 | 0.08 | 0.04 | 0.12 | 0.05 |
| 3 | 174567744 | rs9828728 | 0.629 | -0.971 | 0.617 | 0.131 | 0 | 0.038 | 0.038 | 0 | 0.18 | 0.19 | 0.3 | 0.47 | 0.18 | 0.21 |
| 3 | 174568398 | rs9871227 | 0.206 | -0.974 | 0.659 | 0.129 | -0.004 | 0.055 | 0.055 | 0 | 0.2 | 0.28 | 0.31 | 0.47 | 0.19 | 0.21 |
| 3 | 174579820 | rs16865212 | -0.331 | -0.971 | 0.957 | 0.046 | -0.021 | -0.038 | -0.038 | 0 | 0.14 | 0.36 | 0.09 | 0.04 | 0.13 | 0.05 |
| 3 | 174580571 | rs6767604 | -1.438 | -1.09 | 1.102 | -0.003 | 0.036 | -0.03 | -0.03 | 0 | 0.43 | 0.7 | 0.49 | 0.49 | 0.43 | 0.31 |
| 3 | 174580594 | rs6801168 | -0.46 | -1.082 | 1.098 | 0.002 | -0.004 | 0.013 | 0.013 | 0 | 0.3 | 0.33 | 0.4 | 0.45 | 0.3 | 0.27 |
| 3 | 174581888 | rs6445184 | -1.964 | -1.168 | 0.543 | -0.455 | -0.111 | 0.091 | 0.091 | 0 | 0.65 | 0.89 | 0.68 | 0.83 | 0.63 | 0.67 |
| 3 | 174582723 | rs16865252 | -0.144 | -1.829 | 0.984 | -0.372 | -0.022 | -0.035 | -0.035 | 0 | 0.13 | 0.28 | 0.09 | 0.04 | 0.13 | 0.05 |
| 3 | 174585719 | rs10513726 | -0.441 | -1.849 | 0.941 | -0.381 | -0.023 | -0.04 | -0.04 | 0 | 0.14 | 0.38 | 0.11 | 0.04 | 0.13 | 0.05 |
| 3 | 174586440 | rs2861851 | -0.586 | -1.778 | 0.494 | -0.245 | -0.003 | 0.004 | 0.004 | 0 | 0.3 | 0.31 | 0.39 | 0.32 | 0.3 | 0.26 |
| 3 | 174592237 | rs12696366 | 0.658 | -1.745 | 0.033 | -0.121 | 0 | -0.019 | -0.019 | 0 | 0.16 | 0.1 | 0.31 | 0.31 | 0.16 | 0.18 |
| 3 | 174596166 | rs3849522 | 0.167 | -1.575 | -0.161 | -0.218 | -0.009 | -0.012 | -0.012 | 0 | 0.11 | 0.1 | 0.08 | 0.04 | 0.13 | 0.05 |
| 3 | 174608936 | rs7623634 | -0.855 | -1.535 | -0.186 | -0.147 | -0.005 | 0 | 0 | 0 | 0.35 | 0.6 | 0.42 | 0.32 | 0.3 | 0.28 |
| 3 | 174627921 | rs940059 | -1.298 | -0.877 | -1.161 | -0.258 | -0.19 | 0.105 | 0.105 | 0 | 0.74 | 0.97 | 0.72 | 0.94 | 0.63 | 0.76 |
| 3 | 174628523 | rs4527385 | -0.769 | -1.305 | 0.092 | 0.651 | -0.012 | 0.022 | 0.022 | 0 | 0.24 | 0.3 | 0.34 | 0.34 | 0.18 | 0.21 |
| 3 | 174646603 | rs7615452 | -0.382 | -0.943 | -0.565 | 0.766 | 0 | 0.001 | 0.001 | 0 | 0.21 | 0.25 | 0.35 | 0.35 | 0.18 | 0.23 |
| 3 | 174667446 | rs6801807 | 0.176 | -1.173 | -0.831 | 0.525 | 0.011 | -0.036 | -0.036 | 0.129 | 0.7 | 0.73 | 0.53 | 0.52 | 0.78 | 0.58 |
| 3 | 174669007 | rs9874488 | -0.881 | -1.478 | -0.95 | 0.052 | 0.086 | 0.146 | 0.146 | 0 | 0.48 | 0.7 | 0.59 | 0.86 | 0.33 | 0.62 |
| 3 | 174672849 | rs11708020 | 0.21 | -1.274 | -1.057 | 0.411 | 0.026 | -0.038 | -0.038 | 3.353 | 0.66 | 0.78 | 0.51 | 0.48 | 0.76 | 0.64 |
| 3 | 174681247 | rs10513729 | 0.278 | -0.997 | -0.761 | -0.414 | 0.012 | 0.046 | 0.046 | 0 | 0.49 | 0.36 | 0.4 | 0.14 | 0.56 | 0.36 |
| 3 | 174683126 | rs9845780 | 0.109 | -1.056 | -1.096 | -0.639 | 0.013 | 0.18 | 0.18 | 1.01 | 0.49 | 0.73 | 0.39 | 0.11 | 0.53 | 0.31 |
| 3 | 174684649 | rs4425273 | -0.139 | -1.814 | -1.536 | -0.097 | 0.034 | -0.009 | -0.009 | 0 | 0.51 | 0.92 | 0.47 | 0.53 | 0.57 | 0.4 |
| 3 | 174685535 | rs13080899 | 1.197 | -2.302 | -1.497 | 0.074 | -0.068 | -0.011 | -0.011 | 0 | 0.31 | 0.11 | 0.36 | 0.25 | 0.52 | 0.28 |
| 3 | 174690419 | rs11719400 | -0.581 | -2.36 | -1.245 | 0.097 | 0.066 | 0.013 | 0.013 | 1.112 | 0.53 | 0.85 | 0.49 | 0.57 | 0.61 | 0.45 |
| 3 | 174694718 | rs7625517 | -0.183 | -2.427 | -0.255 | 0.202 | -0.001 | -0.001 | -0.001 | 0 | 0.44 | 0.48 | 0.42 | 0.38 | 0.55 | 0.32 |
| 3 | 174697959 | rs13083861 | 1.253 | -2.252 | -0.701 | 0.162 | -0.109 | -0.023 | -0.023 | 0 | 0.29 | 0.07 | 0.34 | 0.25 | 0.48 | 0.26 |
| 3 | 174708659 | rs9813316 | -1.454 | -3.597 | 0.138 | -0.188 | 0.107 | 0.017 | 0.017 | 1.045 | 0.43 | 0.78 | 0.28 | 0.42 | 0.16 | 0.31 |
| 3 | 174710700 | rs6806492 | -1.146 | -3.639 | 1.096 | -0.148 | -0.053 | -0.003 | -0.003 | 2.868 | 0.38 | 0.5 | 0.24 | 0.4 | 0.15 | 0.27 |
| 3 | 174717183 | rs6445188 | -0.388 | -3.741 | 1.488 | -0.189 | -0.087 | -0.099 | -0.099 | 0 | 0.23 | 0.54 | 0.2 | 0.1 | 0.08 | 0.11 |
| 3 | 174719405 | rs7615084 | -1.355 | -3.648 | 1.015 | -0.262 | 0.081 | 0.01 | 0.01 | 2.257 | 0.4 | 0.76 | 0.25 | 0.4 | 0.15 | 0.23 |
| 3 | 174724681 | rs4348006 | -1.581 | -3.644 | 0.896 | -0.241 | 0.162 | 0.02 | 0.02 | 1.237 | 0.43 | 0.82 | 0.28 | 0.41 | 0.16 | 0.27 |
| 3 | 174726543 | rs6786297 | -0.88 | -3.594 | 0.924 | -0.264 | -0.097 | -0.018 | -0.018 | 0 | 0.33 | 0.5 | 0.22 | 0.28 | 0.11 | 0.22 |
| 3 | 174732012 | rs9863905 | -0.083 | -3.334 | 0.721 | -0.344 | -0.001 | 0 | 0 | 0.012 | 0.1 | 0.06 | 0.04 | 0.12 | 0.07 | 0.05 |
| 3 | 174733081 | rs9868994 | -1.452 | -3.169 | 0.478 | -0.243 | 0.221 | 0.015 | 0.015 | 2.499 | 0.44 | 0.9 | 0.35 | 0.41 | 0.17 | 0.27 |
| 3 | 174737429 | rs9863038 | -0.743 | -2.624 | 1.071 | -0.426 | -0.038 | -0.058 | -0.058 | 4.684 | 0.29 | 0.35 | 0.1 | 0.11 | 0.13 | 0.11 |
| 3 | 174737795 | rs9825659 | -0.876 | -2.635 | 1.325 | -0.434 | -0.08 | -0.116 | -0.116 | 0 | 0.3 | 0.45 | 0.11 | 0.11 | 0.13 | 0.11 |
| 3 | 174739787 | rs4555530 | 1.098 | -2.699 | 0.991 | -0.465 | -0.002 | -0.016 | -0.016 | 0 | 0.11 | 0.13 | 0.03 | 0 | 0.04 | 0.03 |
| 3 | 174740103 | rs9878181 | -1.426 | -2.897 | 1.333 | -0.365 | 0.013 | 0.002 | 0.002 | 3.172 | 0.36 | 0.34 | 0.3 | 0.38 | 0.17 | 0.23 |
| 3 | 174740389 | rs9878945 | -0.415 | -2.846 | 1.339 | -0.371 | -0.026 | 0.012 | 0.012 | 0.284 | 0.2 | 0.3 | 0.24 | 0.28 | 0.08 | 0.17 |
| 3 | 174741628 | rs7653217 | 0.029 | -2.663 | 1.195 | -0.263 | -0.017 | 0.062 | 0.062 | 0 | 0.09 | 0.17 | 0.21 | 0.27 | 0.05 | 0.14 |
| 3 | 174744836 | rs9845338 | -0.49 | -2.553 | 0.889 | -0.152 | 0.033 | 0.013 | 0.013 | 0.433 | 0.17 | 0.04 | 0.06 | 0.14 | 0.09 | 0.05 |
| 3 | 174747326 | rs13318274 | -0.681 | -2.577 | 0.949 | -0.163 | 0.018 | 0.01 | 0.01 | 0.255 | 0.18 | 0.09 | 0.06 | 0.14 | 0.09 | 0.05 |
| 3 | 174757260 | rs9837420 | -0.915 | -2.686 | 0.806 | -0.346 | 0.035 | -0.023 | -0.023 | 1.251 | 0.3 | 0.16 | 0.26 | 0.41 | 0.21 | 0.24 |
| 3 | 174764910 | rs9851386 | -1.184 | -2.665 | 0.807 | -0.392 | 0.035 | -0.019 | -0.019 | 0 | 0.33 | 0.18 | 0.33 | 0.41 | 0.25 | 0.27 |
| 3 | 174767354 | rs6788065 | -0.463 | -2.661 | 1.396 | -0.375 | -0.041 | 0.038 | 0.038 | 0 | 0.18 | 0.31 | 0.21 | 0.26 | 0.12 | 0.21 |
| 3 | 174767848 | rs7630646 | -1.807 | -2.828 | 0.888 | -0.465 | 0.089 | 0.034 | 0.034 | 2.012 | 0.51 | 0.7 | 0.4 | 0.41 | 0.3 | 0.34 |
| 3 | 174778214 | rs9854799 | -0.426 | -2.461 | 0.947 | -0.332 | 0.013 | 0.008 | 0.008 | 0.094 | 0.18 | 0.1 | 0.09 | 0.15 | 0.12 | 0.1 |
| 3 | 174788812 | rs9880874 | -1.499 | -2.477 | 1.117 | -0.302 | 0.045 | 0.006 | 0.006 | 1.2 | 0.36 | 0.71 | 0.34 | 0.36 | 0.23 | 0.27 |
| 3 | 174790334 | rs6773248 | 0.045 | -2.035 | 1.251 | -0.46 | 0.005 | -0.006 | -0.006 | 0.446 | 0.13 | 0.09 | 0.16 | 0.22 | 0.09 | 0.17 |
| 3 | 174791319 | rs11928561 | -0.682 | -2.008 | 1.107 | -0.448 | -0.05 | 0.028 | 0.028 | 0.038 | 0.18 | 0.52 | 0.24 | 0.22 | 0.11 | 0.2 |
| 3 | 174798056 | rs4452337 | -0.67 | -0.77 | 0.266 | -0.543 | 0.011 | 0.05 | 0.05 | 0 | 0.45 | 0.76 | 0.51 | 0.31 | 0.43 | 0.38 |

| | | | | | | | | | | | | | | | | |
|---|---|---|---|---|---|---|---|---|---|---|---|---|---|---|---|---|
| 3 | 174800760 | rs9823658 | 0.24 | -0.117 | 0.067 | -0.704 | 0.017 | -0.008 | -0.008 | 0 | 0.28 | 0.21 | 0.26 | 0.33 | 0.23 | 0.26 |
| 3 | 174807929 | rs1463524 | -1.451 | -0.36 | 0.575 | -0.522 | 0.003 | -0.005 | -0.005 | 0 | 0.54 | 0.7 | 0.61 | 0.54 | 0.56 | 0.5 |
| 3 | 174808332 | rs9866323 | 0.884 | -0.388 | 0.456 | -0.508 | 0 | -0.001 | -0.001 | 0.984 | 0.21 | 0.24 | 0.23 | 0.33 | 0.16 | 0.23 |
| 3 | 174812658 | rs6791132 | -0.109 | -0.211 | 0.744 | -0.598 | -0.006 | -0.001 | -0.001 | 0 | 0.4 | 0.4 | 0.51 | 0.44 | 0.46 | 0.44 |
| 3 | 174814738 | rs9823911 | 0.466 | 0.116 | 0.357 | -0.634 | -0.005 | -0.012 | -0.012 | 0 | 0.33 | 0.29 | 0.42 | 0.44 | 0.39 | 0.4 |
| 3 | 174814920 | rs9836841 | 0.387 | -0.192 | 0.509 | -0.481 | -0.017 | -0.009 | -0.009 | 0 | 0.37 | 0.29 | 0.45 | 0.44 | 0.44 | 0.41 |
| 3 | 174816927 | rs1381130 | -1.095 | -0.239 | 0.669 | -0.306 | -0.014 | -0.024 | -0.024 | 0 | 0.59 | 0.82 | 0.63 | 0.61 | 0.58 | 0.49 |
| 3 | 174819996 | rs6443282 | -1.121 | -0.71 | 0.84 | -0.189 | -0.037 | -0.035 | -0.035 | 0 | 0.22 | 0.46 | 0.17 | 0.17 | 0.14 | 0.11 |
| 3 | 174823729 | rs9290531 | -1.019 | -0.658 | 0.467 | -0.185 | -0.025 | -0.016 | -0.016 | 0 | 0.61 | 0.74 | 0.58 | 0.61 | 0.53 | 0.55 |
| 3 | 174824444 | rs10049218 | -0.222 | -0.667 | 0.526 | -0.183 | 0.003 | 0.004 | 0.004 | 0.016 | 0.14 | 0.1 | 0.08 | 0.11 | 0.11 | 0.08 |
| 3 | 174824539 | rs4894477 | 0.063 | -0.612 | 0.419 | -0.098 | -0.002 | 0.001 | 0.001 | 0 | 0.46 | 0.52 | 0.49 | 0.51 | 0.41 | 0.47 |
| 3 | 174827967 | rs6782183 | 0.659 | -0.413 | -0.113 | -0.142 | -0.008 | -0.014 | -0.014 | 0 | 0.35 | 0.25 | 0.41 | 0.43 | 0.39 | 0.43 |
| 3 | 174829687 | rs9847129 | -0.32 | 0.403 | -0.508 | 0.085 | -0.054 | -0.05 | -0.05 | 0 | 0.63 | 0.87 | 0.59 | 0.61 | 0.56 | 0.58 |
| 3 | 174831994 | rs9824182 | -1.066 | 0.248 | 0.166 | 0.004 | -0.024 | -0.044 | -0.044 | 0 | 0.08 | 0.21 | 0.04 | 0 | 0.03 | 0.02 |
| 3 | 174852459 | rs9874963 | -1.133 | 0.831 | 0.22 | -0.09 | 0.031 | -0.066 | -0.066 | 0 | 0.11 | 0.26 | 0.16 | 0 | 0.12 | 0.13 |
| 3 | 174854012 | rs6799562 | 0.019 | 0.674 | 0.184 | -0.44 | -0.008 | -0.02 | -0.02 | 0 | 0.42 | 0.45 | 0.34 | 0.29 | 0.36 | 0.47 |
| 3 | 174861035 | rs6792698 | -0.039 | 0.134 | 0.648 | -0.88 | -0.019 | -0.05 | -0.05 | 1.428 | 0.39 | 0.48 | 0.3 | 0.21 | 0.28 | 0.36 |
| 3 | 174861684 | rs7651836 | -0.028 | 0.082 | 0.728 | -0.898 | -0.004 | -0.017 | -0.017 | 0.031 | 0.39 | 0.36 | 0.29 | 0.21 | 0.28 | 0.34 |
| 3 | 174864481 | rs1381136 | -0.2 | -0.414 | 1.325 | -0.215 | -0.003 | -0.01 | -0.01 | 0.229 | 0.44 | 0.51 | 0.49 | 0.38 | 0.43 | 0.46 |
| 3 | 174868463 | rs3866331 | 0.655 | -0.415 | 1.278 | -0.161 | -0.017 | 0.059 | 0.059 | 0 | 0.33 | 0.12 | 0.3 | 0.22 | 0.33 | 0.32 |
| 3 | 174869139 | rs936480 | 0.572 | -0.43 | 1.311 | -0.168 | -0.003 | 0.045 | 0.045 | 0 | 0.33 | 0.19 | 0.3 | 0.21 | 0.33 | 0.33 |
| 3 | 174872038 | rs13061641 | 0.964 | -0.07 | 1.203 | -0.59 | -0.005 | -0.003 | -0.003 | 0.125 | 0.77 | 0.36 | 0.73 | 0.75 | 0.78 | 0.78 |
| 3 | 174872316 | rs11706404 | 0.151 | -0.067 | 1.315 | -0.581 | -0.002 | -0.009 | -0.009 | 0 | 0.43 | 0.48 | 0.49 | 0.37 | 0.43 | 0.45 |
| 3 | 174875397 | rs9819816 | -0.872 | 0.004 | 1.184 | -0.592 | -0.032 | -0.008 | -0.008 | 0 | 0.08 | 0.19 | 0.04 | 0.06 | 0.02 | 0.09 |
| 3 | 174876464 | rs10470502 | -1.111 | -0.021 | 1.197 | -0.593 | -0.015 | 0.07 | 0.07 | 0.014 | 0.11 | 0.3 | 0.2 | 0.23 | 0.1 | 0.18 |
| 3 | 174879055 | rs9858268 | -1.674 | -0.033 | 1.177 | -0.548 | -0.036 | 0.046 | 0.046 | 0.008 | 0.15 | 0.35 | 0.22 | 0.23 | 0.11 | 0.18 |
| 3 | 174879330 | rs9985272 | -0.241 | -0.003 | 1.146 | -0.536 | 0.001 | 0.033 | 0.033 | 0 | 0.07 | 0.16 | 0.17 | 0.16 | 0.09 | 0.09 |
| 3 | 174880072 | rs1461272 | -0.26 | -0.061 | 1.093 | -0.329 | -0.001 | -0.004 | -0.004 | 0 | 0.52 | 0.49 | 0.63 | 0.74 | 0.54 | 0.6 |
| 3 | 174884006 | rs11922187 | 1.11 | 0.038 | 0.839 | -0.305 | -0.033 | 0.04 | 0.04 | 0 | 0.33 | 0.07 | 0.29 | 0.24 | 0.34 | 0.33 |
| 3 | 174895840 | rs2035101 | 1.046 | 0.762 | 0.752 | -0.265 | -0.039 | -0.08 | -0.08 | 1.14 | 0.36 | 0.09 | 0.39 | 0.51 | 0.36 | 0.41 |
| 3 | 174898957 | rs3979497 | 0.688 | 0.614 | 1.101 | -0.104 | -0.002 | -0.003 | -0.003 | 0.058 | 0.77 | 0.56 | 0.77 | 0.76 | 0.78 | 0.77 |
| 3 | 174901394 | rs7648439 | -0.359 | 0.606 | 1.1 | -0.113 | 0 | -0.034 | -0.034 | 0 | 0.84 | 0.73 | 0.92 | 0.9 | 0.86 | 0.87 |
| 3 | 174904407 | rs2861999 | 0.71 | 0.314 | 0.251 | -0.206 | -0.039 | -0.037 | -0.037 | 3.62 | 0.41 | 0.16 | 0.43 | 0.5 | 0.45 | 0.4 |
| 3 | 174905811 | rs9849745 | -0.281 | 0.28 | -0.099 | -0.045 | 0.004 | 0.029 | 0.029 | 0 | 0.07 | 0.14 | 0.15 | 0.14 | 0.09 | 0.09 |
| 3 | 174909952 | rs7615453 | 0.747 | 0.26 | -0.778 | -0.114 | -0.018 | 0.027 | 0.027 | 0 | 0.33 | 0.06 | 0.3 | 0.26 | 0.32 | 0.35 |
| 3 | 174926485 | rs4243408 | -0.288 | 0.521 | -1.057 | 0.305 | -0.018 | -0.062 | -0.062 | 0 | 0.63 | 0.87 | 0.75 | 0.51 | 0.64 | 0.48 |
| 3 | 174926597 | rs4312664 | -0.317 | 0.517 | -1.089 | 0.302 | -0.021 | -0.07 | -0.07 | 1.946 | 0.63 | 0.89 | 0.75 | 0.51 | 0.64 | 0.48 |
| 3 | 174929045 | rs7641680 | 0.109 | 0.443 | -1.03 | 0.274 | -0.014 | 0.053 | 0.053 | 0.059 | 0.08 | 0.16 | 0.14 | 0.2 | 0.06 | 0.07 |
| 3 | 174930740 | rs6804034 | -0.782 | 0.384 | -1.605 | 0.808 | -0.079 | -0.116 | -0.116 | 0 | 0.69 | 0.97 | 0.78 | 0.69 | 0.68 | 0.62 |
| 3 | 174931730 | rs4894701 | 0.441 | 0.222 | -1.354 | 0.336 | -0.006 | 0.136 | 0.136 | 0.224 | 0.46 | 0.17 | 0.42 | 0.29 | 0.55 | 0.41 |
| 3 | 174933353 | rs6443290 | -0.393 | 0.186 | -2.223 | 0.332 | 0.025 | 0 | 0 | 0 | 0.48 | 0.71 | 0.55 | 0.56 | 0.38 | 0.42 |
| 3 | 174933478 | rs10936827 | 0.816 | 0.111 | -2.479 | 0.275 | -0.041 | 0.042 | 0.042 | 0.13 | 0.44 | 0.13 | 0.31 | 0.3 | 0.54 | 0.44 |
| 3 | 174935547 | rs1381122 | -0.858 | 0.124 | -2.353 | 0.262 | 0.009 | 0.025 | 0.025 | 0 | 0.91 | 0.77 | 0.84 | 0.87 | 0.91 | 0.86 |
| 3 | 174936391 | rs6443291 | 0.084 | -0.077 | -2.08 | 0.272 | -0.035 | -0.012 | -0.012 | 1.65 | 0.3 | 0.5 | 0.22 | 0.3 | 0.18 | 0.27 |
| 3 | 174939222 | rs6765162 | -0.907 | -0.039 | -2.14 | 0.262 | 0.025 | -0.003 | -0.003 | 0 | 0.48 | 0.68 | 0.55 | 0.56 | 0.37 | 0.42 |
| 3 | 174940266 | rs6768672 | 0.621 | 0.284 | -1.938 | 0.515 | -0.022 | -0.042 | -0.042 | 0 | 0.16 | 0.07 | 0.31 | 0.25 | 0.19 | 0.15 |
| 3 | 174942578 | rs4894702 | -0.884 | 0.289 | -1.824 | 0.346 | 0.014 | 0.047 | 0.047 | 0 | 0.91 | 0.76 | 0.84 | 0.86 | 0.91 | 0.86 |
| 3 | 174942871 | rs11915650 | 1.6 | 0.542 | -2.126 | 0.067 | -0.02 | 0.036 | 0.036 | 0 | 0.13 | 0.03 | 0.06 | 0 | 0.16 | 0.02 |
| 3 | 174943598 | rs1461250 | -0.938 | 0.729 | -1.896 | 0.356 | 0.002 | 0.035 | 0.035 | 0 | 0.92 | 0.81 | 0.85 | 0.85 | 0.91 | 0.86 |
| 3 | 174961387 | rs1463525 | 1.095 | 0.667 | -1.666 | 0.21 | -0.004 | -0.003 | -0.003 | 0.379 | 0.36 | 0.35 | 0.24 | 0.34 | 0.39 | 0.42 |
| 3 | 174962279 | rs1463526 | 0.885 | 0.746 | -1.766 | 0.266 | -0.047 | -0.053 | -0.053 | 0 | 0.12 | 0.01 | 0.3 | 0.2 | 0.18 | 0.15 |
| 3 | 174966522 | rs7619752 | -0.924 | 0.771 | -1.645 | 0.236 | 0.018 | -0.004 | -0.004 | 0 | 0.85 | 0.75 | 0.8 | 0.87 | 0.81 | 0.85 |
| 3 | 174968553 | rs7622037 | 1.176 | 0.681 | -2.085 | 0.385 | -0.005 | -0.004 | -0.004 | 0 | 0.34 | 0.32 | 0.28 | 0.32 | 0.35 | 0.42 |
| 3 | 174971516 | rs12633712 | -0.575 | 0.304 | -2.206 | 0.388 | 0.012 | -0.067 | -0.067 | 0 | 0.41 | 0.17 | 0.52 | 0.54 | 0.39 | 0.39 |
| 3 | 174972810 | rs1461249 | 0.496 | 0.605 | -1.886 | 0.805 | 0.014 | -0.039 | -0.039 | 0 | 0.21 | 0.1 | 0.19 | 0.33 | 0.17 | 0.19 |
| 3 | 174978751 | rs17533447 | -0.051 | 1.149 | -1.465 | 0.852 | -0.05 | -0.048 | -0.048 | 0 | 0.09 | 0.02 | 0.12 | 0.16 | 0.16 | 0.13 |
| 3 | 174980057 | rs1461248 | -0.732 | 1.23 | -1.386 | 0.897 | -0.001 | 0.032 | 0.032 | 3.403 | 0.51 | 0.63 | 0.62 | 0.62 | 0.49 | 0.7 |
| 3 | 174985106 | rs6443300 | -0.957 | 1.062 | -0.697 | 0.625 | 0.042 | 0.027 | 0.027 | 1.239 | 0.16 | 0.42 | 0.32 | 0.18 | 0.21 | 0.14 |
| 3 | 174996509 | rs13067238 | 0.116 | 0.829 | -1.568 | 0.55 | -0.129 | -0.086 | -0.086 | 0.084 | 0.28 | 0.03 | 0.3 | 0.3 | 0.4 | 0.44 |
| 3 | 175001029 | rs9879950 | 0.411 | 0.439 | -1.638 | 0.604 | 0.057 | -0.012 | -0.012 | 0 | 0.56 | 0.83 | 0.49 | 0.47 | 0.35 | 0.36 |
| 3 | 175002708 | rs6809621 | -0.21 | 0.643 | -1.583 | 0.637 | -0.017 | -0.011 | -0.011 | 0 | 0.85 | 0.9 | 0.82 | 0.79 | 0.77 | 0.84 |
| 3 | 175006190 | rs6443301 | -0.713 | 0.173 | -1.257 | 0.477 | 0.07 | -0.02 | -0.02 | 1.598 | 0.49 | 0.84 | 0.54 | 0.49 | 0.62 | 0.61 |
| 3 | 175008948 | rs6799976 | -0.426 | -0.622 | -0.798 | -0.085 | -0.001 | -0.007 | -0.007 | 2.886 | 0.39 | 0.57 | 0.27 | 0.21 | 0.37 | 0.16 |
| 3 | 175012620 | rs13059425 | 0.593 | -0.585 | -0.538 | -0.138 | -0.033 | -0.027 | -0.027 | 0.072 | 0.35 | 0.26 | 0.45 | 0.44 | 0.49 | 0.53 |
| 3 | 175015466 | rs7630477 | 0.503 | -0.989 | 0.072 | -0.132 | -0.047 | -0.159 | -0.159 | 1.986 | 0.24 | 0.43 | 0.15 | 0 | 0.17 | 0.04 |
| 3 | 175019533 | rs12630370 | -0.401 | -0.734 | -0.205 | 0.314 | 0.013 | 0.145 | 0.145 | 0.571 | 0.05 | 0.22 | 0.05 | 0.14 | 0.04 | 0.03 |
| 3 | 175020834 | rs9844160 | -0.627 | -0.85 | -0.658 | 0.27 | 0.039 | -0.107 | -0.107 | 0 | 0.73 | 0.94 | 0.79 | 0.55 | 0.82 | 0.68 |
| 3 | 175021928 | rs7623764 | -0.931 | -0.631 | 0.058 | 0.307 | -0.008 | 0.152 | 0.152 | 0.431 | 0.08 | 0.26 | 0.14 | 0.19 | 0.04 | 0.04 |
| 3 | 175027803 | rs7625843 | -0.422 | -0.399 | -0.438 | -0.032 | 0.014 | 0.06 | 0.06 | 0 | 0.54 | 0.7 | 0.46 | 0.69 | 0.43 | 0.44 |

| | | | | | | | | | | | | | | | | |
|---|---|---|---|---|---|---|---|---|---|---|---|---|---|---|---|---|
| 3 | 175042776 | rs4894713 | 0.36 | -0.437 | -1.133 | 0.088 | -0.003 | -0.019 | -0.019 | 0 | 0.17 | 0.08 | 0.2 | 0.22 | 0.22 | 0.14 |
| 3 | 175043656 | rs13065290 | 0.181 | -0.718 | -0.895 | -0.049 | 0.014 | 0.022 | 0.022 | 1.012 | 0.2 | 0.13 | 0.17 | 0 | 0.17 | 0.08 |
| 3 | 175050468 | rs2161044 | -0.49 | -0.317 | -0.328 | 0.895 | -0.034 | 0.059 | 0.059 | 0 | 0.3 | 0.44 | 0.24 | 0.51 | 0.17 | 0.3 |
| 3 | 175055095 | rs9821455 | -1.268 | 0.263 | -0.415 | 0.625 | 0 | -0.007 | -0.007 | 0.001 | 0.07 | 0.08 | 0.15 | 0.03 | 0.1 | 0.1 |
| 3 | 175056030 | rs10490874 | -0.931 | -0.035 | -0.955 | 1.222 | -0.009 | -0.072 | -0.072 | 1.381 | 0.11 | 0.01 | 0.1 | 0.2 | 0.1 | 0.08 |
| 3 | 175057188 | rs16825471 | -0.672 | 0.433 | 0.208 | 0.999 | 0.076 | -0.004 | -0.004 | 0 | 0.06 | 0.27 | 0.06 | 0 | 0.06 | 0.04 |
| 3 | 175059171 | rs7629234 | -1.766 | -0.289 | 0.919 | 0.833 | -0.008 | -0.065 | -0.065 | 0 | 0.13 | 0.58 | 0.09 | 0.07 | 0.07 | 0.08 |
| 3 | 175061871 | rs6769077 | 0.138 | 0.332 | 0.239 | 0.057 | -0.049 | -0.023 | -0.023 | 0.025 | 0.29 | 0.21 | 0.56 | 0.63 | 0.46 | 0.54 |
| 3 | 175064233 | rs16825489 | 0.094 | 0.543 | 0.664 | -0.004 | 0.01 | 0.005 | 0.005 | 0 | 0.29 | 0.33 | 0.55 | 0.65 | 0.42 | 0.54 |
| 3 | 175064809 | rs10936845 | -1.008 | 0.503 | 1.051 | -0.046 | 0.003 | -0.048 | -0.048 | 1.127 | 0.12 | 0.33 | 0.07 | 0.08 | 0.08 | 0.06 |
| 3 | 175066203 | rs6807589 | -1.306 | 1.216 | 0.965 | 0.573 | 0.008 | 0.015 | 0.015 | 0.055 | 0.33 | 0.64 | 0.51 | 0.56 | 0.41 | 0.36 |
| 3 | 175067922 | rs16825525 | 0.054 | 1.408 | 0.866 | 0.762 | -0.001 | 0 | 0 | 0.138 | 0.07 | 0.06 | 0.16 | 0.15 | 0.1 | 0.14 |
| 3 | 175068143 | rs16825528 | -0.026 | 1.699 | 0.739 | 1.07 | -0.105 | -0.121 | -0.121 | 0.637 | 0.19 | 0.04 | 0.39 | 0.39 | 0.33 | 0.3 |
| 3 | 175068218 | rs16825529 | -1.526 | 1.57 | 1.331 | 0.881 | 0.017 | 0.013 | 0.013 | 0 | 0.36 | 0.49 | 0.52 | 0.45 | 0.52 | 0.38 |
| 3 | 175068537 | rs6774623 | -2.027 | 1.54 | 1.493 | 0.926 | -0.003 | -0.004 | -0.004 | 0 | 0.45 | 0.61 | 0.69 | 0.7 | 0.62 | 0.55 |
| 3 | 175069717 | rs11915209 | -1.249 | 1.496 | 1.65 | 0.939 | -0.005 | 0.041 | 0.041 | 0 | 0.13 | 0.31 | 0.2 | 0.29 | 0.12 | 0.18 |
| 3 | 175075109 | rs6771725 | 0.129 | 1.78 | 1.539 | 0.869 | -0.001 | 0.01 | 0.01 | 0 | 0.19 | 0.19 | 0.37 | 0.48 | 0.29 | 0.27 |
| 3 | 175077686 | rs4894485 | -2.098 | 1.678 | 1.847 | 0.814 | 0.018 | 0.023 | 0.023 | 0 | 0.45 | 0.69 | 0.7 | 0.69 | 0.62 | 0.52 |
| 3 | 175087884 | rs3915774 | -1.151 | 1.653 | 1.586 | 0.72 | -0.008 | 0.003 | 0.003 | 0.204 | 0.11 | 0.08 | 0.1 | 0.01 | 0.2 | 0.08 |
| 3 | 175090662 | rs7631033 | -1.094 | 0.964 | 1.701 | 0.715 | -0.025 | -0.031 | -0.031 | 0 | 0.09 | 0.28 | 0.05 | 0.04 | 0.04 | 0.04 |
| 3 | 175094357 | rs9869663 | -1.632 | 1.322 | 1.924 | 1.032 | 0.008 | 0.007 | 0.007 | 0 | 0.34 | 0.59 | 0.59 | 0.62 | 0.42 | 0.45 |
| 3 | 175096546 | rs9290550 | -0.992 | 1.354 | 1.913 | 1.076 | 0.019 | 0.087 | 0.087 | 0 | 0.25 | 0.4 | 0.42 | 0.52 | 0.3 | 0.31 |
| 3 | 175102338 | rs6793266 | -1.196 | 1.681 | 1.627 | 1.095 | 0.026 | 0.047 | 0.047 | 0 | 0.27 | 0.36 | 0.55 | 0.6 | 0.38 | 0.47 |
| 3 | 175109345 | rs7643142 | -0.304 | 1.794 | 1.312 | 1.125 | 0.021 | 0.028 | 0.028 | 0 | 0.08 | 0.14 | 0.17 | 0.15 | 0.13 | 0.16 |
| 3 | 175113985 | rs12490133 | -0.839 | 0.678 | 0.621 | 0.919 | -0.001 | -0.002 | -0.002 | 7.261 | 0.27 | 0.25 | 0.42 | 0.46 | 0.33 | 0.34 |
| 3 | 175115541 | rs12495889 | -0.816 | 0.706 | 0.328 | 0.958 | -0.019 | -0.059 | -0.059 | 0.034 | 0.25 | 0.14 | 0.4 | 0.45 | 0.32 | 0.34 |
| 3 | 175115709 | rs2193848 | -0.027 | 0.57 | 0.328 | 1.052 | -0.008 | -0.037 | -0.037 | 0 | 0.18 | 0.09 | 0.35 | 0.44 | 0.22 | 0.27 |
| 3 | 175116207 | rs6774177 | -0.433 | -0.39 | 0.326 | 0.408 | -0.004 | -0.002 | -0.002 | 2.927 | 0.2 | 0.25 | 0.12 | 0.21 | 0.16 | 0.21 |
| 3 | 175118132 | rs1862077 | 1.297 | -0.605 | 0.136 | 0.301 | 0.022 | 0.01 | 0.01 | 0 | 0.55 | 0.32 | 0.55 | 0.59 | 0.62 | 0.62 |
| 3 | 175118665 | rs2193849 | 0.796 | -0.672 | 0.303 | 0.296 | -0.005 | 0.017 | 0.017 | 0 | 0.83 | 0.9 | 0.83 | 0.88 | 0.81 | 0.79 |
| 3 | 175124340 | rs9843194 | 1.439 | -0.71 | 1.34 | 0.341 | 0.032 | 0.015 | 0.015 | 0.614 | 0.87 | 0.69 | 0.83 | 0.85 | 0.82 | 0.8 |
| 3 | 175127141 | rs13082925 | 2.159 | -0.857 | 1.398 | 0.27 | 0.002 | -0.001 | -0.001 | 0 | 0.66 | 0.39 | 0.61 | 0.67 | 0.69 | 0.67 |
| 3 | 175128075 | rs1073231 | -0.373 | -0.7 | 1.773 | 0.283 | 0.001 | -0.015 | -0.015 | 0 | 0.19 | 0.13 | 0.09 | 0.03 | 0.13 | 0.12 |
| 3 | 175136656 | rs6773706 | 1.251 | -0.627 | 2.141 | 0.27 | 0.005 | -0.012 | -0.012 | 0.484 | 0.81 | 0.79 | 0.79 | 0.88 | 0.76 | 0.74 |
| 3 | 175139842 | rs9815671 | 2.013 | -0.744 | 2.079 | 0.244 | 0 | -0.002 | -0.002 | 0.137 | 0.65 | 0.39 | 0.62 | 0.66 | 0.66 | 0.66 |
| 3 | 175174602 | rs9818285 | 2.058 | -0.519 | 2.661 | 0.614 | 0.117 | 0.235 | 0.235 | 0.793 | 0.88 | 0.57 | 0.71 | 0.74 | 0.81 | 0.81 |
| 3 | 175179451 | rs2902108 | 1.847 | -0.598 | 2.64 | 0.558 | 0.011 | 0.077 | 0.077 | 0 | 0.77 | 0.59 | 0.68 | 0.67 | 0.75 | 0.77 |
| 3 | 175180353 | rs10460885 | -2.954 | -0.561 | 2.902 | 0.665 | 0.056 | 0.414 | 0.414 | 3.739 | 0.06 | 0.36 | 0.16 | 0.24 | 0.03 | 0.04 |
| 3 | 175182072 | rs6789345 | -1.048 | -0.542 | 2.891 | 0.639 | -0.043 | 0.097 | 0.097 | 0.024 | 0.17 | 0.49 | 0.19 | 0.22 | 0.09 | 0.05 |
| 3 | 175185060 | rs9839245 | 2.112 | -0.961 | 2.355 | 0.687 | 0.026 | -0.046 | -0.046 | 0 | 0.61 | 0.24 | 0.56 | 0.55 | 0.66 | 0.52 |
| 3 | 175186191 | rs7433056 | 2.086 | -1.004 | 2.217 | 0.44 | 0.01 | 0.009 | 0.009 | 0 | 0.6 | 0.41 | 0.67 | 0.69 | 0.65 | 0.52 |
| 3 | 175187978 | rs2161042 | 1.723 | -0.16 | 2.138 | 0.559 | -0.007 | -0.006 | -0.006 | 0 | 0.73 | 0.61 | 0.71 | 0.76 | 0.72 | 0.56 |
| 3 | 175194867 | rs9290557 | 0.353 | 0.273 | 1.689 | 0.382 | 0.031 | -0.003 | -0.003 | 0 | 0.87 | 0.81 | 0.81 | 0.89 | 0.81 | 0.88 |
| 3 | 175200278 | rs4894720 | 2.33 | 0.121 | 1.967 | 0.285 | 0.002 | -0.012 | -0.012 | 0 | 0.73 | 0.29 | 0.69 | 0.89 | 0.69 | 0.73 |
| 3 | 175204418 | rs6763187 | -2.288 | 0.321 | 2.195 | 0.656 | -0.002 | 0.002 | 0.002 | 0.558 | 0.31 | 0.7 | 0.37 | 0.34 | 0.26 | 0.35 |
| 3 | 175205148 | rs7646041 | 0.892 | 0.361 | 2.231 | 0.676 | 0.004 | 0.038 | 0.038 | 0 | 0.83 | 0.79 | 0.69 | 0.72 | 0.82 | 0.71 |
| 3 | 175207539 | rs9816719 | 1.697 | 0.418 | 2.028 | 0.662 | 0.052 | -0.019 | -0.019 | 0 | 0.91 | 0.84 | 0.83 | 0.97 | 0.8 | 0.75 |
| 3 | 175209406 | rs7619984 | 0.073 | 0.382 | 2.23 | 0.697 | -0.026 | 0.002 | 0.002 | 0 | 0.11 | 0.19 | 0.09 | 0.13 | 0.07 | 0.17 |
| 3 | 175215959 | rs2098570 | 0.152 | 0.464 | 1.577 | 0.543 | -0.003 | 0.001 | 0.001 | 0 | 0.1 | 0.2 | 0.09 | 0.09 | 0.08 | 0.02 |
| 3 | 175216853 | rs4894489 | 1.64 | 0.573 | 1.136 | 0.415 | -0.011 | -0.035 | -0.035 | 0.686 | 0.66 | 0.2 | 0.52 | 0.63 | 0.63 | 0.56 |
| 3 | 175218006 | rs6443305 | -1.28 | 0.299 | 1.003 | 0.448 | -0.001 | 0 | 0 | 0.608 | 0.31 | 0.67 | 0.37 | 0.35 | 0.25 | 0.34 |
| 3 | 175224459 | rs7628569 | 1.173 | 0.99 | 1.19 | 0.001 | 0.038 | -0.008 | -0.008 | 0 | 0.94 | 0.89 | 0.89 | 0.96 | 0.84 | 0.87 |
| 3 | 175228372 | rs13067631 | -1.734 | 0.8 | 1.276 | -0.022 | 0.004 | 0.003 | 0.003 | 0 | 0.09 | 0.15 | 0.16 | 0.08 | 0.13 | 0.14 |
| 3 | 175233654 | rs2112044 | 2.303 | 0.77 | 1.65 | -0.101 | 0.002 | -0.004 | -0.004 | 0 | 0.78 | 0.27 | 0.71 | 0.86 | 0.78 | 0.76 |
| 3 | 175238672 | rs12494093 | -1.932 | 0.753 | 2.161 | -0.176 | 0.057 | 0.028 | 0.028 | 0.095 | 0.12 | 0.47 | 0.2 | 0.08 | 0.13 | 0.14 |
| 3 | 175242041 | rs10513733 | -0.498 | 0.786 | 1.985 | -0.143 | 0.013 | -0.033 | -0.033 | 0 | 0.08 | 0.22 | 0.06 | 0.03 | 0.06 | 0.01 |
| 3 | 175245827 | rs10804857 | 1.796 | 0.707 | 1.517 | -0.146 | 0.004 | -0.011 | -0.011 | 0 | 0.8 | 0.28 | 0.72 | 0.88 | 0.82 | 0.77 |
| 3 | 175259933 | rs6797470 | 1.449 | 1.088 | 1.419 | -0.382 | 0.02 | -0.039 | -0.039 | 0 | 0.74 | 0.38 | 0.69 | 0.87 | 0.72 | 0.82 |
| 3 | 175263907 | rs6773768 | 1.351 | 1.047 | 0.397 | -0.62 | -0.07 | -0.176 | -0.176 | 0 | 0.25 | 0.02 | 0.34 | 0.6 | 0.29 | 0.44 |
| 3 | 175267563 | rs7638295 | -1.608 | 0.916 | 1.488 | -1.732 | 0.007 | -0.107 | -0.107 | 1.676 | 0.23 | 0.44 | 0.26 | 0.03 | 0.22 | 0.09 |
| 3 | 175281578 | rs16825963 | -0.645 | 0.822 | 1.348 | -1.565 | -0.006 | 0.002 | 0.002 | 0 | 0.14 | 0.1 | 0.11 | 0 | 0.15 | 0.06 |
| 3 | 175282048 | rs4243412 | 1.731 | 0.757 | 1.808 | -1.574 | -0.001 | -0.001 | -0.001 | 0 | 0.77 | 0.28 | 0.71 | 0.84 | 0.77 | 0.83 |
| 3 | 175295076 | rs6784858 | -1.351 | 0.761 | 2.46 | -1.613 | -0.001 | -0.113 | -0.113 | 0 | 0.2 | 0.59 | 0.25 | 0.03 | 0.17 | 0.08 |
| 3 | 175296589 | rs6791525 | -1.365 | 0.763 | 2.461 | -1.612 | -0.003 | -0.107 | -0.107 | 0 | 0.2 | 0.62 | 0.25 | 0.03 | 0.17 | 0.08 |
| 3 | 175326982 | rs10936861 | 1.435 | 0.467 | 1.994 | -1.751 | -0.001 | -0.008 | -0.008 | 0 | 0.74 | 0.32 | 0.7 | 0.85 | 0.77 | 0.82 |
| 3 | 175328953 | rs7648032 | 1.334 | 0.474 | 2.181 | -1.748 | 0 | -0.048 | -0.048 | 3.292 | 0.75 | 0.48 | 0.72 | 0.85 | 0.78 | 0.82 |
| 3 | 175334302 | rs6443312 | -0.628 | -0.359 | 1.748 | -2.102 | 0.008 | -0.117 | -0.117 | 0 | 0.17 | 0.34 | 0.13 | 0 | 0.16 | 0.06 |
| 3 | 175338777 | rs4605557 | 1.342 | -0.644 | 1.229 | -2.136 | -0.021 | -0.05 | -0.05 | 0 | 0.71 | 0.47 | 0.82 | 0.86 | 0.8 | 0.82 |
| 3 | 175344146 | rs12489573 | 0.378 | -0.766 | 0.671 | -1.952 | -0.045 | 0.072 | 0.072 | 0 | 0.13 | 0.01 | 0.18 | 0.01 | 0.15 | 0.07 |

| | | | | | | | | | | | | | | | | |
|---|---|---|---|---|---|---|---|---|---|---|---|---|---|---|---|---|
| 3 | 175349612 | rs9681019 | -0.47 | -1.047 | 1.587 | -1.368 | 0.01 | -0.008 | -0.008 | 0.091 | 0.08 | 0.05 | 0.02 | 0.13 | 0.02 | 0.11 |
| 3 | 175389857 | rs12107747 | -1.895 | -1.111 | 1.977 | -1.543 | -0.041 | -0.02 | -0.02 | 0 | 0.15 | 0.68 | 0.1 | 0.12 | 0.05 | 0.15 |
| 3 | 175403065 | rs16826153 | -0.31 | -1.083 | 1.747 | -1.511 | -0.003 | -0.001 | -0.001 | 0 | 0.08 | 0.06 | 0.03 | 0.12 | 0.02 | 0.14 |
| 3 | 175403905 | rs9833254 | -1.954 | -1.318 | 1.377 | -1.692 | 0.079 | 0.095 | 0.095 | 0 | 0.32 | 0.84 | 0.24 | 0.16 | 0.17 | 0.23 |
| 3 | 175410177 | rs1889261 | -1.753 | -1.24 | 1.814 | -1.414 | -0.045 | -0.013 | -0.013 | 0 | 0.16 | 0.66 | 0.15 | 0.15 | 0.07 | 0.17 |
| 3 | 175415175 | rs4558765 | -1.404 | -1.281 | 1.602 | -1.644 | -0.011 | -0.025 | -0.025 | 0 | 0.28 | 0.64 | 0.24 | 0.15 | 0.22 | 0.24 |
| 3 | 175422241 | rs4383533 | -1.082 | -1.006 | 1.82 | -1.657 | -0.004 | -0.017 | -0.017 | 0.406 | 0.23 | 0.68 | 0.23 | 0.12 | 0.21 | 0.17 |
| 3 | 175425449 | rs6784051 | -0.301 | -0.882 | 1.892 | -1.644 | 0.024 | -0.063 | -0.063 | 0.196 | 0.17 | 0.45 | 0.14 | 0.07 | 0.18 | 0.13 |
| 3 | 175450586 | rs16826200 | -0.671 | -0.941 | 1.425 | -1.688 | -0.012 | -0.088 | -0.088 | 0 | 0.31 | 0.33 | 0.17 | 0.01 | 0.23 | 0.08 |
| 3 | 175458558 | rs6791612 | -0.962 | -1.042 | 1.551 | -1.735 | -0.028 | -0.114 | -0.114 | 0.326 | 0.37 | 0.51 | 0.27 | 0.05 | 0.26 | 0.15 |
| 3 | 175472118 | rs9876798 | -0.059 | -0.949 | 0.894 | -1.647 | -0.034 | -0.122 | -0.122 | 0 | 0.21 | 0.35 | 0.12 | 0 | 0.15 | 0.04 |
| 3 | 175481669 | rs2902212 | -0.026 | -0.943 | 0.252 | -1.644 | 0.001 | -0.034 | -0.034 | 0 | 0.21 | 0.2 | 0.12 | 0 | 0.15 | 0.04 |
| 3 | 175488587 | rs1931156 | -0.526 | -0.367 | -0.189 | -1.314 | 0.105 | 0.257 | 0.257 | 0.913 | 0.46 | 0.85 | 0.33 | 0.08 | 0.29 | 0.27 |
| 3 | 175496866 | rs3114967 | -1.378 | 0.8 | 0.661 | 0.229 | 0.015 | 0.007 | 0.007 | 0 | 0.37 | 0.46 | 0.47 | 0.41 | 0.44 | 0.58 |
| 3 | 175497466 | rs1931164 | -1.291 | 0.457 | 0.222 | -0.277 | -0.001 | -0.013 | -0.013 | 0 | 0.25 | 0.67 | 0.34 | 0.08 | 0.24 | 0.18 |
| 3 | 175507453 | rs3114972 | -1.06 | 0.722 | 0.295 | -0.079 | 0.019 | -0.069 | -0.069 | 0 | 0.23 | 0.51 | 0.22 | 0.09 | 0.22 | 0.25 |
| 3 | 175510469 | rs7609959 | -1.772 | 0.443 | 1.066 | 0.16 | 0.051 | 0.088 | 0.088 | 0 | 0.06 | 0.38 | 0.05 | 0.04 | 0.03 | 0.01 |
| 3 | 175510677 | rs2184207 | -1.082 | 0.957 | -0.36 | 0.419 | -0.124 | -0.062 | -0.062 | 0.153 | 0.39 | 0.92 | 0.48 | 0.4 | 0.49 | 0.5 |
| 3 | 175512408 | rs6805025 | -1.149 | 0.319 | -0.271 | 0.745 | 0.019 | -0.026 | -0.026 | 4.252 | 0.31 | 0.8 | 0.36 | 0.45 | 0.23 | 0.38 |
| 3 | 175513789 | rs9880439 | -1.215 | 0.199 | -0.187 | 0.931 | 0.022 | -0.025 | -0.025 | 1.125 | 0.31 | 0.79 | 0.38 | 0.43 | 0.21 | 0.4 |
| 3 | 175515266 | rs6790940 | -1.073 | 0.152 | 0.709 | 0.808 | 0.009 | -0.002 | -0.002 | 0 | 0.12 | 0.26 | 0.25 | 0.13 | 0.11 | 0.26 |
| 3 | 175515423 | rs6791089 | -1.087 | 0.151 | 0.7 | 0.8 | 0.042 | 0.075 | 0.075 | 0.096 | 0.06 | 0.26 | 0.24 | 0.13 | 0.06 | 0.19 |
| 3 | 175516157 | rs9849649 | -0.498 | 0.101 | 1.504 | 0.918 | -0.033 | 0.064 | 0.064 | 1.632 | 0.18 | 0.29 | 0.12 | 0.34 | 0.1 | 0.14 |
| 3 | 175520736 | rs12107944 | -1.359 | 0.14 | 1.649 | 0.891 | 0.086 | 0.128 | 0.128 | 0.443 | 0.07 | 0.44 | 0.25 | 0.12 | 0.06 | 0.2 |
| 3 | 175521509 | rs7638113 | -1.883 | 0.167 | 1.629 | 0.87 | 0.066 | 0.047 | 0.047 | 0.287 | 0.1 | 0.44 | 0.28 | 0.12 | 0.13 | 0.21 |
| 3 | 175525659 | rs1931169 | -2.16 | 0.151 | 1.544 | 0.92 | 0.02 | -0.027 | -0.027 | 0 | 0.34 | 0.79 | 0.41 | 0.47 | 0.28 | 0.37 |
| 3 | 175529750 | rs9871961 | -0.566 | 0.129 | 1.724 | 0.931 | -0.059 | 0.116 | 0.116 | 2.477 | 0.18 | 0.5 | 0.13 | 0.32 | 0.1 | 0.09 |
| 3 | 175530097 | rs4386512 | -1.883 | 0.146 | 1.563 | 0.911 | 0.027 | 0.021 | 0.021 | 0.105 | 0.1 | 0.23 | 0.27 | 0.13 | 0.13 | 0.21 |
| 3 | 175539275 | rs9840501 | -0.469 | -0.163 | 1.504 | 1.013 | -0.036 | 0.064 | 0.064 | 2.19 | 0.17 | 0.31 | 0.11 | 0.3 | 0.09 | 0.09 |
| 3 | 175541402 | rs6801642 | -1.626 | -0.181 | 1.188 | 0.933 | 0.012 | -0.016 | -0.016 | 0 | 0.3 | 0.77 | 0.37 | 0.43 | 0.21 | 0.34 |
| 3 | 175545048 | rs6790597 | -1.546 | -0.636 | 0.528 | 0.365 | 0.082 | 0.154 | 0.154 | 0 | 0.49 | 0.77 | 0.53 | 0.8 | 0.34 | 0.71 |
| 3 | 175548952 | rs9882704 | -1.372 | -0.603 | 0.504 | 0.389 | 0.066 | 0.117 | 0.117 | 0 | 0.46 | 0.86 | 0.48 | 0.77 | 0.32 | 0.67 |
| 3 | 175555183 | rs10936871 | -0.633 | -1.15 | 1.112 | 0.049 | -0.011 | -0.12 | -0.12 | 0 | 0.18 | 0.37 | 0.12 | 0 | 0.11 | 0.06 |
| 3 | 175555304 | rs9869315 | -1.812 | -1.268 | 0.771 | 0.224 | 0.006 | 0.005 | 0.005 | 0.073 | 0.29 | 0.75 | 0.2 | 0.12 | 0.19 | 0.2 |
| 3 | 175566986 | rs9917624 | -0.882 | -1.247 | 0.838 | 0.564 | -0.018 | 0.044 | 0.044 | 0 | 0.12 | 0.34 | 0.08 | 0.14 | 0.08 | 0.14 |
| 3 | 175573756 | rs12633710 | -0.451 | -1.456 | 0.273 | 0.711 | 0.031 | -0.113 | -0.113 | 0.005 | 0.18 | 0.03 | 0.28 | 0.62 | 0.12 | 0.44 |
| 3 | 175576447 | rs6794349 | -0.907 | -1.168 | 0.809 | 0.661 | -0.024 | 0.026 | 0.026 | 0 | 0.11 | 0.26 | 0.07 | 0.14 | 0.07 | 0.14 |
| 3 | 175586997 | rs12493932 | -0.257 | -2.344 | -0.271 | 0.25 | 0.103 | -0.145 | -0.145 | 0 | 0.24 | 0.08 | 0.33 | 0.62 | 0.13 | 0.29 |
| 3 | 175593960 | rs4243416 | -1.284 | -2.404 | 0.216 | -0.208 | 0.054 | 0.018 | 0.018 | 6.88 | 0.46 | 0.37 | 0.46 | 0.7 | 0.28 | 0.48 |
| 3 | 175619643 | rs11921736 | -1.234 | -2.358 | 0.494 | -0.264 | -0.014 | -0.06 | -0.06 | 0 | 0.29 | 0.32 | 0.17 | 0.07 | 0.2 | 0.23 |
| 3 | 175630821 | rs13091384 | 0.752 | -2.122 | 0.575 | -0.161 | -0.019 | 0.047 | 0.047 | 0 | 0.1 | 0.02 | 0.07 | 0.01 | 0.09 | 0.08 |
| 3 | 175632046 | rs9809859 | -1.789 | -1.475 | 0.534 | -0.107 | -0.014 | 0.059 | 0.059 | 3.237 | 0.6 | 0.86 | 0.58 | 0.81 | 0.41 | 0.63 |
| 3 | 175632101 | rs9847111 | -1.287 | -1.501 | 0.744 | -0.093 | 0.059 | 0.028 | 0.028 | 4.938 | 0.52 | 0.6 | 0.51 | 0.73 | 0.32 | 0.5 |
| 3 | 175640593 | rs9862314 | -1.722 | -1.365 | 0.544 | -0.05 | -0.015 | 0.058 | 0.058 | 3.282 | 0.6 | 0.85 | 0.58 | 0.81 | 0.42 | 0.64 |
| 3 | 175651713 | rs7617330 | 0.025 | -1.223 | -0.138 | -0.098 | -0.009 | -0.048 | -0.048 | 0 | 0.21 | 0.26 | 0.12 | 0.07 | 0.15 | 0.18 |
| 3 | 175653894 | rs7635524 | -0.185 | -1.244 | -0.142 | -0.108 | -0.014 | -0.057 | -0.057 | 0 | 0.23 | 0.27 | 0.12 | 0.07 | 0.15 | 0.19 |
| 3 | 175656506 | rs9815577 | 0.083 | -0.278 | -0.74 | 0.387 | 0.04 | 0.027 | 0.027 | 0 | 0.06 | 0.1 | 0.06 | 0.09 | 0.09 | 0.13 |
| 3 | 175658506 | rs9884045 | -0.952 | 0.049 | -0.862 | 0.325 | -0.001 | 0.02 | 0.02 | 2.752 | 0.59 | 0.8 | 0.56 | 0.74 | 0.43 | 0.55 |
| 3 | 175669896 | rs9823496 | 0.405 | 0.068 | -0.634 | 0.582 | -0.004 | -0.006 | -0.006 | 0 | 0.54 | 0.52 | 0.66 | 0.62 | 0.62 | 0.55 |
| 3 | 175670278 | rs9824424 | -0.882 | -0.361 | -0.049 | 1.174 | -0.017 | 0.065 | 0.065 | 0 | 0.12 | 0.28 | 0.11 | 0.26 | 0.07 | 0.19 |
| 3 | 175672207 | rs7614793 | -0.476 | 0.044 | -0.193 | 1.42 | -0.001 | 0.003 | 0.003 | 0.504 | 0.19 | 0.24 | 0.27 | 0.34 | 0.25 | 0.18 |
| 3 | 175681864 | rs9859162 | -0.756 | -0.097 | -0.575 | 0.409 | 0.051 | 0.013 | 0.013 | 0 | 0.57 | 0.84 | 0.61 | 0.38 | 0.68 | 0.49 |
| 3 | 175683589 | rs6801541 | 0.273 | -0.173 | -0.38 | 0.501 | 0.001 | 0.008 | 0.008 | 0.644 | 0.2 | 0.24 | 0.28 | 0.33 | 0.25 | 0.18 |
| 3 | 175693993 | rs2862403 | 0.384 | -0.172 | -0.398 | 0.532 | -0.008 | -0.017 | -0.017 | 0 | 0.19 | 0.17 | 0.27 | 0.29 | 0.25 | 0.17 |
| 3 | 175705041 | rs4496497 | 1.851 | -0.404 | -0.382 | 0.317 | 0 | -0.006 | -0.006 | 0 | 0.07 | 0.09 | 0.03 | 0.01 | 0.03 | 0.11 |
| 3 | 175708175 | rs2862404 | -1.116 | -0.967 | -0.066 | 0.491 | 0.019 | -0.005 | -0.005 | 1.286 | 0.64 | 0.75 | 0.78 | 0.58 | 0.75 | 0.57 |
| 3 | 175718927 | rs9841807 | -0.056 | -0.887 | 0.064 | 0.408 | -0.002 | 0.014 | 0.014 | 0.049 | 0.27 | 0.44 | 0.32 | 0.34 | 0.28 | 0.28 |
| 3 | 175757340 | rs6787411 | 0.588 | -0.427 | 0.173 | 0.039 | 0.055 | -0.108 | -0.108 | 0 | 0.29 | 0.37 | 0.31 | 0.04 | 0.45 | 0.21 |
| 3 | 175767480 | rs9882330 | -0.171 | -1.495 | 0.553 | 0.861 | -0.03 | 0.041 | 0.041 | 0.159 | 0.22 | 0.48 | 0.22 | 0.36 | 0.15 | 0.26 |
| 3 | 175775312 | rs9821038 | -1.196 | -2.077 | -0.065 | 0.901 | -0.011 | -0.063 | -0.063 | 0.813 | 0.86 | 0.98 | 0.88 | 0.81 | 0.86 | 0.85 |
| 3 | 175790003 | rs7653056 | 0.317 | -2.224 | 0.001 | 0.887 | -0.018 | 0.034 | 0.034 | 0 | 0.2 | 0.37 | 0.25 | 0.34 | 0.15 | 0.28 |
| 3 | 175793476 | rs7640642 | 0.317 | -2.224 | 0.001 | 0.887 | -0.018 | 0.034 | 0.034 | 0 | 0.2 | 0.37 | 0.25 | 0.34 | 0.15 | 0.28 |
| 3 | 175802031 | rs9869872 | 1.183 | -2.243 | 0 | 0.918 | 0 | 0.004 | 0.004 | 0 | 0.13 | 0.16 | 0.21 | 0.33 | 0.13 | 0.14 |
| 3 | 175808125 | rs9847632 | 1.183 | -2.243 | 0 | 0.918 | 0 | 0.004 | 0.004 | 0 | 0.13 | 0.16 | 0.22 | 0.33 | 0.13 | 0.14 |
| 3 | 175821406 | rs4577477 | -0.44 | -2.225 | 0.153 | 0.92 | -0.019 | 0.023 | 0.023 | 0 | 0.35 | 0.51 | 0.39 | 0.54 | 0.3 | 0.43 |
| 3 | 175831874 | rs12497666 | 0.838 | -2.749 | -0.009 | 0.829 | 0.018 | -0.007 | -0.007 | 0 | 0.26 | 0.14 | 0.19 | 0.3 | 0.12 | 0.3 |
| 3 | 175846905 | rs5001165 | 0.489 | -2.177 | -0.23 | 0.832 | 0.009 | -0.012 | -0.012 | 0 | 0.21 | 0.17 | 0.25 | 0.34 | 0.15 | 0.29 |
| 3 | 175859004 | rs7609811 | -0.35 | -1.939 | -0.486 | 0.893 | -0.013 | 0.003 | 0.003 | 0 | 0.38 | 0.51 | 0.4 | 0.47 | 0.3 | 0.43 |
| 3 | 175861931 | rs6808187 | -0.059 | -2.006 | -0.689 | 0.957 | 0.008 | -0.003 | -0.003 | 1.743 | 0.31 | 0.32 | 0.23 | 0.33 | 0.16 | 0.26 |

| | | | | | | | | | | | | | | | | |
|---|---|---|---|---|---|---|---|---|---|---|---|---|---|---|---|---|
| 3 | 175862678 | rs2902242 | -0.517 | -1.917 | -0.303 | 1.2 | 0.01 | -0.016 | -0.016 | 0 | 0.26 | 0.74 | 0.33 | 0.49 | 0.16 | 0.3 |
| 3 | 175865817 | rs1921467 | 0.37 | -1.699 | -0.33 | 1.089 | 0 | -0.039 | -0.039 | 0.01 | 0.16 | 0.07 | 0.24 | 0.32 | 0.15 | 0.1 |
| 3 | 175870797 | rs13316219 | 0.658 | -2 | -0.196 | 0.768 | -0.012 | -0.007 | -0.007 | 0 | 0.09 | 0.14 | 0.12 | 0.07 | 0.05 | 0.07 |
| 3 | 175874225 | rs6796072 | -1.149 | -2.471 | 0.476 | 0.786 | -0.009 | 0.003 | 0.003 | 5.46 | 0.38 | 0.54 | 0.39 | 0.51 | 0.21 | 0.39 |
| 3 | 175893082 | rs13061380 | -1.183 | -2.164 | -0.229 | 0.327 | -0.061 | 0.003 | 0.003 | 2.526 | 0.65 | 0.83 | 0.6 | 0.76 | 0.41 | 0.75 |
| 3 | 175893784 | rs16826704 | 0.848 | -2.181 | -0.028 | 0.268 | -0.018 | -0.013 | -0.013 | 0.082 | 0.1 | 0.18 | 0.12 | 0.07 | 0.05 | 0.07 |
| 3 | 175895736 | rs12495724 | 0.297 | -2.412 | 0.1 | 0.328 | 0.063 | 0.004 | 0.004 | 0 | 0.34 | 0.17 | 0.21 | 0.35 | 0.21 | 0.56 |
| 3 | 175896348 | rs12496526 | 0.274 | -2.007 | 0.056 | 0.25 | 0.04 | 0.011 | 0.011 | 0 | 0.36 | 0.18 | 0.25 | 0.35 | 0.27 | 0.57 |
| 3 | 175901580 | rs13078340 | -0.677 | -2.028 | 0.435 | 0.275 | 0.003 | 0.015 | 0.015 | 0.966 | 0.55 | 0.36 | 0.5 | 0.69 | 0.42 | 0.67 |
| 3 | 175905626 | rs7618770 | 1.121 | -2.092 | 0.695 | 0.26 | -0.034 | 0.014 | 0.014 | 1.164 | 0.34 | 0.27 | 0.4 | 0.25 | 0.53 | 0.26 |
| 3 | 175907342 | rs2067613 | 0.096 | -2.157 | 0.849 | 0.265 | 0.011 | 0.006 | 0.006 | 0 | 0.53 | 0.57 | 0.61 | 0.57 | 0.68 | 0.36 |
| 3 | 175920884 | rs4894505 | 0.136 | -2.302 | 0.823 | 0.245 | 0.003 | 0.001 | 0.001 | 0 | 0.55 | 0.58 | 0.61 | 0.56 | 0.68 | 0.37 |
| 3 | 175927451 | rs11923600 | 0.76 | -2.279 | 0.885 | 0.245 | -0.001 | 0.019 | 0.019 | 0.209 | 0.17 | 0.21 | 0.2 | 0.32 | 0.14 | 0.1 |
| 3 | 175929674 | rs1921476 | 1.064 | -2.284 | 0.798 | 0.133 | -0.028 | 0.019 | 0.019 | 0.771 | 0.37 | 0.28 | 0.4 | 0.27 | 0.54 | 0.26 |
| 3 | 175931891 | rs6443331 | -0.97 | -2.024 | 0.921 | 0.163 | 0 | -0.009 | -0.009 | 0 | 0.89 | 0.85 | 0.83 | 0.92 | 0.88 | 0.92 |
| 3 | 175934613 | rs7615573 | 0.686 | -2.035 | 0.883 | 0.233 | -0.001 | 0.016 | 0.016 | 0.201 | 0.18 | 0.21 | 0.23 | 0.34 | 0.15 | 0.1 |
| 3 | 175937783 | rs10936879 | -0.488 | -2.095 | 0.921 | 0.162 | -0.005 | -0.004 | -0.004 | 0 | 0.52 | 0.5 | 0.43 | 0.64 | 0.34 | 0.65 |
| 3 | 175939204 | rs17689331 | 0.749 | -1.731 | 1.102 | 0.241 | -0.031 | 0.039 | 0.039 | 0 | 0.43 | 0.27 | 0.38 | 0.29 | 0.54 | 0.43 |
| 3 | 175946326 | rs7645789 | -0.195 | -1.69 | 0.811 | 0.296 | -0.006 | -0.002 | -0.002 | 0 | 0.43 | 0.48 | 0.44 | 0.61 | 0.33 | 0.47 |
| 3 | 175954186 | rs7609918 | -0.195 | -1.69 | 0.811 | 0.296 | -0.006 | -0.002 | -0.002 | 0 | 0.43 | 0.48 | 0.44 | 0.61 | 0.33 | 0.47 |
| 3 | 175957859 | rs7615876 | 1.141 | -1.596 | 0.794 | 0.278 | -0.007 | 0.001 | 0.001 | 0 | 0.24 | 0.27 | 0.18 | 0.27 | 0.18 | 0.35 |
| 3 | 175965350 | rs1471695 | 0.003 | -1.152 | 0.551 | 0.372 | 0 | 0.002 | 0.002 | 0 | 0.46 | 0.43 | 0.46 | 0.62 | 0.39 | 0.48 |
| 3 | 175969241 | rs17689750 | 1.189 | -1.155 | 0.504 | 0.379 | 0.001 | 0.002 | 0.002 | 0 | 0.27 | 0.2 | 0.19 | 0.27 | 0.24 | 0.36 |
| 3 | 175972919 | rs12489622 | 0.659 | -1.144 | 0.52 | 0.371 | 0 | -0.005 | -0.005 | 0 | 0.18 | 0.16 | 0.26 | 0.34 | 0.15 | 0.12 |
| 3 | 175979713 | rs11719850 | 0.03 | -1.152 | 0.493 | 0.353 | 0.001 | 0.002 | 0.002 | 0 | 0.46 | 0.41 | 0.45 | 0.61 | 0.39 | 0.48 |
| 3 | 175992530 | rs4544626 | 0.699 | -1.13 | 0.424 | 0.38 | -0.001 | -0.021 | -0.021 | 0 | 0.18 | 0.13 | 0.26 | 0.35 | 0.15 | 0.12 |
| 3 | 176004686 | rs16826837 | 0.559 | -1.115 | 0.221 | 0.386 | 0 | -0.007 | -0.007 | 0 | 0.18 | 0.14 | 0.26 | 0.35 | 0.15 | 0.12 |
| 3 | 176014233 | rs2046350 | 0.562 | -1.088 | 0.152 | 0.301 | 0.002 | -0.01 | -0.01 | 0 | 0.42 | 0.5 | 0.37 | 0.31 | 0.49 | 0.42 |
| 3 | 176014495 | rs2046352 | -0.633 | -0.864 | -0.094 | 0.158 | 0 | 0.001 | 0.001 | 0 | 0.88 | 0.9 | 0.83 | 0.91 | 0.88 | 0.9 |
| 3 | 176022286 | rs953068 | 0.142 | -1.2 | -0.094 | 0.333 | 0.009 | 0.026 | 0.026 | 0 | 0.5 | 0.59 | 0.48 | 0.37 | 0.58 | 0.44 |
| 3 | 176022635 | rs951161 | -0.517 | -1.072 | -0.38 | 0.36 | -0.007 | 0.007 | 0.007 | 0.888 | 0.31 | 0.67 | 0.33 | 0.44 | 0.21 | 0.29 |
| 3 | 176022917 | rs5008411 | -0.552 | -0.697 | -0.667 | 0.712 | 0.005 | 0.009 | 0.009 | 0 | 0.69 | 0.77 | 0.76 | 0.75 | 0.73 | 0.62 |
| 3 | 176025504 | rs1032077 | 0.187 | -0.511 | -1.396 | 0.81 | 0.004 | -0.004 | -0.004 | 0 | 0.18 | 0.2 | 0.26 | 0.4 | 0.11 | 0.15 |
| 3 | 176030096 | rs6799767 | 0.002 | 0.159 | -1.56 | 0.4 | 0.014 | 0.023 | 0.023 | 0 | 0.49 | 0.4 | 0.49 | 0.35 | 0.62 | 0.47 |
| 3 | 176031564 | rs9873527 | -0.232 | 0.706 | -2.275 | 0.465 | -0.074 | 0.001 | 0.001 | 0 | 0.28 | 0.05 | 0.24 | 0.24 | 0.3 | 0.44 |
| 3 | 176031687 | rs9835842 | -0.13 | -0.094 | -2.408 | 0.542 | 0.033 | -0.019 | -0.019 | 0 | 0.19 | 0.05 | 0.13 | 0.21 | 0.16 | 0.15 |
| 3 | 176032666 | rs10936885 | -0.4 | 0.531 | -2.349 | 0.396 | -0.022 | 0.025 | 0.025 | 0 | 0.09 | 0.03 | 0.16 | 0.08 | 0.12 | 0.07 |
| 3 | 176033450 | rs1483630 | -0.584 | -0.038 | -2.806 | 0.126 | 0.048 | 0.082 | 0.082 | 1.769 | 0.55 | 0.8 | 0.73 | 0.75 | 0.59 | 0.68 |
| 3 | 176034991 | rs4894763 | 1.086 | -0.336 | -2.792 | 0.379 | -0.094 | -0.098 | -0.098 | 0 | 0.27 | 0.06 | 0.5 | 0.54 | 0.41 | 0.46 |
| 3 | 176035490 | rs1156636 | -1.03 | -0.413 | -2.796 | 0.364 | 0.048 | 0.085 | 0.085 | 1.575 | 0.55 | 0.8 | 0.73 | 0.75 | 0.59 | 0.68 |
| 3 | 176048311 | rs12489250 | 0.098 | -0.409 | -1.682 | 0.335 | -0.01 | -0.01 | -0.01 | 0 | 0.37 | 0.31 | 0.67 | 0.62 | 0.53 | 0.54 |
| 3 | 176051150 | rs2128849 | 1.074 | -0.322 | -1.759 | 0.728 | -0.106 | -0.095 | -0.095 | 0 | 0.27 | 0.05 | 0.43 | 0.4 | 0.41 | 0.46 |
| 3 | 176052011 | rs13065682 | 0.378 | -0.367 | -1.362 | 0.532 | -0.015 | 0.025 | 0.025 | 0 | 0.09 | 0.03 | 0.16 | 0.06 | 0.12 | 0.07 |
| 3 | 176055763 | rs9841755 | 0.271 | -0.504 | -1.188 | 0.538 | -0.044 | -0.003 | -0.003 | 0 | 0.17 | 0.48 | 0.06 | 0.13 | 0.06 | 0.15 |
| 3 | 176062509 | rs7650171 | 0.168 | -0.5 | -0.926 | 0.54 | -0.045 | -0.009 | -0.009 | 0 | 0.18 | 0.52 | 0.06 | 0.13 | 0.07 | 0.15 |
| 3 | 176082414 | rs9853201 | 0.645 | -0.478 | -0.19 | 0.563 | -0.02 | -0.005 | -0.005 | 0.193 | 0.1 | 0.2 | 0.03 | 0.09 | 0.04 | 0.02 |
| 3 | 176087269 | rs301230 | 0.381 | -1.098 | -0.35 | 0.293 | -0.011 | -0.015 | -0.015 | 0 | 0.4 | 0.44 | 0.25 | 0.23 | 0.36 | 0.28 |
| 3 | 176097014 | rs1564731 | 1.63 | -0.884 | -0.42 | 0.467 | -0.074 | -0.062 | -0.062 | 0 | 0.27 | 0.1 | 0.43 | 0.4 | 0.4 | 0.49 |
| 3 | 176116524 | rs1483944 | -0.969 | -0.976 | -0.144 | 0.267 | 0.004 | -0.022 | -0.022 | 0 | 0.9 | 0.86 | 0.83 | 0.95 | 0.88 | 0.93 |
| 3 | 176126533 | rs301181 | -0.92 | -1.016 | -0.18 | 0.249 | 0 | -0.007 | -0.007 | 0 | 0.9 | 0.91 | 0.84 | 0.96 | 0.89 | 0.93 |
| 3 | 176128315 | rs9875303 | 0.277 | -1.01 | -0.204 | 0.182 | -0.009 | -0.015 | -0.015 | 0.102 | 0.41 | 0.45 | 0.25 | 0.23 | 0.36 | 0.3 |
| 3 | 176131530 | rs301178 | 0.3 | -1.094 | -0.116 | 0.288 | -0.027 | -0.023 | -0.023 | 0 | 0.67 | 0.53 | 0.69 | 0.66 | 0.76 | 0.78 |
| 3 | 176132164 | rs301177 | 0.316 | -1.108 | -0.122 | 0.282 | -0.027 | -0.024 | -0.024 | 0 | 0.67 | 0.53 | 0.69 | 0.66 | 0.76 | 0.78 |
| 3 | 176135457 | rs301172 | -0.309 | -1.159 | -0.254 | -0.058 | -0.027 | -0.022 | -0.022 | 0.009 | 0.31 | 0.54 | 0.24 | 0.31 | 0.28 | 0.21 |
| 3 | 176138839 | rs301191 | 1.535 | -1.112 | -0.178 | -0.065 | -0.003 | -0.002 | -0.002 | 0.023 | 0.07 | 0.06 | 0.09 | 0 | 0.14 | 0.05 |
| 3 | 176149354 | rs6778967 | 0.774 | -1.454 | -0.088 | -0.236 | -0.005 | -0.009 | -0.009 | 0 | 0.59 | 0.44 | 0.62 | 0.63 | 0.62 | 0.74 |
| 3 | 176157272 | rs1483940 | 0.241 | -1.55 | 0.109 | -0.28 | -0.054 | 0 | 0 | 0.94 | 0.24 | 0.42 | 0.14 | 0.31 | 0.13 | 0.15 |
| 3 | 176159192 | rs7651271 | 0.901 | -1.336 | 0.082 | -0.216 | 0.016 | 0.015 | 0.015 | 0.136 | 0.33 | 0.28 | 0.16 | 0.2 | 0.23 | 0.21 |
| 3 | 176172315 | rs7619454 | 0.784 | -0.642 | 0.178 | -0.135 | -0.009 | -0.019 | -0.019 | 0 | 0.59 | 0.52 | 0.62 | 0.63 | 0.62 | 0.74 |
| 3 | 176181980 | rs2862476 | 0.514 | -0.372 | 0.662 | -0.107 | -0.001 | -0.121 | -0.121 | 0.115 | 0.81 | 0.49 | 0.81 | 0.96 | 0.78 | 0.89 |
| 3 | 176194786 | rs1483927 | 0.101 | -0.31 | 0.56 | 0.086 | -0.002 | -0.02 | -0.02 | 0 | 0.87 | 0.58 | 0.9 | 0.94 | 0.9 | 0.94 |
| 3 | 176203825 | rs9832479 | 0.679 | -0.632 | -0.551 | -0.059 | 0.021 | 0.017 | 0.017 | 0.193 | 0.22 | 0.1 | 0.13 | 0.17 | 0.16 | 0.17 |
| 3 | 176206226 | rs7628033 | 0.329 | 0.576 | -0.496 | 0.318 | -0.119 | -0.097 | -0.097 | 0 | 0.36 | 0.07 | 0.52 | 0.47 | 0.45 | 0.53 |
| 3 | 176212508 | rs1156688 | 0.826 | 1.116 | 0.248 | 0.324 | -0.028 | 0.09 | 0.09 | 2.497 | 0.31 | 0.07 | 0.18 | 0.14 | 0.26 | 0.19 |
| 3 | 176216767 | rs9838609 | 0.031 | 1.4 | 0.592 | 0.626 | -0.001 | -0.003 | -0.003 | 0 | 0.38 | 0.47 | 0.36 | 0.38 | 0.36 | 0.28 |
| 3 | 176225642 | rs7645298 | 0.561 | 1.275 | 0.718 | 0.786 | -0.003 | 0.001 | 0.001 | 2.511 | 0.4 | 0.41 | 0.37 | 0.43 | 0.36 | 0.58 |
| 3 | 176233955 | rs1454149 | -0.814 | 1.252 | 0.593 | 0.635 | -0.024 | 0.034 | 0.034 | 0 | 0.78 | 0.9 | 0.83 | 0.89 | 0.72 | 0.87 |
| 3 | 176237200 | rs1351800 | 0.617 | 1.234 | 0.622 | 0.702 | -0.002 | 0 | 0 | 1.635 | 0.42 | 0.44 | 0.45 | 0.48 | 0.38 | 0.59 |
| 3 | 176243614 | rs17620274 | 0.788 | 1.251 | -0.493 | -0.786 | 0.011 | 0.045 | 0.045 | 2.225 | 0.39 | 0.24 | 0.48 | 0.76 | 0.36 | 0.61 |

| | | | | | | | | | | | | | | | | |
|---|---|---|---|---|---|---|---|---|---|---|---|---|---|---|---|---|
| 3 | 176246528 | rs9818162 | 0.703 | 1.351 | -1.339 | -0.783 | 0.01 | 0.075 | 0.075 | 2.075 | 0.32 | 0.06 | 0.19 | 0.18 | 0.23 | 0.19 |
| 3 | 176261513 | rs1377820 | 0.467 | 1.08 | -0.915 | -0.478 | -0.008 | -0.033 | -0.033 | 0.032 | 0.41 | 0.5 | 0.52 | 0.79 | 0.39 | 0.61 |
| 3 | 176267268 | rs1349837 | 0.116 | 1.394 | -1.162 | -0.522 | -0.023 | 0.042 | 0.042 | 0 | 0.78 | 0.89 | 0.84 | 0.98 | 0.74 | 0.88 |
| 3 | 176271598 | rs936030 | 0.255 | 0.962 | -0.669 | 0.004 | -0.001 | -0.004 | -0.004 | 0 | 0.57 | 0.34 | 0.45 | 0.54 | 0.55 | 0.57 |
| 3 | 176289744 | rs6443357 | 0.048 | 0.817 | 0.526 | -0.21 | 0.007 | -0.048 | -0.048 | 0 | 0.79 | 0.64 | 0.85 | 0.9 | 0.78 | 0.71 |
| 3 | 176294153 | rs2862481 | 0.948 | 0.832 | 0.551 | -0.115 | 0.002 | -0.006 | -0.006 | 0.761 | 0.69 | 0.32 | 0.68 | 0.76 | 0.71 | 0.64 |
| 3 | 176312533 | rs9290596 | -0.996 | 0.544 | 0.862 | 0.272 | -0.002 | 0.173 | 0.173 | 0 | 0.06 | 0.19 | 0.09 | 0.19 | 0.02 | 0.17 |
| 3 | 176313261 | rs9848490 | -1.192 | 0.545 | 0.892 | 0.271 | 0.007 | 0.195 | 0.195 | 0 | 0.06 | 0.23 | 0.1 | 0.19 | 0.02 | 0.16 |
| 3 | 176318239 | rs1377827 | -1.513 | 0.394 | 0.735 | 0.151 | -0.004 | 0.1 | 0.1 | 0 | 0.16 | 0.43 | 0.18 | 0.21 | 0.1 | 0.24 |
| 3 | 176328274 | rs1545542 | -0.905 | 0.386 | 0.538 | -0.076 | -0.003 | 0.044 | 0.044 | 0.958 | 0.31 | 0.57 | 0.41 | 0.49 | 0.37 | 0.45 |
| 3 | 176337924 | rs9290599 | -1.239 | 0.185 | 0.266 | -0.383 | 0 | -0.024 | -0.024 | 0 | 0.64 | 0.79 | 0.66 | 0.54 | 0.68 | 0.64 |
| 3 | 176352763 | rs838209 | -0.281 | 0.934 | 0.351 | -0.343 | -0.002 | -0.002 | -0.002 | 0 | 0.25 | 0.28 | 0.37 | 0.32 | 0.34 | 0.17 |
| 3 | 176372607 | rs1027183 | 1.441 | 0.397 | 0.221 | -0.436 | 0.054 | -0.04 | -0.04 | 0 | 0.41 | 0.19 | 0.37 | 0.53 | 0.29 | 0.45 |
| 3 | 176382135 | rs4857788 | -0.447 | 1.071 | 0.384 | -0.009 | -0.041 | -0.038 | -0.038 | 0.68 | 0.84 | 0.91 | 0.69 | 0.69 | 0.68 | 0.86 |
| 3 | 176382268 | rs7610013 | 0.637 | 1.137 | 0.428 | 0.019 | -0.002 | -0.057 | -0.057 | 7.94 | 0.12 | 0.21 | 0.08 | 0 | 0.09 | 0.01 |
| 3 | 176382972 | rs9815008 | -0.834 | 1.117 | 0.529 | 0.007 | -0.003 | 0.05 | 0.05 | 0.215 | 0.08 | 0.23 | 0.04 | 0.1 | 0.06 | 0.22 |
| 3 | 176383197 | rs1490049 | -0.834 | 1.117 | 0.416 | 0.007 | 0.007 | 0.069 | 0.069 | 0.215 | 0.08 | 0.25 | 0.05 | 0.1 | 0.06 | 0.22 |
| 3 | 176384090 | rs4857654 | 0.803 | 1.437 | -0.314 | -0.179 | 0.016 | 0 | 0 | 0 | 0.68 | 0.6 | 0.57 | 0.66 | 0.55 | 0.78 |
| 3 | 176387145 | rs7647703 | -0.343 | 0.668 | -0.522 | -0.498 | -0.001 | -0.019 | -0.019 | 0 | 0.17 | 0.2 | 0.11 | 0.03 | 0.14 | 0.07 |
| 3 | 176407064 | rs1490051 | -0.353 | 0.96 | -0.467 | -0.187 | 0.001 | -0.014 | -0.014 | 0.853 | 0.19 | 0.18 | 0.16 | 0.05 | 0.22 | 0.12 |
| 3 | 176409064 | rs9880529 | 0.619 | 0.302 | -0.084 | 0.138 | -0.015 | 0.003 | 0.003 | 0 | 0.81 | 0.77 | 0.85 | 0.8 | 0.87 | 0.78 |
| 3 | 176412862 | rs6772784 | -0.747 | -0.225 | -0.036 | 0.389 | -0.01 | 0.039 | 0.039 | 1.304 | 0.14 | 0.56 | 0.14 | 0.14 | 0.11 | 0.19 |
| 3 | 176417144 | rs1490070 | 0.15 | -0.269 | 0.097 | 0.374 | 0.011 | -0.006 | -0.006 | 0 | 0.82 | 0.9 | 0.86 | 0.8 | 0.87 | 0.78 |
| 3 | 176418794 | rs9826338 | -0.433 | -0.281 | 1.2 | 0.352 | -0.004 | 0.021 | 0.021 | 0 | 0.11 | 0.22 | 0.12 | 0.14 | 0.1 | 0.19 |
| 3 | 176423123 | rs4857658 | -0.981 | -0.301 | 1.646 | 0.343 | -0.008 | 0.033 | 0.033 | 0.404 | 0.13 | 0.36 | 0.13 | 0.14 | 0.11 | 0.19 |
| 3 | 176430561 | rs12494343 | 0.968 | -0.009 | 0.592 | 0.554 | -0.046 | -0.051 | -0.051 | 2.571 | 0.17 | 0.04 | 0.25 | 0.22 | 0.25 | 0.15 |
| 3 | 176430786 | rs977943 | 0.914 | -0.009 | 0.706 | 0.554 | -0.01 | -0.012 | -0.012 | 0 | 0.17 | 0.11 | 0.26 | 0.22 | 0.25 | 0.15 |
| 3 | 176456234 | rs9839536 | -0.838 | -0.02 | 0.679 | 0.544 | 0 | 0.002 | 0.002 | 0 | 0.33 | 0.64 | 0.39 | 0.36 | 0.35 | 0.35 |
| 3 | 176462187 | rs6802030 | -0.607 | -0.019 | 0.937 | 0.546 | 0.007 | 0.034 | 0.034 | 0 | 0.31 | 0.53 | 0.39 | 0.36 | 0.35 | 0.35 |
| 3 | 176464937 | rs7632805 | 0.306 | -0.272 | 0.136 | 0.338 | 0.011 | -0.002 | -0.002 | 0 | 0.58 | 0.75 | 0.58 | 0.58 | 0.59 | 0.6 |
| 3 | 176468053 | rs6804392 | -0.279 | -0.239 | 0.282 | 0.414 | 0.002 | 0.058 | 0.058 | 0 | 0.14 | 0.43 | 0.13 | 0.2 | 0.06 | 0.2 |
| 3 | 176471332 | rs1490054 | -0.13 | -0.268 | -0.673 | 0.498 | -0.034 | -0.021 | -0.021 | 0 | 0.34 | 0.1 | 0.38 | 0.28 | 0.4 | 0.25 |
| 3 | 176477852 | rs901932 | 0.896 | 0.281 | -0.862 | 0.509 | 0.025 | 0.024 | 0.024 | 1.783 | 0.41 | 0.15 | 0.34 | 0.34 | 0.35 | 0.45 |
| 3 | 176479883 | rs6778657 | -0.129 | 0.937 | -0.389 | 0.433 | 0.013 | -0.013 | -0.013 | 0 | 0.6 | 0.73 | 0.56 | 0.43 | 0.61 | 0.57 |
| 3 | 176480903 | rs6782137 | 0.26 | 0.949 | -0.444 | 0.828 | 0.019 | -0.016 | -0.016 | 2.313 | 0.36 | 0.23 | 0.37 | 0.45 | 0.3 | 0.33 |
| 3 | 176481047 | rs11920836 | -0.026 | 1.038 | 0.538 | 0.558 | 0.038 | -0.012 | -0.012 | 0 | 0.07 | 0.29 | 0.04 | 0 | 0.06 | 0.03 |
| 3 | 176489072 | rs9874890 | 1.134 | 0.079 | 0.166 | -0.105 | -0.046 | -0.084 | -0.084 | 0 | 0.16 | 0.22 | 0.07 | 0.01 | 0.05 | 0.03 |
| 3 | 176489547 | rs12493292 | 0.572 | 0.368 | 0.375 | -0.072 | 0 | -0.002 | -0.002 | 2.023 | 0.66 | 0.3 | 0.73 | 0.83 | 0.62 | 0.77 |
| 3 | 176498235 | rs12487119 | 0.56 | 0.415 | 0.371 | -0.089 | 0 | -0.007 | -0.007 | 0 | 0.66 | 0.31 | 0.73 | 0.88 | 0.63 | 0.77 |
| 3 | 176501869 | rs1490047 | -0.559 | 0.419 | 0.584 | -0.052 | -0.03 | -0.044 | -0.044 | 0 | 0.77 | 0.63 | 0.85 | 0.91 | 0.84 | 0.87 |
| 3 | 176505740 | rs4323017 | 0.007 | 0.427 | 0.651 | -0.01 | 0.053 | 0.107 | 0.107 | 0.071 | 0.94 | 0.74 | 0.92 | 0.91 | 0.92 | 0.9 |
| 3 | 176507084 | rs9829134 | 0.583 | -0.054 | 0.436 | 0.079 | -0.026 | 0.034 | 0.034 | 0.109 | 0.1 | 0.29 | 0.06 | 0.15 | 0.05 | 0.03 |
| 3 | 176507415 | rs9867221 | -0.562 | 0.172 | 0.648 | -0.124 | 0.001 | -0.019 | -0.019 | 0 | 0.34 | 0.63 | 0.26 | 0.15 | 0.32 | 0.22 |
| 3 | 176515103 | rs9843890 | 0.08 | 0.053 | 0.696 | -0.094 | 0.022 | -0.111 | -0.111 | 0 | 0.25 | 0.57 | 0.17 | 0.03 | 0.23 | 0.08 |
| 3 | 176516795 | rs9850321 | 0.08 | 0.053 | 0.696 | -0.094 | 0.022 | -0.111 | -0.111 | 0 | 0.25 | 0.57 | 0.17 | 0.03 | 0.23 | 0.08 |
| 3 | 176524770 | rs2862632 | -1.162 | -0.297 | 0.28 | -0.186 | 0.019 | -0.022 | -0.022 | 0 | 0.91 | 0.98 | 0.94 | 0.91 | 0.92 | 0.89 |
| 3 | 176525654 | rs6762255 | 0.16 | -0.284 | 0.467 | -0.2 | 0.001 | -0.055 | -0.055 | 0 | 0.67 | 0.56 | 0.78 | 0.88 | 0.69 | 0.8 |
| 3 | 176530374 | rs1906658 | 0.539 | -0.293 | 0.392 | -0.182 | -0.004 | -0.033 | -0.033 | 0 | 0.62 | 0.49 | 0.71 | 0.79 | 0.66 | 0.71 |
| 3 | 176537330 | rs13081803 | 1.581 | -0.596 | 0.008 | -0.241 | 0.021 | 0.019 | 0.019 | 3.537 | 0.17 | 0.08 | 0.07 | 0 | 0.08 | 0.03 |
| 3 | 176544103 | rs2862636 | 0.512 | -0.571 | 0.561 | -0.121 | -0.061 | -0.004 | -0.004 | 0 | 0.83 | 0.5 | 0.79 | 0.85 | 0.89 | 0.79 |
| 3 | 176556131 | rs12486909 | 0.157 | -0.411 | 0.451 | -0.376 | 0 | -0.09 | -0.09 | 0.153 | 0.71 | 0.59 | 0.77 | 0.91 | 0.77 | 0.81 |
| 3 | 176556670 | rs9874964 | 1.181 | -0.287 | 0.481 | -0.273 | 0.066 | -0.029 | -0.029 | 0.38 | 0.88 | 0.69 | 0.83 | 0.91 | 0.84 | 0.83 |
| 3 | 176561166 | rs6772982 | 0.722 | -0.21 | -0.19 | -0.049 | 0.02 | -0.065 | -0.065 | 0 | 0.36 | 0.19 | 0.37 | 0.54 | 0.31 | 0.37 |
| 3 | 176563562 | rs7628387 | 0.122 | 0.054 | -0.073 | 0.095 | 0.047 | -0.007 | -0.007 | 0 | 0.13 | 0.35 | 0.13 | 0.11 | 0.17 | 0.17 |
| 3 | 176568103 | rs6803362 | -0.716 | -0.068 | -0.347 | 0.224 | 0.002 | 0.002 | 0.002 | 0 | 0.93 | 0.95 | 0.9 | 0.92 | 0.95 | 0.9 |
| 3 | 176571410 | rs993040 | -0.636 | -0.314 | -0.562 | 0.29 | -0.069 | -0.053 | -0.053 | 0.131 | 0.65 | 0.88 | 0.53 | 0.55 | 0.59 | 0.5 |
| 3 | 176573023 | rs13097506 | 1.508 | -0.057 | -0.476 | 0.449 | -0.051 | -0.054 | -0.054 | 0 | 0.23 | 0.12 | 0.31 | 0.3 | 0.38 | 0.31 |
| 3 | 176573734 | rs2902280 | 0.794 | -0.231 | 0.321 | 0.563 | -0.027 | 0.009 | 0.009 | 0 | 0.09 | 0.31 | 0.08 | 0.07 | 0.05 | 0.1 |
| 3 | 176575760 | rs4626108 | 0.416 | -0.055 | 0.044 | 0.603 | -0.044 | 0.003 | 0.003 | 0 | 0.86 | 0.71 | 0.91 | 0.86 | 0.92 | 0.94 |
| 3 | 176590693 | rs12630639 | 0.376 | 0.554 | -0.747 | 0.754 | -0.119 | -0.055 | -0.055 | 0 | 0.29 | 0.07 | 0.42 | 0.36 | 0.44 | 0.43 |
| 3 | 176592004 | rs4555533 | 0.773 | 0.594 | -0.88 | 0.654 | -0.067 | -0.019 | -0.019 | 0 | 0.25 | 0.01 | 0.22 | 0.22 | 0.27 | 0.32 |
| 3 | 176594897 | rs6794098 | 0.241 | 0.172 | -0.281 | 0.504 | -0.001 | -0.017 | -0.017 | 0 | 0.6 | 0.7 | 0.53 | 0.51 | 0.6 | 0.6 |
| 3 | 176595334 | rs11924000 | 0.724 | -0.167 | -0.089 | 0.163 | 0.024 | -0.125 | -0.125 | 0.483 | 0.21 | 0.39 | 0.13 | 0.03 | 0.17 | 0.11 |
| 3 | 176600522 | rs9647367 | 1.022 | -0.183 | -0.36 | 0.231 | 0.004 | -0.024 | -0.024 | 0.003 | 0.4 | 0.32 | 0.4 | 0.48 | 0.43 | 0.48 |
| 3 | 176614990 | rs9879244 | -0.207 | -0.671 | -0.242 | 0.126 | -0.004 | 0 | 0 | 0 | 0.87 | 0.88 | 0.93 | 0.88 | 0.93 | 0.92 |
| 3 | 176615819 | rs13097413 | 1.102 | -0.569 | -0.207 | 0.183 | -0.009 | -0.022 | -0.022 | 0.092 | 0.4 | 0.25 | 0.34 | 0.45 | 0.45 | 0.48 |
| 3 | 176616271 | rs2130014 | 0.314 | -0.514 | 0.251 | 0.135 | 0.001 | -0.001 | -0.001 | 0 | 0.42 | 0.4 | 0.53 | 0.4 | 0.38 | 0.35 |
| 3 | 176617426 | rs6808937 | -0.25 | -0.481 | 0.19 | 0.141 | 0 | 0 | 0 | 0 | 0.87 | 0.9 | 0.93 | 0.88 | 0.93 | 0.92 |
| 3 | 176621577 | rs6807139 | -0.19 | -0.402 | -0.045 | 0.182 | 0.136 | 0.018 | 0.018 | 0 | 0.06 | 0.21 | 0.06 | 0.03 | 0.1 | 0.09 |

| | | | | | | | | | | | | | | | | |
|---|---|---|---|---|---|---|---|---|---|---|---|---|---|---|---|---|
| 3 | 176621874 | rs7629105 | -0.16 | -0.445 | -0.17 | 0.141 | -0.053 | -0.056 | -0.056 | 0 | 0.72 | 0.62 | 0.87 | 0.88 | 0.88 | 0.91 |
| 3 | 176622316 | rs972131 | 2.395 | -0.652 | -0.798 | 0.135 | -0.122 | -0.107 | -0.107 | 0.135 | 0.23 | 0.02 | 0.25 | 0.31 | 0.39 | 0.22 |
| 3 | 176623079 | rs1387161 | 1.236 | -0.519 | -0.69 | 0.21 | -0.11 | -0.153 | -0.153 | 0.766 | 0.25 | 0.02 | 0.46 | 0.4 | 0.33 | 0.33 |
| 3 | 176627032 | rs1488891 | 0.184 | -0.802 | -0.384 | -0.106 | -0.021 | -0.06 | -0.06 | 0.016 | 0.66 | 0.51 | 0.82 | 0.85 | 0.78 | 0.82 |
| 3 | 176634914 | rs6443406 | -0.052 | -0.95 | -0.776 | 0.302 | -0.006 | -0.005 | -0.005 | 0.308 | 0.31 | 0.62 | 0.14 | 0.24 | 0.19 | 0.28 |
| 3 | 176638068 | rs12635195 | 1.19 | -0.81 | -0.988 | 0.456 | -0.11 | -0.165 | -0.165 | 0 | 0.26 | 0.02 | 0.44 | 0.43 | 0.33 | 0.35 |
| 3 | 176639894 | rs4490374 | -0.359 | -0.263 | -0.552 | 0.586 | -0.011 | -0.035 | -0.035 | 0 | 0.93 | 0.85 | 0.94 | 1 | 0.96 | 0.99 |
| 3 | 176644781 | rs4362729 | 2.12 | -0.373 | -0.663 | 0.547 | -0.106 | -0.061 | -0.061 | 0 | 0.25 | 0.03 | 0.27 | 0.27 | 0.39 | 0.24 |
| 3 | 176647139 | rs4274761 | -0.185 | -0.298 | 0.743 | 0.529 | -0.038 | -0.096 | -0.096 | 0 | 0.79 | 0.69 | 0.88 | 1 | 0.91 | 0.98 |
| 3 | 176647385 | rs6784900 | -0.49 | -0.145 | 0.583 | 0.619 | 0.06 | 0.054 | 0.054 | 0 | 0.14 | 0.41 | 0.08 | 0.17 | 0.13 | 0.15 |
| 3 | 176647549 | rs4857664 | 0.455 | -0.363 | 0.252 | 0.607 | -0.01 | -0.031 | -0.031 | 0 | 0.93 | 0.87 | 0.94 | 1 | 0.96 | 0.99 |
| 3 | 176651806 | rs6783389 | 0.409 | -0.177 | -0.034 | 0.803 | -0.023 | -0.019 | -0.019 | 0.992 | 0.16 | 0.21 | 0.09 | 0.07 | 0.05 | 0.12 |
| 3 | 176659412 | rs7634696 | -0.156 | 0.626 | -0.027 | 1.249 | 0.005 | 0.003 | 0.003 | 0 | 0.3 | 0.31 | 0.32 | 0.3 | 0.34 | 0.41 |
| 3 | 176659521 | rs6443419 | 0.734 | 0.806 | 0.127 | 1.764 | 0.005 | 0.033 | 0.033 | 0 | 0.87 | 0.86 | 0.72 | 0.69 | 0.81 | 0.85 |
| 3 | 176667289 | rs936698 | -0.919 | 0.135 | 0.371 | 1.247 | -0.037 | 0.006 | 0.006 | 0.68 | 0.25 | 0.41 | 0.34 | 0.25 | 0.13 | 0.11 |
| 3 | 176672414 | rs10936921 | 0.166 | 0.038 | 0.308 | 1.128 | 0 | -0.005 | -0.005 | 0 | 0.15 | 0.17 | 0.05 | 0.04 | 0.05 | 0.02 |
| 3 | 176673732 | rs9942133 | 0.763 | 0.462 | 0.531 | 1.88 | 0.021 | 0.19 | 0.19 | 0 | 0.94 | 0.84 | 0.88 | 0.65 | 0.91 | 0.95 |
| 3 | 176675286 | rs10936922 | 1.051 | 0.17 | 0.528 | 1.5 | 0.039 | 0.003 | 0.003 | 0.055 | 0.57 | 0.31 | 0.49 | 0.4 | 0.69 | 0.77 |
| 3 | 176680084 | rs6443423 | 1.492 | 0.734 | 0.261 | 1.375 | -0.023 | 0.088 | 0.088 | 0 | 0.44 | 0.19 | 0.36 | 0.28 | 0.45 | 0.45 |
| 3 | 176683434 | rs7644128 | -0.079 | 1.489 | 0.224 | 1.54 | 0.009 | 0.008 | 0.008 | 0.66 | 0.23 | 0.21 | 0.32 | 0.19 | 0.14 | 0.12 |
| 3 | 176688497 | rs3921860 | -0.531 | 2.322 | 0.386 | 1.813 | 0.001 | 0 | 0 | 1.87 | 0.13 | 0.11 | 0.13 | 0.1 | 0.25 | 0.32 |
| 3 | 176688676 | rs2200251 | 0.647 | 2.286 | 0.499 | 1.844 | 0.038 | 0.248 | 0.248 | 0 | 0.94 | 0.8 | 0.88 | 0.68 | 0.91 | 0.96 |
| 3 | 176691296 | rs6803575 | 0.605 | 2.135 | 0.183 | 1.6 | 0.05 | 0.089 | 0.089 | 0.003 | 0.81 | 0.71 | 0.75 | 0.58 | 0.67 | 0.64 |
| 3 | 176695772 | rs12486786 | -1.316 | 1.975 | 0.481 | 1.087 | 0.088 | 0.031 | 0.031 | 0.04 | 0.09 | 0.27 | 0.15 | 0.1 | 0.15 | 0.43 |
| 3 | 176703201 | rs11916879 | -1.368 | 1.95 | 0.601 | 1.101 | 0.071 | 0.024 | 0.024 | 0.048 | 0.09 | 0.24 | 0.15 | 0.11 | 0.15 | 0.44 |
| 3 | 176703372 | rs11924728 | -2.137 | 1.555 | 1.043 | 0.879 | 0.045 | -0.012 | -0.012 | 0.428 | 0.14 | 0.38 | 0.16 | 0.11 | 0.16 | 0.46 |
| 3 | 176705746 | rs12489936 | -1.248 | 1.517 | 1.341 | 0.862 | 0.003 | 0 | 0 | 0.017 | 0.09 | 0.12 | 0.14 | 0.11 | 0.15 | 0.44 |
| 3 | 176715918 | rs1905499 | -0.641 | 1.854 | 0.508 | 0.892 | 0.003 | 0.024 | 0.024 | 0.537 | 0.31 | 0.44 | 0.44 | 0.56 | 0.33 | 0.1 |
| 3 | 176720780 | rs1905496 | -0.659 | 1.834 | 0.499 | 0.895 | 0.003 | 0.022 | 0.022 | 0.392 | 0.31 | 0.44 | 0.44 | 0.57 | 0.33 | 0.1 |
| 3 | 176728407 | rs1344201 | -0.179 | 2.009 | 0.641 | 1.13 | 0.017 | 0.011 | 0.011 | 0 | 0.09 | 0.21 | 0.19 | 0.13 | 0.11 | 0.02 |
| 3 | 176738798 | rs6983 | -0.629 | 1.961 | 0.859 | 1.121 | -0.001 | -0.041 | -0.041 | 0 | 0.22 | 0.15 | 0.23 | 0.46 | 0.2 | 0.08 |
| 3 | 176739663 | rs1130272 | -1.598 | 1.392 | 0.719 | 0.357 | -0.005 | -0.005 | -0.005 | 0 | 0.42 | 0.5 | 0.51 | 0.57 | 0.4 | 0.21 |
| 3 | 176750426 | rs9290610 | -2.44 | 0.768 | 0.886 | 0.007 | -0.091 | 0.098 | 0.098 | 0 | 0.56 | 0.96 | 0.69 | 0.72 | 0.57 | 0.65 |
| 3 | 176757832 | rs2129017 | 0.007 | 0.782 | 0.966 | 0.025 | -0.002 | 0.067 | 0.067 | 0.011 | 0.22 | 0.3 | 0.24 | 0.44 | 0.2 | 0.08 |
| 3 | 176758477 | rs1905501 | -0.899 | 0.76 | 0.977 | 0.012 | -0.011 | 0.027 | 0.027 | 0 | 0.34 | 0.42 | 0.32 | 0.45 | 0.31 | 0.19 |
| 3 | 176761002 | rs6795240 | 0.418 | 0.678 | 0.919 | -0.025 | 0 | -0.019 | -0.019 | 0.395 | 0.13 | 0.12 | 0.09 | 0 | 0.1 | 0.12 |
| 3 | 176765269 | rs3749234 | 1.103 | 0.603 | 0.868 | -0.053 | -0.053 | -0.116 | -0.116 | 0 | 0.16 | 0.02 | 0.22 | 0.44 | 0.2 | 0.06 |
| 3 | 176770400 | rs2862644 | 0.553 | -0.002 | 0.536 | -0.469 | -0.003 | 0.012 | 0.012 | 0 | 0.09 | 0.07 | 0.1 | 0.05 | 0.08 | 0.2 |
| 3 | 176785767 | rs10936928 | 0.157 | -0.018 | 0.554 | -0.46 | -0.003 | 0.096 | 0.096 | 0.006 | 0.23 | 0.36 | 0.24 | 0.44 | 0.2 | 0.08 |
| 3 | 176787698 | rs10936930 | -1.083 | -0.019 | 0.524 | -0.476 | -0.006 | -0.007 | -0.007 | 0 | 0.41 | 0.48 | 0.51 | 0.57 | 0.4 | 0.21 |
| 3 | 176792166 | rs1681655 | -2.694 | -0.567 | -0.604 | -1.314 | 0.012 | 0.019 | 0.019 | 1.455 | 0.82 | 0.98 | 0.93 | 0.9 | 0.91 | 0.97 |
| 3 | 176799074 | rs6775004 | 0.888 | -0.497 | -0.493 | -1.248 | -0.001 | -0.028 | -0.028 | 0 | 0.22 | 0.22 | 0.23 | 0.44 | 0.2 | 0.08 |
| 3 | 176800388 | rs1201288 | 0.407 | -0.513 | -0.51 | -1.291 | -0.006 | 0.01 | 0.01 | 0 | 0.14 | 0.11 | 0.14 | 0.1 | 0.16 | 0.44 |
| 3 | 176802500 | rs11915541 | 1.486 | -0.572 | -0.457 | -1.264 | -0.011 | -0.054 | -0.054 | 0.097 | 0.12 | 0.15 | 0.09 | 0 | 0.1 | 0.12 |
| 3 | 176803648 | rs9858226 | 0.108 | -0.556 | -0.548 | -1.211 | -0.001 | -0.006 | -0.006 | 0 | 0.29 | 0.31 | 0.41 | 0.56 | 0.3 | 0.09 |
| 3 | 176816531 | rs2046380 | 1.828 | -0.358 | -0.564 | -0.995 | -0.004 | -0.008 | -0.008 | 0 | 0.07 | 0.04 | 0.19 | 0.13 | 0.08 | 0.02 |
| 3 | 176818583 | rs13315476 | 0.941 | -0.412 | -0.287 | -1.025 | 0.003 | 0.015 | 0.015 | 0 | 0.1 | 0.13 | 0.21 | 0.18 | 0.09 | 0.02 |
| 3 | 176822676 | rs1201289 | -1.22 | -0.603 | -0.263 | -1.245 | -0.012 | -0.031 | -0.031 | 0 | 0.41 | 0.55 | 0.45 | 0.29 | 0.37 | 0.62 |
| 3 | 176824755 | rs1681657 | -1.667 | -0.479 | -0.156 | -0.799 | -0.006 | 0.016 | 0.016 | 0 | 0.58 | 0.64 | 0.67 | 0.73 | 0.56 | 0.69 |
| 3 | 176827927 | rs1209035 | 1.333 | -0.396 | -0.195 | -0.785 | 0 | -0.001 | -0.001 | 0 | 0.06 | 0.08 | 0.05 | 0.06 | 0.1 | 0.25 |
| 3 | 176830302 | rs1201296 | -1.15 | -0.592 | -0.267 | -0.911 | -0.001 | 0.01 | 0.01 | 0 | 0.49 | 0.57 | 0.63 | 0.72 | 0.49 | 0.59 |
| 3 | 176835597 | rs6785871 | -0.253 | -0.599 | -0.501 | -0.872 | -0.011 | -0.046 | -0.046 | 0 | 0.69 | 0.79 | 0.69 | 0.51 | 0.67 | 0.62 |
| 3 | 176839656 | rs7638794 | 1.874 | -0.65 | -0.113 | -0.905 | -0.093 | 0.067 | 0.067 | 0 | 0.4 | 0.1 | 0.31 | 0.27 | 0.44 | 0.32 |
| 3 | 176859444 | rs7426456 | -0.192 | -0.609 | -0.132 | -0.828 | -0.011 | -0.04 | -0.04 | 0 | 0.69 | 0.78 | 0.69 | 0.49 | 0.66 | 0.61 |
| 3 | 176861144 | rs1881977 | -0.592 | -0.55 | -0.172 | -0.844 | -0.004 | -0.016 | -0.016 | 0.016 | 0.86 | 0.84 | 0.9 | 0.94 | 0.84 | 0.65 |
| 3 | 176863563 | rs2902287 | 2.347 | -0.265 | -0.346 | -0.993 | -0.059 | 0.056 | 0.056 | 0.657 | 0.19 | 0.03 | 0.14 | 0.08 | 0.17 | 0.05 |
| 3 | 176865688 | rs12489569 | 1.688 | -0.368 | 0.244 | -0.842 | 0.005 | 0.076 | 0.076 | 0 | 0.43 | 0.26 | 0.39 | 0.24 | 0.53 | 0.33 |
| 3 | 176867736 | rs6774257 | 0.456 | -0.523 | 0.084 | -0.731 | 0.016 | 0.02 | 0.02 | 1.083 | 0.77 | 0.84 | 0.87 | 0.93 | 0.81 | 0.59 |
| 3 | 176868053 | rs7632766 | 0.1 | -0.292 | -0.024 | -0.411 | -0.007 | -0.08 | -0.08 | 0.25 | 0.26 | 0.14 | 0.26 | 0.47 | 0.24 | 0.13 |
| 3 | 176868116 | rs7621281 | -0.204 | -0.078 | 0.327 | -0.379 | 0.013 | 0.026 | 0.026 | 0.025 | 0.28 | 0.47 | 0.44 | 0.66 | 0.29 | 0.08 |
| 3 | 176868978 | rs6778192 | 0.288 | -0.176 | -0.224 | 0.073 | 0.017 | 0.027 | 0.027 | 0 | 0.5 | 0.39 | 0.42 | 0.28 | 0.43 | 0.53 |
| 3 | 176869498 | rs6443429 | -0.224 | -0.691 | 0.399 | 0.014 | -0.025 | -0.05 | -0.05 | 0.057 | 0.64 | 0.56 | 0.6 | 0.73 | 0.7 | 0.54 |
| 3 | 176879005 | rs1920133 | 2.57 | -0.459 | 0.206 | 0.645 | 0.008 | 0.091 | 0.091 | 4.912 | 0.05 | 0.03 | 0.09 | 0.47 | 0.02 | 0.04 |
| 3 | 176887272 | rs6786351 | -0.16 | -0.428 | 0.213 | 0.548 | 0.037 | 0.09 | 0.09 | 0 | 0.55 | 0.66 | 0.54 | 0.72 | 0.61 | 0.44 |
| 3 | 176910577 | rs12490455 | 0.123 | 0.293 | -0.356 | 0.071 | 0.021 | 0.033 | 0.033 | 0 | 0.48 | 0.32 | 0.42 | 0.25 | 0.59 | 0.4 |
| 3 | 176925740 | rs13077541 | 0.158 | 0.317 | -1.054 | 0.24 | -0.007 | 0.072 | 0.072 | 0 | 0.43 | 0.21 | 0.39 | 0.25 | 0.54 | 0.37 |
| 3 | 176929742 | rs7611527 | -0.285 | -0.332 | -0.451 | 0.4 | 0.019 | 0.012 | 0.012 | 6.605 | 0.52 | 0.56 | 0.64 | 0.65 | 0.68 | 0.59 |
| 3 | 176933598 | rs4857827 | 0.249 | -0.225 | 0.316 | 0.803 | -0.013 | 0.2 | 0.2 | 0 | 0.06 | 0.19 | 0.08 | 0.24 | 0.03 | 0.14 |
| 3 | 176940408 | rs3923106 | -0.542 | 0.054 | 0.625 | 0.922 | -0.001 | 0.03 | 0.03 | 0 | 0.47 | 0.61 | 0.54 | 0.66 | 0.5 | 0.58 |

| | | | | | | | | | | | | | | | | |
|---|---|---|---|---|---|---|---|---|---|---|---|---|---|---|---|---|
| 3 | 176944249 | rs9821966 | 0.537 | 0.118 | 0.224 | 0.956 | -0.013 | 0.196 | 0.196 | 0 | 0.06 | 0.19 | 0.08 | 0.24 | 0.03 | 0.14 |
| 3 | 176948553 | rs4857674 | -0.142 | 0.107 | 0.288 | 0.935 | 0 | -0.012 | -0.012 | 0 | 0.46 | 0.49 | 0.53 | 0.66 | 0.5 | 0.55 |
| 3 | 176952068 | rs12486925 | 0.114 | 0.391 | -0.564 | 0.643 | -0.025 | -0.01 | -0.01 | 0 | 0.39 | 0.28 | 0.45 | 0.42 | 0.47 | 0.41 |
| 3 | 176954641 | rs9290616 | 0.201 | 0.429 | -0.842 | 0.66 | -0.048 | -0.027 | -0.027 | 0 | 0.38 | 0.18 | 0.44 | 0.42 | 0.47 | 0.41 |
| 3 | 176980658 | rs9864317 | 0.499 | 0.241 | -0.54 | 0.154 | 0.001 | -0.006 | -0.006 | 0.072 | 0.4 | 0.4 | 0.59 | 0.64 | 0.51 | 0.37 |
| 3 | 176981307 | rs6775864 | 0.364 | 0.211 | -0.553 | 0.113 | -0.001 | -0.006 | -0.006 | 0.068 | 0.43 | 0.43 | 0.61 | 0.64 | 0.56 | 0.38 |
| 3 | 176990782 | rs6443443 | -0.203 | -0.049 | -0.76 | 0.156 | 0.053 | 0.068 | 0.068 | 0 | 0.54 | 0.76 | 0.67 | 0.63 | 0.61 | 0.53 |
| 3 | 177038879 | rs6788853 | 0.625 | -0.839 | -0.548 | -0.347 | -0.008 | 0.101 | 0.101 | 0 | 0.57 | 0.26 | 0.67 | 0.83 | 0.59 | 0.47 |
| 3 | 177057139 | rs7431825 | 0.325 | -0.88 | -0.96 | -0.406 | -0.025 | 0.129 | 0.129 | 0 | 0.61 | 0.82 | 0.71 | 0.85 | 0.63 | 0.44 |
| 3 | 177063272 | rs6773879 | 0.096 | -0.684 | -1.407 | -0.276 | -0.082 | 0.175 | 0.175 | 0 | 0.61 | 0.91 | 0.73 | 0.85 | 0.63 | 0.42 |
| 3 | 177064814 | rs9871240 | 0.324 | -0.701 | -1.388 | -0.267 | -0.088 | 0.219 | 0.219 | 0 | 0.57 | 0.91 | 0.71 | 0.85 | 0.58 | 0.41 |
| 3 | 177066783 | rs9836057 | -0.091 | -0.614 | -1.109 | -0.178 | -0.011 | 0.126 | 0.126 | 0 | 0.66 | 0.83 | 0.77 | 0.92 | 0.68 | 0.67 |
| 3 | 177067339 | rs9840651 | -0.129 | -0.786 | -1.022 | 0.247 | -0.007 | 0 | 0 | 0 | 0.21 | 0.24 | 0.13 | 0.2 | 0.1 | 0.11 |
| 3 | 177074926 | rs6443448 | 0.002 | -0.179 | -1.747 | 0.519 | -0.071 | 0.013 | 0.013 | 0.286 | 0.66 | 0.89 | 0.51 | 0.68 | 0.6 | 0.58 |
| 3 | 177082829 | rs9873211 | -0.379 | -0.005 | -1.458 | 0.489 | -0.051 | -0.021 | -0.021 | 0 | 0.69 | 0.88 | 0.52 | 0.68 | 0.64 | 0.65 |
| 3 | 177085723 | rs9861140 | 0.412 | 0.491 | -1.363 | 0.472 | 0 | 0 | 0 | 0.056 | 0.42 | 0.5 | 0.35 | 0.43 | 0.49 | 0.35 |
| 3 | 177092297 | rs7432880 | -0.153 | 1.615 | -1.607 | 1.324 | -0.101 | -0.081 | -0.081 | 0.112 | 0.76 | 0.96 | 0.72 | 0.69 | 0.68 | 0.7 |
| 3 | 177095072 | rs12629106 | -0.91 | 1.058 | -0.362 | 1.248 | 0.002 | 0.002 | 0.002 | 0 | 0.14 | 0.13 | 0.1 | 0.17 | 0.16 | 0.07 |
| 3 | 177113056 | rs9850957 | -1.322 | 0.897 | -0.175 | 1.502 | 0.001 | 0.052 | 0.052 | 0.392 | 0.24 | 0.63 | 0.19 | 0.47 | 0.23 | 0.35 |
| 3 | 177113635 | rs9813987 | -1.473 | 0.592 | 0.424 | 1.821 | 0.023 | 0.246 | 0.246 | 0.632 | 0.13 | 0.51 | 0.11 | 0.35 | 0.09 | 0.28 |
| 3 | 177114488 | rs7374825 | -0.147 | 0.465 | -1.026 | 1.286 | -0.067 | 0.014 | 0.014 | 0 | 0.62 | 0.92 | 0.63 | 0.75 | 0.57 | 0.7 |
| 3 | 177119143 | rs6804202 | -0.543 | 0.523 | -0.732 | 1.1 | 0.003 | 0.014 | 0.014 | 0 | 0.7 | 0.69 | 0.62 | 0.48 | 0.66 | 0.78 |
| 3 | 177120303 | rs7639599 | -0.527 | 0.142 | -0.768 | 0.734 | -0.008 | 0.01 | 0.01 | 0 | 0.81 | 0.9 | 0.85 | 0.82 | 0.8 | 0.81 |
| 3 | 177125620 | rs6766374 | 0.089 | 0.771 | -0.614 | 0.757 | 0.01 | 0.114 | 0.114 | 0 | 0.15 | 0.23 | 0.22 | 0.51 | 0.15 | 0.23 |
| 3 | 177127074 | rs9968207 | 0.302 | 0.93 | -0.17 | 0.197 | 0.049 | -0.068 | -0.068 | 0.35 | 0.12 | 0.39 | 0.09 | 0 | 0.13 | 0.03 |
| 3 | 177127453 | rs9968208 | -0.624 | 1.013 | -0.064 | 0.049 | 0.02 | -0.138 | -0.138 | 0.089 | 0.22 | 0.55 | 0.2 | 0 | 0.28 | 0.07 |
| 3 | 177133282 | rs869122 | 2.139 | 0.993 | -0.617 | -0.095 | -0.08 | -0.067 | -0.067 | 0 | 0.3 | 0.02 | 0.37 | 0.28 | 0.29 | 0.29 |
| 3 | 177138752 | rs2047564 | 0.754 | 1.492 | -0.466 | -0.127 | -0.038 | 0.048 | 0.048 | 0 | 0.11 | 0.03 | 0.06 | 0 | 0.17 | 0.03 |
| 3 | 177142736 | rs7645227 | 0.28 | 1.808 | 0.375 | 0.1 | -0.007 | -0.031 | -0.031 | 0 | 0.62 | 0.51 | 0.71 | 0.76 | 0.67 | 0.43 |
| 3 | 177150899 | rs4857688 | 0.128 | 1.925 | 0.579 | 0.227 | 0.02 | 0.224 | 0.224 | 0 | 0.12 | 0.31 | 0.3 | 0.5 | 0.19 | 0.12 |
| 3 | 177152632 | rs1995243 | 1.044 | 1.692 | 0.339 | -0.019 | 0.002 | 0.055 | 0.055 | 0 | 0.48 | 0.6 | 0.69 | 0.81 | 0.49 | 0.44 |
| 3 | 177155921 | rs9825638 | 0.452 | 1.5 | 0.4 | 0.052 | 0.009 | 0.05 | 0.05 | 0 | 0.6 | 0.71 | 0.76 | 0.81 | 0.65 | 0.48 |
| 3 | 177157160 | rs6783019 | -0.471 | 1.467 | 0.035 | 0.009 | -0.035 | 0.093 | 0.093 | 0.69 | 0.74 | 0.96 | 0.85 | 0.88 | 0.75 | 0.84 |
| 3 | 177164062 | rs11922932 | -0.42 | 1.463 | 0.056 | 0.007 | -0.01 | 0.08 | 0.08 | 0.687 | 0.74 | 0.92 | 0.84 | 0.89 | 0.75 | 0.84 |
| 3 | 177165928 | rs13314733 | -0.3 | 1.333 | 0.311 | 0.149 | -0.011 | -0.013 | -0.013 | 0 | 0.14 | 0.25 | 0.09 | 0.07 | 0.1 | 0.36 |
| 3 | 177166139 | rs4353846 | 1.172 | 1.4 | -0.1 | 0.214 | -0.009 | 0.025 | 0.025 | 0 | 0.42 | 0.32 | 0.5 | 0.35 | 0.46 | 0.4 |
| 3 | 177167376 | rs1875107 | 0.071 | 1.372 | -1.829 | 0.095 | -0.004 | 0.138 | 0.138 | 0.036 | 0.66 | 0.92 | 0.83 | 0.87 | 0.69 | 0.8 |
| 3 | 177171101 | rs4539982 | 0.778 | 1.205 | -1.664 | 0.447 | -0.024 | -0.03 | -0.03 | 0.215 | 0.27 | 0.2 | 0.52 | 0.31 | 0.35 | 0.22 |
| 3 | 177174689 | rs10936943 | -0.125 | 1.099 | -1.201 | 0.429 | 0.012 | 0.031 | 0.031 | 0.315 | 0.34 | 0.45 | 0.42 | 0.46 | 0.35 | 0.49 |
| 3 | 177175323 | rs9758492 | -1.163 | 0.59 | -1.96 | 0.241 | -0.01 | -0.005 | -0.005 | 0.82 | 0.7 | 0.94 | 0.72 | 0.75 | 0.73 | 0.78 |
| 3 | 177176516 | rs6799682 | 0.794 | 0.523 | -0.957 | 0.143 | -0.018 | -0.02 | -0.02 | 1.538 | 0.25 | 0.44 | 0.19 | 0.28 | 0.21 | 0.23 |
| 3 | 177177991 | rs7632844 | 0.666 | 0.537 | -0.945 | 0.149 | -0.017 | -0.019 | -0.019 | 0 | 0.25 | 0.42 | 0.19 | 0.28 | 0.21 | 0.23 |
| 3 | 177183874 | rs4857692 | 0.85 | 0.879 | -1.306 | -0.027 | -0.026 | 0.057 | 0.057 | 0 | 0.12 | 0.04 | 0.1 | 0 | 0.15 | 0.06 |
| 3 | 177189575 | rs9880546 | -0.238 | 0.709 | -1.152 | 0.306 | -0.001 | 0.006 | 0.006 | 0 | 0.24 | 0.3 | 0.26 | 0.39 | 0.24 | 0.33 |
| 3 | 177191797 | rs9853442 | -1.486 | 0.493 | -1.497 | 0.048 | -0.019 | -0.058 | -0.058 | 1.656 | 0.62 | 0.87 | 0.57 | 0.6 | 0.62 | 0.63 |
| 3 | 177198248 | rs17631673 | 1.576 | 0.606 | -1.339 | -0.141 | -0.057 | 0.073 | 0.073 | 0 | 0.13 | 0.03 | 0.11 | 0 | 0.18 | 0.07 |
| 3 | 177199626 | rs9830263 | -0.416 | 0.178 | -0.74 | -0.008 | -0.006 | 0.009 | 0.009 | 0 | 0.33 | 0.43 | 0.32 | 0.44 | 0.32 | 0.42 |
| 3 | 177206690 | rs4857693 | -0.465 | 0.127 | -0.191 | -0.061 | -0.006 | 0.017 | 0.017 | 0.096 | 0.3 | 0.61 | 0.32 | 0.44 | 0.28 | 0.36 |
| 3 | 177211328 | rs6794095 | -0.817 | 0.472 | -0.171 | -0.358 | -0.027 | 0.076 | 0.076 | 0.006 | 0.47 | 0.79 | 0.38 | 0.38 | 0.47 | 0.46 |
| 3 | 177212302 | rs9881791 | -0.621 | 0.633 | 0.405 | -0.039 | 0.023 | 0.035 | 0.035 | 0.219 | 0.14 | 0.4 | 0.11 | 0.23 | 0.14 | 0.2 |
| 3 | 177214656 | rs13091554 | 0.624 | 0.688 | -0.084 | -0.06 | 0.004 | 0.011 | 0.011 | 0 | 0.09 | 0.2 | 0.08 | 0.19 | 0.09 | 0.17 |
| 3 | 177224309 | rs10804867 | 0.085 | 0.844 | -0.428 | 0.121 | -0.032 | -0.053 | -0.053 | 0 | 0.16 | 0.12 | 0.27 | 0.53 | 0.25 | 0.26 |
| 3 | 177226791 | rs7637681 | 0.963 | 0.349 | -0.587 | -0.097 | 0.018 | -0.036 | -0.036 | 0.005 | 0.14 | 0.03 | 0.07 | 0.23 | 0.1 | 0.22 |
| 3 | 177228979 | rs1353899 | 0.78 | 0.965 | -0.395 | -0.427 | 0.012 | 0.018 | 0.018 | 0 | 0.12 | 0.11 | 0.3 | 0.23 | 0.19 | 0.21 |
| 3 | 177229309 | rs9878375 | -0.857 | 0.387 | -0.42 | -0.939 | 0.062 | 0.095 | 0.095 | 0 | 0.6 | 0.75 | 0.78 | 0.76 | 0.71 | 0.62 |
| 3 | 177231614 | rs4857702 | 0.49 | 0.604 | 0.119 | -0.708 | 0.027 | 0.084 | 0.084 | 0 | 0.19 | 0.3 | 0.29 | 0.52 | 0.25 | 0.26 |
| 3 | 177243737 | rs1499880 | -0.123 | 0.303 | -0.141 | -0.746 | -0.003 | -0.002 | -0.002 | 0.427 | 0.48 | 0.57 | 0.38 | 0.49 | 0.49 | 0.44 |
| 3 | 177248741 | rs1499881 | -1.281 | -0.882 | -0.683 | -0.868 | 0.014 | -0.004 | -0.004 | 0.156 | 0.73 | 0.75 | 0.85 | 0.99 | 0.81 | 0.83 |
| 3 | 177254210 | rs2863009 | 0.023 | -0.797 | -1.016 | -0.814 | 0.06 | 0.081 | 0.081 | 0 | 0.5 | 0.35 | 0.7 | 0.76 | 0.66 | 0.52 |
| 3 | 177262543 | rs13070316 | 0.569 | 0.017 | -0.47 | 0.267 | 0.024 | -0.024 | -0.024 | 0.236 | 0.15 | 0.07 | 0.07 | 0.26 | 0.11 | 0.2 |
| 3 | 177264970 | rs3851413 | -0.117 | 0.345 | -0.047 | 0.219 | 0.023 | 0.339 | 0.339 | 0 | 0.1 | 0.3 | 0.12 | 0.4 | 0.08 | 0.17 |
| 3 | 177266750 | rs10470510 | -0.212 | 0.682 | -0.513 | 0.108 | -0.002 | 0.001 | 0.001 | 0 | 0.47 | 0.47 | 0.62 | 0.58 | 0.49 | 0.58 |
| 3 | 177282809 | rs17632548 | -0.786 | 0.489 | -0.44 | -0.052 | -0.005 | -0.026 | -0.026 | 0 | 0.1 | 0.09 | 0.1 | 0.02 | 0.12 | 0.07 |
| 3 | 177286946 | rs7627379 | -0.854 | 0.72 | 0.169 | -0.103 | 0.002 | 0.021 | 0.021 | 0.015 | 0.34 | 0.57 | 0.49 | 0.54 | 0.38 | 0.44 |
| 3 | 177287843 | rs11707350 | 0.768 | 0.589 | 0.026 | -0.165 | -0.088 | -0.153 | -0.153 | 0 | 0.19 | 0.01 | 0.35 | 0.52 | 0.26 | 0.35 |
| 3 | 177292543 | rs493795 | -0.566 | 0.751 | -0.411 | -0.801 | -0.011 | 0.019 | 0.019 | 0.781 | 0.8 | 0.88 | 0.82 | 0.94 | 0.75 | 0.75 |
| 3 | 177295215 | rs552305 | 1.292 | 0.652 | -0.326 | -0.878 | 0.008 | -0.02 | -0.02 | 0 | 0.46 | 0.13 | 0.33 | 0.4 | 0.4 | 0.34 |
| 3 | 177297824 | rs6801849 | -0.402 | 0.75 | 0.231 | -0.634 | 0.003 | 0.043 | 0.043 | 0.056 | 0.33 | 0.55 | 0.49 | 0.54 | 0.38 | 0.43 |
| 3 | 177303610 | rs2133595 | -0.405 | 0.72 | 0.217 | -0.647 | 0.002 | 0.039 | 0.039 | 0.238 | 0.33 | 0.55 | 0.49 | 0.54 | 0.38 | 0.43 |

| | | | | | | | | | | | | | | | | |
|---|---|---|---|---|---|---|---|---|---|---|---|---|---|---|---|---|
| 3 | 177306883 | rs11923361 | 0.179 | 0.747 | 0.094 | -0.64 | -0.007 | -0.021 | -0.021 | 0 | 0.1 | 0.09 | 0.1 | 0.02 | 0.1 | 0.07 |
| 3 | 177308158 | rs6769488 | 0.673 | 0.879 | 0.733 | -0.448 | 0.002 | 0.018 | 0.018 | 0 | 0.22 | 0.25 | 0.37 | 0.53 | 0.28 | 0.36 |
| 3 | 177312876 | rs4857716 | -0.963 | 0.548 | 0.536 | -0.585 | 0.017 | 0.029 | 0.029 | 0.774 | 0.51 | 0.63 | 0.64 | 0.67 | 0.59 | 0.65 |
| 3 | 177316085 | rs4241484 | -0.965 | 0.545 | 0.58 | -0.566 | 0.017 | 0.04 | 0.04 | 0.805 | 0.51 | 0.63 | 0.65 | 0.73 | 0.59 | 0.66 |
| 3 | 177328829 | rs6804861 | 0.671 | 0.723 | 0.171 | -0.486 | -0.004 | -0.005 | -0.005 | 0.456 | 0.26 | 0.25 | 0.39 | 0.56 | 0.31 | 0.42 |
| 3 | 177335573 | rs17633480 | -0.461 | 0.743 | 0.019 | -0.541 | -0.002 | 0.006 | 0.006 | 0 | 0.24 | 0.18 | 0.24 | 0.12 | 0.29 | 0.24 |
| 3 | 177336402 | rs13083260 | 0.466 | 0.848 | -0.136 | -0.279 | -0.026 | -0.053 | -0.053 | 0 | 0.26 | 0.14 | 0.41 | 0.58 | 0.31 | 0.44 |
| 3 | 177338280 | rs2242401 | 0.445 | 0.834 | -0.144 | -0.274 | -0.021 | -0.046 | -0.046 | 1.022 | 0.26 | 0.16 | 0.41 | 0.58 | 0.31 | 0.43 |
| 3 | 177346889 | rs17633881 | 0.376 | -1.3 | -0.07 | -1.682 | 0.035 | 0.031 | 0.031 | 3.029 | 0.2 | 0.13 | 0.09 | 0 | 0.11 | 0.05 |
| 3 | 177347155 | rs2133600 | 0.325 | -1.352 | -0.375 | -1.68 | 0.008 | 0 | 0 | 0 | 0.68 | 0.72 | 0.73 | 0.75 | 0.74 | 0.84 |
| 3 | 177351280 | rs6443478 | -0.591 | -1.301 | -0.502 | -1.659 | -0.033 | -0.067 | -0.067 | 0.724 | 0.89 | 0.99 | 0.84 | 0.75 | 0.86 | 0.89 |
| 3 | 177351845 | rs937508 | 0.071 | -1.463 | 0.073 | -1.998 | -0.001 | -0.063 | -0.063 | 0.444 | 0.16 | 0.29 | 0.12 | 0.05 | 0.12 | 0.18 |
| 3 | 177359599 | rs10513752 | 0.687 | -1.38 | -0.278 | -1.843 | 0.015 | -0.008 | -0.008 | 0 | 0.2 | 0.16 | 0.1 | 0 | 0.11 | 0.04 |
| 3 | 177363666 | rs13074870 | 0.694 | -1.422 | -0.317 | -2.177 | 0.006 | 0.016 | 0.016 | 2.31 | 0.46 | 0.36 | 0.57 | 0.85 | 0.55 | 0.75 |
| 3 | 177374031 | rs6789946 | 0.11 | -1.076 | -0.194 | -1.927 | -0.005 | -0.058 | -0.058 | 0 | 0.66 | 0.51 | 0.66 | 0.85 | 0.67 | 0.79 |
| 3 | 177374604 | rs16828425 | 0.541 | -0.529 | 0.012 | -1.654 | -0.001 | 0.011 | 0.011 | 0 | 0.08 | 0.08 | 0.11 | 0.16 | 0.11 | 0.14 |
| 3 | 177389109 | rs12487312 | 0.396 | -0.604 | 0.072 | -1.919 | -0.003 | -0.001 | -0.001 | 0 | 0.13 | 0.12 | 0.18 | 0.19 | 0.19 | 0.14 |
| 3 | 177390314 | rs10513750 | -0.616 | -1.02 | 0.38 | -2.292 | 0.002 | -0.018 | -0.018 | 0 | 0.41 | 0.53 | 0.38 | 0.24 | 0.41 | 0.23 |
| 3 | 177402994 | rs6793629 | -0.784 | -1.15 | 0.338 | -2.356 | 0.004 | 0.056 | 0.056 | 1.004 | 0.46 | 0.75 | 0.4 | 0.24 | 0.42 | 0.23 |
| 3 | 177408279 | rs6768558 | 0.436 | -1.143 | 0.609 | -2.454 | -0.002 | -0.085 | -0.085 | 0 | 0.25 | 0.34 | 0.19 | 0.04 | 0.21 | 0.08 |
| 3 | 177412574 | rs10513748 | -1.585 | -1.375 | 0.359 | -2.602 | -0.051 | 0.059 | 0.059 | 9.247 | 0.6 | 0.87 | 0.45 | 0.21 | 0.5 | 0.29 |
| 3 | 177413613 | rs16828497 | -0.412 | -0.609 | 0.707 | -2.049 | -0.01 | -0.017 | -0.017 | 0.056 | 0.07 | 0.31 | 0.03 | 0 | 0.01 | 0.01 |
| 3 | 177421455 | rs17635597 | 0.461 | -0.167 | -0.118 | -1.845 | -0.003 | -0.006 | -0.006 | 0 | 0.15 | 0.11 | 0.19 | 0.18 | 0.19 | 0.16 |
| 3 | 177431683 | rs4857725 | -1.059 | -0.16 | -0.27 | -1.859 | -0.004 | 0.076 | 0.076 | 0 | 0.49 | 0.73 | 0.43 | 0.21 | 0.51 | 0.28 |
| 3 | 177433299 | rs6782388 | -0.175 | -0.16 | -0.08 | -1.897 | 0.001 | -0.116 | -0.116 | 0 | 0.29 | 0.41 | 0.2 | 0.03 | 0.26 | 0.08 |
| 3 | 177433541 | rs6772134 | -0.725 | -0.487 | -0.613 | -1.655 | 0.002 | 0.067 | 0.067 | 0.027 | 0.48 | 0.73 | 0.42 | 0.21 | 0.46 | 0.28 |
| 3 | 177435527 | rs1499806 | -0.378 | -0.453 | -0.386 | -1.849 | 0.003 | -0.085 | -0.085 | 0 | 0.2 | 0.28 | 0.14 | 0.04 | 0.18 | 0.1 |
| 3 | 177435752 | rs1499807 | -0.105 | -0.343 | -0.435 | -1.703 | -0.001 | -0.048 | -0.048 | 0.918 | 0.22 | 0.21 | 0.14 | 0.03 | 0.2 | 0.13 |
| 3 | 177443968 | rs6788878 | -0.192 | -0.331 | -0.485 | -1.732 | 0.005 | -0.003 | -0.003 | 0 | 0.29 | 0.21 | 0.23 | 0.04 | 0.27 | 0.16 |
| 3 | 177444417 | rs7634440 | -0.312 | -0.192 | -0.327 | -1.783 | -0.005 | -0.125 | -0.125 | 0.14 | 0.33 | 0.41 | 0.29 | 0.04 | 0.33 | 0.17 |
| 3 | 177452007 | rs4857620 | 0.043 | -1.05 | 0.028 | -1.995 | -0.048 | -0.065 | -0.065 | 0 | 0.87 | 0.72 | 0.81 | 0.98 | 0.92 | 0.84 |
| 3 | 177459881 | rs9824206 | 0.724 | -0.921 | -0.439 | -2.345 | -0.039 | 0.052 | 0.052 | 0.919 | 0.35 | 0.2 | 0.3 | 0.83 | 0.39 | 0.63 |
| 3 | 177467569 | rs4857735 | 0.896 | -0.966 | -0.33 | -2.636 | 0.007 | -0.01 | -0.01 | 0.491 | 0.17 | 0.25 | 0.25 | 0.08 | 0.24 | 0.07 |
| 3 | 177469219 | rs9824506 | 0.955 | -0.787 | -0.703 | -2.502 | -0.066 | 0.064 | 0.064 | 0.65 | 0.34 | 0.12 | 0.28 | 0.83 | 0.38 | 0.6 |
| 3 | 177470593 | rs6443496 | -0.57 | -0.632 | -0.805 | -2.389 | -0.032 | -0.046 | -0.046 | 0.105 | 0.87 | 0.78 | 0.81 | 0.98 | 0.92 | 0.84 |
| 3 | 177474472 | rs7634430 | -0.352 | -0.637 | -0.952 | -2.405 | -0.008 | -0.029 | -0.029 | 0.103 | 0.86 | 0.83 | 0.8 | 0.98 | 0.89 | 0.83 |
| 3 | 177477967 | rs7641352 | 0.992 | -0.578 | -1.498 | -2.476 | 0 | 0.019 | 0.019 | 0 | 0.22 | 0.14 | 0.2 | 0.07 | 0.19 | 0.12 |
| 3 | 177490268 | rs1106387 | 1.179 | -0.284 | -1.759 | -2.232 | -0.099 | -0.053 | -0.053 | 0 | 0.21 | 0.04 | 0.2 | 0.75 | 0.29 | 0.53 |
| 3 | 177490311 | rs1106388 | 1.179 | -0.241 | -1.759 | -2.232 | -0.097 | -0.053 | -0.053 | 0 | 0.21 | 0.04 | 0.2 | 0.75 | 0.29 | 0.53 |
| 3 | 177494187 | rs9825912 | 0.65 | -0.622 | -1.31 | -2.287 | 0.007 | 0.009 | 0.009 | 0 | 0.1 | 0.09 | 0.09 | 0.03 | 0.09 | 0.06 |
| 3 | 177495631 | rs1369569 | -0.205 | -0.377 | -1.16 | -2.121 | 0.003 | -0.006 | -0.006 | 0.06 | 0.66 | 0.72 | 0.7 | 0.93 | 0.68 | 0.83 |
| 3 | 177502714 | rs4857631 | -0.312 | -0.416 | -1.494 | -1.169 | -0.03 | -0.054 | -0.054 | 2.93 | 0.86 | 0.99 | 0.83 | 0.31 | 0.86 | 0.64 |
| 3 | 177507406 | rs9811323 | -0.236 | -1.267 | -1.405 | -1.95 | 0.001 | -0.007 | -0.007 | 0 | 0.32 | 0.62 | 0.36 | 0.17 | 0.24 | 0.24 |
| 3 | 177519305 | rs9809387 | 0.204 | -0.899 | -1.906 | -2.353 | -0.007 | 0.046 | 0.046 | 0 | 0.65 | 0.74 | 0.7 | 0.28 | 0.67 | 0.47 |
| 3 | 177520768 | rs4857746 | 0.846 | -1.236 | -2.184 | -2.504 | 0.04 | 0.101 | 0.101 | 0 | 0.2 | 0.05 | 0.19 | 0.07 | 0.13 | 0.15 |
| 3 | 177524783 | rs17574276 | 1.155 | -1.225 | -2.354 | -2.646 | -0.06 | 0.201 | 0.201 | 0.317 | 0.4 | 0.1 | 0.33 | 0.12 | 0.52 | 0.3 |
| 3 | 177527363 | rs9861373 | 1.806 | -1.071 | -2.433 | -2.586 | -0.128 | 0.165 | 0.165 | 0 | 0.33 | 0.05 | 0.29 | 0.12 | 0.45 | 0.24 |
| 3 | 177527638 | rs13075805 | 0.712 | -1.175 | -2.312 | -2.178 | 0.042 | 0.076 | 0.076 | 0 | 0.21 | 0.06 | 0.19 | 0.09 | 0.13 | 0.17 |
| 3 | 177528689 | rs6802821 | 0.469 | -0.337 | -1.735 | -1.596 | 0.008 | 0.012 | 0.012 | 0.045 | 0.15 | 0.15 | 0.09 | 0.03 | 0.13 | 0.09 |
| 3 | 177529463 | rs1347447 | -0.097 | 0.079 | -1.462 | -1.002 | -0.01 | -0.006 | -0.006 | 0 | 0.8 | 0.86 | 0.8 | 0.32 | 0.8 | 0.56 |
| 3 | 177531847 | rs4857747 | -0.631 | -0.956 | -1.318 | -0.652 | 0.027 | 0.023 | 0.023 | 0.089 | 0.3 | 0.77 | 0.35 | 0.2 | 0.19 | 0.25 |
| 3 | 177533260 | rs13068858 | 0.727 | -0.49 | -1.364 | -0.391 | -0.025 | -0.016 | -0.016 | 0 | 0.32 | 0.24 | 0.36 | 0.38 | 0.39 | 0.22 |
| 3 | 177535748 | rs2164349 | -0.37 | -1.091 | -1.631 | -0.35 | -0.042 | -0.088 | -0.088 | 0.015 | 0.71 | 0.92 | 0.79 | 0.55 | 0.7 | 0.59 |
| 3 | 177535841 | rs12633177 | 1.086 | -1.281 | -1.773 | -0.377 | -0.142 | -0.052 | -0.052 | 0 | 0.34 | 0.08 | 0.32 | 0.35 | 0.44 | 0.28 |
| 3 | 177542852 | rs7641262 | -0.622 | -1.497 | -1.911 | -0.68 | -0.003 | -0.013 | -0.013 | 1.045 | 0.23 | 0.73 | 0.34 | 0.12 | 0.23 | 0.17 |
| 3 | 177551779 | rs1561030 | 0.66 | -1.634 | -1.207 | -0.551 | -0.001 | 0.014 | 0.014 | 0 | 0.58 | 0.3 | 0.59 | 0.38 | 0.6 | 0.46 |
| 3 | 177552252 | rs1002240 | 1.565 | -1.789 | -0.999 | -0.316 | -0.159 | -0.057 | -0.057 | 0 | 0.36 | 0.02 | 0.32 | 0.34 | 0.48 | 0.3 |
| 3 | 177552702 | rs9825805 | -0.823 | -0.617 | 0.317 | 0.797 | 0.006 | 0.051 | 0.051 | 0.029 | 0.15 | 0.55 | 0.2 | 0.18 | 0.15 | 0.12 |
| 3 | 177562035 | rs11922738 | 0.762 | -0.462 | -0.106 | 0.737 | 0.011 | -0.005 | -0.005 | 0 | 0.63 | 0.25 | 0.61 | 0.39 | 0.66 | 0.53 |
| 3 | 177590561 | rs7614917 | -1.388 | -0.358 | 0.924 | 0.536 | -0.005 | -0.008 | -0.008 | 0 | 0.08 | 0.32 | 0.11 | 0.02 | 0.04 | 0.07 |
| 3 | 177591074 | rs10936959 | 1.127 | -0.021 | 0.666 | 1.042 | -0.002 | 0.058 | 0.058 | 0 | 0.74 | 0.62 | 0.7 | 0.5 | 0.75 | 0.7 |
| 3 | 177594938 | rs1435649 | 1.354 | -0.171 | 1.075 | 0.93 | 0.012 | 0.055 | 0.055 | 0 | 0.71 | 0.47 | 0.67 | 0.44 | 0.69 | 0.54 |
| 3 | 177598482 | rs7643184 | 1.398 | -0.13 | 1.531 | 0.957 | -0.005 | 0.054 | 0.054 | 0 | 0.71 | 0.43 | 0.63 | 0.44 | 0.73 | 0.68 |
| 3 | 177603519 | rs7626364 | -2.17 | -0.369 | 1.432 | 0.794 | -0.005 | -0.003 | -0.003 | 0 | 0.14 | 0.75 | 0.23 | 0.1 | 0.11 | 0.12 |
| 3 | 177607087 | rs1362826 | -1.62 | -0.658 | 1.33 | 0.743 | -0.01 | 0.026 | 0.026 | 0 | 0.35 | 0.59 | 0.38 | 0.46 | 0.3 | 0.44 |
| 3 | 177614691 | rs6443513 | -2.021 | -0.73 | 0.716 | 0.594 | 0.049 | 0.008 | 0.008 | 0.054 | 0.42 | 0.85 | 0.46 | 0.6 | 0.34 | 0.55 |
| 3 | 177632888 | rs12486411 | -1.072 | -0.827 | 0.826 | 0.765 | 0.01 | -0.068 | -0.068 | 0 | 0.27 | 0.12 | 0.3 | 0.53 | 0.25 | 0.41 |
| 3 | 177636693 | rs1421163 | -0.847 | -0.803 | 0.885 | 0.618 | 0.002 | -0.033 | -0.033 | 0.414 | 0.22 | 0.13 | 0.28 | 0.54 | 0.21 | 0.3 |
| 3 | 177639827 | rs10212574 | -1.483 | -0.734 | 1.463 | 0.554 | 0.024 | -0.002 | -0.002 | 0 | 0.09 | 0.42 | 0.12 | 0.04 | 0.05 | 0.07 |

| | | | | | | | | | | | | | | | | |
|---|---|---|---|---|---|---|---|---|---|---|---|---|---|---|---|---|
| 3 | 177685168 | rs6781923 | -2.064 | -0.83 | 0.972 | 0.297 | 0.058 | 0.029 | 0.029 | 0 | 0.46 | 0.77 | 0.47 | 0.59 | 0.35 | 0.57 |
| 3 | 177701293 | rs9830203 | -2.286 | -1.027 | 0.993 | 0.329 | 0.1 | 0.017 | 0.017 | 0 | 0.44 | 0.86 | 0.44 | 0.59 | 0.29 | 0.56 |
| 3 | 177705068 | rs6443523 | -1.762 | -1.08 | 1.023 | 0.332 | 0.028 | -0.007 | -0.007 | 0 | 0.36 | 0.72 | 0.35 | 0.58 | 0.2 | 0.51 |
| 3 | 177706945 | rs6792736 | -1.333 | -1.078 | 1.027 | 0.291 | -0.033 | -0.077 | -0.077 | 0.314 | 0.14 | 0.28 | 0.09 | 0.01 | 0.09 | 0.2 |
| 3 | 177707225 | rs6779930 | -2.291 | -1.054 | 0.744 | 0.314 | 0.094 | -0.026 | -0.026 | 0 | 0.43 | 0.94 | 0.42 | 0.58 | 0.27 | 0.55 |
| 3 | 177710225 | rs2160836 | -2.291 | -1.048 | 0.751 | 0.314 | 0.094 | -0.018 | -0.018 | 0 | 0.43 | 0.93 | 0.41 | 0.58 | 0.27 | 0.55 |
| 3 | 177712676 | rs9817294 | -1.768 | 0.115 | 0.983 | 0.003 | 0.005 | -0.042 | -0.042 | 0 | 0.2 | 0.76 | 0.2 | 0.01 | 0.27 | 0.12 |
| 3 | 177720393 | rs6788525 | -0.926 | -0.748 | 1.429 | 0.105 | 0.007 | -0.065 | -0.065 | 0 | 0.12 | 0.26 | 0.11 | 0 | 0.15 | 0.1 |
| 3 | 177729343 | rs7645340 | -1.215 | -0.726 | 1.151 | 0.12 | -0.002 | -0.1 | -0.1 | 0 | 0.16 | 0.37 | 0.12 | 0.01 | 0.15 | 0.23 |
| 3 | 177735032 | rs2160835 | -1.77 | -0.56 | -0.057 | 0.333 | -0.012 | 0.004 | 0.004 | 0.086 | 0.44 | 0.96 | 0.45 | 0.61 | 0.34 | 0.57 |
| 3 | 177743488 | rs4611248 | -0.711 | 0.057 | 0.409 | 1.221 | 0.02 | 0.181 | 0.181 | 0.446 | 0.07 | 0.12 | 0.11 | 0.47 | 0.09 | 0.3 |
| 3 | 177744154 | rs4420316 | -1.246 | -0.285 | 0.463 | 0.553 | 0 | 0.005 | 0.005 | 0 | 0.29 | 0.29 | 0.28 | 0.55 | 0.3 | 0.46 |
| 3 | 177744450 | rs6786229 | -1.41 | -0.639 | 0.818 | 0.143 | 0 | -0.016 | -0.016 | 0.006 | 0.35 | 0.45 | 0.5 | 0.69 | 0.38 | 0.55 |
| 3 | 177744507 | rs1362830 | -1.374 | -0.998 | 0.174 | 0.037 | 0.017 | 0.04 | 0.04 | 0 | 0.42 | 0.7 | 0.58 | 0.69 | 0.39 | 0.57 |
| 3 | 177761920 | rs1982822 | -0.53 | -1.191 | 0.096 | -0.135 | 0.013 | -0.005 | -0.005 | 3.423 | 0.31 | 0.23 | 0.33 | 0.61 | 0.23 | 0.4 |
| 3 | 177763155 | rs9851864 | -1.16 | -1.402 | -0.173 | -0.281 | 0.068 | 0.022 | 0.022 | 0.487 | 0.38 | 0.84 | 0.48 | 0.64 | 0.25 | 0.55 |
| 3 | 177767236 | rs2902414 | -0.352 | -1.483 | -0.221 | -0.271 | 0.024 | -0.059 | -0.059 | 0.441 | 0.28 | 0.04 | 0.32 | 0.62 | 0.23 | 0.39 |
| 3 | 177775345 | rs1489630 | -1.554 | -1.373 | 0.154 | -0.581 | 0.033 | 0.115 | 0.115 | 0.272 | 0.43 | 0.82 | 0.53 | 0.75 | 0.38 | 0.53 |
| 3 | 177790822 | rs12490085 | -1.56 | -1.481 | 0.259 | -0.575 | 0.041 | 0.091 | 0.091 | 0.109 | 0.43 | 0.77 | 0.53 | 0.75 | 0.38 | 0.53 |
| 3 | 177802197 | rs6785366 | -1.079 | -1.016 | 0.727 | -0.543 | 0.008 | -0.048 | -0.048 | 0 | 0.16 | 0.25 | 0.15 | 0.05 | 0.21 | 0.12 |
| 3 | 177816610 | rs13101116 | -0.087 | -1.523 | 0.348 | -0.505 | 0.032 | -0.023 | -0.023 | 0.207 | 0.25 | 0.08 | 0.34 | 0.66 | 0.19 | 0.38 |
| 3 | 177823987 | rs1489623 | -1.333 | -1.318 | 1.246 | -1.563 | 0 | -0.006 | -0.006 | 0 | 0.48 | 0.5 | 0.54 | 0.76 | 0.44 | 0.64 |
| 3 | 177826047 | rs7628308 | -1.232 | -1.305 | 1.033 | -1.554 | 0.013 | 0.081 | 0.081 | 0 | 0.43 | 0.24 | 0.47 | 0.73 | 0.42 | 0.5 |
| 3 | 177837221 | rs13098361 | 0.177 | -1.285 | 1.052 | -1.493 | 0.028 | -0.03 | -0.03 | 0 | 0.25 | 0.09 | 0.32 | 0.67 | 0.19 | 0.38 |
| 3 | 177838441 | rs1565327 | -1.07 | -1.317 | 0.992 | -1.636 | 0.011 | 0.089 | 0.089 | 0 | 0.47 | 0.26 | 0.47 | 0.72 | 0.46 | 0.51 |
| 3 | 177863108 | rs2902413 | 0.397 | -0.931 | 1.127 | -1.619 | -0.009 | 0 | 0 | 0 | 0.78 | 0.77 | 0.83 | 0.94 | 0.71 | 0.89 |
| 3 | 177864536 | rs2863110 | 0.191 | -1.136 | 0.932 | -1.586 | 0.019 | -0.015 | -0.015 | 0 | 0.26 | 0.21 | 0.35 | 0.66 | 0.18 | 0.39 |
| 3 | 177865313 | rs923456 | 1.306 | -1.187 | 0.735 | -1.589 | -0.014 | 0.071 | 0.071 | 0 | 0.47 | 0.24 | 0.45 | 0.26 | 0.52 | 0.46 |
| 3 | 177869073 | rs16829162 | 0.787 | -1.106 | 0.222 | -1.456 | -0.006 | -0.108 | -0.108 | 0 | 0.18 | 0.01 | 0.27 | 0.65 | 0.14 | 0.36 |
| 3 | 177871232 | rs7642011 | 1.24 | -1.329 | 1.213 | -1.841 | -0.009 | 0.046 | 0.046 | 0 | 0.44 | 0.36 | 0.44 | 0.22 | 0.47 | 0.41 |
| 3 | 177873533 | rs1489616 | 1.243 | -1.322 | 1.393 | -1.84 | -0.006 | 0.048 | 0.048 | 0 | 0.46 | 0.62 | 0.46 | 0.22 | 0.47 | 0.41 |
| 3 | 177876600 | rs7637967 | 0.561 | -1.31 | 1.432 | -1.921 | -0.01 | 0.1 | 0.1 | 0 | 0.55 | 0.78 | 0.51 | 0.24 | 0.53 | 0.45 |
| 3 | 177887326 | rs10936964 | 0.723 | -1.304 | 1.484 | -1.818 | -0.002 | 0.035 | 0.035 | 0 | 0.54 | 0.66 | 0.5 | 0.24 | 0.52 | 0.43 |
| 3 | 177896888 | rs1906888 | 0.734 | -1.308 | 1.458 | -1.856 | -0.002 | 0.009 | 0.009 | 0 | 0.54 | 0.55 | 0.53 | 0.28 | 0.54 | 0.51 |
| 3 | 177899353 | rs7620308 | 0.064 | -1.287 | 1.325 | -1.838 | -0.024 | 0.016 | 0.016 | 0 | 0.63 | 0.81 | 0.58 | 0.32 | 0.6 | 0.55 |
| 3 | 177912894 | rs9290640 | 0.189 | -1.274 | 0.57 | -1.833 | -0.024 | 0.033 | 0.033 | 0 | 0.62 | 0.79 | 0.58 | 0.28 | 0.59 | 0.53 |
| 3 | 177917303 | rs6781744 | 1.505 | -1.038 | 0.478 | -1.845 | -0.006 | 0.039 | 0.039 | 0 | 0.42 | 0.32 | 0.39 | 0.24 | 0.42 | 0.39 |
| 3 | 177920039 | rs3924997 | 1.069 | -0.991 | 0.507 | -1.68 | -0.001 | 0.051 | 0.051 | 0 | 0.48 | 0.36 | 0.42 | 0.28 | 0.46 | 0.38 |
| 3 | 177932934 | rs6762634 | 0.934 | -0.97 | 0.326 | -1.71 | 0.001 | 0.008 | 0.008 | 0 | 0.5 | 0.51 | 0.45 | 0.32 | 0.49 | 0.4 |
| 3 | 177944933 | rs6810087 | 1.27 | -1.075 | 0.309 | -1.735 | -0.002 | 0.016 | 0.016 | 0 | 0.46 | 0.63 | 0.44 | 0.31 | 0.46 | 0.41 |
| 3 | 177949454 | rs4568149 | -0.887 | -1.085 | 0.265 | -1.862 | -0.008 | 0.005 | 0.005 | 0 | 0.88 | 0.91 | 0.88 | 0.91 | 0.78 | 0.9 |
| 3 | 177961323 | rs6780470 | -0.005 | -1.166 | 0.314 | -1.822 | -0.018 | -0.004 | -0.004 | 0 | 0.63 | 0.82 | 0.59 | 0.35 | 0.62 | 0.57 |
| 3 | 177963171 | rs12492103 | 0.698 | -1.182 | 0.388 | -1.821 | -0.002 | 0.031 | 0.031 | 0 | 0.53 | 0.66 | 0.47 | 0.26 | 0.51 | 0.43 |
| 3 | 177970767 | rs6808802 | 0.893 | -1.174 | 0.424 | -1.82 | 0 | 0.007 | 0.007 | 0 | 0.51 | 0.57 | 0.45 | 0.24 | 0.51 | 0.32 |
| 3 | 177990186 | rs7629595 | 1.376 | -1.129 | 0.335 | -1.824 | 0 | -0.006 | -0.006 | 0 | 0.43 | 0.41 | 0.41 | 0.24 | 0.46 | 0.31 |
| 3 | 178007676 | rs17578740 | -0.904 | -0.817 | 0.368 | -1.803 | 0.003 | 0.015 | 0.015 | 0 | 0.21 | 0.14 | 0.14 | 0.1 | 0.18 | 0.16 |
| 3 | 178008970 | rs4857765 | 1.097 | -0.667 | -0.054 | -1.933 | 0.004 | 0.044 | 0.044 | 0 | 0.21 | 0.4 | 0.26 | 0.29 | 0.25 | 0.16 |
| 3 | 178022371 | rs9854383 | 0.857 | -1.35 | -0.215 | -1.583 | -0.008 | 0.074 | 0.074 | 0 | 0.49 | 0.31 | 0.53 | 0.32 | 0.53 | 0.46 |
| 3 | 178022390 | rs6791314 | -0.064 | -1.285 | -0.057 | -1.439 | -0.019 | -0.036 | -0.036 | 0.1 | 0.64 | 0.81 | 0.66 | 0.42 | 0.63 | 0.57 |
| 3 | 178052017 | rs1961019 | 1.088 | -1.287 | -0.18 | -1.435 | 0.009 | 0.015 | 0.015 | 0.893 | 0.46 | 0.36 | 0.58 | 0.42 | 0.57 | 0.54 |
| 3 | 178072978 | rs1822174 | -0.624 | -1.485 | -0.284 | -1.337 | -0.015 | 0 | 0 | 0.904 | 0.41 | 0.5 | 0.32 | 0.53 | 0.22 | 0.36 |
| 3 | 178076337 | rs1583919 | 0.438 | -0.959 | -0.211 | -1.032 | -0.046 | 0.078 | 0.078 | 0 | 0.25 | 0.37 | 0.28 | 0.53 | 0.16 | 0.34 |
| 3 | 178086048 | rs2634152 | 1.429 | -0.847 | -0.583 | -0.994 | -0.026 | 0.029 | 0.029 | 0 | 0.42 | 0.3 | 0.5 | 0.31 | 0.5 | 0.52 |
| 3 | 178091627 | rs1837788 | 0 | -1.362 | -0.502 | -0.436 | -0.018 | 0.019 | 0.019 | 0 | 0.24 | 0.27 | 0.26 | 0.57 | 0.15 | 0.3 |
| 3 | 178097463 | rs7650856 | -0.183 | -0.757 | -0.819 | -0.028 | -0.074 | 0.014 | 0.014 | 0 | 0.36 | 0.45 | 0.31 | 0.57 | 0.18 | 0.31 |
| 3 | 178105341 | rs282875 | 0.307 | 0.994 | -1.031 | -0.183 | -0.016 | 0.01 | 0.01 | 0 | 0.88 | 0.91 | 0.89 | 0.97 | 0.79 | 0.92 |
| 3 | 178107949 | rs7629696 | -0.381 | 0.765 | -1.187 | 0.063 | -0.006 | -0.024 | -0.024 | 1.473 | 0.33 | 0.26 | 0.43 | 0.47 | 0.4 | 0.35 |
| 3 | 178118676 | rs11718301 | -0.699 | 0.276 | -0.342 | -0.396 | -0.034 | -0.027 | -0.027 | 0.058 | 0.12 | 0.27 | 0.08 | 0.09 | 0.07 | 0.17 |
| 3 | 178119369 | rs282861 | 1.14 | -0.296 | -0.557 | -0.852 | 0.019 | -0.011 | -0.011 | 0 | 0.29 | 0.34 | 0.38 | 0.29 | 0.38 | 0.34 |
| 3 | 178122321 | rs4857768 | 0.111 | -0.419 | -0.772 | -0.623 | -0.023 | -0.041 | -0.041 | 0 | 0.32 | 0.12 | 0.38 | 0.55 | 0.36 | 0.27 |
| 3 | 178122486 | rs444537 | -0.904 | -0.732 | -0.855 | -0.775 | 0.004 | 0.011 | 0.011 | 0 | 0.84 | 0.89 | 0.88 | 0.96 | 0.86 | 0.82 |
| 3 | 178124484 | rs100608 | -0.08 | -0.728 | -0.935 | -0.617 | -0.051 | -0.045 | -0.045 | 0.193 | 0.66 | 0.89 | 0.59 | 0.46 | 0.6 | 0.71 |
| 3 | 178126001 | rs13100119 | 0.877 | -0.93 | -0.893 | -0.744 | 0.027 | 0.026 | 0.026 | 0 | 0.11 | 0.04 | 0.07 | 0 | 0.07 | 0.01 |
| 3 | 178126597 | rs6782287 | 0.535 | -0.973 | -0.767 | -0.763 | 0.008 | -0.006 | -0.006 | 0.703 | 0.13 | 0.07 | 0.08 | 0 | 0.07 | 0.01 |
| 3 | 178127824 | rs1812958 | -0.904 | -1.149 | 0.034 | -0.81 | -0.043 | -0.047 | -0.047 | 0 | 0.25 | 0.55 | 0.19 | 0.12 | 0.19 | 0.23 |
| 3 | 178135091 | rs369516 | -0.407 | -1.394 | 0.068 | -0.88 | 0 | 0 | 0 | 0 | 0.69 | 0.69 | 0.6 | 0.46 | 0.66 | 0.67 |
| 3 | 178148233 | rs9868938 | -0.949 | -1.345 | 0.17 | -0.837 | 0.085 | 0.086 | 0.086 | 2.543 | 0.07 | 0.35 | 0.06 | 0.06 | 0.06 | 0.12 |
| 3 | 178155754 | rs282877 | 1.008 | -1.357 | -0.234 | -0.869 | -0.046 | 0.123 | 0.123 | 0 | 0.45 | 0.2 | 0.42 | 0.34 | 0.47 | 0.44 |
| 3 | 178159452 | rs282880 | 1.044 | -1.271 | -0.434 | -0.824 | -0.008 | 0.051 | 0.051 | 0 | 0.46 | 0.35 | 0.44 | 0.34 | 0.47 | 0.43 |

| | | | | | | | | | | | | | | | | |
|---|---|---|---|---|---|---|---|---|---|---|---|---|---|---|---|---|
| 3 | 178159539 | rs282881 | -0.089 | -0.504 | -0.809 | -0.447 | -0.028 | -0.05 | -0.05 | 0 | 0.71 | 0.89 | 0.66 | 0.46 | 0.67 | 0.68 |
| 3 | 178167962 | rs282887 | 0.212 | -0.48 | -0.826 | -0.514 | 0.01 | -0.002 | -0.002 | 0 | 0.66 | 0.62 | 0.61 | 0.43 | 0.61 | 0.68 |
| 3 | 178171044 | rs17581727 | 0.645 | 0 | -1.521 | -0.455 | -0.015 | 0.029 | 0.029 | 0 | 0.11 | 0.03 | 0.07 | 0.02 | 0.12 | 0.08 |
| 3 | 178173268 | rs11710597 | 0.29 | -0.266 | -2.094 | 0.05 | -0.07 | -0.021 | -0.021 | 0 | 0.42 | 0.14 | 0.44 | 0.49 | 0.47 | 0.55 |
| 3 | 178174699 | rs2291478 | -0.075 | 0.457 | -1.989 | 0.275 | -0.006 | 0.008 | 0.008 | 0 | 0.45 | 0.25 | 0.41 | 0.38 | 0.47 | 0.46 |
| 3 | 178176128 | rs9290646 | -0.233 | 0.692 | -2.134 | 0.197 | 0.039 | -0.009 | -0.009 | 0 | 0.52 | 0.21 | 0.65 | 0.44 | 0.61 | 0.54 |
| 3 | 178183705 | rs2902423 | -1.734 | 0.484 | -2.571 | 0.276 | -0.079 | -0.065 | -0.065 | 0 | 0.84 | 0.98 | 0.8 | 0.64 | 0.77 | 0.86 |
| 3 | 178195183 | rs3816843 | 0.467 | -0.025 | -2.083 | -0.219 | -0.01 | -0.008 | -0.008 | 0 | 0.4 | 0.34 | 0.56 | 0.46 | 0.49 | 0.51 |
| 3 | 178196694 | rs1016911 | 1.189 | -0.59 | -1.543 | -0.374 | -0.034 | 0.089 | 0.089 | 0.095 | 0.1 | 0.26 | 0.05 | 0.14 | 0.07 | 0.22 |
| 3 | 178198831 | rs6769947 | 1.189 | -0.59 | -1.543 | -0.374 | -0.034 | 0.089 | 0.089 | 0.095 | 0.1 | 0.25 | 0.05 | 0.14 | 0.07 | 0.22 |
| 3 | 178200226 | rs9854298 | -0.052 | -0.936 | -1.82 | -0.628 | 0.004 | -0.002 | -0.002 | 0.436 | 0.58 | 0.39 | 0.67 | 0.5 | 0.64 | 0.58 |
| 3 | 178200399 | rs12493097 | 1.188 | -0.974 | -1.788 | -0.648 | -0.105 | -0.089 | -0.089 | 0 | 0.38 | 0.11 | 0.53 | 0.46 | 0.49 | 0.51 |
| 3 | 178205810 | rs10936968 | 2.084 | -0.899 | -2.205 | -0.714 | -0.07 | -0.11 | -0.11 | 0 | 0.15 | 0.01 | 0.28 | 0.31 | 0.25 | 0.22 |
| 3 | 178207097 | rs11711450 | -0.168 | -1.188 | -1.921 | -0.727 | 0.004 | -0.006 | -0.006 | 3.729 | 0.58 | 0.39 | 0.67 | 0.45 | 0.64 | 0.58 |
| 3 | 178209053 | rs17582581 | 1.802 | -1.133 | -2.039 | -0.739 | 0.001 | 0.081 | 0.081 | 0 | 0.19 | 0.04 | 0.11 | 0.04 | 0.15 | 0.07 |
| 3 | 178210300 | rs12492876 | 0.737 | -1.033 | -1.965 | -0.672 | -0.094 | -0.079 | -0.079 | 0 | 0.38 | 0.11 | 0.53 | 0.41 | 0.49 | 0.52 |
| 3 | 178223239 | rs6768143 | -2.047 | -0.902 | -1.707 | -0.536 | -0.025 | -0.028 | -0.028 | 0 | 0.82 | 0.85 | 0.76 | 0.59 | 0.74 | 0.84 |
| 3 | 178225884 | rs11929122 | 1.327 | -0.92 | -1.466 | -0.398 | -0.027 | 0.093 | 0.093 | 0.209 | 0.1 | 0.26 | 0.05 | 0.16 | 0.06 | 0.22 |
| 3 | 178229935 | rs11918167 | 0.211 | -0.833 | -2.04 | -0.514 | -0.017 | -0.012 | -0.012 | 0.187 | 0.31 | 0.55 | 0.18 | 0.2 | 0.22 | 0.29 |
| 3 | 178230172 | rs7642169 | 1.52 | -0.422 | -2.517 | -0.537 | -0.019 | 0.092 | 0.092 | 0 | 0.19 | 0.03 | 0.11 | 0.04 | 0.16 | 0.07 |
| 3 | 178231482 | rs1441881 | 0.195 | -0.566 | -2.471 | -0.463 | 0.017 | -0.014 | -0.014 | 0.059 | 0.49 | 0.1 | 0.61 | 0.39 | 0.59 | 0.53 |
| 3 | 178238319 | rs1441883 | 1.407 | -0.449 | -2.171 | -0.652 | 0.016 | 0.038 | 0.038 | 0.049 | 0.2 | 0.06 | 0.12 | 0.16 | 0.16 | 0.08 |
| 3 | 178239558 | rs13321110 | 1.525 | -0.252 | -2.279 | -0.986 | -0.001 | 0.027 | 0.027 | 0 | 0.19 | 0.14 | 0.2 | 0.06 | 0.15 | 0.15 |
| 3 | 178246465 | rs4857772 | -1.674 | -0.393 | -1.798 | -0.996 | 0.042 | 0.01 | 0.01 | 0 | 0.81 | 0.62 | 0.76 | 0.74 | 0.74 | 0.83 |
| 3 | 178256957 | rs17590394 | 0.723 | -0.434 | -1.779 | -1.002 | 0.021 | -0.006 | -0.006 | 0 | 0.13 | 0.33 | 0.12 | 0.1 | 0.16 | 0.31 |
| 3 | 178264581 | rs1477779 | 1.358 | -0.513 | -2.181 | -0.987 | -0.022 | -0.129 | -0.129 | 0.023 | 0.27 | 0.05 | 0.31 | 0.42 | 0.27 | 0.29 |
| 3 | 178267549 | rs9837011 | 1.193 | -0.702 | -2.227 | -1.285 | 0.004 | 0.076 | 0.076 | 0 | 0.26 | 0.12 | 0.22 | 0.07 | 0.2 | 0.29 |
| 3 | 178272167 | rs34278351 | 0.71 | -0.757 | -2.057 | -1.252 | 0.015 | 0.016 | 0.016 | 0.271 | 0.32 | 0.73 | 0.26 | 0.29 | 0.28 | 0.26 |
| 3 | 178276886 | rs10936973 | 0.772 | -0.555 | -1.996 | -1.166 | 0.024 | 0.075 | 0.075 | 0 | 0.48 | 0.1 | 0.43 | 0.59 | 0.43 | 0.39 |
| 3 | 178280455 | rs16829756 | 1.028 | -0.395 | -1.507 | -1.125 | 0.071 | 0.093 | 0.093 | 0.271 | 0.09 | 0.39 | 0.11 | 0.13 | 0.12 | 0.17 |
| 3 | 178286940 | rs9848437 | 0.131 | -0.44 | -1.869 | -1.164 | -0.001 | -0.005 | -0.005 | 0 | 0.34 | 0.61 | 0.36 | 0.33 | 0.3 | 0.44 |
| 3 | 178290081 | rs11918114 | 0.198 | -0.469 | -1.876 | -1.177 | -0.001 | -0.018 | -0.018 | 0.008 | 0.33 | 0.5 | 0.35 | 0.33 | 0.3 | 0.44 |
| 3 | 178292353 | rs9290656 | 0.866 | -0.567 | -2.486 | -1.187 | 0.004 | -0.078 | -0.078 | 0 | 0.29 | 0.13 | 0.33 | 0.4 | 0.29 | 0.34 |
| 3 | 178310836 | rs7636991 | 1.872 | -0.985 | -2.426 | -1.3 | 0.034 | 0.084 | 0.084 | 0.089 | 0.22 | 0.07 | 0.09 | 0.04 | 0.17 | 0.07 |
| 3 | 178311942 | rs12497665 | 1.395 | -1 | -2.401 | -1.304 | -0.016 | -0.045 | -0.045 | 0 | 0.24 | 0.35 | 0.12 | 0.05 | 0.17 | 0.09 |
| 3 | 178319253 | rs1477773 | -0.225 | -0.676 | -2.552 | -1.09 | -0.034 | 0.089 | 0.089 | 0.461 | 0.56 | 0.83 | 0.46 | 0.36 | 0.47 | 0.53 |
| 3 | 178337631 | rs9290658 | -0.824 | -0.178 | -2.631 | -1.064 | -0.056 | -0.059 | -0.059 | 0 | 0.61 | 0.85 | 0.6 | 0.58 | 0.6 | 0.61 |
| 3 | 178341582 | rs2054399 | -1.099 | -0.591 | -2.872 | -1.304 | -0.041 | -0.067 | -0.067 | 0 | 0.72 | 0.88 | 0.64 | 0.56 | 0.62 | 0.71 |
| 3 | 178343683 | rs6802120 | 1.868 | -0.445 | -2.727 | -1.326 | 0.002 | -0.002 | -0.002 | 0 | 0.17 | 0.19 | 0.21 | 0.23 | 0.14 | 0.17 |
| 3 | 178353935 | rs13095759 | 1.81 | -0.446 | -2.844 | -1.437 | 0.012 | 0.13 | 0.13 | 1.137 | 0.25 | 0.05 | 0.1 | 0.02 | 0.17 | 0.13 |
| 3 | 178355720 | rs1382044 | -1.223 | -0.388 | -2.881 | -1.418 | -0.058 | -0.089 | -0.089 | 0 | 0.72 | 0.89 | 0.63 | 0.56 | 0.62 | 0.71 |
| 3 | 178356269 | rs6443552 | 1.352 | -0.634 | -2.632 | -1.519 | 0.086 | 0.075 | 0.075 | 0.244 | 0.08 | 0.42 | 0.09 | 0.1 | 0.1 | 0.08 |
| 3 | 178357197 | rs9847335 | -0.497 | -0.603 | -2.721 | -1.444 | -0.016 | 0.06 | 0.06 | 0 | 0.54 | 0.66 | 0.42 | 0.33 | 0.48 | 0.56 |
| 3 | 178364921 | rs6789584 | 0.637 | -0.893 | -2.709 | -1.483 | 0.036 | 0.092 | 0.092 | 0.522 | 0.41 | 0.23 | 0.31 | 0.21 | 0.31 | 0.3 |
| 3 | 178365792 | rs10755134 | 1.023 | -0.704 | -2.643 | -1.447 | 0.046 | 0.115 | 0.115 | 2.689 | 0.4 | 0.26 | 0.2 | 0.07 | 0.29 | 0.17 |
| 3 | 178365840 | rs7615444 | 0.526 | -0.652 | -2.674 | -1.318 | 0.012 | 0.026 | 0.026 | 0.472 | 0.28 | 0.48 | 0.28 | 0.36 | 0.3 | 0.41 |
| 3 | 178371354 | rs9856750 | -0.475 | -0.685 | -2.669 | -1.298 | 0.012 | 0.002 | 0.002 | 0.021 | 0.45 | 0.69 | 0.49 | 0.55 | 0.43 | 0.57 |
| 3 | 178372037 | rs9861190 | -1.691 | -0.041 | -2.266 | -0.904 | -0.091 | -0.113 | -0.113 | 0.02 | 0.86 | 0.99 | 0.68 | 0.61 | 0.72 | 0.76 |
| 3 | 178374095 | rs9871964 | 0.664 | -0.721 | -1.761 | -0.854 | -0.011 | -0.016 | -0.016 | 0 | 0.21 | 0.29 | 0.28 | 0.2 | 0.18 | 0.2 |
| 3 | 178374124 | rs7429015 | -1.831 | -0.395 | -1.967 | -0.811 | -0.128 | -0.173 | -0.173 | 0.252 | 0.78 | 0.98 | 0.62 | 0.6 | 0.67 | 0.7 |
| 3 | 178378283 | rs6798240 | -1.182 | -0.776 | -1.576 | -1.031 | 0.001 | 0.002 | 0.002 | 0 | 0.55 | 0.46 | 0.44 | 0.52 | 0.46 | 0.5 |
| 3 | 178381419 | rs7433956 | 0.74 | -0.873 | -1.573 | -1.115 | 0.012 | 0.002 | 0.002 | 0 | 0.21 | 0.21 | 0.09 | 0.17 | 0.13 | 0.18 |
| 3 | 178386451 | rs11921195 | 0.075 | -0.733 | -1.576 | -1.101 | 0.021 | 0.011 | 0.011 | 0 | 0.35 | 0.33 | 0.16 | 0.28 | 0.24 | 0.26 |
| 3 | 178397422 | rs6792781 | 0.354 | -0.449 | -1.342 | -0.918 | 0.006 | -0.082 | -0.082 | 0 | 0.27 | 0.49 | 0.2 | 0.12 | 0.28 | 0.23 |
| 3 | 178397892 | rs11920669 | -2.211 | -0.47 | -1.397 | -0.902 | -0.05 | -0.08 | -0.08 | 0 | 0.76 | 0.88 | 0.6 | 0.61 | 0.67 | 0.68 |
| 3 | 178399551 | rs13100616 | 0.349 | -0.463 | -1.39 | -0.903 | 0.008 | -0.079 | -0.079 | 0 | 0.28 | 0.5 | 0.21 | 0.12 | 0.28 | 0.23 |
| 3 | 178402265 | rs6802875 | -0.371 | -0.947 | -1.33 | -0.829 | 0.046 | 0.018 | 0.018 | 0.071 | 0.48 | 0.39 | 0.39 | 0.5 | 0.39 | 0.46 |
| 3 | 178416735 | rs13067125 | 0.513 | -0.643 | -0.294 | -0.771 | -0.008 | -0.079 | -0.079 | 1.79 | 0.22 | 0.39 | 0.14 | 0.07 | 0.21 | 0.17 |
| 3 | 178430105 | rs12487889 | -0.042 | -0.441 | -0.718 | -0.497 | 0.02 | 0.055 | 0.055 | 0 | 0.27 | 0.13 | 0.32 | 0.22 | 0.27 | 0.2 |
| 3 | 178430587 | rs9825195 | -0.366 | -0.429 | -0.642 | -0.417 | 0.029 | -0.001 | -0.001 | 0 | 0.33 | 0.26 | 0.33 | 0.39 | 0.29 | 0.36 |
| 3 | 178446068 | rs6443569 | -0.719 | -0.082 | -0.093 | -0.852 | -0.002 | -0.004 | -0.004 | 1.963 | 0.41 | 0.61 | 0.48 | 0.4 | 0.44 | 0.51 |
| 3 | 178449388 | rs7622551 | -0.347 | -0.074 | 0.446 | -0.887 | -0.016 | -0.004 | -0.004 | 0 | 0.34 | 0.48 | 0.4 | 0.37 | 0.32 | 0.4 |
| 3 | 178449906 | rs6443571 | -2.056 | -0.134 | 0.111 | -1.171 | -0.107 | -0.12 | -0.12 | 0.035 | 0.73 | 0.96 | 0.63 | 0.49 | 0.65 | 0.71 |
| 3 | 178452941 | rs7612729 | 0.903 | 0.181 | 0.423 | -0.794 | 0.035 | -0.016 | -0.016 | 0 | 0.07 | 0.08 | 0.06 | 0.04 | 0.12 | 0.1 |
| 3 | 178453925 | rs7615682 | 0.631 | 0.067 | 0.507 | -0.854 | 0.002 | -0.001 | -0.001 | 0 | 0.2 | 0.19 | 0.17 | 0.23 | 0.14 | 0.23 |
| 3 | 178459665 | rs9839376 | 0.522 | 0.095 | 0.408 | -0.842 | -0.02 | 0 | 0 | 0 | 0.21 | 0.31 | 0.18 | 0.23 | 0.14 | 0.23 |
| 3 | 178461771 | rs7624046 | -0.949 | 0.139 | -0.215 | -0.815 | -0.002 | -0.003 | -0.003 | 0 | 0.41 | 0.6 | 0.48 | 0.41 | 0.44 | 0.51 |
| 3 | 178483476 | rs7613104 | -0.368 | -0.04 | -0.278 | -0.785 | -0.013 | -0.015 | -0.015 | 0 | 0.29 | 0.61 | 0.26 | 0.27 | 0.25 | 0.33 |
| 3 | 178495858 | rs12634309 | 0.795 | 0.057 | -0.848 | -0.866 | -0.024 | -0.019 | -0.019 | 0 | 0.12 | 0.02 | 0.22 | 0.14 | 0.18 | 0.16 |

| | | | | | | | | | | | | | | | | |
|---|---|---|---|---|---|---|---|---|---|---|---|---|---|---|---|---|
| 3 | 178520951 | rs12696465 | 0.247 | 0.292 | -0.216 | -1.039 | -0.016 | -0.009 | -0.009 | 0.004 | 0.21 | 0.3 | 0.18 | 0.15 | 0.15 | 0.22 |
| 3 | 178522077 | rs4370006 | -0.577 | 0.421 | -0.705 | -0.881 | 0.035 | 0.09 | 0.09 | 0 | 0.49 | 0.73 | 0.36 | 0.27 | 0.42 | 0.48 |
| 3 | 178539486 | rs6807583 | -0.294 | 0.477 | -0.955 | -1.087 | -0.005 | -0.025 | -0.025 | 1.966 | 0.4 | 0.42 | 0.31 | 0.05 | 0.44 | 0.27 |
| 3 | 178540316 | rs11914342 | -0.586 | 0.285 | -1.137 | -0.986 | -0.034 | -0.003 | -0.003 | 0 | 0.23 | 0.56 | 0.22 | 0.17 | 0.16 | 0.25 |
| 3 | 178542817 | rs6809208 | -0.65 | 0.505 | -1.437 | -0.933 | 0.002 | 0.078 | 0.078 | 1.208 | 0.38 | 0.84 | 0.39 | 0.19 | 0.32 | 0.39 |
| 3 | 178543156 | rs9844536 | -0.631 | 0.505 | -1.422 | -0.933 | 0.005 | 0.076 | 0.076 | 0.81 | 0.38 | 0.84 | 0.39 | 0.19 | 0.32 | 0.39 |
| 3 | 178544317 | rs4624543 | -1.216 | 0.581 | -1.679 | -0.94 | -0.064 | 0.037 | 0.037 | 0.498 | 0.62 | 0.93 | 0.52 | 0.23 | 0.58 | 0.53 |
| 3 | 178545276 | rs9844737 | -1.216 | 0.581 | -1.679 | -0.94 | -0.064 | 0.037 | 0.037 | 0.498 | 0.62 | 0.93 | 0.52 | 0.23 | 0.58 | 0.53 |
| 3 | 178551573 | rs9877109 | -0.332 | 0.36 | -1.393 | -1.059 | -0.019 | -0.036 | -0.036 | 0 | 0.1 | 0.15 | 0.05 | 0.01 | 0.04 | 0.02 |
| 3 | 178552671 | rs13100117 | 0.449 | 0.719 | -1.109 | -0.838 | -0.004 | 0.004 | 0.004 | 0 | 0.25 | 0.23 | 0.14 | 0.03 | 0.26 | 0.13 |
| 3 | 178556483 | rs6443582 | -0.531 | 0.528 | -0.739 | -0.376 | -0.004 | -0.013 | -0.013 | 0 | 0.32 | 0.57 | 0.34 | 0.18 | 0.28 | 0.36 |
| 3 | 178569163 | rs6443589 | 0.242 | 0.975 | -0.576 | 0.139 | -0.001 | 0 | 0 | 0 | 0.62 | 0.59 | 0.68 | 0.66 | 0.59 | 0.78 |
| 3 | 178575038 | rs11929266 | -0.809 | 0.954 | -0.207 | -0.327 | -0.006 | -0.03 | -0.03 | 0.638 | 0.13 | 0.15 | 0.05 | 0 | 0.06 | 0.05 |
| 3 | 178585678 | rs9832653 | 0.79 | 1.08 | 0.608 | -0.12 | -0.006 | 0.005 | 0.005 | 0.011 | 0.53 | 0.39 | 0.59 | 0.62 | 0.54 | 0.71 |
| 3 | 178588348 | rs6770789 | -1.313 | 0.294 | 0.842 | -0.305 | -0.025 | -0.025 | -0.025 | 0 | 0.54 | 0.76 | 0.47 | 0.41 | 0.46 | 0.36 |
| 3 | 178610090 | rs10433478 | -0.992 | 0.219 | 1.067 | -0.577 | 0.008 | 0.019 | 0.019 | 0 | 0.35 | 0.45 | 0.36 | 0.38 | 0.35 | 0.29 |
| 3 | 178617251 | rs13089659 | 0.906 | 0.3 | 0.855 | -0.551 | -0.04 | 0.064 | 0.064 | 0 | 0.13 | 0.03 | 0.08 | 0.03 | 0.15 | 0.12 |
| 3 | 178633207 | rs11916133 | -1.339 | 0.171 | 1.183 | -0.367 | -0.002 | 0.004 | 0.004 | 2.491 | 0.38 | 0.6 | 0.37 | 0.42 | 0.31 | 0.22 |
| 3 | 178635292 | rs11715386 | -0.097 | 0.253 | 1.501 | -0.313 | 0.264 | 0.591 | 0.591 | 0 | 0.05 | 0.32 | 0.21 | 0.42 | 0.13 | 0.12 |
| 3 | 178639454 | rs10936984 | -1.666 | 0.123 | 1.174 | -0.246 | -0.001 | 0.009 | 0.009 | 0.325 | 0.51 | 0.63 | 0.46 | 0.38 | 0.46 | 0.34 |
| 3 | 178662309 | rs4955803 | 0.544 | 1.11 | 0.735 | -0.234 | 0.06 | 0.038 | 0.038 | 0 | 0.06 | 0.2 | 0.18 | 0.22 | 0.15 | 0.22 |
| 3 | 178665472 | rs6762374 | 0.361 | 1.073 | 0.769 | -0.339 | 0.064 | 0.003 | 0.003 | 0 | 0.06 | 0.2 | 0.18 | 0.14 | 0.14 | 0.22 |
| 3 | 178682459 | rs9862902 | 0.011 | 0.927 | 0.437 | -0.531 | 0.016 | 0.037 | 0.037 | 0 | 0.75 | 0.67 | 0.68 | 0.55 | 0.69 | 0.8 |
| 3 | 178687926 | rs4621371 | 2.02 | 0.396 | -0.609 | -0.98 | -0.027 | -0.07 | -0.07 | 0 | 0.32 | 0.03 | 0.31 | 0.34 | 0.31 | 0.37 |
| 3 | 178710585 | rs6787571 | 1.454 | 0.35 | -0.53 | -1.014 | -0.004 | 0.018 | 0.018 | 0 | 0.44 | 0.16 | 0.38 | 0.35 | 0.41 | 0.42 |
| 3 | 178713099 | rs13072825 | 0.897 | 0.409 | -0.612 | -1.844 | -0.038 | 0.098 | 0.098 | 0 | 0.19 | 0.03 | 0.12 | 0.03 | 0.18 | 0.14 |
| 3 | 178719335 | rs4417911 | -0.833 | 0.719 | -0.663 | -1.862 | -0.075 | 0.067 | 0.067 | 0.05 | 0.81 | 0.96 | 0.84 | 0.94 | 0.78 | 0.84 |
| 3 | 178738432 | rs6790867 | -0.744 | 0.517 | -0.549 | -1.903 | -0.059 | 0.072 | 0.072 | 0.033 | 0.8 | 0.93 | 0.84 | 0.94 | 0.77 | 0.82 |
| 3 | 178752778 | rs9842103 | -0.479 | 0.266 | 0.133 | -2.002 | -0.065 | -0.084 | -0.084 | 0 | 0.26 | 0.4 | 0.12 | 0.06 | 0.13 | 0.16 |
| 3 | 178755116 | rs6769215 | 0.284 | 0.428 | 0.229 | -2.123 | 0.02 | -0.052 | -0.052 | 0.016 | 0.69 | 0.6 | 0.61 | 0.8 | 0.62 | 0.62 |
| 3 | 178765726 | rs4401404 | 0.309 | 0.39 | 0.182 | -2.119 | 0.002 | -0.018 | -0.018 | 0 | 0.69 | 0.64 | 0.61 | 0.8 | 0.62 | 0.62 |
| 3 | 178782021 | rs4955793 | 1.806 | 0.28 | -0.077 | -2.181 | 0.005 | 0.002 | 0.002 | 0.136 | 0.12 | 0.12 | 0.06 | 0 | 0.1 | 0.01 |
| 3 | 178784789 | rs4416420 | 0.911 | 0.159 | -0.025 | -2.207 | 0.008 | 0.008 | 0.008 | 0.068 | 0.11 | 0.07 | 0.05 | 0 | 0.08 | 0.02 |
| 3 | 178785549 | rs6772028 | -0.792 | 0.191 | -0.068 | -2.17 | -0.079 | 0.069 | 0.069 | 0.004 | 0.8 | 0.97 | 0.84 | 0.94 | 0.77 | 0.82 |
| 3 | 178787131 | rs4955806 | -0.377 | 0.081 | 0.186 | -2.294 | 0.013 | -0.04 | -0.04 | 0 | 0.12 | 0.25 | 0.21 | 0.12 | 0.16 | 0.26 |
| 3 | 178794797 | rs6804195 | 1.717 | 0.142 | 0.334 | -2.348 | -0.022 | -0.057 | -0.057 | 0 | 0.12 | 0.19 | 0.07 | 0 | 0.08 | 0.01 |
| 3 | 178800879 | rs10936990 | 0.827 | -0.34 | -0.274 | -2.62 | -0.001 | -0.033 | -0.033 | 0 | 0.53 | 0.47 | 0.59 | 0.84 | 0.58 | 0.63 |
| 3 | 178833722 | rs1976765 | 0.302 | -0.272 | -0.128 | -2.542 | -0.022 | -0.015 | -0.015 | 2.874 | 0.11 | 0.22 | 0.14 | 0.1 | 0.06 | 0.08 |
| 3 | 178835186 | rs9814424 | -0.066 | -0.281 | -0.131 | -2.546 | -0.023 | -0.017 | -0.017 | 0 | 0.13 | 0.22 | 0.14 | 0.11 | 0.06 | 0.24 |
| 3 | 178844004 | rs9844482 | -0.836 | -0.201 | -0.248 | -2.584 | -0.032 | -0.098 | -0.098 | 0.32 | 0.3 | 0.52 | 0.28 | 0.11 | 0.24 | 0.27 |
| 3 | 178854075 | rs7611081 | -0.689 | -0.217 | -0.247 | -2.583 | -0.054 | -0.094 | -0.094 | 0 | 0.29 | 0.47 | 0.24 | 0.11 | 0.19 | 0.26 |
| 3 | 178871906 | rs11919383 | -0.683 | -0.207 | -0.241 | -2.579 | -0.054 | -0.094 | -0.094 | 0 | 0.29 | 0.48 | 0.25 | 0.11 | 0.19 | 0.26 |
| 3 | 178872355 | rs11706842 | 0.516 | -0.214 | -0.213 | -2.581 | -0.009 | -0.005 | -0.005 | 0.313 | 0.11 | 0.14 | 0.11 | 0.1 | 0.06 | 0.09 |
| 3 | 178881270 | rs7635669 | -0.449 | -0.181 | -0.216 | -2.563 | -0.053 | -0.101 | -0.101 | 0 | 0.28 | 0.48 | 0.25 | 0.09 | 0.19 | 0.1 |
| 3 | 178893029 | rs7651265 | -0.085 | -0.176 | -0.216 | -2.56 | -0.004 | -0.006 | -0.006 | 0 | 0.14 | 0.18 | 0.15 | 0.1 | 0.11 | 0.09 |
| 3 | 178897674 | rs6443624 | -0.754 | -0.157 | -0.236 | -2.561 | -0.032 | -0.098 | -0.098 | 0.007 | 0.3 | 0.52 | 0.28 | 0.11 | 0.24 | 0.27 |
| 3 | 178903264 | rs2677760 | -2.268 | -0.235 | -0.174 | -2.55 | -0.028 | 0.049 | 0.049 | 2.066 | 0.62 | 0.79 | 0.51 | 0.21 | 0.51 | 0.47 |
| 3 | 178907849 | rs9866361 | -0.454 | -0.368 | -0.47 | -2.54 | -0.063 | -0.091 | -0.091 | 0 | 0.28 | 0.48 | 0.2 | 0.11 | 0.18 | 0.23 |
| 3 | 178910350 | rs17592582 | 1.648 | -0.391 | -0.449 | -2.542 | 0.035 | 0.028 | 0.028 | 3.108 | 0.13 | 0.06 | 0.07 | 0.05 | 0.04 | 0.04 |
| 3 | 178911534 | rs2677770 | 1.077 | -0.346 | -0.342 | -2.538 | -0.036 | 0.037 | 0.037 | 0.171 | 0.18 | 0.07 | 0.15 | 0.06 | 0.23 | 0.16 |
| 3 | 178921158 | rs7623154 | -0.384 | -0.595 | -0.44 | -2.472 | -0.033 | -0.063 | -0.063 | 0 | 0.28 | 0.44 | 0.25 | 0.1 | 0.19 | 0.11 |
| 3 | 178923807 | rs2677764 | 2.03 | -0.459 | -0.516 | -2.572 | -0.034 | -0.016 | -0.016 | 0 | 0.06 | 0 | 0.08 | 0.06 | 0.09 | 0.03 |
| 3 | 178941323 | rs13320527 | 2.057 | -0.429 | -0.451 | -2.553 | -0.021 | -0.056 | -0.056 | 0 | 0.12 | 0.22 | 0.08 | 0 | 0.08 | 0.01 |
| 3 | 178947325 | rs3729692 | 1.252 | -0.359 | -0.598 | -2.521 | 0.004 | 0.005 | 0.005 | 0 | 0.06 | 0.04 | 0.05 | 0 | 0.06 | 0.01 |
| 3 | 178950297 | rs1607237 | -2.929 | -0.342 | -0.695 | -2.49 | -0.134 | -0.033 | -0.033 | 0 | 0.72 | 0.97 | 0.58 | 0.27 | 0.58 | 0.62 |
| 3 | 178952507 | rs9838117 | -1.162 | -0.68 | -0.422 | -2.365 | -0.048 | -0.067 | -0.067 | 12.813 | 0.42 | 0.54 | 0.33 | 0.15 | 0.24 | 0.3 |
| 3 | 178962897 | rs6799756 | 0.284 | -0.157 | -0.433 | -1.965 | -0.013 | -0.007 | -0.007 | 0 | 0.1 | 0.17 | 0.12 | 0.1 | 0.06 | 0.09 |
| 3 | 178968634 | rs7645550 | -0.838 | 0.053 | -0.549 | -2.126 | 0.008 | -0.069 | -0.069 | 0.089 | 0.34 | 0.42 | 0.31 | 0.12 | 0.37 | 0.32 |
| 3 | 178969107 | rs1183319 | -1.12 | 0.035 | -1.022 | -2.05 | 0.004 | 0.16 | 0.16 | 0 | 0.5 | 0.73 | 0.44 | 0.13 | 0.54 | 0.5 |
| 3 | 178970010 | rs1170673 | -0.031 | -0.217 | -0.44 | -1.621 | -0.007 | -0.005 | -0.005 | 0 | 0.77 | 0.76 | 0.83 | 0.86 | 0.87 | 0.86 |
| 3 | 178975408 | rs10804876 | 1.477 | 0.708 | -0.354 | -1.553 | -0.003 | 0.197 | 0.197 | 0 | 0.52 | 0.22 | 0.63 | 0.89 | 0.51 | 0.65 |
| 3 | 178976699 | rs7648422 | 1.256 | -0.139 | 0.107 | -1.451 | -0.024 | -0.053 | -0.053 | 0.046 | 0.12 | 0.2 | 0.06 | 0 | 0.07 | 0.01 |
| 3 | 178977074 | rs10936996 | -0.403 | 0.073 | 0.025 | -1.35 | -0.026 | -0.053 | -0.053 | 0 | 0.13 | 0.22 | 0.05 | 0.01 | 0.06 | 0.15 |
| 3 | 178982493 | rs4854984 | -1.123 | 0.482 | -0.282 | -1.449 | 0.03 | 0.072 | 0.072 | 0 | 0.4 | 0.74 | 0.33 | 0.06 | 0.32 | 0.28 |
| 3 | 178982869 | rs1468923 | -1.144 | 0.461 | -0.282 | -1.433 | 0.032 | 0.069 | 0.069 | 0 | 0.41 | 0.74 | 0.35 | 0.07 | 0.32 | 0.28 |
| 3 | 178985197 | rs10936997 | 1.699 | 0.816 | -0.653 | -0.866 | -0.073 | 0.005 | 0.005 | 0 | 0.44 | 0.09 | 0.56 | 0.58 | 0.47 | 0.39 |
| 3 | 178988564 | rs6781173 | -0.678 | 0.473 | 0.17 | -0.91 | 0.04 | 0.012 | 0.012 | 1.044 | 0.18 | 0.53 | 0.17 | 0.16 | 0.18 | 0.22 |
| 3 | 178988701 | rs1861803 | 1.668 | 0.491 | -0.164 | -0.88 | -0.075 | 0.01 | 0.01 | 0 | 0.43 | 0.09 | 0.55 | 0.59 | 0.45 | 0.4 |
| 3 | 178991951 | rs13076233 | -0.089 | 0.928 | 0.365 | -0.559 | 0.003 | 0.004 | 0.004 | 0 | 0.14 | 0.09 | 0.13 | 0.09 | 0.13 | 0.23 |

| | | | | | | | | | | | | | | | | |
|---|---|---|---|---|---|---|---|---|---|---|---|---|---|---|---|---|
| 3 | 178995657 | rs9813206 | 0.57 | 1.705 | 0.78 | -0.157 | 0.058 | 0.052 | 0.052 | 0 | 0.93 | 0.87 | 0.86 | 0.82 | 0.8 | 0.86 |
| 3 | 178996626 | rs9856476 | 0.378 | 1.677 | 0.95 | -0.207 | 0.012 | -0.027 | -0.027 | 0 | 0.06 | 0.15 | 0.04 | 0 | 0.05 | 0.01 |
| 3 | 178997778 | rs4854991 | 0.84 | 1.768 | 0.504 | -0.027 | 0.09 | 0.082 | 0.082 | 0 | 0.92 | 0.81 | 0.86 | 0.82 | 0.81 | 0.86 |
| 3 | 179005976 | rs6443636 | 0.878 | 1.326 | 0.151 | -0.303 | -0.006 | 0.024 | 0.024 | 0 | 0.53 | 0.3 | 0.6 | 0.58 | 0.5 | 0.51 |
| 3 | 179010761 | rs6807729 | 0.808 | 1.32 | 0.204 | -0.313 | -0.006 | 0.018 | 0.018 | 0 | 0.53 | 0.37 | 0.6 | 0.58 | 0.5 | 0.49 |
| 3 | 179013860 | rs7628077 | -0.454 | 1.57 | 0.581 | -0.335 | 0.002 | -0.008 | -0.008 | 0.12 | 0.38 | 0.35 | 0.29 | 0.19 | 0.28 | 0.37 |
| 3 | 179015775 | rs6776229 | 0.642 | 1.626 | 0.564 | -0.297 | 0.146 | 0.142 | 0.142 | 0 | 0.92 | 0.74 | 0.9 | 0.77 | 0.79 | 0.86 |
| 3 | 179017949 | rs9834223 | -0.563 | 0.71 | 0.184 | -0.672 | -0.003 | -0.018 | -0.018 | 0.009 | 0.38 | 0.36 | 0.3 | 0.19 | 0.28 | 0.37 |
| 3 | 179022115 | rs7636818 | -0.153 | 1.933 | 0.354 | -0.627 | 0.1 | -0.064 | -0.064 | 0 | 0.89 | 0.81 | 0.84 | 1 | 0.78 | 0.93 |
| 3 | 179022528 | rs16848314 | 1.241 | 1.141 | -0.245 | -0.251 | -0.017 | -0.063 | -0.063 | 0 | 0.28 | 0.08 | 0.38 | 0.4 | 0.34 | 0.33 |
| 3 | 179040177 | rs2032700 | 1.134 | 1.135 | -0.135 | -0.513 | -0.06 | 0.002 | 0.002 | 2.11 | 0.11 | 0.02 | 0.09 | 0.07 | 0.17 | 0.06 |
| 3 | 179041975 | rs7610174 | 1.57 | 1.115 | -0.305 | -0.388 | -0.053 | -0.111 | -0.111 | 0 | 0.27 | 0.03 | 0.4 | 0.37 | 0.34 | 0.31 |
| 3 | 179045493 | rs6786409 | -0.716 | 0.005 | -0.197 | -0.686 | -0.006 | -0.011 | -0.011 | 0.344 | 0.68 | 0.79 | 0.79 | 0.67 | 0.69 | 0.73 |
| 3 | 179050721 | rs12491673 | 2.02 | -0.256 | -0.588 | -0.846 | -0.001 | 0.038 | 0.038 | 0 | 0.08 | 0.01 | 0.12 | 0 | 0.1 | 0.11 |
| 3 | 179056135 | rs10460887 | 1.606 | -0.415 | -0.36 | -0.891 | 0.011 | 0.005 | 0.005 | 0.001 | 0.23 | 0.14 | 0.23 | 0.22 | 0.18 | 0.21 |
| 3 | 179058502 | rs6805876 | -0.085 | -0.359 | -0.329 | -0.878 | 0 | 0.003 | 0.003 | 0 | 0.55 | 0.68 | 0.69 | 0.59 | 0.52 | 0.52 |
| 3 | 179070168 | rs2287312 | 1.498 | -0.458 | -0.265 | -0.851 | 0.007 | 0.004 | 0.004 | 0.008 | 0.22 | 0.15 | 0.23 | 0.21 | 0.18 | 0.22 |
| 3 | 179074615 | rs9822116 | -0.604 | -0.475 | -0.183 | -0.966 | 0.008 | 0.127 | 0.127 | 0 | 0.49 | 0.8 | 0.5 | 0.29 | 0.46 | 0.53 |
| 3 | 179076879 | rs6779649 | 0.471 | -0.502 | -0.068 | -0.973 | 0.045 | 0.041 | 0.041 | 1.609 | 0.06 | 0.53 | 0.06 | 0 | 0.01 | 0 |
| 3 | 179084046 | rs7355960 | 0.471 | -0.502 | -0.068 | -0.973 | 0.045 | 0.041 | 0.041 | 1.609 | 0.06 | 0.53 | 0.06 | 0 | 0.01 | 0 |
| 3 | 179085791 | rs7356002 | 0.471 | -0.502 | -0.068 | -0.973 | 0.045 | 0.041 | 0.041 | 1.609 | 0.06 | 0.53 | 0.06 | 0 | 0.01 | 0 |
| 3 | 179090772 | rs2111534 | -0.634 | -0.517 | -0.131 | -0.985 | 0.01 | 0.13 | 0.13 | 0 | 0.49 | 0.8 | 0.5 | 0.29 | 0.46 | 0.53 |
| 3 | 179106847 | rs10049286 | 0.468 | -0.522 | -0.119 | -0.985 | 0.002 | 0.002 | 0.002 | 0.741 | 0.3 | 0.7 | 0.29 | 0.22 | 0.19 | 0.36 |
| 3 | 179108917 | rs13083369 | 0.742 | -0.472 | -0.134 | -0.975 | -0.034 | 0.042 | 0.042 | 0.087 | 0.19 | 0.1 | 0.21 | 0.08 | 0.26 | 0.17 |
| 3 | 179115380 | rs10937003 | 2.527 | -0.451 | 0.084 | -0.727 | -0.031 | -0.089 | -0.089 | 0.068 | 0.2 | 0.05 | 0.27 | 0.36 | 0.23 | 0.2 |
| 3 | 179118397 | rs11714353 | 2.451 | -0.526 | -0.164 | -0.651 | -0.025 | -0.079 | -0.079 | 0 | 0.19 | 0.05 | 0.27 | 0.37 | 0.23 | 0.2 |
| 3 | 179118998 | rs3774225 | 1.522 | -0.523 | -0.107 | -0.647 | 0.011 | 0.004 | 0.004 | 0 | 0.22 | 0.13 | 0.23 | 0.23 | 0.18 | 0.22 |
| 3 | 179124814 | rs12634184 | 2.619 | -0.453 | -0.141 | -0.586 | -0.043 | -0.102 | -0.102 | 0.237 | 0.19 | 0.03 | 0.26 | 0.36 | 0.23 | 0.2 |
| 3 | 179137273 | rs1362650 | 0.475 | -0.389 | -0.015 | -0.657 | 0.003 | 0 | 0 | 0 | 0.41 | 0.45 | 0.51 | 0.44 | 0.5 | 0.5 |
| 3 | 179143941 | rs1078749 | 1.852 | -0.945 | -0.437 | -0.919 | 0.059 | 0.061 | 0.061 | 3.129 | 0.13 | 0.02 | 0.05 | 0 | 0.04 | 0.01 |
| 3 | 179146844 | rs9844549 | -0.099 | -0.729 | -0.17 | -0.895 | 0.014 | -0.092 | -0.092 | 0.965 | 0.26 | 0.5 | 0.27 | 0.08 | 0.27 | 0.22 |
| 3 | 179147171 | rs2024497 | -0.501 | -0.542 | -0.931 | -0.802 | -0.01 | -0.02 | -0.02 | 0.858 | 0.68 | 0.82 | 0.77 | 0.68 | 0.68 | 0.73 |
| 3 | 179156647 | rs12488540 | 0.587 | -0.228 | -0.824 | -0.708 | -0.043 | 0.054 | 0.054 | 0 | 0.19 | 0.08 | 0.21 | 0.08 | 0.26 | 0.15 |
| 3 | 179162316 | rs2287208 | 0.484 | -0.634 | -0.643 | -0.68 | -0.003 | 0.007 | 0.007 | 0 | 0.47 | 0.63 | 0.53 | 0.44 | 0.52 | 0.38 |
| 3 | 179172979 | rs7612445 | -0.391 | -1.377 | -0.818 | -0.854 | -0.008 | -0.009 | -0.009 | 0 | 0.35 | 0.39 | 0.26 | 0.17 | 0.17 | 0.28 |
| 3 | 179174789 | rs7618251 | 0.77 | -1.182 | -0.817 | -0.888 | -0.04 | -0.059 | -0.059 | 0 | 0.15 | 0.18 | 0.08 | 0 | 0.04 | 0.02 |
| 3 | 179176081 | rs7621827 | 1.67 | -0.578 | -1.384 | -0.523 | 0.006 | -0.098 | -0.098 | 0 | 0.26 | 0.06 | 0.26 | 0.41 | 0.25 | 0.25 |
| 3 | 179178245 | rs9850338 | -0.915 | 0.882 | -0.876 | -0.63 | -0.007 | -0.016 | -0.016 | 0.034 | 0.37 | 0.46 | 0.24 | 0.26 | 0.27 | 0.28 |
| 3 | 179180132 | rs13074744 | 2.401 | 1.396 | -0.997 | -0.654 | 0.011 | -0.05 | -0.05 | 0.696 | 0.34 | 0.07 | 0.31 | 0.42 | 0.31 | 0.4 |
| 3 | 179180153 | rs4855077 | -0.194 | 2.389 | 0.394 | -0.22 | -0.045 | 0.045 | 0.045 | 0 | 0.9 | 0.98 | 0.82 | 1 | 0.77 | 0.85 |
| 3 | 179180937 | rs7428403 | -1.239 | 2.133 | 0.481 | -0.18 | -0.009 | -0.02 | -0.02 | 0 | 0.37 | 0.47 | 0.24 | 0.26 | 0.27 | 0.28 |
| 3 | 179189543 | rs7638443 | 0.184 | 2.301 | 0.446 | -0.255 | -0.028 | 0.051 | 0.051 | 0 | 0.75 | 0.84 | 0.72 | 0.98 | 0.69 | 0.75 |
| 3 | 179191682 | rs7622921 | 0.816 | 2.349 | 0.523 | -0.208 | 0.063 | -0.009 | -0.009 | 0.307 | 0.06 | 0.2 | 0.07 | 0.06 | 0.11 | 0.06 |
| 3 | 179191821 | rs7623033 | 0.211 | 2.244 | 0.641 | -0.2 | 0.001 | -0.061 | -0.061 | 0.376 | 0.75 | 0.72 | 0.71 | 0.98 | 0.68 | 0.75 |
| 3 | 179207592 | rs6807740 | -0.325 | 1.655 | 0.73 | 0.089 | -0.045 | 0.035 | 0.035 | 0.249 | 0.15 | 0.25 | 0.08 | 0.21 | 0.05 | 0.12 |
| 3 | 179221407 | rs4855081 | 1.907 | 1.838 | 0.284 | 0.182 | 0.022 | 0.014 | 0.014 | 0 | 0.46 | 0.3 | 0.34 | 0.56 | 0.4 | 0.51 |
| 3 | 179228364 | rs2339802 | 1.396 | 2.134 | 0.504 | 0.17 | -0.001 | 0.018 | 0.018 | 0.622 | 0.48 | 0.3 | 0.53 | 0.41 | 0.5 | 0.44 |
| 3 | 179247185 | rs10513760 | 0.022 | 1.515 | 0.373 | -0.47 | 0.006 | -0.007 | -0.007 | 0.492 | 0.15 | 0.13 | 0.06 | 0.02 | 0.06 | 0.09 |
| 3 | 179247616 | rs713177 | -0.138 | 1.991 | 0.556 | -0.136 | 0.045 | 0.066 | 0.066 | 1.404 | 0.11 | 0.22 | 0.33 | 0.29 | 0.18 | 0.12 |
| 3 | 179250724 | rs1345445 | 1.156 | 1.514 | 0.804 | -0.636 | 0.002 | -0.058 | -0.058 | 0 | 0.74 | 0.72 | 0.79 | 0.99 | 0.64 | 0.71 |
| 3 | 179252263 | rs9832757 | -1.289 | 0.758 | 0.925 | -0.337 | 0.002 | -0.011 | -0.011 | 0 | 0.16 | 0.16 | 0.11 | 0.01 | 0.16 | 0.03 |
| 3 | 179252544 | rs6779664 | 0.54 | 0.814 | 1.231 | 0.057 | -0.041 | 0.05 | 0.05 | 0 | 0.64 | 0.77 | 0.57 | 0.73 | 0.51 | 0.63 |
| 3 | 179256722 | rs2077675 | 0.543 | 0.374 | 1.893 | -0.181 | 0.008 | -0.06 | -0.06 | 0.152 | 0.61 | 0.48 | 0.64 | 0.87 | 0.62 | 0.71 |
| 3 | 179272481 | rs7630511 | -0.192 | 0.305 | 1.546 | -0.083 | -0.058 | 0.086 | 0.086 | 0.181 | 0.71 | 0.89 | 0.73 | 0.91 | 0.65 | 0.84 |
| 3 | 179278768 | rs12492220 | -0.224 | 0.304 | 1.54 | -0.083 | -0.061 | 0.087 | 0.087 | 0.116 | 0.71 | 0.89 | 0.73 | 0.91 | 0.65 | 0.84 |
| 3 | 179288279 | rs10513761 | 0.055 | 0.254 | 1.461 | 0.149 | 0 | -0.003 | -0.003 | 0 | 0.06 | 0.06 | 0.15 | 0.1 | 0.08 | 0.18 |
| 3 | 179288496 | rs4855090 | -0.119 | 0.254 | 1.483 | 0.149 | 0.002 | 0.039 | 0.039 | 0 | 0.06 | 0.1 | 0.15 | 0.1 | 0.08 | 0.18 |
| 3 | 179289095 | rs9861172 | -0.19 | 0.22 | 1.718 | 0.121 | 0 | -0.018 | -0.018 | 0 | 0.64 | 0.63 | 0.68 | 0.89 | 0.62 | 0.71 |
| 3 | 179289429 | rs6790272 | 0.824 | 0.204 | 1.702 | 0.291 | -0.064 | -0.005 | -0.005 | 0 | 0.52 | 0.09 | 0.49 | 0.62 | 0.53 | 0.47 |
| 3 | 179290093 | rs4855092 | 0.025 | 0.302 | 1.702 | 0.307 | 0.004 | 0.043 | 0.043 | 0 | 0.06 | 0.1 | 0.15 | 0.1 | 0.08 | 0.18 |
| 3 | 179306775 | rs10513762 | 0.018 | 0.306 | 1.635 | 0.315 | 0.004 | 0.043 | 0.043 | 0 | 0.06 | 0.11 | 0.15 | 0.1 | 0.08 | 0.18 |
| 3 | 179322385 | rs2339844 | 0.002 | 0.354 | 1.574 | 0.285 | 0.002 | 0.039 | 0.039 | 0 | 0.06 | 0.1 | 0.15 | 0.1 | 0.08 | 0.19 |
| 3 | 179327082 | rs9829395 | -0.156 | 0.355 | 1.579 | 0.278 | 0.001 | -0.024 | -0.024 | 0 | 0.64 | 0.63 | 0.68 | 0.89 | 0.62 | 0.71 |
| 3 | 179329015 | rs4147787 | -0.019 | 0.401 | 1.344 | 0.309 | 0.002 | 0.039 | 0.039 | 0 | 0.06 | 0.11 | 0.15 | 0.1 | 0.08 | 0.19 |
| 3 | 179331178 | rs4147788 | 0.483 | 0.398 | 1.361 | 0.299 | -0.001 | -0.003 | -0.003 | 0.239 | 0.57 | 0.45 | 0.53 | 0.79 | 0.54 | 0.53 |
| 3 | 179347189 | rs10937007 | 0.377 | 0.863 | -0.622 | 0.017 | -0.016 | 0.035 | 0.035 | 0.284 | 0.59 | 0.33 | 0.65 | 0.74 | 0.6 | 0.65 |
| 3 | 179363840 | rs12494466 | -1.301 | 0.554 | -0.308 | 0.779 | -0.01 | 0.055 | 0.055 | 0 | 0.07 | 0.18 | 0.03 | 0.17 | 0.03 | 0.05 |
| 3 | 179376292 | rs9874079 | -0.215 | 0.696 | -0.654 | -0.499 | 0.003 | 0.015 | 0.015 | 0 | 0.49 | 0.41 | 0.37 | 0.13 | 0.46 | 0.34 |
| 3 | 179382523 | rs6775913 | 0.318 | -0.118 | -1.617 | -0.732 | 0.03 | 0.032 | 0.032 | 0.104 | 0.73 | 0.92 | 0.85 | 0.85 | 0.75 | 0.81 |

| | | | | | | | | | | | | | | | |
|---|---|---|---|---|---|---|---|---|---|---|---|---|---|---|---|
| 3 | 179383908 | rs4855102 | 0.607 | 0.39 | -2.319 | -1.047 | 0.006 | 0.054 | 0.054 | 0.785 | 0.46 | 0.79 | 0.59 | 0.64 | 0.4 | 0.58 |
| 3 | 179385000 | rs6443664 | 0.042 | -0.519 | -1.461 | -1.357 | 0.013 | 0.015 | 0.015 | 0 | 0.48 | 0.57 | 0.56 | 0.31 | 0.63 | 0.48 |
| 3 | 179386744 | rs10513763 | 0.289 | -1.124 | -1.417 | -0.935 | -0.039 | -0.014 | -0.014 | 0 | 0.18 | 0.22 | 0.05 | 0.14 | 0.06 | 0.13 |
| 3 | 179389656 | rs12633396 | 0.719 | -0.85 | -1.581 | -0.84 | 0.009 | -0.108 | -0.108 | 0 | 0.23 | 0.05 | 0.29 | 0.39 | 0.23 | 0.27 |
| 3 | 179391320 | rs4854945 | 0.514 | -0.819 | -1.287 | -1.31 | 0.001 | -0.046 | -0.046 | 0 | 0.38 | 0.58 | 0.31 | 0.08 | 0.41 | 0.31 |
| 3 | 179394123 | rs4855104 | 1.41 | -0.499 | -1.051 | -0.991 | -0.009 | -0.008 | -0.008 | 0.063 | 0.14 | 0.12 | 0.26 | 0.26 | 0.21 | 0.2 |
| 3 | 179399570 | rs13080037 | 0.573 | -0.249 | -0.843 | -1.273 | -0.005 | 0.064 | 0.064 | 0 | 0.47 | 0.3 | 0.62 | 0.66 | 0.55 | 0.47 |
| 3 | 179400345 | rs7629853 | 1.397 | 0.761 | -0.283 | -0.324 | -0.004 | 0.003 | 0.003 | 0 | 0.16 | 0.15 | 0.15 | 0.08 | 0.2 | 0.17 |
| 3 | 179417976 | rs4854948 | -1.046 | 0.725 | 0.443 | -0.222 | 0.023 | 0.142 | 0.142 | 0 | 0.92 | 0.77 | 0.94 | 0.77 | 0.94 | 0.87 |
| 3 | 179420390 | rs2300772 | -0.164 | 0.493 | 0.39 | -0.447 | -0.002 | -0.062 | -0.062 | 0 | 0.33 | 0.56 | 0.26 | 0.14 | 0.32 | 0.29 |
| 3 | 179420666 | rs2300769 | 1.102 | 0.634 | 0.169 | -0.439 | -0.059 | -0.119 | -0.119 | 1.234 | 0.43 | 0.05 | 0.38 | 0.38 | 0.44 | 0.39 |
| 3 | 179421547 | rs2268941 | 1.608 | 0.724 | 0.326 | -0.439 | -0.013 | 0.026 | 0.026 | 0 | 0.19 | 0.14 | 0.15 | 0.09 | 0.23 | 0.2 |
| 3 | 179424410 | rs17788187 | 2.09 | 0.697 | 0.433 | -0.408 | -0.005 | -0.005 | -0.005 | 0 | 0.14 | 0.12 | 0.28 | 0.26 | 0.18 | 0.19 |
| 3 | 179425147 | rs2284887 | 0.128 | 0.603 | 0.447 | -0.336 | 0.007 | 0.001 | 0.001 | 0 | 0.71 | 0.58 | 0.78 | 0.69 | 0.7 | 0.66 |
| 3 | 179430941 | rs800368 | -1.221 | 0.698 | 0.301 | -0.263 | 0.026 | 0.125 | 0.125 | 0 | 0.93 | 0.81 | 0.94 | 0.79 | 0.93 | 0.86 |
| 3 | 179431962 | rs16830755 | 0.859 | 0.486 | 0.173 | -0.356 | -0.023 | -0.039 | -0.039 | 0 | 0.46 | 0.14 | 0.43 | 0.42 | 0.5 | 0.46 |
| 3 | 179436298 | rs4854951 | 0.195 | 0.384 | 0.236 | -0.728 | -0.018 | -0.011 | -0.011 | 0 | 0.27 | 0.45 | 0.36 | 0.3 | 0.2 | 0.2 |
| 3 | 179439466 | rs2119898 | 1.008 | 0.638 | -0.605 | -0.578 | -0.095 | -0.123 | -0.123 | 0.423 | 0.43 | 0.04 | 0.4 | 0.45 | 0.47 | 0.42 |
| 3 | 179441421 | rs4346543 | 0.973 | 0.587 | -0.551 | -0.574 | -0.059 | -0.074 | -0.074 | 0 | 0.43 | 0.08 | 0.41 | 0.45 | 0.48 | 0.42 |
| 3 | 179443431 | rs2111391 | -1.171 | 0.741 | -0.376 | -0.579 | -0.013 | 0.041 | 0.041 | 0 | 0.77 | 0.83 | 0.85 | 0.93 | 0.74 | 0.76 |
| 3 | 179448383 | rs3213761 | -0.054 | 0.607 | 0.033 | -0.534 | 0.005 | -0.015 | -0.015 | 0 | 0.7 | 0.64 | 0.8 | 0.76 | 0.68 | 0.62 |
| 3 | 179451466 | rs9824503 | 1.166 | 0.527 | 0.14 | -0.489 | 0.019 | 0.158 | 0.158 | 0 | 0.06 | 0.2 | 0.04 | 0.17 | 0.03 | 0.14 |
| 3 | 179452888 | rs1025864 | -0.912 | 0.585 | -0.071 | -0.568 | -0.027 | 0.075 | 0.075 | 0 | 0.75 | 0.83 | 0.84 | 0.93 | 0.71 | 0.74 |
| 3 | 179455611 | rs2268933 | 0.095 | 0.424 | 0.111 | -0.607 | -0.001 | -0.019 | -0.019 | 0 | 0.63 | 0.53 | 0.72 | 0.85 | 0.64 | 0.68 |
| 3 | 179456888 | rs2261441 | 1.324 | -0.2 | -0.282 | -0.383 | -0.077 | -0.165 | -0.165 | 0.496 | 0.4 | 0.04 | 0.37 | 0.43 | 0.42 | 0.35 |
| 3 | 179460825 | rs2268929 | -0.229 | -0.216 | 0.071 | -0.46 | -0.002 | 0.002 | 0.002 | 2.37 | 0.61 | 0.32 | 0.48 | 0.65 | 0.47 | 0.5 |
| 3 | 179465758 | rs9862823 | 0.058 | 0.069 | -0.083 | -0.232 | 0.009 | 0.027 | 0.027 | 0 | 0.51 | 0.52 | 0.45 | 0.63 | 0.44 | 0.47 |
| 3 | 179469995 | rs2239639 | 1.271 | 0.406 | -1.207 | -0.153 | -0.019 | -0.182 | -0.182 | 0.152 | 0.36 | 0.02 | 0.3 | 0.44 | 0.33 | 0.31 |
| 3 | 179472674 | rs2287240 | 0.195 | 0.456 | -0.762 | -0.248 | -0.025 | 0.122 | 0.122 | 0.041 | 0.52 | 0.18 | 0.46 | 0.64 | 0.5 | 0.54 |
| 3 | 179490367 | rs2284884 | 0.108 | 1.146 | -0.102 | -0.135 | -0.043 | -0.038 | -0.038 | 0.03 | 0.15 | 0.08 | 0.16 | 0.22 | 0.19 | 0.22 |
| 3 | 179498965 | rs759596 | 0.264 | 0.738 | 0.675 | -0.754 | -0.017 | -0.108 | -0.108 | 0 | 0.17 | 0.49 | 0.17 | 0.05 | 0.13 | 0.07 |
| 3 | 179507761 | rs9813117 | -1.213 | 0.412 | 0.511 | -0.869 | -0.004 | 0.052 | 0.052 | 1.169 | 0.37 | 0.69 | 0.34 | 0.16 | 0.32 | 0.28 |
| 3 | 179514386 | rs9869557 | -1.38 | 0.925 | -0.087 | -0.98 | -0.031 | 0.011 | 0.011 | 0 | 0.86 | 0.94 | 0.83 | 0.92 | 0.78 | 0.77 |
| 3 | 179517234 | rs13091139 | -0.15 | 0.879 | 0.534 | -0.976 | 0.066 | -0.001 | -0.001 | 0 | 0.22 | 0.39 | 0.23 | 0.19 | 0.26 | 0.28 |
| 3 | 179517671 | rs3774227 | -0.623 | 0.802 | 0.412 | -1.008 | 0.012 | -0.031 | -0.031 | 0 | 0.31 | 0.44 | 0.3 | 0.23 | 0.28 | 0.29 |
| 3 | 179521935 | rs1442495 | 0.835 | 0.897 | 0.411 | -0.96 | 0.002 | 0 | 0 | 0 | 0.16 | 0.16 | 0.17 | 0.19 | 0.18 | 0.23 |
| 3 | 179524975 | rs4855113 | 1.558 | 0.767 | 0.218 | -1.038 | 0.02 | 0.066 | 0.066 | 0 | 0.49 | 0.32 | 0.5 | 0.69 | 0.5 | 0.48 |
| 3 | 179530595 | rs6809820 | -0.916 | 0.566 | 0.02 | -1.03 | -0.015 | 0.037 | 0.037 | 0 | 0.8 | 0.91 | 0.82 | 0.92 | 0.78 | 0.77 |
| 3 | 179531062 | rs9845643 | 0.316 | 0.821 | 0.264 | -0.853 | 0.077 | 0.007 | 0.007 | 0 | 0.18 | 0.39 | 0.21 | 0.16 | 0.19 | 0.21 |
| 3 | 179545261 | rs10937011 | 0.414 | 0.397 | -0.118 | -0.803 | -0.003 | 0.02 | 0.02 | 0.101 | 0.52 | 0.44 | 0.54 | 0.72 | 0.58 | 0.54 |
| 3 | 179561502 | rs7643653 | 0.937 | 0.703 | -0.647 | -1.171 | -0.004 | 0.002 | 0.002 | 1.249 | 0.16 | 0.16 | 0.23 | 0.05 | 0.23 | 0.22 |
| 3 | 179567416 | rs2056596 | 0.472 | 0.263 | -0.779 | -1.406 | 0.013 | 0.042 | 0.042 | 0 | 0.41 | 0.73 | 0.45 | 0.75 | 0.41 | 0.45 |
| 3 | 179568847 | rs7651639 | 0.475 | 0.23 | -0.519 | -1.38 | 0.001 | 0.002 | 0.002 | 0 | 0.4 | 0.64 | 0.45 | 0.72 | 0.41 | 0.45 |
| 3 | 179568940 | rs7640024 | -0.247 | 0.063 | -0.693 | -1.201 | -0.034 | 0.11 | 0.11 | 0.136 | 0.57 | 0.89 | 0.69 | 0.79 | 0.63 | 0.68 |
| 3 | 179572539 | rs6808678 | 0.01 | -0.029 | -0.432 | -1.095 | -0.006 | 0.072 | 0.072 | 0 | 0.52 | 0.68 | 0.62 | 0.76 | 0.58 | 0.55 |
| 3 | 179579536 | rs6443674 | 0.039 | -0.312 | -0.061 | -0.936 | 0.008 | 0.022 | 0.022 | 0 | 0.53 | 0.58 | 0.54 | 0.73 | 0.46 | 0.62 |
| 3 | 179592240 | rs2339914 | 0.136 | -0.387 | 0.266 | -1.208 | 0.018 | 0.056 | 0.056 | 0 | 0.51 | 0.4 | 0.54 | 0.79 | 0.46 | 0.61 |
| 3 | 179598716 | rs3774245 | 1.49 | -0.261 | -0.083 | -1.28 | -0.012 | -0.025 | -0.025 | 2.107 | 0.08 | 0 | 0.04 | 0.05 | 0.06 | 0.13 |
| 3 | 179606442 | rs9290684 | -1.253 | -0.005 | -0.013 | -1.344 | -0.003 | 0 | 0 | 0 | 0.92 | 0.94 | 0.82 | 0.94 | 0.9 | 0.97 |
| 3 | 179614457 | rs3821796 | 1.023 | -0.542 | -0.317 | -1.385 | 0.017 | 0.012 | 0.012 | 4.412 | 0.26 | 0.22 | 0.2 | 0.19 | 0.15 | 0.34 |
| 3 | 179627093 | rs9822592 | 1.007 | -0.314 | -0.046 | -1.529 | 0.02 | -0.054 | -0.054 | 0 | 0.11 | 0.15 | 0.07 | 0 | 0.13 | 0.09 |
| 3 | 179629514 | rs7636571 | 0.126 | -0.141 | -0.026 | -1.448 | -0.005 | 0.029 | 0.029 | 0.013 | 0.55 | 0.36 | 0.42 | 0.31 | 0.47 | 0.61 |
| 3 | 179632209 | rs1404271 | 1.171 | -0.153 | 0.043 | -1.369 | -0.004 | 0.02 | 0.02 | 0.059 | 0.3 | 0.24 | 0.22 | 0.13 | 0.32 | 0.29 |
| 3 | 179635808 | rs11708154 | 1.19 | -0.128 | -0.035 | -1.362 | -0.024 | 0.054 | 0.054 | 0.063 | 0.29 | 0.13 | 0.21 | 0.13 | 0.32 | 0.29 |
| 3 | 179637810 | rs13083997 | 1.046 | -0.186 | -0.184 | -1.319 | 0.04 | 0.041 | 0.041 | 5.837 | 0.24 | 0.12 | 0.2 | 0.16 | 0.15 | 0.22 |
| 3 | 179638781 | rs7650575 | -1.642 | -0.089 | -0.052 | -1.281 | 0.003 | -0.004 | -0.004 | 0 | 0.93 | 0.89 | 0.83 | 0.95 | 0.91 | 0.98 |
| 3 | 179641429 | rs12632198 | 0.319 | 0.003 | 0.068 | -1.208 | 0.008 | 0.006 | 0.006 | 0.274 | 0.37 | 0.54 | 0.41 | 0.58 | 0.43 | 0.35 |
| 3 | 179642067 | rs1524511 | -0.715 | -0.021 | 0.051 | -1.263 | 0.04 | 0.037 | 0.037 | 5.764 | 0.68 | 0.87 | 0.64 | 0.76 | 0.76 | 0.65 |
| 3 | 179642221 | rs9836757 | 0.997 | -0.011 | 0.057 | -1.261 | 0 | -0.007 | -0.007 | 0 | 0.31 | 0.33 | 0.22 | 0.17 | 0.33 | 0.3 |
| 3 | 179646264 | rs6794543 | -0.761 | 0.004 | -0.404 | -1.184 | 0.036 | 0.057 | 0.057 | 6.4 | 0.68 | 0.87 | 0.64 | 0.8 | 0.76 | 0.75 |
| 3 | 179660640 | rs7652323 | 1.162 | -0.006 | -0.323 | -1.037 | 0 | 0.021 | 0.021 | 0.159 | 0.25 | 0.31 | 0.35 | 0.59 | 0.29 | 0.26 |
| 3 | 179661318 | rs7630877 | 0.621 | 0.234 | -0.779 | -1.309 | -0.001 | 0.001 | 0.001 | 0 | 0.31 | 0.32 | 0.22 | 0.17 | 0.32 | 0.3 |
| 3 | 179664065 | rs11917103 | -0.044 | 0.226 | -0.8 | -1.367 | 0.042 | -0.084 | -0.084 | 0 | 0.15 | 0.3 | 0.1 | 0 | 0.18 | 0.1 |
| 3 | 179664102 | rs9842133 | 0.509 | -0.128 | -1.119 | -1.486 | -0.005 | -0.01 | -0.01 | 0 | 0.34 | 0.57 | 0.38 | 0.67 | 0.33 | 0.4 |
| 3 | 179666018 | rs7623941 | -0.093 | -0.128 | -1.666 | -1.335 | -0.031 | 0.092 | 0.092 | 1.44 | 0.45 | 0.81 | 0.48 | 0.68 | 0.5 | 0.48 |
| 3 | 179670217 | rs7642327 | -0.675 | 0.246 | -1.374 | -1.529 | -0.027 | 0.074 | 0.074 | 0.089 | 0.71 | 0.88 | 0.67 | 0.89 | 0.65 | 0.84 |
| 3 | 179671825 | rs6804380 | -0.671 | 0.26 | -1.369 | -1.523 | -0.027 | 0.074 | 0.074 | 0.126 | 0.71 | 0.87 | 0.67 | 0.89 | 0.65 | 0.85 |
| 3 | 179672337 | rs9856937 | -0.726 | 0.285 | -1.367 | -1.524 | -0.035 | 0.087 | 0.087 | 0.077 | 0.71 | 0.9 | 0.67 | 0.89 | 0.65 | 0.84 |
| 3 | 179672836 | rs1524512 | -0.675 | 0.289 | -1.366 | -1.519 | -0.027 | 0.074 | 0.074 | 0.126 | 0.71 | 0.87 | 0.67 | 0.89 | 0.65 | 0.84 |

| | | | | | | | | | | | | | | | | |
|---|---|---|---|---|---|---|---|---|---|---|---|---|---|---|---|---|
| 3 | 179675554 | rs9825224 | 0.101 | 0.136 | -1.199 | -1.651 | -0.006 | 0.01 | 0.01 | 0.215 | 0.68 | 0.69 | 0.77 | 0.94 | 0.69 | 0.82 |
| 3 | 179678135 | rs6782786 | 0.035 | 0.158 | -0.995 | -1.644 | -0.013 | 0.029 | 0.029 | 0 | 0.71 | 0.83 | 0.87 | 0.95 | 0.71 | 0.82 |
| 3 | 179678562 | rs4855129 | 0.265 | 0.057 | -0.857 | -1.725 | 0.03 | 0.145 | 0.145 | 0 | 0.48 | 0.3 | 0.66 | 0.88 | 0.44 | 0.68 |
| 3 | 179684793 | rs1534024 | 0.017 | 0.218 | -0.976 | -1.338 | -0.045 | 0.092 | 0.092 | 0 | 0.74 | 0.92 | 0.88 | 0.95 | 0.71 | 0.83 |
| 3 | 179688604 | rs2031043 | -0.64 | -0.358 | -0.196 | -0.254 | 0.011 | 0.074 | 0.074 | 0 | 0.21 | 0.3 | 0.29 | 0.53 | 0.2 | 0.25 |
| 3 | 179689264 | rs3774256 | -0.55 | -0.353 | -0.066 | -0.251 | 0.02 | 0.117 | 0.117 | 0 | 0.2 | 0.33 | 0.29 | 0.53 | 0.2 | 0.25 |
| 3 | 179691697 | rs12488363 | -0.527 | -0.584 | -0.364 | -0.543 | 0.006 | -0.004 | -0.004 | 1.468 | 0.38 | 0.65 | 0.4 | 0.36 | 0.32 | 0.4 |
| 3 | 179698241 | rs956395 | -0.9 | -0.761 | -0.358 | -0.525 | -0.005 | 0.005 | 0.005 | 0.825 | 0.35 | 0.31 | 0.57 | 0.74 | 0.4 | 0.41 |
| 3 | 179701679 | rs12492673 | 0.381 | -0.667 | -0.437 | -0.555 | -0.016 | -0.046 | -0.046 | 0 | 0.19 | 0.04 | 0.28 | 0.19 | 0.2 | 0.17 |
| 3 | 179706448 | rs1522116 | -1.763 | -0.767 | -0.401 | -0.63 | -0.016 | 0.011 | 0.011 | 0.006 | 0.49 | 0.84 | 0.46 | 0.36 | 0.44 | 0.49 |
| 3 | 179706629 | rs1522115 | -0.601 | -0.789 | -0.21 | -0.778 | -0.064 | -0.032 | -0.032 | 0 | 0.23 | 0.51 | 0.11 | 0.15 | 0.13 | 0.24 |
| 3 | 179709928 | rs9824299 | -0.529 | -0.64 | -0.183 | -0.665 | 0.004 | -0.004 | -0.004 | 0 | 0.3 | 0.34 | 0.36 | 0.2 | 0.32 | 0.25 |
| 3 | 179711169 | rs6764772 | -1.701 | -0.139 | 0.099 | -1.076 | -0.038 | 0.072 | 0.072 | 0.479 | 0.33 | 0.9 | 0.24 | 0.18 | 0.33 | 0.36 |
| 3 | 179728337 | rs2339932 | -1.332 | -0.37 | 0.461 | -0.76 | 0 | 0.015 | 0.015 | 0.701 | 0.27 | 0.78 | 0.18 | 0.18 | 0.24 | 0.33 |
| 3 | 179731684 | rs6800148 | -0.823 | -0.367 | 1.127 | -0.767 | -0.043 | -0.013 | -0.013 | 0 | 0.2 | 0.37 | 0.12 | 0.15 | 0.15 | 0.25 |
| 3 | 179733753 | rs9816641 | -1.144 | -0.376 | 1.482 | -0.769 | 0.013 | -0.01 | -0.01 | 0 | 0.21 | 0.56 | 0.15 | 0.17 | 0.21 | 0.29 |
| 3 | 179738924 | rs6443686 | -0.381 | -0.328 | 1.402 | -0.81 | 0.172 | 0 | 0 | 1.355 | 0.09 | 0.47 | 0.07 | 0.03 | 0.11 | 0.09 |
| 3 | 179746438 | rs9828621 | -0.507 | -0.366 | 1.296 | -0.781 | -0.035 | 0.015 | 0.015 | 0 | 0.15 | 0.37 | 0.09 | 0.13 | 0.1 | 0.2 |
| 3 | 179751306 | rs2204095 | -1.549 | -0.349 | 0.635 | -0.789 | 0 | 0.003 | 0.003 | 0.137 | 0.27 | 0.7 | 0.18 | 0.16 | 0.26 | 0.33 |
| 3 | 179755981 | rs6797922 | -1.116 | -0.277 | 0.535 | -0.732 | -0.002 | 0.004 | 0.004 | 0.582 | 0.22 | 0.75 | 0.16 | 0.16 | 0.22 | 0.29 |
| 3 | 179777060 | rs9853277 | -0.316 | -0.443 | 0.575 | -0.639 | -0.016 | -0.045 | -0.045 | 0 | 0.09 | 0.27 | 0.05 | 0 | 0.05 | 0.06 |
| 3 | 179777387 | rs1568554 | -1.398 | -0.51 | 0.532 | -0.83 | 0.001 | -0.012 | -0.012 | 0 | 0.38 | 0.6 | 0.56 | 0.77 | 0.36 | 0.49 |
| 3 | 179778368 | rs6806193 | -0.98 | -0.553 | 0.488 | -0.796 | 0.074 | -0.088 | -0.088 | 0 | 0.16 | 0.5 | 0.12 | 0.03 | 0.16 | 0.15 |
| 3 | 179779589 | rs7611246 | -0.149 | -1.521 | 0.131 | -0.488 | -0.013 | -0.043 | -0.043 | 0.105 | 0.08 | 0.27 | 0.05 | 0 | 0.05 | 0.06 |
| 3 | 179785100 | rs10937016 | -0.23 | -1.337 | -0.644 | -0.388 | -0.002 | -0.062 | -0.062 | 0 | 0.22 | 0.05 | 0.43 | 0.74 | 0.18 | 0.33 |
| 3 | 179786726 | rs1980275 | -0.894 | -0.963 | -0.381 | -0.736 | 0.05 | 0.081 | 0.081 | 0.593 | 0.47 | 0.7 | 0.59 | 0.88 | 0.34 | 0.61 |
| 3 | 179797647 | rs9809186 | 0.8 | -0.757 | -0.069 | 0.18 | -0.017 | 0.012 | 0.012 | 0.023 | 0.64 | 0.6 | 0.67 | 0.53 | 0.76 | 0.49 |
| 3 | 179810898 | rs1608071 | 1.424 | -0.822 | -0.021 | -0.251 | 0.021 | 0.011 | 0.011 | 0.138 | 0.51 | 0.4 | 0.52 | 0.53 | 0.65 | 0.42 |
| 3 | 179815191 | rs10513767 | -0.13 | -0.962 | 0.199 | -0.401 | 0.001 | -0.051 | -0.051 | 0.792 | 0.09 | 0.22 | 0.07 | 0.01 | 0.05 | 0.13 |
| 3 | 179823887 | rs1522117 | -0.915 | -1.056 | 0.431 | -0.455 | -0.006 | -0.083 | -0.083 | 0.987 | 0.12 | 0.44 | 0.07 | 0.01 | 0.06 | 0.13 |
| 3 | 179827381 | rs1522118 | 0.659 | -1.112 | 0.384 | -0.43 | 0.017 | -0.024 | -0.024 | 0.272 | 0.63 | 0.78 | 0.6 | 0.54 | 0.71 | 0.55 |
| 3 | 179831733 | rs7652876 | -2.076 | -1.053 | 0.567 | -0.03 | 0.007 | -0.005 | -0.005 | 1.698 | 0.39 | 0.65 | 0.44 | 0.47 | 0.27 | 0.52 |
| 3 | 179844161 | rs12633907 | -0.853 | -0.965 | 0.432 | 0.071 | -0.007 | -0.047 | -0.047 | 0 | 0.24 | 0.07 | 0.34 | 0.36 | 0.22 | 0.37 |
| 3 | 179846858 | rs4622902 | -1.409 | -0.82 | 0.973 | 0.106 | -0.013 | -0.089 | -0.089 | 0.243 | 0.13 | 0.56 | 0.09 | 0.01 | 0.06 | 0.14 |
| 3 | 179858044 | rs10513770 | -0.256 | -0.778 | 0.707 | 0.129 | 0.002 | -0.047 | -0.047 | 0 | 0.08 | 0.19 | 0.05 | 0.01 | 0.06 | 0.14 |
| 3 | 179873285 | rs1404743 | -0.985 | -0.899 | 0.608 | 0.231 | 0.007 | -0.009 | -0.009 | 0 | 0.26 | 0.22 | 0.32 | 0.38 | 0.2 | 0.35 |
| 3 | 179891121 | rs12487650 | -1.258 | -0.809 | 0.471 | 0.319 | 0 | -0.001 | -0.001 | 0 | 0.31 | 0.3 | 0.36 | 0.4 | 0.26 | 0.5 |
| 3 | 179909578 | rs11711435 | -0.51 | -0.712 | 0.584 | 0.237 | -0.004 | -0.045 | -0.045 | 0 | 0.1 | 0.22 | 0.07 | 0.01 | 0.08 | 0.17 |
| 3 | 179919078 | rs4854964 | -1.737 | -0.847 | 0.969 | 0.268 | -0.01 | -0.001 | -0.001 | 0 | 0.34 | 0.59 | 0.39 | 0.4 | 0.27 | 0.5 |
| 3 | 179934309 | rs10937022 | -0.469 | -0.908 | 0.916 | 0.604 | -0.031 | -0.152 | -0.152 | 0 | 0.2 | 0.03 | 0.34 | 0.52 | 0.19 | 0.35 |
| 3 | 179983857 | rs7643438 | -1.563 | -0.69 | 1.06 | 0.556 | -0.007 | 0 | 0 | 0 | 0.33 | 0.42 | 0.42 | 0.39 | 0.32 | 0.5 |
| 3 | 179996119 | rs6780998 | -1.638 | -0.699 | 1.072 | 0.494 | -0.015 | 0.036 | 0.036 | 0 | 0.36 | 0.51 | 0.51 | 0.52 | 0.33 | 0.53 |
| 3 | 180018650 | rs16831374 | -0.201 | -0.557 | 0.561 | 0.384 | -0.01 | -0.023 | -0.023 | 0 | 0.11 | 0.22 | 0.07 | 0.02 | 0.08 | 0.13 |
| 3 | 180032301 | rs1198109 | 0.606 | 0.009 | 0.273 | 0.27 | -0.016 | 0.011 | 0.011 | 0 | 0.1 | 0.07 | 0.05 | 0.1 | 0.13 | 0.11 |
| 3 | 180035517 | rs16831406 | -0.782 | -0.365 | 0.196 | 0.274 | -0.029 | -0.083 | -0.083 | 0 | 0.26 | 0.04 | 0.35 | 0.38 | 0.24 | 0.37 |
| 3 | 180037039 | rs11928576 | -0.745 | -0.58 | 0.506 | 0.03 | 0.011 | 0.016 | 0.016 | 0 | 0.15 | 0.09 | 0.07 | 0.03 | 0.1 | 0.14 |
| 3 | 180042078 | rs9867478 | -0.819 | -0.568 | 0.554 | -0.109 | -0.011 | -0.053 | -0.053 | 0 | 0.15 | 0.18 | 0.08 | 0.01 | 0.1 | 0.12 |
| 3 | 180045406 | rs6786124 | 0.245 | -0.503 | 0.671 | -0.036 | 0.017 | 0.005 | 0.005 | 0 | 0.05 | 0.25 | 0.04 | 0 | 0.01 | 0 |
| 3 | 180046054 | rs10212210 | 0.297 | -0.501 | 0.67 | -0.035 | 0.016 | 0.004 | 0.004 | 0 | 0.05 | 0.24 | 0.04 | 0 | 0.01 | 0 |
| 3 | 180066007 | rs9817579 | 0.464 | -0.542 | 0.87 | -0.065 | 0.011 | -0.009 | -0.009 | 0.071 | 0.05 | 0.23 | 0.04 | 0 | 0.02 | 0 |
| 3 | 180067592 | rs4854973 | -1.044 | -0.562 | 0.576 | 0 | -0.028 | -0.074 | -0.074 | 0 | 0.28 | 0.07 | 0.38 | 0.4 | 0.3 | 0.47 |
| 3 | 180071986 | rs1402227 | -1.742 | -0.58 | 0.628 | -0.009 | -0.014 | -0.001 | -0.001 | 0 | 0.37 | 0.47 | 0.44 | 0.4 | 0.32 | 0.47 |
| 3 | 180073549 | rs1464359 | -1.679 | -0.581 | 0.653 | -0.009 | 0.004 | -0.001 | -0.001 | 0 | 0.37 | 0.34 | 0.42 | 0.4 | 0.32 | 0.47 |
| 3 | 180076400 | rs16831521 | 0.124 | -0.465 | 0.382 | 0.046 | 0.005 | -0.001 | -0.001 | 0 | 0.11 | 0.07 | 0.05 | 0 | 0.08 | 0.11 |
| 3 | 180076602 | rs6792185 | -0.159 | -0.369 | 0.386 | 0.202 | -0.055 | -0.115 | -0.115 | 2.906 | 0.18 | 0.02 | 0.33 | 0.4 | 0.21 | 0.35 |
| 3 | 180103221 | rs6443694 | 0.101 | -0.452 | 0.753 | 0.045 | -0.007 | -0.026 | -0.026 | 0 | 0.06 | 0.22 | 0.04 | 0 | 0.02 | 0 |
| 3 | 180104190 | rs7612449 | -1.573 | -0.689 | 1.382 | -0.068 | -0.06 | -0.093 | -0.093 | 0 | 0.14 | 0.6 | 0.08 | 0 | 0.02 | 0.01 |
| 3 | 180114551 | rs11924799 | -0.049 | -0.61 | 1.371 | -0.021 | 0.018 | 0.004 | 0.004 | 0.579 | 0.07 | 0.35 | 0.04 | 0 | 0.01 | 0 |
| 3 | 180128288 | rs12493613 | -0.37 | -0.462 | 0.839 | 0.149 | -0.018 | -0.025 | -0.025 | 0 | 0.1 | 0.05 | 0.2 | 0.35 | 0.15 | 0.31 |
| 3 | 180135368 | rs10513772 | -1.074 | -0.543 | 0.849 | -0.18 | -0.023 | -0.106 | -0.106 | 0 | 0.25 | 0.65 | 0.21 | 0 | 0.14 | 0.07 |
| 3 | 180147892 | rs7634356 | -2.639 | -0.827 | 0.258 | -0.106 | -0.124 | -0.009 | -0.009 | 0.007 | 0.62 | 0.95 | 0.58 | 0.69 | 0.51 | 0.7 |
| 3 | 180171589 | rs9823620 | -2.584 | -0.75 | 0.42 | -0.074 | -0.04 | 0.07 | 0.07 | 0 | 0.58 | 0.86 | 0.61 | 0.78 | 0.48 | 0.69 |
| 3 | 180174467 | rs9290705 | 1.667 | -0.777 | 0.497 | -0.119 | -0.001 | -0.009 | -0.009 | 0 | 0.1 | 0.08 | 0.11 | 0 | 0.1 | 0.06 |
| 3 | 180182024 | rs1497891 | -0.265 | -0.862 | 0.619 | -0.19 | 0.001 | -0.04 | -0.04 | 0 | 0.23 | 0.23 | 0.17 | 0 | 0.19 | 0.17 |
| 3 | 180189540 | rs2338578 | -1.53 | -0.886 | 0.538 | -0.206 | 0.024 | -0.004 | -0.004 | 0.365 | 0.37 | 0.72 | 0.26 | 0 | 0.23 | 0.18 |
| 3 | 180196951 | rs1009336 | 1.827 | -0.862 | 0.755 | -0.047 | -0.005 | -0.016 | -0.016 | 2.721 | 0.07 | 0.06 | 0.12 | 0.21 | 0.06 | 0.08 |
| 3 | 180199917 | rs16831657 | 0.15 | -0.816 | 0.707 | -0.261 | -0.017 | -0.033 | -0.033 | 0 | 0.1 | 0.05 | 0.2 | 0.37 | 0.14 | 0.3 |
| 3 | 180199945 | rs11916013 | 0.26 | -0.838 | 0.721 | -0.727 | 0.017 | 0.032 | 0.032 | 0 | 0.12 | 0.06 | 0.05 | 0 | 0.08 | 0.11 |
| 3 | 180234393 | rs7633440 | -1.568 | -0.758 | 0.736 | -0.729 | 0.02 | -0.013 | -0.013 | 0.094 | 0.37 | 0.71 | 0.26 | 0 | 0.23 | 0.18 |

| | | | | | | | | | | | | | | | | |
|---|---|---|---|---|---|---|---|---|---|---|---|---|---|---|---|---|
| 3 | 180268361 | rs13097806 | 1.92 | -0.217 | 0.141 | -1.089 | -0.061 | 0.08 | 0.08 | 0 | 0.13 | 0.01 | 0.07 | 0 | 0.17 | 0.04 |
| 3 | 180284747 | rs12497999 | 2.33 | -0.669 | 0.854 | -0.457 | -0.081 | 0.068 | 0.068 | 0 | 0.39 | 0.07 | 0.39 | 0.22 | 0.49 | 0.3 |
| 3 | 180291037 | rs11718627 | 0.006 | -0.638 | 0.674 | -0.462 | 0.01 | 0.008 | 0.008 | 0 | 0.12 | 0.07 | 0.05 | 0 | 0.08 | 0.11 |
| 3 | 180293187 | rs7636793 | -0.949 | -0.64 | 0.665 | -0.469 | -0.039 | -0.162 | -0.162 | 0 | 0.28 | 0.53 | 0.2 | 0 | 0.17 | 0.17 |
| 3 | 180304277 | rs12496529 | 2.144 | -0.479 | 0.187 | -0.384 | -0.12 | 0.062 | 0.062 | 0 | 0.39 | 0.04 | 0.39 | 0.22 | 0.49 | 0.3 |
| 3 | 180319981 | rs1532485 | 0.687 | -0.034 | 1.235 | -0.138 | -0.001 | -0.009 | -0.009 | 0 | 0.1 | 0.08 | 0.12 | 0 | 0.1 | 0.06 |
| 3 | 180325487 | rs9858483 | 0.687 | -0.035 | 1.276 | -0.138 | -0.001 | -0.009 | -0.009 | 0.131 | 0.1 | 0.08 | 0.12 | 0 | 0.1 | 0.06 |
| 3 | 180325754 | rs7645817 | -1.189 | 0.039 | 0.933 | -0.097 | -0.007 | -0.073 | -0.073 | 0 | 0.27 | 0.68 | 0.21 | 0 | 0.17 | 0.17 |
| 3 | 180347917 | rs6798564 | -0.175 | 0.174 | -0.172 | -0.153 | -0.004 | -0.018 | -0.018 | 0 | 0.17 | 0.14 | 0.21 | 0 | 0.21 | 0.1 |
| 3 | 180348965 | rs869889 | -0.369 | 0.148 | -0.295 | 0.107 | -0.014 | -0.053 | -0.053 | 0 | 0.1 | 0.02 | 0.13 | 0.32 | 0.11 | 0.31 |
| 3 | 180365956 | rs6769457 | -1.456 | 0.173 | 0.283 | -0.752 | 0.01 | -0.007 | -0.007 | 0 | 0.31 | 0.74 | 0.29 | 0 | 0.25 | 0.17 |
| 3 | 180366067 | rs2338436 | -1.24 | 0.044 | 0.672 | 0.09 | -0.033 | -0.087 | -0.087 | 2.594 | 0.13 | 0.46 | 0.08 | 0 | 0.06 | 0.07 |
| 3 | 180377071 | rs1356606 | -1.471 | 1.07 | 1.02 | 1.195 | 0.054 | 0.165 | 0.165 | 0 | 0.14 | 0.31 | 0.26 | 0.35 | 0.18 | 0.21 |
| 3 | 180386213 | rs13097883 | -0.635 | 1.111 | 1.234 | 1.142 | 0.005 | -0.024 | -0.024 | 0 | 0.06 | 0.09 | 0.05 | 0 | 0.09 | 0.01 |
| 3 | 180396974 | rs2292896 | -2.183 | 1.047 | 1.304 | 1.176 | 0.044 | 0.137 | 0.137 | 0 | 0.14 | 0.29 | 0.26 | 0.35 | 0.18 | 0.21 |
| 3 | 180399599 | rs12485400 | -2.363 | 1.046 | 1.328 | 1.204 | 0.071 | 0.205 | 0.205 | 0 | 0.14 | 0.44 | 0.27 | 0.35 | 0.18 | 0.21 |
| 3 | 180402098 | rs1474112 | -2.34 | 1.098 | 1.678 | 1.197 | 0.077 | 0.199 | 0.199 | 0 | 0.14 | 0.42 | 0.28 | 0.35 | 0.19 | 0.22 |
| 3 | 180433381 | rs7628371 | 0.05 | 0.79 | 0.442 | 0.28 | -0.001 | -0.001 | -0.001 | 0 | 0.47 | 0.46 | 0.51 | 0.52 | 0.51 | 0.65 |
| 3 | 180435217 | rs2090578 | -0.303 | 0.794 | 0.42 | 0.296 | 0.005 | 0.001 | 0.001 | 0 | 0.51 | 0.66 | 0.53 | 0.52 | 0.52 | 0.66 |
| 3 | 180437487 | rs1401337 | -0.409 | 0.789 | 0.445 | 0.292 | -0.017 | -0.035 | -0.035 | 0 | 0.52 | 0.87 | 0.55 | 0.52 | 0.53 | 0.68 |
| 3 | 180438462 | rs7627545 | -0.233 | 0.791 | 0.417 | 0.296 | 0.004 | 0.002 | 0.002 | 0 | 0.51 | 0.57 | 0.52 | 0.52 | 0.52 | 0.66 |
| 3 | 180449589 | rs6779680 | 0.153 | 0.757 | 0.134 | 0.453 | -0.023 | 0.061 | 0.061 | 0 | 0.27 | 0.44 | 0.26 | 0.42 | 0.19 | 0.35 |
| 3 | 180449758 | rs11714847 | 0.057 | 1.482 | -0.104 | 0.288 | -0.068 | -0.017 | -0.017 | 3.032 | 0.14 | 0 | 0.11 | 0.12 | 0.19 | 0.17 |
| 3 | 180480576 | rs9838353 | 0.637 | 1.212 | 0.669 | 0.207 | -0.013 | 0.055 | 0.055 | 0 | 0.12 | 0.14 | 0.16 | 0.42 | 0.04 | 0.25 |
| 3 | 180488885 | rs4621329 | -0.317 | 1.321 | 0.342 | 0.267 | -0.001 | 0 | 0 | 0 | 0.28 | 0.7 | 0.29 | 0.42 | 0.19 | 0.35 |
| 3 | 180499783 | rs9827893 | -0.81 | 1.382 | -0.104 | 0.261 | 0.01 | -0.062 | -0.062 | 0 | 0.36 | 0.89 | 0.36 | 0.42 | 0.28 | 0.39 |
| 3 | 180513363 | rs6773765 | -0.422 | 1.212 | -0.329 | 0.334 | 0.019 | -0.031 | -0.031 | 0.899 | 0.29 | 0.81 | 0.31 | 0.42 | 0.2 | 0.36 |
| 3 | 180520800 | rs6795421 | -0.508 | 1.271 | -0.734 | 0.214 | 0.046 | -0.06 | -0.06 | 0 | 0.23 | 0.45 | 0.2 | 0.12 | 0.27 | 0.19 |
| 3 | 180521624 | rs6443721 | -0.966 | 0.974 | -1.76 | 0.515 | -0.123 | -0.06 | -0.06 | 0 | 0.51 | 0.97 | 0.49 | 0.58 | 0.46 | 0.56 |
| 3 | 180523951 | rs16832027 | -0.485 | 0.132 | -1.115 | 1.212 | 0.007 | -0.008 | -0.008 | 0.194 | 0.11 | 0.06 | 0.04 | 0.2 | 0.07 | 0.17 |
| 3 | 180524108 | rs12635682 | -1.279 | -0.631 | -0.62 | 1.741 | -0.032 | 0.025 | 0.025 | 0 | 0.31 | 0.43 | 0.33 | 0.48 | 0.16 | 0.35 |
| 3 | 180530195 | rs16832041 | -0.197 | -0.925 | -0.77 | 1.875 | 0.013 | -0.016 | -0.016 | 0.002 | 0.12 | 0.05 | 0.08 | 0.25 | 0.06 | 0.15 |
| 3 | 180530726 | rs1949547 | -0.402 | -2.429 | -0.738 | 1.493 | -0.111 | 0.049 | 0.049 | 3.613 | 0.3 | 0.53 | 0.22 | 0.35 | 0.08 | 0.28 |
| 3 | 180533251 | rs9838229 | -0.134 | -2.185 | -1.657 | 1.224 | -0.034 | 0.056 | 0.056 | 0 | 0.76 | 0.98 | 0.82 | 0.83 | 0.74 | 0.72 |
| 3 | 180544880 | rs12634070 | 1.222 | -1.189 | -1.79 | 1.448 | -0.037 | -0.031 | -0.031 | 0 | 0.06 | 0.01 | 0.1 | 0.22 | 0.19 | 0.17 |
| 3 | 180559330 | rs13067678 | 1.595 | -1.391 | -1.796 | 0.968 | -0.064 | -0.148 | -0.148 | 0.896 | 0.38 | 0.08 | 0.48 | 0.5 | 0.56 | 0.42 |
| 3 | 180570320 | rs6763026 | -0.167 | -1.546 | -1.215 | 1.033 | 0.024 | -0.01 | -0.01 | 0.14 | 0.33 | 0.7 | 0.3 | 0.34 | 0.15 | 0.29 |
| 3 | 180615583 | rs34133935 | 0.273 | -1.434 | -0.976 | 0.821 | 0.001 | 0.039 | 0.039 | 0 | 0.22 | 0.07 | 0.15 | 0.17 | 0.21 | 0.22 |
| 3 | 180627303 | rs496250 | 0.166 | -1.583 | -0.617 | 0.904 | -0.088 | 0.022 | 0.022 | 4.039 | 0.33 | 0.48 | 0.24 | 0.35 | 0.13 | 0.3 |
| 3 | 180633485 | rs1805564 | 0.226 | -1.507 | -0.556 | 0.878 | 0.002 | 0.04 | 0.04 | 0 | 0.22 | 0.07 | 0.15 | 0.16 | 0.21 | 0.26 |
| 3 | 180650962 | rs536127 | -1.7 | -1.439 | -0.484 | 0.909 | 0.018 | -0.073 | -0.073 | 0 | 0.59 | 0.89 | 0.49 | 0.51 | 0.4 | 0.53 |
| 3 | 180713021 | rs13073459 | 0.211 | -1.343 | -0.359 | 0.744 | -0.004 | 0.048 | 0.048 | 0 | 0.22 | 0.07 | 0.15 | 0.16 | 0.21 | 0.28 |
| 3 | 180713903 | rs7620793 | 0.206 | -1.458 | -0.326 | 0.933 | -0.079 | 0.024 | 0.024 | 4.534 | 0.32 | 0.52 | 0.19 | 0.33 | 0.13 | 0.29 |
| 3 | 180714307 | rs9810292 | -1.445 | -1.332 | -0.438 | 0.942 | 0.059 | -0.063 | -0.063 | 0 | 0.57 | 0.87 | 0.38 | 0.49 | 0.35 | 0.57 |
| 3 | 180733341 | rs6443732 | -0.864 | -1.573 | -0.365 | 0.744 | 0.023 | 0.021 | 0.021 | 0.188 | 0.35 | 0.7 | 0.29 | 0.25 | 0.22 | 0.4 |
| 3 | 180736253 | rs10049181 | -0.272 | -1.615 | -0.219 | 0.727 | -0.005 | -0.023 | -0.023 | 0 | 0.24 | 0.27 | 0.18 | 0.16 | 0.22 | 0.28 |
| 3 | 180737641 | rs34293605 | 0.529 | -1.614 | -0.234 | 0.702 | -0.024 | 0.051 | 0.051 | 0 | 0.21 | 0.03 | 0.15 | 0.16 | 0.2 | 0.28 |
| 3 | 180751651 | rs4855011 | -2.481 | -1.151 | -0.424 | 0.796 | -0.13 | 0.02 | 0.02 | 0.433 | 0.66 | 0.98 | 0.63 | 0.77 | 0.63 | 0.72 |
| 3 | 180751808 | rs6783677 | 1.72 | -1.354 | -0.187 | 0.652 | -0.05 | 0.056 | 0.056 | 0.56 | 0.11 | 0.43 | 0.11 | 0.09 | 0.01 | 0.12 |
| 3 | 180752138 | rs13096176 | 0.531 | -1.343 | -0.193 | 0.591 | -0.028 | 0.048 | 0.048 | 0 | 0.21 | 0.03 | 0.15 | 0.15 | 0.2 | 0.25 |
| 3 | 180752365 | rs16832272 | 1.856 | -1.234 | -0.195 | 0.705 | -0.03 | 0.003 | 0.003 | 0.13 | 0.05 | 0.01 | 0.17 | 0.29 | 0.17 | 0.17 |
| 3 | 180765861 | rs12630851 | 1.512 | -2.334 | 0.052 | -0.006 | -0.049 | 0.091 | 0.091 | 0.671 | 0.11 | 0.43 | 0.11 | 0.12 | 0.01 | 0.12 |
| 3 | 180778763 | rs1806190 | -0.498 | -1.925 | -0.246 | 0.135 | -0.005 | -0.024 | -0.024 | 0 | 0.24 | 0.27 | 0.18 | 0.16 | 0.21 | 0.28 |
| 3 | 180797921 | rs12635178 | 1.178 | -1.859 | -0.283 | 0.151 | -0.021 | -0.03 | -0.03 | 0 | 0.06 | 0 | 0.08 | 0.14 | 0.09 | 0.13 |
| 3 | 180799879 | rs4855015 | -0.138 | -1.826 | -0.209 | 0.171 | -0.003 | -0.03 | -0.03 | 0 | 0.23 | 0.26 | 0.17 | 0.15 | 0.21 | 0.28 |
| 3 | 180831977 | rs2543163 | -1.375 | -1.975 | -0.468 | 0.117 | 0.053 | 0.102 | 0.102 | 1.802 | 0.56 | 0.82 | 0.37 | 0.27 | 0.35 | 0.45 |
| 3 | 180832739 | rs6801189 | -0.054 | -1.529 | -0.521 | 0.246 | -0.005 | 0.05 | 0.05 | 0 | 0.22 | 0.09 | 0.16 | 0.14 | 0.2 | 0.28 |
| 3 | 180872987 | rs13071279 | -0.06 | -1.462 | -0.742 | 0.277 | -0.004 | 0.048 | 0.048 | 0 | 0.22 | 0.09 | 0.16 | 0.14 | 0.2 | 0.25 |
| 3 | 180874238 | rs13100468 | -0.06 | -1.462 | -0.741 | 0.277 | -0.002 | 0.044 | 0.044 | 0 | 0.22 | 0.1 | 0.16 | 0.14 | 0.2 | 0.25 |
| 3 | 180890956 | rs2718783 | -0.141 | -1.898 | -0.394 | 0.279 | -0.087 | -0.059 | -0.059 | 0.142 | 0.68 | 0.5 | 0.8 | 0.86 | 0.86 | 0.8 |
| 3 | 180896383 | rs2567668 | -0.149 | -1.902 | -0.337 | 0.296 | -0.091 | -0.059 | -0.059 | 0 | 0.68 | 0.51 | 0.8 | 0.86 | 0.86 | 0.8 |
| 3 | 180902019 | rs2567665 | 1.228 | -1.991 | -0.436 | 0.166 | 0.023 | 0.033 | 0.033 | 1.536 | 0.45 | 0.35 | 0.63 | 0.72 | 0.65 | 0.56 |
| 3 | 180904082 | rs11716918 | 0.106 | -1.968 | -0.464 | 0.164 | -0.006 | 0.053 | 0.053 | 0 | 0.22 | 0.11 | 0.16 | 0.14 | 0.21 | 0.24 |
| 3 | 180913191 | rs2718784 | -0.274 | -1.994 | -0.413 | 0.153 | -0.087 | -0.056 | -0.056 | 0.254 | 0.68 | 0.51 | 0.79 | 0.85 | 0.86 | 0.8 |
| 3 | 180932119 | rs2718790 | 0.137 | -2.091 | 0.092 | 0.089 | 0.128 | 0.095 | 0.095 | 7.077 | 0.64 | 0.15 | 0.75 | 0.86 | 0.85 | 0.8 |
| 3 | 180952891 | rs2718786 | 1.432 | -2.011 | -0.243 | 0.648 | -0.051 | -0.154 | -0.154 | 0 | 0.4 | 0.03 | 0.57 | 0.53 | 0.62 | 0.46 |
| 3 | 180981244 | rs4339108 | 0.583 | -2.331 | 0.142 | 0.307 | -0.073 | 0.026 | 0.026 | 2.278 | 0.14 | 0.62 | 0.08 | 0.13 | 0.02 | 0.12 |
| 3 | 180991951 | rs2008306 | 1.787 | -2.345 | 0.441 | 0.3 | -0.043 | 0.055 | 0.055 | 0 | 0.09 | 0.21 | 0.04 | 0.13 | 0.01 | 0.12 |
| 3 | 180993339 | rs2176612 | 1.084 | -2.52 | -0.088 | 0.169 | 0.012 | -0.015 | -0.015 | 2.457 | 0.21 | 0.15 | 0.09 | 0 | 0.12 | 0.08 |

| | | | | | | | | | | | | | | | |
|---|---|---|---|---|---|---|---|---|---|---|---|---|---|---|---|
| 3 | 180996959 | rs7619173 | -1.628 | -2.482 | -0.372 | 0.191 | 0.029 | 0.107 | 0.107 | 2.494 | 0.57 | 0.88 | 0.33 | 0.27 | 0.36 | 0.44 |
| 3 | 180998071 | rs7617548 | 0.183 | -2.208 | -0.297 | 0.252 | -0.033 | 0.068 | 0.068 | 0 | 0.22 | 0.03 | 0.16 | 0.14 | 0.21 | 0.24 |
| 3 | 181020115 | rs2139181 | 0.042 | -1.793 | -0.789 | 0.08 | 0.02 | 0.014 | 0.014 | 0 | 0.53 | 0.31 | 0.49 | 0.54 | 0.6 | 0.51 |
| 3 | 181023400 | rs7651261 | -0.222 | -2.063 | -1.167 | -0.083 | 0.122 | 0.097 | 0.097 | 5.977 | 0.41 | 0.85 | 0.25 | 0.27 | 0.24 | 0.28 |
| 3 | 181033842 | rs6778227 | 0.048 | -1.989 | -0.553 | 0.027 | 0.002 | 0.001 | 0.001 | 0 | 0.54 | 0.54 | 0.51 | 0.54 | 0.59 | 0.5 |
| 3 | 181045873 | rs2879464 | -0.497 | -1.792 | -1.093 | 0.137 | -0.126 | -0.005 | -0.005 | 0 | 0.64 | 0.97 | 0.57 | 0.68 | 0.62 | 0.62 |
| 3 | 181055378 | rs1518933 | 0.694 | -1.622 | -0.344 | 0.49 | -0.069 | -0.02 | -0.02 | 0.302 | 0.35 | 0.14 | 0.37 | 0.36 | 0.52 | 0.37 |
| 3 | 181073854 | rs6807049 | -1.599 | -1.796 | -0.309 | 0.391 | -0.038 | 0.074 | 0.074 | 0 | 0.76 | 0.9 | 0.71 | 0.93 | 0.72 | 0.83 |
| 3 | 181074751 | rs1518929 | -0.22 | -1.961 | 0.037 | 0.196 | -0.006 | -0.001 | -0.001 | 0.177 | 0.59 | 0.46 | 0.49 | 0.38 | 0.68 | 0.53 |
| 3 | 181101544 | rs6782970 | 0.711 | -1.948 | 0.075 | -0.04 | -0.01 | -0.065 | -0.065 | 0 | 0.25 | 0.25 | 0.11 | 0 | 0.16 | 0.1 |
| 3 | 181126962 | rs4855026 | 1.11 | -1.983 | -0.027 | -0.026 | 0.021 | 0.008 | 0.008 | 3.865 | 0.23 | 0.16 | 0.1 | 0 | 0.14 | 0.1 |
| 3 | 181166738 | rs2127238 | 0.874 | -1.947 | 0.023 | 0.184 | -0.077 | -0.009 | -0.009 | 0.067 | 0.34 | 0.13 | 0.37 | 0.32 | 0.51 | 0.36 |
| 3 | 181169949 | rs1479162 | -1.494 | -1.902 | 0.066 | 0.448 | -0.044 | -0.028 | -0.028 | 0 | 0.75 | 0.9 | 0.71 | 0.72 | 0.71 | 0.75 |
| 3 | 181180667 | rs9825132 | -0.286 | -1.704 | 0.272 | 0.123 | -0.004 | -0.006 | -0.006 | 0 | 0.59 | 0.48 | 0.49 | 0.32 | 0.66 | 0.52 |
| 3 | 181182392 | rs9873013 | -0.632 | -1.657 | 0.288 | 0.158 | -0.007 | 0.009 | 0.009 | 0 | 0.63 | 0.54 | 0.62 | 0.45 | 0.69 | 0.52 |
| 3 | 181183727 | rs7373337 | 0.611 | -1.635 | 0.303 | 0.102 | -0.013 | -0.069 | -0.069 | 0 | 0.25 | 0.32 | 0.11 | 0 | 0.15 | 0.1 |
| 3 | 181195182 | rs7611361 | 0.71 | -1.596 | 0.051 | 0.097 | -0.002 | -0.052 | -0.052 | 0 | 0.25 | 0.29 | 0.11 | 0 | 0.16 | 0.11 |
| 3 | 181204102 | rs2199718 | 0.399 | -1.118 | -0.298 | 0.502 | -0.046 | -0.027 | -0.027 | 0 | 0.34 | 0.19 | 0.48 | 0.44 | 0.5 | 0.4 |
| 3 | 181208850 | rs4859120 | -0.167 | -1.32 | -0.085 | 0.158 | 0.009 | 0 | 0 | 0 | 0.69 | 0.73 | 0.77 | 0.69 | 0.74 | 0.7 |
| 3 | 181212727 | rs6801673 | 0.488 | -1.992 | 0.261 | 0.344 | -0.006 | -0.001 | -0.001 | 0 | 0.17 | 0.18 | 0.1 | 0.24 | 0.06 | 0.2 |
| 3 | 181218016 | rs1817424 | -1.279 | -1.959 | -0.065 | 0.398 | 0.036 | -0.038 | -0.038 | 0 | 0.57 | 0.82 | 0.46 | 0.57 | 0.35 | 0.6 |
| 3 | 181222668 | rs16832533 | -1.058 | -1.866 | -0.428 | 0.443 | 0.026 | -0.003 | -0.003 | 0 | 0.53 | 0.39 | 0.4 | 0.57 | 0.33 | 0.59 |
| 3 | 181229833 | rs746058 | -0.315 | -0.703 | -0.692 | 0.081 | 0.04 | 0.019 | 0.019 | 0.003 | 0.35 | 0.14 | 0.3 | 0.33 | 0.26 | 0.38 |
| 3 | 181247797 | rs766743 | -0.62 | -0.601 | -0.607 | 0.108 | -0.02 | -0.02 | -0.02 | 0 | 0.39 | 0.54 | 0.35 | 0.33 | 0.29 | 0.4 |
| 3 | 181254051 | rs10513779 | -0.455 | 0.303 | -1.323 | 0.488 | 0.007 | 0.022 | 0.022 | 0 | 0.34 | 0.17 | 0.34 | 0.32 | 0.29 | 0.32 |
| 3 | 181264122 | rs6776298 | -0.875 | 0.705 | -0.551 | 1.032 | 0 | 0.024 | 0.024 | 0 | 0.14 | 0.21 | 0.17 | 0.34 | 0.13 | 0.21 |
| 3 | 181278376 | rs6443745 | -0.616 | 0.572 | -0.572 | 1.028 | 0.003 | -0.011 | -0.011 | 0 | 0.13 | 0.13 | 0.15 | 0.32 | 0.1 | 0.25 |
| 3 | 181278496 | rs6443746 | -1.031 | 0.542 | -0.167 | 0.999 | -0.002 | 0.068 | 0.068 | 0 | 0.15 | 0.32 | 0.18 | 0.33 | 0.11 | 0.25 |
| 3 | 181310591 | rs9854461 | -0.569 | 0.616 | -1.002 | 0.949 | 0.033 | 0.004 | 0.004 | 0.964 | 0.51 | 0.81 | 0.48 | 0.46 | 0.42 | 0.28 |
| 3 | 181320396 | rs6414490 | -0.815 | 0.897 | -1.252 | 1.078 | -0.136 | -0.064 | -0.064 | 0 | 0.55 | 0.98 | 0.52 | 0.61 | 0.48 | 0.46 |
| 3 | 181321265 | rs6443749 | -0.699 | 0.834 | -0.75 | 1.193 | -0.038 | -0.041 | -0.041 | 0 | 0.53 | 0.9 | 0.51 | 0.51 | 0.48 | 0.44 |
| 3 | 181324494 | rs9842957 | 0.063 | 0.419 | 0.208 | 1.168 | 0.134 | 0.211 | 0.211 | 0 | 0.07 | 0.32 | 0.11 | 0.17 | 0.12 | 0.06 |
| 3 | 181334964 | rs6443751 | -0.588 | 0.007 | 0.351 | 0.756 | 0.032 | 0.021 | 0.021 | 3.656 | 0.48 | 0.63 | 0.39 | 0.41 | 0.34 | 0.39 |
| 3 | 181340141 | rs13086529 | -1.187 | -0.178 | 1.123 | 0.975 | 0.04 | 0.006 | 0.006 | 0 | 0.18 | 0.48 | 0.23 | 0.15 | 0.19 | 0.11 |
| 3 | 181342415 | rs9845058 | 0.237 | -0.196 | 1.253 | 1.258 | -0.031 | 0.006 | 0.006 | 4.173 | 0.29 | 0.42 | 0.21 | 0.32 | 0.2 | 0.27 |
| 3 | 181346790 | rs9838604 | -0.883 | -0.168 | 0.954 | 1.182 | 0.043 | -0.008 | -0.008 | 0.001 | 0.46 | 0.7 | 0.4 | 0.47 | 0.37 | 0.42 |
| 3 | 181372001 | rs6784538 | -0.097 | -0.076 | 1.078 | 1.191 | -0.041 | -0.007 | -0.007 | 0 | 0.31 | 0.6 | 0.23 | 0.28 | 0.21 | 0.27 |
| 3 | 181378692 | rs4855035 | -1.494 | 1.096 | 1.711 | 1.629 | 0.082 | 0.171 | 0.171 | 0.061 | 0.11 | 0.59 | 0.25 | 0.21 | 0.14 | 0.1 |
| 3 | 181402480 | rs9844617 | -2.232 | 1.023 | 1.44 | 1.337 | -0.002 | -0.002 | -0.002 | 0.067 | 0.19 | 0.78 | 0.17 | 0.17 | 0.18 | 0.11 |
| 3 | 181413035 | rs7632216 | -1.041 | 1.373 | 1.882 | 1.074 | 0.087 | -0.058 | -0.058 | 1.186 | 0.1 | 0.55 | 0.13 | 0 | 0.13 | 0.03 |
| 3 | 181422854 | rs4434184 | -1.843 | 1.302 | 1.338 | 1.331 | 0.002 | 0.009 | 0.009 | 1.085 | 0.16 | 0.78 | 0.16 | 0.22 | 0.16 | 0.07 |
| 3 | 181426554 | rs35788479 | -0.942 | 1.293 | 1.338 | 0.79 | 0.129 | -0.03 | -0.03 | 0 | 0.09 | 0.4 | 0.11 | 0 | 0.13 | 0.03 |
| 3 | 181435148 | rs4575941 | -0.464 | 0.999 | -0.033 | 1.243 | 0.003 | 0.04 | 0.04 | 0.001 | 0.23 | 0.32 | 0.47 | 0.6 | 0.25 | 0.33 |
| 3 | 181455519 | rs7653630 | -0.254 | 1.551 | -1.402 | 0.326 | -0.136 | 0.048 | 0.048 | 0 | 0.75 | 0.98 | 0.85 | 0.87 | 0.63 | 0.66 |
| 3 | 181455914 | rs7633815 | 0.174 | 1.324 | -0.788 | 0.97 | -0.025 | -0.002 | -0.002 | 0 | 0.68 | 0.8 | 0.78 | 0.64 | 0.59 | 0.51 |
| 3 | 181457601 | rs6765739 | 0.261 | 0.069 | -1.032 | 1.056 | -0.013 | -0.01 | -0.01 | 0 | 0.34 | 0.19 | 0.37 | 0.35 | 0.36 | 0.21 |
| 3 | 181460587 | rs7610679 | -0.677 | -0.105 | -1.077 | 0.913 | 0.012 | 0.065 | 0.065 | 0 | 0.5 | 0.67 | 0.71 | 0.73 | 0.56 | 0.53 |
| 3 | 181466810 | rs4855037 | -0.737 | -0.697 | -0.682 | 0.983 | 0.002 | 0.006 | 0.006 | 0.655 | 0.43 | 0.59 | 0.59 | 0.76 | 0.41 | 0.51 |
| 3 | 181466976 | rs1558798 | -0.32 | -1.055 | -0.622 | 1.293 | 0.02 | 0.073 | 0.073 | 0.896 | 0.3 | 0.5 | 0.45 | 0.5 | 0.36 | 0.41 |
| 3 | 181467235 | rs12489879 | 0.425 | -1.007 | -0.726 | 0.654 | 0.02 | -0.024 | -0.024 | 0 | 0.12 | 0.07 | 0.13 | 0.26 | 0.06 | 0.11 |
| 3 | 181475226 | rs6793883 | -0.519 | -0.594 | -0.476 | 0.689 | 0.008 | -0.047 | -0.047 | 0.123 | 0.29 | 0.5 | 0.22 | 0.18 | 0.32 | 0.32 |
| 3 | 181482803 | rs9868748 | -0.521 | -1.001 | -0.461 | 0.86 | -0.018 | 0.092 | 0.092 | 0 | 0.22 | 0.34 | 0.36 | 0.48 | 0.19 | 0.18 |
| 3 | 181484270 | rs9840545 | 0.42 | -1.016 | -0.396 | 0.825 | -0.019 | -0.058 | -0.058 | 0.405 | 0.16 | 0.21 | 0.1 | 0.04 | 0.12 | 0.06 |
| 3 | 181490205 | rs16832650 | -0.051 | -0.85 | -0.443 | 0.856 | 0.004 | 0.159 | 0.159 | 0 | 0.06 | 0.1 | 0.26 | 0.44 | 0.07 | 0.13 |
| 3 | 181492405 | rs11923857 | -1.035 | -1.141 | -0.659 | 0.358 | -0.019 | -0.014 | -0.014 | 0 | 0.4 | 0.42 | 0.49 | 0.74 | 0.26 | 0.34 |
| 3 | 181498753 | rs17540772 | 0.598 | -0.393 | -0.869 | 0.485 | 0.006 | 0.01 | 0.01 | 0 | 0.09 | 0.09 | 0.06 | 0 | 0.08 | 0.01 |
| 3 | 181503845 | rs11714692 | 0.227 | -0.388 | -0.635 | 0.744 | -0.003 | 0.019 | 0.019 | 0 | 0.11 | 0.11 | 0.11 | 0.27 | 0.06 | 0.11 |
| 3 | 181504459 | rs6773639 | -0.099 | -0.364 | -0.085 | 0.217 | -0.009 | -0.076 | -0.076 | 0 | 0.14 | 0.33 | 0.1 | 0.04 | 0.11 | 0.14 |
| 3 | 181504668 | rs10513780 | 0.383 | 0.416 | -0.334 | 0.04 | 0.015 | -0.047 | -0.047 | 0 | 0.13 | 0.22 | 0.09 | 0.03 | 0.12 | 0.12 |
| 3 | 181509166 | rs7618962 | -0.412 | 0.592 | -0.476 | 0.556 | 0.052 | 0.279 | 0.279 | 0.016 | 0.16 | 0.42 | 0.38 | 0.49 | 0.2 | 0.24 |
| 3 | 181510576 | rs2339463 | -0.685 | 0.48 | -1.632 | -0.537 | -0.021 | 0.212 | 0.212 | 0.235 | 0.57 | 0.88 | 0.75 | 0.88 | 0.57 | 0.62 |
| 3 | 181511375 | rs17613182 | 0.146 | 0.18 | -0.926 | -0.303 | 0 | -0.002 | -0.002 | 0.192 | 0.56 | 0.57 | 0.54 | 0.61 | 0.48 | 0.67 |
| 3 | 181511904 | rs12638589 | -0.535 | -0.042 | -1.056 | -0.748 | 0.018 | 0.048 | 0.048 | 0 | 0.37 | 0.33 | 0.35 | 0.2 | 0.35 | 0.35 |
| 3 | 181515848 | rs13315015 | 0.636 | -0.841 | -0.832 | -0.343 | 0.001 | -0.002 | -0.002 | 0 | 0.34 | 0.29 | 0.32 | 0.56 | 0.31 | 0.44 |
| 3 | 181515948 | rs2339461 | -0.797 | -1.054 | -0.603 | -0.275 | 0.051 | 0.043 | 0.043 | 0.188 | 0.17 | 0.08 | 0.18 | 0.06 | 0.13 | 0.23 |
| 3 | 181522020 | rs833272 | -0.947 | -0.923 | -0.221 | -0.218 | -0.045 | -0.042 | -0.042 | 0.848 | 0.19 | 0.22 | 0.18 | 0.07 | 0.14 | 0.26 |
| 3 | 181524517 | rs2339460 | -0.874 | -0.132 | -0.226 | 0.406 | 0.002 | -0.041 | -0.041 | 0.006 | 0.15 | 0.25 | 0.24 | 0.04 | 0.17 | 0.2 |
| 3 | 181531075 | rs855296 | -0.464 | -0.229 | -0.785 | 0.455 | 0 | 0.084 | 0.084 | 0.051 | 0.21 | 0.05 | 0.1 | 0.09 | 0.15 | 0.12 |
| 3 | 181556123 | rs6763608 | -0.49 | -0.697 | -0.939 | -0.205 | 0.037 | 0.035 | 0.035 | 0 | 0.22 | 0.13 | 0.07 | 0.02 | 0.09 | 0.08 |

| | | | | | | | | | | | | | | | | |
|---|---|---|---|---|---|---|---|---|---|---|---|---|---|---|---|---|
| 3 | 181573945 | rs11922688 | -0.159 | 1.146 | -1.168 | -0.179 | -0.03 | 0.019 | 0.019 | 0 | 0.14 | 0.09 | 0.26 | 0.05 | 0.32 | 0.08 |
| 3 | 181579709 | rs12489180 | 0.75 | -0.374 | -1.8 | -0.238 | -0.034 | 0.056 | 0.056 | 1.292 | 0.38 | 0.05 | 0.3 | 0.26 | 0.36 | 0.41 |
| 3 | 181584903 | rs9834123 | -0.002 | -1.689 | -0.976 | 0.388 | 0.013 | -0.018 | -0.018 | 0 | 0.75 | 0.75 | 0.76 | 0.49 | 0.89 | 0.76 |
| 3 | 181586376 | rs10937067 | -0.376 | -0.838 | -0.859 | 0.78 | -0.015 | 0.079 | 0.079 | 2.994 | 0.09 | 0.08 | 0.09 | 0.56 | 0.01 | 0.12 |
| 3 | 181588762 | rs833282 | -0.386 | 0.075 | -1.24 | -0.618 | -0.153 | 0.129 | 0.129 | 0 | 0.25 | 0.03 | 0.38 | 0.04 | 0.55 | 0.23 |
| 3 | 181602745 | rs9877982 | 0.157 | -1.506 | -0.352 | -0.495 | -0.003 | -0.022 | -0.022 | 0 | 0.53 | 0.46 | 0.44 | 0.87 | 0.25 | 0.59 |
| 3 | 181607662 | rs4855052 | 0.211 | -1.571 | -0.188 | -0.524 | 0.109 | 0.145 | 0.145 | 0 | 0.52 | 0.27 | 0.43 | 0.87 | 0.25 | 0.59 |
| 3 | 181609741 | rs4855054 | -0.215 | -0.978 | -0.464 | -0.871 | -0.038 | 0.136 | 0.136 | 0.843 | 0.67 | 0.91 | 0.6 | 0.97 | 0.39 | 0.75 |
| 3 | 181617660 | rs6783518 | -0.401 | -1.892 | -0.293 | -0.168 | 0.011 | -0.008 | -0.008 | 0 | 0.23 | 0.17 | 0.23 | 0.28 | 0.14 | 0.2 |
| 3 | 181617927 | rs13096015 | 0.562 | -2.132 | -0.593 | -0.397 | -0.078 | 0.12 | 0.12 | 0 | 0.3 | 0.08 | 0.47 | 0.1 | 0.61 | 0.3 |
| 3 | 181629223 | rs9834564 | -0.626 | -2.167 | 0.313 | -0.312 | -0.021 | -0.006 | -0.006 | 0 | 0.56 | 0.51 | 0.73 | 0.38 | 0.76 | 0.53 |
| 3 | 181631615 | rs7623553 | -0.737 | -2.088 | -0.069 | -0.278 | 0.021 | 0.005 | 0.005 | 0.582 | 0.58 | 0.65 | 0.74 | 0.38 | 0.76 | 0.54 |
| 3 | 181637995 | rs7614481 | -0.682 | -2.129 | -0.176 | 0.161 | -0.059 | 0.06 | 0.06 | 0.31 | 0.3 | 0.48 | 0.2 | 0.43 | 0.16 | 0.24 |
| 3 | 181638250 | rs4855056 | 0.049 | -1.371 | -0.526 | -0.397 | 0.004 | 0.002 | 0.002 | 0 | 0.5 | 0.42 | 0.49 | 0.51 | 0.54 | 0.42 |
| 3 | 181638448 | rs4855057 | -0.187 | -0.985 | -0.607 | -0.284 | 0.025 | 0.01 | 0.01 | 0.444 | 0.54 | 0.75 | 0.52 | 0.56 | 0.61 | 0.57 |
| 3 | 181658257 | rs16832803 | 0.736 | -0.913 | -0.387 | -0.528 | -0.029 | -0.061 | -0.061 | 0.773 | 0.12 | 0.21 | 0.06 | 0 | 0.04 | 0 |
| 3 | 181659359 | rs1392340 | 0.199 | -0.226 | -0.53 | -0.753 | -0.008 | 0.007 | 0.007 | 0.014 | 0.64 | 0.49 | 0.59 | 0.6 | 0.64 | 0.58 |
| 3 | 181665698 | rs7646383 | -0.372 | -0.306 | -0.441 | -0.924 | 0.008 | -0.018 | -0.018 | 0.26 | 0.74 | 0.81 | 0.68 | 0.64 | 0.74 | 0.65 |
| 3 | 181669056 | rs11720370 | 1.516 | -0.369 | -0.628 | -0.953 | -0.022 | -0.043 | -0.043 | 1.549 | 0.38 | 0.19 | 0.37 | 0.53 | 0.41 | 0.37 |
| 3 | 181685198 | rs4855065 | 0.541 | 0.06 | -0.122 | -0.283 | -0.012 | 0.003 | 0.003 | 0.042 | 0.64 | 0.57 | 0.6 | 0.63 | 0.66 | 0.57 |
| 3 | 181689647 | rs13088708 | 0.211 | 0.09 | -0.408 | -0.403 | -0.001 | 0.026 | 0.026 | 0 | 0.08 | 0.04 | 0.05 | 0.01 | 0.07 | 0.08 |
| 3 | 181700562 | rs9810038 | -0.248 | 0.07 | -0.275 | -0.292 | -0.006 | 0.011 | 0.011 | 0.154 | 0.74 | 0.69 | 0.65 | 0.66 | 0.75 | 0.65 |
| 3 | 181705738 | rs4854927 | -0.23 | 0.698 | -0.827 | -0.133 | -0.048 | 0.012 | 0.012 | 0.663 | 0.15 | 0.05 | 0.18 | 0.12 | 0.23 | 0.23 |
| 3 | 181727995 | rs4855069 | 0.895 | 0.308 | -1.13 | 0.203 | 0 | -0.013 | -0.013 | 0 | 0.54 | 0.27 | 0.39 | 0.48 | 0.49 | 0.4 |
| 3 | 181734542 | rs4854932 | 0.361 | 0.598 | -0.629 | 0.544 | 0.002 | 0.043 | 0.043 | 0.359 | 0.73 | 0.44 | 0.58 | 0.62 | 0.71 | 0.66 |
| 3 | 181747300 | rs17188166 | -0.346 | 1.021 | -1.296 | 0.161 | -0.089 | 0.061 | 0.061 | 0 | 0.2 | 0.03 | 0.19 | 0.05 | 0.3 | 0.15 |
| 3 | 181752619 | rs1561072 | -0.211 | -0.07 | -1.297 | 0.089 | -0.002 | 0 | 0 | 2.105 | 0.23 | 0.78 | 0.18 | 0.2 | 0.1 | 0.11 |
| 3 | 181785515 | rs11706418 | 0.238 | 0.026 | -1.239 | 0.131 | -0.089 | 0.059 | 0.059 | 0 | 0.2 | 0.03 | 0.19 | 0.05 | 0.3 | 0.15 |
| 3 | 181801136 | rs1435959 | -1.402 | -0.079 | -1.641 | 0.534 | -0.024 | -0.021 | -0.021 | 0.348 | 0.54 | 0.8 | 0.51 | 0.47 | 0.51 | 0.4 |
| 3 | 181855149 | rs16832893 | 0.136 | -0.171 | 0.173 | 0.334 | -0.019 | 0.012 | 0.012 | 0 | 0.19 | 0.24 | 0.14 | 0.23 | 0.15 | 0.14 |
| 3 | 181908937 | rs6443778 | 0.07 | 0.115 | -0.244 | 0.339 | -0.009 | 0.006 | 0.006 | 0 | 0.2 | 0.25 | 0.15 | 0.23 | 0.17 | 0.14 |
| 3 | 181932928 | rs6788528 | -0.416 | 0.492 | -0.149 | 0.124 | 0.013 | -0.053 | -0.053 | 0 | 0.25 | 0.36 | 0.23 | 0.06 | 0.29 | 0.14 |
| 3 | 181935178 | rs9853234 | -0.787 | -0.309 | -0.813 | 0.539 | 0.05 | -0.023 | -0.023 | 0 | 0.41 | 0.74 | 0.29 | 0.42 | 0.28 | 0.31 |
| 3 | 181938437 | rs7611110 | -1.016 | -0.618 | -1.241 | 0.748 | 0.072 | 0.081 | 0.081 | 0 | 0.48 | 0.87 | 0.33 | 0.64 | 0.31 | 0.41 |
| 3 | 181938870 | rs7635994 | -0.686 | -0.876 | -0.716 | 0.735 | 0.016 | 0.002 | 0.002 | 0 | 0.26 | 0.72 | 0.13 | 0.24 | 0.13 | 0.19 |
| 3 | 181941060 | rs4264727 | 0.527 | -0.863 | -0.761 | 0.725 | -0.002 | 0.111 | 0.111 | 0.067 | 0.07 | 0.07 | 0.13 | 0.4 | 0.06 | 0.15 |
| 3 | 181954164 | rs9850886 | -1.223 | -0.932 | -0.867 | 0.469 | 0.046 | -0.019 | -0.019 | 0.017 | 0.33 | 0.84 | 0.26 | 0.64 | 0.19 | 0.35 |
| 3 | 181955695 | rs7639661 | 0.285 | -0.823 | 0.255 | 0.347 | -0.011 | -0.097 | -0.097 | 0.193 | 0.16 | 0.34 | 0.06 | 0.02 | 0.1 | 0.07 |
| 3 | 181958418 | rs6443784 | -1.947 | -0.524 | -0.841 | 0.004 | -0.161 | 0.088 | 0.088 | 0 | 0.65 | 0.94 | 0.51 | 0.83 | 0.47 | 0.64 |
| 3 | 181969294 | rs7636148 | 1.102 | -0.743 | -0.307 | 0.168 | 0.007 | 0.014 | 0.014 | 0 | 0.11 | 0.06 | 0.03 | 0 | 0.08 | 0.01 |
| 3 | 181975033 | rs6787138 | 0.256 | -0.874 | -0.111 | 0.023 | 0.022 | 0.028 | 0.028 | 0.058 | 0.27 | 0.13 | 0.1 | 0 | 0.19 | 0.08 |
| 3 | 181975303 | rs4507242 | -0.785 | -0.862 | 0.185 | 0.588 | 0.059 | 0.366 | 0.366 | 1.514 | 0.06 | 0.34 | 0.05 | 0.33 | 0.03 | 0.14 |
| 3 | 181979272 | rs9290737 | -0.92 | -0.842 | -0.09 | 0.53 | 0.054 | 0.151 | 0.151 | 0 | 0.45 | 0.88 | 0.47 | 0.76 | 0.32 | 0.43 |
| 3 | 181988782 | rs12634003 | 1.034 | -0.95 | 0.005 | 0.668 | 0.012 | -0.018 | -0.018 | 0.225 | 0.17 | 0.17 | 0.09 | 0.32 | 0.13 | 0.16 |
| 3 | 182002353 | rs1982808 | -0.42 | -1 | 0.927 | 0.553 | 0.056 | 0.14 | 0.14 | 0 | 0.05 | 0.3 | 0.18 | 0.06 | 0.01 | 0.01 |
| 3 | 182004287 | rs4859198 | -1.239 | -1.009 | 0.489 | 0.414 | -0.004 | -0.011 | -0.011 | 0.328 | 0.45 | 0.51 | 0.46 | 0.74 | 0.33 | 0.37 |
| 3 | 182010044 | rs11916677 | -0.75 | -0.912 | 0.607 | 0.468 | 0.07 | 0.163 | 0.163 | 0.969 | 0.06 | 0.49 | 0.19 | 0.06 | 0.01 | 0.01 |
| 3 | 182023431 | rs12633943 | 0.274 | -0.873 | -0.133 | 0.651 | -0.044 | 0.063 | 0.063 | 0 | 0.23 | 0.68 | 0.28 | 0.41 | 0.12 | 0.18 |
| 3 | 182050012 | rs17217692 | 0.024 | -0.806 | -0.471 | 0.507 | -0.016 | -0.038 | -0.038 | 0 | 0.22 | 0.08 | 0.2 | 0.32 | 0.22 | 0.19 |
| 3 | 182055057 | rs7624385 | -0.123 | -0.439 | -0.455 | 0.797 | -0.014 | -0.012 | -0.012 | 0 | 0.4 | 0.57 | 0.31 | 0.68 | 0.33 | 0.36 |
| 3 | 182080081 | rs16833068 | 0.263 | -0.476 | -1.012 | 0.724 | -0.014 | -0.036 | -0.036 | 0 | 0.08 | 0.02 | 0.15 | 0.32 | 0.09 | 0.17 |
| 3 | 182096311 | rs9836621 | -1.578 | -0.019 | -1.177 | -0.045 | -0.099 | 0.181 | 0.181 | 0.07 | 0.64 | 0.93 | 0.59 | 0.91 | 0.52 | 0.65 |
| 3 | 182132052 | rs17826716 | -0.705 | -0.436 | -0.972 | 0.295 | 0.052 | 0.043 | 0.043 | 0 | 0.53 | 0.76 | 0.36 | 0.66 | 0.37 | 0.52 |
| 3 | 182132383 | rs16833118 | -0.02 | -0.421 | -1.073 | 0.301 | -0.01 | -0.006 | -0.006 | 0.652 | 0.43 | 0.56 | 0.32 | 0.65 | 0.34 | 0.39 |
| 3 | 182135821 | rs7640799 | -0.537 | -0.503 | -1.371 | 0.282 | -0.019 | -0.026 | -0.026 | 0 | 0.08 | 0.09 | 0.03 | 0 | 0.03 | 0.12 |
| 3 | 182153406 | rs13074509 | 0.537 | -0.102 | -2.034 | 0.208 | -0.084 | 0.127 | 0.127 | 0 | 0.31 | 0.03 | 0.32 | 0.1 | 0.33 | 0.39 |
| 3 | 182153436 | rs17217699 | -1.019 | -0.507 | -0.617 | 1.605 | 0.004 | -0.015 | -0.015 | 0 | 0.1 | 0.04 | 0.15 | 0.3 | 0.08 | 0.15 |
| 3 | 182207825 | rs9877496 | -0.982 | -1.003 | -0.747 | 0.527 | -0.023 | 0.264 | 0.264 | 2.699 | 0.55 | 0.93 | 0.62 | 0.89 | 0.41 | 0.75 |
| 3 | 182211107 | rs9880110 | -0.785 | -1.843 | -0.134 | 0.976 | 0.076 | 0.082 | 0.082 | 0.28 | 0.48 | 0.71 | 0.56 | 0.76 | 0.32 | 0.65 |
| 3 | 182222315 | rs1457578 | -1.404 | -1.632 | 1.052 | 1.547 | -0.034 | 0.033 | 0.033 | 0 | 0.2 | 0.36 | 0.21 | 0.28 | 0.11 | 0.29 |
| 3 | 182222834 | rs9816312 | -1.616 | -1.702 | 1.107 | 1.554 | -0.049 | 0.028 | 0.028 | 0 | 0.25 | 0.65 | 0.26 | 0.36 | 0.13 | 0.33 |
| 3 | 182232391 | rs2710932 | 0.654 | -1.613 | 1.065 | 1.06 | -0.009 | -0.03 | -0.03 | 1.728 | 0.85 | 0.8 | 0.93 | 0.99 | 0.94 | 0.84 |
| 3 | 182238748 | rs13079695 | 1.594 | -1.891 | 1.397 | 1.33 | -0.071 | -0.015 | -0.015 | 0 | 0.76 | 0.32 | 0.88 | 0.78 | 0.93 | 0.78 |
| 3 | 182245072 | rs1457567 | 1.484 | -1.818 | 1.394 | 1.004 | -0.075 | -0.055 | -0.055 | 0 | 0.77 | 0.31 | 0.88 | 0.87 | 0.93 | 0.83 |
| 3 | 182256685 | rs2339722 | -1.827 | -0.531 | 1.453 | 1.785 | 0.002 | -0.006 | -0.006 | 0 | 0.32 | 0.75 | 0.42 | 0.4 | 0.34 | 0.39 |
| 3 | 182265580 | rs4859226 | -1.627 | -0.577 | 1.916 | 1.878 | 0.012 | -0.078 | -0.078 | 0.264 | 0.28 | 0.43 | 0.21 | 0.14 | 0.32 | 0.24 |
| 3 | 182286102 | rs4859231 | -1.8 | -0.563 | 1.596 | 1.866 | -0.002 | 0.015 | 0.015 | 0 | 0.31 | 0.53 | 0.41 | 0.4 | 0.34 | 0.39 |
| 3 | 182287145 | rs9813493 | -2.086 | -0.589 | 0.629 | 1.853 | 0.006 | -0.012 | -0.012 | 0 | 0.34 | 0.83 | 0.43 | 0.4 | 0.34 | 0.39 |
| 3 | 182301473 | rs9822514 | -0.894 | -0.432 | 0.35 | 1.572 | -0.001 | -0.002 | -0.002 | 0 | 0.18 | 0.13 | 0.14 | 0.13 | 0.22 | 0.22 |

| | | | | | | | | | | | | | | | | |
|---|---|---|---|---|---|---|---|---|---|---|---|---|---|---|---|---|
| 3 | 182311332 | rs4859234 | -1.931 | -0.464 | -0.416 | 1.82 | 0.002 | -0.048 | -0.048 | 0 | 0.33 | 0.84 | 0.33 | 0.5 | 0.34 | 0.38 |
| 3 | 182311870 | rs6797190 | -1.212 | -0.441 | -0.365 | 1.777 | -0.018 | 0.102 | 0.102 | 12.702 | 0.06 | 0.1 | 0.11 | 0.38 | 0.02 | 0.17 |
| 3 | 182312152 | rs6772756 | -1.534 | -0.285 | -0.677 | 1.669 | -0.002 | -0.033 | -0.033 | 0 | 0.32 | 0.81 | 0.33 | 0.48 | 0.34 | 0.38 |
| 3 | 182314029 | rs2879560 | -1.286 | -0.238 | -0.528 | 1.452 | 0.001 | 0.003 | 0.003 | 0.591 | 0.24 | 0.83 | 0.19 | 0.24 | 0.21 | 0.21 |
| 3 | 182315570 | rs2607953 | -1.3 | -0.546 | -0.875 | 2.115 | 0.007 | -0.089 | -0.089 | 5.101 | 0.15 | 0.03 | 0.38 | 0.45 | 0.14 | 0.23 |
| 3 | 182321461 | rs3843881 | 1.386 | -0.695 | -0.882 | 1.55 | 0.026 | 0.123 | 0.123 | 0 | 0.49 | 0.26 | 0.29 | 0.15 | 0.54 | 0.34 |
| 3 | 182324992 | rs7643238 | 1.584 | -1.206 | -0.077 | 1.317 | 0.014 | 0.009 | 0.009 | 0 | 0.6 | 0.65 | 0.45 | 0.25 | 0.72 | 0.44 |
| 3 | 182326726 | rs1463444 | 1.143 | -1.058 | 0.214 | 1.504 | 0.01 | -0.02 | -0.02 | 0 | 0.75 | 0.78 | 0.58 | 0.48 | 0.8 | 0.67 |
| 3 | 182332485 | rs1463443 | 0.881 | -1.347 | 0.165 | 1.749 | 0.05 | -0.129 | -0.129 | 0 | 0.82 | 0.98 | 0.78 | 0.53 | 0.89 | 0.74 |
| 3 | 182333358 | rs6784627 | -2.131 | -0.686 | 1.382 | 1.718 | 0.109 | 0.024 | 0.024 | 0 | 0.1 | 0.38 | 0.13 | 0.11 | 0.15 | 0.13 |
| 3 | 182347019 | rs9862173 | -1.712 | -0.857 | 1.944 | 1.89 | -0.027 | 0.091 | 0.091 | 0.017 | 0.11 | 0.24 | 0.12 | 0.25 | 0.07 | 0.2 |
| 3 | 182353074 | rs2315505 | -2.129 | -0.753 | 2.338 | 1.843 | 0.007 | 0.012 | 0.012 | 0.008 | 0.27 | 0.61 | 0.3 | 0.38 | 0.29 | 0.32 |
| 3 | 182353090 | rs6780863 | -0.927 | -1.039 | 2.562 | 1.947 | 0.002 | -0.012 | -0.012 | 0 | 0.16 | 0.13 | 0.17 | 0.26 | 0.14 | 0.19 |
| 3 | 182353819 | rs7617785 | -2.054 | -0.985 | 2.783 | 1.915 | 0.055 | 0.012 | 0.012 | 0 | 0.07 | 0.35 | 0.08 | 0.06 | 0.07 | 0.06 |
| 3 | 182355770 | rs7355912 | -2.422 | -1.538 | 2.547 | 1.988 | 0.006 | 0.006 | 0.006 | 0 | 0.36 | 0.64 | 0.44 | 0.82 | 0.28 | 0.42 |
| 3 | 182364788 | rs12485872 | -0.967 | -1.366 | 1.996 | 1.473 | 0.004 | -0.039 | -0.039 | 0.157 | 0.18 | 0.07 | 0.16 | 0.25 | 0.14 | 0.19 |
| 3 | 182366158 | rs6443818 | -1.671 | -1.355 | 1.982 | 1.485 | -0.01 | 0.008 | 0.008 | 0.476 | 0.25 | 0.42 | 0.24 | 0.31 | 0.21 | 0.25 |
| 3 | 182373879 | rs6767825 | -2.19 | -0.854 | 1.941 | 1.306 | 0.001 | -0.003 | -0.003 | 0 | 0.32 | 0.68 | 0.4 | 0.45 | 0.28 | 0.32 |
| 3 | 182374458 | rs6804316 | -2.179 | -0.087 | 2.243 | 0.36 | 0 | -0.012 | -0.012 | 2.864 | 0.23 | 0.75 | 0.21 | 0.11 | 0.23 | 0.13 |
| 3 | 182375715 | rs12637956 | -1.638 | -0.062 | 2.137 | 0.328 | -0.007 | -0.003 | -0.003 | 0.517 | 0.07 | 0.06 | 0.08 | 0.1 | 0.12 | 0.07 |
| 3 | 182382262 | rs2700862 | 2.543 | -0.122 | 2.41 | 0.253 | 0.03 | -0.017 | -0.017 | 0.007 | 0.86 | 0.64 | 0.87 | 0.89 | 0.84 | 0.83 |
| 3 | 182390083 | rs2700845 | 1.804 | -0.101 | 1.976 | 0.264 | 0.012 | -0.092 | -0.092 | 5.39 | 0.76 | 0.51 | 0.79 | 0.89 | 0.74 | 0.8 |
| 3 | 182392033 | rs2700837 | 1.529 | 0.134 | 1.699 | 0.45 | 0 | -0.016 | -0.016 | 0 | 0.78 | 0.24 | 0.79 | 0.89 | 0.78 | 0.83 |
| 3 | 182392108 | rs2717302 | 1.019 | -0.092 | 1.44 | 0.294 | -0.014 | -0.12 | -0.12 | 0.526 | 0.82 | 0.48 | 0.83 | 0.99 | 0.86 | 0.88 |
| 3 | 182394127 | rs7617305 | 0.942 | 1.028 | 0.584 | 0.33 | 0.082 | -0.084 | -0.084 | 0 | 0.73 | 0.52 | 0.55 | 0.86 | 0.53 | 0.81 |
| 3 | 182429032 | rs11920872 | -1.472 | 0.429 | 0.843 | 0.75 | 0.004 | 0.076 | 0.076 | 0 | 0.09 | 0.29 | 0.07 | 0.13 | 0.05 | 0.11 |
| 3 | 182429315 | rs11916914 | -1.442 | 0.426 | 0.848 | 0.75 | 0.008 | 0.077 | 0.077 | 0 | 0.08 | 0.29 | 0.07 | 0.13 | 0.05 | 0.11 |
| 3 | 182432644 | rs6795829 | 0.203 | 1.137 | -0.238 | 0.19 | -0.032 | 0.026 | 0.026 | 0 | 0.65 | 0.75 | 0.61 | 0.8 | 0.53 | 0.73 |
| 3 | 182433424 | rs12633691 | -0.224 | -0.099 | -0.553 | 0.176 | -0.02 | 0.011 | 0.011 | 0 | 0.3 | 0.61 | 0.26 | 0.62 | 0.2 | 0.33 |
| 3 | 182434708 | rs6765224 | 1.06 | 0.109 | -0.617 | -0.138 | -0.002 | 0.005 | 0.005 | 0 | 0.36 | 0.37 | 0.38 | 0.19 | 0.34 | 0.45 |
| 3 | 182440135 | rs2700868 | -0.508 | 0.203 | -0.219 | 0.659 | 0.011 | 0.051 | 0.051 | 0.382 | 0.16 | 0.21 | 0.22 | 0.31 | 0.18 | 0.17 |
| 3 | 182446674 | rs3107664 | -1.147 | 0.266 | -0.523 | 0.563 | 0.036 | 0.012 | 0.012 | 0 | 0.41 | 0.83 | 0.35 | 0.64 | 0.33 | 0.32 |
| 3 | 182450613 | rs1523060 | -0.858 | -0.378 | 0.023 | 0.698 | -0.001 | 0.009 | 0.009 | 0 | 0.31 | 0.35 | 0.25 | 0.35 | 0.27 | 0.24 |
| 3 | 182451327 | rs2687803 | -1.481 | -0.561 | 0.296 | 0.659 | 0.019 | 0.006 | 0.006 | 0 | 0.45 | 0.59 | 0.31 | 0.43 | 0.35 | 0.39 |
| 3 | 182453765 | rs16833435 | 1.343 | -0.314 | -0.093 | 0.563 | 0 | 0.005 | 0.005 | 0 | 0.13 | 0.12 | 0.2 | 0.2 | 0.2 | 0.12 |
| 3 | 182454798 | rs16833441 | -0.434 | -0.41 | 0.124 | 0.586 | -0.019 | 0.006 | 0.006 | 0 | 0.27 | 0.4 | 0.27 | 0.28 | 0.29 | 0.27 |
| 3 | 182456625 | rs6793157 | 0.022 | -0.411 | 0.126 | 0.573 | -0.045 | -0.043 | -0.043 | 2.903 | 0.14 | 0.25 | 0.07 | 0.08 | 0.1 | 0.15 |
| 3 | 182456918 | rs2700842 | -1.207 | -0.368 | 0.035 | 0.585 | 0.003 | 0.009 | 0.009 | 0 | 0.49 | 0.45 | 0.45 | 0.57 | 0.49 | 0.41 |
| 3 | 182457097 | rs2717297 | -2.031 | -0.329 | -0.432 | 0.595 | -0.087 | 0.012 | 0.012 | 0 | 0.59 | 0.88 | 0.53 | 0.63 | 0.55 | 0.51 |
| 3 | 182457367 | rs1879990 | -0.649 | -0.334 | -0.435 | 0.574 | 0.036 | 0 | 0 | 0.611 | 0.36 | 0.82 | 0.33 | 0.33 | 0.33 | 0.36 |
| 3 | 182457759 | rs1879989 | -0.34 | -0.231 | -0.337 | 0.635 | -0.035 | -0.078 | -0.078 | 0 | 0.23 | 0.06 | 0.19 | 0.29 | 0.22 | 0.15 |
| 3 | 182467550 | rs13089627 | 1.363 | -0.486 | -0.577 | 0.417 | -0.084 | 0.012 | 0.012 | 0.696 | 0.4 | 0.06 | 0.42 | 0.31 | 0.43 | 0.49 |
| 3 | 182470146 | rs1115037 | 1.383 | -0.502 | -0.562 | 0.41 | -0.082 | 0.013 | 0.013 | 0.701 | 0.4 | 0.06 | 0.42 | 0.31 | 0.43 | 0.49 |
| 3 | 182471086 | rs2717274 | -1.643 | -0.454 | -0.442 | 0.24 | -0.002 | -0.002 | -0.002 | 0 | 0.87 | 0.91 | 0.9 | 0.99 | 0.9 | 0.94 |
| 3 | 182471372 | rs2717272 | -0.984 | -0.366 | -0.053 | 0.528 | 0.005 | 0.024 | 0.024 | 0.043 | 0.45 | 0.69 | 0.47 | 0.58 | 0.46 | 0.38 |
| 3 | 182471928 | rs7640210 | 0.068 | -0.283 | 0.395 | 0.65 | -0.001 | 0.006 | 0.006 | 0 | 0.26 | 0.31 | 0.27 | 0.28 | 0.28 | 0.25 |
| 3 | 182474593 | rs940072 | -0.763 | -0.549 | 0.583 | 0.857 | 0.008 | 0.096 | 0.096 | 0 | 0.18 | 0.38 | 0.19 | 0.28 | 0.16 | 0.13 |
| 3 | 182478533 | rs2717294 | -1.327 | -1.481 | 0.438 | 0.705 | -0.014 | 0.012 | 0.012 | 0 | 0.36 | 0.63 | 0.34 | 0.45 | 0.27 | 0.3 |
| 3 | 182482857 | rs1403476 | -1.048 | -2.046 | 0.329 | 0.498 | 0.017 | -0.041 | -0.041 | 1.728 | 0.32 | 0.15 | 0.32 | 0.32 | 0.26 | 0.19 |
| 3 | 182488961 | rs1879986 | 1.145 | -1.999 | 0.497 | 0.549 | -0.001 | 0.002 | 0.002 | 1.104 | 0.66 | 0.63 | 0.67 | 0.59 | 0.73 | 0.73 |
| 3 | 182491415 | rs7629226 | 0.382 | -2.084 | 0.614 | 0.565 | -0.002 | 0.012 | 0.012 | 1.263 | 0.15 | 0.32 | 0.11 | 0.18 | 0.11 | 0.14 |
| 3 | 182494774 | rs9868197 | 1.195 | -2.072 | 0.562 | 0.573 | -0.005 | 0.006 | 0.006 | 1.751 | 0.66 | 0.65 | 0.68 | 0.56 | 0.74 | 0.73 |
| 3 | 182496141 | rs4859242 | 0.323 | -2.025 | 0.375 | 0.547 | 0.019 | -0.068 | -0.068 | 0.503 | 0.8 | 0.92 | 0.77 | 0.74 | 0.82 | 0.87 |
| 3 | 182497683 | rs6788905 | -0.572 | -2.113 | 0.652 | 0.14 | -0.043 | -0.066 | -0.066 | 0 | 0.18 | 0.46 | 0.1 | 0.15 | 0.1 | 0.09 |
| 3 | 182510867 | rs6791488 | 0.525 | -1.984 | 0.649 | 0.123 | 0 | -0.042 | -0.042 | 0 | 0.14 | 0.28 | 0.1 | 0.04 | 0.09 | 0.05 |
| 3 | 182513714 | rs10937101 | 1.916 | -2.468 | 0.756 | 0.195 | 0.08 | 0.038 | 0.038 | 1.368 | 0.51 | 0.29 | 0.61 | 0.42 | 0.68 | 0.65 |
| 3 | 182521127 | rs12696497 | 1.431 | -2.465 | 0.849 | 0.238 | -0.016 | 0.004 | 0.004 | 0 | 0.65 | 0.63 | 0.71 | 0.44 | 0.77 | 0.7 |
| 3 | 182547256 | rs2049283 | 1.653 | -2.53 | 0.76 | 0.192 | 0.023 | -0.009 | -0.009 | 0 | 0.63 | 0.38 | 0.68 | 0.44 | 0.76 | 0.69 |
| 3 | 182580276 | rs622205 | 0.321 | -2.599 | 0.705 | 0.198 | 0.032 | -0.097 | -0.097 | 0 | 0.8 | 0.92 | 0.78 | 0.61 | 0.84 | 0.8 |
| 3 | 182595815 | rs492310 | 1.359 | -2.498 | 0.719 | 0.148 | -0.051 | 0.021 | 0.021 | 0 | 0.64 | 0.5 | 0.69 | 0.46 | 0.77 | 0.7 |
| 3 | 182608551 | rs7650693 | 0.754 | -2.415 | 0.672 | 0.105 | -0.003 | -0.023 | -0.023 | 0 | 0.14 | 0.22 | 0.1 | 0.04 | 0.09 | 0.05 |
| 3 | 182613683 | rs10937103 | -1.454 | -2.543 | 0.786 | 0.096 | -0.01 | 0.002 | 0.002 | 0 | 0.36 | 0.37 | 0.29 | 0.54 | 0.24 | 0.3 |
| 3 | 182637840 | rs16833596 | 0.481 | -2.343 | 0.691 | 0.076 | -0.006 | -0.056 | -0.056 | 0 | 0.14 | 0.27 | 0.1 | 0.02 | 0.09 | 0.04 |
| 3 | 182640615 | rs1133034 | -0.241 | -2.501 | 0.779 | 0.097 | -0.035 | -0.013 | -0.013 | 0 | 0.15 | 0.25 | 0.06 | 0.18 | 0.07 | 0.13 |
| 3 | 182640753 | rs1133035 | 0.636 | -2.366 | 0.848 | -0.106 | 0.006 | -0.062 | -0.062 | 0 | 0.14 | 0.22 | 0.12 | 0.02 | 0.11 | 0.04 |
| 3 | 182640877 | rs11710878 | -1.553 | -2.706 | 0.87 | 0.422 | -0.009 | 0 | 0 | 0 | 0.35 | 0.37 | 0.29 | 0.56 | 0.24 | 0.34 |
| 3 | 182650847 | rs2089612 | 1.054 | -2.419 | 0.566 | -0.109 | -0.003 | -0.011 | -0.011 | 0 | 0.55 | 0.44 | 0.63 | 0.24 | 0.65 | 0.53 |
| 3 | 182659953 | rs9878775 | 0.566 | -2.628 | 1.016 | -0.085 | 0.014 | 0.013 | 0.013 | 0.001 | 0.67 | 0.73 | 0.71 | 0.27 | 0.74 | 0.61 |
| 3 | 182663516 | rs1509229 | 0.626 | -2.641 | 0.504 | -0.08 | 0.008 | 0.008 | 0.008 | 0.018 | 0.67 | 0.67 | 0.7 | 0.27 | 0.74 | 0.59 |

| | | | | | | | | | | | | | | | | |
|---|---|---|---|---|---|---|---|---|---|---|---|---|---|---|---|---|
| 3 | 182664547 | rs4859259 | 1.845 | -2.795 | 0.081 | -0.184 | 0.182 | 0.126 | 0.126 | 0.311 | 0.55 | 0.04 | 0.63 | 0.24 | 0.76 | 0.56 |
| 3 | 182669780 | rs6786608 | 1.893 | -2.674 | 0.221 | -0.059 | 0.096 | 0.187 | 0.187 | 0.154 | 0.51 | 0.14 | 0.61 | 0.24 | 0.65 | 0.53 |
| 3 | 182671158 | rs9814216 | 0.799 | -2.527 | -0.132 | 0.01 | 0.005 | 0.005 | 0.005 | 0.024 | 0.67 | 0.68 | 0.7 | 0.27 | 0.74 | 0.59 |
| 3 | 182685440 | rs4859148 | 1.455 | -2.392 | -0.433 | -0.006 | 0.042 | 0.079 | 0.079 | 0 | 0.55 | 0.66 | 0.64 | 0.24 | 0.65 | 0.53 |
| 3 | 182720206 | rs9869993 | 0.051 | -1.664 | -0.346 | 0.199 | 0.011 | 0.001 | 0.001 | 0.539 | 0.12 | 0.07 | 0.03 | 0 | 0.07 | 0.04 |
| 3 | 182727097 | rs9841498 | -1.576 | -1.984 | -0.515 | 0.91 | 0.04 | -0.083 | -0.083 | 0 | 0.4 | 0.86 | 0.22 | 0.49 | 0.28 | 0.37 |
| 3 | 182729293 | rs7635560 | -0.348 | -2.112 | 0.261 | 0.681 | -0.04 | -0.07 | -0.07 | 0 | 0.15 | 0.23 | 0.03 | 0 | 0.07 | 0.04 |
| 3 | 182731930 | rs1547910 | -0.271 | -2.033 | 0.234 | 0.697 | -0.025 | -0.059 | -0.059 | 0 | 0.15 | 0.21 | 0.04 | 0 | 0.07 | 0.04 |
| 3 | 182732230 | rs9290751 | -0.575 | -2.288 | 0.529 | 1.321 | -0.005 | -0.031 | -0.031 | 0 | 0.23 | 0.17 | 0.3 | 0.57 | 0.2 | 0.31 |
| 3 | 182736304 | rs11917378 | -0.08 | -2.278 | 0.487 | 1.309 | 0.028 | 0.022 | 0.022 | 2.62 | 0.86 | 0.92 | 0.97 | 1 | 0.93 | 0.96 |
| 3 | 182743267 | rs7640612 | 2.043 | -2.376 | 0.374 | 1.342 | 0.024 | 0.107 | 0.107 | 0.278 | 0.55 | 0.05 | 0.44 | 0.23 | 0.7 | 0.52 |
| 3 | 182755209 | rs2270968 | 1.694 | -2.394 | 0.717 | 1.415 | 0.039 | 0.036 | 0.036 | 0.223 | 0.6 | 0.19 | 0.63 | 0.33 | 0.73 | 0.57 |
| 3 | 182763580 | rs1502762 | 1.532 | -2.384 | 0.814 | 1.5 | 0.036 | -0.05 | -0.05 | 0 | 0.65 | 0.19 | 0.68 | 0.5 | 0.79 | 0.65 |
| 3 | 182778941 | rs13078931 | 1.976 | -2.349 | -0.138 | 1.554 | 0.033 | -0.069 | -0.069 | 0 | 0.6 | 0.05 | 0.5 | 0.37 | 0.75 | 0.59 |
| 3 | 182802874 | rs9822789 | -0.251 | -2.09 | 0.024 | 1.377 | 0 | 0.015 | 0.015 | 0 | 0.18 | 0.2 | 0.25 | 0.4 | 0.14 | 0.25 |
| 3 | 182815090 | rs4859156 | 1.865 | -2.057 | -0.524 | 0.986 | 0.042 | -0.028 | -0.028 | 0 | 0.5 | 0.08 | 0.41 | 0.44 | 0.63 | 0.57 |
| 3 | 182826316 | rs7618830 | 0.682 | -1.993 | -0.047 | 0.929 | 0.019 | 0.002 | 0.002 | 2.638 | 0.73 | 0.78 | 0.7 | 0.72 | 0.83 | 0.81 |
| 3 | 182831612 | rs13098857 | 1.18 | -2.034 | 0.095 | 0.971 | 0.024 | -0.001 | -0.001 | 0.635 | 0.68 | 0.32 | 0.49 | 0.65 | 0.79 | 0.74 |
| 3 | 182831688 | rs11716740 | -0.255 | -1.776 | -0.096 | 1.299 | -0.028 | -0.017 | -0.017 | 1.088 | 0.17 | 0.05 | 0.07 | 0.19 | 0.15 | 0.17 |
| 3 | 182832506 | rs2134428 | 0.519 | -1.732 | 0.233 | 1.331 | -0.006 | 0.035 | 0.035 | 0 | 0.9 | 0.89 | 0.8 | 0.73 | 0.92 | 0.8 |
| 3 | 182846387 | rs500193 | 0.562 | -1.764 | 1.187 | 1.346 | -0.01 | 0.107 | 0.107 | 0 | 0.89 | 0.78 | 0.78 | 0.72 | 0.93 | 0.81 |
| 3 | 182849564 | rs3772715 | 1.186 | -1.851 | 1.092 | 1.334 | 0.006 | 0.037 | 0.037 | 0 | 0.75 | 0.71 | 0.72 | 0.54 | 0.78 | 0.7 |
| 3 | 182851441 | rs4358320 | -0.317 | -2.062 | 1.119 | 1.247 | -0.018 | -0.017 | -0.017 | 2.303 | 0.12 | 0.5 | 0.21 | 0.06 | 0.05 | 0.06 |
| 3 | 182853212 | rs3736551 | 1.256 | -2.321 | 0.891 | 1.139 | -0.044 | 0.03 | 0.03 | 0.17 | 0.75 | 0.39 | 0.58 | 0.66 | 0.88 | 0.75 |
| 3 | 182853670 | rs482912 | 1.88 | -2.338 | 0.788 | 1.107 | 0.02 | -0.051 | -0.051 | 0 | 0.61 | 0.22 | 0.5 | 0.47 | 0.72 | 0.65 |
| 3 | 182858461 | rs831256 | 1.806 | -1.879 | 0.701 | 1.008 | 0.002 | -0.013 | -0.013 | 0.062 | 0.6 | 0.36 | 0.48 | 0.47 | 0.67 | 0.64 |
| 3 | 182859353 | rs1393586 | 0.271 | -1.686 | 0.234 | 0.74 | -0.032 | -0.019 | -0.019 | 0 | 0.14 | 0.06 | 0.07 | 0.19 | 0.15 | 0.1 |
| 3 | 182861174 | rs7639990 | -1.171 | -1.745 | 0.42 | 0.703 | -0.001 | -0.009 | -0.009 | 0 | 0.32 | 0.59 | 0.32 | 0.25 | 0.26 | 0.19 |
| 3 | 182862976 | rs10513791 | 0.546 | -1.421 | 0.408 | 0.972 | 0 | -0.013 | -0.013 | 0.554 | 0.1 | 0.05 | 0.05 | 0.17 | 0.08 | 0.12 |
| 3 | 182867869 | rs605105 | 1.147 | -1.511 | 0.542 | 1.1 | -0.017 | 0.019 | 0.019 | 0 | 0.77 | 0.64 | 0.81 | 0.73 | 0.83 | 0.77 |
| 3 | 182869818 | rs683395 | 0.505 | -1.483 | 0.205 | 1.091 | -0.011 | 0.002 | 0.002 | 0.008 | 0.87 | 0.81 | 0.88 | 0.9 | 0.91 | 0.88 |
| 3 | 182870436 | rs514636 | 0.505 | -1.483 | 0.21 | 1.091 | -0.011 | 0.002 | 0.002 | 0.008 | 0.87 | 0.81 | 0.88 | 0.9 | 0.91 | 0.88 |
| 3 | 182871962 | rs653316 | 0.947 | -1.488 | 0.075 | 1.102 | -0.001 | 0.001 | 0.001 | 0 | 0.78 | 0.78 | 0.83 | 0.72 | 0.83 | 0.78 |
| 3 | 182872318 | rs2280563 | 0.654 | -1.344 | -0.258 | 1.064 | 0 | -0.012 | -0.012 | 0 | 0.09 | 0.05 | 0.05 | 0.19 | 0.08 | 0.1 |
| 3 | 182877292 | rs658672 | 1.322 | -1.527 | 0.065 | 1.396 | -0.005 | 0.024 | 0.024 | 2.562 | 0.77 | 0.29 | 0.54 | 0.57 | 0.81 | 0.74 |
| 3 | 182878123 | rs17803349 | 0.571 | -1.646 | -0.33 | 1.159 | 0.006 | 0.012 | 0.012 | 0.008 | 0.12 | 0.04 | 0.05 | 0.12 | 0.09 | 0.09 |
| 3 | 182878897 | rs1965484 | 1.504 | -1.238 | -0.898 | 0.713 | -0.032 | -0.016 | -0.016 | 0.643 | 0.42 | 0.11 | 0.4 | 0.39 | 0.57 | 0.58 |
| 3 | 182880111 | rs6443856 | -0.049 | -0.802 | 0.087 | 0.944 | -0.001 | 0 | 0 | 0 | 0.88 | 0.84 | 0.88 | 0.91 | 0.9 | 0.85 |
| 3 | 182889001 | rs521981 | -0.015 | -0.795 | 0.37 | 0.924 | -0.004 | 0 | 0 | 0 | 0.86 | 0.72 | 0.86 | 0.89 | 0.9 | 0.81 |
| 3 | 182896412 | rs6771207 | -0.343 | -0.106 | 0.498 | 1.251 | 0.061 | 0.104 | 0.104 | 0 | 0.08 | 0.1 | 0.35 | 0.3 | 0.17 | 0.1 |
| 3 | 182903072 | rs11708313 | -0.313 | -0.097 | 0.545 | 1.274 | 0.069 | 0.114 | 0.114 | 0 | 0.08 | 0.11 | 0.35 | 0.29 | 0.17 | 0.1 |
| 3 | 182905808 | rs472795 | -0.682 | -0.445 | 0.588 | 0.351 | -0.001 | -0.027 | -0.027 | 0 | 0.88 | 0.86 | 0.95 | 1 | 0.91 | 0.84 |
| 3 | 182907865 | rs534333 | 0.34 | -0.679 | 0.817 | 0.522 | 0.044 | -0.013 | -0.013 | 0.106 | 0.76 | 0.4 | 0.64 | 0.84 | 0.75 | 0.71 |
| 3 | 182912282 | rs684174 | -0.589 | -1.219 | 0.406 | 0.181 | 0.016 | 0.015 | 0.015 | 6.327 | 0.54 | 0.33 | 0.43 | 0.39 | 0.48 | 0.51 |
| 3 | 182921803 | rs4630966 | 0.985 | -2.082 | 0.025 | -0.256 | -0.013 | -0.055 | -0.055 | 0 | 0.21 | 0.04 | 0.19 | 0.28 | 0.24 | 0.41 |
| 3 | 182942402 | rs12489712 | 0.574 | -1.786 | 0.569 | -0.278 | 0.058 | 0.071 | 0.071 | 7.461 | 0.48 | 0.3 | 0.63 | 0.72 | 0.62 | 0.58 |
| 3 | 182942671 | rs13066057 | -0.124 | -1.961 | 0.278 | -0.372 | 0.081 | 0.086 | 0.086 | 4.99 | 0.43 | 0.12 | 0.29 | 0.28 | 0.34 | 0.42 |
| 3 | 182944282 | rs4859159 | 0.074 | -1.869 | 0.202 | -0.322 | -0.011 | -0.03 | -0.03 | 0.092 | 0.94 | 0.8 | 0.96 | 1 | 0.96 | 0.99 |
| 3 | 182946304 | rs2272115 | 0.778 | -1.739 | -0.333 | -0.266 | 0.057 | 0.079 | 0.079 | 8.447 | 0.45 | 0.22 | 0.62 | 0.71 | 0.61 | 0.53 |
| 3 | 182949977 | rs940056 | -0.112 | -1.095 | 0.187 | 0.502 | 0.038 | 0.031 | 0.031 | 0.183 | 0.32 | 0.11 | 0.28 | 0.25 | 0.27 | 0.37 |
| 3 | 182967435 | rs9883461 | -1.916 | -0.967 | 1.486 | 1.096 | -0.068 | 0.01 | 0.01 | 5.929 | 0.12 | 0.65 | 0.07 | 0.15 | 0.01 | 0.14 |
| 3 | 182970805 | rs6809841 | -1.59 | -0.744 | 1.708 | 0.946 | -0.004 | 0.008 | 0.008 | 0.705 | 0.09 | 0.4 | 0.07 | 0.06 | 0.03 | 0.15 |
| 3 | 182973255 | rs6793163 | -1.651 | -0.752 | 1.615 | 0.943 | -0.026 | -0.015 | -0.015 | 0.011 | 0.11 | 0.5 | 0.08 | 0.06 | 0.03 | 0.14 |
| 3 | 182981159 | rs10049125 | -1.213 | -0.403 | 1.465 | 1.651 | -0.006 | 0.089 | 0.089 | 0.07 | 0.23 | 0.58 | 0.21 | 0.54 | 0.18 | 0.39 |
| 3 | 183003626 | rs11924380 | -0.198 | -0.485 | 1.607 | 1.554 | -0.008 | 0 | 0 | 0 | 0.05 | 0.04 | 0.01 | 0.05 | 0.01 | 0.02 |
| 3 | 183005320 | rs6807075 | -1.396 | -0.694 | 2.186 | 1.819 | -0.012 | 0.054 | 0.054 | 0.038 | 0.25 | 0.63 | 0.23 | 0.55 | 0.18 | 0.42 |
| 3 | 183006918 | rs3732602 | -0.269 | -0.596 | 1.765 | 1.444 | -0.008 | 0 | 0 | 0 | 0.05 | 0.04 | 0.01 | 0.05 | 0.01 | 0.02 |
| 3 | 183017385 | rs884910 | -0.798 | -0.915 | 2.064 | 1.535 | -0.057 | 0.012 | 0.012 | 4.304 | 0.12 | 0.25 | 0.02 | 0.15 | 0.02 | 0.02 |
| 3 | 183020592 | rs7634425 | -0.946 | -1.144 | 1.899 | 1.791 | -0.02 | 0.023 | 0.023 | 0.003 | 0.33 | 0.43 | 0.22 | 0.51 | 0.23 | 0.4 |
| 3 | 183024333 | rs1496802 | 0.615 | -0.735 | 1.045 | 1.234 | -0.007 | 0.064 | 0.064 | 0 | 0.31 | 0.17 | 0.33 | 0.1 | 0.3 | 0.2 |
| 3 | 183027542 | rs7639705 | -0.308 | -0.557 | 1.817 | 1.864 | -0.002 | 0.05 | 0.05 | 0 | 0.24 | 0.42 | 0.24 | 0.36 | 0.22 | 0.38 |
| 3 | 183029331 | rs3732595 | 0.135 | -0.479 | 1.015 | 1.899 | -0.007 | -0.037 | -0.037 | 0 | 0.21 | 0.12 | 0.19 | 0.36 | 0.21 | 0.37 |
| 3 | 183031713 | rs6443863 | -1.891 | -0.332 | 1.754 | 2.229 | 0.001 | -0.004 | -0.004 | 0 | 0.19 | 0.21 | 0.28 | 0.36 | 0.21 | 0.21 |
| 3 | 183035175 | rs10513792 | -1.468 | -0.557 | 2.196 | 1.609 | -0.018 | 0.004 | 0.004 | 0.376 | 0.16 | 0.32 | 0.11 | 0.18 | 0.15 | 0.07 |
| 3 | 183036526 | rs13434301 | -2.606 | -0.652 | 2.631 | 1.726 | -0.012 | 0.106 | 0.106 | 0.861 | 0.17 | 0.59 | 0.19 | 0.29 | 0.18 | 0.19 |
| 3 | 183040637 | rs6773123 | -0.87 | -0.592 | 2.476 | 1.72 | 0.041 | 0.011 | 0.011 | 0 | 0.07 | 0.24 | 0.09 | 0.07 | 0.13 | 0.05 |
| 3 | 183043851 | rs6770912 | -2.221 | -0.605 | 2.495 | 1.719 | 0.01 | 0.138 | 0.138 | 0.108 | 0.15 | 0.41 | 0.17 | 0.3 | 0.18 | 0.19 |
| 3 | 183050387 | rs10937123 | -1.628 | -0.621 | 2.1 | 1.748 | -0.049 | 0.097 | 0.097 | 2.783 | 0.15 | 0.31 | 0.1 | 0.32 | 0.06 | 0.17 |
| 3 | 183051675 | rs6780072 | -1.552 | -0.534 | 1.406 | 1.768 | -0.032 | 0.026 | 0.026 | 2.826 | 0.21 | 0.61 | 0.17 | 0.28 | 0.15 | 0.19 |

| | | | | | | | | | | | | | | | | |
|---|---|---|---|---|---|---|---|---|---|---|---|---|---|---|---|---|
| 3 | 183054273 | rs9868599 | -1.83 | -0.521 | 1.028 | 1.693 | 0.021 | -0.028 | -0.028 | 0 | 0.27 | 0.8 | 0.22 | 0.41 | 0.2 | 0.22 |
| 3 | 183075106 | rs7620788 | -1.795 | -0.386 | 0.998 | 1.492 | -0.015 | 0.034 | 0.034 | 3.296 | 0.25 | 0.68 | 0.2 | 0.43 | 0.19 | 0.23 |
| 3 | 183076891 | rs9811069 | -1.541 | -0.621 | 1.187 | 1.472 | -0.038 | 0.076 | 0.076 | 1.88 | 0.22 | 0.63 | 0.18 | 0.39 | 0.17 | 0.22 |
| 3 | 183080524 | rs6786692 | -1.812 | -0.682 | 1.269 | 1.18 | -0.029 | 0.025 | 0.025 | 3.083 | 0.27 | 0.67 | 0.2 | 0.41 | 0.17 | 0.24 |
| 3 | 183085464 | rs6791370 | -1.654 | -0.69 | 1.251 | 1.229 | -0.039 | 0.022 | 0.022 | 1.558 | 0.26 | 0.64 | 0.19 | 0.35 | 0.16 | 0.19 |
| 3 | 183095918 | rs10937128 | -1.418 | -0.331 | 0.613 | 1.295 | -0.007 | 0.045 | 0.045 | 0 | 0.08 | 0.17 | 0.07 | 0.16 | 0.08 | 0.13 |
| 3 | 183099294 | rs4859178 | 1.041 | -0.644 | 0.058 | 0.285 | -0.009 | -0.009 | -0.009 | 0.044 | 0.63 | 0.58 | 0.8 | 0.75 | 0.76 | 0.75 |
| 3 | 183103260 | rs2136233 | 0.175 | -0.324 | -0.292 | 0.025 | -0.006 | -0.013 | -0.013 | 0 | 0.84 | 0.83 | 0.9 | 0.94 | 0.93 | 0.8 |
| 3 | 183110863 | rs879713 | -0.816 | 0.224 | -0.938 | -0.216 | -0.013 | 0.103 | 0.103 | 0 | 0.52 | 0.73 | 0.49 | 0.31 | 0.46 | 0.48 |
| 3 | 183112220 | rs16857430 | -0.5 | 0.439 | -0.832 | -0.431 | 0.019 | 0.022 | 0.022 | 0 | 0.39 | 0.6 | 0.39 | 0.22 | 0.27 | 0.37 |
| 3 | 183125354 | rs12491399 | -0.559 | 0.226 | -1.011 | -0.65 | -0.064 | 0.104 | 0.104 | 0 | 0.31 | 0.07 | 0.25 | 0.14 | 0.33 | 0.29 |
| 3 | 183127212 | rs7652892 | -0.619 | 0.143 | 0.069 | -0.753 | 0.018 | -0.022 | -0.022 | 4.561 | 0.14 | 0.27 | 0.23 | 0.15 | 0.16 | 0.15 |
| 3 | 183141028 | rs11709177 | -2.053 | 0.222 | 0.115 | -0.832 | -0.011 | 0.115 | 0.115 | 0 | 0.48 | 0.7 | 0.35 | 0.23 | 0.43 | 0.39 |
| 3 | 183141152 | rs16857451 | -0.636 | 0.233 | 0.438 | -0.883 | 0.016 | 0.01 | 0.01 | 0 | 0.06 | 0.09 | 0.04 | 0.09 | 0.08 | 0.07 |
| 3 | 183142544 | rs4859188 | -2.115 | 0.508 | 0.397 | -0.806 | -0.001 | 0.116 | 0.116 | 0 | 0.48 | 0.7 | 0.36 | 0.24 | 0.43 | 0.39 |
| 3 | 183144470 | rs16857470 | -2.084 | 0.058 | 1.216 | -0.697 | 0.101 | 0.069 | 0.069 | 0.103 | 0.08 | 0.33 | 0.07 | 0.09 | 0.08 | 0.04 |
| 3 | 183147036 | rs4859190 | -1.518 | 0.403 | 1.276 | -0.748 | -0.002 | -0.009 | -0.009 | 0.898 | 0.33 | 0.6 | 0.28 | 0.21 | 0.28 | 0.28 |
| 3 | 183151660 | rs10937129 | -0.682 | 0.067 | 1.22 | -0.706 | 0.033 | 0.007 | 0.007 | 0 | 0.24 | 0.05 | 0.32 | 0.19 | 0.17 | 0.25 |
| 3 | 183152681 | rs1509012 | 0.574 | -0.03 | 0.991 | -1.111 | 0.001 | 0.006 | 0.006 | 0 | 0.78 | 0.86 | 0.9 | 0.98 | 0.81 | 0.84 |
| 3 | 183161510 | rs6805188 | -1.157 | -0.115 | 1.049 | -0.895 | -0.005 | 0.001 | 0.001 | 0 | 0.32 | 0.32 | 0.4 | 0.38 | 0.26 | 0.37 |
| 3 | 183182376 | rs3732592 | 1.114 | 0.095 | 0.685 | -0.73 | 0.034 | 0.104 | 0.104 | 0 | 0.55 | 0.88 | 0.56 | 0.68 | 0.63 | 0.52 |
| 3 | 183182461 | rs3732591 | -2.004 | 0.149 | 0.879 | -0.651 | 0.066 | 0.044 | 0.044 | 0 | 0.1 | 0.34 | 0.06 | 0.09 | 0.1 | 0.05 |
| 3 | 183183776 | rs6809726 | 1.811 | 0.149 | 0.646 | -0.669 | -0.005 | 0.032 | 0.032 | 0 | 0.46 | 0.61 | 0.51 | 0.68 | 0.54 | 0.49 |
| 3 | 183193375 | rs9828957 | 1.067 | -0.141 | -0.109 | -0.987 | -0.012 | 0.058 | 0.058 | 0 | 0.65 | 0.85 | 0.69 | 0.84 | 0.66 | 0.72 |
| 3 | 183194506 | rs11919592 | -0.24 | -0.389 | 0.164 | -0.885 | -0.013 | -0.008 | -0.008 | 0 | 0.17 | 0.25 | 0.11 | 0.14 | 0.13 | 0.21 |
| 3 | 183195708 | rs2686423 | -0.006 | -0.178 | -0.321 | -0.75 | -0.006 | -0.02 | -0.02 | 0 | 0.9 | 1 | 0.98 | 0.93 | 0.93 | 0.93 |
| 3 | 183195991 | rs2971459 | -0.006 | -0.178 | -0.321 | -0.75 | -0.006 | -0.02 | -0.02 | 0 | 0.9 | 1 | 0.98 | 0.93 | 0.93 | 0.93 |
| 3 | 183196803 | rs869023 | 1.463 | -0.204 | -0.305 | -0.769 | -0.012 | 0.061 | 0.061 | 0 | 0.65 | 0.87 | 0.69 | 0.84 | 0.66 | 0.72 |
| 3 | 183197823 | rs6793484 | 0.769 | -0.205 | -0.303 | -0.982 | -0.012 | 0.009 | 0.009 | 0 | 0.86 | 0.91 | 0.94 | 0.95 | 0.81 | 0.91 |
| 3 | 183198269 | rs7638522 | 1.432 | -0.219 | -0.295 | -0.976 | -0.008 | 0.089 | 0.089 | 0 | 0.65 | 0.87 | 0.69 | 0.86 | 0.67 | 0.72 |
| 3 | 183200368 | rs2686431 | 1.459 | 0.12 | -0.023 | -0.657 | 0.003 | -0.004 | -0.004 | 0 | 0.78 | 0.7 | 0.88 | 0.8 | 0.76 | 0.73 |
| 3 | 183203289 | rs11923573 | -0.785 | 0.227 | 0.428 | -0.614 | 0.002 | -0.05 | -0.05 | 8.054 | 0.11 | 0.32 | 0.06 | 0.02 | 0.06 | 0.12 |
| 3 | 183203603 | rs872426 | 1.041 | 0.124 | -1.049 | -0.61 | -0.034 | -0.025 | -0.025 | 0.601 | 0.3 | 0.21 | 0.48 | 0.61 | 0.4 | 0.41 |
| 3 | 183204923 | rs17543340 | -0.41 | 0.214 | -1.44 | -0.591 | 0 | -0.018 | -0.018 | 0.328 | 0.12 | 0.11 | 0.05 | 0.05 | 0.12 | 0.01 |
| 3 | 183204997 | rs906721 | -0.303 | -0.388 | -1.84 | -0.523 | -0.027 | 0.166 | 0.166 | 3.02 | 0.57 | 0.97 | 0.64 | 0.85 | 0.59 | 0.62 |
| 3 | 183207072 | rs16857672 | -0.53 | -0.521 | -0.312 | -0.429 | -0.052 | -0.068 | -0.068 | 1.005 | 0.11 | 0.3 | 0.04 | 0 | 0.02 | 0.01 |
| 3 | 183207297 | rs1553693 | -0.32 | 0.054 | -1.582 | -0.049 | -0.055 | 0.016 | 0.016 | 0.55 | 0.58 | 0.96 | 0.54 | 0.74 | 0.59 | 0.7 |
| 3 | 183208737 | rs7644322 | -0.147 | 0.648 | -0.642 | -0.644 | -0.011 | 0.009 | 0.009 | 0.162 | 0.64 | 0.69 | 0.53 | 0.8 | 0.47 | 0.65 |
| 3 | 183210972 | rs1001190 | 0.635 | 0.314 | -0.177 | -1.005 | -0.013 | -0.024 | -0.024 | 0.166 | 0.45 | 0.52 | 0.45 | 0.79 | 0.31 | 0.51 |
| 3 | 183210991 | rs1979983 | 0.61 | 0.271 | -0.108 | -1.016 | 0.005 | 0.005 | 0.005 | 2.423 | 0.45 | 0.58 | 0.45 | 0.79 | 0.3 | 0.5 |
| 3 | 183211906 | rs1520101 | -0.85 | 0.277 | 0.024 | -1.078 | -0.083 | 0.032 | 0.032 | 0.313 | 0.69 | 0.87 | 0.58 | 0.8 | 0.48 | 0.67 |
| 3 | 183212026 | rs2256061 | -0.346 | -0.09 | 0.055 | -1.001 | 0.001 | -0.026 | -0.026 | 0 | 0.23 | 0.24 | 0.12 | 0.01 | 0.18 | 0.16 |
| 3 | 183215287 | rs2971458 | -0.582 | -0.181 | 0.045 | -0.516 | -0.006 | -0.004 | -0.004 | 0 | 0.26 | 0.36 | 0.16 | 0.22 | 0.18 | 0.24 |
| 3 | 183226378 | rs7637137 | 0.221 | -0.152 | 0.38 | -0.706 | 0.013 | 0.007 | 0.007 | 0.205 | 0.52 | 0.44 | 0.4 | 0.6 | 0.31 | 0.54 |
| 3 | 183234142 | rs7623174 | -0.1 | 0.481 | 0.417 | -0.467 | 0 | 0 | 0 | 0 | 0.38 | 0.41 | 0.49 | 0.34 | 0.58 | 0.43 |
| 3 | 183234383 | rs10804891 | 0.133 | 0.477 | 0.169 | -0.427 | -0.041 | 0.088 | 0.088 | 0.265 | 0.35 | 0.16 | 0.44 | 0.16 | 0.58 | 0.37 |
| 3 | 183236106 | rs12496193 | -0.086 | 0.482 | 0.173 | -0.427 | -0.007 | -0.001 | -0.001 | 1.458 | 0.38 | 0.34 | 0.48 | 0.34 | 0.58 | 0.43 |
| 3 | 183248484 | rs2056324 | 0.223 | 0.61 | -0.063 | -0.746 | -0.058 | 0.077 | 0.077 | 0.221 | 0.33 | 0.11 | 0.45 | 0.17 | 0.58 | 0.36 |
| 3 | 183259865 | rs6800237 | 0.479 | 0.546 | 0.424 | -0.466 | -0.011 | 0.025 | 0.025 | 0.471 | 0.78 | 0.83 | 0.85 | 0.99 | 0.77 | 0.81 |
| 3 | 183286534 | rs4859121 | 0.427 | 0.507 | 0.329 | -0.121 | -0.001 | 0 | 0 | 1.467 | 0.63 | 0.41 | 0.59 | 0.69 | 0.64 | 0.59 |
| 3 | 183292676 | rs4859211 | 0.881 | -0.089 | 0.075 | -0.685 | -0.02 | 0.075 | 0.075 | 0 | 0.43 | 0.27 | 0.5 | 0.82 | 0.43 | 0.57 |
| 3 | 183293954 | rs11718362 | 1.074 | -0.161 | -0.082 | -0.732 | -0.058 | 0.118 | 0.118 | 0.014 | 0.41 | 0.12 | 0.47 | 0.82 | 0.42 | 0.57 |
| 3 | 183295453 | rs9290759 | 0.99 | -0.001 | 0.097 | -0.633 | -0.018 | 0.067 | 0.067 | 0.231 | 0.42 | 0.28 | 0.49 | 0.79 | 0.42 | 0.57 |
| 3 | 183297187 | rs6807272 | 0.488 | -0.581 | 0.186 | -0.571 | 0 | -0.022 | -0.022 | 0.258 | 0.72 | 0.73 | 0.7 | 0.96 | 0.7 | 0.8 |
| 3 | 183301774 | rs9824873 | -1.167 | -0.431 | 0.333 | 0.149 | -0.02 | -0.007 | -0.007 | 0 | 0.32 | 0.59 | 0.29 | 0.3 | 0.24 | 0.27 |
| 3 | 183306876 | rs9856481 | -1.582 | 0.103 | 0.871 | 0.693 | -0.002 | 0.124 | 0.124 | 0 | 0.13 | 0.44 | 0.17 | 0.24 | 0.08 | 0.1 |
| 3 | 183309763 | rs11706582 | -1.076 | 0.274 | 0.857 | 0.298 | 0.047 | -0.032 | -0.032 | 0.709 | 0.09 | 0.38 | 0.04 | 0 | 0.05 | 0.02 |
| 3 | 183312587 | rs7611339 | -1.249 | 0.426 | 0.559 | 0.931 | 0.005 | 0.075 | 0.075 | 0 | 0.14 | 0.28 | 0.19 | 0.28 | 0.16 | 0.15 |
| 3 | 183320754 | rs2313214 | -1.555 | 0.328 | 1.792 | 0.927 | 0.027 | 0.208 | 0.208 | 0 | 0.14 | 0.48 | 0.18 | 0.29 | 0.14 | 0.13 |
| 3 | 183335081 | rs10155052 | -2.03 | 0.195 | 1.737 | 1.245 | -0.004 | 0.174 | 0.174 | 0 | 0.21 | 0.52 | 0.22 | 0.42 | 0.2 | 0.16 |
| 3 | 183335322 | rs9290763 | -0.499 | 0.192 | 1.618 | 1.263 | -0.001 | 0.003 | 0.003 | 0 | 0.06 | 0.07 | 0.04 | 0.11 | 0.05 | 0.04 |
| 3 | 183339096 | rs6799350 | -2.425 | 0.214 | 2.021 | 1.238 | 0.002 | -0.006 | -0.006 | 0 | 0.23 | 0.76 | 0.24 | 0.42 | 0.2 | 0.16 |
| 3 | 183352334 | rs6777400 | -2.562 | 0.001 | 2.581 | 1.365 | 0.014 | -0.053 | -0.053 | 1.439 | 0.29 | 0.83 | 0.3 | 0.42 | 0.26 | 0.17 |
| 3 | 183368891 | rs2293031 | -2.281 | -0.056 | 2.571 | 1.404 | -0.008 | 0.079 | 0.079 | 0 | 0.27 | 0.52 | 0.29 | 0.44 | 0.27 | 0.18 |
| 3 | 183380035 | rs830179 | -2.42 | -0.2 | 2.164 | 0.985 | -0.006 | 0.03 | 0.03 | 2.006 | 0.34 | 0.52 | 0.3 | 0.44 | 0.32 | 0.19 |
| 3 | 183384674 | rs9844222 | -1.935 | -0.186 | 2.291 | 1.022 | -0.007 | 0.11 | 0.11 | 0 | 0.27 | 0.44 | 0.28 | 0.44 | 0.26 | 0.18 |
| 3 | 183386204 | rs9825921 | -2.38 | -0.188 | 2.299 | 1.03 | 0.015 | -0.053 | -0.053 | 0 | 0.3 | 0.82 | 0.3 | 0.43 | 0.27 | 0.18 |
| 3 | 183393037 | rs17621407 | 0.67 | 0.328 | 1.338 | 0.297 | -0.045 | 0.077 | 0.077 | 0 | 0.14 | 0.02 | 0.1 | 0.02 | 0.19 | 0.2 |
| 3 | 183394462 | rs6807489 | -1.424 | 0.29 | 1.551 | 0.44 | 0.003 | 0.037 | 0.037 | 0.27 | 0.24 | 0.35 | 0.25 | 0.29 | 0.26 | 0.15 |

| | | | | | | | | | | | | | | | |
|---|---|---|---|---|---|---|---|---|---|---|---|---|---|---|---|
| 3 | 183397507 | rs9856997 | -1.562 | 0.312 | 1.432 | 0.397 | -0.008 | 0.09 | 0.09 | 0 | 0.27 | 0.5 | 0.28 | 0.44 | 0.26 | 0.18 |
| 3 | 183398071 | rs10937148 | -1.579 | 0.312 | 1.427 | 0.397 | -0.008 | 0.082 | 0.082 | 0 | 0.27 | 0.51 | 0.28 | 0.44 | 0.27 | 0.18 |
| 3 | 183416341 | rs12630496 | -1.444 | 0.225 | 0.904 | 0.408 | -0.001 | -0.003 | -0.003 | 0.093 | 0.32 | 0.66 | 0.45 | 0.32 | 0.34 | 0.3 |
| 3 | 183421149 | rs7619431 | 0.411 | 0.139 | 0.699 | 0.501 | -0.009 | 0.016 | 0.016 | 0 | 0.14 | 0.08 | 0.1 | 0.04 | 0.19 | 0.19 |
| 3 | 183432983 | rs3736535 | -0.12 | -0.061 | 0.969 | 0.675 | 0.001 | 0.036 | 0.036 | 3.016 | 0.13 | 0.14 | 0.33 | 0.29 | 0.14 | 0.11 |
| 3 | 183435754 | rs262985 | 1.362 | -1.201 | -0.053 | 0.491 | 0.031 | 0.009 | 0.009 | 0 | 0.15 | 0.05 | 0.06 | 0.14 | 0.06 | 0.14 |
| 3 | 183443634 | rs262974 | -0.838 | 0.282 | -0.088 | 0.044 | -0.031 | 0.022 | 0.022 | 0.001 | 0.8 | 0.88 | 0.81 | 0.96 | 0.66 | 0.7 |
| 3 | 183452125 | rs262982 | 1.215 | -0.065 | 0.116 | 0.587 | -0.004 | 0.021 | 0.021 | 4.398 | 0.58 | 0.36 | 0.48 | 0.66 | 0.45 | 0.54 |
| 3 | 183473203 | rs262996 | 1.346 | -0.255 | 0.03 | 0.537 | -0.009 | 0.01 | 0.01 | 2.585 | 0.6 | 0.36 | 0.48 | 0.66 | 0.46 | 0.54 |
| 3 | 183476685 | rs262993 | 1.52 | -0.326 | 0.015 | 0.592 | -0.024 | -0.014 | -0.014 | 0 | 0.58 | 0.26 | 0.45 | 0.52 | 0.45 | 0.51 |
| 3 | 183477126 | rs262992 | 1.344 | -0.323 | 0.023 | 0.592 | -0.026 | 0.035 | 0.035 | 2.548 | 0.59 | 0.27 | 0.47 | 0.66 | 0.45 | 0.54 |
| 3 | 183481246 | rs6443904 | -0.501 | -0.303 | 0.352 | 0.563 | 0.008 | 0.082 | 0.082 | 0.918 | 0.16 | 0.3 | 0.3 | 0.3 | 0.18 | 0.15 |
| 3 | 183486117 | rs262956 | -1.127 | -0.011 | -0.023 | 0.487 | -0.056 | 0.06 | 0.06 | 0.127 | 0.81 | 0.92 | 0.81 | 0.96 | 0.66 | 0.67 |
| 3 | 183491045 | rs262963 | 2.445 | -0.823 | -0.583 | 0.253 | 0.001 | -0.015 | -0.015 | 0 | 0.09 | 0.01 | 0.05 | 0.09 | 0.06 | 0.05 |
| 3 | 183521603 | rs2278968 | 1.536 | -0.902 | -1.501 | 0.093 | -0.064 | -0.147 | -0.147 | 0 | 0.41 | 0.01 | 0.38 | 0.44 | 0.36 | 0.46 |
| 3 | 183528442 | rs263024 | 0.745 | -0.382 | -0.697 | 0.402 | -0.005 | -0.004 | -0.004 | 0.062 | 0.59 | 0.38 | 0.41 | 0.51 | 0.47 | 0.51 |
| 3 | 183533376 | rs2582093 | 0.831 | -0.064 | -0.899 | 0.626 | 0.033 | 0.03 | 0.03 | 2.373 | 0.5 | 0.41 | 0.43 | 0.65 | 0.35 | 0.5 |
| 3 | 183534170 | rs1061594 | 1.049 | 0.048 | -0.204 | 1.178 | -0.032 | -0.016 | -0.016 | 0 | 0.59 | 0.25 | 0.46 | 0.52 | 0.47 | 0.51 |
| 3 | 183534538 | rs1061593 | 1.049 | 0.048 | -0.204 | 1.178 | -0.032 | -0.016 | -0.016 | 0 | 0.59 | 0.25 | 0.46 | 0.53 | 0.47 | 0.51 |
| 3 | 183534976 | rs2255015 | 1.083 | 0.067 | -0.268 | 1.313 | -0.017 | -0.004 | -0.004 | 0 | 0.58 | 0.33 | 0.4 | 0.43 | 0.47 | 0.46 |
| 3 | 183535763 | rs897604 | 0.784 | 0.229 | -0.181 | 0.863 | -0.001 | 0 | 0 | 0 | 0.63 | 0.38 | 0.52 | 0.62 | 0.53 | 0.55 |
| 3 | 183537759 | rs2606228 | -0.942 | 0.319 | -0.606 | 0.744 | -0.111 | 0.06 | 0.06 | 1.223 | 0.83 | 0.97 | 0.79 | 0.95 | 0.68 | 0.67 |
| 3 | 183545750 | rs1378909 | -1.109 | 0.118 | -0.259 | 1.103 | 0 | 0.025 | 0.025 | 0 | 0.15 | 0.15 | 0.26 | 0.43 | 0.18 | 0.17 |
| 3 | 183546083 | rs263004 | 1.28 | 0.136 | 0.166 | 0.996 | -0.007 | -0.007 | -0.007 | 0.003 | 0.63 | 0.34 | 0.53 | 0.52 | 0.57 | 0.55 |
| 3 | 183546965 | rs263003 | -0.368 | 0.19 | 0.347 | 0.877 | 0.032 | -0.101 | -0.101 | 0 | 0.79 | 0.6 | 0.8 | 0.95 | 0.74 | 0.72 |
| 3 | 183551123 | rs262998 | -0.383 | 0.249 | 0.362 | 0.892 | 0.034 | -0.105 | -0.105 | 0.948 | 0.79 | 0.58 | 0.81 | 0.95 | 0.74 | 0.72 |
| 3 | 183555074 | rs9866941 | 0.912 | 0.173 | 0.338 | 1.109 | 0.003 | 0.033 | 0.033 | 0.559 | 0.68 | 0.48 | 0.58 | 0.58 | 0.66 | 0.56 |
| 3 | 183566942 | rs9844777 | -1.114 | 0.383 | 0.795 | 1.362 | 0.024 | -0.064 | -0.064 | 2.243 | 0.82 | 0.76 | 0.84 | 0.95 | 0.76 | 0.74 |
| 3 | 183572790 | rs12107202 | -0.983 | 0.473 | 0.816 | 1.483 | 0.029 | 0.042 | 0.042 | 0 | 0.12 | 0.22 | 0.13 | 0.26 | 0.19 | 0.13 |
| 3 | 183575770 | rs12631031 | 0.958 | -0.657 | 0.55 | 1.6 | -0.015 | -0.035 | -0.035 | 0 | 0.08 | 0.03 | 0.23 | 0.28 | 0.14 | 0.17 |
| 3 | 183577904 | rs6809370 | 1.2 | -0.665 | 0.63 | 1.503 | 0.015 | 0.013 | 0.013 | 0.754 | 0.61 | 0.5 | 0.5 | 0.41 | 0.48 | 0.4 |
| 3 | 183585541 | rs1402000 | -0.689 | -0.669 | 0.63 | 1.526 | 0.034 | -0.021 | -0.021 | 0.69 | 0.79 | 0.75 | 0.8 | 0.86 | 0.68 | 0.71 |
| 3 | 183585883 | rs953419 | -0.75 | -0.606 | 0.439 | 1.485 | -0.002 | 0.048 | 0.048 | 0 | 0.18 | 0.26 | 0.3 | 0.46 | 0.2 | 0.33 |
| 3 | 183586005 | rs3749446 | -0.718 | -0.602 | 0.43 | 1.483 | 0.036 | -0.027 | -0.027 | 0 | 0.8 | 0.75 | 0.8 | 0.87 | 0.68 | 0.72 |
| 3 | 183592567 | rs6776999 | 0.315 | -0.607 | 0.16 | 1.478 | 0.012 | 0.03 | 0.03 | 0.346 | 0.71 | 0.65 | 0.58 | 0.49 | 0.61 | 0.44 |
| 3 | 183596184 | rs7644746 | 0.682 | -0.582 | 0.162 | 1.497 | 0.003 | 0.004 | 0.004 | 0.36 | 0.65 | 0.63 | 0.54 | 0.49 | 0.55 | 0.42 |
| 3 | 183607661 | rs6792309 | -1.447 | -0.395 | 0.382 | 1.8 | 0.015 | 0.105 | 0.105 | 0 | 0.23 | 0.57 | 0.36 | 0.54 | 0.28 | 0.34 |
| 3 | 183621991 | rs17683012 | -0.699 | -0.369 | 0.123 | 1.614 | 0.031 | 0.145 | 0.145 | 0.089 | 0.16 | 0.39 | 0.3 | 0.39 | 0.2 | 0.22 |
| 3 | 183624010 | rs16858210 | -1.309 | -0.399 | -0.056 | 1.623 | 0.018 | 0.139 | 0.139 | 0.076 | 0.23 | 0.42 | 0.34 | 0.55 | 0.28 | 0.36 |
| 3 | 183627089 | rs872013 | 0.72 | -0.487 | -0.718 | 1.336 | -0.034 | 0.041 | 0.041 | 0 | 0.37 | 0.19 | 0.25 | 0.26 | 0.38 | 0.39 |
| 3 | 183628585 | rs11716770 | 0.309 | -0.188 | -0.663 | 1.468 | -0.022 | -0.056 | -0.056 | 0.014 | 0.1 | 0.03 | 0.22 | 0.31 | 0.12 | 0.15 |
| 3 | 183632423 | rs7613247 | -1.279 | -0.313 | -0.102 | 1.011 | 0.029 | -0.011 | -0.011 | 0 | 0.12 | 0.59 | 0.1 | 0.07 | 0.1 | 0.07 |
| 3 | 183634879 | rs1533682 | -0.514 | -0.566 | -0.472 | 1.046 | 0.01 | 0.141 | 0.141 | 0 | 0.55 | 0.82 | 0.6 | 0.85 | 0.57 | 0.58 |
| 3 | 183635488 | rs1533683 | -0.102 | -0.547 | -0.382 | 1.228 | -0.001 | 0.06 | 0.06 | 0 | 0.48 | 0.7 | 0.54 | 0.68 | 0.49 | 0.45 |
| 3 | 183635768 | rs1000002 | 0.345 | -0.62 | -0.217 | 1.517 | -0.074 | -0.06 | -0.06 | 1.121 | 0.43 | 0.13 | 0.45 | 0.51 | 0.48 | 0.44 |
| 3 | 183635815 | rs1533684 | -0.661 | -0.668 | -0.195 | 1.534 | 0.013 | 0.144 | 0.144 | 0 | 0.55 | 0.82 | 0.6 | 0.85 | 0.57 | 0.58 |
| 3 | 183635997 | rs2313217 | -0.661 | -0.668 | -0.196 | 1.534 | 0.01 | 0.142 | 0.142 | 0 | 0.55 | 0.82 | 0.6 | 0.85 | 0.57 | 0.58 |
| 3 | 183636195 | rs6768149 | 0.353 | -0.693 | -0.196 | 1.535 | -0.077 | -0.06 | -0.06 | 4.49 | 0.43 | 0.13 | 0.45 | 0.51 | 0.48 | 0.44 |
| 3 | 183636273 | rs2872249 | -0.658 | -0.688 | -0.182 | 1.533 | 0.007 | 0.142 | 0.142 | 0 | 0.55 | 0.82 | 0.6 | 0.85 | 0.57 | 0.58 |
| 3 | 183637722 | rs3805114 | 0.993 | -0.687 | -0.216 | 1.535 | 0.004 | 0.019 | 0.019 | 0 | 0.07 | 0.09 | 0.05 | 0.16 | 0.08 | 0.14 |
| 3 | 183637845 | rs562 | -0.131 | -0.699 | -0.016 | 1.532 | -0.002 | 0.004 | 0.004 | 0 | 0.48 | 0.69 | 0.5 | 0.51 | 0.48 | 0.45 |
| 3 | 183640865 | rs7646621 | 0.026 | -0.663 | 0.091 | 1.536 | -0.005 | 0 | 0 | 0 | 0.47 | 0.63 | 0.49 | 0.51 | 0.48 | 0.44 |
| 3 | 183642303 | rs8180093 | 1.281 | -0.703 | 0.069 | 1.535 | -0.015 | -0.025 | -0.025 | 2.081 | 0.06 | 0.01 | 0.04 | 0.16 | 0.08 | 0.13 |
| 3 | 183642794 | rs1402002 | -0.663 | -0.704 | 0.074 | 1.531 | 0.012 | 0.124 | 0.124 | 0 | 0.55 | 0.78 | 0.6 | 0.84 | 0.57 | 0.58 |
| 3 | 183645662 | rs3805112 | 0.297 | -0.706 | 0.083 | 1.532 | -0.036 | -0.024 | -0.024 | 0.535 | 0.43 | 0.21 | 0.46 | 0.51 | 0.48 | 0.44 |
| 3 | 183649870 | rs4148593 | 0.415 | -0.699 | 0.006 | 1.514 | -0.002 | 0 | 0 | 0 | 0.4 | 0.58 | 0.32 | 0.4 | 0.4 | 0.39 |
| 3 | 183650330 | rs2037379 | -0.755 | -0.755 | 0.016 | 1.511 | 0.01 | 0.139 | 0.139 | 0 | 0.55 | 0.82 | 0.6 | 0.84 | 0.57 | 0.58 |
| 3 | 183652736 | rs9858774 | -0.728 | -0.815 | 0.009 | 1.51 | 0.01 | 0.136 | 0.136 | 0 | 0.55 | 0.81 | 0.6 | 0.84 | 0.57 | 0.58 |
| 3 | 183653718 | rs4912515 | 0.389 | -0.792 | 0.022 | 1.518 | -0.002 | 0.001 | 0.001 | 0 | 0.4 | 0.58 | 0.32 | 0.4 | 0.39 | 0.39 |
| 3 | 183653937 | rs2139559 | 1.335 | -0.787 | 0.026 | 1.515 | -0.034 | 0.014 | 0.014 | 0 | 0.31 | 0.14 | 0.23 | 0.24 | 0.31 | 0.25 |
| 3 | 183660585 | rs3749442 | -0.311 | -0.77 | 0.024 | 1.467 | 0.004 | 0.056 | 0.056 | 0.016 | 0.15 | 0.19 | 0.27 | 0.44 | 0.18 | 0.19 |
| 3 | 183663173 | rs9812777 | 0.647 | -0.774 | 0.026 | 1.535 | -0.003 | 0.054 | 0.054 | 0.725 | 0.1 | 0.14 | 0.23 | 0.35 | 0.1 | 0.14 |
| 3 | 183663833 | rs2292998 | -0.733 | -0.796 | 0.008 | 1.542 | 0.007 | 0.149 | 0.149 | 0 | 0.55 | 0.81 | 0.6 | 0.85 | 0.57 | 0.57 |
| 3 | 183664678 | rs9838667 | -1.025 | -0.774 | 0.008 | 1.545 | 0.005 | 0.043 | 0.043 | 0 | 0.21 | 0.27 | 0.32 | 0.6 | 0.25 | 0.32 |
| 3 | 183665742 | rs2280392 | 1.183 | -0.643 | 0.399 | 1.946 | -0.013 | -0.025 | -0.025 | 0 | 0.06 | 0.01 | 0.04 | 0.16 | 0.08 | 0.14 |
| 3 | 183666862 | rs1464322 | 1.25 | -0.584 | 0.601 | 1.954 | -0.102 | -0.063 | -0.063 | 0 | 0.29 | 0.02 | 0.22 | 0.26 | 0.31 | 0.25 |
| 3 | 183670642 | rs4148585 | -1.034 | -0.786 | 0.611 | 1.959 | 0.006 | 0.156 | 0.156 | 0 | 0.55 | 0.82 | 0.6 | 0.86 | 0.57 | 0.57 |
| 3 | 183670672 | rs3817404 | 1.004 | -0.759 | 0.613 | 1.961 | -0.01 | 0.006 | 0.006 | 0 | 0.31 | 0.21 | 0.25 | 0.26 | 0.32 | 0.25 |
| 3 | 183672280 | rs2139564 | -0.578 | -0.822 | 0.611 | 1.968 | 0.004 | 0.056 | 0.056 | 0.016 | 0.15 | 0.19 | 0.27 | 0.44 | 0.18 | 0.19 |

| | | | | | | | | | | | | | | | |
|---|---|---|---|---|---|---|---|---|---|---|---|---|---|---|---|
| 3 | 183679532 | rs6443924 | -0.99 | -0.849 | 0.614 | 1.956 | 0.011 | 0.147 | 0.147 | 0 | 0.55 | 0.79 | 0.6 | 0.86 | 0.57 | 0.57 |
| 3 | 183681056 | rs2292999 | 1.419 | -0.847 | 0.612 | 1.962 | -0.009 | -0.023 | -0.023 | 0 | 0.05 | 0.01 | 0.04 | 0.16 | 0.07 | 0.13 |
| 3 | 183682695 | rs2271936 | 1.253 | -0.855 | 0.605 | 1.965 | -0.111 | -0.063 | -0.063 | 0.661 | 0.29 | 0.02 | 0.23 | 0.26 | 0.32 | 0.25 |
| 3 | 183682897 | rs1879257 | -0.438 | -0.856 | 0.615 | 1.965 | 0.004 | 0.048 | 0.048 | 0.062 | 0.15 | 0.19 | 0.27 | 0.44 | 0.19 | 0.19 |
| 3 | 183683938 | rs3792583 | -0.438 | -0.849 | 0.62 | 1.978 | 0.004 | 0.048 | 0.048 | 0.062 | 0.15 | 0.19 | 0.27 | 0.44 | 0.19 | 0.19 |
| 3 | 183685165 | rs4148581 | 1.253 | -0.849 | 0.619 | 1.978 | -0.111 | -0.063 | -0.063 | 3.492 | 0.29 | 0.02 | 0.23 | 0.26 | 0.32 | 0.25 |
| 3 | 183685183 | rs4148580 | 0.624 | -0.851 | 0.621 | 1.978 | -0.011 | -0.052 | -0.052 | 0 | 0.1 | 0.02 | 0.22 | 0.35 | 0.11 | 0.14 |
| 3 | 183685249 | rs4148579 | -1.013 | -0.851 | 0.622 | 1.977 | 0.007 | 0.155 | 0.155 | 0 | 0.55 | 0.81 | 0.6 | 0.86 | 0.57 | 0.57 |
| 3 | 183685534 | rs939336 | -1.004 | -0.851 | 0.621 | 1.981 | 0.011 | 0.15 | 0.15 | 0 | 0.55 | 0.79 | 0.6 | 0.86 | 0.57 | 0.57 |
| 3 | 183685893 | rs6800217 | -1.004 | -0.851 | 0.621 | 1.968 | 0.011 | 0.147 | 0.147 | 0 | 0.55 | 0.79 | 0.6 | 0.86 | 0.57 | 0.57 |
| 3 | 183688627 | rs9861983 | -0.449 | -0.851 | 0.622 | 1.968 | 0.004 | 0.048 | 0.048 | 0.062 | 0.15 | 0.2 | 0.28 | 0.44 | 0.19 | 0.19 |
| 3 | 183689138 | rs869417 | -0.994 | -0.851 | 0.621 | 1.968 | 0.011 | 0.148 | 0.148 | 0 | 0.55 | 0.79 | 0.6 | 0.86 | 0.57 | 0.57 |
| 3 | 183690243 | rs869335 | 1.421 | -0.84 | 0.623 | 1.964 | -0.016 | -0.031 | -0.031 | 0 | 0.05 | 0 | 0.04 | 0.16 | 0.07 | 0.12 |
| 3 | 183691451 | rs3828469 | -0.452 | -0.808 | 0.636 | 1.743 | 0.004 | 0.048 | 0.048 | 0.062 | 0.15 | 0.19 | 0.27 | 0.44 | 0.19 | 0.19 |
| 3 | 183691725 | rs3805107 | 1.572 | -0.808 | 0.636 | 1.743 | -0.017 | -0.03 | -0.03 | 3.534 | 0.05 | 0 | 0.04 | 0.16 | 0.07 | 0.12 |
| 3 | 183694602 | rs3792582 | 1.26 | -0.808 | 0.635 | 1.743 | -0.111 | -0.056 | -0.056 | 0.725 | 0.29 | 0.02 | 0.23 | 0.25 | 0.32 | 0.25 |
| 3 | 183695870 | rs3792581 | -0.454 | -0.807 | 0.638 | 1.714 | 0.004 | 0.049 | 0.049 | 0.062 | 0.15 | 0.19 | 0.27 | 0.44 | 0.19 | 0.19 |
| 3 | 183696077 | rs4148578 | -0.454 | -0.807 | 0.638 | 1.714 | 0.004 | 0.049 | 0.049 | 0.062 | 0.15 | 0.19 | 0.27 | 0.44 | 0.19 | 0.19 |
| 3 | 183696402 | rs1132776 | -1.107 | -0.805 | 0.645 | 1.708 | 0.015 | 0.124 | 0.124 | 0 | 0.55 | 0.75 | 0.59 | 0.85 | 0.57 | 0.57 |
| 3 | 183698219 | rs6775518 | -0.459 | -0.8 | 0.647 | 1.719 | 0.004 | 0.049 | 0.049 | 0.062 | 0.15 | 0.19 | 0.27 | 0.44 | 0.19 | 0.19 |
| 3 | 183698341 | rs4912471 | 1.258 | -0.804 | 0.635 | 1.685 | -0.111 | -0.056 | -0.056 | 0.452 | 0.29 | 0.02 | 0.23 | 0.25 | 0.32 | 0.25 |
| 3 | 183698943 | rs6802849 | -1.025 | -0.808 | 0.643 | 1.684 | 0.006 | 0.149 | 0.149 | 0 | 0.55 | 0.82 | 0.6 | 0.85 | 0.57 | 0.57 |
| 3 | 183699516 | rs7636910 | 0.47 | -0.808 | 0.658 | 1.685 | -0.017 | -0.011 | -0.011 | 0 | 0.37 | 0.22 | 0.29 | 0.42 | 0.39 | 0.38 |
| 3 | 183699781 | rs2872247 | 1.26 | -0.808 | 0.656 | 1.685 | -0.111 | -0.056 | -0.056 | 2.386 | 0.29 | 0.02 | 0.23 | 0.25 | 0.32 | 0.25 |
| 3 | 183700211 | rs2293001 | 0.147 | -0.792 | 0.26 | 1.683 | 0 | -0.001 | -0.001 | 0 | 0.4 | 0.55 | 0.31 | 0.42 | 0.39 | 0.38 |
| 3 | 183700928 | rs2313212 | -0.98 | -0.792 | 0.259 | 1.682 | 0.011 | 0.142 | 0.142 | 0 | 0.55 | 0.79 | 0.6 | 0.85 | 0.57 | 0.57 |
| 3 | 183701147 | rs939337 | -0.989 | -0.792 | 0.259 | 1.682 | 0.007 | 0.149 | 0.149 | 0 | 0.55 | 0.81 | 0.6 | 0.85 | 0.57 | 0.57 |
| 3 | 183702089 | rs3749440 | 0.147 | -0.792 | 0.259 | 1.683 | 0 | -0.001 | -0.001 | 0 | 0.4 | 0.55 | 0.31 | 0.42 | 0.39 | 0.38 |
| 3 | 183702275 | rs4148575 | -1.085 | -0.792 | 0.259 | 1.678 | 0.015 | 0.124 | 0.124 | 0 | 0.55 | 0.75 | 0.6 | 0.85 | 0.57 | 0.57 |
| 3 | 183704068 | rs939338 | -0.987 | -0.79 | 0.248 | 1.679 | 0.007 | 0.149 | 0.149 | 0 | 0.55 | 0.81 | 0.61 | 0.85 | 0.57 | 0.57 |
| 3 | 183706630 | rs6443926 | -1.094 | -0.789 | 0.243 | 1.68 | 0.015 | 0.124 | 0.124 | 0 | 0.55 | 0.75 | 0.59 | 0.85 | 0.57 | 0.57 |
| 3 | 183708327 | rs10937157 | -0.481 | -0.788 | 0.245 | 1.685 | 0.005 | 0.06 | 0.06 | 0.052 | 0.16 | 0.22 | 0.28 | 0.44 | 0.19 | 0.19 |
| 3 | 183708439 | rs10937158 | -0.948 | -0.789 | 0.234 | 1.658 | 0.014 | 0.132 | 0.132 | 0 | 0.55 | 0.77 | 0.59 | 0.85 | 0.57 | 0.57 |
| 3 | 183714063 | rs16858328 | 0.528 | -0.786 | 0.24 | 1.68 | -0.001 | 0.031 | 0.031 | 0.729 | 0.1 | 0.11 | 0.22 | 0.35 | 0.11 | 0.14 |
| 3 | 183714467 | rs4148573 | 1.103 | -0.801 | 0.223 | 1.671 | -0.008 | -0.024 | -0.024 | 0.025 | 0.06 | 0.01 | 0.04 | 0.16 | 0.07 | 0.12 |
| 3 | 183716604 | rs6773404 | -0.344 | -0.753 | 0.271 | 1.688 | 0.003 | 0.233 | 0.233 | 0 | 0.13 | 0.47 | 0.25 | 0.35 | 0.11 | 0.14 |
| 3 | 183719175 | rs9840067 | 0.396 | -0.705 | 0.178 | 1.705 | -0.002 | 0.047 | 0.047 | 0.866 | 0.11 | 0.14 | 0.23 | 0.35 | 0.11 | 0.14 |
| 3 | 183720576 | rs4148571 | -0.99 | -0.721 | 0.135 | 1.687 | 0.009 | 0.13 | 0.13 | 0 | 0.55 | 0.78 | 0.59 | 0.85 | 0.57 | 0.56 |
| 3 | 183720745 | rs4148570 | 0.413 | -0.597 | 0.159 | 1.742 | -0.011 | -0.054 | -0.054 | 0 | 0.1 | 0.02 | 0.22 | 0.35 | 0.11 | 0.14 |
| 3 | 183722779 | rs4148569 | 1.479 | -0.596 | 0.148 | 1.72 | -0.01 | -0.022 | -0.022 | 0 | 0.05 | 0.01 | 0.04 | 0.16 | 0.07 | 0.12 |
| 3 | 183723021 | rs4148568 | 0.092 | -0.632 | 0.162 | 1.899 | 0.003 | -0.016 | -0.016 | 0.294 | 0.11 | 0.09 | 0.2 | 0.24 | 0.11 | 0.13 |
| 3 | 183723075 | rs4148567 | -1.044 | -0.625 | 0.103 | 1.126 | 0.008 | 0.131 | 0.131 | 0 | 0.54 | 0.78 | 0.59 | 0.85 | 0.57 | 0.57 |
| 3 | 183724072 | rs2292997 | 0.668 | -0.604 | 0.147 | 1.164 | -0.003 | -0.032 | -0.032 | 0 | 0.1 | 0.06 | 0.23 | 0.36 | 0.11 | 0.14 |
| 3 | 183725982 | rs4148564 | -1.035 | -0.621 | 0.168 | 1.144 | 0.01 | 0.096 | 0.096 | 0 | 0.56 | 0.73 | 0.59 | 0.85 | 0.57 | 0.56 |
| 3 | 183726218 | rs4148562 | 0.714 | -0.636 | 0.14 | 1.14 | -0.009 | -0.009 | -0.009 | 0 | 0.05 | 0.01 | 0.04 | 0.08 | 0.08 | 0.05 |
| 3 | 183727455 | rs4148559 | 1.47 | -0.643 | 0.137 | 1.135 | -0.014 | -0.031 | -0.031 | 0.001 | 0.05 | 0 | 0.04 | 0.16 | 0.06 | 0.12 |
| 3 | 183729111 | rs7624838 | 0.458 | -0.656 | 0.206 | 1.11 | -0.01 | -0.007 | -0.007 | 0 | 0.36 | 0.25 | 0.29 | 0.42 | 0.38 | 0.38 |
| 3 | 183729487 | rs6443928 | -1.065 | -0.654 | 0.23 | 1.111 | 0.012 | 0.075 | 0.075 | 0 | 0.54 | 0.67 | 0.58 | 0.85 | 0.57 | 0.56 |
| 3 | 183730295 | rs4074672 | 0.431 | -0.627 | -0.417 | 1.123 | -0.01 | -0.007 | -0.007 | 0.076 | 0.36 | 0.25 | 0.29 | 0.42 | 0.38 | 0.38 |
| 3 | 183733784 | rs4148557 | -0.964 | -0.498 | -0.696 | 1.14 | 0.012 | 0.081 | 0.081 | 0 | 0.55 | 0.68 | 0.59 | 0.85 | 0.57 | 0.56 |
| 3 | 183741134 | rs7613237 | -0.037 | -0.456 | -0.78 | 1.176 | 0.027 | 0.254 | 0.254 | 0 | 0.12 | 0.47 | 0.25 | 0.36 | 0.11 | 0.12 |
| 3 | 183745663 | rs939335 | -0.6 | -0.4 | -1.036 | 1.091 | -0.01 | 0.116 | 0.116 | 0 | 0.51 | 0.78 | 0.56 | 0.79 | 0.5 | 0.53 |
| 3 | 183749731 | rs939334 | 0.527 | -0.279 | -1.025 | 1.026 | 0.004 | -0.015 | -0.015 | 0 | 0.35 | 0.49 | 0.29 | 0.28 | 0.33 | 0.29 |
| 3 | 183752301 | rs10937159 | -0.377 | -0.3 | -1.353 | 0.994 | 0 | 0.056 | 0.056 | 0 | 0.52 | 0.31 | 0.58 | 0.69 | 0.51 | 0.46 |
| 3 | 183752773 | rs12493550 | 0.881 | -0.34 | -1.149 | 0.552 | 0.005 | 0.006 | 0.006 | 0 | 0.11 | 0.04 | 0.11 | 0.04 | 0.08 | 0.05 |
| 3 | 183753337 | rs7622660 | -0.867 | -0.157 | -0.806 | 0.693 | -0.021 | 0.102 | 0.102 | 0 | 0.68 | 0.88 | 0.71 | 0.92 | 0.65 | 0.63 |
| 3 | 183754029 | rs6792482 | 0.164 | -0.108 | -0.8 | 0.759 | -0.009 | -0.007 | -0.007 | 0 | 0.52 | 0.71 | 0.46 | 0.48 | 0.47 | 0.47 |
| 3 | 183758506 | rs7621975 | -0.943 | -0.046 | 0.32 | 1.453 | 0.006 | 0.095 | 0.095 | 0 | 0.19 | 0.65 | 0.3 | 0.46 | 0.18 | 0.15 |
| 3 | 183761125 | rs10937161 | -0.51 | 0.269 | -0.683 | 1.272 | 0 | 0.006 | 0.006 | 1.094 | 0.23 | 0.22 | 0.27 | 0.56 | 0.2 | 0.24 |
| 3 | 183771986 | rs7648564 | 0.008 | 0.71 | -1.483 | 0.574 | -0.057 | -0.018 | -0.018 | 0 | 0.66 | 0.88 | 0.7 | 0.67 | 0.61 | 0.43 |
| 3 | 183773771 | rs6762938 | 0.576 | 0.016 | -0.759 | 0.024 | -0.001 | 0.002 | 0.002 | 0 | 0.44 | 0.4 | 0.54 | 0.64 | 0.47 | 0.36 |
| 3 | 183774762 | rs6766410 | 0.598 | 0.095 | -0.694 | 0.119 | -0.001 | 0.002 | 0.002 | 0 | 0.44 | 0.39 | 0.53 | 0.65 | 0.46 | 0.33 |
| 3 | 183778298 | rs6807670 | -0.219 | 0.001 | -0.795 | -0.152 | 0.024 | -0.018 | -0.018 | 0 | 0.69 | 0.5 | 0.73 | 0.76 | 0.68 | 0.51 |
| 3 | 183786537 | rs12492217 | 0.514 | 0.033 | -0.846 | -0.214 | 0 | -0.011 | -0.011 | 0 | 0.55 | 0.49 | 0.68 | 0.75 | 0.57 | 0.42 |
| 3 | 183786678 | rs9869582 | 0.035 | -0.03 | -0.712 | -0.216 | 0.013 | 0.015 | 0.015 | 2.587 | 0.18 | 0.14 | 0.14 | 0.03 | 0.12 | 0.1 |
| 3 | 183797963 | rs4912521 | -0.102 | 0.09 | -0.546 | -0.263 | -0.028 | -0.079 | -0.079 | 0 | 0.19 | 0.38 | 0.16 | 0.03 | 0.14 | 0.15 |
| 3 | 183800463 | rs7648737 | -0.34 | 0.213 | -0.969 | -0.593 | -0.065 | 0.028 | 0.028 | 2.012 | 0.67 | 0.83 | 0.65 | 0.76 | 0.62 | 0.5 |
| 3 | 183800537 | rs7628671 | 0.523 | 0.169 | -0.869 | -0.568 | -0.014 | -0.053 | -0.053 | 0.046 | 0.17 | 0.25 | 0.09 | 0.02 | 0.14 | 0.14 |
| 3 | 183815439 | rs6765267 | 0.646 | 0.247 | -0.367 | -0.583 | 0 | 0.009 | 0.009 | 0 | 0.49 | 0.47 | 0.52 | 0.74 | 0.49 | 0.36 |

| | | | | | | | | | | | | | | | | |
|---|---|---|---|---|---|---|---|---|---|---|---|---|---|---|---|---|
| 3 | 183818416 | rs7627615 | -0.048 | -0.054 | -1.264 | -1.16 | -0.051 | 0.052 | 0.052 | 0.747 | 0.64 | 0.93 | 0.68 | 0.77 | 0.62 | 0.59 |
| 3 | 183819155 | rs7432211 | -1.005 | -0.441 | -0.998 | -0.916 | 0.04 | 0.053 | 0.053 | 0 | 0.49 | 0.33 | 0.45 | 0.34 | 0.43 | 0.52 |
| 3 | 183821694 | rs12634774 | -0.721 | -0.417 | -1.38 | -0.981 | 0 | 0.091 | 0.091 | 0.036 | 0.43 | 0.07 | 0.34 | 0.25 | 0.4 | 0.49 |
| 3 | 183830393 | rs11718245 | 0.253 | -0.236 | -1.16 | -0.943 | -0.019 | 0.063 | 0.063 | 0.805 | 0.15 | 0.06 | 0.1 | 0.03 | 0.17 | 0.21 |
| 3 | 183832187 | rs6443953 | -0.206 | -0.574 | -1.206 | -0.845 | 0.011 | -0.007 | -0.007 | 3.165 | 0.25 | 0.18 | 0.29 | 0.25 | 0.21 | 0.23 |
| 3 | 183834822 | rs1490497 | 1.221 | -0.492 | -1.138 | -0.881 | -0.013 | -0.031 | -0.031 | 1.036 | 0.06 | 0.12 | 0.02 | 0 | 0.04 | 0.01 |
| 3 | 183843135 | rs6806377 | -1.174 | -0.614 | -1.301 | -1.089 | -0.03 | 0.017 | 0.017 | 0 | 0.55 | 0.73 | 0.53 | 0.39 | 0.53 | 0.47 |
| 3 | 183844312 | rs9860004 | -0.355 | -0.547 | -0.611 | -0.885 | -0.026 | 0.041 | 0.041 | 0 | 0.29 | 0.43 | 0.32 | 0.38 | 0.24 | 0.28 |
| 3 | 183851740 | rs11718304 | -0.341 | -0.568 | -0.557 | -0.878 | -0.028 | 0.039 | 0.039 | 0 | 0.29 | 0.43 | 0.32 | 0.38 | 0.23 | 0.28 |
| 3 | 183852861 | rs2292611 | -0.319 | -0.348 | -0.584 | -0.785 | -0.021 | 0.048 | 0.048 | 0 | 0.29 | 0.43 | 0.32 | 0.39 | 0.24 | 0.28 |
| 3 | 183861243 | rs843358 | -0.086 | 0.747 | -0.856 | -0.455 | 0.01 | 0.033 | 0.033 | 0 | 0.27 | 0.33 | 0.36 | 0.53 | 0.3 | 0.44 |
| 3 | 183866486 | rs4912474 | -0.245 | 0.764 | -0.861 | -0.504 | 0.017 | 0.039 | 0.039 | 0 | 0.29 | 0.4 | 0.26 | 0.54 | 0.31 | 0.44 |
| 3 | 183867622 | rs16858473 | 2.136 | 0.546 | -0.844 | -0.673 | -0.007 | -0.019 | -0.019 | 0.711 | 0.05 | 0.09 | 0.02 | 0 | 0.03 | 0.01 |
| 3 | 183876512 | rs1969253 | -1.19 | 0.435 | -1.038 | -0.788 | -0.001 | 0 | 0 | 0 | 0.47 | 0.56 | 0.37 | 0.54 | 0.47 | 0.6 |
| 3 | 183886945 | rs2175525 | 1.406 | 0.242 | -0.815 | -0.947 | 0 | -0.061 | -0.061 | 0.166 | 0.1 | 0.31 | 0.06 | 0 | 0.07 | 0.01 |
| 3 | 183890868 | rs11919795 | -0.605 | 0.193 | -1.14 | -1.056 | -0.029 | -0.171 | -0.171 | 0.119 | 0.31 | 0.53 | 0.18 | 0.01 | 0.23 | 0.19 |
| 3 | 183893519 | rs843346 | 1.411 | 0.762 | -1.69 | -0.191 | -0.073 | -0.114 | -0.114 | 0 | 0.16 | 0.01 | 0.16 | 0.32 | 0.22 | 0.39 |
| 3 | 183896641 | rs2231216 | -0.54 | 0.846 | -1.773 | -0.398 | -0.074 | -0.039 | -0.039 | 0 | 0.41 | 0.18 | 0.38 | 0.57 | 0.47 | 0.57 |
| 3 | 183900338 | rs843367 | -1.014 | 0.684 | -1.212 | -0.438 | 0.003 | 0.019 | 0.019 | 0 | 0.5 | 0.6 | 0.45 | 0.6 | 0.48 | 0.58 |
| 3 | 183900836 | rs843364 | -1.014 | 0.684 | -1.212 | -0.438 | 0.003 | 0.019 | 0.019 | 0 | 0.5 | 0.61 | 0.45 | 0.6 | 0.48 | 0.58 |
| 3 | 183902801 | rs6443954 | -1.025 | 0.492 | -1.211 | -0.436 | 0.003 | 0.019 | 0.019 | 0 | 0.5 | 0.61 | 0.45 | 0.6 | 0.48 | 0.58 |
| 3 | 183904049 | rs11539876 | 2.246 | -0.074 | -2.287 | -0.575 | 0.019 | -0.072 | -0.072 | 0 | 0.22 | 0.02 | 0.33 | 0.31 | 0.19 | 0.1 |
| 3 | 183907879 | rs17774065 | 1.725 | 0.697 | -1.977 | -0.427 | 0.043 | -0.061 | -0.061 | 0.277 | 0.35 | 0.03 | 0.37 | 0.35 | 0.28 | 0.26 |
| 3 | 183919950 | rs7371826 | 0.576 | 0.926 | -1.678 | 0.224 | -0.03 | -0.075 | -0.075 | 0.689 | 0.26 | 0.16 | 0.3 | 0.49 | 0.31 | 0.43 |
| 3 | 183933235 | rs10446348 | 0.667 | 0.624 | -1.663 | -0.192 | -0.021 | 0.08 | 0.08 | 0.012 | 0.14 | 0.02 | 0.08 | 0.01 | 0.14 | 0.15 |
| 3 | 183936680 | rs2668193 | 0.782 | 0.727 | -1.304 | 0.053 | 0.001 | 0.014 | 0.014 | 0 | 0.48 | 0.34 | 0.53 | 0.59 | 0.48 | 0.36 |
| 3 | 183936947 | rs9837978 | -0.466 | 0.222 | -0.753 | 0.029 | -0.044 | -0.045 | -0.045 | 0 | 0.21 | 0.54 | 0.22 | 0.16 | 0.16 | 0.17 |
| 3 | 183938418 | rs9882147 | -0.918 | 0.094 | -0.787 | 0.077 | -0.04 | -0.026 | -0.026 | 0 | 0.27 | 0.48 | 0.22 | 0.29 | 0.19 | 0.21 |
| 3 | 183942187 | rs1709678 | -1.209 | 0.488 | -0.828 | 0.039 | -0.021 | -0.028 | -0.028 | 0 | 0.3 | 0.5 | 0.25 | 0.32 | 0.26 | 0.23 |
| 3 | 183943141 | rs1709679 | -0.281 | 0.513 | -0.831 | 0.035 | -0.009 | 0.021 | 0.021 | 0.235 | 0.2 | 0.09 | 0.18 | 0.23 | 0.23 | 0.22 |
| 3 | 183953972 | rs1709657 | -0.284 | 0.46 | -0.772 | 0.028 | -0.001 | 0.019 | 0.019 | 0.167 | 0.21 | 0.14 | 0.19 | 0.24 | 0.23 | 0.23 |
| 3 | 183957917 | rs16848529 | 1.357 | 0.437 | -0.667 | 0.063 | -0.011 | 0.004 | 0.004 | 0 | 0.06 | 0.12 | 0.04 | 0.08 | 0.03 | 0.01 |
| 3 | 183976103 | rs11546878 | 0.915 | 1.371 | -0.876 | 0.072 | -0.047 | -0.016 | -0.016 | 0 | 0.07 | 0.01 | 0.11 | 0.07 | 0.18 | 0.11 |
| 3 | 183976108 | rs902415 | -0.76 | 1.136 | -0.678 | 0.043 | -0.01 | -0.005 | -0.005 | 0 | 0.25 | 0.48 | 0.25 | 0.34 | 0.25 | 0.23 |
| 3 | 183976241 | rs6767237 | 0.377 | 1.151 | -0.685 | 0.027 | -0.001 | -0.031 | -0.031 | 0.022 | 0.17 | 0.21 | 0.12 | 0.02 | 0.15 | 0.15 |
| 3 | 183977194 | rs9824 | 1.737 | 0.762 | -1.613 | -0.09 | 0.005 | -0.154 | -0.154 | 4.643 | 0.3 | 0.03 | 0.36 | 0.38 | 0.23 | 0.22 |
| 3 | 183982887 | rs1021376 | -0.466 | 0.988 | -1.049 | 0.284 | -0.005 | -0.001 | -0.001 | 0 | 0.24 | 0.44 | 0.25 | 0.34 | 0.24 | 0.23 |
| 3 | 183995341 | rs1001817 | 0.759 | 1.13 | -1.244 | -0.015 | 0.003 | 0.006 | 0.006 | 1.455 | 0.59 | 0.53 | 0.6 | 0.42 | 0.53 | 0.36 |
| 3 | 183996068 | rs3752904 | 0.767 | 1.182 | -1.294 | 0.007 | 0 | 0 | 0 | 0 | 0.6 | 0.59 | 0.61 | 0.42 | 0.53 | 0.36 |
| 3 | 183997735 | rs865809 | -1.53 | 0.847 | -1.732 | 0.067 | -0.059 | -0.08 | -0.08 | 0.028 | 0.86 | 0.97 | 0.83 | 0.68 | 0.79 | 0.62 |
| 3 | 184003194 | rs1687230 | 0.05 | 0.763 | -1.682 | 0.047 | -0.041 | -0.036 | -0.036 | 0.011 | 0.68 | 0.89 | 0.72 | 0.66 | 0.63 | 0.49 |
| 3 | 184010230 | rs3892286 | 0.527 | 0.498 | -1.792 | -0.545 | -0.033 | 0.148 | 0.148 | 0 | 0.27 | 0.01 | 0.13 | 0.05 | 0.22 | 0.28 |
| 3 | 184011561 | rs2376524 | -1.641 | 0.591 | -1.859 | -0.147 | -0.059 | -0.081 | -0.081 | 0 | 0.86 | 0.97 | 0.8 | 0.65 | 0.77 | 0.67 |
| 3 | 184013551 | rs4912533 | 1.136 | 0.461 | -1.492 | -0.295 | 0.015 | 0.004 | 0.004 | 1.232 | 0.44 | 0.6 | 0.51 | 0.37 | 0.27 | 0.24 |
| 3 | 184016599 | rs4912534 | 1.172 | 0.502 | -1.474 | -0.288 | 0.021 | 0.007 | 0.007 | 1.426 | 0.45 | 0.61 | 0.51 | 0.38 | 0.27 | 0.24 |
| 3 | 184016694 | rs4912535 | 1.004 | 0.481 | -1.374 | -0.298 | 0.133 | 0.028 | 0.028 | 2.134 | 0.47 | 0.81 | 0.52 | 0.38 | 0.27 | 0.24 |
| 3 | 184018528 | rs12488922 | 1.172 | 0.486 | -1.416 | -0.295 | 0.023 | 0.007 | 0.007 | 1.556 | 0.45 | 0.61 | 0.51 | 0.38 | 0.27 | 0.24 |
| 3 | 184021628 | rs9883929 | 1.801 | -0.055 | -1.26 | -0.564 | -0.032 | -0.07 | -0.07 | 0 | 0.19 | 0.22 | 0.05 | 0.02 | 0.08 | 0.05 |
| 3 | 184029232 | rs10804892 | -1.739 | 0.869 | -1.783 | -0.1 | -0.049 | -0.065 | -0.065 | 0.386 | 0.87 | 0.96 | 0.8 | 0.66 | 0.77 | 0.67 |
| 3 | 184030387 | rs34881379 | -0.638 | 0.891 | -1.881 | -0.188 | -0.102 | 0.103 | 0.103 | 0.009 | 0.39 | 0.12 | 0.26 | 0.29 | 0.47 | 0.43 |
| 3 | 184038969 | rs4912537 | 0.953 | 1.05 | -1.752 | -0.084 | 0.003 | -0.002 | -0.002 | 0 | 0.11 | 0.14 | 0.14 | 0.24 | 0.25 | 0.14 |
| 3 | 184039666 | rs2178403 | -1.234 | 0.86 | -2.038 | -0.044 | -0.087 | -0.119 | -0.119 | 0.859 | 0.84 | 0.98 | 0.8 | 0.65 | 0.75 | 0.66 |
| 3 | 184046470 | rs2230571 | 0.484 | 0.239 | -2.227 | -0.383 | -0.043 | 0.143 | 0.143 | 0 | 0.26 | 0.02 | 0.14 | 0.05 | 0.23 | 0.27 |
| 3 | 184058581 | rs13322510 | 0.875 | -0.003 | -1.617 | -0.103 | -0.002 | -0.001 | -0.001 | 0 | 0.4 | 0.4 | 0.46 | 0.36 | 0.25 | 0.18 |
| 3 | 184061387 | rs4912539 | 0.041 | 0.29 | -2.45 | -0.312 | -0.086 | 0.16 | 0.16 | 0 | 0.28 | 0.02 | 0.19 | 0.05 | 0.3 | 0.28 |
| 3 | 184063791 | rs9944 | -1.08 | 0.576 | -2.129 | 0.667 | -0.053 | -0.08 | -0.08 | 0 | 0.86 | 0.98 | 0.79 | 0.71 | 0.79 | 0.64 |
| 3 | 184064916 | rs2293603 | 1.435 | -0.159 | -2.223 | 0.344 | 0.027 | -0.027 | -0.027 | 0.438 | 0.2 | 0.12 | 0.34 | 0.31 | 0.11 | 0.09 |
| 3 | 184065064 | rs6770808 | -1.271 | 0.011 | -2.355 | 0.49 | -0.053 | -0.09 | -0.09 | 0 | 0.86 | 0.98 | 0.82 | 0.63 | 0.79 | 0.64 |
| 3 | 184066615 | rs16858720 | 1.153 | -1.446 | -2.126 | 0.456 | 0.051 | -0.069 | -0.069 | 1.838 | 0.21 | 0.05 | 0.35 | 0.33 | 0.13 | 0.11 |
| 3 | 184075047 | rs2228291 | 0.025 | -1.419 | -1.31 | 0.31 | -0.03 | 0.007 | 0.007 | 0 | 0.38 | 0.47 | 0.44 | 0.37 | 0.24 | 0.24 |
| 3 | 184077557 | rs13099401 | -1.329 | -0.254 | -1.81 | 0.591 | -0.125 | -0.144 | -0.144 | 0.016 | 0.82 | 0.98 | 0.71 | 0.49 | 0.64 | 0.56 |
| 3 | 184084846 | rs28467468 | 0.049 | -0.739 | -1.463 | 0.084 | -0.008 | 0.074 | 0.074 | 0.609 | 0.56 | 0.69 | 0.3 | 0.17 | 0.44 | 0.35 |
| 3 | 184086969 | rs4912486 | -0.437 | -0.614 | -1.468 | 0.213 | -0.038 | 0.042 | 0.042 | 0 | 0.64 | 0.75 | 0.37 | 0.2 | 0.52 | 0.46 |
| 3 | 184087736 | rs9850030 | -0.409 | -0.572 | -1.268 | 0.241 | -0.011 | 0.026 | 0.026 | 0 | 0.1 | 0.21 | 0.03 | 0.13 | 0.07 | 0.05 |
| 3 | 184090266 | rs6141 | -0.38 | -0.436 | -1.741 | 0.235 | -0.191 | -0.166 | -0.166 | 1.692 | 0.72 | 0.98 | 0.66 | 0.42 | 0.55 | 0.49 |
| 3 | 184091149 | rs6142 | 0.096 | -0.92 | -1.238 | -0.761 | 0.025 | 0.061 | 0.061 | 0 | 0.49 | 0.6 | 0.31 | 0.06 | 0.39 | 0.4 |
| 3 | 184092481 | rs10513797 | -0.721 | -0.518 | -0.29 | 0.293 | -0.01 | 0.038 | 0.038 | 0 | 0.11 | 0.25 | 0.13 | 0.12 | 0.08 | 0.06 |
| 3 | 184093527 | rs2280740 | -0.468 | -0.334 | -0.345 | 0.397 | -0.002 | 0.025 | 0.025 | 0 | 0.07 | 0.13 | 0.15 | 0.12 | 0.09 | 0.06 |
| 3 | 184094183 | rs956732 | -0.195 | -0.224 | -0.091 | 0.511 | 0.003 | 0.004 | 0.004 | 0.174 | 0.64 | 0.61 | 0.64 | 0.39 | 0.5 | 0.39 |

| | | | | | | | | | | | | | | | | |
|---|---|---|---|---|---|---|---|---|---|---|---|---|---|---|---|---|
| 3 | 184096578 | rs885838 | -0.679 | 0.253 | -0.318 | 0.526 | 0.001 | -0.003 | -0.003 | 1.736 | 0.54 | 0.53 | 0.6 | 0.36 | 0.48 | 0.43 |
| 3 | 184097085 | rs2855306 | 0.358 | 0.328 | -0.342 | 0.398 | 0.014 | 0.063 | 0.063 | 0 | 0.41 | 0.25 | 0.2 | 0.06 | 0.33 | 0.2 |
| 3 | 184099206 | rs3749229 | -1.489 | 0.62 | 0.168 | 1.628 | 0 | 0.056 | 0.056 | 0 | 0.06 | 0.12 | 0.21 | 0.19 | 0.07 | 0.14 |
| 3 | 184102396 | rs16858778 | -0.936 | 0.49 | -0.178 | 1.641 | 0.004 | 0.01 | 0.01 | 0 | 0.05 | 0.02 | 0.25 | 0.2 | 0.05 | 0.15 |
| 3 | 184103903 | rs16858780 | -2.07 | 0.479 | -0.099 | 1.674 | 0 | 0.114 | 0.114 | 0 | 0.16 | 0.34 | 0.41 | 0.31 | 0.15 | 0.25 |
| 3 | 184106493 | rs2293606 | -1.12 | 0.445 | -0.447 | 1.044 | 0 | 0.054 | 0.054 | 0 | 0.07 | 0.16 | 0.13 | 0.11 | 0.06 | 0.08 |
| 3 | 184125634 | rs4912542 | -0.135 | 0.712 | -1.143 | 0.812 | -0.028 | -0.027 | -0.027 | 0 | 0.64 | 0.85 | 0.7 | 0.41 | 0.6 | 0.52 |
| 3 | 184129172 | rs10937167 | 0.328 | 0.335 | -0.615 | 0.154 | -0.001 | -0.039 | -0.039 | 0 | 0.34 | 0.45 | 0.32 | 0.19 | 0.34 | 0.23 |
| 3 | 184131031 | rs10937169 | -0.136 | 0.159 | -0.817 | 0.203 | -0.002 | -0.003 | -0.003 | 0.793 | 0.42 | 0.46 | 0.53 | 0.35 | 0.44 | 0.3 |
| 3 | 184133839 | rs1533679 | 0.224 | 0.181 | -0.47 | 0.136 | -0.011 | -0.089 | -0.089 | 0 | 0.21 | 0.39 | 0.16 | 0.06 | 0.16 | 0.22 |
| 3 | 184137229 | rs4640598 | 0.273 | 0.131 | -0.317 | 0.157 | -0.002 | -0.029 | -0.029 | 0 | 0.19 | 0.25 | 0.15 | 0.07 | 0.16 | 0.21 |
| 3 | 184137656 | rs2376515 | -0.479 | 0.177 | -0.011 | 0.146 | -0.005 | -0.042 | -0.042 | 0 | 0.3 | 0.5 | 0.39 | 0.22 | 0.3 | 0.28 |
| 3 | 184140711 | rs6796172 | -0.796 | 0.217 | -1.11 | 0.326 | -0.123 | -0.13 | -0.13 | 0 | 0.64 | 0.98 | 0.68 | 0.39 | 0.59 | 0.51 |
| 3 | 184141428 | rs12493455 | 0.351 | -0.222 | -1.375 | -0.178 | -0.018 | 0.008 | 0.008 | 0 | 0.27 | 0.09 | 0.38 | 0.21 | 0.25 | 0.29 |
| 3 | 184144646 | rs7630982 | 0.295 | -0.481 | -0.357 | -0.411 | -0.047 | -0.129 | -0.129 | 0 | 0.2 | 0.48 | 0.09 | 0.02 | 0.11 | 0.15 |
| 3 | 184146786 | rs7637387 | 0.038 | -0.508 | 0.137 | -0.453 | 0.002 | -0.019 | -0.019 | 1.871 | 0.31 | 0.35 | 0.46 | 0.21 | 0.33 | 0.31 |
| 3 | 184150007 | rs9858198 | 0.499 | -0.513 | 0.102 | -0.369 | 0.015 | 0.053 | 0.053 | 0 | 0.52 | 0.32 | 0.51 | 0.77 | 0.61 | 0.56 |
| 3 | 184155408 | rs7429010 | -0.712 | -0.216 | -0.017 | -0.265 | 0.004 | -0.004 | -0.004 | 0 | 0.44 | 0.49 | 0.6 | 0.37 | 0.5 | 0.38 |
| 3 | 184157645 | rs4371543 | 0.955 | -0.246 | -0.171 | -0.334 | -0.006 | 0.128 | 0.128 | 0.01 | 0.48 | 0.11 | 0.44 | 0.75 | 0.58 | 0.54 |
| 3 | 184157882 | rs6806701 | -0.02 | -0.292 | 0.068 | -0.362 | -0.038 | -0.105 | -0.105 | 0.074 | 0.76 | 0.42 | 0.85 | 0.98 | 0.86 | 0.85 |
| 3 | 184157976 | rs11927393 | 0.324 | -0.279 | -0.228 | -0.316 | -0.05 | -0.101 | -0.101 | 0 | 0.2 | 0.4 | 0.09 | 0.02 | 0.1 | 0.14 |
| 3 | 184161735 | rs4912490 | 0.518 | 0.163 | -1.053 | 0.113 | 0.006 | 0.105 | 0.105 | 0.083 | 0.48 | 0.11 | 0.44 | 0.72 | 0.57 | 0.54 |
| 3 | 184164617 | rs7638033 | 0.573 | 0.235 | -0.974 | 0.225 | -0.025 | -0.042 | -0.042 | 0 | 0.39 | 0.15 | 0.26 | 0.59 | 0.4 | 0.5 |
| 3 | 184165344 | rs4558788 | 0.779 | 0.08 | -1.277 | 0.23 | -0.091 | -0.11 | -0.11 | 0 | 0.36 | 0.06 | 0.29 | 0.59 | 0.43 | 0.48 |
| 3 | 184170962 | rs6443968 | -0.184 | 0.074 | -1.002 | 0.656 | 0 | -0.001 | -0.001 | 0 | 0.84 | 0.83 | 0.79 | 0.73 | 0.83 | 0.82 |
| 3 | 184170992 | rs13069321 | 0.268 | 0.499 | -1.402 | 0.45 | -0.003 | 0.001 | 0.001 | 0 | 0.13 | 0.14 | 0.18 | 0.09 | 0.2 | 0.06 |
| 3 | 184171422 | rs6772307 | -0.598 | 0.23 | -1.254 | -0.117 | 0 | -0.014 | -0.014 | 0 | 0.42 | 0.58 | 0.43 | 0.17 | 0.36 | 0.32 |
| 3 | 184173466 | rs4912555 | -0.797 | 0.155 | -1.306 | 0.155 | 0.005 | 0.003 | 0.003 | 0 | 0.48 | 0.58 | 0.56 | 0.41 | 0.42 | 0.43 |
| 3 | 184175082 | rs13062189 | 0.891 | 0.216 | -1.274 | 0.15 | -0.002 | 0 | 0 | 0 | 0.12 | 0.14 | 0.16 | 0.09 | 0.19 | 0.06 |
| 3 | 184177368 | rs6414508 | -0.484 | -0.024 | -1.378 | 0.096 | -0.015 | -0.046 | -0.046 | 0 | 0.8 | 0.62 | 0.82 | 0.89 | 0.83 | 0.85 |
| 3 | 184178374 | rs7429036 | 1.552 | 0.079 | -1.87 | 0.169 | -0.12 | -0.134 | -0.134 | 1.915 | 0.33 | 0.04 | 0.26 | 0.47 | 0.41 | 0.42 |
| 3 | 184178412 | rs7429078 | 0.258 | -0.11 | -1.588 | -0.496 | -0.006 | 0.02 | 0.02 | 0 | 0.31 | 0.25 | 0.25 | 0.11 | 0.33 | 0.2 |
| 3 | 184179187 | rs13086359 | 1.166 | 0.479 | -1.922 | 0.068 | -0.116 | -0.041 | -0.041 | 0.019 | 0.3 | 0.03 | 0.25 | 0.26 | 0.39 | 0.38 |
| 3 | 184179757 | rs6797449 | -1.397 | 0.444 | -1.025 | -0.099 | 0.003 | -0.003 | -0.003 | 0 | 0.9 | 0.87 | 0.78 | 0.91 | 0.85 | 0.87 |
| 3 | 184181808 | rs9844738 | -0.358 | 0.259 | -0.972 | -0.569 | -0.001 | -0.019 | -0.019 | 0.004 | 0.3 | 0.27 | 0.25 | 0.06 | 0.26 | 0.07 |
| 3 | 184184235 | rs6798973 | 0.993 | 0.399 | -1.008 | -0.638 | -0.005 | 0.004 | 0.004 | 0 | 0.11 | 0.09 | 0.15 | 0.02 | 0.16 | 0.03 |
| 3 | 184185085 | rs12696521 | -1.27 | 0.245 | -1.421 | -1.052 | -0.024 | 0.023 | 0.023 | 0 | 0.71 | 0.83 | 0.79 | 0.78 | 0.64 | 0.72 |
| 3 | 184186015 | rs9841832 | 0.932 | -0.117 | -1.303 | -1.252 | 0.024 | -0.048 | -0.048 | 0 | 0.28 | 0.09 | 0.3 | 0.55 | 0.23 | 0.52 |
| 3 | 184186619 | rs6808271 | 1.703 | -0.18 | -1.208 | -1.527 | 0.007 | 0.049 | 0.049 | 0.169 | 0.17 | 0.06 | 0.09 | 0.05 | 0.1 | 0.05 |
| 3 | 184188338 | rs6777514 | 1.355 | -0.284 | -0.96 | -1.624 | -0.005 | -0.027 | -0.027 | 1.737 | 0.2 | 0.16 | 0.11 | 0.05 | 0.11 | 0.07 |
| 3 | 184188503 | rs6777648 | 0.534 | -0.286 | -0.593 | -1.668 | -0.002 | -0.023 | -0.023 | 0.728 | 0.34 | 0.36 | 0.38 | 0.14 | 0.28 | 0.2 |
| 3 | 184193055 | rs9823910 | 0.086 | -0.369 | -0.67 | -1.7 | -0.003 | -0.003 | -0.003 | 0 | 0.41 | 0.51 | 0.37 | 0.69 | 0.27 | 0.45 |
| 3 | 184193482 | rs3932485 | -0.5 | -0.285 | -0.758 | -1.709 | 0.019 | 0.065 | 0.065 | 0 | 0.52 | 0.62 | 0.47 | 0.68 | 0.43 | 0.48 |
| 3 | 184198443 | rs9822959 | -1.907 | -0.244 | -0.763 | -1.715 | -0.009 | 0.004 | 0.004 | 0.421 | 0.75 | 0.78 | 0.81 | 0.77 | 0.69 | 0.75 |
| 3 | 184198475 | rs9840462 | -0.146 | -0.352 | -0.463 | -1.684 | 0.001 | -0.044 | -0.044 | 0 | 0.37 | 0.49 | 0.45 | 0.09 | 0.42 | 0.3 |
| 3 | 184203031 | rs3892035 | 1.901 | -0.101 | -0.712 | -1.374 | -0.005 | 0.005 | 0.005 | 0 | 0.13 | 0.16 | 0.27 | 0.09 | 0.17 | 0.13 |
| 3 | 184211984 | rs13096649 | 0.166 | -0.083 | -0.648 | -1.524 | -0.001 | 0.01 | 0.01 | 0 | 0.48 | 0.41 | 0.43 | 0.81 | 0.46 | 0.63 |
| 3 | 184213807 | rs11709926 | 0.772 | 0.033 | -0.811 | -1.181 | -0.007 | 0.011 | 0.011 | 0 | 0.25 | 0.28 | 0.42 | 0.15 | 0.34 | 0.17 |
| 3 | 184214116 | rs4912567 | 0.662 | 0.069 | -0.799 | -1.165 | -0.007 | 0.01 | 0.01 | 0 | 0.25 | 0.28 | 0.42 | 0.15 | 0.34 | 0.16 |
| 3 | 184216307 | rs4912494 | 0.189 | -0.36 | -0.826 | -0.906 | 0.003 | 0.001 | 0.001 | 0.026 | 0.47 | 0.54 | 0.61 | 0.55 | 0.36 | 0.43 |
| 3 | 184217526 | rs9821450 | -0.265 | -0.406 | -0.807 | -1.594 | -0.004 | -0.016 | -0.016 | 1.172 | 0.47 | 0.42 | 0.28 | 0.32 | 0.39 | 0.49 |
| 3 | 184217677 | rs11916995 | -0.195 | 0.23 | -0.839 | -1.387 | 0.004 | 0.018 | 0.018 | 0 | 0.26 | 0.17 | 0.15 | 0.24 | 0.26 | 0.42 |
| 3 | 184221680 | rs7433081 | 0.533 | 0.526 | 0.022 | -1.559 | 0.005 | -0.101 | -0.101 | 0 | 0.85 | 0.68 | 0.92 | 0.98 | 0.87 | 0.84 |
| 3 | 184223392 | rs4912574 | 1.453 | 0.385 | -0.06 | -2.039 | 0 | 0.013 | 0.013 | 0 | 0.48 | 0.39 | 0.69 | 0.91 | 0.52 | 0.59 |
| 3 | 184223454 | rs6770142 | 0.566 | 0.483 | -0.304 | -2.076 | 0.023 | -0.091 | -0.091 | 0.681 | 0.72 | 0.69 | 0.81 | 0.97 | 0.65 | 0.75 |
| 3 | 184232464 | rs6807172 | -0.555 | -0.247 | 0.074 | -1.52 | 0.026 | 0.03 | 0.03 | 0 | 0.23 | 0.19 | 0.11 | 0.05 | 0.14 | 0.16 |
| 3 | 184242854 | rs6443978 | 0.778 | -0.271 | 0.346 | -1.227 | -0.086 | -0.116 | -0.116 | 0.157 | 0.76 | 0.56 | 0.92 | 0.93 | 0.91 | 0.85 |
| 3 | 184263673 | rs11720282 | -0.071 | 0.894 | 0.082 | -1.008 | -0.016 | 0.022 | 0.022 | 0 | 0.19 | 0.12 | 0.12 | 0.07 | 0.24 | 0.26 |
| 3 | 184264927 | rs9860156 | -1.371 | 0.75 | 0.226 | -0.984 | 0.005 | 0.023 | 0.023 | 0.098 | 0.49 | 0.59 | 0.34 | 0.23 | 0.53 | 0.45 |
| 3 | 184266407 | rs9290790 | -0.243 | 0.836 | 0.048 | -0.957 | 0.001 | -0.006 | -0.006 | 0 | 0.23 | 0.23 | 0.17 | 0.16 | 0.24 | 0.12 |
| 3 | 184268194 | rs10804894 | 0.702 | 0.883 | -0.068 | -0.965 | 0.003 | -0.015 | -0.015 | 0 | 0.15 | 0.07 | 0.09 | 0.14 | 0.14 | 0.09 |
| 3 | 184271242 | rs9854599 | -1.199 | 0.517 | 0.147 | -0.866 | 0 | -0.003 | -0.003 | 1.157 | 0.49 | 0.47 | 0.31 | 0.24 | 0.54 | 0.45 |
| 3 | 184278103 | rs11713341 | -0.338 | 0.593 | -0.201 | -0.779 | -0.026 | 0.043 | 0.043 | 0 | 0.35 | 0.19 | 0.19 | 0.22 | 0.38 | 0.34 |
| 3 | 184278371 | rs11709414 | -0.444 | 0.595 | -0.2 | -0.778 | -0.026 | 0.042 | 0.042 | 0 | 0.35 | 0.19 | 0.19 | 0.22 | 0.38 | 0.34 |
| 3 | 184284279 | rs9647371 | -0.336 | 0.658 | 0.122 | -0.922 | 0.009 | -0.075 | -0.075 | 0.361 | 0.17 | 0.2 | 0.12 | 0 | 0.18 | 0.03 |
| 3 | 184289982 | rs7640108 | -0.593 | 0.624 | 0.093 | -0.424 | -0.007 | -0.01 | -0.01 | 0 | 0.41 | 0.34 | 0.26 | 0.51 | 0.45 | 0.44 |
| 3 | 184290348 | rs9862375 | 0.484 | 0.286 | 0.269 | -0.764 | 0.01 | -0.034 | -0.034 | 0.966 | 0.08 | 0.09 | 0.07 | 0 | 0.11 | 0.01 |
| 3 | 184293610 | rs9823034 | 0.515 | 0.378 | 0.031 | -0.711 | -0.007 | 0.001 | 0.001 | 0 | 0.15 | 0.1 | 0.08 | 0.16 | 0.16 | 0.18 |
| 3 | 184293769 | rs7652597 | -1.036 | 0.381 | 0.194 | -0.676 | -0.001 | -0.011 | -0.011 | 0 | 0.4 | 0.56 | 0.25 | 0.17 | 0.36 | 0.25 |

| | | | | | | | | | | | | | | | | |
|---|---|---|---|---|---|---|---|---|---|---|---|---|---|---|---|---|
| 3 | 184302634 | rs7652280 | -0.003 | 0.497 | -0.69 | -0.396 | -0.001 | 0.018 | 0.018 | 0 | 0.23 | 0.13 | 0.11 | 0.16 | 0.19 | 0.18 |
| 3 | 184306559 | rs7620519 | -0.098 | 0.452 | -0.733 | -0.394 | 0 | 0.006 | 0.006 | 0 | 0.24 | 0.16 | 0.12 | 0.16 | 0.19 | 0.18 |
| 3 | 184312106 | rs4582031 | -0.06 | 0.759 | -0.972 | -0.163 | 0.01 | 0.086 | 0.086 | 0 | 0.55 | 0.29 | 0.54 | 0.81 | 0.6 | 0.74 |
| 3 | 184322950 | rs10937179 | 0.216 | 0.717 | -1.751 | 0.837 | -0.037 | -0.087 | -0.087 | 0 | 0.72 | 0.91 | 0.72 | 0.62 | 0.77 | 0.76 |
| 3 | 184326485 | rs9877051 | 0.456 | 0.735 | -1.93 | 1.254 | 0.001 | -0.061 | -0.061 | 0.144 | 0.69 | 0.82 | 0.74 | 0.53 | 0.71 | 0.72 |
| 3 | 184328020 | rs9849690 | 0.266 | 0.545 | -1.527 | 1.252 | -0.004 | -0.003 | -0.003 | 0 | 0.45 | 0.43 | 0.38 | 0.53 | 0.52 | 0.38 |
| 3 | 184331242 | rs4074869 | 0.343 | 0.059 | -1.155 | 1.304 | -0.015 | -0.016 | -0.016 | 0 | 0.34 | 0.24 | 0.29 | 0.45 | 0.55 | 0.3 |
| 3 | 184331446 | rs4074871 | -0.201 | -0.428 | -0.909 | 1.127 | -0.002 | 0.006 | 0.006 | 0 | 0.19 | 0.24 | 0.36 | 0.25 | 0.16 | 0.38 |
| 3 | 184335641 | rs6770501 | -0.929 | -0.543 | -0.881 | 1.025 | 0.092 | 0.105 | 0.105 | 0 | 0.58 | 0.89 | 0.7 | 0.7 | 0.71 | 0.67 |
| 3 | 184337818 | rs13085530 | -0.875 | -0.322 | -0.72 | 1.125 | 0.028 | 0.031 | 0.031 | 0 | 0.55 | 0.61 | 0.68 | 0.7 | 0.72 | 0.68 |
| 3 | 184339757 | rs13097965 | 0.575 | -0.21 | -0.825 | 1.231 | -0.017 | -0.018 | -0.018 | 0 | 0.34 | 0.24 | 0.29 | 0.45 | 0.55 | 0.29 |
| 3 | 184341513 | rs10937181 | -0.021 | 0.072 | -0.743 | 1.505 | -0.022 | -0.029 | -0.029 | 0 | 0.06 | 0.01 | 0.06 | 0.07 | 0.06 | 0.21 |
| 3 | 184343037 | rs7623163 | -1.015 | 0.07 | -0.65 | 1.447 | -0.002 | 0.035 | 0.035 | 0 | 0.19 | 0.3 | 0.36 | 0.24 | 0.16 | 0.38 |
| 3 | 184348577 | rs4687113 | -0.634 | 0.191 | -0.898 | 1.532 | 0.001 | -0.002 | -0.002 | 0.071 | 0.15 | 0.11 | 0.43 | 0.29 | 0.15 | 0.14 |
| 3 | 184355123 | rs4074988 | -0.961 | 0.6 | -0.627 | 0.777 | 0.002 | -0.034 | -0.034 | 0 | 0.41 | 0.45 | 0.67 | 0.73 | 0.45 | 0.54 |
| 3 | 184359176 | rs12493138 | -0.757 | 0.039 | -0.975 | 0.809 | 0.054 | 0.024 | 0.024 | 0.98 | 0.49 | 0.35 | 0.72 | 0.62 | 0.67 | 0.47 |
| 3 | 184359263 | rs4284958 | -0.764 | 0.039 | -0.818 | 0.809 | 0.038 | 0.017 | 0.017 | 0.9 | 0.49 | 0.39 | 0.72 | 0.62 | 0.67 | 0.47 |
| 3 | 184366735 | rs10937184 | 0.897 | 0.105 | -0.924 | 0.427 | 0.004 | 0.081 | 0.081 | 0 | 0.44 | 0.22 | 0.22 | 0.26 | 0.43 | 0.36 |
| 3 | 184377320 | rs4640536 | -1.582 | 0.255 | -1.249 | 0.663 | 0 | -0.003 | -0.003 | 0.297 | 0.86 | 0.8 | 0.92 | 0.97 | 0.89 | 0.92 |
| 3 | 184377986 | rs4475017 | -1.027 | 0.39 | -1.3 | 0.769 | -0.001 | 0.007 | 0.007 | 0 | 0.82 | 0.79 | 0.9 | 0.93 | 0.86 | 0.75 |
| 3 | 184382792 | rs9846698 | -0.579 | 0.729 | -1.303 | 1.098 | -0.076 | -0.085 | -0.085 | 0 | 0.32 | 0.14 | 0.61 | 0.43 | 0.39 | 0.36 |
| 3 | 184391573 | rs9865980 | -0.205 | 0.716 | -1.308 | 1.182 | -0.064 | -0.062 | -0.062 | 0 | 0.28 | 0.12 | 0.58 | 0.37 | 0.37 | 0.2 |
| 3 | 184397940 | rs4299462 | -0.848 | 0.376 | -1.237 | 0.859 | -0.05 | -0.023 | -0.023 | 0 | 0.41 | 0.21 | 0.65 | 0.56 | 0.52 | 0.46 |
| 3 | 184402122 | rs9290796 | -0.989 | -0.168 | -1.028 | 1.109 | -0.032 | -0.089 | -0.089 | 0 | 0.33 | 0.15 | 0.54 | 0.53 | 0.37 | 0.43 |
| 3 | 184402715 | rs4687146 | -1.119 | -1.098 | -0.417 | 1.495 | 0.001 | 0.005 | 0.005 | 0 | 0.87 | 0.84 | 0.93 | 0.98 | 0.9 | 0.93 |
| 3 | 184406682 | rs3934720 | -0.789 | -1.133 | -0.563 | 1.688 | 0.017 | -0.041 | -0.041 | 0 | 0.21 | 0.12 | 0.43 | 0.43 | 0.15 | 0.4 |
| 3 | 184406997 | rs6443987 | 0.942 | -1.15 | -0.345 | 1.55 | 0.017 | -0.009 | -0.009 | 0.379 | 0.64 | 0.72 | 0.49 | 0.43 | 0.72 | 0.51 |
| 3 | 184409778 | rs10937185 | -1.056 | -1.046 | -0.48 | 1.861 | 0.009 | -0.074 | -0.074 | 0 | 0.13 | 0.04 | 0.12 | 0.3 | 0.14 | 0.4 |
| 3 | 184410071 | rs4621309 | 1.172 | -1.211 | 0.42 | 1.264 | -0.037 | -0.042 | -0.042 | 0 | 0.7 | 0.44 | 0.82 | 0.8 | 0.79 | 0.67 |
| 3 | 184410718 | rs4686574 | 0.88 | -1.232 | 0.687 | 1.499 | -0.045 | -0.002 | -0.002 | 0.396 | 0.73 | 0.56 | 0.86 | 0.77 | 0.82 | 0.67 |
| 3 | 184412473 | rs7631034 | -0.092 | -1.306 | 0.829 | 1.629 | -0.039 | 0.069 | 0.069 | 0 | 0.78 | 0.57 | 0.68 | 0.68 | 0.85 | 0.88 |
| 3 | 184416009 | rs6443991 | -0.661 | -1.218 | -0.02 | 1.422 | 0.025 | -0.029 | -0.029 | 0 | 0.82 | 0.89 | 0.71 | 0.78 | 0.88 | 0.89 |
| 3 | 184424852 | rs13084650 | 0.387 | -1.144 | 0.351 | 1.154 | -0.035 | 0.008 | 0.008 | 0 | 0.64 | 0.48 | 0.5 | 0.44 | 0.76 | 0.57 |
| 3 | 184426945 | rs10937186 | 0.34 | -1.159 | 0.475 | 1.148 | -0.016 | 0.005 | 0.005 | 0 | 0.65 | 0.63 | 0.51 | 0.44 | 0.76 | 0.57 |
| 3 | 184428035 | rs10212515 | 0.514 | -1.071 | -0.106 | 1.172 | 0.018 | -0.002 | -0.002 | 0 | 0.6 | 0.39 | 0.48 | 0.44 | 0.73 | 0.57 |
| 3 | 184437480 | rs4687478 | -0.067 | -0.064 | -0.546 | 1.064 | -0.002 | -0.003 | -0.003 | 0.16 | 0.65 | 0.35 | 0.51 | 0.49 | 0.59 | 0.63 |
| 3 | 184440848 | rs9850765 | 0.256 | -0.708 | -0.918 | 0.798 | 0.009 | 0.039 | 0.039 | 0 | 0.54 | 0.26 | 0.4 | 0.34 | 0.52 | 0.62 |
| 3 | 184445284 | rs9844571 | 0.196 | -0.851 | -0.391 | 1.032 | 0 | -0.001 | -0.001 | 0.641 | 0.58 | 0.43 | 0.51 | 0.46 | 0.53 | 0.67 |
| 3 | 184446069 | rs4686687 | 0.523 | -1.185 | -0.609 | 0.917 | 0.019 | 0.048 | 0.048 | 0 | 0.5 | 0.36 | 0.44 | 0.39 | 0.48 | 0.5 |
| 3 | 184452573 | rs4686690 | -1.32 | -0.834 | -1.182 | 0.649 | -0.012 | -0.021 | -0.021 | 2.258 | 0.84 | 0.91 | 0.84 | 0.79 | 0.86 | 0.81 |
| 3 | 184455035 | rs9831802 | 0.179 | -0.982 | -0.444 | 1.028 | -0.007 | 0.007 | 0.007 | 0.015 | 0.54 | 0.68 | 0.66 | 0.53 | 0.46 | 0.49 |
| 3 | 184459667 | rs11923588 | -0.262 | -1.004 | 0.068 | 1.09 | -0.04 | 0.044 | 0.044 | 0 | 0.11 | 0.27 | 0.16 | 0.2 | 0.06 | 0.22 |
| 3 | 184460101 | rs16859163 | 0.014 | -0.98 | 0.207 | 1.11 | -0.074 | 0.12 | 0.12 | 0.886 | 0.16 | 0.39 | 0.28 | 0.27 | 0.08 | 0.22 |
| 3 | 184461586 | rs6804092 | 0.043 | -0.768 | -0.329 | 0.647 | 0 | 0.01 | 0.01 | 0 | 0.56 | 0.61 | 0.45 | 0.43 | 0.51 | 0.58 |
| 3 | 184462940 | rs11719284 | -0.173 | -0.106 | -0.528 | 0.325 | 0 | 0.002 | 0.002 | 0 | 0.19 | 0.16 | 0.11 | 0.1 | 0.2 | 0.15 |
| 3 | 184463505 | rs9875166 | 1.305 | -0.359 | -0.796 | 0.291 | -0.069 | 0.121 | 0.121 | 0.125 | 0.31 | 0.11 | 0.18 | 0.05 | 0.33 | 0.25 |
| 3 | 184465325 | rs6766579 | 0.059 | -0.397 | -0.526 | 0.772 | -0.006 | 0 | 0 | 0 | 0.41 | 0.56 | 0.38 | 0.56 | 0.26 | 0.47 |
| 3 | 184466359 | rs2377100 | -0.055 | 0.521 | -0.488 | 1.337 | 0.024 | 0.051 | 0.051 | 0 | 0.86 | 0.79 | 0.65 | 0.7 | 0.76 | 0.84 |
| 3 | 184467803 | rs4330241 | -0.702 | 0.959 | -0.809 | 1.331 | 0 | -0.002 | -0.002 | 0 | 0.88 | 0.88 | 0.8 | 0.82 | 0.77 | 0.86 |
| 3 | 184468603 | rs4381915 | 0.325 | 0.798 | -0.861 | 0.788 | 0 | 0.005 | 0.005 | 0 | 0.15 | 0.11 | 0.23 | 0.11 | 0.19 | 0.14 |
| 3 | 184468677 | rs4392394 | 0.35 | 0.807 | -0.909 | 0.792 | 0 | 0.007 | 0.007 | 0 | 0.15 | 0.09 | 0.23 | 0.11 | 0.19 | 0.14 |
| 3 | 184469113 | rs6786156 | 0.19 | 0.589 | -1.165 | 1.039 | -0.004 | -0.004 | -0.004 | 0 | 0.56 | 0.69 | 0.57 | 0.5 | 0.56 | 0.59 |
| 3 | 184470330 | rs13100880 | 0.184 | 0.741 | -1.095 | 0.815 | -0.004 | 0.024 | 0.024 | 0 | 0.57 | 0.69 | 0.5 | 0.4 | 0.56 | 0.6 |
| 3 | 184474305 | rs9829962 | -0.414 | 1.541 | -1.115 | 1.122 | -0.012 | -0.004 | -0.004 | 0 | 0.87 | 0.9 | 0.78 | 0.66 | 0.75 | 0.83 |
| 3 | 184474826 | rs4309706 | -0.054 | 0.772 | -1.153 | 0.316 | 0.021 | -0.007 | -0.007 | 0.039 | 0.41 | 0.19 | 0.35 | 0.37 | 0.37 | 0.44 |
| 3 | 184477755 | rs9818534 | -0.221 | 0.964 | -0.59 | 0.099 | -0.001 | -0.001 | -0.001 | 0 | 0.43 | 0.49 | 0.37 | 0.42 | 0.45 | 0.44 |
| 3 | 184485263 | rs12632166 | -0.431 | 0.343 | -0.441 | 0.223 | -0.022 | 0.018 | 0.018 | 2.673 | 0.12 | 0.2 | 0.08 | 0.15 | 0.06 | 0.14 |
| 3 | 184485548 | rs2377102 | 0.812 | 0.27 | -0.556 | -0.082 | -0.007 | -0.002 | -0.002 | 0 | 0.24 | 0.13 | 0.22 | 0.24 | 0.27 | 0.29 |
| 3 | 184485570 | rs2377101 | -0.297 | 0.299 | -0.505 | -0.085 | 0 | -0.001 | -0.001 | 0 | 0.37 | 0.37 | 0.32 | 0.4 | 0.32 | 0.44 |
| 3 | 184486297 | rs17818937 | 0.521 | 0.234 | -0.824 | -0.243 | 0.002 | 0.028 | 0.028 | 0 | 0.12 | 0.07 | 0.04 | 0.02 | 0.07 | 0.06 |
| 3 | 184488916 | rs10513798 | -0.619 | 0.265 | -0.523 | -0.066 | -0.017 | 0.04 | 0.04 | 1.684 | 0.14 | 0.27 | 0.11 | 0.17 | 0.08 | 0.17 |
| 3 | 184490958 | rs4686722 | 0.267 | 0.285 | -1.052 | -0.692 | 0.003 | 0.026 | 0.026 | 0.012 | 0.31 | 0.13 | 0.29 | 0.26 | 0.31 | 0.36 |
| 3 | 184491252 | rs12152460 | -0.103 | 0.227 | -0.666 | -0.808 | -0.024 | -0.078 | -0.078 | 0 | 0.26 | 0.35 | 0.07 | 0.04 | 0.16 | 0.16 |
| 3 | 184493177 | rs4686726 | 0.07 | 0.815 | -1.455 | -0.682 | -0.004 | 0.132 | 0.132 | 0 | 0.49 | 0.3 | 0.41 | 0.3 | 0.55 | 0.49 |
| 3 | 184497063 | rs6443994 | -0.196 | 0.909 | -1.135 | -0.594 | 0.003 | 0.081 | 0.081 | 0 | 0.49 | 0.65 | 0.38 | 0.16 | 0.48 | 0.36 |
| 3 | 184499277 | rs4686407 | -0.071 | 1.041 | -1.057 | 0.198 | -0.015 | -0.01 | -0.01 | 0 | 0.43 | 0.36 | 0.48 | 0.5 | 0.5 | 0.52 |
| 3 | 184499620 | rs2377110 | 0.163 | 0.886 | -1.106 | 0.586 | -0.071 | -0.045 | -0.045 | 0 | 0.38 | 0.14 | 0.57 | 0.52 | 0.49 | 0.49 |
| 3 | 184504435 | rs2377112 | -0.075 | 1.001 | -0.603 | 0.53 | -0.006 | 0.008 | 0.008 | 0 | 0.45 | 0.6 | 0.53 | 0.42 | 0.49 | 0.41 |
| 3 | 184506578 | rs6771739 | 0.611 | 1.01 | -1.224 | 0.534 | -0.014 | 0.031 | 0.031 | 0 | 0.35 | 0.15 | 0.44 | 0.31 | 0.39 | 0.34 |

| | | | | | | | | | | | | | | | | |
|---|---|---|---|---|---|---|---|---|---|---|---|---|---|---|---|---|
| 3 | 184508101 | rs7627535 | -0.339 | 0.895 | -1.643 | 0.433 | -0.009 | 0.004 | 0.004 | 0 | 0.52 | 0.84 | 0.59 | 0.45 | 0.55 | 0.49 |
| 3 | 184514613 | rs4686760 | 0.45 | 0.556 | -1.704 | 0.066 | -0.027 | 0.085 | 0.085 | 0.006 | 0.45 | 0.19 | 0.53 | 0.3 | 0.48 | 0.44 |
| 3 | 184515299 | rs4686762 | -0.075 | 0.354 | -1.83 | 0.13 | -0.069 | -0.056 | -0.056 | 0 | 0.59 | 0.84 | 0.55 | 0.43 | 0.54 | 0.52 |
| 3 | 184517445 | rs767848 | -0.244 | -0.306 | -1.098 | -0.665 | 0.004 | 0.013 | 0.013 | 0.13 | 0.31 | 0.26 | 0.14 | 0.1 | 0.24 | 0.26 |
| 3 | 184518380 | rs6786934 | -1.73 | -0.319 | 0.55 | 0.259 | 0.067 | 0.097 | 0.097 | 0 | 0.07 | 0.39 | 0.04 | 0.05 | 0.04 | 0.04 |
| 3 | 184520938 | rs9290800 | 1.025 | -0.823 | -0.114 | 0.073 | 0.037 | 0.027 | 0.027 | 0 | 0.83 | 0.94 | 0.86 | 0.89 | 0.9 | 0.86 |
| 3 | 184523185 | rs10937190 | -1.501 | -0.743 | 0.714 | 0.353 | -0.005 | -0.001 | -0.001 | 0 | 0.3 | 0.36 | 0.41 | 0.7 | 0.33 | 0.51 |
| 3 | 184525971 | rs2377115 | -0.956 | -0.65 | 0.876 | 0.326 | 0.013 | -0.016 | -0.016 | 0 | 0.24 | 0.16 | 0.35 | 0.66 | 0.24 | 0.49 |
| 3 | 184526974 | rs11711955 | -1.012 | -0.658 | 0.876 | 0.315 | -0.004 | -0.016 | -0.016 | 0 | 0.25 | 0.16 | 0.38 | 0.67 | 0.32 | 0.51 |
| 3 | 184533531 | rs6443997 | -2.338 | -0.59 | 1.701 | 0.291 | -0.033 | -0.059 | -0.059 | 0 | 0.15 | 0.6 | 0.11 | 0.05 | 0.06 | 0.08 |
| 3 | 184549539 | rs10513799 | -0.373 | -0.627 | 1.755 | 0.375 | 0.004 | -0.039 | -0.039 | 0.277 | 0.19 | 0.11 | 0.32 | 0.61 | 0.23 | 0.46 |
| 3 | 184550501 | rs9830734 | -2.1 | -0.646 | 1.681 | 0.332 | 0.022 | 0.006 | 0.006 | 6.963 | 0.37 | 0.71 | 0.45 | 0.67 | 0.32 | 0.55 |
| 3 | 184573555 | rs6443999 | -1.994 | -0.755 | 1.837 | 0.46 | 0.006 | 0.001 | 0.001 | 4.472 | 0.36 | 0.68 | 0.46 | 0.66 | 0.36 | 0.55 |
| 3 | 184575831 | rs724273 | -2.054 | -0.756 | 1.828 | 0.462 | 0.001 | 0.001 | 0.001 | 5.113 | 0.37 | 0.66 | 0.47 | 0.66 | 0.39 | 0.56 |
| 3 | 184589984 | rs6779667 | -2.138 | -0.72 | 2.334 | 0.549 | 0.041 | -0.047 | -0.047 | 0.015 | 0.14 | 0.42 | 0.13 | 0.05 | 0.14 | 0.1 |
| 3 | 184607480 | rs1000270 | -0.408 | -0.728 | 2.279 | 0.567 | -0.001 | -0.047 | -0.047 | 0 | 0.2 | 0.1 | 0.32 | 0.61 | 0.24 | 0.46 |
| 3 | 184633247 | rs4686879 | 1.826 | -0.823 | 2.484 | 0.545 | 0 | 0 | 0 | 0 | 0.65 | 0.36 | 0.54 | 0.34 | 0.63 | 0.46 |
| 3 | 184647413 | rs4643688 | 0.987 | -0.765 | 2.566 | 0.557 | 0.061 | 0.011 | 0.011 | 0.293 | 0.92 | 0.73 | 0.91 | 0.95 | 0.9 | 0.93 |
| 3 | 184665304 | rs4422281 | 0.391 | -0.803 | 2.175 | 0.64 | 0.003 | -0.039 | -0.039 | 0 | 0.78 | 0.86 | 0.68 | 0.38 | 0.76 | 0.53 |
| 3 | 184682214 | rs6799328 | 0.656 | -0.618 | 2.109 | 0.593 | -0.009 | -0.047 | -0.047 | 0 | 0.76 | 0.86 | 0.65 | 0.39 | 0.7 | 0.51 |
| 3 | 184684029 | rs721432 | -0.553 | -0.832 | 2.09 | 0.384 | 0.002 | 0.002 | 0.002 | 0 | 0.12 | 0.09 | 0.11 | 0.06 | 0.07 | 0.13 |
| 3 | 184691216 | rs6444007 | -2.518 | -0.754 | 2.396 | 0.425 | -0.009 | -0.074 | -0.074 | 0 | 0.22 | 0.37 | 0.16 | 0.07 | 0.18 | 0.2 |
| 3 | 184700426 | rs11555405 | -0.391 | -0.802 | 2.488 | 0.842 | -0.067 | -0.165 | -0.165 | 2.696 | 0.22 | 0.02 | 0.33 | 0.51 | 0.29 | 0.39 |
| 3 | 184724719 | rs4546129 | -1.836 | -0.724 | 2.608 | 0.825 | -0.005 | -0.057 | -0.057 | 0.129 | 0.18 | 0.32 | 0.14 | 0.06 | 0.17 | 0.15 |
| 3 | 184756694 | rs9869435 | -2.718 | -0.671 | 2.173 | 0.856 | -0.008 | -0.006 | -0.006 | 0 | 0.18 | 0.68 | 0.17 | 0.16 | 0.16 | 0.2 |
| 3 | 184760449 | rs1011147 | -2.068 | -0.667 | 2.242 | 0.866 | -0.003 | 0.008 | 0.008 | 2.242 | 0.15 | 0.61 | 0.15 | 0.15 | 0.14 | 0.2 |
| 3 | 184762324 | rs6765837 | -2.467 | -0.659 | 2.17 | 0.805 | -0.001 | 0.047 | 0.047 | 3.735 | 0.47 | 0.74 | 0.54 | 0.68 | 0.49 | 0.69 |
| 3 | 184765449 | rs1509504 | -0.741 | -0.27 | 2.472 | 1.137 | -0.011 | -0.05 | -0.05 | 0 | 0.23 | 0.14 | 0.35 | 0.61 | 0.3 | 0.49 |
| 3 | 184766301 | rs3821750 | -1.823 | -0.274 | 2.592 | 1.135 | 0.044 | -0.044 | -0.044 | 0 | 0.13 | 0.41 | 0.14 | 0.05 | 0.14 | 0.11 |
| 3 | 184769941 | rs2271258 | -2.049 | -0.273 | 2.553 | 1.135 | 0.003 | -0.001 | -0.001 | 0.973 | 0.37 | 0.55 | 0.49 | 0.67 | 0.44 | 0.6 |
| 3 | 184782202 | rs9755247 | -0.816 | 0.045 | 2.034 | 1.313 | -0.013 | -0.054 | -0.054 | 0.348 | 0.23 | 0.12 | 0.35 | 0.61 | 0.3 | 0.49 |
| 3 | 184792444 | rs4687016 | -1.036 | 0.174 | 2.428 | 1.157 | 0.043 | 0.006 | 0.006 | 0 | 0.07 | 0.21 | 0.07 | 0.05 | 0.09 | 0.07 |
| 3 | 184797711 | rs2377257 | 1.299 | -0.072 | 2.703 | 1.095 | -0.022 | -0.037 | -0.037 | 0 | 0.86 | 0.79 | 0.89 | 0.92 | 0.94 | 0.86 |
| 3 | 184797765 | rs1043435 | -2.668 | -0.137 | 3.176 | 1.141 | 0.045 | 0.062 | 0.062 | 1.608 | 0.13 | 0.47 | 0.14 | 0.15 | 0.15 | 0.19 |
| 3 | 184798122 | rs6444028 | -0.703 | -0.118 | 3.162 | 1.088 | 0.032 | 0.003 | 0.003 | 0 | 0.06 | 0.16 | 0.07 | 0.05 | 0.08 | 0.06 |
| 3 | 184798828 | rs6773358 | -0.703 | -0.095 | 3.131 | 1.111 | 0.032 | 0.003 | 0.003 | 0 | 0.06 | 0.16 | 0.07 | 0.05 | 0.08 | 0.06 |
| 3 | 184799288 | rs876453 | 1.221 | -0.127 | 3.051 | 1.11 | -0.018 | -0.032 | -0.032 | 0 | 0.87 | 0.81 | 0.9 | 0.92 | 0.94 | 0.86 |
| 3 | 184806706 | rs7644823 | -1.408 | -0.005 | 2.948 | 1.124 | 0.092 | 0.003 | 0.003 | 0.033 | 0.09 | 0.29 | 0.11 | 0.05 | 0.15 | 0.08 |
| 3 | 184808281 | rs11925854 | -1.578 | -0.095 | 2.598 | 1.038 | -0.005 | 0.005 | 0.005 | 0.262 | 0.16 | 0.16 | 0.16 | 0.06 | 0.23 | 0.22 |
| 3 | 184809896 | rs9857572 | 2.559 | -0.124 | 2.654 | 1.101 | -0.006 | -0.018 | -0.018 | 0.102 | 0.71 | 0.39 | 0.71 | 0.78 | 0.7 | 0.61 |
| 3 | 184812416 | rs13321350 | 2.74 | -0.24 | 2.137 | 1.312 | -0.017 | 0.059 | 0.059 | 0.272 | 0.48 | 0.21 | 0.4 | 0.32 | 0.48 | 0.3 |
| 3 | 184814444 | rs6770129 | -1.37 | 0.12 | 2.433 | 1.717 | 0.003 | 0.011 | 0.011 | 0.318 | 0.27 | 0.3 | 0.34 | 0.51 | 0.36 | 0.41 |
| 3 | 184814484 | rs6794937 | -2.433 | 0.358 | 2.57 | 1.733 | 0.03 | 0.006 | 0.006 | 3.332 | 0.16 | 0.64 | 0.15 | 0.14 | 0.19 | 0.22 |
| 3 | 184816854 | rs4686509 | -0.326 | -0.672 | 2.41 | 1.977 | -0.004 | 0.007 | 0.007 | 0 | 0.08 | 0.06 | 0.21 | 0.22 | 0.11 | 0.13 |
| 3 | 184817478 | rs10937204 | -0.093 | -0.736 | 2.346 | 2.022 | -0.004 | -0.068 | -0.068 | 0 | 0.18 | 0.08 | 0.29 | 0.41 | 0.2 | 0.24 |
| 3 | 184836082 | rs4687046 | 0.335 | -0.764 | 2.399 | 1.94 | 0.003 | 0.004 | 0.004 | 0 | 0.93 | 0.93 | 0.91 | 0.94 | 0.95 | 0.95 |
| 3 | 184838045 | rs7619914 | 0.651 | -0.76 | 2.057 | 1.941 | 0.004 | 0.005 | 0.005 | 0 | 0.92 | 0.93 | 0.9 | 0.94 | 0.95 | 0.95 |
| 3 | 184844707 | rs13074297 | -0.519 | -0.781 | 2.046 | 1.94 | 0.006 | -0.007 | -0.007 | 0 | 0.09 | 0.08 | 0.06 | 0.1 | 0.08 | 0.19 |
| 3 | 184845544 | rs16859665 | -1.284 | -0.798 | 2.086 | 1.9 | 0.014 | -0.046 | -0.046 | 0 | 0.29 | 0.22 | 0.36 | 0.52 | 0.29 | 0.44 |
| 3 | 184853053 | rs9878804 | 2.564 | -0.721 | 1.59 | 1.888 | 0.035 | 0.196 | 0.196 | 2.148 | 0.5 | 0.2 | 0.31 | 0.2 | 0.54 | 0.37 |
| 3 | 184853342 | rs9841506 | 1.517 | -0.175 | 1.236 | 1.983 | 0 | 0.006 | 0.006 | 0 | 0.83 | 0.83 | 0.69 | 0.72 | 0.83 | 0.82 |
| 3 | 184857970 | rs4687083 | -0.874 | -0.074 | 1.105 | 1.979 | 0.006 | 0.035 | 0.035 | 0 | 0.1 | 0.11 | 0.21 | 0.22 | 0.12 | 0.14 |
| 3 | 184872794 | rs11714089 | -0.277 | -0.08 | 0.75 | 2.037 | -0.013 | -0.074 | -0.074 | 0 | 0.17 | 0.06 | 0.22 | 0.34 | 0.21 | 0.2 |
| 3 | 184875746 | rs6800626 | 1.743 | -0.34 | 1.898 | 2.17 | -0.01 | -0.007 | -0.007 | 0.112 | 0.74 | 0.53 | 0.71 | 0.79 | 0.77 | 0.68 |
| 3 | 184876783 | rs7647854 | 1.14 | -0.219 | 1.891 | 2.273 | 0.011 | 0.034 | 0.034 | 0 | 0.82 | 0.69 | 0.78 | 0.81 | 0.82 | 0.8 |
| 3 | 184886137 | rs16859708 | -2.401 | -0.446 | 2.513 | 2.487 | -0.038 | 0.142 | 0.142 | 0.846 | 0.19 | 0.42 | 0.22 | 0.35 | 0.13 | 0.24 |
| 3 | 184889938 | rs16859736 | -1.341 | -0.435 | 2.27 | 2.542 | 0.002 | 0.033 | 0.033 | 0 | 0.08 | 0.08 | 0.11 | 0.3 | 0.07 | 0.12 |
| 3 | 184895463 | rs12629404 | -2.934 | -0.462 | 2.718 | 2.506 | -0.034 | 0.127 | 0.127 | 0.651 | 0.2 | 0.44 | 0.24 | 0.35 | 0.14 | 0.23 |
| 3 | 184897801 | rs6804537 | -1.879 | -0.429 | 3.002 | 2.518 | -0.006 | 0.184 | 0.184 | 0 | 0.09 | 0.24 | 0.13 | 0.3 | 0.07 | 0.12 |
| 3 | 184906201 | rs6783938 | -2.373 | -0.447 | 3.134 | 2.62 | -0.011 | 0.122 | 0.122 | 2.759 | 0.14 | 0.49 | 0.22 | 0.22 | 0.09 | 0.19 |
| 3 | 184910133 | rs11919970 | -2.047 | -0.443 | 3.009 | 2.559 | 0.009 | -0.005 | -0.005 | 0.399 | 0.12 | 0.39 | 0.16 | 0.07 | 0.08 | 0.12 |
| 3 | 184911151 | rs3732933 | -0.996 | -0.438 | 2.781 | 2.563 | -0.01 | -0.009 | -0.009 | 0.288 | 0.08 | 0.02 | 0.12 | 0.07 | 0.07 | 0.12 |
| 3 | 184917949 | rs6788796 | -1.983 | -0.416 | 2.865 | 2.581 | -0.005 | 0.003 | 0.003 | 2.321 | 0.1 | 0.26 | 0.14 | 0.07 | 0.06 | 0.07 |
| 3 | 184925384 | rs721814 | 1.507 | -0.547 | 2.723 | 2.452 | -0.032 | 0.143 | 0.143 | 0 | 0.78 | 0.66 | 0.78 | 0.48 | 0.85 | 0.71 |
| 3 | 184930238 | rs2269228 | 0.833 | -0.546 | 2.56 | 2.462 | 0.007 | -0.084 | -0.084 | 0 | 0.87 | 1 | 0.9 | 0.7 | 0.91 | 0.81 |
| 3 | 184931013 | rs16859825 | -1.646 | -0.575 | 2.683 | 2.443 | -0.006 | -0.002 | -0.002 | 0 | 0.1 | 0.26 | 0.13 | 0.06 | 0.06 | 0.07 |
| 3 | 184940498 | rs6779662 | 0.784 | -0.578 | 2.669 | 2.454 | -0.009 | 0.027 | 0.027 | 0 | 0.87 | 0.89 | 0.89 | 0.7 | 0.91 | 0.82 |
| 3 | 184941725 | rs3774051 | 2.356 | -0.56 | 2.493 | 2.21 | -0.007 | 0.006 | 0.006 | 0 | 0.69 | 0.39 | 0.64 | 0.42 | 0.8 | 0.64 |
| 3 | 184947422 | rs719393 | -1.942 | 0.003 | 2.126 | 1.583 | -0.006 | -0.002 | -0.002 | 0 | 0.1 | 0.28 | 0.14 | 0.06 | 0.06 | 0.07 |

| | | | | | | | | | | | | | | | | |
|---|---|---|---|---|---|---|---|---|---|---|---|---|---|---|---|---|
| 3 | 184980371 | rs6795830 | -1.836 | 0.093 | 1.975 | 1.59 | -0.007 | -0.003 | -0.003 | 0.049 | 0.09 | 0.26 | 0.14 | 0.06 | 0.06 | 0.06 |
| 3 | 184998296 | rs12491575 | -0.226 | 0.067 | 1.396 | 1.678 | -0.02 | -0.024 | -0.024 | 0 | 0.07 | 0 | 0.07 | 0.07 | 0.08 | 0.12 |
| 3 | 185001311 | rs7429119 | 1.561 | 0.043 | 2.063 | 1.956 | -0.009 | -0.002 | -0.002 | 0 | 0.8 | 0.72 | 0.78 | 0.79 | 0.84 | 0.75 |
| 3 | 185025889 | rs6779106 | 2.139 | -0.039 | 2.241 | 1.939 | -0.012 | 0.036 | 0.036 | 0.441 | 0.75 | 0.47 | 0.68 | 0.57 | 0.79 | 0.64 |
| 3 | 185036664 | rs7432310 | 1.534 | -0.04 | 2.28 | 1.938 | -0.016 | -0.006 | -0.006 | 0 | 0.79 | 0.66 | 0.77 | 0.78 | 0.84 | 0.75 |
| 3 | 185042323 | rs12639437 | -0.6 | -0.022 | 2.004 | 1.916 | -0.019 | -0.032 | -0.032 | 0.677 | 0.11 | 0.01 | 0.09 | 0.16 | 0.1 | 0.19 |
| 3 | 185043280 | rs7633909 | -0.918 | 0.415 | 2.202 | 1.902 | 0 | 0 | 0 | 0.046 | 0.12 | 0.16 | 0.15 | 0.11 | 0.18 | 0.19 |
| 3 | 185048314 | rs7432658 | 2.038 | 0.243 | 2.321 | 1.925 | -0.012 | 0.054 | 0.054 | 0.345 | 0.74 | 0.43 | 0.61 | 0.51 | 0.79 | 0.64 |
| 3 | 185057627 | rs4687248 | 1.467 | 0.224 | 1.924 | 1.815 | -0.018 | 0.013 | 0.013 | 0 | 0.79 | 0.62 | 0.7 | 0.72 | 0.84 | 0.75 |
| 3 | 185069658 | rs6444054 | 1.528 | 0.275 | 1.852 | 1.875 | -0.012 | 0.027 | 0.027 | 0.218 | 0.79 | 0.36 | 0.78 | 0.62 | 0.84 | 0.7 |
| 3 | 185079487 | rs6804066 | 0.867 | 0.156 | 1.984 | 1.757 | -0.017 | -0.012 | -0.012 | 0.292 | 0.84 | 0.65 | 0.88 | 0.84 | 0.89 | 0.81 |
| 3 | 185097580 | rs7428983 | -0.788 | 0.371 | 1.507 | 1.816 | 0 | 0.002 | 0.002 | 0 | 0.05 | 0.08 | 0.17 | 0.13 | 0.05 | 0.05 |
| 3 | 185099352 | rs9866735 | -2.037 | 0.434 | 1.562 | 1.834 | 0.019 | 0.061 | 0.061 | 0.017 | 0.22 | 0.37 | 0.42 | 0.42 | 0.28 | 0.37 |
| 3 | 185110520 | rs6777516 | -0.947 | 0.342 | 1.611 | 1.657 | 0.013 | 0.027 | 0.027 | 0 | 0.11 | 0.21 | 0.19 | 0.15 | 0.13 | 0.18 |
| 3 | 185115396 | rs7651004 | 0.823 | -0.05 | 1.614 | 1.331 | -0.006 | -0.024 | -0.024 | 0 | 0.91 | 0.94 | 0.97 | 1 | 0.95 | 0.99 |
| 3 | 185120842 | rs7428082 | 1.935 | -0.091 | 1.195 | 1.175 | -0.003 | 0.008 | 0.008 | 0.029 | 0.58 | 0.53 | 0.54 | 0.46 | 0.62 | 0.46 |
| 3 | 185123416 | rs6803944 | -1.647 | -0.126 | 1.391 | 1.456 | -0.005 | 0.134 | 0.134 | 1.42 | 0.17 | 0.45 | 0.26 | 0.34 | 0.14 | 0.27 |
| 3 | 185124607 | rs9842230 | 0.801 | -0.162 | 1.105 | 1.402 | -0.038 | -0.041 | -0.041 | 0 | 0.83 | 0.98 | 0.86 | 0.8 | 0.84 | 0.74 |
| 3 | 185128219 | rs6444068 | 0.781 | -0.162 | 1.105 | 1.402 | -0.038 | -0.042 | -0.042 | 0 | 0.83 | 0.98 | 0.86 | 0.8 | 0.84 | 0.74 |
| 3 | 185141857 | rs7629245 | -0.944 | -0.138 | 1.092 | 1.41 | -0.033 | -0.051 | -0.051 | 0.007 | 0.16 | 0.01 | 0.14 | 0.22 | 0.16 | 0.25 |
| 3 | 185146702 | rs3732577 | -0.931 | -0.13 | 1.096 | 1.414 | -0.034 | -0.051 | -0.051 | 12.478 | 0.16 | 0.01 | 0.14 | 0.22 | 0.16 | 0.25 |
| 3 | 185148605 | rs7646052 | -2.474 | -0.267 | 1.334 | 1.407 | -0.003 | 0.026 | 0.026 | 0.323 | 0.34 | 0.42 | 0.41 | 0.55 | 0.32 | 0.52 |
| 3 | 185151885 | rs1017502 | 1.09 | -0.362 | 1.871 | 1.421 | -0.013 | 0.136 | 0.136 | 1.642 | 0.84 | 0.47 | 0.73 | 0.68 | 0.89 | 0.74 |
| 3 | 185152113 | rs1017501 | 2.565 | -0.366 | 1.848 | 1.428 | -0.004 | 0.007 | 0.007 | 2.917 | 0.65 | 0.39 | 0.58 | 0.45 | 0.73 | 0.49 |
| 3 | 185155482 | rs2304815 | -1.11 | -0.375 | 1.824 | 1.468 | -0.016 | -0.033 | -0.033 | 0.839 | 0.17 | 0.05 | 0.15 | 0.24 | 0.16 | 0.26 |
| 3 | 185158409 | rs894276 | 0.671 | -0.386 | 1.969 | 1.456 | -0.004 | 0.014 | 0.014 | 0.097 | 0.87 | 0.59 | 0.81 | 0.9 | 0.91 | 0.83 |
| 3 | 185162426 | rs1992454 | 2.109 | -0.271 | 1.805 | 1.358 | -0.005 | 0.008 | 0.008 | 2.818 | 0.65 | 0.4 | 0.59 | 0.45 | 0.73 | 0.48 |
| 3 | 185175405 | rs9868878 | 0.929 | -0.355 | 1.96 | 1.092 | -0.027 | 0.083 | 0.083 | 0.703 | 0.79 | 0.53 | 0.72 | 0.66 | 0.85 | 0.75 |
| 3 | 185175570 | rs9832000 | 1.225 | -0.331 | 1.678 | 1.096 | -0.023 | 0.027 | 0.027 | 3.131 | 0.77 | 0.4 | 0.71 | 0.66 | 0.85 | 0.75 |
| 3 | 185180812 | rs9878704 | 0.823 | -0.653 | 1.65 | 0.967 | -0.064 | 0.058 | 0.058 | 0.52 | 0.79 | 0.54 | 0.75 | 0.68 | 0.87 | 0.74 |
| 3 | 185199555 | rs13060295 | -0.581 | -0.554 | 1.844 | 1.64 | -0.003 | 0.06 | 0.06 | 0.11 | 0.1 | 0.14 | 0.18 | 0.37 | 0.09 | 0.28 |
| 3 | 185200783 | rs3755647 | 0.725 | -0.317 | 1.147 | 0.825 | -0.001 | -0.002 | -0.002 | 0 | 0.48 | 0.47 | 0.45 | 0.3 | 0.52 | 0.34 |
| 3 | 185201338 | rs13091808 | -0.243 | -0.321 | 1.161 | 0.836 | -0.007 | 0.09 | 0.09 | 0 | 0.11 | 0.17 | 0.17 | 0.37 | 0.09 | 0.27 |
| 3 | 185206717 | rs1054843 | 0.642 | -0.264 | 1.169 | 0.716 | 0 | 0.004 | 0.004 | 0 | 0.48 | 0.46 | 0.45 | 0.31 | 0.51 | 0.36 |
| 3 | 185218275 | rs16860081 | 0.576 | -0.531 | 1.063 | 0.48 | -0.001 | 0 | 0 | 0 | 0.39 | 0.39 | 0.42 | 0.28 | 0.46 | 0.29 |
| 3 | 185218549 | rs7633364 | 0.586 | -0.518 | 1.079 | 0.471 | 0.007 | -0.004 | -0.004 | 0 | 0.39 | 0.41 | 0.42 | 0.3 | 0.46 | 0.29 |
| 3 | 185223193 | rs6444074 | 0.609 | -0.629 | 1.11 | 0.455 | -0.01 | 0.004 | 0.004 | 0 | 0.39 | 0.36 | 0.39 | 0.3 | 0.45 | 0.25 |
| 3 | 185224119 | rs1065096 | 0.502 | -0.645 | 1.148 | 0.448 | -0.003 | 0.001 | 0.001 | 0 | 0.4 | 0.39 | 0.39 | 0.29 | 0.45 | 0.25 |
| 3 | 185226396 | rs6788865 | 0.47 | -0.626 | 1.157 | 0.446 | -0.001 | 0 | 0 | 0 | 0.4 | 0.4 | 0.4 | 0.29 | 0.45 | 0.25 |
| 3 | 185229757 | rs12233604 | 0.394 | -0.647 | 1.165 | 0.426 | 0.004 | -0.006 | -0.006 | 3.084 | 0.42 | 0.47 | 0.4 | 0.29 | 0.45 | 0.25 |
| 3 | 185234543 | rs9790230 | -0.143 | -0.71 | 1.082 | 0.247 | -0.001 | -0.004 | -0.004 | 0 | 0.54 | 0.45 | 0.54 | 0.72 | 0.54 | 0.59 |
| 3 | 185235838 | rs2030579 | -1.459 | -0.174 | 1.094 | 0.171 | -0.01 | -0.025 | -0.025 | 0.531 | 0.07 | 0.25 | 0.06 | 0.01 | 0.04 | 0.01 |
| 3 | 185244569 | rs602149 | 0.334 | -0.073 | 0.621 | 0.232 | -0.012 | -0.021 | -0.021 | 0 | 0.94 | 0.78 | 0.95 | 0.99 | 0.96 | 0.98 |
| 3 | 185251001 | rs612304 | 0.473 | -0.058 | 0.34 | 0.239 | -0.021 | -0.03 | -0.03 | 3.178 | 0.94 | 0.8 | 0.95 | 0.99 | 0.96 | 0.98 |
| 3 | 185258933 | rs4626118 | 0.421 | 0.574 | 0.017 | 0.444 | 0.002 | -0.001 | -0.001 | 0.121 | 0.35 | 0.35 | 0.34 | 0.28 | 0.41 | 0.24 |
| 3 | 185295704 | rs17217776 | -0.237 | 0.556 | -0.491 | 0.996 | 0.035 | -0.085 | -0.085 | 0 | 0.15 | 0.01 | 0.08 | 0.3 | 0.08 | 0.4 |
| 3 | 185296602 | rs11918734 | 0.08 | 0.321 | -0.549 | -0.032 | -0.004 | -0.063 | -0.063 | 0.305 | 0.29 | 0.36 | 0.25 | 0.06 | 0.25 | 0.19 |
| 3 | 185302098 | rs13095912 | -0.901 | 0.385 | -0.39 | 0.037 | -0.001 | -0.001 | -0.001 | 0 | 0.44 | 0.45 | 0.34 | 0.36 | 0.32 | 0.59 |
| 3 | 185302885 | rs10937226 | -0.917 | 0.453 | -0.401 | 0.039 | -0.005 | -0.005 | -0.005 | 0 | 0.44 | 0.46 | 0.34 | 0.36 | 0.33 | 0.59 |
| 3 | 185331165 | rs6762208 | -0.969 | 0.362 | 0.048 | 0.055 | -0.001 | -0.003 | -0.003 | 1.487 | 0.45 | 0.51 | 0.34 | 0.32 | 0.32 | 0.59 |
| 3 | 185348351 | rs4687477 | -0.901 | 0.028 | 0.439 | 0.4 | 0.014 | 0.001 | 0.001 | 0 | 0.49 | 0.44 | 0.45 | 0.48 | 0.37 | 0.62 |
| 3 | 185348402 | rs3087964 | -0.679 | 0.007 | 0.659 | 0.34 | 0.035 | 0.041 | 0.041 | 0 | 0.44 | 0.33 | 0.33 | 0.32 | 0.32 | 0.58 |
| 3 | 185350706 | rs7638853 | -0.713 | -0.048 | 0.729 | 0.325 | 0.025 | 0.03 | 0.03 | 0 | 0.44 | 0.37 | 0.34 | 0.31 | 0.32 | 0.59 |
| 3 | 185353235 | rs7613002 | 0.212 | -0.099 | 0.806 | 0.22 | -0.005 | -0.06 | -0.06 | 0 | 0.29 | 0.33 | 0.25 | 0.06 | 0.25 | 0.19 |
| 3 | 185356176 | rs9878208 | 0.34 | -0.374 | 0.588 | 0.153 | 0.004 | -0.006 | -0.006 | 0.206 | 0.11 | 0.09 | 0.07 | 0.24 | 0.05 | 0.39 |
| 3 | 185357260 | rs9826022 | 0.34 | -0.374 | 0.588 | 0.203 | 0.004 | -0.005 | -0.005 | 0 | 0.11 | 0.1 | 0.07 | 0.24 | 0.05 | 0.39 |
| 3 | 185357531 | rs9864104 | -1.131 | -0.919 | 0.99 | 0.193 | -0.072 | 0.161 | 0.161 | 0 | 0.21 | 0.52 | 0.25 | 0.47 | 0.16 | 0.43 |
| 3 | 185421647 | rs750753 | -0.447 | -1.007 | 0.109 | -1.021 | 0 | 0 | 0 | 0 | 0.44 | 0.48 | 0.31 | 0.3 | 0.38 | 0.54 |
| 3 | 185425976 | rs7609665 | -1.67 | -1.107 | -0.065 | -1.225 | 0.004 | 0 | 0 | 0 | 0.74 | 0.78 | 0.47 | 0.35 | 0.62 | 0.74 |
| 3 | 185431283 | rs17826758 | 1.787 | -0.953 | -0.114 | -1.002 | -0.023 | 0.083 | 0.083 | 0 | 0.2 | 0.07 | 0.13 | 0.02 | 0.2 | 0.11 |
| 3 | 185445144 | rs6791275 | -0.587 | -0.986 | 0.185 | -1.148 | 0.001 | 0.007 | 0.007 | 0 | 0.48 | 0.56 | 0.35 | 0.36 | 0.44 | 0.58 |
| 3 | 185445842 | rs2290066 | -1.057 | -1.212 | 0.494 | -0.903 | -0.032 | -0.001 | -0.001 | 0 | 0.29 | 0.52 | 0.23 | 0.34 | 0.25 | 0.48 |
| 3 | 185447038 | rs12635769 | 0.85 | -0.953 | 0.366 | -0.997 | 0.02 | -0.009 | -0.009 | 0 | 0.1 | 0.21 | 0.08 | 0.08 | 0.1 | 0.04 |
| 3 | 185454844 | rs9850770 | -1.137 | -0.956 | 0.199 | -0.773 | -0.023 | 0.001 | 0.001 | 0.544 | 0.23 | 0.63 | 0.19 | 0.21 | 0.17 | 0.34 |
| 3 | 185464488 | rs13076013 | 0.915 | -0.968 | -0.068 | -0.948 | -0.007 | -0.047 | -0.047 | 0 | 0.19 | 0.19 | 0.09 | 0 | 0.16 | 0.12 |
| 3 | 185473065 | rs6784185 | -0.279 | -0.634 | 0.04 | -0.868 | -0.001 | 0.006 | 0.006 | 0 | 0.2 | 0.26 | 0.16 | 0.21 | 0.2 | 0.34 |
| 3 | 185482618 | rs6763887 | -1.386 | -0.283 | -0.728 | -0.825 | -0.059 | -0.051 | -0.051 | 0.202 | 0.73 | 0.91 | 0.52 | 0.4 | 0.66 | 0.69 |
| 3 | 185488882 | rs16860216 | -0.638 | -0.542 | 0.298 | -0.707 | 0.009 | -0.03 | -0.03 | 0 | 0.24 | 0.48 | 0.21 | 0.16 | 0.24 | 0.28 |
| 3 | 185490240 | rs13073992 | -0.595 | -0.504 | 0.399 | -0.711 | 0.011 | -0.024 | -0.024 | 0 | 0.23 | 0.43 | 0.2 | 0.16 | 0.26 | 0.28 |

| | | | | | | | | | | | | | | | | |
|---|---|---|---|---|---|---|---|---|---|---|---|---|---|---|---|---|
| 3 | 185501238 | rs6808808 | 0.069 | -0.677 | 0.374 | -0.747 | -0.001 | 0.042 | 0.042 | 0.003 | 0.07 | 0.19 | 0.08 | 0.09 | 0.03 | 0.11 |
| 3 | 185505787 | rs4481184 | -1.755 | -0.839 | 0.352 | -0.906 | 0.012 | 0.023 | 0.023 | 2.725 | 0.37 | 0.7 | 0.29 | 0.25 | 0.3 | 0.46 |
| 3 | 185510018 | rs7619232 | 0.534 | -0.931 | 0.311 | -0.925 | -0.011 | 0.007 | 0.007 | 0 | 0.06 | 0.1 | 0.07 | 0.09 | 0.03 | 0.11 |
| 3 | 185510613 | rs7633675 | -2.011 | -0.935 | 0.309 | -0.93 | 0.033 | 0.054 | 0.054 | 3.745 | 0.39 | 0.84 | 0.3 | 0.26 | 0.3 | 0.46 |
| 3 | 185510884 | rs16860234 | -0.986 | -0.899 | 0.317 | -0.92 | -0.005 | -0.044 | -0.044 | 0 | 0.3 | 0.49 | 0.21 | 0.16 | 0.27 | 0.34 |
| 3 | 185511687 | rs4402960 | -1.51 | -0.883 | 0.156 | -0.915 | -0.009 | -0.022 | -0.022 | 0.028 | 0.35 | 0.57 | 0.28 | 0.25 | 0.3 | 0.46 |
| 3 | 185520342 | rs10428126 | -1.596 | -0.931 | 0.134 | -0.911 | -0.007 | -0.014 | -0.014 | 0 | 0.36 | 0.62 | 0.29 | 0.25 | 0.3 | 0.46 |
| 3 | 185520732 | rs6767577 | -0.395 | -0.959 | 0.246 | -0.93 | 0.004 | -0.011 | -0.011 | 0 | 0.23 | 0.3 | 0.18 | 0.15 | 0.25 | 0.29 |
| 3 | 185529080 | rs1470579 | -1.477 | -0.747 | -1.433 | -0.852 | 0.033 | 0.054 | 0.054 | 3.72 | 0.39 | 0.85 | 0.3 | 0.26 | 0.3 | 0.46 |
| 3 | 185536223 | rs6444082 | -0.834 | -0.666 | -1.37 | -0.81 | -0.003 | -0.004 | -0.004 | 0.349 | 0.23 | 0.78 | 0.26 | 0.25 | 0.24 | 0.38 |
| 3 | 185544309 | rs11705701 | -1.036 | -0.228 | -1.738 | -0.758 | 0.005 | 0.134 | 0.134 | 1.942 | 0.49 | 0.88 | 0.35 | 0.24 | 0.43 | 0.49 |
| 3 | 185562292 | rs4634163 | 0.137 | 0.035 | -1.137 | -0.549 | -0.008 | -0.025 | -0.025 | 0 | 0.57 | 0.54 | 0.71 | 0.8 | 0.63 | 0.58 |
| 3 | 185565723 | rs6776536 | 0.105 | 0.772 | -1.576 | 0.004 | 0.013 | -0.002 | -0.002 | 0 | 0.58 | 0.82 | 0.51 | 0.33 | 0.45 | 0.41 |
| 3 | 185574131 | rs12639517 | -0.57 | 1.256 | -0.713 | -0.024 | -0.006 | -0.005 | -0.005 | 0 | 0.33 | 0.51 | 0.45 | 0.68 | 0.34 | 0.37 |
| 3 | 185574682 | rs12631770 | -1.007 | 1.879 | -0.643 | -0.057 | -0.004 | 0.006 | 0.006 | 1.399 | 0.41 | 0.64 | 0.54 | 0.68 | 0.49 | 0.48 |
| 3 | 185579020 | rs4686704 | -0.571 | 1.315 | -0.489 | 0.144 | -0.004 | -0.005 | -0.005 | 0.736 | 0.33 | 0.55 | 0.45 | 0.67 | 0.35 | 0.37 |
| 3 | 185592046 | rs741604 | -0.922 | 0.788 | -0.378 | 0.058 | 0.008 | -0.046 | -0.046 | 1.897 | 0.09 | 0.27 | 0.1 | 0.03 | 0.08 | 0.06 |
| 3 | 185597788 | rs6809601 | -0.879 | 0.927 | -0.816 | 0.04 | 0.017 | 0.025 | 0.025 | 0.934 | 0.45 | 0.59 | 0.56 | 0.66 | 0.65 | 0.52 |
| 3 | 185598500 | rs10460894 | 0.474 | 0.808 | -1.015 | -0.038 | -0.004 | 0.038 | 0.038 | 0 | 0.37 | 0.22 | 0.41 | 0.3 | 0.35 | 0.41 |
| 3 | 185598552 | rs12107951 | 0.467 | 0.8 | -0.998 | -0.031 | -0.002 | 0.029 | 0.029 | 0 | 0.37 | 0.26 | 0.42 | 0.3 | 0.35 | 0.42 |
| 3 | 185608957 | rs16860296 | 0.299 | 0.948 | -0.689 | 0.458 | 0.004 | 0.022 | 0.022 | 0 | 0.06 | 0.09 | 0.03 | 0.12 | 0.06 | 0.07 |
| 3 | 185610274 | rs17196046 | 0.208 | 0.928 | -0.67 | 0.266 | -0.003 | 0.074 | 0.074 | 0.219 | 0.32 | 0.14 | 0.19 | 0.1 | 0.3 | 0.24 |
| 3 | 185613272 | rs6782205 | -0.846 | -0.416 | 0.145 | 1.189 | 0.052 | 0.414 | 0.414 | 2.575 | 0.07 | 0.41 | 0.17 | 0.33 | 0.05 | 0.12 |
| 3 | 185616564 | rs6770735 | -0.179 | -0.603 | -0.368 | 0.626 | 0.002 | 0.002 | 0.002 | 0 | 0.42 | 0.48 | 0.55 | 0.64 | 0.32 | 0.59 |
| 3 | 185618266 | rs4686399 | -0.638 | -0.604 | -0.36 | 0.579 | 0.002 | 0.006 | 0.006 | 0.085 | 0.68 | 0.78 | 0.76 | 0.75 | 0.69 | 0.81 |
| 3 | 185621829 | rs6444087 | -0.552 | -1.221 | -0.222 | 0.519 | -0.004 | -0.001 | -0.001 | 0 | 0.67 | 0.7 | 0.7 | 0.69 | 0.73 | 0.79 |
| 3 | 185629299 | rs4686709 | -0.286 | -1.02 | 0.298 | 0.93 | 0.041 | 0.253 | 0.253 | 0.613 | 0.05 | 0.22 | 0.2 | 0.22 | 0.06 | 0.13 |
| 3 | 185631793 | rs6761961 | -0.305 | -1.303 | -0.045 | 0.642 | 0.03 | 0.032 | 0.032 | 2.559 | 0.48 | 0.72 | 0.62 | 0.62 | 0.57 | 0.69 |
| 3 | 185633136 | rs4686710 | -0.747 | -1.287 | 0.16 | 0.987 | -0.003 | 0.189 | 0.189 | 4.542 | 0.2 | 0.46 | 0.4 | 0.47 | 0.16 | 0.48 |
| 3 | 185643454 | rs2551372 | 0.543 | -1.184 | -0.21 | 0.943 | -0.039 | -0.075 | -0.075 | 0 | 0.12 | 0 | 0.19 | 0.25 | 0.1 | 0.34 |
| 3 | 185652184 | rs1862903 | -0.865 | -1.276 | -0.26 | 0.879 | 0.031 | -0.028 | -0.028 | 0 | 0.68 | 0.88 | 0.72 | 0.67 | 0.74 | 0.79 |
| 3 | 185654924 | rs2251407 | -0.404 | -0.633 | 0.229 | 1.141 | 0 | 0.003 | 0.003 | 1.067 | 0.14 | 0.19 | 0.17 | 0.25 | 0.1 | 0.37 |
| 3 | 185664946 | rs4686400 | -1.286 | -0.678 | 0.593 | 1.039 | 0.072 | 0.282 | 0.282 | 0.947 | 0.07 | 0.46 | 0.2 | 0.16 | 0.06 | 0.15 |
| 3 | 185676350 | rs6444089 | -0.073 | -0.95 | 0.139 | 0.73 | 0 | -0.001 | -0.001 | 0.905 | 0.42 | 0.41 | 0.42 | 0.61 | 0.54 | 0.6 |
| 3 | 185678110 | rs7356015 | 0.111 | -1.968 | 0.03 | 0.323 | 0.031 | 0.025 | 0.025 | 0 | 0.44 | 0.39 | 0.35 | 0.27 | 0.34 | 0.25 |
| 3 | 185680124 | rs6787684 | 0.306 | -2.133 | -0.105 | 0.265 | 0.003 | 0.002 | 0.002 | 1.044 | 0.5 | 0.48 | 0.51 | 0.48 | 0.53 | 0.41 |
| 3 | 185680144 | rs10937233 | 1.05 | -2.043 | -0.407 | 0.323 | 0.003 | 0.018 | 0.018 | 0 | 0.12 | 0.14 | 0.17 | 0.29 | 0.13 | 0.32 |
| 3 | 185682129 | rs9840713 | -1.185 | -1.741 | -0.326 | -0.007 | 0.019 | 0.008 | 0.008 | 0 | 0.86 | 0.93 | 0.92 | 1 | 0.92 | 0.86 |
| 3 | 185683829 | rs4563434 | 0.239 | -1.649 | -0.091 | 0.092 | -0.001 | 0 | 0 | 0 | 0.27 | 0.26 | 0.23 | 0.23 | 0.25 | 0.19 |
| 3 | 185686741 | rs10049246 | -1.32 | -1.602 | -0.3 | 0.192 | -0.054 | 0.005 | 0.005 | 0 | 0.75 | 0.86 | 0.78 | 0.77 | 0.58 | 0.78 |
| 3 | 185688779 | rs9845685 | 0.59 | -1.476 | -0.356 | -0.055 | -0.001 | 0 | 0 | 0 | 0.22 | 0.2 | 0.22 | 0.23 | 0.23 | 0.11 |
| 3 | 185693840 | rs13065531 | 1.082 | -1.35 | -0.694 | -0.186 | 0.011 | 0.009 | 0.009 | 0 | 0.13 | 0.07 | 0.09 | 0 | 0.1 | 0.17 |
| 3 | 185696444 | rs10937235 | -0.372 | -1.226 | -0.405 | 0.144 | -0.04 | -0.1 | -0.1 | 0 | 0.78 | 0.64 | 0.87 | 0.95 | 0.82 | 0.87 |
| 3 | 185699585 | rs4686718 | -0.784 | -1.068 | -0.433 | 0.228 | -0.03 | -0.097 | -0.097 | 0.203 | 0.81 | 0.62 | 0.87 | 0.95 | 0.83 | 0.93 |
| 3 | 185704236 | rs4686719 | 0.519 | -1.055 | -0.566 | 0.414 | -0.004 | 0.001 | 0.001 | 0 | 0.45 | 0.49 | 0.44 | 0.51 | 0.56 | 0.46 |
| 3 | 185704345 | rs6768604 | 0.399 | -0.901 | -0.944 | 0.422 | 0.004 | -0.116 | -0.116 | 0 | 0.35 | 0.11 | 0.43 | 0.42 | 0.27 | 0.47 |
| 3 | 185707842 | rs6763645 | -0.512 | -0.725 | -0.844 | 0.533 | -0.026 | 0.056 | 0.056 | 0 | 0.77 | 0.65 | 0.77 | 0.66 | 0.79 | 0.88 |
| 3 | 185708275 | rs11718099 | -0.09 | 0.138 | -0.621 | -0.004 | 0.004 | -0.023 | -0.023 | 0 | 0.42 | 0.48 | 0.34 | 0.28 | 0.51 | 0.39 |
| 3 | 185730227 | rs7614436 | -0.288 | 0.589 | -0.595 | -0.112 | 0.008 | -0.012 | -0.012 | 0.097 | 0.36 | 0.39 | 0.32 | 0.17 | 0.49 | 0.26 |
| 3 | 185734921 | rs6771203 | -0.096 | 1.142 | -0.686 | -0.246 | -0.014 | 0.001 | 0.001 | 0.156 | 0.22 | 0.22 | 0.22 | 0.16 | 0.33 | 0.12 |
| 3 | 185746890 | rs9867846 | -0.575 | -0.216 | -0.471 | -0.21 | -0.006 | 0 | 0 | 0 | 0.58 | 0.46 | 0.6 | 0.35 | 0.79 | 0.57 |
| 3 | 185752767 | rs9820527 | 0.07 | -0.254 | -0.38 | -0.324 | 0.048 | 0.038 | 0.038 | 0 | 0.46 | 0.32 | 0.51 | 0.3 | 0.66 | 0.45 |
| 3 | 185757744 | rs9818739 | 0.958 | 0.492 | -0.813 | -0.286 | -0.051 | -0.026 | -0.026 | 0 | 0.08 | 0.01 | 0.15 | 0.09 | 0.2 | 0.08 |
| 3 | 185761846 | rs9844271 | -1.01 | -0.448 | -0.43 | -0.179 | -0.001 | -0.019 | -0.019 | 0 | 0.86 | 0.86 | 0.91 | 0.94 | 0.86 | 0.87 |
| 3 | 185763219 | rs4461451 | -0.999 | -1.084 | -0.269 | -0.289 | 0.087 | -0.044 | -0.044 | 13.205 | 0.67 | 0.86 | 0.66 | 0.37 | 0.85 | 0.64 |
| 3 | 185763566 | rs6780296 | -0.228 | -1.044 | -0.19 | -0.279 | 0.041 | 0.015 | 0.015 | 0 | 0.54 | 0.67 | 0.57 | 0.32 | 0.72 | 0.51 |
| 3 | 185785996 | rs9831938 | 0.192 | -0.415 | 0.752 | 0.757 | -0.024 | 0.199 | 0.199 | 0.004 | 0.79 | 0.52 | 0.68 | 0.48 | 0.84 | 0.77 |
| 3 | 185786406 | rs4234588 | -1.024 | -0.39 | 0.442 | 0.21 | 0 | -0.01 | -0.01 | 0 | 0.87 | 0.9 | 0.92 | 0.94 | 0.87 | 0.87 |
| 3 | 185795855 | rs12638263 | -0.66 | -0.428 | 0.719 | 0.342 | -0.008 | 0.041 | 0.041 | 0.247 | 0.24 | 0.38 | 0.37 | 0.62 | 0.19 | 0.29 |
| 3 | 185801628 | rs7433760 | -1.33 | -0.885 | 0.217 | 0.026 | 0.017 | 0.022 | 0.022 | 0 | 0.45 | 0.6 | 0.51 | 0.77 | 0.32 | 0.56 |
| 3 | 185802823 | rs10937240 | -1.096 | 0.045 | 0.704 | 0.769 | -0.015 | 0.074 | 0.074 | 0.288 | 0.25 | 0.51 | 0.39 | 0.61 | 0.19 | 0.27 |
| 3 | 185811495 | rs6444106 | -1.078 | 0.061 | 0.591 | 0.841 | -0.014 | 0.106 | 0.106 | 0.471 | 0.24 | 0.49 | 0.39 | 0.55 | 0.19 | 0.27 |
| 3 | 185822353 | rs10513801 | 0.407 | 0.169 | 0.125 | 0.556 | -0.015 | 0.063 | 0.063 | 2.653 | 0.14 | 0.01 | 0.08 | 0.06 | 0.14 | 0.12 |
| 3 | 185830514 | rs4686733 | 0.704 | 0.2 | 0.337 | 0.754 | 0.016 | 0.271 | 0.271 | 0.721 | 0.91 | 0.65 | 0.69 | 0.46 | 0.94 | 0.84 |
| 3 | 185833759 | rs7635103 | 0.502 | 0.291 | -0.065 | 0.628 | -0.01 | 0.043 | 0.043 | 1.056 | 0.71 | 0.46 | 0.57 | 0.4 | 0.76 | 0.66 |
| 3 | 185842523 | rs11718073 | -0.46 | 0.255 | -0.649 | 0.487 | 0.026 | 0.001 | 0.001 | 0.012 | 0.49 | 0.59 | 0.39 | 0.55 | 0.58 | 0.56 |
| 3 | 185845583 | rs11924144 | -0.434 | 0.885 | -0.507 | 0.435 | 0 | 0.002 | 0.002 | 0 | 0.44 | 0.54 | 0.34 | 0.53 | 0.52 | 0.51 |
| 3 | 185847441 | rs16860471 | 0.293 | 0.677 | -0.383 | -0.205 | 0.002 | -0.01 | -0.01 | 0 | 0.17 | 0.12 | 0.11 | 0.01 | 0.17 | 0.17 |
| 3 | 185848147 | rs16848688 | 0.38 | 0.037 | -1.122 | -0.263 | -0.026 | -0.062 | -0.062 | 2.438 | 0.16 | 0.01 | 0.17 | 0.2 | 0.13 | 0.3 |

| | | | | | | | | | | | | | | | | |
|---|---|---|---|---|---|---|---|---|---|---|---|---|---|---|---|---|
| 3 | 185848681 | rs4686734 | 0.431 | -0.04 | -0.282 | -0.86 | 0 | -0.033 | -0.033 | 0 | 0.74 | 0.55 | 0.82 | 0.76 | 0.76 | 0.75 |
| 3 | 185849219 | rs9877031 | -0.859 | -0.551 | -0.23 | -1.557 | 0.003 | -0.033 | -0.033 | 0 | 0.29 | 0.72 | 0.19 | 0.06 | 0.2 | 0.29 |
| 3 | 185849390 | rs9877630 | -0.64 | -0.918 | -0.184 | -1.531 | -0.011 | -0.056 | -0.056 | 0 | 0.26 | 0.68 | 0.16 | 0.06 | 0.17 | 0.27 |
| 3 | 185853075 | rs6783586 | -0.567 | -1.159 | -0.202 | -1.842 | -0.004 | -0.013 | -0.013 | 0 | 0.46 | 0.53 | 0.33 | 0.23 | 0.37 | 0.57 |
| 3 | 185856797 | rs11706414 | -0.722 | -1.303 | -0.403 | -2.053 | 0.002 | -0.009 | -0.009 | 0 | 0.23 | 0.75 | 0.13 | 0.05 | 0.2 | 0.25 |
| 3 | 185867283 | rs6444109 | 0.062 | -1.427 | -0.482 | -2.434 | 0.023 | 0.044 | 0.044 | 0 | 0.56 | 0.67 | 0.75 | 0.91 | 0.63 | 0.66 |
| 3 | 185867391 | rs2575 | 1.106 | -1.375 | -0.817 | -2.538 | -0.153 | 0.021 | 0.021 | 0.071 | 0.3 | 0.06 | 0.54 | 0.75 | 0.45 | 0.33 |
| 3 | 185867637 | rs3732940 | 1.066 | -1.398 | -0.641 | -2.506 | 0.015 | 0.009 | 0.009 | 0 | 0.2 | 0.1 | 0.17 | 0.17 | 0.16 | 0.3 |
| 3 | 185877382 | rs7627157 | -0.075 | -1.344 | -0.475 | -2.502 | 0.018 | 0.019 | 0.019 | 0 | 0.55 | 0.66 | 0.76 | 0.91 | 0.63 | 0.66 |
| 3 | 185878587 | rs4686740 | 1.129 | -1.121 | -0.66 | -2.455 | -0.143 | 0.005 | 0.005 | 0 | 0.28 | 0.07 | 0.54 | 0.75 | 0.45 | 0.34 |
| 3 | 185883562 | rs1400362 | -0.456 | -1.375 | -0.277 | -2.535 | -0.001 | -0.04 | -0.04 | 0 | 0.69 | 0.68 | 0.84 | 0.92 | 0.71 | 0.83 |
| 3 | 185890427 | rs3774055 | 0.473 | -1.048 | -0.215 | -2.153 | -0.035 | -0.085 | -0.085 | 0 | 0.16 | 0.24 | 0.09 | 0.01 | 0.1 | 0.17 |
| 3 | 185894484 | rs9815931 | -0.255 | -0.486 | -0.357 | -2.145 | -0.015 | 0.022 | 0.022 | 0.367 | 0.67 | 0.73 | 0.47 | 0.22 | 0.64 | 0.65 |
| 3 | 185894919 | rs2090394 | 0.371 | -1.07 | 0.056 | -2.078 | -0.008 | 0.011 | 0.011 | 0.852 | 0.39 | 0.33 | 0.65 | 0.81 | 0.5 | 0.44 |
| 3 | 185896787 | rs4686742 | -0.481 | -1.195 | -0.165 | -1.996 | -0.015 | 0.109 | 0.109 | 0.212 | 0.71 | 0.9 | 0.86 | 0.93 | 0.69 | 0.59 |
| 3 | 185900713 | rs1400360 | 0.417 | -1.305 | 0.023 | -1.721 | -0.051 | 0.02 | 0.02 | 0 | 0.21 | 0.34 | 0.18 | 0.23 | 0.1 | 0.15 |
| 3 | 185901333 | rs4686743 | 0.262 | -1.31 | -0.749 | -1.723 | -0.091 | 0.045 | 0.045 | 0.001 | 0.22 | 0.43 | 0.18 | 0.23 | 0.09 | 0.15 |
| 3 | 185905876 | rs13087397 | 0.933 | -0.985 | -2.231 | -1.665 | -0.096 | 0.016 | 0.016 | 1.32 | 0.43 | 0.06 | 0.65 | 0.7 | 0.57 | 0.43 |
| 3 | 185909651 | rs2284842 | 0.679 | -0.335 | -1.935 | -1.639 | 0.035 | 0.102 | 0.102 | 0.105 | 0.49 | 0.67 | 0.73 | 0.74 | 0.59 | 0.44 |
| 3 | 185911823 | rs6762646 | 0.359 | 0.741 | -1.7 | -2.415 | -0.032 | 0.104 | 0.104 | 0.803 | 0.71 | 0.89 | 0.87 | 0.93 | 0.68 | 0.6 |
| 3 | 185926701 | rs875571 | 0.144 | 0.6 | -1.213 | -2.509 | 0 | -0.065 | -0.065 | 0 | 0.55 | 0.52 | 0.69 | 0.94 | 0.56 | 0.64 |
| 3 | 185929467 | rs7619111 | -0.722 | -0.195 | -1.442 | -2.985 | 0 | -0.044 | -0.044 | 0 | 0.28 | 0.29 | 0.14 | 0.01 | 0.2 | 0.07 |
| 3 | 185930711 | rs16860553 | -0.863 | -0.298 | -1.331 | -3.054 | 0.017 | 0.016 | 0.016 | 3.547 | 0.29 | 0.21 | 0.14 | 0.01 | 0.2 | 0.07 |
| 3 | 185934593 | rs17295554 | -0.725 | -0.328 | -1.368 | -2.774 | -0.005 | -0.031 | -0.031 | 1.435 | 0.16 | 0.13 | 0.06 | 0 | 0.09 | 0.05 |
| 3 | 185939759 | rs12492576 | 0.133 | 0.24 | -1.364 | -2.681 | -0.015 | 0.069 | 0.069 | 1.11 | 0.43 | 0.26 | 0.61 | 0.85 | 0.55 | 0.62 |
| 3 | 185944837 | rs2284838 | -0.291 | 0.433 | -1.134 | -2.498 | -0.007 | 0 | 0 | 0 | 0.34 | 0.28 | 0.26 | 0.07 | 0.36 | 0.27 |
| 3 | 185951648 | rs4686753 | 0.436 | -0.165 | -0.758 | -2.151 | 0.005 | -0.033 | -0.033 | 0.008 | 0.67 | 0.29 | 0.81 | 0.98 | 0.8 | 0.9 |
| 3 | 185953613 | rs9968130 | 1.11 | -0.155 | 0.413 | -1.798 | 0.039 | -0.012 | -0.012 | 0.23 | 0.92 | 0.64 | 0.94 | 1 | 0.96 | 0.99 |
| 3 | 185955511 | rs12497361 | 1.094 | -0.68 | -0.325 | -2.415 | 0.061 | 0.178 | 0.178 | 0.153 | 0.62 | 0.14 | 0.78 | 0.94 | 0.78 | 0.82 |
| 3 | 185965264 | rs6793168 | 0.577 | -0.515 | 1.301 | -2.285 | 0.028 | -0.011 | -0.011 | 0.139 | 0.94 | 0.71 | 0.95 | 1 | 0.96 | 0.99 |
| 3 | 185965546 | rs1878297 | 1.578 | -0.826 | 1.583 | -2.45 | -0.043 | -0.105 | -0.105 | 0 | 0.85 | 0.45 | 0.9 | 1 | 0.92 | 0.96 |
| 3 | 185969052 | rs6801129 | -1.557 | -1.01 | 1.903 | -2.403 | -0.028 | -0.069 | -0.069 | 0 | 0.24 | 0.4 | 0.13 | 0.08 | 0.16 | 0.15 |
| 3 | 185972670 | rs7623321 | -1.744 | -0.8 | 1.738 | -2.714 | -0.012 | -0.076 | -0.076 | 0.009 | 0.29 | 0.48 | 0.15 | 0.09 | 0.21 | 0.14 |
| 3 | 185973696 | rs9816818 | -0.974 | -0.88 | 1.804 | -2.709 | -0.016 | -0.073 | -0.073 | 0 | 0.14 | 0.26 | 0.09 | 0 | 0.09 | 0.06 |
| 3 | 185975789 | rs13315397 | -0.496 | -0.865 | 1.48 | -2.675 | 0.003 | 0.005 | 0.005 | 0 | 0.12 | 0.07 | 0.07 | 0 | 0.09 | 0.06 |
| 3 | 185976029 | rs918220 | 1.954 | -1.03 | 1.484 | -2.173 | 0.03 | 0.078 | 0.078 | 0.857 | 0.63 | 0.23 | 0.8 | 0.9 | 0.77 | 0.85 |
| 3 | 185976248 | rs2268837 | -0.514 | -0.953 | 1.346 | -1.972 | -0.001 | -0.003 | -0.003 | 0 | 0.17 | 0.17 | 0.05 | 0.09 | 0.1 | 0.09 |
| 3 | 185977310 | rs2268835 | -1.221 | -0.981 | 1.434 | -1.965 | -0.023 | -0.038 | -0.038 | 0 | 0.24 | 0.31 | 0.11 | 0.12 | 0.14 | 0.13 |
| 3 | 185980589 | rs2268833 | -0.368 | -0.054 | 0.697 | -1.191 | 0.011 | 0.013 | 0.013 | 0 | 0.19 | 0.29 | 0.27 | 0.27 | 0.25 | 0.32 |
| 3 | 185982955 | rs3815620 | 0.809 | -0.847 | 0.47 | -1.702 | -0.001 | -0.002 | -0.002 | 0 | 0.6 | 0.41 | 0.63 | 0.71 | 0.61 | 0.5 |
| 3 | 185982972 | rs3815619 | 1.313 | -0.95 | 0.178 | -1.787 | -0.019 | 0.011 | 0.011 | 0 | 0.47 | 0.16 | 0.42 | 0.61 | 0.52 | 0.46 |
| 3 | 185984803 | rs1014695 | 0.165 | -1.141 | 0.556 | -1.625 | -0.07 | -0.106 | -0.106 | 0 | 0.78 | 0.62 | 0.88 | 0.98 | 0.86 | 0.83 |
| 3 | 185990096 | rs2193587 | -0.154 | -1.094 | -0.101 | -1.294 | 0.008 | -0.015 | -0.015 | 5.072 | 0.78 | 0.92 | 0.84 | 0.74 | 0.8 | 0.72 |
| 3 | 185994734 | rs6788108 | -0.319 | -0.973 | -0.511 | -1.71 | -0.016 | 0.052 | 0.052 | 0 | 0.77 | 0.9 | 0.75 | 0.89 | 0.81 | 0.82 |
| 3 | 185995196 | rs1541614 | -0.328 | -0.446 | 0.138 | -0.757 | -0.004 | 0.085 | 0.085 | 0 | 0.29 | 0.51 | 0.26 | 0.49 | 0.35 | 0.37 |
| 3 | 185996345 | rs9834264 | 0.352 | -0.489 | -0.161 | -0.858 | -0.032 | -0.074 | -0.074 | 0 | 0.13 | 0.29 | 0.07 | 0 | 0.07 | 0.04 |
| 3 | 185998884 | rs2239626 | -0.808 | -0.339 | -1.172 | -0.606 | 0.066 | 0.027 | 0.027 | 0.968 | 0.4 | 0.81 | 0.28 | 0.29 | 0.28 | 0.33 |
| 3 | 186005592 | rs3819863 | 0.52 | 0.061 | -0.727 | -0.345 | 0.009 | 0.019 | 0.019 | 0 | 0.5 | 0.34 | 0.54 | 0.58 | 0.57 | 0.51 |
| 3 | 186006165 | rs1558909 | 0.295 | 0.282 | -0.961 | -0.196 | 0.006 | 0.035 | 0.035 | 0 | 0.51 | 0.58 | 0.56 | 0.8 | 0.51 | 0.61 |
| 3 | 186006307 | rs1004589 | -0.453 | 0.086 | -0.275 | 0.285 | 0.025 | 0.091 | 0.091 | 0 | 0.1 | 0.49 | 0.09 | 0.14 | 0.1 | 0.09 |
| 3 | 186006566 | rs710443 | 0.13 | 0.536 | -1.094 | 0.397 | 0.001 | 0.003 | 0.003 | 0 | 0.64 | 0.69 | 0.7 | 0.71 | 0.7 | 0.68 |
| 3 | 186013603 | rs9870928 | 0.106 | 0.868 | -1.157 | 0.658 | -0.007 | -0.016 | -0.016 | 0 | 0.27 | 0.2 | 0.35 | 0.39 | 0.3 | 0.34 |
| 3 | 186014444 | rs6804820 | -0.219 | -0.156 | -1.099 | -0.444 | 0.007 | 0.025 | 0.025 | 0 | 0.31 | 0.26 | 0.2 | 0.14 | 0.27 | 0.31 |
| 3 | 186015283 | rs3819860 | 0.364 | -0.633 | -0.47 | -0.732 | 0.016 | -0.007 | -0.007 | 0 | 0.54 | 0.35 | 0.51 | 0.5 | 0.58 | 0.51 |
| 3 | 186015714 | rs6762330 | 0.156 | -0.732 | -0.164 | -0.794 | 0.002 | -0.008 | -0.008 | 0.119 | 0.58 | 0.63 | 0.55 | 0.5 | 0.58 | 0.51 |
| 3 | 186018203 | rs9868652 | 0.023 | -0.858 | -0.188 | -1.026 | -0.004 | -0.048 | -0.048 | 0.033 | 0.28 | 0.3 | 0.23 | 0.04 | 0.22 | 0.22 |
| 3 | 186022133 | rs2268819 | -0.591 | -0.622 | -0.314 | -0.828 | -0.005 | -0.025 | -0.025 | 0 | 0.77 | 0.84 | 0.69 | 0.57 | 0.71 | 0.68 |
| 3 | 186025431 | rs2239624 | 0.581 | -0.62 | -0.372 | -0.814 | 0.015 | 0.009 | 0.009 | 0 | 0.53 | 0.33 | 0.49 | 0.54 | 0.56 | 0.5 |
| 3 | 186030397 | rs2268817 | -0.055 | -0.479 | -0.037 | -0.454 | 0.006 | -0.013 | -0.013 | 0 | 0.43 | 0.3 | 0.41 | 0.49 | 0.42 | 0.46 |
| 3 | 186033242 | rs3819851 | -0.355 | -0.397 | 0.287 | -0.351 | -0.007 | 0.002 | 0.002 | 0 | 0.63 | 0.41 | 0.61 | 0.58 | 0.66 | 0.55 |
| 3 | 186048931 | rs6767474 | -1.391 | -0.278 | 0.554 | -0.596 | 0.008 | 0.019 | 0.019 | 1.099 | 0.36 | 0.67 | 0.2 | 0.18 | 0.28 | 0.16 |
| 3 | 186054149 | rs1981783 | -1.179 | 0.045 | -0.256 | -0.455 | 0.001 | 0.008 | 0.008 | 0.262 | 0.32 | 0.29 | 0.17 | 0.18 | 0.28 | 0.16 |
| 3 | 186067505 | rs17196108 | -1.038 | -0.08 | -0.539 | -0.172 | 0.01 | 0.054 | 0.054 | 0.194 | 0.16 | 0.1 | 0.09 | 0.06 | 0.13 | 0.08 |
| 3 | 186079990 | rs7617588 | -1.41 | -0.157 | 0.211 | -0.186 | -0.028 | -0.094 | -0.094 | 0 | 0.19 | 0.54 | 0.13 | 0.07 | 0.14 | 0.08 |
| 3 | 186082149 | rs6789403 | -1.545 | -0.177 | 1.027 | -0.173 | 0.02 | -0.027 | -0.027 | 0.064 | 0.13 | 0.32 | 0.09 | 0.08 | 0.14 | 0.07 |
| 3 | 186083611 | rs7618021 | -0.523 | -0.468 | 0.663 | -0.013 | -0.013 | -0.007 | -0.007 | 0 | 0.43 | 0.49 | 0.42 | 0.58 | 0.34 | 0.42 |
| 3 | 186086283 | rs4608682 | -0.648 | -0.46 | 0.178 | -0.432 | 0.011 | 0.136 | 0.136 | 0 | 0.53 | 0.81 | 0.57 | 0.75 | 0.48 | 0.58 |
| 3 | 186086878 | rs10804905 | 0.77 | -0.542 | 0.047 | 0.181 | 0.01 | -0.081 | -0.081 | 0 | 0.28 | 0.13 | 0.24 | 0.52 | 0.27 | 0.31 |
| 3 | 186087169 | rs7628096 | -0.294 | -0.607 | 0.433 | 0.144 | -0.001 | -0.001 | -0.001 | 0 | 0.45 | 0.6 | 0.41 | 0.55 | 0.43 | 0.4 |

| | | | | | | | | | | | | | | | | |
|---|---|---|---|---|---|---|---|---|---|---|---|---|---|---|---|---|
| 3 | 186087279 | rs7639756 | 0.464 | 0.362 | -0.208 | 0.423 | -0.029 | -0.059 | -0.059 | 0 | 0.29 | 0.15 | 0.27 | 0.53 | 0.33 | 0.32 |
| 3 | 186092574 | rs9839753 | 0.257 | 0.472 | 0.007 | 0.421 | 0.002 | 0.001 | 0.001 | 0 | 0.48 | 0.45 | 0.33 | 0.4 | 0.42 | 0.37 |
| 3 | 186094684 | rs7432284 | -0.684 | 0.616 | 0.088 | 0.575 | 0.001 | 0.047 | 0.047 | 0 | 0.08 | 0.16 | 0.13 | 0.23 | 0.08 | 0.18 |
| 3 | 186100848 | rs6776796 | -0.818 | 0.47 | 0.397 | -0.14 | 0.011 | -0.084 | -0.084 | 0 | 0.14 | 0.59 | 0.11 | 0.02 | 0.14 | 0.09 |
| 3 | 186101425 | rs9821408 | -0.667 | 0.359 | -1.072 | -0.423 | 0.039 | 0.197 | 0.197 | 0.131 | 0.46 | 0.88 | 0.33 | 0.24 | 0.41 | 0.32 |
| 3 | 186107980 | rs7628341 | -1.066 | 1.073 | 0.251 | 0.235 | 0.09 | 0.138 | 0.138 | 0.656 | 0.11 | 0.41 | 0.19 | 0.2 | 0.16 | 0.05 |
| 3 | 186108018 | rs7628509 | -1.029 | 1.061 | 0.259 | 0.267 | 0.087 | 0.09 | 0.09 | 0.963 | 0.1 | 0.4 | 0.19 | 0.13 | 0.15 | 0.05 |
| 3 | 186117187 | rs4234594 | 0.187 | 1.249 | -0.126 | -0.017 | -0.002 | 0.013 | 0.013 | 0 | 0.78 | 0.82 | 0.72 | 0.83 | 0.84 | 0.66 |
| 3 | 186126310 | rs4583642 | 0.957 | 1.084 | -0.12 | 0.024 | 0.003 | -0.001 | -0.001 | 0.031 | 0.67 | 0.35 | 0.53 | 0.64 | 0.68 | 0.61 |
| 3 | 186129391 | rs6794653 | 0.42 | 0.618 | -0.647 | -0.228 | 0.008 | 0.002 | 0.002 | 0.635 | 0.86 | 0.92 | 0.72 | 0.88 | 0.86 | 0.79 |
| 3 | 186134871 | rs7635507 | 0.832 | -0.122 | -0.474 | -1.319 | -0.042 | -0.173 | -0.173 | 0 | 0.27 | 0.46 | 0.17 | 0 | 0.19 | 0.08 |
| 3 | 186135861 | rs11926434 | 0.832 | -0.09 | -0.474 | -1.319 | -0.04 | -0.173 | -0.173 | 0.052 | 0.27 | 0.46 | 0.17 | 0 | 0.19 | 0.08 |
| 3 | 186143744 | rs6444127 | -0.817 | 0.123 | -1.455 | -1.28 | -0.092 | 0.019 | 0.019 | 0.346 | 0.68 | 0.92 | 0.44 | 0.29 | 0.56 | 0.28 |
| 3 | 186158483 | rs11916012 | -0.002 | 0.207 | -1.261 | -1.413 | -0.015 | 0.138 | 0.138 | 0.688 | 0.57 | 0.79 | 0.38 | 0.14 | 0.45 | 0.19 |
| 3 | 186166038 | rs7650276 | -0.259 | 0.142 | -1.487 | -1.337 | -0.073 | 0.081 | 0.081 | 0.372 | 0.6 | 0.91 | 0.42 | 0.3 | 0.53 | 0.23 |
| 3 | 186170632 | rs12629868 | -1.076 | -0.39 | -1.005 | -1.43 | 0.006 | 0.017 | 0.017 | 0 | 0.45 | 0.56 | 0.25 | 0.31 | 0.35 | 0.21 |
| 3 | 186174630 | rs3935586 | -0.071 | -0.538 | -0.47 | -1.525 | -0.017 | 0 | 0 | 0 | 0.28 | 0.18 | 0.17 | 0.3 | 0.3 | 0.14 |
| 3 | 186177610 | rs6798365 | -0.557 | -0.646 | -0.097 | -1.676 | -0.01 | 0.017 | 0.017 | 0 | 0.62 | 0.68 | 0.35 | 0.3 | 0.52 | 0.26 |
| 3 | 186184328 | rs6771063 | 1.263 | -0.451 | -0.21 | -1.308 | 0.004 | 0.016 | 0.016 | 0 | 0.49 | 0.43 | 0.5 | 0.6 | 0.51 | 0.56 |
| 3 | 186188610 | rs9290833 | 0.484 | -0.521 | -0.116 | -1.384 | 0 | -0.021 | -0.021 | 0 | 0.66 | 0.67 | 0.62 | 0.82 | 0.66 | 0.71 |
| 3 | 186191665 | rs7632097 | 0.379 | -0.503 | -0.646 | -1.401 | -0.018 | 0.073 | 0.073 | 0.728 | 0.42 | 0.23 | 0.56 | 0.79 | 0.5 | 0.6 |
| 3 | 186191696 | rs7610178 | 0.918 | -0.55 | -0.612 | -1.542 | -0.009 | 0.034 | 0.034 | 1.208 | 0.1 | 0.05 | 0.09 | 0.01 | 0.15 | 0.06 |
| 3 | 186193569 | rs4374519 | -0.124 | -0.427 | -0.769 | -1.533 | -0.028 | -0.037 | -0.037 | 0.189 | 0.88 | 0.95 | 0.8 | 0.74 | 0.82 | 0.8 |
| 3 | 186194398 | rs6770219 | 0.645 | -0.667 | -0.667 | -1.432 | 0.002 | -0.009 | -0.009 | 0 | 0.38 | 0.67 | 0.19 | 0.1 | 0.25 | 0.16 |
| 3 | 186195323 | rs9845510 | 0.235 | -0.942 | -0.751 | -1.461 | -0.017 | 0.045 | 0.045 | 0.462 | 0.48 | 0.14 | 0.47 | 0.63 | 0.45 | 0.64 |
| 3 | 186204424 | rs4490348 | 0.752 | -0.975 | -0.742 | -1.447 | -0.038 | -0.003 | -0.003 | 0 | 0.39 | 0.13 | 0.45 | 0.66 | 0.41 | 0.6 |
| 3 | 186205356 | rs9681406 | 1.564 | -1.026 | -0.344 | -1.381 | -0.022 | -0.14 | -0.14 | 0.005 | 0.27 | 0.45 | 0.19 | 0.01 | 0.19 | 0.11 |
| 3 | 186205729 | rs9844915 | 0.647 | -1.024 | -0.435 | -1.354 | -0.026 | -0.027 | -0.027 | 0.102 | 0.76 | 0.87 | 0.69 | 0.66 | 0.7 | 0.66 |
| 3 | 186205930 | rs9866421 | 0.894 | -1.007 | -0.39 | -1.265 | -0.019 | -0.021 | -0.021 | 0.367 | 0.39 | 0.22 | 0.45 | 0.57 | 0.46 | 0.48 |
| 3 | 186208478 | rs9878039 | 1.015 | -0.514 | -0.287 | -1.115 | -0.005 | 0.005 | 0.005 | 0 | 0.1 | 0.04 | 0.1 | 0.09 | 0.1 | 0.13 |
| 3 | 186209848 | rs9870051 | -0.454 | -0.46 | -0.301 | -1.129 | -0.015 | 0.042 | 0.042 | 0.003 | 0.18 | 0.07 | 0.19 | 0.1 | 0.22 | 0.18 |
| 3 | 186212977 | rs7642903 | 0.186 | -0.627 | 0.205 | -0.775 | -0.003 | -0.023 | -0.023 | 0 | 0.38 | 0.46 | 0.39 | 0.25 | 0.35 | 0.28 |
| 3 | 186213784 | rs11918665 | -0.787 | -0.476 | 0.105 | -0.761 | -0.041 | 0.001 | 0.001 | 0.704 | 0.92 | 0.98 | 0.89 | 0.94 | 0.88 | 0.89 |
| 3 | 186214151 | rs9834116 | 1.08 | -0.556 | 0.233 | -0.705 | -0.013 | -0.015 | -0.015 | 0 | 0.42 | 0.25 | 0.44 | 0.58 | 0.47 | 0.52 |
| 3 | 186222970 | rs1868143 | 0.466 | -0.79 | -0.017 | -0.48 | -0.004 | -0.005 | -0.005 | 0 | 0.83 | 0.93 | 0.87 | 0.86 | 0.84 | 0.83 |
| 3 | 186223224 | rs3941831 | 0.202 | -0.742 | 0.314 | -0.882 | 0.004 | -0.042 | -0.042 | 0.443 | 0.6 | 0.62 | 0.71 | 0.97 | 0.72 | 0.73 |
| 3 | 186225120 | rs13089682 | 0.863 | -0.573 | -0.554 | -0.527 | -0.016 | 0.018 | 0.018 | 0.145 | 0.45 | 0.25 | 0.53 | 0.58 | 0.58 | 0.57 |
| 3 | 186226983 | rs6804256 | 0.928 | -0.489 | -0.888 | -0.331 | 0.006 | -0.019 | -0.019 | 0 | 0.22 | 0.18 | 0.12 | 0 | 0.14 | 0.07 |
| 3 | 186228287 | rs6778486 | -0.028 | -0.092 | -0.881 | 0.064 | -0.001 | -0.002 | -0.002 | 0 | 0.58 | 0.57 | 0.67 | 0.68 | 0.68 | 0.64 |
| 3 | 186230003 | rs745588 | -0.766 | 0.348 | -1.198 | 0.434 | -0.005 | -0.038 | -0.038 | 0.143 | 0.24 | 0.11 | 0.23 | 0.32 | 0.2 | 0.28 |
| 3 | 186235907 | rs1530641 | 0.531 | 0.374 | -0.424 | -0.44 | -0.011 | -0.04 | -0.04 | 0.674 | 0.69 | 0.45 | 0.69 | 0.77 | 0.68 | 0.65 |
| 3 | 186238966 | rs2377869 | 0.411 | 0.991 | -0.65 | -0.628 | -0.029 | 0.038 | 0.038 | 0.115 | 0.14 | 0.05 | 0.07 | 0.06 | 0.15 | 0.09 |
| 3 | 186247373 | rs4686428 | -0.148 | 0.806 | -0.176 | -0.741 | 0.021 | -0.006 | -0.006 | 0.704 | 0.9 | 0.87 | 0.87 | 0.89 | 0.78 | 0.87 |
| 3 | 186252771 | rs2889756 | 0.064 | 0.553 | 0.401 | -0.792 | -0.036 | -0.017 | -0.017 | 1.808 | 0.17 | 0.25 | 0.16 | 0.13 | 0.1 | 0.2 |
| 3 | 186254832 | rs1447670 | -0.608 | 0.621 | 0.7 | -1.013 | -0.053 | -0.057 | -0.057 | 0 | 0.22 | 0.4 | 0.22 | 0.13 | 0.16 | 0.24 |
| 3 | 186256133 | rs1374808 | 2.063 | 0.384 | 0.566 | -1.206 | -0.104 | 0.139 | 0.139 | 0.612 | 0.46 | 0.02 | 0.54 | 0.71 | 0.45 | 0.52 |
| 3 | 186260740 | rs10937252 | 2.063 | 0.384 | 0.566 | -1.206 | -0.104 | 0.139 | 0.139 | 0.612 | 0.46 | 0.02 | 0.54 | 0.71 | 0.45 | 0.52 |
| 3 | 186285706 | rs12330837 | -0.903 | 0.333 | 1.435 | -0.694 | 0 | -0.048 | -0.048 | 0.02 | 0.08 | 0.2 | 0.06 | 0 | 0.08 | 0.06 |
| 3 | 186287649 | rs12330139 | -1.479 | 0.344 | 1.378 | -0.647 | -0.068 | -0.069 | -0.069 | 0.175 | 0.28 | 0.56 | 0.24 | 0.15 | 0.19 | 0.26 |
| 3 | 186287980 | rs12330397 | -1.479 | 0.344 | 1.378 | -0.647 | -0.068 | -0.069 | -0.069 | 0.175 | 0.28 | 0.56 | 0.24 | 0.15 | 0.19 | 0.26 |
| 3 | 186288565 | rs3806714 | 1.323 | 0.567 | 1.475 | -0.655 | 0.119 | 0.035 | 0.035 | 0.023 | 0.91 | 0.61 | 0.87 | 0.89 | 0.84 | 0.9 |
| 3 | 186289322 | rs6776042 | 0.953 | 0.563 | 1.486 | -0.624 | 0.047 | 0.008 | 0.008 | 0 | 0.92 | 0.8 | 0.89 | 0.89 | 0.84 | 0.9 |
| 3 | 186296304 | rs2121752 | 2.208 | 0.458 | 1.292 | -0.751 | -0.001 | 0.112 | 0.112 | 5.035 | 0.63 | 0.03 | 0.63 | 0.79 | 0.65 | 0.65 |
| 3 | 186299107 | rs11921733 | -0.034 | 0.56 | 1.479 | -0.692 | -0.033 | -0.007 | -0.007 | 0.508 | 0.15 | 0.25 | 0.16 | 0.15 | 0.1 | 0.21 |
| 3 | 186301703 | rs8147 | -1.533 | 0.565 | 1.491 | -0.691 | -0.069 | -0.07 | -0.07 | 0.936 | 0.27 | 0.55 | 0.24 | 0.15 | 0.19 | 0.26 |
| 3 | 186310374 | rs6763361 | 0.655 | 0.613 | 1.367 | -0.687 | -0.004 | -0.085 | -0.085 | 0 | 0.8 | 0.68 | 0.84 | 1 | 0.77 | 0.77 |
| 3 | 186317857 | rs4686432 | -1.535 | 0.789 | 1.522 | -0.622 | -0.025 | -0.046 | -0.046 | 0 | 0.24 | 0.38 | 0.21 | 0.14 | 0.16 | 0.2 |
| 3 | 186321513 | rs9842063 | -1.137 | 0.848 | 1.582 | -0.665 | 0.023 | -0.059 | -0.059 | 0.19 | 0.1 | 0.23 | 0.08 | 0 | 0.11 | 0.06 |
| 3 | 186328121 | rs6788635 | 0.498 | 1.717 | 0.987 | -0.26 | 0.157 | 0.095 | 0.095 | 0 | 0.9 | 0.74 | 0.64 | 0.84 | 0.76 | 0.83 |
| 3 | 186330462 | rs2077119 | 2.31 | 1.372 | 0.558 | -0.441 | -0.093 | -0.004 | -0.004 | 0.065 | 0.51 | 0.03 | 0.35 | 0.59 | 0.45 | 0.51 |
| 3 | 186332571 | rs2593813 | 0.818 | 1.634 | 0.769 | -0.374 | 0.09 | 0.069 | 0.069 | 0.021 | 0.8 | 0.64 | 0.57 | 0.72 | 0.67 | 0.77 |
| 3 | 186335941 | rs2070633 | 1.818 | 1.535 | 0.62 | -0.416 | -0.015 | 0.007 | 0.007 | 3.829 | 0.6 | 0.34 | 0.44 | 0.62 | 0.54 | 0.63 |
| 3 | 186337713 | rs4917 | 0.483 | 1.539 | 0.634 | -0.438 | 0.031 | 0.018 | 0.018 | 0.021 | 0.82 | 0.79 | 0.6 | 0.75 | 0.68 | 0.82 |
| 3 | 186337827 | rs2518136 | 1.947 | 1.541 | 0.639 | -0.452 | -0.02 | 0.014 | 0.014 | 4.853 | 0.58 | 0.31 | 0.43 | 0.62 | 0.52 | 0.58 |
| 3 | 186338135 | rs1900618 | 0.967 | 2.064 | 0.919 | -0.12 | 0.05 | 0.031 | 0.031 | 0.048 | 0.81 | 0.71 | 0.59 | 0.75 | 0.68 | 0.82 |
| 3 | 186338382 | rs4918 | 0.967 | 2.064 | 0.919 | -0.12 | 0.05 | 0.031 | 0.031 | 0.048 | 0.81 | 0.71 | 0.59 | 0.75 | 0.68 | 0.82 |
| 3 | 186339617 | rs4488820 | -0.884 | 2.068 | 0.737 | -0.214 | -0.016 | -0.046 | -0.046 | 0 | 0.09 | 0.23 | 0.08 | 0 | 0.1 | 0.07 |
| 3 | 186342060 | rs13073106 | 0.769 | 2.259 | -0.076 | -0.042 | 0.028 | 0.013 | 0.013 | 0 | 0.83 | 0.81 | 0.6 | 0.76 | 0.67 | 0.82 |
| 3 | 186345900 | rs9872086 | -1.119 | 0.96 | -0.311 | -0.499 | -0.031 | -0.14 | -0.14 | 0 | 0.27 | 0.53 | 0.19 | 0.02 | 0.24 | 0.14 |

| | | | | | | | | | | | | | | | | |
|---|---|---|---|---|---|---|---|---|---|---|---|---|---|---|---|---|
| 3 | 186348831 | rs11717166 | 1.206 | 0.963 | -1.624 | -0.119 | -0.006 | -0.046 | -0.046 | 0 | 0.46 | 0.05 | 0.35 | 0.54 | 0.36 | 0.46 |
| 3 | 186348873 | rs4686791 | -0.311 | 1.204 | -1.27 | -0.125 | -0.004 | -0.05 | -0.05 | 0 | 0.26 | 0.41 | 0.31 | 0.14 | 0.28 | 0.33 |
| 3 | 186349412 | rs13084035 | 0.623 | 0.393 | -2.173 | 0.179 | -0.004 | -0.04 | -0.04 | 0 | 0.47 | 0.05 | 0.36 | 0.54 | 0.4 | 0.5 |
| 3 | 186357538 | rs3755838 | -0.622 | 0.048 | -1.126 | 0.236 | -0.018 | -0.008 | -0.008 | 0 | 0.09 | 0.12 | 0.05 | 0.03 | 0.04 | 0.04 |
| 3 | 186358366 | rs1047115 | -0.712 | 0.027 | -1.076 | 0.164 | 0 | -0.011 | -0.011 | 1.394 | 0.12 | 0.17 | 0.09 | 0.12 | 0.12 | 0.05 |
| 3 | 186359162 | rs2292994 | -0.455 | -0.269 | -1.086 | 0.154 | -0.022 | -0.013 | -0.013 | 0 | 0.1 | 0.12 | 0.06 | 0.1 | 0.04 | 0.04 |
| 3 | 186360409 | rs3733159 | -0.804 | -0.764 | -1.093 | 0.45 | 0.031 | -0.017 | -0.017 | 0 | 0.41 | 0.64 | 0.4 | 0.51 | 0.3 | 0.37 |
| 3 | 186370333 | rs1131364 | -1.089 | -1.346 | -0.915 | 0.4 | -0.016 | 0.055 | 0.055 | 0.697 | 0.6 | 0.8 | 0.59 | 0.75 | 0.46 | 0.62 |
| 3 | 186370807 | rs3817400 | -0.153 | -1.69 | -0.707 | 0.474 | -0.014 | 0.002 | 0.002 | 0.786 | 0.35 | 0.41 | 0.32 | 0.35 | 0.28 | 0.35 |
| 3 | 186374671 | rs1868154 | -1.457 | -1.444 | -0.384 | 0.462 | -0.002 | 0.047 | 0.047 | 1.09 | 0.59 | 0.68 | 0.57 | 0.76 | 0.45 | 0.62 |
| 3 | 186381578 | rs9824398 | 0.556 | -1.469 | -0.398 | 0.458 | 0.004 | 0.006 | 0.006 | 0.482 | 0.52 | 0.48 | 0.51 | 0.3 | 0.59 | 0.45 |
| 3 | 186383180 | rs9814412 | -0.003 | -1.777 | -0.625 | 0.607 | 0.017 | -0.018 | -0.018 | 2.445 | 0.28 | 0.19 | 0.28 | 0.34 | 0.26 | 0.31 |
| 3 | 186387299 | rs16860974 | 0.644 | -1.668 | -0.697 | 0.604 | -0.031 | -0.116 | -0.116 | 0 | 0.19 | 0.03 | 0.18 | 0.38 | 0.15 | 0.27 |
| 3 | 186388028 | rs2555515 | 1.436 | -1.266 | -1.201 | 0.115 | -0.159 | 0.026 | 0.026 | 2.461 | 0.3 | 0.02 | 0.3 | 0.15 | 0.4 | 0.34 |
| 3 | 186389559 | rs10770 | 0.942 | -1.415 | -0.637 | 0.441 | -0.021 | -0.023 | -0.023 | 0 | 0.17 | 0.2 | 0.11 | 0.09 | 0.12 | 0.13 |
| 3 | 186390627 | rs9898 | -0.692 | -1.474 | -0.603 | 0.412 | 0.052 | 0.006 | 0.006 | 0.382 | 0.49 | 0.63 | 0.4 | 0.57 | 0.33 | 0.46 |
| 3 | 186395436 | rs1042445 | -0.178 | -0.736 | -0.508 | 0.708 | -0.007 | 0.002 | 0.002 | 0 | 0.25 | 0.24 | 0.28 | 0.28 | 0.25 | 0.33 |
| 3 | 186397474 | rs6444156 | -0.269 | -0.788 | -0.64 | 0.704 | 0.006 | -0.002 | -0.002 | 0 | 0.27 | 0.23 | 0.29 | 0.29 | 0.27 | 0.36 |
| 3 | 186398690 | rs4686794 | 0.509 | -0.587 | -0.492 | 0.446 | 0 | 0.001 | 0.001 | 0 | 0.46 | 0.52 | 0.52 | 0.62 | 0.43 | 0.44 |
| 3 | 186399024 | rs16861008 | 0.574 | -0.71 | -0.363 | 0.153 | 0.035 | 0.034 | 0.034 | 0 | 0.14 | 0.02 | 0.04 | 0.06 | 0.07 | 0.1 |
| 3 | 186420170 | rs7646517 | -0.513 | 0.25 | 0.394 | -0.074 | 0.062 | -0.071 | -0.071 | 0 | 0.13 | 0.35 | 0.18 | 0.02 | 0.13 | 0.15 |
| 3 | 186427340 | rs7624355 | 0.076 | 0.404 | 0.158 | 0.483 | 0.006 | -0.024 | -0.024 | 0.052 | 0.2 | 0.16 | 0.17 | 0.4 | 0.18 | 0.24 |
| 3 | 186428061 | rs10513803 | 0.944 | 0.542 | 0.211 | 0.305 | 0.008 | 0.017 | 0.017 | 0 | 0.6 | 0.54 | 0.57 | 0.52 | 0.56 | 0.54 |
| 3 | 186431432 | rs11918289 | -0.86 | 0.474 | 0.334 | 0.331 | 0.007 | 0.035 | 0.035 | 0 | 0.33 | 0.49 | 0.35 | 0.46 | 0.34 | 0.34 |
| 3 | 186433046 | rs11927941 | 0.301 | 0.024 | -0.241 | -0.028 | -0.003 | -0.013 | -0.013 | 0.008 | 0.37 | 0.47 | 0.31 | 0.34 | 0.32 | 0.45 |
| 3 | 186435370 | rs1050274 | 0.195 | 0.005 | -0.274 | -0.077 | -0.006 | -0.015 | -0.015 | 0 | 0.39 | 0.47 | 0.31 | 0.34 | 0.32 | 0.46 |
| 3 | 186436398 | rs1851665 | 0.201 | 0.008 | -0.13 | -0.076 | -0.001 | -0.001 | -0.001 | 0 | 0.39 | 0.4 | 0.3 | 0.34 | 0.32 | 0.46 |
| 3 | 186437944 | rs5029980 | 0.967 | 0.186 | -0.099 | -0.055 | -0.013 | 0.013 | 0.013 | 0.81 | 0.09 | 0.06 | 0.15 | 0.06 | 0.13 | 0.13 |
| 3 | 186438935 | rs1624230 | -0.592 | -0.084 | -0.231 | -0.08 | 0 | 0 | 0 | 0.024 | 0.53 | 0.49 | 0.36 | 0.42 | 0.4 | 0.63 |
| 3 | 186440243 | rs1469859 | 1.602 | 0.005 | -0.298 | -0.08 | -0.026 | 0.029 | 0.029 | 0 | 0.3 | 0.16 | 0.39 | 0.21 | 0.34 | 0.2 |
| 3 | 186440559 | rs5029999 | 1.555 | -0.041 | -0.276 | -0.065 | 0.023 | -0.014 | -0.014 | 0 | 0.27 | 0.13 | 0.14 | 0.28 | 0.2 | 0.33 |
| 3 | 186443018 | rs1656922 | -0.231 | -0.006 | -0.075 | -0.09 | -0.001 | 0.013 | 0.013 | 0 | 0.56 | 0.62 | 0.61 | 0.36 | 0.54 | 0.5 |
| 3 | 186444644 | rs5030023 | 1.363 | -0.041 | -0.439 | -0.122 | 0.03 | 0.002 | 0.002 | 0 | 0.3 | 0.13 | 0.15 | 0.28 | 0.22 | 0.34 |
| 3 | 186447047 | rs266723 | -0.417 | -0.244 | -0.362 | -0.046 | 0.001 | -0.005 | -0.005 | 0.072 | 0.57 | 0.39 | 0.38 | 0.41 | 0.45 | 0.61 |
| 3 | 186449916 | rs1648700 | 0.192 | -0.182 | -0.347 | -0.034 | -0.002 | -0.009 | -0.009 | 0 | 0.43 | 0.39 | 0.33 | 0.35 | 0.38 | 0.5 |
| 3 | 186451236 | rs4686799 | -0.737 | 0.352 | -0.631 | 0.303 | 0.01 | 0.048 | 0.048 | 0 | 0.86 | 0.8 | 0.79 | 0.64 | 0.78 | 0.85 |
| 3 | 186452917 | rs5030060 | 1.253 | 0.217 | -0.717 | -0.006 | -0.096 | -0.041 | -0.041 | 0.589 | 0.23 | 0.02 | 0.36 | 0.21 | 0.32 | 0.19 |
| 3 | 186454180 | rs5030062 | 0.381 | 0.159 | -0.515 | -0.042 | -0.011 | 0.011 | 0.011 | 0 | 0.33 | 0.31 | 0.42 | 0.29 | 0.39 | 0.33 |
| 3 | 186457332 | rs5030115 | 0.825 | 0.219 | -0.636 | 0.013 | 0 | 0.003 | 0.003 | 0 | 0.06 | 0.05 | 0.02 | 0.06 | 0.05 | 0.13 |
| 3 | 186459927 | rs710446 | -0.037 | 0.031 | -0.204 | -0.109 | 0.006 | -0.028 | -0.028 | 0.253 | 0.38 | 0.52 | 0.47 | 0.3 | 0.42 | 0.36 |
| 3 | 186460877 | rs5030091 | -0.429 | 0.071 | -0.341 | 0.185 | 0.004 | 0.005 | 0.005 | 0 | 0.54 | 0.45 | 0.37 | 0.41 | 0.43 | 0.65 |
| 3 | 186461181 | rs2062632 | -0.096 | 0.578 | -1.076 | 0.026 | -0.041 | 0.029 | 0.029 | 0 | 0.22 | 0.11 | 0.23 | 0.15 | 0.25 | 0.2 |
| 3 | 186463343 | rs1972703 | 0.445 | 0.559 | -1.073 | -0.032 | -0.012 | 0.016 | 0.016 | 0 | 0.14 | 0.07 | 0.18 | 0.1 | 0.15 | 0.21 |
| 3 | 186464055 | rs6807774 | 0.461 | 0.841 | -1.048 | -0.084 | -0.012 | 0.034 | 0.034 | 0 | 0.39 | 0.26 | 0.42 | 0.32 | 0.48 | 0.33 |
| 3 | 186464107 | rs6796803 | -0.386 | 0.442 | -0.912 | 0.103 | 0.003 | 0.016 | 0.016 | 0.153 | 0.82 | 0.8 | 0.83 | 0.69 | 0.75 | 0.82 |
| 3 | 186466252 | rs1656966 | 0.434 | 0.483 | -0.889 | 0.167 | 0.004 | 0.064 | 0.064 | 0 | 0.13 | 0.2 | 0.12 | 0.28 | 0.15 | 0.15 |
| 3 | 186466858 | rs822368 | -0.032 | 0.111 | -0.763 | 0.228 | -0.003 | -0.003 | -0.003 | 0 | 0.53 | 0.48 | 0.54 | 0.6 | 0.64 | 0.48 |
| 3 | 186471186 | rs822373 | 0.81 | 0.67 | -0.922 | 0.419 | -0.01 | 0.016 | 0.016 | 0 | 0.3 | 0.16 | 0.36 | 0.3 | 0.36 | 0.27 |
| 3 | 186472919 | rs266745 | -0.102 | 0.922 | -1.035 | 0.478 | 0.012 | -0.017 | -0.017 | 0 | 0.79 | 0.73 | 0.73 | 0.87 | 0.75 | 0.76 |
| 3 | 186475791 | rs266743 | 0.225 | 1.082 | -1.262 | 0.691 | 0.013 | 0.078 | 0.078 | 1.22 | 0.78 | 0.67 | 0.71 | 0.59 | 0.78 | 0.73 |
| 3 | 186475845 | rs266742 | 0.438 | 1.027 | -2.032 | 0.322 | -0.079 | -0.03 | -0.03 | 0 | 0.27 | 0.04 | 0.33 | 0.29 | 0.34 | 0.24 |
| 3 | 186480123 | rs822355 | -0.347 | 0.89 | -1.564 | 0.133 | -0.009 | 0.036 | 0.036 | 0 | 0.7 | 0.8 | 0.59 | 0.85 | 0.66 | 0.79 |
| 3 | 186487601 | rs12639287 | 0.156 | 0.495 | -1.78 | 0.753 | 0.013 | -0.074 | -0.074 | 0 | 0.25 | 0.09 | 0.25 | 0.4 | 0.22 | 0.26 |
| 3 | 186494422 | rs185554 | -0.257 | 0.57 | -1.472 | 0.498 | -0.002 | -0.008 | -0.008 | 0 | 0.4 | 0.3 | 0.48 | 0.4 | 0.47 | 0.44 |
| 3 | 186501648 | rs266719 | -0.726 | 1.329 | -1.799 | 0.151 | -0.063 | 0.021 | 0.021 | 0.003 | 0.87 | 0.96 | 0.8 | 0.9 | 0.79 | 0.78 |
| 3 | 186503435 | rs1426810 | -0.37 | 0.619 | -1.737 | -0.037 | -0.056 | 0.033 | 0.033 | 0 | 0.69 | 0.82 | 0.7 | 0.75 | 0.64 | 0.64 |
| 3 | 186505900 | rs1354091 | 0.92 | 0.672 | -1.851 | -0.186 | -0.007 | -0.107 | -0.107 | 1.055 | 0.25 | 0.09 | 0.36 | 0.43 | 0.28 | 0.34 |
| 3 | 186507757 | rs1047148 | 1.916 | 0.612 | -1.902 | -0.237 | -0.03 | -0.018 | -0.018 | 0 | 0.16 | 0.01 | 0.07 | 0.14 | 0.13 | 0.14 |
| 3 | 186507922 | rs2066500 | 0.495 | 1.03 | -1.266 | 0.122 | -0.007 | -0.108 | -0.108 | 0.069 | 0.25 | 0.08 | 0.36 | 0.43 | 0.28 | 0.34 |
| 3 | 186509517 | rs187868 | -0.712 | 0.815 | -0.51 | 0.045 | 0 | 0.002 | 0.002 | 0 | 0.42 | 0.44 | 0.49 | 0.5 | 0.49 | 0.51 |
| 3 | 186516428 | rs843991 | -1.026 | 0.706 | -0.305 | -0.09 | 0.011 | 0.024 | 0.024 | 0 | 0.48 | 0.63 | 0.52 | 0.56 | 0.52 | 0.53 |
| 3 | 186519136 | rs1648703 | 0.559 | 1.154 | -0.5 | -0.192 | 0.025 | -0.082 | -0.082 | 0 | 0.15 | 0.24 | 0.11 | 0 | 0.2 | 0.11 |
| 3 | 186519141 | rs1656941 | 0.559 | 1.154 | -0.5 | -0.192 | 0.025 | -0.082 | -0.082 | 0 | 0.15 | 0.24 | 0.11 | 0 | 0.2 | 0.11 |
| 3 | 186534114 | rs13094303 | -0.252 | 1.298 | -1.137 | 0.058 | -0.021 | -0.069 | -0.069 | 1.276 | 0.39 | 0.14 | 0.44 | 0.43 | 0.46 | 0.44 |
| 3 | 186548565 | rs6810075 | -0.442 | 1.763 | -0.296 | 0.539 | 0.003 | 0.006 | 0.006 | 0 | 0.32 | 0.32 | 0.43 | 0.45 | 0.4 | 0.4 |
| 3 | 186549695 | rs10937273 | 1.094 | 1.628 | -0.579 | 0.479 | 0.076 | 0.076 | 0.076 | 0.011 | 0.5 | 0.14 | 0.33 | 0.4 | 0.39 | 0.4 |
| 3 | 186554292 | rs864265 | -0.064 | 1.652 | 0.309 | 0.397 | 0.012 | 0.003 | 0.003 | 0 | 0.91 | 0.88 | 0.86 | 0.88 | 0.87 | 0.78 |
| 3 | 186556037 | rs822387 | -0.353 | 1.563 | 0.113 | 0.391 | 0.05 | -0.022 | -0.022 | 0 | 0.09 | 0.35 | 0.12 | 0.04 | 0.08 | 0.01 |
| 3 | 186566728 | rs822394 | -0.315 | 1.465 | -0.54 | 0.711 | -0.046 | -0.019 | -0.019 | 0 | 0.89 | 0.99 | 0.84 | 0.88 | 0.82 | 0.8 |

| | | | | | | | | | | | | | | | | |
|---|---|---|---|---|---|---|---|---|---|---|---|---|---|---|---|---|
| 3 | 186566877 | rs822396 | -0.024 | 1.453 | -0.17 | 0.699 | 0.009 | -0.006 | -0.006 | 0.569 | 0.87 | 0.82 | 0.81 | 0.88 | 0.82 | 0.8 |
| 3 | 186568799 | rs7627128 | -0.183 | 1.564 | -0.394 | 1.171 | -0.002 | 0.002 | 0.002 | 0 | 0.15 | 0.11 | 0.2 | 0.25 | 0.19 | 0.17 |
| 3 | 186571123 | rs1501299 | 0.206 | 1.109 | -0.391 | 1.06 | -0.002 | 0.001 | 0.001 | 0.169 | 0.32 | 0.39 | 0.3 | 0.29 | 0.28 | 0.21 |
| 3 | 186571486 | rs3821799 | 0.661 | 0.822 | -0.454 | 0.981 | -0.003 | 0.01 | 0.01 | 0.014 | 0.46 | 0.39 | 0.48 | 0.39 | 0.53 | 0.54 |
| 3 | 186571814 | rs3774262 | 0.835 | 0.685 | -0.324 | 1.032 | 0.014 | -0.06 | -0.06 | 0 | 0.15 | 0.04 | 0.18 | 0.3 | 0.13 | 0.15 |
| 3 | 186573705 | rs6773957 | 0.477 | 0.102 | -0.723 | 1.084 | 0.019 | 0.02 | 0.02 | 0 | 0.5 | 0.39 | 0.49 | 0.41 | 0.59 | 0.64 |
| 3 | 186574075 | rs1063537 | 0.741 | 0.103 | -0.727 | 1.089 | -0.002 | -0.081 | -0.081 | 0 | 0.15 | 0.01 | 0.18 | 0.3 | 0.13 | 0.15 |
| 3 | 186574183 | rs1063538 | 0.39 | 0.232 | -0.643 | 1.024 | 0.019 | 0.02 | 0.02 | 0 | 0.5 | 0.39 | 0.49 | 0.41 | 0.59 | 0.64 |
| 3 | 186579744 | rs6444175 | 0.054 | 0.577 | -1.037 | 0.649 | 0 | 0 | 0 | 0 | 0.32 | 0.32 | 0.3 | 0.28 | 0.28 | 0.25 |
| 3 | 186597161 | rs9865432 | 0.352 | -0.119 | -1.283 | 0.419 | 0.059 | 0.079 | 0.079 | 0.107 | 0.57 | 0.24 | 0.47 | 0.27 | 0.65 | 0.53 |
| 3 | 186599003 | rs7611851 | -0.242 | -0.463 | -0.453 | 1.083 | -0.046 | 0.091 | 0.091 | 0.747 | 0.17 | 0.5 | 0.18 | 0.26 | 0.11 | 0.21 |
| 3 | 186600420 | rs2117986 | 0.756 | -0.541 | -0.589 | 0.984 | 0.049 | 0.055 | 0.055 | 0.049 | 0.58 | 0.27 | 0.48 | 0.28 | 0.66 | 0.53 |
| 3 | 186606285 | rs9821232 | -0.284 | 0.613 | -0.388 | 0.25 | 0.013 | -0.018 | -0.018 | 0 | 0.27 | 0.3 | 0.26 | 0.1 | 0.43 | 0.3 |
| 3 | 186609682 | rs9820270 | 0.462 | -0.116 | -0.539 | 0.449 | -0.003 | 0.006 | 0.006 | 0 | 0.29 | 0.31 | 0.26 | 0.42 | 0.27 | 0.3 |
| 3 | 186612467 | rs9853541 | -0.613 | 0.069 | -0.538 | 0.125 | 0 | -0.004 | -0.004 | 0.203 | 0.69 | 0.67 | 0.67 | 0.78 | 0.65 | 0.61 |
| 3 | 186613364 | rs11716002 | 1.129 | -0.014 | -0.391 | 0.184 | 0.016 | -0.049 | -0.049 | 0 | 0.28 | 0.17 | 0.31 | 0.41 | 0.26 | 0.31 |
| 3 | 186614478 | rs7610110 | 2.44 | 0.326 | -0.601 | 0.205 | -0.027 | -0.031 | -0.031 | 0.022 | 0.07 | 0 | 0.07 | 0.09 | 0.05 | 0.02 |
| 3 | 186614800 | rs3872720 | 0.896 | 0.364 | -0.21 | 0.064 | 0.005 | 0.004 | 0.004 | 0 | 0.24 | 0.17 | 0.25 | 0.24 | 0.23 | 0.21 |
| 3 | 186616305 | rs7648806 | 1.19 | 0.413 | -0.093 | 0.027 | 0.072 | -0.007 | -0.007 | 0 | 0.11 | 0.29 | 0.07 | 0.05 | 0.13 | 0.07 |
| 3 | 186619943 | rs4686443 | 0.962 | 0.26 | -0.277 | 0.007 | 0.001 | -0.005 | -0.005 | 0 | 0.33 | 0.32 | 0.39 | 0.5 | 0.32 | 0.33 |
| 3 | 186630007 | rs6767800 | -0.188 | -0.142 | -0.371 | -0.104 | -0.002 | 0 | 0 | 0 | 0.63 | 0.66 | 0.64 | 0.61 | 0.74 | 0.6 |
| 3 | 186634074 | rs9876387 | 0.584 | 0.329 | -0.545 | -0.194 | -0.034 | 0.035 | 0.035 | 0 | 0.25 | 0.13 | 0.17 | 0.11 | 0.3 | 0.27 |
| 3 | 186635984 | rs7645316 | 0.964 | 0.536 | -0.844 | -0.159 | -0.003 | 0.027 | 0.027 | 0 | 0.37 | 0.45 | 0.4 | 0.5 | 0.34 | 0.33 |
| 3 | 186638219 | rs6801612 | 0.838 | 0.237 | -1.119 | -0.197 | 0.008 | -0.003 | -0.003 | 0 | 0.23 | 0.15 | 0.22 | 0.24 | 0.2 | 0.21 |
| 3 | 186647652 | rs9814673 | 0.537 | 0.657 | -0.94 | -0.26 | -0.053 | 0.068 | 0.068 | 0.939 | 0.24 | 0.08 | 0.17 | 0.08 | 0.29 | 0.28 |
| 3 | 186648117 | rs1468906 | -0.072 | 0.446 | -0.817 | -0.334 | 0.008 | 0.006 | 0.006 | 0 | 0.38 | 0.34 | 0.4 | 0.39 | 0.35 | 0.38 |
| 3 | 186650790 | rs10937275 | -0.724 | 0.713 | -0.881 | -0.333 | 0.021 | -0.029 | -0.029 | 0 | 0.94 | 0.87 | 0.92 | 1 | 0.89 | 0.99 |
| 3 | 186650942 | rs12493118 | -1.037 | 0.702 | -0.958 | -0.376 | -0.006 | 0.013 | 0.013 | 0 | 0.82 | 0.86 | 0.81 | 0.96 | 0.74 | 0.78 |
| 3 | 186651000 | rs7647396 | -1.026 | 0.715 | -0.952 | -0.37 | -0.006 | 0.014 | 0.014 | 0 | 0.82 | 0.86 | 0.81 | 0.96 | 0.74 | 0.78 |
| 3 | 186656019 | rs2889789 | 0.206 | 0.031 | -0.671 | 0.024 | 0.009 | 0.029 | 0.029 | 0.043 | 0.47 | 0.6 | 0.43 | 0.27 | 0.4 | 0.42 |
| 3 | 186656113 | rs10513807 | 0.279 | -0.002 | -0.201 | 0.077 | 0 | 0.009 | 0.009 | 0 | 0.5 | 0.42 | 0.6 | 0.59 | 0.53 | 0.58 |
| 3 | 186656535 | rs3864105 | -0.241 | 0.4 | -0.589 | 0.459 | -0.006 | -0.005 | -0.005 | 0 | 0.66 | 0.67 | 0.69 | 0.63 | 0.6 | 0.69 |
| 3 | 186664728 | rs16861317 | 0.426 | -0.385 | -0.318 | 0.75 | 0.008 | -0.027 | -0.027 | 0 | 0.16 | 0.07 | 0.22 | 0.24 | 0.13 | 0.25 |
| 3 | 186665645 | rs3887925 | 1.321 | -0.626 | -0.513 | 0.607 | 0.012 | -0.033 | -0.033 | 0 | 0.48 | 0.16 | 0.36 | 0.46 | 0.55 | 0.51 |
| 3 | 186666546 | rs9290841 | -0.232 | -0.561 | -0.145 | 0.651 | -0.003 | 0.01 | 0.01 | 0 | 0.93 | 0.89 | 0.81 | 0.86 | 0.91 | 0.92 |
| 3 | 186668108 | rs16861339 | 0.308 | -0.73 | 0.634 | 0.526 | -0.026 | -0.035 | -0.035 | 1.542 | 0.18 | 0.24 | 0.07 | 0.05 | 0.07 | 0.1 |
| 3 | 186673839 | rs6444186 | -0.391 | -0.817 | 0.846 | 0.536 | -0.054 | -0.048 | -0.048 | 0 | 0.22 | 0.62 | 0.1 | 0.09 | 0.07 | 0.12 |
| 3 | 186675277 | rs7645517 | 0.176 | -0.802 | 1.075 | 0.537 | -0.057 | -0.072 | -0.072 | 0 | 0.18 | 0.39 | 0.08 | 0.05 | 0.07 | 0.1 |
| 3 | 186676455 | rs9799068 | 0.772 | -0.794 | 0.755 | 0.552 | -0.003 | 0.001 | 0.001 | 0.792 | 0.72 | 0.31 | 0.77 | 0.77 | 0.86 | 0.81 |
| 3 | 186677936 | rs6444188 | 0.588 | -0.21 | 0.533 | 0.737 | 0.006 | 0.001 | 0.001 | 0 | 0.55 | 0.46 | 0.5 | 0.57 | 0.63 | 0.58 |
| 3 | 186681272 | rs4686820 | 0.997 | 0.989 | 0.848 | 0.884 | 0.013 | 0.018 | 0.018 | 0 | 0.9 | 0.84 | 0.76 | 0.78 | 0.85 | 0.86 |
| 3 | 186682324 | rs6771438 | -1.157 | 0.579 | 1.116 | 0.627 | 0.066 | 0.052 | 0.052 | 0.098 | 0.07 | 0.31 | 0.12 | 0.05 | 0.11 | 0.03 |
| 3 | 186682612 | rs6809750 | -1.709 | 0.506 | 0.981 | 0.615 | 0.032 | 0.014 | 0.014 | 1.281 | 0.14 | 0.42 | 0.18 | 0.1 | 0.16 | 0.11 |
| 3 | 186685534 | rs9828559 | 0.657 | 0.817 | -0.011 | 0.698 | -0.007 | 0 | 0 | 0 | 0.93 | 0.95 | 0.92 | 0.94 | 0.91 | 0.9 |
| 3 | 186686239 | rs4686823 | 0.977 | 0.776 | 0.171 | 0.68 | 0.006 | -0.004 | -0.004 | 0.849 | 0.92 | 0.89 | 0.92 | 0.94 | 0.91 | 0.9 |
| 3 | 186690614 | rs6766574 | -0.589 | 0.503 | 0.021 | 0.894 | -0.001 | 0.039 | 0.039 | 0.041 | 0.31 | 0.53 | 0.35 | 0.38 | 0.27 | 0.28 |
| 3 | 186691291 | rs11720551 | 0.473 | 0.417 | -0.888 | 0.587 | -0.04 | 0.112 | 0.112 | 0.018 | 0.44 | 0.15 | 0.34 | 0.26 | 0.45 | 0.44 |
| 3 | 186691605 | rs6444190 | -0.31 | 0.935 | -0.051 | 1.059 | 0.025 | 0.008 | 0.008 | 0 | 0.15 | 0.3 | 0.13 | 0.15 | 0.18 | 0.15 |
| 3 | 186694029 | rs3864108 | 0.823 | 0.769 | -0.01 | 1.53 | 0.013 | 0.088 | 0.088 | 0.013 | 0.85 | 0.73 | 0.74 | 0.66 | 0.85 | 0.82 |
| 3 | 186698098 | rs9869132 | -0.38 | 0.295 | 0.005 | 1.521 | -0.013 | 0.05 | 0.05 | 0 | 0.13 | 0.23 | 0.18 | 0.32 | 0.07 | 0.26 |
| 3 | 186698993 | rs4626092 | -0.227 | 0.136 | -0.142 | 1.508 | -0.001 | 0.032 | 0.032 | 0 | 0.17 | 0.16 | 0.22 | 0.44 | 0.18 | 0.16 |
| 3 | 186699610 | rs6444191 | -0.865 | 0.074 | -0.344 | 0.383 | -0.001 | -0.016 | -0.016 | 0.255 | 0.41 | 0.57 | 0.58 | 0.84 | 0.39 | 0.6 |
| 3 | 186708804 | rs759602 | -0.055 | 0.417 | -0.501 | 0.033 | 0 | -0.008 | -0.008 | 0 | 0.78 | 0.81 | 0.85 | 0.94 | 0.8 | 0.89 |
| 3 | 186710637 | rs2019082 | 0.126 | 0.32 | -0.247 | 0.572 | -0.011 | 0.039 | 0.039 | 2.285 | 0.23 | 0.31 | 0.38 | 0.35 | 0.19 | 0.19 |
| 3 | 186710785 | rs3888059 | -0.384 | -0.601 | -0.204 | 0.547 | -0.013 | 0.003 | 0.003 | 0.167 | 0.37 | 0.59 | 0.44 | 0.52 | 0.25 | 0.38 |
| 3 | 186711098 | rs6444192 | -0.279 | 0.203 | -0.891 | 0.082 | -0.001 | 0.001 | 0.001 | 2.335 | 0.44 | 0.39 | 0.58 | 0.46 | 0.45 | 0.34 |
| 3 | 186711608 | rs17718970 | 0.278 | 0.832 | -0.827 | 0.056 | -0.031 | 0.067 | 0.067 | 0.308 | 0.22 | 0.1 | 0.22 | 0.16 | 0.26 | 0.21 |
| 3 | 186718462 | rs11707680 | 0.201 | 1.355 | -0.285 | 0.289 | 0.011 | -0.016 | -0.016 | 0 | 0.15 | 0.2 | 0.17 | 0.14 | 0.18 | 0.13 |
| 3 | 186719374 | rs6808800 | -0.64 | 1.487 | -0.175 | 0.379 | -0.001 | 0.013 | 0.013 | 0 | 0.46 | 0.6 | 0.5 | 0.38 | 0.51 | 0.54 |
| 3 | 186722362 | rs17775791 | 0.716 | 1.126 | 0.474 | 0.453 | -0.02 | -0.001 | -0.001 | 0 | 0.21 | 0.16 | 0.26 | 0.24 | 0.26 | 0.37 |
| 3 | 186722944 | rs7652995 | -1.71 | 1.049 | 0.539 | 0.469 | -0.001 | -0.002 | -0.002 | 0.121 | 0.86 | 0.87 | 0.87 | 0.89 | 0.85 | 0.86 |
| 3 | 186723975 | rs6764279 | 0.085 | 1.061 | 0.643 | 0.459 | 0.002 | -0.006 | -0.006 | 0 | 0.27 | 0.39 | 0.34 | 0.32 | 0.29 | 0.49 |
| 3 | 186724219 | rs6788832 | -0.587 | 1.071 | 0.453 | 0.556 | -0.004 | -0.008 | -0.008 | 0 | 0.37 | 0.58 | 0.41 | 0.39 | 0.36 | 0.51 |
| 3 | 186724553 | rs6791737 | -0.332 | 1.011 | 0.681 | 0.499 | -0.039 | -0.064 | -0.064 | 0.893 | 0.15 | 0.37 | 0.15 | 0.08 | 0.09 | 0.09 |
| 3 | 186727083 | rs7643068 | -1.432 | 1.576 | 0.066 | 0.636 | -0.001 | -0.002 | -0.002 | 0 | 0.87 | 0.87 | 0.88 | 0.89 | 0.85 | 0.86 |
| 3 | 186729365 | rs4686836 | -0.273 | 1.554 | 0.268 | 0.594 | -0.016 | -0.013 | -0.013 | 0 | 0.77 | 0.56 | 0.79 | 0.82 | 0.78 | 0.85 |
| 3 | 186732679 | rs17776120 | -0.106 | 1.559 | -0.457 | 0.944 | -0.063 | -0.06 | -0.06 | 1.517 | 0.28 | 0.14 | 0.42 | 0.4 | 0.37 | 0.41 |
| 3 | 186739677 | rs4686837 | -0.087 | 1.176 | -0.395 | 0.824 | -0.005 | -0.013 | -0.013 | 0 | 0.28 | 0.23 | 0.4 | 0.38 | 0.3 | 0.4 |
| 3 | 186743053 | rs4686838 | -0.706 | 0.709 | -0.775 | 0.658 | 0 | -0.009 | -0.009 | 0 | 0.44 | 0.62 | 0.52 | 0.53 | 0.42 | 0.45 |

| | | | | | | | | | | | | | | | | |
|---|---|---|---|---|---|---|---|---|---|---|---|---|---|---|---|---|
| 3 | 186755882 | rs6444196 | -0.258 | 0.451 | -1.102 | 0.637 | 0.05 | 0.003 | 0.003 | 0.973 | 0.46 | 0.8 | 0.57 | 0.41 | 0.38 | 0.35 |
| 3 | 186756908 | rs2268536 | 0.056 | 0.376 | -0.5 | 0.619 | 0.038 | 0.089 | 0.089 | 1.462 | 0.11 | 0.33 | 0.18 | 0.25 | 0.13 | 0.1 |
| 3 | 186757083 | rs2268535 | -0.017 | 0.007 | 0.197 | 0.466 | -0.028 | -0.032 | -0.032 | 0.652 | 0.35 | 0.56 | 0.38 | 0.17 | 0.25 | 0.26 |
| 3 | 186758583 | rs4234598 | 0.777 | -0.118 | 0.1 | 0.392 | 0.022 | 0.008 | 0.008 | 0 | 0.25 | 0.16 | 0.3 | 0.17 | 0.19 | 0.22 |
| 3 | 186763651 | rs6780016 | -0.219 | 0.138 | -0.048 | 0.428 | 0.057 | 0.052 | 0.052 | 0 | 0.39 | 0.77 | 0.42 | 0.21 | 0.27 | 0.31 |
| 3 | 186766122 | rs6800338 | -0.318 | 0.646 | -0.383 | 0.295 | -0.017 | 0.018 | 0.018 | 0 | 0.6 | 0.76 | 0.7 | 0.69 | 0.56 | 0.63 |
| 3 | 186767945 | rs16861549 | 0.489 | 0.627 | -0.659 | 0.256 | -0.003 | -0.036 | -0.036 | 0.281 | 0.08 | 0.18 | 0.06 | 0 | 0.07 | 0.05 |
| 3 | 186778828 | rs257101 | -0.12 | 1.284 | -1.563 | 0.592 | -0.002 | 0.001 | 0.001 | 0 | 0.83 | 0.84 | 0.75 | 0.83 | 0.8 | 0.85 |
| 3 | 186779864 | rs4012249 | 0.268 | 1.319 | -1.545 | 0.677 | 0.044 | 0.007 | 0.007 | 0 | 0.5 | 0.75 | 0.54 | 0.52 | 0.42 | 0.43 |
| 3 | 186780866 | rs10433485 | -0.368 | 1.134 | -1.743 | 0.857 | -0.047 | -0.043 | -0.043 | 0 | 0.65 | 0.91 | 0.79 | 0.69 | 0.61 | 0.63 |
| 3 | 186784789 | rs736341 | -0.608 | 2.598 | -1.729 | 1.107 | -0.14 | -0.032 | -0.032 | 0.144 | 0.18 | 0.02 | 0.21 | 0.2 | 0.35 | 0.25 |
| 3 | 186791795 | rs9821439 | -0.663 | 1.251 | -1.971 | 0.979 | -0.032 | 0.035 | 0.035 | 1.996 | 0.42 | 0.89 | 0.53 | 0.68 | 0.4 | 0.39 |
| 3 | 186794091 | rs2284750 | -0.012 | 0.776 | -0.686 | 1.937 | -0.029 | 0.155 | 0.155 | 0 | 0.06 | 0.13 | 0.08 | 0.37 | 0.01 | 0.13 |
| 3 | 186794103 | rs2284749 | 0.252 | 0.543 | -0.044 | 1.962 | -0.072 | 0.138 | 0.138 | 0 | 0.21 | 0.43 | 0.32 | 0.42 | 0.12 | 0.13 |
| 3 | 186796053 | rs7559 | 0.217 | 0.568 | -0.08 | 0.8 | -0.038 | -0.008 | -0.008 | 0 | 0.21 | 0.3 | 0.25 | 0.17 | 0.12 | 0.12 |
| 3 | 186798428 | rs4686849 | -0.92 | 0.642 | -0.55 | 0.731 | -0.018 | -0.024 | -0.024 | 0 | 0.4 | 0.81 | 0.47 | 0.43 | 0.39 | 0.37 |
| 3 | 186800321 | rs7634671 | 0.081 | 0.761 | -0.731 | 0.786 | -0.028 | 0.027 | 0.027 | 0 | 0.05 | 0.13 | 0.02 | 0.12 | 0.01 | 0.11 |
| 3 | 186807282 | rs9290844 | -0.546 | 1.26 | -0.736 | 0.692 | 0.021 | 0.001 | 0.001 | 0.248 | 0.33 | 0.5 | 0.43 | 0.31 | 0.38 | 0.27 |
| 3 | 186821199 | rs849755 | -0.223 | 0.533 | -1.279 | 0.144 | -0.065 | -0.005 | -0.005 | 1.161 | 0.66 | 0.82 | 0.58 | 0.68 | 0.56 | 0.75 |
| 3 | 186821570 | rs849754 | 0.653 | 0.127 | -1.316 | -0.105 | -0.014 | -0.02 | -0.02 | 0 | 0.06 | 0.1 | 0.03 | 0 | 0.02 | 0 |
| 3 | 186822991 | rs10490807 | 0.452 | -0.229 | -1.285 | -0.201 | 0.004 | 0.004 | 0.004 | 0 | 0.17 | 0.1 | 0.15 | 0.09 | 0.11 | 0.11 |
| 3 | 186829496 | rs270159 | -1.082 | -0.096 | -1.645 | -0.202 | -0.146 | 0.013 | 0.013 | 0.299 | 0.75 | 0.98 | 0.72 | 0.82 | 0.64 | 0.79 |
| 3 | 186830661 | rs270161 | -0.155 | -0.166 | -1.342 | 0.057 | 0.004 | 0.033 | 0.033 | 1.047 | 0.49 | 0.81 | 0.51 | 0.61 | 0.45 | 0.7 |
| 3 | 186836148 | rs9817493 | 0.623 | 0.365 | -1.009 | 0.362 | -0.095 | -0.109 | -0.109 | 0 | 0.27 | 0.09 | 0.31 | 0.46 | 0.38 | 0.48 |
| 3 | 186838080 | rs6798581 | -1.064 | 0.188 | 0.741 | 0.348 | 0.026 | 0.018 | 0.018 | 0.472 | 0.05 | 0.25 | 0.02 | 0 | 0.01 | 0.05 |
| 3 | 186838657 | rs6801553 | -1.098 | 0.164 | 0.367 | 0.336 | 0.031 | 0.015 | 0.015 | 3.681 | 0.07 | 0.45 | 0.04 | 0 | 0.01 | 0.05 |
| 3 | 186844824 | rs6444200 | -1.859 | 0.199 | -0.104 | 0.64 | 0 | -0.002 | -0.002 | 0 | 0.24 | 0.73 | 0.27 | 0.22 | 0.24 | 0.19 |
| 3 | 186850686 | rs2270354 | -0.345 | 0.276 | -0.182 | 0.667 | -0.009 | -0.008 | -0.008 | 0 | 0.13 | 0.12 | 0.21 | 0.17 | 0.23 | 0.06 |
| 3 | 186862974 | rs6444206 | -2.002 | -0.099 | -0.27 | 0.585 | 0.048 | 0.014 | 0.014 | 2.953 | 0.34 | 0.81 | 0.28 | 0.25 | 0.26 | 0.28 |
| 3 | 186875232 | rs9875174 | -0.485 | 0.216 | -0.259 | 0.338 | 0.004 | -0.001 | -0.001 | 0 | 0.15 | 0.18 | 0.2 | 0.1 | 0.22 | 0.06 |
| 3 | 186880956 | rs270144 | 1.152 | 0.197 | -0.922 | 0.402 | -0.011 | 0.041 | 0.041 | 2.021 | 0.63 | 0.17 | 0.63 | 0.76 | 0.63 | 0.73 |
| 3 | 186882254 | rs270151 | 0.74 | 1.205 | -0.481 | 0.621 | 0.004 | -0.004 | -0.004 | 0 | 0.74 | 0.3 | 0.65 | 0.83 | 0.67 | 0.83 |
| 3 | 186891908 | rs9290845 | -0.537 | 0.471 | -0.913 | 0.811 | 0.006 | 0.01 | 0.01 | 0 | 0.31 | 0.23 | 0.16 | 0.21 | 0.24 | 0.26 |
| 3 | 186902640 | rs6444209 | 0.713 | 0.419 | -0.769 | -0.097 | -0.008 | 0.001 | 0.001 | 0 | 0.73 | 0.81 | 0.87 | 0.98 | 0.76 | 0.83 |
| 3 | 186905512 | rs1521595 | 0.955 | -0.218 | -0.09 | 0.019 | -0.005 | 0.005 | 0.005 | 0 | 0.59 | 0.66 | 0.67 | 0.69 | 0.62 | 0.6 |
| 3 | 186906479 | rs2016011 | 0.583 | 0.053 | -0.065 | 0.058 | -0.005 | 0.004 | 0.004 | 0 | 0.08 | 0.13 | 0.03 | 0.11 | 0.07 | 0.06 |
| 3 | 186910659 | rs6444210 | 1.555 | -0.176 | 0.043 | -0.044 | -0.016 | 0 | 0 | 0 | 0.44 | 0.24 | 0.6 | 0.58 | 0.55 | 0.49 |
| 3 | 186912351 | rs9290846 | 1.03 | -0.173 | 0.065 | -0.048 | 0 | 0.019 | 0.019 | 0 | 0.52 | 0.41 | 0.64 | 0.69 | 0.62 | 0.55 |
| 3 | 186913248 | rs6766251 | 0.557 | -0.175 | 0.085 | -0.032 | -0.008 | 0.007 | 0.007 | 0 | 0.6 | 0.69 | 0.67 | 0.69 | 0.62 | 0.6 |
| 3 | 186914144 | rs11916885 | 1.306 | -0.065 | 0.093 | -0.067 | 0.008 | 0.04 | 0.04 | 0 | 0.1 | 0.02 | 0.03 | 0.01 | 0.06 | 0.03 |
| 3 | 186914603 | rs6772329 | 1.299 | -0.558 | -0.123 | -0.574 | -0.008 | -0.037 | -0.037 | 0 | 0.08 | 0.17 | 0.09 | 0.01 | 0.07 | 0.05 |
| 3 | 186918117 | rs16861709 | 1.152 | -0.594 | 0.091 | -0.599 | 0.005 | 0.072 | 0.072 | 0 | 0.15 | 0.3 | 0.21 | 0.28 | 0.13 | 0.26 |
| 3 | 186918213 | rs11714564 | -0.652 | -0.82 | -0.067 | -0.898 | -0.01 | -0.015 | -0.015 | 0 | 0.2 | 0.25 | 0.1 | 0.12 | 0.17 | 0.19 |
| 3 | 186924035 | rs3852052 | 1.862 | -0.914 | -0.606 | -0.365 | -0.125 | 0.014 | 0.014 | 0.016 | 0.3 | 0.03 | 0.4 | 0.18 | 0.45 | 0.34 |
| 3 | 186928756 | rs3908206 | 1.868 | -0.946 | -0.577 | -0.364 | -0.011 | -0.006 | -0.006 | 0 | 0.1 | 0.03 | 0.05 | 0.12 | 0.11 | 0.05 |
| 3 | 186932041 | rs720511 | 0.984 | -1.101 | -0.625 | -0.025 | -0.072 | 0.045 | 0.045 | 0.19 | 0.46 | 0.07 | 0.58 | 0.68 | 0.6 | 0.6 |
| 3 | 186936778 | rs9854618 | 1.514 | -1.168 | -0.469 | 0.017 | -0.018 | -0.021 | -0.021 | 0 | 0.08 | 0.13 | 0.01 | 0.01 | 0.02 | 0.03 |
| 3 | 186937037 | rs3864095 | -0.059 | -1.22 | -0.312 | 0.025 | -0.058 | 0.072 | 0.072 | 0 | 0.13 | 0.33 | 0.15 | 0.18 | 0.04 | 0.15 |
| 3 | 186937047 | rs3843010 | 0.407 | -1.262 | -0.226 | -0.067 | 0.002 | 0.061 | 0.061 | 0 | 0.53 | 0.25 | 0.63 | 0.69 | 0.61 | 0.62 |
| 3 | 186937200 | rs3852053 | 0.999 | -1.242 | -0.29 | -0.325 | -0.042 | 0.194 | 0.194 | 0.441 | 0.46 | 0.17 | 0.68 | 0.86 | 0.56 | 0.65 |
| 3 | 186937221 | rs3821801 | -0.304 | -1.121 | -0.367 | -0.283 | 0.013 | -0.068 | -0.068 | 0 | 0.64 | 0.54 | 0.79 | 0.87 | 0.65 | 0.77 |
| 3 | 186937276 | rs3864096 | 0.578 | -1.053 | -0.294 | -0.156 | -0.009 | -0.034 | -0.034 | 0 | 0.08 | 0.15 | 0.09 | 0.01 | 0.07 | 0.05 |
| 3 | 186938956 | rs3733001 | -0.52 | -1.039 | -0.497 | -0.153 | -0.019 | -0.008 | -0.008 | 0.124 | 0.35 | 0.56 | 0.41 | 0.69 | 0.24 | 0.4 |
| 3 | 186941429 | rs11720718 | 1.888 | -0.717 | -0.866 | -0.065 | -0.096 | -0.019 | -0.019 | 0 | 0.27 | 0.02 | 0.36 | 0.19 | 0.39 | 0.25 |
| 3 | 186944705 | rs1001073 | -0.161 | -0.646 | -0.387 | -0.382 | -0.059 | 0.089 | 0.089 | 0 | 0.29 | 0.12 | 0.19 | 0.13 | 0.32 | 0.21 |
| 3 | 186945058 | rs12638131 | 0.902 | 0.017 | -0.542 | -0.098 | -0.055 | -0.119 | -0.119 | 0.17 | 0.22 | 0.08 | 0.24 | 0.48 | 0.27 | 0.3 |
| 3 | 186948788 | rs710451 | 1.055 | -0.046 | -0.315 | -0.44 | 0.006 | 0.056 | 0.056 | 0.04 | 0.43 | 0.24 | 0.44 | 0.31 | 0.45 | 0.32 |
| 3 | 186950635 | rs3864097 | 0.501 | 0.095 | -0.021 | -0.28 | -0.024 | -0.031 | -0.031 | 0.069 | 0.25 | 0.17 | 0.36 | 0.66 | 0.29 | 0.39 |
| 3 | 186952415 | rs12489890 | 1.047 | 0.249 | -0.174 | -0.377 | -0.063 | 0.077 | 0.077 | 0 | 0.13 | 0.01 | 0.08 | 0.01 | 0.17 | 0.13 |
| 3 | 186952587 | rs850314 | 2.531 | 0.094 | -0.293 | -0.462 | -0.061 | -0.005 | -0.005 | 0 | 0.26 | 0.04 | 0.33 | 0.19 | 0.33 | 0.24 |
| 3 | 186952588 | rs1109452 | 0.912 | 0.129 | -0.225 | -0.281 | -0.052 | -0.075 | -0.075 | 0 | 0.23 | 0.1 | 0.35 | 0.65 | 0.29 | 0.39 |
| 3 | 186953037 | rs874603 | 0.425 | 0.116 | 0.198 | -0.325 | 0.115 | 0.036 | 0.036 | 0.743 | 0.94 | 0.64 | 0.91 | 0.99 | 0.94 | 0.89 |
| 3 | 186953808 | rs850312 | 2.077 | 0.054 | -0.213 | -0.298 | -0.064 | -0.03 | -0.03 | 0 | 0.29 | 0.03 | 0.35 | 0.2 | 0.35 | 0.24 |
| 3 | 186954324 | rs3774268 | 0.628 | -0.042 | 0.028 | -0.388 | -0.003 | -0.018 | -0.018 | 0 | 0.18 | 0.2 | 0.13 | 0.13 | 0.15 | 0.12 |
| 3 | 186956223 | rs696405 | 1.103 | -0.267 | -1.364 | -0.592 | 0.023 | 0.206 | 0.206 | 0.362 | 0.53 | 0.24 | 0.71 | 0.85 | 0.64 | 0.61 |
| 3 | 186959042 | rs3815623 | -0.315 | -0.105 | -1.054 | -0.424 | -0.017 | 0.038 | 0.038 | 0 | 0.45 | 0.31 | 0.47 | 0.66 | 0.46 | 0.53 |
| 3 | 186960482 | rs710459 | 0.242 | 0.577 | -1.767 | -0.2 | -0.086 | -0.066 | -0.066 | 0 | 0.33 | 0.12 | 0.47 | 0.58 | 0.44 | 0.37 |
| 3 | 186963130 | rs710462 | 0.102 | 0.242 | -0.748 | 0.108 | 0.003 | 0.032 | 0.032 | 0 | 0.51 | 0.35 | 0.56 | 0.38 | 0.54 | 0.3 |
| 3 | 186965268 | rs3774275 | 1.637 | 0.239 | -1.015 | 0.604 | -0.083 | -0.134 | -0.134 | 0.051 | 0.28 | 0.02 | 0.16 | 0.42 | 0.31 | 0.34 |

| | | | | | | | | | | | | | | | | |
|---|---|---|---|---|---|---|---|---|---|---|---|---|---|---|---|---|
| 3 | 186969634 | rs698092 | -1.038 | 0.254 | -0.535 | 0.435 | -0.002 | -0.005 | -0.005 | 0 | 0.4 | 0.47 | 0.46 | 0.42 | 0.38 | 0.37 |
| 3 | 186970808 | rs698094 | -1.784 | 0.256 | -1.139 | 0.49 | -0.011 | -0.043 | -0.043 | 0.11 | 0.71 | 0.89 | 0.85 | 0.66 | 0.72 | 0.64 |
| 3 | 186971296 | rs3105782 | -0.588 | -0.057 | -0.666 | 0.147 | -0.006 | -0.023 | -0.023 | 1.023 | 0.3 | 0.66 | 0.28 | 0.2 | 0.26 | 0.18 |
| 3 | 186976991 | rs4686864 | 1.183 | 0.181 | -0.819 | 0.57 | -0.085 | -0.009 | -0.009 | 1.168 | 0.27 | 0.06 | 0.41 | 0.21 | 0.33 | 0.3 |
| 3 | 186979583 | rs850309 | 0.064 | -0.013 | -0.314 | 0.157 | -0.012 | 0.003 | 0.003 | 0.03 | 0.17 | 0.29 | 0.2 | 0.21 | 0.17 | 0.24 |
| 3 | 186980264 | rs710468 | -0.436 | -0.099 | 0.163 | 0.124 | -0.029 | -0.019 | -0.019 | 0 | 0.19 | 0.51 | 0.19 | 0.18 | 0.17 | 0.25 |
| 3 | 186982208 | rs3774282 | 1.307 | -0.199 | -0.243 | 0.045 | 0.037 | 0.075 | 0.075 | 1.651 | 0.59 | 0.14 | 0.63 | 0.69 | 0.68 | 0.62 |
| 3 | 186984518 | rs710469 | -0.08 | -0.126 | -0.19 | 0.315 | -0.013 | -0.015 | -0.015 | 0 | 0.43 | 0.35 | 0.59 | 0.51 | 0.51 | 0.57 |
| 3 | 186989512 | rs16861812 | 0.472 | -0.087 | 0.253 | 0.304 | 0.003 | -0.01 | -0.01 | 0 | 0.15 | 0.32 | 0.16 | 0.13 | 0.09 | 0.11 |
| 3 | 186990811 | rs12106739 | 2.032 | 0.023 | 0.145 | 0.327 | -0.006 | -0.04 | -0.04 | 0 | 0.08 | 0.2 | 0.06 | 0 | 0.06 | 0.02 |
| 3 | 186993281 | rs698102 | 0.049 | 0.208 | 0.035 | 0.454 | -0.008 | -0.004 | -0.004 | 0 | 0.16 | 0.26 | 0.15 | 0.16 | 0.16 | 0.25 |
| 3 | 186995860 | rs3864098 | 0.565 | 0.198 | 0.156 | 0.497 | -0.013 | -0.06 | -0.06 | 0 | 0.22 | 0.36 | 0.17 | 0.12 | 0.16 | 0.13 |
| 3 | 186997423 | rs698085 | -1.475 | 0.217 | 0.016 | 0.531 | -0.016 | -0.009 | -0.009 | 0 | 0.65 | 0.85 | 0.81 | 0.65 | 0.66 | 0.71 |
| 3 | 186997648 | rs698084 | 0.101 | -0.1 | 0.008 | 0.424 | -0.002 | 0 | 0 | 0 | 0.14 | 0.3 | 0.17 | 0.18 | 0.16 | 0.24 |
| 3 | 187002517 | rs698104 | -0.182 | -0.269 | 0.027 | 0.399 | -0.05 | -0.068 | -0.068 | 0 | 0.76 | 0.5 | 0.82 | 0.88 | 0.86 | 0.81 |
| 3 | 187004183 | rs13094773 | 1.237 | 0.013 | -0.201 | 0.587 | -0.025 | -0.039 | -0.039 | 0 | 0.27 | 0.13 | 0.43 | 0.33 | 0.34 | 0.33 |
| 3 | 187012061 | rs710474 | -1.423 | 0.135 | -0.616 | 0.381 | -0.023 | 0.012 | 0.012 | 4.658 | 0.66 | 0.89 | 0.82 | 0.69 | 0.69 | 0.73 |
| 3 | 187012658 | rs16861903 | -0.06 | 0.018 | -0.379 | 0.218 | -0.05 | -0.068 | -0.068 | 0 | 0.24 | 0.54 | 0.19 | 0.12 | 0.14 | 0.19 |
| 3 | 187017841 | rs4686870 | 0.156 | 0.748 | -0.765 | 0.522 | 0.006 | 0.006 | 0.006 | 0.059 | 0.5 | 0.38 | 0.6 | 0.5 | 0.51 | 0.5 |
| 3 | 187020200 | rs850303 | -0.536 | 1.146 | -0.825 | 0.884 | 0.003 | 0.003 | 0.003 | 0.6 | 0.12 | 0.17 | 0.15 | 0.2 | 0.18 | 0.2 |
| 3 | 187021886 | rs12492885 | 0.357 | 0.663 | -1.258 | 0.6 | 0.004 | 0.01 | 0.01 | 0 | 0.45 | 0.29 | 0.64 | 0.55 | 0.58 | 0.39 |
| 3 | 187023952 | rs710475 | 0.425 | 0.646 | -1.105 | 0.463 | -0.001 | 0 | 0 | 0.11 | 0.32 | 0.32 | 0.34 | 0.62 | 0.3 | 0.45 |
| 3 | 187028051 | rs698108 | 1.07 | 0.139 | -0.237 | 0.68 | 0.004 | -0.001 | -0.001 | 0 | 0.71 | 0.66 | 0.77 | 0.69 | 0.67 | 0.68 |
| 3 | 187036636 | rs12496181 | -0.996 | 0.527 | -0.177 | 0.774 | 0.087 | 0.039 | 0.039 | 0 | 0.07 | 0.21 | 0.13 | 0.1 | 0.16 | 0.17 |
| 3 | 187045616 | rs3914757 | -0.818 | 0.41 | -0.805 | 0.588 | 0.001 | -0.039 | -0.039 | 0 | 0.26 | 0.67 | 0.2 | 0.04 | 0.29 | 0.21 |
| 3 | 187050471 | rs4011965 | -0.7 | 0.719 | -1.148 | 1.67 | -0.061 | -0.072 | -0.072 | 0 | 0.12 | 0.05 | 0.18 | 0.5 | 0.22 | 0.14 |
| 3 | 187053312 | rs882009 | -0.713 | -0.061 | -0.722 | 1.415 | -0.01 | 0.042 | 0.042 | 0 | 0.19 | 0.24 | 0.18 | 0.56 | 0.1 | 0.2 |
| 3 | 187053964 | rs723 | -1.131 | 0.203 | -0.115 | 0.724 | 0.044 | -0.034 | -0.034 | 0 | 0.15 | 0.28 | 0.22 | 0.07 | 0.24 | 0.27 |
| 3 | 187060668 | rs4686872 | -0.046 | -1.188 | -0.082 | 0.727 | -0.017 | 0.038 | 0.038 | 0 | 0.76 | 0.7 | 0.87 | 0.57 | 0.85 | 0.82 |
| 3 | 187069183 | rs1122739 | -0.853 | -1.44 | -0.095 | 0.569 | -0.001 | -0.001 | -0.001 | 0 | 0.25 | 0.27 | 0.29 | 0.35 | 0.31 | 0.31 |
| 3 | 187071300 | rs751396 | 1.012 | -1.78 | -0.08 | 0.467 | -0.004 | -0.006 | -0.006 | 0.012 | 0.45 | 0.34 | 0.59 | 0.51 | 0.51 | 0.48 |
| 3 | 187080751 | rs6773671 | -0.921 | -1.581 | -0.556 | 0.104 | 0.014 | 0.013 | 0.013 | 0.125 | 0.75 | 0.8 | 0.88 | 0.96 | 0.82 | 0.82 |
| 3 | 187084509 | rs6797770 | 0.639 | -1.566 | -0.742 | 0.197 | 0.005 | 0.009 | 0.009 | 0 | 0.31 | 0.28 | 0.14 | 0.17 | 0.23 | 0.31 |
| 3 | 187088926 | rs1533594 | -0.745 | -1.458 | -0.443 | -0.357 | -0.057 | -0.101 | -0.101 | 0 | 0.73 | 0.61 | 0.87 | 0.96 | 0.82 | 0.84 |
| 3 | 187089031 | rs1003995 | 1.603 | -1.445 | -0.669 | -0.279 | -0.125 | -0.143 | -0.143 | 0 | 0.23 | 0.03 | 0.45 | 0.45 | 0.34 | 0.29 |
| 3 | 187089297 | rs9028 | 0.771 | -1.419 | -0.067 | -0.758 | 0.018 | 0.052 | 0.052 | 0.357 | 0.56 | 0.38 | 0.62 | 0.91 | 0.57 | 0.62 |
| 3 | 187091182 | rs9853501 | -0.7 | -0.789 | -0.172 | -0.168 | 0.006 | -0.034 | -0.034 | 0 | 0.16 | 0.19 | 0.24 | 0.05 | 0.24 | 0.21 |
| 3 | 187095439 | rs9846780 | 0.13 | -1.236 | -0.281 | -0.187 | 0.001 | -0.02 | -0.02 | 0 | 0.67 | 0.63 | 0.67 | 0.84 | 0.65 | 0.68 |
| 3 | 187100897 | rs2378246 | 0.435 | -1.259 | -0.648 | 0.515 | -0.013 | 0.005 | 0.005 | 0.709 | 0.06 | 0 | 0.06 | 0.39 | 0.05 | 0.18 |
| 3 | 187101435 | rs1568293 | 0.855 | -1.956 | -0.487 | -0.056 | 0 | 0 | 0 | 0 | 0.38 | 0.32 | 0.46 | 0.66 | 0.45 | 0.41 |
| 3 | 187118321 | rs1518874 | 0.137 | -1.432 | -0.218 | 0.129 | -0.004 | 0.096 | 0.096 | 0 | 0.29 | 0.1 | 0.26 | 0.03 | 0.31 | 0.28 |
| 3 | 187121796 | rs1518872 | 0.172 | -1.413 | -0.177 | 0.132 | -0.001 | -0.035 | -0.035 | 0 | 0.66 | 0.67 | 0.7 | 0.97 | 0.66 | 0.69 |
| 3 | 187122532 | rs13065558 | 0.144 | -1.421 | -0.142 | 0.139 | 0.004 | 0.054 | 0.054 | 0 | 0.3 | 0.2 | 0.27 | 0.05 | 0.31 | 0.28 |
| 3 | 187128567 | rs2679545 | 0.768 | -1.661 | -0.116 | 0.151 | -0.006 | -0.003 | -0.003 | 0 | 0.57 | 0.48 | 0.51 | 0.63 | 0.6 | 0.57 |
| 3 | 187153371 | rs2687923 | -1.06 | -1.683 | -0.093 | -0.026 | 0.009 | 0.011 | 0.011 | 0.285 | 0.44 | 0.65 | 0.5 | 0.34 | 0.4 | 0.45 |
| 3 | 187154182 | rs7629178 | 1.6 | -1.677 | -0.664 | -0.017 | -0.11 | 0.07 | 0.07 | 0.079 | 0.31 | 0.06 | 0.33 | 0.16 | 0.41 | 0.25 |
| 3 | 187158529 | rs7637064 | -1.995 | -0.968 | 0.092 | 0.195 | 0.029 | 0.075 | 0.075 | 0.455 | 0.22 | 0.49 | 0.38 | 0.33 | 0.21 | 0.27 |
| 3 | 187160825 | rs2653315 | -0.904 | -0.896 | -0.215 | 0.2 | 0.037 | -0.039 | -0.039 | 0 | 0.13 | 0.24 | 0.18 | 0.03 | 0.16 | 0.14 |
| 3 | 187161616 | rs4686885 | -0.997 | -1.691 | -0.548 | 0.594 | -0.027 | 0.092 | 0.092 | 0.268 | 0.09 | 0.18 | 0.19 | 0.3 | 0.05 | 0.12 |
| 3 | 187163046 | rs9824450 | -0.066 | -1.52 | -1.085 | 0.185 | 0.021 | 0.123 | 0.123 | 0 | 0.2 | 0.02 | 0.11 | 0.01 | 0.19 | 0.17 |
| 3 | 187163706 | rs7643457 | -1.321 | -0.551 | -0.336 | 1.02 | -0.001 | 0.003 | 0.003 | 0.427 | 0.1 | 0.12 | 0.2 | 0.22 | 0.07 | 0.14 |
| 3 | 187165774 | rs4686880 | -1.11 | -0.442 | 0.008 | 0.939 | 0.016 | 0.205 | 0.205 | 0 | 0.1 | 0.32 | 0.2 | 0.29 | 0.07 | 0.24 |
| 3 | 187166031 | rs9850128 | -0.478 | -0.363 | -0.884 | -0.001 | 0.004 | 0.012 | 0.012 | 0 | 0.16 | 0.16 | 0.11 | 0.01 | 0.18 | 0.11 |
| 3 | 187169895 | rs9833091 | -0.344 | -0.954 | -1.418 | -0.353 | 0.026 | 0.152 | 0.152 | 0.595 | 0.23 | 0.01 | 0.1 | 0 | 0.16 | 0.08 |
| 3 | 187174310 | rs9855124 | -0.716 | -0.451 | -1.125 | -0.212 | 0.008 | 0.005 | 0.005 | 0 | 0.67 | 0.58 | 0.5 | 0.66 | 0.58 | 0.51 |
| 3 | 187196213 | rs17780614 | 0.393 | -0.963 | -1.214 | 0.158 | 0.038 | 0.175 | 0.175 | 0.367 | 0.28 | 0.01 | 0.13 | 0.01 | 0.21 | 0.16 |
| 3 | 187202227 | rs2378265 | -0.596 | -0.804 | -0.129 | 0.489 | -0.016 | -0.013 | -0.013 | 0 | 0.68 | 0.78 | 0.53 | 0.64 | 0.57 | 0.61 |
| 3 | 187203329 | rs12492252 | -0.869 | -0.7 | -0.086 | 0.58 | 0.011 | 0.138 | 0.138 | 0.029 | 0.06 | 0.3 | 0.09 | 0.19 | 0.03 | 0.14 |
| 3 | 187203412 | rs12492283 | -1.061 | -1.06 | 0.135 | 0.336 | -0.045 | 0.214 | 0.214 | 2.217 | 0.14 | 0.42 | 0.19 | 0.61 | 0.06 | 0.31 |
| 3 | 187208183 | rs6807004 | -0.457 | -0.707 | 0.052 | 0.044 | -0.012 | -0.014 | -0.014 | 0.004 | 0.63 | 0.7 | 0.44 | 0.49 | 0.49 | 0.56 |
| 3 | 187218336 | rs9846911 | 0.048 | -0.482 | 0.053 | -0.836 | -0.002 | -0.046 | -0.046 | 0 | 0.55 | 0.57 | 0.33 | 0.06 | 0.46 | 0.38 |
| 3 | 187220633 | rs1597800 | -0.001 | -0.414 | -0.319 | -0.754 | -0.007 | 0.062 | 0.062 | 0 | 0.42 | 0.31 | 0.63 | 0.92 | 0.48 | 0.6 |
| 3 | 187221936 | rs9867503 | -0.462 | 0.009 | -0.122 | -1.145 | -0.013 | 0.068 | 0.068 | 0.141 | 0.61 | 0.72 | 0.8 | 0.88 | 0.6 | 0.66 |
| 3 | 187222367 | rs9830704 | -0.212 | 0.06 | 0.48 | -1.221 | 0.011 | -0.069 | -0.069 | 0 | 0.23 | 0.47 | 0.18 | 0.12 | 0.22 | 0.2 |
| 3 | 187223226 | rs2378269 | 0.695 | -0.014 | 0.236 | -1.336 | -0.031 | -0.022 | -0.022 | 0 | 0.31 | 0.06 | 0.56 | 0.72 | 0.33 | 0.36 |
| 3 | 187225416 | rs6444236 | -0.832 | -0.166 | 0.332 | -1.374 | -0.015 | -0.073 | -0.073 | 0 | 0.3 | 0.51 | 0.22 | 0.16 | 0.26 | 0.32 |
| 3 | 187233157 | rs10937304 | -1.252 | -0.194 | 0.631 | -1.404 | -0.005 | 0.02 | 0.02 | 0.281 | 0.59 | 0.66 | 0.78 | 0.88 | 0.59 | 0.67 |
| 3 | 187235072 | rs12491180 | 0.718 | -0.254 | 0.523 | -1.353 | -0.021 | -0.002 | -0.002 | 0 | 0.29 | 0.12 | 0.57 | 0.73 | 0.34 | 0.36 |
| 3 | 187236486 | rs11922608 | 0.764 | -0.338 | 0.879 | -1.26 | 0.015 | 0.006 | 0.006 | 0 | 0.07 | 0.13 | 0.09 | 0.09 | 0.09 | 0.12 |

| | | | | | | | | | | | | | | | | |
|---|---|---|---|---|---|---|---|---|---|---|---|---|---|---|---|---|
| 3 | 187239645 | rs2553996 | 1.133 | -0.157 | 0.763 | -1.361 | 0.002 | 0.047 | 0.047 | 0.42 | 0.39 | 0.27 | 0.19 | 0.12 | 0.35 | 0.32 |
| 3 | 187242051 | rs2653294 | -0.073 | -0.103 | 1.156 | -0.972 | 0.001 | -0.011 | -0.011 | 0.111 | 0.64 | 0.38 | 0.67 | 0.8 | 0.61 | 0.72 |
| 3 | 187247088 | rs2553991 | 0.331 | -0.07 | 1.32 | -0.971 | 0.035 | -0.063 | -0.063 | 2.459 | 0.78 | 0.44 | 0.77 | 0.89 | 0.75 | 0.79 |
| 3 | 187248459 | rs6774872 | 1.504 | -0.124 | 1.174 | -0.955 | -0.044 | -0.011 | -0.011 | 0 | 0.2 | 0.07 | 0.2 | 0.19 | 0.25 | 0.18 |
| 3 | 187248921 | rs9290852 | 0.89 | 0.16 | 1.585 | -0.787 | 0.036 | -0.062 | -0.062 | 0 | 0.12 | 0.28 | 0.1 | 0.01 | 0.14 | 0.09 |
| 3 | 187251425 | rs2167616 | 0.898 | 0.002 | 1.137 | -0.759 | 0 | 0.045 | 0.045 | 0.99 | 0.47 | 0.25 | 0.54 | 0.65 | 0.47 | 0.48 |
| 3 | 187264516 | rs2653312 | -1.208 | 0.127 | 1.169 | -0.76 | 0.012 | -0.038 | -0.038 | 0 | 0.89 | 0.81 | 0.91 | 0.99 | 0.87 | 0.91 |
| 3 | 187264951 | rs2653309 | 1.095 | 0.235 | 1.425 | -0.753 | 0.028 | 0.012 | 0.012 | 1.658 | 0.73 | 0.43 | 0.75 | 0.71 | 0.71 | 0.66 |
| 3 | 187268087 | rs17726238 | 1.095 | 0.209 | 1.069 | -0.77 | -0.044 | 0.038 | 0.038 | 0 | 0.27 | 0.06 | 0.21 | 0.17 | 0.27 | 0.24 |
| 3 | 187269748 | rs7644608 | 0.065 | 0.227 | 1.343 | -0.739 | 0.001 | -0.005 | -0.005 | 0.125 | 0.37 | 0.39 | 0.35 | 0.31 | 0.39 | 0.36 |
| 3 | 187269839 | rs7644850 | 0.287 | 0.226 | 1.39 | -0.739 | -0.023 | 0.018 | 0.018 | 0 | 0.35 | 0.17 | 0.33 | 0.31 | 0.39 | 0.36 |
| 3 | 187269986 | rs7644965 | -0.319 | 0.252 | 1.379 | -0.738 | -0.001 | -0.007 | -0.007 | 0 | 0.42 | 0.46 | 0.36 | 0.36 | 0.4 | 0.43 |
| 3 | 187277603 | rs1452193 | -0.416 | 0.227 | 1.053 | -0.76 | 0.006 | 0.029 | 0.029 | 0 | 0.46 | 0.35 | 0.4 | 0.36 | 0.47 | 0.43 |
| 3 | 187278108 | rs9809088 | 0.634 | 0.235 | 0.982 | -0.763 | 0.008 | 0.026 | 0.026 | 0 | 0.32 | 0.2 | 0.22 | 0.23 | 0.27 | 0.3 |
| 3 | 187282085 | rs2378274 | -0.387 | 0.144 | 0.974 | -0.71 | 0.015 | 0.05 | 0.05 | 0 | 0.47 | 0.3 | 0.39 | 0.34 | 0.46 | 0.43 |
| 3 | 187283848 | rs2600998 | 1.129 | 0.129 | 1.087 | -0.782 | -0.015 | 0.004 | 0.004 | 0.009 | 0.39 | 0.25 | 0.5 | 0.63 | 0.4 | 0.47 |
| 3 | 187284441 | rs1452188 | 1.079 | 0.193 | 1.17 | -0.754 | -0.003 | 0.007 | 0.007 | 0 | 0.41 | 0.35 | 0.51 | 0.63 | 0.4 | 0.47 |
| 3 | 187289701 | rs2600994 | -0.824 | 0.599 | 0.92 | -0.606 | 0.036 | -0.064 | -0.064 | 0 | 0.88 | 0.75 | 0.9 | 0.99 | 0.86 | 0.9 |
| 3 | 187294715 | rs17727032 | 1.04 | 0.609 | 0.682 | -0.602 | -0.002 | -0.01 | -0.01 | 0 | 0.11 | 0.13 | 0.09 | 0.01 | 0.14 | 0.09 |
| 3 | 187295112 | rs6444239 | -0.046 | 0.223 | 0.646 | -0.726 | -0.006 | 0.005 | 0.005 | 0 | 0.42 | 0.39 | 0.35 | 0.39 | 0.42 | 0.42 |
| 3 | 187301956 | rs13076423 | 1.624 | 0.221 | -0.336 | -0.708 | -0.029 | -0.094 | -0.094 | 0 | 0.33 | 0.12 | 0.47 | 0.55 | 0.35 | 0.4 |
| 3 | 187305068 | rs1559772 | 0.211 | 0.21 | -0.218 | -0.687 | -0.005 | 0.018 | 0.018 | 0 | 0.55 | 0.33 | 0.62 | 0.6 | 0.58 | 0.57 |
| 3 | 187312018 | rs1863392 | -0.371 | 0.236 | -0.2 | -0.569 | 0.003 | 0.019 | 0.019 | 0 | 0.2 | 0.56 | 0.18 | 0.24 | 0.17 | 0.2 |
| 3 | 187315241 | rs1947345 | 1.337 | 0.351 | -0.623 | -0.694 | -0.015 | -0.048 | -0.048 | 0 | 0.34 | 0.18 | 0.48 | 0.58 | 0.37 | 0.37 |
| 3 | 187318035 | rs2053955 | -1.071 | 0.354 | -0.601 | -0.723 | 0 | 0.023 | 0.023 | 0 | 0.39 | 0.65 | 0.31 | 0.23 | 0.38 | 0.39 |
| 3 | 187318908 | rs2059552 | -1.03 | 0.353 | -0.643 | -0.724 | -0.001 | 0.045 | 0.045 | 0 | 0.4 | 0.7 | 0.31 | 0.23 | 0.38 | 0.4 |
| 3 | 187319124 | rs6788371 | -1.081 | 0.374 | -0.75 | -0.707 | 0 | 0.026 | 0.026 | 0 | 0.39 | 0.66 | 0.31 | 0.23 | 0.38 | 0.37 |
| 3 | 187319689 | rs10937312 | 0.512 | 0.342 | -1.295 | -0.709 | -0.009 | -0.002 | -0.002 | 0.187 | 0.12 | 0.03 | 0.1 | 0.09 | 0.12 | 0.14 |
| 3 | 187322302 | rs17795015 | 0.585 | 0.496 | -0.916 | -0.624 | -0.003 | -0.007 | -0.007 | 0.055 | 0.1 | 0.13 | 0.08 | 0.02 | 0.13 | 0.1 |
| 3 | 187322482 | rs10049099 | 0.001 | 0.338 | -0.862 | -0.218 | 0.002 | 0.014 | 0.014 | 0 | 0.27 | 0.16 | 0.21 | 0.18 | 0.25 | 0.23 |
| 3 | 187328475 | rs9833173 | -0.244 | 0.35 | -1.19 | -0.241 | 0.011 | 0.07 | 0.07 | 0 | 0.36 | 0.15 | 0.28 | 0.18 | 0.38 | 0.25 |
| 3 | 187330821 | rs4686894 | 0.142 | 0.007 | -0.755 | -0.001 | -0.017 | -0.017 | -0.017 | 0 | 0.69 | 0.46 | 0.73 | 0.76 | 0.74 | 0.71 |
| 3 | 187331590 | rs7635068 | -0.304 | -0.471 | -0.88 | 0.485 | 0.014 | -0.029 | -0.029 | 0 | 0.25 | 0.19 | 0.42 | 0.45 | 0.17 | 0.37 |
| 3 | 187333377 | rs4686896 | 0.491 | -0.162 | -0.294 | -0.436 | 0 | -0.067 | -0.067 | 0 | 0.77 | 0.52 | 0.81 | 0.87 | 0.77 | 0.81 |
| 3 | 187337208 | rs1863390 | 0.63 | 0.022 | -0.61 | -0.743 | -0.004 | -0.02 | -0.02 | 0 | 0.35 | 0.37 | 0.42 | 0.21 | 0.3 | 0.34 |
| 3 | 187340489 | rs1452182 | 0.488 | 0.245 | -0.756 | -0.665 | -0.039 | 0.041 | 0.041 | 0 | 0.42 | 0.17 | 0.39 | 0.65 | 0.47 | 0.44 |
| 3 | 187344935 | rs9290855 | 0.181 | 0.579 | -0.103 | -0.663 | -0.003 | -0.029 | -0.029 | 0 | 0.12 | 0.16 | 0.08 | 0.03 | 0.12 | 0.1 |
| 3 | 187346363 | rs17728084 | 0.313 | 0.626 | -0.107 | -0.423 | -0.003 | -0.004 | -0.004 | 0 | 0.05 | 0.04 | 0.07 | 0.1 | 0.11 | 0.08 |
| 3 | 187349009 | rs1452201 | -0.298 | 0.184 | 0.077 | -0.853 | 0.003 | 0.05 | 0.05 | 0 | 0.51 | 0.58 | 0.52 | 0.8 | 0.57 | 0.52 |
| 3 | 187372642 | rs9824174 | -0.353 | 0.069 | 0.829 | -0.534 | 0.023 | 0.028 | 0.028 | 0 | 0.08 | 0.2 | 0.09 | 0.14 | 0.12 | 0.03 |
| 3 | 187379755 | rs3864101 | -1.674 | -0.251 | 1.232 | -0.757 | 0 | -0.024 | -0.024 | 0.007 | 0.23 | 0.45 | 0.18 | 0.2 | 0.22 | 0.16 |
| 3 | 187384852 | rs10513818 | 0.299 | -0.623 | 1.342 | -0.884 | -0.006 | -0.009 | -0.009 | 0 | 0.13 | 0.15 | 0.2 | 0.06 | 0.09 | 0.07 |
| 3 | 187386831 | rs4988514 | 0.204 | -0.65 | 1.332 | -0.879 | -0.005 | -0.008 | -0.008 | 0 | 0.13 | 0.15 | 0.2 | 0.06 | 0.09 | 0.07 |
| 3 | 187389636 | rs7624906 | -1.018 | 0.037 | 1.557 | -0.427 | -0.025 | -0.068 | -0.068 | 0 | 0.16 | 0.36 | 0.11 | 0.06 | 0.11 | 0.13 |
| 3 | 187391498 | rs10513819 | 0.195 | 0.085 | 1.192 | -0.408 | -0.01 | -0.011 | -0.011 | 0.919 | 0.07 | 0.06 | 0.05 | 0.14 | 0.1 | 0.03 |
| 3 | 187391918 | rs12638067 | -0.625 | 0.078 | 1.281 | -0.412 | 0.001 | -0.003 | -0.003 | 0 | 0.13 | 0.14 | 0.09 | 0.06 | 0.11 | 0.13 |
| 3 | 187392728 | rs1365113 | 0.15 | 0.155 | 1.131 | -0.373 | -0.01 | -0.011 | -0.011 | 0 | 0.07 | 0.07 | 0.05 | 0.14 | 0.1 | 0.03 |
| 3 | 187395755 | rs9842483 | -1.011 | -0.038 | 1.203 | -0.672 | -0.024 | -0.067 | -0.067 | 0.116 | 0.15 | 0.34 | 0.11 | 0.06 | 0.11 | 0.13 |
| 3 | 187401981 | rs6802392 | -1.053 | -0.05 | 0.894 | -0.769 | -0.01 | -0.064 | -0.064 | 0 | 0.26 | 0.34 | 0.14 | 0.08 | 0.2 | 0.25 |
| 3 | 187402378 | rs6765474 | -2.314 | -0.073 | 1.054 | -0.941 | 0.015 | 0.144 | 0.144 | 1.097 | 0.44 | 0.84 | 0.42 | 0.21 | 0.37 | 0.37 |
| 3 | 187407916 | rs3915052 | -1.332 | 0.463 | 1.042 | -0.611 | 0.018 | -0.056 | -0.056 | 0.116 | 0.18 | 0.5 | 0.28 | 0.13 | 0.17 | 0.12 |
| 3 | 187416089 | rs1851682 | 1.853 | 0.456 | 0.149 | -0.481 | -0.062 | 0.129 | 0.129 | 2.263 | 0.6 | 0.05 | 0.56 | 0.74 | 0.64 | 0.65 |
| 3 | 187416253 | rs1523468 | 1.798 | 0.461 | 0.151 | -0.478 | -0.061 | 0.131 | 0.131 | 2.373 | 0.6 | 0.05 | 0.56 | 0.74 | 0.64 | 0.64 |
| 3 | 187421036 | rs9821619 | -1.622 | 0.481 | 1.002 | -0.48 | 0 | -0.003 | -0.003 | 0.263 | 0.35 | 0.61 | 0.22 | 0.22 | 0.32 | 0.31 |
| 3 | 187421219 | rs7616923 | -0.286 | 0.364 | 1.308 | -0.484 | 0.071 | -0.001 | -0.001 | 0 | 0.1 | 0.32 | 0.1 | 0.08 | 0.12 | 0.04 |
| 3 | 187427720 | rs7633309 | 1.919 | 0.384 | 0.738 | -0.483 | -0.034 | 0.128 | 0.128 | 1.88 | 0.61 | 0.1 | 0.59 | 0.74 | 0.64 | 0.66 |
| 3 | 187438522 | rs4686904 | 1.72 | 0.618 | 0.725 | -0.507 | -0.052 | 0.117 | 0.117 | 2.604 | 0.65 | 0.11 | 0.75 | 0.79 | 0.66 | 0.7 |
| 3 | 187440506 | rs1005099 | 2.112 | 0.544 | 0.708 | -0.511 | -0.067 | 0.144 | 0.144 | 2.51 | 0.61 | 0.07 | 0.59 | 0.74 | 0.62 | 0.66 |
| 3 | 187444210 | rs1523475 | 0.942 | 0.561 | 1.188 | -0.485 | 0.01 | -0.068 | -0.068 | 0.089 | 0.78 | 0.52 | 0.88 | 0.88 | 0.81 | 0.78 |
| 3 | 187447032 | rs1056932 | 1.465 | 0.626 | 0.288 | -0.42 | -0.038 | 0.127 | 0.127 | 2.312 | 0.65 | 0.11 | 0.75 | 0.8 | 0.67 | 0.7 |
| 3 | 187456709 | rs3733017 | -0.095 | 1.259 | 0.577 | 0.438 | 0 | 0.01 | 0.01 | 0 | 0.08 | 0.09 | 0.05 | 0.23 | 0.07 | 0.03 |
| 3 | 187459088 | rs3172469 | -1.213 | 1.272 | 0.527 | 0.412 | 0.089 | 0.036 | 0.036 | 0 | 0.22 | 0.45 | 0.19 | 0.36 | 0.28 | 0.2 |
| 3 | 187460099 | rs3774298 | 1.298 | 1.117 | 0.321 | 0.667 | 0.029 | 0.028 | 0.028 | 0.044 | 0.74 | 0.48 | 0.57 | 0.57 | 0.67 | 0.76 |
| 3 | 187462296 | rs4686467 | 0.032 | 1.293 | -0.72 | 0.662 | -0.048 | 0.01 | 0.01 | 0.684 | 0.66 | 0.77 | 0.67 | 0.65 | 0.54 | 0.62 |
| 3 | 187468973 | rs10937320 | -0.189 | 1.478 | -0.405 | 1.22 | -0.014 | -0.02 | -0.02 | 0 | 0.09 | 0.07 | 0.11 | 0.15 | 0.13 | 0.07 |
| 3 | 187474009 | rs9831920 | -1.037 | 1.224 | 0.306 | 1.15 | 0.017 | 0.022 | 0.022 | 0.019 | 0.13 | 0.33 | 0.14 | 0.16 | 0.15 | 0.2 |
| 3 | 187475080 | rs13079223 | 0.332 | 1.245 | 0.018 | 1.273 | -0.022 | -0.042 | -0.042 | 0 | 0.12 | 0.01 | 0.11 | 0.15 | 0.15 | 0.1 |
| 3 | 187475208 | rs2378341 | -1.496 | 1.387 | 0.111 | 1.093 | -0.04 | 0.037 | 0.037 | 7.91 | 0.47 | 0.86 | 0.59 | 0.58 | 0.46 | 0.65 |
| 3 | 187479288 | rs9811040 | -0.664 | 1.376 | 0.115 | 1.095 | 0.014 | 0.022 | 0.022 | 0.036 | 0.13 | 0.32 | 0.14 | 0.16 | 0.15 | 0.2 |

| | | | | | | | | | | | | | | | | |
|---|---|---|---|---|---|---|---|---|---|---|---|---|---|---|---|---|
| 3 | 187486631 | rs12487326 | -0.261 | 1.299 | 0.241 | 1.503 | -0.03 | 0.003 | 0.003 | 2.298 | 0.29 | 0.08 | 0.21 | 0.2 | 0.28 | 0.4 |
| 3 | 187490116 | rs17797685 | -0.279 | 1.245 | 0.504 | 1.211 | -0.026 | -0.047 | -0.047 | 0 | 0.29 | 0.09 | 0.38 | 0.27 | 0.28 | 0.4 |
| 3 | 187494790 | rs1980301 | -1.025 | 1.227 | 0.52 | 1.029 | 0.009 | -0.016 | -0.016 | 0.374 | 0.39 | 0.81 | 0.5 | 0.54 | 0.33 | 0.57 |
| 3 | 187501080 | rs9871350 | -1.145 | 1.222 | 0.959 | 1.022 | -0.002 | 0.006 | 0.006 | 6.084 | 0.45 | 0.61 | 0.59 | 0.65 | 0.46 | 0.66 |
| 3 | 187502713 | rs7648069 | -1.227 | 1.28 | 0.828 | 0.885 | -0.005 | 0.011 | 0.011 | 4.857 | 0.43 | 0.59 | 0.54 | 0.59 | 0.46 | 0.55 |
| 3 | 187509547 | rs11924583 | -0.812 | 1.037 | 0.935 | 0.927 | -0.005 | 0.026 | 0.026 | 1.845 | 0.34 | 0.52 | 0.4 | 0.46 | 0.3 | 0.44 |
| 3 | 187511001 | rs3821820 | -0.551 | 1.064 | 0.778 | 0.578 | -0.002 | 0.006 | 0.006 | 2.039 | 0.31 | 0.36 | 0.37 | 0.28 | 0.28 | 0.4 |
| 3 | 187514203 | rs7610514 | -1.124 | 0.927 | 0.479 | 0.384 | -0.001 | -0.039 | -0.039 | 0.176 | 0.24 | 0.7 | 0.13 | 0 | 0.15 | 0.13 |
| 3 | 187518999 | rs2035020 | -0.905 | 1.184 | 0.454 | 0.588 | -0.02 | -0.038 | -0.038 | 0 | 0.9 | 0.96 | 0.74 | 0.72 | 0.9 | 0.78 |
| 3 | 187529428 | rs6802646 | -1.406 | 0.897 | 0.831 | -0.29 | -0.007 | -0.053 | -0.053 | 2.386 | 0.23 | 0.69 | 0.13 | 0 | 0.15 | 0.13 |
| 3 | 187540133 | rs11709267 | -0.742 | 1.224 | 0.738 | -0.023 | -0.002 | 0.011 | 0.011 | 0.447 | 0.15 | 0.2 | 0.11 | 0.23 | 0.15 | 0.16 |
| 3 | 187543958 | rs715680 | -0.679 | 1.051 | 0.761 | -0.029 | -0.013 | 0.016 | 0.016 | 0.171 | 0.14 | 0.2 | 0.08 | 0.23 | 0.1 | 0.16 |
| 3 | 187549526 | rs512178 | -1.353 | 1.3 | 0.781 | -0.252 | 0.005 | 0 | 0 | 0 | 0.29 | 0.4 | 0.27 | 0.28 | 0.34 | 0.31 |
| 3 | 187568380 | rs974671 | -1.769 | 1.257 | 1.102 | -0.239 | -0.01 | -0.003 | -0.003 | 0 | 0.34 | 0.78 | 0.33 | 0.32 | 0.38 | 0.32 |
| 3 | 187570173 | rs517116 | -1.454 | 1.307 | 1.005 | -0.45 | 0.018 | 0 | 0 | 0.58 | 0.3 | 0.49 | 0.31 | 0.29 | 0.38 | 0.32 |
| 3 | 187594092 | rs6778044 | -0.818 | 1.507 | 0.868 | -0.519 | 0.034 | -0.071 | -0.071 | 0 | 0.17 | 0.61 | 0.21 | 0.05 | 0.21 | 0.15 |
| 3 | 187595294 | rs6805813 | -0.818 | 1.507 | 0.868 | -0.519 | 0.034 | -0.071 | -0.071 | 0.597 | 0.17 | 0.61 | 0.21 | 0.05 | 0.21 | 0.15 |
| 3 | 187596315 | rs6798156 | -0.836 | 1.717 | 1.002 | -0.595 | 0.036 | -0.056 | -0.056 | 0.546 | 0.19 | 0.34 | 0.21 | 0.05 | 0.26 | 0.2 |
| 3 | 187596744 | rs6788294 | -1.01 | 1.66 | 0.859 | -0.619 | 0.029 | -0.056 | -0.056 | 0.105 | 0.2 | 0.34 | 0.21 | 0.05 | 0.26 | 0.2 |
| 3 | 187598450 | rs6786945 | -1.312 | 1.978 | 0.487 | -0.625 | 0.027 | -0.038 | -0.038 | 0.112 | 0.26 | 0.63 | 0.27 | 0.13 | 0.34 | 0.21 |
| 3 | 187605315 | rs6444265 | -1.396 | 1.206 | 0.641 | -0.527 | 0.054 | -0.058 | -0.058 | 0.851 | 0.16 | 0.59 | 0.32 | 0.08 | 0.19 | 0.03 |
| 3 | 187615000 | rs9822360 | -0.519 | 1.213 | 0.7 | -0.508 | 0.057 | -0.01 | -0.01 | 0.189 | 0.1 | 0.35 | 0.26 | 0.08 | 0.13 | 0.01 |
| 3 | 187615862 | rs523079 | 0.062 | 1.042 | -0.238 | -0.457 | -0.008 | 0.034 | 0.034 | 0 | 0.13 | 0.17 | 0.09 | 0.25 | 0.12 | 0.16 |
| 3 | 187617441 | rs3099226 | -2.022 | 1.151 | -0.925 | -0.618 | -0.073 | 0.012 | 0.012 | 0.176 | 0.43 | 0.9 | 0.57 | 0.38 | 0.52 | 0.33 |
| 3 | 187618140 | rs546343 | -1.864 | 0.925 | -0.698 | -0.445 | 0.003 | 0.014 | 0.014 | 0 | 0.37 | 0.8 | 0.43 | 0.37 | 0.36 | 0.26 |
| 3 | 187620527 | rs1157840 | -1.508 | 0.317 | -0.384 | -0.185 | -0.002 | 0 | 0 | 0 | 0.26 | 0.7 | 0.19 | 0.26 | 0.23 | 0.2 |
| 3 | 187624906 | rs13059988 | -0.448 | 0.438 | -0.953 | 0.288 | 0.01 | -0.027 | -0.027 | 0 | 0.13 | 0.07 | 0.08 | 0.26 | 0.11 | 0.16 |
| 3 | 187629503 | rs1903216 | -0.673 | 0.24 | -1.112 | -0.132 | -0.026 | 0.006 | 0.006 | 0 | 0.58 | 0.79 | 0.57 | 0.44 | 0.51 | 0.4 |
| 3 | 187629577 | rs2611621 | 0.232 | -0.746 | -0.159 | -0.373 | -0.047 | -0.073 | -0.073 | 0 | 0.85 | 0.57 | 0.92 | 0.96 | 0.93 | 0.91 |
| 3 | 187630727 | rs11711595 | -0.315 | -1.33 | -0.656 | -0.74 | -0.006 | -0.004 | -0.004 | 0 | 0.4 | 0.43 | 0.22 | 0.32 | 0.25 | 0.19 |
| 3 | 187630831 | rs11719540 | 0.007 | -1.539 | -1.147 | -0.804 | 0.043 | -0.01 | -0.01 | 0.917 | 0.28 | 0.15 | 0.17 | 0.32 | 0.2 | 0.14 |
| 3 | 187639375 | rs763342 | -0.552 | -1.02 | -1.052 | -0.358 | -0.003 | -0.022 | -0.022 | 0 | 0.34 | 0.27 | 0.31 | 0.51 | 0.35 | 0.25 |
| 3 | 187639657 | rs2611619 | -0.729 | -1.141 | -1.869 | -0.704 | 0.019 | 0.081 | 0.081 | 0 | 0.74 | 0.94 | 0.67 | 0.89 | 0.82 | 0.8 |
| 3 | 187645566 | rs9823495 | 0.376 | -1.189 | -1.684 | -0.396 | 0.005 | -0.025 | -0.025 | 0 | 0.14 | 0.09 | 0.15 | 0.35 | 0.13 | 0.15 |
| 3 | 187649419 | rs6773854 | -0.015 | -1.136 | -1.171 | -0.578 | 0 | -0.024 | -0.024 | 0 | 0.24 | 0.3 | 0.16 | 0.17 | 0.21 | 0.11 |
| 3 | 187655626 | rs7651910 | 0.445 | -1.184 | -1.202 | -0.656 | -0.007 | 0.066 | 0.066 | 0.001 | 0.21 | 0.06 | 0.1 | 0 | 0.19 | 0.07 |
| 3 | 187664598 | rs481145 | -1.052 | -1.759 | -0.53 | -0.952 | 0.004 | -0.002 | -0.002 | 0.066 | 0.46 | 0.62 | 0.5 | 0.53 | 0.38 | 0.28 |
| 3 | 187665490 | rs576072 | -0.564 | -1.794 | -0.316 | -0.817 | -0.042 | 0.046 | 0.046 | 0 | 0.25 | 0.52 | 0.39 | 0.36 | 0.17 | 0.2 |
| 3 | 187667357 | rs2590452 | 0.394 | -1.829 | -0.241 | -0.848 | -0.029 | -0.079 | -0.079 | 0.94 | 0.14 | 0.37 | 0.23 | 0.02 | 0.07 | 0.07 |
| 3 | 187667808 | rs2590453 | -0.516 | -1.815 | -0.508 | -0.826 | -0.048 | 0.051 | 0.051 | 0 | 0.25 | 0.57 | 0.39 | 0.36 | 0.17 | 0.2 |
| 3 | 187668469 | rs2248422 | -0.733 | -1.81 | -0.472 | -0.84 | -0.021 | 0.07 | 0.07 | 0 | 0.27 | 0.62 | 0.41 | 0.53 | 0.2 | 0.21 |
| 3 | 187668721 | rs567713 | 0.082 | -1.819 | -0.459 | -0.88 | -0.048 | -0.108 | -0.108 | 0 | 0.16 | 0.53 | 0.24 | 0.02 | 0.07 | 0.08 |
| 3 | 187673930 | rs2590443 | -1.345 | -1.845 | -0.941 | -0.875 | 0.019 | 0.023 | 0.023 | 0.695 | 0.83 | 0.92 | 0.87 | 0.91 | 0.87 | 0.88 |
| 3 | 187678374 | rs4686469 | 1.507 | -1.77 | -0.951 | -0.74 | -0.005 | -0.03 | -0.03 | 0 | 0.09 | 0.04 | 0.15 | 0.35 | 0.09 | 0.06 |
| 3 | 187678985 | rs568066 | -1.32 | -1.665 | -0.994 | -0.834 | 0.024 | 0.025 | 0.025 | 0 | 0.85 | 0.95 | 0.89 | 0.93 | 0.89 | 0.9 |
| 3 | 187679835 | rs498688 | -1.045 | -1.409 | -0.721 | -0.523 | 0.019 | -0.01 | -0.01 | 0.259 | 0.49 | 0.78 | 0.56 | 0.54 | 0.39 | 0.25 |
| 3 | 187682109 | rs484479 | -0.038 | -1.009 | -0.273 | -0.415 | 0.01 | 0.013 | 0.013 | 0 | 0.3 | 0.37 | 0.3 | 0.36 | 0.3 | 0.17 |
| 3 | 187683685 | rs6777331 | -0.292 | -0.417 | -0.761 | -0.653 | 0.001 | 0.008 | 0.008 | 1.204 | 0.53 | 0.42 | 0.38 | 0.37 | 0.55 | 0.34 |
| 3 | 187687074 | rs9820070 | -0.238 | 0.416 | -0.951 | -0.636 | 0.001 | -0.006 | -0.006 | 0 | 0.39 | 0.41 | 0.31 | 0.34 | 0.39 | 0.22 |
| 3 | 187687890 | rs16862782 | 1.166 | 0.391 | -1.21 | -0.699 | -0.021 | -0.002 | -0.002 | 0 | 0.18 | 0.02 | 0.17 | 0.14 | 0.17 | 0.11 |
| 3 | 187688058 | rs17806690 | 1.526 | 0.484 | -0.946 | -0.715 | -0.001 | -0.061 | -0.061 | 0.213 | 0.18 | 0.15 | 0.11 | 0 | 0.15 | 0.09 |
| 3 | 187688669 | rs1348637 | 0.912 | 0.43 | -1.02 | -0.705 | 0.009 | 0.023 | 0.023 | 0 | 0.19 | 0.06 | 0.18 | 0.14 | 0.17 | 0.12 |
| 3 | 187690288 | rs2030033 | 0.285 | 0.382 | -0.907 | -0.787 | 0.001 | -0.048 | -0.048 | 1.172 | 0.25 | 0.3 | 0.19 | 0.09 | 0.23 | 0.15 |
| 3 | 187691182 | rs17745063 | 1.555 | 0.41 | -1.067 | -0.809 | -0.004 | 0.021 | 0.021 | 0 | 0.18 | 0.08 | 0.1 | 0 | 0.15 | 0.08 |
| 3 | 187695429 | rs17745214 | 1.821 | 0.338 | -1.112 | -0.757 | 0.015 | 0.043 | 0.043 | 0 | 0.13 | 0.03 | 0.1 | 0.05 | 0.09 | 0.06 |
| 3 | 187696799 | rs2030034 | 1.861 | 0.464 | -1.355 | -0.567 | -0.024 | -0.027 | -0.027 | 5.505 | 0.07 | 0.01 | 0.06 | 0.05 | 0.09 | 0.19 |
| 3 | 187698333 | rs13086331 | 0.237 | 0.274 | -1.094 | -0.546 | 0.002 | 0.012 | 0.012 | 0 | 0.15 | 0.25 | 0.08 | 0.22 | 0.15 | 0.12 |
| 3 | 187702211 | rs4686912 | -1.162 | 0.583 | -1.031 | -0.263 | 0.003 | -0.003 | -0.003 | 0 | 0.68 | 0.67 | 0.71 | 0.77 | 0.58 | 0.47 |
| 3 | 187702763 | rs1874751 | 0.247 | 0.644 | -1.112 | -0.161 | -0.002 | -0.008 | -0.008 | 0 | 0.36 | 0.4 | 0.25 | 0.26 | 0.31 | 0.17 |
| 3 | 187710426 | rs9860469 | 0.651 | 0.947 | -1.465 | 0.019 | -0.038 | -0.051 | -0.051 | 0 | 0.26 | 0.14 | 0.26 | 0.4 | 0.32 | 0.19 |
| 3 | 187714053 | rs12330507 | 1.612 | 0.954 | -1.455 | -0.068 | 0 | -0.013 | -0.013 | 0 | 0.18 | 0.12 | 0.14 | 0.25 | 0.18 | 0.09 |
| 3 | 187716886 | rs1553091 | 0.655 | 0.87 | -1.466 | -0.097 | -0.039 | -0.073 | -0.073 | 0 | 0.26 | 0.13 | 0.28 | 0.42 | 0.33 | 0.19 |
| 3 | 187718701 | rs9837670 | 1.936 | 0.737 | -1.436 | -0.205 | 0.016 | 0.025 | 0.025 | 2.553 | 0.16 | 0.1 | 0.12 | 0.04 | 0.11 | 0.07 |
| 3 | 187719473 | rs9843063 | -0.423 | 0.689 | -1.253 | -0.283 | -0.003 | 0.011 | 0.011 | 0.077 | 0.46 | 0.56 | 0.39 | 0.4 | 0.46 | 0.28 |
| 3 | 187723901 | rs1498034 | 1.561 | 0.466 | -1.52 | -0.337 | -0.002 | -0.144 | -0.144 | 0 | 0.2 | 0.02 | 0.37 | 0.43 | 0.17 | 0.29 |
| 3 | 187726113 | rs4686917 | -0.965 | 0.745 | -1.213 | -0.196 | -0.002 | -0.029 | -0.029 | 0 | 0.66 | 0.56 | 0.76 | 0.83 | 0.65 | 0.57 |
| 3 | 187727515 | rs11924318 | 0.387 | 1.127 | -1.301 | 0.189 | -0.023 | -0.032 | -0.032 | 0 | 0.27 | 0.18 | 0.27 | 0.4 | 0.32 | 0.19 |
| 3 | 187737376 | rs2173321 | 1.169 | 0.865 | -1.273 | 0.046 | -0.049 | -0.049 | -0.049 | 0 | 0.09 | 0.01 | 0.12 | 0.16 | 0.13 | 0.11 |
| 3 | 187738766 | rs10937330 | -0.015 | 0.594 | -1.064 | -0.295 | -0.003 | 0.011 | 0.011 | 0 | 0.47 | 0.44 | 0.36 | 0.31 | 0.46 | 0.48 |

| | | | | | | | | | | | | | | | | |
|---|---|---|---|---|---|---|---|---|---|---|---|---|---|---|---|---|
| 3 | 187740523 | rs6808574 | -0.944 | 0.45 | -1.013 | -0.701 | -0.008 | 0.001 | 0.001 | 0 | 0.65 | 0.74 | 0.8 | 0.99 | 0.61 | 0.8 |
| 3 | 187742584 | rs1553092 | 0.639 | -0.016 | -0.314 | -0.256 | -0.018 | 0.083 | 0.083 | 0 | 0.24 | 0.41 | 0.41 | 0.53 | 0.21 | 0.3 |
| 3 | 187751857 | rs1587312 | -1.648 | 0.245 | -0.36 | -0.516 | 0.005 | 0.002 | 0.002 | 0 | 0.81 | 0.89 | 0.89 | 1 | 0.79 | 0.9 |
| 3 | 187753889 | rs16862908 | 0.027 | 0.197 | -0.377 | -0.538 | -0.007 | -0.011 | -0.011 | 0.34 | 0.39 | 0.51 | 0.53 | 0.69 | 0.36 | 0.63 |
| 3 | 187753995 | rs11710227 | 0.607 | 0.189 | -0.514 | -0.562 | -0.054 | 0.029 | 0.029 | 0 | 0.4 | 0.17 | 0.34 | 0.3 | 0.43 | 0.26 |
| 3 | 187754948 | rs2378358 | -1.674 | 0.338 | -0.599 | -0.554 | 0 | 0.033 | 0.033 | 0 | 0.79 | 0.91 | 0.87 | 1 | 0.77 | 0.89 |
| 3 | 187755019 | rs1818599 | -0.337 | 0.163 | -0.185 | -0.046 | -0.009 | -0.014 | -0.014 | 0 | 0.39 | 0.54 | 0.53 | 0.69 | 0.36 | 0.63 |
| 3 | 187756856 | rs9856368 | -0.112 | 0.629 | 0.336 | 0.257 | 0.109 | 0.157 | 0.157 | 0.02 | 0.07 | 0.39 | 0.1 | 0.12 | 0.08 | 0.02 |
| 3 | 187757401 | rs9856948 | 0.511 | 0.021 | -0.512 | 0.381 | 0.035 | 0.037 | 0.037 | 0 | 0.54 | 0.31 | 0.43 | 0.3 | 0.64 | 0.46 |
| 3 | 187758098 | rs13323803 | 1.563 | 0.372 | -0.959 | 0.143 | -0.055 | 0.054 | 0.054 | 4.475 | 0.31 | 0.04 | 0.25 | 0.19 | 0.35 | 0.18 |
| 3 | 187759440 | rs882788 | -0.986 | 0.189 | -0.874 | 0.357 | 0.016 | -0.02 | -0.02 | 0.326 | 0.68 | 0.72 | 0.6 | 0.49 | 0.79 | 0.61 |
| 3 | 187759862 | rs11710971 | 1.562 | 0.58 | -1.253 | 0.189 | -0.027 | 0.029 | 0.029 | 0 | 0.14 | 0.05 | 0.08 | 0 | 0.21 | 0.11 |
| 3 | 187760414 | rs9817429 | 1.182 | 0.43 | -1.007 | 0.661 | -0.055 | -0.008 | -0.008 | 0 | 0.31 | 0.04 | 0.26 | 0.24 | 0.35 | 0.18 |
| 3 | 187763947 | rs17808817 | 1.091 | 0.478 | -0.942 | 0.454 | -0.006 | 0.06 | 0.06 | 4.539 | 0.32 | 0.18 | 0.26 | 0.2 | 0.35 | 0.18 |
| 3 | 187764218 | rs2221070 | -0.411 | 0.41 | -0.299 | 1.068 | -0.011 | -0.013 | -0.013 | 2.481 | 0.13 | 0.1 | 0.16 | 0.16 | 0.16 | 0.3 |
| 3 | 187766215 | rs6444268 | -1.06 | 0.104 | 0.425 | 1.305 | 0.016 | 0.152 | 0.152 | 0.293 | 0.19 | 0.39 | 0.32 | 0.51 | 0.2 | 0.42 |
| 3 | 187767950 | rs7645293 | -0.66 | -0.078 | 0.435 | 1.221 | -0.022 | 0.101 | 0.101 | 1.181 | 0.14 | 0.34 | 0.22 | 0.32 | 0.08 | 0.27 |
| 3 | 187770606 | rs12633719 | 0.084 | -0.553 | -0.282 | 0.333 | 0.037 | -0.019 | -0.019 | 0 | 0.38 | 0.29 | 0.29 | 0.41 | 0.23 | 0.4 |
| 3 | 187771570 | rs12631700 | -0.115 | -0.344 | -0.529 | 0.41 | 0.047 | -0.033 | -0.033 | 0 | 0.43 | 0.29 | 0.35 | 0.46 | 0.32 | 0.42 |
| 3 | 187772141 | rs16862945 | 0.027 | -0.317 | -0.668 | 0.481 | 0.004 | 0.008 | 0.008 | 0.47 | 0.13 | 0.12 | 0.19 | 0.39 | 0.09 | 0.27 |
| 3 | 187773480 | rs12493799 | 0.001 | -0.273 | -0.82 | 0.5 | 0.009 | -0.023 | -0.023 | 0 | 0.13 | 0.08 | 0.19 | 0.39 | 0.09 | 0.27 |
| 3 | 187782348 | rs6783007 | -0.524 | -0.323 | -0.919 | 0.193 | 0.014 | 0.068 | 0.068 | 0 | 0.54 | 0.68 | 0.48 | 0.84 | 0.42 | 0.69 |
| 3 | 187784805 | rs1005984 | 0.04 | 0.123 | -1.146 | 1.051 | -0.002 | -0.054 | -0.054 | 0 | 0.13 | 0.06 | 0.23 | 0.46 | 0.11 | 0.39 |
| 3 | 187789441 | rs17809289 | 0.461 | 0.394 | -0.948 | 0.702 | -0.003 | 0.06 | 0.06 | 0 | 0.11 | 0.02 | 0.19 | 0 | 0.09 | 0.05 |
| 3 | 187793410 | rs9833256 | -0.302 | 0.537 | -0.427 | 0.926 | -0.009 | 0.113 | 0.113 | 0 | 0.12 | 0.25 | 0.08 | 0.31 | 0.08 | 0.18 |
| 3 | 187795657 | rs2037184 | 0.048 | 1.064 | -1.135 | 0.222 | -0.001 | 0.006 | 0.006 | 0 | 0.42 | 0.39 | 0.54 | 0.59 | 0.44 | 0.62 |
| 3 | 187795786 | rs2037183 | -0.028 | 1.273 | -1.607 | 0.22 | 0.062 | 0.03 | 0.03 | 0 | 0.77 | 0.64 | 0.75 | 0.67 | 0.66 | 0.77 |
| 3 | 187796009 | rs13071719 | 0.43 | 1.859 | -2.674 | 0.167 | -0.036 | -0.046 | -0.046 | 0 | 0.08 | 0.01 | 0.12 | 0.13 | 0.11 | 0.17 |
| 3 | 187799940 | rs9821867 | 0.222 | 1.815 | -2.468 | -0.478 | -0.001 | 0.173 | 0.173 | 5.654 | 0.35 | 0.1 | 0.22 | 0.08 | 0.31 | 0.15 |
| 3 | 187800184 | rs2889887 | -0.333 | 1.53 | -2.199 | -0.154 | -0.038 | -0.004 | -0.004 | 0.244 | 0.84 | 0.9 | 0.78 | 0.82 | 0.7 | 0.82 |
| 3 | 187802669 | rs9833769 | 0.033 | 0.096 | -2.076 | -0.339 | 0.046 | 0.155 | 0.155 | 2.226 | 0.29 | 0.06 | 0.13 | 0.01 | 0.19 | 0.11 |
| 3 | 187809352 | rs9816138 | -0.439 | 0.593 | -1.741 | 0.545 | -0.002 | -0.014 | -0.014 | 0.245 | 0.15 | 0.11 | 0.1 | 0.36 | 0.16 | 0.22 |
| 3 | 187809661 | rs3856916 | -0.806 | 0.019 | -1.931 | 0.009 | 0.018 | 0.032 | 0.032 | 0.686 | 0.8 | 0.9 | 0.79 | 0.94 | 0.84 | 0.89 |
| 3 | 187810262 | rs6444269 | 0.133 | 0.666 | -1.475 | 0.471 | -0.001 | -0.003 | -0.003 | 1.115 | 0.45 | 0.54 | 0.52 | 0.33 | 0.42 | 0.48 |
| 3 | 187810765 | rs12630875 | 0.392 | 0.586 | -1.829 | 0.618 | 0 | -0.027 | -0.027 | 0.07 | 0.07 | 0.03 | 0.1 | 0.16 | 0.07 | 0.23 |
| 3 | 187814477 | rs7651169 | -0.25 | 0.392 | -1.301 | 0.374 | -0.009 | -0.001 | -0.001 | 0.187 | 0.29 | 0.37 | 0.22 | 0.29 | 0.24 | 0.38 |
| 3 | 187818463 | rs17753354 | 0.541 | 0.291 | -1.598 | 0.422 | -0.004 | 0.008 | 0.008 | 0 | 0.15 | 0.08 | 0.28 | 0.06 | 0.17 | 0.14 |
| 3 | 187832419 | rs4686933 | -0.283 | 0.097 | -1.61 | 0.631 | 0.013 | -0.071 | -0.071 | 0 | 0.15 | 0.04 | 0.12 | 0.28 | 0.11 | 0.29 |
| 3 | 187838455 | rs344942 | -0.407 | 0.309 | -1.163 | 0.22 | 0 | -0.001 | -0.001 | 0 | 0.6 | 0.65 | 0.68 | 0.77 | 0.62 | 0.8 |
| 3 | 187838663 | rs9810988 | -0.318 | 0.602 | -0.704 | 0.286 | 0.045 | 0.04 | 0.04 | 0 | 0.14 | 0.34 | 0.14 | 0.17 | 0.16 | 0.08 |
| 3 | 187838965 | rs4686934 | -0.46 | 0.735 | -0.673 | 0.483 | 0.031 | 0.033 | 0.033 | 0 | 0.24 | 0.48 | 0.4 | 0.29 | 0.29 | 0.22 |
| 3 | 187839070 | rs189372 | -0.576 | 0.379 | -0.769 | 0.385 | 0.053 | 0.06 | 0.06 | 0 | 0.5 | 0.72 | 0.66 | 0.66 | 0.62 | 0.48 |
| 3 | 187840546 | rs6785081 | -0.155 | 0.477 | -0.161 | 0.339 | 0.044 | 0.03 | 0.03 | 0.249 | 0.1 | 0.38 | 0.12 | 0.04 | 0.09 | 0.04 |
| 3 | 187842721 | rs9852144 | 0.183 | 0.362 | -0.579 | 0.495 | 0.003 | 0.022 | 0.022 | 0 | 0.16 | 0.21 | 0.12 | 0.3 | 0.18 | 0.36 |
| 3 | 187845956 | rs344945 | -1.457 | 0.499 | -1.388 | -0.254 | 0.001 | 0.156 | 0.156 | 0 | 0.72 | 0.96 | 0.84 | 0.97 | 0.76 | 0.81 |
| 3 | 187847027 | rs7638075 | 0.37 | 1.051 | -0.511 | 0.5 | 0.005 | -0.006 | -0.006 | 0.262 | 0.15 | 0.2 | 0.28 | 0.13 | 0.2 | 0.18 |
| 3 | 187853266 | rs3844290 | -0.232 | 0.839 | -0.471 | 0.579 | 0 | -0.003 | -0.003 | 0.351 | 0.29 | 0.33 | 0.36 | 0.34 | 0.31 | 0.23 |
| 3 | 187856109 | rs4686935 | -0.462 | 0.799 | -0.158 | 0.957 | 0.039 | 0.106 | 0.106 | 0 | 0.09 | 0.26 | 0.23 | 0.14 | 0.1 | 0.14 |
| 3 | 187859571 | rs6776574 | 0.225 | 0.602 | -0.43 | 1.043 | 0.002 | -0.004 | -0.004 | 0 | 0.16 | 0.13 | 0.16 | 0.32 | 0.13 | 0.34 |
| 3 | 187865100 | rs13064299 | 0.181 | 0.282 | -0.327 | 0.602 | -0.036 | -0.03 | -0.03 | 0.006 | 0.36 | 0.17 | 0.35 | 0.49 | 0.45 | 0.3 |
| 3 | 187874933 | rs844106 | -0.511 | 0.135 | -0.336 | 0.591 | -0.01 | 0.041 | 0.041 | 0 | 0.47 | 0.28 | 0.48 | 0.8 | 0.53 | 0.61 |
| 3 | 187875552 | rs4234604 | -1.561 | -0.204 | 0.051 | 0.435 | 0.005 | 0.006 | 0.006 | 0 | 0.66 | 0.73 | 0.78 | 0.98 | 0.69 | 0.8 |
| 3 | 187877584 | rs9867814 | 0.99 | -0.034 | -0.077 | 0.723 | -0.001 | 0 | 0 | 0 | 0.22 | 0.17 | 0.2 | 0.23 | 0.3 | 0.15 |
| 3 | 187879004 | rs1711001 | -0.303 | -0.004 | -0.388 | 0.763 | 0.002 | -0.009 | -0.009 | 0 | 0.37 | 0.56 | 0.52 | 0.7 | 0.42 | 0.55 |
| 3 | 187880162 | rs12637043 | -0.001 | -0.314 | -0.575 | 0.893 | -0.016 | 0.137 | 0.137 | 0.28 | 0.14 | 0.32 | 0.32 | 0.47 | 0.12 | 0.37 |
| 3 | 187880447 | rs1679195 | -0.452 | -0.081 | -0.904 | 0.239 | 0.017 | 0.04 | 0.04 | 0 | 0.56 | 0.7 | 0.71 | 0.7 | 0.6 | 0.52 |
| 3 | 187883273 | rs16863057 | -0.363 | -0.56 | -0.893 | 0.511 | -0.015 | 0 | 0 | 0 | 0.09 | 0.11 | 0.02 | 0.1 | 0.01 | 0.07 |
| 3 | 187884853 | rs13098017 | 0.117 | -0.599 | -1.163 | 0.092 | 0.001 | 0.023 | 0.023 | 0.786 | 0.63 | 0.68 | 0.67 | 0.78 | 0.62 | 0.76 |
| 3 | 187887192 | rs9816875 | -0.167 | -1.338 | -1.201 | 0.208 | -0.032 | -0.003 | -0.003 | 1.717 | 0.39 | 0.49 | 0.32 | 0.4 | 0.17 | 0.52 |
| 3 | 187887562 | rs12493257 | 0.928 | -1.464 | -1.435 | 0.075 | -0.099 | -0.067 | -0.067 | 0.168 | 0.32 | 0.11 | 0.38 | 0.39 | 0.48 | 0.32 |
| 3 | 187887694 | rs9817667 | -0.077 | -1.456 | -1.464 | 0.142 | -0.031 | -0.004 | -0.004 | 0 | 0.39 | 0.49 | 0.32 | 0.39 | 0.17 | 0.51 |
| 3 | 187889723 | rs1007308 | -0.169 | -1.226 | -1.621 | 0.018 | -0.007 | -0.001 | -0.001 | 0 | 0.43 | 0.49 | 0.34 | 0.39 | 0.22 | 0.56 |
| 3 | 187890435 | rs9989991 | 0.136 | -1.172 | -1.701 | -0.095 | -0.019 | -0.007 | -0.007 | 0 | 0.39 | 0.32 | 0.42 | 0.4 | 0.49 | 0.29 |
| 3 | 187891687 | rs3930402 | -0.568 | -1.091 | -1.829 | -0.517 | -0.025 | 0.03 | 0.03 | 3.192 | 0.69 | 0.8 | 0.72 | 0.8 | 0.64 | 0.79 |
| 3 | 187891855 | rs344961 | -0.044 | -1.412 | -1.271 | -0.236 | -0.016 | -0.004 | -0.004 | 0 | 0.37 | 0.41 | 0.44 | 0.7 | 0.27 | 0.67 |
| 3 | 187896912 | rs13096262 | 0.727 | -0.952 | -1.99 | -0.432 | 0.003 | 0.187 | 0.187 | 0 | 0.5 | 0.23 | 0.67 | 0.86 | 0.56 | 0.59 |
| 3 | 187897424 | rs4577458 | 1.566 | -0.999 | -2.092 | -0.071 | -0.032 | -0.033 | -0.033 | 0 | 0.33 | 0.19 | 0.42 | 0.46 | 0.44 | 0.23 |
| 3 | 187898005 | rs344953 | 0.488 | -1.063 | -1.923 | -0.166 | -0.034 | -0.038 | -0.038 | 0 | 0.11 | 0.17 | 0.04 | 0.01 | 0.02 | 0.05 |
| 3 | 187898593 | rs344952 | 0.839 | -0.763 | -1.801 | 0.066 | -0.001 | 0 | 0 | 0 | 0.45 | 0.36 | 0.46 | 0.47 | 0.47 | 0.28 |

| | | | | | | | | | | | | | | | | |
|---|---|---|---|---|---|---|---|---|---|---|---|---|---|---|---|---|
| 3 | 187902463 | rs13084334 | 0.128 | -1.162 | -1.46 | 0.282 | 0 | -0.01 | -0.01 | 0 | 0.68 | 0.33 | 0.77 | 0.91 | 0.72 | 0.78 |
| 3 | 187902932 | rs2222103 | 0.95 | -1.298 | -1.378 | 0.502 | 0.008 | -0.112 | -0.112 | 0 | 0.15 | 0.02 | 0.24 | 0.4 | 0.11 | 0.35 |
| 3 | 187908259 | rs1710997 | 0.064 | -1.056 | -1.372 | 0.195 | -0.002 | -0.051 | -0.051 | 0 | 0.7 | 0.41 | 0.81 | 0.91 | 0.73 | 0.77 |
| 3 | 187909210 | rs9851426 | -0.215 | -1.458 | -1.538 | 0.156 | -0.033 | -0.071 | -0.071 | 0.169 | 0.75 | 0.42 | 0.88 | 0.93 | 0.86 | 0.81 |
| 3 | 187909753 | rs1679182 | -0.172 | -1.545 | -1.499 | 0.331 | 0.025 | -0.052 | -0.052 | 0 | 0.22 | 0.13 | 0.27 | 0.5 | 0.13 | 0.39 |
| 3 | 187910011 | rs9838692 | 0.899 | -1.597 | -1.682 | 0.312 | 0.048 | 0.081 | 0.081 | 0 | 0.51 | 0.27 | 0.41 | 0.32 | 0.6 | 0.31 |
| 3 | 187912292 | rs4686938 | -0.932 | -0.68 | -1.69 | 1.143 | 0.042 | -0.1 | -0.1 | 0 | 0.23 | 0.07 | 0.34 | 0.53 | 0.16 | 0.51 |
| 3 | 187915210 | rs4572756 | -0.963 | -0.689 | -1.684 | 1.133 | 0.038 | -0.074 | -0.074 | 0.675 | 0.24 | 0.11 | 0.34 | 0.53 | 0.16 | 0.51 |
| 3 | 187924902 | rs7645376 | -0.635 | -0.183 | -1.174 | 0.51 | 0.006 | 0 | 0 | 0 | 0.09 | 0.09 | 0.1 | 0.05 | 0.06 | 0.16 |
| 3 | 187935634 | rs6785028 | 0.582 | 0.709 | -1.078 | 0.549 | -0.002 | -0.001 | -0.001 | 0 | 0.86 | 0.83 | 0.78 | 0.84 | 0.75 | 0.89 |
| 3 | 187936974 | rs7633866 | -0.498 | -0.713 | -1.136 | 0.192 | 0.008 | 0.004 | 0.004 | 0 | 0.43 | 0.57 | 0.52 | 0.59 | 0.26 | 0.58 |
| 3 | 187944924 | rs1975991 | -0.062 | -0.137 | -1.608 | 0.15 | -0.075 | -0.124 | -0.124 | 0.075 | 0.76 | 0.95 | 0.62 | 0.4 | 0.68 | 0.56 |
| 3 | 187947668 | rs6777135 | 0.575 | -0.126 | -0.952 | -0.145 | 0 | -0.001 | -0.001 | 0 | 0.66 | 0.67 | 0.51 | 0.37 | 0.57 | 0.46 |
| 3 | 187951892 | rs1871517 | 0.778 | -0.235 | -1.079 | -0.801 | -0.017 | 0.032 | 0.032 | 0.203 | 0.18 | 0.02 | 0.13 | 0.06 | 0.21 | 0.21 |
| 3 | 187968192 | rs6809682 | 0.506 | -0.024 | -0.008 | -0.352 | 0.033 | 0.014 | 0.014 | 0 | 0.72 | 0.61 | 0.52 | 0.33 | 0.6 | 0.57 |
| 3 | 187971048 | rs9844029 | 1.378 | -0.341 | 0.014 | -0.571 | 0.003 | 0.143 | 0.143 | 0.218 | 0.48 | 0.18 | 0.34 | 0.23 | 0.44 | 0.34 |
| 3 | 187975289 | rs6809410 | 0.293 | 0.616 | 0.4 | -0.102 | 0.023 | -0.009 | -0.009 | 0 | 0.05 | 0.18 | 0.02 | 0 | 0.03 | 0.03 |
| 3 | 187979733 | rs6784029 | -1.681 | 0.568 | 0.325 | -0.175 | -0.009 | -0.052 | -0.052 | 0.048 | 0.19 | 0.55 | 0.2 | 0.1 | 0.15 | 0.2 |
| 3 | 187985004 | rs715790 | 0.418 | 0.656 | -0.122 | -0.143 | -0.061 | -0.045 | -0.045 | 0 | 0.71 | 0.85 | 0.56 | 0.35 | 0.6 | 0.6 |
| 3 | 187986482 | rs9868086 | -0.556 | 0.642 | 0.146 | -0.148 | 0.008 | 0.033 | 0.033 | 0.269 | 0.07 | 0.23 | 0.11 | 0.09 | 0.08 | 0.15 |
| 3 | 187988025 | rs16848776 | 0.427 | 0.682 | 0.301 | -0.156 | 0.001 | 0.002 | 0.002 | 0 | 0.73 | 0.66 | 0.57 | 0.35 | 0.68 | 0.63 |
| 3 | 187992608 | rs6793337 | -2.059 | 0.687 | 0.185 | -0.175 | 0.011 | -0.054 | -0.054 | 0.006 | 0.22 | 0.53 | 0.18 | 0.1 | 0.19 | 0.21 |
| 3 | 187995491 | rs9830100 | 0.793 | 0.575 | -0.133 | -0.048 | -0.016 | -0.02 | -0.02 | 0 | 0.05 | 0.12 | 0.03 | 0.01 | 0.01 | 0.06 |
| 3 | 188001875 | rs11713973 | 1.082 | 0.665 | -0.379 | -0.06 | -0.004 | 0.012 | 0.012 | 0 | 0.61 | 0.3 | 0.51 | 0.26 | 0.61 | 0.51 |
| 3 | 188003578 | rs6777007 | 1.069 | 0.447 | -0.388 | -0.055 | -0.008 | 0.018 | 0.018 | 0 | 0.6 | 0.3 | 0.48 | 0.26 | 0.58 | 0.51 |
| 3 | 188004254 | rs9840345 | 0.428 | 1.106 | -0.955 | 0.289 | -0.032 | -0.027 | -0.027 | 0 | 0.73 | 0.79 | 0.55 | 0.35 | 0.63 | 0.64 |
| 3 | 188006256 | rs4686945 | 0.415 | 0.632 | -1.164 | 0.379 | -0.114 | -0.07 | -0.07 | 0 | 0.71 | 0.9 | 0.55 | 0.34 | 0.56 | 0.62 |
| 3 | 188011407 | rs4686947 | -0.193 | -0.196 | -0.725 | 0.903 | -0.003 | -0.018 | -0.018 | 0.022 | 0.21 | 0.13 | 0.25 | 0.34 | 0.15 | 0.1 |
| 3 | 188026277 | rs10937344 | -1.386 | -0.133 | 0.08 | 1.263 | 0.001 | 0.011 | 0.011 | 0 | 0.33 | 0.48 | 0.47 | 0.7 | 0.28 | 0.37 |
| 3 | 188029842 | rs6774792 | 0.362 | -0.305 | 0.455 | 1.261 | 0.034 | 0.084 | 0.084 | 8.033 | 0.05 | 0.26 | 0.03 | 0.04 | 0.01 | 0.15 |
| 3 | 188039200 | rs4277678 | -0.328 | -0.206 | 0.344 | 1.528 | 0.033 | 0.291 | 0.291 | 2.421 | 0.06 | 0.2 | 0.18 | 0.35 | 0.04 | 0.2 |
| 3 | 188042029 | rs7631930 | -0.922 | -0.347 | 0.402 | 1.435 | 0.002 | 0.036 | 0.036 | 0.027 | 0.24 | 0.36 | 0.43 | 0.7 | 0.19 | 0.3 |
| 3 | 188045319 | rs4377487 | -1.323 | -0.17 | 0.181 | 1.527 | 0.017 | 0.243 | 0.243 | 1.103 | 0.08 | 0.23 | 0.2 | 0.35 | 0.05 | 0.21 |
| 3 | 188048842 | rs4686952 | -1.798 | -0.506 | 0.017 | 1.35 | 0.027 | 0.032 | 0.032 | 0 | 0.44 | 0.66 | 0.52 | 0.75 | 0.33 | 0.53 |
| 3 | 188050534 | rs12494557 | -0.373 | -0.455 | 0.087 | 1.386 | 0.041 | 0.319 | 0.319 | 2.556 | 0.07 | 0.25 | 0.18 | 0.35 | 0.04 | 0.2 |
| 3 | 188059443 | rs13076750 | -1.662 | -0.861 | -0.016 | 0.534 | 0.033 | 0.008 | 0.008 | 0 | 0.43 | 0.66 | 0.37 | 0.43 | 0.29 | 0.47 |
| 3 | 188061714 | rs4132172 | -1.029 | -0.722 | -0.057 | 0.573 | -0.011 | -0.003 | -0.003 | 0 | 0.35 | 0.45 | 0.33 | 0.39 | 0.29 | 0.3 |
| 3 | 188069330 | rs1403050 | 0.727 | -0.201 | -0.585 | 0.757 | -0.005 | 0.001 | 0.001 | 0.219 | 0.67 | 0.39 | 0.65 | 0.61 | 0.81 | 0.53 |
| 3 | 188070267 | rs9817266 | 0.397 | 0.521 | -0.891 | 0.226 | 0.001 | 0.005 | 0.005 | 0 | 0.57 | 0.38 | 0.49 | 0.28 | 0.61 | 0.56 |
| 3 | 188070292 | rs16863212 | -0.881 | 0.456 | -0.834 | 0.247 | 0 | -0.022 | -0.022 | 1.493 | 0.07 | 0.09 | 0.04 | 0 | 0.07 | 0 |
| 3 | 188070408 | rs17602709 | 0.506 | 0.598 | -1.61 | 0.007 | -0.091 | 0.104 | 0.104 | 1.124 | 0.35 | 0.03 | 0.21 | 0.18 | 0.35 | 0.16 |
| 3 | 188070570 | rs2103025 | -0.941 | 0.459 | -0.253 | 1.365 | -0.001 | 0.161 | 0.161 | 0 | 0.14 | 0.33 | 0.3 | 0.37 | 0.11 | 0.16 |
| 3 | 188070964 | rs2103022 | 0.249 | 0.531 | -0.483 | 1.158 | -0.016 | -0.034 | -0.034 | 0.123 | 0.81 | 0.88 | 0.69 | 0.64 | 0.75 | 0.75 |
| 3 | 188074440 | rs2030525 | -0.336 | 0.432 | -0.541 | 1.115 | -0.016 | -0.059 | -0.059 | 0.955 | 0.18 | 0.08 | 0.25 | 0.36 | 0.21 | 0.25 |
| 3 | 188082285 | rs6790359 | -0.143 | 0.136 | 0.085 | 0.687 | -0.007 | -0.096 | -0.096 | 0.162 | 0.75 | 0.5 | 0.73 | 0.92 | 0.76 | 0.69 |
| 3 | 188087628 | rs9851967 | 1.274 | -0.043 | 0.059 | 0.877 | -0.104 | 0.004 | 0.004 | 4.042 | 0.43 | 0.08 | 0.35 | 0.29 | 0.45 | 0.26 |
| 3 | 188089254 | rs13076312 | -0.571 | -0.197 | 0.805 | 0.843 | -0.003 | 0 | 0 | 0 | 0.43 | 0.37 | 0.54 | 0.66 | 0.48 | 0.56 |
| 3 | 188102716 | rs2889896 | -0.593 | -0.253 | 0.826 | 0.842 | -0.003 | 0 | 0 | 0.759 | 0.43 | 0.32 | 0.54 | 0.66 | 0.49 | 0.5 |
| 3 | 188109002 | rs4686482 | 1.22 | -0.295 | 0.936 | 0.558 | -0.042 | 0.005 | 0.005 | 0.254 | 0.88 | 0.55 | 0.94 | 0.95 | 0.98 | 0.8 |
| 3 | 188114458 | rs12107245 | -1.079 | -0.24 | 1.141 | 0.596 | -0.023 | 0.066 | 0.066 | 0.213 | 0.11 | 0.39 | 0.08 | 0.22 | 0.04 | 0.23 |
| 3 | 188114589 | rs13091753 | -0.226 | -0.17 | 1.026 | 0.691 | -0.003 | 0.002 | 0.002 | 0.27 | 0.38 | 0.29 | 0.48 | 0.41 | 0.44 | 0.44 |
| 3 | 188120643 | rs6785284 | -0.25 | -0.026 | 0.693 | 0.713 | 0 | 0.004 | 0.004 | 0.498 | 0.4 | 0.33 | 0.49 | 0.42 | 0.44 | 0.58 |
| 3 | 188123395 | rs972338 | 0.304 | -0.033 | 0.578 | 0.71 | -0.012 | 0.069 | 0.069 | 0.037 | 0.88 | 0.65 | 0.9 | 0.75 | 0.93 | 0.84 |
| 3 | 188124354 | rs1559810 | -0.502 | 0.273 | 0.269 | 0.601 | -0.012 | 0.004 | 0.004 | 0.518 | 0.39 | 0.21 | 0.48 | 0.42 | 0.44 | 0.55 |
| 3 | 188125341 | rs6444286 | 1.424 | 0.497 | 0.91 | 0.623 | 0.084 | 0.025 | 0.025 | 2.258 | 0.93 | 0.55 | 0.93 | 1 | 0.95 | 0.97 |
| 3 | 188138332 | rs9880787 | 0.566 | 0.543 | 0.265 | 1.074 | 0.012 | 0.127 | 0.127 | 0 | 0.81 | 0.5 | 0.78 | 0.58 | 0.82 | 0.76 |
| 3 | 188147274 | rs9862474 | -0.105 | 0.587 | -0.501 | -0.271 | 0 | -0.044 | -0.044 | 0 | 0.32 | 0.31 | 0.24 | 0.02 | 0.32 | 0.23 |
| 3 | 188152306 | rs9851525 | -0.025 | 0.5 | -0.571 | -0.346 | 0 | -0.047 | -0.047 | 0.082 | 0.27 | 0.3 | 0.22 | 0.01 | 0.3 | 0.22 |
| 3 | 188153208 | rs7431346 | 0.736 | -0.178 | -1.419 | 0.237 | -0.011 | -0.111 | -0.111 | 0.013 | 0.16 | 0 | 0.1 | 0.31 | 0.14 | 0.12 |
| 3 | 188155409 | rs7611312 | 0.19 | 0.295 | -1.08 | -0.195 | 0.009 | -0.093 | -0.093 | 0 | 0.32 | 0.35 | 0.29 | 0.01 | 0.4 | 0.24 |
| 3 | 188158045 | rs11707430 | 0.189 | 0.983 | -1.436 | 0.92 | -0.072 | -0.078 | -0.078 | 0 | 0.13 | 0.01 | 0.31 | 0.3 | 0.22 | 0.14 |
| 3 | 188173145 | rs12492561 | 0.397 | 0.431 | -0.984 | 0.972 | -0.003 | 0.042 | 0.042 | 0.264 | 0.1 | 0.11 | 0.09 | 0.28 | 0.08 | 0.25 |
| 3 | 188181560 | rs3732913 | -0.405 | 0.225 | -0.856 | 0.555 | 0 | 0 | 0 | 0 | 0.4 | 0.59 | 0.47 | 0.54 | 0.42 | 0.43 |
| 3 | 188192294 | rs6444295 | 0.519 | 0.709 | -0.566 | 1.036 | 0.035 | 0.032 | 0.032 | 0 | 0.09 | 0.11 | 0.25 | 0.3 | 0.19 | 0.1 |
| 3 | 188192386 | rs6788663 | -0.328 | -0.076 | -0.473 | 0.684 | 0.003 | 0.003 | 0.003 | 0 | 0.41 | 0.4 | 0.33 | 0.39 | 0.34 | 0.34 |
| 3 | 188197515 | rs919079 | -0.266 | 0 | -0.432 | 0.717 | 0.005 | 0.004 | 0.004 | 0 | 0.4 | 0.39 | 0.32 | 0.39 | 0.34 | 0.33 |
| 3 | 188199624 | rs9830597 | 0.268 | 0.179 | -0.162 | 1.196 | 0.003 | -0.003 | -0.003 | 0 | 0.21 | 0.18 | 0.17 | 0.24 | 0.19 | 0.16 |
| 3 | 188200897 | rs1974968 | 0.16 | 0.406 | 0.329 | 1.495 | -0.014 | 0.028 | 0.028 | 0 | 0.77 | 0.65 | 0.83 | 0.77 | 0.8 | 0.72 |
| 3 | 188203731 | rs6762273 | -1.19 | 0.276 | 0.526 | 1.529 | -0.065 | 0.092 | 0.092 | 6.823 | 0.19 | 0.53 | 0.14 | 0.29 | 0.06 | 0.15 |

| | | | | | | | | | | | | | | | | |
|---|---|---|---|---|---|---|---|---|---|---|---|---|---|---|---|---|
| 3 | 188205217 | rs4686964 | -1.322 | 1.091 | -0.299 | 1.014 | -0.024 | 0.207 | 0.207 | 0 | 0.47 | 0.8 | 0.51 | 0.84 | 0.51 | 0.52 |
| 3 | 188206259 | rs7638476 | -0.923 | 0.208 | 0.472 | 1.337 | -0.061 | 0.098 | 0.098 | 0.516 | 0.2 | 0.49 | 0.17 | 0.32 | 0.1 | 0.18 |
| 3 | 188206842 | rs2162261 | 1.312 | 0.802 | -0.226 | 0.856 | 0 | 0.242 | 0.242 | 0.057 | 0.54 | 0.07 | 0.51 | 0.16 | 0.6 | 0.52 |
| 3 | 188210104 | rs4541397 | 1.182 | 0.782 | 0.882 | 2.062 | -0.023 | 0.273 | 0.273 | 0 | 0.88 | 0.62 | 0.86 | 0.45 | 0.91 | 0.71 |
| 3 | 188211745 | rs2114236 | -0.989 | 1.036 | 0.717 | 1.402 | 0.007 | 0.004 | 0.004 | 5.998 | 0.05 | 0.25 | 0.02 | 0 | 0 | 0.04 |
| 3 | 188224549 | rs7428265 | -1.032 | 1.861 | 0.076 | 1.623 | -0.007 | -0.003 | -0.003 | 0 | 0.11 | 0.07 | 0.14 | 0.1 | 0.11 | 0.13 |
| 3 | 188226359 | rs10937349 | -0.924 | 2.075 | 0.074 | 1.999 | 0.004 | 0.054 | 0.054 | 2.84 | 0.1 | 0.09 | 0.23 | 0.48 | 0.19 | 0.34 |
| 3 | 188229396 | rs9879182 | -1.334 | 1.739 | 0.254 | 1.78 | 0.015 | 0.08 | 0.08 | 1.19 | 0.17 | 0.3 | 0.32 | 0.47 | 0.21 | 0.35 |
| 3 | 188232012 | rs9874528 | -0.686 | 2.091 | -0.185 | 0.633 | -0.013 | 0.011 | 0.011 | 0 | 0.16 | 0.08 | 0.12 | 0.01 | 0.18 | 0.14 |
| 3 | 188232680 | rs16863396 | 0.773 | 2.157 | 0.01 | 0.683 | 0.04 | 0.005 | 0.005 | 0 | 0.7 | 0.51 | 0.58 | 0.34 | 0.66 | 0.57 |
| 3 | 188233404 | rs6807897 | -0.69 | 1.613 | 0.077 | 0.063 | 0.027 | -0.011 | -0.011 | 0 | 0.12 | 0.3 | 0.1 | 0.07 | 0.09 | 0.03 |
| 3 | 188236356 | rs4686969 | 1.122 | 0.946 | -0.519 | -0.188 | 0.021 | 0.034 | 0.034 | 1.211 | 0.71 | 0.23 | 0.79 | 0.82 | 0.78 | 0.71 |
| 3 | 188241115 | rs933843 | -0.425 | 0.028 | -1.197 | -1.216 | 0.037 | 0.083 | 0.083 | 0 | 0.73 | 0.85 | 0.78 | 0.94 | 0.79 | 0.82 |
| 3 | 188241949 | rs890348 | -0.622 | -0.2 | -1.072 | -1.46 | 0.013 | 0.011 | 0.011 | 0 | 0.77 | 0.78 | 0.8 | 0.94 | 0.83 | 0.82 |
| 3 | 188244609 | rs6444300 | -0.649 | -0.217 | -1.075 | -1.451 | 0.022 | 0.028 | 0.028 | 0 | 0.77 | 0.8 | 0.79 | 0.94 | 0.83 | 0.82 |
| 3 | 188246257 | rs11920292 | 0.57 | -0.14 | -0.8 | -2.253 | -0.007 | 0.009 | 0.009 | 0 | 0.43 | 0.35 | 0.5 | 0.92 | 0.46 | 0.52 |
| 3 | 188248624 | rs11709891 | 0.62 | -0.113 | -0.85 | -2.273 | -0.009 | 0.021 | 0.021 | 0 | 0.43 | 0.33 | 0.5 | 0.92 | 0.47 | 0.52 |
| 3 | 188249417 | rs2081550 | 1.095 | -0.74 | -1.052 | -2.69 | -0.086 | 0.103 | 0.103 | 0 | 0.31 | 0.15 | 0.46 | 0.92 | 0.42 | 0.45 |
| 3 | 188260703 | rs1426260 | 0.206 | -0.606 | -0.476 | -2.545 | -0.005 | -0.001 | -0.001 | 0 | 0.44 | 0.59 | 0.53 | 0.92 | 0.47 | 0.52 |
| 3 | 188274288 | rs13088424 | 0.71 | -0.585 | -0.348 | -2.656 | 0.007 | -0.101 | -0.101 | 0.016 | 0.19 | 0.33 | 0.2 | 0.01 | 0.21 | 0.12 |
| 3 | 188279625 | rs6793410 | 0.095 | -0.585 | -0.352 | -2.472 | 0 | -0.115 | -0.115 | 0 | 0.33 | 0.44 | 0.3 | 0.02 | 0.36 | 0.3 |
| 3 | 188280046 | rs13092374 | -1.18 | -0.262 | -0.464 | -2.181 | -0.007 | -0.023 | -0.023 | 0 | 0.75 | 0.72 | 0.79 | 0.94 | 0.82 | 0.83 |
| 3 | 188289697 | rs10937355 | 0.578 | 0.96 | -0.45 | -1.223 | -0.007 | -0.01 | -0.01 | 0.741 | 0.23 | 0.16 | 0.28 | 0.7 | 0.25 | 0.34 |
| 3 | 188290243 | rs2037121 | -1.03 | 0.916 | -0.46 | -1.261 | 0.008 | 0.02 | 0.02 | 3.44 | 0.57 | 0.62 | 0.57 | 0.72 | 0.59 | 0.67 |
| 3 | 188291761 | rs7632060 | -0.503 | 0.891 | -0.471 | -1.375 | 0 | -0.049 | -0.049 | 0.104 | 0.34 | 0.36 | 0.29 | 0.02 | 0.34 | 0.33 |
| 3 | 188292489 | rs10937356 | 0.291 | 1.176 | -1.019 | -1.216 | -0.008 | -0.016 | -0.016 | 0.552 | 0.64 | 0.72 | 0.7 | 0.98 | 0.56 | 0.64 |
| 3 | 188295053 | rs1515424 | 0.311 | 0.639 | -1.205 | -0.205 | -0.008 | -0.014 | -0.014 | 0 | 0.59 | 0.42 | 0.58 | 0.51 | 0.51 | 0.61 |
| 3 | 188298166 | rs1515426 | -0.085 | -0.246 | -1.903 | -0.538 | -0.068 | 0.096 | 0.096 | 0 | 0.38 | 0.08 | 0.43 | 0.27 | 0.52 | 0.34 |
| 3 | 188299793 | rs6444303 | -0.135 | -0.585 | -1.07 | -0.738 | -0.011 | -0.029 | -0.029 | 0.054 | 0.6 | 0.5 | 0.71 | 0.86 | 0.7 | 0.69 |
| 3 | 188302773 | rs4686979 | 0.691 | 0.005 | -0.924 | -0.349 | -0.031 | 0.006 | 0.006 | 0 | 0.41 | 0.54 | 0.34 | 0.47 | 0.25 | 0.46 |
| 3 | 188302851 | rs4686980 | -0.225 | 0.773 | -0.82 | -0.276 | 0.009 | 0.009 | 0.009 | 0 | 0.64 | 0.58 | 0.56 | 0.54 | 0.62 | 0.71 |
| 3 | 188315629 | rs6807988 | -1.079 | 1.966 | -1.172 | -0.749 | -0.036 | 0.049 | 0.049 | 0 | 0.25 | 0.17 | 0.26 | 0.07 | 0.35 | 0.22 |
| 3 | 188320064 | rs9837401 | -0.393 | 2.15 | -1.42 | -0.624 | -0.118 | 0.099 | 0.099 | 0.081 | 0.19 | 0.01 | 0.18 | 0.04 | 0.3 | 0.18 |
| 3 | 188323593 | rs7615767 | -0.733 | 1.593 | -1.013 | -0.693 | -0.006 | -0.026 | -0.026 | 0 | 0.33 | 0.31 | 0.34 | 0.07 | 0.47 | 0.25 |
| 3 | 188327461 | rs1059380 | 0.362 | 1.412 | -1.129 | -0.19 | -0.008 | 0.004 | 0.004 | 0 | 0.61 | 0.73 | 0.56 | 0.44 | 0.57 | 0.58 |
| 3 | 188327683 | rs3732909 | -1.213 | 0.817 | -0.116 | -0.484 | -0.016 | -0.004 | -0.004 | 0 | 0.14 | 0.35 | 0.1 | 0.08 | 0.08 | 0.28 |
| 3 | 188332618 | rs9873800 | -0.12 | 0.835 | -0.318 | -0.906 | 0.005 | 0.017 | 0.017 | 0 | 0.49 | 0.44 | 0.47 | 0.25 | 0.41 | 0.58 |
| 3 | 188336370 | rs4686986 | 0.649 | 0.91 | -0.503 | -1.137 | 0.065 | 0.088 | 0.088 | 3.572 | 0.33 | 0.13 | 0.34 | 0.16 | 0.22 | 0.37 |
| 3 | 188338763 | rs9877898 | 0.296 | 1.227 | -0.228 | -1.815 | 0.013 | -0.019 | -0.019 | 0 | 0.34 | 0.5 | 0.36 | 0.77 | 0.39 | 0.32 |
| 3 | 188341678 | rs4686487 | 0.917 | 0.606 | -0.492 | -1.832 | 0.065 | 0.1 | 0.1 | 0.573 | 0.35 | 0.14 | 0.35 | 0.16 | 0.23 | 0.37 |
| 3 | 188343729 | rs6790135 | -0.794 | 0.689 | -0.528 | -1.814 | -0.016 | -0.111 | -0.111 | 0 | 0.76 | 0.7 | 0.8 | 1 | 0.79 | 0.91 |
| 3 | 188352914 | rs721397 | 0.713 | 0.409 | -1.023 | -1.4 | 0.06 | 0.09 | 0.09 | 3.692 | 0.33 | 0.06 | 0.35 | 0.17 | 0.22 | 0.39 |
| 3 | 188367252 | rs1818553 | -0.309 | 0.884 | -0.471 | -0.677 | -0.009 | -0.011 | -0.011 | 0.734 | 0.17 | 0.13 | 0.24 | 0.76 | 0.25 | 0.18 |
| 3 | 188370583 | rs10937363 | 0.472 | 0.09 | 0.349 | -0.997 | -0.039 | -0.053 | -0.053 | 0.618 | 0.85 | 0.57 | 0.92 | 0.96 | 0.95 | 0.88 |
| 3 | 188378771 | rs12488441 | 0.296 | -0.243 | 0.579 | -1.059 | -0.056 | -0.035 | -0.035 | 0 | 0.87 | 0.66 | 0.94 | 0.95 | 0.97 | 0.89 |
| 3 | 188381903 | rs12485376 | 1.699 | 0.021 | 0.177 | -1.277 | -0.033 | 0.133 | 0.133 | 0 | 0.57 | 0.13 | 0.58 | 0.18 | 0.57 | 0.61 |
| 3 | 188392304 | rs1356095 | -0.166 | 0.089 | 0.554 | -1.158 | -0.04 | -0.024 | -0.024 | 8.244 | 0.89 | 0.72 | 0.95 | 0.96 | 0.98 | 0.89 |
| 3 | 188393053 | rs7646105 | -1.034 | 0.618 | 0.129 | -0.63 | 0.035 | -0.008 | -0.008 | 0 | 0.29 | 0.43 | 0.35 | 0.78 | 0.4 | 0.3 |
| 3 | 188397584 | rs11710948 | -0.66 | -0.144 | -0.316 | -0.97 | -0.013 | -0.061 | -0.061 | 0.716 | 0.19 | 0.19 | 0.1 | 0.01 | 0.15 | 0.05 |
| 3 | 188398655 | rs4686491 | -0.625 | -0.022 | -0.43 | -0.285 | -0.04 | 0.036 | 0.036 | 0 | 0.28 | 0.42 | 0.4 | 0.6 | 0.24 | 0.46 |
| 3 | 188403035 | rs17607619 | -0.468 | 0.969 | -0.677 | -0.245 | -0.005 | 0.003 | 0.003 | 0 | 0.09 | 0.07 | 0.1 | 0.05 | 0.16 | 0.06 |
| 3 | 188405299 | rs9859863 | 0.242 | 0.905 | -1.163 | -0.25 | -0.038 | -0.038 | -0.038 | 0 | 0.11 | 0.04 | 0.19 | 0.2 | 0.18 | 0.19 |
| 3 | 188407867 | rs11719436 | 0.111 | 0.243 | -0.879 | -0.354 | 0.01 | -0.039 | -0.039 | 0.493 | 0.07 | 0.18 | 0.04 | 0 | 0.08 | 0.01 |
| 3 | 188408718 | rs11916786 | 0.723 | -0.111 | -1.437 | -0.51 | 0.013 | 0.009 | 0.009 | 1.521 | 0.64 | 0.36 | 0.55 | 0.3 | 0.68 | 0.55 |
| 3 | 188409087 | rs16863529 | 0.393 | 0.186 | -1.781 | 0.205 | -0.003 | 0.062 | 0.062 | 0 | 0.11 | 0.1 | 0.11 | 0.46 | 0.09 | 0.11 |
| 3 | 188421979 | rs17670991 | 0.322 | 0.606 | -2.118 | 0.292 | -0.026 | -0.068 | -0.068 | 0 | 0.13 | 0.01 | 0.25 | 0.24 | 0.13 | 0.22 |
| 3 | 188430230 | rs1152843 | -0.357 | 1.489 | -2.141 | -0.436 | -0.119 | -0.066 | -0.066 | 0 | 0.18 | 0.03 | 0.3 | 0.26 | 0.34 | 0.3 |
| 3 | 188436575 | rs4686492 | -0.647 | 0.705 | -2.022 | -0.188 | 0.089 | 0.122 | 0.122 | 0.052 | 0.51 | 0.72 | 0.5 | 0.74 | 0.38 | 0.56 |
| 3 | 188449052 | rs925564 | -0.507 | 1.808 | -1.837 | 0.838 | -0.058 | -0.048 | -0.048 | 0 | 0.21 | 0.11 | 0.48 | 0.65 | 0.35 | 0.38 |
| 3 | 188457831 | rs16863564 | -0.394 | 0.999 | -1.764 | 0.655 | 0.047 | 0.058 | 0.058 | 0.991 | 0.13 | 0.03 | 0.04 | 0.02 | 0.04 | 0.14 |
| 3 | 188469617 | rs6762043 | -0.649 | 0.783 | -1.229 | 0.33 | 0.007 | -0.031 | -0.031 | 0 | 0.06 | 0.16 | 0.04 | 0 | 0.07 | 0.02 |
| 3 | 188476787 | rs1398646 | -0.61 | 0.265 | -1.325 | 0.166 | 0.008 | -0.007 | -0.007 | 0 | 0.14 | 0.1 | 0.04 | 0 | 0.04 | 0.14 |
| 3 | 188477590 | rs3817462 | -0.872 | 0.819 | -1.495 | 0.826 | -0.013 | -0.02 | -0.02 | 0.016 | 0.08 | 0.05 | 0.06 | 0.14 | 0.11 | 0.22 |
| 3 | 188479543 | rs13083450 | -0.028 | 0.328 | -2.37 | 0.297 | -0.015 | -0.036 | -0.036 | 0.013 | 0.3 | 0.08 | 0.37 | 0.71 | 0.31 | 0.16 |
| 3 | 188480881 | rs1355720 | -0.523 | 0.531 | -1.917 | -0.712 | 0.001 | -0.043 | -0.043 | 0 | 0.54 | 0.42 | 0.65 | 0.96 | 0.54 | 0.71 |
| 3 | 188485240 | rs1040033 | 0.648 | 0.395 | -1.567 | -0.344 | 0.016 | -0.031 | -0.031 | 0.849 | 0.28 | 0.17 | 0.34 | 0.65 | 0.22 | 0.13 |
| 3 | 188498272 | rs11920167 | 1.061 | 0.585 | -1.403 | -1.216 | -0.02 | 0.057 | 0.057 | 0 | 0.24 | 0.13 | 0.18 | 0.01 | 0.3 | 0.12 |
| 3 | 188500658 | rs3846183 | -1.931 | 0.919 | -0.603 | -1.959 | 0.002 | -0.02 | -0.02 | 0 | 0.64 | 0.69 | 0.75 | 0.92 | 0.66 | 0.83 |
| 3 | 188507780 | rs12495383 | -0.573 | 0.961 | -0.152 | -2.181 | -0.026 | -0.025 | -0.025 | 0 | 0.33 | 0.38 | 0.46 | 0.78 | 0.19 | 0.45 |

| | | | | | | | | | | | | | | | | |
|---|---|---|---|---|---|---|---|---|---|---|---|---|---|---|---|---|
| 3 | 188508886 | rs3898140 | -2.221 | 1.267 | -0.17 | -2.195 | 0.006 | 0.005 | 0.005 | 0 | 0.63 | 0.7 | 0.76 | 0.92 | 0.64 | 0.83 |
| 3 | 188510214 | rs6810359 | -0.439 | 1.66 | -0.212 | -2.087 | -0.001 | -0.007 | -0.007 | 0 | 0.07 | 0.07 | 0.05 | 0.01 | 0.07 | 0.07 |
| 3 | 188520252 | rs3856919 | -2.142 | 1.478 | -0.043 | -2.179 | 0.024 | 0.098 | 0.098 | 0.01 | 0.61 | 0.72 | 0.75 | 0.87 | 0.64 | 0.81 |
| 3 | 188539861 | rs4686997 | -0.287 | 1.369 | 0.593 | -1.939 | 0.022 | 0.024 | 0.024 | 0 | 0.28 | 0.18 | 0.46 | 0.81 | 0.19 | 0.43 |
| 3 | 188552852 | rs6444332 | -0.757 | 1.515 | 0.78 | -1.932 | -0.031 | -0.062 | -0.062 | 0.498 | 0.29 | 0.38 | 0.53 | 0.85 | 0.22 | 0.5 |
| 3 | 188555882 | rs4686998 | -1.689 | 1.8 | 0.799 | -1.656 | -0.007 | 0.083 | 0.083 | 0 | 0.46 | 0.63 | 0.68 | 0.85 | 0.52 | 0.63 |
| 3 | 188561017 | rs3914270 | -2.582 | 1.816 | 0.647 | -1.722 | 0.018 | 0.127 | 0.127 | 0.018 | 0.55 | 0.72 | 0.73 | 0.87 | 0.6 | 0.76 |
| 3 | 188566306 | rs9861473 | -0.829 | 1.996 | 1.02 | -1.341 | 0.099 | -0.003 | -0.003 | 0 | 0.1 | 0.25 | 0.14 | 0.05 | 0.21 | 0.17 |
| 3 | 188574366 | rs3915462 | -1.916 | 1.947 | 0.914 | -1.532 | -0.002 | 0.005 | 0.005 | 0 | 0.46 | 0.53 | 0.67 | 0.85 | 0.52 | 0.63 |
| 3 | 188574996 | rs1513181 | -0.206 | 1.405 | 0.562 | -1.387 | 0 | 0 | 0 | 0 | 0.19 | 0.17 | 0.46 | 0.81 | 0.15 | 0.4 |
| 3 | 188581424 | rs4525856 | -0.302 | 1.423 | 0.511 | -1.363 | 0.003 | 0.002 | 0.002 | 0 | 0.2 | 0.16 | 0.46 | 0.81 | 0.17 | 0.41 |
| 3 | 188584679 | rs13067593 | 0.002 | 1.668 | 0.482 | -1.414 | -0.002 | 0.018 | 0.018 | 0.154 | 0.17 | 0.07 | 0.08 | 0 | 0.16 | 0.05 |
| 3 | 188586447 | rs12632821 | 0.593 | 1.965 | 0.643 | -1.269 | -0.062 | 0.061 | 0.061 | 0 | 0.75 | 0.9 | 0.85 | 1 | 0.65 | 0.84 |
| 3 | 188589606 | rs3796280 | 1.875 | -0.143 | 1.18 | -0.621 | -0.004 | 0.033 | 0.033 | 0.015 | 0.47 | 0.33 | 0.4 | 0.38 | 0.53 | 0.43 |
| 3 | 188590446 | rs1136644 | 1.94 | -0.156 | 1.023 | -0.733 | -0.011 | 0.008 | 0.008 | 0.051 | 0.47 | 0.26 | 0.39 | 0.43 | 0.53 | 0.43 |
| 3 | 188590650 | rs2278096 | -0.101 | -0.048 | 0.834 | -0.625 | -0.012 | 0.089 | 0.089 | 0 | 0.22 | 0.33 | 0.39 | 0.53 | 0.16 | 0.37 |
| 3 | 188599369 | rs4686496 | 1.845 | -0.044 | 0.896 | -0.573 | -0.006 | 0.006 | 0.006 | 0 | 0.49 | 0.29 | 0.4 | 0.46 | 0.55 | 0.49 |
| 3 | 188600300 | rs6793523 | -0.038 | -0.064 | 0.181 | -0.666 | -0.01 | 0.083 | 0.083 | 0 | 0.22 | 0.34 | 0.4 | 0.49 | 0.15 | 0.39 |
| 3 | 188602952 | rs3796283 | 0.644 | -0.212 | 0.136 | -0.32 | -0.001 | 0.001 | 0.001 | 0 | 0.54 | 0.5 | 0.45 | 0.5 | 0.59 | 0.5 |
| 3 | 188605426 | rs9290883 | 1.035 | -0.289 | 0.127 | -0.305 | -0.001 | -0.001 | -0.001 | 0.037 | 0.51 | 0.44 | 0.43 | 0.5 | 0.55 | 0.47 |
| 3 | 188609999 | rs16863691 | 0.65 | -0.916 | 0.269 | 0.074 | -0.001 | 0.002 | 0.002 | 0 | 0.58 | 0.56 | 0.5 | 0.5 | 0.65 | 0.55 |
| 3 | 188621549 | rs9290884 | 0.971 | -1.802 | 0.659 | -0.404 | -0.029 | -0.117 | -0.117 | 0 | 0.65 | 0.53 | 0.79 | 0.96 | 0.78 | 0.72 |
| 3 | 188625888 | rs6797432 | -0.815 | -1.982 | 0.375 | -0.465 | -0.011 | -0.071 | -0.071 | 0 | 0.26 | 0.29 | 0.09 | 0 | 0.15 | 0.08 |
| 3 | 188638678 | rs10937377 | 0.821 | -2.265 | -0.535 | 0.014 | 0.124 | 0.225 | 0.225 | 0.721 | 0.63 | 0.11 | 0.78 | 0.93 | 0.85 | 0.76 |
| 3 | 188639324 | rs4687006 | 0.972 | -2.242 | -0.7 | 0.299 | 0.116 | 0.094 | 0.094 | 0.765 | 0.61 | 0.06 | 0.76 | 0.75 | 0.84 | 0.67 |
| 3 | 188653768 | rs13060586 | 1.729 | -1.569 | 0.782 | 1.048 | -0.036 | 0.031 | 0.031 | 2.006 | 0.89 | 0.29 | 0.88 | 0.88 | 0.97 | 0.91 |
| 3 | 188657297 | rs16863752 | -1.613 | -1.497 | 1.51 | 1.014 | 0.017 | 0.12 | 0.12 | 0.379 | 0.05 | 0.25 | 0.06 | 0.1 | 0.01 | 0.08 |
| 3 | 188673378 | rs6444341 | -1.885 | -1.784 | 1.781 | 0.997 | -0.011 | 0.042 | 0.042 | 0.029 | 0.06 | 0.21 | 0.03 | 0.09 | 0.01 | 0.08 |
| 3 | 188675732 | rs4687010 | 2.562 | -1.999 | 2.076 | 0.952 | -0.026 | -0.001 | -0.001 | 0 | 0.85 | 0.17 | 0.88 | 0.84 | 0.98 | 0.86 |
| 3 | 188682282 | rs16863812 | -3.05 | -2.066 | 2.348 | 0.93 | -0.066 | -0.001 | -0.001 | 0.072 | 0.14 | 0.68 | 0.1 | 0.16 | 0.02 | 0.14 |
| 3 | 188683430 | rs2378458 | -0.194 | -2.592 | 1.293 | 1.184 | -0.028 | -0.112 | -0.112 | 0.212 | 0.28 | 0.02 | 0.37 | 0.37 | 0.27 | 0.45 |
| 3 | 188689987 | rs16863823 | -2.333 | -2.241 | 2.158 | 1.04 | -0.062 | 0.061 | 0.061 | 1.05 | 0.12 | 0.5 | 0.07 | 0.16 | 0.02 | 0.13 |
| 3 | 188694749 | rs710493 | 2.625 | -2.125 | 1.569 | 1.186 | 0.003 | -0.002 | -0.002 | 0 | 0.86 | 0.16 | 0.89 | 0.82 | 0.98 | 0.83 |
| 3 | 188696071 | rs10490883 | -2.126 | -1.981 | 2.062 | 1.06 | -0.004 | 0.092 | 0.092 | 0.804 | 0.08 | 0.41 | 0.07 | 0.1 | 0.01 | 0.09 |
| 3 | 188699808 | rs13092581 | -0.356 | -1.89 | 0.549 | 1.373 | -0.007 | -0.121 | -0.121 | 0 | 0.26 | 0.01 | 0.36 | 0.4 | 0.22 | 0.4 |
| 3 | 188741418 | rs6787392 | -2.049 | -1.232 | 1.681 | 1.101 | -0.014 | 0.052 | 0.052 | 0 | 0.07 | 0.23 | 0.04 | 0.12 | 0.02 | 0.1 |
| 3 | 188757737 | rs710511 | 2.582 | -1.239 | 2.42 | 1 | 0 | 0.051 | 0.051 | 0.177 | 0.91 | 0.51 | 0.93 | 0.9 | 0.95 | 0.89 |
| 3 | 188758301 | rs710512 | 2.448 | -1.244 | 2.53 | 0.998 | 0.012 | 0.063 | 0.063 | 0 | 0.91 | 0.6 | 0.93 | 0.89 | 0.96 | 0.9 |
| 3 | 188766239 | rs9840231 | -2.323 | -1.245 | 2.64 | 1.065 | 0.01 | 0.069 | 0.069 | 0 | 0.07 | 0.25 | 0.11 | 0.12 | 0.04 | 0.1 |
| 3 | 188781109 | rs3914986 | -2.791 | -1.249 | 2.626 | 1.081 | -0.002 | 0.071 | 0.071 | 0.199 | 0.09 | 0.47 | 0.12 | 0.12 | 0.05 | 0.11 |
| 3 | 188791804 | rs6780254 | -2.192 | -1.154 | 1.91 | 1.006 | -0.007 | 0.002 | 0.002 | 0 | 0.06 | 0.17 | 0.11 | 0.08 | 0.06 | 0.08 |
| 3 | 188825770 | rs2378469 | -1.131 | -1.919 | 1.488 | 1.359 | -0.019 | 0.013 | 0.013 | 0.061 | 0.33 | 0.59 | 0.48 | 0.49 | 0.24 | 0.43 |
| 3 | 188829698 | rs2378472 | -1.25 | -1.705 | 1.305 | 1.324 | 0.001 | -0.002 | -0.002 | 0 | 0.34 | 0.65 | 0.49 | 0.49 | 0.26 | 0.46 |
| 3 | 188860551 | rs9815495 | -0.549 | -1.614 | 0.838 | 1.379 | -0.004 | 0.012 | 0.012 | 0.093 | 0.35 | 0.35 | 0.51 | 0.41 | 0.31 | 0.44 |
| 3 | 188868715 | rs1052437 | -0.514 | -1.775 | 0.998 | 1.521 | 0.001 | 0 | 0 | 0 | 0.33 | 0.29 | 0.48 | 0.28 | 0.32 | 0.43 |
| 3 | 188868753 | rs1052439 | -0.647 | -1.839 | 1.386 | 1.492 | -0.006 | -0.004 | -0.004 | 5.514 | 0.34 | 0.37 | 0.49 | 0.28 | 0.32 | 0.45 |
| 3 | 188878471 | rs6766978 | -0.828 | -1.865 | 2.326 | 1.195 | -0.011 | -0.022 | -0.022 | 0.115 | 0.27 | 0.7 | 0.32 | 0.07 | 0.16 | 0.22 |
| 3 | 188879064 | rs7616752 | -1.642 | -2.037 | 1.683 | 1.039 | -0.008 | 0.014 | 0.014 | 1.395 | 0.48 | 0.92 | 0.57 | 0.36 | 0.42 | 0.62 |
| 3 | 188881482 | rs9290889 | -0.653 | -1.893 | 1.776 | 1.088 | -0.006 | -0.007 | -0.007 | 0 | 0.22 | 0.17 | 0.25 | 0.29 | 0.26 | 0.39 |
| 3 | 188884425 | rs16864045 | -0.093 | -2.05 | 1.158 | 1.439 | 0.007 | -0.011 | -0.011 | 0.042 | 0.09 | 0.05 | 0.05 | 0.19 | 0.08 | 0.06 |
| 3 | 188886628 | rs4687025 | -1.418 | -2.299 | 1.294 | 0.977 | -0.06 | -0.005 | -0.005 | 0 | 0.29 | 0.58 | 0.3 | 0.24 | 0.16 | 0.3 |
| 3 | 188890300 | rs9834801 | -0.719 | -1.554 | 1.215 | 0.874 | 0.001 | -0.001 | -0.001 | 0 | 0.36 | 0.35 | 0.43 | 0.38 | 0.37 | 0.52 |
| 3 | 188898647 | rs4686505 | -2.656 | -1.765 | 1.234 | 0.835 | -0.062 | -0.03 | -0.03 | 2.076 | 0.63 | 0.87 | 0.64 | 0.62 | 0.58 | 0.73 |
| 3 | 188909868 | rs4686507 | -1.924 | -1.775 | 1.316 | 0.842 | -0.021 | 0.002 | 0.002 | 3.436 | 0.53 | 0.8 | 0.59 | 0.45 | 0.49 | 0.63 |
| 3 | 188911398 | rs1584576 | 0.123 | -1.846 | 1.183 | 0.854 | -0.015 | -0.016 | -0.016 | 0 | 0.22 | 0.15 | 0.24 | 0.4 | 0.31 | 0.39 |
| 3 | 188916053 | rs6765588 | -2.803 | -1.804 | 0.957 | 0.762 | -0.126 | -0.078 | -0.078 | 2.811 | 0.64 | 0.95 | 0.64 | 0.63 | 0.58 | 0.73 |
| 3 | 188918464 | rs7635429 | -0.373 | -2.052 | 1.606 | 0.425 | -0.027 | -0.027 | -0.027 | 0 | 0.16 | 0.43 | 0.15 | 0.07 | 0.12 | 0.12 |
| 3 | 188919056 | rs16864083 | -0.024 | -2.059 | 1.539 | 0.422 | -0.015 | -0.012 | -0.012 | 0.118 | 0.14 | 0.22 | 0.13 | 0.07 | 0.11 | 0.12 |
| 3 | 188923369 | rs6783147 | -1.624 | -2.072 | 1.15 | 0.463 | -0.014 | -0.001 | -0.001 | 0 | 0.29 | 0.65 | 0.21 | 0.25 | 0.21 | 0.2 |
| 3 | 188940512 | rs16864109 | 0.251 | -1.546 | -0.132 | 0.509 | 0.031 | 0.035 | 0.035 | 0 | 0.13 | 0.08 | 0.18 | 0.01 | 0.06 | 0.11 |
| 3 | 188941449 | rs16864111 | -0.533 | -1.006 | -0.667 | 0.929 | 0.006 | -0.053 | -0.053 | 0 | 0.11 | 0.01 | 0.04 | 0.18 | 0.08 | 0.08 |
| 3 | 188946496 | rs4687034 | -0.592 | -0.348 | -0.77 | 0.316 | -0.018 | -0.018 | -0.018 | 0 | 0.34 | 0.41 | 0.33 | 0.3 | 0.23 | 0.49 |
| 3 | 188949286 | rs9820421 | -0.643 | 1.035 | -1.136 | 0.151 | -0.015 | 0.018 | 0.018 | 0 | 0.17 | 0.12 | 0.15 | 0.03 | 0.22 | 0.09 |
| 3 | 188954611 | rs6777676 | 0.167 | 1.047 | -2.045 | 0.234 | -0.163 | -0.031 | -0.031 | 0 | 0.67 | 0.95 | 0.54 | 0.76 | 0.52 | 0.53 |
| 3 | 188956297 | rs7627629 | 0.744 | 0.302 | -1.173 | 0.359 | 0.006 | -0.001 | -0.001 | 0 | 0.46 | 0.63 | 0.41 | 0.58 | 0.38 | 0.27 |
| 3 | 188956787 | rs7640523 | 0.357 | -0.394 | -0.825 | -0.37 | 0 | -0.045 | -0.045 | 0.057 | 0.09 | 0.33 | 0.05 | 0 | 0.05 | 0 |
| 3 | 188965222 | rs9831552 | -1.072 | -0.348 | -1.136 | -0.348 | 0.016 | 0.055 | 0.055 | 0 | 0.64 | 0.8 | 0.59 | 0.26 | 0.68 | 0.61 |
| 3 | 188969148 | rs6777323 | -0.836 | -0.15 | -0.325 | 0.288 | 0.002 | -0.003 | -0.003 | 0 | 0.21 | 0.22 | 0.12 | 0.24 | 0.19 | 0.22 |
| 3 | 188976010 | rs6801108 | -1.418 | -0.243 | -0.439 | 0.119 | 0.018 | 0 | 0 | 0.737 | 0.26 | 0.77 | 0.19 | 0.26 | 0.21 | 0.27 |

| | | | | | | | | | | | | | | | | |
|---|---|---|---|---|---|---|---|---|---|---|---|---|---|---|---|---|
| 3 | 189000601 | rs2176038 | -1.222 | -0.248 | -0.117 | 0.116 | -0.009 | 0.001 | 0.001 | 1.104 | 0.25 | 0.64 | 0.18 | 0.26 | 0.2 | 0.27 |
| 3 | 189004818 | rs9682650 | 0.236 | -0.122 | -0.39 | 0.192 | -0.017 | -0.032 | -0.032 | 0 | 0.09 | 0.01 | 0.05 | 0.14 | 0.13 | 0.14 |
| 3 | 189010687 | rs9875124 | -1.124 | -0.105 | -0.58 | -0.275 | 0.043 | 0.069 | 0.069 | 0 | 0.52 | 0.72 | 0.43 | 0.29 | 0.58 | 0.47 |
| 3 | 189010993 | rs1877935 | 0.591 | 0.056 | -0.138 | -0.412 | -0.021 | -0.011 | -0.011 | 0 | 0.27 | 0.19 | 0.22 | 0 | 0.36 | 0.19 |
| 3 | 189014219 | rs6769057 | -0.449 | 0.007 | -0.091 | -0.321 | -0.032 | -0.01 | -0.01 | 0 | 0.16 | 0.62 | 0.12 | 0.11 | 0.08 | 0.12 |
| 3 | 189020311 | rs7640560 | 0.593 | 0.106 | 0.098 | -0.554 | -0.002 | -0.009 | -0.009 | 0 | 0.09 | 0.05 | 0.09 | 0.18 | 0.13 | 0.15 |
| 3 | 189027921 | rs9820403 | 0.048 | -0.182 | 0.124 | -0.363 | -0.023 | -0.068 | -0.068 | 1.096 | 0.12 | 0.26 | 0.04 | 0 | 0.06 | 0.05 |
| 3 | 189029220 | rs6784306 | -0.279 | -0.004 | -0.561 | -0.087 | 0.01 | 0.046 | 0.046 | 0 | 0.49 | 0.58 | 0.36 | 0.18 | 0.55 | 0.39 |
| 3 | 189032004 | rs9822201 | -0.523 | -0.071 | -1.298 | -0.035 | 0.035 | 0.137 | 0.137 | 0 | 0.52 | 0.75 | 0.4 | 0.21 | 0.56 | 0.46 |
| 3 | 189034821 | rs10513835 | -0.882 | 0.344 | -0.919 | 0.503 | -0.006 | 0.001 | 0.001 | 0 | 0.14 | 0.1 | 0.12 | 0.08 | 0.12 | 0.16 |
| 3 | 189037348 | rs9820955 | -1.147 | -0.503 | -0.928 | 0.89 | -0.04 | 0.036 | 0.036 | 0 | 0.2 | 0.35 | 0.18 | 0.28 | 0.09 | 0.31 |
| 3 | 189038648 | rs1055152 | -0.754 | -0.328 | -2.239 | 1.04 | 0.063 | 0 | 0 | 0 | 0.47 | 0.88 | 0.54 | 0.62 | 0.35 | 0.61 |
| 3 | 189039611 | rs9851974 | -0.904 | 0.178 | -1.282 | 1.358 | 0.003 | 0.009 | 0.009 | 0.051 | 0.1 | 0.1 | 0.16 | 0.3 | 0.06 | 0.19 |
| 3 | 189044503 | rs6793853 | -0.282 | 0.312 | -1.814 | 0.508 | 0.045 | 0.133 | 0.133 | 2.211 | 0.52 | 0.77 | 0.57 | 0.8 | 0.58 | 0.56 |
| 3 | 189044675 | rs4686510 | 0.21 | 0.475 | -1.655 | 0.959 | 0.01 | -0.031 | -0.031 | 0 | 0.65 | 0.74 | 0.64 | 0.53 | 0.68 | 0.61 |
| 3 | 189045268 | rs9872797 | 0.281 | 0.939 | -1.193 | 0.632 | 0.003 | -0.002 | -0.002 | 0 | 0.59 | 0.5 | 0.33 | 0.4 | 0.45 | 0.56 |
| 3 | 189045487 | rs1606789 | 0.619 | 0.925 | -0.696 | 0.361 | -0.008 | -0.045 | -0.045 | 0 | 0.14 | 0.22 | 0.11 | 0.01 | 0.14 | 0.04 |
| 3 | 189047108 | rs16848815 | -0.171 | 1.486 | -0.652 | 0.781 | -0.004 | -0.003 | -0.003 | 0 | 0.1 | 0.1 | 0.2 | 0.16 | 0.14 | 0.09 |
| 3 | 189048860 | rs2063795 | 0.948 | 1.223 | -1.106 | 0.58 | -0.003 | -0.053 | -0.053 | 2.834 | 0.19 | 0.06 | 0.09 | 0.31 | 0.18 | 0.26 |
| 3 | 189049768 | rs9857394 | -0.686 | 1.492 | 0.004 | 0.812 | 0.001 | 0.016 | 0.016 | 0 | 0.36 | 0.41 | 0.39 | 0.46 | 0.4 | 0.47 |
| 3 | 189050664 | rs13076888 | -0.152 | 1.058 | -0.193 | 0.36 | -0.043 | -0.026 | -0.026 | 1.589 | 0.1 | 0.02 | 0.19 | 0.14 | 0.12 | 0.05 |
| 3 | 189052464 | rs10513837 | -0.587 | 0.415 | -0.02 | 0.249 | 0 | -0.034 | -0.034 | 0 | 0.26 | 0.28 | 0.2 | 0.01 | 0.25 | 0.16 |
| 3 | 189054068 | rs13099998 | -0.193 | 0.56 | -0.356 | 0.415 | -0.004 | -0.01 | -0.01 | 0.478 | 0.09 | 0.07 | 0.06 | 0.17 | 0.09 | 0.06 |
| 3 | 189057350 | rs1399615 | -1.803 | -0.092 | -0.394 | 0.557 | 0.059 | -0.005 | -0.005 | 0.087 | 0.49 | 0.9 | 0.36 | 0.44 | 0.37 | 0.36 |
| 3 | 189059421 | rs16864260 | -0.381 | 0.144 | -0.127 | 0.853 | -0.001 | -0.001 | -0.001 | 0.006 | 0.09 | 0.08 | 0.07 | 0.15 | 0.09 | 0.09 |
| 3 | 189061076 | rs13067334 | -1.05 | -0.048 | 0.22 | 0.796 | -0.004 | 0.034 | 0.034 | 0.095 | 0.13 | 0.27 | 0.09 | 0.19 | 0.1 | 0.1 |
| 3 | 189062649 | rs1022304 | -1.89 | -0.094 | 0.543 | 0.79 | 0.011 | -0.004 | -0.004 | 0 | 0.41 | 0.58 | 0.29 | 0.48 | 0.31 | 0.31 |
| 3 | 189065057 | rs16864263 | -1.519 | 0.002 | 0.742 | 0.856 | 0.013 | 0.223 | 0.223 | 0.356 | 0.09 | 0.35 | 0.08 | 0.32 | 0.06 | 0.1 |
| 3 | 189065156 | rs13059863 | -1.691 | -0.096 | 0.55 | 0.836 | -0.013 | 0.022 | 0.022 | 0.03 | 0.33 | 0.54 | 0.25 | 0.52 | 0.26 | 0.25 |
| 3 | 189069410 | rs1399613 | -0.55 | 0.335 | -0.138 | 0.398 | 0.012 | -0.028 | -0.028 | 0.359 | 0.16 | 0.08 | 0.14 | 0.29 | 0.13 | 0.18 |
| 3 | 189075327 | rs6797146 | -1.325 | 0.639 | -0.154 | 0.407 | -0.009 | -0.01 | -0.01 | 0 | 0.09 | 0.09 | 0.21 | 0.33 | 0.11 | 0.24 |
| 3 | 189075833 | rs6775310 | -0.944 | 0.567 | -0.175 | -0.516 | 0.011 | 0.051 | 0.051 | 0 | 0.21 | 0.12 | 0.15 | 0.03 | 0.17 | 0.11 |
| 3 | 189077920 | rs4687042 | -2.173 | 0.929 | 0.044 | -0.518 | 0.002 | -0.013 | -0.013 | 0.552 | 0.37 | 0.42 | 0.39 | 0.21 | 0.39 | 0.37 |
| 3 | 189080480 | rs3846201 | -1.075 | 0.951 | -0.191 | -0.438 | -0.009 | -0.014 | -0.014 | 0 | 0.11 | 0.12 | 0.2 | 0.19 | 0.12 | 0.27 |
| 3 | 189082892 | rs2360692 | -1.091 | 0.995 | -0.175 | -0.5 | 0 | -0.001 | -0.001 | 0 | 0.22 | 0.23 | 0.2 | 0.19 | 0.23 | 0.28 |
| 3 | 189084461 | rs7630774 | -0.326 | 0.307 | 0.104 | -0.327 | -0.003 | 0.013 | 0.013 | 0 | 0.06 | 0.07 | 0.16 | 0.03 | 0.06 | 0.09 |
| 3 | 189084917 | rs13073419 | -0.814 | 0.332 | 0.176 | -0.366 | 0.007 | -0.026 | -0.026 | 0.257 | 0.2 | 0.33 | 0.15 | 0.16 | 0.2 | 0.24 |
| 3 | 189086515 | rs7626770 | -1.223 | 0.317 | 0.151 | -0.323 | 0.006 | -0.04 | -0.04 | 2.407 | 0.27 | 0.48 | 0.32 | 0.2 | 0.26 | 0.34 |
| 3 | 189088573 | rs182009 | 0.915 | 0.343 | -0.094 | -0.75 | 0.007 | 0.017 | 0.017 | 0 | 0.56 | 0.35 | 0.62 | 0.83 | 0.64 | 0.55 |
| 3 | 189101446 | rs264146 | 0.702 | 0.328 | 0.229 | -0.711 | -0.002 | -0.064 | -0.064 | 0.038 | 0.65 | 0.43 | 0.65 | 0.84 | 0.7 | 0.62 |
| 3 | 189106781 | rs16864324 | -0.096 | 0.192 | -0.1 | -0.063 | 0.013 | -0.038 | -0.038 | 1.531 | 0.08 | 0.02 | 0.02 | 0.17 | 0.03 | 0.06 |
| 3 | 189107451 | rs260555 | 0.794 | 0.122 | 0.519 | -0.329 | 0 | -0.023 | -0.023 | 0 | 0.63 | 0.43 | 0.6 | 0.79 | 0.64 | 0.65 |
| 3 | 189107467 | rs260556 | -0.376 | 0.109 | 1.115 | -0.474 | 0.001 | -0.059 | -0.059 | 1.227 | 0.81 | 0.68 | 0.86 | 0.93 | 0.84 | 0.81 |
| 3 | 189110132 | rs6778546 | -1.096 | 0.159 | 0.939 | -0.449 | -0.001 | -0.043 | -0.043 | 0 | 0.21 | 0.29 | 0.26 | 0.13 | 0.21 | 0.18 |
| 3 | 189112946 | rs260560 | 1.623 | 0.145 | 0.683 | -0.347 | 0.008 | 0.025 | 0.025 | 0 | 0.54 | 0.36 | 0.57 | 0.63 | 0.61 | 0.59 |
| 3 | 189119792 | rs12152518 | 0.617 | 0.183 | 0.436 | 0.039 | -0.004 | 0.01 | 0.01 | 9.457 | 0.09 | 0.04 | 0.17 | 0.07 | 0.09 | 0.12 |
| 3 | 189120128 | rs7639459 | -0.899 | 0.155 | 0.471 | 0.041 | 0.011 | -0.028 | -0.028 | 0.268 | 0.17 | 0.12 | 0.19 | 0.24 | 0.12 | 0.18 |
| 3 | 189121959 | rs13085813 | -0.091 | 1.013 | 0.436 | -0.452 | -0.029 | 0.021 | 0.021 | 5.628 | 0.19 | 0.1 | 0.24 | 0.09 | 0.27 | 0.19 |
| 3 | 189125964 | rs260579 | -0.251 | 0.758 | 0.663 | -0.745 | 0.006 | -0.014 | -0.014 | 0 | 0.81 | 0.8 | 0.88 | 1 | 0.84 | 0.94 |
| 3 | 189127707 | rs260580 | 1.711 | 0.553 | 0.973 | -0.697 | 0.003 | 0.126 | 0.126 | 0.031 | 0.51 | 0.24 | 0.53 | 0.74 | 0.48 | 0.55 |
| 3 | 189131433 | rs11707115 | 0.142 | 0.684 | 1.039 | -0.701 | -0.033 | 0.023 | 0.023 | 5.457 | 0.19 | 0.1 | 0.24 | 0.09 | 0.27 | 0.21 |
| 3 | 189133577 | rs6775595 | -0.498 | 0.681 | 1.362 | -0.708 | 0.076 | -0.023 | -0.023 | 1.402 | 0.23 | 0.46 | 0.3 | 0.18 | 0.32 | 0.36 |
| 3 | 189134681 | rs260581 | 1.28 | 0.684 | 1.102 | -0.663 | -0.018 | 0.067 | 0.067 | 0.625 | 0.57 | 0.31 | 0.58 | 0.8 | 0.52 | 0.57 |
| 3 | 189148605 | rs155069 | 0.718 | 0.417 | 1.034 | -0.691 | 0.002 | -0.016 | -0.016 | 0.154 | 0.62 | 0.37 | 0.58 | 0.85 | 0.55 | 0.75 |
| 3 | 189150107 | rs13073685 | -0.031 | 0.542 | 0.583 | -0.596 | 0.016 | -0.006 | -0.006 | 0 | 0.2 | 0.27 | 0.29 | 0.15 | 0.29 | 0.2 |
| 3 | 189159352 | rs6771859 | -1.026 | 0.405 | -0.081 | -0.453 | -0.009 | 0.077 | 0.077 | 0 | 0.47 | 0.7 | 0.49 | 0.25 | 0.52 | 0.42 |
| 3 | 189164574 | rs17491469 | 0.359 | 0.515 | -0.069 | -0.42 | -0.002 | 0.003 | 0.003 | 0 | 0.13 | 0.12 | 0.1 | 0.02 | 0.16 | 0.03 |
| 3 | 189165194 | rs3846206 | 0.08 | 0.17 | -0.071 | -0.429 | -0.003 | -0.001 | -0.001 | 0.563 | 0.82 | 0.76 | 0.63 | 0.87 | 0.86 | 0.71 |
| 3 | 189165356 | rs3909572 | 0.059 | 0.174 | -0.095 | -0.427 | 0 | 0 | 0 | 0 | 0.82 | 0.81 | 0.63 | 0.87 | 0.86 | 0.71 |
| 3 | 189170062 | rs9834913 | 1.011 | -0.333 | -0.364 | -0.568 | -0.008 | 0.004 | 0.004 | 0.42 | 0.4 | 0.5 | 0.25 | 0.62 | 0.36 | 0.25 |
| 3 | 189170348 | rs9855126 | 1.074 | -0.196 | -0.737 | -0.527 | 0.007 | 0.019 | 0.019 | 0 | 0.45 | 0.25 | 0.27 | 0.66 | 0.42 | 0.44 |
| 3 | 189172796 | rs4687045 | 0.443 | 0.342 | -0.956 | -0.458 | -0.055 | 0.008 | 0.008 | 0.849 | 0.19 | 0.06 | 0.25 | 0.15 | 0.31 | 0.24 |
| 3 | 189190614 | rs11706039 | 0.838 | 0.179 | -1.349 | -0.663 | -0.088 | 0.034 | 0.034 | 0.461 | 0.25 | 0.03 | 0.2 | 0.15 | 0.34 | 0.26 |
| 3 | 189194905 | rs1152857 | 0.436 | -0.484 | -0.958 | -0.559 | -0.003 | -0.003 | -0.003 | 0 | 0.16 | 0.07 | 0.3 | 0.1 | 0.16 | 0.22 |
| 3 | 189195811 | rs6765410 | -0.171 | -0.606 | -0.31 | -0.542 | -0.013 | 0.004 | 0.004 | 0 | 0.81 | 0.68 | 0.66 | 0.85 | 0.83 | 0.77 |
| 3 | 189199930 | rs7630698 | 0.523 | -0.809 | -0.271 | -0.79 | -0.007 | 0.065 | 0.065 | 0.644 | 0.43 | 0.29 | 0.43 | 0.16 | 0.56 | 0.37 |
| 3 | 189203007 | rs9850625 | -0.54 | -0.647 | -0.325 | -0.74 | -0.002 | 0 | 0 | 0.062 | 0.81 | 0.79 | 0.67 | 0.86 | 0.83 | 0.77 |
| 3 | 189206601 | rs1869112 | 0.336 | -0.482 | -0.433 | -0.485 | -0.012 | -0.002 | -0.002 | 0 | 0.37 | 0.42 | 0.24 | 0.7 | 0.26 | 0.4 |
| 3 | 189216167 | rs2600846 | 0.952 | -0.144 | -0.507 | -0.079 | 0.005 | 0.027 | 0.027 | 0.034 | 0.2 | 0.24 | 0.32 | 0.4 | 0.23 | 0.19 |

| | | | | | | | | | | | | | | | | |
|---|---|---|---|---|---|---|---|---|---|---|---|---|---|---|---|---|
| 3 | 189219130 | rs2014894 | -0.169 | -0.473 | -1.042 | -0.469 | 0.015 | 0 | 0 | 0.382 | 0.42 | 0.35 | 0.31 | 0.46 | 0.36 | 0.51 |
| 3 | 189221832 | rs9809592 | 0.041 | -0.712 | -1.287 | -0.714 | 0.003 | -0.01 | -0.01 | 0 | 0.56 | 0.26 | 0.53 | 0.57 | 0.58 | 0.72 |
| 3 | 189221904 | rs17493690 | 0.86 | -0.317 | -1.455 | -1.071 | -0.112 | 0.114 | 0.114 | 0.703 | 0.32 | 0.05 | 0.33 | 0.12 | 0.41 | 0.3 |
| 3 | 189223024 | rs2552362 | -0.025 | -0.523 | -0.397 | -0.505 | -0.012 | -0.089 | -0.089 | 0.617 | 0.8 | 0.73 | 0.89 | 1 | 0.81 | 0.92 |
| 3 | 189227476 | rs2046693 | -0.074 | -0.854 | -0.474 | -0.02 | -0.035 | 0.025 | 0.025 | 0 | 0.62 | 0.51 | 0.66 | 0.44 | 0.77 | 0.62 |
| 3 | 189229935 | rs3856927 | 1.272 | -0.322 | -1.139 | -0.64 | -0.101 | 0.144 | 0.144 | 0 | 0.22 | 0.02 | 0.32 | 0.01 | 0.27 | 0.13 |
| 3 | 189230204 | rs1364593 | -0.86 | -0.61 | -0.91 | -0.602 | -0.021 | 0.029 | 0.029 | 0.079 | 0.66 | 0.56 | 0.67 | 0.44 | 0.78 | 0.64 |
| 3 | 189230306 | rs6785685 | -0.296 | -0.722 | -0.353 | -0.667 | 0.024 | -0.016 | -0.016 | 0 | 0.6 | 0.34 | 0.64 | 0.44 | 0.77 | 0.61 |
| 3 | 189230922 | rs6444380 | 0.045 | -0.708 | -0.299 | -0.793 | 0.01 | 0.03 | 0.03 | 0.064 | 0.36 | 0.45 | 0.31 | 0.42 | 0.45 | 0.47 |
| 3 | 189232641 | rs10513838 | -0.182 | -0.762 | -0.305 | -0.715 | -0.015 | -0.02 | -0.02 | 0 | 0.11 | 0.13 | 0.05 | 0.04 | 0.06 | 0.08 |
| 3 | 189236003 | rs9849231 | -1.318 | -0.948 | 0 | -0.356 | 0.002 | 0.002 | 0.002 | 0.152 | 0.7 | 0.71 | 0.75 | 0.74 | 0.81 | 0.72 |
| 3 | 189237651 | rs6769971 | 0.489 | -0.969 | 0.079 | -0.332 | 0.005 | -0.02 | -0.02 | 0 | 0.33 | 0.44 | 0.43 | 0.28 | 0.33 | 0.19 |
| 3 | 189239521 | rs1895663 | -0.394 | -1.09 | -0.007 | -0.308 | 0.026 | -0.012 | -0.012 | 0 | 0.61 | 0.34 | 0.66 | 0.48 | 0.79 | 0.65 |
| 3 | 189245214 | rs7617035 | 0.175 | -1.318 | -0.026 | -0.504 | 0.001 | 0.017 | 0.017 | 0.201 | 0.52 | 0.41 | 0.51 | 0.69 | 0.54 | 0.66 |
| 3 | 189245356 | rs16864503 | 1.049 | -1.296 | -0.069 | -0.501 | -0.004 | -0.009 | -0.009 | 0 | 0.08 | 0.06 | 0.03 | 0.01 | 0.04 | 0.06 |
| 3 | 189245545 | rs6797709 | 0.287 | -1.119 | -0.15 | -0.373 | 0.005 | -0.001 | -0.001 | 0.981 | 0.49 | 0.33 | 0.47 | 0.43 | 0.53 | 0.61 |
| 3 | 189245593 | rs17429138 | 1.378 | -0.901 | -0.572 | -0.266 | -0.109 | 0.03 | 0.03 | 1.275 | 0.21 | 0.02 | 0.22 | 0.09 | 0.37 | 0.35 |
| 3 | 189246758 | rs6776619 | 0.253 | -0.772 | -0.22 | -0.234 | -0.002 | -0.021 | -0.021 | 7.246 | 0.4 | 0.5 | 0.47 | 0.33 | 0.37 | 0.27 |
| 3 | 189260625 | rs1364596 | 0.109 | -0.68 | -0.505 | -0.355 | -0.01 | 0.005 | 0.005 | 0.752 | 0.34 | 0.27 | 0.32 | 0.67 | 0.46 | 0.49 |
| 3 | 189261856 | rs16864553 | 1.662 | -1.089 | -0.737 | -0.596 | 0.057 | 0.087 | 0.087 | 1.092 | 0.2 | 0.06 | 0.18 | 0.02 | 0.09 | 0.14 |
| 3 | 189284257 | rs11720501 | 1.728 | -1.284 | -0.9 | -0.807 | -0.09 | 0.157 | 0.157 | 0.281 | 0.25 | 0.04 | 0.33 | 0.03 | 0.3 | 0.17 |
| 3 | 189286891 | rs7615734 | -0.378 | -1.455 | -0.24 | -0.716 | -0.022 | -0.015 | -0.015 | 0 | 0.66 | 0.56 | 0.71 | 0.73 | 0.81 | 0.73 |
| 3 | 189287175 | rs16864619 | 1.854 | -1.378 | -0.425 | -0.69 | 0.067 | 0.082 | 0.082 | 1.556 | 0.19 | 0.05 | 0.17 | 0.02 | 0.07 | 0.14 |
| 3 | 189296353 | rs1920244 | -0.749 | -1.411 | -0.081 | -0.607 | -0.019 | 0 | 0 | 0.359 | 0.7 | 0.57 | 0.72 | 0.69 | 0.83 | 0.7 |
| 3 | 189298408 | rs3856776 | 1.282 | -0.85 | -0.203 | -0.809 | -0.068 | 0.103 | 0.103 | 0 | 0.32 | 0.17 | 0.47 | 0.1 | 0.51 | 0.28 |
| 3 | 189301594 | rs2141610 | -0.848 | -1.269 | -0.024 | -0.769 | -0.035 | -0.009 | -0.009 | 0 | 0.71 | 0.54 | 0.73 | 0.73 | 0.84 | 0.73 |
| 3 | 189303711 | rs2378502 | -0.681 | -1.017 | 0.018 | -0.732 | 0.001 | -0.009 | -0.009 | 0 | 0.45 | 0.43 | 0.37 | 0.67 | 0.53 | 0.53 |
| 3 | 189303939 | rs1920246 | -0.796 | -0.963 | 0.008 | -0.698 | 0.052 | 0.147 | 0.147 | 0 | 0.89 | 0.63 | 0.88 | 0.72 | 0.89 | 0.84 |
| 3 | 189304472 | rs6787246 | -0.477 | -0.965 | 0.046 | -0.682 | -0.009 | 0.02 | 0.02 | 2.245 | 0.42 | 0.36 | 0.37 | 0.71 | 0.53 | 0.53 |
| 3 | 189320167 | rs6792756 | 0.454 | -1.18 | 0.069 | -0.229 | -0.018 | 0.205 | 0.205 | 0 | 0.13 | 0.32 | 0.1 | 0.37 | 0.07 | 0.17 |
| 3 | 189323685 | rs6444389 | -0.788 | -1.231 | -0.081 | -0.398 | 0.008 | -0.001 | -0.001 | 0 | 0.47 | 0.34 | 0.43 | 0.95 | 0.6 | 0.67 |
| 3 | 189330551 | rs9882796 | -0.16 | -1.61 | 0.079 | -0.359 | -0.013 | 0.001 | 0.001 | 0 | 0.61 | 0.38 | 0.71 | 0.63 | 0.81 | 0.66 |
| 3 | 189331804 | rs9812901 | 0.592 | -1.226 | 0.131 | -0.184 | -0.011 | -0.011 | -0.011 | 0 | 0.27 | 0.21 | 0.27 | 0.37 | 0.39 | 0.32 |
| 3 | 189333335 | rs1920286 | 0.552 | -1.274 | 0.073 | -0.242 | -0.009 | -0.011 | -0.011 | 0 | 0.2 | 0.17 | 0.27 | 0.42 | 0.35 | 0.32 |
| 3 | 189333966 | rs1920285 | 0.976 | -1.381 | 0.008 | -0.133 | -0.009 | 0.052 | 0.052 | 2.204 | 0.59 | 0.22 | 0.62 | 0.26 | 0.53 | 0.52 |
| 3 | 189336473 | rs4687076 | -0.437 | -1.324 | 0.034 | 0.131 | 0.016 | 0.361 | 0.361 | 0 | 0.88 | 0.5 | 0.91 | 0.6 | 0.93 | 0.83 |
| 3 | 189340384 | rs13085076 | 1.546 | -1.273 | -0.093 | 0.121 | 0 | 0.068 | 0.068 | 0 | 0.08 | 0.08 | 0.05 | 0.32 | 0.07 | 0.14 |
| 3 | 189341616 | rs16864664 | 1.427 | -1.261 | -0.23 | -0.216 | -0.004 | -0.017 | -0.017 | 0 | 0.06 | 0.06 | 0.03 | 0 | 0.05 | 0.03 |
| 3 | 189343424 | rs7610017 | -0.491 | -1.317 | -0.316 | -0.227 | -0.013 | -0.04 | -0.04 | 0 | 0.91 | 0.63 | 0.94 | 1 | 0.95 | 0.97 |
| 3 | 189344011 | rs4398410 | 0.625 | -1.223 | -0.349 | -0.156 | -0.001 | -0.001 | -0.001 | 0 | 0.28 | 0.26 | 0.27 | 0.3 | 0.38 | 0.28 |
| 3 | 189344149 | rs6790258 | 1.199 | -1.178 | -0.377 | -0.037 | 0.002 | 0.176 | 0.176 | 0 | 0.09 | 0.18 | 0.06 | 0.4 | 0.07 | 0.17 |
| 3 | 189348411 | rs7631358 | 0.839 | -1.171 | -0.364 | -0.184 | 0.001 | 0.161 | 0.161 | 0 | 0.11 | 0.19 | 0.08 | 0.44 | 0.09 | 0.2 |
| 3 | 189356941 | rs7636839 | -0.207 | -1.105 | -1.047 | -0.308 | -0.01 | -0.01 | -0.01 | 0 | 0.39 | 0.28 | 0.36 | 0.52 | 0.49 | 0.47 |
| 3 | 189375694 | rs9830662 | 0.243 | -1.133 | -1.127 | -0.337 | -0.023 | 0.072 | 0.072 | 2.168 | 0.59 | 0.21 | 0.62 | 0.27 | 0.52 | 0.51 |
| 3 | 189379493 | rs1920273 | 1.064 | -1.138 | -1.428 | -0.36 | 0.015 | 0.263 | 0.263 | 1.043 | 0.47 | 0.17 | 0.54 | 0.06 | 0.42 | 0.33 |
| 3 | 189383183 | rs10937405 | 0.218 | -1.145 | -1.078 | 0.062 | -0.005 | 0.001 | 0.001 | 0.016 | 0.33 | 0.27 | 0.29 | 0.31 | 0.43 | 0.32 |
| 3 | 189383430 | rs4600802 | 0.029 | -0.331 | -0.734 | 0.291 | 0.009 | 0.321 | 0.321 | 0 | 0.87 | 0.46 | 0.9 | 0.58 | 0.91 | 0.8 |
| 3 | 189389626 | rs1920280 | 0.314 | -0.011 | -1.607 | -0.399 | -0.065 | 0.086 | 0.086 | 0.242 | 0.36 | 0.14 | 0.45 | 0.23 | 0.41 | 0.34 |
| 3 | 189391823 | rs9812619 | 0.264 | 0.039 | -1.585 | -0.377 | -0.064 | 0.092 | 0.092 | 0.409 | 0.37 | 0.14 | 0.45 | 0.23 | 0.41 | 0.34 |
| 3 | 189394372 | rs7649341 | 0.156 | 0.376 | -1.734 | -0.043 | 0.004 | 0.029 | 0.029 | 0.357 | 0.47 | 0.2 | 0.5 | 0.57 | 0.52 | 0.49 |
| 3 | 189394427 | rs13080141 | 1.972 | 0.171 | -1.844 | 0.294 | -0.003 | 0.003 | 0.003 | 0 | 0.06 | 0.04 | 0.01 | 0.14 | 0.01 | 0.09 |
| 3 | 189395680 | rs13097576 | 0.452 | 0.727 | -2.171 | -0.161 | -0.007 | -0.072 | -0.072 | 0 | 0.28 | 0.09 | 0.26 | 0.57 | 0.27 | 0.33 |
| 3 | 189400282 | rs2378510 | 0.956 | 0.222 | -1.616 | -0.521 | -0.025 | -0.054 | -0.054 | 0.224 | 0.31 | 0.5 | 0.34 | 0.14 | 0.25 | 0.31 |
| 3 | 189408144 | rs1950084 | 0.813 | 0.393 | -1.813 | -0.478 | 0 | -0.021 | -0.021 | 0 | 0.14 | 0.07 | 0.07 | 0.26 | 0.12 | 0.17 |
| 3 | 189409196 | rs2889918 | 1.085 | 0.356 | -1.84 | -0.298 | -0.145 | -0.029 | -0.029 | 0 | 0.29 | 0.02 | 0.3 | 0.25 | 0.37 | 0.3 |
| 3 | 189409341 | rs7618742 | 1.291 | -0.17 | -0.991 | -0.06 | 0.002 | 0.001 | 0.001 | 0 | 0.22 | 0.17 | 0.27 | 0.07 | 0.15 | 0.14 |
| 3 | 189411830 | rs17445954 | 1.901 | -0.346 | -1.234 | -0.093 | 0.02 | 0.038 | 0.038 | 0 | 0.17 | 0.05 | 0.13 | 0.06 | 0.13 | 0.08 |
| 3 | 189416115 | rs1920238 | 0.617 | -0.055 | -1.238 | -0.024 | -0.029 | -0.069 | -0.069 | 0 | 0.28 | 0.1 | 0.25 | 0.57 | 0.3 | 0.33 |
| 3 | 189417208 | rs10513840 | 0.568 | -0.225 | -0.683 | -0.173 | -0.059 | -0.072 | -0.072 | 0 | 0.28 | 0.52 | 0.31 | 0.07 | 0.17 | 0.15 |
| 3 | 189421319 | rs4575879 | -0.533 | 0.615 | -0.582 | 0.547 | 0.005 | -0.009 | -0.009 | 0.599 | 0.38 | 0.43 | 0.34 | 0.64 | 0.38 | 0.49 |
| 3 | 189424634 | rs9839197 | 0.937 | 0.626 | -1.09 | 0.564 | -0.003 | 0.001 | 0.001 | 0 | 0.06 | 0.04 | 0.01 | 0.13 | 0.01 | 0.09 |
| 3 | 189432432 | rs11915751 | 0.166 | 0.922 | -1.065 | 0.631 | 0.002 | 0.016 | 0.016 | 0.405 | 0.64 | 0.51 | 0.64 | 0.59 | 0.67 | 0.63 |
| 3 | 189435199 | rs13095039 | -0.089 | 1.335 | -1.134 | 0.742 | -0.01 | -0.007 | -0.007 | 0 | 0.18 | 0.14 | 0.3 | 0.25 | 0.27 | 0.35 |
| 3 | 189437794 | rs9815203 | 0.278 | 1.208 | -1.353 | 0.721 | -0.052 | -0.032 | -0.032 | 1.89 | 0.19 | 0.06 | 0.29 | 0.25 | 0.25 | 0.37 |
| 3 | 189438979 | rs4687090 | -0.128 | 1.187 | -1.089 | 0.432 | 0.005 | 0.002 | 0.002 | 0.3 | 0.58 | 0.5 | 0.56 | 0.49 | 0.57 | 0.48 |
| 3 | 189441242 | rs11711203 | 0.637 | 0.929 | -1.867 | 0.208 | -0.036 | -0.056 | -0.056 | 0 | 0.27 | 0.05 | 0.27 | 0.35 | 0.31 | 0.32 |
| 3 | 189443562 | rs9942139 | -0.42 | 0.97 | -1.605 | 0.016 | -0.003 | 0.005 | 0.005 | 0.105 | 0.58 | 0.55 | 0.67 | 0.58 | 0.64 | 0.51 |
| 3 | 189448545 | rs4687091 | -0.195 | 0.788 | -0.786 | 0.592 | -0.001 | 0.026 | 0.026 | 0 | 0.32 | 0.5 | 0.48 | 0.5 | 0.35 | 0.34 |
| 3 | 189451290 | rs4607088 | -0.362 | 0.791 | -0.843 | 0.643 | -0.003 | 0.009 | 0.009 | 0 | 0.38 | 0.55 | 0.51 | 0.5 | 0.38 | 0.39 |

| | | | | | | | | | | | | | | | | |
|---|---|---|---|---|---|---|---|---|---|---|---|---|---|---|---|---|
| 3 | 189453482 | rs6777607 | 0.034 | 0.784 | -0.831 | 0.469 | -0.002 | 0.014 | 0.014 | 0 | 0.4 | 0.66 | 0.43 | 0.28 | 0.43 | 0.21 |
| 3 | 189457674 | rs6800495 | 0.479 | 0.73 | -0.477 | 0.335 | 0.001 | 0.001 | 0.001 | 0 | 0.29 | 0.33 | 0.3 | 0.23 | 0.3 | 0.18 |
| 3 | 189462592 | rs6444395 | -0.43 | 0.865 | -0.797 | 0.347 | -0.027 | 0.043 | 0.043 | 0 | 0.66 | 0.8 | 0.7 | 0.78 | 0.57 | 0.65 |
| 3 | 189471639 | rs6794551 | 0.109 | 0.515 | -0.802 | 0.707 | 0.017 | 0.026 | 0.026 | 0.017 | 0.51 | 0.43 | 0.47 | 0.61 | 0.45 | 0.4 |
| 3 | 189481564 | rs6797860 | -0.832 | 0.592 | -0.498 | 1.007 | -0.01 | 0.018 | 0.018 | 0 | 0.31 | 0.36 | 0.29 | 0.44 | 0.3 | 0.39 |
| 3 | 189483649 | rs9871604 | 0.531 | 0.67 | -0.657 | 0.863 | -0.006 | 0.059 | 0.059 | 0 | 0.91 | 0.77 | 0.92 | 0.86 | 0.96 | 0.92 |
| 3 | 189484508 | rs11708278 | -0.479 | 1.184 | -1.552 | 0.499 | -0.07 | 0.032 | 0.032 | 0 | 0.21 | 0.02 | 0.15 | 0.13 | 0.25 | 0.26 |
| 3 | 189489850 | rs9824061 | -0.806 | 0.988 | -0.78 | 0.532 | 0.012 | -0.027 | -0.027 | 0 | 0.07 | 0.15 | 0.12 | 0.05 | 0.09 | 0.08 |
| 3 | 189491376 | rs9830137 | -0.727 | 1.291 | -1.158 | 0.695 | -0.044 | -0.026 | -0.026 | 0 | 0.08 | 0.01 | 0.17 | 0.05 | 0.12 | 0.12 |
| 3 | 189498809 | rs12630584 | -0.414 | 0.91 | -1.041 | 1.217 | 0.005 | -0.002 | -0.002 | 0 | 0.12 | 0.1 | 0.1 | 0.32 | 0.1 | 0.2 |
| 3 | 189498828 | rs17505775 | 0.804 | 0.872 | -2.15 | 0.085 | -0.068 | 0.156 | 0.156 | 0.385 | 0.23 | 0.02 | 0.14 | 0 | 0.26 | 0.07 |
| 3 | 189500067 | rs7650285 | 0.197 | 0.876 | -1.865 | 0.179 | -0.002 | 0.023 | 0.023 | 0.69 | 0.12 | 0.12 | 0.1 | 0.32 | 0.1 | 0.23 |
| 3 | 189500313 | rs9849766 | 0.616 | 0.645 | -1.291 | -0.264 | -0.001 | -0.017 | -0.017 | 0.417 | 0.37 | 0.41 | 0.21 | 0.15 | 0.32 | 0.32 |
| 3 | 189506531 | rs1464117 | -0.281 | 0.596 | -0.729 | -0.304 | -0.004 | 0.033 | 0.033 | 0.018 | 0.19 | 0.2 | 0.15 | 0.44 | 0.15 | 0.28 |
| 3 | 189508471 | rs9854771 | 0.185 | 0.515 | -0.475 | -0.779 | -0.007 | 0.018 | 0.018 | 0 | 0.33 | 0.25 | 0.46 | 0.14 | 0.39 | 0.25 |
| 3 | 189521319 | rs17506395 | 1.247 | 0.542 | -0.881 | -0.917 | -0.034 | 0.055 | 0.055 | 0.299 | 0.2 | 0.07 | 0.15 | 0 | 0.28 | 0.07 |
| 3 | 189522258 | rs7619549 | 0.782 | -0.049 | -0.584 | -0.711 | -0.008 | 0.075 | 0.075 | 0 | 0.47 | 0.3 | 0.42 | 0.19 | 0.5 | 0.32 |
| 3 | 189528852 | rs4687094 | 0.599 | 0.07 | -0.501 | -0.22 | -0.005 | -0.035 | -0.035 | 0 | 0.59 | 0.49 | 0.67 | 0.2 | 0.61 | 0.51 |
| 3 | 189529969 | rs16864809 | -0.57 | -0.024 | -0.81 | -0.292 | -0.001 | -0.003 | -0.003 | 0.025 | 0.32 | 0.31 | 0.18 | 0.28 | 0.25 | 0.34 |
| 3 | 189530734 | rs16864811 | -0.456 | -0.271 | -0.956 | 0.243 | 0.009 | -0.002 | -0.002 | 0 | 0.21 | 0.14 | 0.13 | 0.27 | 0.12 | 0.19 |
| 3 | 189534583 | rs13075375 | 0.055 | 0.604 | -0.909 | 0.369 | 0.035 | 0.029 | 0.029 | 0 | 0.68 | 0.49 | 0.76 | 0.62 | 0.63 | 0.58 |
| 3 | 189534621 | rs11720810 | 0.254 | 0.304 | -0.958 | 0.285 | -0.001 | -0.001 | -0.001 | 0.058 | 0.65 | 0.38 | 0.75 | 0.62 | 0.62 | 0.58 |
| 3 | 189536660 | rs9881595 | 0.742 | -0.157 | -1.498 | -0.48 | 0.004 | 0.037 | 0.037 | 0 | 0.48 | 0.18 | 0.65 | 0.62 | 0.46 | 0.58 |
| 3 | 189539320 | rs9879356 | -0.151 | -0.443 | -2.066 | -0.568 | -0.001 | -0.033 | -0.033 | 0 | 0.41 | 0.26 | 0.33 | 0.51 | 0.38 | 0.28 |
| 3 | 189548461 | rs17447411 | 1.01 | -0.443 | -2.142 | -0.592 | 0.024 | 0.066 | 0.066 | 0 | 0.25 | 0.11 | 0.41 | 0.09 | 0.23 | 0.35 |
| 3 | 189548862 | rs2037128 | 0.92 | -0.197 | -2.248 | -0.527 | -0.041 | 0.086 | 0.086 | 0 | 0.14 | 0.02 | 0.08 | 0 | 0.16 | 0.05 |
| 3 | 189551420 | rs7616437 | 0.3 | 0.25 | -2.255 | -0.487 | -0.001 | -0.018 | -0.018 | 0 | 0.21 | 0.2 | 0.13 | 0.03 | 0.24 | 0.09 |
| 3 | 189552371 | rs7619556 | 0.21 | 0.103 | -1.935 | -0.107 | -0.016 | -0.014 | -0.014 | 0.671 | 0.45 | 0.27 | 0.37 | 0.55 | 0.46 | 0.31 |
| 3 | 189553287 | rs17447607 | 0.207 | 0.708 | -1.926 | 0.21 | -0.09 | -0.03 | -0.03 | 0 | 0.22 | 0.02 | 0.12 | 0.19 | 0.25 | 0.16 |
| 3 | 189559757 | rs2138244 | 0.288 | 0.181 | -1.253 | 0.363 | -0.013 | 0.002 | 0.002 | 0 | 0.68 | 0.42 | 0.75 | 0.64 | 0.76 | 0.66 |
| 3 | 189560617 | rs13326685 | -0.23 | -0.19 | -1.628 | -0.139 | -0.006 | 0.036 | 0.036 | 0 | 0.46 | 0.38 | 0.52 | 0.33 | 0.49 | 0.43 |
| 3 | 189563530 | rs9817981 | -0.226 | -0.271 | -1.828 | 0.302 | 0.037 | 0.011 | 0.011 | 0 | 0.51 | 0.71 | 0.32 | 0.51 | 0.41 | 0.53 |
| 3 | 189565689 | rs7624251 | -0.902 | 0.193 | -2.086 | 0.406 | 0.023 | -0.025 | -0.025 | 0 | 0.78 | 0.91 | 0.82 | 0.79 | 0.84 | 0.87 |
| 3 | 189570630 | rs13059768 | 0.399 | 0.693 | -1.965 | 0.587 | -0.009 | -0.018 | -0.018 | 5.298 | 0.22 | 0.2 | 0.44 | 0.32 | 0.29 | 0.34 |
| 3 | 189572668 | rs11719667 | 0.602 | 0.39 | -1.982 | 0.034 | 0.004 | 0.051 | 0.051 | 0 | 0.3 | 0.2 | 0.2 | 0.2 | 0.29 | 0.35 |
| 3 | 189579511 | rs17448036 | -0.244 | 0.204 | -1.828 | 0.005 | -0.031 | -0.045 | -0.045 | 1.552 | 0.63 | 0.78 | 0.48 | 0.53 | 0.59 | 0.53 |
| 3 | 189580420 | rs6790791 | 1.686 | -0.137 | -1.813 | 0.012 | -0.033 | -0.01 | -0.01 | 0 | 0.22 | 0.63 | 0.11 | 0.18 | 0.11 | 0.17 |
| 3 | 189583298 | rs3773926 | 2.293 | -0.318 | -0.869 | -0.064 | -0.044 | -0.002 | -0.002 | 0.395 | 0.17 | 0.26 | 0.08 | 0.15 | 0.1 | 0.14 |
| 3 | 189583705 | rs3773928 | 0.469 | -0.212 | -1.049 | -0.079 | 0.013 | 0.014 | 0.014 | 0.045 | 0.47 | 0.66 | 0.36 | 0.38 | 0.43 | 0.3 |
| 3 | 189586030 | rs1399773 | 0.411 | -0.147 | -1.326 | -0.107 | 0.039 | 0.037 | 0.037 | 0.775 | 0.48 | 0.72 | 0.37 | 0.35 | 0.42 | 0.28 |
| 3 | 189587091 | rs6789961 | 0.453 | -0.355 | -1.444 | -0.555 | 0.016 | 0.017 | 0.017 | 0.182 | 0.36 | 0.66 | 0.2 | 0.15 | 0.22 | 0.19 |
| 3 | 189587274 | rs6790167 | -0.591 | -0.263 | -1.448 | -0.519 | -0.021 | -0.011 | -0.011 | 0 | 0.63 | 0.7 | 0.43 | 0.39 | 0.51 | 0.41 |
| 3 | 189588376 | rs1515489 | 1.583 | 0.273 | -0.987 | -0.237 | -0.015 | 0.002 | 0.002 | 0.076 | 0.14 | 0.03 | 0.11 | 0.17 | 0.16 | 0.22 |
| 3 | 189592202 | rs9812089 | 0.079 | 0.303 | -0.952 | -0.228 | -0.009 | -0.002 | -0.002 | 0.031 | 0.59 | 0.66 | 0.43 | 0.36 | 0.56 | 0.36 |
| 3 | 189600160 | rs12107036 | -0.619 | 0.315 | -0.747 | -0.18 | 0.008 | 0.057 | 0.057 | 0 | 0.48 | 0.28 | 0.42 | 0.38 | 0.54 | 0.39 |
| 3 | 189611194 | rs9681004 | 1.025 | 0.383 | -0.535 | -0.22 | -0.001 | -0.003 | -0.003 | 0 | 0.39 | 0.28 | 0.3 | 0.35 | 0.4 | 0.34 |
| 3 | 189615854 | rs11709791 | 1.962 | 0.381 | -0.446 | -0.128 | -0.002 | -0.002 | -0.002 | 0 | 0.26 | 0.23 | 0.15 | 0.16 | 0.21 | 0.2 |
| 3 | 189618496 | rs7628595 | 0.398 | 0.471 | -0.504 | -0.08 | 0.009 | -0.004 | -0.004 | 0 | 0.26 | 0.38 | 0.26 | 0.21 | 0.28 | 0.17 |
| 3 | 189628072 | rs7642848 | -0.195 | 0.567 | -1.046 | -0.052 | -0.028 | -0.015 | -0.015 | 0 | 0.61 | 0.72 | 0.46 | 0.37 | 0.58 | 0.39 |
| 3 | 189631932 | rs4479569 | 0.4 | 0.369 | -1.134 | 0.27 | -0.028 | -0.008 | -0.008 | 0 | 0.23 | 0.15 | 0.22 | 0.19 | 0.28 | 0.14 |
| 3 | 189645933 | rs710521 | 0.38 | 0.387 | -1.111 | 0.281 | -0.023 | -0.005 | -0.005 | 0 | 0.23 | 0.19 | 0.22 | 0.2 | 0.28 | 0.14 |
| 3 | 189647882 | rs12696598 | 1.549 | 0.276 | -0.914 | 0.126 | 0 | 0 | 0 | 0 | 0.29 | 0.27 | 0.42 | 0.19 | 0.32 | 0.29 |
| 3 | 189647906 | rs710525 | 0.245 | 0.41 | -0.774 | -0.126 | -0.002 | 0.001 | 0.001 | 0 | 0.24 | 0.23 | 0.18 | 0.18 | 0.26 | 0.2 |
| 3 | 189651639 | rs710527 | -1.033 | 0.352 | -1.028 | 0.199 | -0.034 | -0.013 | -0.013 | 0.516 | 0.91 | 0.99 | 0.87 | 0.94 | 0.87 | 0.81 |
| 3 | 189655039 | rs6806309 | 0.884 | -0.028 | -0.683 | 0.105 | 0 | 0.003 | 0.003 | 0 | 0.61 | 0.4 | 0.69 | 0.69 | 0.66 | 0.49 |
| 3 | 189659181 | rs698109 | 0.618 | -0.052 | -0.986 | 0.379 | -0.022 | -0.008 | -0.008 | 0.016 | 0.55 | 0.85 | 0.64 | 0.61 | 0.54 | 0.51 |
| 3 | 189660500 | rs4687111 | -0.164 | 0.073 | -0.57 | 0.414 | 0.011 | -0.007 | -0.007 | 0 | 0.74 | 0.66 | 0.75 | 0.75 | 0.71 | 0.65 |
| 3 | 189660809 | rs724438 | 0.925 | -0.031 | -0.505 | 0.428 | -0.003 | 0 | 0 | 0.549 | 0.47 | 0.54 | 0.56 | 0.49 | 0.42 | 0.52 |
| 3 | 189662409 | rs696408 | 0.703 | 0.352 | -0.271 | 0.466 | 0.013 | -0.008 | -0.008 | 0.192 | 0.17 | 0.33 | 0.11 | 0.18 | 0.17 | 0.17 |
| 3 | 189663385 | rs4687112 | 1.154 | -0.284 | -0.435 | 0.562 | -0.001 | -0.032 | -0.032 | 1.579 | 0.35 | 0.27 | 0.49 | 0.43 | 0.37 | 0.32 |
| 3 | 189664617 | rs7641791 | 1.061 | -0.098 | 0.606 | 0.645 | 0.004 | 0.004 | 0.004 | 0 | 0.62 | 0.39 | 0.63 | 0.49 | 0.59 | 0.56 |
| 3 | 189672468 | rs696066 | -0.031 | -0.044 | 0.275 | 0.513 | -0.029 | 0.024 | 0.024 | 0 | 0.9 | 0.99 | 0.89 | 0.97 | 0.9 | 0.81 |
| 3 | 189672558 | rs696067 | 0.03 | 0.027 | 0.737 | 0.789 | -0.028 | 0.019 | 0.019 | 0 | 0.2 | 0.13 | 0.16 | 0.17 | 0.29 | 0.2 |
| 3 | 189680945 | rs837682 | 0.01 | 0.398 | 0.027 | 0.057 | -0.026 | 0.017 | 0.017 | 0 | 0.2 | 0.13 | 0.16 | 0.17 | 0.28 | 0.2 |
| 3 | 189692565 | rs1985690 | 0.623 | -0.308 | 0.132 | -0.427 | -0.003 | 0.045 | 0.045 | 0 | 0.51 | 0.41 | 0.62 | 0.8 | 0.5 | 0.55 |
| 3 | 189693880 | rs13072268 | 0.644 | -0.316 | 0.139 | -0.43 | -0.002 | 0.057 | 0.057 | 0 | 0.51 | 0.4 | 0.61 | 0.8 | 0.5 | 0.55 |
| 3 | 189697935 | rs837767 | -0.186 | -0.347 | 0.586 | -0.512 | -0.02 | -0.074 | -0.074 | 0.046 | 0.73 | 0.6 | 0.77 | 0.93 | 0.78 | 0.77 |
| 3 | 189698779 | rs710552 | -1.391 | -0.466 | 0.452 | -0.518 | 0.017 | 0.024 | 0.024 | 0 | 0.84 | 0.9 | 0.86 | 0.93 | 0.88 | 0.8 |
| 3 | 189701445 | rs710555 | -0.367 | -0.566 | 0.463 | -0.201 | 0.003 | -0.003 | -0.003 | 0 | 0.42 | 0.42 | 0.47 | 0.42 | 0.47 | 0.42 |

| | | | | | | | | | | | | | | | | |
|---|---|---|---|---|---|---|---|---|---|---|---|---|---|---|---|---|
| 3 | 189706124 | rs1040293 | -0.6 | -0.844 | 0.366 | -0.297 | -0.005 | -0.034 | -0.034 | 0.896 | 0.73 | 0.67 | 0.78 | 0.93 | 0.77 | 0.77 |
| 3 | 189708251 | rs9851100 | 1.532 | -0.432 | 0.174 | -0.144 | -0.018 | -0.061 | -0.061 | 0 | 0.29 | 0.15 | 0.31 | 0.49 | 0.3 | 0.34 |
| 3 | 189712010 | rs1719600 | 0.611 | -0.191 | 0.284 | 0.128 | -0.052 | 0.021 | 0.021 | 0 | 0.18 | 0.05 | 0.16 | 0.16 | 0.28 | 0.2 |
| 3 | 189712525 | rs2222442 | 0.313 | -0.195 | 0.346 | 0.127 | -0.017 | 0.012 | 0.012 | 0 | 0.19 | 0.13 | 0.17 | 0.16 | 0.28 | 0.22 |
| 3 | 189715295 | rs710563 | -0.568 | -0.27 | 0.404 | 0.102 | 0.013 | 0.001 | 0.001 | 0 | 0.4 | 0.46 | 0.47 | 0.45 | 0.47 | 0.42 |
| 3 | 189732606 | rs1018343 | 0.282 | 0.035 | 0.046 | -0.184 | -0.024 | 0.022 | 0.022 | 0 | 0.19 | 0.13 | 0.15 | 0.15 | 0.27 | 0.18 |
| 3 | 189733204 | rs710576 | 0.556 | -0.041 | 0.498 | 0.318 | -0.005 | 0.038 | 0.038 | 0 | 0.21 | 0.29 | 0.36 | 0.36 | 0.19 | 0.21 |
| 3 | 189744621 | rs967119 | 0.868 | 0.162 | 0.868 | 0.368 | -0.001 | 0.01 | 0.01 | 0 | 0.29 | 0.31 | 0.28 | 0.42 | 0.26 | 0.35 |
| 3 | 189748884 | rs1349036 | 1.234 | 0.285 | 1.441 | 0.597 | 0.001 | 0.002 | 0.002 | 0 | 0.62 | 0.29 | 0.75 | 0.64 | 0.67 | 0.71 |
| 3 | 189749583 | rs1719625 | -0.092 | 0.224 | 1.732 | 0.523 | 0.031 | -0.015 | -0.015 | 0.459 | 0.84 | 0.8 | 0.86 | 0.85 | 0.78 | 0.79 |
| 3 | 189751821 | rs754981 | 1.552 | 0.234 | 1.468 | 0.536 | -0.001 | 0.001 | 0.001 | 0 | 0.62 | 0.21 | 0.73 | 0.63 | 0.67 | 0.7 |
| 3 | 189755231 | rs1666390 | -0.145 | 0.296 | 1.2 | 0.55 | -0.007 | 0.004 | 0.004 | 0 | 0.85 | 0.87 | 0.86 | 0.85 | 0.78 | 0.8 |
| 3 | 189758098 | rs2594173 | 1.439 | 0.328 | 0.997 | 0.567 | -0.013 | -0.009 | -0.009 | 0 | 0.62 | 0.18 | 0.74 | 0.63 | 0.67 | 0.7 |
| 3 | 189764268 | rs2594174 | -0.018 | 0.598 | 1.062 | 0.469 | 0.004 | -0.002 | -0.002 | 0.112 | 0.84 | 0.86 | 0.86 | 0.87 | 0.77 | 0.8 |
| 3 | 189768681 | rs1447939 | -0.03 | 0.606 | 1.013 | 0.472 | -0.011 | 0.007 | 0.007 | 0 | 0.85 | 0.89 | 0.88 | 0.87 | 0.77 | 0.8 |
| 3 | 189771201 | rs10937417 | -1.352 | 0.399 | 1.509 | 0.524 | 0.002 | -0.005 | -0.005 | 0.09 | 0.36 | 0.55 | 0.24 | 0.31 | 0.34 | 0.28 |
| 3 | 189775242 | rs13096237 | -1.15 | 0.293 | 1.393 | 0.376 | -0.014 | 0.044 | 0.044 | 0.077 | 0.32 | 0.29 | 0.18 | 0.09 | 0.32 | 0.25 |
| 3 | 189778883 | rs1526031 | -0.99 | 0.347 | 1.52 | 0.56 | 0.04 | 0.067 | 0.067 | 0 | 0.2 | 0.3 | 0.12 | 0.34 | 0.21 | 0.25 |
| 3 | 189781195 | rs7636581 | -0.312 | 0.343 | 1.771 | 0.578 | -0.029 | -0.09 | -0.09 | 0 | 0.18 | 0.4 | 0.12 | 0 | 0.11 | 0.05 |
| 3 | 189788068 | rs11917702 | -0.024 | 0.349 | 1.662 | 0.582 | -0.021 | -0.071 | -0.071 | 3.977 | 0.17 | 0.36 | 0.11 | 0 | 0.11 | 0.03 |
| 3 | 189794491 | rs6789122 | -0.165 | 0.583 | 1.202 | 0.797 | -0.062 | 0.007 | 0.007 | 0 | 0.12 | 0.02 | 0.09 | 0.09 | 0.19 | 0.18 |
| 3 | 189795575 | rs10937420 | -0.918 | 0.572 | 1.347 | 0.867 | 0.071 | 0.114 | 0.114 | 0 | 0.16 | 0.32 | 0.12 | 0.29 | 0.19 | 0.22 |
| 3 | 189801991 | rs7630486 | -1.676 | 0.169 | 1 | 0.991 | 0.064 | 0.371 | 0.371 | 0.037 | 0.05 | 0.37 | 0.04 | 0.2 | 0.01 | 0.06 |
| 3 | 189809163 | rs710580 | -0.963 | 0.465 | -0.34 | 0.312 | -0.006 | 0.081 | 0.081 | 1.976 | 0.39 | 0.78 | 0.27 | 0.19 | 0.39 | 0.26 |
| 3 | 189814339 | rs3106807 | -0.315 | 0.366 | -0.772 | 0.682 | -0.021 | -0.032 | -0.032 | 0 | 0.9 | 0.99 | 0.73 | 0.89 | 0.87 | 0.79 |
| 3 | 189816756 | rs710586 | -0.676 | 0.473 | -0.429 | 0.674 | -0.002 | -0.002 | -0.002 | 0.628 | 0.15 | 0.18 | 0.11 | 0.19 | 0.15 | 0.17 |
| 3 | 189822654 | rs710589 | -1.499 | 0.726 | -0.598 | 0.895 | -0.025 | -0.001 | -0.001 | 0 | 0.41 | 0.85 | 0.32 | 0.4 | 0.38 | 0.24 |
| 3 | 189823398 | rs16865070 | 0.188 | 0.899 | 0.097 | 0.457 | 0.031 | -0.03 | -0.03 | 0 | 0.06 | 0.15 | 0.04 | 0 | 0.05 | 0.02 |
| 3 | 189824114 | rs710591 | -0.194 | 0.698 | 0.014 | 0.582 | -0.005 | -0.003 | -0.003 | 0 | 0.13 | 0.13 | 0.1 | 0.14 | 0.14 | 0.13 |
| 3 | 189824163 | rs13083331 | 0.238 | 0.64 | 0.177 | 0.529 | 0 | -0.028 | -0.028 | 0 | 0.15 | 0.14 | 0.08 | 0 | 0.16 | 0.03 |
| 3 | 189830038 | rs838692 | -1.132 | 0.583 | 0.247 | 0.638 | -0.001 | -0.027 | -0.027 | 0 | 0.33 | 0.4 | 0.21 | 0.16 | 0.33 | 0.2 |
| 3 | 189835951 | rs710595 | -0.511 | -0.254 | -0.496 | 1.077 | 0.001 | -0.002 | -0.002 | 0 | 0.25 | 0.21 | 0.23 | 0.43 | 0.18 | 0.37 |
| 3 | 189839297 | rs938513 | -0.425 | 0.071 | -0.739 | 0.83 | -0.006 | -0.003 | -0.003 | 0.556 | 0.39 | 0.5 | 0.31 | 0.33 | 0.31 | 0.31 |
| 3 | 189840887 | rs838699 | -0.761 | 0.148 | -1.428 | 1.303 | -0.123 | 0.089 | 0.089 | 3.871 | 0.62 | 0.96 | 0.62 | 0.77 | 0.52 | 0.52 |
| 3 | 189844961 | rs2675414 | -0.648 | 0.222 | -1.286 | 1.118 | -0.048 | -0.048 | -0.048 | 0 | 0.6 | 0.86 | 0.62 | 0.51 | 0.52 | 0.45 |
| 3 | 189845608 | rs7612998 | -0.475 | -0.335 | -0.488 | 1.374 | -0.038 | 0.057 | 0.057 | 0 | 0.15 | 0.19 | 0.23 | 0.34 | 0.06 | 0.25 |
| 3 | 189848916 | rs838269 | -0.226 | -0.288 | -0.855 | 0.931 | 0.014 | -0.009 | -0.009 | 0 | 0.81 | 0.9 | 0.86 | 0.8 | 0.85 | 0.86 |
| 3 | 189849198 | rs838268 | -0.06 | 0.062 | -1.045 | 0.415 | -0.001 | -0.001 | -0.001 | 0.065 | 0.31 | 0.65 | 0.36 | 0.62 | 0.33 | 0.4 |
| 3 | 189849259 | rs17424551 | 0.951 | -0.022 | -1.085 | 0.276 | -0.034 | 0.282 | 0.282 | 0 | 0.49 | 0.17 | 0.39 | 0.09 | 0.5 | 0.4 |
| 3 | 189857353 | rs9818813 | 0.381 | -0.039 | -0.387 | 0.544 | -0.014 | -0.001 | -0.001 | 0 | 0.15 | 0.24 | 0.09 | 0.13 | 0.08 | 0.27 |
| 3 | 189858976 | rs501443 | -0.085 | -0.106 | 0.194 | 0.559 | 0.036 | 0.115 | 0.115 | 0 | 0.81 | 0.69 | 0.63 | 0.38 | 0.73 | 0.72 |
| 3 | 189861477 | rs2944385 | -0.617 | -0.057 | 0.192 | 0.578 | 0.066 | 0.22 | 0.22 | 0 | 0.84 | 0.67 | 0.64 | 0.48 | 0.74 | 0.87 |
| 3 | 189869596 | rs12330208 | 0.549 | -0.09 | 0.142 | 0.509 | 0.006 | 0 | 0 | 0 | 0.15 | 0.13 | 0.09 | 0.13 | 0.1 | 0.29 |
| 3 | 189871715 | rs537205 | 1.115 | -0.174 | 0.195 | 0.448 | 0.009 | -0.005 | -0.005 | 0 | 0.18 | 0.15 | 0.15 | 0.2 | 0.15 | 0.14 |
| 3 | 189875918 | rs573533 | -0.733 | -0.152 | 0.157 | 0.426 | 0.034 | 0.124 | 0.124 | 0 | 0.84 | 0.75 | 0.64 | 0.47 | 0.73 | 0.86 |
| 3 | 189876198 | rs10513848 | 1.852 | 0.362 | -0.28 | -0.077 | 0.023 | 0.09 | 0.09 | 0 | 0.33 | 0.18 | 0.23 | 0.05 | 0.3 | 0.22 |
| 3 | 189876421 | rs568904 | 0.271 | 0.36 | -0.27 | -0.067 | 0.009 | -0.006 | -0.006 | 0.059 | 0.68 | 0.62 | 0.55 | 0.29 | 0.63 | 0.51 |
| 3 | 189877157 | rs2667706 | 1.198 | 0.484 | -0.365 | 0 | -0.014 | 0.008 | 0.008 | 0 | 0.2 | 0.26 | 0.16 | 0.24 | 0.15 | 0.22 |
| 3 | 189877822 | rs3098262 | -0.847 | 0.474 | -0.379 | 0.018 | 0 | 0.004 | 0.004 | 0 | 0.51 | 0.58 | 0.51 | 0.45 | 0.46 | 0.64 |
| 3 | 189880471 | rs17491071 | 0.776 | 1.365 | -0.169 | 0.707 | -0.003 | 0.002 | 0.002 | 0 | 0.15 | 0.13 | 0.24 | 0.12 | 0.2 | 0.21 |
| 3 | 189884801 | rs9831273 | -0.14 | 1.228 | -0.258 | 0.799 | -0.001 | 0.001 | 0.001 | 0.123 | 0.31 | 0.32 | 0.33 | 0.26 | 0.3 | 0.51 |
| 3 | 189884869 | rs536435 | 0.5 | 0.782 | -0.773 | 0.449 | 0.003 | 0.012 | 0.012 | 0.476 | 0.53 | 0.44 | 0.41 | 0.25 | 0.45 | 0.36 |
| 3 | 189903318 | rs554956 | 0.588 | 0.903 | -1.015 | 1.196 | -0.012 | -0.024 | -0.024 | 0.695 | 0.88 | 0.93 | 0.84 | 0.53 | 0.87 | 0.78 |
| 3 | 189906773 | rs505974 | -1.095 | 1.025 | -1.057 | 0.876 | -0.012 | -0.028 | -0.028 | 0.851 | 0.23 | 0.17 | 0.26 | 0.49 | 0.31 | 0.3 |
| 3 | 189908762 | rs552805 | -0.099 | 0.653 | -1.046 | 0.476 | 0.001 | -0.01 | -0.01 | 0 | 0.88 | 0.92 | 0.77 | 0.82 | 0.91 | 0.69 |
| 3 | 189914180 | rs977265 | 0.341 | 0.043 | -0.87 | -0.395 | 0.003 | -0.002 | -0.002 | 0.427 | 0.38 | 0.62 | 0.23 | 0.2 | 0.29 | 0.16 |
| 3 | 189916813 | rs563134 | -0.629 | 0.613 | -0.717 | -0.117 | 0.019 | 0.075 | 0.075 | 0.259 | 0.22 | 0.36 | 0.35 | 0.52 | 0.25 | 0.29 |
| 3 | 189932145 | rs1523270 | 1.904 | -0.078 | -0.98 | -0.506 | -0.007 | 0.036 | 0.036 | 0 | 0.18 | 0.07 | 0.1 | 0.13 | 0.19 | 0.11 |
| 3 | 189933794 | rs9290925 | 0.265 | 0.191 | -0.646 | -0.213 | 0.04 | 0.237 | 0.237 | 0 | 0.08 | 0.25 | 0.2 | 0.42 | 0.12 | 0.1 |
| 3 | 189945500 | rs9856741 | -0.616 | -0.086 | 0.103 | -0.362 | 0.032 | 0.295 | 0.295 | 0.066 | 0.14 | 0.46 | 0.22 | 0.42 | 0.13 | 0.1 |
| 3 | 189946080 | rs17426472 | 1.541 | -0.017 | -0.411 | -0.798 | -0.021 | 0.115 | 0.115 | 0 | 0.3 | 0.04 | 0.23 | 0.1 | 0.28 | 0.19 |
| 3 | 189948611 | rs6805413 | 1.106 | -0.043 | -0.315 | -0.771 | 0.003 | -0.004 | -0.004 | 0 | 0.06 | 0.26 | 0.02 | 0 | 0.01 | 0 |
| 3 | 189949098 | rs1436659 | -0.029 | -0.115 | -0.184 | -0.852 | 0.016 | -0.004 | -0.004 | 0 | 0.43 | 0.67 | 0.42 | 0.62 | 0.41 | 0.26 |
| 3 | 189952141 | rs13091722 | 2.453 | -0.155 | -0.179 | -0.878 | -0.002 | 0.03 | 0.03 | 0 | 0.19 | 0.09 | 0.11 | 0.13 | 0.19 | 0.11 |
| 3 | 189954280 | rs493186 | -0.895 | -0.146 | -0.21 | -0.877 | -0.026 | 0.049 | 0.049 | 0.403 | 0.57 | 0.83 | 0.57 | 0.72 | 0.54 | 0.45 |
| 3 | 189956100 | rs9811462 | 1.587 | -0.136 | -0.126 | -0.875 | -0.026 | -0.023 | -0.023 | 0 | 0.13 | 0.21 | 0.11 | 0.06 | 0.09 | 0.05 |
| 3 | 189956461 | rs9811787 | 0.943 | -0.107 | -0.099 | -0.866 | 0.002 | 0.037 | 0.037 | 0 | 0.3 | 0.38 | 0.3 | 0.55 | 0.31 | 0.21 |
| 3 | 189969372 | rs9852867 | 0.585 | 0.357 | -0.158 | -0.656 | -0.001 | -0.018 | -0.018 | 1.86 | 0.42 | 0.47 | 0.37 | 0.25 | 0.41 | 0.24 |
| 3 | 189971757 | rs4571226 | 1.028 | 0.431 | -0.128 | -0.581 | -0.043 | 0.004 | 0.004 | 0 | 0.1 | 0.03 | 0.15 | 0.06 | 0.13 | 0.08 |

| | | | | | | | | | | | | | | | | |
|---|---|---|---|---|---|---|---|---|---|---|---|---|---|---|---|---|
| 3 | 189977046 | rs1996903 | 0.329 | 0.328 | 0.011 | -0.486 | 0.026 | 0.209 | 0.209 | 0 | 0.11 | 0.27 | 0.19 | 0.43 | 0.13 | 0.1 |
| 3 | 189998942 | rs1436652 | -0.999 | 0.712 | -0.806 | -0.673 | -0.003 | -0.002 | -0.002 | 0 | 0.87 | 0.92 | 0.81 | 0.82 | 0.82 | 0.64 |
| 3 | 190014705 | rs9878318 | -0.306 | 0.198 | -0.617 | -0.533 | -0.007 | 0.033 | 0.033 | 0 | 0.53 | 0.73 | 0.52 | 0.67 | 0.54 | 0.34 |
| 3 | 190015700 | rs6776530 | -1.216 | 0.283 | -0.701 | -0.807 | -0.051 | 0.028 | 0.028 | 0 | 0.71 | 0.91 | 0.73 | 0.82 | 0.7 | 0.56 |
| 3 | 190021448 | rs7632915 | -0.602 | 0.113 | -0.78 | -0.656 | -0.03 | 0.049 | 0.049 | 0 | 0.6 | 0.79 | 0.6 | 0.74 | 0.6 | 0.4 |
| 3 | 190023552 | rs8298 | -1.107 | 0.518 | -0.842 | -0.589 | -0.106 | 0.052 | 0.052 | 0.22 | 0.67 | 0.94 | 0.61 | 0.8 | 0.6 | 0.54 |
| 3 | 190023837 | rs8798 | -1.102 | 0.528 | -0.846 | -0.572 | -0.084 | 0.052 | 0.052 | 0.219 | 0.67 | 0.91 | 0.61 | 0.8 | 0.6 | 0.54 |
| 3 | 190024057 | rs1364722 | -1.041 | 0.568 | -0.836 | -0.591 | -0.068 | 0.058 | 0.058 | 0 | 0.66 | 0.86 | 0.61 | 0.8 | 0.6 | 0.54 |
| 3 | 190024457 | rs9290928 | -1.041 | 0.477 | -0.836 | -0.591 | -0.068 | 0.058 | 0.058 | 0 | 0.66 | 0.86 | 0.61 | 0.8 | 0.6 | 0.54 |
| 3 | 190026654 | rs17501010 | -0.332 | 0.392 | -0.391 | -0.434 | -0.003 | -0.044 | -0.044 | 0 | 0.18 | 0.25 | 0.16 | 0.05 | 0.13 | 0.09 |
| 3 | 190030526 | rs6776378 | 0.283 | 0.331 | -0.54 | -0.197 | 0.013 | 0.015 | 0.015 | 0 | 0.34 | 0.32 | 0.23 | 0.19 | 0.26 | 0.28 |
| 3 | 190030680 | rs9869263 | 0.438 | 0.906 | -0.142 | 0.617 | 0 | 0.024 | 0.024 | 0 | 0.85 | 0.75 | 0.8 | 0.82 | 0.86 | 0.84 |
| 3 | 190030989 | rs10513846 | -0.714 | 1.281 | -0.448 | 1.038 | -0.006 | 0.051 | 0.051 | 0.042 | 0.08 | 0.05 | 0.15 | 0.46 | 0.16 | 0.23 |
| 3 | 190032594 | rs6809685 | -1.191 | 0.884 | -0.123 | 0.026 | 0.012 | -0.049 | -0.049 | 0.453 | 0.15 | 0.28 | 0.17 | 0.05 | 0.14 | 0.17 |
| 3 | 190033808 | rs9848283 | -0.389 | 0.805 | -1.009 | -0.72 | -0.06 | 0.19 | 0.19 | 0 | 0.6 | 0.9 | 0.67 | 0.93 | 0.56 | 0.65 |
| 3 | 190036418 | rs3774021 | -0.368 | 0.359 | -0.477 | -0.232 | -0.004 | 0.042 | 0.042 | 0 | 0.1 | 0.18 | 0.17 | 0.23 | 0.08 | 0.18 |
| 3 | 190037414 | rs3774019 | 0.341 | 0.393 | -0.031 | -0.454 | 0.012 | 0.032 | 0.032 | 0 | 0.21 | 0.25 | 0.28 | 0.57 | 0.3 | 0.17 |
| 3 | 190044017 | rs6781278 | -0.796 | 0.287 | -0.086 | -0.614 | -0.029 | 0.086 | 0.086 | 0 | 0.57 | 0.78 | 0.58 | 0.77 | 0.55 | 0.55 |
| 3 | 190046808 | rs16865373 | 0.6 | 0.231 | -0.055 | -0.58 | -0.005 | 0 | 0 | 0 | 0.07 | 0.06 | 0.07 | 0.07 | 0.04 | 0.08 |
| 3 | 190049397 | rs9876372 | 1.878 | -0.109 | 0.345 | -0.765 | 0.01 | 0.01 | 0.01 | 0.176 | 0.2 | 0.19 | 0.15 | 0.08 | 0.1 | 0.15 |
| 3 | 190053776 | rs7641462 | 0.431 | -0.124 | 0.527 | -0.793 | 0.006 | 0.015 | 0.015 | 0 | 0.23 | 0.25 | 0.28 | 0.61 | 0.32 | 0.21 |
| 3 | 190054083 | rs3909578 | 1.861 | -0.23 | 0.543 | -0.817 | 0 | 0 | 0 | 0 | 0.2 | 0.21 | 0.15 | 0.08 | 0.1 | 0.15 |
| 3 | 190055093 | rs6789805 | 0.457 | -0.156 | 0.46 | -0.733 | -0.008 | 0.005 | 0.005 | 0 | 0.07 | 0.15 | 0.08 | 0.07 | 0.04 | 0.08 |
| 3 | 190056322 | rs6780419 | -0.314 | -0.123 | 0.375 | -0.728 | 0.01 | 0.003 | 0.003 | 0 | 0.3 | 0.4 | 0.4 | 0.68 | 0.36 | 0.29 |
| 3 | 190063620 | rs3846102 | -1.22 | 0.026 | -0.196 | -0.759 | -0.1 | 0.231 | 0.231 | 0.027 | 0.6 | 0.91 | 0.67 | 0.93 | 0.58 | 0.64 |
| 3 | 190065672 | rs12696601 | 0.184 | 0.222 | 0.291 | -0.274 | 0 | 0.002 | 0.002 | 0.073 | 0.22 | 0.2 | 0.32 | 0.61 | 0.32 | 0.21 |
| 3 | 190070819 | rs13340153 | -0.235 | -0.103 | 0.757 | -0.333 | 0.03 | -0.012 | -0.012 | 0.128 | 0.07 | 0.25 | 0.08 | 0.02 | 0.09 | 0.13 |
| 3 | 190074689 | rs9290930 | 1.094 | -0.465 | -0.112 | -1.013 | -0.017 | 0.234 | 0.234 | 0.005 | 0.5 | 0.21 | 0.37 | 0.16 | 0.47 | 0.35 |
| 3 | 190075367 | rs2113892 | -0.766 | -0.451 | 0.167 | -0.73 | -0.006 | -0.047 | -0.047 | 0 | 0.89 | 0.84 | 0.9 | 1 | 0.94 | 0.87 |
| 3 | 190077120 | rs16865412 | -0.083 | -0.421 | 0.029 | -0.148 | 0 | 0.121 | 0.121 | 0 | 0.12 | 0.17 | 0.22 | 0.48 | 0.14 | 0.18 |
| 3 | 190079780 | rs1593314 | 0.4 | -0.46 | 0.076 | -0.173 | -0.002 | 0.044 | 0.044 | 0.481 | 0.79 | 0.75 | 0.69 | 0.52 | 0.81 | 0.69 |
| 3 | 190084208 | rs4553958 | -1.181 | -0.337 | -0.288 | -0.409 | -0.008 | 0.035 | 0.035 | 1.253 | 0.92 | 0.98 | 0.91 | 1 | 0.94 | 0.87 |
| 3 | 190087933 | rs1425104 | -0.145 | -0.376 | -0.262 | -0.402 | 0 | 0.017 | 0.017 | 1.752 | 0.87 | 0.89 | 0.85 | 0.95 | 0.9 | 0.82 |
| 3 | 190090661 | rs876234 | -0.95 | -0.91 | -0.634 | -0.617 | 0.05 | 0.075 | 0.075 | 0 | 0.73 | 0.88 | 0.82 | 0.87 | 0.85 | 0.72 |
| 3 | 190091308 | rs1425108 | 1.298 | -0.483 | -0.362 | -0.128 | -0.006 | -0.009 | -0.009 | 0 | 0.15 | 0.13 | 0.17 | 0.33 | 0.21 | 0.16 |
| 3 | 190094024 | rs10804925 | 0.112 | -0.592 | -0.486 | -0.343 | 0.011 | -0.009 | -0.009 | 0.008 | 0.47 | 0.67 | 0.55 | 0.48 | 0.53 | 0.47 |
| 3 | 190094453 | rs10937430 | 1.933 | -0.12 | -0.661 | -0.502 | -0.049 | 0.108 | 0.108 | 0 | 0.24 | 0.06 | 0.15 | 0.02 | 0.23 | 0.15 |
| 3 | 190094864 | rs1425109 | -0.744 | -0.14 | -0.29 | -0.469 | 0.058 | 0.098 | 0.098 | 0 | 0.63 | 0.8 | 0.73 | 0.82 | 0.74 | 0.64 |
| 3 | 190100842 | rs1425113 | 0.226 | -0.099 | 0.143 | -0.516 | -0.013 | 0.183 | 0.183 | 0 | 0.15 | 0.33 | 0.28 | 0.45 | 0.14 | 0.2 |
| 3 | 190104123 | rs6788908 | -1.292 | -0.074 | 0.178 | -0.444 | 0.056 | 0.101 | 0.101 | 0 | 0.68 | 0.85 | 0.79 | 0.86 | 0.8 | 0.69 |
| 3 | 190104303 | rs6764454 | 1.319 | -0.286 | 0.31 | -0.368 | -0.008 | -0.016 | -0.016 | 0 | 0.16 | 0.13 | 0.2 | 0.36 | 0.22 | 0.17 |
| 3 | 190104855 | rs1425115 | -0.027 | -0.288 | 0.33 | -0.375 | 0.065 | 0.088 | 0.088 | 0 | 0.24 | 0.42 | 0.33 | 0.38 | 0.38 | 0.29 |
| 3 | 190105451 | rs13080849 | 1.595 | -0.257 | 0.129 | -0.333 | -0.009 | 0 | 0 | 0.376 | 0.14 | 0.11 | 0.13 | 0.13 | 0.21 | 0.08 |
| 3 | 190119679 | rs11927924 | 1.663 | -0.332 | -0.471 | -0.544 | 0.046 | 0.048 | 0.048 | 0 | 0.23 | 0.12 | 0.11 | 0.13 | 0.14 | 0.14 |
| 3 | 190142205 | rs7642120 | -0.862 | -0.356 | -0.486 | -0.467 | -0.004 | -0.023 | -0.023 | 0 | 0.25 | 0.67 | 0.22 | 0.09 | 0.25 | 0.31 |
| 3 | 190144608 | rs2049672 | -0.806 | -0.282 | -0.681 | -0.388 | -0.023 | -0.042 | -0.042 | 0 | 0.4 | 0.79 | 0.42 | 0.45 | 0.47 | 0.48 |
| 3 | 190147799 | rs1425093 | -0.634 | -0.419 | -0.597 | -0.23 | 0.032 | -0.088 | -0.088 | 0.715 | 0.63 | 0.86 | 0.57 | 0.47 | 0.69 | 0.62 |
| 3 | 190154740 | rs2378570 | -1.144 | -0.207 | -0.642 | 0.035 | -0.051 | -0.098 | -0.098 | 0 | 0.86 | 0.98 | 0.67 | 0.64 | 0.85 | 0.79 |
| 3 | 190161365 | rs6791513 | -0.131 | -0.132 | -0.164 | -0.125 | 0.085 | -0.017 | -0.017 | 0 | 0.06 | 0.19 | 0.15 | 0.02 | 0.16 | 0.11 |
| 3 | 190161842 | rs1820338 | 0.315 | -0.44 | -0.139 | -0.228 | 0.004 | 0.003 | 0.003 | 0.057 | 0.45 | 0.42 | 0.52 | 0.36 | 0.61 | 0.44 |
| 3 | 190168529 | rs6797395 | -0.192 | -0.482 | -0.084 | -0.221 | 0.057 | 0.074 | 0.074 | 0.113 | 0.46 | 0.67 | 0.32 | 0.3 | 0.4 | 0.45 |
| 3 | 190169443 | rs16865457 | 1.808 | -0.511 | -0.029 | -0.239 | 0.026 | 0.025 | 0.025 | 0.115 | 0.23 | 0.17 | 0.1 | 0.14 | 0.15 | 0.17 |
| 3 | 190169644 | rs11922741 | 0.442 | -0.539 | -0.03 | -0.252 | -0.009 | -0.036 | -0.036 | 0.657 | 0.34 | 0.36 | 0.28 | 0.17 | 0.34 | 0.34 |
| 3 | 190175151 | rs1024935 | 1.528 | -0.454 | -0.227 | -0.218 | 0 | -0.001 | -0.001 | 0 | 0.24 | 0.26 | 0.11 | 0.14 | 0.16 | 0.17 |
| 3 | 190176528 | rs6763260 | -0.211 | -0.433 | -0.156 | -0.193 | 0.046 | -0.019 | -0.019 | 0 | 0.55 | 0.75 | 0.56 | 0.5 | 0.65 | 0.55 |
| 3 | 190179974 | rs13095380 | 1.554 | -0.384 | -0.251 | -0.204 | -0.001 | 0.005 | 0.005 | 0.182 | 0.16 | 0.15 | 0.2 | 0.32 | 0.21 | 0.21 |
| 3 | 190180242 | rs3943979 | 0.075 | -0.557 | -0.341 | -0.182 | 0 | -0.001 | -0.001 | 0 | 0.49 | 0.5 | 0.48 | 0.51 | 0.57 | 0.55 |
| 3 | 190183099 | rs9819973 | 0.448 | -0.544 | -0.549 | -0.935 | 0 | 0.009 | 0.009 | 0 | 0.34 | 0.36 | 0.21 | 0.14 | 0.27 | 0.23 |
| 3 | 190188644 | rs11916404 | 1.351 | -0.119 | -1.431 | -1.139 | -0.071 | 0.126 | 0.126 | 0 | 0.22 | 0.07 | 0.16 | 0.01 | 0.22 | 0.12 |
| 3 | 190194568 | rs1008800 | 0.642 | -0.702 | -0.808 | -0.538 | 0.025 | 0.028 | 0.028 | 0.122 | 0.23 | 0.18 | 0.12 | 0.14 | 0.16 | 0.17 |
| 3 | 190212706 | rs9864293 | -0.403 | 0.035 | -0.528 | -0.194 | -0.005 | -0.005 | -0.005 | 0 | 0.47 | 0.51 | 0.42 | 0.39 | 0.59 | 0.49 |
| 3 | 190216491 | rs9857115 | -0.161 | 0.393 | -0.948 | -0.091 | -0.007 | 0.01 | 0.01 | 0 | 0.39 | 0.26 | 0.37 | 0.37 | 0.55 | 0.4 |
| 3 | 190220784 | rs13316596 | -0.052 | 0.579 | -1.478 | -0.039 | -0.128 | -0.039 | -0.039 | 0.71 | 0.38 | 0.07 | 0.29 | 0.38 | 0.48 | 0.38 |
| 3 | 190221825 | rs2042865 | -0.504 | 0.226 | -1.145 | 0.227 | -0.013 | -0.006 | -0.006 | 0 | 0.57 | 0.54 | 0.45 | 0.38 | 0.66 | 0.51 |
| 3 | 190222954 | rs766442 | -0.21 | 0.817 | -1.519 | 0.358 | -0.119 | 0.031 | 0.031 | 0 | 0.23 | 0.03 | 0.19 | 0.14 | 0.31 | 0.22 |
| 3 | 190234304 | rs6800609 | -0.883 | -0.291 | -0.366 | 1.313 | -0.05 | 0.203 | 0.203 | 0.016 | 0.12 | 0.28 | 0.13 | 0.46 | 0.05 | 0.16 |
| 3 | 190235788 | rs3773997 | -0.336 | -0.895 | -0.222 | 0.442 | -0.006 | 0.001 | 0.001 | 0.555 | 0.34 | 0.35 | 0.24 | 0.33 | 0.28 | 0.19 |
| 3 | 190237101 | rs11719243 | -0.336 | -0.895 | -0.222 | 0.442 | -0.006 | 0.001 | 0.001 | 0.555 | 0.34 | 0.35 | 0.24 | 0.33 | 0.28 | 0.19 |
| 3 | 190237351 | rs9290935 | -0.856 | -0.836 | -0.086 | 0.443 | 0.082 | -0.007 | -0.007 | 0 | 0.43 | 0.76 | 0.32 | 0.45 | 0.3 | 0.23 |

| | | | | | | | | | | | | | | | | |
|---|---|---|---|---|---|---|---|---|---|---|---|---|---|---|---|---|
| 3 | 190239148 | rs10212103 | -0.634 | -0.759 | 0.01 | 0.477 | 0.011 | -0.003 | -0.003 | 0.976 | 0.41 | 0.6 | 0.3 | 0.45 | 0.3 | 0.23 |
| 3 | 190253665 | rs3773992 | -1.269 | -0.193 | 0.1 | 1.167 | -0.009 | 0.086 | 0.086 | 0 | 0.12 | 0.2 | 0.1 | 0.36 | 0.06 | 0.11 |
| 3 | 190255192 | rs9876152 | -1.226 | -0.185 | 0.019 | 1.171 | -0.005 | 0.056 | 0.056 | 0 | 0.11 | 0.17 | 0.1 | 0.35 | 0.06 | 0.11 |
| 3 | 190257179 | rs9849030 | -1.726 | -0.278 | 0.486 | 1.357 | -0.018 | 0.162 | 0.162 | 0 | 0.16 | 0.34 | 0.15 | 0.42 | 0.09 | 0.16 |
| 3 | 190260037 | rs2361832 | 1.674 | 0.068 | 0.398 | 1.106 | -0.004 | 0.099 | 0.099 | 0.134 | 0.76 | 0.57 | 0.73 | 0.57 | 0.81 | 0.71 |
| 3 | 190260163 | rs997534 | 1.22 | 0.387 | 0.02 | 1.138 | 0.003 | 0.025 | 0.025 | 0 | 0.83 | 0.78 | 0.77 | 0.65 | 0.84 | 0.77 |
| 3 | 190262052 | rs3773990 | -1.35 | 0.591 | -0.016 | 1.509 | 0.018 | 0.095 | 0.095 | 0 | 0.06 | 0.12 | 0.07 | 0.27 | 0.06 | 0.15 |
| 3 | 190262799 | rs2193880 | 0.543 | 0.171 | -1.233 | 0.658 | -0.064 | 0.017 | 0.017 | 1.855 | 0.47 | 0.12 | 0.58 | 0.6 | 0.57 | 0.53 |
| 3 | 190264270 | rs4687148 | -0.462 | -0.106 | -0.887 | 0.414 | -0.004 | -0.037 | -0.037 | 0.058 | 0.09 | 0.09 | 0.11 | 0 | 0.08 | 0.1 |
| 3 | 190265429 | rs3773989 | 0.487 | 0.279 | -1.328 | 0.301 | -0.075 | -0.009 | -0.009 | 0 | 0.37 | 0.14 | 0.52 | 0.6 | 0.49 | 0.47 |
| 3 | 190271177 | rs2080794 | 0.448 | 0.726 | -0.494 | 0.516 | 0.033 | 0.062 | 0.062 | 0.035 | 0.76 | 0.56 | 0.8 | 0.68 | 0.72 | 0.74 |
| 3 | 190273818 | rs4686554 | 0.583 | 0.494 | -0.163 | 0.448 | 0.036 | 0.047 | 0.047 | 0.01 | 0.75 | 0.49 | 0.79 | 0.69 | 0.71 | 0.78 |
| 3 | 190275545 | rs16865597 | 0.272 | -0.164 | -0.372 | 0.295 | -0.031 | -0.009 | -0.009 | 0 | 0.17 | 0.32 | 0.13 | 0.14 | 0.1 | 0.17 |
| 3 | 190278915 | rs3773986 | 0.641 | 0 | -0.363 | 0.556 | 0.002 | -0.009 | -0.009 | 0.487 | 0.09 | 0.09 | 0.08 | 0.12 | 0.1 | 0.11 |
| 3 | 190280052 | rs3773982 | 0.264 | 0.07 | -0.241 | 0.529 | 0.046 | 0.033 | 0.033 | 3.011 | 0.14 | 0.37 | 0.12 | 0.19 | 0.16 | 0.12 |
| 3 | 190280884 | rs4687150 | 1.214 | -0.356 | -0.694 | 0.202 | 0.017 | 0.057 | 0.057 | 0 | 0.59 | 0.19 | 0.66 | 0.67 | 0.65 | 0.66 |
| 3 | 190281376 | rs9883249 | 1.612 | -0.371 | -0.748 | 0.191 | 0.008 | -0.009 | -0.009 | 0 | 0.49 | 0.11 | 0.57 | 0.55 | 0.54 | 0.55 |
| 3 | 190282186 | rs6764286 | 0.958 | -0.668 | -0.457 | 0.035 | 0.002 | -0.034 | -0.034 | 0.011 | 0.06 | 0.14 | 0.1 | 0 | 0.06 | 0.05 |
| 3 | 190282714 | rs2241343 | -0.741 | -0.959 | -0.548 | -0.053 | -0.068 | -0.052 | -0.052 | 0 | 0.25 | 0.45 | 0.2 | 0.14 | 0.14 | 0.23 |
| 3 | 190283007 | rs16865627 | -0.125 | -0.535 | -0.903 | -0.102 | 0 | -0.02 | -0.02 | 0.043 | 0.13 | 0.05 | 0.1 | 0.12 | 0.13 | 0.12 |
| 3 | 190284586 | rs3773980 | 0.334 | -0.445 | -0.765 | -0.05 | 0.003 | -0.013 | -0.013 | 0 | 0.23 | 0.19 | 0.36 | 0.55 | 0.23 | 0.32 |
| 3 | 190285822 | rs9877268 | 1.905 | 0.129 | -1.056 | -0.669 | -0.034 | 0.105 | 0.105 | 0 | 0.28 | 0.1 | 0.25 | 0 | 0.31 | 0.18 |
| 3 | 190286886 | rs3773978 | -0.306 | -0.062 | -0.433 | -0.443 | -0.01 | 0.049 | 0.049 | 0 | 0.24 | 0.3 | 0.22 | 0.36 | 0.21 | 0.26 |
| 3 | 190287285 | rs3773976 | 0.027 | 0.397 | -0.396 | -0.347 | 0.006 | 0.066 | 0.066 | 0 | 0.07 | 0.11 | 0.14 | 0.24 | 0.1 | 0.11 |
| 3 | 190292028 | rs3773971 | 0.101 | 0.583 | -0.35 | -0.428 | 0.03 | 0.035 | 0.035 | 0 | 0.84 | 0.8 | 0.8 | 0.75 | 0.77 | 0.83 |
| 3 | 190292102 | rs3773970 | 0.627 | -0.216 | -0.245 | -0.467 | 0.004 | 0.058 | 0.058 | 0.003 | 0.11 | 0.18 | 0.13 | 0.19 | 0.12 | 0.14 |
| 3 | 190293555 | rs10937439 | -1.185 | -0.113 | -0.274 | -0.912 | 0.013 | -0.15 | -0.15 | 0.014 | 0.65 | 0.46 | 0.67 | 1 | 0.61 | 0.81 |
| 3 | 190294188 | rs10937441 | -1.661 | 0.185 | -1.094 | -0.784 | -0.065 | 0.113 | 0.113 | 0 | 0.7 | 0.89 | 0.72 | 1 | 0.62 | 0.81 |
| 3 | 190297083 | rs1559018 | -0.331 | -0.147 | -0.496 | 0.153 | -0.002 | 0.028 | 0.028 | 0 | 0.31 | 0.36 | 0.43 | 0.53 | 0.33 | 0.32 |
| 3 | 190301872 | rs9862847 | 0.082 | -0.313 | -0.811 | -0.097 | -0.001 | -0.03 | -0.03 | 0 | 0.08 | 0.1 | 0.04 | 0 | 0.07 | 0.07 |
| 3 | 190303805 | rs6444435 | -0.317 | -0.777 | -1.022 | -0.41 | 0.027 | 0.031 | 0.031 | 0 | 0.47 | 0.43 | 0.5 | 0.81 | 0.35 | 0.63 |
| 3 | 190305018 | rs3773967 | -0.376 | -0.278 | -0.989 | 0.348 | -0.002 | -0.038 | -0.038 | 0 | 0.15 | 0.13 | 0.26 | 0.46 | 0.18 | 0.19 |
| 3 | 190307448 | rs3773963 | 0.113 | -0.096 | -1.128 | 0.192 | -0.039 | -0.097 | -0.097 | 0 | 0.16 | 0.02 | 0.24 | 0.46 | 0.19 | 0.22 |
| 3 | 190308527 | rs7628250 | 0.034 | 0.225 | -1.362 | -0.028 | -0.023 | -0.012 | -0.012 | 0.772 | 0.79 | 0.92 | 0.74 | 0.86 | 0.77 | 0.79 |
| 3 | 190312433 | rs7652346 | 0.787 | 0.275 | -1.046 | -0.227 | -0.012 | 0.021 | 0.021 | 0 | 0.69 | 0.82 | 0.69 | 0.83 | 0.67 | 0.68 |
| 3 | 190313507 | rs2193873 | 1.358 | 0.289 | -0.493 | -0.231 | 0.005 | 0.023 | 0.023 | 0 | 0.51 | 0.58 | 0.43 | 0.36 | 0.48 | 0.46 |
| 3 | 190314720 | rs3773953 | -0.394 | 0.147 | -0.133 | -0.048 | 0.002 | -0.019 | -0.019 | 0 | 0.1 | 0.12 | 0.05 | 0.02 | 0.1 | 0.11 |
| 3 | 190323875 | rs716984 | 1.348 | -0.135 | 0.153 | -0.06 | 0.003 | 0.036 | 0.036 | 0.669 | 0.63 | 0.22 | 0.63 | 0.82 | 0.67 | 0.72 |
| 3 | 190330777 | rs759783 | 1.209 | -0.201 | 0.202 | 0.005 | -0.011 | -0.023 | -0.023 | 0.975 | 0.64 | 0.5 | 0.64 | 0.81 | 0.69 | 0.7 |
| 3 | 190333876 | rs9290939 | -0.003 | -0.214 | 0.3 | -0.002 | -0.011 | -0.031 | -0.031 | 0 | 0.11 | 0.18 | 0.08 | 0.03 | 0.08 | 0.1 |
| 3 | 190340177 | rs9813227 | -0.55 | 0.126 | -0.247 | 0.053 | -0.057 | 0.063 | 0.063 | 0.012 | 0.89 | 0.99 | 0.91 | 1 | 0.82 | 0.94 |
| 3 | 190341241 | rs4687160 | 0.96 | 0.072 | -0.177 | 0.057 | -0.009 | -0.02 | -0.02 | 0 | 0.64 | 0.49 | 0.64 | 0.81 | 0.68 | 0.69 |
| 3 | 190342166 | rs2193876 | 0.987 | 0.088 | -0.39 | 0.064 | -0.014 | -0.029 | -0.029 | 1.122 | 0.63 | 0.45 | 0.63 | 0.81 | 0.68 | 0.69 |
| 3 | 190347523 | rs1024946 | 1.073 | -0.099 | -0.327 | 0.096 | 0.003 | 0.007 | 0.007 | 0.064 | 0.62 | 0.35 | 0.63 | 0.82 | 0.68 | 0.69 |
| 3 | 190347732 | rs1024949 | 0.211 | -0.899 | -0.654 | 0.758 | -0.037 | -0.106 | -0.106 | 0.303 | 0.14 | 0 | 0.11 | 0.51 | 0.12 | 0.19 |
| 3 | 190350461 | rs7626795 | -1.249 | -0.998 | 0.149 | 0.22 | -0.066 | -0.01 | -0.01 | 0.032 | 0.25 | 0.5 | 0.27 | 0.17 | 0.13 | 0.25 |
| 3 | 190361497 | rs9831803 | -1.193 | -0.801 | 0.14 | -0.199 | -0.039 | -0.096 | -0.096 | 0.48 | 0.26 | 0.52 | 0.13 | 0.07 | 0.16 | 0.21 |
| 3 | 190363116 | rs7650510 | -1.262 | -0.262 | 0.505 | -0.423 | 0.024 | -0.107 | -0.107 | 0 | 0.21 | 0.33 | 0.13 | 0.03 | 0.24 | 0.2 |
| 3 | 190370764 | rs11915384 | -2.167 | -0.921 | 1.335 | 0.539 | -0.054 | 0.209 | 0.209 | 0.387 | 0.16 | 0.46 | 0.13 | 0.33 | 0.06 | 0.12 |
| 3 | 190413415 | rs6798694 | -2.397 | -0.956 | 1.768 | 0.555 | -0.041 | 0.144 | 0.144 | 0.89 | 0.15 | 0.55 | 0.13 | 0.31 | 0.06 | 0.12 |
| 3 | 190414518 | rs4687167 | -0.044 | -1.013 | 1.214 | 0.014 | -0.024 | -0.05 | -0.05 | 0.16 | 0.22 | 0.14 | 0.42 | 0.62 | 0.28 | 0.49 |
| 3 | 190421204 | rs17514402 | 0.744 | -0.882 | 1.302 | 0.014 | -0.01 | 0.016 | 0.016 | 0 | 0.05 | 0.01 | 0.04 | 0 | 0.06 | 0.04 |
| 3 | 190433597 | rs9841126 | -1.125 | -1.491 | 0.968 | -0.402 | -0.001 | 0 | 0 | 2.598 | 0.45 | 0.36 | 0.5 | 0.6 | 0.45 | 0.66 |
| 3 | 190442316 | rs16848932 | -0.556 | -1.527 | 0.909 | -0.049 | 0 | 0.034 | 0.034 | 0 | 0.06 | 0.05 | 0.07 | 0.28 | 0.02 | 0.07 |
| 3 | 190443615 | rs17514723 | -0.372 | -1.632 | 0.831 | -0.222 | 0.014 | 0.032 | 0.032 | 0 | 0.16 | 0.07 | 0.06 | 0.06 | 0.14 | 0.19 |
| 3 | 190446487 | rs9873487 | -1.227 | -1.783 | 0.694 | -0.039 | 0.011 | -0.002 | -0.002 | 0 | 0.4 | 0.86 | 0.5 | 0.65 | 0.32 | 0.55 |
| 3 | 190447440 | rs9858730 | -0.834 | -1.758 | 1.071 | -0.061 | 0.008 | -0.015 | -0.015 | 0.049 | 0.36 | 0.67 | 0.44 | 0.57 | 0.25 | 0.51 |
| 3 | 190458193 | rs16865790 | -1.382 | -1.756 | 1.358 | -0.122 | -0.002 | -0.006 | -0.006 | 0.147 | 0.44 | 0.51 | 0.45 | 0.66 | 0.4 | 0.65 |
| 3 | 190468049 | rs4687170 | -0.876 | -1.758 | 1.478 | -0.112 | 0 | -0.006 | -0.006 | 0 | 0.34 | 0.65 | 0.43 | 0.59 | 0.3 | 0.51 |
| 3 | 190470973 | rs6444457 | -2.857 | -1.961 | 1.034 | -0.448 | -0.165 | 0.186 | 0.186 | 0 | 0.66 | 0.98 | 0.65 | 0.95 | 0.53 | 0.84 |
| 3 | 190477598 | rs9858906 | -1.583 | -1.592 | 1.592 | -0.226 | 0.003 | 0.022 | 0.022 | 0 | 0.49 | 0.62 | 0.52 | 0.64 | 0.46 | 0.67 |
| 3 | 190484483 | rs9858191 | -0.214 | -1.422 | 1.593 | -0.035 | 0.01 | 0.055 | 0.055 | 0 | 0.27 | 0.4 | 0.39 | 0.55 | 0.28 | 0.46 |
| 3 | 190498709 | rs4687174 | -0.87 | -1.21 | 1.693 | 0.082 | -0.001 | -0.002 | -0.002 | 0 | 0.45 | 0.5 | 0.45 | 0.61 | 0.42 | 0.63 |
| 3 | 190500306 | rs7617445 | -0.958 | -1.275 | 1.608 | 0.096 | 0 | 0.001 | 0.001 | 0 | 0.46 | 0.47 | 0.46 | 0.6 | 0.45 | 0.65 |
| 3 | 190512235 | rs13315893 | -2.112 | -0.989 | 2.245 | 0.625 | 0.021 | 0.028 | 0.028 | 0.316 | 0.07 | 0.37 | 0.05 | 0.02 | 0.02 | 0.05 |
| 3 | 190513856 | rs6775728 | -2.466 | -0.98 | 2.141 | 0.726 | 0.025 | 0.031 | 0.031 | 0.03 | 0.08 | 0.35 | 0.06 | 0.05 | 0.05 | 0.06 |
| 3 | 190514985 | rs13079048 | -0.675 | -0.853 | 1.561 | 0.873 | -0.011 | 0.083 | 0.083 | 0 | 0.18 | 0.23 | 0.17 | 0.44 | 0.13 | 0.35 |
| 3 | 190515906 | rs13062230 | -0.641 | -0.85 | 1.563 | 0.875 | -0.011 | 0.086 | 0.086 | 0 | 0.18 | 0.23 | 0.17 | 0.44 | 0.13 | 0.35 |
| 3 | 190519325 | rs13061181 | -0.641 | -0.85 | 1.563 | 0.875 | -0.011 | 0.086 | 0.086 | 0 | 0.18 | 0.23 | 0.17 | 0.44 | 0.13 | 0.35 |

| | | | | | | | | | | | | | | | | |
|---|---|---|---|---|---|---|---|---|---|---|---|---|---|---|---|---|
| 3 | 190520710 | rs7649452 | -2.351 | -0.885 | 1.314 | 0.805 | 0.032 | -0.028 | -0.028 | 2.271 | 0.47 | 0.89 | 0.39 | 0.57 | 0.37 | 0.57 |
| 3 | 190522579 | rs16865892 | -0.825 | -0.76 | 1.577 | 0.791 | 0.001 | 0.001 | 0.001 | 0 | 0.17 | 0.17 | 0.09 | 0.09 | 0.14 | 0.18 |
| 3 | 190525516 | rs9870207 | -1.946 | -0.801 | 1.685 | 0.722 | 0.017 | -0.026 | -0.026 | 1.358 | 0.39 | 0.71 | 0.34 | 0.53 | 0.33 | 0.55 |
| 3 | 190528576 | rs16865914 | -0.776 | -0.852 | 1.292 | 0.845 | 0.001 | -0.001 | -0.001 | 0 | 0.16 | 0.17 | 0.09 | 0.13 | 0.14 | 0.18 |
| 3 | 190536777 | rs9863612 | 0.333 | -0.703 | 1.192 | 0.952 | 0.014 | 0.069 | 0.069 | 0 | 0.15 | 0.2 | 0.25 | 0.36 | 0.17 | 0.3 |
| 3 | 190537545 | rs9830737 | -1.988 | -0.617 | 0.307 | 0.874 | 0.038 | -0.009 | -0.009 | 0 | 0.49 | 0.88 | 0.43 | 0.58 | 0.38 | 0.61 |
| 3 | 190543141 | rs10513861 | -0.281 | -0.491 | 0.322 | 0.731 | 0.009 | -0.027 | -0.027 | 0 | 0.05 | 0.17 | 0.05 | 0 | 0.05 | 0.03 |
| 3 | 190546144 | rs9290953 | -1.93 | -0.534 | 0.312 | 0.779 | 0.047 | -0.014 | -0.014 | 1.33 | 0.43 | 0.75 | 0.39 | 0.56 | 0.34 | 0.59 |
| 3 | 190550565 | rs4234618 | 0.009 | -0.223 | 0.471 | 0.986 | 0.024 | 0.134 | 0.134 | 0 | 0.17 | 0.37 | 0.26 | 0.37 | 0.18 | 0.34 |
| 3 | 190554485 | rs6444462 | -0.354 | -0.105 | -0.215 | 1.342 | 0 | 0.034 | 0.034 | 0.194 | 0.28 | 0.48 | 0.33 | 0.4 | 0.29 | 0.49 |
| 3 | 190561016 | rs16865970 | -1.097 | 0.21 | -0.549 | 1.288 | 0.014 | -0.015 | -0.015 | 0 | 0.13 | 0.12 | 0.05 | 0.16 | 0.05 | 0.08 |
| 3 | 190579558 | rs6766389 | -0.823 | 1.597 | -0.188 | 1.326 | 0.035 | 0.038 | 0.038 | 0 | 0.17 | 0.29 | 0.2 | 0.2 | 0.23 | 0.13 |
| 3 | 190581161 | rs9682599 | -0.676 | 1.562 | 0.088 | 0.775 | 0.022 | -0.017 | -0.017 | 3.055 | 0.14 | 0.28 | 0.11 | 0.09 | 0.14 | 0.17 |
| 3 | 190583269 | rs6764542 | -1.109 | 1.49 | -0.099 | 0.97 | 0.027 | -0.004 | -0.004 | 0 | 0.3 | 0.56 | 0.34 | 0.68 | 0.3 | 0.43 |
| 3 | 190583583 | rs9812368 | -1.235 | 1.503 | -0.049 | 0.952 | 0.023 | -0.001 | -0.001 | 0 | 0.29 | 0.59 | 0.34 | 0.67 | 0.29 | 0.41 |
| 3 | 190583928 | rs7636699 | -0.068 | 1.059 | -0.588 | 0.263 | -0.003 | 0.02 | 0.02 | 0 | 0.15 | 0.09 | 0.23 | 0.03 | 0.17 | 0.15 |
| 3 | 190586181 | rs10937447 | -1.009 | 0.823 | 0.293 | 1.4 | 0.022 | 0.231 | 0.231 | 0 | 0.1 | 0.27 | 0.16 | 0.42 | 0.07 | 0.26 |
| 3 | 190586978 | rs7624080 | -1.113 | -0.332 | -0.362 | 0.727 | 0.042 | -0.049 | -0.049 | 0 | 0.3 | 0.82 | 0.27 | 0.43 | 0.16 | 0.32 |
| 3 | 190588344 | rs12634559 | -0.869 | -0.934 | -1.136 | 0.159 | 0.108 | 0.082 | 0.082 | 0.001 | 0.52 | 0.88 | 0.46 | 0.62 | 0.33 | 0.53 |
| 3 | 190591090 | rs9863733 | 0.129 | -0.862 | -0.485 | 0.021 | 0.003 | 0.003 | 0.003 | 0.053 | 0.27 | 0.25 | 0.28 | 0.2 | 0.16 | 0.2 |
| 3 | 190592125 | rs9868955 | -0.514 | 0.054 | 0.412 | 0.427 | -0.003 | 0.001 | 0.001 | 0.105 | 0.42 | 0.56 | 0.4 | 0.36 | 0.56 | 0.31 |
| 3 | 190593084 | rs17518905 | -0.538 | 0.049 | 0.453 | 0.425 | -0.007 | 0.004 | 0.004 | 0.072 | 0.42 | 0.6 | 0.4 | 0.36 | 0.56 | 0.31 |
| 3 | 190595760 | rs13323015 | -0.282 | 0.115 | 0.458 | 0.467 | 0.001 | -0.002 | -0.002 | 0 | 0.38 | 0.58 | 0.37 | 0.34 | 0.47 | 0.26 |
| 3 | 190602087 | rs9818981 | -1.254 | 0.081 | 0.796 | 0.403 | 0.01 | -0.036 | -0.036 | 1.186 | 0.11 | 0.42 | 0.06 | 0.02 | 0.09 | 0.05 |
| 3 | 190621019 | rs1501600 | -0.011 | 0.22 | 1.041 | 0.45 | 0.004 | -0.002 | -0.002 | 0 | 0.36 | 0.4 | 0.36 | 0.34 | 0.45 | 0.25 |
| 3 | 190632775 | rs9868693 | -0.369 | 0.3 | 1.058 | 0.627 | 0.014 | -0.001 | -0.001 | 0.114 | 0.37 | 0.45 | 0.35 | 0.36 | 0.49 | 0.3 |
| 3 | 190657163 | rs1316356 | 0.521 | 0.347 | 0.714 | 0.534 | -0.029 | -0.02 | -0.02 | 0 | 0.27 | 0.14 | 0.26 | 0.33 | 0.4 | 0.22 |
| 3 | 190671328 | rs9812571 | -0.425 | 0.13 | 0.622 | -0.316 | -0.003 | 0.033 | 0.033 | 0 | 0.46 | 0.61 | 0.49 | 0.84 | 0.54 | 0.59 |
| 3 | 190682313 | rs9880180 | -1.827 | -0.046 | 1.688 | 0.441 | -0.009 | 0.018 | 0.018 | 0.076 | 0.2 | 0.48 | 0.21 | 0.23 | 0.16 | 0.3 |
| 3 | 190687002 | rs9860559 | -1.096 | -0.125 | 1.77 | 0.541 | -0.006 | 0.088 | 0.088 | 0 | 0.08 | 0.16 | 0.09 | 0.23 | 0.04 | 0.19 |
| 3 | 190687732 | rs6444475 | -2.878 | -0.252 | 2.335 | 0.459 | -0.013 | 0.001 | 0.001 | 0 | 0.24 | 0.65 | 0.23 | 0.28 | 0.18 | 0.39 |
| 3 | 190688176 | rs7650354 | -1.931 | -0.227 | 2.429 | 0.469 | -0.01 | 0.07 | 0.07 | 0 | 0.12 | 0.26 | 0.14 | 0.23 | 0.08 | 0.23 |
| 3 | 190689247 | rs1501602 | -1.991 | -0.227 | 2.511 | 0.432 | -0.012 | -0.097 | -0.097 | 0 | 0.14 | 0.38 | 0.1 | 0.02 | 0.11 | 0.14 |
| 3 | 190690194 | rs7644897 | -2.226 | -0.227 | 2.499 | 0.433 | -0.014 | -0.095 | -0.095 | 0 | 0.16 | 0.46 | 0.12 | 0.05 | 0.12 | 0.22 |
| 3 | 190700956 | rs13314422 | -1.146 | -0.211 | 2.255 | 0.453 | -0.009 | -0.021 | -0.021 | 0 | 0.07 | 0.13 | 0.04 | 0.04 | 0.03 | 0.16 |
| 3 | 190714498 | rs6774178 | -0.987 | -0.19 | 2.202 | 0.498 | 0.006 | 0.098 | 0.098 | 0 | 0.1 | 0.25 | 0.13 | 0.23 | 0.1 | 0.17 |
| 3 | 190714587 | rs9864764 | -1.591 | -0.025 | 1.51 | 0.168 | 0.01 | -0.021 | -0.021 | 0 | 0.08 | 0.26 | 0.06 | 0.04 | 0.04 | 0.16 |
| 3 | 190722423 | rs6807826 | -1.627 | -0.053 | 1.012 | 0.133 | -0.015 | -0.069 | -0.069 | 0.163 | 0.13 | 0.33 | 0.09 | 0.02 | 0.08 | 0.13 |
| 3 | 190737794 | rs9754606 | 0.017 | 0.522 | -0.15 | 0.453 | -0.018 | -0.007 | -0.007 | 0 | 0.63 | 0.78 | 0.46 | 0.45 | 0.56 | 0.49 |
| 3 | 190752382 | rs6783375 | -0.329 | 0.606 | -0.309 | 0.85 | -0.093 | -0.138 | -0.138 | 0 | 0.69 | 0.96 | 0.51 | 0.53 | 0.61 | 0.61 |
| 3 | 190758923 | rs11922651 | 0.437 | -0.159 | 0.154 | 0.264 | -0.016 | -0.013 | -0.013 | 0 | 0.21 | 0.24 | 0.09 | 0.13 | 0.12 | 0.22 |
| 3 | 190769705 | rs1510900 | 0.447 | -0.613 | 0.897 | 0.618 | 0.015 | 0.026 | 0.026 | 0 | 0.38 | 0.51 | 0.46 | 0.49 | 0.45 | 0.29 |
| 3 | 190775418 | rs7613308 | 0.316 | -0.661 | 0.901 | 0.58 | -0.003 | -0.005 | -0.005 | 0.136 | 0.4 | 0.66 | 0.47 | 0.49 | 0.45 | 0.29 |
| 3 | 190782523 | rs2048417 | 0.107 | -0.708 | 0.914 | 0.583 | -0.018 | -0.042 | -0.042 | 0.475 | 0.42 | 0.8 | 0.45 | 0.48 | 0.45 | 0.31 |
| 3 | 190783307 | rs6775178 | 0.704 | -0.825 | 0.869 | 0.53 | 0.012 | 0.009 | 0.009 | 0 | 0.2 | 0.16 | 0.07 | 0.13 | 0.12 | 0.21 |
| 3 | 190785207 | rs6769238 | 0.092 | -0.952 | 1.154 | 0.472 | -0.023 | -0.027 | -0.027 | 0.052 | 0.24 | 0.33 | 0.12 | 0.13 | 0.16 | 0.29 |
| 3 | 190786415 | rs2048415 | -0.902 | -0.89 | 1.077 | 0.476 | -0.018 | -0.002 | -0.002 | 0 | 0.64 | 0.78 | 0.52 | 0.61 | 0.6 | 0.58 |
| 3 | 190787990 | rs13078649 | -0.026 | -0.923 | 1.171 | 0.433 | -0.048 | -0.054 | -0.054 | 0 | 0.26 | 0.49 | 0.14 | 0.13 | 0.16 | 0.29 |
| 3 | 190791994 | rs6794276 | 0.891 | -0.812 | 1.113 | -0.045 | -0.011 | -0.006 | -0.006 | 0 | 0.16 | 0.13 | 0.13 | 0 | 0.26 | 0.16 |
| 3 | 190801723 | rs2063421 | -0.251 | -1.005 | 1.234 | 0.031 | 0.055 | -0.028 | -0.028 | 0.441 | 0.43 | 0.77 | 0.37 | 0.49 | 0.28 | 0.3 |
| 3 | 190804380 | rs13058800 | 0.098 | -0.962 | 1.354 | 0.017 | -0.002 | 0 | 0 | 0.018 | 0.28 | 0.28 | 0.13 | 0.25 | 0.19 | 0.33 |
| 3 | 190809722 | rs1355365 | 0.036 | -0.683 | 1.36 | -0.172 | 0.052 | -0.118 | -0.118 | 0.384 | 0.21 | 0.34 | 0.18 | 0 | 0.32 | 0.23 |
| 3 | 190811250 | rs9290973 | 0.995 | -0.727 | 1.333 | 0.087 | -0.007 | -0.001 | -0.001 | 0 | 0.21 | 0.22 | 0.08 | 0.16 | 0.12 | 0.23 |
| 3 | 190812290 | rs9814786 | -0.579 | -0.686 | 1.04 | 0.13 | -0.002 | 0.032 | 0.032 | 0.908 | 0.43 | 0.6 | 0.27 | 0.16 | 0.44 | 0.46 |
| 3 | 190818456 | rs1195356 | -1.328 | -0.676 | -0.075 | 0.117 | -0.043 | 0.01 | 0.01 | 0 | 0.65 | 0.91 | 0.55 | 0.64 | 0.62 | 0.6 |
| 3 | 190821307 | rs1195361 | -1.287 | -0.616 | -0.076 | 0.115 | -0.039 | 0.012 | 0.012 | 0 | 0.65 | 0.91 | 0.55 | 0.63 | 0.63 | 0.6 |
| 3 | 190830562 | rs4441706 | 0.662 | -0.48 | -0.191 | 0.015 | -0.087 | 0.095 | 0.095 | 0.236 | 0.15 | 0.01 | 0.13 | 0 | 0.25 | 0.16 |
| 3 | 190842139 | rs10804931 | 0.601 | -0.352 | -0.04 | 0.343 | 0.018 | 0.069 | 0.069 | 0 | 0.47 | 0.13 | 0.53 | 0.39 | 0.59 | 0.53 |
| 3 | 190844007 | rs10937458 | 0.272 | -0.745 | 0.227 | 0.117 | 0.032 | 0.028 | 0.028 | 0 | 0.57 | 0.26 | 0.6 | 0.39 | 0.68 | 0.64 |
| 3 | 190845905 | rs12497814 | 0.895 | -0.694 | 0.209 | 0.117 | 0.025 | 0.078 | 0.078 | 0 | 0.48 | 0.19 | 0.54 | 0.39 | 0.59 | 0.54 |
| 3 | 190847809 | rs9851126 | -0.934 | -0.908 | 0.959 | 0.444 | -0.002 | 0.004 | 0.004 | 0.457 | 0.79 | 0.77 | 0.81 | 0.62 | 0.83 | 0.88 |
| 3 | 190847898 | rs13068656 | 0.578 | -0.905 | 0.867 | 0.386 | 0.04 | 0.032 | 0.032 | 0 | 0.57 | 0.24 | 0.6 | 0.4 | 0.68 | 0.64 |
| 3 | 190848369 | rs9855788 | -0.29 | -0.937 | 1.352 | 0.633 | -0.011 | 0.068 | 0.068 | 0 | 0.23 | 0.28 | 0.22 | 0.41 | 0.18 | 0.3 |
| 3 | 190853429 | rs9882405 | -0.883 | -0.988 | 0.846 | 0.574 | 0.003 | 0.002 | 0.002 | 0.229 | 0.42 | 0.41 | 0.38 | 0.44 | 0.46 | 0.49 |
| 3 | 190856753 | rs2363961 | 0.905 | -0.62 | 0.643 | 0.639 | -0.007 | 0.016 | 0.016 | 0 | 0.4 | 0.31 | 0.45 | 0.36 | 0.45 | 0.44 |
| 3 | 190859781 | rs1551424 | -0.016 | -0.636 | 0.784 | 0.699 | -0.002 | 0 | 0 | 0 | 0.55 | 0.44 | 0.58 | 0.58 | 0.65 | 0.6 |
| 3 | 190861768 | rs7641416 | 0.823 | -0.465 | 0.34 | 0.579 | -0.01 | 0.023 | 0.023 | 0 | 0.4 | 0.26 | 0.44 | 0.36 | 0.45 | 0.44 |
| 3 | 190863542 | rs9870155 | -0.913 | -0.182 | -0.158 | 0.78 | 0.012 | -0.001 | -0.001 | 0 | 0.4 | 0.43 | 0.32 | 0.36 | 0.44 | 0.49 |
| 3 | 190865991 | rs2603658 | -1.255 | -1.183 | -0.93 | 0.811 | 0.091 | -0.003 | -0.003 | 0 | 0.74 | 0.97 | 0.84 | 0.74 | 0.88 | 0.85 |

| | | | | | | | | | | | | | | | | |
|---|---|---|---|---|---|---|---|---|---|---|---|---|---|---|---|---|
| 3 | 190873984 | rs2363970 | -0.938 | -0.526 | -1.2 | 0.907 | -0.05 | -0.004 | -0.004 | 0 | 0.56 | 0.95 | 0.47 | 0.65 | 0.42 | 0.52 |
| 3 | 190877111 | rs16866197 | 0.264 | -1.022 | -0.075 | 1.275 | -0.084 | 0.025 | 0.025 | 6.167 | 0.35 | 0.54 | 0.27 | 0.47 | 0.15 | 0.19 |
| 3 | 190880134 | rs11922372 | -0.607 | -0.812 | -0.2 | 1.061 | 0.033 | 0.005 | 0.005 | 6.415 | 0.44 | 0.71 | 0.37 | 0.6 | 0.26 | 0.42 |
| 3 | 190881904 | rs6793890 | 0.541 | -1.004 | -0.09 | 0.821 | -0.008 | 0 | 0 | 4.094 | 0.3 | 0.28 | 0.18 | 0.26 | 0.12 | 0.15 |
| 3 | 190884998 | rs9826780 | 0.859 | 0.196 | -0.672 | 0.582 | -0.038 | 0.047 | 0.047 | 0.142 | 0.15 | 0.05 | 0.12 | 0.04 | 0.2 | 0.07 |
| 3 | 190886678 | rs7619900 | -0.599 | -0.008 | -0.116 | 1.051 | 0.007 | -0.003 | -0.003 | 0.204 | 0.55 | 0.63 | 0.42 | 0.59 | 0.4 | 0.47 |
| 3 | 190891939 | rs13100749 | -0.414 | -0.047 | 0.131 | 1.251 | 0.007 | 0.009 | 0.009 | 0 | 0.53 | 0.46 | 0.35 | 0.4 | 0.38 | 0.42 |
| 3 | 190892500 | rs6780671 | -0.626 | -0.08 | 0.256 | 1.205 | 0.009 | -0.004 | -0.004 | 0.724 | 0.56 | 0.67 | 0.42 | 0.58 | 0.39 | 0.47 |
| 3 | 190895026 | rs17463141 | 1.378 | 0.708 | -0.509 | 1.208 | -0.043 | -0.053 | -0.053 | 0 | 0.11 | 0.02 | 0.11 | 0.19 | 0.2 | 0.25 |
| 3 | 190913446 | rs7615048 | -0.481 | 0.262 | -0.086 | 0.92 | 0.001 | 0.003 | 0.003 | 0.007 | 0.54 | 0.57 | 0.36 | 0.38 | 0.39 | 0.42 |
| 3 | 190928594 | rs9842080 | -0.082 | 0.292 | -0.77 | 0.36 | 0.019 | 0.015 | 0.015 | 1.212 | 0.46 | 0.66 | 0.25 | 0.1 | 0.3 | 0.23 |
| 3 | 190941728 | rs9883559 | 1.034 | -0.018 | -0.932 | 0.353 | -0.038 | 0.042 | 0.042 | 0.244 | 0.14 | 0.04 | 0.1 | 0.04 | 0.17 | 0.07 |
| 3 | 190945633 | rs6801419 | 0.031 | 0.266 | -0.759 | 0.746 | 0.034 | 0.037 | 0.037 | 0.453 | 0.48 | 0.67 | 0.3 | 0.3 | 0.3 | 0.28 |
| 3 | 190953635 | rs6762027 | -0.861 | 0.641 | -0.856 | 1.073 | -0.017 | -0.018 | -0.018 | 0 | 0.81 | 0.91 | 0.82 | 0.79 | 0.76 | 0.78 |
| 3 | 190955098 | rs9862736 | 0.479 | 0.231 | -0.328 | 1.414 | -0.014 | 0.04 | 0.04 | 1.326 | 0.71 | 0.37 | 0.66 | 0.53 | 0.73 | 0.71 |
| 3 | 190956807 | rs9869038 | 0.923 | 0.14 | -1.694 | 1.165 | -0.095 | -0.024 | -0.024 | 0 | 0.46 | 0.05 | 0.37 | 0.39 | 0.46 | 0.36 |
| 3 | 190958369 | rs12494322 | 0.49 | 0.357 | -1.669 | 0.935 | -0.046 | -0.075 | -0.075 | 0 | 0.26 | 0.06 | 0.35 | 0.37 | 0.28 | 0.41 |
| 3 | 190961156 | rs9857505 | 0.417 | 0.245 | -1.59 | 0.863 | -0.002 | 0.176 | 0.176 | 2.829 | 0.53 | 0.81 | 0.37 | 0.19 | 0.45 | 0.32 |
| 3 | 190961658 | rs16866360 | 0.416 | 0.416 | -1.731 | 0.948 | -0.067 | -0.075 | -0.075 | 0.048 | 0.25 | 0.06 | 0.36 | 0.38 | 0.29 | 0.39 |
| 3 | 190965972 | rs9855647 | 0.904 | -0.013 | -1.684 | 0.367 | -0.063 | 0.228 | 0.228 | 0.485 | 0.44 | 0.09 | 0.28 | 0.15 | 0.44 | 0.31 |
| 3 | 190967779 | rs2034771 | 0.251 | 0.111 | -1.171 | 0.662 | -0.043 | 0.006 | 0.006 | 5.354 | 0.24 | 0.09 | 0.35 | 0.18 | 0.27 | 0.35 |
| 3 | 190968964 | rs9833453 | -0.506 | 0.195 | -1.367 | 0.723 | -0.04 | 0.046 | 0.046 | 1.51 | 0.37 | 0.8 | 0.49 | 0.22 | 0.34 | 0.45 |
| 3 | 190973854 | rs9822737 | -0.145 | 0.509 | -0.579 | 0.753 | 0.031 | 0.074 | 0.074 | 0.002 | 0.31 | 0.43 | 0.5 | 0.41 | 0.32 | 0.47 |
| 3 | 190976054 | rs6444521 | -0.751 | 0.571 | -1.073 | 0.681 | -0.007 | -0.032 | -0.032 | 0 | 0.8 | 0.9 | 0.83 | 0.67 | 0.76 | 0.78 |
| 3 | 190978515 | rs9816545 | 0.198 | 0.636 | -1.075 | -0.232 | -0.016 | 0.131 | 0.131 | 0 | 0.31 | 0.09 | 0.21 | 0.06 | 0.35 | 0.22 |
| 3 | 190981868 | rs7637404 | -0.674 | 0.751 | -0.796 | -0.426 | 0.004 | 0.032 | 0.032 | 0 | 0.45 | 0.32 | 0.45 | 0.1 | 0.54 | 0.42 |
| 3 | 190982476 | rs7637969 | 0.131 | 0.876 | -0.993 | -0.346 | -0.014 | 0.011 | 0.011 | 1.247 | 0.17 | 0.1 | 0.24 | 0.04 | 0.21 | 0.24 |
| 3 | 190987067 | rs17465071 | -0.095 | 0.928 | -0.769 | -0.934 | 0.006 | -0.025 | -0.025 | 0 | 0.2 | 0.21 | 0.28 | 0.04 | 0.25 | 0.28 |
| 3 | 190987344 | rs9856076 | -0.335 | 0.238 | -0.601 | -0.481 | 0.016 | 0.019 | 0.019 | 0 | 0.49 | 0.44 | 0.48 | 0.1 | 0.59 | 0.47 |
| 3 | 190991384 | rs9857697 | -0.338 | 0.261 | -0.592 | -0.471 | 0.016 | 0.02 | 0.02 | 0 | 0.5 | 0.44 | 0.48 | 0.1 | 0.58 | 0.47 |
| 3 | 190994143 | rs4399929 | -0.751 | 0.204 | -0.191 | 0.347 | -0.013 | 0.008 | 0.008 | 0.056 | 0.28 | 0.56 | 0.3 | 0.3 | 0.26 | 0.28 |
| 3 | 190996107 | rs3816495 | -1.433 | -0.637 | 0.633 | 0.373 | -0.074 | 0.045 | 0.045 | 0.087 | 0.14 | 0.38 | 0.16 | 0.14 | 0.05 | 0.14 |
| 3 | 190996409 | rs16866431 | -1.433 | -0.637 | 0.633 | 0.375 | -0.074 | 0.038 | 0.038 | 0.087 | 0.14 | 0.38 | 0.16 | 0.13 | 0.05 | 0.13 |
| 3 | 191013346 | rs13067368 | -0.332 | -0.535 | 0.358 | 0.521 | -0.004 | 0.002 | 0.002 | 0 | 0.38 | 0.25 | 0.4 | 0.64 | 0.44 | 0.46 |
| 3 | 191014928 | rs1435066 | -1.199 | -0.179 | 0.283 | 0.044 | 0.004 | 0.005 | 0.005 | 0.017 | 0.6 | 0.68 | 0.64 | 0.64 | 0.65 | 0.59 |
| 3 | 191019603 | rs6796119 | -0.886 | -0.167 | 0.263 | 0.04 | 0.009 | 0.005 | 0.005 | 0.698 | 0.57 | 0.39 | 0.65 | 0.64 | 0.66 | 0.63 |
| 3 | 191020382 | rs293862 | -0.633 | -0.106 | 0.105 | 0.156 | -0.017 | -0.027 | -0.027 | 0.498 | 0.66 | 0.76 | 0.65 | 0.5 | 0.57 | 0.62 |
| 3 | 191020976 | rs150503 | -0.111 | -0.086 | 0.057 | 0.121 | -0.005 | 0 | 0 | 0 | 0.61 | 0.72 | 0.58 | 0.36 | 0.55 | 0.51 |
| 3 | 191031960 | rs10937476 | -0.818 | -0.329 | 0.574 | 0.248 | -0.005 | -0.04 | -0.04 | 1.647 | 0.26 | 0.44 | 0.31 | 0.13 | 0.23 | 0.25 |
| 3 | 191033815 | rs12495302 | -0.166 | -0.526 | -0.005 | 0.723 | 0.026 | -0.004 | -0.004 | 0 | 0.05 | 0 | 0.06 | 0.22 | 0.03 | 0.12 |
| 3 | 191040174 | rs4677722 | -1.117 | -0.276 | -0.095 | 0.39 | -0.015 | 0.057 | 0.057 | 0.287 | 0.54 | 0.73 | 0.56 | 0.67 | 0.5 | 0.58 |
| 3 | 191043229 | rs6779253 | 0.407 | -0.166 | 0.24 | 0.465 | -0.033 | 0.065 | 0.065 | 2.074 | 0.27 | 0.46 | 0.22 | 0.46 | 0.17 | 0.25 |
| 3 | 191043993 | rs13319838 | 0.243 | -0.068 | -0.092 | 0.166 | -0.015 | -0.004 | -0.004 | 0 | 0.19 | 0.16 | 0.26 | 0 | 0.28 | 0.15 |
| 3 | 191045551 | rs4677628 | -1.414 | -0.242 | -0.528 | 0.033 | -0.089 | 0.178 | 0.178 | 0.568 | 0.62 | 0.89 | 0.7 | 0.86 | 0.58 | 0.73 |
| 3 | 191046330 | rs6789787 | 0.181 | -0.751 | 0.025 | 0.649 | -0.038 | 0.109 | 0.109 | 1.381 | 0.25 | 0.47 | 0.31 | 0.54 | 0.16 | 0.3 |
| 3 | 191048462 | rs10937477 | -0.973 | -0.778 | -0.162 | 0.075 | 0.04 | 0.077 | 0.077 | 0.137 | 0.42 | 0.75 | 0.47 | 0.91 | 0.28 | 0.58 |
| 3 | 191053196 | rs293864 | 1.634 | -0.551 | -0.719 | -0.117 | -0.038 | 0.019 | 0.019 | 0.019 | 0.09 | 0.01 | 0.08 | 0.05 | 0.12 | 0.07 |
| 3 | 191059322 | rs1473869 | 1.499 | -0.695 | -0.235 | 0.517 | 0.001 | -0.001 | -0.001 | 0 | 0.09 | 0.09 | 0.07 | 0.26 | 0.08 | 0.09 |
| 3 | 191060954 | rs1473868 | 0.137 | -0.808 | 0.061 | 0.611 | -0.008 | 0.063 | 0.063 | 0 | 0.17 | 0.21 | 0.18 | 0.47 | 0.15 | 0.24 |
| 3 | 191072060 | rs448133 | -0.963 | -0.744 | -0.56 | 0.253 | 0.025 | 0.039 | 0.039 | 0 | 0.4 | 0.67 | 0.44 | 0.92 | 0.28 | 0.58 |
| 3 | 191075012 | rs1520203 | -0.466 | -0.385 | -0.488 | 0.49 | 0.011 | -0.022 | -0.022 | 0 | 0.31 | 0.4 | 0.35 | 0.26 | 0.36 | 0.27 |
| 3 | 191081344 | rs293801 | -0.35 | -0.425 | -0.282 | 0.935 | -0.039 | 0.088 | 0.088 | 0.476 | 0.25 | 0.54 | 0.25 | 0.44 | 0.17 | 0.3 |
| 3 | 191081486 | rs9868286 | 0.681 | -0.236 | -0.536 | 0.672 | -0.042 | 0.038 | 0.038 | 0.501 | 0.14 | 0.08 | 0.25 | 0 | 0.26 | 0.14 |
| 3 | 191087259 | rs4677726 | -0.43 | -0.366 | -0.473 | 0.744 | 0.013 | -0.006 | -0.006 | 0.003 | 0.25 | 0.31 | 0.33 | 0.25 | 0.33 | 0.22 |
| 3 | 191088902 | rs510014 | 1.508 | -0.391 | -0.656 | 0.993 | -0.015 | -0.028 | -0.028 | 0 | 0.06 | 0.01 | 0.09 | 0.22 | 0.08 | 0.17 |
| 3 | 191091320 | rs1435061 | -1.054 | -0.492 | -0.424 | 0.921 | -0.009 | -0.001 | -0.001 | 0.815 | 0.41 | 0.72 | 0.47 | 0.47 | 0.41 | 0.35 |
| 3 | 191093053 | rs2028572 | -1.054 | -0.492 | -0.424 | 0.921 | -0.009 | -0.001 | -0.001 | 0 | 0.41 | 0.72 | 0.47 | 0.47 | 0.41 | 0.35 |
| 3 | 191093080 | rs2028573 | 0.996 | -0.502 | -0.326 | 0.919 | -0.041 | 0.043 | 0.043 | 2.537 | 0.17 | 0.47 | 0.15 | 0.22 | 0.08 | 0.13 |
| 3 | 191093310 | rs4677728 | -1.051 | -0.503 | -0.388 | 0.914 | -0.008 | 0 | 0 | 0 | 0.41 | 0.72 | 0.47 | 0.47 | 0.41 | 0.35 |
| 3 | 191097966 | rs293813 | -1.638 | -0.49 | -0.422 | 0.872 | -0.03 | 0.053 | 0.053 | 0.815 | 0.49 | 0.85 | 0.58 | 0.69 | 0.5 | 0.52 |
| 3 | 191099468 | rs432085 | -2.143 | -0.644 | -0.497 | 0.786 | -0.048 | 0.256 | 0.256 | 0.459 | 0.57 | 0.91 | 0.7 | 0.91 | 0.54 | 0.71 |
| 3 | 191103490 | rs547195 | 1.464 | -0.632 | -0.494 | 0.792 | -0.029 | 0.057 | 0.057 | 0.685 | 0.5 | 0.15 | 0.42 | 0.31 | 0.51 | 0.48 |
| 3 | 191111160 | rs9841435 | -0.481 | -0.639 | -0.416 | 0.782 | 0 | -0.003 | -0.003 | 1.57 | 0.32 | 0.67 | 0.42 | 0.22 | 0.34 | 0.26 |
| 3 | 191112226 | rs6774730 | 1.021 | -0.61 | -0.39 | 0.811 | -0.043 | 0.039 | 0.039 | 2.494 | 0.17 | 0.47 | 0.15 | 0.22 | 0.08 | 0.13 |
| 3 | 191113239 | rs167151 | -0.414 | -0.607 | -0.436 | 0.814 | 0 | 0.013 | 0.013 | 0 | 0.83 | 0.83 | 0.84 | 0.54 | 0.84 | 0.74 |
| 3 | 191114266 | rs2293377 | -1.041 | -0.444 | -0.379 | 0.922 | -0.001 | -0.003 | -0.003 | 0 | 0.39 | 0.59 | 0.52 | 0.45 | 0.38 | 0.45 |
| 3 | 191117867 | rs9812544 | -0.391 | -0.29 | -0.65 | 0.935 | -0.072 | -0.071 | -0.071 | 0 | 0.22 | 0.08 | 0.36 | 0.23 | 0.31 | 0.32 |
| 3 | 191124232 | rs293855 | -1.369 | -1.12 | -0.143 | 1.482 | -0.015 | -0.025 | -0.025 | 0 | 0.94 | 0.99 | 0.91 | 0.78 | 0.92 | 0.83 |
| 3 | 191128441 | rs293861 | 0.695 | -1.236 | -0.012 | 1.449 | -0.012 | 0.009 | 0.009 | 2.026 | 0.68 | 0.43 | 0.7 | 0.32 | 0.77 | 0.62 |

| | | | | | | | | | | | | | | | | |
|---|---|---|---|---|---|---|---|---|---|---|---|---|---|---|---|---|
| 3 | 191129438 | rs440266 | 0.838 | -1.242 | -0.082 | 1.446 | 0.01 | -0.007 | -0.007 | 0.038 | 0.67 | 0.32 | 0.68 | 0.32 | 0.77 | 0.62 |
| 3 | 191132167 | rs293845 | -1.04 | -0.913 | -0.115 | 1.315 | -0.015 | -0.025 | -0.025 | 0 | 0.94 | 0.99 | 0.91 | 0.78 | 0.92 | 0.83 |
| 3 | 191135558 | rs7618303 | -0.576 | -1.045 | 0.098 | 1.202 | -0.044 | 0.132 | 0.132 | 0.171 | 0.24 | 0.42 | 0.27 | 0.45 | 0.12 | 0.32 |
| 3 | 191143631 | rs569767 | 0.162 | -1.006 | 0.435 | 0.97 | -0.04 | 0.027 | 0.027 | 0 | 0.86 | 0.65 | 0.86 | 0.79 | 0.92 | 0.85 |
| 3 | 191144599 | rs9290987 | -0.692 | -0.982 | 1.056 | 0.656 | 0.027 | -0.043 | -0.043 | 0 | 0.1 | 0.34 | 0.08 | 0 | 0.1 | 0.09 |
| 3 | 191145049 | rs696352 | 1.495 | -1.345 | 0.519 | -0.127 | -0.005 | 0.064 | 0.064 | 0 | 0.42 | 0.3 | 0.43 | 0.07 | 0.51 | 0.3 |
| 3 | 191147527 | rs16822489 | 0.817 | -1.44 | 0.134 | 0.798 | -0.003 | -0.016 | -0.016 | 0.016 | 0.06 | 0.01 | 0.13 | 0.24 | 0.06 | 0.14 |
| 3 | 191147791 | rs389062 | -0.05 | -1.64 | 0.688 | 0.812 | -0.069 | -0.029 | -0.029 | 0.029 | 0.78 | 0.42 | 0.86 | 0.82 | 0.89 | 0.83 |
| 3 | 191150836 | rs293820 | -0.328 | -1.629 | 0.697 | 0.844 | -0.067 | 0.03 | 0.03 | 0.358 | 0.8 | 0.47 | 0.82 | 0.74 | 0.89 | 0.82 |
| 3 | 191154590 | rs13100475 | 0.203 | -1.2 | 0.069 | 0.893 | 0.001 | -0.042 | -0.042 | 0.31 | 0.23 | 0.14 | 0.22 | 0.44 | 0.22 | 0.3 |
| 3 | 191159716 | rs12633934 | 0.575 | -1.075 | -0.004 | 0.963 | -0.017 | -0.019 | -0.019 | 0.011 | 0.2 | 0.09 | 0.15 | 0.27 | 0.25 | 0.18 |
| 3 | 191169024 | rs167149 | 1.736 | -0.831 | 0.079 | 0.305 | -0.028 | 0.046 | 0.046 | 0 | 0.36 | 0.25 | 0.37 | 0.06 | 0.44 | 0.26 |
| 3 | 191171150 | rs406821 | 0.692 | -0.96 | 0.334 | 0.548 | -0.008 | 0.002 | 0.002 | 0 | 0.64 | 0.56 | 0.6 | 0.42 | 0.72 | 0.59 |
| 3 | 191172809 | rs430975 | -0.726 | -0.991 | -0.298 | 0.687 | 0 | -0.046 | -0.046 | 0.654 | 0.83 | 0.92 | 0.85 | 0.69 | 0.85 | 0.79 |
| 3 | 191173649 | rs550947 | 0.411 | -1.028 | -0.232 | 0.666 | 0.004 | -0.002 | -0.002 | 0 | 0.72 | 0.74 | 0.75 | 0.69 | 0.77 | 0.73 |
| 3 | 191186292 | rs430271 | -1.925 | -0.902 | -0.331 | 0.75 | -0.017 | -0.017 | -0.017 | 0 | 0.91 | 0.99 | 0.89 | 0.84 | 0.91 | 0.87 |
| 3 | 191186349 | rs7356085 | 0.275 | -0.909 | -0.217 | 0.74 | 0.009 | -0.014 | -0.014 | 0 | 0.28 | 0.17 | 0.22 | 0.36 | 0.29 | 0.33 |
| 3 | 191189965 | rs2886242 | 0.026 | -0.575 | -0.454 | 1.148 | -0.015 | -0.097 | -0.097 | 0 | 0.17 | 0.02 | 0.2 | 0.27 | 0.13 | 0.22 |
| 3 | 191191370 | rs430388 | -0.231 | -0.266 | -0.089 | 1.107 | 0 | 0.015 | 0.015 | 0 | 0.85 | 0.77 | 0.83 | 0.69 | 0.86 | 0.8 |
| 3 | 191207636 | rs4234621 | -0.543 | -0.524 | -0.196 | 0.803 | 0.035 | 0.055 | 0.055 | 0 | 0.45 | 0.7 | 0.5 | 0.68 | 0.37 | 0.44 |
| 3 | 191213432 | rs2090721 | 0.05 | 0.411 | -0.697 | 0.377 | -0.079 | 0.178 | 0.178 | 0 | 0.61 | 0.87 | 0.65 | 0.8 | 0.52 | 0.57 |
| 3 | 191213468 | rs4687218 | -0.404 | 0.469 | 0.529 | 1.125 | -0.004 | 0.007 | 0.007 | 0 | 0.39 | 0.62 | 0.5 | 0.37 | 0.42 | 0.3 |
| 3 | 191216000 | rs7642724 | -1.352 | 0.491 | 0.739 | 2.242 | 0.025 | 0.469 | 0.469 | 0 | 0.07 | 0.33 | 0.2 | 0.36 | 0.04 | 0.09 |
| 3 | 191227296 | rs11922996 | 0.674 | 1.054 | -0.867 | 1.395 | -0.035 | -0.034 | -0.034 | 0.47 | 0.61 | 0.28 | 0.45 | 0.48 | 0.55 | 0.6 |
| 3 | 191228990 | rs1016618 | -0.84 | 1.146 | -0.379 | 1.547 | -0.015 | 0.001 | 0.001 | 0 | 0.07 | 0.01 | 0.17 | 0.33 | 0.08 | 0.1 |
| 3 | 191232772 | rs9820708 | 0.175 | 1.107 | -0.143 | 1.296 | -0.006 | 0.001 | 0.001 | 0 | 0.73 | 0.75 | 0.67 | 0.84 | 0.64 | 0.73 |
| 3 | 191246141 | rs2365028 | -0.981 | 0.724 | -0.412 | 1.702 | 0.019 | 0.008 | 0.008 | 0 | 0.07 | 0.01 | 0.16 | 0.39 | 0.04 | 0.12 |
| 3 | 191254431 | rs872581 | 0.938 | 0.822 | -0.003 | 1.432 | -0.017 | 0.004 | 0.004 | 1.137 | 0.59 | 0.29 | 0.46 | 0.39 | 0.55 | 0.53 |
| 3 | 191256891 | rs6794065 | 0.609 | 0.88 | -0.03 | 1.453 | -0.012 | -0.012 | -0.012 | 0 | 0.68 | 0.72 | 0.55 | 0.41 | 0.63 | 0.62 |
| 3 | 191257864 | rs7647049 | -0.091 | 1.014 | -0.125 | 1.571 | -0.068 | -0.008 | -0.008 | 0.467 | 0.8 | 0.9 | 0.78 | 0.78 | 0.69 | 0.79 |
| 3 | 191258074 | rs7625165 | 1.112 | 0.808 | 1.049 | 1.901 | 0.026 | -0.014 | -0.014 | 0 | 0.67 | 0.54 | 0.6 | 0.29 | 0.62 | 0.52 |
| 3 | 191262206 | rs11924324 | -0.077 | 0.119 | 0.899 | 0.964 | -0.003 | -0.023 | -0.023 | 0.014 | 0.3 | 0.41 | 0.31 | 0.17 | 0.28 | 0.26 |
| 3 | 191267203 | rs1405037 | 0.032 | -0.465 | 0.593 | 0.703 | 0.01 | 0.016 | 0.016 | 0 | 0.13 | 0.09 | 0.07 | 0.05 | 0.08 | 0.15 |
| 3 | 191267342 | rs4686577 | 0.399 | -0.288 | 0.764 | 0.305 | 0.021 | 0.051 | 0.051 | 0 | 0.41 | 0.26 | 0.35 | 0.08 | 0.37 | 0.27 |
| 3 | 191268184 | rs1525923 | 0.412 | -0.3 | 0.759 | 0.3 | 0.021 | 0.05 | 0.05 | 0 | 0.4 | 0.26 | 0.35 | 0.08 | 0.37 | 0.27 |
| 3 | 191272414 | rs2012405 | -0.829 | -0.158 | 0.756 | 0.506 | 0.014 | -0.004 | -0.004 | 0 | 0.74 | 0.69 | 0.76 | 0.73 | 0.69 | 0.68 |
| 3 | 191274891 | rs7629043 | -0.706 | -0.228 | 0.879 | 0.481 | 0.029 | 0.11 | 0.11 | 0 | 0.49 | 0.71 | 0.51 | 0.78 | 0.41 | 0.59 |
| 3 | 191275468 | rs6444557 | 0.359 | -0.137 | 1.076 | 0.613 | -0.002 | -0.007 | -0.007 | 0 | 0.33 | 0.62 | 0.34 | 0.18 | 0.3 | 0.33 |
| 3 | 191286527 | rs6766478 | -1.228 | 0.564 | 0.859 | 0.756 | -0.106 | -0.007 | -0.007 | 0 | 0.8 | 0.99 | 0.8 | 0.81 | 0.74 | 0.72 |
| 3 | 191291423 | rs10937502 | -0.6 | 0.416 | 1.278 | 0.524 | -0.026 | -0.029 | -0.029 | 0.516 | 0.75 | 0.84 | 0.72 | 0.34 | 0.7 | 0.63 |
| 3 | 191300252 | rs7434135 | -0.305 | 0.389 | 1.171 | 0.53 | 0.018 | 0.023 | 0.023 | 0.02 | 0.72 | 0.62 | 0.71 | 0.34 | 0.7 | 0.62 |
| 3 | 191302048 | rs11709456 | -0.095 | 0.471 | 1.052 | 0.343 | 0.006 | 0 | 0 | 0.09 | 0.7 | 0.41 | 0.69 | 0.28 | 0.7 | 0.58 |
| 3 | 191305238 | rs7613979 | -0.149 | 0.469 | 1.131 | 0.342 | 0.012 | 0.001 | 0.001 | 0.033 | 0.71 | 0.47 | 0.69 | 0.28 | 0.7 | 0.58 |
| 3 | 191312423 | rs7629648 | 0.021 | 0.392 | 1.036 | 0.087 | -0.013 | 0.041 | 0.041 | 0 | 0.38 | 0.27 | 0.39 | 0.26 | 0.39 | 0.44 |
| 3 | 191314079 | rs4686582 | 0.393 | 0.459 | 0.978 | 0.175 | -0.009 | 0.05 | 0.05 | 0 | 0.59 | 0.33 | 0.62 | 0.22 | 0.62 | 0.44 |
| 3 | 191317215 | rs1949751 | 0.342 | 0.472 | 0.94 | 0.181 | -0.007 | 0.033 | 0.033 | 0 | 0.6 | 0.38 | 0.62 | 0.23 | 0.62 | 0.45 |
| 3 | 191318381 | rs7610040 | -0.027 | 0.356 | 0.897 | 0.192 | -0.014 | -0.028 | -0.028 | 0 | 0.13 | 0.25 | 0.09 | 0.05 | 0.08 | 0.13 |
| 3 | 191320278 | rs9840268 | 1.013 | 0.279 | -0.41 | -0.054 | -0.015 | 0.031 | 0.031 | 1.468 | 0.1 | 0.04 | 0.16 | 0 | 0.13 | 0.09 |
| 3 | 191320935 | rs7618931 | 0.747 | 0.341 | -0.74 | -0.043 | -0.027 | 0.003 | 0.003 | 0.012 | 0.23 | 0.11 | 0.27 | 0.24 | 0.26 | 0.27 |
| 3 | 191320988 | rs9825318 | 0.292 | 0.361 | -0.11 | 0.366 | -0.002 | -0.007 | -0.007 | 10.708 | 0.13 | 0.19 | 0.16 | 0.07 | 0.13 | 0.09 |
| 3 | 191323202 | rs4686584 | -1.096 | 0.487 | -0.968 | 0.443 | -0.118 | -0.067 | -0.067 | 0.018 | 0.75 | 0.99 | 0.81 | 0.68 | 0.71 | 0.74 |
| 3 | 191323429 | rs1523477 | 1.754 | -0.258 | -1.152 | -0.374 | -0.002 | 0.153 | 0.153 | 3.297 | 0.24 | 0.03 | 0.17 | 0.01 | 0.21 | 0.05 |
| 3 | 191323910 | rs1523478 | 1.615 | -0.287 | -0.593 | -0.3 | 0.009 | 0.008 | 0.008 | 2.267 | 0.26 | 0.21 | 0.2 | 0.04 | 0.21 | 0.08 |
| 3 | 191331505 | rs10937504 | -0.583 | -0.282 | 0.75 | 0.379 | -0.028 | -0.107 | -0.107 | 1.132 | 0.19 | 0.5 | 0.22 | 0.03 | 0.15 | 0.13 |
| 3 | 191342386 | rs2037578 | -1.532 | -0.23 | 0.906 | 0.801 | -0.004 | 0.054 | 0.054 | 1.194 | 0.28 | 0.53 | 0.32 | 0.31 | 0.26 | 0.31 |
| 3 | 191342471 | rs9843703 | 1.025 | -0.222 | 0.849 | 0.917 | -0.005 | -0.004 | -0.004 | 0 | 0.06 | 0.04 | 0.05 | 0.07 | 0.07 | 0.13 |
| 3 | 191343146 | rs2037576 | -1.569 | -0.233 | 1.043 | 0.899 | -0.002 | 0.037 | 0.037 | 1.313 | 0.28 | 0.47 | 0.32 | 0.3 | 0.26 | 0.31 |
| 3 | 191344113 | rs9816423 | 0.884 | -0.308 | 1.059 | 0.795 | -0.003 | -0.043 | -0.043 | 0 | 0.1 | 0.18 | 0.16 | 0 | 0.12 | 0.07 |
| 3 | 191348064 | rs13059172 | 0.659 | -0.277 | 0.976 | 0.835 | -0.01 | -0.028 | -0.028 | 0 | 0.06 | 0.1 | 0.03 | 0 | 0.03 | 0.02 |
| 3 | 191348677 | rs12491931 | -1.562 | -0.326 | 1.127 | 0.928 | -0.027 | 0.209 | 0.209 | 0.311 | 0.13 | 0.55 | 0.13 | 0.24 | 0.07 | 0.1 |
| 3 | 191351574 | rs11713378 | 0.773 | -0.311 | 0.781 | 0.923 | 0.001 | 0.001 | 0.001 | 1.015 | 0.11 | 0.08 | 0.16 | 0.09 | 0.12 | 0.13 |
| 3 | 191361882 | rs13068846 | -0.696 | -0.337 | 0.82 | 0.908 | -0.017 | -0.011 | -0.011 | 0 | 0.09 | 0.31 | 0.06 | 0.04 | 0.03 | 0.05 |
| 3 | 191362353 | rs1916631 | -0.666 | -0.331 | 0.823 | 0.901 | -0.016 | -0.018 | -0.018 | 0 | 0.09 | 0.31 | 0.06 | 0.03 | 0.03 | 0.05 |
| 3 | 191365134 | rs9843456 | -1.836 | -0.261 | 0.277 | 1.014 | 0.001 | -0.01 | -0.01 | 0 | 0.31 | 0.73 | 0.35 | 0.39 | 0.27 | 0.36 |
| 3 | 191366468 | rs1916636 | -1.8 | -0.243 | 0.304 | 1.023 | 0.001 | -0.007 | -0.007 | 0 | 0.31 | 0.72 | 0.35 | 0.39 | 0.26 | 0.35 |
| 3 | 191366512 | rs1916637 | -0.946 | -0.367 | 0.308 | 0.903 | -0.03 | -0.034 | -0.034 | 0.114 | 0.22 | 0.66 | 0.24 | 0.12 | 0.15 | 0.18 |
| 3 | 191370717 | rs1818869 | -1.602 | 0.124 | -0.661 | 0.638 | -0.036 | 0.13 | 0.13 | 0 | 0.79 | 0.99 | 0.9 | 0.98 | 0.82 | 0.84 |
| 3 | 191390350 | rs13100354 | 0.249 | 0.221 | 0.43 | 1.349 | -0.011 | -0.034 | -0.034 | 0 | 0.06 | 0.15 | 0.04 | 0 | 0.03 | 0.02 |
| 3 | 191401028 | rs9837138 | -1.403 | 0.132 | 0.715 | 1.536 | -0.025 | 0.041 | 0.041 | 1.09 | 0.25 | 0.64 | 0.29 | 0.31 | 0.19 | 0.24 |

| | | | | | | | | | | | | | | | | |
|---|---|---|---|---|---|---|---|---|---|---|---|---|---|---|---|---|
| 3 | 191401581 | rs9290998 | 0.05 | 0.361 | 0.497 | 1.022 | 0.003 | 0.006 | 0.006 | 0.733 | 0.13 | 0.33 | 0.18 | 0.09 | 0.12 | 0.12 |
| 3 | 191410032 | rs13324976 | 0.479 | 0.655 | -0.003 | 1.17 | -0.004 | -0.004 | -0.004 | 0 | 0.06 | 0.04 | 0.05 | 0.09 | 0.09 | 0.12 |
| 3 | 191411106 | rs4687252 | -0.647 | 0.319 | 0.127 | 0.888 | -0.024 | -0.045 | -0.045 | 0 | 0.19 | 0.48 | 0.21 | 0.09 | 0.15 | 0.15 |
| 3 | 191413314 | rs1083698 | -1.173 | 0.376 | -1.116 | 1.031 | -0.011 | 0.035 | 0.035 | 1.306 | 0.48 | 0.86 | 0.55 | 0.59 | 0.42 | 0.62 |
| 3 | 191416890 | rs779504 | 0.095 | 0.984 | -1.219 | 1.332 | -0.071 | -0.09 | -0.09 | 0 | 0.19 | 0.03 | 0.24 | 0.31 | 0.28 | 0.26 |
| 3 | 191423639 | rs1083696 | -1.455 | 0.72 | -0.497 | 1.151 | -0.004 | 0.003 | 0.003 | 0 | 0.41 | 0.61 | 0.5 | 0.58 | 0.43 | 0.5 |
| 3 | 191429026 | rs3106816 | -0.557 | 0.717 | -0.003 | 1.249 | -0.01 | -0.019 | -0.019 | 0 | 0.29 | 0.25 | 0.31 | 0.41 | 0.35 | 0.38 |
| 3 | 191432232 | rs9816458 | -1.756 | 0.423 | 0.069 | 1.281 | -0.002 | 0.017 | 0.017 | 0.498 | 0.27 | 0.64 | 0.41 | 0.51 | 0.24 | 0.36 |
| 3 | 191432503 | rs11707389 | -0.609 | 0.509 | 0.432 | 1.393 | 0 | 0.068 | 0.068 | 0 | 0.14 | 0.22 | 0.24 | 0.38 | 0.17 | 0.27 |
| 3 | 191433799 | rs7619568 | -0.528 | 0.52 | 0.221 | 1.268 | 0.039 | 0.078 | 0.078 | 0 | 0.07 | 0.23 | 0.15 | 0.13 | 0.06 | 0.06 |
| 3 | 191456220 | rs709086 | 0.709 | 0.594 | -0.039 | 1.367 | -0.061 | -0.076 | -0.076 | 0 | 0.18 | 0.05 | 0.25 | 0.32 | 0.28 | 0.28 |
| 3 | 191460727 | rs9844345 | 0.323 | 0.456 | -0.163 | 1.201 | -0.013 | 0.05 | 0.05 | 0 | 0.09 | 0.01 | 0.15 | 0 | 0.11 | 0.04 |
| 3 | 191465461 | rs9842612 | -0.81 | 0.439 | 0.133 | 1.391 | 0 | -0.004 | -0.004 | 5.854 | 0.15 | 0.13 | 0.21 | 0.35 | 0.18 | 0.23 |
| 3 | 191465981 | rs1516493 | -1.225 | 0.366 | 0.354 | 1.359 | -0.015 | 0.077 | 0.077 | 0.29 | 0.18 | 0.4 | 0.25 | 0.35 | 0.18 | 0.23 |
| 3 | 191470965 | rs709102 | -0.725 | 0.318 | 0.287 | 1.257 | 0.036 | 0.059 | 0.059 | 0 | 0.31 | 0.45 | 0.37 | 0.42 | 0.37 | 0.44 |
| 3 | 191478656 | rs709113 | -1.683 | 0.432 | 0.201 | 1.171 | -0.002 | 0.005 | 0.005 | 0 | 0.44 | 0.57 | 0.56 | 0.76 | 0.54 | 0.62 |
| 3 | 191498409 | rs709125 | 0.65 | 0.762 | -0.298 | 1.395 | -0.059 | -0.064 | -0.064 | 0 | 0.18 | 0.05 | 0.25 | 0.27 | 0.28 | 0.26 |
| 3 | 191499500 | rs1516496 | 0.474 | 0.668 | -0.005 | 1.194 | 0.002 | 0.001 | 0.001 | 0.018 | 0.06 | 0.06 | 0.06 | 0.09 | 0.08 | 0.13 |
| 3 | 191506171 | rs6803696 | 0.709 | 0.692 | -0.059 | 1.204 | 0.01 | 0.008 | 0.008 | 0.049 | 0.05 | 0.07 | 0.06 | 0.09 | 0.08 | 0.13 |
| 3 | 191512568 | rs11711997 | -0.08 | 0.241 | -0.145 | 0.94 | 0.007 | 0.016 | 0.016 | 0.081 | 0.09 | 0.05 | 0.14 | 0 | 0.12 | 0.08 |
| 3 | 191516568 | rs1083661 | -0.239 | 0.458 | -0.221 | 1.453 | -0.086 | -0.077 | -0.077 | 0 | 0.25 | 0.07 | 0.31 | 0.32 | 0.35 | 0.34 |
| 3 | 191517245 | rs779264 | -0.623 | 0.244 | 0.562 | 1.517 | 0.013 | 0.018 | 0.018 | 0 | 0.3 | 0.37 | 0.34 | 0.38 | 0.37 | 0.37 |
| 3 | 191518473 | rs11929015 | -0.67 | -0.106 | 0.896 | 1.651 | 0.002 | 0.183 | 0.183 | 0.172 | 0.08 | 0.35 | 0.09 | 0.28 | 0.03 | 0.14 |
| 3 | 191537433 | rs13062679 | 0.276 | -0.131 | 1.102 | 1.285 | 0.004 | -0.015 | -0.015 | 0.455 | 0.05 | 0.21 | 0.03 | 0 | 0.02 | 0.01 |
| 3 | 191575240 | rs1083673 | -0.451 | 0.057 | -0.116 | 1.379 | -0.048 | -0.02 | -0.02 | 0 | 0.31 | 0.19 | 0.42 | 0.64 | 0.45 | 0.53 |
| 3 | 191597705 | rs6444590 | 0.829 | -0.231 | -0.497 | 0.772 | 0.001 | -0.001 | -0.001 | 0.148 | 0.3 | 0.27 | 0.3 | 0.12 | 0.26 | 0.25 |
| 3 | 191608214 | rs9816055 | -0.197 | -0.657 | -0.46 | 0.604 | -0.015 | -0.009 | -0.009 | 0.052 | 0.66 | 0.76 | 0.61 | 0.63 | 0.55 | 0.49 |
| 3 | 191624783 | rs6444592 | 0.919 | -0.67 | -0.432 | 0.589 | 0 | 0 | 0 | 0 | 0.31 | 0.3 | 0.36 | 0.33 | 0.36 | 0.33 |
| 3 | 191625177 | rs4497982 | 0.109 | -0.262 | -0.609 | 0.706 | -0.011 | -0.007 | -0.007 | 0.09 | 0.66 | 0.72 | 0.61 | 0.63 | 0.61 | 0.49 |
| 3 | 191628429 | rs7621589 | 0.176 | -0.197 | -0.818 | 0.378 | -0.003 | -0.006 | -0.006 | 0 | 0.68 | 0.73 | 0.59 | 0.41 | 0.62 | 0.46 |
| 3 | 191629821 | rs13087722 | 1.273 | -0.525 | -1.359 | -0.278 | -0.084 | 0.163 | 0.163 | 0.062 | 0.29 | 0.07 | 0.3 | 0.01 | 0.35 | 0.19 |
| 3 | 191630044 | rs9838431 | -0.405 | -0.113 | -1.347 | 0.321 | 0.015 | -0.006 | -0.006 | 0 | 0.7 | 0.75 | 0.72 | 0.68 | 0.72 | 0.56 |
| 3 | 191637415 | rs6806006 | -0.113 | -0.3 | -1.276 | 0.191 | 0.006 | -0.003 | -0.003 | 0 | 0.68 | 0.71 | 0.6 | 0.4 | 0.68 | 0.46 |
| 3 | 191640921 | rs4299432 | 2.51 | -0.581 | -1.115 | 0.123 | -0.011 | -0.003 | -0.003 | 0 | 0.21 | 0.14 | 0.18 | 0.26 | 0.22 | 0.12 |
| 3 | 191644758 | rs7613485 | 0.143 | -0.743 | -0.56 | 0.167 | -0.013 | -0.009 | -0.009 | 0.244 | 0.64 | 0.54 | 0.59 | 0.38 | 0.67 | 0.46 |
| 3 | 191650264 | rs9851745 | 1.705 | -0.572 | -1 | -0.075 | 0.005 | -0.044 | -0.044 | 0 | 0.1 | 0.02 | 0.08 | 0.19 | 0.09 | 0.13 |
| 3 | 191655531 | rs4443117 | 0.827 | -0.587 | -1.144 | -0.18 | -0.059 | -0.09 | -0.09 | 0 | 0.15 | 0.01 | 0.17 | 0.3 | 0.21 | 0.18 |
| 3 | 191657940 | rs9291002 | -0.325 | -0.501 | -0.569 | -0.444 | 0.001 | -0.019 | -0.019 | 0.192 | 0.47 | 0.54 | 0.42 | 0.14 | 0.44 | 0.33 |
| 3 | 191659343 | rs13086796 | 0.813 | -0.379 | -0.825 | -0.438 | -0.092 | 0.189 | 0.189 | 0.109 | 0.29 | 0.04 | 0.29 | 0.01 | 0.33 | 0.19 |
| 3 | 191671970 | rs1499840 | 0.174 | -0.73 | 0.058 | 0.117 | 0.006 | 0.106 | 0.106 | 0 | 0.11 | 0.3 | 0.19 | 0.25 | 0.07 | 0.14 |
| 3 | 191675760 | rs12494587 | 0.229 | -0.636 | -0.205 | 0.027 | 0.002 | 0.009 | 0.009 | 0 | 0.32 | 0.27 | 0.29 | 0.29 | 0.31 | 0.41 |
| 3 | 191679473 | rs3863984 | -2.989 | -0.524 | -0.136 | 0.238 | -0.002 | -0.03 | -0.03 | 0.759 | 0.82 | 0.91 | 0.87 | 0.69 | 0.82 | 0.87 |
| 3 | 191682693 | rs2366568 | -1.178 | -0.541 | -0.105 | 0.045 | -0.001 | 0.042 | 0.042 | 0.319 | 0.74 | 0.5 | 0.69 | 0.41 | 0.77 | 0.74 |
| 3 | 191684827 | rs2669973 | -1.268 | -0.826 | -0.153 | -0.344 | -0.009 | 0.015 | 0.015 | 0.783 | 0.69 | 0.47 | 0.67 | 0.46 | 0.75 | 0.74 |
| 3 | 191687812 | rs9861268 | -0.281 | -0.797 | -0.216 | -0.592 | -0.005 | 0.019 | 0.019 | 0.154 | 0.46 | 0.56 | 0.47 | 0.35 | 0.49 | 0.43 |
| 3 | 191694215 | rs4687267 | 0.509 | -0.751 | -0.355 | -0.52 | -0.002 | -0.004 | -0.004 | 3.816 | 0.28 | 0.24 | 0.28 | 0.34 | 0.3 | 0.37 |
| 3 | 191708375 | rs11915453 | 1.733 | -0.789 | -0.816 | -0.602 | -0.042 | 0.05 | 0.05 | 0 | 0.22 | 0.09 | 0.27 | 0.05 | 0.33 | 0.17 |
| 3 | 191711064 | rs6764696 | -2.788 | -0.834 | -0.867 | -0.601 | 0.007 | -0.043 | -0.043 | 0 | 0.77 | 0.89 | 0.85 | 0.71 | 0.8 | 0.83 |
| 3 | 191719292 | rs1392204 | 0.975 | -0.896 | -0.725 | -0.667 | 0.007 | 0.058 | 0.058 | 0.814 | 0.12 | 0.21 | 0.13 | 0.31 | 0.11 | 0.18 |
| 3 | 191720301 | rs13062497 | -1.22 | -0.888 | -0.758 | -0.671 | 0.001 | -0.002 | -0.002 | 0 | 0.66 | 0.69 | 0.72 | 0.41 | 0.68 | 0.65 |
| 3 | 191724304 | rs2649802 | -2.31 | -0.591 | -1.356 | -0.55 | 0.006 | -0.053 | -0.053 | 0 | 0.77 | 0.92 | 0.86 | 0.71 | 0.8 | 0.83 |
| 3 | 191724654 | rs7627247 | 1.398 | -0.164 | -1.324 | -0.611 | -0.051 | 0.063 | 0.063 | 0 | 0.22 | 0.06 | 0.27 | 0.04 | 0.33 | 0.17 |
| 3 | 191726598 | rs884298 | 0.027 | -0.389 | -0.699 | -0.288 | -0.001 | -0.007 | -0.007 | 0.406 | 0.28 | 0.26 | 0.28 | 0.34 | 0.29 | 0.37 |
| 3 | 191728386 | rs2649803 | -2.088 | -0.496 | -0.972 | -0.191 | 0.009 | -0.099 | -0.099 | 0 | 0.78 | 0.99 | 0.88 | 0.71 | 0.84 | 0.89 |
| 3 | 191729056 | rs13084806 | -0.09 | -0.626 | -0.453 | -0.412 | -0.005 | 0.011 | 0.011 | 0 | 0.36 | 0.51 | 0.47 | 0.46 | 0.39 | 0.53 |
| 3 | 191735799 | rs13090097 | 2.11 | -0.005 | -0.645 | -0.474 | -0.08 | 0.047 | 0.047 | 0 | 0.14 | 0.02 | 0.2 | 0.04 | 0.22 | 0.12 |
| 3 | 191736737 | rs1491523 | -0.9 | -0.311 | 0.119 | -0.47 | 0.005 | 0.006 | 0.006 | 0 | 0.52 | 0.58 | 0.55 | 0.47 | 0.48 | 0.62 |
| 3 | 191738026 | rs2087698 | -1.579 | -0.325 | 0.038 | -0.444 | -0.054 | -0.032 | -0.032 | 0.058 | 0.63 | 0.88 | 0.66 | 0.68 | 0.62 | 0.76 |
| 3 | 191739586 | rs6796558 | -0.034 | -1.029 | 0.436 | -1.18 | 0.028 | 0.019 | 0.019 | 0.566 | 0.06 | 0.33 | 0.04 | 0 | 0.02 | 0 |
| 3 | 191760451 | rs1491520 | 0.798 | -0.95 | 0.258 | -1.111 | 0.036 | 0.079 | 0.079 | 0 | 0.11 | 0.27 | 0.1 | 0.24 | 0.14 | 0.15 |
| 3 | 191761351 | rs7629822 | -1.422 | -1.229 | -0.105 | -1.422 | -0.001 | 0.065 | 0.065 | 0.856 | 0.52 | 0.94 | 0.5 | 0.35 | 0.42 | 0.51 |
| 3 | 191765223 | rs6773456 | -1.606 | -1.225 | -0.114 | -1.42 | -0.041 | 0.023 | 0.023 | 1.231 | 0.54 | 0.95 | 0.57 | 0.38 | 0.45 | 0.55 |
| 3 | 191769485 | rs6769307 | -0.747 | -1.519 | 0.102 | -1.566 | 0.039 | 0.046 | 0.046 | 0 | 0.39 | 0.67 | 0.39 | 0.11 | 0.28 | 0.35 |
| 3 | 191775123 | rs10755147 | 0.282 | -1.558 | 0.144 | -1.567 | -0.005 | -0.021 | -0.021 | 0 | 0.26 | 0.23 | 0.31 | 0.11 | 0.23 | 0.25 |
| 3 | 191778698 | rs6784780 | 1.102 | -1.417 | 0.304 | -1.574 | -0.017 | -0.044 | -0.044 | 0 | 0.12 | 0.15 | 0.14 | 0.01 | 0.09 | 0.1 |
| 3 | 191780532 | rs12053855 | 0.519 | -1.441 | 0.15 | -1.571 | -0.004 | -0.04 | -0.04 | 0 | 0.62 | 0.44 | 0.69 | 0.91 | 0.72 | 0.65 |
| 3 | 191783287 | rs9291016 | 0.503 | -1.232 | 0.131 | -1.318 | 0.002 | 0.002 | 0.002 | 0 | 0.63 | 0.69 | 0.69 | 0.66 | 0.68 | 0.63 |
| 3 | 191797614 | rs6444628 | -0.212 | -1.3 | 0.151 | -1.468 | -0.012 | -0.045 | -0.045 | 0.074 | 0.75 | 0.76 | 0.86 | 0.91 | 0.8 | 0.76 |
| 3 | 191802191 | rs10804940 | 0.96 | -0.685 | -0.611 | -1.014 | 0.077 | 0.108 | 0.108 | 0 | 0.59 | 0.21 | 0.69 | 0.78 | 0.68 | 0.54 |

| | | | | | | | | | | | | | | | | |
|---|---|---|---|---|---|---|---|---|---|---|---|---|---|---|---|---|
| 3 | 191803483 | rs9878232 | 1.043 | -0.789 | -0.807 | -1.052 | 0.003 | 0.002 | 0.002 | 0 | 0.09 | 0.03 | 0.04 | 0.1 | 0.11 | 0.16 |
| 3 | 191805160 | rs1602517 | 1.014 | -0.749 | -0.719 | -1.038 | 0.085 | 0.121 | 0.121 | 0 | 0.58 | 0.19 | 0.69 | 0.78 | 0.68 | 0.54 |
| 3 | 191817026 | rs10937527 | 1.179 | -0.243 | -0.883 | -1.154 | -0.019 | 0.148 | 0.148 | 0 | 0.47 | 0.19 | 0.55 | 0.77 | 0.46 | 0.45 |
| 3 | 191817287 | rs10937530 | 0.957 | -0.282 | -0.624 | -1.172 | -0.003 | 0.072 | 0.072 | 0.038 | 0.49 | 0.34 | 0.58 | 0.77 | 0.47 | 0.45 |
| 3 | 191818642 | rs4585146 | 1.863 | -0.295 | -1.036 | -1.053 | -0.008 | -0.029 | -0.029 | 1.244 | 0.36 | 0.11 | 0.34 | 0.52 | 0.35 | 0.29 |
| 3 | 191820205 | rs2951798 | 0.65 | -0.507 | -0.572 | -1.348 | 0.007 | 0.002 | 0.002 | 0 | 0.1 | 0 | 0.05 | 0.1 | 0.1 | 0.16 |
| 3 | 191821726 | rs13090037 | -0.148 | -0.418 | -0.297 | -1.341 | 0.011 | -0.035 | -0.035 | 0.024 | 0.27 | 0.37 | 0.29 | 0.14 | 0.33 | 0.28 |
| 3 | 191828984 | rs9867755 | 0.546 | 0.175 | -0.47 | -1.008 | -0.003 | -0.013 | -0.013 | 0 | 0.1 | 0.09 | 0.05 | 0.01 | 0.05 | 0.09 |
| 3 | 191833747 | rs1907541 | 0.912 | 0.459 | -0.661 | -0.869 | -0.03 | 0.026 | 0.026 | 0 | 0.14 | 0.08 | 0.21 | 0.04 | 0.22 | 0.12 |
| 3 | 191839031 | rs1380990 | 0.599 | -0.741 | -0.159 | -0.377 | -0.04 | -0.094 | -0.094 | 0 | 0.73 | 0.55 | 0.83 | 0.9 | 0.78 | 0.72 |
| 3 | 191844140 | rs870808 | -0.901 | -0.608 | -0.385 | 0.262 | 0.007 | 0 | 0 | 0 | 0.35 | 0.38 | 0.36 | 0.35 | 0.3 | 0.5 |
| 3 | 191845112 | rs4686599 | 1.068 | -0.631 | 0.563 | 0.223 | 0.018 | 0.136 | 0.136 | 1.911 | 0.87 | 0.49 | 0.77 | 0.74 | 0.92 | 0.74 |
| 3 | 191847799 | rs6779599 | 0.365 | -0.821 | 0.43 | 0.091 | 0.001 | 0.007 | 0.007 | 0 | 0.14 | 0.12 | 0.1 | 0.13 | 0.16 | 0.21 |
| 3 | 191852283 | rs12497223 | 1.626 | -0.564 | -0.042 | 0.287 | -0.025 | 0 | 0 | 0 | 0.39 | 0.2 | 0.36 | 0.36 | 0.46 | 0.26 |
| 3 | 191852657 | rs9879329 | -0.708 | -0.734 | 0.412 | 0.244 | -0.022 | 0.031 | 0.031 | 0 | 0.47 | 0.2 | 0.41 | 0.38 | 0.47 | 0.51 |
| 3 | 191853822 | rs9809647 | 0.032 | -0.67 | 0.366 | 0.231 | -0.001 | 0.009 | 0.009 | 0.359 | 0.69 | 0.35 | 0.59 | 0.5 | 0.74 | 0.62 |
| 3 | 191853836 | rs6775772 | 1.846 | -0.637 | 0.294 | 0.242 | -0.042 | -0.056 | -0.056 | 0.167 | 0.17 | 0.03 | 0.18 | 0.23 | 0.18 | 0.15 |
| 3 | 191855182 | rs13101203 | 0.994 | -0.704 | 0.259 | 0.214 | 0 | 0.01 | 0.01 | 0 | 0.14 | 0.08 | 0.1 | 0.13 | 0.16 | 0.21 |
| 3 | 191855497 | rs972459 | 1.941 | -0.52 | 0.257 | 0.112 | -0.067 | -0.045 | -0.045 | 0 | 0.37 | 0.06 | 0.31 | 0.35 | 0.41 | 0.23 |
| 3 | 191856805 | rs6774732 | 0.68 | -0.458 | 0.459 | 0.145 | 0.006 | 0.044 | 0.044 | 0 | 0.16 | 0.04 | 0.13 | 0.02 | 0.14 | 0.15 |
| 3 | 191860535 | rs1460924 | -0.188 | -0.461 | 0.554 | 0.236 | 0.01 | 0.067 | 0.067 | 0.383 | 0.72 | 0.44 | 0.6 | 0.56 | 0.75 | 0.65 |
| 3 | 191861056 | rs11720096 | 1.704 | -0.467 | 0.908 | 0.256 | -0.004 | -0.005 | -0.005 | 0 | 0.18 | 0.17 | 0.19 | 0.19 | 0.18 | 0.14 |
| 3 | 191861115 | rs10049459 | -0.191 | -0.464 | 0.811 | 0.238 | 0.01 | 0.065 | 0.065 | 0.383 | 0.72 | 0.44 | 0.6 | 0.56 | 0.75 | 0.65 |
| 3 | 191862675 | rs9809505 | 1.04 | -0.675 | 0.361 | 0.044 | -0.012 | 0.096 | 0.096 | 0.296 | 0.48 | 0.25 | 0.42 | 0.34 | 0.58 | 0.47 |
| 3 | 191862817 | rs9829249 | 0.063 | -0.577 | 0.352 | 0.131 | 0 | 0 | 0 | 0 | 0.65 | 0.4 | 0.55 | 0.38 | 0.72 | 0.62 |
| 3 | 191865260 | rs1380992 | 1.223 | -0.436 | 0.161 | 0.192 | 0.008 | -0.007 | -0.007 | 0 | 0.63 | 0.37 | 0.56 | 0.51 | 0.72 | 0.55 |
| 3 | 191866779 | rs9877993 | -0.086 | -0.353 | 0.224 | 0.268 | 0 | 0.01 | 0.01 | 0 | 0.39 | 0.33 | 0.37 | 0.24 | 0.37 | 0.33 |
| 3 | 191873224 | rs10937534 | 0.466 | -0.393 | 0.035 | -0.093 | 0.009 | -0.013 | -0.013 | 0.523 | 0.78 | 0.67 | 0.8 | 0.74 | 0.73 | 0.68 |
| 3 | 191883836 | rs6800252 | -0.774 | -0.226 | -0.018 | -0.185 | 0.027 | 0.106 | 0.106 | 0 | 0.5 | 0.69 | 0.52 | 0.83 | 0.37 | 0.62 |
| 3 | 191894055 | rs1460925 | -0.656 | -0.65 | -0.052 | 0.205 | -0.009 | 0.006 | 0.006 | 0 | 0.08 | 0.12 | 0.06 | 0.11 | 0.07 | 0.06 |
| 3 | 191897218 | rs975121 | -1.206 | -0.875 | -0.796 | -0.468 | -0.165 | 0.185 | 0.185 | 0 | 0.64 | 0.97 | 0.63 | 0.95 | 0.48 | 0.74 |
| 3 | 191897771 | rs11721107 | -1.038 | -1.353 | -0.03 | 0.2 | -0.016 | -0.007 | -0.007 | 0 | 0.08 | 0 | 0.1 | 0.07 | 0.08 | 0.28 |
| 3 | 191901623 | rs9848565 | -1.789 | -1.432 | 0.755 | 0.258 | -0.031 | 0.015 | 0.015 | 0.111 | 0.2 | 0.47 | 0.21 | 0.24 | 0.16 | 0.31 |
| 3 | 191932318 | rs4687295 | -0.603 | -1.394 | 0.507 | -0.224 | 0.016 | 0.094 | 0.094 | 0.071 | 0.38 | 0.23 | 0.38 | 0.89 | 0.33 | 0.56 |
| 3 | 191935121 | rs9869920 | -0.749 | -1.458 | 0.554 | -0.253 | 0.021 | 0.092 | 0.092 | 0.009 | 0.39 | 0.24 | 0.4 | 0.9 | 0.34 | 0.56 |
| 3 | 191939237 | rs6790664 | -1.092 | -1.55 | 1.525 | -0.334 | 0.018 | 0.006 | 0.006 | 0 | 0.46 | 0.61 | 0.47 | 0.99 | 0.39 | 0.66 |
| 3 | 191966701 | rs940901 | -1.136 | -1.548 | 1.818 | -0.333 | 0.016 | 0.026 | 0.026 | 0 | 0.45 | 0.36 | 0.45 | 0.94 | 0.38 | 0.65 |
| 3 | 191989376 | rs4687309 | -0.579 | -1.515 | 1.864 | -0.311 | 0.012 | 0.047 | 0.047 | 0.219 | 0.39 | 0.29 | 0.39 | 0.9 | 0.33 | 0.56 |
| 3 | 191989642 | rs1812757 | -0.742 | -1.522 | 2.002 | -0.314 | 0.004 | -0.004 | -0.004 | 0.534 | 0.4 | 0.36 | 0.41 | 0.91 | 0.34 | 0.56 |
| 3 | 191990575 | rs7633010 | -0.572 | -1.506 | 1.839 | -0.306 | 0.011 | 0.043 | 0.043 | 0.191 | 0.38 | 0.29 | 0.39 | 0.9 | 0.33 | 0.56 |
| 3 | 191996202 | rs903043 | -0.677 | -1.419 | 1.461 | -0.271 | -0.004 | -0.042 | -0.042 | 0 | 0.41 | 0.42 | 0.41 | 0.91 | 0.33 | 0.56 |
| 3 | 191996779 | rs1522271 | -0.829 | -1.346 | 0.854 | -0.344 | 0.017 | 0.015 | 0.015 | 0.03 | 0.46 | 0.59 | 0.46 | 0.94 | 0.38 | 0.65 |
| 3 | 192020055 | rs2049212 | -0.522 | -1.629 | 0.86 | 0.852 | -0.003 | 0 | 0 | 0.056 | 0.16 | 0.18 | 0.13 | 0.15 | 0.08 | 0.21 |
| 3 | 192028670 | rs7641260 | -1.713 | -1.883 | 0.689 | 1.024 | -0.011 | 0.003 | 0.003 | 0.565 | 0.3 | 0.68 | 0.25 | 0.38 | 0.16 | 0.43 |
| 3 | 192038154 | rs7631206 | -1.083 | -1.613 | 0.538 | 1.033 | -0.03 | 0.026 | 0.026 | 0.12 | 0.25 | 0.66 | 0.27 | 0.39 | 0.15 | 0.31 |
| 3 | 192042183 | rs9850123 | -0.677 | -1.58 | 0.63 | 1.045 | -0.062 | 0.101 | 0.101 | 0 | 0.22 | 0.45 | 0.24 | 0.35 | 0.14 | 0.31 |
| 3 | 192045982 | rs4687321 | -0.906 | -1.419 | -0.322 | 1.034 | 0.003 | -0.006 | -0.006 | 0 | 0.31 | 0.7 | 0.29 | 0.42 | 0.18 | 0.48 |
| 3 | 192066029 | rs923820 | -0.674 | 0.194 | -0.599 | 0.975 | -0.02 | -0.012 | -0.012 | 0 | 0.17 | 0.14 | 0.29 | 0.23 | 0.25 | 0.16 |
| 3 | 192067566 | rs1994855 | -0.232 | -0.629 | -1.791 | 0.093 | 0.136 | 0.083 | 0.083 | 0 | 0.45 | 0.81 | 0.43 | 0.7 | 0.24 | 0.59 |
| 3 | 192068227 | rs10937543 | -0.138 | -0.648 | -1.165 | 0.085 | 0.02 | 0.007 | 0.007 | 0 | 0.18 | 0.28 | 0.31 | 0.23 | 0.25 | 0.16 |
| 3 | 192070275 | rs12487104 | -0.714 | -1.342 | -1.128 | 0.12 | 0.012 | 0.122 | 0.122 | 0 | 0.46 | 0.73 | 0.58 | 0.84 | 0.39 | 0.6 |
| 3 | 192070587 | rs4602335 | 0.052 | -1.401 | -1.115 | 0.115 | -0.018 | 0.007 | 0.007 | 0 | 0.31 | 0.61 | 0.32 | 0.61 | 0.18 | 0.45 |
| 3 | 192077217 | rs4687326 | 0.407 | -1.358 | -0.098 | 0.06 | -0.021 | -0.012 | -0.012 | 1.398 | 0.17 | 0.12 | 0.27 | 0.23 | 0.23 | 0.15 |
| 3 | 192078887 | rs11711838 | -0.583 | -1.445 | -0.065 | 0.027 | -0.019 | -0.009 | -0.009 | 0.018 | 0.28 | 0.18 | 0.36 | 0.32 | 0.29 | 0.29 |
| 3 | 192079832 | rs6784434 | -0.569 | -1.59 | -0.119 | 0.342 | -0.02 | -0.002 | -0.002 | 0 | 0.38 | 0.6 | 0.4 | 0.61 | 0.19 | 0.57 |
| 3 | 192079951 | rs9864008 | 0.288 | -1.567 | -0.059 | 0.376 | -0.055 | 0.078 | 0.078 | 0 | 0.26 | 0.4 | 0.29 | 0.53 | 0.12 | 0.42 |
| 3 | 192086091 | rs6444637 | -1.124 | -1.578 | 0.107 | 0.886 | 0.005 | 0.013 | 0.013 | 0 | 0.37 | 0.44 | 0.48 | 0.51 | 0.39 | 0.46 |
| 3 | 192087751 | rs2366659 | -1.575 | -1.417 | -0.463 | 0.391 | -0.13 | 0.234 | 0.234 | 0 | 0.61 | 0.93 | 0.7 | 0.93 | 0.47 | 0.75 |
| 3 | 192088525 | rs9859577 | -1.06 | -1.137 | 0.361 | 1.013 | -0.032 | 0.059 | 0.059 | 0 | 0.21 | 0.47 | 0.23 | 0.26 | 0.16 | 0.3 |
| 3 | 192089178 | rs6804526 | -1.153 | -0.364 | 0.369 | 1.138 | 0.007 | 0.042 | 0.042 | 0 | 0.17 | 0.38 | 0.33 | 0.2 | 0.16 | 0.23 |
| 3 | 192092640 | rs9881054 | -1.003 | -0.326 | 0.477 | 1.089 | 0.017 | 0.036 | 0.036 | 0 | 0.14 | 0.44 | 0.32 | 0.12 | 0.12 | 0.22 |
| 3 | 192093296 | rs9810385 | -1.27 | -0.48 | 0.655 | 2.077 | 0.02 | 0.322 | 0.322 | 0 | 0.08 | 0.37 | 0.14 | 0.38 | 0.06 | 0.08 |
| 3 | 192093611 | rs4429577 | -0.996 | -0.097 | -0.859 | 1.692 | -0.018 | -0.037 | -0.037 | 0 | 0.42 | 0.91 | 0.36 | 0.44 | 0.36 | 0.46 |
| 3 | 192093975 | rs4453795 | -0.283 | 0.167 | -1.219 | 1.306 | 0.027 | 0.026 | 0.026 | 0 | 0.47 | 0.64 | 0.38 | 0.39 | 0.42 | 0.5 |
| 3 | 192095111 | rs4299433 | -0.291 | 1.17 | -1.417 | 1.301 | -0.018 | -0.021 | -0.021 | 0 | 0.12 | 0.08 | 0.19 | 0.34 | 0.14 | 0.24 |
| 3 | 192105055 | rs1525910 | -0.612 | 1.113 | -1.396 | 1.036 | -0.026 | -0.029 | -0.029 | 0 | 0.52 | 0.8 | 0.47 | 0.52 | 0.48 | 0.47 |
| 3 | 192119973 | rs2686464 | 0.37 | 0.976 | -0.643 | 0.949 | 0.001 | -0.003 | -0.003 | 0 | 0.71 | 0.77 | 0.66 | 0.74 | 0.71 | 0.73 |
| 3 | 192126757 | rs3109189 | 1.048 | 0.663 | -1.128 | 0.711 | -0.006 | 0.031 | 0.031 | 0 | 0.26 | 0.1 | 0.28 | 0.19 | 0.25 | 0.28 |
| 3 | 192126987 | rs884309 | 0.147 | 0.601 | -0.532 | 1.037 | 0 | 0.028 | 0.028 | 0 | 0.12 | 0.13 | 0.13 | 0.45 | 0.09 | 0.17 |

| | | | | | | | | | | | | | | | | |
|---|---|---|---|---|---|---|---|---|---|---|---|---|---|---|---|---|
| 3 | 192135606 | rs12488767 | -0.245 | 0.766 | 0.015 | 0.351 | -0.001 | -0.002 | -0.002 | 0 | 0.35 | 0.31 | 0.43 | 0.32 | 0.4 | 0.39 |
| 3 | 192145679 | rs9836854 | -0.816 | 0.525 | -0.01 | 0.487 | 0.006 | 0.007 | 0.007 | 0 | 0.55 | 0.71 | 0.55 | 0.38 | 0.58 | 0.61 |
| 3 | 192149956 | rs3892203 | -0.827 | 0.582 | -0.019 | 0.332 | 0.014 | -0.002 | -0.002 | 0 | 0.55 | 0.74 | 0.59 | 0.39 | 0.59 | 0.57 |
| 3 | 192150817 | rs12107377 | -0.635 | 0.637 | 0.056 | 0.335 | 0.008 | 0.019 | 0.019 | 0 | 0.51 | 0.68 | 0.56 | 0.4 | 0.53 | 0.56 |
| 3 | 192155879 | rs6779518 | -0.755 | 0.711 | 0.245 | 0.339 | 0.025 | 0.011 | 0.011 | 1.15 | 0.51 | 0.6 | 0.51 | 0.4 | 0.58 | 0.54 |
| 3 | 192157175 | rs6807480 | -0.6 | 0.968 | 0.705 | -0.168 | 0.01 | -0.04 | -0.04 | 0 | 0.3 | 0.33 | 0.34 | 0.12 | 0.36 | 0.32 |
| 3 | 192159654 | rs1501619 | -1.902 | 0.369 | 1.69 | 0.264 | 0.068 | 0.064 | 0.064 | 0.657 | 0.05 | 0.5 | 0.04 | 0 | 0.01 | 0 |
| 3 | 192163051 | rs7653308 | -1.149 | 0.531 | 1.627 | 0.345 | -0.005 | -0.008 | -0.008 | 0 | 0.16 | 0.74 | 0.14 | 0.2 | 0.12 | 0.22 |
| 3 | 192198358 | rs2366676 | -1.047 | 0.548 | 1.771 | 0.36 | -0.004 | -0.008 | -0.008 | 8.456 | 0.16 | 0.73 | 0.13 | 0.2 | 0.12 | 0.21 |
| 3 | 192223737 | rs4687336 | 1.07 | 0.529 | 1.533 | 0.342 | 0.023 | 0.015 | 0.015 | 0 | 0.82 | 0.61 | 0.61 | 0.86 | 0.81 | 0.78 |
| 3 | 192227290 | rs12152403 | 0.882 | 0.556 | 1.686 | 0.444 | 0.033 | 0.062 | 0.062 | 0 | 0.91 | 0.72 | 0.81 | 0.9 | 0.89 | 0.86 |
| 3 | 192229661 | rs4686610 | 1.998 | 0.527 | 1.967 | 0.414 | 0.012 | 0.023 | 0.023 | 0 | 0.81 | 0.49 | 0.59 | 0.82 | 0.81 | 0.77 |
| 3 | 192234311 | rs9832414 | 1.451 | 0.48 | 2.143 | 0.466 | 0.016 | 0.037 | 0.037 | 0 | 0.88 | 0.46 | 0.81 | 0.88 | 0.88 | 0.84 |
| 3 | 192238709 | rs9843820 | 1.555 | 0.418 | 1.726 | 0.458 | 0.01 | 0.036 | 0.036 | 0.043 | 0.87 | 0.49 | 0.82 | 0.88 | 0.88 | 0.85 |
| 3 | 192249620 | rs4312609 | -0.266 | 0.11 | 0.06 | -0.163 | 0.006 | 0.13 | 0.13 | 0.007 | 0.55 | 0.77 | 0.64 | 0.87 | 0.6 | 0.71 |
| 3 | 192251697 | rs11705857 | -0.299 | 0.092 | -0.023 | -0.021 | -0.012 | 0.075 | 0.075 | 0 | 0.55 | 0.8 | 0.57 | 0.77 | 0.58 | 0.6 |
| 3 | 192257991 | rs9855262 | 0.427 | -0.122 | -0.003 | -0.268 | 0.024 | 0.047 | 0.047 | 0 | 0.37 | 0.2 | 0.22 | 0.14 | 0.29 | 0.28 |
| 3 | 192263633 | rs3845944 | 0.139 | -0.064 | -0.03 | -0.134 | -0.004 | -0.004 | -0.004 | 0 | 0.71 | 0.81 | 0.66 | 0.61 | 0.7 | 0.61 |
| 3 | 192266586 | rs1393989 | 1.26 | -0.1 | -0.231 | -0.176 | 0.022 | 0.096 | 0.096 | 0 | 0.32 | 0.15 | 0.12 | 0.04 | 0.24 | 0.21 |
| 3 | 192273198 | rs2654703 | 0.593 | 0.165 | 0.652 | 0.14 | 0.029 | -0.059 | -0.059 | 0 | 0.86 | 0.66 | 0.76 | 0.93 | 0.88 | 0.91 |
| 3 | 192275595 | rs2134635 | -0.068 | 0.141 | 0.434 | 0.332 | -0.005 | -0.002 | -0.002 | 0 | 0.36 | 0.42 | 0.5 | 0.56 | 0.44 | 0.39 |
| 3 | 192282528 | rs6802533 | 0.352 | 0.743 | 0.073 | 0.872 | 0.022 | 0.029 | 0.029 | 0 | 0.71 | 0.67 | 0.46 | 0.49 | 0.58 | 0.72 |
| 3 | 192287263 | rs1532684 | 0.086 | 0.89 | -0.67 | 0.925 | -0.04 | -0.047 | -0.047 | 0.39 | 0.73 | 0.84 | 0.49 | 0.5 | 0.59 | 0.73 |
| 3 | 192291771 | rs2062829 | 0.133 | 0.448 | -0.863 | 0.363 | -0.002 | 0.006 | 0.006 | 0.314 | 0.29 | 0.19 | 0.2 | 0.29 | 0.27 | 0.35 |
| 3 | 192291867 | rs781414 | -0.097 | -0.075 | -0.336 | 0.168 | 0.021 | 0.007 | 0.007 | 0 | 0.48 | 0.56 | 0.44 | 0.61 | 0.62 | 0.52 |
| 3 | 192293753 | rs781415 | 0.374 | -0.021 | -0.113 | 0.219 | -0.01 | 0.067 | 0.067 | 0.062 | 0.8 | 0.65 | 0.57 | 0.67 | 0.84 | 0.71 |
| 3 | 192298277 | rs781422 | -0.428 | 0.307 | -0.599 | -1.019 | 0.004 | 0.001 | 0.001 | 0 | 0.71 | 0.79 | 0.73 | 0.97 | 0.72 | 0.75 |
| 3 | 192302970 | rs3897920 | 0.817 | 0.465 | -0.613 | -1.028 | -0.001 | 0.011 | 0.011 | 0.182 | 0.24 | 0.21 | 0.34 | 0.52 | 0.25 | 0.37 |
| 3 | 192305208 | rs781425 | 0.852 | 0.711 | -0.92 | -1.213 | -0.115 | -0.087 | -0.087 | 0 | 0.26 | 0.01 | 0.24 | 0.36 | 0.31 | 0.17 |
| 3 | 192309520 | rs7609933 | 0.359 | 0.622 | -0.631 | -1.261 | -0.001 | 0 | 0 | 0 | 0.14 | 0.16 | 0.1 | 0.1 | 0.16 | 0.13 |
| 3 | 192311220 | rs2701595 | 0.965 | 0.644 | -0.653 | -1.269 | -0.089 | -0.071 | -0.071 | 0 | 0.26 | 0.04 | 0.24 | 0.36 | 0.3 | 0.17 |
| 3 | 192312270 | rs2654694 | -0.536 | 0.741 | -0.541 | -1.247 | -0.002 | 0.001 | 0.001 | 0.074 | 0.46 | 0.61 | 0.4 | 0.46 | 0.46 | 0.33 |
| 3 | 192315067 | rs2708309 | 0.427 | 0.793 | -0.531 | -1.194 | 0.005 | -0.002 | -0.002 | 0.536 | 0.3 | 0.43 | 0.3 | 0.36 | 0.3 | 0.19 |
| 3 | 192320179 | rs905566 | -0.212 | 1.122 | -0.977 | -1.284 | -0.008 | 0 | 0 | 0 | 0.41 | 0.39 | 0.36 | 0.46 | 0.46 | 0.3 |
| 3 | 192325362 | rs7610664 | 0.017 | 1.215 | -1.109 | -1.664 | -0.008 | -0.002 | -0.002 | 0 | 0.12 | 0.09 | 0.06 | 0.01 | 0.15 | 0.08 |
| 3 | 192327238 | rs1997439 | -0.072 | 0.607 | -1.115 | -1.194 | 0.003 | -0.011 | -0.011 | 0 | 0.71 | 0.78 | 0.74 | 0.97 | 0.7 | 0.77 |
| 3 | 192330494 | rs2708305 | -0.043 | 0.671 | -1.124 | -1.138 | 0.003 | -0.034 | -0.034 | 0.339 | 0.68 | 0.69 | 0.71 | 0.97 | 0.69 | 0.75 |
| 3 | 192337670 | rs2701588 | -0.843 | 1.016 | -1.186 | -0.565 | 0 | -0.014 | -0.014 | 0 | 0.14 | 0.13 | 0.14 | 0 | 0.12 | 0.1 |
| 3 | 192338861 | rs1922910 | 0.249 | 1.741 | -1.878 | -0.243 | -0.109 | -0.061 | -0.061 | 0 | 0.79 | 0.99 | 0.78 | 0.76 | 0.67 | 0.69 |
| 3 | 192341051 | rs2062832 | 0.249 | 1.741 | -1.878 | -0.243 | -0.109 | -0.061 | -0.061 | 0 | 0.79 | 0.99 | 0.78 | 0.75 | 0.67 | 0.69 |
| 3 | 192357901 | rs6803287 | 0.523 | 1.453 | -1.22 | -0.279 | 0.013 | -0.019 | -0.019 | 0.335 | 0.6 | 0.51 | 0.42 | 0.67 | 0.61 | 0.63 |
| 3 | 192375440 | rs9813078 | 0.438 | 1.307 | -0.519 | -0.406 | 0.017 | -0.018 | -0.018 | 2.576 | 0.62 | 0.48 | 0.45 | 0.67 | 0.61 | 0.63 |
| 3 | 192376137 | rs6798763 | 0.347 | 1.298 | -0.254 | -0.41 | -0.006 | 0.003 | 0.003 | 0.032 | 0.63 | 0.63 | 0.47 | 0.67 | 0.61 | 0.63 |
| 3 | 192379973 | rs1995472 | 0.66 | 0.636 | -0.445 | -0.378 | 0 | -0.002 | -0.002 | 0 | 0.45 | 0.48 | 0.39 | 0.53 | 0.52 | 0.42 |
| 3 | 192388678 | rs13068005 | 0.249 | 1.457 | -0.814 | -0.7 | -0.08 | 0.062 | 0.062 | 0.236 | 0.75 | 0.91 | 0.59 | 0.85 | 0.61 | 0.7 |
| 3 | 192390750 | rs880774 | -0.702 | 1.481 | 0.072 | -0.806 | 0.083 | -0.046 | -0.046 | 0 | 0.11 | 0.35 | 0.1 | 0.02 | 0.12 | 0.17 |
| 3 | 192395419 | rs1948883 | -1.197 | 1.404 | -0.009 | -0.934 | 0.053 | -0.085 | -0.085 | 0.278 | 0.15 | 0.36 | 0.14 | 0.02 | 0.19 | 0.19 |
| 3 | 192399612 | rs12696657 | 1.381 | 0.795 | 0.113 | -0.88 | 0.013 | 0.02 | 0.02 | 0.677 | 0.57 | 0.37 | 0.34 | 0.61 | 0.44 | 0.46 |
| 3 | 192407268 | rs1605083 | 0.46 | 0.777 | 0.913 | -0.776 | 0.006 | -0.009 | -0.009 | 0 | 0.69 | 0.72 | 0.54 | 0.79 | 0.55 | 0.59 |
| 3 | 192424298 | rs9859403 | 1.358 | 0.587 | 0.795 | -0.723 | -0.007 | 0.03 | 0.03 | 0.668 | 0.58 | 0.33 | 0.37 | 0.63 | 0.5 | 0.63 |
| 3 | 192426058 | rs6801152 | -0.616 | 0.861 | 0.565 | -0.757 | -0.072 | 0.033 | 0.033 | 0.335 | 0.81 | 0.96 | 0.62 | 0.85 | 0.67 | 0.78 |
| 3 | 192432100 | rs13069056 | -0.833 | 0.419 | 0.859 | 0.098 | 0.028 | 0.119 | 0.119 | 0 | 0.16 | 0.29 | 0.16 | 0.39 | 0.19 | 0.09 |
| 3 | 192434058 | rs1553560 | -1.001 | 0.419 | 0.825 | 0.092 | 0.031 | 0.129 | 0.129 | 0 | 0.17 | 0.33 | 0.18 | 0.39 | 0.2 | 0.1 |
| 3 | 192434415 | rs1995574 | 0.285 | 0.356 | 0.774 | 0.151 | 0.014 | 0.192 | 0.192 | 0 | 0.05 | 0.2 | 0.13 | 0.22 | 0.01 | 0.1 |
| 3 | 192435920 | rs13081379 | -0.079 | 0.374 | 0.387 | 0.142 | -0.059 | -0.071 | -0.071 | 0 | 0.11 | 0.01 | 0.11 | 0.39 | 0.18 | 0.09 |
| 3 | 192438594 | rs11920769 | -0.767 | -0.066 | -0.038 | -0.373 | -0.063 | 0.064 | 0.064 | 0.698 | 0.75 | 0.9 | 0.62 | 0.85 | 0.67 | 0.79 |
| 3 | 192440779 | rs7614462 | -1.571 | 0.287 | 0.365 | -0.09 | 0.001 | -0.003 | -0.003 | 0 | 0.27 | 0.7 | 0.33 | 0.61 | 0.22 | 0.2 |
| 3 | 192446037 | rs13089551 | -1.318 | 0.319 | 0.386 | -0.068 | -0.001 | 0.046 | 0.046 | 0.015 | 0.24 | 0.57 | 0.32 | 0.61 | 0.21 | 0.21 |
| 3 | 192451718 | rs2202479 | -0.802 | -0.422 | -0.586 | 0.05 | -0.031 | 0.121 | 0.121 | 0 | 0.63 | 0.87 | 0.59 | 0.82 | 0.62 | 0.77 |
| 3 | 192453666 | rs10460907 | 0.859 | -0.371 | -0.493 | 0.117 | 0.005 | 0.034 | 0.034 | 0 | 0.42 | 0.32 | 0.29 | 0.25 | 0.41 | 0.57 |
| 3 | 192456208 | rs11921051 | 1.115 | -0.401 | -0.995 | 0.152 | -0.02 | 0.072 | 0.072 | 0.365 | 0.38 | 0.14 | 0.27 | 0.23 | 0.41 | 0.56 |
| 3 | 192465696 | rs6768136 | -0.905 | -0.273 | -1.085 | 0.277 | -0.043 | 0.111 | 0.111 | 0 | 0.65 | 0.87 | 0.59 | 0.83 | 0.62 | 0.78 |
| 3 | 192470103 | rs1505449 | -0.867 | -0.227 | -1.076 | 0.285 | -0.039 | 0.115 | 0.115 | 0 | 0.64 | 0.87 | 0.59 | 0.83 | 0.62 | 0.77 |
| 3 | 192490238 | rs4687358 | -0.87 | -0.264 | -1.231 | 0.33 | -0.052 | -0.091 | -0.091 | 0.021 | 0.62 | 0.92 | 0.7 | 0.46 | 0.62 | 0.64 |
| 3 | 192501056 | rs1505440 | -0.917 | -0.195 | -0.802 | 0.315 | 0.016 | -0.003 | -0.003 | 0 | 0.24 | 0.53 | 0.29 | 0.23 | 0.2 | 0.16 |
| 3 | 192504337 | rs4687360 | 0.626 | -0.153 | -1.491 | 0.039 | -0.044 | 0.059 | 0.059 | 0.412 | 0.35 | 0.09 | 0.26 | 0.22 | 0.41 | 0.48 |
| 3 | 192515557 | rs1061860 | -0.3 | -0.129 | -1.229 | -0.02 | -0.077 | -0.048 | -0.048 | 0 | 0.57 | 0.86 | 0.43 | 0.56 | 0.56 | 0.57 |
| 3 | 192531202 | rs2986 | -0.283 | -0.24 | -0.17 | -0.064 | -0.001 | -0.001 | -0.001 | 0.166 | 0.57 | 0.61 | 0.42 | 0.56 | 0.56 | 0.61 |
| 3 | 192536466 | rs9826763 | 0.626 | -0.478 | 0.073 | -0.537 | -0.01 | -0.015 | -0.015 | 0 | 0.38 | 0.29 | 0.32 | 0.59 | 0.42 | 0.56 |

| | | | | | | | | | | | | | | | | |
|---|---|---|---|---|---|---|---|---|---|---|---|---|---|---|---|---|
| 3 | 192538160 | rs6444661 | -0.45 | -0.238 | -0.077 | -0.411 | -0.037 | -0.013 | -0.013 | 0 | 0.57 | 0.79 | 0.44 | 0.59 | 0.54 | 0.59 |
| 3 | 192546196 | rs5026725 | -0.824 | 0.125 | -0.283 | -0.696 | -0.125 | -0.009 | -0.009 | 0 | 0.64 | 0.94 | 0.54 | 0.74 | 0.57 | 0.71 |
| 3 | 192555298 | rs9291033 | -0.528 | -0.378 | 0.106 | -0.767 | 0.003 | -0.009 | -0.009 | 0 | 0.6 | 0.58 | 0.48 | 0.78 | 0.52 | 0.69 |
| 3 | 192562429 | rs9870609 | 0.134 | -0.284 | 0.026 | -0.671 | 0.02 | 0.017 | 0.017 | 0 | 0.48 | 0.62 | 0.36 | 0.6 | 0.41 | 0.58 |
| 3 | 192576014 | rs9812883 | -0.175 | -0.156 | -0.436 | -0.695 | -0.002 | 0.008 | 0.008 | 0 | 0.6 | 0.63 | 0.46 | 0.78 | 0.48 | 0.66 |
| 3 | 192587703 | rs6783864 | -0.592 | 0.055 | -1.326 | -0.632 | -0.081 | 0.025 | 0.025 | 0 | 0.64 | 0.83 | 0.5 | 0.77 | 0.52 | 0.71 |
| 3 | 192591103 | rs13059477 | -0.628 | 0.099 | -1.558 | -0.612 | -0.135 | 0.001 | 0.001 | 0 | 0.65 | 0.89 | 0.5 | 0.77 | 0.53 | 0.71 |
| 3 | 192592587 | rs2133865 | 0.53 | -0.163 | -1.267 | -0.489 | 0 | -0.004 | -0.004 | 3.851 | 0.35 | 0.29 | 0.29 | 0.56 | 0.37 | 0.43 |
| 3 | 192595307 | rs2367129 | -0.719 | 0.011 | -1.337 | -0.973 | -0.001 | -0.047 | -0.047 | 0.186 | 0.29 | 0.25 | 0.17 | 0 | 0.25 | 0.17 |
| 3 | 192596365 | rs7621131 | -0.234 | -0.167 | -1.441 | -0.046 | -0.011 | -0.002 | -0.002 | 0 | 0.18 | 0.74 | 0.12 | 0.2 | 0.09 | 0.22 |
| 3 | 192599336 | rs6444670 | 0.12 | 1.145 | -2.036 | -0.176 | -0.181 | 0.018 | 0.018 | 0 | 0.65 | 0.96 | 0.54 | 0.82 | 0.6 | 0.73 |
| 3 | 192614308 | rs4455277 | 0.076 | 0.677 | -0.774 | -0.706 | 0 | 0 | 0 | 0 | 0.49 | 0.52 | 0.55 | 0.33 | 0.52 | 0.55 |
| 3 | 192625190 | rs9869202 | 0.121 | 0.089 | -0.867 | -0.897 | 0.067 | 0.142 | 0.142 | 0.012 | 0.51 | 0.11 | 0.54 | 0.26 | 0.59 | 0.52 |
| 3 | 192629447 | rs1948721 | -0.231 | -0.006 | -0.573 | -0.939 | 0.024 | 0.022 | 0.022 | 0.037 | 0.57 | 0.71 | 0.61 | 0.34 | 0.61 | 0.58 |
| 3 | 192631810 | rs4687368 | 0.211 | 0.776 | -0.722 | -1.158 | -0.001 | 0.199 | 0.199 | 0 | 0.49 | 0.16 | 0.5 | 0.24 | 0.46 | 0.48 |
| 3 | 192637324 | rs769589 | 0.304 | 1.08 | -0.601 | -1.111 | -0.02 | 0.136 | 0.136 | 1.309 | 0.54 | 0.16 | 0.55 | 0.24 | 0.5 | 0.48 |
| 3 | 192647763 | rs769592 | 0.359 | 1.223 | -0.131 | -1.018 | 0.009 | 0.026 | 0.026 | 0 | 0.54 | 0.41 | 0.57 | 0.36 | 0.48 | 0.52 |
| 3 | 192675756 | rs924321 | 0.072 | 1.251 | -0.151 | -0.99 | 0.001 | -0.008 | -0.008 | 0 | 0.32 | 0.2 | 0.28 | 0.61 | 0.31 | 0.39 |
| 3 | 192689067 | rs709139 | -0.531 | 0.408 | 0.003 | -0.889 | -0.003 | 0.013 | 0.013 | 0 | 0.44 | 0.34 | 0.43 | 0.6 | 0.51 | 0.54 |
| 3 | 192689376 | rs6444675 | -0.487 | 0.327 | 0.225 | -0.979 | 0.015 | 0.042 | 0.042 | 0.593 | 0.08 | 0.36 | 0.05 | 0.04 | 0.05 | 0.02 |
| 3 | 192706711 | rs1447840 | -0.036 | 0.225 | -0.043 | -1.049 | -0.028 | 0.003 | 0.003 | 0 | 0.36 | 0.25 | 0.39 | 0.62 | 0.45 | 0.53 |
| 3 | 192709305 | rs1697323 | -1.138 | 0.04 | -1.148 | -1.323 | -0.054 | 0.183 | 0.183 | 0 | 0.58 | 0.95 | 0.61 | 0.84 | 0.62 | 0.71 |
| 3 | 192727562 | rs1021662 | -0.678 | 0.438 | -0.448 | -0.833 | 0.003 | 0.096 | 0.096 | 0 | 0.49 | 0.75 | 0.47 | 0.66 | 0.53 | 0.55 |
| 3 | 192750246 | rs9831317 | -0.907 | 0.98 | -1.224 | -0.495 | -0.014 | 0.159 | 0.159 | 0 | 0.51 | 0.95 | 0.54 | 0.69 | 0.56 | 0.65 |
| 3 | 192752645 | rs1157623 | -1.288 | 0.842 | 0.151 | 0.06 | 0.002 | -0.016 | -0.016 | 0 | 0.14 | 0.35 | 0.16 | 0.08 | 0.13 | 0.12 |
| 3 | 192758191 | rs9283662 | -1.283 | 1.008 | -0.082 | 0.521 | 0.009 | 0.047 | 0.047 | 1.541 | 0.09 | 0.14 | 0.11 | 0.19 | 0.1 | 0.14 |
| 3 | 192759024 | rs9813423 | -0.615 | 1.023 | -0.54 | -0.327 | -0.005 | 0.025 | 0.025 | 0.145 | 0.26 | 0.2 | 0.23 | 0.12 | 0.25 | 0.19 |
| 3 | 192759040 | rs4356766 | -0.087 | 0.242 | -0.479 | -0.363 | -0.044 | -0.039 | -0.039 | 0 | 0.7 | 0.47 | 0.77 | 0.8 | 0.77 | 0.75 |
| 3 | 192761717 | rs2367278 | 0.55 | 1.388 | -1.026 | 0.106 | -0.031 | -0.037 | -0.037 | 0.367 | 0.13 | 0.07 | 0.21 | 0.56 | 0.18 | 0.28 |
| 3 | 192765819 | rs9855714 | 0.068 | 0.836 | -0.705 | -0.464 | 0.009 | 0.037 | 0.037 | 0 | 0.53 | 0.32 | 0.55 | 0.27 | 0.6 | 0.42 |
| 3 | 192772250 | rs9816743 | 0.606 | 1.054 | -0.937 | -0.532 | -0.018 | 0.017 | 0.017 | 0.04 | 0.15 | 0.08 | 0.19 | 0.11 | 0.19 | 0.18 |
| 3 | 192772295 | rs13068101 | 1.513 | 0.867 | -1.433 | -1.03 | -0.079 | -0.021 | -0.021 | 2.857 | 0.16 | 0.01 | 0.26 | 0.16 | 0.27 | 0.12 |
| 3 | 192774090 | rs1021660 | -0.936 | 0.015 | -0.582 | -1.031 | -0.046 | -0.055 | -0.055 | 0.007 | 0.65 | 0.46 | 0.78 | 0.84 | 0.77 | 0.7 |
| 3 | 192784094 | rs9852198 | -0.112 | 0.285 | -0.818 | -1.098 | 0.099 | 0.188 | 0.188 | 3.162 | 0.48 | 0.22 | 0.68 | 0.85 | 0.67 | 0.59 |
| 3 | 192785307 | rs1472459 | -0.914 | 0.111 | -0.083 | -1.209 | -0.031 | -0.087 | -0.087 | 0.117 | 0.66 | 0.5 | 0.79 | 0.91 | 0.74 | 0.78 |
| 3 | 192792970 | rs9834173 | -1.053 | 0.727 | 0.225 | -0.504 | -0.001 | -0.001 | -0.001 | 0.053 | 0.36 | 0.56 | 0.4 | 0.65 | 0.33 | 0.53 |
| 3 | 192797746 | rs6772678 | -0.307 | 1.414 | 0.105 | -0.167 | -0.003 | 0.025 | 0.025 | 0 | 0.2 | 0.2 | 0.22 | 0.57 | 0.17 | 0.3 |
| 3 | 192811289 | rs9810201 | -0.489 | 1.207 | 0.005 | -0.208 | 0.002 | 0.006 | 0.006 | 0.413 | 0.22 | 0.19 | 0.29 | 0.61 | 0.26 | 0.41 |
| 3 | 192815895 | rs6803784 | -1.822 | 0.522 | 0.527 | -0.34 | -0.01 | 0.064 | 0.064 | 0.32 | 0.1 | 0.42 | 0.05 | 0.06 | 0.02 | 0.06 |
| 3 | 192816905 | rs7628342 | -1.599 | 0.542 | 0.617 | -0.332 | -0.011 | 0.05 | 0.05 | 0 | 0.08 | 0.27 | 0.03 | 0.06 | 0.02 | 0.06 |
| 3 | 192835072 | rs2886928 | 0.579 | 0.858 | 0.268 | -0.38 | 0.016 | -0.045 | -0.045 | 0.913 | 0.87 | 0.81 | 0.88 | 0.98 | 0.84 | 0.93 |
| 3 | 192843266 | rs4686626 | -0.071 | 0.387 | -0.05 | -0.479 | -0.011 | -0.022 | -0.022 | 0 | 0.32 | 0.41 | 0.23 | 0.23 | 0.25 | 0.17 |
| 3 | 192848225 | rs922331 | 0.473 | 0.317 | -0.158 | -0.205 | 0.018 | -0.011 | -0.011 | 0.27 | 0.56 | 0.36 | 0.58 | 0.42 | 0.61 | 0.56 |
| 3 | 192850845 | rs9872146 | 0.129 | 0.312 | 0.106 | -0.81 | 0 | -0.003 | -0.003 | 0 | 0.2 | 0.19 | 0.17 | 0.1 | 0.17 | 0.09 |
| 3 | 192850865 | rs9872413 | 0.994 | 0.343 | 0.069 | -0.81 | -0.049 | 0.069 | 0.069 | 0 | 0.15 | 0.03 | 0.1 | 0.02 | 0.17 | 0.09 |
| 3 | 192852792 | rs13327008 | 0.831 | 0.358 | 0.147 | -0.822 | -0.026 | 0.044 | 0.044 | 0 | 0.16 | 0.07 | 0.11 | 0.02 | 0.18 | 0.09 |
| 3 | 192856058 | rs10937566 | -0.768 | 0.309 | 0.286 | -1.111 | 0.009 | -0.126 | -0.126 | 0 | 0.31 | 0.52 | 0.26 | 0.03 | 0.29 | 0.25 |
| 3 | 192864085 | rs6804743 | 0.078 | 0.27 | 0.417 | -1.077 | -0.009 | -0.023 | -0.023 | 0.13 | 0.14 | 0.31 | 0.17 | 0.08 | 0.11 | 0.16 |
| 3 | 192866499 | rs2169524 | -0.988 | 0.251 | 0.245 | -1.123 | -0.028 | 0.051 | 0.051 | 0 | 0.82 | 0.92 | 0.87 | 0.98 | 0.79 | 0.9 |
| 3 | 192870449 | rs9842842 | 0.966 | 0.247 | 0.188 | -1.094 | 0.002 | -0.063 | -0.063 | 0 | 0.66 | 0.48 | 0.69 | 0.9 | 0.68 | 0.74 |
| 3 | 192886861 | rs9837828 | 0.37 | 0.354 | 0.172 | -1.14 | -0.004 | -0.029 | -0.029 | 0.006 | 0.72 | 0.8 | 0.77 | 0.98 | 0.68 | 0.74 |
| 3 | 192893504 | rs4687384 | -0.283 | 0.393 | 0.241 | -1.076 | 0.024 | -0.097 | -0.097 | 3.553 | 0.8 | 0.64 | 0.84 | 0.98 | 0.79 | 0.9 |
| 3 | 192909721 | rs1531464 | 1.047 | -0.338 | 0.217 | -1.012 | -0.018 | -0.069 | -0.069 | 0.4 | 0.66 | 0.48 | 0.7 | 0.9 | 0.72 | 0.75 |
| 3 | 192913538 | rs4686628 | -0.199 | -0.391 | 0.24 | -0.962 | -0.016 | -0.039 | -0.039 | 0 | 0.79 | 0.71 | 0.82 | 0.91 | 0.83 | 0.89 |
| 3 | 192919373 | rs9861073 | 0.918 | 0.044 | -0.138 | -0.8 | 0.003 | -0.018 | -0.018 | 0 | 0.58 | 0.64 | 0.53 | 0.75 | 0.57 | 0.64 |
| 3 | 192937401 | rs7635381 | -0.908 | 0.085 | -0.448 | -0.762 | -0.012 | 0.044 | 0.044 | 0 | 0.85 | 0.95 | 0.91 | 0.98 | 0.84 | 0.9 |
| 3 | 192948186 | rs2276723 | 1.903 | -0.03 | -0.554 | -0.746 | -0.015 | 0.031 | 0.031 | 0 | 0.32 | 0.19 | 0.27 | 0.24 | 0.34 | 0.26 |
| 3 | 192948729 | rs7650724 | -0.265 | 0.021 | -0.831 | -0.853 | -0.064 | 0.077 | 0.077 | 0 | 0.73 | 0.95 | 0.73 | 0.84 | 0.68 | 0.8 |
| 3 | 192949982 | rs7642439 | 1.4 | -0.16 | -0.354 | -0.943 | -0.011 | -0.067 | -0.067 | 0.119 | 0.11 | 0.19 | 0.09 | 0 | 0.07 | 0.12 |
| 3 | 192954665 | rs1462483 | 0.577 | -0.137 | -0.314 | -0.933 | 0 | -0.021 | -0.021 | 0.235 | 0.61 | 0.71 | 0.63 | 0.83 | 0.6 | 0.68 |
| 3 | 192954886 | rs10428211 | 0.602 | -0.141 | -0.301 | -0.935 | 0.001 | -0.025 | -0.025 | 0 | 0.6 | 0.7 | 0.63 | 0.83 | 0.6 | 0.68 |
| 3 | 192967226 | rs9824804 | 1.123 | 0.022 | -0.73 | -0.734 | -0.032 | 0.065 | 0.065 | 0 | 0.44 | 0.18 | 0.44 | 0.36 | 0.49 | 0.37 |
| 3 | 192979624 | rs11720679 | 1.203 | 0.018 | -0.747 | -0.768 | -0.063 | 0.066 | 0.066 | 0 | 0.43 | 0.11 | 0.44 | 0.36 | 0.49 | 0.37 |
| 3 | 192982638 | rs10937575 | -0.562 | -0.133 | -0.666 | -0.775 | 0.037 | 0.003 | 0.003 | 0 | 0.43 | 0.66 | 0.4 | 0.54 | 0.36 | 0.53 |
| 3 | 192983331 | rs10804951 | -1.702 | -0.1 | -0.682 | -0.769 | -0.025 | 0.015 | 0.015 | 0 | 0.93 | 0.99 | 0.93 | 0.98 | 0.86 | 0.92 |
| 3 | 192984366 | rs7611149 | -0.229 | -0.113 | -0.667 | -0.794 | 0.003 | -0.003 | -0.003 | 0 | 0.35 | 0.62 | 0.38 | 0.54 | 0.34 | 0.51 |
| 3 | 192986527 | rs2367474 | 1.145 | -0.033 | -0.881 | -0.786 | -0.06 | 0.063 | 0.063 | 0 | 0.43 | 0.11 | 0.44 | 0.36 | 0.49 | 0.37 |
| 3 | 192988692 | rs2367475 | 2.357 | -0.166 | -0.994 | -0.86 | -0.094 | 0.015 | 0.015 | 0 | 0.28 | 0.03 | 0.26 | 0.22 | 0.34 | 0.26 |
| 3 | 192994543 | rs2271791 | 1.283 | -0.189 | -0.648 | -0.627 | -0.063 | 0.066 | 0.066 | 0 | 0.43 | 0.11 | 0.44 | 0.36 | 0.49 | 0.37 |

| | | | | | | | | | | | | | | | | |
|---|---|---|---|---|---|---|---|---|---|---|---|---|---|---|---|---|
| 3 | 193002227 | rs2886978 | 0.811 | -0.101 | -0.532 | -0.528 | 0.009 | 0.082 | 0.082 | 0.032 | 0.53 | 0.21 | 0.47 | 0.36 | 0.52 | 0.39 |
| 3 | 193008876 | rs2367477 | 1.026 | 0.071 | -0.494 | -0.439 | -0.021 | 0.082 | 0.082 | 0 | 0.46 | 0.18 | 0.46 | 0.36 | 0.49 | 0.39 |
| 3 | 193013810 | rs6794806 | -0.058 | -0.088 | 0.4 | -0.534 | -0.01 | 0.002 | 0.002 | 0 | 0.37 | 0.39 | 0.34 | 0.56 | 0.35 | 0.51 |
| 3 | 193015642 | rs2367478 | -0.928 | 0.032 | 0.732 | -0.444 | 0.016 | -0.03 | -0.03 | 0.04 | 0.89 | 0.84 | 0.9 | 0.98 | 0.85 | 0.91 |
| 3 | 193018778 | rs12630096 | 0.621 | -0.411 | 0.98 | -0.604 | 0.004 | -0.01 | -0.01 | 0.129 | 0.26 | 0.25 | 0.26 | 0.56 | 0.27 | 0.39 |
| 3 | 193019767 | rs7641152 | -0.38 | -0.423 | 0.958 | -0.624 | -0.011 | -0.004 | -0.004 | 0 | 0.36 | 0.54 | 0.39 | 0.62 | 0.33 | 0.51 |
| 3 | 193031926 | rs2280268 | 0.987 | -0.228 | 1.033 | -0.18 | 0.009 | 0.011 | 0.011 | 0.017 | 0.57 | 0.32 | 0.54 | 0.65 | 0.64 | 0.57 |
| 3 | 193037415 | rs2886984 | -0.121 | -0.02 | 1.103 | -0.161 | 0.053 | -0.085 | -0.085 | 0 | 0.82 | 0.7 | 0.77 | 0.98 | 0.76 | 0.77 |
| 3 | 193038596 | rs6444702 | 0.038 | 0.338 | 0.793 | 0.111 | 0.016 | -0.023 | -0.023 | 0 | 0.83 | 0.74 | 0.75 | 0.93 | 0.8 | 0.87 |
| 3 | 193047583 | rs3845934 | -0.376 | 0.369 | 0.671 | 0.013 | 0.053 | -0.048 | -0.048 | 0 | 0.86 | 0.69 | 0.81 | 0.96 | 0.81 | 0.88 |
| 3 | 193047723 | rs3845936 | 0.299 | 0.234 | 0.134 | 0.128 | 0.001 | 0.003 | 0.003 | 0.515 | 0.66 | 0.26 | 0.65 | 0.71 | 0.72 | 0.66 |
| 3 | 193058273 | rs4687401 | 0.226 | 0.305 | -0.118 | 0.244 | -0.004 | -0.036 | -0.036 | 0 | 0.32 | 0.23 | 0.37 | 0.51 | 0.37 | 0.36 |
| 3 | 193065480 | rs13085410 | -0.691 | 0.163 | 0.455 | 0.303 | -0.041 | 0.007 | 0.007 | 1.207 | 0.79 | 0.66 | 0.88 | 0.79 | 0.91 | 0.91 |
| 3 | 193066869 | rs1980263 | 0.635 | 0.142 | 0.388 | 0.201 | 0.001 | -0.011 | -0.011 | 0.213 | 0.43 | 0.39 | 0.44 | 0.22 | 0.48 | 0.38 |
| 3 | 193070674 | rs6784067 | 0.165 | 0.248 | 0.178 | 0.198 | -0.006 | -0.005 | -0.005 | 0.115 | 0.57 | 0.66 | 0.52 | 0.4 | 0.52 | 0.43 |
| 3 | 193077515 | rs4687410 | 0.285 | 0.164 | -0.065 | 0.233 | -0.011 | -0.028 | -0.028 | 0.212 | 0.36 | 0.24 | 0.43 | 0.57 | 0.42 | 0.52 |
| 3 | 193082138 | rs923494 | 1.138 | 0.512 | -0.244 | -0.361 | -0.003 | -0.001 | -0.001 | 1.417 | 0.11 | 0.1 | 0.14 | 0.02 | 0.16 | 0.11 |
| 3 | 193083891 | rs1813604 | -0.048 | -0.231 | -0.177 | -0.135 | 0.003 | 0.02 | 0.02 | 0 | 0.94 | 0.94 | 0.96 | 0.97 | 0.94 | 0.96 |
| 3 | 193090806 | rs9851984 | 0.039 | -0.281 | 0.01 | -0.006 | -0.006 | 0.004 | 0.004 | 0 | 0.72 | 0.62 | 0.71 | 0.42 | 0.72 | 0.72 |
| 3 | 193093702 | rs951016 | 0.126 | -0.723 | -0.183 | -0.069 | 0.02 | 0.052 | 0.052 | 0 | 0.29 | 0.21 | 0.27 | 0.21 | 0.23 | 0.32 |
| 3 | 193096066 | rs1878717 | 0.155 | -0.449 | -0.364 | 0.084 | -0.001 | -0.01 | -0.01 | 0 | 0.64 | 0.63 | 0.68 | 0.76 | 0.71 | 0.64 |
| 3 | 193103028 | rs7632075 | -0.253 | 0.106 | -0.118 | 0.8 | 0.004 | 0.039 | 0.039 | 0.016 | 0.89 | 0.82 | 0.9 | 0.81 | 0.89 | 0.89 |
| 3 | 193105372 | rs7649049 | -0.366 | 0.121 | -0.023 | 0.804 | 0.009 | 0.054 | 0.054 | 0 | 0.89 | 0.83 | 0.9 | 0.8 | 0.89 | 0.89 |
| 3 | 193111865 | rs902192 | 0.562 | -0.189 | -0.205 | 0.594 | -0.013 | -0.026 | -0.026 | 0.089 | 0.18 | 0.25 | 0.25 | 0.1 | 0.16 | 0.13 |
| 3 | 193112022 | rs6762308 | 0.316 | -0.161 | -0.34 | 0.626 | -0.016 | -0.01 | -0.01 | 0.136 | 0.7 | 0.6 | 0.66 | 0.73 | 0.72 | 0.76 |
| 3 | 193114643 | rs4687426 | 1.087 | 0.331 | -0.699 | 0.802 | -0.001 | 0.018 | 0.018 | 0 | 0.5 | 0.44 | 0.46 | 0.44 | 0.49 | 0.63 |
| 3 | 193117009 | rs2280476 | 0.049 | 0.314 | -0.995 | 0.444 | -0.044 | -0.006 | -0.006 | 0 | 0.11 | 0 | 0.04 | 0.08 | 0.1 | 0.04 |
| 3 | 193119305 | rs10439973 | 0.607 | -0.016 | -0.559 | 0.488 | -0.008 | -0.005 | -0.005 | 0 | 0.71 | 0.67 | 0.67 | 0.73 | 0.73 | 0.75 |
| 3 | 193121323 | rs6444717 | 0.152 | 0.052 | -0.742 | 0.767 | 0.001 | -0.006 | -0.006 | 0 | 0.89 | 0.91 | 0.9 | 0.84 | 0.9 | 0.89 |
| 3 | 193121928 | rs7646828 | -0.541 | 0.304 | -0.675 | 0.378 | -0.004 | 0.006 | 0.006 | 0 | 0.23 | 0.26 | 0.16 | 0.21 | 0.2 | 0.09 |
| 3 | 193123303 | rs9823181 | -0.73 | -0.495 | -0.35 | 0.684 | -0.019 | 0.013 | 0.013 | 0 | 0.09 | 0.16 | 0.09 | 0.08 | 0.05 | 0.12 |
| 3 | 193130384 | rs6764121 | -0.525 | -0.373 | -0.746 | 0.678 | 0.054 | 0.006 | 0.006 | 0.348 | 0.48 | 0.73 | 0.49 | 0.55 | 0.39 | 0.27 |
| 3 | 193134759 | rs10446356 | -0.36 | -0.444 | -0.24 | 0.846 | -0.005 | 0.011 | 0.011 | 0.171 | 0.32 | 0.36 | 0.41 | 0.52 | 0.28 | 0.22 |
| 3 | 193136476 | rs7630292 | -1.342 | -0.497 | 0.576 | 1.246 | 0.001 | 0.125 | 0.125 | 0.017 | 0.13 | 0.42 | 0.11 | 0.24 | 0.07 | 0.2 |
| 3 | 193139228 | rs4687428 | -0.128 | -0.613 | 0.318 | 1.069 | -0.002 | -0.003 | -0.003 | 0.018 | 0.43 | 0.48 | 0.53 | 0.3 | 0.58 | 0.47 |
| 3 | 193140348 | rs4368445 | -0.118 | -0.62 | 0.39 | 1.066 | -0.004 | -0.005 | -0.005 | 0 | 0.43 | 0.5 | 0.53 | 0.3 | 0.58 | 0.47 |
| 3 | 193144406 | rs2367612 | 0.393 | -1.225 | -0.04 | 0.966 | -0.015 | 0.04 | 0.04 | 0 | 0.2 | 0.07 | 0.14 | 0.16 | 0.17 | 0.27 |
| 3 | 193149450 | rs971538 | 0.558 | -1.338 | 0.102 | 1.045 | 0.002 | 0.023 | 0.023 | 0.106 | 0.4 | 0.3 | 0.41 | 0.25 | 0.56 | 0.45 |
| 3 | 193149634 | rs4687435 | -0.24 | -1.51 | 0.546 | 1.752 | -0.051 | 0 | 0 | 0.077 | 0.63 | 0.53 | 0.74 | 0.61 | 0.76 | 0.76 |
| 3 | 193150750 | rs7643458 | -0.374 | -1.533 | 0.463 | 1.718 | -0.006 | 0 | 0 | 0 | 0.65 | 0.68 | 0.76 | 0.61 | 0.76 | 0.76 |
| 3 | 193152408 | rs4686638 | -0.821 | -1.688 | 0.362 | 1.896 | 0.065 | -0.032 | -0.032 | 1.495 | 0.72 | 0.99 | 0.84 | 0.8 | 0.81 | 0.79 |
| 3 | 193157924 | rs1489930 | -0.344 | -1.535 | 0.448 | 1.771 | -0.007 | 0 | 0 | 0.339 | 0.65 | 0.68 | 0.76 | 0.61 | 0.76 | 0.76 |
| 3 | 193174782 | rs9879227 | -1.419 | -0.148 | 1.091 | 1.601 | 0.114 | 0.193 | 0.193 | 0 | 0.07 | 0.4 | 0.17 | 0.16 | 0.08 | 0.05 |
| 3 | 193182715 | rs12696666 | -0.944 | 0.848 | 0.964 | 1.29 | 0.032 | -0.011 | -0.011 | 0 | 0.17 | 0.3 | 0.2 | 0.15 | 0.25 | 0.12 |
| 3 | 193183817 | rs3796181 | -1.586 | 0.441 | 1.115 | 1.14 | 0 | -0.011 | -0.011 | 0 | 0.44 | 0.68 | 0.53 | 0.54 | 0.45 | 0.52 |
| 3 | 193187108 | rs12696667 | -0.833 | 0.308 | 1.012 | 1.152 | 0.013 | 0 | 0 | 0.207 | 0.21 | 0.3 | 0.2 | 0.23 | 0.26 | 0.14 |
| 3 | 193187341 | rs6771577 | 0.237 | 0.261 | 1.366 | 1.134 | -0.008 | 0.008 | 0.008 | 0 | 0.22 | 0.22 | 0.31 | 0.3 | 0.19 | 0.36 |
| 3 | 193192795 | rs1017972 | -1.499 | 0.231 | 1.489 | 1.117 | 0.003 | 0.009 | 0.009 | 0 | 0.42 | 0.53 | 0.5 | 0.54 | 0.45 | 0.49 |
| 3 | 193200730 | rs13091394 | -1.64 | 0.191 | 1.58 | 1.099 | -0.001 | -0.007 | -0.007 | 0 | 0.43 | 0.62 | 0.52 | 0.54 | 0.45 | 0.5 |
| 3 | 193203579 | rs7633847 | -1.054 | 0.07 | 1.624 | 1.083 | -0.002 | -0.004 | -0.004 | 0 | 0.36 | 0.36 | 0.47 | 0.47 | 0.39 | 0.46 |
| 3 | 193209178 | rs6788448 | -1.233 | 0.03 | 1.752 | 1.065 | 0 | 0.005 | 0.005 | 0 | 0.38 | 0.53 | 0.49 | 0.47 | 0.39 | 0.46 |
| 3 | 193217412 | rs6803383 | -0.56 | 0.105 | 1.626 | 1.091 | -0.007 | 0.106 | 0.106 | 0 | 0.09 | 0.24 | 0.19 | 0.26 | 0.09 | 0.07 |
| 3 | 193220928 | rs2056367 | -1.231 | 0.041 | 1.981 | 0.99 | -0.007 | -0.012 | -0.012 | 0 | 0.29 | 0.59 | 0.37 | 0.28 | 0.29 | 0.28 |
| 3 | 193223257 | rs13072847 | 0.235 | 0.305 | 1.634 | 0.745 | 0.001 | -0.016 | -0.016 | 0 | 0.16 | 0.2 | 0.12 | 0.11 | 0.16 | 0.06 |
| 3 | 193227545 | rs1489927 | -0.887 | 0.22 | 1.645 | 0.8 | 0 | 0.002 | 0.002 | 0.276 | 0.3 | 0.35 | 0.38 | 0.41 | 0.3 | 0.32 |
| 3 | 193228868 | rs9283663 | -1.935 | 0.209 | 1.542 | 0.785 | -0.002 | 0.002 | 0.002 | 0 | 0.49 | 0.71 | 0.53 | 0.57 | 0.49 | 0.38 |
| 3 | 193233780 | rs13317338 | 0.742 | 0.278 | 1.519 | 0.794 | -0.006 | 0.003 | 0.003 | 0 | 0.13 | 0.08 | 0.13 | 0.13 | 0.16 | 0.05 |
| 3 | 193233926 | rs11918943 | -0.642 | 0.235 | 1.51 | 0.83 | -0.011 | -0.004 | -0.004 | 0 | 0.27 | 0.24 | 0.33 | 0.3 | 0.29 | 0.29 |
| 3 | 193252724 | rs657235 | 0.585 | 0.218 | 1.538 | 0.813 | -0.01 | -0.003 | -0.003 | 0.091 | 0.73 | 0.76 | 0.67 | 0.7 | 0.71 | 0.69 |
| 3 | 193257498 | rs9814777 | 0.826 | 0.339 | 1.423 | 0.817 | -0.01 | 0.002 | 0.002 | 0 | 0.13 | 0.07 | 0.13 | 0.14 | 0.16 | 0.05 |
| 3 | 193276672 | rs281811 | 1.917 | 1.115 | 1.405 | 0.396 | 0.019 | -0.001 | -0.001 | 0.056 | 0.47 | 0.29 | 0.48 | 0.52 | 0.43 | 0.61 |
| 3 | 193277513 | rs13099487 | 0.094 | 1.114 | 1.29 | 0.41 | -0.004 | 0.001 | 0.001 | 0 | 0.17 | 0.13 | 0.13 | 0.16 | 0.18 | 0.06 |
| 3 | 193283519 | rs281808 | 1.03 | 1.113 | 1.376 | 0.442 | -0.023 | 0.004 | 0.004 | 0.051 | 0.61 | 0.69 | 0.57 | 0.59 | 0.55 | 0.66 |
| 3 | 193284624 | rs1872532 | -0.145 | 1.342 | 0.955 | 0.567 | 0.034 | 0.001 | 0.001 | 0 | 0.25 | 0.58 | 0.37 | 0.24 | 0.31 | 0.38 |
| 3 | 193286182 | rs6444728 | -0.672 | 1.347 | 0.812 | 0.593 | -0.003 | -0.002 | -0.002 | 0 | 0.29 | 0.68 | 0.39 | 0.27 | 0.32 | 0.41 |
| 3 | 193288385 | rs189737 | 2.722 | 1.214 | 0.779 | 0.602 | 0.049 | -0.041 | -0.041 | 0 | 0.33 | 0.07 | 0.2 | 0.32 | 0.23 | 0.26 |
| 3 | 193290216 | rs169464 | 0.155 | 1.319 | 0.795 | 0.6 | -0.012 | -0.004 | -0.004 | 0.064 | 0.73 | 0.76 | 0.68 | 0.69 | 0.7 | 0.71 |
| 3 | 193298496 | rs381417 | -0.925 | 1.37 | 0.828 | 0.536 | -0.033 | 0.037 | 0.037 | 0.049 | 0.87 | 0.95 | 0.9 | 0.96 | 0.84 | 0.9 |
| 3 | 193303535 | rs13064790 | -0.374 | 1.321 | 1.115 | 0.559 | 0.012 | -0.007 | -0.007 | 1.913 | 0.34 | 0.4 | 0.43 | 0.3 | 0.43 | 0.39 |

| | | | | | | | | | | | | | | | | |
|---|---|---|---|---|---|---|---|---|---|---|---|---|---|---|---|---|
| 3 | 193311756 | rs281797 | 0.208 | 1.332 | 0.874 | 0.579 | -0.01 | -0.006 | -0.006 | 0 | 0.73 | 0.76 | 0.78 | 0.69 | 0.69 | 0.71 |
| 3 | 193316705 | rs9838374 | 1.132 | 1.42 | 0.987 | 0.6 | 0.024 | 0.036 | 0.036 | 0.065 | 0.15 | 0.27 | 0.29 | 0.2 | 0.19 | 0.32 |
| 3 | 193330022 | rs9828315 | -0.92 | 1.269 | 0.665 | 0.624 | -0.005 | 0.001 | 0.001 | 0 | 0.4 | 0.58 | 0.47 | 0.37 | 0.46 | 0.42 |
| 3 | 193334991 | rs7624750 | -0.92 | 1.269 | 0.665 | 0.624 | -0.005 | 0.001 | 0.001 | 0 | 0.4 | 0.58 | 0.47 | 0.37 | 0.46 | 0.42 |
| 3 | 193358734 | rs100774 | 2.116 | 1.096 | 0.139 | 0.45 | 0.099 | -0.026 | -0.026 | 0 | 0.36 | 0.07 | 0.28 | 0.32 | 0.22 | 0.41 |
| 3 | 193365544 | rs9845217 | -0.685 | 1.2 | 0.545 | 0.449 | 0.018 | -0.005 | -0.005 | 1.235 | 0.39 | 0.46 | 0.46 | 0.37 | 0.46 | 0.43 |
| 3 | 193383943 | rs10937594 | -0.293 | 1.198 | 0.507 | 0.456 | 0.003 | -0.003 | -0.003 | 0.04 | 0.36 | 0.57 | 0.44 | 0.3 | 0.43 | 0.4 |
| 3 | 193386538 | rs414237 | 0.167 | 1.184 | 0.549 | 0.45 | -0.01 | -0.006 | -0.006 | 0 | 0.76 | 0.77 | 0.76 | 0.69 | 0.68 | 0.84 |
| 3 | 193387345 | rs7616448 | -0.118 | 1.178 | 0.657 | 0.468 | 0.02 | -0.011 | -0.011 | 0 | 0.33 | 0.4 | 0.43 | 0.29 | 0.43 | 0.39 |
| 3 | 193389692 | rs366284 | 0.298 | 1.156 | 0.512 | 0.436 | -0.023 | 0 | 0 | 0 | 0.7 | 0.76 | 0.72 | 0.69 | 0.62 | 0.7 |
| 3 | 193401021 | rs4443116 | -0.792 | 1.092 | 0.478 | 0.382 | -0.024 | 0.011 | 0.011 | 0.027 | 0.41 | 0.7 | 0.48 | 0.37 | 0.46 | 0.43 |
| 3 | 193409094 | rs6797542 | -0.284 | 1.189 | 0.194 | 0.48 | -0.007 | 0.002 | 0.002 | 0.074 | 0.81 | 0.8 | 0.83 | 0.8 | 0.71 | 0.88 |
| 3 | 193412672 | rs1061648 | 0.539 | 0.981 | 0.342 | 0.331 | 0.013 | -0.001 | -0.001 | 0 | 0.67 | 0.58 | 0.7 | 0.68 | 0.62 | 0.67 |
| 3 | 193414733 | rs1056392 | 0.123 | 1.167 | 0.551 | 0.304 | 0.014 | 0 | 0 | 0 | 0.27 | 0.33 | 0.39 | 0.28 | 0.34 | 0.37 |
| 3 | 193415750 | rs9831563 | 0.123 | 1.167 | 0.534 | 0.304 | 0.013 | 0 | 0 | 0 | 0.27 | 0.33 | 0.39 | 0.28 | 0.34 | 0.37 |
| 3 | 193417179 | rs7644847 | -0.182 | 0.9 | 0.254 | 0.129 | 0.002 | -0.012 | -0.012 | 0.807 | 0.11 | 0.21 | 0.06 | 0.03 | 0.09 | 0.06 |
| 3 | 193423253 | rs280529 | -0.618 | 0.943 | -0.131 | 0.002 | -0.003 | -0.013 | -0.013 | 0 | 0.29 | 0.36 | 0.23 | 0.26 | 0.26 | 0.17 |
| 3 | 193423644 | rs6781742 | -1.634 | 1.09 | -0.451 | 0.025 | -0.008 | -0.017 | -0.017 | 0 | 0.58 | 0.75 | 0.46 | 0.47 | 0.62 | 0.42 |
| 3 | 193424496 | rs7650473 | 1.147 | 0.89 | -0.659 | -0.113 | -0.049 | 0.049 | 0.049 | 0 | 0.18 | 0.01 | 0.11 | 0.1 | 0.23 | 0.09 |
| 3 | 193425389 | rs280533 | -0.013 | 0.889 | -0.331 | -0.113 | 0.006 | -0.043 | -0.043 | 0 | 0.19 | 0.34 | 0.17 | 0.09 | 0.17 | 0.11 |
| 3 | 193425749 | rs280534 | -1.478 | 0.923 | -0.816 | 0.126 | -0.025 | -0.013 | -0.013 | 0 | 0.56 | 0.74 | 0.4 | 0.58 | 0.55 | 0.44 |
| 3 | 193426056 | rs166812 | -0.949 | 0.841 | -0.703 | -0.316 | 0.005 | 0.023 | 0.023 | 0 | 0.49 | 0.58 | 0.31 | 0.22 | 0.47 | 0.39 |
| 3 | 193427781 | rs280537 | -0.797 | 0.841 | -0.624 | -0.227 | -0.001 | -0.009 | -0.009 | 0.423 | 0.43 | 0.57 | 0.3 | 0.22 | 0.42 | 0.26 |
| 3 | 193437652 | rs280552 | -1.19 | 0.676 | -0.562 | 0.461 | 0.005 | -0.003 | -0.003 | 0.102 | 0.43 | 0.66 | 0.43 | 0.56 | 0.39 | 0.52 |
| 3 | 193444413 | rs280560 | -1.255 | 0.739 | -0.91 | 0.45 | 0.024 | 0 | 0 | 0 | 0.46 | 0.73 | 0.44 | 0.54 | 0.4 | 0.55 |
| 3 | 193449406 | rs7634783 | 0.066 | 0.709 | -1.032 | 0.839 | -0.046 | -0.101 | -0.101 | 1.984 | 0.77 | 0.94 | 0.79 | 0.52 | 0.75 | 0.66 |
| 3 | 193451757 | rs11717055 | 0.777 | 0.848 | -0.736 | 0.8 | -0.002 | -0.007 | -0.007 | 0 | 0.66 | 0.35 | 0.63 | 0.74 | 0.68 | 0.54 |
| 3 | 193453733 | rs6805912 | -0.746 | 0.819 | -0.642 | 1.041 | 0.012 | 0.033 | 0.033 | 0.038 | 0.36 | 0.48 | 0.38 | 0.48 | 0.39 | 0.52 |
| 3 | 193455574 | rs6777024 | -0.958 | 0.303 | -0.878 | 0.964 | 0 | -0.001 | -0.001 | 0.072 | 0.11 | 0.08 | 0.08 | 0.14 | 0.07 | 0.13 |
| 3 | 193464213 | rs9865670 | -0.516 | -0.046 | -1.195 | 0.908 | 0.031 | -0.02 | -0.02 | 0 | 0.41 | 0.77 | 0.39 | 0.5 | 0.32 | 0.47 |
| 3 | 193474957 | rs1568665 | -0.017 | -0.553 | -0.87 | 0.852 | 0.001 | 0.001 | 0.001 | 0 | 0.16 | 0.18 | 0.09 | 0.1 | 0.1 | 0.25 |
| 3 | 193477063 | rs1846808 | -0.249 | -0.45 | 0.092 | 0.911 | -0.009 | 0.067 | 0.067 | 0 | 0.23 | 0.43 | 0.28 | 0.39 | 0.2 | 0.21 |
| 3 | 193485565 | rs904879 | -1.24 | -0.506 | -0.812 | 0.511 | 0.016 | 0.007 | 0.007 | 0 | 0.49 | 0.9 | 0.5 | 0.48 | 0.46 | 0.67 |
| 3 | 193488501 | rs11915891 | -1.319 | -0.485 | 0 | 0.699 | 0.012 | 0.083 | 0.083 | 0 | 0.27 | 0.47 | 0.31 | 0.49 | 0.27 | 0.51 |
| 3 | 193495155 | rs6444746 | 0.018 | -0.521 | -0.278 | 0.872 | -0.006 | 0.024 | 0.024 | 0 | 0.93 | 0.95 | 0.84 | 0.71 | 0.91 | 0.89 |
| 3 | 193495514 | rs9880995 | 0.871 | -1.347 | -0.972 | 0.325 | 0.036 | 0.014 | 0.014 | 0.082 | 0.54 | 0.19 | 0.59 | 0.58 | 0.63 | 0.61 |
| 3 | 193505467 | rs940063 | 1.772 | -1.054 | -1.005 | 0.51 | -0.093 | -0.025 | -0.025 | 1.105 | 0.4 | 0.1 | 0.48 | 0.33 | 0.45 | 0.26 |
| 3 | 193513016 | rs476927 | -0.426 | -1.212 | -0.779 | 0.424 | 0.005 | 0.001 | 0.001 | 0.487 | 0.34 | 0.63 | 0.22 | 0.12 | 0.26 | 0.23 |
| 3 | 193523984 | rs9835579 | 1.47 | -1.119 | -0.846 | 0.612 | -0.035 | 0.011 | 0.011 | 0 | 0.45 | 0.22 | 0.52 | 0.39 | 0.48 | 0.37 |
| 3 | 193527389 | rs533857 | -0.32 | -1.179 | -0.466 | 0.51 | -0.022 | -0.047 | -0.047 | 0.833 | 0.35 | 0.51 | 0.22 | 0.22 | 0.28 | 0.3 |
| 3 | 193529841 | rs10470297 | 1.418 | -1.123 | -0.882 | 0.605 | -0.058 | 0.009 | 0.009 | 0 | 0.43 | 0.15 | 0.5 | 0.36 | 0.48 | 0.32 |
| 3 | 193529913 | rs9683037 | -0.69 | -1.198 | -0.12 | 0.491 | -0.012 | -0.006 | -0.006 | 4.465 | 0.32 | 0.6 | 0.23 | 0.26 | 0.3 | 0.41 |
| 3 | 193533703 | rs9814527 | -0.761 | -0.774 | -0.374 | 0.345 | -0.001 | -0.003 | -0.003 | 0.907 | 0.38 | 0.62 | 0.27 | 0.26 | 0.34 | 0.5 |
| 3 | 193537559 | rs587612 | -0.993 | -0.766 | -0.511 | 0.366 | 0.006 | 0.019 | 0.019 | 0.023 | 0.47 | 0.69 | 0.33 | 0.37 | 0.43 | 0.62 |
| 3 | 193537991 | rs1165640 | -0.109 | -0.805 | -0.495 | 0.66 | 0.001 | 0.097 | 0.097 | 0.057 | 0.92 | 0.81 | 0.85 | 0.71 | 0.93 | 0.92 |
| 3 | 193546594 | rs2030581 | -0.93 | -0.873 | -0.54 | -0.232 | 0 | 0.004 | 0.004 | 0 | 0.44 | 0.6 | 0.33 | 0.37 | 0.42 | 0.59 |
| 3 | 193557558 | rs2513 | 0.727 | -1.296 | -0.31 | -0.33 | -0.037 | -0.027 | -0.027 | 0.043 | 0.87 | 0.74 | 0.91 | 0.85 | 0.91 | 0.89 |
| 3 | 193561684 | rs2651148 | 0.565 | -0.826 | -0.717 | -0.301 | -0.004 | -0.01 | -0.01 | 0 | 0.81 | 0.91 | 0.89 | 0.76 | 0.82 | 0.72 |
| 3 | 193567516 | rs7629215 | 1.239 | -1.06 | -0.881 | -0.572 | 0.035 | 0.068 | 0.068 | 0.019 | 0.49 | 0.12 | 0.57 | 0.59 | 0.56 | 0.41 |
| 3 | 193568108 | rs1797855 | 0.056 | -1.304 | -0.375 | -0.698 | -0.048 | -0.068 | -0.068 | 0 | 0.73 | 0.54 | 0.82 | 0.77 | 0.78 | 0.71 |
| 3 | 193568496 | rs12330159 | 1.371 | -1.248 | -0.379 | -0.703 | 0.049 | 0.068 | 0.068 | 0 | 0.51 | 0.22 | 0.57 | 0.59 | 0.57 | 0.41 |
| 3 | 193576832 | rs9878109 | 0.735 | -1.098 | -0.499 | -0.707 | 0.013 | 0.021 | 0.021 | 0 | 0.57 | 0.34 | 0.64 | 0.58 | 0.62 | 0.46 |
| 3 | 193578770 | rs2246195 | 1.161 | -0.932 | -0.669 | -0.645 | -0.022 | 0.074 | 0.074 | 0 | 0.16 | 0.02 | 0.16 | 0.04 | 0.15 | 0.14 |
| 3 | 193581231 | rs12496695 | 2.163 | -1.052 | -0.767 | -0.728 | -0.104 | -0.1 | -0.1 | 0 | 0.34 | 0.11 | 0.48 | 0.48 | 0.41 | 0.26 |
| 3 | 193581469 | rs2630242 | -0.644 | -1.116 | -0.414 | -0.544 | 0.016 | 0.057 | 0.057 | 0.334 | 0.9 | 0.81 | 0.9 | 0.75 | 0.88 | 0.76 |
| 3 | 193582584 | rs11917777 | -0.271 | -1.214 | -0.313 | -0.61 | -0.044 | -0.064 | -0.064 | 0 | 0.29 | 0.46 | 0.2 | 0.12 | 0.24 | 0.21 |
| 3 | 193583710 | rs604222 | -0.1 | -0.625 | -0.489 | -0.527 | -0.001 | 0.019 | 0.019 | 0 | 0.24 | 0.17 | 0.2 | 0.15 | 0.2 | 0.26 |
| 3 | 193584617 | rs968861 | 0.797 | -0.629 | -0.528 | -0.503 | -0.001 | 0 | 0 | 0 | 0.67 | 0.69 | 0.61 | 0.6 | 0.68 | 0.49 |
| 3 | 193584982 | rs2651158 | -0.895 | -0.633 | -0.582 | -0.515 | 0 | 0 | 0 | 0 | 0.33 | 0.3 | 0.37 | 0.26 | 0.3 | 0.43 |
| 3 | 193585494 | rs2651159 | 0.577 | -0.473 | -0.803 | -0.52 | 0.013 | 0.067 | 0.067 | 0 | 0.21 | 0.11 | 0.17 | 0.04 | 0.15 | 0.14 |
| 3 | 193585931 | rs10439974 | 0.192 | -0.385 | -0.653 | -0.453 | 0.029 | 0.064 | 0.064 | 0 | 0.55 | 0.65 | 0.64 | 0.81 | 0.59 | 0.68 |
| 3 | 193587300 | rs1354037 | -0.358 | 0.173 | -0.542 | 0.558 | -0.02 | -0.02 | -0.02 | 0.894 | 0.09 | 0.03 | 0.19 | 0.1 | 0.13 | 0.17 |
| 3 | 193587435 | rs2630239 | 0.914 | 0.211 | -0.334 | 0.727 | 0.002 | 0.025 | 0.025 | 0 | 0.83 | 0.81 | 0.72 | 0.6 | 0.78 | 0.62 |
| 3 | 193589457 | rs6444753 | 0.879 | -0.095 | -0.295 | 0.088 | -0.01 | -0.012 | -0.012 | 0 | 0.9 | 0.78 | 0.89 | 0.86 | 0.89 | 0.89 |
| 3 | 193591466 | rs1165647 | 0.378 | -0.387 | -0.628 | -0.189 | -0.008 | -0.019 | -0.019 | 0 | 0.65 | 0.41 | 0.75 | 0.73 | 0.67 | 0.64 |
| 3 | 193592218 | rs2630236 | -0.048 | -0.145 | -1.237 | 0.035 | 0.007 | 0.027 | 0.027 | 0 | 0.27 | 0.04 | 0.29 | 0.21 | 0.22 | 0.24 |
| 3 | 193592677 | rs612815 | 0.034 | 0.818 | -0.731 | 0.173 | -0.003 | -0.001 | -0.001 | 0.162 | 0.24 | 0.23 | 0.31 | 0.21 | 0.35 | 0.28 |
| 3 | 193593201 | rs1165649 | -0.899 | 0.55 | -0.65 | 0.417 | -0.004 | 0 | 0 | 0 | 0.77 | 0.8 | 0.71 | 0.66 | 0.8 | 0.71 |
| 3 | 193607758 | rs12107613 | 0.37 | 0.395 | -0.488 | 0.352 | 0.011 | 0.009 | 0.009 | 0 | 0.31 | 0.44 | 0.32 | 0.3 | 0.33 | 0.28 |

| | | | | | | | | | | | | | | | | |
|---|---|---|---|---|---|---|---|---|---|---|---|---|---|---|---|---|
| 3 | 193614072 | rs613893 | -0.693 | 0.188 | -0.798 | 0.43 | 0.023 | -0.026 | -0.026 | 0 | 0.64 | 0.77 | 0.6 | 0.44 | 0.73 | 0.61 |
| 3 | 193618305 | rs1797847 | -0.709 | 0.19 | -0.798 | 0.454 | 0.023 | -0.027 | -0.027 | 0.017 | 0.64 | 0.77 | 0.56 | 0.43 | 0.72 | 0.61 |
| 3 | 193618979 | rs13324043 | 1.037 | 0.443 | -1.167 | 0.528 | -0.009 | -0.031 | -0.031 | 0 | 0.14 | 0.08 | 0.12 | 0.24 | 0.15 | 0.19 |
| 3 | 193619878 | rs614498 | 0.187 | 0.641 | -0.7 | 0.178 | -0.054 | 0.073 | 0.073 | 0 | 0.33 | 0.13 | 0.27 | 0.15 | 0.41 | 0.33 |
| 3 | 193621698 | rs1797858 | 0.59 | 0.782 | -0.879 | 0.274 | -0.089 | 0.039 | 0.039 | 0 | 0.27 | 0.06 | 0.23 | 0.15 | 0.38 | 0.3 |
| 3 | 193639623 | rs13072095 | -0.368 | 0.561 | -0.321 | 0.089 | -0.005 | -0.063 | -0.063 | 0 | 0.67 | 0.44 | 0.76 | 0.89 | 0.68 | 0.7 |
| 3 | 193649040 | rs4686653 | 0.378 | 0.906 | -0.909 | 0.588 | -0.001 | 0.001 | 0.001 | 0 | 0.57 | 0.39 | 0.58 | 0.6 | 0.47 | 0.49 |
| 3 | 193653537 | rs2636914 | 0.171 | 1.322 | -1.749 | 0.796 | -0.014 | -0.012 | -0.012 | 2.586 | 0.88 | 0.86 | 0.83 | 0.68 | 0.8 | 0.78 |
| 3 | 193660447 | rs2630230 | 0.673 | 1.235 | -2.012 | -0.033 | -0.063 | 0.104 | 0.104 | 0 | 0.27 | 0.08 | 0.2 | 0.09 | 0.33 | 0.26 |
| 3 | 193664994 | rs1986907 | 0.464 | 0.521 | -1.965 | 0.23 | 0.005 | 0.001 | 0.001 | 0 | 0.31 | 0.24 | 0.39 | 0.3 | 0.25 | 0.23 |
| 3 | 193667472 | rs6444754 | -0.309 | 1.025 | -2.265 | -0.083 | -0.147 | 0.16 | 0.16 | 0 | 0.64 | 0.95 | 0.7 | 0.9 | 0.52 | 0.65 |
| 3 | 193667833 | rs2636909 | 0.225 | 0.779 | -1.824 | 0.474 | -0.008 | 0.013 | 0.013 | 0.163 | 0.57 | 0.26 | 0.63 | 0.39 | 0.56 | 0.5 |
| 3 | 193668899 | rs1872542 | -0.46 | 0.4 | -1.925 | 0.666 | -0.01 | 0.02 | 0.02 | 0 | 0.49 | 0.23 | 0.49 | 0.43 | 0.48 | 0.36 |
| 3 | 193669436 | rs6771223 | -0.812 | -0.587 | -1.206 | 0.865 | -0.012 | 0.018 | 0.018 | 1.013 | 0.34 | 0.57 | 0.38 | 0.44 | 0.26 | 0.17 |
| 3 | 193672434 | rs4686655 | -1.386 | -1.02 | -1.277 | 0.549 | 0.005 | 0.007 | 0.007 | 0 | 0.5 | 0.48 | 0.52 | 0.69 | 0.38 | 0.59 |
| 3 | 193674581 | rs11914434 | 0.175 | -1.348 | -1.089 | 0.581 | -0.001 | -0.036 | -0.036 | 0 | 0.08 | 0.13 | 0.07 | 0 | 0.07 | 0.02 |
| 3 | 193676362 | rs4687473 | -0.943 | -1.43 | -1.256 | 0.579 | 0.01 | 0.001 | 0.001 | 0 | 0.37 | 0.35 | 0.41 | 0.67 | 0.24 | 0.51 |
| 3 | 193679581 | rs4687474 | -0.86 | -0.04 | -1.469 | 0.573 | 0.003 | -0.037 | -0.037 | 0 | 0.22 | 0.12 | 0.28 | 0.41 | 0.18 | 0.2 |
| 3 | 193682140 | rs1388098 | -0.589 | 0.478 | -1.037 | 0.435 | 0.004 | -0.037 | -0.037 | 0 | 0.07 | 0.18 | 0.06 | 0 | 0.09 | 0.01 |
| 3 | 193693543 | rs9834951 | -0.447 | 0.794 | -1.726 | 0.667 | -0.075 | -0.093 | -0.093 | 0 | 0.2 | 0.05 | 0.15 | 0.28 | 0.22 | 0.2 |
| 3 | 193694298 | rs13319619 | -0.435 | 0.395 | -1.248 | 0.392 | 0.007 | -0.001 | -0.001 | 0 | 0.42 | 0.3 | 0.32 | 0.38 | 0.38 | 0.33 |
| 3 | 193697110 | rs9855415 | -0.962 | 0.427 | -1.714 | 0.086 | -0.074 | 0.109 | 0.109 | 0 | 0.64 | 0.84 | 0.54 | 0.87 | 0.54 | 0.64 |
| 3 | 193697644 | rs2367937 | -0.73 | -0.03 | -1.325 | 0.837 | 0.017 | -0.073 | -0.073 | 0 | 0.15 | 0.07 | 0.14 | 0.47 | 0.11 | 0.29 |
| 3 | 193698518 | rs6799681 | -0.976 | 0.072 | -0.703 | 0.379 | -0.004 | -0.002 | -0.002 | 0.113 | 0.28 | 0.63 | 0.29 | 0.28 | 0.29 | 0.26 |
| 3 | 193699775 | rs7649230 | -1.217 | -0.441 | -0.329 | 0.185 | 0.001 | 0.004 | 0.004 | 0 | 0.44 | 0.58 | 0.48 | 0.68 | 0.38 | 0.57 |
| 3 | 193700313 | rs9871733 | -0.244 | -0.502 | -0.696 | 0.519 | -0.01 | -0.028 | -0.028 | 0.59 | 0.09 | 0.02 | 0.13 | 0.42 | 0.08 | 0.23 |
| 3 | 193703873 | rs6770661 | -1.473 | -0.762 | 0.051 | 0.318 | -0.021 | 0.072 | 0.072 | 0 | 0.2 | 0.35 | 0.27 | 0.51 | 0.12 | 0.4 |
| 3 | 193706323 | rs1472075 | -0.357 | -0.474 | -0.302 | 0.276 | -0.021 | 0.019 | 0.019 | 0.1 | 0.8 | 0.9 | 0.84 | 0.84 | 0.78 | 0.88 |
| 3 | 193713770 | rs6762387 | -1.872 | -0.417 | 1.001 | 0.548 | -0.013 | 0.125 | 0.125 | 0 | 0.24 | 0.44 | 0.25 | 0.45 | 0.21 | 0.34 |
| 3 | 193715150 | rs2054964 | -2.095 | -0.214 | 1.024 | -0.15 | -0.002 | -0.067 | -0.067 | 0 | 0.24 | 0.45 | 0.27 | 0.18 | 0.24 | 0.24 |
| 3 | 193717118 | rs7641641 | -1.008 | -0.101 | 0.754 | -0.171 | 0.01 | -0.072 | -0.072 | 0.181 | 0.13 | 0.26 | 0.11 | 0.01 | 0.15 | 0.08 |
| 3 | 193722330 | rs724767 | -1.152 | -0.482 | 0.522 | 0.053 | -0.012 | -0.012 | -0.012 | 0 | 0.13 | 0.19 | 0.14 | 0.16 | 0.11 | 0.15 |
| 3 | 193724534 | rs4687493 | -1.198 | -0.847 | 0.705 | 0.051 | -0.024 | -0.013 | -0.013 | 0 | 0.22 | 0.39 | 0.26 | 0.26 | 0.18 | 0.2 |
| 3 | 193728370 | rs4687494 | 0.635 | -0.982 | 0.413 | 0.066 | 0.001 | 0.002 | 0.002 | 4.058 | 0.52 | 0.44 | 0.54 | 0.51 | 0.52 | 0.65 |
| 3 | 193730859 | rs9874421 | -0.433 | -1.007 | 0.6 | 0.049 | 0 | 0.005 | 0.005 | 1.661 | 0.68 | 0.77 | 0.76 | 0.72 | 0.68 | 0.81 |
| 3 | 193733031 | rs1078814 | 1.14 | -1.159 | 0.611 | 0.091 | -0.007 | 0.029 | 0.029 | 3.559 | 0.43 | 0.35 | 0.39 | 0.12 | 0.44 | 0.3 |
| 3 | 193735174 | rs4686660 | -0.428 | -0.928 | 0.508 | 0.288 | -0.007 | -0.001 | -0.001 | 0 | 0.15 | 0.33 | 0.22 | 0.21 | 0.16 | 0.17 |
| 3 | 193740233 | rs13320013 | -0.505 | -0.828 | 0.381 | 0.421 | -0.006 | 0.003 | 0.003 | 0.071 | 0.69 | 0.83 | 0.77 | 0.72 | 0.67 | 0.81 |
| 3 | 193748075 | rs7628226 | 0.411 | -0.453 | 0.757 | 0.4 | 0.002 | 0.006 | 0.006 | 0 | 0.51 | 0.49 | 0.57 | 0.3 | 0.57 | 0.52 |
| 3 | 193752149 | rs4687500 | -0.466 | -0.003 | 0.607 | 0.935 | -0.001 | 0.125 | 0.125 | 0 | 0.12 | 0.15 | 0.16 | 0.44 | 0.12 | 0.3 |
| 3 | 193754482 | rs4687504 | -1.062 | 0.096 | 0.141 | 0.53 | -0.001 | 0.132 | 0.132 | 0 | 0.19 | 0.35 | 0.28 | 0.55 | 0.2 | 0.36 |
| 3 | 193757098 | rs4686667 | -0.043 | 0.147 | -0.352 | -0.096 | -0.005 | -0.004 | -0.004 | 0.147 | 0.56 | 0.52 | 0.63 | 0.65 | 0.63 | 0.73 |
| 3 | 193759302 | rs6802592 | -0.714 | 0.635 | -0.151 | -0.054 | 0.005 | -0.038 | -0.038 | 0 | 0.15 | 0.3 | 0.12 | 0.06 | 0.15 | 0.15 |
| 3 | 193760848 | rs4417818 | -1.386 | 0.858 | -0.752 | 0.274 | -0.063 | -0.011 | -0.011 | 0 | 0.12 | 0.01 | 0.19 | 0.14 | 0.23 | 0.18 |
| 3 | 193769266 | rs1490752 | -1.448 | 0.45 | -0.074 | 0.166 | 0.007 | -0.045 | -0.045 | 0 | 0.1 | 0.29 | 0.15 | 0.04 | 0.11 | 0.2 |
| 3 | 193782232 | rs1546050 | 0.035 | 0.656 | -1.094 | 0.084 | 0.005 | 0.093 | 0.093 | 0.845 | 0.52 | 0.25 | 0.53 | 0.68 | 0.57 | 0.63 |
| 3 | 193783178 | rs6444767 | -0.045 | 0.697 | -0.775 | 0.052 | 0 | 0.003 | 0.003 | 0.769 | 0.61 | 0.59 | 0.52 | 0.32 | 0.6 | 0.64 |
| 3 | 193787775 | rs12487368 | -0.064 | 0.547 | -0.919 | 0.631 | -0.006 | -0.015 | -0.015 | 0 | 0.19 | 0.16 | 0.31 | 0.34 | 0.24 | 0.21 |
| 3 | 193792533 | rs9855959 | -0.842 | -0.067 | -0.731 | 0.072 | -0.002 | 0.036 | 0.036 | 0 | 0.64 | 0.76 | 0.76 | 0.91 | 0.68 | 0.81 |
| 3 | 193796498 | rs1873412 | -0.56 | -0.291 | -0.92 | 0.592 | 0.002 | -0.012 | -0.012 | 0 | 0.57 | 0.72 | 0.68 | 0.58 | 0.61 | 0.64 |
| 3 | 193799549 | rs4686671 | -0.891 | -0.431 | -0.764 | 0.695 | 0.026 | -0.03 | -0.03 | 0 | 0.35 | 0.8 | 0.4 | 0.55 | 0.28 | 0.43 |
| 3 | 193801183 | rs7628703 | 0.063 | -0.305 | -0.498 | 0.501 | -0.011 | 0.014 | 0.014 | 0 | 0.22 | 0.46 | 0.26 | 0.21 | 0.19 | 0.23 |
| 3 | 193806588 | rs9872487 | -0.113 | -0.013 | -1.209 | 0.513 | -0.014 | 0.022 | 0.022 | 0 | 0.3 | 0.68 | 0.32 | 0.52 | 0.22 | 0.26 |
| 3 | 193809078 | rs7648783 | -1.101 | 0.474 | -2.04 | 0.291 | -0.052 | 0.096 | 0.096 | 0 | 0.67 | 0.96 | 0.81 | 0.86 | 0.72 | 0.8 |
| 3 | 193811162 | rs10933711 | -0.734 | -0.257 | -1.005 | 0.914 | 0.034 | 0.01 | 0.01 | 0 | 0.41 | 0.74 | 0.44 | 0.65 | 0.31 | 0.45 |
| 3 | 193821569 | rs9848849 | 0.758 | 0.043 | -1.062 | 1.024 | 0.041 | -0.002 | -0.002 | 0 | 0.49 | 0.19 | 0.41 | 0.43 | 0.59 | 0.44 |
| 3 | 193824905 | rs9840796 | -0.412 | -0.327 | -0.313 | 0.926 | -0.035 | -0.045 | -0.045 | 0.032 | 0.85 | 0.78 | 0.89 | 0.94 | 0.95 | 0.88 |
| 3 | 193827215 | rs6787480 | 1.081 | -0.576 | 0.171 | 1.13 | -0.006 | 0.026 | 0.026 | 0 | 0.66 | 0.55 | 0.63 | 0.47 | 0.7 | 0.6 |
| 3 | 193830681 | rs4270456 | -0.39 | -0.392 | -0.086 | 1.223 | 0 | 0.006 | 0.006 | 0 | 0.19 | 0.23 | 0.26 | 0.45 | 0.25 | 0.28 |
| 3 | 193834940 | rs6796696 | 0.952 | -0.257 | 0.107 | 0.923 | -0.028 | -0.039 | -0.039 | 0 | 0.61 | 0.54 | 0.55 | 0.77 | 0.72 | 0.62 |
| 3 | 193839467 | rs6444771 | -1.682 | -0.081 | -0.339 | 1.364 | -0.051 | 0.096 | 0.096 | 0.251 | 0.48 | 0.95 | 0.59 | 0.67 | 0.45 | 0.51 |
| 3 | 193842582 | rs7649147 | -1.178 | -0.019 | 0.688 | 1.544 | -0.001 | 0.005 | 0.005 | 0 | 0.36 | 0.62 | 0.38 | 0.47 | 0.35 | 0.43 |
| 3 | 193843374 | rs7627589 | -1.515 | 0.06 | 1.296 | 1.774 | -0.002 | 0.042 | 0.042 | 1.841 | 0.25 | 0.44 | 0.22 | 0.34 | 0.23 | 0.22 |
| 3 | 193845922 | rs7626182 | -1.487 | 0.072 | 1.711 | 1.485 | 0.02 | -0.051 | -0.051 | 0.004 | 0.14 | 0.4 | 0.23 | 0.07 | 0.17 | 0.14 |
| 3 | 193846565 | rs7639323 | -0.6 | 0.07 | 1.522 | 1.548 | -0.005 | -0.025 | -0.025 | 0 | 0.1 | 0.24 | 0.16 | 0.05 | 0.1 | 0.06 |
| 3 | 193847511 | rs6771393 | -0.445 | 0.103 | 1.511 | 1.683 | -0.006 | -0.001 | -0.001 | 0.052 | 0.17 | 0.23 | 0.25 | 0.17 | 0.15 | 0.18 |
| 3 | 193847627 | rs6444772 | -0.923 | 0.159 | 1.478 | 1.911 | 0.001 | 0.086 | 0.086 | 0.404 | 0.16 | 0.29 | 0.2 | 0.33 | 0.14 | 0.18 |
| 3 | 193851390 | rs6767127 | -1.247 | -0.088 | 1.603 | 1.259 | -0.028 | -0.008 | -0.008 | 0 | 0.12 | 0.39 | 0.09 | 0.1 | 0.1 | 0.09 |
| 3 | 193861242 | rs4686673 | -0.949 | 0.483 | 1.281 | 1.624 | 0.015 | 0.088 | 0.088 | 0 | 0.26 | 0.49 | 0.37 | 0.48 | 0.31 | 0.31 |

| | | | | | | | | | | | | | | | | |
|---|---|---|---|---|---|---|---|---|---|---|---|---|---|---|---|---|
| 3 | 193864097 | rs6805836 | -0.768 | 0.497 | 1.237 | 1.691 | 0.025 | 0.144 | 0.144 | 0 | 0.23 | 0.42 | 0.35 | 0.47 | 0.27 | 0.26 |
| 3 | 193865878 | rs6763995 | -1.424 | 0.725 | 1.454 | 1.175 | 0.003 | -0.039 | -0.039 | 0.111 | 0.19 | 0.41 | 0.14 | 0.15 | 0.17 | 0.21 |
| 3 | 193875909 | rs4380361 | -2.064 | 0.566 | 1.737 | 1.718 | 0.014 | 0.136 | 0.136 | 0 | 0.06 | 0.25 | 0.08 | 0.2 | 0.04 | 0.1 |
| 3 | 193876042 | rs2368046 | -2.776 | 0.522 | 1.834 | 1.648 | 0.009 | 0.106 | 0.106 | 0 | 0.08 | 0.3 | 0.1 | 0.2 | 0.06 | 0.13 |
| 3 | 193878035 | rs9831565 | -2.847 | 0.497 | 1.998 | 1.648 | 0.03 | 0.16 | 0.16 | 0 | 0.09 | 0.41 | 0.11 | 0.2 | 0.06 | 0.13 |
| 3 | 193879515 | rs7652270 | -2.851 | 0.724 | 2.013 | 1.645 | 0.047 | 0.106 | 0.106 | 0.022 | 0.13 | 0.61 | 0.13 | 0.28 | 0.12 | 0.15 |
| 3 | 193884752 | rs11918995 | -2.273 | 0.495 | 1.97 | 1.175 | 0.053 | 0.025 | 0.025 | 0.901 | 0.07 | 0.52 | 0.07 | 0 | 0.02 | 0.04 |
| 3 | 193887269 | rs7632519 | -0.549 | 0.351 | 0.958 | 1.282 | -0.007 | 0.024 | 0.024 | 0 | 0.34 | 0.43 | 0.4 | 0.45 | 0.32 | 0.43 |
| 3 | 193887636 | rs6803664 | -0.964 | 1.512 | 1.334 | 1.15 | 0.105 | 0.019 | 0.019 | 0 | 0.13 | 0.42 | 0.17 | 0.13 | 0.15 | 0.09 |
| 3 | 193891012 | rs11711981 | 0.465 | 1.245 | 0.32 | 1.121 | 0.003 | -0.003 | -0.003 | 0 | 0.6 | 0.47 | 0.34 | 0.36 | 0.43 | 0.44 |
| 3 | 193895196 | rs6779726 | -0.667 | 0.491 | -0.232 | 1.376 | -0.001 | 0.011 | 0.011 | 0 | 0.25 | 0.28 | 0.3 | 0.34 | 0.24 | 0.33 |
| 3 | 193897683 | rs4532075 | 0.792 | 0.681 | -0.973 | 1.311 | 0.013 | 0.012 | 0.012 | 0 | 0.55 | 0.29 | 0.58 | 0.56 | 0.63 | 0.4 |
| 3 | 193899859 | rs2001726 | -0.586 | 0.89 | -1.226 | 1.338 | 0.009 | 0.022 | 0.022 | 0 | 0.21 | 0.29 | 0.35 | 0.39 | 0.24 | 0.4 |
| 3 | 193913552 | rs4074131 | 0.399 | 0.243 | -2.552 | 0.858 | -0.014 | -0.078 | -0.078 | 0.785 | 0.28 | 0.01 | 0.15 | 0.34 | 0.19 | 0.22 |
| 3 | 193916237 | rs6791279 | -0.419 | 0.947 | -2.86 | 0.913 | -0.121 | -0.059 | -0.059 | 0 | 0.78 | 0.98 | 0.8 | 0.82 | 0.64 | 0.79 |
| 3 | 193918902 | rs9858096 | 0.376 | 0.434 | -2.232 | 0.875 | 0.015 | 0.008 | 0.008 | 0.03 | 0.67 | 0.51 | 0.56 | 0.72 | 0.58 | 0.65 |
| 3 | 193921560 | rs4687530 | -0.344 | 0.659 | -3.137 | 0.875 | -0.112 | -0.061 | -0.061 | 0 | 0.77 | 0.98 | 0.8 | 0.81 | 0.64 | 0.79 |
| 3 | 193923004 | rs4594573 | 0.53 | 0.569 | -3.114 | 0.849 | -0.041 | -0.067 | -0.067 | 2.156 | 0.23 | 0.02 | 0.17 | 0.3 | 0.22 | 0.3 |
| 3 | 193929080 | rs2368045 | 0.857 | -0.29 | -3.195 | 0.408 | 0.027 | 0.015 | 0.015 | 0 | 0.23 | 0.06 | 0.13 | 0.25 | 0.13 | 0.16 |
| 3 | 193931183 | rs4078029 | 0.225 | -0.592 | -3.109 | 0.702 | -0.015 | 0.008 | 0.008 | 1.394 | 0.17 | 0.76 | 0.36 | 0.2 | 0.09 | 0.09 |
| 3 | 193931852 | rs4613405 | 0.291 | 0.044 | -3.459 | 0.425 | 0.001 | 0.062 | 0.062 | 0 | 0.35 | 0.07 | 0.21 | 0.27 | 0.3 | 0.37 |
| 3 | 193936094 | rs6774543 | -0.07 | 1.292 | -3.043 | 0.702 | -0.124 | -0.064 | -0.064 | 0 | 0.76 | 0.99 | 0.81 | 0.77 | 0.6 | 0.78 |
| 3 | 193938611 | rs7647261 | -0.398 | 0.847 | -2.738 | 0.698 | -0.03 | -0.046 | -0.046 | 0 | 0.4 | 0.82 | 0.6 | 0.43 | 0.59 | 0.37 |
| 3 | 193944478 | rs6784330 | 0.135 | 0.622 | -2.915 | 0.782 | -0.072 | 0.057 | 0.057 | 0 | 0.45 | 0.12 | 0.4 | 0.28 | 0.53 | 0.32 |
| 3 | 193946147 | rs6766839 | -0.349 | 0.251 | -1.651 | 1.377 | -0.023 | 0.059 | 0.059 | 0 | 0.13 | 0.35 | 0.09 | 0.18 | 0.07 | 0.19 |
| 3 | 193946512 | rs6444779 | -0.574 | 0.183 | -1.695 | 1.308 | -0.032 | 0.028 | 0.028 | 0.363 | 0.15 | 0.34 | 0.09 | 0.18 | 0.08 | 0.19 |
| 3 | 193946534 | rs12495904 | -0.186 | 0.588 | -2.365 | 1.461 | -0.033 | -0.054 | -0.054 | 0 | 0.09 | 0.01 | 0.22 | 0.2 | 0.1 | 0.24 |
| 3 | 193954754 | rs6803149 | 0.42 | -0.037 | -2.531 | 0.739 | 0.018 | 0.082 | 0.082 | 0 | 0.62 | 0.84 | 0.71 | 0.7 | 0.68 | 0.7 |
| 3 | 193955440 | rs4349452 | 0.389 | 0.008 | -2.147 | 1.445 | 0.003 | -0.054 | -0.054 | 0 | 0.9 | 0.99 | 0.89 | 0.78 | 0.94 | 0.88 |
| 3 | 193960464 | rs4687537 | -0.858 | 0.155 | -1.982 | 1.798 | -0.002 | -0.052 | -0.052 | 0.105 | 0.1 | 0.02 | 0.24 | 0.27 | 0.08 | 0.17 |
| 3 | 193963483 | rs4547644 | -0.77 | 0.485 | -1.553 | 0.65 | 0.001 | -0.021 | -0.021 | 0 | 0.44 | 0.45 | 0.64 | 0.79 | 0.44 | 0.6 |
| 3 | 193967428 | rs6778204 | -0.757 | -0.087 | -1.825 | 0.983 | 0.035 | -0.011 | -0.011 | 0 | 0.41 | 0.67 | 0.43 | 0.54 | 0.31 | 0.57 |
| 3 | 193968817 | rs7613544 | -0.433 | -0.128 | -2.094 | 0.521 | 0.011 | 0.082 | 0.082 | 0 | 0.23 | 0.06 | 0.22 | 0.12 | 0.22 | 0.37 |
| 3 | 193969806 | rs9811548 | 0.491 | 0.765 | -1.924 | 0.464 | -0.003 | 0.001 | 0.001 | 0 | 0.3 | 0.31 | 0.31 | 0.24 | 0.34 | 0.21 |
| 3 | 193976071 | rs7613652 | -0.17 | 0.661 | -2.138 | 0.52 | -0.041 | -0.064 | -0.064 | 0 | 0.56 | 0.86 | 0.58 | 0.51 | 0.49 | 0.56 |
| 3 | 193986199 | rs10933727 | 0.168 | 0.497 | -1.348 | -0.328 | 0 | -0.009 | -0.009 | 0 | 0.67 | 0.68 | 0.73 | 0.86 | 0.64 | 0.54 |
| 3 | 193998650 | rs4311173 | -0.7 | 0.209 | -1.023 | 0.216 | 0.009 | -0.022 | -0.022 | 0.201 | 0.34 | 0.74 | 0.3 | 0.51 | 0.28 | 0.42 |
| 3 | 194000979 | rs9862745 | -0.469 | 0.416 | -0.567 | 0.364 | -0.002 | 0.026 | 0.026 | 0 | 0.25 | 0.69 | 0.25 | 0.48 | 0.22 | 0.3 |
| 3 | 194012176 | rs11922293 | -0.491 | 0.727 | -0.739 | -0.359 | 0.012 | 0.016 | 0.016 | 0 | 0.31 | 0.22 | 0.19 | 0.33 | 0.28 | 0.27 |
| 3 | 194018342 | rs4586759 | -0.588 | 1.487 | -0.169 | -0.498 | 0.011 | -0.027 | -0.027 | 0.42 | 0.4 | 0.5 | 0.33 | 0.34 | 0.38 | 0.32 |
| 3 | 194020431 | rs6785704 | -0.496 | 1.832 | 0.056 | -0.218 | 0.008 | -0.016 | -0.016 | 0.756 | 0.1 | 0.13 | 0.13 | 0.01 | 0.15 | 0.04 |
| 3 | 194023689 | rs11928683 | -1.213 | 1.016 | 0.252 | -0.247 | 0.019 | -0.049 | -0.049 | 0 | 0.12 | 0.63 | 0.12 | 0.03 | 0.11 | 0.06 |
| 3 | 194034498 | rs9853035 | -0.905 | 0.695 | 0.1 | 0.007 | -0.006 | -0.052 | -0.052 | 0 | 0.16 | 0.53 | 0.13 | 0.05 | 0.13 | 0.06 |
| 3 | 194057168 | rs12629557 | -0.625 | 0.495 | -0.741 | 0.764 | 0.013 | -0.041 | -0.041 | 0 | 0.08 | 0.01 | 0.09 | 0.36 | 0.04 | 0.14 |
| 3 | 194059425 | rs6437403 | -0.626 | 1.103 | -0.835 | -0.256 | -0.005 | -0.012 | -0.012 | 0 | 0.34 | 0.67 | 0.26 | 0.07 | 0.39 | 0.15 |
| 3 | 194059960 | rs9836980 | -0.119 | 0.657 | -1.256 | 0.124 | 0.007 | -0.021 | -0.021 | 0 | 0.35 | 0.27 | 0.51 | 0.56 | 0.39 | 0.43 |
| 3 | 194060008 | rs6806282 | -0.242 | 0.451 | -1.004 | -0.205 | 0.003 | 0.045 | 0.045 | 0.126 | 0.53 | 0.64 | 0.65 | 0.77 | 0.51 | 0.66 |
| 3 | 194060294 | rs6783012 | -0.079 | -0.67 | -0.334 | -0.166 | -0.007 | -0.001 | -0.001 | 0 | 0.31 | 0.65 | 0.21 | 0.34 | 0.19 | 0.37 |
| 3 | 194060475 | rs6809081 | 0 | -0.292 | 0.164 | -0.178 | 0.008 | 0.01 | 0.01 | 0 | 0.52 | 0.54 | 0.46 | 0.38 | 0.45 | 0.51 |
| 3 | 194061561 | rs6774653 | -0.298 | -0.134 | 0.882 | 0.263 | -0.036 | -0.019 | -0.019 | 0.062 | 0.27 | 0.18 | 0.23 | 0.38 | 0.35 | 0.2 |
| 3 | 194061578 | rs11713634 | 0.183 | -0.129 | 0.887 | 0.204 | -0.009 | -0.004 | -0.004 | 0 | 0.39 | 0.42 | 0.27 | 0.32 | 0.31 | 0.41 |
| 3 | 194061826 | rs11711157 | 0.636 | -0.137 | 0.937 | 0.136 | 0.02 | -0.011 | -0.011 | 0 | 0.34 | 0.17 | 0.25 | 0.32 | 0.31 | 0.4 |
| 3 | 194061907 | rs4974539 | -0.433 | -0.023 | 0.669 | 0.15 | -0.025 | -0.01 | -0.01 | 0.011 | 0.28 | 0.23 | 0.24 | 0.38 | 0.35 | 0.2 |
| 3 | 194062519 | rs3732477 | 0.394 | 0.107 | 0.181 | 0.156 | -0.017 | -0.004 | -0.004 | 0.142 | 0.38 | 0.46 | 0.28 | 0.32 | 0.31 | 0.41 |
| 3 | 194062565 | rs3732476 | -1.042 | -0.106 | 0.083 | -0.87 | -0.004 | -0.003 | -0.003 | 0 | 0.66 | 0.61 | 0.5 | 0.7 | 0.67 | 0.62 |
| 3 | 194064075 | rs7618278 | -0.287 | 0.519 | -0.307 | -0.824 | -0.065 | 0.081 | 0.081 | 0.794 | 0.21 | 0.05 | 0.15 | 0.08 | 0.27 | 0.14 |
| 3 | 194066332 | rs6763011 | -0.474 | 0.363 | -0.085 | -0.795 | 0.046 | -0.07 | -0.07 | 0 | 0.23 | 0.46 | 0.18 | 0.12 | 0.26 | 0.14 |
| 3 | 194067766 | rs9870895 | -0.556 | 0.329 | -0.745 | -0.46 | -0.003 | 0.014 | 0.014 | 0.012 | 0.7 | 0.94 | 0.57 | 0.79 | 0.72 | 0.65 |
| 3 | 194079911 | rs6789870 | -0.931 | 0.884 | 0.492 | -0.33 | 0.035 | -0.025 | -0.025 | 0 | 0.15 | 0.21 | 0.14 | 0.07 | 0.24 | 0.12 |
| 3 | 194081900 | rs923934 | -0.854 | 0.912 | 0.305 | -0.316 | 0.008 | -0.005 | -0.005 | 0 | 0.14 | 0.16 | 0.14 | 0.07 | 0.24 | 0.12 |
| 3 | 194085047 | rs923931 | -0.641 | -0.907 | 0.769 | -0.188 | 0.013 | -0.054 | -0.054 | 0 | 0.1 | 0.26 | 0.07 | 0.01 | 0.1 | 0.07 |
| 3 | 194094582 | rs11916274 | -1.157 | -1.05 | 0.754 | 0.256 | -0.019 | 0.053 | 0.053 | 0 | 0.09 | 0.23 | 0.09 | 0.19 | 0.07 | 0.16 |
| 3 | 194096787 | rs7642341 | -1.453 | -2.149 | 0 | -0.249 | 0.132 | 0.077 | 0.077 | 0 | 0.41 | 0.89 | 0.27 | 0.3 | 0.23 | 0.44 |
| 3 | 194105437 | rs6799698 | -0.482 | -2.3 | 1.061 | -0.334 | -0.039 | -0.098 | -0.098 | 0.064 | 0.19 | 0.34 | 0.09 | 0.01 | 0.11 | 0.13 |
| 3 | 194108758 | rs7626054 | -1.25 | -2.26 | 1.589 | -0.125 | -0.017 | 0.059 | 0.059 | 0 | 0.11 | 0.45 | 0.08 | 0.13 | 0.05 | 0.18 |
| 3 | 194110884 | rs7632356 | -1.42 | -2.525 | 1.416 | -0.313 | -0.059 | -0.076 | -0.076 | 0.336 | 0.32 | 0.47 | 0.16 | 0.13 | 0.18 | 0.3 |
| 3 | 194112249 | rs6791246 | -2.221 | -2.49 | 1.32 | -0.315 | 0.134 | 0.076 | 0.076 | 0.291 | 0.43 | 0.87 | 0.27 | 0.3 | 0.25 | 0.46 |
| 3 | 194114092 | rs9864088 | -1.101 | -2.434 | 1.492 | -0.226 | -0.021 | 0.05 | 0.05 | 0 | 0.11 | 0.3 | 0.11 | 0.17 | 0.07 | 0.16 |
| 3 | 194119108 | rs1538863 | -1.876 | -2.443 | 1.519 | -0.263 | -0.029 | 0.005 | 0.005 | 0 | 0.16 | 0.31 | 0.13 | 0.16 | 0.1 | 0.27 |

| | | | | | | | | | | | | | | | | |
|---|---|---|---|---|---|---|---|---|---|---|---|---|---|---|---|---|
| 3 | 194122207 | rs3755617 | -0.671 | -2.324 | 1.404 | -0.118 | -0.01 | 0.011 | 0.011 | 0 | 0.08 | 0.11 | 0.09 | 0.15 | 0.06 | 0.12 |
| 3 | 194124421 | rs11900 | -1.773 | -2.158 | 1.268 | -0.206 | -0.01 | -0.01 | -0.01 | 0 | 0.34 | 0.63 | 0.23 | 0.21 | 0.21 | 0.42 |
| 3 | 194140256 | rs6784093 | -1.579 | -2.146 | 1.186 | -0.203 | -0.036 | 0.067 | 0.067 | 0.027 | 0.14 | 0.5 | 0.14 | 0.2 | 0.08 | 0.3 |
| 3 | 194140923 | rs6793096 | -1.632 | -2.122 | 1.244 | -0.197 | 0.07 | 0.063 | 0.063 | 2.351 | 0.36 | 0.79 | 0.23 | 0.21 | 0.19 | 0.42 |
| 3 | 194162411 | rs1122426 | -0.72 | -1.976 | 1.277 | -0.077 | -0.012 | 0.17 | 0.17 | 0 | 0.1 | 0.49 | 0.12 | 0.2 | 0.05 | 0.16 |
| 3 | 194172153 | rs953487 | -1.253 | -1.797 | 1.02 | 0.059 | 0.095 | 0.035 | 0.035 | 1.223 | 0.37 | 0.83 | 0.25 | 0.29 | 0.18 | 0.4 |
| 3 | 194172225 | rs953486 | -1.385 | -1.859 | 0.736 | 0.043 | 0.102 | 0.045 | 0.045 | 1.028 | 0.38 | 0.87 | 0.27 | 0.29 | 0.21 | 0.41 |
| 3 | 194184392 | rs6798801 | -1.628 | -1.786 | 0.819 | 0.075 | 0.044 | 0.104 | 0.104 | 0 | 0.07 | 0.35 | 0.05 | 0.07 | 0.02 | 0.02 |
| 3 | 194190426 | rs9874152 | -1.068 | -1.645 | -0.634 | 0.139 | 0.095 | 0.048 | 0.048 | 0.469 | 0.38 | 0.84 | 0.26 | 0.29 | 0.21 | 0.44 |
| 3 | 194199814 | rs4632483 | -0.878 | -1.494 | -0.699 | 0.036 | 0.078 | 0.075 | 0.075 | 1.83 | 0.37 | 0.79 | 0.23 | 0.23 | 0.2 | 0.43 |
| 3 | 194215878 | rs6437379 | -0.926 | -0.767 | -0.272 | 1.346 | 0 | 0.133 | 0.133 | 0 | 0.07 | 0.11 | 0.21 | 0.62 | 0.07 | 0.22 |
| 3 | 194219566 | rs2185478 | -0.456 | -0.452 | -0.222 | 0.493 | 0.002 | -0.004 | -0.004 | 0 | 0.29 | 0.34 | 0.31 | 0.24 | 0.31 | 0.24 |
| 3 | 194219913 | rs2153519 | -1.315 | -1.174 | -0.956 | -0.511 | -0.064 | 0.11 | 0.11 | 0 | 0.66 | 0.87 | 0.64 | 0.88 | 0.55 | 0.69 |
| 3 | 194225655 | rs9824773 | 0.422 | -1.354 | -1.359 | -0.667 | -0.054 | -0.003 | -0.003 | 1.146 | 0.16 | 0.08 | 0.23 | 0.11 | 0.24 | 0.18 |
| 3 | 194226486 | rs10933656 | -0.664 | -1.951 | -0.864 | -0.747 | 0.063 | 0.068 | 0.068 | 0 | 0.47 | 0.7 | 0.39 | 0.76 | 0.31 | 0.49 |
| 3 | 194226525 | rs10933657 | -0.62 | -2.005 | -0.611 | -0.765 | 0.022 | 0.023 | 0.023 | 0 | 0.46 | 0.58 | 0.38 | 0.76 | 0.31 | 0.49 |
| 3 | 194228564 | rs9812355 | -0.975 | -2.4 | -1.185 | 0.42 | 0.067 | -0.087 | -0.087 | 0 | 0.34 | 0.86 | 0.32 | 0.45 | 0.25 | 0.16 |
| 3 | 194229185 | rs11915221 | -0.765 | -2.3 | -0.201 | 1.154 | -0.046 | 0.026 | 0.026 | 0 | 0.25 | 0.36 | 0.1 | 0.35 | 0.15 | 0.14 |
| 3 | 194232753 | rs7627066 | -1.089 | -1.746 | -0.486 | 1.291 | 0.003 | 0.001 | 0.001 | 0 | 0.49 | 0.42 | 0.44 | 0.43 | 0.43 | 0.42 |
| 3 | 194233500 | rs1572837 | -1.55 | -1.71 | -0.435 | 1.084 | -0.001 | -0.006 | -0.006 | 0.006 | 0.57 | 0.42 | 0.52 | 0.65 | 0.48 | 0.51 |
| 3 | 194238390 | rs1807193 | 1.284 | -1.701 | -0.963 | 1.102 | 0.001 | 0.003 | 0.003 | 0 | 0.44 | 0.49 | 0.48 | 0.33 | 0.53 | 0.51 |
| 3 | 194238889 | rs6437384 | 0.486 | -1.386 | -0.709 | 1.286 | -0.02 | 0.018 | 0.018 | 0 | 0.65 | 0.55 | 0.58 | 0.47 | 0.67 | 0.75 |
| 3 | 194244722 | rs9862208 | -0.287 | -0.884 | -1.427 | 1.342 | 0.011 | -0.042 | -0.042 | 0 | 0.14 | 0.05 | 0.17 | 0.28 | 0.1 | 0.13 |
| 3 | 194245931 | rs4493370 | 0.323 | -0.645 | -1.288 | 0.862 | -0.017 | -0.006 | -0.006 | 0 | 0.9 | 0.93 | 0.81 | 0.89 | 0.89 | 0.85 |
| 3 | 194248036 | rs7373647 | 0.659 | -0.708 | -1.522 | 0.583 | 0.019 | 0.18 | 0.18 | 0 | 0.52 | 0.15 | 0.54 | 0.79 | 0.58 | 0.58 |
| 3 | 194266725 | rs1705984 | 0.594 | -0.488 | -1.119 | 1.148 | 0.007 | 0.007 | 0.007 | 0 | 0.83 | 0.81 | 0.72 | 0.76 | 0.79 | 0.74 |
| 3 | 194266844 | rs1705983 | 0.873 | -0.801 | -0.954 | 0.87 | -0.005 | -0.001 | -0.001 | 0.046 | 0.61 | 0.42 | 0.55 | 0.39 | 0.64 | 0.54 |
| 3 | 194273145 | rs12485613 | -0.336 | -0.766 | -1.256 | 0.776 | 0.022 | 0.005 | 0.005 | 0 | 0.19 | 0.11 | 0.16 | 0.18 | 0.09 | 0.22 |
| 3 | 194275435 | rs705298 | 0.852 | -0.798 | -1.179 | 0.839 | 0.018 | 0.003 | 0.003 | 0 | 0.68 | 0.82 | 0.74 | 0.7 | 0.78 | 0.7 |
| 3 | 194289291 | rs1773241 | 0.852 | -0.859 | -0.532 | 1.048 | 0.014 | 0.009 | 0.009 | 0 | 0.66 | 0.5 | 0.41 | 0.38 | 0.64 | 0.5 |
| 3 | 194291478 | rs12695035 | -0.872 | -1.736 | -0.322 | 0.909 | -0.004 | 0.004 | 0.004 | 0 | 0.37 | 0.54 | 0.49 | 0.58 | 0.3 | 0.49 |
| 3 | 194291805 | rs938087 | 1.257 | -1.96 | -0.211 | 1.07 | 0.001 | 0.039 | 0.039 | 0 | 0.5 | 0.31 | 0.44 | 0.41 | 0.58 | 0.37 |
| 3 | 194295696 | rs7620693 | -0.656 | -2.261 | 0.66 | 1.155 | -0.054 | 0.124 | 0.124 | 0 | 0.22 | 0.55 | 0.31 | 0.47 | 0.1 | 0.35 |
| 3 | 194308619 | rs7612496 | -0.805 | -2.477 | 0.356 | 0.923 | -0.013 | 0 | 0 | 0.317 | 0.12 | 0.14 | 0.09 | 0.2 | 0.07 | 0.08 |
| 3 | 194308983 | rs6779853 | -1.209 | -2.528 | 0.669 | 0.796 | -0.014 | -0.004 | -0.004 | 0.088 | 0.3 | 0.37 | 0.24 | 0.23 | 0.18 | 0.33 |
| 3 | 194310529 | rs9846532 | -0.728 | -2.798 | 0.214 | -0.226 | 0.023 | 0.075 | 0.075 | 0 | 0.41 | 0.75 | 0.57 | 0.72 | 0.33 | 0.51 |
| 3 | 194310731 | rs1055161 | -0.623 | -3.029 | 0.523 | -0.587 | 0.005 | 0.02 | 0.02 | 0 | 0.41 | 0.69 | 0.59 | 0.74 | 0.32 | 0.52 |
| 3 | 194311594 | rs7621704 | -0.994 | -3.357 | 0.632 | -1.058 | -0.054 | -0.119 | -0.119 | 0.329 | 0.22 | 0.41 | 0.1 | 0.02 | 0.11 | 0.26 |
| 3 | 194319882 | rs1706016 | 2.291 | -3.416 | -0.32 | -1.433 | -0.122 | 0.05 | 0.05 | 0 | 0.28 | 0.04 | 0.29 | 0.2 | 0.47 | 0.2 |
| 3 | 194324470 | rs2898610 | -0.725 | -3.365 | 0.619 | -1.399 | 0.046 | 0.025 | 0.025 | 0 | 0.06 | 0.4 | 0.06 | 0 | 0.02 | 0.03 |
| 3 | 194324939 | rs6768028 | 0.74 | -3.664 | 0.437 | -1.575 | 0.026 | -0.023 | -0.023 | 0 | 0.3 | 0.39 | 0.52 | 0.75 | 0.31 | 0.47 |
| 3 | 194325010 | rs745168 | 0.751 | -3.662 | 0.129 | -1.574 | -0.001 | 0 | 0 | 0 | 0.3 | 0.3 | 0.5 | 0.75 | 0.31 | 0.47 |
| 3 | 194325383 | rs745170 | 0.729 | -3.663 | 0.132 | -1.574 | 0 | -0.001 | -0.001 | 0 | 0.3 | 0.31 | 0.51 | 0.75 | 0.31 | 0.49 |
| 3 | 194325496 | rs789850 | -0.456 | -3.618 | -0.363 | -1.424 | 0.041 | 0.054 | 0.054 | 0 | 0.85 | 0.96 | 0.95 | 0.97 | 0.92 | 0.96 |
| 3 | 194327098 | rs789852 | -0.234 | -3.6 | -0.369 | -1.507 | 0.041 | 0.065 | 0.065 | 0 | 0.85 | 0.96 | 0.95 | 0.99 | 0.91 | 0.94 |
| 3 | 194327140 | rs813150 | 1.198 | -3.702 | -0.353 | -1.619 | 0.086 | 0.111 | 0.111 | 0.046 | 0.65 | 0.81 | 0.84 | 0.98 | 0.79 | 0.72 |
| 3 | 194328524 | rs789997 | -0.049 | -2.803 | -0.191 | -1.193 | 0.039 | 0.021 | 0.021 | 0 | 0.85 | 0.96 | 0.93 | 0.89 | 0.93 | 0.96 |
| 3 | 194329805 | rs789856 | 0.206 | -2.622 | 0.111 | -1.182 | -0.013 | -0.014 | -0.014 | 0 | 0.82 | 0.76 | 0.9 | 0.89 | 0.9 | 0.94 |
| 3 | 194330250 | rs789857 | -0.071 | -2.595 | 0.019 | -1.169 | 0.013 | 0.013 | 0.013 | 0 | 0.84 | 0.87 | 0.91 | 0.89 | 0.9 | 0.96 |
| 3 | 194334197 | rs1773158 | 0.082 | -2.49 | 0.018 | -1.166 | -0.002 | -0.003 | -0.003 | 0 | 0.83 | 0.79 | 0.89 | 0.9 | 0.88 | 0.93 |
| 3 | 194337901 | rs922282 | -0.119 | -2.18 | -0.058 | -1.165 | 0.002 | 0.011 | 0.011 | 0 | 0.84 | 0.86 | 0.89 | 0.9 | 0.85 | 0.96 |
| 3 | 194339984 | rs3845907 | 1.101 | -2.385 | 0.024 | -1.658 | -0.003 | 0.023 | 0.023 | 0 | 0.44 | 0.31 | 0.33 | 0.15 | 0.56 | 0.39 |
| 3 | 194340143 | rs6768976 | 0.171 | -2.271 | -0.01 | -1.423 | -0.001 | -0.003 | -0.003 | 0 | 0.37 | 0.38 | 0.53 | 0.72 | 0.29 | 0.53 |
| 3 | 194341327 | rs790003 | 1.889 | -1.491 | -0.448 | -1.365 | -0.025 | 0.021 | 0.021 | 0 | 0.3 | 0.21 | 0.27 | 0.13 | 0.47 | 0.23 |
| 3 | 194346803 | rs1773189 | 0.38 | -1.25 | -0.252 | -1.192 | -0.001 | -0.043 | -0.043 | 0 | 0.79 | 0.72 | 0.79 | 0.96 | 0.8 | 0.91 |
| 3 | 194353875 | rs1675955 | 1.209 | -1.086 | -0.297 | -1.249 | -0.004 | 0.009 | 0.009 | 0 | 0.4 | 0.37 | 0.31 | 0.26 | 0.48 | 0.38 |
| 3 | 194354514 | rs6437394 | -0.641 | -1.67 | -0.107 | -0.475 | 0.002 | -0.001 | -0.001 | 0 | 0.38 | 0.62 | 0.35 | 0.56 | 0.31 | 0.43 |
| 3 | 194356026 | rs7649259 | 1.944 | -1.547 | 0.094 | -0.728 | 0.042 | -0.103 | -0.103 | 0 | 0.34 | 0.45 | 0.29 | 0.15 | 0.45 | 0.27 |
| 3 | 194360927 | rs711995 | 2.145 | -1.314 | -0.199 | -0.725 | -0.001 | -0.016 | -0.016 | 1.934 | 0.33 | 0.3 | 0.27 | 0.14 | 0.44 | 0.25 |
| 3 | 194361714 | rs1491 | 0.884 | -1.581 | -0.835 | -0.455 | -0.043 | -0.145 | -0.145 | 0 | 0.21 | 0.02 | 0.21 | 0.43 | 0.22 | 0.24 |
| 3 | 194361863 | rs789990 | -0.037 | -1.37 | -0.564 | -0.4 | 0 | -0.008 | -0.008 | 0 | 0.78 | 0.77 | 0.6 | 0.58 | 0.8 | 0.74 |
| 3 | 194362721 | rs1060827 | 0.493 | -1.413 | -0.638 | 0.109 | 0.002 | -0.003 | -0.003 | 0 | 0.85 | 0.82 | 0.87 | 0.75 | 0.85 | 0.89 |
| 3 | 194366007 | rs2275682 | 1.084 | -1.191 | -0.637 | 0.212 | -0.028 | -0.065 | -0.065 | 0 | 0.15 | 0.04 | 0.17 | 0.42 | 0.17 | 0.24 |
| 3 | 194371496 | rs1773206 | 1.513 | -1.523 | -0.629 | 0.149 | 0.003 | 0.02 | 0.02 | 0 | 0.48 | 0.27 | 0.45 | 0.44 | 0.53 | 0.5 |
| 3 | 194378497 | rs1675951 | 0.962 | -1.404 | -0.346 | -0.059 | 0.014 | 0.025 | 0.025 | 0.031 | 0.63 | 0.34 | 0.63 | 0.87 | 0.7 | 0.75 |
| 3 | 194392152 | rs898820 | 0.738 | -1.376 | -0.274 | -0.045 | -0.022 | -0.065 | -0.065 | 0 | 0.64 | 0.47 | 0.65 | 0.87 | 0.71 | 0.75 |
| 3 | 194398333 | rs1706011 | 0.133 | -1.376 | -0.275 | -0.164 | 0.01 | -0.094 | -0.094 | 0.284 | 0.78 | 0.6 | 0.75 | 0.95 | 0.77 | 0.83 |
| 3 | 194400812 | rs9820108 | -0.955 | -0.969 | -0.547 | -0.297 | 0.008 | 0.011 | 0.011 | 0.064 | 0.48 | 0.58 | 0.51 | 0.81 | 0.39 | 0.62 |
| 3 | 194404941 | rs9846429 | 0.353 | -0.952 | -0.797 | 0.052 | 0 | -0.097 | -0.097 | 0.038 | 0.23 | 0.07 | 0.26 | 0.51 | 0.2 | 0.3 |

| | | | | | | | | | | | | | | | | |
|---|---|---|---|---|---|---|---|---|---|---|---|---|---|---|---|---|
| 3 | 194405888 | rs789859 | 1.916 | -1.103 | -0.493 | 0.054 | -0.003 | -0.038 | -0.038 | 0 | 0.39 | 0.54 | 0.33 | 0.25 | 0.4 | 0.35 |
| 3 | 194406431 | rs810965 | 0.339 | -1.313 | -0.676 | 0.055 | -0.029 | 0.018 | 0.018 | 0 | 0.68 | 0.81 | 0.62 | 0.76 | 0.62 | 0.67 |
| 3 | 194407020 | rs1773218 | 0.34 | -1.39 | -0.676 | 0.068 | -0.029 | 0.02 | 0.02 | 0 | 0.68 | 0.81 | 0.62 | 0.76 | 0.63 | 0.68 |
| 3 | 194407204 | rs813153 | 0.346 | -1.386 | -0.674 | 0.062 | -0.029 | 0.019 | 0.019 | 0 | 0.68 | 0.81 | 0.63 | 0.76 | 0.63 | 0.67 |
| 3 | 194407405 | rs3752798 | 0.378 | -2.174 | -0.674 | 0.282 | -0.018 | -0.13 | -0.13 | 0.088 | 0.23 | 0.05 | 0.26 | 0.51 | 0.2 | 0.3 |
| 3 | 194414147 | rs4677854 | 2.226 | -1.898 | -0.439 | 0.354 | -0.022 | -0.023 | -0.023 | 0 | 0.33 | 0.24 | 0.33 | 0.47 | 0.44 | 0.26 |
| 3 | 194415887 | rs1545721 | -0.73 | -1.864 | -0.288 | -0.396 | -0.004 | 0.003 | 0.003 | 0 | 0.92 | 0.93 | 0.84 | 0.94 | 0.87 | 0.96 |
| 3 | 194417029 | rs898813 | 1.023 | -1.741 | -0.212 | -0.201 | 0.025 | 0.013 | 0.013 | 0 | 0.48 | 0.36 | 0.44 | 0.6 | 0.61 | 0.61 |
| 3 | 194417320 | rs748072 | -0.584 | -1.368 | 0.095 | -0.424 | -0.021 | -0.113 | -0.113 | 0.137 | 0.18 | 0.43 | 0.12 | 0.01 | 0.14 | 0.11 |
| 3 | 194418126 | rs1077729 | 1.268 | -1.093 | -0.483 | -0.055 | -0.026 | 0.023 | 0.023 | 0.028 | 0.32 | 0.22 | 0.28 | 0.25 | 0.43 | 0.29 |
| 3 | 194419810 | rs2169412 | -0.034 | -1.529 | -0.294 | -0.146 | 0.005 | 0.012 | 0.012 | 4.033 | 0.53 | 0.51 | 0.4 | 0.27 | 0.6 | 0.41 |
| 3 | 194422574 | rs6808072 | -0.72 | -2.221 | -0.27 | -0.039 | -0.016 | -0.019 | -0.019 | 1.681 | 0.2 | 0.26 | 0.14 | 0.08 | 0.11 | 0.14 |
| 3 | 194425742 | rs13090351 | 0.62 | -2.511 | -0.046 | 0.31 | -0.013 | -0.03 | -0.03 | 0 | 0.29 | 0.25 | 0.48 | 0.57 | 0.35 | 0.45 |
| 3 | 194426037 | rs12486243 | -0.041 | -3.19 | 0.01 | 0.005 | -0.046 | 0.004 | 0.004 | 0.667 | 0.21 | 0.28 | 0.09 | 0.2 | 0.11 | 0.2 |
| 3 | 194429231 | rs4677697 | 1.848 | -3.39 | -0.004 | -0.18 | -0.019 | 0.011 | 0.011 | 0 | 0.29 | 0.24 | 0.28 | 0.11 | 0.45 | 0.16 |
| 3 | 194440199 | rs2126305 | 0.555 | -3.12 | -0.108 | 0.18 | -0.009 | 0 | 0 | 0.027 | 0.76 | 0.78 | 0.85 | 0.85 | 0.91 | 0.81 |
| 3 | 194440887 | rs12695041 | 0.523 | -2.719 | -0.575 | 0.393 | -0.005 | -0.01 | -0.01 | 2.824 | 0.3 | 0.32 | 0.5 | 0.56 | 0.35 | 0.46 |
| 3 | 194443560 | rs12695042 | 0.354 | -2.658 | -0.694 | 0.376 | 0.002 | 0.009 | 0.009 | 0.748 | 0.32 | 0.39 | 0.5 | 0.57 | 0.35 | 0.46 |
| 3 | 194448231 | rs9854809 | 0.531 | -3.334 | -0.791 | 0.78 | 0.048 | -0.012 | -0.012 | 0.506 | 0.85 | 0.96 | 0.89 | 0.88 | 0.93 | 0.87 |
| 3 | 194461665 | rs4677736 | 0.946 | -2.959 | -0.198 | 1.005 | -0.023 | 0.004 | 0.004 | 0 | 0.65 | 0.37 | 0.79 | 0.59 | 0.82 | 0.66 |
| 3 | 194487957 | rs4677742 | -1.124 | -2.821 | 0.169 | 1.296 | -0.033 | 0.029 | 0.029 | 0 | 0.28 | 0.59 | 0.44 | 0.37 | 0.2 | 0.24 |
| 3 | 194490221 | rs9841777 | -0.173 | -3.183 | -0.04 | 1.527 | -0.001 | 0.004 | 0.004 | 0 | 0.14 | 0.19 | 0.12 | 0.32 | 0.11 | 0.2 |
| 3 | 194491496 | rs7632151 | -0.051 | -3.28 | -0.039 | 1.755 | -0.008 | 0.002 | 0.002 | 0 | 0.08 | 0.03 | 0.29 | 0.25 | 0.09 | 0.08 |
| 3 | 194493973 | rs4677743 | -2.207 | -3.401 | 0.012 | 1.683 | 0.107 | 0.071 | 0.071 | 0 | 0.44 | 0.88 | 0.57 | 0.7 | 0.31 | 0.45 |
| 3 | 194500357 | rs6437415 | -1.497 | -3.322 | 0.372 | 1.683 | 0.021 | -0.008 | -0.008 | 0.025 | 0.35 | 0.74 | 0.28 | 0.44 | 0.22 | 0.36 |
| 3 | 194500372 | rs6437416 | -0.151 | -3.01 | 0.203 | 1.423 | 0.007 | 0.005 | 0.005 | 0 | 0.14 | 0.11 | 0.11 | 0.1 | 0.1 | 0.12 |
| 3 | 194500926 | rs12487771 | -1.151 | -2.368 | 0.241 | 1.67 | -0.001 | 0.017 | 0.017 | 0 | 0.23 | 0.28 | 0.41 | 0.57 | 0.2 | 0.29 |
| 3 | 194506581 | rs6773374 | 1.285 | -2.022 | 0.17 | 1.722 | 0.008 | 0.091 | 0.091 | 0 | 0.86 | 0.76 | 0.68 | 0.62 | 0.87 | 0.82 |
| 3 | 194509192 | rs4497978 | -1.09 | -3.275 | -0.48 | 1.037 | 0.094 | 0.109 | 0.109 | 0 | 0.47 | 0.79 | 0.63 | 0.75 | 0.32 | 0.47 |
| 3 | 194514108 | rs4677746 | -0.729 | -3.252 | -0.031 | 1.323 | 0.02 | 0.13 | 0.13 | 0.553 | 0.12 | 0.25 | 0.42 | 0.39 | 0.15 | 0.16 |
| 3 | 194521981 | rs9874101 | 1.313 | -3.467 | -0.244 | 1.197 | 0.085 | 0.091 | 0.091 | 0 | 0.5 | 0.16 | 0.4 | 0.32 | 0.65 | 0.47 |
| 3 | 194522048 | rs9878967 | 0.861 | -3.74 | 0.174 | 1.32 | -0.001 | 0.045 | 0.045 | 0 | 0.76 | 0.69 | 0.54 | 0.51 | 0.77 | 0.7 |
| 3 | 194524634 | rs11706840 | 0.789 | -3.74 | -0.419 | 1.027 | 0.001 | 0 | 0 | 0 | 0.62 | 0.39 | 0.43 | 0.33 | 0.7 | 0.57 |
| 3 | 194526863 | rs13079588 | 1.634 | -3.924 | -1.243 | 0.603 | -0.087 | 0.092 | 0.092 | 0 | 0.33 | 0.06 | 0.27 | 0.14 | 0.55 | 0.37 |
| 3 | 194529803 | rs4434099 | 0.229 | -3.104 | -0.81 | 0.875 | -0.001 | -0.014 | -0.014 | 0 | 0.14 | 0.11 | 0.34 | 0.29 | 0.14 | 0.18 |
| 3 | 194529917 | rs4677642 | 0.02 | -2.785 | -0.522 | 0.284 | -0.078 | -0.021 | -0.021 | 0.272 | 0.18 | 0.37 | 0.11 | 0.16 | 0.07 | 0.1 |
| 3 | 194532442 | rs4488776 | -0.018 | -2.779 | -0.538 | 0.286 | -0.078 | -0.021 | -0.021 | 0 | 0.18 | 0.38 | 0.11 | 0.16 | 0.07 | 0.13 |
| 3 | 194533470 | rs9822544 | 0.329 | -1.439 | -1.489 | 0.16 | -0.019 | 0.052 | 0.052 | 0 | 0.41 | 0.15 | 0.35 | 0.29 | 0.58 | 0.37 |
| 3 | 194542906 | rs13084613 | 0.31 | -0.221 | -0.945 | -0.314 | 0.003 | -0.095 | -0.095 | 0.571 | 0.68 | 0.56 | 0.74 | 0.9 | 0.66 | 0.68 |
| 3 | 194544290 | rs9860061 | -0.63 | 0.879 | -0.35 | 0.036 | 0.056 | 0.072 | 0.072 | 0.213 | 0.1 | 0.29 | 0.23 | 0.22 | 0.17 | 0.19 |
| 3 | 194545222 | rs11715821 | 0.686 | 1.105 | -1.052 | -0.29 | -0.012 | 0.043 | 0.043 | 0.11 | 0.19 | 0.06 | 0.18 | 0.09 | 0.22 | 0.09 |
| 3 | 194545903 | rs4677644 | 0.194 | 0.491 | -1.383 | -0.29 | -0.002 | 0.013 | 0.013 | 0 | 0.36 | 0.46 | 0.38 | 0.58 | 0.34 | 0.37 |
| 3 | 194551215 | rs4677758 | -1.069 | 0.588 | -1.526 | -0.346 | -0.042 | 0.012 | 0.012 | 0 | 0.83 | 0.96 | 0.74 | 0.8 | 0.79 | 0.78 |
| 3 | 194551660 | rs12637098 | -0.063 | -0.591 | -1.226 | 0.229 | 0.006 | 0.001 | 0.001 | 0 | 0.49 | 0.37 | 0.46 | 0.46 | 0.44 | 0.4 |
| 3 | 194552145 | rs9874252 | 0.977 | -0.703 | -0.916 | 0.278 | -0.031 | 0.007 | 0.007 | 0 | 0.15 | 0.34 | 0.1 | 0.1 | 0.13 | 0.13 |
| 3 | 194553182 | rs4677760 | -0.472 | -0.374 | -0.856 | 0.579 | -0.003 | -0.001 | -0.001 | 0.022 | 0.28 | 0.38 | 0.23 | 0.28 | 0.22 | 0.24 |
| 3 | 194554885 | rs7652843 | 0 | -0.371 | -1.16 | 0.337 | -0.001 | -0.013 | -0.013 | 0.485 | 0.35 | 0.63 | 0.34 | 0.18 | 0.35 | 0.31 |
| 3 | 194557213 | rs2898596 | 0.042 | -0.575 | -1.264 | 0.899 | 0.024 | -0.046 | -0.046 | 0 | 0.15 | 0.02 | 0.1 | 0.22 | 0.07 | 0.08 |
| 3 | 194559473 | rs753628 | -1.144 | -0.236 | -1.134 | 0.956 | -0.008 | -0.023 | -0.023 | 0.038 | 0.39 | 0.82 | 0.54 | 0.66 | 0.39 | 0.42 |
| 3 | 194562734 | rs10933678 | -0.814 | -0.206 | -0.533 | 1.408 | 0 | 0.053 | 0.053 | 0 | 0.09 | 0.17 | 0.27 | 0.24 | 0.06 | 0.06 |
| 3 | 194564931 | rs6764953 | 0.494 | -0.233 | -0.185 | 0.891 | -0.033 | 0.035 | 0.035 | 0 | 0.73 | 0.55 | 0.81 | 0.61 | 0.78 | 0.7 |
| 3 | 194574108 | rs6773029 | -0.123 | 0.236 | -0.246 | 0.518 | 0.003 | -0.032 | -0.032 | 0 | 0.11 | 0.13 | 0.09 | 0 | 0.14 | 0.1 |
| 3 | 194576842 | rs4677766 | -1.621 | -0.16 | 0.342 | 0.667 | -0.006 | -0.024 | -0.024 | 0 | 0.25 | 0.41 | 0.41 | 0.23 | 0.23 | 0.33 |
| 3 | 194576897 | rs6764929 | 0.121 | 0.498 | 0.191 | 0.54 | 0.02 | -0.01 | -0.01 | 1.751 | 0.12 | 0.22 | 0.2 | 0.1 | 0.17 | 0.1 |
| 3 | 194579376 | rs6772240 | 0.485 | 0.467 | 0.089 | 0.621 | 0.001 | 0.003 | 0.003 | 0 | 0.05 | 0.06 | 0.26 | 0.15 | 0.07 | 0.07 |
| 3 | 194580372 | rs1545026 | 0.479 | 0.49 | 0.111 | 0.52 | -0.001 | -0.017 | -0.017 | 0 | 0.1 | 0.1 | 0.08 | 0 | 0.11 | 0.1 |
| 3 | 194581193 | rs7632021 | -1.552 | 0.29 | 0.317 | 0.443 | 0.005 | -0.017 | -0.017 | 0 | 0.37 | 0.55 | 0.6 | 0.32 | 0.38 | 0.42 |
| 3 | 194583085 | rs7638250 | -0.882 | 0.353 | 0.198 | 0.53 | 0 | -0.003 | -0.003 | 0 | 0.22 | 0.36 | 0.4 | 0.25 | 0.21 | 0.32 |
| 3 | 194584206 | rs12695046 | 1.51 | -0.103 | -0.35 | 0.846 | 0.042 | -0.007 | -0.007 | 0 | 0.56 | 0.13 | 0.42 | 0.54 | 0.62 | 0.55 |
| 3 | 194586088 | rs11719165 | 0.739 | 0.436 | -0.321 | 1.046 | -0.005 | -0.022 | -0.022 | 0 | 0.63 | 0.27 | 0.62 | 0.45 | 0.62 | 0.57 |
| 3 | 194597243 | rs7641855 | -1.032 | 0.094 | -0.101 | 1.188 | -0.002 | -0.002 | -0.002 | 0 | 0.39 | 0.62 | 0.57 | 0.56 | 0.39 | 0.51 |
| 3 | 194603569 | rs10933683 | -0.659 | 0.388 | -0.154 | 1.363 | 0.002 | -0.001 | -0.001 | 0.087 | 0.08 | 0.05 | 0.32 | 0.16 | 0.12 | 0.09 |
| 3 | 194612501 | rs7632590 | -0.796 | 0.336 | -0.095 | 1.534 | 0.003 | 0.012 | 0.012 | 0 | 0.08 | 0.07 | 0.32 | 0.24 | 0.12 | 0.13 |
| 3 | 194623370 | rs4352326 | -0.098 | -0.421 | 0.231 | 1.66 | -0.013 | 0.077 | 0.077 | 0 | 0.06 | 0.13 | 0.25 | 0.13 | 0.05 | 0.13 |
| 3 | 194629709 | rs4677776 | -1.559 | -0.505 | 0.825 | 1.631 | -0.01 | 0.225 | 0.225 | 0 | 0.11 | 0.4 | 0.28 | 0.22 | 0.05 | 0.17 |
| 3 | 194634168 | rs4677778 | 1.16 | -0.756 | -0.176 | 1.037 | 0.013 | -0.007 | -0.007 | 0 | 0.42 | 0.28 | 0.67 | 0.52 | 0.61 | 0.4 |
| 3 | 194641346 | rs823048 | 1.195 | -0.726 | -0.177 | 1.049 | 0.009 | -0.007 | -0.007 | 0 | 0.42 | 0.29 | 0.67 | 0.52 | 0.6 | 0.39 |
| 3 | 194648821 | rs726889 | 0.817 | -0.782 | -0.298 | 1.242 | -0.012 | -0.023 | -0.023 | 0.226 | 0.06 | 0.03 | 0.02 | 0.13 | 0.07 | 0.05 |
| 3 | 194649221 | rs4426628 | 0.686 | -1.022 | -0.112 | 1.04 | -0.015 | 0.003 | 0.003 | 0 | 0.56 | 0.48 | 0.74 | 0.59 | 0.68 | 0.56 |

| | | | | | | | | | | | | | | | | |
|---|---|---|---|---|---|---|---|---|---|---|---|---|---|---|---|---|
| 3 | 194651602 | rs6805463 | -0.155 | -1.027 | -0.712 | 0.697 | 0.014 | 0.023 | 0.023 | 0.005 | 0.72 | 0.78 | 0.83 | 0.84 | 0.79 | 0.75 |
| 3 | 194656826 | rs6437437 | 0.582 | -0.057 | -0.974 | 0.927 | 0.026 | -0.006 | -0.006 | 0.272 | 0.43 | 0.29 | 0.71 | 0.59 | 0.61 | 0.49 |
| 3 | 194663349 | rs4311166 | 0.641 | 0.061 | -0.956 | 0.923 | 0.001 | -0.007 | -0.007 | 1.172 | 0.42 | 0.28 | 0.69 | 0.59 | 0.55 | 0.47 |
| 3 | 194666308 | rs2676860 | 0.365 | 0.09 | -1.165 | 0.762 | 0.047 | -0.001 | -0.001 | 0 | 0.51 | 0.3 | 0.51 | 0.47 | 0.56 | 0.56 |
| 3 | 194669375 | rs2568911 | -0.991 | -0.822 | -0.221 | 0.88 | 0 | 0.003 | 0.003 | 0 | 0.39 | 0.45 | 0.44 | 0.46 | 0.39 | 0.52 |
| 3 | 194669416 | rs2676870 | -0.253 | -0.667 | -0.09 | 1.002 | 0.004 | 0.007 | 0.007 | 0 | 0.12 | 0.17 | 0.13 | 0.15 | 0.15 | 0.29 |
| 3 | 194674114 | rs823009 | -0.364 | -0.75 | -0.226 | 0.938 | -0.021 | -0.031 | -0.031 | 0 | 0.77 | 0.88 | 0.71 | 0.68 | 0.75 | 0.76 |
| 3 | 194680000 | rs711993 | 0.955 | -1.122 | -0.066 | 0.887 | 0.01 | 0.007 | 0.007 | 0 | 0.61 | 0.68 | 0.66 | 0.38 | 0.66 | 0.59 |
| 3 | 194681282 | rs1075870 | 1.972 | -1.118 | 0.309 | 0.709 | -0.029 | 0.148 | 0.148 | 0 | 0.44 | 0.21 | 0.44 | 0.02 | 0.49 | 0.28 |
| 3 | 194684216 | rs13062154 | 0.857 | -1.039 | 0.45 | 0.841 | -0.001 | 0.001 | 0.001 | 0 | 0.08 | 0.06 | 0.08 | 0.14 | 0.07 | 0.16 |
| 3 | 194687004 | rs1356358 | -0.327 | -1.084 | 0.475 | 0.861 | -0.013 | 0.022 | 0.022 | 0 | 0.16 | 0.22 | 0.07 | 0.3 | 0.09 | 0.18 |
| 3 | 194687331 | rs6770825 | 2.025 | -1.126 | 0.494 | 0.768 | -0.037 | 0.17 | 0.17 | 0 | 0.44 | 0.18 | 0.44 | 0.02 | 0.48 | 0.28 |
| 3 | 194689413 | rs823025 | -1.116 | -1.132 | 0.528 | 0.972 | -0.023 | 0.049 | 0.049 | 0.02 | 0.21 | 0.32 | 0.12 | 0.34 | 0.16 | 0.28 |
| 3 | 194692862 | rs1356359 | -1.012 | -1.124 | 0.587 | 0.972 | -0.044 | 0.112 | 0.112 | 0 | 0.2 | 0.33 | 0.12 | 0.49 | 0.11 | 0.21 |
| 3 | 194692874 | rs6437444 | 0.306 | -1.172 | 0.44 | 0.934 | -0.009 | 0.039 | 0.039 | 0 | 0.12 | 0.16 | 0.04 | 0.29 | 0.06 | 0.14 |
| 3 | 194695307 | rs4677652 | 1.156 | -1.103 | 0.1 | 0.485 | 0.038 | 0.175 | 0.175 | 0 | 0.51 | 0.19 | 0.53 | 0.19 | 0.57 | 0.45 |
| 3 | 194703666 | rs7622435 | 0.408 | -0.732 | 0.238 | 0.817 | 0.01 | 0.021 | 0.021 | 0 | 0.63 | 0.45 | 0.57 | 0.5 | 0.59 | 0.48 |
| 3 | 194709026 | rs1878428 | 0.289 | -1.282 | 0.27 | 0.464 | 0.013 | 0.001 | 0.001 | 0 | 0.58 | 0.39 | 0.64 | 0.35 | 0.68 | 0.58 |
| 3 | 194709977 | rs2676854 | 0.5 | -1.157 | -0.185 | 0.447 | 0.065 | 0.05 | 0.05 | 0 | 0.53 | 0.23 | 0.6 | 0.33 | 0.63 | 0.56 |
| 3 | 194719238 | rs7623698 | 0.164 | -1.33 | -0.047 | 0.506 | 0.014 | -0.012 | -0.012 | 0 | 0.37 | 0.33 | 0.28 | 0.51 | 0.23 | 0.34 |
| 3 | 194720053 | rs2720917 | 0.377 | -1.114 | -0.003 | 0.617 | -0.013 | 0.031 | 0.031 | 0 | 0.89 | 0.78 | 0.92 | 0.82 | 0.9 | 0.8 |
| 3 | 194722518 | rs2676834 | -0.346 | -0.98 | -0.264 | 0.484 | 0.003 | 0.003 | 0.003 | 0.076 | 0.52 | 0.59 | 0.36 | 0.6 | 0.4 | 0.47 |
| 3 | 194723612 | rs2720912 | -0.104 | -0.967 | -0.036 | 0.432 | 0.004 | -0.019 | -0.019 | 0.06 | 0.87 | 0.68 | 0.9 | 0.88 | 0.89 | 0.85 |
| 3 | 194729050 | rs2676831 | 0.013 | -0.921 | -0.301 | 0.474 | -0.019 | -0.015 | -0.015 | 1.672 | 0.82 | 0.68 | 0.89 | 0.82 | 0.84 | 0.75 |
| 3 | 194733941 | rs878809 | 0.5 | -0.031 | -0.947 | 0.741 | -0.042 | 0.015 | 0.015 | 0.153 | 0.36 | 0.2 | 0.56 | 0.29 | 0.5 | 0.38 |
| 3 | 194737873 | rs2676917 | 0.241 | -0.023 | -1.094 | 0.627 | -0.032 | 0 | 0 | 0.328 | 0.38 | 0.14 | 0.52 | 0.31 | 0.55 | 0.35 |
| 3 | 194740639 | rs2676925 | -0.251 | -0.015 | -0.944 | 0.919 | 0.006 | 0.08 | 0.08 | 0 | 0.1 | 0.12 | 0.04 | 0.34 | 0.11 | 0.1 |
| 3 | 194743015 | rs938712 | 0.019 | -0.106 | -1.315 | 0.558 | -0.045 | -0.042 | -0.042 | 0 | 0.43 | 0.21 | 0.56 | 0.52 | 0.48 | 0.39 |
| 3 | 194752416 | rs2280222 | 0.494 | -0.69 | -0.42 | 0.42 | -0.01 | -0.003 | -0.003 | 0 | 0.72 | 0.68 | 0.71 | 0.69 | 0.79 | 0.84 |
| 3 | 194753890 | rs2720953 | 0.462 | -0.602 | -0.578 | 0.498 | -0.034 | -0.01 | -0.01 | 0 | 0.71 | 0.62 | 0.71 | 0.7 | 0.8 | 0.85 |
| 3 | 194758010 | rs3892715 | -0.269 | 0.109 | -1.018 | 0.535 | 0.013 | 0.04 | 0.04 | 0 | 0.48 | 0.72 | 0.45 | 0.32 | 0.49 | 0.41 |
| 3 | 194759980 | rs11711388 | -0.593 | -0.237 | -0.318 | 0.211 | -0.005 | -0.058 | -0.058 | 0 | 0.12 | 0.58 | 0.06 | 0.03 | 0.07 | 0.06 |
| 3 | 194761024 | rs2676894 | -0.171 | -0.518 | -0.423 | 0.131 | -0.05 | -0.002 | -0.002 | 0.227 | 0.39 | 0.21 | 0.23 | 0.33 | 0.42 | 0.4 |
| 3 | 194763601 | rs2676889 | -0.791 | -0.691 | -0.661 | 0.769 | -0.051 | 0.074 | 0.074 | 0 | 0.49 | 0.9 | 0.39 | 0.65 | 0.46 | 0.56 |
| 3 | 194766665 | rs7639502 | -0.379 | -0.839 | -0.727 | 0.683 | -0.006 | 0.005 | 0.005 | 0 | 0.32 | 0.64 | 0.19 | 0.24 | 0.33 | 0.32 |
| 3 | 194767005 | rs2676879 | -1.428 | -0.96 | -0.68 | 0.606 | -0.03 | 0.045 | 0.045 | 0.267 | 0.57 | 0.86 | 0.42 | 0.68 | 0.5 | 0.63 |
| 3 | 194767128 | rs2720891 | 0.018 | -0.913 | -0.494 | 0.76 | 0.006 | -0.004 | -0.004 | 0 | 0.17 | 0.16 | 0.23 | 0.44 | 0.13 | 0.28 |
| 3 | 194767164 | rs10933693 | -0.787 | -0.227 | 0.301 | 0.327 | 0.008 | 0.097 | 0.097 | 0 | 0.15 | 0.41 | 0.08 | 0.22 | 0.15 | 0.13 |
| 3 | 194767214 | rs12634159 | -1.223 | -0.194 | 0.291 | 0.261 | -0.028 | 0.105 | 0.105 | 0.961 | 0.14 | 0.61 | 0.08 | 0.23 | 0.12 | 0.12 |
| 3 | 194767238 | rs11926488 | -0.046 | -0.103 | 0.151 | -0.308 | -0.006 | -0.048 | -0.048 | 0 | 0.09 | 0.16 | 0.04 | 0 | 0.07 | 0.07 |
| 3 | 194769114 | rs6806968 | 0.271 | -0.012 | -0.197 | -0.24 | -0.069 | 0.09 | 0.09 | 0.361 | 0.21 | 0.06 | 0.1 | 0.01 | 0.21 | 0.19 |
| 3 | 194769289 | rs2720961 | -0.074 | -0.116 | 0.47 | 0.202 | -0.012 | -0.002 | -0.002 | 0 | 0.14 | 0.29 | 0.1 | 0.13 | 0.12 | 0.11 |
| 3 | 194769773 | rs9812839 | 0.258 | -0.473 | -0.3 | -0.11 | 0.003 | 0.056 | 0.056 | 1.558 | 0.39 | 0.28 | 0.17 | 0.09 | 0.28 | 0.31 |
| 3 | 194775336 | rs2720932 | 1.136 | -0.007 | 0.365 | 1.194 | 0.024 | 0.193 | 0.193 | 0 | 0.93 | 0.78 | 0.94 | 0.69 | 0.93 | 0.81 |
| 3 | 194778445 | rs7613935 | -0.142 | 0.371 | 0.441 | 0.346 | 0.011 | -0.08 | -0.08 | 0 | 0.18 | 0.29 | 0.42 | 0.03 | 0.25 | 0.2 |
| 3 | 194780749 | rs3959148 | -0.499 | -0.596 | 0.758 | 1.032 | -0.031 | 0.007 | 0.007 | 0 | 0.26 | 0.34 | 0.12 | 0.34 | 0.17 | 0.17 |
| 3 | 194785443 | rs7627043 | -0.662 | -0.441 | 0.579 | 1.427 | 0.002 | 0.055 | 0.055 | 0 | 0.09 | 0.12 | 0.08 | 0.3 | 0.1 | 0.27 |
| 3 | 194789032 | rs2720930 | -1.473 | -0.472 | 0.473 | 0.85 | 0.052 | 0.182 | 0.182 | 0 | 0.46 | 0.79 | 0.41 | 0.96 | 0.38 | 0.57 |
| 3 | 194791657 | rs2720948 | 0.283 | -0.713 | 0.811 | 1.208 | 0.025 | 0.098 | 0.098 | 0.022 | 0.07 | 0.09 | 0.19 | 0.27 | 0.12 | 0.15 |
| 3 | 194792324 | rs6787005 | -0.919 | -0.84 | 1.25 | 1.168 | -0.058 | 0.005 | 0.005 | 0 | 0.35 | 0.48 | 0.18 | 0.64 | 0.21 | 0.42 |
| 3 | 194792415 | rs6787114 | -1.072 | -0.842 | 1.176 | 1.166 | -0.055 | 0.002 | 0.002 | 0 | 0.36 | 0.52 | 0.17 | 0.64 | 0.2 | 0.42 |
| 3 | 194799732 | rs7653402 | -1.22 | -0.691 | 1.096 | 0.661 | 0.022 | 0.052 | 0.052 | 0 | 0.18 | 0.34 | 0.28 | 0.63 | 0.2 | 0.41 |
| 3 | 194804838 | rs7627506 | -1.663 | -0.734 | 0.329 | 0.142 | 0.081 | 0.226 | 0.226 | 0 | 0.47 | 0.82 | 0.4 | 0.96 | 0.35 | 0.56 |
| 3 | 194807900 | rs10933697 | -0.464 | -1.037 | 0.712 | 0.689 | -0.066 | -0.023 | -0.023 | 0 | 0.22 | 0.35 | 0.09 | 0.19 | 0.1 | 0.11 |
| 3 | 194817484 | rs6437455 | -1.182 | -0.673 | 0.267 | 1.087 | 0.001 | 0 | 0 | 0 | 0.36 | 0.34 | 0.33 | 0.66 | 0.28 | 0.5 |
| 3 | 194820658 | rs9863270 | -0.942 | -0.354 | -0.073 | 1.469 | -0.044 | -0.083 | -0.083 | 0.26 | 0.16 | 0.02 | 0.15 | 0.23 | 0.2 | 0.33 |
| 3 | 194822098 | rs952481 | -1.798 | -0.805 | 0.678 | 1.656 | -0.006 | 0.025 | 0.025 | 0 | 0.23 | 0.28 | 0.21 | 0.43 | 0.2 | 0.42 |
| 3 | 194823070 | rs7635033 | -1.777 | -0.815 | 1.177 | 1.788 | -0.007 | 0.084 | 0.084 | 0 | 0.19 | 0.31 | 0.4 | 0.47 | 0.15 | 0.3 |
| 3 | 194825090 | rs9288754 | 1.767 | -0.387 | 0.946 | 0.566 | 0.01 | 0.041 | 0.041 | 0 | 0.5 | 0.32 | 0.51 | 0.01 | 0.56 | 0.39 |
| 3 | 194826729 | rs6785310 | -1.534 | 0.341 | 1.156 | 0.784 | 0.009 | 0.002 | 0.002 | 0.063 | 0.23 | 0.28 | 0.36 | 0.72 | 0.3 | 0.45 |
| 3 | 194829556 | rs4677660 | 1.086 | 0.366 | 1.21 | 0.88 | 0.024 | 0.048 | 0.048 | 0.795 | 0.77 | 0.65 | 0.7 | 0.38 | 0.71 | 0.62 |
| 3 | 194833940 | rs7652885 | 0.576 | 0.386 | 0.949 | 0.986 | -0.051 | -0.037 | -0.037 | 0.013 | 0.08 | 0.01 | 0.14 | 0.32 | 0.16 | 0.24 |
| 3 | 194834171 | rs903307 | 1.074 | 0.36 | 1.093 | 0.489 | 0.033 | 0.071 | 0.071 | 0.623 | 0.75 | 0.43 | 0.68 | 0.39 | 0.71 | 0.62 |
| 3 | 194836755 | rs6767234 | -0.519 | 0.521 | 0.753 | 0.638 | 0.008 | 0.03 | 0.03 | 0 | 0.12 | 0.16 | 0.16 | 0.26 | 0.14 | 0.2 |
| 3 | 194842322 | rs952586 | -1.002 | 0.288 | 0.794 | 0.672 | 0.055 | 0.288 | 0.288 | 0 | 0.15 | 0.47 | 0.21 | 0.46 | 0.15 | 0.27 |
| 3 | 194846903 | rs4300953 | -1.686 | 0.11 | 0.712 | 0.336 | -0.014 | -0.061 | -0.061 | 0 | 0.38 | 0.5 | 0.41 | 0.83 | 0.29 | 0.51 |
| 3 | 194847752 | rs950256 | -0.512 | 0.107 | 0.592 | 0.355 | -0.009 | -0.116 | -0.116 | 0.918 | 0.23 | 0.06 | 0.24 | 0.58 | 0.18 | 0.31 |
| 3 | 194848766 | rs1874099 | -0.58 | 0.103 | 0.716 | 0.354 | 0.006 | -0.037 | -0.037 | 0 | 0.23 | 0.13 | 0.24 | 0.58 | 0.18 | 0.31 |
| 3 | 194850659 | rs6797160 | -0.832 | -0.053 | -0.048 | -0.585 | 0.002 | -0.021 | -0.021 | 0 | 0.64 | 0.6 | 0.53 | 0.85 | 0.54 | 0.64 |

| | | | | | | | | | | | | | | | | |
|---|---|---|---|---|---|---|---|---|---|---|---|---|---|---|---|---|
| 3 | 194851123 | rs9841564 | 0.74 | -0.112 | 0.663 | 0.344 | -0.008 | -0.002 | -0.002 | 0 | 0.13 | 0.08 | 0.04 | 0.18 | 0.04 | 0.06 |
| 3 | 194851214 | rs7637593 | -0.273 | -0.121 | 0.815 | 0.329 | -0.004 | 0.035 | 0.035 | 0 | 0.14 | 0.2 | 0.16 | 0.27 | 0.11 | 0.2 |
| 3 | 194851700 | rs1552893 | -1.785 | -0.113 | 0.746 | 0.274 | -0.007 | -0.045 | -0.045 | 0 | 0.38 | 0.5 | 0.42 | 0.85 | 0.29 | 0.51 |
| 3 | 194855954 | rs7644208 | -1.602 | -0.204 | 0.95 | 0.638 | 0.027 | 0.026 | 0.026 | 0 | 0.25 | 0.43 | 0.37 | 0.67 | 0.26 | 0.45 |
| 3 | 194866399 | rs9882160 | -1.029 | -0.249 | 0.901 | 0.675 | -0.037 | 0.062 | 0.062 | 0.214 | 0.27 | 0.33 | 0.2 | 0.45 | 0.15 | 0.27 |
| 3 | 194868944 | rs2279634 | -0.923 | -0.239 | 0.483 | 0.633 | -0.003 | 0.016 | 0.016 | 0.099 | 0.25 | 0.29 | 0.26 | 0.58 | 0.19 | 0.31 |
| 3 | 194874292 | rs4677662 | -0.88 | 0.502 | 0.211 | 0.961 | -0.011 | 0.068 | 0.068 | 0 | 0.15 | 0.2 | 0.16 | 0.27 | 0.1 | 0.2 |
| 3 | 194881756 | rs7616009 | -0.263 | 0.889 | -0.122 | 1.069 | -0.004 | 0.034 | 0.034 | 0.338 | 0.1 | 0.08 | 0.19 | 0.39 | 0.15 | 0.26 |
| 3 | 194894133 | rs7653312 | -0.948 | 0.117 | 0.971 | 1.041 | -0.014 | 0.155 | 0.155 | 0 | 0.15 | 0.42 | 0.21 | 0.26 | 0.09 | 0.18 |
| 3 | 194900432 | rs11719455 | 1.008 | 0.201 | 0.03 | 0.319 | -0.066 | -0.079 | -0.079 | 0.146 | 0.44 | 0.13 | 0.28 | 0.48 | 0.44 | 0.47 |
| 3 | 194906614 | rs3796160 | 0.324 | -0.237 | 0.6 | 0.125 | -0.045 | -0.015 | -0.015 | 0 | 0.15 | 0.2 | 0.09 | 0.11 | 0.07 | 0.06 |
| 3 | 194919955 | rs2410806 | -1.225 | -0.24 | 0.691 | 0.095 | -0.054 | 0.054 | 0.054 | 0 | 0.24 | 0.54 | 0.25 | 0.36 | 0.12 | 0.22 |
| 3 | 194924130 | rs9870813 | -0.198 | 0.069 | 0.531 | -0.228 | -0.006 | 0.004 | 0.004 | 0 | 0.68 | 0.74 | 0.53 | 0.93 | 0.58 | 0.72 |
| 3 | 194928138 | rs12495970 | -1.49 | 0.095 | 0.959 | 0.598 | -0.051 | 0.136 | 0.136 | 0 | 0.22 | 0.46 | 0.24 | 0.39 | 0.13 | 0.22 |
| 3 | 194930028 | rs3796161 | -1.29 | 0.096 | 1.279 | 0.012 | -0.03 | 0.029 | 0.029 | 0 | 0.11 | 0.31 | 0.11 | 0.15 | 0.04 | 0.11 |
| 3 | 194934634 | rs9825185 | 0.343 | 0.403 | 0.52 | 0.031 | -0.029 | 0.013 | 0.013 | 0 | 0.91 | 0.95 | 0.93 | 0.94 | 0.86 | 0.85 |
| 3 | 194975720 | rs12486220 | -1.262 | 0.003 | 0.701 | 0.092 | -0.069 | 0.04 | 0.04 | 0 | 0.12 | 0.44 | 0.12 | 0.15 | 0.02 | 0.08 |
| 3 | 194983053 | rs2410840 | 0.038 | 1.09 | 0.106 | -0.019 | 0.001 | 0.001 | 0.001 | 0 | 0.9 | 0.92 | 0.9 | 0.96 | 0.78 | 0.88 |
| 3 | 194984402 | rs752206 | 1.247 | 0.786 | 0.793 | -0.052 | 0.024 | -0.019 | -0.019 | 0 | 0.8 | 0.51 | 0.78 | 0.81 | 0.77 | 0.8 |
| 3 | 194987569 | rs4677666 | -1.14 | 0.997 | 0.718 | -0.384 | 0.001 | -0.02 | -0.02 | 0.272 | 0.28 | 0.34 | 0.35 | 0.83 | 0.22 | 0.51 |
| 3 | 194990363 | rs755202 | 0.712 | 0.958 | 0.662 | -0.288 | 0.024 | -0.032 | -0.032 | 0 | 0.06 | 0 | 0.09 | 0.14 | 0.02 | 0.07 |
| 3 | 194997620 | rs2000052 | -1.142 | 1.069 | 0.948 | -0.383 | 0.004 | -0.02 | -0.02 | 0 | 0.27 | 0.32 | 0.35 | 0.83 | 0.2 | 0.51 |
| 3 | 194999609 | rs3087885 | -0.527 | 1.149 | 0.578 | -0.476 | -0.041 | 0.017 | 0.017 | 0.065 | 0.91 | 0.98 | 0.7 | 0.96 | 0.81 | 0.88 |
| 3 | 195022534 | rs2275519 | -0.184 | 1.089 | 0.702 | -0.465 | -0.001 | -0.001 | -0.001 | 0.051 | 0.19 | 0.19 | 0.26 | 0.69 | 0.19 | 0.41 |
| 3 | 195025889 | rs12494820 | -0.489 | 0.994 | 0.775 | -0.474 | -0.034 | 0.008 | 0.008 | 0 | 0.08 | 0.16 | 0.09 | 0.13 | 0.01 | 0.07 |
| 3 | 195027003 | rs11715578 | 0.382 | 1.167 | 0.605 | 0.493 | -0.034 | -0.015 | -0.015 | 0 | 0.07 | 0 | 0.14 | 0.36 | 0.09 | 0.26 |
| 3 | 195027339 | rs12491026 | -0.321 | 0.94 | 0.666 | -0.456 | 0.021 | -0.007 | -0.007 | 2.016 | 0.08 | 0.05 | 0.08 | 0.13 | 0.01 | 0.07 |
| 3 | 195047912 | rs4677825 | -0.776 | 0.979 | 0.239 | -0.917 | 0.014 | -0.011 | -0.011 | 0 | 0.91 | 0.89 | 0.69 | 0.96 | 0.81 | 0.87 |
| 3 | 195048369 | rs2050809 | -0.807 | 0.972 | 0.263 | -0.92 | 0.002 | -0.002 | -0.002 | 0 | 0.91 | 0.91 | 0.69 | 0.96 | 0.81 | 0.87 |
| 3 | 195050247 | rs2050806 | -1.002 | 1.052 | 0.184 | -0.945 | 0.007 | -0.029 | -0.029 | 0 | 0.28 | 0.41 | 0.36 | 0.82 | 0.22 | 0.5 |
| 3 | 195057001 | rs13082563 | 1.071 | 0.591 | -0.636 | -1.376 | -0.002 | 0.086 | 0.086 | 0.045 | 0.19 | 0.01 | 0.07 | 0.04 | 0.16 | 0.14 |
| 3 | 195078942 | rs6803763 | -0.782 | 1.187 | 0.34 | -0.632 | 0.019 | 0.011 | 0.011 | 0 | 0.24 | 0.38 | 0.28 | 0.69 | 0.21 | 0.42 |
| 3 | 195080491 | rs10439949 | -0.405 | 1.156 | 0.303 | -0.646 | 0.041 | 0.03 | 0.03 | 0.013 | 0.21 | 0.38 | 0.27 | 0.69 | 0.21 | 0.42 |
| 3 | 195103381 | rs6801750 | -1.089 | 1.246 | 0.343 | -0.634 | 0.003 | -0.02 | -0.02 | 0 | 0.29 | 0.38 | 0.36 | 0.82 | 0.22 | 0.5 |
| 3 | 195106167 | rs823305 | 0.731 | 1.119 | -0.22 | -0.502 | 0.018 | -0.011 | -0.011 | 0 | 0.08 | 0.12 | 0.31 | 0.04 | 0.18 | 0.12 |
| 3 | 195166449 | rs9859822 | -1.167 | 0.711 | -0.3 | -0.368 | -0.001 | -0.015 | -0.015 | 0 | 0.31 | 0.35 | 0.36 | 0.84 | 0.25 | 0.54 |
| 3 | 195167114 | rs12630162 | -0.483 | 1.013 | -0.981 | -0.226 | -0.025 | -0.017 | -0.017 | 0 | 0.22 | 0.14 | 0.26 | 0.69 | 0.24 | 0.46 |
| 3 | 195182985 | rs823530 | -0.955 | 0.659 | -0.521 | -0.011 | -0.01 | 0.001 | 0.001 | 0 | 0.14 | 0.29 | 0.38 | 0.16 | 0.13 | 0.15 |
| 3 | 195238559 | rs34950021 | 0.022 | 0.649 | -1.731 | 0.195 | -0.018 | -0.061 | -0.061 | 0.015 | 0.31 | 0.1 | 0.55 | 0.43 | 0.3 | 0.35 |
| 3 | 195240670 | rs823499 | -0.803 | 0.883 | -1.54 | -0.377 | 0.003 | -0.066 | -0.066 | 0.239 | 0.67 | 0.66 | 0.78 | 0.95 | 0.63 | 0.75 |
| 3 | 195242593 | rs1136 | 0.595 | 0.714 | -1.784 | -0.066 | -0.033 | -0.093 | -0.093 | 0 | 0.3 | 0.05 | 0.55 | 0.47 | 0.3 | 0.35 |
| 3 | 195257170 | rs2259661 | -0.565 | 0.898 | -1.187 | -0.599 | 0.008 | -0.105 | -0.105 | 0 | 0.66 | 0.62 | 0.76 | 0.95 | 0.63 | 0.75 |
| 3 | 195258384 | rs34907067 | 0.202 | 0.607 | -1.002 | -0.639 | 0.011 | 0.062 | 0.062 | 0.479 | 0.39 | 0.29 | 0.61 | 0.81 | 0.36 | 0.45 |
| 3 | 195262272 | rs823508 | -1.145 | 0.638 | -0.953 | -0.699 | 0.003 | -0.064 | -0.064 | 0.487 | 0.67 | 0.67 | 0.78 | 0.95 | 0.63 | 0.75 |
| 3 | 195262521 | rs823510 | 1.033 | 0.438 | -0.893 | -0.916 | -0.014 | 0.031 | 0.031 | 0 | 0.19 | 0.16 | 0.12 | 0.02 | 0.22 | 0.15 |
| 3 | 195264241 | rs823513 | -1.333 | 0.428 | -0.893 | -0.932 | 0.003 | -0.064 | -0.064 | 0 | 0.67 | 0.67 | 0.78 | 0.95 | 0.63 | 0.75 |
| 3 | 195272207 | rs1532291 | 1.455 | 0.376 | -1.143 | -0.887 | -0.008 | -0.029 | -0.029 | 0 | 0.06 | 0 | 0.03 | 0.12 | 0.06 | 0.15 |
| 3 | 195273944 | rs1908173 | 0.655 | 0.619 | -1.511 | -0.728 | -0.033 | -0.056 | -0.056 | 0 | 0.38 | 0.05 | 0.57 | 0.44 | 0.35 | 0.39 |
| 3 | 195285633 | rs823504 | -0.324 | 0.239 | -1.087 | -0.402 | -0.006 | -0.013 | -0.013 | 0 | 0.58 | 0.31 | 0.7 | 0.54 | 0.55 | 0.54 |
| 3 | 195295166 | rs4677689 | -0.901 | 0.223 | -1.344 | -1.028 | 0.039 | 0.062 | 0.062 | 0 | 0.83 | 0.9 | 0.92 | 0.99 | 0.89 | 0.87 |
| 3 | 195304653 | rs6605338 | 0.68 | 0.654 | -1.01 | -1.365 | 0.003 | -0.104 | -0.104 | 0 | 0.27 | 0.52 | 0.17 | 0.04 | 0.25 | 0.13 |
| 3 | 195310679 | rs4677695 | -0.511 | 0.722 | -1.634 | -0.514 | -0.01 | -0.051 | -0.051 | 0 | 0.15 | 0.09 | 0.1 | 0.4 | 0.15 | 0.32 |
| 3 | 195477791 | rs2291652 | -0.11 | 0.479 | -1.604 | -1.173 | -0.136 | -0.089 | -0.089 | 0.011 | 0.24 | 0.08 | 0.42 | 0.77 | 0.46 | 0.41 |
| 3 | 195479256 | rs2258447 | -0.055 | -0.475 | -0.758 | -1.088 | -0.038 | 0.185 | 0.185 | 0.065 | 0.87 | 0.65 | 0.93 | 0.82 | 0.93 | 0.82 |
| 3 | 195489009 | rs2246901 | -0.267 | -0.799 | -1.173 | -1.41 | 0.002 | 0.003 | 0.003 | 0 | 0.58 | 0.48 | 0.58 | 0.81 | 0.71 | 0.59 |
| 3 | 195496129 | rs2688492 | 0.222 | -0.769 | -1.512 | -1.615 | -0.005 | 0.001 | 0.001 | 3.698 | 0.7 | 0.4 | 0.6 | 0.81 | 0.76 | 0.66 |
| 3 | 195500758 | rs11928440 | 0.874 | -1.09 | -1.412 | -1.515 | -0.091 | -0.168 | -0.168 | 0 | 0.22 | 0.4 | 0.09 | 0 | 0.12 | 0.04 |
| 3 | 195501149 | rs2259292 | 0.318 | -0.188 | -1.591 | -1.318 | 0.004 | 0.02 | 0.02 | 0 | 0.51 | 0.56 | 0.51 | 0.76 | 0.58 | 0.51 |
| 3 | 195501258 | rs2259331 | -0.37 | -0.144 | -1.654 | -1.388 | -0.018 | -0.007 | -0.007 | 0.023 | 0.61 | 0.85 | 0.58 | 0.77 | 0.65 | 0.56 |
| 3 | 195505417 | rs2259419 | 0.221 | -0.204 | -1.631 | -1.302 | 0.004 | 0.016 | 0.016 | 0 | 0.52 | 0.42 | 0.46 | 0.76 | 0.59 | 0.3 |
| 3 | 195505664 | rs2688513 | -0.796 | -0.096 | -2.096 | -0.804 | -0.011 | -0.102 | -0.102 | 0.551 | 0.82 | 0.99 | 0.66 | 0.81 | 0.84 | 0.54 |
| 3 | 195516305 | rs3749331 | 1.293 | 0.077 | -2.034 | -1.078 | -0.016 | 0.06 | 0.06 | 2.585 | 0.2 | 0.12 | 0.09 | 0.05 | 0.17 | 0.13 |
| 3 | 195516630 | rs3103954 | -0.971 | -0.038 | -2.268 | -1.031 | -0.015 | -0.102 | -0.102 | 0 | 0.82 | 0.99 | 0.69 | 0.82 | 0.84 | 0.69 |
| 3 | 195516878 | rs2177336 | -0.395 | -0.353 | -1.929 | -1.17 | 0.007 | -0.009 | -0.009 | 0 | 0.78 | 0.81 | 0.68 | 0.82 | 0.84 | 0.68 |
| 3 | 195517124 | rs1806440 | 1.572 | -0.452 | -1.876 | -1.244 | -0.006 | 0.038 | 0.038 | 3.363 | 0.21 | 0.17 | 0.1 | 0.05 | 0.17 | 0.13 |
| 3 | 195517258 | rs1106502 | -0.331 | -0.26 | -1.993 | -1.061 | 0.007 | -0.008 | -0.008 | 0 | 0.78 | 0.81 | 0.68 | 0.82 | 0.84 | 0.68 |
| 3 | 195517321 | rs1104760 | -0.324 | -0.256 | -1.991 | -1.059 | 0.007 | -0.009 | -0.009 | 0.127 | 0.78 | 0.81 | 0.68 | 0.82 | 0.84 | 0.68 |
| 3 | 195517553 | rs882605 | -0.324 | -0.256 | -1.991 | -1.059 | 0.007 | -0.009 | -0.009 | 0.003 | 0.78 | 0.81 | 0.69 | 0.82 | 0.84 | 0.68 |
| 3 | 195519857 | rs2688488 | 0.455 | -0.02 | -2.256 | -1.318 | -0.135 | -0.098 | -0.098 | 0.026 | 0.25 | 0.07 | 0.39 | 0.75 | 0.44 | 0.45 |

| | | | | | | | | | | | | | | | | |
|---|---|---|---|---|---|---|---|---|---|---|---|---|---|---|---|---|
| 3 | 195521531 | rs9844491 | 0.265 | -0.377 | -2.187 | -1.332 | -0.118 | -0.069 | -0.069 | 0 | 0.29 | 0.1 | 0.47 | 0.76 | 0.52 | 0.45 |
| 3 | 195527471 | rs2688515 | -0.171 | -0.434 | -1.9 | -1.393 | -0.012 | 0.007 | 0.007 | 0 | 0.38 | 0.38 | 0.52 | 0.78 | 0.54 | 0.52 |
| 3 | 195528226 | rs2641773 | -0.164 | -0.43 | -1.898 | -1.391 | -0.012 | 0.007 | 0.007 | 0 | 0.38 | 0.38 | 0.52 | 0.78 | 0.54 | 0.51 |
| 3 | 195531841 | rs2641772 | -0.212 | 0.27 | -1.681 | -1.003 | -0.001 | 0.029 | 0.029 | 0 | 0.54 | 0.43 | 0.56 | 0.78 | 0.62 | 0.53 |
| 3 | 195536040 | rs842460 | -0.124 | 0.326 | -1.716 | -0.971 | 0.002 | 0.044 | 0.044 | 0 | 0.53 | 0.36 | 0.56 | 0.78 | 0.62 | 0.53 |
| 3 | 195550320 | rs859700 | -0.525 | 0.765 | -1.757 | -0.789 | -0.062 | 0.106 | 0.106 | 3.246 | 0.83 | 0.99 | 0.73 | 0.99 | 0.83 | 0.72 |
| 3 | 195559289 | rs12630378 | -0.541 | 1.733 | -1.133 | -0.178 | 0.042 | 0.052 | 0.052 | 0.247 | 0.1 | 0.16 | 0.18 | 0.15 | 0.2 | 0.11 |
| 3 | 195559939 | rs6800709 | -0.503 | 1.574 | -0.808 | -0.374 | 0.001 | -0.031 | -0.031 | 0 | 0.29 | 0.57 | 0.44 | 0.19 | 0.3 | 0.25 |
| 3 | 195571024 | rs6806773 | 0.267 | 1.114 | -0.996 | -0.133 | -0.001 | -0.003 | -0.003 | 6.608 | 0.81 | 0.78 | 0.87 | 0.84 | 0.76 | 0.82 |
| 3 | 195573324 | rs903196 | 0.111 | 1.167 | -1.436 | -0.222 | -0.023 | 0.058 | 0.058 | 0 | 0.77 | 0.9 | 0.74 | 0.84 | 0.68 | 0.81 |
| 3 | 195577630 | rs9858622 | -0.179 | 0.591 | -1.139 | -0.219 | -0.003 | -0.004 | -0.004 | 0 | 0.42 | 0.4 | 0.52 | 0.67 | 0.32 | 0.45 |
| 3 | 195586589 | rs9844652 | 0.192 | -0.32 | -1.482 | 0.127 | 0.003 | -0.002 | -0.002 | 0 | 0.67 | 0.34 | 0.72 | 0.57 | 0.77 | 0.82 |
| 3 | 195587257 | rs6583249 | 0.049 | -0.119 | -1.317 | 0.147 | -0.005 | 0.001 | 0.001 | 0.574 | 0.31 | 0.34 | 0.23 | 0.38 | 0.25 | 0.23 |
| 3 | 195596907 | rs2278034 | -0.337 | 0.052 | -1.769 | -0.036 | 0.036 | 0.126 | 0.126 | 0.246 | 0.48 | 0.31 | 0.47 | 0.15 | 0.57 | 0.37 |
| 3 | 195602787 | rs7636635 | 0.407 | 0.584 | -1.527 | 0.438 | 0.02 | -0.019 | -0.019 | 0 | 0.19 | 0.67 | 0.18 | 0.12 | 0.24 | 0.15 |
| 3 | 195606770 | rs881753 | -0.737 | 0.403 | -1.906 | 0.548 | -0.011 | 0.109 | 0.109 | 2.551 | 0.57 | 0.82 | 0.49 | 0.22 | 0.53 | 0.34 |
| 3 | 195609808 | rs6771909 | -0.298 | 0.033 | -1.26 | 1.051 | -0.006 | 0.066 | 0.066 | 0.778 | 0.05 | 0.22 | 0.04 | 0.08 | 0.02 | 0.01 |
| 3 | 195615376 | rs3747669 | -0.176 | 0.356 | -2.049 | 0.689 | 0.03 | -0.025 | -0.025 | 0 | 0.41 | 0.67 | 0.42 | 0.46 | 0.29 | 0.29 |
| 3 | 195652708 | rs6783079 | -0.729 | 1.236 | -1.938 | -0.24 | -0.027 | 0.056 | 0.056 | 0.067 | 0.8 | 0.9 | 0.83 | 1 | 0.68 | 0.78 |
| 3 | 195652973 | rs13061255 | 0.483 | 1.482 | -2.443 | -0.469 | -0.06 | -0.004 | -0.004 | 0 | 0.15 | 0.01 | 0.17 | 0.06 | 0.17 | 0.1 |
| 3 | 195656614 | rs9881504 | -0.542 | 1.4 | -2.101 | -0.03 | -0.025 | -0.035 | -0.035 | 0 | 0.77 | 0.85 | 0.58 | 0.37 | 0.65 | 0.64 |
| 3 | 195734915 | rs6797622 | 0.987 | 0.828 | -2.076 | -0.524 | 0.006 | 0.009 | 0.009 | 0.901 | 0.12 | 0.09 | 0.06 | 0.01 | 0.08 | 0.13 |
| 3 | 195738406 | rs4927848 | -0.888 | 1.081 | -2.354 | -0.676 | -0.034 | 0.006 | 0.006 | 0 | 0.78 | 0.89 | 0.71 | 0.77 | 0.66 | 0.67 |
| 3 | 195746112 | rs9837977 | 0.824 | 0.954 | -2.391 | -0.729 | 0.007 | 0.016 | 0.016 | 0.669 | 0.13 | 0.09 | 0.06 | 0.01 | 0.09 | 0.13 |
| 3 | 195750742 | rs9820939 | -0.367 | 0.915 | -2.213 | -1.076 | 0.009 | 0.022 | 0.022 | 0 | 0.31 | 0.24 | 0.24 | 0.06 | 0.28 | 0.23 |
| 3 | 195751630 | rs4927850 | -0.861 | 1.109 | -1.783 | -0.833 | -0.004 | 0.001 | 0.001 | 0 | 0.78 | 0.8 | 0.69 | 0.76 | 0.66 | 0.66 |
| 3 | 195751853 | rs4927851 | -0.852 | 1.112 | -1.782 | -0.832 | -0.004 | 0.001 | 0.001 | 0 | 0.78 | 0.8 | 0.7 | 0.76 | 0.66 | 0.66 |
| 3 | 195762323 | rs9883087 | 0.979 | 0.983 | -1.824 | -1.118 | -0.014 | 0.052 | 0.052 | 0.731 | 0.11 | 0.03 | 0.08 | 0.01 | 0.11 | 0.14 |
| 3 | 195766737 | rs9829686 | 0.118 | 0.706 | -2.05 | -1.406 | -0.003 | 0.149 | 0.149 | 0 | 0.55 | 0.75 | 0.61 | 0.93 | 0.48 | 0.64 |
| 3 | 195776976 | rs406271 | 0.216 | 0.894 | -1.29 | -1.122 | 0.004 | -0.008 | -0.008 | 0 | 0.24 | 0.3 | 0.33 | 0.16 | 0.35 | 0.37 |
| 3 | 195777211 | rs17788379 | 1.378 | 0.901 | -1.597 | -1.367 | -0.024 | 0.042 | 0.042 | 0.67 | 0.09 | 0.02 | 0.07 | 0.01 | 0.09 | 0.12 |
| 3 | 195793979 | rs714602 | 0.86 | 0.703 | -1.919 | -1.961 | -0.03 | 0.118 | 0.118 | 0.036 | 0.33 | 0.12 | 0.29 | 0.06 | 0.36 | 0.21 |
| 3 | 195796049 | rs4927866 | 1.398 | 0.752 | -1.734 | -1.953 | -0.013 | 0.065 | 0.065 | 0 | 0.19 | 0.07 | 0.18 | 0 | 0.26 | 0.17 |
| 3 | 195798258 | rs480760 | -0.823 | -0.169 | -1.232 | -2.108 | -0.048 | -0.062 | -0.062 | 0.351 | 0.89 | 0.66 | 0.94 | 1 | 0.97 | 1 |
| 3 | 195804763 | rs4927868 | 0.114 | 0.081 | -1.357 | -1.954 | -0.001 | 0.074 | 0.074 | 0 | 0.05 | 0.08 | 0.21 | 0.68 | 0.03 | 0.15 |
| 3 | 195815594 | rs12632706 | 0.949 | 0.185 | -1.509 | -2.111 | -0.044 | 0.118 | 0.118 | 0.081 | 0.33 | 0.13 | 0.3 | 0.06 | 0.38 | 0.23 |
| 3 | 195816195 | rs12490036 | -0.103 | 0.144 | -1.341 | -2.171 | -0.006 | 0.141 | 0.141 | 0.107 | 0.53 | 0.26 | 0.38 | 0.23 | 0.53 | 0.53 |
| 3 | 195820274 | rs1466085 | -0.009 | 0.183 | -1.332 | -1.904 | -0.006 | -0.01 | -0.01 | 0.28 | 0.07 | 0.04 | 0.24 | 0.71 | 0.06 | 0.23 |
| 3 | 195827205 | rs9990333 | -0.104 | 0.127 | -1.287 | -2.322 | 0.001 | -0.063 | -0.063 | 0 | 0.51 | 0.45 | 0.3 | 0.01 | 0.47 | 0.39 |
| 3 | 195841644 | rs7432894 | -0.216 | 0.734 | -1.217 | -1.38 | -0.002 | -0.006 | -0.006 | 0 | 0.68 | 0.45 | 0.59 | 0.3 | 0.67 | 0.61 |
| 3 | 195848264 | rs4075688 | -0.23 | 1.078 | -1.86 | -0.966 | 0.014 | 0.099 | 0.099 | 0 | 0.47 | 0.7 | 0.48 | 0.78 | 0.38 | 0.41 |
| 3 | 195851223 | rs9834962 | -0.158 | 1.151 | -1.944 | 0.021 | -0.011 | -0.039 | -0.039 | 1.618 | 0.33 | 0.21 | 0.27 | 0.45 | 0.34 | 0.32 |
| 3 | 195852026 | rs2343130 | -0.489 | 1.205 | -1.888 | 0.308 | 0.009 | 0.003 | 0.003 | 0 | 0.57 | 0.35 | 0.56 | 0.61 | 0.62 | 0.41 |
| 3 | 195854522 | rs11712192 | -0.454 | 1.612 | -1.936 | 1.374 | -0.06 | -0.078 | -0.078 | 0.269 | 0.11 | 0.01 | 0.16 | 0.45 | 0.2 | 0.15 |
| 3 | 195858887 | rs2343129 | -0.256 | 0.715 | -1.606 | 0.692 | 0.006 | -0.002 | -0.002 | 1.341 | 0.29 | 0.23 | 0.41 | 0.37 | 0.24 | 0.29 |
| 3 | 195866289 | rs11185515 | -0.222 | 1.186 | -1.987 | 0.503 | -0.07 | -0.068 | -0.068 | 0 | 0.33 | 0.15 | 0.54 | 0.45 | 0.52 | 0.4 |
| 3 | 195870161 | rs6765113 | -0.69 | 0.389 | -2.048 | 0.233 | -0.066 | -0.048 | -0.048 | 0 | 0.44 | 0.88 | 0.61 | 0.48 | 0.55 | 0.44 |
| 3 | 195871388 | rs4927884 | -0.392 | 0.168 | -1.864 | -0.567 | 0.008 | -0.1 | -0.1 | 0 | 0.23 | 0.65 | 0.19 | 0 | 0.23 | 0.2 |
| 3 | 195877558 | rs4927880 | 0.829 | -1.242 | -1.841 | -0.639 | 0.034 | -0.055 | -0.055 | 0.442 | 0.22 | 0.12 | 0.22 | 0.52 | 0.12 | 0.27 |
| 3 | 195877603 | rs4927879 | -0.213 | -1.485 | -1.463 | -0.716 | 0.012 | -0.009 | -0.009 | 0 | 0.34 | 0.66 | 0.29 | 0.65 | 0.17 | 0.36 |
| 3 | 195887205 | rs9847948 | 0.167 | -0.97 | -2.11 | -1.156 | 0.085 | 0.136 | 0.136 | 3.45 | 0.39 | 0.14 | 0.4 | 0.85 | 0.25 | 0.44 |
| 3 | 195902930 | rs11929092 | 0.592 | 0.046 | -1.825 | -1.112 | -0.056 | 0.095 | 0.095 | 0 | 0.35 | 0.22 | 0.39 | 0 | 0.48 | 0.26 |
| 3 | 195907653 | rs4916483 | 0.976 | -0.11 | -1.908 | -0.497 | 0.024 | -0.157 | -0.157 | 0.11 | 0.22 | 0.01 | 0.22 | 0.5 | 0.13 | 0.32 |
| 3 | 195911680 | rs12631305 | 0.259 | 0.02 | -1.695 | -0.376 | -0.001 | 0.002 | 0.002 | 1.513 | 0.1 | 0.15 | 0.06 | 0.15 | 0.13 | 0.07 |
| 3 | 195919020 | rs3850643 | -0.324 | -0.14 | -1.259 | -0.333 | -0.012 | 0.01 | 0.01 | 0 | 0.69 | 0.5 | 0.69 | 0.66 | 0.75 | 0.76 |
| 3 | 195922427 | rs9850274 | -0.15 | 0.655 | -1.763 | 0.066 | -0.032 | -0.052 | -0.052 | 0 | 0.7 | 0.91 | 0.67 | 0.54 | 0.73 | 0.58 |
| 3 | 195926234 | rs939892 | -0.595 | 1.086 | -1.332 | -0.048 | -0.024 | 0.02 | 0.02 | 0.617 | 0.91 | 0.94 | 0.91 | 1 | 0.88 | 0.88 |
| 3 | 195928711 | rs11185516 | 0.39 | 0.693 | -1.503 | 0.304 | 0.048 | 0.018 | 0.018 | 0 | 0.53 | 0.27 | 0.45 | 0.4 | 0.39 | 0.51 |
| 3 | 195929252 | rs1105977 | 0.664 | 0.25 | -1.311 | -0.123 | 0 | -0.043 | -0.043 | 0 | 0.43 | 0.51 | 0.28 | 0.05 | 0.28 | 0.3 |
| 3 | 195929268 | rs9812036 | 0.462 | 0.206 | -1.345 | 0.19 | -0.01 | -0.009 | -0.009 | 0.983 | 0.05 | 0.05 | 0.03 | 0.06 | 0.06 | 0.04 |
| 3 | 195936382 | rs11719281 | 0.521 | 0.729 | -1.179 | 0.468 | 0.032 | 0.043 | 0.043 | 0 | 0.09 | 0.23 | 0.1 | 0.16 | 0.14 | 0.05 |
| 3 | 195936446 | rs4916502 | 0.625 | 0.781 | -1.246 | 0.352 | 0.004 | 0.009 | 0.009 | 0 | 0.56 | 0.63 | 0.6 | 0.42 | 0.57 | 0.62 |
| 3 | 195942444 | rs3850645 | 0.488 | 0.332 | -1.472 | 0.141 | -0.011 | -0.02 | -0.02 | 3.886 | 0.07 | 0 | 0.03 | 0.08 | 0.05 | 0.05 |
| 3 | 195946535 | rs11185518 | 0.127 | 0.043 | -1.537 | 0.275 | 0.018 | -0.1 | -0.1 | 8.657 | 0.17 | 0.02 | 0.17 | 0.34 | 0.13 | 0.22 |
| 3 | 195955472 | rs1875088 | 0.325 | 0.618 | -1.338 | -0.889 | -0.008 | 0.005 | 0.005 | 1.964 | 0.29 | 0.22 | 0.4 | 0.73 | 0.32 | 0.34 |
| 3 | 195957926 | rs7638797 | 0.191 | 0.496 | -1.064 | -1.017 | -0.001 | 0 | 0 | 1.348 | 0.31 | 0.27 | 0.41 | 0.73 | 0.34 | 0.34 |
| 3 | 195959924 | rs1476331 | -0.57 | 0.3 | -0.415 | -1.241 | -0.001 | -0.035 | -0.035 | 0 | 0.43 | 0.49 | 0.51 | 0.87 | 0.44 | 0.56 |
| 3 | 195961685 | rs1044988 | 0.541 | 0.413 | -0.367 | -1.375 | -0.002 | -0.01 | -0.01 | 0 | 0.19 | 0.22 | 0.15 | 0.13 | 0.21 | 0.22 |
| 3 | 195963910 | rs2342309 | 1.108 | 0.483 | -0.514 | -1.399 | -0.065 | -0.058 | -0.058 | 0 | 0.2 | 0.02 | 0.34 | 0.73 | 0.25 | 0.3 |

| | | | | | | | | | | | | | | | | |
|---|---|---|---|---|---|---|---|---|---|---|---|---|---|---|---|---|
| 3 | 195964298 | rs884534 | -1.126 | 0.772 | -0.629 | -1.781 | -0.036 | 0.032 | 0.032 | 0 | 0.87 | 0.94 | 0.74 | 0.97 | 0.74 | 0.85 |
| 3 | 195964687 | rs712012 | 0.13 | -0.089 | -0.806 | -2.195 | -0.05 | 0.106 | 0.106 | 0.616 | 0.67 | 0.89 | 0.68 | 0.85 | 0.63 | 0.64 |
| 3 | 195968643 | rs1045075 | 0.57 | 0.257 | -0.68 | -1.836 | -0.041 | 0.014 | 0.014 | 0 | 0.34 | 0.1 | 0.45 | 0.69 | 0.4 | 0.44 |
| 3 | 195973244 | rs7639752 | -0.124 | 0.244 | -0.408 | -2.478 | -0.001 | -0.034 | -0.034 | 0 | 0.45 | 0.39 | 0.54 | 0.87 | 0.49 | 0.58 |
| 3 | 195974181 | rs2280525 | 1.488 | 0.246 | -0.413 | -2.443 | 0.001 | 0.002 | 0.002 | 0.008 | 0.13 | 0.08 | 0.1 | 0.08 | 0.13 | 0.19 |
| 3 | 195989417 | rs1948690 | 0.8 | 0.252 | -0.398 | -2.494 | 0.003 | -0.017 | -0.017 | 0.01 | 0.32 | 0.34 | 0.44 | 0.8 | 0.36 | 0.39 |
| 3 | 195990822 | rs12630070 | -0.087 | 0.27 | -0.432 | -2.506 | 0 | -0.064 | -0.064 | 0 | 0.46 | 0.44 | 0.54 | 0.87 | 0.49 | 0.58 |
| 3 | 196005444 | rs12637288 | 0.486 | 0.077 | -0.257 | -2.613 | -0.026 | 0.087 | 0.087 | 3.719 | 0.57 | 0.75 | 0.6 | 0.8 | 0.52 | 0.51 |
| 3 | 196007493 | rs2047551 | 0.358 | 0.298 | -0.522 | -2.289 | 0 | -0.006 | -0.006 | 0 | 0.33 | 0.31 | 0.44 | 0.8 | 0.36 | 0.39 |
| 3 | 196014073 | rs6766480 | -0.433 | 0.237 | -0.555 | -2.297 | 0.001 | -0.001 | -0.001 | 0.001 | 0.46 | 0.4 | 0.48 | 0.8 | 0.43 | 0.52 |
| 3 | 196016340 | rs1357290 | 1.571 | 0.229 | -0.647 | -2.275 | -0.088 | -0.092 | -0.092 | 0.109 | 0.21 | 0.02 | 0.36 | 0.73 | 0.25 | 0.29 |
| 3 | 196019680 | rs7631913 | -1.202 | 0.284 | -0.569 | -2.71 | 0.008 | -0.078 | -0.078 | 0 | 0.59 | 0.46 | 0.58 | 0.87 | 0.55 | 0.71 |
| 3 | 196023385 | rs6778879 | 0.063 | 0.208 | -0.609 | -2.629 | 0.005 | 0.061 | 0.061 | 0.115 | 0.35 | 0.18 | 0.41 | 0.8 | 0.31 | 0.5 |
| 3 | 196038384 | rs11929532 | 0.136 | 0.216 | -0.62 | -2.625 | 0.001 | 0.064 | 0.064 | 0.144 | 0.34 | 0.15 | 0.4 | 0.8 | 0.31 | 0.5 |
| 3 | 196048895 | rs11919443 | 0.105 | 0.17 | -0.503 | -2.576 | -0.01 | -0.023 | -0.023 | 0 | 0.26 | 0.24 | 0.14 | 0.08 | 0.2 | 0.31 |
| 3 | 196051403 | rs2280526 | -1.096 | 0.291 | -0.5 | -2.695 | 0.004 | -0.084 | -0.084 | 0 | 0.59 | 0.44 | 0.6 | 0.87 | 0.57 | 0.72 |
| 3 | 196054004 | rs2276711 | 1.536 | 0.295 | -0.519 | -2.692 | 0.038 | 0.057 | 0.057 | 0 | 0.12 | 0.02 | 0.02 | 0.01 | 0.05 | 0.12 |
| 3 | 196054439 | rs6785339 | -1.036 | 0.434 | -0.433 | -2.633 | 0.004 | -0.084 | -0.084 | 0 | 0.59 | 0.44 | 0.6 | 0.87 | 0.57 | 0.72 |
| 3 | 196058795 | rs12497601 | -1.031 | 0.422 | -0.531 | -2.622 | 0.002 | -0.071 | -0.071 | 0 | 0.58 | 0.43 | 0.6 | 0.87 | 0.57 | 0.72 |
| 3 | 196066569 | rs7636047 | 0.479 | 0.231 | -0.763 | -2.554 | 0.025 | 0.038 | 0.038 | 0 | 0.23 | 0.13 | 0.11 | 0.01 | 0.16 | 0.18 |
| 3 | 196068626 | rs13083065 | 0.896 | 0.339 | -0.2 | -2.221 | 0.004 | -0.016 | -0.016 | 0 | 0.14 | 0.18 | 0.14 | 0.06 | 0.16 | 0.18 |
| 3 | 196072442 | rs4916490 | -0.95 | 0.15 | -0.258 | -2.397 | 0 | -0.059 | -0.059 | 0 | 0.64 | 0.43 | 0.62 | 0.9 | 0.62 | 0.79 |
| 3 | 196078427 | rs6774852 | -1.681 | 1.108 | -0.049 | -1.929 | -0.016 | 0.011 | 0.011 | 0 | 0.95 | 1 | 0.81 | 0.97 | 0.85 | 0.92 |
| 3 | 196078746 | rs9848247 | -0.276 | 0.504 | 0.396 | -1.787 | -0.001 | -0.016 | -0.016 | 0 | 0.45 | 0.39 | 0.56 | 0.85 | 0.53 | 0.5 |
| 3 | 196088464 | rs6764533 | 0.732 | 0.631 | 0.094 | -1.708 | -0.027 | 0.035 | 0.035 | 3.313 | 0.31 | 0.14 | 0.44 | 0.8 | 0.36 | 0.31 |
| 3 | 196096451 | rs9860931 | -0.337 | 0.684 | -0.191 | -1.727 | 0.001 | -0.057 | -0.057 | 0.278 | 0.46 | 0.47 | 0.56 | 0.89 | 0.53 | 0.6 |
| 3 | 196130005 | rs6799210 | 1.073 | 0.789 | -0.568 | -1.7 | -0.005 | 0.014 | 0.014 | 0.211 | 0.13 | 0.06 | 0.11 | 0.05 | 0.17 | 0.2 |
| 3 | 196141661 | rs4459932 | 0.158 | 0.89 | -0.228 | -1.314 | 0.005 | -0.013 | -0.013 | 0.107 | 0.32 | 0.33 | 0.45 | 0.81 | 0.36 | 0.29 |
| 3 | 196158332 | rs7374260 | -0.405 | 0.685 | -0.313 | -1.561 | 0.004 | -0.035 | -0.035 | 0 | 0.43 | 0.54 | 0.47 | 0.85 | 0.47 | 0.49 |
| 3 | 196158773 | rs6583309 | 1.032 | -0.092 | -0.386 | -2.136 | -0.038 | -0.108 | -0.108 | 0.303 | 0.25 | 0.33 | 0.15 | 0.01 | 0.16 | 0.12 |
| 3 | 196167930 | rs12631706 | 0.579 | 0.693 | -0.811 | -2.498 | -0.008 | 0.049 | 0.049 | 1.997 | 0.27 | 0.14 | 0.23 | 0.09 | 0.32 | 0.32 |
| 3 | 196168101 | rs4538407 | 1.495 | 0.042 | -0.759 | -2.494 | 0.028 | 0.101 | 0.101 | 6.492 | 0.19 | 0.02 | 0.12 | 0 | 0.16 | 0.1 |
| 3 | 196170061 | rs9812425 | -0.606 | 0.418 | -0.068 | -2.178 | -0.01 | 0.169 | 0.169 | 4.759 | 0.48 | 0.78 | 0.49 | 0.87 | 0.47 | 0.47 |
| 3 | 196181228 | rs7639524 | -0.606 | 1.325 | 0.44 | -1.501 | 0.013 | -0.004 | -0.004 | 0.463 | 0.1 | 0.15 | 0.07 | 0.1 | 0.1 | 0.06 |
| 3 | 196181269 | rs7648887 | -0.497 | 1.405 | 0.119 | -1.388 | -0.054 | -0.047 | -0.047 | 0 | 0.86 | 0.95 | 0.71 | 0.28 | 0.75 | 0.72 |
| 3 | 196187608 | rs2342371 | -0.514 | 1.429 | 0.355 | -1.303 | -0.008 | -0.008 | -0.008 | 0.536 | 0.85 | 0.89 | 0.66 | 0.29 | 0.71 | 0.72 |
| 3 | 196190893 | rs13094241 | -0.193 | 1.396 | 0.61 | -1.344 | 0.029 | 0.022 | 0.022 | 0.047 | 0.83 | 0.76 | 0.65 | 0.29 | 0.71 | 0.73 |
| 3 | 196199204 | rs3796129 | 1.195 | 0.698 | 0.435 | -2.097 | -0.006 | 0.062 | 0.062 | 3.585 | 0.6 | 0.24 | 0.48 | 0.18 | 0.6 | 0.6 |
| 3 | 196200737 | rs9864836 | -0.485 | 1.165 | -0.054 | -1.869 | -0.077 | -0.031 | -0.031 | 0 | 0.84 | 0.96 | 0.6 | 0.19 | 0.71 | 0.7 |
| 3 | 196210808 | rs7646174 | 0.376 | 1 | 0.166 | -1.823 | -0.023 | 0.028 | 0.028 | 3.903 | 0.71 | 0.8 | 0.53 | 0.17 | 0.62 | 0.6 |
| 3 | 196233795 | rs6778889 | 0.321 | 0.547 | -0.032 | -2.046 | -0.035 | 0.187 | 0.187 | 2.433 | 0.63 | 0.89 | 0.32 | 0.08 | 0.42 | 0.51 |
| 3 | 196245603 | rs11185544 | 0.691 | 0.686 | 0.534 | -2.037 | 0.03 | 0.098 | 0.098 | 1.411 | 0.48 | 0.34 | 0.28 | 0.06 | 0.4 | 0.39 |
| 3 | 196262959 | rs9848622 | 0.399 | 0.903 | -0.118 | -1.845 | 0.014 | 0.17 | 0.17 | 1.536 | 0.54 | 0.78 | 0.35 | 0.12 | 0.43 | 0.48 |
| 3 | 196278366 | rs9647412 | 0.298 | 1.138 | -0.125 | -1.848 | 0.003 | 0.137 | 0.137 | 2.901 | 0.55 | 0.72 | 0.36 | 0.06 | 0.45 | 0.41 |
| 3 | 196303538 | rs11185546 | 0.325 | 1.909 | -0.982 | -0.412 | -0.033 | -0.043 | -0.043 | 0 | 0.12 | 0.05 | 0.32 | 0.61 | 0.23 | 0.22 |
| 3 | 196312124 | rs2014220 | 0.065 | 1.13 | -0.864 | -1.372 | 0.005 | -0.02 | -0.02 | 0.494 | 0.31 | 0.3 | 0.17 | 0.06 | 0.22 | 0.31 |
| 3 | 196315284 | rs11185547 | 0.511 | 1.874 | -0.94 | -0.388 | 0.005 | -0.02 | -0.02 | 0 | 0.88 | 0.83 | 0.9 | 0.98 | 0.84 | 0.83 |
| 3 | 196318855 | rs9843585 | 0.056 | 1.354 | -1.048 | 0.049 | -0.001 | -0.003 | -0.003 | 0 | 0.68 | 0.54 | 0.63 | 0.7 | 0.66 | 0.74 |
| 3 | 196322359 | rs4916524 | -0.283 | 1.695 | -1.912 | -0.354 | -0.052 | 0.061 | 0.061 | 0 | 0.21 | 0.06 | 0.16 | 0.08 | 0.23 | 0.35 |
| 3 | 196328849 | rs9876053 | -0.001 | 1.827 | -2.361 | 0.036 | -0.113 | -0.115 | -0.115 | 0.088 | 0.29 | 0.04 | 0.37 | 0.56 | 0.41 | 0.25 |
| 3 | 196334416 | rs11715619 | -0.097 | 0.82 | -2.298 | -0.53 | 0.009 | 0.03 | 0.03 | 0.113 | 0.23 | 0.79 | 0.24 | 0.09 | 0.19 | 0.18 |
| 3 | 196339852 | rs6798553 | -0.536 | 0.877 | -1.393 | -0.62 | 0.006 | 0.009 | 0.009 | 0.162 | 0.52 | 0.64 | 0.6 | 0.63 | 0.62 | 0.43 |
| 3 | 196342476 | rs4916511 | 0.418 | 0.954 | -1.743 | -0.662 | -0.157 | -0.089 | -0.089 | 0 | 0.32 | 0.04 | 0.38 | 0.66 | 0.46 | 0.3 |
| 3 | 196344162 | rs9824819 | -0.581 | 0.496 | -1.19 | -0.797 | 0.007 | 0.027 | 0.027 | 0 | 0.54 | 0.72 | 0.54 | 0.63 | 0.57 | 0.43 |
| 3 | 196345149 | rs7426655 | 1.281 | 0.149 | -1.723 | -1.167 | 0.01 | 0.023 | 0.023 | 0 | 0.18 | 0.04 | 0.14 | 0.1 | 0.15 | 0.23 |
| 3 | 196346273 | rs4591526 | -0.592 | 0.181 | -1.608 | -1.127 | -0.007 | -0.002 | -0.002 | 0.004 | 0.71 | 0.79 | 0.68 | 0.73 | 0.72 | 0.65 |
| 3 | 196348356 | rs6779465 | 0.003 | -0.046 | -0.769 | -1.285 | -0.001 | -0.003 | -0.003 | 0.099 | 0.61 | 0.6 | 0.63 | 0.7 | 0.65 | 0.59 |
| 3 | 196351215 | rs1385331 | 0.981 | 0.06 | -0.421 | -1.204 | -0.053 | -0.042 | -0.042 | 0 | 0.36 | 0.16 | 0.38 | 0.59 | 0.46 | 0.32 |
| 3 | 196352809 | rs4605582 | 0.102 | -0.115 | 0.043 | -1.223 | -0.014 | -0.03 | -0.03 | 0 | 0.1 | 0.19 | 0.04 | 0.04 | 0.06 | 0.05 |
| 3 | 196359443 | rs1684467 | -0.306 | -0.444 | -0.772 | -1.568 | -0.003 | -0.017 | -0.017 | 0.864 | 0.47 | 0.48 | 0.52 | 0.79 | 0.39 | 0.54 |
| 3 | 196360596 | rs1798630 | -0.457 | -0.164 | -0.533 | -1.24 | 0 | -0.02 | -0.02 | 0 | 0.26 | 0.28 | 0.16 | 0.07 | 0.22 | 0.16 |
| 3 | 196365765 | rs1798624 | 0.202 | -0.199 | -0.643 | -0.888 | 0.014 | 0.026 | 0.026 | 1.134 | 0.55 | 0.35 | 0.57 | 0.7 | 0.64 | 0.5 |
| 3 | 196365919 | rs1718407 | 0.067 | -0.399 | -0.727 | -0.781 | 0.042 | 0.017 | 0.017 | 0 | 0.58 | 0.75 | 0.71 | 0.64 | 0.68 | 0.44 |
| 3 | 196366264 | rs1684446 | -0.329 | 0.719 | -0.475 | -0.938 | 0.023 | -0.055 | -0.055 | 0 | 0.23 | 0.37 | 0.27 | 0.09 | 0.31 | 0.26 |
| 3 | 196367936 | rs7636495 | -0.415 | 0.482 | -1.225 | -0.716 | 0.008 | 0.037 | 0.037 | 0 | 0.15 | 0.07 | 0.12 | 0.1 | 0.14 | 0.12 |
| 3 | 196369163 | rs843522 | -1.514 | 0.538 | -1.406 | -0.221 | 0.009 | -0.016 | -0.016 | 0 | 0.53 | 0.87 | 0.54 | 0.44 | 0.59 | 0.52 |
| 3 | 196372422 | rs7635042 | 0.162 | 0.451 | -1.13 | -0.181 | 0.016 | 0.021 | 0.021 | 0 | 0.11 | 0.05 | 0.03 | 0.08 | 0.07 | 0.08 |
| 3 | 196374156 | rs1212534 | -0.472 | 0.682 | -0.704 | -0.224 | 0.017 | 0.012 | 0.012 | 1.004 | 0.28 | 0.58 | 0.39 | 0.29 | 0.31 | 0.42 |
| 3 | 196375140 | rs6766451 | -0.664 | 0.663 | -0.816 | -0.152 | 0.008 | 0.012 | 0.012 | 0 | 0.25 | 0.22 | 0.1 | 0.12 | 0.21 | 0.2 |

| | | | | | | | | | | | | | | | | |
|---|---|---|---|---|---|---|---|---|---|---|---|---|---|---|---|---|
| 3 | 196377750 | rs1798629 | -0.522 | 1.333 | -0.573 | 0.127 | 0.021 | 0.012 | 0.012 | 0 | 0.27 | 0.36 | 0.32 | 0.28 | 0.33 | 0.37 |
| 3 | 196379916 | rs843520 | -0.438 | 1.129 | -1.07 | 0.184 | -0.013 | -0.015 | -0.015 | 2.875 | 0.47 | 0.73 | 0.43 | 0.53 | 0.55 | 0.56 |
| 3 | 196384865 | rs4912589 | 0.501 | 1.211 | -0.765 | -0.378 | -0.001 | -0.013 | -0.013 | 0 | 0.55 | 0.46 | 0.54 | 0.22 | 0.46 | 0.37 |
| 3 | 196389580 | rs6803735 | -0.514 | 1.624 | -0.805 | -0.331 | -0.001 | -0.001 | -0.001 | 0 | 0.25 | 0.23 | 0.25 | 0.14 | 0.28 | 0.17 |
| 3 | 196393480 | rs1798628 | 0.493 | 1.493 | -0.958 | -0.612 | -0.017 | 0.01 | 0.01 | 0 | 0.86 | 0.92 | 0.92 | 1 | 0.83 | 0.76 |
| 3 | 196401804 | rs13098826 | -0.451 | 0.676 | -0.936 | -0.646 | 0.006 | 0.004 | 0.004 | 0 | 0.12 | 0.08 | 0.07 | 0 | 0.12 | 0.05 |
| 3 | 196407281 | rs9862217 | 0.106 | -0.509 | -0.398 | -0.844 | -0.059 | -0.054 | -0.054 | 0 | 0.7 | 0.47 | 0.76 | 0.16 | 0.83 | 0.59 |
| 3 | 196408487 | rs2686589 | -0.275 | -0.479 | -0.146 | -0.093 | 0.008 | -0.005 | -0.005 | 0 | 0.76 | 0.77 | 0.88 | 0.71 | 0.85 | 0.85 |
| 3 | 196409312 | rs6583169 | -0.199 | -0.482 | 0.024 | -0.094 | -0.013 | 0.005 | 0.005 | 0 | 0.76 | 0.69 | 0.86 | 0.71 | 0.84 | 0.85 |
| 3 | 196412487 | rs843510 | -0.504 | -0.731 | -0.03 | -0.028 | -0.027 | 0.013 | 0.013 | 0 | 0.81 | 0.77 | 0.9 | 0.72 | 0.9 | 0.85 |
| 3 | 196420444 | rs7622437 | 0.092 | 0.059 | -0.181 | -0.554 | 0.009 | 0.024 | 0.024 | 0 | 0.51 | 0.39 | 0.48 | 0.65 | 0.53 | 0.58 |
| 3 | 196426606 | rs12718037 | 0.119 | 0.013 | 0 | -0.569 | 0.014 | 0.039 | 0.039 | 0 | 0.51 | 0.32 | 0.48 | 0.66 | 0.54 | 0.58 |
| 3 | 196429953 | rs13061191 | -0.079 | -0.09 | 0.409 | -0.581 | 0.002 | 0.028 | 0.028 | 0 | 0.56 | 0.66 | 0.51 | 0.71 | 0.55 | 0.65 |
| 3 | 196432167 | rs2030534 | -0.142 | -0.061 | 0.618 | -1.048 | -0.006 | 0.008 | 0.008 | 2.214 | 0.27 | 0.53 | 0.35 | 0.69 | 0.23 | 0.39 |
| 3 | 196433031 | rs9849684 | 0.261 | 0.48 | 0.302 | -1.585 | 0.005 | -0.038 | -0.038 | 0 | 0.29 | 0.3 | 0.2 | 0.04 | 0.32 | 0.25 |
| 3 | 196435534 | rs6776064 | -0.016 | 0.401 | 0.4 | -1.441 | 0.009 | 0.01 | 0.01 | 0 | 0.33 | 0.43 | 0.32 | 0.6 | 0.35 | 0.46 |
| 3 | 196438744 | rs7629566 | -0.182 | 0.658 | 0.391 | -1.391 | 0.023 | 0.25 | 0.25 | 0 | 0.09 | 0.25 | 0.14 | 0.55 | 0.09 | 0.23 |
| 3 | 196442132 | rs1580470 | -0.013 | 0.39 | 0.448 | -1.526 | 0.028 | 0.029 | 0.029 | 0 | 0.31 | 0.52 | 0.43 | 0.53 | 0.35 | 0.46 |
| 3 | 196452093 | rs2049233 | -1.044 | 0.253 | -0.185 | -1.828 | 0.036 | 0.008 | 0.008 | 2.381 | 0.56 | 0.89 | 0.62 | 0.53 | 0.63 | 0.65 |
| 3 | 196452238 | rs2049232 | 0.117 | 0.457 | -0.112 | -1.487 | 0.02 | 0.019 | 0.019 | 0 | 0.31 | 0.45 | 0.42 | 0.53 | 0.35 | 0.47 |
| 3 | 196453467 | rs1568603 | 0.212 | 0.498 | -0.168 | -1.91 | 0.02 | -0.068 | -0.068 | 0.301 | 0.29 | 0.4 | 0.21 | 0.05 | 0.32 | 0.27 |
| 3 | 196462949 | rs4916543 | 0.246 | -0.061 | -0.208 | -1.791 | -0.004 | 0.03 | 0.03 | 0 | 0.21 | 0.12 | 0.16 | 0 | 0.23 | 0.15 |
| 3 | 196466504 | rs1718408 | -0.172 | -0.45 | -0.052 | -1.682 | 0.004 | 0.008 | 0.008 | 0 | 0.71 | 0.67 | 0.66 | 0.2 | 0.76 | 0.53 |
| 3 | 196467338 | rs11711813 | 0.951 | -0.754 | 0.352 | -1.785 | -0.003 | 0.01 | 0.01 | 0 | 0.43 | 0.36 | 0.39 | 0.2 | 0.52 | 0.39 |
| 3 | 196467603 | rs1684470 | -0.164 | -0.769 | 0.275 | -1.503 | 0.072 | 0.058 | 0.058 | 0 | 0.58 | 0.71 | 0.68 | 0.64 | 0.69 | 0.61 |
| 3 | 196467894 | rs1630029 | 0.189 | -0.657 | 0.367 | -1.585 | 0.02 | 0.107 | 0.107 | 0 | 0.53 | 0.67 | 0.55 | 0.22 | 0.57 | 0.45 |
| 3 | 196475119 | rs2686596 | 0.944 | -0.621 | 0.146 | -1.563 | -0.004 | 0.014 | 0.014 | 0 | 0.43 | 0.39 | 0.39 | 0.21 | 0.53 | 0.42 |
| 3 | 196482211 | rs843532 | 1.645 | -0.522 | -0.312 | -1.674 | -0.025 | 0.057 | 0.057 | 0 | 0.11 | 0.01 | 0.07 | 0 | 0.13 | 0.12 |
| 3 | 196493961 | rs6809369 | -0.119 | -0.398 | 0.094 | -1.488 | -0.003 | -0.028 | -0.028 | 0.148 | 0.06 | 0.17 | 0.03 | 0 | 0.03 | 0.04 |
| 3 | 196501413 | rs2089979 | 1.605 | -0.379 | -0.139 | -1.445 | -0.064 | -0.005 | -0.005 | 7.127 | 0.29 | 0.14 | 0.33 | 0.23 | 0.4 | 0.27 |
| 3 | 196506153 | rs9325377 | 1.097 | -1.287 | 0.221 | -2.208 | 0.015 | -0.018 | -0.018 | 0 | 0.35 | 0.39 | 0.6 | 0.67 | 0.51 | 0.46 |
| 3 | 196510533 | rs4916546 | 0.703 | -0.955 | -0.06 | -2.018 | 0.011 | 0.045 | 0.045 | 0 | 0.18 | 0.07 | 0.13 | 0 | 0.18 | 0.12 |
| 3 | 196518026 | rs12497912 | -0.236 | -1.048 | 0.103 | -2.076 | -0.025 | -0.141 | -0.141 | 0 | 0.31 | 0.47 | 0.37 | 0.01 | 0.25 | 0.18 |
| 3 | 196519209 | rs7646247 | -0.531 | -1.033 | -0.478 | -1.818 | 0.008 | -0.005 | -0.005 | 0 | 0.75 | 0.85 | 0.89 | 0.65 | 0.78 | 0.66 |
| 3 | 196538648 | rs4916555 | -1.16 | -0.685 | -1.012 | -1.881 | -0.093 | 0.07 | 0.07 | 0 | 0.63 | 0.91 | 0.59 | 0.28 | 0.5 | 0.58 |
| 3 | 196541252 | rs13078303 | 0.376 | -0.799 | -0.497 | -1.633 | 0.016 | 0 | 0 | 0 | 0.23 | 0.16 | 0.14 | 0.27 | 0.18 | 0.27 |
| 3 | 196541580 | rs11185455 | -0.24 | 0.113 | -0.14 | -1.285 | 0.006 | -0.058 | -0.058 | 0.339 | 0.84 | 0.78 | 0.89 | 1 | 0.83 | 0.86 |
| 3 | 196546374 | rs10446496 | 0.124 | 0.273 | -0.62 | -1.153 | 0.006 | 0.01 | 0.01 | 0.176 | 0.42 | 0.6 | 0.35 | 0.3 | 0.36 | 0.48 |
| 3 | 196546529 | rs10446497 | -0.2 | 0.134 | -0.819 | -1.087 | 0.02 | 0.058 | 0.058 | 0.019 | 0.45 | 0.68 | 0.37 | 0.3 | 0.39 | 0.5 |
| 3 | 196547538 | rs2166799 | -0.491 | 0.307 | -0.592 | -0.786 | -0.016 | 0.071 | 0.071 | 0.228 | 0.54 | 0.71 | 0.57 | 0.32 | 0.52 | 0.62 |
| 3 | 196554020 | rs2084386 | 0.277 | 0.36 | -0.282 | -0.563 | -0.001 | -0.018 | -0.018 | 0 | 0.88 | 0.91 | 0.92 | 1 | 0.87 | 0.9 |
| 3 | 196554952 | rs6789365 | -1.245 | 0.144 | 0.061 | -0.244 | -0.011 | 0.002 | 0.002 | 0 | 0.23 | 0.43 | 0.31 | 0.19 | 0.19 | 0.21 |
| 3 | 196561452 | rs7616401 | -1.176 | 0.123 | 0.421 | -0.518 | -0.01 | -0.046 | -0.046 | 0 | 0.16 | 0.47 | 0.15 | 0.13 | 0.11 | 0.25 |
| 3 | 196576109 | rs4916561 | 0.602 | 0.284 | -0.129 | -1.099 | -0.001 | -0.022 | -0.022 | 0 | 0.57 | 0.58 | 0.69 | 0.86 | 0.65 | 0.54 |
| 3 | 196583207 | rs4916446 | 0.12 | 0.216 | -0.22 | -1.253 | 0.001 | -0.02 | -0.02 | 0 | 0.74 | 0.7 | 0.79 | 0.9 | 0.77 | 0.62 |
| 3 | 196584613 | rs1823234 | -0.209 | -0.018 | -0.421 | -1.3 | 0.001 | -0.006 | -0.006 | 0 | 0.18 | 0.12 | 0.09 | 0.07 | 0.1 | 0.08 |
| 3 | 196593789 | rs4916565 | 0.057 | 0.081 | -0.539 | -1.328 | -0.016 | 0.064 | 0.064 | 0.815 | 0.75 | 0.86 | 0.8 | 0.92 | 0.76 | 0.63 |
| 3 | 196599322 | rs4916567 | -0.324 | -0.077 | -0.231 | -1.671 | -0.012 | -0.011 | -0.011 | 0.114 | 0.35 | 0.6 | 0.46 | 0.73 | 0.3 | 0.33 |
| 3 | 196614372 | rs12639201 | 0.888 | -0.167 | -0.04 | -1.684 | -0.001 | -0.011 | -0.011 | 0.591 | 0.18 | 0.18 | 0.22 | 0.68 | 0.21 | 0.27 |
| 3 | 196616249 | rs7643566 | 0.621 | -0.008 | -0.277 | -1.764 | -0.081 | 0.062 | 0.062 | 0.838 | 0.23 | 0.01 | 0.17 | 0.07 | 0.23 | 0.38 |
| 3 | 196616593 | rs9879020 | -0.156 | -0.151 | -0.039 | -1.849 | -0.005 | -0.005 | -0.005 | 0 | 0.68 | 0.63 | 0.56 | 0.28 | 0.71 | 0.67 |
| 3 | 196617207 | rs11185459 | -0.156 | -0.151 | 0.001 | -1.849 | -0.014 | -0.013 | -0.013 | 0.277 | 0.68 | 0.59 | 0.56 | 0.28 | 0.71 | 0.67 |
| 3 | 196620294 | rs6797560 | 1.417 | -0.014 | -0.391 | -2.562 | -0.003 | -0.026 | -0.026 | 0.267 | 0.25 | 0.28 | 0.12 | 0.17 | 0.23 | 0.23 |
| 3 | 196641105 | rs6583189 | -0.66 | -0.001 | -0.498 | -2.534 | -0.016 | -0.015 | -0.015 | 0 | 0.68 | 0.58 | 0.56 | 0.28 | 0.71 | 0.67 |
| 3 | 196662521 | rs6734 | 0.424 | -0.009 | -0.513 | -2.537 | -0.019 | -0.018 | -0.018 | 0 | 0.32 | 0.44 | 0.44 | 0.73 | 0.29 | 0.33 |
| 3 | 196662696 | rs6733 | 1.421 | 0.012 | -0.583 | -2.571 | -0.066 | 0.092 | 0.092 | 0 | 0.16 | 0.04 | 0.25 | 0 | 0.24 | 0.05 |
| 3 | 196663321 | rs480668 | -1.408 | 0.001 | -0.57 | -2.559 | -0.072 | 0.067 | 0.067 | 0.697 | 0.77 | 0.99 | 0.83 | 0.93 | 0.77 | 0.62 |
| 3 | 196665854 | rs555582 | -0.354 | 0.046 | -0.377 | -2.429 | 0 | -0.015 | -0.015 | 0 | 0.48 | 0.5 | 0.69 | 0.76 | 0.53 | 0.39 |
| 3 | 196668359 | rs548853 | -0.435 | -0.421 | -0.71 | -2.482 | 0.032 | 0.284 | 0.284 | 1.245 | 0.49 | 0.84 | 0.51 | 0.93 | 0.42 | 0.52 |
| 3 | 196669912 | rs335824 | 0.188 | -0.307 | -0.452 | -2.098 | -0.006 | -0.007 | -0.007 | 0 | 0.31 | 0.37 | 0.44 | 0.72 | 0.3 | 0.33 |
| 3 | 196674518 | rs1147238 | -0.192 | -0.586 | -0.077 | -2.487 | -0.046 | -0.146 | -0.146 | 0 | 0.22 | 0.46 | 0.11 | 0 | 0.15 | 0.09 |
| 3 | 196674749 | rs4916589 | 0.092 | -0.54 | -0.141 | -2.396 | 0.013 | 0.066 | 0.066 | 0.287 | 0.56 | 0.68 | 0.57 | 0.77 | 0.6 | 0.5 |
| 3 | 196674879 | rs12636891 | 0.014 | -0.495 | -0.671 | -3.009 | 0 | 0.043 | 0.043 | 0 | 0.48 | 0.6 | 0.61 | 0.76 | 0.49 | 0.51 |
| 3 | 196674916 | rs1147240 | -0.418 | -0.926 | -0.977 | -3.04 | 0.018 | 0.069 | 0.069 | 0.013 | 0.56 | 0.67 | 0.46 | 0.77 | 0.4 | 0.53 |
| 3 | 196676487 | rs746037 | -0.128 | -0.208 | -0.671 | -2.406 | -0.006 | -0.038 | -0.038 | 0 | 0.32 | 0.43 | 0.43 | 0.86 | 0.31 | 0.47 |
| 3 | 196677409 | rs4347982 | -0.103 | -0.086 | -0.753 | -2.401 | -0.003 | -0.018 | -0.018 | 0 | 0.33 | 0.39 | 0.44 | 0.82 | 0.31 | 0.44 |
| 3 | 196684793 | rs556043 | -0.361 | -0.266 | -0.709 | -2.256 | -0.012 | -0.028 | -0.028 | 0 | 0.65 | 0.58 | 0.66 | 0.8 | 0.71 | 0.44 |
| 3 | 196688027 | rs4916595 | 0.46 | 0.695 | -0.943 | -1.957 | -0.001 | 0 | 0 | 0 | 0.22 | 0.23 | 0.38 | 0.78 | 0.33 | 0.23 |
| 3 | 196692717 | rs2568871 | 0.178 | -0.635 | -0.902 | -2.053 | -0.009 | -0.042 | -0.042 | 0 | 0.57 | 0.46 | 0.6 | 0.84 | 0.64 | 0.38 |

| | | | | | | | | | | | | | | | | |
|---|---|---|---|---|---|---|---|---|---|---|---|---|---|---|---|---|
| 3 | 196693337 | rs9823916 | -0.367 | -1.006 | -0.701 | -1.783 | -0.048 | -0.086 | -0.086 | 0 | 0.18 | 0.37 | 0.11 | 0.03 | 0.08 | 0.21 |
| 3 | 196701941 | rs9822086 | -0.448 | -1.089 | -0.75 | -1.694 | -0.064 | -0.078 | -0.078 | 0 | 0.17 | 0.39 | 0.22 | 0.03 | 0.05 | 0.16 |
| 3 | 196710530 | rs4916596 | -0.125 | -0.388 | -1.783 | -1.879 | -0.082 | 0.179 | 0.179 | 0 | 0.4 | 0.16 | 0.51 | 0.87 | 0.47 | 0.36 |
| 3 | 196712778 | rs498738 | 0.214 | -0.387 | -1.9 | -1.94 | -0.062 | 0.112 | 0.112 | 0 | 0.36 | 0.15 | 0.48 | 0.86 | 0.46 | 0.24 |
| 3 | 196717313 | rs569330 | -0.781 | -0.563 | -2.339 | -1.974 | -0.129 | 0.262 | 0.262 | 0 | 0.54 | 0.95 | 0.58 | 0.88 | 0.51 | 0.46 |
| 3 | 196719271 | rs509016 | 0.021 | -0.238 | -1.988 | -1.08 | -0.118 | -0.147 | -0.147 | 1.722 | 0.34 | 0.04 | 0.44 | 0.6 | 0.38 | 0.23 |
| 3 | 196721395 | rs7622580 | -0.185 | 0.055 | -0.761 | -1.78 | -0.004 | 0.021 | 0.021 | 0.14 | 0.63 | 0.65 | 0.64 | 0.87 | 0.61 | 0.39 |
| 3 | 196726039 | rs4916453 | -0.181 | 0.635 | -0.551 | -1.215 | 0.009 | -0.038 | -0.038 | 0 | 0.37 | 0.61 | 0.5 | 0.85 | 0.42 | 0.29 |
| 3 | 196728706 | rs539510 | 0.292 | 0.672 | -0.307 | -1.116 | 0.037 | -0.133 | -0.133 | 0 | 0.76 | 0.54 | 0.7 | 0.94 | 0.71 | 0.72 |
| 3 | 196734973 | rs2280292 | 0.509 | 0.084 | -1.071 | -1.166 | -0.003 | 0.051 | 0.051 | 0.116 | 0.22 | 0.07 | 0.13 | 0.12 | 0.19 | 0.08 |
| 3 | 196735273 | rs7649700 | -0.625 | -0.262 | -0.515 | -0.975 | 0.016 | 0.125 | 0.125 | 1.52 | 0.06 | 0.35 | 0.04 | 0.12 | 0.02 | 0.09 |
| 3 | 196738953 | rs3821716 | 0.599 | -0.172 | -0.762 | -1.005 | -0.008 | 0.059 | 0.059 | 0.072 | 0.22 | 0.07 | 0.13 | 0.12 | 0.19 | 0.08 |
| 3 | 196746629 | rs1056664 | 0.188 | -0.187 | -0.592 | -0.733 | 0.004 | -0.002 | -0.002 | 0.015 | 0.27 | 0.28 | 0.34 | 0.28 | 0.23 | 0.21 |
| 3 | 196747300 | rs12636071 | -0.272 | -0.059 | -0.974 | -1.31 | 0.037 | 0.057 | 0.057 | 0 | 0.49 | 0.45 | 0.62 | 0.86 | 0.65 | 0.44 |
| 3 | 196749408 | rs17129227 | 0.94 | -0.278 | -0.749 | -1.268 | -0.038 | -0.056 | -0.056 | 0.037 | 0.24 | 0.22 | 0.11 | 0.05 | 0.11 | 0.31 |
| 3 | 196750207 | rs890490 | 0.142 | 0.502 | -1.153 | -0.876 | 0.002 | 0.005 | 0.005 | 0 | 0.79 | 0.79 | 0.76 | 0.8 | 0.81 | 0.69 |
| 3 | 196753355 | rs4916598 | -0.067 | 1.343 | -1.175 | -0.542 | -0.036 | -0.024 | -0.024 | 0.058 | 0.22 | 0.13 | 0.46 | 0.66 | 0.46 | 0.2 |
| 3 | 196758824 | rs3755768 | -0.494 | 0.793 | -1.251 | -1.011 | -0.003 | 0.038 | 0.038 | 0.038 | 0.89 | 0.95 | 0.94 | 0.95 | 0.89 | 0.86 |
| 3 | 196767854 | rs17597256 | -0.389 | 1.695 | -1.891 | -1.137 | -0.152 | -0.073 | -0.073 | 0 | 0.23 | 0.03 | 0.42 | 0.72 | 0.42 | 0.29 |
| 3 | 196769797 | rs7753 | -0.024 | 0.391 | -1.683 | -2.019 | 0.001 | -0.001 | -0.001 | 0.004 | 0.61 | 0.44 | 0.64 | 0.8 | 0.65 | 0.62 |
| 3 | 196774356 | rs7638423 | 0.174 | 0.477 | -1.503 | -1.99 | -0.104 | -0.04 | -0.04 | 0.35 | 0.23 | 0.07 | 0.43 | 0.72 | 0.41 | 0.3 |
| 3 | 196778438 | rs7629533 | 1.003 | 0.415 | -1.551 | -2.079 | 0.018 | 0.028 | 0.028 | 0.3 | 0.41 | 0.37 | 0.23 | 0.07 | 0.26 | 0.35 |
| 3 | 196793858 | rs6795358 | 0.034 | 1.127 | -1.138 | -2.248 | -0.004 | -0.028 | -0.028 | 0 | 0.14 | 0.17 | 0.08 | 0.02 | 0.14 | 0.06 |
| 3 | 196813908 | rs3843375 | -0.785 | 1.072 | -1.076 | -2.266 | -0.02 | 0.002 | 0.002 | 0 | 0.27 | 0.24 | 0.44 | 0.74 | 0.43 | 0.36 |
| 3 | 196819024 | rs11706316 | 1.126 | 0.914 | -0.931 | -2.307 | 0.023 | 0.041 | 0.041 | 0.141 | 0.43 | 0.38 | 0.23 | 0.07 | 0.26 | 0.37 |
| 3 | 196837717 | rs7624859 | 0.676 | 0.605 | -1.196 | -2.42 | 0.014 | -0.069 | -0.069 | 1.505 | 0.13 | 0.31 | 0.1 | 0 | 0.09 | 0.02 |
| 3 | 196838486 | rs1356612 | -1.158 | 0.485 | -0.711 | -2.628 | -0.004 | -0.007 | -0.007 | 0 | 0.74 | 0.78 | 0.71 | 0.82 | 0.75 | 0.79 |
| 3 | 196861043 | rs6803606 | 1.054 | 0.529 | -0.841 | -2.641 | 0.028 | 0.063 | 0.063 | 0.204 | 0.42 | 0.35 | 0.22 | 0.07 | 0.26 | 0.37 |
| 3 | 196863988 | rs7612661 | -0.687 | 0.471 | -0.741 | -2.653 | -0.024 | -0.036 | -0.036 | 0 | 0.71 | 0.66 | 0.7 | 0.81 | 0.75 | 0.77 |
| 3 | 196865242 | rs1134986 | 2.592 | 0.485 | -0.895 | -2.6 | 0.038 | 0.104 | 0.104 | 1.662 | 0.28 | 0.01 | 0.12 | 0.07 | 0.14 | 0.33 |
| 3 | 196898609 | rs4916461 | -0.456 | 0.854 | -0.524 | -2.406 | 0.001 | 0 | 0 | 0.09 | 0.26 | 0.27 | 0.44 | 0.73 | 0.43 | 0.35 |
| 3 | 196910160 | rs11928645 | 0.429 | 0.833 | -0.501 | -2.402 | -0.002 | -0.045 | -0.045 | 6.099 | 0.13 | 0.21 | 0.07 | 0.02 | 0.14 | 0.05 |
| 3 | 196912276 | rs11923220 | 0.514 | 0.823 | -0.419 | -2.407 | 0 | 0.012 | 0.012 | 0 | 0.12 | 0.09 | 0.06 | 0.02 | 0.14 | 0.06 |
| 3 | 196921780 | rs9868546 | -0.676 | 0.688 | -0.251 | -2.525 | -0.012 | -0.019 | -0.019 | 0.382 | 0.72 | 0.71 | 0.7 | 0.82 | 0.75 | 0.77 |
| 3 | 196929799 | rs9839886 | 0.112 | 0.779 | -0.353 | -2.423 | -0.007 | -0.019 | -0.019 | 0 | 0.59 | 0.49 | 0.63 | 0.78 | 0.61 | 0.72 |
| 3 | 196931821 | rs7649443 | 0.999 | 0.859 | -0.537 | -2.429 | -0.06 | -0.054 | -0.054 | 1.102 | 0.14 | 0.06 | 0.36 | 0.71 | 0.29 | 0.3 |
| 3 | 196960901 | rs4635680 | 0.121 | 0.702 | -0.457 | -2.451 | 0.005 | 0.006 | 0.006 | 0 | 0.59 | 0.47 | 0.63 | 0.78 | 0.61 | 0.72 |
| 3 | 196962329 | rs6805920 | 0.18 | 1.169 | -0.542 | -2.118 | -0.066 | -0.047 | -0.047 | 0 | 0.17 | 0.07 | 0.4 | 0.71 | 0.35 | 0.34 |
| 3 | 196985596 | rs9843659 | -1.017 | 0.868 | -0.565 | -1.908 | 0 | -0.008 | -0.008 | 0 | 0.38 | 0.64 | 0.54 | 0.71 | 0.53 | 0.41 |
| 3 | 197006700 | rs1741950 | -1.113 | 0.745 | -0.535 | -1.171 | 0.003 | 0.011 | 0.011 | 0 | 0.91 | 0.93 | 0.93 | 0.94 | 0.91 | 0.9 |
| 3 | 197026927 | rs338217 | -0.031 | 1.295 | -0.165 | -0.755 | -0.08 | -0.057 | -0.057 | 0 | 0.83 | 0.94 | 0.63 | 0.28 | 0.66 | 0.64 |
| 3 | 197031608 | rs17129314 | -0.381 | 0.365 | 0.748 | -0.665 | 0.025 | 0.005 | 0.005 | 0.568 | 0.06 | 0.19 | 0.06 | 0.05 | 0.04 | 0.02 |
| 3 | 197034985 | rs6764023 | 0.581 | 0.129 | 0.774 | -0.732 | -0.007 | -0.004 | -0.004 | 0 | 0.2 | 0.18 | 0.22 | 0.14 | 0.14 | 0.09 |
| 3 | 197037067 | rs7620061 | 0.089 | 0.135 | 1.059 | -0.722 | 0.057 | -0.05 | -0.05 | 0.665 | 0.1 | 0.4 | 0.1 | 0 | 0.09 | 0.03 |
| 3 | 197038062 | rs3844276 | 0.939 | -0.11 | 0.705 | -0.549 | -0.005 | -0.001 | -0.001 | 0 | 0.17 | 0.12 | 0.21 | 0.13 | 0.12 | 0.07 |
| 3 | 197043066 | rs13059981 | -0.029 | -0.151 | 0.542 | -0.428 | -0.008 | -0.001 | -0.001 | 0 | 0.24 | 0.22 | 0.25 | 0.16 | 0.19 | 0.13 |
| 3 | 197048060 | rs1321635 | 0.818 | 0.001 | 0.268 | -0.165 | 0.022 | -0.034 | -0.034 | 0 | 0.19 | 0.07 | 0.22 | 0.19 | 0.13 | 0.14 |
| 3 | 197050163 | rs7644221 | -1.014 | 0.061 | 0.226 | -0.496 | 0.013 | 0.046 | 0.046 | 0 | 0.49 | 0.68 | 0.45 | 0.22 | 0.42 | 0.37 |
| 3 | 197052973 | rs6583202 | -1.433 | 0.116 | -0.318 | -0.761 | 0.009 | 0.157 | 0.157 | 0 | 0.61 | 0.89 | 0.77 | 0.88 | 0.61 | 0.67 |
| 3 | 197058402 | rs6797138 | -0.282 | -0.456 | 0.402 | -0.581 | 0.02 | -0.094 | -0.094 | 0.169 | 0.2 | 0.37 | 0.15 | 0.01 | 0.22 | 0.09 |
| 3 | 197060036 | rs4916471 | -0.356 | -0.575 | -0.203 | -0.254 | 0.018 | 0.079 | 0.079 | 0.533 | 0.41 | 0.79 | 0.38 | 0.18 | 0.35 | 0.23 |
| 3 | 197063308 | rs732190 | -1.097 | -0.956 | 0.806 | 0.736 | -0.001 | 0.18 | 0.18 | 0 | 0.09 | 0.44 | 0.06 | 0.19 | 0.06 | 0.13 |
| 3 | 197067678 | rs6785072 | -0.965 | -1.063 | -0.136 | 0.638 | 0.026 | 0.017 | 0.017 | 0.147 | 0.37 | 0.74 | 0.26 | 0.64 | 0.23 | 0.41 |
| 3 | 197067712 | rs6776479 | -0.203 | -1.066 | -0.075 | 0.551 | -0.047 | 0.058 | 0.058 | 0 | 0.17 | 0.36 | 0.09 | 0.24 | 0.05 | 0.24 |
| 3 | 197070959 | rs6777317 | -0.811 | -0.761 | -1.062 | -0.083 | 0.05 | 0.207 | 0.207 | 0.397 | 0.43 | 0.9 | 0.3 | 0.84 | 0.29 | 0.53 |
| 3 | 197075721 | rs768858 | -1.222 | -0.779 | 0.088 | 0.071 | -0.021 | -0.063 | -0.063 | 0 | 0.16 | 0.35 | 0.12 | 0.06 | 0.12 | 0.24 |
| 3 | 197079586 | rs6583203 | -0.854 | -0.728 | 0.143 | -0.492 | -0.001 | -0.102 | -0.102 | 0.049 | 0.32 | 0.37 | 0.51 | 0.05 | 0.32 | 0.3 |
| 3 | 197080008 | rs1018527 | -0.986 | -1.085 | 0.064 | -0.461 | -0.01 | -0.046 | -0.046 | 0 | 0.33 | 0.36 | 0.52 | 0.15 | 0.31 | 0.31 |
| 3 | 197080597 | rs911219 | -0.833 | -1.552 | 0.119 | -0.619 | -0.01 | -0.032 | -0.032 | 0 | 0.29 | 0.32 | 0.5 | 0.14 | 0.26 | 0.34 |
| 3 | 197081796 | rs6779852 | -0.671 | -2.9 | -0.216 | -1.033 | 0.046 | 0.021 | 0.021 | 0 | 0.3 | 0.2 | 0.11 | 0.27 | 0.19 | 0.42 |
| 3 | 197083723 | rs718501 | -1.147 | -2.804 | 0.309 | -1.459 | 0.007 | 0.001 | 0.001 | 0.031 | 0.34 | 0.7 | 0.15 | 0.11 | 0.17 | 0.25 |
| 3 | 197092657 | rs1321632 | -0.626 | -2.269 | 0.565 | -1.286 | -0.043 | -0.045 | -0.045 | 0 | 0.15 | 0.3 | 0.07 | 0.07 | 0.08 | 0.15 |
| 3 | 197093758 | rs6794537 | -1.055 | -2.217 | 0.29 | -1.396 | -0.026 | -0.047 | -0.047 | 0 | 0.25 | 0.62 | 0.13 | 0.12 | 0.19 | 0.24 |
| 3 | 197101909 | rs9325388 | -1.157 | -2.276 | 0.211 | -1.494 | -0.037 | -0.081 | -0.081 | 0 | 0.28 | 0.51 | 0.12 | 0.12 | 0.21 | 0.3 |
| 3 | 197102197 | rs1024217 | 0.667 | -1.816 | -0.781 | -2.162 | -0.013 | 0.096 | 0.096 | 0.61 | 0.25 | 0.07 | 0.14 | 0.08 | 0.2 | 0.22 |
| 3 | 197108353 | rs4132331 | -0.934 | -2.271 | -0.497 | -2.28 | -0.01 | -0.038 | -0.038 | 0 | 0.41 | 0.56 | 0.16 | 0.13 | 0.27 | 0.36 |
| 3 | 197121593 | rs13099091 | -0.295 | -2.29 | -0.595 | -2.209 | -0.008 | -0.018 | -0.018 | 0 | 0.19 | 0.23 | 0.06 | 0.07 | 0.1 | 0.23 |
| 3 | 197126303 | rs7645524 | -1.179 | -2.284 | -0.621 | -2.382 | -0.02 | -0.071 | -0.071 | 0 | 0.32 | 0.43 | 0.16 | 0.13 | 0.27 | 0.35 |
| 3 | 197129945 | rs7621299 | -0.5 | -2.39 | -1.216 | -2.749 | 0.048 | 0.119 | 0.119 | 0 | 0.42 | 0.27 | 0.18 | 0.15 | 0.3 | 0.35 |

| | | | | | | | | | | | | | | | | |
|---|---|---|---|---|---|---|---|---|---|---|---|---|---|---|---|---|
| 3 | 197130745 | rs7644219 | -0.729 | -2.423 | -1.255 | -2.776 | 0.062 | 0.179 | 0.179 | 0 | 0.47 | 0.26 | 0.22 | 0.16 | 0.36 | 0.39 |
| 3 | 197130874 | rs7630584 | -0.826 | -2.546 | -1.144 | -2.754 | -0.017 | -0.042 | -0.042 | 0.725 | 0.23 | 0.61 | 0.17 | 0.1 | 0.19 | 0.16 |
| 3 | 197133986 | rs9843109 | -0.924 | -2.339 | -0.982 | -2.624 | -0.004 | -0.027 | -0.027 | 0 | 0.39 | 0.42 | 0.22 | 0.19 | 0.38 | 0.42 |
| 3 | 197134110 | rs9843268 | 2.239 | -2.166 | -0.941 | -2.598 | -0.005 | 0 | 0 | 0 | 0.06 | 0.03 | 0.03 | 0.05 | 0.1 | 0.05 |
| 3 | 197136335 | rs9840798 | -0.4 | -2.179 | -0.943 | -2.566 | -0.004 | -0.04 | -0.04 | 0 | 0.33 | 0.42 | 0.18 | 0.18 | 0.33 | 0.38 |
| 3 | 197144582 | rs9855729 | -0.8 | -2.036 | -1.058 | -2.52 | -0.005 | -0.037 | -0.037 | 0.361 | 0.39 | 0.53 | 0.22 | 0.19 | 0.38 | 0.42 |
| 3 | 197147144 | rs7620033 | -1.52 | -2.166 | -1.137 | -2.621 | 0 | -0.003 | -0.003 | 0.552 | 0.55 | 0.53 | 0.26 | 0.19 | 0.46 | 0.44 |
| 3 | 197148659 | rs9857634 | -0.593 | -1.783 | -0.978 | -2.69 | -0.002 | -0.053 | -0.053 | 0 | 0.36 | 0.51 | 0.21 | 0.18 | 0.37 | 0.34 |
| 3 | 197148984 | rs9858256 | -0.039 | -1.615 | -1.071 | -2.437 | -0.018 | -0.032 | -0.032 | 0 | 0.41 | 0.45 | 0.21 | 0.19 | 0.27 | 0.34 |
| 3 | 197151491 | rs9868555 | 0.483 | -1.985 | -1.045 | -2.479 | -0.066 | -0.137 | -0.137 | 0.557 | 0.3 | 0.37 | 0.14 | 0.01 | 0.16 | 0.28 |
| 3 | 197151737 | rs6772127 | -0.075 | -0.779 | -1.146 | -2.572 | 0.001 | -0.003 | -0.003 | 0 | 0.41 | 0.46 | 0.28 | 0.34 | 0.45 | 0.24 |
| 3 | 197151801 | rs6784865 | 0.506 | -0.156 | -0.862 | -2.163 | -0.003 | -0.018 | -0.018 | 0 | 0.2 | 0.35 | 0.13 | 0.17 | 0.16 | 0.14 |
| 3 | 197155097 | rs6798107 | -0.461 | 0.196 | -1.344 | -1.787 | -0.034 | -0.014 | -0.014 | 0 | 0.71 | 0.84 | 0.42 | 0.35 | 0.62 | 0.52 |
| 3 | 197158756 | rs13098723 | 0.033 | 0.113 | -0.907 | -1.921 | -0.006 | 0.017 | 0.017 | 0.654 | 0.27 | 0.21 | 0.17 | 0.16 | 0.32 | 0.17 |
| 3 | 197162976 | rs13092636 | -0.958 | -0.277 | -0.456 | -1.981 | 0.001 | -0.003 | -0.003 | 0 | 0.43 | 0.55 | 0.33 | 0.17 | 0.4 | 0.28 |
| 3 | 197166491 | rs1431499 | 0.66 | -0.801 | -0.439 | -1.873 | 0.002 | -0.028 | -0.028 | 0 | 0.2 | 0.18 | 0.09 | 0 | 0.13 | 0.09 |
| 3 | 197168012 | rs4857595 | 0.234 | -0.717 | -0.72 | -1.778 | -0.01 | 0.049 | 0.049 | 0 | 0.45 | 0.2 | 0.49 | 0.61 | 0.4 | 0.51 |
| 3 | 197169747 | rs1431497 | -0.935 | -0.417 | 0.51 | -1.267 | -0.002 | 0.035 | 0.035 | 1.395 | 0.47 | 0.61 | 0.34 | 0.17 | 0.5 | 0.31 |
| 3 | 197170359 | rs2060089 | -0.505 | -0.43 | 0.476 | -1.276 | -0.005 | 0.016 | 0.016 | 0.679 | 0.41 | 0.36 | 0.31 | 0.16 | 0.48 | 0.26 |
| 3 | 197172252 | rs13099300 | -0.694 | -0.315 | -0.275 | -1.1 | -0.003 | 0.067 | 0.067 | 1.246 | 0.47 | 0.63 | 0.34 | 0.17 | 0.5 | 0.31 |
| 3 | 197185470 | rs11185471 | 1.135 | -1.142 | 0.543 | -0.665 | -0.028 | 0.05 | 0.05 | 0.155 | 0.88 | 0.49 | 0.87 | 0.93 | 0.9 | 0.96 |
| 3 | 197192259 | rs2567344 | 1.084 | -1.64 | -0.401 | -1.177 | 0.114 | 0.152 | 0.152 | 0.695 | 0.54 | 0.19 | 0.71 | 0.86 | 0.68 | 0.66 |
| 3 | 197195013 | rs2567346 | 0.931 | -1.898 | -0.535 | -1.32 | 0.124 | 0.212 | 0.212 | 0.101 | 0.6 | 0.12 | 0.78 | 0.95 | 0.79 | 0.67 |
| 3 | 197196497 | rs1963236 | 1.664 | -1.922 | -0.124 | -1.211 | 0.006 | 0 | 0 | 0.307 | 0.79 | 0.2 | 0.83 | 0.87 | 0.85 | 0.85 |
| 3 | 197210354 | rs9813057 | -1.571 | -1.765 | 0.956 | -1.16 | 0.035 | 0.053 | 0.053 | 0 | 0.07 | 0.4 | 0.05 | 0.05 | 0.04 | 0.06 |
| 3 | 197218463 | rs2686079 | -0.44 | -1.623 | 0.556 | -1.282 | 0.014 | 0.027 | 0.027 | 0 | 0.27 | 0.17 | 0.12 | 0.06 | 0.18 | 0.26 |
| 3 | 197232543 | rs6790736 | 1.326 | -1.741 | 1.079 | -1.166 | -0.039 | 0.063 | 0.063 | 0 | 0.85 | 0.45 | 0.85 | 0.88 | 0.88 | 0.84 |
| 3 | 197234819 | rs2060082 | -0.757 | -1.423 | 0.923 | -1.217 | -0.017 | -0.1 | -0.1 | 0 | 0.29 | 0.35 | 0.12 | 0 | 0.18 | 0.22 |
| 3 | 197241877 | rs7616998 | -1.335 | -1.143 | 1.147 | -0.69 | -0.003 | 0.004 | 0.004 | 0 | 0.28 | 0.65 | 0.31 | 0.27 | 0.24 | 0.19 |
| 3 | 197246898 | rs2686065 | -1.474 | -2.233 | 0.511 | -1.086 | 0.058 | 0.018 | 0.018 | 0 | 0.49 | 0.69 | 0.38 | 0.31 | 0.33 | 0.31 |
| 3 | 197252834 | rs7642375 | -0.712 | -2.042 | 1.453 | -1.171 | 0.012 | -0.05 | -0.05 | 0 | 0.21 | 0.4 | 0.23 | 0.1 | 0.19 | 0.13 |
| 3 | 197254231 | rs13095317 | -0.603 | -2.042 | 1.449 | -1.171 | 0.007 | -0.034 | -0.034 | 0 | 0.2 | 0.31 | 0.22 | 0.1 | 0.19 | 0.14 |
| 3 | 197254550 | rs7638381 | -1.753 | -2.39 | 1.017 | -1.34 | 0.099 | 0.073 | 0.073 | 0 | 0.46 | 0.81 | 0.37 | 0.22 | 0.32 | 0.24 |
| 3 | 197256058 | rs11925979 | -0.633 | -2.096 | 1.055 | -1.179 | -0.023 | -0.002 | -0.002 | 0 | 0.23 | 0.27 | 0.13 | 0.12 | 0.13 | 0.1 |
| 3 | 197263293 | rs7649784 | 0.074 | -1.541 | 0.991 | -1.683 | -0.001 | -0.011 | -0.011 | 0 | 0.05 | 0.08 | 0.04 | 0 | 0.06 | 0.05 |
| 3 | 197277127 | rs891668 | -1.175 | -1.361 | 1.068 | -1.995 | 0.02 | -0.071 | -0.071 | 0 | 0.28 | 0.46 | 0.32 | 0.11 | 0.32 | 0.22 |
| 3 | 197281093 | rs1966617 | -0.806 | -0.828 | 1.057 | -1.886 | 0.012 | -0.02 | -0.02 | 0 | 0.23 | 0.3 | 0.26 | 0.12 | 0.26 | 0.18 |
| 3 | 197288629 | rs11185474 | 0.058 | -0.772 | 1.188 | -2.059 | -0.037 | -0.11 | -0.11 | 0.508 | 0.2 | 0.38 | 0.1 | 0.01 | 0.14 | 0.08 |
| 3 | 197290158 | rs2116294 | -1.19 | -0.632 | 0.529 | -2.013 | -0.005 | -0.058 | -0.058 | 0 | 0.4 | 0.48 | 0.31 | 0.11 | 0.35 | 0.23 |
| 3 | 197299181 | rs7616534 | -0.366 | -0.368 | 0.886 | -1.742 | -0.021 | -0.128 | -0.128 | 0.036 | 0.21 | 0.34 | 0.1 | 0.01 | 0.16 | 0.07 |
| 3 | 197300565 | rs7639745 | -0.412 | -0.34 | 0.725 | -1.729 | -0.025 | -0.149 | -0.149 | 0 | 0.21 | 0.38 | 0.1 | 0.01 | 0.16 | 0.07 |
| 3 | 197304279 | rs2686097 | -0.373 | 0.728 | 0.096 | -1.495 | 0.014 | -0.083 | -0.083 | 0 | 0.36 | 0.46 | 0.3 | 0.08 | 0.42 | 0.28 |
| 3 | 197307725 | rs6781945 | -1.003 | 1.027 | 0.41 | -1.075 | 0.008 | -0.056 | -0.056 | 0 | 0.17 | 0.24 | 0.12 | 0.02 | 0.24 | 0.09 |
| 3 | 197310628 | rs11711172 | -0.977 | 1.014 | 0.425 | -1.072 | 0.003 | -0.039 | -0.039 | 0.57 | 0.17 | 0.21 | 0.12 | 0.02 | 0.23 | 0.09 |
| 3 | 197312425 | rs9841534 | 0.698 | 1.225 | 0.461 | -1.342 | -0.001 | 0.006 | 0.006 | 0 | 0.55 | 0.62 | 0.66 | 0.89 | 0.44 | 0.67 |
| 3 | 197332150 | rs4857506 | -1.859 | 0.487 | 2.041 | 0.334 | 0.039 | 0.341 | 0.341 | 0 | 0.09 | 0.33 | 0.12 | 0.33 | 0.07 | 0.17 |
| 3 | 197346566 | rs28703192 | -1.916 | 0.444 | 2.311 | -0.425 | 0.091 | 0.007 | 0.007 | 0 | 0.09 | 0.43 | 0.06 | 0.02 | 0.08 | 0.11 |
| 3 | 197428127 | rs7612230 | -2.158 | 0.401 | 2.089 | -0.432 | 0.027 | -0.029 | -0.029 | 0 | 0.12 | 0.38 | 0.07 | 0.03 | 0.11 | 0.12 |
| 3 | 197436089 | rs4508802 | -2.138 | 0.364 | 1.902 | -0.411 | 0.033 | -0.027 | -0.027 | 0.526 | 0.12 | 0.41 | 0.07 | 0.03 | 0.1 | 0.1 |
| 3 | 197469638 | rs6787318 | 1.21 | -0.486 | 1.424 | -0.469 | -0.023 | -0.02 | -0.02 | 0 | 0.76 | 0.36 | 0.82 | 0.94 | 0.88 | 0.84 |
| 3 | 197564995 | rs11928427 | -1.852 | 0.286 | 1.722 | -0.488 | 0.081 | -0.045 | -0.045 | 1.72 | 0.1 | 0.53 | 0.08 | 0 | 0.08 | 0.07 |
| 4 | 502911 | rs17164861 | -2.381 | 0.898 | 0.878 | 0.289 | -0.034 | 0.034 | 0.034 | 6.434 | 0.19 | 0.56 | 0.14 | 0.19 | 0.14 | 0.2 |
| 4 | 514629 | rs10029104 | -2.103 | 1.505 | 0.667 | 0.557 | 0.013 | 0.098 | 0.098 | 0.118 | 0.15 | 0.41 | 0.09 | 0.23 | 0.13 | 0.18 |
| 4 | 515489 | rs13115344 | -2.107 | 1.489 | 0.712 | 0.556 | 0.002 | 0.066 | 0.066 | 0.181 | 0.15 | 0.38 | 0.08 | 0.21 | 0.12 | 0.13 |
| 4 | 516398 | rs7672868 | -2.165 | 1.497 | 0.817 | 0.561 | -0.012 | 0.015 | 0.015 | 0.013 | 0.14 | 0.25 | 0.09 | 0.19 | 0.11 | 0.14 |
| 4 | 516586 | rs1135945 | -2.491 | 1.983 | 0.848 | 0.537 | 0.067 | 0.057 | 0.057 | 0.059 | 0.16 | 0.36 | 0.13 | 0.22 | 0.21 | 0.19 |
| 4 | 517376 | rs7666226 | -2.199 | 1.44 | 0.889 | 0.586 | 0.017 | 0.103 | 0.103 | 0.216 | 0.14 | 0.4 | 0.1 | 0.22 | 0.13 | 0.18 |
| 4 | 517461 | rs7666425 | -2.199 | 1.44 | 0.889 | 0.586 | 0.017 | 0.103 | 0.103 | 0.216 | 0.14 | 0.4 | 0.1 | 0.22 | 0.13 | 0.18 |
| 4 | 517622 | rs13150531 | -2.181 | 1.439 | 0.888 | 0.586 | 0.019 | 0.106 | 0.106 | 0.203 | 0.14 | 0.4 | 0.1 | 0.22 | 0.13 | 0.18 |
| 4 | 521480 | rs4624610 | -1.375 | 1.478 | 0.763 | 0.553 | 0.002 | 0.053 | 0.053 | 0.113 | 0.09 | 0.16 | 0.06 | 0.18 | 0.1 | 0.12 |
| 4 | 532200 | rs10213007 | -1.945 | 1.603 | 0.664 | 0.577 | 0.03 | 0.096 | 0.096 | 0.332 | 0.13 | 0.33 | 0.1 | 0.19 | 0.12 | 0.15 |
| 4 | 533001 | rs1127410 | -1.99 | 1.505 | 0.643 | 0.566 | 0.031 | 0.11 | 0.11 | 0.446 | 0.13 | 0.36 | 0.1 | 0.22 | 0.13 | 0.15 |
| 4 | 540025 | rs13149565 | -0.003 | 0.991 | -0.181 | 0.565 | -0.123 | -0.004 | -0.004 | 1.938 | 0.33 | 0.02 | 0.23 | 0.24 | 0.37 | 0.26 |
| 4 | 540099 | rs6822767 | -2.348 | 0.502 | 1.109 | 1.093 | 0.053 | 0.117 | 0.117 | 0.607 | 0.14 | 0.41 | 0.11 | 0.22 | 0.15 | 0.16 |
| 4 | 541182 | rs10014828 | -2.22 | 0.239 | 1.144 | 1.128 | 0.04 | 0.128 | 0.128 | 0.474 | 0.13 | 0.41 | 0.1 | 0.22 | 0.13 | 0.16 |
| 4 | 561631 | rs7655590 | -1.877 | 0.33 | 0.827 | 1.15 | 0.053 | 0.102 | 0.102 | 0.149 | 0.13 | 0.41 | 0.1 | 0.17 | 0.13 | 0.14 |
| 4 | 594372 | rs7661376 | -1.592 | -0.057 | 0.606 | 0.946 | -0.006 | 0.059 | 0.059 | 0 | 0.15 | 0.34 | 0.1 | 0.19 | 0.14 | 0.12 |
| 4 | 607202 | rs9884240 | -1.592 | -0.057 | 0.606 | 0.946 | -0.006 | 0.06 | 0.06 | 0 | 0.15 | 0.34 | 0.1 | 0.19 | 0.14 | 0.12 |
| 4 | 624897 | rs10029928 | -1.57 | -0.053 | 0.576 | 0.953 | -0.007 | 0.064 | 0.064 | 0 | 0.15 | 0.34 | 0.1 | 0.2 | 0.14 | 0.12 |

| | | | | | | | | | | | | | | | | |
|---|---|---|---|---|---|---|---|---|---|---|---|---|---|---|---|---|
| 4 | 639670 | rs10031745 | -1.471 | -0.104 | 0.613 | 0.957 | -0.013 | 0.066 | 0.066 | 0 | 0.16 | 0.38 | 0.11 | 0.21 | 0.15 | 0.12 |
| 4 | 644438 | rs10027866 | -1.21 | -0.505 | 0.497 | 0.759 | -0.041 | -0.001 | -0.001 | 0 | 0.24 | 0.39 | 0.12 | 0.21 | 0.16 | 0.15 |
| 4 | 646409 | rs3864130 | -0.954 | -0.511 | 0.468 | 0.591 | -0.024 | -0.073 | -0.073 | 0.005 | 0.23 | 0.31 | 0.1 | 0 | 0.14 | 0.08 |
| 4 | 658089 | rs7436737 | -0.444 | -0.472 | 0.472 | 0.608 | -0.035 | -0.095 | -0.095 | 1.766 | 0.21 | 0.31 | 0.1 | 0 | 0.13 | 0.08 |
| 4 | 663908 | rs28675771 | -0.494 | -0.668 | -0.068 | 0.695 | -0.003 | 0.01 | 0.01 | 0 | 0.65 | 0.43 | 0.64 | 0.46 | 0.66 | 0.62 |
| 4 | 665737 | rs28470912 | -0.578 | -0.864 | 0.652 | 0.706 | 0.015 | 0.001 | 0.001 | 0.462 | 0.19 | 0.16 | 0.09 | 0 | 0.13 | 0.08 |
| 4 | 666212 | rs4463020 | -0.578 | -0.835 | 0.665 | 0.729 | 0.015 | 0.001 | 0.001 | 0.462 | 0.19 | 0.16 | 0.09 | 0 | 0.13 | 0.08 |
| 4 | 680843 | rs7690350 | -1.451 | -0.794 | 0.977 | 0.964 | -0.065 | 0 | 0 | 0 | 0.29 | 0.46 | 0.15 | 0.26 | 0.2 | 0.25 |
| 4 | 681940 | rs9991613 | -0.953 | -0.758 | 0.687 | 0.564 | -0.046 | -0.121 | -0.121 | 0.073 | 0.26 | 0.37 | 0.12 | 0 | 0.16 | 0.1 |
| 4 | 683842 | rs10026699 | -0.731 | -0.367 | 0.333 | 0.817 | 0.011 | -0.014 | -0.014 | 0 | 0.07 | 0.24 | 0.05 | 0.01 | 0.04 | 0.06 |
| 4 | 685993 | rs35466919 | -0.88 | -0.588 | -0.028 | 1.094 | -0.061 | 0.011 | 0.011 | 0.51 | 0.31 | 0.5 | 0.16 | 0.31 | 0.21 | 0.29 |
| 4 | 694613 | rs11730652 | 0.519 | 0.176 | -0.551 | 1.858 | 0 | 0.014 | 0.014 | 0 | 0.83 | 0.84 | 0.72 | 0.52 | 0.86 | 0.76 |
| 4 | 698292 | rs6837180 | 0.28 | 1.45 | -0.95 | 1.658 | -0.009 | -0.009 | -0.009 | 0 | 0.78 | 0.81 | 0.52 | 0.6 | 0.59 | 0.65 |
| 4 | 703813 | rs4690187 | -0.217 | 1.298 | -1.584 | 1.06 | -0.003 | 0.048 | 0.048 | 0 | 0.35 | 0.16 | 0.26 | 0.23 | 0.32 | 0.32 |
| 4 | 712222 | rs4076064 | 0.877 | 0.804 | -1.607 | 0.701 | -0.022 | -0.024 | -0.024 | 0 | 0.34 | 0.54 | 0.2 | 0.18 | 0.22 | 0.23 |
| 4 | 716248 | rs10452329 | 0.341 | 1.032 | -1.686 | 0.866 | 0.036 | 0.035 | 0.035 | 0 | 0.44 | 0.63 | 0.26 | 0.31 | 0.3 | 0.36 |
| 4 | 726322 | rs10034650 | 0.224 | 0.989 | -1.66 | 0.836 | 0.08 | 0.076 | 0.076 | 0.004 | 0.45 | 0.72 | 0.27 | 0.31 | 0.3 | 0.42 |
| 4 | 732261 | rs4690296 | 0.882 | 1.66 | -1.773 | 0.932 | -0.006 | 0.009 | 0.009 | 0 | 0.08 | 0.05 | 0.09 | 0.05 | 0.08 | 0.16 |
| 4 | 734351 | rs11729193 | 0.219 | 1.778 | -1.912 | 0.998 | -0.051 | -0.033 | -0.033 | 1.885 | 0.39 | 0.21 | 0.59 | 0.53 | 0.53 | 0.31 |
| 4 | 734677 | rs747504 | -0.439 | 1.383 | -0.853 | 0.798 | 0.001 | 0.015 | 0.015 | 0 | 0.52 | 0.43 | 0.4 | 0.37 | 0.44 | 0.52 |
| 4 | 735150 | rs4234853 | 1.062 | 1.403 | -0.939 | 0.819 | -0.025 | 0.022 | 0.022 | 0 | 0.3 | 0.07 | 0.23 | 0.21 | 0.27 | 0.25 |
| 4 | 744098 | rs9332466 | 0.863 | 1.712 | -0.492 | 1.083 | 0.062 | 0.107 | 0.107 | 0 | 0.07 | 0.11 | 0.35 | 0.33 | 0.26 | 0.07 |
| 4 | 744972 | rs1531583 | -0.448 | 1.231 | -0.447 | 0.771 | 0.001 | -0.028 | -0.028 | 0.267 | 0.09 | 0.32 | 0.03 | 0.02 | 0.04 | 0.07 |
| 4 | 748447 | rs4690304 | 0.688 | 1.586 | -1.312 | 0.605 | -0.032 | -0.033 | -0.033 | 0 | 0.66 | 0.76 | 0.43 | 0.49 | 0.52 | 0.71 |
| 4 | 753944 | rs4690281 | 0.968 | 1.224 | -1.834 | 0.526 | -0.033 | -0.009 | -0.009 | 2.576 | 0.44 | 0.07 | 0.29 | 0.33 | 0.37 | 0.31 |
| 4 | 755025 | rs2242233 | 1.4 | 1.144 | -1.925 | 0.51 | -0.026 | 0.071 | 0.071 | 1.703 | 0.38 | 0.06 | 0.24 | 0.2 | 0.31 | 0.26 |
| 4 | 755210 | rs2242235 | 0.711 | 1.112 | -1.718 | 0.464 | -0.002 | 0.003 | 0.003 | 0.041 | 0.08 | 0.04 | 0.09 | 0.07 | 0.08 | 0.16 |
| 4 | 756377 | rs4690309 | -0.587 | 1.439 | -1.365 | 0.46 | 0.047 | 0.083 | 0.083 | 0.035 | 0.83 | 0.69 | 0.58 | 0.58 | 0.69 | 0.73 |
| 4 | 756686 | rs6838241 | 0.019 | 1.291 | -0.806 | 0.351 | -0.003 | -0.005 | -0.005 | 0 | 0.31 | 0.29 | 0.25 | 0.23 | 0.27 | 0.33 |
| 4 | 756986 | rs13138362 | 0.012 | 1.305 | -0.96 | 0.371 | -0.018 | -0.027 | -0.027 | 0 | 0.31 | 0.57 | 0.2 | 0.18 | 0.25 | 0.25 |
| 4 | 759709 | rs6810428 | 0.829 | 1.364 | -0.967 | 0.389 | -0.03 | 0.011 | 0.011 | 0 | 0.52 | 0.16 | 0.39 | 0.4 | 0.45 | 0.48 |
| 4 | 760258 | rs7336 | -0.013 | 1.335 | -0.989 | 0.38 | -0.019 | -0.029 | -0.029 | 0 | 0.32 | 0.57 | 0.2 | 0.18 | 0.25 | 0.25 |
| 4 | 760819 | rs4690194 | 1.776 | 1.305 | -1.151 | 0.348 | -0.024 | 0.073 | 0.073 | 1.655 | 0.38 | 0.05 | 0.24 | 0.2 | 0.31 | 0.26 |
| 4 | 763077 | rs1044147 | 0.948 | 1.152 | -0.892 | 0.381 | -0.009 | -0.029 | -0.029 | 0 | 0.06 | 0.01 | 0.05 | 0.12 | 0.06 | 0.05 |
| 4 | 776911 | rs1564279 | -0.729 | 0.97 | -0.867 | 0.051 | -0.001 | -0.024 | -0.024 | 0.364 | 0.34 | 0.61 | 0.22 | 0.08 | 0.26 | 0.32 |
| 4 | 779077 | rs7340883 | 0.803 | 0.91 | -0.66 | 0.013 | -0.017 | -0.072 | -0.072 | 0.013 | 0.17 | 0.23 | 0.09 | 0 | 0.13 | 0.08 |
| 4 | 779394 | rs2242237 | -1.35 | 0.905 | -0.8 | 0.099 | 0.015 | 0.025 | 0.025 | 0 | 0.4 | 0.62 | 0.29 | 0.24 | 0.33 | 0.42 |
| 4 | 779670 | rs6816868 | -0.152 | 0.874 | -0.721 | 0.034 | -0.033 | -0.116 | -0.116 | 0.045 | 0.26 | 0.55 | 0.13 | 0.01 | 0.18 | 0.16 |
| 4 | 785041 | rs3822028 | -0.713 | 0.838 | -0.428 | 0.056 | -0.011 | -0.061 | -0.061 | 0.107 | 0.18 | 0.43 | 0.13 | 0.08 | 0.12 | 0.23 |
| 4 | 786401 | rs2306251 | 1.506 | 1.113 | -1.061 | 0.162 | -0.006 | 0.096 | 0.096 | 0.124 | 0.46 | 0.13 | 0.34 | 0.27 | 0.39 | 0.43 |
| 4 | 790757 | rs11722977 | 0.133 | 1.107 | -0.985 | 0.227 | 0.006 | 0.011 | 0.011 | 2.1 | 0.76 | 0.73 | 0.5 | 0.51 | 0.6 | 0.57 |
| 4 | 793763 | rs10004297 | 0.092 | 1.023 | -1.094 | 0.17 | -0.028 | -0.122 | -0.122 | 0 | 0.25 | 0.53 | 0.13 | 0.01 | 0.18 | 0.16 |
| 4 | 793929 | rs10004503 | 1.655 | 1.209 | -1.157 | 0.31 | -0.005 | -0.004 | -0.004 | 0 | 0.08 | 0.04 | 0.03 | 0.12 | 0.06 | 0.19 |
| 4 | 795890 | rs7677766 | -0.968 | 1.066 | -1.118 | 0.279 | -0.012 | -0.006 | -0.006 | 0.026 | 0.37 | 0.53 | 0.24 | 0.3 | 0.28 | 0.3 |
| 4 | 798511 | rs17165026 | 0.103 | 0.95 | -1.631 | 0.472 | -0.001 | -0.052 | -0.052 | 0 | 0.11 | 0.01 | 0.11 | 0.2 | 0.09 | 0.14 |
| 4 | 798962 | rs17165034 | 1.739 | 0.96 | -1.629 | 0.156 | -0.069 | 0.032 | 0.032 | 1.068 | 0.36 | 0.02 | 0.23 | 0.21 | 0.31 | 0.26 |
| 4 | 809593 | rs9328757 | 1.5 | 1.056 | -0.767 | 0.536 | 0.01 | 0.017 | 0.017 | 3.99 | 0.45 | 0.44 | 0.35 | 0.22 | 0.34 | 0.29 |
| 4 | 809787 | rs9328758 | 0.44 | 1.329 | -0.713 | 0.888 | 0.028 | 0.033 | 0.033 | 0 | 0.64 | 0.56 | 0.52 | 0.49 | 0.52 | 0.51 |
| 4 | 812753 | rs11248043 | -0.593 | 1.62 | -0.821 | 1.5 | -0.002 | -0.006 | -0.006 | 0 | 0.18 | 0.15 | 0.15 | 0.35 | 0.2 | 0.21 |
| 4 | 813726 | rs7376690 | 1.64 | 1.534 | -0.771 | 1.394 | 0.011 | 0.015 | 0.015 | 2.29 | 0.52 | 0.48 | 0.42 | 0.37 | 0.41 | 0.45 |
| 4 | 814080 | rs6832751 | -0.435 | 1.666 | -0.529 | 1.19 | -0.011 | -0.002 | -0.002 | 0.143 | 0.82 | 0.87 | 0.64 | 0.83 | 0.71 | 0.87 |
| 4 | 814719 | rs10155282 | -0.183 | 1.658 | -0.528 | 1.187 | 0.036 | 0.004 | 0.004 | 0 | 0.81 | 0.77 | 0.63 | 0.83 | 0.71 | 0.87 |
| 4 | 821490 | rs936551 | 1.177 | 1.576 | -0.662 | 1.172 | 0.024 | 0.02 | 0.02 | 1.225 | 0.62 | 0.58 | 0.45 | 0.56 | 0.53 | 0.64 |
| 4 | 831181 | rs11736866 | -0.328 | 1.723 | -1.08 | 0.746 | -0.052 | 0.072 | 0.072 | 0 | 0.2 | 0.01 | 0.1 | 0.11 | 0.2 | 0.27 |
| 4 | 832610 | rs4690321 | 1.171 | 1.661 | -0.421 | 1.089 | 0.027 | 0.044 | 0.044 | 0 | 0.72 | 0.44 | 0.57 | 0.64 | 0.62 | 0.75 |
| 4 | 843184 | rs6964 | 0.347 | 0.961 | -1.214 | 0.5 | 0.035 | 0.048 | 0.048 | 2.152 | 0.44 | 0.25 | 0.3 | 0.28 | 0.31 | 0.26 |
| 4 | 843695 | rs8093 | 0.043 | 1.469 | -1.368 | 0.763 | -0.028 | 0.006 | 0.006 | 0 | 0.8 | 0.89 | 0.64 | 0.88 | 0.67 | 0.85 |
| 4 | 848280 | rs4690340 | 1.581 | 1.113 | -1.368 | 0.736 | 0.076 | -0.018 | -0.018 | 0 | 0.33 | 0.14 | 0.22 | 0.31 | 0.16 | 0.24 |
| 4 | 854712 | rs4690339 | 1.456 | 1.091 | -1.333 | 0.725 | 0.069 | -0.008 | -0.008 | 5.271 | 0.33 | 0.15 | 0.22 | 0.32 | 0.16 | 0.32 |
| 4 | 860146 | rs2306248 | 0.633 | 1.189 | -0.694 | 0.851 | 0.053 | 0.072 | 0.072 | 0 | 0.75 | 0.58 | 0.59 | 0.66 | 0.63 | 0.83 |
| 4 | 861628 | rs3733351 | 1.016 | 1.212 | -0.617 | 0.75 | 0.045 | 0.067 | 0.067 | 0 | 0.69 | 0.5 | 0.54 | 0.6 | 0.57 | 0.76 |
| 4 | 862156 | rs2306245 | 0.645 | 1.32 | -0.617 | 0.791 | 0 | -0.009 | -0.009 | 0 | 0.33 | 0.31 | 0.27 | 0.2 | 0.37 | 0.42 |
| 4 | 864126 | rs7690704 | -1.16 | 1.482 | -0.575 | 0.996 | 0.004 | 0.008 | 0.008 | 0 | 0.93 | 0.95 | 0.71 | 0.85 | 0.82 | 0.93 |
| 4 | 867866 | rs764324 | -0.133 | 1.377 | -0.647 | 0.923 | -0.005 | 0 | 0 | 0 | 0.4 | 0.36 | 0.33 | 0.48 | 0.46 | 0.57 |
| 4 | 871675 | rs1564280 | -0.113 | 1.57 | -0.439 | 0.925 | 0.088 | 0.106 | 0.106 | 0 | 0.92 | 0.82 | 0.69 | 0.85 | 0.82 | 0.93 |
| 4 | 875887 | rs2279186 | 1.389 | 1.487 | -0.606 | 0.593 | 0.008 | -0.022 | -0.022 | 0 | 0.2 | 0.21 | 0.2 | 0.1 | 0.26 | 0.21 |
| 4 | 877412 | rs11736181 | 0.283 | 1.292 | -0.841 | 0.686 | -0.027 | -0.077 | -0.077 | 0.819 | 0.17 | 0.07 | 0.12 | 0.37 | 0.18 | 0.33 |
| 4 | 878079 | rs11946340 | -0.402 | 1.272 | -1.419 | 0.384 | -0.008 | -0.004 | -0.004 | 0.429 | 0.37 | 0.65 | 0.54 | 0.33 | 0.39 | 0.5 |
| 4 | 880094 | rs17781378 | -0.942 | 0.878 | -0.219 | 0.509 | 0.001 | -0.001 | -0.001 | 0.16 | 0.46 | 0.53 | 0.39 | 0.55 | 0.51 | 0.68 |

| | | | | | | | | | | | | | | | | |
|---|---|---|---|---|---|---|---|---|---|---|---|---|---|---|---|---|
| 4 | 882643 | rs2279181 | 1.446 | 0.792 | -0.393 | 0.485 | 0.002 | 0.015 | 0.015 | 0 | 0.13 | 0.09 | 0.06 | 0.1 | 0.12 | 0.22 |
| 4 | 886061 | rs3775127 | 2.505 | 0.603 | -0.487 | 0.501 | 0.057 | -0.079 | -0.079 | 0.134 | 0.31 | 0.03 | 0.21 | 0.31 | 0.16 | 0.19 |
| 4 | 888937 | rs11726508 | 0.738 | 0.748 | -0.321 | 0.502 | 0.001 | 0.01 | 0.01 | 0.133 | 0.16 | 0.13 | 0.1 | 0.19 | 0.15 | 0.24 |
| 4 | 891150 | rs9994159 | 1.419 | 0.748 | -0.32 | 0.498 | 0.002 | 0.017 | 0.017 | 0 | 0.13 | 0.09 | 0.06 | 0.1 | 0.12 | 0.22 |
| 4 | 892116 | rs2306252 | 0.576 | 0.741 | -0.325 | 0.503 | 0.003 | 0.01 | 0.01 | 0.124 | 0.17 | 0.13 | 0.1 | 0.2 | 0.15 | 0.32 |
| 4 | 893287 | rs3775122 | 1.626 | 0.777 | -0.374 | 0.483 | -0.025 | 0.019 | 0.019 | 0.124 | 0.12 | 0.01 | 0.05 | 0.1 | 0.12 | 0.22 |
| 4 | 894255 | rs3755963 | 0.98 | 0.84 | -0.145 | 0.467 | 0.009 | -0.023 | -0.023 | 0 | 0.23 | 0.26 | 0.24 | 0.13 | 0.29 | 0.24 |
| 4 | 896486 | rs3822023 | 0.983 | 0.725 | 0.045 | 0.474 | 0.027 | 0.016 | 0.016 | 1.745 | 0.5 | 0.43 | 0.38 | 0.56 | 0.35 | 0.43 |
| 4 | 924280 | rs754758 | 1.325 | 0.439 | 0.522 | 0.66 | 0.027 | 0.002 | 0.002 | 0 | 0.71 | 0.49 | 0.58 | 0.78 | 0.65 | 0.78 |
| 4 | 925149 | rs873785 | 0.578 | 0.601 | 0.471 | 0.688 | 0.009 | -0.011 | -0.011 | 0 | 0.26 | 0.28 | 0.29 | 0.21 | 0.37 | 0.28 |
| 4 | 928684 | rs935965 | 0.227 | 0.879 | 0.641 | 0.895 | 0.039 | 0.019 | 0.019 | 0 | 0.78 | 0.66 | 0.61 | 0.81 | 0.69 | 0.86 |
| 4 | 939883 | rs7687129 | 0.077 | 0.961 | 0.337 | 0.936 | 0.017 | 0.009 | 0.009 | 0 | 0.79 | 0.77 | 0.54 | 0.79 | 0.69 | 0.87 |
| 4 | 941290 | rs6815138 | 0.38 | 0.837 | 0.204 | 0.888 | 0.027 | 0.002 | 0.002 | 0 | 0.29 | 0.47 | 0.31 | 0.36 | 0.39 | 0.41 |
| 4 | 942006 | rs2290403 | 0.937 | 0.913 | 0.096 | 0.636 | 0.028 | -0.036 | -0.036 | 0 | 0.24 | 0.34 | 0.28 | 0.2 | 0.35 | 0.26 |
| 4 | 942439 | rs10516158 | 1.294 | 1.03 | -0.519 | 0.8 | -0.029 | 0.017 | 0.017 | 0.168 | 0.12 | 0.01 | 0.05 | 0.1 | 0.12 | 0.29 |
| 4 | 946974 | rs2290405 | 0.721 | 1.008 | -0.245 | 0.92 | 0.008 | 0.011 | 0.011 | 0 | 0.65 | 0.63 | 0.46 | 0.53 | 0.57 | 0.62 |
| 4 | 951179 | rs1051613 | -0.006 | 0.906 | 0.094 | 0.916 | -0.007 | 0.038 | 0.038 | 0 | 0.46 | 0.6 | 0.4 | 0.27 | 0.53 | 0.5 |
| 4 | 952409 | rs748483 | 1.127 | 0.84 | -0.653 | 0.61 | -0.035 | -0.006 | -0.006 | 0 | 0.12 | 0.01 | 0.06 | 0.11 | 0.12 | 0.23 |
| 4 | 953698 | rs4690326 | 0.024 | 0.856 | -0.316 | 0.983 | 0.001 | -0.002 | -0.002 | 0.283 | 0.46 | 0.51 | 0.62 | 0.5 | 0.52 | 0.53 |
| 4 | 956047 | rs6599390 | -0.797 | 0.83 | -0.615 | 1.005 | 0.001 | -0.03 | -0.03 | 0.253 | 0.63 | 0.73 | 0.71 | 0.61 | 0.65 | 0.79 |
| 4 | 960081 | rs4690324 | -0.514 | 1.531 | -0.987 | 1.328 | -0.007 | 0.001 | 0.001 | 0 | 0.95 | 0.97 | 0.85 | 0.84 | 0.84 | 0.95 |
| 4 | 964359 | rs11248060 | 0.777 | 0.919 | -0.954 | 0.771 | -0.02 | 0.019 | 0.019 | 0.042 | 0.12 | 0.02 | 0.06 | 0.11 | 0.12 | 0.24 |
| 4 | 965779 | rs11724804 | 0.301 | 0.777 | -0.841 | 0.734 | -0.001 | -0.001 | -0.001 | 0.025 | 0.37 | 0.39 | 0.54 | 0.35 | 0.44 | 0.35 |
| 4 | 973036 | rs4583705 | 0.218 | 1.046 | -0.918 | 0.835 | -0.02 | 0.016 | 0.016 | 0.037 | 0.12 | 0.02 | 0.06 | 0.11 | 0.12 | 0.24 |
| 4 | 976598 | rs17783233 | -0.137 | 1.03 | -0.912 | 0.823 | -0.009 | 0.028 | 0.028 | 0.02 | 0.14 | 0.05 | 0.07 | 0.11 | 0.13 | 0.36 |
| 4 | 977022 | rs11736451 | -0.477 | 1.033 | -0.833 | 0.872 | -0.001 | -0.005 | -0.005 | 0 | 0.16 | 0.11 | 0.08 | 0.23 | 0.17 | 0.38 |
| 4 | 989548 | rs11729777 | 0.383 | 1.173 | -0.987 | 0.615 | -0.028 | 0.02 | 0.02 | 0 | 0.11 | 0.04 | 0.06 | 0.11 | 0.13 | 0.23 |
| 4 | 992236 | rs4690222 | 0.555 | 1.336 | -0.523 | 0.569 | 0.025 | -0.007 | -0.007 | 0 | 0.69 | 0.65 | 0.69 | 0.72 | 0.55 | 0.61 |
| 4 | 994414 | rs3755955 | 0.075 | 0.222 | -0.978 | 0.755 | 0.007 | -0.016 | -0.016 | 0 | 0.19 | 0.13 | 0.09 | 0.21 | 0.17 | 0.24 |
| 4 | 995305 | rs6815946 | -0.145 | 0.117 | -0.723 | 0.702 | -0.006 | 0.001 | 0.001 | 0 | 0.22 | 0.25 | 0.11 | 0.21 | 0.17 | 0.31 |
| 4 | 1002125 | rs6855233 | -0.819 | 0.099 | -1.778 | 0.479 | -0.021 | -0.075 | -0.075 | 0 | 0.7 | 0.92 | 0.53 | 0.54 | 0.72 | 0.59 |
| 4 | 1003022 | rs4505759 | 0.495 | 0.279 | -1.266 | -0.013 | 0.013 | -0.008 | -0.008 | 0.041 | 0.25 | 0.31 | 0.28 | 0.16 | 0.3 | 0.08 |
| 4 | 1006924 | rs11734750 | 0.873 | -0.971 | -1.397 | -0.438 | 0.006 | 0.037 | 0.037 | 4.249 | 0.23 | 0.15 | 0.1 | 0.06 | 0.21 | 0.18 |
| 4 | 1009420 | rs34869253 | 0.074 | -0.934 | -1.312 | -0.253 | 0.021 | 0.092 | 0.092 | 0.261 | 0.46 | 0.28 | 0.24 | 0.23 | 0.44 | 0.54 |
| 4 | 1011115 | rs748651 | -0.512 | -0.984 | -1.468 | 0.02 | 0.01 | 0.009 | 0.009 | 0 | 0.62 | 0.57 | 0.31 | 0.46 | 0.49 | 0.55 |
| 4 | 1019011 | rs4647932 | 0.873 | -0.967 | -2.104 | -0.161 | 0.029 | 0.077 | 0.077 | 0.324 | 0.13 | 0.01 | 0.05 | 0.02 | 0.06 | 0.02 |
| 4 | 1019312 | rs4647939 | -0.053 | -1.095 | -1.905 | -0.08 | -0.004 | 0.023 | 0.023 | 0.378 | 0.56 | 0.32 | 0.27 | 0.38 | 0.48 | 0.47 |
| 4 | 1024287 | rs935973 | -0.551 | -1.444 | -2.214 | 0.166 | 0.008 | 0.001 | 0.001 | 0 | 0.36 | 0.64 | 0.22 | 0.35 | 0.29 | 0.15 |
| 4 | 1024752 | rs11722658 | 0.454 | -1.217 | -3.047 | 0.45 | -0.074 | -0.052 | -0.052 | 0.402 | 0.34 | 0.12 | 0.38 | 0.39 | 0.49 | 0.43 |
| 4 | 1026866 | rs10012074 | 0.153 | -0.864 | -2.366 | 0.427 | 0.003 | -0.003 | -0.003 | 0 | 0.18 | 0.06 | 0.1 | 0.21 | 0.18 | 0.11 |
| 4 | 1033685 | rs11941528 | -0.606 | -0.58 | -2.479 | 0.514 | 0.07 | 0.026 | 0.026 | 0.435 | 0.52 | 0.86 | 0.34 | 0.42 | 0.38 | 0.43 |
| 4 | 1045265 | rs935971 | -1.307 | -0.421 | -2.428 | 0.507 | -0.014 | -0.025 | -0.025 | 1.607 | 0.77 | 0.97 | 0.65 | 0.77 | 0.76 | 0.76 |
| 4 | 1052401 | rs2045066 | -1.12 | -0.564 | -2.291 | 0.628 | 0.014 | -0.062 | -0.062 | 0 | 0.76 | 0.92 | 0.65 | 0.65 | 0.76 | 0.75 |
| 4 | 1054159 | rs12645644 | -1.11 | -0.555 | -2.247 | 0.616 | 0.012 | -0.059 | -0.059 | 0 | 0.76 | 0.9 | 0.64 | 0.65 | 0.76 | 0.75 |
| 4 | 1054807 | rs6817989 | 0.457 | -0.854 | -1.232 | 0.196 | 0.069 | -0.056 | -0.056 | 0 | 0.1 | 0.43 | 0.1 | 0 | 0.11 | 0.08 |
| 4 | 1054868 | rs4690213 | -1.37 | -0.831 | -1.281 | 0.223 | 0.012 | -0.059 | -0.059 | 0.01 | 0.76 | 0.9 | 0.64 | 0.65 | 0.76 | 0.75 |
| 4 | 1064557 | rs11732259 | 0.245 | -0.801 | -1.041 | 0.198 | 0.059 | -0.064 | -0.064 | 0 | 0.11 | 0.44 | 0.1 | 0 | 0.11 | 0.07 |
| 4 | 1068819 | rs1010342 | -1.518 | -0.827 | -1.129 | 0.233 | 0.01 | -0.052 | -0.052 | 0 | 0.77 | 0.91 | 0.64 | 0.65 | 0.76 | 0.75 |
| 4 | 1071907 | rs1466216 | 0.67 | -1.616 | -1.62 | 0.307 | -0.077 | -0.063 | -0.063 | 0.623 | 0.36 | 0.11 | 0.37 | 0.31 | 0.49 | 0.44 |
| 4 | 1074009 | rs11732520 | -0.974 | -1.545 | -1.411 | 0.444 | -0.006 | 0 | 0 | 0 | 0.68 | 0.49 | 0.55 | 0.65 | 0.65 | 0.67 |
| 4 | 1078187 | rs1670533 | -1.97 | -1.547 | -1.451 | 0.452 | -0.017 | -0.096 | -0.096 | 0.109 | 0.77 | 0.96 | 0.65 | 0.65 | 0.76 | 0.75 |
| 4 | 1078596 | rs17164229 | 0.592 | -1.899 | -1.329 | 0.229 | 0.033 | -0.02 | -0.02 | 0 | 0.2 | 0.1 | 0.11 | 0.33 | 0.11 | 0.19 |
| 4 | 1087531 | rs615381 | 0.479 | -1.477 | -0.99 | 0.215 | 0.093 | -0.043 | -0.043 | 0 | 0.1 | 0.48 | 0.1 | 0 | 0.11 | 0.07 |
| 4 | 1090625 | rs4045481 | -0.694 | -1.541 | -0.893 | 0.212 | 0.009 | 0.004 | 0.004 | 0.946 | 0.66 | 0.3 | 0.54 | 0.65 | 0.65 | 0.67 |
| 4 | 1095281 | rs3796619 | -0.447 | -1.553 | -1.029 | 0.205 | 0.017 | 0.01 | 0.01 | 0.862 | 0.64 | 0.27 | 0.53 | 0.64 | 0.65 | 0.61 |
| 4 | 1095422 | rs10033163 | 0.862 | -0.7 | 0.418 | 0.431 | 0.043 | -0.036 | -0.036 | 0 | 0.07 | 0.14 | 0.06 | 0 | 0.11 | 0.07 |
| 4 | 1105680 | rs6840347 | -0.146 | -0.727 | 0.518 | 0.433 | 0.048 | -0.029 | -0.029 | 4.854 | 0.09 | 0.49 | 0.07 | 0 | 0.07 | 0.07 |
| 4 | 1111594 | rs6836158 | -0.396 | -0.679 | 0.505 | 0.313 | 0.047 | -0.073 | -0.073 | 0.119 | 0.12 | 0.31 | 0.11 | 0 | 0.13 | 0.1 |
| 4 | 1124038 | rs6599284 | -0.303 | 0.355 | 0.403 | 0.415 | 0.039 | -0.069 | -0.069 | 0 | 0.12 | 0.28 | 0.1 | 0 | 0.13 | 0.1 |
| 4 | 1124678 | rs7659054 | -1.481 | 0.209 | 0.607 | 0.314 | 0.03 | -0.052 | -0.052 | 0 | 0.19 | 0.43 | 0.16 | 0.15 | 0.2 | 0.23 |
| 4 | 1127010 | rs7671502 | 0.407 | 0.891 | 0.18 | 0.512 | -0.055 | -0.054 | -0.054 | 1.909 | 0.22 | 0.11 | 0.29 | 0.28 | 0.34 | 0.34 |
| 4 | 1135350 | rs4974629 | 1.688 | 0.392 | -0.093 | 0.234 | 0.045 | 0.066 | 0.066 | 0.991 | 0.61 | 0.14 | 0.71 | 0.67 | 0.7 | 0.63 |
| 4 | 1136910 | rs10014526 | 0.928 | 0.345 | 0.034 | 0.228 | 0.013 | -0.001 | -0.001 | 0 | 0.15 | 0.12 | 0.09 | 0.19 | 0.1 | 0.15 |
| 4 | 1147407 | rs11937228 | 0.739 | 0.365 | -0.018 | 0.226 | -0.021 | -0.003 | -0.003 | 0 | 0.16 | 0.21 | 0.11 | 0.18 | 0.1 | 0.15 |
| 4 | 1154202 | rs10014245 | 0.035 | 0.368 | -0.012 | 0.223 | 0.039 | -0.079 | -0.079 | 0 | 0.13 | 0.32 | 0.11 | 0 | 0.13 | 0.1 |
| 4 | 1157776 | rs10002437 | 0.311 | 0.464 | 0.114 | 0.275 | 0.032 | -0.068 | -0.068 | 0.027 | 0.12 | 0.26 | 0.1 | 0 | 0.13 | 0.1 |
| 4 | 1158991 | rs17802266 | 1.164 | 0.914 | -0.116 | 0.641 | 0.008 | -0.002 | -0.002 | 0.019 | 0.08 | 0.09 | 0.23 | 0.08 | 0.24 | 0.07 |
| 4 | 1159680 | rs7669287 | -0.572 | 0.287 | -0.04 | 0.521 | 0.022 | -0.039 | -0.039 | 0 | 0.18 | 0.36 | 0.15 | 0.14 | 0.2 | 0.22 |
| 4 | 1165130 | rs11247975 | 1.045 | 0.468 | -0.433 | 0.594 | 0.026 | 0.049 | 0.049 | 0.498 | 0.62 | 0.23 | 0.71 | 0.68 | 0.68 | 0.63 |

| | | | | | | | | | | | | | | | | |
|---|---|---|---|---|---|---|---|---|---|---|---|---|---|---|---|---|
| 4 | 1165516 | rs2242278 | 1.473 | 0.142 | -0.632 | 0.643 | -0.021 | 0.053 | 0.053 | 4.151 | 0.38 | 0.01 | 0.19 | 0.26 | 0.33 | 0.26 |
| 4 | 1192142 | rs6599290 | -0.447 | 0.245 | 0.261 | 0.854 | 0.031 | 0.05 | 0.05 | 0 | 0.1 | 0.29 | 0.09 | 0.17 | 0.1 | 0.17 |
| 4 | 1192243 | rs1250125 | 1.389 | 0.189 | -0.689 | 0.935 | -0.032 | -0.085 | -0.085 | 0.417 | 0.4 | 0.03 | 0.43 | 0.39 | 0.36 | 0.3 |
| 4 | 1193832 | rs900030 | 0.623 | -0.122 | -0.025 | 1.051 | -0.027 | 0.056 | 0.056 | 0.178 | 0.85 | 0.6 | 0.88 | 0.78 | 0.92 | 0.89 |
| 4 | 1203265 | rs1250105 | 1.282 | -0.062 | -0.405 | 0.987 | -0.011 | 0.131 | 0.131 | 1.497 | 0.45 | 0.15 | 0.26 | 0.26 | 0.41 | 0.3 |
| 4 | 1209174 | rs1265923 | -0.412 | -0.158 | 0.245 | 1.145 | -0.042 | 0.035 | 0.035 | 0 | 0.83 | 0.58 | 0.67 | 0.81 | 0.88 | 0.9 |
| 4 | 1224587 | rs1250116 | -0.418 | -0.028 | 0.308 | 1.144 | -0.022 | 0.04 | 0.04 | 0 | 0.83 | 0.56 | 0.67 | 0.82 | 0.86 | 0.91 |
| 4 | 1224648 | rs1250114 | 0.992 | -0.204 | 0.415 | 1.223 | -0.022 | 0.02 | 0.02 | 0.813 | 0.74 | 0.37 | 0.6 | 0.61 | 0.84 | 0.78 |
| 4 | 1224811 | rs4974574 | 0.181 | -0.214 | 0.295 | 1.145 | -0.111 | 0 | 0 | 0 | 0.26 | 0.09 | 0.31 | 0.19 | 0.39 | 0.35 |
| 4 | 1227469 | rs884421 | -0.138 | -0.284 | 0.562 | 1.185 | -0.035 | 0.026 | 0.026 | 0.393 | 0.82 | 0.51 | 0.66 | 0.81 | 0.86 | 0.89 |
| 4 | 1229688 | rs3733340 | 0.339 | 0.039 | 0.494 | 1.098 | -0.025 | 0.064 | 0.064 | 0.077 | 0.07 | 0.13 | 0.06 | 0.2 | 0.02 | 0.11 |
| 4 | 1236840 | rs3755931 | 0.832 | 0.012 | 0.495 | 1.103 | -0.026 | 0.054 | 0.054 | 0.145 | 0.06 | 0.09 | 0.05 | 0.2 | 0.01 | 0.1 |
| 4 | 1237014 | rs3755930 | -0.478 | -0.197 | 0.091 | 0.764 | 0.013 | 0.013 | 0.013 | 0 | 0.3 | 0.17 | 0.2 | 0.22 | 0.27 | 0.32 |
| 4 | 1237968 | rs2335855 | -0.806 | 0.071 | 1.256 | 1.273 | 0.003 | 0.002 | 0.002 | 0.053 | 0.36 | 0.36 | 0.4 | 0.41 | 0.45 | 0.48 |
| 4 | 1240091 | rs730830 | -0.194 | 0.022 | 0.897 | 1.237 | 0.007 | -0.069 | -0.069 | 0.023 | 0.88 | 0.51 | 0.9 | 0.97 | 0.89 | 0.95 |
| 4 | 1243573 | rs1680032 | -0.194 | 0.022 | 0.856 | 1.237 | 0.007 | -0.069 | -0.069 | 0 | 0.88 | 0.51 | 0.9 | 0.97 | 0.89 | 0.95 |
| 4 | 1243617 | rs3755920 | -1.155 | 0.023 | 0.804 | 1.291 | -0.008 | 0.011 | 0.011 | 0.313 | 0.4 | 0.42 | 0.66 | 0.72 | 0.5 | 0.66 |
| 4 | 1244267 | rs2291199 | -0.163 | 0.048 | 0.795 | 1.293 | -0.134 | -0.002 | -0.002 | 0 | 0.29 | 0.08 | 0.33 | 0.21 | 0.43 | 0.39 |
| 4 | 1244416 | rs1732115 | -0.141 | 0.105 | 0.812 | 1.225 | 0.003 | -0.072 | -0.072 | 0 | 0.88 | 0.51 | 0.9 | 0.97 | 0.89 | 0.95 |
| 4 | 1244879 | rs1564508 | -0.783 | 0.168 | 0.682 | 1.339 | -0.021 | -0.021 | -0.021 | 0 | 0.36 | 0.29 | 0.39 | 0.41 | 0.45 | 0.5 |
| 4 | 1285521 | rs11727167 | -0.1 | 0.499 | 0.659 | 0.779 | -0.135 | 0.099 | 0.099 | 0.008 | 0.27 | 0.08 | 0.33 | 0.1 | 0.45 | 0.34 |
| 4 | 1291337 | rs3817604 | 0.31 | 0.399 | 1.393 | 1.002 | 0.023 | -0.051 | -0.051 | 0 | 0.1 | 0.2 | 0.08 | 0.03 | 0.13 | 0.05 |
| 4 | 1304161 | rs17164620 | 0.343 | 0.367 | 1.295 | 1.108 | -0.014 | 0.199 | 0.199 | 0 | 0.05 | 0.09 | 0.28 | 0.42 | 0.03 | 0.17 |
| 4 | 1315183 | rs11933855 | -1.184 | 0.466 | 1.128 | 0.392 | -0.013 | 0.195 | 0.195 | 0.203 | 0.17 | 0.56 | 0.38 | 0.46 | 0.16 | 0.22 |
| 4 | 1320023 | rs28429103 | 0.328 | 0.509 | 0.854 | 0.328 | 0.028 | -0.051 | -0.051 | 0.233 | 0.1 | 0.23 | 0.09 | 0.03 | 0.13 | 0.05 |
| 4 | 1330759 | rs1128427 | -1.562 | 0.735 | 0.646 | 0.223 | -0.032 | -0.012 | -0.012 | 0 | 0.42 | 0.77 | 0.7 | 0.53 | 0.55 | 0.48 |
| 4 | 1330983 | rs13135102 | 0.203 | 1.092 | 0.348 | -0.303 | -0.088 | 0.063 | 0.063 | 0 | 0.21 | 0.07 | 0.29 | 0.1 | 0.38 | 0.23 |
| 4 | 1338618 | rs12647145 | -0.966 | 1.085 | 0.446 | -0.268 | 0.025 | -0.053 | -0.053 | 0.009 | 0.33 | 0.37 | 0.39 | 0.13 | 0.51 | 0.31 |
| 4 | 1339875 | rs7671242 | -0.172 | 1.213 | 0.04 | 0.001 | -0.003 | -0.003 | -0.003 | 0 | 0.22 | 0.21 | 0.31 | 0.1 | 0.37 | 0.23 |
| 4 | 1340765 | rs13141992 | 0.015 | 1.293 | -0.46 | -0.008 | -0.09 | 0.085 | 0.085 | 0 | 0.21 | 0.07 | 0.3 | 0.07 | 0.37 | 0.23 |
| 4 | 1347675 | rs6847965 | -0.081 | 1.414 | -0.299 | -0.015 | -0.01 | 0.023 | 0.023 | 0 | 0.2 | 0.18 | 0.24 | 0.09 | 0.25 | 0.21 |
| 4 | 1348248 | rs12512607 | 0.146 | 1.132 | -0.366 | -0.12 | -0.089 | 0.092 | 0.092 | 0 | 0.22 | 0.07 | 0.29 | 0.07 | 0.36 | 0.22 |
| 4 | 1350186 | rs3806754 | 0.235 | 0.761 | 0.247 | 0.439 | 0.001 | 0.36 | 0.36 | 0 | 0.07 | 0.27 | 0.29 | 0.43 | 0.03 | 0.18 |
| 4 | 1356924 | rs4974609 | -0.872 | 0.451 | 0.066 | -0.438 | 0.013 | -0.017 | -0.017 | 0 | 0.34 | 0.46 | 0.63 | 0.7 | 0.43 | 0.52 |
| 4 | 1363886 | rs10024529 | 0.345 | 0.66 | -0.445 | -0.62 | 0.019 | -0.046 | -0.046 | 0.018 | 0.09 | 0.18 | 0.08 | 0.03 | 0.13 | 0.06 |
| 4 | 1365623 | rs6826415 | 0.239 | 0.785 | -0.843 | -0.57 | -0.092 | 0.08 | 0.08 | 0 | 0.22 | 0.08 | 0.3 | 0.07 | 0.39 | 0.28 |
| 4 | 1365858 | rs6832148 | -1.318 | 0.235 | -1.347 | -1.124 | 0.016 | 0.099 | 0.099 | 1.307 | 0.64 | 0.86 | 0.83 | 0.82 | 0.72 | 0.74 |
| 4 | 1386570 | rs13142985 | -0.083 | 0.602 | -1.498 | -1.004 | -0.06 | 0.098 | 0.098 | 0.547 | 0.27 | 0.13 | 0.3 | 0.06 | 0.39 | 0.25 |
| 4 | 1387780 | rs6599308 | -1.222 | 0.089 | -1.697 | -0.96 | -0.034 | 0.049 | 0.049 | 0 | 0.63 | 0.94 | 0.8 | 0.71 | 0.67 | 0.68 |
| 4 | 1394016 | rs4974620 | -0.845 | 0.266 | -1.396 | -0.849 | 0.018 | 0.04 | 0.04 | 1.654 | 0.57 | 0.76 | 0.79 | 0.7 | 0.65 | 0.68 |
| 4 | 1415698 | rs6814036 | -0.222 | 0.693 | -2.052 | -0.993 | -0.029 | 0.028 | 0.028 | 0 | 0.72 | 0.79 | 0.66 | 0.95 | 0.55 | 0.72 |
| 4 | 1416863 | rs4419429 | 0.182 | 0.486 | -2.305 | -0.495 | -0.036 | 0.042 | 0.042 | 0.921 | 0.38 | 0.01 | 0.18 | 0.25 | 0.3 | 0.32 |
| 4 | 1500753 | rs11721614 | -0.983 | 0.812 | -1.167 | -0.607 | -0.001 | -0.043 | -0.043 | 2.998 | 0.24 | 0.65 | 0.31 | 0.08 | 0.24 | 0.34 |
| 4 | 1507198 | rs12647020 | -0.197 | 0.985 | -1.93 | -0.257 | 0.01 | -0.024 | -0.024 | 0 | 0.62 | 0.68 | 0.67 | 0.48 | 0.63 | 0.54 |
| 4 | 1519322 | rs11947258 | 0.411 | 1.051 | -1.038 | -0.266 | 0.003 | 0.002 | 0.002 | 0.107 | 0.47 | 0.44 | 0.43 | 0.39 | 0.39 | 0.23 |
| 4 | 1523930 | rs4075071 | -0.948 | 1.13 | -0.635 | -0.615 | 0.027 | -0.04 | -0.04 | 0.25 | 0.15 | 0.25 | 0.18 | 0.05 | 0.21 | 0.25 |
| 4 | 1527766 | rs7666926 | -0.797 | 0.351 | -0.922 | -0.562 | 0.008 | 0.019 | 0.019 | 0.035 | 0.33 | 0.67 | 0.36 | 0.18 | 0.25 | 0.37 |
| 4 | 1533976 | rs10029689 | -1.029 | -0.625 | -0.56 | -1.084 | -0.064 | -0.172 | -0.172 | 0.638 | 0.25 | 0.47 | 0.15 | 0.01 | 0.12 | 0.21 |
| 4 | 1539392 | rs4307023 | -0.165 | -0.542 | -1.538 | -0.894 | 0.05 | 0.04 | 0.04 | 0 | 0.51 | 0.22 | 0.44 | 0.57 | 0.4 | 0.61 |
| 4 | 1541224 | rs6822351 | -0.397 | -0.993 | -1.255 | -1.119 | -0.023 | -0.06 | -0.06 | 0 | 0.21 | 0.23 | 0.07 | 0.01 | 0.06 | 0.25 |
| 4 | 1554748 | rs6838024 | 0.777 | -0.337 | -2.049 | -0.496 | -0.05 | -0.072 | -0.072 | 0 | 0.35 | 0.06 | 0.34 | 0.58 | 0.38 | 0.32 |
| 4 | 1563920 | rs10050053 | 0.546 | -0.34 | -2.052 | -0.408 | -0.02 | 0.056 | 0.056 | 0 | 0.43 | 0.09 | 0.36 | 0.63 | 0.41 | 0.38 |
| 4 | 1566164 | rs4374681 | -0.013 | -0.39 | -1.038 | -0.579 | -0.027 | -0.075 | -0.075 | 0 | 0.17 | 0.28 | 0.07 | 0 | 0.07 | 0.2 |
| 4 | 1568595 | rs4865424 | 0.26 | 0.456 | -0.761 | -0.596 | 0.06 | 0.013 | 0.013 | 1.901 | 0.15 | 0.26 | 0.42 | 0.2 | 0.3 | 0.16 |
| 4 | 1573364 | rs4073272 | -0.819 | 0.098 | -0.916 | -0.856 | 0.007 | -0.055 | -0.055 | 0 | 0.34 | 0.47 | 0.44 | 0.15 | 0.41 | 0.36 |
| 4 | 1578967 | rs6813841 | 0.793 | 0.166 | -1.663 | -0.274 | 0 | -0.093 | -0.093 | 0 | 0.2 | 0.06 | 0.24 | 0.56 | 0.22 | 0.4 |
| 4 | 1586036 | rs6833307 | -0.257 | 0.151 | -1.193 | -0.896 | -0.005 | -0.036 | -0.036 | 0 | 0.57 | 0.5 | 0.75 | 0.82 | 0.68 | 0.6 |
| 4 | 1588665 | rs10023523 | 0.165 | 0.604 | -0.789 | -0.312 | 0.028 | -0.002 | -0.002 | 0 | 0.15 | 0.21 | 0.41 | 0.19 | 0.29 | 0.15 |
| 4 | 1598331 | rs4865438 | -0.177 | 0.403 | -1.319 | -0.006 | 0.004 | -0.007 | -0.007 | 0 | 0.33 | 0.28 | 0.3 | 0.5 | 0.29 | 0.42 |
| 4 | 1610872 | rs6858080 | -0.249 | 0.65 | -1.941 | -0.674 | -0.047 | 0.097 | 0.097 | 0 | 0.54 | 0.86 | 0.4 | 0.32 | 0.42 | 0.41 |
| 4 | 1619530 | rs3936194 | -0.456 | -0.15 | -1.62 | -1.49 | -0.017 | -0.007 | -0.007 | 0.477 | 0.15 | 0.09 | 0.35 | 0.19 | 0.26 | 0.13 |
| 4 | 1631795 | rs2166582 | 0.388 | -0.384 | -1.276 | -1.931 | -0.007 | 0.156 | 0.156 | 0 | 0.32 | 0.09 | 0.17 | 0.01 | 0.29 | 0.25 |
| 4 | 1635312 | rs7667125 | -0.479 | -0.245 | -1.238 | -1.814 | 0.013 | 0.002 | 0.002 | 0 | 0.52 | 0.58 | 0.34 | 0.11 | 0.37 | 0.42 |
| 4 | 1647743 | rs4865453 | -0.107 | 0.369 | -1.746 | -1.194 | -0.013 | -0.006 | -0.006 | 0.059 | 0.09 | 0.06 | 0.3 | 0.11 | 0.24 | 0.11 |
| 4 | 1648626 | rs2044507 | -0.716 | -0.268 | -1.661 | -1.086 | 0.012 | 0.037 | 0.037 | 0 | 0.12 | 0.03 | 0.06 | 0.05 | 0.09 | 0.13 |
| 4 | 1684409 | rs6852065 | 0.408 | -0.19 | -2.027 | -1.281 | -0.057 | 0.087 | 0.087 | 0.729 | 0.47 | 0.07 | 0.6 | 0.71 | 0.55 | 0.68 |
| 4 | 1684457 | rs6599397 | 0.1 | -0.398 | -0.907 | -1.851 | -0.008 | -0.055 | -0.055 | 0 | 0.61 | 0.38 | 0.77 | 0.94 | 0.74 | 0.77 |
| 4 | 1693931 | rs3958122 | 0.679 | -0.094 | -0.984 | -1.791 | -0.015 | 0.031 | 0.031 | 0.106 | 0.35 | 0.13 | 0.37 | 0.69 | 0.33 | 0.51 |
| 4 | 1701317 | rs2247341 | 0.679 | -0.094 | -0.964 | -1.791 | -0.014 | 0.031 | 0.031 | 0.106 | 0.35 | 0.13 | 0.37 | 0.69 | 0.33 | 0.51 |

| | | | | | | | | | | | | | | | | |
|---|---|---|---|---|---|---|---|---|---|---|---|---|---|---|---|---|
| 4 | 1707447 | rs798756 | -0.595 | -0.194 | -0.479 | -2.029 | -0.005 | -0.022 | -0.022 | 0.068 | 0.76 | 0.66 | 0.71 | 0.85 | 0.78 | 0.79 |
| 4 | 1720757 | rs798754 | -0.192 | -0.234 | 0.106 | -2.048 | -0.016 | -0.049 | -0.049 | 2.343 | 0.73 | 0.37 | 0.67 | 0.85 | 0.78 | 0.77 |
| 4 | 1721654 | rs2854915 | -0.257 | -0.248 | 0.222 | -2.069 | -0.005 | 0.014 | 0.014 | 0 | 0.39 | 0.34 | 0.32 | 0.16 | 0.44 | 0.29 |
| 4 | 1724579 | rs1374468 | 0.704 | -0.342 | 0.151 | -2.222 | -0.004 | -0.06 | -0.06 | 0.34 | 0.17 | 0.2 | 0.12 | 0.01 | 0.18 | 0.16 |
| 4 | 1728120 | rs2282763 | 0.538 | -0.343 | 0.162 | -2.223 | -0.021 | 0.019 | 0.019 | 0.287 | 0.22 | 0.15 | 0.2 | 0.15 | 0.26 | 0.13 |
| 4 | 1729660 | rs1063742 | 0.55 | -0.332 | 0.114 | -2.218 | -0.024 | 0.02 | 0.02 | 0.277 | 0.22 | 0.13 | 0.2 | 0.15 | 0.26 | 0.13 |
| 4 | 1729988 | rs1063743 | 0.55 | -0.332 | 0.114 | -2.218 | -0.024 | 0.02 | 0.02 | 0.277 | 0.22 | 0.13 | 0.2 | 0.15 | 0.26 | 0.13 |
| 4 | 1730299 | rs798759 | -0.546 | -0.295 | -0.028 | -2.2 | -0.007 | 0.025 | 0.025 | 0 | 0.39 | 0.32 | 0.32 | 0.16 | 0.44 | 0.29 |
| 4 | 1732978 | rs17680881 | 0.561 | -0.289 | -0.137 | -2.195 | -0.024 | 0.02 | 0.02 | 0.258 | 0.22 | 0.13 | 0.2 | 0.15 | 0.26 | 0.13 |
| 4 | 1737262 | rs3752749 | -0.539 | -0.257 | -0.089 | -1.732 | -0.002 | -0.011 | -0.011 | 0 | 0.77 | 0.69 | 0.71 | 0.85 | 0.79 | 0.78 |
| 4 | 1737502 | rs11248073 | 0.828 | -0.117 | -0.864 | -1.664 | -0.021 | 0.023 | 0.023 | 0 | 0.34 | 0.13 | 0.37 | 0.69 | 0.34 | 0.49 |
| 4 | 1742400 | rs4865463 | -0.255 | -0.102 | -0.847 | -1.663 | -0.026 | -0.082 | -0.082 | 0 | 0.73 | 0.46 | 0.69 | 0.85 | 0.78 | 0.77 |
| 4 | 1744252 | rs2403293 | -0.593 | 0.32 | -0.548 | -1.307 | -0.005 | 0.014 | 0.014 | 0 | 0.39 | 0.34 | 0.32 | 0.16 | 0.44 | 0.29 |
| 4 | 1770357 | rs4865468 | 1.178 | 0.033 | -0.203 | -0.922 | -0.03 | -0.066 | -0.066 | 0 | 0.85 | 0.27 | 0.89 | 0.98 | 0.94 | 0.88 |
| 4 | 1785738 | rs1867926 | -0.246 | -0.075 | -0.322 | -1.658 | -0.01 | -0.052 | -0.052 | 0 | 0.45 | 0.46 | 0.31 | 0.11 | 0.34 | 0.3 |
| 4 | 1786702 | rs7668207 | -1.063 | 1.122 | 0.114 | -1.137 | 0.05 | 0.02 | 0.02 | 0.005 | 0.1 | 0.2 | 0.15 | 0.12 | 0.19 | 0.05 |
| 4 | 1797741 | rs3135848 | -0.721 | 0.441 | -0.424 | -1.964 | 0.08 | 0.157 | 0.157 | 0.582 | 0.46 | 0.77 | 0.32 | 0.12 | 0.32 | 0.31 |
| 4 | 1797852 | rs743682 | -1.215 | 0.556 | 0.895 | -1.786 | 0.077 | 0.056 | 0.056 | 0 | 0.09 | 0.39 | 0.14 | 0.06 | 0.09 | 0.06 |
| 4 | 1815990 | rs4865477 | -0.435 | 0.076 | 1.621 | -1.992 | 0.002 | -0.001 | -0.001 | 0 | 0.93 | 0.66 | 0.97 | 0.98 | 0.99 | 0.99 |
| 4 | 1864994 | rs12503340 | 1.218 | -0.513 | 0.785 | -2.726 | 0.081 | 0.127 | 0.127 | 0 | 0.63 | 0.2 | 0.72 | 0.98 | 0.76 | 0.71 |
| 4 | 1866358 | rs4865483 | 0.664 | -0.437 | 1.568 | -2.611 | -0.035 | -0.109 | -0.109 | 2.021 | 0.76 | 0.64 | 0.81 | 0.99 | 0.84 | 0.84 |
| 4 | 1958171 | rs494893 | -0.192 | -0.151 | 1.404 | -2.396 | -0.014 | -0.076 | -0.076 | 0 | 0.84 | 0.79 | 0.83 | 1 | 0.89 | 0.87 |
| 4 | 1986938 | rs3135123 | 0.088 | -0.086 | 0.856 | -2.785 | 0.016 | 0.109 | 0.109 | 0 | 0.46 | 0.73 | 0.47 | 0.91 | 0.45 | 0.6 |
| 4 | 1986972 | rs495367 | -0.685 | 0.473 | 0.739 | -1.542 | 0.093 | 0.088 | 0.088 | 0.127 | 0.2 | 0.48 | 0.41 | 0.29 | 0.31 | 0.23 |
| 4 | 1988193 | rs1077020 | -0.962 | -0.341 | 1.381 | -1.021 | 0.05 | 0.115 | 0.115 | 0 | 0.16 | 0.39 | 0.24 | 0.26 | 0.19 | 0.12 |
| 4 | 1990425 | rs11725821 | 1.029 | -0.418 | 1.146 | -1.012 | 0.027 | -0.039 | -0.039 | 0.143 | 0.32 | 0.1 | 0.23 | 0.64 | 0.28 | 0.51 |
| 4 | 1993985 | rs580602 | -1.601 | -0.27 | 1.707 | -0.022 | -0.001 | -0.006 | -0.006 | 0 | 0.26 | 0.23 | 0.26 | 0.34 | 0.24 | 0.21 |
| 4 | 1994985 | rs550483 | -0.883 | -0.128 | 1.635 | 0.355 | -0.019 | -0.055 | -0.055 | 1.049 | 0.09 | 0.01 | 0.09 | 0.22 | 0.1 | 0.09 |
| 4 | 2054373 | rs7375177 | 1.989 | -0.382 | 2.213 | -0.075 | -0.033 | 0.02 | 0.02 | 0.086 | 0.75 | 0.43 | 0.78 | 0.72 | 0.86 | 0.77 |
| 4 | 2058475 | rs382939 | 1.901 | -0.659 | 2.308 | 0.083 | -0.042 | 0.029 | 0.029 | 0.046 | 0.77 | 0.4 | 0.8 | 0.7 | 0.89 | 0.78 |
| 4 | 2068337 | rs414210 | 2.114 | -0.637 | 2.309 | 0.011 | -0.015 | 0.003 | 0.003 | 1.713 | 0.75 | 0.29 | 0.79 | 0.7 | 0.89 | 0.78 |
| 4 | 2078853 | rs3117813 | 1.909 | -0.514 | 2.222 | 0.091 | -0.035 | 0.019 | 0.019 | 0.59 | 0.77 | 0.35 | 0.8 | 0.7 | 0.89 | 0.77 |
| 4 | 2087202 | rs3135157 | 1.494 | -0.578 | 2.211 | 0.141 | -0.044 | 0.049 | 0.049 | 0 | 0.8 | 0.66 | 0.83 | 0.72 | 0.91 | 0.79 |
| 4 | 2103096 | rs1923775 | 2.488 | -0.637 | 1.981 | 0.079 | 0.016 | 0.008 | 0.008 | 1.092 | 0.65 | 0.17 | 0.72 | 0.71 | 0.72 | 0.64 |
| 4 | 2106836 | rs3135056 | 1.285 | -0.84 | 1.936 | 0.483 | -0.036 | 0.053 | 0.053 | 0.061 | 0.83 | 0.71 | 0.85 | 0.72 | 0.92 | 0.79 |
| 4 | 2139047 | rs554399 | 1.219 | -0.462 | 1.499 | 0.478 | -0.032 | 0.116 | 0.116 | 0 | 0.88 | 0.66 | 0.81 | 0.75 | 0.95 | 0.84 |
| 4 | 2140742 | rs482519 | 1.312 | -0.538 | 1.55 | 0.39 | -0.026 | 0.06 | 0.06 | 0 | 0.85 | 0.76 | 0.83 | 0.66 | 0.93 | 0.85 |
| 4 | 2145720 | rs1745335 | 1.533 | -0.821 | 1.841 | 0.2 | -0.064 | 0.074 | 0.074 | 0.14 | 0.76 | 0.47 | 0.76 | 0.64 | 0.88 | 0.81 |
| 4 | 2160894 | rs10488836 | -0.612 | -0.784 | 1.912 | 0.232 | -0.018 | 0.109 | 0.109 | 0 | 0.12 | 0.26 | 0.17 | 0.34 | 0.07 | 0.14 |
| 4 | 2176454 | rs9328764 | -1.978 | -0.846 | 1.719 | 0.159 | -0.061 | 0.061 | 0.061 | 0.307 | 0.25 | 0.57 | 0.24 | 0.37 | 0.12 | 0.19 |
| 4 | 2181080 | rs6830513 | -1.978 | -0.846 | 1.719 | 0.159 | -0.061 | 0.061 | 0.061 | 0.307 | 0.25 | 0.57 | 0.24 | 0.37 | 0.12 | 0.19 |
| 4 | 2194946 | rs10011549 | -1.886 | -0.868 | 1.796 | 0.127 | -0.049 | 0.038 | 0.038 | 0.692 | 0.25 | 0.6 | 0.25 | 0.37 | 0.12 | 0.19 |
| 4 | 2195024 | rs10018786 | -1.886 | -0.868 | 1.796 | 0.127 | -0.049 | 0.038 | 0.038 | 0.692 | 0.25 | 0.6 | 0.25 | 0.37 | 0.12 | 0.19 |
| 4 | 2210065 | rs2353552 | -0.065 | -0.441 | 1.641 | 0.014 | -0.039 | 0.029 | 0.029 | 0 | 0.07 | 0.01 | 0.07 | 0 | 0.13 | 0.03 |
| 4 | 2214418 | rs6855461 | -2.045 | -0.96 | 2.158 | 0.234 | -0.011 | 0.005 | 0.005 | 2.244 | 0.25 | 0.72 | 0.26 | 0.37 | 0.13 | 0.18 |
| 4 | 2221814 | rs7677712 | -2.087 | -0.973 | 2.187 | 0.217 | -0.034 | 0.029 | 0.029 | 1.251 | 0.25 | 0.64 | 0.25 | 0.38 | 0.12 | 0.19 |
| 4 | 2231600 | rs6851446 | -2.015 | -1.006 | 2.166 | 0.221 | -0.049 | 0.045 | 0.045 | 0.817 | 0.24 | 0.6 | 0.24 | 0.37 | 0.12 | 0.18 |
| 4 | 2233709 | rs11937432 | -0.267 | -1.017 | 2.332 | 0.218 | -0.008 | -0.011 | -0.011 | 4.663 | 0.1 | 0.15 | 0.06 | 0.03 | 0.05 | 0.04 |
| 4 | 2235690 | rs1179 | -2.073 | -1.014 | 2.281 | 0.22 | -0.049 | 0.044 | 0.044 | 0.721 | 0.25 | 0.6 | 0.24 | 0.37 | 0.12 | 0.19 |
| 4 | 2244865 | rs7660498 | -0.414 | -0.904 | 2.274 | 0.296 | -0.022 | -0.027 | -0.027 | 1.331 | 0.11 | 0.25 | 0.07 | 0.03 | 0.05 | 0.04 |
| 4 | 2249111 | rs762832 | -0.72 | -1.201 | 1.743 | 0.049 | -0.03 | 0.035 | 0.035 | 0 | 0.2 | 0.31 | 0.19 | 0.34 | 0.11 | 0.14 |
| 4 | 2250109 | rs880674 | -1.539 | -1.091 | 0.71 | 0.096 | 0.038 | -0.009 | -0.009 | 0.124 | 0.33 | 0.77 | 0.28 | 0.37 | 0.17 | 0.18 |
| 4 | 2250742 | rs3811803 | -1.378 | -1.084 | 0.702 | 0.099 | -0.025 | 0.004 | 0.004 | 0.678 | 0.32 | 0.63 | 0.26 | 0.37 | 0.17 | 0.18 |
| 4 | 2252201 | rs1052682 | -1.837 | -1.187 | 0.289 | -0.001 | 0.101 | -0.002 | -0.002 | 0 | 0.38 | 0.86 | 0.29 | 0.37 | 0.2 | 0.24 |
| 4 | 2256716 | rs2236998 | -1.56 | -1.074 | 0.334 | 0.213 | 0.032 | -0.008 | -0.008 | 0 | 0.34 | 0.75 | 0.28 | 0.38 | 0.17 | 0.19 |
| 4 | 2261720 | rs2182683 | -1.094 | -0.555 | 0.49 | 0.483 | -0.054 | 0.06 | 0.06 | 0.484 | 0.23 | 0.5 | 0.21 | 0.34 | 0.13 | 0.15 |
| 4 | 2275145 | rs736455 | -0.145 | -0.13 | -0.031 | -0.579 | -0.007 | 0.043 | 0.043 | 0.009 | 0.45 | 0.36 | 0.51 | 0.12 | 0.54 | 0.29 |
| 4 | 2275835 | rs2051561 | -0.149 | -0.138 | -0.034 | -0.582 | -0.007 | 0.044 | 0.044 | 0.012 | 0.45 | 0.36 | 0.51 | 0.12 | 0.54 | 0.29 |
| 4 | 2277497 | rs4974666 | -0.648 | -0.729 | 0.061 | -0.388 | -0.03 | 0.06 | 0.06 | 0 | 0.59 | 0.85 | 0.62 | 0.31 | 0.58 | 0.5 |
| 4 | 2282523 | rs17132396 | -0.937 | -0.838 | 1.146 | -0.22 | -0.02 | 0.092 | 0.092 | 0.839 | 0.08 | 0.3 | 0.08 | 0.19 | 0.02 | 0.25 |
| 4 | 2287421 | rs11737692 | -0.647 | -0.751 | 0.364 | -0.476 | -0.038 | 0.035 | 0.035 | 0 | 0.62 | 0.92 | 0.66 | 0.31 | 0.6 | 0.57 |
| 4 | 2294746 | rs7681318 | -0.19 | -0.704 | 0.743 | 0.015 | 0.001 | 0.018 | 0.018 | 0 | 0.56 | 0.56 | 0.59 | 0.21 | 0.58 | 0.4 |
| 4 | 2297550 | rs13105288 | -0.649 | -0.18 | 0.094 | -0.034 | 0.008 | -0.034 | -0.034 | 11.258 | 0.63 | 0.85 | 0.7 | 0.41 | 0.64 | 0.6 |
| 4 | 2297586 | rs13120062 | -0.069 | -0.334 | -0.58 | -0.508 | -0.065 | 0.047 | 0.047 | 0.721 | 0.4 | 0.15 | 0.51 | 0.31 | 0.47 | 0.41 |
| 4 | 2303099 | rs12640733 | 0.49 | 0 | -0.901 | -0.679 | -0.09 | 0.095 | 0.095 | 0 | 0.27 | 0.09 | 0.35 | 0.11 | 0.38 | 0.2 |
| 4 | 2303576 | rs17768393 | -0.206 | -0.099 | -0.364 | -0.403 | -0.007 | 0.014 | 0.014 | 0 | 0.13 | 0.1 | 0.08 | 0.05 | 0.13 | 0.1 |
| 4 | 2304007 | rs2071664 | -0.901 | -0.194 | 0.386 | -0.009 | 0.022 | 0.258 | 0.258 | 0.019 | 0.06 | 0.28 | 0.09 | 0.19 | 0.01 | 0.16 |
| 4 | 2304946 | rs612428 | -0.048 | 0.34 | -0.836 | -0.36 | -0.013 | -0.006 | -0.006 | 0 | 0.87 | 0.93 | 0.82 | 0.84 | 0.83 | 0.87 |
| 4 | 2305634 | rs13126907 | 0.033 | 0.63 | -1.435 | -0.965 | -0.011 | 0.059 | 0.059 | 0.042 | 0.59 | 0.26 | 0.6 | 0.88 | 0.56 | 0.79 |

| | | | | | | | | | | | | | | | | |
|---|---|---|---|---|---|---|---|---|---|---|---|---|---|---|---|---|
| 4 | 2306053 | rs661301 | 0.321 | 0.561 | -1.152 | -0.392 | -0.009 | 0 | 0 | 0 | 0.77 | 0.65 | 0.76 | 0.68 | 0.83 | 0.72 |
| 4 | 2307466 | rs4974648 | 0.127 | 1.282 | -1.587 | -0.232 | -0.054 | -0.004 | -0.004 | 0 | 0.15 | 0.02 | 0.2 | 0.14 | 0.23 | 0.16 |
| 4 | 2307574 | rs743695 | 0.889 | 1.167 | -1.345 | -0.046 | -0.007 | 0.019 | 0.019 | 0 | 0.31 | 0.45 | 0.29 | 0.33 | 0.26 | 0.32 |
| 4 | 2311361 | rs4974674 | -0.494 | 0.969 | -1.402 | -0.819 | 0.003 | -0.058 | -0.058 | 0 | 0.56 | 0.57 | 0.44 | 0.13 | 0.52 | 0.39 |
| 4 | 2316874 | rs4479771 | 0.1 | 0.527 | -1.595 | -0.985 | 0 | -0.04 | -0.04 | 0 | 0.23 | 0.28 | 0.14 | 0 | 0.22 | 0.15 |
| 4 | 2324865 | rs654207 | 0.324 | 0.739 | -1.953 | -0.879 | -0.064 | 0.082 | 0.082 | 0 | 0.29 | 0.06 | 0.3 | 0.14 | 0.33 | 0.26 |
| 4 | 2329295 | rs2097298 | 0.099 | 0.141 | -1.52 | -0.942 | -0.017 | -0.06 | -0.06 | 0.581 | 0.2 | 0.29 | 0.12 | 0 | 0.18 | 0.12 |
| 4 | 2335396 | rs34426652 | -0.573 | 0.588 | -1.561 | -0.172 | 0.005 | -0.003 | -0.003 | 0 | 0.58 | 0.42 | 0.59 | 0.49 | 0.65 | 0.49 |
| 4 | 2336825 | rs2071684 | 0.462 | 0.531 | -1.763 | 0.397 | -0.015 | -0.117 | -0.117 | 0.569 | 0.15 | 0.01 | 0.28 | 0.48 | 0.12 | 0.22 |
| 4 | 2340417 | rs3118617 | -0.555 | 0.522 | -0.998 | 0.086 | -0.002 | -0.003 | -0.003 | 0 | 0.54 | 0.54 | 0.69 | 0.7 | 0.55 | 0.52 |
| 4 | 2343851 | rs2071678 | -0.473 | 1.072 | -0.973 | 0.42 | -0.04 | -0.128 | -0.128 | 0.432 | 0.19 | 0.03 | 0.32 | 0.5 | 0.22 | 0.25 |
| 4 | 2344374 | rs7663204 | 0.148 | 1.71 | -0.531 | 0.251 | 0.184 | 0.007 | 0.007 | 0.086 | 0.05 | 0.25 | 0.08 | 0 | 0.13 | 0.04 |
| 4 | 2344692 | rs6599427 | -0.903 | 1.085 | -0.344 | 0.272 | 0.033 | 0.064 | 0.064 | 0 | 0.27 | 0.56 | 0.44 | 0.57 | 0.35 | 0.29 |
| 4 | 2350919 | rs1179042 | 0.698 | 0.145 | -1.008 | -0.228 | 0.033 | 0.101 | 0.101 | 0 | 0.2 | 0.04 | 0.13 | 0 | 0.12 | 0.13 |
| 4 | 2351350 | rs11731588 | -0.121 | 0.25 | -0.293 | 0.459 | 0.019 | 0.126 | 0.126 | 0 | 0.08 | 0.22 | 0.09 | 0.21 | 0.1 | 0.05 |
| 4 | 2359802 | rs3135127 | 0.376 | 0.157 | -0.432 | 0.553 | 0.009 | -0.001 | -0.001 | 0.395 | 0.56 | 0.34 | 0.68 | 0.6 | 0.56 | 0.52 |
| 4 | 2367744 | rs3135143 | 1.076 | 0.117 | -0.658 | 0.583 | 0.016 | 0.041 | 0.041 | 0 | 0.28 | 0.18 | 0.25 | 0.19 | 0.2 | 0.15 |
| 4 | 2369253 | rs1894433 | -0.47 | 0.077 | -0.297 | 0.609 | -0.017 | -0.025 | -0.025 | 0 | 0.65 | 0.56 | 0.78 | 0.78 | 0.7 | 0.59 |
| 4 | 2371083 | rs10011996 | 1.092 | 0.219 | -0.812 | 0.636 | -0.023 | -0.014 | -0.014 | 0 | 0.1 | 0 | 0.1 | 0.09 | 0.13 | 0.15 |
| 4 | 2385384 | rs4257708 | 0.819 | 0.216 | -0.719 | 0.782 | -0.034 | -0.055 | -0.055 | 0 | 0.29 | 0.15 | 0.44 | 0.51 | 0.37 | 0.38 |
| 4 | 2386411 | rs13129389 | 1.363 | 0.344 | -0.56 | 0.91 | 0.013 | 0.027 | 0.027 | 0 | 0.05 | 0.08 | 0.07 | 0.11 | 0.11 | 0.05 |
| 4 | 2390303 | rs3135093 | 0.495 | -0.324 | -0.493 | 0.603 | -0.004 | -0.004 | -0.004 | 0.58 | 0.61 | 0.38 | 0.77 | 0.7 | 0.68 | 0.57 |
| 4 | 2392839 | rs9994096 | 0.61 | 0.007 | -0.663 | 0.648 | 0 | 0.001 | 0.001 | 0 | 0.11 | 0.11 | 0.12 | 0.09 | 0.14 | 0.14 |
| 4 | 2401347 | rs9996449 | -0.817 | -0.087 | -0.732 | 0.277 | 0.007 | 0.013 | 0.013 | 0 | 0.81 | 0.87 | 0.92 | 0.87 | 0.84 | 0.74 |
| 4 | 2402706 | rs875130 | 0.73 | -0.185 | -0.676 | 0.241 | 0.009 | 0.038 | 0.038 | 0 | 0.32 | 0.19 | 0.26 | 0.14 | 0.27 | 0.19 |
| 4 | 2403365 | rs875133 | -0.32 | -0.045 | -0.526 | 0.611 | 0.007 | 0.013 | 0.013 | 0 | 0.81 | 0.87 | 0.92 | 0.87 | 0.85 | 0.74 |
| 4 | 2406321 | rs3135071 | -0.171 | -0.02 | -0.467 | 0.68 | -0.005 | -0.003 | -0.003 | 0 | 0.85 | 0.82 | 0.95 | 0.88 | 0.91 | 0.78 |
| 4 | 2408397 | rs10009152 | -0.705 | 0.152 | -0.293 | 1.008 | -0.002 | 0.048 | 0.048 | 0 | 0.17 | 0.21 | 0.18 | 0.34 | 0.17 | 0.29 |
| 4 | 2408411 | rs12499021 | 0.547 | 0.664 | -0.943 | 0.819 | -0.033 | -0.071 | -0.071 | 0 | 0.26 | 0.12 | 0.49 | 0.54 | 0.3 | 0.39 |
| 4 | 2428463 | rs2269493 | 0.461 | 0.648 | -0.933 | 0.791 | -0.027 | -0.075 | -0.075 | 0 | 0.27 | 0.13 | 0.49 | 0.54 | 0.3 | 0.4 |
| 4 | 2437290 | rs1203764 | -0.501 | 0.388 | -0.793 | 0.111 | -0.019 | 0.057 | 0.057 | 0.487 | 0.42 | 0.28 | 0.65 | 0.88 | 0.47 | 0.69 |
| 4 | 2446532 | rs3128843 | 0.348 | 0.061 | -0.467 | 0.598 | -0.005 | -0.001 | -0.001 | 0 | 0.72 | 0.72 | 0.81 | 0.72 | 0.78 | 0.67 |
| 4 | 2452087 | rs1203808 | 0.226 | 0.429 | -0.389 | 0.189 | 0.015 | -0.026 | -0.026 | 0.488 | 0.23 | 0.34 | 0.17 | 0.11 | 0.26 | 0.17 |
| 4 | 2458078 | rs9990857 | -0.257 | 0.274 | -0.621 | 0.211 | -0.009 | -0.004 | -0.004 | 0 | 0.26 | 0.28 | 0.19 | 0.26 | 0.22 | 0.33 |
| 4 | 2465304 | rs846252 | 0.41 | 0.738 | -0.446 | 0.648 | 0.044 | 0.021 | 0.021 | 0.115 | 0.71 | 0.55 | 0.78 | 0.69 | 0.61 | 0.78 |
| 4 | 2470613 | rs704347 | -0.771 | 0.887 | -0.814 | 0.49 | -0.007 | -0.002 | -0.002 | 0.002 | 0.85 | 0.88 | 0.89 | 0.84 | 0.84 | 0.88 |
| 4 | 2471236 | rs13128459 | 0.222 | -0.273 | -1.467 | -0.168 | 0.083 | 0.103 | 0.103 | 1.085 | 0.2 | 0.03 | 0.07 | 0.01 | 0.05 | 0.18 |
| 4 | 2471802 | rs704349 | -1.076 | 0.026 | -1.382 | -0.042 | -0.007 | -0.004 | -0.004 | 0.017 | 0.85 | 0.88 | 0.89 | 0.83 | 0.84 | 0.87 |
| 4 | 2471974 | rs4974695 | 1.158 | -0.188 | -2.073 | 0.118 | -0.099 | -0.072 | -0.072 | 2.791 | 0.29 | 0.07 | 0.55 | 0.39 | 0.37 | 0.37 |
| 4 | 2478521 | rs846238 | 1.22 | -0.616 | -2.09 | 0.062 | 0.013 | 0.041 | 0.041 | 0 | 0.23 | 0.07 | 0.16 | 0.18 | 0.26 | 0.25 |
| 4 | 2481490 | rs11248108 | 0.095 | -0.411 | -1.26 | 0.281 | 0.013 | 0.053 | 0.053 | 0 | 0.13 | 0.21 | 0.07 | 0.24 | 0.16 | 0.09 |
| 4 | 2491695 | rs17132647 | 1.792 | 0.449 | -1.528 | 0.216 | -0.03 | -0.013 | -0.013 | 0 | 0.05 | 0 | 0.11 | 0.12 | 0.09 | 0.12 |
| 4 | 2499117 | rs3749467 | 1.277 | -0.057 | -1.334 | 0.228 | -0.078 | -0.038 | -0.038 | 0 | 0.29 | 0.09 | 0.56 | 0.37 | 0.36 | 0.37 |
| 4 | 2502637 | rs1203863 | -0.057 | -0.177 | -1.19 | 0.311 | 0.03 | 0.15 | 0.15 | 0.762 | 0.13 | 0.28 | 0.09 | 0.39 | 0.19 | 0.25 |
| 4 | 2503587 | rs11935873 | 0.775 | -0.43 | -1.34 | -0.039 | 0.003 | -0.001 | -0.001 | 0.099 | 0.08 | 0.04 | 0.03 | 0 | 0.03 | 0.06 |
| 4 | 2504910 | rs1203861 | 1.834 | -0.408 | -1.439 | -0.023 | -0.017 | 0.02 | 0.02 | 0 | 0.18 | 0.01 | 0.13 | 0.16 | 0.23 | 0.21 |
| 4 | 2505091 | rs1203860 | -0.576 | -0.123 | -1.038 | 0.157 | 0 | 0 | 0 | 0 | 0.26 | 0.3 | 0.16 | 0.28 | 0.23 | 0.32 |
| 4 | 2626569 | rs3864117 | 1.428 | 0.724 | -1.224 | -0.519 | -0.086 | -0.099 | -0.099 | 0 | 0.3 | 0.07 | 0.54 | 0.39 | 0.37 | 0.43 |
| 4 | 2631009 | rs17824073 | 0.748 | 0.659 | -1.214 | -0.723 | 0.039 | 0.09 | 0.09 | 0.369 | 0.34 | 0.15 | 0.2 | 0.21 | 0.33 | 0.31 |
| 4 | 2636881 | rs17824295 | 1.471 | 0.696 | -1.145 | -0.713 | 0.016 | 0.032 | 0.032 | 0 | 0.09 | 0.02 | 0.04 | 0.01 | 0.04 | 0.09 |
| 4 | 2658327 | rs2282766 | 1.473 | 0.724 | -1.188 | -0.681 | -0.087 | -0.093 | -0.093 | 0 | 0.31 | 0.07 | 0.54 | 0.39 | 0.37 | 0.43 |
| 4 | 2674271 | rs17825119 | 0.164 | 0.77 | -0.532 | -0.755 | -0.002 | 0.001 | 0.001 | 0.06 | 0.47 | 0.58 | 0.66 | 0.41 | 0.45 | 0.54 |
| 4 | 2680515 | rs7681728 | 1.31 | 0.94 | -0.789 | -0.67 | -0.108 | -0.097 | -0.097 | 4.298 | 0.32 | 0.07 | 0.56 | 0.39 | 0.41 | 0.44 |
| 4 | 2687521 | rs231704 | -0.336 | 0.902 | -0.621 | -0.641 | 0 | 0.004 | 0.004 | 0 | 0.8 | 0.79 | 0.85 | 0.68 | 0.77 | 0.86 |
| 4 | 2689449 | rs1828803 | 1.697 | 0.34 | -0.703 | -0.753 | -0.136 | -0.146 | -0.146 | 0 | 0.27 | 0.02 | 0.53 | 0.38 | 0.36 | 0.42 |
| 4 | 2694385 | rs231707 | -0.535 | 0.679 | -0.277 | -0.569 | 0.003 | 0.027 | 0.027 | 0 | 0.83 | 0.79 | 0.89 | 0.69 | 0.82 | 0.88 |
| 4 | 2697300 | rs4690085 | 0.775 | 0.634 | -0.092 | -0.578 | 0.01 | 0.004 | 0.004 | 1.384 | 0.37 | 0.45 | 0.62 | 0.4 | 0.44 | 0.47 |
| 4 | 2703081 | rs12499320 | 1.525 | 0.644 | -0.149 | -0.61 | -0.062 | -0.075 | -0.075 | 0 | 0.32 | 0.13 | 0.55 | 0.4 | 0.37 | 0.43 |
| 4 | 2703101 | rs231697 | 0.769 | 0.529 | -0.139 | -0.702 | 0.029 | 0.06 | 0.06 | 0.379 | 0.33 | 0.14 | 0.2 | 0.28 | 0.33 | 0.32 |
| 4 | 2707032 | rs4690015 | 1.166 | 0.495 | -0.159 | -0.741 | 0.048 | 0.035 | 0.035 | 0.787 | 0.28 | 0.09 | 0.14 | 0.26 | 0.25 | 0.24 |
| 4 | 2714023 | rs12502642 | 1.214 | 0.654 | 0.099 | -0.856 | -0.03 | -0.028 | -0.028 | 0 | 0.33 | 0.24 | 0.56 | 0.4 | 0.4 | 0.45 |
| 4 | 2717690 | rs2295499 | 0.808 | 0.661 | 0.19 | -0.874 | 0.007 | 0.004 | 0.004 | 1.222 | 0.37 | 0.44 | 0.6 | 0.4 | 0.41 | 0.47 |
| 4 | 2734626 | rs232722 | 0.057 | 0.624 | 0.055 | -0.89 | -0.001 | 0.014 | 0.014 | 0.316 | 0.49 | 0.81 | 0.66 | 0.42 | 0.47 | 0.56 |
| 4 | 2734859 | rs12511418 | 0.66 | 0.726 | 0.001 | -0.865 | -0.033 | -0.015 | -0.015 | 0 | 0.42 | 0.23 | 0.6 | 0.42 | 0.44 | 0.52 |
| 4 | 2743999 | rs232712 | -0.346 | 0.719 | -0.086 | -0.866 | 0.005 | 0.065 | 0.065 | 0 | 0.83 | 0.8 | 0.88 | 0.61 | 0.82 | 0.86 |
| 4 | 2744480 | rs9683949 | 0.62 | 0.64 | -0.208 | -1.026 | -0.036 | -0.003 | -0.003 | 0 | 0.46 | 0.18 | 0.62 | 0.43 | 0.46 | 0.52 |
| 4 | 2744521 | rs17831665 | 1.186 | 0.483 | -0.53 | -1.123 | 0.018 | 0.088 | 0.088 | 0.516 | 0.28 | 0.09 | 0.16 | 0.17 | 0.29 | 0.26 |
| 4 | 2744937 | rs3733226 | 1.216 | 0.476 | -0.517 | -1.005 | 0.02 | 0.033 | 0.033 | 0 | 0.1 | 0.03 | 0.04 | 0.03 | 0.04 | 0.08 |
| 4 | 2747475 | rs2341303 | -0.067 | 0.53 | -0.462 | -1.011 | 0.008 | 0.009 | 0.009 | 0 | 0.52 | 0.65 | 0.68 | 0.43 | 0.5 | 0.55 |

| | | | | | | | | | | | | | | | | |
|---|---|---|---|---|---|---|---|---|---|---|---|---|---|---|---|---|
| 4 | 2748663 | rs4690055 | 0.553 | 0.714 | -0.955 | -0.905 | -0.019 | -0.005 | -0.005 | 0.046 | 0.39 | 0.34 | 0.6 | 0.39 | 0.45 | 0.46 |
| 4 | 2760222 | rs4690060 | 0.502 | 1.023 | -1.358 | -0.681 | -0.062 | -0.012 | -0.012 | 0.178 | 0.46 | 0.09 | 0.62 | 0.41 | 0.45 | 0.52 |
| 4 | 2762679 | rs1268385 | -0.127 | 0.788 | -0.975 | -0.392 | 0.017 | 0.091 | 0.091 | 0 | 0.8 | 0.66 | 0.84 | 0.62 | 0.77 | 0.86 |
| 4 | 2764088 | rs1263492 | -0.078 | 0.634 | -1.682 | -0.51 | 0.026 | 0.085 | 0.085 | 0.194 | 0.34 | 0.17 | 0.2 | 0.19 | 0.33 | 0.31 |
| 4 | 2768630 | rs6600766 | 0.357 | 0.959 | -1.443 | -0.306 | -0.034 | -0.053 | -0.053 | 0.053 | 0.72 | 0.79 | 0.74 | 0.53 | 0.62 | 0.7 |
| 4 | 2770772 | rs12504990 | 0.821 | 0.311 | -1.508 | -0.739 | -0.023 | -0.013 | -0.013 | 0 | 0.41 | 0.27 | 0.53 | 0.46 | 0.47 | 0.46 |
| 4 | 2772530 | rs7695151 | 0.782 | 0.305 | -1.545 | -0.753 | -0.024 | -0.017 | -0.017 | 0 | 0.39 | 0.26 | 0.53 | 0.46 | 0.45 | 0.47 |
| 4 | 2798170 | rs231331 | -0.457 | -0.451 | -1.844 | -1.41 | 0.05 | 0.113 | 0.113 | 2.934 | 0.32 | 0.81 | 0.29 | 0.06 | 0.23 | 0.29 |
| 4 | 2804476 | rs1751848 | 0.684 | 0.396 | -1.144 | -1.137 | 0 | 0 | 0 | 0.345 | 0.48 | 0.47 | 0.55 | 0.49 | 0.48 | 0.52 |
| 4 | 2815256 | rs422734 | -0.673 | 0.481 | -1.55 | -1.019 | -0.099 | -0.128 | -0.128 | 0.016 | 0.85 | 0.98 | 0.81 | 0.57 | 0.72 | 0.8 |
| 4 | 2816300 | rs1263446 | -0.324 | -0.16 | -1.376 | -1.369 | -0.064 | -0.124 | -0.124 | 0.704 | 0.19 | 0.39 | 0.09 | 0.01 | 0.1 | 0.09 |
| 4 | 2819063 | rs1626813 | 1.455 | 0.015 | -1.782 | -1.335 | -0.084 | -0.067 | -0.067 | 0 | 0.39 | 0.11 | 0.49 | 0.46 | 0.44 | 0.51 |
| 4 | 2825660 | rs3755882 | 0.737 | 0.036 | -1.153 | -1.209 | -0.053 | -0.03 | -0.03 | 12.151 | 0.14 | 0.3 | 0.15 | 0.06 | 0.06 | 0.12 |
| 4 | 2826400 | rs3213501 | -1.144 | 0.529 | -1.641 | -0.98 | -0.09 | -0.097 | -0.097 | 0.015 | 0.87 | 0.99 | 0.83 | 0.55 | 0.72 | 0.8 |
| 4 | 2831383 | rs231399 | 0.471 | 0.195 | -1.04 | -1.027 | -0.001 | 0.012 | 0.012 | 0 | 0.51 | 0.35 | 0.52 | 0.47 | 0.49 | 0.55 |
| 4 | 2832727 | rs189139 | 0.895 | 0.41 | -1.583 | -0.913 | -0.034 | -0.089 | -0.089 | 0.429 | 0.27 | 0.11 | 0.35 | 0.44 | 0.3 | 0.31 |
| 4 | 2834554 | rs231395 | -0.718 | -0.049 | -0.845 | -1.282 | -0.028 | -0.086 | -0.086 | 0 | 0.22 | 0.26 | 0.11 | 0.02 | 0.15 | 0.13 |
| 4 | 2836628 | rs231394 | -0.011 | 0.766 | -0.922 | -0.797 | 0.006 | 0.012 | 0.012 | 0 | 0.59 | 0.51 | 0.67 | 0.47 | 0.57 | 0.59 |
| 4 | 2845274 | rs35780153 | 0.281 | 1.289 | -1.173 | -0.893 | -0.036 | 0.053 | 0.053 | 0 | 0.19 | 0.08 | 0.21 | 0.04 | 0.3 | 0.11 |
| 4 | 2850142 | rs12503220 | 0.799 | 0.37 | -1.562 | -0.94 | 0.053 | 0.056 | 0.056 | 0.012 | 0.27 | 0.1 | 0.29 | 0.17 | 0.17 | 0.42 |
| 4 | 2861430 | rs1263360 | -1.108 | 0.904 | -0.689 | -0.342 | -0.008 | -0.001 | -0.001 | 0 | 0.43 | 0.61 | 0.46 | 0.5 | 0.53 | 0.34 |
| 4 | 2862190 | rs10026792 | 0.253 | 1.064 | -0.426 | -0.367 | -0.026 | 0.042 | 0.042 | 0.33 | 0.23 | 0.11 | 0.22 | 0.04 | 0.31 | 0.1 |
| 4 | 2868514 | rs17833172 | 0.058 | 1.045 | -0.447 | -0.376 | -0.02 | 0.053 | 0.053 | 0.196 | 0.25 | 0.12 | 0.22 | 0.04 | 0.31 | 0.1 |
| 4 | 2881256 | rs624833 | -0.034 | 0.935 | 0.289 | -0.463 | 0.021 | -0.122 | -0.122 | 0 | 0.26 | 0.4 | 0.26 | 0.04 | 0.33 | 0.11 |
| 4 | 2882007 | rs1242228 | -0.913 | 0.935 | 0.344 | -0.119 | 0.02 | 0.026 | 0.026 | 0.078 | 0.37 | 0.5 | 0.42 | 0.43 | 0.5 | 0.3 |
| 4 | 2888441 | rs3775068 | -1.63 | 1.04 | 0.099 | -0.396 | 0.017 | 0.011 | 0.011 | 0 | 0.48 | 0.61 | 0.47 | 0.5 | 0.6 | 0.34 |
| 4 | 2906285 | rs2285084 | 0.145 | 1.534 | 0.139 | 0.196 | 0.019 | 0.085 | 0.085 | 0 | 0.11 | 0.17 | 0.18 | 0.45 | 0.21 | 0.2 |
| 4 | 2906707 | rs4961 | 0.484 | 1.55 | 0.22 | 0.249 | -0.032 | -0.006 | -0.006 | 0 | 0.09 | 0.05 | 0.17 | 0.45 | 0.2 | 0.2 |
| 4 | 2916931 | rs16843589 | 0.58 | 1.192 | 0.486 | -0.033 | 0.012 | -0.052 | -0.052 | 1.634 | 0.2 | 0.23 | 0.22 | 0.04 | 0.3 | 0.09 |
| 4 | 2934308 | rs1263349 | 0.353 | 1.087 | 0.228 | 0.389 | 0.011 | 0.081 | 0.081 | 0.508 | 0.84 | 0.77 | 0.81 | 0.54 | 0.8 | 0.8 |
| 4 | 2939018 | rs1263338 | 0.21 | 1.095 | 0.233 | 0.378 | 0.013 | 0.094 | 0.094 | 0.334 | 0.85 | 0.77 | 0.81 | 0.55 | 0.8 | 0.8 |
| 4 | 2941988 | rs2032463 | -0.983 | 0.906 | 0.131 | 0.367 | -0.01 | 0.074 | 0.074 | 0 | 0.43 | 0.73 | 0.43 | 0.22 | 0.44 | 0.22 |
| 4 | 2943361 | rs1054090 | -0.969 | 0.897 | 0.273 | 0.3 | -0.002 | 0.062 | 0.062 | 0 | 0.43 | 0.68 | 0.41 | 0.19 | 0.44 | 0.22 |
| 4 | 2945516 | rs2301376 | -1.161 | 0.961 | 0.22 | 0.489 | -0.036 | 0.016 | 0.016 | 0.015 | 0.93 | 0.99 | 0.89 | 0.61 | 0.86 | 0.83 |
| 4 | 2949878 | rs7665452 | 0.01 | 0.655 | 0.639 | -0.27 | -0.005 | -0.009 | -0.009 | 0 | 0.11 | 0.2 | 0.1 | 0.1 | 0.08 | 0.06 |
| 4 | 2950717 | rs2995797 | -0.068 | 0.629 | 0.858 | -0.308 | 0.007 | -0.042 | -0.042 | 0 | 0.3 | 0.39 | 0.3 | 0.09 | 0.35 | 0.15 |
| 4 | 2951804 | rs2515960 | -1.446 | 0.57 | 0.519 | -0.353 | -0.001 | 0.073 | 0.073 | 0 | 0.45 | 0.68 | 0.42 | 0.19 | 0.44 | 0.22 |
| 4 | 2952506 | rs1063745 | -1.214 | 0.584 | 0.604 | -0.268 | -0.003 | 0.057 | 0.057 | 0 | 0.42 | 0.68 | 0.41 | 0.19 | 0.43 | 0.21 |
| 4 | 2979601 | rs2022485 | -0.631 | 0.592 | 0.649 | -0.321 | 0.004 | -0.064 | -0.064 | 0 | 0.35 | 0.53 | 0.34 | 0.09 | 0.38 | 0.16 |
| 4 | 2982459 | rs3021136 | -0.236 | 0.591 | 0.79 | -0.335 | 0.005 | -0.034 | -0.034 | 0 | 0.3 | 0.37 | 0.3 | 0.09 | 0.35 | 0.15 |
| 4 | 2990375 | rs2051555 | -0.824 | 0.408 | 0.668 | -0.41 | -0.001 | -0.052 | -0.052 | 0 | 0.37 | 0.55 | 0.33 | 0.09 | 0.38 | 0.15 |
| 4 | 2990499 | rs2960306 | -0.905 | 0.414 | 0.666 | -0.407 | -0.002 | -0.057 | -0.057 | 0 | 0.38 | 0.54 | 0.33 | 0.09 | 0.38 | 0.16 |
| 4 | 3006043 | rs1024323 | -1.009 | 0.366 | 0.647 | -0.438 | -0.001 | 0.009 | 0.009 | 0.046 | 0.37 | 0.63 | 0.38 | 0.19 | 0.4 | 0.17 |
| 4 | 3024044 | rs2471325 | -0.232 | 0.42 | 0.512 | -0.397 | -0.006 | 0.008 | 0.008 | 0 | 0.29 | 0.24 | 0.29 | 0.1 | 0.35 | 0.15 |
| 4 | 3024723 | rs1891710 | -0.663 | 0.377 | 0.542 | -0.378 | 0 | -0.005 | -0.005 | 0 | 0.35 | 0.38 | 0.37 | 0.19 | 0.41 | 0.2 |
| 4 | 3027897 | rs2515933 | -1.195 | 0.425 | 0.175 | -0.428 | 0.001 | 0.044 | 0.044 | 0 | 0.45 | 0.62 | 0.41 | 0.19 | 0.44 | 0.2 |
| 4 | 3029588 | rs2857844 | 0.661 | 0.26 | -0.96 | -0.312 | 0.034 | 0.047 | 0.047 | 0.014 | 0.52 | 0.14 | 0.5 | 0.69 | 0.6 | 0.77 |
| 4 | 3029623 | rs7677445 | -0.181 | -0.501 | -0.239 | -0.53 | -0.035 | -0.06 | -0.06 | 0 | 0.11 | 0.21 | 0.04 | 0 | 0.03 | 0.01 |
| 4 | 3030097 | rs3775065 | 1.333 | -0.359 | -0.937 | 0.051 | -0.016 | -0.052 | -0.052 | 0.418 | 0.09 | 0.01 | 0.12 | 0.35 | 0.07 | 0.2 |
| 4 | 3036004 | rs2515930 | 1.431 | -0.328 | -0.527 | -0.259 | -0.011 | -0.035 | -0.035 | 0 | 0.08 | 0.11 | 0.03 | 0 | 0.03 | 0.02 |
| 4 | 3039150 | rs1801058 | -0.738 | -0.189 | -0.835 | -0.286 | -0.069 | -0.092 | -0.092 | 0 | 0.75 | 0.91 | 0.64 | 0.53 | 0.57 | 0.73 |
| 4 | 3042902 | rs2471343 | 0.661 | -0.432 | -0.58 | -0.421 | -0.016 | -0.036 | -0.036 | 0.076 | 0.11 | 0.14 | 0.04 | 0 | 0.04 | 0.02 |
| 4 | 3046853 | rs2857847 | 1.013 | -0.543 | -0.405 | -0.535 | -0.001 | 0.042 | 0.042 | 0 | 0.51 | 0.31 | 0.48 | 0.43 | 0.45 | 0.68 |
| 4 | 3058583 | rs1313772 | 0.823 | -0.487 | -0.295 | -0.496 | -0.009 | 0.02 | 0.02 | 0 | 0.54 | 0.34 | 0.49 | 0.44 | 0.46 | 0.71 |
| 4 | 3063817 | rs7694687 | -1.502 | -0.977 | 0.083 | -0.759 | -0.022 | -0.054 | -0.054 | 0 | 0.35 | 0.59 | 0.21 | 0.1 | 0.17 | 0.16 |
| 4 | 3068545 | rs9993542 | -0.195 | -1.047 | 0.305 | -0.779 | -0.033 | -0.087 | -0.087 | 0 | 0.17 | 0.29 | 0.07 | 0.01 | 0.08 | 0.1 |
| 4 | 3074795 | rs762855 | -1.544 | -0.685 | 0.046 | -0.594 | -0.006 | 0 | 0 | 0 | 0.44 | 0.68 | 0.51 | 0.53 | 0.46 | 0.34 |
| 4 | 3084269 | rs7664480 | 0.826 | -0.635 | 0.157 | -0.509 | -0.012 | -0.04 | -0.04 | 0 | 0.08 | 0.21 | 0.04 | 0 | 0.04 | 0.01 |
| 4 | 3107664 | rs7693317 | 1.298 | -0.617 | 0.046 | -0.503 | -0.01 | -0.036 | -0.036 | 0.309 | 0.07 | 0.15 | 0.03 | 0 | 0.04 | 0.01 |
| 4 | 3109442 | rs10015979 | 0.769 | -1.095 | -0.891 | -0.973 | -0.09 | -0.103 | -0.103 | 2.235 | 0.37 | 0.09 | 0.36 | 0.48 | 0.38 | 0.55 |
| 4 | 3132436 | rs363084 | 1.961 | -0.874 | -0.182 | -0.906 | -0.01 | -0.042 | -0.042 | 0.579 | 0.08 | 0.16 | 0.04 | 0.01 | 0.03 | 0.09 |
| 4 | 3148276 | rs6855981 | -0.183 | -0.955 | -0.252 | -0.775 | -0.014 | -0.015 | -0.015 | 0 | 0.29 | 0.19 | 0.35 | 0.36 | 0.38 | 0.23 |
| 4 | 3151813 | rs11731237 | 1.231 | -1.428 | -0.719 | -1.067 | -0.087 | -0.11 | -0.11 | 0.034 | 0.27 | 0.08 | 0.28 | 0.49 | 0.34 | 0.44 |
| 4 | 3156344 | rs363101 | -0.455 | -1.834 | -0.497 | -1.361 | -0.071 | -0.158 | -0.158 | 0.008 | 0.3 | 0.5 | 0.12 | 0.01 | 0.11 | 0.11 |
| 4 | 3160630 | rs6839274 | -0.307 | -1.929 | -0.42 | -1.407 | -0.06 | -0.146 | -0.146 | 0 | 0.25 | 0.41 | 0.11 | 0.01 | 0.1 | 0.1 |
| 4 | 3174256 | rs363098 | -0.285 | -1.933 | -0.422 | -1.41 | -0.059 | -0.146 | -0.146 | 0.447 | 0.24 | 0.41 | 0.11 | 0.01 | 0.1 | 0.1 |
| 4 | 3177259 | rs363097 | -0.383 | -1.98 | -0.331 | -1.431 | -0.074 | -0.174 | -0.174 | 0.001 | 0.26 | 0.48 | 0.12 | 0.01 | 0.11 | 0.11 |
| 4 | 3180021 | rs363096 | -0.859 | -2.16 | -0.281 | -1.551 | -0.015 | 0 | 0 | 0 | 0.61 | 0.68 | 0.54 | 0.38 | 0.59 | 0.43 |
| 4 | 3186244 | rs2298969 | -1.43 | -1.839 | -0.018 | -1.257 | 0.002 | 0.009 | 0.009 | 0 | 0.52 | 0.68 | 0.52 | 0.51 | 0.49 | 0.46 |

| | | | | | | | | | | | | | | | | |
|---|---|---|---|---|---|---|---|---|---|---|---|---|---|---|---|---|
| 4 | 3186993 | rs10488840 | 0.223 | -1.369 | 0.318 | -1.12 | -0.004 | -0.044 | -0.044 | 0 | 0.2 | 0.2 | 0.09 | 0.01 | 0.1 | 0.1 |
| 4 | 3189547 | rs363125 | -0.67 | -1.369 | 0.31 | -1.117 | -0.075 | -0.175 | -0.175 | 0 | 0.26 | 0.48 | 0.12 | 0.01 | 0.11 | 0.11 |
| 4 | 3190533 | rs16844026 | -0.222 | -1.355 | 0.076 | -1.111 | -0.025 | -0.087 | -0.087 | 0 | 0.23 | 0.27 | 0.1 | 0.01 | 0.1 | 0.1 |
| 4 | 3196029 | rs363092 | 0.735 | -1.273 | -0.194 | -1.01 | -0.008 | 0 | 0 | 0 | 0.62 | 0.33 | 0.57 | 0.64 | 0.58 | 0.69 |
| 4 | 3203578 | rs363089 | 2.88 | -1.112 | -0.359 | -0.958 | -0.005 | -0.027 | -0.027 | 0 | 0.06 | 0.1 | 0.03 | 0.01 | 0.03 | 0.09 |
| 4 | 3215835 | rs362331 | -0.995 | -1.014 | -0.615 | -0.827 | -0.001 | -0.001 | -0.001 | 0 | 0.42 | 0.59 | 0.43 | 0.39 | 0.43 | 0.32 |
| 4 | 3227419 | rs362273 | 0.811 | -0.316 | -0.592 | -0.333 | 0.014 | 0.022 | 0.022 | 0 | 0.13 | 0.12 | 0.3 | 0.34 | 0.32 | 0.2 |
| 4 | 3231661 | rs2276881 | 1.678 | -0.951 | -0.955 | -0.561 | -0.027 | -0.012 | -0.012 | 0 | 0.07 | 0 | 0.19 | 0.36 | 0.06 | 0.16 |
| 4 | 3234980 | rs362272 | 1.6 | -0.739 | -0.736 | -0.894 | -0.057 | -0.054 | -0.054 | 0.357 | 0.09 | 0.01 | 0.29 | 0.32 | 0.31 | 0.21 |
| 4 | 3235589 | rs362313 | 1.534 | -1.259 | -0.851 | -1.452 | 0.009 | -0.003 | -0.003 | 0 | 0.17 | 0.11 | 0.1 | 0 | 0.16 | 0.11 |
| 4 | 3242100 | rs362306 | 1.549 | -0.64 | -0.723 | -0.948 | -0.05 | -0.041 | -0.041 | 0 | 0.09 | 0.01 | 0.29 | 0.34 | 0.31 | 0.21 |
| 4 | 3242307 | rs362303 | -0.259 | -1.238 | -0.355 | -1.287 | -0.045 | -0.136 | -0.136 | 0 | 0.23 | 0.37 | 0.1 | 0.01 | 0.11 | 0.11 |
| 4 | 3252852 | rs3129322 | -0.395 | -0.09 | -0.608 | -0.51 | 0.04 | 0.051 | 0.051 | 0 | 0.21 | 0.32 | 0.33 | 0.37 | 0.37 | 0.26 |
| 4 | 3263776 | rs3095081 | 0.734 | -0.055 | -1.129 | -0.421 | -0.103 | -0.095 | -0.095 | 0 | 0.14 | 0.02 | 0.31 | 0.34 | 0.38 | 0.25 |
| 4 | 3267668 | rs2798224 | 0.407 | -0.913 | -1.244 | -0.888 | -0.038 | -0.025 | -0.025 | 3.361 | 0.36 | 0.18 | 0.44 | 0.38 | 0.58 | 0.38 |
| 4 | 3270348 | rs16844106 | 0.709 | -1.124 | -1.042 | -0.99 | -0.011 | -0.037 | -0.037 | 0 | 0.09 | 0.13 | 0.04 | 0.01 | 0.04 | 0.07 |
| 4 | 3270488 | rs2798221 | 0.328 | -1.072 | -0.968 | -0.995 | -0.001 | 0 | 0 | 0.738 | 0.4 | 0.33 | 0.47 | 0.38 | 0.61 | 0.39 |
| 4 | 3283422 | rs7658462 | 0.496 | -0.979 | -1.019 | -0.816 | 0 | 0.001 | 0.001 | 0 | 0.17 | 0.15 | 0.16 | 0.15 | 0.17 | 0.2 |
| 4 | 3284560 | rs12506821 | 0.812 | -1.237 | -0.899 | -0.705 | -0.002 | -0.002 | -0.002 | 0 | 0.13 | 0.15 | 0.13 | 0.15 | 0.12 | 0.2 |
| 4 | 3284751 | rs6446731 | -0.401 | -1.513 | -0.834 | -0.778 | 0 | -0.001 | -0.001 | 0.302 | 0.42 | 0.55 | 0.45 | 0.59 | 0.28 | 0.51 |
| 4 | 3287655 | rs3129317 | 0.29 | -1.326 | -0.836 | -0.853 | 0.03 | 0.077 | 0.077 | 0 | 0.4 | 0.24 | 0.26 | 0.17 | 0.34 | 0.4 |
| 4 | 3291694 | rs10022193 | 0.234 | -1.438 | -0.592 | -0.679 | -0.053 | 0.059 | 0.059 | 0 | 0.31 | 0.43 | 0.31 | 0.45 | 0.17 | 0.33 |
| 4 | 3298800 | rs2051559 | 0.735 | -0.865 | -0.42 | -0.37 | 0.002 | 0 | 0 | 0 | 0.13 | 0.09 | 0.13 | 0.15 | 0.12 | 0.2 |
| 4 | 3300783 | rs13129052 | -0.544 | -0.386 | -0.103 | -0.235 | 0.002 | -0.026 | -0.026 | 1.514 | 0.08 | 0.19 | 0.04 | 0.02 | 0.06 | 0.09 |
| 4 | 3314966 | rs2749786 | 1.094 | -0.572 | -0.438 | -0.165 | 0.014 | -0.039 | -0.039 | 0.218 | 0.33 | 0.08 | 0.28 | 0.34 | 0.28 | 0.34 |
| 4 | 3318413 | rs2236052 | -1.224 | -0.469 | -0.564 | -0.149 | -0.141 | -0.119 | -0.119 | 0 | 0.8 | 0.98 | 0.64 | 0.66 | 0.59 | 0.75 |
| 4 | 3323767 | rs2857861 | 0.103 | -0.561 | -0.218 | -0.207 | 0.047 | 0.034 | 0.034 | 0.042 | 0.69 | 0.45 | 0.55 | 0.58 | 0.53 | 0.66 |
| 4 | 3339038 | rs7668784 | 0.738 | -0.616 | -0.143 | -0.244 | -0.002 | -0.001 | -0.001 | 2.49 | 0.31 | 0.3 | 0.26 | 0.26 | 0.22 | 0.33 |
| 4 | 3342137 | rs7688715 | 0.528 | -0.559 | -0.626 | -0.249 | -0.019 | -0.051 | -0.051 | 0 | 0.09 | 0.22 | 0.04 | 0 | 0.04 | 0 |
| 4 | 3346617 | rs2749781 | -1.069 | -0.057 | -1.007 | 0.097 | -0.141 | -0.123 | -0.123 | 0 | 0.8 | 0.98 | 0.64 | 0.65 | 0.59 | 0.77 |
| 4 | 3350248 | rs2749779 | 0.735 | -0.041 | -0.858 | 0.061 | 0.009 | 0.001 | 0.001 | 0.061 | 0.35 | 0.32 | 0.3 | 0.34 | 0.28 | 0.35 |
| 4 | 3369099 | rs16844212 | 0.167 | 0.014 | -0.953 | 0.09 | -0.029 | -0.034 | -0.034 | 0.046 | 0.39 | 0.48 | 0.31 | 0.26 | 0.26 | 0.32 |
| 4 | 3374539 | rs12640186 | 0.633 | 0.065 | -1.149 | 0.197 | 0.021 | -0.002 | -0.002 | 0 | 0.34 | 0.21 | 0.29 | 0.34 | 0.29 | 0.35 |
| 4 | 3375722 | rs6855619 | -0.47 | -0.049 | -0.874 | -0.273 | 0.008 | 0.009 | 0.009 | 0 | 0.44 | 0.57 | 0.33 | 0.29 | 0.3 | 0.41 |
| 4 | 3384844 | rs1005187 | -0.403 | -0.32 | -0.848 | -0.646 | 0.002 | -0.014 | -0.014 | 3.156 | 0.5 | 0.49 | 0.67 | 0.91 | 0.66 | 0.54 |
| 4 | 3385540 | rs11730455 | -0.374 | -1.257 | -0.925 | -0.731 | -0.04 | -0.01 | -0.01 | 0 | 0.34 | 0.43 | 0.27 | 0.25 | 0.16 | 0.48 |
| 4 | 3387148 | rs16844249 | -0.17 | -1.242 | -0.857 | -0.884 | 0.005 | 0 | 0 | 0 | 0.15 | 0.11 | 0.12 | 0.12 | 0.11 | 0.14 |
| 4 | 3390900 | rs12649466 | -0.848 | 0.437 | -0.914 | -1.281 | -0.04 | 0.044 | 0.044 | 0 | 0.37 | 0.17 | 0.33 | 0.21 | 0.37 | 0.52 |
| 4 | 3396963 | rs11732375 | -0.5 | -0.279 | -0.406 | -0.572 | 0.028 | 0.018 | 0.018 | 0 | 0.35 | 0.27 | 0.27 | 0.2 | 0.15 | 0.44 |
| 4 | 3405165 | rs6852186 | -1.018 | -0.299 | -0.05 | -0.653 | 0.001 | -0.017 | -0.017 | 2.1 | 0.47 | 0.45 | 0.4 | 0.2 | 0.43 | 0.53 |
| 4 | 3407694 | rs1962767 | -1.245 | -0.276 | -0.189 | -0.373 | 0 | -0.002 | -0.002 | 0 | 0.55 | 0.52 | 0.49 | 0.53 | 0.52 | 0.65 |
| 4 | 3409024 | rs16844280 | 0.762 | -0.4 | -0.71 | -0.428 | -0.009 | 0.001 | 0.001 | 1.061 | 0.14 | 0.11 | 0.07 | 0.1 | 0.04 | 0.14 |
| 4 | 3409359 | rs1730768 | -2.004 | -0.186 | -1.096 | -0.458 | 0.002 | 0.137 | 0.137 | 0.452 | 0.78 | 0.97 | 0.89 | 1 | 0.79 | 0.88 |
| 4 | 3409700 | rs1406674 | -0.89 | -0.028 | -1.015 | -0.346 | -0.013 | -0.021 | -0.021 | 0 | 0.56 | 0.74 | 0.49 | 0.5 | 0.51 | 0.64 |
| 4 | 3414654 | rs10027926 | 1.456 | 0.41 | -0.657 | 0.001 | -0.017 | 0.006 | 0.006 | 0 | 0.05 | 0.01 | 0.08 | 0.29 | 0.09 | 0.1 |
| 4 | 3415378 | rs16844301 | 0.812 | 0.402 | -0.635 | -0.028 | -0.011 | -0.04 | -0.04 | 0 | 0.22 | 0.2 | 0.41 | 0.49 | 0.27 | 0.25 |
| 4 | 3419840 | rs12641989 | 0.493 | 0.294 | -0.279 | 0.21 | 0.017 | 0.101 | 0.101 | 0 | 0.06 | 0.08 | 0.09 | 0.29 | 0.09 | 0.1 |
| 4 | 3429856 | rs2269497 | 1.673 | -0.044 | -0.583 | -0.107 | 0.01 | -0.013 | -0.013 | 0 | 0.06 | 0.02 | 0.08 | 0.05 | 0.04 | 0.04 |
| 4 | 3430862 | rs1203112 | 0.817 | 0.136 | -0.092 | -0.519 | -0.002 | -0.055 | -0.055 | 0 | 0.83 | 0.46 | 0.89 | 0.91 | 0.84 | 0.78 |
| 4 | 3434885 | rs6818397 | -0.16 | -0.303 | -1.009 | -0.602 | 0.039 | -0.028 | -0.028 | 0 | 0.42 | 0.28 | 0.59 | 0.5 | 0.59 | 0.61 |
| 4 | 3436588 | rs13116176 | 0.354 | 0.586 | -1.073 | -0.485 | -0.034 | -0.031 | -0.031 | 0 | 0.28 | 0.24 | 0.38 | 0.45 | 0.43 | 0.29 |
| 4 | 3438643 | rs16844364 | 0.155 | -0.779 | -2.136 | -0.926 | 0.112 | -0.085 | -0.085 | 0.359 | 0.35 | 0.1 | 0.26 | 0.45 | 0.22 | 0.32 |
| 4 | 3444503 | rs2073505 | 0.676 | -1.078 | -1.924 | -0.898 | 0.002 | 0.005 | 0.005 | 1.476 | 0.19 | 0.12 | 0.11 | 0.09 | 0.08 | 0.16 |
| 4 | 3445745 | rs4690097 | -0.996 | -0.538 | -1.882 | -1.211 | 0 | -0.054 | -0.054 | 0 | 0.7 | 0.66 | 0.71 | 0.92 | 0.72 | 0.66 |
| 4 | 3446091 | rs3748034 | 1.965 | 0.253 | -2.49 | -0.447 | -0.013 | -0.063 | -0.063 | 0 | 0.11 | 0.02 | 0.11 | 0.31 | 0.14 | 0.13 |
| 4 | 3446513 | rs17805201 | 1.378 | 1.12 | -2.504 | -0.718 | -0.023 | 0.014 | 0.014 | 0 | 0.1 | 0.06 | 0.08 | 0.01 | 0.21 | 0.1 |
| 4 | 3447156 | rs4690098 | 2.158 | 0.197 | -2.593 | -0.46 | -0.072 | -0.136 | -0.136 | 0 | 0.17 | 0.01 | 0.25 | 0.45 | 0.23 | 0.21 |
| 4 | 3447243 | rs16844383 | 0.353 | -0.585 | -1.759 | -0.276 | 0.008 | -0.059 | -0.059 | 0 | 0.16 | 0.08 | 0.2 | 0.37 | 0.17 | 0.16 |
| 4 | 3449652 | rs16844401 | 1.349 | -0.481 | -1.702 | -0.514 | 0.014 | 0.002 | 0.002 | 0.523 | 0.1 | 0.02 | 0.09 | 0.09 | 0.07 | 0.13 |
| 4 | 3449886 | rs28468427 | -0.312 | -0.694 | -1.184 | -0.511 | -0.013 | 0.011 | 0.011 | 0 | 0.28 | 0.29 | 0.33 | 0.36 | 0.23 | 0.18 |
| 4 | 3451109 | rs2498323 | 1.59 | -0.133 | -1.011 | -0.456 | 0.002 | -0.009 | -0.009 | 0 | 0.06 | 0.09 | 0.09 | 0.05 | 0.08 | 0.03 |
| 4 | 3454304 | rs1203111 | -0.55 | -0.188 | -1.602 | -0.659 | -0.04 | 0.035 | 0.035 | 0 | 0.57 | 0.84 | 0.61 | 0.61 | 0.49 | 0.61 |
| 4 | 3454454 | rs4690099 | 0.942 | -0.392 | -1.202 | -0.575 | -0.004 | -0.059 | -0.059 | 0 | 0.23 | 0.13 | 0.26 | 0.47 | 0.25 | 0.25 |
| 4 | 3463452 | rs1203104 | -0.409 | -0.444 | -0.599 | -0.664 | 0.01 | -0.009 | -0.009 | 0 | 0.43 | 0.63 | 0.4 | 0.52 | 0.38 | 0.31 |
| 4 | 3467519 | rs10022329 | -0.033 | -0.845 | 0.188 | -1.745 | -0.053 | -0.107 | -0.107 | 0 | 0.22 | 0.47 | 0.16 | 0.07 | 0.12 | 0.07 |
| 4 | 3469443 | rs10937927 | -0.819 | -0.621 | 0.001 | -1.774 | -0.004 | 0.011 | 0.011 | 0.277 | 0.45 | 0.57 | 0.37 | 0.37 | 0.46 | 0.32 |
| 4 | 3469458 | rs10937928 | 0.446 | -1.332 | 0.627 | -1.574 | -0.025 | -0.078 | -0.078 | 0 | 0.17 | 0.31 | 0.11 | 0.06 | 0.11 | 0.04 |
| 4 | 3471754 | rs16844418 | 0.309 | -1.308 | 0.489 | -1.565 | -0.022 | -0.085 | -0.085 | 1.058 | 0.19 | 0.36 | 0.13 | 0.03 | 0.13 | 0.04 |
| 4 | 3473139 | rs6831256 | -1.091 | -1.184 | -0.26 | -1.52 | 0.037 | 0.046 | 0.046 | 0 | 0.51 | 0.67 | 0.54 | 0.38 | 0.4 | 0.49 |

| | | | | | | | | | | | | | | | | |
|---|---|---|---|---|---|---|---|---|---|---|---|---|---|---|---|---|
| 4 | 3477299 | rs6842419 | 1.299 | -0.996 | -0.674 | -1.352 | -0.007 | 0.017 | 0.017 | 0 | 0.11 | 0.14 | 0.14 | 0.22 | 0.11 | 0.08 |
| 4 | 3478323 | rs2699426 | -1.034 | -0.776 | -1.008 | -1.294 | 0.051 | 0.076 | 0.076 | 0 | 0.53 | 0.86 | 0.57 | 0.38 | 0.39 | 0.48 |
| 4 | 3485786 | rs1881803 | -1.029 | -0.457 | -0.688 | -1.669 | 0.004 | -0.022 | -0.022 | 0.417 | 0.67 | 0.62 | 0.7 | 0.92 | 0.55 | 0.74 |
| 4 | 3493215 | rs13146709 | 1.96 | -0.813 | -1.328 | -1.402 | -0.031 | -0.092 | -0.092 | 0 | 0.14 | 0.03 | 0.13 | 0.52 | 0.21 | 0.23 |
| 4 | 3494055 | rs2699416 | -2.062 | -0.967 | -0.99 | -1.573 | 0.102 | -0.066 | -0.066 | 0 | 0.76 | 0.58 | 0.8 | 0.91 | 0.67 | 0.8 |
| 4 | 3494600 | rs6811423 | 2.144 | -0.978 | -1.006 | -1.527 | -0.043 | -0.092 | -0.092 | 0.236 | 0.15 | 0.03 | 0.14 | 0.5 | 0.23 | 0.23 |
| 4 | 3494898 | rs6850908 | 2.103 | -0.981 | -0.998 | -1.527 | -0.039 | -0.089 | -0.089 | 0.328 | 0.15 | 0.04 | 0.14 | 0.5 | 0.22 | 0.22 |
| 4 | 3494956 | rs16844464 | 1.509 | -1.293 | -0.973 | -1.667 | -0.016 | -0.009 | -0.009 | 1.4 | 0.1 | 0.16 | 0.08 | 0.04 | 0.03 | 0.07 |
| 4 | 3495950 | rs2699414 | 0.987 | -1.384 | -1.15 | -1.711 | 0.031 | 0.005 | 0.005 | 0 | 0.21 | 0.09 | 0.21 | 0.24 | 0.1 | 0.14 |
| 4 | 3496058 | rs2252054 | 1 | -1.385 | -1.145 | -1.709 | 0.028 | 0.007 | 0.007 | 0 | 0.21 | 0.1 | 0.22 | 0.24 | 0.1 | 0.14 |
| 4 | 3496110 | rs10937930 | 0.919 | -1.422 | -1.149 | -1.831 | 0.011 | 0.02 | 0.02 | 0 | 0.25 | 0.19 | 0.33 | 0.16 | 0.22 | 0.39 |
| 4 | 3496150 | rs1054662 | -1.283 | -0.854 | -0.938 | -1.629 | 0.018 | -0.063 | -0.063 | 0.001 | 0.64 | 0.45 | 0.69 | 0.91 | 0.54 | 0.76 |
| 4 | 3496940 | rs13136935 | 1.76 | -0.753 | -1.131 | -1.592 | -0.009 | 0.022 | 0.022 | 0 | 0.12 | 0.06 | 0.11 | 0 | 0.13 | 0.05 |
| 4 | 3505448 | rs3468 | 0.37 | -0.823 | -1.16 | -1.583 | -0.019 | -0.058 | -0.058 | 0 | 0.29 | 0.16 | 0.25 | 0.51 | 0.35 | 0.27 |
| 4 | 3510514 | rs3184 | -2.153 | -0.495 | -1.101 | -1.416 | 0.068 | -0.051 | -0.051 | 0 | 0.76 | 0.6 | 0.8 | 0.91 | 0.67 | 0.8 |
| 4 | 3514539 | rs1049574 | 0.96 | -0.649 | -0.873 | -0.603 | -0.001 | 0.018 | 0.018 | 0 | 0.18 | 0.2 | 0.15 | 0.51 | 0.23 | 0.22 |
| 4 | 3514695 | rs6789 | -1.393 | -0.712 | -1.196 | -0.469 | 0.017 | 0.009 | 0.009 | 0.387 | 0.39 | 0.29 | 0.52 | 0.41 | 0.28 | 0.51 |
| 4 | 3525057 | rs1730775 | -0.8 | -0.304 | -0.842 | -0.038 | 0.013 | -0.004 | -0.004 | 0 | 0.22 | 0.16 | 0.27 | 0.28 | 0.11 | 0.18 |
| 4 | 3592160 | rs28585439 | -1.325 | 0.854 | -0.637 | -0.386 | 0.006 | -0.089 | -0.089 | 0.121 | 0.22 | 0.54 | 0.22 | 0.05 | 0.22 | 0.12 |
| 4 | 3602109 | rs6856283 | -0.985 | 0.951 | -1.266 | -0.488 | 0.003 | -0.061 | -0.061 | 0 | 0.26 | 0.65 | 0.29 | 0.02 | 0.29 | 0.27 |
| 4 | 3606875 | rs16844952 | -0.533 | 0.626 | -1.76 | 0.551 | 0.027 | -0.042 | -0.042 | 0 | 0.31 | 0.22 | 0.21 | 0.42 | 0.26 | 0.2 |
| 4 | 3609390 | rs1810606 | -0.463 | 1.124 | -1.688 | 0.386 | 0.003 | 0.052 | 0.052 | 0 | 0.48 | 0.22 | 0.41 | 0.63 | 0.47 | 0.43 |
| 4 | 3632194 | rs17628728 | -0.981 | 1.733 | -1.7 | 0.228 | -0.043 | 0.032 | 0.032 | 0 | 0.24 | 0.11 | 0.25 | 0.14 | 0.33 | 0.21 |
| 4 | 3637536 | rs7673350 | 1.053 | 1.955 | -1.285 | -0.343 | -0.001 | -0.018 | -0.018 | 0 | 0.73 | 0.72 | 0.76 | 0.9 | 0.64 | 0.83 |
| 4 | 3643599 | rs177802 | 0.763 | 2.003 | -1.163 | -0.359 | -0.025 | 0.035 | 0.035 | 0.041 | 0.77 | 0.85 | 0.78 | 0.9 | 0.66 | 0.84 |
| 4 | 3644957 | rs177798 | 1.604 | 2.115 | -0.935 | -0.666 | 0 | -0.025 | -0.025 | 0 | 0.57 | 0.56 | 0.6 | 0.77 | 0.46 | 0.55 |
| 4 | 3650826 | rs17202883 | -0.229 | 2.028 | -0.857 | -0.707 | -0.016 | -0.005 | -0.005 | 0 | 0.14 | 0.06 | 0.15 | 0.13 | 0.19 | 0.29 |
| 4 | 3651101 | rs177789 | 1.756 | 1.139 | -1.058 | 0 | -0.002 | 0.001 | 0.001 | 0 | 0.43 | 0.45 | 0.38 | 0.48 | 0.36 | 0.44 |
| 4 | 3652592 | rs4916611 | 0.583 | 1.081 | -0.881 | 0.027 | -0.001 | 0.011 | 0.011 | 0 | 0.13 | 0.12 | 0.22 | 0.27 | 0.09 | 0.1 |
| 4 | 3656013 | rs177780 | 1.728 | 0.744 | -0.429 | -0.236 | 0.005 | -0.012 | -0.012 | 0.047 | 0.4 | 0.3 | 0.37 | 0.49 | 0.35 | 0.45 |
| 4 | 3656174 | rs12644079 | -1.022 | 0.706 | -0.326 | -0.22 | -0.016 | 0.038 | 0.038 | 0 | 0.32 | 0.47 | 0.38 | 0.4 | 0.29 | 0.38 |
| 4 | 3658324 | rs2180259 | -0.42 | 0.937 | -0.401 | -0.271 | 0.009 | -0.006 | -0.006 | 0 | 0.07 | 0.12 | 0.06 | 0.05 | 0.11 | 0.07 |
| 4 | 3659574 | rs11731847 | 1.976 | 0.999 | -0.295 | -0.243 | 0.005 | -0.008 | -0.008 | 0 | 0.4 | 0.32 | 0.37 | 0.49 | 0.35 | 0.43 |
| 4 | 3661134 | rs17203086 | -0.438 | 1.108 | -0.48 | -0.321 | -0.002 | -0.001 | -0.001 | 0 | 0.14 | 0.14 | 0.16 | 0.13 | 0.18 | 0.27 |
| 4 | 3663463 | rs16844725 | -0.475 | 1.132 | -0.527 | -0.392 | 0.001 | -0.007 | -0.007 | 0.403 | 0.14 | 0.16 | 0.15 | 0.08 | 0.16 | 0.28 |
| 4 | 3664206 | rs177769 | 0.045 | 1.442 | -0.788 | -0.339 | -0.115 | 0.102 | 0.102 | 0 | 0.75 | 0.98 | 0.74 | 0.88 | 0.65 | 0.82 |
| 4 | 3666471 | rs1884412 | 0.681 | 1.01 | -0.489 | -0.515 | -0.009 | -0.038 | -0.038 | 0.244 | 0.57 | 0.53 | 0.4 | 0.21 | 0.43 | 0.3 |
| 4 | 3668328 | rs16844747 | -0.852 | 0.603 | -0.128 | 0.925 | -0.001 | 0.068 | 0.068 | 0 | 0.13 | 0.19 | 0.22 | 0.39 | 0.14 | 0.26 |
| 4 | 3669171 | rs177766 | 0.913 | -0.323 | 0.338 | 0.578 | -0.011 | -0.037 | -0.037 | 0.044 | 0.63 | 0.43 | 0.46 | 0.15 | 0.7 | 0.51 |
| 4 | 3679491 | rs6827536 | -1.648 | -0.267 | 1.099 | 1.604 | -0.015 | 0.022 | 0.022 | 4.057 | 0.19 | 0.71 | 0.17 | 0.39 | 0.11 | 0.31 |
| 4 | 3683269 | rs2880892 | -1.226 | -0.195 | 1.286 | 1.646 | -0.039 | 0.177 | 0.177 | 1.356 | 0.17 | 0.55 | 0.17 | 0.4 | 0.11 | 0.31 |
| 4 | 3688660 | rs16844775 | -1.136 | 0.083 | 0.477 | 1.468 | -0.01 | 0.039 | 0.039 | 0.259 | 0.31 | 0.57 | 0.19 | 0.41 | 0.25 | 0.33 |
| 4 | 3691191 | rs10805007 | -1.013 | 0.393 | -0.381 | 1.473 | -0.008 | 0.034 | 0.034 | 0.031 | 0.51 | 0.79 | 0.56 | 0.57 | 0.44 | 0.68 |
| 4 | 3692118 | rs11930561 | -1.118 | 0.2 | -0.023 | 1.52 | -0.005 | 0.084 | 0.084 | 0.009 | 0.46 | 0.77 | 0.58 | 0.69 | 0.43 | 0.61 |
| 4 | 3702481 | rs4916608 | -1.035 | 0.518 | 0.646 | 1.925 | 0.074 | 0.527 | 0.527 | 2.686 | 0.05 | 0.39 | 0.25 | 0.37 | 0 | 0.11 |
| 4 | 3703725 | rs885797 | -0.747 | 0.678 | -0.24 | 1.083 | 0 | -0.011 | -0.011 | 0 | 0.41 | 0.53 | 0.52 | 0.68 | 0.41 | 0.57 |
| 4 | 3709475 | rs929006 | 0.305 | 0.612 | -0.842 | 1.105 | -0.031 | -0.053 | -0.053 | 0 | 0.78 | 0.91 | 0.64 | 0.68 | 0.78 | 0.73 |
| 4 | 3710982 | rs2748789 | 0.383 | 0.579 | -0.771 | 1.128 | -0.02 | -0.042 | -0.042 | 0 | 0.77 | 0.88 | 0.68 | 0.67 | 0.78 | 0.73 |
| 4 | 3711721 | rs6837039 | 0.552 | 0.492 | -0.745 | 0.715 | 0.005 | -0.012 | -0.012 | 0 | 0.37 | 0.32 | 0.29 | 0.54 | 0.37 | 0.3 |
| 4 | 3712082 | rs2857969 | 0.678 | 0.538 | -0.897 | 0.407 | -0.037 | 0.089 | 0.089 | 0 | 0.36 | 0.18 | 0.31 | 0.14 | 0.39 | 0.41 |
| 4 | 3715551 | rs2748784 | 1.041 | 0.367 | -0.996 | 0.36 | -0.094 | 0.081 | 0.081 | 0 | 0.32 | 0.06 | 0.29 | 0.12 | 0.36 | 0.27 |
| 4 | 3718394 | rs16844810 | 0.056 | 0.401 | -0.995 | 0.588 | 0.018 | -0.055 | -0.055 | 0.163 | 0.09 | 0.01 | 0.21 | 0.24 | 0.05 | 0.17 |
| 4 | 3719380 | rs11938629 | 0.664 | 0.416 | -0.685 | 0.836 | 0 | -0.003 | -0.003 | 1.188 | 0.37 | 0.34 | 0.31 | 0.47 | 0.4 | 0.31 |
| 4 | 3721269 | rs7377501 | -0.047 | 0.467 | -1.49 | 0.862 | 0.004 | 0.003 | 0.003 | 0 | 0.45 | 0.24 | 0.51 | 0.36 | 0.41 | 0.53 |
| 4 | 3721299 | rs7375509 | 0.511 | 0.983 | -1.678 | 0.114 | -0.057 | 0.092 | 0.092 | 0 | 0.34 | 0.12 | 0.29 | 0.13 | 0.36 | 0.36 |
| 4 | 3722407 | rs2748763 | 0.245 | 1.065 | -0.706 | 0.539 | -0.011 | -0.005 | -0.005 | 0.115 | 0.84 | 0.88 | 0.86 | 0.84 | 0.82 | 0.82 |
| 4 | 3729652 | rs2092898 | -1.168 | 0.718 | -0.052 | 0.201 | 0 | -0.047 | -0.047 | 0 | 0.22 | 0.58 | 0.14 | 0.14 | 0.2 | 0.2 |
| 4 | 3734895 | rs1894441 | -1.191 | 0.693 | 0.19 | 0.179 | -0.009 | -0.042 | -0.042 | 0.465 | 0.22 | 0.64 | 0.14 | 0.14 | 0.19 | 0.2 |
| 4 | 3737883 | rs2857962 | -1.057 | 0.814 | 0.527 | 0.109 | 0.06 | -0.031 | -0.031 | 0 | 0.08 | 0.34 | 0.07 | 0 | 0.07 | 0.04 |
| 4 | 3738828 | rs6851496 | -0.246 | 0.841 | -0.184 | 0.046 | -0.012 | 0.011 | 0.011 | 0.313 | 0.06 | 0.04 | 0.04 | 0.04 | 0.08 | 0.05 |
| 4 | 3740612 | rs2857960 | 1.19 | 0.37 | -0.983 | 0.104 | 0.014 | -0.015 | -0.015 | 0 | 0.48 | 0.3 | 0.46 | 0.41 | 0.56 | 0.4 |
| 4 | 3741038 | rs2857958 | 1.004 | 0.465 | -1.177 | -0.23 | 0.023 | -0.01 | -0.01 | 0.334 | 0.48 | 0.37 | 0.48 | 0.41 | 0.56 | 0.47 |
| 4 | 3742998 | rs445275 | -1.736 | 0.244 | -1.47 | -0.226 | -0.08 | -0.073 | -0.073 | 0 | 0.62 | 0.82 | 0.49 | 0.6 | 0.57 | 0.64 |
| 4 | 3747842 | rs4498196 | -1.32 | 0.076 | -1.082 | -0.25 | -0.004 | -0.027 | -0.027 | 0 | 0.41 | 0.34 | 0.46 | 0.56 | 0.42 | 0.51 |
| 4 | 3750155 | rs4916612 | -0.173 | -0.342 | -1.14 | -0.05 | 0.018 | -0.013 | -0.013 | 0 | 0.21 | 0.16 | 0.28 | 0.3 | 0.11 | 0.28 |
| 4 | 3753417 | rs10013535 | -1.661 | -0.131 | -0.929 | -0.899 | 0.034 | 0.05 | 0.05 | 0 | 0.51 | 0.69 | 0.36 | 0.34 | 0.6 | 0.46 |
| 4 | 3754818 | rs9291156 | -2.285 | -0.145 | 0.018 | -0.306 | 0 | 0.135 | 0.135 | 0.329 | 0.51 | 0.78 | 0.62 | 0.76 | 0.49 | 0.52 |
| 4 | 3754853 | rs9291157 | -0.435 | 0.085 | 0.692 | -0.597 | 0.041 | -0.081 | -0.081 | 0.681 | 0.14 | 0.48 | 0.24 | 0.01 | 0.1 | 0.04 |
| 4 | 3760093 | rs3935655 | -2.063 | 0.034 | 0.058 | -0.58 | -0.018 | 0.088 | 0.088 | 0 | 0.43 | 0.79 | 0.45 | 0.33 | 0.42 | 0.44 |

| | | | | | | | | | | | | | | | | |
|---|---|---|---|---|---|---|---|---|---|---|---|---|---|---|---|---|
| 4 | 3762004 | rs7677352 | -1.055 | -0.489 | 0.596 | -0.358 | -0.026 | -0.054 | -0.054 | 0 | 0.26 | 0.57 | 0.3 | 0.16 | 0.15 | 0.15 |
| 4 | 3765755 | rs13118771 | 0.132 | -0.405 | 0.156 | -0.017 | 0.014 | -0.019 | -0.019 | 0 | 0.85 | 0.89 | 0.79 | 0.62 | 0.9 | 0.77 |
| 4 | 3767394 | rs9714431 | -0.222 | -0.368 | -0.064 | 0.008 | 0.022 | -0.048 | -0.048 | 0.563 | 0.86 | 0.95 | 0.79 | 0.62 | 0.91 | 0.77 |
| 4 | 3775685 | rs7681159 | 0.576 | -0.178 | -0.149 | -0.094 | -0.008 | 0.005 | 0.005 | 0.699 | 0.13 | 0.12 | 0.11 | 0.16 | 0.21 | 0.29 |
| 4 | 3775957 | rs13134185 | -0.262 | -0.592 | -0.376 | 0.092 | 0.033 | -0.042 | -0.042 | 0 | 0.85 | 0.91 | 0.78 | 0.65 | 0.91 | 0.74 |
| 4 | 3780072 | rs6822427 | 1.217 | -0.13 | -0.242 | -0.179 | 0.001 | 0.038 | 0.038 | 0.384 | 0.44 | 0.25 | 0.29 | 0.28 | 0.46 | 0.27 |
| 4 | 3787202 | rs7435195 | 0.399 | 0.38 | -0.584 | -0.826 | -0.042 | -0.013 | -0.013 | 0 | 0.08 | 0.02 | 0.23 | 0.06 | 0.13 | 0.04 |
| 4 | 3791661 | rs12501853 | -1.035 | 0.155 | -0.282 | -1.082 | -0.025 | -0.072 | -0.072 | 0.039 | 0.3 | 0.41 | 0.3 | 0.16 | 0.19 | 0.15 |
| 4 | 3792442 | rs7440077 | -0.422 | 0.028 | -0.932 | -0.595 | 0.002 | 0.048 | 0.048 | 0 | 0.46 | 0.57 | 0.46 | 0.3 | 0.44 | 0.54 |
| 4 | 3793990 | rs28622001 | -0.67 | 0.118 | -1.499 | -0.531 | -0.036 | 0.21 | 0.21 | 0.578 | 0.49 | 0.89 | 0.49 | 0.3 | 0.45 | 0.54 |
| 4 | 3796387 | rs28610504 | -0.729 | 0.055 | -1.455 | -0.541 | -0.012 | 0.226 | 0.226 | 1.182 | 0.5 | 0.88 | 0.49 | 0.29 | 0.43 | 0.53 |
| 4 | 3804286 | rs28605619 | -0.406 | 0.717 | -0.806 | -0.457 | -0.001 | -0.02 | -0.02 | 0 | 0.4 | 0.56 | 0.46 | 0.22 | 0.44 | 0.48 |
| 4 | 3813929 | rs28612860 | 0.439 | 0.436 | -0.632 | 0.833 | -0.008 | 0.087 | 0.087 | 0 | 0.86 | 0.81 | 0.8 | 0.58 | 0.88 | 0.71 |
| 4 | 3824221 | rs28590539 | -0.201 | 0.732 | -0.909 | -0.52 | 0.002 | -0.024 | -0.024 | 0.328 | 0.47 | 0.55 | 0.5 | 0.05 | 0.45 | 0.18 |
| 4 | 3845104 | rs28687658 | 0.248 | 0.684 | -1.059 | 0.275 | 0 | -0.003 | -0.003 | 0 | 0.6 | 0.35 | 0.46 | 0.65 | 0.65 | 0.55 |
| 4 | 3856471 | rs28366830 | -0.196 | 0.472 | -1.587 | 0.701 | -0.026 | 0.002 | 0.002 | 0.12 | 0.57 | 0.83 | 0.57 | 0.64 | 0.5 | 0.48 |
| 4 | 3874052 | rs28523451 | -0.564 | -0.396 | -1.065 | 1.105 | 0.03 | -0.025 | -0.025 | 0.053 | 0.28 | 0.79 | 0.16 | 0.37 | 0.14 | 0.24 |
| 4 | 3923728 | rs3889790 | -0.136 | 0.501 | -0.677 | 0.518 | 0.002 | 0.008 | 0.008 | 0.375 | 0.45 | 0.42 | 0.5 | 0.29 | 0.42 | 0.35 |
| 4 | 3942867 | rs6422714 | -0.227 | 1.11 | -1.786 | 0.539 | -0.061 | 0.165 | 0.165 | 6.649 | 0.43 | 0.12 | 0.46 | 0.19 | 0.44 | 0.21 |
| 4 | 3943984 | rs7695071 | -0.244 | 1.091 | -1.718 | 0.529 | -0.052 | 0.16 | 0.16 | 7.108 | 0.44 | 0.13 | 0.48 | 0.19 | 0.44 | 0.21 |
| 4 | 4039718 | rs13103428 | -0.028 | 1.135 | -1.97 | 0.1 | 0.019 | 0.095 | 0.095 | 0 | 0.51 | 0.68 | 0.63 | 0.28 | 0.44 | 0.47 |
| 4 | 4167133 | rs7679962 | -0.605 | 1.581 | -2.127 | -0.266 | -0.038 | 0.074 | 0.074 | 0 | 0.18 | 0.03 | 0.09 | 0.03 | 0.2 | 0.14 |
| 4 | 4183931 | rs2920250 | -0.313 | 1.438 | -2.332 | -0.013 | 0.043 | 0.046 | 0.046 | 0 | 0.53 | 0.19 | 0.48 | 0.66 | 0.57 | 0.69 |
| 4 | 4185610 | rs6840829 | -0.132 | 1.357 | -1.701 | -0.06 | 0.039 | 0.05 | 0.05 | 0 | 0.52 | 0.25 | 0.47 | 0.65 | 0.56 | 0.64 |
| 4 | 4206953 | rs2916422 | 0.449 | 1.773 | -0.93 | 0.609 | -0.007 | -0.007 | -0.007 | 0.009 | 0.48 | 0.54 | 0.29 | 0.35 | 0.4 | 0.52 |
| 4 | 4207366 | rs1991917 | -0.75 | 0.914 | -1.231 | 0.864 | -0.047 | -0.092 | -0.092 | 0 | 0.77 | 0.96 | 0.75 | 0.71 | 0.72 | 0.88 |
| 4 | 4208861 | rs1991915 | 0.14 | 0.808 | -0.652 | 0.265 | -0.002 | -0.001 | -0.001 | 0 | 0.65 | 0.7 | 0.39 | 0.37 | 0.59 | 0.61 |
| 4 | 4209796 | rs2916424 | -0.241 | 0.605 | -0.635 | 0.313 | -0.009 | -0.008 | -0.008 | 2.637 | 0.7 | 0.79 | 0.54 | 0.39 | 0.63 | 0.69 |
| 4 | 4219590 | rs2920227 | -0.566 | 0.264 | -0.416 | 0.209 | -0.001 | -0.011 | -0.011 | 0 | 0.08 | 0.08 | 0.04 | 0.01 | 0.07 | 0.13 |
| 4 | 4238315 | rs2920244 | -0.952 | -0.4 | -0.065 | 0.574 | -0.044 | 0.038 | 0.038 | 0 | 0.27 | 0.56 | 0.41 | 0.28 | 0.16 | 0.34 |
| 4 | 4247059 | rs3846232 | -0.942 | -0.22 | -0.474 | 0.514 | -0.039 | -0.072 | -0.072 | 0 | 0.3 | 0.16 | 0.33 | 0.42 | 0.34 | 0.42 |
| 4 | 4249300 | rs2916468 | -1.122 | -0.105 | -0.047 | 0.57 | 0 | 0.022 | 0.022 | 0 | 0.36 | 0.53 | 0.5 | 0.42 | 0.34 | 0.48 |
| 4 | 4249620 | rs2916464 | -1.11 | -0.095 | -0.043 | 0.575 | 0 | 0.023 | 0.023 | 0 | 0.36 | 0.53 | 0.5 | 0.42 | 0.34 | 0.48 |
| 4 | 4252956 | rs2980098 | -0.902 | 0.037 | 0.128 | 0.681 | 0.017 | 0.037 | 0.037 | 0 | 0.28 | 0.57 | 0.5 | 0.38 | 0.3 | 0.44 |
| 4 | 4253074 | rs2916457 | -1.127 | -0.6 | 0.031 | 0.959 | -0.012 | -0.034 | -0.034 | 0 | 0.9 | 0.97 | 0.83 | 0.77 | 0.86 | 0.89 |
| 4 | 4254798 | rs2916456 | 0.658 | -0.547 | 0.186 | 0.822 | 0.008 | 0.105 | 0.105 | 0 | 0.76 | 0.54 | 0.57 | 0.58 | 0.75 | 0.62 |
| 4 | 4258887 | rs2980091 | 1.234 | -0.607 | -0.335 | 0.418 | -0.002 | 0.001 | 0.001 | 0 | 0.42 | 0.39 | 0.49 | 0.44 | 0.55 | 0.33 |
| 4 | 4259052 | rs11936688 | -0.119 | -0.281 | -0.104 | 0.889 | 0.001 | 0.017 | 0.017 | 0 | 0.12 | 0.17 | 0.23 | 0.19 | 0.1 | 0.25 |
| 4 | 4259085 | rs11943160 | -0.73 | -0.345 | -0.136 | 0.794 | -0.013 | 0.01 | 0.01 | 0.074 | 0.29 | 0.35 | 0.27 | 0.32 | 0.18 | 0.43 |
| 4 | 4263087 | rs10937817 | -0.89 | -0.516 | -0.417 | 0.623 | -0.017 | -0.007 | -0.007 | 0 | 0.33 | 0.42 | 0.31 | 0.27 | 0.22 | 0.43 |
| 4 | 4263579 | rs2920228 | 0.378 | 0.399 | -0.58 | 0.819 | -0.004 | 0.033 | 0.033 | 0 | 0.69 | 0.55 | 0.55 | 0.53 | 0.71 | 0.6 |
| 4 | 4267129 | rs6835291 | -0.899 | 0.205 | -0.748 | 0.301 | 0.001 | -0.019 | -0.019 | 0 | 0.14 | 0.16 | 0.07 | 0.01 | 0.11 | 0.08 |
| 4 | 4267929 | rs1865328 | -0.731 | 1.228 | -0.567 | 0.935 | 0.032 | 0.029 | 0.029 | 0 | 0.26 | 0.28 | 0.38 | 0.51 | 0.43 | 0.32 |
| 4 | 4270726 | rs7671899 | -1.165 | 0.604 | 0.586 | 0.535 | 0.026 | -0.072 | -0.072 | 0 | 0.12 | 0.43 | 0.11 | 0 | 0.08 | 0.02 |
| 4 | 4275287 | rs2272740 | -1.794 | -0.115 | 0.336 | 1.203 | -0.011 | 0.025 | 0.025 | 0.248 | 0.25 | 0.68 | 0.22 | 0.48 | 0.19 | 0.38 |
| 4 | 4277736 | rs4689210 | 1.33 | -0.519 | 0.336 | 1.136 | -0.021 | -0.041 | -0.041 | 0 | 0.78 | 0.36 | 0.85 | 0.96 | 0.86 | 0.89 |
| 4 | 4279395 | rs3733425 | 1.427 | -0.525 | 0.335 | 1.348 | -0.014 | -0.009 | -0.009 | 0.19 | 0.77 | 0.34 | 0.84 | 0.83 | 0.85 | 0.84 |
| 4 | 4285756 | rs2980213 | -1.384 | 0.345 | 0.298 | 1.5 | 0.003 | 0.092 | 0.092 | 0 | 0.26 | 0.47 | 0.35 | 0.55 | 0.29 | 0.33 |
| 4 | 4285813 | rs2980212 | 0.65 | 0.213 | -0.064 | 1.071 | -0.003 | -0.002 | -0.002 | 0 | 0.75 | 0.69 | 0.84 | 0.78 | 0.8 | 0.76 |
| 4 | 4291205 | rs2980155 | -1.08 | 0.049 | 0.078 | 1.265 | 0.004 | 0.092 | 0.092 | 0 | 0.26 | 0.47 | 0.35 | 0.55 | 0.29 | 0.33 |
| 4 | 4291517 | rs2980156 | -1.15 | 0.028 | 0.073 | 1.254 | 0 | 0.065 | 0.065 | 0 | 0.26 | 0.53 | 0.35 | 0.55 | 0.29 | 0.33 |
| 4 | 4296665 | rs4689241 | 2.093 | -0.025 | -0.194 | 1.134 | 0.023 | 0.209 | 0.209 | 0 | 0.5 | 0.11 | 0.51 | 0.2 | 0.55 | 0.45 |
| 4 | 4297389 | rs2980164 | 1.251 | -0.046 | 0 | 1.134 | -0.02 | -0.008 | -0.008 | 0 | 0.73 | 0.38 | 0.82 | 0.77 | 0.83 | 0.78 |
| 4 | 4298599 | rs2920174 | 1.251 | -0.046 | 0 | 1.134 | -0.02 | -0.008 | -0.008 | 0 | 0.73 | 0.38 | 0.83 | 0.77 | 0.83 | 0.78 |
| 4 | 4300855 | rs2920253 | -0.306 | -0.015 | -0.034 | 1.338 | -0.027 | -0.023 | -0.023 | 0.836 | 0.19 | 0.09 | 0.29 | 0.25 | 0.27 | 0.16 |
| 4 | 4304522 | rs2920217 | -0.341 | -0.011 | -0.029 | 1.355 | -0.024 | -0.024 | -0.024 | 0 | 0.19 | 0.1 | 0.29 | 0.25 | 0.27 | 0.16 |
| 4 | 4308513 | rs894486 | 1.419 | -0.05 | 0.055 | 1.391 | 0.002 | -0.019 | -0.019 | 0.662 | 0.61 | 0.66 | 0.59 | 0.52 | 0.58 | 0.62 |
| 4 | 4316736 | rs2980181 | -2.041 | -0.063 | 0.098 | 1.366 | -0.005 | -0.026 | -0.026 | 0 | 0.42 | 0.49 | 0.44 | 0.8 | 0.43 | 0.55 |
| 4 | 4320468 | rs6839561 | -0.377 | 0.135 | -0.22 | 1.398 | -0.021 | -0.027 | -0.027 | 0 | 0.2 | 0.09 | 0.28 | 0.24 | 0.26 | 0.16 |
| 4 | 4323865 | rs894484 | -1.796 | 0.181 | -0.191 | 1.329 | 0.017 | -0.004 | -0.004 | 0 | 0.46 | 0.67 | 0.45 | 0.48 | 0.43 | 0.38 |
| 4 | 4327912 | rs1020265 | 1.721 | 0.478 | -0.596 | 0.86 | 0.05 | 0.213 | 0.213 | 0 | 0.5 | 0.12 | 0.51 | 0.2 | 0.55 | 0.45 |
| 4 | 4333418 | rs1031095 | -1.754 | 0.745 | -0.052 | 1.159 | 0.005 | 0.002 | 0.002 | 0 | 0.44 | 0.56 | 0.44 | 0.61 | 0.45 | 0.36 |
| 4 | 4335824 | rs2980206 | -0.006 | 1.065 | -0.452 | 1.239 | -0.043 | -0.068 | -0.068 | 0 | 0.81 | 0.92 | 0.72 | 0.63 | 0.7 | 0.81 |
| 4 | 4336008 | rs2980207 | -0.96 | 0.999 | -0.591 | 1.08 | 0.009 | 0.02 | 0.02 | 0.035 | 0.21 | 0.11 | 0.16 | 0.16 | 0.19 | 0.21 |
| 4 | 4337286 | rs11735439 | -0.336 | 1.156 | -0.456 | 1.161 | -0.058 | -0.085 | -0.085 | 0 | 0.19 | 0.05 | 0.28 | 0.37 | 0.3 | 0.19 |
| 4 | 4343240 | rs4688956 | 0.567 | 1.045 | -0.177 | 0.801 | -0.024 | -0.03 | -0.03 | 0 | 0.67 | 0.79 | 0.64 | 0.61 | 0.55 | 0.7 |
| 4 | 4343261 | rs4689307 | 1.251 | 0.902 | 0.271 | 0.539 | 0.003 | 0.002 | 0.002 | 0.015 | 0.59 | 0.4 | 0.54 | 0.47 | 0.49 | 0.61 |
| 4 | 4346065 | rs4689314 | -0.4 | 0.927 | 0.223 | 0.272 | 0.003 | -0.02 | -0.02 | 0.167 | 0.94 | 0.91 | 0.95 | 1 | 0.93 | 0.98 |
| 4 | 4346237 | rs10027109 | 1.865 | 1.149 | -0.099 | 0.146 | 0.014 | 0.009 | 0.009 | 0 | 0.47 | 0.34 | 0.48 | 0.5 | 0.41 | 0.57 |

| | | | | | | | | | | | | | | | | |
|---|---|---|---|---|---|---|---|---|---|---|---|---|---|---|---|---|
| 4 | 4346427 | rs3981 | -0.057 | 1.326 | 0.005 | 0.213 | -0.033 | -0.049 | -0.049 | 0 | 0.27 | 0.15 | 0.35 | 0.44 | 0.38 | 0.28 |
| 4 | 4346538 | rs16835627 | 1.241 | 1.373 | 0.058 | 0.236 | 0 | 0.001 | 0.001 | 0 | 0.57 | 0.64 | 0.54 | 0.55 | 0.46 | 0.64 |
| 4 | 4347505 | rs4689316 | 0.604 | 0.827 | 0.526 | 0.238 | 0.01 | -0.054 | -0.054 | 0 | 0.85 | 0.8 | 0.89 | 1 | 0.84 | 0.92 |
| 4 | 4349205 | rs7668915 | 1.246 | 0.765 | 0.302 | 0.255 | 0.02 | -0.046 | -0.046 | 0.113 | 0.89 | 0.7 | 0.89 | 0.95 | 0.88 | 0.92 |
| 4 | 4351730 | rs6843595 | 0.971 | 0.714 | 0.321 | 0.264 | 0.022 | -0.099 | -0.099 | 0 | 0.8 | 0.65 | 0.84 | 0.95 | 0.79 | 0.85 |
| 4 | 4354778 | rs7677584 | -1.353 | 0.624 | 0.441 | 0.053 | 0.005 | 0.099 | 0.099 | 0.022 | 0.16 | 0.45 | 0.18 | 0.33 | 0.15 | 0.14 |
| 4 | 4355380 | rs6446632 | 1.174 | 0.432 | 0.252 | 0.082 | 0.003 | -0.009 | -0.009 | 0.007 | 0.72 | 0.43 | 0.76 | 0.72 | 0.75 | 0.84 |
| 4 | 4355774 | rs13146316 | 0.888 | 0.462 | -0.351 | 0.175 | 0.005 | -0.01 | -0.01 | 0.018 | 0.7 | 0.45 | 0.73 | 0.67 | 0.7 | 0.78 |
| 4 | 4360381 | rs7674400 | -0.38 | 0.421 | -0.943 | -0.348 | -0.008 | 0.005 | 0.005 | 0.932 | 0.35 | 0.62 | 0.58 | 0.71 | 0.47 | 0.3 |
| 4 | 4360662 | rs7698300 | 0.41 | 0.485 | -0.682 | -0.656 | -0.002 | -0.037 | -0.037 | 2.641 | 0.68 | 0.66 | 0.72 | 0.96 | 0.66 | 0.71 |
| 4 | 4362223 | rs10937869 | -0.504 | 0.131 | -0.219 | 0.06 | -0.024 | -0.036 | -0.036 | 3.865 | 0.09 | 0.21 | 0.07 | 0.03 | 0.06 | 0.09 |
| 4 | 4362337 | rs4279148 | -1.444 | 0.335 | -0.001 | 0.601 | -0.012 | 0.083 | 0.083 | 0 | 0.2 | 0.48 | 0.16 | 0.31 | 0.18 | 0.15 |
| 4 | 4363642 | rs10018486 | 0.188 | 0.416 | -0.802 | 0.457 | -0.003 | 0.03 | 0.03 | 0.233 | 0.47 | 0.32 | 0.36 | 0.37 | 0.47 | 0.51 |
| 4 | 4365066 | rs4689340 | -0.13 | 0.256 | -1.019 | 0.314 | -0.003 | 0.041 | 0.041 | 0 | 0.54 | 0.27 | 0.47 | 0.37 | 0.53 | 0.51 |
| 4 | 4366380 | rs4478124 | 0.304 | -0.112 | -1.075 | 0.348 | -0.012 | -0.005 | -0.005 | 3.295 | 0.62 | 0.64 | 0.63 | 0.31 | 0.68 | 0.51 |
| 4 | 4366999 | rs7690552 | 0.04 | -0.261 | -0.81 | 0.413 | -0.01 | -0.023 | -0.023 | 0 | 0.39 | 0.6 | 0.52 | 0.26 | 0.33 | 0.31 |
| 4 | 4369400 | rs4330304 | -0.396 | 0.66 | -0.274 | 0.857 | 0.023 | 0.051 | 0.051 | 0.052 | 0.83 | 0.78 | 0.85 | 0.62 | 0.77 | 0.71 |
| 4 | 4370684 | rs12639864 | -0.175 | 0.559 | -0.21 | 0.761 | 0 | -0.001 | -0.001 | 0 | 0.58 | 0.62 | 0.54 | 0.62 | 0.52 | 0.69 |
| 4 | 4371097 | rs7664767 | 0.256 | 0.554 | -0.582 | 0.492 | 0.028 | -0.019 | -0.019 | 0.001 | 0.16 | 0.33 | 0.15 | 0.13 | 0.19 | 0.08 |
| 4 | 4372439 | rs12512085 | -0.275 | 0.508 | -0.581 | 0.468 | 0.005 | -0.012 | -0.012 | 0 | 0.63 | 0.59 | 0.6 | 0.76 | 0.63 | 0.73 |
| 4 | 4376170 | rs4494977 | -0.926 | 0.835 | -1.479 | 0.222 | -0.093 | 0.102 | 0.102 | 0 | 0.75 | 0.96 | 0.66 | 0.94 | 0.68 | 0.79 |
| 4 | 4376744 | rs6816148 | -0.008 | -0.083 | -0.909 | 0.778 | -0.002 | 0 | 0 | 0 | 0.21 | 0.2 | 0.22 | 0.29 | 0.16 | 0.22 |
| 4 | 4381429 | rs6825630 | -0.839 | 0.107 | -0.859 | 0.905 | 0.004 | -0.007 | -0.007 | 0 | 0.27 | 0.73 | 0.27 | 0.37 | 0.21 | 0.25 |
| 4 | 4387560 | rs4305464 | -1.134 | 0.183 | -0.751 | 0.577 | 0.009 | -0.003 | -0.003 | 0.014 | 0.36 | 0.72 | 0.4 | 0.62 | 0.34 | 0.26 |
| 4 | 4388017 | rs4689412 | 0.842 | 0.268 | -0.389 | 0.214 | -0.001 | 0.002 | 0.002 | 0 | 0.61 | 0.54 | 0.6 | 0.57 | 0.52 | 0.45 |
| 4 | 4392357 | rs4689416 | 0.017 | 0.214 | -0.434 | -0.224 | -0.002 | 0.005 | 0.005 | 0 | 0.68 | 0.76 | 0.75 | 0.67 | 0.65 | 0.52 |
| 4 | 4393068 | rs3792686 | 1.265 | 0.173 | -0.423 | -0.252 | -0.01 | 0 | 0 | 0 | 0.47 | 0.28 | 0.41 | 0.47 | 0.43 | 0.35 |
| 4 | 4394618 | rs10017186 | -1.084 | 0.529 | 0.362 | -0.639 | 0.001 | -0.01 | -0.01 | 0.327 | 0.22 | 0.32 | 0.35 | 0.2 | 0.26 | 0.19 |
| 4 | 4397537 | rs4555567 | 1.474 | -0.051 | 0.441 | -0.88 | -0.01 | -0.115 | -0.115 | 0 | 0.71 | 0.37 | 0.77 | 0.98 | 0.72 | 0.8 |
| 4 | 4402199 | rs4402980 | 1.422 | -0.243 | 0.314 | -0.811 | 0.017 | 0.024 | 0.024 | 1.927 | 0.67 | 0.21 | 0.78 | 0.94 | 0.74 | 0.81 |
| 4 | 4404843 | rs4234746 | 0.123 | -0.158 | 0.363 | -0.701 | 0 | 0.001 | 0.001 | 0 | 0.82 | 0.82 | 0.86 | 0.95 | 0.82 | 0.87 |
| 4 | 4410007 | rs11726155 | -1.773 | -0.297 | 0.534 | -0.775 | 0.004 | 0.001 | 0.001 | 0.149 | 0.29 | 0.76 | 0.21 | 0.07 | 0.2 | 0.16 |
| 4 | 4419611 | rs10002237 | 0.372 | -0.184 | 0.402 | -0.723 | 0.004 | 0.004 | 0.004 | 0 | 0.11 | 0.12 | 0.1 | 0.04 | 0.12 | 0.11 |
| 4 | 4425620 | rs4689519 | 0.143 | 0.108 | 0.147 | -0.581 | -0.031 | -0.048 | -0.048 | 0 | 0.15 | 0.01 | 0.2 | 0.22 | 0.18 | 0.18 |
| 4 | 4427343 | rs3935096 | -0.983 | 0.025 | 0.814 | -0.856 | -0.001 | -0.036 | -0.036 | 0 | 0.1 | 0.28 | 0.07 | 0.03 | 0.08 | 0.06 |
| 4 | 4436436 | rs6825715 | -2.108 | -0.021 | 1.012 | -0.936 | 0.035 | 0.085 | 0.085 | 2.334 | 0.32 | 0.86 | 0.24 | 0.07 | 0.23 | 0.18 |
| 4 | 4442734 | rs11734030 | 0.346 | 0.026 | 0.986 | -0.908 | -0.002 | 0.007 | 0.007 | 0 | 0.13 | 0.1 | 0.12 | 0.05 | 0.15 | 0.11 |
| 4 | 4445315 | rs6446647 | -2.085 | 0.027 | 1.146 | -0.936 | 0.007 | 0.009 | 0.009 | 2.821 | 0.31 | 0.77 | 0.23 | 0.11 | 0.23 | 0.18 |
| 4 | 4445875 | rs7698589 | -1.313 | 0.062 | 1.376 | -0.953 | 0.002 | -0.052 | -0.052 | 0.229 | 0.12 | 0.35 | 0.08 | 0.03 | 0.09 | 0.06 |
| 4 | 4474034 | rs9990629 | 0.142 | 0.087 | 1.183 | -0.933 | -0.001 | 0.001 | 0.001 | 0 | 0.14 | 0.18 | 0.13 | 0.05 | 0.15 | 0.12 |
| 4 | 4487383 | rs6446652 | -2.199 | 0.103 | 1.222 | -0.987 | 0.013 | 0.02 | 0.02 | 2.252 | 0.32 | 0.79 | 0.22 | 0.07 | 0.23 | 0.18 |
| 4 | 4487610 | rs11937935 | 1.566 | 0.041 | 1.437 | -0.999 | -0.016 | -0.023 | -0.023 | 0 | 0.06 | 0.19 | 0.01 | 0 | 0 | 0 |
| 4 | 4490280 | rs11940730 | -2.198 | 0.035 | 1.483 | -1.009 | -0.032 | -0.096 | -0.096 | 0.081 | 0.17 | 0.61 | 0.09 | 0.03 | 0.07 | 0.06 |
| 4 | 4491724 | rs6811336 | 0.943 | -0.063 | 1.408 | -0.925 | -0.017 | -0.024 | -0.024 | 0.004 | 0.07 | 0.27 | 0.02 | 0 | 0 | 0 |
| 4 | 4498594 | rs6838568 | 0.693 | 0.306 | 0.508 | -0.717 | -0.005 | -0.015 | -0.015 | 0.087 | 0.19 | 0.11 | 0.13 | 0.34 | 0.18 | 0.35 |
| 4 | 4520962 | rs9654059 | -0.254 | 0.253 | 0.581 | -0.869 | -0.003 | -0.01 | -0.01 | 0 | 0.18 | 0.22 | 0.16 | 0.06 | 0.19 | 0.15 |
| 4 | 4526248 | rs12644497 | -0.923 | 0.164 | 0.489 | -0.893 | -0.003 | -0.002 | -0.002 | 0.789 | 0.4 | 0.41 | 0.38 | 0.39 | 0.36 | 0.51 |
| 4 | 4533974 | rs7684307 | -0.497 | 0.159 | 0.462 | -0.86 | -0.021 | -0.048 | -0.048 | 0.109 | 0.21 | 0.36 | 0.21 | 0.06 | 0.19 | 0.17 |
| 4 | 4534579 | rs28695605 | 0.072 | 0.189 | 0.039 | -0.81 | 0.015 | 0.025 | 0.025 | 0.902 | 0.17 | 0.1 | 0.16 | 0.06 | 0.17 | 0.17 |
| 4 | 4550632 | rs6829211 | -0.609 | 0.23 | -0.002 | -1.044 | -0.007 | -0.013 | -0.013 | 1.082 | 0.39 | 0.58 | 0.32 | 0.32 | 0.3 | 0.42 |
| 4 | 4555497 | rs7689784 | 0.744 | 0.407 | -0.58 | -1.225 | -0.01 | -0.02 | -0.02 | 0 | 0.22 | 0.13 | 0.37 | 0.61 | 0.31 | 0.33 |
| 4 | 4557259 | rs12646052 | 0.94 | 0.393 | -0.74 | -1.222 | 0 | -0.011 | -0.011 | 0 | 0.22 | 0.11 | 0.22 | 0.32 | 0.18 | 0.36 |
| 4 | 4559234 | rs1345041 | -1.343 | 0.607 | -0.629 | -1.368 | -0.002 | 0.002 | 0.002 | 0.569 | 0.69 | 0.72 | 0.83 | 0.97 | 0.72 | 0.85 |
| 4 | 4560776 | rs4689885 | -1.377 | 0.625 | -0.743 | -1.359 | -0.003 | 0.025 | 0.025 | 0.53 | 0.69 | 0.77 | 0.83 | 0.97 | 0.72 | 0.85 |
| 4 | 4562693 | rs16835890 | 0.644 | 0.695 | -0.97 | -0.878 | 0 | -0.013 | -0.013 | 0 | 0.22 | 0.09 | 0.22 | 0.32 | 0.18 | 0.35 |
| 4 | 4562916 | rs10016411 | -0.296 | 0.913 | -1.026 | -1.316 | 0 | -0.007 | -0.007 | 0 | 0.19 | 0.18 | 0.21 | 0.04 | 0.21 | 0.17 |
| 4 | 4565610 | rs11728302 | -0.269 | 1.238 | -1.115 | -1.077 | 0.001 | -0.084 | -0.084 | 0 | 0.34 | 0.62 | 0.29 | 0.05 | 0.29 | 0.29 |
| 4 | 4569460 | rs4689888 | -0.164 | 1.058 | -1.909 | -0.588 | -0.065 | 0.12 | 0.12 | 0.128 | 0.42 | 0.15 | 0.57 | 0.78 | 0.55 | 0.57 |
| 4 | 4579042 | rs4689174 | -0.35 | 0.803 | -2.076 | 0.071 | -0.013 | -0.048 | -0.048 | 2.65 | 0.28 | 0.06 | 0.28 | 0.38 | 0.3 | 0.3 |
| 4 | 4585470 | rs13142209 | -0.301 | 1.331 | -1.549 | -0.069 | 0.004 | -0.014 | -0.014 | 0.883 | 0.25 | 0.36 | 0.16 | 0.14 | 0.25 | 0.25 |
| 4 | 4586438 | rs10428352 | -0.525 | 0.74 | -1.344 | 0.045 | -0.003 | -0.008 | -0.008 | 0 | 0.61 | 0.67 | 0.5 | 0.52 | 0.58 | 0.73 |
| 4 | 4593893 | rs6446665 | -0.351 | 0.614 | -1.222 | -0.01 | 0.028 | 0.021 | 0.021 | 2.031 | 0.5 | 0.43 | 0.6 | 0.6 | 0.55 | 0.45 |
| 4 | 4594732 | rs4234835 | -0.04 | 0.764 | -2.161 | -0.173 | -0.061 | 0.018 | 0.018 | 2.351 | 0.66 | 0.97 | 0.79 | 0.79 | 0.6 | 0.6 |
| 4 | 4594743 | rs4689893 | 0.841 | 0.266 | -2.02 | 0.007 | -0.01 | -0.013 | -0.013 | 0 | 0.14 | 0.12 | 0.32 | 0.15 | 0.16 | 0.15 |
| 4 | 4595807 | rs751262 | -0.179 | 0.213 | -1.675 | -0.032 | 0.014 | 0.013 | 0.013 | 0 | 0.63 | 0.75 | 0.57 | 0.8 | 0.66 | 0.61 |
| 4 | 4602813 | rs12509575 | 0.449 | 0.684 | -1.734 | 0.485 | -0.017 | -0.031 | -0.031 | 0 | 0.15 | 0.07 | 0.32 | 0.15 | 0.15 | 0.15 |
| 4 | 4602905 | rs11733672 | -0.147 | 0.802 | -1.494 | 0.48 | -0.029 | -0.02 | -0.02 | 0.051 | 0.14 | 0.08 | 0.11 | 0.17 | 0.19 | 0.15 |
| 4 | 4610628 | rs2033958 | 0.605 | 0.657 | -1.484 | 0.586 | -0.029 | -0.046 | -0.046 | 0 | 0.18 | 0.05 | 0.2 | 0.34 | 0.23 | 0.27 |
| 4 | 4614187 | rs6853673 | 0.6 | 0.632 | -1.451 | 0.563 | -0.023 | -0.035 | -0.035 | 0 | 0.18 | 0.1 | 0.2 | 0.34 | 0.23 | 0.27 |

| | | | | | | | | | | | | | | | | |
|---|---|---|---|---|---|---|---|---|---|---|---|---|---|---|---|---|
| 4 | 4615708 | rs1837711 | -0.849 | 0.175 | -1.21 | 0.388 | -0.014 | -0.008 | -0.008 | 3.794 | 0.4 | 0.71 | 0.43 | 0.61 | 0.46 | 0.56 |
| 4 | 4629166 | rs1510798 | 0.625 | 0.395 | -1.434 | 0.683 | -0.024 | -0.037 | -0.037 | 0 | 0.19 | 0.11 | 0.21 | 0.33 | 0.24 | 0.27 |
| 4 | 4631235 | rs2102528 | -0.698 | -0.09 | -1.937 | 0.234 | 0.027 | 0.057 | 0.057 | 0.003 | 0.52 | 0.34 | 0.67 | 0.69 | 0.58 | 0.59 |
| 4 | 4638578 | rs17706055 | 0.492 | -0.059 | -1.957 | 0.276 | 0 | 0 | 0 | 0 | 0.22 | 0.24 | 0.34 | 0.2 | 0.19 | 0.2 |
| 4 | 4645158 | rs6840368 | 0.173 | -0.128 | -1.959 | 0.181 | 0.003 | 0.004 | 0.004 | 0 | 0.29 | 0.26 | 0.37 | 0.21 | 0.24 | 0.23 |
| 4 | 4651063 | rs4689911 | -0.277 | 0.05 | -2.779 | 0.047 | -0.02 | 0.063 | 0.063 | 0.018 | 0.48 | 0.85 | 0.65 | 0.64 | 0.49 | 0.63 |
| 4 | 4653004 | rs6830734 | -0.484 | 0.517 | -2.486 | 0.269 | -0.015 | -0.019 | -0.019 | 3.409 | 0.34 | 0.32 | 0.37 | 0.48 | 0.43 | 0.43 |
| 4 | 4653127 | rs6824817 | -0.115 | 0.562 | -2.37 | 0.169 | -0.001 | -0.001 | -0.001 | 0 | 0.16 | 0.2 | 0.17 | 0.16 | 0.2 | 0.17 |
| 4 | 4673698 | rs7694788 | -1.01 | 0.28 | -2.735 | -0.219 | 0.028 | 0.118 | 0.118 | 0.096 | 0.79 | 0.99 | 0.92 | 0.98 | 0.85 | 0.85 |
| 4 | 4675257 | rs13140219 | 0.62 | 0.242 | -2.907 | -0.25 | -0.01 | 0.001 | 0.001 | 0.157 | 0.17 | 0.11 | 0.14 | 0.17 | 0.18 | 0.14 |
| 4 | 4675296 | rs6446675 | 0.056 | 0.219 | -2.92 | -0.256 | 0.02 | 0.113 | 0.113 | 0 | 0.62 | 0.88 | 0.78 | 0.81 | 0.67 | 0.71 |
| 4 | 4678511 | rs4552410 | -0.747 | 0.891 | -2.745 | 0.218 | -0.02 | -0.019 | -0.019 | 0 | 0.59 | 0.86 | 0.71 | 0.6 | 0.58 | 0.57 |
| 4 | 4682289 | rs13149758 | -0.175 | 1.009 | -1.841 | 0.369 | -0.034 | -0.007 | -0.007 | 0 | 0.16 | 0.05 | 0.13 | 0.16 | 0.17 | 0.09 |
| 4 | 4686072 | rs17707114 | 0.18 | 1.009 | -1.761 | 0.127 | -0.012 | 0.044 | 0.044 | 0 | 0.16 | 0.09 | 0.1 | 0.08 | 0.16 | 0.1 |
| 4 | 4687339 | rs4458410 | -1.232 | 1.324 | -1.329 | 0.205 | 0.004 | 0.025 | 0.025 | 0.012 | 0.45 | 0.67 | 0.54 | 0.69 | 0.57 | 0.53 |
| 4 | 4689987 | rs10937886 | 0.049 | 1.645 | -0.567 | 0.866 | 0.04 | 0.136 | 0.136 | 0 | 0.14 | 0.27 | 0.16 | 0.4 | 0.24 | 0.29 |
| 4 | 4696955 | rs11732485 | -0.466 | 1.494 | -0.596 | 0.816 | 0.021 | 0.043 | 0.043 | 0 | 0.2 | 0.27 | 0.25 | 0.4 | 0.35 | 0.32 |
| 4 | 4697613 | rs17713046 | -0.056 | 1.563 | -0.419 | 0.811 | 0 | 0.005 | 0.005 | 0 | 0.24 | 0.29 | 0.27 | 0.34 | 0.24 | 0.23 |
| 4 | 4700606 | rs7690052 | -0.198 | 0.664 | -0.253 | 0.487 | -0.033 | 0.029 | 0.029 | 0.138 | 0.12 | 0.29 | 0.06 | 0.15 | 0.06 | 0.12 |
| 4 | 4704014 | rs4476553 | -0.894 | 0.895 | -0.853 | 0.657 | -0.01 | 0.007 | 0.007 | 0.84 | 0.44 | 0.79 | 0.51 | 0.56 | 0.53 | 0.42 |
| 4 | 4711863 | rs12503646 | 0.765 | 0.472 | -1.43 | -0.119 | 0.008 | 0.091 | 0.091 | 0.052 | 0.25 | 0.05 | 0.27 | 0.05 | 0.21 | 0.16 |
| 4 | 4716184 | rs12648481 | 0.317 | 0.809 | -1.054 | 0.627 | -0.013 | -0.039 | -0.039 | 0.037 | 0.29 | 0.24 | 0.3 | 0.52 | 0.33 | 0.36 |
| 4 | 4716521 | rs17713587 | 1.461 | 0.851 | -1.217 | 0.748 | -0.053 | 0.006 | 0.006 | 0 | 0.18 | 0.01 | 0.1 | 0.14 | 0.19 | 0.11 |
| 4 | 4719494 | rs1000579 | 0.185 | 0.466 | -0.398 | 0.67 | -0.004 | 0.001 | 0.001 | 0 | 0.55 | 0.47 | 0.61 | 0.54 | 0.41 | 0.51 |
| 4 | 4720455 | rs10004542 | 0.64 | 0.117 | -0.557 | 0.714 | 0.059 | 0.128 | 0.128 | 1.071 | 0.46 | 0.1 | 0.41 | 0.25 | 0.34 | 0.35 |
| 4 | 4721078 | rs1000578 | 1.635 | 0.337 | -0.573 | 0.678 | -0.002 | 0.03 | 0.03 | 0 | 0.06 | 0.05 | 0.09 | 0.26 | 0.14 | 0.19 |
| 4 | 4721690 | rs12644897 | 0.954 | -0.166 | -0.464 | 0.412 | -0.026 | 0.025 | 0.025 | 0 | 0.22 | 0.3 | 0.27 | 0.36 | 0.12 | 0.24 |
| 4 | 4721965 | rs10013273 | -1.958 | 0.476 | -0.804 | 0.126 | -0.056 | 0.032 | 0.032 | 0 | 0.78 | 0.97 | 0.88 | 0.85 | 0.75 | 0.85 |
| 4 | 4723984 | rs11724809 | -0.218 | -0.164 | -0.718 | -0.267 | -0.003 | -0.007 | -0.007 | 0.266 | 0.35 | 0.33 | 0.36 | 0.08 | 0.4 | 0.29 |
| 4 | 4724676 | rs13130037 | 1.125 | -0.015 | -0.916 | -0.221 | 0.03 | -0.007 | -0.007 | 0 | 0.11 | 0.18 | 0.16 | 0.06 | 0.2 | 0.15 |
| 4 | 4727223 | rs10009773 | -1.152 | 0.106 | -0.542 | 0.012 | -0.014 | -0.014 | -0.014 | 0 | 0.7 | 0.75 | 0.78 | 0.61 | 0.62 | 0.65 |
| 4 | 4727507 | rs10005569 | 0.022 | 0.748 | -0.403 | 0.208 | -0.001 | 0.064 | 0.064 | 0 | 0.22 | 0.08 | 0.2 | 0.03 | 0.2 | 0.15 |
| 4 | 4737134 | rs4689928 | -0.286 | 0.718 | -0.997 | 0.552 | -0.037 | -0.006 | -0.006 | 0 | 0.24 | 0.12 | 0.28 | 0.26 | 0.33 | 0.32 |
| 4 | 4739301 | rs4381992 | -0.097 | 0.63 | -1.029 | 0.599 | -0.005 | 0.011 | 0.011 | 0 | 0.22 | 0.17 | 0.28 | 0.16 | 0.25 | 0.2 |
| 4 | 4739625 | rs4689929 | 0.222 | 0.835 | -1.488 | 0.871 | -0.054 | -0.105 | -0.105 | 0 | 0.25 | 0.1 | 0.27 | 0.51 | 0.28 | 0.31 |
| 4 | 4739687 | rs13149006 | -0.332 | 0.501 | -0.322 | 0.69 | 0.095 | 0.047 | 0.047 | 0 | 0.11 | 0.42 | 0.11 | 0.07 | 0.14 | 0.16 |
| 4 | 4744037 | rs10516168 | 0.399 | 0.599 | -0.26 | 0.733 | 0.029 | 0.009 | 0.009 | 0 | 0.07 | 0.11 | 0.08 | 0.06 | 0.12 | 0.14 |
| 4 | 4744593 | rs6822612 | -0.606 | 0.532 | -0.206 | 0.498 | 0.001 | 0.01 | 0.01 | 0.046 | 0.51 | 0.58 | 0.55 | 0.68 | 0.5 | 0.58 |
| 4 | 4745726 | rs2172445 | -0.542 | 0.517 | -0.259 | 0.568 | 0.001 | 0.028 | 0.028 | 0 | 0.52 | 0.42 | 0.55 | 0.68 | 0.5 | 0.57 |
| 4 | 4748816 | rs4574345 | 0.46 | 0.656 | -0.889 | 0.818 | -0.034 | -0.076 | -0.076 | 0 | 0.26 | 0.1 | 0.26 | 0.45 | 0.25 | 0.34 |
| 4 | 4749898 | rs767564 | -0.228 | 0.497 | -0.586 | 0.622 | 0.006 | -0.005 | -0.005 | 0.005 | 0.15 | 0.27 | 0.23 | 0.12 | 0.16 | 0.24 |
| 4 | 4751272 | rs7679624 | -1.613 | 0.47 | -1.201 | 0.621 | -0.019 | 0.037 | 0.037 | 0 | 0.7 | 0.88 | 0.81 | 0.82 | 0.67 | 0.83 |
| 4 | 4754129 | rs16836215 | 0.128 | 0.462 | -0.685 | 1.008 | -0.034 | -0.017 | -0.017 | 0 | 0.12 | 0.02 | 0.13 | 0.17 | 0.17 | 0.16 |
| 4 | 4761862 | rs1318947 | -0.997 | 0.636 | -0.079 | 1.349 | 0.018 | 0.03 | 0.03 | 0 | 0.24 | 0.38 | 0.36 | 0.4 | 0.32 | 0.24 |
| 4 | 4762672 | rs6854227 | -1.994 | 0.421 | 0.054 | 1.262 | -0.009 | -0.004 | -0.004 | 0 | 0.39 | 0.74 | 0.37 | 0.41 | 0.41 | 0.42 |
| 4 | 4764026 | rs1039906 | -0.459 | 0.596 | -0.149 | 1.068 | -0.005 | 0.007 | 0.007 | 0 | 0.22 | 0.2 | 0.23 | 0.21 | 0.25 | 0.21 |
| 4 | 4767695 | rs6830371 | -1.066 | 0.452 | 0.093 | 1.128 | 0.018 | 0.065 | 0.065 | 0.279 | 0.31 | 0.54 | 0.32 | 0.54 | 0.36 | 0.33 |
| 4 | 4774402 | rs12186237 | -0.528 | 0.426 | 0.055 | 0.891 | 0.013 | 0.077 | 0.077 | 0 | 0.08 | 0.24 | 0.08 | 0.14 | 0.07 | 0.14 |
| 4 | 4784580 | rs2933568 | -1.563 | 0.26 | -1.18 | 0.182 | -0.102 | 0.185 | 0.185 | 0 | 0.65 | 0.98 | 0.68 | 0.94 | 0.66 | 0.79 |
| 4 | 4792249 | rs10937891 | 1.614 | 0.022 | -0.789 | 0.525 | -0.003 | 0.029 | 0.029 | 0.303 | 0.11 | 0.14 | 0.06 | 0.24 | 0.11 | 0.15 |
| 4 | 4794814 | rs7681595 | 0.074 | -0.303 | -0.064 | 0.34 | -0.042 | 0.056 | 0.056 | 1.116 | 0.19 | 0.52 | 0.12 | 0.24 | 0.14 | 0.16 |
| 4 | 4799692 | rs2933572 | -1.772 | -0.264 | -0.066 | 0.111 | 0.009 | 0.108 | 0.108 | 0 | 0.62 | 0.83 | 0.67 | 0.86 | 0.69 | 0.75 |
| 4 | 4802946 | rs2933573 | -0.737 | -0.392 | -0.165 | 0.255 | 0.009 | -0.006 | -0.006 | 0 | 0.4 | 0.65 | 0.41 | 0.48 | 0.31 | 0.38 |
| 4 | 4803675 | rs2087865 | -1.567 | -0.345 | -0.125 | 0.117 | 0.005 | -0.028 | -0.028 | 6.043 | 0.54 | 0.44 | 0.6 | 0.92 | 0.69 | 0.74 |
| 4 | 4805454 | rs13117653 | -0.852 | -0.51 | -0.214 | 0.276 | 0 | -0.021 | -0.021 | 0 | 0.33 | 0.67 | 0.24 | 0.45 | 0.33 | 0.39 |
| 4 | 4807203 | rs16836288 | 0.483 | -0.522 | -0.134 | 0.267 | -0.031 | -0.007 | -0.007 | 0.006 | 0.11 | 0.39 | 0.09 | 0.06 | 0.04 | 0.05 |
| 4 | 4809353 | rs4689941 | -1.259 | -0.446 | -0.51 | 0.306 | -0.013 | 0.02 | 0.02 | 0.071 | 0.43 | 0.31 | 0.54 | 0.7 | 0.54 | 0.6 |
| 4 | 4809607 | rs6832194 | -1.309 | -0.43 | -0.675 | 0.35 | 0.007 | -0.03 | -0.03 | 0 | 0.82 | 0.87 | 0.71 | 0.73 | 0.84 | 0.79 |
| 4 | 4810491 | rs10937893 | 0.039 | -0.366 | -0.674 | 0.315 | -0.015 | -0.056 | -0.056 | 0 | 0.68 | 0.82 | 0.57 | 0.54 | 0.65 | 0.64 |
| 4 | 4810993 | rs4689942 | -2.831 | -0.396 | -0.936 | 0.172 | -0.089 | 0.132 | 0.132 | 0 | 0.69 | 0.98 | 0.69 | 0.91 | 0.69 | 0.77 |
| 4 | 4814852 | rs17659785 | -1.768 | -0.421 | -0.784 | 0.22 | -0.024 | 0.076 | 0.076 | 0 | 0.57 | 0.82 | 0.59 | 0.73 | 0.5 | 0.56 |
| 4 | 4818925 | rs1907989 | -0.58 | -0.24 | -0.928 | 0.335 | 0 | -0.019 | -0.019 | 1.108 | 0.34 | 0.3 | 0.46 | 0.53 | 0.36 | 0.4 |
| 4 | 4819835 | rs17716572 | 1.462 | 0.133 | -1.233 | 0.406 | -0.04 | 0.041 | 0.041 | 0 | 0.07 | 0 | 0.07 | 0 | 0.14 | 0.04 |
| 4 | 4820679 | rs749286 | 0.234 | 0.214 | -1.057 | 0.613 | -0.031 | -0.026 | -0.026 | 0 | 0.1 | 0.05 | 0.1 | 0.19 | 0.19 | 0.21 |
| 4 | 4820813 | rs4689946 | 1.174 | -0.223 | -0.886 | 0.613 | -0.026 | -0.063 | -0.063 | 0 | 0.13 | 0.05 | 0.26 | 0.23 | 0.15 | 0.2 |
| 4 | 4822309 | rs10212960 | -0.478 | -0.459 | -0.309 | 0.304 | -0.053 | 0.006 | 0.006 | 0 | 0.25 | 0.52 | 0.13 | 0.21 | 0.15 | 0.17 |
| 4 | 4827883 | rs9291153 | 1.253 | 0.274 | -0.933 | 0.697 | -0.002 | 0.047 | 0.047 | 0.425 | 0.11 | 0.13 | 0.05 | 0.22 | 0.11 | 0.13 |
| 4 | 4837058 | rs7666443 | -0.524 | 0.301 | -1.146 | 0.121 | -0.011 | -0.023 | -0.023 | 0 | 0.41 | 0.72 | 0.4 | 0.52 | 0.47 | 0.49 |
| 4 | 4845777 | rs13121578 | 1.316 | 0.26 | -1.842 | -0.63 | -0.106 | 0.177 | 0.177 | 0 | 0.29 | 0.03 | 0.29 | 0.03 | 0.31 | 0.19 |

| | | | | | | | | | | | | | | | | |
|---|---|---|---|---|---|---|---|---|---|---|---|---|---|---|---|---|
| 4 | 4846142 | rs12640329 | -0.87 | 0.028 | -1.289 | -0.566 | 0.001 | -0.011 | -0.011 | 0.529 | 0.46 | 0.48 | 0.49 | 0.72 | 0.53 | 0.53 |
| 4 | 4848349 | rs4597767 | -0.626 | 0.209 | -1.629 | -0.692 | -0.009 | 0.056 | 0.056 | 0 | 0.43 | 0.28 | 0.46 | 0.87 | 0.47 | 0.58 |
| 4 | 4849402 | rs3733397 | -1.468 | 0.198 | -1.44 | -0.776 | -0.002 | -0.091 | -0.091 | 0 | 0.61 | 0.51 | 0.63 | 0.91 | 0.66 | 0.78 |
| 4 | 4851329 | rs4435686 | 0.519 | 0.236 | -2.174 | -0.542 | -0.006 | -0.01 | -0.01 | 0.521 | 0.22 | 0.16 | 0.28 | 0.62 | 0.3 | 0.36 |
| 4 | 4851979 | rs6819546 | 0.531 | 0.241 | -2.188 | -0.593 | 0.018 | -0.005 | -0.005 | 0 | 0.25 | 0.16 | 0.21 | 0.26 | 0.21 | 0.36 |
| 4 | 4852354 | rs884690 | 0.209 | 0.681 | -2.24 | -0.299 | -0.007 | -0.01 | -0.01 | 0 | 0.22 | 0.16 | 0.27 | 0.62 | 0.29 | 0.36 |
| 4 | 4852867 | rs4348044 | -1.114 | 1.141 | -2.441 | -0.372 | -0.122 | 0.123 | 0.123 | 0 | 0.67 | 0.98 | 0.82 | 0.88 | 0.63 | 0.83 |
| 4 | 4860138 | rs3821947 | -0.756 | 0.545 | -2.281 | -0.115 | -0.017 | -0.038 | -0.038 | 0 | 0.54 | 0.84 | 0.67 | 0.55 | 0.49 | 0.56 |
| 4 | 4860402 | rs3821949 | -0.351 | -0.628 | -1.271 | -0.131 | -0.011 | 0.095 | 0.095 | 0 | 0.23 | 0.54 | 0.34 | 0.42 | 0.21 | 0.17 |
| 4 | 4863745 | rs3775261 | -0.227 | -0.932 | -0.96 | -0.379 | 0.001 | -0.007 | -0.007 | 0.844 | 0.26 | 0.25 | 0.49 | 0.6 | 0.29 | 0.28 |
| 4 | 4864381 | rs1042484 | -0.539 | -0.974 | -0.917 | -0.708 | -0.002 | 0.072 | 0.072 | 0.098 | 0.27 | 0.13 | 0.18 | 0.06 | 0.22 | 0.16 |
| 4 | 4867322 | rs4464513 | -1.35 | -0.982 | -0.672 | -0.63 | -0.026 | 0.019 | 0.019 | 0 | 0.6 | 0.76 | 0.68 | 0.66 | 0.53 | 0.5 |
| 4 | 4872247 | rs4473606 | -1.835 | -0.99 | -0.544 | -0.626 | -0.018 | 0.014 | 0.014 | 0 | 0.67 | 0.75 | 0.64 | 0.72 | 0.59 | 0.55 |
| 4 | 4880597 | rs12501827 | 0.226 | -0.728 | -0.217 | -0.848 | 0 | -0.023 | -0.023 | 0.48 | 0.12 | 0.15 | 0.09 | 0 | 0.07 | 0.05 |
| 4 | 4883409 | rs12500028 | -0.962 | -0.992 | 0.215 | -1.096 | 0.001 | -0.01 | -0.01 | 0 | 0.28 | 0.24 | 0.19 | 0.07 | 0.2 | 0.33 |
| 4 | 4883459 | rs12498543 | -0.116 | -0.978 | -0.007 | -1.089 | 0.015 | 0.041 | 0.041 | 0 | 0.2 | 0.11 | 0.12 | 0 | 0.13 | 0.16 |
| 4 | 4889621 | rs4280680 | -0.718 | -1.107 | 0.205 | -1.061 | 0.009 | 0.006 | 0.006 | 0 | 0.32 | 0.26 | 0.17 | 0.12 | 0.25 | 0.27 |
| 4 | 4890733 | rs16836361 | 0.922 | -1.062 | -0.101 | -0.824 | -0.022 | -0.04 | -0.04 | 0 | 0.16 | 0.08 | 0.33 | 0.59 | 0.24 | 0.25 |
| 4 | 4891148 | rs6817411 | -1.515 | -1.092 | 0.034 | -0.883 | 0.013 | 0.116 | 0.116 | 0 | 0.49 | 0.7 | 0.56 | 0.77 | 0.5 | 0.58 |
| 4 | 4895024 | rs4689185 | 0.724 | -0.903 | -0.013 | -0.777 | 0.001 | 0.002 | 0.002 | 0 | 0.11 | 0.1 | 0.08 | 0 | 0.07 | 0.05 |
| 4 | 4895897 | rs2968686 | -0.546 | -1.141 | 0.284 | -0.889 | -0.018 | -0.049 | -0.049 | 0.286 | 0.27 | 0.4 | 0.17 | 0.1 | 0.18 | 0.2 |
| 4 | 4896233 | rs13118392 | -0.224 | -1.119 | -0.182 | -0.878 | 0.029 | 0.071 | 0.071 | 0 | 0.25 | 0.13 | 0.14 | 0.1 | 0.18 | 0.2 |
| 4 | 4905782 | rs4566595 | -2.343 | -1.132 | -0.237 | -0.781 | -0.021 | 0.031 | 0.031 | 0 | 0.64 | 0.85 | 0.66 | 0.71 | 0.63 | 0.69 |
| 4 | 4905825 | rs4488884 | 0.615 | -0.891 | -0.439 | -0.514 | -0.013 | -0.025 | -0.025 | 0 | 0.17 | 0.13 | 0.34 | 0.53 | 0.26 | 0.22 |
| 4 | 4908020 | rs4689961 | 0 | -0.874 | -0.537 | -0.635 | 0.014 | 0.025 | 0.025 | 1.521 | 0.17 | 0.13 | 0.11 | 0.1 | 0.1 | 0.1 |
| 4 | 4908281 | rs11932240 | 0.83 | -0.545 | -0.417 | -0.405 | 0.003 | 0.004 | 0.004 | 0 | 0.07 | 0.1 | 0.07 | 0.07 | 0.07 | 0.17 |
| 4 | 4911642 | rs6446700 | -0.854 | -0.607 | -0.507 | -0.562 | 0.001 | 0 | 0 | 0.052 | 0.34 | 0.34 | 0.51 | 0.56 | 0.47 | 0.36 |
| 4 | 4912942 | rs4593079 | -1.478 | -0.749 | -0.728 | -0.61 | 0.007 | 0.017 | 0.017 | 0.092 | 0.45 | 0.35 | 0.37 | 0.34 | 0.43 | 0.47 |
| 4 | 4917971 | rs4639019 | -0.956 | 0.127 | -0.077 | -0.122 | -0.016 | 0.02 | 0.02 | 0.152 | 0.3 | 0.56 | 0.3 | 0.39 | 0.27 | 0.42 |
| 4 | 4923296 | rs736405 | 0.552 | 0.529 | -0.474 | 0.118 | 0.003 | 0.057 | 0.057 | 0 | 0.07 | 0.08 | 0.09 | 0.23 | 0.09 | 0.16 |
| 4 | 4932825 | rs7678151 | -1.139 | 0.441 | -0.759 | -0.675 | -0.011 | 0.041 | 0.041 | 0 | 0.65 | 0.77 | 0.77 | 0.91 | 0.62 | 0.75 |
| 4 | 4935309 | rs10516169 | -0.597 | -0.155 | -0.517 | 0.16 | -0.005 | -0.021 | -0.021 | 0.158 | 0.4 | 0.56 | 0.47 | 0.63 | 0.39 | 0.37 |
| 4 | 4937762 | rs6446716 | -0.689 | -0.357 | -0.908 | 0.179 | 0.037 | 0.098 | 0.098 | 0.528 | 0.52 | 0.74 | 0.63 | 0.76 | 0.44 | 0.5 |
| 4 | 4939423 | rs11946368 | 0.397 | -0.881 | -1.004 | 0.295 | 0.025 | 0.014 | 0.014 | 2.199 | 0.15 | 0.08 | 0.12 | 0.09 | 0.1 | 0.13 |
| 4 | 4942150 | rs1010776 | -0.435 | -1.168 | -1.012 | 0.84 | 0.08 | -0.028 | -0.028 | 0 | 0.32 | 0.14 | 0.26 | 0.34 | 0.2 | 0.31 |
| 4 | 4946111 | rs4234841 | 0.598 | -1.185 | 0.379 | 1.035 | 0 | 0.003 | 0.003 | 0 | 0.12 | 0.12 | 0.26 | 0.23 | 0.18 | 0.09 |
| 4 | 4949339 | rs4330303 | -2.292 | -0.758 | -0.292 | 0.546 | -0.1 | 0.011 | 0.011 | 0.197 | 0.68 | 0.98 | 0.78 | 0.79 | 0.7 | 0.68 |
| 4 | 4950210 | rs2968680 | -0.728 | -1.299 | 0.174 | 0.873 | -0.055 | 0.029 | 0.029 | 0 | 0.37 | 0.52 | 0.35 | 0.54 | 0.25 | 0.41 |
| 4 | 4952982 | rs4689190 | -0.156 | -1.231 | 0.255 | 0.766 | 0.065 | 0.208 | 0.208 | 0 | 0.17 | 0.47 | 0.34 | 0.42 | 0.24 | 0.18 |
| 4 | 4965186 | rs4689965 | -1.297 | -1.241 | 0.005 | 0.536 | 0.046 | 0.006 | 0.006 | 0.667 | 0.51 | 0.74 | 0.52 | 0.52 | 0.43 | 0.43 |
| 4 | 4970920 | rs11734790 | -0.718 | -1.264 | -0.53 | 0.204 | -0.039 | -0.025 | -0.025 | 0 | 0.34 | 0.43 | 0.2 | 0.2 | 0.22 | 0.34 |
| 4 | 4972091 | rs10223021 | -0.723 | -1.243 | -0.531 | 0.214 | -0.061 | -0.042 | -0.042 | 0.612 | 0.36 | 0.48 | 0.21 | 0.2 | 0.22 | 0.34 |
| 4 | 4974559 | rs2968706 | -0.411 | -0.917 | -1.002 | 0.161 | -0.03 | -0.033 | -0.033 | 0 | 0.28 | 0.36 | 0.17 | 0.1 | 0.18 | 0.24 |
| 4 | 4984192 | rs4689967 | 0.517 | -0.5 | -1.894 | 0.114 | 0.068 | 0.114 | 0.114 | 0 | 0.51 | 0.24 | 0.48 | 0.3 | 0.61 | 0.55 |
| 4 | 4985810 | rs4689968 | -0.097 | -0.019 | -2.349 | 0.167 | -0.056 | -0.105 | -0.105 | 0 | 0.21 | 0.03 | 0.35 | 0.32 | 0.28 | 0.23 |
| 4 | 4988914 | rs11726928 | 0.338 | 0.009 | -2.285 | -0.914 | -0.089 | 0.167 | 0.167 | 0 | 0.3 | 0.02 | 0.27 | 0.1 | 0.28 | 0.36 |
| 4 | 4991380 | rs10022680 | -0.193 | -0.392 | -1.918 | -1.232 | 0.013 | 0.05 | 0.05 | 2.444 | 0.68 | 0.79 | 0.77 | 0.94 | 0.72 | 0.71 |
| 4 | 4995876 | rs4522817 | 0.296 | -0.287 | -1.454 | -0.85 | 0.009 | -0.007 | -0.007 | 0.293 | 0.46 | 0.35 | 0.47 | 0.45 | 0.55 | 0.39 |
| 4 | 5005171 | rs13124969 | -0.132 | -0.323 | -1.219 | -0.77 | 0.001 | -0.018 | -0.018 | 0.314 | 0.17 | 0.15 | 0.09 | 0 | 0.16 | 0.14 |
| 4 | 5005218 | rs4072934 | -0.439 | -0.484 | -1.078 | -0.999 | 0.005 | -0.017 | -0.017 | 0.14 | 0.78 | 0.8 | 0.71 | 0.57 | 0.83 | 0.64 |
| 4 | 5005698 | rs4392464 | 0.86 | -0.513 | -0.72 | -0.997 | -0.013 | 0.006 | 0.006 | 0 | 0.59 | 0.49 | 0.62 | 0.52 | 0.67 | 0.5 |
| 4 | 5007377 | rs7659395 | -0.694 | -0.104 | -0.608 | -1.481 | -0.006 | -0.039 | -0.039 | 0 | 0.18 | 0.23 | 0.1 | 0.05 | 0.16 | 0.15 |
| 4 | 5010866 | rs9999422 | -0.194 | 0.786 | -1.127 | -1.404 | -0.02 | 0.028 | 0.028 | 0 | 0.15 | 0.09 | 0.2 | 0.01 | 0.2 | 0.06 |
| 4 | 5014223 | rs6810546 | 0.099 | -0.321 | -0.636 | -1.596 | -0.049 | -0.116 | -0.116 | 0.009 | 0.69 | 0.45 | 0.8 | 0.93 | 0.8 | 0.7 |
| 4 | 5015167 | rs6446309 | 0.182 | 0.473 | -0.404 | -0.729 | 0.032 | 0.173 | 0.173 | 0 | 0.8 | 0.58 | 0.71 | 0.54 | 0.79 | 0.67 |
| 4 | 5015486 | rs10011142 | -0.638 | 0.21 | -1.193 | -1.743 | -0.021 | 0.082 | 0.082 | 0 | 0.18 | 0.04 | 0.06 | 0 | 0.14 | 0.16 |
| 4 | 5022496 | rs4689977 | -0.114 | -0.043 | -0.744 | -1.598 | -0.007 | -0.1 | -0.1 | 0 | 0.79 | 0.68 | 0.89 | 1 | 0.86 | 0.87 |
| 4 | 5023112 | rs10937615 | 0.257 | -0.25 | -1.713 | -1.681 | 0.024 | 0.052 | 0.052 | 0.171 | 0.64 | 0.23 | 0.78 | 0.96 | 0.78 | 0.74 |
| 4 | 5043674 | rs11732574 | -0.32 | -0.513 | -1.407 | -0.858 | 0.043 | 0.07 | 0.07 | 0.12 | 0.33 | 0.16 | 0.18 | 0.11 | 0.26 | 0.24 |
| 4 | 5055774 | rs9799796 | -0.527 | -0.483 | -0.894 | 0.276 | -0.03 | 0.034 | 0.034 | 0 | 0.32 | 0.4 | 0.32 | 0.5 | 0.23 | 0.31 |
| 4 | 5057311 | rs10009026 | 0.518 | 0.389 | -1.391 | -0.129 | -0.034 | 0.144 | 0.144 | 0 | 0.45 | 0.17 | 0.36 | 0.2 | 0.46 | 0.21 |
| 4 | 5059668 | rs11727656 | -0.483 | 0.165 | -1.056 | 0.346 | -0.008 | -0.006 | -0.006 | 0.504 | 0.36 | 0.3 | 0.33 | 0.61 | 0.44 | 0.52 |
| 4 | 5060425 | rs7692195 | -0.63 | 0.146 | -0.592 | -0.213 | 0.001 | 0.004 | 0.004 | 0.097 | 0.28 | 0.54 | 0.24 | 0.28 | 0.27 | 0.14 |
| 4 | 5067789 | rs1868047 | -0.332 | 0.493 | -0.67 | -0.508 | -0.008 | 0.007 | 0.007 | 0 | 0.38 | 0.42 | 0.4 | 0.46 | 0.35 | 0.64 |
| 4 | 5071688 | rs1982349 | 0.306 | 0.564 | -1.243 | -1.648 | -0.042 | -0.035 | -0.035 | 0 | 0.14 | 0.05 | 0.18 | 0.17 | 0.21 | 0.13 |
| 4 | 5073415 | rs11940060 | 1.214 | 0.236 | -1.084 | -1.926 | -0.001 | 0.086 | 0.086 | 0 | 0.28 | 0.05 | 0.23 | 0.12 | 0.24 | 0.15 |
| 4 | 5074816 | rs6847646 | -0.898 | 0.588 | -0.07 | -1.748 | 0.025 | -0.057 | -0.057 | 0 | 0.85 | 0.74 | 0.83 | 0.99 | 0.79 | 0.96 |
| 4 | 5077162 | rs10028432 | 0.745 | 0.541 | -0.044 | -1.742 | 0.012 | -0.034 | -0.034 | 0 | 0.14 | 0.15 | 0.15 | 0.02 | 0.21 | 0.05 |
| 4 | 5077822 | rs17732017 | 0.935 | 0.553 | -0.031 | -1.74 | -0.007 | -0.006 | -0.006 | 1.005 | 0.13 | 0.08 | 0.15 | 0.02 | 0.21 | 0.04 |

| | | | | | | | | | | | | | | | | |
|---|---|---|---|---|---|---|---|---|---|---|---|---|---|---|---|---|
| 4 | 5078646 | rs13150536 | 0.859 | 0.549 | 0.008 | -1.742 | 0.019 | -0.044 | -0.044 | 0 | 0.13 | 0.16 | 0.16 | 0.02 | 0.21 | 0.05 |
| 4 | 5078789 | rs10937620 | -1.363 | 0.184 | -0.275 | -2.013 | 0.004 | 0.044 | 0.044 | 0.405 | 0.62 | 0.77 | 0.69 | 0.84 | 0.63 | 0.72 |
| 4 | 5080052 | rs16836608 | 0.3 | -0.013 | 0.033 | -1.555 | 0.016 | 0.008 | 0.008 | 0 | 0.15 | 0.1 | 0.06 | 0 | 0.08 | 0.15 |
| 4 | 5085210 | rs976682 | -0.01 | 0.246 | 0.154 | -1.215 | -0.003 | 0.024 | 0.024 | 0.102 | 0.15 | 0.32 | 0.22 | 0.16 | 0.14 | 0.16 |
| 4 | 5086219 | rs6822181 | 0.799 | 0.569 | 0.133 | -0.918 | -0.003 | 0.103 | 0.103 | 2.108 | 0.39 | 0.15 | 0.27 | 0.19 | 0.38 | 0.29 |
| 4 | 5087553 | rs1447284 | -0.845 | 0.287 | 0.12 | -1.07 | -0.001 | -0.01 | -0.01 | 0 | 0.47 | 0.5 | 0.45 | 0.19 | 0.46 | 0.35 |
| 4 | 5093523 | rs7693293 | -0.697 | 0.408 | -0.389 | -0.927 | -0.001 | 0.032 | 0.032 | 3.136 | 0.44 | 0.39 | 0.42 | 0.18 | 0.43 | 0.34 |
| 4 | 5112978 | rs12507259 | 0.42 | 0.383 | -0.128 | -0.82 | -0.023 | -0.175 | -0.175 | 0.15 | 0.73 | 0.57 | 0.8 | 1 | 0.79 | 0.95 |
| 4 | 5115426 | rs16836654 | -0.733 | 0.316 | -0.192 | -0.482 | -0.011 | -0.033 | -0.033 | 0 | 0.28 | 0.33 | 0.32 | 0.18 | 0.25 | 0.32 |
| 4 | 5116247 | rs12509816 | -1.066 | 0.305 | -0.171 | 0.484 | 0 | -0.014 | -0.014 | 0 | 0.3 | 0.19 | 0.34 | 0.61 | 0.28 | 0.39 |
| 4 | 5120597 | rs12649783 | -0.579 | 0.43 | -0.521 | 1.197 | -0.031 | -0.109 | -0.109 | 0.031 | 0.22 | 0.05 | 0.29 | 0.4 | 0.24 | 0.39 |
| 4 | 5125048 | rs7660270 | -0.83 | 0.628 | -0.298 | 0.6 | -0.01 | 0.017 | 0.017 | 0 | 0.42 | 0.39 | 0.62 | 0.67 | 0.55 | 0.52 |
| 4 | 5127546 | rs747224 | -1.269 | 0.247 | -0.398 | 0.816 | -0.017 | -0.013 | -0.013 | 0 | 0.45 | 0.72 | 0.62 | 0.46 | 0.52 | 0.41 |
| 4 | 5128511 | rs12651470 | 0.012 | 0.273 | 0.2 | 0.944 | 0.058 | 0.018 | 0.018 | 0 | 0.14 | 0.25 | 0.15 | 0.13 | 0.25 | 0.17 |
| 4 | 5128641 | rs11726208 | -0.734 | -0.216 | 0.135 | 0.577 | -0.014 | -0.014 | -0.014 | 0 | 0.17 | 0.26 | 0.15 | 0.15 | 0.14 | 0.1 |
| 4 | 5135979 | rs10011769 | 1.372 | -0.306 | 0.044 | 0.568 | 0.022 | 0.002 | 0.002 | 0 | 0.71 | 0.23 | 0.53 | 0.69 | 0.76 | 0.78 |
| 4 | 5136130 | rs6856163 | 0.587 | -0.156 | 0.27 | 0.199 | -0.021 | -0.026 | -0.026 | 0 | 0.82 | 0.68 | 0.85 | 0.86 | 0.86 | 0.9 |
| 4 | 5137364 | rs4689989 | 0.363 | 0.075 | -0.592 | 0.305 | -0.015 | -0.015 | -0.015 | 0 | 0.8 | 0.31 | 0.82 | 0.85 | 0.86 | 0.9 |
| 4 | 5142212 | rs12511127 | 0.265 | 0.736 | -2.477 | 0.5 | -0.123 | 0.054 | 0.054 | 0 | 0.48 | 0.03 | 0.33 | 0.36 | 0.48 | 0.5 |
| 4 | 5145806 | rs10012571 | -0.099 | 0.746 | -2.337 | 0.312 | -0.024 | 0.045 | 0.045 | 0 | 0.46 | 0.26 | 0.47 | 0.31 | 0.47 | 0.45 |
| 4 | 5146091 | rs1562425 | 0.522 | 0.525 | -2.723 | 0.041 | -0.065 | -0.004 | -0.004 | 0 | 0.29 | 0.11 | 0.21 | 0.26 | 0.4 | 0.3 |
| 4 | 5146683 | rs1562427 | 0.498 | 0.493 | -2.738 | 0.026 | -0.065 | -0.003 | -0.003 | 0 | 0.3 | 0.11 | 0.21 | 0.26 | 0.4 | 0.3 |
| 4 | 5154673 | rs7672067 | -0.522 | 0.091 | -3.465 | -0.852 | 0 | 0.101 | 0.101 | 0 | 0.74 | 0.92 | 0.8 | 0.93 | 0.77 | 0.81 |
| 4 | 5158436 | rs1542203 | -0.298 | -0.586 | -2.509 | -0.878 | -0.005 | -0.041 | -0.041 | 0 | 0.33 | 0.58 | 0.42 | 0.12 | 0.28 | 0.18 |
| 4 | 5159654 | rs7683724 | -0.819 | 0.027 | -2.614 | -0.478 | -0.007 | -0.016 | -0.016 | 0 | 0.63 | 0.74 | 0.69 | 0.52 | 0.59 | 0.53 |
| 4 | 5160771 | rs998984 | 0.945 | -0.238 | -2.913 | -0.675 | -0.06 | 0.044 | 0.044 | 0 | 0.19 | 0.03 | 0.13 | 0.07 | 0.22 | 0.16 |
| 4 | 5162389 | rs13128441 | -1.003 | -0.532 | -2.11 | -0.584 | -0.001 | -0.006 | -0.006 | 0.005 | 0.75 | 0.76 | 0.78 | 0.91 | 0.74 | 0.84 |
| 4 | 5162594 | rs13121814 | 1.309 | -0.347 | -2.23 | -0.521 | 0 | -0.021 | -0.021 | 0.173 | 0.27 | 0.18 | 0.2 | 0.32 | 0.27 | 0.27 |
| 4 | 5163447 | rs1530609 | -0.251 | -0.653 | -1.518 | -0.584 | -0.001 | -0.001 | -0.001 | 0 | 0.45 | 0.54 | 0.4 | 0.54 | 0.47 | 0.55 |
| 4 | 5167734 | rs3806723 | 1.791 | -0.505 | -1.253 | -0.553 | -0.005 | -0.035 | -0.035 | 0 | 0.2 | 0.1 | 0.15 | 0.3 | 0.22 | 0.24 |
| 4 | 5170284 | rs3749566 | 1.863 | -0.469 | -1.329 | -0.551 | -0.044 | -0.088 | -0.088 | 1.032 | 0.19 | 0.03 | 0.14 | 0.3 | 0.21 | 0.22 |
| 4 | 5170549 | rs10022435 | -0.142 | -0.777 | -0.696 | -0.682 | -0.01 | 0.001 | 0.001 | 0 | 0.38 | 0.53 | 0.5 | 0.39 | 0.3 | 0.3 |
| 4 | 5182102 | rs7654310 | 1.365 | -0.773 | -0.473 | -0.694 | 0 | 0.002 | 0.002 | 0 | 0.24 | 0.23 | 0.36 | 0.37 | 0.24 | 0.26 |
| 4 | 5182934 | rs2369670 | -0.382 | -1.051 | -0.349 | -0.906 | -0.01 | 0.007 | 0.007 | 0 | 0.46 | 0.34 | 0.4 | 0.56 | 0.49 | 0.5 |
| 4 | 5185925 | rs13115405 | -1.021 | -1.086 | -0.203 | -0.924 | -0.013 | -0.059 | -0.059 | 0 | 0.71 | 0.44 | 0.77 | 0.93 | 0.75 | 0.78 |
| 4 | 5187398 | rs13123841 | 0.164 | -1.101 | -0.225 | -0.943 | -0.054 | -0.054 | -0.054 | 3.974 | 0.4 | 0.2 | 0.31 | 0.52 | 0.46 | 0.47 |
| 4 | 5187537 | rs12650617 | 0.652 | -1.465 | -0.488 | -1.126 | 0 | -0.007 | -0.007 | 0 | 0.23 | 0.28 | 0.24 | 0.35 | 0.23 | 0.23 |
| 4 | 5188454 | rs10937629 | -1.027 | -1.911 | -0.178 | -1.114 | -0.009 | -0.043 | -0.043 | 0 | 0.7 | 0.61 | 0.78 | 0.92 | 0.74 | 0.77 |
| 4 | 5192533 | rs11730793 | 1.717 | -1.416 | -0.583 | -0.492 | -0.041 | -0.052 | -0.052 | 0 | 0.17 | 0.05 | 0.17 | 0.39 | 0.26 | 0.3 |
| 4 | 5195094 | rs12506939 | 0.483 | -1.566 | -0.738 | -0.623 | -0.053 | -0.097 | -0.097 | 3.354 | 0.29 | 0.09 | 0.25 | 0.48 | 0.35 | 0.43 |
| 4 | 5196546 | rs12504675 | 0.319 | -1.561 | -0.608 | -0.342 | -0.037 | -0.089 | -0.089 | 0 | 0.32 | 0.13 | 0.27 | 0.51 | 0.36 | 0.44 |
| 4 | 5210283 | rs989707 | 0.464 | -1.879 | -0.611 | -0.608 | 0 | 0 | 0 | 1.292 | 0.42 | 0.43 | 0.52 | 0.44 | 0.49 | 0.37 |
| 4 | 5210567 | rs989708 | -1.427 | -1.808 | -0.53 | -0.678 | 0 | -0.022 | -0.022 | 0 | 0.91 | 0.91 | 0.95 | 0.99 | 0.9 | 0.91 |
| 4 | 5211645 | rs17734102 | 2.558 | -1.733 | -0.895 | -0.576 | -0.028 | -0.007 | -0.007 | 0.837 | 0.15 | 0.01 | 0.12 | 0.09 | 0.15 | 0.06 |
| 4 | 5216320 | rs1562429 | 0.735 | -1.78 | -0.759 | -0.641 | 0.081 | 0.086 | 0.086 | 0 | 0.51 | 0.28 | 0.52 | 0.81 | 0.73 | 0.6 |
| 4 | 5216885 | rs11734549 | 0.799 | -1.775 | -0.831 | -0.606 | 0.094 | 0.096 | 0.096 | 0 | 0.49 | 0.24 | 0.5 | 0.8 | 0.72 | 0.6 |
| 4 | 5218136 | rs11731545 | 0.496 | -1.637 | -0.703 | -0.648 | 0.051 | 0.092 | 0.092 | 0.487 | 0.35 | 0.18 | 0.38 | 0.08 | 0.22 | 0.12 |
| 4 | 5219619 | rs16836902 | -0.442 | -0.329 | 0.66 | 0.248 | -0.001 | -0.072 | -0.072 | 0.456 | 0.11 | 0.32 | 0.26 | 0 | 0.05 | 0.04 |
| 4 | 5221578 | rs1812310 | 0.224 | -0.211 | -0.313 | 0.266 | -0.005 | -0.004 | -0.004 | 0 | 0.84 | 0.88 | 0.72 | 0.99 | 0.84 | 0.87 |
| 4 | 5228210 | rs13123245 | 1.236 | -0.059 | -0.075 | 0.738 | -0.09 | -0.083 | -0.083 | 0.844 | 0.15 | 0.04 | 0.13 | 0.31 | 0.3 | 0.24 |
| 4 | 5229884 | rs985222 | -0.447 | -0.36 | 0.348 | 0.767 | 0 | 0.006 | 0.006 | 0 | 0.42 | 0.41 | 0.57 | 0.74 | 0.58 | 0.56 |
| 4 | 5231140 | rs11728257 | 0.499 | -0.127 | 0.356 | 1.009 | -0.027 | -0.048 | -0.048 | 0.02 | 0.16 | 0.05 | 0.19 | 0.38 | 0.23 | 0.29 |
| 4 | 5232548 | rs2369498 | -0.101 | -0.875 | 1.205 | 0.524 | 0.001 | 0.004 | 0.004 | 0 | 0.75 | 0.79 | 0.85 | 0.9 | 0.78 | 0.75 |
| 4 | 5235815 | rs4688922 | 2.065 | -0.808 | 2.558 | 1.434 | -0.031 | 0.172 | 0.172 | 1.077 | 0.89 | 0.53 | 0.9 | 0.81 | 0.96 | 0.89 |
| 4 | 5237152 | rs10000042 | -1.691 | -0.882 | 1.732 | 1.494 | 0.004 | -0.009 | -0.009 | 0 | 0.09 | 0.08 | 0.23 | 0.18 | 0.07 | 0.11 |
| 4 | 5237192 | rs11729112 | 3.323 | -1.155 | 1.602 | 1.407 | 0.104 | -0.083 | -0.083 | 1.065 | 0.7 | 0.1 | 0.62 | 0.64 | 0.86 | 0.75 |
| 4 | 5241571 | rs6817353 | -2.986 | -1.032 | 1.931 | 1.449 | 0 | -0.005 | -0.005 | 0 | 0.2 | 0.76 | 0.15 | 0.24 | 0.07 | 0.15 |
| 4 | 5249944 | rs10001177 | -1.516 | -0.923 | 1.786 | 1.007 | -0.016 | 0.006 | 0.006 | 0 | 0.07 | 0.17 | 0.03 | 0.11 | 0.04 | 0.05 |
| 4 | 5257364 | rs6446345 | -2.546 | -0.97 | 2.514 | 0.948 | -0.033 | 0.084 | 0.084 | 5.666 | 0.1 | 0.43 | 0.07 | 0.1 | 0.01 | 0.08 |
| 4 | 5265305 | rs9985541 | -1.88 | -1.058 | 2.182 | 0.917 | 0.009 | 0 | 0 | 0 | 0.09 | 0.13 | 0.03 | 0.1 | 0.05 | 0.11 |
| 4 | 5279796 | rs7655639 | -2.701 | -1.058 | 2.584 | 0.928 | -0.028 | 0.12 | 0.12 | 0.426 | 0.1 | 0.46 | 0.09 | 0.15 | 0.03 | 0.14 |
| 4 | 5288159 | rs7669364 | -2.293 | -0.915 | 2.728 | 0.856 | -0.004 | -0.014 | -0.014 | 0.333 | 0.1 | 0.17 | 0.06 | 0.06 | 0.1 | 0.11 |
| 4 | 5309317 | rs6849493 | -2.803 | -0.86 | 3.419 | 0.843 | -0.014 | 0.008 | 0.008 | 0.547 | 0.1 | 0.38 | 0.07 | 0.05 | 0.04 | 0.06 |
| 4 | 5310171 | rs6829618 | -3.567 | -0.873 | 3.376 | 0.836 | -0.024 | -0.027 | -0.027 | 0.014 | 0.18 | 0.6 | 0.12 | 0.12 | 0.14 | 0.16 |
| 4 | 5324465 | rs10031235 | -3.714 | -0.934 | 3.971 | 0.806 | -0.038 | -0.044 | -0.044 | 0 | 0.22 | 0.36 | 0.29 | 0.12 | 0.15 | 0.19 |
| 4 | 5324491 | rs6853800 | -1.429 | -1.002 | 3.901 | 0.854 | -0.022 | 0.036 | 0.036 | 0 | 0.07 | 0.2 | 0.05 | 0.13 | 0.03 | 0.09 |
| 4 | 5324579 | rs6853980 | -4.202 | -1.016 | 3.984 | 0.845 | 0.05 | 0.011 | 0.011 | 0.057 | 0.31 | 0.82 | 0.36 | 0.25 | 0.19 | 0.28 |
| 4 | 5326726 | rs10028484 | -3.86 | -1.067 | 4.332 | 0.842 | -0.065 | -0.036 | -0.036 | 1.095 | 0.23 | 0.58 | 0.27 | 0.15 | 0.14 | 0.16 |
| 4 | 5327821 | rs1393861 | -2.373 | -1.053 | 4.067 | 0.882 | -0.013 | 0 | 0 | 0 | 0.12 | 0.28 | 0.05 | 0.11 | 0.1 | 0.13 |

| | | | | | | | | | | | | | | | | |
|---|---|---|---|---|---|---|---|---|---|---|---|---|---|---|---|---|
| 4 | 5332558 | rs878544 | -3.458 | -1.07 | 4.122 | 0.881 | -0.072 | 0.005 | 0.005 | 0.039 | 0.18 | 0.5 | 0.3 | 0.15 | 0.06 | 0.14 |
| 4 | 5333091 | rs16837029 | -1.386 | -1.05 | 3.866 | 0.886 | -0.025 | -0.014 | -0.014 | 0 | 0.07 | 0.22 | 0.02 | 0.04 | 0.03 | 0.05 |
| 4 | 5341817 | rs4689212 | -3.8 | -0.891 | 3.427 | 0.881 | 0.037 | 0.003 | 0.003 | 0.101 | 0.3 | 0.82 | 0.35 | 0.26 | 0.18 | 0.27 |
| 4 | 5347818 | rs10028303 | -2.383 | -0.777 | 3.411 | 0.764 | -0.028 | -0.025 | -0.025 | 0 | 0.14 | 0.33 | 0.07 | 0.09 | 0.1 | 0.13 |
| 4 | 5365859 | rs11727433 | -3.631 | -0.925 | 3.431 | 0.688 | -0.023 | -0.012 | -0.012 | 0 | 0.32 | 0.68 | 0.38 | 0.23 | 0.2 | 0.3 |
| 4 | 5383127 | rs16837061 | -3.187 | -0.625 | 3.213 | 0.777 | -0.021 | -0.01 | -0.01 | 0.018 | 0.28 | 0.63 | 0.39 | 0.23 | 0.21 | 0.25 |
| 4 | 5383242 | rs11930775 | -2.741 | -0.626 | 3.259 | 0.775 | -0.015 | -0.039 | -0.039 | 0 | 0.23 | 0.51 | 0.16 | 0.13 | 0.19 | 0.18 |
| 4 | 5384256 | rs10937641 | -3.022 | -0.635 | 3.198 | 0.855 | -0.019 | -0.016 | -0.016 | 0 | 0.27 | 0.49 | 0.18 | 0.2 | 0.23 | 0.25 |
| 4 | 5389231 | rs195140 | 2.02 | -0.628 | 2.773 | 0.865 | -0.015 | -0.008 | -0.008 | 0 | 0.82 | 0.61 | 0.89 | 0.87 | 0.86 | 0.85 |
| 4 | 5390637 | rs9291090 | -1.882 | -0.481 | 2.2 | 1.082 | 0.012 | -0.016 | -0.016 | 0 | 0.08 | 0.04 | 0.23 | 0.1 | 0.03 | 0.12 |
| 4 | 5391783 | rs195134 | 2.946 | -0.598 | 2.538 | 0.94 | -0.059 | -0.023 | -0.023 | 0 | 0.74 | 0.41 | 0.64 | 0.83 | 0.89 | 0.8 |
| 4 | 5394914 | rs6840421 | -2.173 | -0.521 | 2.673 | 0.999 | -0.032 | 0.032 | 0.032 | 1.622 | 0.17 | 0.38 | 0.32 | 0.16 | 0.08 | 0.1 |
| 4 | 5395071 | rs11726374 | -0.068 | -0.492 | 2.351 | 0.975 | -0.014 | 0.037 | 0.037 | 0 | 0.07 | 0.01 | 0.06 | 0 | 0.06 | 0.03 |
| 4 | 5395967 | rs739968 | -1.345 | -0.466 | 2.396 | 1.023 | -0.027 | -0.024 | -0.024 | 0 | 0.2 | 0.26 | 0.3 | 0.09 | 0.09 | 0.16 |
| 4 | 5396855 | rs195129 | 1.55 | -0.347 | 1.628 | 1.147 | 0.01 | -0.001 | -0.001 | 0 | 0.66 | 0.27 | 0.57 | 0.7 | 0.8 | 0.69 |
| 4 | 5401326 | rs2654498 | 1.036 | 1.176 | 0.578 | 1.327 | 0.008 | 0.011 | 0.011 | 0 | 0.82 | 0.75 | 0.58 | 0.72 | 0.76 | 0.73 |
| 4 | 5403545 | rs2009157 | -0.961 | 0.452 | 1.426 | 1.01 | -0.008 | 0.018 | 0.018 | 0 | 0.1 | 0.33 | 0.06 | 0.07 | 0.04 | 0.12 |
| 4 | 5405805 | rs10010358 | -2.429 | 1.194 | 1.774 | 0.906 | 0.015 | -0.013 | -0.013 | 1.938 | 0.2 | 0.71 | 0.43 | 0.16 | 0.25 | 0.16 |
| 4 | 5417222 | rs195103 | 1.371 | 1.288 | 1.865 | 0.939 | 0 | 0 | 0 | 0 | 0.89 | 0.89 | 0.67 | 0.9 | 0.8 | 0.86 |
| 4 | 5418485 | rs2301857 | -1.476 | 1.062 | 1.84 | 1.837 | -0.001 | 0.021 | 0.021 | 0 | 0.12 | 0.09 | 0.15 | 0.44 | 0.12 | 0.23 |
| 4 | 5419685 | rs16837174 | -1.912 | 0.987 | 2.096 | 1.868 | -0.003 | 0.051 | 0.051 | 0 | 0.14 | 0.18 | 0.17 | 0.52 | 0.13 | 0.28 |
| 4 | 5426224 | rs2654506 | 2.116 | 0.951 | 2.58 | 1.811 | 0.011 | 0.07 | 0.07 | 0 | 0.77 | 0.67 | 0.51 | 0.44 | 0.72 | 0.62 |
| 4 | 5427691 | rs4017782 | -1.538 | 0.788 | 2.6 | 1.84 | 0.006 | 0.211 | 0.211 | 0.221 | 0.11 | 0.3 | 0.15 | 0.45 | 0.09 | 0.23 |
| 4 | 5427944 | rs13141339 | -1.492 | 0.786 | 2.563 | 1.839 | -0.001 | 0.154 | 0.154 | 0.137 | 0.11 | 0.24 | 0.15 | 0.45 | 0.09 | 0.22 |
| 4 | 5436590 | rs2369705 | -2.101 | 0.676 | 3.189 | 1.804 | 0.017 | 0.241 | 0.241 | 0.797 | 0.12 | 0.33 | 0.18 | 0.5 | 0.11 | 0.23 |
| 4 | 5437760 | rs4689226 | -0.291 | -0.448 | 1.984 | 0.99 | -0.003 | 0 | 0 | 0.905 | 0.32 | 0.32 | 0.25 | 0.7 | 0.2 | 0.47 |
| 4 | 5438328 | rs3774836 | -0.802 | -0.386 | 1.853 | 1.195 | -0.002 | 0.101 | 0.101 | 0 | 0.1 | 0.18 | 0.15 | 0.43 | 0.08 | 0.22 |
| 4 | 5439424 | rs4689236 | -0.429 | -0.391 | 1.81 | 1.124 | 0.017 | -0.004 | -0.004 | 0 | 0.33 | 0.26 | 0.26 | 0.69 | 0.23 | 0.48 |
| 4 | 5440224 | rs6827240 | 1.605 | -0.039 | 1.452 | 0.149 | 0.038 | -0.101 | -0.101 | 1.787 | 0.31 | 0.41 | 0.31 | 0.07 | 0.45 | 0.16 |
| 4 | 5448429 | rs3733182 | -0.771 | -0.26 | 1.288 | 0.261 | 0.002 | 0.002 | 0.002 | 0 | 0.16 | 0.14 | 0.17 | 0.54 | 0.11 | 0.32 |
| 4 | 5450124 | rs3733180 | -0.525 | -0.254 | 1.195 | 0.266 | 0 | -0.073 | -0.073 | 0 | 0.15 | 0.06 | 0.15 | 0.51 | 0.11 | 0.32 |
| 4 | 5451856 | rs739965 | 1.215 | -0.336 | 1.189 | 0.232 | 0.041 | -0.031 | -0.031 | 0.527 | 0.24 | 0.11 | 0.12 | 0.27 | 0.15 | 0.29 |
| 4 | 5452736 | rs1001465 | 0.07 | -0.164 | 1.289 | 0.117 | -0.021 | 0.017 | 0.017 | 0 | 0.72 | 0.85 | 0.58 | 0.83 | 0.67 | 0.76 |
| 4 | 5454049 | rs3774828 | 0.928 | -0.21 | 1.262 | 0.116 | -0.023 | -0.001 | -0.001 | 0 | 0.07 | 0.16 | 0.02 | 0.06 | 0.02 | 0.1 |
| 4 | 5454829 | rs11732645 | 1.96 | -0.201 | 1.357 | 0.095 | 0.001 | -0.05 | -0.05 | 0.726 | 0.3 | 0.28 | 0.26 | 0.02 | 0.4 | 0.14 |
| 4 | 5457295 | rs3821925 | 0.862 | -0.192 | 1.37 | 0.147 | -0.004 | -0.014 | -0.014 | 0 | 0.62 | 0.52 | 0.53 | 0.72 | 0.64 | 0.66 |
| 4 | 5460027 | rs6446363 | 0.955 | -0.225 | 1.175 | 0.086 | -0.004 | -0.005 | -0.005 | 0 | 0.61 | 0.65 | 0.43 | 0.4 | 0.58 | 0.55 |
| 4 | 5460088 | rs16837305 | 1.647 | -0.161 | 0.627 | 0.261 | -0.008 | -0.001 | -0.001 | 0.028 | 0.06 | 0.09 | 0.03 | 0.06 | 0.02 | 0.1 |
| 4 | 5461058 | rs3774820 | 1.36 | -0.251 | 0.555 | 0.168 | 0.036 | -0.052 | -0.052 | 0 | 0.23 | 0.1 | 0.12 | 0.33 | 0.14 | 0.31 |
| 4 | 5463014 | rs1034748 | -0.241 | -0.32 | 0.549 | 0.185 | 0.037 | 0.043 | 0.043 | 0 | 0.37 | 0.24 | 0.29 | 0.81 | 0.26 | 0.63 |
| 4 | 5463535 | rs3774817 | 1.213 | -0.39 | 0.411 | 0.264 | 0.018 | -0.022 | -0.022 | 0 | 0.23 | 0.16 | 0.13 | 0.33 | 0.15 | 0.32 |
| 4 | 5466902 | rs16837322 | 0.299 | -0.17 | 0.041 | 0.68 | 0.001 | -0.004 | -0.004 | 0 | 0.05 | 0.01 | 0.08 | 0.32 | 0.05 | 0.17 |
| 4 | 5468231 | rs3815189 | -0.264 | -0.096 | -0.031 | 0.263 | -0.049 | -0.024 | -0.024 | 0 | 0.15 | 0.05 | 0.39 | 0.14 | 0.23 | 0.14 |
| 4 | 5469218 | rs7661234 | 0.046 | -0.556 | 0.126 | 0.37 | -0.052 | -0.018 | -0.018 | 0 | 0.71 | 0.89 | 0.53 | 0.78 | 0.66 | 0.78 |
| 4 | 5469810 | rs2301855 | 0.656 | -0.568 | 0.617 | -0.138 | -0.006 | -0.003 | -0.003 | 0 | 0.38 | 0.31 | 0.29 | 0.1 | 0.45 | 0.24 |
| 4 | 5476176 | rs28607918 | -0.475 | 0.03 | 0.717 | 0.596 | -0.021 | 0.001 | 0.001 | 0.645 | 0.1 | 0.32 | 0.04 | 0.07 | 0.04 | 0.08 |
| 4 | 5480337 | rs3774811 | -0.67 | 0.113 | -0.515 | 0.616 | -0.087 | -0.078 | -0.078 | 0.888 | 0.37 | 0.09 | 0.57 | 0.47 | 0.46 | 0.5 |
| 4 | 5499335 | rs739970 | -0.961 | -0.186 | 0.652 | 0.563 | -0.028 | -0.015 | -0.015 | 0 | 0.11 | 0.34 | 0.04 | 0.07 | 0.04 | 0.07 |
| 4 | 5503503 | rs10000488 | -1.227 | 0.295 | 0.067 | 1.185 | -0.001 | -0.001 | -0.001 | 0 | 0.19 | 0.21 | 0.32 | 0.31 | 0.23 | 0.17 |
| 4 | 5504274 | rs16837352 | -0.893 | -0.007 | -0.177 | 0.477 | -0.023 | -0.048 | -0.048 | 0 | 0.19 | 0.04 | 0.13 | 0.28 | 0.2 | 0.3 |
| 4 | 5508665 | rs10032518 | -0.232 | -0.45 | 0.834 | 0.52 | -0.01 | -0.061 | -0.061 | 0 | 0.14 | 0.04 | 0.23 | 0.6 | 0.14 | 0.27 |
| 4 | 5516179 | rs1468309 | -0.729 | -0.502 | 1.441 | 0.399 | -0.005 | 0.022 | 0.022 | 0 | 0.16 | 0.22 | 0.16 | 0.25 | 0.14 | 0.29 |
| 4 | 5516502 | rs10020769 | -0.554 | -0.521 | 1.392 | 0.436 | 0 | 0.021 | 0.021 | 0 | 0.15 | 0.17 | 0.24 | 0.59 | 0.14 | 0.25 |
| 4 | 5527249 | rs2267654 | -1.119 | -0.619 | 1.572 | 0.307 | -0.016 | 0.031 | 0.031 | 0.603 | 0.24 | 0.31 | 0.3 | 0.63 | 0.18 | 0.26 |
| 4 | 5532985 | rs12641040 | -1.189 | -0.738 | 1.683 | -0.017 | -0.009 | 0.003 | 0.003 | 0.231 | 0.26 | 0.31 | 0.29 | 0.68 | 0.19 | 0.29 |
| 4 | 5536310 | rs16837389 | -0.497 | -0.706 | 1.498 | 0.005 | 0.013 | -0.064 | -0.064 | 0.008 | 0.21 | 0.09 | 0.26 | 0.61 | 0.17 | 0.25 |
| 4 | 5541650 | rs12505562 | -1.077 | -0.772 | 1.911 | -0.151 | -0.012 | 0.063 | 0.063 | 0 | 0.14 | 0.34 | 0.14 | 0.24 | 0.12 | 0.32 |
| 4 | 5543143 | rs3846230 | -1.942 | -0.849 | 1.503 | -0.611 | 0.035 | 0.071 | 0.071 | 1.25 | 0.43 | 0.69 | 0.44 | 0.9 | 0.33 | 0.65 |
| 4 | 5550882 | rs10937654 | -1.424 | -0.914 | 1.347 | -0.822 | 0.009 | 0.003 | 0.003 | 0 | 0.36 | 0.31 | 0.4 | 0.93 | 0.32 | 0.64 |
| 4 | 5554134 | rs13138291 | 1.516 | -0.915 | 0.489 | -1.559 | -0.062 | 0.104 | 0.104 | 0.004 | 0.18 | 0.01 | 0.14 | 0.01 | 0.28 | 0.14 |
| 4 | 5563196 | rs13113867 | 0.99 | -1.036 | 1.038 | -1.665 | 0.01 | -0.001 | -0.001 | 0 | 0.58 | 0.41 | 0.55 | 0.07 | 0.64 | 0.36 |
| 4 | 5570221 | rs12511039 | -0.515 | -0.967 | 0.858 | -1.621 | 0.013 | 0.077 | 0.077 | 0 | 0.32 | 0.16 | 0.37 | 0.88 | 0.31 | 0.56 |
| 4 | 5575751 | rs4689258 | -1.506 | -1.042 | 0.88 | -1.656 | 0.055 | 0.13 | 0.13 | 0.52 | 0.44 | 0.74 | 0.45 | 0.87 | 0.35 | 0.56 |
| 4 | 5576140 | rs7670299 | -1.574 | -1.131 | 0.879 | -1.655 | 0.052 | 0.127 | 0.127 | 0.346 | 0.45 | 0.73 | 0.45 | 0.87 | 0.36 | 0.57 |
| 4 | 5587093 | rs9995842 | -1.221 | -1.16 | 0.638 | -1.662 | -0.003 | -0.029 | -0.029 | 0 | 0.42 | 0.54 | 0.43 | 0.87 | 0.36 | 0.57 |
| 4 | 5591448 | rs13109991 | 0.537 | -1.043 | 0.161 | -1.263 | -0.034 | 0.027 | 0.027 | 0 | 0.3 | 0.12 | 0.25 | 0.25 | 0.43 | 0.47 |
| 4 | 5592348 | rs4334691 | 0.008 | -1.097 | 0.176 | -1.263 | 0.02 | -0.076 | -0.076 | 0 | 0.21 | 0.06 | 0.25 | 0.63 | 0.16 | 0.24 |
| 4 | 5594201 | rs749086 | 0.004 | -0.953 | 0.201 | -1.193 | 0.008 | -0.025 | -0.025 | 0 | 0.19 | 0.14 | 0.25 | 0.63 | 0.15 | 0.23 |
| 4 | 5594969 | rs4333126 | -0.56 | -0.833 | 0.242 | -1.198 | -0.028 | 0.195 | 0.195 | 0.814 | 0.48 | 0.19 | 0.5 | 0.87 | 0.57 | 0.7 |

| | | | | | | | | | | | | | | | |
|---|---|---|---|---|---|---|---|---|---|---|---|---|---|---|---|
| 4 | 5618307 | rs7684204 | -0.873 | -0.753 | 1.134 | -0.37 | -0.004 | 0.015 | 0.015 | 0 | 0.24 | 0.31 | 0.31 | 0.61 | 0.2 | 0.26 |
| 4 | 5619332 | rs4429676 | -0.589 | -0.895 | 1.354 | -0.43 | 0.029 | -0.016 | -0.016 | 0 | 0.08 | 0.34 | 0.07 | 0 | 0.04 | 0.01 |
| 4 | 5621673 | rs9993134 | -0.219 | -0.893 | 1.284 | -0.429 | 0.028 | -0.013 | -0.013 | 0 | 0.07 | 0.27 | 0.05 | 0 | 0.04 | 0.01 |
| 4 | 5622042 | rs13131951 | 1.544 | -0.925 | 1.173 | -0.449 | 0.049 | 0.007 | 0.007 | 0.43 | 0.63 | 0.13 | 0.61 | 0.32 | 0.74 | 0.68 |
| 4 | 5624670 | rs730469 | -1.811 | -0.871 | 1.174 | -0.489 | 0.044 | -0.025 | -0.025 | 0 | 0.37 | 0.88 | 0.4 | 0.62 | 0.26 | 0.28 |
| 4 | 5629298 | rs1078758 | -1.12 | -0.738 | 0.691 | -0.43 | -0.018 | 0.027 | 0.027 | 0.128 | 0.28 | 0.46 | 0.38 | 0.62 | 0.22 | 0.26 |
| 4 | 5631673 | rs7689275 | -1.773 | -0.662 | 0.541 | -0.442 | 0.036 | -0.012 | -0.012 | 0 | 0.37 | 0.86 | 0.45 | 0.63 | 0.27 | 0.28 |
| 4 | 5636212 | rs880291 | -1.48 | -0.56 | 1.459 | -0.595 | 0.016 | 0.013 | 0.013 | 4.086 | 0.06 | 0.38 | 0.03 | 0 | 0 | 0 |
| 4 | 5638652 | rs4353849 | -1.774 | -0.523 | 0.355 | -0.582 | -0.008 | -0.031 | -0.031 | 0.321 | 0.14 | 0.8 | 0.09 | 0 | 0.05 | 0.01 |
| 4 | 5638799 | rs10001971 | -1.796 | -0.522 | 0.291 | -0.582 | -0.012 | -0.036 | -0.036 | 0.575 | 0.15 | 0.78 | 0.09 | 0 | 0.05 | 0.01 |
| 4 | 5640595 | rs6850210 | -1.304 | 0.08 | 0.397 | -0.426 | -0.006 | -0.096 | -0.096 | 0 | 0.14 | 0.31 | 0.11 | 0 | 0.11 | 0.06 |
| 4 | 5641507 | rs747971 | -0.095 | 0.478 | -0.303 | -0.08 | 0.003 | 0.019 | 0.019 | 0.088 | 0.5 | 0.39 | 0.53 | 0.55 | 0.45 | 0.28 |
| 4 | 5641753 | rs10025465 | -0.27 | 0.588 | -0.344 | 0.061 | 0.001 | 0.019 | 0.019 | 0.119 | 0.49 | 0.53 | 0.43 | 0.43 | 0.54 | 0.63 |
| 4 | 5642161 | rs751841 | -0.092 | 0.718 | -0.798 | 0.34 | 0.013 | 0.006 | 0.006 | 0 | 0.53 | 0.63 | 0.56 | 0.48 | 0.62 | 0.58 |
| 4 | 5643287 | rs6833786 | 0.237 | 0.869 | -1.104 | 0.557 | -0.005 | -0.015 | -0.015 | 0 | 0.67 | 0.75 | 0.6 | 0.61 | 0.66 | 0.55 |
| 4 | 5647018 | rs10024750 | 0.387 | 0.862 | -1.013 | 0.414 | 0.011 | 0.017 | 0.017 | 0 | 0.52 | 0.3 | 0.5 | 0.61 | 0.58 | 0.41 |
| 4 | 5656508 | rs10804964 | 0.095 | 0.673 | -0.525 | 0.303 | -0.025 | -0.028 | -0.028 | 0 | 0.68 | 0.87 | 0.6 | 0.63 | 0.65 | 0.55 |
| 4 | 5658004 | rs11735503 | 1.321 | 0.684 | -0.824 | -0.193 | -0.009 | -0.031 | -0.031 | 0 | 0.25 | 0.15 | 0.3 | 0.42 | 0.27 | 0.28 |
| 4 | 5658554 | rs6842115 | 0.903 | 0.463 | -0.177 | -0.497 | 0.006 | -0.024 | -0.024 | 0 | 0.14 | 0.22 | 0.09 | 0.07 | 0.15 | 0.1 |
| 4 | 5658891 | rs6829204 | -1.772 | 0.365 | -0.37 | -0.521 | -0.011 | -0.033 | -0.033 | 0 | 0.83 | 0.95 | 0.7 | 0.76 | 0.83 | 0.69 |
| 4 | 5660264 | rs4688945 | -0.532 | 0.401 | -0.366 | -0.548 | -0.012 | -0.009 | -0.009 | 0.845 | 0.75 | 0.92 | 0.68 | 0.76 | 0.76 | 0.57 |
| 4 | 5665201 | rs4689273 | 1.364 | 0.399 | -0.357 | -0.679 | -0.008 | -0.047 | -0.047 | 0.073 | 0.41 | 0.15 | 0.42 | 0.5 | 0.43 | 0.35 |
| 4 | 5667561 | rs4298088 | -0.057 | 0.437 | -0.234 | -0.689 | -0.005 | 0 | 0 | 0 | 0.67 | 0.74 | 0.61 | 0.69 | 0.63 | 0.48 |
| 4 | 5667671 | rs4524330 | 0.507 | 0.393 | 0.075 | -0.684 | 0.005 | -0.002 | -0.002 | 0 | 0.28 | 0.33 | 0.21 | 0.27 | 0.31 | 0.21 |
| 4 | 5668688 | rs11730653 | -0.274 | 0.213 | 0.202 | -0.694 | -0.008 | -0.008 | -0.008 | 0 | 0.71 | 0.57 | 0.64 | 0.76 | 0.75 | 0.59 |
| 4 | 5677101 | rs13146518 | 0.293 | 0.492 | -0.095 | -0.607 | 0.025 | -0.012 | -0.012 | 0.155 | 0.3 | 0.52 | 0.23 | 0.27 | 0.32 | 0.23 |
| 4 | 5680311 | rs6446389 | -0.642 | 0.637 | -0.676 | -0.462 | -0.007 | -0.005 | -0.005 | 0 | 0.78 | 0.88 | 0.68 | 0.77 | 0.79 | 0.63 |
| 4 | 5682189 | rs4689276 | -0.648 | 0.737 | -0.718 | -0.484 | -0.003 | -0.007 | -0.007 | 0.07 | 0.86 | 0.87 | 0.8 | 0.77 | 0.86 | 0.66 |
| 4 | 5682811 | rs10937658 | 1.331 | 0.517 | -1.038 | -0.104 | -0.043 | -0.067 | -0.067 | 0.597 | 0.23 | 0.05 | 0.32 | 0.33 | 0.28 | 0.24 |
| 4 | 5683395 | rs6837414 | -1.731 | 0.477 | -0.85 | -0.039 | -0.006 | 0.009 | 0.009 | 0 | 0.89 | 0.98 | 0.87 | 0.95 | 0.92 | 0.89 |
| 4 | 5686924 | rs957727 | 0.196 | 0.392 | -0.399 | -0.155 | 0.005 | -0.013 | -0.013 | 0.143 | 0.66 | 0.52 | 0.68 | 0.75 | 0.69 | 0.69 |
| 4 | 5686967 | rs957726 | 0.187 | 0.39 | -0.4 | -0.156 | 0.005 | -0.013 | -0.013 | 0.226 | 0.66 | 0.52 | 0.68 | 0.75 | 0.69 | 0.69 |
| 4 | 5687281 | rs960055 | 0.909 | 0.396 | -0.12 | -0.188 | 0.027 | -0.041 | -0.041 | 0 | 0.12 | 0.14 | 0.08 | 0.02 | 0.17 | 0.13 |
| 4 | 5688643 | rs7684327 | 1.549 | 0.231 | -0.545 | -0.108 | -0.001 | 0.002 | 0.002 | 0.017 | 0.11 | 0.25 | 0.09 | 0.12 | 0.12 | 0.02 |
| 4 | 5688669 | rs10937660 | 0.065 | 0.221 | -1.114 | -0.019 | -0.008 | 0.017 | 0.017 | 0 | 0.55 | 0.25 | 0.59 | 0.63 | 0.58 | 0.67 |
| 4 | 5690902 | rs4689278 | 0.429 | 0.601 | -1.397 | -0.02 | -0.006 | -0.035 | -0.035 | 0 | 0.2 | 0.19 | 0.29 | 0.4 | 0.21 | 0.26 |
| 4 | 5691199 | rs10937661 | -0.315 | 0.557 | -1.398 | -0.09 | -0.005 | 0.029 | 0.029 | 0 | 0.89 | 0.98 | 0.95 | 0.96 | 0.9 | 0.94 |
| 4 | 5691320 | rs10937662 | -0.937 | 0.572 | -1.263 | -0.02 | -0.008 | 0.006 | 0.006 | 0.205 | 0.9 | 0.98 | 0.88 | 0.94 | 0.92 | 0.88 |
| 4 | 5694773 | rs6852113 | 0.201 | 0.359 | -1.02 | -0.145 | 0.029 | 0.009 | 0.009 | 0.044 | 0.48 | 0.63 | 0.42 | 0.41 | 0.37 | 0.43 |
| 4 | 5699097 | rs13126806 | 1.174 | 1.078 | -1.057 | -0.099 | 0.026 | -0.006 | -0.006 | 0.06 | 0.17 | 0.26 | 0.16 | 0.19 | 0.24 | 0.14 |
| 4 | 5703630 | rs11725145 | 1.061 | 0.479 | -1.212 | -0.03 | -0.001 | -0.01 | -0.01 | 0.805 | 0.06 | 0.06 | 0.03 | 0 | 0.1 | 0.06 |
| 4 | 5711368 | rs3774854 | -0.099 | -0.779 | -1.574 | 0.124 | 0.058 | -0.022 | -0.022 | 0.031 | 0.45 | 0.1 | 0.48 | 0.55 | 0.39 | 0.48 |
| 4 | 5717867 | rs7681283 | -0.53 | -1.168 | -1.673 | -0.083 | 0.018 | 0.056 | 0.056 | 0.135 | 0.72 | 0.93 | 0.87 | 0.88 | 0.77 | 0.8 |
| 4 | 5721021 | rs2291157 | 0.952 | -1.335 | -1.424 | 0.205 | 0.023 | -0.033 | -0.033 | 0 | 0.15 | 0.03 | 0.13 | 0.16 | 0.1 | 0.14 |
| 4 | 5721930 | rs1383178 | -0.426 | -0.898 | -0.344 | 0.426 | 0.014 | 0.017 | 0.017 | 1.688 | 0.16 | 0.49 | 0.25 | 0.19 | 0.15 | 0.22 |
| 4 | 5723145 | rs881686 | 1.197 | -0.778 | -0.709 | 0.428 | 0.015 | 0.029 | 0.029 | 0 | 0.48 | 0.35 | 0.38 | 0.35 | 0.54 | 0.35 |
| 4 | 5731518 | rs899688 | 0.196 | -0.681 | -1.19 | 0.803 | -0.017 | -0.067 | -0.067 | 0 | 0.1 | 0.01 | 0.26 | 0.31 | 0.12 | 0.25 |
| 4 | 5732978 | rs1871587 | 0.338 | -0.481 | -0.915 | 0.901 | 0.023 | 0.044 | 0.044 | 0 | 0.05 | 0.16 | 0.07 | 0.08 | 0.05 | 0.12 |
| 4 | 5735520 | rs2291154 | 0.107 | -0.469 | -1.595 | 0.764 | -0.001 | -0.017 | -0.017 | 0 | 0.08 | 0.04 | 0.05 | 0.1 | 0.06 | 0.05 |
| 4 | 5735715 | rs13130333 | -0.398 | -0.382 | -1.456 | 0.734 | 0.025 | -0.044 | -0.044 | 0 | 0.26 | 0.1 | 0.31 | 0.31 | 0.24 | 0.34 |
| 4 | 5742400 | rs10440166 | -0.478 | -0.44 | -1.007 | 0.582 | -0.015 | 0 | 0 | 0 | 0.46 | 0.51 | 0.27 | 0.44 | 0.35 | 0.45 |
| 4 | 5743053 | rs7674034 | -0.397 | -0.481 | -1.045 | 0.431 | 0.004 | -0.002 | -0.002 | 1.02 | 0.45 | 0.58 | 0.43 | 0.5 | 0.34 | 0.5 |
| 4 | 5743509 | rs6446393 | 0.866 | -0.407 | -1.395 | -0.088 | -0.01 | 0.044 | 0.044 | 0 | 0.92 | 1 | 0.98 | 1 | 0.94 | 0.9 |
| 4 | 5743512 | rs6414624 | 0.719 | -0.702 | -0.945 | -0.61 | -0.028 | -0.095 | -0.095 | 2.342 | 0.68 | 0.5 | 0.78 | 0.9 | 0.79 | 0.83 |
| 4 | 5743781 | rs1480169 | -0.731 | 0.125 | -1.55 | -0.694 | -0.007 | 0.055 | 0.055 | 0 | 0.25 | 0.13 | 0.15 | 0.11 | 0.24 | 0.14 |
| 4 | 5745304 | rs17689430 | 0.519 | 0.31 | -1.91 | -0.908 | -0.014 | 0.027 | 0.027 | 6.459 | 0.12 | 0.03 | 0.06 | 0.06 | 0.11 | 0.01 |
| 4 | 5745392 | rs994812 | -0.069 | 0.845 | -1.387 | -0.88 | 0.007 | -0.021 | -0.021 | 0 | 0.26 | 0.58 | 0.16 | 0.19 | 0.25 | 0.27 |
| 4 | 5745720 | rs1871584 | -0.557 | 1.094 | -0.987 | -1.218 | -0.001 | -0.006 | -0.006 | 0 | 0.47 | 0.48 | 0.44 | 0.35 | 0.52 | 0.43 |
| 4 | 5746620 | rs1351357 | -0.901 | 0.961 | -1.045 | -1.253 | -0.002 | -0.003 | -0.003 | 0 | 0.54 | 0.48 | 0.49 | 0.35 | 0.61 | 0.56 |
| 4 | 5747072 | rs2286343 | 1.227 | 0.826 | -0.981 | -1.186 | 0.011 | 0.003 | 0.003 | 0.257 | 0.37 | 0.28 | 0.45 | 0.65 | 0.32 | 0.39 |
| 4 | 5749784 | rs4689312 | -0.061 | 0.361 | -1.106 | -0.608 | -0.005 | -0.004 | -0.004 | 0 | 0.41 | 0.49 | 0.31 | 0.61 | 0.36 | 0.53 |
| 4 | 5749904 | rs4688963 | -0.061 | 0.361 | -1.106 | -0.608 | -0.005 | -0.004 | -0.004 | 0 | 0.41 | 0.49 | 0.31 | 0.62 | 0.36 | 0.53 |
| 4 | 5751253 | rs6813121 | 1.04 | 0.448 | -0.743 | -0.557 | 0.019 | 0.033 | 0.033 | 0.221 | 0.17 | 0.37 | 0.09 | 0.19 | 0.16 | 0.22 |
| 4 | 5752654 | rs3774861 | 0.278 | 0.258 | -0.773 | -0.727 | 0.001 | 0.011 | 0.011 | 0 | 0.52 | 0.44 | 0.58 | 0.62 | 0.56 | 0.52 |
| 4 | 5752753 | rs10011837 | 1.688 | 0.375 | -0.768 | -0.992 | -0.016 | 0.039 | 0.039 | 0 | 0.07 | 0 | 0.04 | 0 | 0.08 | 0.09 |
| 4 | 5753053 | rs899694 | 0.916 | 0.315 | -0.781 | -0.628 | -0.018 | -0.003 | -0.003 | 0 | 0.29 | 0.21 | 0.44 | 0.32 | 0.38 | 0.19 |
| 4 | 5753111 | rs899693 | 0.728 | 0.226 | -0.663 | -0.621 | -0.001 | 0.014 | 0.014 | 0 | 0.39 | 0.49 | 0.25 | 0.49 | 0.33 | 0.51 |
| 4 | 5753249 | rs899692 | 1.126 | 0.287 | -0.987 | -0.57 | -0.062 | -0.028 | -0.028 | 0 | 0.28 | 0.1 | 0.43 | 0.32 | 0.38 | 0.2 |
| 4 | 5753354 | rs2035304 | -0.026 | 0.273 | -0.965 | -0.573 | 0.007 | -0.013 | -0.013 | 0.025 | 0.49 | 0.75 | 0.31 | 0.5 | 0.41 | 0.6 |

| | | | | | | | | | | | | | | | | |
|---|---|---|---|---|---|---|---|---|---|---|---|---|---|---|---|---|
| 4 | 5754115 | rs3774867 | 2.125 | 0.528 | -1.227 | -0.57 | -0.023 | -0.046 | -0.046 | 0 | 0.08 | 0.01 | 0.19 | 0.18 | 0.1 | 0.1 |
| 4 | 5754544 | rs899691 | 0.596 | 0.367 | -0.937 | -0.575 | -0.064 | 0.012 | 0.012 | 0 | 0.36 | 0.16 | 0.47 | 0.32 | 0.47 | 0.28 |
| 4 | 5755542 | rs2302075 | -0.704 | 0.306 | -0.61 | -0.315 | -0.025 | -0.018 | -0.018 | 0.993 | 0.78 | 0.99 | 0.73 | 0.83 | 0.79 | 0.8 |
| 4 | 5756967 | rs883994 | 0.019 | 0.254 | -0.349 | -0.34 | -0.019 | -0.021 | -0.021 | 0.166 | 0.25 | 0.4 | 0.31 | 0.17 | 0.22 | 0.2 |
| 4 | 5757469 | rs899690 | -0.583 | 0.282 | -0.344 | -0.331 | 0.007 | 0.008 | 0.008 | 0 | 0.77 | 0.87 | 0.72 | 0.82 | 0.79 | 0.8 |
| 4 | 5758869 | rs735172 | -0.394 | 0.452 | -0.584 | -0.239 | 0.006 | 0.013 | 0.013 | 0 | 0.31 | 0.3 | 0.33 | 0.18 | 0.3 | 0.29 |
| 4 | 5761366 | rs11939527 | 0.995 | 0.692 | -0.589 | 0.062 | -0.001 | 0.023 | 0.023 | 0 | 0.25 | 0.27 | 0.18 | 0.5 | 0.23 | 0.41 |
| 4 | 5762161 | rs6855229 | 0.006 | 0.606 | -0.467 | 0.188 | -0.002 | 0.001 | 0.001 | 0 | 0.18 | 0.23 | 0.23 | 0.17 | 0.16 | 0.18 |
| 4 | 5770734 | rs17747287 | -0.098 | 0.732 | -0.783 | 0.246 | -0.002 | 0 | 0 | 0 | 0.51 | 0.48 | 0.69 | 0.52 | 0.52 | 0.4 |
| 4 | 5771295 | rs4689308 | -0.24 | 0.822 | -0.852 | 0.034 | 0.002 | 0.029 | 0.029 | 0.414 | 0.49 | 0.46 | 0.58 | 0.32 | 0.49 | 0.28 |
| 4 | 5773055 | rs6446395 | 0.006 | 0.804 | -0.884 | 0.274 | 0.004 | -0.009 | -0.009 | 0 | 0.52 | 0.78 | 0.73 | 0.53 | 0.54 | 0.44 |
| 4 | 5774747 | rs3856953 | -1.064 | 0.749 | -0.709 | 0.492 | 0 | 0.013 | 0.013 | 0 | 0.95 | 0.94 | 0.98 | 0.87 | 0.95 | 0.94 |
| 4 | 5778783 | rs7680147 | 0.256 | 0.594 | -0.911 | 0.026 | 0 | -0.016 | -0.016 | 0 | 0.11 | 0.11 | 0.07 | 0 | 0.09 | 0.02 |
| 4 | 5784934 | rs4689304 | -0.075 | 0.309 | -0.753 | 0.109 | 0.005 | 0.035 | 0.035 | 0.085 | 0.45 | 0.67 | 0.58 | 0.33 | 0.47 | 0.26 |
| 4 | 5785442 | rs1383180 | 1.004 | 0.299 | -0.705 | 0.141 | -0.018 | -0.003 | -0.003 | 0 | 0.3 | 0.18 | 0.47 | 0.33 | 0.4 | 0.22 |
| 4 | 5793156 | rs7678015 | -0.427 | 0.21 | -0.334 | -0.086 | 0 | -0.001 | -0.001 | 0 | 0.52 | 0.41 | 0.44 | 0.54 | 0.44 | 0.61 |
| 4 | 5794481 | rs17691258 | 0.641 | 0.128 | -0.481 | -0.192 | -0.006 | 0.009 | 0.009 | 0 | 0.1 | 0.09 | 0.18 | 0.04 | 0.16 | 0.07 |
| 4 | 5794904 | rs6826372 | 0.322 | 0.112 | -0.271 | -0.2 | 0.012 | -0.025 | -0.025 | 0 | 0.11 | 0.2 | 0.2 | 0.04 | 0.16 | 0.07 |
| 4 | 5798627 | rs2291152 | -0.839 | 0.086 | -0.389 | -0.332 | -0.013 | -0.108 | -0.108 | 0.251 | 0.31 | 0.41 | 0.32 | 0.03 | 0.29 | 0.14 |
| 4 | 5803903 | rs2306399 | 0.737 | 0.61 | -0.951 | 0.021 | -0.048 | -0.05 | -0.05 | 0.629 | 0.32 | 0.15 | 0.3 | 0.53 | 0.33 | 0.55 |
| 4 | 5804830 | rs3774876 | 1.449 | 0.658 | -1.144 | 0.118 | -0.047 | -0.065 | -0.065 | 0 | 0.1 | 0 | 0.17 | 0.21 | 0.12 | 0.22 |
| 4 | 5806366 | rs1552057 | 0.088 | 0.872 | -0.704 | 0.157 | -0.012 | -0.016 | -0.016 | 0 | 0.34 | 0.22 | 0.26 | 0.37 | 0.3 | 0.42 |
| 4 | 5809023 | rs7654977 | 1.2 | 0.84 | -0.818 | 0.149 | -0.035 | -0.065 | -0.065 | 0.1 | 0.22 | 0.07 | 0.17 | 0.34 | 0.2 | 0.35 |
| 4 | 5809187 | rs7680768 | -0.587 | 0.749 | -0.727 | 0.089 | -0.001 | 0.003 | 0.003 | 0 | 0.42 | 0.37 | 0.37 | 0.37 | 0.43 | 0.46 |
| 4 | 5812195 | rs2279250 | -0.967 | 0.618 | -0.824 | -0.088 | 0 | 0.001 | 0.001 | 0 | 0.72 | 0.71 | 0.65 | 0.57 | 0.69 | 0.83 |
| 4 | 5812566 | rs2279247 | -0.899 | 1.018 | -0.712 | 0.113 | -0.005 | -0.061 | -0.061 | 0 | 0.2 | 0.3 | 0.2 | 0.03 | 0.23 | 0.11 |
| 4 | 5812974 | rs16837692 | 0.216 | 1.066 | -0.577 | 0.474 | 0 | -0.003 | -0.003 | 0 | 0.1 | 0.12 | 0.11 | 0.03 | 0.13 | 0.05 |
| 4 | 5815381 | rs3733186 | -0.16 | 1.115 | -0.685 | 0.497 | -0.003 | 0.012 | 0.012 | 0 | 0.42 | 0.36 | 0.36 | 0.33 | 0.43 | 0.46 |
| 4 | 5815635 | rs3733189 | -0.303 | 1.377 | -0.631 | 0.818 | -0.004 | 0.014 | 0.014 | 0 | 0.09 | 0.06 | 0.11 | 0.03 | 0.14 | 0.05 |
| 4 | 5815929 | rs3755847 | -0.352 | 1.378 | -0.634 | 0.819 | -0.001 | 0.006 | 0.006 | 0 | 0.09 | 0.08 | 0.12 | 0.03 | 0.14 | 0.06 |
| 4 | 5816938 | rs6813104 | 0.036 | 1.333 | -0.453 | 1.116 | -0.003 | -0.003 | -0.003 | 0.126 | 0.78 | 0.82 | 0.69 | 0.58 | 0.72 | 0.85 |
| 4 | 5819154 | rs16837696 | -0.635 | 0.785 | -0.215 | 1.064 | 0 | 0.006 | 0.006 | 0 | 0.16 | 0.16 | 0.21 | 0.31 | 0.17 | 0.27 |
| 4 | 5819381 | rs11944616 | -0.857 | 0.358 | 0.803 | 0.337 | -0.028 | -0.091 | -0.091 | 0 | 0.14 | 0.35 | 0.06 | 0.01 | 0.06 | 0.09 |
| 4 | 5819967 | rs10937680 | -0.054 | 0.886 | 0.568 | -0.11 | 0.021 | -0.079 | -0.079 | 3.728 | 0.83 | 0.75 | 0.86 | 1 | 0.79 | 0.9 |
| 4 | 5820761 | rs10937681 | 0.966 | 0.542 | 0.907 | 0.496 | 0.003 | 0.004 | 0.004 | 0 | 0.64 | 0.47 | 0.61 | 0.61 | 0.64 | 0.64 |
| 4 | 5822738 | rs1057052 | -1.173 | 0.134 | 1.205 | 0.764 | -0.032 | -0.003 | -0.003 | 0.342 | 0.12 | 0.19 | 0.04 | 0.1 | 0.05 | 0.17 |
| 4 | 5826597 | rs3774883 | -2.469 | 0.043 | 1.088 | 1.182 | 0.035 | -0.022 | -0.022 | 0.289 | 0.24 | 0.87 | 0.37 | 0.26 | 0.16 | 0.27 |
| 4 | 5827079 | rs2286286 | -2.198 | 0.081 | 1.807 | 0.869 | -0.028 | -0.03 | -0.03 | 0 | 0.22 | 0.64 | 0.18 | 0.13 | 0.16 | 0.22 |
| 4 | 5830414 | rs6818391 | -0.951 | 0.024 | 1.337 | 0.494 | -0.008 | 0.042 | 0.042 | 0 | 0.92 | 0.99 | 0.97 | 1 | 0.94 | 0.93 |
| 4 | 5831521 | rs6820236 | -1.833 | 0.163 | 2.388 | 0.747 | -0.019 | -0.109 | -0.109 | 0 | 0.15 | 0.56 | 0.1 | 0 | 0.11 | 0.1 |
| 4 | 5831665 | rs16837714 | -1.085 | 0.162 | 2.275 | 0.756 | -0.036 | -0.074 | -0.074 | 1.284 | 0.12 | 0.27 | 0.04 | 0 | 0.06 | 0.09 |
| 4 | 5833625 | rs11723228 | -2.29 | 0.809 | 1.685 | 0.481 | -0.024 | 0.074 | 0.074 | 0.937 | 0.33 | 0.86 | 0.3 | 0.16 | 0.35 | 0.35 |
| 4 | 5836702 | rs3755849 | -0.678 | 0.88 | 1.875 | 0.68 | 0.044 | 0 | 0 | 0.028 | 0.14 | 0.31 | 0.17 | 0.14 | 0.19 | 0.22 |
| 4 | 5837325 | rs3821937 | -0.863 | 0.778 | 1.778 | 0.617 | 0.025 | -0.008 | -0.008 | 0 | 0.15 | 0.29 | 0.17 | 0.13 | 0.19 | 0.2 |
| 4 | 5838837 | rs7654446 | -1.892 | 0.855 | 1.526 | 1.134 | -0.001 | 0.005 | 0.005 | 0.176 | 0.31 | 0.63 | 0.31 | 0.58 | 0.29 | 0.4 |
| 4 | 5840887 | rs2240492 | -1.745 | 1.196 | 1.987 | 1.297 | 0.049 | 0.429 | 0.429 | 0 | 0.07 | 0.3 | 0.12 | 0.36 | 0.06 | 0.15 |
| 4 | 5841649 | rs13144677 | -1.528 | 1.452 | 1.709 | 1.116 | 0.065 | 0.089 | 0.089 | 0.833 | 0.21 | 0.58 | 0.27 | 0.44 | 0.32 | 0.27 |
| 4 | 5842856 | rs3774885 | -1.267 | 1.534 | 1.202 | 0.207 | 0.087 | -0.132 | -0.132 | 1.405 | 0.23 | 0.43 | 0.19 | 0.03 | 0.32 | 0.16 |
| 4 | 5843328 | rs2286280 | -1.147 | 1.377 | 0.592 | 0.655 | 0.06 | 0.039 | 0.039 | 0.105 | 0.25 | 0.54 | 0.28 | 0.33 | 0.35 | 0.27 |
| 4 | 5847493 | rs1979427 | 0.238 | 0.132 | -0.323 | -0.964 | 0.035 | 0.067 | 0.067 | 0 | 0.6 | 0.33 | 0.51 | 0.87 | 0.74 | 0.69 |
| 4 | 5848928 | rs3885409 | -0.786 | 0.473 | -0.122 | -1.271 | 0.008 | -0.025 | -0.025 | 0 | 0.25 | 0.38 | 0.26 | 0.16 | 0.29 | 0.26 |
| 4 | 5850343 | rs11945849 | -0.571 | 0.64 | -0.057 | -1.392 | -0.001 | -0.08 | -0.08 | 0.185 | 0.16 | 0.25 | 0.09 | 0 | 0.17 | 0.08 |
| 4 | 5851343 | rs2276877 | 1.404 | 0.527 | -0.516 | -1.422 | -0.156 | -0.034 | -0.034 | 0 | 0.36 | 0.02 | 0.25 | 0.69 | 0.45 | 0.43 |
| 4 | 5852138 | rs984576 | 0.117 | 0.422 | -0.346 | -1.752 | 0.075 | 0.211 | 0.211 | 0.007 | 0.49 | 0.2 | 0.44 | 0.86 | 0.62 | 0.64 |
| 4 | 5856509 | rs6848864 | -0.705 | 0.064 | 0.052 | -1.691 | 0.001 | -0.005 | -0.005 | 0.296 | 0.11 | 0.13 | 0.07 | 0.06 | 0.11 | 0.12 |
| 4 | 5859510 | rs10030434 | -0.147 | -0.377 | 0.379 | -2.095 | 0.015 | 0.027 | 0.027 | 0 | 0.7 | 0.75 | 0.61 | 0.8 | 0.75 | 0.68 |
| 4 | 5860906 | rs7356507 | 1.929 | -0.382 | 0.051 | -2.169 | -0.146 | 0.063 | 0.063 | 0.13 | 0.42 | 0.02 | 0.42 | 0.76 | 0.53 | 0.48 |
| 4 | 5865088 | rs13146086 | 1.144 | -0.725 | 0.772 | -2.244 | 0.036 | 0.079 | 0.079 | 0 | 0.49 | 0.33 | 0.47 | 0.75 | 0.58 | 0.52 |
| 4 | 5865361 | rs7691078 | 0.398 | -0.843 | 1.159 | -2.327 | -0.014 | -0.031 | -0.031 | 0.707 | 0.62 | 0.43 | 0.56 | 0.79 | 0.72 | 0.65 |
| 4 | 5866819 | rs6838020 | -0.145 | -0.882 | 1.094 | -2.327 | -0.008 | -0.012 | -0.012 | 0 | 0.66 | 0.59 | 0.59 | 0.8 | 0.75 | 0.69 |
| 4 | 5866947 | rs6446401 | -1.71 | -1.175 | 0.863 | -2.504 | -0.014 | -0.059 | -0.059 | 0.09 | 0.3 | 0.57 | 0.17 | 0.05 | 0.23 | 0.22 |
| 4 | 5871485 | rs12508543 | 0.791 | -1.038 | 0.764 | -2.472 | 0.009 | 0.031 | 0.031 | 0.021 | 0.54 | 0.34 | 0.48 | 0.76 | 0.6 | 0.51 |
| 4 | 5873648 | rs6823027 | -0.156 | -1.047 | 0.847 | -2.425 | -0.032 | -0.092 | -0.092 | 0 | 0.15 | 0.27 | 0.07 | 0 | 0.1 | 0.04 |
| 4 | 5874579 | rs9790594 | 0.075 | -1.475 | 0.612 | -2.696 | -0.021 | -0.05 | -0.05 | 0.077 | 0.73 | 0.68 | 0.75 | 0.92 | 0.78 | 0.84 |
| 4 | 5882084 | rs4292278 | -0.788 | -1.578 | 0.812 | -2.674 | -0.05 | -0.093 | -0.093 | 0.055 | 0.17 | 0.37 | 0.08 | 0.01 | 0.06 | 0.08 |
| 4 | 5882848 | rs3774889 | 0.26 | -1.433 | 1.007 | -2.633 | 0.003 | 0.005 | 0.005 | 0 | 0.17 | 0.12 | 0.23 | 0.1 | 0.2 | 0.16 |
| 4 | 5887388 | rs3774893 | 1.235 | -1.332 | 0.824 | -2.625 | 0.002 | 0.004 | 0.004 | 0 | 0.56 | 0.45 | 0.5 | 0.75 | 0.59 | 0.61 |
| 4 | 5889251 | rs3755851 | 0.457 | -1.298 | 0.822 | -2.628 | -0.024 | -0.046 | -0.046 | 0 | 0.65 | 0.55 | 0.55 | 0.8 | 0.68 | 0.72 |
| 4 | 5895798 | rs10004248 | 0.28 | -1.076 | 0.779 | -2.571 | -0.001 | -0.005 | -0.005 | 0.021 | 0.67 | 0.67 | 0.54 | 0.81 | 0.67 | 0.74 |

| | | | | | | | | | | | | | | | | |
|---|---|---|---|---|---|---|---|---|---|---|---|---|---|---|---|---|
| 4 | 5896702 | rs6812766 | -1.152 | -1.062 | 0.244 | -2.707 | -0.006 | 0.023 | 0.023 | 0 | 0.84 | 0.9 | 0.81 | 0.9 | 0.85 | 0.84 |
| 4 | 5901291 | rs4456913 | 2.12 | -0.928 | 0.116 | -2.628 | -0.178 | 0.051 | 0.051 | 0 | 0.38 | 0.02 | 0.39 | 0.77 | 0.5 | 0.51 |
| 4 | 5901873 | rs10020979 | 0.916 | -0.807 | 0.436 | -2.539 | 0.009 | 0.085 | 0.085 | 0 | 0.55 | 0.27 | 0.44 | 0.78 | 0.57 | 0.58 |
| 4 | 5902274 | rs6446405 | 0.451 | -0.69 | 0.396 | -2.448 | -0.002 | -0.012 | -0.012 | 0 | 0.62 | 0.41 | 0.51 | 0.82 | 0.66 | 0.69 |
| 4 | 5910469 | rs13146468 | 1.544 | -0.428 | -0.43 | -2.166 | -0.17 | 0.039 | 0.039 | 0 | 0.39 | 0.02 | 0.4 | 0.77 | 0.5 | 0.53 |
| 4 | 5914110 | rs6446407 | 0.604 | -0.067 | 0.688 | -2 | -0.011 | -0.044 | -0.044 | 4.579 | 0.9 | 0.81 | 0.96 | 0.99 | 0.95 | 0.91 |
| 4 | 5919709 | rs4689294 | -1.466 | -0.038 | 1.135 | -2.111 | -0.021 | -0.106 | -0.106 | 0 | 0.15 | 0.47 | 0.09 | 0.01 | 0.07 | 0.08 |
| 4 | 5920389 | rs16837803 | -2.16 | 0.131 | 0.998 | -2.012 | 0.056 | -0.045 | -0.045 | 0 | 0.1 | 0.39 | 0.05 | 0 | 0.07 | 0.09 |
| 4 | 5924289 | rs4425309 | -0.273 | 0.059 | 0.246 | -2.258 | -0.003 | 0.015 | 0.015 | 0 | 0.78 | 0.84 | 0.76 | 0.85 | 0.76 | 0.78 |
| 4 | 5924458 | rs11729343 | 0.05 | -0.081 | 0.388 | -2.609 | 0 | -0.005 | -0.005 | 0 | 0.17 | 0.17 | 0.26 | 0.06 | 0.22 | 0.14 |
| 4 | 5924629 | rs4443226 | -1.551 | -0.173 | 0.07 | -2.826 | 0.006 | 0.072 | 0.072 | 0 | 0.36 | 0.75 | 0.37 | 0.07 | 0.35 | 0.27 |
| 4 | 5924794 | rs12506399 | 2.133 | -0.17 | -0.024 | -2.774 | -0.065 | 0.115 | 0.115 | 0 | 0.42 | 0.1 | 0.39 | 0.79 | 0.41 | 0.51 |
| 4 | 5926312 | rs12505016 | 1.801 | -0.175 | 0.235 | -2.778 | -0.036 | 0.138 | 0.138 | 0 | 0.45 | 0.14 | 0.41 | 0.79 | 0.47 | 0.56 |
| 4 | 5926551 | rs7675169 | 1.679 | -0.205 | 0.655 | -2.753 | -0.006 | 0.051 | 0.051 | 0 | 0.46 | 0.32 | 0.42 | 0.79 | 0.44 | 0.52 |
| 4 | 5927071 | rs9999689 | -1.192 | -0.017 | 0.66 | -2.599 | -0.004 | -0.006 | -0.006 | 0.742 | 0.86 | 0.84 | 0.82 | 0.9 | 0.87 | 0.88 |
| 4 | 5927355 | rs9999966 | 1.233 | 0.087 | 0.433 | -2.528 | -0.001 | 0 | 0 | 0 | 0.08 | 0.08 | 0.06 | 0.05 | 0.11 | 0.1 |
| 4 | 5928187 | rs10016978 | 1.017 | -0.12 | 0.34 | -2.766 | 0.002 | 0.011 | 0.011 | 0 | 0.49 | 0.43 | 0.42 | 0.79 | 0.44 | 0.56 |
| 4 | 5931208 | rs6840908 | -1.019 | 0.017 | -0.388 | -2.852 | -0.007 | 0.014 | 0.014 | 0 | 0.39 | 0.31 | 0.4 | 0.11 | 0.45 | 0.31 |
| 4 | 5931713 | rs6857538 | -0.508 | 0.221 | -0.751 | -2.71 | 0.034 | -0.082 | -0.082 | 0 | 0.27 | 0.5 | 0.36 | 0.1 | 0.35 | 0.23 |
| 4 | 5932608 | rs10937695 | 1.437 | 0.328 | -1.817 | -2.48 | -0.068 | 0.078 | 0.078 | 0 | 0.37 | 0.04 | 0.35 | 0.79 | 0.36 | 0.49 |
| 4 | 5934814 | rs6446410 | 0.135 | 0.655 | -1.209 | -2.478 | -0.035 | 0.03 | 0.03 | 0 | 0.77 | 0.86 | 0.6 | 0.88 | 0.65 | 0.78 |
| 4 | 5935406 | rs11727977 | -0.524 | 0.654 | -0.67 | -2.386 | -0.001 | -0.024 | -0.024 | 0.51 | 0.13 | 0.15 | 0.07 | 0.02 | 0.11 | 0.16 |
| 4 | 5936570 | rs17751160 | -0.497 | 0.863 | -0.883 | -2.81 | 0.007 | 0.076 | 0.076 | 0 | 0.55 | 0.67 | 0.59 | 0.13 | 0.58 | 0.41 |
| 4 | 5936875 | rs11945109 | -1.111 | 0.765 | -0.262 | -2.899 | 0.012 | 0.021 | 0.021 | 0 | 0.65 | 0.72 | 0.69 | 0.19 | 0.75 | 0.51 |
| 4 | 5939053 | rs4539987 | -1.161 | 0.826 | -0.075 | -2.911 | 0.024 | -0.034 | -0.034 | 0 | 0.17 | 0.57 | 0.16 | 0.06 | 0.18 | 0.15 |
| 4 | 5939889 | rs11725173 | 0.23 | 0.681 | -0.451 | -3.249 | -0.031 | 0.207 | 0.207 | 0 | 0.45 | 0.17 | 0.51 | 0.12 | 0.55 | 0.38 |
| 4 | 5941155 | rs16837872 | -1.029 | 0.735 | -0.39 | -3.164 | 0.002 | -0.078 | -0.078 | 0 | 0.2 | 0.57 | 0.17 | 0.04 | 0.21 | 0.13 |
| 4 | 5943692 | rs17694785 | -0.352 | 0.743 | 0.023 | -3.195 | 0.007 | 0.017 | 0.017 | 0 | 0.51 | 0.47 | 0.56 | 0.12 | 0.57 | 0.44 |
| 4 | 5945106 | rs7682973 | 0.068 | 0.695 | -0.138 | -3.28 | -0.025 | 0.047 | 0.047 | 0 | 0.29 | 0.24 | 0.39 | 0.07 | 0.35 | 0.2 |
| 4 | 5948793 | rs9684740 | -0.339 | 0.59 | 0.14 | -3.473 | -0.004 | 0 | 0 | 0 | 0.34 | 0.3 | 0.2 | 0.05 | 0.37 | 0.32 |
| 4 | 5949558 | rs10005242 | -1.575 | 0.453 | 0.375 | -3.541 | 0.01 | 0.094 | 0.094 | 0 | 0.66 | 0.83 | 0.62 | 0.13 | 0.69 | 0.52 |
| 4 | 5954520 | rs4431167 | -1.113 | 0.434 | 1.027 | -3.534 | -0.007 | -0.05 | -0.05 | 0.1 | 0.62 | 0.6 | 0.6 | 0.13 | 0.68 | 0.52 |
| 4 | 5955591 | rs7680750 | -0.3 | 0.471 | 1.062 | -3.429 | 0.002 | -0.114 | -0.114 | 0 | 0.34 | 0.45 | 0.41 | 0.09 | 0.33 | 0.22 |
| 4 | 5956975 | rs6855903 | -2.015 | 0.167 | 1.063 | -3.304 | -0.042 | -0.087 | -0.087 | 0 | 0.13 | 0.47 | 0.07 | 0.01 | 0.04 | 0.08 |
| 4 | 5958489 | rs16837906 | -1.174 | 0.268 | 1.028 | -3.266 | 0.013 | -0.019 | -0.019 | 0 | 0.06 | 0.21 | 0.02 | 0 | 0.03 | 0.06 |
| 4 | 5960352 | rs3733190 | 0.394 | 0.37 | 0.768 | -3.495 | -0.013 | 0.048 | 0.048 | 0 | 0.26 | 0.19 | 0.17 | 0.03 | 0.29 | 0.24 |
| 4 | 5961995 | rs13107008 | -0.421 | 0.417 | 0.085 | -3.484 | 0.034 | 0.077 | 0.077 | 0 | 0.52 | 0.38 | 0.53 | 0.15 | 0.63 | 0.47 |
| 4 | 5964345 | rs6446414 | -1.17 | 0.317 | 0.037 | -3.401 | 0.017 | 0.078 | 0.078 | 0 | 0.62 | 0.75 | 0.62 | 0.15 | 0.68 | 0.5 |
| 4 | 5966801 | rs6839295 | 0.398 | 0.313 | -0.192 | -3.399 | 0.008 | -0.033 | -0.033 | 0 | 0.28 | 0.3 | 0.39 | 0.11 | 0.35 | 0.25 |
| 4 | 5967316 | rs878145 | -1.022 | 0.01 | -0.627 | -2.926 | 0.012 | 0.117 | 0.117 | 0 | 0.65 | 0.86 | 0.63 | 0.15 | 0.69 | 0.55 |
| 4 | 5969640 | rs13124429 | -0.116 | 0.147 | -0.823 | -2.843 | 0.045 | 0.097 | 0.097 | 0 | 0.53 | 0.32 | 0.54 | 0.14 | 0.64 | 0.46 |
| 4 | 5970508 | rs7674007 | -0.455 | 0.48 | -1.571 | -2.758 | 0.001 | 0.192 | 0.192 | 1.484 | 0.62 | 0.91 | 0.47 | 0.12 | 0.66 | 0.44 |
| 4 | 5970798 | rs6813176 | 0.474 | 0.149 | -1.439 | -2.376 | 0.009 | 0.085 | 0.085 | 0.093 | 0.26 | 0.08 | 0.45 | 0.87 | 0.22 | 0.5 |
| 4 | 5971082 | rs9996280 | -0.614 | 0.416 | -0.569 | -2.36 | -0.014 | -0.156 | -0.156 | 1.326 | 0.28 | 0.5 | 0.17 | 0.01 | 0.22 | 0.16 |
| 4 | 5975174 | rs16837957 | -0.211 | 0.357 | -0.774 | -1.905 | -0.001 | -0.01 | -0.01 | 0.616 | 0.51 | 0.49 | 0.51 | 0.7 | 0.46 | 0.63 |
| 4 | 5975538 | rs16837960 | -0.386 | 0.314 | -0.473 | -1.904 | 0.005 | 0.048 | 0.048 | 0 | 0.5 | 0.41 | 0.49 | 0.7 | 0.45 | 0.6 |
| 4 | 5977553 | rs4689290 | 0.904 | 0.142 | -0.596 | -2.131 | 0 | -0.008 | -0.008 | 0 | 0.22 | 0.17 | 0.32 | 0.66 | 0.22 | 0.37 |
| 4 | 5980110 | rs10433662 | 0.878 | 0.117 | -0.572 | -2.144 | -0.011 | -0.04 | -0.04 | 0 | 0.22 | 0.09 | 0.31 | 0.67 | 0.22 | 0.38 |
| 4 | 5981004 | rs11941221 | -0.289 | 0.065 | -0.497 | -2.27 | 0.002 | -0.089 | -0.089 | 0 | 0.21 | 0.31 | 0.15 | 0.02 | 0.19 | 0.17 |
| 4 | 5982108 | rs6821688 | -0.554 | -0.326 | -1.029 | -2.982 | 0.005 | 0.007 | 0.007 | 0 | 0.32 | 0.69 | 0.33 | 0.25 | 0.25 | 0.25 |
| 4 | 5983053 | rs6846882 | -0.051 | -0.4 | -0.387 | -2.963 | 0.021 | -0.065 | -0.065 | 0.781 | 0.11 | 0.41 | 0.06 | 0 | 0.07 | 0.07 |
| 4 | 5983751 | rs7672438 | -1.028 | -0.372 | -0.904 | -3.029 | -0.032 | 0.135 | 0.135 | 0 | 0.6 | 0.83 | 0.7 | 0.91 | 0.56 | 0.71 |
| 4 | 5984952 | rs4689288 | -1.374 | -0.807 | -1.203 | -3.007 | -0.113 | 0.257 | 0.257 | 0.004 | 0.59 | 0.9 | 0.67 | 0.92 | 0.51 | 0.62 |
| 4 | 5985002 | rs4688950 | 1.6 | -0.556 | -0.814 | -2.571 | -0.001 | 0.009 | 0.009 | 0 | 0.21 | 0.23 | 0.32 | 0.65 | 0.23 | 0.31 |
| 4 | 5988237 | rs10014397 | -0.46 | -0.693 | -0.438 | -2.764 | -0.002 | -0.07 | -0.07 | 0.319 | 0.46 | 0.51 | 0.59 | 0.93 | 0.46 | 0.55 |
| 4 | 5990110 | rs4689287 | 0.467 | -0.902 | -0.72 | -2.274 | -0.001 | 0 | 0 | 0 | 0.29 | 0.21 | 0.37 | 0.66 | 0.32 | 0.36 |
| 4 | 5992086 | rs747391 | -0.647 | -0.662 | -0.917 | -1.963 | 0.001 | -0.002 | -0.002 | 0 | 0.17 | 0.17 | 0.17 | 0.23 | 0.1 | 0.17 |
| 4 | 5995255 | rs16838017 | 0.211 | -0.463 | -1.104 | -2.107 | 0.001 | -0.02 | -0.02 | 0 | 0.13 | 0.15 | 0.05 | 0 | 0.07 | 0.04 |
| 4 | 5997514 | rs6857506 | 0.171 | -0.283 | -1.217 | -2.085 | 0.003 | -0.049 | -0.049 | 0 | 0.32 | 0.31 | 0.16 | 0.01 | 0.25 | 0.2 |
| 4 | 6000691 | rs3887346 | -0.823 | 0.615 | -1.921 | -1.212 | -0.122 | 0.048 | 0.048 | 0 | 0.74 | 0.97 | 0.72 | 0.86 | 0.62 | 0.73 |
| 4 | 6002830 | rs7657266 | -0.731 | 0.148 | -0.665 | -0.761 | -0.006 | 0.013 | 0.013 | 0 | 0.18 | 0.29 | 0.21 | 0.23 | 0.15 | 0.2 |
| 4 | 6018866 | rs6838861 | 0.102 | 0.203 | -1.123 | -0.146 | 0.001 | -0.084 | -0.084 | 0 | 0.25 | 0.07 | 0.18 | 0.63 | 0.18 | 0.42 |
| 4 | 6018891 | rs4689318 | -0.366 | -0.025 | -0.782 | -1.269 | -0.003 | -0.062 | -0.062 | 0 | 0.72 | 0.64 | 0.81 | 0.99 | 0.74 | 0.85 |
| 4 | 6024426 | rs878948 | 0.464 | -0.137 | -0.402 | -1.241 | -0.009 | 0.006 | 0.006 | 0 | 0.36 | 0.3 | 0.41 | 0.25 | 0.47 | 0.32 |
| 4 | 6026058 | rs2285986 | -0.186 | -0.675 | -0.181 | -1.375 | 0.012 | 0.014 | 0.014 | 0 | 0.59 | 0.61 | 0.58 | 0.25 | 0.74 | 0.5 |
| 4 | 6026877 | rs13119847 | 0.369 | -0.278 | -0.348 | -1.224 | 0.005 | -0.026 | -0.026 | 0.405 | 0.33 | 0.18 | 0.21 | 0.62 | 0.25 | 0.45 |
| 4 | 6028838 | rs4689322 | 0.49 | 0.302 | 0.003 | -1.417 | -0.016 | 0 | 0 | 0 | 0.29 | 0.22 | 0.38 | 0.26 | 0.44 | 0.3 |
| 4 | 6032178 | rs13105242 | -0.046 | 0.049 | 0.478 | -1.652 | 0.019 | 0.036 | 0.036 | 0.458 | 0.49 | 0.46 | 0.52 | 0.3 | 0.66 | 0.46 |
| 4 | 6032456 | rs10026454 | -0.517 | -0.238 | 0.558 | -1.786 | 0.017 | 0.034 | 0.034 | 0.177 | 0.6 | 0.82 | 0.64 | 0.31 | 0.71 | 0.55 |

| | | | | | | | | | | | | | | | | |
|---|---|---|---|---|---|---|---|---|---|---|---|---|---|---|---|---|
| 4 | 6032801 | rs6813726 | -0.797 | -0.683 | 1.4 | -2.434 | 0.002 | -0.072 | -0.072 | 0 | 0.12 | 0.28 | 0.09 | 0 | 0.09 | 0.08 |
| 4 | 6033657 | rs765831 | -1.646 | -0.562 | 1.282 | -2.649 | 0.003 | 0.183 | 0.183 | 0 | 0.43 | 0.81 | 0.39 | 0.01 | 0.44 | 0.3 |
| 4 | 6034039 | rs6819485 | -0.914 | -0.609 | 1.395 | -2.559 | -0.004 | -0.118 | -0.118 | 0 | 0.33 | 0.53 | 0.31 | 0.01 | 0.37 | 0.23 |
| 4 | 6042732 | rs7665309 | -2.252 | -0.98 | 1.886 | -2.133 | -0.028 | -0.079 | -0.079 | 0 | 0.12 | 0.44 | 0.07 | 0 | 0.05 | 0.08 |
| 4 | 6046867 | rs6446445 | -0.982 | -0.978 | 1.844 | -2.181 | -0.01 | -0.096 | -0.096 | 0 | 0.13 | 0.51 | 0.09 | 0 | 0.07 | 0.08 |
| 4 | 6047757 | rs6446446 | -2.318 | -1.027 | 1.772 | -2.254 | -0.005 | -0.059 | -0.059 | 0 | 0.23 | 0.75 | 0.16 | 0 | 0.13 | 0.17 |
| 4 | 6049030 | rs6815511 | -0.374 | -1.03 | 1.522 | -2.214 | 0.002 | -0.069 | -0.069 | 0 | 0.12 | 0.26 | 0.07 | 0 | 0.08 | 0.09 |
| 4 | 6058701 | rs16838138 | -1.713 | -0.876 | 0.277 | -2.186 | 0.014 | 0.009 | 0.009 | 0.775 | 0.23 | 0.85 | 0.16 | 0 | 0.14 | 0.18 |
| 4 | 6058732 | rs7438838 | 0.444 | -0.876 | -0.13 | -2.327 | -0.032 | 0.109 | 0.109 | 0 | 0.21 | 0.02 | 0.23 | 0.06 | 0.26 | 0.15 |
| 4 | 6059226 | rs6814350 | -1.431 | -0.533 | 0.526 | -1.889 | -0.025 | -0.111 | -0.111 | 0.773 | 0.21 | 0.61 | 0.13 | 0 | 0.11 | 0.16 |
| 4 | 6062639 | rs6821383 | -1.279 | -0.549 | 0.149 | -1.839 | -0.037 | -0.136 | -0.136 | 0.017 | 0.21 | 0.62 | 0.12 | 0 | 0.11 | 0.15 |
| 4 | 6065430 | rs16838164 | -1.193 | -0.209 | -0.668 | -1.299 | 0.032 | -0.013 | -0.013 | 0 | 0.05 | 0.2 | 0.04 | 0 | 0.04 | 0.03 |
| 4 | 6068267 | rs17766335 | -0.526 | -0.728 | -2.112 | -1.778 | 0.056 | 0.164 | 0.164 | 0.02 | 0.38 | 0.14 | 0.26 | 0.09 | 0.33 | 0.34 |
| 4 | 6068495 | rs11731133 | -0.429 | -0.972 | -2.49 | -1.531 | 0.04 | 0.148 | 0.148 | 0.093 | 0.35 | 0.06 | 0.25 | 0.14 | 0.3 | 0.26 |
| 4 | 6071895 | rs3889723 | 0.527 | -0.717 | -2.609 | -1.084 | -0.038 | 0.076 | 0.076 | 0.075 | 0.41 | 0.08 | 0.41 | 0.22 | 0.63 | 0.47 |
| 4 | 6072923 | rs6842183 | 0.333 | -0.886 | -2.674 | -0.903 | 0.02 | -0.01 | -0.01 | 0 | 0.45 | 0.08 | 0.43 | 0.38 | 0.67 | 0.49 |
| 4 | 6075470 | rs3901458 | 0.302 | -0.389 | -2.149 | -0.544 | -0.013 | -0.03 | -0.03 | 0 | 0.22 | 0.14 | 0.44 | 0.59 | 0.26 | 0.36 |
| 4 | 6080817 | rs6446452 | -0.517 | -1.179 | -1.28 | -1.179 | -0.067 | -0.127 | -0.127 | 0.032 | 0.3 | 0.57 | 0.12 | 0.02 | 0.1 | 0.12 |
| 4 | 6085431 | rs4689333 | 0.151 | -0.131 | -1.868 | -1.062 | -0.008 | 0.133 | 0.133 | 0 | 0.49 | 0.23 | 0.63 | 0.7 | 0.51 | 0.54 |
| 4 | 6086752 | rs10516179 | -0.694 | 0.474 | -1.724 | -1.136 | 0.016 | -0.018 | -0.018 | 0 | 0.05 | 0.08 | 0.06 | 0.01 | 0.12 | 0.04 |
| 4 | 6088956 | rs4689336 | 0.402 | 0.317 | -1.892 | -0.644 | -0.004 | 0.068 | 0.068 | 0 | 0.5 | 0.24 | 0.62 | 0.6 | 0.51 | 0.55 |
| 4 | 6096087 | rs17710575 | -0.098 | -0.115 | -2.128 | -0.777 | -0.017 | 0.039 | 0.039 | 0.065 | 0.33 | 0.05 | 0.22 | 0.25 | 0.33 | 0.4 |
| 4 | 6104378 | rs12507895 | 0.644 | 0.106 | -2.242 | -0.719 | -0.114 | 0.015 | 0.015 | 0 | 0.37 | 0.08 | 0.66 | 0.71 | 0.5 | 0.49 |
| 4 | 6106821 | rs7680061 | 0.052 | -0.162 | -1.303 | -1.231 | 0.007 | -0.12 | -0.12 | 3.842 | 0.67 | 0.44 | 0.79 | 0.98 | 0.67 | 0.78 |
| 4 | 6109614 | rs7673767 | -0.693 | -0.301 | -0.945 | -0.89 | 0.001 | 0.004 | 0.004 | 0 | 0.42 | 0.63 | 0.25 | 0.41 | 0.39 | 0.32 |
| 4 | 6113559 | rs11731332 | 0.893 | -0.074 | -0.838 | -0.972 | -0.011 | 0.019 | 0.019 | 0 | 0.4 | 0.26 | 0.68 | 0.69 | 0.52 | 0.56 |
| 4 | 6115394 | rs12186252 | -0.69 | 0.118 | -0.793 | -1.306 | -0.014 | 0.012 | 0.012 | 0 | 0.29 | 0.18 | 0.18 | 0.23 | 0.3 | 0.3 |
| 4 | 6116097 | rs10212632 | -1.048 | 0.809 | -0.182 | -2.028 | 0.027 | -0.15 | -0.15 | 0.056 | 0.27 | 0.42 | 0.29 | 0.01 | 0.29 | 0.19 |
| 4 | 6116708 | rs7694562 | 0.829 | 0.759 | -0.126 | -1.938 | 0.004 | -0.113 | -0.113 | 0.22 | 0.64 | 0.38 | 0.65 | 0.96 | 0.63 | 0.77 |
| 4 | 6122058 | rs11732318 | 0.987 | 0.655 | -0.627 | -1.321 | -0.008 | -0.021 | -0.021 | 0 | 0.19 | 0.13 | 0.42 | 0.55 | 0.24 | 0.46 |
| 4 | 6122135 | rs7664611 | 0.072 | 0.562 | -0.576 | -1.646 | -0.002 | 0.001 | 0.001 | 0 | 0.4 | 0.38 | 0.37 | 0.41 | 0.49 | 0.35 |
| 4 | 6126549 | rs16838298 | 0.488 | 0.393 | -0.397 | -1.463 | 0.003 | -0.055 | -0.055 | 0.663 | 0.18 | 0.2 | 0.11 | 0.01 | 0.18 | 0.07 |
| 4 | 6131812 | rs4580721 | -0.92 | 0.424 | -0.71 | -1.398 | -0.003 | -0.006 | -0.006 | 0.023 | 0.69 | 0.67 | 0.39 | 0.42 | 0.62 | 0.53 |
| 4 | 6135717 | rs4516797 | 0.642 | 0.359 | -0.755 | -1.401 | -0.005 | -0.012 | -0.012 | 0 | 0.29 | 0.29 | 0.52 | 0.55 | 0.37 | 0.47 |
| 4 | 6136653 | rs4689343 | -0.696 | 0.591 | -0.716 | -1.462 | 0.007 | -0.047 | -0.047 | 0 | 0.79 | 0.75 | 0.85 | 0.97 | 0.75 | 0.88 |
| 4 | 6138066 | rs7687079 | -0.335 | 0.265 | -1.252 | -1.405 | 0.007 | -0.001 | -0.001 | 0.097 | 0.58 | 0.58 | 0.66 | 0.57 | 0.56 | 0.63 |
| 4 | 6139210 | rs13115520 | 0.018 | 0.098 | -1.762 | -0.894 | 0.025 | 0.106 | 0.106 | 0.475 | 0.23 | 0.07 | 0.11 | 0.02 | 0.19 | 0.14 |
| 4 | 6140737 | rs4512066 | 0.087 | 0.454 | -1.32 | -0.442 | -0.004 | 0.007 | 0.007 | 0 | 0.45 | 0.36 | 0.63 | 0.58 | 0.56 | 0.57 |
| 4 | 6143608 | rs10017549 | 0.406 | 0.004 | -2.073 | -0.872 | -0.078 | 0.103 | 0.103 | 0.155 | 0.43 | 0.03 | 0.57 | 0.72 | 0.52 | 0.52 |
| 4 | 6147486 | rs7691044 | 0.553 | -0.148 | -0.585 | -1.028 | 0.031 | -0.022 | -0.022 | 0 | 0.92 | 0.46 | 0.93 | 1 | 0.96 | 0.93 |
| 4 | 6151065 | rs4398594 | -0.9 | -0.439 | -0.946 | -1.253 | 0.037 | 0.131 | 0.131 | 0.106 | 0.48 | 0.76 | 0.32 | 0.21 | 0.44 | 0.41 |
| 4 | 6154886 | rs7680135 | -1.269 | -1.022 | -0.493 | -1.333 | -0.026 | -0.051 | -0.051 | 0.068 | 0.21 | 0.32 | 0.23 | 0.02 | 0.16 | 0.17 |
| 4 | 6156083 | rs4522943 | -1.044 | -1.061 | -1.113 | -1.441 | 0.028 | 0.11 | 0.11 | 0 | 0.2 | 0.02 | 0.14 | 0.01 | 0.17 | 0.18 |
| 4 | 6161431 | rs4389634 | 0.921 | -0.74 | -0.81 | -1.19 | 0.031 | 0.088 | 0.088 | 0.396 | 0.58 | 0.18 | 0.65 | 0.77 | 0.62 | 0.67 |
| 4 | 6166456 | rs6824225 | -0.038 | -0.687 | -0.872 | -0.858 | -0.024 | -0.04 | -0.04 | 0.121 | 0.22 | 0.07 | 0.14 | 0.28 | 0.26 | 0.16 |
| 4 | 6168155 | rs6446469 | 0.77 | -0.696 | -0.274 | -0.979 | -0.001 | -0.006 | -0.006 | 0 | 0.66 | 0.63 | 0.79 | 0.78 | 0.65 | 0.75 |
| 4 | 6173521 | rs17773390 | -0.569 | -0.023 | -0.963 | -1.01 | -0.078 | -0.002 | -0.002 | 1.946 | 0.26 | 0.05 | 0.23 | 0.2 | 0.32 | 0.27 |
| 4 | 6175398 | rs4688970 | -0.756 | 0.005 | -0.648 | -0.852 | -0.039 | -0.014 | -0.014 | 0 | 0.28 | 0.12 | 0.25 | 0.26 | 0.35 | 0.28 |
| 4 | 6177056 | rs17773514 | -0.744 | 0.073 | -0.099 | -1.253 | -0.001 | 0 | 0 | 0.195 | 0.25 | 0.25 | 0.18 | 0.21 | 0.21 | 0.3 |
| 4 | 6177135 | rs4689355 | -1.765 | 0.137 | -0.065 | -1.429 | -0.008 | 0.01 | 0.01 | 0 | 0.38 | 0.39 | 0.33 | 0.21 | 0.43 | 0.42 |
| 4 | 6183184 | rs10937705 | 1.013 | -0.044 | -0.271 | -1.479 | 0.01 | -0.132 | -0.132 | 0.422 | 0.77 | 0.42 | 0.81 | 0.98 | 0.75 | 0.8 |
| 4 | 6185816 | rs885420 | 1.3 | -0.03 | -1.811 | -1.518 | -0.084 | 0.092 | 0.092 | 0 | 0.43 | 0.07 | 0.46 | 0.76 | 0.44 | 0.49 |
| 4 | 6191051 | rs1877762 | -0.619 | -0.015 | -1.058 | -0.702 | -0.042 | -0.03 | -0.03 | 0.281 | 0.24 | 0.04 | 0.15 | 0.2 | 0.21 | 0.27 |
| 4 | 6194483 | rs11943701 | 0.242 | 0.354 | -0.077 | -0.655 | 0.038 | -0.076 | -0.076 | 0 | 0.88 | 0.55 | 0.92 | 1 | 0.91 | 0.91 |
| 4 | 6195962 | rs10470721 | 0.717 | 0.596 | -1.322 | 0.008 | -0.021 | 0.006 | 0.006 | 0.043 | 0.53 | 0.19 | 0.59 | 0.61 | 0.51 | 0.53 |
| 4 | 6196421 | rs11728313 | -0.496 | 0.695 | -0.764 | 0.273 | -0.001 | 0.002 | 0.002 | 0 | 0.1 | 0.05 | 0.07 | 0.09 | 0.12 | 0.1 |
| 4 | 6199980 | rs4688973 | 0.977 | 0.413 | -0.345 | 0.211 | -0.013 | -0.037 | -0.037 | 0 | 0.87 | 0.78 | 0.76 | 0.98 | 0.91 | 0.87 |
| 4 | 6200070 | rs7377975 | 0.399 | 0.47 | -0.401 | 0.57 | 0.002 | 0.005 | 0.005 | 0.008 | 0.74 | 0.68 | 0.77 | 0.72 | 0.77 | 0.69 |
| 4 | 6200768 | rs16838428 | -1.297 | 0.465 | -0.67 | 1.207 | -0.003 | -0.02 | -0.02 | 0.233 | 0.13 | 0.1 | 0.22 | 0.31 | 0.15 | 0.25 |
| 4 | 6213622 | rs4274923 | 0.626 | -0.104 | -0.196 | 0.907 | -0.018 | 0.01 | 0.01 | 0.011 | 0.66 | 0.5 | 0.61 | 0.69 | 0.75 | 0.58 |
| 4 | 6218099 | rs6858345 | 0.363 | 0.129 | -0.121 | 1.033 | 0 | 0 | 0 | 1.708 | 0.45 | 0.58 | 0.49 | 0.45 | 0.5 | 0.41 |
| 4 | 6221614 | rs908017 | -0.366 | -0.001 | -0.25 | 1.113 | -0.004 | -0.05 | -0.05 | 1.991 | 0.1 | 0.01 | 0.18 | 0.24 | 0.09 | 0.22 |
| 4 | 6227628 | rs4602567 | -1.24 | -0.23 | 0.284 | 0.646 | -0.041 | 0.022 | 0.022 | 0 | 0.11 | 0.24 | 0.06 | 0.11 | 0.06 | 0.01 |
| 4 | 6233890 | rs4688978 | -0.34 | -0.032 | 0.022 | 0.115 | -0.033 | -0.04 | -0.04 | 0 | 0.67 | 0.78 | 0.61 | 0.54 | 0.56 | 0.71 |
| 4 | 6235553 | rs12510751 | 0.46 | -0.074 | -0.246 | -0.402 | 0 | -0.059 | -0.059 | 0 | 0.27 | 0.23 | 0.16 | 0.01 | 0.26 | 0.31 |
| 4 | 6236514 | rs7376535 | -0.672 | -0.105 | -0.407 | -0.14 | 0.008 | 0.014 | 0.014 | 0.352 | 0.68 | 0.64 | 0.61 | 0.53 | 0.61 | 0.71 |
| 4 | 6237973 | rs13103694 | 0.001 | -0.282 | -0.847 | 0.168 | 0.022 | -0.086 | -0.086 | 0 | 0.33 | 0.11 | 0.38 | 0.6 | 0.3 | 0.38 |
| 4 | 6245217 | rs6446477 | -0.967 | -0.243 | 0.373 | -0.331 | 0.016 | -0.023 | -0.023 | 0 | 0.08 | 0.36 | 0.06 | 0 | 0.03 | 0.1 |
| 4 | 6250182 | rs885030 | -1.518 | -0.275 | 0.353 | -0.571 | -0.016 | -0.103 | -0.103 | 1.012 | 0.16 | 0.59 | 0.17 | 0 | 0.14 | 0.14 |

| | | | | | | | | | | | | | | | | |
|---|---|---|---|---|---|---|---|---|---|---|---|---|---|---|---|---|
| 4 | 6256130 | rs7680068 | -0.355 | -0.639 | -0.486 | -0.618 | 0.039 | 0.005 | 0.005 | 0.082 | 0.51 | 0.66 | 0.54 | 0.54 | 0.43 | 0.45 |
| 4 | 6256489 | rs9993703 | -0.256 | -1.478 | -0.243 | -1.455 | -0.043 | -0.05 | -0.05 | 0 | 0.31 | 0.51 | 0.23 | 0.16 | 0.22 | 0.25 |
| 4 | 6256629 | rs4361456 | -1.257 | -0.769 | -1.484 | -1.908 | -0.039 | 0.104 | 0.104 | 3.582 | 0.7 | 0.89 | 0.81 | 0.99 | 0.68 | 0.76 |
| 4 | 6257188 | rs4689381 | 0.408 | -0.624 | -1.727 | -1.79 | -0.171 | 0.126 | 0.126 | 0 | 0.39 | 0.03 | 0.55 | 0.83 | 0.47 | 0.52 |
| 4 | 6259236 | rs11934878 | -0.29 | -1.032 | -0.274 | -1.057 | -0.043 | -0.038 | -0.038 | 0 | 0.27 | 0.46 | 0.24 | 0.17 | 0.2 | 0.19 |
| 4 | 6261776 | rs3898050 | 0.018 | -0.944 | -0.738 | -0.959 | 0.022 | 0.013 | 0.013 | 3.454 | 0.2 | 0.12 | 0.19 | 0.12 | 0.14 | 0.1 |
| 4 | 6267001 | rs4688982 | 0.174 | -0.834 | -0.548 | -1.89 | 0.026 | 0.014 | 0.014 | 2.152 | 0.52 | 0.43 | 0.65 | 0.91 | 0.61 | 0.67 |
| 4 | 6271071 | rs13107806 | 0.111 | -0.965 | -0.487 | -1.954 | 0.013 | -0.027 | -0.027 | 0 | 0.53 | 0.47 | 0.67 | 0.91 | 0.61 | 0.67 |
| 4 | 6275735 | rs17718958 | 0.888 | -1.288 | -1.054 | -2.319 | -0.025 | 0.082 | 0.082 | 5.478 | 0.15 | 0.04 | 0.16 | 0 | 0.18 | 0.06 |
| 4 | 6276581 | rs10002743 | -0.278 | -1.713 | -0.432 | -2.587 | -0.007 | -0.051 | -0.051 | 0 | 0.81 | 0.78 | 0.87 | 0.91 | 0.84 | 0.81 |
| 4 | 6280449 | rs4689391 | 0.849 | -1.711 | 0.152 | -2.495 | -0.003 | -0.093 | -0.093 | 0 | 0.53 | 0.51 | 0.67 | 0.9 | 0.61 | 0.65 |
| 4 | 6281961 | rs752854 | -0.222 | -1.788 | 0.023 | -2.582 | 0.005 | 0.038 | 0.038 | 4.668 | 0.69 | 0.81 | 0.77 | 1 | 0.68 | 0.84 |
| 4 | 6284633 | rs10001190 | 0.15 | -1.804 | 0.03 | -2.593 | 0.014 | 0.041 | 0.041 | 5.061 | 0.64 | 0.79 | 0.76 | 1 | 0.66 | 0.81 |
| 4 | 6289986 | rs4458523 | 0.806 | -1.844 | 0.252 | -2.615 | 0.008 | -0.028 | -0.028 | 0 | 0.58 | 0.58 | 0.69 | 0.9 | 0.63 | 0.67 |
| 4 | 6292915 | rs10010131 | 0.512 | -1.701 | 0.287 | -2.622 | 0.01 | 0.024 | 0.024 | 4.645 | 0.6 | 0.69 | 0.7 | 0.9 | 0.63 | 0.72 |
| 4 | 6293350 | rs10012946 | 0.522 | -1.697 | 0.287 | -2.622 | 0.011 | 0.024 | 0.024 | 4.679 | 0.6 | 0.69 | 0.7 | 0.9 | 0.63 | 0.72 |
| 4 | 6293967 | rs4416547 | 0.512 | -1.697 | 0.287 | -2.622 | 0.011 | 0.024 | 0.024 | 4.645 | 0.6 | 0.69 | 0.7 | 0.9 | 0.63 | 0.72 |
| 4 | 6298375 | rs4481292 | 0.944 | -1.689 | 0.276 | -2.662 | 0.003 | -0.065 | -0.065 | 0 | 0.55 | 0.53 | 0.68 | 0.9 | 0.62 | 0.64 |
| 4 | 6299940 | rs3821943 | 1.7 | -2.094 | 0.68 | -2.691 | 0.01 | 0.026 | 0.026 | 0.834 | 0.46 | 0.34 | 0.58 | 0.81 | 0.57 | 0.58 |
| 4 | 6302707 | rs1801206 | 1.06 | -2.004 | 0.664 | -2.728 | -0.002 | -0.088 | -0.088 | 1.557 | 0.53 | 0.45 | 0.67 | 0.9 | 0.62 | 0.64 |
| 4 | 6303247 | rs2230719 | 1.444 | -2.068 | 0.565 | -2.738 | 0.006 | -0.009 | -0.009 | 0 | 0.12 | 0.07 | 0.04 | 0 | 0.05 | 0.06 |
| 4 | 6303354 | rs734312 | 1.918 | -2.301 | 0.381 | -2.821 | -0.054 | 0.214 | 0.214 | 2.77 | 0.43 | 0.03 | 0.55 | 0.81 | 0.56 | 0.56 |
| 4 | 6303955 | rs1046314 | 0.908 | -2.076 | 0.675 | -2.786 | -0.004 | -0.098 | -0.098 | 0 | 0.54 | 0.49 | 0.68 | 0.9 | 0.62 | 0.64 |
| 4 | 6304242 | rs1046317 | -0.342 | -1.98 | 0.383 | -2.637 | -0.004 | -0.001 | -0.001 | 0 | 0.73 | 0.8 | 0.75 | 0.91 | 0.69 | 0.76 |
| 4 | 6304448 | rs1046322 | -0.044 | -1.635 | 0.537 | -2.268 | 0.036 | 0.038 | 0.038 | 0 | 0.18 | 0.07 | 0.08 | 0.06 | 0.1 | 0.12 |
| 4 | 6312175 | rs7658531 | 0.288 | -1.345 | 0.774 | -2.15 | 0.004 | -0.016 | -0.016 | 1.856 | 0.2 | 0.19 | 0.07 | 0.09 | 0.1 | 0.16 |
| 4 | 6312976 | rs9986109 | 1.293 | -0.626 | 0.419 | -1.109 | -0.014 | -0.022 | -0.022 | 2.508 | 0.35 | 0.27 | 0.38 | 0.5 | 0.39 | 0.4 |
| 4 | 6315954 | rs10804976 | 0.303 | -0.583 | 0.493 | -1.655 | 0.004 | -0.063 | -0.063 | 0 | 0.5 | 0.5 | 0.65 | 0.91 | 0.57 | 0.69 |
| 4 | 6324246 | rs7697730 | -0.243 | -0.057 | 0.138 | -1.473 | 0.013 | -0.102 | -0.102 | 0.114 | 0.66 | 0.5 | 0.67 | 0.94 | 0.59 | 0.76 |
| 4 | 6324647 | rs6446490 | 0.771 | -0.101 | 0.108 | -1.518 | -0.004 | 0.004 | 0.004 | 0 | 0.45 | 0.35 | 0.6 | 0.85 | 0.49 | 0.59 |
| 4 | 6324785 | rs3796398 | 0.558 | -0.051 | -0.168 | -1.483 | 0 | -0.059 | -0.059 | 1.347 | 0.47 | 0.47 | 0.61 | 0.85 | 0.49 | 0.6 |
| 4 | 6325086 | rs3796403 | 0.334 | 0.606 | 0.452 | -0.728 | 0.016 | -0.097 | -0.097 | 0.242 | 0.65 | 0.43 | 0.66 | 0.94 | 0.58 | 0.74 |
| 4 | 6333126 | rs4689404 | -1.469 | 0.695 | 0.677 | 0.641 | -0.003 | -0.005 | -0.005 | 0 | 0.27 | 0.74 | 0.32 | 0.3 | 0.29 | 0.26 |
| 4 | 6341740 | rs4689407 | 1.074 | 0.742 | 0.643 | 1.143 | -0.002 | -0.03 | -0.03 | 0 | 0.74 | 0.74 | 0.65 | 0.46 | 0.73 | 0.65 |
| 4 | 6342213 | rs4689408 | -0.509 | 0.773 | 0.623 | 1.327 | 0 | 0 | 0 | 0.049 | 0.18 | 0.21 | 0.17 | 0.18 | 0.19 | 0.2 |
| 4 | 6342359 | rs7676409 | 1.527 | 0.824 | 0.273 | 1.228 | 0.008 | -0.029 | -0.029 | 0.038 | 0.61 | 0.24 | 0.54 | 0.43 | 0.65 | 0.56 |
| 4 | 6344204 | rs4689410 | -1.46 | 1.666 | 0.391 | 0.857 | 0.007 | 0.002 | 0.002 | 0.068 | 0.28 | 0.66 | 0.31 | 0.32 | 0.32 | 0.28 |
| 4 | 6344237 | rs4689411 | -1.187 | 1.45 | 0.51 | 0.872 | 0.015 | 0.029 | 0.029 | 0.002 | 0.26 | 0.59 | 0.29 | 0.31 | 0.27 | 0.24 |
| 4 | 6348191 | rs6856061 | 1.016 | 1.082 | 0.682 | 0.188 | 0.032 | -0.09 | -0.09 | 0 | 0.84 | 0.55 | 0.88 | 0.96 | 0.86 | 0.86 |
| 4 | 6351958 | rs4386675 | 0.862 | 0.649 | -0.2 | 0.535 | 0.023 | -0.031 | -0.031 | 0 | 0.63 | 0.16 | 0.62 | 0.68 | 0.7 | 0.75 |
| 4 | 6356513 | rs11730177 | 1.079 | 0.776 | -0.133 | 0.481 | 0.002 | -0.015 | -0.015 | 3.929 | 0.61 | 0.16 | 0.59 | 0.7 | 0.67 | 0.71 |
| 4 | 6361632 | rs4689415 | -0.986 | 0.578 | 0.828 | 0.381 | 0.001 | 0 | 0 | 0 | 0.29 | 0.32 | 0.33 | 0.27 | 0.31 | 0.27 |
| 4 | 6362566 | rs10446500 | -0.426 | 0.414 | 0.604 | 0.352 | -0.014 | -0.021 | -0.021 | 0 | 0.21 | 0.12 | 0.28 | 0.25 | 0.27 | 0.22 |
| 4 | 6363608 | rs13122929 | 0.962 | 0.397 | 0.694 | 0.137 | -0.051 | -0.106 | -0.106 | 1.127 | 0.83 | 0.35 | 0.91 | 1 | 0.95 | 0.95 |
| 4 | 6364311 | rs13124487 | 1.265 | 0.692 | -0.121 | 0.321 | -0.018 | 0 | 0 | 1.732 | 0.6 | 0.13 | 0.58 | 0.67 | 0.63 | 0.7 |
| 4 | 6366210 | rs4689421 | 1.454 | 0.155 | 0.383 | -0.001 | -0.028 | -0.072 | -0.072 | 1.372 | 0.86 | 0.73 | 0.92 | 1 | 0.95 | 0.94 |
| 4 | 6367467 | rs6831971 | 0.374 | 0.088 | -0.39 | 0.238 | 0.005 | -0.001 | -0.001 | 0 | 0.72 | 0.81 | 0.72 | 0.67 | 0.77 | 0.71 |
| 4 | 6369731 | rs4596320 | 0.921 | -0.807 | 0.538 | -0.38 | -0.033 | -0.105 | -0.105 | 0 | 0.84 | 0.38 | 0.87 | 1 | 0.9 | 0.94 |
| 4 | 6375269 | rs16838658 | -1.248 | -1.643 | 1.496 | 0.269 | -0.084 | 0.171 | 0.171 | 0 | 0.21 | 0.45 | 0.32 | 0.34 | 0.13 | 0.18 |
| 4 | 6375986 | rs4585379 | -1.051 | -1.633 | 1.255 | 0.284 | -0.055 | 0.116 | 0.116 | 0 | 0.17 | 0.31 | 0.29 | 0.34 | 0.08 | 0.15 |
| 4 | 6376273 | rs6446495 | -0.51 | -2.108 | 0.623 | -0.595 | -0.029 | -0.121 | -0.121 | 2.757 | 0.28 | 0.33 | 0.14 | 0.01 | 0.19 | 0.24 |
| 4 | 6377309 | rs4689424 | 0.617 | -2.176 | 0.06 | -0.634 | -0.023 | -0.123 | -0.123 | 0 | 0.66 | 0.45 | 0.78 | 0.99 | 0.74 | 0.75 |
| 4 | 6381323 | rs4327561 | 0.179 | -2.204 | -0.672 | -0.171 | -0.001 | 0.002 | 0.002 | 0 | 0.78 | 0.71 | 0.63 | 0.63 | 0.84 | 0.72 |
| 4 | 6383792 | rs4688999 | 1.042 | -1.286 | -1.346 | -0.498 | -0.12 | 0.004 | 0.004 | 0.127 | 0.39 | 0.07 | 0.57 | 0.68 | 0.51 | 0.46 |
| 4 | 6383965 | rs11936718 | 0.697 | -1.568 | -0.946 | -0.505 | -0.033 | 0.034 | 0.034 | 0 | 0.18 | 0.31 | 0.17 | 0.25 | 0.13 | 0.18 |
| 4 | 6384823 | rs6851340 | 0.519 | -1.2 | -0.828 | -0.451 | -0.033 | 0.032 | 0.032 | 0 | 0.25 | 0.47 | 0.17 | 0.31 | 0.18 | 0.27 |
| 4 | 6386039 | rs16838698 | 0.352 | -1.197 | -1.974 | -0.484 | 0.079 | 0.077 | 0.077 | 0.566 | 0.54 | 0.23 | 0.72 | 0.7 | 0.69 | 0.56 |
| 4 | 6389642 | rs755403 | -0.727 | -1.26 | -2.539 | -0.08 | 0.02 | 0.003 | 0.003 | 0.676 | 0.51 | 0.85 | 0.45 | 0.44 | 0.4 | 0.52 |
| 4 | 6391045 | rs872858 | 1.052 | 0.571 | -2.4 | 0.837 | -0.049 | -0.051 | -0.051 | 0 | 0.08 | 0.01 | 0.21 | 0.24 | 0.18 | 0.19 |
| 4 | 6391735 | rs4270639 | -0.406 | -0.49 | -2.12 | 0.469 | 0.001 | 0.017 | 0.017 | 0 | 0.31 | 0.11 | 0.29 | 0.33 | 0.27 | 0.41 |
| 4 | 6392342 | rs2240268 | -0.916 | -0.321 | -1.625 | 0.335 | -0.002 | -0.003 | -0.003 | 0 | 0.58 | 0.45 | 0.41 | 0.49 | 0.45 | 0.63 |
| 4 | 6392879 | rs2240264 | -0.553 | 0.281 | -1.01 | 0.426 | 0.003 | 0.002 | 0.002 | 0 | 0.34 | 0.36 | 0.24 | 0.25 | 0.29 | 0.26 |
| 4 | 6401102 | rs7671165 | -0.242 | 0.719 | -1.088 | 0.514 | -0.013 | -0.012 | -0.012 | 0 | 0.27 | 0.14 | 0.17 | 0.7 | 0.28 | 0.3 |
| 4 | 6402732 | rs7438559 | -1.065 | 0.632 | -0.593 | -0.06 | -0.002 | 0.039 | 0.039 | 0.847 | 0.43 | 0.39 | 0.51 | 0.88 | 0.46 | 0.64 |
| 4 | 6404531 | rs7655407 | 0.342 | 0.734 | -0.718 | 0.551 | 0.003 | -0.069 | -0.069 | 0.531 | 0.26 | 0.08 | 0.15 | 0.52 | 0.23 | 0.3 |
| 4 | 6406411 | rs6857081 | -1.339 | 0.49 | -0.014 | -0.421 | -0.003 | 0.114 | 0.114 | 0.199 | 0.49 | 0.7 | 0.56 | 0.88 | 0.52 | 0.72 |
| 4 | 6407801 | rs7377185 | -0.666 | 0.229 | 0.488 | -0.365 | 0.001 | 0.098 | 0.098 | 1.55 | 0.17 | 0.49 | 0.4 | 0.28 | 0.18 | 0.4 |
| 4 | 6410449 | rs6853134 | 0.301 | 0.385 | 0.347 | -0.306 | 0.007 | 0.012 | 0.012 | 1.797 | 0.12 | 0.19 | 0.33 | 0.17 | 0.17 | 0.35 |
| 4 | 6410746 | rs7434469 | -0.636 | 0.261 | 0.395 | -0.545 | 0 | -0.018 | -0.018 | 0 | 0.44 | 0.42 | 0.52 | 0.87 | 0.43 | 0.67 |

| | | | | | | | | | | | | | | | | |
|---|---|---|---|---|---|---|---|---|---|---|---|---|---|---|---|---|
| 4 | 6411979 | rs10007735 | -0.476 | 0.371 | 0.38 | -0.168 | 0 | -0.014 | -0.014 | 0 | 0.42 | 0.53 | 0.29 | 0.71 | 0.39 | 0.4 |
| 4 | 6414398 | rs13434456 | -0.059 | 0.334 | 0.345 | -0.167 | -0.001 | -0.004 | -0.004 | 0 | 0.23 | 0.23 | 0.45 | 0.18 | 0.26 | 0.3 |
| 4 | 6416437 | rs11724328 | -0.551 | -0.269 | 0.276 | -0.52 | 0 | -0.001 | -0.001 | 0 | 0.31 | 0.27 | 0.47 | 0.18 | 0.28 | 0.33 |
| 4 | 6417028 | rs7658616 | 0.858 | -0.397 | 0.483 | -0.6 | -0.006 | 0.067 | 0.067 | 0 | 0.47 | 0.35 | 0.27 | 0.21 | 0.47 | 0.36 |
| 4 | 6417785 | rs16838780 | -0.315 | -0.936 | 0.662 | -0.831 | 0.007 | -0.004 | -0.004 | 2.474 | 0.31 | 0.25 | 0.27 | 0.61 | 0.27 | 0.34 |
| 4 | 6422476 | rs878281 | 0.289 | -0.805 | 1.208 | -0.478 | -0.03 | -0.027 | -0.027 | 2.477 | 0.65 | 0.54 | 0.51 | 0.76 | 0.72 | 0.66 |
| 4 | 6423060 | rs11734695 | 0.382 | -0.82 | 1.235 | -0.167 | -0.01 | -0.004 | -0.004 | 0 | 0.64 | 0.44 | 0.45 | 0.73 | 0.72 | 0.65 |
| 4 | 6425308 | rs17778826 | 0.565 | -0.698 | 1.023 | 0.101 | -0.015 | -0.035 | -0.035 | 0.182 | 0.23 | 0.07 | 0.49 | 0.22 | 0.26 | 0.29 |
| 4 | 6427070 | rs4345249 | 0.964 | -1.019 | 0.875 | 0.419 | 0.052 | 0.161 | 0.161 | 0.821 | 0.53 | 0.19 | 0.31 | 0.24 | 0.58 | 0.42 |
| 4 | 6427757 | rs4469149 | -0.362 | -0.997 | 1.295 | 0.639 | 0.001 | 0.156 | 0.156 | 0 | 0.9 | 0.81 | 0.88 | 0.56 | 0.88 | 0.76 |
| 4 | 6428296 | rs4689003 | 1.328 | -1.011 | 1.329 | 0.597 | -0.025 | 0.049 | 0.049 | 0.234 | 0.76 | 0.32 | 0.81 | 0.46 | 0.84 | 0.69 |
| 4 | 6429992 | rs4689443 | -0.527 | -0.717 | 1.409 | 0.825 | -0.001 | -0.001 | -0.001 | 0 | 0.7 | 0.73 | 0.48 | 0.74 | 0.72 | 0.68 |
| 4 | 6433905 | rs11724762 | 0.169 | -0.671 | 1.604 | 1.102 | 0.009 | -0.004 | -0.004 | 0.037 | 0.29 | 0.16 | 0.44 | 0.26 | 0.28 | 0.3 |
| 4 | 6435495 | rs4689005 | -0.086 | -0.565 | 1.482 | 1.263 | 0.003 | 0.14 | 0.14 | 0 | 0.05 | 0.06 | 0.14 | 0.43 | 0.1 | 0.22 |
| 4 | 6435926 | rs12510934 | -0.003 | -0.705 | 1.955 | 1.097 | -0.02 | 0 | 0 | 0.001 | 0.65 | 0.44 | 0.46 | 0.7 | 0.71 | 0.69 |
| 4 | 6441415 | rs17722973 | 0.384 | -0.647 | 1.329 | 0.729 | 0.019 | 0.037 | 0.037 | 2.423 | 0.59 | 0.25 | 0.36 | 0.33 | 0.61 | 0.48 |
| 4 | 6442931 | rs6813956 | 0.233 | -0.671 | 1.252 | 0.786 | -0.002 | -0.002 | -0.002 | 0 | 0.62 | 0.37 | 0.38 | 0.34 | 0.64 | 0.48 |
| 4 | 6447500 | rs12640531 | 0.596 | 0.2 | 0.196 | 0.282 | -0.046 | 0.067 | 0.067 | 0.734 | 0.35 | 0.1 | 0.2 | 0.23 | 0.32 | 0.29 |
| 4 | 6447607 | rs9291117 | 0.772 | 0.485 | 0.272 | 0.283 | 0.006 | -0.013 | -0.013 | 0.082 | 0.1 | 0.17 | 0.06 | 0.09 | 0.13 | 0.09 |
| 4 | 6449514 | rs17779675 | 0.919 | 0.308 | 0.319 | 0.095 | -0.007 | 0.06 | 0.06 | 0.092 | 0.37 | 0.16 | 0.2 | 0.22 | 0.34 | 0.31 |
| 4 | 6450950 | rs16838853 | -0.149 | 0.228 | 0.385 | 0.529 | 0 | 0.026 | 0.026 | 0 | 0.15 | 0.19 | 0.23 | 0.43 | 0.19 | 0.29 |
| 4 | 6451080 | rs10014123 | 0.873 | 0.254 | 0.352 | 0.412 | 0.001 | -0.012 | -0.012 | 0 | 0.1 | 0.15 | 0.07 | 0.05 | 0.12 | 0.08 |
| 4 | 6454012 | rs12500979 | 0.378 | 0.296 | 0.217 | 0.444 | -0.033 | 0.092 | 0.092 | 1.033 | 0.45 | 0.17 | 0.28 | 0.29 | 0.46 | 0.38 |
| 4 | 6454335 | rs10029948 | 0.997 | 0.346 | 0.698 | 0.547 | -0.014 | -0.029 | -0.029 | 0.434 | 0.31 | 0.37 | 0.36 | 0.2 | 0.27 | 0.32 |
| 4 | 6456871 | rs6813305 | 0.802 | 0.423 | 0.212 | 0.731 | -0.012 | 0.049 | 0.049 | 0 | 0.05 | 0.01 | 0.11 | 0.37 | 0.07 | 0.21 |
| 4 | 6460539 | rs4404624 | -0.639 | 0.81 | -0.038 | 0.528 | -0.034 | -0.062 | -0.062 | 2.321 | 0.22 | 0.12 | 0.23 | 0.43 | 0.31 | 0.35 |
| 4 | 6465791 | rs7664714 | -0.804 | 0.646 | 0.157 | 0.45 | 0.009 | 0.027 | 0.027 | 1.925 | 0.59 | 0.35 | 0.46 | 0.72 | 0.64 | 0.62 |
| 4 | 6465928 | rs7664961 | 0.428 | 0.641 | 0.008 | 0.519 | -0.031 | 0.049 | 0.049 | 1.056 | 0.44 | 0.17 | 0.27 | 0.35 | 0.45 | 0.36 |
| 4 | 6471246 | rs10937735 | -0.444 | 0.625 | 0.112 | 0.367 | 0 | 0.011 | 0.011 | 0.123 | 0.6 | 0.32 | 0.45 | 0.63 | 0.58 | 0.59 |
| 4 | 6479693 | rs4234745 | -0.862 | 0.451 | 0.014 | 0.176 | 0.001 | -0.009 | -0.009 | 0 | 0.23 | 0.23 | 0.23 | 0.62 | 0.2 | 0.33 |
| 4 | 6480469 | rs11938118 | 0.215 | 0.828 | 0.722 | -0.458 | 0.011 | -0.144 | -0.144 | 0 | 0.81 | 0.54 | 0.86 | 1 | 0.84 | 0.91 |
| 4 | 6481781 | rs4331842 | -0.943 | 0.866 | 0.727 | -0.448 | 0.019 | 0.027 | 0.027 | 0 | 0.6 | 0.63 | 0.43 | 0.32 | 0.65 | 0.49 |
| 4 | 6487726 | rs13117055 | -0.677 | 0.992 | 0.678 | -0.422 | 0.005 | 0.007 | 0.007 | 0 | 0.57 | 0.56 | 0.42 | 0.38 | 0.61 | 0.45 |
| 4 | 6490789 | rs4689465 | -0.255 | 0.834 | 0.595 | -0.391 | 0 | -0.003 | -0.003 | 0.568 | 0.49 | 0.43 | 0.37 | 0.33 | 0.51 | 0.38 |
| 4 | 6499037 | rs7654321 | 1.457 | 0.811 | -0.366 | -0.666 | -0.031 | 0.063 | 0.063 | 0 | 0.3 | 0.09 | 0.2 | 0.22 | 0.32 | 0.28 |
| 4 | 6507841 | rs4689469 | 0.667 | 0.787 | -0.402 | -0.645 | 0.029 | 0.077 | 0.077 | 0.079 | 0.38 | 0.23 | 0.55 | 0.72 | 0.33 | 0.51 |
| 4 | 6509412 | rs10937739 | 0.178 | 0.778 | -0.336 | -0.659 | -0.018 | 0.141 | 0.141 | 0 | 0.47 | 0.27 | 0.63 | 0.78 | 0.49 | 0.61 |
| 4 | 6515067 | rs4623093 | -0.427 | 0.823 | -0.119 | -0.51 | 0.005 | -0.074 | -0.074 | 1.922 | 0.87 | 0.73 | 0.89 | 1 | 0.9 | 0.93 |
| 4 | 6517066 | rs4565150 | 0.264 | 0.954 | -0.253 | -0.52 | 0 | -0.005 | -0.005 | 4.344 | 0.47 | 0.48 | 0.65 | 0.71 | 0.45 | 0.56 |
| 4 | 6518046 | rs3934547 | -0.551 | 0.732 | -0.71 | 0.445 | 0.033 | 0.114 | 0.114 | 2.333 | 0.12 | 0.59 | 0.22 | 0.17 | 0.1 | 0.07 |
| 4 | 6518453 | rs6820874 | 0.419 | 0.539 | -1.599 | -0.013 | 0 | 0.021 | 0.021 | 0 | 0.25 | 0.04 | 0.13 | 0.23 | 0.22 | 0.23 |
| 4 | 6523276 | rs4075006 | 0.474 | 0.688 | -1.727 | -0.049 | -0.075 | -0.013 | -0.013 | 0 | 0.3 | 0.03 | 0.33 | 0.22 | 0.38 | 0.22 |
| 4 | 6523395 | rs4075007 | -0.323 | 0.413 | -0.812 | -0.007 | -0.001 | 0 | 0 | 0 | 0.47 | 0.56 | 0.59 | 0.35 | 0.53 | 0.41 |
| 4 | 6527362 | rs7688267 | 0.19 | 0.611 | -0.548 | 0.122 | 0.018 | 0.037 | 0.037 | 0 | 0.53 | 0.34 | 0.53 | 0.29 | 0.61 | 0.4 |
| 4 | 6532065 | rs3934625 | -0.599 | 0.019 | -0.052 | -0.099 | -0.019 | -0.02 | -0.02 | 0 | 0.19 | 0.34 | 0.22 | 0.12 | 0.12 | 0.15 |
| 4 | 6533061 | rs4689472 | -0.162 | 0.079 | -0.63 | -0.116 | -0.041 | 0.025 | 0.025 | 0 | 0.22 | 0.06 | 0.27 | 0.13 | 0.25 | 0.24 |
| 4 | 6540153 | rs6850064 | -0.686 | -0.201 | -0.449 | -0.334 | -0.003 | 0.005 | 0.005 | 0.181 | 0.34 | 0.27 | 0.28 | 0.18 | 0.35 | 0.37 |
| 4 | 6541749 | rs4689477 | -0.163 | -0.648 | -0.322 | -0.228 | -0.077 | 0.171 | 0.171 | 1.439 | 0.78 | 0.48 | 0.72 | 0.49 | 0.89 | 0.81 |
| 4 | 6542401 | rs4610412 | 0.423 | -0.165 | -0.74 | -0.087 | 0.003 | -0.014 | -0.014 | 0.43 | 0.21 | 0.2 | 0.22 | 0.07 | 0.26 | 0.19 |
| 4 | 6544606 | rs10030485 | -1.058 | -0.319 | -1.045 | -0.117 | 0.007 | 0.075 | 0.075 | 0.032 | 0.5 | 0.25 | 0.51 | 0.7 | 0.46 | 0.59 |
| 4 | 6546433 | rs9654056 | -0.414 | -0.756 | -0.931 | 0.477 | -0.041 | -0.038 | -0.038 | 0 | 0.18 | 0.31 | 0.07 | 0.12 | 0.1 | 0.21 |
| 4 | 6547317 | rs11737849 | -0.24 | -0.081 | -1.667 | 0.978 | 0.027 | -0.088 | -0.088 | 1.685 | 0.15 | 0.03 | 0.23 | 0.54 | 0.11 | 0.2 |
| 4 | 6550849 | rs4358467 | -0.3 | 0.082 | -1.984 | 0.343 | 0.016 | 0.016 | 0.016 | 2.116 | 0.31 | 0.1 | 0.16 | 0.27 | 0.24 | 0.25 |
| 4 | 6552170 | rs10937742 | -0.571 | 0.883 | -2.046 | 0.226 | -0.051 | 0.036 | 0.036 | 0.275 | 0.34 | 0.11 | 0.46 | 0.75 | 0.42 | 0.52 |
| 4 | 6560931 | rs11731327 | -0.726 | 0.667 | -1.104 | 0.54 | -0.004 | 0.009 | 0.009 | 0 | 0.35 | 0.36 | 0.45 | 0.74 | 0.39 | 0.5 |
| 4 | 6570032 | rs4689021 | 0.347 | 0.714 | -0.528 | 0.163 | 0.009 | -0.028 | -0.028 | 0.68 | 0.82 | 0.53 | 0.84 | 0.87 | 0.85 | 0.83 |
| 4 | 6575960 | rs11732212 | -0.226 | 0.854 | -1.377 | 0.278 | -0.094 | -0.17 | -0.17 | 0.048 | 0.3 | 0.03 | 0.34 | 0.52 | 0.33 | 0.34 |
| 4 | 6593117 | rs6847759 | -0.487 | 0.652 | -0.593 | 0.228 | 0.004 | -0.012 | -0.012 | 0.109 | 0.4 | 0.38 | 0.37 | 0.47 | 0.37 | 0.43 |
| 4 | 6593437 | rs12507418 | 0.345 | 0.749 | -0.131 | -0.244 | -0.003 | -0.008 | -0.008 | 0.01 | 0.46 | 0.49 | 0.31 | 0.24 | 0.39 | 0.34 |
| 4 | 6594947 | rs2301796 | -0.361 | 0.942 | 0.19 | -0.093 | -0.021 | 0.001 | 0.001 | 0 | 0.42 | 0.18 | 0.48 | 0.61 | 0.45 | 0.54 |
| 4 | 6596360 | rs2301795 | 0.335 | 0.929 | 0.398 | -0.09 | 0 | 0 | 0 | 0.066 | 0.54 | 0.44 | 0.48 | 0.39 | 0.54 | 0.45 |
| 4 | 6600012 | rs2301790 | -0.388 | 0.873 | 0.594 | -0.117 | 0.002 | -0.002 | -0.002 | 0 | 0.42 | 0.44 | 0.51 | 0.61 | 0.45 | 0.51 |
| 4 | 6602941 | rs6446513 | 0.57 | 0.676 | 0.796 | -0.136 | -0.019 | -0.061 | -0.061 | 0.795 | 0.89 | 0.79 | 0.95 | 1 | 0.95 | 0.91 |
| 4 | 6607830 | rs7659958 | -1.287 | 1.12 | 0.579 | 0.327 | 0.031 | 0 | 0 | 0 | 0.14 | 0.24 | 0.28 | 0.14 | 0.18 | 0.18 |
| 4 | 6608856 | rs6446514 | -0.654 | 0.304 | 0.461 | 0.509 | -0.012 | -0.007 | -0.007 | 0.101 | 0.57 | 0.51 | 0.69 | 0.64 | 0.6 | 0.69 |
| 4 | 6610668 | rs11721430 | -0.601 | 0.548 | 0.596 | 1.003 | 0.007 | 0.013 | 0.013 | 0 | 0.06 | 0.07 | 0.12 | 0.13 | 0.1 | 0.08 |
| 4 | 6616948 | rs10049503 | 1.611 | 0.432 | 0.877 | 1.427 | 0.016 | 0.143 | 0.143 | 0.131 | 0.87 | 0.75 | 0.65 | 0.52 | 0.84 | 0.81 |
| 4 | 6619052 | rs4689488 | 1.944 | -0.237 | 1.577 | 0.682 | -0.012 | -0.022 | -0.022 | 0 | 0.91 | 0.43 | 0.93 | 0.99 | 0.98 | 0.86 |
| 4 | 6619204 | rs4689490 | 2.063 | -0.792 | 1.232 | 0.423 | -0.006 | -0.015 | -0.015 | 0 | 0.7 | 0.28 | 0.84 | 0.9 | 0.86 | 0.73 |

| | | | | | | | | | | | | | | | | |
|---|---|---|---|---|---|---|---|---|---|---|---|---|---|---|---|---|
| 4 | 6619378 | rs7698546 | -0.493 | -0.767 | 0.883 | 0.424 | -0.001 | -0.003 | -0.003 | 0 | 0.13 | 0.11 | 0.07 | 0.08 | 0.1 | 0.09 |
| 4 | 6620710 | rs3733604 | 0.462 | -0.758 | 1.016 | 0.428 | -0.019 | -0.031 | -0.031 | 0 | 0.85 | 0.73 | 0.91 | 0.92 | 0.9 | 0.9 |
| 4 | 6623332 | rs3088248 | 1.478 | -0.683 | 0.322 | 0.481 | 0.047 | 0.052 | 0.052 | 1.193 | 0.7 | 0.18 | 0.84 | 0.9 | 0.86 | 0.73 |
| 4 | 6627084 | rs9790604 | -1.609 | -0.182 | 0.499 | 1.281 | -0.003 | 0.076 | 0.076 | 0 | 0.15 | 0.19 | 0.24 | 0.45 | 0.17 | 0.25 |
| 4 | 6628640 | rs4689022 | 2.044 | -0.494 | 0.893 | 1.156 | -0.052 | 0.052 | 0.052 | 0.803 | 0.75 | 0.37 | 0.77 | 0.6 | 0.88 | 0.71 |
| 4 | 6633831 | rs12645336 | -0.663 | -0.083 | 0.496 | 1.255 | -0.019 | -0.027 | -0.027 | 0 | 0.08 | 0.01 | 0.11 | 0.34 | 0.11 | 0.07 |
| 4 | 6640962 | rs6446517 | -1.949 | -0.126 | 0.641 | 1.219 | 0.002 | -0.007 | -0.007 | 0.876 | 0.19 | 0.14 | 0.24 | 0.42 | 0.18 | 0.29 |
| 4 | 6644327 | rs7660424 | -1.216 | -0.055 | 0.588 | 1.119 | 0.062 | 0.005 | 0.005 | 0.436 | 0.05 | 0.23 | 0.05 | 0 | 0.04 | 0.02 |
| 4 | 6645597 | rs11736897 | 1.831 | 0.173 | -0.977 | 0.55 | -0.005 | -0.011 | -0.011 | 0.671 | 0.52 | 0.11 | 0.6 | 0.55 | 0.63 | 0.46 |
| 4 | 6646220 | rs4416546 | 0.918 | 0.048 | -0.559 | -0.103 | 0.019 | 0.081 | 0.081 | 0.128 | 0.7 | 0.2 | 0.83 | 0.98 | 0.8 | 0.74 |
| 4 | 6647324 | rs10804983 | 0.922 | -0.031 | -0.553 | -0.08 | 0.036 | 0.087 | 0.087 | 0.228 | 0.7 | 0.19 | 0.84 | 0.98 | 0.83 | 0.74 |
| 4 | 6649619 | rs4689024 | 1.473 | 0.109 | -0.816 | 0.161 | -0.006 | 0.165 | 0.165 | 1.207 | 0.57 | 0.09 | 0.61 | 0.78 | 0.67 | 0.65 |
| 4 | 6650566 | rs6810446 | -1.321 | 0.158 | 0.329 | 0.538 | -0.007 | 0.002 | 0.002 | 0 | 0.14 | 0.25 | 0.18 | 0.17 | 0.13 | 0.16 |
| 4 | 6663319 | rs4689498 | 0.676 | 0.31 | 0.08 | 0.472 | -0.001 | -0.001 | -0.001 | 0 | 0.46 | 0.45 | 0.46 | 0.39 | 0.5 | 0.56 |
| 4 | 6670315 | rs2285864 | -0.585 | 0.646 | -0.092 | 0.717 | 0 | 0 | 0 | 0 | 0.14 | 0.17 | 0.19 | 0.19 | 0.16 | 0.15 |
| 4 | 6671959 | rs724858 | 1.261 | 0.456 | -0.243 | 0.58 | -0.055 | 0.134 | 0.134 | 0 | 0.37 | 0.15 | 0.42 | 0.16 | 0.47 | 0.39 |
| 4 | 6673462 | rs4689506 | -0.592 | 0.117 | -0.053 | 0.512 | 0 | -0.006 | -0.006 | 0.645 | 0.69 | 0.73 | 0.69 | 0.59 | 0.68 | 0.75 |
| 4 | 6680577 | rs4689510 | 0.554 | 0.38 | -0.303 | 0.415 | 0.001 | 0.026 | 0.026 | 0 | 0.45 | 0.33 | 0.52 | 0.39 | 0.53 | 0.58 |
| 4 | 6687113 | rs4689516 | 0.745 | 0.331 | -0.32 | -0.06 | 0.002 | 0.001 | 0.001 | 0 | 0.62 | 0.37 | 0.55 | 0.2 | 0.69 | 0.44 |
| 4 | 6687545 | rs11935439 | -1.205 | 0.603 | -0.538 | -0.003 | 0.011 | -0.008 | -0.008 | 0.464 | 0.4 | 0.52 | 0.53 | 0.67 | 0.45 | 0.5 |
| 4 | 6688852 | rs12500833 | -0.863 | 0.301 | -0.788 | 0.429 | 0.062 | 0.005 | 0.005 | 0.035 | 0.49 | 0.75 | 0.51 | 0.59 | 0.35 | 0.51 |
| 4 | 6694778 | rs4645287 | 0.163 | -0.133 | -1.216 | 0.376 | 0.028 | -0.046 | -0.046 | 0.304 | 0.59 | 0.19 | 0.55 | 0.5 | 0.62 | 0.61 |
| 4 | 6697766 | rs3822262 | -0.698 | 0.041 | -1.123 | 0.426 | 0.004 | -0.011 | -0.011 | 0 | 0.28 | 0.25 | 0.34 | 0.64 | 0.25 | 0.35 |
| 4 | 6701203 | rs4493618 | 1.198 | -0.011 | -1.653 | -0.066 | -0.128 | 0.151 | 0.151 | 0.353 | 0.32 | 0.07 | 0.31 | 0.11 | 0.49 | 0.29 |
| 4 | 6708515 | rs10025855 | 0.547 | 0.101 | -0.953 | 0.612 | -0.009 | -0.018 | -0.018 | 0 | 0.11 | 0.05 | 0.09 | 0.16 | 0.14 | 0.13 |
| 4 | 6709345 | rs4689523 | -0.009 | -0.173 | -0.153 | 0.778 | -0.029 | 0.166 | 0.166 | 0.056 | 0.15 | 0.29 | 0.34 | 0.49 | 0.08 | 0.25 |
| 4 | 6718775 | rs3172604 | 0.174 | 0.576 | -0.232 | 0.595 | 0 | 0.004 | 0.004 | 2.032 | 0.62 | 0.45 | 0.71 | 0.55 | 0.64 | 0.62 |
| 4 | 6732660 | rs16839178 | -0.288 | 1.189 | -0.122 | 0.469 | 0.021 | -0.019 | -0.019 | 0 | 0.23 | 0.28 | 0.38 | 0.2 | 0.26 | 0.3 |
| 4 | 6733588 | rs4393997 | -0.77 | 0.965 | 0.247 | 1.048 | 0.009 | 0.043 | 0.043 | 0.218 | 0.28 | 0.58 | 0.48 | 0.55 | 0.29 | 0.37 |
| 4 | 6733659 | rs4420983 | 0.248 | 0.708 | -0.971 | 0.348 | 0.015 | 0.02 | 0.02 | 0.91 | 0.3 | 0.14 | 0.14 | 0.26 | 0.24 | 0.35 |
| 4 | 6734367 | rs4689044 | -0.066 | 1.231 | -0.49 | 0.398 | 0.011 | -0.012 | -0.012 | 0 | 0.21 | 0.26 | 0.38 | 0.17 | 0.27 | 0.28 |
| 4 | 6736365 | rs7661864 | -0.678 | 0.766 | -0.72 | 0.479 | -0.011 | -0.019 | -0.019 | 0 | 0.35 | 0.54 | 0.3 | 0.27 | 0.28 | 0.31 |
| 4 | 6736751 | rs4689534 | -0.252 | 0.914 | -1.312 | 0.776 | -0.003 | -0.061 | -0.061 | 0.887 | 0.39 | 0.17 | 0.36 | 0.46 | 0.37 | 0.35 |
| 4 | 6738883 | rs4689536 | 0.314 | 0.745 | -0.887 | 0.698 | -0.001 | 0 | 0 | 0.07 | 0.23 | 0.22 | 0.25 | 0.28 | 0.2 | 0.26 |
| 4 | 6746425 | rs4543114 | -0.016 | 0.783 | -0.751 | 0.484 | -0.009 | -0.008 | -0.008 | 0 | 0.61 | 0.69 | 0.58 | 0.6 | 0.55 | 0.6 |
| 4 | 6754570 | rs4689542 | 0.338 | 0.392 | -0.109 | 0.266 | -0.001 | -0.016 | -0.016 | 0 | 0.93 | 0.8 | 0.92 | 0.98 | 0.94 | 0.95 |
| 4 | 6755324 | rs4302466 | -0.055 | 0.121 | -0.571 | -0.066 | 0.037 | 0.082 | 0.082 | 3.317 | 0.46 | 0.17 | 0.32 | 0.25 | 0.4 | 0.28 |
| 4 | 6755936 | rs10516190 | 0.49 | -0.285 | -0.848 | -0.357 | 0.003 | -0.009 | -0.009 | 0.459 | 0.18 | 0.11 | 0.19 | 0.23 | 0.16 | 0.12 |
| 4 | 6757184 | rs11935889 | -0.854 | -0.481 | -0.063 | -0.549 | -0.001 | 0.006 | 0.006 | 0 | 0.58 | 0.55 | 0.38 | 0.31 | 0.43 | 0.4 |
| 4 | 6762107 | rs2068344 | -1.417 | -0.055 | -0.34 | -0.443 | -0.05 | -0.066 | -0.066 | 0 | 0.68 | 0.85 | 0.62 | 0.53 | 0.56 | 0.6 |
| 4 | 6763115 | rs4689050 | 1.06 | -0.647 | -0.146 | -0.079 | 0.002 | -0.048 | -0.048 | 0 | 0.15 | 0.04 | 0.23 | 0.3 | 0.12 | 0.12 |
| 4 | 6766870 | rs7681358 | -0.404 | -0.723 | 0.473 | -0.174 | 0.013 | 0.203 | 0.203 | 0.517 | 0.1 | 0.51 | 0.16 | 0.2 | 0.05 | 0.19 |
| 4 | 6767336 | rs4689545 | -0.608 | -0.687 | -0.091 | -0.231 | -0.001 | -0.001 | -0.001 | 0 | 0.52 | 0.47 | 0.45 | 0.52 | 0.42 | 0.49 |
| 4 | 6767551 | rs4689547 | 0.682 | -0.634 | -0.625 | -0.373 | 0.002 | -0.009 | -0.009 | 1.746 | 0.16 | 0.1 | 0.17 | 0.29 | 0.12 | 0.11 |
| 4 | 6768043 | rs6446542 | -0.016 | -0.631 | -0.381 | -1.066 | -0.015 | -0.078 | -0.078 | 3.239 | 0.21 | 0.26 | 0.09 | 0 | 0.14 | 0.05 |
| 4 | 6771474 | rs6845618 | -0.707 | -0.521 | -1.216 | -1.103 | 0.036 | 0.017 | 0.017 | 0.162 | 0.38 | 0.74 | 0.21 | 0.01 | 0.26 | 0.24 |
| 4 | 6776212 | rs9995180 | -0.859 | 0.151 | -2.141 | -0.348 | -0.065 | -0.022 | -0.022 | 0 | 0.55 | 0.89 | 0.59 | 0.6 | 0.49 | 0.54 |
| 4 | 6777024 | rs6824277 | -0.418 | -0.403 | -0.923 | -0.992 | -0.005 | -0.085 | -0.085 | 0.995 | 0.16 | 0.37 | 0.17 | 0.04 | 0.11 | 0.14 |
| 4 | 6777164 | rs6854931 | -0.941 | -0.402 | -1.71 | -1.085 | 0.056 | 0.229 | 0.229 | 0.915 | 0.4 | 0.85 | 0.3 | 0.04 | 0.28 | 0.26 |
| 4 | 6817928 | rs4689552 | -0.444 | 0.039 | -0.384 | 0.223 | -0.011 | 0.052 | 0.052 | 0 | 0.14 | 0.28 | 0.34 | 0.32 | 0.1 | 0.28 |
| 4 | 6833794 | rs6833118 | -0.41 | 0.014 | -0.332 | 0.293 | -0.002 | 0.012 | 0.012 | 0 | 0.88 | 0.84 | 0.69 | 0.7 | 0.92 | 0.75 |
| 4 | 6836662 | rs6848312 | -0.598 | 0.536 | -0.303 | -0.046 | -0.007 | 0.002 | 0.002 | 0 | 0.08 | 0.06 | 0.05 | 0.01 | 0.15 | 0.12 |
| 4 | 6852114 | rs17728979 | 0.435 | 0.337 | -0.607 | -0.409 | -0.002 | 0.07 | 0.07 | 0.424 | 0.3 | 0.15 | 0.18 | 0.09 | 0.28 | 0.21 |
| 4 | 6854060 | rs11723013 | 1.288 | 0.227 | -0.562 | -0.544 | -0.098 | 0.229 | 0.229 | 0.908 | 0.33 | 0.03 | 0.26 | 0.02 | 0.42 | 0.32 |
| 4 | 6855767 | rs6845985 | 0.373 | 0.209 | -0.025 | -0.151 | -0.034 | 0.093 | 0.093 | 0 | 0.76 | 0.54 | 0.59 | 0.53 | 0.84 | 0.7 |
| 4 | 6857278 | rs6836941 | 0.855 | 0.161 | -0.004 | -0.233 | -0.002 | -0.023 | -0.023 | 0 | 0.15 | 0.08 | 0.19 | 0.57 | 0.14 | 0.13 |
| 4 | 6862131 | rs10006395 | 1.38 | 0.092 | -0.039 | -0.263 | -0.003 | 0.076 | 0.076 | 0.128 | 0.23 | 0.07 | 0.13 | 0.06 | 0.19 | 0.17 |
| 4 | 6868155 | rs10022630 | -0.145 | 0.126 | 0.066 | -0.007 | -0.018 | 0.09 | 0.09 | 0.534 | 0.38 | 0.19 | 0.22 | 0.1 | 0.4 | 0.32 |
| 4 | 6868452 | rs10022837 | 0.706 | 0.092 | 0.318 | 0.021 | -0.014 | 0.01 | 0.01 | 0.695 | 0.46 | 0.31 | 0.45 | 0.57 | 0.53 | 0.41 |
| 4 | 6868495 | rs17796902 | -0.155 | 0.187 | 0.254 | -0.147 | -0.019 | 0.094 | 0.094 | 0.506 | 0.38 | 0.18 | 0.22 | 0.1 | 0.4 | 0.33 |
| 4 | 6876569 | rs13115900 | 0.866 | 0.492 | 0.206 | -0.175 | -0.02 | 0.052 | 0.052 | 0.085 | 0.27 | 0.14 | 0.17 | 0.07 | 0.3 | 0.27 |
| 4 | 6877902 | rs6828754 | 0.392 | 0.611 | 0.445 | 0.282 | -0.014 | 0.02 | 0.02 | 0.65 | 0.45 | 0.24 | 0.37 | 0.38 | 0.5 | 0.39 |
| 4 | 6883012 | rs3733606 | 0.676 | 0.393 | 0.153 | -0.302 | -0.007 | -0.051 | -0.051 | 1.563 | 0.15 | 0.06 | 0.16 | 0.51 | 0.15 | 0.14 |
| 4 | 6883514 | rs3822268 | -0.726 | -0.047 | 0.328 | 0.102 | -0.001 | 0.022 | 0.022 | 0 | 0.86 | 0.81 | 0.66 | 0.64 | 0.9 | 0.74 |
| 4 | 6885815 | rs6828784 | -1.417 | -0.024 | 0.243 | -0.338 | 0.04 | -0.056 | -0.056 | 0.008 | 0.11 | 0.35 | 0.07 | 0.03 | 0.1 | 0.15 |
| 4 | 6886650 | rs7694173 | -0.418 | 0.216 | 0.127 | -0.217 | -0.002 | 0.006 | 0.006 | 0 | 0.36 | 0.28 | 0.2 | 0.09 | 0.33 | 0.27 |
| 4 | 6886830 | rs10001636 | -0.456 | 0.3 | -0.121 | -0.166 | -0.007 | 0.041 | 0.041 | 0 | 0.35 | 0.22 | 0.18 | 0.09 | 0.33 | 0.29 |
| 4 | 6889967 | rs9999836 | -1.144 | 0.442 | 0.44 | 0.934 | 0.049 | -0.036 | -0.036 | 1.772 | 0.09 | 0.19 | 0.07 | 0.02 | 0.1 | 0.08 |
| 4 | 6893681 | rs3901368 | -0.836 | 0.051 | -0.262 | 1.531 | -0.032 | 0.148 | 0.148 | 0 | 0.21 | 0.45 | 0.32 | 0.59 | 0.15 | 0.16 |

| | | | | | | | | | | | | | | | | |
|---|---|---|---|---|---|---|---|---|---|---|---|---|---|---|---|---|
| 4 | 6903159 | rs6446548 | 0.698 | 0.036 | -1.07 | 1.521 | 0.019 | -0.034 | -0.034 | 0 | 0.88 | 0.97 | 0.95 | 0.72 | 0.94 | 0.86 |
| 4 | 6904074 | rs6826453 | -0.492 | 0.161 | -1.492 | 0.637 | 0.006 | 0.046 | 0.046 | 0.212 | 0.3 | 0.21 | 0.34 | 0.13 | 0.29 | 0.16 |
| 4 | 6907052 | rs10020189 | -0.253 | 0.103 | -1.513 | 0.517 | 0.002 | 0 | 0 | 0 | 0.28 | 0.26 | 0.21 | 0.2 | 0.24 | 0.28 |
| 4 | 6907365 | rs28396694 | 0.108 | -0.068 | -1.514 | -0.063 | -0.012 | -0.046 | -0.046 | 0 | 0.21 | 0.25 | 0.12 | 0.01 | 0.11 | 0.07 |
| 4 | 6908390 | rs4342189 | -0.482 | 0.264 | -2.094 | 0.042 | 0.038 | 0.093 | 0.093 | 0 | 0.51 | 0.83 | 0.49 | 0.35 | 0.42 | 0.34 |
| 4 | 6913707 | rs4432749 | 0.205 | 0.471 | -1.823 | -0.381 | -0.023 | 0.021 | 0.021 | 0.826 | 0.38 | 0.3 | 0.45 | 0.71 | 0.52 | 0.62 |
| 4 | 6915342 | rs11934364 | 0.168 | 0.066 | -1.934 | -1.021 | 0.047 | 0.106 | 0.106 | 1.902 | 0.33 | 0.12 | 0.16 | 0.03 | 0.19 | 0.16 |
| 4 | 6916263 | rs7688690 | -0.924 | -0.336 | -0.637 | -1.218 | -0.001 | -0.009 | -0.009 | 0 | 0.53 | 0.52 | 0.51 | 0.27 | 0.46 | 0.36 |
| 4 | 6918679 | rs10937762 | 1.174 | -0.312 | -0.387 | -1.339 | -0.021 | 0.015 | 0.015 | 0.018 | 0.4 | 0.32 | 0.41 | 0.7 | 0.47 | 0.64 |
| 4 | 6919661 | rs10804990 | 0.353 | -0.454 | -0.392 | -1.306 | 0.007 | 0.017 | 0.017 | 0 | 0.62 | 0.33 | 0.52 | 0.72 | 0.62 | 0.74 |
| 4 | 6919674 | rs7668673 | 0.125 | -0.491 | -0.133 | -1.403 | 0.001 | 0.001 | 0.001 | 0 | 0.24 | 0.2 | 0.27 | 0.23 | 0.27 | 0.18 |
| 4 | 6924241 | rs6446552 | -0.006 | -0.625 | -0.092 | -1.35 | 0.002 | 0 | 0 | 0 | 0.27 | 0.21 | 0.34 | 0.26 | 0.27 | 0.2 |
| 4 | 6930968 | rs4323102 | 1.165 | -0.612 | -0.018 | -1.324 | 0.009 | 0.012 | 0.012 | 0 | 0.47 | 0.45 | 0.49 | 0.71 | 0.62 | 0.64 |
| 4 | 6937984 | rs9291123 | 1.144 | -0.657 | -0.008 | -1.311 | 0.01 | 0.013 | 0.013 | 0.005 | 0.47 | 0.45 | 0.51 | 0.71 | 0.62 | 0.64 |
| 4 | 6941375 | rs4234788 | -0.589 | -0.704 | 0.139 | -1.257 | -0.002 | -0.008 | -0.008 | 0.014 | 0.78 | 0.79 | 0.88 | 0.97 | 0.91 | 0.84 |
| 4 | 6956134 | rs11732898 | 0.481 | -0.534 | 0.445 | -1.349 | -0.028 | -0.089 | -0.089 | 0 | 0.7 | 0.55 | 0.8 | 0.97 | 0.82 | 0.81 |
| 4 | 6957297 | rs4234791 | -1.177 | -0.594 | 0.486 | -1.366 | 0.025 | 0.025 | 0.025 | 0.001 | 0.81 | 0.88 | 0.9 | 0.98 | 0.92 | 0.86 |
| 4 | 6957332 | rs41410847 | -1.03 | -0.582 | 0.521 | -1.361 | 0.009 | -0.051 | -0.051 | 0.379 | 0.11 | 0.28 | 0.09 | 0 | 0.1 | 0.06 |
| 4 | 6961589 | rs4689568 | 0.221 | -0.594 | 0.571 | -1.354 | -0.035 | -0.095 | -0.095 | 0.487 | 0.72 | 0.55 | 0.81 | 0.97 | 0.84 | 0.83 |
| 4 | 6969919 | rs6831368 | -1.329 | -0.669 | 0.516 | -1.485 | 0.006 | 0.013 | 0.013 | 0 | 0.42 | 0.63 | 0.45 | 0.28 | 0.35 | 0.26 |
| 4 | 6983677 | rs10015163 | 1.945 | -0.655 | 0.522 | -1.471 | -0.018 | 0.007 | 0.007 | 0 | 0.37 | 0.27 | 0.44 | 0.7 | 0.54 | 0.56 |
| 4 | 6988677 | rs1048009 | -0.649 | -0.636 | 0.562 | -1.465 | 0.02 | -0.045 | -0.045 | 0 | 0.1 | 0.39 | 0.08 | 0 | 0.08 | 0.06 |
| 4 | 6996129 | rs3892490 | -0.733 | -0.793 | 0.543 | -1.574 | -0.021 | -0.071 | -0.071 | 0.016 | 0.33 | 0.42 | 0.19 | 0.03 | 0.18 | 0.22 |
| 4 | 7002344 | rs17742611 | 2.177 | -0.786 | 0.506 | -1.364 | -0.041 | -0.006 | -0.006 | 2.35 | 0.36 | 0.22 | 0.43 | 0.66 | 0.51 | 0.55 |
| 4 | 7006541 | rs2301819 | 0.217 | -0.801 | 0.502 | -1.412 | 0.009 | -0.004 | -0.004 | 1.053 | 0.26 | 0.2 | 0.15 | 0.03 | 0.12 | 0.18 |
| 4 | 7008078 | rs10013211 | 2.052 | -0.727 | 0.496 | -1.358 | -0.025 | 0.007 | 0.007 | 0 | 0.36 | 0.26 | 0.43 | 0.7 | 0.51 | 0.56 |
| 4 | 7018195 | rs11732301 | -0.427 | -0.768 | 0.474 | -1.398 | -0.014 | -0.004 | -0.004 | 0 | 0.32 | 0.46 | 0.39 | 0.32 | 0.28 | 0.25 |
| 4 | 7021365 | rs12650264 | 1.931 | -0.631 | 0.542 | -1.342 | 0.004 | 0 | 0 | 0 | 0.36 | 0.37 | 0.43 | 0.64 | 0.5 | 0.54 |
| 4 | 7023479 | rs6854776 | -0.905 | -0.664 | 0.49 | -1.38 | 0.037 | 0.062 | 0.062 | 0 | 0.78 | 0.89 | 0.88 | 0.98 | 0.87 | 0.86 |
| 4 | 7025909 | rs10516182 | -0.146 | -0.711 | 0.713 | -1.412 | 0.004 | 0 | 0 | 0 | 0.32 | 0.26 | 0.36 | 0.34 | 0.28 | 0.26 |
| 4 | 7027041 | rs6834102 | -0.868 | -0.671 | 0.614 | -1.397 | 0.011 | 0.004 | 0.004 | 0 | 0.78 | 0.85 | 0.87 | 0.98 | 0.87 | 0.86 |
| 4 | 7032180 | rs11546450 | -0.618 | -0.479 | 0.476 | -1.248 | -0.002 | -0.032 | -0.032 | 0 | 0.1 | 0.22 | 0.08 | 0.01 | 0.09 | 0.05 |
| 4 | 7033671 | rs12119 | 1.536 | -0.476 | 0.484 | -1.232 | -0.025 | 0.005 | 0.005 | 0 | 0.38 | 0.27 | 0.42 | 0.66 | 0.51 | 0.56 |
| 4 | 7035637 | rs6852613 | -0.066 | -0.447 | 0.691 | -0.993 | 0.016 | -0.016 | -0.016 | 0 | 0.06 | 0.17 | 0.06 | 0.01 | 0.08 | 0.03 |
| 4 | 7039197 | rs3857175 | 0.615 | -0.575 | 0.542 | -1.156 | -0.032 | -0.094 | -0.094 | 0.51 | 0.68 | 0.52 | 0.75 | 0.91 | 0.8 | 0.78 |
| 4 | 7039326 | rs10937779 | 1.871 | -0.536 | 0.569 | -1.083 | -0.024 | 0.014 | 0.014 | 0 | 0.4 | 0.2 | 0.44 | 0.64 | 0.57 | 0.52 |
| 4 | 7039345 | rs3857176 | 0.559 | -0.07 | 0.806 | -0.724 | -0.032 | -0.099 | -0.099 | 0 | 0.72 | 0.54 | 0.76 | 0.92 | 0.81 | 0.8 |
| 4 | 7055253 | rs868489 | -1.215 | 0.437 | 0.734 | -0.38 | 0.005 | 0 | 0 | 0 | 0.27 | 0.22 | 0.29 | 0.26 | 0.24 | 0.23 |
| 4 | 7061378 | rs3796908 | 0.989 | 0.483 | 1.171 | -0.234 | -0.01 | 0.012 | 0.012 | 0 | 0.76 | 0.64 | 0.71 | 0.73 | 0.77 | 0.8 |
| 4 | 7063524 | rs6836129 | -0.956 | 0.715 | 0.841 | -0.214 | -0.003 | -0.002 | -0.002 | 0 | 0.23 | 0.2 | 0.33 | 0.26 | 0.29 | 0.19 |
| 4 | 7069804 | rs3822270 | 1.14 | 0.268 | 0.796 | 0 | -0.014 | -0.004 | -0.004 | 0 | 0.72 | 0.45 | 0.68 | 0.74 | 0.76 | 0.8 |
| 4 | 7074583 | rs4256224 | 0.973 | 0.325 | 0.706 | 0.072 | -0.014 | 0.006 | 0.006 | 0 | 0.73 | 0.47 | 0.69 | 0.74 | 0.76 | 0.8 |
| 4 | 7084835 | rs907526 | 1.143 | 0.303 | 0.611 | 0.31 | -0.007 | 0.001 | 0.001 | 0 | 0.63 | 0.45 | 0.62 | 0.6 | 0.71 | 0.72 |
| 4 | 7084858 | rs3857183 | 1.276 | 0.107 | 0.139 | 0.514 | 0.032 | 0.028 | 0.028 | 0 | 0.3 | 0.18 | 0.27 | 0.21 | 0.2 | 0.37 |
| 4 | 7084985 | rs3857184 | 0.753 | 0.172 | 0.569 | 0.483 | -0.011 | 0.003 | 0.003 | 0 | 0.63 | 0.46 | 0.61 | 0.59 | 0.72 | 0.69 |
| 4 | 7086930 | rs6446566 | -0.908 | -0.004 | 0.674 | 0.652 | -0.025 | 0.07 | 0.07 | 0 | 0.22 | 0.39 | 0.26 | 0.38 | 0.19 | 0.17 |
| 4 | 7093834 | rs11734660 | 0.972 | 0.566 | 0.184 | 0.875 | -0.066 | -0.051 | -0.051 | 4.236 | 0.3 | 0.15 | 0.35 | 0.33 | 0.48 | 0.34 |
| 4 | 7097498 | rs12642809 | -0.73 | 0.503 | 0.822 | 1.419 | 0 | 0.007 | 0.007 | 0 | 0.14 | 0.17 | 0.19 | 0.39 | 0.15 | 0.14 |
| 4 | 7102597 | rs4689599 | -1.652 | 0.458 | 1.063 | 1.343 | -0.003 | 0.018 | 0.018 | 0 | 0.27 | 0.36 | 0.33 | 0.45 | 0.25 | 0.25 |
| 4 | 7109083 | rs4082843 | -1.388 | 0.272 | 1.111 | 1.355 | -0.022 | 0.102 | 0.102 | 0 | 0.2 | 0.51 | 0.33 | 0.44 | 0.19 | 0.21 |
| 4 | 7109830 | rs28401383 | -2.058 | 0.389 | 0.85 | 1.395 | 0.001 | -0.001 | -0.001 | 0 | 0.37 | 0.63 | 0.47 | 0.63 | 0.33 | 0.36 |
| 4 | 7112676 | rs4082846 | -1.835 | 0.502 | 1.069 | 1.162 | -0.015 | 0.018 | 0.018 | 0.228 | 0.28 | 0.61 | 0.35 | 0.43 | 0.26 | 0.25 |
| 4 | 7113986 | rs7378101 | -0.277 | 0.52 | 1.051 | 0.995 | -0.002 | -0.033 | -0.033 | 0 | 0.13 | 0.14 | 0.13 | 0.02 | 0.1 | 0.08 |
| 4 | 7118527 | rs11938388 | -0.682 | 0.513 | 1.235 | 0.974 | 0.01 | -0.085 | -0.085 | 0 | 0.15 | 0.38 | 0.17 | 0.02 | 0.14 | 0.08 |
| 4 | 7120761 | rs12647765 | -0.945 | 0.512 | 1.059 | 1.531 | -0.02 | 0.139 | 0.139 | 0.158 | 0.11 | 0.25 | 0.17 | 0.42 | 0.11 | 0.13 |
| 4 | 7126063 | rs10027697 | -0.068 | 1.108 | 0.576 | 1.115 | 0.025 | 0.015 | 0.015 | 0 | 0.32 | 0.4 | 0.35 | 0.33 | 0.45 | 0.38 |
| 4 | 7128805 | rs4689603 | -0.24 | 1.573 | 0.255 | 0.979 | -0.034 | -0.037 | -0.037 | 0 | 0.3 | 0.21 | 0.29 | 0.48 | 0.41 | 0.37 |
| 4 | 7129001 | rs16843076 | -0.27 | 0.786 | 0.325 | 0.463 | -0.034 | -0.063 | -0.063 | 0 | 0.12 | 0.25 | 0.04 | 0.01 | 0.04 | 0.05 |
| 4 | 7129556 | rs4689604 | -0.68 | 1.498 | -0.442 | 0.779 | 0.002 | 0.031 | 0.031 | 0.004 | 0.34 | 0.53 | 0.27 | 0.4 | 0.3 | 0.34 |
| 4 | 7132365 | rs2359009 | -0.735 | 1.16 | -1.21 | 0.655 | 0 | -0.002 | -0.002 | 0 | 0.27 | 0.3 | 0.39 | 0.48 | 0.33 | 0.33 |
| 4 | 7133905 | rs10155062 | -0.145 | 1.213 | -1.267 | 0.65 | 0.013 | 0.002 | 0.002 | 0.065 | 0.29 | 0.6 | 0.3 | 0.27 | 0.33 | 0.26 |
| 4 | 7134206 | rs17807577 | 0.774 | 1 | -1.275 | 0.763 | -0.002 | 0.024 | 0.024 | 0.049 | 0.1 | 0.03 | 0.05 | 0.02 | 0.12 | 0.1 |
| 4 | 7135912 | rs10804992 | -0.697 | 1.067 | -1.09 | 1.2 | 0 | 0 | 0 | 0 | 0.16 | 0.18 | 0.28 | 0.43 | 0.21 | 0.23 |
| 4 | 7137119 | rs4689608 | -0.747 | 1.227 | -1.616 | 1.259 | -0.052 | -0.101 | -0.101 | 0.308 | 0.15 | 0.03 | 0.27 | 0.5 | 0.24 | 0.25 |
| 4 | 7137140 | rs13127331 | 0.239 | 0.714 | -1.961 | 0.526 | 0.002 | 0.077 | 0.077 | 0 | 0.19 | 0.04 | 0.06 | 0.01 | 0.16 | 0.17 |
| 4 | 7137336 | rs4689609 | -0.705 | 0.794 | -1.747 | 0.807 | 0.048 | 0.098 | 0.098 | 0 | 0.48 | 0.75 | 0.5 | 0.76 | 0.38 | 0.63 |
| 4 | 7137521 | rs16839667 | -0.323 | 1.184 | -1.356 | 1.575 | -0.023 | -0.049 | -0.049 | 0 | 0.11 | 0.04 | 0.22 | 0.42 | 0.17 | 0.24 |
| 4 | 7137631 | rs9762239 | -1.093 | 1.389 | -1.108 | 1.094 | -0.015 | -0.011 | -0.011 | 0 | 0.22 | 0.09 | 0.14 | 0.19 | 0.27 | 0.17 |
| 4 | 7137993 | rs7665697 | -1.111 | 2.033 | -0.725 | 1.47 | 0.021 | 0.031 | 0.031 | 0 | 0.16 | 0.19 | 0.17 | 0.25 | 0.29 | 0.11 |

| | | | | | | | | | | | | | | | | |
|---|---|---|---|---|---|---|---|---|---|---|---|---|---|---|---|---|
| 4 | 7142427 | rs7656556 | -0.875 | 1.727 | -0.806 | 1.135 | -0.007 | -0.002 | -0.002 | 0 | 0.26 | 0.24 | 0.34 | 0.33 | 0.35 | 0.4 |
| 4 | 7142868 | rs7687509 | -0.895 | 1.004 | -0.897 | 0.897 | -0.003 | 0.005 | 0.005 | 0.194 | 0.53 | 0.71 | 0.5 | 0.48 | 0.47 | 0.61 |
| 4 | 7145249 | rs4689069 | -0.41 | 2.117 | -0.461 | 1.204 | 0.097 | 0.019 | 0.019 | 0 | 0.19 | 0.37 | 0.39 | 0.3 | 0.35 | 0.37 |
| 4 | 7146033 | rs4689071 | -0.725 | 1.018 | -1.323 | 0.82 | 0.033 | 0.041 | 0.041 | 0.018 | 0.54 | 0.88 | 0.61 | 0.68 | 0.61 | 0.73 |
| 4 | 7149407 | rs6814125 | -0.107 | 1.154 | -1.03 | 0.596 | -0.002 | 0.001 | 0.001 | 0 | 0.26 | 0.23 | 0.31 | 0.17 | 0.44 | 0.2 |
| 4 | 7151434 | rs12233740 | 0.399 | 0.636 | -1.737 | 0.24 | 0.006 | 0.054 | 0.054 | 2.307 | 0.16 | 0.03 | 0.05 | 0.1 | 0.13 | 0.13 |
| 4 | 7151708 | rs10021205 | -0.946 | 0.216 | -1.237 | -0.015 | 0.009 | 0.034 | 0.034 | 0 | 0.48 | 0.42 | 0.39 | 0.27 | 0.57 | 0.41 |
| 4 | 7151812 | rs6836687 | -1.102 | -0.062 | -1.388 | 0.471 | 0.01 | 0.003 | 0.003 | 0.588 | 0.63 | 0.91 | 0.69 | 0.71 | 0.66 | 0.72 |
| 4 | 7151930 | rs6810509 | -0.294 | -0.288 | -1.035 | 0.622 | -0.03 | 0.063 | 0.063 | 0.139 | 0.31 | 0.49 | 0.34 | 0.53 | 0.21 | 0.46 |
| 4 | 7154743 | rs3915879 | -0.627 | -0.091 | -1.855 | 0.276 | -0.064 | 0.011 | 0.011 | 0.503 | 0.72 | 0.97 | 0.79 | 0.82 | 0.7 | 0.69 |
| 4 | 7157382 | rs12508556 | -0.752 | -0.949 | -0.451 | 0.674 | -0.022 | -0.014 | -0.014 | 0 | 0.37 | 0.55 | 0.24 | 0.29 | 0.24 | 0.22 |
| 4 | 7163405 | rs4689074 | 0.607 | -0.97 | -0.078 | 0.935 | -0.013 | 0.039 | 0.039 | 0 | 0.7 | 0.64 | 0.73 | 0.57 | 0.76 | 0.77 |
| 4 | 7163426 | rs12505331 | 0.113 | -1.007 | -0.678 | 1.288 | -0.001 | -0.102 | -0.102 | 0.746 | 0.14 | 0.01 | 0.27 | 0.42 | 0.08 | 0.36 |
| 4 | 7165415 | rs4689620 | -1.214 | -1.117 | 0.056 | 0.882 | -0.027 | 0.001 | 0.001 | 0.044 | 0.29 | 0.4 | 0.4 | 0.7 | 0.17 | 0.46 |
| 4 | 7168998 | rs4566655 | -0.445 | -0.861 | -0.099 | 0.875 | 0.005 | 0.01 | 0.01 | 1.337 | 0.8 | 0.89 | 0.85 | 0.86 | 0.84 | 0.89 |
| 4 | 7170562 | rs9991298 | 0.821 | -0.998 | -0.125 | 1.144 | 0.036 | -0.049 | -0.049 | 0 | 0.63 | 0.79 | 0.58 | 0.47 | 0.75 | 0.52 |
| 4 | 7172750 | rs3982 | 0.166 | -1.001 | 0.174 | 1.44 | 0 | 0.038 | 0.038 | 0.251 | 0.13 | 0.1 | 0.26 | 0.43 | 0.11 | 0.42 |
| 4 | 7172827 | rs4689623 | -0.86 | -0.983 | 0.61 | 1.133 | -0.017 | 0.009 | 0.009 | 2.916 | 0.35 | 0.63 | 0.27 | 0.44 | 0.21 | 0.21 |
| 4 | 7174993 | rs9999241 | -1.558 | -1.084 | 0.416 | 0.92 | 0.047 | 0.105 | 0.105 | 1.364 | 0.46 | 0.71 | 0.53 | 0.87 | 0.3 | 0.6 |
| 4 | 7178278 | rs4689078 | -1.049 | -0.997 | 0.549 | 0.874 | 0.015 | 0.005 | 0.005 | 0.517 | 0.46 | 0.39 | 0.49 | 0.57 | 0.37 | 0.55 |
| 4 | 7183984 | rs7692512 | -0.517 | -1.013 | 0.157 | 0.922 | -0.022 | 0.019 | 0.019 | 3.474 | 0.33 | 0.6 | 0.23 | 0.44 | 0.19 | 0.21 |
| 4 | 7184370 | rs6851506 | -0.227 | -0.96 | -0.666 | 0.947 | -0.027 | -0.137 | -0.137 | 0 | 0.23 | 0.03 | 0.37 | 0.43 | 0.24 | 0.44 |
| 4 | 7186257 | rs10017600 | -1.42 | -0.75 | -0.366 | 0.627 | -0.005 | 0.135 | 0.135 | 0.527 | 0.56 | 0.79 | 0.61 | 0.87 | 0.43 | 0.66 |
| 4 | 7190015 | rs4689626 | -0.017 | -0.681 | -0.325 | 1.109 | 0.025 | 0.016 | 0.016 | 3.198 | 0.26 | 0.17 | 0.12 | 0.17 | 0.16 | 0.18 |
| 4 | 7195054 | rs4689630 | -1.128 | 0.136 | 0.476 | 1.834 | 0.013 | 0.154 | 0.154 | 0.869 | 0.24 | 0.47 | 0.35 | 0.45 | 0.24 | 0.23 |
| 4 | 7202789 | rs6834325 | -0.902 | 0.145 | 0.283 | 1.968 | 0 | 0.012 | 0.012 | 0.548 | 0.18 | 0.21 | 0.28 | 0.39 | 0.15 | 0.17 |
| 4 | 7203659 | rs4689081 | -1.006 | -0.151 | 0.439 | 0.817 | -0.002 | -0.021 | -0.021 | 0.887 | 0.65 | 0.64 | 0.6 | 0.92 | 0.44 | 0.73 |
| 4 | 7204174 | rs2240200 | -0.66 | -0.231 | 0.946 | 1.147 | 0.042 | 0.14 | 0.14 | 0.017 | 0.07 | 0.25 | 0.17 | 0.25 | 0.1 | 0.09 |
| 4 | 7204493 | rs3815181 | 0.324 | -0.813 | 0.452 | 1.032 | 0.005 | -0.002 | -0.002 | 1.443 | 0.24 | 0.22 | 0.13 | 0.25 | 0.13 | 0.19 |
| 4 | 7206401 | rs951449 | -1.366 | -0.713 | 0.121 | 0.586 | -0.056 | 0.122 | 0.122 | 0 | 0.65 | 0.83 | 0.6 | 0.93 | 0.46 | 0.58 |
| 4 | 7207480 | rs4461527 | -1.776 | -0.608 | 0.048 | 0.651 | -0.082 | 0.11 | 0.11 | 0 | 0.69 | 0.86 | 0.61 | 0.93 | 0.47 | 0.75 |
| 4 | 7208680 | rs10937790 | -0.527 | -0.612 | 0.68 | 1.234 | -0.023 | 0.03 | 0.03 | 0 | 0.35 | 0.43 | 0.31 | 0.44 | 0.24 | 0.48 |
| 4 | 7210808 | rs870860 | -0.437 | -0.246 | 1.245 | 1.653 | 0.059 | 0.301 | 0.301 | 0 | 0.07 | 0.36 | 0.07 | 0.22 | 0.05 | 0.07 |
| 4 | 7211197 | rs4689637 | -0.216 | 0.149 | 1.049 | 1.144 | -0.008 | 0.079 | 0.079 | 0.081 | 0.67 | 0.82 | 0.68 | 0.82 | 0.71 | 0.74 |
| 4 | 7211739 | rs12512052 | 1.701 | 0.365 | 0.929 | 0.698 | -0.016 | 0.028 | 0.028 | 0 | 0.28 | 0.18 | 0.31 | 0.01 | 0.38 | 0.34 |
| 4 | 7215211 | rs13143421 | -0.997 | 0.157 | 1.359 | 1.326 | 0.04 | 0.267 | 0.267 | 0.007 | 0.13 | 0.39 | 0.09 | 0.32 | 0.12 | 0.15 |
| 4 | 7215322 | rs4689642 | -0.406 | 0.056 | 1.111 | 1.463 | -0.006 | 0.014 | 0.014 | 0 | 0.31 | 0.51 | 0.24 | 0.53 | 0.29 | 0.31 |
| 4 | 7218072 | rs12152570 | 1.812 | 0.115 | 0.91 | 1.256 | -0.018 | 0.041 | 0.041 | 0 | 0.3 | 0.18 | 0.32 | 0.01 | 0.4 | 0.34 |
| 4 | 7219907 | rs4234797 | 0.178 | 0.307 | 0.969 | 1.538 | -0.012 | 0.052 | 0.052 | 0 | 0.6 | 0.71 | 0.64 | 0.72 | 0.63 | 0.62 |
| 4 | 7220548 | rs16839840 | 0.224 | 0.26 | 1.564 | 2.057 | -0.015 | 0.022 | 0.022 | 0 | 0.2 | 0.3 | 0.16 | 0.2 | 0.17 | 0.13 |
| 4 | 7221071 | rs12646898 | -0.381 | 0.278 | 1.327 | 1.946 | -0.015 | 0.113 | 0.113 | 0 | 0.25 | 0.41 | 0.2 | 0.45 | 0.22 | 0.19 |
| 4 | 7224235 | rs12641883 | -1.133 | 0.374 | 0.616 | 1.574 | 0.026 | 0.076 | 0.076 | 0.778 | 0.08 | 0.23 | 0.05 | 0.11 | 0.07 | 0.08 |
| 4 | 7226751 | rs4689090 | -0.374 | 0.351 | 0.48 | 1.418 | -0.001 | -0.002 | -0.002 | 0.901 | 0.25 | 0.24 | 0.17 | 0.21 | 0.24 | 0.17 |
| 4 | 7239621 | rs4689093 | -0.463 | 0.111 | 0.361 | 1.322 | -0.007 | 0.004 | 0.004 | 0.259 | 0.37 | 0.35 | 0.22 | 0.49 | 0.26 | 0.24 |
| 4 | 7241780 | rs11931042 | -1.03 | 0.415 | 0.218 | 0.796 | 0.038 | -0.076 | -0.076 | 1.066 | 0.12 | 0.32 | 0.13 | 0 | 0.13 | 0.02 |
| 4 | 7250020 | rs2008121 | -0.224 | -0.074 | -0.438 | 1.262 | 0.045 | -0.045 | -0.045 | 0.689 | 0.34 | 0.15 | 0.2 | 0.5 | 0.26 | 0.25 |
| 4 | 7250232 | rs2008103 | 0.228 | 0.353 | -0.202 | 1.024 | -0.004 | 0.007 | 0.007 | 0 | 0.78 | 0.8 | 0.8 | 0.79 | 0.79 | 0.78 |
| 4 | 7253260 | rs905010 | -1.29 | 0.157 | 0.017 | 1.51 | -0.01 | 0.057 | 0.057 | 0 | 0.18 | 0.35 | 0.15 | 0.26 | 0.12 | 0.18 |
| 4 | 7256620 | rs9291133 | 0.739 | 0.293 | -0.36 | 1.183 | -0.004 | 0.011 | 0.011 | 0 | 0.87 | 0.78 | 0.76 | 0.81 | 0.87 | 0.85 |
| 4 | 7257160 | rs7686843 | -0.28 | 0.383 | -0.682 | 1.279 | -0.001 | 0 | 0 | 0.089 | 0.4 | 0.36 | 0.31 | 0.37 | 0.33 | 0.31 |
| 4 | 7258075 | rs10025225 | 0.259 | 0.488 | -1.162 | 1.279 | -0.01 | -0.026 | -0.026 | 0.156 | 0.89 | 0.93 | 0.77 | 0.8 | 0.87 | 0.87 |
| 4 | 7262170 | rs904989 | -0.744 | 0.37 | -1.28 | 0.816 | 0.071 | 0.054 | 0.054 | 1.763 | 0.49 | 0.71 | 0.41 | 0.61 | 0.37 | 0.46 |
| 4 | 7265635 | rs867350 | -0.606 | 0.588 | -1.51 | 0.191 | 0.003 | 0.143 | 0.143 | 0.134 | 0.47 | 0.71 | 0.38 | 0.2 | 0.44 | 0.36 |
| 4 | 7265945 | rs6845989 | -0.494 | 0.724 | -1.182 | 0.431 | -0.004 | 0.028 | 0.028 | 0.019 | 0.48 | 0.62 | 0.45 | 0.39 | 0.43 | 0.4 |
| 4 | 7266856 | rs6826089 | -1.035 | -0.18 | -1.311 | 1.031 | 0.035 | -0.059 | -0.059 | 0.444 | 0.23 | 0.79 | 0.23 | 0.47 | 0.14 | 0.27 |
| 4 | 7271603 | rs6446590 | -0.277 | 0.374 | -0.746 | 0.462 | -0.005 | -0.023 | -0.023 | 0 | 0.26 | 0.31 | 0.15 | 0.27 | 0.25 | 0.32 |
| 4 | 7279707 | rs6831675 | -0.054 | 0.618 | -0.653 | 0.518 | -0.008 | 0.008 | 0.008 | 0.526 | 0.61 | 0.54 | 0.76 | 0.46 | 0.69 | 0.58 |
| 4 | 7280033 | rs4689677 | 0.551 | 0.95 | -0.587 | 0.678 | -0.001 | 0.036 | 0.036 | 0 | 0.05 | 0.05 | 0.29 | 0.19 | 0.07 | 0.06 |
| 4 | 7280324 | rs6814708 | 0.282 | 1.422 | -0.808 | 0.104 | 0.019 | -0.053 | -0.053 | 0 | 0.31 | 0.42 | 0.29 | 0.07 | 0.39 | 0.17 |
| 4 | 7282332 | rs6857447 | 0.507 | 1.433 | -0.517 | 0.156 | 0.022 | -0.021 | -0.021 | 0 | 0.17 | 0.2 | 0.19 | 0.05 | 0.26 | 0.12 |
| 4 | 7282506 | rs17755381 | 0.461 | 1.406 | -0.194 | 0.149 | 0.048 | -0.035 | -0.035 | 1.25 | 0.17 | 0.27 | 0.2 | 0.05 | 0.26 | 0.13 |
| 4 | 7288703 | rs3857193 | 0.076 | 0.572 | 0.066 | 1.13 | -0.001 | 0 | 0 | 0.704 | 0.61 | 0.64 | 0.77 | 0.43 | 0.67 | 0.56 |
| 4 | 7289113 | rs4689682 | -0.475 | 0.726 | -0.171 | 1.395 | 0.008 | 0.041 | 0.041 | 1.277 | 0.76 | 0.72 | 0.82 | 0.59 | 0.75 | 0.69 |
| 4 | 7291363 | rs11946827 | 0.651 | 0.822 | 0.188 | 1.43 | -0.012 | 0.042 | 0.042 | 0.185 | 0.69 | 0.55 | 0.79 | 0.44 | 0.78 | 0.66 |
| 4 | 7292894 | rs3846424 | 0.509 | 1.159 | -0.397 | 1.188 | 0.003 | 0.001 | 0.001 | 0 | 0.73 | 0.74 | 0.87 | 0.7 | 0.77 | 0.73 |
| 4 | 7295154 | rs7656453 | 0.623 | 0.887 | -0.719 | 1.244 | 0 | -0.026 | -0.026 | 0 | 0.1 | 0.03 | 0.24 | 0.16 | 0.08 | 0.07 |
| 4 | 7297506 | rs12498790 | 0.455 | 1.039 | -0.293 | 1.28 | 0.01 | 0.03 | 0.03 | 0.125 | 0.78 | 0.72 | 0.85 | 0.67 | 0.81 | 0.72 |
| 4 | 7304566 | rs4689687 | -0.614 | 1.052 | -1.018 | 0.855 | -0.065 | -0.024 | -0.024 | 0 | 0.23 | 0.06 | 0.38 | 0.21 | 0.27 | 0.18 |
| 4 | 7309924 | rs952588 | 0.267 | 0.381 | -0.433 | 0.699 | -0.036 | 0.054 | 0.054 | 5.39 | 0.77 | 0.4 | 0.81 | 0.38 | 0.86 | 0.77 |

| | | | | | | | | | | | | | | | | |
|---|---|---|---|---|---|---|---|---|---|---|---|---|---|---|---|---|
| 4 | 7312054 | rs3930439 | 0.821 | 0.606 | -1.078 | 0.067 | -0.023 | 0.049 | 0.049 | 0 | 0.24 | 0.11 | 0.18 | 0.03 | 0.27 | 0.11 |
| 4 | 7312145 | rs11942816 | -0.131 | 0.717 | -0.931 | 0.309 | 0.01 | 0.034 | 0.034 | 2.721 | 0.56 | 0.25 | 0.66 | 0.36 | 0.61 | 0.67 |
| 4 | 7315904 | rs10446501 | 0.739 | 0.749 | -0.594 | 0.139 | -0.006 | -0.043 | -0.043 | 0 | 0.33 | 0.33 | 0.28 | 0.14 | 0.33 | 0.49 |
| 4 | 7321511 | rs3916118 | -0.373 | 0.603 | -0.995 | 0.724 | 0.004 | 0.001 | 0.001 | 1.331 | 0.56 | 0.57 | 0.6 | 0.59 | 0.61 | 0.72 |
| 4 | 7322423 | rs1104939 | -0.952 | 0.824 | -1.645 | 0.835 | -0.035 | 0.063 | 0.063 | 0.389 | 0.72 | 0.95 | 0.74 | 0.81 | 0.7 | 0.55 |
| 4 | 7325002 | rs4689705 | -0.975 | 0.844 | -1.636 | 0.836 | -0.046 | 0.06 | 0.06 | 0.397 | 0.72 | 0.96 | 0.77 | 0.81 | 0.69 | 0.6 |
| 4 | 7326199 | rs4689707 | -0.623 | 0.955 | -0.807 | 1.574 | -0.016 | -0.036 | -0.036 | 0 | 0.1 | 0.03 | 0.22 | 0.21 | 0.13 | 0.06 |
| 4 | 7328221 | rs11946683 | -0.982 | 0.873 | -0.184 | 1.513 | 0.031 | 0.107 | 0.107 | 0.766 | 0.25 | 0.52 | 0.34 | 0.4 | 0.29 | 0.22 |
| 4 | 7329324 | rs6840423 | -0.687 | 0.032 | 0.237 | 1.434 | -0.001 | 0.019 | 0.019 | 0 | 0.12 | 0.24 | 0.16 | 0.13 | 0.11 | 0.08 |
| 4 | 7329426 | rs3864202 | -1.762 | -0.745 | -0.366 | 1.384 | 0.095 | 0.122 | 0.122 | 2.326 | 0.41 | 0.92 | 0.48 | 0.81 | 0.29 | 0.4 |
| 4 | 7329446 | rs6823860 | -1.257 | -0.404 | 0.528 | 1.811 | 0.026 | 0.239 | 0.239 | 1.608 | 0.07 | 0.36 | 0.08 | 0.28 | 0.05 | 0.15 |
| 4 | 7330107 | rs3892224 | -1.096 | -0.298 | 0.151 | 1.34 | -0.016 | 0.01 | 0.01 | 0 | 0.26 | 0.36 | 0.31 | 0.33 | 0.23 | 0.18 |
| 4 | 7330676 | rs16840053 | -1.065 | -0.278 | 0.064 | 0.919 | -0.044 | -0.014 | -0.014 | 6.232 | 0.2 | 0.45 | 0.19 | 0.13 | 0.14 | 0.09 |
| 4 | 7333152 | rs7657904 | -0.181 | -0.266 | -0.505 | 0.289 | 0 | -0.001 | -0.001 | 0.007 | 0.45 | 0.42 | 0.42 | 0.73 | 0.37 | 0.47 |
| 4 | 7334424 | rs3936616 | -0.39 | -0.826 | -0.272 | 0.24 | -0.031 | -0.018 | -0.018 | 0.042 | 0.12 | 0.23 | 0.08 | 0.06 | 0.05 | 0.09 |
| 4 | 7334778 | rs4689710 | -1.398 | -0.517 | -0.704 | 0.075 | -0.002 | 0.006 | 0.006 | 0 | 0.47 | 0.41 | 0.38 | 0.52 | 0.48 | 0.37 |
| 4 | 7335580 | rs7689486 | -0.717 | -0.754 | -0.737 | 0.146 | 0 | -0.018 | -0.018 | 0 | 0.27 | 0.32 | 0.18 | 0.11 | 0.25 | 0.21 |
| 4 | 7335948 | rs16840065 | 0.757 | -0.715 | -0.911 | 0.263 | -0.015 | -0.082 | -0.082 | 0 | 0.12 | 0.02 | 0.16 | 0.4 | 0.16 | 0.11 |
| 4 | 7338619 | rs3907647 | -1.976 | -1.006 | -0.802 | 0.512 | 0.001 | 0 | 0 | 0 | 0.5 | 0.53 | 0.39 | 0.43 | 0.46 | 0.4 |
| 4 | 7338852 | rs7667847 | -0.228 | -1.025 | -0.815 | 0.453 | -0.053 | -0.033 | -0.033 | 0 | 0.22 | 0.37 | 0.14 | 0.11 | 0.11 | 0.16 |
| 4 | 7339378 | rs10005640 | -1.241 | -0.712 | -0.598 | 0.424 | 0.004 | 0.007 | 0.007 | 0 | 0.51 | 0.57 | 0.4 | 0.56 | 0.46 | 0.41 |
| 4 | 7339720 | rs16840080 | -1.377 | -0.714 | -0.628 | 0.423 | 0.004 | 0.007 | 0.007 | 0 | 0.51 | 0.58 | 0.4 | 0.56 | 0.46 | 0.42 |
| 4 | 7341243 | rs10011842 | -0.448 | -0.618 | -0.759 | 0.447 | -0.007 | 0.004 | 0.004 | 0 | 0.33 | 0.35 | 0.29 | 0.39 | 0.27 | 0.25 |
| 4 | 7347203 | rs13105398 | -0.067 | -0.428 | -0.875 | 0.437 | -0.014 | 0.012 | 0.012 | 0.788 | 0.25 | 0.35 | 0.27 | 0.32 | 0.21 | 0.16 |
| 4 | 7348593 | rs11724125 | -0.123 | 0.053 | -1.648 | 0.34 | -0.042 | 0.078 | 0.078 | 0 | 0.19 | 0.06 | 0.12 | 0.04 | 0.23 | 0.18 |
| 4 | 7350677 | rs11734491 | -0.016 | 0.134 | -1.618 | 0.389 | -0.019 | -0.006 | -0.006 | 0 | 0.18 | 0.11 | 0.12 | 0.18 | 0.22 | 0.2 |
| 4 | 7352477 | rs7681365 | -0.746 | -0.233 | -1.818 | 0.093 | 0.015 | 0.005 | 0.005 | 1.17 | 0.49 | 0.66 | 0.4 | 0.52 | 0.44 | 0.37 |
| 4 | 7353052 | rs13110208 | 0.149 | -0.169 | -2.345 | -0.247 | -0.036 | 0.088 | 0.088 | 0 | 0.39 | 0.14 | 0.34 | 0.21 | 0.54 | 0.53 |
| 4 | 7362054 | rs3916114 | -0.771 | -0.484 | -2.105 | -0.102 | 0.025 | 0.049 | 0.049 | 0 | 0.24 | 0.12 | 0.27 | 0.12 | 0.16 | 0.22 |
| 4 | 7362647 | rs17757615 | -0.551 | -0.416 | -2.088 | 0.196 | -0.007 | -0.016 | -0.016 | 0.478 | 0.09 | 0.01 | 0.08 | 0.07 | 0.06 | 0.05 |
| 4 | 7363007 | rs10937807 | 0.211 | -0.442 | -1.769 | 0.149 | -0.012 | -0.01 | -0.01 | 0 | 0.61 | 0.56 | 0.53 | 0.28 | 0.65 | 0.49 |
| 4 | 7363952 | rs17814167 | -0.1 | 0.507 | -2.137 | 0.324 | -0.097 | 0.047 | 0.047 | 0 | 0.3 | 0.12 | 0.4 | 0.16 | 0.47 | 0.28 |
| 4 | 7366947 | rs12500910 | -1.017 | 0.074 | -1.562 | 1.115 | 0.016 | -0.045 | -0.045 | 0.277 | 0.12 | 0.03 | 0.14 | 0.25 | 0.11 | 0.11 |
| 4 | 7369230 | rs36022449 | -0.979 | -0.081 | -0.523 | 1.348 | -0.017 | 0.099 | 0.099 | 0 | 0.18 | 0.4 | 0.15 | 0.29 | 0.19 | 0.3 |
| 4 | 7372806 | rs13126941 | 0.889 | 0.221 | -1.127 | 0.652 | -0.07 | 0.145 | 0.145 | 0.821 | 0.44 | 0.11 | 0.41 | 0.18 | 0.45 | 0.34 |
| 4 | 7373836 | rs2878610 | 0.662 | 0.338 | -0.104 | 1.109 | -0.006 | 0.003 | 0.003 | 1.578 | 0.62 | 0.46 | 0.58 | 0.38 | 0.59 | 0.52 |
| 4 | 7373948 | rs3864206 | 0.238 | 0.268 | -0.578 | 0.786 | 0.011 | 0.02 | 0.02 | 0.031 | 0.68 | 0.74 | 0.67 | 0.94 | 0.71 | 0.7 |
| 4 | 7376694 | rs3864210 | 0.35 | 0.297 | -0.557 | 0.81 | 0.01 | 0.019 | 0.019 | 0 | 0.67 | 0.72 | 0.66 | 0.94 | 0.69 | 0.68 |
| 4 | 7380808 | rs3864218 | 0.533 | 0.699 | -0.247 | 1.24 | -0.004 | 0.044 | 0.044 | 0 | 0.68 | 0.55 | 0.63 | 0.44 | 0.66 | 0.64 |
| 4 | 7381431 | rs3864220 | -1.235 | 0.711 | -0.304 | 1.853 | 0.01 | 0.194 | 0.194 | 0 | 0.1 | 0.22 | 0.15 | 0.57 | 0.13 | 0.2 |
| 4 | 7385168 | rs3905843 | 0.692 | 0.751 | -1.042 | 1.554 | -0.006 | -0.071 | -0.071 | 0.844 | 0.79 | 0.9 | 0.89 | 0.43 | 0.85 | 0.68 |
| 4 | 7390415 | rs4689719 | -0.817 | 0.908 | -0.862 | 1.084 | -0.002 | -0.002 | -0.002 | 0.023 | 0.3 | 0.26 | 0.34 | 0.52 | 0.29 | 0.36 |
| 4 | 7390448 | rs4689721 | -0.817 | 0.908 | -0.862 | 1.084 | -0.002 | -0.002 | -0.002 | 0 | 0.3 | 0.26 | 0.34 | 0.52 | 0.29 | 0.36 |
| 4 | 7390715 | rs13142618 | -0.819 | 0.91 | -0.839 | 1.081 | -0.014 | -0.031 | -0.031 | 0 | 0.28 | 0.2 | 0.33 | 0.52 | 0.29 | 0.36 |
| 4 | 7391471 | rs12641352 | 0.77 | 0.509 | -1.832 | 0.189 | 0.045 | 0.112 | 0.112 | 1.19 | 0.35 | 0.14 | 0.19 | 0.09 | 0.33 | 0.24 |
| 4 | 7393898 | rs6851949 | -0.407 | 0.756 | -1.84 | 0.425 | -0.059 | -0.122 | -0.122 | 0.16 | 0.29 | 0.1 | 0.33 | 0.5 | 0.29 | 0.39 |
| 4 | 7395823 | rs7356467 | -0.575 | 1.486 | -0.996 | 0.009 | 0.058 | -0.066 | -0.066 | 0 | 0.15 | 0.28 | 0.21 | 0.03 | 0.28 | 0.15 |
| 4 | 7399054 | rs9994834 | -0.763 | 1.493 | -0.998 | 0.475 | 0.019 | 0.001 | 0.001 | 0 | 0.12 | 0.14 | 0.23 | 0.11 | 0.2 | 0.15 |
| 4 | 7399225 | rs9997426 | -1.295 | 1.501 | -0.777 | 0.472 | 0.029 | 0.003 | 0.003 | 0 | 0.13 | 0.19 | 0.3 | 0.13 | 0.25 | 0.16 |
| 4 | 7399604 | rs16840236 | 1.201 | 1.384 | -1.031 | 0.742 | -0.011 | -0.036 | -0.036 | 0 | 0.06 | 0 | 0.02 | 0.14 | 0.04 | 0.15 |
| 4 | 7400117 | rs6827958 | -1.63 | 1.084 | -0.478 | 0.654 | -0.02 | 0.108 | 0.108 | 1.478 | 0.23 | 0.43 | 0.32 | 0.47 | 0.17 | 0.33 |
| 4 | 7401208 | rs4689726 | 1.752 | 0.839 | -1.3 | -0.19 | -0.019 | 0.241 | 0.241 | 2.272 | 0.36 | 0.03 | 0.26 | 0.05 | 0.35 | 0.27 |
| 4 | 7401468 | rs6830500 | 0.822 | 0.861 | -1.068 | -0.003 | 0.007 | 0.185 | 0.185 | 0.699 | 0.51 | 0.15 | 0.44 | 0.2 | 0.59 | 0.53 |
| 4 | 7414226 | rs6446597 | -0.39 | 0.933 | -0.396 | 0.43 | -0.018 | 0.014 | 0.014 | 0.214 | 0.3 | 0.46 | 0.5 | 0.63 | 0.21 | 0.27 |
| 4 | 7415038 | rs4558873 | 0.602 | 1.564 | -0.543 | 0.667 | -0.005 | -0.001 | -0.001 | 1.441 | 0.75 | 0.79 | 0.8 | 0.7 | 0.64 | 0.57 |
| 4 | 7419276 | rs6837589 | 0.546 | 0.931 | -0.424 | 0.851 | -0.015 | 0.01 | 0.01 | 0.017 | 0.81 | 0.77 | 0.83 | 0.82 | 0.83 | 0.84 |
| 4 | 7419553 | rs4689739 | 1.513 | 2.575 | -0.615 | 1.291 | 0.023 | 0.022 | 0.022 | 0 | 0.72 | 0.65 | 0.57 | 0.42 | 0.6 | 0.55 |
| 4 | 7419924 | rs10022359 | -0.131 | 2.474 | -0.534 | 0.951 | -0.013 | 0.024 | 0.024 | 0 | 0.22 | 0.31 | 0.2 | 0.29 | 0.17 | 0.25 |
| 4 | 7420184 | rs13105690 | 0.806 | 2.428 | -1.094 | 1.051 | 0.016 | -0.024 | -0.024 | 3.637 | 0.66 | 0.76 | 0.69 | 0.54 | 0.73 | 0.62 |
| 4 | 7420575 | rs7682286 | 1.863 | 3.047 | -0.64 | 1.478 | 0.023 | 0.005 | 0.005 | 0 | 0.71 | 0.64 | 0.52 | 0.34 | 0.56 | 0.52 |
| 4 | 7420785 | rs4292336 | 0.1 | 2.829 | -0.608 | 1.464 | -0.006 | 0.013 | 0.013 | 0 | 0.83 | 0.82 | 0.84 | 0.82 | 0.84 | 0.84 |
| 4 | 7421007 | rs7663695 | 0.93 | 2.689 | -0.864 | 1.027 | -0.024 | -0.019 | -0.019 | 0.338 | 0.08 | 0.04 | 0.14 | 0.09 | 0.14 | 0.06 |
| 4 | 7422453 | rs7660356 | 1.319 | 2.396 | -0.761 | 0.797 | -0.005 | 0.002 | 0.002 | 0 | 0.64 | 0.63 | 0.44 | 0.33 | 0.47 | 0.48 |
| 4 | 7423842 | rs11731502 | -0.183 | 2.085 | -0.681 | 0.768 | -0.001 | 0.012 | 0.012 | 0 | 0.19 | 0.13 | 0.11 | 0.06 | 0.23 | 0.12 |
| 4 | 7423851 | rs7693497 | -0.83 | 2.374 | -0.543 | 1.472 | 0.042 | 0.123 | 0.123 | 0 | 0.08 | 0.2 | 0.19 | 0.32 | 0.12 | 0.25 |
| 4 | 7424945 | rs9999662 | -0.968 | 1.821 | -0.898 | 0.793 | 0.011 | -0.004 | -0.004 | 0.017 | 0.48 | 0.66 | 0.42 | 0.52 | 0.5 | 0.37 |
| 4 | 7426937 | rs12508214 | 0.953 | 1.748 | -0.319 | 0.847 | 0.009 | 0.005 | 0.005 | 0 | 0.54 | 0.43 | 0.38 | 0.47 | 0.35 | 0.44 |
| 4 | 7427899 | rs4025904 | -0.647 | 2.013 | -0.257 | 1.309 | -0.003 | 0.003 | 0.003 | 0 | 0.06 | 0.04 | 0.07 | 0.22 | 0.08 | 0.23 |
| 4 | 7428355 | rs10023533 | 0.531 | 2.406 | 0.164 | 1.246 | 0.08 | 0.095 | 0.095 | 0 | 0.91 | 0.74 | 0.8 | 0.83 | 0.84 | 0.96 |

| | | | | | | | | | | | | | | | | |
|---|---|---|---|---|---|---|---|---|---|---|---|---|---|---|---|---|
| 4 | 7431266 | rs4458454 | 0.76 | 2.404 | -0.108 | 0.779 | 0 | -0.001 | -0.001 | 0 | 0.9 | 0.87 | 0.87 | 0.94 | 0.84 | 0.79 |
| 4 | 7432402 | rs6854437 | 0.269 | 2.022 | -0.33 | 0.51 | -0.008 | -0.023 | -0.023 | 0 | 0.37 | 0.23 | 0.29 | 0.44 | 0.34 | 0.37 |
| 4 | 7434017 | rs10033032 | -0.003 | 1.85 | -0.155 | 0.604 | 0.017 | 0.005 | 0.005 | 0 | 0.08 | 0.09 | 0.11 | 0.11 | 0.18 | 0.07 |
| 4 | 7434607 | rs10014338 | -0.003 | 1.85 | -0.155 | 0.602 | 0.017 | 0.004 | 0.004 | 0 | 0.08 | 0.09 | 0.11 | 0.11 | 0.18 | 0.07 |
| 4 | 7435721 | rs6850206 | 1.759 | 1.081 | -0.229 | 0.183 | 0.07 | 0.115 | 0.115 | 4.052 | 0.31 | 0.1 | 0.14 | 0.08 | 0.19 | 0.14 |
| 4 | 7437059 | rs9997955 | 0.504 | 1.238 | 0.294 | 0.35 | 0.015 | 0.067 | 0.067 | 0 | 0.79 | 0.66 | 0.63 | 0.58 | 0.74 | 0.77 |
| 4 | 7438352 | rs7660079 | -0.825 | 1.231 | 0.241 | 0.17 | -0.005 | -0.021 | -0.021 | 0 | 0.58 | 0.57 | 0.73 | 0.81 | 0.64 | 0.82 |
| 4 | 7439376 | rs13435654 | 1.693 | 1.019 | -0.408 | 0.283 | 0.042 | 0.048 | 0.048 | 5.93 | 0.26 | 0.13 | 0.12 | 0.11 | 0.17 | 0.14 |
| 4 | 7440206 | rs13117409 | 1.043 | 1.476 | -0.997 | 0.298 | -0.083 | -0.134 | -0.134 | 0 | 0.29 | 0.02 | 0.23 | 0.42 | 0.28 | 0.36 |
| 4 | 7441323 | rs4234805 | -0.378 | 1.681 | -0.16 | 0.845 | 0.098 | 0.071 | 0.071 | 0 | 0.91 | 0.77 | 0.85 | 0.88 | 0.83 | 0.97 |
| 4 | 7442126 | rs4689748 | 0.052 | 1.343 | -0.611 | 0.457 | 0 | 0.004 | 0.004 | 1.448 | 0.49 | 0.5 | 0.45 | 0.46 | 0.51 | 0.6 |
| 4 | 7445048 | rs4356913 | 0.88 | 2.378 | 0.195 | 1.13 | 0.079 | 0.125 | 0.125 | 0.965 | 0.76 | 0.59 | 0.55 | 0.55 | 0.67 | 0.72 |
| 4 | 7448581 | rs4443286 | -1.311 | 1.807 | -0.201 | 0.739 | 0.011 | 0.007 | 0.007 | 0.75 | 0.19 | 0.22 | 0.4 | 0.24 | 0.25 | 0.4 |
| 4 | 7459344 | rs7694518 | 1.043 | 1.368 | 0.042 | 0.309 | 0.004 | -0.016 | -0.016 | 2.078 | 0.81 | 0.67 | 0.78 | 0.88 | 0.82 | 0.77 |
| 4 | 7460233 | rs4639074 | 1.508 | 1.302 | -0.144 | 0.632 | -0.002 | 0.012 | 0.012 | 0 | 0.87 | 0.84 | 0.88 | 0.82 | 0.9 | 0.73 |
| 4 | 7460789 | rs4380522 | 0.768 | 0.857 | -0.572 | -0.573 | -0.018 | 0.129 | 0.129 | 0.02 | 0.56 | 0.29 | 0.48 | 0.08 | 0.57 | 0.44 |
| 4 | 7463304 | rs4599414 | 0.145 | 0.777 | -0.589 | -0.632 | -0.057 | 0.146 | 0.146 | 0 | 0.63 | 0.87 | 0.55 | 0.09 | 0.57 | 0.35 |
| 4 | 7465277 | rs7654086 | 0.604 | 0.735 | -0.457 | -0.691 | -0.011 | 0.077 | 0.077 | 0 | 0.58 | 0.71 | 0.53 | 0.08 | 0.56 | 0.35 |
| 4 | 7466466 | rs4411993 | 0.524 | 0.664 | -0.654 | -0.337 | -0.005 | -0.052 | -0.052 | 0 | 0.88 | 0.96 | 0.9 | 0.76 | 0.88 | 0.72 |
| 4 | 7466980 | rs4613570 | 1.54 | 0.483 | -0.037 | -0.671 | 0.002 | -0.056 | -0.056 | 0 | 0.4 | 0.42 | 0.3 | 0.01 | 0.38 | 0.18 |
| 4 | 7467316 | rs4689126 | 1.401 | 0.489 | 0.517 | 0.005 | -0.011 | 0.002 | 0.002 | 0 | 0.68 | 0.57 | 0.69 | 0.72 | 0.74 | 0.57 |
| 4 | 7473205 | rs4495038 | -0.595 | 0.865 | 0.525 | -0.193 | 0.006 | -0.043 | -0.043 | 0 | 0.17 | 0.2 | 0.23 | 0.03 | 0.18 | 0.17 |
| 4 | 7475424 | rs7683874 | -1.296 | 0.592 | 0.832 | 0.441 | 0.012 | 0.183 | 0.183 | 0.077 | 0.08 | 0.26 | 0.2 | 0.24 | 0.07 | 0.26 |
| 4 | 7478631 | rs16840358 | -1.258 | 0.545 | 0.704 | 0.417 | -0.006 | 0.118 | 0.118 | 0 | 0.1 | 0.22 | 0.2 | 0.24 | 0.07 | 0.26 |
| 4 | 7480400 | rs10937822 | -0.341 | 0.613 | 0.325 | 0.505 | -0.002 | -0.005 | -0.005 | 1.333 | 0.07 | 0.04 | 0.17 | 0.24 | 0.06 | 0.25 |
| 4 | 7480422 | rs10937823 | -0.341 | 0.613 | 0.325 | 0.505 | -0.002 | -0.005 | -0.005 | 0 | 0.07 | 0.04 | 0.17 | 0.24 | 0.06 | 0.25 |
| 4 | 7481564 | rs13353636 | -1.832 | 0.505 | 0.701 | 0.346 | -0.006 | 0.078 | 0.078 | 0 | 0.16 | 0.3 | 0.25 | 0.29 | 0.14 | 0.27 |
| 4 | 7483346 | rs11930924 | -0.433 | 0.588 | 0.583 | 0.175 | 0.001 | -0.005 | -0.005 | 0 | 0.07 | 0.13 | 0.05 | 0.05 | 0.07 | 0.02 |
| 4 | 7489235 | rs11737430 | -0.667 | 0.829 | 0.856 | 0.64 | 0 | 0.127 | 0.127 | 0 | 0.08 | 0.18 | 0.19 | 0.26 | 0.07 | 0.25 |
| 4 | 7489534 | rs4234818 | -0.268 | 0.85 | 0.712 | -0.24 | -0.041 | -0.001 | -0.001 | 0 | 0.38 | 0.21 | 0.48 | 0.69 | 0.45 | 0.54 |
| 4 | 7491757 | rs10032900 | 0.882 | 0.817 | 0.836 | -0.495 | -0.005 | -0.085 | -0.085 | 0 | 0.47 | 0.48 | 0.3 | 0.05 | 0.45 | 0.21 |
| 4 | 7492414 | rs6820154 | -0.463 | 0.749 | 0.786 | -0.268 | 0.009 | 0.096 | 0.096 | 0 | 0.09 | 0.19 | 0.19 | 0.26 | 0.09 | 0.25 |
| 4 | 7495063 | rs12506309 | -0.172 | 0.356 | 0.461 | -1.064 | -0.012 | 0.219 | 0.219 | 0.165 | 0.46 | 0.14 | 0.64 | 0.85 | 0.57 | 0.77 |
| 4 | 7495484 | rs6857793 | -0.815 | 0.225 | 1.114 | -1.164 | -0.003 | -0.049 | -0.049 | 0.1 | 0.55 | 0.49 | 0.72 | 0.89 | 0.63 | 0.78 |
| 4 | 7496857 | rs4689131 | -1.56 | -0.001 | 1.091 | -1.73 | 0.034 | 0.11 | 0.11 | 0 | 0.64 | 0.81 | 0.79 | 0.99 | 0.69 | 0.86 |
| 4 | 7498864 | rs9647418 | 0.715 | 0.075 | 1.513 | -1.545 | -0.013 | -0.017 | -0.017 | 0.065 | 0.91 | 0.61 | 0.94 | 0.99 | 0.99 | 1 |
| 4 | 7501447 | rs7692314 | -1.86 | 0.126 | 1.264 | -1.481 | -0.018 | -0.002 | -0.002 | 1.502 | 0.13 | 0.53 | 0.11 | 0.06 | 0.08 | 0.03 |
| 4 | 7508309 | rs6446605 | -1.219 | 0.12 | 1.39 | -1.459 | 0.033 | 0.012 | 0.012 | 3.309 | 0.1 | 0.42 | 0.09 | 0.05 | 0.06 | 0.02 |
| 4 | 7508446 | rs4689767 | -0.414 | 0.152 | 1.091 | -1.606 | 0.054 | 0.268 | 0.268 | 0.107 | 0.49 | 0.17 | 0.65 | 0.92 | 0.59 | 0.84 |
| 4 | 7509758 | rs10000185 | -2.362 | 0.287 | 1.335 | -1.733 | 0.022 | -0.069 | -0.069 | 0.659 | 0.11 | 0.55 | 0.1 | 0.01 | 0.08 | 0.03 |
| 4 | 7513983 | rs13128906 | 0.238 | 0.729 | 0.902 | -1.701 | -0.002 | 0 | 0 | 0.105 | 0.27 | 0.31 | 0.32 | 0.78 | 0.31 | 0.42 |
| 4 | 7515705 | rs12650698 | 0.409 | 0.694 | 0.735 | -1.69 | -0.048 | -0.04 | -0.04 | 0 | 0.25 | 0.1 | 0.31 | 0.73 | 0.29 | 0.41 |
| 4 | 7517179 | rs4487342 | -1.23 | 0.604 | 1.062 | -2.068 | 0.013 | -0.003 | -0.003 | 0 | 0.58 | 0.66 | 0.73 | 0.96 | 0.62 | 0.83 |
| 4 | 7521802 | rs13117257 | 0.896 | 0.417 | 0.732 | -2.162 | -0.004 | -0.088 | -0.088 | 0 | 0.39 | 0.39 | 0.24 | 0.02 | 0.33 | 0.11 |
| 4 | 7522239 | rs4689773 | -0.266 | 0.508 | 0.299 | -1.996 | -0.001 | -0.002 | -0.002 | 0.263 | 0.29 | 0.15 | 0.36 | 0.22 | 0.3 | 0.37 |
| 4 | 7523273 | rs744356 | -1.48 | 0.32 | 0.44 | -2.073 | 0.009 | -0.035 | -0.035 | 0 | 0.63 | 0.66 | 0.8 | 0.99 | 0.7 | 0.9 |
| 4 | 7523364 | rs744355 | -0.902 | 0.712 | 0.435 | -1.84 | 0 | -0.001 | -0.001 | 0 | 0.34 | 0.34 | 0.43 | 0.25 | 0.36 | 0.41 |
| 4 | 7525739 | rs6840716 | 1.275 | -0.238 | 0.085 | -2.6 | -0.023 | -0.075 | -0.075 | 0 | 0.77 | 0.62 | 0.83 | 0.9 | 0.8 | 0.76 |
| 4 | 7529367 | rs4429728 | -0.326 | -0.008 | -0.453 | -2.869 | -0.005 | 0.056 | 0.056 | 0.009 | 0.48 | 0.62 | 0.52 | 0.92 | 0.5 | 0.64 |
| 4 | 7533041 | rs7694823 | 0.977 | 0.133 | -0.092 | -2.341 | 0.004 | -0.048 | -0.048 | 0.412 | 0.92 | 0.81 | 0.89 | 0.99 | 0.9 | 0.93 |
| 4 | 7533234 | rs10007474 | -0.583 | 0.149 | -0.069 | -2.281 | 0.007 | -0.03 | -0.03 | 0 | 0.06 | 0.15 | 0.09 | 0.01 | 0.07 | 0.06 |
| 4 | 7533674 | rs10021708 | -0.331 | 0.058 | -0.131 | -2.318 | -0.015 | -0.035 | -0.035 | 0 | 0.07 | 0.11 | 0.09 | 0.02 | 0.06 | 0.09 |
| 4 | 7534603 | rs6812745 | 0.577 | -0.352 | -0.537 | -3.027 | 0.002 | -0.039 | -0.039 | 0 | 0.24 | 0.21 | 0.23 | 0 | 0.19 | 0.13 |
| 4 | 7535214 | rs6446607 | -0.712 | -0.128 | -1.192 | -3.055 | 0.075 | 0.266 | 0.266 | 0 | 0.51 | 0.81 | 0.45 | 0.1 | 0.38 | 0.34 |
| 4 | 7535574 | rs16840471 | -0.348 | 0.459 | -0.472 | -2.326 | -0.009 | -0.026 | -0.026 | 0 | 0.12 | 0.19 | 0.06 | 0.07 | 0.08 | 0.09 |
| 4 | 7536746 | rs7696497 | 0.107 | 1.136 | -1.044 | -2.157 | -0.083 | 0.1 | 0.1 | 2.27 | 0.28 | 0.1 | 0.4 | 0.89 | 0.39 | 0.6 |
| 4 | 7544610 | rs10028107 | -0.087 | 0.707 | -0.152 | -1.777 | -0.002 | -0.032 | -0.032 | 0 | 0.09 | 0.13 | 0.07 | 0 | 0.09 | 0.01 |
| 4 | 7546598 | rs4689138 | 0.695 | 0.799 | 0.543 | -2.028 | 0.027 | -0.056 | -0.056 | 0 | 0.78 | 0.51 | 0.78 | 0.88 | 0.78 | 0.79 |
| 4 | 7547490 | rs10470724 | 0.772 | 0.805 | 1.349 | -2.093 | 0.02 | -0.028 | -0.028 | 0 | 0.93 | 0.75 | 0.9 | 0.99 | 0.96 | 0.9 |
| 4 | 7547547 | rs11721353 | 1.908 | 0.686 | 1.625 | -2.155 | -0.002 | 0.036 | 0.036 | 0.226 | 0.67 | 0.25 | 0.7 | 0.88 | 0.69 | 0.78 |
| 4 | 7549473 | rs11937027 | 0.682 | 0.8 | 2.41 | -2.109 | 0.019 | -0.032 | -0.032 | 0 | 0.83 | 0.57 | 0.86 | 0.89 | 0.83 | 0.89 |
| 4 | 7551651 | rs882266 | 1.81 | 0.814 | 2.176 | -2.128 | 0.006 | -0.032 | -0.032 | 0.053 | 0.75 | 0.34 | 0.77 | 0.89 | 0.76 | 0.79 |
| 4 | 7554721 | rs4637403 | 0.143 | 0.901 | 1.885 | -2.007 | -0.011 | 0.003 | 0.003 | 0 | 0.12 | 0.06 | 0.12 | 0.1 | 0.14 | 0.1 |
| 4 | 7564758 | rs16840505 | -1.36 | 0.866 | 2.113 | -2.009 | 0.027 | -0.046 | -0.046 | 0 | 0.2 | 0.38 | 0.19 | 0.1 | 0.21 | 0.15 |
| 4 | 7565734 | rs4290892 | -2.005 | 0.972 | 1.811 | -2.071 | 0.012 | -0.068 | -0.068 | 0.303 | 0.32 | 0.56 | 0.29 | 0.11 | 0.34 | 0.24 |
| 4 | 7570358 | rs7667548 | 0.643 | 0.925 | 1.495 | -1.905 | 0 | -0.08 | -0.08 | 0.587 | 0.88 | 0.64 | 0.9 | 1 | 0.9 | 0.95 |
| 4 | 7572718 | rs11942379 | -0.951 | 0.856 | 0.956 | -2 | -0.03 | 0.042 | 0.042 | 0 | 0.25 | 0.19 | 0.22 | 0.1 | 0.31 | 0.15 |
| 4 | 7573488 | rs4689781 | 1.05 | 0.911 | 0.199 | -2.429 | -0.018 | -0.007 | -0.007 | 0 | 0.26 | 0.1 | 0.39 | 0.72 | 0.3 | 0.48 |
| 4 | 7573950 | rs6858110 | -0.322 | 0.356 | 0.739 | -2.537 | -0.04 | -0.05 | -0.05 | 0 | 0.81 | 0.71 | 0.87 | 0.93 | 0.91 | 0.89 |

| | | | | | | | | | | | | | | | | |
|---|---|---|---|---|---|---|---|---|---|---|---|---|---|---|---|---|
| 4 | 7576024 | rs13144333 | 0.184 | 0.559 | 0.886 | -2.518 | -0.009 | 0.01 | 0.01 | 0 | 0.22 | 0.2 | 0.24 | 0.1 | 0.3 | 0.15 |
| 4 | 7576612 | rs7440772 | -0.2 | 0.65 | 0.966 | -1.994 | -0.016 | -0.027 | -0.027 | 0.271 | 0.19 | 0.27 | 0.13 | 0.12 | 0.13 | 0.14 |
| 4 | 7584560 | rs4689786 | 1.311 | 0.639 | 0.755 | -1.974 | -0.036 | -0.039 | -0.039 | 0.048 | 0.26 | 0.1 | 0.38 | 0.69 | 0.31 | 0.49 |
| 4 | 7584767 | rs10012347 | 0.162 | 0.6 | 0.951 | -2.175 | -0.006 | 0.136 | 0.136 | 0 | 0.49 | 0.3 | 0.43 | 0.13 | 0.53 | 0.32 |
| 4 | 7586924 | rs6835799 | 0.373 | 0.246 | 0.969 | -1.972 | -0.032 | -0.004 | -0.004 | 0.039 | 0.16 | 0.18 | 0.09 | 0.13 | 0.05 | 0.07 |
| 4 | 7588936 | rs4689789 | 1.16 | 0.567 | 0.699 | -1.717 | -0.009 | -0.018 | -0.018 | 0 | 0.26 | 0.19 | 0.37 | 0.66 | 0.31 | 0.51 |
| 4 | 7590048 | rs11735301 | -1.027 | 0.573 | 0.908 | -1.701 | 0.005 | 0.004 | 0.004 | 0 | 0.72 | 0.66 | 0.6 | 0.29 | 0.69 | 0.44 |
| 4 | 7592068 | rs4594727 | 1.167 | 0.569 | 0.422 | -1.673 | -0.016 | -0.03 | -0.03 | 0.061 | 0.25 | 0.13 | 0.38 | 0.69 | 0.29 | 0.5 |
| 4 | 7597452 | rs734526 | 0.338 | 0.497 | 1.012 | -1.854 | -0.002 | 0 | 0 | 0.536 | 0.33 | 0.5 | 0.44 | 0.66 | 0.34 | 0.52 |
| 4 | 7597712 | rs11933482 | 0.198 | 0.955 | 0.991 | -2.449 | -0.011 | 0.015 | 0.015 | 0 | 0.22 | 0.2 | 0.22 | 0.09 | 0.29 | 0.17 |
| 4 | 7600936 | rs7667970 | -0.092 | 0.727 | 1.033 | -2.395 | -0.003 | -0.101 | -0.101 | 0 | 0.37 | 0.46 | 0.27 | 0.04 | 0.35 | 0.21 |
| 4 | 7607757 | rs757240 | 0.404 | 0.791 | 1.059 | -2.32 | 0 | -0.07 | -0.07 | 0.05 | 0.32 | 0.39 | 0.22 | 0.04 | 0.31 | 0.19 |
| 4 | 7607924 | rs12503735 | 0.55 | 0.878 | 0.556 | -2.27 | -0.003 | 0.088 | 0.088 | 0 | 0.29 | 0.14 | 0.2 | 0.04 | 0.29 | 0.19 |
| 4 | 7612515 | rs16840640 | 0.9 | 1.009 | 0.785 | -2.072 | 0.019 | -0.042 | -0.042 | 0 | 0.08 | 0.2 | 0.08 | 0 | 0.1 | 0.01 |
| 4 | 7612772 | rs10021084 | -0.353 | 0.997 | 0.474 | -2.201 | 0.005 | -0.029 | -0.029 | 0 | 0.23 | 0.27 | 0.2 | 0.09 | 0.24 | 0.17 |
| 4 | 7615377 | rs16840670 | 0.073 | 0.877 | 0.059 | -2.335 | 0.005 | -0.098 | -0.098 | 3.105 | 0.39 | 0.41 | 0.32 | 0.05 | 0.42 | 0.2 |
| 4 | 7619653 | rs13145735 | 0.066 | 0.793 | 0.056 | -1.791 | 0.02 | -0.006 | -0.006 | 0.985 | 0.95 | 0.84 | 0.93 | 0.97 | 0.96 | 0.9 |
| 4 | 7620682 | rs10937843 | -0.356 | 0.794 | -0.601 | -1.773 | -0.003 | 0.023 | 0.023 | 0 | 0.21 | 0.15 | 0.18 | 0.09 | 0.2 | 0.17 |
| 4 | 7625979 | rs10937844 | -0.131 | 0.233 | -0.839 | -1.981 | 0.01 | 0.013 | 0.013 | 0.924 | 0.19 | 0.13 | 0.15 | 0.12 | 0.11 | 0.08 |
| 4 | 7626136 | rs4689806 | -0.522 | 0.17 | -0.827 | -2.292 | -0.02 | -0.063 | -0.063 | 0 | 0.77 | 0.68 | 0.84 | 0.96 | 0.8 | 0.84 |
| 4 | 7628406 | rs2884800 | 0.405 | 0.743 | -1.471 | -2.098 | -0.021 | -0.004 | -0.004 | 0 | 0.24 | 0.17 | 0.31 | 0.74 | 0.29 | 0.47 |
| 4 | 7629251 | rs886372 | -0.587 | 0.163 | -1.229 | -2.658 | -0.016 | -0.082 | -0.082 | 0.307 | 0.77 | 0.64 | 0.85 | 0.95 | 0.81 | 0.84 |
| 4 | 7629288 | rs757239 | -0.205 | 0.147 | -1.296 | -2.589 | -0.012 | -0.084 | -0.084 | 0 | 0.7 | 0.62 | 0.8 | 0.95 | 0.74 | 0.83 |
| 4 | 7632223 | rs4689808 | 0.764 | 0.679 | -1.665 | -1.962 | -0.049 | -0.036 | -0.036 | 0.004 | 0.19 | 0.07 | 0.34 | 0.74 | 0.24 | 0.48 |
| 4 | 7637297 | rs4689148 | 0.279 | 0.074 | -1.296 | -2.665 | 0.01 | 0.008 | 0.008 | 0 | 0.51 | 0.61 | 0.68 | 0.95 | 0.56 | 0.7 |
| 4 | 7639906 | rs2158276 | 0.122 | -0.428 | -1.289 | -1.877 | 0.017 | 0.054 | 0.054 | 0 | 0.77 | 0.9 | 0.86 | 0.95 | 0.79 | 0.83 |
| 4 | 7645289 | rs2158275 | 0.322 | -0.284 | -1.295 | -1.651 | 0.043 | 0.263 | 0.263 | 2.711 | 0.54 | 0.13 | 0.72 | 0.93 | 0.64 | 0.79 |
| 4 | 7646256 | rs11932646 | -1.47 | -0.266 | 0.112 | -0.883 | -0.021 | -0.074 | -0.074 | 0 | 0.15 | 0.73 | 0.11 | 0 | 0.07 | 0.05 |
| 4 | 7654022 | rs886366 | 1.271 | -0.317 | 0.144 | -0.872 | -0.023 | -0.101 | -0.101 | 0 | 0.83 | 0.34 | 0.89 | 1 | 0.9 | 0.92 |
| 4 | 7655855 | rs2285789 | 0.839 | -0.139 | -0.773 | -0.552 | 0.024 | 0.164 | 0.164 | 3.039 | 0.61 | 0.09 | 0.67 | 0.89 | 0.69 | 0.6 |
| 4 | 7660146 | rs11737898 | 0.187 | -0.402 | -0.013 | 0.027 | -0.025 | 0.017 | 0.017 | 0.546 | 0.7 | 0.39 | 0.64 | 0.68 | 0.76 | 0.6 |
| 4 | 7661649 | rs1158073 | -1.267 | -0.149 | 0.528 | -0.132 | 0.009 | -0.019 | -0.019 | 0 | 0.08 | 0.33 | 0.05 | 0 | 0.05 | 0.03 |
| 4 | 7665945 | rs6810814 | 0.714 | -0.081 | 0.13 | -0.231 | 0.003 | -0.058 | -0.058 | 0 | 0.81 | 0.59 | 0.75 | 0.91 | 0.8 | 0.86 |
| 4 | 7667486 | rs2214459 | 0.416 | -0.685 | -1.355 | -0.386 | 0.124 | 0.137 | 0.137 | 3.201 | 0.51 | 0.15 | 0.69 | 0.66 | 0.69 | 0.74 |
| 4 | 7669798 | rs10488964 | -0.284 | -0.458 | -1.32 | -0.628 | 0.007 | 0.074 | 0.074 | 3.32 | 0.23 | 0.11 | 0.1 | 0 | 0.16 | 0.05 |
| 4 | 7672424 | rs6816649 | 0.55 | -0.599 | -1.017 | 0.259 | 0.001 | -0.002 | -0.002 | 1.213 | 0.73 | 0.78 | 0.81 | 0.51 | 0.82 | 0.57 |
| 4 | 7672785 | rs11724048 | 0.389 | -0.174 | -0.538 | 0.723 | 0.002 | 0.147 | 0.147 | 0 | 0.77 | 0.58 | 0.71 | 0.53 | 0.77 | 0.64 |
| 4 | 7680962 | rs10013239 | 0.238 | 0.36 | -0.823 | 0.939 | -0.001 | 0.003 | 0.003 | 0 | 0.65 | 0.65 | 0.55 | 0.57 | 0.67 | 0.52 |
| 4 | 7683417 | rs7697323 | 0.376 | 0.744 | -1.047 | 0.436 | -0.011 | -0.06 | -0.06 | 0.792 | 0.43 | 0.48 | 0.33 | 0.16 | 0.37 | 0.35 |
| 4 | 7684641 | rs4689820 | -0.183 | 0.963 | -1.18 | 0.739 | 0 | -0.001 | -0.001 | 0 | 0.62 | 0.63 | 0.64 | 0.7 | 0.62 | 0.52 |
| 4 | 7685529 | rs2079169 | 0.207 | 0.773 | -1.984 | 0.735 | -0.014 | -0.037 | -0.037 | 0 | 0.37 | 0.15 | 0.37 | 0.45 | 0.38 | 0.3 |
| 4 | 7685758 | rs2359633 | 1.17 | 1.082 | -1.783 | 1.028 | -0.017 | -0.033 | -0.033 | 0 | 0.17 | 0.12 | 0.3 | 0.39 | 0.23 | 0.17 |
| 4 | 7685801 | rs2884743 | -0.599 | 0.704 | -1.35 | 0.672 | 0.006 | 0.007 | 0.007 | 0 | 0.85 | 0.84 | 0.8 | 0.75 | 0.8 | 0.81 |
| 4 | 7687588 | rs16840826 | 0.796 | 0.451 | -1.438 | 0.924 | -0.012 | -0.032 | -0.032 | 0.777 | 0.06 | 0.01 | 0.06 | 0.13 | 0.04 | 0.13 |
| 4 | 7687652 | rs1006467 | 0.745 | 0.794 | -1.679 | 0.775 | -0.047 | -0.012 | -0.012 | 0.295 | 0.2 | 0.08 | 0.21 | 0.25 | 0.31 | 0.18 |
| 4 | 7689152 | rs886378 | -0.761 | 0.68 | -1.419 | -0.253 | 0.002 | -0.015 | -0.015 | 0 | 0.62 | 0.62 | 0.72 | 0.91 | 0.63 | 0.75 |
| 4 | 7690403 | rs886376 | -1.575 | 0.566 | -1.72 | -0.361 | -0.001 | -0.014 | -0.014 | 0 | 0.77 | 0.81 | 0.81 | 0.91 | 0.76 | 0.78 |
| 4 | 7690857 | rs17381732 | -0.686 | 1.08 | -1.278 | 0.349 | -0.001 | 0 | 0 | 0 | 0.45 | 0.38 | 0.53 | 0.45 | 0.41 | 0.57 |
| 4 | 7693469 | rs4689825 | -0.41 | 1.044 | -1.157 | -0.062 | -0.001 | 0 | 0 | 0 | 0.36 | 0.31 | 0.39 | 0.3 | 0.34 | 0.49 |
| 4 | 7694472 | rs10023670 | -0.037 | 0.98 | -1.884 | -0.243 | -0.004 | 0.119 | 0.119 | 0.174 | 0.51 | 0.75 | 0.53 | 0.71 | 0.53 | 0.39 |
| 4 | 7698316 | rs2057723 | -0.605 | 1.033 | -1.673 | -0.319 | 0.012 | 0.052 | 0.052 | 0 | 0.64 | 0.76 | 0.77 | 0.91 | 0.68 | 0.74 |
| 4 | 7700712 | rs6845903 | -0.213 | 1.409 | -1.296 | 0.283 | -0.001 | 0 | 0 | 0.019 | 0.35 | 0.63 | 0.49 | 0.71 | 0.43 | 0.4 |
| 4 | 7704795 | rs2359630 | -0.668 | 1.658 | -1.269 | 0.993 | -0.007 | -0.008 | -0.008 | 0 | 0.21 | 0.17 | 0.24 | 0.31 | 0.3 | 0.29 |
| 4 | 7709880 | rs12500759 | -0.142 | 1.617 | -1.552 | 1.041 | -0.035 | -0.043 | -0.043 | 1.805 | 0.24 | 0.12 | 0.26 | 0.36 | 0.29 | 0.25 |
| 4 | 7710640 | rs766008 | -0.13 | 0.454 | -0.789 | 0.428 | -0.018 | -0.053 | -0.053 | 1.445 | 0.75 | 0.62 | 0.81 | 0.89 | 0.78 | 0.82 |
| 4 | 7712266 | rs2359610 | 0.06 | 0.444 | -1.113 | 0.806 | -0.002 | -0.022 | -0.022 | 0 | 0.81 | 0.9 | 0.88 | 0.72 | 0.81 | 0.72 |
| 4 | 7714212 | rs4689832 | 0.628 | -0.359 | -0.895 | 0.539 | 0.017 | 0.008 | 0.008 | 0 | 0.58 | 0.28 | 0.68 | 0.61 | 0.63 | 0.65 |
| 4 | 7714490 | rs4689833 | 0.121 | -0.416 | -0.961 | 0.067 | 0.024 | 0.031 | 0.031 | 0 | 0.71 | 0.79 | 0.8 | 1 | 0.73 | 0.92 |
| 4 | 7715181 | rs11726260 | 0.882 | -0.128 | -1.583 | 0.565 | -0.114 | 0 | 0 | 0.223 | 0.3 | 0.09 | 0.31 | 0.27 | 0.39 | 0.27 |
| 4 | 7716932 | rs6816604 | -1.355 | -0.461 | -0.536 | 0.966 | -0.011 | -0.053 | -0.053 | 0 | 0.91 | 0.84 | 0.96 | 1 | 0.96 | 0.98 |
| 4 | 7717012 | rs2285781 | 0.669 | -0.603 | -0.972 | 0.709 | 0.022 | 0.118 | 0.118 | 0 | 0.46 | 0.13 | 0.42 | 0.26 | 0.57 | 0.33 |
| 4 | 7717629 | rs16840900 | 0.588 | -0.875 | -1.323 | 0.962 | -0.04 | -0.17 | -0.17 | 0.092 | 0.23 | 0.01 | 0.33 | 0.52 | 0.24 | 0.43 |
| 4 | 7718534 | rs4689835 | -0.328 | -1.143 | -0.515 | 0.491 | -0.06 | -0.076 | -0.076 | 0 | 0.79 | 0.52 | 0.82 | 0.88 | 0.87 | 0.82 |
| 4 | 7718802 | rs4689836 | 0.302 | -0.801 | -1.027 | 0.72 | -0.043 | -0.089 | -0.089 | 0 | 0.25 | 0.04 | 0.35 | 0.62 | 0.28 | 0.47 |
| 4 | 7720077 | rs2285780 | 0.492 | -0.46 | -0.377 | 0.741 | -0.02 | -0.009 | -0.009 | 0 | 0.11 | 0.27 | 0.09 | 0.1 | 0.08 | 0.17 |
| 4 | 7720834 | rs4689838 | 0.309 | 0.022 | -1.046 | 0.328 | -0.073 | 0.061 | 0.061 | 0 | 0.34 | 0.17 | 0.35 | 0.21 | 0.49 | 0.22 |
| 4 | 7722458 | rs2097769 | -0.235 | -0.168 | -0.218 | 0.894 | -0.015 | 0.034 | 0.034 | 0 | 0.1 | 0.25 | 0.05 | 0.14 | 0.06 | 0.15 |
| 4 | 7726847 | rs2285777 | -0.057 | -0.029 | -0.435 | 0.792 | 0 | -0.008 | -0.008 | 0 | 0.36 | 0.36 | 0.28 | 0.44 | 0.42 | 0.44 |
| 4 | 7729704 | rs7679804 | -0.405 | 0.641 | -0.772 | 0.446 | -0.02 | 0.039 | 0.039 | 0.174 | 0.74 | 0.84 | 0.86 | 0.85 | 0.75 | 0.81 |

| | | | | | | | | | | | | | | | | |
|---|---|---|---|---|---|---|---|---|---|---|---|---|---|---|---|---|
| 4 | 7730392 | rs4689157 | 1.301 | 0.543 | -0.73 | 0.437 | 0.004 | 0.003 | 0.003 | 2.384 | 0.19 | 0.15 | 0.13 | 0.17 | 0.15 | 0.15 |
| 4 | 7730560 | rs17466657 | -0.035 | 0.587 | -0.87 | 0.347 | -0.093 | -0.061 | -0.061 | 0 | 0.32 | 0.08 | 0.51 | 0.36 | 0.37 | 0.39 |
| 4 | 7732805 | rs6821770 | 0.709 | 0.494 | -0.562 | 0.434 | 0 | -0.003 | -0.003 | 0.129 | 0.19 | 0.19 | 0.41 | 0.3 | 0.17 | 0.26 |
| 4 | 7733843 | rs12233824 | -0.533 | 0.436 | 0.106 | 0.368 | 0.001 | 0.002 | 0.002 | 0 | 0.54 | 0.41 | 0.68 | 0.58 | 0.52 | 0.57 |
| 4 | 7736455 | rs6843811 | -0.433 | 0.424 | 0.15 | 0.385 | 0.018 | 0.002 | 0.002 | 0 | 0.52 | 0.39 | 0.63 | 0.44 | 0.53 | 0.51 |
| 4 | 7736890 | rs12499617 | 0.221 | 0.695 | -0.51 | 0.312 | -0.016 | -0.022 | -0.022 | 0.014 | 0.41 | 0.22 | 0.53 | 0.42 | 0.39 | 0.49 |
| 4 | 7737137 | rs4689845 | 1.35 | 0.143 | -0.496 | 0.345 | 0.007 | 0.003 | 0.003 | 0.194 | 0.16 | 0.12 | 0.12 | 0.17 | 0.14 | 0.15 |
| 4 | 7738369 | rs886374 | -1.035 | 0.367 | -0.659 | 0.728 | -0.003 | 0.004 | 0.004 | 0 | 0.71 | 0.67 | 0.78 | 0.69 | 0.7 | 0.72 |
| 4 | 7738730 | rs6840685 | -0.72 | 0.819 | -0.998 | 0.701 | -0.009 | -0.024 | -0.024 | 0 | 0.45 | 0.23 | 0.57 | 0.49 | 0.49 | 0.54 |
| 4 | 7743135 | rs17382228 | -0.375 | 1.275 | -1.506 | 0.924 | -0.107 | -0.137 | -0.137 | 0 | 0.24 | 0.05 | 0.48 | 0.42 | 0.34 | 0.31 |
| 4 | 7743283 | rs13442 | -0.624 | 1.013 | -0.966 | 0.566 | -0.003 | -0.006 | -0.006 | 0 | 0.38 | 0.34 | 0.46 | 0.43 | 0.39 | 0.41 |
| 4 | 7744367 | rs12498937 | -0.222 | 0.572 | -0.557 | 0.127 | -0.022 | -0.038 | -0.038 | 0 | 0.84 | 0.56 | 0.9 | 0.91 | 0.88 | 0.84 |
| 4 | 7746390 | rs2359608 | 0.443 | 0.511 | -0.806 | -0.057 | -0.022 | -0.054 | -0.054 | 0 | 0.72 | 0.44 | 0.82 | 0.86 | 0.78 | 0.74 |
| 4 | 7746820 | rs4689160 | 0.484 | 0.786 | -1.122 | 0.037 | 0.001 | -0.025 | -0.025 | 0 | 0.58 | 0.48 | 0.74 | 0.75 | 0.58 | 0.57 |
| 4 | 7751894 | rs4689848 | 0.577 | 0.792 | -1.695 | 0.39 | -0.048 | -0.088 | -0.088 | 0.687 | 0.33 | 0.07 | 0.42 | 0.41 | 0.33 | 0.37 |
| 4 | 7752091 | rs17466832 | 0.984 | 1.066 | -1.805 | -0.066 | -0.069 | 0.007 | 0.007 | 0 | 0.16 | 0.02 | 0.08 | 0.11 | 0.21 | 0.19 |
| 4 | 7752580 | rs4689851 | -0.206 | 0.708 | -0.915 | -0.292 | -0.001 | -0.043 | -0.043 | 0 | 0.6 | 0.51 | 0.76 | 0.8 | 0.58 | 0.7 |
| 4 | 7753589 | rs9968354 | -0.591 | 0.705 | -0.873 | -0.043 | 0.001 | 0.009 | 0.009 | 0 | 0.5 | 0.55 | 0.31 | 0.31 | 0.55 | 0.44 |
| 4 | 7755379 | rs17466860 | 0.557 | 0.934 | -1.012 | 0.256 | -0.021 | 0.006 | 0.006 | 0 | 0.17 | 0.13 | 0.12 | 0.13 | 0.24 | 0.19 |
| 4 | 7756195 | rs12501350 | -0.106 | 0.607 | -1.088 | 0.582 | -0.029 | -0.045 | -0.045 | 0.812 | 0.26 | 0.13 | 0.29 | 0.36 | 0.32 | 0.37 |
| 4 | 7757650 | rs17382358 | 0.153 | 0.764 | -1.123 | 0.244 | -0.007 | 0.018 | 0.018 | 0 | 0.33 | 0.24 | 0.35 | 0.3 | 0.38 | 0.28 |
| 4 | 7759804 | rs11729332 | 0.884 | 0.813 | -1.087 | 0.185 | -0.028 | 0.067 | 0.067 | 0 | 0.25 | 0.09 | 0.17 | 0.11 | 0.31 | 0.11 |
| 4 | 7761114 | rs2285771 | 0.991 | 0.325 | -0.751 | 0.302 | -0.024 | 0.157 | 0.157 | 0 | 0.16 | 0.31 | 0.16 | 0.47 | 0.1 | 0.26 |
| 4 | 7761211 | rs2285770 | 1.32 | 0.292 | -0.753 | 0.288 | 0.013 | -0.047 | -0.047 | 0 | 0.14 | 0.05 | 0.14 | 0.47 | 0.1 | 0.26 |
| 4 | 7761286 | rs2285769 | 1.243 | 0.272 | -0.763 | 0.271 | 0.014 | -0.05 | -0.05 | 0.148 | 0.15 | 0.05 | 0.14 | 0.47 | 0.1 | 0.26 |
| 4 | 7762502 | rs11736215 | 0.887 | 0.157 | -0.536 | -0.143 | -0.034 | 0.134 | 0.134 | 0 | 0.39 | 0.14 | 0.37 | 0.12 | 0.45 | 0.2 |
| 4 | 7762848 | rs2285766 | 1.57 | 0.138 | -0.369 | 0.098 | 0.013 | -0.048 | -0.048 | 0 | 0.14 | 0.05 | 0.15 | 0.47 | 0.1 | 0.26 |
| 4 | 7763161 | rs2285764 | -0.307 | 0.167 | -0.409 | -0.086 | -0.007 | 0.011 | 0.011 | 0 | 0.76 | 0.56 | 0.64 | 0.71 | 0.79 | 0.68 |
| 4 | 7763304 | rs2285763 | 1.723 | 0.181 | -0.354 | -0.089 | 0 | 0.003 | 0.003 | 0 | 0.09 | 0.09 | 0.19 | 0.2 | 0.1 | 0.18 |
| 4 | 7763761 | rs2269850 | -0.23 | 0.155 | -0.351 | -0.102 | -0.007 | 0.011 | 0.011 | 0 | 0.76 | 0.56 | 0.64 | 0.71 | 0.79 | 0.68 |
| 4 | 7764148 | rs2269849 | 1.685 | 0.108 | -0.382 | -0.051 | 0.013 | -0.048 | -0.048 | 0 | 0.14 | 0.05 | 0.14 | 0.47 | 0.1 | 0.26 |
| 4 | 7764821 | rs2107182 | -1.204 | 0.146 | -0.17 | -0.138 | -0.006 | -0.029 | -0.029 | 0.698 | 0.84 | 0.66 | 0.82 | 0.91 | 0.87 | 0.85 |
| 4 | 7765193 | rs2240054 | 1.161 | 0.185 | -0.237 | 0.467 | -0.028 | 0.155 | 0.155 | 0 | 0.17 | 0.37 | 0.17 | 0.39 | 0.1 | 0.26 |
| 4 | 7767318 | rs16841112 | 1.601 | 0.233 | -0.677 | 0.487 | 0.013 | -0.047 | -0.047 | 0 | 0.14 | 0.05 | 0.14 | 0.38 | 0.09 | 0.26 |
| 4 | 7769493 | rs13127069 | 0.963 | 0.253 | -0.997 | 0.24 | -0.156 | 0.077 | 0.077 | 0 | 0.43 | 0.05 | 0.24 | 0.22 | 0.54 | 0.34 |
| 4 | 7770723 | rs2285761 | 0.641 | 0.273 | -0.525 | 0.345 | -0.003 | 0.017 | 0.017 | 0 | 0.23 | 0.26 | 0.34 | 0.66 | 0.18 | 0.43 |
| 4 | 7772175 | rs4234829 | -1.735 | 0.12 | -0.251 | -0.174 | -0.011 | -0.008 | -0.008 | 0 | 0.88 | 0.79 | 0.87 | 0.91 | 0.94 | 0.86 |
| 4 | 7772599 | rs1557816 | -0.472 | 0.092 | -0.076 | -0.181 | 0.005 | 0.016 | 0.016 | 0 | 0.41 | 0.61 | 0.6 | 0.75 | 0.34 | 0.53 |
| 4 | 7774352 | rs6828005 | 0.303 | 0.16 | -0.109 | -0.132 | -0.004 | 0.002 | 0.002 | 0 | 0.55 | 0.3 | 0.53 | 0.53 | 0.55 | 0.41 |
| 4 | 7774610 | rs2276967 | 1.799 | 0.304 | -0.253 | 0.427 | 0.009 | -0.028 | -0.028 | 0 | 0.13 | 0.05 | 0.14 | 0.41 | 0.09 | 0.23 |
| 4 | 7777057 | rs4689859 | 0.313 | 0.277 | -0.183 | 0.125 | -0.002 | 0.01 | 0.01 | 0 | 0.25 | 0.26 | 0.29 | 0.55 | 0.2 | 0.39 |
| 4 | 7780344 | rs2269878 | 1.645 | 0.322 | -0.377 | 0.25 | 0.005 | 0.008 | 0.008 | 0 | 0.14 | 0.1 | 0.15 | 0.46 | 0.09 | 0.23 |
| 4 | 7782434 | rs16841209 | 1.345 | 0.332 | -0.242 | 0.223 | 0.003 | 0.03 | 0.03 | 0 | 0.1 | 0.16 | 0.19 | 0.23 | 0.1 | 0.2 |
| 4 | 7783286 | rs10516189 | 1.666 | 0.304 | -0.383 | 0.222 | 0.005 | 0.007 | 0.007 | 0 | 0.14 | 0.1 | 0.15 | 0.46 | 0.09 | 0.24 |
| 4 | 7787049 | rs13128495 | 1.034 | 0.27 | -0.46 | -0.125 | -0.017 | 0.101 | 0.101 | 0 | 0.4 | 0.19 | 0.37 | 0.07 | 0.45 | 0.17 |
| 4 | 7792662 | rs877707 | 1.569 | 0.293 | -0.267 | 0.056 | -0.001 | -0.002 | -0.002 | 0 | 0.09 | 0.09 | 0.19 | 0.27 | 0.1 | 0.2 |
| 4 | 7794001 | rs10022249 | 1.786 | 0.299 | -0.337 | -0.206 | 0 | -0.06 | -0.06 | 2.91 | 0.15 | 0.18 | 0.22 | 0.01 | 0.14 | 0.09 |
| 4 | 7796922 | rs10937862 | 1.756 | 0.304 | -0.348 | -0.204 | 0 | -0.045 | -0.045 | 6.173 | 0.15 | 0.16 | 0.22 | 0.01 | 0.14 | 0.09 |
| 4 | 7800699 | rs756255 | 0.724 | 0.293 | -0.122 | -0.251 | 0.01 | -0.023 | -0.023 | 0 | 0.15 | 0.31 | 0.18 | 0.09 | 0.15 | 0.16 |
| 4 | 7801581 | rs4689866 | -0.244 | 0.213 | -0.415 | -0.264 | 0 | 0.001 | 0.001 | 0 | 0.38 | 0.36 | 0.56 | 0.74 | 0.33 | 0.53 |
| 4 | 7802292 | rs11728778 | 2.019 | 0.222 | -0.53 | -0.35 | -0.076 | 0.094 | 0.094 | 0 | 0.25 | 0.01 | 0.15 | 0.06 | 0.3 | 0.08 |
| 4 | 7802686 | rs6821044 | 0.494 | 0.22 | -0.384 | -0.366 | 0.004 | -0.048 | -0.048 | 0.377 | 0.42 | 0.52 | 0.35 | 0.15 | 0.48 | 0.25 |
| 4 | 7804265 | rs7376483 | 0.758 | 0.236 | -0.54 | -0.211 | 0.005 | -0.011 | -0.011 | 0 | 0.22 | 0.18 | 0.34 | 0.73 | 0.18 | 0.43 |
| 4 | 7807834 | rs1106248 | 0.488 | 0.052 | -0.879 | -0.375 | -0.025 | 0.085 | 0.085 | 0 | 0.44 | 0.26 | 0.55 | 0.28 | 0.51 | 0.34 |
| 4 | 7809042 | rs4689868 | -1.143 | 0.137 | -1.051 | -0.29 | -0.089 | 0.203 | 0.203 | 0 | 0.54 | 0.88 | 0.76 | 0.83 | 0.48 | 0.69 |
| 4 | 7809472 | rs4689869 | -0.548 | 0.093 | -1.034 | -0.253 | 0.013 | 0.004 | 0.004 | 0.043 | 0.45 | 0.83 | 0.57 | 0.56 | 0.37 | 0.49 |
| 4 | 7809979 | rs7693234 | 0.215 | 0.167 | -0.847 | -0.188 | -0.001 | -0.001 | -0.001 | 0 | 0.55 | 0.56 | 0.56 | 0.6 | 0.52 | 0.44 |
| 4 | 7810027 | rs6846245 | 0.37 | 0.21 | -0.894 | -0.584 | 0.003 | -0.004 | -0.004 | 0.384 | 0.16 | 0.19 | 0.14 | 0.14 | 0.16 | 0.22 |
| 4 | 7812674 | rs6446622 | 0.013 | 0.234 | -0.933 | -1.182 | 0.003 | -0.035 | -0.035 | 0.034 | 0.63 | 0.53 | 0.63 | 0.82 | 0.63 | 0.66 |
| 4 | 7813170 | rs885262 | -0.413 | 0.457 | -1.124 | -1.352 | 0.003 | -0.029 | -0.029 | 0.758 | 0.36 | 0.54 | 0.5 | 0.31 | 0.38 | 0.36 |
| 4 | 7815054 | rs6844380 | -0.133 | 0.662 | -1.175 | -0.989 | -0.05 | -0.007 | -0.007 | 0 | 0.66 | 0.85 | 0.72 | 0.61 | 0.63 | 0.52 |
| 4 | 7822186 | rs4234831 | -1.184 | 0.704 | -1.156 | -1.085 | -0.026 | 0.033 | 0.033 | 0 | 0.79 | 0.92 | 0.91 | 0.88 | 0.78 | 0.77 |
| 4 | 7831463 | rs7685914 | 0.589 | 0.471 | -1.063 | -1.311 | -0.001 | -0.025 | -0.025 | 1.7 | 0.56 | 0.53 | 0.54 | 0.11 | 0.56 | 0.32 |
| 4 | 7831977 | rs6446629 | -1.113 | 0.678 | -1.15 | -1.285 | -0.087 | 0.012 | 0.012 | 0 | 0.79 | 0.97 | 0.91 | 0.88 | 0.75 | 0.77 |
| 4 | 7840696 | rs4296679 | -0.011 | 0.486 | -0.911 | -1.134 | -0.002 | 0 | 0 | 0.464 | 0.63 | 0.61 | 0.72 | 0.62 | 0.62 | 0.51 |
| 4 | 7841487 | rs4580651 | -0.459 | 0.47 | -0.889 | -1.131 | -0.086 | -0.011 | -0.011 | 0.179 | 0.67 | 0.9 | 0.76 | 0.64 | 0.62 | 0.53 |
| 4 | 7841833 | rs4689166 | 0.004 | 0.163 | -0.952 | -1.237 | 0.008 | 0.003 | 0.003 | 0 | 0.44 | 0.58 | 0.6 | 0.54 | 0.39 | 0.43 |
| 4 | 7847468 | rs4473655 | 0.242 | 0.066 | 0.253 | -0.961 | -0.014 | -0.057 | -0.057 | 0.431 | 0.82 | 0.72 | 0.72 | 0.93 | 0.83 | 0.83 |
| 4 | 7852851 | rs4361385 | 0.953 | 0.274 | 0.009 | -0.993 | -0.064 | -0.029 | -0.029 | 1.251 | 0.15 | 0.03 | 0.07 | 0.13 | 0.16 | 0.11 |

| | | | | | | | | | | | | | | | | |
|---|---|---|---|---|---|---|---|---|---|---|---|---|---|---|---|---|
| 4 | 7853160 | rs4619890 | 2.084 | -0.026 | 0.001 | -1.175 | -0.041 | 0.163 | 0.163 | 3.36 | 0.47 | 0.14 | 0.4 | 0.3 | 0.53 | 0.38 |
| 4 | 7853361 | rs4413405 | 0.273 | -0.064 | 0.224 | -1.183 | -0.031 | -0.071 | -0.071 | 1.66 | 0.77 | 0.69 | 0.72 | 0.93 | 0.82 | 0.77 |
| 4 | 7865324 | rs28738908 | -0.151 | 0.055 | 0.622 | -1.123 | 0.025 | -0.016 | -0.016 | 1.199 | 0.22 | 0.42 | 0.15 | 0.16 | 0.24 | 0.2 |
| 4 | 7866500 | rs7441783 | 0.507 | 0.036 | 0.369 | -1.185 | 0.011 | -0.003 | -0.003 | 0 | 0.69 | 0.43 | 0.68 | 0.37 | 0.69 | 0.61 |
| 4 | 7879440 | rs28622214 | 2.306 | 0.201 | -0.146 | -1.283 | -0.075 | 0.205 | 0.205 | 4.55 | 0.37 | 0.11 | 0.25 | 0.05 | 0.44 | 0.15 |
| 4 | 7894887 | rs17770177 | 1.077 | 0.252 | -0.137 | -1.239 | -0.035 | -0.005 | -0.005 | 0 | 0.15 | 0.04 | 0.07 | 0.13 | 0.15 | 0.11 |
| 4 | 7896866 | rs17770213 | 2.286 | 0.106 | 0.076 | -1.268 | -0.049 | 0.16 | 0.16 | 4.245 | 0.37 | 0.15 | 0.26 | 0.05 | 0.44 | 0.16 |
| 4 | 7902003 | rs4478172 | -0.844 | -0.35 | 0.905 | -1.038 | -0.024 | 0.1 | 0.1 | 0.001 | 0.25 | 0.43 | 0.37 | 0.49 | 0.23 | 0.32 |
| 4 | 7904431 | rs6831381 | 0.484 | -0.086 | 1.032 | -0.974 | -0.028 | -0.023 | -0.023 | 0 | 0.66 | 0.63 | 0.57 | 0.67 | 0.74 | 0.56 |
| 4 | 7906178 | rs6824410 | -1.317 | -0.014 | 0.676 | -0.951 | 0.012 | 0.018 | 0.018 | 0.289 | 0.4 | 0.62 | 0.5 | 0.36 | 0.35 | 0.54 |
| 4 | 7913961 | rs13127935 | -1.322 | 0.047 | 0.507 | -0.921 | -0.008 | -0.013 | -0.013 | 0 | 0.39 | 0.56 | 0.48 | 0.35 | 0.35 | 0.54 |
| 4 | 7917806 | rs6838291 | -0.156 | 0.054 | 0.238 | -0.771 | -0.02 | -0.044 | -0.044 | 0 | 0.21 | 0.26 | 0.27 | 0.07 | 0.16 | 0.21 |
| 4 | 7923973 | rs10049998 | -1.012 | 0.638 | -0.404 | -0.441 | -0.044 | -0.024 | -0.024 | 0 | 0.85 | 0.97 | 0.93 | 0.87 | 0.84 | 0.88 |
| 4 | 7929041 | rs9991239 | -0.504 | 0.429 | 0.118 | -0.158 | 0.005 | 0.125 | 0.125 | 0.277 | 0.12 | 0.21 | 0.22 | 0.49 | 0.14 | 0.29 |
| 4 | 7931635 | rs1344223 | -0.537 | 0.261 | 0.109 | -0.925 | -0.017 | -0.088 | -0.088 | 0.896 | 0.31 | 0.46 | 0.38 | 0.1 | 0.3 | 0.3 |
| 4 | 7931725 | rs2385914 | -0.55 | 0.259 | 0.096 | -0.927 | -0.022 | -0.108 | -0.108 | 0.351 | 0.31 | 0.49 | 0.39 | 0.1 | 0.3 | 0.3 |
| 4 | 7939194 | rs4696687 | -0.538 | 0.297 | 0.249 | -0.655 | -0.011 | 0.014 | 0.014 | 0.191 | 0.24 | 0.4 | 0.35 | 0.72 | 0.17 | 0.42 |
| 4 | 7943106 | rs6447872 | 0.978 | 0.644 | -0.007 | -0.839 | 0.009 | -0.054 | -0.054 | 0 | 0.63 | 0.52 | 0.59 | 0.14 | 0.67 | 0.44 |
| 4 | 7947266 | rs2891926 | 0.044 | 0.636 | 0.043 | -0.795 | 0 | 0.003 | 0.003 | 0.021 | 0.21 | 0.2 | 0.32 | 0.71 | 0.18 | 0.43 |
| 4 | 7951137 | rs4696833 | 0.606 | 0.708 | 0.006 | -0.718 | 0.027 | -0.002 | -0.002 | 0 | 0.72 | 0.66 | 0.76 | 0.66 | 0.74 | 0.65 |
| 4 | 7951542 | rs7685634 | -0.846 | 0.738 | -0.279 | -0.697 | -0.044 | -0.012 | -0.012 | 0 | 0.86 | 0.97 | 0.95 | 0.86 | 0.84 | 0.88 |
| 4 | 7953583 | rs10023401 | 0.184 | 0.602 | 0.009 | -0.714 | 0.023 | 0.018 | 0.018 | 0.177 | 0.14 | 0.37 | 0.19 | 0.21 | 0.1 | 0.22 |
| 4 | 7955488 | rs11732351 | 1.486 | 0.498 | -1.081 | -1.078 | -0.05 | 0.208 | 0.208 | 0.838 | 0.35 | 0.13 | 0.25 | 0.03 | 0.44 | 0.18 |
| 4 | 7959029 | rs6824093 | 0.144 | 0.353 | -0.549 | -0.11 | -0.001 | 0.014 | 0.014 | 0 | 0.18 | 0.15 | 0.23 | 0.47 | 0.19 | 0.23 |
| 4 | 7959980 | rs12512108 | 0.681 | 0.233 | -0.511 | -0.071 | 0.001 | -0.005 | -0.005 | 0 | 0.11 | 0.08 | 0.14 | 0.19 | 0.08 | 0.24 |
| 4 | 7962856 | rs7356321 | 0.58 | 0.019 | -0.207 | -0.496 | 0 | 0.008 | 0.008 | 0.014 | 0.46 | 0.55 | 0.35 | 0.3 | 0.5 | 0.34 |
| 4 | 7966812 | rs4696709 | -1.319 | -0.457 | 0.033 | -0.16 | -0.039 | 0.013 | 0.013 | 0.387 | 0.24 | 0.38 | 0.3 | 0.3 | 0.15 | 0.29 |
| 4 | 7967683 | rs4696886 | -0.896 | -0.395 | -0.176 | -0.008 | -0.024 | 0.022 | 0.022 | 0 | 0.17 | 0.24 | 0.16 | 0.3 | 0.08 | 0.18 |
| 4 | 7967875 | rs2292735 | -0.794 | -0.453 | -0.153 | 0.023 | -0.036 | 0.038 | 0.038 | 0 | 0.16 | 0.25 | 0.16 | 0.31 | 0.07 | 0.18 |
| 4 | 7980298 | rs1317285 | 0.303 | -0.209 | -0.431 | 0.177 | -0.011 | 0.085 | 0.085 | 0 | 0.15 | 0.24 | 0.25 | 0.5 | 0.12 | 0.15 |
| 4 | 7980652 | rs7699715 | 1.152 | -0.087 | -1.101 | -0.873 | -0.004 | 0.068 | 0.068 | 0 | 0.33 | 0.25 | 0.2 | 0.06 | 0.32 | 0.19 |
| 4 | 7985977 | rs16841523 | -0.045 | -0.084 | -0.952 | -0.379 | -0.022 | 0.119 | 0.119 | 1.751 | 0.22 | 0.59 | 0.35 | 0.38 | 0.19 | 0.23 |
| 4 | 7986860 | rs7677869 | -0.01 | -0.064 | -1.321 | -0.71 | -0.01 | 0.025 | 0.025 | 0.057 | 0.65 | 0.64 | 0.6 | 0.45 | 0.66 | 0.5 |
| 4 | 7987764 | rs16841530 | 0.417 | -0.194 | -0.898 | -0.266 | -0.01 | 0.152 | 0.152 | 0 | 0.12 | 0.25 | 0.07 | 0.35 | 0.1 | 0.12 |
| 4 | 7991587 | rs4696662 | -0.217 | 0.054 | -0.751 | -0.085 | -0.043 | 0.163 | 0.163 | 0.942 | 0.2 | 0.53 | 0.31 | 0.38 | 0.15 | 0.19 |
| 4 | 7992632 | rs4696726 | -0.993 | -0.059 | -1.002 | -0.232 | -0.009 | 0.016 | 0.016 | 0.368 | 0.38 | 0.56 | 0.38 | 0.42 | 0.29 | 0.26 |
| 4 | 7992749 | rs13129405 | 0.892 | 0.397 | -1.623 | -0.113 | -0.061 | 0.08 | 0.08 | 0 | 0.14 | 0.04 | 0.07 | 0 | 0.15 | 0.07 |
| 4 | 7993976 | rs4696663 | 0.089 | -0.141 | -1.084 | 0.374 | -0.022 | 0.094 | 0.094 | 0 | 0.17 | 0.23 | 0.26 | 0.37 | 0.13 | 0.17 |
| 4 | 7997642 | rs17772026 | 0.016 | 0.43 | -2.032 | 0.153 | -0.088 | 0.027 | 0.027 | 0.347 | 0.36 | 0.12 | 0.39 | 0.3 | 0.41 | 0.47 |
| 4 | 8003927 | rs11729198 | 0.22 | 0.463 | -1.395 | 0.017 | 0.001 | 0.017 | 0.017 | 0 | 0.33 | 0.21 | 0.18 | 0.26 | 0.34 | 0.2 |
| 4 | 8008264 | rs4696664 | -0.084 | 0.613 | -1.554 | -0.16 | -0.048 | 0.091 | 0.091 | 0 | 0.61 | 0.82 | 0.62 | 0.8 | 0.55 | 0.67 |
| 4 | 8009737 | rs16841552 | -0.239 | 0.257 | -1.262 | 0.94 | -0.004 | 0.061 | 0.061 | 0.011 | 0.08 | 0.13 | 0.19 | 0.31 | 0.06 | 0.16 |
| 4 | 8009759 | rs17180256 | -0.759 | -0.32 | -1.925 | 0.414 | 0.036 | 0.082 | 0.082 | 0 | 0.63 | 0.86 | 0.71 | 0.8 | 0.72 | 0.72 |
| 4 | 8010112 | rs13125250 | -1.364 | -0.522 | -2.127 | 0.225 | 0.021 | 0.003 | 0.003 | 0 | 0.74 | 0.93 | 0.77 | 0.8 | 0.81 | 0.78 |
| 4 | 8011696 | rs1317368 | 0.42 | -0.533 | -1.674 | 0.223 | -0.007 | -0.043 | -0.043 | 0.01 | 0.13 | 0.21 | 0.05 | 0 | 0.08 | 0.03 |
| 4 | 8012167 | rs16841559 | -0.222 | -0.249 | -1.202 | 0.251 | -0.008 | 0.066 | 0.066 | 0.037 | 0.3 | 0.4 | 0.43 | 0.55 | 0.33 | 0.37 |
| 4 | 8015421 | rs4696729 | -1.049 | -0.262 | -1.175 | 0.249 | 0.007 | 0.024 | 0.024 | 0 | 0.43 | 0.6 | 0.49 | 0.7 | 0.43 | 0.55 |
| 4 | 8017124 | rs9990943 | -1.559 | -0.349 | -1.445 | 0.216 | -0.003 | -0.088 | -0.088 | 0.122 | 0.73 | 0.94 | 0.72 | 0.67 | 0.77 | 0.63 |
| 4 | 8019447 | rs4696666 | 0.201 | -0.209 | -0.866 | 0.48 | -0.014 | 0.025 | 0.025 | 0 | 0.17 | 0.27 | 0.12 | 0.26 | 0.13 | 0.26 |
| 4 | 8020502 | rs11932811 | 0.538 | 0.097 | -0.772 | 0.446 | 0.071 | 0.198 | 0.198 | 0.016 | 0.1 | 0.24 | 0.29 | 0.3 | 0.16 | 0.12 |
| 4 | 8021592 | rs13131680 | -0.288 | -0.159 | -0.95 | 0.245 | -0.002 | 0 | 0 | 3.652 | 0.39 | 0.36 | 0.52 | 0.41 | 0.54 | 0.33 |
| 4 | 8026176 | rs11729381 | 0.296 | -0.273 | -1.312 | -0.045 | 0.03 | 0.058 | 0.058 | 0 | 0.14 | 0.05 | 0.1 | 0.03 | 0.11 | 0.12 |
| 4 | 8026238 | rs11726230 | 0.136 | -0.315 | -1.255 | -0.087 | 0.021 | 0.028 | 0.028 | 0 | 0.17 | 0.15 | 0.13 | 0.03 | 0.12 | 0.15 |
| 4 | 8027403 | rs6833644 | 0.271 | 0.464 | -0.727 | 0.606 | 0.039 | 0.163 | 0.163 | 0.045 | 0.09 | 0.2 | 0.2 | 0.32 | 0.14 | 0.12 |
| 4 | 8028937 | rs17795172 | 0.694 | 0.379 | -1.111 | 0.204 | -0.074 | 0.044 | 0.044 | 0 | 0.22 | 0.12 | 0.22 | 0.13 | 0.33 | 0.15 |
| 4 | 8031590 | rs11942146 | -0.89 | 0.001 | -0.769 | 0.65 | -0.005 | 0.025 | 0.025 | 0.48 | 0.23 | 0.25 | 0.26 | 0.61 | 0.23 | 0.29 |
| 4 | 8035102 | rs4696734 | -0.147 | 0.57 | -0.898 | 0.299 | -0.002 | -0.015 | -0.015 | 0.004 | 0.38 | 0.48 | 0.36 | 0.26 | 0.33 | 0.34 |
| 4 | 8045492 | rs4696738 | 0.046 | 1.047 | -0.885 | 0.914 | 0.004 | 0.058 | 0.058 | 0 | 0.11 | 0.12 | 0.29 | 0.34 | 0.14 | 0.09 |
| 4 | 8046433 | rs6830813 | 0.567 | 0.666 | -0.952 | 0.102 | 0.001 | -0.148 | -0.148 | 0 | 0.35 | 0.47 | 0.18 | 0 | 0.31 | 0.18 |
| 4 | 8047973 | rs2385916 | -1.161 | 1.173 | -1.333 | 0.395 | -0.082 | 0.008 | 0.008 | 0.404 | 0.8 | 0.97 | 0.79 | 0.78 | 0.72 | 0.68 |
| 4 | 8050056 | rs1574414 | 0.126 | 0.89 | -1.188 | 0.467 | 0.009 | 0.067 | 0.067 | 0 | 0.1 | 0.12 | 0.28 | 0.3 | 0.13 | 0.08 |
| 4 | 8050343 | rs10029338 | 0.039 | 0.702 | -1.407 | 0.344 | -0.002 | 0.021 | 0.021 | 0.024 | 0.26 | 0.3 | 0.32 | 0.34 | 0.27 | 0.42 |
| 4 | 8051014 | rs1320061 | -0.88 | 0.965 | -1.581 | -0.094 | -0.04 | 0.028 | 0.028 | 0 | 0.83 | 0.97 | 0.88 | 0.84 | 0.81 | 0.8 |
| 4 | 8056846 | rs7661897 | -0.287 | 0.778 | -0.727 | -0.288 | -0.004 | -0.079 | -0.079 | 0.006 | 0.25 | 0.36 | 0.25 | 0.12 | 0.24 | 0.28 |
| 4 | 8060637 | rs1320058 | 0.21 | 0.918 | -1.486 | -1.121 | -0.019 | 0.081 | 0.081 | 0 | 0.35 | 0.15 | 0.45 | 0.8 | 0.37 | 0.46 |
| 4 | 8062202 | rs4696669 | -0.775 | 0.994 | -0.88 | -1.318 | -0.003 | -0.085 | -0.085 | 0 | 0.63 | 0.47 | 0.69 | 0.92 | 0.63 | 0.83 |
| 4 | 8062830 | rs16841624 | 0.827 | 1.119 | -1.116 | -0.203 | -0.004 | 0.04 | 0.04 | 0 | 0.05 | 0.03 | 0.22 | 0.31 | 0.15 | 0.13 |
| 4 | 8063501 | rs6845573 | 0.121 | 0.587 | -1.003 | -0.766 | -0.001 | -0.023 | -0.023 | 0 | 0.28 | 0.28 | 0.25 | 0.1 | 0.26 | 0.35 |
| 4 | 8066573 | rs6850368 | 0.123 | 0.37 | -1.93 | -1.33 | -0.008 | 0.146 | 0.146 | 0 | 0.31 | 0.06 | 0.23 | 0.09 | 0.27 | 0.39 |

| | | | | | | | | | | | | | | | | |
|---|---|---|---|---|---|---|---|---|---|---|---|---|---|---|---|---|
| 4 | 8070793 | rs6829649 | 0.601 | 0.04 | -1.255 | -1.332 | -0.01 | -0.023 | -0.023 | 0 | 0.19 | 0.23 | 0.08 | 0.15 | 0.13 | 0.19 |
| 4 | 8071136 | rs13132381 | 1.483 | -0.17 | -1.577 | -0.741 | -0.046 | -0.075 | -0.075 | 0 | 0.15 | 0.03 | 0.1 | 0.19 | 0.19 | 0.15 |
| 4 | 8072684 | rs9991680 | -0.616 | -0.188 | -1.164 | -0.944 | -0.021 | 0.069 | 0.069 | 0.253 | 0.44 | 0.73 | 0.58 | 0.74 | 0.48 | 0.47 |
| 4 | 8074362 | rs7660009 | 0.831 | -0.39 | -1.251 | -0.838 | -0.015 | -0.027 | -0.027 | 0 | 0.21 | 0.23 | 0.07 | 0.12 | 0.09 | 0.18 |
| 4 | 8075134 | rs6842276 | 0.335 | -0.121 | -1.522 | -0.894 | 0.003 | 0.012 | 0.012 | 0 | 0.46 | 0.34 | 0.37 | 0.24 | 0.44 | 0.47 |
| 4 | 8078157 | rs1573843 | 1.084 | 0.054 | -1.257 | -0.497 | -0.016 | -0.041 | -0.041 | 0 | 0.2 | 0.25 | 0.16 | 0.1 | 0.16 | 0.1 |
| 4 | 8080481 | rs6856548 | 0.282 | 0.516 | -1.208 | 0.105 | 0.001 | 0.044 | 0.044 | 0 | 0.51 | 0.35 | 0.36 | 0.37 | 0.46 | 0.53 |
| 4 | 8080573 | rs6826670 | -0.033 | 1.031 | -0.707 | 0.873 | 0.036 | 0.132 | 0.132 | 0 | 0.74 | 0.47 | 0.5 | 0.57 | 0.69 | 0.68 |
| 4 | 8083563 | rs17178920 | -0.111 | 1.133 | -1.601 | 0.559 | -0.053 | 0.016 | 0.016 | 0 | 0.15 | 0.04 | 0.15 | 0.14 | 0.22 | 0.27 |
| 4 | 8084560 | rs6447849 | -0.237 | 0.348 | -2.217 | 0.724 | -0.084 | -0.037 | -0.037 | 0 | 0.42 | 0.1 | 0.3 | 0.4 | 0.49 | 0.47 |
| 4 | 8085174 | rs11724908 | 0.909 | -0.077 | -2.389 | 0.145 | -0.044 | 0.163 | 0.163 | 1.685 | 0.32 | 0.04 | 0.17 | 0.06 | 0.3 | 0.21 |
| 4 | 8086173 | rs11731965 | 0.362 | -0.237 | -1.805 | 0.668 | 0.008 | 0.003 | 0.003 | 0.815 | 0.39 | 0.25 | 0.49 | 0.38 | 0.35 | 0.33 |
| 4 | 8086655 | rs6817164 | 0.315 | 0.046 | -1.826 | 0.478 | -0.016 | 0.008 | 0.008 | 0 | 0.26 | 0.21 | 0.19 | 0.2 | 0.3 | 0.17 |
| 4 | 8089598 | rs2292739 | 0.969 | 0.212 | -1.622 | 0.253 | 0.015 | 0.127 | 0.127 | 6.761 | 0.38 | 0.17 | 0.2 | 0.07 | 0.32 | 0.14 |
| 4 | 8097469 | rs7674677 | 0.545 | 0.746 | -1.498 | 0.945 | -0.039 | -0.036 | -0.036 | 0 | 0.2 | 0.12 | 0.28 | 0.33 | 0.29 | 0.16 |
| 4 | 8100171 | rs12500841 | 0.032 | 0.476 | -1.579 | 1.062 | -0.019 | -0.053 | -0.053 | 0.411 | 0.25 | 0.12 | 0.28 | 0.36 | 0.29 | 0.48 |
| 4 | 8100375 | rs4464550 | -0.203 | 0.556 | -0.405 | 1.236 | 0.035 | 0.308 | 0.308 | 1.402 | 0.07 | 0.44 | 0.21 | 0.26 | 0.03 | 0.11 |
| 4 | 8103323 | rs17795478 | 0.915 | 0.731 | -1.08 | 0.923 | -0.017 | -0.008 | -0.008 | 0 | 0.57 | 0.3 | 0.48 | 0.53 | 0.47 | 0.58 |
| 4 | 8104173 | rs12647620 | 0.535 | 0.603 | -0.951 | 1.006 | -0.001 | -0.003 | -0.003 | 0 | 0.64 | 0.33 | 0.56 | 0.62 | 0.66 | 0.68 |
| 4 | 8105010 | rs12511735 | -0.091 | 1.14 | -1.533 | 0.629 | -0.124 | -0.027 | -0.027 | 0.054 | 0.23 | 0.03 | 0.22 | 0.17 | 0.33 | 0.19 |
| 4 | 8105320 | rs16841721 | -0.575 | 0.746 | -1.009 | 0.562 | -0.001 | -0.007 | -0.007 | 0 | 0.06 | 0.05 | 0.07 | 0 | 0.07 | 0.03 |
| 4 | 8106249 | rs4696758 | -0.258 | 0.631 | -0.853 | 0.502 | -0.001 | 0 | 0 | 0 | 0.46 | 0.51 | 0.61 | 0.47 | 0.52 | 0.3 |
| 4 | 8108027 | rs12108540 | -0.079 | 0.712 | -0.883 | 0.309 | 0.003 | -0.002 | -0.002 | 0.146 | 0.78 | 0.72 | 0.81 | 0.75 | 0.77 | 0.55 |
| 4 | 8110782 | rs1878519 | -0.327 | 1.034 | -0.894 | 0.214 | -0.006 | 0.012 | 0.012 | 0 | 0.46 | 0.33 | 0.35 | 0.4 | 0.45 | 0.55 |
| 4 | 8111117 | rs1105616 | -0.83 | 0.963 | -0.792 | 0.004 | -0.008 | -0.006 | -0.006 | 3.547 | 0.73 | 0.68 | 0.82 | 0.83 | 0.81 | 0.63 |
| 4 | 8114525 | rs17181205 | 0.25 | 0.915 | -1.031 | -0.167 | 0.006 | -0.02 | -0.02 | 0 | 0.16 | 0.19 | 0.21 | 0.1 | 0.24 | 0.05 |
| 4 | 8116202 | rs2138826 | -0.291 | 0.884 | -0.9 | -0.182 | 0.044 | -0.078 | -0.078 | 1.048 | 0.83 | 0.55 | 0.76 | 0.9 | 0.76 | 0.95 |
| 4 | 8116432 | rs1517523 | 0.623 | 1.008 | -0.872 | -0.231 | 0.08 | -0.091 | -0.091 | 0.023 | 0.76 | 0.44 | 0.68 | 0.84 | 0.61 | 0.81 |
| 4 | 8117671 | rs1125307 | 0.272 | 0.797 | -1.078 | -0.196 | -0.028 | 0.013 | 0.013 | 0 | 0.44 | 0.26 | 0.3 | 0.38 | 0.45 | 0.54 |
| 4 | 8118328 | rs11736271 | 0.143 | 0.827 | -1.066 | -0.187 | -0.01 | -0.003 | -0.003 | 0 | 0.46 | 0.44 | 0.6 | 0.55 | 0.39 | 0.3 |
| 4 | 8118561 | rs10938692 | -0.368 | 0.793 | -1.086 | -0.116 | -0.005 | 0 | 0 | 0 | 0.57 | 0.48 | 0.52 | 0.45 | 0.66 | 0.59 |
| 4 | 8119937 | rs1517525 | 1.039 | 0.659 | -1.563 | 0.129 | 0.027 | -0.01 | -0.01 | 0.146 | 0.32 | 0.19 | 0.3 | 0.34 | 0.17 | 0.22 |
| 4 | 8120120 | rs13122226 | 0.321 | 0.859 | -1.535 | 0.088 | 0.038 | -0.011 | -0.011 | 1.339 | 0.38 | 0.19 | 0.36 | 0.42 | 0.29 | 0.37 |
| 4 | 8120666 | rs4696763 | 0.27 | 1.666 | -1.537 | 0.198 | 0.054 | 0.019 | 0.019 | 0 | 0.76 | 0.51 | 0.59 | 0.67 | 0.63 | 0.76 |
| 4 | 8121799 | rs17181631 | -0.066 | 2.068 | -2.326 | -0.296 | -0.086 | 0.022 | 0.022 | 0 | 0.19 | 0.03 | 0.14 | 0.14 | 0.27 | 0.16 |
| 4 | 8122496 | rs11936149 | -0.671 | 1.241 | -2.165 | -0.159 | -0.071 | 0.033 | 0.033 | 0.741 | 0.33 | 0.2 | 0.28 | 0.29 | 0.44 | 0.4 |
| 4 | 8125788 | rs4696765 | -0.1 | 1.169 | -1.953 | -0.246 | 0.002 | -0.015 | -0.015 | 0 | 0.46 | 0.61 | 0.41 | 0.25 | 0.5 | 0.45 |
| 4 | 8127714 | rs1109531 | -0.739 | 1.337 | -2.427 | -0.049 | -0.046 | 0.046 | 0.046 | 0 | 0.21 | 0.05 | 0.23 | 0.11 | 0.24 | 0.29 |
| 4 | 8131414 | rs4696766 | -0.288 | 1.461 | -2.34 | 0.442 | -0.166 | -0.142 | -0.142 | 0.436 | 0.25 | 0.02 | 0.32 | 0.45 | 0.4 | 0.27 |
| 4 | 8138645 | rs1878521 | -0.681 | 0.428 | -1.688 | -0.071 | 0 | 0.005 | 0.005 | 0 | 0.11 | 0.08 | 0.06 | 0.06 | 0.1 | 0.13 |
| 4 | 8139958 | rs4422405 | -0.717 | -0.308 | -1.187 | -0.573 | -0.019 | -0.053 | -0.053 | 0.006 | 0.33 | 0.57 | 0.16 | 0.07 | 0.19 | 0.28 |
| 4 | 8142672 | rs16841848 | -0.016 | -0.242 | -1.09 | -0.45 | -0.051 | -0.115 | -0.115 | 1.397 | 0.23 | 0.34 | 0.09 | 0.01 | 0.11 | 0.16 |
| 4 | 8148159 | rs4696769 | 0.206 | 0.576 | -1.553 | 0.434 | 0.009 | -0.004 | -0.004 | 0 | 0.56 | 0.2 | 0.57 | 0.52 | 0.56 | 0.42 |
| 4 | 8154534 | rs12506374 | -1.146 | 0.439 | -1.214 | 0.77 | 0.006 | -0.006 | -0.006 | 0.018 | 0.2 | 0.16 | 0.13 | 0.28 | 0.16 | 0.21 |
| 4 | 8156982 | rs6816775 | -0.912 | 1.536 | -0.392 | 0.551 | 0.079 | -0.007 | -0.007 | 0 | 0.11 | 0.23 | 0.12 | 0.1 | 0.2 | 0.12 |
| 4 | 8157710 | rs7660199 | -1.1 | 2.116 | -0.438 | 1.016 | -0.011 | -0.012 | -0.012 | 0 | 0.26 | 0.21 | 0.26 | 0.4 | 0.36 | 0.3 |
| 4 | 8158352 | rs1533508 | -1.571 | 1.362 | 0.053 | 0.619 | 0.007 | 0.01 | 0.01 | 0 | 0.37 | 0.59 | 0.33 | 0.45 | 0.44 | 0.35 |
| 4 | 8171401 | rs10938694 | 1.714 | 0.463 | 0.072 | 0.688 | 0.009 | -0.025 | -0.025 | 0.398 | 0.45 | 0.2 | 0.53 | 0.43 | 0.42 | 0.4 |
| 4 | 8177703 | rs13151241 | -0.795 | 1.129 | 0.319 | 0.754 | -0.001 | -0.007 | -0.007 | 0 | 0.21 | 0.2 | 0.23 | 0.34 | 0.19 | 0.2 |
| 4 | 8178177 | rs13152324 | -1.32 | 1.144 | 0.52 | 0.71 | 0.013 | 0.013 | 0.013 | 0 | 0.25 | 0.36 | 0.29 | 0.34 | 0.29 | 0.23 |
| 4 | 8189780 | rs753386 | 0.926 | 0.764 | 0.594 | 0.5 | -0.004 | 0.022 | 0.022 | 0 | 0.77 | 0.63 | 0.76 | 0.69 | 0.79 | 0.83 |
| 4 | 8190219 | rs1078701 | -0.588 | 0.684 | 0.398 | 0.414 | -0.017 | -0.031 | -0.031 | 3.411 | 0.93 | 0.86 | 0.98 | 1 | 0.98 | 1 |
| 4 | 8199833 | rs1678263 | 2.056 | 0.484 | -0.496 | 0.371 | -0.078 | -0.125 | -0.125 | 0 | 0.37 | 0.02 | 0.35 | 0.41 | 0.35 | 0.39 |
| 4 | 8201805 | rs753959 | 0.135 | 0.829 | -0.308 | 0.559 | -0.051 | -0.07 | -0.07 | 0.759 | 0.14 | 0.02 | 0.19 | 0.21 | 0.18 | 0.14 |
| 4 | 8202211 | rs1281121 | 0.41 | 1.012 | 0.438 | 0.43 | 0.052 | -0.09 | -0.09 | 0 | 0.86 | 0.61 | 0.89 | 1 | 0.84 | 0.98 |
| 4 | 8202841 | rs1298472 | 0.453 | 0.933 | 0.536 | 0.416 | 0.045 | -0.109 | -0.109 | 0 | 0.85 | 0.53 | 0.87 | 1 | 0.83 | 0.98 |
| 4 | 8204854 | rs7677412 | -0.576 | 1.457 | 0.319 | 0.812 | -0.009 | -0.014 | -0.014 | 1.184 | 0.17 | 0.15 | 0.23 | 0.22 | 0.21 | 0.15 |
| 4 | 8207083 | rs2677605 | 1.402 | 1.407 | 0.407 | 0.923 | -0.001 | -0.001 | -0.001 | 0 | 0.54 | 0.55 | 0.47 | 0.3 | 0.53 | 0.4 |
| 4 | 8207810 | rs922521 | 1.596 | 1.345 | 0.089 | 0.958 | 0.041 | 0.057 | 0.057 | 0.5 | 0.52 | 0.39 | 0.41 | 0.29 | 0.37 | 0.39 |
| 4 | 8209383 | rs1281132 | 0.567 | 0.825 | 0.051 | 1.299 | -0.042 | 0.049 | 0.049 | 0 | 0.8 | 0.68 | 0.79 | 0.71 | 0.87 | 0.74 |
| 4 | 8211624 | rs1712316 | 0.768 | 0.835 | -0.24 | 1.053 | -0.045 | 0.049 | 0.049 | 0 | 0.79 | 0.63 | 0.76 | 0.71 | 0.87 | 0.73 |
| 4 | 8212524 | rs13152394 | -0.792 | 1.037 | -0.79 | 0.648 | -0.054 | -0.093 | -0.093 | 0 | 0.19 | 0.02 | 0.15 | 0.27 | 0.2 | 0.19 |
| 4 | 8214636 | rs1712287 | 0.869 | 0.908 | 0.023 | 0.071 | 0.004 | -0.086 | -0.086 | 0 | 0.86 | 0.71 | 0.91 | 1 | 0.88 | 0.89 |
| 4 | 8220400 | rs1298145 | -0.585 | 1.197 | -1.32 | -0.171 | -0.015 | 0.068 | 0.068 | 0 | 0.32 | 0.16 | 0.23 | 0.23 | 0.35 | 0.31 |
| 4 | 8222157 | rs7660416 | -0.474 | 1.192 | -1.044 | 0.098 | 0.004 | 0.07 | 0.07 | 0.323 | 0.26 | 0.53 | 0.48 | 0.53 | 0.28 | 0.39 |
| 4 | 8226488 | rs1281142 | 0.086 | 1.146 | -1.551 | -0.226 | -0.056 | 0.046 | 0.046 | 0 | 0.25 | 0.03 | 0.14 | 0.21 | 0.25 | 0.29 |
| 4 | 8230456 | rs1281152 | 0.605 | 1.211 | -0.859 | 0.169 | 0.005 | 0.026 | 0.026 | 0 | 0.69 | 0.43 | 0.49 | 0.5 | 0.65 | 0.66 |
| 4 | 8233341 | rs3756193 | 0.06 | 1.153 | -1.564 | -0.245 | -0.028 | 0.038 | 0.038 | 0 | 0.36 | 0.19 | 0.31 | 0.29 | 0.44 | 0.34 |
| 4 | 8234381 | rs3775905 | 0.352 | 1.354 | -1.554 | -0.153 | -0.042 | 0.038 | 0.038 | 0 | 0.32 | 0.15 | 0.3 | 0.24 | 0.4 | 0.27 |

| | | | | | | | | | | | | | | | | |
|---|---|---|---|---|---|---|---|---|---|---|---|---|---|---|---|---|
| 4 | 8254784 | rs1678336 | -0.712 | 0.928 | -1.637 | -0.32 | -0.069 | 0.04 | 0.04 | 0 | 0.69 | 0.88 | 0.75 | 0.78 | 0.55 | 0.8 |
| 4 | 8258650 | rs12186177 | -0.271 | 0.515 | -1.235 | 0.159 | 0 | 0.016 | 0.016 | 0.006 | 0.21 | 0.23 | 0.46 | 0.47 | 0.23 | 0.24 |
| 4 | 8267897 | rs4455399 | -0.002 | 0.349 | -0.873 | 0.05 | 0.001 | 0.026 | 0.026 | 0 | 0.78 | 0.73 | 0.54 | 0.5 | 0.8 | 0.68 |
| 4 | 8270320 | rs6833227 | 0.593 | -0.053 | -0.857 | -0.457 | 0.024 | 0.036 | 0.036 | 0.106 | 0.4 | 0.28 | 0.23 | 0.31 | 0.32 | 0.51 |
| 4 | 8280040 | rs12639755 | 0.473 | 0.182 | -0.993 | -0.442 | -0.055 | 0.012 | 0.012 | 0 | 0.27 | 0.11 | 0.23 | 0.24 | 0.42 | 0.2 |
| 4 | 8287563 | rs7690786 | -0.249 | -0.464 | -0.865 | -0.7 | -0.04 | 0.063 | 0.063 | 0 | 0.63 | 0.73 | 0.66 | 0.81 | 0.51 | 0.72 |
| 4 | 8291434 | rs13140502 | 1.38 | -1.111 | -0.965 | -0.595 | 0.022 | -0.018 | -0.018 | 0 | 0.26 | 0.15 | 0.16 | 0.3 | 0.19 | 0.38 |
| 4 | 8291918 | rs13132047 | -0.594 | -0.949 | -0.887 | -1.088 | -0.083 | 0.105 | 0.105 | 0 | 0.64 | 0.87 | 0.63 | 0.83 | 0.52 | 0.75 |
| 4 | 8292827 | rs13132711 | -0.358 | -1.043 | -0.344 | -1.128 | -0.005 | 0.01 | 0.01 | 0.847 | 0.6 | 0.65 | 0.61 | 0.83 | 0.52 | 0.75 |
| 4 | 8293356 | rs4696793 | 0.679 | -1.056 | -0.726 | -1.122 | -0.002 | 0.031 | 0.031 | 0.736 | 0.43 | 0.33 | 0.54 | 0.72 | 0.44 | 0.54 |
| 4 | 8293974 | rs4235258 | -0.687 | -0.408 | -1.461 | -0.844 | -0.118 | 0.068 | 0.068 | 0 | 0.67 | 0.91 | 0.64 | 0.83 | 0.53 | 0.73 |
| 4 | 8299162 | rs4583740 | 0.766 | -0.213 | -0.824 | -0.616 | 0.026 | 0.027 | 0.027 | 0.018 | 0.35 | 0.28 | 0.22 | 0.16 | 0.23 | 0.39 |
| 4 | 8303557 | rs4696796 | -0.678 | 0.125 | -0.643 | -0.49 | 0.014 | -0.034 | -0.034 | 0 | 0.64 | 0.58 | 0.69 | 0.86 | 0.53 | 0.79 |
| 4 | 8304097 | rs4696798 | 0.243 | 0.183 | -1.002 | -0.488 | -0.04 | -0.075 | -0.075 | 0 | 0.24 | 0.09 | 0.42 | 0.64 | 0.28 | 0.36 |
| 4 | 8304137 | rs13144236 | 0.676 | 0.265 | -1.053 | -0.441 | 0.024 | 0.03 | 0.03 | 0.525 | 0.37 | 0.27 | 0.25 | 0.17 | 0.26 | 0.41 |
| 4 | 8306091 | rs4696802 | -0.744 | 1.118 | -1.041 | 0.442 | -0.006 | -0.015 | -0.015 | 0 | 0.27 | 0.22 | 0.43 | 0.64 | 0.29 | 0.38 |
| 4 | 8310257 | rs6447857 | 0.905 | 1.048 | -0.774 | -0.279 | 0.001 | -0.023 | -0.023 | 0.413 | 0.71 | 0.65 | 0.76 | 0.84 | 0.7 | 0.68 |
| 4 | 8311043 | rs6818156 | 1.098 | 0.834 | -0.592 | -0.318 | 0.001 | -0.016 | -0.016 | 0.595 | 0.72 | 0.69 | 0.77 | 0.8 | 0.72 | 0.67 |
| 4 | 8312324 | rs4629434 | 0.824 | 0.711 | -0.628 | -0.346 | -0.002 | -0.019 | -0.019 | 0 | 0.77 | 0.7 | 0.86 | 0.8 | 0.8 | 0.7 |
| 4 | 8313595 | rs10023502 | 0.906 | 0.295 | -0.667 | -0.373 | -0.036 | -0.025 | -0.025 | 0 | 0.76 | 0.66 | 0.87 | 0.8 | 0.85 | 0.72 |
| 4 | 8315938 | rs6823378 | 0.812 | 0.272 | -0.65 | -0.404 | -0.005 | -0.02 | -0.02 | 0.658 | 0.87 | 0.86 | 0.95 | 0.98 | 0.94 | 0.92 |
| 4 | 8316128 | rs4696679 | 1.451 | 0.447 | -0.602 | -0.404 | -0.029 | -0.033 | -0.033 | 0 | 0.71 | 0.62 | 0.85 | 0.8 | 0.82 | 0.72 |
| 4 | 8320126 | rs12640734 | 0.196 | 0.78 | -1.519 | -0.646 | -0.081 | 0.05 | 0.05 | 0.09 | 0.29 | 0.1 | 0.32 | 0.16 | 0.39 | 0.23 |
| 4 | 8321215 | rs4696811 | 0.464 | 0.439 | -0.847 | -0.868 | -0.004 | -0.005 | -0.005 | 3.11 | 0.7 | 0.38 | 0.82 | 0.8 | 0.84 | 0.74 |
| 4 | 8321456 | rs6447860 | 0.395 | 0.46 | -0.459 | -0.773 | -0.021 | 0.006 | 0.006 | 0 | 0.15 | 0.24 | 0.1 | 0.18 | 0.11 | 0.18 |
| 4 | 8321709 | rs4696812 | 0.865 | 0.593 | -0.755 | -0.564 | -0.013 | 0.018 | 0.018 | 0 | 0.41 | 0.28 | 0.53 | 0.65 | 0.45 | 0.52 |
| 4 | 8326441 | rs1678330 | -0.506 | 1.005 | -0.378 | -0.743 | 0.001 | -0.003 | -0.003 | 0 | 0.28 | 0.32 | 0.31 | 0.24 | 0.31 | 0.31 |
| 4 | 8328353 | rs940141 | -0.581 | 0.986 | 0.089 | -0.759 | 0.01 | -0.019 | -0.019 | 0 | 0.28 | 0.38 | 0.32 | 0.24 | 0.31 | 0.31 |
| 4 | 8329912 | rs13144781 | -0.86 | 1.088 | 0.064 | -0.79 | -0.021 | -0.008 | -0.008 | 0 | 0.82 | 0.89 | 0.75 | 0.77 | 0.76 | 0.79 |
| 4 | 8331883 | rs754739 | -1.163 | 0.711 | -0.334 | -0.815 | -0.001 | 0.012 | 0.012 | 0.023 | 0.71 | 0.68 | 0.57 | 0.48 | 0.72 | 0.61 |
| 4 | 8331956 | rs754740 | 0.551 | 0.4 | -0.136 | -0.999 | 0.002 | 0.03 | 0.03 | 1.917 | 0.36 | 0.23 | 0.22 | 0.24 | 0.36 | 0.3 |
| 4 | 8333694 | rs1720011 | 1.061 | 0.459 | -0.122 | -0.931 | -0.039 | -0.013 | -0.013 | 0 | 0.18 | 0.08 | 0.24 | 0.23 | 0.25 | 0.21 |
| 4 | 8334382 | rs4696816 | 1.088 | 0.523 | 0.024 | -0.963 | -0.004 | -0.012 | -0.012 | 0.235 | 0.29 | 0.25 | 0.44 | 0.51 | 0.32 | 0.38 |
| 4 | 8336320 | rs2046027 | 0.25 | 0.448 | 0.033 | -0.912 | -0.003 | -0.015 | -0.015 | 0 | 0.37 | 0.45 | 0.25 | 0.25 | 0.37 | 0.28 |
| 4 | 8342853 | rs827018 | 0.801 | 0.694 | 0.018 | -0.78 | 0.003 | 0 | 0 | 1.806 | 0.18 | 0.23 | 0.25 | 0.21 | 0.26 | 0.21 |
| 4 | 8350786 | rs7682386 | -0.196 | 0.614 | 0.083 | -1.013 | -0.002 | 0.006 | 0.006 | 0.043 | 0.46 | 0.48 | 0.4 | 0.26 | 0.47 | 0.34 |
| 4 | 8353829 | rs1678272 | 0.927 | 0.364 | -0.434 | -0.795 | -0.01 | -0.024 | -0.024 | 0.004 | 0.15 | 0.07 | 0.09 | 0.24 | 0.2 | 0.2 |
| 4 | 8354600 | rs4696820 | 0.104 | -0.221 | -0.432 | -1.172 | -0.02 | -0.003 | -0.003 | 0 | 0.57 | 0.67 | 0.62 | 0.54 | 0.51 | 0.51 |
| 4 | 8355890 | rs4696821 | 1.106 | -0.422 | -0.649 | -1.219 | -0.031 | -0.102 | -0.102 | 0.382 | 0.27 | 0.1 | 0.44 | 0.48 | 0.31 | 0.36 |
| 4 | 8356092 | rs11933783 | 0.694 | -1.07 | -0.387 | -1.536 | -0.031 | -0.049 | -0.049 | 0 | 0.09 | 0.27 | 0.03 | 0 | 0.02 | 0.08 |
| 4 | 8356253 | rs13103277 | 0.435 | -1.094 | -0.796 | -1.515 | 0.007 | 0.075 | 0.075 | 0 | 0.27 | 0.15 | 0.16 | 0.05 | 0.24 | 0.14 |
| 4 | 8357645 | rs13129831 | -0.568 | -0.873 | -0.491 | -1.379 | -0.035 | -0.107 | -0.107 | 0 | 0.4 | 0.52 | 0.2 | 0.05 | 0.25 | 0.22 |
| 4 | 8367799 | rs899491 | 1.355 | -0.843 | -1.142 | -1.073 | 0.003 | 0.032 | 0.032 | 0.248 | 0.41 | 0.29 | 0.61 | 0.73 | 0.59 | 0.51 |
| 4 | 8368433 | rs7809 | -0.531 | -0.801 | -0.729 | -1.023 | 0.005 | 0.001 | 0.001 | 1.458 | 0.33 | 0.28 | 0.15 | 0.27 | 0.27 | 0.3 |
| 4 | 8368604 | rs708244 | -0.713 | -0.814 | -0.818 | -1.097 | 0.012 | 0.008 | 0.008 | 0.519 | 0.37 | 0.28 | 0.16 | 0.27 | 0.29 | 0.3 |
| 4 | 8371554 | rs735812 | -0.342 | -0.966 | -0.836 | -1.223 | -0.04 | -0.11 | -0.11 | 0.139 | 0.26 | 0.46 | 0.13 | 0.05 | 0.16 | 0.15 |
| 4 | 8373088 | rs747580 | -2.166 | -0.395 | -1.305 | -0.956 | -0.057 | -0.117 | -0.117 | 0 | 0.71 | 0.9 | 0.51 | 0.5 | 0.61 | 0.66 |
| 4 | 8373832 | rs747582 | -1.158 | -0.215 | -0.747 | -0.803 | -0.002 | 0.007 | 0.007 | 0 | 0.57 | 0.54 | 0.43 | 0.5 | 0.55 | 0.56 |
| 4 | 8377058 | rs11735859 | 1.059 | -0.167 | -0.863 | -0.559 | -0.003 | -0.019 | -0.019 | 0 | 0.14 | 0.12 | 0.08 | 0.24 | 0.17 | 0.17 |
| 4 | 8378959 | rs3796751 | -0.124 | -0.028 | -0.661 | -0.351 | 0 | 0.005 | 0.005 | 0.011 | 0.2 | 0.13 | 0.16 | 0.14 | 0.15 | 0.08 |
| 4 | 8380149 | rs4481223 | -0.76 | -0.238 | -0.051 | -0.536 | -0.018 | -0.041 | -0.041 | 0 | 0.26 | 0.6 | 0.15 | 0.12 | 0.17 | 0.1 |
| 4 | 8383899 | rs12504709 | 0.37 | -0.411 | -0.159 | -0.859 | -0.011 | -0.038 | -0.038 | 1.074 | 0.08 | 0.13 | 0.03 | 0 | 0.05 | 0.07 |
| 4 | 8386107 | rs7697154 | -1.797 | -0.436 | -0.229 | -0.869 | 0.026 | 0.065 | 0.065 | 0.037 | 0.34 | 0.78 | 0.19 | 0.11 | 0.23 | 0.18 |
| 4 | 8389187 | rs7696306 | -0.647 | -0.404 | 0.276 | -0.741 | -0.007 | -0.019 | -0.019 | 0.006 | 0.25 | 0.38 | 0.19 | 0.24 | 0.21 | 0.3 |
| 4 | 8390646 | rs1568692 | -2.151 | -0.431 | 0.287 | -0.769 | 0.025 | 0.072 | 0.072 | 0 | 0.38 | 0.77 | 0.24 | 0.24 | 0.29 | 0.39 |
| 4 | 8394463 | rs11934642 | 0.062 | -0.523 | 0.631 | -0.768 | -0.016 | -0.007 | -0.007 | 0.008 | 0.1 | 0.17 | 0.09 | 0.13 | 0.07 | 0.22 |
| 4 | 8397625 | rs6849528 | -0.532 | -0.509 | 0.664 | -0.779 | -0.011 | -0.096 | -0.096 | 0 | 0.14 | 0.39 | 0.05 | 0 | 0.09 | 0.1 |
| 4 | 8397686 | rs6810854 | 1.981 | -0.48 | 0.859 | -0.747 | 0.023 | -0.017 | -0.017 | 0 | 0.05 | 0.21 | 0.02 | 0 | 0.04 | 0.02 |
| 4 | 8403036 | rs4543095 | -0.821 | -0.486 | 0.729 | -0.775 | -0.027 | -0.019 | -0.019 | 1.043 | 0.24 | 0.37 | 0.17 | 0.23 | 0.17 | 0.29 |
| 4 | 8404346 | rs1880025 | 0.623 | -0.416 | -0.071 | -0.741 | -0.042 | -0.043 | -0.043 | 3.327 | 0.35 | 0.14 | 0.35 | 0.42 | 0.42 | 0.47 |
| 4 | 8409237 | rs10016221 | -0.463 | -0.434 | 0.107 | -0.705 | -0.008 | -0.005 | -0.005 | 0 | 0.24 | 0.26 | 0.16 | 0.23 | 0.16 | 0.28 |
| 4 | 8416976 | rs3775889 | 1.901 | -0.206 | -0.29 | -0.596 | -0.045 | -0.086 | -0.086 | 0 | 0.18 | 0.03 | 0.22 | 0.32 | 0.23 | 0.24 |
| 4 | 8418801 | rs11736150 | -0.416 | -0.594 | 0.121 | -0.685 | -0.011 | -0.011 | -0.011 | 0 | 0.12 | 0.16 | 0.1 | 0.12 | 0.09 | 0.22 |
| 4 | 8421345 | rs2688228 | 0.632 | -0.619 | 0.119 | -0.725 | -0.01 | -0.014 | -0.014 | 0.898 | 0.36 | 0.23 | 0.34 | 0.42 | 0.41 | 0.47 |
| 4 | 8424831 | rs2140336 | -0.604 | -0.694 | 0.165 | -0.775 | -0.001 | 0.002 | 0.002 | 0 | 0.49 | 0.51 | 0.4 | 0.43 | 0.51 | 0.57 |
| 4 | 8429817 | rs2631731 | -3.243 | -0.622 | -0.108 | -0.722 | -0.029 | -0.073 | -0.073 | 0.366 | 0.75 | 0.9 | 0.59 | 0.66 | 0.72 | 0.86 |
| 4 | 8458394 | rs13129077 | 1.732 | -0.96 | 0.057 | -0.755 | 0.011 | -0.006 | -0.006 | 0 | 0.11 | 0.02 | 0.05 | 0.11 | 0.07 | 0.06 |
| 4 | 8461517 | rs2631749 | 0.144 | -1.111 | 0.18 | -0.827 | 0.001 | 0.001 | 0.001 | 0.014 | 0.49 | 0.7 | 0.55 | 0.47 | 0.48 | 0.32 |
| 4 | 8467375 | rs10009833 | -0.074 | -0.817 | 0.291 | -0.723 | -0.013 | -0.006 | -0.006 | 0 | 0.11 | 0.16 | 0.1 | 0.13 | 0.08 | 0.22 |

| | | | | | | | | | | | | | | | | |
|---|---|---|---|---|---|---|---|---|---|---|---|---|---|---|---|---|
| 4 | 8473050 | rs2631768 | -2.297 | -0.852 | 0.164 | -0.603 | -0.009 | -0.003 | -0.003 | 0 | 0.7 | 0.71 | 0.75 | 0.76 | 0.64 | 0.83 |
| 4 | 8476118 | rs2631735 | -3.478 | -0.89 | 0.156 | -0.616 | -0.039 | -0.029 | -0.029 | 0 | 0.76 | 0.87 | 0.77 | 0.76 | 0.68 | 0.85 |
| 4 | 8479754 | rs2631738 | -0.659 | -0.919 | 0.196 | -1.202 | 0.003 | -0.001 | -0.001 | 0 | 0.5 | 0.37 | 0.45 | 0.55 | 0.5 | 0.69 |
| 4 | 8490934 | rs2631746 | -2.879 | -0.943 | -0.223 | -1.267 | -0.039 | -0.077 | -0.077 | 0.077 | 0.7 | 0.9 | 0.57 | 0.65 | 0.67 | 0.8 |
| 4 | 8498456 | rs10001657 | -0.703 | -0.706 | 0.112 | -0.943 | -0.023 | -0.019 | -0.019 | 0.215 | 0.14 | 0.22 | 0.11 | 0.12 | 0.09 | 0.22 |
| 4 | 8503281 | rs13111630 | 0.326 | -0.344 | 0.311 | -0.788 | 0 | 0.004 | 0.004 | 0 | 0.15 | 0.15 | 0.07 | 0.11 | 0.13 | 0.1 |
| 4 | 8503359 | rs1949733 | -2.265 | -0.066 | 0.45 | -0.594 | -0.029 | -0.069 | -0.069 | 0.833 | 0.74 | 0.89 | 0.59 | 0.66 | 0.71 | 0.87 |
| 4 | 8512889 | rs3115397 | -0.207 | -0.136 | 0.747 | -0.718 | -0.003 | -0.022 | -0.022 | 0.32 | 0.82 | 0.79 | 0.83 | 0.92 | 0.84 | 0.73 |
| 4 | 8516605 | rs3103075 | 0.385 | -0.019 | 0.718 | -0.677 | 0 | -0.024 | -0.024 | 0 | 0.08 | 0.11 | 0.05 | 0 | 0.08 | 0.05 |
| 4 | 8518896 | rs7665475 | 0.912 | 0.003 | 0.771 | -0.57 | 0 | -0.001 | -0.001 | 0.154 | 0.32 | 0.34 | 0.3 | 0.43 | 0.35 | 0.44 |
| 4 | 8522221 | rs13127781 | 0.317 | -0.068 | 0.507 | -0.566 | 0.013 | -0.024 | -0.024 | 0 | 0.09 | 0.16 | 0.09 | 0.06 | 0.13 | 0.23 |
| 4 | 8524767 | rs3103078 | 0.334 | -0.36 | 0.329 | -0.413 | -0.03 | -0.022 | -0.022 | 0 | 0.44 | 0.21 | 0.5 | 0.48 | 0.45 | 0.28 |
| 4 | 8525665 | rs7679494 | -0.216 | -0.233 | 0.669 | -0.476 | -0.039 | -0.07 | -0.07 | 1.912 | 0.83 | 0.72 | 0.9 | 0.98 | 0.92 | 0.93 |
| 4 | 8527461 | rs10461081 | 0.286 | -0.024 | 0.091 | -0.296 | 0.002 | 0.002 | 0.002 | 0.039 | 0.12 | 0.17 | 0.09 | 0.13 | 0.12 | 0.11 |
| 4 | 8529240 | rs4696710 | 1.366 | -0.395 | -0.014 | -0.606 | 0.028 | 0.062 | 0.062 | 0.18 | 0.61 | 0.21 | 0.76 | 0.84 | 0.75 | 0.84 |
| 4 | 8529812 | rs13140128 | 0.605 | -0.471 | 0 | -0.539 | -0.007 | -0.016 | -0.016 | 0 | 0.37 | 0.3 | 0.46 | 0.39 | 0.39 | 0.19 |
| 4 | 8534283 | rs6850810 | -1.279 | -0.558 | 0.077 | -0.491 | -0.042 | -0.034 | -0.034 | 0.987 | 0.87 | 0.64 | 0.9 | 0.95 | 0.94 | 0.95 |
| 4 | 8536360 | rs3103082 | -0.673 | -0.194 | 0.138 | -0.318 | -0.014 | -0.023 | -0.023 | 0 | 0.63 | 0.73 | 0.52 | 0.61 | 0.6 | 0.81 |
| 4 | 8538007 | rs3115401 | -0.671 | -0.261 | 0.332 | 0.007 | -0.038 | -0.002 | -0.002 | 0.004 | 0.78 | 0.63 | 0.86 | 0.83 | 0.88 | 0.88 |
| 4 | 8538981 | rs3103091 | 0.444 | -0.26 | 0.3 | -0.153 | -0.033 | -0.019 | -0.019 | 0 | 0.22 | 0.35 | 0.14 | 0.15 | 0.12 | 0.12 |
| 4 | 8541869 | rs6447879 | -0.115 | 0.108 | -0.356 | -0.062 | -0.031 | 0.015 | 0.015 | 0 | 0.63 | 0.78 | 0.62 | 0.65 | 0.5 | 0.48 |
| 4 | 8546816 | rs3103086 | -0.2 | 0.805 | -0.356 | 0.307 | 0.069 | 0.025 | 0.025 | 0 | 0.31 | 0.42 | 0.4 | 0.46 | 0.47 | 0.46 |
| 4 | 8549845 | rs10805209 | -0.053 | 0.645 | -1.238 | 0.599 | -0.031 | -0.034 | -0.034 | 0 | 0.31 | 0.1 | 0.33 | 0.46 | 0.4 | 0.44 |
| 4 | 8554894 | rs4696842 | -0.205 | 0.561 | -0.861 | 0.73 | 0.015 | -0.022 | -0.022 | 0.035 | 0.44 | 0.26 | 0.45 | 0.49 | 0.36 | 0.54 |
| 4 | 8556609 | rs7682449 | -1.293 | 0.43 | 0.294 | 0.773 | 0 | 0.053 | 0.053 | 0.415 | 0.12 | 0.41 | 0.06 | 0.19 | 0.05 | 0.13 |
| 4 | 8557865 | rs3115378 | 0.528 | 0.513 | 0.054 | 0.849 | -0.002 | 0.021 | 0.021 | 0 | 0.65 | 0.53 | 0.51 | 0.48 | 0.62 | 0.62 |
| 4 | 8559576 | rs11937563 | -1.214 | 0.199 | 0.503 | 1.217 | -0.035 | 0.269 | 0.269 | 2.939 | 0.12 | 0.52 | 0.3 | 0.35 | 0.08 | 0.14 |
| 4 | 8562668 | rs3103074 | -0.143 | 0.335 | -0.273 | 0.521 | -0.04 | 0.032 | 0.032 | 0.211 | 0.21 | 0.08 | 0.16 | 0.17 | 0.27 | 0.24 |
| 4 | 8566213 | rs4358385 | 0.302 | 1.719 | -0.812 | 0.519 | -0.059 | -0.038 | -0.038 | 0 | 0.09 | 0.02 | 0.24 | 0.15 | 0.24 | 0.13 |
| 4 | 8567305 | rs13108246 | -0.751 | 0.875 | -1.008 | -0.212 | -0.006 | 0.098 | 0.098 | 0 | 0.36 | 0.23 | 0.22 | 0.16 | 0.35 | 0.29 |
| 4 | 8570973 | rs11735274 | -0.304 | 0.954 | -1.403 | -0.374 | -0.084 | 0.087 | 0.087 | 0.515 | 0.28 | 0.02 | 0.16 | 0.12 | 0.3 | 0.25 |
| 4 | 8574490 | rs3115395 | -1.064 | 0.463 | -0.622 | -0.578 | 0.005 | 0.007 | 0.007 | 0 | 0.47 | 0.36 | 0.51 | 0.83 | 0.48 | 0.58 |
| 4 | 8579149 | rs6838103 | -0.153 | 0.766 | -0.94 | -0.918 | 0.043 | -0.038 | -0.038 | 0 | 0.78 | 0.66 | 0.68 | 0.88 | 0.69 | 0.78 |
| 4 | 8579193 | rs6838127 | -0.278 | 0.904 | -1.103 | -0.887 | 0.144 | -0.019 | -0.019 | 1.184 | 0.84 | 0.59 | 0.69 | 0.88 | 0.72 | 0.79 |
| 4 | 8579758 | rs1282 | 1.214 | 0.205 | -2.384 | -1.22 | 0.022 | 0.027 | 0.027 | 0 | 0.27 | 0.18 | 0.14 | 0.02 | 0.16 | 0.17 |
| 4 | 8580952 | rs2270269 | 0.613 | 0.325 | -2.114 | -0.387 | -0.026 | 0.188 | 0.188 | 0.716 | 0.52 | 0.19 | 0.33 | 0.14 | 0.54 | 0.44 |
| 4 | 8584879 | rs4235272 | 0.444 | 0.9 | -1.74 | 0.392 | 0.002 | 0.052 | 0.052 | 0 | 0.46 | 0.24 | 0.55 | 0.42 | 0.47 | 0.47 |
| 4 | 8586739 | rs7696507 | 0.437 | 1.774 | -1.478 | 0.749 | -0.106 | -0.015 | -0.015 | 0.487 | 0.8 | 0.94 | 0.81 | 0.79 | 0.68 | 0.81 |
| 4 | 8589023 | rs9685931 | -0.846 | 0.745 | -0.781 | 0.692 | -0.005 | -0.022 | -0.022 | 0 | 0.11 | 0.2 | 0.05 | 0.04 | 0.11 | 0.12 |
| 4 | 8589263 | rs9799717 | -1.678 | 0.4 | -0.051 | 0.526 | -0.032 | -0.067 | -0.067 | 0.005 | 0.17 | 0.45 | 0.08 | 0.05 | 0.13 | 0.2 |
| 4 | 8589714 | rs9799378 | -1.762 | 0.352 | -0.052 | 1.174 | -0.035 | 0.048 | 0.048 | 0.121 | 0.18 | 0.44 | 0.07 | 0.21 | 0.15 | 0.28 |
| 4 | 8590511 | rs4696853 | 1.544 | 0.351 | -0.36 | 0.647 | -0.021 | -0.053 | -0.053 | 0 | 0.71 | 0.55 | 0.86 | 0.91 | 0.74 | 0.66 |
| 4 | 8596478 | rs10938726 | -0.092 | 1.09 | -0.716 | 0.8 | -0.054 | -0.015 | -0.015 | 0 | 0.13 | 0.03 | 0.19 | 0.16 | 0.23 | 0.09 |
| 4 | 8597140 | rs4696855 | 1.074 | 0.788 | -0.338 | 1.177 | -0.02 | 0.008 | 0.008 | 2.753 | 0.73 | 0.55 | 0.88 | 0.7 | 0.79 | 0.69 |
| 4 | 8604153 | rs3822202 | 1.48 | 0.534 | -0.061 | 1.031 | 0.001 | -0.004 | -0.004 | 0 | 0.58 | 0.42 | 0.67 | 0.65 | 0.56 | 0.64 |
| 4 | 8606485 | rs6834153 | -0.638 | 0.435 | 0.095 | 1.033 | 0.012 | 0.075 | 0.075 | 0.031 | 0.05 | 0.19 | 0.02 | 0.13 | 0.02 | 0.04 |
| 4 | 8606863 | rs6838212 | -1.275 | 0.071 | -0.087 | 0.876 | -0.061 | -0.001 | -0.001 | 0.06 | 0.19 | 0.42 | 0.1 | 0.2 | 0.11 | 0.27 |
| 4 | 8608634 | rs2302580 | 2.029 | 0.275 | -0.551 | 1.027 | -0.068 | -0.015 | -0.015 | 0 | 0.44 | 0.1 | 0.58 | 0.42 | 0.46 | 0.46 |
| 4 | 8620464 | rs2302569 | -0.091 | 0.163 | -1.102 | 1.293 | 0.017 | -0.039 | -0.039 | 0 | 0.15 | 0.08 | 0.07 | 0.31 | 0.12 | 0.24 |
| 4 | 8621386 | rs1050464 | 1.218 | 0.339 | -1.252 | 1.047 | -0.003 | 0.012 | 0.012 | 2.214 | 0.48 | 0.19 | 0.62 | 0.44 | 0.48 | 0.51 |
| 4 | 9996890 | rs13111638 | -1.111 | -1.272 | -0.422 | -0.454 | -0.009 | 0.012 | 0.012 | 0 | 0.83 | 0.85 | 0.74 | 0.98 | 0.83 | 0.84 |
| 4 | 9997060 | rs4547795 | 0.643 | -1.268 | -0.337 | -0.447 | 0.021 | -0.038 | -0.038 | 1.474 | 0.19 | 0.39 | 0.13 | 0.1 | 0.22 | 0.03 |
| 4 | 9998493 | rs13113918 | -0.169 | -1.262 | -0.334 | -0.468 | -0.001 | -0.027 | -0.027 | 0.018 | 0.78 | 0.78 | 0.72 | 0.97 | 0.8 | 0.81 |
| 4 | 10000236 | rs7696536 | 0.582 | -1.103 | -0.341 | -0.362 | 0.021 | -0.038 | -0.038 | 0 | 0.19 | 0.39 | 0.13 | 0.1 | 0.22 | 0.03 |
| 4 | 10001861 | rs1014290 | 0.353 | -1.198 | -0.388 | -0.225 | 0.002 | -0.004 | -0.004 | 0 | 0.68 | 0.71 | 0.6 | 0.59 | 0.76 | 0.69 |
| 4 | 10004832 | rs10023068 | 0.773 | -1.263 | -0.54 | -0.345 | -0.03 | 0.012 | 0.012 | 0 | 0.64 | 0.59 | 0.63 | 0.51 | 0.81 | 0.65 |
| 4 | 10005435 | rs6853437 | 0.476 | -1.141 | -0.472 | -0.278 | -0.032 | 0.035 | 0.035 | 0 | 0.69 | 0.59 | 0.64 | 0.51 | 0.82 | 0.65 |
| 4 | 10024303 | rs12509955 | 0.169 | -1.185 | -0.544 | -0.643 | -0.031 | -0.158 | -0.158 | 0.07 | 0.7 | 0.53 | 0.69 | 0.98 | 0.8 | 0.83 |
| 4 | 10027542 | rs6820230 | -0.538 | -0.906 | -0.508 | -0.679 | 0.018 | -0.058 | -0.058 | 5.728 | 0.24 | 0.55 | 0.22 | 0.1 | 0.3 | 0.07 |
| 4 | 10036140 | rs10006397 | 0.249 | -1.058 | -0.704 | -0.552 | 0.033 | -0.027 | -0.027 | 0 | 0.66 | 0.75 | 0.67 | 0.51 | 0.82 | 0.67 |
| 4 | 10038112 | rs10012779 | 0.52 | -1.141 | -0.47 | -0.485 | -0.015 | -0.086 | -0.086 | 0 | 0.13 | 0.4 | 0.11 | 0 | 0.09 | 0.02 |
| 4 | 10062326 | rs6834555 | 0.492 | -1.102 | -1.086 | -0.376 | 0.023 | -0.016 | -0.016 | 0.972 | 0.65 | 0.74 | 0.55 | 0.52 | 0.79 | 0.66 |
| 4 | 10062849 | rs6820756 | 0.467 | -1.1 | -1.085 | -0.375 | 0.023 | -0.016 | -0.016 | 0.037 | 0.65 | 0.74 | 0.55 | 0.52 | 0.79 | 0.66 |
| 4 | 10099340 | rs13441 | -0.13 | -1.031 | -1.791 | -0.576 | -0.008 | 0.013 | 0.013 | 0 | 0.68 | 0.39 | 0.71 | 0.65 | 0.71 | 0.5 |
| 4 | 10104670 | rs717615 | -0.933 | -1.029 | -1.86 | -0.64 | -0.012 | 0.002 | 0.002 | 0 | 0.46 | 0.3 | 0.57 | 0.5 | 0.48 | 0.39 |
| 4 | 10105739 | rs3756224 | -0.725 | -1.137 | -2.509 | -0.446 | 0.013 | -0.026 | -0.026 | 0 | 0.38 | 0.18 | 0.37 | 0.54 | 0.32 | 0.55 |
| 4 | 10105797 | rs3756223 | -0.796 | -1.192 | -2.147 | -0.431 | 0.001 | 0.003 | 0.003 | 0 | 0.52 | 0.38 | 0.48 | 0.51 | 0.46 | 0.41 |
| 4 | 10115065 | rs4604059 | 0.728 | -0.944 | -1.884 | -0.299 | -0.001 | 0.016 | 0.016 | 0 | 0.48 | 0.36 | 0.48 | 0.64 | 0.54 | 0.37 |
| 4 | 10120609 | rs12374320 | 0.863 | -1.049 | -2.054 | -0.328 | 0.057 | 0.098 | 0.098 | 0.298 | 0.48 | 0.21 | 0.55 | 0.34 | 0.61 | 0.39 |

| | | | | | | | | | | | | | | | | |
|---|---|---|---|---|---|---|---|---|---|---|---|---|---|---|---|---|
| 4 | 10125808 | rs7699512 | -0.498 | -1.088 | -1.966 | -0.22 | -0.008 | -0.002 | -0.002 | 0.821 | 0.43 | 0.33 | 0.57 | 0.49 | 0.46 | 0.39 |
| 4 | 10125874 | rs7699671 | 0.288 | -1.085 | -2.011 | -0.275 | -0.032 | 0.037 | 0.037 | 2.065 | 0.3 | 0.21 | 0.39 | 0.2 | 0.39 | 0.26 |
| 4 | 10127141 | rs4467562 | 0.515 | -1.287 | -1.243 | -0.162 | -0.004 | -0.001 | -0.001 | 0 | 0.45 | 0.42 | 0.36 | 0.44 | 0.36 | 0.28 |
| 4 | 10131932 | rs12501508 | -0.757 | -0.616 | -0.946 | 0.226 | 0.055 | 0.228 | 0.228 | 0 | 0.86 | 0.62 | 0.78 | 0.59 | 0.82 | 0.59 |
| 4 | 10138746 | rs11732828 | 0.54 | -0.674 | -0.991 | -0.448 | 0.01 | 0.048 | 0.048 | 0.018 | 0.49 | 0.4 | 0.45 | 0.17 | 0.45 | 0.29 |
| 4 | 10145890 | rs10024447 | 0.711 | -0.725 | -0.766 | -0.289 | 0.001 | 0.011 | 0.011 | 0 | 0.26 | 0.33 | 0.25 | 0.36 | 0.27 | 0.49 |
| 4 | 10146654 | rs4697715 | 0.71 | -0.727 | -0.848 | -0.389 | 0.006 | 0.065 | 0.065 | 0 | 0.5 | 0.38 | 0.44 | 0.17 | 0.48 | 0.3 |
| 4 | 10148828 | rs6813385 | -0.218 | -0.626 | -0.701 | -0.338 | 0.009 | 0.008 | 0.008 | 0.24 | 0.37 | 0.43 | 0.36 | 0.53 | 0.43 | 0.56 |
| 4 | 10149595 | rs4697936 | 0.758 | -0.677 | -0.761 | -0.387 | 0.009 | 0.049 | 0.049 | 0 | 0.48 | 0.38 | 0.44 | 0.17 | 0.45 | 0.3 |
| 4 | 10154554 | rs6814532 | -1.305 | 0.047 | -0.512 | 0.112 | 0.005 | 0.036 | 0.036 | 0 | 0.84 | 0.76 | 0.79 | 0.69 | 0.84 | 0.84 |
| 4 | 10156321 | rs11929718 | -0.79 | -0.911 | -0.426 | -0.176 | -0.007 | 0.003 | 0.003 | 0.242 | 0.61 | 0.61 | 0.64 | 0.56 | 0.65 | 0.69 |
| 4 | 10165779 | rs16894270 | 1.059 | -0.981 | -0.6 | -0.436 | 0.005 | -0.003 | -0.003 | 0.585 | 0.29 | 0.36 | 0.28 | 0.24 | 0.3 | 0.38 |
| 4 | 10168849 | rs1001216 | 0.879 | -0.997 | -0.564 | -0.436 | -0.002 | 0.002 | 0.002 | 0 | 0.17 | 0.19 | 0.26 | 0.23 | 0.12 | 0.27 |
| 4 | 10172479 | rs6835689 | -0.043 | -0.809 | -0.49 | -0.547 | 0.034 | -0.033 | -0.033 | 0 | 0.27 | 0.39 | 0.33 | 0.2 | 0.34 | 0.2 |
| 4 | 10173019 | rs4697952 | -0.492 | -0.852 | -0.529 | -0.573 | 0.006 | 0.004 | 0.004 | 1.836 | 0.53 | 0.59 | 0.57 | 0.57 | 0.58 | 0.68 |
| 4 | 10173696 | rs10034405 | 0.665 | -0.787 | -0.401 | -0.583 | -0.012 | -0.069 | -0.069 | 4.387 | 0.12 | 0.25 | 0.1 | 0 | 0.1 | 0.02 |
| 4 | 10179769 | rs6449342 | -0.646 | -0.66 | -0.5 | -0.393 | 0.01 | 0.001 | 0.001 | 1.988 | 0.54 | 0.63 | 0.64 | 0.55 | 0.66 | 0.7 |
| 4 | 10180823 | rs6811287 | 0.253 | -0.575 | -0.821 | -0.403 | -0.008 | -0.02 | -0.02 | 0 | 0.42 | 0.34 | 0.51 | 0.44 | 0.42 | 0.66 |
| 4 | 10183108 | rs887733 | -0.048 | -0.659 | -0.785 | -0.309 | -0.021 | -0.018 | -0.018 | 0 | 0.3 | 0.25 | 0.42 | 0.43 | 0.42 | 0.33 |
| 4 | 10188326 | rs956312 | -0.194 | -0.819 | -0.591 | -0.283 | 0.031 | 0.008 | 0.008 | 2.759 | 0.53 | 0.42 | 0.62 | 0.56 | 0.66 | 0.68 |
| 4 | 10202997 | rs2868420 | -0.337 | -0.828 | -0.519 | -0.292 | -0.008 | -0.001 | -0.001 | 0.378 | 0.55 | 0.52 | 0.62 | 0.57 | 0.66 | 0.68 |
| 4 | 10209251 | rs11947517 | -0.586 | -0.927 | -0.415 | -0.333 | 0.023 | -0.002 | -0.002 | 0.696 | 0.58 | 0.72 | 0.64 | 0.58 | 0.67 | 0.71 |
| 4 | 10240663 | rs11734783 | 0.943 | -0.907 | -0.422 | -0.375 | -0.002 | -0.005 | -0.005 | 0 | 0.16 | 0.24 | 0.25 | 0.15 | 0.16 | 0.15 |
| 4 | 10241132 | rs917821 | 0.109 | -1.163 | -0.431 | -0.336 | -0.084 | 0.06 | 0.06 | 0 | 0.78 | 0.54 | 0.79 | 0.69 | 0.91 | 0.85 |
| 4 | 10245087 | rs1978274 | 0.907 | -1.114 | -0.605 | -0.446 | -0.003 | -0.009 | -0.009 | 0.211 | 0.22 | 0.24 | 0.29 | 0.15 | 0.2 | 0.14 |
| 4 | 10258968 | rs4697986 | 0.273 | -0.791 | -1.183 | -0.222 | 0.037 | 0.006 | 0.006 | 1.208 | 0.52 | 0.67 | 0.62 | 0.54 | 0.64 | 0.68 |
| 4 | 10259959 | rs4697988 | 0.047 | -0.908 | -0.924 | -0.111 | -0.024 | 0.002 | 0.002 | 0.473 | 0.73 | 0.7 | 0.8 | 0.69 | 0.89 | 0.85 |
| 4 | 10272591 | rs7676442 | 0.106 | -0.45 | -1.01 | -0.052 | -0.001 | 0.003 | 0.003 | 0 | 0.67 | 0.68 | 0.7 | 0.52 | 0.68 | 0.74 |
| 4 | 10273244 | rs10489072 | -0.388 | -0.44 | -1.026 | -0.048 | 0.001 | 0.006 | 0.006 | 0 | 0.75 | 0.75 | 0.72 | 0.52 | 0.69 | 0.74 |
| 4 | 10273329 | rs10939814 | -0.214 | -0.433 | -1.034 | -0.072 | 0.004 | 0.012 | 0.012 | 0 | 0.74 | 0.72 | 0.71 | 0.58 | 0.69 | 0.81 |
| 4 | 10275470 | rs1860910 | 0.935 | -0.566 | -1.077 | -0.229 | 0 | -0.001 | -0.001 | 3.136 | 0.51 | 0.46 | 0.43 | 0.44 | 0.48 | 0.67 |
| 4 | 10276467 | rs12510549 | 1.256 | -0.676 | -1.066 | -0.195 | -0.003 | 0.012 | 0.012 | 0.064 | 0.19 | 0.13 | 0.28 | 0.15 | 0.2 | 0.13 |
| 4 | 10281141 | rs734662 | -1.477 | -0.473 | -0.955 | -0.004 | -0.02 | 0.039 | 0.039 | 0 | 0.84 | 0.79 | 0.82 | 0.69 | 0.91 | 0.85 |
| 4 | 10283627 | rs4698005 | -1.636 | -0.322 | -1.063 | 0.053 | 0.004 | -0.006 | -0.006 | 0 | 0.85 | 0.87 | 0.82 | 0.7 | 0.91 | 0.88 |
| 4 | 10310495 | rs9991653 | 0.758 | -0.024 | -1.115 | -0.136 | 0.03 | 0.021 | 0.021 | 0 | 0.33 | 0.44 | 0.36 | 0.41 | 0.45 | 0.55 |
| 4 | 10313042 | rs714086 | 1.945 | 0.007 | -1.367 | -0.026 | -0.066 | -0.017 | -0.017 | 0.057 | 0.19 | 0.02 | 0.16 | 0.12 | 0.21 | 0.29 |
| 4 | 10313460 | rs6853389 | 1.42 | 0.128 | -0.945 | -0.068 | 0.002 | 0.011 | 0.011 | 0 | 0.24 | 0.33 | 0.24 | 0.23 | 0.23 | 0.42 |
| 4 | 10318433 | rs2024276 | -0.794 | -0.232 | -0.996 | -0.036 | -0.005 | 0.004 | 0.004 | 0 | 0.78 | 0.8 | 0.81 | 0.6 | 0.9 | 0.84 |
| 4 | 10323935 | rs993173 | -0.223 | -0.345 | -1.138 | -0.24 | 0.012 | -0.027 | -0.027 | 0 | 0.38 | 0.49 | 0.42 | 0.32 | 0.43 | 0.27 |
| 4 | 10324160 | rs9291683 | 1.092 | -0.243 | -1.2 | -0.179 | -0.015 | 0.02 | 0.02 | 0 | 0.42 | 0.33 | 0.39 | 0.28 | 0.48 | 0.56 |
| 4 | 10335524 | rs2192090 | -1.721 | -0.528 | -1.217 | -0.095 | 0.007 | -0.004 | -0.004 | 0 | 0.85 | 0.87 | 0.82 | 0.61 | 0.93 | 0.86 |
| 4 | 10342723 | rs10938761 | -1.571 | -0.432 | -1.171 | -0.047 | 0.016 | -0.023 | -0.023 | 0 | 0.83 | 0.89 | 0.82 | 0.61 | 0.92 | 0.85 |
| 4 | 10344431 | rs7654423 | 0.817 | -0.561 | -0.952 | -0.304 | -0.038 | -0.057 | -0.057 | 5.942 | 0.1 | 0.22 | 0.02 | 0 | 0.02 | 0 |
| 4 | 10365093 | rs4235361 | -1.711 | -0.448 | -1.336 | -0.211 | 0.014 | -0.023 | -0.023 | 0 | 0.83 | 0.89 | 0.82 | 0.61 | 0.92 | 0.85 |
| 4 | 10375328 | rs10938772 | 1.404 | -0.223 | -1.365 | -0.271 | -0.011 | 0.021 | 0.021 | 0.061 | 0.19 | 0.1 | 0.28 | 0.14 | 0.21 | 0.17 |
| 4 | 10378602 | rs4698043 | 0.321 | -0.399 | -1.166 | -0.542 | 0 | 0 | 0 | 0 | 0.51 | 0.53 | 0.63 | 0.55 | 0.64 | 0.69 |
| 4 | 10383005 | rs7677806 | 0.365 | -0.396 | -1.154 | -0.541 | -0.001 | 0 | 0 | 0 | 0.5 | 0.52 | 0.62 | 0.55 | 0.64 | 0.7 |
| 4 | 10388313 | rs9990427 | -1.818 | -0.163 | -0.992 | -0.248 | 0.004 | -0.004 | -0.004 | 0 | 0.84 | 0.87 | 0.82 | 0.69 | 0.92 | 0.86 |
| 4 | 10388610 | rs9990701 | 0.116 | -0.198 | -0.929 | -0.303 | -0.001 | 0 | 0 | 1.067 | 0.57 | 0.57 | 0.64 | 0.56 | 0.65 | 0.69 |
| 4 | 10404512 | rs7654258 | -1.812 | -0.17 | -1.262 | -0.148 | 0.009 | -0.013 | -0.013 | 2.674 | 0.84 | 0.89 | 0.82 | 0.72 | 0.92 | 0.87 |
| 4 | 10407572 | rs4643800 | 0.861 | -0.1 | -1.317 | -0.318 | 0.019 | 0.059 | 0.059 | 0 | 0.25 | 0.13 | 0.28 | 0.03 | 0.21 | 0.12 |
| 4 | 10408757 | rs7697246 | -0.708 | 0.036 | -1.589 | -0.229 | -0.01 | -0.046 | -0.046 | 0 | 0.64 | 0.83 | 0.64 | 0.56 | 0.66 | 0.7 |
| 4 | 10409841 | rs7656072 | 0.368 | 0.092 | -1.417 | -0.221 | 0.027 | 0.024 | 0.024 | 0 | 0.36 | 0.49 | 0.36 | 0.53 | 0.46 | 0.58 |
| 4 | 10411638 | rs3923393 | 1.176 | 0.187 | -1.434 | 0.185 | -0.002 | 0.057 | 0.057 | 0 | 0.24 | 0.39 | 0.24 | 0.35 | 0.22 | 0.43 |
| 4 | 10413168 | rs4610325 | 0.159 | 0.078 | -1.764 | 0.512 | -0.013 | -0.044 | -0.044 | 0 | 0.65 | 0.85 | 0.65 | 0.57 | 0.66 | 0.71 |
| 4 | 10413908 | rs7661365 | -0.661 | -0.381 | -0.914 | 1.014 | 0.008 | -0.012 | -0.012 | 0.009 | 0.84 | 0.89 | 0.83 | 0.73 | 0.92 | 0.87 |
| 4 | 10419465 | rs7683253 | -1.061 | -0.067 | -0.4 | 0.935 | -0.006 | 0.016 | 0.016 | 0 | 0.19 | 0.4 | 0.17 | 0.2 | 0.15 | 0.15 |
| 4 | 10424323 | rs4552452 | 1.362 | -0.483 | -0.014 | 1.242 | 0.01 | -0.013 | -0.013 | 1.556 | 0.75 | 0.21 | 0.77 | 0.63 | 0.83 | 0.72 |
| 4 | 10432402 | rs6811292 | 0.731 | -0.523 | 0.193 | 1.236 | 0.041 | 0.195 | 0.195 | 3.876 | 0.94 | 0.64 | 0.96 | 0.84 | 1 | 0.91 |
| 4 | 10435344 | rs4273464 | 1.194 | -0.955 | -0.624 | 1.279 | 0 | -0.028 | -0.028 | 1.513 | 0.24 | 0.04 | 0.27 | 0.19 | 0.2 | 0.23 |
| 4 | 10437391 | rs10003864 | 0.83 | -0.855 | -1.335 | 0.992 | -0.089 | 0.04 | 0.04 | 0.655 | 0.29 | 0.07 | 0.23 | 0.23 | 0.41 | 0.25 |
| 4 | 10440704 | rs35565862 | 0.842 | -0.874 | -1.341 | 0.985 | -0.086 | 0.042 | 0.042 | 0.744 | 0.29 | 0.07 | 0.23 | 0.23 | 0.4 | 0.25 |
| 4 | 10443384 | rs4422413 | 1.235 | -0.871 | -1.342 | 0.987 | -0.085 | 0.021 | 0.021 | 0.019 | 0.26 | 0.07 | 0.22 | 0.23 | 0.39 | 0.23 |
| 4 | 10444650 | rs3217 | 1.235 | -0.871 | -1.342 | 0.993 | -0.085 | 0.021 | 0.021 | 0.019 | 0.26 | 0.07 | 0.22 | 0.23 | 0.39 | 0.23 |
| 4 | 10446906 | rs10016022 | -1.742 | -1.074 | -1.165 | 0.925 | -0.007 | -0.002 | -0.002 | 5.067 | 0.79 | 0.89 | 0.72 | 0.79 | 0.76 | 0.78 |
| 4 | 10447640 | rs10016702 | -0.255 | -1.052 | -0.797 | 0.988 | 0.031 | 0.03 | 0.03 | 2.14 | 0.13 | 0.43 | 0.12 | 0.17 | 0.09 | 0.15 |
| 4 | 10447848 | rs10021506 | -0.031 | -1.034 | -0.768 | 1.028 | 0.021 | 0.162 | 0.162 | 4.039 | 0.06 | 0.32 | 0.04 | 0.16 | 0 | 0.12 |
| 4 | 10462106 | rs4339203 | 0.566 | -1.059 | -0.834 | 0.958 | -0.003 | 0.002 | 0.002 | 0.089 | 0.34 | 0.37 | 0.37 | 0.4 | 0.26 | 0.4 |
| 4 | 10475280 | rs4333176 | -0.572 | -1.335 | -0.945 | 0.977 | 0.039 | 0.029 | 0.029 | 0.065 | 0.47 | 0.74 | 0.48 | 0.57 | 0.37 | 0.52 |

| | | | | | | | | | | | | | | | | |
|---|---|---|---|---|---|---|---|---|---|---|---|---|---|---|---|---|
| 4 | 10484987 | rs6835433 | -0.706 | -1.4 | -0.829 | 1.125 | -0.059 | 0.119 | 0.119 | 9.326 | 0.13 | 0.68 | 0.12 | 0.36 | 0.05 | 0.24 |
| 4 | 10485577 | rs6836689 | -0.73 | -1.399 | -0.828 | 1.125 | -0.06 | 0.116 | 0.116 | 9.597 | 0.13 | 0.68 | 0.12 | 0.36 | 0.05 | 0.24 |
| 4 | 10486083 | rs10440278 | -1.838 | -1.198 | -0.92 | 0.963 | 0.006 | -0.011 | -0.011 | 0 | 0.76 | 0.93 | 0.72 | 0.8 | 0.79 | 0.75 |
| 4 | 10490278 | rs10488945 | -0.76 | -1.166 | -0.723 | 0.947 | -0.036 | 0 | 0 | 0 | 0.65 | 0.59 | 0.67 | 0.62 | 0.78 | 0.59 |
| 4 | 10491719 | rs13127079 | -0.86 | -1.166 | -0.7 | 0.948 | -0.04 | 0.004 | 0.004 | 0 | 0.66 | 0.59 | 0.67 | 0.62 | 0.79 | 0.6 |
| 4 | 10500431 | rs17467273 | 1.306 | -0.816 | -0.818 | 0.976 | -0.049 | -0.003 | -0.003 | 0 | 0.2 | 0.08 | 0.2 | 0.19 | 0.37 | 0.2 |
| 4 | 10501373 | rs6834691 | 1.754 | -1.451 | -0.979 | 0.955 | 0.03 | -0.017 | -0.017 | 0.19 | 0.3 | 0.03 | 0.29 | 0.24 | 0.22 | 0.26 |
| 4 | 10501625 | rs13109939 | 1.321 | -1.554 | -1.011 | 0.998 | -0.015 | 0.007 | 0.007 | 0 | 0.23 | 0.08 | 0.28 | 0.2 | 0.25 | 0.22 |
| 4 | 10502334 | rs5000857 | -0.953 | -1.6 | -0.508 | 0.997 | 0.02 | -0.01 | -0.01 | 0 | 0.58 | 0.65 | 0.61 | 0.54 | 0.71 | 0.6 |
| 4 | 10503752 | rs10488947 | 0.222 | -1.54 | -0.342 | 1.024 | 0.012 | 0.201 | 0.201 | 1.835 | 0.08 | 0.32 | 0.09 | 0.23 | 0.06 | 0.11 |
| 4 | 10506599 | rs6833095 | 1.119 | -1.41 | -0.55 | 1.004 | 0.011 | -0.025 | -0.025 | 0.046 | 0.49 | 0.08 | 0.5 | 0.44 | 0.6 | 0.46 |
| 4 | 10507984 | rs887112 | 0.159 | -1.331 | -0.327 | 0.439 | 0.033 | 0.042 | 0.042 | 1.233 | 0.11 | 0.46 | 0.11 | 0.14 | 0.06 | 0.15 |
| 4 | 10510881 | rs13109005 | 0.395 | -1.298 | -0.464 | 0.51 | 0.013 | 0.01 | 0.01 | 0 | 0.58 | 0.26 | 0.53 | 0.56 | 0.64 | 0.55 |
| 4 | 10511483 | rs1004327 | -0.479 | -1.755 | 0.076 | 0.751 | -0.011 | 0.005 | 0.005 | 0 | 0.69 | 0.5 | 0.65 | 0.65 | 0.7 | 0.67 |
| 4 | 10511996 | rs1004329 | 1.543 | -1.84 | -0.389 | 0.74 | 0.046 | -0.114 | -0.114 | 0.777 | 0.31 | 0.04 | 0.28 | 0.36 | 0.23 | 0.27 |
| 4 | 10513072 | rs13125670 | 0.491 | -1.583 | -0.24 | 0.741 | -0.001 | 0 | 0 | 0.287 | 0.28 | 0.27 | 0.33 | 0.27 | 0.45 | 0.34 |
| 4 | 10518525 | rs10034180 | 0.128 | -1.71 | -0.196 | 0.771 | 0 | 0.008 | 0.008 | 0 | 0.58 | 0.4 | 0.59 | 0.63 | 0.47 | 0.59 |
| 4 | 10520693 | rs13125086 | 1.268 | -1.092 | 0.076 | 1.132 | 0.007 | -0.032 | -0.032 | 0 | 0.12 | 0.01 | 0.03 | 0.14 | 0.08 | 0.08 |
| 4 | 10528226 | rs12641877 | 0.128 | -0.91 | -0.088 | 0.938 | 0.023 | 0.003 | 0.003 | 0 | 0.62 | 0.34 | 0.64 | 0.59 | 0.71 | 0.54 |
| 4 | 10545768 | rs6856070 | -0.399 | -0.328 | -0.348 | 1.057 | -0.019 | 0.007 | 0.007 | 0 | 0.07 | 0.12 | 0.03 | 0.14 | 0.03 | 0.12 |
| 4 | 10547343 | rs13112750 | -0.144 | -1.032 | -0.502 | 1.14 | -0.012 | 0.009 | 0.009 | 0 | 0.67 | 0.67 | 0.58 | 0.57 | 0.77 | 0.7 |
| 4 | 10549891 | rs4698066 | 0.92 | -1.09 | -0.778 | 1.036 | 0.029 | -0.006 | -0.006 | 0 | 0.09 | 0.02 | 0.16 | 0.07 | 0.05 | 0.13 |
| 4 | 10550193 | rs2159245 | 0.073 | -0.436 | -1.009 | 0.657 | -0.001 | -0.019 | -0.019 | 0 | 0.36 | 0.47 | 0.31 | 0.29 | 0.32 | 0.37 |
| 4 | 10551500 | rs10938845 | 0.33 | 0.061 | -1.496 | 0.858 | -0.085 | -0.029 | -0.029 | 0 | 0.26 | 0.11 | 0.34 | 0.31 | 0.52 | 0.36 |
| 4 | 10551560 | rs6848730 | 0.326 | -0.721 | -1.046 | 0.821 | -0.003 | -0.003 | -0.003 | 0 | 0.26 | 0.29 | 0.22 | 0.19 | 0.24 | 0.24 |
| 4 | 10557861 | rs1974584 | 0.624 | -1.291 | -0.587 | 0.683 | -0.075 | -0.076 | -0.076 | 0 | 0.27 | 0.13 | 0.34 | 0.39 | 0.53 | 0.35 |
| 4 | 10559041 | rs4698069 | 0.39 | -1.428 | -0.397 | 0.714 | -0.015 | -0.002 | -0.002 | 0 | 0.47 | 0.47 | 0.44 | 0.44 | 0.25 | 0.43 |
| 4 | 10572867 | rs17467553 | 1.574 | -1.399 | -0.52 | 0.542 | 0 | 0.012 | 0.012 | 0 | 0.06 | 0.03 | 0.04 | 0.06 | 0.06 | 0.04 |
| 4 | 10574056 | rs17383040 | 0.85 | -1.423 | -0.416 | 0.555 | -0.012 | -0.035 | -0.035 | 0.164 | 0.19 | 0.29 | 0.17 | 0.08 | 0.16 | 0.19 |
| 4 | 10575214 | rs6448095 | -0.176 | -1.431 | -0.394 | 0.574 | -0.031 | -0.081 | -0.081 | 0 | 0.73 | 0.54 | 0.77 | 0.85 | 0.79 | 0.78 |
| 4 | 10577980 | rs12643937 | 1.442 | -1.047 | -0.754 | 0.695 | -0.023 | -0.028 | -0.028 | 0 | 0.07 | 0.02 | 0.06 | 0.17 | 0.14 | 0.11 |
| 4 | 10581872 | rs10030578 | 0.151 | -1.626 | -1.024 | 0.511 | -0.038 | -0.069 | -0.069 | 0.033 | 0.39 | 0.14 | 0.47 | 0.45 | 0.58 | 0.41 |
| 4 | 10589553 | rs11934521 | 0.752 | -1.199 | -0.555 | 0.475 | -0.01 | -0.034 | -0.034 | 1.112 | 0.18 | 0.27 | 0.17 | 0.08 | 0.16 | 0.19 |
| 4 | 10591448 | rs10488949 | 1.432 | -0.836 | -0.878 | 0.668 | -0.073 | -0.056 | -0.056 | 0 | 0.19 | 0.07 | 0.27 | 0.24 | 0.36 | 0.25 |
| 4 | 10594679 | rs1990272 | -0.093 | -1.308 | -0.196 | 0.155 | -0.032 | -0.112 | -0.112 | 0 | 0.73 | 0.54 | 0.77 | 0.92 | 0.79 | 0.75 |
| 4 | 10597423 | rs6853762 | -0.131 | -1.114 | 0.041 | 0.438 | -0.028 | -0.036 | -0.036 | 0 | 0.66 | 0.58 | 0.75 | 0.76 | 0.75 | 0.72 |
| 4 | 10601918 | rs4697760 | 1.086 | -1.203 | -0.149 | 0.423 | -0.003 | -0.004 | -0.004 | 0 | 0.19 | 0.16 | 0.17 | 0.21 | 0.19 | 0.22 |
| 4 | 10605821 | rs2531185 | 1.04 | -1.121 | -0.197 | 0.418 | -0.049 | -0.044 | -0.044 | 0 | 0.25 | 0.16 | 0.26 | 0.38 | 0.45 | 0.33 |
| 4 | 10609118 | rs2079685 | -0.343 | -1.314 | 0.186 | 0.072 | -0.017 | -0.024 | -0.024 | 0 | 0.65 | 0.45 | 0.74 | 0.75 | 0.75 | 0.71 |
| 4 | 10621516 | rs17383444 | 0.989 | -1.349 | 0.313 | 0.072 | -0.007 | -0.008 | -0.008 | 0 | 0.23 | 0.26 | 0.17 | 0.18 | 0.18 | 0.2 |
| 4 | 10626314 | rs2720363 | -0.506 | -0.693 | 0.26 | 0.381 | 0.004 | 0.001 | 0.001 | 0.02 | 0.58 | 0.43 | 0.74 | 0.76 | 0.74 | 0.71 |
| 4 | 10649456 | rs12650571 | -0.228 | -0.318 | 0.023 | 0.485 | -0.008 | -0.007 | -0.007 | 0 | 0.34 | 0.4 | 0.42 | 0.29 | 0.27 | 0.35 |
| 4 | 10650693 | rs10516207 | 1.454 | -0.327 | -0.3 | 0.476 | -0.006 | -0.022 | -0.022 | 0.236 | 0.11 | 0.01 | 0.2 | 0.09 | 0.07 | 0.09 |
| 4 | 10651891 | rs11723092 | 0.328 | -0.212 | -0.109 | 0.562 | -0.029 | -0.026 | -0.026 | 0.401 | 0.31 | 0.25 | 0.31 | 0.46 | 0.49 | 0.44 |
| 4 | 10654346 | rs35699234 | -0.49 | -0.282 | 0.024 | 0.525 | -0.002 | -0.004 | -0.004 | 0 | 0.4 | 0.58 | 0.52 | 0.3 | 0.3 | 0.36 |
| 4 | 10655218 | rs12233843 | 0.04 | 0.047 | -0.214 | 0.642 | -0.042 | -0.047 | -0.047 | 0 | 0.33 | 0.23 | 0.39 | 0.49 | 0.53 | 0.45 |
| 4 | 10658129 | rs13119277 | 0.173 | -0.01 | -0.174 | 0.61 | -0.045 | -0.039 | -0.039 | 0 | 0.31 | 0.23 | 0.31 | 0.46 | 0.5 | 0.44 |
| 4 | 10666272 | rs6448177 | 0.204 | 0.037 | -0.045 | 0.553 | -0.009 | -0.009 | -0.009 | 0 | 0.32 | 0.27 | 0.4 | 0.52 | 0.49 | 0.43 |
| 4 | 10666909 | rs13144136 | -0.215 | -0.183 | 0.174 | 0.541 | -0.001 | 0 | 0 | 0 | 0.37 | 0.39 | 0.43 | 0.29 | 0.31 | 0.38 |
| 4 | 10667476 | rs12502030 | -1.564 | -0.169 | 0.159 | 0.359 | -0.022 | -0.01 | -0.01 | 0 | 0.7 | 0.66 | 0.88 | 0.8 | 0.85 | 0.85 |
| 4 | 10669522 | rs13136927 | -0.086 | 0.074 | -0.052 | 0.688 | -0.018 | -0.005 | -0.005 | 0 | 0.35 | 0.3 | 0.45 | 0.35 | 0.54 | 0.47 |
| 4 | 10670046 | rs13115026 | 1.373 | 0.554 | -0.024 | 0.819 | 0.012 | 0.021 | 0.021 | 0 | 0.18 | 0.22 | 0.29 | 0.3 | 0.29 | 0.26 |
| 4 | 10674007 | rs4582144 | -0.279 | 0.388 | -0.057 | 0.77 | -0.002 | -0.014 | -0.014 | 0.125 | 0.35 | 0.28 | 0.41 | 0.45 | 0.31 | 0.37 |
| 4 | 10674772 | rs17268650 | 0.72 | 0.397 | -0.133 | 0.806 | -0.003 | 0 | 0 | 0 | 0.2 | 0.16 | 0.09 | 0.18 | 0.19 | 0.22 |
| 4 | 10675030 | rs4697777 | -0.727 | 0.704 | -0.061 | 0.911 | 0 | 0 | 0 | 0 | 0.46 | 0.49 | 0.55 | 0.5 | 0.5 | 0.55 |
| 4 | 10679925 | rs10033332 | 0.102 | 0.716 | -0.027 | 0.928 | -0.001 | 0.011 | 0.011 | 0 | 0.36 | 0.4 | 0.43 | 0.45 | 0.31 | 0.37 |
| 4 | 10688637 | rs6827843 | -0.667 | 0.246 | -0.085 | 0.608 | 0 | 0.003 | 0.003 | 0 | 0.5 | 0.46 | 0.57 | 0.37 | 0.51 | 0.56 |
| 4 | 10689292 | rs4550919 | -0.478 | 0.306 | -0.063 | 0.63 | 0.003 | -0.009 | -0.009 | 1.933 | 0.44 | 0.45 | 0.53 | 0.37 | 0.45 | 0.53 |
| 4 | 10690043 | rs12510590 | 0.005 | 0.429 | -0.508 | 0.871 | 0.007 | 0.002 | 0.002 | 0 | 0.29 | 0.32 | 0.42 | 0.33 | 0.48 | 0.43 |
| 4 | 10691942 | rs13147855 | 0.954 | 0.593 | -1.143 | 0.753 | -0.073 | -0.037 | -0.037 | 0 | 0.17 | 0.03 | 0.12 | 0.18 | 0.24 | 0.15 |
| 4 | 10692788 | rs11721652 | -0.423 | 0.7 | -0.381 | 1.017 | 0.029 | 0.006 | 0.006 | 0 | 0.3 | 0.35 | 0.42 | 0.31 | 0.48 | 0.4 |
| 4 | 10693222 | rs6448182 | -0.578 | 0.376 | -0.729 | 1.406 | -0.03 | 0.003 | 0.003 | 0.249 | 0.7 | 0.66 | 0.87 | 0.65 | 0.84 | 0.78 |
| 4 | 10694991 | rs11725929 | -0.075 | 0.959 | -0.857 | 1.152 | -0.011 | -0.01 | -0.01 | 0 | 0.19 | 0.15 | 0.3 | 0.26 | 0.32 | 0.22 |
| 4 | 10698265 | rs4697648 | -0.01 | -0.449 | -0.69 | 0.752 | -0.031 | -0.006 | -0.006 | 0.051 | 0.73 | 0.68 | 0.88 | 0.78 | 0.87 | 0.8 |
| 4 | 10702156 | rs6819946 | 0.339 | 0.333 | -1.067 | 0.866 | -0.028 | -0.015 | -0.015 | 0 | 0.6 | 0.22 | 0.55 | 0.55 | 0.55 | 0.52 |
| 4 | 10702568 | rs11726285 | -0.238 | 0.546 | -0.004 | 0.935 | 0.055 | -0.014 | -0.014 | 0 | 0.22 | 0.4 | 0.33 | 0.18 | 0.29 | 0.17 |
| 4 | 10704867 | rs4697781 | 0.675 | 0.445 | 0.936 | 1.085 | 0.035 | 0.004 | 0.004 | 0 | 0.72 | 0.52 | 0.65 | 0.72 | 0.67 | 0.77 |
| 4 | 10706422 | rs4697650 | -0.551 | 0.421 | 0.938 | 1.091 | -0.005 | 0.017 | 0.017 | 0 | 0.95 | 0.87 | 0.98 | 0.91 | 0.96 | 0.94 |
| 4 | 10707508 | rs4549371 | 0.534 | 0.649 | 1.009 | 1.069 | 0.002 | 0.014 | 0.014 | 0 | 0.92 | 0.82 | 0.96 | 0.89 | 0.93 | 0.92 |

| | | | | | | | | | | | | | | | | |
|---|---|---|---|---|---|---|---|---|---|---|---|---|---|---|---|---|
| 4 | 10709812 | rs11733408 | 0.78 | 0.485 | 1.014 | 0.988 | -0.005 | -0.001 | -0.001 | 0.238 | 0.8 | 0.7 | 0.88 | 0.78 | 0.82 | 0.69 |
| 4 | 10711468 | rs4452425 | 0.351 | 0.48 | 1.002 | 0.989 | 0 | 0 | 0 | 1.732 | 0.12 | 0.13 | 0.08 | 0.11 | 0.1 | 0.24 |
| 4 | 10713674 | rs9291444 | 1.896 | 0.305 | 0.94 | 0.637 | -0.01 | 0.01 | 0.01 | 1.864 | 0.51 | 0.14 | 0.5 | 0.55 | 0.5 | 0.48 |
| 4 | 10714457 | rs4561924 | 0.836 | 0.415 | 0.905 | 0.664 | -0.001 | -0.004 | -0.004 | 0 | 0.78 | 0.7 | 0.85 | 0.78 | 0.77 | 0.67 |
| 4 | 10714656 | rs4368598 | 0.827 | 0.428 | 0.902 | 0.661 | -0.002 | -0.004 | -0.004 | 0 | 0.78 | 0.7 | 0.86 | 0.78 | 0.78 | 0.68 |
| 4 | 10726853 | rs16872571 | -1.108 | 0.447 | 0.925 | 0.873 | 0.012 | 0.009 | 0.009 | 2.438 | 0.38 | 0.39 | 0.4 | 0.49 | 0.46 | 0.24 |
| 4 | 10727697 | rs13116306 | 1.338 | 0.391 | 0.481 | 0.597 | 0.037 | 0.039 | 0.039 | 2.481 | 0.5 | 0.22 | 0.48 | 0.56 | 0.43 | 0.47 |
| 4 | 10728396 | rs11736462 | 1.742 | 0.082 | -0.336 | 0.646 | -0.036 | -0.056 | -0.056 | 0 | 0.18 | 0.07 | 0.28 | 0.38 | 0.29 | 0.08 |
| 4 | 10728759 | rs4697653 | 0.73 | 0.685 | -0.09 | 0.492 | 0.005 | 0.06 | 0.06 | 0.944 | 0.53 | 0.27 | 0.64 | 0.68 | 0.45 | 0.51 |
| 4 | 10730199 | rs13130534 | 0.791 | 0.736 | -0.116 | 0.632 | 0.019 | 0.048 | 0.048 | 1.458 | 0.52 | 0.27 | 0.64 | 0.58 | 0.44 | 0.47 |
| 4 | 10740267 | rs12644305 | 0.128 | 1.049 | 0.653 | 0.867 | 0.002 | 0.02 | 0.02 | 0.061 | 0.18 | 0.12 | 0.12 | 0.11 | 0.16 | 0.26 |
| 4 | 10744548 | rs12650576 | -0.23 | 1.046 | 0.882 | 0.859 | -0.015 | -0.066 | -0.066 | 2.701 | 0.21 | 0.42 | 0.14 | 0.1 | 0.16 | 0.24 |
| 4 | 10749071 | rs10007887 | -0.692 | 1.188 | 1.067 | 0.95 | 0.006 | -0.011 | -0.011 | 0 | 0.25 | 0.48 | 0.17 | 0.3 | 0.25 | 0.3 |
| 4 | 10751743 | rs4420973 | -1.139 | 1.15 | 1.19 | 0.934 | -0.001 | 0 | 0 | 0 | 0.29 | 0.65 | 0.2 | 0.35 | 0.28 | 0.35 |
| 4 | 10753416 | rs16872862 | 0.144 | 1.1 | 1.21 | 0.931 | -0.001 | -0.003 | -0.003 | 0 | 0.19 | 0.18 | 0.12 | 0.11 | 0.16 | 0.24 |
| 4 | 10792729 | rs12651622 | -0.27 | 1.062 | 1.095 | 0.972 | -0.012 | -0.022 | -0.022 | 0.291 | 0.22 | 0.44 | 0.15 | 0.15 | 0.18 | 0.26 |
| 4 | 10804019 | rs10938980 | -0.953 | 1.133 | 0.736 | 0.952 | -0.006 | 0.009 | 0.009 | 1.054 | 0.28 | 0.6 | 0.19 | 0.36 | 0.26 | 0.35 |
| 4 | 10831659 | rs7691429 | 0.332 | 0.766 | 0.431 | 0.36 | -0.004 | 0.015 | 0.015 | 0.259 | 0.54 | 0.38 | 0.53 | 0.32 | 0.48 | 0.55 |
| 4 | 10831736 | rs13141536 | -0.17 | 0.74 | 0.535 | 0.365 | -0.037 | 0.002 | 0.002 | 0 | 0.41 | 0.27 | 0.43 | 0.61 | 0.49 | 0.41 |
| 4 | 10833701 | rs12696865 | 1.276 | 0.771 | 0.459 | 0.364 | -0.079 | -0.101 | -0.101 | 0 | 0.21 | 0.06 | 0.31 | 0.55 | 0.35 | 0.15 |
| 4 | 10834255 | rs1872090 | 1.251 | 0.773 | 0.461 | 0.379 | -0.076 | -0.099 | -0.099 | 0 | 0.21 | 0.06 | 0.31 | 0.54 | 0.35 | 0.15 |
| 4 | 10835165 | rs2869756 | -0.015 | 0.725 | 0.504 | 0.328 | -0.03 | -0.009 | -0.009 | 0.063 | 0.38 | 0.26 | 0.4 | 0.62 | 0.47 | 0.39 |
| 4 | 10836045 | rs16873628 | 0.226 | 0.762 | 0.588 | 0.378 | -0.004 | 0.015 | 0.015 | 0.229 | 0.54 | 0.38 | 0.53 | 0.32 | 0.48 | 0.55 |
| 4 | 10845207 | rs4257647 | 0.746 | 0.397 | 0.212 | -0.392 | 0.034 | 0.039 | 0.039 | 0 | 0.33 | 0.22 | 0.16 | 0.16 | 0.23 | 0.37 |
| 4 | 10847303 | rs7673890 | 0.313 | 0.994 | 0.192 | -0.132 | 0.02 | 0.041 | 0.041 | 0.519 | 0.52 | 0.37 | 0.37 | 0.3 | 0.43 | 0.56 |
| 4 | 10857992 | rs2869749 | 0.039 | 0.846 | -0.243 | 0.026 | -0.04 | 0.021 | 0.021 | 1.885 | 0.4 | 0.26 | 0.54 | 0.63 | 0.5 | 0.33 |
| 4 | 10867056 | rs4074743 | -0.023 | 0.217 | -0.154 | -0.031 | 0 | 0.068 | 0.068 | 1.891 | 0.42 | 0.27 | 0.59 | 0.72 | 0.56 | 0.36 |
| 4 | 10871655 | rs10516213 | 0.848 | 0.174 | -0.118 | -0.265 | 0.001 | -0.001 | -0.001 | 0 | 0.16 | 0.16 | 0.11 | 0.06 | 0.15 | 0.18 |
| 4 | 10872167 | rs4697799 | -0.17 | -0.025 | 0.143 | -0.285 | -0.028 | -0.081 | -0.081 | 1.456 | 0.78 | 0.48 | 0.83 | 0.89 | 0.83 | 0.79 |
| 4 | 10874361 | rs7667727 | 0.346 | -0.291 | -0.544 | -0.138 | -0.04 | 0.055 | 0.055 | 0 | 0.15 | 0.37 | 0.18 | 0.22 | 0.09 | 0.19 |
| 4 | 10874933 | rs13107834 | 1.445 | -0.222 | -0.697 | -0.123 | -0.118 | -0.127 | -0.127 | 0 | 0.23 | 0.06 | 0.46 | 0.48 | 0.41 | 0.16 |
| 4 | 10877223 | rs7698615 | 0.418 | -0.257 | -0.686 | -0.133 | -0.049 | 0.056 | 0.056 | 0 | 0.15 | 0.37 | 0.11 | 0.22 | 0.08 | 0.19 |
| 4 | 10886524 | rs16874378 | 0.377 | -0.119 | -1.132 | -0.113 | 0.056 | 0.129 | 0.129 | 0.318 | 0.43 | 0.19 | 0.2 | 0.19 | 0.31 | 0.43 |
| 4 | 10886810 | rs11735551 | -0.135 | -0.154 | -0.973 | -0.161 | 0.013 | 0.01 | 0.01 | 1.153 | 0.69 | 0.28 | 0.76 | 0.73 | 0.75 | 0.63 |
| 4 | 10892111 | rs4697806 | -0.008 | -0.186 | -0.897 | -0.111 | 0.015 | 0.006 | 0.006 | 0.751 | 0.66 | 0.27 | 0.66 | 0.67 | 0.71 | 0.58 |
| 4 | 10902159 | rs959233 | -0.238 | 0.29 | -1.047 | 0.018 | -0.012 | 0.012 | 0.012 | 0 | 0.57 | 0.7 | 0.42 | 0.4 | 0.49 | 0.55 |
| 4 | 10903005 | rs7660096 | 0.327 | -0.17 | -1.039 | 0.118 | 0.026 | 0.018 | 0.018 | 0 | 0.25 | 0.19 | 0.12 | 0.14 | 0.13 | 0.31 |
| 4 | 10904684 | rs2904192 | 0.327 | -0.17 | -1.039 | 0.118 | 0.026 | 0.018 | 0.018 | 0 | 0.25 | 0.19 | 0.12 | 0.14 | 0.13 | 0.31 |
| 4 | 10905127 | rs1105503 | 0.862 | 0.213 | -0.959 | 0.266 | -0.005 | -0.001 | -0.001 | 0 | 0.3 | 0.35 | 0.18 | 0.29 | 0.25 | 0.3 |
| 4 | 10909225 | rs10028469 | 0.202 | 0.428 | -1.136 | 0.723 | -0.025 | -0.008 | -0.008 | 0 | 0.2 | 0.26 | 0.12 | 0.14 | 0.12 | 0.29 |
| 4 | 10910291 | rs1384837 | 0.155 | 1.028 | -1.083 | 0.574 | 0.002 | 0.011 | 0.011 | 0 | 0.51 | 0.57 | 0.43 | 0.4 | 0.51 | 0.55 |
| 4 | 10916915 | rs12511310 | -0.428 | 1.377 | -1.431 | 0.764 | -0.04 | -0.017 | -0.017 | 0 | 0.32 | 0.19 | 0.4 | 0.32 | 0.44 | 0.17 |
| 4 | 10920805 | rs6818760 | -0.148 | 0.549 | -1.141 | 0.532 | -0.007 | 0.036 | 0.036 | 0.382 | 0.84 | 0.83 | 0.82 | 0.41 | 0.88 | 0.57 |
| 4 | 10931542 | rs17280854 | 0.633 | 0.206 | -2.105 | -0.597 | -0.057 | 0.24 | 0.24 | 0.475 | 0.45 | 0.11 | 0.37 | 0.13 | 0.5 | 0.39 |
| 4 | 10931588 | rs11935551 | 0.494 | 0.211 | -2.055 | -0.549 | 0 | 0.005 | 0.005 | 0.56 | 0.16 | 0.13 | 0.18 | 0.6 | 0.12 | 0.44 |
| 4 | 10931705 | rs4697808 | 0.821 | 0.119 | -2.251 | -0.805 | -0.059 | 0.046 | 0.046 | 0 | 0.12 | 0.03 | 0.27 | 0.05 | 0.25 | 0.06 |
| 4 | 10939039 | rs12643516 | -0.375 | -0.595 | -1.954 | -1.151 | -0.025 | 0.008 | 0.008 | 0 | 0.27 | 0.29 | 0.36 | 0.61 | 0.14 | 0.46 |
| 4 | 10942197 | rs12642366 | 0.713 | -0.642 | -1.912 | -1.091 | -0.001 | 0.008 | 0.008 | 0.825 | 0.17 | 0.13 | 0.19 | 0.6 | 0.12 | 0.44 |
| 4 | 10943926 | rs2869768 | 0.043 | -0.646 | -0.81 | -1.191 | -0.003 | -0.023 | -0.023 | 0 | 0.64 | 0.48 | 0.57 | 0.73 | 0.62 | 0.82 |
| 4 | 10945449 | rs7660235 | 0.476 | -0.455 | -1.073 | -1.373 | 0.003 | 0.088 | 0.088 | 0.077 | 0.53 | 0.24 | 0.49 | 0.69 | 0.51 | 0.72 |
| 4 | 10949825 | rs13124128 | -0.579 | -0.578 | -0.547 | -1.461 | -0.008 | -0.013 | -0.013 | 0 | 0.69 | 0.56 | 0.79 | 0.75 | 0.74 | 0.72 |
| 4 | 10950989 | rs10222862 | 1.498 | -0.642 | -0.739 | -1.489 | -0.104 | 0.101 | 0.101 | 0.226 | 0.31 | 0.09 | 0.35 | 0.15 | 0.39 | 0.31 |
| 4 | 10951093 | rs12510358 | -0.612 | -0.607 | -0.569 | -1.358 | -0.009 | -0.003 | -0.003 | 0 | 0.38 | 0.47 | 0.44 | 0.59 | 0.35 | 0.41 |
| 4 | 10951486 | rs13132998 | -0.612 | -0.607 | -0.492 | -1.358 | -0.007 | -0.003 | -0.003 | 0.789 | 0.38 | 0.48 | 0.44 | 0.59 | 0.35 | 0.41 |
| 4 | 10952541 | rs1466529 | 0.625 | -0.59 | -0.293 | -1.12 | -0.005 | 0.004 | 0.004 | 0 | 0.21 | 0.16 | 0.27 | 0.55 | 0.1 | 0.36 |
| 4 | 10953205 | rs6821267 | 0.099 | -0.36 | -0.377 | -1.284 | -0.006 | 0.023 | 0.023 | 0 | 0.56 | 0.56 | 0.66 | 0.7 | 0.49 | 0.67 |
| 4 | 10961185 | rs7683621 | -0.729 | -0.341 | 0.136 | -1.053 | -0.009 | 0.056 | 0.056 | 0 | 0.79 | 0.69 | 0.7 | 0.43 | 0.82 | 0.64 |
| 4 | 10971423 | rs10009111 | -0.928 | -0.45 | -0.177 | -1.67 | -0.016 | 0.177 | 0.177 | 4.421 | 0.55 | 0.79 | 0.63 | 0.95 | 0.52 | 0.77 |
| 4 | 10972640 | rs7675535 | 0.963 | -0.468 | 0.119 | -1.43 | -0.002 | 0.017 | 0.017 | 0 | 0.2 | 0.2 | 0.3 | 0.54 | 0.18 | 0.38 |
| 4 | 10972886 | rs7662952 | 0.877 | -0.487 | 0.255 | -1.732 | -0.008 | 0.035 | 0.035 | 0 | 0.21 | 0.33 | 0.31 | 0.64 | 0.18 | 0.37 |
| 4 | 10973261 | rs10516217 | 1.805 | -0.619 | -0.185 | -1.807 | -0.13 | 0.124 | 0.124 | 0 | 0.23 | 0.02 | 0.28 | 0.04 | 0.37 | 0.17 |
| 4 | 10975416 | rs10516218 | 0.093 | -0.402 | 0.102 | -1.737 | -0.001 | -0.008 | -0.008 | 0 | 0.22 | 0.26 | 0.22 | 0.17 | 0.22 | 0.16 |
| 4 | 10975428 | rs16876186 | 1.292 | -0.413 | 0.125 | -1.741 | 0.01 | 0 | 0 | 0 | 0.12 | 0.19 | 0.1 | 0.13 | 0.13 | 0.22 |
| 4 | 10975896 | rs2128866 | -0.264 | -0.443 | 0.135 | -1.762 | 0.01 | 0.047 | 0.047 | 0 | 0.53 | 0.64 | 0.49 | 0.78 | 0.41 | 0.66 |
| 4 | 10980181 | rs13143997 | 1.55 | -0.452 | -0.033 | -1.773 | -0.053 | 0.094 | 0.094 | 0.528 | 0.26 | 0.14 | 0.3 | 0.04 | 0.38 | 0.18 |
| 4 | 10981163 | rs10012274 | -0.833 | -0.205 | -0.059 | -1.528 | 0.01 | -0.016 | -0.016 | 0 | 0.35 | 0.48 | 0.33 | 0.31 | 0.35 | 0.39 |
| 4 | 10986574 | rs16876402 | -0.166 | -0.115 | -0.162 | -1.259 | 0.002 | -0.005 | -0.005 | 0 | 0.22 | 0.25 | 0.25 | 0.19 | 0.24 | 0.16 |
| 4 | 10989008 | rs2046359 | -0.186 | -0.337 | -0.145 | -0.713 | -0.023 | 0.024 | 0.024 | 0.398 | 0.74 | 0.42 | 0.67 | 0.36 | 0.82 | 0.62 |
| 4 | 11011598 | rs875394 | 0.651 | -0.248 | -0.779 | -0.883 | 0.027 | 0.112 | 0.112 | 1.514 | 0.18 | 0.02 | 0.07 | 0.01 | 0.11 | 0.06 |

| | | | | | | | | | | | | | | | | |
|---|---|---|---|---|---|---|---|---|---|---|---|---|---|---|---|---|
| 4 | 11016733 | rs9291483 | 0.132 | -0.042 | -0.279 | -0.644 | -0.003 | -0.003 | -0.003 | 0.391 | 0.33 | 0.61 | 0.4 | 0.71 | 0.27 | 0.56 |
| 4 | 11019347 | rs6448447 | -0.673 | 0.212 | -0.695 | -0.589 | -0.039 | -0.047 | -0.047 | 0 | 0.92 | 0.99 | 0.92 | 0.85 | 0.89 | 0.9 |
| 4 | 11022799 | rs4373155 | -0.428 | 0.118 | -0.231 | -0.757 | -0.021 | -0.025 | -0.025 | 0 | 0.33 | 0.45 | 0.32 | 0.2 | 0.26 | 0.22 |
| 4 | 11025131 | rs6846529 | 0.257 | 0.117 | -0.286 | -0.744 | -0.008 | -0.012 | -0.012 | 0 | 0.65 | 0.6 | 0.67 | 0.8 | 0.72 | 0.78 |
| 4 | 11038268 | rs17218817 | 0.756 | 0.177 | -0.588 | -0.871 | 0.003 | 0.123 | 0.123 | 0.304 | 0.22 | 0.02 | 0.2 | 0.04 | 0.17 | 0.12 |
| 4 | 11040290 | rs2904297 | -0.132 | 0.335 | -0.232 | -0.543 | -0.007 | -0.009 | -0.009 | 0 | 0.66 | 0.63 | 0.67 | 0.78 | 0.72 | 0.77 |
| 4 | 11044254 | rs6820330 | -0.023 | 0.113 | -0.104 | -0.026 | -0.004 | -0.001 | -0.001 | 0.188 | 0.32 | 0.59 | 0.39 | 0.69 | 0.27 | 0.56 |
| 4 | 11049844 | rs1110365 | 0.097 | 0.137 | -0.077 | 0.232 | -0.008 | 0.035 | 0.035 | 1.355 | 0.31 | 0.47 | 0.38 | 0.61 | 0.26 | 0.56 |
| 4 | 11050130 | rs1110364 | -0.72 | -0.152 | -0.567 | -0.288 | -0.021 | 0.044 | 0.044 | 0 | 0.57 | 0.82 | 0.61 | 0.68 | 0.44 | 0.68 |
| 4 | 11050420 | rs1039682 | -0.105 | -0.289 | 0.026 | -0.302 | -0.028 | -0.077 | -0.077 | 0 | 0.26 | 0.36 | 0.24 | 0.06 | 0.18 | 0.13 |
| 4 | 11050853 | rs7694991 | 0.115 | 0.432 | 0.146 | 0.019 | 0.02 | -0.034 | -0.034 | 0 | 0.89 | 0.76 | 0.82 | 0.95 | 0.86 | 0.97 |
| 4 | 11053537 | rs6846243 | -1.045 | 0.073 | 0.179 | 0.719 | -0.003 | 0.006 | 0.006 | 1.72 | 0.33 | 0.66 | 0.27 | 0.57 | 0.32 | 0.31 |
| 4 | 11054542 | rs16877315 | -0.585 | -0.067 | 0.752 | 1.023 | -0.034 | -0.026 | -0.026 | 0 | 0.22 | 0.48 | 0.14 | 0.2 | 0.17 | 0.25 |
| 4 | 11055432 | rs2063056 | -1.586 | -0.21 | 0.694 | 1.117 | -0.017 | 0.013 | 0.013 | 1.009 | 0.3 | 0.6 | 0.38 | 0.32 | 0.26 | 0.2 |
| 4 | 11059843 | rs4345174 | -1.898 | -0.392 | 0.454 | 1.205 | 0.031 | 0.037 | 0.037 | 0.531 | 0.41 | 0.87 | 0.56 | 0.7 | 0.34 | 0.45 |
| 4 | 11060009 | rs7698375 | -1.066 | -0.44 | 1.41 | 1.384 | -0.031 | 0.083 | 0.083 | 0 | 0.26 | 0.58 | 0.37 | 0.51 | 0.22 | 0.28 |
| 4 | 11069107 | rs10003771 | -0.452 | -0.495 | 1.647 | 1.541 | -0.043 | 0.101 | 0.101 | 0.133 | 0.19 | 0.45 | 0.32 | 0.27 | 0.15 | 0.16 |
| 4 | 11069265 | rs7697986 | -0.579 | -0.435 | 1.536 | 1.543 | 0.015 | 0.017 | 0.017 | 0 | 0.13 | 0.29 | 0.19 | 0.16 | 0.12 | 0.16 |
| 4 | 11071463 | rs2202592 | -1.505 | -0.307 | 0.748 | 1.15 | -0.039 | 0.027 | 0.027 | 1.479 | 0.88 | 0.99 | 0.92 | 0.97 | 0.86 | 0.82 |
| 4 | 11076268 | rs7662758 | -2.242 | -0.722 | 1.569 | 1.461 | 0.027 | 0.019 | 0.019 | 0.361 | 0.4 | 0.76 | 0.53 | 0.65 | 0.33 | 0.46 |
| 4 | 11077316 | rs4697834 | -0.943 | -0.724 | 1.76 | 1.146 | 0.01 | 0.014 | 0.014 | 0.016 | 0.16 | 0.32 | 0.2 | 0.16 | 0.13 | 0.17 |
| 4 | 11080161 | rs10516219 | 0.43 | -0.777 | 2.068 | 1.096 | -0.007 | -0.045 | -0.045 | 0 | 0.14 | 0.19 | 0.07 | 0 | 0.1 | 0.07 |
| 4 | 11081033 | rs4697836 | -0.887 | -0.758 | 1.963 | 1.112 | 0.011 | 0.002 | 0.002 | 0.083 | 0.16 | 0.32 | 0.2 | 0.15 | 0.13 | 0.17 |
| 4 | 11085160 | rs6822701 | 0.046 | -0.761 | 1.952 | 1.13 | -0.004 | 0.045 | 0.045 | 0.002 | 0.06 | 0.11 | 0.04 | 0.22 | 0.05 | 0.13 |
| 4 | 11087189 | rs6811589 | 0.274 | -0.777 | 1.997 | 1.114 | -0.023 | -0.09 | -0.09 | 0.189 | 0.15 | 0.31 | 0.08 | 0 | 0.1 | 0.07 |
| 4 | 11087495 | rs7655566 | -0.012 | -0.649 | 1.873 | 1.236 | -0.004 | 0.039 | 0.039 | 0.002 | 0.06 | 0.11 | 0.04 | 0.22 | 0.05 | 0.12 |
| 4 | 11088906 | rs16877810 | -0.105 | -0.648 | 1.872 | 1.236 | -0.003 | 0.036 | 0.036 | 0 | 0.06 | 0.11 | 0.04 | 0.22 | 0.05 | 0.13 |
| 4 | 11091396 | rs6837846 | -2.016 | -0.679 | 1.905 | 1.239 | 0.002 | 0.071 | 0.071 | 0.645 | 0.22 | 0.43 | 0.25 | 0.37 | 0.19 | 0.29 |
| 4 | 11092841 | rs4697839 | -0.973 | -0.639 | 1.764 | 1.138 | 0.011 | 0.002 | 0.002 | 0.083 | 0.16 | 0.32 | 0.2 | 0.15 | 0.13 | 0.17 |
| 4 | 11099374 | rs12501933 | 0.365 | -0.628 | 1.351 | 1.103 | -0.031 | 0.047 | 0.047 | 0 | 0.09 | 0.01 | 0.06 | 0.02 | 0.1 | 0.05 |
| 4 | 11099518 | rs6448494 | -2.339 | -0.827 | 1.508 | 1.284 | 0.027 | 0.052 | 0.052 | 0.59 | 0.4 | 0.76 | 0.54 | 0.71 | 0.33 | 0.47 |
| 4 | 11100735 | rs16878079 | 0.321 | -0.776 | 1.669 | 1.215 | 0.003 | -0.034 | -0.034 | 0.116 | 0.1 | 0.11 | 0.07 | 0.02 | 0.09 | 0.05 |
| 4 | 11100788 | rs16878083 | -0.877 | -0.773 | 1.64 | 1.358 | -0.007 | 0.028 | 0.028 | 0 | 0.27 | 0.31 | 0.35 | 0.42 | 0.24 | 0.23 |
| 4 | 11112475 | rs8180283 | -2.146 | -0.793 | 1.111 | 1.3 | 0.034 | 0.094 | 0.094 | 0.653 | 0.41 | 0.79 | 0.55 | 0.79 | 0.34 | 0.47 |
| 4 | 11112491 | rs11944172 | -0.863 | -0.854 | 1.404 | 1.23 | 0.063 | 0.177 | 0.177 | 3.157 | 0.06 | 0.37 | 0.13 | 0.12 | 0.02 | 0.11 |
| 4 | 11115865 | rs16878196 | 0.06 | -0.799 | 0.664 | 1.363 | -0.003 | 0.055 | 0.055 | 0.008 | 0.06 | 0.09 | 0.05 | 0.27 | 0.05 | 0.13 |
| 4 | 11116063 | rs7694486 | -1.495 | -0.742 | 0.612 | 1.099 | -0.003 | 0 | 0 | 0 | 0.35 | 0.35 | 0.4 | 0.61 | 0.3 | 0.35 |
| 4 | 11117827 | rs9291494 | -1.884 | -0.844 | -0.507 | 0.544 | 0.007 | 0.126 | 0.126 | 0.773 | 0.46 | 0.93 | 0.58 | 0.73 | 0.36 | 0.49 |
| 4 | 11119349 | rs1506794 | -0.291 | -0.822 | -0.129 | 0.279 | -0.014 | 0.008 | 0.008 | 2.572 | 0.39 | 0.49 | 0.48 | 0.49 | 0.35 | 0.33 |
| 4 | 11120491 | rs1506791 | -1.664 | -0.94 | -0.605 | 0.154 | -0.068 | 0.099 | 0.099 | 1.988 | 0.61 | 0.84 | 0.62 | 0.76 | 0.54 | 0.52 |
| 4 | 11127123 | rs10034047 | -0.519 | -0.869 | -0.423 | 0.2 | 0.006 | -0.01 | -0.01 | 0 | 0.18 | 0.08 | 0.12 | 0.3 | 0.16 | 0.18 |
| 4 | 11127820 | rs16878277 | -0.431 | -0.803 | -0.076 | 0.257 | -0.006 | -0.009 | -0.009 | 1.73 | 0.2 | 0.23 | 0.1 | 0.06 | 0.1 | 0.14 |
| 4 | 11132123 | rs16878282 | 0.337 | -0.342 | -0.216 | 0.595 | 0.001 | 0.046 | 0.046 | 0 | 0.05 | 0.06 | 0.04 | 0.26 | 0.07 | 0.08 |
| 4 | 11133145 | rs4697674 | 0.409 | -0.196 | -0.218 | 0.395 | -0.003 | -0.005 | -0.005 | 0.168 | 0.05 | 0.03 | 0.06 | 0 | 0.08 | 0.05 |
| 4 | 11142798 | rs13152779 | 1.503 | -0.239 | -0.332 | -0.055 | -0.057 | 0.044 | 0.044 | 0 | 0.13 | 0.01 | 0.08 | 0.03 | 0.18 | 0.11 |
| 4 | 11156010 | rs12509327 | -0.812 | -0.361 | 0.151 | -0.129 | 0.022 | -0.066 | -0.066 | 2.109 | 0.11 | 0.23 | 0.1 | 0 | 0.12 | 0.11 |
| 4 | 11162918 | rs1876265 | -0.346 | -0.868 | 0.47 | 0.214 | -0.009 | 0.022 | 0.022 | 0.338 | 0.25 | 0.3 | 0.42 | 0.43 | 0.17 | 0.34 |
| 4 | 11173785 | rs6448544 | -2.299 | -1.059 | 0.553 | 0.099 | 0.044 | 0.089 | 0.089 | 0.101 | 0.45 | 0.82 | 0.57 | 0.69 | 0.36 | 0.53 |
| 4 | 11175752 | rs7659670 | -0.381 | -0.831 | 0.865 | 0.253 | 0.008 | -0.04 | -0.04 | 0 | 0.1 | 0.12 | 0.08 | 0 | 0.12 | 0.11 |
| 4 | 11177208 | rs9291505 | 1.314 | -0.989 | 0.753 | 0.166 | -0.002 | -0.033 | -0.033 | 0 | 0.13 | 0.11 | 0.09 | 0 | 0.09 | 0.05 |
| 4 | 11179846 | rs1399334 | -1.26 | -0.706 | 0.886 | 0.376 | -0.003 | 0.05 | 0.05 | 0.323 | 0.54 | 0.66 | 0.63 | 0.7 | 0.54 | 0.54 |
| 4 | 11179967 | rs1399336 | -1.049 | -0.767 | 0.891 | 0.376 | 0.014 | 0.05 | 0.05 | 1.227 | 0.49 | 0.63 | 0.57 | 0.7 | 0.45 | 0.5 |
| 4 | 11182041 | rs10034712 | 0.646 | -0.458 | 0.447 | 0.016 | 0.001 | 0.05 | 0.05 | 0 | 0.05 | 0.06 | 0.04 | 0.26 | 0.07 | 0.08 |
| 4 | 11186450 | rs1506798 | -0.525 | -0.64 | 0.002 | -0.411 | 0.007 | 0.022 | 0.022 | 1.88 | 0.52 | 0.5 | 0.61 | 0.69 | 0.45 | 0.54 |
| 4 | 11190557 | rs908107 | -1.103 | -0.521 | -0.05 | -0.237 | 0.006 | -0.014 | -0.014 | 0.065 | 0.62 | 0.63 | 0.69 | 0.7 | 0.56 | 0.65 |
| 4 | 11191470 | rs6835872 | 0.058 | -0.603 | -0.019 | 0.373 | -0.009 | 0.03 | 0.03 | 0 | 0.15 | 0.2 | 0.32 | 0.57 | 0.12 | 0.25 |
| 4 | 11192848 | rs1606724 | -0.757 | -0.146 | -0.07 | 0.284 | 0.014 | -0.026 | -0.026 | 0 | 0.63 | 0.61 | 0.67 | 0.7 | 0.51 | 0.65 |
| 4 | 11199352 | rs16878773 | 1.031 | -0.256 | 0.029 | 1.307 | 0.024 | -0.012 | -0.012 | 1.257 | 0.08 | 0.03 | 0.23 | 0.2 | 0.04 | 0.02 |
| 4 | 11200248 | rs7681223 | -0.58 | -0.318 | 0.776 | 1.151 | 0.016 | 0.048 | 0.048 | 0.444 | 0.08 | 0.3 | 0.14 | 0.1 | 0.04 | 0.12 |
| 4 | 11201074 | rs9683691 | -1.373 | -0.588 | 1.202 | 1.101 | -0.024 | 0.027 | 0.027 | 0.362 | 0.17 | 0.5 | 0.21 | 0.28 | 0.09 | 0.25 |
| 4 | 11202360 | rs13146400 | -1.39 | -0.592 | 1.2 | 1.099 | -0.024 | 0.023 | 0.023 | 0.503 | 0.17 | 0.5 | 0.21 | 0.27 | 0.09 | 0.25 |
| 4 | 11208907 | rs16878957 | 0.161 | -0.59 | 1.093 | 1.104 | -0.002 | 0.02 | 0.02 | 0 | 0.1 | 0.19 | 0.07 | 0.18 | 0.06 | 0.14 |
| 4 | 11209409 | rs16878962 | -1.706 | -0.604 | 0.725 | 1.068 | -0.026 | 0 | 0 | 2.877 | 0.21 | 0.6 | 0.22 | 0.28 | 0.1 | 0.26 |
| 4 | 11216238 | rs10009253 | -1.599 | -0.467 | 0.681 | 1.366 | 0.022 | -0.033 | -0.033 | 1.891 | 0.22 | 0.78 | 0.42 | 0.47 | 0.11 | 0.17 |
| 4 | 11216471 | rs10031075 | -0.652 | -0.47 | 1.028 | 1.382 | -0.079 | 0.123 | 0.123 | 2.632 | 0.15 | 0.42 | 0.27 | 0.19 | 0.05 | 0.02 |
| 4 | 11221352 | rs4697681 | 0.542 | 0.311 | 0.614 | 1.335 | -0.004 | 0.025 | 0.025 | 0 | 0.73 | 0.58 | 0.54 | 0.47 | 0.71 | 0.67 |
| 4 | 11224411 | rs1003271 | -0.167 | 0.441 | 0.127 | 0.936 | -0.045 | 0.015 | 0.015 | 0 | 0.66 | 0.81 | 0.71 | 0.67 | 0.55 | 0.67 |
| 4 | 11227687 | rs1355967 | 0.732 | 0.49 | 0.47 | 0.879 | -0.001 | 0 | 0 | 0.076 | 0.54 | 0.44 | 0.42 | 0.46 | 0.49 | 0.61 |
| 4 | 11229468 | rs12511580 | -0.3 | 0.718 | 0.453 | 0.978 | -0.013 | 0.025 | 0.025 | 0 | 0.7 | 0.83 | 0.84 | 0.78 | 0.67 | 0.71 |

| | | | | | | | | | | | | | | | | |
|---|---|---|---|---|---|---|---|---|---|---|---|---|---|---|---|---|
| 4 | 11229690 | rs1949275 | 0.35 | 0.516 | 0.571 | 0.931 | 0.046 | 0.189 | 0.189 | 0 | 0.07 | 0.29 | 0.16 | 0.24 | 0.05 | 0.13 |
| 4 | 11231335 | rs1355966 | 0.598 | 0.584 | 0.576 | 0.813 | -0.007 | 0.038 | 0.038 | 0 | 0.54 | 0.36 | 0.46 | 0.27 | 0.48 | 0.6 |
| 4 | 11232290 | rs11726519 | -0.304 | 0.605 | 0.468 | 0.882 | -0.006 | -0.005 | -0.005 | 0 | 0.66 | 0.67 | 0.74 | 0.52 | 0.55 | 0.64 |
| 4 | 11232758 | rs965001 | 1.418 | 0.444 | -0.144 | 0.628 | -0.044 | 0.167 | 0.167 | 0 | 0.44 | 0.16 | 0.3 | 0.19 | 0.42 | 0.43 |
| 4 | 11233074 | rs11727553 | 0.15 | 0.911 | 0.054 | 0.717 | 0.004 | 0.037 | 0.037 | 0 | 0.18 | 0.25 | 0.36 | 0.25 | 0.22 | 0.08 |
| 4 | 11234427 | rs12651349 | -0.338 | 1.001 | 0.067 | 0.672 | 0 | 0.077 | 0.077 | 0.292 | 0.29 | 0.51 | 0.34 | 0.57 | 0.31 | 0.5 |
| 4 | 11235402 | rs16879350 | 1.15 | 1.03 | 0.041 | 0.773 | 0 | 0.001 | 0.001 | 0 | 0.17 | 0.2 | 0.17 | 0.3 | 0.23 | 0.31 |
| 4 | 11235941 | rs10034384 | -0.435 | 0.841 | -0.139 | 0.75 | 0.016 | 0.008 | 0.008 | 0 | 0.34 | 0.44 | 0.52 | 0.54 | 0.45 | 0.39 |
| 4 | 11236408 | rs4697859 | 0.165 | 0.734 | -0.038 | 0.821 | -0.013 | 0.118 | 0.118 | 0.153 | 0.12 | 0.35 | 0.18 | 0.27 | 0.08 | 0.19 |
| 4 | 11244792 | rs1548184 | -1.087 | 0.849 | -0.294 | 0.564 | -0.039 | 0.036 | 0.036 | 0 | 0.31 | 0.42 | 0.47 | 0.51 | 0.19 | 0.28 |
| 4 | 11251244 | rs2671878 | 0.685 | 0.917 | -0.121 | 0.725 | -0.05 | 0.101 | 0.101 | 0 | 0.75 | 0.52 | 0.55 | 0.49 | 0.84 | 0.82 |
| 4 | 11252591 | rs6840284 | 1.345 | 0.597 | -0.432 | 0.599 | -0.016 | 0.181 | 0.181 | 0 | 0.44 | 0.15 | 0.23 | 0.19 | 0.42 | 0.36 |
| 4 | 11258260 | rs782758 | 0.186 | 0.872 | -0.08 | 0.973 | -0.053 | 0.095 | 0.095 | 0 | 0.79 | 0.63 | 0.59 | 0.49 | 0.87 | 0.83 |
| 4 | 11264808 | rs782754 | -0.353 | 1.077 | -0.573 | 1.095 | 0.094 | 0.045 | 0.045 | 0.234 | 0.82 | 0.49 | 0.69 | 0.74 | 0.73 | 0.78 |
| 4 | 11266157 | rs782756 | 0.646 | 1.195 | -0.63 | 0.979 | 0.016 | 0.029 | 0.029 | 0 | 0.69 | 0.47 | 0.43 | 0.49 | 0.64 | 0.75 |
| 4 | 11267586 | rs1533338 | 0.601 | 0.59 | -0.546 | 1.115 | -0.002 | 0.002 | 0.002 | 0 | 0.56 | 0.45 | 0.35 | 0.49 | 0.48 | 0.73 |
| 4 | 11273101 | rs6830548 | 0.108 | 0.405 | -0.285 | 1.694 | -0.005 | 0.012 | 0.012 | 0 | 0.11 | 0.14 | 0.06 | 0.16 | 0.08 | 0.02 |
| 4 | 11275048 | rs12643282 | -0.161 | 0.828 | -0.256 | 1.655 | -0.004 | -0.022 | -0.022 | 0 | 0.21 | 0.17 | 0.38 | 0.44 | 0.25 | 0.35 |
| 4 | 11276716 | rs6852625 | 0.869 | 0.372 | 0.229 | 1.001 | -0.004 | 0.061 | 0.061 | 0 | 0.92 | 0.74 | 0.83 | 0.86 | 0.95 | 0.81 |
| 4 | 11285331 | rs1609245 | 0.017 | 0.801 | -0.12 | 1.151 | -0.004 | -0.003 | -0.003 | 0 | 0.19 | 0.17 | 0.41 | 0.37 | 0.32 | 0.36 |
| 4 | 11288642 | rs1528079 | -0.191 | 0.294 | -0.285 | 0.79 | 0.002 | 0.002 | 0.002 | 0.066 | 0.87 | 0.9 | 0.94 | 0.9 | 0.9 | 0.91 |
| 4 | 11289125 | rs7663785 | 0.402 | -0.236 | 0.061 | 0.128 | -0.01 | -0.04 | -0.04 | 0 | 0.23 | 0.33 | 0.11 | 0.08 | 0.18 | 0.29 |
| 4 | 11290673 | rs11737841 | -1.171 | -0.212 | -0.008 | 0.109 | 0 | -0.001 | -0.001 | 0 | 0.89 | 0.92 | 0.95 | 0.9 | 0.9 | 0.95 |
| 4 | 11292775 | rs11945818 | -0.413 | -0.708 | 0.109 | 0.03 | -0.032 | -0.034 | -0.034 | 0 | 0.34 | 0.55 | 0.28 | 0.18 | 0.23 | 0.39 |
| 4 | 11295820 | rs13104553 | 0.925 | -0.612 | -0.094 | -0.258 | -0.003 | 0.036 | 0.036 | 0.265 | 0.46 | 0.34 | 0.55 | 0.71 | 0.53 | 0.52 |
| 4 | 11302032 | rs9997158 | -0.18 | -0.955 | -0.049 | -0.428 | -0.11 | -0.058 | -0.058 | 0 | 0.19 | 0.45 | 0.2 | 0.08 | 0.06 | 0.14 |
| 4 | 11302039 | rs7666445 | -1.012 | -0.965 | -0.139 | -0.379 | 0.014 | -0.002 | -0.002 | 0 | 0.61 | 0.74 | 0.79 | 0.57 | 0.69 | 0.79 |
| 4 | 11302760 | rs6829669 | 0.687 | -0.877 | -0.159 | -0.371 | -0.039 | -0.036 | -0.036 | 0 | 0.33 | 0.22 | 0.48 | 0.49 | 0.49 | 0.6 |
| 4 | 11303176 | rs4697869 | -1.13 | -0.906 | -0.153 | -0.385 | 0.012 | -0.005 | -0.005 | 1.804 | 0.62 | 0.78 | 0.8 | 0.58 | 0.69 | 0.79 |
| 4 | 11303208 | rs12501555 | 0.618 | -0.822 | -0.226 | -0.334 | -0.034 | -0.031 | -0.031 | 0 | 0.34 | 0.25 | 0.49 | 0.49 | 0.49 | 0.6 |
| 4 | 11305539 | rs2903016 | -1.418 | -0.85 | -0.244 | -0.26 | -0.003 | -0.002 | -0.002 | 0 | 0.88 | 0.84 | 0.93 | 0.9 | 0.9 | 0.93 |
| 4 | 11307275 | rs10488971 | 1.83 | -0.891 | -0.3 | -0.292 | -0.007 | -0.03 | -0.03 | 0 | 0.25 | 0.08 | 0.14 | 0.32 | 0.21 | 0.14 |
| 4 | 11307669 | rs6828885 | 1.853 | -0.962 | -0.075 | -0.328 | -0.025 | -0.034 | -0.034 | 0 | 0.18 | 0.1 | 0.4 | 0.38 | 0.31 | 0.35 |
| 4 | 11311393 | rs1029867 | 0.963 | -0.975 | 0.018 | -0.409 | 0.037 | 0.058 | 0.058 | 0.611 | 0.57 | 0.32 | 0.63 | 0.82 | 0.68 | 0.75 |
| 4 | 11315928 | rs1528078 | 0.997 | -0.99 | 0.117 | -0.377 | 0.024 | 0.034 | 0.034 | 0.851 | 0.57 | 0.34 | 0.64 | 0.82 | 0.69 | 0.74 |
| 4 | 11317278 | rs12512670 | 1.903 | -0.947 | 0.103 | -0.337 | -0.001 | 0.006 | 0.006 | 0 | 0.18 | 0.14 | 0.4 | 0.38 | 0.31 | 0.34 |
| 4 | 11320411 | rs17383790 | 1.405 | -0.962 | 0.218 | -0.441 | -0.007 | 0.002 | 0.002 | 0 | 0.15 | 0.11 | 0.09 | 0.11 | 0.16 | 0.25 |
| 4 | 11327966 | rs7699977 | 0.422 | -1.027 | 0.15 | -0.474 | -0.039 | -0.026 | -0.026 | 0 | 0.67 | 0.43 | 0.75 | 0.82 | 0.82 | 0.79 |
| 4 | 11328556 | rs1980381 | 0.369 | -1.069 | 0.309 | -0.509 | -0.058 | -0.033 | -0.033 | 0.008 | 0.16 | 0.31 | 0.18 | 0.08 | 0.05 | 0.15 |
| 4 | 11333844 | rs16880595 | 1.964 | -0.957 | 0.184 | -0.249 | -0.024 | -0.024 | -0.024 | 0 | 0.17 | 0.1 | 0.4 | 0.31 | 0.3 | 0.33 |
| 4 | 11337983 | rs981667 | -1.683 | -1.031 | -0.404 | -0.249 | 0.002 | 0.001 | 0.001 | 0 | 0.89 | 0.93 | 0.95 | 0.91 | 0.92 | 0.94 |
| 4 | 11341565 | rs1406077 | 0.967 | -1.035 | -0.418 | -0.255 | -0.022 | 0.033 | 0.033 | 0.752 | 0.35 | 0.15 | 0.25 | 0.32 | 0.35 | 0.19 |
| 4 | 11344212 | rs7676615 | 1.043 | -1.062 | -0.202 | -0.251 | -0.041 | -0.069 | -0.069 | 0 | 0.11 | 0.31 | 0.05 | 0 | 0.04 | 0.05 |
| 4 | 11352031 | rs917684 | 1.732 | -0.934 | -0.557 | -0.145 | -0.01 | -0.077 | -0.077 | 0 | 0.26 | 0.09 | 0.15 | 0.42 | 0.23 | 0.19 |
| 4 | 11357481 | rs1558402 | -0.424 | -0.911 | -0.509 | -0.204 | -0.004 | -0.012 | -0.012 | 0.152 | 0.53 | 0.72 | 0.69 | 0.49 | 0.55 | 0.7 |
| 4 | 11364771 | rs13112974 | 1.654 | -0.858 | -0.845 | -0.705 | -0.013 | -0.026 | -0.026 | 0 | 0.25 | 0.07 | 0.14 | 0.33 | 0.22 | 0.17 |
| 4 | 11366016 | rs12641904 | -0.2 | -0.873 | -0.873 | -1.138 | -0.001 | -0.015 | -0.015 | 6.935 | 0.78 | 0.78 | 0.85 | 0.91 | 0.79 | 0.87 |
| 4 | 11372956 | rs2109603 | 0.908 | -0.597 | -0.799 | -0.851 | -0.003 | -0.009 | -0.009 | 0 | 0.1 | 0.07 | 0.09 | 0 | 0.12 | 0.05 |
| 4 | 11380054 | rs725432 | -0.336 | -0.735 | -0.712 | -0.816 | -0.053 | -0.028 | -0.028 | 0 | 0.2 | 0.32 | 0.19 | 0.1 | 0.1 | 0.11 |
| 4 | 11390069 | rs11731587 | 0.107 | -0.159 | -0.919 | -0.563 | -0.015 | 0.02 | 0.02 | 0 | 0.24 | 0.17 | 0.19 | 0.08 | 0.29 | 0.27 |
| 4 | 11390743 | rs6854770 | -0.136 | -0.205 | -0.783 | -0.194 | -0.045 | -0.013 | -0.013 | 0 | 0.15 | 0.25 | 0.18 | 0.1 | 0.06 | 0.11 |
| 4 | 11398882 | rs1528081 | -0.741 | 0.652 | -1.366 | 0.205 | -0.002 | -0.002 | -0.002 | 0 | 0.91 | 0.93 | 0.98 | 0.94 | 0.96 | 0.95 |
| 4 | 11401087 | rs1047389 | 0.323 | 0.636 | -1.373 | 0.154 | 0.004 | -0.004 | -0.004 | 0 | 0.22 | 0.32 | 0.19 | 0.19 | 0.2 | 0.32 |
| 4 | 11402508 | rs4305508 | -0.588 | 0.63 | -1.531 | 0.278 | -0.002 | -0.002 | -0.002 | 0 | 0.91 | 0.93 | 0.98 | 0.94 | 0.96 | 0.95 |
| 4 | 11403704 | rs1406076 | 0.17 | 1.071 | -1.682 | 0.585 | -0.078 | -0.071 | -0.071 | 1.721 | 0.27 | 0.04 | 0.28 | 0.29 | 0.33 | 0.21 |
| 4 | 11408585 | rs16881446 | -0.753 | 1.215 | -0.695 | 0.35 | 0.004 | -0.005 | -0.005 | 0 | 0.25 | 0.3 | 0.22 | 0.18 | 0.26 | 0.34 |
| 4 | 11412087 | rs9999523 | -1.243 | 1.486 | -0.531 | 0.053 | 0.02 | -0.052 | -0.052 | 0 | 0.1 | 0.2 | 0.1 | 0 | 0.13 | 0.06 |
| 4 | 11412625 | rs16881495 | -0.474 | 1.645 | -1.383 | 0.636 | -0.025 | -0.11 | -0.11 | 0 | 0.22 | 0.09 | 0.16 | 0.47 | 0.22 | 0.29 |
| 4 | 11414037 | rs224463 | 0.404 | 1.348 | -1.139 | -0.244 | -0.022 | -0.011 | -0.011 | 0 | 0.39 | 0.52 | 0.48 | 0.37 | 0.28 | 0.32 |
| 4 | 11415299 | rs17423205 | 0.38 | 2.289 | -1.213 | -0.508 | -0.009 | 0.013 | 0.013 | 0 | 0.17 | 0.12 | 0.15 | 0.13 | 0.2 | 0.22 |
| 4 | 11416758 | rs17488994 | 0.51 | 2.252 | -1.226 | -0.522 | -0.018 | 0.016 | 0.016 | 0 | 0.16 | 0.09 | 0.14 | 0.13 | 0.2 | 0.22 |
| 4 | 11418534 | rs16881554 | 0.858 | 2.82 | -1.268 | -0.728 | -0.001 | -0.015 | -0.015 | 0 | 0.07 | 0.07 | 0.02 | 0 | 0.05 | 0.04 |
| 4 | 11419617 | rs13135853 | 0.072 | 3.008 | -1.311 | -0.756 | -0.033 | 0.031 | 0.031 | 0.098 | 0.2 | 0.09 | 0.2 | 0.13 | 0.3 | 0.24 |
| 4 | 11420619 | rs2109600 | -0.225 | 2.309 | -0.73 | -1.014 | -0.009 | -0.006 | -0.006 | 0 | 0.41 | 0.3 | 0.27 | 0.5 | 0.43 | 0.4 |
| 4 | 11421969 | rs16881636 | 1.158 | 1.482 | -0.86 | -1.58 | -0.016 | -0.099 | -0.099 | 0 | 0.16 | 0.32 | 0.1 | 0 | 0.11 | 0.13 |
| 4 | 11424898 | rs7660531 | 0.737 | 1.816 | -0.849 | -1.495 | -0.002 | -0.004 | -0.004 | 0 | 0.33 | 0.31 | 0.25 | 0.46 | 0.37 | 0.3 |
| 4 | 11425012 | rs7660733 | -0.021 | 1.63 | -1.089 | -1.407 | 0.028 | 0.004 | 0.004 | 0 | 0.49 | 0.73 | 0.34 | 0.46 | 0.39 | 0.45 |
| 4 | 11425821 | rs12054490 | -0.098 | 1.548 | -1.194 | -1.4 | -0.012 | -0.02 | -0.02 | 0 | 0.55 | 0.77 | 0.34 | 0.46 | 0.41 | 0.49 |
| 4 | 11432165 | rs224489 | 0.865 | 0.931 | -1.318 | -0.824 | 0.004 | -0.112 | -0.112 | 0 | 0.2 | 0.04 | 0.24 | 0.44 | 0.14 | 0.26 |

| | | | | | | | | | | | | | | | | |
|---|---|---|---|---|---|---|---|---|---|---|---|---|---|---|---|---|
| 4 | 11435516 | rs11936371 | 0.493 | 1.193 | -0.966 | -1.128 | -0.001 | -0.036 | -0.036 | 0.169 | 0.18 | 0.19 | 0.12 | 0 | 0.19 | 0.1 |
| 4 | 11439946 | rs224484 | 0.311 | 1.394 | -0.838 | -1.199 | -0.003 | 0.004 | 0.004 | 3.972 | 0.4 | 0.45 | 0.59 | 0.55 | 0.34 | 0.47 |
| 4 | 11441634 | rs224481 | -0.099 | 1.27 | -1.182 | -0.779 | -0.015 | -0.022 | -0.022 | 0 | 0.61 | 0.76 | 0.75 | 0.57 | 0.59 | 0.59 |
| 4 | 11448182 | rs411035 | 0.299 | 0.865 | -0.616 | -0.525 | 0.011 | -0.042 | -0.042 | 0.058 | 0.86 | 0.72 | 0.88 | 0.95 | 0.83 | 0.89 |
| 4 | 11449199 | rs17424739 | 0.04 | 1.057 | -1.202 | -0.576 | -0.048 | 0.053 | 0.053 | 0.006 | 0.17 | 0.06 | 0.15 | 0.09 | 0.25 | 0.21 |
| 4 | 11449278 | rs17490471 | 0.155 | 0.978 | -1.106 | -0.583 | -0.027 | 0.047 | 0.047 | 0 | 0.17 | 0.07 | 0.21 | 0.09 | 0.22 | 0.2 |
| 4 | 11455518 | rs402575 | 1.042 | 0.183 | -1.127 | -0.729 | 0.014 | 0.015 | 0.015 | 0.854 | 0.38 | 0.23 | 0.22 | 0.33 | 0.34 | 0.25 |
| 4 | 11459738 | rs224474 | 0.145 | -0.254 | -0.608 | -0.923 | 0.034 | 0.044 | 0.044 | 1.07 | 0.49 | 0.65 | 0.3 | 0.33 | 0.39 | 0.3 |
| 4 | 11460840 | rs16882139 | 0.512 | -0.063 | -0.021 | -0.476 | -0.017 | 0.02 | 0.02 | 0.533 | 0.09 | 0.25 | 0.05 | 0.07 | 0.04 | 0.06 |
| 4 | 11461954 | rs224470 | -0.565 | 0.163 | -0.333 | -0.439 | -0.003 | -0.003 | -0.003 | 0 | 0.66 | 0.7 | 0.51 | 0.43 | 0.62 | 0.51 |
| 4 | 11463773 | rs224480 | -0.589 | 0.175 | -0.328 | -0.484 | -0.003 | -0.005 | -0.005 | 0 | 0.66 | 0.7 | 0.52 | 0.5 | 0.62 | 0.51 |
| 4 | 11464975 | rs13137776 | 1.147 | 0.192 | -0.899 | -0.38 | -0.099 | -0.158 | -0.158 | 0 | 0.24 | 0.02 | 0.26 | 0.44 | 0.3 | 0.37 |
| 4 | 11467103 | rs7699705 | -0.47 | -0.306 | -0.76 | -0.876 | 0.005 | 0.02 | 0.02 | 0 | 0.59 | 0.61 | 0.67 | 0.84 | 0.64 | 0.74 |
| 4 | 11468244 | rs6822160 | -0.473 | -0.343 | -0.798 | -1.031 | 0.011 | 0.044 | 0.044 | 0 | 0.59 | 0.69 | 0.68 | 0.9 | 0.65 | 0.72 |
| 4 | 11470385 | rs11733131 | 0.585 | 0.023 | -1.095 | -0.744 | -0.091 | -0.065 | -0.065 | 0 | 0.38 | 0.11 | 0.3 | 0.42 | 0.45 | 0.4 |
| 4 | 11470916 | rs737488 | 0.086 | -0.019 | -0.219 | -0.965 | -0.011 | 0.056 | 0.056 | 0.878 | 0.06 | 0.22 | 0.2 | 0.11 | 0.01 | 0.12 |
| 4 | 11474624 | rs10023768 | -1.068 | -0.103 | 0.043 | -1.063 | -0.026 | 0.006 | 0.006 | 1.181 | 0.14 | 0.45 | 0.25 | 0.11 | 0.07 | 0.14 |
| 4 | 11475911 | rs9291554 | -0.49 | 0.077 | -0.206 | -1.128 | 0.002 | 0.066 | 0.066 | 0.993 | 0.28 | 0.54 | 0.36 | 0.45 | 0.27 | 0.31 |
| 4 | 11481304 | rs11735501 | 1.295 | 0.193 | -0.859 | -1.344 | -0.02 | 0.066 | 0.066 | 0 | 0.2 | 0.08 | 0.13 | 0.02 | 0.2 | 0.16 |
| 4 | 11483569 | rs2029799 | -0.192 | 0.095 | -0.655 | -1.307 | -0.037 | 0.023 | 0.023 | 0 | 0.41 | 0.26 | 0.28 | 0.36 | 0.46 | 0.36 |
| 4 | 11487322 | rs12163751 | -1.284 | -0.319 | -0.468 | -1.463 | -0.005 | -0.003 | -0.003 | 0 | 0.71 | 0.67 | 0.59 | 0.79 | 0.82 | 0.75 |
| 4 | 11498145 | rs10004232 | 1.597 | -0.236 | -0.204 | -1.429 | 0.014 | -0.017 | -0.017 | 0 | 0.05 | 0.21 | 0.06 | 0 | 0.05 | 0.01 |
| 4 | 11500166 | rs923997 | -0.141 | -0.319 | -0.299 | -1.563 | 0.071 | 0.104 | 0.104 | 0.294 | 0.57 | 0.24 | 0.5 | 0.79 | 0.71 | 0.66 |
| 4 | 11500214 | rs17236493 | 2.435 | 0.05 | -0.391 | -1.516 | -0.042 | 0.053 | 0.053 | 0 | 0.09 | 0 | 0.08 | 0 | 0.16 | 0.03 |
| 4 | 11503860 | rs1874238 | -0.246 | -0.162 | -0.053 | -1.355 | 0.013 | -0.095 | -0.095 | 0 | 0.13 | 0.43 | 0.09 | 0 | 0.11 | 0.09 |
| 4 | 11506753 | rs13139849 | 1.513 | 0.051 | -0.376 | -1.281 | -0.043 | -0.071 | -0.071 | 0 | 0.18 | 0.07 | 0.23 | 0.44 | 0.26 | 0.32 |
| 4 | 11507052 | rs6847037 | 0.801 | 0.05 | -0.337 | -1.29 | 0.002 | 0.01 | 0.01 | 0 | 0.23 | 0.25 | 0.28 | 0.44 | 0.31 | 0.32 |
| 4 | 11507454 | rs4916640 | 0.772 | 0.048 | -0.366 | -1.222 | 0.013 | 0.027 | 0.027 | 0 | 0.23 | 0.31 | 0.29 | 0.39 | 0.31 | 0.32 |
| 4 | 11511291 | rs2088131 | -0.362 | -0.073 | -0.374 | -1.462 | 0.025 | 0.048 | 0.048 | 0.549 | 0.34 | 0.47 | 0.31 | 0.44 | 0.38 | 0.42 |
| 4 | 11511452 | rs1565924 | -0.182 | -0.231 | -0.776 | -1.638 | 0.007 | 0.003 | 0.003 | 0 | 0.56 | 0.41 | 0.51 | 0.78 | 0.7 | 0.64 |
| 4 | 11513562 | rs11725667 | 1.332 | -0.146 | -0.511 | -1.503 | -0.003 | 0.039 | 0.039 | 0 | 0.21 | 0.11 | 0.14 | 0.02 | 0.19 | 0.16 |
| 4 | 11515688 | rs9993962 | 0.362 | -0.156 | -0.457 | -1.463 | 0.003 | 0.003 | 0.003 | 0 | 0.4 | 0.39 | 0.38 | 0.45 | 0.45 | 0.47 |
| 4 | 11516724 | rs9997161 | -0.278 | -0.235 | 0.019 | -0.916 | 0.011 | 0.012 | 0.012 | 0.009 | 0.66 | 0.65 | 0.55 | 0.78 | 0.76 | 0.73 |
| 4 | 11517436 | rs6848315 | 0.24 | 0.002 | -0.227 | -0.884 | -0.004 | 0.016 | 0.016 | 0.096 | 0.46 | 0.37 | 0.3 | 0.35 | 0.5 | 0.41 |
| 4 | 11519011 | rs10003762 | -1.165 | -0.106 | 0.049 | -1.069 | -0.031 | -0.035 | -0.035 | 0 | 0.16 | 0.57 | 0.27 | 0.12 | 0.09 | 0.14 |
| 4 | 11521906 | rs11729702 | 0.522 | 0.3 | -1.167 | -1.176 | -0.067 | -0.046 | -0.046 | 0 | 0.34 | 0.17 | 0.48 | 0.42 | 0.5 | 0.47 |
| 4 | 11524263 | rs12508213 | -0.188 | 0.259 | -1.551 | -1.131 | 0.011 | 0.006 | 0.006 | 4.683 | 0.53 | 0.86 | 0.62 | 0.42 | 0.59 | 0.55 |
| 4 | 11531314 | rs12647573 | -0.251 | 0.661 | -1.856 | -1.093 | 0.004 | 0.007 | 0.007 | 0 | 0.39 | 0.53 | 0.4 | 0.59 | 0.39 | 0.45 |
| 4 | 11537238 | rs1603186 | -0.38 | 0.486 | -2.004 | -0.802 | 0.023 | 0.016 | 0.016 | 0.749 | 0.71 | 0.79 | 0.86 | 1 | 0.82 | 0.93 |
| 4 | 11538001 | rs10155331 | 0.506 | 0.511 | -1.975 | -0.799 | -0.005 | -0.029 | -0.029 | 0 | 0.31 | 0.22 | 0.31 | 0.6 | 0.3 | 0.44 |
| 4 | 11538947 | rs7687011 | -0.211 | 1.257 | -1.414 | 0.088 | -0.003 | -0.001 | -0.001 | 0 | 0.4 | 0.57 | 0.55 | 0.4 | 0.52 | 0.5 |
| 4 | 11544142 | rs17491203 | 0.061 | 1.38 | -1.575 | 0.075 | 0.001 | -0.007 | -0.007 | 0 | 0.07 | 0.05 | 0.05 | 0.01 | 0.06 | 0.03 |
| 4 | 11546275 | rs956469 | 0.74 | 1.967 | -1.925 | 0.217 | -0.014 | 0.001 | 0.001 | 0 | 0.62 | 0.71 | 0.55 | 0.38 | 0.48 | 0.5 |
| 4 | 11547106 | rs1032502 | -0.206 | 1.385 | -2.085 | 0.35 | -0.023 | -0.113 | -0.113 | 3.166 | 0.76 | 0.94 | 0.69 | 0.47 | 0.74 | 0.62 |
| 4 | 11547681 | rs6846137 | -0.441 | 1.031 | -1.456 | -0.036 | -0.006 | -0.05 | -0.05 | 0.015 | 0.1 | 0.15 | 0.08 | 0.01 | 0.08 | 0.03 |
| 4 | 11547967 | rs756418 | -0.436 | 0.976 | -2.007 | 0.239 | 0 | 0.009 | 0.009 | 0.431 | 0.18 | 0.01 | 0.21 | 0.12 | 0.12 | 0.13 |
| 4 | 11556893 | rs6448784 | 0.434 | 1.841 | -1.576 | 0.567 | -0.008 | -0.002 | -0.002 | 0.243 | 0.82 | 0.84 | 0.78 | 0.83 | 0.75 | 0.84 |
| 4 | 11571164 | rs10049912 | -0.516 | 1.506 | -2.01 | -0.129 | 0.031 | 0.059 | 0.059 | 0 | 0.51 | 0.72 | 0.51 | 0.3 | 0.56 | 0.38 |
| 4 | 11577136 | rs10488991 | -0.679 | 1.396 | -0.709 | -0.311 | 0.051 | -0.115 | -0.115 | 0 | 0.18 | 0.49 | 0.15 | 0.04 | 0.2 | 0.12 |
| 4 | 11594112 | rs2215232 | 0.413 | 1.421 | -1.138 | 0.406 | 0.012 | -0.085 | -0.085 | 0.025 | 0.69 | 0.89 | 0.5 | 0.51 | 0.72 | 0.65 |
| 4 | 11596870 | rs11728997 | -0.091 | 1.248 | -0.443 | 0.42 | -0.012 | 0.008 | 0.008 | 0 | 0.11 | 0.15 | 0.07 | 0.14 | 0.11 | 0.14 |
| 4 | 11600333 | rs2052044 | -0.567 | 1.201 | 0.005 | 0.442 | -0.019 | 0.018 | 0.018 | 0.09 | 0.13 | 0.29 | 0.07 | 0.14 | 0.11 | 0.13 |
| 4 | 11601925 | rs12506708 | 1.864 | 1.244 | 0.364 | 0.216 | -0.007 | 0.036 | 0.036 | 0 | 0.46 | 0.34 | 0.31 | 0.33 | 0.46 | 0.46 |
| 4 | 11605860 | rs17427761 | 1.414 | 1.363 | 0.31 | 0.331 | -0.006 | 0.091 | 0.091 | 0 | 0.51 | 0.81 | 0.36 | 0.36 | 0.46 | 0.47 |
| 4 | 11605925 | rs16884475 | 0.447 | 1.35 | 0.319 | 0.726 | -0.002 | 0.011 | 0.011 | 0 | 0.1 | 0.07 | 0.06 | 0.2 | 0.1 | 0.13 |
| 4 | 11606915 | rs10939361 | 0.82 | 1.348 | 0.297 | 0.516 | 0.019 | 0.03 | 0.03 | 0 | 0.61 | 0.88 | 0.45 | 0.36 | 0.61 | 0.5 |
| 4 | 11609704 | rs6816174 | 0.276 | 1.356 | 0.285 | 0.565 | -0.011 | -0.146 | -0.146 | 0.006 | 0.7 | 0.95 | 0.51 | 0.51 | 0.72 | 0.64 |
| 4 | 11617783 | rs10032590 | 1.605 | 1.393 | 0.431 | 0.529 | 0.007 | 0.065 | 0.065 | 0 | 0.48 | 0.72 | 0.33 | 0.35 | 0.43 | 0.46 |
| 4 | 11619288 | rs6842740 | 0.801 | 1.376 | 0.74 | 0.521 | -0.008 | 0.037 | 0.037 | 0 | 0.59 | 0.88 | 0.43 | 0.37 | 0.59 | 0.5 |
| 4 | 11621121 | rs723563 | 1.353 | 1.482 | 0.712 | 0.508 | 0.005 | 0.018 | 0.018 | 0 | 0.49 | 0.5 | 0.3 | 0.25 | 0.44 | 0.5 |
| 4 | 11621242 | rs3886917 | 0.148 | 1.477 | 0.556 | 0.532 | -0.015 | -0.076 | -0.076 | 0.012 | 0.67 | 0.87 | 0.48 | 0.5 | 0.69 | 0.62 |
| 4 | 11621877 | rs11730207 | 0.472 | 1.469 | 0.569 | 0.536 | -0.008 | 0.013 | 0.013 | 0 | 0.11 | 0.17 | 0.07 | 0.14 | 0.1 | 0.14 |
| 4 | 11623160 | rs11730600 | -1.041 | 1.335 | 0.381 | 0.567 | -0.052 | -0.105 | -0.105 | 0.023 | 0.86 | 0.99 | 0.69 | 0.59 | 0.84 | 0.84 |
| 4 | 11623872 | rs6822439 | 1.305 | 1.223 | 0.588 | 0.453 | 0 | 0.001 | 0.001 | 0 | 0.48 | 0.55 | 0.34 | 0.15 | 0.49 | 0.41 |
| 4 | 11624085 | rs6845061 | 2.029 | 1.148 | 0.367 | 0.434 | 0.002 | -0.002 | -0.002 | 0 | 0.36 | 0.3 | 0.19 | 0.11 | 0.29 | 0.35 |
| 4 | 11625588 | rs9884312 | 0.894 | 1.203 | 0.247 | 0.582 | -0.001 | -0.001 | -0.001 | 0.013 | 0.55 | 0.54 | 0.36 | 0.47 | 0.53 | 0.58 |
| 4 | 11628238 | rs2865155 | 1.076 | 0.914 | 0.007 | 0.377 | 0.035 | 0.169 | 0.169 | 0 | 0.46 | 0.78 | 0.29 | 0.15 | 0.37 | 0.4 |
| 4 | 11630049 | rs6448799 | 0.551 | 0.392 | 0.467 | 1.191 | -0.039 | -0.111 | -0.111 | 0 | 0.72 | 0.92 | 0.49 | 0.54 | 0.68 | 0.63 |
| 4 | 11631446 | rs11728092 | 1.738 | 0.31 | 1.33 | 1.151 | 0.003 | 0 | 0 | 3.356 | 0.57 | 0.37 | 0.36 | 0.4 | 0.58 | 0.57 |

| | | | | | | | | | | | | | | | | |
|---|---|---|---|---|---|---|---|---|---|---|---|---|---|---|---|---|
| 4 | 11633455 | rs13106661 | 1.24 | 0.317 | 1.624 | 1.139 | 0.002 | 0.01 | 0.01 | 0 | 0.62 | 0.53 | 0.4 | 0.4 | 0.6 | 0.58 |
| 4 | 11642534 | rs966240 | 1.19 | 0.411 | 1.386 | 1.153 | -0.028 | -0.027 | -0.027 | 0 | 0.62 | 0.85 | 0.4 | 0.38 | 0.56 | 0.56 |
| 4 | 11651556 | rs1860292 | 1.293 | 0.42 | 1.557 | 1.204 | -0.001 | -0.002 | -0.002 | 0 | 0.61 | 0.64 | 0.4 | 0.39 | 0.6 | 0.58 |
| 4 | 11664714 | rs4397010 | 1.536 | 0.548 | 1.715 | 1.277 | -0.001 | -0.002 | -0.002 | 3.684 | 0.59 | 0.51 | 0.38 | 0.39 | 0.57 | 0.57 |
| 4 | 11674661 | rs7671189 | 0.748 | 0.573 | 1.733 | 1.277 | -0.002 | -0.01 | -0.01 | 0.093 | 0.72 | 0.79 | 0.64 | 0.66 | 0.71 | 0.66 |
| 4 | 11677334 | rs12649160 | -0.75 | 0.602 | 1.552 | 1.359 | 0.001 | 0.028 | 0.028 | 0 | 0.12 | 0.28 | 0.25 | 0.24 | 0.12 | 0.07 |
| 4 | 11692300 | rs2870411 | 0.965 | 0.596 | 1.661 | 1.345 | -0.002 | 0.018 | 0.018 | 0.067 | 0.71 | 0.64 | 0.62 | 0.66 | 0.71 | 0.66 |
| 4 | 11700529 | rs6831780 | -0.382 | 0.573 | 1.382 | 1.342 | -0.034 | -0.094 | -0.094 | 4.815 | 0.86 | 0.99 | 0.85 | 0.71 | 0.85 | 0.87 |
| 4 | 11701193 | rs7696515 | -0.675 | 0.518 | 1.623 | 1.358 | -0.001 | 0.005 | 0.005 | 0 | 0.11 | 0.16 | 0.24 | 0.24 | 0.12 | 0.08 |
| 4 | 11721992 | rs11729868 | 1.408 | 0.5 | 1.593 | 1.3 | 0.001 | -0.007 | -0.007 | 0.328 | 0.59 | 0.36 | 0.39 | 0.42 | 0.59 | 0.59 |
| 4 | 11732810 | rs2279546 | -0.714 | 0.728 | 1.615 | 1.256 | -0.001 | 0.007 | 0.007 | 0 | 0.11 | 0.19 | 0.11 | 0.24 | 0.11 | 0.08 |
| 4 | 11734841 | rs3912887 | 1.408 | 0.424 | 1.762 | 1.216 | 0 | 0 | 0 | 0 | 0.57 | 0.42 | 0.5 | 0.4 | 0.59 | 0.61 |
| 4 | 11743843 | rs6825825 | -0.4 | 0.431 | 1.733 | 1.186 | -0.01 | 0.015 | 0.015 | 0 | 0.09 | 0.15 | 0.09 | 0.24 | 0.05 | 0.08 |
| 4 | 11744388 | rs993835 | 0.029 | 0.449 | 1.389 | 1.195 | -0.05 | -0.052 | -0.052 | 3.54 | 0.7 | 0.9 | 0.62 | 0.65 | 0.65 | 0.67 |
| 4 | 11747298 | rs7654642 | 1.308 | 0.499 | 1.299 | 1.241 | 0.002 | 0 | 0 | 0 | 0.57 | 0.39 | 0.5 | 0.41 | 0.59 | 0.61 |
| 4 | 11754426 | rs1388867 | 0.574 | 0.708 | 0.716 | 0.99 | 0.001 | 0.012 | 0.012 | 0 | 0.7 | 0.65 | 0.61 | 0.64 | 0.7 | 0.67 |
| 4 | 11755497 | rs9291579 | 0.458 | 0.737 | 0.41 | 1.005 | -0.002 | -0.008 | -0.008 | 0.013 | 0.71 | 0.76 | 0.62 | 0.64 | 0.7 | 0.67 |
| 4 | 11759562 | rs7699210 | 0.532 | 0.737 | 0.364 | 0.993 | 0.001 | 0.004 | 0.004 | 0 | 0.7 | 0.68 | 0.62 | 0.64 | 0.7 | 0.67 |
| 4 | 11760145 | rs6845480 | 0.392 | 0.82 | -0.338 | 0.967 | -0.008 | -0.021 | -0.021 | 0 | 0.71 | 0.82 | 0.63 | 0.65 | 0.7 | 0.67 |
| 4 | 11775089 | rs923747 | -1.253 | 0.505 | 0.422 | 0.298 | -0.001 | -0.024 | -0.024 | 0.314 | 0.26 | 0.6 | 0.26 | 0.25 | 0.25 | 0.28 |
| 4 | 11786048 | rs12639854 | -1.229 | 0.407 | 1.731 | 0.392 | -0.011 | -0.008 | -0.008 | 0 | 0.19 | 0.17 | 0.22 | 0.29 | 0.25 | 0.27 |
| 4 | 11792289 | rs12505790 | -2.05 | 0.121 | 1.991 | -0.156 | -0.04 | -0.139 | -0.139 | 0.6 | 0.25 | 0.53 | 0.18 | 0.01 | 0.13 | 0.17 |
| 4 | 11816679 | rs7661409 | -1.304 | 0.23 | 1.883 | -0.131 | -0.008 | -0.064 | -0.064 | 0.113 | 0.17 | 0.2 | 0.15 | 0 | 0.13 | 0.16 |
| 4 | 11825987 | rs1489473 | -1.859 | 0.172 | 1.802 | -0.116 | -0.03 | -0.132 | -0.132 | 0 | 0.18 | 0.54 | 0.17 | 0.01 | 0.11 | 0.17 |
| 4 | 11845784 | rs16886280 | -1.095 | -0.264 | 1.548 | -0.372 | -0.013 | -0.064 | -0.064 | 0 | 0.2 | 0.24 | 0.15 | 0.01 | 0.11 | 0.17 |
| 4 | 11850963 | rs10024612 | -1.657 | 0.655 | 1.44 | -0.619 | -0.005 | 0.021 | 0.021 | 0.232 | 0.38 | 0.68 | 0.43 | 0.13 | 0.39 | 0.34 |
| 4 | 11861566 | rs6448841 | -0.715 | 0.639 | 1.514 | -0.032 | 0.031 | 0.018 | 0.018 | 0.287 | 0.21 | 0.64 | 0.3 | 0.21 | 0.28 | 0.2 |
| 4 | 11864538 | rs17256075 | -0.163 | 0.587 | 1.554 | -0.063 | 0.005 | -0.017 | -0.017 | 0 | 0.17 | 0.13 | 0.14 | 0 | 0.11 | 0.15 |
| 4 | 11865461 | rs993091 | 0.012 | 0.741 | 1.502 | 0.382 | -0.002 | 0 | 0 | 0 | 0.92 | 0.92 | 0.92 | 0.78 | 0.87 | 0.91 |
| 4 | 11865749 | rs12506582 | -0.135 | 0.494 | 1.593 | 0.286 | 0.015 | 0.084 | 0.084 | 0.05 | 0.17 | 0.05 | 0.13 | 0 | 0.11 | 0.15 |
| 4 | 11868097 | rs12645979 | -1.848 | 0.573 | 1.606 | 0.243 | -0.008 | 0.031 | 0.031 | 0.237 | 0.39 | 0.75 | 0.45 | 0.21 | 0.4 | 0.34 |
| 4 | 11868786 | rs2200819 | -1.365 | 0.841 | 1.577 | 0.216 | -0.002 | 0.004 | 0.004 | 0 | 0.32 | 0.33 | 0.37 | 0.21 | 0.33 | 0.32 |
| 4 | 11869456 | rs12510141 | -0.127 | 0.829 | 1.28 | 0.137 | 0.014 | 0.09 | 0.09 | 0.262 | 0.17 | 0.05 | 0.13 | 0 | 0.11 | 0.15 |
| 4 | 11870144 | rs17322411 | -0.106 | 0.84 | 1.292 | 0.187 | -0.037 | -0.033 | -0.033 | 0.929 | 0.15 | 0.1 | 0.22 | 0.19 | 0.22 | 0.17 |
| 4 | 11872010 | rs6840259 | -0.207 | 0.774 | 1.77 | 0.16 | -0.013 | -0.071 | -0.071 | 1.776 | 0.18 | 0.22 | 0.15 | 0 | 0.12 | 0.14 |
| 4 | 11873033 | rs17256627 | -0.141 | 0.772 | 1.83 | 0.16 | -0.016 | -0.015 | -0.015 | 0 | 0.16 | 0.16 | 0.23 | 0.2 | 0.21 | 0.18 |
| 4 | 11873630 | rs6843251 | -2.07 | 0.703 | 2.079 | 0.143 | -0.006 | 0.05 | 0.05 | 0.021 | 0.4 | 0.75 | 0.44 | 0.2 | 0.4 | 0.35 |
| 4 | 11884262 | rs11934095 | -1.188 | 0.726 | 2.188 | 0.158 | 0.05 | 0.012 | 0.012 | 0.068 | 0.22 | 0.55 | 0.29 | 0.2 | 0.27 | 0.21 |
| 4 | 11892822 | rs2866059 | -1.742 | 0.661 | 1.832 | 0.125 | -0.007 | 0.03 | 0.03 | 0 | 0.38 | 0.33 | 0.4 | 0.2 | 0.39 | 0.35 |
| 4 | 11900932 | rs4394018 | -1.001 | 0.638 | 1.846 | 0.117 | 0.011 | -0.036 | -0.036 | 2.183 | 0.07 | 0.21 | 0.06 | 0 | 0.07 | 0.03 |
| 4 | 11901216 | rs6852116 | 0.043 | 0.617 | 1.761 | 0.093 | 0.014 | 0.035 | 0.035 | 0 | 0.17 | 0.07 | 0.13 | 0 | 0.12 | 0.14 |
| 4 | 11904548 | rs17323516 | 1.196 | 0.744 | 1.07 | -0.046 | -0.044 | -0.027 | -0.027 | 1.461 | 0.1 | 0.01 | 0.07 | 0.08 | 0.15 | 0.11 |
| 4 | 11909465 | rs16886377 | -0.824 | 0.706 | 1.333 | -0.21 | 0.038 | -0.023 | -0.023 | 0 | 0.07 | 0.28 | 0.06 | 0 | 0.07 | 0.02 |
| 4 | 11911000 | rs1387459 | -1.121 | 0.226 | 1.004 | -0.614 | -0.003 | -0.022 | -0.022 | 0.072 | 0.41 | 0.53 | 0.42 | 0.23 | 0.38 | 0.34 |
| 4 | 11912625 | rs13111622 | -0.554 | 0.338 | 1.276 | -0.367 | 0.022 | 0.032 | 0.032 | 0.108 | 0.2 | 0.63 | 0.29 | 0.23 | 0.23 | 0.19 |
| 4 | 11913343 | rs17258155 | 0.508 | 0.342 | 1.122 | -0.37 | 0.014 | 0.067 | 0.067 | 0 | 0.17 | 0.07 | 0.13 | 0 | 0.12 | 0.14 |
| 4 | 11917328 | rs10084803 | 0.029 | 0.404 | 1.206 | -0.302 | 0.054 | 0.064 | 0.064 | 0 | 0.17 | 0.4 | 0.25 | 0.23 | 0.23 | 0.18 |
| 4 | 11922712 | rs2171726 | -2.094 | 0.25 | 0.623 | -0.843 | -0.021 | 0.094 | 0.094 | 0 | 0.64 | 0.89 | 0.77 | 0.78 | 0.64 | 0.64 |
| 4 | 11924046 | rs11939914 | -1.917 | 0.366 | 1.268 | -0.713 | 0.004 | 0.106 | 0.106 | 0 | 0.44 | 0.75 | 0.46 | 0.17 | 0.43 | 0.38 |
| 4 | 11926778 | rs11725886 | -0.042 | 0.546 | 1.277 | -0.531 | -0.025 | 0.003 | 0.003 | 0 | 0.18 | 0.08 | 0.24 | 0.16 | 0.25 | 0.21 |
| 4 | 11927652 | rs10939381 | -0.029 | 0.559 | 1.275 | -0.536 | -0.024 | 0.004 | 0.004 | 0 | 0.17 | 0.08 | 0.23 | 0.15 | 0.24 | 0.21 |
| 4 | 11938729 | rs10939384 | -0.136 | 0.348 | 1.783 | -0.644 | -0.028 | -0.114 | -0.114 | 0 | 0.19 | 0.36 | 0.16 | 0 | 0.11 | 0.14 |
| 4 | 11948435 | rs10213500 | -1.207 | 0.292 | 1.895 | -0.672 | -0.009 | -0.064 | -0.064 | 0 | 0.28 | 0.65 | 0.22 | 0.03 | 0.18 | 0.17 |
| 4 | 11948654 | rs10213502 | -1.253 | 0.269 | 1.892 | -0.673 | 0.006 | -0.024 | -0.024 | 0 | 0.28 | 0.71 | 0.23 | 0.03 | 0.18 | 0.17 |
| 4 | 11955722 | rs13149855 | -0.299 | 0.501 | 0.925 | -0.584 | -0.019 | 0.008 | 0.008 | 0 | 0.18 | 0.09 | 0.25 | 0.15 | 0.25 | 0.21 |
| 4 | 11956720 | rs16886480 | -1.283 | 0.332 | 0.906 | -0.724 | -0.001 | -0.04 | -0.04 | 0.982 | 0.27 | 0.67 | 0.22 | 0.03 | 0.18 | 0.17 |
| 4 | 11977042 | rs7696250 | -0.016 | 0.532 | 0.204 | -0.579 | 0.017 | 0.005 | 0.005 | 0 | 0.19 | 0.15 | 0.14 | 0 | 0.12 | 0.12 |
| 4 | 11981872 | rs13133185 | -1.749 | 0.852 | 0.114 | 0.081 | 0 | 0.006 | 0.006 | 0 | 0.5 | 0.49 | 0.46 | 0.51 | 0.54 | 0.45 |
| 4 | 11982994 | rs7666669 | -0.434 | 1.066 | -0.19 | 0.162 | -0.014 | -0.029 | -0.029 | 0 | 0.23 | 0.15 | 0.26 | 0.49 | 0.35 | 0.3 |
| 4 | 11983193 | rs16886509 | -0.59 | 1.107 | -0.215 | 0.187 | -0.002 | -0.003 | -0.003 | 0 | 0.26 | 0.25 | 0.27 | 0.49 | 0.35 | 0.3 |
| 4 | 12000904 | rs1372887 | -1.169 | 1.133 | -0.418 | 0.473 | 0.002 | -0.001 | -0.001 | 0.133 | 0.48 | 0.56 | 0.58 | 0.43 | 0.51 | 0.46 |
| 4 | 12008411 | rs12639801 | -0.374 | 1.527 | -0.748 | 0.696 | -0.05 | -0.077 | -0.077 | 0 | 0.18 | 0.07 | 0.34 | 0.53 | 0.29 | 0.28 |
| 4 | 12014109 | rs10212892 | 0.187 | 1.02 | -0.543 | 0.112 | 0.002 | -0.034 | -0.034 | 0 | 0.22 | 0.18 | 0.16 | 0 | 0.16 | 0.1 |
| 4 | 12017993 | rs6448852 | -1.512 | 1.266 | -0.044 | 0.001 | -0.001 | 0 | 0 | 0 | 0.45 | 0.41 | 0.54 | 0.53 | 0.5 | 0.4 |
| 4 | 12028032 | rs16886566 | 0.076 | 1.28 | -0.058 | 0.055 | -0.071 | -0.1 | -0.1 | 0 | 0.19 | 0.06 | 0.35 | 0.52 | 0.31 | 0.29 |
| 4 | 12036029 | rs11721877 | -0.478 | 1.155 | 0.038 | 0.174 | -0.013 | -0.033 | -0.033 | 0 | 0.26 | 0.32 | 0.18 | 0.08 | 0.2 | 0.17 |
| 4 | 12050985 | rs7672776 | 0.317 | 1.114 | 0.059 | 0.147 | 0.004 | -0.027 | -0.027 | 0.325 | 0.21 | 0.17 | 0.1 | 0 | 0.14 | 0.1 |
| 4 | 12069337 | rs16886607 | -0.489 | 1.125 | 0.017 | 0.161 | -0.013 | -0.03 | -0.03 | 0.527 | 0.26 | 0.3 | 0.14 | 0.1 | 0.19 | 0.17 |
| 4 | 12074137 | rs1025877 | -0.674 | 1.228 | -0.18 | 0.361 | -0.01 | -0.005 | -0.005 | 0 | 0.26 | 0.19 | 0.44 | 0.63 | 0.37 | 0.37 |

| | | | | | | | | | | | | | | | | |
|---|---|---|---|---|---|---|---|---|---|---|---|---|---|---|---|---|
| 4 | 12083991 | rs1479273 | -0.76 | 0.496 | 0.277 | -0.144 | -0.004 | 0.047 | 0.047 | 0.918 | 0.46 | 0.62 | 0.49 | 0.66 | 0.48 | 0.48 |
| 4 | 12088133 | rs12641414 | -0.443 | 0.502 | 0.3 | -0.137 | -0.026 | 0.043 | 0.043 | 1.122 | 0.42 | 0.23 | 0.46 | 0.62 | 0.48 | 0.42 |
| 4 | 12098774 | rs13152023 | -0.9 | -0.05 | -0.176 | -0.705 | 0 | -0.021 | -0.021 | 0 | 0.23 | 0.21 | 0.1 | 0.05 | 0.14 | 0.17 |
| 4 | 12099307 | rs6810659 | -0.077 | 0.049 | -0.149 | -0.899 | -0.018 | -0.089 | -0.089 | 0 | 0.17 | 0.37 | 0.25 | 0.05 | 0.17 | 0.2 |
| 4 | 12108239 | rs4834882 | -0.963 | 0.135 | -0.569 | -0.857 | 0 | 0.01 | 0.01 | 0.05 | 0.59 | 0.6 | 0.71 | 0.68 | 0.65 | 0.61 |
| 4 | 12138827 | rs1382751 | -0.949 | 0.137 | -0.628 | -0.861 | -0.001 | 0.019 | 0.019 | 0.046 | 0.59 | 0.61 | 0.71 | 0.67 | 0.64 | 0.61 |
| 4 | 12143367 | rs1382752 | 0.663 | 0.229 | -1.203 | -0.868 | 0.009 | 0.019 | 0.019 | 0 | 0.12 | 0.06 | 0.2 | 0.05 | 0.12 | 0.15 |
| 4 | 12150124 | rs1479266 | 0.353 | 0.244 | -1.197 | -0.761 | -0.075 | 0.073 | 0.073 | 0.535 | 0.36 | 0.09 | 0.42 | 0.72 | 0.44 | 0.4 |
| 4 | 12154213 | rs10470840 | 0.423 | 0.443 | -1.275 | -0.6 | -0.088 | -0.031 | -0.031 | 0 | 0.35 | 0.07 | 0.43 | 0.59 | 0.45 | 0.41 |
| 4 | 12171491 | rs7670873 | 0.161 | 0.655 | -1.079 | -0.601 | -0.023 | 0.024 | 0.024 | 0 | 0.38 | 0.24 | 0.44 | 0.7 | 0.48 | 0.42 |
| 4 | 12176643 | rs6448866 | -1.054 | 0.756 | -1.302 | -0.596 | 0.04 | 0.039 | 0.039 | 0.333 | 0.61 | 0.71 | 0.72 | 0.66 | 0.73 | 0.62 |
| 4 | 12180791 | rs724317 | 0.29 | 0.863 | -1.252 | -0.561 | 0.028 | -0.059 | -0.059 | 0.625 | 0.17 | 0.44 | 0.25 | 0.07 | 0.18 | 0.13 |
| 4 | 12182361 | rs2127270 | 0.683 | 0.955 | -1.33 | -0.473 | -0.084 | -0.042 | -0.042 | 0 | 0.33 | 0.07 | 0.42 | 0.59 | 0.43 | 0.4 |
| 4 | 12186039 | rs35429502 | 0.947 | 0.958 | -1.323 | -0.733 | -0.08 | -0.011 | -0.011 | 0 | 0.31 | 0.06 | 0.4 | 0.68 | 0.39 | 0.37 |
| 4 | 12186445 | rs10516248 | -0.998 | 0.819 | -1.387 | -0.693 | 0.034 | 0.052 | 0.052 | 0.449 | 0.62 | 0.71 | 0.72 | 0.75 | 0.74 | 0.64 |
| 4 | 12188297 | rs13114953 | -0.801 | 0.979 | -1.542 | -0.613 | 0.003 | -0.114 | -0.114 | 0 | 0.27 | 0.48 | 0.32 | 0.07 | 0.29 | 0.25 |
| 4 | 12196573 | rs12331250 | -0.12 | 1.006 | -2.296 | 0.438 | -0.097 | -0.074 | -0.074 | 0 | 0.42 | 0.09 | 0.45 | 0.52 | 0.48 | 0.43 |
| 4 | 12212637 | rs1982655 | 0.427 | 1.15 | -2.08 | 0.67 | 0.005 | 0.049 | 0.049 | 0 | 0.51 | 0.18 | 0.34 | 0.41 | 0.49 | 0.42 |
| 4 | 12216880 | rs10939415 | 0.14 | 1.597 | -0.919 | 0.992 | 0 | -0.007 | -0.007 | 0 | 0.63 | 0.64 | 0.58 | 0.78 | 0.65 | 0.61 |
| 4 | 12218075 | rs11931935 | 0.88 | 1.583 | -1.107 | 0.957 | 0.003 | 0.055 | 0.055 | 0 | 0.5 | 0.13 | 0.36 | 0.42 | 0.48 | 0.42 |
| 4 | 12218164 | rs13108906 | 0.031 | 1.894 | -0.777 | 1.268 | -0.001 | -0.003 | -0.003 | 0 | 0.74 | 0.76 | 0.86 | 0.81 | 0.8 | 0.65 |
| 4 | 12220243 | rs16886803 | -0.188 | 1.838 | 0.146 | 1.289 | 0.083 | 0.17 | 0.17 | 0.153 | 0.1 | 0.32 | 0.19 | 0.28 | 0.14 | 0.1 |
| 4 | 12223752 | rs10440293 | 0.103 | 1.826 | 0.014 | 1.171 | -0.001 | -0.002 | -0.002 | 0 | 0.73 | 0.76 | 0.85 | 0.81 | 0.79 | 0.65 |
| 4 | 12231149 | rs12505670 | 1.2 | 1.989 | 0.461 | 1.174 | -0.004 | -0.004 | -0.004 | 0.004 | 0.61 | 0.45 | 0.56 | 0.67 | 0.64 | 0.56 |
| 4 | 12242026 | rs2124141 | 0.397 | 1.876 | 0.522 | 1.1 | -0.02 | -0.014 | -0.014 | 0.158 | 0.75 | 0.53 | 0.84 | 0.79 | 0.79 | 0.68 |
| 4 | 12246716 | rs1842160 | 1.148 | 1.983 | 0.554 | 1.182 | -0.001 | -0.003 | -0.003 | 0 | 0.64 | 0.44 | 0.57 | 0.68 | 0.64 | 0.57 |
| 4 | 12256501 | rs1454884 | -0.788 | 2.107 | 0.652 | 1.125 | 0.029 | 0.004 | 0.004 | 0 | 0.15 | 0.32 | 0.31 | 0.13 | 0.18 | 0.11 |
| 4 | 12258122 | rs1454883 | 1.859 | 2.257 | 0.578 | 1.206 | -0.074 | 0.022 | 0.022 | 0 | 0.53 | 0.09 | 0.38 | 0.4 | 0.49 | 0.46 |
| 4 | 12262020 | rs1454867 | 0.926 | 2.186 | 0.585 | 1.16 | -0.001 | -0.003 | -0.003 | 0.006 | 0.69 | 0.33 | 0.69 | 0.53 | 0.68 | 0.61 |
| 4 | 12267885 | rs2061062 | 0.227 | 1.987 | 1.095 | 1.123 | 0.014 | 0.011 | 0.011 | 0 | 0.87 | 0.71 | 0.7 | 0.87 | 0.85 | 0.89 |
| 4 | 12275642 | rs1454870 | 0.068 | 1.951 | 1.653 | 1.218 | -0.008 | 0.011 | 0.011 | 0 | 0.79 | 0.72 | 0.87 | 0.79 | 0.81 | 0.72 |
| 4 | 12285990 | rs2866454 | -0.11 | 1.945 | 1.659 | 1.271 | -0.001 | -0.001 | -0.001 | 0 | 0.13 | 0.13 | 0.28 | 0.13 | 0.15 | 0.11 |
| 4 | 12293571 | rs10019282 | -0.608 | 1.956 | 1.639 | 1.261 | -0.012 | -0.001 | -0.001 | 0 | 0.15 | 0.11 | 0.29 | 0.12 | 0.2 | 0.16 |
| 4 | 12297358 | rs4834885 | -1.073 | 1.813 | 1.613 | 1.205 | 0.002 | 0.109 | 0.109 | 0 | 0.09 | 0.17 | 0.16 | 0.27 | 0.06 | 0.15 |
| 4 | 12297535 | rs12499121 | 0.804 | 2.472 | 1.54 | 1.263 | -0.003 | 0.002 | 0.002 | 0 | 0.77 | 0.71 | 0.85 | 0.81 | 0.79 | 0.72 |
| 4 | 12303661 | rs1869644 | 1.579 | 2.488 | 1.533 | 0.984 | 0.006 | -0.013 | -0.013 | 0.509 | 0.64 | 0.58 | 0.54 | 0.74 | 0.58 | 0.57 |
| 4 | 12305958 | rs10018554 | 0.286 | 2.41 | 1.527 | 0.967 | -0.015 | -0.044 | -0.044 | 0 | 0.81 | 0.72 | 0.88 | 0.91 | 0.85 | 0.74 |
| 4 | 12306268 | rs1454878 | 1.855 | 2.333 | 1.373 | 1.099 | -0.001 | -0.048 | -0.048 | 0 | 0.58 | 0.15 | 0.41 | 0.53 | 0.59 | 0.47 |
| 4 | 12306820 | rs4834974 | 1.613 | 2.326 | 1.549 | 1.093 | 0.002 | -0.012 | -0.012 | 0 | 0.6 | 0.34 | 0.44 | 0.53 | 0.6 | 0.47 |
| 4 | 12309109 | rs7680526 | 1.123 | 2.34 | 1.541 | 0.965 | 0.001 | -0.018 | -0.018 | 0.3 | 0.65 | 0.6 | 0.58 | 0.79 | 0.64 | 0.58 |
| 4 | 12313033 | rs11722471 | 1.711 | 2.33 | 1.336 | 1.066 | 0.001 | -0.015 | -0.015 | 0 | 0.59 | 0.27 | 0.43 | 0.56 | 0.59 | 0.47 |
| 4 | 12317013 | rs1902138 | 1.915 | 2.256 | 1.137 | 1.085 | -0.055 | -0.011 | -0.011 | 3.403 | 0.53 | 0.1 | 0.39 | 0.56 | 0.51 | 0.47 |
| 4 | 12317409 | rs1454882 | 1.403 | 2.284 | 1.155 | 0.901 | 0.006 | -0.015 | -0.015 | 0 | 0.6 | 0.58 | 0.55 | 0.79 | 0.55 | 0.57 |
| 4 | 12320759 | rs7677522 | 1.539 | 2.106 | 0.854 | 0.9 | -0.001 | 0 | 0 | 0 | 0.58 | 0.6 | 0.52 | 0.75 | 0.49 | 0.55 |
| 4 | 12322014 | rs10516252 | 1.436 | 2.068 | 0.329 | 0.313 | -0.017 | 0.041 | 0.041 | 0.009 | 0.64 | 0.23 | 0.43 | 0.73 | 0.63 | 0.48 |
| 4 | 12328063 | rs1378233 | 0.985 | 0.944 | -0.459 | -0.535 | -0.017 | 0.206 | 0.206 | 0.614 | 0.48 | 0.09 | 0.45 | 0.81 | 0.43 | 0.68 |
| 4 | 12344593 | rs16886962 | -0.855 | 0.833 | 0.121 | -0.619 | -0.017 | -0.018 | -0.018 | 0 | 0.05 | 0.25 | 0.04 | 0 | 0.01 | 0.04 |
| 4 | 12349114 | rs12643322 | 1.059 | 0.783 | 0.27 | -0.598 | -0.008 | 0.198 | 0.198 | 0.818 | 0.49 | 0.09 | 0.45 | 0.78 | 0.43 | 0.69 |
| 4 | 12356550 | rs6822254 | -1.876 | 0.77 | 0.813 | -0.599 | 0.008 | 0.005 | 0.005 | 5.023 | 0.07 | 0.39 | 0.04 | 0.01 | 0.01 | 0.04 |
| 4 | 12359809 | rs17356154 | -0.11 | 1.006 | 0.557 | -0.552 | -0.041 | -0.038 | -0.038 | 0 | 0.1 | 0.01 | 0.21 | 0.12 | 0.14 | 0.1 |
| 4 | 12361202 | rs6448887 | 0.554 | 0.697 | 1.315 | -0.477 | 0.008 | 0.02 | 0.02 | 0 | 0.53 | 0.65 | 0.46 | 0.7 | 0.41 | 0.65 |
| 4 | 12374586 | rs1454888 | 0.481 | 0.879 | 1.14 | -0.382 | -0.005 | 0.01 | 0.01 | 0 | 0.61 | 0.43 | 0.46 | 0.65 | 0.42 | 0.7 |
| 4 | 12374820 | rs1454889 | 0.26 | 0.877 | 1.225 | -0.384 | 0.011 | -0.01 | -0.01 | 0 | 0.62 | 0.6 | 0.47 | 0.65 | 0.42 | 0.7 |
| 4 | 12375057 | rs10516249 | 0.366 | 0.668 | 1.104 | -0.508 | 0 | -0.012 | -0.012 | 0 | 0.6 | 0.62 | 0.46 | 0.67 | 0.41 | 0.69 |
| 4 | 12375400 | rs9283696 | 0.254 | 0.831 | 1.227 | -0.379 | 0.011 | -0.011 | -0.011 | 0 | 0.62 | 0.6 | 0.47 | 0.65 | 0.43 | 0.7 |
| 4 | 12375716 | rs9291588 | 0.847 | 0.802 | 1.211 | -0.396 | -0.014 | 0.078 | 0.078 | 0.513 | 0.55 | 0.11 | 0.42 | 0.68 | 0.42 | 0.67 |
| 4 | 12378886 | rs2866457 | 0.289 | 0.982 | 1.163 | -0.31 | 0.007 | -0.012 | -0.012 | 0.366 | 0.62 | 0.63 | 0.48 | 0.69 | 0.43 | 0.7 |
| 4 | 12381591 | rs6414745 | -0.921 | 0.924 | 0.915 | -0.304 | -0.012 | 0.03 | 0.03 | 0 | 0.42 | 0.71 | 0.55 | 0.37 | 0.58 | 0.31 |
| 4 | 12383947 | rs4377605 | -0.689 | 1.015 | 0.773 | -0.495 | 0.006 | -0.017 | -0.017 | 0 | 0.38 | 0.42 | 0.53 | 0.3 | 0.57 | 0.3 |
| 4 | 12395247 | rs11942175 | -0.874 | -0.046 | -0.18 | -0.826 | 0.003 | 0.162 | 0.162 | 0.065 | 0.5 | 0.79 | 0.56 | 0.26 | 0.55 | 0.34 |
| 4 | 12420636 | rs13435388 | -0.205 | -0.178 | 0.221 | -0.809 | -0.007 | 0.014 | 0.014 | 0 | 0.43 | 0.57 | 0.5 | 0.25 | 0.5 | 0.24 |
| 4 | 12420960 | rs6448889 | -2.544 | -0.518 | 1.202 | -0.74 | -0.005 | -0.003 | -0.003 | 0 | 0.08 | 0.33 | 0.06 | 0.04 | 0.03 | 0.05 |
| 4 | 12421862 | rs7656574 | -2.311 | -0.437 | 0.868 | -0.695 | -0.004 | 0.003 | 0.003 | 1.821 | 0.05 | 0.18 | 0.04 | 0.03 | 0.03 | 0.05 |
| 4 | 12436544 | rs4447880 | -2.156 | -0.546 | 0.959 | -0.747 | -0.014 | -0.026 | -0.026 | 0.238 | 0.08 | 0.27 | 0.04 | 0.02 | 0.03 | 0.05 |
| 4 | 12440385 | rs7673955 | -2.58 | -0.646 | 0.9 | -0.785 | -0.051 | -0.054 | -0.054 | 0.715 | 0.11 | 0.65 | 0.07 | 0.02 | 0.02 | 0.04 |
| 4 | 12454605 | rs6824314 | -1.283 | -0.89 | 0.455 | -0.78 | -0.033 | -0.011 | -0.011 | 0.01 | 0.2 | 0.69 | 0.12 | 0.17 | 0.1 | 0.07 |
| 4 | 12470500 | rs6812909 | -1.835 | -0.854 | 0.117 | -1.009 | 0.02 | 0.019 | 0.019 | 1.873 | 0.32 | 0.73 | 0.17 | 0.2 | 0.2 | 0.14 |
| 4 | 12471651 | rs6819663 | -2.065 | -0.92 | 0.015 | -1.127 | 0.028 | 0.032 | 0.032 | 0.303 | 0.34 | 0.74 | 0.18 | 0.2 | 0.22 | 0.15 |
| 4 | 12526377 | rs2219943 | -1.342 | -1.002 | 0.347 | -1.327 | 0.01 | 0.013 | 0.013 | 0.466 | 0.34 | 0.32 | 0.15 | 0.18 | 0.23 | 0.16 |

| | | | | | | | | | | | | | | | | |
|---|---|---|---|---|---|---|---|---|---|---|---|---|---|---|---|---|
| 4 | 12526515 | rs1351455 | -1.342 | -1.002 | 0.347 | -1.327 | 0.01 | 0.013 | 0.013 | 0.466 | 0.34 | 0.32 | 0.15 | 0.18 | 0.24 | 0.16 |
| 4 | 12543102 | rs6847440 | -1.727 | -0.975 | 0.388 | -1.077 | 0.017 | 0.009 | 0.009 | 0.326 | 0.37 | 0.67 | 0.2 | 0.25 | 0.24 | 0.16 |
| 4 | 12546096 | rs6448916 | -0.687 | -0.914 | 0.318 | -1.023 | 0.003 | -0.004 | -0.004 | 0 | 0.23 | 0.22 | 0.14 | 0.24 | 0.18 | 0.14 |
| 4 | 12552453 | rs6851677 | -1.885 | -0.993 | 0.339 | -1.09 | 0.039 | 0.021 | 0.021 | 0.485 | 0.38 | 0.73 | 0.21 | 0.25 | 0.24 | 0.16 |
| 4 | 12554693 | rs6448925 | 0.033 | -1.025 | 0.482 | -1.099 | -0.052 | 0.112 | 0.112 | 1.317 | 0.16 | 0.5 | 0.14 | 0.23 | 0.08 | 0.06 |
| 4 | 12557358 | rs16887189 | 0.227 | -1.006 | 0.319 | -1.158 | -0.001 | -0.008 | -0.008 | 0 | 0.11 | 0.13 | 0.04 | 0.01 | 0.1 | 0.07 |
| 4 | 12569163 | rs977629 | -1.997 | -1.022 | 0.28 | -1.156 | 0.04 | 0.025 | 0.025 | 0.392 | 0.38 | 0.73 | 0.21 | 0.25 | 0.24 | 0.16 |
| 4 | 12570464 | rs10939438 | -1.997 | -1.022 | 0.291 | -1.156 | 0.042 | 0.025 | 0.025 | 0.392 | 0.38 | 0.73 | 0.21 | 0.25 | 0.24 | 0.16 |
| 4 | 12573378 | rs6825629 | -1.33 | -1.028 | 0.355 | -1.208 | -0.026 | -0.112 | -0.112 | 0 | 0.26 | 0.59 | 0.11 | 0.02 | 0.16 | 0.09 |
| 4 | 12578444 | rs10029959 | -1.347 | -0.983 | 0.348 | -1.19 | -0.036 | -0.152 | -0.152 | 0 | 0.25 | 0.46 | 0.1 | 0.02 | 0.16 | 0.09 |
| 4 | 12579381 | rs17360757 | 0.285 | -0.846 | -0.288 | -1.121 | -0.01 | 0.034 | 0.034 | 0.264 | 0.1 | 0.03 | 0.03 | 0.02 | 0.1 | 0.07 |
| 4 | 12579382 | rs17360778 | 0.341 | -0.748 | -0.204 | -0.959 | 0.005 | 0.064 | 0.064 | 1.312 | 0.11 | 0.02 | 0.03 | 0 | 0.06 | 0.02 |
| 4 | 12585408 | rs6857588 | -0.809 | -0.692 | -0.171 | -0.908 | 0.001 | -0.002 | -0.002 | 0 | 0.24 | 0.25 | 0.15 | 0.24 | 0.15 | 0.09 |
| 4 | 12588094 | rs6830435 | -0.474 | -0.103 | -0.9 | -1.486 | -0.03 | 0.079 | 0.079 | 0 | 0.25 | 0.14 | 0.18 | 0.01 | 0.32 | 0.14 |
| 4 | 12593370 | rs17295743 | 0.248 | -0.199 | -0.931 | -1.563 | -0.035 | 0.065 | 0.065 | 0 | 0.17 | 0.04 | 0.14 | 0 | 0.26 | 0.11 |
| 4 | 12593395 | rs4571345 | -0.624 | -0.24 | -0.348 | -1.504 | -0.049 | 0.017 | 0.017 | 4.692 | 0.27 | 0.49 | 0.16 | 0.29 | 0.15 | 0.1 |
| 4 | 12595799 | rs6448932 | 0.298 | -0.221 | -0.308 | -1.464 | -0.041 | 0.116 | 0.116 | 3.431 | 0.16 | 0.38 | 0.12 | 0.29 | 0.09 | 0.07 |
| 4 | 12596193 | rs6448933 | -1.5 | -0.163 | -0.334 | -1.459 | -0.002 | -0.006 | -0.006 | 1.287 | 0.43 | 0.53 | 0.29 | 0.29 | 0.41 | 0.21 |
| 4 | 12599135 | rs7664245 | -1.113 | 0.013 | -0.658 | -1.962 | -0.018 | 0.111 | 0.111 | 0 | 0.38 | 0.24 | 0.22 | 0.02 | 0.42 | 0.21 |
| 4 | 12606923 | rs7680227 | -0.803 | 0.291 | -0.289 | -1.952 | 0.005 | -0.064 | -0.064 | 0 | 0.3 | 0.3 | 0.21 | 0.03 | 0.41 | 0.18 |
| 4 | 12607338 | rs7659827 | 0.351 | -0.025 | -0.371 | -1.79 | -0.033 | 0.038 | 0.038 | 1.89 | 0.22 | 0.38 | 0.27 | 0.29 | 0.15 | 0.13 |
| 4 | 12615274 | rs4834846 | -0.115 | -0.071 | -0.418 | -1.838 | -0.034 | 0.029 | 0.029 | 0.232 | 0.39 | 0.11 | 0.39 | 0.3 | 0.46 | 0.25 |
| 4 | 12627162 | rs1545817 | -1.09 | -0.108 | -0.205 | -1.817 | -0.001 | 0.002 | 0.002 | 0 | 0.56 | 0.42 | 0.46 | 0.32 | 0.6 | 0.33 |
| 4 | 12633072 | rs6839361 | -0.739 | 0.025 | -0.111 | -1.721 | 0 | 0.1 | 0.1 | 0.195 | 0.51 | 0.19 | 0.44 | 0.32 | 0.59 | 0.34 |
| 4 | 12634511 | rs6822083 | -0.475 | 0.044 | -0.135 | -1.728 | -0.063 | 0.063 | 0.063 | 0.295 | 0.45 | 0.11 | 0.42 | 0.32 | 0.53 | 0.3 |
| 4 | 12636408 | rs13141338 | 0.55 | 0.073 | 0.128 | -1.694 | 0.016 | -0.038 | -0.038 | 0 | 0.07 | 0.17 | 0.03 | 0 | 0.06 | 0.04 |
| 4 | 12642861 | rs7687042 | 0.433 | 0.086 | -0.51 | -2.158 | 0 | -0.003 | -0.003 | 0 | 0.35 | 0.56 | 0.45 | 0.68 | 0.33 | 0.66 |
| 4 | 12662979 | rs17363850 | 0.041 | 0.046 | -0.699 | -2.26 | -0.052 | 0.078 | 0.078 | 0.106 | 0.46 | 0.12 | 0.42 | 0.31 | 0.53 | 0.3 |
| 4 | 12663829 | rs11946420 | 0.751 | 0.128 | -0.498 | -2.121 | 0.008 | -0.04 | -0.04 | 0 | 0.07 | 0.15 | 0.03 | 0 | 0.06 | 0.04 |
| 4 | 12664089 | rs10939444 | 0.473 | 0.12 | -0.463 | -2.129 | 0.005 | -0.004 | -0.004 | 0 | 0.34 | 0.47 | 0.44 | 0.68 | 0.33 | 0.66 |
| 4 | 12671583 | rs10516260 | -0.247 | 0.206 | -0.251 | -1.862 | 0.001 | -0.036 | -0.036 | 0 | 0.14 | 0.16 | 0.06 | 0.02 | 0.13 | 0.1 |
| 4 | 12673985 | rs11724617 | -0.421 | 0.205 | -0.244 | -1.863 | 0.009 | -0.072 | -0.072 | 0.149 | 0.15 | 0.28 | 0.06 | 0.02 | 0.13 | 0.1 |
| 4 | 12678480 | rs1402033 | 1.011 | 0.28 | -0.513 | -1.983 | -0.001 | -0.001 | -0.001 | 0 | 0.35 | 0.33 | 0.44 | 0.65 | 0.33 | 0.65 |
| 4 | 12679774 | rs2222310 | -1.716 | 0.201 | -0.682 | -2.209 | -0.017 | 0.058 | 0.058 | 0.006 | 0.88 | 0.96 | 0.93 | 1 | 0.89 | 0.99 |
| 4 | 12679993 | rs13151965 | 0.089 | 0.262 | -0.596 | -2.167 | -0.001 | 0 | 0 | 0.041 | 0.49 | 0.5 | 0.5 | 0.67 | 0.46 | 0.75 |
| 4 | 12682297 | rs7690844 | 1.156 | 0.148 | -0.428 | -1.836 | -0.001 | 0.002 | 0.002 | 0 | 0.07 | 0.09 | 0.04 | 0.02 | 0.07 | 0.08 |
| 4 | 12683518 | rs13123654 | 0.777 | 0.135 | -0.44 | -1.819 | 0.005 | -0.034 | -0.034 | 0 | 0.07 | 0.08 | 0.02 | 0 | 0.06 | 0.02 |
| 4 | 12690183 | rs17299409 | 0.105 | 0.313 | -0.566 | -1.887 | -0.052 | 0.072 | 0.072 | 0 | 0.2 | 0.09 | 0.18 | 0.02 | 0.38 | 0.14 |
| 4 | 12697432 | rs1520296 | -0.294 | 0.264 | -0.367 | -1.666 | -0.061 | 0.033 | 0.033 | 0 | 0.42 | 0.19 | 0.41 | 0.32 | 0.54 | 0.31 |
| 4 | 12707180 | rs2903351 | -0.28 | 0.195 | 0.12 | -1.442 | -0.022 | -0.052 | -0.052 | 2.041 | 0.86 | 0.74 | 0.91 | 0.98 | 0.93 | 0.96 |
| 4 | 12723676 | rs10031132 | 0.483 | 0.203 | -0.239 | -1.453 | -0.003 | -0.005 | -0.005 | 0 | 0.53 | 0.55 | 0.55 | 0.67 | 0.53 | 0.74 |
| 4 | 12723707 | rs7663013 | 0.339 | 0.23 | -1.325 | -1.4 | -0.027 | -0.027 | -0.027 | 1.574 | 0.32 | 0.1 | 0.36 | 0.31 | 0.4 | 0.23 |
| 4 | 12725164 | rs11728601 | 0.236 | 0.606 | -1.136 | -0.975 | -0.033 | 0.055 | 0.055 | 2.083 | 0.47 | 0.21 | 0.5 | 0.67 | 0.47 | 0.69 |
| 4 | 12733389 | rs13136382 | -0.91 | 0.46 | -0.033 | -0.926 | 0.008 | -0.067 | -0.067 | 0 | 0.13 | 0.57 | 0.1 | 0.02 | 0.09 | 0.09 |
| 4 | 12736121 | rs9291598 | -0.093 | 0.072 | -0.322 | -1.171 | 0.009 | -0.011 | -0.011 | 0.441 | 0.3 | 0.21 | 0.27 | 0.34 | 0.22 | 0.28 |
| 4 | 12736715 | rs9884719 | 0.888 | -0.065 | -0.444 | -1.275 | -0.012 | 0.023 | 0.023 | 0.033 | 0.16 | 0.17 | 0.22 | 0.32 | 0.1 | 0.18 |
| 4 | 12741313 | rs10022602 | -1.364 | -0.06 | -0.487 | -1.294 | 0.031 | 0.01 | 0.01 | 0 | 0.42 | 0.76 | 0.37 | 0.36 | 0.31 | 0.36 |
| 4 | 12757442 | rs7692324 | 0.513 | -0.09 | 0.22 | -1.322 | -0.015 | -0.023 | -0.023 | 0 | 0.08 | 0.19 | 0.05 | 0.02 | 0.02 | 0.06 |
| 4 | 12758617 | rs2903395 | 1.096 | -0.207 | 0.28 | -1.23 | -0.022 | 0.039 | 0.039 | 0.698 | 0.15 | 0.18 | 0.24 | 0.31 | 0.08 | 0.17 |
| 4 | 12763137 | rs2866811 | 1.485 | 0.258 | -0.768 | -1.966 | -0.109 | -0.121 | -0.121 | 5.995 | 0.28 | 0.02 | 0.35 | 0.65 | 0.32 | 0.56 |
| 4 | 12773994 | rs16887552 | -0.765 | 0.082 | -0.232 | -1.596 | -0.005 | -0.078 | -0.078 | 0 | 0.27 | 0.59 | 0.16 | 0.05 | 0.2 | 0.19 |
| 4 | 12776391 | rs6810977 | -0.17 | 0.086 | -0.084 | -1.585 | -0.007 | -0.038 | -0.038 | 5.374 | 0.12 | 0.28 | 0.08 | 0.03 | 0.08 | 0.1 |
| 4 | 12776895 | rs6448952 | -1.245 | 0.097 | -0.3 | -1.38 | 0.051 | -0.015 | -0.015 | 0.26 | 0.43 | 0.79 | 0.34 | 0.38 | 0.29 | 0.37 |
| 4 | 12778849 | rs17366799 | 0.982 | 0.123 | 0.137 | -1.571 | 0.014 | 0.007 | 0.007 | 0 | 0.54 | 0.31 | 0.61 | 0.62 | 0.64 | 0.62 |
| 4 | 12782701 | rs6857474 | -0.779 | 0.036 | 0.123 | -1.29 | -0.013 | 0.034 | 0.034 | 0 | 0.78 | 0.7 | 0.75 | 0.64 | 0.84 | 0.81 |
| 4 | 12789828 | rs11731395 | 1.128 | -0.034 | -0.079 | -1.287 | 0.006 | -0.003 | -0.003 | 0 | 0.18 | 0.14 | 0.17 | 0.37 | 0.08 | 0.18 |
| 4 | 12790058 | rs12507532 | 0.383 | 0.145 | -0.349 | -1.23 | 0.036 | -0.037 | -0.037 | 0 | 0.65 | 0.22 | 0.65 | 0.61 | 0.77 | 0.7 |
| 4 | 12791282 | rs1402037 | 0.858 | 0.307 | -0.634 | -1.015 | 0.012 | -0.04 | -0.04 | 0 | 0.52 | 0.14 | 0.6 | 0.53 | 0.64 | 0.61 |
| 4 | 12791971 | rs11728626 | -0.197 | 0.442 | -0.112 | -0.912 | 0.016 | -0.027 | -0.027 | 0.142 | 0.05 | 0.11 | 0.03 | 0 | 0.06 | 0.05 |
| 4 | 12793263 | rs17301479 | 0.435 | 0.307 | -0.658 | -1.148 | -0.012 | 0.093 | 0.093 | 0 | 0.19 | 0.02 | 0.14 | 0.02 | 0.14 | 0.15 |
| 4 | 12795135 | rs13105543 | 0.488 | 0.343 | -0.425 | -0.872 | -0.04 | -0.032 | -0.032 | 0 | 0.39 | 0.21 | 0.42 | 0.59 | 0.52 | 0.6 |
| 4 | 12797555 | rs11728489 | 0.22 | 0.347 | -0.263 | -0.864 | -0.001 | -0.025 | -0.025 | 0.018 | 0.06 | 0.09 | 0.02 | 0 | 0.04 | 0.01 |
| 4 | 12802055 | rs807423 | 0.371 | 0.087 | 0.194 | -0.911 | 0.005 | -0.007 | -0.007 | 0 | 0.4 | 0.3 | 0.39 | 0.36 | 0.32 | 0.32 |
| 4 | 12803879 | rs6849099 | -0.024 | 0.102 | 0.254 | -0.885 | 0.007 | -0.001 | -0.001 | 0 | 0.44 | 0.21 | 0.4 | 0.36 | 0.36 | 0.33 |
| 4 | 12808091 | rs807418 | -0.015 | 0.109 | 0.229 | -0.868 | 0.001 | 0.001 | 0.001 | 0.233 | 0.45 | 0.39 | 0.42 | 0.36 | 0.36 | 0.31 |
| 4 | 12809589 | rs16887595 | 1.07 | 0.297 | -0.163 | -1.028 | -0.003 | -0.042 | -0.042 | 0 | 0.31 | 0.17 | 0.3 | 0.59 | 0.3 | 0.54 |
| 4 | 12811827 | rs7662165 | 1.295 | 0.121 | -0.232 | -0.83 | -0.003 | -0.025 | -0.025 | 0 | 0.3 | 0.17 | 0.32 | 0.32 | 0.26 | 0.26 |
| 4 | 12817157 | rs2540818 | 0.88 | 0.06 | 0.075 | -0.894 | -0.025 | -0.028 | -0.028 | 0 | 0.15 | 0.26 | 0.16 | 0.08 | 0.12 | 0.06 |
| 4 | 12819018 | rs2540821 | 0.112 | 0.078 | -0.063 | -0.879 | 0.015 | -0.005 | -0.005 | 4.358 | 0.46 | 0.34 | 0.48 | 0.4 | 0.39 | 0.33 |

| | | | | | | | | | | | | | | | | |
|---|---|---|---|---|---|---|---|---|---|---|---|---|---|---|---|---|
| 4 | 12827562 | rs2540823 | 0.831 | -0.372 | -0.449 | -0.818 | -0.002 | -0.002 | -0.002 | 0 | 0.14 | 0.14 | 0.15 | 0.08 | 0.12 | 0.07 |
| 4 | 12827751 | rs11723612 | 0.966 | -0.181 | -0.625 | -0.892 | -0.049 | -0.113 | -0.113 | 0 | 0.33 | 0.03 | 0.32 | 0.31 | 0.28 | 0.3 |
| 4 | 12828247 | rs2540825 | 0.831 | -0.146 | -0.43 | -0.903 | -0.016 | -0.019 | -0.019 | 0.086 | 0.15 | 0.18 | 0.16 | 0.08 | 0.12 | 0.07 |
| 4 | 12838429 | rs2540834 | 0.003 | -0.081 | -0.083 | -0.756 | 0.022 | 0.01 | 0.01 | 0 | 0.5 | 0.38 | 0.49 | 0.38 | 0.39 | 0.36 |
| 4 | 12839087 | rs2711547 | 0.304 | -0.019 | -0.459 | -0.711 | 0.05 | -0.031 | -0.031 | 8.67 | 0.43 | 0.13 | 0.45 | 0.37 | 0.36 | 0.35 |
| 4 | 12847075 | rs2540841 | 0.133 | 0.16 | -0.424 | -0.757 | 0.047 | -0.021 | -0.021 | 0.139 | 0.47 | 0.15 | 0.43 | 0.39 | 0.39 | 0.33 |
| 4 | 12859756 | rs2110898 | -1.279 | 0.306 | 0.053 | -0.735 | 0.004 | 0.071 | 0.071 | 0 | 0.72 | 0.55 | 0.64 | 0.41 | 0.72 | 0.44 |
| 4 | 12862476 | rs10939459 | 1.18 | 0.315 | -0.079 | -0.733 | -0.003 | -0.014 | -0.014 | 0 | 0.28 | 0.27 | 0.34 | 0.59 | 0.28 | 0.56 |
| 4 | 12875351 | rs17515558 | -0.076 | 0.442 | -0.305 | -0.775 | 0.012 | -0.064 | -0.064 | 0.049 | 0.19 | 0.24 | 0.18 | 0 | 0.3 | 0.11 |
| 4 | 12876257 | rs9291601 | 1.187 | 0.161 | -0.82 | -0.359 | -0.017 | -0.132 | -0.132 | 0 | 0.3 | 0.04 | 0.29 | 0.38 | 0.25 | 0.25 |
| 4 | 12881391 | rs12502445 | 1.137 | 0.363 | -0.788 | -0.472 | -0.009 | -0.088 | -0.088 | 2.253 | 0.3 | 0.04 | 0.28 | 0.32 | 0.26 | 0.26 |
| 4 | 12888409 | rs223914 | -0.081 | -0.319 | -0.733 | 0.012 | 0.01 | -0.015 | -0.015 | 0 | 0.16 | 0.11 | 0.15 | 0.21 | 0.15 | 0.1 |
| 4 | 12888995 | rs7697105 | 1.094 | -0.27 | -0.358 | -0.299 | -0.002 | -0.016 | -0.016 | 0 | 0.57 | 0.54 | 0.66 | 0.79 | 0.54 | 0.74 |
| 4 | 12889196 | rs7697521 | 1.188 | -0.022 | -0.205 | -0.192 | 0.004 | 0.012 | 0.012 | 0.109 | 0.25 | 0.34 | 0.29 | 0.45 | 0.26 | 0.52 |
| 4 | 12894741 | rs12648564 | -0.129 | 0.051 | -0.399 | -0.627 | 0.011 | -0.029 | -0.029 | 0 | 0.19 | 0.24 | 0.14 | 0.06 | 0.25 | 0.18 |
| 4 | 12897698 | rs981269 | 1.569 | -0.06 | -0.805 | -0.431 | -0.014 | -0.026 | -0.026 | 0.319 | 0.23 | 0.2 | 0.29 | 0.41 | 0.27 | 0.53 |
| 4 | 12902074 | rs16887775 | 0.904 | -0.19 | -1.156 | -0.38 | 0.009 | -0.039 | -0.039 | 0 | 0.39 | 0.18 | 0.39 | 0.44 | 0.36 | 0.23 |
| 4 | 12914467 | rs190110 | 0.803 | 0.098 | -0.83 | -0.246 | 0.005 | -0.028 | -0.028 | 0.209 | 0.07 | 0.02 | 0.09 | 0.1 | 0.04 | 0.05 |
| 4 | 12914709 | rs7675207 | 1.575 | -0.024 | -0.659 | -0.432 | 0 | 0.019 | 0.019 | 1.695 | 0.61 | 0.65 | 0.68 | 0.83 | 0.56 | 0.74 |
| 4 | 12919993 | rs1513935 | 0.825 | -0.601 | -0.629 | -0.454 | -0.012 | -0.021 | -0.021 | 0.025 | 0.75 | 0.72 | 0.79 | 0.9 | 0.81 | 0.9 |
| 4 | 12924466 | rs1513955 | 1.471 | -0.525 | -0.491 | -0.51 | -0.013 | -0.02 | -0.02 | 0 | 0.64 | 0.62 | 0.77 | 0.83 | 0.74 | 0.76 |
| 4 | 12928197 | rs4834855 | 2.045 | -0.533 | -0.45 | -0.567 | 0.005 | 0.007 | 0.007 | 3.299 | 0.56 | 0.56 | 0.68 | 0.83 | 0.62 | 0.74 |
| 4 | 12931444 | rs670400 | 0.311 | -0.537 | -0.446 | -0.563 | -0.004 | -0.04 | -0.04 | 1.072 | 0.81 | 0.71 | 0.87 | 0.93 | 0.83 | 0.84 |
| 4 | 12931901 | rs672282 | 0.018 | -0.506 | -0.68 | -0.569 | 0.001 | -0.003 | -0.003 | 0.031 | 0.12 | 0.08 | 0.11 | 0.1 | 0.08 | 0.07 |
| 4 | 12934281 | rs223997 | -1.752 | -0.814 | -1.091 | -0.403 | -0.009 | -0.016 | -0.016 | 0 | 0.73 | 0.85 | 0.67 | 0.67 | 0.75 | 0.79 |
| 4 | 12935474 | rs6811193 | 1.869 | -0.684 | -1.01 | -0.298 | -0.001 | -0.014 | -0.014 | 0 | 0.06 | 0.08 | 0.06 | 0 | 0.09 | 0.02 |
| 4 | 12938532 | rs170369 | 0.592 | -0.7 | -1.04 | -0.289 | -0.001 | -0.025 | -0.025 | 0 | 0.1 | 0.02 | 0.11 | 0.1 | 0.08 | 0.06 |
| 4 | 12938795 | rs7679576 | 0.003 | -0.536 | -0.851 | -0.207 | 0.001 | 0.007 | 0.007 | 0.15 | 0.49 | 0.73 | 0.48 | 0.57 | 0.56 | 0.68 |
| 4 | 12939091 | rs1154862 | 1.831 | -0.583 | -0.632 | -0.229 | -0.028 | -0.05 | -0.05 | 0.049 | 0.28 | 0.23 | 0.35 | 0.5 | 0.37 | 0.52 |
| 4 | 12940751 | rs6850802 | -0.185 | -0.591 | -0.608 | -0.328 | -0.002 | -0.029 | -0.029 | 0.078 | 0.15 | 0.42 | 0.07 | 0.07 | 0.1 | 0.14 |
| 4 | 12942258 | rs728171 | 1.05 | -0.28 | -0.833 | -0.009 | -0.027 | -0.007 | -0.007 | 0 | 0.33 | 0.27 | 0.41 | 0.39 | 0.45 | 0.53 |
| 4 | 12960502 | rs10489033 | -0.011 | -0.371 | -0.798 | -0.121 | -0.003 | -0.003 | -0.003 | 0 | 0.15 | 0.16 | 0.15 | 0 | 0.2 | 0.05 |
| 4 | 12960713 | rs16887898 | 1.626 | -0.374 | -0.797 | -0.113 | 0 | -0.01 | -0.01 | 0 | 0.06 | 0.08 | 0.06 | 0 | 0.09 | 0.02 |
| 4 | 12962308 | rs223975 | 0.953 | -1.002 | -1.216 | -0.356 | 0.054 | -0.004 | -0.004 | 0 | 0.17 | 0.02 | 0.07 | 0.1 | 0.07 | 0.09 |
| 4 | 12962421 | rs223976 | -0.646 | -0.929 | -1.366 | -0.278 | 0.009 | 0.031 | 0.031 | 0.947 | 0.51 | 0.27 | 0.52 | 0.39 | 0.47 | 0.36 |
| 4 | 12963574 | rs763318 | 1.219 | -1.003 | -1.191 | -0.257 | -0.014 | -0.009 | -0.009 | 0 | 0.41 | 0.6 | 0.46 | 0.5 | 0.48 | 0.62 |
| 4 | 12965300 | rs7676689 | 2.142 | -0.923 | -1.132 | -0.185 | -0.033 | -0.034 | -0.034 | 0 | 0.28 | 0.19 | 0.39 | 0.43 | 0.36 | 0.51 |
| 4 | 12965868 | rs4108065 | 1.646 | -0.565 | -1.171 | -0.03 | -0.057 | -0.082 | -0.082 | 0 | 0.19 | 0.01 | 0.29 | 0.29 | 0.2 | 0.22 |
| 4 | 12967160 | rs223982 | -1.3 | -0.56 | -0.963 | -0.093 | -0.018 | -0.011 | -0.011 | 0 | 0.9 | 0.81 | 0.97 | 0.93 | 0.94 | 0.95 |
| 4 | 12972256 | rs223989 | -0.781 | -0.529 | -1.192 | 0.073 | 0.01 | 0.036 | 0.036 | 3.183 | 0.52 | 0.25 | 0.57 | 0.61 | 0.44 | 0.38 |
| 4 | 12993951 | rs2867020 | -1.204 | -0.558 | -1.427 | -0.11 | 0.043 | -0.015 | -0.015 | 0 | 0.84 | 0.97 | 0.93 | 0.89 | 0.93 | 0.91 |
| 4 | 12994201 | rs2867021 | 0.954 | -0.527 | -1.256 | -0.13 | -0.002 | -0.001 | -0.001 | 0 | 0.43 | 0.44 | 0.49 | 0.61 | 0.52 | 0.61 |
| 4 | 12997151 | rs4834856 | 1.321 | -0.048 | -1.989 | 0.133 | -0.049 | -0.094 | -0.094 | 1.333 | 0.19 | 0.02 | 0.31 | 0.29 | 0.2 | 0.23 |
| 4 | 13000803 | rs10516262 | 0.299 | -0.204 | -1.708 | 0.026 | -0.004 | -0.004 | -0.004 | 0.056 | 0.15 | 0.17 | 0.1 | 0 | 0.18 | 0.07 |
| 4 | 13001452 | rs1828760 | -0.919 | -0.224 | -1.774 | 0.255 | 0.052 | -0.009 | -0.009 | 0 | 0.83 | 0.98 | 0.94 | 0.89 | 0.93 | 0.87 |
| 4 | 13002341 | rs2673423 | 0.891 | 0.551 | -1.412 | 0.749 | -0.027 | -0.071 | -0.071 | 0 | 0.19 | 0.06 | 0.32 | 0.4 | 0.23 | 0.24 |
| 4 | 13002905 | rs1040057 | 0.231 | 0.967 | -0.693 | 0.514 | 0.008 | 0.01 | 0.01 | 5.191 | 0.35 | 0.55 | 0.42 | 0.61 | 0.4 | 0.55 |
| 4 | 13003758 | rs1040056 | -0.014 | 0.806 | -0.637 | 0.431 | 0.014 | 0.025 | 0.025 | 0 | 0.5 | 0.58 | 0.45 | 0.36 | 0.56 | 0.58 |
| 4 | 13003905 | rs2673422 | -0.127 | 0.67 | -1.023 | 0.524 | 0.061 | 0.024 | 0.024 | 0 | 0.51 | 0.73 | 0.55 | 0.57 | 0.42 | 0.39 |
| 4 | 13005874 | rs1913704 | 0.266 | 0.988 | -0.822 | 0.732 | -0.004 | -0.016 | -0.016 | 0 | 0.9 | 0.93 | 0.96 | 0.86 | 0.89 | 0.86 |
| 4 | 13006337 | rs1040055 | 0.886 | 0.973 | -1.276 | 0.647 | 0.02 | -0.048 | -0.048 | 0 | 0.37 | 0.15 | 0.54 | 0.43 | 0.37 | 0.34 |
| 4 | 13009987 | rs2056116 | 0.452 | 1.378 | -0.53 | 0.392 | -0.01 | 0.006 | 0.006 | 0 | 0.36 | 0.27 | 0.25 | 0.28 | 0.37 | 0.43 |
| 4 | 13010694 | rs2621243 | -0.017 | 1.032 | -0.214 | 0.14 | -0.03 | -0.078 | -0.078 | 0.062 | 0.14 | 0.29 | 0.06 | 0 | 0.06 | 0.02 |
| 4 | 13020235 | rs1947850 | 1.341 | 1.154 | -0.435 | 0.344 | 0.021 | -0.056 | -0.056 | 0 | 0.36 | 0.1 | 0.51 | 0.43 | 0.35 | 0.33 |
| 4 | 13026562 | rs7679886 | -0.324 | 1.442 | -0.108 | 0.243 | -0.004 | 0.002 | 0.002 | 0 | 0.44 | 0.56 | 0.34 | 0.43 | 0.4 | 0.5 |
| 4 | 13026792 | rs1513947 | 0.085 | 1.37 | -0.147 | 0.352 | -0.004 | -0.007 | -0.007 | 0 | 0.84 | 0.86 | 0.91 | 0.85 | 0.82 | 0.83 |
| 4 | 13027245 | rs1513948 | 0.124 | 1.354 | -0.155 | 0.344 | -0.004 | -0.007 | -0.007 | 0.836 | 0.84 | 0.86 | 0.91 | 0.85 | 0.82 | 0.83 |
| 4 | 13028973 | rs1399019 | 0.527 | 1.37 | 0.102 | 0.343 | 0.01 | -0.005 | -0.005 | 0.93 | 0.1 | 0.18 | 0.11 | 0.14 | 0.12 | 0.09 |
| 4 | 13030876 | rs4834860 | 0.364 | 1.327 | 0.264 | 0.315 | -0.001 | 0 | 0 | 0 | 0.54 | 0.47 | 0.63 | 0.58 | 0.48 | 0.43 |
| 4 | 13036900 | rs1318883 | -0.843 | 1.259 | -0.109 | 0.043 | -0.023 | -0.045 | -0.045 | 0.659 | 0.89 | 0.97 | 0.96 | 0.84 | 0.86 | 0.84 |
| 4 | 13037163 | rs981561 | 0.987 | 1.165 | 0.366 | 0.293 | 0.006 | 0.007 | 0.007 | 0 | 0.45 | 0.44 | 0.6 | 0.45 | 0.41 | 0.42 |
| 4 | 13037938 | rs1973655 | 1.545 | 1.235 | 0.317 | 0.321 | 0.031 | 0.033 | 0.033 | 0 | 0.38 | 0.25 | 0.5 | 0.32 | 0.31 | 0.32 |
| 4 | 13038094 | rs1947851 | 0.434 | 1.328 | 0.342 | 0.332 | -0.012 | 0.005 | 0.005 | 0 | 0.78 | 0.85 | 0.89 | 0.84 | 0.76 | 0.82 |
| 4 | 13047066 | rs2673420 | 0.912 | 1.371 | 0.643 | 0.366 | 0.017 | 0.009 | 0.009 | 0 | 0.77 | 0.58 | 0.81 | 0.71 | 0.72 | 0.74 |
| 4 | 13048129 | rs1515407 | -1 | 1.4 | 0.225 | 0.369 | -0.05 | -0.019 | -0.019 | 1.576 | 0.48 | 0.84 | 0.41 | 0.52 | 0.45 | 0.49 |
| 4 | 13048594 | rs7678073 | 0.643 | 1.433 | 0.411 | 0.403 | 0 | -0.001 | -0.001 | 10.945 | 0.09 | 0.16 | 0.08 | 0.14 | 0.07 | 0.09 |
| 4 | 13049139 | rs1904794 | -0.214 | 1.469 | 0.125 | 0.429 | -0.016 | -0.015 | -0.015 | 0 | 0.86 | 0.89 | 0.9 | 0.84 | 0.79 | 0.83 |
| 4 | 13052713 | rs1913699 | -0.883 | 1.415 | 0.164 | 0.381 | -0.029 | -0.007 | -0.007 | 0.434 | 0.47 | 0.8 | 0.39 | 0.52 | 0.45 | 0.49 |
| 4 | 13056251 | rs2127657 | 0.119 | 1.571 | 0.617 | 0.3 | 0.001 | 0.006 | 0.006 | 0.014 | 0.36 | 0.6 | 0.34 | 0.52 | 0.34 | 0.47 |

| | | | | | | | | | | | | | | | | |
|---|---|---|---|---|---|---|---|---|---|---|---|---|---|---|---|---|
| 4 | 13056521 | rs2903472 | 1.577 | 1.382 | 0.832 | 0.28 | -0.02 | 0.03 | 0.03 | 3.833 | 0.64 | 0.22 | 0.71 | 0.7 | 0.61 | 0.72 |
| 4 | 13057870 | rs481266 | -0.986 | 1.435 | 0.538 | 0.297 | -0.002 | -0.055 | -0.055 | 0.027 | 0.89 | 0.82 | 0.93 | 1 | 0.89 | 0.98 |
| 4 | 13067251 | rs2127660 | 0.696 | 1.871 | 0.597 | 0.606 | 0.02 | 0.068 | 0.068 | 0.07 | 0.27 | 0.35 | 0.22 | 0.41 | 0.28 | 0.41 |
| 4 | 13070446 | rs474076 | 0.462 | 1.465 | 0.533 | 0.482 | 0.084 | 0.057 | 0.057 | 0.927 | 0.9 | 0.57 | 0.89 | 0.86 | 0.87 | 0.95 |
| 4 | 13072361 | rs4834919 | 0.4 | 1.196 | 0.066 | 0.314 | 0.032 | 0.033 | 0.033 | 0.321 | 0.52 | 0.38 | 0.59 | 0.46 | 0.4 | 0.43 |
| 4 | 13073411 | rs560164 | 1.109 | 1.138 | 0.434 | 0.418 | 0.045 | 0.023 | 0.023 | 1.501 | 0.88 | 0.59 | 0.89 | 0.85 | 0.85 | 0.93 |
| 4 | 13073618 | rs566737 | -0.145 | 1.787 | 0.211 | 0.555 | 0.013 | 0.041 | 0.041 | 0 | 0.64 | 0.55 | 0.71 | 0.56 | 0.61 | 0.54 |
| 4 | 13074010 | rs481012 | -0.841 | 1.585 | -0.015 | 0.25 | -0.004 | 0.005 | 0.005 | 0.804 | 0.35 | 0.32 | 0.31 | 0.36 | 0.35 | 0.22 |
| 4 | 13075100 | rs16888112 | -0.596 | 1.225 | 0.137 | 0.116 | -0.008 | 0.001 | 0.001 | 1.081 | 0.31 | 0.41 | 0.3 | 0.6 | 0.27 | 0.51 |
| 4 | 13077746 | rs16888117 | -0.314 | 1.352 | -0.05 | 0.474 | -0.005 | -0.01 | -0.01 | 0 | 0.28 | 0.28 | 0.25 | 0.5 | 0.3 | 0.5 |
| 4 | 13080111 | rs6848051 | -0.087 | 1.103 | 0.02 | 0.087 | -0.001 | -0.007 | -0.007 | 0 | 0.18 | 0.28 | 0.21 | 0.19 | 0.18 | 0.16 |
| 4 | 13080633 | rs481213 | 1.14 | 0.561 | -0.035 | -0.036 | -0.046 | -0.005 | -0.005 | 0 | 0.11 | 0.01 | 0.06 | 0.09 | 0.15 | 0.09 |
| 4 | 13081396 | rs10939464 | 0.411 | 0.357 | -0.154 | -0.353 | 0 | -0.067 | -0.067 | 0.362 | 0.7 | 0.44 | 0.79 | 0.91 | 0.7 | 0.86 |
| 4 | 13082236 | rs496099 | 0.882 | 0.684 | 0.33 | -0.204 | 0.017 | -0.02 | -0.02 | 0 | 0.12 | 0.22 | 0.08 | 0.09 | 0.15 | 0.09 |
| 4 | 13082344 | rs497117 | -0.503 | 0.666 | 0.096 | -0.144 | 0.125 | -0.002 | -0.002 | 0.118 | 0.94 | 0.75 | 0.91 | 1 | 0.87 | 0.97 |
| 4 | 13092297 | rs11945548 | 0.446 | 0.64 | -0.314 | -0.097 | -0.004 | 0.012 | 0.012 | 0 | 0.14 | 0.22 | 0.18 | 0.2 | 0.12 | 0.15 |
| 4 | 13093944 | rs4458470 | -0.415 | 0.706 | -0.622 | -0.138 | 0.066 | -0.049 | -0.049 | 0 | 0.83 | 0.66 | 0.85 | 0.92 | 0.73 | 0.88 |
| 4 | 13094261 | rs1021873 | 0.845 | 0.849 | -1.631 | 0.018 | -0.072 | -0.033 | -0.033 | 0 | 0.43 | 0.06 | 0.52 | 0.52 | 0.44 | 0.63 |
| 4 | 13098882 | rs512244 | -1.08 | 0.776 | -1.013 | 0.023 | 0.037 | -0.027 | -0.027 | 0.023 | 0.95 | 0.88 | 0.91 | 1 | 0.87 | 0.95 |
| 4 | 13102351 | rs13122273 | 0.297 | 0.748 | -1.547 | 0.23 | -0.06 | -0.081 | -0.081 | 0 | 0.18 | 0.02 | 0.16 | 0.25 | 0.2 | 0.36 |
| 4 | 13102477 | rs9291606 | 0.672 | 0.526 | -0.645 | -0.046 | -0.003 | -0.005 | -0.005 | 0.027 | 0.66 | 0.4 | 0.77 | 0.69 | 0.68 | 0.72 |
| 4 | 13106197 | rs559283 | 0.339 | 0.382 | -0.923 | -0.063 | 0.007 | -0.005 | -0.005 | 0 | 0.59 | 0.66 | 0.82 | 0.51 | 0.7 | 0.71 |
| 4 | 13107182 | rs524129 | -0.359 | 0.48 | -1.239 | -0.136 | 0 | 0.05 | 0.05 | 0 | 0.76 | 0.66 | 0.79 | 0.52 | 0.74 | 0.72 |
| 4 | 13116181 | rs4834921 | 1.07 | 0.284 | -1.687 | 0.143 | -0.053 | -0.058 | -0.058 | 0 | 0.11 | 0 | 0.2 | 0.22 | 0.15 | 0.1 |
| 4 | 13118169 | rs525216 | 0.63 | 0.353 | -1.265 | 0.092 | 0.002 | 0.001 | 0.001 | 0.045 | 0.52 | 0.43 | 0.71 | 0.53 | 0.56 | 0.65 |
| 4 | 13120238 | rs510171 | -0.43 | 0.563 | -1.605 | 0.158 | 0.004 | 0.006 | 0.006 | 0 | 0.82 | 0.84 | 0.91 | 0.91 | 0.83 | 0.83 |
| 4 | 13124794 | rs568118 | -0.271 | 0.954 | -1.768 | 0.467 | -0.024 | -0.016 | -0.016 | 2.107 | 0.38 | 0.11 | 0.45 | 0.33 | 0.36 | 0.52 |
| 4 | 13127303 | rs6841445 | -0.347 | 0.528 | -1.403 | -0.074 | 0.024 | 0.041 | 0.041 | 0 | 0.41 | 0.29 | 0.33 | 0.28 | 0.34 | 0.47 |
| 4 | 13130251 | rs536983 | 0.187 | 0.511 | -0.804 | -0.804 | 0.008 | -0.004 | -0.004 | 0 | 0.49 | 0.55 | 0.74 | 0.86 | 0.65 | 0.54 |
| 4 | 13131206 | rs11935085 | -0.211 | 0.496 | -0.515 | -0.91 | 0.049 | -0.03 | -0.03 | 1.331 | 0.08 | 0.21 | 0.07 | 0.01 | 0.08 | 0.11 |
| 4 | 13140134 | rs519639 | 0.558 | 0.589 | -0.119 | -0.92 | -0.002 | -0.02 | -0.02 | 0.618 | 0.44 | 0.45 | 0.57 | 0.72 | 0.6 | 0.48 |
| 4 | 13147622 | rs490325 | -0.855 | 0.2 | 0.177 | -1.059 | 0.013 | 0.021 | 0.021 | 0 | 0.57 | 0.65 | 0.44 | 0.33 | 0.35 | 0.57 |
| 4 | 13149122 | rs503506 | -0.32 | -0.286 | 0.078 | -1.313 | -0.015 | -0.032 | -0.032 | 0.955 | 0.43 | 0.56 | 0.23 | 0.16 | 0.25 | 0.36 |
| 4 | 13167596 | rs562920 | -0.949 | -0.035 | 0.025 | -1.603 | -0.004 | -0.043 | -0.043 | 0 | 0.57 | 0.54 | 0.49 | 0.16 | 0.39 | 0.55 |
| 4 | 13168944 | rs3101937 | -1.216 | -0.135 | 0.075 | -1.878 | 0.022 | 0.048 | 0.048 | 0 | 0.48 | 0.6 | 0.48 | 0.17 | 0.37 | 0.5 |
| 4 | 13171640 | rs16888291 | -1.301 | 0.359 | 0.019 | -2.122 | 0.01 | -0.055 | -0.055 | 0.046 | 0.14 | 0.25 | 0.1 | 0.01 | 0.14 | 0.17 |
| 4 | 13182254 | rs10013514 | -0.481 | 0.248 | -0.344 | -2.19 | -0.003 | 0.005 | 0.005 | 0 | 0.1 | 0.1 | 0.07 | 0.01 | 0.11 | 0.14 |
| 4 | 13182756 | rs7685975 | -0.872 | -0.087 | -0.382 | -2.446 | 0.002 | -0.032 | -0.032 | 0 | 0.36 | 0.35 | 0.22 | 0.01 | 0.31 | 0.48 |
| 4 | 13184283 | rs10012046 | -0.74 | -0.106 | -0.411 | -2.39 | 0.002 | -0.031 | -0.031 | 0 | 0.12 | 0.18 | 0.08 | 0.01 | 0.11 | 0.16 |
| 4 | 13193295 | rs870835 | 0.832 | -0.166 | -0.306 | -2.411 | 0.004 | 0.013 | 0.013 | 0 | 0.19 | 0.12 | 0.12 | 0 | 0.18 | 0.28 |
| 4 | 13201728 | rs17315056 | 1.095 | -0.057 | -0.332 | -2.406 | -0.026 | 0.042 | 0.042 | 6.369 | 0.08 | 0.01 | 0.11 | 0 | 0.08 | 0.04 |
| 4 | 13213672 | rs10008923 | -0.807 | -0.066 | -0.294 | -2.427 | 0.002 | -0.035 | -0.035 | 0 | 0.12 | 0.22 | 0.08 | 0.01 | 0.11 | 0.16 |
| 4 | 13220338 | rs4574432 | -0.35 | -0.101 | -0.291 | -2.439 | 0.01 | -0.004 | -0.004 | 0 | 0.29 | 0.2 | 0.21 | 0.01 | 0.23 | 0.34 |
| 4 | 13221390 | rs6838271 | 0.492 | -0.205 | -0.287 | -2.416 | 0.005 | -0.028 | -0.028 | 1.503 | 0.21 | 0.2 | 0.1 | 0 | 0.15 | 0.3 |
| 4 | 13242797 | rs733048 | 0.956 | -0.042 | -0.755 | -1.99 | -0.004 | -0.057 | -0.057 | 0.185 | 0.24 | 0.13 | 0.26 | 0.34 | 0.25 | 0.21 |
| 4 | 13245646 | rs6847194 | -1.169 | -0.389 | -0.664 | -2.534 | 0.008 | -0.027 | -0.027 | 1.033 | 0.42 | 0.41 | 0.28 | 0.01 | 0.33 | 0.5 |
| 4 | 13257662 | rs16888414 | -0.081 | -0.239 | -0.64 | -2.389 | 0 | -0.008 | -0.008 | 0 | 0.1 | 0.15 | 0.06 | 0.01 | 0.09 | 0.15 |
| 4 | 13261188 | rs16888421 | -0.358 | -0.245 | -0.642 | -2.392 | 0 | -0.006 | -0.006 | 0.029 | 0.11 | 0.15 | 0.06 | 0.01 | 0.09 | 0.15 |
| 4 | 13264141 | rs16888428 | 0.625 | -0.254 | -0.644 | -2.413 | 0.006 | -0.016 | -0.016 | 0 | 0.21 | 0.17 | 0.11 | 0 | 0.17 | 0.31 |
| 4 | 13265092 | rs6852153 | 0.374 | 0.138 | -0.006 | -1.716 | 0.002 | -0.029 | -0.029 | 0 | 0.9 | 0.8 | 0.93 | 0.99 | 0.91 | 0.86 |
| 4 | 13265492 | rs11729455 | 0.805 | 0.318 | -0.381 | -1.735 | -0.026 | -0.027 | -0.027 | 0 | 0.1 | 0.01 | 0.06 | 0.11 | 0.11 | 0.1 |
| 4 | 13265681 | rs10022758 | 1.41 | 0.274 | 0.185 | -1.754 | -0.005 | -0.06 | -0.06 | 0 | 0.79 | 0.69 | 0.83 | 0.99 | 0.84 | 0.78 |
| 4 | 13266612 | rs7654928 | 1.728 | 0.336 | 0.096 | -1.716 | -0.026 | -0.124 | -0.124 | 0.407 | 0.73 | 0.51 | 0.79 | 0.99 | 0.82 | 0.71 |
| 4 | 13269293 | rs16888449 | -0.735 | 0.374 | -0.057 | -1.677 | 0.003 | 0.022 | 0.022 | 0 | 0.14 | 0.12 | 0.07 | 0.01 | 0.11 | 0.16 |
| 4 | 13269649 | rs16888451 | -0.949 | 0.366 | -0.084 | -1.691 | 0.005 | 0.028 | 0.028 | 0 | 0.15 | 0.12 | 0.07 | 0.01 | 0.11 | 0.17 |
| 4 | 13276652 | rs16888468 | -1.82 | 0.469 | -0.224 | -1.622 | -0.011 | -0.114 | -0.114 | 0 | 0.3 | 0.46 | 0.24 | 0.01 | 0.23 | 0.32 |
| 4 | 13280762 | rs1014335 | 0.647 | 0.691 | -0.989 | -1.17 | -0.001 | -0.05 | -0.05 | 0.28 | 0.75 | 0.69 | 0.84 | 0.98 | 0.74 | 0.74 |
| 4 | 13297604 | rs3213705 | -0.844 | 0.247 | -1.166 | -0.583 | -0.013 | -0.003 | -0.003 | 0 | 0.12 | 0.2 | 0.09 | 0.11 | 0.08 | 0.17 |
| 4 | 13302267 | rs6851825 | -2.058 | 0.063 | -0.748 | -0.645 | -0.019 | -0.032 | -0.032 | 0 | 0.22 | 0.37 | 0.18 | 0.12 | 0.16 | 0.21 |
| 4 | 13303269 | rs16888511 | -1.066 | -0.054 | -0.763 | -0.541 | -0.023 | -0.006 | -0.006 | 0.025 | 0.13 | 0.27 | 0.09 | 0.12 | 0.08 | 0.17 |
| 4 | 13359047 | rs10016671 | 0.04 | -0.23 | -1.068 | -0.694 | -0.004 | -0.01 | -0.01 | 0 | 0.06 | 0.08 | 0.01 | 0 | 0.01 | 0.01 |
| 4 | 13360630 | rs11947530 | -0.977 | -0.734 | -1.385 | -0.763 | 0.011 | 0.003 | 0.003 | 0 | 0.17 | 0.19 | 0.09 | 0.18 | 0.09 | 0.17 |
| 4 | 13370308 | rs10003958 | -1.222 | -0.694 | -1.358 | -0.743 | -0.031 | -0.048 | -0.048 | 0 | 0.09 | 0.26 | 0.04 | 0 | 0.03 | 0.01 |
| 4 | 13377286 | rs2109590 | 1.362 | -0.614 | -2.575 | -0.997 | -0.11 | -0.051 | -0.051 | 0 | 0.41 | 0.07 | 0.63 | 0.57 | 0.53 | 0.48 |
| 4 | 13391279 | rs10939477 | -0.299 | -0.918 | -2.161 | -1.138 | 0.006 | 0.004 | 0.004 | 0.051 | 0.19 | 0.14 | 0.09 | 0.18 | 0.1 | 0.31 |
| 4 | 13393883 | rs10001262 | 0.658 | -0.831 | -2.32 | -1.055 | -0.02 | -0.028 | -0.028 | 0 | 0.07 | 0.14 | 0.02 | 0 | 0.01 | 0.01 |
| 4 | 13439808 | rs10805318 | 1.244 | -0.549 | -2.58 | -1.248 | -0.133 | -0.081 | -0.081 | 0.112 | 0.36 | 0.06 | 0.57 | 0.56 | 0.48 | 0.4 |
| 4 | 13459953 | rs4518326 | -0.2 | -0.434 | -2.284 | -1.34 | -0.024 | -0.042 | -0.042 | 0 | 0.77 | 0.44 | 0.87 | 0.92 | 0.87 | 0.76 |
| 4 | 13502412 | rs10489095 | 1.28 | 0.338 | -2.342 | -1.192 | -0.005 | 0.04 | 0.04 | 0 | 0.08 | 0 | 0.04 | 0 | 0.07 | 0.05 |

| | | | | | | | | | | | | | | | | |
|---|---|---|---|---|---|---|---|---|---|---|---|---|---|---|---|---|
| 4 | 13526386 | rs2075050 | -0.5 | 0.403 | -2.525 | -0.577 | -0.001 | -0.008 | -0.008 | 0 | 0.62 | 0.43 | 0.48 | 0.6 | 0.63 | 0.52 |
| 4 | 13530824 | rs10034452 | -0.567 | -0.113 | -1.86 | -1.06 | -0.022 | -0.018 | -0.018 | 0.836 | 0.22 | 0.38 | 0.12 | 0.18 | 0.15 | 0.19 |
| 4 | 13534408 | rs1157946 | -0.117 | 0.044 | -2.558 | -1.6 | -0.012 | 0.193 | 0.193 | 0 | 0.29 | 0.04 | 0.18 | 0.01 | 0.27 | 0.16 |
| 4 | 13538956 | rs10939485 | 0.562 | -0.137 | -2.615 | -1.955 | 0.019 | 0.124 | 0.124 | 0.202 | 0.72 | 0.14 | 0.83 | 0.96 | 0.81 | 0.81 |
| 4 | 13539373 | rs758973 | 0.398 | -0.098 | -2.56 | -1.875 | 0.007 | 0.1 | 0.1 | 0.559 | 0.75 | 0.14 | 0.85 | 0.96 | 0.83 | 0.86 |
| 4 | 13540518 | rs12501307 | 0.398 | -0.098 | -2.533 | -1.875 | 0.008 | 0.096 | 0.096 | 0.559 | 0.75 | 0.15 | 0.85 | 0.96 | 0.83 | 0.85 |
| 4 | 13542036 | rs1476211 | 0.398 | -0.098 | -2.534 | -1.875 | 0.007 | 0.1 | 0.1 | 0.559 | 0.75 | 0.14 | 0.85 | 0.96 | 0.83 | 0.86 |
| 4 | 13542674 | rs3822226 | -0.259 | -0.03 | -0.923 | -1.91 | 0.002 | -0.114 | -0.114 | 0.331 | 0.2 | 0.4 | 0.1 | 0.04 | 0.15 | 0.14 |
| 4 | 13555006 | rs1379086 | 0.34 | -0.239 | -1.76 | -2.681 | 0.033 | 0.217 | 0.217 | 0 | 0.55 | 0.22 | 0.64 | 0.97 | 0.65 | 0.67 |
| 4 | 13559103 | rs13105197 | 0.473 | -0.258 | -1.795 | -2.716 | 0.04 | 0.235 | 0.235 | 0 | 0.53 | 0.22 | 0.64 | 0.97 | 0.65 | 0.67 |
| 4 | 13561643 | rs4698247 | 0.746 | 1.304 | -1.986 | -2.471 | -0.028 | -0.024 | -0.024 | 0.741 | 0.07 | 0.03 | 0.14 | 0.13 | 0.21 | 0.12 |
| 4 | 13566458 | rs12651430 | -0.548 | -0.484 | -1.697 | -2.46 | -0.001 | 0 | 0 | 0.594 | 0.56 | 0.47 | 0.65 | 0.69 | 0.68 | 0.6 |
| 4 | 13578682 | rs7695814 | 0.802 | -0.239 | 0.1 | -2.992 | 0.058 | 0.004 | 0.004 | 0 | 0.05 | 0.18 | 0.03 | 0.01 | 0.04 | 0.05 |
| 4 | 13584108 | rs17745075 | 0.65 | -0.229 | -0.493 | -2.993 | -0.001 | -0.044 | -0.044 | 0 | 0.2 | 0.16 | 0.21 | 0 | 0.16 | 0.18 |
| 4 | 13595559 | rs1013300 | 0.628 | -0.282 | -0.796 | -3.09 | 0.021 | 0.342 | 0.342 | 0 | 0.53 | 0.12 | 0.63 | 0.97 | 0.65 | 0.68 |
| 4 | 13604182 | rs17745676 | 0.722 | -0.315 | -0.715 | -3.046 | 0 | -0.038 | -0.038 | 0 | 0.19 | 0.14 | 0.21 | 0 | 0.15 | 0.18 |
| 4 | 13615174 | rs2035820 | 0.722 | -0.311 | -0.717 | -3.045 | -0.003 | -0.049 | -0.049 | 0 | 0.19 | 0.16 | 0.21 | 0 | 0.15 | 0.18 |
| 4 | 13623200 | rs11726392 | 0.605 | -0.278 | -0.757 | -3.087 | 0.022 | 0.319 | 0.319 | 0 | 0.54 | 0.13 | 0.63 | 0.97 | 0.66 | 0.68 |
| 4 | 13625648 | rs4235372 | -0.418 | -0.281 | -0.743 | -3.089 | -0.002 | -0.055 | -0.055 | 0 | 0.73 | 0.29 | 0.84 | 0.97 | 0.81 | 0.86 |
| 4 | 13625902 | rs4698090 | 0.521 | -0.215 | -0.922 | -3.025 | 0.022 | 0.319 | 0.319 | 0 | 0.54 | 0.13 | 0.63 | 0.97 | 0.66 | 0.68 |
| 4 | 13643437 | rs16888892 | 1.334 | 0.215 | -0.396 | -2.302 | -0.02 | 0.011 | 0.011 | 0 | 0.08 | 0.02 | 0.06 | 0.06 | 0.1 | 0.03 |
| 4 | 13677430 | rs1388300 | 0.388 | -0.077 | -0.005 | -2.326 | 0.006 | -0.026 | -0.026 | 0.276 | 0.68 | 0.28 | 0.83 | 0.98 | 0.78 | 0.79 |
| 4 | 13681606 | rs10017022 | -0.621 | -0.293 | 0.269 | -2.33 | -0.016 | -0.104 | -0.104 | 0 | 0.29 | 0.56 | 0.15 | 0.02 | 0.21 | 0.21 |
| 4 | 13689343 | rs2058579 | 0.515 | -0.436 | 0.029 | -2.73 | 0.059 | 0.103 | 0.103 | 1.108 | 0.56 | 0.27 | 0.67 | 0.97 | 0.69 | 0.65 |
| 4 | 13692300 | rs1491262 | 0.068 | -0.568 | 0.244 | -2.821 | -0.021 | -0.122 | -0.122 | 0 | 0.26 | 0.38 | 0.23 | 0.02 | 0.2 | 0.25 |
| 4 | 13697961 | rs10939499 | 0.846 | -0.539 | -0.498 | -2.82 | 0.096 | 0.359 | 0.359 | 0.869 | 0.53 | 0.1 | 0.65 | 0.97 | 0.69 | 0.65 |
| 4 | 13699931 | rs6850309 | -0.579 | -0.372 | 0.455 | -2.298 | -0.037 | -0.111 | -0.111 | 0.063 | 0.84 | 0.6 | 0.91 | 0.99 | 0.92 | 0.93 |
| 4 | 13703081 | rs1491267 | -0.266 | 0.258 | 0.066 | -2.325 | 0 | -0.048 | -0.048 | 0 | 0.82 | 0.74 | 0.86 | 0.99 | 0.82 | 0.9 |
| 4 | 13706858 | rs11727013 | 0.763 | 0.652 | -0.253 | -2.59 | -0.035 | 0.057 | 0.057 | 1.714 | 0.21 | 0.08 | 0.21 | 0.01 | 0.25 | 0.28 |
| 4 | 13708614 | rs16888984 | 0.479 | 0.237 | -0.415 | -2.233 | -0.016 | 0.009 | 0.009 | 1.123 | 0.24 | 0.28 | 0.21 | 0.28 | 0.19 | 0.17 |
| 4 | 13710530 | rs6835289 | 0.375 | 0.38 | -0.392 | -2.24 | 0.006 | 0.008 | 0.008 | 0.895 | 0.26 | 0.3 | 0.26 | 0.28 | 0.3 | 0.18 |
| 4 | 13712206 | rs13141860 | 1.139 | 0.341 | -0.316 | -2.246 | -0.02 | 0.023 | 0.023 | 1.106 | 0.21 | 0.13 | 0.21 | 0.01 | 0.25 | 0.28 |
| 4 | 13713538 | rs10025528 | 0.227 | 0.491 | -0.378 | -2.276 | 0.008 | 0.004 | 0.004 | 0 | 0.28 | 0.32 | 0.29 | 0.28 | 0.33 | 0.21 |
| 4 | 13715903 | rs875373 | -0.73 | 0.223 | -0.386 | -2.367 | 0.038 | 0.068 | 0.068 | 0 | 0.54 | 0.72 | 0.53 | 0.29 | 0.61 | 0.54 |
| 4 | 13717262 | rs10489085 | 1.58 | 0.168 | -0.576 | -2.431 | -0.062 | -0.105 | -0.105 | 0 | 0.27 | 0.02 | 0.35 | 0.68 | 0.29 | 0.38 |
| 4 | 13717471 | rs13120921 | -0.477 | 0.401 | -0.36 | -2.298 | 0.006 | 0.007 | 0.007 | 0 | 0.49 | 0.44 | 0.51 | 0.29 | 0.59 | 0.49 |
| 4 | 13719489 | rs10084768 | 0.121 | 0.675 | 0.081 | -1.682 | -0.011 | 0.007 | 0.007 | 1.091 | 0.26 | 0.29 | 0.22 | 0.28 | 0.22 | 0.19 |
| 4 | 13732086 | rs16889099 | 0.957 | 0.603 | -0.017 | -1.942 | -0.003 | 0.001 | 0.001 | 0 | 0.12 | 0.1 | 0.09 | 0.12 | 0.09 | 0.12 |
| 4 | 13733371 | rs9994682 | -0.49 | 0.919 | -0.115 | -1.928 | 0.001 | -0.003 | -0.003 | 0.082 | 0.61 | 0.62 | 0.67 | 0.97 | 0.65 | 0.65 |
| 4 | 13735895 | rs7688193 | 1.165 | 0.977 | -0.869 | -1.8 | -0.058 | 0.14 | 0.14 | 5.807 | 0.3 | 0.07 | 0.26 | 0.03 | 0.32 | 0.33 |
| 4 | 13737915 | rs759261 | -0.079 | 1.21 | -0.698 | -1.479 | -0.01 | 0.035 | 0.035 | 0 | 0.4 | 0.36 | 0.52 | 0.81 | 0.45 | 0.51 |
| 4 | 13755011 | rs2270279 | 0.494 | 0.694 | -0.593 | -0.989 | -0.01 | 0.002 | 0.002 | 1.155 | 0.2 | 0.23 | 0.16 | 0.19 | 0.17 | 0.09 |
| 4 | 13760844 | rs16889152 | 0.925 | 0.859 | -0.332 | -0.811 | 0.007 | -0.003 | -0.003 | 0.313 | 0.1 | 0.12 | 0.04 | 0 | 0.04 | 0.06 |
| 4 | 13761641 | rs3931694 | 0.221 | 0.801 | -0.128 | -0.601 | -0.021 | 0.096 | 0.096 | 0.011 | 0.16 | 0.24 | 0.34 | 0.6 | 0.15 | 0.3 |
| 4 | 13761712 | rs3931695 | -0.297 | 1.112 | -0.37 | -0.835 | 0.006 | 0.035 | 0.035 | 0.042 | 0.48 | 0.56 | 0.68 | 0.71 | 0.54 | 0.71 |
| 4 | 13767656 | rs741217 | -1.052 | 1.219 | -0.544 | -0.753 | -0.007 | -0.003 | -0.003 | 0 | 0.9 | 0.9 | 0.92 | 0.87 | 0.88 | 0.87 |
| 4 | 13780176 | rs1491250 | 0.487 | 1.626 | -1.188 | -0.511 | -0.012 | -0.099 | -0.099 | 3.129 | 0.15 | 0.02 | 0.32 | 0.61 | 0.17 | 0.28 |
| 4 | 13782594 | rs1472092 | -0.67 | 1.731 | -0.608 | -0.934 | 0.001 | 0.009 | 0.009 | 0.006 | 0.8 | 0.77 | 0.88 | 0.87 | 0.83 | 0.83 |
| 4 | 13783235 | rs1158808 | -0.321 | 1.853 | -0.397 | -0.459 | 0.003 | 0.007 | 0.007 | 3.007 | 0.49 | 0.46 | 0.56 | 0.77 | 0.49 | 0.39 |
| 4 | 13785225 | rs1008218 | -0.218 | 1.536 | -0.971 | -0.399 | 0 | 0.048 | 0.048 | 3.673 | 0.52 | 0.55 | 0.61 | 0.77 | 0.57 | 0.39 |
| 4 | 13788219 | rs7441080 | -0.775 | 1.278 | -0.923 | 0.455 | 0.005 | -0.004 | -0.004 | 1.172 | 0.36 | 0.63 | 0.43 | 0.45 | 0.31 | 0.5 |
| 4 | 13788316 | rs1019271 | 0.493 | 1.528 | -0.816 | 0.434 | -0.007 | -0.02 | -0.02 | 0 | 0.69 | 0.34 | 0.58 | 0.53 | 0.62 | 0.67 |
| 4 | 13797533 | rs1016029 | -0.321 | 0.649 | -0.426 | -0.456 | -0.018 | -0.024 | -0.024 | 0 | 0.39 | 0.58 | 0.32 | 0.21 | 0.26 | 0.36 |
| 4 | 13798314 | rs716424 | -0.807 | 1.348 | -0.64 | -0.796 | -0.017 | 0.045 | 0.045 | 0 | 0.62 | 0.75 | 0.61 | 0.22 | 0.57 | 0.57 |
| 4 | 13798796 | rs16889224 | 0.057 | 0.558 | 0.017 | -0.15 | -0.012 | -0.034 | -0.034 | 0 | 0.24 | 0.28 | 0.1 | 0.04 | 0.11 | 0.17 |
| 4 | 13799411 | rs17754182 | 0.225 | 0.58 | -0.052 | -0.151 | 0.018 | 0.004 | 0.004 | 0.08 | 0.23 | 0.2 | 0.09 | 0.02 | 0.11 | 0.11 |
| 4 | 13799857 | rs10939508 | -0.302 | 1.055 | -0.223 | 0.16 | -0.027 | -0.036 | -0.036 | 0 | 0.39 | 0.5 | 0.32 | 0.21 | 0.27 | 0.36 |
| 4 | 13807335 | rs7684636 | -0.344 | 1.02 | -0.875 | 0.027 | -0.012 | -0.028 | -0.028 | 0 | 0.39 | 0.58 | 0.3 | 0.17 | 0.26 | 0.36 |
| 4 | 13808360 | rs10489088 | -1.016 | 1.915 | -1.114 | 1.144 | -0.033 | -0.031 | -0.031 | 0.356 | 0.07 | 0.02 | 0.08 | 0.2 | 0.14 | 0.13 |
| 4 | 13809978 | rs10012159 | -0.437 | 1.034 | -0.957 | 0.949 | 0.008 | 0.017 | 0.017 | 0.326 | 0.51 | 0.65 | 0.35 | 0.64 | 0.46 | 0.41 |
| 4 | 13812406 | rs10007245 | -0.498 | 1.927 | -0.816 | 1.34 | 0.008 | 0.011 | 0.011 | 0 | 0.23 | 0.25 | 0.25 | 0.4 | 0.35 | 0.3 |
| 4 | 13815155 | rs10939516 | -0.534 | 1.992 | -1.055 | 1.041 | 0.012 | 0.001 | 0.001 | 0 | 0.07 | 0.09 | 0.17 | 0.12 | 0.17 | 0.09 |
| 4 | 13818881 | rs7697386 | 0.383 | 1.069 | -1.568 | 0.962 | 0.043 | 0.007 | 0.007 | 0 | 0.53 | 0.22 | 0.58 | 0.58 | 0.56 | 0.54 |
| 4 | 13820617 | rs7660348 | -0.332 | 1.155 | -1.55 | 0.605 | -0.001 | 0.006 | 0.006 | 0 | 0.68 | 0.69 | 0.59 | 0.78 | 0.72 | 0.83 |
| 4 | 13820801 | rs9784575 | -0.236 | 0.079 | -1.059 | 0.183 | 0.018 | 0.018 | 0.018 | 0 | 0.48 | 0.43 | 0.4 | 0.27 | 0.42 | 0.4 |
| 4 | 13824622 | rs7686779 | 0.015 | 0.298 | -1.095 | 0.083 | -0.013 | 0.003 | 0.003 | 0 | 0.13 | 0.07 | 0.19 | 0.1 | 0.15 | 0.11 |
| 4 | 13824647 | rs4395500 | -0.35 | 0.444 | -1.421 | 0.375 | -0.026 | -0.058 | -0.058 | 0.224 | 0.16 | 0.09 | 0.14 | 0.25 | 0.21 | 0.29 |
| 4 | 13825610 | rs887924 | -1.195 | 0.245 | -1.176 | 0.318 | -0.015 | -0.029 | -0.029 | 0 | 0.32 | 0.2 | 0.34 | 0.37 | 0.36 | 0.4 |
| 4 | 13831792 | rs13151645 | 1.006 | 0.031 | 0.036 | -0.048 | 0.006 | -0.001 | -0.001 | 0 | 0.67 | 0.38 | 0.62 | 0.7 | 0.62 | 0.59 |

| | | | | | | | | | | | | | | | | |
|---|---|---|---|---|---|---|---|---|---|---|---|---|---|---|---|---|
| 4 | 13833161 | rs887925 | 0.921 | -0.142 | 0.402 | -0.104 | -0.024 | 0.028 | 0.028 | 0 | 0.88 | 0.74 | 0.93 | 0.84 | 0.92 | 0.87 |
| 4 | 13841321 | rs2160120 | 1.787 | -0.289 | 1.421 | -0.222 | -0.001 | 0.01 | 0.01 | 0.045 | 0.6 | 0.3 | 0.62 | 0.71 | 0.62 | 0.59 |
| 4 | 13855897 | rs4698178 | 0.623 | -0.252 | 1.439 | -0.492 | -0.038 | -0.068 | -0.068 | 0.018 | 0.74 | 0.5 | 0.73 | 0.89 | 0.82 | 0.75 |
| 4 | 13861249 | rs1861135 | 1.319 | -0.077 | 1.473 | -0.497 | 0.059 | -0.041 | -0.041 | 0.29 | 0.86 | 0.64 | 0.87 | 0.94 | 0.8 | 0.94 |
| 4 | 13867556 | rs7669912 | 0.699 | -0.083 | 1.514 | -0.507 | -0.005 | -0.026 | -0.026 | 0 | 0.08 | 0.1 | 0.04 | 0 | 0.05 | 0.13 |
| 4 | 13870359 | rs11724465 | -0.737 | -0.215 | 1.766 | -0.427 | -0.01 | 0.008 | 0.008 | 0.132 | 0.19 | 0.25 | 0.3 | 0.21 | 0.16 | 0.23 |
| 4 | 13872720 | rs7690022 | -0.745 | -0.288 | 1.721 | -0.473 | 0.012 | 0.012 | 0.012 | 0.092 | 0.19 | 0.16 | 0.15 | 0.09 | 0.07 | 0.29 |
| 4 | 13874500 | rs6824694 | 0.535 | -0.278 | 1.904 | -0.481 | -0.009 | -0.008 | -0.008 | 0 | 0.8 | 0.75 | 0.83 | 0.92 | 0.93 | 0.71 |
| 4 | 13877759 | rs2867578 | -1.799 | -0.24 | 1.893 | -0.465 | 0.025 | 0.012 | 0.012 | 0.039 | 0.24 | 0.34 | 0.38 | 0.25 | 0.28 | 0.27 |
| 4 | 13882785 | rs1157531 | 1.78 | -0.37 | 1.806 | -0.579 | -0.008 | -0.038 | -0.038 | 0.664 | 0.67 | 0.4 | 0.72 | 0.87 | 0.77 | 0.66 |
| 4 | 13883763 | rs807764 | 1.28 | -0.34 | 1.769 | -0.563 | -0.023 | -0.004 | -0.004 | 2.663 | 0.77 | 0.51 | 0.64 | 0.82 | 0.82 | 0.76 |
| 4 | 13887264 | rs10005317 | 1.314 | -0.539 | 1.402 | -0.615 | 0.006 | 0.025 | 0.025 | 0 | 0.07 | 0.01 | 0.04 | 0 | 0.05 | 0.13 |
| 4 | 13887783 | rs1397623 | 2.446 | -0.788 | 1.496 | -0.615 | -0.001 | -0.004 | -0.004 | 0 | 0.67 | 0.37 | 0.58 | 0.8 | 0.71 | 0.69 |
| 4 | 13892169 | rs2058582 | 2.13 | -0.777 | 1.454 | -0.56 | 0.001 | -0.006 | -0.006 | 0 | 0.72 | 0.35 | 0.56 | 0.8 | 0.72 | 0.69 |
| 4 | 13894966 | rs1553853 | 2.299 | -0.823 | 1.428 | -0.625 | 0.001 | 0.017 | 0.017 | 0.225 | 0.64 | 0.31 | 0.52 | 0.77 | 0.67 | 0.55 |
| 4 | 13896514 | rs1606082 | 2.75 | -1.091 | 1.311 | -0.771 | 0.013 | 0.043 | 0.043 | 0 | 0.58 | 0.33 | 0.52 | 0.79 | 0.66 | 0.55 |
| 4 | 13897193 | rs1606083 | 0.269 | -1.676 | 1.179 | -0.581 | 0.023 | 0.001 | 0.001 | 0.247 | 0.1 | 0.06 | 0.12 | 0.08 | 0.02 | 0.21 |
| 4 | 13902266 | rs807780 | -0.429 | -1.455 | 1.387 | -0.562 | -0.009 | -0.061 | -0.061 | 0.115 | 0.89 | 0.78 | 0.91 | 1 | 0.91 | 0.86 |
| 4 | 13904303 | rs16889409 | -0.121 | -1.387 | 1.33 | -0.466 | -0.001 | -0.002 | -0.002 | 0 | 0.07 | 0.06 | 0.18 | 0.09 | 0.1 | 0.03 |
| 4 | 13905871 | rs16889425 | -0.323 | -1.792 | 1.265 | -0.776 | 0.039 | 0.026 | 0.026 | 0 | 0.15 | 0.05 | 0.14 | 0.08 | 0.04 | 0.2 |
| 4 | 13906669 | rs807783 | -2.761 | -2.089 | 1.529 | -0.98 | -0.022 | -0.032 | -0.032 | 0 | 0.4 | 0.57 | 0.41 | 0.15 | 0.24 | 0.37 |
| 4 | 13908785 | rs4453935 | -0.047 | -2.06 | 1.416 | -0.955 | 0.042 | 0.026 | 0.026 | 0 | 0.14 | 0.04 | 0.13 | 0.08 | 0.04 | 0.2 |
| 4 | 13914373 | rs10805321 | -1.141 | -2.274 | 1.155 | -1.213 | 0.049 | 0.071 | 0.071 | 0.109 | 0.28 | 0.13 | 0.2 | 0.18 | 0.16 | 0.28 |
| 4 | 13914459 | rs10939525 | -0.323 | -2.28 | 1.139 | -1.195 | 0.045 | 0.039 | 0.039 | 0.258 | 0.2 | 0.1 | 0.15 | 0.08 | 0.05 | 0.2 |
| 4 | 13915477 | rs4698655 | 2.624 | -2.163 | 1.215 | -1.17 | 0.024 | 0.079 | 0.079 | 0 | 0.47 | 0.3 | 0.5 | 0.71 | 0.58 | 0.53 |
| 4 | 13920982 | rs505703 | -0.963 | -2.267 | 1.262 | -1.296 | 0 | 0.001 | 0.001 | 0 | 0.25 | 0.23 | 0.33 | 0.13 | 0.16 | 0.23 |
| 4 | 13921238 | rs13112792 | 2.485 | -2.168 | 1.452 | -1.277 | 0.041 | 0.121 | 0.121 | 0.219 | 0.49 | 0.25 | 0.51 | 0.72 | 0.57 | 0.51 |
| 4 | 13922335 | rs527313 | -1.415 | -2.059 | 1.63 | -1.182 | -0.017 | -0.013 | -0.013 | 0.729 | 0.28 | 0.34 | 0.35 | 0.23 | 0.21 | 0.24 |
| 4 | 13925186 | rs578319 | 1.441 | -1.821 | 1.638 | -1.045 | 0.055 | 0.063 | 0.063 | 0.093 | 0.89 | 0.67 | 0.83 | 0.89 | 0.88 | 0.75 |
| 4 | 13928993 | rs579137 | 0.163 | -1.784 | 1.541 | -1.046 | 0.031 | -0.002 | -0.002 | 0 | 0.94 | 0.9 | 0.96 | 0.96 | 0.9 | 0.87 |
| 4 | 13929751 | rs551592 | -1.718 | -1.993 | 1.856 | -1.17 | -0.058 | -0.066 | -0.066 | 0.329 | 0.19 | 0.43 | 0.19 | 0.08 | 0.11 | 0.15 |
| 4 | 13930615 | rs11939384 | -0.409 | -1.932 | 1.902 | -1.126 | 0.004 | 0.062 | 0.062 | 0 | 0.06 | 0.22 | 0.12 | 0.06 | 0.01 | 0.11 |
| 4 | 13930991 | rs10016970 | 1.029 | -2.028 | 1.969 | -1.238 | -0.003 | -0.043 | -0.043 | 0 | 0.11 | 0.15 | 0.08 | 0 | 0.11 | 0.13 |
| 4 | 13931019 | rs11940421 | -1.091 | -2.042 | 1.895 | -1.137 | -0.011 | 0.095 | 0.095 | 0 | 0.07 | 0.26 | 0.12 | 0.14 | 0.02 | 0.12 |
| 4 | 13932083 | rs564771 | 1.369 | -2.048 | 1.62 | -1.22 | -0.01 | -0.002 | -0.002 | 0.518 | 0.71 | 0.61 | 0.64 | 0.74 | 0.74 | 0.64 |
| 4 | 13936707 | rs552056 | 0.28 | -1.921 | 1.649 | -1.179 | 0.058 | 0.016 | 0.016 | 0.071 | 0.9 | 0.69 | 0.84 | 0.94 | 0.89 | 0.88 |
| 4 | 13940961 | rs7691637 | 0.949 | -1.872 | 1.244 | -1.154 | 0.003 | -0.03 | -0.03 | 0.219 | 0.06 | 0.11 | 0.04 | 0.02 | 0.07 | 0.04 |
| 4 | 13951077 | rs971079 | 2.331 | -2.011 | 1.071 | -1.212 | 0.025 | 0.099 | 0.099 | 0 | 0.48 | 0.26 | 0.5 | 0.72 | 0.53 | 0.47 |
| 4 | 13953276 | rs536993 | -0.085 | -1.951 | 1.186 | -1.119 | 0.008 | -0.001 | -0.001 | 0.176 | 0.19 | 0.17 | 0.16 | 0.17 | 0.15 | 0.24 |
| 4 | 13953713 | rs2120555 | 1.395 | -2.129 | 1.292 | -1.284 | 0.007 | 0.005 | 0.005 | 0.04 | 0.65 | 0.66 | 0.68 | 0.76 | 0.71 | 0.58 |
| 4 | 13954615 | rs11734382 | 0.314 | -2.099 | 1.231 | -1.26 | 0.019 | 0.022 | 0.022 | 0 | 0.18 | 0.05 | 0.1 | 0.09 | 0.14 | 0.23 |
| 4 | 13955163 | rs971048 | 2.183 | -2.144 | 1.373 | -1.289 | 0.035 | 0.117 | 0.117 | 0.23 | 0.49 | 0.24 | 0.56 | 0.73 | 0.56 | 0.42 |
| 4 | 13963977 | rs504227 | 1.181 | -2.108 | 1.396 | -1.27 | -0.01 | -0.035 | -0.035 | 0.695 | 0.72 | 0.58 | 0.79 | 0.83 | 0.75 | 0.66 |
| 4 | 13966851 | rs524675 | 0.111 | -2.045 | 1.38 | -1.249 | 0.009 | 0.027 | 0.027 | 0 | 0.22 | 0.13 | 0.21 | 0.09 | 0.19 | 0.24 |
| 4 | 13975619 | rs1444152 | 1.893 | -2.098 | 1.457 | -1.266 | 0.001 | 0.003 | 0.003 | 0.642 | 0.52 | 0.43 | 0.58 | 0.75 | 0.55 | 0.46 |
| 4 | 13977266 | rs541373 | -0.315 | -1.99 | 1.385 | -1.209 | 0.012 | 0.021 | 0.021 | 0 | 0.89 | 0.82 | 0.88 | 0.86 | 0.86 | 0.81 |
| 4 | 13988911 | rs535932 | 0.233 | -2.141 | 1.274 | -1.308 | 0.002 | 0.027 | 0.027 | 0 | 0.21 | 0.14 | 0.2 | 0.08 | 0.2 | 0.21 |
| 4 | 13992268 | rs10516270 | 0.157 | -1.961 | 1.379 | -0.998 | -0.004 | 0.055 | 0.055 | 0 | 0.1 | 0.19 | 0.14 | 0.17 | 0.12 | 0.2 |
| 4 | 13992929 | rs476133 | -1.186 | -2.246 | 1.126 | -1.307 | 0.001 | -0.006 | -0.006 | 0 | 0.36 | 0.37 | 0.29 | 0.1 | 0.31 | 0.33 |
| 4 | 13992974 | rs16889528 | 0.4 | -1.851 | 0.937 | -1.171 | 0.014 | 0.025 | 0.025 | 0.047 | 0.13 | 0.07 | 0.08 | 0 | 0.11 | 0.12 |
| 4 | 13996896 | rs576093 | -1.217 | -2.047 | 0.92 | -0.959 | 0.023 | 0.003 | 0.003 | 0 | 0.53 | 0.63 | 0.44 | 0.37 | 0.44 | 0.56 |
| 4 | 14001392 | rs10008925 | -0.504 | -1.949 | 0.969 | -0.952 | 0.007 | -0.006 | -0.006 | 0 | 0.21 | 0.19 | 0.09 | 0.04 | 0.13 | 0.14 |
| 4 | 14017269 | rs492134 | 1.018 | -2.113 | 0.82 | -1.024 | 0.018 | -0.002 | -0.002 | 0 | 0.45 | 0.32 | 0.55 | 0.63 | 0.56 | 0.42 |
| 4 | 14024946 | rs794095 | -0.392 | -2.062 | 0.366 | -1.086 | -0.011 | 0.005 | 0.005 | 0 | 0.64 | 0.6 | 0.65 | 0.69 | 0.68 | 0.55 |
| 4 | 14026281 | rs518592 | 0.841 | -1.492 | -0.625 | -0.806 | 0.023 | -0.01 | -0.01 | 0 | 0.43 | 0.25 | 0.54 | 0.64 | 0.55 | 0.41 |
| 4 | 14028747 | rs522578 | -0.03 | -1.092 | -1.179 | -0.621 | 0.074 | 0.02 | 0.02 | 0.181 | 0.53 | 0.24 | 0.62 | 0.68 | 0.67 | 0.52 |
| 4 | 14033010 | rs13121292 | 0.301 | -0.192 | -1.374 | -0.205 | -0.051 | -0.021 | -0.021 | 0 | 0.37 | 0.17 | 0.52 | 0.67 | 0.48 | 0.37 |
| 4 | 14037453 | rs4082824 | 0.213 | 0.218 | -1.137 | -0.926 | 0 | 0.034 | 0.034 | 0.12 | 0.54 | 0.42 | 0.77 | 0.83 | 0.56 | 0.52 |
| 4 | 14037934 | rs1877147 | -0.062 | -0.317 | -0.677 | -1.27 | 0.027 | -0.063 | -0.063 | 0.794 | 0.67 | 0.58 | 0.8 | 0.87 | 0.6 | 0.7 |
| 4 | 14053263 | rs4698240 | 0.827 | -0.557 | -0.263 | -2.071 | -0.026 | -0.114 | -0.114 | 0 | 0.65 | 0.46 | 0.87 | 0.93 | 0.8 | 0.74 |
| 4 | 14056930 | rs3857159 | 1.472 | -0.521 | 0.156 | -1.991 | -0.005 | 0.009 | 0.009 | 0 | 0.43 | 0.41 | 0.71 | 0.82 | 0.5 | 0.51 |
| 4 | 14060248 | rs7682816 | -0.931 | -0.369 | 0.316 | -1.841 | 0.001 | -0.019 | -0.019 | 0 | 0.9 | 0.89 | 0.97 | 0.98 | 0.9 | 0.93 |
| 4 | 14062571 | rs11736290 | 0.484 | -0.586 | 0.64 | -2.041 | 0.01 | -0.031 | -0.031 | 0 | 0.66 | 0.42 | 0.79 | 0.82 | 0.63 | 0.66 |
| 4 | 14063649 | rs4698241 | -0.968 | -0.581 | 0.738 | -2.051 | 0.004 | -0.028 | -0.028 | 0 | 0.9 | 0.84 | 0.97 | 0.98 | 0.9 | 0.93 |
| 4 | 14064059 | rs10516272 | 0.139 | -0.594 | 0.743 | -2.062 | 0.003 | -0.006 | -0.006 | 0 | 0.23 | 0.22 | 0.18 | 0.17 | 0.3 | 0.31 |
| 4 | 14077835 | rs9917971 | 1.637 | -0.93 | 0.673 | -2.221 | -0.015 | 0.059 | 0.059 | 0 | 0.44 | 0.32 | 0.73 | 0.81 | 0.53 | 0.55 |
| 4 | 14082952 | rs10028728 | 0.657 | -0.942 | 0.606 | -2.193 | -0.008 | 0.005 | 0.005 | 0.418 | 0.19 | 0.15 | 0.14 | 0.12 | 0.27 | 0.21 |
| 4 | 14092353 | rs4089881 | -1.175 | -0.943 | 0.619 | -2.192 | -0.009 | -0.048 | -0.048 | 0 | 0.44 | 0.5 | 0.24 | 0.12 | 0.37 | 0.32 |
| 4 | 14098935 | rs12499108 | 1.232 | -1.01 | 0.534 | -2.35 | 0.016 | 0.206 | 0.206 | 1.275 | 0.5 | 0.22 | 0.73 | 0.83 | 0.59 | 0.63 |

| | | | | | | | | | | | | | | | | |
|---|---|---|---|---|---|---|---|---|---|---|---|---|---|---|---|---|
| 4 | 14099183 | rs4608810 | 0.284 | -0.935 | 0.64 | -1.638 | -0.016 | 0.009 | 0.009 | 0.69 | 0.7 | 0.75 | 0.84 | 0.77 | 0.64 | 0.72 |
| 4 | 14099571 | rs3915263 | 0.37 | -1.452 | 0.296 | -2.237 | -0.004 | 0.003 | 0.003 | 0 | 0.63 | 0.66 | 0.82 | 0.88 | 0.64 | 0.71 |
| 4 | 14102069 | rs3846386 | 0.074 | -1.462 | 0.313 | -1.846 | -0.012 | -0.043 | -0.043 | 0.153 | 0.68 | 0.56 | 0.77 | 0.85 | 0.72 | 0.77 |
| 4 | 14104524 | rs3915267 | 0.991 | -1.443 | 0.147 | -2.108 | -0.005 | -0.076 | -0.076 | 0 | 0.61 | 0.54 | 0.8 | 0.88 | 0.69 | 0.8 |
| 4 | 14110597 | rs9291621 | 0.434 | -1.498 | 0.389 | -2.105 | -0.026 | -0.067 | -0.067 | 0.842 | 0.7 | 0.45 | 0.82 | 0.91 | 0.79 | 0.86 |
| 4 | 14112550 | rs11730712 | 2.046 | -1.267 | -0.818 | -2.33 | -0.026 | -0.005 | -0.005 | 0 | 0.14 | 0.01 | 0.1 | 0.07 | 0.16 | 0.05 |
| 4 | 14112933 | rs2204618 | -0.988 | -1.185 | -0.366 | -2.234 | -0.006 | -0.018 | -0.018 | 0 | 0.83 | 0.74 | 0.88 | 0.91 | 0.85 | 0.88 |
| 4 | 14117046 | rs3846395 | 1.346 | -1.12 | -0.748 | -2.172 | -0.021 | -0.008 | -0.008 | 0 | 0.11 | 0.01 | 0.09 | 0.11 | 0.11 | 0.18 |
| 4 | 14117252 | rs17402733 | 1.447 | -1.213 | -0.424 | -2.323 | -0.018 | 0.071 | 0.071 | 0 | 0.42 | 0.3 | 0.68 | 0.78 | 0.56 | 0.55 |
| 4 | 14123352 | rs3846396 | 0.378 | -1.151 | -0.406 | -2.172 | 0.009 | 0.018 | 0.018 | 0 | 0.19 | 0.09 | 0.13 | 0.11 | 0.15 | 0.18 |
| 4 | 14125088 | rs4610328 | 0.466 | -0.544 | -0.061 | -2.446 | 0.025 | -0.055 | -0.055 | 0 | 0.19 | 0.34 | 0.16 | 0.06 | 0.21 | 0.18 |
| 4 | 14127483 | rs3846401 | 0.575 | -0.826 | -0.494 | -2.632 | -0.001 | -0.007 | -0.007 | 0 | 0.58 | 0.6 | 0.73 | 0.82 | 0.63 | 0.63 |
| 4 | 14128820 | rs17403552 | 0.36 | -0.79 | -0.639 | -2.585 | 0.016 | 0.046 | 0.046 | 0 | 0.23 | 0.1 | 0.14 | 0.12 | 0.17 | 0.2 |
| 4 | 14131925 | rs3846407 | 1.421 | -0.73 | -0.679 | -2.656 | -0.004 | 0.052 | 0.052 | 0.001 | 0.42 | 0.31 | 0.65 | 0.79 | 0.49 | 0.54 |
| 4 | 14135068 | rs1834514 | -1.93 | -0.676 | -0.725 | -2.597 | -0.014 | 0.011 | 0.011 | 0 | 0.93 | 0.97 | 0.97 | 0.97 | 0.89 | 0.97 |
| 4 | 14135172 | rs1426119 | -1.364 | -0.855 | -0.598 | -2.666 | -0.009 | -0.016 | -0.016 | 0 | 0.92 | 0.93 | 0.98 | 1 | 0.97 | 0.95 |
| 4 | 14137784 | rs1426123 | 0.389 | -1.032 | -0.615 | -2.818 | 0.01 | 0.057 | 0.057 | 0.173 | 0.65 | 0.76 | 0.8 | 0.85 | 0.7 | 0.72 |
| 4 | 14138415 | rs7670043 | 0.119 | -0.949 | -0.713 | -3.023 | 0.016 | -0.137 | -0.137 | 0.065 | 0.29 | 0.47 | 0.2 | 0.03 | 0.3 | 0.27 |
| 4 | 14139678 | rs3893808 | 0.866 | -0.49 | -0.828 | -3.123 | 0.007 | 0.008 | 0.008 | 0 | 0.49 | 0.38 | 0.71 | 0.94 | 0.52 | 0.64 |
| 4 | 14140211 | rs3846413 | -0.11 | -0.46 | -0.765 | -3.078 | 0.016 | -0.125 | -0.125 | 0 | 0.29 | 0.43 | 0.19 | 0.04 | 0.3 | 0.26 |
| 4 | 14141243 | rs16889750 | 1.285 | -0.373 | -0.851 | -3.022 | 0.013 | 0.004 | 0.004 | 0.258 | 0.14 | 0.06 | 0.1 | 0.13 | 0.13 | 0.18 |
| 4 | 14143365 | rs2912200 | 1.274 | -0.227 | -0.77 | -3.093 | 0.021 | -0.078 | -0.078 | 0 | 0.18 | 0.23 | 0.16 | 0.01 | 0.23 | 0.18 |
| 4 | 14146572 | rs17345360 | 1.61 | -0.427 | -0.904 | -2.983 | -0.001 | -0.013 | -0.013 | 0.57 | 0.12 | 0.02 | 0.1 | 0.13 | 0.13 | 0.18 |
| 4 | 14148374 | rs4698716 | 1.845 | -0.406 | -0.804 | -3.044 | 0 | 0.016 | 0.016 | 0 | 0.36 | 0.29 | 0.61 | 0.82 | 0.39 | 0.46 |
| 4 | 14152469 | rs34855554 | -0.385 | -0.434 | -0.991 | -3.052 | 0.014 | 0.055 | 0.055 | 0.438 | 0.32 | 0.21 | 0.18 | 0.15 | 0.28 | 0.24 |
| 4 | 14152937 | rs3846415 | -0.763 | -0.584 | -1.129 | -3.134 | 0.034 | 0.188 | 0.188 | 0 | 0.51 | 0.22 | 0.29 | 0.17 | 0.55 | 0.44 |
| 4 | 14153819 | rs11937314 | -0.469 | -0.414 | -1.049 | -3.038 | 0.018 | 0.087 | 0.087 | 0.138 | 0.34 | 0.16 | 0.18 | 0.16 | 0.28 | 0.28 |
| 4 | 14156515 | rs10939534 | 1.532 | -0.301 | -0.701 | -3.016 | 0.007 | -0.043 | -0.043 | 0 | 0.13 | 0.21 | 0.08 | 0.03 | 0.11 | 0.02 |
| 4 | 14157416 | rs6818005 | 0.508 | -0.32 | -0.655 | -3.118 | -0.011 | -0.002 | -0.002 | 0 | 0.29 | 0.18 | 0.21 | 0.01 | 0.31 | 0.22 |
| 4 | 14158633 | rs17346022 | 1.719 | -0.345 | -0.771 | -3.133 | -0.002 | 0.005 | 0.005 | 0.275 | 0.14 | 0.06 | 0.1 | 0.13 | 0.15 | 0.19 |
| 4 | 14158921 | rs6830592 | 0.483 | -0.438 | -0.73 | -3.192 | 0 | -0.106 | -0.106 | 0 | 0.55 | 0.5 | 0.71 | 0.96 | 0.58 | 0.69 |
| 4 | 14161125 | rs3909661 | -0.575 | -0.512 | -0.782 | -3.118 | 0.019 | 0.03 | 0.03 | 0.037 | 0.33 | 0.27 | 0.17 | 0.15 | 0.25 | 0.34 |
| 4 | 14163726 | rs3111819 | -0.487 | -0.263 | -0.754 | -3.094 | -0.035 | 0.063 | 0.063 | 0.017 | 0.68 | 0.83 | 0.74 | 0.92 | 0.6 | 0.67 |
| 4 | 14164082 | rs10024326 | -0.095 | -0.342 | -0.634 | -2.804 | -0.009 | -0.055 | -0.055 | 0.37 | 0.18 | 0.32 | 0.1 | 0.09 | 0.16 | 0.16 |
| 4 | 14164509 | rs3111820 | 0.158 | -0.334 | -0.843 | -2.206 | -0.035 | 0.079 | 0.079 | 0 | 0.57 | 0.76 | 0.67 | 0.69 | 0.54 | 0.53 |
| 4 | 14165975 | rs3096592 | 2.007 | -0.266 | -0.759 | -2.064 | 0 | -0.001 | -0.001 | 4.703 | 0.31 | 0.28 | 0.52 | 0.58 | 0.33 | 0.24 |
| 4 | 14167059 | rs2903626 | -1.013 | -0.385 | -0.623 | -1.941 | -0.018 | -0.029 | -0.029 | 0 | 0.37 | 0.52 | 0.22 | 0.33 | 0.27 | 0.39 |
| 4 | 14179198 | rs3111816 | -1.109 | 0.088 | -0.976 | -1.55 | -0.013 | -0.006 | -0.006 | 0 | 0.78 | 0.85 | 0.82 | 0.96 | 0.65 | 0.81 |
| 4 | 14183204 | rs2004908 | 1.382 | -0.256 | -1.044 | -1.515 | 0.008 | 0.015 | 0.015 | 0 | 0.33 | 0.2 | 0.51 | 0.63 | 0.35 | 0.35 |
| 4 | 14185184 | rs6846915 | 0.413 | -0.293 | -0.887 | -1.615 | 0.013 | 0.019 | 0.019 | 0 | 0.49 | 0.55 | 0.62 | 0.88 | 0.43 | 0.55 |
| 4 | 14188496 | rs3096587 | -1.817 | 0.033 | -0.916 | -1.573 | -0.047 | 0.045 | 0.045 | 0 | 0.82 | 0.94 | 0.84 | 0.99 | 0.72 | 0.84 |
| 4 | 14198023 | rs12512931 | 1.282 | -0.269 | -0.613 | -1.498 | 0.004 | -0.001 | -0.001 | 0 | 0.13 | 0.1 | 0.09 | 0.25 | 0.08 | 0.2 |
| 4 | 14198860 | rs1549457 | -0.069 | -0.306 | -0.538 | -1.538 | -0.002 | -0.017 | -0.017 | 0 | 0.3 | 0.34 | 0.18 | 0.11 | 0.24 | 0.27 |
| 4 | 14204807 | rs890299 | -0.087 | -0.301 | -0.649 | -1.542 | -0.005 | -0.038 | -0.038 | 0 | 0.3 | 0.41 | 0.19 | 0.11 | 0.25 | 0.27 |
| 4 | 14216574 | rs11724569 | 2.393 | -0.481 | -1.622 | -1.411 | -0.014 | -0.016 | -0.016 | 0.065 | 0.15 | 0.01 | 0.09 | 0.09 | 0.14 | 0.05 |
| 4 | 14217009 | rs2969168 | -0.028 | -0.341 | -1.335 | -1.444 | 0.003 | 0.005 | 0.005 | 0 | 0.33 | 0.72 | 0.22 | 0.11 | 0.25 | 0.27 |
| 4 | 14221055 | rs16889857 | 1.016 | -0.238 | -1.476 | -1.319 | -0.025 | -0.028 | -0.028 | 0 | 0.09 | 0.03 | 0.08 | 0.28 | 0.13 | 0.21 |
| 4 | 14222240 | rs3111828 | 0.015 | -0.467 | -1.455 | -1.489 | 0.004 | -0.047 | -0.047 | 0 | 0.58 | 0.51 | 0.67 | 0.71 | 0.51 | 0.61 |
| 4 | 14222539 | rs3096566 | 0.806 | -0.489 | -1.398 | -1.447 | -0.008 | -0.016 | -0.016 | 0 | 0.25 | 0.32 | 0.16 | 0.08 | 0.17 | 0.25 |
| 4 | 14222634 | rs10516275 | 0.521 | -0.262 | -1.658 | -1.143 | 0.012 | 0.037 | 0.037 | 0 | 0.38 | 0.2 | 0.53 | 0.63 | 0.38 | 0.37 |
| 4 | 14234630 | rs10939537 | 1.905 | -0.147 | -1.382 | -0.08 | 0.006 | -0.027 | -0.027 | 0 | 0.06 | 0.01 | 0.07 | 0.18 | 0.06 | 0.13 |
| 4 | 14245329 | rs16889917 | 0.487 | -0.291 | -1.49 | -0.124 | 0.011 | -0.042 | -0.042 | 0 | 0.2 | 0.09 | 0.17 | 0.27 | 0.16 | 0.22 |
| 4 | 14245558 | rs10025267 | -1.198 | -0.125 | -1.859 | -0.297 | -0.026 | -0.026 | -0.026 | 0 | 0.47 | 0.82 | 0.43 | 0.56 | 0.5 | 0.62 |
| 4 | 14250959 | rs10470841 | 0.314 | -0.142 | -1.461 | -0.349 | 0.009 | -0.027 | -0.027 | 0.429 | 0.21 | 0.49 | 0.16 | 0.26 | 0.19 | 0.28 |
| 4 | 14255076 | rs7691835 | -1.09 | -0.34 | -1.823 | -0.518 | 0.013 | 0.005 | 0.005 | 0.073 | 0.5 | 0.28 | 0.36 | 0.52 | 0.42 | 0.51 |
| 4 | 14255866 | rs12645353 | 1.919 | -0.15 | -1.53 | 0.089 | -0.021 | -0.021 | -0.021 | 0 | 0.06 | 0 | 0.13 | 0.25 | 0.07 | 0.07 |
| 4 | 14257631 | rs3096584 | -1.317 | -0.123 | -1.414 | -0.777 | -0.012 | 0.103 | 0.103 | 0.643 | 0.47 | 0.68 | 0.34 | 0.27 | 0.47 | 0.47 |
| 4 | 14259503 | rs2052711 | -0.265 | -0.016 | -1.4 | -0.749 | -0.045 | 0.062 | 0.062 | 0 | 0.33 | 0.17 | 0.25 | 0.23 | 0.42 | 0.4 |
| 4 | 14263549 | rs3096581 | -1.746 | -0.07 | -1.525 | -0.688 | -0.008 | 0.061 | 0.061 | 0.496 | 0.63 | 0.86 | 0.6 | 0.84 | 0.62 | 0.71 |
| 4 | 14267241 | rs13108994 | 0.436 | 0.19 | -1.139 | -0.249 | 0.004 | -0.016 | -0.016 | 0 | 0.21 | 0.17 | 0.28 | 0.56 | 0.17 | 0.23 |
| 4 | 14267496 | rs3096579 | -0.522 | 0.343 | -1.084 | -0.188 | 0.007 | -0.002 | -0.002 | 0 | 0.36 | 0.67 | 0.37 | 0.59 | 0.22 | 0.31 |
| 4 | 14268049 | rs13148052 | 0.623 | 0.987 | -0.685 | -0.255 | -0.002 | -0.006 | -0.006 | 0 | 0.11 | 0.07 | 0.12 | 0.03 | 0.2 | 0.19 |
| 4 | 14269174 | rs890300 | 0.002 | 1.071 | -0.881 | -0.214 | 0.087 | -0.094 | -0.094 | 0 | 0.15 | 0.45 | 0.15 | 0.03 | 0.21 | 0.17 |
| 4 | 14269562 | rs17415839 | 0.556 | 0.578 | -1.711 | 0.561 | -0.01 | -0.041 | -0.041 | 0 | 0.08 | 0.01 | 0.06 | 0.1 | 0.05 | 0.05 |
| 4 | 14270358 | rs12499689 | -0.711 | 0.842 | -1.319 | 0.401 | -0.039 | 0.005 | 0.005 | 0 | 0.18 | 0.04 | 0.11 | 0.22 | 0.18 | 0.22 |
| 4 | 14272582 | rs16889949 | -0.88 | 0.939 | -0.433 | 0.455 | 0.03 | -0.008 | -0.008 | 0 | 0.13 | 0.28 | 0.08 | 0.09 | 0.14 | 0.05 |
| 4 | 14273807 | rs16889955 | -0.058 | 1.39 | -0.474 | 0.867 | 0.004 | 0.01 | 0.01 | 0 | 0.14 | 0.13 | 0.17 | 0.28 | 0.15 | 0.14 |
| 4 | 14275327 | rs3111813 | -0.653 | 2.048 | -1.148 | 0.296 | -0.042 | 0.047 | 0.047 | 0 | 0.9 | 0.97 | 0.86 | 0.99 | 0.8 | 0.84 |
| 4 | 14277183 | rs1991268 | -0.652 | 2.072 | -1.104 | 0.276 | -0.03 | 0.029 | 0.029 | 0 | 0.9 | 0.95 | 0.86 | 0.99 | 0.8 | 0.85 |

| | | | | | | | | | | | | | | | | |
|---|---|---|---|---|---|---|---|---|---|---|---|---|---|---|---|---|
| 4 | 14279189 | rs1426110 | 1.084 | 1.144 | -1.143 | 0.309 | 0.015 | 0.01 | 0.01 | 1.058 | 0.48 | 0.42 | 0.46 | 0.37 | 0.39 | 0.38 |
| 4 | 14280282 | rs16890027 | 0.698 | 0.685 | -1.453 | 0.189 | -0.054 | -0.041 | -0.041 | 0 | 0.16 | 0.03 | 0.1 | 0.21 | 0.2 | 0.14 |
| 4 | 14280826 | rs6826155 | 1.084 | 0.133 | -1.682 | -0.057 | -0.006 | -0.002 | -0.002 | 0 | 0.26 | 0.27 | 0.12 | 0.14 | 0.16 | 0.13 |
| 4 | 14282131 | rs10013659 | -0.549 | 0.312 | -1.783 | 0.199 | -0.001 | 0.013 | 0.013 | 1.39 | 0.75 | 0.74 | 0.79 | 0.85 | 0.69 | 0.72 |
| 4 | 14284343 | rs16890060 | 0.805 | 0.766 | -1.534 | 0.587 | -0.001 | -0.009 | -0.009 | 0 | 0.05 | 0.02 | 0.09 | 0.06 | 0.05 | 0.12 |
| 4 | 14287766 | rs13131008 | 0.214 | 0.825 | -1.751 | 0.51 | -0.13 | -0.165 | -0.165 | 0.322 | 0.26 | 0.03 | 0.44 | 0.42 | 0.34 | 0.39 |
| 4 | 14294745 | rs1544874 | -0.196 | 0.672 | -0.275 | 0.01 | 0.048 | -0.017 | -0.017 | 0 | 0.1 | 0.33 | 0.21 | 0.04 | 0.12 | 0.06 |
| 4 | 14296300 | rs1426107 | -0.446 | -0.178 | -0.754 | -0.14 | 0.059 | 0.005 | 0.005 | 0 | 0.18 | 0.06 | 0.13 | 0.16 | 0.11 | 0.29 |
| 4 | 14296613 | rs2079949 | 1.367 | 0.562 | -0.27 | -0.413 | 0.007 | 0.048 | 0.048 | 0 | 0.57 | 0.29 | 0.49 | 0.78 | 0.53 | 0.46 |
| 4 | 14302221 | rs16890073 | 0.589 | 0.403 | 0.177 | -0.099 | 0.044 | -0.015 | -0.015 | 0 | 0.08 | 0.28 | 0.19 | 0.03 | 0.13 | 0.04 |
| 4 | 14306031 | rs2554377 | -1.519 | 0.043 | 0.231 | -0.136 | 0.001 | 0.001 | 0.001 | 0 | 0.3 | 0.73 | 0.37 | 0.22 | 0.26 | 0.37 |
| 4 | 14312034 | rs4698250 | 1.298 | 0.349 | -0.322 | 0.134 | -0.029 | -0.018 | -0.018 | 0.058 | 0.42 | 0.14 | 0.33 | 0.4 | 0.41 | 0.31 |
| 4 | 14312197 | rs2903643 | -1.193 | 0.384 | 0.103 | -0.092 | 0.012 | -0.058 | -0.058 | 0.086 | 0.8 | 0.75 | 0.83 | 0.98 | 0.72 | 0.8 |
| 4 | 14312437 | rs10488976 | 0.972 | 0.361 | 0.213 | 0.01 | 0.015 | -0.017 | -0.017 | 0 | 0.09 | 0.23 | 0.2 | 0.03 | 0.13 | 0.04 |
| 4 | 14314054 | rs11733030 | 0.853 | 0.296 | 0.38 | 0.099 | 0.005 | -0.004 | -0.004 | 0 | 0.09 | 0.2 | 0.2 | 0.06 | 0.13 | 0.08 |
| 4 | 14317116 | rs6449012 | 0.633 | 0.084 | 0.527 | 0.202 | -0.003 | -0.007 | -0.007 | 0 | 0.57 | 0.54 | 0.59 | 0.6 | 0.56 | 0.48 |
| 4 | 14320086 | rs758877 | 0.535 | 0.045 | 0.779 | 0.555 | -0.016 | 0.024 | 0.024 | 0.134 | 0.65 | 0.54 | 0.51 | 0.53 | 0.64 | 0.54 |
| 4 | 14323343 | rs4698728 | 0.323 | 0.058 | 0.429 | 0.692 | 0.025 | -0.027 | -0.027 | 0.193 | 0.11 | 0.05 | 0.17 | 0.32 | 0.07 | 0.25 |
| 4 | 14324623 | rs10805326 | 0.268 | 0.042 | 0.604 | 0.457 | -0.001 | -0.007 | -0.007 | 0.022 | 0.72 | 0.34 | 0.8 | 0.88 | 0.72 | 0.65 |
| 4 | 14327348 | rs4698729 | -0.674 | 0.07 | 0.659 | 0.411 | -0.009 | 0.02 | 0.02 | 0 | 0.79 | 0.85 | 0.84 | 0.96 | 0.74 | 0.75 |
| 4 | 14331472 | rs17464021 | 1.447 | -0.033 | 0.381 | 0.477 | -0.103 | -0.037 | -0.037 | 1.467 | 0.38 | 0.06 | 0.35 | 0.46 | 0.42 | 0.22 |
| 4 | 14338342 | rs4698732 | 0.907 | 0.338 | 0.355 | 0.729 | 0.013 | -0.031 | -0.031 | 0 | 0.59 | 0.27 | 0.57 | 0.46 | 0.62 | 0.39 |
| 4 | 14340073 | rs4140926 | 0.837 | 0.235 | 0.42 | 0.671 | 0.011 | -0.017 | -0.017 | 0 | 0.61 | 0.35 | 0.63 | 0.54 | 0.64 | 0.4 |
| 4 | 14345605 | rs10022773 | 0.142 | 1.174 | 0.44 | 0.487 | 0.033 | -0.041 | -0.041 | 0 | 0.74 | 0.65 | 0.8 | 0.88 | 0.66 | 0.65 |
| 4 | 14350981 | rs9996349 | -0.661 | 0.974 | 0.477 | 0.82 | -0.004 | -0.012 | -0.012 | 0 | 0.41 | 0.49 | 0.44 | 0.67 | 0.42 | 0.52 |
| 4 | 14355389 | rs2109678 | 0.388 | 0.779 | 0.264 | 0.618 | -0.015 | -0.005 | -0.005 | 0 | 0.4 | 0.48 | 0.37 | 0.36 | 0.26 | 0.32 |
| 4 | 14358260 | rs10805327 | -0.772 | 0.885 | 0.256 | 0.705 | -0.022 | 0.086 | 0.086 | 0 | 0.65 | 0.88 | 0.66 | 0.91 | 0.64 | 0.7 |
| 4 | 14359888 | rs10155152 | 0.854 | 0.735 | 0.365 | 0.818 | -0.027 | -0.032 | -0.032 | 2.32 | 0.36 | 0.51 | 0.24 | 0.25 | 0.25 | 0.24 |
| 4 | 14362571 | rs6816401 | 0.66 | 0.766 | 0.092 | 0.831 | 0.007 | -0.003 | -0.003 | 0 | 0.4 | 0.66 | 0.37 | 0.42 | 0.32 | 0.25 |
| 4 | 14365576 | rs11932589 | 0.006 | 0.467 | 0.075 | 1.208 | -0.017 | -0.089 | -0.089 | 0.161 | 0.28 | 0.08 | 0.24 | 0.46 | 0.34 | 0.19 |
| 4 | 14367070 | rs758878 | -1.328 | 0.343 | 0.124 | 0.784 | 0.005 | 0.122 | 0.122 | 0 | 0.45 | 0.79 | 0.52 | 0.75 | 0.46 | 0.53 |
| 4 | 14368628 | rs4698737 | -0.006 | 0.144 | 0.041 | 0.68 | 0.001 | -0.001 | -0.001 | 0 | 0.09 | 0.07 | 0.13 | 0.11 | 0.05 | 0.26 |
| 4 | 14369076 | rs6449020 | 1.276 | 0.189 | -0.061 | 0.79 | -0.004 | 0.01 | 0.01 | 0.612 | 0.15 | 0.12 | 0.15 | 0.19 | 0.12 | 0.14 |
| 4 | 14370747 | rs16890321 | 0.35 | 0.439 | -0.352 | 0.763 | 0.016 | 0.028 | 0.028 | 0 | 0.17 | 0.25 | 0.15 | 0.34 | 0.24 | 0.11 |
| 4 | 14373371 | rs17466384 | -0.464 | 0.595 | -0.885 | 0.449 | 0.017 | 0.012 | 0.012 | 0 | 0.43 | 0.64 | 0.39 | 0.63 | 0.39 | 0.51 |
| 4 | 14384751 | rs6839912 | -0.745 | 0.996 | -1.336 | 0.132 | -0.026 | 0.078 | 0.078 | 0 | 0.77 | 0.91 | 0.73 | 0.99 | 0.73 | 0.78 |
| 4 | 14385809 | rs1860673 | -0.081 | 0.743 | -1.267 | 0.227 | -0.021 | 0.056 | 0.056 | 2.705 | 0.32 | 0.47 | 0.31 | 0.41 | 0.26 | 0.35 |
| 4 | 14388846 | rs10488980 | 0.782 | 0.71 | -1.369 | 0.333 | -0.001 | 0 | 0 | 0 | 0.13 | 0.12 | 0.12 | 0.21 | 0.12 | 0.14 |
| 4 | 14389355 | rs1013284 | 0.676 | 0.598 | -1.788 | 0.498 | -0.001 | 0.001 | 0.001 | 0.381 | 0.36 | 0.3 | 0.36 | 0.3 | 0.35 | 0.38 |
| 4 | 14391806 | rs16890379 | 1 | 0.696 | -1.918 | 0.626 | -0.001 | 0.007 | 0.007 | 0 | 0.1 | 0.12 | 0.11 | 0.31 | 0.12 | 0.06 |
| 4 | 14392330 | rs2191684 | 0.076 | 0.499 | -1.796 | 0.441 | -0.059 | 0.111 | 0.111 | 0.372 | 0.25 | 0.52 | 0.2 | 0.38 | 0.17 | 0.27 |
| 4 | 14393864 | rs17468022 | -0.177 | 1.019 | -1.598 | 0.879 | -0.004 | -0.003 | -0.003 | 0 | 0.44 | 0.4 | 0.47 | 0.59 | 0.47 | 0.42 |
| 4 | 14394452 | rs7654792 | -0.49 | 0.941 | -1.99 | 0.988 | -0.051 | -0.105 | -0.105 | 0 | 0.34 | 0.15 | 0.46 | 0.43 | 0.44 | 0.34 |
| 4 | 14397827 | rs6449024 | 0.604 | 0.623 | -1.263 | 0.906 | -0.016 | 0.016 | 0.016 | 1.169 | 0.61 | 0.29 | 0.47 | 0.3 | 0.55 | 0.56 |
| 4 | 14403377 | rs2109673 | -0.632 | 0.622 | -1.07 | 0.96 | 0.004 | 0.002 | 0.002 | 0.616 | 0.33 | 0.64 | 0.47 | 0.66 | 0.37 | 0.4 |
| 4 | 14420921 | rs2109674 | -0.207 | 0.898 | -1.323 | 1.152 | -0.068 | -0.026 | -0.026 | 0 | 0.22 | 0.03 | 0.3 | 0.2 | 0.3 | 0.28 |
| 4 | 14428542 | rs218769 | 0.232 | 0.787 | -0.971 | 1.049 | -0.001 | -0.006 | -0.006 | 0.643 | 0.93 | 0.91 | 0.92 | 0.96 | 0.91 | 0.94 |
| 4 | 14433990 | rs7693669 | -0.673 | 0.357 | -1.199 | 0.796 | 0.033 | 0.076 | 0.076 | 0 | 0.5 | 0.2 | 0.38 | 0.37 | 0.42 | 0.53 |
| 4 | 14437458 | rs218775 | 0.644 | 0.066 | -1.342 | 0.611 | 0.012 | 0.029 | 0.029 | 0 | 0.56 | 0.74 | 0.58 | 0.64 | 0.56 | 0.48 |
| 4 | 14438997 | rs6824715 | -1.034 | -0.122 | -0.317 | 0.66 | 0.003 | -0.002 | -0.002 | 0 | 0.45 | 0.58 | 0.47 | 0.46 | 0.41 | 0.57 |
| 4 | 14442252 | rs1875355 | -1.245 | -0.237 | -0.029 | 0.676 | 0 | 0 | 0 | 4.677 | 0.48 | 0.52 | 0.52 | 0.55 | 0.46 | 0.63 |
| 4 | 14442433 | rs6449031 | -0.329 | -0.232 | 0.293 | 0.664 | 0.063 | 0.108 | 0.108 | 0 | 0.06 | 0.26 | 0.12 | 0.14 | 0.05 | 0.13 |
| 4 | 14443769 | rs7700111 | 0.746 | -0.385 | 0.108 | 0.574 | -0.009 | -0.017 | -0.017 | 0 | 0.16 | 0.14 | 0.03 | 0.02 | 0.05 | 0.22 |
| 4 | 14453452 | rs16890501 | 0.675 | -0.189 | 0.018 | 0.757 | -0.036 | -0.046 | -0.046 | 2.802 | 0.2 | 0.07 | 0.31 | 0.28 | 0.28 | 0.19 |
| 4 | 14462605 | rs1501159 | -0.206 | -0.173 | 0.33 | 0.784 | 0.003 | 0.013 | 0.013 | 0 | 0.06 | 0.07 | 0.11 | 0.14 | 0.06 | 0.13 |
| 4 | 14472634 | rs1909917 | 0.64 | -0.222 | 0.482 | 0.738 | -0.001 | 0 | 0 | 0.277 | 0.2 | 0.19 | 0.29 | 0.3 | 0.28 | 0.21 |
| 4 | 14497339 | rs13152405 | -1.335 | -0.261 | 0.755 | 0.685 | 0.019 | 0.044 | 0.044 | 0 | 0.12 | 0.2 | 0.19 | 0.25 | 0.15 | 0.2 |
| 4 | 14500927 | rs13130965 | -0.5 | -0.303 | 0.686 | 0.543 | 0.021 | 0.036 | 0.036 | 0 | 0.3 | 0.38 | 0.48 | 0.53 | 0.41 | 0.35 |
| 4 | 14505552 | rs10939557 | -1.095 | -0.39 | 0.878 | 0.415 | 0 | 0.003 | 0.003 | 0 | 0.46 | 0.63 | 0.52 | 0.55 | 0.46 | 0.6 |
| 4 | 14511712 | rs16890598 | -1.424 | -0.822 | 1.688 | -0.13 | -0.011 | -0.03 | -0.03 | 0 | 0.28 | 0.42 | 0.24 | 0.17 | 0.2 | 0.31 |
| 4 | 14522601 | rs16890604 | -0.175 | -0.822 | 1.527 | -0.102 | -0.002 | -0.001 | -0.001 | 0.069 | 0.07 | 0.06 | 0.13 | 0.09 | 0.07 | 0.09 |
| 4 | 14526801 | rs17351936 | 0.051 | -0.669 | 1.436 | -0.1 | -0.032 | -0.015 | -0.015 | 0 | 0.05 | 0 | 0.07 | 0.07 | 0.11 | 0.08 |
| 4 | 14533433 | rs1553278 | -1.216 | -1.059 | 1.709 | -0.183 | -0.023 | -0.041 | -0.041 | 0.701 | 0.89 | 0.89 | 0.98 | 1 | 0.98 | 0.83 |
| 4 | 14539140 | rs905043 | -0.911 | -1.138 | 2.088 | -0.213 | -0.001 | -0.004 | -0.004 | 2.798 | 0.06 | 0.24 | 0.04 | 0 | 0.02 | 0.01 |
| 4 | 14539436 | rs747978 | -2.141 | -1.196 | 2.034 | -0.239 | -0.003 | -0.007 | -0.007 | 0.013 | 0.38 | 0.59 | 0.35 | 0.28 | 0.32 | 0.36 |
| 4 | 14552520 | rs1501180 | -0.711 | -1.11 | 1.876 | -0.301 | -0.005 | -0.014 | -0.014 | 0 | 0.05 | 0.1 | 0.05 | 0.02 | 0.05 | 0.04 |
| 4 | 14556388 | rs16868209 | -1.671 | -1.125 | 1.997 | -0.317 | 0.005 | -0.007 | -0.007 | 0 | 0.17 | 0.21 | 0.24 | 0.16 | 0.22 | 0.2 |
| 4 | 14557065 | rs6823611 | -1.535 | -1.125 | 1.962 | -0.317 | -0.007 | 0.005 | 0.005 | 0 | 0.17 | 0.13 | 0.23 | 0.16 | 0.22 | 0.2 |
| 4 | 14557480 | rs1875358 | 2.272 | -1.247 | 2.106 | -0.349 | 0.06 | 0.112 | 0.112 | 0 | 0.51 | 0.28 | 0.63 | 0.73 | 0.66 | 0.46 |

| | | | | | | | | | | | | | | | | |
|---|---|---|---|---|---|---|---|---|---|---|---|---|---|---|---|---|
| 4 | 14558941 | rs16890645 | -0.621 | -1.072 | 2.166 | -0.239 | -0.028 | -0.017 | -0.017 | 0 | 0.2 | 0.3 | 0.1 | 0.11 | 0.1 | 0.16 |
| 4 | 14561973 | rs16890647 | -0.632 | -1.063 | 1.929 | -0.237 | -0.004 | -0.002 | -0.002 | 0 | 0.19 | 0.2 | 0.09 | 0.11 | 0.1 | 0.15 |
| 4 | 14562019 | rs12639674 | 1.409 | -0.919 | 1.979 | -0.188 | 0.008 | -0.008 | -0.008 | 0 | 0.83 | 0.81 | 0.76 | 0.84 | 0.78 | 0.79 |
| 4 | 14564502 | rs1392908 | 1.484 | -1.003 | 1.999 | -0.193 | 0.003 | -0.008 | -0.008 | 0 | 0.86 | 0.83 | 0.79 | 0.87 | 0.83 | 0.84 |
| 4 | 14572063 | rs16890684 | -0.998 | -0.98 | 2.112 | -0.182 | 0.029 | -0.001 | -0.001 | 0 | 0.11 | 0.2 | 0.19 | 0.13 | 0.15 | 0.15 |
| 4 | 14574597 | rs10012584 | 0.261 | -1.139 | 2.001 | -0.355 | 0.011 | -0.002 | -0.002 | 0 | 0.13 | 0.09 | 0.04 | 0 | 0.05 | 0.18 |
| 4 | 14577711 | rs10029194 | 1.064 | -1.185 | 2.204 | -0.37 | -0.021 | -0.023 | -0.023 | 0 | 0.94 | 0.8 | 0.96 | 1 | 0.98 | 0.99 |
| 4 | 14578377 | rs16890724 | -1.247 | -1.185 | 2.086 | -0.37 | -0.031 | -0.033 | -0.033 | 0 | 0.06 | 0.16 | 0.04 | 0 | 0.02 | 0.01 |
| 4 | 14582702 | rs16890738 | -1.463 | -1.113 | 1.86 | -0.35 | -0.004 | -0.028 | -0.028 | 0 | 0.24 | 0.27 | 0.23 | 0.13 | 0.2 | 0.33 |
| 4 | 14592876 | rs6847363 | 1.047 | -1.19 | 1.608 | -0.476 | -0.051 | -0.104 | -0.104 | 0 | 0.77 | 0.42 | 0.88 | 1 | 0.88 | 0.78 |
| 4 | 14594724 | rs1501154 | -0.212 | -1.138 | 1.432 | -0.449 | -0.009 | -0.042 | -0.042 | 0 | 0.85 | 0.87 | 0.94 | 1 | 0.9 | 0.79 |
| 4 | 14597832 | rs1501157 | 1.63 | -1.246 | 1.586 | -0.514 | -0.014 | -0.027 | -0.027 | 0.004 | 0.82 | 0.35 | 0.76 | 0.88 | 0.85 | 0.83 |
| 4 | 14598431 | rs1501158 | 2.333 | -1.57 | 1.227 | -0.444 | 0.1 | 0.135 | 0.135 | 0 | 0.53 | 0.23 | 0.65 | 0.76 | 0.68 | 0.48 |
| 4 | 14602461 | rs1501170 | 1.847 | -1.538 | 1.397 | -0.629 | 0.044 | 0.089 | 0.089 | 0 | 0.64 | 0.26 | 0.68 | 0.88 | 0.73 | 0.63 |
| 4 | 14602568 | rs17367774 | -0.769 | -1.369 | 1.234 | -0.526 | -0.013 | 0 | 0 | 0 | 0.1 | 0.05 | 0.17 | 0.12 | 0.13 | 0.17 |
| 4 | 14602883 | rs17367859 | -0.769 | -1.369 | 1.234 | -0.526 | -0.013 | 0 | 0 | 0 | 0.1 | 0.05 | 0.18 | 0.12 | 0.13 | 0.17 |
| 4 | 14609901 | rs6837615 | -0.05 | -1.468 | 1.345 | -0.6 | 0.008 | -0.008 | -0.008 | 0 | 0.83 | 0.88 | 0.93 | 1 | 0.87 | 0.81 |
| 4 | 14614441 | rs12641168 | 0.683 | -1.522 | 1.888 | -0.628 | -0.036 | -0.107 | -0.107 | 0 | 0.79 | 0.69 | 0.89 | 1 | 0.85 | 0.81 |
| 4 | 14617465 | rs9996860 | -0.748 | -1.526 | 2.272 | -0.63 | -0.042 | -0.13 | -0.13 | 1.875 | 0.2 | 0.38 | 0.09 | 0 | 0.13 | 0.2 |
| 4 | 14620966 | rs6852076 | 1.76 | -1.455 | 2.156 | -0.522 | 0.013 | 0.001 | 0.001 | 0 | 0.89 | 0.68 | 0.78 | 0.88 | 0.89 | 0.9 |
| 4 | 14626431 | rs16890883 | -1.765 | -1.493 | 1.903 | -0.537 | -0.029 | -0.086 | -0.086 | 1.165 | 0.32 | 0.38 | 0.26 | 0.12 | 0.27 | 0.37 |
| 4 | 14649304 | rs17371992 | -1.147 | -1.217 | 1.428 | -0.447 | -0.026 | 0.005 | 0.005 | 0 | 0.1 | 0.04 | 0.16 | 0.09 | 0.17 | 0.23 |
| 4 | 14650646 | rs4550921 | 0.636 | -1.476 | 1.716 | -0.57 | -0.089 | -0.164 | -0.164 | 0 | 0.78 | 0.6 | 0.9 | 1 | 0.88 | 0.75 |
| 4 | 14660073 | rs764759 | -1.684 | -1.055 | 1.837 | -0.157 | 0.039 | 0.003 | 0.003 | 0 | 0.12 | 0.23 | 0.2 | 0.1 | 0.21 | 0.22 |
| 4 | 14660437 | rs956518 | 2.29 | -1.713 | 1.889 | -0.519 | 0.074 | 0.132 | 0.132 | 0 | 0.58 | 0.24 | 0.7 | 0.89 | 0.7 | 0.54 |
| 4 | 14671493 | rs10001851 | -1.736 | -1.025 | 1.898 | 0.016 | 0.014 | 0.012 | 0.012 | 0 | 0.12 | 0.23 | 0.1 | 0.11 | 0.17 | 0.21 |
| 4 | 14673461 | rs4698079 | 1.665 | -0.716 | 1.998 | 0.272 | 0.029 | 0.041 | 0.041 | 0.785 | 0.81 | 0.5 | 0.84 | 0.84 | 0.8 | 0.83 |
| 4 | 14678186 | rs3891377 | 1.575 | -0.801 | 0.653 | 0.557 | 0.027 | 0.001 | 0.001 | 0 | 0.51 | 0.27 | 0.65 | 0.49 | 0.54 | 0.4 |
| 4 | 14689222 | rs7679234 | 1.487 | -0.646 | 0.642 | 0.888 | 0.033 | 0.024 | 0.024 | 0.235 | 0.55 | 0.27 | 0.68 | 0.61 | 0.61 | 0.48 |
| 4 | 14691708 | rs6856233 | -0.584 | -0.932 | 1.108 | 0.619 | -0.028 | -0.053 | -0.053 | 0 | 0.2 | 0.26 | 0.09 | 0.04 | 0.12 | 0.22 |
| 4 | 14695432 | rs11932201 | -0.91 | -0.908 | 0.859 | 1.065 | -0.011 | -0.004 | -0.004 | 1.142 | 0.28 | 0.31 | 0.3 | 0.26 | 0.19 | 0.32 |
| 4 | 14699227 | rs4698281 | 0.233 | -0.818 | 1.15 | 1.455 | 0.005 | 0.083 | 0.083 | 0 | 0.82 | 0.62 | 0.89 | 0.68 | 0.82 | 0.79 |
| 4 | 14700515 | rs4327484 | 2.151 | -0.941 | 0.825 | 1.383 | 0.034 | 0.003 | 0.003 | 0.138 | 0.5 | 0.29 | 0.52 | 0.43 | 0.57 | 0.45 |
| 4 | 14706991 | rs4293781 | -1.266 | -0.884 | 0.78 | 1.306 | 0.002 | 0.003 | 0.003 | 0.241 | 0.32 | 0.33 | 0.37 | 0.25 | 0.26 | 0.34 |
| 4 | 14727209 | rs4515159 | 0.72 | -0.635 | 0.77 | 1.322 | 0.009 | 0.019 | 0.019 | 0 | 0.77 | 0.63 | 0.82 | 0.68 | 0.76 | 0.76 |
| 4 | 14733772 | rs10016726 | 1.585 | -0.824 | 0.13 | 0.912 | 0.012 | 0.068 | 0.068 | 0.022 | 0.46 | 0.22 | 0.46 | 0.28 | 0.55 | 0.42 |
| 4 | 14741456 | rs6814769 | -0.549 | -0.961 | 0.258 | 0.927 | -0.005 | 0.042 | 0.042 | 0 | 0.15 | 0.06 | 0.08 | 0.06 | 0.14 | 0.21 |
| 4 | 14743209 | rs11735459 | 2.242 | -0.828 | 0.675 | 1.108 | 0.016 | 0.02 | 0.02 | 1.45 | 0.51 | 0.61 | 0.54 | 0.38 | 0.66 | 0.41 |
| 4 | 14744996 | rs4464561 | 1.145 | -0.81 | 0.611 | 1.061 | -0.053 | 0.034 | 0.034 | 0.371 | 0.68 | 0.59 | 0.67 | 0.46 | 0.83 | 0.65 |
| 4 | 14745825 | rs11724609 | 2.325 | -0.82 | 0.724 | 1.062 | 0.018 | 0.018 | 0.018 | 0.895 | 0.5 | 0.46 | 0.53 | 0.39 | 0.66 | 0.41 |
| 4 | 14751518 | rs4577572 | 1.619 | -1.107 | 1.075 | 1.391 | -0.034 | 0.028 | 0.028 | 0.608 | 0.66 | 0.58 | 0.61 | 0.46 | 0.79 | 0.62 |
| 4 | 14764460 | rs6846158 | -2.005 | -1.161 | 0.929 | 1.16 | -0.012 | 0.036 | 0.036 | 0 | 0.28 | 0.55 | 0.45 | 0.61 | 0.22 | 0.42 |
| 4 | 14765943 | rs7670595 | 2.404 | -1.232 | 0.942 | 1.117 | 0.064 | -0.007 | -0.007 | 0 | 0.61 | 0.22 | 0.51 | 0.39 | 0.75 | 0.55 |
| 4 | 14769762 | rs4568238 | -1.456 | -1.298 | 1.191 | 1.421 | -0.002 | 0.267 | 0.267 | 0 | 0.08 | 0.27 | 0.22 | 0.28 | 0.04 | 0.1 |
| 4 | 14771119 | rs4364258 | -1.381 | -1.308 | 1.199 | 1.427 | 0.001 | 0.017 | 0.017 | 0 | 0.19 | 0.28 | 0.22 | 0.35 | 0.18 | 0.33 |
| 4 | 14771349 | rs4342182 | 2.504 | -1.306 | 1.091 | 1.355 | 0.073 | -0.009 | -0.009 | 0 | 0.6 | 0.19 | 0.5 | 0.37 | 0.74 | 0.53 |
| 4 | 14783158 | rs6829622 | 2.059 | -1.349 | 0.256 | 1.277 | -0.023 | 0.006 | 0.006 | 4.853 | 0.73 | 0.38 | 0.65 | 0.73 | 0.87 | 0.85 |
| 4 | 14783489 | rs6812089 | -1.097 | -1.552 | 0.333 | 1.422 | 0.036 | 0.425 | 0.425 | 2.45 | 0.07 | 0.46 | 0.22 | 0.27 | 0.02 | 0.07 |
| 4 | 14790478 | rs12502866 | 2.102 | -1.56 | 0.151 | 1.41 | 0.12 | 0.004 | 0.004 | 0.045 | 0.6 | 0.18 | 0.56 | 0.35 | 0.78 | 0.55 |
| 4 | 14803794 | rs4298137 | 1.844 | -1.337 | 0.311 | 1.534 | 0.025 | -0.008 | -0.008 | 0.11 | 0.65 | 0.37 | 0.6 | 0.41 | 0.8 | 0.59 |
| 4 | 14804417 | rs13121851 | -0.842 | -0.384 | 0.386 | 1.88 | 0.001 | -0.001 | -0.001 | 0 | 0.18 | 0.17 | 0.16 | 0.36 | 0.15 | 0.32 |
| 4 | 14805560 | rs11723660 | 1.337 | -0.23 | -0.073 | 1.418 | -0.001 | -0.001 | -0.001 | 0.045 | 0.72 | 0.38 | 0.6 | 0.4 | 0.8 | 0.57 |
| 4 | 14806347 | rs10011351 | -0.619 | 0.502 | -0.253 | 0.715 | 0.001 | 0.054 | 0.054 | 1.369 | 0.32 | 0.14 | 0.21 | 0.17 | 0.26 | 0.32 |
| 4 | 14811750 | rs11731529 | -1.448 | 0.905 | -0.114 | 0.913 | -0.008 | -0.006 | -0.006 | 0.39 | 0.53 | 0.76 | 0.5 | 0.47 | 0.48 | 0.55 |
| 4 | 14811962 | rs13145996 | 0.006 | 0.866 | -0.214 | 0.871 | 0.026 | 0.083 | 0.083 | 0.538 | 0.31 | 0.11 | 0.15 | 0.14 | 0.23 | 0.32 |
| 4 | 14812561 | rs6853723 | 0.067 | 0.872 | -0.331 | 0.874 | 0.018 | 0.095 | 0.095 | 0.673 | 0.31 | 0.07 | 0.14 | 0.14 | 0.23 | 0.32 |
| 4 | 14812935 | rs4698295 | -0.566 | 1.03 | 0.13 | 1.02 | 0.066 | 0.166 | 0.166 | 0.277 | 0.15 | 0.34 | 0.28 | 0.33 | 0.22 | 0.18 |
| 4 | 14818982 | rs7658940 | -0.888 | 1.05 | -0.635 | 1.029 | -0.004 | 0.002 | 0.002 | 2.264 | 0.53 | 0.72 | 0.5 | 0.49 | 0.48 | 0.56 |
| 4 | 14832006 | rs4569748 | 0.909 | 1.434 | 0.308 | 1.583 | 0.09 | 0.228 | 0.228 | 4.857 | 0.83 | 0.45 | 0.58 | 0.62 | 0.78 | 0.84 |
| 4 | 14840148 | rs7695275 | -0.98 | 1.246 | -0.915 | 0.736 | -0.079 | 0.067 | 0.067 | 0.259 | 0.6 | 0.91 | 0.73 | 0.72 | 0.5 | 0.63 |
| 4 | 14844405 | rs4235370 | -0.15 | 0.866 | -0.691 | 0.937 | 0.001 | 0.004 | 0.004 | 0 | 0.28 | 0.29 | 0.27 | 0.38 | 0.3 | 0.27 |
| 4 | 14853337 | rs1826705 | -1.227 | 1.434 | -1.068 | 1.1 | -0.067 | 0.033 | 0.033 | 0.575 | 0.65 | 0.93 | 0.77 | 0.75 | 0.61 | 0.72 |
| 4 | 14853491 | rs1316686 | -1.507 | 1.491 | 0.34 | 1.338 | 0.066 | 0.096 | 0.096 | 0.092 | 0.18 | 0.53 | 0.22 | 0.32 | 0.23 | 0.23 |
| 4 | 14860842 | rs16891104 | -1.113 | 0.826 | -0.069 | 0.75 | -0.003 | -0.01 | -0.01 | 0 | 0.21 | 0.26 | 0.25 | 0.12 | 0.21 | 0.25 |
| 4 | 14862873 | rs16891108 | -1.053 | 0.553 | -0.432 | 0.936 | 0.005 | -0.011 | -0.011 | 0 | 0.28 | 0.32 | 0.33 | 0.46 | 0.25 | 0.42 |
| 4 | 14863713 | rs4698299 | -0.465 | 0.613 | -0.47 | 0.987 | 0 | 0.001 | 0.001 | 0 | 0.23 | 0.25 | 0.27 | 0.46 | 0.21 | 0.36 |
| 4 | 14864422 | rs6848922 | -1.301 | 0.814 | -0.154 | 0.839 | 0 | -0.001 | -0.001 | 0 | 0.43 | 0.49 | 0.53 | 0.54 | 0.44 | 0.54 |
| 4 | 14866685 | rs6850085 | -0.774 | 0.966 | -0.353 | 0.87 | -0.011 | -0.005 | -0.005 | 0.164 | 0.17 | 0.14 | 0.12 | 0.2 | 0.18 | 0.16 |
| 4 | 14871034 | rs28597787 | 0.95 | 0.035 | -0.791 | 0.575 | 0.015 | -0.031 | -0.031 | 0 | 0.22 | 0.13 | 0.37 | 0.55 | 0.18 | 0.16 |

| | | | | | | | | | | | | | | | | |
|---|---|---|---|---|---|---|---|---|---|---|---|---|---|---|---|---|
| 4 | 14872639 | rs11726071 | -0.828 | -0.151 | -0.005 | 0.225 | 0.006 | -0.058 | -0.058 | 0.024 | 0.26 | 0.38 | 0.17 | 0.09 | 0.25 | 0.27 |
| 4 | 14873480 | rs11944971 | 0.279 | -0.211 | 0.318 | 0.291 | -0.003 | -0.028 | -0.028 | 0 | 0.09 | 0.2 | 0.06 | 0.03 | 0.08 | 0.12 |
| 4 | 14874104 | rs923983 | 1.142 | -0.161 | 0.226 | 0.388 | 0.015 | -0.03 | -0.03 | 0 | 0.22 | 0.13 | 0.37 | 0.54 | 0.18 | 0.16 |
| 4 | 14874657 | rs12646676 | 0.472 | -0.388 | -0.021 | 0.163 | -0.007 | 0.031 | 0.031 | 0 | 0.06 | 0.09 | 0.06 | 0.15 | 0.03 | 0.02 |
| 4 | 14874745 | rs13138689 | -1.574 | -0.255 | -0.266 | -0.36 | 0.005 | -0.015 | -0.015 | 0 | 0.69 | 0.68 | 0.77 | 0.83 | 0.61 | 0.59 |
| 4 | 14876427 | rs6858687 | -0.769 | -0.109 | -0.251 | -0.537 | -0.004 | 0.02 | 0.02 | 0.782 | 0.4 | 0.37 | 0.33 | 0.14 | 0.4 | 0.39 |
| 4 | 14879196 | rs6848585 | 0.188 | -0.118 | -0.194 | -0.028 | -0.009 | 0.028 | 0.028 | 0 | 0.06 | 0.11 | 0.06 | 0.15 | 0.02 | 0.02 |
| 4 | 14886330 | rs9990522 | -0.28 | -0.244 | -0.062 | -0.172 | 0.009 | 0.01 | 0.01 | 0.291 | 0.53 | 0.61 | 0.45 | 0.37 | 0.43 | 0.37 |
| 4 | 14888787 | rs9884305 | -0.991 | -0.389 | 0.023 | -0.36 | -0.022 | -0.008 | -0.008 | 2.209 | 0.65 | 0.77 | 0.51 | 0.39 | 0.49 | 0.47 |
| 4 | 14892595 | rs1494951 | 0.532 | -0.386 | -0.055 | -0.294 | 0.004 | 0.001 | 0.001 | 0.048 | 0.11 | 0.08 | 0.05 | 0.01 | 0.06 | 0.11 |
| 4 | 14895945 | rs16891135 | 0.775 | -0.211 | 0.024 | -0.104 | 0 | 0.008 | 0.008 | 0 | 0.05 | 0.05 | 0.07 | 0.12 | 0.05 | 0.03 |
| 4 | 14902808 | rs16891148 | 0.831 | -0.209 | 0.014 | -0.103 | 0 | 0.007 | 0.007 | 0 | 0.05 | 0.05 | 0.07 | 0.12 | 0.05 | 0.03 |
| 4 | 14903605 | rs750703 | 0.322 | -0.553 | 0.199 | -0.466 | 0.03 | 0.034 | 0.034 | 0 | 0.42 | 0.3 | 0.33 | 0.25 | 0.33 | 0.33 |
| 4 | 14904847 | rs11931562 | 0.163 | -0.51 | 0.188 | -0.445 | 0.013 | 0.014 | 0.014 | 0 | 0.45 | 0.41 | 0.35 | 0.25 | 0.33 | 0.33 |
| 4 | 14906935 | rs6815830 | 0.29 | -0.02 | 0.295 | -0.51 | 0.001 | -0.049 | -0.049 | 0 | 0.14 | 0.24 | 0.08 | 0.01 | 0.12 | 0.1 |
| 4 | 14911059 | rs7680772 | -1.031 | -0.291 | 0.344 | -0.877 | -0.006 | -0.096 | -0.096 | 4.45 | 0.27 | 0.42 | 0.15 | 0.03 | 0.22 | 0.27 |
| 4 | 14911772 | rs10213507 | 0.327 | 0.28 | 0.16 | -1.368 | 0.063 | -0.038 | -0.038 | 0.042 | 0.22 | 0.3 | 0.32 | 0.13 | 0.35 | 0.17 |
| 4 | 14918627 | rs1994945 | 0.661 | 0.109 | 0.098 | -1.399 | 0.003 | 0.015 | 0.015 | 0 | 0.08 | 0.08 | 0.04 | 0.02 | 0.06 | 0.15 |
| 4 | 14919063 | rs10022899 | -1.478 | 0.088 | 0.076 | -1.41 | 0.01 | 0.006 | 0.006 | 0 | 0.47 | 0.58 | 0.46 | 0.16 | 0.56 | 0.44 |
| 4 | 14921247 | rs13119988 | 0.211 | 0.697 | -0.263 | -1.076 | -0.032 | 0.022 | 0.022 | 1.387 | 0.22 | 0.16 | 0.32 | 0.13 | 0.35 | 0.18 |
| 4 | 14926053 | rs6449092 | -1.345 | -0.01 | -0.467 | -0.815 | 0.027 | 0.028 | 0.028 | 0 | 0.58 | 0.67 | 0.61 | 0.38 | 0.63 | 0.62 |
| 4 | 14926218 | rs6857060 | 0.441 | -0.189 | -0.794 | -0.925 | 0.004 | 0.019 | 0.019 | 0.93 | 0.56 | 0.65 | 0.54 | 0.28 | 0.57 | 0.44 |
| 4 | 14927008 | rs12646389 | -0.804 | 0.004 | -1.039 | -0.753 | 0.053 | 0.094 | 0.094 | 0.601 | 0.58 | 0.83 | 0.69 | 0.71 | 0.71 | 0.66 |
| 4 | 14927179 | rs12640213 | 0.776 | 0.031 | -0.59 | -0.165 | 0.027 | 0.038 | 0.038 | 0 | 0.23 | 0.3 | 0.48 | 0.4 | 0.35 | 0.22 |
| 4 | 14929433 | rs12498517 | 0.952 | -0.051 | -0.493 | -0.143 | -0.008 | -0.008 | -0.008 | 0 | 0.29 | 0.36 | 0.17 | 0.25 | 0.28 | 0.4 |
| 4 | 14929874 | rs7656729 | -0.198 | -0.106 | -0.317 | -0.17 | 0.018 | 0.024 | 0.024 | 0 | 0.08 | 0.34 | 0.06 | 0.06 | 0.07 | 0.03 |
| 4 | 14931835 | rs10031277 | -0.659 | -0.079 | -0.545 | -0.172 | 0.048 | 0.081 | 0.081 | 0.47 | 0.52 | 0.74 | 0.66 | 0.65 | 0.63 | 0.62 |
| 4 | 14934107 | rs11722368 | 0.731 | 0.034 | -0.501 | -0.068 | 0.016 | 0.023 | 0.023 | 0 | 0.22 | 0.25 | 0.48 | 0.39 | 0.35 | 0.22 |
| 4 | 14944898 | rs7685796 | -2.01 | -0.36 | -1.049 | -0.191 | 0.08 | 0.108 | 0.108 | 2.16 | 0.73 | 0.97 | 0.88 | 0.9 | 0.84 | 0.85 |
| 4 | 14948831 | rs1494957 | -0.618 | 0.771 | 0.261 | -0.054 | 0.065 | 0.068 | 0.068 | 0 | 0.3 | 0.51 | 0.52 | 0.42 | 0.4 | 0.36 |
| 4 | 14962496 | rs6449093 | 0.116 | 0.363 | 0.608 | -0.646 | -0.013 | 0.035 | 0.035 | 0.143 | 0.12 | 0.32 | 0.08 | 0.15 | 0.1 | 0.25 |
| 4 | 14973588 | rs11941349 | 0.288 | 0.372 | 0.502 | -0.62 | -0.009 | -0.003 | -0.003 | 0.064 | 0.05 | 0.02 | 0.03 | 0.06 | 0.08 | 0.04 |
| 4 | 14981989 | rs7661252 | -0.336 | 0.315 | 0.757 | -0.599 | 0.015 | 0.126 | 0.126 | 0.108 | 0.05 | 0.26 | 0.1 | 0.15 | 0.02 | 0.21 |
| 4 | 14984117 | rs7673375 | -0.235 | 0.325 | 0.723 | -0.594 | 0.023 | 0.141 | 0.141 | 0.199 | 0.05 | 0.26 | 0.1 | 0.15 | 0.02 | 0.21 |
| 4 | 14990889 | rs13117049 | -0.253 | 0.552 | 0.759 | -0.948 | -0.002 | 0.001 | 0.001 | 0 | 0.32 | 0.33 | 0.6 | 0.73 | 0.52 | 0.26 |
| 4 | 15028262 | rs13140166 | -0.027 | 0.615 | 0.765 | -0.933 | -0.016 | 0.009 | 0.009 | 0 | 0.3 | 0.23 | 0.59 | 0.73 | 0.52 | 0.26 |
| 4 | 15031397 | rs4453931 | 1.597 | 0.474 | 0.768 | -1.051 | 0.002 | 0.052 | 0.052 | 0.564 | 0.85 | 0.59 | 0.84 | 0.77 | 0.86 | 0.6 |
| 4 | 15038411 | rs2604573 | -1.064 | 0.636 | 0.7 | -0.998 | 0.017 | 0.103 | 0.103 | 0.038 | 0.1 | 0.31 | 0.14 | 0.21 | 0.1 | 0.25 |
| 4 | 15045896 | rs2604578 | -1.101 | 0.726 | 0.706 | -0.858 | -0.002 | -0.004 | -0.004 | 0 | 0.38 | 0.42 | 0.5 | 0.38 | 0.59 | 0.41 |
| 4 | 15051670 | rs17450086 | -0.355 | 0.56 | 0.356 | -1.006 | -0.092 | 0.078 | 0.078 | 0.027 | 0.33 | 0.07 | 0.38 | 0.18 | 0.52 | 0.23 |
| 4 | 15067580 | rs3749570 | -0.607 | 1.02 | 0.71 | -0.67 | -0.041 | 0.039 | 0.039 | 0 | 0.33 | 0.12 | 0.4 | 0.23 | 0.57 | 0.2 |
| 4 | 15079533 | rs2604536 | -0.936 | 0.953 | 1.25 | -0.495 | 0.026 | 0.019 | 0.019 | 0 | 0.07 | 0.26 | 0.11 | 0.07 | 0.05 | 0.1 |
| 4 | 15089689 | rs2604547 | -0.737 | 0.796 | 1.202 | -0.436 | 0.038 | 0.176 | 0.176 | 1.113 | 0.06 | 0.33 | 0.11 | 0.15 | 0.02 | 0.21 |
| 4 | 15101916 | rs11729197 | -0.083 | 0.543 | 0.79 | -0.765 | -0.033 | 0.083 | 0.083 | 0 | 0.35 | 0.17 | 0.38 | 0.15 | 0.53 | 0.17 |
| 4 | 15102929 | rs12646901 | -0.585 | 0.536 | 0.87 | -0.694 | 0.017 | 0.055 | 0.055 | 0 | 0.39 | 0.22 | 0.62 | 0.75 | 0.59 | 0.33 |
| 4 | 15104690 | rs7687891 | -1.414 | 0.626 | 1.329 | -0.529 | -0.024 | 0.092 | 0.092 | 3.286 | 0.09 | 0.37 | 0.06 | 0.16 | 0.05 | 0.23 |
| 4 | 15112646 | rs1378343 | -1.834 | 0.48 | 1.144 | -0.731 | -0.04 | 0.063 | 0.063 | 1.988 | 0.16 | 0.57 | 0.16 | 0.22 | 0.12 | 0.29 |
| 4 | 15112737 | rs2034797 | -0.393 | 0.442 | 0.846 | -0.826 | 0.009 | 0.044 | 0.044 | 0 | 0.37 | 0.21 | 0.62 | 0.77 | 0.57 | 0.37 |
| 4 | 15112878 | rs2034796 | -0.526 | 0.531 | 0.895 | -0.901 | 0.007 | 0.034 | 0.034 | 0 | 0.39 | 0.25 | 0.62 | 0.73 | 0.57 | 0.37 |
| 4 | 15117192 | rs1378342 | -0.725 | 0.771 | 0.825 | -0.731 | 0.028 | 0.241 | 0.241 | 0 | 0.06 | 0.19 | 0.25 | 0.56 | 0.06 | 0.19 |
| 4 | 15118979 | rs1378341 | -1.643 | 0.665 | 0.83 | -0.784 | -0.036 | 0.07 | 0.07 | 2 | 0.16 | 0.54 | 0.15 | 0.26 | 0.12 | 0.29 |
| 4 | 15120374 | rs2702546 | -0.782 | 0.851 | 0.646 | -0.693 | 0.051 | 0.31 | 0.31 | 0 | 0.06 | 0.23 | 0.25 | 0.56 | 0.06 | 0.19 |
| 4 | 15127630 | rs11721627 | -1.009 | 0.783 | 0.472 | -0.81 | -0.001 | 0.003 | 0.003 | 0 | 0.44 | 0.57 | 0.46 | 0.4 | 0.57 | 0.38 |
| 4 | 15128633 | rs1455207 | -1.718 | 0.782 | -0.049 | -0.798 | 0.068 | 0.015 | 0.015 | 0.687 | 0.54 | 0.88 | 0.57 | 0.43 | 0.66 | 0.63 |
| 4 | 15131114 | rs16891402 | -0.118 | 1.257 | 0.231 | -0.631 | -0.019 | 0.036 | 0.036 | 0 | 0.12 | 0.22 | 0.08 | 0.24 | 0.12 | 0.21 |
| 4 | 15134596 | rs13136309 | 0.286 | 1.774 | -0.541 | -0.713 | -0.125 | 0.15 | 0.15 | 0 | 0.25 | 0.02 | 0.28 | 0.01 | 0.39 | 0.11 |
| 4 | 15143489 | rs17452586 | -0.221 | 1.445 | -0.197 | -0.455 | -0.023 | 0.02 | 0.02 | 0 | 0.33 | 0.28 | 0.34 | 0.18 | 0.46 | 0.24 |
| 4 | 15145180 | rs2604562 | -0.05 | 1.008 | -0.409 | -0.205 | -0.016 | 0.081 | 0.081 | 0 | 0.86 | 0.66 | 0.91 | 0.78 | 0.87 | 0.78 |
| 4 | 15146875 | rs13146533 | 0.889 | 0.91 | -1.179 | -0.367 | 0.058 | -0.037 | -0.037 | 0.033 | 0.4 | 0.12 | 0.44 | 0.56 | 0.29 | 0.27 |
| 4 | 15147403 | rs2702534 | -0.676 | 1.598 | -0.318 | -0.746 | 0.014 | -0.096 | -0.096 | 0 | 0.29 | 0.35 | 0.36 | 0.01 | 0.36 | 0.19 |
| 4 | 15148917 | rs16891473 | -0.707 | 1.262 | -0.37 | -0.256 | 0.003 | -0.01 | -0.01 | 0.102 | 0.08 | 0.11 | 0.05 | 0.04 | 0.08 | 0.08 |
| 4 | 15149977 | rs16891486 | -0.46 | 1.033 | -0.414 | -0.182 | 0.004 | -0.001 | -0.001 | 0.338 | 0.08 | 0.06 | 0.05 | 0.11 | 0.05 | 0.06 |
| 4 | 15151976 | rs930830 | -0.382 | 0.871 | -0.271 | -0.384 | -0.002 | -0.017 | -0.017 | 0 | 0.32 | 0.44 | 0.44 | 0.18 | 0.29 | 0.34 |
| 4 | 15154795 | rs1455202 | -1.664 | 1.454 | 0.064 | -0.321 | 0.006 | -0.01 | -0.01 | 0 | 0.27 | 0.37 | 0.31 | 0.81 | 0.3 | 0.38 |
| 4 | 15167548 | rs1817186 | -0.229 | 1.449 | -0.131 | 0.409 | -0.001 | 0.052 | 0.052 | 0 | 0.1 | 0.11 | 0.18 | 0.6 | 0.07 | 0.1 |
| 4 | 15175036 | rs974179 | -0.099 | 1.559 | -0.115 | 0.399 | -0.007 | -0.001 | -0.001 | 0.182 | 0.06 | 0.05 | 0.06 | 0.17 | 0.11 | 0.15 |
| 4 | 15175539 | rs10155296 | -1.467 | 1.505 | -0.014 | 0.298 | -0.005 | 0.013 | 0.013 | 0 | 0.18 | 0.27 | 0.26 | 0.77 | 0.19 | 0.25 |
| 4 | 15181885 | rs10014445 | -1.566 | 1.612 | -0.098 | 0.097 | 0.001 | -0.018 | -0.018 | 0 | 0.28 | 0.33 | 0.33 | 0.87 | 0.33 | 0.36 |
| 4 | 15184903 | rs10021046 | -0.041 | 1.362 | 0.02 | 0.339 | 0.002 | 0.011 | 0.011 | 0.036 | 0.14 | 0.08 | 0.33 | 0.08 | 0.13 | 0.21 |

| | | | | | | | | | | | | | | | | |
|---|---|---|---|---|---|---|---|---|---|---|---|---|---|---|---|---|
| 4 | 15193122 | rs10805344 | -0.862 | 1.33 | 0.573 | 0.89 | 0.053 | 0.337 | 0.337 | 0 | 0.13 | 0.39 | 0.22 | 0.54 | 0.14 | 0.17 |
| 4 | 15197936 | rs10008462 | 0.158 | 1.131 | -0.012 | 0.746 | 0.005 | -0.002 | -0.002 | 2.421 | 0.08 | 0.06 | 0.05 | 0.12 | 0.06 | 0.05 |
| 4 | 15200699 | rs10516285 | 0.218 | 1.234 | -0.116 | 0.714 | -0.038 | -0.026 | -0.026 | 2.074 | 0.11 | 0.02 | 0.07 | 0.1 | 0.15 | 0.11 |
| 4 | 15204833 | rs4309834 | -1.74 | 0.877 | 0.131 | 0.8 | 0 | 0.023 | 0.023 | 0 | 0.36 | 0.71 | 0.35 | 0.77 | 0.34 | 0.5 |
| 4 | 15206338 | rs10516284 | -1.124 | 0.888 | 0.367 | 0.968 | -0.011 | 0.05 | 0.05 | 0 | 0.22 | 0.31 | 0.26 | 0.55 | 0.18 | 0.2 |
| 4 | 15210204 | rs1869696 | -1.179 | 0.886 | 0.389 | 0.967 | -0.013 | 0.052 | 0.052 | 0 | 0.23 | 0.32 | 0.26 | 0.55 | 0.18 | 0.2 |
| 4 | 15210390 | rs16891600 | -0.231 | 0.943 | 0.35 | 1.086 | 0.005 | 0.234 | 0.234 | 0 | 0.06 | 0.14 | 0.19 | 0.48 | 0.07 | 0.07 |
| 4 | 15219683 | rs4698323 | -0.217 | 0.845 | 0.019 | 0.666 | 0.006 | -0.004 | -0.004 | 0 | 0.09 | 0.06 | 0.35 | 0.1 | 0.06 | 0.15 |
| 4 | 15219710 | rs9991734 | -0.555 | 0.979 | 0.182 | 0.687 | -0.005 | -0.012 | -0.012 | 0 | 0.17 | 0.18 | 0.07 | 0.06 | 0.1 | 0.14 |
| 4 | 15223639 | rs6449117 | 0.793 | 1.394 | 0.283 | 1.082 | 0.053 | 0.069 | 0.069 | 0 | 0.86 | 0.64 | 0.9 | 0.77 | 0.84 | 0.67 |
| 4 | 15233866 | rs4074082 | -0.947 | 0.424 | -0.053 | 1.366 | -0.034 | 0 | 0 | 0.488 | 0.15 | 0.21 | 0.29 | 0.13 | 0.07 | 0.13 |
| 4 | 15239767 | rs6813890 | -0.867 | 0.279 | 0.687 | 1.372 | -0.033 | 0.179 | 0.179 | 0.018 | 0.16 | 0.32 | 0.08 | 0.57 | 0.1 | 0.08 |
| 4 | 15243220 | rs6811795 | 1.548 | 0.414 | 0.142 | 0.493 | 0 | -0.018 | -0.018 | 0.35 | 0.48 | 0.49 | 0.52 | 0.13 | 0.41 | 0.35 |
| 4 | 15246897 | rs10011591 | 0.984 | 0.95 | 0.123 | 0.694 | 0.036 | -0.005 | -0.005 | 0 | 0.68 | 0.57 | 0.63 | 0.29 | 0.59 | 0.65 |
| 4 | 15252649 | rs12646586 | -1.466 | 0.558 | -0.124 | 1.02 | 0.024 | -0.003 | -0.003 | 0 | 0.36 | 0.43 | 0.39 | 0.65 | 0.45 | 0.38 |
| 4 | 15254268 | rs13144510 | -1.76 | 0.576 | -0.091 | 0.874 | -0.001 | -0.026 | -0.026 | 0 | 0.44 | 0.48 | 0.45 | 0.74 | 0.53 | 0.52 |
| 4 | 15254870 | rs4698099 | -2.459 | 0.463 | -0.297 | 0.747 | 0 | 0.141 | 0.141 | 0.12 | 0.58 | 0.87 | 0.51 | 0.8 | 0.6 | 0.68 |
| 4 | 15255882 | rs13143217 | -2.459 | 0.463 | -0.309 | 0.747 | -0.002 | 0.141 | 0.141 | 0.12 | 0.58 | 0.87 | 0.51 | 0.81 | 0.6 | 0.68 |
| 4 | 15258995 | rs12641060 | -0.276 | 1.183 | -0.49 | 1.329 | -0.004 | -0.112 | -0.112 | 0 | 0.22 | 0.06 | 0.1 | 0.54 | 0.17 | 0.2 |
| 4 | 15295105 | rs35315148 | -0.669 | 1.754 | 0.056 | 2.069 | 0.007 | 0.006 | 0.006 | 0 | 0.16 | 0.16 | 0.32 | 0.31 | 0.34 | 0.26 |
| 4 | 15315490 | rs16891689 | -0.728 | 0.59 | 0.039 | 1.463 | 0.009 | 0.002 | 0.002 | 0 | 0.11 | 0.09 | 0.03 | 0.07 | 0.05 | 0.16 |
| 4 | 15336287 | rs16891741 | -1.268 | 0.682 | 0.185 | 1.511 | -0.024 | 0.112 | 0.112 | 0.2 | 0.13 | 0.34 | 0.3 | 0.26 | 0.08 | 0.25 |
| 4 | 15353889 | rs13139715 | -1.03 | 1.403 | -0.551 | 1.19 | -0.007 | -0.024 | -0.024 | 0 | 0.34 | 0.73 | 0.57 | 0.48 | 0.36 | 0.46 |
| 4 | 15369280 | rs16891785 | -0.225 | 1.238 | -1.297 | 0.932 | -0.001 | -0.068 | -0.068 | 0.512 | 0.18 | 0.07 | 0.13 | 0.41 | 0.16 | 0.09 |
| 4 | 15375984 | rs4303974 | -0.008 | 1.494 | -1.323 | 1.165 | -0.031 | -0.031 | -0.031 | 0 | 0.19 | 0.11 | 0.28 | 0.28 | 0.27 | 0.25 |
| 4 | 15380517 | rs6449134 | -1.581 | 1.087 | -1.382 | 0.552 | -0.008 | 0.206 | 0.206 | 0 | 0.53 | 0.83 | 0.69 | 0.9 | 0.53 | 0.55 |
| 4 | 15409103 | rs6449138 | -0.419 | 1.102 | -0.836 | 1.101 | -0.003 | -0.005 | -0.005 | 0.328 | 0.3 | 0.3 | 0.49 | 0.52 | 0.37 | 0.37 |
| 4 | 15420679 | rs2192355 | -0.076 | 1.022 | -0.795 | 1.033 | -0.048 | -0.063 | -0.063 | 0 | 0.26 | 0.08 | 0.25 | 0.36 | 0.33 | 0.28 |
| 4 | 15424591 | rs2192356 | 0.646 | 0.315 | -1.273 | 0.471 | 0.044 | -0.049 | -0.049 | 1.901 | 0.59 | 0.15 | 0.45 | 0.44 | 0.67 | 0.6 |
| 4 | 15430626 | rs13116208 | 0.709 | 0.257 | -1.199 | 0.445 | 0.048 | -0.064 | -0.064 | 0.047 | 0.58 | 0.13 | 0.46 | 0.43 | 0.67 | 0.6 |
| 4 | 15438851 | rs1029973 | 0.51 | -0.065 | -1.301 | 0.421 | 0.022 | -0.072 | -0.072 | 0 | 0.18 | 0.04 | 0.12 | 0.32 | 0.13 | 0.09 |
| 4 | 15441980 | rs17469634 | 2.25 | -0.056 | -1.435 | 0.242 | -0.094 | 0.122 | 0.122 | 0 | 0.31 | 0.02 | 0.18 | 0.08 | 0.34 | 0.32 |
| 4 | 15445533 | rs2314590 | 1.925 | -0.021 | -1.405 | 0.236 | -0.019 | 0.127 | 0.127 | 0 | 0.33 | 0.08 | 0.2 | 0.08 | 0.34 | 0.32 |
| 4 | 15453870 | rs2160046 | -0.038 | -0.094 | -1.151 | 0.421 | -0.014 | -0.019 | -0.019 | 0.15 | 0.31 | 0.3 | 0.49 | 0.52 | 0.39 | 0.36 |
| 4 | 15460334 | rs722235 | 0.221 | -0.241 | -0.637 | 0.291 | -0.108 | -0.105 | -0.105 | 0.211 | 0.32 | 0.04 | 0.28 | 0.37 | 0.39 | 0.29 |
| 4 | 15460758 | rs13123050 | 0.155 | 0.243 | -0.704 | 0.083 | -0.004 | 0.044 | 0.044 | 0 | 0.29 | 0.46 | 0.27 | 0.37 | 0.29 | 0.28 |
| 4 | 15467035 | rs719993 | 0.551 | -0.24 | -0.95 | -0.076 | -0.01 | 0.009 | 0.009 | 0 | 0.72 | 0.68 | 0.79 | 0.61 | 0.81 | 0.7 |
| 4 | 15474996 | rs11728800 | 2.1 | -0.265 | -0.993 | -0.148 | -0.114 | 0.119 | 0.119 | 0.417 | 0.31 | 0.02 | 0.24 | 0.08 | 0.37 | 0.28 |
| 4 | 15480225 | rs9993580 | -0.638 | -0.069 | -0.803 | -0.088 | -0.007 | -0.05 | -0.05 | 0 | 0.84 | 0.81 | 0.84 | 0.97 | 0.87 | 0.76 |
| 4 | 15481976 | rs1861048 | 0.131 | -0.124 | -0.713 | 0.195 | 0.027 | 0.018 | 0.018 | 1.369 | 0.5 | 0.72 | 0.54 | 0.57 | 0.45 | 0.41 |
| 4 | 15482322 | rs1861049 | -0.582 | 0.177 | -0.452 | 0.031 | 0.068 | 0.213 | 0.213 | 1.613 | 0.07 | 0.49 | 0.25 | 0.17 | 0.04 | 0.12 |
| 4 | 15487432 | rs4698112 | 0.747 | 0.317 | -0.922 | 0.091 | 0.001 | 0.016 | 0.016 | 0.164 | 0.41 | 0.62 | 0.51 | 0.24 | 0.42 | 0.4 |
| 4 | 15493755 | rs4698390 | -0.796 | 0.405 | -1.154 | 0.65 | -0.008 | 0 | 0 | 0 | 0.1 | 0.06 | 0.32 | 0.33 | 0.13 | 0.16 |
| 4 | 15496310 | rs6811203 | -1.305 | 0.191 | -0.882 | 0.618 | 0.004 | 0.046 | 0.046 | 0 | 0.2 | 0.28 | 0.41 | 0.38 | 0.26 | 0.27 |
| 4 | 15499251 | rs2160043 | -1.28 | -0.017 | -0.986 | 0.408 | 0 | 0 | 0 | 0 | 0.47 | 0.52 | 0.54 | 0.52 | 0.48 | 0.52 |
| 4 | 15501503 | rs7682982 | 1.346 | -0.233 | -0.898 | 0.637 | 0.016 | 0.005 | 0.005 | 0.3 | 0.08 | 0.04 | 0.03 | 0.35 | 0.02 | 0.02 |
| 4 | 15507032 | rs1154578 | -2.12 | -0.348 | -0.86 | 0.371 | -0.011 | -0.017 | -0.017 | 0 | 0.38 | 0.57 | 0.48 | 0.75 | 0.34 | 0.39 |
| 4 | 15509291 | rs767092 | -0.42 | -0.296 | -0.818 | 0.475 | -0.008 | 0.01 | 0.01 | 0 | 0.09 | 0.08 | 0.33 | 0.34 | 0.14 | 0.18 |
| 4 | 15515791 | rs7686668 | -2.016 | -0.341 | -0.524 | 0.457 | -0.003 | -0.003 | -0.003 | 0 | 0.37 | 0.61 | 0.45 | 0.73 | 0.31 | 0.38 |
| 4 | 15516389 | rs2286976 | 1.434 | -0.261 | -0.651 | 0.567 | 0.016 | 0.004 | 0.004 | 0.106 | 0.08 | 0.04 | 0.03 | 0.35 | 0.03 | 0.02 |
| 4 | 15522209 | rs16892107 | -0.613 | -0.11 | -0.596 | 0.481 | 0.002 | 0.035 | 0.035 | 0 | 0.11 | 0.13 | 0.35 | 0.36 | 0.15 | 0.18 |
| 4 | 15522282 | rs13131119 | 0.563 | -0.112 | -0.488 | 0.193 | 0.006 | -0.038 | -0.038 | 0 | 0.1 | 0.19 | 0.07 | 0.04 | 0.12 | 0.11 |
| 4 | 15527808 | rs7675761 | 1.997 | -0.126 | -0.816 | 0.062 | -0.032 | 0.101 | 0.101 | 0.026 | 0.32 | 0.14 | 0.27 | 0.07 | 0.38 | 0.25 |
| 4 | 15539498 | rs1861044 | 0.859 | -0.202 | -0.636 | 0.254 | 0 | 0.001 | 0.001 | 0 | 0.48 | 0.47 | 0.59 | 0.34 | 0.56 | 0.51 |
| 4 | 15541609 | rs6812708 | -0.743 | -0.157 | -0.64 | 0.212 | -0.001 | 0.001 | 0.001 | 0.58 | 0.71 | 0.72 | 0.74 | 0.8 | 0.73 | 0.67 |
| 4 | 15545410 | rs9918079 | 0.859 | 0.162 | -0.512 | 0.183 | 0.001 | 0.024 | 0.024 | 0 | 0.28 | 0.19 | 0.12 | 0.12 | 0.22 | 0.22 |
| 4 | 15548415 | rs12508186 | 1.869 | 0.453 | -0.76 | -0.02 | -0.004 | 0.039 | 0.039 | 0 | 0.22 | 0.11 | 0.19 | 0.07 | 0.25 | 0.18 |
| 4 | 15548716 | rs7697314 | 0.353 | 0.415 | -0.414 | 0.201 | 0.001 | -0.007 | -0.007 | 0 | 0.39 | 0.36 | 0.39 | 0.51 | 0.43 | 0.42 |
| 4 | 15549778 | rs16892141 | 1.335 | 0.41 | -0.378 | 0.247 | -0.007 | 0.082 | 0.082 | 0 | 0.08 | 0.09 | 0.03 | 0.4 | 0.02 | 0.1 |
| 4 | 15552295 | rs2041672 | 1.737 | 0.519 | -0.382 | -0.089 | 0.01 | -0.022 | -0.022 | 0 | 0.08 | 0.12 | 0.09 | 0.03 | 0.16 | 0.14 |
| 4 | 15557442 | rs13144428 | 1.737 | 0.498 | -0.395 | -0.089 | 0.009 | -0.023 | -0.023 | 0 | 0.08 | 0.13 | 0.09 | 0.03 | 0.16 | 0.14 |
| 4 | 15565758 | rs11932730 | -1.237 | 0.396 | -0.202 | -0.077 | -0.017 | 0.044 | 0.044 | 0 | 0.68 | 0.6 | 0.69 | 0.53 | 0.73 | 0.73 |
| 4 | 15569146 | rs13116304 | -1.32 | 0.36 | -0.2 | -0.089 | -0.014 | 0.044 | 0.044 | 0 | 0.69 | 0.61 | 0.77 | 0.53 | 0.73 | 0.73 |
| 4 | 15577587 | rs12649803 | -0.549 | 0.401 | -0.45 | -0.097 | -0.003 | 0.012 | 0.012 | 0 | 0.61 | 0.47 | 0.68 | 0.49 | 0.57 | 0.58 |
| 4 | 15580754 | rs12500587 | 0.686 | 0.371 | -0.521 | -0.233 | 0.012 | 0.063 | 0.063 | 1.308 | 0.42 | 0.29 | 0.29 | 0.19 | 0.34 | 0.38 |
| 4 | 15586225 | rs4267734 | -1.62 | 0.378 | -0.478 | -0.156 | 0.057 | -0.032 | -0.032 | 0 | 0.9 | 0.68 | 0.9 | 0.96 | 0.84 | 0.86 |
| 4 | 15594898 | rs11728037 | 2.073 | 0.234 | -0.678 | -0.232 | -0.061 | 0.045 | 0.045 | 0 | 0.27 | 0.01 | 0.14 | 0.13 | 0.23 | 0.22 |
| 4 | 15600799 | rs4270584 | 0.631 | 0.419 | -0.065 | 0.14 | 0.004 | 0.008 | 0.008 | 0.663 | 0.45 | 0.4 | 0.34 | 0.39 | 0.37 | 0.49 |
| 4 | 15606431 | rs10214 | 0.637 | 0.39 | -0.043 | 0.148 | 0.004 | 0.006 | 0.006 | 0.755 | 0.45 | 0.42 | 0.34 | 0.39 | 0.36 | 0.49 |

| | | | | | | | | | | | | | | | | |
|---|---|---|---|---|---|---|---|---|---|---|---|---|---|---|---|---|
| 4 | 15623955 | rs10008376 | -0.883 | 0.384 | -0.072 | 0.145 | 0.004 | 0.007 | 0.007 | 0.767 | 0.56 | 0.6 | 0.66 | 0.61 | 0.64 | 0.51 |
| 4 | 15624571 | rs10008975 | -0.875 | 0.407 | -0.062 | 0.156 | 0.005 | 0.007 | 0.007 | 0.788 | 0.55 | 0.6 | 0.66 | 0.61 | 0.64 | 0.51 |
| 4 | 15624770 | rs7438356 | -0.819 | 0.353 | -0.076 | 0.13 | 0.006 | 0.008 | 0.008 | 1.723 | 0.54 | 0.59 | 0.66 | 0.61 | 0.64 | 0.51 |
| 4 | 15626475 | rs6414764 | 1.066 | 0.477 | -0.061 | 0.14 | 0.031 | -0.028 | -0.028 | 0 | 0.18 | 0.32 | 0.18 | 0.09 | 0.25 | 0.19 |
| 4 | 15630737 | rs6826058 | 1.066 | 0.477 | -0.061 | 0.14 | 0.031 | -0.028 | -0.028 | 0.015 | 0.18 | 0.32 | 0.18 | 0.09 | 0.25 | 0.19 |
| 4 | 15639171 | rs6827406 | -0.267 | 0.465 | -0.087 | 0.108 | 0.031 | -0.049 | -0.049 | 0 | 0.26 | 0.38 | 0.3 | 0.12 | 0.37 | 0.24 |
| 4 | 15646794 | rs13133213 | -0.006 | 0.594 | 0.018 | 0.657 | 0.003 | 0.001 | 0.001 | 0.066 | 0.4 | 0.4 | 0.5 | 0.35 | 0.51 | 0.37 |
| 4 | 15658481 | rs10939630 | 0.929 | 0.528 | 0.051 | 0.647 | -0.017 | 0.027 | 0.027 | 0 | 0.09 | 0.13 | 0.04 | 0.23 | 0.02 | 0.09 |
| 4 | 15659649 | rs9994518 | -0.235 | 0.637 | 0.072 | 0.505 | 0.031 | -0.037 | -0.037 | 0.004 | 0.24 | 0.33 | 0.3 | 0.12 | 0.37 | 0.23 |
| 4 | 15684234 | rs12640192 | 0.734 | 0.446 | 0.008 | 0.529 | 0.012 | -0.039 | -0.039 | 0 | 0.31 | 0.22 | 0.36 | 0.47 | 0.27 | 0.29 |
| 4 | 15692058 | rs6815254 | 0.669 | 0.667 | 0.069 | 0.514 | 0.051 | -0.03 | -0.03 | 0.042 | 0.18 | 0.32 | 0.18 | 0.09 | 0.29 | 0.19 |
| 4 | 15692954 | rs6449161 | -0.921 | 0.617 | 0.068 | 0.52 | 0.016 | 0.012 | 0.012 | 0.891 | 0.56 | 0.62 | 0.67 | 0.61 | 0.68 | 0.52 |
| 4 | 15699005 | rs4583752 | 0.555 | 0.382 | -0.141 | 0.7 | 0.011 | -0.04 | -0.04 | 1.047 | 0.31 | 0.22 | 0.36 | 0.47 | 0.28 | 0.29 |
| 4 | 15705039 | rs2302467 | -0.478 | 0.417 | -0.032 | 0.647 | 0.006 | 0.046 | 0.046 | 0 | 0.44 | 0.31 | 0.54 | 0.73 | 0.45 | 0.51 |
| 4 | 15709192 | rs2302465 | 0.004 | 0.411 | 0.738 | 1.161 | -0.035 | 0.021 | 0.021 | 0 | 0.06 | 0.01 | 0.12 | 0.05 | 0.12 | 0.03 |
| 4 | 15709252 | rs2302464 | 0.448 | 0.391 | 0.766 | 1.369 | 0.015 | -0.019 | -0.019 | 2.505 | 0.07 | 0.03 | 0.03 | 0.22 | 0.02 | 0.1 |
| 4 | 15712787 | rs4541502 | 1.957 | 0.364 | 1.006 | 1.146 | -0.005 | -0.006 | -0.006 | 0 | 0.58 | 0.39 | 0.59 | 0.44 | 0.53 | 0.57 |
| 4 | 15713003 | rs16892260 | -1.455 | 0.386 | 0.838 | 1.196 | -0.001 | 0.043 | 0.043 | 0 | 0.17 | 0.25 | 0.21 | 0.48 | 0.15 | 0.22 |
| 4 | 15717321 | rs3213710 | 1.959 | 0.175 | 0.628 | 1.082 | -0.024 | -0.02 | -0.02 | 0.002 | 0.56 | 0.23 | 0.57 | 0.43 | 0.53 | 0.56 |
| 4 | 15722573 | rs10001565 | 0.156 | -0.003 | 1.427 | 0.783 | -0.008 | -0.016 | -0.016 | 0 | 0.14 | 0.41 | 0.11 | 0.05 | 0.1 | 0.05 |
| 4 | 15723526 | rs9942275 | -0.555 | 0.097 | 1.448 | 1.129 | -0.024 | 0.176 | 0.176 | 0 | 0.1 | 0.33 | 0.08 | 0.24 | 0.03 | 0.1 |
| 4 | 15724443 | rs7688526 | 0.087 | 0.161 | 1.297 | 1.155 | -0.017 | 0.112 | 0.112 | 0 | 0.07 | 0.16 | 0.06 | 0.23 | 0.02 | 0.09 |
| 4 | 15725766 | rs11931532 | -0.981 | 0.068 | 1.276 | 1.143 | -0.045 | 0.224 | 0.224 | 0 | 0.12 | 0.3 | 0.12 | 0.5 | 0.03 | 0.18 |
| 4 | 15728896 | rs4588454 | -0.369 | 0.239 | 1.023 | 1.057 | 0.041 | -0.014 | -0.014 | 2.413 | 0.06 | 0.16 | 0.12 | 0.06 | 0.12 | 0.03 |
| 4 | 15733454 | rs1058212 | -0.631 | 0.128 | 0.849 | 1.109 | 0.001 | 0.043 | 0.043 | 0 | 0.11 | 0.12 | 0.1 | 0.5 | 0.03 | 0.18 |
| 4 | 15737348 | rs4698412 | 2.254 | 0.152 | 0.514 | 0.85 | -0.039 | -0.003 | -0.003 | 0 | 0.57 | 0.1 | 0.55 | 0.38 | 0.56 | 0.59 |
| 4 | 15737937 | rs4538475 | -2.079 | 0.087 | 0.94 | 0.925 | -0.019 | 0.114 | 0.114 | 0 | 0.2 | 0.35 | 0.23 | 0.56 | 0.16 | 0.21 |
| 4 | 15738253 | rs11934811 | -1.203 | 0.1 | 0.975 | 1.043 | 0.007 | 0.062 | 0.062 | 0 | 0.1 | 0.19 | 0.17 | 0.32 | 0.14 | 0.11 |
| 4 | 15738370 | rs6835705 | -1.22 | 0.073 | 0.772 | 0.908 | -0.032 | 0.093 | 0.093 | 0 | 0.14 | 0.19 | 0.11 | 0.5 | 0.03 | 0.18 |
| 4 | 15738795 | rs4698119 | 0.898 | 0.221 | 0.8 | 0.863 | -0.001 | -0.019 | -0.019 | 0 | 0.69 | 0.77 | 0.72 | 0.44 | 0.7 | 0.62 |
| 4 | 15738973 | rs4698414 | -0.697 | 0.46 | 0.923 | 0.622 | 0.017 | -0.036 | -0.036 | 0 | 0.9 | 0.91 | 0.89 | 0.99 | 0.81 | 0.85 |
| 4 | 15739498 | rs997250 | -0.448 | 0.389 | 1.143 | 0.758 | 0.024 | -0.028 | -0.028 | 0 | 0.17 | 0.36 | 0.17 | 0.1 | 0.21 | 0.08 |
| 4 | 15740292 | rs3815466 | -1.231 | 0.298 | 1.124 | 0.936 | 0.002 | 0.147 | 0.147 | 0 | 0.12 | 0.39 | 0.13 | 0.32 | 0.12 | 0.11 |
| 4 | 15742716 | rs1110258 | 1.258 | 0.283 | 0.031 | 0.354 | -0.051 | -0.077 | -0.077 | 0.003 | 0.23 | 0.02 | 0.36 | 0.22 | 0.24 | 0.23 |
| 4 | 15742986 | rs6852450 | 1.488 | 0.528 | 0.606 | 0.518 | -0.013 | 0.001 | 0.001 | 0 | 0.6 | 0.23 | 0.61 | 0.37 | 0.58 | 0.57 |
| 4 | 15743391 | rs4698415 | -0.277 | 0.593 | 0.967 | 0.501 | 0.067 | -0.005 | -0.005 | 3.927 | 0.87 | 0.67 | 0.85 | 0.93 | 0.8 | 0.81 |
| 4 | 15743573 | rs10019008 | -0.252 | 0.264 | 1.004 | 0.845 | -0.005 | 0.07 | 0.07 | 0 | 0.2 | 0.38 | 0.19 | 0.39 | 0.19 | 0.13 |
| 4 | 15744528 | rs11933202 | -1.191 | 0.281 | 0.849 | 0.797 | -0.025 | 0.101 | 0.101 | 0 | 0.27 | 0.51 | 0.25 | 0.54 | 0.21 | 0.24 |
| 4 | 15753687 | rs6837105 | 0.115 | 0.048 | 0.086 | 0.616 | -0.004 | 0.021 | 0.021 | 0 | 0.07 | 0.08 | 0.06 | 0.15 | 0.03 | 0.08 |
| 4 | 15755905 | rs868763 | -1.047 | -0.03 | -0.052 | 0.418 | 0.008 | -0.008 | -0.008 | 0 | 0.3 | 0.69 | 0.27 | 0.6 | 0.2 | 0.23 |
| 4 | 15758931 | rs12645519 | 0.372 | 0.614 | -0.207 | -0.328 | -0.004 | 0.005 | 0.005 | 0 | 0.08 | 0.08 | 0.06 | 0.05 | 0.12 | 0.07 |
| 4 | 15767752 | rs10002171 | -0.057 | 0.341 | -0.099 | -0.635 | -0.001 | -0.001 | -0.001 | 0 | 0.43 | 0.38 | 0.32 | 0.38 | 0.35 | 0.43 |
| 4 | 15773008 | rs11945730 | -0.3 | 0.392 | 0.222 | -0.485 | 0.025 | 0.078 | 0.078 | 0 | 0.12 | 0.27 | 0.11 | 0.24 | 0.12 | 0.14 |
| 4 | 15773936 | rs16892407 | -0.216 | 0.157 | 0.239 | -0.631 | 0.007 | 0.009 | 0.009 | 0 | 0.4 | 0.65 | 0.32 | 0.38 | 0.32 | 0.41 |
| 4 | 15776181 | rs6831585 | -1.078 | 0.303 | -0.257 | -0.93 | -0.029 | 0.023 | 0.023 | 0.017 | 0.54 | 0.88 | 0.45 | 0.41 | 0.55 | 0.57 |
| 4 | 15791464 | rs7666953 | -1.444 | 0.365 | -0.119 | -0.973 | -0.019 | 0.031 | 0.031 | 0.432 | 0.31 | 0.77 | 0.27 | 0.25 | 0.36 | 0.3 |
| 4 | 15810779 | rs16892418 | -0.02 | 0.315 | 0.37 | -0.722 | -0.006 | 0.06 | 0.06 | 0.618 | 0.11 | 0.26 | 0.09 | 0.17 | 0.05 | 0.09 |
| 4 | 15813607 | rs6836946 | -0.247 | 0.268 | 0.179 | -0.791 | 0.004 | 0.007 | 0.007 | 0 | 0.39 | 0.67 | 0.33 | 0.38 | 0.31 | 0.43 |
| 4 | 15814621 | rs7655635 | 1.678 | 0.362 | -0.382 | -0.746 | -0.024 | 0.028 | 0.028 | 0 | 0.21 | 0.05 | 0.18 | 0.15 | 0.18 | 0.24 |
| 4 | 15814814 | rs6449195 | -0.51 | 0.364 | -0.224 | -0.74 | 0.003 | 0.061 | 0.061 | 0 | 0.13 | 0.31 | 0.1 | 0.23 | 0.11 | 0.14 |
| 4 | 15818391 | rs3796868 | 1.662 | 0.374 | -0.403 | -0.773 | -0.024 | 0.028 | 0.028 | 0 | 0.21 | 0.04 | 0.18 | 0.15 | 0.18 | 0.26 |
| 4 | 15818548 | rs3796867 | -0.624 | -0.032 | 0.069 | -0.255 | 0.102 | 0.017 | 0.017 | 0 | 0.08 | 0.44 | 0.07 | 0.06 | 0.09 | 0.06 |
| 4 | 15835365 | rs3796866 | 0.389 | -0.409 | -0.609 | -0.121 | -0.006 | 0.022 | 0.022 | 0 | 0.08 | 0.09 | 0.06 | 0.17 | 0.03 | 0.08 |
| 4 | 15835844 | rs1800051 | 0.389 | -0.409 | -0.609 | -0.121 | -0.006 | 0.022 | 0.022 | 0 | 0.08 | 0.09 | 0.06 | 0.17 | 0.03 | 0.08 |
| 4 | 15840839 | rs10805347 | -0.34 | -0.46 | -1.052 | -0.105 | -0.048 | -0.1 | -0.1 | 0 | 0.7 | 0.96 | 0.57 | 0.45 | 0.7 | 0.59 |
| 4 | 15850330 | rs1803404 | 0.243 | -0.085 | -0.355 | -0.068 | -0.009 | -0.037 | -0.037 | 1.377 | 0.07 | 0.12 | 0.04 | 0 | 0.03 | 0.01 |
| 4 | 15850685 | rs1130169 | -0.143 | -0.151 | -0.596 | -0.199 | 0.002 | -0.02 | -0.02 | 2.21 | 0.42 | 0.54 | 0.42 | 0.29 | 0.53 | 0.46 |
| 4 | 15851968 | rs13137313 | -0.183 | -0.348 | -1.207 | 0.312 | -0.017 | -0.146 | -0.146 | 0.013 | 0.76 | 0.97 | 0.61 | 0.6 | 0.75 | 0.65 |
| 4 | 15852104 | rs17476066 | 0.408 | -0.585 | -1.497 | -0.339 | 0.004 | 0.067 | 0.067 | 2.061 | 0.31 | 0.12 | 0.2 | 0.22 | 0.28 | 0.21 |
| 4 | 15854725 | rs3733593 | 0.704 | -0.448 | -0.436 | -0.414 | 0.011 | -0.019 | -0.019 | 0.087 | 0.37 | 0.49 | 0.34 | 0.3 | 0.44 | 0.38 |
| 4 | 15856799 | rs13136270 | 0.867 | -0.569 | -0.501 | -0.456 | -0.043 | 0.032 | 0.032 | 1.16 | 0.35 | 0.22 | 0.32 | 0.3 | 0.44 | 0.38 |
| 4 | 15866814 | rs9998059 | 0.352 | -0.583 | -0.337 | -0.221 | -0.029 | 0.042 | 0.042 | 1.539 | 0.28 | 0.11 | 0.18 | 0.25 | 0.28 | 0.25 |
| 4 | 15874104 | rs2531166 | 0.063 | -0.968 | -0.017 | -0.092 | -0.024 | 0.036 | 0.036 | 0 | 0.72 | 0.64 | 0.67 | 0.57 | 0.81 | 0.65 |
| 4 | 15881412 | rs2532083 | -0.251 | -0.663 | 0.363 | -0.273 | 0.01 | 0.02 | 0.02 | 2.005 | 0.37 | 0.69 | 0.21 | 0.2 | 0.24 | 0.24 |
| 4 | 15883763 | rs758327 | 0.287 | -0.681 | -0.069 | -0.257 | 0.012 | 0.026 | 0.026 | 2.774 | 0.32 | 0.24 | 0.18 | 0.21 | 0.23 | 0.16 |
| 4 | 15895791 | rs2191003 | 0.733 | -0.787 | -0.483 | -0.067 | -0.087 | 0.008 | 0.008 | 1.953 | 0.4 | 0.11 | 0.55 | 0.39 | 0.57 | 0.44 |
| 4 | 15896088 | rs6845349 | 0.048 | -0.637 | 0.196 | 0.379 | 0.002 | 0.172 | 0.172 | 0.471 | 0.79 | 0.54 | 0.77 | 0.6 | 0.82 | 0.61 |
| 4 | 15896341 | rs727488 | -0.734 | -0.417 | -0.913 | 0.506 | -0.018 | -0.097 | -0.097 | 0 | 0.84 | 0.98 | 0.81 | 0.64 | 0.84 | 0.71 |
| 4 | 15907133 | rs11944611 | -0.213 | -0.071 | -0.726 | -0.19 | 0.016 | 0.026 | 0.026 | 0.108 | 0.55 | 0.71 | 0.63 | 0.4 | 0.59 | 0.55 |

| | | | | | | | | | | | | | | | | |
|---|---|---|---|---|---|---|---|---|---|---|---|---|---|---|---|---|
| 4 | 15910385 | rs2532070 | -0.087 | 0.216 | -0.891 | -0.005 | 0.005 | 0.016 | 0.016 | 1.012 | 0.47 | 0.6 | 0.59 | 0.34 | 0.58 | 0.44 |
| 4 | 15914006 | rs2531149 | -0.396 | 0.716 | -1.089 | 0.131 | -0.002 | 0.001 | 0.001 | 0 | 0.47 | 0.48 | 0.51 | 0.48 | 0.41 | 0.56 |
| 4 | 15914034 | rs1966374 | -0.088 | 0.36 | -1.497 | 0.147 | 0.005 | -0.013 | -0.013 | 0.48 | 0.17 | 0.1 | 0.2 | 0.4 | 0.15 | 0.24 |
| 4 | 15914801 | rs12498177 | 0.062 | 0.577 | -1.807 | -0.761 | -0.024 | 0.041 | 0.041 | 0 | 0.24 | 0.1 | 0.15 | 0.17 | 0.23 | 0.17 |
| 4 | 15915036 | rs4698424 | 0.029 | 0.766 | -1.895 | -0.527 | -0.127 | 0.094 | 0.094 | 0.071 | 0.42 | 0.06 | 0.63 | 0.7 | 0.53 | 0.65 |
| 4 | 15921471 | rs11723486 | -0.31 | 0.748 | -1.159 | -0.479 | 0.001 | 0.061 | 0.061 | 0.083 | 0.51 | 0.48 | 0.56 | 0.7 | 0.56 | 0.69 |
| 4 | 15922502 | rs16892587 | -1.039 | 0.691 | -0.398 | -0.111 | 0.001 | -0.052 | -0.052 | 3.328 | 0.09 | 0.23 | 0.09 | 0 | 0.09 | 0.06 |
| 4 | 15930961 | rs876320 | 0.839 | 0.556 | -0.339 | 0.417 | -0.009 | 0.018 | 0.018 | 0.175 | 0.71 | 0.59 | 0.71 | 0.58 | 0.77 | 0.73 |
| 4 | 15934405 | rs11730943 | 0.043 | 0.221 | -0.478 | -0.506 | 0.01 | 0.021 | 0.021 | 1.361 | 0.49 | 0.45 | 0.52 | 0.55 | 0.55 | 0.33 |
| 4 | 15934723 | rs2532098 | 0.693 | 0.208 | -0.396 | -0.527 | -0.004 | -0.076 | -0.076 | 0 | 0.33 | 0.3 | 0.26 | 0.05 | 0.32 | 0.49 |
| 4 | 15935268 | rs12503130 | 0.726 | 0.369 | -0.765 | -0.333 | -0.052 | -0.012 | -0.012 | 0.053 | 0.38 | 0.21 | 0.47 | 0.54 | 0.47 | 0.26 |
| 4 | 15935497 | rs12503191 | -0.179 | 0.377 | -0.766 | -0.139 | 0.013 | 0.16 | 0.16 | 0.576 | 0.54 | 0.24 | 0.64 | 0.77 | 0.57 | 0.42 |
| 4 | 15935760 | rs2532095 | 0.482 | 0.436 | -0.517 | 0.556 | -0.004 | 0.022 | 0.022 | 0 | 0.81 | 0.77 | 0.78 | 0.59 | 0.87 | 0.82 |
| 4 | 15936638 | rs12499618 | -0.063 | 0.56 | -0.767 | 0.57 | -0.007 | -0.05 | -0.05 | 0.284 | 0.14 | 0.02 | 0.17 | 0.2 | 0.1 | 0.15 |
| 4 | 15938392 | rs16892645 | -0.022 | 0.407 | -1.048 | 0.295 | -0.008 | -0.048 | -0.048 | 0 | 0.14 | 0.02 | 0.17 | 0.2 | 0.1 | 0.15 |
| 4 | 15942802 | rs3822253 | 0.06 | 0.635 | -0.73 | 0.421 | 0 | 0 | 0 | 0 | 0.39 | 0.38 | 0.39 | 0.51 | 0.46 | 0.22 |
| 4 | 15942832 | rs1558198 | -0.419 | -0.295 | -0.64 | -0.212 | 0.001 | -0.021 | -0.021 | 0.004 | 0.63 | 0.68 | 0.67 | 0.94 | 0.67 | 0.54 |
| 4 | 15943140 | rs10939642 | 0.49 | -0.235 | -0.98 | 0.217 | 0.043 | 0.072 | 0.072 | 1.221 | 0.53 | 0.2 | 0.55 | 0.66 | 0.64 | 0.61 |
| 4 | 15947493 | rs10939644 | -0.868 | -0.304 | -0.509 | 0.171 | 0.002 | -0.003 | -0.003 | 0 | 0.87 | 0.89 | 0.87 | 0.92 | 0.85 | 0.82 |
| 4 | 15949531 | rs9968434 | 0.647 | -0.689 | -0.599 | 0.783 | -0.034 | -0.035 | -0.035 | 0.295 | 0.42 | 0.19 | 0.31 | 0.5 | 0.47 | 0.2 |
| 4 | 15949551 | rs9968276 | 0.265 | 0.3 | -0.834 | 0.965 | 0 | -0.001 | -0.001 | 0.15 | 0.19 | 0.16 | 0.16 | 0.33 | 0.22 | 0.14 |
| 4 | 15950731 | rs12509859 | 0.564 | 0.14 | -0.688 | 0.93 | -0.034 | -0.037 | -0.037 | 0 | 0.42 | 0.19 | 0.31 | 0.49 | 0.47 | 0.2 |
| 4 | 15956364 | rs6835120 | 0.942 | 0.039 | -0.56 | 0.996 | 0.017 | -0.016 | -0.016 | 0 | 0.29 | 0.13 | 0.43 | 0.3 | 0.27 | 0.52 |
| 4 | 15957568 | rs10755216 | -0.722 | 0.137 | 0.291 | 0.967 | 0.032 | -0.026 | -0.026 | 0 | 0.84 | 0.58 | 0.83 | 0.9 | 0.85 | 0.81 |
| 4 | 15957763 | rs4698429 | -0.334 | 0.12 | 0.235 | 0.968 | 0.014 | 0.005 | 0.005 | 1.356 | 0.82 | 0.38 | 0.8 | 0.79 | 0.84 | 0.81 |
| 4 | 15961555 | rs4698431 | 0.364 | -0.065 | -0.586 | 0.93 | 0.008 | 0.04 | 0.04 | 0 | 0.38 | 0.18 | 0.51 | 0.29 | 0.36 | 0.56 |
| 4 | 15963673 | rs2286461 | -0.339 | 0.007 | -0.336 | 0.98 | 0 | 0.036 | 0.036 | 0.908 | 0.52 | 0.31 | 0.65 | 0.44 | 0.52 | 0.74 |
| 4 | 15964040 | rs2286460 | -0.678 | 0.036 | -0.339 | 0.836 | -0.001 | 0 | 0 | 0 | 0.55 | 0.43 | 0.67 | 0.49 | 0.53 | 0.78 |
| 4 | 15964485 | rs758329 | 0.564 | 0.07 | -0.771 | 0.833 | -0.025 | -0.023 | -0.023 | 0 | 0.43 | 0.22 | 0.3 | 0.5 | 0.47 | 0.22 |
| 4 | 15967777 | rs12506181 | 0.234 | 0.323 | -1.202 | 0.9 | -0.04 | -0.028 | -0.028 | 0 | 0.24 | 0.04 | 0.34 | 0.27 | 0.28 | 0.24 |
| 4 | 15968574 | rs4698434 | -0.321 | 0.235 | -1.085 | 0.914 | -0.007 | 0.018 | 0.018 | 0.053 | 0.54 | 0.34 | 0.66 | 0.6 | 0.54 | 0.76 |
| 4 | 15970349 | rs2240688 | 0.367 | 0.696 | -1.577 | 0.592 | -0.032 | -0.033 | -0.033 | 0 | 0.22 | 0.05 | 0.32 | 0.27 | 0.26 | 0.24 |
| 4 | 15972549 | rs3857147 | 0.024 | 0.932 | -1.089 | 0.176 | 0.016 | -0.081 | -0.081 | 0 | 0.88 | 0.7 | 0.9 | 1 | 0.9 | 0.93 |
| 4 | 15972905 | rs11735976 | 0.689 | 1.033 | -2.086 | 0.622 | -0.103 | -0.006 | -0.006 | 0 | 0.24 | 0.02 | 0.15 | 0.19 | 0.28 | 0.14 |
| 4 | 15975839 | rs7669462 | -0.252 | 1.07 | -1.979 | 0.328 | -0.065 | 0.02 | 0.02 | 3.378 | 0.79 | 0.94 | 0.74 | 0.84 | 0.72 | 0.68 |
| 4 | 15979621 | rs7690707 | 0.443 | 1.335 | -1.775 | 0.203 | -0.006 | 0.008 | 0.008 | 0 | 0.24 | 0.22 | 0.3 | 0.19 | 0.25 | 0.25 |
| 4 | 15980975 | rs11725522 | 0.407 | 0.796 | -2.225 | -0.002 | -0.073 | 0.074 | 0.074 | 0 | 0.25 | 0.01 | 0.25 | 0.11 | 0.28 | 0.3 |
| 4 | 15981332 | rs9685492 | -0.74 | 0.543 | -1.298 | 0.419 | 0.003 | -0.013 | -0.013 | 0.015 | 0.39 | 0.46 | 0.36 | 0.32 | 0.41 | 0.36 |
| 4 | 15982480 | rs12649852 | 1.105 | -0.137 | -1.286 | 0.057 | 0.002 | 0.006 | 0.006 | 2.212 | 0.32 | 0.24 | 0.33 | 0.25 | 0.27 | 0.26 |
| 4 | 15983317 | rs10428324 | -0.872 | -0.13 | -1.324 | 0.031 | 0.005 | 0.072 | 0.072 | 0.384 | 0.73 | 0.48 | 0.66 | 0.45 | 0.69 | 0.63 |
| 4 | 15983335 | rs7692432 | 1.44 | -0.093 | -0.732 | 0.802 | 0.024 | 0.069 | 0.069 | 0 | 0.11 | 0.27 | 0.14 | 0.21 | 0.15 | 0.13 |
| 4 | 15983618 | rs10428501 | 0.226 | -0.147 | -0.716 | 0.749 | 0.01 | -0.006 | -0.006 | 0.757 | 0.35 | 0.43 | 0.33 | 0.32 | 0.37 | 0.36 |
| 4 | 15988239 | rs1990271 | 1.064 | -0.166 | -0.852 | 1.202 | 0.001 | -0.002 | -0.002 | 0 | 0.17 | 0.12 | 0.2 | 0.32 | 0.17 | 0.25 |
| 4 | 15991015 | rs17477869 | 1.46 | -0.241 | -0.863 | 1 | 0.002 | 0.119 | 0.119 | 3.865 | 0.32 | 0.08 | 0.31 | 0.14 | 0.27 | 0.22 |
| 4 | 15991275 | rs2240685 | -1.046 | -0.32 | -0.426 | 1.104 | -0.004 | 0.001 | 0.001 | 1.885 | 0.62 | 0.58 | 0.57 | 0.53 | 0.69 | 0.67 |
| 4 | 15995314 | rs717176 | 0.234 | -0.306 | -0.437 | 0.946 | 0.001 | 0.006 | 0.006 | 0.009 | 0.49 | 0.32 | 0.47 | 0.5 | 0.48 | 0.43 |
| 4 | 15998470 | rs6449211 | -0.206 | -0.217 | -0.189 | 1.34 | 0.008 | 0.003 | 0.003 | 0.574 | 0.48 | 0.43 | 0.46 | 0.47 | 0.56 | 0.51 |
| 4 | 16003151 | rs13141429 | 0.459 | -0.062 | -0.559 | 1.004 | -0.108 | 0.118 | 0.118 | 1.48 | 0.42 | 0.04 | 0.33 | 0.18 | 0.46 | 0.42 |
| 4 | 16005061 | rs6825026 | 0.688 | -0.294 | -0.306 | 0.61 | -0.008 | -0.032 | -0.032 | 0 | 0.28 | 0.32 | 0.31 | 0.13 | 0.17 | 0.19 |
| 4 | 16005699 | rs3796849 | 0.862 | 0.323 | -0.193 | 1.033 | -0.001 | -0.002 | -0.002 | 3.401 | 0.12 | 0.11 | 0.11 | 0.1 | 0.13 | 0.14 |
| 4 | 16011211 | rs2313734 | -0.026 | 0.401 | -0.334 | 0.999 | 0.004 | 0.002 | 0.002 | 0.762 | 0.5 | 0.56 | 0.53 | 0.54 | 0.45 | 0.5 |
| 4 | 16018244 | rs17387327 | 0.473 | 0.4 | -0.121 | 0.914 | -0.003 | -0.037 | -0.037 | 0 | 0.11 | 0.14 | 0.1 | 0 | 0.11 | 0.1 |
| 4 | 16018282 | rs2677789 | 0.641 | 0.394 | -0.08 | 1.513 | 0.003 | 0.08 | 0.08 | 0 | 0.94 | 0.85 | 0.8 | 0.67 | 0.94 | 0.81 |
| 4 | 16020162 | rs2286455 | -0.162 | 0.119 | -0.415 | 1.478 | 0.032 | -0.041 | -0.041 | 0 | 0.15 | 0.03 | 0.16 | 0.24 | 0.09 | 0.21 |
| 4 | 16020664 | rs1517598 | -0.454 | 0.985 | 0.272 | 1.007 | 0.027 | -0.046 | -0.046 | 0 | 0.09 | 0.21 | 0.08 | 0 | 0.13 | 0.05 |
| 4 | 16024136 | rs2677791 | -0.85 | 0.999 | 0.375 | 1.17 | 0.039 | 0.003 | 0.003 | 0 | 0.1 | 0.25 | 0.09 | 0.07 | 0.14 | 0.06 |
| 4 | 16024565 | rs2240681 | 0.295 | 0.781 | 0.458 | 1.301 | 0 | 0.007 | 0.007 | 0 | 0.84 | 0.76 | 0.72 | 0.73 | 0.84 | 0.71 |
| 4 | 16026458 | rs1517599 | 1.676 | 0.802 | 0.593 | 1.796 | 0.064 | 0.097 | 0.097 | 0 | 0.85 | 0.6 | 0.71 | 0.66 | 0.8 | 0.75 |
| 4 | 16026983 | rs7663999 | -0.724 | 0.174 | 0.244 | 1.444 | -0.024 | -0.006 | -0.006 | 0 | 0.12 | 0.19 | 0.09 | 0.08 | 0.07 | 0.05 |
| 4 | 16031448 | rs884718 | 1.636 | 0.606 | 0.522 | 2.047 | 0.028 | 0.157 | 0.157 | 0 | 0.8 | 0.64 | 0.53 | 0.49 | 0.76 | 0.66 |
| 4 | 16035996 | rs6816182 | 0.832 | 0.342 | 0.078 | 2.162 | -0.045 | -0.065 | -0.065 | 0 | 0.9 | 0.99 | 0.65 | 0.71 | 0.88 | 0.73 |
| 4 | 16038752 | rs950946 | -0.318 | 0.285 | -0.417 | 1.829 | -0.021 | -0.103 | -0.103 | 0 | 0.21 | 0.01 | 0.11 | 0.32 | 0.16 | 0.18 |
| 4 | 16038800 | rs1400773 | 0.989 | 0.363 | 0.08 | 1.871 | 0.026 | 0.059 | 0.059 | 0 | 0.84 | 0.62 | 0.68 | 0.75 | 0.84 | 0.67 |
| 4 | 16041487 | rs6849988 | 1.045 | 0.339 | 0.106 | 1.488 | 0 | -0.003 | -0.003 | 0 | 0.63 | 0.61 | 0.63 | 0.48 | 0.69 | 0.47 |
| 4 | 16046575 | rs11733193 | 0.343 | 0.237 | 0.353 | 0.858 | -0.006 | -0.05 | -0.05 | 0 | 0.8 | 0.62 | 0.9 | 0.94 | 0.86 | 0.85 |
| 4 | 16048413 | rs12511202 | -0.931 | 0.302 | -0.079 | 1.213 | -0.014 | -0.045 | -0.045 | 0 | 0.09 | 0.03 | 0.13 | 0.18 | 0.09 | 0.18 |
| 4 | 16054926 | rs6823548 | -1.173 | 0.246 | 0.297 | 1.337 | -0.001 | 0.056 | 0.056 | 0 | 0.1 | 0.16 | 0.16 | 0.23 | 0.1 | 0.18 |
| 4 | 16057694 | rs13137175 | 0.164 | 0.077 | -0.269 | 1.156 | -0.053 | -0.035 | -0.035 | 0 | 0.21 | 0.02 | 0.15 | 0.24 | 0.19 | 0.19 |
| 4 | 16063055 | rs1554243 | 0.63 | -0.287 | 0.166 | 0.909 | -0.018 | -0.011 | -0.011 | 0 | 0.67 | 0.5 | 0.71 | 0.76 | 0.75 | 0.71 |

| | | | | | | | | | | | | | | | | |
|---|---|---|---|---|---|---|---|---|---|---|---|---|---|---|---|---|
| 4 | 16064027 | rs10516295 | 0.163 | -0.354 | 0.354 | 0.87 | -0.001 | -0.003 | -0.003 | 0 | 0.75 | 0.74 | 0.74 | 0.76 | 0.79 | 0.72 |
| 4 | 16064648 | rs13150823 | 0.732 | -0.367 | 0.372 | 0.914 | -0.008 | 0.006 | 0.006 | 0 | 0.67 | 0.58 | 0.69 | 0.7 | 0.75 | 0.71 |
| 4 | 16067266 | rs3857152 | 1.145 | -0.51 | 0.595 | 0.907 | -0.011 | 0.002 | 0.002 | 0.258 | 0.46 | 0.3 | 0.52 | 0.46 | 0.57 | 0.47 |
| 4 | 16072487 | rs1040209 | 0.641 | -0.373 | 0.676 | 0.913 | -0.001 | 0 | 0 | 0 | 0.72 | 0.7 | 0.74 | 0.75 | 0.79 | 0.76 |
| 4 | 16073092 | rs884227 | 0.817 | -0.344 | 0.954 | 0.874 | -0.02 | 0.021 | 0.021 | 0 | 0.69 | 0.43 | 0.69 | 0.69 | 0.79 | 0.78 |
| 4 | 16073221 | rs908645 | 0.145 | -0.236 | 0.676 | 0.718 | -0.003 | -0.001 | -0.001 | 0 | 0.78 | 0.73 | 0.77 | 0.8 | 0.87 | 0.83 |
| 4 | 16076784 | rs10010994 | -0.159 | -0.109 | 0.732 | 0.654 | -0.003 | 0.019 | 0.019 | 0 | 0.79 | 0.7 | 0.76 | 0.75 | 0.86 | 0.79 |
| 4 | 16085850 | rs7656857 | -0.055 | -0.117 | -0.139 | 0.504 | -0.031 | -0.003 | -0.003 | 0.062 | 0.68 | 0.46 | 0.71 | 0.71 | 0.81 | 0.76 |
| 4 | 16087856 | rs1004861 | 1.005 | 0.003 | -0.317 | 0.553 | -0.006 | -0.001 | -0.001 | 0 | 0.42 | 0.26 | 0.57 | 0.41 | 0.52 | 0.52 |
| 4 | 16104248 | rs1115917 | -0.235 | 0.161 | -0.498 | 0.629 | -0.025 | -0.011 | -0.011 | 0 | 0.64 | 0.53 | 0.7 | 0.73 | 0.79 | 0.76 |
| 4 | 16105431 | rs16893029 | 0.321 | 0.27 | -1.185 | 0.685 | -0.092 | -0.066 | -0.066 | 0 | 0.22 | 0.02 | 0.11 | 0.29 | 0.24 | 0.21 |
| 4 | 16107125 | rs1554245 | 0.453 | -0.129 | -0.482 | 0.655 | 0.007 | 0.004 | 0.004 | 0 | 0.61 | 0.36 | 0.68 | 0.7 | 0.74 | 0.72 |
| 4 | 16113889 | rs1712493 | -0.091 | 0.421 | -0.722 | 0.508 | -0.001 | -0.001 | -0.001 | 0 | 0.56 | 0.61 | 0.43 | 0.53 | 0.46 | 0.51 |
| 4 | 16121755 | rs317881 | 0.274 | 0.15 | -0.732 | 0.394 | 0.001 | -0.006 | -0.006 | 0 | 0.19 | 0.17 | 0.22 | 0.25 | 0.17 | 0.18 |
| 4 | 16128075 | rs317892 | -0.423 | -0.345 | -0.587 | 0.101 | -0.03 | -0.037 | -0.037 | 0 | 0.32 | 0.48 | 0.2 | 0.22 | 0.21 | 0.27 |
| 4 | 16134623 | rs2120236 | 0.542 | -0.276 | -0.247 | 0.127 | 0.001 | -0.015 | -0.015 | 0.017 | 0.22 | 0.13 | 0.14 | 0.25 | 0.23 | 0.22 |
| 4 | 16135475 | rs317900 | -0.157 | -0.713 | -0.333 | -0.481 | -0.004 | -0.003 | -0.003 | 0 | 0.37 | 0.39 | 0.24 | 0.29 | 0.33 | 0.28 |
| 4 | 16138165 | rs317903 | 0.196 | -0.089 | -0.705 | -0.3 | -0.002 | 0.006 | 0.006 | 0 | 0.54 | 0.42 | 0.42 | 0.39 | 0.51 | 0.47 |
| 4 | 16138797 | rs11729630 | 0.556 | 0.053 | -0.822 | -0.13 | -0.003 | -0.03 | -0.03 | 0.267 | 0.74 | 0.18 | 0.57 | 0.67 | 0.78 | 0.72 |
| 4 | 16145588 | rs11730743 | 0.856 | 0.047 | -1.174 | -0.266 | -0.068 | 0.012 | 0.012 | 1.858 | 0.41 | 0.06 | 0.36 | 0.4 | 0.44 | 0.41 |
| 4 | 16146149 | rs12647911 | 1.141 | 0.495 | -0.963 | -0.083 | -0.056 | -0.081 | -0.081 | 0 | 0.19 | 0.04 | 0.14 | 0.28 | 0.25 | 0.23 |
| 4 | 16156273 | rs317866 | -0.319 | 0.564 | -0.403 | 0.007 | 0.044 | 0.093 | 0.093 | 0 | 0.86 | 0.73 | 0.63 | 0.71 | 0.8 | 0.77 |
| 4 | 16156964 | rs16893097 | 1.148 | 0.045 | -0.387 | -0.124 | 0.009 | 0.007 | 0.007 | 0 | 0.18 | 0.15 | 0.07 | 0.05 | 0.09 | 0.13 |
| 4 | 16159734 | rs10034143 | 0.646 | -0.005 | -0.238 | -0.151 | -0.053 | -0.073 | -0.073 | 0.179 | 0.22 | 0.36 | 0.1 | 0.05 | 0.09 | 0.13 |
| 4 | 16161137 | rs16893119 | 1.311 | 0.089 | -0.645 | -0.222 | -0.036 | -0.072 | -0.072 | 0.006 | 0.16 | 0.03 | 0.13 | 0.29 | 0.21 | 0.22 |
| 4 | 16163209 | rs13634 | -0.484 | 0.317 | -0.408 | -0.225 | 0.033 | 0.054 | 0.054 | 0 | 0.84 | 0.74 | 0.62 | 0.71 | 0.78 | 0.77 |
| 4 | 16163221 | rs9946 | 0.871 | 0.237 | -0.234 | -0.274 | -0.069 | -0.095 | -0.095 | 1.886 | 0.21 | 0.45 | 0.1 | 0.05 | 0.09 | 0.13 |
| 4 | 16163486 | rs10516299 | -0.32 | 0.199 | -0.281 | -0.219 | 0.008 | 0.002 | 0.002 | 1.665 | 0.39 | 0.65 | 0.27 | 0.34 | 0.33 | 0.35 |
| 4 | 16164230 | rs10516298 | 1.597 | 0.225 | -0.282 | -0.202 | -0.038 | -0.067 | -0.067 | 0 | 0.15 | 0.03 | 0.13 | 0.29 | 0.21 | 0.22 |
| 4 | 16164489 | rs7359 | 0.049 | -0.182 | -0.521 | -0.418 | -0.04 | -0.082 | -0.082 | 0.596 | 0.34 | 0.41 | 0.14 | 0.06 | 0.16 | 0.25 |
| 4 | 16166404 | rs10516297 | 1.354 | 0.164 | -0.332 | -0.161 | -0.054 | -0.017 | -0.017 | 0.048 | 0.32 | 0.07 | 0.32 | 0.36 | 0.38 | 0.28 |
| 4 | 16183180 | rs317874 | -0.235 | 0.119 | -0.427 | -0.191 | 0.021 | 0.024 | 0.024 | 0.168 | 0.48 | 0.34 | 0.28 | 0.31 | 0.4 | 0.46 |
| 4 | 16185525 | rs16893162 | 0.531 | 0.298 | -0.452 | -0.221 | 0.014 | -0.034 | -0.034 | 0 | 0.1 | 0.24 | 0.09 | 0.02 | 0.09 | 0.1 |
| 4 | 16204829 | rs10017532 | 0.04 | 0.308 | -0.457 | -0.192 | 0.009 | 0.016 | 0.016 | 1.683 | 0.49 | 0.57 | 0.34 | 0.4 | 0.44 | 0.43 |
| 4 | 16232330 | rs35141398 | 0.823 | 1.253 | -1.35 | 0.222 | -0.023 | 0.043 | 0.043 | 2.809 | 0.28 | 0.1 | 0.21 | 0.24 | 0.25 | 0.19 |
| 4 | 16234265 | rs203434 | 0.589 | 1.907 | -0.398 | 0.929 | 0.03 | 0.035 | 0.035 | 0 | 0.82 | 0.74 | 0.62 | 0.72 | 0.74 | 0.76 |
| 4 | 16242215 | rs203438 | 0.167 | 1.706 | -0.517 | 0.954 | 0.004 | 0.004 | 0.004 | 0.095 | 0.79 | 0.78 | 0.74 | 0.74 | 0.71 | 0.72 |
| 4 | 16255566 | rs203446 | 0.3 | 1.124 | -0.643 | 0.732 | 0.003 | 0.002 | 0.002 | 0 | 0.12 | 0.07 | 0.11 | 0.1 | 0.12 | 0.19 |
| 4 | 16257677 | rs6822469 | -0.405 | 1.017 | -0.077 | 0.78 | 0.001 | 0.08 | 0.08 | 0 | 0.23 | 0.42 | 0.26 | 0.46 | 0.24 | 0.28 |
| 4 | 16258670 | rs2215637 | -0.951 | 0.651 | -0.573 | 0.108 | -0.039 | -0.038 | -0.038 | 0 | 0.57 | 0.8 | 0.51 | 0.49 | 0.53 | 0.54 |
| 4 | 16260060 | rs11728731 | -0.067 | 0.811 | -0.296 | 0.13 | -0.006 | -0.062 | -0.062 | 0 | 0.38 | 0.5 | 0.29 | 0.08 | 0.33 | 0.29 |
| 4 | 16261094 | rs12502603 | 1.266 | 0.871 | -0.552 | 0.214 | -0.009 | -0.01 | -0.01 | 0 | 0.09 | 0.05 | 0.11 | 0.26 | 0.13 | 0.19 |
| 4 | 16268263 | rs17559158 | 0.022 | 0.621 | -0.361 | -0.276 | -0.005 | -0.079 | -0.079 | 0 | 0.37 | 0.36 | 0.28 | 0.03 | 0.32 | 0.27 |
| 4 | 16272456 | rs16893213 | -0.979 | 0.608 | -0.095 | -0.04 | -0.004 | -0.019 | -0.019 | 0 | 0.3 | 0.62 | 0.32 | 0.22 | 0.28 | 0.24 |
| 4 | 16277491 | rs6851785 | 1.561 | 0.52 | 0.005 | -0.035 | 0.002 | -0.003 | -0.003 | 0 | 0.1 | 0.1 | 0.08 | 0.11 | 0.1 | 0.04 |
| 4 | 16281681 | rs11721706 | 0.604 | 0.532 | 0.388 | -0.087 | 0.008 | -0.034 | -0.034 | 0 | 0.14 | 0.23 | 0.16 | 0.06 | 0.14 | 0.16 |
| 4 | 16284487 | rs6449223 | 0.329 | 1.468 | 0.167 | 0.292 | -0.047 | -0.02 | -0.02 | 0 | 0.73 | 0.8 | 0.59 | 0.38 | 0.6 | 0.66 |
| 4 | 16292409 | rs10489066 | 1.167 | 1.401 | 0.408 | 0.303 | -0.009 | 0 | 0 | 0.282 | 0.12 | 0.29 | 0.1 | 0.11 | 0.1 | 0.05 |
| 4 | 16293543 | rs2041540 | 0.543 | 1.462 | 0.407 | 0.289 | 0.008 | -0.05 | -0.05 | 3.006 | 0.15 | 0.31 | 0.2 | 0.04 | 0.14 | 0.16 |
| 4 | 16295798 | rs887649 | -0.339 | 1.458 | 0.427 | 0.296 | -0.006 | -0.038 | -0.038 | 0 | 0.26 | 0.42 | 0.28 | 0.16 | 0.24 | 0.21 |
| 4 | 16296121 | rs887648 | -1.482 | 1.765 | 0.254 | 0.673 | -0.026 | -0.004 | -0.004 | 0 | 0.95 | 0.99 | 0.78 | 0.76 | 0.85 | 0.87 |
| 4 | 16304314 | rs7660122 | 1.385 | 0.968 | 0.194 | 0.055 | -0.001 | 0 | 0 | 0 | 0.12 | 0.1 | 0.1 | 0.12 | 0.11 | 0.19 |
| 4 | 16318066 | rs975777 | -0.203 | 1.016 | 0.279 | 0.028 | -0.007 | -0.038 | -0.038 | 0 | 0.26 | 0.53 | 0.28 | 0.19 | 0.24 | 0.23 |
| 4 | 16334352 | rs2191958 | 0.591 | 0.699 | 0.245 | -0.26 | -0.004 | -0.012 | -0.012 | 0 | 0.32 | 0.36 | 0.22 | 0.16 | 0.26 | 0.21 |
| 4 | 16336013 | rs2041538 | -0.248 | 0.737 | 0.103 | -0.226 | 0.006 | 0.011 | 0.011 | 0 | 0.45 | 0.58 | 0.37 | 0.17 | 0.4 | 0.34 |
| 4 | 16352116 | rs17562392 | 1.53 | 0.791 | -0.138 | -0.198 | -0.001 | -0.001 | -0.001 | 0.28 | 0.12 | 0.1 | 0.1 | 0.12 | 0.12 | 0.2 |
| 4 | 16353313 | rs13131868 | 2.383 | 0.861 | -0.313 | -0.162 | -0.008 | 0.031 | 0.031 | 0.068 | 0.23 | 0.06 | 0.15 | 0.17 | 0.18 | 0.15 |
| 4 | 16354757 | rs4321631 | -0.745 | 1.081 | -0.181 | 0.061 | -0.066 | -0.076 | -0.076 | 0 | 0.83 | 0.93 | 0.65 | 0.49 | 0.69 | 0.71 |
| 4 | 16362803 | rs16893390 | 1.263 | 0.789 | 0.229 | -0.062 | -0.023 | -0.005 | -0.005 | 0 | 0.12 | 0.18 | 0.1 | 0.12 | 0.11 | 0.2 |
| 4 | 16363818 | rs16893392 | 1.215 | 0.788 | 0.223 | -0.063 | -0.023 | -0.006 | -0.006 | 0 | 0.12 | 0.19 | 0.1 | 0.12 | 0.11 | 0.21 |
| 4 | 16365531 | rs16893404 | 0.993 | 0.839 | 0.44 | -0.133 | 0.013 | -0.06 | -0.06 | 0 | 0.13 | 0.26 | 0.16 | 0.01 | 0.14 | 0.14 |
| 4 | 16375164 | rs11933865 | -0.069 | 0.764 | 0.33 | -0.148 | -0.004 | -0.034 | -0.034 | 0 | 0.25 | 0.46 | 0.28 | 0.19 | 0.25 | 0.22 |
| 4 | 16386449 | rs1005391 | 1.232 | 0.649 | 0.072 | -0.066 | -0.011 | -0.003 | -0.003 | 0.061 | 0.12 | 0.15 | 0.1 | 0.13 | 0.11 | 0.21 |
| 4 | 16390191 | rs17563866 | 1.399 | 0.601 | -0.195 | -0.245 | 0.016 | 0.081 | 0.081 | 0 | 0.19 | 0.05 | 0.1 | 0.01 | 0.15 | 0.12 |
| 4 | 16390974 | rs16893441 | 1.622 | 0.884 | -0.001 | -0.015 | 0.004 | 0 | 0 | 0 | 0.11 | 0.11 | 0.07 | 0.11 | 0.1 | 0.06 |
| 4 | 16396682 | rs6812305 | 0.215 | 0.944 | -0.144 | -0.003 | -0.002 | 0.009 | 0.009 | 1.155 | 0.31 | 0.24 | 0.26 | 0.17 | 0.34 | 0.33 |
| 4 | 16397925 | rs7662668 | -0.849 | 1.253 | -0.149 | 0.511 | 0.022 | 0.044 | 0.044 | 0 | 0.92 | 0.92 | 0.72 | 0.74 | 0.8 | 0.88 |
| 4 | 16410294 | rs1501139 | 0.089 | 0.435 | -0.108 | 0.235 | 0.002 | 0.002 | 0.002 | 0 | 0.32 | 0.31 | 0.25 | 0.27 | 0.27 | 0.35 |
| 4 | 16410913 | rs4698482 | 2.161 | 0.398 | -0.072 | 0.221 | 0.013 | 0.013 | 0.013 | 0 | 0.26 | 0.18 | 0.15 | 0.21 | 0.18 | 0.16 |

| | | | | | | | | | | | | | | | | |
|---|---|---|---|---|---|---|---|---|---|---|---|---|---|---|---|---|
| 4 | 16411704 | rs7674517 | 0.352 | 0.532 | -0.051 | 0.276 | -0.002 | 0.005 | 0.005 | 3.597 | 0.58 | 0.51 | 0.4 | 0.53 | 0.46 | 0.53 |
| 4 | 16412051 | rs7657627 | -0.144 | 0.771 | -0.318 | 0.051 | 0.004 | -0.054 | -0.054 | 0 | 0.28 | 0.35 | 0.2 | 0.12 | 0.26 | 0.19 |
| 4 | 16413099 | rs10025710 | 1.379 | 0.598 | -0.364 | 0.175 | 0.001 | 0.01 | 0.01 | 0 | 0.46 | 0.4 | 0.24 | 0.19 | 0.39 | 0.3 |
| 4 | 16414889 | rs6449244 | 1.206 | 0.685 | -0.852 | 0.229 | 0.008 | 0.074 | 0.074 | 0 | 0.48 | 0.63 | 0.27 | 0.19 | 0.39 | 0.3 |
| 4 | 16418399 | rs4698486 | 2.201 | 0.699 | -0.883 | 0.229 | 0.007 | -0.004 | -0.004 | 0 | 0.09 | 0.06 | 0.13 | 0.11 | 0.09 | 0.05 |
| 4 | 16419601 | rs7687486 | -0.521 | 0.683 | -1.302 | 0.407 | 0.024 | -0.007 | -0.007 | 0 | 0.36 | 0.27 | 0.32 | 0.41 | 0.29 | 0.46 |
| 4 | 16419618 | rs6449249 | -0.671 | 0.693 | -1.302 | 0.442 | -0.08 | -0.084 | -0.084 | 0.049 | 0.84 | 0.93 | 0.6 | 0.67 | 0.69 | 0.79 |
| 4 | 16422288 | rs2036361 | 1.002 | 0.588 | -1.122 | 0.144 | 0.011 | 0.087 | 0.087 | 0 | 0.48 | 0.66 | 0.27 | 0.26 | 0.4 | 0.33 |
| 4 | 16427211 | rs13147951 | 0.798 | 1.188 | -0.582 | 0.338 | -0.01 | 0.006 | 0.006 | 0 | 0.64 | 0.67 | 0.35 | 0.33 | 0.49 | 0.49 |
| 4 | 16427592 | rs2313547 | 0.78 | 1.212 | -0.697 | 0.35 | -0.019 | 0.01 | 0.01 | 0 | 0.64 | 0.71 | 0.35 | 0.33 | 0.49 | 0.49 |
| 4 | 16428830 | rs12503110 | 0.105 | 1.014 | -0.565 | 0.483 | 0.005 | -0.01 | -0.01 | 6.251 | 0.11 | 0.07 | 0.11 | 0.18 | 0.11 | 0.19 |
| 4 | 16433770 | rs1392898 | -0.061 | 1.411 | -0.123 | 0.262 | 0.071 | -0.008 | -0.008 | 0 | 0.05 | 0.18 | 0.06 | 0 | 0.07 | 0.02 |
| 4 | 16447313 | rs4698149 | 0.097 | 1.308 | -0.567 | 0.337 | 0.003 | -0.006 | -0.006 | 0 | 0.65 | 0.69 | 0.62 | 0.77 | 0.62 | 0.74 |
| 4 | 16451719 | rs16893497 | 0.204 | 1.088 | -0.305 | -0.013 | -0.005 | -0.028 | -0.028 | 1.183 | 0.07 | 0.11 | 0.03 | 0 | 0.07 | 0.06 |
| 4 | 16460247 | rs10020270 | -1.66 | 1.367 | -0.488 | 0.395 | -0.036 | -0.063 | -0.063 | 0 | 0.87 | 0.98 | 0.75 | 0.74 | 0.83 | 0.85 |
| 4 | 16462094 | rs7440228 | -0.072 | 1.106 | -0.327 | 0.088 | -0.006 | 0.068 | 0.068 | 0 | 0.48 | 0.38 | 0.5 | 0.3 | 0.52 | 0.43 |
| 4 | 16462250 | rs10026610 | 0.279 | 0.967 | -0.07 | 0.172 | 0.009 | 0.091 | 0.091 | 0.78 | 0.09 | 0.47 | 0.08 | 0.16 | 0.04 | 0.04 |
| 4 | 16468136 | rs884092 | 0.971 | 1.192 | -0.254 | -0.103 | 0.001 | -0.029 | -0.029 | 0 | 0.25 | 0.26 | 0.24 | 0.1 | 0.25 | 0.15 |
| 4 | 16469668 | rs6830747 | 0.302 | 1.041 | -0.111 | 0.246 | 0 | 0 | 0 | 0 | 0.26 | 0.31 | 0.28 | 0.22 | 0.27 | 0.29 |
| 4 | 16469713 | rs6449254 | -0.742 | 1.293 | -0.131 | 0.822 | 0.024 | 0.142 | 0.142 | 0.24 | 0.84 | 0.69 | 0.72 | 0.61 | 0.83 | 0.83 |
| 4 | 16476442 | rs1120373 | -1.654 | 1.067 | -0.606 | 0.685 | -0.033 | -0.077 | -0.077 | 0 | 0.85 | 0.97 | 0.73 | 0.71 | 0.79 | 0.83 |
| 4 | 16478276 | rs905041 | 1.113 | 0.84 | -1.002 | 0.283 | 0.022 | 0.096 | 0.096 | 0 | 0.25 | 0.06 | 0.14 | 0.1 | 0.19 | 0.32 |
| 4 | 16480357 | rs6816070 | -0.377 | 0.845 | -0.838 | 0.374 | 0.005 | -0.042 | -0.042 | 0 | 0.59 | 0.85 | 0.58 | 0.48 | 0.59 | 0.49 |
| 4 | 16481679 | rs7672573 | 0.095 | 0.875 | -0.595 | 0.286 | -0.008 | -0.012 | -0.012 | 0 | 0.38 | 0.42 | 0.24 | 0.27 | 0.32 | 0.43 |
| 4 | 16487781 | rs1354079 | -2.108 | 0.886 | -0.706 | 0.445 | -0.031 | -0.077 | -0.077 | 0.208 | 0.85 | 0.98 | 0.73 | 0.7 | 0.8 | 0.84 |
| 4 | 16491284 | rs1501140 | -0.14 | 0.828 | -0.586 | 0.387 | -0.002 | 0.009 | 0.009 | 0.194 | 0.69 | 0.61 | 0.62 | 0.41 | 0.68 | 0.69 |
| 4 | 16492812 | rs1392896 | -0.435 | 0.941 | -0.966 | 0.33 | 0.003 | 0.014 | 0.014 | 0 | 0.65 | 0.46 | 0.5 | 0.53 | 0.63 | 0.65 |
| 4 | 16494951 | rs1501141 | -2.085 | 1.038 | -1.512 | 0.445 | -0.03 | -0.079 | -0.079 | 7.602 | 0.84 | 0.98 | 0.72 | 0.71 | 0.79 | 0.83 |
| 4 | 16498825 | rs16893547 | 1.365 | 1.133 | -1.39 | 0.16 | -0.025 | -0.042 | -0.042 | 0 | 0.07 | 0.01 | 0.04 | 0.17 | 0.09 | 0.1 |
| 4 | 16499833 | rs7698804 | 1.155 | 1.067 | -1.257 | -0.241 | 0 | 0.002 | 0.002 | 0 | 0.23 | 0.21 | 0.26 | 0.14 | 0.21 | 0.17 |
| 4 | 16511887 | rs16893565 | 0.392 | 0.88 | -1.217 | 0.009 | -0.003 | 0.008 | 0.008 | 0 | 0.17 | 0.19 | 0.19 | 0.06 | 0.17 | 0.17 |
| 4 | 16512436 | rs3792679 | 0.545 | 0.711 | -1.423 | 0.24 | -0.004 | 0 | 0 | 0 | 0.39 | 0.35 | 0.38 | 0.39 | 0.41 | 0.32 |
| 4 | 16513748 | rs6824949 | -0.72 | 0.588 | -1.264 | 0.43 | 0.014 | 0.063 | 0.063 | 0 | 0.26 | 0.09 | 0.15 | 0.18 | 0.22 | 0.33 |
| 4 | 16518215 | rs13126663 | -0.466 | 0.587 | -1.242 | 0.572 | 0.012 | 0.052 | 0.052 | 0 | 0.23 | 0.06 | 0.13 | 0.16 | 0.2 | 0.31 |
| 4 | 16521723 | rs6843498 | -0.918 | 0.958 | -0.575 | 1.127 | -0.001 | 0.003 | 0.003 | 0 | 0.31 | 0.32 | 0.29 | 0.38 | 0.35 | 0.38 |
| 4 | 16522293 | rs12510434 | -0.049 | 0.787 | -0.125 | 1.58 | 0.001 | 0.004 | 0.004 | 1.768 | 0.14 | 0.13 | 0.14 | 0.28 | 0.1 | 0.12 |
| 4 | 16524379 | rs11732672 | -0.073 | 1.381 | -0.412 | 1.535 | -0.061 | -0.034 | -0.034 | 0.158 | 0.79 | 0.88 | 0.6 | 0.7 | 0.72 | 0.72 |
| 4 | 16526736 | rs4698491 | -0.082 | 1.469 | -0.363 | 1.497 | -0.043 | -0.019 | -0.019 | 0 | 0.79 | 0.87 | 0.62 | 0.71 | 0.72 | 0.73 |
| 4 | 16527611 | rs10020336 | 1.404 | 1.685 | 0.361 | 1.444 | -0.002 | -0.008 | -0.008 | 0 | 0.57 | 0.4 | 0.34 | 0.34 | 0.45 | 0.44 |
| 4 | 16530079 | rs10019178 | -2.517 | 0.586 | 0.845 | 1.909 | 0.004 | 0.017 | 0.017 | 0.037 | 0.41 | 0.62 | 0.36 | 0.76 | 0.39 | 0.58 |
| 4 | 16531401 | rs10805350 | -1.09 | 0.825 | 0.693 | 2.036 | -0.022 | 0.078 | 0.078 | 0.023 | 0.22 | 0.26 | 0.2 | 0.5 | 0.18 | 0.29 |
| 4 | 16531505 | rs10805351 | -2.157 | 0.874 | 0.726 | 1.976 | 0.001 | 0.008 | 0.008 | 0 | 0.34 | 0.47 | 0.33 | 0.6 | 0.37 | 0.48 |
| 4 | 16533376 | rs13138387 | -1.057 | 1.562 | 0.773 | 1.594 | -0.002 | 0.002 | 0.002 | 0 | 0.1 | 0.06 | 0.08 | 0.02 | 0.1 | 0.17 |
| 4 | 16537510 | rs16893606 | -0.999 | 1.347 | 0.796 | 1.671 | -0.009 | 0.076 | 0.076 | 0 | 0.1 | 0.18 | 0.06 | 0.23 | 0.07 | 0.19 |
| 4 | 16540004 | rs11947427 | -1.536 | 1.734 | 0.194 | 1.414 | 0.019 | -0.004 | -0.004 | 0.128 | 0.1 | 0.23 | 0.1 | 0.1 | 0.16 | 0.18 |
| 4 | 16546672 | rs13110049 | -1.71 | 1.503 | 0.461 | 1.781 | 0.014 | 0.114 | 0.114 | 0.368 | 0.21 | 0.47 | 0.29 | 0.39 | 0.2 | 0.41 |
| 4 | 16546995 | rs6819114 | -1.35 | 1.277 | 0.45 | 1.789 | 0.001 | 0 | 0 | 0 | 0.1 | 0.08 | 0.07 | 0.12 | 0.07 | 0.19 |
| 4 | 16549115 | rs6855048 | -2.417 | 1.119 | 0.561 | 2.26 | -0.019 | 0.086 | 0.086 | 2.334 | 0.22 | 0.62 | 0.22 | 0.59 | 0.16 | 0.36 |
| 4 | 16550094 | rs13133023 | -2.342 | 1.255 | 0.51 | 2.289 | -0.011 | 0.071 | 0.071 | 0 | 0.2 | 0.26 | 0.19 | 0.6 | 0.15 | 0.36 |
| 4 | 16551411 | rs1501145 | 1.698 | 1.252 | 0.577 | 2.295 | -0.008 | 0.047 | 0.047 | 0.11 | 0.82 | 0.8 | 0.84 | 0.41 | 0.86 | 0.64 |
| 4 | 16551576 | rs13113158 | 0.29 | 1.359 | -0.317 | 1.79 | -0.057 | 0.072 | 0.072 | 0 | 0.26 | 0.04 | 0.16 | 0.07 | 0.3 | 0.3 |
| 4 | 16551836 | rs12233679 | -0.693 | 1.107 | -0.035 | 2.126 | -0.013 | 0.128 | 0.128 | 0 | 0.08 | 0.12 | 0.11 | 0.45 | 0.04 | 0.18 |
| 4 | 16553922 | rs16893622 | -2.029 | 1.233 | -0.095 | 2.331 | -0.025 | -0.058 | -0.058 | 3.8 | 0.16 | 0.05 | 0.13 | 0.33 | 0.18 | 0.27 |
| 4 | 16557143 | rs2055418 | 1.626 | 1.14 | 0.053 | 2.328 | -0.026 | 0.1 | 0.1 | 0 | 0.83 | 0.75 | 0.8 | 0.37 | 0.9 | 0.65 |
| 4 | 16565404 | rs12508513 | -0.448 | 1.423 | -0.318 | 2.031 | -0.001 | 0.001 | 0.001 | 0 | 0.06 | 0.04 | 0.03 | 0.22 | 0.05 | 0.09 |
| 4 | 16572822 | rs11725152 | -0.283 | 1.876 | -0.614 | 1.538 | -0.128 | 0.152 | 0.152 | 1.162 | 0.36 | 0.08 | 0.26 | 0.08 | 0.5 | 0.37 |
| 4 | 16575046 | rs1392895 | -1.463 | 1.696 | 0.39 | 2.442 | 0.002 | 0.037 | 0.037 | 0 | 0.07 | 0.09 | 0.13 | 0.26 | 0.09 | 0.1 |
| 4 | 16578179 | rs4698494 | -1.885 | 1.492 | 0.867 | 2.478 | -0.02 | 0.275 | 0.275 | 1.571 | 0.13 | 0.43 | 0.17 | 0.46 | 0.08 | 0.2 |
| 4 | 16581810 | rs13120039 | -1.342 | 1.542 | 0.586 | 2.254 | 0 | -0.001 | -0.001 | 0 | 0.17 | 0.21 | 0.18 | 0.3 | 0.2 | 0.21 |
| 4 | 16585917 | rs4072813 | -1.405 | 2.309 | 0.834 | 2.063 | 0.03 | -0.018 | -0.018 | 0 | 0.06 | 0.15 | 0.06 | 0.01 | 0.1 | 0.1 |
| 4 | 16586122 | rs13109710 | -1.108 | 2.336 | 0.816 | 2.101 | 0 | 0.001 | 0.001 | 0 | 0.09 | 0.09 | 0.07 | 0.11 | 0.1 | 0.14 |
| 4 | 16588729 | rs10755217 | -2.33 | 2.627 | 0.876 | 2.166 | -0.002 | -0.001 | -0.001 | 0.03 | 0.23 | 0.3 | 0.24 | 0.34 | 0.35 | 0.37 |
| 4 | 16590588 | rs11945652 | -1.893 | 2.65 | 0.863 | 2.166 | 0 | -0.007 | -0.007 | 1.123 | 0.19 | 0.2 | 0.19 | 0.34 | 0.24 | 0.3 |
| 4 | 16591598 | rs2135442 | 2.609 | 2.543 | 1.717 | 2.328 | -0.002 | 0.005 | 0.005 | 3.862 | 0.7 | 0.32 | 0.71 | 0.55 | 0.66 | 0.6 |
| 4 | 16594879 | rs10017860 | -0.866 | 3.622 | 1.594 | 2.651 | 0.06 | 0.161 | 0.161 | 0.781 | 0.14 | 0.26 | 0.33 | 0.44 | 0.19 | 0.29 |
| 4 | 16598510 | rs2173074 | -1.42 | 3.66 | 1.188 | 2.432 | -0.001 | -0.002 | -0.002 | 0 | 0.18 | 0.21 | 0.19 | 0.34 | 0.24 | 0.27 |
| 4 | 16599234 | rs1496747 | 2.234 | 3.908 | 1.44 | 2.81 | 0.034 | 0.008 | 0.008 | 2.74 | 0.68 | 0.51 | 0.54 | 0.35 | 0.53 | 0.48 |
| 4 | 16601009 | rs11931315 | -2.397 | 3.478 | 1.014 | 2.868 | -0.02 | -0.021 | -0.021 | 0 | 0.29 | 0.3 | 0.41 | 0.62 | 0.39 | 0.48 |
| 4 | 16604857 | rs16893678 | -0.244 | 3.561 | 0.881 | 2.964 | -0.054 | -0.053 | -0.053 | 0.348 | 0.09 | 0.01 | 0.21 | 0.26 | 0.13 | 0.16 |

| | | | | | | | | | | | | | | | | |
|---|---|---|---|---|---|---|---|---|---|---|---|---|---|---|---|---|
| 4 | 16605178 | rs17490415 | -1.804 | 3.228 | 0.957 | 2.547 | -0.023 | -0.019 | -0.019 | 0.022 | 0.29 | 0.27 | 0.41 | 0.67 | 0.4 | 0.44 |
| 4 | 16608497 | rs13118746 | -1.83 | 3.106 | 0.959 | 2.514 | -0.021 | -0.019 | -0.019 | 0.749 | 0.29 | 0.27 | 0.41 | 0.67 | 0.38 | 0.43 |
| 4 | 16610295 | rs33948812 | -1.348 | 2.994 | 0.672 | 2.484 | -0.019 | -0.035 | -0.035 | 0 | 0.26 | 0.24 | 0.38 | 0.59 | 0.35 | 0.4 |
| 4 | 16624209 | rs1496750 | -2.045 | 2.982 | 0.64 | 2.4 | -0.012 | -0.003 | -0.003 | 0 | 0.35 | 0.38 | 0.45 | 0.69 | 0.45 | 0.55 |
| 4 | 16637779 | rs12506426 | -1.524 | 3.046 | 0.561 | 2.455 | -0.008 | -0.011 | -0.011 | 0 | 0.25 | 0.28 | 0.37 | 0.59 | 0.35 | 0.41 |
| 4 | 16638981 | rs7693848 | 1.964 | 2.69 | 0.866 | 2.342 | 0.083 | 0.16 | 0.16 | 0 | 0.83 | 0.58 | 0.73 | 0.56 | 0.7 | 0.71 |
| 4 | 16647943 | rs3763967 | -2.213 | 2.52 | 0.368 | 2.419 | 0.004 | 0.05 | 0.05 | 0 | 0.08 | 0.09 | 0.15 | 0.3 | 0.09 | 0.21 |
| 4 | 16648246 | rs3763969 | -1.441 | 2.467 | 0.523 | 2.351 | 0.003 | 0.003 | 0.003 | 0 | 0.1 | 0.15 | 0.11 | 0.12 | 0.17 | 0.14 |
| 4 | 16650090 | rs1566066 | -1.955 | 2.326 | 0.297 | 2.192 | 0.043 | 0.156 | 0.156 | 0 | 0.09 | 0.21 | 0.16 | 0.35 | 0.11 | 0.2 |
| 4 | 16656578 | rs13110156 | -0.395 | 1.258 | -0.936 | 0.643 | 0.003 | 0.114 | 0.114 | 0.004 | 0.19 | 0.01 | 0.07 | 0.02 | 0.17 | 0.14 |
| 4 | 16666332 | rs1027784 | 0.297 | 1.583 | -0.338 | 1.066 | 0.011 | 0.072 | 0.072 | 1.226 | 0.71 | 0.56 | 0.61 | 0.45 | 0.63 | 0.6 |
| 4 | 16668043 | rs6449262 | -0.291 | 1.72 | -0.695 | 0.873 | 0.002 | 0.013 | 0.013 | 0 | 0.84 | 0.79 | 0.91 | 0.72 | 0.85 | 0.77 |
| 4 | 16671394 | rs10939683 | -0.63 | 1.565 | -0.483 | 0.846 | -0.003 | -0.014 | -0.014 | 0 | 0.34 | 0.67 | 0.43 | 0.58 | 0.41 | 0.47 |
| 4 | 16674801 | rs6851110 | 0.061 | 1.178 | -0.749 | -0.051 | -0.001 | -0.003 | -0.003 | 0.055 | 0.54 | 0.46 | 0.62 | 0.83 | 0.54 | 0.68 |
| 4 | 16677019 | rs17492017 | -0.785 | 1.153 | -0.671 | 0.721 | 0.002 | 0.012 | 0.012 | 0 | 0.17 | 0.19 | 0.17 | 0.34 | 0.19 | 0.28 |
| 4 | 16677557 | rs10027272 | 0.754 | 1.216 | -0.584 | 0.499 | -0.001 | -0.001 | -0.001 | 0 | 0.56 | 0.55 | 0.65 | 0.48 | 0.56 | 0.57 |
| 4 | 16680651 | rs6848653 | -0.249 | 1.349 | -0.227 | 0.861 | 0 | 0.035 | 0.035 | 0 | 0.18 | 0.2 | 0.29 | 0.37 | 0.19 | 0.19 |
| 4 | 16680736 | rs7661729 | 0.56 | 1.382 | -0.198 | 0.667 | 0 | -0.001 | -0.001 | 0 | 0.55 | 0.46 | 0.53 | 0.45 | 0.57 | 0.66 |
| 4 | 16681217 | rs1395291 | -0.629 | 1.473 | 0.091 | 0.763 | -0.005 | 0.043 | 0.043 | 0 | 0.75 | 0.88 | 0.86 | 0.88 | 0.75 | 0.87 |
| 4 | 16681543 | rs7692454 | -0.11 | 1.317 | 0.359 | 1.431 | 0.018 | 0 | 0 | 0.559 | 0.1 | 0.18 | 0.07 | 0.1 | 0.14 | 0.21 |
| 4 | 16694863 | rs16893797 | -0.426 | 0.927 | 0.911 | 1.979 | 0.007 | 0.121 | 0.121 | 0 | 0.05 | 0.12 | 0.2 | 0.19 | 0.07 | 0.07 |
| 4 | 16694900 | rs13110423 | -0.17 | 0.914 | 0.636 | 1.9 | -0.004 | -0.008 | -0.008 | 0.022 | 0.3 | 0.32 | 0.37 | 0.61 | 0.39 | 0.34 |
| 4 | 16698184 | rs283029 | -0.22 | 0.839 | 0.644 | 1.71 | -0.002 | -0.003 | -0.003 | 2.177 | 0.31 | 0.34 | 0.34 | 0.65 | 0.39 | 0.34 |
| 4 | 16713075 | rs157619 | -0.891 | 0.821 | 0.768 | 2.062 | 0.006 | -0.051 | -0.051 | 0.013 | 0.37 | 0.55 | 0.55 | 0.78 | 0.46 | 0.4 |
| 4 | 16717423 | rs283038 | -0.297 | 0.871 | 0.718 | 2.114 | -0.01 | -0.021 | -0.021 | 0.051 | 0.31 | 0.23 | 0.45 | 0.4 | 0.41 | 0.33 |
| 4 | 16719156 | rs17428815 | -0.854 | 0.867 | 0.724 | 2.074 | -0.002 | -0.001 | -0.001 | 0 | 0.36 | 0.34 | 0.54 | 0.78 | 0.46 | 0.4 |
| 4 | 16723640 | rs1514644 | -0.577 | 0.834 | 0.613 | 2.058 | -0.002 | -0.004 | -0.004 | 0.279 | 0.33 | 0.29 | 0.46 | 0.43 | 0.42 | 0.35 |
| 4 | 16726650 | rs16893890 | -0.108 | 0.906 | 0.465 | 1.944 | 0.029 | 0.112 | 0.112 | 0 | 0.06 | 0.16 | 0.21 | 0.13 | 0.08 | 0.08 |
| 4 | 16736065 | rs533825 | -1.059 | 0.879 | -0.063 | 1.634 | 0.003 | -0.023 | -0.023 | 0.27 | 0.39 | 0.57 | 0.55 | 0.76 | 0.47 | 0.42 |
| 4 | 16739648 | rs283019 | 0.535 | 0.837 | -0.077 | 1.722 | -0.073 | -0.064 | -0.064 | 0 | 0.23 | 0.03 | 0.22 | 0.26 | 0.31 | 0.23 |
| 4 | 16740168 | rs283018 | -0.407 | 0.608 | 0.186 | 1.269 | 0.003 | -0.02 | -0.02 | 0.488 | 0.41 | 0.57 | 0.59 | 0.77 | 0.52 | 0.44 |
| 4 | 16744084 | rs883086 | -0.469 | 0.336 | 0.541 | 1.574 | -0.019 | 0.115 | 0.115 | 0 | 0.24 | 0.04 | 0.17 | 0.06 | 0.19 | 0.1 |
| 4 | 16747687 | rs1546182 | -1.939 | 0.185 | 1.94 | 2.862 | 0.074 | 0.438 | 0.438 | 0.132 | 0.08 | 0.46 | 0.14 | 0.37 | 0.09 | 0.09 |
| 4 | 16753012 | rs157630 | 2.055 | -0.214 | 2.032 | 2.536 | -0.059 | 0.104 | 0.104 | 1.783 | 0.83 | 0.39 | 0.63 | 0.6 | 0.89 | 0.91 |
| 4 | 16753863 | rs283027 | 2.254 | -0.078 | 2.11 | 2.621 | -0.056 | 0.242 | 0.242 | 4.303 | 0.87 | 0.45 | 0.64 | 0.6 | 0.91 | 0.92 |
| 4 | 16773323 | rs207685 | 1.404 | 0.353 | 2.053 | 2.803 | 0.059 | 0.454 | 0.454 | 0 | 0.94 | 0.65 | 0.7 | 0.66 | 0.94 | 0.92 |
| 4 | 16786858 | rs207701 | 1.261 | 0.412 | 1.374 | 2.512 | 0.023 | 0.387 | 0.387 | 0 | 0.93 | 0.72 | 0.73 | 0.59 | 0.93 | 0.93 |
| 4 | 16789915 | rs17757703 | -0.994 | 0.783 | 0.158 | 1.423 | -0.004 | -0.01 | -0.01 | 0.014 | 0.1 | 0.08 | 0.12 | 0.15 | 0.15 | 0.19 |
| 4 | 16823900 | rs123043 | 0.273 | 0.824 | 0.783 | 1.054 | 0.003 | 0.028 | 0.028 | 0 | 0.68 | 0.53 | 0.7 | 0.68 | 0.67 | 0.48 |
| 4 | 16828794 | rs207703 | 0.512 | 0.793 | 1.087 | 1.068 | 0.053 | 0.153 | 0.153 | 1.13 | 0.95 | 0.71 | 0.95 | 0.89 | 0.97 | 0.97 |
| 4 | 16837966 | rs629782 | 0.405 | 1.034 | 0.479 | 0.208 | -0.008 | 0.024 | 0.024 | 3.053 | 0.2 | 0.12 | 0.09 | 0.1 | 0.17 | 0.14 |
| 4 | 16839420 | rs158260 | -0.057 | 0.964 | 0.581 | 0.503 | 0 | 0.013 | 0.013 | 0 | 0.68 | 0.6 | 0.72 | 0.65 | 0.68 | 0.5 |
| 4 | 16842716 | rs12503223 | 0.295 | 1.141 | -0.12 | -0.557 | -0.028 | 0.029 | 0.029 | 0 | 0.15 | 0.06 | 0.23 | 0 | 0.26 | 0.19 |
| 4 | 16851024 | rs207713 | 0.262 | 0.799 | 0.129 | -0.735 | -0.014 | 0.055 | 0.055 | 0.235 | 0.57 | 0.65 | 0.53 | 0.76 | 0.46 | 0.32 |
| 4 | 16855730 | rs13135819 | -0.618 | 0.56 | 0.048 | -1.082 | 0.009 | 0.12 | 0.12 | 0.32 | 0.5 | 0.71 | 0.34 | 0.28 | 0.41 | 0.62 |
| 4 | 16856564 | rs6839117 | 0.636 | 0.67 | 0.608 | -1.093 | -0.001 | -0.003 | -0.003 | 0 | 0.25 | 0.31 | 0.23 | 0.19 | 0.24 | 0.47 |
| 4 | 16861512 | rs6829935 | -0.264 | 0.687 | 0.741 | -1.118 | 0.004 | -0.015 | -0.015 | 0 | 0.79 | 0.8 | 0.74 | 0.92 | 0.7 | 0.76 |
| 4 | 16865939 | rs13113385 | -1.131 | 0.479 | 0.733 | -1.21 | -0.003 | 0.071 | 0.071 | 6.274 | 0.78 | 0.93 | 0.9 | 0.91 | 0.82 | 0.85 |
| 4 | 16867757 | rs11734663 | 0.049 | 0.761 | 0.874 | -1.097 | -0.032 | 0.076 | 0.076 | 0.931 | 0.6 | 0.75 | 0.64 | 0.83 | 0.53 | 0.62 |
| 4 | 16871315 | rs13104451 | 0.847 | 0.737 | 1.112 | -0.755 | 0.017 | -0.003 | -0.003 | 0 | 0.31 | 0.23 | 0.42 | 0.65 | 0.28 | 0.14 |
| 4 | 16873743 | rs10939693 | 1.22 | 0.674 | 0.901 | -0.77 | 0.012 | -0.034 | -0.034 | 0 | 0.27 | 0.13 | 0.39 | 0.66 | 0.26 | 0.12 |
| 4 | 16892585 | rs1483009 | -1.35 | 0.702 | 1.268 | -1.28 | 0.003 | -0.021 | -0.021 | 0.047 | 0.36 | 0.36 | 0.39 | 0.19 | 0.44 | 0.38 |
| 4 | 16893893 | rs1483012 | -1.726 | 0.678 | 1.228 | -1.292 | -0.018 | 0.11 | 0.11 | 0 | 0.4 | 0.75 | 0.44 | 0.18 | 0.45 | 0.38 |
| 4 | 16899946 | rs7690911 | -0.586 | 0.827 | 1.088 | -1.311 | -0.024 | -0.011 | -0.011 | 0.04 | 0.14 | 0.26 | 0.12 | 0.1 | 0.08 | 0.11 |
| 4 | 16906388 | rs992704 | -1.319 | 0.91 | 1.085 | -1.374 | -0.001 | -0.029 | -0.029 | 0 | 0.3 | 0.26 | 0.31 | 0.08 | 0.36 | 0.23 |
| 4 | 16913766 | rs6816423 | 0.251 | 0.925 | 0.866 | -1.335 | -0.012 | 0.022 | 0.022 | 0 | 0.11 | 0.06 | 0.06 | 0 | 0.11 | 0.06 |
| 4 | 16914239 | rs13126845 | -1.849 | 0.879 | 1.021 | -1.329 | -0.005 | 0.008 | 0.008 | 0 | 0.36 | 0.35 | 0.39 | 0.19 | 0.45 | 0.38 |
| 4 | 16914874 | rs2108937 | -0.462 | 0.909 | 0.807 | -1.339 | -0.017 | 0.024 | 0.024 | 0 | 0.2 | 0.13 | 0.26 | 0.08 | 0.31 | 0.22 |
| 4 | 16915245 | rs13129387 | -1.131 | 0.893 | 0.768 | -1.339 | -0.037 | 0.051 | 0.051 | 0 | 0.28 | 0.16 | 0.3 | 0.09 | 0.4 | 0.26 |
| 4 | 16922182 | rs4698165 | -1.261 | 0.73 | 1.206 | -1.173 | 0.009 | -0.038 | -0.038 | 0 | 0.27 | 0.55 | 0.34 | 0.18 | 0.33 | 0.3 |
| 4 | 16929643 | rs725858 | -1.67 | 0.691 | 1.255 | -1.205 | 0 | -0.013 | -0.013 | 0.053 | 0.34 | 0.6 | 0.39 | 0.19 | 0.38 | 0.3 |
| 4 | 16932945 | rs2872565 | -0.758 | 0.504 | 1.374 | -1.043 | 0.01 | 0.008 | 0.008 | 0 | 0.06 | 0.19 | 0.09 | 0.08 | 0.08 | 0.04 |
| 4 | 16935552 | rs917494 | -1.201 | 0.28 | 1.376 | -1.185 | -0.017 | -0.037 | -0.037 | 0 | 0.15 | 0.25 | 0.12 | 0.08 | 0.12 | 0.05 |
| 4 | 16940151 | rs16894200 | -1.745 | 0.22 | 1.652 | -1.136 | -0.026 | 0.069 | 0.069 | 0.324 | 0.07 | 0.23 | 0.08 | 0.11 | 0.02 | 0.07 |
| 4 | 16942990 | rs1949292 | -0.556 | 0.308 | 1.491 | -1.128 | 0.005 | 0.001 | 0.001 | 0 | 0.06 | 0.16 | 0.08 | 0.08 | 0.07 | 0.04 |
| 4 | 16985245 | rs6414774 | -2.425 | 0.335 | 1.451 | -1.32 | -0.03 | -0.034 | -0.034 | 2.9 | 0.24 | 0.44 | 0.2 | 0.2 | 0.19 | 0.19 |
| 4 | 16992028 | rs4698527 | 0.094 | -0.039 | 0.562 | -1.861 | -0.01 | -0.021 | -0.021 | 0 | 0.45 | 0.5 | 0.3 | 0.27 | 0.33 | 0.54 |
| 4 | 16993463 | rs4698528 | -1.212 | -0.053 | 0.688 | -1.817 | -0.01 | 0.089 | 0.089 | 0 | 0.09 | 0.25 | 0.13 | 0.22 | 0.08 | 0.1 |
| 4 | 16995023 | rs13109226 | -0.162 | -0.033 | 0.645 | -1.799 | -0.009 | -0.024 | -0.024 | 0 | 0.46 | 0.49 | 0.3 | 0.26 | 0.34 | 0.55 |

| | | | | | | | | | | | | | | | | |
|---|---|---|---|---|---|---|---|---|---|---|---|---|---|---|---|---|
| 4 | 16995752 | rs4698529 | -1.399 | -0.053 | 0.655 | -1.792 | -0.017 | 0.042 | 0.042 | 0 | 0.1 | 0.25 | 0.13 | 0.19 | 0.08 | 0.1 |
| 4 | 16997327 | rs2314597 | 0.128 | -0.043 | 0.498 | -1.765 | -0.012 | -0.034 | -0.034 | 0 | 0.45 | 0.49 | 0.3 | 0.24 | 0.33 | 0.54 |
| 4 | 17016263 | rs6858060 | -0.48 | -0.004 | 0.449 | -1.295 | -0.001 | 0.036 | 0.036 | 0.122 | 0.57 | 0.72 | 0.41 | 0.32 | 0.43 | 0.57 |
| 4 | 17025691 | rs2102702 | -1.773 | 0.604 | 0.069 | -1.43 | -0.051 | -0.043 | -0.043 | 0.035 | 0.72 | 0.94 | 0.59 | 0.33 | 0.67 | 0.76 |
| 4 | 17035562 | rs11730753 | 0.738 | 0.227 | 0.24 | -1.345 | 0.006 | 0.1 | 0.1 | 0 | 0.4 | 0.21 | 0.26 | 0.15 | 0.34 | 0.49 |
| 4 | 17036244 | rs6449275 | 0.747 | 0.23 | 0.259 | -1.409 | 0.006 | 0.108 | 0.108 | 0 | 0.4 | 0.21 | 0.26 | 0.12 | 0.34 | 0.49 |
| 4 | 17039677 | rs13109476 | 0.429 | 0.33 | 0.474 | -1.351 | 0.018 | 0.039 | 0.039 | 0.259 | 0.46 | 0.42 | 0.34 | 0.31 | 0.36 | 0.57 |
| 4 | 17041687 | rs11724970 | 0.138 | 0.031 | 0.022 | -1.822 | 0.002 | 0.001 | 0.001 | 0 | 0.37 | 0.37 | 0.29 | 0.22 | 0.3 | 0.44 |
| 4 | 17042063 | rs1514989 | -0.479 | 0.047 | -0.066 | -1.887 | 0.017 | 0.096 | 0.096 | 0.537 | 0.55 | 0.75 | 0.4 | 0.22 | 0.42 | 0.58 |
| 4 | 17043787 | rs13353571 | 1.85 | 0.03 | 0.027 | -1.8 | -0.013 | -0.092 | -0.092 | 0.586 | 0.17 | 0.3 | 0.1 | 0 | 0.11 | 0.14 |
| 4 | 17044466 | rs16894247 | 1.015 | 0.088 | -0.005 | -1.837 | 0.013 | 0.017 | 0.017 | 0 | 0.27 | 0.2 | 0.23 | 0.21 | 0.24 | 0.43 |
| 4 | 17044760 | rs13107569 | -0.049 | 0.276 | 0.036 | -1.559 | 0.003 | 0.002 | 0.002 | 0.852 | 0.49 | 0.48 | 0.37 | 0.22 | 0.39 | 0.62 |
| 4 | 17045368 | rs4426772 | -0.575 | 0.581 | -0.178 | -1.08 | -0.029 | 0.02 | 0.02 | 0.027 | 0.59 | 0.78 | 0.43 | 0.33 | 0.48 | 0.64 |
| 4 | 17047470 | rs16894256 | 1.413 | 0.576 | -0.041 | -1.017 | 0.005 | -0.014 | -0.014 | 2.133 | 0.27 | 0.25 | 0.3 | 0.34 | 0.26 | 0.44 |
| 4 | 17047686 | rs1356028 | -0.541 | 0.675 | -0.361 | -1.099 | 0.001 | 0.017 | 0.017 | 0.608 | 0.51 | 0.66 | 0.48 | 0.43 | 0.53 | 0.6 |
| 4 | 17050660 | rs10001817 | 1.337 | 0.799 | -0.427 | -0.906 | -0.004 | 0.009 | 0.009 | 1.61 | 0.28 | 0.37 | 0.32 | 0.34 | 0.28 | 0.45 |
| 4 | 17051156 | rs11929846 | 0.578 | 1.246 | -0.998 | -0.417 | -0.01 | -0.048 | -0.048 | 0 | 0.18 | 0.07 | 0.32 | 0.56 | 0.21 | 0.2 |
| 4 | 17054262 | rs16894263 | 0.206 | 1.219 | -0.861 | -0.729 | 0.022 | -0.026 | -0.026 | 0 | 0.14 | 0.23 | 0.12 | 0.09 | 0.2 | 0.14 |
| 4 | 17054716 | rs13143173 | 0.776 | 0.617 | -1.179 | -0.213 | 0.009 | 0.006 | 0.006 | 2.605 | 0.17 | 0.12 | 0.11 | 0 | 0.12 | 0.16 |
| 4 | 17060545 | rs828164 | -0.263 | 0.819 | -1.273 | 0.969 | -0.031 | -0.068 | -0.068 | 0 | 0.19 | 0.06 | 0.3 | 0.35 | 0.23 | 0.13 |
| 4 | 17062952 | rs828154 | 0.147 | 0.431 | -0.34 | 1.647 | -0.003 | 0.002 | 0.002 | 1.475 | 0.66 | 0.36 | 0.56 | 0.38 | 0.72 | 0.62 |
| 4 | 17063564 | rs828152 | 0.42 | 0.382 | -0.195 | 1.965 | -0.004 | 0.005 | 0.005 | 0 | 0.86 | 0.81 | 0.7 | 0.56 | 0.85 | 0.81 |
| 4 | 17064024 | rs16894291 | 1.008 | 0.147 | 0.226 | 1.991 | -0.023 | -0.009 | -0.009 | 0.59 | 0.19 | 0.27 | 0.12 | 0.18 | 0.13 | 0.2 |
| 4 | 17070502 | rs1464091 | -0.263 | 0.552 | -0.653 | 2.027 | -0.056 | -0.096 | -0.096 | 0 | 0.16 | 0.05 | 0.33 | 0.45 | 0.19 | 0.13 |
| 4 | 17091852 | rs16894314 | 0.462 | 0.253 | -0.604 | 1.303 | -0.008 | -0.031 | -0.031 | 0 | 0.12 | 0.17 | 0.08 | 0.02 | 0.07 | 0.06 |
| 4 | 17092275 | rs16894315 | 0.476 | 0.252 | -0.604 | 1.303 | -0.008 | -0.031 | -0.031 | 0.162 | 0.12 | 0.17 | 0.08 | 0.02 | 0.07 | 0.06 |
| 4 | 17095691 | rs12647067 | -0.927 | 0.233 | -0.438 | 1.503 | -0.024 | -0.012 | -0.012 | 0 | 0.24 | 0.2 | 0.23 | 0.27 | 0.3 | 0.21 |
| 4 | 17104059 | rs13121153 | 0.498 | -0.285 | -0.302 | 1.4 | -0.006 | 0.036 | 0.036 | 0.632 | 0.48 | 0.66 | 0.6 | 0.35 | 0.55 | 0.53 |
| 4 | 17105434 | rs2056394 | 1.626 | -0.345 | -0.239 | 1.491 | 0.002 | -0.002 | -0.002 | 0.854 | 0.34 | 0.34 | 0.44 | 0.27 | 0.44 | 0.39 |
| 4 | 17106331 | rs2056393 | 1.448 | -0.335 | -0.217 | 1.463 | 0.008 | -0.009 | -0.009 | 0.107 | 0.36 | 0.4 | 0.5 | 0.29 | 0.45 | 0.39 |
| 4 | 17112493 | rs12640561 | -0.229 | -0.298 | -0.586 | 1.599 | -0.097 | -0.021 | -0.021 | 0.307 | 0.24 | 0.05 | 0.18 | 0.2 | 0.27 | 0.14 |
| 4 | 17115201 | rs10000500 | 1.359 | -0.355 | -0.448 | 1.63 | -0.011 | 0.006 | 0.006 | 4.053 | 0.2 | 0.2 | 0.14 | 0.26 | 0.14 | 0.17 |
| 4 | 17115730 | rs1522077 | 1.978 | -0.029 | -0.844 | 1.496 | -0.092 | -0.033 | -0.033 | 0 | 0.27 | 0.01 | 0.22 | 0.21 | 0.36 | 0.35 |
| 4 | 17123821 | rs675984 | -0.602 | -0.233 | -0.054 | 1.825 | 0.008 | -0.011 | -0.011 | 0 | 0.91 | 0.95 | 0.92 | 0.8 | 0.95 | 0.84 |
| 4 | 17126696 | rs9291654 | 0.05 | -0.071 | 0.072 | 1.697 | 0 | -0.004 | -0.004 | 0.499 | 0.08 | 0.12 | 0.08 | 0.06 | 0.1 | 0.1 |
| 4 | 17133221 | rs12645194 | 0.266 | 0.219 | -0.334 | 1.975 | -0.018 | -0.012 | -0.012 | 0.014 | 0.09 | 0.08 | 0.31 | 0.28 | 0.16 | 0.12 |
| 4 | 17133297 | rs9291655 | 0.953 | -0.052 | -0.297 | 1.903 | 0.01 | -0.003 | -0.003 | 0 | 0.19 | 0.14 | 0.18 | 0.22 | 0.12 | 0.15 |
| 4 | 17136795 | rs970428 | -0.24 | -0.06 | -0.286 | 1.88 | -0.035 | -0.045 | -0.045 | 0 | 0.17 | 0.09 | 0.35 | 0.29 | 0.2 | 0.13 |
| 4 | 17137284 | rs629542 | 1.218 | -0.106 | -0.042 | 1.857 | 0.016 | 0.028 | 0.028 | 0 | 0.73 | 0.59 | 0.57 | 0.64 | 0.71 | 0.74 |
| 4 | 17145268 | rs10489116 | 0.388 | -0.242 | -0.094 | 1.881 | 0.017 | -0.01 | -0.01 | 0 | 0.14 | 0.09 | 0.1 | 0.2 | 0.07 | 0.18 |
| 4 | 17145515 | rs492959 | 1.441 | -0.223 | -0.088 | 1.886 | 0 | 0.002 | 0.002 | 0 | 0.6 | 0.58 | 0.47 | 0.44 | 0.64 | 0.57 |
| 4 | 17147731 | rs596777 | 0.899 | -0.266 | -0.089 | 1.922 | -0.028 | -0.015 | -0.015 | 0 | 0.68 | 0.58 | 0.55 | 0.72 | 0.77 | 0.65 |
| 4 | 17148643 | rs16894476 | 0.989 | -0.321 | 0.03 | 1.999 | 0.005 | -0.001 | -0.001 | 0 | 0.2 | 0.15 | 0.19 | 0.22 | 0.13 | 0.15 |
| 4 | 17149944 | rs627206 | 0.472 | -0.347 | -0.047 | 1.931 | -0.02 | -0.007 | -0.007 | 0.084 | 0.77 | 0.72 | 0.82 | 0.78 | 0.85 | 0.71 |
| 4 | 17156183 | rs504670 | 1.208 | -0.295 | -0.043 | 1.673 | 0.01 | 0.015 | 0.015 | 0.427 | 0.66 | 0.53 | 0.5 | 0.4 | 0.69 | 0.58 |
| 4 | 17165360 | rs667932 | -0.354 | -0.221 | -0.047 | 1.69 | 0.002 | 0.006 | 0.006 | 0.588 | 0.91 | 0.79 | 0.93 | 0.92 | 0.92 | 0.88 |
| 4 | 17167808 | rs566672 | 0.665 | -0.185 | -0.087 | 1.833 | 0.063 | 0.08 | 0.08 | 0 | 0.81 | 0.63 | 0.61 | 0.62 | 0.77 | 0.76 |
| 4 | 17168001 | rs16894543 | 0.254 | -0.212 | -0.2 | 1.745 | 0.025 | -0.034 | -0.034 | 0 | 0.15 | 0.07 | 0.11 | 0.23 | 0.09 | 0.22 |
| 4 | 17168826 | rs16868330 | 0.913 | -0.132 | -0.154 | 1.766 | -0.054 | -0.051 | -0.051 | 0.573 | 0.08 | 0.01 | 0.1 | 0.23 | 0.13 | 0.09 |
| 4 | 17178058 | rs1522075 | -0.38 | -1.158 | 0.062 | 1.713 | 0.034 | -0.014 | -0.014 | 0 | 0.34 | 0.24 | 0.28 | 0.39 | 0.2 | 0.33 |
| 4 | 17181579 | rs608897 | -0.259 | -1.104 | 0.138 | 1.704 | 0 | -0.003 | -0.003 | 0.061 | 0.91 | 0.73 | 0.93 | 0.94 | 0.93 | 0.9 |
| 4 | 17187440 | rs574427 | 0.715 | -1.037 | 0.001 | 1.748 | 0.02 | -0.003 | -0.003 | 0 | 0.55 | 0.43 | 0.64 | 0.55 | 0.72 | 0.57 |
| 4 | 17188778 | rs7669359 | 0.625 | -0.988 | -0.048 | 1.786 | -0.03 | -0.029 | -0.029 | 0.124 | 0.09 | 0.06 | 0.16 | 0.29 | 0.16 | 0.11 |
| 4 | 17188787 | rs7667911 | 0.855 | -1.13 | 0.014 | 1.101 | 0 | 0 | 0 | 0.91 | 0.2 | 0.16 | 0.18 | 0.18 | 0.13 | 0.13 |
| 4 | 17194641 | rs6449278 | 1.09 | -1.042 | -0.165 | 1.094 | -0.008 | 0.04 | 0.04 | 0.086 | 0.46 | 0.35 | 0.47 | 0.24 | 0.56 | 0.43 |
| 4 | 17196209 | rs721166 | 0.83 | -0.635 | -0.02 | 0.916 | -0.007 | -0.003 | -0.003 | 0 | 0.2 | 0.19 | 0.12 | 0.19 | 0.15 | 0.14 |
| 4 | 17200948 | rs13142734 | 0.169 | -0.672 | -0.566 | 0.863 | -0.051 | -0.081 | -0.081 | 0 | 0.27 | 0.1 | 0.28 | 0.36 | 0.26 | 0.27 |
| 4 | 17201248 | rs567404 | -0.104 | -0.722 | -0.418 | 0.848 | 0 | -0.01 | -0.01 | 0 | 0.7 | 0.7 | 0.71 | 0.62 | 0.74 | 0.72 |
| 4 | 17205620 | rs16894648 | 0.211 | -0.672 | -0.502 | 0.822 | -0.055 | -0.082 | -0.082 | 0 | 0.26 | 0.07 | 0.28 | 0.36 | 0.26 | 0.25 |
| 4 | 17209596 | rs12640889 | 0.293 | -0.61 | -0.326 | 0.835 | -0.013 | 0.095 | 0.095 | 0 | 0.16 | 0.24 | 0.17 | 0.39 | 0.11 | 0.22 |
| 4 | 17210786 | rs6837748 | 0.094 | -0.368 | 0.024 | 0.031 | -0.001 | 0.055 | 0.055 | 0 | 0.41 | 0.32 | 0.46 | 0.74 | 0.37 | 0.49 |
| 4 | 17223625 | rs965572 | 0.151 | -0.359 | -0.007 | 0.068 | -0.001 | 0.015 | 0.015 | 0 | 0.39 | 0.28 | 0.43 | 0.65 | 0.36 | 0.48 |
| 4 | 17224160 | rs501015 | -0.043 | -0.391 | 0.058 | 0.053 | -0.002 | -0.002 | -0.002 | 0 | 0.56 | 0.5 | 0.56 | 0.35 | 0.64 | 0.52 |
| 4 | 17224345 | rs10025799 | 0.007 | -0.241 | -0.176 | -0.453 | -0.002 | 0.055 | 0.055 | 0 | 0.4 | 0.28 | 0.45 | 0.74 | 0.37 | 0.48 |
| 4 | 17228017 | rs509994 | -0.099 | 0.522 | 0.03 | -0.016 | 0.01 | -0.026 | -0.026 | 0 | 0.35 | 0.38 | 0.44 | 0.16 | 0.48 | 0.35 |
| 4 | 17240787 | rs9995238 | -1.39 | 0.05 | 0.441 | 0.785 | -0.034 | 0.074 | 0.074 | 0.513 | 0.16 | 0.56 | 0.16 | 0.27 | 0.11 | 0.16 |
| 4 | 17242936 | rs1402949 | -1.683 | 0.008 | 1.319 | 0.496 | 0.002 | 0.012 | 0.012 | 1.639 | 0.1 | 0.41 | 0.05 | 0.1 | 0.05 | 0.07 |
| 4 | 17248161 | rs16894737 | -0.703 | 0.066 | 1.564 | -0.018 | 0.023 | -0.068 | -0.068 | 1.495 | 0.11 | 0.25 | 0.06 | 0 | 0.1 | 0.04 |
| 4 | 17255650 | rs584272 | -0.634 | -0.045 | 0.471 | -1.112 | -0.021 | 0.114 | 0.114 | 0 | 0.56 | 0.74 | 0.37 | 0.15 | 0.37 | 0.4 |

| | | | | | | | | | | | | | | | | |
|---|---|---|---|---|---|---|---|---|---|---|---|---|---|---|---|---|
| 4 | 17256096 | rs16894739 | -0.493 | 0.034 | 0.734 | -1.066 | -0.005 | -0.001 | -0.001 | 0 | 0.08 | 0.12 | 0.03 | 0.1 | 0.04 | 0.07 |
| 4 | 17257605 | rs613801 | 0.369 | -0.009 | 0.819 | -1.042 | -0.037 | 0.152 | 0.152 | 3.089 | 0.39 | 0.17 | 0.29 | 0.07 | 0.34 | 0.34 |
| 4 | 17259908 | rs144246 | -0.119 | -0.026 | 0.968 | -0.968 | -0.002 | 0.058 | 0.058 | 1.044 | 0.46 | 0.36 | 0.32 | 0.17 | 0.35 | 0.4 |
| 4 | 17261697 | rs673248 | -0.246 | -0.022 | 0.915 | -0.958 | 0 | 0.016 | 0.016 | 1.005 | 0.47 | 0.45 | 0.33 | 0.18 | 0.36 | 0.4 |
| 4 | 17268118 | rs531934 | -0.102 | -0.227 | 0.814 | -1.252 | -0.001 | -0.055 | -0.055 | 0 | 0.43 | 0.38 | 0.3 | 0.09 | 0.34 | 0.31 |
| 4 | 17268148 | rs12509299 | -0.16 | -0.251 | 0.096 | -0.653 | -0.012 | -0.114 | -0.114 | 0 | 0.15 | 0.01 | 0.18 | 0.59 | 0.1 | 0.21 |
| 4 | 17269405 | rs649405 | 0.445 | 0.136 | 0.538 | -0.855 | -0.015 | 0.101 | 0.101 | 0.956 | 0.37 | 0.19 | 0.29 | 0.09 | 0.32 | 0.31 |
| 4 | 17272182 | rs7356497 | -0.237 | 0.494 | 0.487 | -0.736 | 0 | 0.015 | 0.015 | 0.016 | 0.48 | 0.54 | 0.35 | 0.14 | 0.38 | 0.4 |
| 4 | 17273603 | rs12498656 | -0.274 | 0.544 | 0.06 | -0.706 | 0.003 | 0.046 | 0.046 | 0.028 | 0.48 | 0.58 | 0.36 | 0.14 | 0.38 | 0.4 |
| 4 | 17275696 | rs4585300 | 0.387 | 1.364 | 0.068 | -0.064 | -0.001 | -0.001 | -0.001 | 0 | 0.8 | 0.79 | 0.59 | 0.26 | 0.77 | 0.67 |
| 4 | 17276304 | rs12507442 | 1.08 | 1.89 | -0.247 | -0.548 | -0.023 | 0.09 | 0.09 | 0 | 0.57 | 0.26 | 0.36 | 0.26 | 0.56 | 0.51 |
| 4 | 17279997 | rs1981693 | 0.227 | 2.178 | -0.617 | -0.417 | 0.016 | 0.004 | 0.004 | 0.125 | 0.81 | 0.74 | 0.57 | 0.26 | 0.74 | 0.68 |
| 4 | 17280522 | rs6847269 | -0.747 | 1.704 | -0.673 | -0.312 | -0.01 | 0.004 | 0.004 | 0 | 0.25 | 0.23 | 0.23 | 0.05 | 0.25 | 0.2 |
| 4 | 17291007 | rs7663121 | -0.304 | 1.965 | -0.769 | 0.929 | -0.01 | -0.003 | -0.003 | 0.244 | 0.11 | 0.07 | 0.12 | 0 | 0.23 | 0.05 |
| 4 | 17291396 | rs4698560 | 0.123 | 1.092 | -0.789 | 0.602 | -0.006 | -0.003 | -0.003 | 0 | 0.65 | 0.46 | 0.64 | 0.64 | 0.68 | 0.65 |
| 4 | 17293238 | rs7694269 | 0.18 | 1.077 | -0.862 | 1.014 | -0.001 | 0 | 0 | 0.13 | 0.64 | 0.42 | 0.61 | 0.42 | 0.68 | 0.65 |
| 4 | 17294785 | rs6810740 | -0.597 | 0.739 | -1.053 | 1.254 | 0.006 | -0.033 | -0.033 | 0 | 0.21 | 0.17 | 0.21 | 0.31 | 0.15 | 0.16 |
| 4 | 17298007 | rs7690593 | 0.914 | 0.781 | -0.956 | 1.311 | 0.012 | 0.056 | 0.056 | 1.84 | 0.57 | 0.2 | 0.48 | 0.31 | 0.56 | 0.62 |
| 4 | 17298445 | rs7692897 | 0.53 | 0.126 | -0.168 | 1.287 | -0.039 | 0.087 | 0.087 | 0.913 | 0.83 | 0.66 | 0.82 | 0.7 | 0.92 | 0.84 |
| 4 | 17301096 | rs16894806 | 0.333 | 0.294 | -0.756 | 1.388 | -0.024 | -0.095 | -0.095 | 0 | 0.13 | 0.02 | 0.09 | 0.38 | 0.14 | 0.18 |
| 4 | 17318278 | rs6843133 | 1.039 | 0.39 | -0.37 | 1.263 | 0.019 | 0.016 | 0.016 | 3.977 | 0.63 | 0.19 | 0.62 | 0.34 | 0.61 | 0.7 |
| 4 | 17321714 | rs730308 | 0.898 | 0.427 | -0.293 | 1.278 | 0.009 | 0.008 | 0.008 | 2.949 | 0.64 | 0.34 | 0.63 | 0.34 | 0.61 | 0.7 |
| 4 | 17330381 | rs2872795 | 0.985 | 0.484 | -0.252 | 2.052 | 0 | -0.086 | -0.086 | 0.859 | 0.74 | 0.14 | 0.63 | 0.56 | 0.72 | 0.83 |
| 4 | 17330760 | rs13107521 | 1.573 | 0.696 | -0.751 | 1.695 | -0.052 | 0.244 | 0.244 | 2.433 | 0.53 | 0.09 | 0.51 | 0.17 | 0.5 | 0.62 |
| 4 | 17331894 | rs6823069 | -0.041 | 0.682 | 0.602 | 1.718 | -0.011 | 0.049 | 0.049 | 3.021 | 0.17 | 0.39 | 0.24 | 0.17 | 0.2 | 0.06 |
| 4 | 17335027 | rs16894897 | -0.992 | 0.788 | 1.008 | 1.777 | 0 | 0.016 | 0.016 | 0 | 0.24 | 0.34 | 0.16 | 0.4 | 0.24 | 0.27 |
| 4 | 17336925 | rs10003191 | 1.104 | 0.798 | 0.991 | 1.774 | 0.025 | 0.048 | 0.048 | 0.219 | 0.73 | 0.5 | 0.77 | 0.33 | 0.7 | 0.69 |
| 4 | 17344492 | rs16894917 | 0.057 | 0.772 | 0.429 | 1.838 | -0.048 | -0.099 | -0.099 | 0 | 0.17 | 0.02 | 0.11 | 0.38 | 0.2 | 0.2 |
| 4 | 17348363 | rs10000081 | 0.819 | 0.664 | 1.329 | 1.785 | 0.001 | 0.131 | 0.131 | 3.338 | 0.76 | 0.51 | 0.66 | 0.57 | 0.73 | 0.88 |
| 4 | 17349030 | rs13130097 | 0.062 | 0.72 | 1.452 | 1.876 | 0.023 | 0.187 | 0.187 | 2.047 | 0.82 | 0.63 | 0.69 | 0.57 | 0.74 | 0.9 |
| 4 | 17361114 | rs16894953 | -0.666 | 0.792 | 1.399 | 1.774 | 0 | -0.001 | -0.001 | 0 | 0.22 | 0.25 | 0.27 | 0.38 | 0.23 | 0.22 |
| 4 | 17361909 | rs2314958 | -0.481 | 0.806 | 1.328 | 1.779 | -0.006 | -0.035 | -0.035 | 0 | 0.21 | 0.11 | 0.26 | 0.38 | 0.23 | 0.24 |
| 4 | 17373412 | rs6449287 | -0.092 | 0.769 | 1.503 | 1.728 | 0.068 | 0.296 | 0.296 | 0.065 | 0.84 | 0.5 | 0.69 | 0.57 | 0.74 | 0.88 |
| 4 | 17379583 | rs1901168 | 0.784 | 0.552 | 0.112 | 1.148 | 0.007 | 0.011 | 0.011 | 0 | 0.54 | 0.42 | 0.47 | 0.37 | 0.6 | 0.43 |
| 4 | 17381717 | rs4698581 | -0.489 | 0.843 | -0.686 | 0.91 | -0.065 | -0.057 | -0.057 | 0 | 0.22 | 0.02 | 0.25 | 0.29 | 0.25 | 0.25 |
| 4 | 17385551 | rs2122574 | 0.292 | 0.554 | -0.075 | 0.437 | -0.002 | -0.012 | -0.012 | 0 | 0.73 | 0.42 | 0.58 | 0.76 | 0.7 | 0.83 |
| 4 | 17401363 | rs1868737 | -0.406 | 0.763 | -0.056 | 0.039 | 0.04 | 0.109 | 0.109 | 0.55 | 0.5 | 0.69 | 0.39 | 0.23 | 0.56 | 0.48 |
| 4 | 17402150 | rs10516312 | 0.56 | 1.137 | -0.084 | -0.331 | 0 | -0.016 | -0.016 | 0.104 | 0.23 | 0.24 | 0.12 | 0.12 | 0.2 | 0.28 |
| 4 | 17405513 | rs10012313 | 0.348 | 1.016 | -0.49 | -0.354 | 0.023 | -0.037 | -0.037 | 0 | 0.26 | 0.37 | 0.26 | 0.11 | 0.35 | 0.21 |
| 4 | 17408416 | rs1528604 | -0.47 | 0.585 | -1.244 | 0.15 | 0.077 | 0.01 | 0.01 | 0.907 | 0.53 | 0.82 | 0.62 | 0.41 | 0.63 | 0.52 |
| 4 | 17419691 | rs6856880 | 0.146 | 0.539 | -0.897 | 0.561 | 0.018 | -0.064 | -0.064 | 0.073 | 0.24 | 0.15 | 0.29 | 0.59 | 0.19 | 0.28 |
| 4 | 17431869 | rs7681641 | -0.152 | 0.858 | 0.779 | 0.187 | 0 | 0 | 0 | 0.659 | 0.71 | 0.37 | 0.61 | 0.7 | 0.71 | 0.66 |
| 4 | 17432275 | rs7682455 | -0.152 | 0.858 | 0.735 | 0.187 | 0 | 0 | 0 | 0.659 | 0.71 | 0.37 | 0.61 | 0.7 | 0.71 | 0.66 |
| 4 | 17435958 | rs2697684 | 0.442 | 0.733 | 0.844 | -0.098 | 0.001 | 0.029 | 0.029 | 0 | 0.24 | 0.16 | 0.17 | 0.13 | 0.22 | 0.3 |
| 4 | 17439346 | rs7671329 | 0.905 | 0.73 | 0.792 | -0.406 | 0 | -0.027 | -0.027 | 0 | 0.41 | 0.39 | 0.32 | 0.13 | 0.46 | 0.32 |
| 4 | 17448903 | rs2518652 | -1.051 | 0.754 | 0.966 | -0.338 | -0.002 | -0.007 | -0.007 | 1.226 | 0.58 | 0.62 | 0.68 | 0.86 | 0.54 | 0.69 |
| 4 | 17449199 | rs2697695 | -0.271 | 0.762 | 0.804 | -0.26 | 0.024 | 0.098 | 0.098 | 0 | 0.47 | 0.28 | 0.45 | 0.74 | 0.41 | 0.57 |
| 4 | 17450023 | rs1852748 | -0.086 | 0.76 | 0.849 | -0.215 | 0.062 | 0.053 | 0.053 | 0 | 0.89 | 0.67 | 0.77 | 0.88 | 0.86 | 0.88 |
| 4 | 17451760 | rs6855040 | 0.82 | 0.848 | 0.296 | -0.331 | -0.004 | 0.012 | 0.012 | 0 | 0.41 | 0.33 | 0.32 | 0.15 | 0.47 | 0.31 |
| 4 | 17453405 | rs2597810 | 0.086 | 0.923 | -0.081 | 0.155 | 0.001 | 0.025 | 0.025 | 0 | 0.23 | 0.16 | 0.16 | 0.14 | 0.21 | 0.31 |
| 4 | 17465704 | rs2053882 | 0.007 | 0.938 | -0.036 | 0.698 | 0.001 | 0.001 | 0.001 | 0 | 0.22 | 0.22 | 0.17 | 0.29 | 0.22 | 0.26 |
| 4 | 17467715 | rs2518641 | -0.762 | 0.618 | -0.088 | 0.725 | -0.009 | 0.045 | 0.045 | 0 | 0.24 | 0.27 | 0.24 | 0.47 | 0.22 | 0.28 |
| 4 | 17468047 | rs13112606 | 0.33 | 1.246 | -0.341 | 0.407 | -0.01 | 0.017 | 0.017 | 0 | 0.19 | 0.15 | 0.28 | 0.14 | 0.2 | 0.21 |
| 4 | 17469817 | rs10003746 | -0.022 | 0.841 | -0.244 | -0.423 | -0.003 | -0.103 | -0.103 | 0 | 0.56 | 0.47 | 0.77 | 0.97 | 0.64 | 0.62 |
| 4 | 17471978 | rs10007905 | 0.947 | 1.546 | -0.262 | -0.054 | 0.103 | 0.072 | 0.072 | 0.018 | 0.05 | 0.18 | 0.11 | 0.11 | 0.17 | 0.09 |
| 4 | 17472734 | rs2597749 | -0.333 | 1.065 | -0.579 | -0.185 | 0.061 | -0.025 | -0.025 | 0 | 0.62 | 0.28 | 0.68 | 0.51 | 0.75 | 0.68 |
| 4 | 17475541 | rs2597751 | -0.379 | 1.122 | -0.027 | 0.12 | 0.003 | 0.02 | 0.02 | 0 | 0.53 | 0.66 | 0.46 | 0.56 | 0.55 | 0.56 |
| 4 | 17478268 | rs4698599 | -0.719 | 1.147 | -0.03 | -0.231 | 0.031 | -0.027 | -0.027 | 0 | 0.33 | 0.45 | 0.43 | 0.25 | 0.44 | 0.3 |
| 4 | 17479402 | rs13128422 | 1.123 | 0.985 | -0.185 | -0.164 | -0.011 | -0.001 | -0.001 | 0 | 0.11 | 0.05 | 0.2 | 0.14 | 0.13 | 0.15 |
| 4 | 17483682 | rs13138275 | -0.003 | 1.046 | 0.011 | -0.227 | 0.002 | -0.001 | -0.001 | 1.26 | 0.15 | 0.18 | 0.26 | 0.18 | 0.2 | 0.22 |
| 4 | 17484602 | rs2597754 | 0.841 | 0.809 | -0.237 | -0.702 | 0.008 | 0.129 | 0.129 | 0 | 0.34 | 0.1 | 0.15 | 0.02 | 0.28 | 0.38 |
| 4 | 17485081 | rs2597756 | 0.017 | 1.029 | 0.193 | -0.632 | 0.014 | -0.029 | -0.029 | 0 | 0.69 | 0.46 | 0.54 | 0.76 | 0.69 | 0.68 |
| 4 | 17488064 | rs1031326 | 0.133 | 1.009 | 0.144 | -0.628 | 0.032 | -0.015 | -0.015 | 0 | 0.71 | 0.57 | 0.52 | 0.71 | 0.67 | 0.65 |
| 4 | 17488072 | rs699459 | -0.049 | 1.017 | 0.143 | -0.628 | 0.026 | -0.032 | -0.032 | 0 | 0.72 | 0.57 | 0.56 | 0.76 | 0.71 | 0.69 |
| 4 | 17488132 | rs10604 | 0.814 | 0.909 | 0.137 | -0.768 | 0.037 | 0.125 | 0.125 | 0 | 0.42 | 0.22 | 0.19 | 0.02 | 0.33 | 0.4 |
| 4 | 17488286 | rs699460 | -0.125 | 0.907 | 0.163 | -0.739 | 0.026 | -0.032 | -0.032 | 0 | 0.72 | 0.57 | 0.56 | 0.76 | 0.71 | 0.68 |
| 4 | 17488346 | rs1049601 | 0.356 | 0.864 | 0.201 | -0.458 | 0.004 | 0.426 | 0.426 | 0 | 0.06 | 0.27 | 0.13 | 0.43 | 0.02 | 0.09 |
| 4 | 17488351 | rs1049600 | 0.356 | 0.864 | 0.201 | -0.458 | 0.004 | 0.426 | 0.426 | 0 | 0.06 | 0.27 | 0.13 | 0.43 | 0.02 | 0.09 |
| 4 | 17492843 | rs744731 | 1.412 | 0.91 | -0.227 | -0.488 | -0.046 | -0.011 | -0.011 | 0 | 0.23 | 0.09 | 0.2 | 0.26 | 0.31 | 0.16 |

| | | | | | | | | | | | | | | | | |
|---|---|---|---|---|---|---|---|---|---|---|---|---|---|---|---|---|
| 4 | 17494643 | rs12512471 | 0.721 | 0.866 | -0.155 | -0.342 | 0.026 | 0.507 | 0.507 | 0 | 0.05 | 0.24 | 0.13 | 0.43 | 0.02 | 0.09 |
| 4 | 17500905 | rs11722315 | 1.366 | 0.984 | -1.137 | -1.047 | -0.022 | 0.005 | 0.005 | 0.162 | 0.14 | 0.06 | 0.2 | 0.11 | 0.16 | 0.13 |
| 4 | 17503382 | rs2597775 | 0.362 | 0.632 | -1.082 | -1.322 | 0.027 | 0.103 | 0.103 | 0 | 0.41 | 0.22 | 0.19 | 0.02 | 0.33 | 0.4 |
| 4 | 17510236 | rs17458406 | 0.803 | 1.2 | -0.932 | -0.515 | -0.013 | 0.004 | 0.004 | 0 | 0.1 | 0.05 | 0.2 | 0.1 | 0.13 | 0.18 |
| 4 | 17516691 | rs2597783 | 0.2 | 0.781 | -1.303 | -0.485 | 0.023 | -0.045 | -0.045 | 0.065 | 0.36 | 0.22 | 0.44 | 0.47 | 0.31 | 0.32 |
| 4 | 17518920 | rs7661303 | 0.556 | 0.717 | -1.125 | -0.773 | -0.023 | -0.117 | -0.117 | 0.727 | 0.31 | 0.45 | 0.18 | 0.1 | 0.24 | 0.29 |
| 4 | 17519867 | rs4698603 | 1.279 | 0.941 | -1.134 | -0.946 | -0.034 | -0.054 | -0.054 | 0.189 | 0.23 | 0.1 | 0.21 | 0.33 | 0.31 | 0.21 |
| 4 | 17526682 | rs10020773 | 0.434 | 0.41 | -0.831 | -1.155 | -0.02 | -0.161 | -0.161 | 0.312 | 0.3 | 0.45 | 0.18 | 0.01 | 0.24 | 0.25 |
| 4 | 17528463 | rs2597791 | 0.413 | 0.443 | -0.872 | -0.396 | -0.011 | -0.032 | -0.032 | 1.396 | 0.09 | 0.12 | 0.03 | 0 | 0.03 | 0.06 |
| 4 | 17533530 | rs1609899 | 0.314 | 0.46 | -0.804 | -0.394 | -0.02 | -0.052 | -0.052 | 0 | 0.1 | 0.15 | 0.04 | 0 | 0.04 | 0.06 |
| 4 | 17537159 | rs2518649 | -0.312 | 0.638 | -0.921 | -0.283 | -0.001 | 0.002 | 0.002 | 0 | 0.46 | 0.37 | 0.42 | 0.41 | 0.5 | 0.43 |
| 4 | 17546381 | rs6843189 | -0.346 | 0.403 | -1.559 | -0.119 | -0.005 | 0.115 | 0.115 | 0 | 0.51 | 0.32 | 0.56 | 0.8 | 0.55 | 0.6 |
| 4 | 17550349 | rs2109520 | -0.343 | 0.505 | -1.11 | 0.246 | -0.011 | -0.012 | -0.012 | 0 | 0.31 | 0.27 | 0.26 | 0.35 | 0.38 | 0.49 |
| 4 | 17561139 | rs6828925 | 0.559 | -0.255 | -0.639 | -0.414 | 0.03 | 0.09 | 0.09 | 0.041 | 0.57 | 0.32 | 0.7 | 0.81 | 0.65 | 0.42 |
| 4 | 17582051 | rs3796812 | 0.572 | -0.424 | -0.532 | -0.568 | -0.009 | 0.175 | 0.175 | 0.008 | 0.61 | 0.15 | 0.68 | 0.85 | 0.65 | 0.55 |
| 4 | 17586703 | rs3733576 | 0.057 | -0.67 | 0.258 | -0.687 | -0.017 | 0.073 | 0.073 | 0.017 | 0.68 | 0.84 | 0.76 | 0.9 | 0.66 | 0.63 |
| 4 | 17597549 | rs6812675 | 0.065 | -0.74 | 1.899 | -0.728 | 0.003 | -0.059 | -0.059 | 0 | 0.69 | 0.67 | 0.76 | 0.9 | 0.67 | 0.63 |
| 4 | 17608620 | rs4698628 | 0.407 | -0.774 | 2.239 | -0.735 | 0 | -0.037 | -0.037 | 0 | 0.64 | 0.4 | 0.7 | 0.86 | 0.65 | 0.54 |
| 4 | 17617111 | rs4642241 | 0.882 | -0.86 | 2.219 | -0.747 | -0.007 | 0.187 | 0.187 | 0 | 0.59 | 0.15 | 0.66 | 0.85 | 0.63 | 0.54 |
| 4 | 17618713 | rs1006722 | 0.858 | -0.862 | 2.578 | -0.748 | 0.005 | 0.099 | 0.099 | 0 | 0.6 | 0.24 | 0.67 | 0.85 | 0.63 | 0.54 |
| 4 | 17625658 | rs555 | 0.868 | -0.862 | 2.579 | -0.766 | 0.004 | 0.117 | 0.117 | 0 | 0.6 | 0.21 | 0.67 | 0.85 | 0.63 | 0.54 |
| 4 | 17643848 | rs2286771 | 0.872 | -0.85 | 2.686 | -0.754 | -0.004 | 0.179 | 0.179 | 0 | 0.59 | 0.17 | 0.66 | 0.85 | 0.62 | 0.54 |
| 4 | 17650758 | rs1558377 | 0.596 | -0.851 | 2.701 | -0.754 | -0.001 | -0.077 | -0.077 | 0.223 | 0.62 | 0.5 | 0.69 | 0.86 | 0.62 | 0.57 |
| 4 | 17652450 | rs10805358 | 0.854 | -0.839 | 2.638 | -0.748 | -0.004 | 0.178 | 0.178 | 0 | 0.59 | 0.17 | 0.66 | 0.85 | 0.62 | 0.54 |
| 4 | 17660082 | rs1860596 | 0.729 | -0.832 | 2.539 | -0.745 | -0.002 | -0.041 | -0.041 | 0 | 0.61 | 0.4 | 0.68 | 0.85 | 0.63 | 0.55 |
| 4 | 17667766 | rs11934968 | -0.304 | -0.908 | 2.242 | -0.817 | -0.022 | 0.053 | 0.053 | 0 | 0.29 | 0.19 | 0.24 | 0.1 | 0.35 | 0.38 |
| 4 | 17682218 | rs10009541 | -0.652 | -0.859 | 2.532 | -0.526 | 0.007 | -0.111 | -0.111 | 1.632 | 0.34 | 0.47 | 0.27 | 0.1 | 0.36 | 0.38 |
| 4 | 17682834 | rs1008375 | 0.677 | -0.843 | 2.241 | -0.479 | 0.002 | 0.024 | 0.024 | 0 | 0.61 | 0.3 | 0.69 | 0.81 | 0.63 | 0.57 |
| 4 | 17683811 | rs11722037 | 0.798 | -0.821 | 1.566 | -0.468 | 0.006 | 0.084 | 0.084 | 0 | 0.6 | 0.24 | 0.68 | 0.81 | 0.63 | 0.54 |
| 4 | 17684951 | rs10939746 | 0.725 | -0.799 | 1.536 | -0.458 | 0.004 | 0.045 | 0.045 | 0 | 0.61 | 0.31 | 0.68 | 0.81 | 0.63 | 0.54 |
| 4 | 17691047 | rs11947857 | -0.225 | -0.439 | 1.303 | -0.279 | -0.013 | 0.02 | 0.02 | 0.126 | 0.3 | 0.21 | 0.28 | 0.11 | 0.36 | 0.41 |
| 4 | 17692782 | rs4698649 | 0.271 | -0.473 | 1.517 | -0.124 | 0.014 | -0.078 | -0.078 | 0.002 | 0.66 | 0.49 | 0.7 | 0.85 | 0.63 | 0.56 |
| 4 | 17696434 | rs6825100 | 0.975 | -0.238 | 0.786 | 0.112 | -0.093 | 0.082 | 0.082 | 1.714 | 0.38 | 0.07 | 0.3 | 0.28 | 0.41 | 0.26 |
| 4 | 17713717 | rs11735608 | 0.144 | 0.069 | 1.691 | 0.943 | 0.001 | 0.121 | 0.121 | 0 | 0.82 | 0.57 | 0.62 | 0.41 | 0.79 | 0.75 |
| 4 | 17727941 | rs4631043 | -0.4 | 0.091 | 1.455 | 0.585 | 0 | -0.001 | -0.001 | 0 | 0.34 | 0.3 | 0.26 | 0.21 | 0.29 | 0.35 |
| 4 | 17728356 | rs13148666 | -0.225 | 0.133 | 1.247 | 0.578 | 0.002 | 0.034 | 0.034 | 1.249 | 0.32 | 0.22 | 0.23 | 0.14 | 0.29 | 0.32 |
| 4 | 17747023 | rs6844932 | -0.125 | 0.5 | 0.193 | 1.154 | 0.006 | 0.118 | 0.118 | 0 | 0.84 | 0.68 | 0.62 | 0.48 | 0.8 | 0.79 |
| 4 | 17765622 | rs11730953 | 0.73 | 0.571 | -0.804 | 0.292 | -0.069 | 0.231 | 0.231 | 3.887 | 0.46 | 0.09 | 0.37 | 0.18 | 0.49 | 0.4 |
| 4 | 17769238 | rs17583557 | 0.748 | 0.606 | -0.873 | 0.295 | -0.09 | 0.233 | 0.233 | 3.947 | 0.45 | 0.07 | 0.37 | 0.18 | 0.49 | 0.39 |
| 4 | 17780932 | rs7682632 | -1.002 | 1.852 | 0.206 | 1.566 | 0.095 | 0.141 | 0.141 | 0 | 0.12 | 0.21 | 0.35 | 0.36 | 0.25 | 0.14 |
| 4 | 17781209 | rs16895707 | -0.753 | 0.795 | 0.359 | 1.207 | -0.008 | 0.001 | 0.001 | 0 | 0.25 | 0.26 | 0.18 | 0.31 | 0.2 | 0.31 |
| 4 | 17782264 | rs740672 | -0.885 | 0.876 | 0.363 | 1.223 | -0.003 | 0 | 0 | 0 | 0.27 | 0.25 | 0.21 | 0.31 | 0.22 | 0.32 |
| 4 | 17789420 | rs16895737 | -0.873 | 0.412 | 0.601 | 1.132 | 0.018 | 0.013 | 0.013 | 0 | 0.22 | 0.16 | 0.13 | 0.16 | 0.1 | 0.27 |
| 4 | 17797455 | rs7672919 | -1.6 | 0.234 | 1.482 | 1.259 | -0.073 | -0.003 | -0.003 | 0 | 0.28 | 0.53 | 0.17 | 0.26 | 0.12 | 0.36 |
| 4 | 17797966 | rs7678436 | -1.632 | 0.22 | 1.475 | 1.462 | -0.073 | 0.02 | 0.02 | 0 | 0.28 | 0.55 | 0.18 | 0.32 | 0.12 | 0.41 |
| 4 | 17809538 | rs2286540 | -0.85 | 0.25 | 1.464 | 1.379 | 0.018 | 0.013 | 0.013 | 0 | 0.22 | 0.16 | 0.13 | 0.16 | 0.1 | 0.25 |
| 4 | 17811914 | rs7667864 | 1.257 | 1.33 | 1.623 | 1.563 | 0.118 | 0.198 | 0.198 | 0 | 0.86 | 0.7 | 0.63 | 0.55 | 0.71 | 0.85 |
| 4 | 17812615 | rs2074974 | 1.311 | 1.365 | 1.995 | 1.375 | 0.129 | 0.217 | 0.217 | 0 | 0.86 | 0.69 | 0.63 | 0.54 | 0.71 | 0.87 |
| 4 | 17813711 | rs1395371 | -0.561 | 1.574 | 1.838 | 1.363 | -0.03 | 0.026 | 0.026 | 0 | 0.07 | 0.03 | 0.28 | 0.46 | 0.2 | 0.08 |
| 4 | 17846522 | rs16895877 | -0.361 | 1.282 | 1.968 | 1.303 | 0.022 | 0.008 | 0.008 | 0 | 0.17 | 0.1 | 0.12 | 0.16 | 0.09 | 0.26 |
| 4 | 17847281 | rs16895878 | -0.343 | 1.283 | 1.946 | 1.304 | 0.026 | 0.009 | 0.009 | 0 | 0.17 | 0.09 | 0.12 | 0.16 | 0.09 | 0.26 |
| 4 | 17854055 | rs6842303 | 1.072 | 1.338 | 2.274 | 1.319 | 0.099 | 0.176 | 0.176 | 0 | 0.88 | 0.76 | 0.66 | 0.54 | 0.72 | 0.87 |
| 4 | 17854768 | rs16895895 | -0.367 | 1.114 | 1.829 | 1.271 | 0.026 | 0.009 | 0.009 | 0 | 0.17 | 0.09 | 0.12 | 0.16 | 0.09 | 0.26 |
| 4 | 17919811 | rs925098 | 0.86 | 1.705 | 1.936 | 1.344 | 0.135 | 0.21 | 0.21 | 0 | 0.84 | 0.64 | 0.64 | 0.53 | 0.7 | 0.86 |
| 4 | 17957354 | rs7684221 | -1.131 | 1.424 | 1.809 | 1.173 | -0.065 | -0.047 | -0.047 | 3.38 | 0.25 | 0.46 | 0.17 | 0.16 | 0.13 | 0.3 |
| 4 | 17970655 | rs6853216 | -1.031 | 1.417 | 1.803 | 1.169 | -0.081 | -0.059 | -0.059 | 0 | 0.25 | 0.52 | 0.18 | 0.16 | 0.13 | 0.3 |
| 4 | 17992332 | rs16896210 | -1.124 | 1.406 | 1.791 | 1.164 | -0.076 | -0.054 | -0.054 | 0 | 0.25 | 0.48 | 0.18 | 0.16 | 0.13 | 0.3 |
| 4 | 17993410 | rs724577 | 0.654 | 1.566 | 1.594 | 1.245 | 0.117 | 0.19 | 0.19 | 0 | 0.85 | 0.67 | 0.65 | 0.53 | 0.71 | 0.87 |
| 4 | 18004014 | rs13116230 | -0.987 | 0.745 | 1.376 | 0.715 | -0.045 | -0.027 | -0.027 | 1.974 | 0.23 | 0.4 | 0.16 | 0.16 | 0.11 | 0.29 |
| 4 | 18011864 | rs11936089 | -0.967 | 0.756 | 1.358 | 0.72 | -0.044 | -0.026 | -0.026 | 0 | 0.23 | 0.39 | 0.15 | 0.16 | 0.1 | 0.13 |
| 4 | 18035250 | rs2061455 | -1.14 | 0.833 | 1.043 | 0.806 | -0.061 | -0.035 | -0.035 | 0.465 | 0.23 | 0.44 | 0.18 | 0.17 | 0.13 | 0.27 |
| 4 | 18049335 | rs1459331 | 0.776 | 2.032 | 0.876 | 1.476 | 0.034 | 0.07 | 0.07 | 0 | 0.84 | 0.76 | 0.65 | 0.59 | 0.69 | 0.89 |
| 4 | 18073943 | rs4698667 | -1.535 | 1.826 | 0.739 | 1.299 | 0.006 | 0.046 | 0.046 | 0 | 0.19 | 0.25 | 0.32 | 0.41 | 0.21 | 0.14 |
| 4 | 18079517 | rs9291662 | -2.216 | 1.606 | 1.425 | 1.315 | 0 | 0.039 | 0.039 | 0.956 | 0.25 | 0.68 | 0.37 | 0.5 | 0.22 | 0.21 |
| 4 | 18111823 | rs10004929 | -1.538 | 1.594 | 1.996 | 1.362 | 0.01 | 0.054 | 0.054 | 0 | 0.2 | 0.28 | 0.33 | 0.4 | 0.21 | 0.15 |
| 4 | 18124727 | rs6854314 | -1.847 | 1.676 | 2.217 | 1.286 | 0.021 | 0.113 | 0.113 | 0.686 | 0.23 | 0.55 | 0.36 | 0.41 | 0.22 | 0.15 |
| 4 | 18124909 | rs6854692 | -1.883 | 1.274 | 1.841 | 1.058 | 0 | 0 | 0 | 0.043 | 0.33 | 0.67 | 0.54 | 0.57 | 0.27 | 0.23 |
| 4 | 18131245 | rs1503872 | 1.354 | 1.499 | 0.634 | 0.442 | 0.09 | 0.154 | 0.154 | 3.863 | 0.48 | 0.15 | 0.35 | 0.27 | 0.29 | 0.55 |
| 4 | 18135757 | rs7687236 | -0.38 | 1.834 | 0.441 | 0.585 | -0.113 | -0.115 | -0.115 | 1.608 | 0.75 | 0.98 | 0.73 | 0.65 | 0.55 | 0.74 |

| | | | | | | | | | | | | | | | | |
|---|---|---|---|---|---|---|---|---|---|---|---|---|---|---|---|---|
| 4 | 18140181 | rs7684006 | 1.482 | 1.695 | 0.981 | 0.433 | -0.002 | -0.012 | -0.012 | 0 | 0.53 | 0.44 | 0.44 | 0.26 | 0.34 | 0.56 |
| 4 | 18147151 | rs11934728 | -1.462 | 1.616 | 1.256 | 0.478 | 0.02 | -0.065 | -0.065 | 0 | 0.11 | 0.54 | 0.1 | 0.01 | 0.07 | 0.07 |
| 4 | 18162104 | rs2320289 | -0.654 | 1.764 | 1.051 | 0.714 | 0.017 | 0.076 | 0.076 | 0 | 0.15 | 0.21 | 0.27 | 0.46 | 0.2 | 0.17 |
| 4 | 18169817 | rs1847401 | -1.844 | 1.607 | 1.35 | 0.603 | 0.036 | 0.032 | 0.032 | 4.081 | 0.06 | 0.45 | 0.06 | 0.01 | 0.01 | 0 |
| 4 | 18179501 | rs2320467 | -0.169 | 2.369 | 1.004 | 0.635 | -0.039 | -0.006 | -0.006 | 1.283 | 0.89 | 0.94 | 0.88 | 0.9 | 0.71 | 0.86 |
| 4 | 18181274 | rs4698221 | 1.49 | 2.328 | 0.895 | 0.679 | -0.002 | 0 | 0 | 0.946 | 0.65 | 0.33 | 0.69 | 0.68 | 0.51 | 0.68 |
| 4 | 18181391 | rs16896557 | 0.146 | 2.284 | 0.57 | 0.757 | -0.003 | 0.005 | 0.005 | 0.711 | 0.16 | 0.14 | 0.11 | 0.15 | 0.18 | 0.17 |
| 4 | 18191999 | rs6449361 | 0.656 | 1.915 | 0.749 | 0.531 | 0.01 | -0.003 | -0.003 | 0 | 0.8 | 0.75 | 0.84 | 0.85 | 0.69 | 0.85 |
| 4 | 18193446 | rs13150576 | -0.441 | 2.164 | 0.8 | 0.681 | -0.028 | -0.041 | -0.041 | 3.779 | 0.22 | 0.2 | 0.34 | 0.54 | 0.31 | 0.24 |
| 4 | 18194063 | rs6449362 | -1.458 | 2.128 | 1.105 | 0.626 | 0.053 | 0.031 | 0.031 | 0.46 | 0.29 | 0.47 | 0.47 | 0.64 | 0.38 | 0.34 |
| 4 | 18195120 | rs6856758 | -0.778 | 2.079 | 0.873 | 0.826 | 0.01 | 0.028 | 0.028 | 0.947 | 0.23 | 0.32 | 0.35 | 0.52 | 0.31 | 0.23 |
| 4 | 18202615 | rs1394084 | 0.321 | 1.952 | 0.635 | 0.79 | -0.009 | -0.007 | -0.007 | 0 | 0.82 | 0.85 | 0.85 | 0.81 | 0.71 | 0.85 |
| 4 | 18203435 | rs13106632 | -0.756 | 2.109 | 0.498 | 0.812 | -0.01 | -0.025 | -0.025 | 0.368 | 0.25 | 0.25 | 0.36 | 0.5 | 0.29 | 0.25 |
| 4 | 18207538 | rs1503884 | 1.665 | 2.118 | 0.469 | 0.669 | -0.003 | 0.029 | 0.029 | 0.032 | 0.55 | 0.33 | 0.46 | 0.31 | 0.42 | 0.6 |
| 4 | 18208997 | rs4698673 | 0.316 | 2.146 | 0.241 | 0.822 | -0.022 | -0.005 | -0.005 | 0.322 | 0.84 | 0.91 | 0.85 | 0.84 | 0.7 | 0.85 |
| 4 | 18218019 | rs1503888 | 0.046 | 2.161 | 0.165 | 1.001 | -0.051 | -0.039 | -0.039 | 0 | 0.85 | 0.96 | 0.86 | 0.82 | 0.71 | 0.86 |
| 4 | 18218684 | rs10009303 | 0.587 | 2.101 | 0.235 | 0.972 | -0.024 | -0.012 | -0.012 | 0.518 | 0.82 | 0.91 | 0.84 | 0.81 | 0.69 | 0.85 |
| 4 | 18220536 | rs6841071 | 0.174 | 1.321 | 0.23 | 0.595 | -0.058 | -0.075 | -0.075 | 0 | 0.23 | 0.48 | 0.12 | 0.12 | 0.12 | 0.22 |
| 4 | 18243314 | rs6811915 | -0.337 | 1.492 | 0.029 | 0.67 | -0.051 | -0.023 | -0.023 | 0 | 0.84 | 0.96 | 0.85 | 0.83 | 0.7 | 0.86 |
| 4 | 18246425 | rs17527941 | 1.492 | 1.691 | 0.033 | 0.433 | 0.016 | 0.048 | 0.048 | 0 | 0.5 | 0.36 | 0.45 | 0.27 | 0.39 | 0.51 |
| 4 | 18250003 | rs10488994 | 0.584 | 1.639 | -0.117 | 0.352 | -0.018 | -0.024 | -0.024 | 0 | 0.21 | 0.3 | 0.11 | 0.12 | 0.12 | 0.23 |
| 4 | 18255647 | rs4356908 | 1.185 | 1.677 | -0.232 | 0.314 | 0.035 | 0.17 | 0.17 | 0.275 | 0.45 | 0.14 | 0.4 | 0.15 | 0.34 | 0.35 |
| 4 | 18264644 | rs2192521 | 0.203 | 1.396 | -0.643 | 0.484 | -0.008 | 0.031 | 0.031 | 0 | 0.43 | 0.69 | 0.44 | 0.35 | 0.37 | 0.5 |
| 4 | 18272243 | rs2110497 | -0.985 | 1.779 | -0.358 | 0.994 | 0.013 | 0.012 | 0.012 | 0 | 0.94 | 0.95 | 0.94 | 0.94 | 0.87 | 0.95 |
| 4 | 18277716 | rs9291670 | -0.057 | 1.498 | 0.067 | 1.015 | 0.014 | 0.046 | 0.046 | 3.32 | 0.06 | 0.2 | 0.05 | 0.13 | 0.04 | 0.16 |
| 4 | 18280561 | rs1558588 | 1.173 | 1.51 | -0.072 | 0.758 | 0.006 | 0.024 | 0.024 | 0 | 0.13 | 0.15 | 0.23 | 0.22 | 0.11 | 0.11 |
| 4 | 18281475 | rs10488992 | 0.971 | 1.352 | -0.416 | 0.65 | -0.019 | 0.003 | 0.003 | 2.143 | 0.18 | 0.04 | 0.11 | 0.14 | 0.2 | 0.11 |
| 4 | 18282476 | rs4505820 | 0.195 | 1.162 | -0.26 | 0.566 | -0.009 | -0.019 | -0.019 | 0 | 0.31 | 0.21 | 0.28 | 0.38 | 0.32 | 0.23 |
| 4 | 18293144 | rs13147572 | 1.083 | 1.365 | -0.559 | 0.522 | 0.02 | 0.015 | 0.015 | 0.073 | 0.46 | 0.2 | 0.35 | 0.43 | 0.41 | 0.49 |
| 4 | 18293651 | rs16896868 | -0.935 | 0.898 | 0.166 | 0.729 | -0.001 | -0.005 | -0.005 | 0 | 0.43 | 0.6 | 0.45 | 0.53 | 0.41 | 0.42 |
| 4 | 18296221 | rs11730060 | -0.345 | 0.769 | 0.777 | 0.868 | 0.022 | 0.064 | 0.064 | 2.59 | 0.05 | 0.19 | 0.05 | 0.13 | 0.05 | 0.16 |
| 4 | 18315750 | rs16896934 | -0.377 | 0.95 | -0.128 | 0.939 | 0.006 | 0.016 | 0.016 | 0 | 0.15 | 0.16 | 0.2 | 0.25 | 0.18 | 0.16 |
| 4 | 18315992 | rs4140904 | 0.658 | 0.221 | -0.212 | 0.803 | -0.029 | -0.001 | -0.001 | 0 | 0.41 | 0.51 | 0.19 | 0.42 | 0.24 | 0.44 |
| 4 | 18321092 | rs741207 | -0.079 | 0.176 | -0.006 | 0.765 | 0.001 | 0.046 | 0.046 | 3.193 | 0.53 | 0.79 | 0.4 | 0.62 | 0.4 | 0.56 |
| 4 | 18321896 | rs7693245 | -0.683 | -0.303 | -0.454 | 0.469 | -0.088 | -0.032 | -0.032 | 0.104 | 0.67 | 0.86 | 0.45 | 0.63 | 0.46 | 0.68 |
| 4 | 18325391 | rs6849596 | 0.41 | -0.308 | -0.297 | 0.479 | 0.003 | 0.01 | 0.01 | 3.051 | 0.53 | 0.6 | 0.38 | 0.62 | 0.4 | 0.56 |
| 4 | 18377649 | rs11733969 | 1.528 | -0.406 | -0.217 | 0.361 | 0.039 | -0.021 | -0.021 | 0 | 0.41 | 0.22 | 0.19 | 0.43 | 0.29 | 0.44 |
| 4 | 18381481 | rs2160107 | 0.767 | -0.029 | -0.271 | 0.698 | -0.005 | 0.001 | 0.001 | 0.022 | 0.13 | 0.1 | 0.19 | 0.29 | 0.16 | 0.13 |
| 4 | 18382212 | rs1395099 | -0.598 | -0.024 | -0.356 | 0.703 | -0.084 | -0.031 | -0.031 | 0.32 | 0.73 | 0.87 | 0.49 | 0.64 | 0.52 | 0.7 |
| 4 | 18396021 | rs6833761 | 0.4 | -0.121 | 0.108 | 0.838 | 0.007 | 0.035 | 0.035 | 0 | 0.14 | 0.16 | 0.2 | 0.3 | 0.17 | 0.15 |
| 4 | 18407127 | rs10019574 | -1.141 | -0.51 | 0.133 | 0.675 | 0.025 | -0.03 | -0.03 | 0 | 0.33 | 0.74 | 0.31 | 0.38 | 0.23 | 0.35 |
| 4 | 18408920 | rs6843156 | -0.388 | -0.501 | 0.543 | 0.68 | -0.006 | 0.015 | 0.015 | 0 | 0.27 | 0.32 | 0.26 | 0.33 | 0.21 | 0.32 |
| 4 | 18417138 | rs4698690 | 1.161 | -0.522 | 0.244 | 0.705 | 0.021 | -0.004 | -0.004 | 0.52 | 0.12 | 0.07 | 0.04 | 0.13 | 0.05 | 0.18 |
| 4 | 18432045 | rs976812 | -1.232 | -0.626 | 0.282 | 0.544 | 0.037 | 0.041 | 0.041 | 0.798 | 0.36 | 0.76 | 0.32 | 0.21 | 0.24 | 0.24 |
| 4 | 18433540 | rs13114254 | 1.516 | -0.333 | -0.067 | 0.608 | 0.028 | 0.056 | 0.056 | 0.094 | 0.39 | 0.14 | 0.17 | 0.26 | 0.28 | 0.35 |
| 4 | 18437923 | rs12646743 | 1.011 | -0.325 | 0.047 | 0.6 | -0.006 | -0.006 | -0.006 | 3.278 | 0.13 | 0.1 | 0.2 | 0.2 | 0.17 | 0.12 |
| 4 | 18442072 | rs13140382 | 1.395 | -0.423 | 0.077 | 0.527 | 0.027 | 0.045 | 0.045 | 0.136 | 0.39 | 0.22 | 0.18 | 0.26 | 0.28 | 0.36 |
| 4 | 18448350 | rs10022178 | 0.934 | -0.525 | 0.265 | 0.375 | 0.018 | 0.015 | 0.015 | 0.017 | 0.46 | 0.6 | 0.23 | 0.32 | 0.3 | 0.37 |
| 4 | 18453713 | rs13136984 | 1.001 | -0.319 | 0.152 | 0.353 | 0.023 | 0.026 | 0.026 | 0 | 0.44 | 0.36 | 0.22 | 0.32 | 0.35 | 0.39 |
| 4 | 18461495 | rs10034918 | -1.135 | 0.053 | -0.281 | 0.391 | -0.097 | -0.087 | -0.087 | 0.131 | 0.84 | 0.96 | 0.55 | 0.66 | 0.63 | 0.81 |
| 4 | 18465884 | rs11728685 | 1.711 | -0.255 | -0.163 | 0.366 | 0.008 | 0.015 | 0.015 | 0.251 | 0.38 | 0.19 | 0.18 | 0.31 | 0.32 | 0.36 |
| 4 | 18473310 | rs1354676 | 0.035 | -0.16 | -0.103 | 0.423 | -0.032 | -0.035 | -0.035 | 0.106 | 0.72 | 0.82 | 0.48 | 0.53 | 0.56 | 0.6 |
| 4 | 18481635 | rs10516323 | 0.705 | -0.297 | -0.03 | 0.353 | 0.006 | 0.014 | 0.014 | 0 | 0.59 | 0.72 | 0.28 | 0.33 | 0.39 | 0.48 |
| 4 | 18482287 | rs16897089 | 0.855 | -0.291 | 0.224 | 0.372 | 0.008 | 0.005 | 0.005 | 4.791 | 0.06 | 0.29 | 0.04 | 0 | 0.01 | 0 |
| 4 | 18502355 | rs11936950 | 1.696 | -0.288 | 0.109 | 0.369 | 0.008 | 0.013 | 0.013 | 0.063 | 0.39 | 0.29 | 0.19 | 0.31 | 0.32 | 0.36 |
| 4 | 18511288 | rs1477892 | -1.374 | 0.535 | -0.064 | 0.478 | -0.044 | -0.065 | -0.065 | 0 | 0.9 | 0.99 | 0.74 | 0.69 | 0.83 | 0.92 |
| 4 | 18515483 | rs1477886 | -0.021 | 0.039 | 0.226 | 0.502 | 0.001 | 0.001 | 0.001 | 0 | 0.7 | 0.69 | 0.46 | 0.53 | 0.54 | 0.57 |
| 4 | 18553345 | rs13128839 | 1.824 | -0.468 | -0.609 | -0.262 | -0.048 | 0.131 | 0.131 | 3.013 | 0.23 | 0.04 | 0.12 | 0.01 | 0.23 | 0.17 |
| 4 | 18569229 | rs12507985 | 0.975 | -0.487 | -0.642 | -0.01 | -0.021 | -0.029 | -0.029 | 0 | 0.14 | 0.08 | 0.27 | 0.22 | 0.17 | 0.14 |
| 4 | 18572708 | rs2643431 | 0.619 | -0.573 | -0.698 | 0.001 | -0.084 | -0.032 | -0.032 | 0 | 0.24 | 0.09 | 0.42 | 0.33 | 0.43 | 0.3 |
| 4 | 18575715 | rs4235396 | -0.69 | -0.991 | -0.156 | 0.063 | 0.001 | 0 | 0 | 0.174 | 0.57 | 0.47 | 0.78 | 0.63 | 0.67 | 0.67 |
| 4 | 18581701 | rs922388 | 1.116 | -1.21 | 0.6 | 0.683 | 0.004 | -0.004 | -0.004 | 0 | 0.16 | 0.14 | 0.07 | 0.35 | 0.11 | 0.17 |
| 4 | 18596590 | rs6835867 | -1.267 | -0.514 | 1.255 | 0.662 | -0.025 | 0.053 | 0.053 | 0.482 | 0.28 | 0.42 | 0.45 | 0.43 | 0.2 | 0.37 |
| 4 | 18605502 | rs10021749 | -0.685 | -0.521 | 0.776 | 0.658 | 0.018 | -0.067 | -0.067 | 0.233 | 0.23 | 0.15 | 0.4 | 0.43 | 0.18 | 0.37 |
| 4 | 18609772 | rs11734672 | 1.171 | -0.528 | 0.613 | 0.656 | 0.035 | -0.042 | -0.042 | 0 | 0.67 | 0.23 | 0.52 | 0.57 | 0.81 | 0.63 |
| 4 | 18612043 | rs7696154 | 0.635 | -0.303 | 0.133 | 0.433 | 0.007 | -0.003 | -0.003 | 0 | 0.1 | 0.11 | 0.04 | 0 | 0.04 | 0.06 |
| 4 | 18617189 | rs16897260 | -0.304 | -0.062 | 0.147 | 0.61 | 0.004 | 0.006 | 0.006 | 0.086 | 0.06 | 0.1 | 0.07 | 0.11 | 0.08 | 0.1 |
| 4 | 18617775 | rs6449399 | 1.379 | -0.264 | 0.453 | 0.672 | -0.022 | 0.013 | 0.013 | 0 | 0.64 | 0.43 | 0.38 | 0.46 | 0.73 | 0.56 |
| 4 | 18629525 | rs4698702 | 0.058 | -0.292 | 0.259 | 0.624 | 0 | 0.018 | 0.018 | 0.124 | 0.14 | 0.12 | 0.46 | 0.43 | 0.14 | 0.28 |

| | | | | | | | | | | | | | | | | |
|---|---|---|---|---|---|---|---|---|---|---|---|---|---|---|---|---|
| 4 | 18638021 | rs2320979 | 0.922 | -0.288 | 0.305 | 0.588 | -0.011 | 0.012 | 0.012 | 0 | 0.68 | 0.56 | 0.42 | 0.46 | 0.74 | 0.56 |
| 4 | 18665579 | rs4235399 | 0.534 | -0.62 | 0.684 | 0.487 | -0.062 | -0.045 | -0.045 | 0.536 | 0.81 | 0.4 | 0.89 | 0.9 | 0.94 | 0.91 |
| 4 | 18668957 | rs4398521 | 1.233 | -0.26 | 0.368 | 0.485 | 0.028 | -0.025 | -0.025 | 0.141 | 0.64 | 0.17 | 0.48 | 0.46 | 0.73 | 0.61 |
| 4 | 18670019 | rs12505952 | 0.332 | -0.354 | 0.774 | 0.37 | -0.086 | -0.064 | -0.064 | 2.455 | 0.79 | 0.46 | 0.89 | 0.9 | 0.94 | 0.9 |
| 4 | 18682975 | rs12501690 | -0.076 | -0.317 | 0.716 | 0.322 | 0.001 | -0.023 | -0.023 | 0 | 0.16 | 0.14 | 0.07 | 0 | 0.08 | 0.06 |
| 4 | 18694002 | rs938841 | 0.954 | -0.252 | 0.96 | 0.406 | -0.008 | 0.038 | 0.038 | 1.191 | 0.68 | 0.52 | 0.51 | 0.47 | 0.73 | 0.62 |
| 4 | 18695770 | rs13118998 | -0.288 | -0.093 | 0.55 | 0.502 | -0.022 | -0.039 | -0.039 | 0.04 | 0.84 | 0.9 | 0.59 | 0.57 | 0.8 | 0.72 |
| 4 | 18696592 | rs1546658 | -0.847 | -0.043 | 0.441 | 0.666 | -0.063 | -0.086 | -0.086 | 0 | 0.88 | 0.99 | 0.63 | 0.68 | 0.84 | 0.79 |
| 4 | 18701138 | rs7656740 | 1.541 | -0.244 | -0.093 | -0.031 | -0.048 | -0.038 | -0.038 | 0.198 | 0.38 | 0.11 | 0.3 | 0.45 | 0.44 | 0.39 |
| 4 | 18704754 | rs16897436 | 0.012 | -0.156 | 0.617 | 0.287 | -0.082 | -0.13 | -0.13 | 1.773 | 0.18 | 0.45 | 0.09 | 0 | 0.06 | 0.06 |
| 4 | 18709859 | rs7656712 | -0.232 | -0.104 | 0.06 | 0.332 | -0.042 | -0.028 | -0.028 | 1.275 | 0.2 | 0.71 | 0.11 | 0.1 | 0.06 | 0.11 |
| 4 | 18722405 | rs1878644 | -0.076 | -0.083 | -0.172 | 0.263 | -0.054 | -0.106 | -0.106 | 0 | 0.76 | 0.91 | 0.54 | 0.57 | 0.72 | 0.72 |
| 4 | 18725182 | rs1400938 | -0.792 | -0.062 | -0.387 | 0.203 | -0.084 | -0.108 | -0.108 | 0 | 0.82 | 0.96 | 0.57 | 0.69 | 0.76 | 0.81 |
| 4 | 18725802 | rs2321533 | 1.021 | 0.002 | -0.252 | -0.315 | 0.031 | -0.004 | -0.004 | 0 | 0.56 | 0.21 | 0.42 | 0.47 | 0.66 | 0.6 |
| 4 | 18737869 | rs1517986 | -0.151 | 0.272 | -0.188 | -0.315 | -0.016 | -0.099 | -0.099 | 0 | 0.23 | 0.31 | 0.15 | 0.01 | 0.16 | 0.12 |
| 4 | 18748499 | rs11732784 | 0.348 | 0.499 | -0.662 | 0.087 | -0.003 | 0.021 | 0.021 | 0 | 0.51 | 0.39 | 0.38 | 0.47 | 0.56 | 0.55 |
| 4 | 18752731 | rs1400931 | 0.238 | 0.412 | -0.473 | -0.151 | 0.004 | -0.026 | -0.026 | 0 | 0.81 | 0.74 | 0.86 | 0.84 | 0.83 | 0.86 |
| 4 | 18752760 | rs16897498 | 0.042 | -0.304 | -1.253 | -0.43 | 0.033 | 0.038 | 0.038 | 0.349 | 0.29 | 0.13 | 0.2 | 0.22 | 0.18 | 0.41 |
| 4 | 18753186 | rs2203758 | 0.042 | 0.43 | -0.999 | -0.104 | 0 | -0.011 | -0.011 | 0 | 0.58 | 0.7 | 0.5 | 0.52 | 0.46 | 0.58 |
| 4 | 18754324 | rs4697637 | 0.709 | 1.057 | -1.384 | -0.238 | 0.013 | 0.03 | 0.03 | 0 | 0.23 | 0.23 | 0.36 | 0.48 | 0.35 | 0.25 |
| 4 | 18756789 | rs1464189 | 0.514 | 0.597 | -1.189 | -0.259 | 0.014 | 0.012 | 0.012 | 0 | 0.49 | 0.6 | 0.38 | 0.45 | 0.39 | 0.39 |
| 4 | 18757927 | rs16897505 | 1.147 | 1 | -1.132 | 0.019 | -0.001 | 0.01 | 0.01 | 0.827 | 0.08 | 0.13 | 0.12 | 0.07 | 0.06 | 0.19 |
| 4 | 18773006 | rs1400935 | 0.243 | 1.086 | -1.504 | -0.091 | -0.037 | -0.04 | -0.04 | 0.084 | 0.07 | 0 | 0.13 | 0.21 | 0.09 | 0.13 |
| 4 | 18776097 | rs1607309 | 0.142 | 0.618 | -1.699 | -0.649 | -0.011 | 0.15 | 0.15 | 0 | 0.71 | 0.94 | 0.84 | 1 | 0.76 | 0.81 |
| 4 | 18779280 | rs10024472 | -0.479 | 0.756 | -0.693 | -0.044 | 0.038 | 0.075 | 0.075 | 0 | 0.09 | 0.25 | 0.14 | 0.22 | 0.09 | 0.14 |
| 4 | 18779862 | rs1517987 | -0.675 | 0.256 | -0.44 | -0.363 | -0.001 | 0.004 | 0.004 | 0 | 0.3 | 0.29 | 0.25 | 0.26 | 0.25 | 0.2 |
| 4 | 18786401 | rs10027565 | -0.196 | 0.295 | -0.679 | -0.04 | -0.006 | -0.001 | -0.001 | 0 | 0.55 | 0.52 | 0.58 | 0.46 | 0.62 | 0.52 |
| 4 | 18787232 | rs2321454 | -0.319 | 0.343 | -0.286 | 0.505 | 0.061 | 0.046 | 0.046 | 0.082 | 0.61 | 0.79 | 0.82 | 0.69 | 0.73 | 0.56 |
| 4 | 18789372 | rs1568234 | 0.383 | 0.754 | -0.248 | 0.406 | 0.026 | 0.005 | 0.005 | 0 | 0.14 | 0.26 | 0.22 | 0.16 | 0.18 | 0.13 |
| 4 | 18794118 | rs9291679 | 0.599 | 0.81 | -0.403 | 0.476 | -0.01 | 0.025 | 0.025 | 0 | 0.41 | 0.29 | 0.35 | 0.31 | 0.43 | 0.39 |
| 4 | 18794800 | rs16897583 | 1.213 | 0.537 | -0.547 | 0.523 | 0.001 | -0.002 | -0.002 | 0 | 0.18 | 0.15 | 0.1 | 0.2 | 0.12 | 0.19 |
| 4 | 18800428 | rs4566649 | -0.265 | 1.076 | -0.596 | 0.073 | -0.009 | 0.03 | 0.03 | 0 | 0.36 | 0.27 | 0.31 | 0.18 | 0.34 | 0.31 |
| 4 | 18800595 | rs13147495 | 1.192 | 1.803 | -0.893 | 0.317 | -0.066 | 0.001 | 0.001 | 0 | 0.11 | 0.01 | 0.21 | 0.08 | 0.18 | 0.08 |
| 4 | 18801755 | rs1356472 | -1.178 | 1.419 | -1.002 | 0.66 | -0.022 | 0.038 | 0.038 | 0 | 0.72 | 0.84 | 0.78 | 0.81 | 0.73 | 0.79 |
| 4 | 18803401 | rs10516330 | -0.185 | 1.642 | -0.501 | 1.184 | -0.002 | 0.021 | 0.021 | 0.203 | 0.09 | 0.11 | 0.18 | 0.24 | 0.22 | 0.17 |
| 4 | 18803428 | rs16897611 | 0.368 | 1.245 | -0.457 | 0.528 | -0.01 | 0.013 | 0.013 | 0 | 0.1 | 0.18 | 0.09 | 0.12 | 0.06 | 0.04 |
| 4 | 18812552 | rs1356468 | -1.507 | 1.629 | -0.926 | 0.631 | -0.012 | -0.004 | -0.004 | 0.006 | 0.84 | 0.85 | 0.73 | 0.81 | 0.78 | 0.83 |
| 4 | 18813683 | rs12640123 | -1.278 | 1.592 | -0.663 | 0.62 | 0.008 | 0 | 0 | 0 | 0.83 | 0.76 | 0.72 | 0.82 | 0.78 | 0.83 |
| 4 | 18835874 | rs6449425 | -1.165 | 2.386 | -1.081 | 0.877 | -0.03 | -0.014 | -0.014 | 0 | 0.79 | 0.86 | 0.69 | 0.74 | 0.72 | 0.72 |
| 4 | 18837642 | rs16897644 | 0.836 | 1.902 | -0.682 | 1.171 | 0.005 | 0.074 | 0.074 | 0 | 0.13 | 0.21 | 0.18 | 0.24 | 0.14 | 0.04 |
| 4 | 18839839 | rs4147043 | -0.128 | 1.236 | -1.486 | 0.875 | -0.018 | 0.008 | 0.008 | 0 | 0.56 | 0.69 | 0.51 | 0.56 | 0.51 | 0.52 |
| 4 | 18840158 | rs4147042 | 0.266 | 1.134 | -1.508 | 0.361 | 0.026 | 0.052 | 0.052 | 0 | 0.41 | 0.24 | 0.36 | 0.32 | 0.36 | 0.48 |
| 4 | 18841225 | rs7699469 | 0.592 | 0.948 | -1.354 | 0.521 | 0 | 0.019 | 0.019 | 0.363 | 0.42 | 0.23 | 0.41 | 0.39 | 0.42 | 0.47 |
| 4 | 18841427 | rs4697158 | 0.541 | 0.286 | -1.194 | 0.783 | 0.033 | 0.021 | 0.021 | 0 | 0.51 | 0.13 | 0.58 | 0.47 | 0.58 | 0.57 |
| 4 | 18843657 | rs1518002 | 1.192 | -0.109 | -0.412 | 0.519 | -0.002 | 0.002 | 0.002 | 0 | 0.35 | 0.32 | 0.33 | 0.28 | 0.41 | 0.38 |
| 4 | 18851987 | rs938840 | -0.382 | -0.143 | -0.2 | 0.61 | 0.008 | 0.027 | 0.027 | 0.012 | 0.7 | 0.84 | 0.75 | 0.83 | 0.69 | 0.74 |
| 4 | 18852745 | rs16897709 | 0.135 | -0.106 | -0.43 | 0.725 | -0.013 | 0 | 0 | 0 | 0.25 | 0.2 | 0.29 | 0.24 | 0.25 | 0.25 |
| 4 | 18866169 | rs6845621 | 0.982 | -0.291 | -0.416 | 0.481 | 0 | 0 | 0 | 0.041 | 0.39 | 0.41 | 0.43 | 0.57 | 0.44 | 0.46 |
| 4 | 18869343 | rs6819510 | -0.331 | 0.804 | -0.609 | 0.792 | -0.011 | 0.027 | 0.027 | 0 | 0.25 | 0.2 | 0.28 | 0.18 | 0.24 | 0.25 |
| 4 | 18878845 | rs6832325 | -0.568 | 0.533 | -0.558 | 0.985 | 0.004 | -0.009 | -0.009 | 0.016 | 0.68 | 0.75 | 0.61 | 0.6 | 0.68 | 0.77 |
| 4 | 18889088 | rs7676193 | 1.239 | 0.566 | -0.613 | 0.954 | 0.008 | 0.026 | 0.026 | 0.453 | 0.34 | 0.44 | 0.31 | 0.37 | 0.38 | 0.39 |
| 4 | 18894117 | rs4254763 | 0.924 | 0.579 | -0.814 | 0.884 | -0.07 | 0.021 | 0.021 | 0 | 0.12 | 0.01 | 0.09 | 0.09 | 0.19 | 0.16 |
| 4 | 18894787 | rs4444827 | -0.505 | 0.322 | -0.41 | 1.218 | 0.003 | 0.004 | 0.004 | 0 | 0.71 | 0.74 | 0.7 | 0.86 | 0.71 | 0.85 |
| 4 | 18900428 | rs12508082 | 0.477 | 0.74 | -0.586 | 1.379 | -0.021 | 0.035 | 0.035 | 0 | 0.29 | 0.18 | 0.28 | 0.24 | 0.3 | 0.34 |
| 4 | 18908518 | rs6449435 | -0.713 | 0.758 | -0.577 | 1.387 | -0.022 | 0.034 | 0.034 | 0 | 0.71 | 0.82 | 0.72 | 0.76 | 0.7 | 0.66 |
| 4 | 18910602 | rs2215762 | 0.829 | 0.678 | -0.581 | 1.472 | 0.002 | 0 | 0 | 1.11 | 0.33 | 0.35 | 0.3 | 0.32 | 0.38 | 0.39 |
| 4 | 18921383 | rs207291 | 0.232 | 0.48 | -0.757 | 0.834 | -0.008 | -0.019 | -0.019 | 0.048 | 0.33 | 0.36 | 0.31 | 0.12 | 0.29 | 0.15 |
| 4 | 18933156 | rs10489055 | 0.523 | 0.697 | -0.871 | 0.847 | 0.006 | -0.008 | -0.008 | 0.147 | 0.36 | 0.42 | 0.4 | 0.64 | 0.4 | 0.48 |
| 4 | 18936647 | rs1452556 | 0.533 | 0.672 | -0.908 | 0.295 | 0.007 | 0.012 | 0.012 | 0.444 | 0.35 | 0.39 | 0.32 | 0.41 | 0.4 | 0.43 |
| 4 | 18937840 | rs207283 | 0.126 | 1.535 | -1.053 | 1.083 | -0.02 | -0.035 | -0.035 | 0 | 0.7 | 0.8 | 0.64 | 0.51 | 0.69 | 0.6 |
| 4 | 18944656 | rs7664024 | -0.759 | 1.521 | -0.459 | 0.75 | 0.024 | -0.006 | -0.006 | 0 | 0.15 | 0.25 | 0.15 | 0.14 | 0.19 | 0.19 |
| 4 | 18954655 | rs2601573 | -0.739 | 0.424 | -0.347 | -0.106 | -0.028 | -0.065 | -0.065 | 1.24 | 0.16 | 0.22 | 0.06 | 0.01 | 0.06 | 0.15 |
| 4 | 18958075 | rs1452559 | 0.652 | 0.662 | -0.472 | 0.292 | 0.041 | 0.018 | 0.018 | 0.543 | 0.51 | 0.21 | 0.6 | 0.52 | 0.63 | 0.3 |
| 4 | 18960220 | rs2537942 | 0.179 | 0.641 | -0.374 | 0.324 | -0.017 | -0.049 | -0.049 | 0.019 | 0.15 | 0.18 | 0.05 | 0.01 | 0.06 | 0.15 |
| 4 | 18961510 | rs1452565 | -0.232 | 0.62 | 0.162 | 0.339 | -0.052 | -0.093 | -0.093 | 0.055 | 0.17 | 0.34 | 0.08 | 0.04 | 0.06 | 0.17 |
| 4 | 18968306 | rs1349670 | -0.056 | 0.618 | 0.207 | 0.338 | -0.036 | -0.077 | -0.077 | 0.291 | 0.16 | 0.26 | 0.06 | 0.01 | 0.06 | 0.15 |
| 4 | 18983905 | rs2601593 | -0.131 | 0.618 | 0.21 | 0.338 | -0.043 | -0.079 | -0.079 | 0 | 0.16 | 0.28 | 0.07 | 0.04 | 0.06 | 0.18 |
| 4 | 18985469 | rs2537914 | -0.457 | 0.647 | 0.295 | 0.145 | 0.004 | 0.023 | 0.023 | 0.692 | 0.72 | 0.65 | 0.71 | 0.56 | 0.73 | 0.56 |
| 4 | 18990266 | rs207397 | -1.08 | 0.616 | -0.092 | 0.038 | -0.048 | 0.011 | 0.011 | 0 | 0.87 | 0.97 | 0.91 | 0.88 | 0.84 | 0.81 |

| | | | | | | | | | | | | | | | | |
|---|---|---|---|---|---|---|---|---|---|---|---|---|---|---|---|---|
| 4 | 19005988 | rs207363 | -0.774 | 0.188 | 0.738 | -0.136 | -0.014 | -0.017 | -0.017 | 0 | 0.09 | 0.58 | 0.08 | 0.03 | 0.02 | 0.02 |
| 4 | 19020766 | rs16897980 | -0.015 | 0.002 | 0.567 | -0.35 | 0.005 | 0.027 | 0.027 | 0.032 | 0.11 | 0.07 | 0.04 | 0.01 | 0.08 | 0.17 |
| 4 | 19024888 | rs207348 | -1.534 | 0 | 0.571 | -0.351 | -0.036 | -0.091 | -0.091 | 0.305 | 0.2 | 0.66 | 0.12 | 0.04 | 0.1 | 0.19 |
| 4 | 19035608 | rs10001989 | 0.027 | -0.012 | 0.925 | -0.359 | 0.017 | 0.026 | 0.026 | 0.022 | 0.11 | 0.07 | 0.04 | 0.01 | 0.06 | 0.17 |
| 4 | 19036305 | rs207343 | -1.466 | -0.01 | 0.889 | -0.357 | -0.051 | -0.091 | -0.091 | 5.307 | 0.2 | 0.66 | 0.12 | 0.04 | 0.08 | 0.18 |
| 4 | 19038807 | rs207342 | -0.041 | -0.083 | 0.524 | -0.041 | -0.029 | -0.075 | -0.075 | 1.513 | 0.74 | 0.89 | 0.73 | 0.56 | 0.75 | 0.59 |
| 4 | 19040579 | rs12644264 | 1.61 | -0.102 | 0.367 | -0.111 | 0 | -0.013 | -0.013 | 0.011 | 0.48 | 0.17 | 0.57 | 0.51 | 0.6 | 0.35 |
| 4 | 19047617 | rs16898029 | -0.94 | -0.098 | 0.844 | -0.266 | -0.038 | -0.087 | -0.087 | 0.668 | 0.17 | 0.36 | 0.08 | 0.01 | 0.08 | 0.22 |
| 4 | 19056475 | rs16898037 | -0.792 | 0.111 | 0.105 | -0.569 | 0.005 | 0.021 | 0.021 | 0 | 0.24 | 0.17 | 0.17 | 0.09 | 0.25 | 0.18 |
| 4 | 19058245 | rs207332 | -0.079 | -0.366 | -0.108 | -0.278 | -0.002 | 0.004 | 0.004 | 0 | 0.58 | 0.65 | 0.58 | 0.41 | 0.61 | 0.4 |
| 4 | 19058288 | rs16898044 | 0.054 | -0.372 | -0.101 | -0.293 | 0.003 | -0.002 | -0.002 | 0 | 0.19 | 0.16 | 0.14 | 0.05 | 0.21 | 0.09 |
| 4 | 19059824 | rs207331 | 0.134 | -0.304 | -0.247 | -0.083 | -0.001 | 0.007 | 0.007 | 0 | 0.58 | 0.65 | 0.57 | 0.38 | 0.61 | 0.38 |
| 4 | 19075878 | rs207311 | 0.374 | -0.097 | -1.11 | 0.268 | -0.003 | -0.012 | -0.012 | 0 | 0.61 | 0.25 | 0.75 | 0.62 | 0.66 | 0.71 |
| 4 | 19087520 | rs182289 | -0.422 | 0.25 | -0.358 | -0.34 | -0.012 | -0.059 | -0.059 | 0.67 | 0.18 | 0.37 | 0.14 | 0.06 | 0.16 | 0.16 |
| 4 | 19092157 | rs207305 | -0.536 | -0.175 | -0.672 | -0.903 | 0.004 | 0.056 | 0.056 | 0.285 | 0.68 | 0.83 | 0.83 | 0.87 | 0.71 | 0.78 |
| 4 | 19093088 | rs16898064 | 1.372 | -0.145 | -0.104 | -1.156 | -0.007 | -0.057 | -0.057 | 0 | 0.1 | 0.17 | 0.05 | 0 | 0.05 | 0.01 |
| 4 | 19095959 | rs207301 | -0.062 | -0.6 | -0.07 | -1.374 | 0.014 | 0.017 | 0.017 | 0 | 0.67 | 0.76 | 0.84 | 0.97 | 0.74 | 0.75 |
| 4 | 19096163 | rs207300 | -0.136 | -0.601 | -0.082 | -1.374 | 0.019 | 0.042 | 0.042 | 0 | 0.68 | 0.82 | 0.85 | 0.97 | 0.74 | 0.75 |
| 4 | 19101375 | rs10020700 | -0.21 | 0.106 | 0.002 | -1.132 | 0.003 | -0.017 | -0.017 | 0 | 0.23 | 0.26 | 0.17 | 0.13 | 0.23 | 0.23 |
| 4 | 19111493 | rs12649695 | 0.088 | 0.198 | -0.103 | -0.444 | -0.003 | -0.002 | -0.002 | 0 | 0.06 | 0.05 | 0.03 | 0.22 | 0.05 | 0.07 |
| 4 | 19116730 | rs6815911 | -0.128 | 0.208 | -0.092 | -0.443 | 0.003 | -0.017 | -0.017 | 0 | 0.23 | 0.25 | 0.17 | 0.13 | 0.23 | 0.23 |
| 4 | 19118335 | rs11931960 | -0.259 | 0.156 | -0.118 | -0.468 | 0.002 | -0.016 | -0.016 | 0 | 0.24 | 0.25 | 0.17 | 0.13 | 0.23 | 0.23 |
| 4 | 19119813 | rs17495360 | -0.054 | -0.222 | -0.108 | -0.732 | 0.009 | 0.006 | 0.006 | 0 | 0.33 | 0.23 | 0.18 | 0.25 | 0.28 | 0.27 |
| 4 | 19120064 | rs17495401 | 0.036 | -0.059 | -0.066 | -0.63 | 0.003 | -0.017 | -0.017 | 2.642 | 0.23 | 0.25 | 0.17 | 0.13 | 0.23 | 0.23 |
| 4 | 19122442 | rs10461114 | 1.333 | -0.078 | -0.112 | -0.653 | -0.056 | -0.016 | -0.016 | 0 | 0.13 | 0.03 | 0.12 | 0.13 | 0.18 | 0.21 |
| 4 | 19129715 | rs13125998 | -0.086 | -0.079 | -0.065 | -0.65 | 0.003 | -0.017 | -0.017 | 0.097 | 0.23 | 0.25 | 0.17 | 0.13 | 0.23 | 0.23 |
| 4 | 19133988 | rs9992304 | -0.035 | -0.005 | -0.047 | -0.654 | 0.002 | -0.016 | -0.016 | 0 | 0.23 | 0.25 | 0.17 | 0.13 | 0.22 | 0.23 |
| 4 | 19136574 | rs2169705 | -0.035 | -0.005 | -0.06 | -0.654 | 0.002 | -0.017 | -0.017 | 0 | 0.23 | 0.26 | 0.17 | 0.13 | 0.22 | 0.23 |
| 4 | 19150192 | rs7671224 | -1.039 | -0.071 | -0.054 | -0.693 | -0.001 | 0.006 | 0.006 | 0 | 0.52 | 0.43 | 0.33 | 0.38 | 0.49 | 0.49 |
| 4 | 19150738 | rs5013969 | -1.173 | -0.072 | -0.024 | -0.686 | -0.001 | -0.001 | -0.001 | 0 | 0.53 | 0.49 | 0.34 | 0.38 | 0.5 | 0.49 |
| 4 | 19167507 | rs13139953 | -0.667 | -0.079 | -0.094 | -0.708 | 0 | 0.004 | 0.004 | 0 | 0.28 | 0.31 | 0.19 | 0.37 | 0.26 | 0.29 |
| 4 | 19174530 | rs12374356 | 0.927 | -0.083 | -0.128 | -0.741 | 0.001 | 0.006 | 0.006 | 0 | 0.23 | 0.14 | 0.13 | 0 | 0.23 | 0.18 |
| 4 | 19186822 | rs11944202 | 1.329 | -0.009 | -0.137 | -0.649 | -0.019 | 0 | 0 | 0 | 0.13 | 0.07 | 0.12 | 0.13 | 0.17 | 0.21 |
| 4 | 19189389 | rs1478051 | -2.273 | -0.043 | 0.074 | -0.625 | -0.011 | -0.054 | -0.054 | 0.013 | 0.9 | 0.8 | 0.95 | 1 | 0.95 | 0.99 |
| 4 | 19193583 | rs1478049 | 0.959 | -0.074 | 0.014 | -0.644 | -0.004 | 0.002 | 0.002 | 0 | 0.43 | 0.4 | 0.62 | 0.62 | 0.45 | 0.49 |
| 4 | 19201819 | rs12506297 | -0.791 | -0.084 | -0.041 | -0.57 | -0.001 | -0.002 | -0.002 | 1.341 | 0.53 | 0.5 | 0.33 | 0.37 | 0.5 | 0.49 |
| 4 | 19239803 | rs1382155 | 0.345 | 0.29 | -0.601 | -0.403 | -0.036 | 0.002 | 0.002 | 0.053 | 0.22 | 0.07 | 0.18 | 0.21 | 0.24 | 0.24 |
| 4 | 19253659 | rs414289 | -1.533 | 0.121 | -0.649 | -0.508 | 0.001 | 0.032 | 0.032 | 0 | 0.78 | 0.89 | 0.89 | 0.95 | 0.79 | 0.81 |
| 4 | 19259307 | rs7668345 | 0.538 | 0.245 | -0.705 | -0.546 | -0.051 | 0.009 | 0.009 | 0.06 | 0.22 | 0.06 | 0.18 | 0.18 | 0.24 | 0.24 |
| 4 | 19259854 | rs312357 | 1.381 | 0.158 | -0.359 | -0.417 | 0.005 | 0.015 | 0.015 | 0 | 0.36 | 0.44 | 0.58 | 0.54 | 0.4 | 0.42 |
| 4 | 19262251 | rs312353 | 0.946 | 0.16 | -0.488 | -0.394 | 0.002 | 0.006 | 0.006 | 0 | 0.41 | 0.57 | 0.63 | 0.54 | 0.47 | 0.44 |
| 4 | 19262896 | rs1026681 | 2.3 | 0.151 | -0.392 | -0.355 | -0.008 | -0.037 | -0.037 | 0 | 0.08 | 0.11 | 0.04 | 0 | 0.05 | 0.01 |
| 4 | 19267283 | rs16898171 | 0.005 | 0.178 | -0.454 | -0.303 | -0.023 | 0.035 | 0.035 | 0 | 0.16 | 0.26 | 0.08 | 0.21 | 0.08 | 0.12 |
| 4 | 19275229 | rs312368 | 0.968 | 0.384 | -0.443 | -0.3 | 0.012 | 0.028 | 0.028 | 0 | 0.37 | 0.52 | 0.6 | 0.56 | 0.41 | 0.43 |
| 4 | 19302457 | rs17564753 | -0.201 | 0.558 | -0.773 | -0.477 | -0.008 | 0.022 | 0.022 | 0 | 0.13 | 0.06 | 0.18 | 0.03 | 0.16 | 0.18 |
| 4 | 19306711 | rs312330 | 0.115 | 0.524 | -0.872 | -0.562 | -0.008 | 0.072 | 0.072 | 0 | 0.66 | 0.83 | 0.7 | 0.91 | 0.63 | 0.63 |
| 4 | 19326717 | rs2322090 | -0.368 | 0.678 | -1.066 | -0.387 | -0.001 | 0.008 | 0.008 | 0.384 | 0.76 | 0.71 | 0.77 | 0.74 | 0.75 | 0.77 |
| 4 | 19333182 | rs1010743 | 0.78 | 0.61 | -1.137 | -0.516 | 0 | -0.003 | -0.003 | 0 | 0.61 | 0.57 | 0.64 | 0.72 | 0.57 | 0.63 |
| 4 | 19335464 | rs1490874 | 0.334 | 0.63 | -1.093 | -0.457 | 0 | 0.129 | 0.129 | 0 | 0.08 | 0.26 | 0.14 | 0.2 | 0.06 | 0.07 |
| 4 | 19368572 | rs12642807 | -0.306 | 0.687 | -1.156 | -0.6 | 0.002 | -0.008 | -0.008 | 0.005 | 0.13 | 0.13 | 0.13 | 0.03 | 0.19 | 0.13 |
| 4 | 19381302 | rs1552660 | -2.206 | 0.67 | -1.164 | -0.661 | -0.026 | 0.044 | 0.044 | 0.424 | 0.84 | 0.95 | 0.91 | 0.94 | 0.82 | 0.84 |
| 4 | 19406545 | rs13123977 | -0.417 | 0.64 | -1.05 | -0.624 | -0.001 | 0.011 | 0.011 | 0.393 | 0.75 | 0.66 | 0.76 | 0.74 | 0.75 | 0.77 |
| 4 | 19407186 | rs13102773 | 1.044 | 0.675 | -1.183 | -0.7 | 0 | -0.006 | -0.006 | 0 | 0.55 | 0.51 | 0.6 | 0.73 | 0.51 | 0.62 |
| 4 | 19410177 | rs10023012 | 0.538 | 0.594 | -1.165 | -0.677 | -0.004 | 0.008 | 0.008 | 0 | 0.14 | 0.19 | 0.17 | 0.2 | 0.12 | 0.09 |
| 4 | 19414766 | rs7696764 | 0.783 | 0.611 | -1.165 | -0.263 | -0.001 | -0.001 | -0.001 | 0 | 0.58 | 0.56 | 0.66 | 0.73 | 0.61 | 0.63 |
| 4 | 19424787 | rs10010555 | 0.951 | 0.715 | -1.133 | -0.149 | -0.005 | 0.063 | 0.063 | 0 | 0.07 | 0.14 | 0.13 | 0.2 | 0.06 | 0.07 |
| 4 | 19425386 | rs6829074 | -0.144 | 0.644 | -1.176 | -0.227 | -0.001 | -0.002 | -0.002 | 0 | 0.73 | 0.75 | 0.82 | 0.92 | 0.72 | 0.71 |
| 4 | 19428544 | rs4074143 | 1.192 | 0.817 | -1.435 | 0.013 | -0.023 | -0.037 | -0.037 | 0.181 | 0.18 | 0.05 | 0.17 | 0.22 | 0.18 | 0.24 |
| 4 | 19445650 | rs10009055 | 1.006 | 0.741 | -1.408 | 0.049 | -0.006 | 0.061 | 0.061 | 0 | 0.07 | 0.14 | 0.13 | 0.21 | 0.06 | 0.07 |
| 4 | 19452605 | rs12233736 | 0.743 | 0.805 | -1.419 | 0.007 | -0.028 | -0.013 | -0.013 | 0.012 | 0.21 | 0.11 | 0.23 | 0.22 | 0.29 | 0.24 |
| 4 | 19463717 | rs10004437 | 0.84 | 0.736 | -1.21 | 0.124 | -0.006 | 0.059 | 0.059 | 0 | 0.07 | 0.14 | 0.13 | 0.21 | 0.06 | 0.07 |
| 4 | 19484765 | rs7437307 | -0.349 | 0.641 | -1.425 | -0.031 | -0.062 | 0.023 | 0.023 | 0 | 0.64 | 0.83 | 0.67 | 0.72 | 0.52 | 0.61 |
| 4 | 19487028 | rs7683977 | -0.874 | 0.513 | -1.292 | -0.101 | -0.014 | 0.025 | 0.025 | 0 | 0.19 | 0.31 | 0.23 | 0.23 | 0.15 | 0.21 |
| 4 | 19488871 | rs10011402 | -0.953 | 0.471 | -1.313 | -0.169 | -0.016 | -0.001 | -0.001 | 0 | 0.26 | 0.38 | 0.27 | 0.23 | 0.2 | 0.22 |
| 4 | 19492256 | rs4525972 | 1.231 | 0.459 | -1.304 | -0.149 | -0.012 | 0.016 | 0.016 | 0 | 0.38 | 0.45 | 0.41 | 0.49 | 0.32 | 0.38 |
| 4 | 19496230 | rs10017144 | 1.81 | 0.482 | -1.184 | -0.123 | 0.017 | -0.02 | -0.02 | 0 | 0.06 | 0.19 | 0.08 | 0.02 | 0.06 | 0.02 |
| 4 | 19497475 | rs4346638 | -1.386 | 0.529 | -1.25 | -0.159 | -0.002 | -0.01 | -0.01 | 0 | 0.29 | 0.44 | 0.33 | 0.23 | 0.31 | 0.23 |
| 4 | 19508543 | rs6842946 | 0.013 | 0.556 | -1.263 | -0.123 | 0.009 | 0.039 | 0.039 | 0 | 0.1 | 0.17 | 0.19 | 0.21 | 0.17 | 0.08 |
| 4 | 19511507 | rs6837423 | -0.328 | 0.535 | -1.278 | -0.135 | 0.001 | 0.005 | 0.005 | 0 | 0.17 | 0.26 | 0.23 | 0.21 | 0.23 | 0.09 |

| | | | | | | | | | | | | | | | | |
|---|---|---|---|---|---|---|---|---|---|---|---|---|---|---|---|---|
| 4 | 19512213 | rs10006185 | -0.658 | 0.537 | -1.399 | -0.2 | -0.043 | 0.005 | 0.005 | 0 | 0.67 | 0.88 | 0.74 | 0.71 | 0.63 | 0.61 |
| 4 | 19517296 | rs12507186 | 0.816 | 0.518 | -1.264 | -0.163 | 0.001 | 0.038 | 0.038 | 0 | 0.1 | 0.03 | 0.06 | 0.01 | 0.08 | 0.13 |
| 4 | 19525991 | rs7700114 | -0.196 | 0.518 | -1.22 | -0.164 | -0.007 | -0.05 | -0.05 | 0 | 0.12 | 0.19 | 0.1 | 0.02 | 0.09 | 0.14 |
| 4 | 19551260 | rs4478186 | -0.586 | 0.499 | -1.411 | -0.131 | -0.04 | 0.006 | 0.006 | 0 | 0.67 | 0.86 | 0.73 | 0.71 | 0.63 | 0.62 |
| 4 | 19570941 | rs6850501 | 0.948 | 0.262 | -1.989 | -0.65 | -0.037 | 0.068 | 0.068 | 0 | 0.3 | 0.45 | 0.41 | 0.49 | 0.3 | 0.35 |
| 4 | 19571122 | rs13120547 | -0.415 | 0.249 | -1.977 | -0.638 | -0.005 | 0.015 | 0.015 | 0 | 0.37 | 0.33 | 0.31 | 0.23 | 0.33 | 0.26 |
| 4 | 19573855 | rs9996345 | -3.365 | 0.308 | -2.225 | -0.765 | -0.022 | 0.027 | 0.027 | 0.391 | 0.85 | 0.94 | 0.91 | 0.93 | 0.81 | 0.84 |
| 4 | 19574923 | rs10027409 | -0.952 | 0.319 | -2.222 | -0.751 | -0.03 | 0.015 | 0.015 | 0 | 0.67 | 0.82 | 0.73 | 0.72 | 0.63 | 0.62 |
| 4 | 19576080 | rs4295256 | 1.868 | 0.399 | -2.138 | -0.768 | -0.056 | -0.054 | -0.054 | 0 | 0.17 | 0.03 | 0.17 | 0.21 | 0.18 | 0.22 |
| 4 | 19576271 | rs12511971 | -0.569 | 0.405 | -1.951 | -0.729 | -0.005 | 0.015 | 0.015 | 0 | 0.37 | 0.33 | 0.31 | 0.23 | 0.33 | 0.26 |
| 4 | 19576975 | rs10000908 | 2.347 | 0.442 | -1.818 | -0.679 | -0.009 | -0.045 | -0.045 | 1.818 | 0.08 | 0.18 | 0.06 | 0 | 0.06 | 0.01 |
| 4 | 19584331 | rs17508080 | 1.569 | 0.582 | -1.925 | -0.627 | -0.068 | 0.087 | 0.087 | 0.04 | 0.16 | 0.03 | 0.1 | 0.01 | 0.18 | 0.06 |
| 4 | 19589352 | rs11937925 | 1.437 | 0.396 | -1.683 | -0.351 | -0.005 | 0.069 | 0.069 | 0 | 0.07 | 0.12 | 0.13 | 0.22 | 0.06 | 0.07 |
| 4 | 19607089 | rs10033656 | -0.355 | 0.307 | -1.856 | -0.504 | 0 | -0.002 | -0.002 | 0 | 0.36 | 0.4 | 0.37 | 0.43 | 0.3 | 0.32 |
| 4 | 19609939 | rs3846382 | -0.743 | 0.383 | -1.99 | -0.684 | 0 | 0.001 | 0.001 | 0.643 | 0.5 | 0.51 | 0.46 | 0.45 | 0.42 | 0.38 |
| 4 | 19616531 | rs3904756 | 1.974 | -0.097 | -1.634 | -0.553 | 0.007 | -0.005 | -0.005 | 0 | 0.16 | 0.06 | 0.23 | 0.21 | 0.19 | 0.24 |
| 4 | 19616649 | rs10025939 | 0.547 | 0.05 | -1.202 | -0.487 | 0.005 | -0.062 | -0.062 | 0.143 | 0.17 | 0.24 | 0.12 | 0.02 | 0.12 | 0.08 |
| 4 | 19617286 | rs7682588 | 1.279 | 0.088 | -1.3 | -0.468 | -0.042 | 0.067 | 0.067 | 0 | 0.13 | 0.03 | 0.07 | 0.01 | 0.12 | 0.06 |
| 4 | 19629309 | rs3846385 | -0.382 | 0.01 | -1.254 | -0.415 | -0.004 | -0.008 | -0.008 | 0.032 | 0.3 | 0.39 | 0.28 | 0.23 | 0.23 | 0.16 |
| 4 | 19633338 | rs3846388 | 1.23 | -0.027 | -1.167 | -0.291 | -0.006 | 0.013 | 0.013 | 0 | 0.14 | 0.17 | 0.17 | 0.21 | 0.11 | 0.08 |
| 4 | 19643571 | rs11939712 | -1.124 | 0.009 | -1.421 | -0.501 | -0.001 | -0.001 | -0.001 | 0 | 0.53 | 0.51 | 0.46 | 0.44 | 0.44 | 0.38 |
| 4 | 19684918 | rs17570168 | 1.537 | 0.323 | -1.599 | -0.788 | -0.042 | 0.08 | 0.08 | 0 | 0.16 | 0.03 | 0.09 | 0.01 | 0.16 | 0.06 |
| 4 | 19696428 | rs3842911 | 1.612 | 0.271 | -1.525 | -0.617 | -0.028 | -0.055 | -0.055 | 0 | 0.14 | 0.02 | 0.14 | 0.2 | 0.14 | 0.2 |
| 4 | 19717625 | rs9291405 | 1.039 | 0.185 | -1.123 | -0.843 | -0.023 | -0.062 | -0.062 | 0 | 0.17 | 0.26 | 0.2 | 0.02 | 0.11 | 0.08 |
| 4 | 19724641 | rs4696937 | -1.076 | 0.723 | -0.603 | -0.745 | 0.06 | -0.038 | -0.038 | 0 | 0.93 | 0.85 | 0.93 | 1 | 0.88 | 0.89 |
| 4 | 19740879 | rs10029490 | -0.127 | 0.38 | -0.656 | -0.899 | 0 | -0.019 | -0.019 | 0.237 | 0.31 | 0.28 | 0.3 | 0.03 | 0.27 | 0.15 |
| 4 | 19751585 | rs3935613 | 1.339 | 0.491 | -0.749 | -0.818 | -0.002 | -0.024 | -0.024 | 0 | 0.16 | 0.17 | 0.19 | 0.02 | 0.11 | 0.08 |
| 4 | 19753174 | rs13137813 | -0.888 | 0.522 | -0.875 | -0.187 | 0 | 0.003 | 0.003 | 1.066 | 0.55 | 0.45 | 0.58 | 0.41 | 0.46 | 0.38 |
| 4 | 19763176 | rs4525973 | 1.087 | 0.455 | -0.986 | -0.21 | 0.013 | -0.05 | -0.05 | 0 | 0.19 | 0.09 | 0.27 | 0.37 | 0.19 | 0.22 |
| 4 | 19798055 | rs13145150 | -0.547 | 0.411 | -0.859 | -0.203 | 0.009 | 0.011 | 0.011 | 1.975 | 0.5 | 0.48 | 0.57 | 0.42 | 0.43 | 0.33 |
| 4 | 19800481 | rs10938771 | 1.105 | 0.447 | -1.089 | -0.107 | 0.014 | -0.029 | -0.029 | 0 | 0.14 | 0.1 | 0.27 | 0.4 | 0.15 | 0.18 |
| 4 | 19810878 | rs4380519 | -1.105 | 1.139 | -0.729 | -0.169 | -0.01 | -0.008 | -0.008 | 0 | 0.63 | 0.78 | 0.66 | 0.63 | 0.59 | 0.5 |
| 4 | 19810968 | rs4315783 | -0.313 | 0.556 | -0.771 | 0.091 | -0.002 | -0.002 | -0.002 | 0 | 0.51 | 0.55 | 0.57 | 0.63 | 0.43 | 0.34 |
| 4 | 19821565 | rs7658418 | 0.205 | 0.828 | -0.983 | -0.302 | -0.005 | 0.012 | 0.012 | 0.453 | 0.27 | 0.25 | 0.22 | 0.13 | 0.29 | 0.31 |
| 4 | 19827258 | rs6833662 | 0.764 | 0.747 | -0.958 | -0.4 | -0.007 | -0.004 | -0.004 | 0 | 0.26 | 0.23 | 0.38 | 0.75 | 0.3 | 0.42 |
| 4 | 19831621 | rs6447929 | 0.055 | -0.093 | -0.425 | -0.287 | -0.028 | 0.018 | 0.018 | 1.221 | 0.67 | 0.52 | 0.73 | 0.66 | 0.71 | 0.56 |
| 4 | 19832995 | rs4261984 | 0.556 | -0.239 | -0.617 | -0.462 | -0.002 | -0.001 | -0.001 | 4.72 | 0.56 | 0.49 | 0.6 | 0.65 | 0.6 | 0.47 |
| 4 | 19834012 | rs6833091 | 1.286 | -0.147 | -0.531 | -0.3 | -0.02 | 0.046 | 0.046 | 0 | 0.05 | 0.17 | 0.03 | 0.11 | 0.02 | 0.11 |
| 4 | 19840731 | rs6844265 | 2.235 | 0.179 | -0.698 | -0.188 | -0.004 | 0.001 | 0.001 | 0 | 0.15 | 0.19 | 0.16 | 0.21 | 0.19 | 0.23 |
| 4 | 19842636 | rs6852734 | 0.687 | 0.133 | -0.715 | -0.253 | 0.004 | 0.008 | 0.008 | 0 | 0.21 | 0.39 | 0.19 | 0.32 | 0.22 | 0.34 |
| 4 | 19847106 | rs4516715 | 0.091 | 0.1 | -1.257 | -0.421 | 0.02 | 0.043 | 0.043 | 0.944 | 0.59 | 0.8 | 0.66 | 0.76 | 0.61 | 0.65 |
| 4 | 19857997 | rs12649350 | 0.161 | 0.079 | -1.296 | -0.447 | 0.007 | 0 | 0 | 0.85 | 0.3 | 0.18 | 0.22 | 0.23 | 0.29 | 0.3 |
| 4 | 19872318 | rs7693487 | 1.73 | 0.048 | -1.736 | -0.5 | -0.094 | -0.149 | -0.149 | 0.014 | 0.32 | 0.04 | 0.36 | 0.43 | 0.32 | 0.23 |
| 4 | 19873071 | rs10516339 | 1.933 | 0.113 | -1.757 | -0.522 | -0.076 | -0.152 | -0.152 | 0 | 0.28 | 0.04 | 0.33 | 0.44 | 0.27 | 0.22 |
| 4 | 19875509 | rs4404540 | 0.566 | 0.032 | -1.142 | -0.483 | 0.007 | 0.024 | 0.024 | 2.727 | 0.17 | 0.1 | 0.19 | 0.01 | 0.17 | 0.1 |
| 4 | 19896225 | rs6858624 | -0.497 | -0.85 | -0.94 | -0.376 | 0.011 | -0.005 | -0.005 | 0 | 0.59 | 0.51 | 0.56 | 0.55 | 0.51 | 0.35 |
| 4 | 19899630 | rs11733327 | 1.588 | -0.914 | -1.585 | -0.426 | -0.052 | -0.158 | -0.158 | 1.176 | 0.25 | 0.03 | 0.22 | 0.43 | 0.21 | 0.2 |
| 4 | 19901649 | rs12512109 | -0.499 | -0.933 | -1.365 | -0.451 | -0.013 | 0.008 | 0.008 | 1.459 | 0.58 | 0.36 | 0.55 | 0.55 | 0.5 | 0.35 |
| 4 | 19906275 | rs11735138 | 1.497 | -0.905 | -1.533 | -0.408 | -0.046 | -0.153 | -0.153 | 3.792 | 0.26 | 0.05 | 0.24 | 0.44 | 0.21 | 0.22 |
| 4 | 19915956 | rs41532447 | -0.394 | -0.886 | -1.268 | -0.326 | -0.014 | 0.007 | 0.007 | 1.801 | 0.58 | 0.41 | 0.55 | 0.55 | 0.49 | 0.35 |
| 4 | 19933069 | rs4464568 | 0.654 | -1.259 | -1.171 | -0.395 | -0.006 | 0.007 | 0.007 | 0.678 | 0.1 | 0.16 | 0.08 | 0.11 | 0.07 | 0.02 |
| 4 | 19942443 | rs12650682 | 0.605 | -1.294 | -1.069 | -0.404 | -0.011 | 0.012 | 0.012 | 0.254 | 0.1 | 0.16 | 0.09 | 0.12 | 0.07 | 0.03 |
| 4 | 19971235 | rs13135681 | -0.393 | -1.374 | -1.319 | -0.649 | 0 | -0.001 | -0.001 | 0 | 0.67 | 0.41 | 0.64 | 0.64 | 0.66 | 0.49 |
| 4 | 19972465 | rs4549380 | 0.845 | -1.424 | -1.582 | -0.72 | 0.001 | 0.01 | 0.01 | 3.252 | 0.16 | 0.12 | 0.19 | 0.01 | 0.17 | 0.06 |
| 4 | 19974862 | rs10516343 | -2.149 | -1.53 | -1.503 | -0.649 | -0.019 | 0.003 | 0.003 | 0.956 | 0.83 | 0.91 | 0.8 | 0.81 | 0.75 | 0.78 |
| 4 | 19980071 | rs4128232 | 1.314 | -1.553 | -1.559 | -0.377 | -0.117 | -0.132 | -0.132 | 0 | 0.36 | 0.06 | 0.34 | 0.46 | 0.37 | 0.39 |
| 4 | 19983104 | rs4626202 | 0.616 | -1.53 | -1.534 | -0.436 | -0.048 | 0.077 | 0.077 | 1.904 | 0.46 | 0.2 | 0.42 | 0.63 | 0.43 | 0.42 |
| 4 | 19992481 | rs2629722 | -1.935 | -1.329 | -0.903 | -0.006 | -0.071 | -0.04 | -0.04 | 0 | 0.84 | 0.98 | 0.73 | 0.81 | 0.76 | 0.78 |
| 4 | 19998503 | rs2247321 | 0.222 | -1.131 | -0.62 | 0.232 | -0.004 | 0.028 | 0.028 | 0.354 | 0.1 | 0.15 | 0.09 | 0.2 | 0.06 | 0.02 |
| 4 | 20011236 | rs991645 | -1.072 | -1.196 | -0.56 | 0.203 | -0.01 | 0.013 | 0.013 | 5.25 | 0.26 | 0.37 | 0.18 | 0.35 | 0.2 | 0.25 |
| 4 | 20020240 | rs2629715 | -0.708 | -1.233 | -0.564 | 0.089 | -0.004 | -0.017 | -0.017 | 1.065 | 0.69 | 0.45 | 0.57 | 0.78 | 0.69 | 0.63 |
| 4 | 20025385 | rs7684055 | -0.031 | -1.198 | -0.555 | 0.137 | -0.009 | 0.003 | 0.003 | 0.619 | 0.11 | 0.27 | 0.07 | 0.13 | 0.06 | 0.04 |
| 4 | 20025539 | rs13107783 | 1.096 | -1.16 | -0.904 | 0.14 | -0.128 | -0.066 | -0.066 | 0.33 | 0.43 | 0.06 | 0.42 | 0.51 | 0.49 | 0.39 |
| 4 | 20032963 | rs2708987 | -1.037 | -0.92 | -0.519 | -0.059 | -0.001 | -0.001 | -0.001 | 5.383 | 0.25 | 0.29 | 0.15 | 0.28 | 0.2 | 0.24 |
| 4 | 20043516 | rs13148770 | 0.222 | -0.47 | -1.024 | -0.338 | 0.001 | 0 | 0 | 0.005 | 0.49 | 0.25 | 0.39 | 0.47 | 0.47 | 0.54 |
| 4 | 20050184 | rs1872469 | -0.042 | -0.272 | -1.408 | -0.684 | 0.005 | 0.005 | 0.005 | 0 | 0.19 | 0.15 | 0.21 | 0.08 | 0.17 | 0.16 |
| 4 | 20050733 | rs901123 | -0.14 | -0.172 | -1.395 | -0.574 | 0 | -0.001 | -0.001 | 0 | 0.39 | 0.53 | 0.43 | 0.37 | 0.37 | 0.32 |
| 4 | 20053908 | rs1032133 | 1.605 | -0.122 | -1.038 | 0.436 | 0 | 0.011 | 0.011 | 2.448 | 0.52 | 0.52 | 0.52 | 0.36 | 0.52 | 0.48 |
| 4 | 20054356 | rs6845174 | -0.713 | -0.288 | -1.024 | 0.464 | -0.015 | -0.005 | -0.005 | 0 | 0.2 | 0.3 | 0.22 | 0.12 | 0.15 | 0.17 |

| | | | | | | | | | | | | | | | | |
|---|---|---|---|---|---|---|---|---|---|---|---|---|---|---|---|---|
| 4 | 20063063 | rs10018142 | -0.341 | 0.425 | -1.232 | 0.57 | 0.013 | 0.006 | 0.006 | 2.156 | 0.13 | 0.07 | 0.05 | 0.14 | 0.08 | 0.18 |
| 4 | 20064263 | rs2874743 | -0.15 | 0.534 | -1.276 | 0.608 | -0.025 | -0.029 | -0.029 | 0 | 0.82 | 0.85 | 0.76 | 0.61 | 0.71 | 0.79 |
| 4 | 20069071 | rs28461290 | -0.383 | 0.796 | -1.18 | 0.689 | 0.01 | 0.005 | 0.005 | 2.363 | 0.13 | 0.08 | 0.05 | 0.14 | 0.08 | 0.18 |
| 4 | 20078019 | rs961521 | 0.232 | 0.956 | -1.058 | 0.495 | -0.01 | -0.024 | -0.024 | 0 | 0.64 | 0.7 | 0.71 | 0.45 | 0.62 | 0.61 |
| 4 | 20088468 | rs10938787 | 0.313 | 0.552 | -0.688 | -0.39 | -0.009 | -0.046 | -0.046 | 0 | 0.7 | 0.39 | 0.74 | 0.88 | 0.73 | 0.79 |
| 4 | 20089011 | rs4642249 | 0.25 | 0.363 | -0.186 | -0.63 | 0.007 | -0.043 | -0.043 | 0 | 0.91 | 0.79 | 0.93 | 1 | 0.89 | 0.94 |
| 4 | 20097540 | rs2322346 | 1.097 | 0.183 | 0.527 | -0.857 | -0.02 | -0.123 | -0.123 | 0 | 0.77 | 0.57 | 0.84 | 0.96 | 0.83 | 0.83 |
| 4 | 20100285 | rs4697140 | 1.102 | 0.456 | 0.63 | -1.358 | 0.02 | -0.061 | -0.061 | 0 | 0.84 | 0.61 | 0.84 | 0.95 | 0.85 | 0.92 |
| 4 | 20108488 | rs6852889 | -2.32 | 0.083 | 0.585 | -1.599 | -0.003 | -0.045 | -0.045 | 0.005 | 0.36 | 0.62 | 0.28 | 0.1 | 0.3 | 0.2 |
| 4 | 20111944 | rs6855958 | -1.151 | 0.06 | 0.837 | -1.592 | -0.01 | -0.061 | -0.061 | 0 | 0.2 | 0.23 | 0.12 | 0.04 | 0.16 | 0.13 |
| 4 | 20112399 | rs13123715 | -1.124 | 0.06 | 0.833 | -1.591 | -0.02 | -0.091 | -0.091 | 0 | 0.21 | 0.32 | 0.13 | 0.04 | 0.16 | 0.13 |
| 4 | 20117416 | rs1872467 | -1.401 | -0.053 | 1.142 | -1.625 | 0.003 | -0.002 | -0.002 | 0 | 0.23 | 0.24 | 0.18 | 0.1 | 0.17 | 0.15 |
| 4 | 20118073 | rs10516346 | -1.83 | -0.06 | 1.152 | -1.628 | -0.026 | -0.061 | -0.061 | 0 | 0.27 | 0.37 | 0.19 | 0.1 | 0.18 | 0.15 |
| 4 | 20123954 | rs16869206 | -1.351 | -0.094 | 1.136 | -1.643 | 0.008 | 0.011 | 0.011 | 0.03 | 0.22 | 0.2 | 0.19 | 0.1 | 0.17 | 0.15 |
| 4 | 20126912 | rs16869213 | -0.529 | -0.041 | 1.156 | -1.637 | 0.038 | -0.056 | -0.056 | 0 | 0.1 | 0.21 | 0.08 | 0 | 0.12 | 0.06 |
| 4 | 20127041 | rs6826653 | 1.394 | -0.022 | 0.355 | -2.593 | -0.042 | 0.091 | 0.091 | 0 | 0.31 | 0.13 | 0.21 | 0.13 | 0.37 | 0.29 |
| 4 | 20141256 | rs12510334 | -1.535 | -0.181 | 0.119 | -2.944 | -0.003 | -0.049 | -0.049 | 0 | 0.38 | 0.59 | 0.29 | 0.13 | 0.35 | 0.34 |
| 4 | 20168391 | rs1436294 | 1.021 | 0.137 | -0.794 | -3.622 | 0 | -0.015 | -0.015 | 0 | 0.55 | 0.41 | 0.68 | 0.83 | 0.61 | 0.57 |
| 4 | 20169556 | rs13117394 | 0.506 | 0.195 | -0.765 | -3.595 | 0.002 | 0.002 | 0.002 | 1.922 | 0.14 | 0.17 | 0.11 | 0.09 | 0.08 | 0.16 |
| 4 | 20171488 | rs7659298 | 0.506 | 0.195 | -0.765 | -3.595 | 0.002 | 0.002 | 0.002 | 1.922 | 0.14 | 0.17 | 0.11 | 0.09 | 0.08 | 0.16 |
| 4 | 20176829 | rs10027583 | 0.666 | 0.208 | -0.744 | -3.609 | 0.05 | -0.063 | -0.063 | 0 | 0.1 | 0.27 | 0.08 | 0 | 0.12 | 0.06 |
| 4 | 20190068 | rs17598758 | 0.807 | 0.188 | -0.787 | -3.591 | 0.005 | -0.003 | -0.003 | 0 | 0.13 | 0.02 | 0.09 | 0.1 | 0.08 | 0.16 |
| 4 | 20190972 | rs587668 | 0.031 | 0.016 | -0.889 | -3.711 | 0.016 | -0.168 | -0.168 | 0 | 0.75 | 0.52 | 0.79 | 0.96 | 0.72 | 0.82 |
| 4 | 20193573 | rs12507342 | 0.597 | 0.014 | -0.935 | -3.7 | 0.016 | 0.014 | 0.014 | 1.184 | 0.14 | 0.1 | 0.1 | 0.09 | 0.08 | 0.16 |
| 4 | 20206183 | rs1165446 | 1.215 | 0.022 | -0.985 | -3.728 | 0.009 | 0.036 | 0.036 | 1.382 | 0.55 | 0.33 | 0.68 | 0.83 | 0.61 | 0.57 |
| 4 | 20206215 | rs685064 | -1.283 | 0.109 | -0.948 | -3.572 | 0.001 | -0.007 | -0.007 | 0.057 | 0.85 | 0.87 | 0.89 | 0.96 | 0.84 | 0.88 |
| 4 | 20211471 | rs534860 | -1.168 | 0.094 | -0.945 | -3.579 | 0.002 | -0.003 | -0.003 | 1.23 | 0.85 | 0.86 | 0.89 | 0.96 | 0.84 | 0.88 |
| 4 | 20213740 | rs812652 | 1.137 | 0.13 | -0.891 | -3.865 | -0.006 | 0.113 | 0.113 | 0 | 0.51 | 0.3 | 0.64 | 0.84 | 0.54 | 0.45 |
| 4 | 20214277 | rs556208 | 0.104 | 0.211 | -0.758 | -3.79 | 0.003 | 0.009 | 0.009 | 0 | 0.69 | 0.7 | 0.77 | 0.88 | 0.71 | 0.61 |
| 4 | 20216543 | rs1923841 | 0.674 | 0.235 | -0.968 | -3.756 | -0.023 | 0.03 | 0.03 | 0.013 | 0.16 | 0.05 | 0.12 | 0.08 | 0.15 | 0.27 |
| 4 | 20229030 | rs10516353 | 0.695 | 0.236 | -0.968 | -3.756 | -0.023 | 0.03 | 0.03 | 0.004 | 0.15 | 0.05 | 0.12 | 0.08 | 0.15 | 0.27 |
| 4 | 20230757 | rs509217 | 0.726 | 0.244 | -0.97 | -3.714 | -0.009 | 0.055 | 0.055 | 0 | 0.58 | 0.3 | 0.66 | 0.87 | 0.57 | 0.55 |
| 4 | 20235828 | rs544236 | -0.889 | 0.108 | -0.76 | -3.363 | -0.002 | -0.033 | -0.033 | 0 | 0.85 | 0.82 | 0.89 | 0.96 | 0.84 | 0.88 |
| 4 | 20242529 | rs559767 | -0.288 | 0.421 | -1.033 | -3.37 | -0.007 | 0.162 | 0.162 | 0.033 | 0.62 | 0.85 | 0.77 | 0.87 | 0.61 | 0.73 |
| 4 | 20244146 | rs507134 | -1.258 | 0.42 | -0.449 | -2.863 | 0.001 | -0.019 | -0.019 | 0 | 0.92 | 0.9 | 0.93 | 1 | 0.88 | 0.96 |
| 4 | 20249840 | rs640501 | 0.882 | 0.406 | -0.392 | -2.65 | -0.002 | -0.035 | -0.035 | 0.019 | 0.7 | 0.41 | 0.69 | 0.82 | 0.72 | 0.71 |
| 4 | 20251117 | rs614158 | 0.621 | 0.407 | -0.504 | -2.651 | -0.001 | -0.009 | -0.009 | 0 | 0.7 | 0.34 | 0.69 | 0.82 | 0.72 | 0.71 |
| 4 | 20255306 | rs7655084 | -0.12 | 0.534 | -0.348 | -2.686 | -0.014 | 0.048 | 0.048 | 0.078 | 0.75 | 0.84 | 0.79 | 0.95 | 0.72 | 0.79 |
| 4 | 20262768 | rs1323067 | 0.503 | 0.469 | -0.191 | -2.688 | -0.007 | 0.029 | 0.029 | 0 | 0.66 | 0.7 | 0.68 | 0.79 | 0.66 | 0.67 |
| 4 | 20263058 | rs1323068 | 1.386 | 0.618 | -0.182 | -2.948 | 0.003 | -0.001 | -0.001 | 0 | 0.27 | 0.32 | 0.46 | 0.67 | 0.31 | 0.49 |
| 4 | 20267915 | rs715213 | 1.907 | 0.628 | -0.337 | -2.815 | 0 | 0.032 | 0.032 | 0.485 | 0.05 | 0.14 | 0.02 | 0.07 | 0.03 | 0.01 |
| 4 | 20271773 | rs587032 | 0.965 | 0.7 | -0.785 | -2.847 | -0.011 | 0.034 | 0.034 | 1.553 | 0.16 | 0.09 | 0.14 | 0.05 | 0.15 | 0.15 |
| 4 | 20283069 | rs17601458 | 1.535 | 0.512 | -0.918 | -2.79 | -0.05 | 0.148 | 0.148 | 0 | 0.26 | 0.07 | 0.15 | 0.03 | 0.28 | 0.08 |
| 4 | 20289373 | rs7680945 | 0.956 | 0.782 | -0.954 | -1.651 | -0.005 | 0.003 | 0.003 | 0 | 0.26 | 0.29 | 0.44 | 0.66 | 0.25 | 0.37 |
| 4 | 20292239 | rs593943 | -1.463 | 1.177 | -1.982 | -1.455 | -0.012 | 0.021 | 0.021 | 0 | 0.91 | 0.96 | 0.92 | 1 | 0.87 | 0.95 |
| 4 | 20303108 | rs12511950 | 1.165 | 0.879 | -1.736 | -1.14 | -0.007 | -0.054 | -0.054 | 0 | 0.21 | 0.11 | 0.4 | 0.59 | 0.23 | 0.35 |
| 4 | 20309742 | rs1323062 | -0.459 | 0.681 | -1.738 | -1.261 | -0.013 | 0.067 | 0.067 | 0 | 0.66 | 0.77 | 0.74 | 0.92 | 0.67 | 0.73 |
| 4 | 20310173 | rs13434430 | -0.48 | 0.692 | -1.752 | -1.261 | -0.016 | 0.069 | 0.069 | 0 | 0.66 | 0.78 | 0.74 | 0.92 | 0.67 | 0.73 |
| 4 | 20310276 | rs13435197 | -0.2 | 0.609 | -1.789 | -1.072 | -0.003 | 0.005 | 0.005 | 0 | 0.61 | 0.61 | 0.68 | 0.87 | 0.6 | 0.7 |
| 4 | 20318837 | rs12646142 | 0.157 | 0.588 | -1.882 | 0.214 | -0.003 | -0.052 | -0.052 | 0 | 0.21 | 0.11 | 0.41 | 0.59 | 0.22 | 0.35 |
| 4 | 20322789 | rs9993087 | -0.482 | 0.089 | -2.031 | -0.592 | 0.042 | 0.095 | 0.095 | 0 | 0.27 | 0.16 | 0.12 | 0.08 | 0.2 | 0.35 |
| 4 | 20322913 | rs4696953 | 0.329 | 0.295 | -2.405 | -1.37 | -0.011 | 0.126 | 0.126 | 0.941 | 0.54 | 0.26 | 0.66 | 0.72 | 0.6 | 0.49 |
| 4 | 20327010 | rs10938796 | 0.318 | 0.294 | -3.164 | -1.233 | 0.004 | -0.01 | -0.01 | 0 | 0.55 | 0.23 | 0.46 | 0.56 | 0.57 | 0.57 |
| 4 | 20333722 | rs12503652 | 0.707 | 0.063 | -2.504 | -1.224 | -0.012 | -0.016 | -0.016 | 0 | 0.12 | 0.19 | 0.12 | 0.06 | 0.08 | 0.09 |
| 4 | 20334919 | rs12502618 | -1.49 | 0.282 | -2.379 | -1.123 | -0.009 | -0.013 | -0.013 | 0 | 0.92 | 0.94 | 0.76 | 0.82 | 0.86 | 0.88 |
| 4 | 20347939 | rs6447955 | -1.39 | 0.261 | -2.179 | -1.133 | 0.001 | 0.009 | 0.009 | 0 | 0.92 | 0.9 | 0.76 | 0.82 | 0.86 | 0.85 |
| 4 | 20362716 | rs4406013 | 1.864 | 0.166 | -2.903 | -1.303 | -0.047 | -0.144 | -0.144 | 0 | 0.18 | 0.01 | 0.19 | 0.45 | 0.19 | 0.3 |
| 4 | 20370143 | rs5016268 | -0.329 | 0.154 | -3.373 | -1.785 | -0.032 | 0.035 | 0.035 | 0.055 | 0.59 | 0.8 | 0.49 | 0.33 | 0.57 | 0.42 |
| 4 | 20377167 | rs4403045 | -1.23 | 0.265 | -3.381 | -1.805 | -0.023 | -0.103 | -0.103 | 1.3 | 0.68 | 0.86 | 0.73 | 0.52 | 0.71 | 0.54 |
| 4 | 20380257 | rs9884539 | 1.407 | 0.15 | -3.001 | -1.807 | -0.009 | 0 | 0 | 0 | 0.12 | 0.18 | 0.15 | 0.13 | 0.09 | 0.1 |
| 4 | 20386044 | rs4552454 | -2.017 | 0.272 | -3.049 | -1.749 | -0.002 | 0.012 | 0.012 | 0 | 0.87 | 0.88 | 0.93 | 0.97 | 0.9 | 0.84 |
| 4 | 20391615 | rs13142826 | 1.222 | 0.248 | -2.66 | -1.483 | -0.006 | -0.014 | -0.014 | 0.026 | 0.08 | 0.06 | 0.24 | 0.19 | 0.13 | 0.12 |
| 4 | 20405032 | rs9996157 | 0.477 | 0.245 | -2.495 | -1.576 | -0.021 | 0.059 | 0.059 | 0 | 0.23 | 0.14 | 0.19 | 0.08 | 0.27 | 0.08 |
| 4 | 20425588 | rs17609702 | 0.462 | 0.244 | -2.503 | -1.576 | -0.031 | 0.078 | 0.078 | 0 | 0.23 | 0.12 | 0.19 | 0.07 | 0.27 | 0.08 |
| 4 | 20444781 | rs1473080 | 0.428 | 0.271 | -2.301 | -1.555 | -0.012 | 0.04 | 0.04 | 0 | 0.24 | 0.2 | 0.2 | 0.08 | 0.27 | 0.11 |
| 4 | 20447040 | rs884770 | -0.431 | 0.264 | -1.44 | -1.629 | -0.001 | -0.044 | -0.044 | 1.312 | 0.21 | 0.36 | 0.19 | 0.13 | 0.21 | 0.17 |
| 4 | 20449733 | rs16869609 | 0.73 | 0.882 | -1.237 | -1.254 | 0.024 | -0.021 | -0.021 | 0 | 0.07 | 0.08 | 0.04 | 0.01 | 0.11 | 0.06 |
| 4 | 20454884 | rs16869615 | 0.879 | 0.907 | -1.337 | -1.245 | 0.012 | -0.022 | -0.022 | 1.361 | 0.06 | 0.05 | 0.03 | 0 | 0.11 | 0.06 |
| 4 | 20455955 | rs9990844 | 0.168 | 0.822 | -1.442 | -1.145 | -0.004 | 0.007 | 0.007 | 0 | 0.08 | 0.09 | 0.09 | 0.04 | 0.1 | 0.05 |

| | | | | | | | | | | | | | | | | |
|---|---|---|---|---|---|---|---|---|---|---|---|---|---|---|---|---|
| 4 | 20458688 | rs11733205 | -0.614 | 0.942 | -1.492 | -0.916 | -0.063 | -0.027 | -0.027 | 0 | 0.61 | 0.87 | 0.66 | 0.38 | 0.62 | 0.4 |
| 4 | 20459118 | rs11942897 | -0.363 | 1.183 | -1.086 | -1.083 | 0 | 0.036 | 0.036 | 0.183 | 0.47 | 0.57 | 0.4 | 0.2 | 0.48 | 0.28 |
| 4 | 20467767 | rs1033111 | -0.541 | 1.42 | -0.967 | 0.136 | -0.009 | -0.011 | -0.011 | 0 | 0.41 | 0.44 | 0.72 | 0.47 | 0.57 | 0.36 |
| 4 | 20473278 | rs11936122 | 0.334 | 2.1 | -1.368 | -0.392 | -0.047 | 0.005 | 0.005 | 0.074 | 0.17 | 0.07 | 0.22 | 0.09 | 0.24 | 0.18 |
| 4 | 20476800 | rs625614 | -0.163 | 1.925 | -0.594 | -0.206 | -0.002 | 0.001 | 0.001 | 0 | 0.87 | 0.87 | 0.87 | 0.93 | 0.78 | 0.88 |
| 4 | 20479620 | rs16869643 | -0.522 | 0.535 | -0.638 | -0.345 | 0.002 | -0.027 | -0.027 | 0 | 0.26 | 0.17 | 0.09 | 0.05 | 0.18 | 0.23 |
| 4 | 20487711 | rs10019416 | -0.794 | 1.08 | 0.048 | 0.666 | -0.008 | 0.034 | 0.034 | 0 | 0.21 | 0.4 | 0.2 | 0.3 | 0.2 | 0.13 |
| 4 | 20488464 | rs484614 | 0.498 | 1.16 | -0.602 | 0.299 | 0.004 | -0.041 | -0.041 | 0 | 0.38 | 0.18 | 0.4 | 0.44 | 0.33 | 0.31 |
| 4 | 20489479 | rs3756147 | -0.515 | 1.627 | -0.691 | -0.063 | 0.001 | 0.001 | 0.001 | 0 | 0.36 | 0.64 | 0.32 | 0.46 | 0.4 | 0.39 |
| 4 | 20497277 | rs522501 | -0.659 | 1.571 | -0.741 | -0.136 | 0 | 0.045 | 0.045 | 0 | 0.74 | 0.83 | 0.74 | 0.95 | 0.73 | 0.7 |
| 4 | 20518603 | rs6825654 | -0.668 | 1.402 | -0.198 | 0.046 | -0.004 | 0.005 | 0.005 | 0 | 0.36 | 0.57 | 0.39 | 0.46 | 0.34 | 0.37 |
| 4 | 20520993 | rs519813 | -0.823 | 1.425 | -0.423 | -0.385 | 0 | -0.024 | -0.024 | 0 | 0.88 | 0.89 | 0.92 | 0.99 | 0.88 | 0.95 |
| 4 | 20524715 | rs16869706 | 0.309 | 1.175 | -0.039 | -0.126 | 0.005 | 0.014 | 0.014 | 0.099 | 0.18 | 0.14 | 0.11 | 0.14 | 0.15 | 0.1 |
| 4 | 20527864 | rs7690660 | -0.329 | 1.148 | 0.348 | 0.045 | 0.026 | -0.049 | -0.049 | 0 | 0.17 | 0.43 | 0.19 | 0.13 | 0.18 | 0.16 |
| 4 | 20529670 | rs3775815 | -0.693 | 1.068 | 0.02 | 0.302 | 0.004 | -0.015 | -0.015 | 0 | 0.18 | 0.17 | 0.09 | 0.31 | 0.16 | 0.25 |
| 4 | 20530374 | rs9992559 | 0.872 | 0.866 | -0.411 | -0.898 | 0.016 | 0.033 | 0.033 | 3.235 | 0.3 | 0.16 | 0.16 | 0 | 0.24 | 0.06 |
| 4 | 20530981 | rs2168801 | 0.483 | 1.2 | -0.325 | -0.174 | 0.007 | 0.018 | 0.018 | 0.021 | 0.41 | 0.23 | 0.37 | 0.33 | 0.35 | 0.14 |
| 4 | 20533014 | rs2322557 | 0.925 | 1.234 | -0.31 | -0.079 | -0.006 | 0.004 | 0.004 | 0 | 0.1 | 0.07 | 0.22 | 0.33 | 0.11 | 0.08 |
| 4 | 20533034 | rs632152 | -0.813 | 1.475 | -0.613 | -0.318 | -0.04 | 0.031 | 0.031 | 0 | 0.89 | 0.98 | 0.87 | 0.97 | 0.87 | 0.87 |
| 4 | 20534467 | rs6816854 | 0.04 | 1.218 | 0.073 | -0.351 | 0.006 | -0.008 | -0.008 | 3.632 | 0.59 | 0.49 | 0.48 | 0.67 | 0.52 | 0.46 |
| 4 | 20535876 | rs520535 | 0.882 | 1.057 | -0.133 | -0.381 | 0.009 | 0.012 | 0.012 | 0.813 | 0.44 | 0.35 | 0.24 | 0.24 | 0.34 | 0.34 |
| 4 | 20537784 | rs12512830 | 1.358 | 1.269 | -0.58 | -0.054 | -0.001 | -0.009 | -0.009 | 0 | 0.07 | 0.03 | 0.05 | 0.2 | 0.08 | 0.15 |
| 4 | 20540540 | rs3775820 | 1.088 | 1.267 | -0.404 | 0.096 | 0.029 | 0.039 | 0.039 | 1.482 | 0.4 | 0.25 | 0.22 | 0.25 | 0.33 | 0.33 |
| 4 | 20542485 | rs3775823 | 1.378 | 1.52 | -0.699 | 0.188 | -0.02 | 0.016 | 0.016 | 1.523 | 0.15 | 0.1 | 0.28 | 0.1 | 0.25 | 0.21 |
| 4 | 20543709 | rs16869739 | 0.846 | 1.3 | -0.526 | 0.776 | -0.012 | -0.013 | -0.013 | 0.14 | 0.1 | 0.04 | 0.22 | 0.34 | 0.11 | 0.09 |
| 4 | 20543817 | rs675465 | 0.584 | 1.074 | -0.112 | 0.587 | -0.003 | -0.002 | -0.002 | 1.039 | 0.7 | 0.71 | 0.75 | 0.68 | 0.7 | 0.65 |
| 4 | 20544184 | rs7690492 | -0.756 | 0.559 | 0.139 | 0.244 | -0.032 | 0.024 | 0.024 | 0 | 0.3 | 0.49 | 0.18 | 0.37 | 0.23 | 0.38 |
| 4 | 20544416 | rs7669094 | -0.547 | 0.725 | -0.418 | -0.085 | -0.002 | 0.002 | 0.002 | 1.39 | 0.26 | 0.23 | 0.16 | 0.3 | 0.25 | 0.34 |
| 4 | 20544912 | rs3775824 | -0.01 | 1.248 | -0.793 | -0.265 | -0.023 | -0.032 | -0.032 | 4.528 | 0.19 | 0.08 | 0.15 | 0.26 | 0.22 | 0.28 |
| 4 | 20545314 | rs3775827 | 0.915 | 1.264 | -0.163 | -0.777 | -0.007 | -0.017 | -0.017 | 1.008 | 0.77 | 0.61 | 0.88 | 0.79 | 0.79 | 0.66 |
| 4 | 20548976 | rs11936468 | -0.393 | 1.053 | -0.12 | -1.045 | 0 | 0.058 | 0.058 | 0.28 | 0.6 | 0.72 | 0.8 | 0.84 | 0.6 | 0.7 |
| 4 | 20552510 | rs17542486 | 0.777 | 1.215 | -0.06 | -1.051 | -0.005 | 0.003 | 0.003 | 0 | 0.08 | 0.06 | 0.18 | 0 | 0.09 | 0.03 |
| 4 | 20552853 | rs4697169 | 0.688 | 1.358 | 0.25 | -1.174 | 0.003 | 0.003 | 0.003 | 0 | 0.38 | 0.39 | 0.5 | 0.78 | 0.29 | 0.38 |
| 4 | 20563108 | rs10516357 | -0.709 | 1.639 | 0.296 | -1.144 | 0.015 | -0.037 | -0.037 | 0 | 0.17 | 0.21 | 0.27 | 0.01 | 0.29 | 0.13 |
| 4 | 20565861 | rs506358 | 0.461 | 1.472 | 0.205 | -1.205 | -0.012 | -0.032 | -0.032 | 0 | 0.42 | 0.54 | 0.52 | 0.81 | 0.3 | 0.46 |
| 4 | 20579372 | rs7666974 | -0.813 | 1.534 | -0.273 | -1.226 | -0.069 | 0.269 | 0.269 | 0.177 | 0.61 | 0.94 | 0.83 | 0.97 | 0.61 | 0.67 |
| 4 | 20585883 | rs2168802 | -1.151 | 1.626 | 0.075 | -1.034 | 0.084 | 0.028 | 0.028 | 0 | 0.14 | 0.37 | 0.14 | 0.18 | 0.22 | 0.27 |
| 4 | 20599591 | rs9992598 | -0.55 | 1.373 | -0.288 | -1.323 | -0.005 | 0.082 | 0.082 | 0.042 | 0.54 | 0.68 | 0.8 | 0.97 | 0.58 | 0.67 |
| 4 | 20599796 | rs12507257 | -0.148 | 1.484 | -0.4 | -0.851 | 0.027 | 0.008 | 0.008 | 0.066 | 0.15 | 0.25 | 0.26 | 0.17 | 0.25 | 0.21 |
| 4 | 20601311 | rs10010466 | -0.083 | 0.775 | -0.591 | -0.377 | 0.003 | 0.003 | 0.003 | 0 | 0.39 | 0.34 | 0.5 | 0.73 | 0.32 | 0.31 |
| 4 | 20605594 | rs2322559 | -0.382 | 1.102 | -0.749 | -0.242 | -0.001 | 0 | 0 | 0 | 0.57 | 0.58 | 0.75 | 0.67 | 0.55 | 0.61 |
| 4 | 20608975 | rs12511349 | 0.461 | 1.342 | -1.175 | -0.487 | -0.031 | 0.046 | 0.046 | 0 | 0.08 | 0.05 | 0.17 | 0 | 0.09 | 0.07 |
| 4 | 20611629 | rs3733510 | 0.478 | 0.636 | -1.26 | -0.107 | 0.018 | -0.012 | -0.012 | 0 | 0.1 | 0.04 | 0.14 | 0.12 | 0.04 | 0.05 |
| 4 | 20612056 | rs9993853 | 0.148 | 1.028 | -0.791 | -0.301 | 0.014 | 0.074 | 0.074 | 0 | 0.28 | 0.35 | 0.34 | 0.49 | 0.31 | 0.35 |
| 4 | 20621690 | rs8663 | -0.241 | 0.884 | -0.901 | -0.471 | 0.005 | -0.004 | -0.004 | 0 | 0.21 | 0.16 | 0.14 | 0.03 | 0.15 | 0.12 |
| 4 | 20623976 | rs11930632 | -0.43 | 1.149 | -1.12 | -0.003 | -0.013 | -0.001 | -0.001 | 0 | 0.31 | 0.31 | 0.48 | 0.31 | 0.37 | 0.24 |
| 4 | 20627764 | rs16869812 | -0.718 | 1.278 | -0.842 | 0.123 | 0.002 | 0.007 | 0.007 | 0 | 0.52 | 0.58 | 0.63 | 0.38 | 0.52 | 0.35 |
| 4 | 20628259 | rs936359 | -0.613 | 1.122 | -0.974 | 0.192 | 0.001 | 0.012 | 0.012 | 0 | 0.51 | 0.63 | 0.62 | 0.41 | 0.49 | 0.42 |
| 4 | 20637946 | rs1465522 | 0.046 | 1.048 | -1.213 | 0.043 | -0.006 | -0.001 | -0.001 | 0.012 | 0.38 | 0.28 | 0.36 | 0.6 | 0.42 | 0.51 |
| 4 | 20640647 | rs6824850 | 0.409 | 0.577 | -0.995 | -0.644 | -0.019 | -0.043 | -0.043 | 0.006 | 0.35 | 0.38 | 0.24 | 0.19 | 0.3 | 0.32 |
| 4 | 20660406 | rs10024417 | 0.207 | 0.412 | -1.305 | -1.063 | -0.025 | -0.141 | -0.141 | 1.785 | 0.2 | 0.36 | 0.14 | 0 | 0.16 | 0.23 |
| 4 | 20663866 | rs967942 | -0.828 | 0.83 | -1.727 | -0.603 | -0.007 | -0.011 | -0.011 | 0 | 0.75 | 0.82 | 0.75 | 0.69 | 0.72 | 0.7 |
| 4 | 20674741 | rs10805218 | 0.9 | 0.651 | -1.62 | -0.65 | -0.013 | -0.004 | -0.004 | 0.422 | 0.11 | 0.08 | 0.15 | 0.09 | 0.11 | 0.11 |
| 4 | 20674761 | rs10755201 | -0.776 | 0.682 | -1.576 | -0.597 | -0.019 | -0.01 | -0.01 | 0 | 0.74 | 0.83 | 0.74 | 0.69 | 0.7 | 0.69 |
| 4 | 20685983 | rs2195033 | 2.092 | 0.669 | -1.347 | -0.453 | 0 | 0.018 | 0.018 | 0 | 0.17 | 0.12 | 0.3 | 0.41 | 0.2 | 0.2 |
| 4 | 20687802 | rs1365580 | 0.819 | 0.682 | -0.919 | -0.561 | 0.014 | -0.004 | -0.004 | 4.909 | 0.11 | 0.17 | 0.15 | 0.08 | 0.12 | 0.07 |
| 4 | 20692096 | rs11736062 | -0.056 | 0.596 | -0.983 | -0.724 | -0.005 | -0.002 | -0.002 | 0 | 0.63 | 0.6 | 0.6 | 0.59 | 0.59 | 0.57 |
| 4 | 20692580 | rs16869853 | 1.216 | 0.421 | -0.731 | -0.852 | -0.003 | -0.003 | -0.003 | 0.002 | 0.39 | 0.3 | 0.44 | 0.58 | 0.42 | 0.35 |
| 4 | 20696980 | rs10938801 | -0.164 | 0.348 | -0.685 | -1.005 | -0.008 | -0.039 | -0.039 | 0 | 0.45 | 0.47 | 0.29 | 0.17 | 0.39 | 0.37 |
| 4 | 20714719 | rs3764967 | 1.672 | 0.382 | -0.705 | -0.9 | -0.004 | 0.03 | 0.03 | 2.335 | 0.22 | 0.12 | 0.14 | 0.16 | 0.23 | 0.15 |
| 4 | 20717173 | rs11721944 | 0.104 | 0.263 | -0.727 | -0.972 | -0.002 | 0.013 | 0.013 | 0.119 | 0.19 | 0.14 | 0.2 | 0.08 | 0.18 | 0.1 |
| 4 | 20717561 | rs11722085 | 0.407 | 0.237 | -1.056 | -1.116 | -0.032 | 0.105 | 0.105 | 0.163 | 0.17 | 0.03 | 0.09 | 0 | 0.15 | 0.09 |
| 4 | 20727464 | rs9291412 | 0.902 | 0.195 | -0.655 | -0.785 | -0.042 | -0.1 | -0.1 | 0.077 | 0.15 | 0.35 | 0.11 | 0 | 0.1 | 0.2 |
| 4 | 20730601 | rs6811030 | 0.212 | 0.359 | -0.684 | -0.74 | -0.014 | -0.046 | -0.046 | 0 | 0.37 | 0.47 | 0.25 | 0.18 | 0.33 | 0.39 |
| 4 | 20730780 | rs16869915 | 1.368 | 0.344 | -0.62 | -0.791 | -0.025 | -0.073 | -0.073 | 0.683 | 0.13 | 0.22 | 0.1 | 0 | 0.1 | 0.2 |
| 4 | 20731047 | rs2288308 | 0.225 | 0.321 | -0.645 | -0.771 | -0.014 | -0.055 | -0.055 | 0 | 0.37 | 0.47 | 0.25 | 0.17 | 0.33 | 0.34 |
| 4 | 20731332 | rs12640448 | 1.338 | 0.33 | -0.642 | -0.772 | -0.002 | 0.019 | 0.019 | 3.149 | 0.24 | 0.2 | 0.16 | 0.16 | 0.24 | 0.15 |
| 4 | 20732334 | rs7688805 | 1.284 | 0.392 | -0.628 | -0.857 | -0.033 | -0.076 | -0.076 | 0.136 | 0.13 | 0.25 | 0.09 | 0 | 0.08 | 0.2 |
| 4 | 20736742 | rs6447978 | 0.543 | 0.805 | -0.732 | -0.497 | -0.013 | -0.073 | -0.073 | 0 | 0.38 | 0.47 | 0.25 | 0.11 | 0.33 | 0.34 |

| | | | | | | | | | | | | | | | | |
|---|---|---|---|---|---|---|---|---|---|---|---|---|---|---|---|---|
| 4 | 20746450 | rs10084802 | 0.494 | 0.866 | -0.776 | -0.546 | -0.007 | -0.063 | -0.063 | 0 | 0.4 | 0.47 | 0.26 | 0.11 | 0.36 | 0.36 |
| 4 | 20749299 | rs10938803 | -0.553 | 0.861 | -0.71 | -0.539 | -0.018 | 0.042 | 0.042 | 0 | 0.59 | 0.71 | 0.46 | 0.26 | 0.52 | 0.49 |
| 4 | 20751278 | rs2322688 | 0.76 | 0.98 | -0.844 | -0.519 | -0.013 | -0.049 | -0.049 | 0.167 | 0.38 | 0.47 | 0.25 | 0.17 | 0.33 | 0.39 |
| 4 | 20755240 | rs7689421 | -0.114 | 1.199 | -0.558 | -0.14 | 0.006 | -0.026 | -0.026 | 0 | 0.58 | 0.6 | 0.44 | 0.21 | 0.51 | 0.49 |
| 4 | 20760089 | rs4697177 | -0.142 | 1.227 | -1.13 | 0.336 | -0.024 | -0.026 | -0.026 | 0 | 0.76 | 0.84 | 0.75 | 0.62 | 0.72 | 0.7 |
| 4 | 20765867 | rs7657542 | -0.83 | 0.706 | -0.537 | 0.632 | 0.001 | 0.061 | 0.061 | 1.297 | 0.2 | 0.28 | 0.24 | 0.31 | 0.2 | 0.17 |
| 4 | 20767337 | rs7686877 | -0.54 | 0.232 | -0.4 | 0.377 | -0.022 | 0.003 | 0.003 | 2.78 | 0.24 | 0.42 | 0.14 | 0.23 | 0.16 | 0.31 |
| 4 | 20767378 | rs16869940 | 0.119 | 0.05 | -1.379 | 0.408 | -0.004 | -0.081 | -0.081 | 5.391 | 0.18 | 0.03 | 0.37 | 0.28 | 0.14 | 0.24 |
| 4 | 20769558 | rs13129008 | -0.082 | 0.256 | -1.352 | -0.065 | -0.001 | 0 | 0 | 0 | 0.46 | 0.44 | 0.53 | 0.69 | 0.52 | 0.5 |
| 4 | 20770444 | rs16869949 | 2.228 | -0.128 | -1.455 | 0.028 | 0.001 | 0.004 | 0.004 | 1.778 | 0.1 | 0.11 | 0.04 | 0.19 | 0.04 | 0.14 |
| 4 | 20771062 | rs3857162 | -0.71 | -0.256 | -1.522 | -0.006 | -0.002 | 0.015 | 0.015 | 0 | 0.53 | 0.44 | 0.66 | 0.66 | 0.56 | 0.57 |
| 4 | 20773916 | rs13137029 | 2.886 | -0.19 | -1.416 | 0.12 | -0.006 | 0.082 | 0.082 | 0.088 | 0.07 | 0.11 | 0.04 | 0.19 | 0.04 | 0.09 |
| 4 | 20776516 | rs10938805 | -1.016 | -0.233 | -1.486 | -0.269 | 0.012 | 0.08 | 0.08 | 5.307 | 0.54 | 0.76 | 0.64 | 0.67 | 0.49 | 0.72 |
| 4 | 20780035 | rs16869959 | 1.836 | 0.064 | -1.721 | 0.005 | -0.056 | -0.053 | -0.053 | 0 | 0.1 | 0.01 | 0.12 | 0.19 | 0.23 | 0.08 |
| 4 | 20796706 | rs6843196 | -0.543 | -0.097 | -1.454 | -0.101 | -0.006 | -0.011 | -0.011 | 0 | 0.41 | 0.46 | 0.59 | 0.65 | 0.44 | 0.52 |
| 4 | 20798129 | rs9998730 | -0.582 | -0.095 | -1.456 | -0.103 | -0.004 | -0.011 | -0.011 | 0 | 0.42 | 0.46 | 0.59 | 0.65 | 0.45 | 0.52 |
| 4 | 20801649 | rs4697184 | -1.313 | 0.14 | -1.767 | -0.294 | -0.034 | 0.012 | 0.012 | 0 | 0.87 | 0.99 | 0.85 | 0.94 | 0.88 | 0.93 |
| 4 | 20805765 | rs16869987 | 0.127 | 0.049 | -1.566 | -0.157 | 0.003 | -0.019 | -0.019 | 0.167 | 0.33 | 0.28 | 0.54 | 0.47 | 0.4 | 0.37 |
| 4 | 20806697 | rs12641748 | 2.591 | 0.164 | -1.848 | -0.191 | -0.008 | -0.02 | -0.02 | 0.972 | 0.08 | 0 | 0.02 | 0.1 | 0.06 | 0.1 |
| 4 | 20807977 | rs16870017 | 2.558 | 0 | -1.464 | 0.099 | -0.011 | 0.015 | 0.015 | 0 | 0.11 | 0.2 | 0.05 | 0.14 | 0.05 | 0.21 |
| 4 | 20809704 | rs1863632 | -0.012 | -0.094 | -1.471 | 0.086 | 0.001 | -0.006 | -0.006 | 0.016 | 0.37 | 0.34 | 0.57 | 0.47 | 0.44 | 0.39 |
| 4 | 20814133 | rs16870076 | 2.405 | 0.044 | -1.621 | 0.085 | -0.008 | 0.008 | 0.008 | 0 | 0.12 | 0.21 | 0.07 | 0.14 | 0.06 | 0.21 |
| 4 | 20814178 | rs9998323 | -0.444 | 0.453 | -1.78 | -0.073 | -0.004 | 0.024 | 0.024 | 2.735 | 0.56 | 0.67 | 0.67 | 0.67 | 0.53 | 0.72 |
| 4 | 20814216 | rs9998327 | 0.728 | 0.786 | -2.071 | 0.39 | -0.043 | -0.029 | -0.029 | 0.157 | 0.22 | 0.05 | 0.45 | 0.26 | 0.3 | 0.27 |
| 4 | 20815387 | rs1873530 | -1.306 | 0.931 | -1.526 | 0.346 | -0.028 | -0.007 | -0.007 | 0 | 0.79 | 0.92 | 0.9 | 0.73 | 0.75 | 0.83 |
| 4 | 20820088 | rs12501548 | -0.551 | 0.625 | -1.298 | 0.224 | 0 | 0.007 | 0.007 | 0.244 | 0.44 | 0.32 | 0.65 | 0.37 | 0.47 | 0.48 |
| 4 | 20820808 | rs6838129 | -0.005 | 0.206 | -1.128 | 0.344 | 0.003 | 0.002 | 0.002 | 0 | 0.63 | 0.72 | 0.47 | 0.65 | 0.61 | 0.64 |
| 4 | 20821110 | rs11734471 | -0.262 | -0.083 | -1.046 | 0.496 | -0.009 | 0.006 | 0.006 | 0 | 0.39 | 0.6 | 0.28 | 0.42 | 0.33 | 0.56 |
| 4 | 20827366 | rs10013103 | 0.338 | -0.331 | -1.076 | -0.288 | 0.01 | 0.104 | 0.104 | 0 | 0.47 | 0.25 | 0.36 | 0.2 | 0.52 | 0.48 |
| 4 | 20828809 | rs976376 | -0.425 | -0.207 | -0.418 | 0.11 | 0.007 | 0.07 | 0.07 | 0 | 0.81 | 0.72 | 0.72 | 0.59 | 0.76 | 0.85 |
| 4 | 20831331 | rs12649113 | 1.245 | -0.228 | -0.894 | -0.317 | 0.043 | 0.03 | 0.03 | 0 | 0.32 | 0.21 | 0.11 | 0.19 | 0.17 | 0.39 |
| 4 | 20832402 | rs17555380 | 0.702 | -0.144 | -0.831 | -0.338 | 0.024 | 0.06 | 0.06 | 0 | 0.42 | 0.27 | 0.2 | 0.21 | 0.29 | 0.45 |
| 4 | 20834721 | rs902665 | -0.468 | 0.031 | -0.603 | -0.336 | 0.009 | 0.019 | 0.019 | 0 | 0.52 | 0.55 | 0.61 | 0.37 | 0.6 | 0.46 |
| 4 | 20836574 | rs10025134 | 0.469 | 0.224 | -0.607 | -0.121 | -0.022 | 0.032 | 0.032 | 0 | 0.31 | 0.25 | 0.34 | 0.16 | 0.46 | 0.39 |
| 4 | 20836911 | rs2130920 | -1.229 | 0.228 | -0.743 | 0.008 | -0.002 | 0.007 | 0.007 | 0.06 | 0.85 | 0.84 | 0.74 | 0.58 | 0.78 | 0.84 |
| 4 | 20838476 | rs11932876 | 1.047 | 0.231 | -0.594 | -0.021 | 0.005 | 0.011 | 0.011 | 0 | 0.41 | 0.35 | 0.2 | 0.19 | 0.28 | 0.39 |
| 4 | 20843206 | rs10001877 | 0.09 | 0.752 | -0.982 | -0.304 | -0.007 | 0.002 | 0.002 | 0 | 0.3 | 0.28 | 0.32 | 0.16 | 0.44 | 0.37 |
| 4 | 20851050 | rs4697192 | -0.179 | 1.173 | -1.344 | 0.187 | -0.046 | -0.07 | -0.07 | 1.04 | 0.17 | 0.07 | 0.39 | 0.43 | 0.25 | 0.34 |
| 4 | 20852166 | rs3765121 | -0.442 | 1.192 | -1.6 | -0.14 | -0.071 | -0.037 | -0.037 | 1.438 | 0.33 | 0.12 | 0.48 | 0.43 | 0.43 | 0.4 |
| 4 | 20852244 | rs3765122 | -0.442 | 1.192 | -1.6 | -0.14 | -0.071 | -0.037 | -0.037 | 3.108 | 0.33 | 0.12 | 0.48 | 0.43 | 0.43 | 0.4 |
| 4 | 20857804 | rs16870149 | 1.183 | 0.875 | -1.679 | -0.281 | 0.018 | -0.001 | -0.001 | 0 | 0.27 | 0.2 | 0.15 | 0.21 | 0.19 | 0.37 |
| 4 | 20861179 | rs9291414 | -0.667 | 1.416 | -1.136 | -0.12 | 0.012 | 0.058 | 0.058 | 0 | 0.8 | 0.66 | 0.8 | 0.63 | 0.73 | 0.84 |
| 4 | 20861514 | rs17556698 | 0.303 | 1.004 | -1.087 | -0.729 | 0.009 | 0.038 | 0.038 | 0 | 0.54 | 0.67 | 0.38 | 0.26 | 0.39 | 0.46 |
| 4 | 20864229 | rs17556817 | -0.342 | 1.027 | -1.055 | -0.527 | -0.061 | -0.03 | -0.03 | 0.436 | 0.7 | 0.83 | 0.44 | 0.27 | 0.53 | 0.5 |
| 4 | 20866065 | rs6835474 | -0.349 | 1.002 | -0.529 | -0.777 | 0.011 | 0.012 | 0.012 | 0 | 0.07 | 0.14 | 0.13 | 0.09 | 0.07 | 0.14 |
| 4 | 20875314 | rs17632612 | 1.203 | 0.516 | -0.823 | -0.991 | 0.003 | -0.001 | -0.001 | 0 | 0.11 | 0.09 | 0.04 | 0.01 | 0.07 | 0.24 |
| 4 | 20875574 | rs17632634 | -0.055 | 0.643 | -0.67 | -0.686 | -0.003 | 0.003 | 0.003 | 0 | 0.19 | 0.11 | 0.11 | 0.01 | 0.22 | 0.09 |
| 4 | 20885254 | rs10516368 | 0.301 | 0.556 | -0.1 | -0.033 | 0.01 | 0.062 | 0.062 | 0 | 0.94 | 0.75 | 0.97 | 0.94 | 0.99 | 0.98 |
| 4 | 20891419 | rs7679910 | 0.539 | 0.778 | -0.739 | -0.059 | -0.023 | -0.01 | -0.01 | 2.743 | 0.08 | 0.01 | 0.17 | 0.1 | 0.13 | 0.1 |
| 4 | 20892799 | rs13143622 | 0.624 | 0.399 | -1.09 | -0.523 | 0.002 | 0.016 | 0.016 | 0 | 0.12 | 0.02 | 0.06 | 0.06 | 0.09 | 0.04 |
| 4 | 20894093 | rs1352796 | -0.901 | 0.577 | -1.096 | -0.719 | -0.001 | 0.001 | 0.001 | 0 | 0.84 | 0.87 | 0.83 | 0.84 | 0.85 | 0.68 |
| 4 | 20895013 | rs7670834 | 0.239 | 0.709 | -1.073 | -0.517 | -0.019 | -0.026 | -0.026 | 0 | 0.28 | 0.23 | 0.46 | 0.56 | 0.43 | 0.27 |
| 4 | 20896318 | rs7677781 | -0.015 | 0.523 | -0.589 | -0.769 | -0.004 | -0.036 | -0.036 | 2.784 | 0.67 | 0.65 | 0.76 | 0.83 | 0.75 | 0.65 |
| 4 | 20898220 | rs1565534 | 0.312 | 0.137 | -0.778 | -1.081 | 0.01 | 0.044 | 0.044 | 0 | 0.5 | 0.41 | 0.69 | 0.82 | 0.6 | 0.4 |
| 4 | 20908415 | rs1491370 | 0.633 | 0.365 | -1.209 | -0.846 | -0.097 | -0.135 | -0.135 | 0.07 | 0.26 | 0.07 | 0.44 | 0.49 | 0.42 | 0.24 |
| 4 | 20914984 | rs6843757 | -0.661 | 0.32 | -0.843 | -0.865 | -0.001 | -0.03 | -0.03 | 0 | 0.78 | 0.68 | 0.81 | 0.83 | 0.84 | 0.67 |
| 4 | 20923261 | rs12500896 | 0.291 | 0.436 | -1.318 | -0.676 | -0.103 | -0.151 | -0.151 | 1.934 | 0.27 | 0.07 | 0.45 | 0.49 | 0.42 | 0.23 |
| 4 | 20926006 | rs6849543 | 1.681 | 0.429 | -1.238 | -0.835 | -0.009 | 0.016 | 0.016 | 0 | 0.12 | 0.1 | 0.05 | 0.01 | 0.12 | 0.22 |
| 4 | 20926819 | rs1434957 | 1.263 | 0.398 | -0.887 | -0.537 | -0.009 | -0.005 | -0.005 | 0.184 | 0.35 | 0.38 | 0.3 | 0.28 | 0.29 | 0.38 |
| 4 | 20930535 | rs1114663 | 0.378 | 0.598 | -1.123 | -0.345 | -0.104 | -0.15 | -0.15 | 0 | 0.25 | 0.09 | 0.44 | 0.49 | 0.41 | 0.23 |
| 4 | 20939779 | rs4697198 | 0.368 | 0.342 | -0.663 | -0.58 | -0.012 | -0.036 | -0.036 | 0 | 0.6 | 0.46 | 0.75 | 0.77 | 0.71 | 0.6 |
| 4 | 20946907 | rs1434954 | 0.399 | 0.152 | -0.633 | -0.69 | -0.034 | -0.054 | -0.054 | 0.237 | 0.18 | 0.32 | 0.07 | 0.06 | 0.1 | 0.05 |
| 4 | 20948059 | rs6830607 | 0.795 | 0.704 | -0.843 | -0.774 | 0 | 0.017 | 0.017 | 0 | 0.51 | 0.43 | 0.63 | 0.74 | 0.5 | 0.53 |
| 4 | 20948497 | rs6448000 | 0.318 | 1.015 | -1.37 | -0.299 | -0.024 | -0.113 | -0.113 | 0.22 | 0.17 | 0.06 | 0.35 | 0.47 | 0.21 | 0.14 |
| 4 | 20949917 | rs1434950 | -0.066 | 0.977 | -0.861 | -0.646 | 0.001 | -0.013 | -0.013 | 4.948 | 0.78 | 0.75 | 0.81 | 0.82 | 0.81 | 0.65 |
| 4 | 20951955 | rs7677635 | 0.258 | 0.975 | -0.841 | -0.647 | -0.002 | -0.015 | -0.015 | 1.585 | 0.75 | 0.72 | 0.8 | 0.82 | 0.8 | 0.64 |
| 4 | 20952556 | rs6811024 | 0.689 | 0.976 | -0.918 | -0.639 | -0.016 | 0.034 | 0.034 | 0 | 0.66 | 0.74 | 0.7 | 0.78 | 0.59 | 0.56 |
| 4 | 20959057 | rs6821566 | 0.637 | 1.301 | -1.139 | -0.352 | -0.047 | 0.013 | 0.013 | 1.918 | 0.73 | 0.8 | 0.71 | 0.72 | 0.61 | 0.6 |
| 4 | 20960640 | rs1434943 | 0.186 | 0.987 | -0.777 | -0.251 | 0.012 | -0.006 | -0.006 | 0 | 0.81 | 0.69 | 0.83 | 0.82 | 0.82 | 0.69 |

| | | | | | | | | | | | | | | | | |
|---|---|---|---|---|---|---|---|---|---|---|---|---|---|---|---|---|
| 4 | 20972919 | rs17573138 | 0.505 | 0.489 | -1.272 | -0.554 | 0.021 | 0.037 | 0.037 | 0 | 0.14 | 0.04 | 0.04 | 0 | 0.1 | 0.06 |
| 4 | 20985138 | rs6841954 | -0.213 | 0.596 | -1.364 | -0.218 | 0.011 | 0.018 | 0.018 | 3.819 | 0.45 | 0.67 | 0.57 | 0.38 | 0.41 | 0.31 |
| 4 | 20987443 | rs17635673 | 0.336 | 0.836 | -2.045 | -0.274 | -0.145 | -0.008 | -0.008 | 0 | 0.23 | 0.02 | 0.27 | 0.18 | 0.38 | 0.3 |
| 4 | 20995935 | rs7675477 | 1.42 | 1.202 | -1.541 | -0.032 | 0.031 | 0.005 | 0.005 | 0.022 | 0.06 | 0.11 | 0.11 | 0.06 | 0.12 | 0.08 |
| 4 | 21000867 | rs12640604 | 0.097 | 0.848 | -1.251 | 0.007 | -0.001 | -0.001 | -0.001 | 0.056 | 0.43 | 0.57 | 0.55 | 0.38 | 0.4 | 0.28 |
| 4 | 21001815 | rs12504474 | 0.592 | 0.845 | -1.864 | 0.37 | -0.01 | -0.068 | -0.068 | 0 | 0.26 | 0.14 | 0.26 | 0.31 | 0.22 | 0.31 |
| 4 | 21002294 | rs6825511 | -0.374 | 0.737 | -1.393 | 0.399 | 0.001 | -0.004 | -0.004 | 0 | 0.69 | 0.68 | 0.74 | 0.66 | 0.64 | 0.6 |
| 4 | 21005251 | rs10034991 | -0.104 | 0.811 | -1.312 | 0 | 0.005 | -0.003 | -0.003 | 0 | 0.55 | 0.65 | 0.57 | 0.45 | 0.53 | 0.45 |
| 4 | 21007589 | rs10938823 | -0.853 | 0.821 | -1.619 | -0.315 | -0.039 | 0.066 | 0.066 | 0 | 0.54 | 0.79 | 0.49 | 0.4 | 0.52 | 0.58 |
| 4 | 21010644 | rs9631707 | -0.166 | 0.598 | -0.956 | -0.407 | -0.025 | -0.025 | -0.025 | 0 | 0.35 | 0.47 | 0.34 | 0.26 | 0.26 | 0.32 |
| 4 | 21011157 | rs12644707 | 1.666 | 0.58 | -1.162 | -0.335 | 0.001 | 0 | 0 | 0 | 0.06 | 0.03 | 0.09 | 0.06 | 0.06 | 0.08 |
| 4 | 21019900 | rs4348088 | -0.476 | 0.77 | -1.163 | -0.428 | 0.007 | 0.011 | 0.011 | 0 | 0.41 | 0.57 | 0.39 | 0.29 | 0.32 | 0.43 |
| 4 | 21021688 | rs13133807 | 1.499 | 0.731 | -1.575 | -0.398 | -0.049 | -0.03 | -0.03 | 2.6 | 0.12 | 0.01 | 0.12 | 0.07 | 0.12 | 0.15 |
| 4 | 21022508 | rs4389575 | -1.152 | 0.9 | -1.059 | -0.717 | 0.004 | 0.054 | 0.054 | 0 | 0.49 | 0.68 | 0.47 | 0.42 | 0.5 | 0.56 |
| 4 | 21026460 | rs6825103 | -0.082 | 0.287 | -0.768 | -0.673 | 0.019 | 0.029 | 0.029 | 0.006 | 0.19 | 0.1 | 0.13 | 0.06 | 0.14 | 0.13 |
| 4 | 21028953 | rs6819439 | -1.204 | 0.276 | -0.178 | -0.81 | -0.011 | -0.014 | -0.014 | 0 | 0.4 | 0.55 | 0.32 | 0.32 | 0.36 | 0.38 |
| 4 | 21033853 | rs6853609 | -1.366 | 0.219 | -0.148 | -0.838 | 0.063 | 0.077 | 0.077 | 0 | 0.42 | 0.77 | 0.34 | 0.32 | 0.36 | 0.38 |
| 4 | 21041975 | rs10938827 | 0.057 | 0.177 | 0.176 | -0.984 | -0.006 | 0 | 0 | 0 | 0.21 | 0.26 | 0.18 | 0.22 | 0.21 | 0.12 |
| 4 | 21042894 | rs6855710 | -0.73 | 0.098 | 0.228 | -1.059 | -0.022 | -0.023 | -0.023 | 0.58 | 0.31 | 0.4 | 0.21 | 0.23 | 0.28 | 0.17 |
| 4 | 21052369 | rs10033946 | 0.843 | 0.09 | 0.643 | -1.093 | 0.043 | -0.027 | -0.027 | 0.302 | 0.08 | 0.25 | 0.11 | 0 | 0.08 | 0.14 |
| 4 | 21057559 | rs6854888 | -1.658 | 0.036 | 0.673 | -1.099 | 0.04 | 0.066 | 0.066 | 0 | 0.4 | 0.71 | 0.34 | 0.25 | 0.36 | 0.31 |
| 4 | 21079749 | rs12510687 | -0.674 | 0.09 | 0.549 | -1.071 | 0.019 | 0.021 | 0.021 | 0.01 | 0.31 | 0.23 | 0.2 | 0.24 | 0.28 | 0.17 |
| 4 | 21085330 | rs6817687 | -0.665 | 0.243 | 0.356 | -1.144 | -0.005 | -0.004 | -0.004 | 0 | 0.26 | 0.28 | 0.17 | 0.24 | 0.23 | 0.15 |
| 4 | 21089763 | rs6448011 | -0.243 | 0.685 | 0.188 | -0.641 | -0.002 | 0.008 | 0.008 | 0.171 | 0.2 | 0.64 | 0.31 | 0.14 | 0.21 | 0.22 |
| 4 | 21096145 | rs720853 | -0.262 | 0.777 | 0.08 | -0.529 | 0.011 | 0.003 | 0.003 | 0 | 0.22 | 0.14 | 0.15 | 0.22 | 0.2 | 0.14 |
| 4 | 21104154 | rs1368645 | 0.457 | 0.833 | -0.076 | -0.463 | 0.008 | -0.047 | -0.047 | 0.009 | 0.11 | 0.01 | 0.05 | 0.18 | 0.11 | 0.06 |
| 4 | 21112046 | rs7696887 | -0.287 | 1.029 | 0.433 | -0.573 | 0.068 | 0.068 | 0.068 | 0.634 | 0.17 | 0.48 | 0.28 | 0.17 | 0.23 | 0.22 |
| 4 | 21113871 | rs6846038 | 0.666 | 1.204 | -0.194 | -1.127 | -0.069 | 0.099 | 0.099 | 0 | 0.27 | 0.12 | 0.28 | 0.11 | 0.33 | 0.27 |
| 4 | 21114825 | rs11944192 | 0.214 | 1.619 | 0.187 | -0.767 | 0.036 | 0.103 | 0.103 | 0.259 | 0.15 | 0.47 | 0.26 | 0.16 | 0.18 | 0.22 |
| 4 | 21115516 | rs16870400 | 1.453 | 1.621 | 0.307 | -0.777 | 0.006 | -0.045 | -0.045 | 0 | 0.08 | 0.23 | 0.09 | 0 | 0.12 | 0.04 |
| 4 | 21116890 | rs16870421 | 0.677 | 1.779 | 0.189 | -0.617 | 0.063 | 0.126 | 0.126 | 0 | 0.08 | 0.18 | 0.19 | 0.15 | 0.11 | 0.18 |
| 4 | 21118914 | rs5027019 | 1.082 | 2.092 | -0.323 | -0.758 | -0.054 | 0.033 | 0.033 | 0 | 0.15 | 0.08 | 0.13 | 0.08 | 0.24 | 0.2 |
| 4 | 21125935 | rs7687191 | -1.58 | 1.623 | -0.381 | -0.733 | 0.099 | -0.042 | -0.042 | 0 | 0.55 | 0.82 | 0.57 | 0.49 | 0.68 | 0.56 |
| 4 | 21138108 | rs6448023 | -1.615 | 1.429 | -0.308 | -0.613 | 0.043 | -0.039 | -0.039 | 0 | 0.55 | 0.78 | 0.56 | 0.49 | 0.63 | 0.56 |
| 4 | 21138383 | rs1433482 | -1.615 | 1.429 | -0.308 | -0.613 | 0.043 | -0.039 | -0.039 | 1.185 | 0.55 | 0.78 | 0.56 | 0.49 | 0.63 | 0.56 |
| 4 | 21140045 | rs10011670 | -0.842 | 1.448 | -0.056 | -0.65 | -0.015 | 0.011 | 0.011 | 1.203 | 0.41 | 0.66 | 0.44 | 0.3 | 0.5 | 0.46 |
| 4 | 21141832 | rs10007205 | 0.674 | 1.366 | 0.445 | -0.597 | 0.045 | 0.012 | 0.012 | 0 | 0.06 | 0.22 | 0.07 | 0.01 | 0.03 | 0.15 |
| 4 | 21151856 | rs17446667 | 1.22 | 1.379 | 0.272 | -0.591 | 0.003 | -0.019 | -0.019 | 0 | 0.11 | 0.04 | 0.1 | 0.19 | 0.11 | 0.09 |
| 4 | 21155716 | rs17506318 | 1.543 | 1.965 | 0.162 | -0.663 | -0.085 | 0.015 | 0.015 | 0 | 0.15 | 0.01 | 0.12 | 0.11 | 0.22 | 0.2 |
| 4 | 21155998 | rs4580648 | 0.704 | 1.575 | 0.442 | -0.777 | -0.003 | -0.016 | -0.016 | 0 | 0.17 | 0.26 | 0.13 | 0.03 | 0.18 | 0.08 |
| 4 | 21156962 | rs876478 | 0.084 | 1.638 | 0.385 | -0.739 | 0.076 | 0.036 | 0.036 | 0 | 0.25 | 0.43 | 0.31 | 0.27 | 0.3 | 0.38 |
| 4 | 21159965 | rs7667000 | 0.644 | 1.445 | 0.042 | -0.704 | 0.005 | 0.007 | 0.007 | 0.258 | 0.17 | 0.18 | 0.12 | 0.03 | 0.18 | 0.08 |
| 4 | 21160249 | rs1347240 | 0.18 | 1.462 | -0.053 | -0.668 | -0.003 | -0.046 | -0.046 | 1.981 | 0.2 | 0.29 | 0.14 | 0.03 | 0.23 | 0.09 |
| 4 | 21175619 | rs10516374 | 0.747 | 1.438 | -0.665 | -0.461 | -0.004 | -0.016 | -0.016 | 0.016 | 0.12 | 0.05 | 0.11 | 0.19 | 0.12 | 0.09 |
| 4 | 21179758 | rs2113966 | 0.758 | 1.461 | -0.652 | -0.458 | -0.005 | -0.015 | -0.015 | 0 | 0.12 | 0.06 | 0.12 | 0.19 | 0.12 | 0.09 |
| 4 | 21180995 | rs1319225 | -0.904 | 1.468 | -1.035 | -0.313 | 0.038 | -0.048 | -0.048 | 0 | 0.56 | 0.83 | 0.58 | 0.51 | 0.64 | 0.56 |
| 4 | 21183041 | rs7655488 | -1.545 | 0.573 | -1.125 | -0.301 | 0.085 | -0.113 | -0.113 | 0 | 0.69 | 0.93 | 0.75 | 0.52 | 0.78 | 0.63 |
| 4 | 21188231 | rs10013055 | -0.579 | 0.496 | -0.356 | -0.341 | -0.007 | -0.016 | -0.016 | 0 | 0.34 | 0.64 | 0.27 | 0.24 | 0.33 | 0.18 |
| 4 | 21191990 | rs1364841 | -0.242 | 0.803 | -0.028 | -0.061 | 0.052 | 0.012 | 0.012 | 0 | 0.83 | 0.7 | 0.86 | 0.77 | 0.8 | 0.76 |
| 4 | 21198867 | rs13117927 | 0.116 | 0.236 | -0.733 | -0.266 | -0.008 | 0.106 | 0.106 | 0 | 0.33 | 0.08 | 0.23 | 0.16 | 0.27 | 0.44 |
| 4 | 21200024 | rs6448033 | -0.825 | -0.071 | -0.405 | -0.213 | 0.045 | 0.076 | 0.076 | 0.919 | 0.44 | 0.76 | 0.33 | 0.32 | 0.34 | 0.51 |
| 4 | 21215907 | rs6448037 | -1.153 | 0.174 | -0.568 | 0.112 | 0.005 | -0.037 | -0.037 | 0 | 0.49 | 0.87 | 0.47 | 0.49 | 0.41 | 0.53 |
| 4 | 21220921 | rs7672506 | -0.968 | 0.161 | -0.239 | -0.023 | 0.014 | 0.047 | 0.047 | 0.072 | 0.47 | 0.7 | 0.44 | 0.34 | 0.41 | 0.53 |
| 4 | 21236883 | rs11733487 | -1.619 | 0.138 | -0.858 | -0.369 | -0.046 | -0.147 | -0.147 | 0 | 0.68 | 0.95 | 0.76 | 0.49 | 0.68 | 0.64 |
| 4 | 21244175 | rs6833463 | 0.543 | 0.064 | -0.52 | -0.245 | -0.005 | 0.098 | 0.098 | 0 | 0.32 | 0.09 | 0.25 | 0.16 | 0.27 | 0.44 |
| 4 | 21247995 | rs6823638 | -1.251 | 0.154 | -0.074 | -0.077 | -0.013 | 0.045 | 0.045 | 0.274 | 0.16 | 0.57 | 0.23 | 0.2 | 0.12 | 0.26 |
| 4 | 21254536 | rs1967398 | -1.543 | -0.08 | -0.318 | -0.104 | -0.021 | 0.026 | 0.026 | 0.693 | 0.19 | 0.73 | 0.28 | 0.32 | 0.11 | 0.29 |
| 4 | 21255650 | rs1425333 | -1.364 | -0.084 | -0.319 | -0.085 | -0.022 | 0 | 0 | 0 | 0.18 | 0.73 | 0.24 | 0.18 | 0.09 | 0.26 |
| 4 | 21256272 | rs4697213 | 0.462 | -0.303 | -0.629 | -0.11 | 0.003 | 0.036 | 0.036 | 0 | 0.38 | 0.19 | 0.27 | 0.29 | 0.34 | 0.33 |
| 4 | 21257521 | rs6827486 | 0.447 | -0.299 | -0.451 | -0.193 | 0.005 | 0.052 | 0.052 | 0 | 0.38 | 0.25 | 0.25 | 0.17 | 0.32 | 0.31 |
| 4 | 21260528 | rs10018523 | -0.484 | -0.257 | -0.049 | -0.128 | 0.002 | 0.074 | 0.074 | 0.001 | 0.14 | 0.47 | 0.21 | 0.16 | 0.09 | 0.22 |
| 4 | 21261499 | rs12648055 | 0.55 | -0.187 | -0.568 | -0.102 | -0.008 | -0.029 | -0.029 | 0.333 | 0.1 | 0.05 | 0.18 | 0.16 | 0.09 | 0.22 |
| 4 | 21263330 | rs11722994 | 0.543 | -0.237 | -0.6 | -0.178 | 0.002 | 0.065 | 0.065 | 0 | 0.37 | 0.2 | 0.23 | 0.16 | 0.32 | 0.3 |
| 4 | 21265036 | rs1863289 | -1.685 | -0.084 | -1.432 | 0.342 | -0.067 | -0.143 | -0.143 | 0 | 0.78 | 0.98 | 0.87 | 0.58 | 0.74 | 0.74 |
| 4 | 21267346 | rs16870654 | -0.414 | -0.673 | -0.5 | 0.418 | -0.018 | 0.081 | 0.081 | 0 | 0.15 | 0.36 | 0.23 | 0.31 | 0.11 | 0.25 |
| 4 | 21275159 | rs2322883 | 0.378 | -0.58 | -0.726 | 0.323 | 0.008 | 0.051 | 0.051 | 0 | 0.34 | 0.23 | 0.25 | 0.17 | 0.27 | 0.33 |
| 4 | 21276080 | rs901113 | 0.345 | -0.581 | -0.727 | 0.361 | 0.008 | 0.019 | 0.019 | 0.002 | 0.34 | 0.24 | 0.25 | 0.28 | 0.27 | 0.34 |
| 4 | 21281970 | rs6850985 | 0.027 | -0.324 | -0.943 | 0.558 | 0.007 | 0.068 | 0.068 | 0 | 0.46 | 0.18 | 0.43 | 0.33 | 0.38 | 0.52 |
| 4 | 21286777 | rs993611 | -0.237 | -0.264 | -1.226 | 0.451 | 0.002 | 0 | 0 | 0 | 0.53 | 0.6 | 0.48 | 0.47 | 0.4 | 0.57 |

| | | | | | | | | | | | | | | | | |
|---|---|---|---|---|---|---|---|---|---|---|---|---|---|---|---|---|
| 4 | 21292213 | rs4696973 | -0.565 | 0.051 | -2.495 | 0.56 | -0.113 | -0.094 | -0.094 | 0.041 | 0.61 | 0.92 | 0.6 | 0.51 | 0.47 | 0.61 |
| 4 | 21295315 | rs13111323 | -0.17 | -0.232 | -2.24 | -0.134 | 0.005 | 0.117 | 0.117 | 0 | 0.34 | 0.07 | 0.36 | 0.18 | 0.3 | 0.32 |
| 4 | 21296926 | rs6858452 | -0.25 | -0.395 | -2.107 | 0.139 | -0.034 | -0.029 | -0.029 | 0 | 0.6 | 0.72 | 0.62 | 0.55 | 0.53 | 0.57 |
| 4 | 21300220 | rs16870742 | -0.132 | -0.907 | -1.205 | 0.058 | -0.018 | 0.012 | 0.012 | 0 | 0.12 | 0.26 | 0.08 | 0.1 | 0.09 | 0.09 |
| 4 | 21300290 | rs1425339 | -1.339 | -1.066 | -1.448 | 0.084 | -0.019 | -0.014 | -0.014 | 0.65 | 0.7 | 0.72 | 0.72 | 0.59 | 0.59 | 0.67 |
| 4 | 21301546 | rs2007762 | 0.915 | -1.019 | -1.505 | 0.045 | 0.003 | -0.015 | -0.015 | 0 | 0.1 | 0.01 | 0.11 | 0.06 | 0.08 | 0.1 |
| 4 | 21303220 | rs16870748 | -0.17 | -1.17 | -1.647 | 0.071 | -0.019 | -0.013 | -0.013 | 2.904 | 0.56 | 0.7 | 0.48 | 0.5 | 0.49 | 0.41 |
| 4 | 21306003 | rs16870755 | 1.893 | -1.283 | -1.973 | 0.107 | -0.01 | -0.028 | -0.028 | 0 | 0.07 | 0.01 | 0.04 | 0.11 | 0.07 | 0.03 |
| 4 | 21306185 | rs13101486 | 1.056 | -1.245 | -1.917 | 0.163 | 0.02 | -0.016 | -0.016 | 0 | 0.1 | 0 | 0.1 | 0.06 | 0.07 | 0.1 |
| 4 | 21307467 | rs11726974 | 0.519 | -1.607 | -1.675 | 0.075 | -0.024 | 0.053 | 0.053 | 0 | 0.38 | 0.14 | 0.38 | 0.3 | 0.4 | 0.26 |
| 4 | 21310634 | rs11734360 | 0.709 | -1.11 | -1.095 | 0.128 | 0.001 | 0.001 | 0.001 | 0 | 0.11 | 0.11 | 0.12 | 0.06 | 0.08 | 0.1 |
| 4 | 21314419 | rs16870778 | -0.016 | -1.015 | -0.776 | 0.069 | -0.004 | -0.049 | -0.049 | 0 | 0.1 | 0.14 | 0.07 | 0 | 0.09 | 0.03 |
| 4 | 21324191 | rs17520130 | 0.037 | -0.606 | -1.002 | 0.259 | 0.036 | 0.021 | 0.021 | 0 | 0.5 | 0.63 | 0.47 | 0.42 | 0.42 | 0.45 |
| 4 | 21333411 | rs1364836 | -1.26 | 0.013 | -1.068 | 0.325 | -0.063 | -0.065 | -0.065 | 1.945 | 0.77 | 0.9 | 0.77 | 0.6 | 0.67 | 0.67 |
| 4 | 21334285 | rs16870831 | -0.112 | -0.365 | -0.38 | 0.115 | -0.003 | -0.007 | -0.007 | 1.02 | 0.37 | 0.37 | 0.33 | 0.28 | 0.33 | 0.28 |
| 4 | 21336070 | rs16870839 | -0.106 | -0.553 | -0.312 | 0.12 | -0.011 | -0.013 | -0.013 | 0 | 0.36 | 0.36 | 0.26 | 0.19 | 0.28 | 0.29 |
| 4 | 21344139 | rs16870878 | 0.202 | -0.265 | -0.622 | 0.596 | -0.003 | -0.007 | -0.007 | 0 | 0.11 | 0.07 | 0.09 | 0.14 | 0.13 | 0.07 |
| 4 | 21348980 | rs17520919 | -0.296 | -0.032 | -0.798 | 0.62 | -0.001 | -0.002 | -0.002 | 0 | 0.1 | 0.11 | 0.15 | 0.13 | 0.08 | 0.15 |
| 4 | 21351260 | rs7661530 | -1.337 | -0.133 | -1.437 | 0.16 | -0.087 | -0.104 | -0.104 | 0 | 0.74 | 0.9 | 0.77 | 0.59 | 0.64 | 0.54 |
| 4 | 21351470 | rs17521122 | 0.991 | -0.376 | -1.372 | -0.372 | -0.002 | 0.022 | 0.022 | 0 | 0.14 | 0.06 | 0.07 | 0.01 | 0.13 | 0.07 |
| 4 | 21361701 | rs12501359 | 0.697 | -0.53 | -1.063 | -0.101 | 0.026 | 0.153 | 0.153 | 0.048 | 0.32 | 0.13 | 0.32 | 0.02 | 0.29 | 0.13 |
| 4 | 21368339 | rs2323001 | 0.38 | -0.12 | -0.744 | 0.554 | 0.003 | -0.002 | -0.002 | 4.081 | 0.88 | 0.88 | 0.76 | 0.69 | 0.92 | 0.79 |
| 4 | 21382332 | rs12641447 | 0.304 | -0.347 | -0.302 | 0.266 | 0.008 | 0.011 | 0.011 | 0 | 0.57 | 0.41 | 0.44 | 0.66 | 0.62 | 0.63 |
| 4 | 21387660 | rs1390262 | 0.651 | -0.517 | 0.104 | 0.336 | -0.037 | 0.14 | 0.14 | 1.089 | 0.8 | 0.52 | 0.71 | 0.59 | 0.87 | 0.71 |
| 4 | 21389678 | rs1908552 | 0.173 | -0.536 | -0.137 | 0.387 | -0.002 | 0 | 0 | 0.085 | 0.85 | 0.82 | 0.84 | 0.86 | 0.88 | 0.83 |
| 4 | 21393616 | rs1495509 | 1.216 | -0.362 | -0.877 | 0.228 | -0.057 | -0.089 | -0.089 | 0 | 0.28 | 0.1 | 0.23 | 0.46 | 0.33 | 0.43 |
| 4 | 21393750 | rs13113329 | 1.043 | -0.314 | -0.474 | 0.153 | 0 | 0.08 | 0.08 | 0 | 0.25 | 0.1 | 0.19 | 0 | 0.26 | 0.09 |
| 4 | 21401617 | rs10012593 | -0.686 | -0.124 | -0.684 | 0.362 | -0.071 | -0.085 | -0.085 | 0.097 | 0.62 | 0.89 | 0.65 | 0.45 | 0.6 | 0.44 |
| 4 | 21407759 | rs6448050 | -0.125 | 0.073 | -0.771 | 0.558 | 0.007 | 0.002 | 0.002 | 0 | 0.35 | 0.66 | 0.37 | 0.32 | 0.31 | 0.25 |
| 4 | 21415503 | rs1874341 | -0.222 | 0.553 | -1.153 | 0.679 | 0.003 | 0.034 | 0.034 | 0 | 0.53 | 0.7 | 0.55 | 0.65 | 0.47 | 0.61 |
| 4 | 21424434 | rs2322963 | 0.191 | 0.696 | -1.429 | 0.219 | -0.041 | 0.029 | 0.029 | 0.374 | 0.24 | 0.14 | 0.23 | 0 | 0.36 | 0.17 |
| 4 | 21425057 | rs10022151 | -0.391 | -0.351 | -1.146 | 0.638 | -0.049 | 0.031 | 0.031 | 0 | 0.35 | 0.49 | 0.38 | 0.58 | 0.2 | 0.31 |
| 4 | 21438786 | rs11730849 | 1.007 | 0.227 | -1.5 | 0.367 | -0.004 | -0.03 | -0.03 | 0 | 0.13 | 0.05 | 0.1 | 0.2 | 0.1 | 0.1 |
| 4 | 21452250 | rs1476279 | -0.158 | 0.38 | -1.205 | -0.388 | -0.035 | 0.09 | 0.09 | 2.239 | 0.42 | 0.2 | 0.29 | 0.01 | 0.48 | 0.31 |
| 4 | 21453446 | rs168583 | -0.557 | 0.24 | -0.961 | -0.102 | 0 | -0.003 | -0.003 | 0.879 | 0.53 | 0.54 | 0.47 | 0.37 | 0.45 | 0.57 |
| 4 | 21455396 | rs358574 | -0.94 | 0.308 | -0.594 | 0.467 | 0.061 | -0.063 | -0.063 | 2.492 | 0.62 | 0.87 | 0.66 | 0.49 | 0.69 | 0.58 |
| 4 | 21455877 | rs1460485 | 0.334 | 0.218 | -0.089 | 0.201 | -0.008 | 0.148 | 0.148 | 0 | 0.45 | 0.22 | 0.35 | 0.17 | 0.54 | 0.34 |
| 4 | 21456023 | rs358573 | 1.148 | 0.259 | 0.666 | 0.373 | 0.02 | 0.006 | 0.006 | 0 | 0.1 | 0.22 | 0.26 | 0.16 | 0.13 | 0.14 |
| 4 | 21461121 | rs358569 | 0.814 | 0.259 | 0.682 | 0.375 | 0.052 | 0.024 | 0.024 | 0.068 | 0.12 | 0.36 | 0.27 | 0.16 | 0.13 | 0.14 |
| 4 | 21471568 | rs6855589 | -0.662 | 0.423 | 0.668 | 0.348 | 0.029 | -0.018 | -0.018 | 0 | 0.6 | 0.72 | 0.64 | 0.47 | 0.66 | 0.57 |
| 4 | 21472214 | rs17523841 | 1.153 | 0.568 | 0.58 | 0.219 | -0.006 | 0.012 | 0.012 | 0 | 0.16 | 0.07 | 0.12 | 0.11 | 0.15 | 0.18 |
| 4 | 21472843 | rs1460491 | 0.227 | 0.52 | 0.647 | 0.154 | 0.003 | 0.035 | 0.035 | 0 | 0.21 | 0.41 | 0.3 | 0.32 | 0.2 | 0.26 |
| 4 | 21475938 | rs17523974 | 1.606 | 0.674 | 0.57 | 0.437 | -0.009 | 0.008 | 0.008 | 0 | 0.22 | 0.16 | 0.21 | 0.04 | 0.31 | 0.13 |
| 4 | 21476990 | rs358592 | -1.541 | 0.67 | 0.573 | 0.438 | 0.036 | -0.008 | -0.008 | 0 | 0.61 | 0.83 | 0.67 | 0.61 | 0.66 | 0.63 |
| 4 | 21477084 | rs1460492 | -0.313 | 0.671 | 0.592 | 0.449 | 0 | 0.002 | 0.002 | 0 | 0.37 | 0.48 | 0.42 | 0.43 | 0.35 | 0.44 |
| 4 | 21477096 | rs358591 | 1.05 | 1.033 | 0.679 | -0.008 | 0.027 | 0 | 0 | 0 | 0.24 | 0.34 | 0.26 | 0.18 | 0.31 | 0.19 |
| 4 | 21489423 | rs4314271 | -0.086 | 0.64 | 0.38 | -0.093 | -0.012 | -0.045 | -0.045 | 0 | 0.27 | 0.51 | 0.37 | 0.19 | 0.22 | 0.3 |
| 4 | 21490361 | rs1565947 | -0.379 | 0.464 | 0.071 | -0.007 | 0.013 | 0.013 | 0.013 | 0 | 0.5 | 0.57 | 0.59 | 0.37 | 0.59 | 0.49 |
| 4 | 21492445 | rs6830109 | 1.71 | 0.6 | -0.354 | -0.2 | -0.021 | 0.014 | 0.014 | 0 | 0.2 | 0.11 | 0.18 | 0 | 0.28 | 0.1 |
| 4 | 21494694 | rs1013568 | 0.423 | 0.308 | 0.072 | 0.314 | -0.023 | 0.041 | 0.041 | 0 | 0.22 | 0.15 | 0.31 | 0.15 | 0.23 | 0.26 |
| 4 | 21506851 | rs358558 | -0.673 | 0.043 | 0.009 | 0.522 | -0.002 | -0.011 | -0.011 | 0.63 | 0.36 | 0.42 | 0.35 | 0.52 | 0.36 | 0.31 |
| 4 | 21511850 | rs17467572 | 1.681 | -0.087 | -0.163 | 0.393 | -0.037 | -0.037 | -0.037 | 0 | 0.17 | 0.02 | 0.14 | 0.15 | 0.18 | 0.12 |
| 4 | 21514249 | rs6810518 | -0.239 | 0.506 | 0.161 | 0.654 | 0.018 | 0.008 | 0.008 | 0 | 0.27 | 0.36 | 0.3 | 0.35 | 0.32 | 0.22 |
| 4 | 21520250 | rs2279674 | -0.095 | 0.625 | 0.156 | 0.708 | 0.032 | 0.012 | 0.012 | 0 | 0.23 | 0.32 | 0.28 | 0.32 | 0.29 | 0.17 |
| 4 | 21526226 | rs1460494 | 0.065 | 0.399 | 0.048 | 0.524 | 0.01 | -0.014 | -0.014 | 0 | 0.22 | 0.31 | 0.34 | 0.17 | 0.24 | 0.31 |
| 4 | 21539014 | rs11734403 | 0.836 | -0.402 | -0.746 | 0.336 | 0.024 | 0.022 | 0.022 | 0 | 0.25 | 0.04 | 0.15 | 0.15 | 0.22 | 0.17 |
| 4 | 21546217 | rs870368 | 0.508 | -0.005 | -0.623 | 0.462 | 0.033 | 0.098 | 0.098 | 0.016 | 0.35 | 0.05 | 0.2 | 0.15 | 0.29 | 0.23 |
| 4 | 21546537 | rs1817324 | -0.324 | 0.079 | -0.082 | 0.453 | 0.07 | 0.046 | 0.046 | 1.241 | 0.48 | 0.3 | 0.28 | 0.34 | 0.37 | 0.53 |
| 4 | 21549276 | rs13115047 | -0.497 | 0.047 | -0.132 | 0.414 | 0.026 | 0.003 | 0.003 | 0.697 | 0.54 | 0.35 | 0.5 | 0.52 | 0.4 | 0.66 |
| 4 | 21558828 | rs2199286 | -0.472 | 0.261 | 0.145 | 0.144 | 0.004 | 0.006 | 0.006 | 0 | 0.5 | 0.45 | 0.34 | 0.63 | 0.41 | 0.38 |
| 4 | 21562013 | rs13119355 | 1.035 | 0.099 | 0.072 | -0.132 | 0.006 | -0.018 | -0.018 | 0.153 | 0.19 | 0.09 | 0.07 | 0.01 | 0.11 | 0.15 |
| 4 | 21562391 | rs358840 | 0.277 | 0.176 | 0.159 | -0.452 | -0.002 | 0.008 | 0.008 | 0 | 0.25 | 0.3 | 0.44 | 0.33 | 0.25 | 0.42 |
| 4 | 21565222 | rs358839 | 0.747 | 0.13 | 0.191 | -0.503 | -0.001 | 0.033 | 0.033 | 0 | 0.2 | 0.27 | 0.41 | 0.42 | 0.22 | 0.37 |
| 4 | 21565822 | rs1460475 | 0.64 | 0.274 | 0.185 | -0.113 | 0.022 | -0.049 | -0.049 | 0 | 0.27 | 0.12 | 0.13 | 0.32 | 0.24 | 0.23 |
| 4 | 21566484 | rs6448057 | -0.523 | 0.425 | 0.106 | -0.246 | -0.001 | -0.001 | -0.001 | 0 | 0.55 | 0.46 | 0.35 | 0.54 | 0.44 | 0.44 |
| 4 | 21569545 | rs16871465 | -0.46 | 0.427 | 0.15 | -0.273 | 0.002 | -0.001 | -0.001 | 0 | 0.56 | 0.62 | 0.4 | 0.56 | 0.54 | 0.45 |
| 4 | 21571087 | rs1604805 | 0.923 | 0.181 | -0.223 | 0.033 | 0.006 | -0.013 | -0.013 | 0.023 | 0.1 | 0.01 | 0.06 | 0.05 | 0.08 | 0.09 |
| 4 | 21571755 | rs17483325 | 1.269 | 0.455 | 0.085 | 0.09 | 0.087 | -0.027 | -0.027 | 0.029 | 0.05 | 0.17 | 0.07 | 0.01 | 0.13 | 0.06 |
| 4 | 21572845 | rs402802 | -0.435 | 0.063 | 0.069 | 0.064 | 0 | 0.011 | 0.011 | 0 | 0.54 | 0.56 | 0.42 | 0.65 | 0.54 | 0.64 |

| | | | | | | | | | | | | | | | | |
|---|---|---|---|---|---|---|---|---|---|---|---|---|---|---|---|---|
| 4 | 21575623 | rs1966048 | 1.24 | -0.074 | 0.359 | -1.822 | -0.019 | 0.067 | 0.067 | 0 | 0.2 | 0.3 | 0.36 | 0.59 | 0.16 | 0.22 |
| 4 | 21581191 | rs7657068 | 0.99 | 0.051 | -0.164 | -1.931 | -0.016 | 0.021 | 0.021 | 0 | 0.17 | 0.07 | 0.13 | 0.11 | 0.2 | 0.24 |
| 4 | 21582320 | rs6843837 | 1.034 | -0.083 | -0.028 | -1.966 | -0.009 | -0.016 | -0.016 | 0 | 0.06 | 0.15 | 0.02 | 0.03 | 0.04 | 0.13 |
| 4 | 21590684 | rs11722935 | 0.644 | -0.115 | 0.064 | -1.998 | 0.011 | -0.004 | -0.004 | 0 | 0.29 | 0.26 | 0.42 | 0.64 | 0.25 | 0.31 |
| 4 | 21601372 | rs17557643 | 1.177 | 0.008 | -0.118 | -2.073 | -0.026 | 0.024 | 0.024 | 0 | 0.16 | 0.05 | 0.12 | 0.09 | 0.21 | 0.24 |
| 4 | 21602926 | rs1827593 | -2.3 | -0.077 | -0.265 | -2.24 | -0.001 | 0.024 | 0.024 | 0 | 0.82 | 0.89 | 0.86 | 1 | 0.81 | 0.9 |
| 4 | 21607597 | rs3913663 | -0.969 | -0.042 | -0.25 | -2.216 | -0.013 | 0.078 | 0.078 | 0 | 0.65 | 0.79 | 0.74 | 0.9 | 0.61 | 0.63 |
| 4 | 21614184 | rs7676653 | -0.148 | -0.089 | -0.141 | -2.222 | 0.003 | 0.001 | 0.001 | 0 | 0.48 | 0.46 | 0.54 | 0.74 | 0.46 | 0.6 |
| 4 | 21614663 | rs6826086 | 0.586 | -0.484 | -0.085 | -2.197 | 0 | 0 | 0 | 0 | 0.23 | 0.21 | 0.2 | 0.23 | 0.18 | 0.16 |
| 4 | 21614825 | rs12647718 | 1.703 | -0.487 | -0.098 | -2.083 | 0 | -0.087 | -0.087 | 0 | 0.18 | 0.06 | 0.33 | 0.52 | 0.18 | 0.21 |
| 4 | 21622448 | rs9291428 | 0.419 | 0.172 | 0.409 | -2.347 | 0.028 | -0.026 | -0.026 | 0 | 0.23 | 0.27 | 0.19 | 0.11 | 0.33 | 0.33 |
| 4 | 21622793 | rs2340 | -2.267 | 0.164 | 0.299 | -2.388 | -0.046 | 0.109 | 0.109 | 0 | 0.83 | 0.98 | 0.86 | 1 | 0.81 | 0.9 |
| 4 | 21624110 | rs12502842 | 0.221 | 0.157 | 0.412 | -2.34 | 0.001 | 0.002 | 0.002 | 0 | 0.24 | 0.21 | 0.21 | 0.22 | 0.19 | 0.16 |
| 4 | 21627809 | rs2323070 | -1.151 | 0.168 | 0.439 | -2.343 | 0.03 | 0.074 | 0.074 | 0 | 0.49 | 0.66 | 0.42 | 0.33 | 0.52 | 0.5 |
| 4 | 21628375 | rs10516395 | 0.135 | 0.115 | 0.392 | -2.286 | -0.006 | -0.01 | -0.01 | 0 | 0.25 | 0.29 | 0.23 | 0.23 | 0.19 | 0.16 |
| 4 | 21629865 | rs1039549 | -0.218 | 0.105 | 0.157 | -2.296 | -0.016 | -0.027 | -0.027 | 0 | 0.28 | 0.4 | 0.25 | 0.22 | 0.2 | 0.16 |
| 4 | 21632162 | rs6448063 | 0.213 | 0.745 | -0.173 | -2.616 | -0.001 | -0.025 | -0.025 | 1.11 | 0.55 | 0.53 | 0.64 | 0.87 | 0.45 | 0.47 |
| 4 | 21633160 | rs6832022 | 0.665 | 1.472 | -0.261 | -1.982 | -0.035 | -0.01 | -0.01 | 0 | 0.06 | 0.01 | 0.04 | 0.05 | 0.09 | 0.06 |
| 4 | 21634922 | rs1503979 | -0.564 | 1.14 | -0.162 | -1.952 | 0.01 | -0.007 | -0.007 | 0 | 0.29 | 0.29 | 0.23 | 0.16 | 0.43 | 0.4 |
| 4 | 21647862 | rs10014328 | 0.633 | 0.197 | 0.268 | -2.038 | -0.05 | -0.127 | -0.127 | 8.149 | 0.27 | 0.38 | 0.53 | 0.91 | 0.21 | 0.34 |
| 4 | 21651620 | rs1503990 | -0.327 | 0.428 | 0.468 | -2.477 | -0.001 | -0.08 | -0.08 | 0.139 | 0.46 | 0.5 | 0.27 | 0.08 | 0.44 | 0.45 |
| 4 | 21651850 | rs6817879 | -1.394 | 0.481 | 0.43 | -2.388 | -0.05 | 0.063 | 0.063 | 0 | 0.77 | 0.88 | 0.81 | 0.9 | 0.68 | 0.8 |
| 4 | 21655717 | rs1026523 | 0.906 | 0.445 | 0.62 | -2.295 | -0.02 | -0.058 | -0.058 | 0.077 | 0.27 | 0.33 | 0.52 | 0.9 | 0.21 | 0.37 |
| 4 | 21655863 | rs1026522 | 0.629 | 0.455 | 0.685 | -2.293 | -0.036 | -0.117 | -0.117 | 0.022 | 0.3 | 0.43 | 0.55 | 0.9 | 0.24 | 0.38 |
| 4 | 21658285 | rs7666642 | 0.446 | 0.723 | 0.442 | -2.162 | -0.038 | -0.063 | -0.063 | 0.016 | 0.35 | 0.46 | 0.55 | 0.81 | 0.24 | 0.36 |
| 4 | 21658543 | rs1824371 | 0.727 | 0.712 | 0.439 | -2.092 | -0.052 | -0.029 | -0.029 | 0 | 0.32 | 0.46 | 0.53 | 0.71 | 0.21 | 0.34 |
| 4 | 21661079 | rs6843682 | 0.749 | 0.84 | 0.137 | -2.226 | -0.002 | -0.002 | -0.002 | 0 | 0.09 | 0.13 | 0.08 | 0.03 | 0.13 | 0.09 |
| 4 | 21663995 | rs4697227 | -0.243 | 0.781 | 0.012 | -2.082 | -0.007 | 0.049 | 0.049 | 0 | 0.57 | 0.63 | 0.69 | 0.96 | 0.51 | 0.67 |
| 4 | 21671519 | rs7671181 | 0.842 | 0.863 | -0.19 | -2.008 | -0.004 | -0.004 | -0.004 | 0 | 0.32 | 0.29 | 0.53 | 0.74 | 0.24 | 0.37 |
| 4 | 21671589 | rs13145193 | 0.472 | 0.853 | -0.158 | -2.092 | 0.003 | -0.056 | -0.056 | 0 | 0.23 | 0.27 | 0.15 | 0.07 | 0.27 | 0.29 |
| 4 | 21673431 | rs4697230 | 0.163 | 0.239 | -0.397 | -1.249 | -0.004 | -0.017 | -0.017 | 0 | 0.32 | 0.28 | 0.53 | 0.83 | 0.24 | 0.37 |
| 4 | 21689430 | rs1354390 | -0.853 | 0.457 | 0.124 | -0.188 | -0.015 | -0.044 | -0.044 | 0 | 0.24 | 0.5 | 0.15 | 0.19 | 0.17 | 0.17 |
| 4 | 21701301 | rs2167246 | -1.121 | 0.224 | 0.718 | -0.236 | 0.034 | 0.019 | 0.019 | 0.211 | 0.06 | 0.34 | 0.05 | 0 | 0.01 | 0.01 |
| 4 | 21707600 | rs1459282 | -0.309 | 0.341 | -0.609 | -0.306 | -0.043 | -0.016 | -0.016 | 0.068 | 0.65 | 0.9 | 0.55 | 0.31 | 0.61 | 0.41 |
| 4 | 21708295 | rs1459283 | -0.309 | 0.341 | -0.609 | -0.306 | -0.043 | -0.016 | -0.016 | 0 | 0.65 | 0.9 | 0.55 | 0.31 | 0.61 | 0.41 |
| 4 | 21712050 | rs1503976 | 0.479 | -0.068 | -1.387 | -0.959 | -0.128 | 0.008 | 0.008 | 0 | 0.26 | 0.03 | 0.21 | 0.17 | 0.4 | 0.19 |
| 4 | 21715034 | rs6850480 | -0.712 | 0.043 | -1.301 | -0.856 | -0.035 | -0.015 | -0.015 | 0.043 | 0.63 | 0.91 | 0.55 | 0.32 | 0.6 | 0.4 |
| 4 | 21719296 | rs9884923 | -0.046 | -0.027 | -1.771 | -1.404 | 0.004 | 0.143 | 0.143 | 0.235 | 0.53 | 0.77 | 0.65 | 0.75 | 0.37 | 0.67 |
| 4 | 21726861 | rs11721958 | 0.702 | 0.002 | -2.034 | -1.693 | -0.01 | 0.042 | 0.042 | 0.742 | 0.14 | 0.08 | 0.12 | 0 | 0.13 | 0.06 |
| 4 | 21734047 | rs12509958 | 0.901 | -0.577 | -1.779 | -1.82 | 0.009 | 0.017 | 0.017 | 0.655 | 0.38 | 0.24 | 0.54 | 0.66 | 0.28 | 0.65 |
| 4 | 21735319 | rs7697475 | 0.606 | -0.671 | -1.679 | -1.864 | 0.032 | 0.044 | 0.044 | 0.344 | 0.42 | 0.7 | 0.59 | 0.66 | 0.29 | 0.65 |
| 4 | 21738299 | rs1450730 | -1.46 | -0.782 | -1.749 | -1.911 | -0.048 | -0.039 | -0.039 | 0 | 0.65 | 0.95 | 0.48 | 0.35 | 0.73 | 0.38 |
| 4 | 21751771 | rs6838195 | 1.009 | -0.764 | -1.629 | -1.929 | -0.001 | -0.015 | -0.015 | 1.571 | 0.14 | 0.12 | 0.12 | 0 | 0.13 | 0.06 |
| 4 | 21755399 | rs1380270 | -0.472 | -0.781 | -1.615 | -1.886 | -0.028 | 0.18 | 0.18 | 0.403 | 0.52 | 0.88 | 0.39 | 0.25 | 0.62 | 0.29 |
| 4 | 21764761 | rs717531 | -1.4 | -0.784 | -1.458 | -1.857 | -0.028 | -0.017 | -0.017 | 0 | 0.65 | 0.93 | 0.48 | 0.35 | 0.73 | 0.39 |
| 4 | 21769360 | rs16871885 | -0.338 | -0.759 | -1.274 | -1.839 | 0.007 | 0 | 0 | 0 | 0.06 | 0.6 | 0.06 | 0 | 0 | 0 |
| 4 | 21770028 | rs12646630 | -0.09 | -0.76 | -1.325 | -1.854 | 0.001 | 0.102 | 0.102 | 0.06 | 0.46 | 0.27 | 0.33 | 0.25 | 0.61 | 0.29 |
| 4 | 21793217 | rs10021705 | -1.124 | -0.615 | -0.352 | -1.504 | 0.024 | 0.071 | 0.071 | 2.477 | 0.08 | 0.53 | 0.08 | 0.09 | 0.04 | 0.07 |
| 4 | 21794660 | rs7696044 | 1.264 | -0.734 | -0.46 | -1.578 | 0.001 | 0 | 0 | 0.896 | 0.77 | 0.27 | 0.85 | 0.81 | 0.87 | 0.87 |
| 4 | 21796070 | rs4631029 | 0.239 | -0.725 | -0.436 | -1.574 | -0.007 | -0.005 | -0.005 | 6.754 | 0.86 | 0.8 | 0.93 | 0.9 | 0.9 | 0.94 |
| 4 | 21798563 | rs10516402 | 0.524 | -0.771 | -0.704 | -1.575 | -0.063 | 0.072 | 0.072 | 0.473 | 0.29 | 0.13 | 0.21 | 0.18 | 0.45 | 0.25 |
| 4 | 21820620 | rs10034770 | 0.85 | -0.811 | -0.73 | -1.617 | -0.007 | 0.087 | 0.087 | 0.02 | 0.49 | 0.21 | 0.63 | 0.63 | 0.41 | 0.59 |
| 4 | 21825726 | rs4377576 | -0.254 | -0.822 | -0.634 | -1.625 | 0.003 | -0.013 | -0.013 | 0 | 0.37 | 0.54 | 0.28 | 0.27 | 0.5 | 0.35 |
| 4 | 21833931 | rs2137650 | 0.11 | -0.794 | -0.625 | -1.605 | -0.012 | -0.01 | -0.01 | 0 | 0.86 | 0.75 | 0.92 | 0.9 | 0.91 | 0.94 |
| 4 | 21842795 | rs957196 | 0.608 | -0.842 | -0.671 | -1.642 | -0.027 | 0.141 | 0.141 | 0 | 0.52 | 0.25 | 0.66 | 0.71 | 0.44 | 0.65 |
| 4 | 21848082 | rs966045 | 0.2 | -0.832 | -0.753 | -1.655 | -0.065 | 0.045 | 0.045 | 0.284 | 0.33 | 0.13 | 0.23 | 0.26 | 0.49 | 0.32 |
| 4 | 21849857 | rs1398835 | -0.084 | -0.807 | -0.785 | -1.639 | -0.009 | -0.006 | -0.006 | 0 | 0.87 | 0.81 | 0.92 | 0.9 | 0.9 | 0.94 |
| 4 | 21850305 | rs1513572 | 0.978 | -0.81 | -0.833 | -1.64 | -0.083 | 0.054 | 0.054 | 0.119 | 0.47 | 0.09 | 0.63 | 0.63 | 0.41 | 0.59 |
| 4 | 21852757 | rs13136049 | -0.477 | -0.772 | -0.715 | -1.576 | -0.013 | -0.012 | -0.012 | 0.256 | 0.15 | 0.23 | 0.08 | 0.1 | 0.1 | 0.06 |
| 4 | 21866646 | rs4697244 | 0.962 | -0.757 | -1.007 | -1.633 | -0.101 | 0.037 | 0.037 | 0.28 | 0.47 | 0.07 | 0.62 | 0.62 | 0.41 | 0.58 |
| 4 | 21870314 | rs6839725 | -0.025 | -0.709 | -0.823 | -1.628 | -0.003 | -0.002 | -0.002 | 0.089 | 0.13 | 0.12 | 0.08 | 0.11 | 0.09 | 0.07 |
| 4 | 21871986 | rs12504827 | 0.358 | -0.729 | -0.641 | -1.631 | 0.013 | -0.031 | -0.031 | 0 | 0.31 | 0.33 | 0.23 | 0.23 | 0.45 | 0.25 |
| 4 | 21877250 | rs10009391 | 0.813 | -0.686 | -1.194 | -1.492 | -0.1 | 0.052 | 0.052 | 0.605 | 0.48 | 0.07 | 0.63 | 0.62 | 0.42 | 0.58 |
| 4 | 21883570 | rs10516407 | -0.385 | 0.099 | -0.852 | -1.317 | -0.027 | -0.007 | -0.007 | 0 | 0.14 | 0.03 | 0.12 | 0.13 | 0.18 | 0.08 |
| 4 | 21887265 | rs4697247 | 0.366 | -0.132 | -1.044 | -1.471 | -0.009 | 0.027 | 0.027 | 0.042 | 0.69 | 0.21 | 0.79 | 0.71 | 0.68 | 0.69 |
| 4 | 21895517 | rs10000092 | -0.212 | -0.033 | -0.73 | -1.405 | 0.022 | -0.007 | -0.007 | 0 | 0.26 | 0.41 | 0.17 | 0.28 | 0.31 | 0.26 |
| 4 | 21910553 | rs4277766 | 0.332 | 0.06 | -0.688 | -1.257 | -0.001 | -0.009 | -0.009 | 0.076 | 0.83 | 0.79 | 0.87 | 0.91 | 0.82 | 0.92 |
| 4 | 21911866 | rs2874977 | -0.831 | 0.499 | -0.715 | -0.784 | 0.025 | 0.017 | 0.017 | 0.688 | 0.6 | 0.66 | 0.54 | 0.33 | 0.74 | 0.44 |
| 4 | 21913523 | rs9995524 | 1.218 | 0.592 | -1.155 | -0.612 | 0.006 | -0.088 | -0.088 | 0.054 | 0.33 | 0.1 | 0.41 | 0.58 | 0.23 | 0.46 |

| | | | | | | | | | | | | | | | | |
|---|---|---|---|---|---|---|---|---|---|---|---|---|---|---|---|---|
| 4 | 21916439 | rs1398837 | 1.218 | 0.592 | -1.155 | -0.59 | 0.006 | -0.075 | -0.075 | 0 | 0.33 | 0.1 | 0.41 | 0.59 | 0.23 | 0.46 |
| 4 | 21920543 | rs727633 | 1.186 | 0.588 | -1.059 | -0.605 | 0.021 | -0.065 | -0.065 | 0 | 0.34 | 0.12 | 0.41 | 0.58 | 0.23 | 0.46 |
| 4 | 21921234 | rs2203274 | -1.181 | 0.422 | -0.538 | -0.708 | 0.009 | 0.034 | 0.034 | 3.987 | 0.1 | 0.48 | 0.08 | 0.09 | 0.04 | 0.12 |
| 4 | 21922674 | rs10938866 | 1.253 | 0.44 | -0.514 | -0.669 | 0.022 | -0.062 | -0.062 | 0 | 0.34 | 0.12 | 0.41 | 0.58 | 0.23 | 0.46 |
| 4 | 21929301 | rs1913330 | -0.666 | 1.034 | -0.985 | -1.117 | 0 | -0.022 | -0.022 | 0.171 | 0.67 | 0.63 | 0.67 | 0.9 | 0.6 | 0.87 |
| 4 | 21931195 | rs4697248 | -0.8 | 1.143 | -0.946 | -1.026 | 0.008 | -0.016 | -0.016 | 0 | 0.34 | 0.5 | 0.25 | 0.31 | 0.35 | 0.39 |
| 4 | 21938803 | rs4697249 | 0.51 | 1.211 | -1.328 | -1.067 | -0.035 | 0.1 | 0.1 | 0 | 0.31 | 0.14 | 0.31 | 0.1 | 0.4 | 0.13 |
| 4 | 21940697 | rs4697250 | 0.1 | 1.405 | -0.925 | -0.604 | -0.125 | -0.016 | -0.016 | 0 | 0.25 | 0.03 | 0.19 | 0.26 | 0.34 | 0.36 |
| 4 | 21962153 | rs4696996 | 1.019 | 1.281 | -0.001 | -0.772 | -0.001 | -0.004 | -0.004 | 0 | 0.58 | 0.43 | 0.63 | 0.72 | 0.53 | 0.58 |
| 4 | 21965464 | rs9291431 | 0.003 | 1.848 | 0.264 | -0.766 | 0.126 | 0.027 | 0.027 | 0.067 | 0.07 | 0.27 | 0.09 | 0.02 | 0.1 | 0.05 |
| 4 | 21971493 | rs11940932 | -1.402 | 1.699 | 0.272 | -0.886 | -0.001 | -0.002 | -0.002 | 0 | 0.39 | 0.38 | 0.36 | 0.29 | 0.46 | 0.41 |
| 4 | 21972367 | rs4697253 | -0.696 | 1.708 | -0.215 | -0.886 | -0.036 | 0.063 | 0.063 | 0 | 0.33 | 0.11 | 0.28 | 0.26 | 0.36 | 0.36 |
| 4 | 21973217 | rs2007195 | 1.672 | 1.625 | -0.266 | -0.845 | -0.082 | 0.086 | 0.086 | 0 | 0.54 | 0.12 | 0.59 | 0.66 | 0.53 | 0.53 |
| 4 | 21978266 | rs1533278 | 1.227 | 1.537 | 0.091 | -0.471 | 0.026 | 0.002 | 0.002 | 0.071 | 0.67 | 0.14 | 0.73 | 0.67 | 0.72 | 0.63 |
| 4 | 21979317 | rs4696998 | 1.556 | 1.369 | 0.096 | -0.444 | -0.036 | 0.021 | 0.021 | 0 | 0.63 | 0.13 | 0.66 | 0.67 | 0.65 | 0.57 |
| 4 | 21981053 | rs6827204 | -0.032 | 0.607 | 0.352 | -0.345 | -0.01 | 0.014 | 0.014 | 0 | 0.77 | 0.87 | 0.81 | 0.76 | 0.74 | 0.68 |
| 4 | 21986877 | rs6448104 | 1.359 | 0.583 | 0.278 | -0.045 | 0.032 | -0.037 | -0.037 | 0 | 0.66 | 0.13 | 0.69 | 0.61 | 0.72 | 0.61 |
| 4 | 22007483 | rs7691155 | 1.19 | 0.749 | 0.555 | -0.076 | 0.022 | 0.016 | 0.016 | 0.146 | 0.71 | 0.2 | 0.72 | 0.74 | 0.78 | 0.64 |
| 4 | 22013294 | rs4697259 | -0.663 | 0.835 | 1.298 | -0.027 | -0.002 | -0.047 | -0.047 | 0 | 0.24 | 0.42 | 0.19 | 0.19 | 0.21 | 0.32 |
| 4 | 22028574 | rs6838353 | 0.032 | 0.828 | 1.276 | -0.031 | -0.014 | 0.046 | 0.046 | 0 | 0.19 | 0.09 | 0.17 | 0.15 | 0.2 | 0.29 |
| 4 | 22030668 | rs2175778 | 0.998 | 0.81 | 1.413 | 0.012 | 0.001 | 0 | 0 | 0 | 0.73 | 0.31 | 0.73 | 0.74 | 0.79 | 0.65 |
| 4 | 22034546 | rs11933737 | 0.946 | 0.81 | 1.46 | 0.014 | -0.005 | -0.011 | -0.011 | 0.305 | 0.74 | 0.38 | 0.78 | 0.79 | 0.79 | 0.67 |
| 4 | 22044372 | rs13106072 | 1.102 | 0.805 | 1.462 | 0.011 | 0.01 | 0.005 | 0.005 | 0 | 0.73 | 0.25 | 0.73 | 0.74 | 0.79 | 0.65 |
| 4 | 22048637 | rs7695992 | -0.164 | 0.831 | 1.473 | 0.016 | -0.001 | -0.011 | -0.011 | 0.176 | 0.8 | 0.8 | 0.82 | 0.83 | 0.8 | 0.7 |
| 4 | 22052817 | rs12510345 | 0.912 | 0.848 | 1.209 | 0.054 | -0.006 | -0.002 | -0.002 | 0 | 0.74 | 0.36 | 0.74 | 0.73 | 0.79 | 0.65 |
| 4 | 22057269 | rs10433802 | -0.035 | 0.83 | 1.18 | 0.001 | -0.004 | 0.012 | 0.012 | 0 | 0.19 | 0.07 | 0.16 | 0.2 | 0.19 | 0.24 |
| 4 | 22060315 | rs10031901 | -2.027 | 0.893 | 1.24 | -0.027 | 0.032 | 0.074 | 0.074 | 6.67 | 0.07 | 0.56 | 0.05 | 0.06 | 0.03 | 0.04 |
| 4 | 22061205 | rs17526294 | -0.015 | 0.885 | 1.22 | -0.033 | -0.004 | 0.012 | 0.012 | 0 | 0.19 | 0.07 | 0.16 | 0.2 | 0.19 | 0.24 |
| 4 | 22062457 | rs6816789 | -2.025 | 0.884 | 1.24 | -0.031 | 0.025 | 0.065 | 0.065 | 6.67 | 0.07 | 0.58 | 0.05 | 0.06 | 0.03 | 0.04 |
| 4 | 22064524 | rs6448111 | -1.154 | 0.89 | 1.228 | -0.033 | -0.009 | -0.006 | -0.006 | 0 | 0.27 | 0.63 | 0.21 | 0.26 | 0.22 | 0.29 |
| 4 | 22068673 | rs4697263 | -0.052 | 0.893 | 1.154 | -0.033 | -0.004 | 0.013 | 0.013 | 0 | 0.19 | 0.07 | 0.16 | 0.2 | 0.19 | 0.24 |
| 4 | 22069379 | rs6448112 | -1.101 | 0.905 | 1.161 | -0.025 | -0.005 | -0.004 | -0.004 | 0 | 0.27 | 0.64 | 0.21 | 0.26 | 0.22 | 0.29 |
| 4 | 22073768 | rs7656406 | -1.311 | 0.625 | 0.966 | 0.067 | 0.022 | 0.007 | 0.007 | 0 | 0.31 | 0.75 | 0.36 | 0.32 | 0.24 | 0.42 |
| 4 | 22079213 | rs6448117 | -1.414 | 0.696 | 1.647 | 0.088 | 0.06 | 0.057 | 0.057 | 0.014 | 0.05 | 0.18 | 0.04 | 0.05 | 0.04 | 0.14 |
| 4 | 22079733 | rs11931022 | -1.414 | 0.684 | 1.647 | 0.088 | 0.041 | 0.057 | 0.057 | 0.014 | 0.05 | 0.18 | 0.04 | 0.05 | 0.04 | 0.14 |
| 4 | 22081816 | rs6831346 | -1.233 | 0.688 | 1.852 | 0.106 | 0.019 | 0.006 | 0.006 | 0 | 0.31 | 0.75 | 0.36 | 0.32 | 0.24 | 0.41 |
| 4 | 22087470 | rs13123835 | -1.093 | 0.698 | 1.871 | 0.109 | -0.017 | -0.015 | -0.015 | 0 | 0.3 | 0.56 | 0.29 | 0.27 | 0.24 | 0.38 |
| 4 | 22090575 | rs6851636 | -1.29 | 0.721 | 1.593 | 0.104 | 0.014 | 0.007 | 0.007 | 0 | 0.31 | 0.75 | 0.37 | 0.32 | 0.25 | 0.41 |
| 4 | 22090754 | rs6834054 | -1.274 | 0.725 | 1.577 | 0.197 | 0.018 | -0.008 | -0.008 | 0 | 0.31 | 0.76 | 0.37 | 0.34 | 0.25 | 0.41 |
| 4 | 22091384 | rs6448119 | -0.683 | 0.795 | 1.504 | 0.199 | 0.002 | 0.002 | 0.002 | 0 | 0.27 | 0.24 | 0.27 | 0.27 | 0.24 | 0.37 |
| 4 | 22098163 | rs6820902 | -1.232 | 0.716 | 1.709 | 0.237 | 0.069 | 0.071 | 0.071 | 0.421 | 0.05 | 0.2 | 0.05 | 0.04 | 0.05 | 0.07 |
| 4 | 22106866 | rs4590011 | -0.965 | 0.487 | 1.65 | 0.271 | 0.008 | 0 | 0 | 0 | 0.31 | 0.64 | 0.34 | 0.34 | 0.25 | 0.41 |
| 4 | 22107318 | rs10019171 | -0.319 | 0.446 | 1.36 | 0.195 | -0.003 | -0.001 | -0.001 | 0 | 0.25 | 0.28 | 0.24 | 0.26 | 0.2 | 0.32 |
| 4 | 22109508 | rs6837880 | -0.364 | 0.488 | 1.357 | 0.193 | -0.001 | -0.002 | -0.002 | 0 | 0.26 | 0.28 | 0.24 | 0.24 | 0.21 | 0.32 |
| 4 | 22111275 | rs4697267 | -0.364 | 0.488 | 1.357 | 0.193 | -0.001 | -0.001 | -0.001 | 0 | 0.26 | 0.28 | 0.24 | 0.26 | 0.21 | 0.32 |
| 4 | 22114968 | rs16872408 | -0.395 | 0.429 | 1.358 | 0.194 | -0.021 | -0.017 | -0.017 | 0.228 | 0.26 | 0.37 | 0.24 | 0.24 | 0.2 | 0.32 |
| 4 | 22124060 | rs17528275 | -0.215 | 0.449 | 0.758 | 0.197 | 0.025 | -0.003 | -0.003 | 0 | 0.24 | 0.08 | 0.23 | 0.25 | 0.2 | 0.31 |
| 4 | 22135324 | rs16872414 | 0.04 | 0.51 | 0.768 | 0 | -0.007 | -0.026 | -0.026 | 0 | 0.11 | 0.2 | 0.07 | 0.04 | 0.07 | 0.13 |
| 4 | 22161298 | rs10033940 | -1.384 | 0.281 | 1.409 | 0.292 | -0.009 | -0.004 | -0.004 | 2.038 | 0.35 | 0.44 | 0.35 | 0.36 | 0.3 | 0.5 |
| 4 | 22163667 | rs11728484 | -0.202 | 0.319 | 1.182 | 0.335 | 0.001 | -0.003 | -0.003 | 0 | 0.24 | 0.21 | 0.27 | 0.32 | 0.22 | 0.37 |
| 4 | 22164317 | rs11933151 | -0.207 | 0.317 | 1.155 | 0.57 | -0.002 | 0.005 | 0.005 | 0 | 0.24 | 0.23 | 0.27 | 0.37 | 0.22 | 0.37 |
| 4 | 22165174 | rs9291436 | -0.635 | -0.029 | 0.352 | 0.311 | 0.037 | 0.018 | 0.018 | 0 | 0.48 | 0.72 | 0.45 | 0.45 | 0.37 | 0.54 |
| 4 | 22171998 | rs7688470 | -1.261 | 0.262 | 0.337 | 0.218 | -0.013 | -0.014 | -0.014 | 0 | 0.24 | 0.62 | 0.26 | 0.18 | 0.18 | 0.24 |
| 4 | 22180997 | rs10938878 | -1.174 | 0.414 | -0.47 | 1.106 | -0.017 | -0.128 | -0.128 | 0 | 0.77 | 0.98 | 0.84 | 0.65 | 0.76 | 0.92 |
| 4 | 22189941 | rs17588445 | 0.993 | 1.07 | -0.508 | 1.072 | 0.017 | 0.009 | 0.009 | 0.008 | 0.11 | 0.12 | 0.23 | 0.15 | 0.16 | 0.3 |
| 4 | 22192762 | rs6448132 | 0.227 | 0.793 | -0.654 | 0.898 | -0.001 | -0.008 | -0.008 | 0 | 0.11 | 0.16 | 0.1 | 0.07 | 0.08 | 0.13 |
| 4 | 22202451 | rs4697270 | 1.274 | 0.758 | -0.791 | 0.752 | -0.052 | 0.078 | 0.078 | 0.275 | 0.42 | 0.05 | 0.34 | 0.28 | 0.39 | 0.4 |
| 4 | 22205496 | rs10022457 | -0.096 | 0.749 | 0.112 | 1.058 | -0.01 | -0.007 | -0.007 | 0 | 0.11 | 0.21 | 0.06 | 0.08 | 0.05 | 0.11 |
| 4 | 22220505 | rs9997871 | -0.894 | 0.819 | 0.24 | 1.305 | 0.002 | 0.066 | 0.066 | 0.207 | 0.14 | 0.53 | 0.11 | 0.18 | 0.08 | 0.15 |
| 4 | 22220925 | rs4235294 | -0.527 | 0.88 | -0.599 | 1.312 | -0.011 | -0.024 | -0.024 | 0.054 | 0.35 | 0.33 | 0.43 | 0.46 | 0.43 | 0.39 |
| 4 | 22222487 | rs11730240 | 0.376 | -0.013 | -0.559 | 1.625 | 0.028 | -0.008 | -0.008 | 0.071 | 0.25 | 0.24 | 0.1 | 0.31 | 0.15 | 0.09 |
| 4 | 22222814 | rs11727303 | 0.33 | -0.016 | -0.451 | 1.624 | 0.008 | -0.003 | -0.003 | 0.109 | 0.25 | 0.3 | 0.11 | 0.31 | 0.15 | 0.09 |
| 4 | 22235324 | rs10938882 | -0.028 | -0.028 | -0.251 | 1.506 | -0.043 | 0.036 | 0.036 | 0 | 0.28 | 0.44 | 0.21 | 0.4 | 0.16 | 0.2 |
| 4 | 22235796 | rs4697273 | 0.576 | 0.501 | -0.328 | 1.355 | -0.044 | -0.027 | -0.027 | 0 | 0.16 | 0.06 | 0.26 | 0.2 | 0.22 | 0.42 |
| 4 | 22237340 | rs4697275 | -1.486 | 0.388 | -0.391 | 0.908 | 0.003 | -0.017 | -0.017 | 0 | 0.56 | 0.54 | 0.53 | 0.67 | 0.47 | 0.73 |
| 4 | 22240923 | rs4697003 | -0.11 | 0.734 | -0.916 | 0.904 | -0.013 | 0.013 | 0.013 | 1.192 | 0.22 | 0.19 | 0.3 | 0.11 | 0.25 | 0.46 |
| 4 | 22246755 | rs11944669 | -1.442 | -0.346 | -1.038 | 1.137 | 0.045 | -0.005 | -0.005 | 6.41 | 0.5 | 0.79 | 0.34 | 0.54 | 0.38 | 0.29 |
| 4 | 22256034 | rs11940485 | -1.078 | -0.618 | -1.08 | 1.059 | 0.053 | 0.067 | 0.067 | 1.082 | 0.5 | 0.79 | 0.44 | 0.61 | 0.37 | 0.37 |
| 4 | 22264837 | rs10805233 | 0.01 | -1.1 | -0.594 | 0.95 | 0 | -0.003 | -0.003 | 0 | 0.25 | 0.21 | 0.19 | 0.33 | 0.15 | 0.08 |

| | | | | | | | | | | | | | | | | |
|---|---|---|---|---|---|---|---|---|---|---|---|---|---|---|---|---|
| 4 | 22266680 | rs4285070 | 0.008 | -1.098 | -0.608 | 0.951 | 0.002 | -0.004 | -0.004 | 0 | 0.25 | 0.19 | 0.19 | 0.33 | 0.15 | 0.08 |
| 4 | 22277576 | rs4697283 | 0.057 | -0.596 | -0.616 | 0.334 | -0.028 | 0.005 | 0.005 | 0.002 | 0.19 | 0.04 | 0.24 | 0.13 | 0.18 | 0.37 |
| 4 | 22293243 | rs1500453 | -0.031 | -0.765 | -0.165 | 0.239 | 0.003 | -0.023 | -0.023 | 0 | 0.22 | 0.15 | 0.34 | 0.33 | 0.19 | 0.35 |
| 4 | 22297899 | rs7659139 | -0.207 | -0.614 | 0.403 | 0.042 | 0.081 | -0.052 | -0.052 | 0.169 | 0.1 | 0.27 | 0.1 | 0 | 0.13 | 0.09 |
| 4 | 22300272 | rs1500463 | -1.136 | -0.691 | 0.487 | 0.151 | 0.005 | 0.013 | 0.013 | 0 | 0.33 | 0.59 | 0.5 | 0.39 | 0.34 | 0.53 |
| 4 | 22302352 | rs1392592 | -2.192 | -0.927 | -0.343 | -0.071 | -0.134 | -0.146 | -0.146 | 0 | 0.66 | 0.96 | 0.66 | 0.52 | 0.51 | 0.67 |
| 4 | 22303196 | rs6448143 | -0.471 | -0.904 | 0.224 | -0.405 | 0.084 | -0.066 | -0.066 | 0.732 | 0.12 | 0.41 | 0.13 | 0.02 | 0.14 | 0.09 |
| 4 | 22304073 | rs10938886 | -0.908 | -0.958 | -0.566 | -0.421 | 0.05 | 0.028 | 0.028 | 0 | 0.45 | 0.16 | 0.44 | 0.37 | 0.35 | 0.45 |
| 4 | 22316708 | rs6448144 | -0.996 | -0.46 | -0.371 | -0.746 | 0.015 | 0.069 | 0.069 | 0.834 | 0.38 | 0.71 | 0.25 | 0.05 | 0.28 | 0.19 |
| 4 | 22321784 | rs11929787 | -0.292 | -1.16 | 0.165 | -0.394 | 0.019 | -0.092 | -0.092 | 0 | 0.14 | 0.28 | 0.13 | 0.01 | 0.11 | 0.1 |
| 4 | 22322240 | rs4697007 | -1.765 | -1.602 | -0.38 | -0.3 | -0.076 | -0.087 | -0.087 | 2.146 | 0.59 | 0.9 | 0.59 | 0.44 | 0.42 | 0.66 |
| 4 | 22344500 | rs7674804 | 0.372 | -2.03 | 0.385 | 0.331 | -0.027 | 0.07 | 0.07 | 0.052 | 0.72 | 0.53 | 0.62 | 0.61 | 0.79 | 0.52 |
| 4 | 22344738 | rs16872561 | -0.844 | -2.055 | 0.372 | 0.331 | -0.072 | -0.059 | -0.059 | 0 | 0.2 | 0.42 | 0.13 | 0.08 | 0.1 | 0.08 |
| 4 | 22349886 | rs1983021 | 1.408 | -1.943 | -0.701 | 0.392 | 0.102 | -0.085 | -0.085 | 0.635 | 0.53 | 0.09 | 0.5 | 0.52 | 0.7 | 0.43 |
| 4 | 22370669 | rs955785 | -0.795 | -0.787 | -0.44 | 0.834 | 0.006 | -0.023 | -0.023 | 0 | 0.25 | 0.19 | 0.31 | 0.32 | 0.21 | 0.38 |
| 4 | 22390167 | rs9002 | -0.64 | -0.958 | -0.401 | -0.02 | -0.004 | -0.001 | -0.001 | 0 | 0.27 | 0.3 | 0.23 | 0.23 | 0.2 | 0.36 |
| 4 | 22391761 | rs6855769 | -1.215 | -0.82 | -1.047 | -0.111 | 0.055 | -0.032 | -0.032 | 0.015 | 0.39 | 0.88 | 0.4 | 0.36 | 0.28 | 0.47 |
| 4 | 22396289 | rs4697296 | 1.171 | -1.1 | -0.245 | -0.018 | -0.036 | -0.026 | -0.026 | 0 | 0.77 | 0.34 | 0.77 | 0.89 | 0.88 | 0.8 |
| 4 | 22397135 | rs1846731 | 1.208 | -1.158 | 0.559 | -0.057 | -0.009 | -0.024 | -0.024 | 2.607 | 0.86 | 0.76 | 0.83 | 0.96 | 0.89 | 0.83 |
| 4 | 22401728 | rs4697297 | 2.293 | -0.763 | 1.273 | 0.043 | -0.003 | 0.071 | 0.071 | 0 | 0.73 | 0.47 | 0.64 | 0.64 | 0.77 | 0.66 |
| 4 | 22414023 | rs10002842 | -1.422 | -0.59 | 1.579 | -0.044 | -0.004 | -0.003 | -0.003 | 0 | 0.14 | 0.15 | 0.17 | 0.02 | 0.19 | 0.16 |
| 4 | 22416738 | rs6821181 | -2.403 | -1.141 | 1.832 | -0.108 | -0.029 | 0.13 | 0.13 | 0 | 0.18 | 0.5 | 0.21 | 0.3 | 0.11 | 0.19 |
| 4 | 22428329 | rs10805234 | 2.209 | -1.856 | 1.538 | -0.469 | 0.035 | -0.008 | -0.008 | 0 | 0.63 | 0.3 | 0.61 | 0.58 | 0.78 | 0.6 |
| 4 | 22430507 | rs13135121 | 2.263 | -1.911 | 1.65 | -0.701 | -0.023 | -0.005 | -0.005 | 0 | 0.61 | 0.44 | 0.67 | 0.62 | 0.75 | 0.59 |
| 4 | 22433659 | rs13110697 | -0.022 | -1.941 | 1.523 | -0.706 | 0.01 | -0.008 | -0.008 | 0 | 0.06 | 0.03 | 0.07 | 0.2 | 0.03 | 0.14 |
| 4 | 22435993 | rs6448155 | 2.87 | -1.997 | 1.659 | -0.738 | 0.076 | 0.018 | 0.018 | 0 | 0.59 | 0.24 | 0.64 | 0.62 | 0.75 | 0.59 |
| 4 | 22436684 | rs16872644 | -0.111 | -1.954 | 1.544 | -0.753 | 0.001 | 0.004 | 0.004 | 0 | 0.06 | 0.07 | 0.01 | 0.04 | 0.01 | 0.02 |
| 4 | 22439599 | rs16872651 | -2.23 | -1.774 | 1.716 | -0.777 | 0.011 | 0.012 | 0.012 | 0.035 | 0.22 | 0.21 | 0.15 | 0.12 | 0.14 | 0.14 |
| 4 | 22440261 | rs17532048 | -0.75 | -1.767 | 1.694 | -0.778 | -0.001 | 0.01 | 0.01 | 0 | 0.1 | 0.08 | 0.06 | 0 | 0.1 | 0.06 |
| 4 | 22441466 | rs1463000 | 2.335 | -1.833 | 1.847 | -0.736 | -0.046 | -0.042 | -0.042 | 0 | 0.75 | 0.59 | 0.82 | 0.82 | 0.86 | 0.82 |
| 4 | 22447508 | rs7657579 | 1.582 | -1.826 | 1.601 | -0.596 | -0.008 | 0.009 | 0.009 | 0 | 0.79 | 0.78 | 0.78 | 0.71 | 0.88 | 0.71 |
| 4 | 22450122 | rs28726245 | -1.284 | -1.821 | 1.588 | -0.596 | 0.001 | -0.004 | -0.004 | 3.071 | 0.16 | 0.14 | 0.16 | 0.21 | 0.11 | 0.22 |
| 4 | 22451781 | rs1564138 | 1.484 | -1.745 | 1.585 | -0.591 | -0.012 | 0.013 | 0.013 | 0 | 0.79 | 0.75 | 0.77 | 0.72 | 0.87 | 0.7 |
| 4 | 22453260 | rs964549 | 2.064 | -1.755 | 1.52 | -0.574 | 0.007 | -0.005 | -0.005 | 0 | 0.69 | 0.33 | 0.71 | 0.62 | 0.84 | 0.65 |
| 4 | 22462730 | rs16872686 | -0.541 | -1.442 | 1.214 | -0.42 | -0.012 | 0.087 | 0.087 | 0.429 | 0.07 | 0.16 | 0.07 | 0.19 | 0.03 | 0.15 |
| 4 | 22469890 | rs16872711 | -0.919 | -1.672 | 0.835 | -0.721 | 0.036 | 0.04 | 0.04 | 3.409 | 0.15 | 0.08 | 0.04 | 0.04 | 0.06 | 0.03 |
| 4 | 22470618 | rs6851006 | -0.769 | -1.387 | 0.927 | -0.864 | 0.024 | -0.028 | -0.028 | 0 | 0.06 | 0.12 | 0.06 | 0 | 0.09 | 0.05 |
| 4 | 22470931 | rs7697959 | 2.149 | -1.689 | 0.784 | -1.313 | 0.006 | 0.012 | 0.012 | 0.448 | 0.46 | 0.35 | 0.53 | 0.69 | 0.62 | 0.56 |
| 4 | 22472251 | rs17599392 | -0.427 | -1.582 | 0.722 | -1.26 | 0.024 | -0.028 | -0.028 | 0 | 0.06 | 0.13 | 0.06 | 0 | 0.09 | 0.05 |
| 4 | 22479781 | rs13150142 | 1.741 | -1.823 | 0.605 | -1.263 | 0.005 | 0.005 | 0.005 | 0.816 | 0.51 | 0.41 | 0.59 | 0.69 | 0.71 | 0.62 |
| 4 | 22480252 | rs4235298 | -0.096 | -1.794 | 0.045 | -1.457 | 0 | 0.004 | 0.004 | 0.09 | 0.88 | 0.83 | 0.86 | 0.99 | 0.91 | 0.91 |
| 4 | 22484933 | rs16872734 | -0.614 | -1.696 | -0.069 | -1.327 | 0.023 | 0.02 | 0.02 | 3.89 | 0.15 | 0.1 | 0.04 | 0.04 | 0.06 | 0.03 |
| 4 | 22486548 | rs11730156 | -0.482 | -1.557 | -0.122 | -0.854 | 0.035 | -0.039 | -0.039 | 0 | 0.2 | 0.09 | 0.13 | 0.23 | 0.14 | 0.22 |
| 4 | 22487956 | rs6826101 | 0.239 | -1.55 | -0.257 | -1.479 | -0.003 | -0.055 | -0.055 | 0 | 0.68 | 0.41 | 0.7 | 0.9 | 0.73 | 0.72 |
| 4 | 22493757 | rs1841823 | 1.715 | -1.579 | -0.599 | -1.492 | -0.01 | -0.004 | -0.004 | 0 | 0.42 | 0.25 | 0.5 | 0.62 | 0.57 | 0.5 |
| 4 | 22493834 | rs6818597 | -0.07 | -1.578 | -1.146 | -1.331 | 0.026 | -0.072 | -0.072 | 0 | 0.19 | 0.08 | 0.18 | 0.25 | 0.15 | 0.21 |
| 4 | 22496882 | rs11728370 | 0.303 | -1.489 | -1.062 | -1.331 | 0.03 | -0.067 | -0.067 | 0 | 0.16 | 0.07 | 0.17 | 0.34 | 0.12 | 0.24 |
| 4 | 22501043 | rs6448161 | -0.804 | -1.379 | -0.98 | -1.435 | -0.001 | -0.061 | -0.061 | 3.301 | 0.25 | 0.36 | 0.23 | 0 | 0.2 | 0.21 |
| 4 | 22508257 | rs10516410 | -0.407 | -1.302 | -0.793 | -1.25 | -0.004 | -0.064 | -0.064 | 0 | 0.2 | 0.35 | 0.22 | 0.05 | 0.17 | 0.2 |
| 4 | 22509714 | rs7697048 | 0.995 | -0.662 | -1.1 | -0.806 | -0.076 | -0.045 | -0.045 | 0.061 | 0.38 | 0.16 | 0.45 | 0.59 | 0.52 | 0.45 |
| 4 | 22519313 | rs1381915 | 0.719 | -0.757 | -0.612 | -0.242 | -0.023 | -0.004 | -0.004 | 0 | 0.57 | 0.53 | 0.6 | 0.61 | 0.68 | 0.52 |
| 4 | 22521890 | rs2086010 | 0.28 | -0.96 | -0.925 | 0.02 | 0.037 | -0.025 | -0.025 | 0 | 0.79 | 0.87 | 0.87 | 0.72 | 0.86 | 0.81 |
| 4 | 22530016 | rs6838206 | 0.883 | -1.099 | 0.219 | -0.093 | 0.01 | 0.015 | 0.015 | 0.104 | 0.9 | 0.75 | 0.93 | 0.9 | 0.94 | 0.92 |
| 4 | 22531986 | rs1120044 | 1.58 | -1.229 | 0.539 | 0.613 | 0.002 | 0.158 | 0.158 | 0.302 | 0.88 | 0.65 | 0.88 | 0.76 | 0.93 | 0.87 |
| 4 | 22536001 | rs10017558 | -0.326 | -1.842 | 0.578 | 0.868 | -0.022 | -0.003 | -0.003 | 0 | 0.19 | 0.28 | 0.29 | 0.17 | 0.09 | 0.11 |
| 4 | 22537258 | rs13143909 | 1.094 | -1.363 | -0.141 | 0.416 | -0.02 | -0.044 | -0.044 | 0.484 | 0.37 | 0.18 | 0.27 | 0.5 | 0.37 | 0.47 |
| 4 | 22537387 | rs12163757 | 0.812 | -1.373 | -0.219 | 0.385 | -0.103 | 0.037 | 0.037 | 0 | 0.33 | 0.16 | 0.36 | 0.19 | 0.49 | 0.29 |
| 4 | 22538841 | rs10026861 | -0.196 | -1.355 | -0.215 | 0.401 | 0.001 | -0.008 | -0.008 | 0 | 0.58 | 0.57 | 0.56 | 0.71 | 0.45 | 0.63 |
| 4 | 22538956 | rs10027019 | -0.212 | -1.374 | -0.224 | 0.391 | 0.002 | -0.008 | -0.008 | 0 | 0.58 | 0.57 | 0.56 | 0.71 | 0.45 | 0.63 |
| 4 | 22555317 | rs2323624 | 1.097 | -1.365 | -0.294 | 0.259 | 0 | -0.021 | -0.021 | 0 | 0.35 | 0.23 | 0.25 | 0.54 | 0.31 | 0.46 |
| 4 | 22556784 | rs11722502 | -0.438 | -1.47 | -0.262 | 0.325 | -0.002 | -0.011 | -0.011 | 0.794 | 0.6 | 0.64 | 0.61 | 0.86 | 0.47 | 0.68 |
| 4 | 22559043 | rs10446635 | 0.776 | -1.473 | -0.247 | 0.321 | -0.002 | -0.007 | -0.007 | 0 | 0.39 | 0.31 | 0.28 | 0.54 | 0.37 | 0.52 |
| 4 | 22579261 | rs2323549 | -0.471 | -1.409 | -0.288 | 0.219 | -0.001 | -0.017 | -0.017 | 0 | 0.61 | 0.64 | 0.61 | 0.88 | 0.47 | 0.68 |
| 4 | 22581486 | rs6813399 | 1.189 | -1.422 | -0.317 | 0.195 | -0.152 | 0.164 | 0.164 | 0 | 0.3 | 0.09 | 0.32 | 0.05 | 0.47 | 0.24 |
| 4 | 22585550 | rs2581963 | -0.446 | -1.431 | -0.142 | 0.34 | 0.023 | 0.019 | 0.019 | 0.146 | 0.91 | 0.73 | 0.93 | 0.92 | 0.94 | 0.92 |
| 4 | 22585721 | rs13152115 | 0.328 | -1.326 | -0.5 | 0.485 | 0.022 | 0.003 | 0.003 | 0.112 | 0.19 | 0.17 | 0.28 | 0.19 | 0.09 | 0.11 |
| 4 | 22591935 | rs13120856 | -0.644 | -1.404 | -0.139 | 0.386 | -0.046 | 0.031 | 0.031 | 0.181 | 0.27 | 0.63 | 0.39 | 0.41 | 0.13 | 0.23 |
| 4 | 22592929 | rs6858626 | 0.453 | -1.056 | -0.565 | -0.104 | -0.057 | 0.106 | 0.106 | 0 | 0.37 | 0.2 | 0.36 | 0.07 | 0.52 | 0.3 |
| 4 | 22595658 | rs11944771 | -0.387 | -1.48 | -0.339 | 0.258 | 0.036 | 0.024 | 0.024 | 0.024 | 0.21 | 0.16 | 0.19 | 0.15 | 0.08 | 0.09 |

| | | | | | | | | | | | | | | | | |
|---|---|---|---|---|---|---|---|---|---|---|---|---|---|---|---|---|
| 4 | 22605059 | rs1824864 | -0.314 | -0.845 | -0.163 | 0.613 | 0.006 | 0.021 | 0.021 | 0 | 0.21 | 0.24 | 0.21 | 0.58 | 0.3 | 0.26 |
| 4 | 22613279 | rs2581987 | -1.919 | -1.288 | -0.349 | -0.042 | -0.041 | 0.127 | 0.127 | 0.42 | 0.59 | 0.78 | 0.63 | 0.88 | 0.56 | 0.73 |
| 4 | 22613860 | rs1842098 | 0.181 | -1.202 | -0.286 | 0.093 | -0.001 | 0.011 | 0.011 | 0 | 0.2 | 0.2 | 0.22 | 0.55 | 0.27 | 0.26 |
| 4 | 22614302 | rs7654229 | -0.199 | -1.56 | 0.008 | -0.115 | -0.003 | -0.005 | -0.005 | 0 | 0.18 | 0.16 | 0.09 | 0.13 | 0.15 | 0.19 |
| 4 | 22614378 | rs2618940 | -2.499 | -1.597 | -0.182 | -0.225 | -0.044 | 0.118 | 0.118 | 0.102 | 0.6 | 0.78 | 0.63 | 0.88 | 0.56 | 0.74 |
| 4 | 22625571 | rs2581994 | 0.554 | -1.497 | -0.085 | 0.306 | -0.06 | -0.115 | -0.115 | 0 | 0.15 | 0.02 | 0.21 | 0.44 | 0.24 | 0.19 |
| 4 | 22627981 | rs2581997 | -1.153 | -1.657 | 0.804 | -0.123 | 0.001 | -0.016 | -0.016 | 0 | 0.46 | 0.57 | 0.51 | 0.85 | 0.41 | 0.58 |
| 4 | 22631389 | rs2439468 | -1.917 | -1.623 | 0.584 | -0.108 | 0.001 | 0.058 | 0.058 | 0.011 | 0.54 | 0.59 | 0.6 | 0.87 | 0.52 | 0.68 |
| 4 | 22646725 | rs2618947 | -2.009 | -1.411 | 0.881 | -0.026 | 0.003 | 0.183 | 0.183 | 0.184 | 0.54 | 0.76 | 0.6 | 0.88 | 0.5 | 0.58 |
| 4 | 22648713 | rs16872878 | 0.486 | -1.5 | 1.233 | 0.088 | -0.027 | 0.034 | 0.034 | 0 | 0.1 | 0.27 | 0.14 | 0.11 | 0.05 | 0.04 |
| 4 | 22656105 | rs2245395 | -0.563 | -1.523 | 1.308 | -0.075 | -0.013 | 0.061 | 0.061 | 0 | 0.34 | 0.12 | 0.36 | 0.77 | 0.35 | 0.45 |
| 4 | 22659580 | rs2618954 | -1.395 | -1.518 | 1.552 | -0.073 | -0.005 | -0.039 | -0.039 | 0 | 0.43 | 0.49 | 0.46 | 0.8 | 0.44 | 0.52 |
| 4 | 22661136 | rs12507337 | 0.859 | -1.5 | 1.512 | -0.054 | -0.015 | 0.004 | 0.004 | 0 | 0.09 | 0.14 | 0.13 | 0.09 | 0.05 | 0.04 |
| 4 | 22661376 | rs2258846 | -1.412 | -1.494 | 1.51 | -0.083 | -0.008 | -0.055 | -0.055 | 2.67 | 0.46 | 0.52 | 0.5 | 0.86 | 0.41 | 0.51 |
| 4 | 22671199 | rs2581966 | -1.966 | -1.443 | 1.105 | -0.08 | 0.009 | 0.052 | 0.052 | 0.024 | 0.53 | 0.63 | 0.59 | 0.88 | 0.5 | 0.59 |
| 4 | 22683706 | rs12501294 | 0.399 | -1.321 | 0.435 | 0.113 | 0.004 | 0.045 | 0.045 | 0 | 0.65 | 0.81 | 0.63 | 0.25 | 0.65 | 0.55 |
| 4 | 22685868 | rs6856346 | -0.162 | -0.831 | 1.002 | 0.497 | -0.032 | 0.029 | 0.029 | 0.027 | 0.11 | 0.36 | 0.15 | 0.11 | 0.06 | 0.05 |
| 4 | 22687741 | rs2009731 | 1.018 | -0.564 | 0.453 | 0.74 | 0.012 | 0.009 | 0.009 | 0.098 | 0.6 | 0.35 | 0.58 | 0.43 | 0.6 | 0.62 |
| 4 | 22689850 | rs12506103 | -0.211 | -0.295 | 0.617 | 0.764 | 0.006 | 0.007 | 0.007 | 0 | 0.11 | 0.13 | 0.18 | 0.14 | 0.17 | 0.09 |
| 4 | 22691505 | rs10028144 | -1.779 | -0.398 | 0.985 | 0.704 | 0.011 | -0.013 | -0.013 | 0 | 0.21 | 0.57 | 0.29 | 0.16 | 0.26 | 0.16 |
| 4 | 22693754 | rs7682224 | -2.186 | -0.407 | 0.855 | 0.741 | -0.002 | -0.003 | -0.003 | 0 | 0.28 | 0.57 | 0.36 | 0.27 | 0.35 | 0.24 |
| 4 | 22696560 | rs10004988 | -0.809 | -0.435 | 0.858 | 0.823 | 0 | 0.005 | 0.005 | 0 | 0.14 | 0.1 | 0.15 | 0.13 | 0.18 | 0.15 |
| 4 | 22699639 | rs9991992 | -0.809 | -0.426 | 0.919 | 0.828 | 0 | 0.004 | 0.004 | 0 | 0.14 | 0.11 | 0.15 | 0.13 | 0.18 | 0.15 |
| 4 | 22714805 | rs17539365 | -1.579 | -0.287 | 0.746 | 0.558 | -0.012 | 0.023 | 0.023 | 0 | 0.32 | 0.17 | 0.41 | 0.26 | 0.4 | 0.24 |
| 4 | 22715030 | rs16873068 | -1.791 | -0.32 | 0.971 | 0.541 | -0.004 | -0.029 | -0.029 | 0 | 0.35 | 0.48 | 0.43 | 0.26 | 0.4 | 0.24 |
| 4 | 22715639 | rs1963996 | -0.381 | 0.176 | 0.998 | 0.77 | -0.014 | -0.08 | -0.08 | 0 | 0.76 | 0.87 | 0.84 | 0.59 | 0.78 | 0.7 |
| 4 | 22717410 | rs1485829 | -0.753 | 0.319 | 1.225 | 0.558 | -0.003 | 0.001 | 0.001 | 0 | 0.17 | 0.13 | 0.26 | 0.14 | 0.22 | 0.1 |
| 4 | 22757756 | rs17540321 | -1.372 | -0.047 | -0.656 | 0.56 | 0.002 | 0.009 | 0.009 | 0 | 0.31 | 0.24 | 0.32 | 0.27 | 0.31 | 0.18 |
| 4 | 22770299 | rs1385760 | -1.355 | -0.086 | -0.493 | 0.211 | -0.016 | 0.062 | 0.062 | 0 | 0.28 | 0.17 | 0.21 | 0.13 | 0.32 | 0.14 |
| 4 | 22774617 | rs4103518 | -1.459 | -0.096 | -0.497 | 0.206 | -0.008 | 0.042 | 0.042 | 0 | 0.29 | 0.21 | 0.31 | 0.16 | 0.32 | 0.15 |
| 4 | 22786034 | rs10000431 | -1.289 | 0.265 | -0.346 | 0.581 | -0.003 | 0.006 | 0.006 | 0 | 0.14 | 0.1 | 0.15 | 0.12 | 0.17 | 0.1 |
| 4 | 22793164 | rs6854336 | -1.273 | 0.275 | -0.341 | 0.537 | -0.004 | 0.008 | 0.008 | 0 | 0.14 | 0.1 | 0.14 | 0.11 | 0.17 | 0.1 |
| 4 | 22793514 | rs358255 | -0.029 | 0.222 | -0.413 | 0.492 | -0.008 | 0.008 | 0.008 | 0.489 | 0.84 | 0.85 | 0.89 | 0.88 | 0.82 | 0.94 |
| 4 | 22793528 | rs183084 | 1.812 | 0.208 | -0.209 | 0.486 | 0 | -0.01 | -0.01 | 0 | 0.68 | 0.56 | 0.63 | 0.73 | 0.65 | 0.83 |
| 4 | 22794601 | rs358254 | 0.461 | -0.002 | -0.392 | 0.639 | 0 | -0.001 | -0.001 | 0 | 0.83 | 0.73 | 0.78 | 0.84 | 0.82 | 0.93 |
| 4 | 22803174 | rs370843 | 0.478 | 0.074 | -0.633 | 0.699 | -0.005 | 0.001 | 0.001 | 0.711 | 0.79 | 0.71 | 0.75 | 0.77 | 0.83 | 0.91 |
| 4 | 22812879 | rs358238 | 0.191 | -0.074 | -0.846 | 0.697 | 0.015 | -0.014 | -0.014 | 0 | 0.81 | 0.95 | 0.89 | 0.8 | 0.84 | 0.92 |
| 4 | 22819747 | rs358233 | 1.101 | -0.045 | -1.157 | 0.717 | -0.013 | 0.143 | 0.143 | 0.068 | 0.48 | 0.2 | 0.43 | 0.3 | 0.54 | 0.54 |
| 4 | 22824021 | rs3099557 | 0.86 | -0.243 | -0.56 | 0.79 | -0.03 | 0.008 | 0.008 | 0.273 | 0.77 | 0.64 | 0.74 | 0.75 | 0.84 | 0.9 |
| 4 | 22832447 | rs430976 | 0.632 | -0.007 | -1.335 | 0.528 | -0.014 | 0.014 | 0.014 | 1.942 | 0.81 | 0.98 | 0.93 | 0.86 | 0.83 | 0.93 |
| 4 | 22832682 | rs439863 | -0.309 | 0.092 | -1.378 | 0.411 | -0.008 | 0.004 | 0.004 | 0.219 | 0.36 | 0.31 | 0.36 | 0.67 | 0.36 | 0.4 |
| 4 | 22834211 | rs418932 | -0.979 | -0.754 | -0.017 | 0.68 | -0.053 | 0.051 | 0.051 | 0.015 | 0.1 | 0.44 | 0.09 | 0.12 | 0.02 | 0.03 |
| 4 | 22843458 | rs412218 | 0.452 | -0.877 | -0.653 | 0.667 | -0.028 | -0.04 | -0.04 | 1.057 | 0.74 | 0.89 | 0.6 | 0.33 | 0.74 | 0.6 |
| 4 | 22844392 | rs403543 | -0.533 | -0.867 | -0.624 | 0.675 | -0.053 | 0.051 | 0.051 | 0.063 | 0.1 | 0.44 | 0.09 | 0.12 | 0.02 | 0.03 |
| 4 | 22850202 | rs11946006 | 1.209 | -0.892 | -0.611 | 0.56 | -0.003 | -0.008 | -0.008 | 0 | 0.65 | 0.44 | 0.52 | 0.2 | 0.72 | 0.57 |
| 4 | 22860785 | rs358224 | 0.503 | -0.838 | -0.33 | 0.715 | -0.019 | -0.023 | -0.023 | 0.059 | 0.76 | 0.87 | 0.59 | 0.28 | 0.76 | 0.61 |
| 4 | 22871367 | rs921096 | -0.587 | -0.953 | -0.285 | 1.851 | -0.006 | -0.07 | -0.07 | 0 | 0.16 | 0.09 | 0.32 | 0.43 | 0.18 | 0.17 |
| 4 | 22878904 | rs6840245 | -1.822 | -1.157 | 0.438 | 1.311 | -0.008 | 0 | 0 | 0.975 | 0.29 | 0.63 | 0.4 | 0.7 | 0.2 | 0.3 |
| 4 | 22901347 | rs718137 | -1.746 | -1.021 | 1.091 | 1.324 | -0.047 | 0.007 | 0.007 | 0 | 0.27 | 0.56 | 0.31 | 0.69 | 0.17 | 0.29 |
| 4 | 22915626 | rs4697350 | -1.913 | -1.087 | 0.96 | 1.237 | -0.008 | -0.001 | -0.001 | 0 | 0.3 | 0.65 | 0.41 | 0.7 | 0.19 | 0.31 |
| 4 | 22918094 | rs4697351 | -1.107 | -1.101 | 0.793 | 1.436 | 0.019 | -0.049 | -0.049 | 0 | 0.19 | 0.12 | 0.34 | 0.62 | 0.16 | 0.25 |
| 4 | 22934689 | rs2323700 | -1.4 | -0.056 | 1.172 | 0.691 | 0.029 | 0.008 | 0.008 | 0 | 0.13 | 0.35 | 0.07 | 0.08 | 0.12 | 0.09 |
| 4 | 22937281 | rs2044512 | -1.415 | -0.054 | 1.166 | 0.64 | 0.029 | -0.008 | -0.008 | 0.04 | 0.13 | 0.34 | 0.07 | 0.07 | 0.12 | 0.09 |
| 4 | 22946063 | rs6448180 | -1.251 | -0.186 | 1.438 | 0.743 | 0.021 | 0.049 | 0.049 | 0.047 | 0.09 | 0.31 | 0.07 | 0.06 | 0.09 | 0.06 |
| 4 | 22962472 | rs13113121 | -1.432 | -0.66 | 1.49 | 0.456 | -0.063 | -0.015 | -0.015 | 0 | 0.29 | 0.5 | 0.45 | 0.75 | 0.18 | 0.25 |
| 4 | 23002014 | rs10938926 | -0.989 | -0.55 | 1.314 | 0.488 | 0.001 | -0.008 | -0.008 | 0.926 | 0.19 | 0.18 | 0.43 | 0.73 | 0.18 | 0.25 |
| 4 | 23018704 | rs7664442 | -0.823 | -0.721 | 1.292 | 0.399 | -0.002 | -0.001 | -0.001 | 0 | 0.1 | 0.1 | 0.05 | 0.06 | 0.08 | 0.04 |
| 4 | 23032220 | rs4697365 | -0.978 | -0.493 | 1.377 | 0.382 | 0.004 | 0.024 | 0.024 | 0 | 0.18 | 0.26 | 0.45 | 0.73 | 0.2 | 0.25 |
| 4 | 23038516 | rs35570804 | 1.886 | -0.509 | 0.013 | -0.864 | -0.067 | 0.108 | 0.108 | 0 | 0.22 | 0.01 | 0.12 | 0.03 | 0.21 | 0.33 |
| 4 | 23040791 | rs7660606 | -0.883 | -0.013 | 0.688 | -0.334 | -0.004 | 0.041 | 0.041 | 0 | 0.22 | 0.39 | 0.46 | 0.73 | 0.19 | 0.26 |
| 4 | 23085015 | rs11944694 | -1.506 | 0.068 | 0.503 | -0.638 | -0.01 | -0.03 | -0.03 | 0.155 | 0.32 | 0.56 | 0.52 | 0.78 | 0.29 | 0.31 |
| 4 | 23160661 | rs11733138 | -1.019 | -0.312 | 0.413 | -0.283 | 0.025 | 0.031 | 0.031 | 0 | 0.53 | 0.66 | 0.61 | 0.67 | 0.65 | 0.48 |
| 4 | 23169854 | rs7676830 | -0.751 | 0.477 | 1.198 | 0.632 | 0.019 | 0.087 | 0.087 | 0 | 0.18 | 0.26 | 0.36 | 0.53 | 0.25 | 0.21 |
| 4 | 23173926 | rs4498120 | 1.406 | 0.282 | 1.336 | 0.251 | -0.01 | -0.029 | -0.029 | 0 | 0.86 | 0.66 | 0.84 | 0.93 | 0.87 | 0.84 |
| 4 | 23175107 | rs10805240 | 1.297 | 0.271 | 1.402 | 0.302 | -0.011 | -0.024 | -0.024 | 0 | 0.87 | 0.65 | 0.85 | 0.93 | 0.87 | 0.85 |
| 4 | 23194506 | rs6830403 | 0.643 | 0.639 | 1.279 | 0.59 | -0.009 | -0.009 | -0.009 | 0 | 0.76 | 0.79 | 0.59 | 0.42 | 0.62 | 0.64 |
| 4 | 23209159 | rs13127424 | 1.326 | 0.439 | 1.526 | 0.447 | -0.014 | -0.006 | -0.006 | 0 | 0.63 | 0.21 | 0.52 | 0.39 | 0.59 | 0.61 |
| 4 | 23220263 | rs7668474 | 0.92 | 0.72 | 1.842 | 0.697 | 0.022 | 0.036 | 0.036 | 0.093 | 0.72 | 0.38 | 0.57 | 0.46 | 0.65 | 0.65 |
| 4 | 23235331 | rs20414 | 0.79 | 0.976 | 0.413 | 0.799 | -0.003 | -0.004 | -0.004 | 0 | 0.71 | 0.7 | 0.6 | 0.45 | 0.64 | 0.63 |

| | | | | | | | | | | | | | | | | |
|---|---|---|---|---|---|---|---|---|---|---|---|---|---|---|---|---|
| 4 | 23266728 | rs10938932 | -0.985 | 1.015 | 1.064 | 0.779 | 0.003 | 0.016 | 0.016 | 0.014 | 0.24 | 0.54 | 0.38 | 0.31 | 0.22 | 0.27 |
| 4 | 23267131 | rs16873417 | -0.488 | 0.968 | 1.519 | 0.763 | -0.004 | -0.005 | -0.005 | 0.412 | 0.21 | 0.18 | 0.33 | 0.3 | 0.22 | 0.27 |
| 4 | 23288742 | rs7658635 | 0.821 | 1.299 | 1.542 | 0.437 | 0.067 | -0.055 | -0.055 | 0 | 0.86 | 0.49 | 0.83 | 0.98 | 0.81 | 0.94 |
| 4 | 23289721 | rs10050029 | 1.375 | 1.374 | 1.447 | 0.456 | -0.022 | 0.02 | 0.02 | 0.038 | 0.65 | 0.26 | 0.49 | 0.69 | 0.6 | 0.67 |
| 4 | 23293043 | rs12503311 | -0.381 | 1.385 | 1.471 | 0.488 | -0.003 | -0.003 | -0.003 | 0 | 0.21 | 0.2 | 0.35 | 0.29 | 0.21 | 0.25 |
| 4 | 23296159 | rs10938939 | -0.583 | 1.356 | 1.604 | 0.475 | -0.001 | 0 | 0 | 0 | 0.22 | 0.29 | 0.36 | 0.29 | 0.2 | 0.27 |
| 4 | 23307693 | rs12507992 | -0.42 | 1.343 | 1.139 | 1.038 | -0.003 | -0.046 | -0.046 | 0.018 | 0.21 | 0.17 | 0.36 | 0.44 | 0.2 | 0.3 |
| 4 | 23307920 | rs2036713 | 1.224 | 1.362 | 0.94 | 1.028 | -0.015 | -0.024 | -0.024 | 0 | 0.65 | 0.28 | 0.48 | 0.54 | 0.61 | 0.64 |
| 4 | 23317124 | rs2175069 | 1.165 | 1.361 | 0.915 | 1.046 | -0.005 | -0.008 | -0.008 | 4.234 | 0.65 | 0.34 | 0.49 | 0.53 | 0.61 | 0.63 |
| 4 | 23320940 | rs16873495 | -0.113 | 1.265 | 0.531 | 1.268 | -0.001 | -0.021 | -0.021 | 1.712 | 0.16 | 0.15 | 0.33 | 0.47 | 0.16 | 0.24 |
| 4 | 23323428 | rs11731411 | -0.204 | 1.146 | -0.091 | 0.792 | 0.011 | 0.012 | 0.012 | 0 | 0.25 | 0.15 | 0.11 | 0.18 | 0.22 | 0.13 |
| 4 | 23324095 | rs1858113 | 1.085 | 1.147 | 0.177 | 0.464 | -0.019 | 0.016 | 0.016 | 0 | 0.58 | 0.31 | 0.48 | 0.59 | 0.53 | 0.51 |
| 4 | 23325040 | rs882348 | -0.105 | 0.883 | -0.039 | 0.903 | -0.002 | 0.003 | 0.003 | 0 | 0.12 | 0.11 | 0.11 | 0.19 | 0.11 | 0.18 |
| 4 | 23333141 | rs10517011 | -0.248 | 0.859 | 0.108 | 0.509 | -0.001 | -0.006 | -0.006 | 0 | 0.23 | 0.26 | 0.32 | 0.18 | 0.24 | 0.2 |
| 4 | 23338785 | rs2940777 | -0.207 | 0.896 | 0.12 | 0.554 | -0.001 | -0.005 | -0.005 | 0 | 0.23 | 0.25 | 0.31 | 0.17 | 0.24 | 0.19 |
| 4 | 23340149 | rs11728319 | 1.237 | 1.166 | 0.239 | 0.092 | 0.007 | 0.122 | 0.122 | 0 | 0.51 | 0.28 | 0.49 | 0.71 | 0.48 | 0.61 |
| 4 | 23340766 | rs6448201 | 0.559 | 0.946 | 0.775 | -0.064 | 0.002 | -0.031 | -0.031 | 0 | 0.74 | 0.76 | 0.82 | 0.92 | 0.72 | 0.8 |
| 4 | 23343798 | rs1509247 | 0.736 | 0.771 | 1.021 | 0.162 | 0.003 | 0.04 | 0.04 | 0 | 0.52 | 0.38 | 0.54 | 0.71 | 0.51 | 0.59 |
| 4 | 23346349 | rs13151220 | 0.162 | 0.785 | 0.895 | 0.083 | 0.001 | -0.002 | -0.002 | 0.14 | 0.1 | 0.09 | 0.06 | 0.03 | 0.08 | 0.08 |
| 4 | 23348610 | rs2940776 | -0.821 | 0.98 | 0.892 | -0.046 | -0.008 | -0.011 | -0.011 | 0.636 | 0.82 | 0.86 | 0.87 | 1 | 0.76 | 0.86 |
| 4 | 23350299 | rs12509827 | 0.442 | 1.147 | 0.449 | 0.098 | -0.044 | -0.018 | -0.018 | 0 | 0.08 | 0.01 | 0.12 | 0.08 | 0.14 | 0.07 |
| 4 | 23351305 | rs10005577 | -0.607 | 0.588 | 0.876 | 0.584 | -0.018 | 0.076 | 0.076 | 0.035 | 0.14 | 0.31 | 0.14 | 0.24 | 0.08 | 0.15 |
| 4 | 23351883 | rs7690639 | 0.48 | 0.601 | 0.436 | 0.583 | 0.005 | 0.013 | 0.013 | 0 | 0.26 | 0.09 | 0.29 | 0.24 | 0.24 | 0.25 |
| 4 | 23352821 | rs10938942 | 1.628 | 0.539 | 0.447 | 0.542 | -0.066 | -0.076 | -0.076 | 0.142 | 0.41 | 0.12 | 0.42 | 0.46 | 0.44 | 0.47 |
| 4 | 23354371 | rs7682902 | 0.234 | 0.748 | 0.595 | 0.705 | -0.001 | 0.013 | 0.013 | 0 | 0.26 | 0.09 | 0.29 | 0.25 | 0.26 | 0.24 |
| 4 | 23356724 | rs7666498 | 0.245 | 0.765 | 0.603 | 0.713 | -0.001 | 0.013 | 0.013 | 0 | 0.26 | 0.09 | 0.29 | 0.24 | 0.26 | 0.24 |
| 4 | 23386583 | rs169998 | -0.31 | 0.743 | 1.142 | 0.486 | -0.003 | -0.032 | -0.032 | 0 | 0.24 | 0.59 | 0.21 | 0.14 | 0.24 | 0.2 |
| 4 | 23387315 | rs215478 | 0.662 | 0.73 | 1.38 | 0.46 | 0.003 | -0.047 | -0.047 | 0 | 0.18 | 0.21 | 0.13 | 0 | 0.23 | 0.14 |
| 4 | 23388399 | rs215475 | -0.561 | 0.922 | 0.674 | 0.594 | -0.008 | -0.018 | -0.018 | 0.829 | 0.75 | 0.9 | 0.69 | 0.72 | 0.73 | 0.76 |
| 4 | 23390149 | rs976892 | 0.889 | 0.859 | 0.734 | 0.587 | 0.011 | 0.029 | 0.029 | 0 | 0.5 | 0.31 | 0.49 | 0.57 | 0.51 | 0.55 |
| 4 | 23390161 | rs17539073 | 0.519 | 0.858 | 0.673 | 0.584 | -0.001 | -0.001 | -0.001 | 0 | 0.08 | 0.09 | 0.05 | 0.07 | 0.08 | 0.07 |
| 4 | 23394710 | rs4697034 | 1.299 | 1.054 | 0.628 | 0.59 | -0.079 | -0.085 | -0.085 | 0 | 0.44 | 0.12 | 0.45 | 0.51 | 0.47 | 0.47 |
| 4 | 23416412 | rs2159518 | 1.395 | 0.854 | 0.619 | 0.598 | -0.016 | -0.031 | -0.031 | 0 | 0.42 | 0.22 | 0.43 | 0.5 | 0.43 | 0.47 |
| 4 | 23424060 | rs215410 | -0.642 | 0.712 | 0.66 | 0.579 | -0.004 | -0.02 | -0.02 | 2.384 | 0.33 | 0.62 | 0.25 | 0.21 | 0.31 | 0.28 |
| 4 | 23424293 | rs215408 | -0.575 | 0.773 | -0.149 | 0.556 | 0.003 | -0.022 | -0.022 | 0.078 | 0.75 | 0.93 | 0.7 | 0.71 | 0.76 | 0.77 |
| 4 | 23430768 | rs1605813 | 1.219 | 0.929 | -0.587 | 0.684 | -0.071 | -0.079 | -0.079 | 0 | 0.41 | 0.14 | 0.43 | 0.5 | 0.45 | 0.48 |
| 4 | 23438665 | rs215446 | 0.304 | 0.72 | -0.336 | 0.733 | 0.016 | 0.003 | 0.003 | 0 | 0.65 | 0.8 | 0.66 | 0.64 | 0.68 | 0.69 |
| 4 | 23451237 | rs6817487 | -0.551 | 0.986 | 0.265 | 0.611 | 0.022 | -0.036 | -0.036 | 0.071 | 0.13 | 0.27 | 0.14 | 0.07 | 0.17 | 0.05 |
| 4 | 23451900 | rs215428 | 1.213 | 0.733 | 0.396 | 0.647 | 0.006 | -0.011 | -0.011 | 0 | 0.64 | 0.47 | 0.54 | 0.62 | 0.59 | 0.68 |
| 4 | 23454910 | rs17613288 | 1.277 | 0.492 | 0.251 | 0.581 | 0.004 | 0.005 | 0.005 | 1.026 | 0.19 | 0.11 | 0.06 | 0.1 | 0.14 | 0.23 |
| 4 | 23456531 | rs4697389 | -0.504 | 0.494 | 0.301 | 0.309 | -0.009 | -0.032 | -0.032 | 0.968 | 0.79 | 0.89 | 0.69 | 0.69 | 0.76 | 0.77 |
| 4 | 23456789 | rs4697390 | 0.08 | 0.5 | 0.334 | 0.283 | 0.023 | -0.037 | -0.037 | 0 | 0.13 | 0.28 | 0.14 | 0.06 | 0.17 | 0.05 |
| 4 | 23462730 | rs13129559 | 1.364 | 0.433 | 0.365 | 0.857 | -0.039 | 0.075 | 0.075 | 0.311 | 0.42 | 0.16 | 0.29 | 0.33 | 0.45 | 0.43 |
| 4 | 23466369 | rs1304559 | -0.438 | 0.518 | 0.374 | 0.927 | -0.001 | 0 | 0 | 0 | 0.78 | 0.79 | 0.69 | 0.69 | 0.77 | 0.78 |
| 4 | 23473748 | rs723394 | -0.363 | 0.494 | 0.065 | 0.907 | 0 | -0.017 | -0.017 | 0 | 0.78 | 0.85 | 0.69 | 0.68 | 0.76 | 0.71 |
| 4 | 23473797 | rs723393 | -0.363 | 0.494 | 0.065 | 0.907 | 0 | -0.017 | -0.017 | 3.038 | 0.78 | 0.85 | 0.69 | 0.68 | 0.76 | 0.71 |
| 4 | 23474365 | rs12331728 | 1.295 | 0.483 | 0.135 | 0.895 | -0.013 | 0.05 | 0.05 | 0.317 | 0.43 | 0.29 | 0.3 | 0.34 | 0.46 | 0.43 |
| 4 | 23474539 | rs6829223 | -0.343 | 0.496 | 0.112 | 0.912 | 0.001 | -0.017 | -0.017 | 0 | 0.78 | 0.85 | 0.69 | 0.68 | 0.76 | 0.72 |
| 4 | 23476755 | rs887471 | -0.366 | 0.455 | -0.031 | 0.916 | 0.003 | -0.013 | -0.013 | 0 | 0.78 | 0.84 | 0.69 | 0.69 | 0.77 | 0.77 |
| 4 | 23477482 | rs12498626 | 0.741 | 0.34 | -0.032 | 0.974 | -0.019 | 0.043 | 0.043 | 0.119 | 0.23 | 0.29 | 0.25 | 0.29 | 0.13 | 0.26 |
| 4 | 23478842 | rs11728280 | 1.007 | 0.827 | -0.339 | 0.895 | -0.001 | -0.001 | -0.001 | 0 | 0.65 | 0.63 | 0.56 | 0.62 | 0.6 | 0.68 |
| 4 | 23480613 | rs16873729 | -0.225 | 0.685 | -0.973 | 1.273 | 0.008 | -0.049 | -0.049 | 0.967 | 0.14 | 0.04 | 0.13 | 0.26 | 0.14 | 0.14 |
| 4 | 23485012 | rs17613848 | 0.418 | 1.244 | -1.273 | 0.918 | -0.07 | -0.039 | -0.039 | 0 | 0.14 | 0.03 | 0.12 | 0.17 | 0.22 | 0.16 |
| 4 | 23485264 | rs6857002 | 0.078 | 0.735 | -0.579 | 0.039 | -0.013 | -0.107 | -0.107 | 0 | 0.32 | 0.45 | 0.3 | 0.07 | 0.26 | 0.34 |
| 4 | 23485419 | rs6831833 | 0.603 | 0.535 | -0.247 | 0.283 | -0.013 | 0.011 | 0.011 | 0 | 0.07 | 0.03 | 0.17 | 0.03 | 0.1 | 0.08 |
| 4 | 23487104 | rs756516 | 0.074 | 0.248 | -0.023 | 0.567 | -0.02 | 0.014 | 0.014 | 0 | 0.1 | 0.14 | 0.09 | 0.18 | 0.06 | 0.08 |
| 4 | 23488892 | rs12505641 | -1.242 | 0.009 | 0.675 | 0.769 | -0.031 | 0.031 | 0.031 | 0 | 0.21 | 0.56 | 0.48 | 0.32 | 0.17 | 0.19 |
| 4 | 23495538 | rs4317189 | -0.86 | 0.588 | 0.71 | 0.461 | -0.012 | 0.005 | 0.005 | 0 | 0.67 | 0.76 | 0.78 | 0.68 | 0.65 | 0.61 |
| 4 | 23520711 | rs17554625 | 0.332 | 0.502 | 0.845 | 0.493 | 0.006 | -0.009 | -0.009 | 0.563 | 0.07 | 0.13 | 0.23 | 0.06 | 0.08 | 0.08 |
| 4 | 23532789 | rs1003561 | 0.696 | 0.525 | 0.7 | 0.141 | 0.006 | -0.007 | -0.007 | 0 | 0.07 | 0.08 | 0.22 | 0.06 | 0.08 | 0.08 |
| 4 | 23539244 | rs10517015 | 0.696 | 0.525 | 0.7 | 0.141 | 0.006 | -0.007 | -0.007 | 0 | 0.07 | 0.08 | 0.22 | 0.06 | 0.08 | 0.08 |
| 4 | 23546394 | rs2323996 | 0.808 | 0.525 | 0.333 | 0.108 | -0.059 | 0.054 | 0.054 | 0 | 0.31 | 0.1 | 0.18 | 0.19 | 0.35 | 0.37 |
| 4 | 23555237 | rs10021542 | 0.106 | 0.151 | 0.423 | 0.34 | -0.004 | -0.011 | -0.011 | 0 | 0.29 | 0.24 | 0.36 | 0.6 | 0.34 | 0.32 |
| 4 | 23560044 | rs10517018 | 0.517 | 0.087 | -0.418 | -0.598 | -0.002 | -0.031 | -0.031 | 0 | 0.17 | 0.24 | 0.09 | 0 | 0.07 | 0.04 |
| 4 | 23561662 | rs950950 | -0.08 | 0.749 | -0.342 | -0.054 | 0.031 | 0.207 | 0.207 | 0 | 0.51 | 0.84 | 0.55 | 0.84 | 0.43 | 0.59 |
| 4 | 23563103 | rs4331766 | 0.008 | 0.956 | -0.217 | 0.14 | 0.025 | 0.046 | 0.046 | 0 | 0.5 | 0.62 | 0.44 | 0.74 | 0.42 | 0.47 |
| 4 | 23566334 | rs10017437 | -0.402 | 0.918 | -0.079 | -0.108 | 0.015 | -0.066 | -0.066 | 0.077 | 0.14 | 0.26 | 0.1 | 0.02 | 0.14 | 0.09 |
| 4 | 23572152 | rs13103252 | -1.106 | 0.996 | -0.401 | -0.152 | -0.009 | 0.067 | 0.067 | 0 | 0.54 | 0.84 | 0.5 | 0.33 | 0.54 | 0.54 |
| 4 | 23574876 | rs11727130 | -0.501 | 1.3 | -0.407 | -0.02 | 0.015 | -0.009 | -0.009 | 0 | 0.15 | 0.23 | 0.12 | 0.13 | 0.2 | 0.12 |

| | | | | | | | | | | | | | | | | |
|---|---|---|---|---|---|---|---|---|---|---|---|---|---|---|---|---|
| 4 | 23575883 | rs10009448 | 0.002 | 0.758 | -0.395 | 0.488 | 0.019 | 0.09 | 0.09 | 0 | 0.06 | 0.18 | 0.03 | 0.12 | 0.03 | 0.06 |
| 4 | 23581156 | rs10470807 | -0.15 | 1.033 | -1.241 | -0.08 | -0.013 | 0.068 | 0.068 | 0 | 0.34 | 0.21 | 0.34 | 0.11 | 0.35 | 0.3 |
| 4 | 23587465 | rs10027997 | -0.058 | 0.417 | -1.576 | 0.556 | 0.029 | -0.066 | -0.066 | 4.965 | 0.65 | 0.83 | 0.8 | 0.48 | 0.7 | 0.51 |
| 4 | 23591442 | rs4697039 | 0.339 | 0.026 | -1.279 | 0.063 | 0 | -0.001 | -0.001 | 0 | 0.29 | 0.27 | 0.36 | 0.15 | 0.35 | 0.28 |
| 4 | 23600766 | rs17632146 | 0.516 | -0.013 | -1.367 | 0.034 | -0.133 | 0.023 | 0.023 | 0.342 | 0.27 | 0.04 | 0.34 | 0.15 | 0.35 | 0.27 |
| 4 | 23607172 | rs6810541 | -0.827 | 0.197 | -0.718 | -0.221 | 0.067 | -0.034 | -0.034 | 7.598 | 0.92 | 0.86 | 0.92 | 0.98 | 0.85 | 0.9 |
| 4 | 23613573 | rs7693965 | -0.441 | 0.17 | -0.053 | -0.237 | 0.135 | -0.028 | -0.028 | 0 | 0.91 | 0.73 | 0.9 | 0.98 | 0.85 | 0.91 |
| 4 | 23615048 | rs13121107 | 0.292 | -0.258 | 0.09 | -0.813 | 0.008 | 0.001 | 0.001 | 4.027 | 0.5 | 0.51 | 0.49 | 0.82 | 0.4 | 0.6 |
| 4 | 23616145 | rs12108313 | 1.041 | -0.831 | 0.116 | -0.257 | 0.01 | -0.011 | -0.011 | 0 | 0.35 | 0.37 | 0.33 | 0.57 | 0.33 | 0.5 |
| 4 | 23622531 | rs13121520 | 0.659 | -0.84 | 0.099 | -0.145 | -0.016 | 0.019 | 0.019 | 0 | 0.13 | 0.14 | 0.16 | 0.21 | 0.06 | 0.09 |
| 4 | 23626018 | rs10001239 | 0.053 | -0.875 | 0.108 | 0.032 | 0.004 | 0.004 | 0.004 | 3.803 | 0.49 | 0.51 | 0.49 | 0.7 | 0.39 | 0.53 |
| 4 | 23631034 | rs6845167 | 0.909 | -0.872 | 0.107 | 0.032 | 0.014 | -0.012 | -0.012 | 0 | 0.36 | 0.37 | 0.33 | 0.57 | 0.33 | 0.5 |
| 4 | 23631474 | rs868356 | 0.099 | -0.866 | 0.107 | 0.031 | -0.01 | 0.034 | 0.034 | 4.576 | 0.43 | 0.35 | 0.43 | 0.19 | 0.46 | 0.31 |
| 4 | 23631863 | rs13118539 | 0.659 | -0.839 | 0.086 | 0.043 | -0.018 | 0.019 | 0.019 | 0 | 0.13 | 0.14 | 0.15 | 0.21 | 0.06 | 0.09 |
| 4 | 23632143 | rs16873952 | 0.322 | -0.844 | 0.268 | 0.017 | 0 | -0.004 | -0.004 | 0 | 0.17 | 0.18 | 0.08 | 0.09 | 0.12 | 0.11 |
| 4 | 23638099 | rs13123614 | 1.403 | -0.849 | 0.159 | 0.017 | -0.056 | 0.015 | 0.015 | 0.923 | 0.27 | 0.13 | 0.35 | 0.18 | 0.35 | 0.27 |
| 4 | 23642370 | rs1565535 | 1.008 | -0.866 | 0.238 | 0.033 | 0.016 | -0.02 | -0.02 | 0 | 0.35 | 0.36 | 0.33 | 0.51 | 0.32 | 0.44 |
| 4 | 23663235 | rs2035827 | 1.15 | -0.859 | 0.254 | 0.163 | -0.001 | -0.004 | -0.004 | 0 | 0.62 | 0.5 | 0.68 | 0.67 | 0.67 | 0.71 |
| 4 | 23669223 | rs306050 | 0.607 | -0.767 | 0.207 | 0.286 | 0.017 | -0.018 | -0.018 | 0 | 0.13 | 0.08 | 0.15 | 0.21 | 0.06 | 0.09 |
| 4 | 23683303 | rs10008808 | 0.617 | -0.756 | 0.233 | 0.301 | 0.095 | 0.02 | 0.02 | 0 | 0.09 | 0.23 | 0.09 | 0.11 | 0.15 | 0.15 |
| 4 | 23686819 | rs2054996 | 0.165 | -0.758 | 0.064 | 0.274 | 0.015 | 0.058 | 0.058 | 0 | 0.78 | 0.6 | 0.75 | 0.68 | 0.79 | 0.76 |
| 4 | 23688573 | rs11736033 | 1.259 | -0.8 | -0.136 | 0.153 | -0.067 | 0.044 | 0.044 | 0.86 | 0.26 | 0.07 | 0.34 | 0.15 | 0.34 | 0.25 |
| 4 | 23702558 | rs1491377 | 0.326 | -1.334 | -0.108 | 0.396 | 0.016 | -0.001 | -0.001 | 0 | 0.46 | 0.43 | 0.34 | 0.45 | 0.4 | 0.5 |
| 4 | 23705256 | rs10938956 | 0.907 | -1.423 | -0.039 | 0.359 | -0.019 | 0.016 | 0.016 | 0.319 | 0.33 | 0.23 | 0.42 | 0.21 | 0.48 | 0.33 |
| 4 | 23708557 | rs1491393 | -0.472 | -1.1 | 0.205 | 0.563 | 0.025 | -0.046 | -0.046 | 1.464 | 0.87 | 0.97 | 0.85 | 0.82 | 0.91 | 0.92 |
| 4 | 23712340 | rs13434906 | 1.004 | -1.217 | 0.264 | 0.464 | 0 | -0.04 | -0.04 | 0 | 0.1 | 0.12 | 0.06 | 0.01 | 0.08 | 0.04 |
| 4 | 23712830 | rs2130926 | 0.761 | -1.118 | -0.003 | 0.629 | -0.039 | 0.053 | 0.053 | 0 | 0.79 | 0.65 | 0.76 | 0.65 | 0.87 | 0.84 |
| 4 | 23714895 | rs1021400 | 0.735 | -1.116 | 0.003 | 0.63 | -0.039 | 0.056 | 0.056 | 0 | 0.79 | 0.65 | 0.76 | 0.65 | 0.87 | 0.84 |
| 4 | 23719003 | rs10010697 | 0.044 | -0.86 | -0.332 | 0.71 | 0.002 | -0.001 | -0.001 | 0.501 | 0.86 | 0.87 | 0.84 | 0.74 | 0.92 | 0.86 |
| 4 | 23724215 | rs17572286 | -0.15 | -0.387 | -0.181 | 0.403 | -0.025 | 0.01 | 0.01 | 0 | 0.39 | 0.27 | 0.55 | 0.33 | 0.49 | 0.33 |
| 4 | 23732218 | rs10517023 | -0.351 | -0.77 | -0.088 | 0.352 | 0.004 | 0 | 0 | 0.099 | 0.64 | 0.71 | 0.79 | 0.63 | 0.71 | 0.51 |
| 4 | 23734719 | rs1491389 | 0.414 | -0.692 | 0.188 | 0.633 | 0.014 | 0.016 | 0.016 | 0 | 0.31 | 0.6 | 0.57 | 0.4 | 0.41 | 0.33 |
| 4 | 23736348 | rs1352802 | 1.406 | -0.652 | -0.279 | 0.672 | -0.037 | -0.022 | -0.022 | 0.192 | 0.23 | 0.14 | 0.45 | 0.25 | 0.35 | 0.17 |
| 4 | 23736573 | rs13107733 | 1.216 | -0.781 | -0.311 | 0.576 | 0.001 | -0.002 | -0.002 | 0 | 0.2 | 0.19 | 0.12 | 0.34 | 0.13 | 0.38 |
| 4 | 23738906 | rs7656180 | 1.614 | -0.777 | -0.184 | 0.591 | -0.023 | -0.014 | -0.014 | 0 | 0.22 | 0.13 | 0.45 | 0.23 | 0.31 | 0.17 |
| 4 | 23739736 | rs1027228 | -0.544 | -0.809 | -0.157 | 0.542 | 0.003 | -0.085 | -0.085 | 0 | 0.71 | 0.65 | 0.84 | 0.96 | 0.76 | 0.82 |
| 4 | 23739796 | rs1565537 | 1.164 | -0.683 | -0.586 | 0.648 | -0.001 | 0.001 | 0.001 | 0 | 0.2 | 0.19 | 0.12 | 0.34 | 0.13 | 0.38 |
| 4 | 23743450 | rs16874072 | 0.356 | -0.68 | -0.584 | 0.257 | 0.003 | 0 | 0 | 0 | 0.05 | 0.09 | 0.04 | 0.06 | 0.06 | 0.1 |
| 4 | 23744268 | rs963209 | 1.567 | -0.612 | -0.96 | 0.443 | 0.007 | 0.001 | 0.001 | 0 | 0.2 | 0.08 | 0.17 | 0.2 | 0.22 | 0.14 |
| 4 | 23746582 | rs13142156 | 1.406 | -0.486 | -0.885 | 0.509 | -0.023 | -0.004 | -0.004 | 0 | 0.22 | 0.13 | 0.45 | 0.2 | 0.31 | 0.17 |
| 4 | 23746704 | rs1491384 | 0.267 | -0.513 | -1.401 | 0.497 | 0.038 | 0.009 | 0.009 | 0 | 0.42 | 0.28 | 0.29 | 0.59 | 0.36 | 0.52 |
| 4 | 23759262 | rs974083 | -0.043 | -0.177 | -0.747 | 0.073 | 0.002 | 0.007 | 0.007 | 0 | 0.59 | 0.6 | 0.68 | 0.73 | 0.61 | 0.71 |
| 4 | 23759328 | rs13122003 | 1.188 | -0.142 | -0.795 | 0.078 | -0.011 | 0.007 | 0.007 | 0 | 0.4 | 0.22 | 0.51 | 0.58 | 0.41 | 0.56 |
| 4 | 23759713 | rs4697418 | 1.157 | -0.143 | -0.795 | 0.079 | -0.01 | 0.052 | 0.052 | 0 | 0.41 | 0.22 | 0.55 | 0.67 | 0.42 | 0.56 |
| 4 | 23759757 | rs1491379 | -0.782 | -0.439 | -0.693 | 0.746 | 0.025 | -0.029 | -0.029 | 0 | 0.19 | 0.41 | 0.17 | 0.15 | 0.19 | 0.16 |
| 4 | 23767480 | rs16874121 | -0.492 | -0.251 | -1.443 | 0.975 | 0.012 | 0.02 | 0.02 | 0 | 0.24 | 0.16 | 0.19 | 0.2 | 0.27 | 0.14 |
| 4 | 23770218 | rs885395 | 1.253 | 0.036 | -0.906 | 1.469 | -0.002 | 0.004 | 0.004 | 1.848 | 0.86 | 0.21 | 0.72 | 0.74 | 0.93 | 0.87 |
| 4 | 23781649 | rs2932978 | -0.793 | 0.243 | -1.196 | 1.031 | 0.014 | 0.01 | 0.01 | 0 | 0.38 | 0.71 | 0.5 | 0.31 | 0.34 | 0.38 |
| 4 | 23788341 | rs2970882 | -0.747 | 0.454 | -0.062 | 1.796 | 0.004 | 0.022 | 0.022 | 0 | 0.32 | 0.48 | 0.42 | 0.35 | 0.33 | 0.41 |
| 4 | 23794252 | rs2279525 | -0.577 | 0.39 | 0.589 | 1.66 | -0.001 | -0.031 | -0.031 | 0.987 | 0.28 | 0.43 | 0.33 | 0.2 | 0.28 | 0.28 |
| 4 | 23797000 | rs6821591 | 0.238 | 0.454 | 0.295 | 1.381 | -0.004 | 0.009 | 0.009 | 0.165 | 0.58 | 0.62 | 0.47 | 0.69 | 0.53 | 0.65 |
| 4 | 23798818 | rs768695 | 0.306 | 0.441 | 0.76 | 1.425 | 0.005 | -0.017 | -0.017 | 2.917 | 0.57 | 0.51 | 0.46 | 0.69 | 0.53 | 0.65 |
| 4 | 23801187 | rs12650562 | 0.459 | 0.396 | 0.739 | 1.408 | 0 | 0 | 0 | 0.38 | 0.54 | 0.54 | 0.38 | 0.52 | 0.48 | 0.5 |
| 4 | 23805486 | rs2932965 | -0.643 | 0.243 | 1.219 | 1.224 | -0.016 | -0.006 | -0.006 | 0.033 | 0.81 | 0.87 | 0.88 | 0.78 | 0.81 | 0.87 |
| 4 | 23806778 | rs1873532 | 1.708 | -0.37 | 1.162 | 0.971 | -0.012 | 0.008 | 0.008 | 3.454 | 0.41 | 0.23 | 0.31 | 0.44 | 0.43 | 0.34 |
| 4 | 23815522 | rs3755863 | 1.724 | -0.352 | 1.168 | 0.979 | -0.012 | 0.008 | 0.008 | 3.371 | 0.4 | 0.24 | 0.33 | 0.45 | 0.43 | 0.34 |
| 4 | 23815662 | rs8192678 | 2.204 | -0.672 | 0.858 | 0.531 | -0.074 | -0.084 | -0.084 | 0 | 0.29 | 0.05 | 0.26 | 0.44 | 0.36 | 0.29 |
| 4 | 23815924 | rs2970847 | -1.568 | -0.221 | 1.098 | 0.823 | -0.069 | -0.059 | -0.059 | 0 | 0.85 | 0.99 | 0.9 | 0.78 | 0.82 | 0.87 |
| 4 | 23817027 | rs2970848 | -0.211 | -0.232 | 1.298 | 0.808 | -0.008 | 0.018 | 0.018 | 0.347 | 0.28 | 0.49 | 0.52 | 0.26 | 0.33 | 0.36 |
| 4 | 23818884 | rs2932971 | 0.648 | -0.213 | 1.129 | 0.763 | 0 | -0.005 | -0.005 | 0.409 | 0.21 | 0.22 | 0.38 | 0.08 | 0.26 | 0.2 |
| 4 | 23822328 | rs2932976 | 0.604 | -0.121 | 0.935 | 0.821 | -0.007 | 0.054 | 0.054 | 0.586 | 0.2 | 0.07 | 0.37 | 0.08 | 0.26 | 0.2 |
| 4 | 23823519 | rs2970853 | 0.293 | 0.193 | 1.275 | 1.187 | 0.005 | 0.005 | 0.005 | 0 | 0.75 | 0.7 | 0.83 | 0.72 | 0.77 | 0.79 |
| 4 | 23828708 | rs3774909 | 0.21 | 0.225 | 0.996 | 1.202 | 0.007 | 0.026 | 0.026 | 0 | 0.06 | 0.02 | 0.03 | 0 | 0.04 | 0.04 |
| 4 | 23828934 | rs3774908 | -0.722 | 0.306 | 1.036 | 1.267 | 0.012 | 0.069 | 0.069 | 0 | 0.06 | 0.07 | 0.12 | 0.19 | 0.09 | 0.16 |
| 4 | 23829314 | rs1388332 | -0.746 | 0.306 | 1.012 | 1.268 | 0.023 | 0.093 | 0.093 | 0 | 0.06 | 0.1 | 0.13 | 0.19 | 0.09 | 0.16 |
| 4 | 23836653 | rs6448226 | 1.011 | 0.302 | 0.893 | 0.977 | 0.008 | 0.011 | 0.011 | 0 | 0.62 | 0.59 | 0.7 | 0.53 | 0.66 | 0.54 |
| 4 | 23840088 | rs10938964 | 0.716 | 0.275 | 0.525 | 1.108 | 0.007 | 0.021 | 0.021 | 0 | 0.66 | 0.59 | 0.73 | 0.53 | 0.7 | 0.56 |
| 4 | 23853011 | rs7665116 | -1.408 | 0.082 | 0.493 | 1.124 | -0.032 | 0.063 | 0.063 | 0 | 0.23 | 0.55 | 0.19 | 0.27 | 0.14 | 0.33 |
| 4 | 23855974 | rs7437482 | -1.413 | 1.646 | 0.681 | 1.337 | 0.077 | 0.029 | 0.029 | 0 | 0.22 | 0.39 | 0.2 | 0.26 | 0.37 | 0.27 |

| | | | | | | | | | | | | | | | | |
|---|---|---|---|---|---|---|---|---|---|---|---|---|---|---|---|---|
| 4 | 23863873 | rs4452416 | -0.673 | 2.158 | 0.976 | 1.475 | 0.07 | 0.051 | 0.051 | 0 | 0.07 | 0.14 | 0.1 | 0.12 | 0.14 | 0.05 |
| 4 | 23864412 | rs4235308 | -1.222 | 1.481 | -0.713 | 1.108 | 0.011 | 0 | 0 | 0 | 0.49 | 0.36 | 0.41 | 0.46 | 0.42 | 0.46 |
| 4 | 23864809 | rs4697426 | 0.274 | 1.455 | -0.899 | 1.112 | 0.009 | 0.023 | 0.023 | 0.377 | 0.3 | 0.03 | 0.25 | 0.21 | 0.24 | 0.22 |
| 4 | 23866016 | rs7656250 | -2.087 | 1.373 | -1.079 | 1.058 | -0.007 | -0.06 | -0.06 | 0.022 | 0.69 | 0.92 | 0.69 | 0.63 | 0.74 | 0.68 |
| 4 | 23866339 | rs10213440 | -0.616 | 1.393 | -0.862 | 0.933 | 0.025 | 0.04 | 0.04 | 0 | 0.16 | 0.26 | 0.14 | 0.22 | 0.19 | 0.14 |
| 4 | 23871021 | rs9996943 | -0.907 | 0.83 | -0.478 | 0.92 | 0.018 | -0.005 | -0.005 | 0 | 0.53 | 0.66 | 0.48 | 0.49 | 0.61 | 0.52 |
| 4 | 23875311 | rs6838600 | 0.798 | 0.199 | -0.286 | 0.385 | -0.025 | -0.017 | -0.017 | 1.628 | 0.6 | 0.48 | 0.58 | 0.68 | 0.74 | 0.71 |
| 4 | 23875675 | rs4361373 | -0.495 | 0.225 | -0.783 | 0.277 | -0.024 | -0.025 | -0.025 | 0 | 0.85 | 0.96 | 0.69 | 0.83 | 0.84 | 0.89 |
| 4 | 23880985 | rs17637318 | 0.959 | 1.095 | -0.901 | -0.251 | -0.009 | 0.01 | 0.01 | 0.291 | 0.11 | 0.09 | 0.13 | 0.03 | 0.3 | 0.19 |
| 4 | 23881977 | rs16874276 | 0.863 | 0.541 | -0.851 | -0.281 | 0.006 | -0.007 | -0.007 | 0 | 0.1 | 0.09 | 0.04 | 0 | 0.06 | 0.04 |
| 4 | 23883796 | rs4469064 | 0.388 | 0.413 | -0.336 | 0.045 | -0.038 | 0.012 | 0.012 | 0 | 0.12 | 0.27 | 0.06 | 0.12 | 0.07 | 0.08 |
| 4 | 23884711 | rs3796407 | 1.373 | 0.012 | -1.091 | -0.466 | -0.067 | 0.095 | 0.095 | 0 | 0.26 | 0.03 | 0.18 | 0.07 | 0.3 | 0.1 |
| 4 | 23885568 | rs12500214 | 1.519 | -0.02 | -1.233 | -0.404 | -0.039 | 0.036 | 0.036 | 0 | 0.24 | 0.03 | 0.15 | 0.13 | 0.25 | 0.1 |
| 4 | 23886323 | rs2946385 | 0.036 | -0.339 | -0.968 | -0.46 | -0.001 | -0.002 | -0.002 | 0 | 0.54 | 0.49 | 0.29 | 0.34 | 0.41 | 0.32 |
| 4 | 23889813 | rs2970872 | -0.066 | -0.228 | -1.169 | -0.421 | -0.001 | 0.073 | 0.073 | 0 | 0.56 | 0.71 | 0.42 | 0.21 | 0.56 | 0.38 |
| 4 | 23890782 | rs3774902 | 0.897 | -0.659 | -1.285 | 0.298 | 0.039 | -0.055 | -0.055 | 0 | 0.15 | 0.08 | 0.18 | 0.39 | 0.06 | 0.2 |
| 4 | 23893017 | rs2970870 | 0.238 | -0.347 | -0.925 | 0.034 | -0.015 | 0.037 | 0.037 | 0 | 0.54 | 0.72 | 0.36 | 0.6 | 0.47 | 0.38 |
| 4 | 23895413 | rs1878949 | 0.213 | -0.488 | -0.749 | -0.152 | -0.066 | 0.078 | 0.078 | 0 | 0.24 | 0.49 | 0.17 | 0.38 | 0.14 | 0.22 |
| 4 | 23896474 | rs7657071 | 1.139 | -0.642 | -0.817 | -0.303 | -0.005 | 0.018 | 0.018 | 0 | 0.36 | 0.3 | 0.22 | 0.1 | 0.35 | 0.25 |
| 4 | 23897796 | rs2970863 | -1.331 | -0.326 | -1.116 | -0.339 | 0.002 | -0.159 | -0.159 | 3.686 | 0.75 | 0.96 | 0.54 | 0.51 | 0.79 | 0.69 |
| 4 | 23898430 | rs4568225 | 1.367 | -0.218 | -1.113 | -0.384 | -0.093 | 0.06 | 0.06 | 0.87 | 0.14 | 0.02 | 0.13 | 0.03 | 0.3 | 0.22 |
| 4 | 23900959 | rs2946390 | 0.08 | -0.648 | -0.616 | -0.468 | -0.003 | -0.031 | -0.031 | 0 | 0.54 | 0.51 | 0.39 | 0.13 | 0.68 | 0.48 |
| 4 | 23901275 | rs2970860 | 1.136 | -0.505 | -0.971 | -0.412 | 0.017 | -0.024 | -0.024 | 0.622 | 0.16 | 0.19 | 0.16 | 0.03 | 0.31 | 0.22 |
| 4 | 23901422 | rs10030083 | 0.542 | -1.17 | -1.157 | 0.01 | -0.032 | 0.053 | 0.053 | 0 | 0.21 | 0.33 | 0.13 | 0.38 | 0.11 | 0.22 |
| 4 | 23903867 | rs2946392 | -1.085 | -0.501 | -1.657 | 0.119 | 0.009 | -0.101 | -0.101 | 0 | 0.8 | 0.99 | 0.58 | 0.67 | 0.87 | 0.78 |
| 4 | 23907727 | rs17576576 | 0.897 | 0.217 | -1.201 | -0.347 | 0.037 | -0.061 | -0.061 | 0.491 | 0.23 | 0.31 | 0.2 | 0.03 | 0.42 | 0.19 |
| 4 | 23907928 | rs2946394 | 0.283 | -0.253 | -1.485 | -0.43 | 0.003 | -0.057 | -0.057 | 0 | 0.34 | 0.63 | 0.31 | 0.04 | 0.51 | 0.39 |
| 4 | 23909565 | rs957615 | 0.649 | -0.337 | -1.888 | 0.355 | -0.008 | 0.016 | 0.016 | 0 | 0.2 | 0.2 | 0.13 | 0.38 | 0.12 | 0.29 |
| 4 | 23913579 | rs2946399 | -0.761 | 0.158 | -2.314 | 0.794 | -0.026 | -0.064 | -0.064 | 0 | 0.86 | 0.99 | 0.66 | 0.81 | 0.88 | 0.89 |
| 4 | 23921453 | rs2946403 | 0.094 | 0.11 | -2.244 | 0.548 | -0.021 | -0.064 | -0.064 | 0 | 0.61 | 0.9 | 0.44 | 0.52 | 0.63 | 0.72 |
| 4 | 23925233 | rs2946405 | -0.174 | 0.6 | -1.772 | 1.079 | -0.042 | -0.092 | -0.092 | 0 | 0.78 | 0.95 | 0.66 | 0.69 | 0.75 | 0.8 |
| 4 | 23935053 | rs4697428 | 1.227 | 0.296 | -0.034 | 1.004 | 0.034 | 0.123 | 0.123 | 0 | 0.91 | 0.53 | 0.93 | 0.86 | 0.92 | 0.96 |
| 4 | 23937074 | rs4697429 | -0.295 | 0.224 | -0.39 | 1.106 | -0.012 | -0.101 | -0.101 | 0.32 | 0.21 | 0.07 | 0.37 | 0.34 | 0.21 | 0.17 |
| 4 | 23955370 | rs10017711 | 0.592 | 0.197 | -0.594 | 0.549 | -0.042 | 0.061 | 0.061 | 0.093 | 0.34 | 0.2 | 0.25 | 0.1 | 0.45 | 0.35 |
| 4 | 23957254 | rs10517026 | 0.883 | -0.147 | -0.759 | 1.054 | 0.003 | -0.036 | -0.036 | 0.557 | 0.16 | 0.07 | 0.19 | 0.3 | 0.15 | 0.2 |
| 4 | 23959089 | rs10938973 | 1.342 | 0.175 | -0.329 | 0.946 | -0.023 | 0.039 | 0.039 | 0.114 | 0.31 | 0.2 | 0.23 | 0.1 | 0.38 | 0.33 |
| 4 | 23962031 | rs9291455 | -1.423 | 0.087 | -0.152 | 1.214 | -0.003 | 0.132 | 0.132 | 0 | 0.81 | 0.54 | 0.66 | 0.72 | 0.84 | 0.78 |
| 4 | 23962061 | rs13111174 | -0.173 | 0.017 | -0.06 | 1.152 | 0.012 | 0.02 | 0.02 | 0 | 0.44 | 0.24 | 0.39 | 0.48 | 0.39 | 0.42 |
| 4 | 23962961 | rs10020457 | 1.412 | 0.069 | -0.132 | 0.991 | -0.021 | 0.039 | 0.039 | 0.017 | 0.3 | 0.19 | 0.21 | 0.1 | 0.36 | 0.32 |
| 4 | 23966042 | rs2324237 | 1.856 | 0.013 | -0.288 | 1.196 | -0.042 | -0.064 | -0.064 | 0 | 0.15 | 0.02 | 0.3 | 0.18 | 0.16 | 0.15 |
| 4 | 23968295 | rs10517031 | 0.879 | 0.003 | -0.258 | 1.183 | -0.002 | -0.053 | -0.053 | 0.787 | 0.16 | 0.05 | 0.19 | 0.3 | 0.15 | 0.2 |
| 4 | 23972257 | rs10470808 | -0.217 | 0.01 | 0.093 | 1.178 | 0.013 | 0.016 | 0.016 | 0 | 0.46 | 0.31 | 0.4 | 0.49 | 0.4 | 0.41 |
| 4 | 23972751 | rs2324233 | 0.277 | -0.06 | 0.443 | 1.208 | -0.01 | 0.066 | 0.066 | 0 | 0.24 | 0.36 | 0.36 | 0.41 | 0.23 | 0.26 |
| 4 | 23976060 | rs1316862 | -0.5 | 0.028 | 0.304 | 1.007 | 0.014 | -0.02 | -0.02 | 0.866 | 0.62 | 0.57 | 0.71 | 0.79 | 0.49 | 0.71 |
| 4 | 23977778 | rs1316861 | 0.158 | -0.042 | 0.351 | 0.984 | 0.002 | 0.002 | 0.002 | 0 | 0.51 | 0.46 | 0.63 | 0.28 | 0.62 | 0.5 |
| 4 | 23986967 | rs1714638 | 0.814 | -0.012 | 0.331 | 0.947 | 0.028 | -0.054 | -0.054 | 6.175 | 0.33 | 0.46 | 0.26 | 0.19 | 0.43 | 0.26 |
| 4 | 23990989 | rs609893 | 1.332 | -0.197 | 0.211 | 0.916 | 0.022 | 0.043 | 0.043 | 0 | 0.25 | 0.13 | 0.17 | 0.18 | 0.17 | 0.2 |
| 4 | 23991100 | rs16874486 | 2.046 | -0.177 | 0.13 | 0.985 | -0.043 | -0.085 | -0.085 | 4.832 | 0.15 | 0.02 | 0.35 | 0.21 | 0.16 | 0.15 |
| 4 | 23994155 | rs6833157 | 0.235 | -0.186 | 0.237 | 1.065 | -0.001 | -0.002 | -0.002 | 0 | 0.22 | 0.22 | 0.23 | 0.32 | 0.23 | 0.21 |
| 4 | 23996375 | rs16874509 | 1.107 | -0.326 | 0.035 | 1.073 | -0.004 | -0.046 | -0.046 | 0.103 | 0.2 | 0.07 | 0.32 | 0.22 | 0.17 | 0.27 |
| 4 | 23996637 | rs597082 | 0.33 | -0.308 | 0.1 | 1.039 | -0.002 | 0.004 | 0.004 | 0 | 0.91 | 0.88 | 0.95 | 0.84 | 0.93 | 0.86 |
| 4 | 23998787 | rs637282 | 1.229 | -0.288 | 0.115 | 0.982 | 0.016 | 0.028 | 0.028 | 0 | 0.25 | 0.17 | 0.18 | 0.19 | 0.16 | 0.2 |
| 4 | 24000480 | rs10026136 | 1.17 | -0.293 | 0.122 | 0.977 | 0.016 | 0 | 0 | 0.154 | 0.26 | 0.17 | 0.18 | 0.26 | 0.16 | 0.26 |
| 4 | 24001226 | rs668672 | 1.144 | -0.293 | 0.111 | 0.949 | 0.018 | -0.001 | -0.001 | 0 | 0.26 | 0.17 | 0.15 | 0.26 | 0.17 | 0.26 |
| 4 | 24002981 | rs6858449 | 1.074 | 0.17 | 0.325 | 0.798 | 0 | -0.034 | -0.034 | 0 | 0.29 | 0.31 | 0.23 | 0.05 | 0.4 | 0.24 |
| 4 | 24009437 | rs16874564 | 0.46 | 0.149 | 0.347 | 0.931 | 0.012 | 0.05 | 0.05 | 0 | 0.2 | 0.38 | 0.27 | 0.31 | 0.19 | 0.19 |
| 4 | 24010844 | rs609501 | -0.715 | 0.092 | 0.172 | 0.953 | 0.009 | -0.074 | -0.074 | 0 | 0.76 | 0.86 | 0.66 | 0.56 | 0.77 | 0.65 |
| 4 | 24011608 | rs7672552 | -0.1 | 0.101 | 0.164 | 0.901 | 0.03 | 0.069 | 0.069 | 0 | 0.48 | 0.71 | 0.45 | 0.37 | 0.58 | 0.43 |
| 4 | 24014210 | rs13142815 | -0.091 | 0.2 | 0.251 | 0.93 | 0.011 | 0.017 | 0.017 | 0 | 0.23 | 0.31 | 0.28 | 0.34 | 0.27 | 0.21 |
| 4 | 24016473 | rs2191598 | -0.278 | 0.087 | -0.156 | 0.989 | 0.037 | -0.018 | -0.018 | 0.521 | 0.53 | 0.79 | 0.46 | 0.47 | 0.6 | 0.6 |
| 4 | 24026356 | rs694917 | 0.821 | 0.252 | 0.155 | 1.087 | -0.006 | 0.005 | 0.005 | 0 | 0.2 | 0.23 | 0.33 | 0.21 | 0.17 | 0.25 |
| 4 | 24026481 | rs648674 | -0.457 | 0.29 | 0.134 | 1.05 | -0.014 | -0.017 | -0.017 | 1.836 | 0.38 | 0.25 | 0.59 | 0.53 | 0.43 | 0.37 |
| 4 | 24028621 | rs630902 | -0.153 | 0.063 | 0.202 | 1.121 | -0.002 | -0.021 | -0.021 | 0 | 0.35 | 0.24 | 0.56 | 0.53 | 0.35 | 0.35 |
| 4 | 24030761 | rs6858568 | -0.19 | 0.049 | 0.247 | 1.112 | -0.001 | -0.024 | -0.024 | 0.121 | 0.36 | 0.23 | 0.55 | 0.53 | 0.36 | 0.35 |
| 4 | 24034708 | rs10008679 | 0.888 | -0.397 | 0.259 | 1.169 | 0.01 | 0.072 | 0.072 | 0 | 0.25 | 0.05 | 0.14 | 0.17 | 0.17 | 0.21 |
| 4 | 24035754 | rs618283 | 1.242 | -0.475 | 0.462 | 1.525 | 0.001 | -0.006 | -0.006 | 0.011 | 0.17 | 0.13 | 0.29 | 0.2 | 0.16 | 0.16 |
| 4 | 24040555 | rs10025406 | 0.684 | -0.573 | 0.412 | 1.3 | 0.047 | 0.072 | 0.072 | 0 | 0.48 | 0.33 | 0.37 | 0.35 | 0.6 | 0.45 |
| 4 | 24041100 | rs650356 | -0.257 | -0.395 | 0.049 | 1.027 | 0.006 | -0.002 | -0.002 | 0 | 0.72 | 0.73 | 0.78 | 0.66 | 0.8 | 0.69 |
| 4 | 24041241 | rs6820543 | 0.964 | -0.111 | -0.426 | 0.785 | 0.027 | 0.046 | 0.046 | 0 | 0.21 | 0.06 | 0.15 | 0.17 | 0.13 | 0.2 |

| | | | | | | | | | | | | | | | | |
|---|---|---|---|---|---|---|---|---|---|---|---|---|---|---|---|---|
| 4 | 24047613 | rs4697435 | 0.247 | 0.126 | 0.018 | 1.01 | 0.026 | 0.02 | 0.02 | 0 | 0.38 | 0.19 | 0.45 | 0.35 | 0.29 | 0.35 |
| 4 | 24048711 | rs625365 | -0.056 | 0.281 | 0.537 | 1.271 | 0.004 | 0.069 | 0.069 | 0 | 0.93 | 0.85 | 0.94 | 0.84 | 0.91 | 0.96 |
| 4 | 24055017 | rs626610 | -0.017 | -0.142 | 0.894 | 1.179 | 0.038 | -0.008 | -0.008 | 0.065 | 0.54 | 0.66 | 0.55 | 0.5 | 0.64 | 0.62 |
| 4 | 24057548 | rs584374 | 0.378 | 0.811 | 0.806 | 0.817 | 0.132 | 0.012 | 0.012 | 0 | 0.15 | 0.47 | 0.17 | 0.1 | 0.23 | 0.11 |
| 4 | 24060940 | rs647536 | 0.354 | 0.857 | 0.878 | 0.76 | 0.005 | 0.004 | 0.004 | 0 | 0.4 | 0.32 | 0.37 | 0.37 | 0.37 | 0.45 |
| 4 | 24064404 | rs16874916 | 0.321 | 1.049 | 0.908 | 0.715 | 0.152 | -0.105 | -0.105 | 0 | 0.15 | 0.38 | 0.17 | 0 | 0.28 | 0.09 |
| 4 | 24069770 | rs3815420 | -0.116 | 0.608 | 0.499 | 0.851 | 0 | 0.002 | 0.002 | 0 | 0.89 | 0.89 | 0.95 | 0.82 | 0.9 | 0.87 |
| 4 | 24072063 | rs2215488 | -0.756 | 0.613 | 0.466 | 0.854 | 0 | -0.01 | -0.01 | 0.201 | 0.93 | 0.97 | 0.98 | 0.82 | 0.93 | 0.88 |
| 4 | 24074065 | rs10938978 | 1.02 | 0.47 | 0.363 | 0.75 | 0.032 | 0.002 | 0.002 | 0 | 0.33 | 0.1 | 0.39 | 0.33 | 0.24 | 0.3 |
| 4 | 24092877 | rs16874990 | -0.655 | 0.541 | 1.112 | 0.856 | -0.017 | 0.159 | 0.159 | 0 | 0.14 | 0.45 | 0.06 | 0.25 | 0.08 | 0.16 |
| 4 | 24105609 | rs17188642 | 0.547 | 0.863 | 1.171 | 0.618 | 0.045 | -0.056 | -0.056 | 0 | 0.14 | 0.18 | 0.16 | 0 | 0.28 | 0.1 |
| 4 | 24114572 | rs6848087 | -0.732 | 0.349 | 0.823 | 0.569 | -0.005 | -0.046 | -0.046 | 0 | 0.7 | 0.65 | 0.7 | 0.89 | 0.67 | 0.66 |
| 4 | 24115074 | rs13109143 | -0.685 | 0.077 | 0.416 | 0.764 | -0.006 | 0.025 | 0.025 | 0 | 0.25 | 0.36 | 0.25 | 0.59 | 0.21 | 0.29 |
| 4 | 24118165 | rs7680610 | -0.26 | 0.051 | 0.098 | 0.728 | 0.013 | -0.043 | -0.043 | 0 | 0.63 | 0.82 | 0.64 | 0.57 | 0.63 | 0.56 |
| 4 | 24125887 | rs10489098 | -0.799 | -0.035 | -0.17 | 0.158 | 0.027 | 0.036 | 0.036 | 0 | 0.61 | 0.73 | 0.67 | 0.77 | 0.74 | 0.59 |
| 4 | 24128287 | rs2215492 | -0.469 | 0.232 | -0.323 | 0.435 | 0.019 | 0.009 | 0.009 | 0 | 0.52 | 0.63 | 0.63 | 0.58 | 0.7 | 0.48 |
| 4 | 24130517 | rs12644722 | 0.3 | 0.691 | -0.431 | 0.681 | 0.001 | -0.017 | -0.017 | 0 | 0.06 | 0.02 | 0.01 | 0.17 | 0.04 | 0.12 |
| 4 | 24133750 | rs929530 | 0.697 | 1.548 | -0.023 | 0.232 | 0.08 | -0.057 | -0.057 | 0.85 | 0.1 | 0.2 | 0.12 | 0 | 0.22 | 0.04 |
| 4 | 24134431 | rs12512863 | -0.739 | 1.081 | -0.201 | 0.331 | -0.001 | 0 | 0 | 0 | 0.49 | 0.51 | 0.61 | 0.52 | 0.66 | 0.44 |
| 4 | 24135578 | rs16875100 | -0.06 | 0.849 | -0.439 | -0.048 | -0.007 | 0.02 | 0.02 | 0.077 | 0.26 | 0.2 | 0.13 | 0.08 | 0.27 | 0.17 |
| 4 | 24138139 | rs7663527 | -0.561 | 0.636 | -0.166 | 0.583 | 0 | -0.001 | -0.001 | 0 | 0.32 | 0.65 | 0.4 | 0.33 | 0.23 | 0.35 |
| 4 | 24141403 | rs6841821 | -0.652 | 0.595 | 0.122 | 0.564 | 0.009 | -0.002 | -0.002 | 0 | 0.33 | 0.68 | 0.42 | 0.33 | 0.23 | 0.36 |
| 4 | 24144512 | rs1705098 | -0.89 | 0.596 | 0.881 | 0.589 | -0.026 | -0.039 | -0.039 | 0 | 0.35 | 0.49 | 0.39 | 0.25 | 0.28 | 0.37 |
| 4 | 24150414 | rs13110084 | 0.033 | 0.575 | 1.18 | 0.483 | 0.006 | -0.017 | -0.017 | 0 | 0.07 | 0.09 | 0.04 | 0.02 | 0.09 | 0.06 |
| 4 | 24151604 | rs1267016 | -1.076 | 0.51 | 1.191 | 0.578 | 0.02 | -0.005 | -0.005 | 0.071 | 0.34 | 0.7 | 0.42 | 0.35 | 0.23 | 0.36 |
| 4 | 24154002 | rs1267012 | -1.024 | 1.006 | 0.779 | 1.062 | -0.045 | -0.052 | -0.052 | 0.772 | 0.91 | 0.99 | 0.88 | 0.71 | 0.81 | 0.89 |
| 4 | 24156429 | rs10213261 | -1.12 | -0.377 | 1.961 | 1.098 | 0.042 | -0.02 | -0.02 | 0 | 0.09 | 0.26 | 0.05 | 0.03 | 0.09 | 0.06 |
| 4 | 24171888 | rs10517034 | -0.853 | -0.353 | 1.381 | 1.214 | 0.011 | 0.001 | 0.001 | 0 | 0.23 | 0.19 | 0.34 | 0.24 | 0.17 | 0.3 |
| 4 | 24176596 | rs6839666 | -1.789 | -0.342 | 1.597 | 1.404 | -0.02 | 0.035 | 0.035 | 0.046 | 0.31 | 0.54 | 0.41 | 0.41 | 0.25 | 0.35 |
| 4 | 24178227 | rs16875239 | -0.287 | -0.306 | 1.121 | 1.471 | 0.034 | -0.032 | -0.032 | 0 | 0.18 | 0.02 | 0.28 | 0.22 | 0.12 | 0.28 |
| 4 | 24190844 | rs7676251 | -1.816 | 0.496 | 2.193 | 1.11 | 0.042 | -0.022 | -0.022 | 0 | 0.09 | 0.24 | 0.06 | 0.03 | 0.11 | 0.07 |
| 4 | 24193409 | rs6822382 | 2.396 | 0.408 | 2.646 | 1.225 | 0.004 | 0.078 | 0.078 | 3.666 | 0.83 | 0.35 | 0.82 | 0.75 | 0.84 | 0.84 |
| 4 | 24195161 | rs6848086 | -1.446 | 0.498 | 2.653 | 1.202 | 0.045 | 0.008 | 0.008 | 0 | 0.07 | 0.24 | 0.05 | 0.03 | 0.05 | 0.04 |
| 4 | 24195219 | rs1877089 | 1.555 | 0.469 | 2.591 | 1.204 | 0.01 | -0.006 | -0.006 | 0.008 | 0.9 | 0.8 | 0.89 | 0.94 | 0.89 | 0.88 |
| 4 | 24198653 | rs12647637 | -0.612 | 0.506 | 2.577 | 1.227 | 0.005 | 0.007 | 0.007 | 0 | 0.06 | 0.09 | 0.07 | 0.06 | 0.08 | 0.11 |
| 4 | 24199845 | rs1397490 | 2.763 | 0.488 | 2.59 | 1.263 | 0.022 | 0.099 | 0.099 | 3.149 | 0.82 | 0.42 | 0.8 | 0.74 | 0.83 | 0.84 |
| 4 | 24205824 | rs13131294 | 2.602 | 0.496 | 2.558 | 1.248 | 0.041 | 0.042 | 0.042 | 2.029 | 0.83 | 0.51 | 0.82 | 0.83 | 0.83 | 0.84 |
| 4 | 24207765 | rs1995917 | 1.754 | 0.52 | 2.486 | 1.255 | 0.03 | 0.007 | 0.007 | 0.005 | 0.92 | 0.71 | 0.95 | 0.96 | 0.95 | 0.98 |
| 4 | 24212477 | rs9684046 | 2.675 | 0.499 | 2.457 | 1.277 | 0.022 | 0.102 | 0.102 | 3.133 | 0.82 | 0.43 | 0.8 | 0.74 | 0.83 | 0.83 |
| 4 | 24213123 | rs6837980 | -1.964 | 0.498 | 2.498 | 1.094 | 0.081 | 0.172 | 0.172 | 0.366 | 0.08 | 0.43 | 0.09 | 0.12 | 0.06 | 0.04 |
| 4 | 24215638 | rs4697051 | 1.731 | 0.584 | 1.705 | 1.029 | 0.039 | -0.014 | -0.014 | 0 | 0.9 | 0.68 | 0.93 | 0.96 | 0.91 | 0.96 |
| 4 | 24223553 | rs4697052 | 1.271 | 0.504 | 1.085 | 1.177 | 0.033 | 0.146 | 0.146 | 1.4 | 0.89 | 0.63 | 0.86 | 0.84 | 0.89 | 0.89 |
| 4 | 24225918 | rs6820131 | 0.121 | -0.055 | -0.118 | 0.782 | 0.06 | 0.003 | 0.003 | 0.059 | 0.51 | 0.25 | 0.44 | 0.51 | 0.6 | 0.53 |
| 4 | 24229043 | rs10016733 | 0.149 | -0.276 | 0.072 | 0.737 | 0.035 | 0.032 | 0.032 | 0 | 0.5 | 0.37 | 0.48 | 0.64 | 0.61 | 0.53 |
| 4 | 24230323 | rs6448251 | -0.013 | -0.341 | 0.064 | 0.751 | 0.04 | 0.009 | 0.009 | 0 | 0.49 | 0.36 | 0.46 | 0.46 | 0.6 | 0.51 |
| 4 | 24232043 | rs6813754 | -0.356 | -0.644 | 0.297 | 0.512 | 0.002 | -0.024 | -0.024 | 0.104 | 0.15 | 0.18 | 0.18 | 0.05 | 0.18 | 0.13 |
| 4 | 24232169 | rs10938983 | 0.961 | -0.724 | 0.582 | 0.532 | 0.021 | 0.048 | 0.048 | 0 | 0.05 | 0.17 | 0.08 | 0.06 | 0.05 | 0.12 |
| 4 | 24233053 | rs4235311 | 0.705 | -0.762 | 0.567 | 0.436 | 0.01 | -0.09 | -0.09 | 0 | 0.79 | 0.58 | 0.77 | 0.95 | 0.78 | 0.87 |
| 4 | 24233227 | rs4235312 | 0.473 | -0.724 | 0.299 | 0.609 | 0.041 | 0.01 | 0.01 | 0.007 | 0.5 | 0.36 | 0.45 | 0.47 | 0.6 | 0.51 |
| 4 | 24234074 | rs17641468 | 0.697 | 0.148 | -0.255 | -0.099 | -0.024 | 0.187 | 0.187 | 0 | 0.46 | 0.17 | 0.34 | 0.17 | 0.48 | 0.39 |
| 4 | 24235281 | rs1510784 | 0.587 | -0.091 | 0.048 | 0.012 | 0.011 | -0.049 | -0.049 | 1.635 | 0.83 | 0.45 | 0.81 | 0.94 | 0.84 | 0.79 |
| 4 | 24237854 | rs10016565 | 0.652 | 0.191 | -0.321 | 0.114 | -0.025 | 0.006 | 0.006 | 0 | 0.47 | 0.17 | 0.38 | 0.44 | 0.47 | 0.4 |
| 4 | 24238470 | rs7662477 | 0.361 | 0.275 | -0.071 | 0.061 | 0.019 | -0.006 | -0.006 | 0 | 0.52 | 0.28 | 0.46 | 0.5 | 0.52 | 0.56 |
| 4 | 24238868 | rs6448252 | 0.732 | 0.287 | -0.195 | -0.016 | -0.027 | 0.189 | 0.189 | 0 | 0.46 | 0.17 | 0.36 | 0.17 | 0.47 | 0.38 |
| 4 | 24240273 | rs10023214 | 0.773 | 0.321 | -0.236 | -0.115 | -0.017 | -0.012 | -0.012 | 0 | 0.2 | 0.14 | 0.15 | 0.01 | 0.29 | 0.15 |
| 4 | 24241504 | rs2324421 | 0.194 | 0.345 | -0.115 | 0.042 | 0.011 | -0.026 | -0.026 | 0 | 0.36 | 0.26 | 0.43 | 0.49 | 0.37 | 0.44 |
| 4 | 24242223 | rs7666134 | 0.572 | 0.183 | 0.068 | -0.008 | -0.006 | -0.047 | -0.047 | 0.988 | 0.1 | 0.15 | 0.05 | 0 | 0.07 | 0.05 |
| 4 | 24245365 | rs7673666 | 0.456 | 0.114 | -0.724 | -0.066 | -0.015 | 0.043 | 0.043 | 0 | 0.29 | 0.37 | 0.33 | 0.44 | 0.27 | 0.4 |
| 4 | 24246136 | rs7679088 | -0.606 | 0.086 | -0.771 | -0.167 | 0.016 | -0.009 | -0.009 | 0 | 0.36 | 0.39 | 0.3 | 0.31 | 0.46 | 0.25 |
| 4 | 24246892 | rs7663508 | 0.902 | -0.118 | -0.693 | -0.07 | 0.038 | -0.017 | -0.017 | 0 | 0.36 | 0.27 | 0.3 | 0.47 | 0.26 | 0.3 |
| 4 | 24253157 | rs12642909 | 0.643 | 0.011 | -0.726 | -0.005 | -0.003 | 0.033 | 0.033 | 0 | 0.09 | 0.09 | 0.06 | 0.3 | 0.07 | 0.06 |
| 4 | 24255842 | rs6448254 | -0.4 | 0.14 | -0.791 | -0.144 | -0.011 | 0 | 0 | 0.103 | 0.32 | 0.25 | 0.27 | 0.31 | 0.47 | 0.24 |
| 4 | 24256292 | rs7688742 | -1.842 | 0.014 | -0.839 | -0.217 | 0.048 | 0.092 | 0.092 | 0 | 0.7 | 0.82 | 0.72 | 0.85 | 0.78 | 0.76 |
| 4 | 24256496 | rs7356453 | 0.797 | -0.022 | -0.805 | -0.116 | -0.048 | 0.073 | 0.073 | 0 | 0.27 | 0.4 | 0.3 | 0.43 | 0.2 | 0.28 |
| 4 | 24259055 | rs1995919 | 0.497 | 0.291 | -1.301 | -0.099 | -0.008 | 0.003 | 0.003 | 0 | 0.23 | 0.17 | 0.22 | 0.16 | 0.39 | 0.19 |
| 4 | 24262089 | rs17581643 | -0.199 | 0.278 | -1.297 | -0.095 | -0.042 | 0.033 | 0.033 | 0.202 | 0.31 | 0.17 | 0.26 | 0.16 | 0.45 | 0.24 |
| 4 | 24263196 | rs1969192 | -0.65 | 0.193 | -1.3 | -0.098 | -0.019 | 0.065 | 0.065 | 0 | 0.36 | 0.17 | 0.32 | 0.16 | 0.51 | 0.25 |
| 4 | 24263853 | rs6852227 | 1.016 | 0.245 | -1.24 | -0.153 | -0.007 | 0.012 | 0.012 | 0 | 0.09 | 0.14 | 0.05 | 0.14 | 0.06 | 0.06 |
| 4 | 24264014 | rs10938987 | -1.325 | 0.099 | -1.324 | -0.166 | 0.013 | 0.002 | 0.002 | 0 | 0.45 | 0.36 | 0.4 | 0.4 | 0.58 | 0.47 |

| | | | | | | | | | | | | | | | | |
|---|---|---|---|---|---|---|---|---|---|---|---|---|---|---|---|---|
| 4 | 24264580 | rs16875458 | 0.488 | 0.188 | -1.438 | -0.209 | -0.01 | -0.016 | -0.016 | 0 | 0.11 | 0.04 | 0.14 | 0.11 | 0.12 | 0.23 |
| 4 | 24264901 | rs17581877 | 0.201 | 0.171 | -1.384 | -0.242 | -0.012 | -0.026 | -0.026 | 0 | 0.25 | 0.19 | 0.21 | 0.01 | 0.4 | 0.18 |
| 4 | 24267149 | rs17648420 | -0.726 | 0.142 | -1.411 | -0.162 | -0.002 | -0.001 | -0.001 | 0 | 0.37 | 0.33 | 0.36 | 0.35 | 0.52 | 0.42 |
| 4 | 24267644 | rs16875481 | 2.093 | -0.018 | -1.693 | -0.214 | 0.031 | 0.015 | 0.015 | 0 | 0.26 | 0.03 | 0.22 | 0.21 | 0.18 | 0.25 |
| 4 | 24268934 | rs1511358 | -0.333 | -0.167 | -1.53 | -0.166 | -0.005 | 0 | 0 | 1.859 | 0.38 | 0.53 | 0.47 | 0.55 | 0.32 | 0.5 |
| 4 | 24269622 | rs4697446 | 0.9 | -0.072 | -1.494 | -0.121 | 0.03 | 0.019 | 0.019 | 0 | 0.53 | 0.6 | 0.56 | 0.64 | 0.39 | 0.53 |
| 4 | 24271684 | rs11945373 | 0.719 | 0.011 | -1.394 | -0.095 | -0.044 | 0.101 | 0.101 | 0 | 0.26 | 0.45 | 0.33 | 0.44 | 0.2 | 0.28 |
| 4 | 24271794 | rs11931856 | -0.19 | 0.384 | -1.582 | -0.412 | 0.027 | -0.02 | -0.02 | 0.137 | 0.73 | 0.65 | 0.74 | 0.84 | 0.57 | 0.82 |
| 4 | 24275114 | rs4697447 | -0.634 | 0.654 | -1.758 | -0.543 | 0.002 | -0.031 | -0.031 | 0 | 0.88 | 0.83 | 0.89 | 1 | 0.87 | 0.98 |
| 4 | 24276706 | rs7685185 | 0.48 | 0.738 | -1.856 | -0.529 | -0.018 | -0.007 | -0.007 | 0 | 0.21 | 0.13 | 0.2 | 0.01 | 0.35 | 0.17 |
| 4 | 24277125 | rs17582609 | 1.093 | 0.109 | -2.005 | 0.099 | 0.034 | 0.04 | 0.04 | 0 | 0.27 | 0.06 | 0.25 | 0.21 | 0.2 | 0.25 |
| 4 | 24280482 | rs7659588 | 0.644 | 0.548 | -1.034 | 0.66 | -0.001 | -0.001 | -0.001 | 0.703 | 0.78 | 0.75 | 0.84 | 0.76 | 0.8 | 0.92 |
| 4 | 24283000 | rs6448255 | 0.478 | 0.47 | -1.129 | 0.923 | 0.002 | -0.004 | -0.004 | 0.019 | 0.65 | 0.36 | 0.65 | 0.42 | 0.71 | 0.71 |
| 4 | 24298140 | rs12646616 | 0.166 | 0.562 | -1.535 | 0.771 | -0.018 | 0.025 | 0.025 | 4.98 | 0.29 | 0.1 | 0.21 | 0.25 | 0.29 | 0.36 |
| 4 | 24302081 | rs17582833 | 0.172 | 0.548 | -1.369 | 0.769 | -0.01 | 0.027 | 0.027 | 5.127 | 0.29 | 0.13 | 0.21 | 0.25 | 0.29 | 0.34 |
| 4 | 24302785 | rs1849098 | 0.522 | 0.529 | -1.352 | 0.461 | 0.002 | 0.018 | 0.018 | 0 | 0.28 | 0.23 | 0.31 | 0.17 | 0.27 | 0.22 |
| 4 | 24303151 | rs1464008 | 0.308 | 0.369 | -0.544 | 0.708 | -0.007 | -0.013 | -0.013 | 0 | 0.3 | 0.35 | 0.32 | 0.22 | 0.28 | 0.33 |
| 4 | 24305034 | rs1949112 | -0.233 | 0.328 | -0.068 | 1.292 | -0.006 | -0.003 | -0.003 | 0 | 0.68 | 0.73 | 0.66 | 0.63 | 0.63 | 0.72 |
| 4 | 24307070 | rs7685773 | -0.273 | 0.326 | -0.069 | 1.291 | -0.006 | -0.003 | -0.003 | 0 | 0.68 | 0.73 | 0.67 | 0.63 | 0.64 | 0.73 |
| 4 | 24310076 | rs6815672 | -0.341 | 0.251 | 0.003 | 1.57 | -0.056 | 0.046 | 0.046 | 0.027 | 0.7 | 0.91 | 0.76 | 0.79 | 0.65 | 0.73 |
| 4 | 24315694 | rs2063409 | -1.102 | 0.273 | 0.141 | 1.215 | -0.01 | -0.028 | -0.028 | 0 | 0.29 | 0.48 | 0.17 | 0.22 | 0.25 | 0.3 |
| 4 | 24316984 | rs6448258 | 1.646 | 0.302 | -0.174 | 1.233 | -0.024 | -0.051 | -0.051 | 2.889 | 0.33 | 0.12 | 0.39 | 0.37 | 0.37 | 0.32 |
| 4 | 24317057 | rs9998315 | -0.862 | 0.401 | 0.239 | 1.23 | -0.004 | -0.014 | -0.014 | 0 | 0.28 | 0.36 | 0.16 | 0.23 | 0.25 | 0.3 |
| 4 | 24317449 | rs6827997 | -1.473 | 0.396 | 0.36 | 1.256 | -0.003 | 0.011 | 0.011 | 1.246 | 0.41 | 0.67 | 0.27 | 0.36 | 0.41 | 0.49 |
| 4 | 24322900 | rs9994454 | -1.595 | 0.488 | 0.357 | 1.197 | -0.008 | -0.002 | -0.002 | 3.77 | 0.42 | 0.68 | 0.36 | 0.48 | 0.44 | 0.5 |
| 4 | 24324261 | rs9685879 | -1.492 | 0.487 | 0.267 | 1.2 | 0.005 | -0.01 | -0.01 | 0 | 0.42 | 0.55 | 0.36 | 0.41 | 0.44 | 0.48 |
| 4 | 24335226 | rs16835 | 1.027 | 0.663 | 0.615 | 0.968 | 0.01 | 0.02 | 0.02 | 1.589 | 0.76 | 0.39 | 0.75 | 0.7 | 0.74 | 0.73 |
| 4 | 24340511 | rs11938023 | 0.06 | 0.551 | 0.421 | 0.661 | 0.001 | -0.017 | -0.017 | 0 | 0.16 | 0.18 | 0.13 | 0.08 | 0.18 | 0.19 |
| 4 | 24342273 | rs17649407 | -0.492 | 0.594 | 0.287 | 0.76 | -0.013 | 0.017 | 0.017 | 1.507 | 0.18 | 0.14 | 0.11 | 0.13 | 0.22 | 0.24 |
| 4 | 24353666 | rs4529036 | -0.32 | 0.67 | 0.281 | 0.788 | -0.019 | 0.014 | 0.014 | 0.115 | 0.16 | 0.14 | 0.12 | 0.13 | 0.22 | 0.23 |
| 4 | 24358657 | rs10024907 | 0.388 | 0.731 | 0.254 | 0.773 | -0.009 | -0.004 | -0.004 | 0 | 0.08 | 0.07 | 0.07 | 0.12 | 0.17 | 0.16 |
| 4 | 24358882 | rs10002552 | -1.49 | 0.57 | 0.457 | 0.629 | 0.004 | -0.014 | -0.014 | 0 | 0.33 | 0.36 | 0.25 | 0.23 | 0.4 | 0.42 |
| 4 | 24364497 | rs17650027 | 1.12 | -0.21 | 0.245 | 0.592 | 0.011 | 0.045 | 0.045 | 0.516 | 0.24 | 0.12 | 0.23 | 0.11 | 0.2 | 0.2 |
| 4 | 24368837 | rs10805250 | 0.352 | -0.226 | 0.188 | 0.582 | 0 | -0.01 | -0.01 | 3.047 | 0.16 | 0.16 | 0.13 | 0.1 | 0.17 | 0.18 |
| 4 | 24369452 | rs10007635 | -1.405 | -0.093 | 0.337 | 0.686 | 0.003 | -0.016 | -0.016 | 1.142 | 0.33 | 0.43 | 0.29 | 0.33 | 0.37 | 0.43 |
| 4 | 24373046 | rs6848313 | 1.131 | 0.061 | 0.107 | 0.848 | -0.037 | -0.057 | -0.057 | 1.476 | 0.39 | 0.24 | 0.38 | 0.46 | 0.43 | 0.36 |
| 4 | 24377777 | rs10018759 | -0.618 | 0.142 | 0.132 | 0.735 | 0.002 | -0.008 | -0.008 | 1.514 | 0.16 | 0.2 | 0.15 | 0.13 | 0.19 | 0.23 |
| 4 | 24378477 | rs3857105 | 0.165 | 0.242 | 0.131 | 0.738 | 0.003 | 0.006 | 0.006 | 2.217 | 0.51 | 0.53 | 0.59 | 0.7 | 0.59 | 0.56 |
| 4 | 24379666 | rs3857106 | -0.607 | 0.494 | 0.042 | 0.929 | 0.032 | 0.067 | 0.067 | 0.001 | 0.14 | 0.25 | 0.21 | 0.29 | 0.2 | 0.26 |
| 4 | 24384208 | rs17591101 | 0.837 | 0.35 | -0.782 | 0.79 | 0.017 | 0.063 | 0.063 | 0.164 | 0.28 | 0.12 | 0.27 | 0.13 | 0.22 | 0.21 |
| 4 | 24392437 | rs16875599 | 1.147 | 0.464 | -0.936 | 0.654 | -0.033 | -0.081 | -0.081 | 0 | 0.37 | 0.13 | 0.36 | 0.53 | 0.38 | 0.36 |
| 4 | 24393041 | rs4697455 | 0.705 | 0.373 | -0.664 | 0.37 | 0.016 | 0.064 | 0.064 | 0.189 | 0.28 | 0.11 | 0.26 | 0.13 | 0.22 | 0.21 |
| 4 | 24397676 | rs11721883 | -0.379 | 0.564 | 0.016 | 0.494 | 0.003 | -0.054 | -0.054 | 0 | 0.17 | 0.55 | 0.15 | 0.09 | 0.16 | 0.17 |
| 4 | 24403899 | rs6854638 | 0.579 | 0.497 | -0.65 | 0.91 | -0.025 | -0.027 | -0.027 | 0 | 0.32 | 0.1 | 0.38 | 0.36 | 0.36 | 0.24 |
| 4 | 24410336 | rs4443265 | -0.19 | 0.492 | -0.218 | 0.757 | 0.014 | 0.008 | 0.008 | 4.564 | 0.49 | 0.54 | 0.53 | 0.53 | 0.43 | 0.35 |
| 4 | 24410439 | rs4406005 | -1.254 | 0.709 | -0.344 | 0.633 | 0.001 | 0.022 | 0.022 | 0.452 | 0.69 | 0.76 | 0.79 | 0.78 | 0.72 | 0.62 |
| 4 | 24413937 | rs7680809 | 1.048 | 0.173 | 0.159 | 0.801 | 0.003 | -0.014 | -0.014 | 0.538 | 0.32 | 0.21 | 0.38 | 0.38 | 0.32 | 0.19 |
| 4 | 24415276 | rs6858097 | -1.212 | 0.409 | 0.275 | 0.818 | 0.018 | 0.023 | 0.023 | 0.427 | 0.68 | 0.74 | 0.77 | 0.79 | 0.75 | 0.61 |
| 4 | 24415312 | rs6835454 | -1.171 | 0.397 | 0.327 | 0.812 | 0.007 | 0.009 | 0.009 | 0.352 | 0.68 | 0.68 | 0.76 | 0.79 | 0.75 | 0.61 |
| 4 | 24416173 | rs744790 | -1.382 | 1.07 | 0.939 | 1.255 | -0.031 | 0.089 | 0.089 | 0.133 | 0.18 | 0.4 | 0.22 | 0.25 | 0.11 | 0.16 |
| 4 | 24422187 | rs11938561 | -0.741 | 1.359 | 0.525 | 0.871 | 0.015 | -0.006 | -0.006 | 0 | 0.21 | 0.27 | 0.18 | 0.19 | 0.33 | 0.21 |
| 4 | 24424154 | rs7690462 | -1.821 | 1.331 | 0.573 | 0.951 | 0.023 | 0.008 | 0.008 | 0 | 0.36 | 0.54 | 0.37 | 0.4 | 0.42 | 0.36 |
| 4 | 24429852 | rs4697458 | -0.708 | 1.582 | 0.084 | 1.04 | 0.056 | -0.002 | -0.002 | 0 | 0.21 | 0.32 | 0.19 | 0.22 | 0.33 | 0.2 |
| 4 | 24433341 | rs17651313 | -0.526 | 1.136 | -0.176 | 1.154 | -0.004 | -0.011 | -0.011 | 0 | 0.09 | 0.12 | 0.05 | 0.05 | 0.06 | 0.04 |
| 4 | 24435780 | rs16875639 | -1.583 | 1.144 | -0.598 | 1.238 | -0.006 | 0.007 | 0.007 | 0 | 0.15 | 0.22 | 0.2 | 0.18 | 0.09 | 0.16 |
| 4 | 24436905 | rs16875643 | -0.667 | 1.305 | -0.785 | 1.355 | -0.004 | 0.014 | 0.014 | 0 | 0.07 | 0.1 | 0.15 | 0.13 | 0.03 | 0.11 |
| 4 | 24438499 | rs5015586 | -0.653 | 2.091 | -0.844 | 1.145 | 0.04 | 0.01 | 0.01 | 5.25 | 0.13 | 0.15 | 0.14 | 0.2 | 0.28 | 0.13 |
| 4 | 24438686 | rs6817993 | 0.186 | 1.222 | -1.722 | 0.868 | 0.02 | 0.034 | 0.034 | 6.251 | 0.65 | 0.17 | 0.65 | 0.72 | 0.72 | 0.68 |
| 4 | 24439865 | rs11723961 | -0.09 | 1.354 | -1.634 | 0.892 | 0.01 | 0.032 | 0.032 | 0 | 0.39 | 0.15 | 0.23 | 0.27 | 0.32 | 0.36 |
| 4 | 24443965 | rs16875658 | -0.305 | 1.98 | -1.361 | 1.613 | -0.05 | -0.05 | -0.05 | 0.374 | 0.08 | 0.01 | 0.21 | 0.22 | 0.19 | 0.08 |
| 4 | 24444785 | rs6823107 | 1.007 | 1.592 | -0.698 | 1.335 | 0.013 | 0.053 | 0.053 | 0 | 0.72 | 0.55 | 0.58 | 0.41 | 0.66 | 0.74 |
| 4 | 24448584 | rs17651758 | -1.366 | 1.723 | -0.815 | 0.631 | -0.024 | 0.013 | 0.013 | 0 | 0.3 | 0.22 | 0.22 | 0.27 | 0.37 | 0.22 |
| 4 | 24448620 | rs17592374 | -1.115 | 1.71 | -0.874 | 0.595 | -0.069 | -0.02 | -0.02 | 0 | 0.23 | 0.08 | 0.15 | 0.27 | 0.34 | 0.22 |
| 4 | 24451107 | rs10027557 | -0.488 | 1.636 | -0.549 | 0.494 | -0.012 | -0.002 | -0.002 | 0 | 0.19 | 0.42 | 0.22 | 0.16 | 0.18 | 0.19 |
| 4 | 24451810 | rs10018808 | 0.471 | 1.652 | -0.363 | 0.518 | -0.012 | 0.065 | 0.065 | 0 | 0.1 | 0.32 | 0.14 | 0.12 | 0.06 | 0.12 |
| 4 | 24454502 | rs10025642 | -1.091 | 1.705 | -0.524 | 0.538 | -0.027 | -0.035 | -0.035 | 0 | 0.34 | 0.57 | 0.31 | 0.18 | 0.22 | 0.24 |
| 4 | 24455932 | rs10026526 | -2.023 | 1.726 | -0.522 | 0.584 | 0.003 | 0.018 | 0.018 | 0.19 | 0.49 | 0.65 | 0.43 | 0.43 | 0.52 | 0.4 |
| 4 | 24456779 | rs13112452 | -0.225 | 1.821 | -0.552 | 0.626 | -0.044 | -0.033 | -0.033 | 0 | 0.13 | 0.05 | 0.11 | 0.25 | 0.28 | 0.15 |
| 4 | 24462075 | rs13435868 | -1.611 | 1.479 | -0.181 | 0.17 | 0.004 | 0.008 | 0.008 | 0.037 | 0.4 | 0.64 | 0.35 | 0.24 | 0.31 | 0.29 |

| | | | | | | | | | | | | | | | | |
|---|---|---|---|---|---|---|---|---|---|---|---|---|---|---|---|---|
| 4 | 24467089 | rs11737023 | -1.376 | 1.95 | -0.136 | 0.013 | -0.011 | 0.02 | 0.02 | 0 | 0.22 | 0.18 | 0.16 | 0.07 | 0.21 | 0.15 |
| 4 | 24470143 | rs3811738 | -0.713 | 1.625 | -0.258 | 0.13 | 0.013 | 0.025 | 0.025 | 0 | 0.52 | 0.41 | 0.51 | 0.39 | 0.55 | 0.6 |
| 4 | 24470847 | rs3828483 | -0.704 | 1.641 | -0.29 | 0.138 | 0.012 | 0.022 | 0.022 | 0 | 0.52 | 0.42 | 0.52 | 0.39 | 0.55 | 0.6 |
| 4 | 24480956 | rs12506649 | 0.883 | 1.44 | 0.399 | 0.513 | -0.003 | 0.026 | 0.026 | 0 | 0.51 | 0.36 | 0.49 | 0.61 | 0.49 | 0.48 |
| 4 | 24483311 | rs4697062 | 0.539 | 0.962 | 0.735 | 0.676 | 0 | 0.004 | 0.004 | 0.223 | 0.58 | 0.55 | 0.53 | 0.68 | 0.56 | 0.55 |
| 4 | 24488420 | rs16875707 | -0.039 | 1.106 | -0.114 | 0.488 | 0 | 0.004 | 0.004 | 0 | 0.64 | 0.54 | 0.58 | 0.6 | 0.59 | 0.52 |
| 4 | 24489441 | rs7656078 | 0.312 | 1.385 | -1.071 | 0.707 | 0.008 | 0.01 | 0.01 | 0.455 | 0.56 | 0.3 | 0.51 | 0.6 | 0.55 | 0.51 |
| 4 | 24514957 | rs12651262 | -0.179 | 1.33 | -0.917 | 1.129 | 0 | -0.018 | -0.018 | 0 | 0.25 | 0.19 | 0.28 | 0.38 | 0.23 | 0.09 |
| 4 | 24521431 | rs6448278 | 0.408 | 1.45 | -2.301 | 0.593 | -0.094 | 0.005 | 0.005 | 0.031 | 0.3 | 0.02 | 0.14 | 0.24 | 0.3 | 0.32 |
| 4 | 24526581 | rs10517052 | -0.501 | 1.154 | -1.334 | 0.514 | -0.001 | 0.001 | 0.001 | 0 | 0.58 | 0.38 | 0.49 | 0.67 | 0.57 | 0.46 |
| 4 | 24534687 | rs3828558 | 0.57 | 1.517 | -0.27 | 0.015 | 0.007 | 0.035 | 0.035 | 0 | 0.13 | 0.18 | 0.33 | 0.3 | 0.21 | 0.34 |
| 4 | 24546752 | rs2048026 | -0.394 | 1.313 | 0.307 | -0.092 | 0.013 | 0.011 | 0.011 | 0 | 0.38 | 0.29 | 0.45 | 0.31 | 0.35 | 0.5 |
| 4 | 24548678 | rs7690565 | 1.138 | 1.328 | 0.288 | -0.092 | -0.1 | 0.012 | 0.012 | 0.059 | 0.3 | 0.02 | 0.14 | 0.24 | 0.31 | 0.32 |
| 4 | 24554084 | rs12509760 | -0.364 | 1.262 | 0.519 | -0.06 | 0.008 | 0.008 | 0.008 | 0 | 0.37 | 0.33 | 0.45 | 0.31 | 0.35 | 0.5 |
| 4 | 24570155 | rs13115594 | 0.603 | 1.222 | 0.783 | -0.08 | -0.012 | -0.007 | -0.007 | 0 | 0.57 | 0.46 | 0.48 | 0.62 | 0.61 | 0.44 |
| 4 | 24584445 | rs6812715 | 0.983 | 1.081 | 0.721 | -0.023 | -0.003 | 0.019 | 0.019 | 1.359 | 0.28 | 0.3 | 0.33 | 0.43 | 0.25 | 0.13 |
| 4 | 24593495 | rs10939001 | 1.244 | 1.282 | 0.648 | -0.024 | -0.008 | -0.061 | -0.061 | 0 | 0.31 | 0.14 | 0.31 | 0.39 | 0.26 | 0.14 |
| 4 | 24593935 | rs9994148 | -1.148 | 1.52 | 0.664 | -0.03 | -0.015 | 0.022 | 0.022 | 0 | 0.71 | 0.86 | 0.87 | 0.71 | 0.71 | 0.68 |
| 4 | 24604616 | rs3857113 | 0.581 | 1.621 | 0.853 | -0.027 | -0.007 | -0.013 | -0.013 | 0 | 0.13 | 0.13 | 0.34 | 0.28 | 0.23 | 0.35 |
| 4 | 24605273 | rs13113294 | 0.07 | 1.295 | 0.623 | -0.183 | -0.007 | 0.005 | 0.005 | 0.229 | 0.69 | 0.59 | 0.55 | 0.67 | 0.68 | 0.56 |
| 4 | 24610039 | rs10033408 | 0.7 | 1.282 | 0.669 | -0.187 | 0 | 0.002 | 0.002 | 0.64 | 0.4 | 0.39 | 0.38 | 0.42 | 0.36 | 0.22 |
| 4 | 24618440 | rs6825842 | 0.102 | 1.251 | 0.51 | -0.106 | -0.011 | -0.002 | -0.002 | 0 | 0.69 | 0.54 | 0.61 | 0.69 | 0.69 | 0.57 |
| 4 | 24624177 | rs12640588 | 0.832 | 1.268 | 0.359 | -0.102 | -0.005 | 0.002 | 0.002 | 1.006 | 0.39 | 0.38 | 0.34 | 0.39 | 0.28 | 0.2 |
| 4 | 24624240 | rs13137535 | 1.055 | 1.38 | -0.282 | 0.011 | -0.079 | -0.019 | -0.019 | 0 | 0.27 | 0.03 | 0.19 | 0.27 | 0.3 | 0.32 |
| 4 | 24626903 | rs1395221 | 0.55 | 1.094 | -0.248 | 0.025 | -0.079 | -0.002 | -0.002 | 0 | 0.3 | 0.07 | 0.27 | 0.31 | 0.41 | 0.36 |
| 4 | 24646772 | rs12501699 | 0.216 | 0.747 | -0.207 | -0.11 | -0.001 | 0.012 | 0.012 | 1.784 | 0.75 | 0.69 | 0.52 | 0.48 | 0.71 | 0.42 |
| 4 | 24650004 | rs7695254 | -0.14 | 1.201 | 0.131 | -0.67 | 0.003 | -0.062 | -0.062 | 0 | 0.34 | 0.6 | 0.22 | 0.08 | 0.38 | 0.22 |
| 4 | 24651750 | rs4312745 | 1.055 | 0.887 | 0.544 | -0.241 | -0.039 | -0.077 | -0.077 | 0.516 | 0.82 | 0.68 | 0.71 | 0.89 | 0.89 | 0.84 |
| 4 | 24652142 | rs10939004 | 0.682 | 1.033 | 0.555 | -0.076 | -0.005 | -0.014 | -0.014 | 0 | 0.88 | 0.58 | 0.75 | 0.89 | 0.93 | 0.85 |
| 4 | 24653384 | rs7693909 | 0.292 | 1.861 | -0.516 | 0.677 | -0.057 | -0.066 | -0.066 | 0 | 0.84 | 0.93 | 0.56 | 0.63 | 0.66 | 0.63 |
| 4 | 24672521 | rs4295245 | -0.494 | 1.879 | -0.621 | 0.692 | -0.065 | -0.078 | -0.078 | 0 | 0.15 | 0.05 | 0.36 | 0.35 | 0.32 | 0.42 |
| 4 | 24673420 | rs12507254 | -0.834 | 1.528 | -1.294 | 0.167 | 0.045 | 0.002 | 0.002 | 0 | 0.47 | 0.27 | 0.53 | 0.44 | 0.63 | 0.57 |
| 4 | 24676205 | rs4499689 | -0.399 | 0.882 | -0.679 | 0.26 | -0.025 | 0.099 | 0.099 | 0 | 0.14 | 0.25 | 0.25 | 0.36 | 0.11 | 0.28 |
| 4 | 24676608 | rs4351004 | 0.055 | 1.099 | -1.197 | -0.284 | -0.03 | 0.098 | 0.098 | 0 | 0.3 | 0.13 | 0.16 | 0.09 | 0.29 | 0.15 |
| 4 | 24686179 | rs13101306 | 0.554 | 1.002 | -0.358 | -0.162 | 0.013 | -0.014 | -0.014 | 0.677 | 0.62 | 0.48 | 0.36 | 0.34 | 0.55 | 0.3 |
| 4 | 24688574 | rs11736386 | 0.478 | 0.962 | -0.915 | -0.234 | -0.028 | 0.014 | 0.014 | 0.179 | 0.16 | 0.04 | 0.07 | 0.14 | 0.18 | 0.08 |
| 4 | 24691463 | rs16875929 | -0.221 | 1.161 | -0.515 | 0.427 | -0.062 | -0.099 | -0.099 | 2.036 | 0.25 | 0.1 | 0.47 | 0.45 | 0.34 | 0.52 |
| 4 | 24710623 | rs2080014 | -0.528 | 1.137 | -0.323 | -0.184 | -0.062 | -0.02 | -0.02 | 0.917 | 0.32 | 0.16 | 0.59 | 0.6 | 0.42 | 0.66 |
| 4 | 24710866 | rs10013569 | -0.619 | 1.168 | 0.235 | -0.172 | 0.011 | -0.003 | -0.003 | 0.464 | 0.33 | 0.37 | 0.62 | 0.66 | 0.42 | 0.67 |
| 4 | 24728480 | rs7666030 | -0.058 | 1.004 | -0.159 | -0.073 | 0.011 | -0.024 | -0.024 | 0 | 0.27 | 0.19 | 0.39 | 0.59 | 0.23 | 0.6 |
| 4 | 24736258 | rs1476880 | -0.493 | 0.38 | -0.879 | -0.44 | 0 | 0.018 | 0.018 | 5.696 | 0.55 | 0.74 | 0.6 | 0.62 | 0.56 | 0.73 |
| 4 | 24740958 | rs12500612 | 0.233 | 0.557 | -0.26 | -0.505 | 0.006 | 0.045 | 0.045 | 3.054 | 0.19 | 0.08 | 0.09 | 0 | 0.16 | 0.09 |
| 4 | 24750063 | rs6851578 | 0.005 | 0.845 | -1.487 | -0.322 | -0.012 | -0.045 | -0.045 | 0 | 0.26 | 0.67 | 0.17 | 0.08 | 0.17 | 0.12 |
| 4 | 24750388 | rs6448303 | 0.71 | 1.446 | -1.518 | 0.515 | -0.031 | 0.007 | 0.007 | 4.203 | 0.75 | 0.91 | 0.81 | 0.77 | 0.71 | 0.85 |
| 4 | 24754107 | rs758951 | 0.379 | 0.933 | -1.694 | 0.546 | -0.042 | -0.019 | -0.019 | 0 | 0.8 | 0.99 | 0.89 | 0.81 | 0.81 | 0.89 |
| 4 | 24760492 | rs13104450 | 0.454 | 1.139 | -1.269 | 0.853 | -0.012 | -0.004 | -0.004 | 0 | 0.65 | 0.68 | 0.6 | 0.65 | 0.59 | 0.6 |
| 4 | 24762301 | rs1014262 | 1.294 | 0.519 | -0.619 | 1.078 | -0.008 | 0.006 | 0.006 | 0.447 | 0.86 | 0.85 | 0.78 | 0.88 | 0.9 | 0.77 |
| 4 | 24765819 | rs7435399 | -0.682 | 0.248 | -0.83 | 1.125 | 0 | 0.002 | 0.002 | 0 | 0.39 | 0.71 | 0.3 | 0.32 | 0.38 | 0.24 |
| 4 | 24773476 | rs13134000 | 0.013 | 0.337 | -0.888 | 1.241 | -0.007 | -0.025 | -0.025 | 1.331 | 0.07 | 0.02 | 0.08 | 0.08 | 0.06 | 0.16 |
| 4 | 24776982 | rs12501287 | 1.436 | 0.204 | -0.93 | 1.528 | -0.063 | 0.015 | 0.015 | 0 | 0.48 | 0.08 | 0.48 | 0.39 | 0.53 | 0.49 |
| 4 | 24779506 | rs13138705 | -0.717 | -0.096 | -0.875 | 1.449 | -0.118 | -0.119 | -0.119 | 0 | 0.34 | 0.1 | 0.29 | 0.51 | 0.41 | 0.42 |
| 4 | 24785911 | rs800421 | 0.229 | -0.998 | -0.753 | 1.431 | 0.016 | -0.034 | -0.034 | 0 | 0.68 | 0.81 | 0.79 | 0.54 | 0.72 | 0.72 |
| 4 | 24786904 | rs13434451 | -1.049 | -0.935 | 0.412 | 1.407 | -0.03 | 0.009 | 0.009 | 0 | 0.16 | 0.28 | 0.23 | 0.12 | 0.09 | 0.19 |
| 4 | 24791146 | rs12649653 | -0.612 | -0.908 | 0.039 | 1.439 | -0.041 | -0.097 | -0.097 | 0 | 0.33 | 0.09 | 0.26 | 0.35 | 0.31 | 0.38 |
| 4 | 24792175 | rs6849330 | 0.085 | -0.575 | 0.355 | 1.069 | -0.016 | 0.034 | 0.034 | 1.394 | 0.09 | 0.37 | 0.17 | 0.04 | 0.05 | 0.03 |
| 4 | 24794797 | rs2284659 | 0.943 | -0.176 | -0.504 | 0.975 | 0.061 | 0.043 | 0.043 | 0 | 0.48 | 0.1 | 0.52 | 0.38 | 0.62 | 0.51 |
| 4 | 24796568 | rs8192287 | -0.129 | -0.13 | -0.489 | 1.076 | -0.017 | -0.037 | -0.037 | 0 | 0.07 | 0.01 | 0.07 | 0.09 | 0.06 | 0.17 |
| 4 | 24796803 | rs699473 | 0.471 | 0.118 | -0.195 | 0.702 | 0.079 | 0.063 | 0.063 | 0 | 0.49 | 0.25 | 0.53 | 0.37 | 0.63 | 0.52 |
| 4 | 24804722 | rs17552548 | -1.052 | 0.624 | -0.359 | 0.746 | -0.019 | 0.034 | 0.034 | 0 | 0.06 | 0 | 0.01 | 0 | 0.06 | 0.05 |
| 4 | 24807287 | rs758946 | 0.806 | 0.305 | -0.242 | 0.958 | 0.003 | -0.002 | -0.002 | 0 | 0.88 | 0.93 | 0.93 | 0.91 | 0.91 | 0.93 |
| 4 | 24809884 | rs1019229 | 1.039 | 0.386 | -0.276 | 0.931 | -0.001 | -0.001 | -0.001 | 0 | 0.87 | 0.93 | 0.92 | 0.91 | 0.88 | 0.9 |
| 4 | 24818097 | rs800452 | 0.882 | 0.453 | -0.119 | 1.079 | -0.001 | 0 | 0 | 0 | 0.58 | 0.41 | 0.53 | 0.53 | 0.58 | 0.61 |
| 4 | 24828799 | rs800456 | 0.184 | 0.786 | -0.033 | 1.648 | -0.013 | -0.029 | -0.029 | 0 | 0.86 | 0.94 | 0.85 | 0.71 | 0.84 | 0.89 |
| 4 | 24829427 | rs800457 | 0.253 | 0.761 | 0.365 | 1.642 | 0 | 0.011 | 0.011 | 0.236 | 0.86 | 0.9 | 0.86 | 0.73 | 0.84 | 0.9 |
| 4 | 24831450 | rs16876184 | -0.36 | 0.701 | 0.867 | 1.535 | -0.005 | -0.035 | -0.035 | 0 | 0.1 | 0.16 | 0.06 | 0 | 0.08 | 0.07 |
| 4 | 24832724 | rs16876189 | -0.578 | 0.672 | 0.818 | 1.473 | -0.004 | -0.035 | -0.035 | 0 | 0.11 | 0.16 | 0.06 | 0 | 0.09 | 0.08 |
| 4 | 24836416 | rs6838713 | -0.437 | 0.399 | 1.853 | 1.309 | 0.011 | 0.025 | 0.025 | 0 | 0.18 | 0.29 | 0.26 | 0.18 | 0.17 | 0.25 |
| 4 | 24836619 | rs16876194 | -0.201 | 0.288 | 1.783 | 1.305 | -0.005 | -0.037 | -0.037 | 0 | 0.1 | 0.16 | 0.06 | 0 | 0.08 | 0.07 |
| 4 | 24842936 | rs12510049 | -0.291 | 0.317 | 1.98 | 1.343 | 0.005 | 0.018 | 0.018 | 0 | 0.17 | 0.2 | 0.26 | 0.2 | 0.17 | 0.25 |

| | | | | | | | | | | | | | | | | |
|---|---|---|---|---|---|---|---|---|---|---|---|---|---|---|---|---|
| 4 | 24846501 | rs800460 | 2.158 | 0.236 | 1.923 | 1.299 | -0.019 | 0.006 | 0.006 | 0 | 0.54 | 0.18 | 0.51 | 0.53 | 0.58 | 0.56 |
| 4 | 24856909 | rs800464 | 1.231 | 0.254 | 2.065 | 1.344 | -0.001 | 0.002 | 0.002 | 0 | 0.75 | 0.75 | 0.84 | 0.89 | 0.76 | 0.84 |
| 4 | 24860242 | rs17625018 | -0.398 | 0.244 | 1.965 | 1.315 | 0.002 | -0.008 | -0.008 | 0 | 0.1 | 0.08 | 0.06 | 0 | 0.09 | 0.08 |
| 4 | 24860527 | rs800478 | 1.287 | 0.254 | 2.002 | 1.383 | 0 | 0.001 | 0.001 | 0 | 0.76 | 0.76 | 0.85 | 0.88 | 0.78 | 0.84 |
| 4 | 24871204 | rs800379 | 0.1 | 0.256 | 1.994 | 1.401 | 0 | 0 | 0 | 0 | 0.85 | 0.83 | 0.9 | 0.88 | 0.85 | 0.92 |
| 4 | 24871999 | rs800383 | -0.06 | 0.268 | 1.867 | 1.407 | -0.017 | -0.012 | -0.012 | 0.247 | 0.13 | 0.05 | 0.09 | 0.12 | 0.15 | 0.08 |
| 4 | 24873604 | rs13131918 | 2.593 | -0.336 | 1.318 | 1.429 | -0.05 | 0.072 | 0.072 | 0 | 0.44 | 0.11 | 0.46 | 0.4 | 0.49 | 0.34 |
| 4 | 24881852 | rs941265 | 2.279 | -0.281 | 1.323 | 1.422 | -0.006 | 0.083 | 0.083 | 0 | 0.54 | 0.17 | 0.54 | 0.41 | 0.59 | 0.51 |
| 4 | 24885284 | rs10517056 | -1.189 | -0.329 | 1.403 | 1.47 | -0.014 | -0.023 | -0.023 | 0 | 0.3 | 0.21 | 0.38 | 0.31 | 0.32 | 0.33 |
| 4 | 24885567 | rs1527354 | 1.576 | -0.28 | 0.84 | 1.066 | -0.009 | 0.071 | 0.071 | 0 | 0.55 | 0.26 | 0.49 | 0.39 | 0.58 | 0.42 |
| 4 | 24885755 | rs1527353 | 1.994 | -0.168 | 0.585 | 1.208 | -0.029 | 0.087 | 0.087 | 0 | 0.45 | 0.17 | 0.43 | 0.38 | 0.49 | 0.34 |
| 4 | 24886099 | rs4697489 | 1.994 | -0.168 | 0.585 | 1.221 | -0.029 | 0.104 | 0.104 | 0 | 0.45 | 0.17 | 0.43 | 0.37 | 0.49 | 0.34 |
| 4 | 24897032 | rs10939012 | -1.111 | -0.649 | 0.878 | 1.398 | 0.009 | 0.007 | 0.007 | 0 | 0.19 | 0.11 | 0.16 | 0.15 | 0.15 | 0.25 |
| 4 | 24899586 | rs4697493 | -1.035 | -0.687 | 1.089 | 1.406 | -0.002 | -0.007 | -0.007 | 0 | 0.18 | 0.18 | 0.24 | 0.09 | 0.13 | 0.23 |
| 4 | 24900112 | rs9993370 | 1.984 | -0.65 | 1.381 | 1.791 | 0.001 | 0.003 | 0.003 | 0 | 0.5 | 0.43 | 0.49 | 0.58 | 0.57 | 0.42 |
| 4 | 24901466 | rs4053966 | 1.066 | -0.923 | 1.769 | 2.073 | -0.008 | 0.038 | 0.038 | 0.451 | 0.77 | 0.38 | 0.71 | 0.67 | 0.81 | 0.71 |
| 4 | 24902583 | rs4697075 | -0.711 | -0.949 | 1.637 | 2.024 | -0.005 | -0.05 | -0.05 | 0 | 0.2 | 0.1 | 0.14 | 0.23 | 0.17 | 0.18 |
| 4 | 24908601 | rs4697076 | 0.093 | -0.914 | 1.767 | 2.031 | -0.009 | 0.083 | 0.083 | 0 | 0.84 | 0.64 | 0.81 | 0.77 | 0.87 | 0.81 |
| 4 | 24924432 | rs11721701 | 2.889 | -0.909 | 1.237 | 1.792 | -0.118 | 0.001 | 0.001 | 0 | 0.43 | 0.07 | 0.37 | 0.41 | 0.52 | 0.4 |
| 4 | 24924497 | rs7680589 | -0.386 | -0.87 | 1.339 | 1.712 | -0.008 | -0.045 | -0.045 | 0.033 | 0.17 | 0.28 | 0.14 | 0.05 | 0.13 | 0.18 |
| 4 | 24932206 | rs1527359 | 2.551 | -0.844 | 1.042 | 1.739 | -0.087 | 0.009 | 0.009 | 0 | 0.45 | 0.1 | 0.49 | 0.44 | 0.52 | 0.44 |
| 4 | 24936925 | rs11931422 | -1.659 | -0.798 | 1.099 | 1.641 | -0.002 | -0.013 | -0.013 | 0 | 0.33 | 0.58 | 0.3 | 0.16 | 0.27 | 0.36 |
| 4 | 24938446 | rs10008999 | -1.979 | -0.754 | 0.491 | 1.779 | 0.015 | -0.073 | -0.073 | 0 | 0.39 | 0.83 | 0.4 | 0.44 | 0.3 | 0.48 |
| 4 | 24939037 | rs12645335 | -0.835 | -0.74 | 0.548 | 1.783 | -0.024 | 0.098 | 0.098 | 0 | 0.18 | 0.44 | 0.18 | 0.28 | 0.12 | 0.21 |
| 4 | 24940002 | rs13105803 | -0.97 | -0.359 | 0.799 | 1.404 | 0.003 | -0.014 | -0.014 | 0 | 0.21 | 0.37 | 0.22 | 0.16 | 0.18 | 0.27 |
| 4 | 24949675 | rs2167955 | 0.211 | -0.594 | -0.02 | 1.373 | -0.01 | -0.018 | -0.018 | 0 | 0.84 | 0.91 | 0.89 | 0.88 | 0.82 | 0.92 |
| 4 | 24955907 | rs4235320 | 1.351 | -0.99 | 0.325 | 1.184 | 0.017 | 0.029 | 0.029 | 0.348 | 0.59 | 0.32 | 0.49 | 0.17 | 0.66 | 0.43 |
| 4 | 24956062 | rs4697503 | -1.025 | -1.128 | 0.293 | 1.726 | -0.038 | 0.045 | 0.045 | 0 | 0.19 | 0.37 | 0.15 | 0.28 | 0.11 | 0.32 |
| 4 | 24957653 | rs1453815 | 0.627 | -0.504 | -0.239 | 1.913 | 0 | -0.006 | -0.006 | 0.115 | 0.78 | 0.78 | 0.68 | 0.51 | 0.76 | 0.78 |
| 4 | 24978429 | rs13126958 | -1.013 | -0.477 | -0.526 | 1.703 | 0 | -0.023 | -0.023 | 0 | 0.16 | 0.13 | 0.13 | 0.22 | 0.13 | 0.34 |
| 4 | 24981812 | rs764496 | -0.72 | -1.432 | -0.178 | 1.214 | -0.056 | -0.026 | -0.026 | 0 | 0.14 | 0.22 | 0.05 | 0.06 | 0.04 | 0.14 |
| 4 | 24983923 | rs7668800 | -1.712 | -1.687 | -0.019 | 1.711 | -0.008 | 0.031 | 0.031 | 0 | 0.28 | 0.63 | 0.37 | 0.51 | 0.25 | 0.48 |
| 4 | 24985937 | rs4697516 | -1.582 | -1.656 | 0.229 | 1.746 | -0.008 | 0.078 | 0.078 | 0 | 0.26 | 0.57 | 0.35 | 0.5 | 0.23 | 0.42 |
| 4 | 24991254 | rs13130781 | -0.55 | -1.545 | -0.245 | 1.739 | -0.045 | -0.073 | -0.073 | 0 | 0.15 | 0.02 | 0.07 | 0.21 | 0.15 | 0.3 |
| 4 | 24991618 | rs4317191 | -1.726 | -1.721 | 0.475 | 1.718 | -0.089 | 0.094 | 0.094 | 0.158 | 0.24 | 0.52 | 0.31 | 0.41 | 0.13 | 0.31 |
| 4 | 24992827 | rs4697082 | -0.953 | -1.232 | 0.783 | 1.9 | -0.013 | 0.176 | 0.176 | 2.545 | 0.1 | 0.28 | 0.26 | 0.34 | 0.09 | 0.19 |
| 4 | 24994457 | rs2098300 | -1.157 | -0.514 | 0.39 | 1.614 | 0 | 0 | 0 | 0 | 0.37 | 0.66 | 0.41 | 0.57 | 0.36 | 0.52 |
| 4 | 24996698 | rs13135230 | -1.034 | -0.946 | 0.017 | 1.777 | -0.025 | -0.086 | -0.086 | 0 | 0.28 | 0.11 | 0.21 | 0.43 | 0.26 | 0.38 |
| 4 | 24999299 | rs17406878 | -0.599 | -0.271 | 0.242 | 2.105 | -0.001 | -0.029 | -0.029 | 0 | 0.17 | 0.04 | 0.07 | 0.18 | 0.13 | 0.28 |
| 4 | 24999316 | rs16876537 | -0.308 | -0.846 | 0.158 | 2.089 | -0.017 | 0.045 | 0.045 | 0 | 0.13 | 0.18 | 0.18 | 0.38 | 0.06 | 0.25 |
| 4 | 25000053 | rs4697519 | -1.746 | -0.814 | 0.421 | 2.115 | 0.006 | 0.031 | 0.031 | 0.545 | 0.28 | 0.38 | 0.48 | 0.62 | 0.3 | 0.37 |
| 4 | 25000555 | rs7659172 | 0.298 | -0.812 | 0.403 | 2.112 | -0.047 | -0.064 | -0.064 | 0.02 | 0.11 | 0.01 | 0.14 | 0.23 | 0.13 | 0.09 |
| 4 | 25001685 | rs7670138 | -2.563 | -0.796 | 0.925 | 2.057 | 0 | -0.006 | -0.006 | 0 | 0.46 | 0.47 | 0.55 | 0.8 | 0.42 | 0.65 |
| 4 | 25002119 | rs6853990 | -2.747 | -0.996 | 1.191 | 1.944 | -0.003 | 0.062 | 0.062 | 0 | 0.53 | 0.65 | 0.63 | 0.92 | 0.46 | 0.7 |
| 4 | 25003503 | rs8235 | -3.22 | -1.026 | 1.105 | 1.93 | -0.028 | 0.189 | 0.189 | 0.039 | 0.55 | 0.81 | 0.64 | 0.91 | 0.46 | 0.7 |
| 4 | 25004004 | rs6841999 | -0.457 | -0.967 | 1.086 | 2.016 | -0.014 | 0.066 | 0.066 | 0 | 0.07 | 0.2 | 0.07 | 0.12 | 0.04 | 0.05 |
| 4 | 25004757 | rs3796786 | -0.175 | -0.961 | 1.074 | 2.019 | -0.01 | 0.071 | 0.071 | 0 | 0.07 | 0.18 | 0.07 | 0.12 | 0.04 | 0.05 |
| 4 | 25004778 | rs3796787 | -0.145 | -0.961 | 1.074 | 2.019 | -0.01 | 0.073 | 0.073 | 0 | 0.07 | 0.18 | 0.07 | 0.12 | 0.04 | 0.05 |
| 4 | 25004971 | rs2232030 | 0.283 | -0.871 | 0.942 | 2.044 | -0.047 | -0.065 | -0.065 | 0 | 0.11 | 0.01 | 0.14 | 0.23 | 0.13 | 0.09 |
| 4 | 25005023 | rs2232029 | -0.783 | -0.888 | 1.025 | 1.912 | -0.002 | 0.05 | 0.05 | 0.595 | 0.17 | 0.24 | 0.33 | 0.39 | 0.16 | 0.26 |
| 4 | 25009491 | rs4235326 | 1.362 | -0.131 | 0.237 | 1.392 | 0.031 | 0 | 0 | 3.153 | 0.48 | 0.38 | 0.41 | 0.46 | 0.36 | 0.4 |
| 4 | 25011001 | rs11733058 | 1.728 | -0.119 | 0.042 | 1.383 | 0.091 | 0.071 | 0.071 | 0.072 | 0.43 | 0.15 | 0.33 | 0.31 | 0.32 | 0.37 |
| 4 | 25013576 | rs1000946 | 1.246 | 0.066 | 0.382 | 1.637 | 0.008 | 0.005 | 0.005 | 1.258 | 0.61 | 0.42 | 0.38 | 0.42 | 0.47 | 0.63 |
| 4 | 25014793 | rs2110453 | -1.193 | 0.431 | 0.106 | 1.219 | -0.047 | 0.04 | 0.04 | 0.138 | 0.87 | 0.95 | 0.75 | 0.98 | 0.68 | 0.93 |
| 4 | 25017735 | rs10488953 | 0.132 | 0.336 | 0.3 | 1.298 | 0.004 | 0.026 | 0.026 | 0 | 0.12 | 0.18 | 0.21 | 0.23 | 0.13 | 0.09 |
| 4 | 25018625 | rs4697525 | 0.898 | 0.178 | 0.422 | 0.945 | 0.04 | 0.036 | 0.036 | 0.018 | 0.72 | 0.64 | 0.51 | 0.58 | 0.54 | 0.79 |
| 4 | 25019263 | rs2324642 | -1.675 | -0.152 | 0.681 | 1.113 | -0.007 | 0.006 | 0.006 | 0 | 0.27 | 0.62 | 0.35 | 0.69 | 0.22 | 0.28 |
| 4 | 25020194 | rs10022509 | -0.503 | -0.111 | 0.465 | 1.152 | -0.007 | 0.096 | 0.096 | 0 | 0.14 | 0.34 | 0.12 | 0.29 | 0.09 | 0.14 |
| 4 | 25022408 | rs6448323 | -1.488 | -0.196 | 0.01 | 0.194 | 0.003 | 0 | 0 | 0 | 0.29 | 0.75 | 0.37 | 0.69 | 0.22 | 0.28 |
| 4 | 25023893 | rs4560399 | -1.05 | 0.271 | -0.263 | -0.327 | -0.004 | 0.077 | 0.077 | 1.029 | 0.4 | 0.81 | 0.49 | 0.76 | 0.35 | 0.46 |
| 4 | 25024935 | rs16876650 | 1.193 | 1.028 | -0.27 | -0.808 | 0.014 | 0.055 | 0.055 | 0 | 0.35 | 0.23 | 0.25 | 0.06 | 0.31 | 0.27 |
| 4 | 25025389 | rs6843764 | -0.572 | 1.299 | -0.686 | -0.71 | 0.005 | 0.002 | 0.002 | 0 | 0.49 | 0.54 | 0.52 | 0.91 | 0.42 | 0.66 |
| 4 | 25028158 | rs4428268 | -0.131 | 1.886 | -0.525 | 0.252 | -0.002 | -0.018 | -0.018 | 0.266 | 0.86 | 0.85 | 0.78 | 1 | 0.75 | 0.94 |
| 4 | 25029817 | rs757637 | 0.525 | 2.002 | -0.246 | 0.561 | 0.083 | -0.008 | -0.008 | 0 | 0.81 | 0.62 | 0.71 | 0.83 | 0.66 | 0.89 |
| 4 | 25033008 | rs736831 | 0.118 | 1.52 | -1.375 | 0.559 | -0.041 | 0.037 | 0.037 | 0 | 0.57 | 0.86 | 0.57 | 0.67 | 0.42 | 0.6 |
| 4 | 25036142 | rs10939032 | -0.338 | 0.822 | -1.485 | 0.43 | -0.005 | -0.012 | -0.012 | 0 | 0.71 | 0.32 | 0.68 | 0.73 | 0.78 | 0.57 |
| 4 | 25036320 | rs7676843 | 0.502 | 0.945 | -1.536 | 0.404 | -0.033 | 0.036 | 0.036 | 0 | 0.44 | 0.14 | 0.44 | 0.33 | 0.59 | 0.4 |
| 4 | 25040676 | rs4235328 | 0.053 | 0.826 | -1.474 | 0.469 | 0.024 | -0.019 | -0.019 | 0.124 | 0.62 | 0.15 | 0.58 | 0.38 | 0.73 | 0.64 |
| 4 | 25041068 | rs10488952 | 0.944 | 0.742 | -1.474 | 0.971 | 0.005 | -0.009 | -0.009 | 0 | 0.24 | 0.18 | 0.2 | 0.23 | 0.18 | 0.12 |

| | | | | | | | | | | | | | | | | |
|---|---|---|---|---|---|---|---|---|---|---|---|---|---|---|---|---|
| 4 | 25041513 | rs2324649 | -0.323 | 0.86 | -1.818 | 0.761 | 0.023 | -0.049 | -0.049 | 0 | 0.64 | 0.17 | 0.62 | 0.54 | 0.75 | 0.67 |
| 4 | 25048536 | rs13124493 | 0.255 | 0.941 | -1.699 | 0.602 | -0.041 | 0.047 | 0.047 | 0 | 0.44 | 0.13 | 0.39 | 0.34 | 0.47 | 0.39 |
| 4 | 25053498 | rs11931069 | -0.092 | 0.864 | -0.541 | 1.23 | -0.014 | 0.06 | 0.06 | 0 | 0.18 | 0.27 | 0.24 | 0.37 | 0.16 | 0.17 |
| 4 | 25056765 | rs6856359 | -0.261 | 0.665 | -1.041 | 0.878 | -0.001 | -0.002 | -0.002 | 1.495 | 0.57 | 0.41 | 0.59 | 0.64 | 0.6 | 0.45 |
| 4 | 25058232 | rs759245 | 0.645 | 0.653 | -1.055 | 0.883 | -0.042 | 0.06 | 0.06 | 0 | 0.39 | 0.15 | 0.35 | 0.26 | 0.44 | 0.28 |
| 4 | 25060758 | rs6448329 | 0.092 | 0.66 | -0.887 | 0.847 | 0.002 | -0.006 | -0.006 | 0.086 | 0.42 | 0.48 | 0.44 | 0.35 | 0.44 | 0.53 |
| 4 | 25066309 | rs7654693 | 0.211 | 0.703 | -1.04 | 0.807 | -0.022 | -0.052 | -0.052 | 0 | 0.39 | 0.46 | 0.29 | 0.14 | 0.29 | 0.51 |
| 4 | 25066802 | rs12502737 | -0.258 | 1.103 | -1.542 | 0.958 | 0.026 | -0.013 | -0.013 | 0 | 0.56 | 0.7 | 0.5 | 0.53 | 0.42 | 0.67 |
| 4 | 25067218 | rs6817894 | 0.704 | 1.256 | -1.796 | 0.979 | -0.008 | 0.006 | 0.006 | 0.033 | 0.92 | 0.93 | 0.77 | 0.73 | 0.83 | 0.93 |
| 4 | 25067377 | rs6841005 | 0.607 | 0.933 | -2.122 | 0.447 | -0.053 | 0.118 | 0.118 | 0 | 0.32 | 0.09 | 0.28 | 0.17 | 0.34 | 0.23 |
| 4 | 25067465 | rs4697534 | 0.324 | 0.788 | -1.398 | 1.272 | 0.004 | -0.035 | -0.035 | 0.047 | 0.82 | 0.88 | 0.86 | 0.73 | 0.84 | 0.86 |
| 4 | 25068062 | rs13116533 | -0.553 | 0.577 | -1.258 | 0.489 | 0.008 | 0.004 | 0.004 | 0 | 0.24 | 0.21 | 0.31 | 0.76 | 0.25 | 0.36 |
| 4 | 25068446 | rs2098302 | 0.497 | 0.79 | -1.547 | 0.163 | -0.015 | 0.094 | 0.094 | 0 | 0.54 | 0.28 | 0.33 | 0.24 | 0.5 | 0.4 |
| 4 | 25072651 | rs2324654 | -0.571 | 0.467 | -0.93 | -0.482 | -0.003 | -0.062 | -0.062 | 0.379 | 0.69 | 0.68 | 0.77 | 0.98 | 0.69 | 0.7 |
| 4 | 25081118 | rs7693334 | 0.226 | 0.746 | -1.319 | 0.078 | -0.084 | -0.14 | -0.14 | 0.029 | 0.22 | 0.03 | 0.34 | 0.39 | 0.25 | 0.26 |
| 4 | 25083127 | rs16876761 | -0.148 | 0.343 | -0.683 | 0.142 | -0.012 | 0.034 | 0.034 | 0.468 | 0.06 | 0.21 | 0.06 | 0.08 | 0.01 | 0.08 |
| 4 | 25087708 | rs7682064 | 0.687 | 0.451 | -0.671 | -0.025 | -0.006 | -0.066 | -0.066 | 4.674 | 0.28 | 0.32 | 0.22 | 0.04 | 0.27 | 0.31 |
| 4 | 25088919 | rs6846989 | -0.927 | 0.671 | -0.771 | 0.192 | 0.005 | 0.019 | 0.019 | 0 | 0.65 | 0.55 | 0.69 | 0.52 | 0.62 | 0.71 |
| 4 | 25102070 | rs16876811 | -0.595 | 0.727 | -0.664 | 0.584 | -0.002 | 0.002 | 0.002 | 0 | 0.06 | 0.06 | 0.07 | 0.11 | 0.05 | 0.06 |
| 4 | 25107115 | rs759243 | -0.929 | 0.658 | -0.691 | 0.597 | -0.001 | 0 | 0 | 0 | 0.39 | 0.58 | 0.48 | 0.48 | 0.36 | 0.47 |
| 4 | 25113048 | rs16876855 | -1.547 | 0.431 | -0.832 | 0.539 | 0.001 | 0.012 | 0.012 | 0.12 | 0.1 | 0.04 | 0.1 | 0.08 | 0.08 | 0.12 |
| 4 | 25114103 | rs16876859 | -1.082 | 0.476 | -1.02 | 0.898 | -0.01 | -0.027 | -0.027 | 0.783 | 0.22 | 0.12 | 0.24 | 0.26 | 0.19 | 0.25 |
| 4 | 25114664 | rs10939038 | 1.168 | 0.006 | -1.566 | 0.427 | -0.06 | 0.078 | 0.078 | 0 | 0.34 | 0.09 | 0.35 | 0.17 | 0.41 | 0.28 |
| 4 | 25130283 | rs16876882 | -0.407 | -0.794 | 0.172 | 1.05 | 0.001 | 0.008 | 0.008 | 0 | 0.11 | 0.07 | 0.08 | 0.01 | 0.11 | 0.09 |
| 4 | 25133669 | rs3756208 | 0.238 | -0.925 | 0.518 | 0.992 | -0.016 | 0.003 | 0.003 | 0 | 0.59 | 0.47 | 0.62 | 0.38 | 0.66 | 0.62 |
| 4 | 25135095 | rs10213282 | -1.619 | -1.106 | 0.876 | 1.14 | 0.001 | -0.011 | -0.011 | 0.539 | 0.79 | 0.82 | 0.83 | 0.65 | 0.81 | 0.85 |
| 4 | 25146307 | rs2302565 | 0.583 | -1.108 | 0.878 | 1.141 | -0.015 | 0.016 | 0.016 | 0 | 0.6 | 0.48 | 0.69 | 0.49 | 0.66 | 0.66 |
| 4 | 25148136 | rs3796795 | -1.197 | -1.295 | 0.903 | 1.134 | -0.018 | 0.007 | 0.007 | 0 | 0.16 | 0.32 | 0.13 | 0.16 | 0.13 | 0.15 |
| 4 | 25163672 | rs11937742 | 1.092 | -1.271 | 0.897 | 1.107 | 0.001 | 0.017 | 0.017 | 0 | 0.45 | 0.38 | 0.51 | 0.28 | 0.56 | 0.42 |
| 4 | 25168417 | rs13139513 | 1.655 | -1.261 | 0.901 | 1.077 | -0.06 | 0.088 | 0.088 | 0 | 0.39 | 0.14 | 0.39 | 0.18 | 0.5 | 0.32 |
| 4 | 25168491 | rs6845806 | -0.523 | -1.129 | 1.08 | 1.211 | -0.012 | 0.07 | 0.07 | 0 | 0.06 | 0.15 | 0.04 | 0.15 | 0.03 | 0.07 |
| 4 | 25174251 | rs6838393 | 0.032 | -1.111 | 0.971 | 1.152 | -0.007 | 0.002 | 0.002 | 0.362 | 0.66 | 0.58 | 0.72 | 0.64 | 0.69 | 0.74 |
| 4 | 25176971 | rs10018167 | -1.353 | -1.258 | 1.023 | 1.389 | 0.003 | -0.011 | -0.011 | 0 | 0.79 | 0.83 | 0.83 | 0.66 | 0.81 | 0.86 |
| 4 | 25200584 | rs1909475 | -0.075 | -1.131 | 1.115 | 1.403 | -0.012 | 0.045 | 0.045 | 0.534 | 0.67 | 0.48 | 0.66 | 0.53 | 0.7 | 0.68 |
| 4 | 25204724 | rs12505283 | 0.4 | -1.187 | 1.041 | 1.359 | 0.019 | 0.001 | 0.001 | 0.665 | 0.63 | 0.27 | 0.57 | 0.35 | 0.69 | 0.63 |
| 4 | 25226538 | rs10007349 | 0.887 | -1.171 | 0.904 | 1.429 | 0 | 0.008 | 0.008 | 0 | 0.07 | 0.12 | 0.11 | 0.11 | 0.06 | 0.13 |
| 4 | 25227917 | rs7654358 | 1.29 | -1.457 | 0.83 | 1.226 | 0.034 | 0.09 | 0.09 | 0 | 0.46 | 0.2 | 0.51 | 0.27 | 0.61 | 0.44 |
| 4 | 25237391 | rs313548 | -0.443 | -1.465 | 0.996 | 1.347 | -0.017 | 0.025 | 0.025 | 0.361 | 0.72 | 0.56 | 0.77 | 0.66 | 0.76 | 0.78 |
| 4 | 25253995 | rs313567 | -1.431 | -1.739 | 1.012 | 1.407 | 0.002 | -0.003 | -0.003 | 0 | 0.8 | 0.8 | 0.82 | 0.68 | 0.84 | 0.82 |
| 4 | 25255841 | rs10006479 | 1.26 | -1.814 | 1.104 | 1.339 | 0.026 | 0.028 | 0.028 | 0.079 | 0.48 | 0.32 | 0.46 | 0.36 | 0.6 | 0.42 |
| 4 | 25268962 | rs313543 | -0.823 | -1.834 | 1.382 | 1.352 | -0.036 | 0.054 | 0.054 | 1.021 | 0.76 | 0.58 | 0.71 | 0.34 | 0.82 | 0.73 |
| 4 | 25270011 | rs3115232 | 0.972 | -1.858 | 1.23 | 1.293 | -0.002 | 0.069 | 0.069 | 0 | 0.15 | 0.03 | 0.11 | 0.03 | 0.11 | 0.19 |
| 4 | 25272880 | rs16876963 | -0.05 | -1.851 | 1.391 | 1.292 | 0.011 | -0.025 | -0.025 | 1.218 | 0.06 | 0.15 | 0.05 | 0 | 0.06 | 0.03 |
| 4 | 25273241 | rs7696265 | 1.061 | -1.845 | 1.026 | 1.303 | 0 | 0.002 | 0.002 | 0 | 0.07 | 0.1 | 0.11 | 0.1 | 0.06 | 0.15 |
| 4 | 25278988 | rs3796780 | 1.494 | -1.812 | 0.998 | 1.342 | -0.002 | 0.122 | 0.122 | 0.175 | 0.44 | 0.14 | 0.43 | 0.19 | 0.58 | 0.36 |
| 4 | 25279320 | rs6834255 | 1.166 | -1.78 | 0.926 | 1.422 | 0.006 | -0.01 | -0.01 | 0 | 0.2 | 0.17 | 0.18 | 0.32 | 0.16 | 0.18 |
| 4 | 25281948 | rs12505006 | 2.088 | -2.097 | 0.731 | 1.21 | -0.121 | 0.046 | 0.046 | 0.207 | 0.36 | 0.05 | 0.39 | 0.19 | 0.55 | 0.3 |
| 4 | 25282410 | rs4370127 | -0.281 | -1.603 | 1.078 | 1.371 | 0.006 | 0.031 | 0.031 | 0 | 0.78 | 0.87 | 0.78 | 0.86 | 0.83 | 0.67 |
| 4 | 25285913 | rs12649921 | 1.086 | -1.573 | 1.023 | 1.398 | 0.001 | -0.002 | -0.002 | 0 | 0.2 | 0.2 | 0.18 | 0.32 | 0.16 | 0.18 |
| 4 | 25294239 | rs6448346 | -1.451 | -1.297 | 1.177 | 1.209 | 0.02 | 0.139 | 0.139 | 0.548 | 0.1 | 0.46 | 0.08 | 0.17 | 0.07 | 0.07 |
| 4 | 25296348 | rs7672293 | 0.782 | -1.249 | 0.588 | 1.199 | -0.011 | -0.006 | -0.006 | 0 | 0.67 | 0.39 | 0.69 | 0.69 | 0.76 | 0.58 |
| 4 | 25309815 | rs13133104 | 0.23 | -0.579 | 0.092 | 1.478 | 0.004 | 0.053 | 0.053 | 0 | 0.09 | 0.17 | 0.14 | 0.35 | 0.1 | 0.07 |
| 4 | 25310060 | rs11726511 | -1.715 | -1.502 | 0.477 | 1.716 | -0.032 | 0.019 | 0.019 | 0 | 0.26 | 0.39 | 0.2 | 0.31 | 0.16 | 0.29 |
| 4 | 25310351 | rs316807 | 1.097 | -1.509 | 0.314 | 1.73 | 0.013 | -0.022 | -0.022 | 0 | 0.67 | 0.75 | 0.66 | 0.47 | 0.78 | 0.53 |
| 4 | 25310628 | rs16876992 | -0.255 | -1.663 | 0.367 | 1.82 | -0.017 | 0.002 | 0.002 | 1.393 | 0.28 | 0.63 | 0.25 | 0.28 | 0.21 | 0.23 |
| 4 | 25312242 | rs1533131 | 0.121 | -1.627 | 0.314 | 1.817 | 0.024 | -0.001 | -0.001 | 2.266 | 0.76 | 0.87 | 0.84 | 0.72 | 0.85 | 0.65 |
| 4 | 25312357 | rs1533132 | 0.171 | -1.639 | 0.308 | 1.811 | 0.026 | 0 | 0 | 0 | 0.76 | 0.87 | 0.84 | 0.72 | 0.85 | 0.65 |
| 4 | 25313994 | rs16877005 | -0.745 | -1.624 | 0.329 | 1.883 | -0.012 | 0.055 | 0.055 | 0.104 | 0.09 | 0.12 | 0.1 | 0.35 | 0.05 | 0.14 |
| 4 | 25316815 | rs12508686 | 1.496 | -1.696 | 0.43 | 1.759 | 0.036 | 0.057 | 0.057 | 0 | 0.48 | 0.31 | 0.52 | 0.34 | 0.61 | 0.41 |
| 4 | 25317277 | rs1553876 | 2.218 | -1.69 | 0.398 | 1.747 | -0.026 | 0.095 | 0.095 | 0 | 0.4 | 0.2 | 0.43 | 0.18 | 0.56 | 0.3 |
| 4 | 25318984 | rs4697567 | -0.869 | -1.709 | 0.536 | 1.795 | -0.021 | 0.012 | 0.012 | 0 | 0.14 | 0.24 | 0.13 | 0.11 | 0.09 | 0.17 |
| 4 | 25319698 | rs10939051 | 1.537 | -1.735 | 0.533 | 1.796 | 0.037 | 0.058 | 0.058 | 0 | 0.48 | 0.31 | 0.52 | 0.34 | 0.61 | 0.41 |
| 4 | 25320465 | rs10031354 | 1.537 | -1.735 | 0.533 | 1.796 | 0.037 | 0.058 | 0.058 | 0 | 0.48 | 0.3 | 0.52 | 0.34 | 0.61 | 0.41 |
| 4 | 25324742 | rs1963656 | 2.162 | -1.753 | 0.524 | 1.79 | -0.04 | 0.096 | 0.096 | 0 | 0.4 | 0.15 | 0.42 | 0.18 | 0.56 | 0.3 |
| 4 | 25324899 | rs12642624 | 1.292 | -1.606 | 0.592 | 1.938 | -0.031 | 0.092 | 0.092 | 1.812 | 0.17 | 0.46 | 0.18 | 0.25 | 0.13 | 0.15 |
| 4 | 25325264 | rs13128106 | -0.467 | -1.519 | 0.386 | 1.989 | -0.001 | 0.004 | 0.004 | 0 | 0.09 | 0.05 | 0.06 | 0.05 | 0.09 | 0.1 |
| 4 | 25327288 | rs2667295 | -1.2 | -1.519 | 0.386 | 1.99 | -0.021 | -0.052 | -0.052 | 4.642 | 0.84 | 0.99 | 0.88 | 0.75 | 0.89 | 0.77 |
| 4 | 25342463 | rs316783 | 0.578 | -1.518 | 0.386 | 1.972 | -0.013 | -0.037 | -0.037 | 0 | 0.75 | 0.94 | 0.82 | 0.7 | 0.81 | 0.68 |
| 4 | 25344591 | rs16877042 | 1.106 | -1.493 | 0.432 | 1.958 | -0.001 | 0.008 | 0.008 | 0 | 0.07 | 0.07 | 0.1 | 0.1 | 0.07 | 0.12 |

| | | | | | | | | | | | | | | | | |
|---|---|---|---|---|---|---|---|---|---|---|---|---|---|---|---|---|
| 4 | 25347771 | rs13114588 | -0.488 | -1.487 | 0.407 | 1.889 | -0.001 | 0.004 | 0.004 | 0 | 0.09 | 0.06 | 0.06 | 0.05 | 0.09 | 0.13 |
| 4 | 25349921 | rs2875425 | 0.162 | -1.45 | 0.448 | 1.984 | -0.007 | 0.088 | 0.088 | 0 | 0.1 | 0.17 | 0.19 | 0.25 | 0.07 | 0.13 |
| 4 | 25352640 | rs10009504 | -1.077 | -1.39 | 0.276 | 1.834 | -0.021 | -0.052 | -0.052 | 0.375 | 0.84 | 0.99 | 0.88 | 0.75 | 0.89 | 0.8 |
| 4 | 25357164 | rs316776 | -0.327 | -1.356 | 0.419 | 1.798 | 0.012 | -0.017 | -0.017 | 0 | 0.81 | 0.9 | 0.86 | 0.73 | 0.88 | 0.79 |
| 4 | 25362101 | rs184360 | -0.355 | -1.392 | 0.947 | 1.75 | -0.035 | 0.014 | 0.014 | 0.857 | 0.27 | 0.48 | 0.24 | 0.29 | 0.22 | 0.26 |
| 4 | 25367546 | rs315674 | 1.552 | -1.374 | 0.955 | 1.759 | 0.006 | -0.01 | -0.01 | 0.246 | 0.16 | 0.13 | 0.14 | 0.25 | 0.12 | 0.16 |
| 4 | 25371815 | rs315688 | 0.959 | -1.374 | 1.026 | 1.759 | -0.037 | 0.06 | 0.06 | 2.259 | 0.18 | 0.39 | 0.18 | 0.25 | 0.13 | 0.14 |
| 4 | 25382783 | rs2324761 | -0.303 | -1.406 | 0.755 | 1.773 | 0.01 | -0.006 | -0.006 | 0.115 | 0.81 | 0.87 | 0.87 | 0.76 | 0.87 | 0.76 |
| 4 | 25386559 | rs10027814 | 2.066 | -1.427 | 0.589 | 1.57 | -0.023 | 0.074 | 0.074 | 0 | 0.4 | 0.2 | 0.48 | 0.18 | 0.55 | 0.31 |
| 4 | 25391584 | rs186408 | -1.485 | -1.523 | 0.499 | 1.722 | -0.038 | -0.078 | -0.078 | 0 | 0.88 | 0.99 | 0.89 | 0.67 | 0.89 | 0.8 |
| 4 | 25394785 | rs315669 | 0.985 | -1.544 | 0.728 | 1.753 | -0.001 | 0.001 | 0.001 | 2.6 | 0.2 | 0.24 | 0.21 | 0.22 | 0.19 | 0.12 |
| 4 | 25397459 | rs13152467 | -0.189 | -1.536 | 0.738 | 1.728 | 0.001 | 0.001 | 0.001 | 0 | 0.08 | 0.07 | 0.04 | 0.04 | 0.03 | 0.12 |
| 4 | 25400468 | rs10032586 | -0.24 | -1.514 | 0.609 | 1.769 | -0.008 | -0.003 | -0.003 | 1.181 | 0.08 | 0.1 | 0.05 | 0.06 | 0.04 | 0.12 |
| 4 | 25401863 | rs13109196 | 0.699 | -1.462 | 0.567 | 1.816 | 0.002 | -0.013 | -0.013 | 0 | 0.06 | 0 | 0.02 | 0.05 | 0.03 | 0.09 |
| 4 | 25402066 | rs12506959 | 0.904 | -1.414 | 0.811 | 1.868 | 0.003 | 0.003 | 0.003 | 0 | 0.54 | 0.41 | 0.54 | 0.36 | 0.62 | 0.52 |
| 4 | 25407356 | rs3756202 | 1.22 | -1.4 | 0.891 | 1.869 | -0.001 | -0.002 | -0.002 | 0 | 0.19 | 0.22 | 0.13 | 0.17 | 0.18 | 0.11 |
| 4 | 25415106 | rs1993602 | 2.367 | -1.365 | 0.851 | 1.839 | -0.055 | 0.106 | 0.106 | 0 | 0.41 | 0.12 | 0.46 | 0.2 | 0.55 | 0.31 |
| 4 | 25417244 | rs3816587 | 1.664 | -1.315 | 0.835 | 1.865 | 0.001 | 0.003 | 0.003 | 0 | 0.49 | 0.46 | 0.52 | 0.31 | 0.6 | 0.42 |
| 4 | 25417371 | rs10517073 | 1.652 | -1.306 | 0.712 | 1.87 | 0.019 | 0.046 | 0.046 | 0 | 0.5 | 0.57 | 0.54 | 0.3 | 0.6 | 0.42 |
| 4 | 25419283 | rs9174 | 1.336 | -1.306 | 0.702 | 1.882 | 0.024 | 0.059 | 0.059 | 0 | 0.5 | 0.62 | 0.55 | 0.31 | 0.6 | 0.42 |
| 4 | 25427554 | rs4697095 | 0.008 | -1.345 | 0.44 | 1.965 | 0.006 | -0.051 | -0.051 | 0 | 0.26 | 0.16 | 0.2 | 0.42 | 0.25 | 0.28 |
| 4 | 25429854 | rs1996180 | -0.745 | -1.307 | 0.494 | 1.771 | 0.008 | -0.007 | -0.007 | 0 | 0.36 | 0.39 | 0.31 | 0.37 | 0.28 | 0.35 |
| 4 | 25432828 | rs13131313 | -0.242 | -1.202 | 0.226 | 1.783 | 0.012 | -0.017 | -0.017 | 0 | 0.17 | 0.13 | 0.21 | 0.18 | 0.11 | 0.26 |
| 4 | 25438518 | rs6837837 | -1.096 | -0.86 | -0.443 | 1.986 | -0.026 | -0.068 | -0.068 | 0 | 0.88 | 0.98 | 0.89 | 0.67 | 0.9 | 0.79 |
| 4 | 25440525 | rs6852016 | -1.206 | -0.962 | -0.346 | 1.847 | 0.008 | 0.001 | 0.001 | 0 | 0.44 | 0.66 | 0.37 | 0.38 | 0.33 | 0.45 |
| 4 | 25440832 | rs7698526 | 1.023 | -0.729 | -0.61 | 1.523 | -0.002 | -0.001 | -0.001 | 0 | 0.17 | 0.14 | 0.09 | 0.18 | 0.17 | 0.09 |
| 4 | 25444762 | rs10008104 | -0.46 | -0.607 | -0.772 | 1.471 | -0.008 | -0.001 | -0.001 | 0 | 0.43 | 0.57 | 0.34 | 0.39 | 0.34 | 0.48 |
| 4 | 25446100 | rs6448356 | -0.08 | 0.118 | -0.907 | 1.909 | -0.026 | -0.05 | -0.05 | 0 | 0.88 | 0.96 | 0.87 | 0.63 | 0.88 | 0.79 |
| 4 | 25455404 | rs922054 | 0.173 | -0.338 | -0.942 | 0.972 | 0.023 | 0.073 | 0.073 | 0 | 0.58 | 0.23 | 0.57 | 0.21 | 0.65 | 0.52 |
| 4 | 25457501 | rs1996182 | 0.014 | -0.46 | -0.32 | 0.659 | -0.001 | -0.017 | -0.017 | 0 | 0.47 | 0.57 | 0.44 | 0.17 | 0.5 | 0.36 |
| 4 | 25457661 | rs1996183 | -0.848 | -0.818 | -0.132 | 0.893 | -0.012 | -0.001 | -0.001 | 0 | 0.2 | 0.26 | 0.13 | 0.18 | 0.18 | 0.14 |
| 4 | 25464527 | rs12507905 | -0.733 | -0.83 | -0.033 | 1.346 | -0.021 | 0.088 | 0.088 | 0 | 0.13 | 0.32 | 0.29 | 0.23 | 0.12 | 0.16 |
| 4 | 25464878 | rs3851436 | 0.79 | -0.814 | -0.247 | 1.074 | 0 | -0.073 | -0.073 | 0 | 0.38 | 0.35 | 0.24 | 0.02 | 0.4 | 0.29 |
| 4 | 25465790 | rs13114364 | 0.188 | -0.798 | -0.611 | 1.386 | -0.028 | -0.029 | -0.029 | 0.004 | 0.41 | 0.35 | 0.44 | 0.52 | 0.45 | 0.35 |
| 4 | 25466614 | rs13116199 | 0.217 | -0.121 | -0.753 | 1.198 | -0.013 | -0.006 | -0.006 | 0.496 | 0.41 | 0.33 | 0.64 | 0.51 | 0.48 | 0.38 |
| 4 | 25467149 | rs1399240 | 0.481 | -0.335 | -0.642 | 1.726 | 0.008 | -0.011 | -0.011 | 0 | 0.88 | 0.93 | 0.95 | 0.69 | 0.95 | 0.81 |
| 4 | 25475192 | rs12650434 | 0.913 | -0.358 | -0.229 | 1.667 | -0.017 | 0.026 | 0.026 | 0.01 | 0.81 | 0.63 | 0.73 | 0.68 | 0.83 | 0.77 |
| 4 | 25475510 | rs959018 | 0.838 | -0.41 | -0.178 | 1.775 | -0.013 | 0.011 | 0.011 | 0 | 0.93 | 0.92 | 0.94 | 0.85 | 0.96 | 0.89 |
| 4 | 25486335 | rs4697098 | 1.277 | -0.709 | 0.11 | 1.758 | -0.002 | 0 | 0 | 0 | 0.55 | 0.44 | 0.5 | 0.43 | 0.63 | 0.55 |
| 4 | 25500479 | rs6851586 | 0.077 | -0.731 | 0.423 | 1.816 | 0.017 | 0.086 | 0.086 | 0.05 | 0.06 | 0.15 | 0.12 | 0.21 | 0.06 | 0.05 |
| 4 | 25502383 | rs989263 | 0.972 | -0.799 | 0.462 | 1.773 | -0.011 | -0.022 | -0.022 | 0 | 0.61 | 0.5 | 0.61 | 0.74 | 0.69 | 0.6 |
| 4 | 25505735 | rs6854268 | 1.049 | -0.786 | 0.47 | 1.785 | -0.009 | -0.006 | -0.006 | 0 | 0.61 | 0.52 | 0.61 | 0.63 | 0.69 | 0.6 |
| 4 | 25505932 | rs6831932 | 1.515 | -0.686 | 0.733 | 1.871 | 0.028 | 0.012 | 0.012 | 0.008 | 0.51 | 0.28 | 0.47 | 0.38 | 0.58 | 0.51 |
| 4 | 25507698 | rs6448365 | 1.497 | -0.223 | 1.025 | 2.226 | 0.004 | 0.091 | 0.091 | 0.234 | 0.9 | 0.71 | 0.85 | 0.78 | 0.9 | 0.85 |
| 4 | 25514402 | rs4053860 | -2.065 | -0.276 | 1.246 | 2.249 | 0.001 | 0.072 | 0.072 | 0.697 | 0.2 | 0.38 | 0.35 | 0.38 | 0.21 | 0.23 |
| 4 | 25531234 | rs6448367 | -0.442 | -0.268 | 1.094 | 2.201 | 0.001 | 0.028 | 0.028 | 0 | 0.1 | 0.15 | 0.21 | 0.3 | 0.11 | 0.09 |
| 4 | 25533554 | rs4697584 | -1.9 | -0.313 | 1.289 | 2.177 | -0.013 | 0.077 | 0.077 | 0.234 | 0.18 | 0.34 | 0.27 | 0.33 | 0.15 | 0.19 |
| 4 | 25536386 | rs10018150 | -0.473 | 0.017 | 0.85 | 1.94 | 0.001 | 0.003 | 0.003 | 0 | 0.09 | 0.13 | 0.2 | 0.15 | 0.11 | 0.09 |
| 4 | 25542206 | rs12331861 | -0.623 | 0.07 | 1.082 | 1.935 | 0.012 | 0.016 | 0.016 | 0 | 0.08 | 0.13 | 0.18 | 0.14 | 0.1 | 0.1 |
| 4 | 25543152 | rs871476 | 0.721 | -0.574 | 1.339 | 2.039 | -0.017 | 0.027 | 0.027 | 0 | 0.82 | 0.66 | 0.77 | 0.79 | 0.86 | 0.71 |
| 4 | 25546127 | rs2176838 | 0.54 | -0.423 | 1.345 | 1.993 | 0.022 | -0.008 | -0.008 | 0.507 | 0.83 | 0.67 | 0.77 | 0.86 | 0.83 | 0.74 |
| 4 | 25554332 | rs9991078 | -0.578 | -0.31 | 0.96 | 2.092 | 0 | 0.015 | 0.015 | 0 | 0.07 | 0.08 | 0.19 | 0.19 | 0.1 | 0.11 |
| 4 | 25557047 | rs7686265 | -1.997 | -0.706 | 1.423 | 2.447 | -0.039 | 0.318 | 0.318 | 0.681 | 0.15 | 0.46 | 0.28 | 0.55 | 0.11 | 0.18 |
| 4 | 25558594 | rs11937512 | -1.558 | -0.662 | 0.78 | 2.285 | -0.001 | 0.015 | 0.015 | 0.05 | 0.07 | 0.06 | 0.11 | 0.21 | 0.08 | 0.06 |
| 4 | 25562241 | rs1980239 | 0.577 | -0.084 | 0.016 | 1.588 | -0.034 | -0.002 | -0.002 | 0 | 0.74 | 0.9 | 0.75 | 0.77 | 0.69 | 0.85 |
| 4 | 25562386 | rs6830401 | -0.681 | 0.074 | 0.394 | 1.363 | 0.005 | 0.001 | 0.001 | 0 | 0.2 | 0.22 | 0.18 | 0.2 | 0.22 | 0.31 |
| 4 | 25565070 | rs6813987 | -0.203 | -0.164 | 0.176 | 1.133 | -0.02 | 0.035 | 0.035 | 0.095 | 0.89 | 0.97 | 0.94 | 0.98 | 0.89 | 0.95 |
| 4 | 25575537 | rs7689639 | -1.665 | -1.092 | -0.519 | 1.548 | 0.099 | -0.18 | -0.18 | 0 | 0.35 | 0.94 | 0.36 | 0.53 | 0.19 | 0.52 |
| 4 | 25576330 | rs13147366 | -1.284 | -1.051 | 0.281 | 1.383 | -0.045 | 0.094 | 0.094 | 0 | 0.16 | 0.41 | 0.21 | 0.26 | 0.09 | 0.29 |
| 4 | 25577184 | rs10013819 | -0.968 | -0.557 | 0.319 | 0.87 | -0.045 | 0 | 0 | 0 | 0.18 | 0.43 | 0.14 | 0.18 | 0.13 | 0.18 |
| 4 | 25583481 | rs6830679 | -0.086 | -0.028 | -0.602 | 0.38 | -0.05 | 0.018 | 0.018 | 0 | 0.86 | 0.97 | 0.84 | 0.86 | 0.85 | 0.93 |
| 4 | 25584201 | rs2308974 | 0.968 | -0.037 | 0.37 | 0.173 | 0.028 | -0.076 | -0.076 | 0 | 0.78 | 0.56 | 0.77 | 0.86 | 0.78 | 0.93 |
| 4 | 25587743 | rs4697106 | -0.258 | 0.281 | -1.004 | 0.133 | 0.08 | 0.018 | 0.018 | 0.874 | 0.42 | 0.77 | 0.35 | 0.29 | 0.29 | 0.48 |
| 4 | 25589180 | rs6448380 | -0.348 | 0.23 | -0.907 | 0.076 | 0.065 | -0.007 | -0.007 | 0 | 0.43 | 0.7 | 0.36 | 0.38 | 0.3 | 0.48 |
| 4 | 25591895 | rs6448381 | 0.722 | 0.002 | -0.631 | 0.262 | 0.004 | 0.039 | 0.039 | 0.029 | 0.73 | 0.41 | 0.65 | 0.55 | 0.73 | 0.7 |
| 4 | 25592650 | rs16877296 | 1.032 | 0.82 | -1.058 | 0.139 | -0.022 | -0.172 | -0.172 | 1.375 | 0.32 | 0.05 | 0.36 | 0.48 | 0.29 | 0.41 |
| 4 | 25594437 | rs10939067 | 0.54 | 0.825 | -0.98 | 0.133 | 0.008 | -0.048 | -0.048 | 0 | 0.37 | 0.2 | 0.41 | 0.48 | 0.35 | 0.43 |
| 4 | 25594556 | rs7664681 | -0.351 | 0.622 | -1.158 | 0.461 | 0.011 | 0.015 | 0.015 | 0 | 0.52 | 0.77 | 0.47 | 0.38 | 0.55 | 0.51 |
| 4 | 25615723 | rs939356 | -0.586 | 0.037 | -0.807 | 0.522 | 0.008 | -0.006 | -0.006 | 1.991 | 0.38 | 0.63 | 0.29 | 0.47 | 0.28 | 0.39 |

| | | | | | | | | | | | | | | | | |
|---|---|---|---|---|---|---|---|---|---|---|---|---|---|---|---|---|
| 4 | 25616103 | rs9992129 | -0.128 | -0.245 | -0.31 | 0.476 | -0.046 | -0.07 | -0.07 | 0 | 0.22 | 0.44 | 0.12 | 0.06 | 0.16 | 0.1 |
| 4 | 25620658 | rs6821985 | -0.759 | 0.423 | -0.514 | 0.202 | 0.006 | 0.024 | 0.024 | 0 | 0.27 | 0.34 | 0.26 | 0.43 | 0.26 | 0.39 |
| 4 | 25625150 | rs7671379 | -0.109 | 0.408 | -0.548 | 0.213 | 0.001 | -0.021 | -0.021 | 0 | 0.1 | 0.08 | 0.08 | 0 | 0.13 | 0.11 |
| 4 | 25627026 | rs12499526 | -0.872 | 0.302 | -0.413 | 0.252 | 0.004 | 0.026 | 0.026 | 0 | 0.47 | 0.77 | 0.35 | 0.45 | 0.44 | 0.55 |
| 4 | 25628002 | rs12504139 | -0.438 | -0.331 | -0.013 | 0.128 | 0.002 | -0.001 | -0.001 | 0 | 0.47 | 0.44 | 0.45 | 0.22 | 0.38 | 0.48 |
| 4 | 25629255 | rs7676077 | 0.979 | -0.729 | 0.677 | 0.893 | 0.003 | -0.014 | -0.014 | 0 | 0.69 | 0.53 | 0.61 | 0.23 | 0.68 | 0.76 |
| 4 | 25638736 | rs10517077 | -1.468 | -0.973 | 0.394 | 1.303 | 0.007 | -0.006 | -0.006 | 0.667 | 0.34 | 0.65 | 0.33 | 0.41 | 0.16 | 0.26 |
| 4 | 25641870 | rs4327471 | 0.387 | -0.471 | 0.003 | 1.474 | -0.087 | -0.095 | -0.095 | 0.159 | 0.86 | 0.99 | 0.85 | 0.66 | 0.76 | 0.88 |
| 4 | 25645165 | rs4580639 | 1.677 | -0.491 | 0.368 | 1.434 | 0.02 | 0.05 | 0.05 | 5.536 | 0.53 | 0.29 | 0.53 | 0.3 | 0.6 | 0.63 |
| 4 | 25654566 | rs6448388 | -2.059 | -1.302 | 0.481 | 1.798 | -0.051 | 0.016 | 0.016 | 3.122 | 0.35 | 0.49 | 0.36 | 0.64 | 0.2 | 0.31 |
| 4 | 25656404 | rs11721800 | -1.067 | -1.233 | 0.142 | 2.144 | -0.045 | 0.048 | 0.048 | 5.034 | 0.16 | 0.3 | 0.17 | 0.32 | 0.06 | 0.14 |
| 4 | 25671043 | rs3815464 | -1.431 | -0.934 | -0.142 | 1.988 | -0.005 | 0.034 | 0.034 | 0 | 0.14 | 0.17 | 0.2 | 0.6 | 0.04 | 0.17 |
| 4 | 25674194 | rs3796777 | -0.92 | -0.835 | -0.093 | 1.605 | -0.029 | 0.008 | 0.008 | 0.196 | 0.26 | 0.36 | 0.18 | 0.34 | 0.15 | 0.17 |
| 4 | 25675125 | rs2240997 | -1.844 | -0.175 | 0.147 | 1.72 | 0.009 | 0.086 | 0.086 | 0 | 0.12 | 0.34 | 0.21 | 0.27 | 0.1 | 0.09 |
| 4 | 25675203 | rs2240998 | -1.697 | 0.164 | -0.288 | 1.819 | 0.011 | 0.08 | 0.08 | 0.016 | 0.05 | 0.08 | 0.16 | 0.31 | 0.09 | 0.08 |
| 4 | 25676982 | rs12505556 | -0.699 | -0.322 | -0.48 | 1.328 | 0.005 | 0.002 | 0.002 | 0 | 0.17 | 0.12 | 0.27 | 0.58 | 0.13 | 0.17 |
| 4 | 25678396 | rs3733545 | -1.041 | -0.173 | -0.037 | 1.104 | -0.001 | 0.107 | 0.107 | 0 | 0.2 | 0.31 | 0.31 | 0.58 | 0.2 | 0.2 |
| 4 | 25678659 | rs4374611 | -0.083 | -0.572 | -1.181 | -0.181 | -0.015 | 0.138 | 0.138 | 0 | 0.79 | 0.98 | 0.88 | 1 | 0.84 | 0.81 |
| 4 | 25679209 | rs4697598 | 0.577 | -0.294 | -1.368 | -0.326 | -0.074 | -0.03 | -0.03 | 0 | 0.31 | 0.11 | 0.37 | 0.3 | 0.46 | 0.44 |
| 4 | 25681990 | rs10005751 | -0.742 | -1.37 | 0.373 | -1.57 | -0.003 | -0.025 | -0.025 | 0.116 | 0.32 | 0.34 | 0.28 | 0.11 | 0.24 | 0.23 |
| 4 | 25702694 | rs9654104 | -0.421 | -1.6 | 0.653 | -1.613 | -0.001 | -0.03 | -0.03 | 0.038 | 0.16 | 0.15 | 0.11 | 0 | 0.12 | 0.11 |
| 4 | 25705533 | rs868838 | -0.537 | -1.695 | 0.404 | -1.663 | 0.021 | 0.082 | 0.082 | 0 | 0.17 | 0.04 | 0.09 | 0 | 0.12 | 0.11 |
| 4 | 25705912 | rs7692656 | -1.609 | -1.787 | 0.454 | -1.847 | 0.017 | 0.006 | 0.006 | 0 | 0.42 | 0.64 | 0.35 | 0.13 | 0.26 | 0.25 |
| 4 | 25707422 | rs13152663 | -1.627 | -1.795 | 0.483 | -1.866 | 0.025 | 0.023 | 0.023 | 0 | 0.42 | 0.67 | 0.35 | 0.14 | 0.26 | 0.25 |
| 4 | 25721489 | rs16877508 | -0.985 | -0.97 | 0.242 | -1.138 | 0.013 | 0.015 | 0.015 | 0 | 0.21 | 0.13 | 0.2 | 0.13 | 0.12 | 0.14 |
| 4 | 25729059 | rs4328901 | -1.217 | -0.639 | 0.552 | -1.019 | -0.028 | -0.05 | -0.05 | 0 | 0.22 | 0.55 | 0.28 | 0.13 | 0.16 | 0.1 |
| 4 | 25730546 | rs11721812 | -0.81 | -0.264 | 0.657 | -1.078 | 0.003 | -0.052 | -0.052 | 0.361 | 0.15 | 0.36 | 0.08 | 0.05 | 0.11 | 0.12 |
| 4 | 25734246 | rs11935381 | -1.164 | -0.036 | 0.436 | -1.331 | -0.014 | -0.148 | -0.148 | 0.089 | 0.24 | 0.52 | 0.17 | 0.01 | 0.21 | 0.16 |
| 4 | 25740206 | rs4395489 | 0.191 | 0.202 | -0.212 | -0.984 | -0.009 | 0.014 | 0.014 | 0.102 | 0.39 | 0.32 | 0.48 | 0.66 | 0.47 | 0.38 |
| 4 | 25744431 | rs6823673 | 0.38 | 0.293 | 0.219 | -1.173 | 0.003 | -0.012 | -0.012 | 0 | 0.73 | 0.7 | 0.66 | 0.79 | 0.72 | 0.71 |
| 4 | 25745088 | rs7684489 | -0.248 | 0.187 | 0.294 | -1.301 | -0.004 | 0.036 | 0.036 | 0 | 0.15 | 0.33 | 0.25 | 0.21 | 0.12 | 0.19 |
| 4 | 25745489 | rs4697608 | 0.916 | 0.257 | 0.334 | -1.321 | 0.01 | -0.023 | -0.023 | 0 | 0.66 | 0.46 | 0.62 | 0.69 | 0.66 | 0.6 |
| 4 | 25749303 | rs7701 | 0.61 | -0.34 | -0.679 | -1.548 | -0.046 | 0.051 | 0.051 | 0 | 0.39 | 0.15 | 0.44 | 0.69 | 0.45 | 0.45 |
| 4 | 25750849 | rs4697610 | 0.567 | -0.435 | -0.236 | -1.593 | -0.018 | -0.029 | -0.029 | 0 | 0.33 | 0.42 | 0.28 | 0.23 | 0.24 | 0.2 |
| 4 | 25758940 | rs4697613 | -0.442 | -0.217 | -0.307 | -1.678 | 0.013 | -0.042 | -0.042 | 0 | 0.87 | 0.85 | 0.9 | 1 | 0.81 | 0.87 |
| 4 | 25758995 | rs4697614 | 0.874 | -0.521 | -0.451 | -1.686 | 0.016 | 0.052 | 0.052 | 0 | 0.31 | 0.09 | 0.25 | 0.24 | 0.22 | 0.24 |
| 4 | 25760828 | rs2276953 | -0.121 | -0.733 | -0.255 | -1.73 | 0.025 | 0.057 | 0.057 | 0 | 0.44 | 0.34 | 0.4 | 0.27 | 0.32 | 0.36 |
| 4 | 25761406 | rs11731086 | -0.451 | -0.698 | -0.301 | -1.541 | 0.021 | 0.038 | 0.038 | 0 | 0.49 | 0.39 | 0.48 | 0.38 | 0.38 | 0.44 |
| 4 | 25765527 | rs6810985 | -0.077 | -0.471 | -1.337 | -1.099 | -0.005 | 0.001 | 0.001 | 0 | 0.78 | 0.84 | 0.83 | 0.81 | 0.75 | 0.73 |
| 4 | 25769327 | rs2290579 | 0.859 | -0.337 | -1.599 | -0.953 | 0.009 | 0.046 | 0.046 | 0 | 0.31 | 0.09 | 0.25 | 0.24 | 0.24 | 0.25 |
| 4 | 25771115 | rs4697617 | 0.271 | 0.24 | -1.196 | -0.649 | -0.061 | -0.01 | -0.01 | 0 | 0.77 | 0.95 | 0.84 | 0.8 | 0.74 | 0.73 |
| 4 | 25771973 | rs6448402 | 0.307 | 0.479 | -0.968 | -0.436 | -0.073 | 0.062 | 0.062 | 0 | 0.63 | 0.86 | 0.56 | 0.23 | 0.51 | 0.45 |
| 4 | 25772477 | rs13128580 | -0.518 | 0.794 | -0.257 | -0.378 | 0.027 | 0.02 | 0.02 | 2.246 | 0.76 | 0.76 | 0.65 | 0.38 | 0.66 | 0.68 |
| 4 | 25775034 | rs6819016 | 0.208 | 0.717 | -0.342 | -0.52 | -0.006 | -0.008 | -0.008 | 0 | 0.12 | 0.16 | 0.11 | 0.1 | 0.11 | 0.21 |
| 4 | 25775618 | rs4697618 | 0.353 | 0.262 | -0.718 | -1.174 | 0.012 | -0.032 | -0.032 | 0.426 | 0.35 | 0.3 | 0.17 | 0 | 0.28 | 0.25 |
| 4 | 25779985 | rs2309286 | 0.198 | 0.076 | -0.806 | -1.072 | 0.003 | 0.002 | 0.002 | 0 | 0.41 | 0.61 | 0.51 | 0.38 | 0.33 | 0.42 |
| 4 | 25780891 | rs4697619 | -1.026 | 0.304 | -0.884 | -0.946 | -0.099 | -0.07 | -0.07 | 0 | 0.77 | 0.93 | 0.7 | 0.38 | 0.63 | 0.68 |
| 4 | 25784732 | rs7667794 | 0.644 | 0.256 | -0.383 | -0.821 | -0.008 | -0.002 | -0.002 | 0 | 0.28 | 0.29 | 0.39 | 0.28 | 0.22 | 0.21 |
| 4 | 25784979 | rs9997403 | -0.341 | 0.561 | -0.567 | -0.713 | -0.002 | -0.065 | -0.065 | 0 | 0.3 | 0.28 | 0.14 | 0 | 0.21 | 0.24 |
| 4 | 25803830 | rs3733543 | 0.4 | 1.685 | -0.38 | 0.519 | 0.001 | 0.001 | 0.001 | 1.057 | 0.79 | 0.77 | 0.73 | 0.37 | 0.67 | 0.75 |
| 4 | 25813070 | rs936232 | 0.241 | 0.707 | -0.154 | -0.237 | 0.043 | 0.072 | 0.072 | 0 | 0.45 | 0.69 | 0.43 | 0.25 | 0.34 | 0.31 |
| 4 | 25813119 | rs6448405 | -0.21 | 1.636 | -0.482 | 0.514 | -0.014 | 0.003 | 0.003 | 0 | 0.87 | 0.9 | 0.86 | 0.57 | 0.71 | 0.83 |
| 4 | 25816048 | rs10000609 | -0.658 | 0.359 | -0.479 | 0.409 | -0.027 | 0.018 | 0.018 | 3.193 | 0.15 | 0.77 | 0.23 | 0.27 | 0.03 | 0.15 |
| 4 | 25818473 | rs1378952 | -0.065 | 0.298 | -0.609 | 0.44 | 0.099 | 0.015 | 0.015 | 0.139 | 0.48 | 0.82 | 0.59 | 0.49 | 0.34 | 0.42 |
| 4 | 25818674 | rs729317 | 1.258 | 0.304 | -0.626 | 0.434 | -0.041 | 0.11 | 0.11 | 0.404 | 0.33 | 0.02 | 0.35 | 0.2 | 0.31 | 0.26 |
| 4 | 25822281 | rs10939083 | -0.768 | 0.262 | 0.666 | 0.726 | -0.027 | 0.023 | 0.023 | 3.038 | 0.15 | 0.77 | 0.23 | 0.29 | 0.03 | 0.15 |
| 4 | 25823624 | rs7676030 | 0.213 | 0.351 | 0.365 | 0.408 | 0.015 | -0.012 | -0.012 | 0 | 0.08 | 0.06 | 0.02 | 0.08 | 0.03 | 0.09 |
| 4 | 25837310 | rs6847559 | -0.477 | 0.743 | -0.163 | 0.007 | 0.019 | 0.07 | 0.07 | 0.141 | 0.39 | 0.76 | 0.37 | 0.12 | 0.32 | 0.33 |
| 4 | 25838526 | rs17681098 | 1.264 | 0.637 | -0.598 | 0.335 | 0.002 | 0.046 | 0.046 | 2.039 | 0.38 | 0.15 | 0.45 | 0.32 | 0.36 | 0.3 |
| 4 | 25846377 | rs6448409 | -0.492 | 0.672 | -0.055 | 0.141 | -0.041 | -0.091 | -0.091 | 1.847 | 0.34 | 0.48 | 0.28 | 0.11 | 0.25 | 0.29 |
| 4 | 25861359 | rs6842695 | -0.376 | 0.107 | -0.835 | -0.089 | 0.013 | 0.002 | 0.002 | 0.682 | 0.23 | 0.81 | 0.17 | 0.01 | 0.13 | 0.15 |
| 4 | 25863375 | rs6831024 | 0.337 | 0.709 | -0.942 | 0.37 | -0.003 | -0.003 | -0.003 | 0 | 0.65 | 0.32 | 0.63 | 0.54 | 0.52 | 0.64 |
| 4 | 25867118 | rs13140429 | -0.176 | 0.162 | -1.278 | -0.04 | 0.032 | 0.028 | 0.028 | 0 | 0.17 | 0.1 | 0.07 | 0 | 0.09 | 0.14 |
| 4 | 25871101 | rs731284 | 0.75 | 0.582 | -2.185 | -0.032 | 0.002 | 0.098 | 0.098 | 0.433 | 0.56 | 0.23 | 0.53 | 0.74 | 0.44 | 0.42 |
| 4 | 25874058 | rs6820540 | 0.292 | 0.941 | -1.976 | -0.087 | 0 | -0.001 | -0.001 | 0 | 0.72 | 0.33 | 0.73 | 0.75 | 0.63 | 0.58 |
| 4 | 25875261 | rs1545606 | 1.238 | 0.343 | -2.639 | 0.267 | -0.076 | 0.014 | 0.014 | 0.043 | 0.47 | 0.04 | 0.52 | 0.35 | 0.43 | 0.4 |
| 4 | 25880334 | rs13118224 | 0.729 | 0.277 | -2.207 | 0.429 | -0.038 | 0.024 | 0.024 | 0 | 0.61 | 0.25 | 0.69 | 0.64 | 0.53 | 0.52 |
| 4 | 25881877 | rs7681279 | -1.091 | -0.067 | -2.071 | 0.388 | -0.018 | 0 | 0 | 0.686 | 0.32 | 0.43 | 0.23 | 0.29 | 0.24 | 0.27 |
| 4 | 25887058 | rs12642988 | 0.385 | -0.268 | -2.17 | 0.401 | -0.003 | 0.009 | 0.009 | 0.497 | 0.17 | 0.23 | 0.16 | 0.29 | 0.12 | 0.19 |

| | | | | | | | | | | | | | | | | |
|---|---|---|---|---|---|---|---|---|---|---|---|---|---|---|---|---|
| 4 | 25895258 | rs12512962 | 0.034 | -0.147 | -2.089 | 0.122 | 0.02 | 0.076 | 0.076 | 0.012 | 0.13 | 0.01 | 0.04 | 0 | 0.1 | 0.08 |
| 4 | 25908919 | rs6818008 | -0.013 | 0.087 | -2.003 | 0.128 | 0.067 | -0.006 | -0.006 | 0 | 0.72 | 0.51 | 0.73 | 0.71 | 0.6 | 0.63 |
| 4 | 25911176 | rs9647441 | -0.989 | 0.488 | -1.923 | 0.515 | -0.027 | -0.022 | -0.022 | 0 | 0.31 | 0.2 | 0.24 | 0.31 | 0.35 | 0.36 |
| 4 | 25913155 | rs878552 | 1.614 | 0.416 | -2.025 | 0.424 | 0.003 | 0.017 | 0.017 | 0.063 | 0.51 | 0.55 | 0.62 | 0.42 | 0.51 | 0.41 |
| 4 | 25915858 | rs2305670 | 0.179 | 0.441 | -2.054 | 0.484 | -0.031 | -0.039 | -0.039 | 0 | 0.19 | 0.13 | 0.19 | 0.32 | 0.24 | 0.3 |
| 4 | 25919109 | rs936234 | -0.962 | 0.348 | -2.076 | 0.539 | -0.022 | -0.012 | -0.012 | 0 | 0.32 | 0.2 | 0.4 | 0.61 | 0.34 | 0.42 |
| 4 | 25919478 | rs6828696 | -0.866 | 0.374 | -1.969 | 0.685 | -0.013 | -0.015 | -0.015 | 0.143 | 0.2 | 0.12 | 0.26 | 0.31 | 0.23 | 0.2 |
| 4 | 25924426 | rs1123943 | 0.729 | 0.242 | -2.211 | 0.513 | 0.036 | -0.026 | -0.026 | 0 | 0.54 | 0.84 | 0.66 | 0.43 | 0.6 | 0.46 |
| 4 | 25924857 | rs2168523 | 1.76 | 0.23 | -2.227 | 0.387 | 0 | -0.015 | -0.015 | 0 | 0.4 | 0.5 | 0.38 | 0.14 | 0.41 | 0.29 |
| 4 | 25930928 | rs14553 | 0.77 | 0.368 | -2.414 | 0.396 | -0.039 | -0.036 | -0.036 | 0 | 0.1 | 0.03 | 0.26 | 0.31 | 0.17 | 0.17 |
| 4 | 25931325 | rs554 | 0.325 | 0.099 | -2.444 | 0.363 | 0.02 | 0.005 | 0.005 | 0 | 0.73 | 0.79 | 0.82 | 0.74 | 0.77 | 0.71 |
| 4 | 25933737 | rs16877751 | 0.585 | 0.244 | -2.342 | 0.587 | -0.004 | -0.097 | -0.097 | 0.02 | 0.19 | 0.04 | 0.17 | 0.31 | 0.17 | 0.26 |
| 4 | 25940635 | rs2309291 | 0.989 | 0.362 | -2.184 | 0.563 | -0.013 | -0.089 | -0.089 | 0.35 | 0.21 | 0.02 | 0.18 | 0.25 | 0.19 | 0.33 |
| 4 | 25942460 | rs6448417 | -0.908 | 0.287 | -2.349 | 0.561 | -0.014 | -0.059 | -0.059 | 0 | 0.87 | 0.97 | 0.88 | 0.77 | 0.89 | 0.82 |
| 4 | 25942592 | rs1456435 | 1.099 | 0.337 | -2.333 | 0.585 | -0.017 | -0.089 | -0.089 | 0 | 0.21 | 0.02 | 0.18 | 0.25 | 0.19 | 0.33 |
| 4 | 25942751 | rs7654585 | 0.479 | 0.329 | -1.867 | 0.479 | 0.042 | -0.004 | -0.004 | 0 | 0.53 | 0.6 | 0.63 | 0.5 | 0.6 | 0.43 |
| 4 | 25950606 | rs10939086 | 1.373 | 0.47 | -1.974 | 0.416 | 0.003 | 0.05 | 0.05 | 0 | 0.41 | 0.6 | 0.38 | 0.13 | 0.43 | 0.26 |
| 4 | 25950625 | rs10939087 | 1.316 | 0.518 | -1.926 | 0.44 | -0.002 | -0.07 | -0.07 | 0 | 0.4 | 0.4 | 0.36 | 0.12 | 0.43 | 0.26 |
| 4 | 25951539 | rs4692188 | 0.042 | 0.805 | -1.574 | 0.855 | -0.07 | -0.077 | -0.077 | 0.969 | 0.12 | 0.02 | 0.26 | 0.37 | 0.17 | 0.17 |
| 4 | 25957151 | rs4692198 | -0.257 | 0.496 | -1.438 | 0.805 | -0.039 | -0.068 | -0.068 | 0 | 0.24 | 0.11 | 0.29 | 0.38 | 0.25 | 0.23 |
| 4 | 25958994 | rs17627409 | -0.439 | 0.412 | -1.017 | 0.354 | 0 | 0.004 | 0.004 | 0 | 0.5 | 0.57 | 0.52 | 0.64 | 0.46 | 0.55 |
| 4 | 25961788 | rs6820447 | -0.169 | 0.641 | -1.396 | 0.505 | -0.004 | 0.033 | 0.033 | 0 | 0.46 | 0.36 | 0.48 | 0.64 | 0.44 | 0.52 |
| 4 | 25967254 | rs7440308 | 0.305 | 0.375 | -1.031 | 0.794 | -0.01 | 0.001 | 0.001 | 0 | 0.84 | 0.79 | 0.86 | 0.86 | 0.86 | 0.89 |
| 4 | 25968931 | rs2045329 | -0.827 | 0.489 | -1.882 | 1.209 | -0.041 | -0.109 | -0.109 | 0 | 0.27 | 0.12 | 0.31 | 0.39 | 0.29 | 0.36 |
| 4 | 25969256 | rs7676237 | -1.283 | 0.113 | -1.979 | 1.231 | 0.011 | -0.061 | -0.061 | 0 | 0.33 | 0.8 | 0.4 | 0.5 | 0.33 | 0.38 |
| 4 | 25969618 | rs2198844 | -0.863 | 0.115 | -1.928 | 1.416 | -0.009 | -0.054 | -0.054 | 0 | 0.2 | 0.12 | 0.21 | 0.34 | 0.22 | 0.2 |
| 4 | 25970163 | rs936235 | -0.821 | 0.077 | -1.926 | 1.419 | -0.008 | -0.053 | -0.053 | 0 | 0.19 | 0.12 | 0.21 | 0.34 | 0.22 | 0.2 |
| 4 | 25970325 | rs936236 | -1.498 | 0.139 | -1.529 | 1.381 | -0.001 | 0 | 0 | 0.175 | 0.36 | 0.66 | 0.39 | 0.57 | 0.34 | 0.44 |
| 4 | 25975111 | rs16877848 | 0.434 | 0.452 | -2.335 | 1.271 | -0.022 | -0.037 | -0.037 | 0 | 0.14 | 0.05 | 0.24 | 0.32 | 0.19 | 0.21 |
| 4 | 25976544 | rs10006741 | 0.477 | 0.174 | -2.46 | 0.544 | 0.014 | 0.085 | 0.085 | 1.822 | 0.53 | 0.14 | 0.46 | 0.3 | 0.56 | 0.52 |
| 4 | 25983887 | rs11728123 | 0.806 | -0.056 | -2.089 | 0.85 | 0.043 | 0.07 | 0.07 | 4.586 | 0.62 | 0.16 | 0.58 | 0.33 | 0.69 | 0.48 |
| 4 | 25984065 | rs11724348 | 0.571 | 0.784 | -1.57 | 1.584 | -0.023 | -0.023 | -0.023 | 0 | 0.88 | 0.91 | 0.77 | 0.61 | 0.84 | 0.81 |
| 4 | 25987596 | rs10939088 | -0.804 | 0.401 | -1.309 | 0.921 | 0.002 | 0.003 | 0.003 | 0 | 0.19 | 0.15 | 0.09 | 0.09 | 0.15 | 0.11 |
| 4 | 25987977 | rs6841503 | 1.137 | 0.139 | -1.293 | 0.849 | 0.007 | 0.002 | 0.002 | 0 | 0.66 | 0.69 | 0.78 | 0.74 | 0.72 | 0.65 |
| 4 | 25988674 | rs4234990 | -0.119 | 0.137 | -1.231 | 0.811 | 0.001 | -0.001 | -0.001 | 0 | 0.84 | 0.83 | 0.86 | 0.82 | 0.86 | 0.75 |
| 4 | 25991628 | rs4406006 | -0.98 | 0.393 | -0.907 | 0.403 | 0 | 0.014 | 0.014 | 0.018 | 0.41 | 0.54 | 0.54 | 0.74 | 0.44 | 0.6 |
| 4 | 25992329 | rs5029078 | 0.662 | 0.356 | -1.029 | 0.474 | 0.015 | -0.015 | -0.015 | 0 | 0.15 | 0.05 | 0.05 | 0.09 | 0.11 | 0.08 |
| 4 | 25993373 | rs6828245 | 1.478 | 0.491 | -0.856 | 0.288 | -0.025 | 0.108 | 0.108 | 0 | 0.45 | 0.25 | 0.39 | 0.25 | 0.47 | 0.28 |
| 4 | 26000595 | rs6840416 | 0.102 | 0.506 | 0.253 | 0.25 | 0 | -0.103 | -0.103 | 0 | 0.16 | 0.36 | 0.11 | 0.02 | 0.13 | 0.12 |
| 4 | 26005161 | rs6837551 | 0.488 | 0.405 | 0.606 | 0.55 | 0.004 | 0.006 | 0.006 | 0 | 0.63 | 0.68 | 0.5 | 0.33 | 0.63 | 0.47 |
| 4 | 26016156 | rs12647908 | 0.175 | 0.407 | 0.702 | 0.667 | -0.004 | 0.003 | 0.003 | 0 | 0.17 | 0.13 | 0.43 | 0.61 | 0.26 | 0.44 |
| 4 | 26017406 | rs1027998 | -0.778 | 0.059 | 1.082 | 0.395 | -0.007 | -0.024 | -0.024 | 0 | 0.29 | 0.35 | 0.15 | 0.11 | 0.24 | 0.18 |
| 4 | 26018690 | rs6847589 | 0.143 | -0.049 | 0.649 | -0.36 | -0.012 | 0.013 | 0.013 | 0 | 0.42 | 0.24 | 0.55 | 0.66 | 0.46 | 0.64 |
| 4 | 26021208 | rs17685537 | 1.902 | -0.063 | 0.695 | -0.362 | -0.037 | 0.006 | 0.006 | 0 | 0.25 | 0.09 | 0.28 | 0.23 | 0.3 | 0.2 |
| 4 | 26021310 | rs1512104 | -0.373 | -0.122 | 0.845 | -0.367 | -0.036 | -0.062 | -0.062 | 0.651 | 0.29 | 0.48 | 0.16 | 0.11 | 0.22 | 0.18 |
| 4 | 26025073 | rs4263391 | -0.754 | -0.084 | 0.985 | -0.287 | 0.027 | 0.005 | 0.005 | 0 | 0.95 | 0.74 | 0.97 | 1 | 0.98 | 0.98 |
| 4 | 26025323 | rs4287995 | 0.101 | -0.103 | 0.981 | -0.294 | 0.022 | -0.005 | -0.005 | 0 | 0.11 | 0.33 | 0.1 | 0.05 | 0.1 | 0.15 |
| 4 | 26025487 | rs12499714 | 0.207 | -0.105 | 0.843 | -0.176 | -0.001 | 0.002 | 0.002 | 0 | 0.51 | 0.52 | 0.4 | 0.38 | 0.49 | 0.4 |
| 4 | 26025756 | rs4269167 | 0.344 | 0.018 | 0.422 | -0.394 | 0.011 | 0.13 | 0.13 | 0 | 0.44 | 0.17 | 0.7 | 0.86 | 0.59 | 0.64 |
| 4 | 26026106 | rs1567648 | 0.089 | 0.182 | 0.508 | -0.118 | -0.024 | -0.013 | -0.013 | 0 | 0.21 | 0.16 | 0.35 | 0.22 | 0.33 | 0.42 |
| 4 | 26027797 | rs9884830 | 1 | -0.077 | 0.668 | -0.62 | 0.001 | -0.008 | -0.008 | 0 | 0.2 | 0.17 | 0.12 | 0 | 0.17 | 0.1 |
| 4 | 26028187 | rs9884387 | -1.156 | -0.023 | 0.924 | -0.567 | 0.034 | -0.038 | -0.038 | 0 | 0.7 | 0.5 | 0.71 | 0.8 | 0.65 | 0.77 |
| 4 | 26028805 | rs6846440 | -0.532 | -0.157 | 0.829 | -0.607 | -0.001 | 0.003 | 0.003 | 0.293 | 0.63 | 0.35 | 0.69 | 0.81 | 0.63 | 0.74 |
| 4 | 26031528 | rs12502605 | 0.153 | -0.264 | 0.73 | -0.683 | -0.014 | -0.057 | -0.057 | 0 | 0.27 | 0.33 | 0.14 | 0.04 | 0.2 | 0.2 |
| 4 | 26032228 | rs1033118 | -0.012 | -0.246 | 0.916 | -0.466 | -0.016 | 0.031 | 0.031 | 0 | 0.08 | 0.26 | 0.03 | 0.04 | 0.02 | 0.1 |
| 4 | 26034030 | rs9790578 | -0.28 | -0.136 | 0.889 | -0.489 | 0.01 | 0.103 | 0.103 | 0 | 0.45 | 0.28 | 0.58 | 0.8 | 0.45 | 0.65 |
| 4 | 26037602 | rs4692317 | -0.702 | 0.009 | 0.503 | -0.506 | 0.003 | -0.002 | -0.002 | 1.039 | 0.53 | 0.54 | 0.61 | 0.85 | 0.47 | 0.75 |
| 4 | 26047288 | rs11736287 | 0.992 | -0.216 | 0.735 | -0.161 | -0.015 | 0.023 | 0.023 | 1.048 | 0.87 | 0.44 | 0.84 | 0.89 | 0.91 | 0.83 |
| 4 | 26048141 | rs2137231 | 0.7 | -0.235 | 0.451 | -0.194 | 0.009 | -0.005 | -0.005 | 0 | 0.89 | 0.64 | 0.9 | 0.95 | 0.91 | 0.82 |
| 4 | 26050047 | rs11731522 | 0.476 | -0.078 | -0.525 | 0.131 | -0.065 | -0.086 | -0.086 | 0 | 0.17 | 0.01 | 0.29 | 0.33 | 0.2 | 0.28 |
| 4 | 26054418 | rs17697268 | 1.106 | 0.295 | -0.352 | -0.863 | -0.104 | 0.043 | 0.043 | 0.194 | 0.27 | 0.04 | 0.23 | 0.17 | 0.34 | 0.2 |
| 4 | 26056021 | rs10155365 | -0.002 | 0.258 | -0.065 | -0.907 | 0.012 | 0.09 | 0.09 | 0 | 0.53 | 0.34 | 0.36 | 0.17 | 0.55 | 0.3 |
| 4 | 26057402 | rs13126784 | 0.256 | 0.467 | -0.547 | -0.739 | -0.024 | 0.045 | 0.045 | 0 | 0.38 | 0.12 | 0.48 | 0.72 | 0.38 | 0.52 |
| 4 | 26060048 | rs10032780 | 0.207 | 0.502 | -0.228 | -0.587 | 0.013 | -0.061 | -0.061 | 0.288 | 0.35 | 0.47 | 0.28 | 0.17 | 0.37 | 0.21 |
| 4 | 26068018 | rs1105912 | -0.196 | 0.558 | -0.853 | -0.366 | -0.023 | 0.025 | 0.025 | 0 | 0.59 | 0.83 | 0.48 | 0.33 | 0.57 | 0.44 |
| 4 | 26072585 | rs4692346 | -0.32 | 0.999 | -0.828 | -0.479 | 0.003 | -0.001 | -0.001 | 0 | 0.83 | 0.77 | 0.87 | 0.88 | 0.78 | 0.78 |
| 4 | 26079129 | rs4558863 | 0.853 | 0.588 | 0.039 | -1.046 | 0.002 | -0.041 | -0.041 | 0 | 0.25 | 0.29 | 0.16 | 0 | 0.19 | 0.1 |
| 4 | 26079164 | rs2048507 | 0.872 | 0.581 | 0.043 | -1.05 | 0.009 | -0.008 | -0.008 | 0.018 | 0.24 | 0.25 | 0.15 | 0 | 0.19 | 0.1 |
| 4 | 26079216 | rs17697588 | 1.402 | 0.556 | -0.198 | -0.863 | -0.063 | -0.045 | -0.045 | 0 | 0.13 | 0.01 | 0.1 | 0.17 | 0.17 | 0.17 |

| | | | | | | | | | | | | | | | | |
|---|---|---|---|---|---|---|---|---|---|---|---|---|---|---|---|---|
| 4 | 26081846 | rs907496 | 0.021 | 0.56 | 0.136 | -1.195 | 0 | 0.005 | 0.005 | 0 | 0.59 | 0.51 | 0.62 | 0.72 | 0.57 | 0.67 |
| 4 | 26083058 | rs6818978 | -0.421 | 0.751 | 0.278 | -1.541 | 0.014 | 0.014 | 0.014 | 0 | 0.83 | 0.69 | 0.83 | 0.81 | 0.79 | 0.75 |
| 4 | 26085511 | rs10517086 | 0.719 | 0.986 | 0.47 | -1.481 | 0.005 | -0.077 | -0.077 | 0.069 | 0.25 | 0.28 | 0.21 | 0 | 0.3 | 0.11 |
| 4 | 26085546 | rs10517087 | 0.674 | 1.135 | -0.373 | -1.816 | -0.018 | 0.088 | 0.088 | 0 | 0.17 | 0.04 | 0.13 | 0 | 0.16 | 0.15 |
| 4 | 26092005 | rs4692369 | 0.687 | 1.098 | 0.032 | -1.491 | 0.02 | 0.064 | 0.064 | 0.083 | 0.39 | 0.16 | 0.44 | 0.75 | 0.32 | 0.44 |
| 4 | 26094755 | rs6827904 | 0.192 | 1.188 | 0.04 | -1.523 | -0.003 | 0.004 | 0.004 | 0 | 0.44 | 0.62 | 0.37 | 0.01 | 0.47 | 0.26 |
| 4 | 26100786 | rs4692073 | -0.572 | 0.655 | 0.27 | -1.128 | 0 | 0 | 0 | 0 | 0.94 | 0.94 | 0.98 | 0.94 | 0.92 | 0.96 |
| 4 | 26112846 | rs1877384 | -0.29 | 0.765 | 0.311 | -1.263 | 0.021 | 0.073 | 0.073 | 0 | 0.52 | 0.63 | 0.52 | 0.17 | 0.61 | 0.43 |
| 4 | 26132361 | rs4692386 | 0.539 | 1.322 | -0.044 | -0.423 | 0.026 | -0.061 | -0.061 | 0.028 | 0.67 | 0.51 | 0.71 | 0.79 | 0.64 | 0.47 |
| 4 | 26133708 | rs10939096 | 0.695 | 1.481 | 0.383 | -0.708 | 0.032 | -0.088 | -0.088 | 0.525 | 0.8 | 0.67 | 0.8 | 0.98 | 0.77 | 0.77 |
| 4 | 26136773 | rs7682322 | -0.219 | 1.328 | -0.004 | -0.051 | 0.001 | 0 | 0 | 0 | 0.46 | 0.53 | 0.49 | 0.59 | 0.49 | 0.37 |
| 4 | 26139677 | rs907498 | 0.554 | 1.104 | 0.452 | -0.419 | 0.09 | -0.049 | -0.049 | 0.274 | 0.88 | 0.68 | 0.84 | 0.98 | 0.81 | 0.84 |
| 4 | 26142837 | rs10033468 | -0.246 | 0.661 | 0.214 | -0.104 | -0.016 | 0.026 | 0.026 | 0 | 0.43 | 0.34 | 0.44 | 0.33 | 0.53 | 0.3 |
| 4 | 26147138 | rs13140653 | 0.727 | 0.379 | -0.624 | -0.044 | -0.073 | -0.13 | -0.13 | 0.704 | 0.42 | 0.03 | 0.41 | 0.5 | 0.38 | 0.42 |
| 4 | 26151057 | rs4692399 | 0.144 | 0.501 | -0.107 | -0.01 | -0.017 | 0.014 | 0.014 | 0 | 0.83 | 0.48 | 0.86 | 0.83 | 0.9 | 0.79 |
| 4 | 26155360 | rs1581089 | 0.718 | 0.584 | -0.975 | -0.192 | -0.124 | 0.042 | 0.042 | 0 | 0.29 | 0.02 | 0.32 | 0.16 | 0.39 | 0.15 |
| 4 | 26160310 | rs12650853 | -0.02 | 0.644 | -0.531 | -0.063 | -0.005 | 0.007 | 0.007 | 0 | 0.42 | 0.41 | 0.44 | 0.36 | 0.51 | 0.37 |
| 4 | 26163108 | rs13115567 | 1.047 | 0.612 | -1.095 | -0.27 | -0.06 | -0.149 | -0.149 | 0.065 | 0.36 | 0.03 | 0.39 | 0.5 | 0.34 | 0.43 |
| 4 | 26163751 | rs7697053 | -0.342 | 0.775 | -0.855 | -0.218 | 0.001 | 0.008 | 0.008 | 0 | 0.47 | 0.46 | 0.47 | 0.4 | 0.56 | 0.4 |
| 4 | 26166250 | rs13150492 | 0.326 | 0.793 | -0.873 | 0.08 | -0.069 | -0.106 | -0.106 | 0.326 | 0.38 | 0.05 | 0.36 | 0.5 | 0.39 | 0.44 |
| 4 | 26167227 | rs9291482 | -0.421 | 0.859 | -0.806 | 0.023 | -0.001 | 0 | 0 | 0 | 0.55 | 0.59 | 0.5 | 0.65 | 0.52 | 0.67 |
| 4 | 26172899 | rs16878154 | 0.084 | 0.78 | -1.019 | -0.174 | 0 | 0.018 | 0.018 | 0 | 0.16 | 0.43 | 0.15 | 0.16 | 0.14 | 0.25 |
| 4 | 26175955 | rs1512105 | 0.353 | 0.95 | -2.394 | -0.079 | -0.1 | 0.075 | 0.075 | 0 | 0.4 | 0.12 | 0.46 | 0.24 | 0.47 | 0.22 |
| 4 | 26177372 | rs6824647 | -0.887 | 0.902 | -1.076 | 0.07 | -0.002 | -0.003 | -0.003 | 0 | 0.22 | 0.7 | 0.19 | 0.17 | 0.18 | 0.26 |
| 4 | 26192444 | rs6817712 | 0.351 | 1.5 | -1.634 | 0.168 | -0.064 | -0.016 | -0.016 | 0 | 0.75 | 0.95 | 0.68 | 0.79 | 0.74 | 0.84 |
| 4 | 26197873 | rs10017756 | 0.835 | 1.329 | -1.154 | -0.016 | -0.008 | 0.01 | 0.01 | 0 | 0.67 | 0.77 | 0.57 | 0.73 | 0.68 | 0.64 |
| 4 | 26205938 | rs4552442 | 1.001 | 2.008 | -0.497 | -0.035 | 0.049 | 0.098 | 0.098 | 0.916 | 0.85 | 0.67 | 0.67 | 0.75 | 0.81 | 0.7 |
| 4 | 26208843 | rs17823089 | -1.067 | 2.247 | -0.814 | -0.482 | -0.044 | 0.023 | 0.023 | 0.238 | 0.13 | 0.02 | 0.14 | 0.06 | 0.16 | 0.21 |
| 4 | 26213763 | rs11735620 | 1.426 | 1.97 | -0.351 | -0.75 | 0.009 | -0.055 | -0.055 | 1.359 | 0.59 | 0.53 | 0.57 | 0.87 | 0.53 | 0.61 |
| 4 | 26218025 | rs10939100 | 1.719 | 1.893 | -0.131 | -0.901 | 0.002 | -0.024 | -0.024 | 0 | 0.69 | 0.71 | 0.71 | 0.91 | 0.64 | 0.63 |
| 4 | 26219793 | rs7692708 | 1.194 | 2.106 | -0.102 | -0.905 | -0.017 | 0.018 | 0.018 | 0.798 | 0.77 | 0.84 | 0.82 | 0.93 | 0.7 | 0.65 |
| 4 | 26220299 | rs16878186 | -1.151 | 1.758 | 0.088 | -0.692 | 0.008 | -0.04 | -0.04 | 0 | 0.08 | 0.24 | 0.11 | 0.02 | 0.07 | 0.01 |
| 4 | 26229299 | rs7685350 | -1.081 | 1.664 | -0.266 | -0.649 | -0.024 | 0.015 | 0.015 | 0 | 0.11 | 0.06 | 0.12 | 0.06 | 0.18 | 0.2 |
| 4 | 26230407 | rs6850487 | -0.536 | 1.585 | -0.012 | -0.499 | -0.008 | -0.001 | -0.001 | 0 | 0.07 | 0.14 | 0.1 | 0.08 | 0.05 | 0.07 |
| 4 | 26237157 | rs1006802 | -2.28 | 2.067 | 0.778 | -0.558 | 0.048 | -0.051 | -0.051 | 0 | 0.22 | 0.47 | 0.21 | 0.15 | 0.28 | 0.29 |
| 4 | 26259369 | rs10028140 | 0.721 | 0.773 | 0.683 | -1.034 | -0.059 | -0.106 | -0.106 | 0 | 0.82 | 0.56 | 0.9 | 0.99 | 0.93 | 0.95 |
| 4 | 26273910 | rs7655052 | 0.659 | -0.246 | -0.179 | -0.731 | 0.002 | 0.011 | 0.011 | 0 | 0.54 | 0.43 | 0.67 | 0.36 | 0.56 | 0.43 |
| 4 | 26275139 | rs11933748 | 0.058 | -0.365 | 0.677 | -0.305 | 0.05 | 0.122 | 0.122 | 0 | 0.06 | 0.27 | 0.06 | 0.11 | 0.07 | 0.02 |
| 4 | 26275336 | rs6831303 | -0.165 | -0.174 | 0.289 | -0.269 | -0.057 | 0.013 | 0.013 | 0 | 0.9 | 0.99 | 0.92 | 0.95 | 0.82 | 0.82 |
| 4 | 26280575 | rs13109468 | 1.446 | -0.945 | -0.499 | -0.87 | -0.113 | -0.083 | -0.083 | 0 | 0.28 | 0.07 | 0.49 | 0.33 | 0.39 | 0.35 |
| 4 | 26292555 | rs2871198 | 0.707 | -1.348 | 0.506 | -0.665 | 0.016 | 0.009 | 0.009 | 0 | 0.55 | 0.72 | 0.67 | 0.37 | 0.59 | 0.41 |
| 4 | 26299125 | rs1877209 | 0.263 | -1.126 | 1.19 | -0.477 | -0.04 | -0.005 | -0.005 | 0 | 0.13 | 0.34 | 0.09 | 0.11 | 0.08 | 0.03 |
| 4 | 26300494 | rs2271384 | 0.76 | -1.091 | 0.776 | -0.215 | -0.006 | -0.03 | -0.03 | 0 | 0.59 | 0.77 | 0.72 | 0.44 | 0.59 | 0.4 |
| 4 | 26308792 | rs17644283 | 0.938 | -0.943 | 0.835 | 0.433 | 0.03 | 0.07 | 0.07 | 0 | 0.54 | 0.76 | 0.73 | 0.69 | 0.61 | 0.45 |
| 4 | 26321906 | rs3822223 | -0.456 | -0.922 | 1.273 | 0.418 | 0.091 | 0.406 | 0.406 | 0 | 0.06 | 0.33 | 0.08 | 0.36 | 0.09 | 0.06 |
| 4 | 26337615 | rs13128399 | 1.619 | -0.901 | 1.013 | 0.41 | 0.001 | 0.002 | 0.002 | 0 | 0.49 | 0.48 | 0.63 | 0.36 | 0.53 | 0.44 |
| 4 | 26342538 | rs6821126 | -0.277 | 0.065 | 1.253 | 0.535 | -0.023 | 0.026 | 0.026 | 0 | 0.91 | 0.97 | 0.87 | 0.97 | 0.82 | 0.88 |
| 4 | 26348730 | rs7695991 | 0.057 | -0.228 | 1.236 | 0.402 | 0.017 | -0.005 | -0.005 | 0 | 0.07 | 0.1 | 0.03 | 0.07 | 0.09 | 0.14 |
| 4 | 26408827 | rs3109836 | 0.452 | -0.364 | 1.1 | 0.347 | 0.012 | 0.034 | 0.034 | 0 | 0.73 | 0.8 | 0.9 | 0.88 | 0.83 | 0.72 |
| 4 | 26416113 | rs2788864 | 1.48 | -0.352 | 0.98 | 0.319 | 0 | -0.062 | -0.062 | 0 | 0.63 | 0.58 | 0.75 | 0.85 | 0.64 | 0.61 |
| 4 | 26431386 | rs946346 | -0.606 | -0.374 | 0.47 | 0.305 | 0.045 | 0.08 | 0.08 | 0 | 0.8 | 0.95 | 0.94 | 0.95 | 0.91 | 0.87 |
| 4 | 26439223 | rs10805262 | 1.613 | -0.468 | 0.556 | 0.643 | 0.016 | 0.017 | 0.017 | 0 | 0.54 | 0.25 | 0.65 | 0.52 | 0.56 | 0.56 |
| 4 | 26439486 | rs10517099 | -0.536 | -0.414 | 0.667 | 0.729 | -0.011 | 0.156 | 0.156 | 0 | 0.08 | 0.2 | 0.09 | 0.33 | 0.08 | 0.05 |
| 4 | 26440988 | rs6833606 | -0.536 | -0.414 | 0.667 | 0.729 | -0.011 | 0.156 | 0.156 | 0 | 0.08 | 0.2 | 0.09 | 0.33 | 0.08 | 0.05 |
| 4 | 26451326 | rs6818128 | 1.391 | -0.304 | 0.364 | 0.815 | 0.002 | 0.007 | 0.007 | 0 | 0.57 | 0.39 | 0.7 | 0.56 | 0.59 | 0.6 |
| 4 | 26451697 | rs13116206 | 1.539 | -0.301 | 0.27 | 0.816 | 0.008 | 0.016 | 0.016 | 0 | 0.56 | 0.37 | 0.69 | 0.56 | 0.59 | 0.57 |
| 4 | 26456971 | rs17578817 | 1.492 | -0.073 | -0.413 | 0.952 | -0.056 | -0.066 | -0.066 | 0.071 | 0.54 | 0.1 | 0.64 | 0.46 | 0.52 | 0.57 |
| 4 | 26457168 | rs3109848 | 0.217 | -0.105 | 0.091 | 0.859 | -0.009 | -0.031 | -0.031 | 0.009 | 0.74 | 0.61 | 0.88 | 0.84 | 0.82 | 0.79 |
| 4 | 26457487 | rs7691160 | 0.445 | -0.1 | -0.133 | 0.862 | -0.016 | -0.048 | -0.048 | 0 | 0.73 | 0.39 | 0.84 | 0.84 | 0.81 | 0.79 |
| 4 | 26460991 | rs6826952 | 0.362 | 0.041 | -0.711 | 0.864 | 0.005 | -0.015 | -0.015 | 0 | 0.09 | 0.1 | 0.12 | 0.04 | 0.18 | 0.12 |
| 4 | 26461373 | rs13144195 | 1.424 | -0.03 | -1.092 | 0.991 | -0.015 | 0.025 | 0.025 | 0 | 0.58 | 0.18 | 0.68 | 0.6 | 0.59 | 0.64 |
| 4 | 26464646 | rs2788853 | -1.785 | 0.143 | -0.421 | 1.253 | -0.008 | -0.011 | -0.011 | 0 | 0.25 | 0.85 | 0.25 | 0.28 | 0.27 | 0.2 |
| 4 | 26466861 | rs6854815 | -1.265 | 0.081 | 0.103 | 1.48 | -0.003 | 0.126 | 0.126 | 0 | 0.14 | 0.43 | 0.07 | 0.25 | 0.08 | 0.09 |
| 4 | 26468733 | rs716075 | 0.068 | -0.148 | -1.223 | 1.668 | -0.008 | -0.117 | -0.117 | 1.312 | 0.83 | 0.97 | 0.87 | 0.49 | 0.88 | 0.84 |
| 4 | 26468820 | rs716073 | -0.336 | 0.364 | -0.626 | 1.537 | 0.012 | 0.015 | 0.015 | 0 | 0.07 | 0.15 | 0.07 | 0.11 | 0.1 | 0.07 |
| 4 | 26468947 | rs716074 | 1.015 | 0.518 | -1.283 | 1.395 | -0.064 | -0.073 | -0.073 | 0 | 0.65 | 0.15 | 0.66 | 0.42 | 0.59 | 0.68 |
| 4 | 26473220 | rs3113032 | -0.706 | -0.347 | -1.057 | 1.263 | 0.067 | -0.045 | -0.045 | 0.751 | 0.36 | 0.77 | 0.29 | 0.58 | 0.26 | 0.29 |
| 4 | 26480145 | rs967413 | -0.897 | 0.066 | -0.416 | 0.825 | -0.002 | 0.06 | 0.06 | 0 | 0.44 | 0.36 | 0.35 | 0.26 | 0.46 | 0.51 |
| 4 | 26482021 | rs2854030 | -0.46 | -0.286 | 0.284 | 1.098 | -0.016 | -0.004 | -0.004 | 0.006 | 0.34 | 0.41 | 0.27 | 0.35 | 0.29 | 0.41 |

| | | | | | | | | | | | | | | | | |
|---|---|---|---|---|---|---|---|---|---|---|---|---|---|---|---|---|
| 4 | 26486909 | rs7665027 | -0.504 | -0.329 | 0.727 | 1.314 | -0.009 | -0.002 | -0.002 | 0 | 0.19 | 0.2 | 0.14 | 0.23 | 0.14 | 0.32 |
| 4 | 26489057 | rs2725307 | -1.243 | -0.454 | 0.793 | 1.414 | -0.047 | 0.056 | 0.056 | 0.532 | 0.31 | 0.54 | 0.21 | 0.56 | 0.22 | 0.46 |
| 4 | 26493209 | rs2854038 | -1.556 | 0.03 | 0.786 | 1.176 | -0.003 | 0.032 | 0.032 | 0 | 0.23 | 0.35 | 0.23 | 0.35 | 0.23 | 0.26 |
| 4 | 26498513 | rs869902 | -0.441 | 0.111 | 0.737 | 1.152 | -0.009 | 0.065 | 0.065 | 0 | 0.1 | 0.18 | 0.05 | 0.27 | 0.07 | 0.14 |
| 4 | 26499696 | rs10031898 | -1.289 | -0.211 | 1.425 | 0.997 | -0.012 | -0.041 | -0.041 | 0 | 0.28 | 0.5 | 0.32 | 0.22 | 0.27 | 0.43 |
| 4 | 26501114 | rs11733591 | 0.092 | -0.193 | 1.419 | 1.062 | -0.004 | 0.021 | 0.021 | 0 | 0.1 | 0.16 | 0.06 | 0.19 | 0.07 | 0.09 |
| 4 | 26501372 | rs11725410 | -1.941 | -0.182 | 1.347 | 1.06 | 0.007 | -0.001 | -0.001 | 2.152 | 0.38 | 0.66 | 0.39 | 0.41 | 0.34 | 0.51 |
| 4 | 26504157 | rs2130250 | -1.963 | -0.182 | 1.339 | 1.061 | 0.008 | -0.014 | -0.014 | 0 | 0.38 | 0.66 | 0.38 | 0.49 | 0.34 | 0.57 |
| 4 | 26504723 | rs7667060 | -0.189 | -0.7 | 1.899 | 1.077 | -0.027 | 0.003 | 0.003 | 4.01 | 0.15 | 0.23 | 0.15 | 0.19 | 0.11 | 0.31 |
| 4 | 26504936 | rs2130248 | -0.718 | -0.808 | 1.833 | 0.928 | 0.04 | 0.025 | 0.025 | 0 | 0.11 | 0.37 | 0.16 | 0.09 | 0.1 | 0.13 |
| 4 | 26512016 | rs13115937 | 1.536 | -0.86 | 1.062 | 0.755 | 0.049 | -0.015 | -0.015 | 0 | 0.61 | 0.21 | 0.62 | 0.56 | 0.72 | 0.38 |
| 4 | 26522349 | rs902095 | 1.386 | -0.539 | 1.545 | 0.549 | -0.016 | 0 | 0 | 0 | 0.75 | 0.32 | 0.74 | 0.76 | 0.83 | 0.7 |
| 4 | 26522786 | rs902096 | 1.377 | -0.534 | 1.547 | 0.552 | -0.016 | 0.001 | 0.001 | 0 | 0.75 | 0.32 | 0.74 | 0.76 | 0.83 | 0.7 |
| 4 | 26530421 | rs13135516 | 0.94 | -0.284 | 2.077 | 0.229 | -0.007 | -0.031 | -0.031 | 0 | 0.83 | 0.29 | 0.78 | 0.91 | 0.85 | 0.78 |
| 4 | 26530798 | rs4692540 | 0.751 | -0.261 | 2.376 | 0.224 | 0.017 | -0.074 | -0.074 | 0 | 0.84 | 0.43 | 0.79 | 0.95 | 0.85 | 0.8 |
| 4 | 26533333 | rs1387542 | 1.188 | -0.342 | 2.272 | 0.151 | -0.004 | -0.016 | -0.016 | 0 | 0.79 | 0.27 | 0.77 | 0.92 | 0.83 | 0.76 |
| 4 | 26570153 | rs3899075 | 1.751 | -0.295 | 3.018 | 0.415 | 0.064 | 0.114 | 0.114 | 4.888 | 0.95 | 0.48 | 0.96 | 0.96 | 1 | 0.98 |
| 4 | 26573764 | rs2014308 | -0.115 | -0.191 | 2.306 | 0.211 | 0.012 | 0.006 | 0.006 | 0 | 0.8 | 0.5 | 0.87 | 0.76 | 0.83 | 0.92 |
| 4 | 26585311 | rs10805265 | 1.143 | -0.324 | 2.189 | 0.26 | -0.036 | 0.025 | 0.025 | 7.366 | 0.93 | 0.31 | 0.94 | 0.93 | 1 | 0.99 |
| 4 | 26585881 | rs4692549 | -0.564 | -0.718 | 1.121 | -0.243 | -0.032 | 0.026 | 0.026 | 0 | 0.4 | 0.17 | 0.28 | 0.36 | 0.39 | 0.4 |
| 4 | 26591253 | rs12650361 | -0.819 | -0.85 | 0.851 | -0.49 | -0.036 | 0.068 | 0.068 | 0 | 0.47 | 0.17 | 0.31 | 0.36 | 0.45 | 0.43 |
| 4 | 26625185 | rs260958 | -0.812 | -0.857 | 1.443 | -0.514 | -0.064 | 0.062 | 0.062 | 0 | 0.46 | 0.11 | 0.31 | 0.37 | 0.45 | 0.43 |
| 4 | 26672973 | rs1558722 | 0.96 | -0.863 | 1.15 | -0.691 | -0.059 | 0.061 | 0.061 | 0 | 0.13 | 0.01 | 0.16 | 0.03 | 0.17 | 0.26 |
| 4 | 26756855 | rs1129230 | -0.687 | -0.8 | -0.365 | -0.462 | -0.033 | 0.064 | 0.064 | 0 | 0.46 | 0.18 | 0.31 | 0.37 | 0.45 | 0.44 |
| 4 | 26780582 | rs5024581 | 0.536 | -0.523 | -0.833 | -0.061 | -0.005 | 0.013 | 0.013 | 0 | 0.51 | 0.16 | 0.66 | 0.56 | 0.58 | 0.56 |
| 4 | 26793681 | rs4428270 | 0.094 | -0.056 | -1.131 | -0.08 | -0.053 | -0.033 | -0.033 | 0 | 0.38 | 0.13 | 0.47 | 0.55 | 0.42 | 0.51 |
| 4 | 26809177 | rs4261979 | -0.791 | 0.952 | -0.882 | 0.427 | -0.023 | -0.035 | -0.035 | 0.008 | 0.84 | 0.91 | 0.91 | 0.68 | 0.82 | 0.93 |
| 4 | 26817610 | rs4299571 | 0.243 | 0.903 | -0.836 | 0.127 | 0.012 | 0.127 | 0.127 | 0 | 0.44 | 0.23 | 0.4 | 0.15 | 0.42 | 0.45 |
| 4 | 26822377 | rs7692829 | -0.176 | 0.78 | -0.159 | 0.925 | 0.011 | 0.009 | 0.009 | 3.199 | 0.37 | 0.58 | 0.52 | 0.38 | 0.42 | 0.47 |
| 4 | 26826309 | rs7676115 | -0.295 | 1.047 | -0.076 | 1.203 | 0.075 | 0.194 | 0.194 | 0 | 0.06 | 0.38 | 0.05 | 0.15 | 0.05 | 0.08 |
| 4 | 26828438 | rs6821586 | 0.279 | 1.112 | -1.297 | 1.188 | -0.02 | -0.011 | -0.011 | 0 | 0.28 | 0.08 | 0.3 | 0.24 | 0.3 | 0.35 |
| 4 | 26831827 | rs6448475 | -0.141 | 0.869 | -1.168 | 1.016 | -0.004 | 0.01 | 0.01 | 0 | 0.49 | 0.76 | 0.53 | 0.47 | 0.53 | 0.43 |
| 4 | 26835952 | rs6839242 | 0.53 | 0.66 | -0.348 | 1.301 | -0.012 | 0.044 | 0.044 | 0 | 0.74 | 0.54 | 0.74 | 0.64 | 0.79 | 0.82 |
| 4 | 26843494 | rs3924178 | 0.487 | 0.784 | -0.62 | 1.13 | -0.012 | 0.043 | 0.043 | 0 | 0.53 | 0.27 | 0.4 | 0.42 | 0.5 | 0.44 |
| 4 | 26844469 | rs7688662 | -0.632 | 1.2 | -0.022 | 1.57 | 0.014 | -0.004 | -0.004 | 0 | 0.42 | 0.58 | 0.39 | 0.4 | 0.34 | 0.31 |
| 4 | 26852438 | rs7660174 | -0.463 | 1.148 | 1.014 | 1.812 | -0.015 | -0.009 | -0.009 | 0.091 | 0.4 | 0.45 | 0.34 | 0.28 | 0.31 | 0.26 |
| 4 | 26861656 | rs9790789 | 0.948 | 1.023 | 1.045 | 1.685 | -0.043 | 0.008 | 0.008 | 0 | 0.56 | 0.14 | 0.41 | 0.42 | 0.53 | 0.49 |
| 4 | 26872460 | rs6833053 | 0.248 | -0.065 | 1.127 | 1.237 | -0.001 | 0.055 | 0.055 | 0 | 0.22 | 0.37 | 0.25 | 0.41 | 0.21 | 0.33 |
| 4 | 26880755 | rs13150500 | 0.482 | -0.469 | 1.253 | 1.225 | 0.007 | 0.001 | 0.001 | 0 | 0.12 | 0.11 | 0.04 | 0.08 | 0.06 | 0.1 |
| 4 | 26881854 | rs6815719 | 0.153 | -0.499 | 1.497 | 1.17 | -0.011 | 0.085 | 0.085 | 0 | 0.24 | 0.44 | 0.27 | 0.41 | 0.21 | 0.33 |
| 4 | 26894656 | rs10022599 | -0.525 | -0.537 | 1.492 | 1.156 | -0.014 | -0.004 | -0.004 | 0.001 | 0.36 | 0.44 | 0.3 | 0.28 | 0.28 | 0.23 |
| 4 | 26902361 | rs10023256 | -0.568 | -0.483 | 1.541 | 1.153 | 0.001 | 0.001 | 0.001 | 0.156 | 0.45 | 0.51 | 0.5 | 0.61 | 0.43 | 0.45 |
| 4 | 26911000 | rs12642608 | -0.323 | -0.751 | 0.869 | 0.941 | -0.004 | -0.015 | -0.015 | 0 | 0.38 | 0.24 | 0.46 | 0.61 | 0.4 | 0.39 |
| 4 | 26938136 | rs7692953 | -1.61 | -0.671 | 0.444 | 0.946 | -0.117 | 0.035 | 0.035 | 0 | 0.59 | 0.92 | 0.59 | 0.71 | 0.51 | 0.56 |
| 4 | 26945149 | rs7658174 | -1.382 | -0.82 | 0.533 | 0.96 | -0.013 | 0.046 | 0.046 | 0 | 0.57 | 0.74 | 0.53 | 0.72 | 0.49 | 0.55 |
| 4 | 26950265 | rs10030655 | -0.206 | -0.808 | 0.34 | 0.882 | 0.031 | -0.007 | -0.007 | 1.883 | 0.33 | 0.21 | 0.27 | 0.3 | 0.27 | 0.22 |
| 4 | 26962873 | rs10032186 | 0.368 | -0.823 | 0.334 | 1.019 | 0.001 | -0.002 | -0.002 | 0.166 | 0.23 | 0.17 | 0.32 | 0.33 | 0.22 | 0.12 |
| 4 | 26966277 | rs16878655 | 0.434 | -0.442 | 0.438 | 0.878 | -0.005 | -0.019 | -0.019 | 0 | 0.16 | 0.07 | 0.17 | 0.21 | 0.18 | 0.25 |
| 4 | 26970376 | rs10433805 | 0.094 | -0.57 | 0.564 | 0.849 | -0.004 | 0 | 0 | 0 | 0.35 | 0.42 | 0.36 | 0.64 | 0.31 | 0.29 |
| 4 | 27000995 | rs6822297 | 1.305 | -0.614 | -0.038 | 0.79 | -0.055 | 0.099 | 0.099 | 0 | 0.38 | 0.15 | 0.39 | 0.16 | 0.47 | 0.4 |
| 4 | 27005235 | rs4692137 | 1.276 | -0.577 | -0.184 | 0.887 | -0.048 | -0.02 | -0.02 | 0 | 0.38 | 0.15 | 0.45 | 0.4 | 0.49 | 0.44 |
| 4 | 27011357 | rs6855865 | -0.443 | -0.143 | -0.81 | 1.145 | 0.023 | -0.006 | -0.006 | 0 | 0.39 | 0.35 | 0.31 | 0.39 | 0.3 | 0.3 |
| 4 | 27016468 | rs6830950 | 0.217 | -0.032 | -0.947 | 0.184 | 0.015 | -0.032 | -0.032 | 0 | 0.32 | 0.15 | 0.34 | 0.63 | 0.28 | 0.28 |
| 4 | 27023347 | rs6853163 | 0.861 | 0.061 | -0.985 | 0.282 | -0.008 | -0.024 | -0.024 | 0 | 0.07 | 0.08 | 0.02 | 0 | 0.04 | 0.06 |
| 4 | 27034672 | rs9998590 | -0.17 | 0.405 | -1.203 | 0.581 | 0.015 | -0.003 | -0.003 | 0.237 | 0.66 | 0.68 | 0.63 | 0.37 | 0.71 | 0.73 |
| 4 | 27045049 | rs16878720 | -0.531 | 0.868 | -1 | 1.029 | -0.009 | -0.021 | -0.021 | 0 | 0.15 | 0.06 | 0.17 | 0.22 | 0.16 | 0.24 |
| 4 | 27056744 | rs16878739 | -1.112 | 1.133 | -0.893 | 0.669 | -0.013 | 0.02 | 0.02 | 0 | 0.09 | 0.09 | 0.08 | 0.01 | 0.13 | 0.05 |
| 4 | 27060775 | rs7678869 | 0.491 | 1.018 | -0.567 | 0.614 | -0.001 | 0.039 | 0.039 | 0 | 0.56 | 0.31 | 0.64 | 0.6 | 0.59 | 0.49 |
| 4 | 27067027 | rs4692020 | 0.563 | 0.676 | -0.057 | 0.488 | -0.016 | 0.014 | 0.014 | 0.022 | 0.85 | 0.63 | 0.86 | 0.81 | 0.91 | 0.69 |
| 4 | 27070339 | rs17570945 | 1.516 | 0.787 | 0.723 | 0.598 | -0.014 | 0.087 | 0.087 | 0 | 0.37 | 0.21 | 0.34 | 0.1 | 0.37 | 0.29 |
| 4 | 27078376 | rs6448497 | 0.401 | 0.836 | 0.775 | 0.542 | -0.006 | 0.009 | 0.009 | 0 | 0.86 | 0.74 | 0.88 | 0.81 | 0.9 | 0.68 |
| 4 | 27079446 | rs7682298 | 1.232 | 0.986 | 0.577 | 1.519 | -0.005 | -0.013 | -0.013 | 0 | 0.38 | 0.24 | 0.37 | 0.39 | 0.37 | 0.32 |
| 4 | 27083248 | rs741135 | 0.434 | 0.877 | 0.075 | 1.302 | 0.015 | 0.061 | 0.061 | 0 | 0.49 | 0.25 | 0.64 | 0.6 | 0.57 | 0.4 |
| 4 | 27087177 | rs16878829 | -0.777 | 0.546 | 0.453 | 1.702 | 0.032 | 0.248 | 0.248 | 0.126 | 0.09 | 0.3 | 0.13 | 0.36 | 0.11 | 0.14 |
| 4 | 27087570 | rs4692163 | 1.382 | 0.77 | -0.138 | 1.239 | -0.02 | 0.223 | 0.223 | 0 | 0.36 | 0.04 | 0.3 | 0.03 | 0.33 | 0.26 |
| 4 | 27087761 | rs4692164 | 0.642 | 0.861 | 0.049 | 1.257 | 0.008 | 0.004 | 0.004 | 0 | 0.45 | 0.41 | 0.43 | 0.39 | 0.44 | 0.4 |
| 4 | 27096261 | rs7677573 | -0.475 | 0.574 | -0.189 | 0.92 | -0.066 | 0.141 | 0.141 | 0 | 0.63 | 0.87 | 0.79 | 0.91 | 0.57 | 0.7 |
| 4 | 27097098 | rs12503097 | 1.253 | 0.145 | 0.06 | 1.099 | 0.033 | 0.159 | 0.159 | 0 | 0.38 | 0.16 | 0.31 | 0.04 | 0.31 | 0.31 |
| 4 | 27097696 | rs2075093 | -0.373 | 0.153 | 0.039 | 1.098 | -0.031 | 0.188 | 0.188 | 0 | 0.58 | 0.86 | 0.73 | 0.92 | 0.47 | 0.64 |

| | | | | | | | | | | | | | | | | |
|---|---|---|---|---|---|---|---|---|---|---|---|---|---|---|---|---|
| 4 | 27102429 | rs2041627 | -0.59 | 0.17 | 0.354 | 1.449 | -0.036 | 0.129 | 0.129 | 0 | 0.13 | 0.34 | 0.26 | 0.26 | 0.06 | 0.06 |
| 4 | 27103530 | rs6835095 | 0.265 | 0.191 | -0.174 | 1.458 | 0.011 | 0.01 | 0.01 | 0 | 0.51 | 0.58 | 0.59 | 0.31 | 0.37 | 0.36 |
| 4 | 27115301 | rs13113425 | 1.091 | 0.354 | -1.054 | 1.2 | 0.014 | 0.2 | 0.2 | 0.295 | 0.38 | 0.07 | 0.31 | 0.04 | 0.33 | 0.3 |
| 4 | 27122947 | rs969949 | 0.209 | 0.743 | -1.282 | 1.529 | 0.037 | 0.158 | 0.158 | 0.094 | 0.52 | 0.77 | 0.49 | 0.2 | 0.4 | 0.55 |
| 4 | 27124674 | rs4692169 | 0.149 | 0.827 | -1.241 | 1.549 | 0.032 | 0.022 | 0.022 | 0.104 | 0.53 | 0.74 | 0.53 | 0.6 | 0.4 | 0.59 |
| 4 | 27125382 | rs7685249 | -0.511 | 1.344 | -1.005 | 1.511 | -0.012 | -0.01 | -0.01 | 0.015 | 0.06 | 0.03 | 0.09 | 0.17 | 0.08 | 0.26 |
| 4 | 27127678 | rs6813524 | -0.302 | 0.877 | -2.022 | 1.099 | 0.005 | -0.004 | -0.004 | 0 | 0.51 | 0.89 | 0.45 | 0.53 | 0.4 | 0.46 |
| 4 | 27132182 | rs17589537 | -0.441 | 0.78 | -1.771 | 0.854 | -0.007 | 0.001 | 0.001 | 0.06 | 0.16 | 0.15 | 0.17 | 0.12 | 0.17 | 0.33 |
| 4 | 27134618 | rs10939150 | -1.209 | -0.146 | -1.946 | 0.724 | 0.085 | 0.224 | 0.224 | 0 | 0.51 | 0.82 | 0.59 | 0.82 | 0.32 | 0.56 |
| 4 | 27139247 | rs6853864 | 1.156 | -0.485 | -0.681 | 1.273 | -0.034 | 0.005 | 0.005 | 0.387 | 0.22 | 0.26 | 0.22 | 0.26 | 0.12 | 0.09 |
| 4 | 27148758 | rs16878895 | 0.709 | -0.459 | -0.189 | 1.317 | 0.021 | 0.126 | 0.126 | 0 | 0.05 | 0.11 | 0.15 | 0.24 | 0.05 | 0.05 |
| 4 | 27148768 | rs13105110 | 0.289 | -0.381 | -0.408 | 1.393 | -0.021 | -0.011 | -0.011 | 0 | 0.16 | 0.1 | 0.16 | 0.16 | 0.17 | 0.34 |
| 4 | 27157313 | rs1013532 | -0.139 | -0.408 | -0.017 | 1.414 | -0.029 | 0.184 | 0.184 | 0 | 0.9 | 0.8 | 0.81 | 0.58 | 0.97 | 0.84 |
| 4 | 27166535 | rs6448500 | 0.899 | -0.418 | 0.062 | 1.321 | -0.001 | 0.014 | 0.014 | 0.721 | 0.75 | 0.7 | 0.65 | 0.41 | 0.8 | 0.5 |
| 4 | 27172128 | rs7668273 | -1.308 | -0.32 | -0.341 | 0.77 | 0.067 | 0.236 | 0.236 | 0 | 0.51 | 0.85 | 0.6 | 0.82 | 0.33 | 0.59 |
| 4 | 27175502 | rs13101823 | -0.666 | 0.592 | 0.484 | 1.019 | 0.059 | 0.005 | 0.005 | 0.425 | 0.26 | 0.38 | 0.31 | 0.27 | 0.42 | 0.43 |
| 4 | 27180604 | rs1488274 | -0.432 | 0.856 | -0.072 | 0.516 | -0.061 | 0.058 | 0.058 | 0 | 0.26 | 0.14 | 0.24 | 0.07 | 0.42 | 0.23 |
| 4 | 27180841 | rs993967 | -1.44 | 0.921 | -0.106 | 0.78 | -0.003 | 0.008 | 0.008 | 0 | 0.4 | 0.67 | 0.56 | 0.72 | 0.53 | 0.29 |
| 4 | 27186835 | rs1982810 | 0.729 | 0.349 | -0.234 | 0.686 | 0.004 | -0.003 | -0.003 | 0 | 0.4 | 0.38 | 0.31 | 0.48 | 0.27 | 0.42 |
| 4 | 27197786 | rs1488290 | 0.112 | 0.483 | -0.331 | 0.517 | 0.049 | -0.022 | -0.022 | 0.08 | 0.67 | 0.5 | 0.51 | 0.72 | 0.5 | 0.78 |
| 4 | 27202594 | rs6826441 | -0.986 | 0.331 | 0.143 | 0.557 | -0.028 | 0.008 | 0.008 | 0 | 0.57 | 0.5 | 0.67 | 0.51 | 0.72 | 0.59 |
| 4 | 27209559 | rs1488268 | 1.003 | 0.327 | 0.226 | 0.556 | 0.024 | -0.013 | -0.013 | 0 | 0.41 | 0.36 | 0.31 | 0.49 | 0.27 | 0.41 |
| 4 | 27209596 | rs13130950 | 0.28 | 0.351 | 0.176 | 0.404 | -0.029 | 0.022 | 0.022 | 0 | 0.23 | 0.17 | 0.22 | 0.07 | 0.41 | 0.19 |
| 4 | 27211040 | rs2046701 | 0.894 | 0.344 | -0.044 | 0.412 | 0.002 | 0.002 | 0.002 | 0 | 0.26 | 0.12 | 0.2 | 0.23 | 0.23 | 0.36 |
| 4 | 27211999 | rs13127219 | 0.179 | 0.583 | -0.046 | 0.522 | -0.056 | 0.048 | 0.048 | 0 | 0.23 | 0.12 | 0.21 | 0.07 | 0.41 | 0.19 |
| 4 | 27216501 | rs4639059 | -1.232 | 0.718 | -0.131 | 0.623 | -0.004 | 0.007 | 0.007 | 0 | 0.39 | 0.45 | 0.59 | 0.44 | 0.6 | 0.41 |
| 4 | 27216662 | rs901832 | -0.876 | 0.753 | -0.164 | 0.67 | 0.019 | 0.021 | 0.021 | 0 | 0.34 | 0.42 | 0.58 | 0.44 | 0.54 | 0.39 |
| 4 | 27218939 | rs17591786 | 0.198 | 1.215 | -1.04 | 0.227 | -0.051 | 0.05 | 0.05 | 0.14 | 0.23 | 0.12 | 0.2 | 0.08 | 0.38 | 0.18 |
| 4 | 27219806 | rs11724172 | 0.189 | 1.282 | -1.098 | 0.602 | -0.068 | 0.058 | 0.058 | 0.109 | 0.23 | 0.1 | 0.21 | 0.1 | 0.38 | 0.17 |
| 4 | 27223375 | rs13142026 | 0.734 | 1.838 | -1.284 | -0.082 | -0.062 | 0.069 | 0.069 | 9.393 | 0.12 | 0.02 | 0.09 | 0.01 | 0.17 | 0.1 |
| 4 | 27225759 | rs12511504 | 1.684 | 1.296 | -1.736 | -0.197 | -0.028 | 0.016 | 0.016 | 0 | 0.19 | 0.01 | 0.15 | 0.09 | 0.17 | 0.15 |
| 4 | 27229288 | rs2029319 | 0.999 | 1.59 | -1.587 | -0.042 | -0.012 | -0.013 | -0.013 | 0 | 0.06 | 0.02 | 0.11 | 0.15 | 0.09 | 0.24 |
| 4 | 27230197 | rs13118370 | 1.232 | 1.198 | -1.503 | -0.79 | -0.003 | 0.033 | 0.033 | 2.768 | 0.27 | 0.11 | 0.13 | 0.01 | 0.25 | 0.28 |
| 4 | 27235976 | rs7673254 | 0.379 | 1.655 | -1.473 | -0.152 | -0.007 | 0.015 | 0.015 | 0 | 0.86 | 0.91 | 0.65 | 0.46 | 0.78 | 0.72 |
| 4 | 27241002 | rs10015926 | 0.771 | 1.03 | -1.662 | -0.126 | -0.005 | -0.004 | -0.004 | 0.951 | 0.13 | 0.09 | 0.19 | 0.27 | 0.21 | 0.09 |
| 4 | 27252907 | rs9291493 | 0.774 | 1.032 | -1.661 | -0.192 | -0.005 | -0.002 | -0.002 | 0 | 0.13 | 0.09 | 0.19 | 0.29 | 0.21 | 0.09 |
| 4 | 27254091 | rs7356206 | -0.089 | 0.992 | -1.611 | -0.187 | 0.013 | 0.024 | 0.024 | 0.009 | 0.19 | 0.24 | 0.22 | 0.29 | 0.28 | 0.11 |
| 4 | 27256570 | rs6448506 | 0.503 | 0.267 | -1.842 | -0.391 | 0.025 | 0.059 | 0.059 | 0 | 0.19 | 0.06 | 0.1 | 0.06 | 0.13 | 0.19 |
| 4 | 27261129 | rs1488298 | -0.065 | 0.776 | -1.47 | -0.091 | 0 | 0.003 | 0.003 | 0 | 0.59 | 0.41 | 0.49 | 0.45 | 0.59 | 0.56 |
| 4 | 27261443 | rs1488299 | -0.769 | 0.848 | -2.162 | -0.198 | -0.033 | 0.174 | 0.174 | 0 | 0.42 | 0.19 | 0.69 | 0.94 | 0.54 | 0.57 |
| 4 | 27262518 | rs16879070 | -0.426 | 0.995 | -2.298 | 0.267 | -0.118 | -0.121 | -0.121 | 1.762 | 0.36 | 0.09 | 0.47 | 0.43 | 0.47 | 0.31 |
| 4 | 27263045 | rs5002502 | -0.002 | 1.388 | -1.593 | 0.532 | -0.017 | -0.043 | -0.043 | 3.254 | 0.65 | 0.85 | 0.61 | 0.49 | 0.61 | 0.6 |
| 4 | 27264542 | rs6854240 | 0.185 | 1.416 | -1.304 | -0.397 | -0.014 | 0.027 | 0.027 | 0.932 | 0.62 | 0.75 | 0.38 | 0.06 | 0.53 | 0.55 |
| 4 | 27268342 | rs748057 | -0.303 | 1.714 | -1.459 | 0.17 | -0.074 | 0.027 | 0.027 | 0.231 | 0.36 | 0.14 | 0.43 | 0.65 | 0.49 | 0.5 |
| 4 | 27269408 | rs1488281 | 0.455 | 1.574 | -1.021 | 0.503 | 0 | 0.01 | 0.01 | 0 | 0.92 | 0.9 | 0.84 | 0.77 | 0.89 | 0.74 |
| 4 | 27276248 | rs10939154 | 0.072 | 0.638 | -0.431 | -0.216 | -0.016 | -0.053 | -0.053 | 0 | 0.16 | 0.19 | 0.13 | 0 | 0.09 | 0.14 |
| 4 | 27280902 | rs6848074 | 1.268 | 0.608 | 0.251 | -0.351 | -0.026 | -0.031 | -0.031 | 0 | 0.72 | 0.41 | 0.69 | 0.77 | 0.78 | 0.62 |
| 4 | 27287137 | rs2029321 | 0.214 | 0.532 | 0.279 | -0.332 | -0.047 | -0.131 | -0.131 | 0.254 | 0.79 | 0.42 | 0.84 | 1 | 0.89 | 0.85 |
| 4 | 27288616 | rs6812821 | 0.853 | 0.495 | 0.32 | -0.316 | -0.048 | -0.026 | -0.026 | 0 | 0.75 | 0.43 | 0.73 | 0.77 | 0.83 | 0.67 |
| 4 | 27292819 | rs1488275 | 0.077 | 0.592 | 0.423 | -0.463 | -0.044 | -0.137 | -0.137 | 0.814 | 0.81 | 0.45 | 0.84 | 1 | 0.89 | 0.84 |
| 4 | 27294467 | rs6854189 | 0.188 | 0.622 | 0.251 | -0.448 | 0.001 | -0.026 | -0.026 | 0.035 | 0.15 | 0.16 | 0.13 | 0 | 0.11 | 0.15 |
| 4 | 27296405 | rs16879148 | -0.35 | 0.594 | 0.486 | -0.535 | -0.017 | -0.085 | -0.085 | 0 | 0.18 | 0.29 | 0.16 | 0 | 0.13 | 0.16 |
| 4 | 27298022 | rs10489014 | 0.453 | 0.957 | -0.308 | -0.787 | -0.044 | 0.05 | 0.05 | 0 | 0.12 | 0.02 | 0.1 | 0.04 | 0.16 | 0.17 |
| 4 | 27298305 | rs6821695 | -0.278 | 0.967 | 0.093 | -0.239 | -0.001 | 0.013 | 0.013 | 0 | 0.1 | 0.11 | 0.14 | 0.36 | 0.1 | 0.29 |
| 4 | 27310241 | rs2309558 | 0.427 | 0.811 | 0.501 | -0.254 | 0.005 | 0.128 | 0.128 | 0 | 0.88 | 0.68 | 0.8 | 0.66 | 0.9 | 0.72 |
| 4 | 27311479 | rs6849084 | -0.279 | 0.862 | 0.264 | -0.553 | 0 | -0.042 | -0.042 | 0.04 | 0.23 | 0.26 | 0.24 | 0.01 | 0.25 | 0.18 |
| 4 | 27314726 | rs2192195 | 1.419 | 1.074 | -0.886 | -0.486 | -0.038 | 0.065 | 0.065 | 0.05 | 0.3 | 0.03 | 0.16 | 0.1 | 0.28 | 0.24 |
| 4 | 27323221 | rs13115333 | 0.481 | 1.106 | -0.882 | -0.471 | -0.07 | -0.067 | -0.067 | 0 | 0.4 | 0.04 | 0.36 | 0.44 | 0.42 | 0.53 |
| 4 | 27324215 | rs12644961 | 0.22 | 1.147 | -0.167 | -0.354 | 0.001 | -0.134 | -0.134 | 0.37 | 0.74 | 0.42 | 0.73 | 1 | 0.74 | 0.8 |
| 4 | 27326253 | rs11727052 | 1.225 | 1.111 | 0.255 | 0.276 | 0.001 | 0.061 | 0.061 | 0.955 | 0.81 | 0.44 | 0.73 | 0.65 | 0.84 | 0.65 |
| 4 | 27331116 | rs10021654 | -0.749 | 0.977 | -0.037 | 0.105 | -0.001 | -0.024 | -0.024 | 0 | 0.05 | 0.13 | 0.04 | 0 | 0.03 | 0.13 |
| 4 | 27335003 | rs7662818 | -0.053 | 1.343 | -0.247 | 0 | 0.007 | -0.011 | -0.011 | 0 | 0.08 | 0.09 | 0.07 | 0.04 | 0.12 | 0.14 |
| 4 | 27337756 | rs759097 | 0.739 | 0.792 | 0.137 | 0.858 | 0.003 | 0.284 | 0.284 | 2.375 | 0.9 | 0.49 | 0.85 | 0.81 | 0.95 | 0.85 |
| 4 | 27339904 | rs4692196 | 0.867 | 0.782 | 0.041 | 0.841 | -0.01 | 0.199 | 0.199 | 2.579 | 0.88 | 0.45 | 0.84 | 0.8 | 0.94 | 0.82 |
| 4 | 27342170 | rs6819555 | 0.297 | 0.926 | -1.445 | 0.41 | -0.079 | -0.053 | -0.053 | 0 | 0.42 | 0.03 | 0.37 | 0.44 | 0.43 | 0.51 |
| 4 | 27352006 | rs1000516 | 0.086 | 0.766 | -1.277 | 0.576 | -0.031 | -0.058 | -0.058 | 0.693 | 0.12 | 0.02 | 0.18 | 0.28 | 0.14 | 0.16 |
| 4 | 27355782 | rs9291495 | 0.471 | 0.645 | -1.115 | 0.341 | -0.01 | -0.021 | -0.021 | 0 | 0.82 | 0.9 | 0.75 | 0.71 | 0.76 | 0.84 |
| 4 | 27358898 | rs2215745 | 1.322 | 0.367 | -1.461 | -0.454 | -0.04 | 0.09 | 0.09 | 0.762 | 0.43 | 0.18 | 0.55 | 0.76 | 0.45 | 0.67 |
| 4 | 27360257 | rs7690010 | -0.44 | 1.175 | -0.753 | 0.052 | 0.025 | 0.057 | 0.057 | 0 | 0.86 | 0.78 | 0.8 | 0.76 | 0.85 | 0.84 |

| | | | | | | | | | | | | | | | |
|---|---|---|---|---|---|---|---|---|---|---|---|---|---|---|---|
| 4 | 27360968 | rs13147990 | -0.261 | 1.156 | -0.664 | -0.073 | 0.044 | 0.119 | 0.119 | 0 | 0.86 | 0.7 | 0.8 | 0.72 | 0.85 | 0.83 |
| 4 | 27362740 | rs13134892 | 0.547 | 0.653 | -1.379 | -0.803 | -0.052 | 0.113 | 0.113 | 0 | 0.22 | 0.08 | 0.08 | 0 | 0.25 | 0.15 |
| 4 | 27367496 | rs2871278 | 0.547 | 0.655 | -1.378 | -0.803 | -0.052 | 0.113 | 0.113 | 0 | 0.22 | 0.08 | 0.08 | 0 | 0.25 | 0.15 |
| 4 | 27373502 | rs10489015 | 0.34 | 0.915 | -0.982 | -0.601 | -0.001 | -0.028 | -0.028 | 2.178 | 0.07 | 0.12 | 0.07 | 0 | 0.09 | 0.01 |
| 4 | 27373821 | rs10012661 | -0.469 | 0.997 | -0.736 | -0.263 | 0.002 | 0.019 | 0.019 | 0 | 0.87 | 0.88 | 0.81 | 0.72 | 0.85 | 0.84 |
| 4 | 27374368 | rs6448515 | -0.788 | 0.326 | -0.7 | -1.035 | -0.001 | -0.08 | -0.08 | 2.061 | 0.36 | 0.41 | 0.17 | 0 | 0.35 | 0.18 |
| 4 | 27383785 | rs13107491 | -0.335 | 0.967 | -0.657 | -0.446 | 0.006 | -0.047 | -0.047 | 0.261 | 0.19 | 0.21 | 0.11 | 0 | 0.28 | 0.16 |
| 4 | 27388554 | rs4355380 | -1.201 | -0.422 | -0.441 | 0.193 | 0.005 | -0.002 | -0.002 | 2.662 | 0.28 | 0.68 | 0.4 | 0.66 | 0.17 | 0.24 |
| 4 | 27390449 | rs6448518 | -1.014 | -0.533 | 0.784 | 0.783 | 0.038 | 0.133 | 0.133 | 1.151 | 0.1 | 0.34 | 0.12 | 0.16 | 0.09 | 0.08 |
| 4 | 27406431 | rs210755 | -0.292 | -0.622 | 1.515 | 1.039 | -0.006 | -0.019 | -0.019 | 0.112 | 0.06 | 0.03 | 0.08 | 0.15 | 0.08 | 0.07 |
| 4 | 27409678 | rs4692206 | 1.116 | -0.811 | 0.81 | 0.014 | -0.089 | 0.157 | 0.157 | 0.891 | 0.33 | 0.11 | 0.4 | 0.01 | 0.49 | 0.31 |
| 4 | 27412242 | rs210760 | -0.546 | -0.823 | 1.018 | 0.059 | 0.002 | 0.062 | 0.062 | 0 | 0.08 | 0.16 | 0.1 | 0.15 | 0.08 | 0.08 |
| 4 | 27412694 | rs210761 | -0.132 | -1.241 | 0.771 | -0.192 | 0.012 | 0.012 | 0.012 | 0 | 0.53 | 0.43 | 0.6 | 0.2 | 0.66 | 0.42 |
| 4 | 27432523 | rs210777 | -0.975 | -0.826 | 0.976 | 0.371 | -0.032 | 0.104 | 0.104 | 0.013 | 0.21 | 0.41 | 0.31 | 0.64 | 0.16 | 0.21 |
| 4 | 27438947 | rs210749 | -0.298 | -0.921 | 0.787 | 0.109 | 0.029 | 0.048 | 0.048 | 0 | 0.61 | 0.82 | 0.79 | 0.69 | 0.65 | 0.53 |
| 4 | 27442060 | rs2110177 | -0.886 | -0.85 | 0.736 | 0.162 | 0.011 | 0.015 | 0.015 | 0 | 0.68 | 0.8 | 0.82 | 0.71 | 0.69 | 0.55 |
| 4 | 27450750 | rs210782 | 0.212 | -0.591 | 0.744 | 0.443 | 0.013 | -0.038 | -0.038 | 0 | 0.14 | 0.04 | 0.11 | 0.17 | 0.12 | 0.09 |
| 4 | 27459128 | rs10025989 | -1.402 | -0.247 | 1.2 | 0.678 | 0.008 | 0.191 | 0.191 | 0 | 0.07 | 0.11 | 0.23 | 0.5 | 0.07 | 0.11 |
| 4 | 27483170 | rs6856266 | 0.013 | -0.411 | 0.344 | 0.618 | 0.024 | 0.006 | 0.006 | 0 | 0.49 | 0.68 | 0.65 | 0.48 | 0.53 | 0.44 |
| 4 | 27490495 | rs10461103 | -0.299 | -0.372 | 0.095 | 0.587 | 0.019 | -0.06 | -0.06 | 0 | 0.55 | 0.87 | 0.77 | 0.51 | 0.6 | 0.39 |
| 4 | 27496294 | rs1378788 | 0.41 | 0.196 | 0.241 | 0.359 | -0.006 | 0.147 | 0.147 | 0.172 | 0.44 | 0.74 | 0.41 | 0.02 | 0.42 | 0.33 |
| 4 | 27496410 | rs1993400 | -0.253 | 0.128 | 0.102 | 0.539 | 0.034 | -0.061 | -0.061 | 0.108 | 0.55 | 0.87 | 0.77 | 0.53 | 0.61 | 0.4 |
| 4 | 27509411 | rs12505179 | -0.971 | 0.477 | 0.399 | 0.935 | -0.006 | 0.03 | 0.03 | 0 | 0.1 | 0.11 | 0.35 | 0.54 | 0.16 | 0.07 |
| 4 | 27509621 | rs12505247 | -0.971 | 0.485 | 0.415 | 0.937 | -0.005 | 0.034 | 0.034 | 0 | 0.1 | 0.12 | 0.35 | 0.54 | 0.17 | 0.07 |
| 4 | 27530174 | rs12506982 | -1.039 | 0.445 | 0.803 | 0.937 | 0.013 | 0.115 | 0.115 | 0 | 0.11 | 0.18 | 0.38 | 0.57 | 0.16 | 0.08 |
| 4 | 27534772 | rs1375757 | -0.989 | 0.446 | 0.82 | 0.894 | 0.013 | 0.103 | 0.103 | 0 | 0.11 | 0.16 | 0.38 | 0.58 | 0.16 | 0.08 |
| 4 | 27534917 | rs9992655 | 0.975 | 0.153 | 0.842 | 0.459 | -0.051 | 0.156 | 0.156 | 0 | 0.37 | 0.15 | 0.33 | 0.01 | 0.42 | 0.32 |
| 4 | 27539764 | rs12504241 | -1.292 | 0.095 | 1.404 | 0.438 | 0.017 | 0.165 | 0.165 | 0 | 0.14 | 0.29 | 0.41 | 0.58 | 0.17 | 0.08 |
| 4 | 27542201 | rs16879469 | -1.916 | -0.366 | 1.644 | 0.167 | -0.03 | 0.06 | 0.06 | 0.16 | 0.25 | 0.57 | 0.46 | 0.59 | 0.2 | 0.2 |
| 4 | 27542684 | rs1449756 | -0.622 | -0.358 | 1.556 | 0.179 | -0.007 | -0.006 | -0.006 | 0 | 0.63 | 0.75 | 0.79 | 0.6 | 0.63 | 0.52 |
| 4 | 27548905 | rs10009898 | 1.047 | -0.558 | 1.391 | 0.1 | -0.037 | -0.058 | -0.058 | 0.323 | 0.28 | 0.05 | 0.17 | 0.36 | 0.29 | 0.35 |
| 4 | 27553045 | rs13124025 | -2.044 | -0.524 | 1.407 | 0.08 | -0.033 | 0.033 | 0.033 | 0.38 | 0.26 | 0.61 | 0.38 | 0.59 | 0.19 | 0.21 |
| 4 | 27560693 | rs10506518 | 0.432 | -0.294 | 0.9 | 0.202 | -0.001 | -0.003 | -0.003 | 2.55 | 0.52 | 0.47 | 0.66 | 0.59 | 0.58 | 0.42 |
| 4 | 27570669 | rs16879491 | -0.096 | -0.111 | 0.407 | 0.397 | -0.023 | -0.028 | -0.028 | 1.269 | 0.07 | 0.1 | 0.02 | 0.01 | 0.01 | 0.12 |
| 4 | 27584802 | rs6856564 | 1.178 | 0.312 | 0.619 | 1.32 | -0.003 | 0.001 | 0.001 | 0 | 0.82 | 0.82 | 0.62 | 0.56 | 0.79 | 0.8 |
| 4 | 27600788 | rs4599400 | -2.218 | 0.206 | 0.799 | 1.303 | -0.017 | 0.09 | 0.09 | 0.577 | 0.12 | 0.47 | 0.14 | 0.23 | 0.07 | 0.13 |
| 4 | 27601868 | rs1375761 | 1.165 | 0.289 | 0.841 | 1.12 | -0.02 | -0.053 | -0.053 | 0 | 0.83 | 0.91 | 0.64 | 0.55 | 0.79 | 0.69 |
| 4 | 27602132 | rs16879587 | 0.897 | 0.422 | 0.436 | 0.954 | 0.001 | -0.021 | -0.021 | 0 | 0.49 | 0.46 | 0.33 | 0.08 | 0.44 | 0.47 |
| 4 | 27603144 | rs6847492 | 0.671 | 0.362 | 0.377 | 1.006 | 0.001 | 0.034 | 0.034 | 0 | 0.84 | 0.78 | 0.63 | 0.56 | 0.82 | 0.8 |
| 4 | 27606650 | rs10212920 | 0.667 | 0.426 | 0.387 | 0.417 | 0.002 | -0.005 | -0.005 | 0 | 0.48 | 0.48 | 0.34 | 0.07 | 0.46 | 0.38 |
| 4 | 27614790 | rs7697958 | -1.851 | 0.589 | 0.628 | 1.194 | -0.012 | -0.031 | -0.031 | 0 | 0.07 | 0.19 | 0.03 | 0.01 | 0.04 | 0.12 |
| 4 | 27624165 | rs6448527 | 1.309 | 1.298 | -0.01 | 1.716 | -0.001 | 0.011 | 0.011 | 0 | 0.86 | 0.85 | 0.57 | 0.65 | 0.73 | 0.64 |
| 4 | 27624182 | rs6448528 | 1.435 | 1.266 | -0.042 | 1.649 | -0.013 | -0.012 | -0.012 | 0 | 0.83 | 0.84 | 0.56 | 0.65 | 0.7 | 0.61 |
| 4 | 27624322 | rs6448529 | 0.689 | 1.42 | -0.118 | 1.838 | -0.069 | -0.068 | -0.068 | 1.929 | 0.89 | 0.97 | 0.6 | 0.69 | 0.75 | 0.76 |
| 4 | 27637428 | rs7671071 | 0.498 | 0.98 | -0.079 | 1.893 | -0.061 | -0.063 | -0.063 | 0.02 | 0.89 | 0.97 | 0.62 | 0.67 | 0.77 | 0.76 |
| 4 | 27642019 | rs297282 | -1.444 | 0.376 | 0.043 | 1.714 | -0.001 | -0.001 | -0.001 | 0 | 0.39 | 0.6 | 0.57 | 0.5 | 0.38 | 0.38 |
| 4 | 27655235 | rs454414 | -0.358 | 0.392 | -0.234 | 1.737 | -0.072 | -0.08 | -0.08 | 0 | 0.27 | 0.05 | 0.46 | 0.3 | 0.31 | 0.26 |
| 4 | 27658052 | rs9291503 | 1.178 | 0.454 | -0.155 | 1.63 | 0.012 | -0.019 | -0.019 | 0.236 | 0.59 | 0.29 | 0.49 | 0.53 | 0.64 | 0.61 |
| 4 | 27661645 | rs2034472 | 1.285 | 0.448 | -0.178 | 1.613 | 0.004 | 0.002 | 0.002 | 0 | 0.57 | 0.25 | 0.48 | 0.5 | 0.61 | 0.51 |
| 4 | 27661893 | rs7671299 | -1.619 | 0.093 | 0.873 | 1.889 | -0.04 | 0.188 | 0.188 | 0.435 | 0.14 | 0.49 | 0.19 | 0.22 | 0.06 | 0.12 |
| 4 | 27671803 | rs297291 | 1.413 | 0.272 | 0.732 | 1.925 | -0.006 | 0.151 | 0.151 | 0.743 | 0.95 | 0.69 | 0.88 | 0.84 | 0.99 | 0.91 |
| 4 | 27675410 | rs6846226 | 1.308 | 0.34 | 0.717 | 1.737 | -0.007 | 0.018 | 0.018 | 0 | 0.66 | 0.35 | 0.62 | 0.62 | 0.69 | 0.65 |
| 4 | 27680483 | rs297279 | -0.983 | 0.295 | 0.474 | 1.656 | -0.054 | -0.043 | -0.043 | 0 | 0.27 | 0.08 | 0.29 | 0.25 | 0.3 | 0.29 |
| 4 | 27687397 | rs1449766 | 1.38 | 0.187 | 0.868 | 1.656 | 0.001 | -0.004 | -0.004 | 0 | 0.63 | 0.26 | 0.55 | 0.59 | 0.67 | 0.62 |
| 4 | 27694893 | rs1031311 | 1.177 | 0.295 | 1.068 | 1.709 | -0.004 | 0.033 | 0.033 | 0 | 0.66 | 0.44 | 0.58 | 0.59 | 0.67 | 0.62 |
| 4 | 27704222 | rs6448532 | 1.315 | 0.422 | 1.1 | 1.88 | -0.002 | 0.003 | 0.003 | 0 | 0.62 | 0.35 | 0.53 | 0.46 | 0.59 | 0.56 |
| 4 | 27714566 | rs297308 | -1.333 | 0.575 | 0.584 | 1.203 | -0.025 | 0.08 | 0.08 | 0 | 0.23 | 0.05 | 0.13 | 0.08 | 0.22 | 0.26 |
| 4 | 27715503 | rs297306 | -1.775 | 0.646 | 0.626 | 0.905 | 0.001 | 0.029 | 0.029 | 0 | 0.58 | 0.66 | 0.62 | 0.71 | 0.62 | 0.62 |
| 4 | 27719790 | rs1992981 | -1.227 | 0.706 | 1.268 | 1.384 | 0.008 | 0.222 | 0.222 | 2.181 | 0.11 | 0.2 | 0.2 | 0.42 | 0.13 | 0.13 |
| 4 | 27725689 | rs13126571 | 0.432 | 0.689 | 1.228 | 1.387 | 0.002 | 0.022 | 0.022 | 0 | 0.27 | 0.16 | 0.27 | 0.21 | 0.26 | 0.23 |
| 4 | 27726979 | rs1472691 | -0.02 | 0.703 | 1.295 | 1.397 | 0.009 | 0.023 | 0.023 | 1.701 | 0.31 | 0.21 | 0.29 | 0.21 | 0.29 | 0.24 |
| 4 | 27727259 | rs6841905 | -0.29 | 1.064 | 0.962 | 1.208 | -0.082 | 0.062 | 0.062 | 0.79 | 0.21 | 0.02 | 0.12 | 0.08 | 0.24 | 0.26 |
| 4 | 27735936 | rs10028171 | -0.813 | 0.989 | 1.109 | 1.301 | -0.006 | -0.008 | -0.008 | 0 | 0.38 | 0.52 | 0.49 | 0.63 | 0.39 | 0.35 |
| 4 | 27739598 | rs4450914 | -1.741 | 1.059 | 1.215 | 1.486 | 0.004 | 0.035 | 0.035 | 0.047 | 0.33 | 0.5 | 0.33 | 0.49 | 0.37 | 0.39 |
| 4 | 27742689 | rs13108163 | 0.264 | 0.891 | 0.85 | 1.206 | -0.02 | 0.06 | 0.06 | 0 | 0.34 | 0.11 | 0.32 | 0.22 | 0.34 | 0.27 |
| 4 | 27744290 | rs1503465 | 1.306 | 0.867 | 1.036 | 1.174 | 0.001 | 0.044 | 0.044 | 0.008 | 0.42 | 0.26 | 0.39 | 0.31 | 0.4 | 0.43 |
| 4 | 27744718 | rs4692256 | 0.615 | 0.815 | 1.19 | 0.7 | 0.001 | 0.109 | 0.109 | 0.349 | 0.51 | 0.74 | 0.61 | 0.72 | 0.54 | 0.58 |
| 4 | 27757391 | rs13110484 | 0.329 | 0.752 | 1.331 | 0.684 | -0.062 | 0.058 | 0.058 | 0 | 0.34 | 0.07 | 0.32 | 0.22 | 0.37 | 0.28 |
| 4 | 27757473 | rs17669368 | 0.259 | 0.778 | 1.33 | 0.732 | -0.009 | 0.01 | 0.01 | 0.016 | 0.09 | 0.02 | 0.05 | 0.06 | 0.1 | 0.14 |

| | | | | | | | | | | | | | | | | |
|---|---|---|---|---|---|---|---|---|---|---|---|---|---|---|---|---|
| 4 | 27760141 | rs10014448 | 0.827 | 0.722 | 1.56 | 0.769 | -0.001 | 0.053 | 0.053 | 0.13 | 0.47 | 0.41 | 0.57 | 0.72 | 0.47 | 0.57 |
| 4 | 27763567 | rs7677623 | 0.465 | 0.669 | 1.331 | 0.827 | -0.001 | 0.016 | 0.016 | 0.197 | 0.5 | 0.6 | 0.59 | 0.71 | 0.51 | 0.57 |
| 4 | 27779332 | rs2202258 | -0.786 | 0.634 | 1.207 | 0.782 | -0.005 | 0.085 | 0.085 | 0 | 0.49 | 0.25 | 0.39 | 0.27 | 0.49 | 0.44 |
| 4 | 27782417 | rs7685758 | -1.186 | 0.514 | 1.364 | 1.067 | -0.022 | 0.168 | 0.168 | 0 | 0.14 | 0.34 | 0.22 | 0.5 | 0.14 | 0.16 |
| 4 | 27782716 | rs16879755 | 0.175 | 0.457 | 1.341 | 0.855 | -0.015 | -0.027 | -0.027 | 0 | 0.08 | 0.08 | 0.02 | 0 | 0.04 | 0.05 |
| 4 | 27783448 | rs1503454 | -0.82 | 0.55 | 1.495 | 0.756 | 0.003 | 0.018 | 0.018 | 0 | 0.51 | 0.59 | 0.56 | 0.74 | 0.5 | 0.44 |
| 4 | 27790945 | rs2309628 | 1.813 | 0.487 | 1.48 | 0.67 | -0.037 | 0.073 | 0.073 | 0 | 0.34 | 0.17 | 0.35 | 0.19 | 0.37 | 0.32 |
| 4 | 27792360 | rs13119429 | -0.788 | 0.519 | 1.516 | 0.689 | -0.003 | 0.04 | 0.04 | 0 | 0.52 | 0.44 | 0.55 | 0.74 | 0.52 | 0.49 |
| 4 | 27793744 | rs12640417 | -0.695 | 0.625 | 1.502 | 0.752 | 0.012 | 0.116 | 0.116 | 0 | 0.08 | 0.2 | 0.23 | 0.55 | 0.14 | 0.12 |
| 4 | 27795958 | rs16879791 | -1 | 0.597 | 1.516 | 0.738 | -0.005 | 0.081 | 0.081 | 0 | 0.14 | 0.27 | 0.25 | 0.55 | 0.17 | 0.16 |
| 4 | 27797571 | rs13144861 | 0.251 | 0.71 | 1.218 | 0.619 | -0.005 | 0.074 | 0.074 | 0 | 0.36 | 0.18 | 0.29 | 0.19 | 0.34 | 0.32 |
| 4 | 27805588 | rs7686302 | 0.364 | 1.213 | 0.725 | 0.611 | -0.072 | 0.113 | 0.113 | 0 | 0.37 | 0.11 | 0.3 | 0.2 | 0.39 | 0.33 |
| 4 | 27811805 | rs12643611 | -1.035 | 1.456 | 0.914 | 1.016 | 0.027 | 0.237 | 0.237 | 0.284 | 0.11 | 0.25 | 0.25 | 0.51 | 0.17 | 0.16 |
| 4 | 27812345 | rs13112218 | -0.448 | 1.423 | 0.552 | 0.94 | -0.008 | 0.051 | 0.051 | 0 | 0.54 | 0.34 | 0.38 | 0.24 | 0.5 | 0.51 |
| 4 | 27813390 | rs17671452 | 1.44 | 1.451 | -0.038 | 0.844 | -0.017 | 0.035 | 0.035 | 0 | 0.31 | 0.16 | 0.35 | 0.24 | 0.31 | 0.29 |
| 4 | 27821792 | rs16879832 | 0.382 | 0.852 | -0.007 | 0.448 | 0.002 | -0.025 | -0.025 | 0 | 0.1 | 0.15 | 0.13 | 0.05 | 0.09 | 0.11 |
| 4 | 27824071 | rs7660570 | 1.43 | 1.195 | -0.881 | 0.954 | -0.032 | -0.023 | -0.023 | 0.252 | 0.29 | 0.19 | 0.33 | 0.36 | 0.31 | 0.31 |
| 4 | 27828760 | rs11728477 | 0.731 | 1.248 | -0.645 | 0.55 | -0.008 | 0.019 | 0.019 | 0 | 0.47 | 0.41 | 0.5 | 0.64 | 0.48 | 0.51 |
| 4 | 27828999 | rs1905564 | -0.597 | 1.76 | -0.933 | 1.287 | -0.023 | -0.02 | -0.02 | 0 | 0.08 | 0.02 | 0.08 | 0.21 | 0.13 | 0.06 |
| 4 | 27831637 | rs7686975 | -0.339 | 1.239 | -0.736 | 0.694 | 0 | 0.001 | 0.001 | 0 | 0.27 | 0.28 | 0.28 | 0.19 | 0.3 | 0.3 |
| 4 | 27833499 | rs6832303 | -0.645 | 1.146 | -0.575 | 0.699 | 0.01 | 0.022 | 0.022 | 0 | 0.32 | 0.51 | 0.25 | 0.43 | 0.3 | 0.24 |
| 4 | 27837969 | rs6829231 | -0.474 | 1.168 | -0.17 | 0.751 | 0.002 | -0.037 | -0.037 | 0.69 | 0.27 | 0.38 | 0.17 | 0.17 | 0.21 | 0.21 |
| 4 | 27848155 | rs12640014 | -1.356 | 0.958 | -0.516 | 0.752 | -0.004 | 0.002 | 0.002 | 0 | 0.43 | 0.57 | 0.45 | 0.5 | 0.37 | 0.54 |
| 4 | 27867951 | rs12648098 | 0.676 | 1.224 | -0.519 | 0.826 | -0.034 | -0.006 | -0.006 | 0 | 0.09 | 0.03 | 0.07 | 0.12 | 0.13 | 0.09 |
| 4 | 27871868 | rs6856472 | 0.321 | 1.031 | -0.416 | 0.863 | 0.001 | 0.003 | 0.003 | 0 | 0.11 | 0.12 | 0.26 | 0.34 | 0.14 | 0.35 |
| 4 | 27882142 | rs10939176 | 1.429 | 1.032 | -0.557 | 0.344 | 0.007 | 0.007 | 0.007 | 0.007 | 0.35 | 0.34 | 0.2 | 0.04 | 0.25 | 0.12 |
| 4 | 27896510 | rs6842063 | -1.418 | 0.983 | -0.415 | 0.377 | 0.054 | 0.161 | 0.161 | 0 | 0.89 | 0.68 | 0.87 | 0.74 | 0.83 | 0.86 |
| 4 | 27897437 | rs11736171 | 1.815 | 0.938 | -0.833 | -0.05 | 0.01 | 0.01 | 0.01 | 0.013 | 0.26 | 0.26 | 0.15 | 0.04 | 0.18 | 0.09 |
| 4 | 27902479 | rs6857890 | -0.779 | 1.068 | -0.94 | 0.53 | -0.005 | 0.02 | 0.02 | 0 | 0.46 | 0.32 | 0.42 | 0.39 | 0.44 | 0.5 |
| 4 | 27902507 | rs6857913 | 0.117 | 1.137 | -1.123 | 0.536 | -0.02 | 0.032 | 0.032 | 0 | 0.23 | 0.08 | 0.12 | 0.16 | 0.2 | 0.16 |
| 4 | 27902560 | rs16879958 | -0.012 | 0.91 | -0.73 | 0.979 | 0.001 | -0.002 | -0.002 | 0 | 0.23 | 0.21 | 0.29 | 0.22 | 0.24 | 0.34 |
| 4 | 27905536 | rs6841769 | 0.19 | 1.048 | -1.031 | 0.04 | -0.034 | 0.118 | 0.118 | 0 | 0.43 | 0.27 | 0.27 | 0.18 | 0.38 | 0.26 |
| 4 | 27906829 | rs17616472 | -0.281 | 1.875 | -1.337 | 0.332 | -0.064 | -0.034 | -0.034 | 1.246 | 0.15 | 0.04 | 0.09 | 0.2 | 0.22 | 0.11 |
| 4 | 27907123 | rs1038441 | 0.222 | 1.127 | -1.107 | 0.013 | 0.01 | 0.008 | 0.008 | 0 | 0.46 | 0.56 | 0.62 | 0.59 | 0.42 | 0.68 |
| 4 | 27907530 | rs17616508 | -0.346 | 1.201 | -1.012 | 0.03 | 0.006 | 0.002 | 0.002 | 0 | 0.18 | 0.25 | 0.12 | 0.21 | 0.22 | 0.16 |
| 4 | 27912160 | rs1026947 | 0.952 | 0.628 | -1.557 | 0.456 | 0.007 | -0.085 | -0.085 | 0.069 | 0.13 | 0.01 | 0.26 | 0.28 | 0.12 | 0.25 |
| 4 | 27914446 | rs1292621 | -1.027 | 1.211 | -1.163 | -0.337 | -0.003 | 0.064 | 0.064 | 0 | 0.67 | 0.85 | 0.79 | 0.79 | 0.71 | 0.86 |
| 4 | 27918627 | rs2110510 | -1.147 | 1.276 | -1.248 | -0.326 | -0.01 | 0.041 | 0.041 | 0 | 0.69 | 0.92 | 0.79 | 0.75 | 0.73 | 0.86 |
| 4 | 27924831 | rs4692262 | 0.858 | 1.245 | -1.286 | -0.509 | 0.05 | -0.064 | -0.064 | 0 | 0.34 | 0.11 | 0.48 | 0.56 | 0.28 | 0.56 |
| 4 | 27925154 | rs4692044 | 0.644 | 1.234 | -0.351 | -0.996 | 0.013 | 0.069 | 0.069 | 0 | 0.47 | 0.32 | 0.6 | 0.77 | 0.46 | 0.71 |
| 4 | 27928459 | rs6844406 | 0.731 | 1.279 | -0.623 | -0.993 | -0.055 | -0.039 | -0.039 | 0 | 0.11 | 0.03 | 0.1 | 0.17 | 0.2 | 0.07 |
| 4 | 27931573 | rs10016121 | 0.659 | 1.298 | -0.468 | -1.024 | 0.01 | 0.062 | 0.062 | 0 | 0.48 | 0.34 | 0.6 | 0.77 | 0.48 | 0.73 |
| 4 | 27940513 | rs7684360 | -1.371 | 1.769 | -0.924 | -1.016 | 0.014 | 0.029 | 0.029 | 0 | 0.76 | 0.88 | 0.84 | 0.96 | 0.83 | 0.92 |
| 4 | 27943218 | rs6813435 | -0.619 | 1.784 | -0.696 | -1.003 | 0.029 | 0.005 | 0.005 | 0 | 0.2 | 0.32 | 0.17 | 0.22 | 0.28 | 0.17 |
| 4 | 27943852 | rs756526 | -0.654 | 1.804 | -0.717 | -0.974 | 0.002 | 0.06 | 0.06 | 0 | 0.72 | 0.88 | 0.8 | 0.95 | 0.76 | 0.87 |
| 4 | 27955784 | rs2067760 | -0.565 | 1.826 | -0.838 | -0.915 | 0.022 | 0 | 0 | 0 | 0.22 | 0.38 | 0.18 | 0.23 | 0.27 | 0.16 |
| 4 | 27961533 | rs7659592 | -0.174 | 1.31 | -0.997 | -1.257 | -0.002 | -0.037 | -0.037 | 0.058 | 0.7 | 0.75 | 0.79 | 0.96 | 0.77 | 0.9 |
| 4 | 27961828 | rs7665009 | -0.059 | 1.264 | -1.124 | -1.165 | 0 | 0.022 | 0.022 | 0 | 0.65 | 0.78 | 0.75 | 0.95 | 0.68 | 0.85 |
| 4 | 27966505 | rs9884587 | 0.005 | 1.704 | -1.564 | -0.973 | -0.11 | 0.116 | 0.116 | 0 | 0.82 | 0.98 | 0.81 | 1 | 0.73 | 0.95 |
| 4 | 27971596 | rs2058574 | 0.272 | 1.478 | -1.54 | -0.544 | 0.002 | -0.006 | -0.006 | 0 | 0.11 | 0.09 | 0.18 | 0.19 | 0.14 | 0.15 |
| 4 | 27971792 | rs16880029 | 0.951 | 0.869 | -1.249 | -1.292 | 0.002 | 0.002 | 0.002 | 0 | 0.15 | 0.09 | 0.06 | 0.09 | 0.07 | 0.06 |
| 4 | 27975847 | rs10029165 | 0.691 | 0.906 | -0.702 | -1.473 | -0.001 | 0.012 | 0.012 | 0.007 | 0.54 | 0.44 | 0.66 | 0.72 | 0.52 | 0.77 |
| 4 | 27983025 | rs929620 | 0.218 | 0.833 | -0.412 | -1.401 | 0.006 | 0.003 | 0.003 | 0 | 0.12 | 0.11 | 0.07 | 0.18 | 0.15 | 0.13 |
| 4 | 28000670 | rs11932274 | 0.312 | 0.513 | -0.408 | -1.45 | -0.002 | 0.001 | 0.001 | 0 | 0.11 | 0.1 | 0.06 | 0.17 | 0.1 | 0.11 |
| 4 | 28003277 | rs4692270 | 0.083 | 0.464 | -0.153 | -1.703 | 0.013 | -0.122 | -0.122 | 0 | 0.67 | 0.53 | 0.74 | 0.92 | 0.66 | 0.89 |
| 4 | 28005075 | rs16880055 | 0.437 | 0.406 | -0.175 | -1.647 | -0.003 | 0.001 | 0.001 | 0 | 0.11 | 0.1 | 0.06 | 0.17 | 0.09 | 0.11 |
| 4 | 28016367 | rs1125825 | -0.519 | 0.242 | 0.714 | -1.759 | -0.047 | -0.034 | -0.034 | 5.447 | 0.3 | 0.5 | 0.15 | 0.24 | 0.19 | 0.17 |
| 4 | 28017651 | rs1405674 | 0.994 | 0.366 | 0.343 | -1.737 | -0.001 | 0.135 | 0.135 | 0 | 0.53 | 0.25 | 0.67 | 0.76 | 0.57 | 0.77 |
| 4 | 28024351 | rs1527339 | 0.421 | 0.3 | 0.745 | -1.718 | 0.008 | -0.069 | -0.069 | 0 | 0.65 | 0.47 | 0.73 | 0.86 | 0.65 | 0.85 |
| 4 | 28032948 | rs1358311 | 0.371 | 0.296 | 0.751 | -1.72 | 0.009 | -0.072 | -0.072 | 0 | 0.66 | 0.48 | 0.74 | 0.86 | 0.65 | 0.85 |
| 4 | 28036809 | rs4692272 | 0.769 | 0.218 | 1.143 | -1.716 | -0.026 | -0.019 | -0.019 | 0 | 0.17 | 0.22 | 0.08 | 0.14 | 0.11 | 0.09 |
| 4 | 28046364 | rs2110506 | 0.232 | -0.159 | 0.577 | -2.308 | 0.02 | 0.065 | 0.065 | 0 | 0.34 | 0.19 | 0.32 | 0.19 | 0.31 | 0.42 |
| 4 | 28052469 | rs725070 | 1.134 | -0.15 | 0.62 | -2.271 | -0.03 | -0.022 | -0.022 | 0.642 | 0.17 | 0.25 | 0.08 | 0.14 | 0.11 | 0.09 |
| 4 | 28055549 | rs16880200 | 1.083 | -0.169 | 0.603 | -2.28 | -0.029 | -0.022 | -0.022 | 0.062 | 0.18 | 0.23 | 0.08 | 0.14 | 0.11 | 0.09 |
| 4 | 28057728 | rs11726566 | 2.141 | 0.119 | 0.301 | -2.168 | -0.004 | 0.028 | 0.028 | 0 | 0.05 | 0 | 0.01 | 0 | 0.04 | 0.04 |
| 4 | 28067758 | rs13116924 | -0.095 | 0.33 | 0.616 | -2 | 0.052 | 0.095 | 0.095 | 0 | 0.49 | 0.27 | 0.37 | 0.26 | 0.39 | 0.47 |
| 4 | 28071228 | rs1918326 | 1.077 | 0.425 | 0.538 | -2.022 | -0.013 | 0.011 | 0.011 | 0 | 0.3 | 0.18 | 0.39 | 0.67 | 0.32 | 0.43 |
| 4 | 28077002 | rs6850690 | -0.02 | 0.805 | 1.056 | -1.503 | 0.087 | -0.072 | -0.072 | 0.513 | 0.81 | 0.57 | 0.79 | 0.93 | 0.72 | 0.9 |
| 4 | 28087933 | rs13109572 | -0.15 | 0.847 | 0.767 | -1.482 | 0.053 | -0.048 | -0.048 | 0 | 0.81 | 0.66 | 0.79 | 0.93 | 0.72 | 0.9 |

| | | | | | | | | | | | | | | | | |
|---|---|---|---|---|---|---|---|---|---|---|---|---|---|---|---|---|
| 4 | 28090064 | rs10033212 | -0.15 | 0.847 | 0.767 | -1.482 | 0.053 | -0.048 | -0.048 | 0.045 | 0.81 | 0.66 | 0.79 | 0.93 | 0.72 | 0.9 |
| 4 | 28099442 | rs6858327 | 0.645 | 0.601 | 0.808 | -1.55 | -0.009 | -0.076 | -0.076 | 0 | 0.73 | 0.53 | 0.78 | 0.85 | 0.75 | 0.87 |
| 4 | 28101706 | rs10031833 | 0.635 | 0.608 | 0.806 | -1.547 | -0.009 | -0.077 | -0.077 | 1.291 | 0.73 | 0.52 | 0.77 | 0.85 | 0.75 | 0.87 |
| 4 | 28104319 | rs2037766 | 0.277 | 0.431 | 0.623 | -1.679 | 0 | -0.013 | -0.013 | 0 | 0.32 | 0.33 | 0.33 | 0.19 | 0.34 | 0.38 |
| 4 | 28122494 | rs1609116 | 0.659 | 0.336 | 0.746 | -1.828 | -0.001 | -0.008 | -0.008 | 0.398 | 0.44 | 0.42 | 0.47 | 0.78 | 0.45 | 0.53 |
| 4 | 28157447 | rs1527328 | 1.064 | 0.338 | 0.611 | -1.84 | -0.004 | 0.058 | 0.058 | 0 | 0.42 | 0.29 | 0.43 | 0.67 | 0.4 | 0.49 |
| 4 | 28158128 | rs1918317 | -0.486 | 0.359 | 0.623 | -1.834 | 0.006 | 0.035 | 0.035 | 0 | 0.43 | 0.57 | 0.41 | 0.24 | 0.4 | 0.41 |
| 4 | 28158977 | rs1918316 | -0.563 | 0.487 | 0.387 | -1.773 | 0.006 | -0.016 | -0.016 | 0 | 0.84 | 0.76 | 0.83 | 0.91 | 0.81 | 0.91 |
| 4 | 28161145 | rs1534293 | -1.319 | 0.512 | -0.054 | -1.725 | -0.009 | 0.004 | 0.004 | 0 | 0.88 | 0.89 | 0.86 | 1 | 0.85 | 0.95 |
| 4 | 28162665 | rs16880317 | 0.957 | 0.266 | 0.041 | -1.323 | 0.018 | -0.03 | -0.03 | 1.764 | 0.11 | 0.02 | 0.05 | 0.18 | 0.07 | 0.04 |
| 4 | 28170335 | rs13130395 | -0.3 | 0.365 | 0.254 | -1.151 | -0.003 | 0.096 | 0.096 | 0 | 0.39 | 0.17 | 0.35 | 0.22 | 0.39 | 0.39 |
| 4 | 28175258 | rs6835345 | -0.497 | 0.403 | -0.189 | -1.096 | -0.001 | -0.011 | -0.011 | 0.004 | 0.43 | 0.43 | 0.4 | 0.33 | 0.44 | 0.45 |
| 4 | 28182874 | rs4692290 | -0.387 | 0.322 | -0.436 | -0.514 | -0.012 | 0.031 | 0.031 | 0 | 0.57 | 0.72 | 0.5 | 0.78 | 0.53 | 0.53 |
| 4 | 28194516 | rs12642383 | 0.539 | -0.019 | -0.997 | -0.45 | -0.017 | -0.12 | -0.12 | 0 | 0.29 | 0.09 | 0.26 | 0.57 | 0.26 | 0.41 |
| 4 | 28194748 | rs10009033 | -0.858 | -0.3 | -0.346 | -0.574 | -0.003 | 0.011 | 0.011 | 0.464 | 0.54 | 0.66 | 0.46 | 0.72 | 0.5 | 0.51 |
| 4 | 28204105 | rs4318636 | 0.451 | -0.408 | 0.205 | -0.578 | -0.005 | 0.004 | 0.004 | 0 | 0.32 | 0.35 | 0.3 | 0.64 | 0.26 | 0.44 |
| 4 | 28224533 | rs4692293 | -0.019 | -0.276 | -0.064 | -0.681 | -0.004 | 0 | 0 | 0 | 0.36 | 0.64 | 0.32 | 0.64 | 0.27 | 0.44 |
| 4 | 28226334 | rs10517137 | 0.667 | -0.219 | -0.626 | -0.636 | -0.066 | 0.084 | 0.084 | 0 | 0.23 | 0.01 | 0.14 | 0.08 | 0.24 | 0.07 |
| 4 | 28232617 | rs7660232 | -0.333 | 0.022 | 0.073 | -0.346 | 0 | -0.005 | -0.005 | 0 | 0.54 | 0.5 | 0.46 | 0.72 | 0.51 | 0.52 |
| 4 | 28233327 | rs1497691 | -0.333 | 0.022 | 0.073 | -0.346 | 0 | -0.005 | -0.005 | 0 | 0.54 | 0.5 | 0.46 | 0.72 | 0.51 | 0.52 |
| 4 | 28233491 | rs4452420 | -0.333 | 0.022 | 0.073 | -0.346 | 0 | -0.005 | -0.005 | 0 | 0.54 | 0.5 | 0.46 | 0.72 | 0.51 | 0.52 |
| 4 | 28234453 | rs12648254 | -0.305 | 0.017 | 0.101 | -0.349 | -0.007 | 0.025 | 0.025 | 0.038 | 0.55 | 0.59 | 0.47 | 0.72 | 0.51 | 0.52 |
| 4 | 28234938 | rs973586 | -0.357 | 0.072 | -0.219 | -0.322 | 0 | -0.005 | -0.005 | 0 | 0.54 | 0.5 | 0.46 | 0.72 | 0.51 | 0.52 |
| 4 | 28237783 | rs2173279 | -0.357 | 0.072 | -0.219 | -0.322 | 0 | -0.005 | -0.005 | 0 | 0.54 | 0.5 | 0.46 | 0.72 | 0.51 | 0.52 |
| 4 | 28250238 | rs2309717 | -0.477 | -0.459 | -0.414 | -0.025 | -0.008 | 0.006 | 0.006 | 0 | 0.22 | 0.28 | 0.23 | 0.31 | 0.14 | 0.2 |
| 4 | 28251958 | rs11729367 | -0.004 | -0.531 | -0.433 | 0.057 | 0.018 | -0.011 | -0.011 | 0 | 0.18 | 0.14 | 0.2 | 0.22 | 0.13 | 0.19 |
| 4 | 28270157 | rs1497716 | 1.062 | -0.545 | -0.648 | 0.305 | -0.001 | -0.029 | -0.029 | 0 | 0.3 | 0.2 | 0.31 | 0.39 | 0.28 | 0.34 |
| 4 | 28292048 | rs7435189 | -0.339 | -0.796 | -0.194 | 0.234 | -0.051 | 0.069 | 0.069 | 0 | 0.18 | 0.38 | 0.23 | 0.32 | 0.1 | 0.2 |
| 4 | 28305543 | rs7654040 | -0.501 | -0.635 | -0.53 | -0.414 | -0.015 | -0.008 | -0.008 | 1.079 | 0.4 | 0.49 | 0.38 | 0.33 | 0.27 | 0.45 |
| 4 | 28315343 | rs4267724 | 1.308 | -0.092 | -0.854 | -0.33 | -0.123 | -0.039 | -0.039 | 0.02 | 0.19 | 0.01 | 0.15 | 0.16 | 0.33 | 0.11 |
| 4 | 28319636 | rs13137916 | 0.285 | -0.193 | -0.475 | -0.337 | -0.083 | -0.017 | -0.017 | 0 | 0.29 | 0.1 | 0.3 | 0.29 | 0.45 | 0.22 |
| 4 | 28324928 | rs10025742 | 0.128 | -0.362 | -0.186 | -0.342 | -0.028 | 0.001 | 0.001 | 0 | 0.31 | 0.45 | 0.34 | 0.32 | 0.2 | 0.34 |
| 4 | 28345775 | rs2036184 | 0.215 | -0.476 | 0.018 | -0.689 | 0.003 | -0.015 | -0.015 | 0 | 0.58 | 0.53 | 0.69 | 0.7 | 0.47 | 0.66 |
| 4 | 28352417 | rs10517141 | 2.216 | -0.339 | -0.528 | -0.689 | 0.011 | -0.12 | -0.12 | 0 | 0.22 | 0.05 | 0.33 | 0.42 | 0.2 | 0.29 |
| 4 | 28353949 | rs12649612 | 1.137 | -0.325 | -0.415 | -0.778 | 0.006 | 0.022 | 0.022 | 0 | 0.21 | 0.14 | 0.13 | 0.14 | 0.2 | 0.25 |
| 4 | 28360614 | rs10939207 | -0.299 | -0.154 | -0.241 | -0.795 | -0.026 | 0.034 | 0.034 | 0 | 0.41 | 0.29 | 0.28 | 0.3 | 0.51 | 0.37 |
| 4 | 28374425 | rs12645676 | 1.747 | -0.187 | -0.274 | -0.768 | 0.006 | 0.011 | 0.011 | 0.078 | 0.17 | 0.05 | 0.09 | 0.14 | 0.16 | 0.23 |
| 4 | 28378505 | rs16880558 | 0.45 | -0.151 | -0.147 | -0.772 | 0.01 | -0.016 | -0.016 | 0.015 | 0.05 | 0.22 | 0.05 | 0 | 0.04 | 0.01 |
| 4 | 28378795 | rs664330 | -1.249 | -0.271 | -0.224 | -0.93 | -0.057 | 0.015 | 0.015 | 0.597 | 0.78 | 0.93 | 0.81 | 0.84 | 0.66 | 0.87 |
| 4 | 28378913 | rs16880561 | -0.505 | -0.366 | 0.069 | -0.903 | 0.005 | 0.003 | 0.003 | 1.027 | 0.07 | 0.25 | 0.09 | 0.04 | 0.06 | 0.03 |
| 4 | 28379146 | rs7697734 | 1.582 | -0.532 | -0.003 | -0.966 | 0 | 0.055 | 0.055 | 0 | 0.18 | 0.06 | 0.13 | 0.02 | 0.17 | 0.14 |
| 4 | 28379401 | rs1282883 | 0.141 | -0.407 | 0.136 | -0.78 | -0.009 | -0.039 | -0.039 | 0.125 | 0.33 | 0.49 | 0.35 | 0.23 | 0.27 | 0.33 |
| 4 | 28391626 | rs7674497 | 0.8 | -0.271 | 0.088 | -0.591 | -0.017 | 0.006 | 0.006 | 0 | 0.21 | 0.29 | 0.11 | 0.29 | 0.15 | 0.26 |
| 4 | 28405478 | rs1510705 | 0.448 | -0.156 | 0.028 | -0.732 | -0.013 | -0.013 | -0.013 | 0 | 0.57 | 0.4 | 0.68 | 0.49 | 0.5 | 0.59 |
| 4 | 28419100 | rs12502704 | 0.456 | -0.156 | -0.301 | -0.759 | -0.013 | -0.012 | -0.012 | 0 | 0.56 | 0.4 | 0.68 | 0.5 | 0.5 | 0.59 |
| 4 | 28419921 | rs16880624 | 2.15 | -0.158 | -0.58 | -0.583 | 0.007 | -0.12 | -0.12 | 0.039 | 0.21 | 0.04 | 0.31 | 0.44 | 0.19 | 0.25 |
| 4 | 28433243 | rs2250259 | -2.309 | 0.34 | -0.654 | -0.696 | -0.087 | -0.05 | -0.05 | 0.715 | 0.87 | 0.99 | 0.86 | 0.84 | 0.73 | 0.89 |
| 4 | 28440972 | rs3907987 | -0.578 | 0.138 | -0.423 | -0.847 | -0.02 | 0.097 | 0.097 | 0.018 | 0.56 | 0.31 | 0.46 | 0.21 | 0.5 | 0.6 |
| 4 | 28450862 | rs10028579 | 1.669 | -0.04 | 0.05 | -0.705 | -0.016 | 0.052 | 0.052 | 0.062 | 0.24 | 0.3 | 0.33 | 0.44 | 0.21 | 0.25 |
| 4 | 28454474 | rs17685351 | 1.067 | 0.077 | -0.3 | -0.793 | -0.015 | 0.022 | 0.022 | 0.01 | 0.2 | 0.03 | 0.12 | 0.13 | 0.19 | 0.23 |
| 4 | 28460356 | rs7687734 | 0.92 | 0.083 | -0.184 | -0.779 | 0.012 | 0.024 | 0.024 | 0 | 0.21 | 0.1 | 0.12 | 0.15 | 0.19 | 0.26 |
| 4 | 28470784 | rs12642662 | 0.907 | 0.088 | -0.055 | -0.775 | 0.014 | 0.02 | 0.02 | 0 | 0.22 | 0.12 | 0.12 | 0.15 | 0.19 | 0.28 |
| 4 | 28473048 | rs16880691 | 1.049 | 0.773 | 0.527 | -0.688 | -0.011 | -0.001 | -0.001 | 0.156 | 0.65 | 0.29 | 0.61 | 0.63 | 0.58 | 0.78 |
| 4 | 28473524 | rs962625 | 0.874 | 0.92 | 0.238 | -0.808 | 0.001 | -0.016 | -0.016 | 0.059 | 0.15 | 0.09 | 0.15 | 0.02 | 0.28 | 0.12 |
| 4 | 28480415 | rs7656054 | 1.074 | 0.657 | 0.079 | -0.794 | 0.009 | 0.032 | 0.032 | 0 | 0.32 | 0.19 | 0.48 | 0.79 | 0.28 | 0.43 |
| 4 | 28481058 | rs7662460 | 1.929 | 0.636 | 0.015 | -0.506 | -0.005 | -0.129 | -0.129 | 0.157 | 0.23 | 0.07 | 0.34 | 0.5 | 0.2 | 0.35 |
| 4 | 28485201 | rs7653895 | 0.364 | 0.63 | 0.885 | -0.987 | 0.008 | 0.014 | 0.014 | 0 | 0.41 | 0.61 | 0.52 | 0.79 | 0.31 | 0.48 |
| 4 | 28503413 | rs11937136 | -0.198 | 0.645 | 1.281 | -1.009 | 0.005 | -0.031 | -0.031 | 0.982 | 0.07 | 0.26 | 0.03 | 0 | 0.03 | 0.05 |
| 4 | 28513407 | rs11934051 | 0.051 | 0.626 | 1.433 | -1.018 | -0.006 | -0.035 | -0.035 | 0 | 0.06 | 0.17 | 0.03 | 0 | 0.03 | 0.05 |
| 4 | 28525051 | rs292049 | -0.529 | 0.823 | 1.215 | -1.206 | 0.031 | -0.155 | -0.155 | 0.077 | 0.25 | 0.51 | 0.18 | 0.01 | 0.3 | 0.16 |
| 4 | 28534342 | rs292044 | -0.552 | 0.865 | 0.785 | -1.68 | 0.035 | 0.069 | 0.069 | 0 | 0.58 | 0.28 | 0.47 | 0.21 | 0.69 | 0.52 |
| 4 | 28539800 | rs2136818 | -0.24 | 0.843 | 0.626 | -1.56 | -0.001 | -0.028 | -0.028 | 0 | 0.45 | 0.47 | 0.33 | 0.2 | 0.42 | 0.41 |
| 4 | 28544298 | rs292016 | 0.284 | 1.189 | 0.708 | -1.261 | 0 | 0.005 | 0.005 | 0 | 0.53 | 0.41 | 0.57 | 0.56 | 0.54 | 0.53 |
| 4 | 28547108 | rs557255 | 0.722 | 1.32 | 0.089 | -1.449 | -0.087 | 0.087 | 0.087 | 0.902 | 0.16 | 0.01 | 0.15 | 0.01 | 0.27 | 0.09 |
| 4 | 28549589 | rs17698720 | 0.89 | 1.32 | 0.058 | -1.635 | 0.003 | 0.006 | 0.006 | 0 | 0.18 | 0.11 | 0.12 | 0.14 | 0.18 | 0.26 |
| 4 | 28555328 | rs12645493 | 2.394 | 1.244 | -0.128 | -1.696 | 0.004 | -0.12 | -0.12 | 0.123 | 0.3 | 0.04 | 0.39 | 0.56 | 0.25 | 0.39 |
| 4 | 28561990 | rs6448587 | 0.05 | 1.516 | 0.289 | -1.718 | -0.005 | 0.007 | 0.007 | 0 | 0.23 | 0.27 | 0.2 | 0.29 | 0.22 | 0.3 |
| 4 | 28563455 | rs2458614 | 0.776 | 1.645 | 0.002 | -1.669 | -0.087 | 0.087 | 0.087 | 0.902 | 0.16 | 0.01 | 0.15 | 0.01 | 0.27 | 0.09 |
| 4 | 28587729 | rs2943159 | -1.28 | 1.743 | 0.551 | -1.643 | -0.001 | 0.001 | 0.001 | 0 | 0.2 | 0.21 | 0.22 | 0.17 | 0.25 | 0.2 |

| | | | | | | | | | | | | | | | | |
|---|---|---|---|---|---|---|---|---|---|---|---|---|---|---|---|---|
| 4 | 28588964 | rs7655801 | 0.116 | 1.72 | 0.765 | -1.726 | -0.006 | 0.034 | 0.034 | 0 | 0.4 | 0.31 | 0.28 | 0.2 | 0.44 | 0.37 |
| 4 | 28589663 | rs2471712 | -1.914 | 1.7 | 0.875 | -1.713 | 0.006 | -0.001 | -0.001 | 0.385 | 0.64 | 0.69 | 0.51 | 0.37 | 0.69 | 0.57 |
| 4 | 28590366 | rs3114003 | -1.188 | 1.704 | 0.883 | -1.723 | 0.012 | 0.033 | 0.033 | 0 | 0.58 | 0.68 | 0.44 | 0.22 | 0.64 | 0.48 |
| 4 | 28590750 | rs11946058 | -0.625 | 1.664 | 0.975 | -1.696 | 0 | 0.03 | 0.03 | 0 | 0.48 | 0.59 | 0.32 | 0.2 | 0.47 | 0.41 |
| 4 | 28602089 | rs16880825 | 0.265 | 1.618 | 0.875 | -1.558 | -0.003 | -0.006 | -0.006 | 0 | 0.06 | 0.03 | 0.07 | 0.15 | 0.06 | 0.09 |
| 4 | 28603169 | rs11945665 | 0.147 | 1.645 | 1.18 | -1.743 | -0.001 | -0.127 | -0.127 | 1.724 | 0.24 | 0.48 | 0.17 | 0.04 | 0.19 | 0.13 |
| 4 | 28603878 | rs10939219 | -0.084 | 1.626 | 0.919 | -1.77 | -0.027 | 0.047 | 0.047 | 0.01 | 0.66 | 0.79 | 0.73 | 0.82 | 0.55 | 0.65 |
| 4 | 28608116 | rs17644064 | 1.063 | 1.648 | 0.948 | -1.753 | -0.019 | 0.035 | 0.035 | 0 | 0.18 | 0.1 | 0.2 | 0.07 | 0.2 | 0.11 |
| 4 | 28614705 | rs12509357 | 0.285 | 1.604 | 1.26 | -1.795 | -0.003 | 0 | 0 | 0.622 | 0.62 | 0.62 | 0.7 | 0.83 | 0.53 | 0.61 |
| 4 | 28618870 | rs11943453 | 0.807 | 1.687 | 1.248 | -1.812 | -0.037 | 0.017 | 0.017 | 0 | 0.24 | 0.05 | 0.14 | 0.16 | 0.28 | 0.28 |
| 4 | 28634864 | rs16880871 | 0.503 | 1.741 | 1.251 | -1.756 | -0.008 | -0.001 | -0.001 | 0.107 | 0.06 | 0.03 | 0.09 | 0.21 | 0.07 | 0.12 |
| 4 | 28635577 | rs11944357 | 0.105 | 1.705 | 1.289 | -1.805 | -0.001 | -0.022 | -0.022 | 0 | 0.29 | 0.37 | 0.18 | 0.16 | 0.3 | 0.33 |
| 4 | 28637078 | rs7672198 | -0.977 | 1.746 | 1.161 | -1.823 | -0.053 | 0.042 | 0.042 | 0.1 | 0.91 | 0.99 | 0.88 | 0.99 | 0.83 | 0.94 |
| 4 | 28641062 | rs6831370 | 0.357 | 1.756 | 1.214 | -1.828 | -0.002 | -0.024 | -0.024 | 0.022 | 0.06 | 0.14 | 0.03 | 0.01 | 0.03 | 0.05 |
| 4 | 28642726 | rs6844353 | -1.062 | 1.713 | 1.054 | -1.806 | -0.053 | 0.042 | 0.042 | 0.089 | 0.91 | 0.99 | 0.88 | 0.99 | 0.83 | 0.94 |
| 4 | 28648262 | rs4334744 | -0.846 | 1.689 | 1.335 | -1.837 | 0.003 | 0.076 | 0.076 | 0 | 0.49 | 0.66 | 0.32 | 0.19 | 0.47 | 0.42 |
| 4 | 28652911 | rs7695573 | 0.345 | 1.674 | 1.443 | -1.84 | -0.002 | -0.008 | -0.008 | 0 | 0.22 | 0.24 | 0.14 | 0.15 | 0.19 | 0.16 |
| 4 | 28658506 | rs2172455 | 1.579 | 1.684 | 1.325 | -1.85 | -0.008 | 0.019 | 0.019 | 0 | 0.15 | 0.08 | 0.13 | 0.03 | 0.17 | 0.09 |
| 4 | 28659854 | rs6839719 | -1.112 | 1.747 | 1.112 | -1.811 | -0.053 | 0.031 | 0.031 | 0.069 | 0.91 | 0.99 | 0.89 | 0.98 | 0.83 | 0.94 |
| 4 | 28660856 | rs9996729 | 0.052 | 1.738 | 1.205 | -1.845 | 0.052 | -0.039 | -0.039 | 0.059 | 0.87 | 0.73 | 0.8 | 0.95 | 0.79 | 0.92 |
| 4 | 28661714 | rs1493137 | 1.979 | 1.68 | 0.984 | -1.814 | 0.015 | -0.021 | -0.021 | 0 | 0.34 | 0.24 | 0.47 | 0.59 | 0.29 | 0.4 |
| 4 | 28664019 | rs953276 | 2.302 | 1.686 | 0.831 | -1.811 | 0.014 | -0.063 | -0.063 | 0 | 0.33 | 0.13 | 0.46 | 0.59 | 0.29 | 0.4 |
| 4 | 28665838 | rs16880901 | 0.573 | 1.634 | 0.757 | -1.712 | -0.013 | -0.032 | -0.032 | 0 | 0.07 | 0.02 | 0.09 | 0.2 | 0.07 | 0.13 |
| 4 | 28669712 | rs1389169 | 0.863 | 1.616 | 0.815 | -1.714 | -0.001 | -0.002 | -0.002 | 0 | 0.53 | 0.48 | 0.61 | 0.63 | 0.46 | 0.48 |
| 4 | 28687406 | rs7663065 | -0.207 | 1.561 | 0.03 | -1.667 | -0.002 | -0.017 | -0.017 | 0 | 0.33 | 0.19 | 0.25 | 0.36 | 0.34 | 0.41 |
| 4 | 28691851 | rs4692338 | 2.187 | 1.421 | -0.215 | -1.652 | 0.019 | -0.162 | -0.162 | 0.007 | 0.3 | 0.03 | 0.37 | 0.55 | 0.21 | 0.38 |
| 4 | 28692384 | rs11937968 | 0 | 1.563 | 0.306 | -1.526 | 0.002 | -0.018 | -0.018 | 1.125 | 0.08 | 0.07 | 0.1 | 0.21 | 0.06 | 0.13 |
| 4 | 28702554 | rs16880969 | 0.312 | 1.561 | 0.708 | -1.692 | 0.022 | -0.011 | -0.011 | 0 | 0.05 | 0.23 | 0.02 | 0 | 0.02 | 0.05 |
| 4 | 28702690 | rs11945047 | 0.014 | 1.549 | 0.601 | -1.691 | 0.003 | -0.019 | -0.019 | 0 | 0.09 | 0.07 | 0.1 | 0.17 | 0.07 | 0.12 |
| 4 | 28703576 | rs13114499 | 1.844 | 1.591 | 0.369 | -1.79 | 0 | -0.144 | -0.144 | 0 | 0.32 | 0.03 | 0.39 | 0.56 | 0.25 | 0.38 |
| 4 | 28703891 | rs7693029 | 0.618 | 1.767 | 0.644 | -1.821 | -0.016 | 0.008 | 0.008 | 0 | 0.58 | 0.65 | 0.59 | 0.62 | 0.49 | 0.52 |
| 4 | 28710321 | rs7698388 | 1.252 | 1.668 | 0.525 | -1.762 | 0.022 | -0.014 | -0.014 | 0.086 | 0.38 | 0.28 | 0.41 | 0.56 | 0.28 | 0.43 |
| 4 | 28712968 | rs10939225 | -0.12 | 1.791 | 0.39 | -1.938 | -0.027 | 0.031 | 0.031 | 0 | 0.66 | 0.78 | 0.69 | 0.79 | 0.55 | 0.65 |
| 4 | 28718528 | rs9685881 | -0.189 | 1.675 | 0.488 | -1.95 | -0.006 | 0.018 | 0.018 | 0 | 0.43 | 0.57 | 0.37 | 0.26 | 0.49 | 0.39 |
| 4 | 28718982 | rs1389182 | 0.505 | 1.655 | 0.533 | -1.968 | -0.004 | 0.003 | 0.003 | 0 | 0.8 | 0.8 | 0.78 | 0.83 | 0.76 | 0.8 |
| 4 | 28723573 | rs4692342 | 0.938 | 1.259 | 0.642 | -1.82 | 0.019 | -0.005 | -0.005 | 3.285 | 0.38 | 0.3 | 0.42 | 0.58 | 0.27 | 0.41 |
| 4 | 28723854 | rs11722802 | 1.484 | 1.228 | -0.039 | -1.744 | 0.009 | -0.109 | -0.109 | 0 | 0.32 | 0.03 | 0.39 | 0.58 | 0.25 | 0.36 |
| 4 | 28732481 | rs2309883 | 1.509 | 1.319 | 0.069 | -2.534 | 0.046 | 0.033 | 0.033 | 0.107 | 0.33 | 0.04 | 0.4 | 0.73 | 0.27 | 0.34 |
| 4 | 28737332 | rs13143929 | -0.847 | 1.678 | 0.516 | -2.781 | -0.027 | 0.049 | 0.049 | 0 | 0.4 | 0.67 | 0.44 | 0.27 | 0.45 | 0.38 |
| 4 | 28737881 | rs2047033 | 1.243 | 1.805 | 0.984 | -2.853 | 0.009 | -0.015 | -0.015 | 4.233 | 0.18 | 0.18 | 0.22 | 0.12 | 0.25 | 0.17 |
| 4 | 28738088 | rs10001415 | -0.192 | 1.81 | 1.189 | -2.885 | 0.018 | -0.069 | -0.069 | 0.239 | 0.13 | 0.36 | 0.11 | 0.01 | 0.13 | 0.06 |
| 4 | 28738315 | rs2047032 | 1.229 | 1.857 | 0.889 | -2.812 | 0.009 | -0.015 | -0.015 | 0 | 0.18 | 0.18 | 0.22 | 0.12 | 0.25 | 0.17 |
| 4 | 28745882 | rs12508326 | 1.312 | 1.976 | 0.918 | -2.812 | 0.023 | 0.155 | 0.155 | 0 | 0.5 | 0.22 | 0.6 | 0.86 | 0.49 | 0.51 |
| 4 | 28750923 | rs16881041 | 1.165 | 1.976 | 0.917 | -2.748 | 0.006 | -0.013 | -0.013 | 0 | 0.19 | 0.18 | 0.22 | 0.12 | 0.23 | 0.2 |
| 4 | 28752825 | rs9997504 | -0.547 | 2.048 | 1.177 | -2.764 | 0.034 | 0.016 | 0.016 | 0 | 0.11 | 0.25 | 0.14 | 0.13 | 0.11 | 0.14 |
| 4 | 28752932 | rs9997604 | -0.571 | 1.999 | 1.009 | -2.854 | -0.043 | 0.107 | 0.107 | 0 | 0.75 | 0.9 | 0.85 | 1 | 0.72 | 0.78 |
| 4 | 28753519 | rs11946866 | -0.298 | 2.006 | 0.955 | -2.701 | 0.006 | 0.002 | 0.002 | 0 | 0.1 | 0.15 | 0.13 | 0.13 | 0.11 | 0.14 |
| 4 | 28758097 | rs12508012 | 0.154 | 1.912 | 0.987 | -2.762 | 0.01 | -0.007 | -0.007 | 0.068 | 0.28 | 0.28 | 0.32 | 0.25 | 0.31 | 0.34 |
| 4 | 28764584 | rs1353053 | 0.691 | 1.945 | 1.026 | -2.762 | -0.02 | 0.02 | 0.02 | 0 | 0.23 | 0.15 | 0.3 | 0.17 | 0.29 | 0.28 |
| 4 | 28769842 | rs7664880 | 0.59 | 1.947 | 0.99 | -2.761 | 0.007 | -0.011 | -0.011 | 0 | 0.24 | 0.23 | 0.31 | 0.17 | 0.29 | 0.28 |
| 4 | 28776140 | rs1493139 | 1.961 | 1.812 | 0.794 | -2.856 | 0.075 | 0.024 | 0.024 | 0 | 0.31 | 0.09 | 0.39 | 0.74 | 0.24 | 0.36 |
| 4 | 28780231 | rs12512663 | -0.17 | 2.212 | 1.061 | -2.623 | 0.021 | -0.02 | -0.02 | 0.656 | 0.31 | 0.37 | 0.35 | 0.25 | 0.34 | 0.35 |
| 4 | 28790775 | rs7686662 | 1.368 | 2.149 | 0.956 | -2.761 | 0.077 | 0.126 | 0.126 | 0 | 0.38 | 0.14 | 0.49 | 0.84 | 0.31 | 0.44 |
| 4 | 28795213 | rs9991797 | -0.481 | 2.276 | 1.003 | -2.807 | -0.024 | 0.065 | 0.065 | 0 | 0.74 | 0.86 | 0.85 | 1 | 0.72 | 0.78 |
| 4 | 28795758 | rs2131464 | 0.32 | 1.999 | 1.294 | -2.847 | -0.01 | 0 | 0 | 0.055 | 0.65 | 0.67 | 0.76 | 0.98 | 0.6 | 0.7 |
| 4 | 28796401 | rs12499758 | 1.197 | 2.25 | 0.758 | -2.682 | -0.018 | 0.029 | 0.029 | 0 | 0.19 | 0.13 | 0.23 | 0.07 | 0.23 | 0.19 |
| 4 | 28797388 | rs4234998 | 1.908 | 2.21 | 0.543 | -2.66 | 0.087 | 0.063 | 0.063 | 0 | 0.34 | 0.08 | 0.39 | 0.78 | 0.26 | 0.37 |
| 4 | 28815914 | rs1602798 | 0.906 | 2.294 | 0.944 | -2.609 | 0.004 | -0.026 | -0.026 | 0.004 | 0.19 | 0.21 | 0.25 | 0.07 | 0.23 | 0.19 |
| 4 | 28816220 | rs7349599 | 1.248 | 2.137 | 0.951 | -2.513 | -0.031 | 0.013 | 0.013 | 0 | 0.05 | 0 | 0.07 | 0.01 | 0.12 | 0.06 |
| 4 | 28821006 | rs6844183 | 0.812 | 2.003 | 1.169 | -2.57 | 0.011 | -0.079 | -0.079 | 0 | 0.59 | 0.48 | 0.64 | 0.93 | 0.51 | 0.63 |
| 4 | 28826217 | rs1873861 | 1.115 | 1.967 | 1.071 | -2.587 | 0.005 | 0.049 | 0.049 | 0 | 0.54 | 0.35 | 0.62 | 0.85 | 0.48 | 0.57 |
| 4 | 28829163 | rs11942403 | 0.565 | 1.842 | 0.953 | -2.354 | 0.026 | 0.028 | 0.028 | 0 | 0.06 | 0.13 | 0.02 | 0.07 | 0.04 | 0.07 |
| 4 | 28831639 | rs16881070 | 1.411 | 1.783 | 0.635 | -2.616 | 0.102 | 0.124 | 0.124 | 0 | 0.4 | 0.1 | 0.47 | 0.81 | 0.28 | 0.48 |
| 4 | 28832388 | rs6850240 | -0.956 | 2.007 | 0.812 | -2.685 | 0.014 | -0.069 | -0.069 | 0 | 0.8 | 0.75 | 0.87 | 0.99 | 0.75 | 0.87 |
| 4 | 28832545 | rs6850637 | -0.63 | 2.026 | 0.873 | -2.621 | 0.035 | -0.042 | -0.042 | 0 | 0.34 | 0.47 | 0.35 | 0.24 | 0.43 | 0.39 |
| 4 | 28832740 | rs17653404 | 1.476 | 2.107 | 0.343 | -2.609 | -0.027 | 0.012 | 0.012 | 0.104 | 0.05 | 0 | 0.05 | 0.01 | 0.11 | 0.06 |
| 4 | 28832788 | rs6833159 | 0.511 | 2.127 | 0.446 | -2.608 | -0.003 | 0.002 | 0.002 | 0 | 0.23 | 0.24 | 0.28 | 0.16 | 0.29 | 0.26 |
| 4 | 28833274 | rs2029669 | 1.083 | 2.072 | 0.204 | -2.607 | 0.075 | 0.09 | 0.09 | 0 | 0.45 | 0.28 | 0.52 | 0.75 | 0.32 | 0.47 |
| 4 | 28834160 | rs1493164 | 1.214 | 2.228 | -0.361 | -2.495 | -0.035 | 0.054 | 0.054 | 0 | 0.17 | 0.09 | 0.19 | 0.05 | 0.23 | 0.18 |

| | | | | | | | | | | | | | | | | |
|---|---|---|---|---|---|---|---|---|---|---|---|---|---|---|---|---|
| 4 | 28838826 | rs2047028 | -1.068 | 1.456 | -0.177 | -2.251 | -0.01 | 0.038 | 0.038 | 0 | 0.37 | 0.66 | 0.22 | 0.09 | 0.37 | 0.25 |
| 4 | 28841188 | rs11937670 | 0.329 | 1.456 | -0.175 | -2.25 | -0.001 | 0.005 | 0.005 | 0 | 0.68 | 0.35 | 0.82 | 0.92 | 0.74 | 0.8 |
| 4 | 28843468 | rs4441746 | 0.36 | 1.414 | -0.195 | -2.25 | -0.001 | 0.015 | 0.015 | 0 | 0.68 | 0.32 | 0.82 | 0.92 | 0.74 | 0.8 |
| 4 | 28845331 | rs7661449 | 0.013 | 1.409 | -0.148 | -2.27 | -0.036 | 0.048 | 0.048 | 0 | 0.35 | 0.27 | 0.38 | 0.18 | 0.49 | 0.4 |
| 4 | 28850524 | rs7695045 | -0.908 | 1.547 | -0.206 | -2.192 | 0.031 | 0.143 | 0.143 | 0.798 | 0.56 | 0.8 | 0.49 | 0.17 | 0.6 | 0.48 |
| 4 | 28851586 | rs12645808 | 0.066 | 1.495 | -0.343 | -1.606 | -0.013 | 0.015 | 0.015 | 0 | 0.3 | 0.28 | 0.35 | 0.17 | 0.4 | 0.35 |
| 4 | 28863159 | rs7698028 | 0.088 | 1.476 | -0.681 | -1.463 | -0.017 | 0.055 | 0.055 | 0 | 0.23 | 0.13 | 0.12 | 0.02 | 0.27 | 0.12 |
| 4 | 28865191 | rs7438460 | -0.049 | 1.779 | 0.047 | -0.874 | 0.026 | 0.036 | 0.036 | 0 | 0.05 | 0.12 | 0.02 | 0.09 | 0.04 | 0.07 |
| 4 | 28866526 | rs7661966 | 0.17 | 1.533 | 0.228 | -1.157 | 0.017 | -0.042 | -0.042 | 0 | 0.68 | 0.64 | 0.64 | 0.82 | 0.61 | 0.62 |
| 4 | 28872833 | rs10049756 | -1.212 | 0.843 | 0.598 | -0.702 | 0 | -0.016 | -0.016 | 0 | 0.12 | 0.56 | 0.1 | 0.09 | 0.08 | 0.11 |
| 4 | 28887284 | rs7658708 | 0.744 | 0.321 | 0.543 | -1.367 | -0.035 | -0.118 | -0.118 | 0.22 | 0.78 | 0.52 | 0.87 | 0.96 | 0.84 | 0.79 |
| 4 | 28895078 | rs10805281 | 0.13 | 0.791 | -0.539 | -1.982 | -0.027 | 0.042 | 0.042 | 0 | 0.22 | 0.15 | 0.23 | 0.11 | 0.28 | 0.23 |
| 4 | 28901589 | rs12647787 | 1.061 | 0.779 | -1.093 | -2.13 | -0.007 | 0.125 | 0.125 | 1.435 | 0.45 | 0.17 | 0.58 | 0.71 | 0.48 | 0.45 |
| 4 | 28912626 | rs13109384 | 0.838 | 0.778 | -1.709 | -1.96 | -0.015 | 0.015 | 0.015 | 0 | 0.22 | 0.14 | 0.22 | 0.08 | 0.29 | 0.07 |
| 4 | 28920120 | rs1344017 | 0.177 | 0.196 | -1.385 | -1.761 | -0.018 | -0.03 | -0.03 | 0 | 0.79 | 0.44 | 0.85 | 0.83 | 0.87 | 0.8 |
| 4 | 28921523 | rs13112132 | -0.437 | 0.021 | -1.315 | -1.724 | -0.038 | -0.093 | -0.093 | 0 | 0.84 | 0.51 | 0.9 | 0.97 | 0.91 | 0.9 |
| 4 | 28922949 | rs4692065 | 0.461 | 0.33 | -2.061 | -1.366 | 0.027 | -0.093 | -0.093 | 0.028 | 0.28 | 0.05 | 0.36 | 0.6 | 0.25 | 0.44 |
| 4 | 28923633 | rs1402492 | 0.461 | 0.33 | -2.061 | -1.366 | 0.027 | -0.093 | -0.093 | 0.037 | 0.28 | 0.05 | 0.36 | 0.6 | 0.25 | 0.44 |
| 4 | 28928041 | rs1402499 | 0.134 | 1.247 | -1.395 | -1.407 | 0.003 | -0.021 | -0.021 | 0 | 0.72 | 0.69 | 0.67 | 0.88 | 0.69 | 0.8 |
| 4 | 28932131 | rs2222357 | -0.506 | 0.973 | -2.053 | -1.206 | 0.006 | 0.039 | 0.039 | 3.317 | 0.39 | 0.22 | 0.38 | 0.72 | 0.33 | 0.44 |
| 4 | 28933593 | rs12642441 | -0.003 | 0.461 | -1.777 | -1.096 | 0.011 | -0.007 | -0.007 | 0 | 0.24 | 0.19 | 0.36 | 0.75 | 0.2 | 0.35 |
| 4 | 28935317 | rs4326073 | 0.061 | 0.496 | -1.388 | -1.045 | 0.008 | 0.013 | 0.013 | 0 | 0.61 | 0.35 | 0.71 | 0.84 | 0.7 | 0.53 |
| 4 | 28936849 | rs1402504 | -0.226 | 0.265 | -1.16 | -2.365 | -0.001 | -0.026 | -0.026 | 0 | 0.43 | 0.55 | 0.36 | 0.11 | 0.45 | 0.38 |
| 4 | 28940143 | rs1521131 | 0.297 | 0.359 | -1.408 | -2.641 | -0.052 | 0.155 | 0.155 | 0 | 0.3 | 0.06 | 0.28 | 0.04 | 0.35 | 0.22 |
| 4 | 28945208 | rs4639141 | -0.608 | -0.212 | -0.871 | -2.808 | -0.001 | 0.006 | 0.006 | 0 | 0.55 | 0.45 | 0.45 | 0.25 | 0.58 | 0.52 |
| 4 | 28948730 | rs1521123 | -1.077 | -0.591 | -1.359 | -3.208 | -0.06 | 0.007 | 0.007 | 0 | 0.69 | 0.9 | 0.52 | 0.25 | 0.66 | 0.58 |
| 4 | 28949043 | rs12649705 | 0.772 | -0.632 | -1.079 | -3.227 | -0.019 | 0.031 | 0.031 | 0 | 0.11 | 0.17 | 0.07 | 0.14 | 0.06 | 0.14 |
| 4 | 28950110 | rs16881195 | 0.151 | -0.621 | -1.183 | -3.415 | -0.008 | 0.001 | 0.001 | 0 | 0.16 | 0.17 | 0.1 | 0.14 | 0.12 | 0.2 |
| 4 | 28955155 | rs17705257 | 0.241 | -0.801 | 0.131 | -3.484 | 0.001 | -0.01 | -0.01 | 0 | 0.15 | 0.2 | 0.17 | 0.07 | 0.18 | 0.14 |
| 4 | 28955778 | rs16881228 | 1.191 | -0.902 | 0.101 | -3.447 | 0.007 | -0.015 | -0.015 | 3.434 | 0.08 | 0.05 | 0.06 | 0.15 | 0.06 | 0.15 |
| 4 | 28956150 | rs7674882 | -0.731 | -1 | 0.156 | -3.555 | -0.01 | -0.078 | -0.078 | 0 | 0.32 | 0.55 | 0.24 | 0.07 | 0.29 | 0.23 |
| 4 | 28956166 | rs13148843 | 1.676 | -1.085 | 0.065 | -3.646 | -0.044 | 0.148 | 0.148 | 0 | 0.28 | 0.05 | 0.2 | 0.04 | 0.31 | 0.2 |
| 4 | 28956361 | rs7696952 | -0.354 | -1.021 | 0.112 | -3.582 | -0.002 | -0.009 | -0.009 | 0.038 | 0.6 | 0.6 | 0.44 | 0.11 | 0.6 | 0.42 |
| 4 | 28957536 | rs16881249 | 1.544 | -0.986 | 0.088 | -3.544 | -0.015 | -0.056 | -0.056 | 0 | 0.12 | 0.19 | 0.05 | 0 | 0.08 | 0.02 |
| 4 | 28958205 | rs1357033 | -1.226 | -0.939 | 0.077 | -3.519 | -0.028 | 0.005 | 0.005 | 0.011 | 0.7 | 0.81 | 0.51 | 0.26 | 0.66 | 0.57 |
| 4 | 28959562 | rs7670784 | -1.126 | -0.939 | 0.163 | -3.519 | -0.014 | 0.007 | 0.007 | 0 | 0.69 | 0.72 | 0.5 | 0.26 | 0.66 | 0.57 |
| 4 | 28967598 | rs7688399 | 0.247 | -0.942 | 0.183 | -3.568 | 0.009 | 0.029 | 0.029 | 0.172 | 0.47 | 0.6 | 0.29 | 0.05 | 0.43 | 0.29 |
| 4 | 28972033 | rs6448619 | -0.03 | -0.873 | 0.08 | -3.504 | 0 | 0 | 0 | 0 | 0.4 | 0.4 | 0.3 | 0.22 | 0.36 | 0.37 |
| 4 | 28975242 | rs6815902 | -0.234 | -0.885 | -0.046 | -3.506 | 0 | -0.02 | -0.02 | 0 | 0.5 | 0.5 | 0.4 | 0.11 | 0.52 | 0.4 |
| 4 | 28979210 | rs6843077 | -1.298 | -0.841 | -0.015 | -3.44 | -0.025 | 0.003 | 0.003 | 0.224 | 0.7 | 0.79 | 0.51 | 0.26 | 0.66 | 0.57 |
| 4 | 28983128 | rs7664803 | 0.073 | -0.759 | 0.182 | -3.376 | -0.02 | -0.105 | -0.105 | 1.031 | 0.33 | 0.44 | 0.25 | 0.07 | 0.3 | 0.26 |
| 4 | 28990864 | rs11944473 | -0.386 | -0.842 | 0.024 | -3.328 | 0.057 | 0.148 | 0.148 | 0 | 0.45 | 0.74 | 0.3 | 0.07 | 0.37 | 0.23 |
| 4 | 28995299 | rs4285150 | 0.743 | -0.485 | 0.209 | -2.81 | 0.003 | -0.006 | -0.006 | 0 | 0.1 | 0.1 | 0.07 | 0.14 | 0.07 | 0.15 |
| 4 | 28999923 | rs13108377 | 0.272 | -0.215 | 0.618 | -2.435 | 0.004 | 0.023 | 0.023 | 0 | 0.24 | 0.21 | 0.22 | 0.06 | 0.26 | 0.17 |
| 4 | 29004411 | rs11936386 | 0.265 | -0.208 | 0.621 | -2.432 | 0.004 | 0.024 | 0.024 | 0 | 0.25 | 0.21 | 0.22 | 0.06 | 0.26 | 0.17 |
| 4 | 29016593 | rs1402487 | -0.604 | -0.273 | 0.549 | -2.544 | -0.006 | -0.002 | -0.002 | 2.962 | 0.78 | 0.82 | 0.55 | 0.24 | 0.73 | 0.58 |
| 4 | 29018173 | rs2049156 | 1.033 | -0.233 | 0.375 | -2.594 | -0.002 | 0.051 | 0.051 | 0 | 0.44 | 0.31 | 0.3 | 0.17 | 0.42 | 0.29 |
| 4 | 29030161 | rs17655022 | 0.502 | -0.262 | 0.519 | -2.702 | -0.001 | -0.019 | -0.019 | 0 | 0.25 | 0.27 | 0.22 | 0.07 | 0.26 | 0.16 |
| 4 | 29046559 | rs3924180 | 0.3 | -0.053 | 0.423 | -2.684 | 0.024 | -0.036 | -0.036 | 0 | 0.07 | 0.16 | 0.06 | 0 | 0.08 | 0.06 |
| 4 | 29048204 | rs7677192 | 1.052 | -0.033 | 0.211 | -2.512 | 0.008 | 0.044 | 0.044 | 0 | 0.42 | 0.31 | 0.26 | 0.16 | 0.37 | 0.28 |
| 4 | 29066896 | rs7684905 | -0.455 | -0.022 | -0.162 | -2.431 | -0.003 | -0.046 | -0.046 | 3.215 | 0.35 | 0.38 | 0.3 | 0.07 | 0.34 | 0.24 |
| 4 | 29069894 | rs1402481 | 0.923 | -0.005 | -0.193 | -2.359 | 0 | -0.017 | -0.017 | 1.807 | 0.47 | 0.46 | 0.29 | 0.18 | 0.43 | 0.27 |
| 4 | 29081102 | rs4571399 | -0.037 | 0.059 | -0.457 | -2.389 | 0.001 | -0.052 | -0.052 | 0 | 0.66 | 0.7 | 0.7 | 0.99 | 0.67 | 0.76 |
| 4 | 29081919 | rs13103421 | 0.752 | 0.087 | -0.471 | -2.366 | -0.009 | -0.079 | -0.079 | 0 | 0.21 | 0.26 | 0.2 | 0.01 | 0.21 | 0.15 |
| 4 | 29095909 | rs11934543 | 0.478 | 0.305 | -0.374 | -1.974 | -0.007 | 0.077 | 0.077 | 0.075 | 0.44 | 0.26 | 0.27 | 0.24 | 0.43 | 0.28 |
| 4 | 29106713 | rs12644773 | 0.048 | -0.149 | -0.504 | -2.194 | 0.009 | 0.006 | 0.006 | 0 | 0.65 | 0.73 | 0.7 | 0.99 | 0.67 | 0.76 |
| 4 | 29126230 | rs10009272 | 0.169 | 0.145 | -0.815 | -1.966 | -0.014 | -0.143 | -0.143 | 6.262 | 0.59 | 0.48 | 0.67 | 0.99 | 0.63 | 0.74 |
| 4 | 29129308 | rs4401510 | 0.236 | -0.215 | -0.984 | -1.042 | 0.019 | 0.024 | 0.024 | 0 | 0.64 | 0.76 | 0.82 | 0.78 | 0.74 | 0.68 |
| 4 | 29129922 | rs10939244 | 0.029 | -0.884 | -1.377 | -2.228 | 0.018 | 0.01 | 0.01 | 0.155 | 0.65 | 0.72 | 0.81 | 0.96 | 0.68 | 0.75 |
| 4 | 29135014 | rs7694767 | -0.146 | -0.629 | -1.605 | -2.364 | -0.026 | 0.013 | 0.013 | 0 | 0.2 | 0.07 | 0.21 | 0.16 | 0.22 | 0.24 |
| 4 | 29135608 | rs7671430 | -0.349 | -0.739 | -0.761 | -2.478 | 0.001 | -0.004 | -0.004 | 0 | 0.22 | 0.21 | 0.23 | 0.16 | 0.23 | 0.23 |
| 4 | 29141525 | rs6820781 | -1.717 | -0.699 | -0.496 | -2.606 | -0.048 | 0.011 | 0.011 | 0 | 0.68 | 0.85 | 0.53 | 0.26 | 0.61 | 0.66 |
| 4 | 29153921 | rs6448630 | -0.259 | -1.289 | -0.046 | -2.692 | -0.021 | -0.099 | -0.099 | 0 | 0.62 | 0.46 | 0.78 | 0.96 | 0.67 | 0.75 |
| 4 | 29164477 | rs4285149 | -0.498 | -1.405 | 0.377 | -2.758 | 0.01 | -0.021 | -0.021 | 0 | 0.63 | 0.62 | 0.79 | 0.97 | 0.65 | 0.73 |
| 4 | 29195105 | rs1507439 | 1.419 | -1.471 | 0.144 | -2.872 | -0.175 | 0.048 | 0.048 | 0.062 | 0.33 | 0.05 | 0.49 | 0.78 | 0.43 | 0.36 |
| 4 | 29203553 | rs10517155 | -0.719 | -1.591 | 0.378 | -3.044 | 0.003 | -0.155 | -0.155 | 0.765 | 0.78 | 0.54 | 0.84 | 1 | 0.8 | 0.86 |
| 4 | 29203881 | rs16881492 | 1.481 | -1.627 | 0.239 | -3.067 | 0.031 | 0.024 | 0.024 | 0 | 0.18 | 0.14 | 0.07 | 0.01 | 0.15 | 0.14 |
| 4 | 29220153 | rs7696414 | -1.026 | -1.455 | 0.364 | -2.955 | 0.001 | -0.055 | -0.055 | 0 | 0.8 | 0.73 | 0.86 | 1 | 0.8 | 0.86 |
| 4 | 29224491 | rs2019100 | -0.037 | -1.537 | 0.328 | -3.002 | -0.01 | -0.004 | -0.004 | 0 | 0.2 | 0.08 | 0.23 | 0.19 | 0.2 | 0.33 |

| | | | | | | | | | | | | | | | | |
|---|---|---|---|---|---|---|---|---|---|---|---|---|---|---|---|---|
| 4 | 29229619 | rs2117896 | -0.948 | -1.45 | 0.435 | -2.944 | 0.003 | -0.086 | -0.086 | 0 | 0.8 | 0.68 | 0.86 | 1 | 0.8 | 0.86 |
| 4 | 29241920 | rs6824303 | -1.337 | -1.441 | 0.389 | -2.939 | 0 | -0.036 | -0.036 | 0.019 | 0.82 | 0.81 | 0.87 | 1 | 0.8 | 0.86 |
| 4 | 29247786 | rs13121140 | 0.462 | -1.45 | 0.08 | -2.82 | -0.008 | -0.016 | -0.016 | 0 | 0.16 | 0.06 | 0.2 | 0.17 | 0.19 | 0.22 |
| 4 | 29249394 | rs6837434 | -1.24 | -1.59 | 0.113 | -2.883 | 0.103 | 0.143 | 0.143 | 0 | 0.48 | 0.8 | 0.39 | 0.24 | 0.38 | 0.5 |
| 4 | 29275203 | rs11947802 | -0.253 | -1.619 | 0.539 | -2.926 | -0.058 | -0.099 | -0.099 | 0 | 0.3 | 0.55 | 0.18 | 0.07 | 0.18 | 0.29 |
| 4 | 29287401 | rs10033492 | 1.479 | -1.576 | -0.203 | -2.979 | -0.159 | 0.036 | 0.036 | 0.004 | 0.34 | 0.06 | 0.46 | 0.78 | 0.44 | 0.35 |
| 4 | 29289447 | rs12331067 | -0.635 | -1.593 | 0.094 | -2.98 | -0.006 | -0.041 | -0.041 | 0 | 0.07 | 0.18 | 0.05 | 0.02 | 0.04 | 0.12 |
| 4 | 29293875 | rs13148111 | -0.187 | -1.751 | 0.258 | -3.095 | -0.022 | -0.046 | -0.046 | 1.402 | 0.36 | 0.44 | 0.3 | 0.17 | 0.32 | 0.37 |
| 4 | 29294481 | rs7696776 | 0.671 | -1.496 | 0.105 | -2.841 | 0.015 | 0.116 | 0.116 | 0 | 0.57 | 0.21 | 0.57 | 0.79 | 0.6 | 0.62 |
| 4 | 29296845 | rs1369801 | 0.37 | -1.475 | 0.14 | -2.928 | -0.013 | -0.009 | -0.009 | 0.059 | 0.12 | 0.14 | 0.15 | 0.12 | 0.12 | 0.18 |
| 4 | 29299647 | rs17714421 | 1.32 | -1.749 | -0.202 | -3.198 | -0.161 | 0.129 | 0.129 | 0.857 | 0.34 | 0.05 | 0.47 | 0.82 | 0.45 | 0.38 |
| 4 | 29302397 | rs4264889 | 0.051 | -1.544 | 0.265 | -3.072 | 0.047 | 0.094 | 0.094 | 0.061 | 0.55 | 0.69 | 0.73 | 0.99 | 0.64 | 0.67 |
| 4 | 29303608 | rs4588519 | 0.01 | -1.472 | 0.198 | -3.025 | 0.004 | -0.012 | -0.012 | 0.37 | 0.14 | 0.22 | 0.2 | 0.15 | 0.17 | 0.18 |
| 4 | 29306747 | rs4521412 | -0.414 | -1.547 | 0.17 | -3.075 | -0.005 | -0.021 | -0.021 | 1.35 | 0.36 | 0.35 | 0.29 | 0.16 | 0.32 | 0.34 |
| 4 | 29309079 | rs1821687 | 0.086 | -1.13 | 0.064 | -2.763 | 0 | -0.003 | -0.003 | 0.015 | 0.14 | 0.11 | 0.19 | 0.15 | 0.17 | 0.18 |
| 4 | 29309684 | rs2310011 | -0.059 | -1.112 | -0.099 | -2.755 | 0.004 | -0.012 | -0.012 | 0 | 0.14 | 0.22 | 0.2 | 0.15 | 0.17 | 0.18 |
| 4 | 29310397 | rs1436063 | 0.14 | -1.031 | -0.355 | -2.575 | 0.008 | -0.052 | -0.052 | 0 | 0.54 | 0.56 | 0.71 | 0.99 | 0.63 | 0.66 |
| 4 | 29311144 | rs1821685 | 0.14 | -1.031 | -0.355 | -2.575 | 0.008 | -0.052 | -0.052 | 0 | 0.54 | 0.56 | 0.71 | 0.99 | 0.63 | 0.66 |
| 4 | 29314359 | rs10517158 | 0.046 | -0.849 | -0.901 | -2.365 | -0.018 | 0.114 | 0.114 | 0 | 0.24 | 0.07 | 0.18 | 0 | 0.21 | 0.15 |
| 4 | 29314598 | rs13114893 | -0.472 | -1.504 | -0.832 | -1.819 | 0.013 | 0.003 | 0.003 | 0.507 | 0.53 | 0.43 | 0.44 | 0.55 | 0.39 | 0.34 |
| 4 | 29317918 | rs4604115 | -1.628 | -0.917 | -0.688 | -1.434 | 0.114 | -0.041 | -0.041 | 1.516 | 0.8 | 0.68 | 0.63 | 0.88 | 0.64 | 0.82 |
| 4 | 29318283 | rs11939462 | 0.118 | -1.323 | -0.833 | -1.557 | -0.003 | 0.05 | 0.05 | 0 | 0.41 | 0.25 | 0.31 | 0.31 | 0.38 | 0.45 |
| 4 | 29318326 | rs34359528 | 1.554 | -1.268 | -0.825 | -1.571 | -0.011 | 0.1 | 0.1 | 0 | 0.24 | 0.08 | 0.18 | 0 | 0.21 | 0.15 |
| 4 | 29324896 | rs16867887 | -0.605 | -1.681 | -1.003 | -1.713 | 0.01 | -0.03 | -0.03 | 0.2 | 0.64 | 0.67 | 0.56 | 0.86 | 0.52 | 0.62 |
| 4 | 29330467 | rs1836779 | 0.77 | -1.701 | -1.023 | -1.672 | -0.001 | -0.006 | -0.006 | 0.048 | 0.17 | 0.17 | 0.14 | 0.29 | 0.18 | 0.29 |
| 4 | 29331117 | rs1025605 | 1.315 | -1.777 | -0.999 | -1.7 | -0.061 | 0.064 | 0.064 | 0.065 | 0.23 | 0.36 | 0.23 | 0.56 | 0.14 | 0.19 |
| 4 | 29334731 | rs4611992 | -2.189 | -1.683 | -0.645 | -1.801 | 0.025 | -0.008 | -0.008 | 0.326 | 0.81 | 0.85 | 0.64 | 0.89 | 0.64 | 0.82 |
| 4 | 29335025 | rs13103862 | 1.767 | -1.709 | -0.75 | -1.813 | -0.01 | 0.101 | 0.101 | 0 | 0.24 | 0.09 | 0.18 | 0 | 0.21 | 0.15 |
| 4 | 29335563 | rs6448638 | -0.531 | -1.576 | -0.667 | -1.716 | 0.007 | -0.023 | -0.023 | 0 | 0.64 | 0.68 | 0.55 | 0.86 | 0.52 | 0.61 |
| 4 | 29338216 | rs4434324 | -0.367 | -1.797 | -0.298 | -1.783 | 0.009 | -0.021 | -0.021 | 0 | 0.49 | 0.59 | 0.45 | 0.04 | 0.4 | 0.39 |
| 4 | 29339030 | rs6817648 | 0.787 | -1.865 | -0.207 | -1.687 | 0.043 | 0.011 | 0.011 | 0 | 0.46 | 0.41 | 0.4 | 0.56 | 0.34 | 0.32 |
| 4 | 29339512 | rs1436058 | -0.953 | -1.472 | -0.19 | -1.442 | -0.018 | -0.006 | -0.006 | 0 | 0.68 | 0.8 | 0.66 | 0.6 | 0.53 | 0.57 |
| 4 | 29342591 | rs2310014 | 1.516 | -1.614 | -0.362 | -1.349 | -0.007 | 0.013 | 0.013 | 0 | 0.19 | 0.21 | 0.2 | 0.56 | 0.12 | 0.17 |
| 4 | 29347765 | rs6817219 | -0.914 | -0.944 | -0.542 | -1.1 | -0.07 | -0.044 | -0.044 | 0.147 | 0.69 | 0.87 | 0.67 | 0.57 | 0.57 | 0.55 |
| 4 | 29364680 | rs195444 | -0.354 | -1.062 | -0.585 | -1.063 | -0.055 | -0.043 | -0.043 | 0.303 | 0.62 | 0.84 | 0.62 | 0.55 | 0.52 | 0.54 |
| 4 | 29365966 | rs181999 | 0.562 | -1.325 | -0.12 | -1.261 | 0.001 | -0.019 | -0.019 | 0 | 0.4 | 0.36 | 0.4 | 0.02 | 0.38 | 0.37 |
| 4 | 29368526 | rs479245 | -0.832 | -0.856 | -0.074 | -0.941 | -0.073 | -0.047 | -0.047 | 1.432 | 0.69 | 0.87 | 0.65 | 0.57 | 0.56 | 0.55 |
| 4 | 29376155 | rs260930 | 0.738 | -0.599 | -0.344 | -0.872 | -0.035 | 0.032 | 0.032 | 0.439 | 0.1 | 0.01 | 0.19 | 0.02 | 0.13 | 0.22 |
| 4 | 29379799 | rs997412 | 0.941 | -0.756 | 0.098 | -0.597 | 0.006 | -0.009 | -0.009 | 0 | 0.21 | 0.25 | 0.2 | 0.55 | 0.13 | 0.15 |
| 4 | 29382730 | rs988366 | -1.496 | -0.458 | 0.118 | -0.983 | 0.091 | -0.052 | -0.052 | 0 | 0.81 | 0.71 | 0.73 | 0.91 | 0.67 | 0.83 |
| 4 | 29395383 | rs16881736 | 0.968 | -0.852 | -0.161 | -0.565 | 0.013 | -0.019 | -0.019 | 0.298 | 0.21 | 0.21 | 0.19 | 0.59 | 0.14 | 0.14 |
| 4 | 29400203 | rs11936766 | 0.634 | -1.042 | -0.292 | -0.888 | 0.002 | -0.003 | -0.003 | 0.07 | 0.11 | 0.1 | 0.06 | 0.23 | 0.08 | 0.35 |
| 4 | 29404976 | rs166333 | -0.174 | -1.007 | -0.246 | -1.038 | 0.005 | -0.003 | -0.003 | 0.494 | 0.64 | 0.63 | 0.66 | 0.68 | 0.55 | 0.5 |
| 4 | 29407530 | rs260122 | 0.661 | -0.815 | -0.313 | -1.065 | 0.001 | -0.003 | -0.003 | 0 | 0.42 | 0.35 | 0.36 | 0.03 | 0.41 | 0.36 |
| 4 | 29419157 | rs1155595 | -0.664 | -1.346 | -0.203 | -0.45 | -0.037 | -0.019 | -0.019 | 1.22 | 0.32 | 0.4 | 0.36 | 0.8 | 0.21 | 0.5 |
| 4 | 29420040 | rs16881791 | 0.739 | -0.912 | -0.76 | -0.256 | 0.034 | -0.07 | -0.07 | 0 | 0.18 | 0.07 | 0.27 | 0.65 | 0.13 | 0.15 |
| 4 | 29423741 | rs16881804 | 0.739 | -0.912 | -0.76 | -0.277 | 0.034 | -0.068 | -0.068 | 0.477 | 0.18 | 0.07 | 0.27 | 0.65 | 0.14 | 0.15 |
| 4 | 29430546 | rs16881823 | 0.221 | -0.844 | -0.597 | -0.404 | 0.002 | -0.003 | -0.003 | 5.406 | 0.22 | 0.21 | 0.28 | 0.66 | 0.16 | 0.22 |
| 4 | 29441731 | rs2310033 | -0.296 | 0.051 | -0.686 | -1.19 | 0.004 | -0.039 | -0.039 | 0 | 0.61 | 0.53 | 0.57 | 0.8 | 0.58 | 0.68 |
| 4 | 29442576 | rs6811398 | -0.354 | 0.044 | -0.736 | -1.299 | 0.002 | -0.04 | -0.04 | 0 | 0.59 | 0.52 | 0.58 | 0.81 | 0.57 | 0.69 |
| 4 | 29452617 | rs6843590 | -0.177 | -0.586 | -0.611 | -1.374 | -0.002 | -0.004 | -0.004 | 0 | 0.31 | 0.28 | 0.42 | 0.81 | 0.27 | 0.3 |
| 4 | 29458432 | rs13142018 | 0.73 | -0.705 | -1.305 | -1.865 | -0.109 | 0.144 | 0.144 | 0 | 0.41 | 0.05 | 0.39 | 0.15 | 0.54 | 0.6 |
| 4 | 29478879 | rs1980023 | 0.343 | -0.769 | -0.354 | -1.686 | -0.001 | -0.004 | -0.004 | 0.054 | 0.29 | 0.18 | 0.29 | 0.69 | 0.26 | 0.22 |
| 4 | 29489354 | rs4692392 | 1.179 | -0.853 | -0.969 | -2.052 | -0.115 | 0.034 | 0.034 | 0 | 0.19 | 0.03 | 0.37 | 0.08 | 0.4 | 0.27 |
| 4 | 29496936 | rs7666825 | -0.986 | -1.313 | 0.149 | -1.601 | 0.04 | 0.199 | 0.199 | 0 | 0.51 | 0.81 | 0.51 | 0.86 | 0.41 | 0.38 |
| 4 | 29497594 | rs6851037 | 0.457 | -1.207 | 0.49 | -1.457 | -0.033 | -0.03 | -0.03 | 0 | 0.18 | 0.37 | 0.18 | 0.15 | 0.12 | 0.12 |
| 4 | 29502336 | rs7654387 | 1.01 | -1.054 | 0.235 | -1.322 | 0.013 | 0.119 | 0.119 | 0 | 0.48 | 0.25 | 0.5 | 0.17 | 0.57 | 0.61 |
| 4 | 29508260 | rs13106649 | -0.867 | -1.097 | 0.52 | -1.323 | -0.009 | -0.059 | -0.059 | 0 | 0.82 | 0.7 | 0.88 | 0.99 | 0.86 | 0.91 |
| 4 | 29508413 | rs10517167 | -0.493 | -0.877 | 0.246 | -1.107 | -0.011 | -0.037 | -0.037 | 0 | 0.34 | 0.44 | 0.39 | 0.82 | 0.29 | 0.3 |
| 4 | 29508477 | rs10517168 | -0.027 | -0.783 | -0.439 | -1.073 | -0.007 | -0.008 | -0.008 | 0 | 0.28 | 0.13 | 0.28 | 0.7 | 0.27 | 0.22 |
| 4 | 29510895 | rs9991320 | -0.152 | -0.742 | -0.398 | -1.044 | 0.002 | 0.004 | 0.004 | 0 | 0.31 | 0.2 | 0.31 | 0.7 | 0.28 | 0.25 |
| 4 | 29520647 | rs6818222 | -0.158 | -0.737 | -0.416 | -1.042 | 0.002 | 0.004 | 0.004 | 0 | 0.31 | 0.21 | 0.31 | 0.7 | 0.28 | 0.25 |
| 4 | 29524988 | rs10019636 | -0.961 | -0.9 | 0.198 | -0.809 | -0.023 | 0.019 | 0.019 | 0 | 0.2 | 0.3 | 0.22 | 0.74 | 0.15 | 0.24 |
| 4 | 29531282 | rs1837811 | -0.877 | -0.92 | 0.349 | -0.819 | -0.029 | 0.024 | 0.024 | 0 | 0.2 | 0.34 | 0.22 | 0.74 | 0.15 | 0.24 |
| 4 | 29532602 | rs11723936 | 1.323 | -0.972 | 0.436 | -1.097 | 0.009 | 0.013 | 0.013 | 0 | 0.6 | 0.34 | 0.63 | 0.26 | 0.69 | 0.68 |
| 4 | 29538611 | rs13122823 | -0.816 | -0.835 | 0.475 | -0.876 | -0.029 | 0.024 | 0.024 | 0 | 0.2 | 0.35 | 0.22 | 0.74 | 0.14 | 0.24 |
| 4 | 29554873 | rs10019440 | 1.278 | -0.748 | 0.115 | -1.177 | 0.026 | 0.111 | 0.111 | 0 | 0.49 | 0.26 | 0.47 | 0.12 | 0.55 | 0.55 |
| 4 | 29555818 | rs1372309 | -1.166 | -0.544 | -0.245 | -1.134 | -0.034 | 0.11 | 0.11 | 0 | 0.83 | 0.99 | 0.88 | 1 | 0.82 | 0.9 |
| 4 | 29574009 | rs1441671 | -1.083 | -0.562 | 0.019 | -1.143 | 0.002 | 0.032 | 0.032 | 0 | 0.82 | 0.94 | 0.88 | 1 | 0.82 | 0.9 |

| | | | | | | | | | | | | | | | | |
|---|---|---|---|---|---|---|---|---|---|---|---|---|---|---|---|---|
| 4 | 29590146 | rs1441670 | 1.205 | -0.691 | 0.234 | -1.319 | -0.006 | -0.076 | -0.076 | 0 | 0.55 | 0.51 | 0.71 | 0.95 | 0.59 | 0.78 |
| 4 | 29608313 | rs4407458 | -0.895 | -0.566 | 0.351 | -1.214 | 0.002 | 0.045 | 0.045 | 0 | 0.81 | 0.92 | 0.86 | 1 | 0.81 | 0.89 |
| 4 | 29612612 | rs11722750 | 2.088 | -0.598 | 0.598 | -0.919 | -0.008 | 0.026 | 0.026 | 0 | 0.39 | 0.29 | 0.44 | 0.21 | 0.41 | 0.59 |
| 4 | 29612773 | rs10029448 | -0.749 | -0.597 | 0.617 | -0.917 | -0.004 | -0.06 | -0.06 | 0 | 0.8 | 0.81 | 0.85 | 1 | 0.81 | 0.89 |
| 4 | 29618562 | rs7688471 | -0.93 | -0.595 | 0.586 | -0.914 | 0.003 | 0.032 | 0.032 | 0 | 0.81 | 0.91 | 0.86 | 1 | 0.81 | 0.89 |
| 4 | 29620820 | rs11726757 | 0.338 | -0.612 | 0.686 | -0.682 | -0.007 | -0.027 | -0.027 | 0 | 0.72 | 0.7 | 0.82 | 0.89 | 0.74 | 0.79 |
| 4 | 29626865 | rs10002254 | 0.859 | -0.462 | 0.461 | -0.539 | 0 | -0.049 | -0.049 | 0 | 0.16 | 0.21 | 0.11 | 0 | 0.15 | 0.04 |
| 4 | 29636690 | rs10805294 | -0.954 | -0.342 | 0.309 | -0.289 | 0.002 | -0.01 | -0.01 | 0 | 0.82 | 0.87 | 0.87 | 1 | 0.82 | 0.9 |
| 4 | 29638883 | rs10939268 | -1.251 | -0.341 | 0.288 | -0.289 | -0.031 | 0.106 | 0.106 | 0 | 0.83 | 0.99 | 0.89 | 1 | 0.82 | 0.9 |
| 4 | 29649572 | rs16882044 | 0.37 | -0.348 | 0.348 | -0.257 | 0.002 | 0.023 | 0.023 | 0 | 0.16 | 0.22 | 0.2 | 0.69 | 0.18 | 0.16 |
| 4 | 29655893 | rs7671757 | -0.426 | -0.51 | 0.114 | -0.254 | 0.004 | 0.012 | 0.012 | 0 | 0.81 | 0.88 | 0.86 | 1 | 0.81 | 0.89 |
| 4 | 29656702 | rs10024677 | -0.41 | -0.5 | 0.116 | -0.254 | 0.002 | -0.002 | -0.002 | 0 | 0.81 | 0.87 | 0.86 | 1 | 0.81 | 0.89 |
| 4 | 29660543 | rs4692409 | -0.709 | -0.506 | 0.111 | -0.248 | 0.001 | 0 | 0 | 0 | 0.82 | 0.88 | 0.87 | 1 | 0.82 | 0.9 |
| 4 | 29663459 | rs4692410 | -0.877 | -0.512 | -0.055 | -0.244 | -0.03 | 0.109 | 0.109 | 0 | 0.83 | 0.99 | 0.89 | 1 | 0.82 | 0.9 |
| 4 | 29666276 | rs4624623 | -0.53 | -0.491 | 0.02 | -0.236 | 0.003 | 0.013 | 0.013 | 0 | 0.81 | 0.9 | 0.86 | 1 | 0.81 | 0.89 |
| 4 | 29673009 | rs13103435 | 0.506 | -0.524 | -0.05 | 0.45 | 0.009 | -0.033 | -0.033 | 0.325 | 0.72 | 0.79 | 0.75 | 0.36 | 0.73 | 0.76 |
| 4 | 29673019 | rs13103446 | -1 | -0.502 | -0.062 | 0.694 | -0.008 | -0.085 | -0.085 | 1.89 | 0.82 | 0.94 | 0.79 | 0.5 | 0.82 | 0.87 |
| 4 | 29673981 | rs9291530 | -1 | -0.502 | -0.062 | 0.694 | -0.008 | -0.085 | -0.085 | 0 | 0.82 | 0.94 | 0.79 | 0.5 | 0.82 | 0.87 |
| 4 | 29680847 | rs7673510 | 0.866 | -0.477 | -0.127 | 0.678 | -0.018 | -0.023 | -0.023 | 0.188 | 0.12 | 0.09 | 0.1 | 0.15 | 0.15 | 0.11 |
| 4 | 29683641 | rs6846124 | 0.339 | -0.5 | -0.121 | 0.672 | 0.007 | -0.062 | -0.062 | 0 | 0.73 | 0.86 | 0.76 | 0.36 | 0.73 | 0.76 |
| 4 | 29684913 | rs10018343 | -0.965 | -0.513 | 0.273 | 0.661 | 0 | 0.036 | 0.036 | 0.199 | 0.23 | 0.43 | 0.15 | 0.29 | 0.23 | 0.22 |
| 4 | 29685111 | rs16882126 | -0.05 | -0.512 | 0.267 | 0.59 | -0.004 | 0.048 | 0.048 | 0 | 0.11 | 0.29 | 0.05 | 0.14 | 0.09 | 0.11 |
| 4 | 29688391 | rs16882129 | 0.818 | -0.503 | 0.083 | 0.631 | -0.018 | -0.027 | -0.027 | 0.109 | 0.12 | 0.09 | 0.1 | 0.16 | 0.15 | 0.11 |
| 4 | 29689796 | rs10033986 | 1.13 | -0.526 | 0.099 | 0.487 | 0 | 0.057 | 0.057 | 0 | 0.61 | 0.69 | 0.65 | 0.19 | 0.58 | 0.64 |
| 4 | 29690077 | rs10011902 | 1.154 | -0.514 | 0.112 | 0.49 | 0.003 | 0.038 | 0.038 | 0 | 0.6 | 0.64 | 0.65 | 0.2 | 0.58 | 0.65 |
| 4 | 29690129 | rs10939271 | 1.121 | -0.514 | 0.097 | 0.49 | 0.002 | 0.055 | 0.055 | 0 | 0.61 | 0.69 | 0.65 | 0.2 | 0.58 | 0.65 |
| 4 | 29691666 | rs7664033 | 0.055 | -0.522 | 0.255 | 0.341 | -0.009 | 0.014 | 0.014 | 0 | 0.11 | 0.19 | 0.05 | 0.14 | 0.09 | 0.11 |
| 4 | 29693864 | rs13128006 | 1.489 | -0.4 | -0.031 | 0.165 | -0.005 | 0.092 | 0.092 | 0 | 0.46 | 0.66 | 0.5 | 0.15 | 0.46 | 0.61 |
| 4 | 29696261 | rs13104601 | 1.982 | -0.357 | 0.109 | 0.111 | 0 | -0.012 | -0.012 | 0 | 0.42 | 0.36 | 0.46 | 0.15 | 0.43 | 0.61 |
| 4 | 29700316 | rs13134458 | 1.579 | -0.559 | -0.508 | 0.219 | -0.004 | 0.051 | 0.051 | 0 | 0.46 | 0.59 | 0.49 | 0.15 | 0.46 | 0.62 |
| 4 | 29710556 | rs7692836 | 0.226 | -0.409 | -0.445 | 0.466 | 0.01 | 0.004 | 0.004 | 0 | 0.21 | 0.16 | 0.11 | 0.15 | 0.19 | 0.12 |
| 4 | 29712547 | rs16882167 | -1.696 | -0.35 | -0.407 | 0.419 | -0.002 | 0.094 | 0.094 | 0 | 0.53 | 0.32 | 0.5 | 0.85 | 0.52 | 0.37 |
| 4 | 29713447 | rs16882170 | 0.88 | -0.007 | -0.416 | 0.776 | -0.034 | -0.12 | -0.12 | 0 | 0.15 | 0.01 | 0.25 | 0.54 | 0.16 | 0.11 |
| 4 | 29725653 | rs12641649 | 0.349 | -0.073 | -0.301 | 0.735 | -0.002 | -0.004 | -0.004 | 0 | 0.13 | 0.13 | 0.11 | 0.16 | 0.15 | 0.12 |
| 4 | 29726094 | rs12641755 | -0.028 | -0.042 | -0.143 | 0.176 | -0.005 | -0.021 | -0.021 | 0 | 0.29 | 0.19 | 0.36 | 0.7 | 0.31 | 0.22 |
| 4 | 29729748 | rs2310077 | -0.197 | -0.089 | -0.269 | 0.143 | 0.01 | 0.007 | 0.007 | 0 | 0.78 | 0.83 | 0.89 | 0.85 | 0.81 | 0.88 |
| 4 | 29735485 | rs6853827 | 0.232 | 0.332 | -0.048 | 0.132 | 0.003 | 0.017 | 0.017 | 0 | 0.49 | 0.53 | 0.52 | 0.16 | 0.5 | 0.65 |
| 4 | 29739635 | rs2310084 | 0.667 | 0.532 | 0.03 | 0.309 | -0.001 | -0.009 | -0.009 | 0.043 | 0.35 | 0.37 | 0.35 | 0.18 | 0.32 | 0.59 |
| 4 | 29740429 | rs16882206 | -1.021 | 0.26 | 1.046 | 0.782 | 0.041 | 0.001 | 0.001 | 0.085 | 0.1 | 0.38 | 0.06 | 0.05 | 0.07 | 0.02 |
| 4 | 29743040 | rs16882209 | -0.468 | 0.255 | 1.02 | 0.777 | -0.011 | 0.014 | 0.014 | 0.032 | 0.08 | 0.27 | 0.03 | 0.05 | 0.03 | 0.02 |
| 4 | 29743168 | rs17735010 | -1.415 | 0.361 | 0.089 | 1.166 | -0.063 | 0.09 | 0.09 | 0.015 | 0.46 | 0.89 | 0.61 | 0.7 | 0.5 | 0.23 |
| 4 | 29755511 | rs7664238 | -0.473 | 0.303 | 0.18 | 1.062 | 0.008 | -0.004 | -0.004 | 0.077 | 0.34 | 0.39 | 0.53 | 0.68 | 0.43 | 0.21 |
| 4 | 29756029 | rs10020813 | -1.137 | 0.458 | -0.601 | 0.88 | -0.087 | 0.249 | 0.249 | 0.012 | 0.51 | 0.96 | 0.63 | 0.84 | 0.56 | 0.29 |
| 4 | 29763937 | rs10001236 | -1.96 | 0.251 | 0.415 | 2.009 | 0.02 | -0.042 | -0.042 | 0 | 0.43 | 0.9 | 0.51 | 0.58 | 0.38 | 0.29 |
| 4 | 29770254 | rs16882245 | -1.183 | -0.333 | 0.601 | 2.058 | -0.027 | -0.009 | -0.009 | 0 | 0.12 | 0.2 | 0.05 | 0.08 | 0.05 | 0.1 |
| 4 | 29770652 | rs2310091 | -1.815 | -0.196 | -0.191 | 2.04 | 0.054 | -0.05 | -0.05 | 0 | 0.4 | 0.84 | 0.47 | 0.57 | 0.32 | 0.28 |
| 4 | 29773273 | rs6811040 | -1.666 | 0.015 | 0.355 | 2.273 | -0.011 | 0.072 | 0.072 | 0 | 0.13 | 0.23 | 0.08 | 0.36 | 0.08 | 0.12 |
| 4 | 29775082 | rs16882258 | -1.098 | -0.024 | 0.352 | 2.409 | -0.026 | 0.093 | 0.093 | 0 | 0.09 | 0.16 | 0.06 | 0.35 | 0.03 | 0.08 |
| 4 | 29777723 | rs2047870 | -2.052 | -0.176 | 0.092 | 2.171 | 0.031 | -0.055 | -0.055 | 0.871 | 0.38 | 0.86 | 0.27 | 0.54 | 0.28 | 0.21 |
| 4 | 29783037 | rs2062844 | -2.065 | 0.245 | 1.161 | 1.59 | 0.003 | 0.007 | 0.007 | 1.277 | 0.29 | 0.76 | 0.21 | 0.2 | 0.23 | 0.12 |
| 4 | 29796112 | rs7654268 | -0.946 | 0.174 | 1.568 | 1.519 | 0.001 | 0.029 | 0.029 | 0 | 0.14 | 0.07 | 0.08 | 0 | 0.13 | 0.02 |
| 4 | 29801069 | rs1995488 | -2.82 | 0.123 | 1.679 | 1.629 | 0.006 | 0.004 | 0.004 | 1.257 | 0.29 | 0.79 | 0.27 | 0.27 | 0.24 | 0.14 |
| 4 | 29802111 | rs16882322 | -0.952 | 0.117 | 2.772 | 1.645 | 0.04 | 0.142 | 0.142 | 0.932 | 0.06 | 0.28 | 0.03 | 0.11 | 0.03 | 0.05 |
| 4 | 29802167 | rs16882325 | -1.927 | 0.117 | 2.851 | 1.643 | -0.031 | -0.051 | -0.051 | 0 | 0.2 | 0.41 | 0.12 | 0.12 | 0.16 | 0.07 |
| 4 | 29802767 | rs4367136 | -2.676 | 0.135 | 3.134 | 1.648 | 0.061 | 0.06 | 0.06 | 0 | 0.06 | 0.19 | 0.14 | 0.13 | 0.1 | 0.04 |
| 4 | 29804231 | rs11722527 | -2.484 | 0.157 | 3.208 | 1.622 | -0.013 | -0.036 | -0.036 | 0 | 0.23 | 0.59 | 0.15 | 0.14 | 0.2 | 0.09 |
| 4 | 29811570 | rs1503971 | -1.137 | 0.209 | 3.124 | 1.674 | 0.034 | 0.163 | 0.163 | 1.779 | 0.06 | 0.28 | 0.04 | 0.15 | 0.02 | 0.06 |
| 4 | 29817230 | rs16882354 | -1.009 | 0.204 | 3.097 | 1.723 | 0.02 | 0.174 | 0.174 | 2.479 | 0.06 | 0.24 | 0.04 | 0.17 | 0.03 | 0.07 |
| 4 | 29819581 | rs11932609 | -1.009 | 0.204 | 3.097 | 1.723 | 0.02 | 0.174 | 0.174 | 2.479 | 0.06 | 0.24 | 0.04 | 0.17 | 0.03 | 0.07 |
| 4 | 29826189 | rs2202308 | -2.439 | 0.213 | 3.372 | 1.708 | -0.001 | 0.018 | 0.018 | 0.318 | 0.23 | 0.46 | 0.22 | 0.28 | 0.19 | 0.11 |
| 4 | 29837834 | rs1472711 | -2.909 | 0.214 | 3.26 | 1.707 | 0 | -0.001 | -0.001 | 0 | 0.27 | 0.74 | 0.27 | 0.31 | 0.23 | 0.13 |
| 4 | 29838345 | rs11723563 | -1.599 | 0.221 | 3.137 | 1.704 | -0.005 | 0.007 | 0.007 | 0.008 | 0.17 | 0.17 | 0.18 | 0.11 | 0.16 | 0.04 |
| 4 | 29857785 | rs10939282 | -2.908 | 0.179 | 3.122 | 1.72 | 0.002 | -0.001 | -0.001 | 0 | 0.28 | 0.74 | 0.27 | 0.31 | 0.23 | 0.13 |
| 4 | 29869342 | rs1394130 | -2.57 | 0.185 | 3.067 | 1.628 | -0.008 | 0.013 | 0.013 | 0.155 | 0.24 | 0.48 | 0.22 | 0.28 | 0.19 | 0.12 |
| 4 | 29897820 | rs17737828 | -1.659 | 0.167 | 2.222 | 1.667 | -0.008 | 0.016 | 0.016 | 0.013 | 0.17 | 0.12 | 0.17 | 0.11 | 0.17 | 0.05 |
| 4 | 29899300 | rs7672855 | -2.172 | 0.164 | 2.552 | 1.665 | -0.002 | 0.015 | 0.015 | 0.093 | 0.23 | 0.43 | 0.22 | 0.28 | 0.19 | 0.12 |
| 4 | 29903052 | rs7442317 | -2.373 | 0.265 | 2.456 | 1.55 | 0.008 | 0.046 | 0.046 | 0.274 | 0.31 | 0.5 | 0.21 | 0.39 | 0.28 | 0.27 |
| 4 | 29909867 | rs16882465 | -2.28 | 0.283 | 2.867 | 1.677 | 0.002 | 0.006 | 0.006 | 1.647 | 0.19 | 0.29 | 0.13 | 0.2 | 0.16 | 0.1 |
| 4 | 29912308 | rs10001348 | -1.281 | 0.283 | 2.871 | 1.677 | 0.01 | 0.052 | 0.052 | 0 | 0.12 | 0.24 | 0.09 | 0.19 | 0.13 | 0.17 |

| | | | | | | | | | | | | | | | | |
|---|---|---|---|---|---|---|---|---|---|---|---|---|---|---|---|---|
| 4 | 29917438 | rs10023178 | -2.864 | 0.289 | 2.852 | 1.656 | 0.003 | 0.025 | 0.025 | 0.368 | 0.31 | 0.57 | 0.22 | 0.39 | 0.28 | 0.27 |
| 4 | 29929502 | rs1503966 | -2.958 | -0.025 | 2.919 | 1.545 | -0.01 | 0.022 | 0.022 | 0.069 | 0.32 | 0.55 | 0.21 | 0.39 | 0.28 | 0.29 |
| 4 | 29943810 | rs12506164 | -2.588 | -0.27 | 2.781 | 1.457 | 0.017 | 0.003 | 0.003 | 0.145 | 0.24 | 0.16 | 0.11 | 0.2 | 0.17 | 0.24 |
| 4 | 29947072 | rs6846051 | -1.688 | -0.315 | 3.329 | 1.426 | 0 | -0.001 | -0.001 | 0 | 0.15 | 0.12 | 0.09 | 0.2 | 0.11 | 0.2 |
| 4 | 29949591 | rs6837917 | -3.55 | -0.259 | 3.311 | 1.412 | 0.012 | -0.007 | -0.007 | 0 | 0.34 | 0.66 | 0.21 | 0.37 | 0.25 | 0.28 |
| 4 | 29953808 | rs10028915 | -2.715 | -0.229 | 2.97 | 1.437 | -0.016 | 0.021 | 0.021 | 0.339 | 0.28 | 0.35 | 0.15 | 0.37 | 0.19 | 0.28 |
| 4 | 29962266 | rs4692430 | -2.814 | -0.187 | 2.963 | 0.806 | -0.029 | -0.043 | -0.043 | 0 | 0.31 | 0.45 | 0.18 | 0.17 | 0.23 | 0.24 |
| 4 | 29982657 | rs11943991 | -1.77 | -0.363 | 3.079 | 0.935 | 0.022 | -0.008 | -0.008 | 0 | 0.06 | 0.18 | 0.06 | 0.05 | 0.07 | 0.03 |
| 4 | 29983877 | rs4466010 | -2.26 | -0.46 | 3.348 | 0.888 | -0.007 | -0.01 | -0.01 | 0.012 | 0.06 | 0.24 | 0.02 | 0 | 0.01 | 0.04 |
| 4 | 29985899 | rs10939287 | 1.24 | -0.483 | 2.719 | 0.553 | -0.012 | 0.066 | 0.066 | 0 | 0.83 | 0.93 | 0.85 | 0.98 | 0.81 | 0.92 |
| 4 | 29998461 | rs3884638 | 3.854 | -0.649 | 2.575 | 0.77 | 0.008 | -0.005 | -0.005 | 2.055 | 0.56 | 0.06 | 0.62 | 0.61 | 0.62 | 0.66 |
| 4 | 30004497 | rs4593088 | 2.736 | -0.535 | 2.872 | 0.251 | -0.003 | -0.093 | -0.093 | 0 | 0.73 | 0.39 | 0.76 | 0.93 | 0.73 | 0.88 |
| 4 | 30005434 | rs10012732 | 2.846 | -0.5 | 2.773 | 0.268 | 0.01 | 0.021 | 0.021 | 0.315 | 0.68 | 0.26 | 0.74 | 0.93 | 0.73 | 0.84 |
| 4 | 30025294 | rs6840125 | -2.562 | -0.389 | 2.878 | 0.61 | 0.07 | 0.012 | 0.012 | 1.866 | 0.09 | 0.5 | 0.09 | 0.05 | 0.07 | 0.03 |
| 4 | 30055645 | rs4692435 | -1.201 | -0.514 | 2.268 | 0.498 | -0.016 | -0.052 | -0.052 | 0 | 0.12 | 0.25 | 0.08 | 0.02 | 0.08 | 0.06 |
| 4 | 30059039 | rs4377550 | -2.135 | -0.522 | 2.034 | 0.472 | -0.016 | -0.108 | -0.108 | 0.115 | 0.17 | 0.46 | 0.13 | 0.02 | 0.12 | 0.06 |
| 4 | 30063389 | rs12499826 | -2.378 | -0.53 | 1.393 | 0.478 | -0.017 | -0.061 | -0.061 | 0 | 0.21 | 0.66 | 0.16 | 0.08 | 0.14 | 0.09 |
| 4 | 30086484 | rs2310135 | 2.096 | -0.375 | 0.976 | 0.45 | 0.007 | 0.036 | 0.036 | 0 | 0.66 | 0.28 | 0.74 | 0.86 | 0.72 | 0.82 |
| 4 | 30092152 | rs6813280 | 1.398 | -0.587 | 1.455 | 0.766 | 0 | 0.012 | 0.012 | 0.042 | 0.88 | 0.65 | 0.93 | 0.9 | 0.93 | 0.94 |
| 4 | 30102488 | rs2310129 | 1.873 | -0.622 | 0.964 | 0.693 | -0.014 | -0.024 | -0.024 | 0 | 0.78 | 0.3 | 0.87 | 0.88 | 0.88 | 0.86 |
| 4 | 30108394 | rs13120387 | 2.849 | -0.729 | 0.34 | 1.186 | -0.012 | -0.091 | -0.091 | 0 | 0.55 | 0.05 | 0.69 | 0.58 | 0.62 | 0.62 |
| 4 | 30132119 | rs2613180 | -2.099 | -1.021 | 0.86 | 1.075 | -0.015 | -0.017 | -0.017 | 0 | 0.22 | 0.68 | 0.13 | 0.15 | 0.12 | 0.14 |
| 4 | 30134048 | rs2571494 | -2.697 | -1.112 | 0.029 | 1.165 | 0.026 | -0.021 | -0.021 | 0 | 0.39 | 0.93 | 0.24 | 0.31 | 0.28 | 0.33 |
| 4 | 30135529 | rs17688808 | -0.855 | -0.557 | 0.693 | 1.517 | -0.008 | -0.02 | -0.02 | 0 | 0.08 | 0.01 | 0.08 | 0.1 | 0.11 | 0.16 |
| 4 | 30144820 | rs2571483 | -1.619 | -0.506 | 1.23 | 1.425 | 0.009 | 0.045 | 0.045 | 0.086 | 0.1 | 0.34 | 0.06 | 0.12 | 0.07 | 0.04 |
| 4 | 30147719 | rs7689614 | -1.525 | -0.533 | 1.265 | 1.461 | 0.01 | 0.063 | 0.063 | 0.414 | 0.09 | 0.34 | 0.06 | 0.12 | 0.06 | 0.04 |
| 4 | 30156463 | rs1469784 | -1.405 | -0.485 | 0.784 | 1.657 | 0.006 | 0.08 | 0.08 | 0 | 0.05 | 0.14 | 0.03 | 0.16 | 0.05 | 0.19 |
| 4 | 30161847 | rs6836424 | -1.65 | -1.114 | 1.081 | 2.037 | -0.056 | 0.138 | 0.138 | 0 | 0.2 | 0.48 | 0.24 | 0.43 | 0.18 | 0.32 |
| 4 | 30162762 | rs2613167 | -1.731 | -1.456 | -0.21 | 1.504 | 0.056 | -0.064 | -0.064 | 1.048 | 0.41 | 0.91 | 0.4 | 0.56 | 0.34 | 0.37 |
| 4 | 30167692 | rs6815106 | -1.618 | -1.9 | 0.123 | 0.713 | 0.064 | 0.101 | 0.101 | 5.462 | 0.41 | 0.68 | 0.36 | 0.11 | 0.29 | 0.12 |
| 4 | 30168595 | rs16882976 | -1.579 | -1.905 | 1.383 | 1.366 | -0.016 | -0.004 | -0.004 | 0 | 0.37 | 0.58 | 0.46 | 0.32 | 0.29 | 0.2 |
| 4 | 30169493 | rs7691359 | -1.599 | -1.919 | 1.376 | 1.359 | -0.015 | -0.004 | -0.004 | 0 | 0.37 | 0.58 | 0.46 | 0.31 | 0.29 | 0.2 |
| 4 | 30170140 | rs16882984 | -0.353 | -2.069 | 1.803 | 1.241 | 0.011 | -0.011 | -0.011 | 0 | 0.2 | 0.14 | 0.09 | 0 | 0.14 | 0.03 |
| 4 | 30174983 | rs11732479 | -1.584 | -1.927 | 2 | 1.406 | -0.015 | -0.004 | -0.004 | 0 | 0.37 | 0.59 | 0.46 | 0.32 | 0.29 | 0.21 |
| 4 | 30180666 | rs1369184 | -1.288 | -1.928 | 1.889 | 1.324 | -0.027 | -0.039 | -0.039 | 0.002 | 0.33 | 0.37 | 0.32 | 0.17 | 0.26 | 0.06 |
| 4 | 30184038 | rs2256587 | -2.256 | -1.153 | 2.405 | 1.934 | 0.021 | 0.249 | 0.249 | 0.019 | 0.08 | 0.37 | 0.3 | 0.28 | 0.05 | 0.13 |
| 4 | 30193761 | rs976057 | -2.337 | -1.135 | 2.421 | 1.66 | 0.005 | 0.011 | 0.011 | 0.117 | 0.15 | 0.26 | 0.3 | 0.16 | 0.14 | 0.05 |
| 4 | 30198753 | rs2247063 | -2.318 | -0.727 | 2.711 | 1.99 | 0.055 | 0.196 | 0.196 | 0.088 | 0.12 | 0.48 | 0.29 | 0.26 | 0.1 | 0.07 |
| 4 | 30203202 | rs2613184 | -2.895 | -0.812 | 2.759 | 2.215 | 0.029 | 0.113 | 0.113 | 0 | 0.2 | 0.41 | 0.39 | 0.4 | 0.23 | 0.1 |
| 4 | 30205924 | rs1821544 | -1.315 | -0.792 | 2.656 | 2.219 | -0.018 | -0.027 | -0.027 | 0 | 0.1 | 0.04 | 0.12 | 0.14 | 0.13 | 0.05 |
| 4 | 30214893 | rs2871431 | -2.698 | -0.925 | 2.808 | 2.164 | 0.009 | 0.091 | 0.091 | 0 | 0.16 | 0.41 | 0.29 | 0.27 | 0.12 | 0.12 |
| 4 | 30215673 | rs2571468 | -3.782 | -0.928 | 3.011 | 2.384 | -0.001 | 0.043 | 0.043 | 0 | 0.29 | 0.63 | 0.44 | 0.49 | 0.27 | 0.21 |
| 4 | 30216257 | rs7662079 | -1.132 | -0.842 | 2.259 | 2.214 | -0.012 | -0.024 | -0.024 | 0 | 0.06 | 0.11 | 0.02 | 0.01 | 0.02 | 0.05 |
| 4 | 30218577 | rs2571486 | -3.194 | -0.847 | 2.241 | 2.215 | 0.005 | 0.067 | 0.067 | 0 | 0.26 | 0.53 | 0.41 | 0.41 | 0.25 | 0.17 |
| 4 | 30224009 | rs7667202 | -1.765 | -1.015 | 2.125 | 2.206 | 0.023 | 0.172 | 0.172 | 0 | 0.07 | 0.26 | 0.25 | 0.27 | 0.06 | 0.11 |
| 4 | 30231510 | rs3113774 | -1.676 | -0.993 | 1.902 | 1.741 | 0.003 | -0.054 | -0.054 | 0 | 0.09 | 0.23 | 0.05 | 0 | 0.06 | 0.02 |
| 4 | 30235629 | rs2571499 | -2.479 | -0.942 | 1.937 | 2.065 | 0.001 | 0.057 | 0.057 | 0 | 0.15 | 0.3 | 0.22 | 0.37 | 0.16 | 0.11 |
| 4 | 30242957 | rs7668302 | -1.599 | -0.799 | 2.012 | 2.19 | -0.009 | 0.017 | 0.017 | 0 | 0.15 | 0.26 | 0.14 | 0.16 | 0.1 | 0.2 |
| 4 | 30277791 | rs1434827 | -1.722 | -0.749 | 2.143 | 1.853 | 0.026 | 0.184 | 0.184 | 0 | 0.05 | 0.17 | 0.13 | 0.33 | 0.07 | 0.06 |
| 4 | 30283750 | rs3857027 | -3.447 | -0.848 | 2.352 | 1.82 | -0.006 | 0.074 | 0.074 | 0.284 | 0.22 | 0.66 | 0.28 | 0.43 | 0.19 | 0.19 |
| 4 | 30286607 | rs16883227 | -2.73 | -0.865 | 2.358 | 1.81 | -0.014 | -0.03 | -0.03 | 0 | 0.17 | 0.53 | 0.15 | 0.1 | 0.12 | 0.13 |
| 4 | 30295729 | rs1595049 | -1.881 | -0.864 | 2.391 | 1.81 | -0.004 | -0.013 | -0.013 | 2.826 | 0.13 | 0.31 | 0.09 | 0.07 | 0.09 | 0.11 |
| 4 | 30312547 | rs16883282 | -2.976 | -0.841 | 2.331 | 1.796 | -0.027 | -0.052 | -0.052 | 0.198 | 0.21 | 0.53 | 0.15 | 0.1 | 0.13 | 0.15 |
| 4 | 30313458 | rs1435779 | -1.133 | -0.846 | 2.35 | 1.794 | -0.003 | -0.006 | -0.006 | 0 | 0.11 | 0.26 | 0.07 | 0.06 | 0.07 | 0.1 |
| 4 | 30315305 | rs6858072 | -2.623 | -0.847 | 2.125 | 1.802 | -0.028 | -0.053 | -0.053 | 0 | 0.21 | 0.53 | 0.15 | 0.1 | 0.13 | 0.15 |
| 4 | 30324241 | rs12647147 | -1.401 | -0.81 | 1.656 | 1.837 | 0.006 | 0.086 | 0.086 | 0 | 0.06 | 0.12 | 0.16 | 0.36 | 0.09 | 0.08 |
| 4 | 30328822 | rs10049908 | -1.401 | -0.81 | 1.658 | 1.837 | 0.007 | 0.09 | 0.09 | 0 | 0.06 | 0.12 | 0.16 | 0.36 | 0.09 | 0.08 |
| 4 | 30328911 | rs7675877 | -1.544 | -0.821 | 1.665 | 1.824 | 0.002 | -0.01 | -0.01 | 0 | 0.14 | 0.42 | 0.1 | 0.07 | 0.09 | 0.11 |
| 4 | 30334017 | rs7661846 | -1.891 | -0.761 | 1.127 | 1.905 | 0 | 0.016 | 0.016 | 0 | 0.1 | 0.15 | 0.16 | 0.37 | 0.1 | 0.1 |
| 4 | 30346867 | rs16883353 | -2.051 | -0.826 | 1.178 | 1.871 | 0 | 0.085 | 0.085 | 0 | 0.11 | 0.24 | 0.17 | 0.4 | 0.1 | 0.1 |
| 4 | 30361791 | rs9985782 | -2.273 | -1.454 | 0.526 | 1.475 | 0.062 | -0.034 | -0.034 | 0.429 | 0.32 | 0.84 | 0.26 | 0.41 | 0.18 | 0.15 |
| 4 | 30363282 | rs7655653 | -2.258 | -1.14 | 1.322 | 1.769 | -0.041 | 0.192 | 0.192 | 0.507 | 0.16 | 0.52 | 0.17 | 0.35 | 0.09 | 0.07 |
| 4 | 30371605 | rs7698788 | -2.292 | -1.118 | 1.335 | 1.707 | 0.012 | 0.193 | 0.193 | 0.258 | 0.1 | 0.46 | 0.14 | 0.23 | 0.05 | 0.04 |
| 4 | 30384018 | rs12509641 | -1.725 | -1.043 | 1.209 | 1.228 | -0.027 | 0.003 | 0.003 | 0 | 0.11 | 0.29 | 0.08 | 0.1 | 0.06 | 0.04 |
| 4 | 30392084 | rs7695486 | -1.065 | -1.279 | 0.337 | 0.479 | 0.017 | 0.033 | 0.033 | 0 | 0.35 | 0.68 | 0.22 | 0.13 | 0.27 | 0.19 |
| 4 | 30394278 | rs11931286 | -1.165 | -1.306 | 0.383 | 0.466 | 0.035 | 0.082 | 0.082 | 0 | 0.36 | 0.75 | 0.22 | 0.13 | 0.27 | 0.19 |
| 4 | 30395518 | rs7681280 | -1.813 | -0.936 | 0.073 | 1.322 | 0.033 | 0.05 | 0.05 | 0 | 0.46 | 0.88 | 0.45 | 0.62 | 0.36 | 0.4 |
| 4 | 30397570 | rs1435782 | -1.036 | -1.079 | 0.628 | 1.06 | -0.007 | -0.058 | -0.058 | 0.15 | 0.14 | 0.48 | 0.1 | 0.05 | 0.12 | 0.07 |
| 4 | 30403040 | rs4346608 | -0.083 | -0.986 | 0.671 | 1.145 | 0.003 | 0.004 | 0.004 | 0 | 0.22 | 0.22 | 0.11 | 0.07 | 0.15 | 0.12 |

| | | | | | | | | | | | | | | | | |
|---|---|---|---|---|---|---|---|---|---|---|---|---|---|---|---|---|
| 4 | 30407029 | rs1435787 | -2.047 | -1.048 | 1.097 | 1.4 | -0.01 | 0.01 | 0.01 | 0 | 0.39 | 0.53 | 0.38 | 0.47 | 0.33 | 0.38 |
| 4 | 30409883 | rs1369660 | -2.432 | -1.051 | 1.063 | 1.423 | 0.027 | -0.019 | -0.019 | 0.002 | 0.44 | 0.88 | 0.44 | 0.51 | 0.35 | 0.39 |
| 4 | 30412285 | rs6847061 | -0.281 | -1.02 | 1.018 | 1.745 | -0.004 | -0.016 | -0.016 | 0 | 0.23 | 0.28 | 0.19 | 0.11 | 0.17 | 0.13 |
| 4 | 30412796 | rs3846270 | -1.441 | -1.034 | 0.983 | 1.728 | 0 | 0.012 | 0.012 | 0 | 0.16 | 0.18 | 0.18 | 0.36 | 0.16 | 0.22 |
| 4 | 30413369 | rs1836702 | 0.075 | -0.983 | 0.913 | 1.377 | 0.007 | -0.009 | -0.009 | 0.564 | 0.06 | 0.03 | 0.01 | 0.08 | 0.03 | 0.1 |
| 4 | 30416966 | rs12503525 | -0.188 | -0.937 | 1.027 | 1.333 | 0.003 | 0.001 | 0.001 | 0 | 0.22 | 0.22 | 0.11 | 0.07 | 0.15 | 0.12 |
| 4 | 30431211 | rs2871445 | -1.951 | -0.821 | 0.759 | 1.842 | -0.006 | 0.116 | 0.116 | 0.553 | 0.16 | 0.67 | 0.3 | 0.31 | 0.12 | 0.17 |
| 4 | 30431573 | rs6448707 | 1.976 | -0.942 | 0.068 | 1.506 | 0.005 | 0.022 | 0.022 | 0 | 0.54 | 0.11 | 0.54 | 0.43 | 0.62 | 0.58 |
| 4 | 30436488 | rs2203140 | -0.112 | -0.931 | 0.04 | 1.453 | -0.025 | 0.044 | 0.044 | 0 | 0.92 | 0.99 | 0.95 | 1 | 0.88 | 0.97 |
| 4 | 30437096 | rs9291540 | -1.808 | -0.908 | 0.159 | 1.593 | 0.023 | -0.05 | -0.05 | 0 | 0.36 | 0.83 | 0.4 | 0.47 | 0.25 | 0.36 |
| 4 | 30439397 | rs6841092 | -1.088 | -0.919 | 0.202 | 1.764 | 0.011 | -0.049 | -0.049 | 0 | 0.29 | 0.19 | 0.21 | 0.34 | 0.2 | 0.31 |
| 4 | 30439878 | rs4692457 | -0.58 | -1.042 | 0.222 | 1.184 | -0.038 | -0.022 | -0.022 | 0 | 0.18 | 0.64 | 0.21 | 0.12 | 0.1 | 0.13 |
| 4 | 30451158 | rs966770 | -1.462 | -0.607 | 0.335 | 1.408 | -0.016 | 0.065 | 0.065 | 0.899 | 0.12 | 0.65 | 0.2 | 0.28 | 0.07 | 0.08 |
| 4 | 30452567 | rs2164384 | 0.187 | -0.869 | -0.725 | 0.686 | -0.013 | 0.076 | 0.076 | 0 | 0.82 | 0.97 | 0.92 | 0.97 | 0.8 | 0.89 |
| 4 | 30453124 | rs9997877 | 0.507 | -0.602 | -0.152 | 0.839 | 0.023 | -0.015 | -0.015 | 0 | 0.91 | 0.73 | 0.93 | 0.96 | 0.94 | 0.97 |
| 4 | 30453156 | rs12641659 | 1.157 | -0.593 | -1.177 | 1.218 | -0.035 | -0.007 | -0.007 | 4.29 | 0.46 | 0.1 | 0.52 | 0.46 | 0.57 | 0.59 |
| 4 | 30455876 | rs7700123 | -0.691 | -0.247 | -0.636 | 1.312 | 0.006 | 0.048 | 0.048 | 0.103 | 0.33 | 0.52 | 0.32 | 0.42 | 0.31 | 0.53 |
| 4 | 30459400 | rs17763343 | 0.478 | -0.857 | -1.457 | 0.534 | 0.027 | 0.099 | 0.099 | 0 | 0.16 | 0.01 | 0.08 | 0 | 0.14 | 0.12 |
| 4 | 30461539 | rs1512155 | -1.537 | -1.181 | -2.05 | 0.499 | -0.183 | 0.068 | 0.068 | 0.015 | 0.67 | 0.97 | 0.53 | 0.83 | 0.54 | 0.72 |
| 4 | 30466859 | rs10008525 | 0.276 | -1.279 | -1.073 | 1.564 | 0.009 | 0.247 | 0.247 | 0 | 0.9 | 0.71 | 0.88 | 0.63 | 0.94 | 0.85 |
| 4 | 30467399 | rs13143903 | 1.606 | -1.378 | -1.381 | 1.425 | -0.163 | 0.095 | 0.095 | 0 | 0.36 | 0.05 | 0.49 | 0.17 | 0.49 | 0.31 |
| 4 | 30467623 | rs1896180 | 0.416 | -1.601 | -0.612 | 1.503 | -0.002 | -0.002 | -0.002 | 0 | 0.25 | 0.26 | 0.13 | 0.19 | 0.23 | 0.2 |
| 4 | 30471257 | rs1955311 | 0.076 | -1.54 | -0.187 | 1.546 | 0.022 | 0.326 | 0.326 | 0 | 0.89 | 0.61 | 0.88 | 0.66 | 0.94 | 0.85 |
| 4 | 30473099 | rs1980054 | 0.859 | -1.64 | 0.022 | 1.421 | 0.022 | 0.006 | 0.006 | 0 | 0.61 | 0.3 | 0.7 | 0.37 | 0.72 | 0.54 |
| 4 | 30476154 | rs1473094 | -0.539 | -1.593 | 0.188 | 1.713 | 0.015 | 0.175 | 0.175 | 0 | 0.92 | 0.79 | 0.9 | 0.75 | 0.95 | 0.86 |
| 4 | 30478873 | rs4692459 | 0.22 | -1.709 | 0.201 | 1.669 | 0.001 | -0.003 | -0.003 | 0 | 0.68 | 0.7 | 0.69 | 0.5 | 0.75 | 0.55 |
| 4 | 30478965 | rs4692090 | 1.905 | -1.547 | -0.362 | 1.301 | -0.159 | 0.143 | 0.143 | 0 | 0.36 | 0.05 | 0.48 | 0.12 | 0.49 | 0.31 |
| 4 | 30484155 | rs1553923 | 0.231 | -1.479 | -0.107 | 1.404 | 0.01 | -0.006 | -0.006 | 0.39 | 0.23 | 0.09 | 0.21 | 0.25 | 0.2 | 0.31 |
| 4 | 30490490 | rs9291543 | -0.762 | -1.307 | 0.43 | 1.313 | 0.027 | 0.107 | 0.107 | 0.032 | 0.07 | 0.32 | 0.07 | 0.15 | 0.03 | 0.03 |
| 4 | 30508931 | rs1588610 | -0.265 | -1.475 | 0.247 | 1.524 | -0.007 | 0.012 | 0.012 | 0 | 0.69 | 0.7 | 0.79 | 0.6 | 0.75 | 0.66 |
| 4 | 30510223 | rs13147511 | 1.842 | -1.539 | -0.097 | 1.275 | -0.185 | 0.119 | 0.119 | 0 | 0.36 | 0.04 | 0.4 | 0.13 | 0.48 | 0.3 |
| 4 | 30511492 | rs6812829 | -1.02 | -1.418 | 0.444 | 1.381 | 0.06 | 0.107 | 0.107 | 0 | 0.07 | 0.3 | 0.16 | 0.2 | 0.09 | 0.05 |
| 4 | 30533811 | rs1818442 | 0.828 | -1.529 | 0.497 | 1.383 | -0.008 | -0.013 | -0.013 | 0.389 | 0.23 | 0.2 | 0.12 | 0.14 | 0.18 | 0.17 |
| 4 | 30535299 | rs1588614 | 0.527 | -1.238 | 0.707 | 1.338 | 0.012 | 0.001 | 0.001 | 0.125 | 0.3 | 0.2 | 0.14 | 0.08 | 0.24 | 0.2 |
| 4 | 30536779 | rs6846798 | 0.67 | -1.234 | 0.557 | 1.59 | -0.018 | -0.119 | -0.119 | 0.81 | 0.18 | 0.04 | 0.21 | 0.41 | 0.17 | 0.31 |
| 4 | 30536826 | rs2018994 | 0.06 | -1.228 | 0.77 | 1.638 | -0.013 | 0.002 | 0.002 | 0 | 0.61 | 0.57 | 0.76 | 0.61 | 0.71 | 0.66 |
| 4 | 30544014 | rs11727470 | 0.258 | -1.192 | 0.414 | 1.61 | 0.016 | -0.003 | -0.003 | 0 | 0.58 | 0.4 | 0.65 | 0.58 | 0.7 | 0.64 |
| 4 | 30550919 | rs12374418 | 0.651 | -1.117 | 0.24 | 1.686 | -0.019 | -0.119 | -0.119 | 0.111 | 0.18 | 0.04 | 0.21 | 0.41 | 0.17 | 0.3 |
| 4 | 30556489 | rs1512148 | 0.644 | -1.032 | 0.499 | 1.583 | 0.001 | -0.002 | -0.002 | 0 | 0.69 | 0.54 | 0.66 | 0.4 | 0.71 | 0.56 |
| 4 | 30559836 | rs1398181 | 0.725 | -1.065 | 0.432 | 1.574 | -0.005 | 0.003 | 0.003 | 0.018 | 0.71 | 0.61 | 0.69 | 0.43 | 0.77 | 0.56 |
| 4 | 30560475 | rs7678701 | 1.411 | -1.042 | 0.19 | 1.44 | -0.103 | 0.13 | 0.13 | 0 | 0.38 | 0.13 | 0.4 | 0.15 | 0.48 | 0.33 |
| 4 | 30569583 | rs10014617 | -1.221 | -1.088 | 0.117 | 1.606 | -0.043 | -0.098 | -0.098 | 0 | 0.56 | 0.93 | 0.52 | 0.55 | 0.47 | 0.5 |
| 4 | 30572878 | rs7682155 | -0.068 | -1.133 | 0.434 | 1.662 | -0.015 | 0.026 | 0.026 | 0 | 0.22 | 0.31 | 0.21 | 0.3 | 0.18 | 0.28 |
| 4 | 30574424 | rs6847353 | 0.782 | -1.234 | 0.461 | 1.541 | -0.005 | 0 | 0 | 0 | 0.68 | 0.64 | 0.68 | 0.41 | 0.74 | 0.52 |
| 4 | 30575515 | rs1512160 | -1.161 | -1.249 | 0.419 | 1.951 | -0.002 | 0.031 | 0.031 | 0 | 0.93 | 0.94 | 0.95 | 0.74 | 0.94 | 0.85 |
| 4 | 30575592 | rs10028403 | 0.772 | -1.262 | 0.501 | 1.835 | -0.014 | -0.047 | -0.047 | 0.064 | 0.3 | 0.25 | 0.23 | 0.13 | 0.22 | 0.21 |
| 4 | 30576335 | rs10019706 | 0.035 | -1.27 | 0.551 | 1.86 | 0.011 | 0.206 | 0.206 | 0 | 0.89 | 0.7 | 0.88 | 0.69 | 0.93 | 0.84 |
| 4 | 30577159 | rs10517209 | 0.61 | -1.271 | 0.547 | 1.815 | -0.004 | 0.003 | 0.003 | 0 | 0.68 | 0.65 | 0.73 | 0.46 | 0.75 | 0.54 |
| 4 | 30578175 | rs13131444 | 1.341 | -1.261 | 0.538 | 1.73 | -0.011 | 0.018 | 0.018 | 0 | 0.38 | 0.38 | 0.46 | 0.28 | 0.52 | 0.32 |
| 4 | 30579183 | rs7687443 | 1.568 | -1.256 | 0.135 | 1.744 | -0.126 | 0.125 | 0.125 | 0.026 | 0.36 | 0.07 | 0.41 | 0.15 | 0.52 | 0.31 |
| 4 | 30591791 | rs7690439 | 0.649 | -1.172 | 0.218 | 1.689 | -0.029 | -0.034 | -0.034 | 0 | 0.32 | 0.48 | 0.27 | 0.26 | 0.22 | 0.22 |
| 4 | 30597133 | rs10019377 | 0.295 | -1.112 | -0.181 | 1.751 | -0.005 | -0.065 | -0.065 | 0.789 | 0.73 | 0.9 | 0.76 | 0.44 | 0.75 | 0.54 |
| 4 | 30599882 | rs9994236 | -0.31 | -1.091 | -0.106 | 1.77 | 0.007 | 0.139 | 0.139 | 0 | 0.89 | 0.72 | 0.93 | 0.71 | 0.93 | 0.85 |
| 4 | 30610801 | rs10939307 | 0.309 | -1.041 | -0.442 | 1.634 | -0.002 | -0.083 | -0.083 | 0 | 0.2 | 0.06 | 0.19 | 0.29 | 0.19 | 0.32 |
| 4 | 30612026 | rs12374222 | 0.309 | -1.082 | -0.442 | 1.634 | 0.002 | -0.083 | -0.083 | 2.465 | 0.2 | 0.06 | 0.19 | 0.29 | 0.19 | 0.32 |
| 4 | 30613111 | rs13115431 | 1.486 | -0.928 | -0.445 | 1.41 | -0.185 | 0.071 | 0.071 | 0 | 0.35 | 0.03 | 0.42 | 0.17 | 0.46 | 0.31 |
| 4 | 30625276 | rs12511839 | 1.181 | -1.063 | -0.368 | 1.475 | -0.083 | 0.047 | 0.047 | 0 | 0.38 | 0.13 | 0.48 | 0.28 | 0.52 | 0.32 |
| 4 | 30629161 | rs10939310 | -0.498 | -1.155 | -0.079 | 1.656 | 0.043 | 0.027 | 0.027 | 0 | 0.52 | 0.54 | 0.44 | 0.44 | 0.41 | 0.53 |
| 4 | 30629389 | rs7694762 | 1.556 | -1.135 | -0.285 | 1.646 | -0.192 | 0.066 | 0.066 | 0 | 0.35 | 0.03 | 0.42 | 0.17 | 0.47 | 0.32 |
| 4 | 30630382 | rs767478 | 0.792 | -1.265 | -0.04 | 1.581 | -0.035 | -0.093 | -0.093 | 0.953 | 0.31 | 0.31 | 0.24 | 0.14 | 0.22 | 0.21 |
| 4 | 30633334 | rs1906110 | 0.692 | -1.107 | -0.042 | 1.534 | -0.003 | 0.108 | 0.108 | 0 | 0.44 | 0.14 | 0.54 | 0.28 | 0.6 | 0.35 |
| 4 | 30633835 | rs4692473 | -0.206 | -1.06 | -0.029 | 1.291 | -0.014 | -0.004 | -0.004 | 0 | 0.44 | 0.35 | 0.37 | 0.44 | 0.33 | 0.5 |
| 4 | 30639658 | rs10015964 | -0.315 | -0.563 | -0.021 | 0.926 | 0.014 | -0.059 | -0.059 | 0.112 | 0.11 | 0.23 | 0.09 | 0 | 0.14 | 0.04 |
| 4 | 30640272 | rs13109270 | 1.288 | -0.937 | -0.918 | 1.084 | -0.191 | -0.002 | -0.002 | 0.299 | 0.36 | 0.02 | 0.42 | 0.21 | 0.45 | 0.34 |
| 4 | 30641018 | rs13115999 | -1.092 | -0.784 | -0.473 | 1.622 | 0 | 0.039 | 0.039 | 0 | 0.93 | 0.94 | 0.95 | 0.73 | 0.94 | 0.85 |
| 4 | 30645543 | rs6448713 | -0.708 | -0.881 | -0.496 | 1.58 | 0 | 0.015 | 0.015 | 0 | 0.91 | 0.93 | 0.95 | 0.73 | 0.94 | 0.85 |
| 4 | 30645986 | rs6821889 | -0.9 | -1.136 | -0.111 | 1.288 | -0.012 | 0.014 | 0.014 | 0.04 | 0.25 | 0.5 | 0.24 | 0.27 | 0.25 | 0.3 |
| 4 | 30647182 | rs16883928 | -0.836 | -1.196 | 0.563 | 1.253 | -0.016 | 0.018 | 0.018 | 0.048 | 0.24 | 0.48 | 0.23 | 0.27 | 0.23 | 0.31 |
| 4 | 30649819 | rs6848945 | 0.191 | -0.87 | 0.597 | 1.108 | 0.015 | -0.042 | -0.042 | 0 | 0.09 | 0.19 | 0.08 | 0 | 0.12 | 0.04 |

| | | | | | | | | | | | | | | | | |
|---|---|---|---|---|---|---|---|---|---|---|---|---|---|---|---|---|
| 4 | 30652381 | rs1486785 | 0.408 | -0.968 | 0.461 | 1.168 | 0.012 | 0.001 | 0.001 | 0 | 0.49 | 0.35 | 0.57 | 0.46 | 0.55 | 0.6 |
| 4 | 30658535 | rs6820780 | 0.392 | -0.994 | 0.288 | 1.173 | 0.031 | 0.041 | 0.041 | 0 | 0.5 | 0.22 | 0.6 | 0.58 | 0.56 | 0.61 |
| 4 | 30666011 | rs7658553 | 0.206 | -0.929 | 0.342 | 1.081 | 0.02 | -0.047 | -0.047 | 0 | 0.09 | 0.28 | 0.09 | 0.01 | 0.12 | 0.04 |
| 4 | 30668535 | rs13146789 | 1.844 | -1.086 | 0.121 | 1.147 | -0.179 | -0.031 | -0.031 | 0 | 0.33 | 0.03 | 0.42 | 0.22 | 0.44 | 0.34 |
| 4 | 30669244 | rs7654222 | -0.193 | -1.129 | 0.337 | 1.14 | 0.031 | 0.067 | 0.067 | 0.008 | 0.57 | 0.76 | 0.67 | 0.86 | 0.65 | 0.77 |
| 4 | 30669328 | rs747606 | 1.399 | -1.118 | 0.194 | 1.136 | -0.037 | 0.036 | 0.036 | 0 | 0.38 | 0.28 | 0.46 | 0.23 | 0.51 | 0.36 |
| 4 | 30677046 | rs7671654 | -1.982 | -1.134 | 0.161 | 1.053 | -0.001 | 0.016 | 0.016 | 0 | 0.95 | 0.99 | 0.96 | 1 | 0.97 | 0.99 |
| 4 | 30691142 | rs4692481 | 0.223 | -1.145 | 0.229 | 1.147 | 0.01 | -0.024 | -0.024 | 0.141 | 0.16 | 0.12 | 0.18 | 0.36 | 0.13 | 0.26 |
| 4 | 30703243 | rs4270551 | -0.169 | -1.1 | 0.197 | 1.027 | -0.014 | 0.063 | 0.063 | 0 | 0.88 | 0.99 | 0.91 | 1 | 0.9 | 0.96 |
| 4 | 30710951 | rs6855728 | -0.245 | -1.383 | 0.225 | 0.852 | 0.01 | 0.006 | 0.006 | 0 | 0.62 | 0.73 | 0.7 | 0.87 | 0.69 | 0.77 |
| 4 | 30711230 | rs12163915 | -0.12 | -1.366 | 0.372 | 1.055 | -0.014 | 0.04 | 0.04 | 0 | 0.17 | 0.21 | 0.2 | 0.37 | 0.14 | 0.27 |
| 4 | 30718333 | rs9291546 | -0.252 | -1.365 | 0.362 | 0.976 | -0.009 | 0.022 | 0.022 | 0.207 | 0.17 | 0.32 | 0.17 | 0.25 | 0.16 | 0.24 |
| 4 | 30725696 | rs1047012 | 0.928 | -1.35 | 0.131 | 1.202 | 0.002 | -0.064 | -0.064 | 0 | 0.11 | 0.02 | 0.14 | 0.31 | 0.11 | 0.25 |
| 4 | 30725987 | rs977931 | 0.371 | -1.463 | 0.237 | 1.149 | -0.014 | 0.056 | 0.056 | 0 | 0.14 | 0.2 | 0.16 | 0.32 | 0.11 | 0.25 |
| 4 | 30727233 | rs2121658 | 1.002 | -1.676 | 0.148 | 0.99 | 0.009 | 0.001 | 0.001 | 0.06 | 0.44 | 0.25 | 0.51 | 0.54 | 0.54 | 0.52 |
| 4 | 30728507 | rs3775331 | 1.375 | -1.671 | 0.046 | 1.149 | -0.114 | -0.05 | -0.05 | 0 | 0.41 | 0.03 | 0.48 | 0.34 | 0.53 | 0.35 |
| 4 | 30732274 | rs3756025 | 0.591 | -1.642 | 0.136 | 1.094 | -0.012 | 0.07 | 0.07 | 0 | 0.13 | 0.2 | 0.15 | 0.32 | 0.12 | 0.25 |
| 4 | 30736301 | rs13105058 | 0.278 | -1.67 | 0.166 | 1.025 | -0.008 | -0.05 | -0.05 | 1.264 | 0.56 | 0.45 | 0.67 | 0.87 | 0.66 | 0.77 |
| 4 | 30736724 | rs13111136 | 0.796 | -1.582 | 0.267 | 0.971 | 0.003 | 0.007 | 0.007 | 0.012 | 0.46 | 0.39 | 0.53 | 0.6 | 0.55 | 0.52 |
| 4 | 30737146 | rs28420079 | 0.704 | -1.571 | 0.268 | 0.978 | 0.007 | 0.013 | 0.013 | 0 | 0.47 | 0.4 | 0.53 | 0.6 | 0.55 | 0.52 |
| 4 | 30749654 | rs9291547 | -1.115 | -1.359 | 0.045 | 0.626 | -0.043 | -0.063 | -0.063 | 0.02 | 0.56 | 0.93 | 0.55 | 0.51 | 0.48 | 0.49 |
| 4 | 30752187 | rs10003467 | -0.076 | -1.408 | 0.208 | 0.554 | -0.025 | 0.046 | 0.046 | 0 | 0.2 | 0.35 | 0.19 | 0.3 | 0.16 | 0.25 |
| 4 | 30767675 | rs4380480 | -0.842 | -0.81 | 0.049 | 0.541 | 0.021 | 0.016 | 0.016 | 0.004 | 0.52 | 0.77 | 0.48 | 0.61 | 0.46 | 0.63 |
| 4 | 30785307 | rs2121659 | 0.105 | -0.798 | 0.086 | 0.449 | -0.014 | -0.095 | -0.095 | 0 | 0.35 | 0.54 | 0.29 | 0.09 | 0.32 | 0.24 |
| 4 | 30796107 | rs4557224 | -0.961 | -0.739 | 0.007 | 0.513 | -0.016 | 0.012 | 0.012 | 0.084 | 0.54 | 0.78 | 0.5 | 0.61 | 0.49 | 0.62 |
| 4 | 30799850 | rs2310224 | 0.478 | -0.773 | 0.117 | 0.592 | -0.005 | 0.012 | 0.012 | 0.067 | 0.17 | 0.16 | 0.18 | 0.3 | 0.12 | 0.21 |
| 4 | 30807244 | rs11732391 | 0.441 | -0.469 | 0.033 | 0.41 | -0.027 | 0.027 | 0.027 | 0 | 0.08 | 0.02 | 0.07 | 0.01 | 0.13 | 0.05 |
| 4 | 30810421 | rs1348923 | 0.522 | -0.501 | 0.082 | 0.407 | -0.022 | -0.009 | -0.009 | 1.002 | 0.29 | 0.57 | 0.24 | 0.31 | 0.25 | 0.37 |
| 4 | 30810487 | rs6847642 | -1.929 | 0.532 | -0.513 | -0.078 | -0.051 | 0.068 | 0.068 | 0 | 0.88 | 0.99 | 0.89 | 1 | 0.85 | 0.97 |
| 4 | 30811774 | rs6812005 | -1.389 | -0.014 | -0.23 | 0.283 | -0.154 | -0.008 | -0.008 | 0.364 | 0.58 | 0.98 | 0.56 | 0.73 | 0.52 | 0.63 |
| 4 | 30814699 | rs2121656 | 1.033 | -0.09 | -0.042 | 0.268 | -0.023 | -0.093 | -0.093 | 0 | 0.25 | 0.39 | 0.22 | 0.09 | 0.22 | 0.2 |
| 4 | 30815089 | rs13138213 | 0.531 | -0.078 | -0.3 | 0.274 | 0.002 | 0.001 | 0.001 | 0.339 | 0.3 | 0.7 | 0.25 | 0.31 | 0.25 | 0.37 |
| 4 | 30827805 | rs4619848 | -0.969 | 0.117 | -0.612 | 0.295 | -0.108 | -0.087 | -0.087 | 0 | 0.54 | 0.93 | 0.5 | 0.57 | 0.47 | 0.61 |
| 4 | 30828781 | rs4692105 | -0.655 | 0.026 | -0.628 | 0.268 | 0.048 | -0.01 | -0.01 | 0.604 | 0.51 | 0.87 | 0.46 | 0.58 | 0.41 | 0.6 |
| 4 | 30830285 | rs11724608 | 1.22 | 0.036 | -0.543 | 0.372 | -0.02 | 0.021 | 0.021 | 0 | 0.06 | 0.01 | 0.06 | 0 | 0.09 | 0.03 |
| 4 | 30832836 | rs1447373 | -0.854 | -0.046 | -0.217 | 0.347 | -0.003 | 0.089 | 0.089 | 0.392 | 0.28 | 0.45 | 0.28 | 0.48 | 0.24 | 0.41 |
| 4 | 30843533 | rs1345148 | -0.685 | 0.429 | -0.277 | 0.357 | -0.049 | -0.01 | -0.01 | 0 | 0.42 | 0.94 | 0.41 | 0.37 | 0.39 | 0.42 |
| 4 | 30845667 | rs6816969 | -0.551 | 0.496 | -0.211 | 0.397 | -0.014 | 0.004 | 0.004 | 1.397 | 0.39 | 0.82 | 0.39 | 0.37 | 0.38 | 0.41 |
| 4 | 30862363 | rs10517220 | 0.819 | 0.591 | -0.064 | 0.427 | -0.009 | 0.005 | 0.005 | 0 | 0.07 | 0.04 | 0.06 | 0.01 | 0.09 | 0.03 |
| 4 | 30864486 | rs13150968 | 0.812 | 0.594 | 0.005 | 0.302 | 0.003 | -0.005 | -0.005 | 0 | 0.29 | 0.24 | 0.21 | 0.07 | 0.28 | 0.21 |
| 4 | 30866392 | rs10001740 | -0.651 | 0.496 | -0.009 | 0.352 | -0.027 | 0.052 | 0.052 | 1.003 | 0.44 | 0.84 | 0.4 | 0.33 | 0.42 | 0.42 |
| 4 | 30872098 | rs4422383 | -0.278 | 0.529 | -0.008 | 0.197 | 0.003 | -0.029 | -0.029 | 0.225 | 0.39 | 0.44 | 0.34 | 0.23 | 0.41 | 0.26 |
| 4 | 30874445 | rs4269140 | -0.071 | 0.528 | -0.007 | 0.175 | 0.002 | -0.086 | -0.086 | 0.04 | 0.37 | 0.44 | 0.3 | 0.08 | 0.37 | 0.25 |
| 4 | 30893493 | rs11937211 | -0.255 | 0.712 | -0.141 | 0.409 | 0.029 | 0.02 | 0.02 | 0 | 0.1 | 0.21 | 0.13 | 0.16 | 0.13 | 0.04 |
| 4 | 30922078 | rs11735048 | -1.013 | 0.506 | -0.422 | 0.157 | 0.007 | -0.002 | -0.002 | 0.002 | 0.72 | 0.66 | 0.68 | 0.77 | 0.7 | 0.67 |
| 4 | 30943209 | rs4293757 | 1.023 | 0.484 | -0.347 | -0.04 | -0.008 | -0.076 | -0.076 | 0 | 0.31 | 0.35 | 0.22 | 0.06 | 0.27 | 0.21 |
| 4 | 30947714 | rs12508739 | -0.249 | 0.502 | -0.65 | 0.149 | 0 | 0 | 0 | 0 | 0.44 | 0.58 | 0.4 | 0.49 | 0.46 | 0.44 |
| 4 | 30970763 | rs7659873 | -0.634 | 0.552 | -1.148 | 0.178 | -0.089 | -0.05 | -0.05 | 1.227 | 0.64 | 0.86 | 0.61 | 0.6 | 0.55 | 0.65 |
| 4 | 30976208 | rs10027212 | 0.313 | 0.497 | -1.005 | 0.19 | 0.022 | 0.011 | 0.011 | 0.657 | 0.38 | 0.78 | 0.33 | 0.33 | 0.32 | 0.37 |
| 4 | 30977401 | rs6822671 | 0.313 | 0.497 | -1.005 | 0.19 | 0.022 | 0.011 | 0.011 | 0.657 | 0.38 | 0.78 | 0.33 | 0.33 | 0.32 | 0.37 |
| 4 | 30984320 | rs6847898 | 1.126 | 0.191 | -1.069 | 0.147 | 0.015 | 0.004 | 0.004 | 0 | 0.3 | 0.28 | 0.21 | 0.06 | 0.23 | 0.2 |
| 4 | 30991363 | rs13102223 | 0.547 | 0.352 | -2.029 | 0.4 | -0.03 | -0.055 | -0.055 | 0 | 0.23 | 0.02 | 0.24 | 0.26 | 0.22 | 0.3 |
| 4 | 30996285 | rs6832127 | -0.081 | 1.187 | -2.138 | 0.784 | -0.005 | -0.005 | -0.005 | 0 | 0.11 | 0.09 | 0.1 | 0.18 | 0.16 | 0.05 |
| 4 | 30996811 | rs6448730 | -0.455 | 1.143 | -2.181 | 0.462 | -0.119 | -0.03 | -0.03 | 0 | 0.71 | 0.97 | 0.67 | 0.8 | 0.65 | 0.73 |
| 4 | 30997909 | rs10939325 | -0.44 | 1.148 | -2.178 | 0.456 | -0.119 | -0.034 | -0.034 | 0 | 0.71 | 0.98 | 0.67 | 0.8 | 0.65 | 0.73 |
| 4 | 31016990 | rs7696101 | -0.697 | 0.622 | -1.294 | 0.69 | -0.004 | 0.012 | 0.012 | 0.87 | 0.28 | 0.7 | 0.43 | 0.45 | 0.26 | 0.35 |
| 4 | 31025354 | rs10939326 | -0.386 | 0.341 | -1.168 | 0.88 | -0.011 | 0.021 | 0.021 | 2.221 | 0.26 | 0.7 | 0.4 | 0.45 | 0.21 | 0.38 |
| 4 | 31032608 | rs10003973 | 0.016 | 0.762 | -1.758 | 0.662 | -0.025 | -0.02 | -0.02 | 0 | 0.31 | 0.14 | 0.19 | 0.32 | 0.31 | 0.35 |
| 4 | 31037602 | rs6837499 | -0.365 | -0.153 | -1.364 | 0.476 | -0.002 | -0.003 | -0.003 | 0 | 0.23 | 0.24 | 0.41 | 0.21 | 0.21 | 0.39 |
| 4 | 31059015 | rs7661111 | -0.063 | -0.336 | -0.934 | 0.723 | -0.007 | 0.03 | 0.03 | 0 | 0.13 | 0.22 | 0.07 | 0.25 | 0.09 | 0.08 |
| 4 | 31069820 | rs1463851 | -1.307 | -0.622 | -2.011 | -0.145 | -0.137 | -0.081 | -0.081 | 0.015 | 0.67 | 0.94 | 0.63 | 0.47 | 0.52 | 0.68 |
| 4 | 31077545 | rs6820873 | -0.5 | -0.816 | -1.993 | -0.233 | 0.06 | 0.15 | 0.15 | 0.566 | 0.5 | 0.79 | 0.62 | 0.78 | 0.43 | 0.63 |
| 4 | 31080447 | rs3947332 | -1.149 | -0.311 | -1.778 | 0.164 | -0.097 | 0.021 | 0.021 | 0 | 0.69 | 0.87 | 0.69 | 0.78 | 0.54 | 0.79 |
| 4 | 31095678 | rs2221128 | -0.344 | -1.206 | -1.595 | 0.267 | -0.006 | 0.033 | 0.033 | 0 | 0.17 | 0.06 | 0.2 | 0.08 | 0.19 | 0.2 |
| 4 | 31100499 | rs2201945 | -1.045 | -1.44 | -1.442 | 0.753 | 0.012 | -0.002 | -0.002 | 0 | 0.38 | 0.65 | 0.39 | 0.45 | 0.29 | 0.5 |
| 4 | 31104796 | rs10010739 | -0.94 | -1.546 | -1.415 | 0.638 | 0.008 | 0.006 | 0.006 | 0 | 0.37 | 0.65 | 0.49 | 0.77 | 0.29 | 0.55 |
| 4 | 31110337 | rs11935515 | -0.535 | -1.888 | -1.068 | 0.507 | 0.03 | 0.079 | 0.079 | 0 | 0.56 | 0.77 | 0.45 | 0.78 | 0.43 | 0.7 |
| 4 | 31117942 | rs3857021 | -0.791 | -1.76 | -1.376 | 0.573 | -0.045 | 0.068 | 0.068 | 0.84 | 0.58 | 0.95 | 0.47 | 0.79 | 0.44 | 0.71 |
| 4 | 31122429 | rs16884372 | -0.813 | -1.84 | -0.725 | 1.242 | 0.034 | 0.468 | 0.468 | 0.014 | 0.05 | 0.3 | 0.13 | 0.46 | 0.03 | 0.18 |

| | | | | | | | | | | | | | | | | |
|---|---|---|---|---|---|---|---|---|---|---|---|---|---|---|---|---|
| 4 | 31135624 | rs12507539 | -0.824 | -2.015 | -0.448 | 1.15 | 0.045 | 0.505 | 0.505 | 0 | 0.05 | 0.31 | 0.13 | 0.46 | 0.03 | 0.18 |
| 4 | 31148688 | rs16884451 | -0.54 | -2.148 | -0.854 | 0.635 | 0.003 | -0.001 | -0.001 | 0 | 0.54 | 0.54 | 0.42 | 0.58 | 0.44 | 0.62 |
| 4 | 31150900 | rs16884471 | -1.346 | -2.119 | -0.913 | 0.603 | -0.105 | -0.168 | -0.168 | 1.456 | 0.62 | 0.97 | 0.48 | 0.58 | 0.47 | 0.65 |
| 4 | 31151060 | rs16884473 | -0.338 | -1.86 | -0.74 | 0.525 | 0.01 | 0.005 | 0.005 | 0 | 0.5 | 0.52 | 0.36 | 0.52 | 0.42 | 0.57 |
| 4 | 31157943 | rs1463844 | -0.187 | -2.304 | -1.307 | 0.051 | 0.042 | 0.062 | 0.062 | 0 | 0.43 | 0.67 | 0.29 | 0.32 | 0.38 | 0.51 |
| 4 | 31162659 | rs13340304 | 0.106 | -2.922 | -1.353 | -0.47 | 0.009 | -0.043 | -0.043 | 0 | 0.36 | 0.38 | 0.17 | 0 | 0.29 | 0.2 |
| 4 | 31166902 | rs6833999 | -0.744 | -2.55 | -1.283 | -0.164 | -0.008 | -0.027 | -0.027 | 0.451 | 0.05 | 0.16 | 0.02 | 0 | 0.02 | 0.03 |
| 4 | 31170286 | rs11938193 | -0.839 | -2.759 | -1.88 | 0.181 | 0.012 | -0.027 | -0.027 | 0 | 0.57 | 0.9 | 0.41 | 0.48 | 0.45 | 0.64 |
| 4 | 31183845 | rs6448753 | -1.183 | -2.772 | -1.671 | 0.321 | 0.036 | -0.03 | -0.03 | 0.412 | 0.28 | 0.79 | 0.21 | 0.47 | 0.12 | 0.29 |
| 4 | 31187185 | rs2036247 | -0.61 | -2.546 | -0.934 | 0.51 | -0.058 | 0.059 | 0.059 | 0.089 | 0.22 | 0.5 | 0.17 | 0.38 | 0.11 | 0.25 |
| 4 | 31187606 | rs6828314 | -0.383 | -2.46 | -0.903 | 0.619 | -0.065 | 0.019 | 0.019 | 0.517 | 0.19 | 0.45 | 0.15 | 0.23 | 0.08 | 0.22 |
| 4 | 31190168 | rs17779235 | -0.198 | -1.627 | -1.74 | 0.582 | -0.034 | 0.015 | 0.015 | 0.042 | 0.1 | 0.01 | 0.07 | 0.07 | 0.12 | 0.2 |
| 4 | 31196514 | rs6448754 | -0.92 | -1.731 | -1.933 | 0.399 | -0.089 | -0.1 | -0.1 | 0 | 0.67 | 0.91 | 0.42 | 0.53 | 0.51 | 0.67 |
| 4 | 31200828 | rs11930062 | -0.281 | -1.887 | -1.347 | 0.501 | 0.065 | -0.033 | -0.033 | 0 | 0.56 | 0.85 | 0.36 | 0.51 | 0.4 | 0.47 |
| 4 | 31209228 | rs11933756 | -0.997 | -1.687 | -0.183 | 0.146 | -0.039 | -0.046 | -0.046 | 1.954 | 0.24 | 0.48 | 0.18 | 0.13 | 0.14 | 0.12 |
| 4 | 31209875 | rs1463842 | -0.601 | -2.142 | -0.489 | 0.242 | -0.001 | 0 | 0 | 2.04 | 0.65 | 0.67 | 0.81 | 0.49 | 0.75 | 0.45 |
| 4 | 31211978 | rs11937481 | 0.115 | -2.235 | 0.073 | -0.276 | -0.017 | 0.021 | 0.021 | 0.217 | 0.12 | 0.35 | 0.07 | 0.11 | 0.06 | 0.09 |
| 4 | 31216675 | rs6849452 | -1.256 | -2.37 | 0.072 | -0.475 | -0.008 | -0.021 | -0.021 | 0 | 0.34 | 0.44 | 0.25 | 0.13 | 0.29 | 0.19 |
| 4 | 31217251 | rs1499466 | -0.205 | -2.404 | 0.023 | -0.544 | -0.008 | -0.025 | -0.025 | 0 | 0.14 | 0.15 | 0.12 | 0.02 | 0.08 | 0.03 |
| 4 | 31217377 | rs17723773 | 1.129 | -2.27 | -0.05 | -0.448 | -0.017 | 0.052 | 0.052 | 0 | 0.1 | 0.02 | 0.08 | 0 | 0.14 | 0.07 |
| 4 | 31217902 | rs4591553 | -0.146 | -2.458 | 0.225 | -0.429 | 0.009 | 0.017 | 0.017 | 0 | 0.19 | 0.13 | 0.11 | 0.04 | 0.16 | 0.16 |
| 4 | 31218368 | rs12331227 | -1.203 | -2.616 | 0.286 | -0.26 | 0.006 | 0.004 | 0.004 | 0 | 0.58 | 0.68 | 0.34 | 0.28 | 0.38 | 0.43 |
| 4 | 31219796 | rs6821356 | -1.052 | -2.593 | 0.298 | -0.283 | 0.01 | 0.009 | 0.009 | 0 | 0.57 | 0.68 | 0.34 | 0.28 | 0.38 | 0.42 |
| 4 | 31230228 | rs6820490 | 0.67 | -2.522 | -0.142 | -0.246 | 0.039 | 0.021 | 0.021 | 3.086 | 0.24 | 0.16 | 0.08 | 0.13 | 0.12 | 0.22 |
| 4 | 31232183 | rs10001703 | -0.083 | -2.371 | 0.462 | -0.242 | -0.005 | 0.003 | 0.003 | 0 | 0.14 | 0.35 | 0.12 | 0.09 | 0.14 | 0.11 |
| 4 | 31232890 | rs951011 | -1.109 | -2.655 | 0.218 | -0.421 | -0.014 | -0.02 | -0.02 | 0 | 0.34 | 0.35 | 0.23 | 0.15 | 0.24 | 0.21 |
| 4 | 31233979 | rs17724390 | 1.297 | -2.395 | -0.158 | -0.357 | 0.005 | 0.012 | 0.012 | 0 | 0.09 | 0.05 | 0.06 | 0 | 0.11 | 0.09 |
| 4 | 31235824 | rs1392056 | -1.203 | -2.481 | -0.138 | -0.313 | 0.012 | 0.009 | 0.009 | 0 | 0.6 | 0.67 | 0.32 | 0.27 | 0.36 | 0.44 |
| 4 | 31249967 | rs10002340 | 0.104 | -2.439 | -0.48 | -0.219 | 0.095 | 0.048 | 0.048 | 0.347 | 0.33 | 0.12 | 0.09 | 0.18 | 0.15 | 0.32 |
| 4 | 31254765 | rs4321596 | -0.649 | -2.485 | -0.732 | 0.206 | 0.087 | 0.044 | 0.044 | 0.24 | 0.45 | 0.66 | 0.23 | 0.63 | 0.21 | 0.61 |
| 4 | 31254780 | rs4624622 | 0.209 | -1.811 | -0.36 | 0.528 | 0.005 | 0.157 | 0.157 | 0 | 0.07 | 0.13 | 0.15 | 0.45 | 0.07 | 0.3 |
| 4 | 31262043 | rs6811209 | -0.819 | -1.796 | -0.681 | 0.34 | 0.013 | -0.033 | -0.033 | 0 | 0.31 | 0.71 | 0.3 | 0.49 | 0.22 | 0.4 |
| 4 | 31271863 | rs11735188 | 0.286 | -1.513 | -1.123 | 0.575 | 0.009 | 0.136 | 0.136 | 0 | 0.07 | 0.11 | 0.09 | 0.45 | 0.1 | 0.3 |
| 4 | 31277116 | rs6815632 | -0.371 | -1.392 | -1.562 | 0.359 | 0.002 | -0.096 | -0.096 | 1.174 | 0.36 | 0.87 | 0.3 | 0.5 | 0.29 | 0.43 |
| 4 | 31302011 | rs4441721 | -0.815 | -0.401 | -1.202 | 0.32 | -0.183 | -0.07 | -0.07 | 5.658 | 0.72 | 0.97 | 0.44 | 0.65 | 0.5 | 0.75 |
| 4 | 31306116 | rs9291566 | -0.157 | -0.521 | -0.652 | 0.464 | 0.012 | 0.012 | 0.012 | 0 | 0.57 | 0.62 | 0.33 | 0.59 | 0.36 | 0.64 |
| 4 | 31316264 | rs11935585 | -1.323 | -0.286 | -0.232 | -0.033 | -0.012 | 0.04 | 0.04 | 1.078 | 0.09 | 0.25 | 0.05 | 0.19 | 0.05 | 0.14 |
| 4 | 31319406 | rs4303942 | -0.102 | -0.239 | -1.039 | -1.138 | -0.019 | 0.086 | 0.086 | 1.751 | 0.54 | 0.74 | 0.39 | 0.28 | 0.52 | 0.5 |
| 4 | 31322470 | rs1156910 | -0.174 | -0.328 | -0.85 | -1.304 | 0 | 0.003 | 0.003 | 0.289 | 0.4 | 0.36 | 0.38 | 0.69 | 0.36 | 0.45 |
| 4 | 31323469 | rs16884777 | 1.202 | -1.203 | -0.955 | -1.826 | 0.007 | 0.002 | 0.002 | 0 | 0.15 | 0.12 | 0.1 | 0.15 | 0.11 | 0.23 |
| 4 | 31324283 | rs10029975 | 0.406 | -0.954 | -0.98 | -1.854 | 0.001 | 0.002 | 0.002 | 0.274 | 0.39 | 0.35 | 0.37 | 0.69 | 0.33 | 0.45 |
| 4 | 31333616 | rs10025805 | 0.69 | -1.059 | -1.062 | -1.771 | 0.01 | 0 | 0 | 1.413 | 0.36 | 0.26 | 0.35 | 0.59 | 0.31 | 0.39 |
| 4 | 31334007 | rs6820923 | -0.272 | -1.144 | -0.415 | -1.918 | 0.003 | 0 | 0 | 1.164 | 0.56 | 0.61 | 0.49 | 0.85 | 0.44 | 0.68 |
| 4 | 31335371 | rs10517227 | 0.848 | -1.397 | -0.774 | -2.225 | 0.004 | -0.001 | -0.001 | 0 | 0.27 | 0.24 | 0.31 | 0.69 | 0.23 | 0.41 |
| 4 | 31336704 | rs1991343 | -0.819 | -1.427 | -0.629 | -2.326 | 0.023 | 0.023 | 0.023 | 0.043 | 0.82 | 0.93 | 0.71 | 0.87 | 0.84 | 0.82 |
| 4 | 31338387 | rs13128378 | -0.073 | -1.177 | -0.424 | -2.248 | -0.006 | -0.051 | -0.051 | 0 | 0.54 | 0.52 | 0.47 | 0.85 | 0.43 | 0.68 |
| 4 | 31339067 | rs13135059 | 0.755 | -1.255 | -0.482 | -2.317 | 0.044 | -0.115 | -0.115 | 0.389 | 0.3 | 0.38 | 0.24 | 0.02 | 0.41 | 0.14 |
| 4 | 31340010 | rs6848921 | -0.022 | -1.204 | -0.47 | -2.237 | -0.006 | -0.051 | -0.051 | 0 | 0.54 | 0.52 | 0.47 | 0.85 | 0.43 | 0.68 |
| 4 | 31341126 | rs10033651 | -0.022 | -1.204 | -0.47 | -2.231 | -0.006 | -0.047 | -0.047 | 0.886 | 0.54 | 0.52 | 0.47 | 0.84 | 0.43 | 0.68 |
| 4 | 31341384 | rs1426985 | -0.047 | -0.883 | -0.482 | -2.243 | -0.003 | -0.049 | -0.049 | 0.041 | 0.55 | 0.52 | 0.5 | 0.84 | 0.47 | 0.69 |
| 4 | 31345976 | rs7694995 | -0.076 | -1.189 | -0.164 | -2.254 | -0.001 | -0.008 | -0.008 | 0 | 0.46 | 0.43 | 0.35 | 0.28 | 0.53 | 0.44 |
| 4 | 31346154 | rs7695413 | -1.228 | -1.198 | -0.193 | -2.264 | 0.003 | 0.007 | 0.007 | 0.011 | 0.85 | 0.89 | 0.72 | 0.88 | 0.84 | 0.83 |
| 4 | 31354090 | rs4692508 | -0.053 | -1.152 | -0.218 | -2.248 | -0.001 | -0.003 | -0.003 | 0 | 0.45 | 0.43 | 0.35 | 0.27 | 0.53 | 0.44 |
| 4 | 31357836 | rs933954 | 1.123 | -1.018 | -0.565 | -2.02 | -0.004 | 0.001 | 0.001 | 0 | 0.16 | 0.17 | 0.11 | 0.26 | 0.14 | 0.28 |
| 4 | 31358792 | rs16884897 | -0.134 | -0.959 | -0.599 | -2.125 | 0.003 | 0.026 | 0.026 | 0 | 0.54 | 0.4 | 0.47 | 0.85 | 0.46 | 0.69 |
| 4 | 31360789 | rs2081665 | -0.254 | -0.677 | -0.756 | -1.865 | 0.001 | 0.03 | 0.03 | 0.232 | 0.41 | 0.38 | 0.45 | 0.84 | 0.43 | 0.64 |
| 4 | 31364659 | rs4692511 | -1.401 | -0.565 | -0.806 | -1.902 | 0.002 | 0.006 | 0.006 | 0.005 | 0.86 | 0.9 | 0.75 | 0.88 | 0.85 | 0.83 |
| 4 | 31365792 | rs10017667 | 0.533 | -0.497 | -0.839 | -1.831 | -0.004 | -0.006 | -0.006 | 1.635 | 0.26 | 0.28 | 0.35 | 0.61 | 0.28 | 0.37 |
| 4 | 31366504 | rs16884929 | 1.423 | -0.519 | -1.144 | -1.799 | 0.008 | -0.003 | -0.003 | 0.061 | 0.14 | 0.1 | 0.09 | 0.15 | 0.13 | 0.23 |
| 4 | 31367210 | rs6854032 | 0.379 | -0.928 | -0.849 | -1.579 | -0.015 | -0.022 | -0.022 | 0 | 0.26 | 0.21 | 0.33 | 0.6 | 0.28 | 0.37 |
| 4 | 31367687 | rs7677024 | 0.147 | -1.058 | -0.802 | -1.546 | -0.006 | -0.002 | -0.002 | 0.014 | 0.27 | 0.26 | 0.32 | 0.68 | 0.29 | 0.41 |
| 4 | 31368170 | rs11943523 | 0.738 | -1.165 | -0.887 | -1.768 | -0.076 | -0.163 | -0.163 | 0 | 0.34 | 0.48 | 0.18 | 0.01 | 0.14 | 0.1 |
| 4 | 31368529 | rs16884956 | 1.115 | -1.25 | -0.954 | -1.82 | -0.061 | -0.144 | -0.144 | 0 | 0.29 | 0.37 | 0.17 | 0.01 | 0.13 | 0.07 |
| 4 | 31370788 | rs10000543 | -0.894 | -0.709 | -1.653 | -1.548 | 0.007 | -0.006 | -0.006 | 0 | 0.83 | 0.88 | 0.73 | 0.77 | 0.84 | 0.79 |
| 4 | 31375593 | rs13140925 | -1.189 | -0.701 | -1.649 | -1.544 | 0.01 | 0.008 | 0.008 | 0 | 0.84 | 0.91 | 0.74 | 0.85 | 0.86 | 0.83 |
| 4 | 31377722 | rs1834709 | -0.01 | -0.693 | -1.665 | -1.57 | -0.008 | 0.011 | 0.011 | 1.093 | 0.6 | 0.71 | 0.49 | 0.6 | 0.41 | 0.47 |
| 4 | 31377963 | rs1834707 | 0.668 | -0.552 | -1.276 | -1.7 | -0.051 | -0.129 | -0.129 | 0.111 | 0.34 | 0.5 | 0.18 | 0.01 | 0.14 | 0.1 |
| 4 | 31381690 | rs1834705 | 0.11 | 0.49 | -1.59 | -0.904 | -0.068 | 0.022 | 0.022 | 0.013 | 0.23 | 0.12 | 0.23 | 0.17 | 0.43 | 0.31 |
| 4 | 31382580 | rs13140921 | 0.104 | 0.494 | -1.632 | -0.902 | -0.041 | 0.016 | 0.016 | 0.002 | 0.23 | 0.16 | 0.23 | 0.17 | 0.43 | 0.31 |

| | | | | | | | | | | | | | | | |
|---|---|---|---|---|---|---|---|---|---|---|---|---|---|---|---|
| 4 | 31397618 | rs6448771 | 0.131 | 0.153 | -1.382 | 0.031 | -0.004 | -0.001 | -0.001 | 0.133 | 0.7 | 0.86 | 0.48 | 0.3 | 0.68 | 0.54 |
| 4 | 31405546 | rs7689511 | -1.164 | -0.317 | -0.589 | 1.041 | -0.014 | 0.032 | 0.032 | 0 | 0.16 | 0.25 | 0.09 | 0.3 | 0.11 | 0.16 |
| 4 | 31413759 | rs7700198 | -1.188 | -2.173 | -0.377 | 0.091 | 0.013 | 0.006 | 0.006 | 4.114 | 0.35 | 0.28 | 0.16 | 0.08 | 0.2 | 0.22 |
| 4 | 31415607 | rs9684021 | -1.671 | -2.524 | 0.566 | 0.047 | 0.046 | 0.027 | 0.027 | 3.439 | 0.41 | 0.68 | 0.2 | 0.29 | 0.19 | 0.3 |
| 4 | 31417540 | rs16885042 | -0.745 | -2.645 | 1.071 | -0.053 | -0.022 | -0.063 | -0.063 | 0.02 | 0.13 | 0.22 | 0.05 | 0 | 0.07 | 0.06 |
| 4 | 31423493 | rs10517228 | -0.616 | -2.822 | 1.154 | -0.098 | -0.065 | -0.058 | -0.058 | 0 | 0.26 | 0.36 | 0.15 | 0.08 | 0.12 | 0.18 |
| 4 | 31427408 | rs1966546 | -1.487 | -2.615 | 1.299 | 0.031 | -0.054 | 0.024 | 0.024 | 0.102 | 0.17 | 0.4 | 0.07 | 0.21 | 0.07 | 0.12 |
| 4 | 31436863 | rs6825297 | -1.918 | -2.618 | 1.61 | -0.001 | -0.062 | -0.035 | -0.035 | 0 | 0.19 | 0.6 | 0.09 | 0.11 | 0.07 | 0.12 |
| 4 | 31447488 | rs454864 | -0.757 | -3.311 | 0.707 | 0.574 | 0.007 | -0.015 | -0.015 | 0.522 | 0.8 | 0.83 | 0.6 | 0.58 | 0.83 | 0.85 |
| 4 | 31448474 | rs409441 | 1.489 | -3.796 | 1.031 | 0.296 | 0.145 | 0.031 | 0.031 | 0.29 | 0.52 | 0.12 | 0.44 | 0.43 | 0.71 | 0.58 |
| 4 | 31448956 | rs397718 | 1.489 | -3.796 | 1.085 | 0.296 | 0.142 | 0.034 | 0.034 | 0.29 | 0.52 | 0.13 | 0.44 | 0.43 | 0.71 | 0.58 |
| 4 | 31456071 | rs424917 | 0.317 | -3.811 | 1.434 | 0.309 | -0.026 | 0.022 | 0.022 | 0 | 0.67 | 0.56 | 0.55 | 0.58 | 0.77 | 0.79 |
| 4 | 31457293 | rs433149 | 1.448 | -3.765 | 0.947 | 0.313 | 0.146 | 0.036 | 0.036 | 0.256 | 0.52 | 0.12 | 0.43 | 0.42 | 0.71 | 0.58 |
| 4 | 31461728 | rs404532 | 1.264 | -3.847 | 0.913 | 0.323 | 0.123 | 0.009 | 0.009 | 0.657 | 0.55 | 0.12 | 0.44 | 0.42 | 0.72 | 0.61 |
| 4 | 31474310 | rs16885247 | -0.769 | -3.876 | 1.453 | 0.233 | -0.044 | 0.011 | 0.011 | 1.17 | 0.14 | 0.43 | 0.07 | 0.11 | 0.04 | 0.09 |
| 4 | 31480930 | rs401044 | 1.943 | -3.846 | 0.382 | 0.128 | -0.113 | 0.066 | 0.066 | 0 | 0.29 | 0.04 | 0.29 | 0.21 | 0.54 | 0.41 |
| 4 | 31481063 | rs11726010 | -0.699 | -3.65 | 0.744 | 0.16 | 0.007 | 0.006 | 0.006 | 0 | 0.11 | 0.09 | 0.07 | 0.05 | 0.06 | 0.14 |
| 4 | 31489510 | rs2925543 | 0.825 | -3.582 | 0.663 | 0.286 | 0.089 | 0.044 | 0.044 | 0.597 | 0.54 | 0.16 | 0.44 | 0.4 | 0.69 | 0.63 |
| 4 | 31489663 | rs2912616 | 0.815 | -3.547 | 0.919 | 0.168 | -0.007 | -0.001 | -0.001 | 0 | 0.55 | 0.46 | 0.48 | 0.45 | 0.67 | 0.68 |
| 4 | 31490705 | rs446149 | 0.937 | -3.519 | 1.107 | 0.027 | -0.006 | 0 | 0 | 0 | 0.52 | 0.5 | 0.45 | 0.56 | 0.61 | 0.61 |
| 4 | 31494159 | rs452052 | 0.032 | -3.621 | 0.962 | 0.073 | 0.015 | -0.003 | -0.003 | 1.488 | 0.64 | 0.72 | 0.49 | 0.67 | 0.71 | 0.77 |
| 4 | 31496548 | rs4145381 | 0.539 | -3.578 | 1.055 | 0.072 | 0.017 | -0.01 | -0.01 | 0.314 | 0.58 | 0.62 | 0.48 | 0.56 | 0.67 | 0.68 |
| 4 | 31510999 | rs10002573 | -0.73 | -3.541 | 0.842 | 0.095 | 0.058 | -0.005 | -0.005 | 0 | 0.64 | 0.81 | 0.54 | 0.67 | 0.78 | 0.78 |
| 4 | 31513263 | rs6448778 | 1.389 | -3.505 | 0.886 | 0.075 | -0.013 | -0.029 | -0.029 | 0.316 | 0.36 | 0.22 | 0.36 | 0.51 | 0.54 | 0.52 |
| 4 | 31525698 | rs10805304 | -1.069 | -3.611 | 0.687 | 0.055 | 0.068 | 0.053 | 0.053 | 0 | 0.67 | 0.89 | 0.65 | 0.76 | 0.79 | 0.81 |
| 4 | 31531497 | rs1598684 | 1.082 | -3.276 | 0.811 | 0.253 | -0.017 | -0.024 | -0.024 | 0 | 0.37 | 0.15 | 0.45 | 0.6 | 0.54 | 0.53 |
| 4 | 31559379 | rs1379127 | 0.146 | -3.149 | 0.914 | 0.211 | 0.014 | 0.017 | 0.017 | 0 | 0.49 | 0.55 | 0.51 | 0.71 | 0.63 | 0.66 |
| 4 | 31581408 | rs7678026 | 0.603 | -3.055 | 0.803 | 0.318 | 0.004 | 0.004 | 0.004 | 0 | 0.43 | 0.35 | 0.47 | 0.6 | 0.58 | 0.55 |
| 4 | 31590431 | rs7656200 | 0.631 | -3.074 | 0.787 | 0.325 | 0.003 | 0.003 | 0.003 | 0 | 0.43 | 0.35 | 0.47 | 0.59 | 0.57 | 0.56 |
| 4 | 31598534 | rs6830965 | 0.668 | -3.067 | 0.709 | 0.236 | 0.004 | -0.002 | -0.002 | 0 | 0.43 | 0.35 | 0.46 | 0.56 | 0.58 | 0.56 |
| 4 | 31599070 | rs10428460 | -0.255 | -3.237 | 0.82 | 0.148 | -0.023 | -0.002 | -0.002 | 0 | 0.12 | 0.21 | 0.06 | 0.13 | 0.08 | 0.12 |
| 4 | 31613185 | rs7690924 | -0.608 | -3.26 | 0.663 | 0.142 | 0.041 | 0.073 | 0.073 | 0 | 0.62 | 0.9 | 0.59 | 0.75 | 0.71 | 0.74 |
| 4 | 31621421 | rs1397500 | 0.562 | -3.321 | 0.787 | 0.082 | 0.003 | 0.004 | 0.004 | 0 | 0.46 | 0.43 | 0.47 | 0.69 | 0.6 | 0.61 |
| 4 | 31629820 | rs10939353 | -0.634 | -3.409 | 0.729 | 0.101 | 0.051 | 0.076 | 0.076 | 0 | 0.62 | 0.87 | 0.6 | 0.75 | 0.72 | 0.75 |
| 4 | 31641279 | rs11932796 | -0.83 | -3.37 | 0.862 | 0.101 | -0.024 | -0.012 | -0.012 | 1.494 | 0.15 | 0.27 | 0.11 | 0.06 | 0.1 | 0.12 |
| 4 | 31643126 | rs9992305 | 0.903 | -3.339 | 0.868 | 0.145 | 0.003 | -0.003 | -0.003 | 0 | 0.42 | 0.32 | 0.46 | 0.57 | 0.6 | 0.57 |
| 4 | 31660739 | rs11944702 | -2.2 | -3.32 | 0.968 | 0.171 | -0.018 | 0.101 | 0.101 | 0 | 0.09 | 0.27 | 0.07 | 0.19 | 0.05 | 0.12 |
| 4 | 31661814 | rs1510806 | 0.419 | -3.398 | 1.007 | 0.107 | 0.016 | 0.014 | 0.014 | 0 | 0.48 | 0.49 | 0.53 | 0.63 | 0.62 | 0.63 |
| 4 | 31666014 | rs16885609 | -0.056 | -3.189 | 0.782 | -0.045 | -0.003 | -0.024 | -0.024 | 0 | 0.1 | 0.11 | 0.05 | 0 | 0.07 | 0.06 |
| 4 | 31676985 | rs9999931 | -0.033 | -2.963 | 0.159 | -0.104 | 0.003 | 0.003 | 0.003 | 0 | 0.51 | 0.49 | 0.49 | 0.68 | 0.64 | 0.69 |
| 4 | 31678181 | rs4692118 | 0.029 | -2.971 | 0.195 | -0.021 | 0.022 | 0.004 | 0.004 | 0 | 0.49 | 0.39 | 0.48 | 0.56 | 0.64 | 0.63 |
| 4 | 31684540 | rs6831283 | -0.32 | -3.099 | -0.28 | -0.106 | 0.022 | 0.016 | 0.016 | 0 | 0.56 | 0.72 | 0.51 | 0.68 | 0.67 | 0.67 |
| 4 | 31698470 | rs4692526 | -1.004 | -3.191 | -0.617 | -0.183 | 0.039 | 0.036 | 0.036 | 0 | 0.65 | 0.86 | 0.59 | 0.74 | 0.73 | 0.77 |
| 4 | 31698958 | rs16885692 | 0.363 | -2.891 | -0.858 | -0.193 | 0.011 | 0.058 | 0.058 | 0 | 0.1 | 0.01 | 0.05 | 0 | 0.08 | 0.08 |
| 4 | 31700291 | rs16885694 | 0.498 | -2.536 | -0.484 | 0.038 | -0.008 | -0.013 | -0.013 | 0.019 | 0.37 | 0.24 | 0.42 | 0.56 | 0.54 | 0.52 |
| 4 | 31709060 | rs2192531 | -0.64 | -3.024 | -0.589 | -1.12 | -0.049 | -0.122 | -0.122 | 0 | 0.21 | 0.36 | 0.12 | 0.01 | 0.14 | 0.1 |
| 4 | 31709237 | rs12643894 | 1.171 | -2.801 | -1.36 | -1.24 | 0.061 | 0.021 | 0.021 | 0 | 0.47 | 0.09 | 0.48 | 0.6 | 0.59 | 0.59 |
| 4 | 31712529 | rs13133369 | -0.801 | -2.707 | -1.211 | -1.1 | 0.002 | -0.004 | -0.004 | 0 | 0.77 | 0.78 | 0.66 | 0.69 | 0.79 | 0.78 |
| 4 | 31715497 | rs7667509 | -0.359 | -2.396 | -1.26 | -1.031 | -0.024 | -0.054 | -0.054 | 0.501 | 0.14 | 0.47 | 0.08 | 0.08 | 0.09 | 0.11 |
| 4 | 31715825 | rs16885723 | 0.768 | -1.989 | -2.326 | -0.863 | -0.009 | -0.027 | -0.027 | 0 | 0.05 | 0.18 | 0.02 | 0 | 0.03 | 0.01 |
| 4 | 31718284 | rs10489008 | 0.599 | -1.484 | -3.333 | -0.799 | -0.074 | -0.074 | -0.074 | 0 | 0.35 | 0.09 | 0.62 | 0.56 | 0.49 | 0.57 |
| 4 | 31718445 | rs10489007 | 0.527 | -0.69 | -2.538 | -0.267 | -0.014 | 0.036 | 0.036 | 2.232 | 0.17 | 0.01 | 0.07 | 0.05 | 0.14 | 0.07 |
| 4 | 31721265 | rs17595220 | 0.51 | -0.691 | -2.534 | -0.27 | 0.006 | 0.033 | 0.033 | 2.216 | 0.17 | 0.03 | 0.07 | 0.07 | 0.15 | 0.08 |
| 4 | 31739254 | rs9990632 | -0.34 | 0.064 | -2.361 | 0.095 | -0.023 | -0.082 | -0.082 | 0 | 0.38 | 0.1 | 0.66 | 0.49 | 0.57 | 0.51 |
| 4 | 31744052 | rs1157893 | -0.093 | 0.497 | -1.759 | -0.253 | -0.052 | -0.056 | -0.056 | 0.087 | 0.26 | 0.15 | 0.49 | 0.51 | 0.42 | 0.4 |
| 4 | 31744915 | rs10002958 | -0.099 | 0.456 | -1.76 | -0.264 | -0.053 | -0.056 | -0.056 | 0 | 0.26 | 0.15 | 0.49 | 0.51 | 0.42 | 0.4 |
| 4 | 31746895 | rs11946605 | 0.014 | -0.636 | -1.409 | -1.297 | 0.012 | 0.001 | 0.001 | 0 | 0.76 | 0.78 | 0.83 | 0.94 | 0.87 | 0.83 |
| 4 | 31758426 | rs13136356 | 0.014 | -0.865 | -1.476 | -1.222 | 0.012 | 0.003 | 0.003 | 0 | 0.76 | 0.78 | 0.83 | 0.94 | 0.86 | 0.83 |
| 4 | 31766080 | rs17603187 | -0.194 | -0.796 | -1.827 | -1.257 | 0.04 | 0.139 | 0.139 | 0.133 | 0.35 | 0.14 | 0.16 | 0.13 | 0.29 | 0.3 |
| 4 | 31766707 | rs7661647 | -1.138 | -1.641 | -0.521 | -1.356 | -0.072 | -0.063 | -0.063 | 0.85 | 0.16 | 0.41 | 0.09 | 0.07 | 0.06 | 0.16 |
| 4 | 31770464 | rs7682874 | -0.242 | -1.464 | -0.595 | -1.195 | -0.02 | -0.008 | -0.008 | 2.324 | 0.1 | 0.14 | 0.07 | 0.1 | 0.05 | 0.15 |
| 4 | 31778484 | rs965954 | -0.447 | -1.191 | -0.722 | -1.49 | 0 | -0.028 | -0.028 | 0 | 0.12 | 0.12 | 0.07 | 0 | 0.12 | 0.1 |
| 4 | 31779135 | rs887918 | -0.769 | -1.448 | -0.665 | -1.656 | -0.042 | -0.12 | -0.12 | 0.128 | 0.3 | 0.39 | 0.15 | 0.02 | 0.17 | 0.15 |
| 4 | 31780528 | rs17604071 | -0.186 | -0.807 | -1.258 | -1.073 | 0.028 | 0.027 | 0.027 | 0 | 0.14 | 0.07 | 0.05 | 0 | 0.07 | 0.05 |
| 4 | 31784698 | rs988466 | -0.089 | 0.457 | -1.566 | -0.036 | -0.083 | -0.079 | -0.079 | 0.117 | 0.74 | 0.85 | 0.67 | 0.57 | 0.46 | 0.54 |
| 4 | 31785073 | rs2310395 | 0.005 | 0.377 | -1.136 | 0.284 | -0.033 | -0.037 | -0.037 | 2.063 | 0.73 | 0.8 | 0.67 | 0.52 | 0.46 | 0.53 |
| 4 | 31791952 | rs10029539 | -0.313 | 0.641 | -2.21 | -0.003 | -0.046 | 0.039 | 0.039 | 0.622 | 0.4 | 0.2 | 0.43 | 0.61 | 0.5 | 0.43 |
| 4 | 31811755 | rs28493679 | -0.562 | 0.743 | -0.797 | 0.587 | -0.005 | 0.017 | 0.017 | 0 | 0.71 | 0.59 | 0.74 | 0.62 | 0.77 | 0.81 |
| 4 | 31849223 | rs7658061 | -0.608 | 0.59 | -0.302 | 0.405 | -0.002 | 0.069 | 0.069 | 0 | 0.45 | 0.67 | 0.48 | 0.78 | 0.5 | 0.46 |

| | | | | | | | | | | | | | | | | |
|---|---|---|---|---|---|---|---|---|---|---|---|---|---|---|---|---|
| 4 | 31849408 | rs6553979 | -0.604 | 0.592 | -0.214 | 0.406 | -0.002 | 0.047 | 0.047 | 0 | 0.45 | 0.67 | 0.48 | 0.78 | 0.5 | 0.46 |
| 4 | 31855487 | rs6554033 | -0.609 | 0.593 | -0.237 | 0.406 | -0.002 | 0.069 | 0.069 | 0 | 0.45 | 0.69 | 0.48 | 0.78 | 0.5 | 0.46 |
| 4 | 31863889 | rs1523145 | -0.42 | 0.53 | 0.694 | 0.376 | 0.001 | -0.032 | -0.032 | 0 | 0.44 | 0.49 | 0.47 | 0.78 | 0.5 | 0.46 |
| 4 | 31867207 | rs6827011 | 0.95 | -0.489 | 0.124 | -0.163 | 0 | -0.006 | -0.006 | 0 | 0.14 | 0.15 | 0.1 | 0.06 | 0.09 | 0.11 |
| 4 | 31872574 | rs13144359 | 1.727 | -0.285 | 0.213 | 0.276 | 0.015 | 0.041 | 0.041 | 0 | 0.14 | 0.17 | 0.22 | 0.34 | 0.27 | 0.26 |
| 4 | 31890470 | rs7680258 | 0.42 | -0.188 | 0.043 | 0.068 | -0.036 | 0.003 | 0.003 | 0 | 0.38 | 0.27 | 0.39 | 0.57 | 0.42 | 0.39 |
| 4 | 31893113 | rs10007812 | 1.056 | -0.218 | 0.199 | 0.03 | 0.04 | 0.1 | 0.1 | 0 | 0.05 | 0.13 | 0.07 | 0.21 | 0.08 | 0.07 |
| 4 | 31895086 | rs6813096 | 1.38 | -0.366 | -0.125 | -0.096 | 0.009 | -0.009 | -0.009 | 0 | 0.24 | 0.08 | 0.17 | 0.23 | 0.16 | 0.13 |
| 4 | 31901631 | rs10006001 | -0.845 | -0.933 | -0.312 | -0.518 | -0.01 | 0.049 | 0.049 | 0.911 | 0.7 | 0.83 | 0.85 | 0.92 | 0.65 | 0.74 |
| 4 | 31907596 | rs11722381 | 2.053 | -1.063 | -0.486 | -0.515 | 0.008 | -0.011 | -0.011 | 0.912 | 0.22 | 0.12 | 0.17 | 0.23 | 0.15 | 0.13 |
| 4 | 31909356 | rs10027346 | -0.579 | -1.186 | -0.085 | -0.434 | -0.002 | 0.006 | 0.006 | 0.105 | 0.46 | 0.56 | 0.67 | 0.63 | 0.49 | 0.59 |
| 4 | 31923203 | rs2140306 | 1.35 | -1.098 | -0.365 | -0.025 | 0.035 | -0.039 | -0.039 | 0.057 | 0.11 | 0.03 | 0.29 | 0.22 | 0.05 | 0.19 |
| 4 | 31927635 | rs7694314 | 2.078 | -0.873 | -0.24 | 0.033 | 0.006 | 0.022 | 0.022 | 0 | 0.14 | 0.16 | 0.2 | 0.35 | 0.27 | 0.26 |
| 4 | 31930676 | rs7690576 | -0.36 | -1.647 | -0.197 | -0.15 | 0.015 | -0.001 | -0.001 | 0 | 0.26 | 0.2 | 0.42 | 0.29 | 0.17 | 0.29 |
| 4 | 31935746 | rs10517232 | 0.96 | -1.788 | -0.251 | -0.228 | 0.007 | 0.004 | 0.004 | 0 | 0.29 | 0.21 | 0.25 | 0.27 | 0.22 | 0.17 |
| 4 | 31941906 | rs10050320 | -0.493 | -1.509 | 0.043 | -0.356 | -0.001 | 0.002 | 0.002 | 0 | 0.46 | 0.57 | 0.64 | 0.57 | 0.53 | 0.58 |
| 4 | 31944649 | rs4446365 | 0.432 | -1.569 | 0.04 | -0.397 | 0.017 | 0.004 | 0.004 | 0 | 0.16 | 0.1 | 0.34 | 0.13 | 0.11 | 0.22 |
| 4 | 31948000 | rs1581453 | 0.065 | -1.547 | 0.145 | -0.346 | 0.026 | -0.002 | -0.002 | 0 | 0.45 | 0.33 | 0.59 | 0.48 | 0.34 | 0.41 |
| 4 | 31950589 | rs1523171 | -1.273 | -1.405 | 0.094 | -0.377 | 0 | 0.005 | 0.005 | 0.453 | 0.75 | 0.78 | 0.88 | 0.85 | 0.76 | 0.76 |
| 4 | 31951574 | rs1523174 | -1.328 | -1.357 | -0.09 | -0.34 | 0 | 0.002 | 0.002 | 0.854 | 0.76 | 0.78 | 0.88 | 0.85 | 0.76 | 0.76 |
| 4 | 31966758 | rs7683244 | -0.005 | -1.491 | -0.054 | -0.277 | -0.008 | -0.008 | -0.008 | 0 | 0.43 | 0.43 | 0.33 | 0.36 | 0.33 | 0.29 |
| 4 | 31967738 | rs1523160 | -1.173 | -1.577 | -0.148 | -0.278 | -0.001 | -0.008 | -0.008 | 0.614 | 0.74 | 0.74 | 0.89 | 0.84 | 0.76 | 0.76 |
| 4 | 31970683 | rs9994107 | -0.203 | -1.574 | -0.151 | -0.233 | 0.006 | 0.007 | 0.007 | 0.211 | 0.62 | 0.63 | 0.59 | 0.73 | 0.71 | 0.59 |
| 4 | 31987807 | rs1879999 | -0.143 | -1.571 | -0.163 | -0.164 | 0.012 | 0.006 | 0.006 | 0.259 | 0.61 | 0.65 | 0.59 | 0.64 | 0.7 | 0.56 |
| 4 | 31991952 | rs7678075 | 0.692 | -1.438 | -0.148 | -0.436 | 0.006 | 0.012 | 0.012 | 0 | 0.32 | 0.22 | 0.28 | 0.29 | 0.29 | 0.21 |
| 4 | 31996147 | rs1403531 | -0.855 | -1.345 | -0.333 | -0.258 | 0 | 0.003 | 0.003 | 0.251 | 0.73 | 0.76 | 0.89 | 0.77 | 0.74 | 0.76 |
| 4 | 32005439 | rs6553983 | -1.459 | -1.041 | -0.265 | -0.126 | 0.023 | 0.012 | 0.012 | 0 | 0.92 | 0.89 | 0.92 | 0.78 | 0.8 | 0.82 |
| 4 | 32006648 | rs1996886 | 0.739 | -1.441 | -0.639 | -0.323 | 0.021 | -0.012 | -0.012 | 0 | 0.74 | 0.41 | 0.79 | 0.71 | 0.67 | 0.75 |
| 4 | 32017834 | rs2037549 | -0.468 | -1.569 | -0.667 | -0.511 | -0.009 | 0.004 | 0.004 | 0 | 0.94 | 0.94 | 0.91 | 0.94 | 0.86 | 0.91 |
| 4 | 32020581 | rs10016652 | -0.159 | -1.91 | -0.753 | -0.538 | 0.017 | -0.003 | -0.003 | 0 | 0.59 | 0.78 | 0.44 | 0.61 | 0.64 | 0.6 |
| 4 | 32051793 | rs4502723 | 1.51 | -1.893 | -1.133 | -0.349 | -0.07 | -0.05 | -0.05 | 0 | 0.24 | 0.15 | 0.28 | 0.37 | 0.46 | 0.44 |
| 4 | 32071464 | rs6816357 | -0.555 | -1.863 | -1.097 | -0.329 | 0.072 | -0.039 | -0.039 | 4.667 | 0.64 | 0.87 | 0.52 | 0.58 | 0.77 | 0.67 |
| 4 | 32095002 | rs6825247 | -1.206 | -1.462 | -0.487 | -0.225 | 0.016 | 0 | 0 | 0 | 0.86 | 0.92 | 0.68 | 0.9 | 0.93 | 0.84 |
| 4 | 32101805 | rs4568279 | -0.939 | -1.474 | -0.471 | -0.22 | 0.001 | -0.002 | -0.002 | 0 | 0.85 | 0.88 | 0.67 | 0.81 | 0.92 | 0.81 |
| 4 | 32114385 | rs10017595 | -0.161 | -1.744 | -0.41 | -0.392 | -0.017 | -0.004 | -0.004 | 0 | 0.77 | 0.77 | 0.64 | 0.81 | 0.88 | 0.81 |
| 4 | 32125671 | rs6554007 | -0.014 | -1.373 | -0.581 | -0.279 | 0 | -0.001 | -0.001 | 0 | 0.79 | 0.79 | 0.58 | 0.85 | 0.79 | 0.74 |
| 4 | 32129472 | rs6419997 | -1.069 | -1.319 | -0.61 | -0.185 | -0.012 | -0.001 | -0.001 | 0.225 | 0.85 | 0.85 | 0.66 | 0.89 | 0.93 | 0.84 |
| 4 | 32143073 | rs10011864 | 0.63 | -0.868 | -0.862 | -0.043 | 0.034 | 0.023 | 0.023 | 1.638 | 0.44 | 0.16 | 0.42 | 0.61 | 0.61 | 0.58 |
| 4 | 32162911 | rs13131255 | -0.058 | 0.75 | 0.478 | 0.714 | -0.002 | -0.001 | -0.001 | 0 | 0.46 | 0.53 | 0.41 | 0.46 | 0.54 | 0.56 |
| 4 | 32162926 | rs7686517 | -0.515 | 0.556 | 0.529 | 0.51 | 0 | -0.028 | -0.028 | 0 | 0.1 | 0.39 | 0.11 | 0.04 | 0.08 | 0.02 |
| 4 | 32167598 | rs3923313 | -0.405 | 0.513 | 0.761 | 0.572 | 0.017 | 0.007 | 0.007 | 0 | 0.49 | 0.67 | 0.42 | 0.57 | 0.55 | 0.57 |
| 4 | 32174131 | rs1491226 | 0.232 | 0.643 | 0.734 | 0.49 | -0.002 | -0.001 | -0.001 | 0 | 0.94 | 0.94 | 0.93 | 1 | 0.87 | 0.93 |
| 4 | 32186096 | rs4862911 | 0.321 | 0.61 | 0.818 | 0.512 | 0.059 | -0.006 | -0.006 | 0.018 | 0.45 | 0.26 | 0.51 | 0.43 | 0.32 | 0.35 |
| 4 | 32190693 | rs834461 | 0.224 | 0.685 | 0.784 | 0.528 | 0.011 | -0.01 | -0.01 | 0.724 | 0.94 | 0.93 | 0.94 | 1 | 0.88 | 0.93 |
| 4 | 32194466 | rs4862912 | -0.901 | -0.214 | 0.823 | 0.555 | -0.022 | -0.08 | -0.08 | 0 | 0.62 | 0.86 | 0.66 | 0.55 | 0.46 | 0.47 |
| 4 | 32194941 | rs17080898 | 0.133 | -0.216 | 1.355 | 0.548 | 0.006 | -0.016 | -0.016 | 0.01 | 0.07 | 0.25 | 0.09 | 0.04 | 0.07 | 0.02 |
| 4 | 32196347 | rs17080902 | 0.526 | -0.174 | 1.384 | 0.644 | 0.093 | 0.038 | 0.038 | 0 | 0.43 | 0.15 | 0.49 | 0.3 | 0.29 | 0.31 |
| 4 | 32196630 | rs6816384 | -0.41 | -0.334 | 1.429 | 0.619 | -0.001 | 0.021 | 0.021 | 0 | 0.09 | 0.31 | 0.09 | 0.12 | 0.08 | 0.04 |
| 4 | 32200425 | rs7656532 | -0.208 | -0.22 | 0.93 | 0.668 | 0.004 | -0.017 | -0.017 | 0 | 0.05 | 0.07 | 0.04 | 0.01 | 0.05 | 0.07 |
| 4 | 32201551 | rs12509554 | -0.308 | -0.243 | 0.911 | 0.685 | 0.054 | -0.014 | -0.014 | 0 | 0.56 | 0.71 | 0.63 | 0.55 | 0.41 | 0.38 |
| 4 | 32206490 | rs1388281 | -0.757 | -0.186 | 0.699 | 0.742 | -0.003 | -0.045 | -0.045 | 0 | 0.61 | 0.8 | 0.66 | 0.56 | 0.46 | 0.44 |
| 4 | 32222974 | rs7699259 | -0.862 | -0.407 | 0.092 | 0.748 | -0.033 | -0.092 | -0.092 | 0.364 | 0.62 | 0.88 | 0.65 | 0.54 | 0.45 | 0.47 |
| 4 | 32226350 | rs1417055 | -0.119 | -0.821 | 0.587 | 0.716 | 0.054 | 0.045 | 0.045 | 0 | 0.51 | 0.61 | 0.54 | 0.3 | 0.34 | 0.38 |
| 4 | 32227281 | rs17080946 | -0.72 | -0.807 | 0.28 | 0.735 | 0.039 | -0.024 | -0.024 | 0 | 0.58 | 0.86 | 0.62 | 0.42 | 0.39 | 0.42 |
| 4 | 32240417 | rs7681502 | -0.199 | -0.684 | 0.506 | 0.798 | 0.025 | 0.021 | 0.021 | 0 | 0.49 | 0.49 | 0.54 | 0.3 | 0.34 | 0.37 |
| 4 | 32244696 | rs731196 | -0.559 | -0.66 | 0.481 | 0.811 | 0.056 | 0.043 | 0.043 | 0 | 0.56 | 0.79 | 0.61 | 0.33 | 0.39 | 0.39 |
| 4 | 32252137 | rs4596286 | -0.291 | -0.626 | 0.426 | 0.762 | 0.028 | 0.025 | 0.025 | 0 | 0.5 | 0.57 | 0.54 | 0.3 | 0.34 | 0.37 |
| 4 | 32252530 | rs12503077 | -0.299 | -0.72 | 0.25 | 0.872 | -0.012 | 0.099 | 0.099 | 0 | 0.09 | 0.25 | 0.09 | 0.24 | 0.08 | 0.09 |
| 4 | 32259468 | rs9654318 | -0.053 | -0.529 | -0.394 | 0.665 | 0.035 | 0.019 | 0.019 | 0 | 0.44 | 0.33 | 0.51 | 0.31 | 0.32 | 0.33 |
| 4 | 32259822 | rs766436 | 0.019 | -0.624 | -0.193 | 0.714 | 0.032 | 0.015 | 0.015 | 0 | 0.43 | 0.33 | 0.5 | 0.31 | 0.31 | 0.3 |
| 4 | 32260388 | rs10004675 | -0.184 | -0.292 | -0.334 | 0.841 | 0.048 | 0.001 | 0.001 | 0 | 0.51 | 0.58 | 0.62 | 0.49 | 0.41 | 0.4 |
| 4 | 32262563 | rs634360 | -0.863 | 0.054 | -0.179 | 1.018 | -0.002 | 0.001 | 0.001 | 0.04 | 0.43 | 0.45 | 0.37 | 0.42 | 0.44 | 0.48 |
| 4 | 32264997 | rs1670754 | -0.348 | -0.739 | -0.065 | 1.428 | -0.001 | 0.007 | 0.007 | 0 | 0.19 | 0.34 | 0.21 | 0.21 | 0.2 | 0.32 |
| 4 | 32265355 | rs10517241 | -0.185 | -1.394 | -0.086 | 1.516 | 0.017 | 0.063 | 0.063 | 0.218 | 0.23 | 0.09 | 0.13 | 0.09 | 0.17 | 0.15 |
| 4 | 32268382 | rs11734603 | 1.37 | -1.872 | -0.683 | 1.289 | -0.145 | -0.09 | -0.09 | 0.362 | 0.29 | 0.03 | 0.31 | 0.37 | 0.54 | 0.35 |
| 4 | 32269494 | rs10016605 | -1.186 | -1.948 | -0.101 | 1.273 | -0.056 | -0.011 | -0.011 | 0 | 0.71 | 0.6 | 0.64 | 0.79 | 0.85 | 0.82 |
| 4 | 32270703 | rs923669 | -1.202 | -1.951 | 0.087 | 1.21 | -0.069 | -0.1 | -0.1 | 0 | 0.71 | 0.55 | 0.89 | 0.91 | 0.86 | 0.88 |
| 4 | 32271390 | rs4142790 | -1.322 | -1.965 | 0.023 | 1.21 | -0.057 | -0.084 | -0.084 | 0 | 0.72 | 0.58 | 0.89 | 0.91 | 0.86 | 0.88 |
| 4 | 32273315 | rs1472088 | -0.852 | -1.934 | -0.033 | 1.205 | 0.007 | 0.006 | 0.006 | 0 | 0.4 | 0.41 | 0.3 | 0.31 | 0.3 | 0.47 |

| | | | | | | | | | | | | | | | | |
|---|---|---|---|---|---|---|---|---|---|---|---|---|---|---|---|---|
| 4 | 32275756 | rs13139441 | -0.33 | -2.132 | 0.617 | 1.432 | -0.031 | 0.031 | 0.031 | 0 | 0.18 | 0.32 | 0.17 | 0.22 | 0.14 | 0.32 |
| 4 | 32292534 | rs11133199 | -0.638 | -2.004 | 1.006 | 1.355 | -0.039 | -0.027 | -0.027 | 0.966 | 0.72 | 0.61 | 0.88 | 0.84 | 0.85 | 0.88 |
| 4 | 32300373 | rs6824483 | -0.739 | -1.984 | 1.208 | 1.352 | -0.008 | -0.016 | -0.016 | 0 | 0.72 | 0.71 | 0.9 | 0.91 | 0.86 | 0.89 |
| 4 | 32300636 | rs13102286 | 0.635 | -2.009 | 1.101 | 1.412 | -0.007 | 0.022 | 0.022 | 0 | 0.11 | 0.18 | 0.14 | 0.21 | 0.1 | 0.28 |
| 4 | 32304898 | rs2404930 | 0.269 | -2.006 | 0.888 | 1.414 | -0.017 | 0.065 | 0.065 | 0 | 0.12 | 0.3 | 0.15 | 0.21 | 0.11 | 0.28 |
| 4 | 32307437 | rs1116083 | 0.57 | -2.088 | 0.667 | 1.189 | 0.129 | -0.031 | -0.031 | 0.012 | 0.57 | 0.17 | 0.44 | 0.51 | 0.74 | 0.55 |
| 4 | 32313387 | rs10049820 | -0.446 | -1.953 | 0.554 | 1.267 | -0.07 | -0.066 | -0.066 | 0.128 | 0.71 | 0.52 | 0.77 | 0.88 | 0.86 | 0.89 |
| 4 | 32315731 | rs902654 | -0.611 | -1.882 | -0.318 | 1.105 | -0.048 | -0.069 | -0.069 | 0 | 0.73 | 0.62 | 0.89 | 0.91 | 0.86 | 0.89 |
| 4 | 32316663 | rs4314329 | -0.207 | -1.749 | -0.635 | 1.189 | -0.045 | 0.106 | 0.106 | 0 | 0.2 | 0.52 | 0.31 | 0.31 | 0.15 | 0.33 |
| 4 | 32318535 | rs13119336 | 0.081 | -1.06 | -0.308 | 0.612 | -0.009 | -0.022 | -0.022 | 0.015 | 0.71 | 0.69 | 0.88 | 0.92 | 0.84 | 0.89 |
| 4 | 32323374 | rs1377892 | -1.053 | -0.204 | -0.257 | 1.444 | -0.022 | 0.025 | 0.025 | 0 | 0.3 | 0.6 | 0.46 | 0.46 | 0.15 | 0.38 |
| 4 | 32329911 | rs2897006 | -0.928 | -0.324 | 0.669 | 1.747 | -0.01 | 0.1 | 0.1 | 0 | 0.13 | 0.38 | 0.29 | 0.32 | 0.07 | 0.19 |
| 4 | 32342392 | rs2219113 | -1.234 | -0.38 | -0.348 | 1.318 | -0.017 | 0.02 | 0.02 | 0 | 0.55 | 0.93 | 0.7 | 0.66 | 0.42 | 0.57 |
| 4 | 32345711 | rs2061001 | -1.133 | -0.451 | 0.084 | 1.282 | 0.026 | 0.048 | 0.048 | 0 | 0.53 | 0.82 | 0.69 | 0.66 | 0.41 | 0.56 |
| 4 | 32356997 | rs763366 | -1.107 | -0.413 | 0.269 | 1.327 | 0.045 | 0.034 | 0.034 | 0 | 0.52 | 0.79 | 0.51 | 0.61 | 0.41 | 0.52 |
| 4 | 32368307 | rs9654325 | 0.054 | -0.646 | 0.314 | 1.309 | 0.017 | 0.022 | 0.022 | 0 | 0.33 | 0.26 | 0.25 | 0.19 | 0.28 | 0.29 |
| 4 | 32373939 | rs1377896 | -1.378 | -0.676 | 0.803 | 1.323 | 0.056 | 0.314 | 0.314 | 0 | 0.13 | 0.37 | 0.42 | 0.5 | 0.12 | 0.2 |
| 4 | 32380665 | rs1349859 | 0.159 | -0.734 | 0.895 | 1.244 | 0.046 | 0.067 | 0.067 | 0 | 0.32 | 0.21 | 0.26 | 0.2 | 0.28 | 0.3 |
| 4 | 32381251 | rs9998298 | -1.47 | -0.713 | 1.346 | 1.238 | 0.043 | 0.027 | 0.027 | 0 | 0.52 | 0.78 | 0.51 | 0.62 | 0.41 | 0.53 |
| 4 | 32384828 | rs1349860 | -1.448 | -0.711 | 1.347 | 1.238 | 0.045 | 0.028 | 0.028 | 0 | 0.52 | 0.78 | 0.51 | 0.62 | 0.41 | 0.53 |
| 4 | 32398607 | rs1454310 | -0.68 | -0.616 | 1.655 | 1.287 | 0.015 | -0.002 | -0.002 | 0 | 0.41 | 0.41 | 0.44 | 0.62 | 0.36 | 0.45 |
| 4 | 32407960 | rs17313860 | 0.223 | -0.652 | 1.558 | 1.28 | 0.048 | 0.063 | 0.063 | 0 | 0.31 | 0.19 | 0.24 | 0.2 | 0.26 | 0.32 |
| 4 | 32419013 | rs6854744 | -1.555 | -0.716 | 1.816 | 1.359 | 0.047 | 0.052 | 0.052 | 0 | 0.52 | 0.8 | 0.7 | 0.69 | 0.42 | 0.57 |
| 4 | 32425677 | rs4862924 | -1.157 | -0.694 | 2.301 | 1.415 | 0.051 | 0.059 | 0.059 | 0 | 0.07 | 0.21 | 0.29 | 0.11 | 0.08 | 0.1 |
| 4 | 32426634 | rs6850712 | 0.174 | -0.645 | 2.221 | 1.409 | 0.038 | 0.049 | 0.049 | 0 | 0.32 | 0.24 | 0.24 | 0.2 | 0.27 | 0.32 |
| 4 | 32429986 | rs10015890 | 0.226 | -0.641 | 2.116 | 1.42 | 0.047 | 0.069 | 0.069 | 0 | 0.32 | 0.2 | 0.24 | 0.2 | 0.27 | 0.32 |
| 4 | 32432985 | rs733046 | -1.455 | -0.462 | 1.81 | 1.446 | 0.033 | 0.041 | 0.041 | 0 | 0.53 | 0.85 | 0.65 | 0.69 | 0.42 | 0.57 |
| 4 | 32435842 | rs12503381 | -0.135 | -0.457 | 1.837 | 1.471 | -0.001 | 0.081 | 0.081 | 0 | 0.05 | 0.05 | 0.07 | 0.39 | 0.05 | 0.1 |
| 4 | 32437731 | rs11133215 | -1.01 | -0.476 | 1.877 | 1.226 | 0.044 | 0.053 | 0.053 | 0.239 | 0.46 | 0.73 | 0.57 | 0.3 | 0.36 | 0.47 |
| 4 | 32446724 | rs10007799 | -1.218 | -0.378 | 1.777 | 1.349 | 0.032 | 0.284 | 0.284 | 0 | 0.06 | 0.17 | 0.08 | 0.43 | 0.07 | 0.12 |
| 4 | 32449832 | rs4862928 | -0.968 | -0.314 | 1.73 | 1.201 | 0.018 | 0.028 | 0.028 | 0 | 0.47 | 0.58 | 0.53 | 0.2 | 0.39 | 0.49 |
| 4 | 32452973 | rs10007938 | -0.461 | -0.345 | 1.477 | 1.216 | -0.007 | -0.051 | -0.051 | 0 | 0.4 | 0.39 | 0.25 | 0.09 | 0.32 | 0.4 |
| 4 | 32461577 | rs10006044 | -1.337 | 0.837 | 0.989 | 1.315 | -0.034 | 0.028 | 0.028 | 0 | 0.94 | 0.99 | 0.96 | 1 | 0.87 | 0.99 |
| 4 | 32473028 | rs11930703 | -0.806 | 0.619 | 1.276 | 1.415 | -0.007 | 0.038 | 0.038 | 0 | 0.06 | 0.18 | 0.2 | 0.07 | 0.01 | 0.09 |
| 4 | 32477273 | rs4862932 | -1.534 | 0.567 | 0.681 | 0.94 | -0.176 | -0.138 | -0.138 | 0 | 0.7 | 0.97 | 0.69 | 0.62 | 0.57 | 0.62 |
| 4 | 32481587 | rs4298181 | -1.717 | 0.637 | 1.711 | 1.72 | -0.017 | 0.282 | 0.282 | 0 | 0.1 | 0.24 | 0.26 | 0.49 | 0.05 | 0.19 |
| 4 | 32495165 | rs17081424 | -0.391 | 0.77 | 1.707 | 0.979 | -0.007 | -0.068 | -0.068 | 0 | 0.35 | 0.42 | 0.23 | 0.07 | 0.3 | 0.37 |
| 4 | 32496303 | rs11732748 | 0.008 | 0.765 | 1.888 | 0.962 | 0.021 | 0.065 | 0.065 | 0.106 | 0.32 | 0.22 | 0.21 | 0.07 | 0.26 | 0.32 |
| 4 | 32498066 | rs10461348 | -1.493 | 0.58 | 2.315 | 1.642 | -0.031 | 0.157 | 0.157 | 0 | 0.06 | 0.13 | 0.05 | 0.35 | 0.02 | 0.15 |
| 4 | 32501462 | rs961422 | -0.991 | 0.669 | 2.307 | 1.637 | 0.001 | -0.001 | -0.001 | 0 | 0.41 | 0.62 | 0.3 | 0.51 | 0.32 | 0.51 |
| 4 | 32509904 | rs1584155 | -0.448 | 0.71 | 2.317 | 1.626 | 0.003 | -0.002 | -0.002 | 0.318 | 0.36 | 0.39 | 0.25 | 0.43 | 0.27 | 0.48 |
| 4 | 32515265 | rs6554028 | -0.752 | 0.706 | 2.387 | 1.649 | -0.002 | 0 | 0 | 0 | 0.38 | 0.62 | 0.26 | 0.43 | 0.27 | 0.49 |
| 4 | 32521580 | rs7657618 | -2.711 | 0.774 | 2.295 | 1.644 | 0.123 | 0.09 | 0.09 | 0.481 | 0.07 | 0.26 | 0.28 | 0.12 | 0.15 | 0.05 |
| 4 | 32526020 | rs12649655 | -0.477 | 0.688 | 2.272 | 1.654 | 0.004 | -0.005 | -0.005 | 0 | 0.37 | 0.39 | 0.28 | 0.52 | 0.28 | 0.5 |
| 4 | 32527996 | rs12500172 | -1.063 | 0.851 | 2.273 | 1.736 | 0.006 | 0.044 | 0.044 | 0.654 | 0.44 | 0.81 | 0.52 | 0.63 | 0.41 | 0.56 |
| 4 | 32529449 | rs12648469 | -2.128 | 0.874 | 2.419 | 1.793 | 0.066 | 0.3 | 0.3 | 0 | 0.07 | 0.24 | 0.28 | 0.41 | 0.12 | 0.21 |
| 4 | 32530706 | rs11944364 | -1.536 | 0.843 | 2.207 | 1.792 | -0.017 | 0.188 | 0.188 | 0 | 0.06 | 0.17 | 0.04 | 0.39 | 0.03 | 0.17 |
| 4 | 32547580 | rs11941142 | -0.9 | 0.96 | 2.104 | 1.826 | 0.002 | 0.03 | 0.03 | 0.599 | 0.43 | 0.81 | 0.52 | 0.61 | 0.4 | 0.56 |
| 4 | 32577729 | rs7680003 | -0.057 | 0.881 | 1.941 | 1.594 | -0.002 | -0.003 | -0.003 | 0.089 | 0.34 | 0.38 | 0.21 | 0.19 | 0.24 | 0.29 |
| 4 | 32577958 | rs7695359 | -0.544 | 0.97 | 1.468 | 1.606 | 0 | 0 | 0 | 0.05 | 0.39 | 0.62 | 0.5 | 0.23 | 0.4 | 0.35 |
| 4 | 32580240 | rs11735577 | -0.781 | 0.901 | 0.646 | 1.479 | 0.079 | 0.131 | 0.131 | 0.039 | 0.49 | 0.81 | 0.35 | 0.71 | 0.36 | 0.5 |
| 4 | 32580391 | rs13129687 | -0.393 | 1.009 | 0.722 | 1.406 | 0.062 | 0.074 | 0.074 | 0 | 0.46 | 0.74 | 0.31 | 0.7 | 0.33 | 0.5 |
| 4 | 32581594 | rs1373869 | -0.952 | 0.873 | 0.522 | 1.465 | 0.019 | 0.136 | 0.136 | 0.013 | 0.5 | 0.83 | 0.37 | 0.69 | 0.44 | 0.54 |
| 4 | 32590048 | rs1037079 | -1.829 | 0.97 | 2.021 | 2.429 | 0.003 | 0.341 | 0.341 | 0 | 0.08 | 0.26 | 0.13 | 0.55 | 0.05 | 0.26 |
| 4 | 32592691 | rs11936953 | -1.782 | 1.466 | 1.207 | 1.321 | 0.144 | 0.157 | 0.157 | 0 | 0.06 | 0.26 | 0.08 | 0.11 | 0.1 | 0.07 |
| 4 | 32593812 | rs6841893 | -0.964 | 1.515 | 0.57 | 1.531 | -0.001 | -0.003 | -0.003 | 0 | 0.26 | 0.26 | 0.25 | 0.52 | 0.34 | 0.38 |
| 4 | 32594107 | rs993054 | -0.476 | 1.584 | -0.088 | 0.476 | -0.032 | 0.052 | 0.052 | 0.103 | 0.77 | 0.86 | 0.83 | 0.99 | 0.72 | 0.83 |
| 4 | 32596378 | rs11947379 | -0.479 | 1.432 | 0.467 | 0.714 | 0.008 | 0.037 | 0.037 | 0 | 0.24 | 0.27 | 0.25 | 0.52 | 0.32 | 0.31 |
| 4 | 32609134 | rs1348664 | 1.48 | 1.104 | 0.417 | 0.337 | 0.011 | 0.023 | 0.023 | 0 | 0.4 | 0.16 | 0.5 | 0.37 | 0.35 | 0.41 |
| 4 | 32617549 | rs1838626 | 0.68 | 1.415 | 1.124 | 0.276 | 0.002 | 0.002 | 0.002 | 0 | 0.53 | 0.44 | 0.64 | 0.47 | 0.53 | 0.59 |
| 4 | 32623768 | rs4862943 | 0.82 | 1.45 | 1.155 | 0.294 | 0.002 | 0.002 | 0.002 | 0 | 0.52 | 0.43 | 0.57 | 0.46 | 0.41 | 0.52 |
| 4 | 32626153 | rs6840469 | -0.74 | 1.436 | 1.39 | 0.327 | 0.017 | 0.042 | 0.042 | 0.499 | 0.78 | 0.86 | 0.88 | 0.95 | 0.85 | 0.85 |
| 4 | 32628884 | rs755773 | 2.93 | 1.52 | 2.614 | 1.737 | 0.054 | 0.143 | 0.143 | 0.292 | 0.85 | 0.54 | 0.79 | 0.67 | 0.82 | 0.79 |
| 4 | 32633208 | rs9884918 | -2.405 | 1.485 | 2.751 | 1.741 | 0.07 | 0.168 | 0.168 | 0 | 0.09 | 0.32 | 0.16 | 0.29 | 0.14 | 0.1 |
| 4 | 32640520 | rs11133230 | 2.242 | 1.538 | 2.707 | 1.636 | -0.012 | 0.002 | 0.002 | 0 | 0.59 | 0.29 | 0.62 | 0.6 | 0.48 | 0.69 |
| 4 | 32640624 | rs1822923 | 0.874 | 1.64 | 2.544 | 1.288 | -0.046 | 0.085 | 0.085 | 0 | 0.74 | 0.89 | 0.84 | 0.92 | 0.66 | 0.87 |
| 4 | 32653512 | rs13112668 | -1.776 | 1.562 | 2.863 | 1.834 | 0.07 | 0.045 | 0.045 | 1.072 | 0.28 | 0.47 | 0.33 | 0.34 | 0.45 | 0.44 |
| 4 | 32662406 | rs6826047 | -1.72 | 1.431 | 2.897 | 1.844 | 0.004 | 0.005 | 0.005 | 0 | 0.31 | 0.29 | 0.27 | 0.34 | 0.36 | 0.38 |
| 4 | 32682680 | rs6554041 | -2.313 | 1.488 | 3.098 | 2.143 | 0.008 | 0.014 | 0.014 | 1.576 | 0.35 | 0.6 | 0.35 | 0.43 | 0.42 | 0.46 |

| | | | | | | | | | | | | | | | | |
|---|---|---|---|---|---|---|---|---|---|---|---|---|---|---|---|---|
| 4 | 32682696 | rs6842905 | -2.065 | 1.408 | 3.078 | 2.123 | -0.006 | 0.179 | 0.179 | 0.206 | 0.1 | 0.23 | 0.13 | 0.31 | 0.08 | 0.28 |
| 4 | 32686781 | rs6812552 | -1.495 | 1.349 | 3.241 | 2.13 | 0.022 | -0.112 | -0.112 | 0 | 0.27 | 0.44 | 0.19 | 0.07 | 0.32 | 0.14 |
| 4 | 32688362 | rs4547836 | -2.506 | 1.32 | 2.782 | 2.086 | -0.044 | -0.021 | -0.021 | 0 | 0.38 | 0.76 | 0.38 | 0.39 | 0.5 | 0.48 |
| 4 | 32707959 | rs4862896 | -2.83 | 1.349 | 2.784 | 2.122 | -0.044 | -0.054 | -0.054 | 0.573 | 0.39 | 0.8 | 0.38 | 0.5 | 0.5 | 0.5 |
| 4 | 32728359 | rs4862897 | -1.98 | 1.61 | 3.153 | 2.093 | 0.01 | 0.021 | 0.021 | 0.009 | 0.3 | 0.46 | 0.26 | 0.31 | 0.31 | 0.32 |
| 4 | 32729725 | rs4353947 | -0.906 | 1.495 | 3.023 | 2.058 | -0.058 | 0.104 | 0.104 | 0 | 0.22 | 0.04 | 0.13 | 0 | 0.27 | 0.12 |
| 4 | 32738289 | rs6814336 | -2.963 | 1.372 | 3.426 | 2.621 | 0.142 | 0.069 | 0.069 | 0.151 | 0.07 | 0.38 | 0.1 | 0.09 | 0.15 | 0.12 |
| 4 | 32745997 | rs9968428 | -2.963 | 1.372 | 3.411 | 2.621 | 0.12 | 0.054 | 0.054 | 0.151 | 0.07 | 0.36 | 0.1 | 0.09 | 0.15 | 0.12 |
| 4 | 32749519 | rs12715993 | -2.809 | -0.015 | 3.472 | 2.685 | 0.038 | 0.371 | 0.371 | 0.499 | 0.07 | 0.31 | 0.11 | 0.3 | 0.05 | 0.2 |
| 4 | 32773352 | rs4130313 | -3.042 | -0.081 | 3.477 | 2.71 | 0.005 | 0.33 | 0.33 | 0 | 0.08 | 0.3 | 0.11 | 0.34 | 0.04 | 0.2 |
| 4 | 32793497 | rs11940854 | -4.436 | -0.088 | 3.482 | 2.644 | -0.017 | 0.208 | 0.208 | 0.654 | 0.15 | 0.6 | 0.16 | 0.42 | 0.11 | 0.25 |
| 4 | 32798521 | rs4340831 | -4.444 | -0.056 | 3.573 | 2.645 | -0.006 | 0.261 | 0.261 | 0.653 | 0.14 | 0.59 | 0.17 | 0.42 | 0.11 | 0.25 |
| 4 | 32803900 | rs9992839 | -4.122 | -0.133 | 3.392 | 2.57 | 0.048 | 0.361 | 0.361 | 0 | 0.1 | 0.47 | 0.15 | 0.33 | 0.08 | 0.19 |
| 4 | 32813842 | rs13110509 | -2.927 | -0.112 | 3.296 | 2.574 | 0.022 | 0.207 | 0.207 | 0.113 | 0.07 | 0.18 | 0.12 | 0.32 | 0.08 | 0.21 |
| 4 | 32851818 | rs7438388 | -1.716 | -0.39 | 3.015 | 2.522 | -0.001 | 0.001 | 0.001 | 0.213 | 0.21 | 0.28 | 0.26 | 0.46 | 0.26 | 0.3 |
| 4 | 32855573 | rs6812308 | -1.716 | -0.39 | 3.018 | 2.513 | -0.001 | 0.003 | 0.003 | 0.213 | 0.21 | 0.28 | 0.25 | 0.46 | 0.26 | 0.3 |
| 4 | 32862457 | rs7439281 | -2.14 | -0.401 | 3.07 | 2.509 | -0.003 | 0.133 | 0.133 | 0.694 | 0.24 | 0.48 | 0.27 | 0.46 | 0.26 | 0.35 |
| 4 | 32866929 | rs12651213 | -2.383 | -0.613 | 2.584 | 2.375 | -0.015 | 0.08 | 0.08 | 0 | 0.28 | 0.43 | 0.29 | 0.53 | 0.3 | 0.4 |
| 4 | 32877827 | rs9993494 | 1.723 | -0.554 | 2.359 | 2.145 | 0.004 | 0.014 | 0.014 | 0 | 0.75 | 0.7 | 0.72 | 0.67 | 0.72 | 0.7 |
| 4 | 32902806 | rs13123433 | -0.698 | -0.543 | 2.197 | 2.14 | -0.012 | -0.027 | -0.027 | 3.715 | 0.15 | 0.11 | 0.23 | 0.33 | 0.21 | 0.22 |
| 4 | 32931940 | rs7659735 | 0.859 | -0.542 | 2.232 | 2.108 | -0.002 | -0.046 | -0.046 | 0 | 0.9 | 0.84 | 0.94 | 1 | 0.92 | 0.97 |
| 4 | 32932415 | rs6827177 | 1.457 | -0.516 | 2.107 | 2.139 | -0.001 | -0.002 | -0.002 | 0 | 0.75 | 0.75 | 0.72 | 0.67 | 0.71 | 0.73 |
| 4 | 32947309 | rs7697063 | -2.876 | -0.982 | 1.559 | 1.887 | 0.043 | -0.003 | -0.003 | 0 | 0.38 | 0.89 | 0.35 | 0.37 | 0.3 | 0.42 |
| 4 | 32952490 | rs7438285 | -2.854 | -1.464 | 2.5 | 2.356 | -0.042 | 0.031 | 0.031 | 0.973 | 0.21 | 0.71 | 0.23 | 0.33 | 0.1 | 0.32 |
| 4 | 32953000 | rs7667876 | -2.899 | -1.435 | 2.634 | 2.348 | -0.062 | 0.07 | 0.07 | 0.811 | 0.2 | 0.65 | 0.22 | 0.33 | 0.08 | 0.31 |
| 4 | 32958989 | rs9996183 | -2.715 | -1.536 | 2.338 | 1.825 | -0.022 | 0 | 0 | 0 | 0.13 | 0.15 | 0.15 | 0.12 | 0.08 | 0.27 |
| 4 | 32962030 | rs10033556 | -3.645 | -1.66 | 2.039 | 1.996 | 0.091 | -0.113 | -0.113 | 0.008 | 0.34 | 0.94 | 0.32 | 0.47 | 0.19 | 0.39 |
| 4 | 32988370 | rs7437969 | -3.241 | -1.708 | 2.417 | 1.417 | 0.015 | 0.015 | 0.015 | 0.03 | 0.26 | 0.77 | 0.17 | 0.13 | 0.14 | 0.12 |
| 4 | 32997372 | rs16988494 | -2.692 | -1.697 | 3.556 | 1.421 | -0.01 | 0.067 | 0.067 | 0.009 | 0.05 | 0.22 | 0.08 | 0.13 | 0.03 | 0.05 |
| 4 | 32999592 | rs6816693 | -2.691 | -1.703 | 3.968 | 1.344 | -0.033 | -0.132 | -0.132 | 0 | 0.19 | 0.36 | 0.11 | 0.01 | 0.12 | 0.07 |
| 4 | 33000930 | rs1448280 | -2.708 | -1.703 | 3.894 | 1.344 | -0.039 | -0.134 | -0.134 | 0.212 | 0.21 | 0.5 | 0.11 | 0.01 | 0.12 | 0.07 |
| 4 | 33011731 | rs6832554 | -2.461 | -1.698 | 3.616 | 1.347 | -0.021 | -0.098 | -0.098 | 1.565 | 0.18 | 0.3 | 0.1 | 0.01 | 0.12 | 0.07 |
| 4 | 33021328 | rs6531322 | -2.96 | -1.695 | 3.035 | 1.384 | -0.039 | -0.049 | -0.049 | 0 | 0.26 | 0.62 | 0.18 | 0.14 | 0.15 | 0.14 |
| 4 | 33030781 | rs1349084 | -3.103 | -1.742 | 2.874 | 1.423 | 0.038 | -0.015 | -0.015 | 0 | 0.28 | 0.8 | 0.32 | 0.39 | 0.16 | 0.28 |
| 4 | 33034172 | rs6857336 | -2.708 | -1.712 | 3.042 | 1.438 | -0.074 | 0.045 | 0.045 | 0 | 0.25 | 0.48 | 0.27 | 0.39 | 0.15 | 0.28 |
| 4 | 33090656 | rs876637 | -0.915 | -1.313 | 2.364 | 0.903 | 0.005 | -0.059 | -0.059 | 1.055 | 0.1 | 0.3 | 0.06 | 0 | 0.06 | 0.01 |
| 4 | 33098368 | rs16988606 | -1.174 | -1.321 | 2.453 | 0.899 | 0.013 | -0.063 | -0.063 | 0.196 | 0.11 | 0.38 | 0.06 | 0 | 0.06 | 0.01 |
| 4 | 33120543 | rs1584385 | -2.786 | -1.248 | 0.822 | 1.061 | 0.063 | -0.096 | -0.096 | 0.012 | 0.32 | 0.91 | 0.34 | 0.55 | 0.19 | 0.34 |
| 4 | 33152713 | rs3928486 | -0.994 | -0.659 | 1.646 | 0.078 | -0.006 | 0.023 | 0.023 | 0.777 | 0.1 | 0.08 | 0.08 | 0 | 0.09 | 0.01 |
| 4 | 33191579 | rs10856819 | -2.403 | -0.614 | 2.38 | 0.362 | 0.057 | 0.046 | 0.046 | 0.237 | 0.06 | 0.44 | 0.08 | 0 | 0.02 | 0.01 |
| 4 | 33200267 | rs11735365 | -0.002 | -0.627 | 0.894 | 0.34 | 0.005 | -0.017 | -0.017 | 0 | 0.34 | 0.19 | 0.4 | 0.31 | 0.26 | 0.43 |
| 4 | 33200295 | rs11096807 | -0.056 | -0.344 | 1.681 | 0.253 | -0.004 | 0.012 | 0.012 | 0 | 0.55 | 0.36 | 0.54 | 0.37 | 0.41 | 0.55 |
| 4 | 33203015 | rs1373053 | -3.281 | -0.494 | 3.665 | 0.43 | -0.034 | 0.015 | 0.015 | 3.29 | 0.11 | 0.59 | 0.1 | 0.06 | 0.03 | 0.1 |
| 4 | 33203140 | rs10517270 | -3.253 | -0.494 | 3.845 | 0.201 | -0.029 | -0.036 | -0.036 | 0 | 0.09 | 0.36 | 0.06 | 0.01 | 0.02 | 0.1 |
| 4 | 33206192 | rs7680581 | -2.498 | -0.482 | 3.667 | 0.207 | -0.011 | -0.017 | -0.017 | 0 | 0.07 | 0.21 | 0.03 | 0.01 | 0.02 | 0.03 |
| 4 | 33217636 | rs7695326 | -2.539 | -0.452 | 2.963 | 0.222 | -0.024 | -0.035 | -0.035 | 0 | 0.1 | 0.55 | 0.08 | 0.01 | 0.02 | 0.1 |
| 4 | 33225232 | rs2586001 | 1.466 | -0.355 | 2.43 | 0.229 | -0.02 | -0.022 | -0.022 | 0 | 0.79 | 0.69 | 0.74 | 0.88 | 0.87 | 0.75 |
| 4 | 33225866 | rs2687427 | 2.449 | -0.217 | 2.14 | 0.391 | -0.007 | -0.083 | -0.083 | 0 | 0.67 | 0.05 | 0.5 | 0.63 | 0.77 | 0.63 |
| 4 | 33229984 | rs2586005 | 1.915 | -0.237 | 3.007 | 0.317 | -0.039 | -0.056 | -0.056 | 0 | 0.76 | 0.38 | 0.71 | 0.88 | 0.86 | 0.66 |
| 4 | 33235506 | rs2586009 | 1.426 | -0.246 | 3.106 | 0.337 | 0.017 | 0.022 | 0.022 | 0 | 0.8 | 0.85 | 0.76 | 0.88 | 0.86 | 0.67 |
| 4 | 33240041 | rs2687463 | 2.026 | -0.123 | 3.101 | 0.268 | -0.026 | -0.047 | -0.047 | 0.248 | 0.69 | 0.43 | 0.66 | 0.83 | 0.8 | 0.55 |
| 4 | 33247495 | rs2585909 | 2.675 | -0.124 | 2.941 | 0.277 | -0.026 | -0.088 | -0.088 | 0 | 0.61 | 0.04 | 0.46 | 0.59 | 0.71 | 0.52 |
| 4 | 33269322 | rs2687412 | 1.858 | -0.394 | 3.527 | -0.566 | 0.015 | 0.028 | 0.028 | 0 | 0.73 | 0.8 | 0.71 | 0.84 | 0.81 | 0.63 |
| 4 | 33288145 | rs2687386 | 1.991 | -0.414 | 3.48 | -0.572 | 0.006 | 0.007 | 0.007 | 0 | 0.72 | 0.72 | 0.69 | 0.84 | 0.81 | 0.63 |
| 4 | 33304013 | rs2687402 | 1.454 | -0.298 | 3.13 | -0.626 | -0.001 | -0.084 | -0.084 | 0.596 | 0.87 | 0.56 | 0.91 | 0.99 | 0.92 | 0.94 |
| 4 | 33305495 | rs1443207 | -1.832 | -0.123 | 2.678 | -0.498 | -0.016 | -0.008 | -0.008 | 0 | 0.18 | 0.22 | 0.24 | 0.13 | 0.12 | 0.29 |
| 4 | 33308646 | rs2687410 | 2.014 | 0.147 | 2.355 | -0.535 | 0.004 | 0.013 | 0.013 | 0.006 | 0.7 | 0.31 | 0.61 | 0.87 | 0.77 | 0.56 |
| 4 | 33309890 | rs3098933 | 2.036 | 0.119 | 1.994 | -0.033 | -0.013 | -0.027 | -0.027 | 0 | 0.86 | 0.71 | 0.8 | 0.93 | 0.9 | 0.78 |
| 4 | 33318082 | rs883617 | 1.951 | 0.126 | 1.878 | -0.026 | -0.023 | -0.033 | -0.033 | 0 | 0.84 | 0.5 | 0.76 | 0.92 | 0.91 | 0.78 |
| 4 | 33377677 | rs10027353 | 0.921 | 0.259 | 1.267 | 0.034 | 0.002 | -0.004 | -0.004 | 3.952 | 0.9 | 0.93 | 0.94 | 1 | 0.92 | 0.93 |
| 4 | 33378232 | rs1506252 | 1.775 | 0.318 | 1.096 | 0.045 | -0.003 | -0.019 | -0.019 | 0 | 0.85 | 0.63 | 0.79 | 0.93 | 0.88 | 0.77 |
| 4 | 33380852 | rs16988955 | -1.823 | 0.34 | 1.129 | 0.48 | 0.036 | 0.117 | 0.117 | 0.233 | 0.07 | 0.43 | 0.18 | 0.08 | 0.03 | 0.15 |
| 4 | 33439168 | rs6824666 | -1.281 | 0.294 | 0.839 | 0.171 | -0.004 | -0.018 | -0.018 | 0 | 0.17 | 0.22 | 0.15 | 0.09 | 0.12 | 0.12 |
| 4 | 33444407 | rs6827883 | -1.654 | 0.105 | 1.715 | 0.053 | 0.034 | 0.07 | 0.07 | 0 | 0.08 | 0.51 | 0.2 | 0.06 | 0.05 | 0.26 |
| 4 | 33461734 | rs2062990 | -2.042 | -0.084 | 0.479 | -0.66 | -0.005 | 0.157 | 0.157 | 2.14 | 0.44 | 0.95 | 0.49 | 0.2 | 0.35 | 0.5 |
| 4 | 33581520 | rs10025535 | -2.01 | 0.457 | 0.782 | -0.75 | -0.033 | 0.149 | 0.149 | 2.543 | 0.4 | 0.95 | 0.5 | 0.15 | 0.33 | 0.47 |
| 4 | 33582784 | rs6811211 | -1.242 | 0.356 | 1.716 | -0.471 | 0.033 | 0.049 | 0.049 | 0 | 0.07 | 0.35 | 0.17 | 0.06 | 0.05 | 0.16 |
| 4 | 33599429 | rs1598194 | -1.421 | 0.351 | 1.814 | -0.472 | 0.048 | 0.065 | 0.065 | 0 | 0.07 | 0.41 | 0.18 | 0.06 | 0.06 | 0.16 |
| 4 | 33640864 | rs1598193 | -1.651 | 0.22 | 0.515 | -0.483 | 0.038 | 0.079 | 0.079 | 4.598 | 0.36 | 0.89 | 0.49 | 0.17 | 0.27 | 0.46 |

| | | | | | | | | | | | | | | | | |
|---|---|---|---|---|---|---|---|---|---|---|---|---|---|---|---|---|
| 4 | 33641476 | rs10025343 | -2.411 | 0.169 | 1.422 | 0.196 | -0.022 | 0.055 | 0.055 | 0.482 | 0.16 | 0.31 | 0.4 | 0.23 | 0.1 | 0.17 |
| 4 | 33653757 | rs717947 | -2.358 | 1.411 | 3.451 | 0.672 | -0.013 | 0.071 | 0.071 | 0 | 0.15 | 0.24 | 0.16 | 0.35 | 0.1 | 0.14 |
| 4 | 33658459 | rs6830471 | -3.264 | 1.318 | 3.992 | 0.625 | 0.007 | -0.011 | -0.011 | 0.053 | 0.2 | 0.75 | 0.26 | 0.36 | 0.13 | 0.17 |
| 4 | 33695359 | rs1902083 | -3.193 | 1.314 | 4.029 | 0.64 | -0.006 | 0.013 | 0.013 | 1.325 | 0.19 | 0.73 | 0.26 | 0.36 | 0.12 | 0.16 |
| 4 | 33700983 | rs6847997 | -2.703 | 1.318 | 4.15 | 0.642 | -0.001 | 0.171 | 0.171 | 0.047 | 0.16 | 0.37 | 0.22 | 0.36 | 0.12 | 0.16 |
| 4 | 33726506 | rs9997046 | -2.831 | 1.216 | 4.245 | 0.654 | -0.003 | 0.153 | 0.153 | 0 | 0.16 | 0.42 | 0.23 | 0.34 | 0.12 | 0.16 |
| 4 | 33745441 | rs13111438 | -2.946 | 1.221 | 4.318 | 0.657 | -0.013 | 0.098 | 0.098 | 0 | 0.16 | 0.55 | 0.25 | 0.34 | 0.12 | 0.16 |
| 4 | 33747424 | rs7684329 | -2.116 | 1.221 | 4.313 | 0.657 | -0.009 | 0.092 | 0.092 | 0 | 0.13 | 0.25 | 0.16 | 0.34 | 0.1 | 0.14 |
| 4 | 33762750 | rs9306981 | -3.485 | 0.851 | 3.736 | 0.204 | 0.043 | -0.081 | -0.081 | 0.192 | 0.27 | 0.94 | 0.29 | 0.37 | 0.17 | 0.32 |
| 4 | 33766955 | rs12645598 | -1.917 | 0.89 | 3.809 | 0.241 | -0.009 | 0.092 | 0.092 | 0 | 0.13 | 0.25 | 0.16 | 0.34 | 0.1 | 0.14 |
| 4 | 33784564 | rs16989244 | -2.835 | 0.874 | 4.001 | 0.328 | 0.008 | 0.134 | 0.134 | 0 | 0.12 | 0.48 | 0.22 | 0.24 | 0.09 | 0.12 |
| 4 | 33810391 | rs13104973 | -3.054 | 0.87 | 3.916 | 0.328 | -0.008 | 0.163 | 0.163 | 0 | 0.14 | 0.34 | 0.21 | 0.34 | 0.09 | 0.14 |
| 4 | 33910151 | rs13120249 | -1.862 | 0.929 | 3.965 | 0.892 | -0.005 | 0.042 | 0.042 | 0 | 0.08 | 0.12 | 0.13 | 0.2 | 0.07 | 0.08 |
| 4 | 33991474 | rs10002222 | -4.413 | 0.936 | 4.035 | 0.862 | 0.02 | -0.053 | -0.053 | 0 | 0.19 | 0.89 | 0.27 | 0.3 | 0.12 | 0.15 |
| 4 | 34043420 | rs13106616 | -3.637 | 0.945 | 4.125 | 0.866 | -0.002 | 0.113 | 0.113 | 0.051 | 0.16 | 0.39 | 0.23 | 0.3 | 0.12 | 0.16 |
| 4 | 34049908 | rs7698333 | -4.005 | 0.958 | 3.504 | 0.767 | 0.038 | -0.045 | -0.045 | 0 | 0.29 | 0.94 | 0.43 | 0.31 | 0.18 | 0.28 |
| 4 | 34069989 | rs6844408 | -3.755 | 1.302 | 4.114 | 0.943 | 0.048 | 0.1 | 0.1 | 0.024 | 0.08 | 0.29 | 0.18 | 0.18 | 0.08 | 0.1 |
| 4 | 34074079 | rs6852667 | -4.331 | 1.299 | 4.199 | 0.947 | 0.036 | 0.129 | 0.129 | 0.189 | 0.12 | 0.39 | 0.2 | 0.27 | 0.11 | 0.14 |
| 4 | 34110314 | rs13149553 | -3.237 | 1.285 | 4.27 | 1.056 | 0.01 | 0.102 | 0.102 | 0 | 0.09 | 0.21 | 0.16 | 0.28 | 0.08 | 0.11 |
| 4 | 34129758 | rs7439164 | -4.478 | 1.258 | 4.275 | 1.051 | 0.035 | 0.153 | 0.153 | 0.326 | 0.12 | 0.42 | 0.23 | 0.28 | 0.11 | 0.14 |
| 4 | 34135538 | rs13113036 | -4.445 | 1.258 | 4.275 | 1.051 | 0.026 | 0.131 | 0.131 | 0.331 | 0.12 | 0.37 | 0.23 | 0.28 | 0.11 | 0.14 |
| 4 | 34149964 | rs7654957 | -4.584 | 1.257 | 4.286 | 1.05 | 0.033 | 0.155 | 0.155 | 0.33 | 0.13 | 0.45 | 0.23 | 0.28 | 0.11 | 0.15 |
| 4 | 34195693 | rs6531299 | -3.911 | 0.912 | 3.861 | 1.091 | 0.005 | 0.076 | 0.076 | 0 | 0.1 | 0.2 | 0.16 | 0.28 | 0.08 | 0.11 |
| 4 | 34303751 | rs2217015 | -2.197 | 0.855 | 3.729 | 1.093 | 0.004 | 0.057 | 0.057 | 0 | 0.06 | 0.11 | 0.12 | 0.21 | 0.05 | 0.07 |
| 4 | 34323177 | rs16989413 | -3.413 | 0.978 | 3.642 | 1.041 | 0.037 | 0.147 | 0.147 | 0.038 | 0.12 | 0.38 | 0.21 | 0.28 | 0.1 | 0.16 |
| 4 | 34348697 | rs1425503 | -2.932 | 0.931 | 3.62 | 0.561 | 0.055 | -0.038 | -0.038 | 1.215 | 0.12 | 0.46 | 0.18 | 0.06 | 0.1 | 0.28 |
| 4 | 34348739 | rs1425502 | -3.116 | 1.023 | 3.539 | 0.557 | 0.039 | -0.049 | -0.049 | 0.088 | 0.13 | 0.58 | 0.2 | 0.06 | 0.13 | 0.22 |
| 4 | 34356554 | rs6849339 | -3.077 | 1.027 | 3.541 | 0.559 | 0.042 | -0.048 | -0.048 | 0 | 0.13 | 0.58 | 0.2 | 0.06 | 0.13 | 0.22 |
| 4 | 34370370 | rs4859319 | -3.516 | 0.957 | 3.482 | 0.767 | 0.024 | -0.006 | -0.006 | 0.009 | 0.18 | 0.6 | 0.27 | 0.18 | 0.18 | 0.28 |
| 4 | 34373027 | rs10517306 | -3.041 | 0.928 | 3.618 | 0.737 | 0.008 | -0.11 | -0.11 | 0 | 0.14 | 0.5 | 0.11 | 0 | 0.1 | 0.2 |
| 4 | 34397464 | rs7665973 | -3.292 | 0.738 | 3.097 | 0.835 | -0.019 | -0.072 | -0.072 | 1.333 | 0.17 | 0.62 | 0.12 | 0.03 | 0.11 | 0.25 |
| 4 | 34410712 | rs10008623 | -3.211 | 0.727 | 2.446 | 0.981 | -0.057 | -0.083 | -0.083 | 0 | 0.36 | 0.95 | 0.41 | 0.31 | 0.32 | 0.52 |
| 4 | 34413064 | rs6822676 | -3.993 | 0.779 | 3.058 | 0.762 | -0.001 | -0.004 | -0.004 | 0.006 | 0.22 | 0.72 | 0.29 | 0.21 | 0.19 | 0.35 |
| 4 | 34415056 | rs2217013 | -3.98 | 0.787 | 3.062 | 0.766 | -0.001 | -0.004 | -0.004 | 1.319 | 0.22 | 0.72 | 0.29 | 0.21 | 0.19 | 0.35 |
| 4 | 34451311 | rs6855693 | -2.892 | 0.832 | 2.987 | 0.743 | -0.01 | -0.096 | -0.096 | 0.234 | 0.15 | 0.59 | 0.11 | 0 | 0.08 | 0.2 |
| 4 | 34462800 | rs10517312 | -3.455 | 0.876 | 3 | 0.778 | 0.042 | 0.057 | 0.057 | 0 | 0.1 | 0.26 | 0.2 | 0.22 | 0.11 | 0.18 |
| 4 | 34466283 | rs1480978 | -2.917 | 0.745 | 3.019 | 0.782 | -0.016 | -0.061 | -0.061 | 0 | 0.14 | 0.6 | 0.1 | 0.06 | 0.08 | 0.18 |
| 4 | 34473606 | rs16989731 | -2.696 | 0.759 | 2.956 | 0.788 | 0.027 | 0.053 | 0.053 | 0 | 0.06 | 0.15 | 0.1 | 0.17 | 0.09 | 0.14 |
| 4 | 34478767 | rs10011680 | -3.601 | 0.751 | 2.788 | 0.78 | 0.001 | 0.003 | 0.003 | 2.884 | 0.21 | 0.81 | 0.2 | 0.23 | 0.16 | 0.33 |
| 4 | 34484035 | rs11096735 | -3.301 | 0.754 | 2.876 | 0.783 | 0.006 | 0.005 | 0.005 | 0.813 | 0.17 | 0.64 | 0.18 | 0.19 | 0.14 | 0.31 |
| 4 | 34488990 | rs10019000 | -3.508 | 0.718 | 2.909 | 0.781 | -0.001 | 0 | 0 | 0 | 0.2 | 0.76 | 0.2 | 0.23 | 0.16 | 0.33 |
| 4 | 34492968 | rs1904897 | -2.67 | 0.651 | 2.894 | 0.566 | -0.005 | -0.065 | -0.065 | 0 | 0.13 | 0.53 | 0.1 | 0.02 | 0.07 | 0.18 |
| 4 | 34520176 | rs16989912 | -2.456 | 0.635 | 2.905 | 0.506 | 0.009 | -0.048 | -0.048 | 0 | 0.11 | 0.25 | 0.08 | 0.06 | 0.12 | 0.1 |
| 4 | 34534283 | rs1380341 | -3.025 | 0.785 | 2.834 | 0.68 | 0.015 | -0.009 | -0.009 | 0 | 0.21 | 0.58 | 0.23 | 0.22 | 0.24 | 0.33 |
| 4 | 34540177 | rs1380337 | -2.324 | 0.724 | 3.145 | 0.674 | -0.016 | -0.074 | -0.074 | 0 | 0.12 | 0.33 | 0.08 | 0.02 | 0.09 | 0.08 |
| 4 | 34542371 | rs13140884 | -2.206 | 0.785 | 3.332 | 0.675 | -0.011 | -0.067 | -0.067 | 0 | 0.12 | 0.24 | 0.07 | 0.02 | 0.09 | 0.08 |
| 4 | 34544661 | rs1459452 | -2.927 | 0.766 | 3.193 | 0.668 | 0.021 | 0.033 | 0.033 | 0 | 0.15 | 0.43 | 0.17 | 0.18 | 0.15 | 0.24 |
| 4 | 34545919 | rs2360784 | -2.87 | 0.778 | 3.199 | 0.674 | 0.025 | 0.036 | 0.036 | 0 | 0.15 | 0.43 | 0.17 | 0.18 | 0.15 | 0.24 |
| 4 | 34547031 | rs4428312 | -2.87 | 0.778 | 3.21 | 0.674 | 0.023 | 0.034 | 0.034 | 0 | 0.15 | 0.42 | 0.17 | 0.18 | 0.15 | 0.24 |
| 4 | 34553989 | rs1842529 | -2.61 | 1.073 | 2.465 | 0.701 | 0.022 | -0.01 | -0.01 | 0 | 0.06 | 0.14 | 0.18 | 0.06 | 0.09 | 0.09 |
| 4 | 34573079 | rs16989994 | -3.029 | 1.06 | 2.507 | 0.82 | 0.022 | 0.04 | 0.04 | 0 | 0.07 | 0.11 | 0.11 | 0.17 | 0.09 | 0.22 |
| 4 | 34582188 | rs4521368 | -3.519 | 1.066 | 2.528 | 0.77 | 0.062 | 0.112 | 0.112 | 0 | 0.06 | 0.2 | 0.25 | 0.26 | 0.12 | 0.19 |
| 4 | 34595042 | rs1459455 | -3.916 | 1.033 | 2.58 | 0.752 | 0.062 | 0.077 | 0.077 | 0 | 0.1 | 0.27 | 0.29 | 0.26 | 0.16 | 0.25 |
| 4 | 34607713 | rs6846061 | -4.015 | 1.021 | 2.613 | 0.766 | 0.076 | 0.097 | 0.097 | 0 | 0.11 | 0.36 | 0.3 | 0.26 | 0.16 | 0.25 |
| 4 | 34613756 | rs10032489 | -4.028 | 1.011 | 2.607 | 0.791 | 0.076 | 0.101 | 0.101 | 0 | 0.11 | 0.36 | 0.29 | 0.27 | 0.16 | 0.24 |
| 4 | 34618584 | rs2054575 | -2.235 | 1.165 | 2.493 | 0.459 | -0.009 | -0.109 | -0.109 | 0 | 0.16 | 0.52 | 0.11 | 0.02 | 0.15 | 0.09 |
| 4 | 34642826 | rs1599110 | -3.153 | 1.151 | 2.207 | 0.381 | 0.047 | 0.002 | 0.002 | 0 | 0.08 | 0.26 | 0.2 | 0.09 | 0.1 | 0.09 |
| 4 | 34649878 | rs898274 | -3.153 | 1.119 | 2.093 | 0.331 | 0.048 | 0.003 | 0.003 | 0 | 0.08 | 0.26 | 0.2 | 0.09 | 0.11 | 0.09 |
| 4 | 34686182 | rs11935755 | 2.393 | 1.153 | 1.905 | 0.365 | 0.019 | 0.035 | 0.035 | 0 | 0.93 | 0.84 | 0.9 | 0.84 | 0.91 | 0.8 |
| 4 | 34692953 | rs4317222 | 3.007 | 1.142 | 2.021 | 0.472 | 0.042 | 0.067 | 0.067 | 0 | 0.91 | 0.76 | 0.73 | 0.72 | 0.84 | 0.77 |
| 4 | 34693304 | rs13119900 | 2.944 | 1.144 | 2.021 | 0.473 | 0.042 | 0.067 | 0.067 | 0 | 0.91 | 0.77 | 0.74 | 0.72 | 0.84 | 0.77 |
| 4 | 34720226 | rs16990210 | -2.153 | 0.746 | 2 | 0.195 | -0.016 | -0.104 | -0.104 | 0.924 | 0.17 | 0.56 | 0.12 | 0.02 | 0.15 | 0.11 |
| 4 | 34723455 | rs953483 | -1.508 | 0.404 | 1.924 | 0.217 | -0.01 | 0.008 | 0.008 | 0.012 | 0.2 | 0.55 | 0.15 | 0.2 | 0.16 | 0.24 |
| 4 | 34723501 | rs10013111 | -1.844 | 0.186 | 2.02 | 0.187 | -0.016 | 0.027 | 0.027 | 0.021 | 0.14 | 0.32 | 0.11 | 0.17 | 0.08 | 0.14 |
| 4 | 34725270 | rs10019384 | -2.68 | 0.113 | 2.22 | 0.048 | 0.022 | -0.036 | -0.036 | 0 | 0.15 | 0.45 | 0.12 | 0.06 | 0.15 | 0.21 |
| 4 | 34726596 | rs16990230 | -3.644 | 0.171 | 2.127 | 0.002 | 0.024 | -0.053 | -0.053 | 1.168 | 0.2 | 0.52 | 0.29 | 0.16 | 0.22 | 0.25 |
| 4 | 34731172 | rs13434494 | -2.39 | 0.158 | 2.189 | 0.005 | 0.009 | -0.028 | -0.028 | 0 | 0.14 | 0.32 | 0.1 | 0.06 | 0.14 | 0.21 |
| 4 | 34749635 | rs11944699 | -2.921 | 0.159 | 2.157 | 0.002 | 0.008 | -0.05 | -0.05 | 0.022 | 0.16 | 0.51 | 0.13 | 0.06 | 0.14 | 0.21 |
| 4 | 34756057 | rs13106336 | -1.173 | 0.155 | 1.803 | 0.012 | -0.007 | -0.022 | -0.022 | 0 | 0.08 | 0.09 | 0.04 | 0 | 0.06 | 0.05 |

| | | | | | | | | | | | | | | | | |
|---|---|---|---|---|---|---|---|---|---|---|---|---|---|---|---|---|
| 4 | 34763475 | rs10012250 | -2.349 | 0.298 | 1.593 | 0.052 | 0.033 | 0.001 | 0.001 | 0 | 0.06 | 0.2 | 0.18 | 0.1 | 0.08 | 0.04 |
| 4 | 34765595 | rs6849320 | -3.038 | 0.059 | 1.051 | -0.006 | -0.028 | -0.022 | -0.022 | 0 | 0.38 | 0.94 | 0.4 | 0.38 | 0.33 | 0.4 |
| 4 | 34772226 | rs6855391 | 0.355 | 0.243 | 0.465 | -1.096 | 0.001 | -0.084 | -0.084 | 0 | 0.28 | 0.3 | 0.21 | 0.08 | 0.3 | 0.12 |
| 4 | 34772494 | rs16990346 | -0.052 | 0.073 | 0.437 | -1.102 | 0.006 | -0.044 | -0.044 | 3.404 | 0.33 | 0.38 | 0.26 | 0.17 | 0.39 | 0.29 |
| 4 | 34773861 | rs10155317 | -1.373 | -0.119 | -0.138 | -1.029 | -0.104 | 0.018 | 0.018 | 1.49 | 0.55 | 0.94 | 0.41 | 0.39 | 0.49 | 0.48 |
| 4 | 34829761 | rs10019802 | -1.597 | -0.13 | 0.875 | -1.009 | 0.005 | 0.113 | 0.113 | 0.103 | 0.09 | 0.36 | 0.07 | 0.14 | 0.03 | 0.13 |
| 4 | 34831143 | rs7678322 | -0.258 | -0.195 | 0.76 | -1.148 | -0.008 | -0.063 | -0.063 | 0.09 | 0.36 | 0.44 | 0.26 | 0.23 | 0.35 | 0.26 |
| 4 | 34846207 | rs9884148 | 0.069 | 0.11 | 1.048 | -0.881 | -0.015 | -0.016 | -0.016 | 0 | 0.32 | 0.42 | 0.2 | 0.3 | 0.21 | 0.27 |
| 4 | 34852124 | rs2055931 | -0.347 | 0.122 | 1.402 | -0.906 | -0.007 | -0.032 | -0.032 | 0.01 | 0.36 | 0.54 | 0.27 | 0.3 | 0.34 | 0.3 |
| 4 | 34855432 | rs16990420 | -0.856 | -0.082 | 1.026 | -0.982 | -0.001 | 0.192 | 0.192 | 0.11 | 0.42 | 0.85 | 0.3 | 0.22 | 0.39 | 0.34 |
| 4 | 34858160 | rs16990438 | -1.48 | -0.005 | 1.174 | -0.909 | -0.019 | -0.027 | -0.027 | 0 | 0.12 | 0.55 | 0.1 | 0.07 | 0.05 | 0.16 |
| 4 | 34863594 | rs10021358 | -2.015 | 0.013 | 1.157 | -0.914 | 0.02 | -0.05 | -0.05 | 0 | 0.15 | 0.6 | 0.14 | 0.07 | 0.18 | 0.19 |
| 4 | 34867696 | rs7670648 | -1.61 | 0.118 | 0.615 | -1.041 | -0.143 | -0.016 | -0.016 | 0 | 0.58 | 0.94 | 0.4 | 0.45 | 0.51 | 0.5 |
| 4 | 34869949 | rs1706212 | 0.1 | 0.4 | 0.031 | -1.844 | -0.001 | -0.046 | -0.046 | 0 | 0.33 | 0.58 | 0.29 | 0.08 | 0.34 | 0.23 |
| 4 | 34871955 | rs2668134 | 0.456 | 0.41 | 0.223 | -1.839 | 0.003 | -0.039 | -0.039 | 0 | 0.29 | 0.34 | 0.27 | 0.08 | 0.34 | 0.23 |
| 4 | 34872926 | rs1912842 | 1.197 | 0.327 | -0.159 | -1.916 | 0.03 | 0.091 | 0.091 | 0 | 0.29 | 0.06 | 0.16 | 0.22 | 0.18 | 0.26 |
| 4 | 34883393 | rs1617678 | 0.188 | 0.435 | 0.184 | -1.831 | -0.002 | -0.003 | -0.003 | 0 | 0.32 | 0.66 | 0.29 | 0.07 | 0.34 | 0.23 |
| 4 | 34883601 | rs3113627 | 0.147 | 0.418 | 0.176 | -1.84 | -0.001 | -0.001 | -0.001 | 0 | 0.33 | 0.66 | 0.29 | 0.07 | 0.34 | 0.23 |
| 4 | 34900365 | rs9998396 | -0.495 | 0.456 | 0.143 | -1.802 | -0.001 | -0.019 | -0.019 | 0 | 0.68 | 0.37 | 0.7 | 0.93 | 0.66 | 0.77 |
| 4 | 34902615 | rs2124423 | -0.495 | 0.449 | 0.151 | -1.81 | -0.006 | 0.037 | 0.037 | 0 | 0.68 | 0.27 | 0.69 | 0.93 | 0.66 | 0.77 |
| 4 | 34910052 | rs9306919 | -0.489 | 0.442 | 0.028 | -1.814 | -0.001 | -0.019 | -0.019 | 0 | 0.68 | 0.36 | 0.7 | 0.93 | 0.66 | 0.77 |
| 4 | 34913289 | rs11730487 | -0.337 | 0.426 | 0.035 | -1.757 | -0.01 | 0.094 | 0.094 | 0.167 | 0.67 | 0.2 | 0.69 | 0.93 | 0.66 | 0.77 |
| 4 | 34913541 | rs10006196 | -0.483 | 0.429 | -0.008 | -1.756 | -0.001 | -0.014 | -0.014 | 0.471 | 0.68 | 0.36 | 0.7 | 0.93 | 0.66 | 0.77 |
| 4 | 34917769 | rs11736165 | 0.415 | 0.378 | -0.18 | -1.811 | -0.022 | 0.025 | 0.025 | 0 | 0.31 | 0.14 | 0.47 | 0.64 | 0.34 | 0.44 |
| 4 | 34928521 | rs13137372 | 0.571 | 0.311 | -0.266 | -1.775 | 0.015 | 0.073 | 0.073 | 0 | 0.36 | 0.2 | 0.23 | 0.3 | 0.31 | 0.33 |
| 4 | 34941651 | rs2370151 | 0.269 | 0.61 | -0.147 | -1.692 | -0.059 | 0.101 | 0.101 | 0 | 0.6 | 0.18 | 0.6 | 0.77 | 0.48 | 0.6 |
| 4 | 34955378 | rs11729074 | -0.363 | 0.617 | 0.205 | -1.689 | 0.006 | -0.019 | -0.019 | 0.767 | 0.67 | 0.62 | 0.68 | 0.8 | 0.61 | 0.63 |
| 4 | 34964349 | rs1455920 | 0.492 | 0.346 | -0.183 | -1.827 | -0.079 | -0.025 | -0.025 | 0 | 0.31 | 0.02 | 0.45 | 0.67 | 0.31 | 0.47 |
| 4 | 34968968 | rs996913 | -0.297 | 0.239 | 0.089 | -1.821 | -0.011 | 0.038 | 0.038 | 0 | 0.4 | 0.3 | 0.53 | 0.68 | 0.45 | 0.55 |
| 4 | 34969327 | rs2370152 | -0.519 | 0.271 | 0.305 | -1.842 | 0.022 | -0.105 | -0.105 | 0 | 0.7 | 0.58 | 0.71 | 0.94 | 0.63 | 0.78 |
| 4 | 34973964 | rs874936 | 0.928 | -0.007 | 0.181 | -1.808 | -0.05 | -0.019 | -0.019 | 0.103 | 0.28 | 0.38 | 0.17 | 0.31 | 0.16 | 0.2 |
| 4 | 34978685 | rs10020332 | -1.502 | 0.288 | 0.079 | -1.989 | -0.005 | -0.03 | -0.03 | 0 | 0.78 | 0.8 | 0.8 | 1 | 0.7 | 0.9 |
| 4 | 34983045 | rs16990592 | 1.272 | 0.294 | -0.326 | -1.513 | -0.054 | -0.138 | -0.138 | 0 | 0.23 | 0.02 | 0.42 | 0.52 | 0.2 | 0.41 |
| 4 | 34984841 | rs921853 | -0.041 | 0.051 | 0.399 | -1.44 | -0.004 | 0.011 | 0.011 | 0 | 0.42 | 0.41 | 0.57 | 0.59 | 0.45 | 0.57 |
| 4 | 34990357 | rs1350117 | -0.655 | 0.054 | 0.478 | -1.445 | 0.004 | -0.025 | -0.025 | 0.024 | 0.7 | 0.67 | 0.74 | 0.9 | 0.61 | 0.75 |
| 4 | 34992661 | rs16990620 | 0.838 | 0.053 | 0.38 | -1.38 | -0.015 | -0.005 | -0.005 | 0.108 | 0.28 | 0.26 | 0.17 | 0.31 | 0.16 | 0.19 |
| 4 | 35019180 | rs6850734 | 0.012 | 0.131 | 0.19 | -1.336 | -0.004 | 0.008 | 0.008 | 0 | 0.42 | 0.39 | 0.59 | 0.59 | 0.47 | 0.59 |
| 4 | 35026277 | rs6839051 | -0.264 | 0.459 | 0.083 | -1.014 | 0.008 | -0.004 | -0.004 | 0 | 0.67 | 0.62 | 0.7 | 0.65 | 0.62 | 0.66 |
| 4 | 35033164 | rs1350115 | 0.154 | 0.2 | 0.313 | -0.697 | 0.002 | 0.004 | 0.004 | 1.425 | 0.36 | 0.3 | 0.28 | 0.17 | 0.32 | 0.23 |
| 4 | 35033504 | rs10011962 | -0.17 | 0.132 | 0.386 | -0.382 | 0.001 | 0.002 | 0.002 | 0.49 | 0.38 | 0.39 | 0.35 | 0.4 | 0.32 | 0.33 |
| 4 | 35041483 | rs7691503 | 0.229 | 0.017 | 0.611 | -0.526 | -0.017 | -0.078 | -0.078 | 0.23 | 0.34 | 0.44 | 0.27 | 0.13 | 0.27 | 0.26 |
| 4 | 35047017 | rs2054129 | 0.228 | 0.017 | 0.614 | -0.526 | -0.016 | -0.076 | -0.076 | 0 | 0.34 | 0.43 | 0.27 | 0.13 | 0.27 | 0.26 |
| 4 | 35054077 | rs4519816 | 0.155 | 0.005 | 0.578 | -0.544 | -0.014 | -0.044 | -0.044 | 0 | 0.35 | 0.44 | 0.29 | 0.2 | 0.29 | 0.28 |
| 4 | 35065154 | rs12498679 | 0.234 | 0.013 | 0.583 | -0.541 | -0.017 | -0.075 | -0.075 | 0 | 0.34 | 0.43 | 0.27 | 0.13 | 0.27 | 0.26 |
| 4 | 35066604 | rs7689990 | -0.783 | -0.233 | 0.721 | -0.583 | 0.054 | 0.092 | 0.092 | 0.097 | 0.46 | 0.7 | 0.33 | 0.24 | 0.32 | 0.4 |
| 4 | 35068091 | rs11934327 | 0.719 | -0.281 | 0.597 | -0.603 | 0.054 | -0.006 | -0.006 | 0.188 | 0.12 | 0.4 | 0.15 | 0.08 | 0.12 | 0.2 |
| 4 | 35073382 | rs7654262 | -0.209 | -0.22 | 0.571 | -0.614 | -0.003 | -0.015 | -0.015 | 0 | 0.39 | 0.55 | 0.33 | 0.2 | 0.32 | 0.32 |
| 4 | 35073545 | rs4859335 | 0.438 | 0.382 | 0.104 | 0.152 | -0.049 | -0.117 | -0.117 | 1.436 | 0.19 | 0.06 | 0.3 | 0.43 | 0.2 | 0.31 |
| 4 | 35079586 | rs7689165 | -1.307 | 0.043 | 1.128 | -0.105 | -0.007 | 0.032 | 0.032 | 0 | 0.23 | 0.53 | 0.22 | 0.26 | 0.19 | 0.35 |
| 4 | 35098359 | rs7655946 | -0.685 | 0.127 | 0.862 | -0.489 | 0.015 | -0.001 | -0.001 | 0.001 | 0.87 | 0.74 | 0.85 | 0.89 | 0.86 | 0.77 |
| 4 | 35105267 | rs13134303 | 0.857 | -0.013 | 0.725 | -0.705 | 0.067 | 0.151 | 0.151 | 0.317 | 0.51 | 0.18 | 0.63 | 0.75 | 0.63 | 0.57 |
| 4 | 35113694 | rs1480976 | -0.49 | 0.067 | 0.85 | -0.654 | 0.005 | 0.003 | 0.003 | 0 | 0.85 | 0.77 | 0.83 | 0.85 | 0.83 | 0.74 |
| 4 | 35119011 | rs10020480 | 1.305 | 0.071 | 0.687 | -0.686 | 0.015 | 0.019 | 0.019 | 1.648 | 0.22 | 0.11 | 0.12 | 0.05 | 0.16 | 0.06 |
| 4 | 35119704 | rs11943039 | 0.38 | -0.06 | 0.743 | -0.683 | 0.028 | 0.058 | 0.058 | 1.114 | 0.57 | 0.32 | 0.66 | 0.75 | 0.65 | 0.6 |
| 4 | 35127797 | rs10001135 | 0.348 | 0.291 | 0.504 | -0.381 | -0.054 | -0.076 | -0.076 | 0 | 0.29 | 0.46 | 0.17 | 0.14 | 0.18 | 0.16 |
| 4 | 35128162 | rs6823854 | 0.756 | 0.295 | 0.205 | -0.381 | -0.011 | -0.032 | -0.032 | 0 | 0.26 | 0.27 | 0.15 | 0.1 | 0.18 | 0.14 |
| 4 | 35131703 | rs9996939 | 0.611 | 0.372 | -0.797 | -0.336 | -0.026 | -0.043 | -0.043 | 0 | 0.28 | 0.27 | 0.15 | 0.1 | 0.18 | 0.11 |
| 4 | 35149660 | rs1023933 | 0.42 | 1.429 | -1.679 | 0.231 | -0.088 | -0.041 | -0.041 | 0.377 | 0.4 | 0.07 | 0.52 | 0.6 | 0.41 | 0.41 |
| 4 | 35150193 | rs11943194 | -1.249 | 1.735 | -0.282 | 0.603 | 0.015 | 0 | 0 | 0.395 | 0.11 | 0.13 | 0.13 | 0.15 | 0.24 | 0.14 |
| 4 | 35151237 | rs13114204 | -1.313 | 1.046 | -0.194 | 0.475 | 0.016 | -0.019 | -0.019 | 0 | 0.14 | 0.22 | 0.09 | 0.09 | 0.17 | 0.09 |
| 4 | 35156419 | rs2035355 | -0.566 | 0.435 | -0.231 | 0.647 | -0.013 | 0.01 | 0.01 | 0 | 0.33 | 0.42 | 0.26 | 0.41 | 0.33 | 0.33 |
| 4 | 35156806 | rs1994012 | -1.301 | -0.101 | -1.058 | 0.545 | -0.134 | -0.066 | -0.066 | 0 | 0.64 | 0.93 | 0.54 | 0.64 | 0.59 | 0.55 |
| 4 | 35161173 | rs11934913 | 0.16 | -0.129 | -0.424 | 0.526 | 0 | -0.001 | -0.001 | 0 | 0.33 | 0.28 | 0.24 | 0.4 | 0.32 | 0.33 |
| 4 | 35166363 | rs4629470 | -0.042 | -0.377 | -0.502 | 0.465 | -0.025 | -0.003 | -0.003 | 0 | 0.27 | 0.62 | 0.29 | 0.26 | 0.25 | 0.2 |
| 4 | 35166628 | rs1383561 | -0.24 | -0.497 | -0.047 | 0.531 | 0 | 0 | 0 | 0 | 0.27 | 0.3 | 0.25 | 0.23 | 0.26 | 0.21 |
| 4 | 35168725 | rs1480967 | 0.21 | -0.534 | 0.191 | 0.512 | 0 | -0.002 | -0.002 | 0.05 | 0.32 | 0.29 | 0.25 | 0.41 | 0.32 | 0.3 |
| 4 | 35175065 | rs1600683 | 0.219 | -0.576 | -0.122 | 0.505 | -0.013 | 0.016 | 0.016 | 0 | 0.32 | 0.44 | 0.27 | 0.41 | 0.32 | 0.31 |
| 4 | 35175249 | rs2127853 | 0.271 | -0.687 | -0.224 | 0.249 | -0.001 | -0.001 | -0.001 | 0 | 0.18 | 0.22 | 0.2 | 0.2 | 0.2 | 0.18 |
| 4 | 35176533 | rs1480974 | -0.635 | -0.665 | -0.262 | 0.231 | -0.011 | 0.002 | 0.002 | 0.043 | 0.24 | 0.34 | 0.26 | 0.26 | 0.25 | 0.24 |

| | | | | | | | | | | | | | | | | |
|---|---|---|---|---|---|---|---|---|---|---|---|---|---|---|---|---|
| 4 | 35177867 | rs16990845 | 0.147 | -0.607 | -0.684 | 0.2 | 0.005 | 0.006 | 0.006 | 0 | 0.19 | 0.17 | 0.21 | 0.2 | 0.21 | 0.18 |
| 4 | 35183361 | rs11934931 | -0.293 | -1.174 | -0.889 | 0.161 | -0.02 | -0.001 | -0.001 | 0 | 0.23 | 0.7 | 0.17 | 0.18 | 0.09 | 0.08 |
| 4 | 35183698 | rs10015056 | -0.95 | -0.812 | -1.415 | 0.141 | 0.116 | 0.04 | 0.04 | 0.365 | 0.5 | 0.89 | 0.44 | 0.41 | 0.4 | 0.32 |
| 4 | 35195922 | rs2127850 | 0.117 | -0.134 | -1.173 | 0.396 | -0.006 | -0.125 | -0.125 | 0.638 | 0.27 | 0.07 | 0.23 | 0.38 | 0.24 | 0.36 |
| 4 | 35214648 | rs3937773 | -1.213 | 0.228 | -0.834 | -0.005 | 0.036 | -0.006 | -0.006 | 0 | 0.69 | 0.59 | 0.55 | 0.66 | 0.62 | 0.73 |
| 4 | 35218959 | rs6531347 | 0.038 | 0.002 | -0.926 | -0.513 | 0.002 | 0.004 | 0.004 | 0 | 0.28 | 0.24 | 0.21 | 0.18 | 0.28 | 0.26 |
| 4 | 35224429 | rs10034866 | -0.19 | 0.124 | -1.767 | -0.886 | -0.019 | 0.094 | 0.094 | 0 | 0.33 | 0.1 | 0.21 | 0.17 | 0.34 | 0.27 |
| 4 | 35224530 | rs7691687 | -1.219 | 0.128 | -1.877 | -0.967 | -0.057 | 0.071 | 0.071 | 0.143 | 0.58 | 0.86 | 0.45 | 0.29 | 0.51 | 0.51 |
| 4 | 35226451 | rs7687153 | 0.898 | -0.064 | -1.603 | -1.015 | -0.01 | 0.037 | 0.037 | 0 | 0.16 | 0.09 | 0.16 | 0.03 | 0.19 | 0.13 |
| 4 | 35230371 | rs10017113 | 1.37 | -0.15 | -0.901 | -1.114 | 0.039 | -0.03 | -0.03 | 0 | 0.08 | 0.3 | 0.08 | 0 | 0.09 | 0.05 |
| 4 | 35235185 | rs6816663 | 0.84 | -0.423 | -0.633 | -1.309 | -0.008 | -0.005 | -0.005 | 0.476 | 0.16 | 0.23 | 0.14 | 0.14 | 0.15 | 0.11 |
| 4 | 35235257 | rs12643815 | 0.5 | -0.375 | -0.663 | -1.231 | -0.016 | 0.004 | 0.004 | 0.071 | 0.22 | 0.3 | 0.19 | 0.26 | 0.2 | 0.22 |
| 4 | 35236015 | rs10016775 | 1.13 | -0.086 | -0.574 | -0.812 | 0.037 | -0.031 | -0.031 | 0 | 0.08 | 0.3 | 0.07 | 0 | 0.09 | 0.05 |
| 4 | 35237089 | rs13117747 | 0.773 | 0.796 | -1.669 | -0.785 | -0.021 | 0.02 | 0.02 | 0 | 0.36 | 0.19 | 0.45 | 0.69 | 0.35 | 0.46 |
| 4 | 35239619 | rs13140030 | 0.661 | 0.82 | -1.686 | -0.772 | -0.006 | 0.015 | 0.015 | 0 | 0.37 | 0.26 | 0.47 | 0.69 | 0.36 | 0.46 |
| 4 | 35248847 | rs7686175 | -0.42 | 1.264 | -1.837 | -0.707 | -0.058 | 0.108 | 0.108 | 0.15 | 0.81 | 0.98 | 0.9 | 0.99 | 0.82 | 0.86 |
| 4 | 35252743 | rs4615191 | 0.223 | 1.242 | -1.271 | 0.393 | -0.046 | -0.077 | -0.077 | 0 | 0.25 | 0.09 | 0.38 | 0.42 | 0.29 | 0.23 |
| 4 | 35256839 | rs4543137 | -0.108 | 1.145 | -0.406 | -0.404 | 0.012 | -0.098 | -0.098 | 0 | 0.76 | 0.71 | 0.86 | 1 | 0.73 | 0.89 |
| 4 | 35264817 | rs10032114 | 0.186 | 0.555 | 0.716 | 0.415 | -0.019 | 0.03 | 0.03 | 0 | 0.18 | 0.24 | 0.14 | 0.29 | 0.13 | 0.11 |
| 4 | 35268816 | rs11934682 | 0.071 | 0.529 | 0.808 | 0.411 | -0.024 | 0.03 | 0.03 | 0 | 0.18 | 0.28 | 0.15 | 0.28 | 0.13 | 0.11 |
| 4 | 35269594 | rs9306924 | -0.876 | 0.576 | 1.062 | 0.57 | -0.009 | 0.031 | 0.031 | 0 | 0.32 | 0.57 | 0.23 | 0.47 | 0.24 | 0.3 |
| 4 | 35283162 | rs7676113 | 0.302 | 0.175 | 1.873 | 0.506 | 0.005 | -0.035 | -0.035 | 0.143 | 0.94 | 0.83 | 0.93 | 0.99 | 0.92 | 0.96 |
| 4 | 35283461 | rs11096770 | 1.617 | 0.143 | 1.643 | 0.415 | 0.028 | -0.023 | -0.023 | 0.019 | 0.54 | 0.12 | 0.46 | 0.5 | 0.58 | 0.61 |
| 4 | 35295899 | rs6837097 | -0.806 | -0.273 | 1.76 | 0.523 | -0.018 | 0.022 | 0.022 | 0.385 | 0.24 | 0.68 | 0.33 | 0.47 | 0.16 | 0.27 |
| 4 | 35299287 | rs10000487 | -0.005 | -0.052 | 2.331 | 0.185 | -0.028 | 0.052 | 0.052 | 0 | 0.19 | 0.38 | 0.31 | 0.28 | 0.14 | 0.11 |
| 4 | 35301330 | rs16990899 | 0.319 | -0.058 | 2.139 | 0.189 | 0.004 | -0.006 | -0.006 | 0 | 0.17 | 0.14 | 0.27 | 0.27 | 0.14 | 0.1 |
| 4 | 35317313 | rs2375438 | 0.374 | -0.017 | 2.135 | 0.146 | -0.006 | 0.043 | 0.043 | 0 | 0.79 | 0.85 | 0.81 | 0.97 | 0.75 | 0.88 |
| 4 | 35319420 | rs10021051 | -0.692 | -0.151 | 2.755 | 0.782 | -0.027 | 0.028 | 0.028 | 0 | 0.19 | 0.35 | 0.31 | 0.28 | 0.14 | 0.11 |
| 4 | 35328365 | rs9999521 | -0.918 | -0.098 | 2.509 | 0.84 | -0.031 | 0.044 | 0.044 | 0 | 0.21 | 0.63 | 0.32 | 0.37 | 0.14 | 0.13 |
| 4 | 35329616 | rs11934837 | -0.402 | 0.032 | 1.503 | 0.93 | -0.012 | 0.032 | 0.032 | 0 | 0.17 | 0.23 | 0.28 | 0.37 | 0.14 | 0.12 |
| 4 | 35331535 | rs2078217 | -0.319 | 0.094 | 1.398 | 0.96 | -0.018 | 0.066 | 0.066 | 0 | 0.17 | 0.23 | 0.28 | 0.37 | 0.14 | 0.12 |
| 4 | 35331719 | rs4859348 | -0.038 | 0.157 | 0.84 | 0.604 | 0 | 0.001 | 0.001 | 0 | 0.16 | 0.13 | 0.27 | 0.27 | 0.14 | 0.1 |
| 4 | 35333439 | rs13143446 | 0.27 | 0.028 | 0.995 | 0.36 | 0.003 | -0.001 | -0.001 | 0 | 0.74 | 0.79 | 0.69 | 0.72 | 0.77 | 0.82 |
| 4 | 35348509 | rs7662185 | 0.552 | 0.023 | 1.246 | 0.305 | -0.003 | 0.001 | 0.001 | 0 | 0.71 | 0.72 | 0.64 | 0.65 | 0.75 | 0.81 |
| 4 | 35360452 | rs1372156 | 0.752 | 0.149 | 0.926 | 0.226 | 0.039 | 0.022 | 0.022 | 0.192 | 0.57 | 0.23 | 0.56 | 0.62 | 0.65 | 0.73 |
| 4 | 35372098 | rs437943 | -0.988 | 0.187 | 0.663 | 0.244 | 0.041 | 0.021 | 0.021 | 0.197 | 0.42 | 0.83 | 0.44 | 0.38 | 0.34 | 0.27 |
| 4 | 35390613 | rs12640154 | -0.726 | 0.108 | 1.151 | -0.233 | 0.04 | 0.075 | 0.075 | 0 | 0.05 | 0.34 | 0.06 | 0.02 | 0.02 | 0.08 |
| 4 | 35401416 | rs12642609 | -0.748 | 0.118 | 0.616 | -0.225 | 0.014 | 0.051 | 0.051 | 0 | 0.06 | 0.31 | 0.06 | 0.02 | 0.02 | 0.08 |
| 4 | 35404286 | rs317014 | 0.292 | 0.598 | -0.027 | -0.399 | -0.004 | -0.033 | -0.033 | 0 | 0.54 | 0.48 | 0.73 | 0.76 | 0.58 | 0.62 |
| 4 | 35425629 | rs317027 | -1.001 | 0.485 | 0.15 | -0.503 | -0.004 | -0.104 | -0.104 | 0 | 0.18 | 0.32 | 0.13 | 0.02 | 0.16 | 0.17 |
| 4 | 35426994 | rs3098733 | -0.765 | 0.127 | -0.587 | -0.402 | -0.002 | -0.003 | -0.003 | 1.083 | 0.48 | 0.46 | 0.5 | 0.35 | 0.44 | 0.58 |
| 4 | 35429711 | rs16991102 | 0.377 | 0.098 | -0.322 | -0.486 | 0.001 | -0.048 | -0.048 | 0 | 0.1 | 0.18 | 0.07 | 0.01 | 0.11 | 0.1 |
| 4 | 35433331 | rs1588755 | 0.605 | 0.066 | -0.336 | -0.485 | -0.013 | -0.088 | -0.088 | 0.97 | 0.3 | 0.05 | 0.39 | 0.33 | 0.29 | 0.42 |
| 4 | 35437344 | rs317042 | 0.491 | 0.052 | -0.314 | -0.419 | 0.004 | -0.027 | -0.027 | 0 | 0.08 | 0.14 | 0.06 | 0.01 | 0.05 | 0.08 |
| 4 | 35445068 | rs317053 | -1.245 | 0.761 | -0.936 | -0.962 | -0.076 | 0.079 | 0.079 | 0 | 0.87 | 0.98 | 0.91 | 1 | 0.79 | 0.9 |
| 4 | 35454737 | rs13131923 | 0.924 | 0.814 | -0.69 | -0.662 | 0.017 | -0.024 | -0.024 | 0.865 | 0.31 | 0.15 | 0.4 | 0.33 | 0.3 | 0.41 |
| 4 | 35468984 | rs4859360 | 0.919 | 0.687 | -0.316 | -0.724 | 0.017 | -0.023 | -0.023 | 0.069 | 0.31 | 0.15 | 0.4 | 0.33 | 0.3 | 0.42 |
| 4 | 35472712 | rs7681453 | -0.615 | 0.676 | -0.242 | -0.732 | 0 | -0.001 | -0.001 | 0 | 0.49 | 0.48 | 0.54 | 0.36 | 0.46 | 0.59 |
| 4 | 35476300 | rs1022261 | -0.412 | 0.679 | -0.279 | -0.731 | 0.015 | 0.023 | 0.023 | 0 | 0.5 | 0.6 | 0.54 | 0.36 | 0.46 | 0.59 |
| 4 | 35501491 | rs13152634 | 0.942 | 0.702 | -0.489 | -0.699 | -0.016 | -0.075 | -0.075 | 0 | 0.31 | 0.06 | 0.4 | 0.33 | 0.31 | 0.42 |
| 4 | 35506681 | rs6841237 | 1.192 | 0.533 | -0.728 | -0.397 | -0.045 | -0.061 | -0.061 | 0.064 | 0.15 | 0.05 | 0.18 | 0.39 | 0.19 | 0.14 |
| 4 | 35514182 | rs13123633 | 0.524 | -0.275 | -0.231 | -0.472 | 0.003 | 0.003 | 0.003 | 0 | 0.34 | 0.37 | 0.42 | 0.27 | 0.31 | 0.42 |
| 4 | 35528798 | rs7694691 | -0.074 | -0.156 | -0.26 | -0.489 | 0.009 | -0.064 | -0.064 | 0 | 0.85 | 0.73 | 0.88 | 0.98 | 0.85 | 0.83 |
| 4 | 35538439 | rs7694515 | 0.773 | -0.166 | -0.405 | -0.483 | -0.007 | -0.009 | -0.009 | 0.862 | 0.33 | 0.05 | 0.4 | 0.27 | 0.31 | 0.42 |
| 4 | 35554589 | rs10517352 | -0.643 | -0.226 | -0.15 | -0.504 | 0.004 | 0.015 | 0.015 | 0 | 0.53 | 0.59 | 0.54 | 0.3 | 0.47 | 0.62 |
| 4 | 35570040 | rs1863309 | 0.793 | -0.081 | -0.503 | -0.432 | 0.002 | -0.012 | -0.012 | 0.232 | 0.32 | 0.05 | 0.42 | 0.28 | 0.32 | 0.4 |
| 4 | 35583122 | rs11936942 | -0.876 | 0.101 | -0.024 | -0.424 | 0.005 | -0.085 | -0.085 | 0 | 0.18 | 0.34 | 0.14 | 0.02 | 0.16 | 0.21 |
| 4 | 35588460 | rs7656274 | 0.498 | 0.111 | -0.075 | -0.399 | 0.001 | -0.018 | -0.018 | 0 | 0.07 | 0.12 | 0.05 | 0.01 | 0.05 | 0.08 |
| 4 | 35595493 | rs2162098 | -1.196 | 0.083 | -0.095 | -0.439 | 0.002 | -0.118 | -0.118 | 0.011 | 0.19 | 0.47 | 0.14 | 0.02 | 0.16 | 0.21 |
| 4 | 35604709 | rs7667040 | -1.088 | 0.096 | -0.342 | -0.405 | 0 | -0.098 | -0.098 | 0 | 0.18 | 0.43 | 0.14 | 0.02 | 0.16 | 0.22 |
| 4 | 35618590 | rs6836342 | 0.571 | 0.035 | -1.429 | -0.237 | -0.007 | -0.044 | -0.044 | 0 | 0.35 | 0.03 | 0.43 | 0.32 | 0.34 | 0.4 |
| 4 | 35630846 | rs1425460 | -0.364 | 0.246 | -0.443 | -0.148 | 0 | -0.02 | -0.02 | 0 | 0.1 | 0.12 | 0.07 | 0.01 | 0.11 | 0.13 |
| 4 | 35654641 | rs1019641 | -0.83 | 0.062 | -0.261 | -0.119 | -0.011 | -0.032 | -0.032 | 0 | 0.1 | 0.17 | 0.07 | 0.01 | 0.05 | 0.11 |
| 4 | 35668267 | rs12640371 | -1.035 | -0.722 | -0.088 | 0.136 | -0.018 | 0.001 | 0.001 | 0 | 0.24 | 0.32 | 0.09 | 0.3 | 0.12 | 0.33 |
| 4 | 35673557 | rs10517350 | -0.307 | -0.807 | -0.085 | 0.118 | -0.03 | -0.044 | -0.044 | 0 | 0.12 | 0.23 | 0.04 | 0.01 | 0.05 | 0.11 |
| 4 | 35676909 | rs6531383 | -1.397 | -0.992 | -0.587 | -0.586 | 0.003 | 0.018 | 0.018 | 0 | 0.57 | 0.84 | 0.57 | 0.4 | 0.46 | 0.6 |
| 4 | 35689019 | rs10517354 | -0.701 | -0.975 | -0.133 | -0.526 | -0.05 | 0.003 | 0.003 | 0 | 0.25 | 0.45 | 0.11 | 0.3 | 0.15 | 0.34 |
| 4 | 35689166 | rs10517355 | 0.645 | -0.872 | -0.322 | -0.785 | 0.001 | -0.003 | -0.003 | 0 | 0.09 | 0.04 | 0.15 | 0.08 | 0.1 | 0.07 |
| 4 | 35698696 | rs2375563 | -0.49 | -1.055 | 0.279 | -1.032 | -0.011 | -0.069 | -0.069 | 0 | 0.34 | 0.53 | 0.35 | 0.04 | 0.28 | 0.3 |

| | | | | | | | | | | | | | | | | |
|---|---|---|---|---|---|---|---|---|---|---|---|---|---|---|---|---|
| 4 | 35699382 | rs12504424 | 0.032 | -0.674 | 0.04 | -0.303 | -0.012 | -0.044 | -0.044 | 0 | 0.12 | 0.06 | 0.2 | 0.3 | 0.13 | 0.22 |
| 4 | 35700161 | rs6824653 | -1.296 | -0.277 | 0.011 | -0.398 | 0.001 | 0.042 | 0.042 | 0 | 0.52 | 0.65 | 0.57 | 0.34 | 0.46 | 0.52 |
| 4 | 35717920 | rs1596575 | -1.208 | -0.471 | 0.032 | -0.376 | 0.027 | 0.047 | 0.047 | 0 | 0.51 | 0.65 | 0.56 | 0.34 | 0.43 | 0.52 |
| 4 | 35724618 | rs1596577 | -1.231 | -0.482 | 0.038 | -0.35 | 0.025 | 0.043 | 0.043 | 0 | 0.51 | 0.67 | 0.56 | 0.35 | 0.43 | 0.51 |
| 4 | 35726689 | rs1822676 | -0.669 | -0.645 | 0.255 | -0.438 | -0.004 | -0.004 | -0.004 | 0 | 0.38 | 0.43 | 0.52 | 0.34 | 0.37 | 0.39 |
| 4 | 35727026 | rs1444372 | -1.498 | -0.757 | 0.068 | -0.608 | 0.044 | 0.003 | 0.003 | 0 | 0.39 | 0.71 | 0.45 | 0.34 | 0.27 | 0.45 |
| 4 | 35731201 | rs1822674 | -0.49 | -0.674 | 0.236 | -0.514 | -0.003 | 0.036 | 0.036 | 0 | 0.15 | 0.22 | 0.24 | 0.31 | 0.16 | 0.21 |
| 4 | 35733449 | rs1373489 | -1.607 | -0.678 | 0.036 | -0.535 | -0.013 | 0.028 | 0.028 | 0 | 0.54 | 0.88 | 0.6 | 0.36 | 0.46 | 0.5 |
| 4 | 35739721 | rs986240 | -1.076 | -0.701 | 0.098 | -0.545 | -0.033 | 0.019 | 0.019 | 0 | 0.28 | 0.43 | 0.28 | 0.33 | 0.21 | 0.32 |
| 4 | 35740465 | rs11731044 | 0.562 | -0.409 | 0.508 | -0.45 | -0.001 | -0.057 | -0.057 | 5.316 | 0.24 | 0.29 | 0.3 | 0.03 | 0.24 | 0.19 |
| 4 | 35751033 | rs5011112 | 0.118 | -0.38 | 0.233 | -0.425 | -0.007 | -0.13 | -0.13 | 0 | 0.27 | 0.45 | 0.34 | 0.03 | 0.28 | 0.2 |
| 4 | 35765765 | rs2889115 | -1.184 | -0.472 | 0.603 | 0.182 | -0.013 | 0.047 | 0.047 | 0 | 0.16 | 0.27 | 0.19 | 0.31 | 0.14 | 0.22 |
| 4 | 35772231 | rs11947034 | -0.69 | -0.724 | 0.878 | -0.131 | -0.065 | -0.117 | -0.117 | 1.687 | 0.17 | 0.43 | 0.07 | 0.01 | 0.07 | 0.13 |
| 4 | 35782245 | rs987863 | -1.617 | -0.504 | 0.201 | -0.81 | -0.04 | 0.094 | 0.094 | 0 | 0.82 | 0.97 | 0.87 | 1 | 0.76 | 0.91 |
| 4 | 35790610 | rs11096782 | 1.345 | -0.343 | 0.175 | -0.037 | -0.01 | -0.007 | -0.007 | 0 | 0.43 | 0.37 | 0.47 | 0.45 | 0.48 | 0.5 |
| 4 | 35795133 | rs13140299 | 1.438 | -0.586 | 0.4 | 0.032 | -0.004 | -0.015 | -0.015 | 0.279 | 0.11 | 0.1 | 0.03 | 0.01 | 0.05 | 0.11 |
| 4 | 35796678 | rs2375567 | 1.391 | -0.616 | 0.296 | 0.195 | -0.021 | -0.058 | -0.058 | 1.515 | 0.12 | 0.03 | 0.17 | 0.33 | 0.12 | 0.22 |
| 4 | 35801429 | rs1867476 | -2.066 | -0.559 | 0.34 | 0.107 | 0 | 0.008 | 0.008 | 0 | 0.83 | 0.91 | 0.87 | 1 | 0.81 | 0.92 |
| 4 | 35803591 | rs12641153 | 0.36 | -0.613 | 0.569 | 0.408 | -0.015 | 0.058 | 0.058 | 0 | 0.08 | 0.14 | 0.16 | 0.18 | 0.04 | 0.14 |
| 4 | 35806331 | rs6854185 | -0.7 | -0.606 | 0.259 | 0.241 | 0.008 | 0.011 | 0.011 | 0 | 0.74 | 0.85 | 0.74 | 0.79 | 0.72 | 0.85 |
| 4 | 35807078 | rs6835544 | 0.769 | -0.512 | 0.08 | 0.343 | 0.005 | -0.009 | -0.009 | 0 | 0.09 | 0.06 | 0.13 | 0.2 | 0.09 | 0.07 |
| 4 | 35808971 | rs6849048 | 1.579 | -0.44 | -0.063 | 0.212 | -0.02 | 0.038 | 0.038 | 0 | 0.39 | 0.3 | 0.36 | 0.27 | 0.5 | 0.37 |
| 4 | 35812208 | rs11725995 | 1.112 | -0.208 | 0.101 | 0.373 | -0.009 | -0.023 | -0.023 | 0 | 0.11 | 0.12 | 0.02 | 0.01 | 0.05 | 0.11 |
| 4 | 35814428 | rs6851284 | 1.193 | -0.208 | 0.088 | 0.373 | 0.002 | -0.005 | -0.005 | 2.564 | 0.11 | 0.09 | 0.02 | 0.01 | 0.05 | 0.11 |
| 4 | 35814521 | rs1020337 | 1.477 | -0.104 | -0.113 | 0.449 | 0.002 | 0.017 | 0.017 | 0 | 0.46 | 0.3 | 0.47 | 0.41 | 0.55 | 0.41 |
| 4 | 35839822 | rs4833120 | -1.863 | -0.174 | -0.145 | 0.268 | -0.005 | 0.039 | 0.039 | 0 | 0.83 | 0.95 | 0.88 | 1 | 0.81 | 0.92 |
| 4 | 35846086 | rs1368610 | 0.473 | -0.078 | -0.099 | 0.623 | -0.017 | -0.049 | -0.049 | 0 | 0.15 | 0.05 | 0.19 | 0.35 | 0.16 | 0.27 |
| 4 | 35859953 | rs10003970 | 0.305 | -0.107 | 0.094 | 0.65 | -0.026 | -0.019 | -0.019 | 0.296 | 0.63 | 0.5 | 0.68 | 0.76 | 0.72 | 0.68 |
| 4 | 35869643 | rs16991611 | 0.705 | 0.225 | -0.037 | 0.949 | -0.028 | -0.057 | -0.057 | 0.572 | 0.12 | 0.02 | 0.18 | 0.31 | 0.14 | 0.22 |
| 4 | 35873240 | rs1465712 | -0.073 | 0.092 | 0.036 | 0.905 | -0.014 | 0.006 | 0.006 | 0 | 0.74 | 0.63 | 0.7 | 0.77 | 0.77 | 0.79 |
| 4 | 35885873 | rs7676055 | 0.26 | 0.475 | 0.022 | 0.473 | -0.005 | 0.008 | 0.008 | 0 | 0.07 | 0.05 | 0.15 | 0.06 | 0.08 | 0.06 |
| 4 | 35892598 | rs10470868 | 1.242 | 0.722 | -0.178 | 0.386 | -0.006 | 0.001 | 0.001 | 0 | 0.47 | 0.4 | 0.49 | 0.51 | 0.51 | 0.47 |
| 4 | 35903030 | rs749064 | 0.587 | 0.791 | -0.43 | 0.057 | 0 | 0.051 | 0.051 | 1.026 | 0.17 | 0.04 | 0.06 | 0.06 | 0.12 | 0.13 |
| 4 | 35903499 | rs753921 | -1.433 | 0.651 | -0.302 | 0.205 | 0.011 | 0.056 | 0.056 | 0 | 0.36 | 0.26 | 0.26 | 0.13 | 0.29 | 0.31 |
| 4 | 35903624 | rs753920 | 0.13 | 0.718 | -0.33 | 0.229 | -0.019 | 0.066 | 0.066 | 0.819 | 0.2 | 0.05 | 0.09 | 0.07 | 0.18 | 0.15 |
| 4 | 35904938 | rs1433430 | -1.273 | 0.745 | -0.196 | 0.255 | 0.036 | -0.057 | -0.057 | 0 | 0.9 | 0.73 | 0.87 | 1 | 0.88 | 0.98 |
| 4 | 35908009 | rs1714034 | -0.61 | 0.738 | -0.14 | 0.337 | 0.012 | 0.07 | 0.07 | 0 | 0.85 | 0.79 | 0.76 | 0.66 | 0.8 | 0.77 |
| 4 | 35909078 | rs6826926 | 1.003 | 0.69 | -0.165 | 0.139 | -0.005 | 0.009 | 0.009 | 0 | 0.08 | 0.05 | 0.16 | 0.05 | 0.08 | 0.07 |
| 4 | 35911736 | rs1655058 | -0.851 | 0.64 | -0.198 | 0.038 | -0.004 | -0.091 | -0.091 | 3.704 | 0.83 | 0.74 | 0.86 | 1 | 0.84 | 0.91 |
| 4 | 35912580 | rs11734481 | 2.478 | 0.647 | -0.095 | 0.002 | -0.007 | 0.017 | 0.017 | 0 | 0.42 | 0.33 | 0.42 | 0.37 | 0.46 | 0.41 |
| 4 | 35912892 | rs1030700 | -0.412 | 0.662 | -0.05 | -0.231 | -0.001 | -0.014 | -0.014 | 0 | 0.21 | 0.23 | 0.1 | 0.08 | 0.18 | 0.17 |
| 4 | 35914787 | rs1030702 | 1.818 | 0.746 | -0.144 | -0.174 | -0.006 | 0.011 | 0.011 | 0 | 0.09 | 0.04 | 0.05 | 0.06 | 0.09 | 0.06 |
| 4 | 35916584 | rs1836080 | 1.445 | 0.741 | -0.117 | -0.181 | 0.007 | 0.037 | 0.037 | 0 | 0.53 | 0.41 | 0.6 | 0.87 | 0.59 | 0.68 |
| 4 | 35918905 | rs12512900 | 1.259 | 0.546 | -0.275 | -0.165 | 0.005 | -0.002 | -0.002 | 0.608 | 0.55 | 0.4 | 0.62 | 0.88 | 0.64 | 0.69 |
| 4 | 35919345 | rs1714058 | 0.699 | 0.576 | -0.236 | 0.029 | -0.022 | -0.03 | -0.03 | 0 | 0.65 | 0.46 | 0.67 | 0.78 | 0.73 | 0.71 |
| 4 | 35921123 | rs4001124 | 1.268 | 0.762 | -0.09 | 0.116 | 0.001 | 0.001 | 0.001 | 0 | 0.55 | 0.43 | 0.6 | 0.72 | 0.63 | 0.62 |
| 4 | 35930651 | rs4832855 | 0.116 | 0.675 | 0.072 | 0.019 | -0.026 | -0.002 | -0.002 | 0 | 0.2 | 0.34 | 0.09 | 0.15 | 0.12 | 0.16 |
| 4 | 35930794 | rs7668984 | 2.376 | 1.035 | -0.124 | 0.052 | 0.001 | 0.004 | 0.004 | 0.008 | 0.41 | 0.38 | 0.39 | 0.38 | 0.4 | 0.4 |
| 4 | 35933237 | rs7677061 | 0.173 | 0.972 | 0.05 | -0.682 | -0.008 | -0.006 | -0.006 | 0 | 0.75 | 0.62 | 0.7 | 0.81 | 0.78 | 0.77 |
| 4 | 35934885 | rs1433439 | -0.518 | 0.99 | 0.055 | -0.701 | 0.001 | -0.022 | -0.022 | 0.009 | 0.24 | 0.43 | 0.14 | 0.15 | 0.22 | 0.18 |
| 4 | 35935826 | rs16991732 | 0.587 | 1.041 | 0.021 | -0.708 | 0.004 | -0.006 | -0.006 | 0 | 0.18 | 0.16 | 0.07 | 0.01 | 0.12 | 0.11 |
| 4 | 35943816 | rs1714084 | 1.603 | 1.068 | 0.027 | -0.712 | -0.001 | -0.005 | -0.005 | 0 | 0.52 | 0.45 | 0.59 | 0.79 | 0.55 | 0.64 |
| 4 | 35944165 | rs11736705 | -1.049 | 1.11 | 0.075 | -0.644 | 0.058 | 0.11 | 0.11 | 0 | 0.06 | 0.23 | 0.07 | 0.14 | 0.1 | 0.07 |
| 4 | 35958133 | rs1655085 | -0.197 | 1.118 | -0.201 | -0.774 | -0.009 | 0.071 | 0.071 | 0 | 0.78 | 0.91 | 0.73 | 0.93 | 0.78 | 0.8 |
| 4 | 35967246 | rs1655036 | 1.172 | 1.033 | -0.136 | -0.739 | -0.014 | 0.112 | 0.112 | 0.001 | 0.56 | 0.74 | 0.62 | 0.92 | 0.56 | 0.67 |
| 4 | 35976090 | rs17447579 | 1.058 | 1.004 | -0.212 | -0.75 | 0.001 | -0.004 | -0.004 | 1.329 | 0.15 | 0.13 | 0.06 | 0.01 | 0.11 | 0.08 |
| 4 | 35976738 | rs1622680 | 0.316 | 1.053 | -0.251 | -0.7 | -0.038 | 0.1 | 0.1 | 0 | 0.72 | 0.9 | 0.69 | 0.93 | 0.67 | 0.75 |
| 4 | 35992101 | rs9992603 | -1.425 | 0.787 | -0.225 | -0.758 | 0 | 0.044 | 0.044 | 0 | 0.52 | 0.39 | 0.43 | 0.09 | 0.51 | 0.34 |
| 4 | 35995064 | rs12502892 | 1.574 | 0.677 | -0.094 | -0.682 | -0.004 | 0.024 | 0.024 | 0 | 0.44 | 0.34 | 0.52 | 0.73 | 0.46 | 0.57 |
| 4 | 35996495 | rs7439956 | -0.263 | 0.525 | -0.496 | -0.829 | -0.031 | 0.098 | 0.098 | 0 | 0.77 | 0.96 | 0.84 | 0.93 | 0.78 | 0.84 |
| 4 | 36000044 | rs11096793 | 0.058 | 0.664 | -0.325 | -0.821 | -0.001 | -0.015 | -0.015 | 0.226 | 0.76 | 0.77 | 0.81 | 0.93 | 0.74 | 0.85 |
| 4 | 36004792 | rs6815215 | -0.693 | 0.656 | -0.366 | -0.817 | -0.039 | 0.086 | 0.086 | 0 | 0.8 | 0.96 | 0.85 | 0.95 | 0.79 | 0.86 |
| 4 | 36004833 | rs6840079 | -0.176 | 0.868 | -0.384 | -0.699 | -0.002 | 0.005 | 0.005 | 0 | 0.31 | 0.2 | 0.28 | 0.28 | 0.31 | 0.3 |
| 4 | 36005220 | rs13111222 | 0.146 | 0.879 | -0.341 | -0.775 | 0.003 | -0.073 | -0.073 | 0 | 0.73 | 0.6 | 0.77 | 0.93 | 0.71 | 0.85 |
| 4 | 36006393 | rs1510656 | -0.608 | 0.561 | -0.608 | -0.258 | 0 | 0.007 | 0.007 | 0 | 0.31 | 0.19 | 0.28 | 0.28 | 0.31 | 0.33 |
| 4 | 36011362 | rs10010285 | 0.204 | 0.577 | -0.67 | -0.173 | 0.005 | 0.036 | 0.036 | 0 | 0.6 | 0.72 | 0.78 | 0.73 | 0.62 | 0.7 |
| 4 | 36013141 | rs2889174 | -0.429 | 0.141 | -0.353 | 0.109 | -0.002 | -0.023 | -0.023 | 1.17 | 0.34 | 0.72 | 0.51 | 0.46 | 0.37 | 0.42 |
| 4 | 36015016 | rs2175322 | 0.25 | 0.023 | -0.339 | -0.012 | -0.056 | -0.124 | -0.124 | 0.041 | 0.25 | 0.09 | 0.43 | 0.45 | 0.32 | 0.41 |
| 4 | 36015133 | rs1510653 | -0.796 | -0.507 | 0.518 | -0.216 | 0.019 | -0.026 | -0.026 | 0.479 | 0.09 | 0.55 | 0.07 | 0.01 | 0.06 | 0.01 |

| | | | | | | | | | | | | | | | | |
|---|---|---|---|---|---|---|---|---|---|---|---|---|---|---|---|---|
| 4 | 36019457 | rs7675286 | -0.817 | -0.811 | 0.469 | -0.332 | -0.037 | -0.048 | -0.048 | 0 | 0.31 | 0.52 | 0.17 | 0.16 | 0.24 | 0.2 |
| 4 | 36020436 | rs6839195 | -0.401 | -0.826 | 0.599 | -0.361 | -0.027 | -0.032 | -0.032 | 0.035 | 0.19 | 0.28 | 0.09 | 0.09 | 0.11 | 0.13 |
| 4 | 36021366 | rs2175324 | -0.993 | -0.99 | 0.735 | -0.386 | -0.014 | -0.019 | -0.019 | 0 | 0.26 | 0.28 | 0.1 | 0.12 | 0.17 | 0.15 |
| 4 | 36021619 | rs872322 | 0.557 | -1.064 | 1.221 | -0.478 | -0.016 | -0.034 | -0.034 | 0 | 0.75 | 0.66 | 0.87 | 0.85 | 0.79 | 0.78 |
| 4 | 36021678 | rs872321 | 1.478 | -1.081 | 1.35 | -0.416 | 0.015 | 0.022 | 0.022 | 0 | 0.63 | 0.35 | 0.8 | 0.75 | 0.7 | 0.69 |
| 4 | 36022360 | rs13117501 | -0.997 | -1.319 | 0.904 | -0.506 | 0.057 | 0.097 | 0.097 | 0 | 0.31 | 0.11 | 0.11 | 0.1 | 0.22 | 0.19 |
| 4 | 36031024 | rs11096795 | 1.998 | -1.276 | 1.7 | -0.471 | -0.004 | 0.025 | 0.025 | 0.081 | 0.79 | 0.41 | 0.84 | 0.75 | 0.8 | 0.76 |
| 4 | 36031082 | rs11096796 | 1.778 | -1.282 | 1.765 | -0.47 | 0.003 | 0.053 | 0.053 | 0 | 0.82 | 0.53 | 0.86 | 0.76 | 0.83 | 0.8 |
| 4 | 36032942 | rs12508463 | 0.755 | -1.196 | 1.588 | -0.469 | -0.006 | -0.003 | -0.003 | 2.408 | 0.88 | 0.91 | 0.9 | 0.86 | 0.85 | 0.92 |
| 4 | 36046629 | rs6812127 | 1.691 | -1.333 | 1.611 | 0.01 | 0.018 | 0.001 | 0.001 | 0 | 0.59 | 0.33 | 0.69 | 0.6 | 0.68 | 0.66 |
| 4 | 36048773 | rs997572 | 2.025 | -1.46 | 1.363 | -0.042 | 0.064 | -0.035 | -0.035 | 0.027 | 0.56 | 0.06 | 0.67 | 0.6 | 0.67 | 0.65 |
| 4 | 36048959 | rs6851590 | -0.079 | -1.291 | 1.529 | 0.087 | 0.007 | -0.014 | -0.014 | 1.332 | 0.09 | 0.02 | 0.04 | 0.09 | 0.08 | 0.05 |
| 4 | 36052495 | rs2048376 | 1.576 | -1.29 | 1.793 | 0.162 | 0.005 | 0.191 | 0.191 | 0 | 0.83 | 0.56 | 0.71 | 0.61 | 0.83 | 0.78 |
| 4 | 36055637 | rs7676461 | 0.64 | -0.902 | 1.03 | 0.06 | -0.027 | 0.063 | 0.063 | 0 | 0.29 | 0.12 | 0.23 | 0.12 | 0.32 | 0.28 |
| 4 | 36059603 | rs11947828 | 0.665 | -0.713 | 1.186 | 0.213 | 0.079 | 0.302 | 0.302 | 0 | 0.92 | 0.61 | 0.75 | 0.71 | 0.9 | 0.83 |
| 4 | 36060318 | rs13152028 | 0.793 | -0.732 | 0.506 | 0.141 | -0.05 | -0.044 | -0.044 | 0 | 0.5 | 0.12 | 0.44 | 0.49 | 0.51 | 0.54 |
| 4 | 36076892 | rs17493657 | -0.256 | -0.783 | 0.73 | 0.283 | -0.007 | 0.015 | 0.015 | 0 | 0.4 | 0.25 | 0.29 | 0.34 | 0.38 | 0.47 |
| 4 | 36081243 | rs2136806 | 0.704 | -0.896 | 0.268 | 0.194 | 0.039 | 0.122 | 0.122 | 1.623 | 0.19 | 0.01 | 0.04 | 0 | 0.1 | 0.1 |
| 4 | 36084658 | rs16991904 | 1.479 | -0.261 | 0.74 | 0.726 | -0.109 | -0.116 | -0.116 | 0 | 0.3 | 0.06 | 0.4 | 0.45 | 0.41 | 0.36 |
| 4 | 36085412 | rs7670868 | 0.92 | -0.261 | 1.003 | 0.624 | 0.009 | 0.005 | 0.005 | 0 | 0.35 | 0.55 | 0.54 | 0.62 | 0.44 | 0.38 |
| 4 | 36089558 | rs6812542 | -1.799 | 0.114 | 1.323 | 0.958 | -0.023 | 0.196 | 0.196 | 0 | 0.15 | 0.47 | 0.18 | 0.29 | 0.12 | 0.09 |
| 4 | 36101738 | rs13110947 | 1.149 | 0.286 | 1.056 | 1.159 | 0.035 | 0.186 | 0.186 | 0.268 | 0.9 | 0.42 | 0.75 | 0.77 | 0.9 | 0.94 |
| 4 | 36101795 | rs13137847 | 1.149 | 0.286 | 1.05 | 1.159 | 0.036 | 0.189 | 0.189 | 0.268 | 0.9 | 0.42 | 0.74 | 0.77 | 0.9 | 0.94 |
| 4 | 36113030 | rs1011288 | -0.123 | 0.278 | 0.488 | 1.14 | -0.002 | 0.008 | 0.008 | 0 | 0.07 | 0.08 | 0.06 | 0.13 | 0.06 | 0.05 |
| 4 | 36119788 | rs4833107 | 1.223 | 0.366 | 0 | 1.029 | -0.107 | -0.14 | -0.14 | 0 | 0.35 | 0.05 | 0.35 | 0.45 | 0.38 | 0.43 |
| 4 | 36142906 | rs4569781 | 1.199 | 0.256 | -0.419 | 1.022 | -0.104 | -0.093 | -0.093 | 0.332 | 0.36 | 0.09 | 0.36 | 0.45 | 0.41 | 0.43 |
| 4 | 36153128 | rs17516600 | 0.332 | 0.542 | 0.053 | 0.449 | 0.012 | 0.106 | 0.106 | 0.131 | 0.53 | 0.7 | 0.61 | 0.83 | 0.56 | 0.57 |
| 4 | 36154555 | rs7667582 | -0.25 | 0.352 | -0.124 | 0.259 | -0.012 | 0.066 | 0.066 | 0 | 0.67 | 0.83 | 0.67 | 0.83 | 0.65 | 0.61 |
| 4 | 36156910 | rs13109154 | 0.2 | 0.317 | -0.154 | 0.253 | -0.002 | 0.089 | 0.089 | 0 | 0.3 | 0.12 | 0.21 | 0.1 | 0.27 | 0.35 |
| 4 | 36159118 | rs4481252 | -0.322 | 0.411 | -0.164 | 0.357 | -0.003 | 0.023 | 0.023 | 1.12 | 0.83 | 0.89 | 0.82 | 0.92 | 0.81 | 0.92 |
| 4 | 36162923 | rs2271330 | -0.325 | 0.412 | -0.043 | 0.357 | -0.001 | -0.003 | -0.003 | 0 | 0.83 | 0.87 | 0.82 | 0.92 | 0.81 | 0.92 |
| 4 | 36164500 | rs2375743 | -0.302 | 0.41 | -0.03 | 0.378 | 0 | -0.011 | -0.011 | 0 | 0.83 | 0.85 | 0.81 | 0.92 | 0.81 | 0.92 |
| 4 | 36184773 | rs9985895 | -1.028 | 0.134 | -0.028 | 0.341 | -0.004 | -0.033 | -0.033 | 0 | 0.87 | 0.85 | 0.95 | 1 | 0.91 | 0.97 |
| 4 | 36211914 | rs4394022 | -1.027 | 0.136 | -0.21 | 0.34 | 0.006 | -0.005 | -0.005 | 0.268 | 0.87 | 0.9 | 0.95 | 1 | 0.91 | 0.97 |
| 4 | 36213761 | rs6836152 | -0.189 | 0.042 | -0.113 | 0.372 | -0.016 | -0.037 | -0.037 | 0.017 | 0.83 | 0.81 | 0.92 | 0.95 | 0.89 | 0.93 |
| 4 | 36220072 | rs4472157 | 1.259 | 0.108 | -0.602 | 0.699 | -0.106 | -0.171 | -0.171 | 3.534 | 0.38 | 0.04 | 0.34 | 0.49 | 0.4 | 0.46 |
| 4 | 36250842 | rs13103714 | -0.828 | -0.575 | 0.483 | 0.422 | 0 | 0 | 0 | 0.121 | 0.38 | 0.38 | 0.33 | 0.19 | 0.35 | 0.35 |
| 4 | 36260279 | rs2055411 | -1.149 | -0.282 | 1.416 | 0.722 | -0.002 | -0.014 | -0.014 | 0 | 0.31 | 0.63 | 0.35 | 0.16 | 0.28 | 0.29 |
| 4 | 36267669 | rs10049867 | -2.453 | -0.267 | 1.443 | 0.727 | -0.015 | -0.002 | -0.002 | 0 | 0.11 | 0.82 | 0.17 | 0.09 | 0.05 | 0.04 |
| 4 | 36273293 | rs6821079 | -0.705 | -0.146 | 1.32 | 0.82 | 0.006 | -0.033 | -0.033 | 1.927 | 0.27 | 0.39 | 0.32 | 0.14 | 0.27 | 0.28 |
| 4 | 36282235 | rs9306930 | -1.103 | -0.107 | 0.966 | 0.841 | 0.009 | 0.016 | 0.016 | 0.016 | 0.05 | 0.25 | 0.03 | 0.02 | 0.01 | 0.03 |
| 4 | 36284941 | rs16992033 | -0.411 | 0.062 | -0.165 | 1.379 | 0 | 0.021 | 0.021 | 0.028 | 0.07 | 0.03 | 0.1 | 0.24 | 0.01 | 0.1 |
| 4 | 36285929 | rs6824507 | -0.673 | 0.563 | -0.258 | 0.498 | -0.004 | 0.033 | 0.033 | 0 | 0.24 | 0.14 | 0.23 | 0.07 | 0.25 | 0.27 |
| 4 | 36305264 | rs12646644 | -1.109 | 0.147 | -0.965 | 0.962 | 0.029 | -0.038 | -0.038 | 0 | 0.33 | 0.82 | 0.37 | 0.38 | 0.27 | 0.45 |
| 4 | 36310049 | rs12507599 | -0.829 | 0.024 | -0.247 | 0.929 | 0.002 | -0.005 | -0.005 | 0.917 | 0.27 | 0.25 | 0.31 | 0.37 | 0.24 | 0.41 |
| 4 | 36311911 | rs10021650 | -1.267 | 0.037 | -0.077 | 0.921 | 0.016 | -0.016 | -0.016 | 0 | 0.33 | 0.76 | 0.37 | 0.39 | 0.27 | 0.44 |
| 4 | 36312202 | rs10007998 | -1.627 | -0.501 | -0.867 | 0.427 | 0.076 | -0.014 | -0.014 | 0 | 0.48 | 0.89 | 0.49 | 0.41 | 0.38 | 0.48 |
| 4 | 36313238 | rs10025139 | -0.958 | -0.734 | 0.171 | 0.558 | -0.013 | 0.018 | 0.018 | 0 | 0.23 | 0.5 | 0.34 | 0.2 | 0.21 | 0.28 |
| 4 | 36317259 | rs17458763 | -0.559 | -1.102 | -0.176 | 0.516 | 0.016 | -0.017 | -0.017 | 0 | 0.09 | 0.02 | 0.02 | 0.13 | 0.03 | 0.17 |
| 4 | 36317970 | rs9995922 | -0.274 | -0.455 | -0.006 | -0.056 | 0.048 | -0.022 | -0.022 | 0 | 0.26 | 0.53 | 0.38 | 0.22 | 0.36 | 0.25 |
| 4 | 36323014 | rs13131463 | 1.003 | -0.515 | -0.213 | -0.091 | -0.088 | 0.054 | 0.054 | 0 | 0.17 | 0.01 | 0.21 | 0.08 | 0.28 | 0.19 |
| 4 | 36325988 | rs13151413 | -1.877 | -0.192 | -0.359 | -0.237 | 0.001 | -0.067 | -0.067 | 0.196 | 0.61 | 0.86 | 0.75 | 0.48 | 0.65 | 0.69 |
| 4 | 36326105 | rs6832047 | -2.491 | -0.501 | -0.526 | 0.787 | -0.079 | -0.037 | -0.037 | 0 | 0.68 | 0.97 | 0.8 | 0.67 | 0.69 | 0.73 |
| 4 | 36329966 | rs13129319 | -1.847 | -0.289 | -0.09 | 0.494 | 0.011 | -0.026 | -0.026 | 0 | 0.62 | 0.74 | 0.75 | 0.49 | 0.68 | 0.7 |
| 4 | 36330976 | rs7684936 | -2.01 | -0.279 | -0.196 | 0.5 | 0.002 | -0.09 | -0.09 | 0 | 0.64 | 0.88 | 0.77 | 0.49 | 0.68 | 0.7 |
| 4 | 36334157 | rs16992086 | -0.762 | -0.373 | -0.036 | 0.507 | 0.008 | 0.043 | 0.043 | 1.42 | 0.36 | 0.22 | 0.21 | 0.26 | 0.31 | 0.41 |
| 4 | 36334295 | rs12508637 | 0.202 | -0.386 | -0.058 | 0.495 | -0.007 | 0.035 | 0.035 | 0.132 | 0.25 | 0.15 | 0.18 | 0.13 | 0.26 | 0.24 |
| 4 | 36336437 | rs13140371 | -0.246 | -0.016 | 0.261 | 0.635 | 0.003 | -0.002 | -0.002 | 0 | 0.28 | 0.66 | 0.47 | 0.23 | 0.37 | 0.29 |
| 4 | 36337266 | rs7664826 | 0.115 | -0.052 | 0.402 | 0.581 | 0.029 | 0.016 | 0.016 | 0 | 0.06 | 0.35 | 0.04 | 0.01 | 0.04 | 0.03 |
| 4 | 36337470 | rs16992103 | -0.157 | 0.005 | -0.133 | 0.625 | 0.001 | -0.012 | -0.012 | 0.372 | 0.1 | 0.02 | 0.02 | 0.13 | 0.06 | 0.17 |
| 4 | 36340753 | rs9654132 | -0.297 | 0.082 | -0.111 | 0.504 | -0.002 | 0.015 | 0.015 | 0 | 0.24 | 0.18 | 0.19 | 0.13 | 0.26 | 0.22 |
| 4 | 36341751 | rs11936656 | -0.934 | 0.059 | 0.186 | 0.681 | -0.009 | 0.008 | 0.008 | 0 | 0.36 | 0.65 | 0.5 | 0.29 | 0.43 | 0.45 |
| 4 | 36345377 | rs2270880 | 0.389 | -0.02 | 0.229 | 0.93 | -0.045 | -0.005 | -0.005 | 0.351 | 0.14 | 0.01 | 0.06 | 0.12 | 0.17 | 0.2 |
| 4 | 36350375 | rs16992136 | -1.39 | -0.146 | 0.869 | 1.253 | 0.029 | 0.203 | 0.203 | 0.369 | 0.1 | 0.58 | 0.21 | 0.2 | 0.05 | 0.04 |
| 4 | 36350425 | rs13101999 | -0.293 | -0.025 | 0.724 | 1.288 | -0.004 | 0.009 | 0.009 | 0 | 0.11 | 0.12 | 0.03 | 0.15 | 0.08 | 0.17 |
| 4 | 36350641 | rs7699358 | -1.47 | 0.183 | 0.51 | 0.964 | -0.001 | -0.001 | -0.001 | 0 | 0.46 | 0.45 | 0.41 | 0.56 | 0.45 | 0.57 |
| 4 | 36354203 | rs1392891 | 1.533 | 0.186 | 0.419 | 1.008 | -0.004 | 0.089 | 0.089 | 0 | 0.51 | 0.24 | 0.53 | 0.37 | 0.53 | 0.42 |
| 4 | 36356690 | rs7671923 | -0.407 | 0.341 | 0.459 | 0.907 | -0.001 | -0.029 | -0.029 | 0 | 0.81 | 0.77 | 0.86 | 0.99 | 0.79 | 0.8 |
| 4 | 36359605 | rs4833131 | -0.279 | 0.373 | 0.625 | 0.929 | 0.011 | -0.065 | -0.065 | 0 | 0.79 | 0.61 | 0.8 | 0.93 | 0.77 | 0.8 |

| | | | | | | | | | | | | | | | | |
|---|---|---|---|---|---|---|---|---|---|---|---|---|---|---|---|---|
| 4 | 36360969 | rs9990579 | 0.735 | 0.355 | 0.751 | 1.111 | 0.003 | 0.007 | 0.007 | 0 | 0.61 | 0.51 | 0.59 | 0.55 | 0.57 | 0.56 |
| 4 | 36363694 | rs4833132 | 0.727 | 0.358 | 0.753 | 1.113 | 0.003 | 0.007 | 0.007 | 0 | 0.61 | 0.51 | 0.59 | 0.55 | 0.57 | 0.56 |
| 4 | 36369934 | rs6857376 | -0.114 | 0.223 | 0.422 | 1.196 | -0.014 | 0.058 | 0.058 | 0.024 | 0.78 | 0.64 | 0.67 | 0.7 | 0.82 | 0.81 |
| 4 | 36375367 | rs2375870 | 0.294 | 0.186 | -0.198 | 0.987 | -0.015 | -0.043 | -0.043 | 0 | 0.08 | 0 | 0.03 | 0.15 | 0.07 | 0.15 |
| 4 | 36387599 | rs1501114 | 0.238 | 0.209 | 0.055 | 0.831 | -0.001 | -0.006 | -0.006 | 0 | 0.19 | 0.18 | 0.2 | 0.05 | 0.2 | 0.17 |
| 4 | 36390881 | rs6856664 | -0.757 | 0.103 | 0.835 | 1.052 | -0.016 | 0.091 | 0.091 | 0 | 0.15 | 0.26 | 0.12 | 0.34 | 0.09 | 0.19 |
| 4 | 36396090 | rs10003626 | -1.661 | 0.129 | 0.396 | 0.999 | -0.016 | 0.109 | 0.109 | 0 | 0.53 | 0.77 | 0.6 | 0.71 | 0.53 | 0.6 |
| 4 | 36398436 | rs949807 | -0.978 | 0.135 | 0.428 | 1.033 | 0.005 | -0.022 | -0.022 | 0.061 | 0.35 | 0.69 | 0.5 | 0.49 | 0.31 | 0.37 |
| 4 | 36403061 | rs1501116 | -1.847 | 0.041 | -0.136 | 0.638 | -0.049 | 0.03 | 0.03 | 0 | 0.62 | 0.89 | 0.61 | 0.7 | 0.6 | 0.61 |
| 4 | 36405777 | rs924536 | -1.866 | -0.039 | 0.365 | 0.755 | -0.053 | 0.028 | 0.028 | 0 | 0.63 | 0.89 | 0.61 | 0.7 | 0.6 | 0.61 |
| 4 | 36409924 | rs4833134 | -0.823 | -0.074 | 0.549 | 0.686 | -0.006 | -0.045 | -0.045 | 0.003 | 0.42 | 0.51 | 0.28 | 0.17 | 0.39 | 0.39 |
| 4 | 36411073 | rs4527502 | 1.155 | -0.072 | 0.365 | 0.593 | -0.041 | 0.107 | 0.107 | 0.034 | 0.17 | 0.01 | 0.11 | 0 | 0.17 | 0.16 |
| 4 | 36419047 | rs4629469 | -1.372 | 0.047 | 0.574 | 0.742 | 0.005 | -0.003 | -0.003 | 0.742 | 0.42 | 0.61 | 0.31 | 0.57 | 0.4 | 0.44 |
| 4 | 36419298 | rs3925092 | 0.857 | 0.228 | 0.64 | 0.939 | -0.033 | -0.044 | -0.044 | 3.689 | 0.15 | 0.02 | 0.07 | 0.22 | 0.19 | 0.21 |
| 4 | 36435081 | rs7696909 | 0.42 | 0.364 | 0.74 | 0.527 | -0.002 | -0.009 | -0.009 | 0 | 0.31 | 0.32 | 0.15 | 0.16 | 0.27 | 0.23 |
| 4 | 36447541 | rs9884802 | -1.089 | 0.248 | 0.504 | 0.473 | -0.043 | -0.051 | -0.051 | 0 | 0.57 | 0.81 | 0.57 | 0.48 | 0.51 | 0.41 |
| 4 | 36449901 | rs4605678 | -0.9 | 0.239 | 0.497 | 0.398 | -0.009 | 0.012 | 0.012 | 3.762 | 0.55 | 0.62 | 0.39 | 0.36 | 0.49 | 0.41 |
| 4 | 36451424 | rs3949 | -0.488 | 0.235 | 0.586 | 0.351 | 0.001 | 0.004 | 0.004 | 1.363 | 0.49 | 0.49 | 0.35 | 0.35 | 0.45 | 0.37 |
| 4 | 36454804 | rs4355409 | -0.94 | 0.256 | 0.557 | 0.386 | -0.007 | 0.013 | 0.013 | 3.516 | 0.55 | 0.59 | 0.39 | 0.35 | 0.49 | 0.4 |
| 4 | 36460981 | rs4131542 | 0.227 | 0.267 | 0.619 | 0.374 | -0.001 | -0.004 | -0.004 | 0 | 0.24 | 0.29 | 0.24 | 0.19 | 0.22 | 0.18 |
| 4 | 36461320 | rs10001470 | 0.029 | 0.245 | 0.675 | 0.424 | -0.004 | 0.044 | 0.044 | 1.061 | 0.26 | 0.49 | 0.42 | 0.31 | 0.24 | 0.18 |
| 4 | 36464948 | rs7662602 | 0.5 | 0.18 | 0.623 | 0.324 | -0.005 | -0.023 | -0.023 | 0.027 | 0.31 | 0.34 | 0.16 | 0.16 | 0.26 | 0.22 |
| 4 | 36467624 | rs10016490 | 1.102 | 0.183 | 0.693 | 0.327 | -0.005 | -0.025 | -0.025 | 0 | 0.07 | 0.13 | 0.05 | 0 | 0.04 | 0.03 |
| 4 | 36468162 | rs13112410 | 0.917 | 0.186 | 0.714 | 0.37 | 0.011 | 0.082 | 0.082 | 0.143 | 0.19 | 0.34 | 0.37 | 0.32 | 0.2 | 0.15 |
| 4 | 36470307 | rs3922552 | -0.322 | 0.302 | 0.371 | 0.407 | 0.002 | 0.018 | 0.018 | 0 | 0.47 | 0.39 | 0.23 | 0.38 | 0.47 | 0.44 |
| 4 | 36475328 | rs4240253 | -1.995 | 0.298 | 0.225 | 0.428 | -0.011 | -0.012 | -0.012 | 0.282 | 0.73 | 0.86 | 0.64 | 0.7 | 0.71 | 0.62 |
| 4 | 36483601 | rs7670898 | -2.007 | 0.298 | 0.169 | 0.428 | -0.02 | -0.021 | -0.021 | 0 | 0.74 | 0.89 | 0.65 | 0.69 | 0.71 | 0.62 |
| 4 | 36484686 | rs13133201 | 1.283 | 0.226 | 0.247 | 0.212 | -0.007 | 0.025 | 0.025 | 0 | 0.21 | 0.08 | 0.08 | 0.14 | 0.18 | 0.21 |
| 4 | 36517634 | rs13128058 | -0.559 | 0.239 | 0.159 | 0.188 | 0.014 | 0.034 | 0.034 | 0.129 | 0.4 | 0.68 | 0.38 | 0.27 | 0.33 | 0.26 |
| 4 | 36520451 | rs2375953 | -0.531 | 0.278 | -0.312 | -0.031 | 0.003 | 0.099 | 0.099 | 1.19 | 0.48 | 0.33 | 0.27 | 0.16 | 0.46 | 0.35 |
| 4 | 36527853 | rs2375971 | 0.118 | 0.388 | -0.445 | -0.034 | 0.002 | 0.004 | 0.004 | 0 | 0.39 | 0.41 | 0.32 | 0.47 | 0.41 | 0.43 |
| 4 | 36531423 | rs6829630 | 0.949 | 0.49 | -0.818 | -0.182 | -0.05 | 0.095 | 0.095 | 0.117 | 0.15 | 0.01 | 0.11 | 0 | 0.18 | 0.13 |
| 4 | 36534404 | rs6531458 | 0.474 | 0.389 | -0.344 | 0.307 | -0.011 | 0.016 | 0.016 | 0 | 0.12 | 0.21 | 0.15 | 0.21 | 0.09 | 0.02 |
| 4 | 36540033 | rs11730794 | -0.167 | 0.771 | -0.456 | -0.3 | -0.001 | -0.066 | -0.066 | 0 | 0.41 | 0.36 | 0.24 | 0.07 | 0.37 | 0.25 |
| 4 | 36546041 | rs12507648 | -0.638 | 1.121 | -0.408 | 0.866 | 0.012 | 0.079 | 0.079 | 0 | 0.06 | 0.12 | 0.18 | 0.2 | 0.08 | 0.05 |
| 4 | 36547345 | rs7675488 | -0.117 | 0.883 | -1.072 | 0.594 | 0.009 | -0.014 | -0.014 | 0 | 0.39 | 0.26 | 0.43 | 0.42 | 0.36 | 0.33 |
| 4 | 36555185 | rs35570301 | 0.141 | 0.637 | -0.643 | 0.904 | -0.027 | 0.091 | 0.091 | 0 | 0.19 | 0.38 | 0.28 | 0.36 | 0.12 | 0.23 |
| 4 | 36573898 | rs10010198 | -0.132 | 1.28 | -1.254 | 0.604 | -0.054 | -0.026 | -0.026 | 0 | 0.25 | 0.13 | 0.29 | 0.26 | 0.35 | 0.12 |
| 4 | 36583469 | rs10010760 | -0.074 | 1.281 | -1.254 | 0.605 | -0.054 | -0.027 | -0.027 | 0 | 0.25 | 0.13 | 0.29 | 0.26 | 0.35 | 0.12 |
| 4 | 36597275 | rs7675733 | 0.776 | 1.005 | -1.057 | 0.499 | -0.013 | 0.003 | 0.003 | 0 | 0.2 | 0.34 | 0.1 | 0.2 | 0.18 | 0.3 |
| 4 | 36603313 | rs6814457 | 1.717 | 0.868 | -1.425 | 0.373 | -0.041 | 0.039 | 0.039 | 0 | 0.32 | 0.03 | 0.23 | 0.21 | 0.29 | 0.29 |
| 4 | 36616598 | rs11737439 | -0.159 | 0.784 | -0.948 | 0.398 | 0.001 | -0.012 | -0.012 | 0 | 0.72 | 0.52 | 0.72 | 0.75 | 0.7 | 0.52 |
| 4 | 36620780 | rs4576003 | -0.354 | 0.799 | -0.926 | 0.396 | 0.017 | -0.012 | -0.012 | 0 | 0.92 | 0.86 | 0.83 | 0.94 | 0.88 | 0.81 |
| 4 | 36628000 | rs6820713 | -0.899 | 0.762 | -0.725 | 0.399 | 0.011 | 0.001 | 0.001 | 0 | 0.49 | 0.54 | 0.49 | 0.49 | 0.58 | 0.53 |
| 4 | 36631155 | rs10015119 | 0.804 | 0.816 | -0.856 | 0.369 | -0.056 | -0.036 | -0.036 | 0 | 0.22 | 0.1 | 0.27 | 0.26 | 0.33 | 0.12 |
| 4 | 36637447 | rs6820169 | 1.068 | 0.777 | -0.694 | 0.382 | 0.055 | -0.001 | -0.001 | 0 | 0.46 | 0.29 | 0.41 | 0.49 | 0.35 | 0.4 |
| 4 | 36643076 | rs2375997 | 0.048 | 0.892 | -0.172 | 0.682 | -0.027 | -0.012 | -0.012 | 0 | 0.25 | 0.19 | 0.29 | 0.26 | 0.35 | 0.12 |
| 4 | 36650104 | rs6819096 | 1.214 | 0.7 | -0.076 | 0.725 | 0.007 | 0.012 | 0.012 | 0 | 0.52 | 0.38 | 0.35 | 0.41 | 0.46 | 0.59 |
| 4 | 36650196 | rs16992544 | 0.704 | 0.633 | -0.094 | 0.862 | -0.019 | 0.033 | 0.033 | 0 | 0.14 | 0.22 | 0.18 | 0.27 | 0.07 | 0.12 |
| 4 | 36656080 | rs17465676 | 1.14 | 0.795 | 0.056 | 0.874 | 0.01 | 0.065 | 0.065 | 0 | 0.53 | 0.25 | 0.33 | 0.35 | 0.46 | 0.61 |
| 4 | 36656412 | rs4146579 | -0.283 | 1.202 | -0.231 | 0.769 | -0.067 | 0.018 | 0.018 | 0 | 0.85 | 0.98 | 0.84 | 0.92 | 0.8 | 0.88 |
| 4 | 36665729 | rs2259467 | 0.477 | 1.227 | 0.052 | 0.827 | 0 | 0.003 | 0.003 | 0 | 0.71 | 0.63 | 0.66 | 0.65 | 0.72 | 0.76 |
| 4 | 36671810 | rs4610347 | 0.608 | 1.137 | 0.016 | 1.044 | 0.035 | 0.078 | 0.078 | 0 | 0.1 | 0.15 | 0.2 | 0.26 | 0.18 | 0.07 |
| 4 | 36679326 | rs4833153 | 1.279 | -0.095 | -0.624 | 0.494 | 0.012 | 0.012 | 0.012 | 0 | 0.56 | 0.13 | 0.38 | 0.37 | 0.6 | 0.66 |
| 4 | 36684936 | rs2376150 | 1.278 | -0.143 | -0.712 | 0.47 | 0.007 | 0.009 | 0.009 | 0 | 0.54 | 0.09 | 0.37 | 0.37 | 0.6 | 0.66 |
| 4 | 36692005 | rs6842982 | -1.254 | -0.193 | 0.044 | 0.455 | -0.003 | 0.005 | 0.005 | 0 | 0.36 | 0.55 | 0.55 | 0.63 | 0.34 | 0.29 |
| 4 | 36699959 | rs10517382 | -0.081 | 0.008 | -0.801 | 0.68 | -0.046 | -0.034 | -0.034 | 0 | 0.13 | 0.01 | 0.33 | 0.14 | 0.15 | 0.05 |
| 4 | 36724296 | rs16992596 | 0.158 | -0.028 | -0.759 | 0.878 | -0.025 | -0.063 | -0.063 | 0 | 0.1 | 0.01 | 0.12 | 0.45 | 0.12 | 0.09 |
| 4 | 36733188 | rs6835248 | -1.457 | -0.062 | -0.468 | 0.258 | 0.03 | 0.002 | 0.002 | 0 | 0.42 | 0.79 | 0.56 | 0.61 | 0.37 | 0.27 |
| 4 | 36742625 | rs12511017 | -0.615 | -0.04 | 0.019 | 0.072 | -0.002 | -0.01 | -0.01 | 0.661 | 0.31 | 0.35 | 0.41 | 0.15 | 0.27 | 0.11 |
| 4 | 36756697 | rs12650943 | -0.317 | -0.075 | 0.659 | 0.5 | -0.001 | 0.02 | 0.02 | 0 | 0.18 | 0.21 | 0.16 | 0.45 | 0.18 | 0.3 |
| 4 | 36788024 | rs9683431 | -0.093 | -0.248 | 1.233 | 0.245 | -0.046 | -0.05 | -0.05 | 0 | 0.77 | 0.54 | 0.63 | 0.86 | 0.81 | 0.91 |
| 4 | 36792676 | rs1114364 | 1.185 | -0.319 | 0.93 | 0.275 | 0.043 | -0.01 | -0.01 | 0 | 0.58 | 0.16 | 0.47 | 0.41 | 0.63 | 0.62 |
| 4 | 36792841 | rs13118018 | -0.016 | -0.217 | 0.974 | 0.356 | -0.006 | -0.031 | -0.031 | 0.209 | 0.17 | 0.12 | 0.15 | 0.45 | 0.18 | 0.29 |
| 4 | 36811363 | rs4832818 | 1.301 | 0.079 | 0.472 | -0.081 | -0.054 | 0.041 | 0.041 | 0 | 0.47 | 0.11 | 0.36 | 0.37 | 0.45 | 0.55 |
| 4 | 36814622 | rs1380425 | -0.276 | -0.093 | 1.387 | -0.207 | -0.036 | -0.039 | -0.039 | 0 | 0.78 | 0.63 | 0.66 | 0.85 | 0.81 | 0.91 |
| 4 | 36832942 | rs13152060 | 0.052 | -0.164 | 1.868 | -0.242 | -0.043 | -0.047 | -0.047 | 0 | 0.76 | 0.42 | 0.62 | 0.85 | 0.81 | 0.91 |
| 4 | 36835439 | rs12648032 | 0.129 | -0.293 | 1.563 | -0.302 | 0.002 | -0.061 | -0.061 | 0.123 | 0.21 | 0.11 | 0.17 | 0.45 | 0.19 | 0.29 |
| 4 | 36837111 | rs1370459 | -0.888 | -0.341 | 1.815 | -0.402 | -0.04 | -0.105 | -0.105 | 0.107 | 0.81 | 0.64 | 0.9 | 1 | 0.86 | 0.95 |

| | | | | | | | | | | | | | | | | |
|---|---|---|---|---|---|---|---|---|---|---|---|---|---|---|---|---|
| 4 | 36844337 | rs6531484 | 0.126 | -0.356 | 1.885 | -0.402 | -0.022 | -0.024 | -0.024 | 1.057 | 0.75 | 0.33 | 0.62 | 0.85 | 0.81 | 0.92 |
| 4 | 36847609 | rs1870504 | 0.258 | -0.362 | 1.828 | -0.39 | 0.003 | -0.054 | -0.054 | 0 | 0.2 | 0.12 | 0.16 | 0.45 | 0.18 | 0.29 |
| 4 | 36848556 | rs6856563 | 1.357 | -0.586 | 1.566 | -0.691 | 0.029 | 0.308 | 0.308 | 0 | 0.47 | 0.09 | 0.31 | 0.09 | 0.58 | 0.38 |
| 4 | 36853355 | rs1156274 | -0.051 | -0.268 | 1.938 | -0.226 | -0.046 | -0.047 | -0.047 | 0 | 0.77 | 0.51 | 0.64 | 0.85 | 0.81 | 0.91 |
| 4 | 36863788 | rs4527484 | -0.099 | -0.223 | 1.902 | -0.203 | -0.039 | -0.04 | -0.04 | 0 | 0.78 | 0.55 | 0.64 | 0.84 | 0.81 | 0.92 |
| 4 | 36863831 | rs6828250 | 0.728 | -0.264 | 1.722 | -0.329 | 0.028 | 0.045 | 0.045 | 0 | 0.17 | 0.07 | 0.08 | 0 | 0.14 | 0.05 |
| 4 | 36880769 | rs7660024 | 0.404 | -0.254 | 1.72 | -0.325 | -0.019 | -0.068 | -0.068 | 0 | 0.18 | 0.25 | 0.11 | 0 | 0.14 | 0.05 |
| 4 | 36890895 | rs6817693 | 0.288 | -0.109 | 1.741 | 0.06 | -0.007 | 0.226 | 0.226 | 0.062 | 0.1 | 0.32 | 0.16 | 0.3 | 0.05 | 0.24 |
| 4 | 36915898 | rs11096840 | -1.095 | 0.111 | 1.547 | -0.758 | 0.003 | 0.116 | 0.116 | 0 | 0.38 | 0.77 | 0.49 | 0.16 | 0.4 | 0.16 |
| 4 | 36916955 | rs6531493 | -0.552 | 0.155 | 1.633 | -0.796 | -0.012 | -0.133 | -0.133 | 0 | 0.32 | 0.46 | 0.21 | 0.02 | 0.33 | 0.13 |
| 4 | 36919306 | rs1531327 | -0.3 | -0.018 | 1.536 | -0.713 | -0.006 | -0.015 | -0.015 | 0 | 0.24 | 0.24 | 0.35 | 0.14 | 0.21 | 0.08 |
| 4 | 36924014 | rs12506032 | 1.008 | 0.342 | 0.862 | -0.444 | -0.073 | -0.119 | -0.119 | 0 | 0.39 | 0.06 | 0.31 | 0.4 | 0.35 | 0.51 |
| 4 | 36931424 | rs1373475 | 0.747 | 0.69 | 1.036 | -0.414 | 0.003 | -0.037 | -0.037 | 1.462 | 0.1 | 0.12 | 0.09 | 0.01 | 0.14 | 0.09 |
| 4 | 36946521 | rs11722571 | -0.418 | 0.553 | 0.35 | -0.455 | 0.016 | 0.02 | 0.02 | 0 | 0.46 | 0.19 | 0.38 | 0.42 | 0.45 | 0.37 |
| 4 | 36948102 | rs12505300 | 0.542 | 0.591 | 0.859 | -0.588 | -0.005 | 0.01 | 0.01 | 0 | 0.48 | 0.36 | 0.5 | 0.54 | 0.44 | 0.55 |
| 4 | 36948351 | rs4832844 | 0.42 | 0.589 | 1.073 | -0.594 | 0 | 0.006 | 0.006 | 0 | 0.49 | 0.45 | 0.51 | 0.54 | 0.44 | 0.55 |
| 4 | 36949073 | rs1020808 | -0.072 | 0.731 | 1.557 | -0.354 | 0.023 | -0.065 | -0.065 | 0 | 0.87 | 0.59 | 0.86 | 0.97 | 0.85 | 0.9 |
| 4 | 36960436 | rs1584108 | 0.682 | 0.787 | 1.332 | -0.318 | 0.042 | -0.032 | -0.032 | 0.711 | 0.81 | 0.47 | 0.68 | 0.86 | 0.77 | 0.87 |
| 4 | 36979771 | rs1444310 | 0.905 | 1.046 | 1.341 | -0.765 | -0.01 | 0.01 | 0.01 | 0 | 0.75 | 0.76 | 0.66 | 0.9 | 0.67 | 0.86 |
| 4 | 36980142 | rs13145944 | 1.39 | 0.635 | 0.986 | -0.85 | 0.007 | 0.034 | 0.034 | 0.044 | 0.5 | 0.26 | 0.39 | 0.4 | 0.5 | 0.57 |
| 4 | 36982008 | rs2166094 | -0.641 | 0.719 | 0.814 | -0.815 | 0.008 | -0.03 | -0.03 | 0 | 0.33 | 0.19 | 0.33 | 0.51 | 0.34 | 0.37 |
| 4 | 36983307 | rs9306942 | 0.431 | 0.7 | 0.7 | -0.768 | 0.011 | -0.019 | -0.019 | 0.048 | 0.22 | 0.13 | 0.23 | 0.5 | 0.17 | 0.29 |
| 4 | 36991491 | rs2044294 | -0.105 | 0.817 | 0.69 | -0.827 | 0.025 | -0.077 | -0.077 | 0.071 | 0.14 | 0.29 | 0.12 | 0.02 | 0.18 | 0.13 |
| 4 | 36994638 | rs11096855 | -0.302 | 0.712 | 0.267 | -0.626 | -0.038 | 0.014 | 0.014 | 0 | 0.88 | 0.95 | 0.76 | 0.91 | 0.85 | 0.94 |
| 4 | 36998478 | rs2588548 | 0.5 | 0.657 | 0.372 | -0.629 | -0.003 | 0.003 | 0.003 | 0.242 | 0.65 | 0.68 | 0.52 | 0.42 | 0.68 | 0.7 |
| 4 | 37020558 | rs13111292 | 0.354 | 0.533 | 0.24 | -0.522 | -0.006 | 0.023 | 0.023 | 0.102 | 0.21 | 0.16 | 0.23 | 0.48 | 0.16 | 0.24 |
| 4 | 37034349 | rs1885881 | 1.103 | 1.006 | -0.768 | -0.966 | -0.042 | -0.089 | -0.089 | 0.185 | 0.12 | 0.02 | 0.2 | 0.37 | 0.16 | 0.26 |
| 4 | 37036317 | rs17289939 | 0.012 | 0.845 | 0.054 | -0.865 | -0.055 | 0.119 | 0.119 | 0.086 | 0.25 | 0.43 | 0.28 | 0.48 | 0.17 | 0.24 |
| 4 | 37040171 | rs9999604 | -0.351 | 1.594 | 0.349 | -1.292 | 0.036 | -0.083 | -0.083 | 0.455 | 0.13 | 0.29 | 0.12 | 0.01 | 0.18 | 0.14 |
| 4 | 37041426 | rs17420490 | 0.643 | 1.853 | -0.008 | -1.469 | -0.012 | 0.012 | 0.012 | 0.003 | 0.09 | 0.05 | 0.06 | 0.01 | 0.13 | 0.04 |
| 4 | 37044482 | rs11724117 | 0.643 | 1.853 | -0.008 | -1.469 | -0.012 | 0.012 | 0.012 | 0.003 | 0.09 | 0.05 | 0.06 | 0.01 | 0.13 | 0.04 |
| 4 | 37044712 | rs2376322 | -1.052 | 1.847 | 0.141 | -1.504 | 0.047 | -0.127 | -0.127 | 0.864 | 0.22 | 0.39 | 0.19 | 0.02 | 0.31 | 0.13 |
| 4 | 37046166 | rs11725553 | 0.192 | 1.643 | -0.511 | -1.36 | -0.016 | 0.025 | 0.025 | 0 | 0.09 | 0.03 | 0.06 | 0.02 | 0.14 | 0.07 |
| 4 | 37054888 | rs1957748 | -0.829 | 1.644 | -0.533 | -1.434 | -0.021 | 0.021 | 0.021 | 0 | 0.89 | 0.93 | 0.78 | 0.91 | 0.85 | 0.94 |
| 4 | 37061532 | rs11726724 | 0.132 | 1.624 | -0.664 | -1.493 | -0.009 | 0.032 | 0.032 | 0 | 0.11 | 0.04 | 0.06 | 0.01 | 0.14 | 0.09 |
| 4 | 37062345 | rs4832859 | 1.088 | 1.694 | 0.117 | -1.319 | 0.045 | -0.072 | -0.072 | 0.663 | 0.75 | 0.58 | 0.63 | 0.87 | 0.66 | 0.83 |
| 4 | 37063069 | rs6817779 | 0.387 | 1.244 | 0.241 | -1.033 | 0.023 | -0.028 | -0.028 | 0 | 0.85 | 0.69 | 0.75 | 0.88 | 0.85 | 0.94 |
| 4 | 37067592 | rs6825814 | 0.361 | 1.278 | 0.06 | -1.059 | 0.014 | -0.041 | -0.041 | 0 | 0.85 | 0.75 | 0.75 | 0.9 | 0.85 | 0.94 |
| 4 | 37069574 | rs4832868 | 0.309 | 0.607 | -0.146 | -0.951 | -0.013 | -0.017 | -0.017 | 0 | 0.23 | 0.32 | 0.43 | 0.77 | 0.23 | 0.41 |
| 4 | 37070668 | rs1469312 | -0.309 | 0.024 | -0.318 | -1.383 | -0.041 | -0.107 | -0.107 | 0.024 | 0.78 | 0.61 | 0.88 | 0.97 | 0.85 | 0.68 |
| 4 | 37070916 | rs13144248 | 0.415 | 0.586 | -0.827 | -1.571 | -0.092 | 0.044 | 0.044 | 0 | 0.16 | 0.02 | 0.2 | 0.1 | 0.22 | 0.08 |
| 4 | 37072029 | rs1951774 | 0.23 | 0.491 | -0.619 | -1.432 | -0.007 | -0.014 | -0.014 | 0.378 | 0.33 | 0.53 | 0.51 | 0.74 | 0.32 | 0.47 |
| 4 | 37074040 | rs10484209 | 1.684 | 1.017 | -0.683 | -1.279 | -0.069 | 0.105 | 0.105 | 0 | 0.24 | 0.05 | 0.15 | 0.05 | 0.29 | 0.11 |
| 4 | 37075996 | rs10484210 | 0.885 | 1.001 | -0.67 | -1.188 | -0.071 | -0.041 | -0.041 | 1.93 | 0.11 | 0.02 | 0.19 | 0.16 | 0.2 | 0.06 |
| 4 | 37085237 | rs1426414 | 0.184 | 0.83 | -0.325 | -0.957 | -0.013 | 0.042 | 0.042 | 0 | 0.41 | 0.26 | 0.38 | 0.32 | 0.53 | 0.19 |
| 4 | 37089445 | rs9684775 | 1.737 | 0.857 | -0.397 | -0.977 | -0.07 | 0.078 | 0.078 | 0 | 0.25 | 0.05 | 0.15 | 0.08 | 0.29 | 0.11 |
| 4 | 37091905 | rs4289452 | -0.201 | 0.823 | -0.147 | -1.033 | 0.01 | 0.025 | 0.025 | 0 | 0.45 | 0.39 | 0.43 | 0.32 | 0.59 | 0.2 |
| 4 | 37094678 | rs919103 | -0.016 | 0.826 | -0.163 | -1.032 | 0.011 | 0.038 | 0.038 | 0 | 0.43 | 0.31 | 0.4 | 0.32 | 0.59 | 0.2 |
| 4 | 37097381 | rs17826890 | 1.858 | 0.871 | -0.295 | -1.04 | -0.069 | 0.106 | 0.106 | 0 | 0.24 | 0.05 | 0.15 | 0.05 | 0.28 | 0.11 |
| 4 | 37098598 | rs11733992 | -0.623 | 0.488 | -0.09 | -0.175 | 0.076 | 0.083 | 0.083 | 0 | 0.54 | 0.78 | 0.72 | 0.7 | 0.69 | 0.27 |
| 4 | 37099533 | rs1365342 | -0.682 | 0.45 | -0.134 | -0.193 | 0.063 | 0.076 | 0.076 | 0 | 0.55 | 0.91 | 0.73 | 0.7 | 0.69 | 0.27 |
| 4 | 37099612 | rs12507016 | 0.524 | 0.524 | 0.5 | 0.121 | 0.013 | 0.196 | 0.196 | 0 | 0.06 | 0.09 | 0.27 | 0.38 | 0.09 | 0.07 |
| 4 | 37109848 | rs2376183 | -0.584 | 0.261 | -0.042 | -0.15 | 0.082 | 0.11 | 0.11 | 0 | 0.48 | 0.84 | 0.57 | 0.69 | 0.35 | 0.73 |
| 4 | 37111552 | rs12500588 | -0.903 | 0.48 | 0.242 | 0.164 | 0.051 | 0.437 | 0.437 | 0 | 0.1 | 0.45 | 0.3 | 0.38 | 0.1 | 0.07 |
| 4 | 37126314 | rs1972697 | -0.856 | 0.694 | -0.657 | -0.653 | -0.1 | 0.034 | 0.034 | 0.195 | 0.78 | 0.98 | 0.67 | 0.83 | 0.72 | 0.87 |
| 4 | 37133279 | rs10031432 | 0.451 | -0.026 | -0.382 | -0.64 | -0.023 | -0.026 | -0.026 | 0 | 0.14 | 0.22 | 0.07 | 0.03 | 0.06 | 0.11 |
| 4 | 37134174 | rs34855198 | 0.071 | 0.259 | -0.515 | -0.559 | -0.001 | -0.003 | -0.003 | 0 | 0.34 | 0.34 | 0.42 | 0.67 | 0.36 | 0.59 |
| 4 | 37139301 | rs4832877 | 0.903 | 0.115 | -0.758 | -0.024 | 0.02 | -0.045 | -0.045 | 0 | 0.3 | 0.09 | 0.23 | 0.3 | 0.22 | 0.5 |
| 4 | 37140039 | rs4832726 | 0.387 | 0.222 | -0.615 | -0.224 | 0.008 | 0.007 | 0.007 | 1.458 | 0.36 | 0.29 | 0.4 | 0.67 | 0.3 | 0.56 |
| 4 | 37140633 | rs12502592 | -0.393 | 0.961 | -0.814 | -0.106 | -0.064 | 0.057 | 0.057 | 0.296 | 0.75 | 0.87 | 0.64 | 0.83 | 0.66 | 0.85 |
| 4 | 37143147 | rs725292 | -0.343 | 0.96 | -0.753 | -0.106 | -0.039 | 0.04 | 0.04 | 0.309 | 0.75 | 0.83 | 0.63 | 0.83 | 0.66 | 0.85 |
| 4 | 37156548 | rs10484216 | 0.25 | 0.829 | -0.168 | 0.167 | 0.002 | 0.001 | 0.001 | 0.099 | 0.57 | 0.6 | 0.43 | 0.42 | 0.52 | 0.65 |
| 4 | 37163951 | rs2376241 | -0.398 | 0.998 | 0.087 | 0.187 | -0.026 | 0.037 | 0.037 | 0 | 0.7 | 0.63 | 0.73 | 0.62 | 0.75 | 0.78 |
| 4 | 37165240 | rs17548458 | -0.173 | 1.302 | -0.141 | 0.048 | -0.06 | 0.004 | 0.004 | 0.135 | 0.17 | 0.08 | 0.33 | 0.19 | 0.27 | 0.14 |
| 4 | 37167548 | rs1776529 | 0.439 | 1.232 | 0.002 | 0.057 | -0.001 | 0 | 0 | 0 | 0.55 | 0.68 | 0.41 | 0.42 | 0.48 | 0.64 |
| 4 | 37173484 | rs12510063 | 0 | 1.682 | 0.312 | 0.384 | -0.084 | -0.033 | -0.033 | 0 | 0.15 | 0.02 | 0.32 | 0.19 | 0.24 | 0.13 |
| 4 | 37175523 | rs1648404 | 0.445 | 1.295 | 0.098 | -0.035 | 0 | 0 | 0 | 0 | 0.46 | 0.57 | 0.39 | 0.42 | 0.48 | 0.62 |
| 4 | 37184279 | rs1627800 | 0.416 | 1.286 | 0.09 | -0.021 | -0.002 | 0.001 | 0.001 | 0.074 | 0.46 | 0.63 | 0.4 | 0.42 | 0.48 | 0.62 |
| 4 | 37187191 | rs1648395 | -0.473 | 1.295 | -0.277 | -0.237 | 0.012 | 0.007 | 0.007 | 0.517 | 0.46 | 0.72 | 0.64 | 0.59 | 0.56 | 0.6 |

| | | | | | | | | | | | | | | | | |
|---|---|---|---|---|---|---|---|---|---|---|---|---|---|---|---|---|
| 4 | 37187913 | rs1342993 | 0.58 | 1.285 | -0.246 | -0.217 | 0.002 | 0.006 | 0.006 | 0.177 | 0.31 | 0.7 | 0.32 | 0.37 | 0.33 | 0.47 |
| 4 | 37197362 | rs1648392 | -0.761 | 1.264 | -0.067 | -0.306 | 0.008 | 0.003 | 0.003 | 0 | 0.6 | 0.7 | 0.7 | 0.64 | 0.66 | 0.74 |
| 4 | 37198777 | rs1426401 | -0.528 | 1.214 | 0.519 | -0.381 | 0.009 | 0.006 | 0.006 | 0 | 0.56 | 0.45 | 0.66 | 0.64 | 0.64 | 0.74 |
| 4 | 37207259 | rs1418876 | 1.159 | 1.191 | 0.499 | -0.508 | -0.001 | -0.006 | -0.006 | 0.15 | 0.23 | 0.22 | 0.24 | 0.31 | 0.23 | 0.44 |
| 4 | 37210649 | rs11937850 | 0.81 | 0.858 | 0.384 | -0.967 | -0.007 | -0.02 | -0.02 | 0 | 0.22 | 0.34 | 0.14 | 0.12 | 0.19 | 0.16 |
| 4 | 37218366 | rs1435386 | -0.191 | 0.975 | 0.152 | -0.9 | -0.002 | -0.044 | -0.044 | 0.002 | 0.44 | 0.49 | 0.71 | 0.87 | 0.52 | 0.64 |
| 4 | 37232282 | rs16993352 | 0.636 | 1.223 | 0.116 | -0.523 | -0.008 | -0.033 | -0.033 | 0 | 0.22 | 0.39 | 0.15 | 0.12 | 0.19 | 0.18 |
| 4 | 37235854 | rs1776575 | 0.273 | 1.229 | 0.081 | -0.554 | 0.004 | -0.052 | -0.052 | 0 | 0.24 | 0.46 | 0.17 | 0.12 | 0.26 | 0.19 |
| 4 | 37238325 | rs1435383 | -1.247 | 0.79 | -0.958 | -0.877 | 0.038 | 0.092 | 0.092 | 0.548 | 0.73 | 0.92 | 0.88 | 0.99 | 0.79 | 0.8 |
| 4 | 37240192 | rs2211439 | 0.867 | 1.291 | -1.041 | -0.897 | -0.032 | 0.035 | 0.035 | 0 | 0.07 | 0.01 | 0.06 | 0 | 0.09 | 0.02 |
| 4 | 37240580 | rs1023730 | 0.126 | 1.059 | -0.907 | -0.412 | 0.002 | 0.008 | 0.008 | 2.84 | 0.46 | 0.67 | 0.37 | 0.42 | 0.44 | 0.58 |
| 4 | 37244135 | rs1656197 | -0.183 | 1.181 | -0.941 | -0.343 | 0.007 | 0.012 | 0.012 | 3.431 | 0.49 | 0.61 | 0.38 | 0.43 | 0.44 | 0.59 |
| 4 | 37250289 | rs1469807 | -0.986 | 0.99 | -1.657 | -0.168 | 0.024 | 0.136 | 0.136 | 0.782 | 0.71 | 0.95 | 0.86 | 0.99 | 0.75 | 0.79 |
| 4 | 37251630 | rs7669853 | -0.004 | 1.159 | -0.994 | 0.655 | 0.002 | -0.002 | -0.002 | 0 | 0.25 | 0.2 | 0.24 | 0.26 | 0.21 | 0.37 |
| 4 | 37254333 | rs989655 | -0.572 | 1.201 | -1.064 | 0.86 | 0 | -0.001 | -0.001 | 0.341 | 0.4 | 0.63 | 0.57 | 0.58 | 0.43 | 0.55 |
| 4 | 37254584 | rs1618748 | -0.45 | 1.043 | -1.457 | -0.08 | 0 | 0.152 | 0.152 | 0 | 0.54 | 0.82 | 0.73 | 0.91 | 0.53 | 0.62 |
| 4 | 37254768 | rs11932737 | -0.057 | 0.441 | -1.22 | 0.866 | 0.01 | -0.018 | -0.018 | 0 | 0.14 | 0.1 | 0.16 | 0.26 | 0.09 | 0.24 |
| 4 | 37255211 | rs7661725 | 0.013 | 0.376 | -0.93 | 0.307 | 0 | -0.001 | -0.001 | 0 | 0.68 | 0.65 | 0.74 | 0.74 | 0.69 | 0.75 |
| 4 | 37257069 | rs7674779 | -0.508 | 0.438 | -0.407 | -0.111 | -0.001 | -0.035 | -0.035 | 0 | 0.26 | 0.32 | 0.23 | 0.17 | 0.3 | 0.13 |
| 4 | 37258086 | rs10015594 | 0.916 | 0.772 | -0.309 | 0.123 | 0.007 | 0.009 | 0.009 | 0 | 0.07 | 0.13 | 0.04 | 0.07 | 0.09 | 0.05 |
| 4 | 37258713 | rs6531521 | 0.736 | 0.94 | -0.781 | -0.528 | -0.03 | 0.035 | 0.035 | 0 | 0.22 | 0.13 | 0.19 | 0.02 | 0.29 | 0.19 |
| 4 | 37262286 | rs6855364 | 1.111 | 0.648 | -0.495 | 0.02 | 0.003 | 0.008 | 0.008 | 0 | 0.07 | 0.14 | 0.06 | 0.08 | 0.09 | 0.05 |
| 4 | 37264627 | rs4832890 | 0.248 | 0.913 | -0.671 | -0.503 | -0.078 | 0.093 | 0.093 | 0 | 0.27 | 0.15 | 0.22 | 0.02 | 0.4 | 0.2 |
| 4 | 37264831 | rs3910363 | 1.611 | 0.232 | -1.079 | -0.293 | 0 | -0.019 | -0.019 | 1.155 | 0.2 | 0.06 | 0.28 | 0.26 | 0.18 | 0.33 |
| 4 | 37268477 | rs13129949 | 1.299 | -0.137 | -1.346 | -0.517 | 0.017 | 0.032 | 0.032 | 0 | 0.17 | 0.07 | 0.09 | 0.06 | 0.15 | 0.12 |
| 4 | 37269105 | rs1435378 | 0.907 | -0.113 | -1.514 | -0.465 | -0.01 | -0.034 | -0.034 | 0 | 0.28 | 0.14 | 0.33 | 0.33 | 0.27 | 0.38 |
| 4 | 37272433 | rs1594983 | -0.7 | -0.059 | -1.688 | -0.522 | -0.146 | -0.111 | -0.111 | 0 | 0.59 | 0.95 | 0.57 | 0.61 | 0.56 | 0.53 |
| 4 | 37285195 | rs1776560 | -1.333 | -0.283 | -1.321 | -0.217 | 0.002 | -0.117 | -0.117 | 0 | 0.79 | 0.96 | 0.75 | 0.63 | 0.85 | 0.73 |
| 4 | 37291336 | rs1656182 | 0.274 | -0.179 | -0.546 | -0.179 | 0.008 | 0.011 | 0.011 | 0 | 0.52 | 0.54 | 0.49 | 0.38 | 0.59 | 0.56 |
| 4 | 37293830 | rs1656235 | -0.293 | 0.045 | -0.617 | -0.375 | -0.017 | 0.045 | 0.045 | 0 | 0.43 | 0.72 | 0.37 | 0.35 | 0.52 | 0.3 |
| 4 | 37295894 | rs1625910 | -0.368 | 0.305 | 0.148 | -0.433 | 0.012 | -0.069 | -0.069 | 0 | 0.21 | 0.33 | 0.16 | 0.07 | 0.23 | 0.08 |
| 4 | 37297719 | rs7664806 | 0.69 | 0.414 | 0.092 | -0.33 | -0.003 | -0.001 | -0.001 | 0.752 | 0.15 | 0.13 | 0.15 | 0.07 | 0.2 | 0.07 |
| 4 | 37302992 | rs13115793 | 0.685 | 0.028 | 0.347 | -0.138 | 0.001 | 0.005 | 0.005 | 0 | 0.5 | 0.45 | 0.41 | 0.39 | 0.5 | 0.59 |
| 4 | 37319418 | rs1435389 | 1.496 | -0.035 | 0.058 | -0.196 | -0.029 | -0.066 | -0.066 | 0.739 | 0.19 | 0.01 | 0.19 | 0.25 | 0.17 | 0.34 |
| 4 | 37322825 | rs4337743 | 1.205 | -0.03 | 0.11 | -0.373 | -0.058 | 0.125 | 0.125 | 0 | 0.29 | 0.1 | 0.2 | 0.01 | 0.33 | 0.25 |
| 4 | 37328396 | rs17603641 | 0.752 | 0.077 | 0.214 | -0.022 | 0.004 | -0.017 | -0.017 | 0 | 0.11 | 0.08 | 0.07 | 0.18 | 0.12 | 0.08 |
| 4 | 37329512 | rs17575890 | 0.123 | 0.026 | 0.634 | -0.034 | 0.001 | -0.002 | -0.002 | 0 | 0.32 | 0.28 | 0.28 | 0.43 | 0.29 | 0.41 |
| 4 | 37330939 | rs936595 | -1.64 | -0.162 | 0.503 | -0.18 | -0.002 | -0.006 | -0.006 | 0.078 | 0.67 | 0.69 | 0.78 | 0.86 | 0.66 | 0.75 |
| 4 | 37331417 | rs7679428 | 0.544 | 0.074 | 0.564 | -0.041 | -0.001 | -0.012 | -0.012 | 0 | 0.18 | 0.17 | 0.18 | 0.07 | 0.24 | 0.09 |
| 4 | 37335338 | rs7434368 | 1.468 | 0.093 | 0.282 | -0.005 | -0.024 | -0.064 | -0.064 | 0.494 | 0.19 | 0.01 | 0.2 | 0.25 | 0.18 | 0.33 |
| 4 | 37335699 | rs12649785 | 0.967 | 0.14 | 0.499 | 0.067 | 0.001 | 0.001 | 0.001 | 0 | 0.16 | 0.15 | 0.32 | 0.36 | 0.13 | 0.24 |
| 4 | 37335756 | rs12650456 | -0.028 | 0.145 | 0.606 | 0.205 | 0.001 | 0.018 | 0.018 | 0 | 0.36 | 0.44 | 0.52 | 0.44 | 0.37 | 0.34 |
| 4 | 37336553 | rs17603726 | 0.686 | 0.152 | 0.292 | 0.421 | 0.003 | -0.007 | -0.007 | 0.128 | 0.19 | 0.13 | 0.17 | 0.25 | 0.17 | 0.3 |
| 4 | 37336803 | rs17575939 | 0.206 | 0.174 | 0.338 | 0.817 | 0.004 | 0.06 | 0.06 | 0 | 0.19 | 0.34 | 0.31 | 0.3 | 0.18 | 0.26 |
| 4 | 37342620 | rs13340226 | 0.15 | 0.232 | -0.783 | 0.737 | -0.021 | -0.041 | -0.041 | 1.594 | 0.34 | 0.21 | 0.53 | 0.49 | 0.36 | 0.38 |
| 4 | 37348193 | rs11096868 | 0.699 | 0.385 | -1.095 | 0.754 | -0.011 | -0.065 | -0.065 | 0 | 0.28 | 0.13 | 0.42 | 0.41 | 0.26 | 0.31 |
| 4 | 37349537 | rs13120150 | 0.348 | 0.053 | -1.338 | 0.694 | -0.054 | 0.137 | 0.137 | 0 | 0.43 | 0.1 | 0.31 | 0.23 | 0.46 | 0.35 |
| 4 | 37357559 | rs1002363 | 0.809 | 0.213 | -1.14 | 0.409 | -0.023 | -0.045 | -0.045 | 0.217 | 0.15 | 0.01 | 0.09 | 0.19 | 0.14 | 0.24 |
| 4 | 37360983 | rs3892368 | -0.278 | 0.355 | -0.478 | 0.272 | 0.002 | 0.138 | 0.138 | 0 | 0.08 | 0.18 | 0.06 | 0.2 | 0.08 | 0.07 |
| 4 | 37368174 | rs6850878 | -0.034 | 0.416 | -1.27 | -0.659 | -0.02 | 0.131 | 0.131 | 0 | 0.39 | 0.21 | 0.29 | 0.01 | 0.41 | 0.3 |
| 4 | 37369204 | rs10517410 | 0.47 | 0.284 | -0.732 | 0.454 | -0.001 | 0.018 | 0.018 | 0 | 0.05 | 0.09 | 0.04 | 0.1 | 0.06 | 0.04 |
| 4 | 37374046 | rs6823583 | 0.139 | 0.15 | -2.183 | 0.441 | 0.015 | -0.06 | -0.06 | 0 | 0.35 | 0.16 | 0.36 | 0.54 | 0.29 | 0.28 |
| 4 | 37382184 | rs764882 | 0.232 | 0.252 | -2.749 | -0.018 | -0.002 | 0.021 | 0.021 | 0 | 0.12 | 0 | 0.04 | 0.04 | 0.11 | 0.14 |
| 4 | 37385010 | rs17493545 | 0.628 | 0.952 | -2.918 | -0.232 | -0.061 | -0.018 | -0.018 | 0 | 0.16 | 0.01 | 0.18 | 0.13 | 0.19 | 0.24 |
| 4 | 37385086 | rs7375557 | 0.081 | 0.804 | -2.512 | -0.345 | -0.108 | 0.148 | 0.148 | 0.074 | 0.26 | 0.04 | 0.22 | 0.06 | 0.34 | 0.28 |
| 4 | 37386498 | rs17603961 | 0.176 | 0.023 | -2.029 | 0.49 | 0.009 | -0.028 | -0.028 | 0 | 0.09 | 0.03 | 0.28 | 0.22 | 0.08 | 0.06 |
| 4 | 37386993 | rs13131217 | -0.273 | -0.334 | -1.853 | -0.019 | 0.015 | 0.018 | 0.018 | 0 | 0.16 | 0.11 | 0.08 | 0.04 | 0.09 | 0.12 |
| 4 | 37390658 | rs6837863 | -0.86 | -0.114 | -1.503 | -0.042 | 0.005 | 0.031 | 0.031 | 0 | 0.49 | 0.65 | 0.45 | 0.34 | 0.55 | 0.55 |
| 4 | 37391031 | rs10013737 | -0.252 | -0.005 | -1.015 | -0.091 | -0.002 | -0.048 | -0.048 | 0 | 0.35 | 0.46 | 0.25 | 0.16 | 0.34 | 0.36 |
| 4 | 37393292 | rs10013774 | -1.308 | -0.268 | -1.142 | -0.272 | -0.003 | -0.005 | -0.005 | 0 | 0.55 | 0.52 | 0.46 | 0.35 | 0.58 | 0.59 |
| 4 | 37395148 | rs17421729 | 1.469 | -0.081 | -1.084 | -0.237 | -0.061 | -0.062 | -0.062 | 0 | 0.13 | 0.01 | 0.19 | 0.18 | 0.2 | 0.19 |
| 4 | 37395513 | rs6531533 | -1.52 | -0.471 | -1.257 | -0.06 | 0.013 | 0.004 | 0.004 | 0 | 0.75 | 0.61 | 0.6 | 0.72 | 0.73 | 0.79 |
| 4 | 37400306 | rs10028027 | 0.485 | -0.327 | -0.504 | -0.04 | 0.038 | -0.013 | -0.013 | 0 | 0.29 | 0.38 | 0.32 | 0.21 | 0.46 | 0.41 |
| 4 | 37402609 | rs12646961 | 0.38 | -0.686 | -0.092 | 0.357 | -0.042 | 0.01 | 0.01 | 0 | 0.2 | 0.32 | 0.15 | 0.2 | 0.08 | 0.14 |
| 4 | 37403481 | rs1105930 | 0.171 | -0.718 | -0.326 | 0.151 | -0.008 | 0.008 | 0.008 | 0 | 0.23 | 0.27 | 0.18 | 0.32 | 0.17 | 0.22 |
| 4 | 37404064 | rs17493678 | 1.975 | -0.628 | -0.741 | 0.024 | 0.004 | 0.019 | 0.019 | 0 | 0.07 | 0.03 | 0.01 | 0.01 | 0.04 | 0.03 |
| 4 | 37404319 | rs9306948 | 1.842 | -0.352 | -0.577 | -0.01 | -0.012 | 0.009 | 0.009 | 0 | 0.13 | 0.08 | 0.13 | 0.06 | 0.26 | 0.11 |
| 4 | 37406051 | rs10016142 | 1.311 | -0.487 | -0.897 | 0.049 | -0.024 | 0.05 | 0.05 | 1.063 | 0.28 | 0.07 | 0.27 | 0.19 | 0.29 | 0.27 |
| 4 | 37417390 | rs4586959 | -0.959 | -0.369 | -0.561 | 0.073 | -0.01 | 0.034 | 0.034 | 1.022 | 0.59 | 0.71 | 0.59 | 0.75 | 0.46 | 0.61 |

| | | | | | | | | | | | | | | | | |
|---|---|---|---|---|---|---|---|---|---|---|---|---|---|---|---|---|
| 4 | 37417596 | rs6816980 | 1.44 | -0.373 | -0.984 | 0.322 | -0.002 | -0.022 | -0.022 | 0 | 0.16 | 0.1 | 0.16 | 0.31 | 0.15 | 0.19 |
| 4 | 37422817 | rs2928288 | 1.255 | -0.28 | -0.956 | -0.031 | -0.003 | 0.022 | 0.022 | 2.982 | 0.27 | 0.15 | 0.29 | 0.21 | 0.29 | 0.32 |
| 4 | 37427661 | rs10856831 | 1.476 | -0.132 | -1.202 | 0.162 | -0.049 | -0.058 | -0.058 | 2.205 | 0.14 | 0 | 0.21 | 0.16 | 0.19 | 0.22 |
| 4 | 37432416 | rs2973217 | -0.044 | -0.265 | -0.822 | 0.109 | -0.013 | 0.015 | 0.015 | 0.638 | 0.54 | 0.29 | 0.52 | 0.39 | 0.52 | 0.57 |
| 4 | 37445315 | rs4634233 | 1.322 | 0.324 | -0.793 | 0.099 | -0.014 | 0.01 | 0.01 | 0 | 0.15 | 0.12 | 0.17 | 0.06 | 0.28 | 0.14 |
| 4 | 37445885 | rs11724784 | 0.24 | -0.055 | -1.044 | -0.043 | -0.005 | 0.006 | 0.006 | 0 | 0.23 | 0.11 | 0.22 | 0.16 | 0.22 | 0.26 |
| 4 | 37448299 | rs17422084 | 1.216 | -0.074 | -1.087 | -0.078 | 0.006 | 0.002 | 0.002 | 0 | 0.08 | 0.05 | 0.01 | 0 | 0.04 | 0.04 |
| 4 | 37449053 | rs936587 | -0.102 | -0.479 | -0.693 | -0.187 | 0.002 | -0.002 | -0.002 | 0.473 | 0.55 | 0.49 | 0.55 | 0.39 | 0.53 | 0.61 |
| 4 | 37449086 | rs936588 | 1.34 | -0.498 | -0.766 | -0.106 | -0.052 | -0.058 | -0.058 | 0 | 0.15 | 0 | 0.21 | 0.16 | 0.18 | 0.22 |
| 4 | 37449803 | rs10020323 | 0.514 | -0.567 | -0.228 | -0.056 | -0.003 | 0.01 | 0.01 | 0 | 0.26 | 0.25 | 0.26 | 0.52 | 0.18 | 0.23 |
| 4 | 37457601 | rs2928297 | 0.284 | -0.563 | -0.255 | -0.081 | -0.001 | -0.002 | -0.002 | 0.978 | 0.46 | 0.51 | 0.53 | 0.39 | 0.49 | 0.54 |
| 4 | 37466250 | rs2928304 | 0.153 | -0.348 | -0.24 | -0.003 | 0.001 | 0.009 | 0.009 | 0.155 | 0.52 | 0.56 | 0.55 | 0.39 | 0.51 | 0.54 |
| 4 | 37469074 | rs2973235 | -0.697 | -0.458 | -0.605 | 0.22 | 0 | 0 | 0 | 0 | 0.84 | 0.81 | 0.92 | 0.81 | 0.87 | 0.89 |
| 4 | 37469174 | rs2928307 | 0.71 | -0.59 | -0.596 | -0.016 | -0.005 | 0.002 | 0.002 | 0 | 0.42 | 0.47 | 0.51 | 0.58 | 0.36 | 0.49 |
| 4 | 37470428 | rs11096877 | 0.77 | -0.577 | -0.83 | -0.044 | -0.011 | 0.016 | 0.016 | 0 | 0.25 | 0.26 | 0.22 | 0.08 | 0.32 | 0.18 |
| 4 | 37471031 | rs10517414 | -0.005 | -0.547 | -0.776 | -0.01 | -0.044 | 0.043 | 0.043 | 0 | 0.4 | 0.26 | 0.4 | 0.23 | 0.5 | 0.39 |
| 4 | 37471169 | rs2376756 | -0.704 | -0.462 | -0.229 | 0.251 | 0 | 0 | 0 | 1.617 | 0.84 | 0.81 | 0.92 | 0.81 | 0.87 | 0.88 |
| 4 | 37473384 | rs11941570 | 0.824 | -0.441 | -0.322 | 0.265 | 0 | 0 | 0 | 0 | 0.17 | 0.2 | 0.19 | 0.15 | 0.19 | 0.22 |
| 4 | 37476233 | rs1382984 | 0.695 | -0.393 | -0.249 | 0.215 | 0.002 | 0.004 | 0.004 | 0.042 | 0.5 | 0.47 | 0.59 | 0.39 | 0.56 | 0.42 |
| 4 | 37480687 | rs1382979 | 0.041 | -0.381 | 0.145 | 0.226 | -0.022 | 0.024 | 0.024 | 0.178 | 0.68 | 0.43 | 0.67 | 0.57 | 0.74 | 0.54 |
| 4 | 37481131 | rs2928311 | 0.113 | -0.437 | 0.121 | 0.217 | -0.02 | 0.02 | 0.02 | 0.163 | 0.67 | 0.43 | 0.66 | 0.57 | 0.73 | 0.54 |
| 4 | 37486417 | rs952903 | -0.392 | -0.001 | 0.453 | 0.277 | 0.025 | 0.007 | 0.007 | 0 | 0.35 | 0.48 | 0.34 | 0.4 | 0.39 | 0.42 |
| 4 | 37491386 | rs1552021 | -1.436 | -0.207 | 0.397 | 0.095 | 0.001 | -0.002 | -0.002 | 0 | 0.61 | 0.65 | 0.62 | 0.56 | 0.57 | 0.8 |
| 4 | 37503658 | rs2973255 | 0.07 | -0.205 | 0.38 | 0.024 | 0.022 | 0.047 | 0.047 | 0 | 0.32 | 0.45 | 0.36 | 0.45 | 0.35 | 0.42 |
| 4 | 37509330 | rs1600392 | -1.826 | -0.452 | 0.064 | 0.126 | -0.005 | 0.005 | 0.005 | 0.884 | 0.92 | 0.9 | 0.97 | 0.87 | 0.95 | 0.99 |
| 4 | 37513594 | rs2973271 | -0.862 | 0.025 | 0.492 | 0.001 | -0.014 | 0.047 | 0.047 | 0 | 0.47 | 0.17 | 0.35 | 0.39 | 0.48 | 0.46 |
| 4 | 37516608 | rs2928330 | -1.779 | 0.067 | -0.033 | 0.035 | 0.004 | -0.027 | -0.027 | 0 | 0.6 | 0.76 | 0.62 | 0.52 | 0.54 | 0.7 |
| 4 | 37517112 | rs1463575 | -1.044 | 0.118 | -0.332 | -0.149 | 0.001 | 0.011 | 0.011 | 0 | 0.48 | 0.42 | 0.39 | 0.35 | 0.48 | 0.46 |
| 4 | 37518464 | rs17576970 | 0.051 | 0.511 | -0.093 | 0.149 | -0.006 | 0.032 | 0.032 | 0.102 | 0.1 | 0.19 | 0.22 | 0.13 | 0.06 | 0.23 |
| 4 | 37519030 | rs4832912 | 0.555 | 1.058 | -0.212 | -0.436 | 0.051 | 0.168 | 0.168 | 0 | 0.13 | 0.37 | 0.15 | 0.31 | 0.14 | 0.17 |
| 4 | 37523672 | rs6819803 | 0.272 | 1.078 | -0.708 | -0.818 | 0.005 | 0.105 | 0.105 | 0 | 0.18 | 0.03 | 0.06 | 0.01 | 0.12 | 0.07 |
| 4 | 37524887 | rs2380725 | -1.397 | 1.075 | -0.657 | -0.814 | 0 | 0 | 0 | 0 | 0.41 | 0.56 | 0.39 | 0.32 | 0.32 | 0.52 |
| 4 | 37524958 | rs2380727 | -0.566 | 1.089 | -0.64 | -0.797 | 0.014 | 0.043 | 0.043 | 0 | 0.23 | 0.53 | 0.33 | 0.31 | 0.2 | 0.45 |
| 4 | 37526213 | rs2973296 | -0.92 | 1.085 | -0.772 | -0.77 | -0.016 | 0.008 | 0.008 | 0 | 0.39 | 0.67 | 0.51 | 0.35 | 0.42 | 0.65 |
| 4 | 37526293 | rs2939712 | -1.16 | 1.132 | -0.385 | -0.351 | 0.011 | 0.02 | 0.02 | 0 | 0.57 | 0.69 | 0.57 | 0.36 | 0.54 | 0.72 |
| 4 | 37527529 | rs2380734 | -0.468 | 1.637 | 0.26 | 0.666 | 0.053 | 0.129 | 0.129 | 0 | 0.14 | 0.37 | 0.14 | 0.27 | 0.14 | 0.16 |
| 4 | 37536119 | rs2380737 | -0.597 | 1.602 | -0.368 | -0.126 | 0.017 | -0.067 | -0.067 | 0.003 | 0.22 | 0.34 | 0.21 | 0.04 | 0.26 | 0.24 |
| 4 | 37537307 | rs2973285 | 0.097 | 0.976 | -0.889 | 0.153 | 0.021 | 0.007 | 0.007 | 1.388 | 0.58 | 0.25 | 0.62 | 0.55 | 0.67 | 0.72 |
| 4 | 37537569 | rs2939748 | 0.196 | 0.799 | -0.991 | -0.193 | -0.01 | 0.099 | 0.099 | 0 | 0.52 | 0.31 | 0.5 | 0.27 | 0.52 | 0.66 |
| 4 | 37538137 | rs13121029 | -0.275 | 0.893 | -0.654 | -0.597 | 0.028 | -0.045 | -0.045 | 0 | 0.31 | 0.54 | 0.28 | 0.14 | 0.36 | 0.22 |
| 4 | 37542625 | rs10517412 | 0.555 | 0.653 | -0.827 | -0.266 | 0.007 | -0.009 | -0.009 | 0.768 | 0.09 | 0.06 | 0.13 | 0.12 | 0.09 | 0.11 |
| 4 | 37542764 | rs17604934 | 0.458 | 0.07 | -1.421 | 0.15 | 0.013 | -0.096 | -0.096 | 0 | 0.27 | 0.06 | 0.25 | 0.37 | 0.25 | 0.3 |
| 4 | 37543344 | rs2939745 | 0.607 | 0.156 | -1.176 | 0.05 | -0.054 | -0.083 | -0.083 | 0.041 | 0.26 | 0.03 | 0.3 | 0.36 | 0.28 | 0.34 |
| 4 | 37548617 | rs2939739 | -0.101 | 0.151 | -0.668 | -0.032 | 0 | 0.01 | 0.01 | 0.02 | 0.33 | 0.35 | 0.39 | 0.47 | 0.32 | 0.5 |
| 4 | 37555201 | rs7655445 | -0.241 | 0.086 | -1.177 | -0.295 | 0.002 | 0.024 | 0.024 | 0 | 0.7 | 0.78 | 0.74 | 0.75 | 0.71 | 0.72 |
| 4 | 37555279 | rs7679643 | 0.05 | -0.372 | -0.874 | -0.732 | 0.002 | -0.037 | -0.037 | 0 | 0.22 | 0.18 | 0.12 | 0 | 0.17 | 0.16 |
| 4 | 37559226 | rs2939720 | 0.073 | -0.127 | -0.848 | -0.399 | -0.001 | -0.008 | -0.008 | 1.471 | 0.42 | 0.53 | 0.44 | 0.37 | 0.45 | 0.39 |
| 4 | 37561254 | rs11942503 | 0.406 | -0.102 | -1.163 | -0.361 | -0.022 | -0.034 | -0.034 | 0 | 0.23 | 0.19 | 0.41 | 0.48 | 0.34 | 0.29 |
| 4 | 37562007 | rs1876804 | -0.29 | -0.237 | -0.731 | -0.454 | -0.012 | -0.014 | -0.014 | 0 | 0.48 | 0.28 | 0.49 | 0.5 | 0.49 | 0.53 |
| 4 | 37562604 | rs2941002 | 1.052 | -0.334 | -0.522 | -0.787 | -0.001 | 0.004 | 0.004 | 0 | 0.35 | 0.3 | 0.34 | 0.22 | 0.4 | 0.29 |
| 4 | 37565263 | rs11723493 | -0.391 | -0.147 | -1.028 | -0.614 | 0.002 | 0.036 | 0.036 | 0 | 0.51 | 0.37 | 0.55 | 0.64 | 0.5 | 0.63 |
| 4 | 37569593 | rs13148220 | 0.546 | -0.075 | -1.141 | -0.804 | 0.014 | 0.044 | 0.044 | 0 | 0.22 | 0.07 | 0.14 | 0.11 | 0.2 | 0.19 |
| 4 | 37569966 | rs9685958 | 0.832 | 0.144 | -1.005 | -0.735 | 0.008 | -0.036 | -0.036 | 0.929 | 0.17 | 0.22 | 0.14 | 0.02 | 0.27 | 0.13 |
| 4 | 37575251 | rs10025159 | 0.006 | 0.05 | -1.088 | -0.638 | 0 | 0.016 | 0.016 | 0 | 0.3 | 0.22 | 0.22 | 0.25 | 0.28 | 0.38 |
| 4 | 37577791 | rs2380763 | 0.935 | 0.151 | -1.417 | -0.768 | -0.042 | 0.167 | 0.167 | 0 | 0.35 | 0.12 | 0.18 | 0.03 | 0.38 | 0.26 |
| 4 | 37584100 | rs2973236 | 0.629 | 0.362 | -0.773 | 0.202 | -0.069 | 0.07 | 0.07 | 0 | 0.2 | 0.01 | 0.23 | 0.08 | 0.26 | 0.16 |
| 4 | 37584399 | rs6857179 | -0.678 | 0.317 | 0.466 | 0.837 | 0.062 | 0.204 | 0.204 | 2.408 | 0.05 | 0.41 | 0.09 | 0.13 | 0.01 | 0.18 |
| 4 | 37585670 | rs2276944 | 0.601 | 0.695 | -0.528 | 1.003 | -0.001 | 0.016 | 0.016 | 0 | 0.09 | 0.05 | 0.33 | 0.62 | 0.1 | 0.17 |
| 4 | 37590570 | rs2279994 | 0.207 | 0.99 | -1.091 | 0.11 | -0.022 | 0.042 | 0.042 | 0 | 0.21 | 0.12 | 0.17 | 0.08 | 0.26 | 0.15 |
| 4 | 37592128 | rs2973275 | 0.442 | 0.949 | -1.162 | 0.133 | -0.047 | 0.062 | 0.062 | 0.046 | 0.19 | 0.05 | 0.15 | 0.08 | 0.25 | 0.14 |
| 4 | 37593173 | rs13333 | -0.227 | 0.227 | -1.519 | 0.501 | 0.032 | -0.031 | -0.031 | 0 | 0.64 | 0.23 | 0.46 | 0.41 | 0.76 | 0.5 |
| 4 | 37594946 | rs705099 | -0.483 | 0.209 | -1.296 | 0.678 | 0.011 | -0.012 | -0.012 | 0 | 0.66 | 0.34 | 0.64 | 0.54 | 0.77 | 0.69 |
| 4 | 37595898 | rs2939744 | 0.215 | -0.336 | -1.615 | 0.497 | 0.105 | 0.105 | 0.105 | 0 | 0.57 | 0.2 | 0.41 | 0.22 | 0.75 | 0.46 |
| 4 | 37596650 | rs953538 | 0.252 | -0.254 | -1.592 | 0.492 | 0.11 | 0.117 | 0.117 | 0 | 0.55 | 0.19 | 0.39 | 0.22 | 0.73 | 0.46 |
| 4 | 37597591 | rs2973276 | -0.628 | -0.244 | -1.228 | 0.525 | -0.009 | 0.004 | 0.004 | 0 | 0.65 | 0.4 | 0.49 | 0.51 | 0.75 | 0.54 |
| 4 | 37600835 | rs2973278 | -1.322 | -0.388 | -1.142 | 0.261 | 0.032 | 0.043 | 0.043 | 0.135 | 0.74 | 0.86 | 0.88 | 1 | 0.83 | 0.91 |
| 4 | 37604570 | rs2973280 | 0.391 | -0.32 | -1.23 | 0.323 | 0.004 | 0.038 | 0.038 | 0 | 0.46 | 0.19 | 0.41 | 0.36 | 0.42 | 0.49 |
| 4 | 37604982 | rs10050295 | 0.521 | -0.453 | -1.226 | 0.138 | -0.065 | 0.144 | 0.144 | 0 | 0.38 | 0.17 | 0.24 | 0.02 | 0.5 | 0.3 |
| 4 | 37606858 | rs2939736 | 0.109 | -0.202 | -1.094 | 0.108 | -0.036 | 0.1 | 0.1 | 0 | 0.84 | 0.71 | 0.83 | 0.69 | 0.91 | 0.88 |

| | | | | | | | | | | | | | | | | |
|---|---|---|---|---|---|---|---|---|---|---|---|---|---|---|---|---|
| 4 | 37608199 | rs2102336 | -0.312 | -0.204 | -1.18 | 0.063 | -0.004 | 0.04 | 0.04 | 0 | 0.47 | 0.23 | 0.33 | 0.32 | 0.53 | 0.42 |
| 4 | 37609699 | rs12640317 | -0.327 | 0.025 | -1.171 | 0.09 | -0.009 | 0.039 | 0.039 | 0 | 0.15 | 0.16 | 0.15 | 0.3 | 0.1 | 0.13 |
| 4 | 37610768 | rs2939732 | 0.877 | 0.046 | -1.276 | 0.041 | 0.018 | 0.087 | 0.087 | 0.036 | 0.42 | 0.2 | 0.25 | 0.23 | 0.39 | 0.26 |
| 4 | 37612526 | rs12142 | 0.825 | 0.338 | -1.235 | 0.095 | -0.119 | 0.201 | 0.201 | 0 | 0.32 | 0.07 | 0.17 | 0.01 | 0.42 | 0.3 |
| 4 | 37613930 | rs17495047 | 0.807 | 0.329 | -1.24 | 0.09 | -0.119 | 0.203 | 0.203 | 0 | 0.32 | 0.07 | 0.17 | 0.01 | 0.42 | 0.3 |
| 4 | 37615611 | rs17423188 | 1.519 | 0.204 | -1.344 | 0.081 | -0.084 | -0.045 | -0.045 | 0 | 0.21 | 0.01 | 0.13 | 0.18 | 0.26 | 0.19 |
| 4 | 37617389 | rs16993717 | 0.288 | 0.183 | -0.716 | 0.25 | 0.047 | 0.238 | 0.238 | 0 | 0.08 | 0.34 | 0.23 | 0.33 | 0.07 | 0.14 |
| 4 | 37618938 | rs2048257 | 0.872 | 0.021 | -0.921 | 0.165 | 0.028 | -0.015 | -0.015 | 0 | 0.31 | 0.28 | 0.21 | 0.35 | 0.16 | 0.19 |
| 4 | 37631874 | rs2174657 | 1.306 | 0.553 | -1.361 | -0.155 | -0.007 | 0.147 | 0.147 | 0 | 0.22 | 0.01 | 0.09 | 0 | 0.19 | 0.16 |
| 4 | 37633759 | rs17577515 | 1.55 | 0.423 | -1.04 | 0.342 | 0.018 | 0.016 | 0.016 | 0 | 0.05 | 0.01 | 0.07 | 0.28 | 0.03 | 0.12 |
| 4 | 37634192 | rs4482771 | 1.461 | 0.573 | -1.018 | 0.337 | -0.075 | -0.077 | -0.077 | 0.168 | 0.21 | 0.02 | 0.13 | 0.23 | 0.26 | 0.2 |
| 4 | 37636750 | rs2276943 | -0.857 | 0.85 | -0.655 | -0.349 | 0.044 | 0.079 | 0.079 | 0.038 | 0.7 | 0.53 | 0.61 | 0.47 | 0.63 | 0.67 |
| 4 | 37639139 | rs11733542 | 0.2 | 0.969 | -0.688 | -0.351 | -0.001 | 0 | 0 | 0.021 | 0.14 | 0.12 | 0.17 | 0.27 | 0.17 | 0.21 |
| 4 | 37640389 | rs3756143 | -0.294 | 0.928 | -0.567 | -0.459 | 0.023 | 0.092 | 0.092 | 0 | 0.19 | 0.29 | 0.38 | 0.41 | 0.23 | 0.3 |
| 4 | 37640483 | rs6818798 | -1.065 | 0.886 | -0.621 | -0.4 | 0.01 | 0.049 | 0.049 | 1.154 | 0.72 | 0.59 | 0.64 | 0.46 | 0.69 | 0.66 |
| 4 | 37647093 | rs17495288 | 0.453 | 0.572 | -0.54 | -0.57 | 0.003 | -0.086 | -0.086 | 1.618 | 0.26 | 0.31 | 0.15 | 0.01 | 0.28 | 0.28 |
| 4 | 37653622 | rs7690781 | 0.832 | 0.121 | -0.86 | -0.169 | 0.029 | 0.047 | 0.047 | 0 | 0.24 | 0.14 | 0.14 | 0.03 | 0.16 | 0.06 |
| 4 | 37653656 | rs9998333 | 0.085 | 0.725 | -0.164 | 0.77 | 0 | 0 | 0 | 0.168 | 0.65 | 0.41 | 0.45 | 0.33 | 0.65 | 0.59 |
| 4 | 37664677 | rs12710806 | -1.157 | 0.087 | -0.308 | 0.005 | 0.002 | 0.009 | 0.009 | 0 | 0.32 | 0.7 | 0.24 | 0.18 | 0.25 | 0.37 |
| 4 | 37668484 | rs6531568 | -0.873 | 0.072 | 0.5 | -0.025 | 0.02 | -0.004 | -0.004 | 1.835 | 0.19 | 0.32 | 0.16 | 0.13 | 0.2 | 0.35 |
| 4 | 37675941 | rs919006 | 0.834 | -0.022 | 0.847 | 0.012 | -0.02 | 0.007 | 0.007 | 2.723 | 0.9 | 0.81 | 0.89 | 0.91 | 0.97 | 0.79 |
| 4 | 37684896 | rs4407515 | 1.511 | -0.725 | 0.317 | -0.726 | 0.037 | 0.15 | 0.15 | 0 | 0.53 | 0.17 | 0.64 | 0.81 | 0.63 | 0.52 |
| 4 | 37689747 | rs1472951 | 1.199 | -0.753 | 0.159 | -0.947 | 0.04 | 0.088 | 0.088 | 0.595 | 0.61 | 0.18 | 0.65 | 0.82 | 0.69 | 0.62 |
| 4 | 37694948 | rs6817818 | -0.839 | -0.478 | 1.178 | -0.699 | -0.023 | -0.032 | -0.032 | 0.336 | 0.11 | 0.24 | 0.06 | 0.04 | 0.05 | 0.02 |
| 4 | 37695637 | rs10011758 | -1.142 | -0.492 | 1.177 | -0.726 | -0.026 | -0.04 | -0.04 | 0 | 0.12 | 0.29 | 0.06 | 0.04 | 0.05 | 0.02 |
| 4 | 37697032 | rs6831628 | -1.075 | -0.363 | 1.08 | -0.996 | 0.001 | -0.026 | -0.026 | 0.313 | 0.2 | 0.24 | 0.12 | 0.01 | 0.17 | 0.15 |
| 4 | 37698453 | rs2009494 | 1.071 | -0.564 | 1.426 | -0.154 | -0.016 | 0.164 | 0.164 | 0.078 | 0.88 | 0.58 | 0.83 | 0.64 | 0.94 | 0.93 |
| 4 | 37699165 | rs1364830 | 1.351 | -0.714 | 0.926 | -0.396 | 0 | -0.007 | -0.007 | 0.248 | 0.6 | 0.41 | 0.72 | 0.84 | 0.66 | 0.59 |
| 4 | 37701453 | rs17423781 | -0.392 | -0.917 | 0.351 | -0.389 | -0.004 | 0.1 | 0.1 | 0 | 0.17 | 0.02 | 0.05 | 0 | 0.15 | 0.15 |
| 4 | 37704067 | rs7660340 | 1.807 | -1.002 | 0.898 | -0.4 | 0.006 | 0.001 | 0.001 | 0.798 | 0.51 | 0.47 | 0.59 | 0.49 | 0.56 | 0.56 |
| 4 | 37704835 | rs17423816 | 1.532 | -1.065 | 1.197 | -0.386 | 0.012 | 0.033 | 0.033 | 0.025 | 0.54 | 0.62 | 0.65 | 0.64 | 0.58 | 0.66 |
| 4 | 37707375 | rs13115041 | 1.895 | -1.124 | 1.263 | -0.391 | 0.003 | 0.001 | 0.001 | 0 | 0.5 | 0.51 | 0.67 | 0.54 | 0.57 | 0.59 |
| 4 | 37708086 | rs13115042 | -1.756 | -1.185 | 1.189 | -0.469 | 0.036 | 0.077 | 0.077 | 0 | 0.4 | 0.22 | 0.3 | 0.17 | 0.31 | 0.4 |
| 4 | 37711228 | rs13137307 | -0.486 | -1.069 | 0.893 | -0.465 | 0.003 | -0.017 | -0.017 | 0 | 0.2 | 0.18 | 0.13 | 0.01 | 0.15 | 0.11 |
| 4 | 37716343 | rs10028206 | -0.76 | -0.895 | 0.452 | -0.275 | 0.004 | 0.057 | 0.057 | 0 | 0.1 | 0.17 | 0.1 | 0.33 | 0.12 | 0.04 |
| 4 | 37719424 | rs10017065 | -0.966 | -0.934 | 0.481 | -0.257 | 0.004 | -0.006 | -0.006 | 0.247 | 0.25 | 0.17 | 0.16 | 0.34 | 0.27 | 0.2 |
| 4 | 37720580 | rs12505604 | 1.909 | -0.891 | 0.096 | -0.358 | -0.054 | -0.082 | -0.082 | 0.196 | 0.44 | 0.1 | 0.53 | 0.46 | 0.52 | 0.53 |
| 4 | 37725293 | rs7666650 | 2.089 | -0.409 | 1.02 | -0.682 | 0.019 | 0.056 | 0.056 | 0 | 0.56 | 0.29 | 0.6 | 0.69 | 0.6 | 0.68 |
| 4 | 37727622 | rs961082 | -0.067 | -0.097 | 0.859 | -0.562 | -0.022 | 0.032 | 0.032 | 0.028 | 0.12 | 0.07 | 0.1 | 0.01 | 0.16 | 0.1 |
| 4 | 37741826 | rs1469236 | 0.31 | -0.22 | 0.997 | -0.465 | 0.009 | -0.064 | -0.064 | 0 | 0.86 | 0.77 | 0.88 | 0.99 | 0.84 | 0.9 |
| 4 | 37750227 | rs17424009 | -1.329 | -0.344 | 0.933 | -0.538 | 0.01 | 0.038 | 0.038 | 0 | 0.26 | 0.19 | 0.18 | 0.01 | 0.26 | 0.16 |
| 4 | 37752668 | rs2995977 | 1.3 | -0.315 | 0.845 | -0.471 | -0.007 | -0.054 | -0.054 | 0 | 0.7 | 0.55 | 0.68 | 0.89 | 0.71 | 0.72 |
| 4 | 37754104 | rs2995976 | -0.84 | -0.374 | 1.051 | 0.14 | 0.014 | 0.047 | 0.047 | 0 | 0.08 | 0.17 | 0.07 | 0.19 | 0.1 | 0.04 |
| 4 | 37756236 | rs1465496 | 1.171 | -0.429 | 0.641 | 0.037 | -0.027 | -0.023 | -0.023 | 0.001 | 0.83 | 0.69 | 0.79 | 0.89 | 0.87 | 0.82 |
| 4 | 37758324 | rs17605986 | -0.722 | -0.391 | -0.02 | 0.156 | 0.017 | -0.016 | -0.016 | 0 | 0.11 | 0.02 | 0.12 | 0.11 | 0.08 | 0.11 |
| 4 | 37758827 | rs11726893 | 1.396 | -0.6 | 0.776 | 0.061 | -0.028 | 0.034 | 0.034 | 0 | 0.85 | 0.58 | 0.92 | 0.82 | 0.9 | 0.96 |
| 4 | 37759253 | rs10517429 | -1.095 | -0.709 | 0.141 | -0.255 | 0.011 | 0.074 | 0.074 | 0 | 0.13 | 0.04 | 0.06 | 0 | 0.11 | 0.06 |
| 4 | 37760049 | rs17496030 | -1.266 | -0.405 | 0.13 | -0.182 | -0.011 | 0.013 | 0.013 | 0 | 0.14 | 0.05 | 0.15 | 0.11 | 0.14 | 0.17 |
| 4 | 37764357 | rs2995972 | -1.217 | -0.389 | 0.719 | -0.136 | -0.016 | -0.063 | -0.063 | 1.001 | 0.13 | 0.21 | 0.1 | 0.01 | 0.11 | 0.02 |
| 4 | 37766932 | rs4240241 | 1.394 | -0.433 | 1.297 | -0.01 | 0.023 | 0.057 | 0.057 | 2.487 | 0.91 | 0.65 | 0.84 | 0.87 | 0.96 | 0.88 |
| 4 | 37768842 | rs929959 | 1.008 | -0.134 | 0.449 | 0.276 | -0.071 | -0.026 | -0.026 | 0 | 0.3 | 0.12 | 0.38 | 0.31 | 0.41 | 0.3 |
| 4 | 37769060 | rs929958 | 0.139 | -0.406 | 0.974 | 0.055 | -0.012 | 0.024 | 0.024 | 0 | 0.43 | 0.27 | 0.49 | 0.32 | 0.54 | 0.37 |
| 4 | 37769527 | rs919005 | 0.902 | -0.482 | 0.747 | -0.011 | -0.001 | -0.001 | -0.001 | 0 | 0.08 | 0.09 | 0.03 | 0.13 | 0.09 | 0.13 |
| 4 | 37770401 | rs1125117 | -0.092 | -0.493 | 0.846 | -0.065 | 0.026 | 0.017 | 0.017 | 0 | 0.49 | 0.37 | 0.35 | 0.56 | 0.42 | 0.54 |
| 4 | 37771002 | rs1025146 | -0.523 | -0.398 | 1.48 | 0.139 | -0.004 | -0.015 | -0.015 | 0 | 0.94 | 0.78 | 0.96 | 1 | 0.99 | 1 |
| 4 | 37772443 | rs7698331 | 1.28 | -0.407 | 1.565 | 0.131 | 0.003 | -0.023 | -0.023 | 0 | 0.84 | 0.6 | 0.76 | 0.86 | 0.87 | 0.89 |
| 4 | 37773485 | rs4832940 | 1.377 | -0.399 | 1.487 | 0.123 | 0.004 | -0.028 | -0.028 | 0 | 0.84 | 0.55 | 0.75 | 0.86 | 0.86 | 0.89 |
| 4 | 37773849 | rs2911862 | -0.121 | -0.432 | 0.936 | 0.111 | 0.004 | -0.039 | -0.039 | 0 | 0.07 | 0.12 | 0.08 | 0.01 | 0.1 | 0.02 |
| 4 | 37780765 | rs2995965 | 0.588 | -0.694 | 1.172 | 0.316 | -0.02 | 0.037 | 0.037 | 0 | 0.81 | 0.57 | 0.78 | 0.76 | 0.84 | 0.92 |
| 4 | 37785000 | rs4463081 | 0.838 | -1.161 | 0.809 | 0.428 | 0.021 | 0.008 | 0.008 | 0 | 0.32 | 0.22 | 0.26 | 0.23 | 0.21 | 0.43 |
| 4 | 37785525 | rs2911905 | 0.819 | -0.919 | 1.06 | 0.188 | 0.002 | -0.017 | -0.017 | 0 | 0.74 | 0.65 | 0.82 | 0.83 | 0.71 | 0.8 |
| 4 | 37786987 | rs870953 | -0.136 | -0.738 | 1.056 | 0.526 | 0.001 | 0.001 | 0.001 | 0 | 0.51 | 0.5 | 0.54 | 0.59 | 0.44 | 0.7 |
| 4 | 37787117 | rs7692977 | -0.198 | -0.734 | 1.05 | 0.503 | 0 | -0.003 | -0.003 | 0 | 0.51 | 0.5 | 0.56 | 0.73 | 0.46 | 0.79 |
| 4 | 37790639 | rs2911902 | 0.532 | -0.721 | 1.067 | 0.37 | -0.012 | 0.013 | 0.013 | 0 | 0.39 | 0.33 | 0.37 | 0.14 | 0.45 | 0.17 |
| 4 | 37795022 | rs12640538 | 0.244 | -0.93 | 1.46 | 1.298 | -0.012 | 0.04 | 0.04 | 0.516 | 0.06 | 0.08 | 0.03 | 0.15 | 0.03 | 0.08 |
| 4 | 37797027 | rs955290 | 0.987 | -1.037 | 1.487 | 1.266 | -0.027 | 0.036 | 0.036 | 0 | 0.79 | 0.71 | 0.88 | 0.7 | 0.86 | 0.87 |
| 4 | 37797841 | rs1863284 | 0.516 | -0.918 | 1.289 | 1.163 | -0.008 | 0.001 | 0.001 | 0 | 0.85 | 0.79 | 0.91 | 0.87 | 0.87 | 0.94 |
| 4 | 37799735 | rs2995958 | 1.392 | -1.004 | 1.784 | 1.352 | -0.018 | 0.023 | 0.023 | 0 | 0.8 | 0.29 | 0.75 | 0.74 | 0.86 | 0.9 |
| 4 | 37803352 | rs1109971 | 0.297 | -1.337 | 1.38 | 1.345 | 0.089 | -0.034 | -0.034 | 0 | 0.39 | 0.14 | 0.29 | 0.44 | 0.25 | 0.5 |

| | | | | | | | | | | | | | | | | |
|---|---|---|---|---|---|---|---|---|---|---|---|---|---|---|---|---|
| 4 | 37803799 | rs2911874 | 0.355 | -1.343 | 1.377 | 1.472 | 0.085 | 0.068 | 0.068 | 0 | 0.38 | 0.14 | 0.28 | 0.21 | 0.25 | 0.44 |
| 4 | 37805051 | rs2911869 | 1.086 | -1.238 | 1.462 | 1.545 | -0.018 | 0.033 | 0.033 | 0 | 0.81 | 0.29 | 0.76 | 0.73 | 0.86 | 0.91 |
| 4 | 37807815 | rs11096906 | 0.481 | -1.174 | 1.824 | 1.561 | -0.008 | 0.015 | 0.015 | 0 | 0.88 | 0.72 | 0.9 | 0.9 | 0.88 | 0.95 |
| 4 | 37808510 | rs7437855 | -0.029 | -1.287 | 1.788 | 1.539 | 0.005 | -0.001 | -0.001 | 0 | 0.44 | 0.56 | 0.46 | 0.6 | 0.27 | 0.55 |
| 4 | 37830383 | rs2911908 | 0.515 | -1.278 | 1.499 | 1.258 | -0.001 | -0.001 | -0.001 | 0.013 | 0.8 | 0.79 | 0.89 | 0.89 | 0.82 | 0.91 |
| 4 | 37837819 | rs2714555 | -1.374 | -1.395 | 1.101 | 1.608 | 0.095 | -0.006 | -0.006 | 0.008 | 0.68 | 0.91 | 0.7 | 0.74 | 0.81 | 0.61 |
| 4 | 37838237 | rs2608307 | -1.562 | -1.411 | 1.452 | 1.64 | -0.013 | 0.003 | 0.003 | 0.235 | 0.27 | 0.68 | 0.28 | 0.27 | 0.22 | 0.14 |
| 4 | 37840606 | rs3775793 | 0.976 | -1.396 | 1.334 | 1.538 | 0.024 | 0.084 | 0.084 | 0 | 0.41 | 0.19 | 0.4 | 0.22 | 0.59 | 0.4 |
| 4 | 37843161 | rs2714560 | -1.346 | -1.385 | 1.256 | 1.544 | 0.098 | -0.007 | -0.007 | 0 | 0.69 | 0.92 | 0.7 | 0.74 | 0.81 | 0.61 |
| 4 | 37854182 | rs4832960 | 0.445 | -1.151 | 1.641 | 1.459 | 0.014 | 0.009 | 0.009 | 0 | 0.45 | 0.39 | 0.54 | 0.42 | 0.62 | 0.45 |
| 4 | 37856777 | rs4444838 | -0.649 | -0.704 | 0.96 | 1.562 | 0.016 | 0.06 | 0.06 | 0.532 | 0.6 | 0.82 | 0.64 | 0.73 | 0.64 | 0.46 |
| 4 | 37857166 | rs6842360 | -1.102 | -0.445 | 1.154 | 1.9 | 0.113 | 0.068 | 0.068 | 0 | 0.6 | 0.9 | 0.68 | 0.75 | 0.76 | 0.59 |
| 4 | 37858954 | rs6531591 | -0.358 | -0.123 | 1.138 | 1.798 | 0.045 | -0.038 | -0.038 | 0 | 0.48 | 0.89 | 0.59 | 0.61 | 0.6 | 0.51 |
| 4 | 37860967 | rs6844466 | -1.172 | -0.076 | 1.618 | 1.769 | -0.001 | -0.001 | -0.001 | 0 | 0.14 | 0.31 | 0.11 | 0.11 | 0.15 | 0.1 |
| 4 | 37862071 | rs4832962 | -0.371 | 0.035 | 1.63 | 1.874 | 0.005 | 0.009 | 0.009 | 0 | 0.48 | 0.54 | 0.46 | 0.28 | 0.61 | 0.37 |
| 4 | 37862522 | rs6820778 | -0.951 | 0.146 | 0.804 | 1.994 | 0.11 | 0.035 | 0.035 | 0 | 0.58 | 0.9 | 0.65 | 0.71 | 0.73 | 0.59 |
| 4 | 37863385 | rs10569 | -0.234 | 0.349 | 0.958 | 1.131 | 0.013 | 0.02 | 0.02 | 0 | 0.1 | 0.21 | 0.1 | 0.1 | 0.13 | 0.09 |
| 4 | 37868667 | rs11734812 | 0.248 | 0.687 | 1.062 | 0.817 | 0.029 | -0.053 | -0.053 | 0 | 0.37 | 0.42 | 0.38 | 0.22 | 0.48 | 0.27 |
| 4 | 37868805 | rs11722588 | 0.242 | 0.693 | 1.002 | 0.817 | 0.032 | -0.057 | -0.057 | 0 | 0.37 | 0.44 | 0.38 | 0.22 | 0.49 | 0.27 |
| 4 | 37870300 | rs28571127 | -0.58 | 0.904 | 0.228 | 1.015 | 0.021 | 0.021 | 0.021 | 0 | 0.09 | 0.17 | 0.16 | 0.17 | 0.11 | 0.15 |
| 4 | 37882167 | rs6531592 | -0.021 | 0.799 | 0.019 | 0.458 | 0.03 | -0.071 | -0.071 | 0.407 | 0.37 | 0.48 | 0.35 | 0.15 | 0.46 | 0.27 |
| 4 | 37883839 | rs7685175 | -0.095 | 0.555 | 0.116 | 1.126 | 0.001 | -0.004 | -0.004 | 0 | 0.42 | 0.45 | 0.63 | 0.36 | 0.49 | 0.41 |
| 4 | 37884116 | rs9996566 | -0.128 | 0.683 | -0.442 | 1.053 | 0.037 | 0.003 | 0.003 | 0.976 | 0.5 | 0.65 | 0.68 | 0.49 | 0.55 | 0.45 |
| 4 | 37898650 | rs2995926 | -0.813 | 1.162 | -0.394 | 0.531 | 0.002 | -0.044 | -0.044 | 0 | 0.16 | 0.15 | 0.11 | 0 | 0.2 | 0.15 |
| 4 | 37900371 | rs3893296 | -0.245 | 1.215 | -0.347 | 0.29 | 0.001 | -0.006 | -0.006 | 0 | 0.28 | 0.36 | 0.34 | 0.33 | 0.27 | 0.31 |
| 4 | 37900725 | rs17578878 | -0.45 | 1.264 | -0.757 | 1.255 | -0.013 | -0.015 | -0.015 | 0.064 | 0.07 | 0.02 | 0.07 | 0.22 | 0.11 | 0.1 |
| 4 | 37901153 | rs3860064 | -0.496 | 1.066 | -1.251 | 0.967 | -0.008 | 0.04 | 0.04 | 0 | 0.22 | 0.07 | 0.16 | 0.18 | 0.27 | 0.24 |
| 4 | 37903756 | rs2279027 | -2.076 | 0.602 | -0.896 | 0.842 | -0.021 | -0.004 | -0.004 | 0 | 0.58 | 0.88 | 0.59 | 0.66 | 0.61 | 0.69 |
| 4 | 37904082 | rs2279026 | -0.846 | 0.461 | -0.261 | 0.783 | -0.013 | 0.01 | 0.01 | 0 | 0.26 | 0.64 | 0.34 | 0.37 | 0.2 | 0.37 |
| 4 | 37904089 | rs35859249 | 0.539 | 0.473 | -0.302 | 0.792 | -0.004 | -0.023 | -0.023 | 0.005 | 0.08 | 0.01 | 0.06 | 0.12 | 0.1 | 0.14 |
| 4 | 37905601 | rs6531596 | 0.008 | 0.469 | -0.009 | 0.809 | -0.006 | 0.024 | 0.024 | 0.005 | 0.18 | 0.57 | 0.27 | 0.25 | 0.1 | 0.22 |
| 4 | 37906032 | rs2995919 | -1.382 | 0.491 | -0.125 | 0.673 | -0.019 | -0.012 | -0.012 | 0 | 0.39 | 0.19 | 0.3 | 0.41 | 0.51 | 0.47 |
| 4 | 37909495 | rs2925947 | -1.531 | 0.044 | 0.085 | 0.403 | 0.001 | 0.002 | 0.002 | 0 | 0.43 | 0.45 | 0.38 | 0.34 | 0.44 | 0.38 |
| 4 | 37909841 | rs2925948 | -0.519 | 0.352 | -0.52 | 0.523 | -0.025 | -0.041 | -0.041 | 0 | 0.2 | 0.12 | 0.15 | 0.22 | 0.26 | 0.28 |
| 4 | 37910996 | rs2995916 | -1.255 | 0.252 | -0.808 | 0.288 | -0.041 | 0.1 | 0.1 | 0.202 | 0.54 | 0.84 | 0.44 | 0.41 | 0.51 | 0.61 |
| 4 | 37911508 | rs2995915 | 0.544 | 0.207 | -0.925 | 0.08 | -0.008 | 0.028 | 0.028 | 0 | 0.13 | 0.05 | 0.08 | 0.01 | 0.16 | 0.12 |
| 4 | 37911638 | rs2925952 | -1.576 | 0.762 | -0.897 | 0.491 | 0.017 | -0.052 | -0.052 | 0.86 | 0.59 | 0.8 | 0.54 | 0.59 | 0.64 | 0.72 |
| 4 | 37915462 | rs11096913 | 0.371 | 0.505 | -1.658 | -0.471 | 0.001 | 0.069 | 0.069 | 2.087 | 0.24 | 0.03 | 0.21 | 0.12 | 0.24 | 0.16 |
| 4 | 37920053 | rs2925956 | -0.431 | 0.994 | -0.703 | -0.759 | 0 | -0.063 | -0.063 | 0 | 0.36 | 0.51 | 0.22 | 0.17 | 0.37 | 0.4 |
| 4 | 37925978 | rs16994074 | 0.647 | 1.106 | -0.351 | -0.714 | 0.005 | -0.034 | -0.034 | 6.386 | 0.13 | 0.14 | 0.1 | 0.06 | 0.17 | 0.1 |
| 4 | 37926784 | rs11096914 | -1.866 | 1.009 | -1.205 | -1.105 | 0.004 | 0.089 | 0.089 | 0 | 0.73 | 0.85 | 0.8 | 0.97 | 0.78 | 0.8 |
| 4 | 37926978 | rs17497040 | 0.67 | 1.219 | -0.907 | -0.346 | 0.018 | 0.03 | 0.03 | 0 | 0.07 | 0.11 | 0.19 | 0.14 | 0.13 | 0.19 |
| 4 | 37929534 | rs4240242 | -0.634 | 1.185 | -0.567 | -0.463 | -0.001 | 0.03 | 0.03 | 0 | 0.5 | 0.39 | 0.65 | 0.8 | 0.55 | 0.47 |
| 4 | 37930312 | rs4832745 | -0.385 | 1.015 | -0.707 | -0.359 | 0.016 | 0.025 | 0.025 | 0 | 0.43 | 0.28 | 0.45 | 0.65 | 0.42 | 0.28 |
| 4 | 37936849 | rs17579011 | -0.723 | 1.229 | -0.559 | 0.14 | 0.001 | 0.007 | 0.007 | 0.107 | 0.26 | 0.31 | 0.45 | 0.53 | 0.31 | 0.33 |
| 4 | 37938518 | rs10010758 | -0.794 | 1.167 | -0.321 | -0.082 | 0.008 | -0.019 | -0.019 | 0 | 0.28 | 0.41 | 0.27 | 0.21 | 0.28 | 0.28 |
| 4 | 37962133 | rs2290740 | 0.078 | 0.853 | -0.44 | -0.195 | -0.015 | 0.019 | 0.019 | 0 | 0.18 | 0.3 | 0.11 | 0.31 | 0.13 | 0.26 |
| 4 | 37965827 | rs11736130 | -0.842 | 1.165 | -0.9 | -0.173 | -0.017 | 0.005 | 0.005 | 0 | 0.32 | 0.16 | 0.55 | 0.63 | 0.37 | 0.33 |
| 4 | 37967083 | rs10517460 | 0.294 | 0.969 | -0.821 | -0.151 | 0.012 | 0.017 | 0.017 | 1.118 | 0.17 | 0.11 | 0.09 | 0.02 | 0.13 | 0.11 |
| 4 | 37970850 | rs9306955 | -1.254 | 1.403 | -0.847 | 0.005 | -0.017 | -0.012 | -0.012 | 1.63 | 0.63 | 0.74 | 0.47 | 0.49 | 0.57 | 0.6 |
| 4 | 37971275 | rs2381131 | -1.652 | 1.138 | -0.756 | -0.545 | -0.001 | -0.008 | -0.008 | 0.086 | 0.68 | 0.75 | 0.78 | 0.96 | 0.69 | 0.75 |
| 4 | 37973722 | rs6858227 | -0.054 | 0.402 | -0.567 | -0.165 | -0.001 | 0.007 | 0.007 | 0 | 0.31 | 0.26 | 0.32 | 0.31 | 0.29 | 0.34 |
| 4 | 37975020 | rs2890591 | -1.441 | 0.151 | -0.92 | -0.136 | 0.014 | 0.045 | 0.045 | 0 | 0.47 | 0.28 | 0.42 | 0.41 | 0.42 | 0.44 |
| 4 | 37979512 | rs6850410 | -1.724 | 0.068 | -0.317 | -0.222 | 0.001 | -0.079 | -0.079 | 0 | 0.55 | 0.55 | 0.67 | 0.91 | 0.51 | 0.65 |
| 4 | 37989306 | rs10011664 | 0.447 | 0.116 | -0.337 | -0.199 | 0.007 | 0.013 | 0.013 | 0 | 0.28 | 0.23 | 0.32 | 0.28 | 0.26 | 0.28 |
| 4 | 37990581 | rs6841167 | -0.71 | 0.068 | -0.056 | -0.25 | 0 | -0.002 | -0.002 | 0 | 0.37 | 0.35 | 0.57 | 0.84 | 0.39 | 0.57 |
| 4 | 37991472 | rs11943860 | -1.145 | 0.248 | 0.088 | -0.415 | -0.012 | -0.096 | -0.096 | 0 | 0.32 | 0.54 | 0.18 | 0.09 | 0.29 | 0.19 |
| 4 | 37995170 | rs6858606 | -0.727 | 0.202 | 0.038 | -0.385 | -0.011 | -0.063 | -0.063 | 0 | 0.28 | 0.42 | 0.17 | 0.09 | 0.25 | 0.14 |
| 4 | 37996334 | rs1344602 | -1.351 | 0.191 | 0.106 | -0.373 | -0.011 | -0.013 | -0.013 | 0 | 0.34 | 0.47 | 0.4 | 0.63 | 0.34 | 0.38 |
| 4 | 37996827 | rs10084926 | 0.539 | 0.187 | 0.037 | -0.348 | 0.008 | 0.015 | 0.015 | 0 | 0.28 | 0.21 | 0.31 | 0.29 | 0.26 | 0.28 |
| 4 | 37999876 | rs16994133 | 1.262 | 0.18 | 0.109 | -0.384 | -0.001 | -0.064 | -0.064 | 4.871 | 0.11 | 0.31 | 0.07 | 0 | 0.1 | 0.04 |
| 4 | 38006686 | rs7672883 | -1.637 | 0.225 | -0.109 | -0.374 | 0.011 | 0.128 | 0.128 | 0 | 0.49 | 0.73 | 0.65 | 0.87 | 0.52 | 0.62 |
| 4 | 38007559 | rs17579337 | 0.38 | 0.372 | 0.017 | -0.114 | 0.003 | 0.096 | 0.096 | 0 | 0.07 | 0.09 | 0.25 | 0.57 | 0.12 | 0.28 |
| 4 | 38007612 | rs6835205 | -2.238 | 0.342 | -0.095 | -0.333 | -0.016 | 0.024 | 0.024 | 0.052 | 0.66 | 0.79 | 0.73 | 0.93 | 0.64 | 0.71 |
| 4 | 38009782 | rs10517464 | 0.276 | 0.438 | -0.015 | -0.088 | -0.006 | -0.009 | -0.009 | 0 | 0.15 | 0.16 | 0.17 | 0.19 | 0.21 | 0.21 |
| 4 | 38010293 | rs4832980 | -2.559 | 0.429 | -0.102 | -0.427 | -0.006 | 0.025 | 0.025 | 0 | 0.67 | 0.8 | 0.68 | 0.79 | 0.67 | 0.71 |
| 4 | 38011565 | rs16994140 | 1.08 | 0.284 | 0.319 | -0.198 | 0.018 | -0.049 | -0.049 | 0.252 | 0.09 | 0.21 | 0.07 | 0 | 0.12 | 0.01 |
| 4 | 38018181 | rs13142228 | -1.185 | -0.088 | 0.43 | 0.267 | -0.004 | 0.037 | 0.037 | 0 | 0.25 | 0.32 | 0.27 | 0.53 | 0.28 | 0.39 |
| 4 | 38019302 | rs7671645 | -0.82 | -0.225 | 0.557 | 0.231 | -0.003 | 0.038 | 0.038 | 0 | 0.17 | 0.18 | 0.19 | 0.44 | 0.18 | 0.4 |

| | | | | | | | | | | | | | | | | |
|---|---|---|---|---|---|---|---|---|---|---|---|---|---|---|---|---|
| 4 | 38019436 | rs1364951 | 1.38 | -0.049 | 0.417 | 0.131 | -0.006 | 0.005 | 0.005 | 0 | 0.09 | 0.1 | 0.1 | 0.05 | 0.11 | 0.15 |
| 4 | 38026106 | rs933819 | -1.481 | -0.515 | 0.698 | 0.289 | 0.006 | -0.014 | -0.014 | 0 | 0.4 | 0.5 | 0.41 | 0.59 | 0.47 | 0.43 |
| 4 | 38026389 | rs933821 | 0.909 | -0.482 | 0.429 | 0.393 | -0.031 | -0.031 | -0.031 | 1.216 | 0.08 | 0.01 | 0.17 | 0.36 | 0.15 | 0.25 |
| 4 | 38032931 | rs7670756 | -1.26 | -0.518 | 0.601 | 0.108 | 0.001 | 0.018 | 0.018 | 0 | 0.34 | 0.57 | 0.36 | 0.5 | 0.38 | 0.45 |
| 4 | 38046569 | rs10018389 | -2.26 | -0.54 | 0.482 | -0.054 | 0.007 | 0.103 | 0.103 | 0 | 0.5 | 0.79 | 0.46 | 0.63 | 0.5 | 0.58 |
| 4 | 38046709 | rs11096921 | 0.041 | -0.507 | 0.599 | -0.043 | 0.042 | 0.034 | 0.034 | 0 | 0.05 | 0.31 | 0.04 | 0.01 | 0.02 | 0.08 |
| 4 | 38058764 | rs17425670 | 0.693 | -0.465 | 0.359 | -0.036 | -0.006 | 0.014 | 0.014 | 0 | 0.19 | 0.16 | 0.13 | 0.04 | 0.21 | 0.08 |
| 4 | 38060485 | rs1863329 | -1.685 | -0.479 | 0.413 | 0.015 | -0.009 | -0.004 | -0.004 | 0.828 | 0.34 | 0.64 | 0.35 | 0.59 | 0.33 | 0.52 |
| 4 | 38062263 | rs16994165 | 0.735 | -0.693 | 0.307 | 0.037 | -0.01 | -0.002 | -0.002 | 0 | 0.13 | 0.11 | 0.17 | 0.12 | 0.09 | 0.04 |
| 4 | 38073974 | rs17497671 | 2.029 | -0.525 | 0.254 | 0.021 | -0.012 | 0.031 | 0.031 | 0 | 0.12 | 0.05 | 0.09 | 0 | 0.15 | 0.01 |
| 4 | 38079294 | rs933818 | 0.691 | -0.293 | 0.204 | 0.433 | -0.03 | -0.035 | -0.035 | 0.013 | 0.09 | 0.01 | 0.17 | 0.41 | 0.15 | 0.27 |
| 4 | 38079475 | rs10009706 | -1.366 | -0.41 | 0.395 | 0.169 | -0.005 | 0.029 | 0.029 | 0 | 0.28 | 0.48 | 0.29 | 0.57 | 0.28 | 0.45 |
| 4 | 38080507 | rs1425576 | -2.194 | -0.424 | -0.11 | 0.112 | -0.078 | 0.1 | 0.1 | 0.116 | 0.62 | 0.86 | 0.62 | 0.74 | 0.57 | 0.61 |
| 4 | 38086795 | rs1863326 | -0.168 | -0.156 | 0.539 | 1.125 | -0.002 | 0.042 | 0.042 | 0 | 0.12 | 0.13 | 0.18 | 0.42 | 0.17 | 0.32 |
| 4 | 38087105 | rs1863325 | -2.097 | -0.142 | 0.398 | 0.549 | 0.004 | 0.008 | 0.008 | 0 | 0.36 | 0.71 | 0.35 | 0.62 | 0.32 | 0.56 |
| 4 | 38092096 | rs2303421 | -1.286 | -0.127 | 0.213 | 0.198 | 0.002 | 0.003 | 0.003 | 0 | 0.21 | 0.19 | 0.11 | 0.13 | 0.15 | 0.21 |
| 4 | 38097478 | rs10517469 | -1.118 | 0.099 | -0.17 | 0.366 | -0.021 | -0.036 | -0.036 | 0 | 0.19 | 0.42 | 0.1 | 0.11 | 0.14 | 0.19 |
| 4 | 38103016 | rs2114014 | -0.662 | 0.3 | -0.967 | 0.645 | 0.007 | 0.009 | 0.009 | 0 | 0.36 | 0.31 | 0.42 | 0.38 | 0.37 | 0.34 |
| 4 | 38103550 | rs933823 | -1.195 | 0.411 | -1.593 | 0.198 | -0.032 | 0.109 | 0.109 | 0 | 0.67 | 0.88 | 0.8 | 0.84 | 0.71 | 0.65 |
| 4 | 38105161 | rs933822 | -1.63 | 0.413 | -1.669 | 0.345 | -0.042 | 0.04 | 0.04 | 0 | 0.73 | 0.91 | 0.87 | 0.76 | 0.74 | 0.75 |
| 4 | 38106236 | rs3890061 | -0.159 | 0.39 | -1.783 | 0.884 | -0.011 | -0.077 | -0.077 | 7.973 | 0.13 | 0.03 | 0.19 | 0.46 | 0.16 | 0.25 |
| 4 | 38107811 | rs10013783 | -0.993 | 0.209 | -0.913 | 0.674 | -0.004 | -0.007 | -0.007 | 0 | 0.45 | 0.4 | 0.55 | 0.62 | 0.41 | 0.53 |
| 4 | 38109847 | rs1344603 | -1.86 | 0.253 | -1.281 | 0.611 | -0.066 | 0.049 | 0.049 | 0 | 0.73 | 0.93 | 0.86 | 0.77 | 0.72 | 0.75 |
| 4 | 38110620 | rs754886 | -1.236 | 0.284 | -1.186 | 1.42 | -0.015 | -0.032 | -0.032 | 0 | 0.67 | 0.82 | 0.73 | 0.61 | 0.64 | 0.56 |
| 4 | 38122459 | rs645469 | -0.509 | 0.199 | -0.972 | 1.095 | 0.004 | 0.009 | 0.009 | 0 | 0.71 | 0.79 | 0.65 | 0.75 | 0.68 | 0.68 |
| 4 | 38123916 | rs2067557 | 0.337 | 0.049 | -0.342 | 1.081 | -0.002 | 0.019 | 0.019 | 0.054 | 0.36 | 0.43 | 0.37 | 0.36 | 0.37 | 0.33 |
| 4 | 38129949 | rs637495 | -0.311 | 0.162 | -0.816 | 0.906 | 0.008 | 0.021 | 0.021 | 0 | 0.71 | 0.79 | 0.65 | 0.76 | 0.68 | 0.69 |
| 4 | 38131076 | rs13119350 | -0.208 | -0.153 | -0.961 | 0.805 | 0.007 | 0.002 | 0.002 | 0 | 0.12 | 0.12 | 0.26 | 0.14 | 0.11 | 0.09 |
| 4 | 38131255 | rs605073 | 0.992 | -0.093 | -1.157 | 0.833 | -0.021 | 0.01 | 0.01 | 0 | 0.35 | 0.14 | 0.18 | 0.38 | 0.34 | 0.37 |
| 4 | 38132999 | rs633059 | 1.514 | -0.21 | -0.664 | 0.833 | 0.004 | -0.007 | -0.007 | 0 | 0.3 | 0.31 | 0.2 | 0.3 | 0.29 | 0.26 |
| 4 | 38137850 | rs6816462 | 1.525 | -0.062 | -1.365 | 0.594 | -0.058 | 0.093 | 0.093 | 0.297 | 0.22 | 0.05 | 0.21 | 0.04 | 0.25 | 0.12 |
| 4 | 38138856 | rs13110318 | 0.705 | -0.25 | -0.843 | 1.107 | 0.005 | -0.002 | -0.002 | 0 | 0.1 | 0.05 | 0.18 | 0.12 | 0.09 | 0.05 |
| 4 | 38139024 | rs9852 | 0.602 | -0.229 | -0.935 | 1.438 | -0.013 | -0.076 | -0.076 | 0 | 0.11 | 0.01 | 0.09 | 0.3 | 0.11 | 0.22 |
| 4 | 38145373 | rs10021783 | -1.167 | -0.203 | -0.62 | 0.858 | 0.019 | 0.022 | 0.022 | 0 | 0.43 | 0.66 | 0.53 | 0.6 | 0.41 | 0.53 |
| 4 | 38154707 | rs12641222 | -0.048 | -0.061 | -1.086 | 0.659 | -0.01 | 0.009 | 0.009 | 0 | 0.22 | 0.08 | 0.23 | 0.21 | 0.26 | 0.16 |
| 4 | 38155431 | rs12645765 | 1.113 | -0.227 | -1.687 | 0.25 | -0.031 | 0.036 | 0.036 | 0 | 0.15 | 0.07 | 0.2 | 0.04 | 0.23 | 0.1 |
| 4 | 38157510 | rs6814900 | -0.502 | -0.478 | -1.377 | 0.26 | 0.012 | -0.008 | -0.008 | 0 | 0.31 | 0.29 | 0.3 | 0.41 | 0.3 | 0.3 |
| 4 | 38158024 | rs17607861 | -0.452 | -0.355 | -1.635 | -0.597 | 0.012 | 0.045 | 0.045 | 0 | 0.42 | 0.2 | 0.42 | 0.38 | 0.44 | 0.32 |
| 4 | 38158043 | rs12510952 | 0.87 | -0.213 | -1.248 | -0.25 | 0 | 0 | 0 | 0 | 0.07 | 0.05 | 0.05 | 0.16 | 0.07 | 0.05 |
| 4 | 38158347 | rs12503618 | -0.026 | -0.064 | -0.923 | -0.073 | -0.012 | -0.028 | -0.028 | 0.058 | 0.23 | 0.16 | 0.38 | 0.32 | 0.3 | 0.36 |
| 4 | 38162567 | rs6844679 | 0.453 | -0.165 | -0.345 | -0.102 | -0.032 | -0.011 | -0.011 | 0 | 0.24 | 0.38 | 0.14 | 0.24 | 0.17 | 0.22 |
| 4 | 38167985 | rs6849562 | 0.04 | 0.032 | -0.504 | -0.239 | 0.008 | 0.026 | 0.026 | 0 | 0.52 | 0.45 | 0.55 | 0.31 | 0.58 | 0.42 |
| 4 | 38169660 | rs4832749 | -0.497 | 0.22 | -0.471 | -0.196 | -0.002 | -0.004 | -0.004 | 0 | 0.61 | 0.66 | 0.71 | 0.64 | 0.61 | 0.71 |
| 4 | 38170057 | rs907311 | 1.329 | 0.184 | -0.5 | -0.221 | 0.004 | 0.005 | 0.005 | 0 | 0.21 | 0.05 | 0.18 | 0.22 | 0.2 | 0.16 |
| 4 | 38170173 | rs1991231 | -0.176 | 0.368 | -0.718 | -0.369 | 0.002 | -0.002 | -0.002 | 1.034 | 0.4 | 0.61 | 0.52 | 0.56 | 0.42 | 0.52 |
| 4 | 38172840 | rs4832992 | 0.216 | 0.373 | -0.692 | -0.415 | 0.009 | -0.02 | -0.02 | 0 | 0.34 | 0.6 | 0.31 | 0.27 | 0.35 | 0.36 |
| 4 | 38175276 | rs2380964 | 1.091 | 0.618 | -0.899 | -0.401 | -0.032 | 0.049 | 0.049 | 0 | 0.24 | 0.12 | 0.22 | 0.1 | 0.3 | 0.25 |
| 4 | 38176864 | rs6856418 | 1.083 | 0.615 | -0.93 | -0.285 | -0.019 | 0.063 | 0.063 | 0 | 0.2 | 0.05 | 0.19 | 0.08 | 0.19 | 0.19 |
| 4 | 38179244 | rs617615 | 1.11 | 0.871 | -0.167 | 0.234 | 0.002 | -0.007 | -0.007 | 0.268 | 0.17 | 0.19 | 0.11 | 0.14 | 0.19 | 0.13 |
| 4 | 38179948 | rs7436530 | -0.656 | 0.827 | -0.424 | 0.298 | -0.005 | -0.008 | -0.008 | 0.171 | 0.74 | 0.74 | 0.8 | 0.77 | 0.77 | 0.81 |
| 4 | 38179990 | rs11932593 | 0.11 | 0.803 | -0.342 | 0.38 | -0.013 | 0.017 | 0.017 | 0 | 0.65 | 0.58 | 0.56 | 0.49 | 0.69 | 0.65 |
| 4 | 38180161 | rs693978 | 0.093 | 0.803 | -0.37 | 0.381 | -0.005 | 0.005 | 0.005 | 0 | 0.65 | 0.63 | 0.57 | 0.49 | 0.69 | 0.65 |
| 4 | 38181054 | rs6854035 | 0.726 | 0.868 | -0.609 | 0.251 | 0 | 0.037 | 0.037 | 0 | 0.1 | 0.1 | 0.22 | 0.28 | 0.08 | 0.16 |
| 4 | 38187084 | rs586844 | 0.211 | 0.908 | -0.577 | 0.214 | -0.004 | 0.013 | 0.013 | 0 | 0.62 | 0.6 | 0.53 | 0.56 | 0.64 | 0.63 |
| 4 | 38188921 | rs17577389 | 0.635 | 0.936 | -0.806 | 0.362 | -0.035 | -0.083 | -0.083 | 0.173 | 0.12 | 0.01 | 0.25 | 0.28 | 0.13 | 0.22 |
| 4 | 38190035 | rs11737462 | 0.8 | 0.983 | -0.498 | -0.603 | -0.007 | 0.001 | 0.001 | 0 | 0.11 | 0.11 | 0.15 | 0.07 | 0.15 | 0.17 |
| 4 | 38194576 | rs17498347 | 0.549 | 0.631 | -0.25 | -0.286 | -0.011 | 0.05 | 0.05 | 0.001 | 0.44 | 0.27 | 0.4 | 0.27 | 0.46 | 0.48 |
| 4 | 38195371 | rs17580275 | -0.473 | 1.056 | 0.272 | 0.271 | 0.006 | 0.009 | 0.009 | 0.031 | 0.32 | 0.46 | 0.38 | 0.59 | 0.32 | 0.4 |
| 4 | 38197189 | rs2380962 | -0.419 | 1.144 | -0.46 | 0.355 | 0.009 | 0.013 | 0.013 | 0.031 | 0.32 | 0.5 | 0.4 | 0.6 | 0.32 | 0.4 |
| 4 | 38203158 | rs6531613 | -0.222 | 0.923 | -0.641 | -0.398 | 0.001 | -0.017 | -0.017 | 0.042 | 0.51 | 0.49 | 0.62 | 0.84 | 0.59 | 0.64 |
| 4 | 38204133 | rs4832752 | 0.264 | 0.778 | -1.079 | -0.704 | 0.021 | 0.149 | 0.149 | 0 | 0.51 | 0.21 | 0.38 | 0.22 | 0.58 | 0.47 |
| 4 | 38205831 | rs1397816 | 0.553 | 1.027 | -1.443 | -1.082 | -0.005 | 0.042 | 0.042 | 0 | 0.35 | 0.18 | 0.28 | 0.21 | 0.34 | 0.26 |
| 4 | 38211434 | rs4389587 | 0.006 | 1.209 | -0.829 | -0.79 | -0.002 | 0.002 | 0.002 | 0 | 0.76 | 0.82 | 0.85 | 0.94 | 0.71 | 0.78 |
| 4 | 38212624 | rs4554086 | 0.933 | 0.94 | -0.463 | -1.159 | 0.013 | 0.035 | 0.035 | 0 | 0.52 | 0.61 | 0.4 | 0.22 | 0.41 | 0.34 |
| 4 | 38213113 | rs11096935 | 0.488 | 1.201 | -0.199 | -0.184 | 0 | 0.116 | 0.116 | 0.236 | 0.09 | 0.15 | 0.26 | 0.45 | 0.05 | 0.3 |
| 4 | 38213124 | rs11096936 | 0.727 | 1.236 | -0.439 | -0.227 | -0.001 | 0.001 | 0.001 | 0 | 0.59 | 0.62 | 0.66 | 0.67 | 0.46 | 0.64 |
| 4 | 38213331 | rs10012308 | 0.311 | 1.257 | -0.288 | -0.104 | 0.006 | -0.029 | -0.029 | 0 | 0.17 | 0.22 | 0.18 | 0.02 | 0.25 | 0.1 |
| 4 | 38216411 | rs6833738 | 0.319 | 1.464 | -0.078 | 0.013 | 0.048 | 0.081 | 0.081 | 0 | 0.15 | 0.29 | 0.2 | 0.28 | 0.22 | 0.14 |
| 4 | 38225077 | rs7654067 | 0.485 | 1.684 | -0.149 | -0.072 | 0.007 | -0.001 | -0.001 | 0 | 0.67 | 0.62 | 0.52 | 0.29 | 0.61 | 0.44 |

| | | | | | | | | | | | | | | | | |
|---|---|---|---|---|---|---|---|---|---|---|---|---|---|---|---|---|
| 4 | 38228570 | rs6531617 | -0.184 | 1.89 | -0.967 | -0.349 | -0.032 | 0.078 | 0.078 | 0 | 0.75 | 0.9 | 0.88 | 0.99 | 0.68 | 0.77 |
| 4 | 38231635 | rs2055889 | 0.616 | 1.693 | -0.587 | 0.595 | -0.03 | 0.004 | 0.004 | 0.031 | 0.07 | 0.01 | 0.28 | 0.53 | 0.07 | 0.32 |
| 4 | 38233376 | rs10022626 | -0.388 | 1.591 | -0.481 | 0.464 | 0.018 | 0.028 | 0.028 | 0.31 | 0.17 | 0.68 | 0.25 | 0.24 | 0.22 | 0.14 |
| 4 | 38233985 | rs12648491 | 0.349 | 1.23 | -0.799 | 0.879 | -0.03 | 0.003 | 0.003 | 0.13 | 0.07 | 0.01 | 0.27 | 0.53 | 0.07 | 0.3 |
| 4 | 38237981 | rs6857949 | -0.716 | 0.897 | -0.295 | 0.407 | -0.018 | 0.148 | 0.148 | 0.245 | 0.14 | 0.53 | 0.12 | 0.25 | 0.08 | 0.13 |
| 4 | 38238718 | rs4833014 | -0.319 | 1.042 | -1.883 | -0.742 | -0.043 | 0.23 | 0.23 | 0 | 0.43 | 0.08 | 0.34 | 0.08 | 0.59 | 0.43 |
| 4 | 38241220 | rs7695667 | -0.342 | 1.666 | -1.335 | -0.645 | -0.001 | -0.006 | -0.006 | 0 | 0.15 | 0.14 | 0.18 | 0.02 | 0.24 | 0.14 |
| 4 | 38243991 | rs13139983 | 0.502 | 1.283 | -1.803 | -0.949 | -0.029 | 0.165 | 0.165 | 0 | 0.44 | 0.16 | 0.23 | 0.17 | 0.42 | 0.29 |
| 4 | 38246265 | rs16994359 | 0.276 | 0.968 | -1.012 | -0.832 | 0 | 0.019 | 0.019 | 0 | 0.34 | 0.22 | 0.55 | 0.78 | 0.33 | 0.56 |
| 4 | 38251292 | rs4833022 | 0.34 | 0.262 | -0.964 | -1.652 | -0.035 | -0.031 | -0.031 | 0.285 | 0.35 | 0.49 | 0.56 | 0.8 | 0.2 | 0.53 |
| 4 | 38253255 | rs17426830 | 1.42 | 0.445 | -1.119 | -1.794 | -0.055 | 0.134 | 0.134 | 0 | 0.26 | 0.02 | 0.13 | 0.06 | 0.28 | 0.22 |
| 4 | 38254453 | rs907314 | -0.097 | 0.659 | -0.889 | -1.597 | 0.005 | -0.037 | -0.037 | 0 | 0.61 | 0.51 | 0.69 | 0.86 | 0.47 | 0.75 |
| 4 | 38262829 | rs1467093 | -0.853 | 0.906 | -0.724 | -1.512 | -0.005 | 0.004 | 0.004 | 0 | 0.76 | 0.75 | 0.85 | 0.86 | 0.69 | 0.86 |
| 4 | 38270176 | rs1467092 | -0.125 | 0.948 | -1.29 | -2.009 | -0.024 | 0.046 | 0.046 | 0.907 | 0.35 | 0.27 | 0.28 | 0.15 | 0.53 | 0.26 |
| 4 | 38270461 | rs1467091 | -0.465 | 0.896 | -0.974 | -2.035 | 0.015 | -0.058 | -0.058 | 0 | 0.37 | 0.47 | 0.29 | 0.15 | 0.54 | 0.26 |
| 4 | 38270567 | rs16994377 | 1.035 | 0.7 | -1.07 | -2.104 | 0.022 | 0.022 | 0.022 | 0.413 | 0.37 | 0.29 | 0.18 | 0.12 | 0.22 | 0.23 |
| 4 | 38271832 | rs1006658 | -0.53 | 1.636 | -0.886 | -1.615 | -0.03 | 0.047 | 0.047 | 0.055 | 0.35 | 0.27 | 0.28 | 0.16 | 0.53 | 0.25 |
| 4 | 38276064 | rs7660869 | 0.211 | 1.734 | -0.753 | -1.558 | 0.039 | -0.016 | -0.016 | 0 | 0.16 | 0.29 | 0.13 | 0.1 | 0.26 | 0.16 |
| 4 | 38280267 | rs4833028 | 0.367 | 1.43 | -1.81 | -1.33 | -0.093 | 0.067 | 0.067 | 0.062 | 0.22 | 0.01 | 0.17 | 0.1 | 0.29 | 0.14 |
| 4 | 38280712 | rs17580746 | 0.803 | 0.797 | -1.47 | -0.973 | -0.02 | -0.067 | -0.067 | 0 | 0.13 | 0.02 | 0.43 | 0.76 | 0.09 | 0.28 |
| 4 | 38283255 | rs4833029 | 0.559 | 1.334 | -0.894 | -0.899 | -0.001 | -0.01 | -0.01 | 0 | 0.57 | 0.41 | 0.64 | 0.82 | 0.45 | 0.66 |
| 4 | 38287192 | rs9684138 | 0.261 | 1.309 | -1.118 | -0.322 | -0.013 | 0.013 | 0.013 | 0 | 0.6 | 0.68 | 0.58 | 0.67 | 0.52 | 0.72 |
| 4 | 38287333 | rs10517472 | -0.999 | 1.56 | -0.862 | -0.647 | -0.024 | 0.039 | 0.039 | 0 | 0.28 | 0.19 | 0.34 | 0.15 | 0.38 | 0.23 |
| 4 | 38288687 | rs4995512 | 0.137 | 1.465 | -0.382 | -0.689 | 0.002 | -0.032 | -0.032 | 0 | 0.11 | 0.13 | 0.09 | 0 | 0.13 | 0.05 |
| 4 | 38291344 | rs1553879 | 0.645 | 1.498 | 0.053 | -0.668 | 0.045 | -0.071 | -0.071 | 0 | 0.78 | 0.54 | 0.84 | 0.9 | 0.75 | 0.76 |
| 4 | 38298269 | rs6847801 | -0.247 | 1.253 | -0.193 | -0.519 | -0.003 | -0.022 | -0.022 | 0.063 | 0.09 | 0.13 | 0.05 | 0 | 0.08 | 0.11 |
| 4 | 38301690 | rs2890608 | 0.439 | 1.279 | 0.22 | 0.032 | 0.034 | 0.204 | 0.204 | 0 | 0.88 | 0.72 | 0.77 | 0.67 | 0.86 | 0.64 |
| 4 | 38302418 | rs13146334 | -0.991 | 1.123 | -0.495 | -0.552 | -0.004 | -0.047 | -0.047 | 0.039 | 0.46 | 0.61 | 0.37 | 0.11 | 0.39 | 0.35 |
| 4 | 38304013 | rs7676429 | 0.882 | 1.454 | -0.944 | -0.19 | 0.002 | 0.001 | 0.001 | 2.116 | 0.5 | 0.17 | 0.45 | 0.56 | 0.55 | 0.4 |
| 4 | 38308761 | rs991331 | 0.878 | 1.189 | -0.044 | -0.272 | -0.001 | 0 | 0 | 0 | 0.88 | 0.88 | 0.87 | 0.93 | 0.81 | 0.9 |
| 4 | 38314170 | rs1586459 | 0.91 | 1.185 | -0.116 | -0.3 | -0.018 | 0.02 | 0.02 | 0 | 0.88 | 0.93 | 0.87 | 0.96 | 0.82 | 0.9 |
| 4 | 38318354 | rs4833034 | 1.762 | 0.712 | 0.577 | -0.658 | -0.002 | -0.101 | -0.101 | 0 | 0.68 | 0.43 | 0.76 | 0.93 | 0.69 | 0.76 |
| 4 | 38318791 | rs1466614 | 1.535 | 0.922 | 0.865 | -0.44 | 0.012 | -0.061 | -0.061 | 0 | 0.77 | 0.61 | 0.82 | 0.93 | 0.77 | 0.85 |
| 4 | 38321682 | rs7677938 | 0.416 | 0.958 | 0.651 | -0.21 | 0.007 | -0.003 | -0.003 | 0 | 0.91 | 0.91 | 0.87 | 0.94 | 0.83 | 0.91 |
| 4 | 38325036 | rs6856616 | -1.014 | 0.887 | 0.907 | 0.46 | -0.007 | 0.131 | 0.131 | 0 | 0.12 | 0.33 | 0.17 | 0.23 | 0.06 | 0.27 |
| 4 | 38325105 | rs1906297 | 1.441 | 0.814 | 0.698 | 0.392 | 0.009 | 0.028 | 0.028 | 0.361 | 0.75 | 0.43 | 0.69 | 0.7 | 0.76 | 0.61 |
| 4 | 38333458 | rs13101978 | -1.653 | 0.579 | -0.068 | -0.154 | -0.024 | 0.067 | 0.067 | 0.15 | 0.38 | 0.29 | 0.44 | 0.03 | 0.59 | 0.33 |
| 4 | 38334899 | rs17614875 | -0.322 | 0.34 | 0.343 | 0.956 | 0.004 | 0.004 | 0.004 | 0 | 0.07 | 0.06 | 0.17 | 0.24 | 0.04 | 0.18 |
| 4 | 38343533 | rs12502507 | -0.783 | 0.213 | 0.157 | 0.608 | -0.011 | -0.049 | -0.049 | 0.304 | 0.12 | 0.21 | 0.1 | 0 | 0.1 | 0.07 |
| 4 | 38344189 | rs12502727 | -0.907 | 0.195 | 0.251 | 0.599 | -0.016 | -0.062 | -0.062 | 0 | 0.13 | 0.24 | 0.1 | 0 | 0.1 | 0.07 |
| 4 | 38349236 | rs716975 | -1.365 | 0.251 | 0.09 | 1.07 | -0.007 | 0.072 | 0.072 | 0 | 0.14 | 0.22 | 0.26 | 0.46 | 0.11 | 0.13 |
| 4 | 38362065 | rs2381149 | -0.206 | 0.265 | -0.239 | 0.79 | -0.002 | 0.035 | 0.035 | 0 | 0.09 | 0.04 | 0.1 | 0 | 0.1 | 0.06 |
| 4 | 38366090 | rs13133642 | 0.616 | 0.416 | 0.157 | 1.037 | 0.029 | 0.073 | 0.073 | 0 | 0.8 | 0.72 | 0.69 | 0.49 | 0.69 | 0.76 |
| 4 | 38367538 | rs1487624 | 1.902 | 0.112 | 0.26 | 0.906 | -0.035 | 0.091 | 0.091 | 0 | 0.54 | 0.19 | 0.47 | 0.3 | 0.49 | 0.43 |
| 4 | 38370063 | rs6810650 | -1.536 | -0.236 | 0.299 | 0.857 | -0.015 | 0.017 | 0.017 | 0 | 0.32 | 0.61 | 0.36 | 0.41 | 0.22 | 0.47 |
| 4 | 38372849 | rs2046643 | 0.486 | 0.198 | 0.255 | 0.591 | 0.006 | -0.052 | -0.052 | 0 | 0.78 | 0.62 | 0.82 | 0.93 | 0.72 | 0.81 |
| 4 | 38372995 | rs1906292 | 1.334 | 0.083 | -0.044 | 0.979 | 0 | 0.008 | 0.008 | 0 | 0.6 | 0.32 | 0.67 | 0.38 | 0.6 | 0.44 |
| 4 | 38374153 | rs11732737 | 1.876 | -0.18 | -0.907 | 0.828 | -0.166 | 0.264 | 0.264 | 0.652 | 0.42 | 0.02 | 0.4 | 0.08 | 0.49 | 0.24 |
| 4 | 38375537 | rs11722631 | 0.92 | -0.104 | 0.216 | 1.717 | -0.005 | -0.006 | -0.006 | 2.754 | 0.76 | 0.78 | 0.76 | 0.71 | 0.73 | 0.71 |
| 4 | 38376515 | rs6821082 | 0.335 | 0.044 | 0.144 | 1.783 | -0.067 | 0.029 | 0.029 | 0 | 0.81 | 0.96 | 0.85 | 0.91 | 0.75 | 0.82 |
| 4 | 38380708 | rs7668402 | 0.738 | -0.021 | 0.301 | 1.54 | 0.004 | 0.019 | 0.019 | 0 | 0.77 | 0.74 | 0.63 | 0.43 | 0.72 | 0.75 |
| 4 | 38387075 | rs2291433 | 1.273 | -0.289 | 0.495 | 1.389 | -0.001 | 0.008 | 0.008 | 0 | 0.52 | 0.38 | 0.43 | 0.42 | 0.54 | 0.38 |
| 4 | 38391687 | rs1487632 | 0.723 | -0.323 | 1.115 | 1.334 | 0.001 | 0.028 | 0.028 | 0 | 0.64 | 0.58 | 0.59 | 0.5 | 0.63 | 0.49 |
| 4 | 38394714 | rs6854078 | -0.513 | -0.415 | 1.331 | 1.37 | 0.002 | -0.028 | -0.028 | 0 | 0.12 | 0.24 | 0.17 | 0.06 | 0.09 | 0.1 |
| 4 | 38395949 | rs1487631 | 1.851 | -0.491 | 1.261 | 1.261 | -0.004 | 0.005 | 0.005 | 0.029 | 0.47 | 0.29 | 0.41 | 0.44 | 0.52 | 0.36 |
| 4 | 38396739 | rs6531631 | 0.574 | -0.521 | 1.669 | 1.521 | -0.024 | 0.087 | 0.087 | 0 | 0.64 | 0.84 | 0.73 | 0.88 | 0.62 | 0.51 |
| 4 | 38398572 | rs6832648 | 1.002 | -0.521 | 1.85 | 1.519 | 0.001 | 0.008 | 0.008 | 0.02 | 0.6 | 0.6 | 0.59 | 0.5 | 0.61 | 0.47 |
| 4 | 38402111 | rs12511283 | -0.584 | -0.689 | 1.803 | 1.645 | 0.002 | -0.03 | -0.03 | 0 | 0.12 | 0.25 | 0.17 | 0.07 | 0.09 | 0.09 |
| 4 | 38413846 | rs901703 | 0.59 | -0.548 | 0.925 | 1.408 | -0.02 | 0.098 | 0.098 | 0 | 0.67 | 0.87 | 0.76 | 0.9 | 0.67 | 0.53 |
| 4 | 38419843 | rs1487618 | -1.147 | -0.557 | 1.308 | 2.135 | -0.02 | 0.08 | 0.08 | 0.647 | 0.13 | 0.65 | 0.22 | 0.27 | 0.06 | 0.15 |
| 4 | 38425619 | rs10012289 | 1.47 | -0.546 | 0.868 | 1.991 | -0.003 | 0.026 | 0.026 | 0.215 | 0.53 | 0.23 | 0.45 | 0.39 | 0.6 | 0.39 |
| 4 | 38433101 | rs1601758 | -1.963 | -0.906 | 1.325 | 2.218 | 0.009 | -0.009 | -0.009 | 0 | 0.34 | 0.69 | 0.43 | 0.61 | 0.22 | 0.52 |
| 4 | 38433730 | rs7673568 | -1.017 | -0.889 | 1.675 | 2.004 | -0.024 | -0.057 | -0.057 | 0 | 0.2 | 0.39 | 0.14 | 0.09 | 0.15 | 0.16 |
| 4 | 38443544 | rs6830905 | -0.615 | -0.861 | 1.474 | 1.96 | -0.005 | -0.029 | -0.029 | 0 | 0.11 | 0.15 | 0.09 | 0.02 | 0.05 | 0.32 |
| 4 | 38452184 | rs4240245 | -1.245 | -0.892 | 1.528 | 2.368 | -0.019 | 0.153 | 0.153 | 0 | 0.22 | 0.49 | 0.34 | 0.44 | 0.17 | 0.19 |
| 4 | 38458620 | rs1004064 | -0.846 | -0.834 | 1.263 | 2.11 | -0.008 | -0.046 | -0.046 | 0 | 0.12 | 0.18 | 0.1 | 0.02 | 0.06 | 0.34 |
| 4 | 38464493 | rs17615370 | -1.25 | -1.069 | 1.162 | 2.328 | -0.015 | 0.002 | 0.002 | 0 | 0.16 | 0.22 | 0.12 | 0.19 | 0.08 | 0.36 |
| 4 | 38465656 | rs4131726 | -2.177 | -1.267 | 1.147 | 2.309 | 0.042 | 0.023 | 0.023 | 0.493 | 0.36 | 0.71 | 0.26 | 0.27 | 0.22 | 0.51 |
| 4 | 38492765 | rs4832778 | -1.076 | -0.697 | 1.138 | 1.841 | -0.013 | -0.007 | -0.007 | 0.602 | 0.17 | 0.24 | 0.13 | 0.12 | 0.17 | 0.13 |

| | | | | | | | | | | | | | | | | |
|---|---|---|---|---|---|---|---|---|---|---|---|---|---|---|---|---|
| 4 | 38503527 | rs9996673 | -1.333 | -0.79 | 1.436 | 2.077 | 0.032 | 0.129 | 0.129 | 0.082 | 0.06 | 0.28 | 0.03 | 0.2 | 0.02 | 0.09 |
| 4 | 38506777 | rs11723479 | 0.727 | -1.569 | 1.252 | 2.087 | -0.012 | 0.001 | 0.001 | 0 | 0.63 | 0.56 | 0.58 | 0.63 | 0.74 | 0.63 |
| 4 | 38509827 | rs1873515 | -0.99 | -1.704 | 1.19 | 1.831 | 0.02 | 0.105 | 0.105 | 0.327 | 0.06 | 0.25 | 0.03 | 0.19 | 0.02 | 0.09 |
| 4 | 38512602 | rs4833065 | 1.379 | -1.825 | 1.235 | 1.604 | 0.111 | 0.184 | 0.184 | 0 | 0.52 | 0.21 | 0.43 | 0.09 | 0.7 | 0.51 |
| 4 | 38513691 | rs2220634 | -1.784 | -1.731 | 1.18 | 2.055 | -0.038 | 0.045 | 0.045 | 0 | 0.19 | 0.28 | 0.21 | 0.69 | 0.08 | 0.26 |
| 4 | 38514293 | rs17500058 | -0.361 | -1.22 | 0.703 | 1.451 | -0.001 | -0.004 | -0.004 | 0 | 0.25 | 0.26 | 0.32 | 0.4 | 0.24 | 0.5 |
| 4 | 38515141 | rs2609467 | 0.067 | -1.108 | 0.437 | 1.155 | -0.035 | -0.015 | -0.015 | 0 | 0.9 | 0.82 | 0.9 | 0.96 | 0.97 | 0.98 |
| 4 | 38519943 | rs11946196 | -0.182 | -0.994 | 0.333 | 1.443 | -0.005 | 0.046 | 0.046 | 0 | 0.22 | 0.36 | 0.22 | 0.38 | 0.2 | 0.31 |
| 4 | 38523522 | rs2637729 | 0.358 | -0.447 | -0.322 | 0.186 | 0.004 | 0.007 | 0.007 | 0 | 0.53 | 0.44 | 0.42 | 0.19 | 0.57 | 0.54 |
| 4 | 38524098 | rs6814050 | 0.426 | -0.498 | -0.984 | -0.217 | 0.049 | 0.065 | 0.065 | 1.134 | 0.22 | 0.05 | 0.18 | 0.08 | 0.15 | 0.17 |
| 4 | 38524446 | rs7687994 | -0.03 | 0.208 | -0.365 | 0.267 | -0.011 | -0.007 | -0.007 | 0 | 0.15 | 0.11 | 0.16 | 0.01 | 0.24 | 0.13 |
| 4 | 38527901 | rs17615522 | -0.8 | 0.036 | -0.398 | 1.023 | -0.001 | 0.057 | 0.057 | 0 | 0.08 | 0.08 | 0.23 | 0.6 | 0.1 | 0.16 |
| 4 | 38532882 | rs16994596 | -0.79 | -0.092 | 0.103 | 0.711 | 0.003 | -0.023 | -0.023 | 0 | 0.09 | 0.24 | 0.06 | 0.03 | 0.09 | 0.04 |
| 4 | 38533692 | rs7688952 | -0.64 | -0.432 | -0.335 | 0.156 | 0.046 | 0.139 | 0.139 | 1.596 | 0.49 | 0.83 | 0.39 | 0.3 | 0.37 | 0.43 |
| 4 | 38537654 | rs16994601 | 0.515 | -0.547 | -0.202 | 0.088 | -0.002 | 0.013 | 0.013 | 0 | 0.09 | 0.14 | 0.08 | 0.21 | 0.06 | 0.09 |
| 4 | 38537708 | rs923665 | 0.554 | -0.731 | 0.171 | 0.078 | -0.006 | -0.046 | -0.046 | 0.76 | 0.32 | 0.36 | 0.24 | 0.1 | 0.23 | 0.34 |
| 4 | 38537844 | rs16994602 | -0.113 | -0.842 | 0.431 | 0.413 | -0.011 | 0.042 | 0.042 | 0 | 0.11 | 0.26 | 0.09 | 0.21 | 0.06 | 0.09 |
| 4 | 38539272 | rs7675136 | -0.219 | -0.523 | 0.265 | 0.057 | 0 | -0.022 | -0.022 | 0 | 0.51 | 0.52 | 0.38 | 0.13 | 0.51 | 0.47 |
| 4 | 38539297 | rs7658907 | 0.312 | -0.431 | 0.061 | 0.052 | -0.005 | -0.004 | -0.004 | 0 | 0.42 | 0.37 | 0.35 | 0.11 | 0.42 | 0.42 |
| 4 | 38540930 | rs3860065 | -0.031 | -0.053 | -0.313 | 0.721 | -0.002 | 0.089 | 0.089 | 0 | 0.09 | 0.22 | 0.15 | 0.25 | 0.06 | 0.09 |
| 4 | 38548187 | rs2703904 | 0.41 | 0.142 | -0.528 | 0.346 | -0.003 | 0.011 | 0.011 | 0.017 | 0.35 | 0.3 | 0.27 | 0.15 | 0.41 | 0.34 |
| 4 | 38551146 | rs1552684 | 0.299 | 0.172 | -0.939 | 0.302 | -0.008 | 0.066 | 0.066 | 0 | 0.37 | 0.17 | 0.26 | 0.17 | 0.4 | 0.33 |
| 4 | 38562694 | rs6853067 | 0.549 | 0.111 | -0.303 | 0.621 | 0.002 | 0.038 | 0.038 | 0 | 0.84 | 0.71 | 0.75 | 0.73 | 0.84 | 0.66 |
| 4 | 38564339 | rs10026499 | -0.303 | 0.242 | -0.059 | -0.34 | 0.025 | -0.151 | -0.151 | 0 | 0.24 | 0.52 | 0.22 | 0.01 | 0.28 | 0.14 |
| 4 | 38570178 | rs2703887 | -0.861 | -0.439 | -0.509 | 0.796 | 0.042 | -0.046 | -0.046 | 0 | 0.09 | 0.01 | 0.15 | 0.38 | 0.05 | 0.12 |
| 4 | 38570506 | rs2703888 | 0.337 | 0.063 | -0.425 | 0.558 | 0.002 | 0.004 | 0.004 | 0 | 0.83 | 0.83 | 0.78 | 0.74 | 0.82 | 0.6 |
| 4 | 38570708 | rs7654396 | 0.259 | 0.116 | -0.988 | -0.372 | -0.04 | 0.023 | 0.023 | 0 | 0.28 | 0.22 | 0.27 | 0.01 | 0.45 | 0.14 |
| 4 | 38578754 | rs11726368 | 0.808 | -0.077 | -0.316 | 0.197 | 0.003 | 0.122 | 0.122 | 0.166 | 0.81 | 0.44 | 0.71 | 0.48 | 0.85 | 0.76 |
| 4 | 38582859 | rs4833071 | 1.024 | -0.273 | -1.041 | -0.21 | 0.08 | 0.116 | 0.116 | 0.631 | 0.45 | 0.17 | 0.34 | 0.31 | 0.32 | 0.4 |
| 4 | 38585268 | rs4401459 | 1.22 | -0.339 | -1.172 | -0.019 | 0.106 | 0.083 | 0.083 | 1.434 | 0.42 | 0.06 | 0.32 | 0.33 | 0.3 | 0.36 |
| 4 | 38592867 | rs7678850 | 1.02 | 0.18 | -0.894 | 0.095 | -0.047 | -0.01 | -0.01 | 0.703 | 0.72 | 0.84 | 0.6 | 0.4 | 0.54 | 0.61 |
| 4 | 38593723 | rs6811246 | 1.619 | 0.362 | -0.922 | 0.1 | -0.025 | 0.026 | 0.026 | 0.331 | 0.08 | 0.03 | 0.14 | 0 | 0.24 | 0.06 |
| 4 | 38596365 | rs2169541 | -0.271 | -0.414 | -0.778 | 0.458 | -0.012 | -0.065 | -0.065 | 0 | 0.89 | 0.99 | 0.88 | 0.61 | 0.9 | 0.87 |
| 4 | 38596653 | rs16994672 | -0.378 | -0.64 | -0.038 | 0.403 | -0.018 | -0.007 | -0.007 | 0.713 | 0.07 | 0.25 | 0.02 | 0.04 | 0.02 | 0.09 |
| 4 | 38611802 | rs337641 | -0.518 | -0.595 | -0.507 | 0.412 | 0.007 | -0.046 | -0.046 | 0 | 0.73 | 0.83 | 0.77 | 0.59 | 0.76 | 0.79 |
| 4 | 38612357 | rs1962053 | 1.537 | -1.074 | -0.342 | -0.327 | 0.043 | -0.087 | -0.087 | 0 | 0.17 | 0.35 | 0.12 | 0.02 | 0.2 | 0.09 |
| 4 | 38616410 | rs337646 | 0.027 | -1.809 | -0.764 | -0.099 | -0.014 | 0 | 0 | 0 | 0.18 | 0.22 | 0.04 | 0.12 | 0.05 | 0.17 |
| 4 | 38627337 | rs2637713 | -0.56 | -1.851 | -0.798 | -0.177 | -0.006 | 0.001 | 0.001 | 0 | 0.24 | 0.28 | 0.08 | 0.13 | 0.13 | 0.21 |
| 4 | 38644050 | rs1993796 | 0.638 | -1.56 | -0.289 | -0.029 | -0.019 | 0.066 | 0.066 | 0.189 | 0.05 | 0.14 | 0.18 | 0.18 | 0.01 | 0.16 |
| 4 | 38644597 | rs1824652 | 0.385 | -1.663 | -0.393 | -0.295 | 0.008 | -0.013 | -0.013 | 0 | 0.36 | 0.35 | 0.45 | 0.21 | 0.51 | 0.37 |
| 4 | 38654681 | rs4833079 | 0.529 | -1.358 | -0.612 | -0.401 | -0.004 | -0.001 | -0.001 | 0 | 0.14 | 0.11 | 0.2 | 0.02 | 0.36 | 0.11 |
| 4 | 38662059 | rs337609 | -2.471 | -1.711 | -1.071 | -0.238 | -0.029 | -0.145 | -0.145 | 0 | 0.81 | 0.99 | 0.81 | 0.59 | 0.81 | 0.87 |
| 4 | 38664324 | rs1466414 | 0.582 | -1.73 | -0.429 | -0.5 | 0.011 | -0.035 | -0.035 | 0 | 0.26 | 0.42 | 0.32 | 0.25 | 0.27 | 0.31 |
| 4 | 38666972 | rs9522 | 0.653 | -1.731 | -0.587 | -0.663 | 0.005 | -0.012 | -0.012 | 0.012 | 0.31 | 0.39 | 0.42 | 0.15 | 0.44 | 0.24 |
| 4 | 38680186 | rs9992667 | 2.197 | -1.805 | -0.493 | -0.451 | 0.024 | -0.025 | -0.025 | 0.143 | 0.15 | 0.21 | 0.1 | 0.07 | 0.19 | 0.09 |
| 4 | 38688362 | rs16994718 | 0.331 | -2.096 | -0.582 | -0.373 | -0.034 | -0.003 | -0.003 | 0 | 0.28 | 0.36 | 0.28 | 0.21 | 0.16 | 0.21 |
| 4 | 38689042 | rs17428931 | 1.218 | -1.522 | -0.936 | -0.324 | -0.001 | -0.001 | -0.001 | 0 | 0.06 | 0.06 | 0.04 | 0.03 | 0.07 | 0.05 |
| 4 | 38692806 | rs3796533 | 1.024 | -1.843 | -1.396 | -0.458 | 0.057 | 0.033 | 0.033 | 0 | 0.25 | 0.05 | 0.28 | 0.16 | 0.21 | 0.2 |
| 4 | 38692835 | rs1000096 | -0.093 | -1.694 | -1.191 | -0.461 | 0.088 | -0.056 | -0.056 | 0 | 0.23 | 0.47 | 0.34 | 0.19 | 0.33 | 0.41 |
| 4 | 38696934 | rs2045767 | -0.629 | -1.638 | -1.226 | -0.487 | 0.001 | 0.017 | 0.017 | 0 | 0.52 | 0.52 | 0.63 | 0.34 | 0.56 | 0.62 |
| 4 | 38699657 | rs3209570 | 1.167 | -1.624 | -1.105 | -0.525 | 0.173 | -0.108 | -0.108 | 0.829 | 0.18 | 0.42 | 0.2 | 0.03 | 0.33 | 0.25 |
| 4 | 38702437 | rs7147 | -1.279 | -1.796 | -1.507 | -0.516 | 0.025 | -0.056 | -0.056 | 0 | 0.69 | 0.9 | 0.76 | 0.45 | 0.76 | 0.73 |
| 4 | 38702929 | rs17500878 | 0.16 | -1.807 | -1.26 | -0.548 | 0.001 | -0.007 | -0.007 | 0 | 0.27 | 0.32 | 0.16 | 0.17 | 0.29 | 0.2 |
| 4 | 38703734 | rs17582575 | 0.759 | -1.891 | -1.302 | -0.247 | 0.058 | 0.008 | 0.008 | 0 | 0.28 | 0.09 | 0.3 | 0.25 | 0.22 | 0.22 |
| 4 | 38705257 | rs10021686 | 2.588 | -1.127 | -1.119 | -0.18 | -0.001 | -0.016 | -0.016 | 0 | 0.12 | 0.11 | 0.07 | 0.01 | 0.15 | 0.08 |
| 4 | 38711684 | rs13132956 | 1.61 | -1.53 | -1.577 | -0.078 | -0.063 | -0.171 | -0.171 | 0 | 0.24 | 0.02 | 0.21 | 0.4 | 0.2 | 0.26 |
| 4 | 38712167 | rs6813851 | 2.384 | -2.544 | -1.28 | -0.326 | -0.001 | -0.017 | -0.017 | 0 | 0.12 | 0.11 | 0.06 | 0.01 | 0.13 | 0.08 |
| 4 | 38712286 | rs6838905 | 2.363 | -2.548 | -1.244 | -0.328 | -0.001 | -0.026 | -0.026 | 0.576 | 0.12 | 0.13 | 0.06 | 0.01 | 0.13 | 0.08 |
| 4 | 38716483 | rs13149231 | -1.844 | -2.195 | -1.122 | -0.472 | -0.031 | 0.026 | 0.026 | 0 | 0.87 | 0.92 | 0.81 | 0.98 | 0.65 | 0.91 |
| 4 | 38722709 | rs1005761 | -0.01 | -3.51 | -0.457 | -0.457 | -0.049 | -0.005 | -0.005 | 0.241 | 0.13 | 0.32 | 0.05 | 0.08 | 0.05 | 0.22 |
| 4 | 38723613 | rs17616338 | 0.687 | -3.669 | -1.226 | -0.565 | 0.028 | -0.035 | -0.035 | 0 | 0.15 | 0.05 | 0.25 | 0.25 | 0.14 | 0.15 |
| 4 | 38724573 | rs7660102 | 0.464 | -3.603 | -0.903 | -0.626 | -0.008 | -0.01 | -0.01 | 0 | 0.36 | 0.55 | 0.4 | 0.7 | 0.24 | 0.52 |
| 4 | 38728076 | rs7658334 | -1.091 | -3.222 | -1.29 | -0.883 | -0.021 | 0.018 | 0.018 | 0 | 0.67 | 0.78 | 0.7 | 0.98 | 0.44 | 0.83 |
| 4 | 38732464 | rs7660114 | -0.968 | -4.18 | -1.485 | -0.348 | 0.146 | -0.019 | -0.019 | 0 | 0.41 | 0.8 | 0.45 | 0.66 | 0.17 | 0.48 |
| 4 | 38736550 | rs2890620 | -0.795 | -4.485 | -0.938 | -0.387 | 0.061 | 0.233 | 0.233 | 0.001 | 0.57 | 0.84 | 0.64 | 0.93 | 0.34 | 0.68 |
| 4 | 38741445 | rs6842064 | 1.514 | -4.294 | -0.421 | 0.201 | -0.059 | 0.051 | 0.051 | 0 | 0.11 | 0.32 | 0.13 | 0.24 | 0.01 | 0.21 |
| 4 | 38741849 | rs9968502 | 0.506 | -4.285 | -0.57 | 0.284 | -0.093 | 0.158 | 0.158 | 0.015 | 0.16 | 0.46 | 0.15 | 0.38 | 0.03 | 0.24 |
| 4 | 38745883 | rs6838865 | -0.878 | -4.452 | -0.607 | 0.126 | -0.012 | -0.003 | -0.003 | 0 | 0.32 | 0.61 | 0.37 | 0.56 | 0.11 | 0.36 |
| 4 | 38750680 | rs11946541 | 0.412 | -3.916 | -0.416 | -0.111 | -0.092 | 0.095 | 0.095 | 0 | 0.16 | 0.44 | 0.14 | 0.3 | 0.03 | 0.23 |

| | | | | | | | | | | | | | | | | |
|---|---|---|---|---|---|---|---|---|---|---|---|---|---|---|---|---|
| 4 | 38751694 | rs4540058 | -0.496 | -3.626 | -1.156 | -0.417 | 0.048 | 0.152 | 0.152 | 0 | 0.58 | 0.77 | 0.63 | 0.93 | 0.33 | 0.66 |
| 4 | 38752985 | rs4565081 | -0.261 | -2.86 | -0.81 | -0.023 | -0.055 | -0.024 | -0.024 | 0 | 0.36 | 0.4 | 0.46 | 0.67 | 0.19 | 0.36 |
| 4 | 38754189 | rs6824105 | 0.553 | -1.278 | -0.992 | 0.279 | -0.012 | 0.002 | 0.002 | 0.592 | 0.58 | 0.53 | 0.36 | 0.48 | 0.37 | 0.55 |
| 4 | 38769408 | rs4331786 | 0.024 | -2.573 | -0.895 | 0.165 | -0.005 | -0.004 | -0.004 | 0 | 0.58 | 0.5 | 0.42 | 0.58 | 0.4 | 0.54 |
| 4 | 38774785 | rs10776482 | 0.648 | -3.905 | -1.363 | -0.289 | 0.036 | -0.044 | -0.044 | 0 | 0.34 | 0.21 | 0.32 | 0.43 | 0.25 | 0.31 |
| 4 | 38774889 | rs4129009 | 1.758 | -4.453 | -1.683 | -0.214 | -0.037 | -0.125 | -0.125 | 0.072 | 0.2 | 0.01 | 0.14 | 0.27 | 0.21 | 0.15 |
| 4 | 38775040 | rs10776483 | 0.317 | -4.444 | -1.677 | -0.312 | 0.037 | -0.044 | -0.044 | 0 | 0.34 | 0.21 | 0.32 | 0.43 | 0.25 | 0.31 |
| 4 | 38776180 | rs11096956 | 0.317 | -5.219 | -1.539 | -0.506 | 0.037 | -0.044 | -0.044 | 0 | 0.34 | 0.21 | 0.32 | 0.43 | 0.25 | 0.31 |
| 4 | 38776235 | rs11466653 | 1.149 | -5.272 | -1.507 | -0.476 | 0.034 | 0.005 | 0.005 | 0 | 0.12 | 0.03 | 0.16 | 0.11 | 0.04 | 0.16 |
| 4 | 38776303 | rs11466652 | 0.643 | -5.312 | -0.867 | -0.502 | -0.031 | -0.041 | -0.041 | 0 | 0.24 | 0.34 | 0.1 | 0.12 | 0.15 | 0.23 |
| 4 | 38776491 | rs11096957 | -0.84 | -5.303 | -0.866 | -0.497 | -0.007 | -0.005 | -0.005 | 0 | 0.58 | 0.55 | 0.42 | 0.55 | 0.4 | 0.54 |
| 4 | 38777059 | rs10856837 | 0.664 | -5.341 | -0.59 | -0.511 | -0.032 | -0.011 | -0.011 | 0 | 0.13 | 0.2 | 0.18 | 0.11 | 0.04 | 0.16 |
| 4 | 38777236 | rs10856839 | -0.484 | -5.345 | -0.603 | -0.51 | -0.051 | -0.041 | -0.041 | 0 | 0.37 | 0.51 | 0.27 | 0.23 | 0.19 | 0.38 |
| 4 | 38777471 | rs4274855 | 1.543 | -5.169 | -0.647 | -0.238 | -0.036 | -0.149 | -0.149 | 0.585 | 0.21 | 0.01 | 0.14 | 0.32 | 0.21 | 0.15 |
| 4 | 38780721 | rs11466612 | 0.121 | -4.677 | 0.337 | -0.236 | -0.051 | -0.011 | -0.011 | 0 | 0.14 | 0.27 | 0.18 | 0.12 | 0.04 | 0.16 |
| 4 | 38783374 | rs9998350 | 0.008 | -4.631 | 0.353 | -0.371 | -0.115 | -0.07 | -0.07 | 0 | 0.21 | 0.39 | 0.09 | 0.11 | 0.07 | 0.25 |
| 4 | 38783848 | rs11725309 | 1.414 | -4.903 | 0.019 | -0.334 | -0.02 | -0.179 | -0.179 | 0.052 | 0.24 | 0.01 | 0.15 | 0.37 | 0.21 | 0.17 |
| 4 | 38785850 | rs7663239 | 0.156 | -4.909 | 0.074 | -0.342 | -0.117 | -0.072 | -0.072 | 0 | 0.21 | 0.39 | 0.09 | 0.11 | 0.07 | 0.25 |
| 4 | 38787216 | rs12233670 | -0.224 | -4.683 | 0.083 | -0.296 | 0.02 | 0.006 | 0.006 | 0 | 0.46 | 0.54 | 0.24 | 0.48 | 0.28 | 0.41 |
| 4 | 38799710 | rs4833095 | -0.305 | -4.136 | -0.713 | -0.311 | 0.205 | 0.101 | 0.101 | 0.892 | 0.51 | 0.88 | 0.47 | 0.6 | 0.28 | 0.51 |
| 4 | 38802528 | rs5743596 | 1.586 | -3.934 | -0.829 | -0.174 | -0.016 | -0.136 | -0.136 | 0 | 0.22 | 0.01 | 0.12 | 0.28 | 0.18 | 0.15 |
| 4 | 38807654 | rs5743551 | -0.055 | -3.507 | -1.141 | -0.354 | 0.206 | 0.112 | 0.112 | 0.47 | 0.51 | 0.9 | 0.47 | 0.6 | 0.29 | 0.48 |
| 4 | 38812876 | rs17616434 | -0.078 | -3.467 | -1.14 | -0.365 | 0.197 | 0.112 | 0.112 | 0.534 | 0.51 | 0.9 | 0.47 | 0.6 | 0.28 | 0.48 |
| 4 | 38820986 | rs7696175 | -1.224 | -3.426 | -1.305 | -0.303 | -0.187 | 0.044 | 0.044 | 0 | 0.76 | 0.98 | 0.66 | 0.85 | 0.57 | 0.75 |
| 4 | 38829163 | rs5743818 | 0.499 | -2.768 | -0.557 | -0.03 | -0.039 | 0.031 | 0.031 | 0 | 0.1 | 0.04 | 0.09 | 0.01 | 0.24 | 0.08 |
| 4 | 38829832 | rs3775073 | -1.364 | -3.592 | -0.182 | 0.013 | 0.022 | 0.004 | 0.004 | 2.235 | 0.37 | 0.74 | 0.32 | 0.4 | 0.31 | 0.39 |
| 4 | 38831596 | rs1039559 | -1.605 | -3.81 | -0.115 | -0.027 | -0.08 | 0.071 | 0.071 | 0 | 0.66 | 0.88 | 0.49 | 0.73 | 0.54 | 0.57 |
| 4 | 38840162 | rs6833914 | -1.188 | -3.86 | 0.316 | -0.014 | -0.007 | 0.001 | 0.001 | 0.692 | 0.33 | 0.61 | 0.29 | 0.4 | 0.28 | 0.38 |
| 4 | 38846885 | rs7685852 | -0.039 | -3.889 | 0.471 | 0.153 | -0.114 | 0.078 | 0.078 | 0.047 | 0.17 | 0.44 | 0.24 | 0.28 | 0.03 | 0.2 |
| 4 | 38847917 | rs6531673 | -1.204 | -3.884 | 0.274 | 0.091 | -0.052 | 0.08 | 0.08 | 0 | 0.61 | 0.86 | 0.52 | 0.7 | 0.51 | 0.5 |
| 4 | 38857310 | rs17429224 | -0.57 | -3.112 | 0.114 | -0.037 | 0.024 | 0.003 | 0.003 | 0.001 | 0.15 | 0.23 | 0.11 | 0.14 | 0.25 | 0.13 |
| 4 | 38859515 | rs17501134 | 0.041 | -3.034 | 0.111 | -0.039 | 0.013 | -0.001 | -0.001 | 0 | 0.12 | 0.13 | 0.11 | 0.06 | 0.27 | 0.1 |
| 4 | 38873998 | rs1472720 | -0.739 | -2.513 | 0.318 | -0.099 | 0.069 | -0.093 | -0.093 | 0 | 0.18 | 0.4 | 0.14 | 0.01 | 0.27 | 0.11 |
| 4 | 38878194 | rs721653 | -0.914 | -2.633 | 0.193 | -0.088 | -0.001 | -0.001 | -0.001 | 0 | 0.51 | 0.51 | 0.4 | 0.43 | 0.51 | 0.36 |
| 4 | 38879949 | rs11096964 | -0.189 | -2.458 | 0.082 | -0.092 | 0.05 | -0.04 | -0.04 | 0 | 0.16 | 0.32 | 0.21 | 0.09 | 0.25 | 0.12 |
| 4 | 38893736 | rs6851463 | 0.034 | -2.53 | -0.438 | -0.159 | -0.049 | -0.062 | -0.062 | 0 | 0.14 | 0.2 | 0.05 | 0 | 0.06 | 0.09 |
| 4 | 38893843 | rs6851685 | 0.041 | -2.058 | -0.245 | 0.133 | 0.082 | 0.012 | 0.012 | 0 | 0.42 | 0.21 | 0.2 | 0.26 | 0.28 | 0.26 |
| 4 | 38897254 | rs1604834 | -0.206 | -1.544 | -0.073 | 0.199 | 0 | -0.004 | -0.004 | 2.993 | 0.55 | 0.43 | 0.39 | 0.4 | 0.51 | 0.45 |
| 4 | 38901769 | rs12641669 | -0.421 | -1.352 | -0.456 | 0.293 | 0.001 | 0.004 | 0.004 | 0 | 0.58 | 0.58 | 0.41 | 0.41 | 0.51 | 0.45 |
| 4 | 38910362 | rs7676657 | -0.362 | -1.77 | -0.444 | -0.093 | -0.056 | -0.037 | -0.037 | 0 | 0.15 | 0.22 | 0.05 | 0.05 | 0.05 | 0.11 |
| 4 | 38911315 | rs16994921 | -0.616 | -2.283 | -0.592 | -0.374 | -0.08 | -0.068 | -0.068 | 0 | 0.27 | 0.34 | 0.11 | 0.07 | 0.11 | 0.18 |
| 4 | 38912621 | rs7655305 | -0.943 | -1.977 | -1.812 | 0.103 | 0.037 | -0.038 | -0.038 | 0 | 0.58 | 0.82 | 0.36 | 0.58 | 0.38 | 0.57 |
| 4 | 38916914 | rs4240248 | -0.989 | -1.828 | -2.024 | 0.092 | -0.005 | -0.069 | -0.069 | 0 | 0.62 | 0.86 | 0.43 | 0.57 | 0.4 | 0.61 |
| 4 | 38917830 | rs11736617 | -0.294 | -0.857 | -1.536 | 0.713 | 0.025 | -0.093 | -0.093 | 3.731 | 0.2 | 0.08 | 0.2 | 0.42 | 0.12 | 0.23 |
| 4 | 38924275 | rs12509328 | -0.646 | -0.317 | -1.209 | 0.027 | -0.007 | 0.015 | 0.015 | 0.002 | 0.2 | 0.13 | 0.13 | 0.04 | 0.23 | 0.11 |
| 4 | 38924330 | rs10000037 | -0.537 | -0.349 | -0.927 | 0.142 | 0 | -0.006 | -0.006 | 0.051 | 0.2 | 0.16 | 0.14 | 0.06 | 0.23 | 0.17 |
| 4 | 38927230 | rs9999294 | -0.938 | -0.686 | -0.724 | 0.002 | 0.039 | -0.028 | -0.028 | 0 | 0.67 | 0.56 | 0.65 | 0.76 | 0.5 | 0.72 |
| 4 | 38927396 | rs10155189 | -0.94 | -0.425 | -0.659 | -0.023 | 0.031 | -0.025 | -0.025 | 0.009 | 0.67 | 0.6 | 0.66 | 0.73 | 0.51 | 0.72 |
| 4 | 38930433 | rs17429570 | 0.298 | -1.036 | -1.175 | 0.002 | -0.012 | 0.024 | 0.024 | 0 | 0.14 | 0.09 | 0.21 | 0.1 | 0.15 | 0.1 |
| 4 | 38930469 | rs3866197 | -0.985 | -1.385 | -0.923 | -0.215 | -0.013 | 0.024 | 0.024 | 0 | 0.62 | 0.35 | 0.62 | 0.73 | 0.46 | 0.7 |
| 4 | 38931097 | rs13150445 | 0.469 | -1.463 | -0.968 | -0.157 | 0.017 | -0.005 | -0.005 | 0.002 | 0.34 | 0.28 | 0.35 | 0.58 | 0.22 | 0.42 |
| 4 | 38933726 | rs4833106 | -0.097 | -1.199 | -0.958 | 0.611 | 0.096 | 0.026 | 0.026 | 0 | 0.48 | 0.22 | 0.33 | 0.43 | 0.31 | 0.56 |
| 4 | 38933859 | rs17429619 | 0.314 | -1.059 | -1.033 | 0.318 | -0.022 | 0.037 | 0.037 | 0.04 | 0.14 | 0.05 | 0.2 | 0.08 | 0.16 | 0.11 |
| 4 | 38936833 | rs2171778 | -0.023 | -0.558 | -0.477 | 0.219 | 0 | 0.006 | 0.006 | 0.183 | 0.51 | 0.43 | 0.46 | 0.65 | 0.46 | 0.53 |
| 4 | 38937055 | rs12506108 | 0.265 | -0.655 | -0.508 | 0.097 | -0.01 | -0.043 | -0.043 | 0.029 | 0.17 | 0.21 | 0.21 | 0.03 | 0.12 | 0.12 |
| 4 | 38937372 | rs2271031 | -0.986 | -0.471 | -0.643 | 0.206 | -0.084 | -0.037 | -0.037 | 0 | 0.72 | 0.89 | 0.69 | 0.69 | 0.59 | 0.7 |
| 4 | 38940630 | rs6531684 | -0.586 | -0.959 | -0.514 | -0.008 | 0.002 | -0.002 | -0.002 | 0 | 0.33 | 0.66 | 0.16 | 0.14 | 0.28 | 0.3 |
| 4 | 38940833 | rs6812702 | -0.31 | -0.729 | -0.129 | 0.334 | 0.009 | -0.013 | -0.013 | 0 | 0.07 | 0.26 | 0.03 | 0.04 | 0.04 | 0.2 |
| 4 | 38944101 | rs2011590 | -0.441 | -0.659 | -0.536 | 0.647 | -0.013 | -0.034 | -0.034 | 0.69 | 0.44 | 0.21 | 0.35 | 0.38 | 0.44 | 0.46 |
| 4 | 38946448 | rs6840077 | 0.743 | -0.231 | -0.257 | 0.781 | -0.019 | -0.01 | -0.01 | 2.307 | 0.09 | 0.05 | 0.17 | 0.09 | 0.12 | 0.13 |
| 4 | 38952030 | rs7694333 | -0.731 | -0.639 | -0.742 | 1.046 | -0.066 | 0.033 | 0.033 | 0.028 | 0.69 | 0.87 | 0.63 | 0.65 | 0.6 | 0.71 |
| 4 | 38952488 | rs2174285 | -0.614 | -0.695 | -0.785 | 1.48 | -0.078 | -0.065 | -0.065 | 0.248 | 0.71 | 0.9 | 0.64 | 0.55 | 0.6 | 0.71 |
| 4 | 38953128 | rs3806735 | -0.608 | -0.645 | -0.759 | 1.193 | -0.07 | 0.021 | 0.021 | 0 | 0.71 | 0.89 | 0.64 | 0.65 | 0.6 | 0.71 |
| 4 | 38955135 | rs17583377 | 0.592 | -0.095 | 0.212 | 1.482 | 0.006 | -0.002 | -0.002 | 1.756 | 0.22 | 0.2 | 0.25 | 0.21 | 0.17 | 0.2 |
| 4 | 38958404 | rs6833864 | -0.373 | -0.026 | 0.313 | 1.403 | 0.086 | 0.162 | 0.162 | 0.012 | 0.05 | 0.3 | 0.04 | 0.06 | 0.02 | 0.06 |
| 4 | 38960412 | rs17583426 | 0.329 | 0.138 | -0.696 | 1.328 | 0.007 | -0.009 | -0.009 | 0.048 | 0.22 | 0.19 | 0.35 | 0.3 | 0.18 | 0.2 |
| 4 | 38961449 | rs4974985 | 0.564 | 1.058 | -0.635 | 1.303 | -0.001 | -0.001 | -0.001 | 0 | 0.67 | 0.32 | 0.63 | 0.6 | 0.63 | 0.52 |
| 4 | 38965986 | rs6841592 | -0.506 | 1.619 | -1.026 | 0.886 | 0.026 | -0.013 | -0.013 | 0.458 | 0.19 | 0.23 | 0.16 | 0.18 | 0.25 | 0.29 |
| 4 | 38966075 | rs2055576 | -0.311 | 2.086 | -1.524 | 1.163 | -0.039 | -0.046 | -0.046 | 0 | 0.8 | 0.88 | 0.69 | 0.68 | 0.66 | 0.73 |

| | | | | | | | | | | | | | | | | |
|---|---|---|---|---|---|---|---|---|---|---|---|---|---|---|---|---|
| 4 | 38966472 | rs6820989 | -0.195 | 3.012 | -1.342 | 0.686 | 0.017 | -0.007 | -0.007 | 0 | 0.15 | 0.16 | 0.15 | 0.09 | 0.25 | 0.12 |
| 4 | 38967146 | rs1504238 | -0.083 | 2.682 | -1.559 | 0.836 | -0.016 | -0.017 | -0.017 | 0 | 0.1 | 0.06 | 0.17 | 0.17 | 0.13 | 0.13 |
| 4 | 38978171 | rs6814637 | -0.182 | -0.001 | -1.294 | 0.385 | 0.018 | 0.009 | 0.009 | 0 | 0.33 | 0.26 | 0.27 | 0.26 | 0.23 | 0.22 |
| 4 | 38980064 | rs7699651 | 0.199 | -0.192 | -1.588 | 0.209 | 0.044 | 0.046 | 0.046 | 0 | 0.64 | 0.3 | 0.74 | 0.79 | 0.74 | 0.8 |
| 4 | 38982755 | rs6811642 | -0.751 | 0.509 | -1.278 | 0.559 | 0 | 0.001 | 0.001 | 1.022 | 0.43 | 0.72 | 0.53 | 0.49 | 0.43 | 0.39 |
| 4 | 38984852 | rs11946488 | -0.64 | 0.583 | -0.988 | 0.665 | 0.002 | 0.006 | 0.006 | 0.695 | 0.41 | 0.65 | 0.43 | 0.39 | 0.4 | 0.3 |
| 4 | 38988587 | rs17583467 | 0.537 | 1.006 | -1.568 | 0.681 | -0.05 | -0.092 | -0.092 | 0 | 0.31 | 0.09 | 0.26 | 0.39 | 0.29 | 0.35 |
| 4 | 38988621 | rs17583481 | 0.472 | 1.352 | -1.207 | 0.91 | -0.01 | -0.004 | -0.004 | 0 | 0.06 | 0.05 | 0.16 | 0.22 | 0.12 | 0.12 |
| 4 | 38992305 | rs13106655 | -0.379 | 0.896 | -0.579 | 0.286 | -0.01 | -0.013 | -0.013 | 0 | 0.61 | 0.62 | 0.64 | 0.69 | 0.66 | 0.61 |
| 4 | 38994187 | rs4974979 | -0.182 | 0.814 | -1.11 | 0.451 | -0.008 | 0.068 | 0.068 | 0 | 0.48 | 0.26 | 0.46 | 0.65 | 0.5 | 0.53 |
| 4 | 39005601 | rs11096969 | -0.156 | 0.672 | -0.68 | 0.48 | 0 | 0 | 0 | 0.776 | 0.53 | 0.5 | 0.5 | 0.64 | 0.55 | 0.54 |
| 4 | 39008793 | rs7662121 | -0.659 | 0.154 | -1.029 | 0.429 | -0.01 | 0.001 | 0.001 | 0 | 0.45 | 0.49 | 0.34 | 0.42 | 0.34 | 0.51 |
| 4 | 39012630 | rs2163276 | -0.247 | 0.062 | -1.45 | 0.431 | 0.031 | -0.038 | -0.038 | 0 | 0.21 | 0.02 | 0.1 | 0.21 | 0.13 | 0.27 |
| 4 | 39016075 | rs2435530 | -0.17 | 0.722 | -1.084 | 0.204 | -0.006 | 0.008 | 0.008 | 0 | 0.77 | 0.74 | 0.6 | 0.83 | 0.68 | 0.84 |
| 4 | 39019877 | rs2566103 | -0.83 | 0.398 | -0.127 | -0.243 | -0.011 | -0.104 | -0.104 | 0.945 | 0.15 | 0.38 | 0.1 | 0.01 | 0.14 | 0.08 |
| 4 | 39019912 | rs2566104 | 0.007 | 0.026 | -0.613 | -0.232 | 0.003 | -0.023 | -0.023 | 1.936 | 0.58 | 0.8 | 0.67 | 0.51 | 0.58 | 0.46 |
| 4 | 39020194 | rs3860068 | 1.076 | -0.081 | -0.358 | -0.258 | -0.026 | -0.015 | -0.015 | 1.798 | 0.42 | 0.3 | 0.55 | 0.51 | 0.45 | 0.38 |
| 4 | 39027082 | rs12642359 | 1.05 | -0.097 | -0.275 | -0.3 | -0.031 | -0.018 | -0.018 | 0 | 0.43 | 0.26 | 0.56 | 0.52 | 0.45 | 0.38 |
| 4 | 39031979 | rs6821102 | 0.378 | -0.145 | -0.573 | -0.287 | 0.001 | -0.064 | -0.064 | 0 | 0.1 | 0.01 | 0.09 | 0.22 | 0.07 | 0.29 |
| 4 | 39036311 | rs1864495 | -0.223 | -0.126 | -0.413 | -0.382 | -0.016 | -0.118 | -0.118 | 2.317 | 0.16 | 0.47 | 0.11 | 0.02 | 0.14 | 0.08 |
| 4 | 39039258 | rs2711981 | -0.266 | -0.003 | -0.379 | -0.283 | -0.014 | 0.017 | 0.017 | 0 | 0.71 | 0.83 | 0.76 | 0.75 | 0.66 | 0.74 |
| 4 | 39045907 | rs6813885 | -0.115 | -0.371 | -0.093 | -0.274 | -0.007 | -0.026 | -0.026 | 0 | 0.72 | 0.62 | 0.79 | 0.82 | 0.75 | 0.76 |
| 4 | 39055870 | rs2566120 | -0.75 | -0.563 | -0.154 | -0.292 | -0.021 | 0.009 | 0.009 | 0.402 | 0.27 | 0.3 | 0.18 | 0.32 | 0.21 | 0.38 |
| 4 | 39062436 | rs2711943 | -0.423 | -0.596 | -0.245 | -0.312 | 0.001 | 0.004 | 0.004 | 0.328 | 0.73 | 0.73 | 0.79 | 0.83 | 0.75 | 0.77 |
| 4 | 39063614 | rs12500509 | 1.686 | -0.326 | -0.501 | -0.414 | -0.181 | -0.173 | -0.173 | 0 | 0.39 | 0.02 | 0.44 | 0.46 | 0.45 | 0.36 |
| 4 | 39063659 | rs11096974 | 1.333 | -0.462 | -0.327 | -0.438 | -0.006 | -0.025 | -0.025 | 0.013 | 0.44 | 0.23 | 0.6 | 0.5 | 0.54 | 0.38 |
| 4 | 39064162 | rs2711941 | -0.103 | -0.356 | -0.259 | -0.318 | -0.005 | -0.019 | -0.019 | 0 | 0.72 | 0.61 | 0.79 | 0.83 | 0.75 | 0.77 |
| 4 | 39074295 | rs2566134 | -0.123 | -0.353 | -0.259 | -0.316 | -0.005 | -0.021 | -0.021 | 0 | 0.72 | 0.62 | 0.79 | 0.83 | 0.75 | 0.77 |
| 4 | 39081681 | rs2566142 | -0.225 | -0.432 | -0.282 | -0.349 | 0.003 | 0.024 | 0.024 | 0.479 | 0.72 | 0.78 | 0.79 | 0.83 | 0.73 | 0.74 |
| 4 | 39084992 | rs1036044 | -0.11 | -0.422 | -0.154 | -0.344 | -0.003 | -0.016 | -0.016 | 0 | 0.68 | 0.65 | 0.77 | 0.83 | 0.72 | 0.72 |
| 4 | 39116911 | rs3733276 | 0.823 | -0.443 | -0.317 | -0.235 | 0.043 | -0.001 | -0.001 | 0 | 0.48 | 0.34 | 0.6 | 0.5 | 0.58 | 0.53 |
| 4 | 39119753 | rs12499266 | 0.824 | -0.428 | -0.22 | -0.227 | 0.052 | -0.005 | -0.005 | 0 | 0.48 | 0.31 | 0.59 | 0.5 | 0.58 | 0.53 |
| 4 | 39120883 | rs9997089 | 0.803 | -0.428 | -0.248 | -0.227 | 0.045 | 0 | 0 | 0 | 0.48 | 0.34 | 0.6 | 0.5 | 0.58 | 0.53 |
| 4 | 39121414 | rs9990742 | 0.803 | -0.428 | -0.248 | -0.227 | 0.045 | 0 | 0 | 0 | 0.48 | 0.34 | 0.59 | 0.5 | 0.57 | 0.53 |
| 4 | 39122697 | rs3733275 | 0.803 | -0.413 | -0.248 | -0.227 | 0.047 | 0 | 0 | 0 | 0.48 | 0.34 | 0.6 | 0.5 | 0.58 | 0.53 |
| 4 | 39134371 | rs12646251 | 1.244 | -0.455 | -0.246 | -0.241 | 0.01 | -0.078 | -0.078 | 0 | 0.45 | 0.09 | 0.57 | 0.5 | 0.58 | 0.53 |
| 4 | 39135564 | rs10018622 | 0.88 | -0.375 | -0.24 | -0.201 | 0.044 | -0.003 | -0.003 | 0 | 0.48 | 0.33 | 0.59 | 0.5 | 0.58 | 0.53 |
| 4 | 39149491 | rs2044917 | 0.209 | -0.451 | 0.106 | 0.056 | -0.038 | 0.115 | 0.115 | 0.057 | 0.81 | 0.56 | 0.82 | 0.67 | 0.83 | 0.88 |
| 4 | 39151353 | rs2711991 | -0.417 | -0.447 | -0.222 | -0.211 | 0.006 | 0.016 | 0.016 | 0 | 0.33 | 0.29 | 0.24 | 0.17 | 0.32 | 0.29 |
| 4 | 39162457 | rs2711935 | -0.443 | -0.455 | -0.232 | -0.215 | 0.013 | 0.036 | 0.036 | 0 | 0.34 | 0.27 | 0.24 | 0.17 | 0.32 | 0.29 |
| 4 | 39183447 | rs1451820 | -0.19 | -0.485 | 0.493 | -0.026 | -0.002 | 0.011 | 0.011 | 0 | 0.85 | 0.82 | 0.89 | 0.67 | 0.87 | 0.89 |
| 4 | 39207318 | rs17584431 | 0.383 | -0.72 | 0.256 | -0.609 | 0.004 | -0.02 | -0.02 | 0 | 0.19 | 0.01 | 0.15 | 0.15 | 0.16 | 0.2 |
| 4 | 39212010 | rs2381349 | -0.875 | -0.683 | 0.659 | -0.591 | 0.009 | 0.022 | 0.022 | 0 | 0.36 | 0.31 | 0.24 | 0.16 | 0.33 | 0.26 |
| 4 | 39216221 | rs2167494 | -0.833 | -0.682 | 0.66 | -0.59 | 0.008 | 0.02 | 0.02 | 0 | 0.35 | 0.3 | 0.24 | 0.16 | 0.32 | 0.26 |
| 4 | 39229771 | rs11730558 | -0.97 | -0.749 | 1.133 | -0.573 | 0.001 | 0 | 0 | 0 | 0.36 | 0.36 | 0.26 | 0.17 | 0.33 | 0.26 |
| 4 | 39231457 | rs11932351 | -1.087 | -0.763 | 1.616 | -0.58 | -0.022 | -0.052 | -0.052 | 0 | 0.37 | 0.45 | 0.26 | 0.17 | 0.33 | 0.26 |
| 4 | 39263814 | rs6531698 | 0.127 | -0.824 | 1.339 | -0.432 | 0.001 | -0.046 | -0.046 | 0 | 0.13 | 0.03 | 0.09 | 0.21 | 0.12 | 0.08 |
| 4 | 39266464 | rs4975001 | -1.95 | -0.906 | 1.385 | -0.46 | 0.038 | 0.038 | 0.038 | 0.331 | 0.53 | 0.79 | 0.38 | 0.38 | 0.46 | 0.35 |
| 4 | 39271541 | rs3733280 | -0.925 | -1.03 | 1.382 | -0.453 | 0.008 | 0.014 | 0.014 | 0 | 0.36 | 0.27 | 0.24 | 0.26 | 0.34 | 0.27 |
| 4 | 39280655 | rs746678 | -1.756 | -0.946 | 1.258 | -0.457 | 0.04 | 0.043 | 0.043 | 0.077 | 0.52 | 0.77 | 0.38 | 0.38 | 0.46 | 0.35 |
| 4 | 39282303 | rs2381373 | -1.756 | -0.946 | 1.258 | -0.457 | 0.04 | 0.043 | 0.043 | 0.077 | 0.52 | 0.77 | 0.38 | 0.38 | 0.46 | 0.35 |
| 4 | 39286766 | rs2062229 | -1.743 | -0.963 | 1.12 | -0.463 | 0.043 | 0.043 | 0.043 | 0.043 | 0.52 | 0.78 | 0.37 | 0.38 | 0.46 | 0.34 |
| 4 | 39289473 | rs1057807 | -1.875 | -0.968 | 1.105 | -0.466 | 0.034 | 0.035 | 0.035 | 0.99 | 0.53 | 0.84 | 0.38 | 0.39 | 0.46 | 0.35 |
| 4 | 39297200 | rs2306597 | 0.189 | -1.043 | 1.064 | -0.506 | 0.015 | 0.045 | 0.045 | 0 | 0.21 | 0.1 | 0.15 | 0.08 | 0.2 | 0.19 |
| 4 | 39302029 | rs2066786 | 1.755 | -0.865 | 0.877 | -0.401 | -0.001 | 0.018 | 0.018 | 0.791 | 0.45 | 0.06 | 0.61 | 0.63 | 0.55 | 0.66 |
| 4 | 39318706 | rs2066790 | -0.778 | -0.617 | 1.016 | -0.49 | 0.007 | 0.014 | 0.014 | 0 | 0.37 | 0.31 | 0.26 | 0.18 | 0.36 | 0.29 |
| 4 | 39329102 | rs11096992 | -1.636 | -0.646 | 1.075 | -0.502 | 0 | 0.001 | 0.001 | 0 | 0.53 | 0.47 | 0.36 | 0.38 | 0.5 | 0.37 |
| 4 | 39364856 | rs17584703 | -0.654 | 0.032 | 0.805 | -0.459 | -0.003 | -0.009 | -0.009 | 0 | 0.18 | 0.17 | 0.12 | 0.25 | 0.2 | 0.12 |
| 4 | 39366590 | rs6844176 | 1.826 | 0.029 | 0.64 | -0.519 | -0.113 | -0.056 | -0.056 | 0 | 0.4 | 0.06 | 0.57 | 0.58 | 0.44 | 0.6 |
| 4 | 39367654 | rs6851075 | -0.759 | -0.037 | 0.579 | -0.703 | 0.007 | 0.019 | 0.019 | 0 | 0.39 | 0.29 | 0.25 | 0.15 | 0.35 | 0.28 |
| 4 | 39377405 | rs13123782 | 1.394 | 0.098 | 0.612 | -0.479 | -0.051 | -0.016 | -0.016 | 0 | 0.44 | 0.16 | 0.53 | 0.53 | 0.46 | 0.63 |
| 4 | 39377646 | rs2062228 | 1.448 | 0.139 | 0.728 | -0.442 | 0.094 | 0.083 | 0.083 | 2.346 | 0.95 | 0.49 | 0.94 | 1 | 1 | 1 |
| 4 | 39385821 | rs6823497 | 1.361 | -0.032 | 0.342 | -0.571 | 0.005 | 0 | 0 | 0.036 | 0.43 | 0.43 | 0.55 | 0.53 | 0.45 | 0.64 |
| 4 | 39395770 | rs2381378 | -0.376 | -0.294 | -0.208 | -0.622 | 0.008 | -0.014 | -0.014 | 0 | 0.24 | 0.17 | 0.23 | 0.36 | 0.19 | 0.17 |
| 4 | 39405151 | rs13135439 | -1.261 | 0.387 | -0.039 | -0.46 | 0.01 | 0.008 | 0.008 | 0 | 0.52 | 0.37 | 0.44 | 0.47 | 0.57 | 0.35 |
| 4 | 39406987 | rs13149518 | -1.424 | 0.399 | -0.019 | -0.467 | 0.001 | 0 | 0 | 0 | 0.54 | 0.44 | 0.45 | 0.49 | 0.58 | 0.34 |
| 4 | 39414993 | rs11940694 | -1.628 | 0.189 | -0.551 | -0.538 | -0.004 | 0.005 | 0.005 | 0.612 | 0.6 | 0.57 | 0.44 | 0.46 | 0.61 | 0.37 |
| 4 | 39419322 | rs13147477 | -0.126 | 0.402 | -0.934 | -0.347 | -0.014 | -0.05 | -0.05 | 0.718 | 0.25 | 0.12 | 0.28 | 0.35 | 0.27 | 0.18 |
| 4 | 39419409 | rs7674434 | -0.216 | 0.179 | -0.476 | -0.888 | 0.012 | -0.055 | -0.055 | 0 | 0.28 | 0.5 | 0.2 | 0.13 | 0.28 | 0.18 |

| | | | | | | | | | | | | | | | | |
|---|---|---|---|---|---|---|---|---|---|---|---|---|---|---|---|---|
| 4 | 39420869 | rs3860070 | 0.015 | -0.062 | -0.75 | -0.44 | 0.001 | -0.007 | -0.007 | 0 | 0.08 | 0.08 | 0.03 | 0 | 0.07 | 0.01 |
| 4 | 39426809 | rs2608816 | -0.179 | -0.464 | -0.78 | -0.585 | -0.014 | -0.031 | -0.031 | 0.871 | 0.15 | 0.16 | 0.06 | 0.03 | 0.1 | 0.09 |
| 4 | 39428295 | rs17431867 | 0.46 | -0.184 | -1.303 | -0.788 | -0.026 | 0.034 | 0.034 | 0 | 0.18 | 0.07 | 0.11 | 0.11 | 0.21 | 0.11 |
| 4 | 39430969 | rs1979283 | 1.121 | -0.57 | -0.97 | -0.725 | 0.009 | 0.017 | 0.017 | 0.166 | 0.6 | 0.37 | 0.73 | 0.7 | 0.67 | 0.71 |
| 4 | 39431431 | rs2608819 | 0.546 | -0.603 | -0.869 | -0.613 | -0.036 | -0.015 | -0.015 | 0 | 0.77 | 0.41 | 0.85 | 0.81 | 0.88 | 0.81 |
| 4 | 39438149 | rs2687963 | 0.949 | -0.888 | -1.072 | -0.969 | 0.046 | 0.078 | 0.078 | 0.846 | 0.55 | 0.33 | 0.75 | 0.79 | 0.69 | 0.76 |
| 4 | 39439264 | rs4975015 | 0.087 | -1.008 | -1.103 | -1.046 | 0.008 | 0.046 | 0.046 | 0 | 0.26 | 0.17 | 0.1 | 0.05 | 0.2 | 0.15 |
| 4 | 39449595 | rs7674617 | 0.921 | -0.881 | -1.417 | -0.989 | -0.002 | 0.01 | 0.01 | 0 | 0.14 | 0.1 | 0.11 | 0.07 | 0.13 | 0.1 |
| 4 | 39450229 | rs4975017 | 0.974 | -0.615 | -2.073 | -0.523 | -0.035 | -0.042 | -0.042 | 4.654 | 0.19 | 0.12 | 0.4 | 0.38 | 0.32 | 0.51 |
| 4 | 39452290 | rs9995998 | 0.382 | -1.006 | -1.385 | -0.685 | -0.013 | -0.023 | -0.023 | 0 | 0.23 | 0.33 | 0.18 | 0.15 | 0.17 | 0.13 |
| 4 | 39453790 | rs10517522 | 1.169 | -0.976 | -1.387 | -0.671 | -0.045 | -0.054 | -0.054 | 0 | 0.19 | 0.09 | 0.4 | 0.38 | 0.32 | 0.51 |
| 4 | 39455378 | rs2249563 | -0.931 | -1.291 | -1.597 | -0.744 | -0.013 | -0.03 | -0.03 | 0 | 0.63 | 0.53 | 0.81 | 0.72 | 0.68 | 0.82 |
| 4 | 39455742 | rs17510142 | 2.346 | -1.407 | -1.97 | -0.917 | -0.053 | 0.113 | 0.113 | 0 | 0.21 | 0 | 0.1 | 0.02 | 0.2 | 0.1 |
| 4 | 39459154 | rs2608830 | 1.751 | -1.275 | -1.783 | -0.783 | 0.009 | -0.005 | -0.005 | 0 | 0.2 | 0.05 | 0.22 | 0.19 | 0.19 | 0.17 |
| 4 | 39474868 | rs749915 | 0.59 | -1.529 | -1.118 | -0.703 | 0.003 | -0.061 | -0.061 | 0.881 | 0.21 | 0.42 | 0.18 | 0.07 | 0.2 | 0.1 |
| 4 | 39475074 | rs2259073 | 1.979 | -1.427 | -1.206 | -0.68 | 0.018 | -0.016 | -0.016 | 0.269 | 0.16 | 0.05 | 0.2 | 0.19 | 0.17 | 0.16 |
| 4 | 39478164 | rs17618369 | 1.162 | -1.224 | -1.106 | -0.614 | -0.054 | -0.06 | -0.06 | 0 | 0.23 | 0.1 | 0.42 | 0.38 | 0.35 | 0.53 |
| 4 | 39478264 | rs16995300 | 1.581 | -1.229 | -1.081 | -0.619 | 0 | -0.034 | -0.034 | 0.215 | 0.07 | 0.14 | 0.04 | 0 | 0.07 | 0.01 |
| 4 | 39478832 | rs1048140 | -0.021 | -1.093 | -1.278 | -0.474 | -0.009 | -0.012 | -0.012 | 0 | 0.46 | 0.5 | 0.42 | 0.34 | 0.41 | 0.3 |
| 4 | 39482684 | rs2260645 | -0.932 | -0.773 | -1.323 | -0.279 | -0.01 | -0.007 | -0.007 | 0 | 0.69 | 0.62 | 0.85 | 0.72 | 0.76 | 0.83 |
| 4 | 39491521 | rs2687965 | -1.106 | -1.028 | -1.35 | 0.093 | -0.035 | -0.011 | -0.011 | 0 | 0.61 | 0.81 | 0.52 | 0.65 | 0.5 | 0.51 |
| 4 | 39496272 | rs2687964 | 1.013 | -1.03 | -1.633 | 0.098 | 0.015 | 0.037 | 0.037 | 0 | 0.19 | 0.05 | 0.16 | 0.13 | 0.19 | 0.2 |
| 4 | 39500514 | rs1450 | 1.303 | -1.089 | -1.84 | 0.125 | -0.166 | -0.124 | -0.124 | 0.398 | 0.36 | 0.02 | 0.47 | 0.34 | 0.48 | 0.45 |
| 4 | 39501086 | rs1046573 | 0.677 | -1.265 | -1.39 | 0.172 | 0.051 | -0.013 | -0.013 | 0 | 0.59 | 0.26 | 0.69 | 0.54 | 0.71 | 0.74 |
| 4 | 39503196 | rs9995522 | 0.672 | -0.912 | -0.821 | 0.237 | 0.001 | -0.03 | -0.03 | 0 | 0.06 | 0.11 | 0.03 | 0 | 0.07 | 0.01 |
| 4 | 39504305 | rs7667766 | -0.138 | -1.033 | -0.992 | 0.163 | 0 | -0.021 | -0.021 | 0.392 | 0.62 | 0.55 | 0.72 | 0.84 | 0.64 | 0.59 |
| 4 | 39506144 | rs1379819 | 0.266 | -0.735 | -0.796 | 0.888 | 0.1 | 0.495 | 0.495 | 0.196 | 0.06 | 0.37 | 0.08 | 0.26 | 0.04 | 0.04 |
| 4 | 39507457 | rs6832089 | -0.846 | -0.858 | -0.942 | 0.8 | -0.042 | 0.168 | 0.168 | 0.283 | 0.18 | 0.52 | 0.17 | 0.41 | 0.1 | 0.17 |
| 4 | 39515736 | rs10019532 | -0.76 | -0.849 | -0.976 | 0.804 | -0.041 | 0.207 | 0.207 | 0.12 | 0.18 | 0.48 | 0.16 | 0.41 | 0.1 | 0.17 |
| 4 | 39518110 | rs6817264 | -0.795 | -0.661 | -1.411 | 0.182 | 0.001 | 0.005 | 0.005 | 0 | 0.7 | 0.72 | 0.79 | 0.89 | 0.69 | 0.73 |
| 4 | 39526641 | rs13129975 | 1.24 | -0.533 | -1.81 | 0.442 | -0.169 | -0.022 | -0.022 | 1.766 | 0.34 | 0.03 | 0.38 | 0.23 | 0.43 | 0.42 |
| 4 | 39529700 | rs3755899 | -0.135 | -0.116 | -1.025 | 0.194 | 0.004 | -0.007 | -0.007 | 0 | 0.08 | 0.1 | 0.08 | 0.15 | 0.06 | 0.12 |
| 4 | 39531928 | rs2381387 | 0.089 | 0.028 | -0.886 | 0.26 | -0.001 | -0.011 | -0.011 | 2.662 | 0.6 | 0.49 | 0.71 | 0.74 | 0.63 | 0.6 |
| 4 | 39537738 | rs13101951 | 0.304 | -0.006 | -0.87 | 0.396 | -0.002 | -0.009 | -0.009 | 2.48 | 0.57 | 0.46 | 0.7 | 0.7 | 0.61 | 0.58 |
| 4 | 39537794 | rs936380 | 0.326 | 0.042 | -1.177 | 0.271 | 0.001 | 0.091 | 0.091 | 0 | 0.52 | 0.63 | 0.67 | 0.66 | 0.52 | 0.58 |
| 4 | 39538245 | rs7692244 | -0.043 | 0.634 | -1.017 | 0.574 | -0.009 | -0.02 | -0.02 | 0.271 | 0.06 | 0.07 | 0.07 | 0.17 | 0.05 | 0.12 |
| 4 | 39557864 | rs11944799 | -0.065 | 0.584 | -0.296 | 0.983 | 0.115 | 0.254 | 0.254 | 0.743 | 0.05 | 0.39 | 0.08 | 0.1 | 0.03 | 0.02 |
| 4 | 39583823 | rs17618779 | -0.109 | 0.849 | -1.464 | 1.306 | -0.032 | -0.058 | -0.058 | 0 | 0.11 | 0.01 | 0.34 | 0.25 | 0.13 | 0.36 |
| 4 | 39600010 | rs7673664 | -0.415 | 0.597 | -1.857 | 0.832 | -0.034 | 0.034 | 0.034 | 0.182 | 0.36 | 0.06 | 0.49 | 0.34 | 0.36 | 0.64 |
| 4 | 39603132 | rs7658676 | 0.313 | 0.563 | -1.617 | 0.84 | -0.036 | 0.131 | 0.131 | 0 | 0.32 | 0.14 | 0.21 | 0.13 | 0.35 | 0.29 |
| 4 | 39605669 | rs6833494 | 1.299 | 0.32 | -1.811 | 1.111 | -0.073 | 0.01 | 0.01 | 0 | 0.3 | 0.02 | 0.21 | 0.17 | 0.31 | 0.16 |
| 4 | 39610904 | rs900832 | 0.013 | 0.569 | -1.425 | 1.324 | -0.109 | -0.004 | -0.004 | 0 | 0.37 | 0.04 | 0.28 | 0.32 | 0.42 | 0.35 |
| 4 | 39614823 | rs11096999 | 0.193 | 0.471 | -1.381 | 0.885 | -0.001 | -0.03 | -0.03 | 0.01 | 0.09 | 0.03 | 0.32 | 0.24 | 0.09 | 0.35 |
| 4 | 39616548 | rs12512211 | -0.173 | 0.563 | -1.325 | 0.582 | -0.045 | 0.106 | 0.106 | 0 | 0.62 | 0.85 | 0.74 | 0.77 | 0.59 | 0.65 |
| 4 | 39619870 | rs12500234 | -0.173 | 0.563 | -1.325 | 0.582 | -0.045 | 0.106 | 0.106 | 0 | 0.62 | 0.85 | 0.74 | 0.77 | 0.59 | 0.65 |
| 4 | 39637826 | rs977385 | 0.281 | 0.534 | -0.865 | 0.799 | -0.001 | 0.003 | 0.003 | 0 | 0.51 | 0.5 | 0.4 | 0.57 | 0.5 | 0.3 |
| 4 | 39641932 | rs1484321 | -0.099 | 1.135 | -0.854 | 1.03 | 0.062 | 0.036 | 0.036 | 0.056 | 0.19 | 0.53 | 0.26 | 0.17 | 0.25 | 0.08 |
| 4 | 39643536 | rs13104353 | -0.913 | 0.901 | -1.219 | 1.169 | 0.005 | 0.024 | 0.024 | 0.1 | 0.34 | 0.63 | 0.34 | 0.44 | 0.35 | 0.17 |
| 4 | 39648294 | rs12510053 | 1.218 | 1.044 | -1.252 | 1.162 | -0.048 | -0.005 | -0.005 | 0 | 0.12 | 0.02 | 0.16 | 0.1 | 0.19 | 0.06 |
| 4 | 39655457 | rs11941588 | -1.001 | 0.736 | -1.515 | 1.11 | -0.012 | 0.021 | 0.021 | 0 | 0.33 | 0.52 | 0.52 | 0.59 | 0.28 | 0.49 |
| 4 | 39657292 | rs10856848 | -1.155 | 0.902 | -1.628 | 0.714 | 0.014 | 0.032 | 0.032 | 0 | 0.51 | 0.63 | 0.72 | 0.69 | 0.57 | 0.55 |
| 4 | 39671142 | rs1484322 | 0.167 | 0.929 | -1.765 | 0.825 | -0.102 | -0.067 | -0.067 | 8.31 | 0.27 | 0.05 | 0.52 | 0.36 | 0.37 | 0.39 |
| 4 | 39686517 | rs6531736 | 0.113 | 0.66 | -1.99 | 0.486 | 0.005 | -0.019 | -0.019 | 0 | 0.28 | 0.81 | 0.23 | 0.3 | 0.25 | 0.3 |
| 4 | 39694314 | rs4388095 | -1.262 | 0.594 | -1.853 | 0.441 | -0.008 | -0.019 | -0.019 | 0.159 | 0.66 | 0.43 | 0.81 | 0.85 | 0.7 | 0.83 |
| 4 | 39698738 | rs3912392 | -0.361 | 0.632 | -1.892 | 0.479 | -0.106 | -0.061 | -0.061 | 0 | 0.43 | 0.09 | 0.64 | 0.62 | 0.51 | 0.54 |
| 4 | 39698824 | rs3912391 | -0.414 | 0.634 | -1.889 | 0.479 | -0.086 | -0.044 | -0.044 | 0 | 0.43 | 0.12 | 0.65 | 0.62 | 0.5 | 0.54 |
| 4 | 39702089 | rs302941 | 0.776 | 0.725 | -1.663 | 0.458 | 0.111 | 0.156 | 0.156 | 1.612 | 0.07 | 0.51 | 0.08 | 0.07 | 0.06 | 0.02 |
| 4 | 39709947 | rs2054718 | 0.761 | 0.649 | -1.84 | 0.461 | 0.002 | -0.021 | -0.021 | 0 | 0.14 | 0.11 | 0.11 | 0.24 | 0.12 | 0.14 |
| 4 | 39713134 | rs4362831 | -1.907 | 0.665 | -1.918 | 0.365 | -0.001 | 0.09 | 0.09 | 0 | 0.72 | 0.97 | 0.88 | 0.92 | 0.77 | 0.84 |
| 4 | 39737672 | rs159987 | -1.892 | 0.654 | -1.899 | 0.376 | 0.019 | 0.08 | 0.08 | 0 | 0.72 | 0.87 | 0.88 | 0.92 | 0.76 | 0.84 |
| 4 | 39747907 | rs724256 | 1.183 | 0.669 | -1.878 | 0.43 | -0.024 | -0.068 | -0.068 | 0 | 0.13 | 0.01 | 0.1 | 0.24 | 0.12 | 0.14 |
| 4 | 39748278 | rs725856 | 1.117 | 0.67 | -1.873 | 0.445 | -0.005 | -0.038 | -0.038 | 0 | 0.13 | 0.04 | 0.11 | 0.24 | 0.12 | 0.14 |
| 4 | 39748523 | rs10517528 | 1.135 | 0.657 | -1.683 | 0.465 | 0 | 0 | 0 | 0 | 0.21 | 0.18 | 0.15 | 0.22 | 0.2 | 0.29 |
| 4 | 39759989 | rs305827 | -0.533 | 0.584 | -1.503 | 0.491 | 0.009 | 0 | 0 | 0.083 | 0.6 | 0.89 | 0.78 | 0.68 | 0.64 | 0.7 |
| 4 | 39776955 | rs459274 | -0.033 | 0.651 | -1.362 | 0.394 | 0.006 | 0.084 | 0.084 | 0.754 | 0.21 | 0.59 | 0.19 | 0.31 | 0.18 | 0.17 |
| 4 | 39783729 | rs305112 | -1.433 | 0.764 | -1.253 | 0.434 | 0.015 | 0.073 | 0.073 | 0 | 0.73 | 0.92 | 0.88 | 0.92 | 0.76 | 0.84 |
| 4 | 39785276 | rs4263408 | 0.131 | 0.714 | -1.621 | 0.404 | 0.008 | -0.003 | -0.003 | 0 | 0.48 | 0.31 | 0.68 | 0.61 | 0.55 | 0.67 |
| 4 | 39788410 | rs192779 | -0.876 | 0.625 | -2.037 | 0.001 | 0.014 | 0.083 | 0.083 | 0 | 0.72 | 0.93 | 0.88 | 0.92 | 0.76 | 0.84 |
| 4 | 39803506 | rs6816573 | 0.044 | 0.928 | -2.343 | 0.042 | -0.052 | 0.073 | 0.073 | 0.476 | 0.36 | 0.09 | 0.24 | 0.27 | 0.37 | 0.28 |

| | | | | | | | | | | | | | | | | |
|---|---|---|---|---|---|---|---|---|---|---|---|---|---|---|---|---|
| 4 | 39803892 | rs12499832 | 0.275 | 1.042 | -1.524 | 0.553 | 0.002 | -0.007 | -0.007 | 0 | 0.45 | 0.37 | 0.43 | 0.44 | 0.44 | 0.42 |
| 4 | 39806505 | rs11097004 | 0.166 | 1.062 | -1.721 | 0.962 | 0.015 | 0.092 | 0.092 | 0.163 | 0.15 | 0.3 | 0.1 | 0.27 | 0.14 | 0.16 |
| 4 | 39810454 | rs11732002 | 0.347 | 0.975 | -1.917 | 0.808 | -0.048 | 0.109 | 0.109 | 2.303 | 0.44 | 0.06 | 0.29 | 0.32 | 0.43 | 0.41 |
| 4 | 39812247 | rs403611 | -0.424 | 0.981 | -0.958 | 0.923 | 0.06 | 0.139 | 0.139 | 0.077 | 0.9 | 0.8 | 0.73 | 0.71 | 0.86 | 0.82 |
| 4 | 39814064 | rs4974949 | -0.439 | 1.036 | -0.88 | 0.49 | 0.009 | -0.014 | -0.014 | 0 | 0.11 | 0.23 | 0.1 | 0.05 | 0.16 | 0.09 |
| 4 | 39818116 | rs1003212 | -0.598 | 0.944 | -0.371 | 0.756 | 0.098 | 0.2 | 0.2 | 0.019 | 0.9 | 0.67 | 0.73 | 0.72 | 0.87 | 0.83 |
| 4 | 39818522 | rs1003213 | 1.957 | 0.853 | -0.989 | 0.335 | -0.016 | 0.166 | 0.166 | 0 | 0.25 | 0.02 | 0.27 | 0.01 | 0.23 | 0.16 |
| 4 | 39822709 | rs3733297 | 1.739 | 0.845 | -1.22 | 0.097 | -0.011 | 0.217 | 0.217 | 0 | 0.33 | 0.02 | 0.31 | 0.04 | 0.29 | 0.18 |
| 4 | 39822849 | rs305835 | -0.264 | 0.854 | -1.101 | 0.158 | -0.002 | 0.045 | 0.045 | 0.013 | 0.51 | 0.31 | 0.35 | 0.67 | 0.51 | 0.62 |
| 4 | 39830157 | rs460334 | 0.284 | 0.85 | -0.88 | 0.363 | 0.054 | 0.092 | 0.092 | 2.723 | 0.83 | 0.36 | 0.66 | 0.71 | 0.8 | 0.81 |
| 4 | 39836933 | rs7657667 | 1.332 | 0.869 | -1.1 | 0.421 | -0.027 | -0.015 | -0.015 | 0 | 0.05 | 0.01 | 0.03 | 0.05 | 0.09 | 0.06 |
| 4 | 39837128 | rs50296 | -0.256 | 0.831 | -1.091 | 0.358 | -0.034 | 0.085 | 0.085 | 0.593 | 0.48 | 0.17 | 0.34 | 0.67 | 0.51 | 0.62 |
| 4 | 39843114 | rs11723274 | 2.62 | 0.964 | -1.587 | -0.08 | -0.017 | 0.046 | 0.046 | 1.354 | 0.16 | 0.01 | 0.11 | 0.12 | 0.15 | 0.12 |
| 4 | 39854464 | rs17619018 | 0.553 | 0.869 | -1.536 | 0.009 | -0.018 | -0.039 | -0.039 | 0.477 | 0.09 | 0.04 | 0.07 | 0.2 | 0.11 | 0.18 |
| 4 | 39864448 | rs4547806 | 0.364 | 0.826 | -1.525 | -0.052 | -0.051 | 0.109 | 0.109 | 1.204 | 0.36 | 0.05 | 0.24 | 0.24 | 0.36 | 0.37 |
| 4 | 39894988 | rs922749 | 0.922 | 1.117 | -1.554 | -0.014 | -0.017 | -0.069 | -0.069 | 0 | 0.29 | 0.1 | 0.22 | 0.46 | 0.3 | 0.35 |
| 4 | 39921551 | rs6839903 | -0.482 | 1.062 | -1.649 | -0.169 | -0.028 | 0.031 | 0.031 | 0.062 | 0.4 | 0.14 | 0.29 | 0.66 | 0.41 | 0.53 |
| 4 | 39933442 | rs2679780 | 1.426 | 1.116 | -1.523 | -0.044 | -0.049 | 0.016 | 0.016 | 1.524 | 0.25 | 0.03 | 0.19 | 0.23 | 0.25 | 0.28 |
| 4 | 39935865 | rs4440231 | -0.104 | 1.207 | -1.049 | -0.107 | 0.084 | 0.134 | 0.134 | 2.028 | 0.84 | 0.46 | 0.67 | 0.71 | 0.8 | 0.81 |
| 4 | 39943056 | rs2732063 | 0.895 | 1.206 | -1.083 | -0.198 | -0.013 | -0.077 | -0.077 | 0 | 0.13 | 0.22 | 0.06 | 0.01 | 0.11 | 0.1 |
| 4 | 39949502 | rs13105733 | -0.084 | 1.316 | -0.767 | 0.084 | 0.085 | 0.121 | 0.121 | 0.563 | 0.85 | 0.61 | 0.68 | 0.71 | 0.8 | 0.81 |
| 4 | 39984938 | rs2732072 | -0.796 | 1.173 | -0.6 | 0.225 | 0.11 | 0.2 | 0.2 | 0 | 0.9 | 0.65 | 0.72 | 0.72 | 0.86 | 0.82 |
| 4 | 39989369 | rs7677857 | 0.115 | 1.074 | -1.073 | 0.139 | -0.03 | -0.022 | -0.022 | 0 | 0.38 | 0.17 | 0.3 | 0.41 | 0.42 | 0.47 |
| 4 | 39999094 | rs10213194 | 2.04 | 1.028 | -1.108 | -0.003 | -0.004 | 0.043 | 0.043 | 0.278 | 0.21 | 0.11 | 0.18 | 0.12 | 0.24 | 0.13 |
| 4 | 40007284 | rs557980 | 0.372 | 1.069 | -0.969 | 0.079 | 0.016 | 0.057 | 0.057 | 0 | 0.5 | 0.25 | 0.43 | 0.36 | 0.48 | 0.35 |
| 4 | 40022041 | rs6838749 | 0.906 | 1.301 | -0.791 | 0.342 | 0.005 | 0.003 | 0.003 | 0.019 | 0.67 | 0.34 | 0.58 | 0.39 | 0.61 | 0.56 |
| 4 | 40022511 | rs7686844 | 0.906 | 1.301 | -0.791 | 0.342 | 0.005 | 0.003 | 0.003 | 0.019 | 0.67 | 0.34 | 0.58 | 0.39 | 0.61 | 0.57 |
| 4 | 40035491 | rs11097011 | 1.08 | 1.311 | -0.752 | 0.236 | -0.026 | -0.024 | -0.024 | 0.056 | 0.64 | 0.2 | 0.56 | 0.57 | 0.64 | 0.49 |
| 4 | 40035613 | rs7665844 | 0.391 | 1.439 | -0.501 | 0.243 | 0.017 | 0.029 | 0.029 | 0.073 | 0.75 | 0.42 | 0.66 | 0.69 | 0.73 | 0.66 |
| 4 | 40042666 | rs7660948 | 0.503 | 1.457 | -0.801 | 0.221 | -0.023 | -0.01 | -0.01 | 0 | 0.1 | 0.04 | 0.08 | 0.12 | 0.1 | 0.17 |
| 4 | 40045687 | rs17585937 | 0.462 | 1.457 | -0.772 | 0.221 | -0.01 | -0.003 | -0.003 | 0 | 0.1 | 0.06 | 0.08 | 0.12 | 0.1 | 0.17 |
| 4 | 40047752 | rs12503993 | 1.71 | 0.937 | -1.312 | -0.172 | -0.015 | -0.018 | -0.018 | 0 | 0.23 | 0.03 | 0.26 | 0.16 | 0.23 | 0.21 |
| 4 | 40048935 | rs11737408 | 1.274 | 0.798 | -1.273 | -0.342 | -0.004 | 0.138 | 0.138 | 0.013 | 0.32 | 0.09 | 0.25 | 0.07 | 0.3 | 0.17 |
| 4 | 40051064 | rs794016 | -0.705 | 1.042 | -0.259 | 0.16 | 0.07 | -0.025 | -0.025 | 0 | 0.93 | 0.72 | 0.92 | 0.99 | 0.91 | 0.97 |
| 4 | 40054818 | rs6824761 | 0.055 | 0.864 | -0.49 | -0.305 | -0.007 | 0.001 | 0.001 | 0 | 0.56 | 0.53 | 0.6 | 0.49 | 0.56 | 0.64 |
| 4 | 40071832 | rs10017867 | -0.271 | 1.135 | -0.524 | -0.537 | 0.008 | -0.028 | -0.028 | 0 | 0.87 | 0.83 | 0.88 | 0.96 | 0.85 | 0.95 |
| 4 | 40097557 | rs4974961 | 1.086 | 1.19 | -0.577 | -0.71 | -0.009 | -0.009 | -0.009 | 0 | 0.35 | 0.28 | 0.46 | 0.42 | 0.39 | 0.56 |
| 4 | 40097615 | rs813230 | -0.51 | 0.839 | -0.47 | -0.947 | 0.003 | -0.005 | -0.005 | 0.755 | 0.33 | 0.3 | 0.16 | 0.53 | 0.25 | 0.26 |
| 4 | 40103767 | rs17511668 | 2.273 | 0.667 | -0.58 | -0.858 | -0.052 | 0.109 | 0.109 | 0.007 | 0.2 | 0.01 | 0.26 | 0.02 | 0.22 | 0.14 |
| 4 | 40110337 | rs10033188 | 0.696 | 0.499 | -0.106 | -0.568 | -0.017 | 0.054 | 0.054 | 0 | 0.13 | 0.11 | 0.05 | 0.41 | 0.05 | 0.15 |
| 4 | 40120897 | rs1706034 | 0.426 | 0.548 | -0.01 | -0.703 | -0.017 | 0.062 | 0.062 | 0.21 | 0.68 | 0.54 | 0.8 | 0.56 | 0.71 | 0.78 |
| 4 | 40121562 | rs794001 | -0.545 | 1.131 | -0.076 | -0.708 | 0.029 | -0.075 | -0.075 | 0 | 0.82 | 0.68 | 0.85 | 0.97 | 0.76 | 0.93 |
| 4 | 40139719 | rs794009 | -0.564 | 1.118 | -0.075 | -0.701 | 0.036 | -0.082 | -0.082 | 0 | 0.82 | 0.66 | 0.84 | 0.97 | 0.76 | 0.93 |
| 4 | 40151656 | rs807787 | 0.767 | 1.042 | -0.663 | -0.452 | -0.004 | 0.003 | 0.003 | 0 | 0.11 | 0.06 | 0.07 | 0.12 | 0.1 | 0.08 |
| 4 | 40158464 | rs9998717 | 0.565 | 1.219 | -0.569 | -0.455 | 0.03 | 0.029 | 0.029 | 0 | 0.68 | 0.46 | 0.62 | 0.55 | 0.57 | 0.61 |
| 4 | 40162971 | rs6531759 | 0.921 | 0.845 | -0.769 | -0.715 | 0 | 0.001 | 0.001 | 0 | 0.31 | 0.28 | 0.45 | 0.38 | 0.32 | 0.52 |
| 4 | 40165305 | rs12645398 | 0.977 | 0.813 | -1.104 | -0.706 | -0.001 | -0.007 | -0.007 | 0 | 0.3 | 0.23 | 0.44 | 0.39 | 0.31 | 0.51 |
| 4 | 40169424 | rs794018 | 0.565 | 0.832 | -1.243 | -0.203 | -0.026 | -0.103 | -0.103 | 0.238 | 0.74 | 0.87 | 0.84 | 0.57 | 0.7 | 0.8 |
| 4 | 40172933 | rs4974968 | -0.378 | 0.992 | -0.917 | -0.623 | -0.02 | 0.024 | 0.024 | 0 | 0.84 | 0.9 | 0.88 | 0.98 | 0.77 | 0.92 |
| 4 | 40176557 | rs11946808 | 0.359 | 1.091 | -0.877 | -0.551 | 0.003 | -0.013 | -0.013 | 0 | 0.76 | 0.74 | 0.77 | 0.88 | 0.7 | 0.88 |
| 4 | 40187537 | rs1397933 | -0.213 | 0.635 | -1.36 | -0.609 | 0 | 0.024 | 0.024 | 0 | 0.25 | 0.09 | 0.16 | 0.19 | 0.25 | 0.11 |
| 4 | 40187596 | rs1397934 | -1.072 | 0.815 | -1.786 | -0.876 | -0.02 | 0.009 | 0.009 | 0 | 0.81 | 0.86 | 0.82 | 0.96 | 0.7 | 0.89 |
| 4 | 40189325 | rs12501987 | 0.962 | 0.63 | -1.639 | -0.576 | -0.036 | -0.018 | -0.018 | 4.413 | 0.16 | 0.01 | 0.1 | 0.13 | 0.17 | 0.07 |
| 4 | 40190463 | rs6819683 | 0.312 | 0.112 | -0.821 | -0.434 | -0.025 | -0.14 | -0.14 | 0 | 0.34 | 0.48 | 0.19 | 0.03 | 0.22 | 0.31 |
| 4 | 40194572 | rs12643103 | -0.14 | 0.346 | -0.554 | 1.402 | -0.018 | -0.057 | -0.057 | 0 | 0.1 | 0.02 | 0.21 | 0.43 | 0.13 | 0.14 |
| 4 | 40196549 | rs1317577 | -1.087 | 0.627 | 0.046 | 0.856 | 0.005 | 0.055 | 0.055 | 0.01 | 0.11 | 0.24 | 0.24 | 0.26 | 0.09 | 0.35 |
| 4 | 40198369 | rs1996555 | -1.278 | 0.838 | -0.191 | 0.541 | 0.004 | 0.007 | 0.007 | 0.439 | 0.09 | 0.13 | 0.19 | 0.14 | 0.09 | 0.31 |
| 4 | 40198842 | rs1053504 | -1.667 | 0.595 | -0.249 | 0.199 | -0.01 | -0.043 | -0.043 | 1.152 | 0.19 | 0.31 | 0.26 | 0.09 | 0.15 | 0.22 |
| 4 | 40199551 | rs2890726 | -1.345 | 1.042 | -0.653 | 0.62 | -0.008 | -0.004 | -0.004 | 0 | 0.15 | 0.17 | 0.24 | 0.13 | 0.19 | 0.2 |
| 4 | 40205742 | rs4552473 | 0.614 | 1.174 | -0.707 | 0.081 | -0.014 | 0.019 | 0.019 | 1.173 | 0.77 | 0.84 | 0.85 | 0.95 | 0.73 | 0.75 |
| 4 | 40207190 | rs17586314 | -0.333 | 0.904 | -0.766 | 0.959 | -0.002 | 0.025 | 0.025 | 0 | 0.16 | 0.21 | 0.27 | 0.4 | 0.15 | 0.12 |
| 4 | 40211997 | rs2381499 | 0.906 | 0.062 | -0.223 | 1.097 | -0.004 | -0.004 | -0.004 | 1.862 | 0.61 | 0.53 | 0.68 | 0.67 | 0.65 | 0.52 |
| 4 | 40213628 | rs4974974 | 1.335 | 0.146 | -0.173 | 1.451 | 0 | 0 | 0 | 0.187 | 0.52 | 0.53 | 0.6 | 0.6 | 0.56 | 0.45 |
| 4 | 40215687 | rs7671812 | 0.568 | 0.048 | 0.347 | 1.468 | -0.013 | 0.065 | 0.065 | 0.486 | 0.73 | 0.63 | 0.72 | 0.53 | 0.76 | 0.63 |
| 4 | 40217706 | rs2381501 | 1.24 | -0.258 | 0.581 | 1.587 | -0.056 | 0.024 | 0.024 | 1.664 | 0.73 | 0.49 | 0.69 | 0.61 | 0.84 | 0.64 |
| 4 | 40219896 | rs13101295 | -0.247 | -0.277 | 0.433 | 1.477 | 0 | 0 | 0 | 0 | 0.15 | 0.15 | 0.2 | 0.31 | 0.12 | 0.25 |
| 4 | 40220469 | rs6531762 | 1.624 | -0.422 | 0.675 | 1.533 | 0 | -0.001 | -0.001 | 0 | 0.66 | 0.33 | 0.66 | 0.55 | 0.76 | 0.55 |
| 4 | 40221675 | rs1878893 | 1.452 | -0.454 | 0.978 | 1.492 | -0.026 | 0.004 | 0.004 | 0.156 | 0.68 | 0.58 | 0.72 | 0.6 | 0.76 | 0.61 |
| 4 | 40222303 | rs6531763 | 1.778 | -0.518 | 0.797 | 1.623 | -0.018 | -0.015 | -0.015 | 0 | 0.63 | 0.51 | 0.62 | 0.25 | 0.7 | 0.52 |

| | | | | | | | | | | | | | | | | |
|---|---|---|---|---|---|---|---|---|---|---|---|---|---|---|---|---|
| 4 | 40224043 | rs2381502 | -0.491 | -0.558 | 0.328 | 1.28 | 0.005 | 0.003 | 0.003 | 4.151 | 0.88 | 0.93 | 0.97 | 0.93 | 0.96 | 0.89 |
| 4 | 40225912 | rs7677758 | 0.829 | -0.452 | 0.287 | 1.254 | -0.007 | 0.002 | 0.002 | 0.02 | 0.72 | 0.7 | 0.81 | 0.64 | 0.8 | 0.63 |
| 4 | 40227671 | rs11729556 | 1.629 | 0.001 | -0.638 | 0.611 | -0.077 | 0.241 | 0.241 | 0.247 | 0.47 | 0.09 | 0.44 | 0.15 | 0.56 | 0.26 |
| 4 | 40242433 | rs7656219 | -1.445 | -0.54 | 0.265 | 0.803 | -0.035 | -0.03 | -0.03 | 0 | 0.23 | 0.66 | 0.24 | 0.19 | 0.13 | 0.16 |
| 4 | 40244347 | rs13142049 | -0.779 | -0.496 | 0.352 | 0.804 | 0.006 | 0.008 | 0.008 | 0.07 | 0.16 | 0.11 | 0.2 | 0.17 | 0.13 | 0.14 |
| 4 | 40246416 | rs4600925 | 1.952 | -0.299 | -0.346 | 0.872 | 0.05 | 0.087 | 0.087 | 0.008 | 0.51 | 0.19 | 0.36 | 0.27 | 0.59 | 0.55 |
| 4 | 40246463 | rs13130216 | -0.287 | -0.23 | -0.119 | 0.721 | 0.006 | 0.009 | 0.009 | 0 | 0.14 | 0.1 | 0.18 | 0.17 | 0.1 | 0.12 |
| 4 | 40248956 | rs7667815 | -1.623 | -0.384 | 0.668 | 1.153 | -0.051 | 0.205 | 0.205 | 0 | 0.19 | 0.47 | 0.21 | 0.61 | 0.1 | 0.12 |
| 4 | 40255193 | rs7669532 | 0.229 | -0.452 | 0.422 | 1.042 | 0.068 | 0.054 | 0.054 | 0 | 0.8 | 0.94 | 0.94 | 0.82 | 0.93 | 0.9 |
| 4 | 40255411 | rs16995675 | -1.301 | -0.539 | 0.505 | 0.966 | 0.022 | 0.095 | 0.095 | 0.003 | 0.24 | 0.54 | 0.31 | 0.5 | 0.28 | 0.28 |
| 4 | 40255501 | rs2381497 | 1.91 | -0.471 | 0.419 | 0.885 | 0.021 | 0.013 | 0.013 | 0 | 0.56 | 0.34 | 0.63 | 0.33 | 0.65 | 0.63 |
| 4 | 40256245 | rs17512296 | -0.019 | -0.206 | 0.182 | 0.754 | -0.028 | 0.024 | 0.024 | 0 | 0.14 | 0.04 | 0.12 | 0.06 | 0.21 | 0.22 |
| 4 | 40259599 | rs12513003 | 1.655 | 0.485 | 0.229 | 0.71 | 0.012 | 0.04 | 0.04 | 0.005 | 0.51 | 0.39 | 0.42 | 0.28 | 0.48 | 0.56 |
| 4 | 40260379 | rs13105678 | 1.374 | 0.639 | 0.68 | 0.834 | 0 | 0.001 | 0.001 | 0 | 0.6 | 0.42 | 0.66 | 0.54 | 0.63 | 0.57 |
| 4 | 40262582 | rs12643250 | -0.155 | 0.251 | 0.235 | 0.692 | -0.014 | -0.019 | -0.019 | 0 | 0.69 | 0.81 | 0.65 | 0.47 | 0.66 | 0.61 |
| 4 | 40262814 | rs12640505 | 0.513 | 0.239 | 0.328 | 0.692 | -0.01 | -0.001 | -0.001 | 0 | 0.59 | 0.72 | 0.63 | 0.43 | 0.48 | 0.44 |
| 4 | 40265273 | rs28802045 | -0.136 | 0.201 | 0.27 | 0.469 | -0.002 | -0.003 | -0.003 | 1.532 | 0.69 | 0.75 | 0.71 | 0.56 | 0.66 | 0.66 |
| 4 | 40267780 | rs9685160 | 1.125 | 0.194 | 0.057 | 0.222 | 0.019 | 0.158 | 0.158 | 0 | 0.45 | 0.23 | 0.46 | 0.1 | 0.43 | 0.25 |
| 4 | 40274255 | rs10022565 | -0.879 | -0.093 | 0.357 | 0.713 | -0.036 | 0.045 | 0.045 | 0.593 | 0.15 | 0.62 | 0.27 | 0.15 | 0.06 | 0.2 |
| 4 | 40276103 | rs9685080 | -1.211 | -0.05 | 0.112 | 0.733 | -0.002 | 0.006 | 0.006 | 3.068 | 0.29 | 0.68 | 0.36 | 0.35 | 0.26 | 0.42 |
| 4 | 40277346 | rs12512351 | 0.464 | 0.07 | -0.847 | 0.762 | -0.034 | -0.039 | -0.039 | 0 | 0.12 | 0.04 | 0.1 | 0.14 | 0.14 | 0.16 |
| 4 | 40279534 | rs9683756 | -2.003 | 0.01 | -1.105 | 0.598 | -0.11 | 0.023 | 0.023 | 1.372 | 0.57 | 0.94 | 0.53 | 0.65 | 0.48 | 0.76 |
| 4 | 40283419 | rs12186133 | 0.294 | 0.14 | -0.309 | 0.831 | 0.004 | -0.018 | -0.018 | 0.174 | 0.14 | 0.17 | 0.1 | 0.05 | 0.19 | 0.25 |
| 4 | 40284434 | rs12512323 | -1.623 | -0.287 | -0.131 | 1.274 | 0.01 | -0.018 | -0.018 | 0 | 0.32 | 0.74 | 0.34 | 0.39 | 0.23 | 0.32 |
| 4 | 40287396 | rs9994274 | -1.277 | -0.026 | -0.264 | 1.264 | -0.022 | 0.042 | 0.042 | 2.096 | 0.22 | 0.69 | 0.26 | 0.45 | 0.11 | 0.3 |
| 4 | 40291232 | rs9684412 | -0.249 | 0.412 | -0.601 | 0.799 | -0.007 | 0.014 | 0.014 | 0 | 0.44 | 0.6 | 0.4 | 0.42 | 0.5 | 0.48 |
| 4 | 40293158 | rs11932152 | -1.235 | -0.04 | 0.27 | 1.1 | -0.047 | 0.123 | 0.123 | 2.222 | 0.15 | 0.43 | 0.08 | 0.33 | 0.06 | 0.22 |
| 4 | 40303633 | rs6832151 | -0.929 | -0.017 | -0.248 | 0.735 | -0.005 | 0.004 | 0.004 | 0 | 0.68 | 0.68 | 0.74 | 0.62 | 0.72 | 0.83 |
| 4 | 40307533 | rs4444866 | 0.546 | 0.577 | -0.719 | 0.018 | -0.02 | 0.048 | 0.048 | 0 | 0.22 | 0.1 | 0.17 | 0.05 | 0.26 | 0.18 |
| 4 | 40314351 | rs12499947 | 1.304 | 0.305 | -1.104 | 0.193 | -0.038 | -0.062 | -0.062 | 0.228 | 0.3 | 0.06 | 0.27 | 0.32 | 0.32 | 0.19 |
| 4 | 40316298 | rs4618346 | 1.207 | 0.703 | -0.675 | -0.298 | -0.06 | -0.036 | -0.036 | 0 | 0.43 | 0.06 | 0.32 | 0.58 | 0.44 | 0.34 |
| 4 | 40318449 | rs6833121 | -0.79 | 0.833 | 0.145 | -0.594 | 0.009 | -0.048 | -0.048 | 0.737 | 0.26 | 0.5 | 0.37 | 0.16 | 0.25 | 0.23 |
| 4 | 40323397 | rs6818783 | -0.479 | 0.66 | 0.64 | -0.763 | 0.014 | -0.047 | -0.047 | 0 | 0.29 | 0.36 | 0.29 | 0.09 | 0.37 | 0.22 |
| 4 | 40329548 | rs11728461 | 1.512 | 0.414 | 0.376 | -0.706 | -0.024 | 0.196 | 0.196 | 0 | 0.5 | 0.12 | 0.4 | 0.79 | 0.48 | 0.58 |
| 4 | 40330079 | rs7672694 | 0.864 | 0.392 | 0.293 | -0.767 | -0.062 | 0.045 | 0.045 | 0 | 0.16 | 0.01 | 0.18 | 0.05 | 0.22 | 0.17 |
| 4 | 40333521 | rs7673930 | 1.834 | 0.417 | 0.299 | -0.674 | -0.036 | 0.115 | 0.115 | 0 | 0.48 | 0.1 | 0.36 | 0.71 | 0.48 | 0.49 |
| 4 | 40333867 | rs7679386 | 1.649 | 0.417 | 0.411 | -0.732 | -0.007 | 0.19 | 0.19 | 0 | 0.5 | 0.16 | 0.41 | 0.79 | 0.48 | 0.58 |
| 4 | 40334852 | rs6838680 | -0.429 | 0.414 | 0.551 | -0.72 | -0.029 | 0.018 | 0.018 | 0 | 0.22 | 0.14 | 0.32 | 0.16 | 0.25 | 0.2 |
| 4 | 40336927 | rs4469115 | 0.635 | 0.455 | 1.003 | -0.716 | 0.005 | -0.02 | -0.02 | 0 | 0.17 | 0.16 | 0.19 | 0.05 | 0.22 | 0.17 |
| 4 | 40337908 | rs10022491 | -2.119 | 0.426 | 0.777 | -0.617 | -0.071 | -0.008 | -0.008 | 0 | 0.55 | 0.93 | 0.64 | 0.4 | 0.53 | 0.62 |
| 4 | 40342618 | rs10021263 | 0.302 | 0.823 | 1.589 | -0.939 | 0.041 | 0.025 | 0.025 | 0 | 0.09 | 0.15 | 0.11 | 0.14 | 0.16 | 0.16 |
| 4 | 40344395 | rs4861065 | -0.733 | 0.769 | 1.8 | -0.964 | -0.005 | -0.094 | -0.094 | 0 | 0.27 | 0.47 | 0.25 | 0.05 | 0.28 | 0.21 |
| 4 | 40345239 | rs4861307 | 1.984 | 0.689 | 1.469 | -1.027 | -0.011 | 0.168 | 0.168 | 0 | 0.5 | 0.19 | 0.45 | 0.8 | 0.5 | 0.59 |
| 4 | 40350147 | rs10029313 | 0.866 | 0.654 | 1.781 | -1.052 | 0.025 | 0.075 | 0.075 | 0 | 0.64 | 0.74 | 0.59 | 0.94 | 0.67 | 0.78 |
| 4 | 40350651 | rs7669882 | -0.754 | 0.688 | 1.703 | -1.015 | -0.006 | -0.108 | -0.108 | 0 | 0.27 | 0.52 | 0.26 | 0.06 | 0.28 | 0.22 |
| 4 | 40350763 | rs10029872 | 1.772 | 0.794 | 1.227 | -0.941 | -0.002 | 0.11 | 0.11 | 0 | 0.51 | 0.25 | 0.46 | 0.79 | 0.51 | 0.6 |
| 4 | 40356422 | rs10009228 | 0.828 | 0.81 | 1.03 | -0.654 | -0.001 | -0.019 | -0.019 | 0 | 0.76 | 0.64 | 0.68 | 0.85 | 0.78 | 0.79 |
| 4 | 40365870 | rs6850417 | 0.722 | 0.721 | 0.818 | -0.896 | -0.01 | -0.06 | -0.06 | 0 | 0.65 | 0.47 | 0.63 | 0.93 | 0.67 | 0.74 |
| 4 | 40367787 | rs4861350 | 1.213 | 1.065 | 0.335 | -0.832 | -0.002 | 0.05 | 0.05 | 0 | 0.65 | 0.21 | 0.6 | 0.84 | 0.65 | 0.68 |
| 4 | 40368848 | rs10021851 | 1.789 | 1.054 | 0.259 | -0.869 | -0.002 | 0.135 | 0.135 | 0 | 0.49 | 0.16 | 0.47 | 0.78 | 0.45 | 0.5 |
| 4 | 40371287 | rs4861079 | 0.33 | 1.062 | 0.536 | -0.92 | -0.032 | -0.124 | -0.124 | 0 | 0.78 | 0.39 | 0.73 | 1 | 0.82 | 0.91 |
| 4 | 40373000 | rs1492506 | 0.193 | 1.104 | 0.692 | -0.914 | -0.027 | -0.162 | -0.162 | 1.533 | 0.79 | 0.48 | 0.74 | 1 | 0.81 | 0.91 |
| 4 | 40373082 | rs4861085 | 0.295 | 1.484 | 0.576 | -0.374 | -0.033 | 0.024 | 0.024 | 0 | 0.16 | 0.05 | 0.13 | 0.08 | 0.21 | 0.18 |
| 4 | 40373150 | rs1492505 | -0.551 | 1.426 | 0.758 | -0.225 | 0.02 | 0.019 | 0.019 | 0 | 0.14 | 0.19 | 0.14 | 0.16 | 0.17 | 0.24 |
| 4 | 40377691 | rs6447623 | -1.47 | 1.406 | 0.268 | -0.425 | 0 | -0.001 | -0.001 | 0 | 0.2 | 0.76 | 0.18 | 0.15 | 0.15 | 0.19 |
| 4 | 40379061 | rs10026108 | 1.61 | 1.31 | 0.023 | -0.647 | 0.017 | 0.115 | 0.115 | 0.041 | 0.51 | 0.21 | 0.5 | 0.76 | 0.48 | 0.57 |
| 4 | 40379339 | rs4861372 | -0.681 | 1.37 | 0.007 | -0.505 | 0.006 | 0.026 | 0.026 | 0 | 0.88 | 0.95 | 0.83 | 1 | 0.88 | 0.96 |
| 4 | 40380907 | rs902856 | 0.225 | 1.388 | 0.211 | -0.489 | -0.042 | -0.092 | -0.092 | 0.404 | 0.82 | 0.51 | 0.79 | 0.94 | 0.88 | 0.9 |
| 4 | 40383379 | rs1826394 | 1.436 | 1.575 | 0.049 | -0.516 | 0.012 | -0.061 | -0.061 | 0 | 0.65 | 0.44 | 0.63 | 0.85 | 0.62 | 0.75 |
| 4 | 40384525 | rs1955259 | -1.188 | 1.984 | -0.044 | 0.124 | 0.048 | -0.023 | -0.023 | 0.195 | 0.19 | 0.32 | 0.17 | 0.15 | 0.26 | 0.19 |
| 4 | 40387873 | rs17440690 | -0.884 | 1.792 | -0.857 | 0.424 | -0.023 | -0.01 | -0.01 | 0.754 | 0.11 | 0.06 | 0.15 | 0.17 | 0.17 | 0.09 |
| 4 | 40391194 | rs13107860 | -1.103 | 1.573 | -0.635 | 0.494 | 0.001 | 0.009 | 0.009 | 0 | 0.07 | 0.17 | 0.1 | 0.09 | 0.04 | 0.09 |
| 4 | 40391816 | rs17512490 | -1.328 | 2.046 | -1.132 | 0.418 | -0.001 | 0 | 0 | 0.233 | 0.16 | 0.16 | 0.19 | 0.17 | 0.23 | 0.14 |
| 4 | 40393828 | rs13123893 | -0.437 | 1.248 | -2.111 | 0.577 | -0.013 | -0.083 | -0.083 | 0.154 | 0.33 | 0.06 | 0.38 | 0.61 | 0.29 | 0.39 |
| 4 | 40396560 | rs278973 | -0.634 | 1.111 | -2.402 | 0.291 | -0.075 | 0.038 | 0.038 | 0 | 0.64 | 0.84 | 0.65 | 0.76 | 0.55 | 0.69 |
| 4 | 40396697 | rs11731787 | -0.204 | 0.315 | -2.413 | 0.58 | -0.01 | -0.154 | -0.154 | 0 | 0.24 | 0.03 | 0.31 | 0.56 | 0.2 | 0.36 |
| 4 | 40397253 | rs4447895 | 0.534 | -0.059 | -2.255 | 0.203 | 0 | -0.001 | -0.001 | 0 | 0.1 | 0.09 | 0.09 | 0.05 | 0.1 | 0.06 |
| 4 | 40397913 | rs4476635 | -1.025 | -0.34 | -1.629 | -0.124 | 0.027 | 0.036 | 0.036 | 0 | 0.44 | 0.62 | 0.54 | 0.78 | 0.36 | 0.63 |
| 4 | 40401610 | rs6832419 | 0.072 | -0.46 | -0.936 | 0.204 | -0.027 | -0.045 | -0.045 | 0.218 | 0.1 | 0.21 | 0.06 | 0.01 | 0.07 | 0.05 |

| | | | | | | | | | | | | | | | | |
|---|---|---|---|---|---|---|---|---|---|---|---|---|---|---|---|---|
| 4 | 40410479 | rs4861106 | -1.433 | -0.545 | -0.426 | -0.036 | -0.004 | -0.006 | -0.006 | 0 | 0.4 | 0.61 | 0.42 | 0.8 | 0.28 | 0.59 |
| 4 | 40414829 | rs924639 | -0.624 | -0.566 | 0.252 | 0.858 | -0.022 | 0.02 | 0.02 | 0.006 | 0.12 | 0.18 | 0.18 | 0.15 | 0.05 | 0.08 |
| 4 | 40415109 | rs278933 | 0.6 | -0.351 | 0.582 | 1.554 | -0.004 | -0.011 | -0.011 | 0 | 0.8 | 0.85 | 0.77 | 0.6 | 0.75 | 0.68 |
| 4 | 40418670 | rs203273 | 1.862 | -0.486 | 1.122 | 1.617 | -0.008 | -0.018 | -0.018 | 0 | 0.62 | 0.5 | 0.47 | 0.31 | 0.66 | 0.48 |
| 4 | 40419414 | rs10007960 | -2.096 | -0.628 | 1.353 | 2.049 | -0.06 | 0.074 | 0.074 | 0 | 0.28 | 0.48 | 0.43 | 0.55 | 0.15 | 0.31 |
| 4 | 40419876 | rs9790641 | -1.231 | -0.63 | 1.26 | 2.196 | -0.001 | 0.02 | 0.02 | 0.294 | 0.14 | 0.17 | 0.32 | 0.38 | 0.09 | 0.14 |
| 4 | 40420131 | rs12505087 | -0.102 | -0.537 | 1.317 | 1.637 | -0.027 | 0.012 | 0.012 | 0 | 0.11 | 0.18 | 0.1 | 0.12 | 0.05 | 0.08 |
| 4 | 40420708 | rs278941 | 0.821 | -0.38 | 1.34 | 1.251 | -0.007 | -0.004 | -0.004 | 0 | 0.68 | 0.62 | 0.58 | 0.32 | 0.71 | 0.53 |
| 4 | 40420804 | rs17512637 | -0.195 | -0.339 | 0.812 | 1.194 | 0.001 | -0.006 | -0.006 | 0 | 0.07 | 0.08 | 0.14 | 0.06 | 0.06 | 0.07 |
| 4 | 40424213 | rs4362858 | 0.652 | -0.256 | 0.715 | 1.258 | 0 | -0.001 | -0.001 | 0.437 | 0.68 | 0.7 | 0.61 | 0.36 | 0.7 | 0.5 |
| 4 | 40424602 | rs2008438 | 1.184 | -0.1 | 0.177 | 1.047 | 0.008 | 0.022 | 0.022 | 0 | 0.57 | 0.34 | 0.44 | 0.25 | 0.64 | 0.38 |
| 4 | 40426677 | rs278982 | 1.221 | -0.092 | 0.092 | 1.038 | 0.014 | 0.041 | 0.041 | 0 | 0.57 | 0.31 | 0.43 | 0.25 | 0.62 | 0.38 |
| 4 | 40428010 | rs278981 | 0.162 | 0.231 | 0.365 | 1.467 | 0.027 | 0.093 | 0.093 | 0 | 0.79 | 0.67 | 0.78 | 0.58 | 0.78 | 0.68 |
| 4 | 40431413 | rs278979 | -1.814 | -0.433 | 0.496 | 1.688 | -0.027 | 0.096 | 0.096 | 0.005 | 0.19 | 0.63 | 0.33 | 0.52 | 0.12 | 0.15 |
| 4 | 40438576 | rs2307046 | 0.449 | -0.839 | 0.443 | 0.957 | -0.023 | 0.011 | 0.011 | 0.886 | 0.85 | 0.71 | 0.85 | 0.77 | 0.87 | 0.88 |
| 4 | 40443966 | rs10755175 | 1.488 | -1.335 | 0.726 | 0.672 | -0.043 | -0.01 | -0.01 | 0.373 | 0.75 | 0.41 | 0.79 | 0.73 | 0.83 | 0.84 |
| 4 | 40444292 | rs10938155 | -0.242 | -1.119 | -0.121 | 0.287 | -0.001 | -0.001 | -0.001 | 0 | 0.27 | 0.24 | 0.37 | 0.47 | 0.3 | 0.15 |
| 4 | 40445369 | rs13129937 | -0.092 | -1.086 | -0.551 | 0.129 | -0.033 | 0.012 | 0.012 | 0.997 | 0.85 | 0.67 | 0.86 | 0.75 | 0.88 | 0.87 |
| 4 | 40446425 | rs10023957 | -0.416 | -1.048 | -0.887 | -0.109 | 0.003 | 0.002 | 0.002 | 0 | 0.21 | 0.14 | 0.14 | 0 | 0.19 | 0.11 |
| 4 | 40447356 | rs10018317 | 0.571 | -0.69 | -0.62 | 0.401 | 0.011 | 0.027 | 0.027 | 0 | 0.81 | 0.62 | 0.68 | 0.7 | 0.73 | 0.82 |
| 4 | 40451394 | rs7689954 | 1.523 | -0.591 | -1.308 | 0.391 | -0.009 | 0.04 | 0.04 | 2.165 | 0.49 | 0.06 | 0.42 | 0.34 | 0.54 | 0.72 |
| 4 | 40453352 | rs12647748 | -0.784 | 0.021 | -0.601 | 0.422 | 0.02 | -0.021 | -0.021 | 0 | 0.13 | 0.36 | 0.14 | 0.16 | 0.11 | 0.09 |
| 4 | 40457354 | rs7695920 | -0.503 | 0.355 | -1.375 | 0.612 | -0.001 | 0.009 | 0.009 | 0.663 | 0.14 | 0.1 | 0.21 | 0.07 | 0.17 | 0.06 |
| 4 | 40461712 | rs278949 | 0.172 | 0.585 | -2.09 | 0.695 | -0.033 | 0.066 | 0.066 | 0.823 | 0.57 | 0.91 | 0.61 | 0.71 | 0.62 | 0.59 |
| 4 | 40462142 | rs9790441 | -0.887 | 0.75 | -0.855 | 1.782 | 0.025 | 0.255 | 0.255 | 0.065 | 0.07 | 0.12 | 0.13 | 0.39 | 0.07 | 0.04 |
| 4 | 40466547 | rs10000300 | -0.558 | 0.649 | -1.721 | 1.018 | -0.003 | -0.012 | -0.012 | 0 | 0.42 | 0.83 | 0.46 | 0.46 | 0.39 | 0.3 |
| 4 | 40473784 | rs3105209 | -1.087 | 0.199 | -0.811 | 1.199 | -0.034 | 0.058 | 0.058 | 0.224 | 0.21 | 0.58 | 0.25 | 0.37 | 0.15 | 0.22 |
| 4 | 40475295 | rs278962 | -1.399 | 0.933 | -1.535 | 0.975 | -0.032 | -0.06 | -0.06 | 0 | 0.38 | 0.85 | 0.42 | 0.56 | 0.46 | 0.35 |
| 4 | 40477729 | rs16852239 | -0.722 | 0.971 | -0.292 | 0.594 | -0.007 | -0.117 | -0.117 | 0.708 | 0.2 | 0.37 | 0.14 | 0.01 | 0.19 | 0.06 |
| 4 | 40488449 | rs278959 | -0.143 | 1.289 | 0.083 | 0.8 | 0.023 | 0.049 | 0.049 | 0 | 0.16 | 0.26 | 0.22 | 0.29 | 0.25 | 0.17 |
| 4 | 40490465 | rs4861170 | 1.205 | 1.658 | -0.018 | 0.927 | -0.031 | -0.038 | -0.038 | 0 | 0.66 | 0.18 | 0.65 | 0.47 | 0.55 | 0.64 |
| 4 | 40491003 | rs1847118 | 1.233 | 1.514 | 0.641 | 0.787 | 0.01 | 0.011 | 0.011 | 0.712 | 0.68 | 0.34 | 0.62 | 0.52 | 0.53 | 0.71 |
| 4 | 40491187 | rs10517543 | 0.425 | 1.54 | 0.109 | 0.313 | -0.003 | 0 | 0 | 0 | 0.11 | 0.08 | 0.07 | 0 | 0.11 | 0.01 |
| 4 | 40491786 | rs12641621 | 1.18 | 0.716 | -0.558 | 0.323 | 0.013 | -0.006 | -0.006 | 0.02 | 0.32 | 0.21 | 0.29 | 0.39 | 0.28 | 0.33 |
| 4 | 40494616 | rs278986 | -0.615 | 1.28 | -0.446 | 0.554 | 0.028 | -0.043 | -0.043 | 1.501 | 0.21 | 0.34 | 0.18 | 0.09 | 0.29 | 0.12 |
| 4 | 40498421 | rs4504294 | -0.611 | 1.229 | -0.407 | 0.658 | 0.014 | 0.046 | 0.046 | 0 | 0.12 | 0.28 | 0.14 | 0.19 | 0.12 | 0.15 |
| 4 | 40503041 | rs16852276 | -0.33 | 1.416 | -1.034 | 0.688 | 0.018 | 0.001 | 0.001 | 0 | 0.19 | 0.25 | 0.21 | 0.19 | 0.32 | 0.12 |
| 4 | 40506255 | rs6857898 | -0.88 | 0.219 | -1.841 | 0.622 | -0.022 | -0.014 | -0.014 | 2.303 | 0.77 | 0.98 | 0.69 | 0.85 | 0.79 | 0.64 |
| 4 | 40509521 | rs6848632 | -0.439 | -0.281 | -1.297 | 0.82 | -0.007 | 0.014 | 0.014 | 0 | 0.17 | 0.22 | 0.18 | 0.26 | 0.09 | 0.18 |
| 4 | 40515702 | rs364773 | -1.178 | 0.259 | -1.87 | 0.327 | -0.054 | 0.058 | 0.058 | 0.415 | 0.79 | 0.97 | 0.72 | 0.91 | 0.77 | 0.66 |
| 4 | 40519458 | rs430730 | -1.612 | -0.702 | -0.914 | 0.786 | -0.024 | -0.073 | -0.073 | 0 | 0.4 | 0.8 | 0.4 | 0.44 | 0.39 | 0.37 |
| 4 | 40528427 | rs11737664 | -0.839 | -0.816 | -0.229 | 0.121 | 0.012 | -0.1 | -0.1 | 0.286 | 0.25 | 0.42 | 0.17 | 0.04 | 0.26 | 0.19 |
| 4 | 40533297 | rs3114381 | -1.082 | -1.36 | -0.537 | 0.491 | 0.046 | 0.046 | 0.046 | 0.714 | 0.37 | 0.74 | 0.33 | 0.23 | 0.27 | 0.25 |
| 4 | 40535566 | rs4632692 | -1.502 | -1.238 | -0.541 | 0.55 | 0.055 | -0.011 | -0.011 | 0 | 0.47 | 0.77 | 0.41 | 0.47 | 0.38 | 0.43 |
| 4 | 40546244 | rs6847478 | -1.328 | -1.409 | 0.051 | 0.441 | 0.011 | -0.003 | -0.003 | 0 | 0.46 | 0.41 | 0.41 | 0.51 | 0.37 | 0.5 |
| 4 | 40548057 | rs7667552 | -1.286 | -1.663 | 0.811 | 0.364 | -0.056 | -0.009 | -0.009 | 0 | 0.23 | 0.52 | 0.2 | 0.29 | 0.13 | 0.26 |
| 4 | 40549870 | rs6817468 | 0.141 | -1.627 | 0.355 | 0.388 | 0.015 | -0.016 | -0.016 | 0.261 | 0.08 | 0.08 | 0.13 | 0.2 | 0.03 | 0.16 |
| 4 | 40553177 | rs373103 | -1.814 | -1.606 | 0.294 | 0.329 | 0.099 | -0.014 | -0.014 | 0.034 | 0.43 | 0.89 | 0.38 | 0.42 | 0.28 | 0.4 |
| 4 | 40558019 | rs11932270 | -1.097 | -1.21 | 1.131 | 0.012 | -0.031 | -0.083 | -0.083 | 0 | 0.13 | 0.51 | 0.07 | 0 | 0.06 | 0.04 |
| 4 | 40565469 | rs6841429 | -1.682 | -1.187 | 0.495 | 0.078 | -0.03 | -0.02 | -0.02 | 1.833 | 0.25 | 0.61 | 0.22 | 0.15 | 0.18 | 0.16 |
| 4 | 40569727 | rs16852342 | -0.018 | -0.593 | -0.057 | 0.122 | -0.016 | -0.041 | -0.041 | 0 | 0.08 | 0.2 | 0.04 | 0 | 0.05 | 0.02 |
| 4 | 40576393 | rs17513191 | -0.293 | -0.209 | -0.303 | 0.466 | 0.005 | -0.002 | -0.002 | 0.494 | 0.3 | 0.28 | 0.4 | 0.27 | 0.26 | 0.33 |
| 4 | 40577228 | rs10025907 | -1.086 | 0.057 | 0.42 | 0.271 | 0.057 | 0.179 | 0.179 | 5.214 | 0.06 | 0.44 | 0.04 | 0.14 | 0.01 | 0.1 |
| 4 | 40578403 | rs3828531 | -0.871 | 0.809 | -0.968 | 0.126 | -0.011 | 0.025 | 0.025 | 0 | 0.17 | 0.1 | 0.27 | 0.12 | 0.21 | 0.16 |
| 4 | 40589717 | rs6829898 | -0.061 | 0.746 | -0.788 | 0.175 | -0.001 | 0 | 0 | 0 | 0.54 | 0.52 | 0.64 | 0.38 | 0.52 | 0.47 |
| 4 | 40590908 | rs17625986 | -0.295 | 0.728 | -0.119 | -0.297 | -0.001 | -0.001 | -0.001 | 0 | 0.22 | 0.26 | 0.24 | 0.23 | 0.2 | 0.2 |
| 4 | 40594008 | rs4861060 | -0.342 | 0.795 | -0.146 | -0.352 | -0.006 | 0.002 | 0.002 | 0 | 0.22 | 0.19 | 0.27 | 0.23 | 0.25 | 0.2 |
| 4 | 40596439 | rs12644323 | -0.016 | 0.744 | -0.223 | -0.377 | -0.08 | -0.04 | -0.04 | 2.224 | 0.19 | 0.04 | 0.26 | 0.23 | 0.25 | 0.2 |
| 4 | 40598634 | rs1384035 | -0.714 | 0.514 | -0.021 | -0.335 | 0.009 | 0.04 | 0.04 | 0 | 0.25 | 0.56 | 0.33 | 0.36 | 0.26 | 0.3 |
| 4 | 40603055 | rs17587482 | -1.079 | 0.077 | 0.391 | -0.338 | -0.006 | -0.012 | -0.012 | 1.049 | 0.33 | 0.47 | 0.34 | 0.33 | 0.36 | 0.37 |
| 4 | 40607397 | rs6820328 | -0.11 | -0.199 | 0.323 | -0.934 | -0.001 | -0.006 | -0.006 | 0 | 0.5 | 0.47 | 0.42 | 0.14 | 0.5 | 0.34 |
| 4 | 40609305 | rs10938275 | -0.375 | -0.194 | 0.189 | -0.922 | -0.008 | 0.049 | 0.049 | 0.175 | 0.52 | 0.67 | 0.47 | 0.27 | 0.51 | 0.43 |
| 4 | 40610732 | rs11936786 | 1.427 | 0.25 | -0.245 | -1.111 | -0.078 | 0.176 | 0.176 | 0.319 | 0.26 | 0.01 | 0.29 | 0.01 | 0.24 | 0.13 |
| 4 | 40620852 | rs6851689 | 0.513 | 0.243 | -0.164 | -1.123 | 0.006 | 0.102 | 0.102 | 0 | 0.35 | 0.15 | 0.33 | 0.11 | 0.3 | 0.27 |
| 4 | 40624014 | rs13152696 | 0.353 | 0.291 | -0.156 | -0.941 | 0.011 | 0.029 | 0.029 | 0 | 0.37 | 0.16 | 0.37 | 0.26 | 0.31 | 0.36 |
| 4 | 40625135 | rs13148961 | 1.18 | 0.247 | -0.33 | -0.963 | -0.056 | 0.036 | 0.036 | 0.038 | 0.27 | 0.03 | 0.33 | 0.14 | 0.24 | 0.22 |
| 4 | 40631309 | rs7664412 | -1.352 | 0.051 | 0.217 | -0.791 | 0.001 | -0.053 | -0.053 | 0 | 0.73 | 0.86 | 0.76 | 0.5 | 0.71 | 0.65 |
| 4 | 40631476 | rs12498424 | 0.763 | 0.023 | 0.149 | -0.81 | -0.024 | 0.015 | 0.015 | 0 | 0.2 | 0.12 | 0.28 | 0.21 | 0.25 | 0.2 |
| 4 | 40635372 | rs13132427 | -1.225 | 0.026 | 0.234 | -0.786 | 0 | -0.023 | -0.023 | 0.332 | 0.72 | 0.82 | 0.76 | 0.49 | 0.7 | 0.64 |

| | | | | | | | | | | | | | | | | |
|---|---|---|---|---|---|---|---|---|---|---|---|---|---|---|---|---|
| 4 | 40644376 | rs10015679 | -0.425 | 0.112 | -0.107 | -1.069 | -0.018 | -0.016 | -0.016 | 0.593 | 0.31 | 0.44 | 0.34 | 0.29 | 0.27 | 0.34 |
| 4 | 40650498 | rs7679359 | 0.567 | 0.605 | -1.579 | -1.294 | -0.031 | 0.118 | 0.118 | 1.022 | 0.32 | 0.12 | 0.29 | 0.03 | 0.31 | 0.18 |
| 4 | 40651632 | rs1552138 | -0.575 | 0.66 | -1.212 | -0.839 | 0.01 | 0.009 | 0.009 | 0.249 | 0.65 | 0.27 | 0.65 | 0.32 | 0.67 | 0.56 |
| 4 | 40652510 | rs4861297 | -0.26 | 0.879 | -1.092 | -0.289 | -0.02 | 0.025 | 0.025 | 0 | 0.19 | 0.14 | 0.25 | 0.17 | 0.21 | 0.18 |
| 4 | 40663704 | rs7437008 | 0.191 | 0.646 | -1.526 | 0.677 | -0.024 | -0.109 | -0.109 | 0 | 0.78 | 0.94 | 0.88 | 0.62 | 0.79 | 0.74 |
| 4 | 40668113 | rs4861300 | 0.1 | 0.624 | -1.545 | 0.96 | -0.053 | -0.051 | -0.051 | 0.518 | 0.79 | 0.99 | 0.9 | 0.78 | 0.79 | 0.76 |
| 4 | 40670323 | rs6836880 | 0.124 | 0.605 | -1.554 | 0.949 | -0.052 | -0.048 | -0.048 | 0 | 0.79 | 0.99 | 0.9 | 0.78 | 0.79 | 0.75 |
| 4 | 40673993 | rs10017429 | -0.456 | 0.41 | -1.681 | 2.025 | -0.006 | 0.063 | 0.063 | 0 | 0.12 | 0.11 | 0.17 | 0.45 | 0.1 | 0.37 |
| 4 | 40681370 | rs2035385 | 0.428 | 0.609 | -1.012 | 1.022 | -0.002 | 0.003 | 0.003 | 0 | 0.65 | 0.59 | 0.73 | 0.78 | 0.6 | 0.65 |
| 4 | 40683976 | rs16852437 | 0.482 | 0.612 | -0.952 | 0.471 | 0.014 | -0.077 | -0.077 | 0 | 0.15 | 0.39 | 0.16 | 0.01 | 0.18 | 0.1 |
| 4 | 40686808 | rs2035381 | 0.468 | 0.464 | -1.38 | -0.247 | 0.004 | 0.079 | 0.079 | 0 | 0.5 | 0.56 | 0.65 | 0.92 | 0.47 | 0.62 |
| 4 | 40687234 | rs6835691 | 1.142 | 0.482 | -1.272 | -0.203 | 0 | -0.022 | -0.022 | 0 | 0.13 | 0.21 | 0.15 | 0.01 | 0.18 | 0.07 |
| 4 | 40690285 | rs6447325 | -0.333 | 0.739 | -1.181 | 0.198 | -0.02 | -0.05 | -0.05 | 0.017 | 0.73 | 0.85 | 0.52 | 0.65 | 0.7 | 0.53 |
| 4 | 40690330 | rs12498704 | 0.703 | 0.607 | -0.543 | 0.392 | 0 | -0.016 | -0.016 | 9.924 | 0.38 | 0.24 | 0.26 | 0.55 | 0.36 | 0.37 |
| 4 | 40696983 | rs6814199 | -0.199 | 0.677 | -0.269 | 0.596 | 0.012 | 0.051 | 0.051 | 0 | 0.22 | 0.36 | 0.54 | 0.29 | 0.28 | 0.37 |
| 4 | 40698099 | rs7661237 | 0.278 | 0.53 | -0.198 | 0.343 | 0.008 | -0.012 | -0.012 | 0.036 | 0.58 | 0.48 | 0.46 | 0.66 | 0.52 | 0.55 |
| 4 | 40702239 | rs12500580 | 0.128 | 0.329 | -0.681 | -0.398 | -0.003 | 0.22 | 0.22 | 0.285 | 0.5 | 0.77 | 0.67 | 0.89 | 0.5 | 0.63 |
| 4 | 40704872 | rs10000266 | -0.48 | 0.59 | -0.764 | -0.326 | -0.057 | 0.178 | 0.178 | 0.025 | 0.67 | 0.94 | 0.78 | 0.94 | 0.65 | 0.72 |
| 4 | 40708301 | rs6830900 | 0.51 | 0.669 | -0.048 | 0.044 | -0.001 | 0.001 | 0.001 | 0.422 | 0.27 | 0.22 | 0.44 | 0.53 | 0.37 | 0.33 |
| 4 | 40710034 | rs6815557 | -0.424 | 0.497 | -0.009 | -0.048 | 0.007 | -0.017 | -0.017 | 0 | 0.63 | 0.82 | 0.65 | 0.69 | 0.63 | 0.52 |
| 4 | 40712344 | rs10012870 | 0.517 | 0.558 | 0.116 | -0.024 | -0.001 | -0.006 | -0.006 | 0 | 0.34 | 0.36 | 0.18 | 0.16 | 0.26 | 0.18 |
| 4 | 40712481 | rs10008483 | -0.177 | 1.257 | 0.114 | 0.46 | 0.029 | 0.043 | 0.043 | 0.436 | 0.28 | 0.31 | 0.46 | 0.53 | 0.37 | 0.33 |
| 4 | 40722850 | rs2200061 | -0.44 | 1.081 | 0.202 | -0.141 | 0.024 | -0.109 | -0.109 | 0 | 0.18 | 0.39 | 0.12 | 0.01 | 0.2 | 0.04 |
| 4 | 40726020 | rs4516756 | -0.071 | 1.225 | -0.289 | 0.085 | -0.001 | 0 | 0 | 0 | 0.48 | 0.49 | 0.57 | 0.56 | 0.43 | 0.41 |
| 4 | 40730503 | rs790142 | 0.413 | 0.977 | -0.611 | -0.557 | 0.003 | 0.03 | 0.03 | 0 | 0.57 | 0.41 | 0.48 | 0.79 | 0.63 | 0.54 |
| 4 | 40739635 | rs11727143 | 0.392 | 0.936 | -0.535 | -0.797 | -0.002 | 0.025 | 0.025 | 0 | 0.57 | 0.39 | 0.46 | 0.79 | 0.6 | 0.54 |
| 4 | 40746510 | rs17513557 | 0.931 | 0.82 | -0.743 | -0.848 | -0.015 | 0.066 | 0.066 | 0 | 0.14 | 0.04 | 0.11 | 0 | 0.13 | 0.17 |
| 4 | 40754154 | rs1545077 | -0.91 | 0.8 | -0.349 | -0.413 | 0.006 | 0.07 | 0.07 | 0 | 0.81 | 0.8 | 0.86 | 0.49 | 0.75 | 0.61 |
| 4 | 40758525 | rs10029833 | 1.218 | 0.99 | -0.588 | -0.854 | -0.01 | 0.027 | 0.027 | 0 | 0.14 | 0.09 | 0.14 | 0.01 | 0.16 | 0.08 |
| 4 | 40771690 | rs7699447 | 0.96 | 0.585 | -0.382 | -0.868 | -0.013 | 0.018 | 0.018 | 0 | 0.18 | 0.23 | 0.39 | 0.36 | 0.12 | 0.25 |
| 4 | 40773665 | rs10938385 | 0.674 | 0.711 | -0.387 | -1.124 | -0.012 | 0.034 | 0.034 | 0 | 0.3 | 0.25 | 0.18 | 0.11 | 0.32 | 0.08 |
| 4 | 40777147 | rs13106136 | -1.767 | 0.668 | -0.283 | -1.226 | 0 | 0.003 | 0.003 | 0 | 0.8 | 0.83 | 0.87 | 0.52 | 0.77 | 0.6 |
| 4 | 40778162 | rs2437323 | -1.766 | 0.682 | -0.277 | -1.205 | 0.001 | -0.034 | -0.034 | 0.069 | 0.8 | 0.87 | 0.88 | 0.52 | 0.77 | 0.6 |
| 4 | 40779968 | rs2465568 | 0.776 | 0.713 | -0.226 | -1.193 | -0.003 | 0.008 | 0.008 | 0 | 0.3 | 0.3 | 0.19 | 0.12 | 0.32 | 0.07 |
| 4 | 40792007 | rs2437317 | -1.57 | 0.674 | -0.214 | -1.052 | -0.001 | 0.024 | 0.024 | 0 | 0.8 | 0.8 | 0.87 | 0.5 | 0.77 | 0.58 |
| 4 | 40797693 | rs2437309 | -0.353 | 0.638 | -0.33 | -1.2 | -0.009 | 0.044 | 0.044 | 0 | 0.52 | 0.63 | 0.7 | 0.4 | 0.46 | 0.51 |
| 4 | 40810663 | rs4861066 | 1.278 | 0.334 | -0.31 | -1.341 | -0.007 | -0.009 | -0.009 | 0.902 | 0.23 | 0.18 | 0.23 | 0.02 | 0.3 | 0.13 |
| 4 | 40810978 | rs882733 | 1.363 | -0.039 | -0.339 | -1.347 | -0.001 | 0 | 0 | 1.454 | 0.18 | 0.1 | 0.13 | 0 | 0.16 | 0.14 |
| 4 | 40813714 | rs11731472 | 0.157 | -0.076 | -0.296 | -1.332 | -0.003 | 0.034 | 0.034 | 0 | 0.37 | 0.32 | 0.24 | 0.11 | 0.35 | 0.19 |
| 4 | 40815268 | rs13518 | 1.345 | -0.029 | -0.367 | -1.262 | -0.024 | 0.047 | 0.047 | 0 | 0.22 | 0.12 | 0.21 | 0.02 | 0.29 | 0.12 |
| 4 | 40823454 | rs4861067 | -0.66 | -0.219 | -0.323 | -1.036 | -0.007 | -0.008 | -0.008 | 2.684 | 0.56 | 0.64 | 0.64 | 0.49 | 0.46 | 0.46 |
| 4 | 40827214 | rs2341597 | -0.221 | -0.164 | -0.169 | -0.968 | -0.02 | -0.014 | -0.014 | 0.187 | 0.38 | 0.47 | 0.54 | 0.37 | 0.28 | 0.42 |
| 4 | 40832310 | rs3816099 | 0.85 | -0.169 | -0.192 | -0.816 | -0.018 | 0.04 | 0.04 | 0 | 0.17 | 0.33 | 0.3 | 0.27 | 0.12 | 0.25 |
| 4 | 40833954 | rs12501553 | 1.314 | -0.1 | -0.264 | -0.79 | -0.008 | 0.039 | 0.039 | 0 | 0.14 | 0.23 | 0.29 | 0.29 | 0.12 | 0.23 |
| 4 | 40835453 | rs4861318 | 0.981 | -0.2 | -0.309 | -0.774 | -0.009 | 0.048 | 0.048 | 0 | 0.15 | 0.23 | 0.39 | 0.35 | 0.12 | 0.24 |
| 4 | 40836953 | rs7666964 | -0.342 | -0.137 | -0.391 | -0.836 | -0.004 | -0.02 | -0.02 | 0 | 0.59 | 0.55 | 0.46 | 0.14 | 0.63 | 0.31 |
| 4 | 40837929 | rs2437339 | -1.707 | 0.14 | -0.514 | -0.664 | -0.014 | -0.105 | -0.105 | 1.337 | 0.79 | 0.93 | 0.87 | 0.48 | 0.75 | 0.55 |
| 4 | 40838617 | rs17587985 | 1.587 | 0.176 | -0.487 | -0.88 | 0.006 | -0.007 | -0.007 | 0.327 | 0.12 | 0.07 | 0.04 | 0.11 | 0.08 | 0.09 |
| 4 | 40839813 | rs13135802 | 1.344 | 0.298 | -0.494 | -1.028 | -0.026 | 0.049 | 0.049 | 0 | 0.24 | 0.13 | 0.33 | 0.1 | 0.3 | 0.15 |
| 4 | 40842367 | rs765850 | -0.54 | 0.234 | -0.435 | -1.09 | -0.005 | -0.012 | -0.012 | 0 | 0.58 | 0.49 | 0.74 | 0.36 | 0.62 | 0.52 |
| 4 | 40842901 | rs719379 | 0.083 | 0.422 | -1.03 | -1.11 | -0.03 | 0.091 | 0.091 | 0 | 0.41 | 0.21 | 0.45 | 0.1 | 0.45 | 0.28 |
| 4 | 40843288 | rs10517551 | 0.651 | 0.479 | -0.855 | -0.38 | -0.003 | 0.004 | 0.004 | 0 | 0.16 | 0.08 | 0.11 | 0 | 0.15 | 0.12 |
| 4 | 40857194 | rs746860 | 0.167 | 0.699 | -1.237 | -0.089 | 0.006 | 0.01 | 0.01 | 0 | 0.62 | 0.68 | 0.72 | 0.36 | 0.64 | 0.53 |
| 4 | 40860553 | rs2465544 | -0.379 | 0.758 | -1.126 | 0.408 | 0.005 | -0.011 | -0.011 | 0.616 | 0.78 | 0.83 | 0.78 | 0.46 | 0.82 | 0.57 |
| 4 | 40867805 | rs2437320 | -0.015 | 0.856 | -1.326 | 0.456 | -0.005 | -0.049 | -0.049 | 0.017 | 0.65 | 0.82 | 0.66 | 0.48 | 0.66 | 0.39 |
| 4 | 40874516 | rs4861322 | -0.474 | 0.949 | -0.697 | 0.121 | -0.001 | -0.002 | -0.002 | 0 | 0.37 | 0.58 | 0.44 | 0.33 | 0.35 | 0.32 |
| 4 | 40881353 | rs13132027 | 0.766 | 0.944 | -1.366 | -0.158 | 0.016 | -0.08 | -0.08 | 0.005 | 0.3 | 0.12 | 0.29 | 0.54 | 0.27 | 0.52 |
| 4 | 40885144 | rs6817509 | -0.775 | 0.761 | -0.85 | -0.295 | -0.013 | 0.019 | 0.019 | 0 | 0.7 | 0.8 | 0.77 | 0.87 | 0.62 | 0.86 |
| 4 | 40886139 | rs17442189 | -0.569 | 1.404 | -0.518 | -0.626 | -0.01 | -0.003 | -0.003 | 3.143 | 0.2 | 0.15 | 0.37 | 0.27 | 0.28 | 0.24 |
| 4 | 40887991 | rs16852584 | 0.266 | 1.131 | -0.205 | -0.534 | 0 | 0.001 | 0.001 | 0 | 0.07 | 0.07 | 0.1 | 0.12 | 0.07 | 0.02 |
| 4 | 40895018 | rs11737809 | 1.351 | 1.024 | 0.189 | -0.699 | -0.01 | 0.034 | 0.034 | 0 | 0.16 | 0.05 | 0.09 | 0.06 | 0.15 | 0.11 |
| 4 | 40895132 | rs6849533 | 0.426 | 0.97 | 0.34 | -0.75 | 0.015 | 0.024 | 0.024 | 0 | 0.51 | 0.44 | 0.43 | 0.6 | 0.44 | 0.64 |
| 4 | 40902988 | rs13136284 | 1.238 | 0.7 | 0.001 | -0.587 | -0.011 | -0.142 | -0.142 | 0.433 | 0.3 | 0.05 | 0.26 | 0.53 | 0.27 | 0.53 |
| 4 | 40913247 | rs4861330 | -0.576 | 0.995 | 0.718 | -0.736 | 0.008 | -0.024 | -0.024 | 0.004 | 0.4 | 0.47 | 0.51 | 0.28 | 0.43 | 0.34 |
| 4 | 40916975 | rs10019646 | 0.055 | 1.045 | 0.752 | -0.789 | -0.001 | -0.011 | -0.011 | 0.289 | 0.29 | 0.32 | 0.28 | 0.14 | 0.3 | 0.15 |
| 4 | 40917488 | rs13102449 | 1.348 | 0.849 | 0.747 | -0.578 | 0.012 | -0.034 | -0.034 | 0 | 0.31 | 0.24 | 0.28 | 0.52 | 0.28 | 0.52 |
| 4 | 40917734 | rs10938442 | -0.712 | 0.875 | 1.111 | -0.343 | -0.033 | -0.008 | -0.008 | 0 | 0.76 | 0.6 | 0.85 | 0.82 | 0.84 | 0.9 |
| 4 | 40926385 | rs3922617 | -0.155 | 0.969 | 0.965 | -0.378 | -0.014 | -0.007 | -0.007 | 0 | 0.69 | 0.6 | 0.74 | 0.7 | 0.73 | 0.87 |
| 4 | 40928213 | rs7690478 | -0.414 | 1.114 | 1.024 | -0.415 | -0.001 | -0.001 | -0.001 | 0 | 0.72 | 0.75 | 0.78 | 0.82 | 0.72 | 0.85 |

| | | | | | | | | | | | | | | | | |
|---|---|---|---|---|---|---|---|---|---|---|---|---|---|---|---|---|
| 4 | 40928378 | rs7657518 | 0.136 | 0.991 | 0.798 | -0.361 | 0.003 | 0.002 | 0.002 | 0 | 0.07 | 0.1 | 0.11 | 0.11 | 0.09 | 0.02 |
| 4 | 40931712 | rs11733857 | 0.838 | 1.017 | 0.89 | -0.365 | 0.001 | 0.004 | 0.004 | 0.007 | 0.52 | 0.43 | 0.44 | 0.54 | 0.47 | 0.67 |
| 4 | 40933286 | rs4861070 | 0.161 | 1.055 | 0.7 | -0.334 | 0 | -0.003 | -0.003 | 0.064 | 0.6 | 0.6 | 0.56 | 0.66 | 0.58 | 0.69 |
| 4 | 40938623 | rs13125675 | 0.644 | 0.959 | 0.091 | -0.307 | -0.008 | 0.004 | 0.004 | 0 | 0.52 | 0.3 | 0.4 | 0.55 | 0.49 | 0.66 |
| 4 | 40941473 | rs13110219 | 0.756 | 0.822 | 0.22 | -0.373 | 0.005 | -0.002 | -0.002 | 0 | 0.7 | 0.4 | 0.53 | 0.7 | 0.64 | 0.76 |
| 4 | 40949691 | rs4861073 | -0.394 | 0.859 | 0.227 | -0.84 | -0.016 | 0.021 | 0.021 | 0 | 0.36 | 0.23 | 0.41 | 0.19 | 0.47 | 0.27 |
| 4 | 40951828 | rs3935357 | -1.29 | 0.505 | 0.463 | -0.935 | -0.019 | 0.026 | 0.026 | 0 | 0.67 | 0.5 | 0.62 | 0.46 | 0.71 | 0.47 |
| 4 | 40953139 | rs7658815 | 0.744 | 0.434 | 0.402 | -1.125 | 0.006 | 0.012 | 0.012 | 0.022 | 0.33 | 0.27 | 0.2 | 0.17 | 0.26 | 0.21 |
| 4 | 40953707 | rs4861074 | 0.397 | 0.589 | -0.159 | -1.144 | -0.047 | -0.023 | -0.023 | 1.355 | 0.16 | 0.05 | 0.25 | 0.17 | 0.22 | 0.16 |
| 4 | 40956574 | rs7697609 | 0.886 | 0.462 | 0.158 | -1.083 | 0.015 | -0.097 | -0.097 | 0.185 | 0.21 | 0.3 | 0.13 | 0.02 | 0.21 | 0.11 |
| 4 | 40958368 | rs6832948 | 0.058 | 0.562 | -0.33 | -0.747 | 0.001 | 0.002 | 0.002 | 0 | 0.07 | 0.09 | 0.1 | 0.11 | 0.09 | 0.02 |
| 4 | 40965913 | rs4861075 | 0.27 | 0.462 | -0.548 | -0.633 | -0.01 | -0.013 | -0.013 | 2.904 | 0.47 | 0.23 | 0.41 | 0.54 | 0.46 | 0.62 |
| 4 | 40967236 | rs6855534 | 0.236 | 0.493 | -0.741 | -1.188 | -0.006 | -0.005 | -0.005 | 0 | 0.23 | 0.22 | 0.15 | 0 | 0.24 | 0.12 |
| 4 | 40973476 | rs4501254 | 0.308 | 0.288 | -1.093 | -1.117 | 0.009 | 0.02 | 0.02 | 0.012 | 0.31 | 0.19 | 0.24 | 0.25 | 0.28 | 0.27 |
| 4 | 40977934 | rs12511584 | 0.945 | 0.372 | -1.118 | -0.769 | -0.005 | -0.008 | -0.008 | 3.098 | 0.13 | 0.04 | 0.16 | 0.15 | 0.14 | 0.09 |
| 4 | 40982744 | rs1032984 | 0.389 | 0.421 | -0.97 | -1.083 | -0.042 | 0.021 | 0.021 | 0 | 0.38 | 0.2 | 0.46 | 0.7 | 0.44 | 0.55 |
| 4 | 40987461 | rs11735241 | 0.147 | 0.516 | -0.632 | -1.164 | 0.007 | 0.024 | 0.024 | 0 | 0.39 | 0.31 | 0.29 | 0.28 | 0.37 | 0.29 |
| 4 | 40989048 | rs10938478 | 1.224 | 0.597 | -0.593 | -1.167 | 0 | 0.006 | 0.006 | 1.313 | 0.13 | 0.05 | 0.18 | 0.13 | 0.13 | 0.1 |
| 4 | 40989826 | rs1396110 | 1.429 | 0.441 | -0.507 | -1.088 | -0.098 | -0.139 | -0.139 | 0 | 0.26 | 0.05 | 0.34 | 0.54 | 0.35 | 0.43 |
| 4 | 40991885 | rs10024411 | -0.041 | 0.399 | 0.024 | -1.062 | 0.007 | 0.024 | 0.024 | 0 | 0.39 | 0.31 | 0.29 | 0.28 | 0.37 | 0.29 |
| 4 | 41003571 | rs6832976 | -0.116 | 0.427 | -0.221 | -1.009 | -0.005 | -0.027 | -0.027 | 0 | 0.06 | 0.12 | 0.08 | 0 | 0.07 | 0.01 |
| 4 | 41015899 | rs4861358 | 0.319 | 0.239 | -0.015 | -1.217 | 0.002 | -0.006 | -0.006 | 0 | 0.71 | 0.32 | 0.73 | 0.9 | 0.75 | 0.8 |
| 4 | 41020707 | rs6842478 | -0.29 | 0.316 | -0.205 | -1.204 | -0.025 | -0.086 | -0.086 | 0 | 0.76 | 0.57 | 0.77 | 0.9 | 0.77 | 0.81 |
| 4 | 41030604 | rs10033248 | 0.964 | 0.271 | -0.707 | -1.175 | -0.014 | -0.01 | -0.01 | 0 | 0.2 | 0.08 | 0.12 | 0.18 | 0.19 | 0.11 |
| 4 | 41056241 | rs11943259 | 0.419 | 0.655 | -0.535 | -1.272 | -0.004 | -0.025 | -0.025 | 0.096 | 0.23 | 0.2 | 0.2 | 0 | 0.3 | 0.18 |
| 4 | 41061049 | rs6831604 | 1.167 | 0.478 | -0.274 | -1.113 | -0.004 | 0.054 | 0.054 | 0 | 0.26 | 0.4 | 0.3 | 0.54 | 0.24 | 0.4 |
| 4 | 41063203 | rs1472869 | 0.156 | 0.32 | 0.251 | -1.206 | 0.009 | 0.012 | 0.012 | 0 | 0.65 | 0.34 | 0.59 | 0.71 | 0.73 | 0.71 |
| 4 | 41066388 | rs13130182 | 0.058 | 0.722 | -1.093 | -1.012 | 0.002 | 0.006 | 0.006 | 0 | 0.31 | 0.2 | 0.43 | 0.29 | 0.27 | 0.29 |
| 4 | 41070095 | rs10938518 | -0.212 | 0.964 | -1.115 | -0.843 | -0.056 | 0.019 | 0.019 | 2.414 | 0.35 | 0.22 | 0.33 | 0.25 | 0.45 | 0.26 |
| 4 | 41070449 | rs16852774 | 0.904 | 0.618 | -1.252 | -1.01 | 0.024 | 0.056 | 0.056 | 0 | 0.13 | 0.04 | 0.04 | 0 | 0.08 | 0.09 |
| 4 | 41074389 | rs13129535 | 0.646 | 0.645 | -0.857 | -1.036 | 0.022 | 0.06 | 0.06 | 0.099 | 0.51 | 0.65 | 0.62 | 0.72 | 0.46 | 0.6 |
| 4 | 41076288 | rs3098914 | 0.679 | 0.313 | -1.02 | -1.347 | -0.003 | -0.059 | -0.059 | 0.072 | 0.23 | 0.57 | 0.22 | 0.12 | 0.24 | 0.26 |
| 4 | 41076426 | rs7692276 | 1.367 | 0.338 | -1.116 | -1.289 | 0.02 | -0.096 | -0.096 | 0 | 0.28 | 0.06 | 0.34 | 0.6 | 0.22 | 0.34 |
| 4 | 41078183 | rs1508067 | 1.302 | 0.941 | -1.122 | -1.387 | -0.008 | -0.003 | -0.003 | 0 | 0.12 | 0.07 | 0.12 | 0.13 | 0.19 | 0.13 |
| 4 | 41080894 | rs7665863 | 1.327 | 0.904 | -1.119 | -1.439 | -0.017 | -0.003 | -0.003 | 0 | 0.12 | 0.06 | 0.13 | 0.13 | 0.19 | 0.13 |
| 4 | 41081976 | rs17527299 | 0.481 | 1.037 | -1.176 | -1.544 | -0.009 | 0.003 | 0.003 | 2.167 | 0.17 | 0.1 | 0.21 | 0.13 | 0.28 | 0.14 |
| 4 | 41091835 | rs9994615 | 0.348 | 0.885 | -0.929 | -1.625 | -0.011 | 0.003 | 0.003 | 0.136 | 0.31 | 0.3 | 0.25 | 0.25 | 0.35 | 0.25 |
| 4 | 41095181 | rs2661661 | 1.887 | 0.63 | -1.078 | -1.633 | -0.021 | -0.17 | -0.17 | 0 | 0.26 | 0.01 | 0.29 | 0.57 | 0.21 | 0.34 |
| 4 | 41099221 | rs10015784 | -0.332 | 0.814 | -0.961 | -1.832 | -0.007 | 0.004 | 0.004 | 0.681 | 0.37 | 0.39 | 0.35 | 0.28 | 0.45 | 0.3 |
| 4 | 41099333 | rs10031772 | 1.18 | 0.434 | -0.888 | -1.75 | 0 | 0 | 0 | 0.144 | 0.2 | 0.28 | 0.14 | 0.15 | 0.17 | 0.16 |
| 4 | 41104347 | rs7679772 | 1.029 | 0.37 | -0.836 | -1.759 | 0.02 | -0.081 | -0.081 | 0 | 0.22 | 0.5 | 0.22 | 0.1 | 0.26 | 0.26 |
| 4 | 41115429 | rs7655823 | -0.9 | 0.631 | -1.1 | -1.734 | 0.057 | 0.002 | 0.002 | 1.206 | 0.58 | 0.86 | 0.56 | 0.39 | 0.67 | 0.54 |
| 4 | 41122798 | rs10026868 | 1.155 | 0.843 | -1.039 | -1.548 | 0.001 | 0.002 | 0.002 | 0 | 0.19 | 0.23 | 0.13 | 0.12 | 0.16 | 0.12 |
| 4 | 41123834 | rs7656191 | -1.446 | 0.821 | -1.117 | -1.548 | 0.018 | -0.06 | -0.06 | 0 | 0.7 | 0.92 | 0.6 | 0.39 | 0.75 | 0.63 |
| 4 | 41131421 | rs12499130 | -0.632 | 0.754 | -0.36 | -0.736 | -0.014 | 0 | 0 | 0.403 | 0.3 | 0.26 | 0.25 | 0.25 | 0.32 | 0.24 |
| 4 | 41140233 | rs2880169 | 0.639 | 1.063 | -0.745 | -0.589 | -0.062 | -0.03 | -0.03 | 0 | 0.11 | 0.01 | 0.12 | 0.13 | 0.16 | 0.14 |
| 4 | 41144591 | rs2585590 | 0.385 | 0.634 | -0.424 | -0.66 | 0.001 | -0.004 | -0.004 | 0 | 0.58 | 0.56 | 0.65 | 0.77 | 0.56 | 0.72 |
| 4 | 41149316 | rs35125077 | -0.754 | 0.166 | -0.162 | -0.531 | 0.001 | 0.003 | 0.003 | 0 | 0.47 | 0.42 | 0.3 | 0.35 | 0.37 | 0.46 |
| 4 | 41151628 | rs2661665 | -0.425 | -0.028 | 0.538 | -0.617 | 0.001 | -0.018 | -0.018 | 0 | 0.36 | 0.5 | 0.27 | 0.3 | 0.33 | 0.31 |
| 4 | 41153134 | rs7688017 | -0.616 | 0.051 | 0.489 | -0.874 | 0.002 | 0.017 | 0.017 | 0 | 0.32 | 0.68 | 0.24 | 0.16 | 0.29 | 0.2 |
| 4 | 41155960 | rs2341855 | 0.281 | -0.069 | 0.297 | -0.74 | -0.02 | -0.068 | -0.068 | 0 | 0.14 | 0.01 | 0.1 | 0.28 | 0.13 | 0.22 |
| 4 | 41156138 | rs7674456 | -0.412 | -0.015 | 1.187 | -0.752 | 0.007 | -0.043 | -0.043 | 0 | 0.25 | 0.41 | 0.19 | 0.16 | 0.22 | 0.16 |
| 4 | 41158203 | rs6447675 | -0.333 | -0.005 | 1.21 | -0.747 | 0.007 | -0.036 | -0.036 | 1.173 | 0.24 | 0.38 | 0.18 | 0.16 | 0.22 | 0.16 |
| 4 | 41160956 | rs7685881 | -0.245 | 0.076 | 1.225 | -0.75 | -0.007 | -0.027 | -0.027 | 0 | 0.23 | 0.36 | 0.16 | 0.17 | 0.19 | 0.16 |
| 4 | 41175148 | rs7683197 | -0.942 | 0.032 | 1.585 | -0.836 | 0.035 | -0.024 | -0.024 | 0.526 | 0.09 | 0.38 | 0.05 | 0 | 0.07 | 0.04 |
| 4 | 41181161 | rs7659645 | -1.494 | -0.049 | 0.969 | -0.939 | 0.002 | 0.006 | 0.006 | 0 | 0.44 | 0.59 | 0.26 | 0.25 | 0.36 | 0.34 |
| 4 | 41186226 | rs10022718 | -0.959 | -0.027 | 0.87 | -0.923 | 0.002 | 0.007 | 0.007 | 0 | 0.37 | 0.33 | 0.31 | 0.19 | 0.3 | 0.25 |
| 4 | 41193039 | rs17443333 | -0.016 | 0.358 | -0.213 | -0.556 | -0.004 | 0.03 | 0.03 | 0 | 0.19 | 0.09 | 0.1 | 0.1 | 0.15 | 0.15 |
| 4 | 41201819 | rs10021613 | -1.045 | 0.349 | 0.222 | -0.493 | -0.007 | -0.029 | -0.029 | 0.889 | 0.17 | 0.43 | 0.18 | 0.11 | 0.14 | 0.08 |
| 4 | 41202789 | rs11723385 | 0.026 | 0.001 | -0.011 | -0.632 | 0.019 | -0.003 | -0.003 | 0 | 0.22 | 0.12 | 0.07 | 0.22 | 0.11 | 0.26 |
| 4 | 41203125 | rs7683746 | -0.673 | 0.03 | 0.097 | -0.742 | -0.006 | -0.057 | -0.057 | 1.732 | 0.17 | 0.42 | 0.18 | 0.06 | 0.13 | 0.08 |
| 4 | 41204395 | rs6835127 | -1.125 | 0.171 | -0.652 | -0.962 | -0.022 | 0.074 | 0.074 | 0.368 | 0.55 | 0.71 | 0.39 | 0.31 | 0.46 | 0.43 |
| 4 | 41204982 | rs4455451 | 1.268 | 0.154 | -0.619 | -0.972 | 0.023 | 0.027 | 0.027 | 0 | 0.15 | 0.04 | 0.04 | 0.07 | 0.05 | 0.11 |
| 4 | 41205240 | rs4634261 | 1.141 | 0.141 | -0.62 | -0.978 | -0.031 | -0.036 | -0.036 | 0.045 | 0.16 | 0.25 | 0.05 | 0.07 | 0.05 | 0.11 |
| 4 | 41205611 | rs12503156 | 1.276 | 0.307 | -1 | -0.787 | -0.021 | -0.052 | -0.052 | 0.014 | 0.22 | 0.11 | 0.18 | 0.35 | 0.25 | 0.32 |
| 4 | 41205759 | rs6842814 | -0.694 | 0.816 | -0.496 | -1.469 | -0.005 | -0.015 | -0.015 | 0.047 | 0.31 | 0.6 | 0.22 | 0.13 | 0.18 | 0.2 |
| 4 | 41206430 | rs7657549 | 0.025 | 1.012 | -0.415 | -1.363 | -0.007 | -0.017 | -0.017 | 0 | 0.28 | 0.36 | 0.24 | 0.21 | 0.24 | 0.26 |
| 4 | 41207624 | rs17443388 | 1.551 | 1.081 | -0.45 | -1.298 | -0.039 | -0.02 | -0.02 | 0 | 0.16 | 0.01 | 0.08 | 0.17 | 0.15 | 0.21 |
| 4 | 41212540 | rs7662463 | -0.953 | 1.045 | -0.365 | -1.297 | 0.003 | -0.024 | -0.024 | 0 | 0.38 | 0.55 | 0.35 | 0.23 | 0.41 | 0.32 |

| | | | | | | | | | | | | | | | | |
|---|---|---|---|---|---|---|---|---|---|---|---|---|---|---|---|---|
| 4 | 41212997 | rs10003861 | 0.374 | 1.29 | -0.312 | -1.008 | 0.004 | -0.011 | -0.011 | 0.403 | 0.1 | 0.31 | 0.16 | 0.03 | 0.08 | 0.05 |
| 4 | 41218909 | rs7695309 | 0.215 | 1.689 | -1.239 | -1.049 | -0.003 | 0.012 | 0.012 | 0 | 0.24 | 0.16 | 0.25 | 0.07 | 0.24 | 0.23 |
| 4 | 41219792 | rs6811777 | -0.219 | 1.864 | -1.192 | -0.446 | -0.013 | 0.021 | 0.021 | 3.091 | 0.19 | 0.08 | 0.09 | 0.18 | 0.2 | 0.23 |
| 4 | 41220288 | rs17528008 | -0.495 | 1.847 | -1.175 | -0.455 | -0.008 | 0.006 | 0.006 | 1.302 | 0.21 | 0.16 | 0.18 | 0.18 | 0.3 | 0.26 |
| 4 | 41220816 | rs17528036 | 0.154 | 0.891 | -1.635 | -0.24 | -0.006 | 0.027 | 0.027 | 2.692 | 0.17 | 0.04 | 0.08 | 0.13 | 0.16 | 0.2 |
| 4 | 41222917 | rs10938677 | -0.315 | 1.024 | -1.784 | -0.508 | -0.016 | 0.089 | 0.089 | 0 | 0.71 | 0.83 | 0.68 | 0.88 | 0.73 | 0.58 |
| 4 | 41224763 | rs11735673 | 0.077 | 0.868 | -2.446 | 0.439 | -0.025 | -0.11 | -0.11 | 0 | 0.14 | 0.01 | 0.3 | 0.45 | 0.12 | 0.21 |
| 4 | 41225831 | rs13101324 | 0.141 | 0.473 | -1.083 | 0.513 | -0.001 | -0.003 | -0.003 | 0 | 0.54 | 0.45 | 0.33 | 0.31 | 0.58 | 0.33 |
| 4 | 41225946 | rs11941433 | 0.141 | 0.46 | -1.089 | 0.507 | -0.001 | -0.003 | -0.003 | 0 | 0.54 | 0.45 | 0.33 | 0.31 | 0.58 | 0.33 |
| 4 | 41231007 | rs17528085 | 0.53 | 0.618 | -1.415 | -0.15 | -0.031 | 0.123 | 0.123 | 0.785 | 0.34 | 0.1 | 0.21 | 0.09 | 0.36 | 0.16 |
| 4 | 41247773 | rs10019509 | 0.854 | 0.895 | -1.027 | -0.373 | 0.005 | -0.012 | -0.012 | 0 | 0.18 | 0.19 | 0.14 | 0.11 | 0.22 | 0.07 |
| 4 | 41247782 | rs11944436 | 0.661 | 1.229 | -0.983 | -0.152 | -0.008 | -0.01 | -0.01 | 0 | 0.08 | 0.09 | 0.12 | 0.16 | 0.12 | 0.09 |
| 4 | 41254862 | rs7693827 | -1.763 | 0.924 | -0.131 | -0.312 | 0.001 | 0.008 | 0.008 | 0 | 0.6 | 0.62 | 0.63 | 0.8 | 0.61 | 0.47 |
| 4 | 41258978 | rs9321 | 0.679 | 0.718 | -0.031 | -0.171 | -0.01 | -0.014 | -0.014 | 0 | 0.08 | 0.06 | 0.12 | 0.16 | 0.12 | 0.09 |
| 4 | 41266295 | rs3775256 | -0.393 | 0.6 | 0.066 | -0.313 | 0.022 | -0.072 | -0.072 | 0 | 0.25 | 0.13 | 0.31 | 0.54 | 0.18 | 0.34 |
| 4 | 41269193 | rs10517003 | 1.337 | 0.645 | 0.162 | -0.578 | -0.011 | 0.011 | 0.011 | 0 | 0.17 | 0.13 | 0.13 | 0 | 0.21 | 0.05 |
| 4 | 41272188 | rs11728559 | 1.391 | 0.651 | 0.117 | -0.576 | -0.01 | 0.006 | 0.006 | 0 | 0.17 | 0.11 | 0.13 | 0 | 0.21 | 0.04 |
| 4 | 41281031 | rs7349641 | 0.819 | 0.683 | 0.621 | -0.456 | -0.003 | -0.004 | -0.004 | 0 | 0.11 | 0.16 | 0.05 | 0.07 | 0.12 | 0.09 |
| 4 | 41283279 | rs11945108 | 0.641 | 0.717 | 0.481 | -0.379 | -0.01 | -0.013 | -0.013 | 0 | 0.07 | 0.06 | 0.11 | 0.16 | 0.11 | 0.08 |
| 4 | 41291280 | rs4861393 | 0.677 | 0.444 | 0.72 | -0.542 | -0.01 | -0.007 | -0.007 | 0 | 0.12 | 0.15 | 0.04 | 0.09 | 0.06 | 0.1 |
| 4 | 41291573 | rs2219001 | -1.888 | 0.465 | 0.769 | -0.473 | 0.001 | -0.011 | -0.011 | 0 | 0.66 | 0.58 | 0.64 | 0.75 | 0.65 | 0.6 |
| 4 | 41297430 | rs12186184 | -1.939 | 0.034 | 0.759 | -0.533 | -0.047 | 0.026 | 0.026 | 0 | 0.65 | 0.85 | 0.6 | 0.69 | 0.57 | 0.6 |
| 4 | 41299492 | rs6828613 | -0.513 | -0.629 | 1.194 | -0.875 | -0.007 | -0.042 | -0.042 | 0 | 0.12 | 0.17 | 0.05 | 0 | 0.09 | 0.07 |
| 4 | 41305694 | rs11727269 | 0.321 | -0.575 | 1.059 | -0.828 | 0.003 | 0.036 | 0.036 | 0 | 0.23 | 0.15 | 0.19 | 0.13 | 0.23 | 0.12 |
| 4 | 41308656 | rs11724596 | -0.866 | -0.627 | 1.062 | -0.846 | -0.003 | -0.007 | -0.007 | 0.186 | 0.37 | 0.58 | 0.25 | 0.21 | 0.28 | 0.16 |
| 4 | 41319962 | rs11735648 | -1.423 | -0.704 | 1.009 | -0.974 | 0.036 | 0.078 | 0.078 | 0 | 0.46 | 0.72 | 0.32 | 0.24 | 0.38 | 0.26 |
| 4 | 41322070 | rs1377345 | -1.221 | -0.772 | 1.157 | -1.022 | 0.002 | -0.014 | -0.014 | 0 | 0.42 | 0.61 | 0.31 | 0.15 | 0.37 | 0.24 |
| 4 | 41322091 | rs16853063 | 0.222 | -0.783 | 1.16 | -1.023 | -0.028 | 0.002 | 0.002 | 0 | 0.09 | 0.18 | 0.03 | 0.09 | 0.03 | 0.03 |
| 4 | 41322398 | rs6838610 | -0.786 | -0.763 | 1.312 | -1.026 | -0.01 | -0.066 | -0.066 | 1.125 | 0.36 | 0.45 | 0.28 | 0.15 | 0.35 | 0.24 |
| 4 | 41330982 | rs16853108 | -0.738 | -0.734 | 0.963 | -0.998 | 0.002 | -0.008 | -0.008 | 0 | 0.15 | 0.14 | 0.07 | 0.01 | 0.13 | 0.07 |
| 4 | 41335884 | rs16853120 | 0.482 | -0.713 | 1.109 | -0.933 | -0.024 | 0.013 | 0.013 | 0 | 0.08 | 0.2 | 0.03 | 0.09 | 0.03 | 0.03 |
| 4 | 41339317 | rs13128229 | -0.09 | -0.725 | 0.726 | -0.967 | 0 | -0.019 | -0.019 | 0 | 0.09 | 0.12 | 0.05 | 0.01 | 0.11 | 0.07 |
| 4 | 41340526 | rs9637557 | -1.718 | -0.71 | 0.406 | -0.933 | 0.022 | 0.047 | 0.047 | 0 | 0.49 | 0.62 | 0.33 | 0.25 | 0.4 | 0.29 |
| 4 | 41342264 | rs7696120 | -0.917 | -0.734 | 0.478 | -0.942 | -0.038 | -0.039 | -0.039 | 2.095 | 0.19 | 0.42 | 0.11 | 0.12 | 0.14 | 0.12 |
| 4 | 41343706 | rs6447057 | -1.568 | -0.692 | -0.123 | -0.922 | 0.057 | 0.146 | 0.146 | 0 | 0.51 | 0.79 | 0.34 | 0.25 | 0.4 | 0.29 |
| 4 | 41345419 | rs4279250 | -0.641 | -0.035 | -0.38 | -0.567 | 0.008 | 0.054 | 0.054 | 0 | 0.29 | 0.2 | 0.22 | 0.13 | 0.26 | 0.16 |
| 4 | 41346808 | rs11721727 | -0.345 | 0.357 | -0.363 | -0.323 | 0.001 | 0.057 | 0.057 | 0 | 0.25 | 0.14 | 0.2 | 0.13 | 0.24 | 0.17 |
| 4 | 41359624 | rs11728170 | 0.818 | 0.37 | -0.537 | -0.278 | -0.002 | -0.005 | -0.005 | 0 | 0.13 | 0.11 | 0.09 | 0 | 0.13 | 0.03 |
| 4 | 41359965 | rs6834482 | -0.373 | 0.319 | -0.907 | 0.482 | 0.003 | -0.012 | -0.012 | 0.08 | 0.41 | 0.64 | 0.51 | 0.51 | 0.35 | 0.52 |
| 4 | 41361984 | rs7685339 | -0.021 | 0.315 | -1.121 | -0.113 | 0.003 | 0.031 | 0.031 | 0.024 | 0.06 | 0.16 | 0.03 | 0.09 | 0.05 | 0.02 |
| 4 | 41362506 | rs4860991 | -1.401 | 0.464 | -1.807 | -0.117 | -0.112 | -0.149 | -0.149 | 0.289 | 0.6 | 0.96 | 0.44 | 0.54 | 0.57 | 0.68 |
| 4 | 41367089 | rs2068783 | -0.734 | 0.252 | -1.344 | 0.141 | -0.004 | -0.024 | -0.024 | 0 | 0.47 | 0.84 | 0.37 | 0.5 | 0.39 | 0.53 |
| 4 | 41373402 | rs1869286 | -1.211 | -0.372 | -0.637 | -0.318 | 0.01 | 0.036 | 0.036 | 0 | 0.3 | 0.2 | 0.28 | 0.14 | 0.26 | 0.27 |
| 4 | 41376347 | rs1901747 | -0.39 | -0.455 | -0.747 | -0.373 | -0.034 | 0.049 | 0.049 | 0 | 0.2 | 0.02 | 0.23 | 0.07 | 0.2 | 0.17 |
| 4 | 41377619 | rs2342561 | -1.539 | -0.447 | -0.523 | -0.364 | 0.007 | 0.022 | 0.022 | 0 | 0.3 | 0.24 | 0.29 | 0.14 | 0.27 | 0.27 |
| 4 | 41378712 | rs11735207 | -2.022 | -0.749 | -0.018 | -0.329 | 0.058 | 0.074 | 0.074 | 0 | 0.47 | 0.73 | 0.41 | 0.41 | 0.37 | 0.43 |
| 4 | 41406130 | rs13140742 | -0.893 | -0.684 | 0.602 | -0.466 | 0.013 | 0.037 | 0.037 | 0 | 0.23 | 0.14 | 0.23 | 0.06 | 0.21 | 0.22 |
| 4 | 41406368 | rs13142854 | 0.703 | -0.639 | 1.28 | -0.671 | 0.002 | -0.061 | -0.061 | 0 | 0.89 | 0.77 | 0.93 | 0.99 | 0.92 | 0.93 |
| 4 | 41406510 | rs13118239 | -1.427 | -0.734 | 1.826 | -0.649 | -0.029 | -0.037 | -0.037 | 0.592 | 0.28 | 0.47 | 0.3 | 0.17 | 0.23 | 0.43 |
| 4 | 41406966 | rs6447065 | 0.225 | -0.736 | 1.801 | -0.654 | 0.006 | -0.051 | -0.051 | 3.646 | 0.91 | 0.77 | 0.94 | 0.99 | 0.93 | 0.95 |
| 4 | 41407668 | rs6447066 | 1.016 | -0.792 | 1.695 | -0.671 | 0.004 | -0.076 | -0.076 | 0.022 | 0.88 | 0.63 | 0.92 | 0.99 | 0.92 | 0.93 |
| 4 | 41411424 | rs12507919 | 1.939 | -0.756 | 0.956 | -0.636 | 0.045 | 0.132 | 0.132 | 1.617 | 0.59 | 0.14 | 0.62 | 0.82 | 0.69 | 0.52 |
| 4 | 41411891 | rs10938151 | 1.939 | -0.756 | 0.955 | -0.624 | 0.047 | 0.123 | 0.123 | 1.617 | 0.59 | 0.17 | 0.62 | 0.82 | 0.69 | 0.52 |
| 4 | 41421908 | rs4579156 | -0.362 | -0.699 | 0.901 | -0.577 | 0.013 | 0.039 | 0.039 | 0 | 0.18 | 0.07 | 0.21 | 0.07 | 0.16 | 0.21 |
| 4 | 41430052 | rs7675059 | -1.138 | -0.733 | 1.602 | -0.968 | -0.041 | -0.112 | -0.112 | 0 | 0.24 | 0.47 | 0.26 | 0.05 | 0.17 | 0.35 |
| 4 | 41443113 | rs4527518 | -1.37 | -0.719 | 1.581 | -0.945 | -0.006 | -0.016 | -0.016 | 0 | 0.18 | 0.38 | 0.12 | 0.11 | 0.15 | 0.14 |
| 4 | 41447510 | rs4560447 | -1.755 | -1.016 | 1.508 | -0.922 | 0.079 | 0.08 | 0.08 | 8.934 | 0.06 | 0.47 | 0.03 | 0.02 | 0.01 | 0.14 |
| 4 | 41448217 | rs11737555 | -0.707 | -0.937 | 0.526 | -0.911 | -0.002 | -0.016 | -0.016 | 0 | 0.2 | 0.18 | 0.22 | 0.05 | 0.16 | 0.22 |
| 4 | 41448957 | rs7679031 | -0.668 | -0.837 | -0.093 | -0.895 | 0.005 | 0.061 | 0.061 | 0 | 0.19 | 0.07 | 0.22 | 0.05 | 0.18 | 0.23 |
| 4 | 41457735 | rs7665984 | -0.913 | -1.668 | -0.818 | -1.242 | 0.079 | 0.251 | 0.251 | 2.486 | 0.51 | 0.89 | 0.39 | 0.18 | 0.4 | 0.44 |
| 4 | 41457923 | rs7696363 | -1.546 | -1.931 | 0.949 | -0.996 | -0.052 | -0.025 | -0.025 | 0 | 0.19 | 0.51 | 0.11 | 0.15 | 0.14 | 0.14 |
| 4 | 41462412 | rs4507392 | -1.388 | -2.29 | 1.088 | -1.151 | -0.058 | -0.061 | -0.061 | 0.093 | 0.29 | 0.35 | 0.17 | 0.13 | 0.16 | 0.4 |
| 4 | 41464663 | rs10470738 | -0.951 | -2.304 | 0.904 | -1.137 | 0.049 | 0.067 | 0.067 | 0 | 0.24 | 0.11 | 0.14 | 0.06 | 0.14 | 0.26 |
| 4 | 41472234 | rs6852190 | -2.44 | -2.33 | 0.86 | -1.185 | 0.121 | 0.118 | 0.118 | 3.104 | 0.52 | 0.85 | 0.31 | 0.3 | 0.36 | 0.55 |
| 4 | 41485212 | rs4349633 | -2.924 | -2.2 | 1.123 | -0.994 | 0.154 | 0.113 | 0.113 | 0.104 | 0.49 | 0.88 | 0.32 | 0.31 | 0.31 | 0.56 |
| 4 | 41488228 | rs13133735 | -0.989 | -2.224 | 1.361 | -1.01 | 0.017 | 0.016 | 0.016 | 0 | 0.23 | 0.17 | 0.15 | 0.11 | 0.11 | 0.26 |
| 4 | 41490455 | rs7696430 | -1.486 | -1.864 | 1.962 | -0.845 | 0.005 | -0.034 | -0.034 | 0 | 0.16 | 0.35 | 0.11 | 0.07 | 0.16 | 0.19 |
| 4 | 41496140 | rs13140281 | -1.912 | -1.906 | 1.909 | -0.828 | 0.002 | 0.003 | 0.003 | 0 | 0.29 | 0.25 | 0.19 | 0.24 | 0.15 | 0.34 |
| 4 | 41499931 | rs7689372 | -2.557 | -1.793 | 1.816 | -0.758 | 0.066 | 0.032 | 0.032 | 0.021 | 0.43 | 0.62 | 0.27 | 0.31 | 0.25 | 0.53 |

| | | | | | | | | | | | | | | | | |
|---|---|---|---|---|---|---|---|---|---|---|---|---|---|---|---|---|
| 4 | 41513423 | rs4392535 | -1.847 | -1.573 | 1.653 | -0.865 | 0.022 | 0.029 | 0.029 | 0 | 0.41 | 0.56 | 0.31 | 0.17 | 0.28 | 0.44 |
| 4 | 41521194 | rs7671360 | -2.754 | -1.579 | 1.53 | -0.853 | 0.033 | 0.082 | 0.082 | 0.954 | 0.52 | 0.91 | 0.4 | 0.3 | 0.39 | 0.55 |
| 4 | 41522232 | rs12696762 | -1.93 | -1.456 | 1.784 | -0.669 | -0.02 | -0.027 | -0.027 | 0 | 0.39 | 0.48 | 0.22 | 0.17 | 0.25 | 0.43 |
| 4 | 41531404 | rs6447084 | -1.825 | -1.308 | 2.101 | -0.714 | -0.035 | -0.054 | -0.054 | 1.933 | 0.36 | 0.43 | 0.19 | 0.12 | 0.23 | 0.45 |
| 4 | 41532328 | rs6447085 | -2.524 | -1.224 | 1.426 | -0.639 | 0.043 | 0.08 | 0.08 | 0.887 | 0.45 | 0.9 | 0.37 | 0.27 | 0.34 | 0.52 |
| 4 | 41543227 | rs4293818 | -2.038 | -1.385 | 2.592 | -0.531 | -0.062 | 0.007 | 0.007 | 1.047 | 0.13 | 0.39 | 0.08 | 0.08 | 0.04 | 0.18 |
| 4 | 41548580 | rs4455452 | -2.96 | -0.841 | 2.07 | -0.695 | 0.037 | 0.093 | 0.093 | 2.704 | 0.33 | 0.84 | 0.26 | 0.11 | 0.21 | 0.45 |
| 4 | 41551813 | rs6447091 | -1.424 | -0.818 | 2.442 | -0.677 | -0.004 | -0.02 | -0.02 | 0 | 0.12 | 0.18 | 0.06 | 0.04 | 0.08 | 0.18 |
| 4 | 41554364 | rs10007810 | -3.359 | -0.917 | 2.24 | -0.709 | 0.011 | 0.101 | 0.101 | 1.579 | 0.33 | 0.96 | 0.26 | 0.11 | 0.21 | 0.45 |
| 4 | 41556138 | rs7656019 | -3.145 | -0.815 | 2.892 | -0.651 | 0.024 | 0.043 | 0.043 | 0.502 | 0.29 | 0.81 | 0.17 | 0.08 | 0.17 | 0.41 |
| 4 | 41558580 | rs4308395 | -1.763 | -0.886 | 2.943 | -0.652 | -0.006 | -0.026 | -0.026 | 0 | 0.13 | 0.21 | 0.06 | 0.05 | 0.08 | 0.19 |
| 4 | 41572048 | rs6850988 | -2.105 | -0.858 | 3.082 | -0.613 | -0.03 | -0.077 | -0.077 | 0.179 | 0.25 | 0.48 | 0.14 | 0.09 | 0.16 | 0.39 |
| 4 | 41585324 | rs4540083 | -2.609 | -0.874 | 2.621 | -0.621 | 0.049 | 0.096 | 0.096 | 0.565 | 0.29 | 0.88 | 0.18 | 0.09 | 0.17 | 0.4 |
| 4 | 41594774 | rs11729552 | -2.036 | -0.867 | 2.663 | -0.618 | -0.023 | -0.068 | -0.068 | 0 | 0.23 | 0.39 | 0.13 | 0.09 | 0.16 | 0.38 |
| 4 | 41599965 | rs10029589 | -2.311 | -0.776 | 1.386 | -0.618 | 0.048 | 0.085 | 0.085 | 0.709 | 0.31 | 0.88 | 0.18 | 0.11 | 0.18 | 0.41 |
| 4 | 41611575 | rs1105886 | 1.51 | -0.293 | 1.493 | -0.365 | 0.014 | -0.048 | -0.048 | 0 | 0.74 | 0.57 | 0.8 | 0.84 | 0.76 | 0.53 |
| 4 | 41612137 | rs10938153 | 2.089 | -0.355 | 1.046 | -0.249 | -0.037 | -0.009 | -0.009 | 0 | 0.66 | 0.05 | 0.73 | 0.69 | 0.75 | 0.48 |
| 4 | 41613931 | rs10026359 | 0.672 | 0.014 | 1.325 | -0.581 | 0.006 | -0.015 | -0.015 | 0 | 0.81 | 0.66 | 0.89 | 0.84 | 0.85 | 0.6 |
| 4 | 41619643 | rs4860998 | 1.096 | -0.014 | -0.086 | -0.472 | 0 | -0.004 | -0.004 | 0 | 0.72 | 0.36 | 0.77 | 0.84 | 0.76 | 0.53 |
| 4 | 41635034 | rs7657178 | -1.193 | -0.723 | 0.246 | -0.578 | -0.067 | -0.06 | -0.06 | 0.056 | 0.2 | 0.62 | 0.09 | 0.04 | 0.05 | 0.12 |
| 4 | 41639999 | rs6830246 | -0.782 | -0.443 | 0.448 | -1.384 | 0.001 | -0.083 | -0.083 | 0 | 0.27 | 0.34 | 0.16 | 0.03 | 0.26 | 0.34 |
| 4 | 41642833 | rs4861123 | -1.2 | -0.034 | 0.369 | -1.252 | 0.005 | -0.028 | -0.028 | 0.169 | 0.39 | 0.42 | 0.46 | 0.2 | 0.4 | 0.48 |
| 4 | 41645980 | rs7694860 | -1.06 | -0.276 | 0.762 | -1.38 | -0.005 | -0.035 | -0.035 | 0 | 0.07 | 0.19 | 0.03 | 0 | 0.03 | 0.02 |
| 4 | 41652442 | rs16853408 | -1.096 | -0.29 | 0.744 | -1.388 | -0.004 | -0.035 | -0.035 | 0 | 0.07 | 0.18 | 0.03 | 0 | 0.03 | 0.02 |
| 4 | 41656630 | rs6811377 | -1.365 | -0.095 | 0.603 | -1.872 | -0.001 | 0.088 | 0.088 | 0 | 0.44 | 0.72 | 0.38 | 0.14 | 0.42 | 0.45 |
| 4 | 41659486 | rs6821712 | -0.761 | -0.166 | 0.959 | -2.087 | 0.021 | -0.099 | -0.099 | 4.418 | 0.27 | 0.56 | 0.19 | 0.03 | 0.31 | 0.31 |
| 4 | 41659835 | rs17590135 | -0.096 | -0.207 | 0.963 | -2.108 | 0.031 | -0.005 | -0.005 | 0 | 0.13 | 0.04 | 0.15 | 0.11 | 0.07 | 0.13 |
| 4 | 41663002 | rs7349706 | -0.782 | -0.187 | 1.24 | -2.222 | 0.071 | -0.114 | -0.114 | 0 | 0.25 | 0.4 | 0.2 | 0.03 | 0.35 | 0.3 |
| 4 | 41663621 | rs17528897 | -1.913 | -0.873 | 0.4 | -3.025 | -0.012 | 0.032 | 0.032 | 0 | 0.6 | 0.63 | 0.47 | 0.14 | 0.55 | 0.48 |
| 4 | 41664647 | rs2304649 | -0.336 | -0.873 | 0.5 | -3.02 | -0.02 | -0.003 | -0.003 | 0 | 0.05 | 0.17 | 0.02 | 0.02 | 0.01 | 0.03 |
| 4 | 41666173 | rs10006144 | 0.052 | -0.842 | 0.818 | -3.033 | 0.028 | -0.078 | -0.078 | 0 | 0.22 | 0.27 | 0.19 | 0.01 | 0.31 | 0.28 |
| 4 | 41669804 | rs7671106 | -2.068 | -0.885 | 0.745 | -3.012 | -0.03 | 0.086 | 0.086 | 0 | 0.61 | 0.76 | 0.48 | 0.13 | 0.55 | 0.48 |
| 4 | 41673604 | rs11734372 | -2.158 | -0.909 | 0.769 | -3.014 | -0.011 | 0.029 | 0.029 | 0 | 0.62 | 0.69 | 0.48 | 0.14 | 0.57 | 0.49 |
| 4 | 41677213 | rs7677377 | -0.752 | -1.088 | 0.825 | -3.005 | -0.039 | -0.025 | -0.025 | 0 | 0.19 | 0.31 | 0.18 | 0.09 | 0.08 | 0.14 |
| 4 | 41679483 | rs6837759 | -2.087 | -1.044 | 0.789 | -3.011 | -0.01 | 0.047 | 0.047 | 0 | 0.61 | 0.69 | 0.5 | 0.14 | 0.58 | 0.48 |
| 4 | 41681742 | rs2304651 | -0.243 | -1.634 | 0.865 | -2.573 | 0.022 | 0.011 | 0.011 | 0 | 0.15 | 0.13 | 0.15 | 0.09 | 0.07 | 0.13 |
| 4 | 41683212 | rs6816179 | -0.809 | -1.671 | 0.838 | -2.571 | 0.02 | 0.021 | 0.021 | 0.004 | 0.27 | 0.24 | 0.25 | 0.12 | 0.2 | 0.1 |
| 4 | 41687911 | rs2304652 | -1.773 | -1.684 | 0.781 | -2.113 | -0.021 | -0.038 | -0.038 | 2.777 | 0.41 | 0.46 | 0.32 | 0.12 | 0.29 | 0.2 |
| 4 | 41691601 | rs1061597 | 0.113 | -1.616 | 0.538 | -2.148 | 0.006 | 0.116 | 0.116 | 0 | 0.19 | 0.02 | 0.13 | 0 | 0.17 | 0.05 |
| 4 | 41700107 | rs7674006 | -0.057 | -1.672 | 0.866 | -2.087 | -0.026 | -0.018 | -0.018 | 0 | 0.13 | 0.19 | 0.08 | 0.07 | 0.06 | 0.1 |
| 4 | 41700361 | rs1061562 | -1.083 | -1.655 | 0.568 | -2.095 | 0.029 | 0.031 | 0.031 | 0.231 | 0.34 | 0.29 | 0.2 | 0.07 | 0.22 | 0.16 |
| 4 | 41707093 | rs12650313 | -0.985 | -0.925 | 0.454 | -1.587 | -0.038 | -0.007 | -0.007 | 0 | 0.18 | 0.28 | 0.18 | 0.13 | 0.1 | 0.12 |
| 4 | 41710577 | rs1439471 | -1.181 | -0.437 | 0.414 | -1.703 | 0.002 | 0.002 | 0.002 | 0 | 0.39 | 0.64 | 0.35 | 0.21 | 0.31 | 0.29 |
| 4 | 41711689 | rs6817913 | -1.371 | 0.127 | 0.885 | -1.65 | 0.009 | -0.019 | -0.019 | 0.16 | 0.21 | 0.44 | 0.13 | 0.19 | 0.17 | 0.23 |
| 4 | 41712381 | rs6847601 | -2.389 | 0.391 | 1.445 | -2.004 | 0.002 | -0.043 | -0.043 | 0 | 0.34 | 0.6 | 0.24 | 0.1 | 0.27 | 0.41 |
| 4 | 41714272 | rs9990806 | -2.233 | 0.534 | 1.557 | -2.003 | 0 | -0.05 | -0.05 | 0 | 0.32 | 0.58 | 0.2 | 0.1 | 0.26 | 0.4 |
| 4 | 41714558 | rs4861003 | -0.974 | 0.653 | 1.689 | -2.018 | -0.002 | -0.031 | -0.031 | 0 | 0.13 | 0.23 | 0.1 | 0.03 | 0.1 | 0.2 |
| 4 | 41714627 | rs1371301 | -1.174 | 0.471 | 1.689 | -2.166 | 0.023 | -0.06 | -0.06 | 0.018 | 0.19 | 0.36 | 0.11 | 0.06 | 0.17 | 0.21 |
| 4 | 41718139 | rs2043905 | -0.919 | 0.427 | 1.566 | -2.1 | -0.007 | -0.047 | -0.047 | 0 | 0.12 | 0.21 | 0.09 | 0.01 | 0.08 | 0.19 |
| 4 | 41730976 | rs16853557 | -2.523 | 0.442 | 1.823 | -2.147 | -0.004 | -0.078 | -0.078 | 0.002 | 0.24 | 0.55 | 0.19 | 0.1 | 0.18 | 0.27 |
| 4 | 41733493 | rs17529323 | 0.287 | 0.499 | 1.808 | -2.125 | 0.01 | -0.033 | -0.033 | 1.615 | 0.07 | 0.12 | 0.05 | 0 | 0.08 | 0.04 |
| 4 | 41736897 | rs11731281 | -1.644 | 0.47 | 1.957 | -2.121 | 0.014 | -0.128 | -0.128 | 0.05 | 0.21 | 0.44 | 0.18 | 0 | 0.18 | 0.24 |
| 4 | 41737080 | rs11733037 | -0.167 | 0.459 | 1.754 | -2.071 | 0.006 | -0.005 | -0.005 | 0 | 0.11 | 0.12 | 0.05 | 0.07 | 0.1 | 0.17 |
| 4 | 41737309 | rs17444900 | -0.302 | 0.457 | 1.754 | -2.08 | 0.007 | -0.008 | -0.008 | 0 | 0.11 | 0.16 | 0.06 | 0.07 | 0.1 | 0.17 |
| 4 | 41746460 | rs6826373 | -2.898 | 0.565 | 1.374 | -2.176 | -0.004 | 0.069 | 0.069 | 0.259 | 0.39 | 0.76 | 0.26 | 0.18 | 0.33 | 0.43 |
| 4 | 41753130 | rs16853571 | -0.312 | 0.351 | 1.607 | -2.185 | -0.006 | -0.042 | -0.042 | 0 | 0.08 | 0.21 | 0.07 | 0 | 0.05 | 0.17 |
| 4 | 41758803 | rs9283670 | -0.417 | 0.337 | 1.606 | -2.209 | -0.001 | -0.045 | -0.045 | 0.019 | 0.13 | 0.17 | 0.07 | 0 | 0.13 | 0.07 |
| 4 | 41763403 | rs1122992 | -3.135 | 0.306 | 1.666 | -2.251 | 0.007 | 0.063 | 0.063 | 0.114 | 0.39 | 0.73 | 0.26 | 0.17 | 0.31 | 0.44 |
| 4 | 41764707 | rs11729701 | -1.042 | 0.316 | 1.746 | -2.23 | 0.006 | 0.02 | 0.02 | 0 | 0.15 | 0.27 | 0.11 | 0.16 | 0.11 | 0.2 |
| 4 | 41766595 | rs6842806 | -2.303 | 0.314 | 1.753 | -2.233 | 0 | -0.045 | -0.045 | 0 | 0.28 | 0.49 | 0.18 | 0.16 | 0.23 | 0.26 |
| 4 | 41771365 | rs1439475 | -0.617 | 0.329 | 1.691 | -2.25 | 0.009 | -0.016 | -0.016 | 0 | 0.13 | 0.19 | 0.09 | 0.07 | 0.11 | 0.18 |
| 4 | 41774747 | rs5020056 | -0.477 | 0.337 | 1.62 | -2.231 | 0 | -0.051 | -0.051 | 0 | 0.13 | 0.19 | 0.07 | 0 | 0.13 | 0.07 |
| 4 | 41782996 | rs9999953 | -0.83 | 0.384 | 1.639 | -2.199 | 0.004 | -0.06 | -0.06 | 0 | 0.11 | 0.28 | 0.08 | 0.01 | 0.08 | 0.17 |
| 4 | 41786214 | rs1540499 | -2.504 | 0.371 | 1.648 | -2.247 | -0.004 | -0.026 | -0.026 | 0.2 | 0.26 | 0.57 | 0.19 | 0.16 | 0.19 | 0.39 |
| 4 | 41787858 | rs11931088 | -1.558 | 0.416 | 1.519 | -2.161 | 0.016 | 0.012 | 0.012 | 0.05 | 0.17 | 0.32 | 0.11 | 0.16 | 0.14 | 0.21 |
| 4 | 41790191 | rs1439483 | -2.261 | 0.272 | 0.766 | -1.958 | 0.004 | 0.091 | 0.091 | 0.411 | 0.51 | 0.7 | 0.45 | 0.27 | 0.43 | 0.47 |
| 4 | 41790600 | rs9997912 | -2.252 | 0.295 | 0.69 | -2.248 | 0.006 | 0.126 | 0.126 | 0.468 | 0.51 | 0.73 | 0.44 | 0.2 | 0.43 | 0.47 |
| 4 | 41791492 | rs1154598 | -2.536 | 0.404 | 0.576 | -2.42 | 0.007 | 0.114 | 0.114 | 0.088 | 0.49 | 0.65 | 0.43 | 0.11 | 0.45 | 0.46 |
| 4 | 41793898 | rs1898889 | -0.874 | 0.455 | 0.364 | -2.359 | 0.011 | -0.014 | -0.014 | 0 | 0.13 | 0.19 | 0.12 | 0.07 | 0.11 | 0.16 |

| | | | | | | | | | | | | | | | | |
|---|---|---|---|---|---|---|---|---|---|---|---|---|---|---|---|---|
| 4 | 41799127 | rs6822112 | -0.86 | 0.288 | -0.444 | -2.302 | 0.009 | 0.055 | 0.055 | 0.472 | 0.3 | 0.2 | 0.19 | 0.08 | 0.26 | 0.36 |
| 4 | 41801286 | rs1371307 | -0.587 | 0.284 | -0.648 | -2.455 | -0.012 | 0.04 | 0.04 | 0 | 0.19 | 0.08 | 0.12 | 0 | 0.24 | 0.14 |
| 4 | 41805686 | rs6834302 | -0.861 | 0.07 | -0.6 | -2.603 | 0.016 | 0.087 | 0.087 | 0 | 0.41 | 0.25 | 0.32 | 0.16 | 0.39 | 0.32 |
| 4 | 41807737 | rs2043904 | -1.272 | 0.144 | 0.195 | -2.649 | 0.014 | 0.121 | 0.121 | 0 | 0.48 | 0.69 | 0.38 | 0.18 | 0.44 | 0.43 |
| 4 | 41812109 | rs7664468 | -0.705 | 0.089 | 0.752 | -2.425 | 0.022 | 0.033 | 0.033 | 2.09 | 0.46 | 0.64 | 0.25 | 0.08 | 0.33 | 0.28 |
| 4 | 41819608 | rs733778 | -0.427 | 0.215 | 0.532 | -2.366 | 0.025 | 0.197 | 0.197 | 2.879 | 0.42 | 0.2 | 0.23 | 0.08 | 0.33 | 0.29 |
| 4 | 41826336 | rs714226 | -1.599 | 0.649 | 1.443 | -1.322 | -0.027 | -0.087 | -0.087 | 0.115 | 0.15 | 0.7 | 0.11 | 0.01 | 0.07 | 0.12 |
| 4 | 41834610 | rs9998239 | -2.055 | 0.739 | 1.744 | -1.38 | -0.018 | -0.039 | -0.039 | 0 | 0.17 | 0.75 | 0.13 | 0.05 | 0.09 | 0.14 |
| 4 | 41836529 | rs4388122 | -1.745 | 0.724 | 2.045 | -1.388 | -0.03 | -0.096 | -0.096 | 0 | 0.15 | 0.65 | 0.12 | 0.02 | 0.08 | 0.14 |
| 4 | 41855573 | rs6447118 | -2.298 | 0.851 | 1.952 | -1.44 | 0.005 | 0.026 | 0.026 | 1.602 | 0.21 | 0.82 | 0.19 | 0.05 | 0.19 | 0.17 |
| 4 | 41882964 | rs4861149 | -2.163 | 0.49 | 1.825 | -1.559 | 0 | -0.011 | -0.011 | 0.213 | 0.14 | 0.15 | 0.08 | 0.03 | 0.12 | 0.08 |
| 4 | 41883483 | rs9993759 | -2.962 | 0.292 | 2.106 | -1.928 | 0.021 | 0.024 | 0.024 | 1.737 | 0.34 | 0.72 | 0.18 | 0.05 | 0.25 | 0.2 |
| 4 | 41889237 | rs6821891 | -1.346 | 0.28 | 2.532 | -1.916 | -0.032 | -0.137 | -0.137 | 1.715 | 0.17 | 0.47 | 0.1 | 0.02 | 0.12 | 0.12 |
| 4 | 41889667 | rs7660832 | -2.815 | 0.254 | 2.391 | -1.92 | 0.05 | 0.067 | 0.067 | 3.882 | 0.32 | 0.79 | 0.16 | 0.05 | 0.2 | 0.18 |
| 4 | 41892087 | rs10517012 | -1.669 | 0.186 | 2.44 | -1.861 | -0.018 | -0.033 | -0.033 | 0.191 | 0.13 | 0.2 | 0.05 | 0.03 | 0.07 | 0.06 |
| 4 | 41901368 | rs1983138 | -2.307 | 0.275 | 2.497 | -1.888 | 0.002 | -0.042 | -0.042 | 0 | 0.25 | 0.72 | 0.14 | 0.02 | 0.19 | 0.15 |
| 4 | 41933626 | rs4861151 | -2.906 | 0.342 | 2.69 | -1.899 | -0.01 | -0.075 | -0.075 | 6.301 | 0.16 | 0.34 | 0.11 | 0.04 | 0.12 | 0.11 |
| 4 | 41940132 | rs16853775 | -3.402 | 0.338 | 2.766 | -1.941 | -0.022 | -0.114 | -0.114 | 0 | 0.18 | 0.48 | 0.12 | 0.03 | 0.12 | 0.11 |
| 4 | 41956313 | rs6830514 | -1.516 | 0.336 | 2.746 | -1.941 | -0.015 | -0.027 | -0.027 | 0.557 | 0.1 | 0.14 | 0.05 | 0.02 | 0.05 | 0.05 |
| 4 | 41956413 | rs1507086 | 2.928 | 0.317 | 2.586 | -1.959 | 0.018 | 0.273 | 0.273 | 4.101 | 0.62 | 0.06 | 0.75 | 0.95 | 0.74 | 0.78 |
| 4 | 41965136 | rs11942311 | -3.327 | 0.303 | 2.65 | -1.927 | -0.021 | -0.108 | -0.108 | 0.155 | 0.18 | 0.51 | 0.12 | 0.03 | 0.12 | 0.11 |
| 4 | 41977766 | rs2660343 | 2.605 | 0.295 | 2.197 | -1.925 | 0.063 | 0.143 | 0.143 | 3.141 | 0.65 | 0.17 | 0.77 | 0.97 | 0.76 | 0.79 |
| 4 | 41984118 | rs2660320 | 2.472 | 0.288 | 2.097 | -1.919 | 0.06 | 0.134 | 0.134 | 2.385 | 0.66 | 0.15 | 0.8 | 0.96 | 0.78 | 0.82 |
| 4 | 41985101 | rs2660332 | 2.468 | 0.287 | 2.117 | -1.919 | 0.059 | 0.129 | 0.129 | 2.423 | 0.66 | 0.15 | 0.8 | 0.96 | 0.78 | 0.83 |
| 4 | 41985689 | rs2660335 | 2.473 | 0.288 | 2.139 | -1.918 | 0.058 | 0.124 | 0.124 | 2.456 | 0.66 | 0.15 | 0.8 | 0.96 | 0.78 | 0.83 |
| 4 | 41986857 | rs2581439 | 2.387 | 0.286 | 2.26 | -1.919 | 0.029 | 0.029 | 0.029 | 2.336 | 0.67 | 0.25 | 0.8 | 0.96 | 0.78 | 0.83 |
| 4 | 41987892 | rs2581438 | 1.817 | 0.266 | 2.351 | -1.932 | -0.015 | -0.072 | -0.072 | 2.764 | 0.75 | 0.34 | 0.84 | 0.96 | 0.82 | 0.86 |
| 4 | 41992458 | rs16853864 | -1.277 | 0.288 | 2.256 | -1.914 | -0.004 | -0.019 | -0.019 | 3.227 | 0.08 | 0.08 | 0.04 | 0 | 0.04 | 0.04 |
| 4 | 42003671 | rs1047626 | 2.612 | 0.282 | 2.237 | -1.927 | 0.063 | 0.196 | 0.196 | 3.443 | 0.64 | 0.11 | 0.77 | 0.95 | 0.76 | 0.78 |
| 4 | 42006057 | rs2581451 | 2.691 | 0.277 | 2.229 | -1.929 | 0.061 | 0.234 | 0.234 | 3.877 | 0.64 | 0.09 | 0.77 | 0.95 | 0.76 | 0.77 |
| 4 | 42011970 | rs2581441 | 2.662 | 0.276 | 2.229 | -1.929 | 0.059 | 0.231 | 0.231 | 3.674 | 0.64 | 0.09 | 0.77 | 0.95 | 0.76 | 0.77 |
| 4 | 42022464 | rs15857 | 2.398 | 0.378 | 2.309 | -1.886 | 0.038 | 0.072 | 0.072 | 1.999 | 0.68 | 0.19 | 0.8 | 0.96 | 0.78 | 0.8 |
| 4 | 42031437 | rs3827590 | 2.444 | 0.7 | 2.284 | -1.879 | 0.024 | 0.075 | 0.075 | 1.141 | 0.68 | 0.18 | 0.79 | 0.96 | 0.73 | 0.8 |
| 4 | 42032966 | rs9654068 | 2.393 | 0.44 | 2.317 | -1.834 | 0.038 | 0.072 | 0.072 | 1.999 | 0.68 | 0.19 | 0.8 | 0.96 | 0.78 | 0.8 |
| 4 | 42033153 | rs9291209 | 2.393 | 0.44 | 2.317 | -1.816 | 0.038 | 0.073 | 0.073 | 1.999 | 0.68 | 0.19 | 0.8 | 0.96 | 0.78 | 0.8 |
| 4 | 42066378 | rs10938175 | 2.26 | 0.264 | 2.387 | -1.753 | 0.027 | 0.034 | 0.034 | 2.912 | 0.7 | 0.21 | 0.81 | 0.96 | 0.79 | 0.82 |
| 4 | 42074633 | rs4619931 | 2.747 | 0.258 | 2.281 | -1.763 | 0.024 | 0.258 | 0.258 | 5.931 | 0.64 | 0.07 | 0.76 | 0.95 | 0.75 | 0.78 |
| 4 | 42078241 | rs6447123 | -1.867 | 0.254 | 2.643 | -1.769 | -0.042 | -0.116 | -0.116 | 0.078 | 0.22 | 0.59 | 0.15 | 0.04 | 0.15 | 0.15 |
| 4 | 42089177 | rs11051 | 2.63 | 0.283 | 2.461 | -1.75 | 0.021 | 0.26 | 0.26 | 6.006 | 0.64 | 0.05 | 0.76 | 0.95 | 0.75 | 0.78 |
| 4 | 42099799 | rs7655320 | -2.486 | 0.319 | 2.529 | -1.741 | -0.016 | -0.078 | -0.078 | 0 | 0.29 | 0.64 | 0.18 | 0.04 | 0.19 | 0.18 |
| 4 | 42107100 | rs6447127 | -2.679 | 0.253 | 2.414 | -1.748 | 0.045 | 0.103 | 0.103 | 4.99 | 0.32 | 0.84 | 0.22 | 0.04 | 0.21 | 0.21 |
| 4 | 42110719 | rs7697558 | -1.716 | 0.209 | 3.395 | -1.768 | -0.022 | -0.081 | -0.081 | 0.256 | 0.2 | 0.28 | 0.13 | 0.03 | 0.13 | 0.16 |
| 4 | 42112734 | rs7682049 | -3.298 | 0.26 | 3.442 | -1.689 | 0.03 | 0.228 | 0.228 | 6.13 | 0.36 | 0.93 | 0.24 | 0.06 | 0.26 | 0.23 |
| 4 | 42114671 | rs7677472 | -2.401 | 0.258 | 3.897 | -1.69 | -0.041 | -0.118 | -0.118 | 1.582 | 0.26 | 0.56 | 0.17 | 0.05 | 0.18 | 0.19 |
| 4 | 42115810 | rs17446061 | -1.396 | 0.266 | 3.98 | -1.696 | 0.037 | -0.025 | -0.025 | 0 | 0.05 | 0.19 | 0.04 | 0 | 0.06 | 0.03 |
| 4 | 42116116 | rs6847380 | -1.741 | 0.265 | 3.904 | -1.694 | -0.021 | -0.078 | -0.078 | 0.071 | 0.2 | 0.27 | 0.13 | 0.03 | 0.13 | 0.16 |
| 4 | 42117173 | rs4861163 | -2.719 | 0.265 | 4.012 | -1.677 | -0.014 | -0.138 | -0.138 | 2.083 | 0.29 | 0.44 | 0.21 | 0.06 | 0.24 | 0.22 |
| 4 | 42117559 | rs16854020 | -3.545 | 0.281 | 3.724 | -1.654 | 0.009 | -0.082 | -0.082 | 2.162 | 0.15 | 0.54 | 0.1 | 0.03 | 0.12 | 0.06 |
| 4 | 42121223 | rs16854033 | -2.691 | 0.288 | 3.189 | -1.614 | 0.005 | 0.003 | 0.003 | 1.126 | 0.05 | 0.27 | 0.01 | 0 | 0.01 | 0 |
| 4 | 42125621 | rs4132888 | -2.766 | 0.296 | 2.515 | -1.612 | 0.01 | 0.235 | 0.235 | 6.201 | 0.36 | 0.95 | 0.25 | 0.06 | 0.26 | 0.22 |
| 4 | 42129997 | rs6447131 | -2.842 | 0.295 | 2.671 | -1.612 | 0.036 | 0.186 | 0.186 | 6.358 | 0.35 | 0.9 | 0.24 | 0.06 | 0.25 | 0.18 |
| 4 | 42146657 | rs6848386 | -2.648 | 0.424 | 2.839 | -1.585 | 0.023 | 0.09 | 0.09 | 6.546 | 0.33 | 0.82 | 0.24 | 0.06 | 0.24 | 0.17 |
| 4 | 42161427 | rs6447133 | -2.134 | 0.458 | 2.871 | -1.314 | -0.015 | -0.079 | -0.079 | 0 | 0.25 | 0.55 | 0.27 | 0.08 | 0.19 | 0.15 |
| 4 | 42164386 | rs4299622 | -1.492 | 0.631 | 3.062 | -1.191 | -0.005 | -0.046 | -0.046 | 0 | 0.19 | 0.25 | 0.15 | 0.02 | 0.14 | 0.11 |
| 4 | 42165799 | rs13151738 | -1.324 | 0.851 | 2.356 | -1.184 | 0.019 | -0.027 | -0.027 | 0 | 0.07 | 0.24 | 0.03 | 0 | 0.06 | 0.05 |
| 4 | 42174810 | rs7672382 | -1.964 | 0.854 | 2.032 | -1.066 | -0.006 | -0.037 | -0.037 | 0 | 0.32 | 0.63 | 0.32 | 0.09 | 0.25 | 0.21 |
| 4 | 42183596 | rs6827980 | -1.491 | 0.856 | 1.222 | -1.036 | -0.009 | -0.087 | -0.087 | 0.041 | 0.26 | 0.38 | 0.3 | 0.05 | 0.21 | 0.21 |
| 4 | 42193790 | rs6812348 | -1.882 | 0.87 | 1.053 | 0.111 | -0.014 | 0.047 | 0.047 | 0 | 0.11 | 0.23 | 0.12 | 0.2 | 0.06 | 0.06 |
| 4 | 42195808 | rs6852533 | -1.814 | 1.709 | 0.812 | 1.219 | 0.005 | 0.03 | 0.03 | 0 | 0.17 | 0.19 | 0.14 | 0.44 | 0.24 | 0.15 |
| 4 | 42201089 | rs13119304 | -1.28 | 1.471 | 0.589 | 1.405 | -0.006 | -0.04 | -0.04 | 0.02 | 0.11 | 0.05 | 0.06 | 0.28 | 0.12 | 0.06 |
| 4 | 42206574 | rs7674997 | -2.183 | 1.457 | 1.391 | 1.213 | -0.003 | 0.086 | 0.086 | 0 | 0.19 | 0.43 | 0.14 | 0.28 | 0.16 | 0.13 |
| 4 | 42218575 | rs4861168 | -0.959 | 1.505 | 0.737 | 0.978 | -0.002 | 0.005 | 0.005 | 0.027 | 0.44 | 0.67 | 0.36 | 0.58 | 0.43 | 0.49 |
| 4 | 42219146 | rs7667510 | 0.279 | 1.782 | 0.245 | 0.642 | -0.038 | 0.036 | 0.036 | 0 | 0.09 | 0.01 | 0.09 | 0.03 | 0.14 | 0.06 |
| 4 | 42223934 | rs6826969 | -1.297 | 1.604 | 1.61 | 1.059 | -0.019 | 0.096 | 0.096 | 0 | 0.14 | 0.49 | 0.13 | 0.25 | 0.07 | 0.11 |
| 4 | 42224011 | rs17629772 | -0.975 | 1.189 | 1.003 | 0.889 | 0.017 | 0.068 | 0.068 | 1.776 | 0.28 | 0.46 | 0.26 | 0.55 | 0.26 | 0.4 |
| 4 | 42226924 | rs13117610 | -1.073 | 0.817 | 0.698 | 0.652 | -0.013 | -0.041 | -0.041 | 0 | 0.29 | 0.19 | 0.26 | 0.56 | 0.27 | 0.4 |
| 4 | 42230777 | rs7664670 | -1.722 | 0.692 | 1.437 | 0.264 | 0.04 | 0.085 | 0.085 | 0.102 | 0.09 | 0.43 | 0.1 | 0.15 | 0.05 | 0.1 |
| 4 | 42237898 | rs4861175 | -1.145 | 1.144 | 1.621 | 0.021 | 0.058 | -0.068 | -0.068 | 0.128 | 0.13 | 0.41 | 0.12 | 0.02 | 0.17 | 0.08 |
| 4 | 42241036 | rs729466 | -1.442 | 0.789 | 1.282 | -0.476 | 0.003 | 0.01 | 0.01 | 0 | 0.46 | 0.59 | 0.58 | 0.86 | 0.4 | 0.56 |

| | | | | | | | | | | | | | | | | |
|---|---|---|---|---|---|---|---|---|---|---|---|---|---|---|---|---|
| 4 | 42241138 | rs729467 | -0.388 | 0.758 | 1.448 | -0.394 | 0 | -0.001 | -0.001 | 2.789 | 0.31 | 0.29 | 0.38 | 0.66 | 0.28 | 0.41 |
| 4 | 42241485 | rs729179 | -1.696 | 0.809 | 0.955 | -0.41 | 0.006 | 0.024 | 0.024 | 0 | 0.47 | 0.62 | 0.58 | 0.86 | 0.4 | 0.56 |
| 4 | 42243086 | rs989313 | -2.402 | 0.816 | 0.693 | -0.403 | -0.051 | 0.209 | 0.209 | 0.554 | 0.57 | 0.94 | 0.69 | 0.89 | 0.57 | 0.64 |
| 4 | 42249776 | rs9291212 | -0.965 | 0.869 | 1.016 | -0.295 | 0.004 | 0.006 | 0.006 | 0.051 | 0.36 | 0.56 | 0.4 | 0.67 | 0.28 | 0.38 |
| 4 | 42250342 | rs17530831 | 0.136 | 0.552 | -0.597 | -0.933 | -0.042 | 0.071 | 0.071 | 0.211 | 0.31 | 0.14 | 0.35 | 0.22 | 0.4 | 0.28 |
| 4 | 42251630 | rs4861178 | -1.902 | 0.451 | -0.374 | -1.08 | 0 | 0.001 | 0.001 | 0.028 | 0.68 | 0.75 | 0.75 | 0.88 | 0.7 | 0.67 |
| 4 | 42253010 | rs17530845 | 0.737 | 0.414 | -0.569 | -1.06 | -0.019 | 0.103 | 0.103 | 0.002 | 0.23 | 0.04 | 0.26 | 0.05 | 0.23 | 0.2 |
| 4 | 42256594 | rs1542686 | -0.254 | 0.603 | -0.483 | -0.913 | 0 | -0.002 | -0.002 | 1.403 | 0.37 | 0.6 | 0.42 | 0.76 | 0.33 | 0.39 |
| 4 | 42265464 | rs12507844 | -0.773 | 0.895 | 0.016 | -0.616 | -0.023 | 0.049 | 0.049 | 0 | 0.63 | 0.78 | 0.69 | 0.82 | 0.56 | 0.6 |
| 4 | 42276583 | rs2171423 | 0.041 | 1.097 | -0.058 | -0.627 | -0.01 | 0.019 | 0.019 | 0 | 0.71 | 0.68 | 0.68 | 0.85 | 0.67 | 0.57 |
| 4 | 42281042 | rs4861181 | 0.578 | 1.007 | -0.007 | -0.627 | 0.02 | -0.04 | -0.04 | 0.133 | 0.64 | 0.51 | 0.61 | 0.81 | 0.55 | 0.52 |
| 4 | 42291959 | rs6823142 | 0.716 | 1.313 | -0.396 | -0.31 | -0.026 | 0.023 | 0.023 | 0 | 0.67 | 0.71 | 0.62 | 0.76 | 0.58 | 0.55 |
| 4 | 42301184 | rs10026772 | -0.825 | 1.429 | -0.687 | -0.115 | -0.045 | -0.019 | -0.019 | 0 | 0.81 | 0.92 | 0.76 | 0.82 | 0.75 | 0.72 |
| 4 | 42313720 | rs2632695 | 1.087 | 1.132 | 0.056 | -0.164 | 0.004 | -0.012 | -0.012 | 0 | 0.65 | 0.46 | 0.63 | 0.65 | 0.62 | 0.56 |
| 4 | 42314717 | rs2262643 | -0.689 | 1.149 | 0.12 | -0.085 | 0.001 | 0.005 | 0.005 | 0 | 0.81 | 0.79 | 0.8 | 0.71 | 0.79 | 0.74 |
| 4 | 42322946 | rs3103190 | 0.672 | 1.149 | 0.182 | -0.123 | 0.004 | 0.004 | 0.004 | 0 | 0.73 | 0.65 | 0.75 | 0.67 | 0.68 | 0.66 |
| 4 | 42326215 | rs2632689 | 0.704 | 1.136 | 0.031 | -0.309 | 0.002 | -0.002 | -0.002 | 0 | 0.69 | 0.63 | 0.73 | 0.67 | 0.66 | 0.66 |
| 4 | 42326229 | rs2632690 | 0.024 | 1.112 | -0.139 | -0.307 | -0.001 | -0.007 | -0.007 | 0 | 0.76 | 0.87 | 0.79 | 0.71 | 0.76 | 0.73 |
| 4 | 42331579 | rs748360 | 1.306 | 1.005 | 0.12 | 0.066 | 0.001 | 0.005 | 0.005 | 0 | 0.61 | 0.45 | 0.61 | 0.57 | 0.59 | 0.55 |
| 4 | 42336312 | rs1023993 | 0.702 | 0.975 | 0.105 | -0.012 | -0.006 | 0.009 | 0.009 | 0 | 0.67 | 0.47 | 0.65 | 0.62 | 0.69 | 0.63 |
| 4 | 42344051 | rs1565111 | 0.812 | 1.102 | -0.077 | 0.034 | -0.001 | -0.001 | -0.001 | 0.025 | 0.64 | 0.61 | 0.63 | 0.58 | 0.59 | 0.56 |
| 4 | 42357010 | rs3103200 | -0.636 | 1.163 | -0.443 | 0.492 | -0.049 | -0.094 | -0.094 | 0 | 0.86 | 0.96 | 0.78 | 0.62 | 0.8 | 0.79 |
| 4 | 42358088 | rs2719949 | 1.055 | 0.555 | -0.689 | -0.96 | 0.013 | 0.036 | 0.036 | 0.159 | 0.22 | 0.13 | 0.25 | 0.05 | 0.18 | 0.12 |
| 4 | 42363367 | rs10025341 | 0.061 | 0.426 | -0.275 | -1.459 | -0.006 | -0.156 | -0.156 | 0.823 | 0.29 | 0.51 | 0.28 | 0.03 | 0.25 | 0.29 |
| 4 | 42368901 | rs9291216 | 0.063 | 0.352 | -0.175 | -1.469 | 0.001 | -0.158 | -0.158 | 0 | 0.29 | 0.5 | 0.28 | 0.02 | 0.26 | 0.29 |
| 4 | 42373005 | rs6832142 | 0.162 | 0.188 | -0.152 | -1.413 | 0.018 | 0.03 | 0.03 | 0 | 0.51 | 0.42 | 0.53 | 0.66 | 0.62 | 0.54 |
| 4 | 42376356 | rs923309 | 1.068 | 0.208 | -0.159 | -1.45 | -0.016 | -0.006 | -0.006 | 0 | 0.32 | 0.27 | 0.3 | 0.64 | 0.41 | 0.4 |
| 4 | 42376769 | rs3103207 | 1.807 | 0.005 | -0.139 | -1.377 | 0.015 | 0.028 | 0.028 | 0 | 0.22 | 0.13 | 0.23 | 0.05 | 0.16 | 0.1 |
| 4 | 42377390 | rs3109630 | -1.572 | 0.073 | -0.265 | -1.325 | -0.017 | -0.076 | -0.076 | 0 | 0.84 | 0.95 | 0.82 | 0.71 | 0.84 | 0.79 |
| 4 | 42384915 | rs2719947 | 0.228 | 0.058 | -0.179 | -1.421 | -0.01 | -0.071 | -0.071 | 0 | 0.34 | 0.56 | 0.29 | 0.05 | 0.21 | 0.24 |
| 4 | 42389740 | rs6447159 | -0.129 | 0.359 | -0.125 | -1.303 | 0.009 | 0.016 | 0.016 | 0 | 0.53 | 0.57 | 0.53 | 0.68 | 0.61 | 0.61 |
| 4 | 42394408 | rs11735318 | -0.189 | -0.041 | -0.101 | -1.25 | 0.014 | 0.013 | 0.013 | 0 | 0.55 | 0.6 | 0.57 | 0.68 | 0.66 | 0.62 |
| 4 | 42395984 | rs3103218 | 0.35 | 0.044 | -0.206 | -1.419 | -0.001 | -0.01 | -0.01 | 0 | 0.54 | 0.48 | 0.49 | 0.71 | 0.57 | 0.5 |
| 4 | 42397387 | rs7676862 | 0.569 | 0.135 | -0.405 | -1.554 | 0 | 0.098 | 0.098 | 0.084 | 0.37 | 0.16 | 0.45 | 0.04 | 0.37 | 0.24 |
| 4 | 42401416 | rs3103222 | 0.458 | 0.282 | -0.465 | -1.048 | -0.033 | -0.025 | -0.025 | 0 | 0.23 | 0.15 | 0.24 | 0.6 | 0.38 | 0.41 |
| 4 | 42411553 | rs7665824 | 0.988 | -0.149 | -0.127 | -1.431 | -0.002 | -0.033 | -0.033 | 0 | 0.26 | 0.25 | 0.25 | 0.11 | 0.23 | 0.17 |
| 4 | 42417526 | rs4481266 | 0.876 | -0.562 | 0.005 | -1.645 | 0.032 | 0.036 | 0.036 | 0 | 0.28 | 0.19 | 0.21 | 0.03 | 0.17 | 0.16 |
| 4 | 42423197 | rs3935448 | 0.393 | -0.52 | 0.041 | -1.657 | -0.007 | -0.051 | -0.051 | 0 | 0.47 | 0.46 | 0.43 | 0.14 | 0.35 | 0.33 |
| 4 | 42428210 | rs9994372 | 1.098 | -0.497 | 0.155 | -1.598 | -0.004 | -0.029 | -0.029 | 0 | 0.25 | 0.31 | 0.25 | 0.11 | 0.22 | 0.18 |
| 4 | 42431550 | rs7665008 | -0.233 | -0.162 | 0.185 | -1.405 | 0.041 | 0.002 | 0.002 | 0 | 0.68 | 0.5 | 0.66 | 0.37 | 0.56 | 0.62 |
| 4 | 42432089 | rs7688175 | 0.685 | -0.195 | 0.146 | -1.425 | -0.007 | -0.005 | -0.005 | 2.305 | 0.21 | 0.08 | 0.23 | 0.23 | 0.2 | 0.29 |
| 4 | 42433734 | rs7695984 | 0.233 | -0.186 | 0.338 | -1.387 | 0.002 | -0.007 | -0.007 | 0 | 0.52 | 0.48 | 0.51 | 0.21 | 0.4 | 0.4 |
| 4 | 42446079 | rs6855691 | 0.644 | -0.255 | 0.218 | -1.4 | 0.035 | 0.097 | 0.097 | 0.658 | 0.45 | 0.25 | 0.41 | 0.16 | 0.35 | 0.36 |
| 4 | 42449874 | rs7684005 | 0.476 | -0.474 | 0.354 | -1.535 | 0.055 | 0.167 | 0.167 | 1.639 | 0.51 | 0.25 | 0.43 | 0.16 | 0.38 | 0.41 |
| 4 | 42451770 | rs10034082 | -1.166 | -0.467 | 0.852 | -1.581 | 0.057 | -0.014 | -0.014 | 0 | 0.08 | 0.39 | 0.1 | 0.04 | 0.04 | 0.07 |
| 4 | 42453303 | rs13139219 | 0.397 | -0.433 | 0.738 | -1.555 | 0.008 | 0.016 | 0.016 | 0.605 | 0.67 | 0.31 | 0.66 | 0.74 | 0.77 | 0.6 |
| 4 | 42453514 | rs17531575 | 1.012 | -0.449 | 0.738 | -1.581 | 0.038 | 0.19 | 0.19 | 2.719 | 0.45 | 0.17 | 0.41 | 0.16 | 0.36 | 0.33 |
| 4 | 42455837 | rs13103298 | -0.053 | -0.387 | 0.6 | -1.543 | -0.073 | 0.153 | 0.153 | 2.191 | 0.39 | 0.14 | 0.45 | 0.8 | 0.58 | 0.51 |
| 4 | 42466641 | rs12647640 | -0.084 | -0.386 | 0.726 | -1.493 | -0.008 | 0.021 | 0.021 | 1.376 | 0.4 | 0.39 | 0.47 | 0.79 | 0.57 | 0.5 |
| 4 | 42472966 | rs10517027 | 2.13 | -0.651 | 0.458 | -1.704 | 0.021 | 0.1 | 0.1 | 0.327 | 0.21 | 0.03 | 0.18 | 0.05 | 0.14 | 0.16 |
| 4 | 42475352 | rs10000708 | -0.191 | -0.421 | 0.731 | -1.497 | 0.017 | 0.016 | 0.016 | 0.626 | 0.71 | 0.38 | 0.72 | 0.43 | 0.59 | 0.72 |
| 4 | 42481276 | rs6447164 | -0.174 | -0.415 | 0.84 | -1.262 | 0.004 | -0.003 | -0.003 | 0 | 0.76 | 0.66 | 0.69 | 0.73 | 0.79 | 0.68 |
| 4 | 42484396 | rs3935029 | 1.226 | -0.426 | 0.776 | -1.235 | -0.006 | -0.021 | -0.021 | 0.005 | 0.17 | 0.1 | 0.22 | 0.23 | 0.17 | 0.24 |
| 4 | 42490680 | rs9291218 | 1.98 | -0.417 | 1.075 | -1.163 | -0.007 | -0.062 | -0.062 | 0 | 0.89 | 0.35 | 0.88 | 0.96 | 0.96 | 0.92 |
| 4 | 42496713 | rs4861030 | 1.001 | -0.263 | 0.966 | -1.087 | 0.093 | 0.013 | 0.013 | 1.545 | 0.92 | 0.64 | 0.9 | 0.96 | 0.96 | 0.92 |
| 4 | 42497919 | rs10034756 | 0.629 | -0.353 | 0.904 | -1.15 | 0 | -0.081 | -0.081 | 0 | 0.87 | 0.63 | 0.88 | 0.96 | 0.94 | 0.83 |
| 4 | 42499224 | rs4280770 | -0.422 | -0.38 | 0.32 | -1.2 | 0.019 | 0.035 | 0.035 | 0 | 0.5 | 0.38 | 0.51 | 0.33 | 0.35 | 0.59 |
| 4 | 42511242 | rs10517038 | 1.481 | -0.29 | 0.014 | -0.566 | -0.026 | -0.077 | -0.077 | 0.963 | 0.15 | 0.02 | 0.2 | 0.24 | 0.14 | 0.24 |
| 4 | 42511312 | rs6819929 | -0.242 | -0.083 | 0.55 | -0.649 | 0.043 | -0.016 | -0.016 | 0 | 0.06 | 0.2 | 0.09 | 0.04 | 0.05 | 0.08 |
| 4 | 42517211 | rs4526008 | -1.2 | -0.032 | 0.384 | -0.731 | -0.001 | -0.018 | -0.018 | 0.037 | 0.76 | 0.77 | 0.84 | 0.89 | 0.77 | 0.83 |
| 4 | 42552434 | rs4395543 | -0.929 | -0.06 | 0.382 | -0.718 | 0.002 | -0.06 | -0.06 | 0 | 0.75 | 0.61 | 0.81 | 0.88 | 0.76 | 0.83 |
| 4 | 42556540 | rs10805104 | 0.118 | 0.034 | -0.202 | -0.1 | -0.025 | -0.019 | -0.019 | 3.196 | 0.26 | 0.23 | 0.39 | 0.35 | 0.42 | 0.29 |
| 4 | 42559530 | rs17593128 | 0.679 | -0.382 | -0.237 | -0.079 | 0.067 | -0.057 | -0.057 | 0.008 | 0.41 | 0.09 | 0.37 | 0.52 | 0.3 | 0.46 |
| 4 | 42567615 | rs6447170 | -0.355 | -0.266 | 0.353 | -0.171 | 0.046 | -0.024 | -0.024 | 0.373 | 0.91 | 0.74 | 0.94 | 0.96 | 0.92 | 0.92 |
| 4 | 42570934 | rs2061526 | 0.096 | 0.034 | 0.214 | -0.413 | -0.001 | -0.033 | -0.033 | 0 | 0.57 | 0.42 | 0.65 | 0.91 | 0.59 | 0.6 |
| 4 | 42571666 | rs1460354 | 1.485 | -0.004 | -0.707 | -0.317 | 0.036 | -0.023 | -0.023 | 0 | 0.27 | 0.09 | 0.23 | 0.26 | 0.2 | 0.28 |
| 4 | 42574373 | rs12502828 | 0.542 | 0.154 | -0.746 | -0.276 | 0.03 | -0.051 | -0.051 | 0.014 | 0.45 | 0.1 | 0.42 | 0.56 | 0.37 | 0.5 |
| 4 | 42574630 | rs16854520 | 1.821 | 0.195 | -0.841 | -0.271 | -0.005 | -0.02 | -0.02 | 1.111 | 0.05 | 0.08 | 0.03 | 0 | 0.03 | 0.06 |
| 4 | 42576420 | rs2290870 | 1.539 | 0.255 | -0.96 | -0.203 | -0.039 | -0.089 | -0.089 | 0 | 0.18 | 0.01 | 0.19 | 0.31 | 0.17 | 0.22 |

| | | | | | | | | | | | | | | | | |
|---|---|---|---|---|---|---|---|---|---|---|---|---|---|---|---|---|
| 4 | 42576472 | rs13148015 | 2.523 | 0.19 | -0.82 | -0.321 | 0 | 0.111 | 0.111 | 0.226 | 0.17 | 0.01 | 0.08 | 0 | 0.12 | 0.04 |
| 4 | 42577517 | rs2290868 | 1.046 | 0.56 | -0.336 | 0.098 | -0.002 | 0.034 | 0.034 | 0 | 0.09 | 0.07 | 0.21 | 0.35 | 0.22 | 0.1 |
| 4 | 42583029 | rs10013589 | -0.97 | 0.404 | 0.047 | -0.345 | 0.007 | -0.059 | -0.059 | 0 | 0.71 | 0.67 | 0.72 | 0.92 | 0.7 | 0.74 |
| 4 | 42587201 | rs1080248 | 1.295 | 0.465 | -0.124 | -0.313 | -0.046 | 0.116 | 0.116 | 0 | 0.25 | 0.08 | 0.25 | 0.04 | 0.28 | 0.25 |
| 4 | 42593240 | rs3749530 | 0.45 | 0.393 | -0.007 | 0.027 | 0.016 | 0.013 | 0.013 | 0 | 0.13 | 0.18 | 0.14 | 0.2 | 0.23 | 0.13 |
| 4 | 42597060 | rs7672084 | 0.124 | 0.14 | 0.089 | 0.142 | 0.001 | -0.003 | -0.003 | 0.2 | 0.41 | 0.41 | 0.37 | 0.47 | 0.41 | 0.37 |
| 4 | 42607736 | rs9291219 | -0.258 | 0.133 | 0.128 | 0.139 | 0.013 | -0.013 | -0.013 | 0 | 0.58 | 0.37 | 0.62 | 0.52 | 0.59 | 0.63 |
| 4 | 42612831 | rs1870611 | -0.536 | 0.135 | 0.133 | 0.124 | 0.008 | -0.017 | -0.017 | 0.222 | 0.6 | 0.48 | 0.6 | 0.76 | 0.49 | 0.61 |
| 4 | 42622148 | rs10938199 | -1.417 | 0.178 | -0.108 | 0.167 | -0.04 | 0.008 | 0.008 | 0 | 0.72 | 0.65 | 0.77 | 0.72 | 0.83 | 0.76 |
| 4 | 42622525 | rs16854618 | 0.695 | 0.788 | -0.25 | 0.46 | 0.011 | -0.022 | -0.022 | 0 | 0.09 | 0.15 | 0.04 | 0.04 | 0.08 | 0.07 |
| 4 | 42627836 | rs2306004 | 0.022 | 0.77 | -0.275 | 0.437 | -0.009 | -0.024 | -0.024 | 0 | 0.27 | 0.25 | 0.35 | 0.48 | 0.28 | 0.31 |
| 4 | 42629038 | rs16854624 | 0.664 | 0.872 | -0.325 | 0.404 | 0.012 | -0.022 | -0.022 | 2.897 | 0.09 | 0.15 | 0.04 | 0.04 | 0.08 | 0.06 |
| 4 | 42632678 | rs1563849 | 0.678 | 0.996 | -0.547 | 0.521 | -0.005 | -0.006 | -0.006 | 0.101 | 0.11 | 0.09 | 0.13 | 0.19 | 0.24 | 0.13 |
| 4 | 42639186 | rs898500 | 0.59 | 0.736 | -0.528 | 0.423 | -0.001 | 0.002 | 0.002 | 1.035 | 0.57 | 0.58 | 0.51 | 0.32 | 0.47 | 0.51 |
| 4 | 42644312 | rs1460362 | -1.348 | 0.659 | -0.769 | 0.611 | -0.003 | -0.062 | -0.062 | 0 | 0.81 | 0.9 | 0.72 | 0.54 | 0.81 | 0.78 |
| 4 | 42646252 | rs6851867 | -0.347 | 0.689 | -0.617 | 0.49 | 0.084 | 0.04 | 0.04 | 0.19 | 0.21 | 0.52 | 0.22 | 0.26 | 0.29 | 0.24 |
| 4 | 42646677 | rs1380754 | 1.543 | 0.232 | -1.036 | 0.019 | 0 | -0.032 | -0.032 | 0 | 0.26 | 0.21 | 0.15 | 0.03 | 0.27 | 0.23 |
| 4 | 42647802 | rs2612520 | 0.208 | 0.306 | -1.037 | 0.101 | 0 | -0.001 | -0.001 | 0 | 0.59 | 0.38 | 0.5 | 0.28 | 0.5 | 0.53 |
| 4 | 42648398 | rs17448506 | 0.506 | 0.192 | -1.488 | 0.471 | 0.027 | 0.013 | 0.013 | 0 | 0.32 | 0.25 | 0.35 | 0.25 | 0.23 | 0.32 |
| 4 | 42649935 | rs3101246 | -0.332 | 0.667 | -1.311 | 0.407 | 0.03 | 0.071 | 0.071 | 0 | 0.54 | 0.78 | 0.41 | 0.3 | 0.59 | 0.53 |
| 4 | 42652004 | rs7693131 | -0.133 | 0.405 | -0.859 | 0.727 | -0.011 | 0.02 | 0.02 | 0.363 | 0.69 | 0.47 | 0.59 | 0.39 | 0.71 | 0.65 |
| 4 | 42652337 | rs10938202 | 0.134 | 0.516 | -1.185 | 0.303 | -0.001 | -0.005 | -0.005 | 0.045 | 0.1 | 0.13 | 0.06 | 0.04 | 0.07 | 0.13 |
| 4 | 42660891 | rs2575568 | 0.013 | -0.032 | -1.132 | 0.503 | -0.025 | 0.048 | 0.048 | 1.63 | 0.29 | 0.46 | 0.37 | 0.41 | 0.25 | 0.26 |
| 4 | 42661922 | rs2612507 | 0.475 | 0.223 | -1.513 | 0.13 | 0.002 | 0.004 | 0.004 | 0 | 0.71 | 0.72 | 0.8 | 0.8 | 0.73 | 0.71 |
| 4 | 42662283 | rs13133617 | -0.603 | 0.388 | -1.366 | 0.583 | -0.002 | 0.002 | 0.002 | 0 | 0.11 | 0.14 | 0.09 | 0.14 | 0.12 | 0.15 |
| 4 | 42665092 | rs2575551 | -0.431 | 0.083 | -1.973 | 0.373 | 0.072 | -0.018 | -0.018 | 0 | 0.51 | 0.83 | 0.48 | 0.49 | 0.4 | 0.63 |
| 4 | 42673312 | rs12501388 | 0.682 | -0.015 | -2.135 | 0.704 | -0.016 | -0.021 | -0.021 | 0.1 | 0.06 | 0.01 | 0.18 | 0.21 | 0.07 | 0.14 |
| 4 | 42676234 | rs2575484 | -1.371 | 0.404 | -2.688 | 0.055 | -0.005 | -0.001 | -0.001 | 1.539 | 0.83 | 0.9 | 0.91 | 1 | 0.84 | 0.83 |
| 4 | 42683596 | rs882642 | 0.215 | 0.196 | -1.964 | -0.223 | 0.036 | -0.027 | -0.027 | 0 | 0.06 | 0.25 | 0.05 | 0 | 0.08 | 0.01 |
| 4 | 42685758 | rs2575453 | 0.005 | -0.541 | -1.907 | -0.606 | -0.028 | 0.032 | 0.032 | 0.234 | 0.36 | 0.49 | 0.39 | 0.5 | 0.26 | 0.53 |
| 4 | 42690604 | rs1460352 | 1.743 | -0.697 | -1.985 | -0.674 | -0.176 | -0.192 | -0.192 | 0.234 | 0.31 | 0.02 | 0.44 | 0.5 | 0.44 | 0.2 |
| 4 | 42694179 | rs7441772 | 0.119 | -0.669 | -1.842 | -0.653 | 0.027 | 0.055 | 0.055 | 0 | 0.52 | 0.42 | 0.7 | 0.73 | 0.64 | 0.43 |
| 4 | 42697553 | rs2575548 | 1.321 | -0.812 | -1.197 | -0.611 | -0.012 | 0.097 | 0.097 | 0 | 0.11 | 0.35 | 0.11 | 0.14 | 0.06 | 0.08 |
| 4 | 42706128 | rs2612468 | 1.321 | -0.829 | -1.111 | -0.611 | -0.034 | 0.089 | 0.089 | 0 | 0.11 | 0.33 | 0.09 | 0.14 | 0.03 | 0.08 |
| 4 | 42706804 | rs2612467 | 1.321 | -0.829 | -1.092 | -0.611 | -0.034 | 0.087 | 0.087 | 0 | 0.11 | 0.31 | 0.09 | 0.14 | 0.03 | 0.07 |
| 4 | 42707463 | rs2612538 | -1.557 | -0.848 | -1.103 | -0.632 | -0.006 | 0.004 | 0.004 | 0 | 0.83 | 0.9 | 0.91 | 1 | 0.83 | 0.83 |
| 4 | 42708098 | rs3101249 | 1.145 | -0.685 | -0.886 | -0.322 | -0.034 | 0.087 | 0.087 | 0 | 0.11 | 0.31 | 0.09 | 0.13 | 0.03 | 0.07 |
| 4 | 42712959 | rs2345467 | 1.22 | -0.696 | -0.708 | -0.335 | -0.032 | 0.087 | 0.087 | 0 | 0.1 | 0.3 | 0.08 | 0.15 | 0.03 | 0.09 |
| 4 | 42714879 | rs12502362 | 1.311 | -0.784 | -1.186 | -0.782 | -0.078 | -0.033 | -0.033 | 0 | 0.38 | 0.13 | 0.5 | 0.65 | 0.49 | 0.49 |
| 4 | 42718502 | rs2612530 | 0.065 | -0.884 | -1.078 | -0.74 | -0.074 | 0.011 | 0.011 | 0 | 0.18 | 0.41 | 0.1 | 0.16 | 0.08 | 0.18 |
| 4 | 42720868 | rs2085782 | -0.601 | -0.879 | -1.079 | -0.799 | -0.048 | 0.077 | 0.077 | 0 | 0.23 | 0.35 | 0.27 | 0.35 | 0.13 | 0.21 |
| 4 | 42723259 | rs2575509 | -0.563 | -0.716 | -1.077 | -0.697 | -0.057 | 0.04 | 0.04 | 0 | 0.31 | 0.45 | 0.3 | 0.34 | 0.19 | 0.3 |
| 4 | 42726868 | rs2252994 | -1.114 | -0.84 | -1.001 | -0.995 | -0.008 | 0.06 | 0.06 | 0 | 0.74 | 0.86 | 0.86 | 0.99 | 0.76 | 0.8 |
| 4 | 42727694 | rs2575566 | -0.11 | -0.933 | -0.739 | -0.58 | -0.085 | -0.024 | -0.024 | 0 | 0.23 | 0.44 | 0.13 | 0.16 | 0.11 | 0.19 |
| 4 | 42735553 | rs1563850 | -0.384 | -0.754 | -1.403 | -0.4 | -0.05 | -0.026 | -0.026 | 0.009 | 0.28 | 0.56 | 0.18 | 0.18 | 0.2 | 0.2 |
| 4 | 42735847 | rs2575529 | -0.492 | -0.753 | -1.591 | -0.42 | -0.029 | -0.013 | -0.013 | 0 | 0.28 | 0.65 | 0.19 | 0.19 | 0.2 | 0.21 |
| 4 | 42748194 | rs13146994 | 0.989 | -0.574 | -1.908 | -0.478 | -0.056 | -0.031 | -0.031 | 0.086 | 0.4 | 0.13 | 0.52 | 0.63 | 0.54 | 0.49 |
| 4 | 42749176 | rs12330944 | -0.142 | -0.519 | -1.146 | -0.186 | -0.021 | 0.067 | 0.067 | 0.566 | 0.14 | 0.64 | 0.14 | 0.19 | 0.11 | 0.11 |
| 4 | 42753188 | rs2345904 | -0.774 | -0.304 | -1.243 | -0.085 | -0.016 | -0.007 | -0.007 | 0.11 | 0.29 | 0.69 | 0.19 | 0.19 | 0.2 | 0.21 |
| 4 | 42788309 | rs13101579 | -0.042 | 0.432 | -1.643 | 0.568 | -0.012 | -0.05 | -0.05 | 0 | 0.23 | 0.11 | 0.24 | 0.37 | 0.28 | 0.3 |
| 4 | 42790130 | rs12331684 | 0.313 | 1.26 | -1.461 | 0.941 | 0.037 | 0.084 | 0.084 | 0 | 0.08 | 0.16 | 0.2 | 0.25 | 0.13 | 0.09 |
| 4 | 42796127 | rs12650389 | 0.604 | 1.242 | -1.439 | 0.934 | 0.025 | 0.067 | 0.067 | 0 | 0.07 | 0.06 | 0.2 | 0.25 | 0.12 | 0.09 |
| 4 | 42796294 | rs2575513 | 0.309 | 1.749 | -1.222 | 0.681 | 0.06 | 0.085 | 0.085 | 0.329 | 0.14 | 0.55 | 0.23 | 0.23 | 0.18 | 0.13 |
| 4 | 42797274 | rs7697036 | 0.252 | 1.602 | -1.373 | 0.379 | 0.056 | 0.123 | 0.123 | 0 | 0.08 | 0.19 | 0.21 | 0.25 | 0.13 | 0.1 |
| 4 | 42810102 | rs3099914 | -1.43 | 1.135 | -2.074 | -0.108 | 0.011 | 0.148 | 0.148 | 0 | 0.64 | 0.96 | 0.77 | 0.84 | 0.76 | 0.68 |
| 4 | 42813098 | rs17449493 | -0.379 | 1.667 | -1.62 | 0.259 | -0.009 | -0.002 | -0.002 | 0 | 0.12 | 0.07 | 0.16 | 0.14 | 0.19 | 0.16 |
| 4 | 42813800 | rs1562839 | 0.701 | 1.391 | -1.578 | 0.774 | -0.008 | 0.011 | 0.011 | 0.013 | 0.11 | 0.15 | 0.24 | 0.36 | 0.17 | 0.22 |
| 4 | 42816051 | rs1450914 | -0.476 | 0.288 | -1.048 | -0.035 | -0.077 | -0.055 | -0.055 | 0 | 0.8 | 0.57 | 0.92 | 0.89 | 0.88 | 0.82 |
| 4 | 42817301 | rs13130624 | -0.967 | 0.388 | -0.995 | -0.081 | 0.027 | 0.076 | 0.076 | 0 | 0.13 | 0.21 | 0.12 | 0.29 | 0.17 | 0.27 |
| 4 | 42819241 | rs1450917 | 0.405 | 0.323 | -1.299 | -0.307 | -0.014 | -0.008 | -0.008 | 1.075 | 0.3 | 0.28 | 0.52 | 0.6 | 0.44 | 0.39 |
| 4 | 42821511 | rs1584622 | -0.835 | 0.302 | -1.275 | -0.394 | 0.009 | 0.017 | 0.017 | 0 | 0.45 | 0.43 | 0.65 | 0.88 | 0.63 | 0.67 |
| 4 | 42825862 | rs10470752 | 0.24 | 0.132 | -0.852 | -0.263 | -0.047 | -0.051 | -0.051 | 1.224 | 0.76 | 0.56 | 0.85 | 0.85 | 0.83 | 0.79 |
| 4 | 42838348 | rs1562840 | -0.378 | 0.356 | -1.222 | -0.403 | -0.003 | 0.016 | 0.016 | 0.214 | 0.44 | 0.39 | 0.41 | 0.34 | 0.49 | 0.56 |
| 4 | 42839254 | rs4861040 | 0.008 | 0.462 | -0.915 | -0.372 | 0.026 | -0.013 | -0.013 | 0 | 0.9 | 0.56 | 0.91 | 0.94 | 0.91 | 0.94 |
| 4 | 42840063 | rs2122648 | 1.001 | 0.223 | -1.335 | -0.389 | 0.047 | -0.057 | -0.057 | 0 | 0.55 | 0.13 | 0.53 | 0.5 | 0.63 | 0.5 |
| 4 | 42840819 | rs4861041 | 1.011 | 0.165 | -0.861 | -0.208 | -0.03 | -0.013 | -0.013 | 0.113 | 0.81 | 0.44 | 0.89 | 0.82 | 0.84 | 0.74 |
| 4 | 42845691 | rs17594783 | 0.415 | 0.368 | -1.286 | -0.034 | 0.005 | -0.006 | -0.006 | 0.043 | 0.06 | 0.03 | 0.01 | 0.08 | 0.05 | 0.1 |
| 4 | 42847901 | rs12642552 | 1.041 | 0.541 | -1.863 | -0.446 | -0.144 | 0.09 | 0.09 | 0 | 0.31 | 0.04 | 0.33 | 0.16 | 0.37 | 0.3 |
| 4 | 42848697 | rs17534138 | 1.642 | 0.758 | -1.933 | -0.505 | -0.08 | 0.059 | 0.059 | 0 | 0.25 | 0.01 | 0.16 | 0.14 | 0.23 | 0.23 |

| | | | | | | | | | | | | | | | | |
|---|---|---|---|---|---|---|---|---|---|---|---|---|---|---|---|---|
| 4 | 42851723 | rs2881215 | 0.342 | 0.222 | -1.423 | -0.524 | -0.053 | 0.129 | 0.129 | 0 | 0.38 | 0.12 | 0.36 | 0.16 | 0.4 | 0.31 |
| 4 | 42856894 | rs1993059 | 1.393 | 0.051 | -1.51 | -0.434 | -0.052 | -0.019 | -0.019 | 0.001 | 0.26 | 0.09 | 0.37 | 0.27 | 0.36 | 0.15 |
| 4 | 42858453 | rs9985652 | -0.831 | -0.114 | -1.051 | -0.425 | -0.055 | 0.163 | 0.163 | 0.03 | 0.79 | 0.47 | 0.8 | 0.58 | 0.89 | 0.6 |
| 4 | 42858855 | rs12643533 | -0.371 | -0.063 | -1.031 | -0.471 | 0 | 0.005 | 0.005 | 0 | 0.54 | 0.4 | 0.44 | 0.31 | 0.53 | 0.47 |
| 4 | 42869436 | rs13141599 | 1.242 | -0.251 | -1.132 | -0.431 | -0.148 | 0.101 | 0.101 | 0 | 0.33 | 0.04 | 0.34 | 0.15 | 0.39 | 0.3 |
| 4 | 42869937 | rs10023065 | -0.437 | -0.357 | -0.895 | -0.319 | 0 | -0.001 | -0.001 | 0 | 0.45 | 0.46 | 0.47 | 0.42 | 0.51 | 0.29 |
| 4 | 42870369 | rs10033259 | -1.135 | -0.437 | -0.736 | -0.217 | -0.041 | -0.009 | -0.009 | 0.425 | 0.19 | 0.35 | 0.1 | 0.15 | 0.16 | 0.15 |
| 4 | 42873798 | rs716693 | 0.126 | -0.517 | -0.566 | -0.164 | -0.049 | 0.058 | 0.058 | 0 | 0.23 | 0.3 | 0.19 | 0.41 | 0.11 | 0.32 |
| 4 | 42874286 | rs1562837 | 0.232 | -0.534 | -0.627 | -0.209 | -0.033 | 0.014 | 0.014 | 0 | 0.22 | 0.19 | 0.15 | 0.31 | 0.08 | 0.31 |
| 4 | 42877064 | rs11937866 | 0.62 | -0.368 | -0.601 | -0.225 | -0.021 | -0.036 | -0.036 | 0 | 0.07 | 0.09 | 0.03 | 0 | 0.04 | 0.02 |
| 4 | 42877463 | rs7696133 | -1.004 | -0.386 | -0.514 | -0.024 | -0.037 | -0.007 | -0.007 | 0 | 0.16 | 0.37 | 0.1 | 0.16 | 0.11 | 0.07 |
| 4 | 42877802 | rs1562842 | -1.417 | -0.403 | -0.505 | -0.024 | -0.046 | 0.017 | 0.017 | 0 | 0.21 | 0.37 | 0.1 | 0.24 | 0.16 | 0.17 |
| 4 | 42881014 | rs1993062 | 0.516 | -0.383 | -0.494 | -0.01 | 0.007 | 0.051 | 0.051 | 0.11 | 0.08 | 0.24 | 0.06 | 0.16 | 0.07 | 0.04 |
| 4 | 42883195 | rs7655156 | -0.948 | -0.317 | -0.611 | -0.048 | -0.047 | 0.019 | 0.019 | 0 | 0.2 | 0.37 | 0.1 | 0.24 | 0.16 | 0.15 |
| 4 | 42889160 | rs2122651 | -0.777 | 0.017 | -1.031 | -0.288 | -0.004 | 0.007 | 0.007 | 0 | 0.59 | 0.54 | 0.59 | 0.75 | 0.54 | 0.57 |
| 4 | 42900562 | rs10517057 | 1.526 | 0.1 | -2.196 | -0.236 | -0.116 | -0.042 | -0.042 | 1.687 | 0.22 | 0.02 | 0.35 | 0.2 | 0.34 | 0.12 |
| 4 | 42902654 | rs7656477 | 0.004 | -0.222 | -1.772 | -0.361 | -0.011 | 0.131 | 0.131 | 0 | 0.4 | 0.15 | 0.27 | 0.15 | 0.38 | 0.31 |
| 4 | 42902734 | rs10517055 | 0.77 | 0.132 | -1.956 | -0.092 | -0.108 | 0.091 | 0.091 | 0 | 0.31 | 0.06 | 0.24 | 0.15 | 0.34 | 0.27 |
| 4 | 42906100 | rs1450906 | -0.421 | -0.359 | -1.297 | 0.294 | 0.046 | -0.014 | -0.014 | 0.045 | 0.56 | 0.69 | 0.63 | 0.54 | 0.75 | 0.39 |
| 4 | 42907803 | rs10018588 | 0.293 | -0.489 | -0.966 | 0.088 | -0.01 | -0.033 | -0.033 | 0 | 0.06 | 0.1 | 0.02 | 0 | 0.03 | 0.02 |
| 4 | 42908357 | rs2122647 | 0.368 | -0.29 | -1.304 | -0.024 | -0.011 | 0.02 | 0.02 | 0.732 | 0.13 | 0.05 | 0.09 | 0.07 | 0.19 | 0.14 |
| 4 | 42914565 | rs1450911 | 1.348 | -0.55 | -1.212 | -0.006 | 0.003 | 0.014 | 0.014 | 0 | 0.28 | 0.24 | 0.16 | 0.15 | 0.21 | 0.33 |
| 4 | 42919356 | rs1450904 | 1.409 | -0.542 | -1.239 | 0.008 | 0.002 | 0.031 | 0.031 | 0 | 0.28 | 0.21 | 0.16 | 0.15 | 0.22 | 0.33 |
| 4 | 42919640 | rs6830507 | 1.769 | -0.09 | -1.507 | 0.148 | -0.095 | -0.031 | -0.031 | 0 | 0.19 | 0.01 | 0.32 | 0.18 | 0.24 | 0.11 |
| 4 | 42924425 | rs980779 | 1.208 | -0.175 | -1.152 | 0.4 | 0.059 | -0.068 | -0.068 | 0.34 | 0.16 | 0.04 | 0.2 | 0.32 | 0.04 | 0.28 |
| 4 | 42924853 | rs7439983 | -1.026 | 0.487 | -0.686 | 0.255 | 0.04 | 0.002 | 0.002 | 0.493 | 0.31 | 0.54 | 0.3 | 0.35 | 0.48 | 0.26 |
| 4 | 42933496 | rs10517062 | 0.129 | 0.559 | -0.94 | 0.373 | 0.001 | 0.005 | 0.005 | 0 | 0.11 | 0.1 | 0.07 | 0.07 | 0.15 | 0.16 |
| 4 | 42937881 | rs1450932 | 0.253 | 0.421 | -0.596 | 0.226 | 0.002 | -0.016 | -0.016 | 0.266 | 0.44 | 0.51 | 0.34 | 0.29 | 0.45 | 0.41 |
| 4 | 42939381 | rs7667015 | 0.269 | 0.969 | -0.239 | 0.626 | -0.002 | -0.001 | -0.001 | 0 | 0.1 | 0.1 | 0.08 | 0.09 | 0.17 | 0.16 |
| 4 | 42942050 | rs6818868 | -0.201 | 0.929 | -0.215 | 0.476 | 0.106 | 0.034 | 0.034 | 0 | 0.16 | 0.41 | 0.21 | 0.2 | 0.26 | 0.1 |
| 4 | 42959499 | rs9992691 | -0.479 | 0.761 | -0.095 | 0.361 | 0.004 | -0.001 | -0.001 | 0 | 0.05 | 0.18 | 0.02 | 0.03 | 0.03 | 0.07 |
| 4 | 42964453 | rs10938227 | 1.276 | 0.675 | -0.415 | 0.585 | -0.089 | -0.056 | -0.056 | 0 | 0.21 | 0.01 | 0.3 | 0.2 | 0.24 | 0.09 |
| 4 | 42975329 | rs2118030 | -0.519 | 0.782 | 0.2 | 0.779 | 0.037 | 0.057 | 0.057 | 0 | 0.23 | 0.38 | 0.29 | 0.38 | 0.36 | 0.24 |
| 4 | 43007627 | rs1436478 | 1.017 | 0.769 | -0.459 | 0.38 | -0.025 | 0.013 | 0.013 | 0.618 | 0.21 | 0.12 | 0.3 | 0.16 | 0.26 | 0.09 |
| 4 | 43015770 | rs1160755 | -0.909 | 0.328 | -0.245 | -0.012 | -0.001 | 0.018 | 0.018 | 0.781 | 0.46 | 0.6 | 0.56 | 0.78 | 0.52 | 0.6 |
| 4 | 43019669 | rs1545101 | -0.612 | 0.186 | 0.216 | -0.074 | -0.003 | 0.022 | 0.022 | 0.579 | 0.41 | 0.36 | 0.52 | 0.79 | 0.49 | 0.58 |
| 4 | 43025255 | rs10938230 | 1.402 | 0.122 | 0.115 | -0.152 | -0.032 | 0.016 | 0.016 | 0.903 | 0.21 | 0.11 | 0.3 | 0.15 | 0.26 | 0.09 |
| 4 | 43032612 | rs4861050 | -0.633 | 0.094 | 0.152 | -0.075 | -0.003 | 0.022 | 0.022 | 0.47 | 0.41 | 0.36 | 0.51 | 0.79 | 0.49 | 0.58 |
| 4 | 43041018 | rs11736676 | 0.151 | 0.262 | 0.377 | 0.426 | 0.005 | -0.041 | -0.041 | 0.069 | 0.26 | 0.14 | 0.34 | 0.35 | 0.26 | 0.32 |
| 4 | 43042976 | rs4512035 | 0.521 | 0.321 | 0.37 | 0.484 | 0.01 | 0.062 | 0.062 | 0 | 0.15 | 0.21 | 0.17 | 0.44 | 0.23 | 0.26 |
| 4 | 43068604 | rs920152 | -1.129 | 0.247 | 0.462 | 0.065 | -0.001 | -0.004 | -0.004 | 0 | 0.44 | 0.57 | 0.53 | 0.79 | 0.49 | 0.58 |
| 4 | 43074301 | rs7672044 | 0.154 | 0.644 | 0.67 | 0.35 | 0.008 | 0.079 | 0.079 | 0 | 0.16 | 0.21 | 0.17 | 0.49 | 0.23 | 0.28 |
| 4 | 43082956 | rs6850625 | -0.994 | 0.698 | 0.036 | 0.131 | 0.011 | 0.019 | 0.019 | 0 | 0.66 | 0.76 | 0.84 | 0.94 | 0.75 | 0.67 |
| 4 | 43106517 | rs6447224 | -1.138 | 0.71 | -0.187 | 0.148 | 0.018 | 0.047 | 0.047 | 0 | 0.67 | 0.81 | 0.86 | 0.94 | 0.78 | 0.68 |
| 4 | 43112381 | rs6845571 | 0.161 | 0.808 | -0.384 | 0.091 | 0.021 | -0.051 | -0.051 | 0 | 0.11 | 0.18 | 0.14 | 0 | 0.2 | 0.07 |
| 4 | 43112603 | rs12498763 | 0.818 | 0.679 | -1.008 | 0.306 | -0.03 | -0.098 | -0.098 | 1.567 | 0.36 | 0.09 | 0.51 | 0.44 | 0.32 | 0.26 |
| 4 | 43115315 | rs1583540 | -1.155 | 0.391 | -1.011 | -0.159 | 0.032 | 0.083 | 0.083 | 0 | 0.68 | 0.82 | 0.87 | 0.93 | 0.8 | 0.7 |
| 4 | 43124710 | rs1436485 | -1.217 | 0.357 | -1.096 | -0.129 | 0.023 | 0.104 | 0.104 | 0 | 0.69 | 0.85 | 0.86 | 0.93 | 0.78 | 0.7 |
| 4 | 43125020 | rs1436487 | -1.314 | 0.341 | -1.193 | -0.172 | 0.004 | 0.126 | 0.126 | 0 | 0.71 | 0.93 | 0.86 | 0.98 | 0.79 | 0.72 |
| 4 | 43128814 | rs6851917 | 0.65 | 0.598 | -1.274 | 0.206 | -0.123 | -0.133 | -0.133 | 0.088 | 0.37 | 0.03 | 0.58 | 0.54 | 0.42 | 0.25 |
| 4 | 43134215 | rs12503562 | 0.568 | 0.622 | -1.059 | 0.26 | -0.037 | -0.038 | -0.038 | 0 | 0.4 | 0.13 | 0.6 | 0.54 | 0.44 | 0.27 |
| 4 | 43136289 | rs6857139 | -0.178 | 0.766 | -0.433 | 0.383 | 0.046 | -0.018 | -0.018 | 0 | 0.12 | 0.35 | 0.13 | 0.05 | 0.18 | 0.08 |
| 4 | 43140284 | rs4861254 | -0.049 | 0.273 | -0.399 | 0.479 | -0.018 | 0.141 | 0.141 | 0.257 | 0.81 | 0.54 | 0.85 | 0.61 | 0.82 | 0.63 |
| 4 | 43145074 | rs4861052 | 0.007 | 0.374 | -0.545 | 0.511 | 0.009 | 0.003 | 0.003 | 0 | 0.51 | 0.49 | 0.73 | 0.51 | 0.62 | 0.35 |
| 4 | 43147514 | rs10213019 | 0.171 | 0.433 | -0.117 | 0.409 | -0.013 | 0.009 | 0.009 | 0 | 0.81 | 0.58 | 0.82 | 0.87 | 0.86 | 0.9 |
| 4 | 43148953 | rs7688320 | 0.226 | 0.24 | -0.775 | 0.153 | -0.001 | -0.003 | -0.003 | 0 | 0.45 | 0.51 | 0.33 | 0.4 | 0.55 | 0.67 |
| 4 | 43149962 | rs4257684 | 0.451 | 0.33 | -0.758 | 0.13 | 0.003 | -0.015 | -0.015 | 0.111 | 0.67 | 0.29 | 0.71 | 0.59 | 0.69 | 0.62 |
| 4 | 43151137 | rs4535364 | 1.098 | 0.345 | -0.605 | 0.145 | 0.017 | 0.066 | 0.066 | 0 | 0.14 | 0.27 | 0.1 | 0.28 | 0.17 | 0.28 |
| 4 | 43161354 | rs13114756 | 1.078 | 0.409 | -0.649 | 0.145 | 0.025 | 0.079 | 0.079 | 0 | 0.14 | 0.29 | 0.11 | 0.28 | 0.17 | 0.28 |
| 4 | 43162880 | rs12646066 | 0.999 | 0.329 | -0.631 | 0.042 | -0.087 | 0.11 | 0.11 | 0.732 | 0.3 | 0.11 | 0.22 | 0.12 | 0.38 | 0.39 |
| 4 | 43165259 | rs13120255 | 0.95 | 0.218 | -0.581 | 0.092 | 0.021 | -0.102 | -0.102 | 0 | 0.35 | 0.07 | 0.49 | 0.47 | 0.3 | 0.23 |
| 4 | 43180186 | rs6447253 | 0.577 | 0.723 | -0.245 | 0.183 | -0.002 | -0.009 | -0.009 | 0.013 | 0.09 | 0.1 | 0.11 | 0 | 0.21 | 0.12 |
| 4 | 43181231 | rs13113494 | -0.494 | 0.263 | 0.001 | 0.4 | -0.004 | 0.022 | 0.022 | 0.06 | 0.28 | 0.37 | 0.18 | 0.35 | 0.27 | 0.35 |
| 4 | 43191706 | rs4263432 | -0.603 | 0.36 | -0.165 | 0.431 | -0.006 | 0.028 | 0.028 | 0 | 0.29 | 0.4 | 0.18 | 0.35 | 0.27 | 0.35 |
| 4 | 43192298 | rs4270612 | -0.603 | 0.36 | -0.165 | 0.431 | -0.006 | 0.028 | 0.028 | 0 | 0.29 | 0.4 | 0.18 | 0.35 | 0.27 | 0.35 |
| 4 | 43197962 | rs13108021 | 0.99 | 0.315 | -0.82 | 0.349 | -0.093 | 0.026 | 0.026 | 0 | 0.25 | 0.07 | 0.36 | 0.18 | 0.36 | 0.26 |
| 4 | 43199190 | rs12505258 | 0.425 | -0.338 | -0.523 | 0.502 | -0.024 | 0.003 | 0.003 | 0.597 | 0.62 | 0.43 | 0.64 | 0.59 | 0.74 | 0.74 |
| 4 | 43205099 | rs4290920 | 0.346 | -0.328 | -0.48 | 0.507 | -0.037 | 0.008 | 0.008 | 0.451 | 0.63 | 0.51 | 0.64 | 0.58 | 0.74 | 0.73 |
| 4 | 43216821 | rs7666204 | -0.169 | -0.565 | -0.095 | 0.523 | 0.015 | -0.005 | -0.005 | 0 | 0.3 | 0.24 | 0.26 | 0.34 | 0.23 | 0.24 |

| | | | | | | | | | | | | | | | | |
|---|---|---|---|---|---|---|---|---|---|---|---|---|---|---|---|---|
| 4 | 43226893 | rs11932535 | 0.603 | -0.42 | 0.009 | 0.621 | 0.034 | -0.018 | -0.018 | 0 | 0.4 | 0.17 | 0.26 | 0.35 | 0.35 | 0.47 |
| 4 | 43232359 | rs4488964 | 1.12 | -0.367 | 0.182 | 0.605 | -0.042 | 0.026 | 0.026 | 0 | 0.3 | 0.23 | 0.4 | 0.23 | 0.42 | 0.29 |
| 4 | 43232462 | rs12645109 | 0.111 | -0.413 | 0.13 | 0.6 | 0.001 | -0.002 | -0.002 | 0 | 0.26 | 0.23 | 0.25 | 0.33 | 0.23 | 0.22 |
| 4 | 43232759 | rs4447896 | -0.897 | -0.16 | 0.308 | 0.556 | 0.033 | -0.018 | -0.018 | 0.482 | 0.67 | 0.43 | 0.52 | 0.69 | 0.59 | 0.69 |
| 4 | 43246631 | rs1442864 | 0.179 | -0.063 | -0.026 | 0.546 | 0.007 | -0.001 | -0.001 | 0 | 0.2 | 0.19 | 0.23 | 0.14 | 0.27 | 0.21 |
| 4 | 43252905 | rs12500109 | -0.335 | -0.228 | -0.182 | 0.499 | 0.044 | 0.026 | 0.026 | 0 | 0.48 | 0.36 | 0.59 | 0.37 | 0.63 | 0.47 |
| 4 | 43254582 | rs10938268 | 1.401 | -0.119 | -0.191 | 0.47 | -0.017 | 0.012 | 0.012 | 0 | 0.27 | 0.24 | 0.35 | 0.23 | 0.33 | 0.25 |
| 4 | 43254889 | rs7683961 | 0.432 | 0.119 | 0.225 | 1.045 | -0.027 | 0.022 | 0.022 | 0.055 | 0.17 | 0.08 | 0.21 | 0.06 | 0.25 | 0.1 |
| 4 | 43284304 | rs11934444 | 1.344 | -0.917 | 1.014 | 1.281 | -0.066 | 0.034 | 0.034 | 2.464 | 0.71 | 0.42 | 0.62 | 0.34 | 0.84 | 0.67 |
| 4 | 43285567 | rs17595833 | 0.77 | -1.022 | 1.035 | 1.456 | 0.022 | -0.016 | -0.016 | 0 | 0.1 | 0.07 | 0.07 | 0.24 | 0.05 | 0.04 |
| 4 | 43293318 | rs11730763 | 0.909 | -0.936 | 0.979 | 1.447 | -0.021 | 0.011 | 0.011 | 0 | 0.16 | 0.06 | 0.1 | 0.08 | 0.2 | 0.1 |
| 4 | 43296003 | rs1442870 | 0.171 | -0.935 | 1.247 | 1.554 | 0.001 | 0.02 | 0.02 | 0 | 0.13 | 0.12 | 0.21 | 0.34 | 0.08 | 0.27 |
| 4 | 43296814 | rs12505465 | 0.592 | -0.86 | 1.348 | 1.425 | -0.09 | 0.04 | 0.04 | 2.73 | 0.83 | 0.59 | 0.84 | 0.69 | 0.92 | 0.94 |
| 4 | 43305565 | rs17595974 | 0.672 | -0.693 | 1.134 | 1.399 | 0.035 | -0.031 | -0.031 | 0.78 | 0.1 | 0.06 | 0.06 | 0.26 | 0.05 | 0.04 |
| 4 | 43305896 | rs10029778 | -0.396 | -0.708 | 1.277 | 1.519 | 0.017 | -0.02 | -0.02 | 6.493 | 0.22 | 0.4 | 0.21 | 0.15 | 0.21 | 0.13 |
| 4 | 43323916 | rs1442855 | 2.398 | -0.768 | 1 | 1.423 | 0.007 | 0.26 | 0.26 | 0 | 0.49 | 0.1 | 0.48 | 0.15 | 0.6 | 0.42 |
| 4 | 43324563 | rs12510088 | 2.165 | -1.155 | 0.961 | 1.53 | 0.033 | 0.074 | 0.074 | 0 | 0.52 | 0.34 | 0.51 | 0.18 | 0.65 | 0.43 |
| 4 | 43328707 | rs6828560 | 1.42 | -1.115 | 0.984 | 1.568 | -0.002 | -0.002 | -0.002 | 0 | 0.62 | 0.43 | 0.57 | 0.43 | 0.69 | 0.48 |
| 4 | 43331637 | rs4861277 | -1.438 | -1.19 | 0.977 | 1.58 | -0.001 | 0 | 0 | 0 | 0.34 | 0.46 | 0.42 | 0.57 | 0.28 | 0.51 |
| 4 | 43344992 | rs6854231 | 1.994 | -1.813 | 1.316 | 1.985 | -0.033 | -0.004 | -0.004 | 0.281 | 0.67 | 0.37 | 0.63 | 0.25 | 0.83 | 0.57 |
| 4 | 43352577 | rs17596310 | 0.241 | -1.391 | 0.855 | 2.245 | -0.016 | 0.022 | 0.022 | 4.603 | 0.08 | 0.12 | 0.09 | 0.33 | 0.01 | 0.05 |
| 4 | 43353022 | rs920693 | -0.544 | -1.27 | 0.909 | 2.288 | 0.007 | -0.01 | -0.01 | 0 | 0.14 | 0.11 | 0.21 | 0.33 | 0.08 | 0.29 |
| 4 | 43363694 | rs7657677 | -1.372 | -1.28 | 0.997 | 2.285 | -0.006 | 0.01 | 0.01 | 0 | 0.21 | 0.26 | 0.32 | 0.67 | 0.12 | 0.33 |
| 4 | 43394110 | rs671254 | -0.558 | -1.244 | 1.354 | 2.239 | -0.027 | 0.053 | 0.053 | 0.414 | 0.07 | 0.22 | 0.06 | 0.09 | 0.05 | 0.11 |
| 4 | 43410470 | rs599290 | 0.939 | -1.16 | 1.221 | 2.266 | -0.021 | 0.039 | 0.039 | 0 | 0.79 | 0.69 | 0.68 | 0.32 | 0.87 | 0.68 |
| 4 | 43424930 | rs17639240 | 0.35 | -1.156 | 1.247 | 2.524 | -0.025 | 0.196 | 0.196 | 0.232 | 0.07 | 0.21 | 0.11 | 0.38 | 0.02 | 0.05 |
| 4 | 43428426 | rs654313 | 0.485 | -0.694 | 0.932 | 1.883 | 0.004 | -0.055 | -0.055 | 0 | 0.86 | 0.93 | 0.79 | 0.7 | 0.9 | 0.73 |
| 4 | 43438919 | rs6447271 | -0.028 | -0.711 | 0.964 | 1.825 | 0.014 | -0.035 | -0.035 | 0 | 0.89 | 0.93 | 0.82 | 0.73 | 0.93 | 0.75 |
| 4 | 43442648 | rs528582 | 2.142 | -1.002 | 1.817 | 1.81 | 0.002 | -0.005 | -0.005 | 0.088 | 0.75 | 0.21 | 0.64 | 0.61 | 0.87 | 0.67 |
| 4 | 43456817 | rs1631011 | 0.75 | -0.965 | 2.069 | 1.938 | -0.017 | 0.264 | 0.264 | 0 | 0.87 | 0.57 | 0.73 | 0.65 | 0.93 | 0.75 |
| 4 | 43468632 | rs10021165 | 0.268 | -0.9 | 1.172 | 1.459 | 0.006 | -0.024 | -0.024 | 0 | 0.68 | 0.88 | 0.66 | 0.7 | 0.75 | 0.55 |
| 4 | 43481992 | rs6447273 | 2.37 | -0.762 | 1.888 | 2.246 | -0.013 | 0.013 | 0.013 | 0.367 | 0.75 | 0.33 | 0.62 | 0.51 | 0.88 | 0.58 |
| 4 | 43484250 | rs7694215 | 0.686 | -0.608 | 1.598 | 2.191 | 0.008 | -0.023 | -0.023 | 0 | 0.85 | 0.91 | 0.73 | 0.71 | 0.91 | 0.73 |
| 4 | 43502330 | rs473726 | 0.926 | -0.727 | 1.55 | 2.199 | 0.025 | -0.04 | -0.04 | 0 | 0.86 | 0.9 | 0.75 | 0.73 | 0.92 | 0.75 |
| 4 | 43520590 | rs4861284 | 0.564 | -0.756 | 1.533 | 2.357 | 0.012 | -0.01 | -0.01 | 0 | 0.9 | 0.9 | 0.87 | 0.87 | 0.94 | 0.84 |
| 4 | 43523299 | rs13117635 | 2.433 | -0.91 | 1.942 | 2.281 | -0.054 | 0.064 | 0.064 | 0.299 | 0.76 | 0.35 | 0.61 | 0.62 | 0.89 | 0.68 |
| 4 | 43523444 | rs13136012 | 2.835 | -0.961 | 1.969 | 2.354 | -0.02 | 0.012 | 0.012 | 0.369 | 0.75 | 0.31 | 0.61 | 0.54 | 0.9 | 0.62 |
| 4 | 43526808 | rs9992804 | 1.576 | -0.709 | 2.163 | 1.884 | -0.04 | 0.091 | 0.091 | 0 | 0.83 | 0.66 | 0.66 | 0.66 | 0.92 | 0.73 |
| 4 | 43561857 | rs11939659 | 1.985 | -0.724 | 2.145 | 1.873 | -0.06 | 0.12 | 0.12 | 0 | 0.81 | 0.61 | 0.64 | 0.66 | 0.92 | 0.73 |
| 4 | 43564831 | rs6816987 | 2.386 | -0.724 | 2.22 | 1.873 | -0.058 | 0.084 | 0.084 | 0.126 | 0.78 | 0.37 | 0.62 | 0.62 | 0.92 | 0.69 |
| 4 | 43595446 | rs6447283 | 1.059 | -0.678 | 2.056 | 1.86 | 0.016 | -0.023 | -0.023 | 0.019 | 0.86 | 0.89 | 0.75 | 0.75 | 0.93 | 0.75 |
| 4 | 43609761 | rs6813203 | 0.796 | -0.663 | 1.929 | 1.817 | 0.023 | -0.067 | -0.067 | 0 | 0.87 | 0.96 | 0.76 | 0.76 | 0.93 | 0.76 |
| 4 | 43634967 | rs2013357 | 1.921 | -0.685 | 2.107 | 1.757 | -0.05 | 0.032 | 0.032 | 0 | 0.79 | 0.64 | 0.69 | 0.76 | 0.89 | 0.76 |
| 4 | 43635055 | rs985774 | 1.615 | -0.659 | 1.988 | 1.773 | 0.008 | -0.009 | -0.009 | 0.029 | 0.81 | 0.81 | 0.71 | 0.76 | 0.9 | 0.77 |
| 4 | 43656260 | rs12649203 | 2.246 | -0.59 | 2.085 | 1.759 | -0.031 | 0.027 | 0.027 | 0.058 | 0.77 | 0.34 | 0.66 | 0.7 | 0.88 | 0.71 |
| 4 | 43664997 | rs1962872 | 0.984 | -0.596 | 2.004 | 1.773 | 0.022 | -0.032 | -0.032 | 0 | 0.84 | 0.9 | 0.72 | 0.77 | 0.9 | 0.76 |
| 4 | 43666123 | rs13138756 | 2.892 | -0.68 | 2.134 | 1.814 | 0.022 | -0.026 | -0.026 | 0.062 | 0.74 | 0.23 | 0.63 | 0.63 | 0.88 | 0.68 |
| 4 | 43666139 | rs13137462 | 1.76 | -0.596 | 2.056 | 1.558 | -0.023 | 0.022 | 0.022 | 0 | 0.81 | 0.75 | 0.68 | 0.77 | 0.88 | 0.75 |
| 4 | 43680722 | rs1594921 | 0.901 | -0.607 | 1.875 | 1.553 | 0.028 | -0.045 | -0.045 | 0 | 0.83 | 0.92 | 0.73 | 0.77 | 0.9 | 0.77 |
| 4 | 43685461 | rs1648082 | 1.731 | -0.601 | 1.963 | 1.557 | -0.04 | 0.039 | 0.039 | 0 | 0.8 | 0.66 | 0.68 | 0.77 | 0.88 | 0.76 |
| 4 | 43689885 | rs1713517 | -0.375 | 0.252 | 1.347 | 0.798 | 0.036 | -0.049 | -0.049 | 0 | 0.23 | 0.35 | 0.16 | 0.11 | 0.25 | 0.24 |
| 4 | 43702376 | rs7659996 | 2.321 | 0.358 | 0.946 | 0.832 | -0.012 | -0.008 | -0.008 | 0.058 | 0.56 | 0.18 | 0.49 | 0.64 | 0.61 | 0.57 |
| 4 | 43705885 | rs1648078 | 0.299 | 0.506 | 1.047 | 0.807 | 0.083 | 0.073 | 0.073 | 0 | 0.14 | 0.36 | 0.26 | 0.17 | 0.22 | 0.22 |
| 4 | 43714649 | rs1369175 | 0.834 | 0.498 | 1.051 | 0.793 | 0.019 | 0.046 | 0.046 | 0 | 0.74 | 0.51 | 0.61 | 0.74 | 0.72 | 0.66 |
| 4 | 43722400 | rs1550046 | 1.873 | 0.499 | 1.063 | 0.782 | -0.002 | 0.007 | 0.007 | 0.003 | 0.61 | 0.32 | 0.54 | 0.74 | 0.63 | 0.6 |
| 4 | 43731044 | rs7340999 | 1.406 | 0.494 | 1.036 | 0.774 | 0 | 0.003 | 0.003 | 0.001 | 0.7 | 0.33 | 0.58 | 0.73 | 0.69 | 0.65 |
| 4 | 43738519 | rs1834429 | -0.196 | 0.505 | 0.969 | 0.779 | -0.005 | -0.055 | -0.055 | 0.054 | 0.13 | 0.18 | 0.07 | 0 | 0.09 | 0.06 |
| 4 | 43739772 | rs1369177 | 1.653 | 0.481 | 0.666 | 0.845 | -0.006 | 0.019 | 0.019 | 0.019 | 0.6 | 0.21 | 0.53 | 0.74 | 0.63 | 0.6 |
| 4 | 43751098 | rs1583408 | 0.871 | 0.492 | 0.433 | 0.905 | -0.008 | 0.002 | 0.002 | 0 | 0.87 | 0.81 | 0.87 | 0.86 | 0.94 | 0.8 |
| 4 | 43772856 | rs3849587 | 1.83 | 1.193 | 0.234 | 0.802 | 0.011 | 0.007 | 0.007 | 0 | 0.54 | 0.2 | 0.45 | 0.65 | 0.48 | 0.52 |
| 4 | 43777370 | rs1713511 | 0.512 | 1.397 | 0.402 | 0.833 | 0.063 | 0.008 | 0.008 | 0.041 | 0.16 | 0.32 | 0.28 | 0.12 | 0.27 | 0.19 |
| 4 | 43781839 | rs10938307 | 0.914 | 1.24 | -0.025 | 0.83 | -0.01 | -0.002 | -0.002 | 0 | 0.68 | 0.3 | 0.53 | 0.74 | 0.57 | 0.61 |
| 4 | 43786489 | rs2347347 | 1.239 | 1.209 | -0.169 | 0.789 | -0.047 | -0.028 | -0.028 | 0.065 | 0.63 | 0.21 | 0.48 | 0.65 | 0.54 | 0.57 |
| 4 | 43791881 | rs1648095 | -0.03 | 1.005 | -0.241 | 0.768 | 0.01 | -0.009 | -0.009 | 0 | 0.92 | 0.98 | 0.89 | 0.92 | 0.95 | 0.9 |
| 4 | 43792690 | rs10938308 | 1.9 | 0.532 | -0.596 | 0.324 | 0.023 | 0.002 | 0.002 | 0.079 | 0.43 | 0.2 | 0.37 | 0.65 | 0.43 | 0.49 |
| 4 | 43798822 | rs2678759 | 0.146 | 0.276 | -0.626 | 0.254 | 0.022 | 0.017 | 0.017 | 0.12 | 0.17 | 0.09 | 0.12 | 0.1 | 0.09 | 0.05 |
| 4 | 43810738 | rs2613157 | -1.173 | 0.24 | -0.564 | 0.128 | 0.052 | 0.088 | 0.088 | 0 | 0.31 | 0.16 | 0.23 | 0.09 | 0.15 | 0.11 |
| 4 | 43815025 | rs10938310 | 0.992 | 0.237 | -0.199 | 0.175 | -0.027 | 0.061 | 0.061 | 0.746 | 0.83 | 0.51 | 0.75 | 0.79 | 0.93 | 0.8 |
| 4 | 43823868 | rs4695155 | 1.07 | 0.808 | -0.538 | 0.512 | -0.037 | -0.035 | -0.035 | 0.05 | 0.07 | 0.01 | 0.16 | 0.22 | 0.25 | 0.18 |

| | | | | | | | | | | | | | | | | |
|---|---|---|---|---|---|---|---|---|---|---|---|---|---|---|---|---|
| 4 | 43827201 | rs1460627 | 2.098 | 0.052 | -0.192 | -0.203 | -0.01 | 0.018 | 0.018 | 0.258 | 0.46 | 0.25 | 0.36 | 0.64 | 0.54 | 0.54 |
| 4 | 43832158 | rs1460613 | -0.145 | 0.049 | -0.224 | -0.249 | 0.034 | 0.093 | 0.093 | 0 | 0.21 | 0.06 | 0.11 | 0 | 0.12 | 0.07 |
| 4 | 43840138 | rs7675193 | 0.785 | 0.066 | -0.187 | -0.234 | 0.027 | 0.002 | 0.002 | 0.108 | 0.16 | 0.22 | 0.29 | 0.13 | 0.27 | 0.2 |
| 4 | 43840953 | rs2715650 | 1.301 | -0.288 | -0.125 | -0.105 | 0.019 | 0.061 | 0.061 | 0 | 0.7 | 0.89 | 0.69 | 0.83 | 0.82 | 0.81 |
| 4 | 43853032 | rs2722147 | 0.554 | -0.294 | -0.11 | -0.081 | -0.01 | -0.001 | -0.001 | 0.046 | 0.82 | 0.89 | 0.77 | 0.83 | 0.89 | 0.83 |
| 4 | 43853511 | rs1460608 | -0.047 | -0.294 | -0.116 | -0.082 | -0.009 | -0.028 | -0.028 | 0 | 0.91 | 0.94 | 0.8 | 0.83 | 0.94 | 0.88 |
| 4 | 43854629 | rs9790652 | 0.571 | -0.294 | -0.117 | -0.082 | -0.013 | -0.002 | -0.002 | 0 | 0.82 | 0.9 | 0.77 | 0.83 | 0.89 | 0.83 |
| 4 | 43857879 | rs7660461 | -0.026 | -0.222 | 0.21 | -0.055 | -0.058 | -0.1 | -0.1 | 0 | 0.15 | 0.35 | 0.1 | 0 | 0.07 | 0.01 |
| 4 | 43866979 | rs2722136 | -0.743 | -0.257 | 0.204 | -0.072 | -0.087 | -0.133 | -0.133 | 0 | 0.19 | 0.42 | 0.12 | 0 | 0.08 | 0.04 |
| 4 | 43872019 | rs1460614 | 1.529 | -0.233 | 0.125 | -0.027 | -0.06 | -0.04 | -0.04 | 0.152 | 0.71 | 0.5 | 0.67 | 0.81 | 0.85 | 0.79 |
| 4 | 43891003 | rs10008858 | 1.401 | -0.073 | 0.078 | -0.017 | -0.004 | 0.043 | 0.043 | 0.473 | 0.48 | 0.26 | 0.31 | 0.66 | 0.58 | 0.56 |
| 4 | 43900285 | rs898573 | -0.559 | -0.261 | 0.166 | -0.041 | -0.007 | -0.03 | -0.03 | 0 | 0.1 | 0.05 | 0.19 | 0.16 | 0.05 | 0.15 |
| 4 | 43903247 | rs11931045 | -1.125 | -0.311 | 0.307 | 0.02 | 0.002 | 0.008 | 0.008 | 0.002 | 0.08 | 0.37 | 0.05 | 0.04 | 0.04 | 0.08 |
| 4 | 43911686 | rs7679442 | 1.175 | -0.176 | 0.126 | 0.057 | -0.032 | -0.037 | -0.037 | 0 | 0.72 | 0.48 | 0.59 | 0.8 | 0.84 | 0.78 |
| 4 | 43913465 | rs12644632 | -0.112 | 0.262 | -0.144 | 0.285 | 0.016 | -0.02 | -0.02 | 1.108 | 0.65 | 0.46 | 0.6 | 0.78 | 0.58 | 0.62 |
| 4 | 43914603 | rs12513382 | 0.577 | 0.438 | -0.133 | 0.388 | 0.049 | 0.05 | 0.05 | 0 | 0.5 | 0.23 | 0.29 | 0.65 | 0.5 | 0.46 |
| 4 | 43927834 | rs1037855 | -0.038 | 0.562 | -0.051 | 0.212 | 0.022 | -0.04 | -0.04 | 0.01 | 0.6 | 0.5 | 0.59 | 0.8 | 0.56 | 0.67 |
| 4 | 43934098 | rs4426822 | 0.534 | 1.043 | -0.107 | 0.274 | 0.003 | -0.032 | -0.032 | 0 | 0.31 | 0.43 | 0.34 | 0.14 | 0.31 | 0.22 |
| 4 | 43935421 | rs11737698 | -0.008 | 1.047 | -0.097 | 0.309 | 0.003 | -0.041 | -0.041 | 0 | 0.36 | 0.49 | 0.38 | 0.18 | 0.34 | 0.26 |
| 4 | 43941075 | rs11934528 | 0.051 | 0.955 | -0.144 | 0.273 | 0.006 | -0.042 | -0.042 | 0 | 0.36 | 0.52 | 0.44 | 0.17 | 0.35 | 0.33 |
| 4 | 43951966 | rs2881695 | -0.277 | 0.126 | -0.708 | 0.488 | 0.014 | -0.002 | -0.002 | 0 | 0.53 | 0.67 | 0.68 | 0.56 | 0.51 | 0.48 |
| 4 | 43952928 | rs1460639 | 0.44 | 0.075 | -0.847 | 0.459 | 0.001 | -0.072 | -0.072 | 0.093 | 0.41 | 0.12 | 0.28 | 0.44 | 0.46 | 0.46 |
| 4 | 43960180 | rs4385106 | 0.27 | 0.103 | -1.163 | 0.787 | 0.039 | -0.022 | -0.022 | 0 | 0.45 | 0.23 | 0.3 | 0.44 | 0.47 | 0.48 |
| 4 | 43963518 | rs991382 | 0.883 | 0.254 | -0.488 | 0.878 | 0.001 | -0.003 | -0.003 | 0 | 0.9 | 0.86 | 0.73 | 0.9 | 0.94 | 0.88 |
| 4 | 43968054 | rs1460634 | -0.849 | 0.18 | -0.652 | 1.116 | 0.002 | -0.003 | -0.003 | 0 | 0.38 | 0.42 | 0.43 | 0.57 | 0.33 | 0.51 |
| 4 | 43971237 | rs17598761 | -0.943 | 0.355 | -0.51 | 1.59 | 0.012 | 0.034 | 0.034 | 4.413 | 0.13 | 0.17 | 0.2 | 0.36 | 0.19 | 0.29 |
| 4 | 43974714 | rs7700023 | 0.111 | 0.732 | -0.891 | 2.062 | -0.039 | -0.036 | -0.036 | 0 | 0.08 | 0.01 | 0.22 | 0.32 | 0.17 | 0.2 |
| 4 | 43982267 | rs7694541 | -0.161 | -0.233 | -0.783 | 1.503 | 0.016 | 0.019 | 0.019 | 0.042 | 0.18 | 0.1 | 0.19 | 0.09 | 0.13 | 0.2 |
| 4 | 43982530 | rs10010379 | 1.239 | -0.406 | 0.153 | 1.559 | -0.014 | 0.029 | 0.029 | 0 | 0.67 | 0.53 | 0.66 | 0.6 | 0.75 | 0.61 |
| 4 | 43986534 | rs11942913 | -0.364 | -0.355 | 0.279 | 1.307 | -0.005 | -0.006 | -0.006 | 0.896 | 0.19 | 0.21 | 0.21 | 0.09 | 0.13 | 0.2 |
| 4 | 44016954 | rs4323141 | -0.499 | -0.303 | 0.401 | 1.42 | -0.003 | 0.013 | 0.013 | 0.035 | 0.18 | 0.25 | 0.26 | 0.19 | 0.18 | 0.28 |
| 4 | 44029345 | rs4348131 | 0.621 | -0.64 | 0.162 | 1.222 | 0 | -0.007 | -0.007 | 0 | 0.84 | 0.86 | 0.9 | 0.7 | 0.85 | 0.87 |
| 4 | 44041667 | rs4695394 | -0.561 | -1.312 | -0.102 | 0.611 | -0.052 | -0.099 | -0.099 | 0 | 0.28 | 0.47 | 0.26 | 0.06 | 0.18 | 0.18 |
| 4 | 44042710 | rs12331900 | 1.333 | -1.281 | -0.348 | 0.722 | 0.083 | 0.027 | 0.027 | 0.277 | 0.52 | 0.2 | 0.61 | 0.59 | 0.66 | 0.64 |
| 4 | 44050101 | rs6858778 | -1.875 | -1.216 | -0.37 | 0.775 | 0.057 | -0.014 | -0.014 | 0 | 0.49 | 0.82 | 0.45 | 0.49 | 0.39 | 0.46 |
| 4 | 44051144 | rs4540086 | -1.731 | -1.455 | -0.342 | 0.49 | 0.078 | 0.031 | 0.031 | 0.143 | 0.47 | 0.76 | 0.39 | 0.41 | 0.34 | 0.35 |
| 4 | 44091711 | rs7693710 | -0.976 | -1.474 | 0.178 | 0.599 | -0.038 | -0.012 | -0.012 | 0 | 0.28 | 0.32 | 0.25 | 0.29 | 0.18 | 0.24 |
| 4 | 44093499 | rs4695447 | -2.258 | -1.789 | 0.288 | 0.373 | 0.084 | 0.035 | 0.035 | 0.2 | 0.48 | 0.75 | 0.37 | 0.43 | 0.33 | 0.35 |
| 4 | 44100094 | rs7666961 | -0.128 | -1.555 | 0.668 | 0.416 | -0.003 | -0.086 | -0.086 | 0 | 0.13 | 0.39 | 0.12 | 0 | 0.12 | 0.03 |
| 4 | 44106293 | rs12498358 | -0.511 | -1.572 | 0.741 | 0.407 | -0.011 | -0.115 | -0.115 | 0.758 | 0.15 | 0.52 | 0.13 | 0 | 0.12 | 0.03 |
| 4 | 44109192 | rs12511977 | -0.996 | -1.712 | 0.698 | 0.38 | 0.009 | -0.018 | -0.018 | 0 | 0.17 | 0.11 | 0.08 | 0.34 | 0.11 | 0.16 |
| 4 | 44112560 | rs6840648 | 0.622 | -1.736 | 0.658 | 0.372 | -0.003 | -0.005 | -0.005 | 0.512 | 0.14 | 0.1 | 0.17 | 0.07 | 0.11 | 0.16 |
| 4 | 44121146 | rs7689034 | -2.318 | -1.451 | 0.765 | 0.493 | -0.026 | -0.005 | -0.005 | 0 | 0.32 | 0.64 | 0.18 | 0.34 | 0.2 | 0.19 |
| 4 | 44133596 | rs6813741 | -1.186 | -0.722 | 0.522 | 0.89 | -0.002 | 0.036 | 0.036 | 1.269 | 0.09 | 0.08 | 0.03 | 0.34 | 0.05 | 0.15 |
| 4 | 44137398 | rs9996682 | -0.704 | -0.605 | 0.468 | 0.249 | 0 | -0.08 | -0.08 | 0.095 | 0.13 | 0.36 | 0.11 | 0 | 0.12 | 0.03 |
| 4 | 44144150 | rs6853456 | -1.123 | -1.055 | 0.691 | 0.355 | -0.054 | -0.017 | -0.017 | 0 | 0.13 | 0.26 | 0.06 | 0.14 | 0.05 | 0.07 |
| 4 | 44149564 | rs6828553 | -1.756 | -0.693 | 0.545 | -0.455 | -0.021 | -0.092 | -0.092 | 0 | 0.32 | 0.46 | 0.16 | 0.21 | 0.25 | 0.34 |
| 4 | 44152250 | rs7666732 | -1.516 | -0.671 | 0.588 | -0.495 | -0.035 | -0.042 | -0.042 | 0.307 | 0.17 | 0.35 | 0.09 | 0.14 | 0.1 | 0.1 |
| 4 | 44152766 | rs13116771 | -0.372 | -0.657 | 0.611 | -0.501 | -0.004 | 0.015 | 0.015 | 0 | 0.16 | 0.08 | 0.07 | 0.07 | 0.15 | 0.24 |
| 4 | 44167271 | rs1027201 | -2.057 | 0.27 | 0.252 | 0.729 | -0.001 | 0.079 | 0.079 | 0 | 0.23 | 0.38 | 0.31 | 0.49 | 0.25 | 0.25 |
| 4 | 44175533 | rs35939382 | 0.018 | 0.274 | 0.311 | 0.714 | -0.031 | 0.06 | 0.06 | 0 | 0.13 | 0.02 | 0.06 | 0 | 0.15 | 0.18 |
| 4 | 44186433 | rs4694999 | -0.229 | 0.323 | 0.306 | 0.844 | -0.036 | 0.015 | 0.015 | 0 | 0.07 | 0.02 | 0.21 | 0.35 | 0.14 | 0.14 |
| 4 | 44187682 | rs4695701 | -2.491 | -0.248 | 0.781 | 0.116 | 0.021 | 0.018 | 0.018 | 0 | 0.51 | 0.71 | 0.44 | 0.54 | 0.46 | 0.51 |
| 4 | 44196006 | rs6811002 | -1.999 | 0.141 | 1.279 | 0.688 | 0.073 | 0.134 | 0.134 | 0.414 | 0.15 | 0.41 | 0.16 | 0.24 | 0.2 | 0.18 |
| 4 | 44210076 | rs17641486 | -0.779 | 0.458 | 1.29 | 0.724 | 0.002 | -0.003 | -0.003 | 0 | 0.23 | 0.16 | 0.22 | 0.28 | 0.2 | 0.32 |
| 4 | 44218748 | rs10028283 | -2.547 | 0.104 | 0.923 | 0.412 | -0.001 | -0.004 | -0.004 | 0 | 0.43 | 0.57 | 0.42 | 0.52 | 0.44 | 0.5 |
| 4 | 44246039 | rs10938330 | 0.208 | 0.296 | 0.798 | 0.468 | -0.027 | -0.005 | -0.005 | 3.657 | 0.07 | 0.03 | 0.09 | 0.24 | 0.16 | 0.15 |
| 4 | 44260690 | rs11941070 | -1.527 | 0.564 | -0.335 | -0.55 | -0.006 | 0.081 | 0.081 | 1.789 | 0.48 | 0.76 | 0.46 | 0.66 | 0.5 | 0.46 |
| 4 | 44273405 | rs1038767 | 1.809 | 0.506 | -0.4 | -0.563 | 0.022 | -0.026 | -0.026 | 0 | 0.09 | 0.01 | 0.16 | 0.11 | 0.05 | 0.1 |
| 4 | 44276751 | rs10007704 | -0.254 | 0.569 | -0.135 | -0.57 | 0.001 | -0.004 | -0.004 | 0 | 0.36 | 0.4 | 0.38 | 0.65 | 0.39 | 0.36 |
| 4 | 44280236 | rs1948314 | 0.452 | 0.395 | 0.011 | -0.554 | -0.019 | 0.024 | 0.024 | 0.056 | 0.29 | 0.4 | 0.31 | 0.4 | 0.23 | 0.22 |
| 4 | 44293409 | rs10013818 | -0.108 | 0.58 | -0.226 | -0.738 | 0.014 | 0.003 | 0.003 | 0.27 | 0.26 | 0.2 | 0.24 | 0.28 | 0.19 | 0.25 |
| 4 | 44300248 | rs3963342 | 0.635 | 0.673 | -0.15 | -0.617 | 0.003 | 0.008 | 0.008 | 0 | 0.64 | 0.67 | 0.69 | 0.72 | 0.69 | 0.65 |
| 4 | 44308547 | rs1490452 | 1.447 | 0.798 | -0.46 | -0.604 | -0.003 | -0.002 | -0.002 | 1.113 | 0.55 | 0.43 | 0.59 | 0.47 | 0.53 | 0.5 |
| 4 | 44315428 | rs17538903 | 2.922 | 0.798 | -0.518 | -0.629 | -0.055 | 0.107 | 0.107 | 0.787 | 0.26 | 0.05 | 0.28 | 0.07 | 0.3 | 0.27 |
| 4 | 44329589 | rs17599591 | 1.84 | 0.313 | -0.494 | -0.198 | -0.006 | 0.009 | 0.009 | 0.013 | 0.07 | 0.07 | 0.18 | 0.14 | 0.04 | 0.07 |
| 4 | 44334829 | rs17599605 | 0.406 | 0.365 | -0.499 | -0.314 | -0.009 | -0.006 | -0.006 | 0 | 0.14 | 0.1 | 0.26 | 0.39 | 0.19 | 0.23 |
| 4 | 44341530 | rs2703833 | 0.766 | 0.331 | -0.506 | -0.427 | -0.006 | -0.002 | -0.002 | 0.253 | 0.22 | 0.33 | 0.13 | 0.25 | 0.2 | 0.15 |
| 4 | 44348032 | rs10517102 | 1.734 | 0.342 | -0.503 | -0.366 | -0.017 | 0.033 | 0.033 | 0.169 | 0.41 | 0.29 | 0.39 | 0.33 | 0.45 | 0.41 |

| | | | | | | | | | | | | | | | | |
|---|---|---|---|---|---|---|---|---|---|---|---|---|---|---|---|---|
| 4 | 44351013 | rs11732997 | -1.382 | 0.333 | -0.936 | -0.636 | 0.014 | 0.03 | 0.03 | 0 | 0.55 | 0.68 | 0.62 | 0.78 | 0.63 | 0.57 |
| 4 | 44351941 | rs10517103 | 1.283 | 0.551 | -1.605 | -0.483 | -0.045 | 0.044 | 0.044 | 0 | 0.08 | 0.02 | 0.03 | 0 | 0.1 | 0.1 |
| 4 | 44358412 | rs6447330 | 1.1 | 0.486 | -2.05 | -0.071 | -0.01 | 0.002 | 0.002 | 0 | 0.08 | 0.03 | 0.11 | 0.26 | 0.15 | 0.12 |
| 4 | 44363703 | rs13132322 | 1.996 | 0.441 | -2.226 | -0.569 | -0.055 | 0.125 | 0.125 | 0.231 | 0.35 | 0.11 | 0.33 | 0.19 | 0.37 | 0.37 |
| 4 | 44372045 | rs7674464 | -0.699 | 0.448 | -1.598 | -0.303 | -0.004 | 0.027 | 0.027 | 0.032 | 0.3 | 0.26 | 0.27 | 0.09 | 0.27 | 0.28 |
| 4 | 44377494 | rs17653012 | 1.179 | 0.358 | -1.518 | 0.396 | -0.028 | -0.027 | -0.027 | 0.401 | 0.06 | 0.02 | 0.03 | 0.22 | 0.1 | 0.08 |
| 4 | 44381632 | rs16856866 | 0.393 | 0.089 | -1.498 | 0.138 | 0.023 | 0.012 | 0.012 | 0.378 | 0.12 | 0.07 | 0.19 | 0.09 | 0.08 | 0.17 |
| 4 | 44381959 | rs4407541 | -0.18 | -0.302 | -1.297 | -0.181 | -0.003 | -0.002 | -0.002 | 0 | 0.69 | 0.63 | 0.83 | 0.72 | 0.73 | 0.86 |
| 4 | 44400111 | rs17599994 | 0.898 | -0.345 | -1.538 | -0.167 | -0.011 | -0.03 | -0.03 | 0.168 | 0.29 | 0.16 | 0.32 | 0.37 | 0.29 | 0.37 |
| 4 | 44401672 | rs11726514 | 1.708 | 0.062 | -1.247 | -0.066 | -0.038 | -0.026 | -0.026 | 0 | 0.21 | 0.13 | 0.29 | 0.27 | 0.26 | 0.28 |
| 4 | 44402331 | rs17461113 | 0.367 | 0.159 | -0.281 | 0.129 | 0.009 | 0.001 | 0.001 | 0.569 | 0.1 | 0.08 | 0.13 | 0.08 | 0.08 | 0.16 |
| 4 | 44412937 | rs4695718 | -0.453 | 0.1 | -0.095 | 0.092 | -0.039 | 0.072 | 0.072 | 0.061 | 0.8 | 0.53 | 0.86 | 0.72 | 0.86 | 0.88 |
| 4 | 44414165 | rs7664938 | 0.93 | 0.192 | -0.223 | -0.066 | -0.003 | -0.012 | -0.012 | 0 | 0.15 | 0.11 | 0.15 | 0 | 0.2 | 0.04 |
| 4 | 44423496 | rs11931753 | -0.445 | -0.01 | -0.592 | 0.159 | 0.021 | 0.246 | 0.246 | 0.008 | 0.07 | 0.38 | 0.14 | 0.2 | 0.02 | 0.11 |
| 4 | 44424487 | rs4312793 | 1.509 | 0.305 | -0.957 | -0.056 | -0.021 | 0.001 | 0.001 | 0 | 0.51 | 0.22 | 0.56 | 0.51 | 0.52 | 0.64 |
| 4 | 44428788 | rs17461197 | 0.816 | 0.442 | -0.954 | -0.029 | -0.035 | 0.032 | 0.032 | 0 | 0.08 | 0.02 | 0.03 | 0 | 0.12 | 0.1 |
| 4 | 44433864 | rs10938341 | 0.837 | 0.218 | -0.92 | 0.047 | 0.046 | -0.053 | -0.053 | 0.322 | 0.59 | 0.18 | 0.59 | 0.5 | 0.63 | 0.74 |
| 4 | 44435648 | rs16856919 | -0.156 | 0.185 | -0.83 | 0.094 | -0.019 | 0.003 | 0.003 | 0 | 0.23 | 0.51 | 0.29 | 0.27 | 0.22 | 0.14 |
| 4 | 44435685 | rs13144404 | -0.833 | 0.193 | -1.117 | 0.121 | -0.004 | 0.004 | 0.004 | 0 | 0.82 | 0.75 | 0.89 | 0.77 | 0.86 | 0.88 |
| 4 | 44437034 | rs16856929 | 0.103 | 0.22 | -0.976 | 0.157 | 0.061 | 0.381 | 0.381 | 1.57 | 0.06 | 0.39 | 0.14 | 0.26 | 0.01 | 0.04 |
| 4 | 44438741 | rs4695004 | 1.805 | 0.348 | -1.365 | 0.035 | -0.043 | -0.049 | -0.049 | 0.416 | 0.42 | 0.17 | 0.45 | 0.44 | 0.44 | 0.53 |
| 4 | 44443246 | rs4419508 | 0.245 | 0.094 | -1.052 | 0.071 | 0 | -0.041 | -0.041 | 0 | 0.17 | 0.18 | 0.15 | 0 | 0.21 | 0.07 |
| 4 | 44445943 | rs10024393 | -0.153 | 0.159 | -1.288 | 0.66 | -0.001 | 0 | 0 | 2.31 | 0.83 | 0.78 | 0.89 | 0.77 | 0.86 | 0.88 |
| 4 | 44451821 | rs17461326 | -0.497 | -0.038 | -1.646 | 0.373 | 0.067 | 0.079 | 0.079 | 0.02 | 0.2 | 0.02 | 0.16 | 0.06 | 0.11 | 0.13 |
| 4 | 44452305 | rs13133564 | 1.21 | 0.631 | -2.071 | 0.696 | -0.104 | -0.017 | -0.017 | 0 | 0.45 | 0.09 | 0.3 | 0.37 | 0.46 | 0.51 |
| 4 | 44465870 | rs4610370 | -0.31 | 0.982 | -0.744 | 1.344 | 0.024 | 0.138 | 0.138 | 0 | 0.11 | 0.26 | 0.21 | 0.31 | 0.14 | 0.25 |
| 4 | 44481810 | rs7697982 | 0.674 | 1.065 | -0.537 | 1.281 | 0.032 | 0.083 | 0.083 | 0 | 0.79 | 0.47 | 0.74 | 0.6 | 0.76 | 0.81 |
| 4 | 44488196 | rs11731229 | 0.668 | 1.063 | -0.528 | 1.28 | 0.032 | 0.083 | 0.083 | 0 | 0.79 | 0.47 | 0.74 | 0.6 | 0.76 | 0.81 |
| 4 | 44494356 | rs11939023 | 1.719 | 0.849 | -1.249 | 0.814 | -0.054 | 0.057 | 0.057 | 0 | 0.41 | 0.07 | 0.24 | 0.22 | 0.36 | 0.41 |
| 4 | 44528928 | rs1595656 | 0.185 | 0.973 | -0.135 | 0.925 | 0.044 | 0.008 | 0.008 | 0 | 0.86 | 0.64 | 0.88 | 0.88 | 0.85 | 0.9 |
| 4 | 44530236 | rs2118934 | -0.734 | 0.971 | -0.308 | 0.897 | -0.001 | -0.008 | -0.008 | 0 | 0.4 | 0.54 | 0.62 | 0.68 | 0.44 | 0.44 |
| 4 | 44531529 | rs1371238 | 1.205 | 0.837 | -0.627 | 1.101 | 0.029 | -0.097 | -0.097 | 0 | 0.66 | 0.17 | 0.62 | 0.49 | 0.74 | 0.67 |
| 4 | 44532032 | rs6838754 | 1.926 | 0.821 | -0.602 | 1.061 | -0.042 | 0.143 | 0.143 | 0 | 0.46 | 0.11 | 0.26 | 0.2 | 0.41 | 0.46 |
| 4 | 44536354 | rs1439382 | -1.836 | 0.773 | -0.264 | 1.041 | 0.018 | 0.098 | 0.098 | 0 | 0.51 | 0.74 | 0.73 | 0.7 | 0.58 | 0.46 |
| 4 | 44542952 | rs1439389 | -0.494 | 0.735 | 0.04 | 1.063 | 0.049 | 0.044 | 0.044 | 0 | 0.27 | 0.34 | 0.45 | 0.42 | 0.43 | 0.3 |
| 4 | 44545381 | rs11736162 | 2.535 | 0.727 | -0.02 | 1.026 | -0.022 | 0.099 | 0.099 | 0 | 0.42 | 0.09 | 0.24 | 0.22 | 0.36 | 0.42 |
| 4 | 44549846 | rs7684647 | 0.378 | 0.784 | 0.02 | 1.084 | -0.073 | -0.068 | -0.068 | 0.257 | 0.2 | 0.06 | 0.35 | 0.29 | 0.33 | 0.22 |
| 4 | 44553869 | rs10517108 | 0.065 | 0.775 | 0.056 | 1.089 | 0.123 | 0.112 | 0.112 | 0 | 0.07 | 0.27 | 0.09 | 0.12 | 0.11 | 0.08 |
| 4 | 44557599 | rs1371235 | 0.266 | 0.65 | 0.146 | 1.072 | -0.025 | 0.013 | 0.013 | 0.812 | 0.77 | 0.7 | 0.72 | 0.71 | 0.85 | 0.84 |
| 4 | 44560636 | rs1898858 | -1.267 | 0.728 | -0.227 | 0.967 | 0.02 | 0.033 | 0.033 | 0.219 | 0.88 | 0.96 | 0.87 | 0.97 | 0.92 | 0.9 |
| 4 | 44575415 | rs12509197 | 0.064 | 1.024 | -0.386 | 1.036 | -0.047 | -0.043 | -0.043 | 0 | 0.2 | 0.11 | 0.35 | 0.29 | 0.33 | 0.22 |
| 4 | 44580312 | rs1371237 | -1.385 | 1.025 | -0.129 | 0.781 | 0.011 | 0.054 | 0.054 | 0 | 0.51 | 0.63 | 0.61 | 0.67 | 0.56 | 0.5 |
| 4 | 44580804 | rs13140831 | 0.361 | 0.732 | 0.235 | 0.932 | -0.01 | -0.007 | -0.007 | 0 | 0.64 | 0.68 | 0.59 | 0.55 | 0.55 | 0.68 |
| 4 | 44582487 | rs10938353 | 0.53 | 0.692 | 0.885 | 0.98 | 0.041 | 0.001 | 0.001 | 0 | 0.84 | 0.62 | 0.71 | 0.83 | 0.82 | 0.83 |
| 4 | 44592060 | rs13138228 | 1.283 | 0.629 | 1.021 | 1.066 | 0.031 | 0.062 | 0.062 | 0.231 | 0.8 | 0.52 | 0.64 | 0.7 | 0.79 | 0.76 |
| 4 | 44603369 | rs11725120 | 1.696 | 0.495 | 0.513 | 0.772 | 0.051 | 0.055 | 0.055 | 0 | 0.39 | 0.14 | 0.15 | 0.25 | 0.27 | 0.34 |
| 4 | 44607572 | rs17571930 | -0.491 | 0.531 | 0.647 | 0.628 | 0.034 | 0.001 | 0.001 | 0 | 0.14 | 0.25 | 0.28 | 0.17 | 0.18 | 0.19 |
| 4 | 44610348 | rs6844093 | -0.559 | 0.56 | 0.605 | 0.791 | 0.004 | -0.003 | -0.003 | 0 | 0.42 | 0.43 | 0.54 | 0.59 | 0.52 | 0.44 |
| 4 | 44613461 | rs16857230 | 0.023 | 0.613 | 0.271 | 0.731 | -0.03 | -0.02 | -0.02 | 0.255 | 0.62 | 0.72 | 0.52 | 0.57 | 0.49 | 0.65 |
| 4 | 44616283 | rs11736925 | 1.493 | 0.361 | 0.051 | 0.454 | 0.041 | 0.035 | 0.035 | 0 | 0.36 | 0.23 | 0.13 | 0.23 | 0.23 | 0.32 |
| 4 | 44616723 | rs1439365 | 0.337 | 0.462 | 0.301 | 0.437 | 0.034 | -0.009 | -0.009 | 0 | 0.48 | 0.7 | 0.46 | 0.53 | 0.39 | 0.54 |
| 4 | 44628607 | rs1439377 | 0.559 | 0.433 | 0.439 | 0.406 | 0.014 | 0 | 0 | 0 | 0.48 | 0.41 | 0.43 | 0.56 | 0.39 | 0.53 |
| 4 | 44648797 | rs6447363 | -0.717 | 0.515 | 0.813 | 0.533 | 0.004 | 0 | 0 | 0 | 0.51 | 0.46 | 0.67 | 0.52 | 0.66 | 0.53 |
| 4 | 44652030 | rs2348353 | -0.263 | 0.51 | 0.706 | 0.597 | -0.002 | -0.014 | -0.014 | 0 | 0.09 | 0.06 | 0.18 | 0.25 | 0.1 | 0.12 |
| 4 | 44656408 | rs17600990 | -0.118 | 0.426 | 0.692 | 0.572 | -0.002 | -0.012 | -0.012 | 0 | 0.09 | 0.06 | 0.18 | 0.25 | 0.1 | 0.12 |
| 4 | 44660248 | rs12639722 | 1.636 | 0.353 | 0.818 | 0.535 | 0.051 | 0.047 | 0.047 | 0 | 0.37 | 0.2 | 0.13 | 0.23 | 0.24 | 0.35 |
| 4 | 44663178 | rs6826194 | 1.138 | 0.469 | 0.459 | 0.306 | -0.127 | -0.137 | -0.137 | 0 | 0.19 | 0.02 | 0.38 | 0.39 | 0.38 | 0.23 |
| 4 | 44687401 | rs12499960 | -1.818 | -0.09 | 0.903 | 0.366 | 0.02 | 0.019 | 0.019 | 0 | 0.61 | 0.64 | 0.85 | 0.77 | 0.76 | 0.65 |
| 4 | 44701750 | rs6812211 | -0.891 | -0.063 | 0.802 | 0.32 | 0.035 | 0.014 | 0.014 | 0 | 0.48 | 0.29 | 0.52 | 0.4 | 0.6 | 0.46 |
| 4 | 44710346 | rs1547525 | -0.863 | -0.061 | 0.749 | 0.321 | 0.034 | 0.013 | 0.013 | 0 | 0.48 | 0.29 | 0.52 | 0.4 | 0.6 | 0.46 |
| 4 | 44732143 | rs1532194 | -0.82 | -0.071 | 0.602 | 0.309 | 0.033 | 0.011 | 0.011 | 0 | 0.48 | 0.3 | 0.52 | 0.41 | 0.6 | 0.46 |
| 4 | 44770222 | rs2046909 | -0.554 | 0.232 | 0.769 | 0.498 | -0.002 | -0.009 | -0.009 | 1.064 | 0.41 | 0.26 | 0.49 | 0.4 | 0.57 | 0.46 |
| 4 | 44778348 | rs6828324 | -0.786 | 0.289 | 0.617 | 0.631 | 0.018 | 0.076 | 0.076 | 0 | 0.11 | 0.18 | 0.32 | 0.37 | 0.16 | 0.18 |
| 4 | 44785389 | rs1491313 | -0.638 | 0.315 | 0.627 | 0.655 | 0.022 | 0.083 | 0.083 | 0 | 0.1 | 0.19 | 0.32 | 0.37 | 0.15 | 0.18 |
| 4 | 44802221 | rs1491328 | 0.296 | 0.399 | 0.527 | 0.591 | -0.021 | -0.033 | -0.033 | 2.645 | 0.6 | 0.82 | 0.47 | 0.6 | 0.43 | 0.52 |
| 4 | 44803683 | rs6845593 | 0.638 | 0.384 | 0.633 | 0.584 | 0 | -0.001 | -0.001 | 1.131 | 0.57 | 0.57 | 0.45 | 0.6 | 0.43 | 0.52 |
| 4 | 44849345 | rs6816452 | -0.482 | 0.412 | 0.479 | 0.616 | -0.072 | -0.054 | -0.054 | 0 | 0.72 | 0.87 | 0.53 | 0.6 | 0.51 | 0.61 |
| 4 | 44852361 | rs17601614 | -0.646 | 0.353 | 0.539 | 0.663 | 0.06 | 0.152 | 0.152 | 0.08 | 0.07 | 0.29 | 0.05 | 0.15 | 0.08 | 0.03 |
| 4 | 44859443 | rs1497008 | 0.094 | 0.004 | 0.482 | 0.646 | 0.004 | 0.001 | 0.001 | 0 | 0.64 | 0.64 | 0.46 | 0.61 | 0.5 | 0.58 |

| | | | | | | | | | | | | | | | | |
|---|---|---|---|---|---|---|---|---|---|---|---|---|---|---|---|---|
| 4 | 44859711 | rs10517119 | 1.295 | -0.116 | 0.5 | 0.429 | 0.054 | 0.056 | 0.056 | 0 | 0.46 | 0.23 | 0.22 | 0.33 | 0.36 | 0.42 |
| 4 | 44859761 | rs10517120 | 0.584 | -0.059 | 0.426 | 0.387 | -0.003 | -0.003 | -0.003 | 4.845 | 0.53 | 0.52 | 0.39 | 0.46 | 0.42 | 0.49 |
| 4 | 44864639 | rs6835058 | 0.857 | -0.106 | 0.276 | 0.301 | 0.001 | 0 | 0 | 0 | 0.63 | 0.39 | 0.62 | 0.6 | 0.76 | 0.71 |
| 4 | 44868270 | rs4694786 | 0.205 | 0.568 | 0.614 | 0.829 | 0.096 | 0.373 | 0.373 | 0.765 | 0.95 | 0.72 | 0.78 | 0.67 | 0.94 | 0.91 |
| 4 | 44870978 | rs6810444 | 0.185 | 0.641 | 0.286 | 0.825 | -0.024 | -0.014 | -0.014 | 0 | 0.28 | 0.24 | 0.45 | 0.36 | 0.47 | 0.42 |
| 4 | 44871619 | rs2348805 | 0.748 | 0.464 | 0.703 | 1.008 | 0.109 | 0.407 | 0.407 | 1.213 | 0.94 | 0.67 | 0.78 | 0.67 | 0.94 | 0.91 |
| 4 | 44897864 | rs10019381 | -0.227 | 0.058 | -0.674 | -0.227 | 0.012 | -0.011 | -0.011 | 0 | 0.22 | 0.2 | 0.09 | 0 | 0.14 | 0.07 |
| 4 | 44897911 | rs7692626 | 1.161 | -0.031 | -0.608 | -0.331 | 0.013 | 0.022 | 0.022 | 0 | 0.37 | 0.29 | 0.19 | 0.25 | 0.3 | 0.32 |
| 4 | 44898738 | rs4695056 | 0.011 | -0.033 | -0.521 | -0.254 | 0.012 | -0.011 | -0.011 | 0 | 0.22 | 0.2 | 0.09 | 0 | 0.14 | 0.07 |
| 4 | 44904578 | rs7675825 | -0.673 | 0.184 | -0.17 | 0.293 | 0.01 | 0.039 | 0.039 | 0 | 0.19 | 0.3 | 0.27 | 0.33 | 0.2 | 0.23 |
| 4 | 44905275 | rs10938371 | 0.999 | -0.086 | -0.075 | 0.138 | -0.016 | -0.006 | -0.006 | 0.007 | 0.63 | 0.49 | 0.67 | 0.62 | 0.72 | 0.73 |
| 4 | 44906396 | rs1107676 | 1.569 | -0.263 | -0.437 | -0.055 | 0.02 | 0.005 | 0.005 | 0 | 0.33 | 0.26 | 0.15 | 0.26 | 0.24 | 0.23 |
| 4 | 44908179 | rs2348518 | -0.386 | -0.147 | -0.443 | -0.138 | -0.002 | -0.05 | -0.05 | 0 | 0.77 | 0.75 | 0.9 | 1 | 0.85 | 0.92 |
| 4 | 44909241 | rs1496991 | 0.052 | -0.129 | -0.499 | -0.089 | 0.014 | 0.033 | 0.033 | 0 | 0.17 | 0.1 | 0.07 | 0.05 | 0.12 | 0.09 |
| 4 | 44916027 | rs10461071 | 1.803 | 0.712 | -0.924 | 0.098 | -0.088 | -0.077 | -0.077 | 0 | 0.13 | 0.03 | 0.24 | 0.24 | 0.27 | 0.13 |
| 4 | 44918770 | rs11932803 | -0.691 | 0.548 | -0.857 | -0.024 | -0.007 | -0.055 | -0.055 | 0.602 | 0.79 | 0.75 | 0.9 | 1 | 0.86 | 0.93 |
| 4 | 44924735 | rs7661117 | -0.062 | 0.519 | -0.876 | -0.031 | 0.01 | -0.005 | -0.005 | 0 | 0.73 | 0.75 | 0.89 | 0.95 | 0.84 | 0.9 |
| 4 | 44928851 | rs12513234 | 0.197 | 0.63 | -0.836 | -0.054 | -0.068 | -0.026 | -0.026 | 0 | 0.3 | 0.17 | 0.47 | 0.36 | 0.49 | 0.45 |
| 4 | 44935266 | rs1472395 | -2.244 | 0.636 | -0.505 | 0.104 | -0.016 | -0.05 | -0.05 | 0 | 0.88 | 0.83 | 0.96 | 1 | 0.94 | 0.98 |
| 4 | 44939860 | rs9291254 | -0.465 | 0.557 | -0.469 | 0.063 | -0.014 | 0.081 | 0.081 | 0.073 | 0.79 | 0.57 | 0.74 | 0.67 | 0.84 | 0.82 |
| 4 | 44954288 | rs2173109 | 0.189 | 0.506 | -0.567 | 0.017 | -0.013 | -0.001 | -0.001 | 0 | 0.69 | 0.4 | 0.68 | 0.67 | 0.75 | 0.76 |
| 4 | 44954941 | rs2882027 | 0.088 | 0.849 | -0.385 | 0.305 | -0.087 | -0.036 | -0.036 | 0 | 0.3 | 0.12 | 0.52 | 0.36 | 0.5 | 0.5 |
| 4 | 44955488 | rs17573940 | 1.853 | 0.963 | -0.369 | 0.365 | -0.086 | -0.077 | -0.077 | 0 | 0.13 | 0.02 | 0.24 | 0.25 | 0.27 | 0.13 |
| 4 | 44955964 | rs10020374 | -0.092 | 0.26 | -0.029 | 0.392 | 0.039 | 0.191 | 0.191 | 0 | 0.09 | 0.24 | 0.22 | 0.33 | 0.11 | 0.17 |
| 4 | 44964929 | rs1908807 | -0.161 | 0.249 | -0.08 | 0.379 | -0.015 | 0.059 | 0.059 | 0.01 | 0.79 | 0.48 | 0.73 | 0.67 | 0.84 | 0.81 |
| 4 | 44976703 | rs17608495 | 0.213 | 0.307 | -0.184 | 0.513 | 0.036 | 0.162 | 0.162 | 0 | 0.08 | 0.19 | 0.22 | 0.33 | 0.11 | 0.18 |
| 4 | 44982038 | rs10021452 | 0.213 | 0.307 | -0.184 | 0.513 | 0.036 | 0.162 | 0.162 | 0 | 0.08 | 0.19 | 0.21 | 0.33 | 0.11 | 0.18 |
| 4 | 44983208 | rs12645733 | 0.708 | 0.186 | -0.295 | 0.42 | 0.052 | 0.04 | 0.04 | 0 | 0.5 | 0.37 | 0.27 | 0.32 | 0.35 | 0.35 |
| 4 | 44985301 | rs6854974 | 0.649 | 0.355 | -0.688 | 0.329 | 0.012 | 0.008 | 0.008 | 0 | 0.5 | 0.53 | 0.42 | 0.57 | 0.42 | 0.44 |
| 4 | 44989500 | rs4560452 | 0.117 | 0.787 | -1.231 | 0.541 | -0.122 | -0.072 | -0.072 | 0 | 0.29 | 0.09 | 0.41 | 0.38 | 0.48 | 0.47 |
| 4 | 45007208 | rs4695069 | 0.162 | 0.819 | -1.248 | 0.551 | -0.125 | -0.053 | -0.053 | 0.051 | 0.29 | 0.1 | 0.37 | 0.37 | 0.46 | 0.46 |
| 4 | 45015525 | rs10018053 | 0.728 | 0.271 | -0.702 | 0.546 | 0.011 | 0.003 | 0.003 | 0 | 0.41 | 0.38 | 0.26 | 0.31 | 0.34 | 0.26 |
| 4 | 45016808 | rs7666773 | 0.161 | 0.4 | -0.729 | 0.526 | -0.002 | 0.002 | 0.002 | 0 | 0.31 | 0.35 | 0.21 | 0.3 | 0.3 | 0.26 |
| 4 | 45018583 | rs13142647 | -0.084 | 1.192 | -0.285 | 0.926 | 0.057 | 0.115 | 0.115 | 0 | 0.1 | 0.2 | 0.36 | 0.29 | 0.17 | 0.32 |
| 4 | 45025291 | rs10032600 | -0.481 | 1.477 | -0.752 | 0.639 | -0.002 | -0.002 | -0.002 | 0.249 | 0.3 | 0.33 | 0.47 | 0.52 | 0.45 | 0.46 |
| 4 | 45030609 | rs13140100 | 0.579 | 1.588 | -0.744 | 0.365 | 0.016 | 0.01 | 0.01 | 0 | 0.19 | 0.25 | 0.18 | 0.23 | 0.27 | 0.18 |
| 4 | 45031256 | rs10155504 | -0.423 | 1.214 | -0.692 | 0.339 | 0 | 0 | 0 | 0 | 0.32 | 0.35 | 0.52 | 0.49 | 0.46 | 0.53 |
| 4 | 45035197 | rs4305551 | 0.038 | 0.877 | -0.786 | 0.179 | -0.011 | 0.03 | 0.03 | 0 | 0.27 | 0.32 | 0.28 | 0.45 | 0.27 | 0.28 |
| 4 | 45045543 | rs7682029 | -0.43 | 0.609 | -0.397 | -0.128 | 0.002 | 0.002 | 0.002 | 0.054 | 0.66 | 0.62 | 0.6 | 0.46 | 0.56 | 0.61 |
| 4 | 45059561 | rs6844797 | 0.262 | 0.406 | -0.608 | -0.476 | 0.004 | -0.029 | -0.029 | 0.059 | 0.39 | 0.39 | 0.26 | 0.04 | 0.29 | 0.22 |
| 4 | 45068593 | rs4306991 | -0.011 | 0.175 | -0.926 | -0.646 | 0.037 | 0.063 | 0.063 | 0.453 | 0.29 | 0.2 | 0.18 | 0.04 | 0.2 | 0.28 |
| 4 | 45077656 | rs1512320 | -0.294 | 0.164 | -0.987 | -0.651 | 0.046 | 0.101 | 0.101 | 0.265 | 0.71 | 0.86 | 0.83 | 0.96 | 0.8 | 0.72 |
| 4 | 45079676 | rs17608635 | 1.414 | 0.177 | -0.92 | -0.557 | -0.01 | -0.019 | -0.019 | 0 | 0.2 | 0.14 | 0.28 | 0.37 | 0.23 | 0.31 |
| 4 | 45083659 | rs17470661 | 1.074 | 0.376 | -0.929 | -0.562 | 0.04 | 0.065 | 0.065 | 0.729 | 0.21 | 0.09 | 0.15 | 0.04 | 0.15 | 0.19 |
| 4 | 45102488 | rs12640145 | -0.174 | 0.388 | -0.931 | -0.566 | 0.041 | 0.088 | 0.088 | 0.495 | 0.72 | 0.84 | 0.83 | 0.96 | 0.8 | 0.72 |
| 4 | 45117658 | rs16858015 | 1.066 | 0.543 | -0.76 | -0.645 | -0.009 | -0.089 | -0.089 | 3.566 | 0.19 | 0.28 | 0.12 | 0 | 0.17 | 0.04 |
| 4 | 45125825 | rs7689003 | -0.679 | 0.603 | -1.019 | -0.295 | -0.002 | -0.005 | -0.005 | 0 | 0.66 | 0.72 | 0.58 | 0.57 | 0.68 | 0.53 |
| 4 | 45127762 | rs979580 | -1.01 | 0.722 | -0.845 | -0.197 | -0.003 | 0.024 | 0.024 | 0 | 0.93 | 0.98 | 0.98 | 1 | 0.95 | 0.91 |
| 4 | 45133240 | rs7659184 | 1.023 | 0.843 | -0.726 | -0.092 | 0.006 | 0.001 | 0.001 | 0 | 0.23 | 0.27 | 0.18 | 0.23 | 0.3 | 0.19 |
| 4 | 45139673 | rs6845132 | -0.016 | 0.194 | -0.929 | -0.083 | 0.002 | 0.007 | 0.007 | 0 | 0.47 | 0.72 | 0.44 | 0.53 | 0.55 | 0.28 |
| 4 | 45164637 | rs1996023 | 0.367 | 0.17 | -0.746 | -0.181 | 0 | -0.003 | -0.003 | 0 | 0.41 | 0.38 | 0.43 | 0.69 | 0.41 | 0.37 |
| 4 | 45175691 | rs13130484 | 0.647 | -0.039 | -0.837 | -0.399 | -0.027 | 0.026 | 0.026 | 0 | 0.37 | 0.2 | 0.4 | 0.3 | 0.42 | 0.37 |
| 4 | 45175804 | rs16858082 | 0.632 | -0.052 | -0.799 | -0.36 | 0 | -0.002 | -0.002 | 0 | 0.41 | 0.38 | 0.43 | 0.65 | 0.41 | 0.35 |
| 4 | 45179883 | rs12641981 | 0.369 | 0.05 | -1.748 | -0.04 | -0.027 | 0.026 | 0.026 | 0 | 0.37 | 0.2 | 0.4 | 0.3 | 0.42 | 0.37 |
| 4 | 45181744 | rs10027118 | 0.603 | 0.188 | -1.663 | 0.008 | 0.019 | 0.027 | 0.027 | 0.736 | 0.13 | 0.09 | 0.11 | 0.05 | 0.12 | 0.19 |
| 4 | 45182527 | rs10938397 | 0.024 | 0.613 | -1.352 | 0.31 | -0.03 | 0.019 | 0.019 | 0 | 0.36 | 0.2 | 0.4 | 0.3 | 0.42 | 0.37 |
| 4 | 45191856 | rs1858410 | 0.477 | 0.151 | -0.772 | 0.629 | -0.004 | 0 | 0 | 0 | 0.74 | 0.3 | 0.77 | 0.76 | 0.82 | 0.78 |
| 4 | 45192865 | rs12509470 | -0.026 | 0.137 | 0.227 | 0.646 | -0.018 | 0.039 | 0.039 | 0.436 | 0.16 | 0.3 | 0.15 | 0.22 | 0.13 | 0.06 |
| 4 | 45224500 | rs1392615 | 0.792 | 0.409 | -0.28 | 0.327 | -0.014 | 0.055 | 0.055 | 0 | 0.37 | 0.2 | 0.38 | 0.23 | 0.37 | 0.43 |
| 4 | 45230009 | rs348548 | -0.083 | 0.793 | 0.016 | 0.31 | -0.001 | 0.003 | 0.003 | 0 | 0.27 | 0.37 | 0.28 | 0.25 | 0.28 | 0.11 |
| 4 | 45254893 | rs1500493 | 0.469 | 0.663 | 0.16 | 0.42 | 0.006 | 0.003 | 0.003 | 0.904 | 0.65 | 0.27 | 0.68 | 0.67 | 0.67 | 0.76 |
| 4 | 45255852 | rs17637290 | 0.665 | 0.599 | 0.04 | 0.433 | -0.052 | -0.111 | -0.111 | 0 | 0.23 | 0.04 | 0.28 | 0.41 | 0.26 | 0.25 |
| 4 | 45299977 | rs12643061 | 0.319 | 0.505 | 0.384 | 0.111 | 0.006 | 0.009 | 0.009 | 1.23 | 0.67 | 0.3 | 0.73 | 0.8 | 0.74 | 0.78 |
| 4 | 45308763 | rs4267772 | 0.158 | 0.491 | 0.421 | 0.115 | -0.006 | -0.013 | -0.013 | 0 | 0.25 | 0.33 | 0.21 | 0.17 | 0.22 | 0.08 |
| 4 | 45338240 | rs1500488 | 0.05 | 0.636 | 0.13 | 0.058 | -0.008 | -0.036 | -0.036 | 0.396 | 0.68 | 0.39 | 0.71 | 0.83 | 0.72 | 0.85 |
| 4 | 45342647 | rs6858099 | -0.34 | 1.064 | -0.276 | 0.635 | -0.004 | -0.004 | -0.004 | 0 | 0.31 | 0.19 | 0.38 | 0.69 | 0.37 | 0.47 |
| 4 | 45346690 | rs1392612 | 0.645 | 0.893 | 0.183 | 0.574 | 0.002 | 0 | 0 | 1.635 | 0.8 | 0.32 | 0.8 | 0.86 | 0.78 | 0.77 |
| 4 | 45348959 | rs2349125 | -0.097 | 0.423 | 0.029 | 0.598 | 0.033 | 0.014 | 0.014 | 0 | 0.14 | 0.04 | 0.07 | 0.13 | 0.08 | 0.18 |
| 4 | 45362545 | rs6826745 | 1.056 | 0.432 | 0.751 | 1.017 | -0.048 | 0.032 | 0.032 | 0 | 0.76 | 0.56 | 0.84 | 0.67 | 0.82 | 0.62 |

| | | | | | | | | | | | | | | | | |
|---|---|---|---|---|---|---|---|---|---|---|---|---|---|---|---|---|
| 4 | 45374305 | rs1476295 | 1.804 | 1.343 | 1 | 1.017 | -0.007 | -0.038 | -0.038 | 0 | 0.68 | 0.25 | 0.7 | 0.54 | 0.62 | 0.56 |
| 4 | 45374989 | rs12508162 | 1.314 | 1.064 | 0.834 | 1.103 | -0.045 | 0.072 | 0.072 | 0 | 0.8 | 0.56 | 0.86 | 0.67 | 0.86 | 0.63 |
| 4 | 45385709 | rs12503671 | 0.171 | 1.06 | 0.962 | 1.101 | -0.005 | -0.008 | -0.008 | 0 | 0.92 | 0.78 | 0.94 | 0.98 | 0.94 | 0.83 |
| 4 | 45392037 | rs12500516 | 0.171 | 1.06 | 0.962 | 1.101 | -0.005 | -0.008 | -0.008 | 0 | 0.92 | 0.78 | 0.94 | 0.98 | 0.94 | 0.83 |
| 4 | 45401173 | rs13107382 | 1.189 | 1.051 | 1.062 | 1.095 | -0.023 | 0.038 | 0.038 | 0 | 0.82 | 0.68 | 0.86 | 0.67 | 0.86 | 0.64 |
| 4 | 45431010 | rs1486617 | -0.98 | 1.061 | 0.869 | 1.103 | -0.008 | 0.014 | 0.014 | 0.363 | 0.1 | 0.36 | 0.07 | 0.07 | 0.07 | 0.26 |
| 4 | 45441814 | rs10020709 | -1.565 | 1.146 | 0.795 | 1.083 | -0.034 | 0.063 | 0.063 | 0 | 0.2 | 0.43 | 0.15 | 0.34 | 0.17 | 0.38 |
| 4 | 45448030 | rs10028314 | -1.547 | 1.201 | 0.786 | 1.083 | -0.031 | 0.066 | 0.066 | 0 | 0.2 | 0.42 | 0.15 | 0.35 | 0.17 | 0.38 |
| 4 | 45469812 | rs17637825 | 0.441 | 1.678 | 0.086 | 1.304 | -0.036 | -0.048 | -0.048 | 0.908 | 0.12 | 0.02 | 0.16 | 0.39 | 0.18 | 0.16 |
| 4 | 45491047 | rs17595063 | 0.047 | 1.983 | 0.508 | 1.085 | 0.001 | -0.011 | -0.011 | 0 | 0.07 | 0.03 | 0.08 | 0.15 | 0.04 | 0.07 |
| 4 | 45496981 | rs990676 | -0.073 | 2.335 | 0.978 | 1.094 | 0.026 | 0.04 | 0.04 | 0 | 0.88 | 0.78 | 0.89 | 0.86 | 0.86 | 0.94 |
| 4 | 45522585 | rs9992791 | 0.799 | 2.195 | 0.928 | 1.074 | 0.018 | 0.04 | 0.04 | 0 | 0.86 | 0.84 | 0.79 | 0.62 | 0.77 | 0.79 |
| 4 | 45525885 | rs1552419 | 1.602 | 2.416 | 0.401 | 0.737 | -0.008 | -0.01 | -0.01 | 0 | 0.59 | 0.35 | 0.53 | 0.46 | 0.44 | 0.52 |
| 4 | 45556947 | rs840193 | -0.28 | 1.915 | -0.402 | 0.603 | -0.003 | -0.002 | -0.002 | 0.048 | 0.95 | 0.97 | 0.96 | 0.98 | 0.94 | 0.95 |
| 4 | 45588229 | rs1620380 | -0.326 | 1.621 | -0.847 | 0.117 | 0.01 | 0.035 | 0.035 | 0 | 0.88 | 0.91 | 0.95 | 1 | 0.9 | 0.84 |
| 4 | 45612701 | rs6829634 | 0.525 | 2.096 | -1.243 | 0.159 | -0.122 | -0.006 | -0.006 | 0 | 0.75 | 0.96 | 0.78 | 0.85 | 0.66 | 0.82 |
| 4 | 45626526 | rs10004397 | -0.385 | 2.062 | -0.825 | 0.184 | -0.001 | -0.001 | -0.001 | 0.053 | 0.17 | 0.19 | 0.19 | 0.15 | 0.25 | 0.2 |
| 4 | 45638847 | rs2061231 | 0.681 | 0.447 | -1.902 | -0.136 | -0.029 | -0.152 | -0.152 | 0.003 | 0.28 | 0.04 | 0.33 | 0.51 | 0.29 | 0.3 |
| 4 | 45702046 | rs12640483 | -0.617 | 0.225 | -1.16 | -0.559 | -0.018 | -0.056 | -0.056 | 0.255 | 0.84 | 0.8 | 0.95 | 0.99 | 0.91 | 0.82 |
| 4 | 45705727 | rs10015537 | 1.377 | -0.127 | -1.379 | -0.963 | 0.021 | 0.029 | 0.029 | 0 | 0.41 | 0.33 | 0.43 | 0.8 | 0.28 | 0.55 |
| 4 | 45717849 | rs1456507 | -0.745 | 0.098 | -1.286 | -0.761 | 0 | -0.027 | -0.027 | 0 | 0.85 | 0.83 | 0.95 | 1 | 0.9 | 0.94 |
| 4 | 45720024 | rs1456508 | 0.433 | -0.141 | -1.358 | -0.943 | 0.005 | -0.027 | -0.027 | 0 | 0.76 | 0.78 | 0.87 | 1 | 0.79 | 0.9 |
| 4 | 45725524 | rs1598654 | -1.046 | -0.142 | -1.338 | -0.929 | 0.007 | -0.005 | -0.005 | 0.057 | 0.86 | 0.92 | 0.95 | 1 | 0.9 | 0.94 |
| 4 | 45732500 | rs13109781 | -0.104 | -0.185 | -1.314 | -0.877 | -0.004 | -0.059 | -0.059 | 0.311 | 0.11 | 0.2 | 0.09 | 0 | 0.11 | 0.04 |
| 4 | 45737256 | rs957858 | 1.143 | -0.467 | -0.875 | -0.743 | -0.019 | -0.039 | -0.039 | 2.726 | 0.42 | 0.41 | 0.45 | 0.82 | 0.31 | 0.67 |
| 4 | 45745877 | rs1389040 | -0.596 | -0.409 | -1.022 | -0.834 | -0.034 | -0.001 | -0.001 | 0.784 | 0.84 | 0.65 | 0.88 | 0.86 | 0.92 | 0.92 |
| 4 | 45757412 | rs9994814 | -0.588 | -0.226 | -1.057 | -0.852 | 0.02 | 0.139 | 0.139 | 0 | 0.42 | 0.24 | 0.43 | 0.04 | 0.62 | 0.25 |
| 4 | 45792228 | rs6847304 | 0.89 | 0.191 | -0.496 | 0.008 | -0.026 | 0 | 0 | 0 | 0.4 | 0.43 | 0.45 | 0.65 | 0.29 | 0.6 |
| 4 | 45795589 | rs6838450 | 0.943 | 0.133 | -0.529 | 0.28 | -0.036 | -0.064 | -0.064 | 0.148 | 0.38 | 0.39 | 0.33 | 0.16 | 0.27 | 0.47 |
| 4 | 45795895 | rs1492839 | -0.292 | 0.538 | -0.861 | 0.31 | -0.032 | 0.022 | 0.022 | 0 | 0.58 | 0.67 | 0.5 | 0.37 | 0.52 | 0.67 |
| 4 | 45800337 | rs7670127 | -0.423 | 0.733 | -0.748 | -0.287 | -0.028 | 0.058 | 0.058 | 0 | 0.6 | 0.69 | 0.59 | 0.82 | 0.55 | 0.78 |
| 4 | 45811691 | rs988543 | 0.943 | 0.69 | -0.487 | -0.332 | -0.007 | 0.001 | 0.001 | 0 | 0.38 | 0.33 | 0.4 | 0.6 | 0.28 | 0.47 |
| 4 | 45817327 | rs2035906 | -0.375 | 0.794 | -0.629 | -0.351 | -0.027 | 0.055 | 0.055 | 0 | 0.59 | 0.69 | 0.58 | 0.76 | 0.54 | 0.75 |
| 4 | 45839740 | rs6826668 | 0.854 | 0.096 | -0.407 | -0.005 | -0.011 | 0.008 | 0.008 | 0 | 0.34 | 0.31 | 0.39 | 0.58 | 0.27 | 0.46 |
| 4 | 45843672 | rs10032936 | -0.506 | 0.69 | 0.068 | 0.312 | 0.002 | 0.004 | 0.004 | 0.099 | 0.18 | 0.39 | 0.14 | 0.17 | 0.15 | 0.25 |
| 4 | 45851084 | rs1353014 | -0.096 | 1.124 | 0.16 | 0.421 | 0.018 | 0.015 | 0.015 | 0.325 | 0.17 | 0.26 | 0.22 | 0.22 | 0.23 | 0.17 |
| 4 | 45858464 | rs11721867 | -0.262 | 1.066 | 0.422 | 0.392 | 0.027 | 0.022 | 0.022 | 0.451 | 0.18 | 0.35 | 0.23 | 0.22 | 0.24 | 0.17 |
| 4 | 45859948 | rs13136580 | -0.169 | 1.075 | 0.37 | 0.397 | 0.015 | 0.012 | 0.012 | 0.309 | 0.18 | 0.26 | 0.22 | 0.22 | 0.23 | 0.17 |
| 4 | 45863268 | rs2349443 | 0.84 | 0.618 | -0.057 | 0.105 | 0.06 | -0.063 | -0.063 | 0 | 0.38 | 0.16 | 0.34 | 0.54 | 0.28 | 0.46 |
| 4 | 45875599 | rs7692404 | 0.976 | 0.626 | 0.124 | 0.09 | 0.043 | -0.056 | -0.056 | 0 | 0.37 | 0.14 | 0.34 | 0.54 | 0.28 | 0.47 |
| 4 | 45882448 | rs1492837 | -0.919 | 0.567 | 1.047 | 0.007 | -0.025 | 0.01 | 0.01 | 0 | 0.58 | 0.56 | 0.63 | 0.46 | 0.71 | 0.51 |
| 4 | 45896156 | rs1353013 | 0.401 | 1.542 | 1.055 | -0.179 | -0.015 | 0.006 | 0.006 | 0 | 0.25 | 0.19 | 0.26 | 0.03 | 0.34 | 0.18 |
| 4 | 45900199 | rs7690208 | 0.22 | 1.556 | 1.011 | 0.002 | 0.006 | 0.006 | 0.006 | 0.323 | 0.18 | 0.22 | 0.21 | 0.26 | 0.23 | 0.24 |
| 4 | 45923348 | rs9994137 | -0.943 | 1.247 | 1.174 | -0.088 | 0.033 | 0.126 | 0.126 | 0.061 | 0.14 | 0.44 | 0.19 | 0.25 | 0.13 | 0.09 |
| 4 | 45924834 | rs1394346 | -1.056 | 1.243 | 1.213 | -0.09 | 0.112 | 0.224 | 0.224 | 0.064 | 0.08 | 0.33 | 0.18 | 0.25 | 0.12 | 0.09 |
| 4 | 45929172 | rs6817621 | -0.091 | 1.223 | 1.039 | -0.307 | -0.034 | 0.047 | 0.047 | 0 | 0.38 | 0.25 | 0.3 | 0.01 | 0.47 | 0.28 |
| 4 | 45930890 | rs9997632 | -1.237 | 1.665 | 1.561 | 0.662 | 0.045 | 0.149 | 0.149 | 0.002 | 0.14 | 0.46 | 0.19 | 0.25 | 0.13 | 0.09 |
| 4 | 45939413 | rs7668279 | -1.385 | 2.121 | 1.375 | 1.022 | 0.1 | 0.254 | 0.254 | 0.484 | 0.06 | 0.27 | 0.15 | 0.28 | 0.09 | 0.09 |
| 4 | 45956676 | rs7699135 | 0.201 | 1.987 | 0.88 | 1.143 | -0.033 | -0.056 | -0.056 | 4.707 | 0.91 | 1 | 0.85 | 0.83 | 0.89 | 0.91 |
| 4 | 45956780 | rs13127070 | -0.127 | 2.745 | 1.4 | 0.845 | -0.003 | -0.03 | -0.03 | 0 | 0.24 | 0.21 | 0.29 | 0.01 | 0.31 | 0.17 |
| 4 | 45972653 | rs924837 | -0.86 | 2.86 | 1.772 | 0.866 | 0.077 | 0.009 | 0.009 | 0 | 0.28 | 0.55 | 0.41 | 0.29 | 0.41 | 0.24 |
| 4 | 45982196 | rs2089154 | 0.231 | 2.916 | 1.689 | 0.834 | -0.053 | 0.063 | 0.063 | 0 | 0.2 | 0.1 | 0.24 | 0.01 | 0.3 | 0.15 |
| 4 | 45995759 | rs1566854 | -1.062 | 2.614 | 1.566 | 0.881 | 0.019 | -0.003 | -0.003 | 0 | 0.33 | 0.61 | 0.48 | 0.33 | 0.44 | 0.33 |
| 4 | 46025913 | rs10030781 | -0.666 | 2.643 | 1.537 | 0.928 | 0.087 | 0.012 | 0.012 | 0 | 0.28 | 0.53 | 0.41 | 0.33 | 0.4 | 0.31 |
| 4 | 46038274 | rs10938423 | -0.875 | 2.604 | 1.169 | 0.918 | -0.002 | 0 | 0 | 0 | 0.31 | 0.67 | 0.49 | 0.33 | 0.46 | 0.33 |
| 4 | 46039718 | rs7666844 | -0.978 | 2.607 | 1.211 | 0.945 | -0.008 | 0.001 | 0.001 | 0 | 0.33 | 0.67 | 0.49 | 0.33 | 0.46 | 0.33 |
| 4 | 46040738 | rs1497567 | -0.955 | 2.608 | 1.197 | 0.945 | -0.006 | 0.001 | 0.001 | 0 | 0.33 | 0.67 | 0.49 | 0.33 | 0.46 | 0.33 |
| 4 | 46040758 | rs1497568 | -0.955 | 2.608 | 1.197 | 0.945 | -0.008 | 0.001 | 0.001 | 0 | 0.33 | 0.67 | 0.49 | 0.33 | 0.46 | 0.33 |
| 4 | 46042083 | rs7683876 | -0.925 | 2.311 | 1.218 | 0.959 | 0.001 | -0.001 | -0.001 | 0.059 | 0.32 | 0.65 | 0.49 | 0.33 | 0.46 | 0.33 |
| 4 | 46055478 | rs993677 | -0.126 | 2.315 | 1.071 | 1.041 | -0.033 | -0.024 | -0.024 | 0 | 0.94 | 1 | 0.85 | 0.83 | 0.89 | 0.94 |
| 4 | 46072596 | rs1391174 | -1.103 | 2.089 | 1.29 | 0.677 | -0.004 | 0.003 | 0.003 | 0 | 0.35 | 0.66 | 0.56 | 0.33 | 0.55 | 0.33 |
| 4 | 46086060 | rs976156 | -1.17 | 2.061 | 1.212 | 0.682 | -0.008 | 0.007 | 0.007 | 0.015 | 0.36 | 0.72 | 0.56 | 0.33 | 0.55 | 0.33 |
| 4 | 46093351 | rs1497575 | -1.28 | 2.057 | 1.17 | 0.644 | -0.074 | -0.01 | -0.01 | 0 | 0.36 | 0.89 | 0.56 | 0.34 | 0.54 | 0.34 |
| 4 | 46105928 | rs1497559 | -0.945 | 2.112 | 1.67 | 0.674 | 0.037 | -0.002 | -0.002 | 1.465 | 0.32 | 0.54 | 0.48 | 0.33 | 0.52 | 0.3 |
| 4 | 46118134 | rs11942023 | -2.101 | 1.988 | 1.709 | 0.606 | 0.063 | 0.046 | 0.046 | 8.282 | 0.06 | 0.45 | 0.05 | 0 | 0.02 | 0 |
| 4 | 46150637 | rs6842243 | -2.039 | 1.99 | 1.701 | 0.607 | 0.073 | 0.057 | 0.057 | 7.965 | 0.06 | 0.43 | 0.04 | 0 | 0.02 | 0 |
| 4 | 46152332 | rs10033451 | -0.713 | 2.1 | 1.38 | 0.686 | 0.007 | 0 | 0 | 0.213 | 0.28 | 0.34 | 0.41 | 0.27 | 0.5 | 0.29 |
| 4 | 46227847 | rs522636 | 1.108 | 2.1 | 1.115 | 1.125 | -0.067 | -0.041 | -0.041 | 0 | 0.66 | 0.22 | 0.49 | 0.52 | 0.58 | 0.79 |
| 4 | 46238737 | rs478179 | 0.947 | 1.55 | 0.517 | 1.109 | 0.027 | -0.026 | -0.026 | 0 | 0.3 | 0.17 | 0.23 | 0.28 | 0.28 | 0.27 |

| | | | | | | | | | | | | | | | | |
|---|---|---|---|---|---|---|---|---|---|---|---|---|---|---|---|---|
| 4 | 46250605 | rs16859227 | -0.196 | 2.805 | 1.465 | 1.281 | -0.027 | 0.024 | 0.024 | 0 | 0.17 | 0.13 | 0.23 | 0.08 | 0.25 | 0.08 |
| 4 | 46256712 | rs3822051 | 0.09 | 2.868 | 1.418 | 1.293 | -0.084 | 0.031 | 0.031 | 0 | 0.15 | 0.03 | 0.23 | 0.08 | 0.26 | 0.08 |
| 4 | 46256805 | rs534459 | 1.366 | 2.562 | 1.548 | 1.314 | -0.04 | -0.021 | -0.021 | 0.719 | 0.65 | 0.27 | 0.49 | 0.52 | 0.57 | 0.76 |
| 4 | 46263391 | rs526805 | 1.045 | 2.637 | 1.474 | 1.351 | 0.01 | 0.005 | 0.005 | 0 | 0.68 | 0.62 | 0.51 | 0.52 | 0.58 | 0.77 |
| 4 | 46305733 | rs279871 | -1.24 | 2.627 | 1.059 | 1.127 | -0.048 | -0.046 | -0.046 | 0 | 0.3 | 0.24 | 0.47 | 0.53 | 0.43 | 0.23 |
| 4 | 46311447 | rs1808851 | -1.178 | 2.633 | 1.078 | 1.109 | -0.025 | -0.024 | -0.024 | 0 | 0.3 | 0.28 | 0.47 | 0.53 | 0.43 | 0.23 |
| 4 | 46316688 | rs183962 | -1.206 | 2.684 | 0.92 | 1.128 | -0.051 | -0.049 | -0.049 | 0 | 0.29 | 0.23 | 0.47 | 0.53 | 0.43 | 0.23 |
| 4 | 46321863 | rs3775282 | 0.215 | 2.779 | 0.623 | 0.904 | -0.029 | 0.028 | 0.028 | 0 | 0.14 | 0.06 | 0.14 | 0.07 | 0.2 | 0.05 |
| 4 | 46346679 | rs189957 | -1.158 | 2.495 | 0.73 | 0.946 | 0.049 | 0.049 | 0.049 | 0 | 0.34 | 0.48 | 0.5 | 0.57 | 0.45 | 0.33 |
| 4 | 46355479 | rs12647055 | 0.945 | 2.441 | -0.068 | 0.946 | -0.035 | -0.032 | -0.032 | 0 | 0.63 | 0.31 | 0.48 | 0.42 | 0.54 | 0.6 |
| 4 | 46366067 | rs1442060 | -1.409 | 1.882 | -0.158 | 0.722 | -0.005 | 0.001 | 0.001 | 0 | 0.44 | 0.65 | 0.57 | 0.63 | 0.51 | 0.37 |
| 4 | 46368641 | rs3849591 | -0.229 | 2.199 | -0.733 | 0.953 | -0.029 | -0.074 | -0.074 | 0.004 | 0.16 | 0.07 | 0.31 | 0.52 | 0.17 | 0.21 |
| 4 | 46371833 | rs9291283 | -0.213 | 1.978 | -0.106 | 0.355 | 0.032 | -0.054 | -0.054 | 0.009 | 0.18 | 0.3 | 0.2 | 0.07 | 0.29 | 0.06 |
| 4 | 46381221 | rs13152740 | -0.162 | 1.994 | -0.205 | 0.367 | 0.021 | -0.037 | -0.037 | 7.431 | 0.18 | 0.25 | 0.2 | 0.07 | 0.29 | 0.06 |
| 4 | 46400331 | rs1025852 | -1.046 | 1.211 | -0.441 | 0.273 | -0.004 | -0.003 | -0.003 | 0 | 0.54 | 0.58 | 0.71 | 0.74 | 0.67 | 0.67 |
| 4 | 46402152 | rs6847731 | -0.89 | 1.333 | -0.423 | 0.222 | 0.001 | 0 | 0 | 0.016 | 0.46 | 0.59 | 0.66 | 0.74 | 0.6 | 0.65 |
| 4 | 46413251 | rs10461072 | 0.02 | 1.543 | -0.861 | 0.842 | 0.036 | -0.087 | -0.087 | 0 | 0.22 | 0.04 | 0.07 | 0.39 | 0.16 | 0.4 |
| 4 | 46422329 | rs996129 | 0.84 | 2.366 | -0.576 | 0.765 | -0.011 | 0.008 | 0.008 | 0 | 0.15 | 0.14 | 0.18 | 0.07 | 0.27 | 0.06 |
| 4 | 46498640 | rs4293822 | -1.025 | 1.297 | -0.627 | 0.737 | 0.012 | 0.011 | 0.011 | 3.616 | 0.77 | 0.86 | 0.76 | 0.86 | 0.79 | 0.86 |
| 4 | 46541559 | rs10517161 | -0.441 | 1.163 | -0.643 | 0.91 | 0.013 | -0.027 | -0.027 | 0.161 | 0.25 | 0.19 | 0.15 | 0.42 | 0.19 | 0.55 |
| 4 | 46541729 | rs10031022 | -0.448 | 1.113 | -0.568 | 0.886 | 0.003 | -0.005 | -0.005 | 0.027 | 0.26 | 0.27 | 0.16 | 0.42 | 0.19 | 0.55 |
| 4 | 46568165 | rs10938448 | 0.334 | 0.949 | -0.775 | 0.894 | -0.012 | -0.048 | -0.048 | 2.567 | 0.12 | 0.03 | 0.05 | 0.39 | 0.14 | 0.41 |
| 4 | 46576065 | rs12513214 | -0.513 | 0.85 | -0.705 | 0.827 | -0.001 | 0.005 | 0.005 | 0 | 0.21 | 0.22 | 0.14 | 0.42 | 0.16 | 0.46 |
| 4 | 46596239 | rs4541476 | 0.878 | 0.544 | -1.343 | -0.048 | -0.055 | 0.187 | 0.187 | 0 | 0.39 | 0.16 | 0.23 | 0.1 | 0.43 | 0.16 |
| 4 | 46625274 | rs10938449 | -0.627 | 0.426 | -1.252 | 0.045 | -0.002 | -0.003 | -0.003 | 0 | 0.31 | 0.33 | 0.45 | 0.65 | 0.31 | 0.57 |
| 4 | 46627357 | rs10019070 | -0.647 | 0.478 | -0.807 | 0.142 | -0.013 | 0.005 | 0.005 | 4.486 | 0.72 | 0.67 | 0.69 | 0.75 | 0.74 | 0.74 |
| 4 | 46635167 | rs10029664 | -1.551 | 0.632 | -1.231 | 0.331 | -0.016 | -0.028 | -0.028 | 0 | 0.82 | 0.95 | 0.74 | 0.86 | 0.82 | 0.91 |
| 4 | 46683061 | rs10470769 | 0.547 | 0.934 | -0.821 | 0.831 | 0.012 | 0.038 | 0.038 | 0 | 0.06 | 0.07 | 0.33 | 0.27 | 0.14 | 0.14 |
| 4 | 46685463 | rs9291289 | -1.631 | 0.764 | -0.831 | 0.718 | 0.006 | 0.015 | 0.015 | 0.271 | 0.33 | 0.47 | 0.46 | 0.65 | 0.33 | 0.61 |
| 4 | 46690369 | rs4538458 | -0.511 | 0.681 | -1.057 | 0.75 | 0.01 | 0.028 | 0.028 | 0 | 0.14 | 0.24 | 0.37 | 0.27 | 0.15 | 0.14 |
| 4 | 46691120 | rs10024620 | 0.077 | 0.796 | -1.03 | 0.813 | 0.056 | 0.103 | 0.103 | 0 | 0.06 | 0.16 | 0.34 | 0.27 | 0.14 | 0.13 |
| 4 | 46721833 | rs6853207 | -1.537 | 0.723 | -1.198 | 0.703 | -0.001 | -0.002 | -0.002 | 0 | 0.33 | 0.33 | 0.46 | 0.67 | 0.32 | 0.61 |
| 4 | 46734901 | rs9790574 | -0.679 | 1.021 | -1.413 | 0.968 | 0.004 | -0.03 | -0.03 | 2.943 | 0.2 | 0.12 | 0.09 | 0.39 | 0.17 | 0.47 |
| 4 | 46742649 | rs4395475 | 0.446 | 1.218 | -1.459 | 1.035 | -0.006 | 0.029 | 0.029 | 0 | 0.06 | 0.02 | 0.34 | 0.29 | 0.15 | 0.14 |
| 4 | 46750333 | rs11945399 | -0.125 | 0.945 | -1.6 | 1.111 | 0.004 | -0.017 | -0.017 | 0.141 | 0.13 | 0 | 0.04 | 0.11 | 0.08 | 0.18 |
| 4 | 46759314 | rs10938457 | -1.893 | 0.741 | -1.448 | 1.219 | -0.004 | -0.016 | -0.016 | 0.255 | 0.32 | 0.31 | 0.46 | 0.44 | 0.32 | 0.52 |
| 4 | 46789898 | rs4478161 | -2.117 | 0.704 | -1.542 | 1.139 | -0.002 | -0.008 | -0.008 | 0 | 0.37 | 0.36 | 0.47 | 0.43 | 0.36 | 0.57 |
| 4 | 46796836 | rs10938458 | -0.695 | 0.718 | -1.455 | 1.186 | 0.001 | 0 | 0 | 0 | 0.17 | 0.15 | 0.07 | 0.16 | 0.15 | 0.41 |
| 4 | 46805796 | rs4129828 | -1.026 | 0.606 | -1.923 | 0.488 | 0.024 | 0.029 | 0.029 | 5.243 | 0.54 | 0.38 | 0.6 | 0.69 | 0.62 | 0.75 |
| 4 | 46838259 | rs12503660 | -1.601 | 0.872 | -1.703 | 0.777 | -0.001 | -0.012 | -0.012 | 0 | 0.37 | 0.35 | 0.47 | 0.45 | 0.36 | 0.55 |
| 4 | 46868162 | rs2221854 | -0.948 | 1.35 | -1.683 | 0.988 | 0.015 | 0.028 | 0.028 | 0 | 0.22 | 0.32 | 0.44 | 0.35 | 0.3 | 0.37 |
| 4 | 46878105 | rs9291290 | -1.32 | 1.078 | -1.917 | 0.756 | 0.011 | -0.014 | -0.014 | 0.421 | 0.4 | 0.34 | 0.5 | 0.44 | 0.35 | 0.56 |
| 4 | 46878219 | rs9291291 | -0.132 | 1.223 | -1.883 | 0.863 | -0.002 | -0.009 | -0.009 | 0.299 | 0.15 | 0.15 | 0.09 | 0.06 | 0.14 | 0.27 |
| 4 | 46890143 | rs10938462 | 0.315 | 1.286 | -1.933 | 1.025 | -0.014 | 0.02 | 0.02 | 0 | 0.06 | 0.05 | 0.34 | 0.29 | 0.11 | 0.12 |
| 4 | 46898265 | rs7683552 | -1.086 | 1.325 | -1.943 | 0.979 | -0.001 | -0.001 | -0.001 | 0 | 0.21 | 0.24 | 0.44 | 0.35 | 0.24 | 0.39 |
| 4 | 46899077 | rs1512128 | 0.011 | 1.322 | -1.932 | 0.968 | 0 | -0.003 | -0.003 | 0.038 | 0.14 | 0.13 | 0.09 | 0.06 | 0.14 | 0.27 |
| 4 | 46903491 | rs13119670 | 2.108 | 1.481 | -2.489 | 0.419 | -0.02 | 0.11 | 0.11 | 2.098 | 0.18 | 0.01 | 0.08 | 0 | 0.15 | 0.03 |
| 4 | 46919749 | rs11722432 | 1.178 | 1.748 | -2.068 | 0.918 | -0.011 | 0.002 | 0.002 | 1.379 | 0.27 | 0.2 | 0.15 | 0.31 | 0.29 | 0.17 |
| 4 | 46922107 | rs7678338 | 1.178 | 1.748 | -2.065 | 0.918 | -0.01 | 0.002 | 0.002 | 1.379 | 0.27 | 0.21 | 0.15 | 0.31 | 0.29 | 0.17 |
| 4 | 46922688 | rs10033500 | 1.178 | 1.748 | -2.065 | 0.918 | -0.01 | 0.002 | 0.002 | 1.379 | 0.27 | 0.21 | 0.15 | 0.31 | 0.29 | 0.18 |
| 4 | 46922950 | rs9291296 | 1.178 | 1.748 | -2.065 | 0.918 | -0.011 | 0.002 | 0.002 | 1.379 | 0.27 | 0.19 | 0.15 | 0.31 | 0.29 | 0.17 |
| 4 | 46923108 | rs9291297 | 1.156 | 1.737 | -2.071 | 0.913 | -0.01 | 0.002 | 0.002 | 1.492 | 0.27 | 0.21 | 0.15 | 0.31 | 0.29 | 0.17 |
| 4 | 46925813 | rs953380 | 1.181 | 1.738 | -2.071 | 0.913 | -0.01 | 0.002 | 0.002 | 1.379 | 0.27 | 0.21 | 0.15 | 0.31 | 0.29 | 0.17 |
| 4 | 46927124 | rs17599074 | 1.181 | 1.738 | -2.071 | 0.913 | -0.01 | 0.002 | 0.002 | 1.379 | 0.27 | 0.21 | 0.15 | 0.31 | 0.29 | 0.18 |
| 4 | 46927145 | rs1398174 | -0.8 | 1.722 | -2.051 | 0.926 | 0.006 | 0.019 | 0.019 | 1.235 | 0.48 | 0.45 | 0.59 | 0.66 | 0.54 | 0.56 |
| 4 | 46927886 | rs10004905 | -0.8 | 1.722 | -2.051 | 0.926 | 0.006 | 0.019 | 0.019 | 1.235 | 0.48 | 0.45 | 0.59 | 0.66 | 0.54 | 0.56 |
| 4 | 46941018 | rs1028005 | -0.587 | 1.539 | -1.978 | 0.975 | 0.001 | 0.011 | 0.011 | 0 | 0.22 | 0.27 | 0.47 | 0.34 | 0.24 | 0.39 |
| 4 | 46952471 | rs1512130 | -1.639 | 1.528 | -2.123 | 0.632 | -0.008 | -0.016 | -0.016 | 0.255 | 0.67 | 0.53 | 0.69 | 0.76 | 0.68 | 0.77 |
| 4 | 46957305 | rs10015891 | -0.885 | 1.697 | -2.008 | 0.848 | 0.002 | 0.015 | 0.015 | 1.123 | 0.49 | 0.48 | 0.63 | 0.65 | 0.53 | 0.56 |
| 4 | 46961617 | rs2221855 | 0.01 | 1.699 | -2.058 | 0.798 | -0.006 | -0.01 | -0.01 | 0 | 0.38 | 0.29 | 0.53 | 0.6 | 0.39 | 0.29 |
| 4 | 46963222 | rs6844842 | -0.901 | 1.719 | -2.013 | 1.031 | 0.001 | 0.025 | 0.025 | 1.298 | 0.48 | 0.48 | 0.65 | 0.69 | 0.53 | 0.56 |
| 4 | 46967279 | rs2055943 | 0.824 | 1.919 | -1.92 | 1.186 | -0.004 | -0.006 | -0.006 | 0 | 0.27 | 0.2 | 0.14 | 0.34 | 0.28 | 0.17 |
| 4 | 46982826 | rs10517174 | -0.194 | 2.102 | -1.635 | 1.552 | 0.006 | 0.009 | 0.009 | 0 | 0.09 | 0.13 | 0.1 | 0.11 | 0.11 | 0.2 |
| 4 | 46985750 | rs7694035 | -1.271 | 1.66 | -2.069 | 1.13 | 0.011 | 0.043 | 0.043 | 2.382 | 0.72 | 0.93 | 0.79 | 0.82 | 0.77 | 0.76 |
| 4 | 46995653 | rs13151759 | 0.431 | 1.672 | -1.918 | 1.148 | 0.008 | 0.004 | 0.004 | 2.442 | 0.37 | 0.37 | 0.3 | 0.37 | 0.37 | 0.2 |
| 4 | 46997913 | rs2055940 | 1.147 | 1.557 | -2.022 | 1.287 | -0.017 | 0.038 | 0.038 | 0.263 | 0.29 | 0.1 | 0.19 | 0.17 | 0.33 | 0.1 |
| 4 | 46999235 | rs7683042 | 0.42 | 1.741 | -1.993 | 1.229 | 0.018 | 0.021 | 0.021 | 0.282 | 0.43 | 0.27 | 0.32 | 0.38 | 0.41 | 0.21 |
| 4 | 47002378 | rs6447520 | 0.354 | 1.958 | -1.874 | 1.16 | -0.006 | 0.009 | 0.009 | 0.47 | 0.44 | 0.43 | 0.62 | 0.61 | 0.51 | 0.51 |
| 4 | 47003076 | rs1159315 | -0.2 | 1.769 | -1.707 | 1.07 | 0.01 | -0.019 | -0.019 | 0 | 0.42 | 0.43 | 0.35 | 0.34 | 0.47 | 0.31 |

| | | | | | | | | | | | | | | | | |
|---|---|---|---|---|---|---|---|---|---|---|---|---|---|---|---|---|
| 4 | 47004037 | rs1512144 | -0.704 | 1.626 | -1.494 | 1.079 | -0.012 | -0.014 | -0.014 | 0 | 0.57 | 0.46 | 0.57 | 0.66 | 0.61 | 0.62 |
| 4 | 47004305 | rs1512142 | 0.198 | 1.522 | -1.54 | 1.195 | 0.017 | 0.013 | 0.013 | 0.152 | 0.36 | 0.4 | 0.31 | 0.42 | 0.43 | 0.28 |
| 4 | 47006399 | rs13139021 | -0.722 | 1.56 | -1.241 | 1.301 | -0.026 | -0.001 | -0.001 | 0 | 0.62 | 0.5 | 0.47 | 0.62 | 0.69 | 0.72 |
| 4 | 47013138 | rs3934674 | -0.865 | 1.584 | -1.458 | 1.017 | -0.003 | -0.003 | -0.003 | 0.419 | 0.63 | 0.62 | 0.66 | 0.75 | 0.71 | 0.79 |
| 4 | 47018520 | rs4521322 | 0.65 | 1.581 | -1.09 | 1.19 | 0.006 | 0.035 | 0.035 | 0 | 0.87 | 0.72 | 0.89 | 0.8 | 0.91 | 0.91 |
| 4 | 47045377 | rs4317179 | 0.401 | 1.582 | -1.11 | 1.186 | -0.002 | 0.024 | 0.024 | 0 | 0.89 | 0.78 | 0.89 | 0.8 | 0.92 | 0.91 |
| 4 | 47047786 | rs4574382 | -0.349 | 1.485 | -1.35 | 1.287 | 0.007 | -0.008 | -0.008 | 1.119 | 0.47 | 0.67 | 0.35 | 0.52 | 0.51 | 0.46 |
| 4 | 47051185 | rs4315750 | 0.945 | 1.989 | -0.299 | 1.671 | -0.01 | 0.046 | 0.046 | 0 | 0.89 | 0.73 | 0.91 | 0.8 | 0.94 | 0.91 |
| 4 | 47057759 | rs10015366 | 1.242 | 2.239 | -0.335 | 1.818 | 0.04 | 0.079 | 0.079 | 0 | 0.85 | 0.74 | 0.87 | 0.67 | 0.84 | 0.83 |
| 4 | 47064496 | rs4597805 | 0.969 | 1.645 | -0.228 | 1.397 | -0.011 | 0.03 | 0.03 | 0 | 0.89 | 0.76 | 0.91 | 0.82 | 0.94 | 0.91 |
| 4 | 47079408 | rs4694834 | 1.564 | 2.064 | -0.351 | 1.583 | 0.024 | 0.078 | 0.078 | 0 | 0.86 | 0.74 | 0.65 | 0.65 | 0.83 | 0.85 |
| 4 | 47089184 | rs4396968 | 1.042 | 2.193 | -0.339 | 1.569 | 0.032 | 0.012 | 0.012 | 0 | 0.89 | 0.8 | 0.87 | 0.8 | 0.86 | 0.91 |
| 4 | 47097460 | rs7663591 | -0.442 | 2.117 | 0.057 | 1.724 | 0.065 | 0.117 | 0.117 | 0 | 0.07 | 0.24 | 0.09 | 0.08 | 0.07 | 0.18 |
| 4 | 47101223 | rs1866990 | -0.508 | 1.926 | -0.226 | 1.474 | 0.009 | -0.012 | -0.012 | 0.055 | 0.36 | 0.44 | 0.27 | 0.27 | 0.37 | 0.46 |
| 4 | 47102993 | rs10517178 | 1.528 | 1.908 | -0.489 | 0.848 | -0.007 | -0.064 | -0.064 | 0 | 0.42 | 0.38 | 0.3 | 0.04 | 0.36 | 0.31 |
| 4 | 47103115 | rs971354 | 0.653 | 1.914 | -0.87 | 0.928 | -0.007 | -0.032 | -0.032 | 0.959 | 0.79 | 0.83 | 0.78 | 0.46 | 0.75 | 0.82 |
| 4 | 47109360 | rs1372491 | 1.495 | 1.953 | -0.857 | 0.934 | 0.01 | -0.016 | -0.016 | 3.533 | 0.42 | 0.32 | 0.3 | 0.04 | 0.36 | 0.3 |
| 4 | 47132261 | rs12645665 | -0.773 | 2.129 | 0.172 | 1.865 | 0.032 | 0.215 | 0.215 | 0 | 0.13 | 0.26 | 0.13 | 0.35 | 0.17 | 0.16 |
| 4 | 47142218 | rs3906823 | 0.407 | 2.136 | 0.182 | 1.814 | 0 | -0.003 | -0.003 | 0.181 | 0.89 | 0.89 | 0.9 | 0.79 | 0.91 | 0.92 |
| 4 | 47143365 | rs2351299 | -0.874 | 2.175 | 0.232 | 1.841 | 0.035 | 0.207 | 0.207 | 0 | 0.13 | 0.26 | 0.13 | 0.35 | 0.17 | 0.16 |
| 4 | 47150997 | rs1442097 | 1.332 | 1.982 | 0.152 | 1.735 | -0.003 | 0.124 | 0.124 | 0 | 0.77 | 0.51 | 0.79 | 0.46 | 0.77 | 0.8 |
| 4 | 47156545 | rs6827925 | 0.653 | 1.986 | 0.181 | 1.791 | -0.01 | 0 | 0 | 0 | 0.87 | 0.92 | 0.85 | 0.81 | 0.86 | 0.87 |
| 4 | 47164556 | rs17599816 | -0.061 | 2.107 | -0.041 | 1.702 | -0.034 | -0.022 | -0.022 | 0 | 0.11 | 0.01 | 0.06 | 0.15 | 0.1 | 0.2 |
| 4 | 47166781 | rs989808 | 0.464 | 2.29 | 0.032 | 1.748 | -0.021 | -0.001 | -0.001 | 0 | 0.87 | 0.93 | 0.85 | 0.81 | 0.85 | 0.86 |
| 4 | 47167315 | rs11940530 | -0.506 | 2.165 | 0.087 | 1.795 | 0.037 | 0.417 | 0.417 | 0.62 | 0.1 | 0.39 | 0.06 | 0.35 | 0.09 | 0.08 |
| 4 | 47174149 | rs7673824 | -0.067 | 2.183 | 0.117 | 1.846 | 0.037 | 0.342 | 0.342 | 0.212 | 0.08 | 0.26 | 0.05 | 0.33 | 0.09 | 0.08 |
| 4 | 47175700 | rs2119779 | 1.227 | 2.191 | 0.1 | 1.823 | 0.009 | 0.191 | 0.191 | 0 | 0.79 | 0.64 | 0.81 | 0.47 | 0.78 | 0.75 |
| 4 | 47181795 | rs1561776 | 0.326 | 2.199 | -0.069 | 1.639 | -0.003 | -0.006 | -0.006 | 0 | 0.88 | 0.9 | 0.85 | 0.8 | 0.87 | 0.83 |
| 4 | 47203863 | rs10004181 | 1.864 | 2.153 | 0.018 | 1.597 | -0.002 | 0.004 | 0.004 | 0.976 | 0.67 | 0.46 | 0.73 | 0.31 | 0.69 | 0.56 |
| 4 | 47211774 | rs6839998 | 0.843 | 2.002 | -0.187 | 1.248 | 0 | -0.001 | -0.001 | 0.701 | 0.87 | 0.84 | 0.85 | 0.8 | 0.87 | 0.84 |
| 4 | 47222338 | rs1372497 | -1.564 | 2.445 | 0.079 | 0.55 | 0.04 | 0.02 | 0.02 | 0 | 0.24 | 0.34 | 0.28 | 0.28 | 0.33 | 0.16 |
| 4 | 47226204 | rs1372498 | -0.252 | 2.712 | 0.212 | 0.28 | 0.072 | 0.138 | 0.138 | 0.165 | 0.12 | 0.24 | 0.13 | 0.26 | 0.18 | 0.13 |
| 4 | 47226436 | rs1372499 | -1.979 | 2.647 | 0.377 | 0.258 | 0.057 | 0.027 | 0.027 | 0.018 | 0.25 | 0.44 | 0.28 | 0.28 | 0.33 | 0.16 |
| 4 | 47231416 | rs17539445 | -0.579 | 2.707 | 0.132 | 0.284 | -0.02 | 0.034 | 0.034 | 0.966 | 0.12 | 0.07 | 0.15 | 0.02 | 0.15 | 0.02 |
| 4 | 47234200 | rs17539459 | -0.401 | 2.679 | 0.134 | 0.285 | -0.019 | 0.029 | 0.029 | 0.932 | 0.11 | 0.05 | 0.14 | 0.02 | 0.14 | 0.02 |
| 4 | 47243813 | rs7695365 | -1.337 | 2.509 | 0.051 | 0.33 | 0.04 | 0.019 | 0.019 | 0 | 0.24 | 0.35 | 0.28 | 0.28 | 0.33 | 0.16 |
| 4 | 47277298 | rs17653240 | -0.806 | 2.127 | -0.559 | 0.254 | -0.007 | -0.007 | -0.007 | 0.476 | 0.11 | 0.05 | 0.05 | 0.16 | 0.07 | 0.26 |
| 4 | 47284233 | rs17653263 | -0.467 | 2.257 | -0.597 | 0.479 | -0.026 | -0.019 | -0.019 | 0.134 | 0.07 | 0.01 | 0.11 | 0.29 | 0.06 | 0.16 |
| 4 | 47289054 | rs2078610 | -0.715 | 2.118 | -1.002 | -0.667 | 0.017 | 0.102 | 0.102 | 0.342 | 0.56 | 0.73 | 0.68 | 0.98 | 0.59 | 0.73 |
| 4 | 47291135 | rs4396966 | -0.274 | 1.946 | -0.889 | -0.58 | -0.013 | 0.062 | 0.062 | 0 | 0.47 | 0.67 | 0.57 | 0.67 | 0.52 | 0.55 |
| 4 | 47322219 | rs6284 | 0.447 | 2.224 | -0.547 | -0.488 | -0.001 | 0 | 0 | 0 | 0.12 | 0.11 | 0.13 | 0.11 | 0.2 | 0.1 |
| 4 | 47325991 | rs7660843 | 0.057 | 2.117 | -0.335 | -0.732 | -0.002 | 0.002 | 0.002 | 0 | 0.39 | 0.44 | 0.45 | 0.59 | 0.39 | 0.49 |
| 4 | 47347289 | rs7665508 | 0.021 | 1.73 | -0.44 | -1.038 | 0.019 | 0.061 | 0.061 | 0 | 0.47 | 0.32 | 0.39 | 0.31 | 0.41 | 0.48 |
| 4 | 47350872 | rs4588429 | 0.06 | 1.767 | -0.581 | -1.042 | 0.018 | 0.102 | 0.102 | 0 | 0.46 | 0.27 | 0.39 | 0.22 | 0.41 | 0.46 |
| 4 | 47356566 | rs6447550 | -0.488 | 1.483 | -0.294 | -1.814 | 0 | 0.02 | 0.02 | 0 | 0.5 | 0.48 | 0.41 | 0.16 | 0.5 | 0.49 |
| 4 | 47359371 | rs6832172 | 0.646 | 1.481 | -0.487 | -1.96 | -0.031 | 0.243 | 0.243 | 0 | 0.35 | 0.04 | 0.3 | 0.02 | 0.36 | 0.22 |
| 4 | 47365544 | rs6848790 | -0.268 | 1.63 | -0.287 | -2.061 | -0.001 | -0.047 | -0.047 | 0 | 0.46 | 0.44 | 0.4 | 0.02 | 0.49 | 0.35 |
| 4 | 47367125 | rs10517184 | -0.604 | 1.585 | -0.116 | -2.044 | 0 | 0.014 | 0.014 | 0 | 0.53 | 0.57 | 0.45 | 0.16 | 0.54 | 0.53 |
| 4 | 47369065 | rs17461905 | 0.458 | 1.649 | -0.249 | -1.936 | -0.028 | 0.028 | 0.028 | 0 | 0.07 | 0.03 | 0.08 | 0.01 | 0.12 | 0.13 |
| 4 | 47375312 | rs11733050 | -1.031 | 1.685 | -0.067 | -1.943 | 0.051 | -0.007 | -0.007 | 0.052 | 0.17 | 0.31 | 0.26 | 0.15 | 0.25 | 0.33 |
| 4 | 47375627 | rs4695224 | 0.585 | 1.483 | -0.334 | -1.981 | 0.029 | 0.016 | 0.016 | 0 | 0.36 | 0.26 | 0.19 | 0 | 0.28 | 0.19 |
| 4 | 47382920 | rs4695226 | -0.666 | 0.92 | -0.314 | -1.886 | -0.004 | 0.144 | 0.144 | 0.004 | 0.53 | 0.73 | 0.36 | 0.17 | 0.44 | 0.45 |
| 4 | 47390820 | rs6813436 | -0.596 | 0.859 | 0.017 | -1.861 | 0.02 | -0.047 | -0.047 | 0.003 | 0.1 | 0.17 | 0.1 | 0.01 | 0.12 | 0.11 |
| 4 | 47406635 | rs3822113 | -0.822 | 1.117 | -0.362 | -2.03 | -0.015 | 0.161 | 0.161 | 0.016 | 0.55 | 0.79 | 0.33 | 0.16 | 0.5 | 0.43 |
| 4 | 47408709 | rs6289 | -0.087 | 0.613 | -0.38 | -1.91 | 0.007 | 0.044 | 0.044 | 0.1 | 0.39 | 0.31 | 0.19 | 0.02 | 0.31 | 0.19 |
| 4 | 47408838 | rs6290 | -0.964 | 0.325 | -0.107 | -0.89 | 0.005 | 0.036 | 0.036 | 0.049 | 0.07 | 0.18 | 0.05 | 0.14 | 0.07 | 0.15 |
| 4 | 47413522 | rs13106855 | -0.535 | -0.153 | -0.938 | -1.466 | 0.052 | 0.197 | 0.197 | 0 | 0.45 | 0.81 | 0.27 | 0.14 | 0.37 | 0.29 |
| 4 | 47415812 | rs16860198 | -0.101 | -0.487 | -0.101 | -1.731 | -0.005 | -0.031 | -0.031 | 0.041 | 0.29 | 0.34 | 0.18 | 0.14 | 0.28 | 0.24 |
| 4 | 47417586 | rs17600665 | -0.295 | -0.751 | -0.359 | -1.836 | 0.018 | 0.022 | 0.022 | 0.763 | 0.24 | 0.2 | 0.1 | 0.09 | 0.14 | 0.14 |
| 4 | 47421325 | rs10155404 | -1.153 | -0.717 | -0.675 | -1.409 | 0.045 | 0.127 | 0.127 | 0.176 | 0.52 | 0.84 | 0.32 | 0.3 | 0.4 | 0.36 |
| 4 | 47428305 | rs10028945 | 0.271 | -0.696 | -0.222 | -1.521 | -0.001 | 0.001 | 0.001 | 0 | 0.23 | 0.24 | 0.16 | 0.14 | 0.26 | 0.19 |
| 4 | 47430927 | rs13102764 | -1.461 | -1.321 | -0.34 | -1.882 | -0.012 | 0.137 | 0.137 | 3.145 | 0.58 | 0.89 | 0.3 | 0.24 | 0.41 | 0.37 |
| 4 | 47435315 | rs4403019 | 0.485 | -1.266 | -0.15 | -1.818 | 0 | -0.003 | -0.003 | 0 | 0.24 | 0.24 | 0.16 | 0.14 | 0.26 | 0.19 |
| 4 | 47436404 | rs4450898 | -0.766 | -1.478 | 0.545 | -1.931 | -0.012 | -0.026 | -0.026 | 0 | 0.39 | 0.58 | 0.22 | 0.24 | 0.3 | 0.28 |
| 4 | 47436571 | rs10470771 | 0.274 | -1.342 | 0.705 | -1.866 | -0.004 | -0.065 | -0.065 | 0.134 | 0.26 | 0.51 | 0.18 | 0.14 | 0.26 | 0.2 |
| 4 | 47465847 | rs13115740 | 0.437 | -0.611 | 0.554 | -1.467 | 0.016 | 0.003 | 0.003 | 0 | 0.18 | 0.13 | 0.07 | 0 | 0.11 | 0.11 |
| 4 | 47467186 | rs4585276 | -0.406 | -0.607 | 0.643 | -1.464 | -0.053 | -0.129 | -0.129 | 0 | 0.8 | 0.64 | 0.91 | 1 | 0.89 | 0.88 |
| 4 | 47476876 | rs2276 | -0.418 | -0.355 | 0.183 | -1.161 | -0.001 | -0.002 | -0.002 | 0.802 | 0.36 | 0.43 | 0.2 | 0.25 | 0.29 | 0.26 |
| 4 | 47477177 | rs6841768 | -0.463 | 0.421 | -0.615 | -1.172 | 0.02 | 0.091 | 0.091 | 0 | 0.57 | 0.77 | 0.59 | 0.91 | 0.6 | 0.6 |

| | | | | | | | | | | | | | | | | |
|---|---|---|---|---|---|---|---|---|---|---|---|---|---|---|---|---|
| 4 | 47484645 | rs4438736 | -0.251 | 0.857 | -0.576 | -0.487 | -0.004 | 0.001 | 0.001 | 0.223 | 0.83 | 0.87 | 0.89 | 0.81 | 0.85 | 0.89 |
| 4 | 47485111 | rs4508853 | 0.204 | 0.822 | -0.511 | -0.526 | -0.001 | 0.002 | 0.002 | 1.391 | 0.78 | 0.78 | 0.86 | 0.79 | 0.81 | 0.81 |
| 4 | 47503661 | rs6813123 | -0.233 | 1.325 | -0.345 | -0.524 | 0.003 | 0.001 | 0.001 | 0 | 0.3 | 0.36 | 0.39 | 0.66 | 0.31 | 0.33 |
| 4 | 47504161 | rs4235151 | 0.498 | 1.305 | -0.245 | -0.94 | -0.006 | 0.013 | 0.013 | 0 | 0.57 | 0.57 | 0.58 | 0.34 | 0.55 | 0.42 |
| 4 | 47508540 | rs6829033 | 0.563 | 1.123 | -0.436 | -1.507 | -0.001 | -0.013 | -0.013 | 0 | 0.43 | 0.41 | 0.68 | 0.78 | 0.46 | 0.47 |
| 4 | 47511781 | rs10000432 | -0.087 | 0.878 | -0.613 | -1.638 | 0 | -0.011 | -0.011 | 0 | 0.61 | 0.53 | 0.78 | 0.81 | 0.63 | 0.75 |
| 4 | 47526473 | rs6817661 | 0.334 | 0.88 | -0.848 | -1.484 | -0.01 | 0.025 | 0.025 | 0 | 0.34 | 0.31 | 0.39 | 0.81 | 0.37 | 0.42 |
| 4 | 47533601 | rs9999398 | -0.366 | 0.801 | -0.662 | -1.744 | -0.006 | -0.137 | -0.137 | 0.612 | 0.34 | 0.43 | 0.19 | 0.02 | 0.3 | 0.33 |
| 4 | 47533838 | rs7674353 | 0.756 | 0.582 | -1.123 | -1.71 | -0.023 | 0.097 | 0.097 | 0 | 0.18 | 0.05 | 0.1 | 0.02 | 0.17 | 0.28 |
| 4 | 47536566 | rs12374260 | -0.278 | 0.734 | -1.064 | -1.665 | 0.001 | 0.083 | 0.083 | 0 | 0.47 | 0.38 | 0.46 | 0.83 | 0.51 | 0.47 |
| 4 | 47541849 | rs4296657 | 1.091 | 0.595 | -1.078 | -1.349 | -0.008 | 0.052 | 0.052 | 0 | 0.3 | 0.12 | 0.36 | 0.14 | 0.32 | 0.25 |
| 4 | 47543891 | rs4694852 | -0.142 | 0.589 | -0.871 | -0.675 | 0 | 0 | 0 | 0 | 0.68 | 0.65 | 0.64 | 0.28 | 0.69 | 0.64 |
| 4 | 47544012 | rs4694853 | -0.171 | 0.609 | -0.957 | -0.666 | 0 | 0.001 | 0.001 | 0 | 0.68 | 0.66 | 0.64 | 0.28 | 0.69 | 0.64 |
| 4 | 47544947 | rs10938490 | -0.649 | 0.41 | -1.204 | -0.755 | 0.009 | 0.018 | 0.018 | 0 | 0.27 | 0.24 | 0.35 | 0.2 | 0.26 | 0.27 |
| 4 | 47548254 | rs12502323 | -0.445 | 0.138 | -1.19 | -0.838 | 0.001 | 0.002 | 0.002 | 0.707 | 0.23 | 0.24 | 0.31 | 0.2 | 0.22 | 0.26 |
| 4 | 47555196 | rs4695248 | -1.677 | 1.182 | -1.582 | -1.266 | -0.046 | 0.042 | 0.042 | 0.001 | 0.89 | 0.96 | 0.9 | 0.99 | 0.79 | 0.73 |
| 4 | 47558132 | rs4694857 | 0.574 | 0.902 | -1.416 | -1.888 | 0.013 | 0.014 | 0.014 | 0.549 | 0.49 | 0.52 | 0.5 | 0.78 | 0.38 | 0.37 |
| 4 | 47560163 | rs11729411 | 1.255 | 0.419 | -1.086 | -1.501 | 0.014 | -0.004 | -0.004 | 0 | 0.29 | 0.18 | 0.29 | 0.3 | 0.21 | 0.17 |
| 4 | 47560386 | rs4298115 | -0.19 | 0.44 | -1.114 | -1.558 | -0.015 | 0.058 | 0.058 | 3.472 | 0.4 | 0.3 | 0.4 | 0.21 | 0.42 | 0.36 |
| 4 | 47567579 | rs13127414 | -1.768 | 0.788 | -0.833 | -1.43 | -0.008 | -0.007 | -0.007 | 0 | 0.87 | 0.86 | 0.89 | 0.99 | 0.83 | 0.74 |
| 4 | 47568964 | rs6854584 | 1.19 | 0.66 | -0.752 | -1.224 | -0.015 | 0.086 | 0.086 | 0 | 0.18 | 0.26 | 0.19 | 0.5 | 0.15 | 0.2 |
| 4 | 47571914 | rs4694858 | -0.292 | 0.807 | -0.89 | -1.311 | -0.005 | -0.01 | -0.01 | 0.96 | 0.39 | 0.48 | 0.5 | 0.69 | 0.41 | 0.42 |
| 4 | 47573660 | rs10004038 | 1.178 | 0.867 | -1.083 | -1.363 | -0.013 | 0.026 | 0.026 | 0 | 0.17 | 0.13 | 0.1 | 0.01 | 0.19 | 0.09 |
| 4 | 47577455 | rs6840205 | 0.374 | 0.875 | -0.935 | -1.191 | -0.001 | 0.002 | 0.002 | 1.671 | 0.22 | 0.24 | 0.33 | 0.2 | 0.26 | 0.23 |
| 4 | 47578971 | rs16851681 | 0.046 | 0.873 | -0.869 | -1.191 | -0.001 | -0.008 | -0.008 | 0 | 0.24 | 0.37 | 0.35 | 0.2 | 0.26 | 0.22 |
| 4 | 47583291 | rs17572095 | 0.046 | 0.907 | -0.892 | -1.145 | -0.021 | 0.023 | 0.023 | 0.062 | 0.3 | 0.26 | 0.4 | 0.13 | 0.41 | 0.49 |
| 4 | 47584046 | rs1058793 | 0.862 | 0.3 | -0.26 | -0.695 | -0.011 | 0.158 | 0.158 | 0 | 0.18 | 0.33 | 0.25 | 0.57 | 0.16 | 0.16 |
| 4 | 47584682 | rs7694197 | -0.101 | 0.582 | -0.388 | -0.543 | -0.002 | 0.003 | 0.003 | 0 | 0.66 | 0.65 | 0.63 | 0.36 | 0.67 | 0.73 |
| 4 | 47589203 | rs1316874 | 1.884 | 0.422 | -0.519 | -0.209 | -0.001 | 0.019 | 0.019 | 0 | 0.09 | 0.03 | 0.17 | 0.49 | 0.1 | 0.1 |
| 4 | 47589299 | rs2351790 | -1.113 | 0.265 | -0.44 | -0.526 | -0.011 | 0.034 | 0.034 | 0.282 | 0.41 | 0.76 | 0.57 | 0.62 | 0.47 | 0.31 |
| 4 | 47589896 | rs10021442 | -0.159 | 0.253 | -0.122 | -0.533 | 0.039 | -0.027 | -0.027 | 0 | 0.1 | 0.44 | 0.05 | 0 | 0.08 | 0.03 |
| 4 | 47593532 | rs12641970 | 2.051 | 0.274 | -0.131 | -0.497 | 0 | 0.047 | 0.047 | 0 | 0.09 | 0.07 | 0.17 | 0.49 | 0.1 | 0.1 |
| 4 | 47593976 | rs11947521 | -0.491 | 0.264 | -0.093 | -0.531 | 0.011 | -0.03 | -0.03 | 1.809 | 0.29 | 0.4 | 0.38 | 0.12 | 0.36 | 0.21 |
| 4 | 47594220 | rs3811829 | 1.504 | 0.196 | -0.181 | -0.568 | -0.063 | -0.036 | -0.036 | 0 | 0.1 | 0.02 | 0.24 | 0.1 | 0.21 | 0.1 |
| 4 | 47596632 | rs4695253 | -0.366 | 0.11 | -0.107 | -0.444 | -0.001 | -0.003 | -0.003 | 0 | 0.32 | 0.69 | 0.4 | 0.12 | 0.36 | 0.2 |
| 4 | 47597590 | rs3749585 | 1.898 | 0.112 | -0.373 | -0.278 | 0 | 0.047 | 0.047 | 0 | 0.09 | 0.07 | 0.17 | 0.49 | 0.11 | 0.1 |
| 4 | 47602599 | rs12650894 | 0.125 | 0.031 | -0.441 | -0.408 | -0.011 | 0.12 | 0.12 | 0 | 0.2 | 0.43 | 0.26 | 0.63 | 0.19 | 0.2 |
| 4 | 47609664 | rs6832495 | -1.453 | 0.086 | -0.647 | -0.525 | -0.001 | 0.148 | 0.148 | 1.158 | 0.49 | 0.82 | 0.64 | 0.75 | 0.57 | 0.4 |
| 4 | 47614900 | rs4695256 | -0.9 | 0.078 | -0.824 | -0.36 | -0.005 | 0.019 | 0.019 | 0.303 | 0.42 | 0.67 | 0.6 | 0.75 | 0.49 | 0.37 |
| 4 | 47622480 | rs10017276 | 0.421 | 0.189 | -0.823 | -0.389 | -0.002 | 0.001 | 0.001 | 1.365 | 0.22 | 0.25 | 0.35 | 0.12 | 0.3 | 0.17 |
| 4 | 47623275 | rs12331639 | 0.421 | 0.189 | -0.823 | -0.389 | -0.002 | 0.001 | 0.001 | 1.365 | 0.22 | 0.25 | 0.35 | 0.12 | 0.3 | 0.17 |
| 4 | 47631983 | rs13105718 | 0.484 | 0.19 | -0.826 | -0.389 | -0.011 | 0.011 | 0.011 | 1.439 | 0.22 | 0.21 | 0.36 | 0.12 | 0.3 | 0.17 |
| 4 | 47637456 | rs7680097 | 0.244 | -0.418 | -0.73 | 0.068 | 0.002 | 0.006 | 0.006 | 0 | 0.11 | 0.28 | 0.08 | 0.16 | 0.09 | 0.09 |
| 4 | 47667064 | rs11934749 | -0.927 | -0.803 | -2.099 | -0.03 | 0.047 | 0.091 | 0.091 | 0 | 0.78 | 0.99 | 0.88 | 0.97 | 0.88 | 0.8 |
| 4 | 47669751 | rs6815552 | 0.8 | 0.135 | -1.339 | 0.837 | -0.005 | 0.143 | 0.143 | 0 | 0.1 | 0.17 | 0.16 | 0.44 | 0.11 | 0.13 |
| 4 | 47672301 | rs4695267 | -0.731 | -0.115 | -2.119 | 0.561 | 0.072 | 0.01 | 0.01 | 0.047 | 0.61 | 0.95 | 0.76 | 0.65 | 0.73 | 0.61 |
| 4 | 47674150 | rs12643755 | -0.768 | 0.321 | -1.641 | 0.836 | 0.001 | -0.006 | -0.006 | 0 | 0.25 | 0.72 | 0.31 | 0.61 | 0.26 | 0.18 |
| 4 | 47679566 | rs12644230 | 1.115 | 0.354 | -1.339 | 1.054 | -0.022 | -0.048 | -0.048 | 0 | 0.1 | 0.01 | 0.16 | 0.43 | 0.13 | 0.12 |
| 4 | 47682174 | rs10517195 | -0.51 | 0.333 | -1.476 | -0.059 | 0.008 | 0.091 | 0.091 | 0 | 0.4 | 0.25 | 0.52 | 0.13 | 0.58 | 0.46 |
| 4 | 47683429 | rs13118104 | -0.369 | 0.342 | -1.525 | -0.055 | -0.009 | 0.116 | 0.116 | 0 | 0.38 | 0.18 | 0.5 | 0.13 | 0.58 | 0.45 |
| 4 | 47688064 | rs7661217 | -0.576 | 0.429 | -1.138 | -0.011 | 0.004 | 0.01 | 0.01 | 0 | 0.42 | 0.46 | 0.53 | 0.13 | 0.58 | 0.45 |
| 4 | 47691035 | rs10028293 | 0.569 | 1.195 | -1.293 | 0.082 | -0.024 | -0.005 | -0.005 | 0 | 0.15 | 0.15 | 0.3 | 0.13 | 0.25 | 0.15 |
| 4 | 47691710 | rs6447580 | -0.901 | 0.279 | -1.156 | 0.467 | -0.01 | 0 | 0 | 0 | 0.56 | 0.54 | 0.75 | 0.38 | 0.74 | 0.64 |
| 4 | 47705578 | rs10517198 | 0.473 | 0.814 | -1.437 | 0.958 | -0.015 | -0.071 | -0.071 | 0 | 0.13 | 0.05 | 0.22 | 0.27 | 0.15 | 0.19 |
| 4 | 47709613 | rs13122612 | 0.11 | 1.402 | -1.759 | 0.288 | -0.086 | 0.107 | 0.107 | 0 | 0.2 | 0.01 | 0.16 | 0.02 | 0.25 | 0.22 |
| 4 | 47735503 | rs17601663 | 0.051 | 1.779 | -0.448 | 0.679 | 0 | 0.003 | 0.003 | 0 | 0.18 | 0.23 | 0.17 | 0.32 | 0.16 | 0.14 |
| 4 | 47742275 | rs6849926 | 0.303 | 1.938 | -0.529 | 0.503 | -0.065 | 0.08 | 0.08 | 0 | 0.17 | 0.03 | 0.16 | 0.02 | 0.25 | 0.25 |
| 4 | 47745012 | rs10008014 | 0.012 | 1.853 | -0.274 | 0.552 | 0.12 | 0.239 | 0.239 | 0.303 | 0.06 | 0.3 | 0.09 | 0.26 | 0.09 | 0.02 |
| 4 | 47752554 | rs4473628 | -0.289 | 1.754 | -0.238 | 0.527 | 0.034 | -0.001 | -0.001 | 0 | 0.23 | 0.31 | 0.24 | 0.28 | 0.33 | 0.27 |
| 4 | 47816387 | rs12508460 | 0.442 | 1.357 | -1.013 | 0.38 | -0.082 | 0.002 | 0.002 | 0.062 | 0.18 | 0.02 | 0.27 | 0.09 | 0.25 | 0.16 |
| 4 | 47817004 | rs12645164 | 0.639 | 0.942 | -0.388 | 0.809 | -0.003 | 0.048 | 0.048 | 0 | 0.14 | 0.2 | 0.23 | 0.32 | 0.14 | 0.2 |
| 4 | 47833809 | rs1866688 | 0.471 | 0.785 | 0.01 | 0.886 | 0 | 0.013 | 0.013 | 0 | 0.16 | 0.18 | 0.19 | 0.3 | 0.13 | 0.19 |
| 4 | 47837883 | rs893864 | -1.588 | 0.368 | 0.25 | -0.003 | 0.011 | 0.01 | 0.01 | 0 | 0.74 | 0.83 | 0.86 | 0.98 | 0.79 | 0.8 |
| 4 | 47838050 | rs893863 | -1.539 | 0.358 | 0.439 | -0.008 | 0.006 | -0.017 | -0.017 | 0 | 0.73 | 0.79 | 0.86 | 0.98 | 0.79 | 0.8 |
| 4 | 47839929 | rs2289433 | -1.14 | 0.299 | 0.867 | -0.035 | -0.021 | -0.015 | -0.015 | 0 | 0.69 | 0.57 | 0.74 | 0.69 | 0.75 | 0.77 |
| 4 | 47843628 | rs1062858 | 0.035 | 0.596 | 0.621 | -0.315 | 0 | 0 | 0 | 0 | 0.31 | 0.38 | 0.34 | 0.6 | 0.33 | 0.35 |
| 4 | 47845920 | rs1440221 | -0.411 | 0.557 | 0.586 | -0.287 | -0.001 | -0.011 | -0.011 | 0 | 0.35 | 0.44 | 0.46 | 0.89 | 0.37 | 0.38 |
| 4 | 47855999 | rs10025104 | -1.528 | 0.577 | 0.511 | -0.296 | -0.006 | -0.004 | -0.004 | 0.258 | 0.7 | 0.66 | 0.75 | 0.69 | 0.75 | 0.77 |
| 4 | 47868889 | rs2033895 | -1.927 | 0.572 | 0.494 | -0.315 | 0.006 | -0.017 | -0.017 | 0 | 0.73 | 0.79 | 0.87 | 0.98 | 0.79 | 0.8 |

| | | | | | | | | | | | | | | | | |
|---|---|---|---|---|---|---|---|---|---|---|---|---|---|---|---|---|
| 4 | 47887536 | rs2053404 | -1.545 | 0.497 | 0.313 | -0.403 | -0.006 | -0.004 | -0.004 | 0 | 0.7 | 0.66 | 0.75 | 0.69 | 0.75 | 0.77 |
| 4 | 47887991 | rs6818556 | -1.545 | 0.497 | 0.313 | -0.403 | -0.006 | -0.004 | -0.004 | 0 | 0.7 | 0.66 | 0.75 | 0.69 | 0.75 | 0.77 |
| 4 | 47901259 | rs4475122 | 0.433 | 0.471 | 0.357 | -0.444 | -0.032 | 0.132 | 0.132 | 2.133 | 0.37 | 0.14 | 0.4 | 0.09 | 0.42 | 0.43 |
| 4 | 47913127 | rs4695306 | -1.503 | 0.448 | 0.249 | -0.369 | -0.006 | -0.005 | -0.005 | 0 | 0.7 | 0.66 | 0.75 | 0.69 | 0.75 | 0.77 |
| 4 | 47917122 | rs4695309 | -1.737 | 0.443 | 0.238 | -0.426 | 0.001 | -0.04 | -0.04 | 0.02 | 0.73 | 0.76 | 0.86 | 0.98 | 0.79 | 0.81 |
| 4 | 47917441 | rs6833320 | -1.756 | 0.445 | 0.244 | -0.42 | 0.001 | -0.041 | -0.041 | 6.757 | 0.73 | 0.76 | 0.86 | 0.98 | 0.79 | 0.81 |
| 4 | 47937727 | rs9291318 | -1.667 | 0.477 | 0.309 | -0.422 | -0.005 | -0.064 | -0.064 | 1.741 | 0.71 | 0.71 | 0.86 | 0.98 | 0.79 | 0.81 |
| 4 | 47945295 | rs28642966 | 1.367 | 0.524 | 0.126 | -0.291 | -0.002 | 0.037 | 0.037 | 0.082 | 0.13 | 0.18 | 0.19 | 0.3 | 0.13 | 0.2 |
| 4 | 47950256 | rs4695317 | 0.336 | 0.552 | 0.005 | -0.363 | -0.005 | 0.028 | 0.028 | 2.255 | 0.37 | 0.23 | 0.4 | 0.09 | 0.42 | 0.4 |
| 4 | 47955758 | rs17573709 | -0.252 | 0.611 | 0.084 | -0.258 | 0.005 | 0.004 | 0.004 | 0.735 | 0.3 | 0.4 | 0.35 | 0.61 | 0.33 | 0.37 |
| 4 | 47965547 | rs12640473 | -0.428 | 0.42 | -0.074 | -0.444 | 0.042 | 0.038 | 0.038 | 0.94 | 0.54 | 0.26 | 0.55 | 0.35 | 0.63 | 0.59 |
| 4 | 47979063 | rs4695318 | -0.421 | 0.42 | -0.074 | -0.443 | 0.042 | 0.039 | 0.039 | 0.893 | 0.54 | 0.27 | 0.54 | 0.35 | 0.63 | 0.59 |
| 4 | 47986088 | rs167692 | 1.475 | 0.324 | -0.195 | -0.509 | 0.008 | -0.015 | -0.015 | 0 | 0.27 | 0.21 | 0.13 | 0.02 | 0.2 | 0.19 |
| 4 | 47990809 | rs17654757 | 0.675 | 0.568 | -0.363 | -0.202 | -0.075 | -0.074 | -0.074 | 0 | 0.15 | 0.03 | 0.15 | 0.29 | 0.19 | 0.17 |
| 4 | 48018229 | rs321631 | 0.421 | -0.241 | -0.33 | -0.429 | 0.037 | 0.073 | 0.073 | 0 | 0.45 | 0.29 | 0.29 | 0.34 | 0.37 | 0.48 |
| 4 | 48018529 | rs321633 | -1.276 | -0.394 | -0.076 | -0.274 | -0.022 | 0.145 | 0.145 | 0.347 | 0.85 | 0.64 | 0.8 | 0.66 | 0.89 | 0.81 |
| 4 | 48028097 | rs11733284 | 0.474 | -0.371 | -0.144 | -0.393 | 0.045 | 0.076 | 0.076 | 0 | 0.45 | 0.31 | 0.27 | 0.33 | 0.36 | 0.48 |
| 4 | 48036711 | rs10517204 | -1.194 | -0.367 | -0.075 | -0.275 | -0.027 | 0.093 | 0.093 | 0.453 | 0.84 | 0.69 | 0.8 | 0.67 | 0.89 | 0.78 |
| 4 | 48037926 | rs13116684 | 1.295 | -0.492 | -0.231 | -0.423 | 0.028 | 0.028 | 0.028 | 0 | 0.29 | 0.23 | 0.19 | 0.16 | 0.22 | 0.21 |
| 4 | 48038584 | rs10938519 | 1.474 | -0.082 | -0.263 | -0.472 | -0.012 | -0.023 | -0.023 | 0.079 | 0.11 | 0.07 | 0.13 | 0.27 | 0.12 | 0.09 |
| 4 | 48044017 | rs4695329 | 1.003 | -0.074 | -0.232 | -0.516 | 0.046 | -0.064 | -0.064 | 0 | 0.27 | 0.31 | 0.39 | 0.06 | 0.4 | 0.22 |
| 4 | 48047367 | rs4340770 | -0.576 | -0.255 | -0.047 | -0.456 | -0.006 | -0.006 | -0.006 | 0 | 0.6 | 0.58 | 0.43 | 0.6 | 0.49 | 0.55 |
| 4 | 48047591 | rs6812991 | 1.159 | -0.354 | -0.098 | -0.484 | 0.019 | 0.019 | 0.019 | 0 | 0.3 | 0.27 | 0.2 | 0.17 | 0.22 | 0.21 |
| 4 | 48052003 | rs17470528 | 0.756 | -0.04 | -0.433 | -0.335 | -0.051 | 0.043 | 0.043 | 0 | 0.17 | 0 | 0.1 | 0.14 | 0.17 | 0.28 |
| 4 | 48060694 | rs1507925 | -0.279 | 0.136 | -0.009 | -0.252 | -0.001 | 0 | 0 | 0 | 0.48 | 0.54 | 0.59 | 0.43 | 0.53 | 0.45 |
| 4 | 48066864 | rs6839501 | 0.45 | -0.058 | -0.083 | -0.318 | 0.009 | -0.006 | -0.006 | 0 | 0.43 | 0.33 | 0.34 | 0.57 | 0.38 | 0.37 |
| 4 | 48068783 | rs1567217 | 1.304 | -0.083 | -0.304 | -0.328 | 0.028 | 0.045 | 0.045 | 0.117 | 0.3 | 0.02 | 0.19 | 0.18 | 0.27 | 0.29 |
| 4 | 48069725 | rs9996535 | -0.022 | -0.013 | 0.035 | -0.334 | -0.011 | -0.019 | -0.019 | 0 | 0.45 | 0.29 | 0.57 | 0.43 | 0.52 | 0.45 |
| 4 | 48073501 | rs906486 | -0.549 | 0.364 | 0.141 | -0.368 | 0.053 | -0.068 | -0.068 | 0 | 0.89 | 0.64 | 0.92 | 1 | 0.9 | 0.81 |
| 4 | 48078593 | rs6850108 | -0.401 | 0.305 | -0.051 | -0.326 | -0.002 | -0.012 | -0.012 | 0 | 0.65 | 0.36 | 0.57 | 0.93 | 0.59 | 0.79 |
| 4 | 48083021 | rs12647931 | 1.012 | 0.295 | -0.14 | 0.01 | -0.012 | 0.01 | 0.01 | 0 | 0.08 | 0.06 | 0.11 | 0.4 | 0.09 | 0.11 |
| 4 | 48088446 | rs6858664 | 0.612 | 0.245 | 0.209 | -0.282 | 0.085 | 0.081 | 0.081 | 4.995 | 0.95 | 0.53 | 0.95 | 1 | 1 | 1 |
| 4 | 48094275 | rs13118073 | 0.334 | 0.297 | 0.142 | -0.298 | 0.016 | -0.043 | -0.043 | 0.822 | 0.33 | 0.37 | 0.39 | 0.08 | 0.41 | 0.21 |
| 4 | 48100497 | rs6822209 | -0.672 | -0.058 | -0.138 | -0.303 | -0.003 | -0.03 | -0.03 | 0 | 0.38 | 0.52 | 0.42 | 0.76 | 0.33 | 0.49 |
| 4 | 48115264 | rs7658300 | 0.801 | 0.376 | -0.586 | -0.331 | -0.144 | 0.107 | 0.107 | 1.909 | 0.28 | 0.03 | 0.34 | 0.07 | 0.37 | 0.16 |
| 4 | 48126086 | rs2704439 | -1.653 | -0.213 | -0.224 | -0.075 | 0.007 | 0.01 | 0.01 | 0 | 0.74 | 0.79 | 0.82 | 0.89 | 0.75 | 0.75 |
| 4 | 48127262 | rs6837324 | 0.547 | -0.108 | -0.036 | -0.394 | -0.026 | 0.062 | 0.062 | 0.831 | 0.34 | 0.22 | 0.38 | 0.02 | 0.38 | 0.18 |
| 4 | 48132891 | rs885443 | -0.283 | -0.191 | 0.155 | -0.247 | -0.032 | -0.003 | -0.003 | 0 | 0.34 | 0.18 | 0.41 | 0.7 | 0.34 | 0.53 |
| 4 | 48134118 | rs11931912 | 1.033 | -0.201 | 0.071 | -0.042 | -0.027 | -0.055 | -0.055 | 0 | 0.12 | 0.05 | 0.12 | 0.35 | 0.14 | 0.14 |
| 4 | 48138235 | rs3805184 | -1.405 | -0.274 | 0.425 | -0.245 | -0.049 | -0.103 | -0.103 | 0 | 0.71 | 0.55 | 0.83 | 0.91 | 0.76 | 0.75 |
| 4 | 48143201 | rs17470892 | 1.237 | -0.249 | 0.29 | -0.277 | -0.029 | 0.039 | 0.039 | 1.299 | 0.3 | 0.2 | 0.32 | 0.02 | 0.37 | 0.11 |
| 4 | 48146864 | rs996223 | 0.511 | -0.271 | 0.358 | -0.231 | -0.006 | 0.008 | 0.008 | 0.663 | 0.36 | 0.31 | 0.42 | 0.21 | 0.43 | 0.21 |
| 4 | 48147309 | rs17574371 | 1.251 | -0.251 | 0.315 | -0.384 | -0.031 | 0.046 | 0.046 | 1.358 | 0.3 | 0.18 | 0.32 | 0.02 | 0.37 | 0.11 |
| 4 | 48147774 | rs2661548 | -1.711 | -0.313 | 0.192 | -0.364 | 0.02 | 0.031 | 0.031 | 0 | 0.73 | 0.8 | 0.86 | 0.91 | 0.77 | 0.75 |
| 4 | 48161944 | rs2704436 | -2.022 | -0.3 | 0.156 | -0.357 | 0.023 | 0.045 | 0.045 | 0 | 0.75 | 0.83 | 0.86 | 0.91 | 0.77 | 0.75 |
| 4 | 48164537 | rs3792622 | -0.572 | -0.248 | 0.101 | -0.333 | -0.003 | 0.012 | 0.012 | 0 | 0.36 | 0.34 | 0.49 | 0.89 | 0.35 | 0.57 |
| 4 | 48165834 | rs13104849 | -0.58 | -0.222 | -0.101 | -0.287 | -0.009 | 0.045 | 0.045 | 0 | 0.35 | 0.28 | 0.48 | 0.89 | 0.35 | 0.57 |
| 4 | 48172266 | rs2271173 | -0.666 | -0.208 | -0.19 | -0.302 | -0.003 | 0.018 | 0.018 | 0 | 0.37 | 0.34 | 0.49 | 0.89 | 0.36 | 0.57 |
| 4 | 48182433 | rs4312730 | -0.544 | -0.32 | -0.176 | 0.146 | 0.011 | 0.002 | 0.002 | 0 | 0.61 | 0.65 | 0.72 | 0.56 | 0.63 | 0.6 |
| 4 | 48186152 | rs7693779 | 0.101 | -0.219 | 0.002 | 0.532 | -0.014 | 0.255 | 0.255 | 0.448 | 0.14 | 0.49 | 0.26 | 0.36 | 0.1 | 0.24 |
| 4 | 48191491 | rs17655249 | 0.265 | 0.324 | -0.068 | 0.612 | -0.031 | -0.062 | -0.062 | 0 | 0.13 | 0.05 | 0.13 | 0.37 | 0.15 | 0.15 |
| 4 | 48195725 | rs17471142 | -0.494 | 0.388 | 0.16 | -0.167 | 0 | -0.016 | -0.016 | 0 | 0.45 | 0.48 | 0.48 | 0.2 | 0.47 | 0.3 |
| 4 | 48195942 | rs16861112 | -0.257 | 0.427 | 0.436 | -0.328 | -0.022 | -0.113 | -0.113 | 0.313 | 0.17 | 0.38 | 0.1 | 0 | 0.1 | 0.14 |
| 4 | 48197767 | rs2352593 | 0.312 | 0.959 | -0.019 | -0.345 | -0.039 | 0.049 | 0.049 | 0 | 0.22 | 0.11 | 0.31 | 0.05 | 0.3 | 0.1 |
| 4 | 48198986 | rs6829390 | -0.679 | 0.98 | 0.732 | -0.146 | 0.102 | 0.277 | 0.277 | 0.099 | 0.08 | 0.47 | 0.18 | 0.2 | 0.07 | 0.06 |
| 4 | 48199814 | rs2704425 | -1.423 | 0.714 | 0.393 | -0.279 | 0.013 | 0.053 | 0.053 | 0 | 0.48 | 0.66 | 0.56 | 0.27 | 0.47 | 0.33 |
| 4 | 48200584 | rs2457415 | -0.437 | 0.524 | 0.433 | 0.103 | 0.01 | 0.198 | 0.198 | 0 | 0.18 | 0.47 | 0.27 | 0.53 | 0.15 | 0.2 |
| 4 | 48201821 | rs2457414 | -1.761 | 0.539 | 0.298 | 0.06 | 0.021 | -0.03 | -0.03 | 0 | 0.6 | 0.74 | 0.68 | 0.59 | 0.59 | 0.51 |
| 4 | 48204594 | rs2017242 | 0.334 | 0.48 | 0.151 | 0.026 | 0.006 | 0.062 | 0.062 | 0 | 0.17 | 0.07 | 0.07 | 0.02 | 0.1 | 0.2 |
| 4 | 48210653 | rs1912163 | 0.093 | 0.155 | 0.757 | 0.31 | -0.02 | 0.054 | 0.054 | 0 | 0.13 | 0.35 | 0.12 | 0.34 | 0.09 | 0.15 |
| 4 | 48213011 | rs1509657 | -0.697 | 0.368 | 0.594 | 0.179 | 0.013 | 0.032 | 0.032 | 0 | 0.46 | 0.34 | 0.55 | 0.25 | 0.5 | 0.37 |
| 4 | 48226267 | rs4694890 | -0.599 | 0.397 | 0.251 | 0.402 | 0 | 0 | 0 | 0 | 0.45 | 0.38 | 0.53 | 0.3 | 0.48 | 0.36 |
| 4 | 48227782 | rs2704423 | -0.76 | 0.051 | -0.075 | 0.613 | 0.01 | 0.004 | 0.004 | 0.237 | 0.53 | 0.58 | 0.52 | 0.48 | 0.61 | 0.51 |
| 4 | 48228266 | rs2704421 | -0.891 | 0.268 | -0.138 | 0.597 | 0.064 | -0.017 | -0.017 | 0 | 0.55 | 0.76 | 0.66 | 0.58 | 0.63 | 0.53 |
| 4 | 48229886 | rs2664017 | -0.831 | 1.473 | -0.011 | 0.642 | 0.07 | 0.044 | 0.044 | 0.017 | 0.21 | 0.56 | 0.32 | 0.28 | 0.25 | 0.21 |
| 4 | 48248258 | rs12643804 | -0.114 | 0.794 | -0.166 | 0.928 | 0.005 | 0.005 | 0.005 | 0.009 | 0.44 | 0.35 | 0.47 | 0.42 | 0.42 | 0.49 |
| 4 | 48260644 | rs1472971 | -0.397 | 0.623 | -0.171 | 0.781 | 0 | 0.001 | 0.001 | 0 | 0.51 | 0.53 | 0.65 | 0.65 | 0.52 | 0.57 |
| 4 | 48263941 | rs10938526 | 0.497 | 0.627 | -0.319 | 0.783 | -0.033 | 0.059 | 0.059 | 0 | 0.46 | 0.18 | 0.32 | 0.35 | 0.48 | 0.43 |
| 4 | 48269985 | rs9994779 | 0.238 | 0.719 | -0.466 | 0.927 | 0.003 | 0.053 | 0.053 | 0.217 | 0.4 | 0.24 | 0.36 | 0.26 | 0.39 | 0.3 |

| | | | | | | | | | | | | | | | | |
|---|---|---|---|---|---|---|---|---|---|---|---|---|---|---|---|---|
| 4 | 48296382 | rs380016 | 0.658 | 0.981 | -0.678 | 0.813 | -0.008 | 0.034 | 0.034 | 0 | 0.49 | 0.08 | 0.37 | 0.39 | 0.54 | 0.45 |
| 4 | 48312390 | rs10014431 | 0.322 | 1.004 | -0.86 | 0.696 | 0.017 | 0.048 | 0.048 | 0.115 | 0.36 | 0.24 | 0.29 | 0.22 | 0.33 | 0.32 |
| 4 | 48313005 | rs6814786 | 0.319 | 1.061 | -0.939 | 0.638 | -0.105 | -0.016 | -0.016 | 0.853 | 0.18 | 0.01 | 0.22 | 0.19 | 0.23 | 0.18 |
| 4 | 48318390 | rs4599384 | -0.862 | 0.917 | -0.618 | 0.893 | 0.003 | 0.009 | 0.009 | 0 | 0.48 | 0.55 | 0.59 | 0.64 | 0.46 | 0.56 |
| 4 | 48332641 | rs4695372 | -0.193 | 0.931 | -0.984 | 0.598 | -0.02 | 0.062 | 0.062 | 0 | 0.38 | 0.23 | 0.37 | 0.23 | 0.4 | 0.27 |
| 4 | 48336985 | rs309872 | 0.663 | 0.394 | -0.857 | 0.604 | 0.016 | 0.009 | 0.009 | 0 | 0.55 | 0.35 | 0.51 | 0.59 | 0.59 | 0.5 |
| 4 | 53339047 | rs1459822 | 0.735 | 0.629 | 1.214 | 0.59 | 0 | 0.011 | 0.011 | 0 | 0.69 | 0.44 | 0.44 | 0.35 | 0.7 | 0.55 |
| 4 | 53383129 | rs346901 | 0.6 | 0.654 | 1.028 | 0.459 | -0.001 | -0.001 | -0.001 | 0.527 | 0.72 | 0.74 | 0.47 | 0.34 | 0.7 | 0.54 |
| 4 | 53414666 | rs10034992 | 1.129 | 0.821 | 0.166 | 0.646 | 0.001 | -0.022 | -0.022 | 0.547 | 0.71 | 0.68 | 0.47 | 0.15 | 0.7 | 0.51 |
| 4 | 53434611 | rs346011 | -1.644 | 0.389 | 0.571 | 0.855 | 0.009 | 0.039 | 0.039 | 0.417 | 0.17 | 0.41 | 0.13 | 0.16 | 0.15 | 0.14 |
| 4 | 53453449 | rs17051578 | -0.991 | 0.536 | 0.538 | 1.007 | 0.015 | 0.04 | 0.04 | 0 | 0.14 | 0.27 | 0.12 | 0.17 | 0.13 | 0.11 |
| 4 | 53454738 | rs346005 | -2.651 | 0.547 | 0.404 | 1.425 | 0.013 | -0.048 | -0.048 | 0 | 0.39 | 0.82 | 0.53 | 0.44 | 0.32 | 0.33 |
| 4 | 53456416 | rs7668895 | -0.965 | 0.97 | 0.41 | 1.705 | 0.012 | 0.033 | 0.033 | 0 | 0.13 | 0.2 | 0.12 | 0.17 | 0.13 | 0.11 |
| 4 | 53459620 | rs3733506 | -1.016 | 0.968 | 0.416 | 1.704 | 0.011 | 0.03 | 0.03 | 0 | 0.13 | 0.2 | 0.12 | 0.17 | 0.13 | 0.11 |
| 4 | 53463566 | rs17051586 | -1.627 | 0.971 | 0.433 | 1.706 | 0.01 | 0.038 | 0.038 | 0.005 | 0.15 | 0.34 | 0.13 | 0.17 | 0.13 | 0.12 |
| 4 | 53468479 | rs2244291 | -0.395 | 1.05 | 0.461 | 1.75 | -0.039 | -0.054 | -0.054 | 0 | 0.09 | 0.01 | 0.15 | 0.21 | 0.09 | 0.17 |
| 4 | 53472171 | rs7680727 | -1.089 | 0.991 | 0.829 | 1.36 | 0.007 | -0.001 | -0.001 | 0 | 0.12 | 0.17 | 0.11 | 0.12 | 0.13 | 0.09 |
| 4 | 53523725 | rs10014556 | -1.907 | 0.986 | 0.832 | 1.373 | 0.013 | 0.041 | 0.041 | 0.004 | 0.15 | 0.35 | 0.13 | 0.17 | 0.13 | 0.12 |
| 4 | 53549643 | rs10517266 | -1.037 | 1.141 | 0.432 | 1.451 | 0.006 | 0 | 0 | 0 | 0.12 | 0.16 | 0.11 | 0.12 | 0.13 | 0.1 |
| 4 | 53578009 | rs10028122 | -1.408 | 1.162 | 0.152 | 1.391 | 0.007 | 0.024 | 0.024 | 0.071 | 0.13 | 0.23 | 0.12 | 0.18 | 0.13 | 0.11 |
| 4 | 53582065 | rs7677282 | -1.526 | 1.092 | 0.132 | 1.374 | 0.002 | 0.019 | 0.019 | 0.003 | 0.14 | 0.25 | 0.12 | 0.18 | 0.13 | 0.11 |
| 4 | 53604587 | rs6840875 | -1.226 | 1.093 | -0.044 | 1.411 | 0.001 | 0.013 | 0.013 | 0 | 0.13 | 0.15 | 0.16 | 0.18 | 0.12 | 0.11 |
| 4 | 53619367 | rs4865408 | -1.345 | 1.108 | -0.047 | 1.389 | 0 | 0.014 | 0.014 | 0.009 | 0.14 | 0.17 | 0.17 | 0.2 | 0.13 | 0.09 |
| 4 | 53623737 | rs10009096 | -1.363 | 1.141 | -0.059 | 1.406 | 0.001 | 0.022 | 0.022 | 0 | 0.13 | 0.19 | 0.17 | 0.2 | 0.12 | 0.09 |
| 4 | 53627606 | rs6838531 | -1.541 | 1.717 | -0.399 | 1.375 | -0.022 | -0.026 | -0.026 | 0 | 0.18 | 0.12 | 0.24 | 0.24 | 0.24 | 0.14 |
| 4 | 53629126 | rs4865412 | -1.518 | 1.692 | 0.038 | 1.491 | 0.015 | 0.085 | 0.085 | 0 | 0.08 | 0.16 | 0.14 | 0.21 | 0.08 | 0.1 |
| 4 | 53646007 | rs10000595 | -1.93 | 1.678 | 0.075 | 1.882 | 0.021 | 0.144 | 0.144 | 0.041 | 0.08 | 0.21 | 0.17 | 0.28 | 0.09 | 0.12 |
| 4 | 53647521 | rs17476009 | 1.185 | 1.663 | -1.011 | 1.091 | -0.055 | 0.185 | 0.185 | 0.007 | 0.42 | 0.05 | 0.45 | 0.13 | 0.35 | 0.33 |
| 4 | 53675834 | rs1384847 | -0.919 | 1.912 | -0.333 | 1.335 | 0 | 0 | 0 | 0 | 0.07 | 0.05 | 0.02 | 0 | 0.08 | 0.1 |
| 4 | 53680696 | rs730247 | -1.73 | 1.855 | 0.638 | 1.695 | 0.052 | 0.162 | 0.162 | 0 | 0.17 | 0.41 | 0.18 | 0.35 | 0.23 | 0.27 |
| 4 | 53681594 | rs4864676 | -2.704 | 1.851 | 0.636 | 1.701 | 0.033 | 0.093 | 0.093 | 0 | 0.23 | 0.45 | 0.2 | 0.36 | 0.28 | 0.37 |
| 4 | 53700565 | rs7667301 | -0.475 | 1.852 | 1.117 | 1.991 | 0.015 | 0.057 | 0.057 | 1.564 | 0.1 | 0.19 | 0.18 | 0.21 | 0.1 | 0.11 |
| 4 | 53710828 | rs6838718 | -1.746 | 1.895 | 1.062 | 1.962 | 0.002 | -0.008 | -0.008 | 0 | 0.17 | 0.25 | 0.14 | 0.1 | 0.2 | 0.3 |
| 4 | 53711634 | rs17698672 | -0.367 | 1.897 | 1.037 | 1.953 | -0.038 | 0.049 | 0.049 | 0 | 0.1 | 0.01 | 0.08 | 0.01 | 0.13 | 0.14 |
| 4 | 53712071 | rs9993034 | -0.718 | 1.916 | 1.299 | 2.072 | 0.067 | 0.267 | 0.267 | 0.019 | 0.05 | 0.21 | 0.07 | 0.27 | 0.08 | 0.05 |
| 4 | 53712304 | rs7662817 | -1.088 | 1.907 | 1.301 | 2.054 | 0.012 | -0.01 | -0.01 | 0 | 0.06 | 0.21 | 0.04 | 0.02 | 0.07 | 0.13 |
| 4 | 53718375 | rs10026162 | -2.451 | 1.922 | 1.266 | 2.045 | 0.036 | 0.109 | 0.109 | 0 | 0.23 | 0.48 | 0.2 | 0.37 | 0.28 | 0.35 |
| 4 | 53724744 | rs6818178 | -1.687 | 1.945 | 1.223 | 1.923 | 0.005 | -0.015 | -0.015 | 0.134 | 0.17 | 0.25 | 0.14 | 0.08 | 0.2 | 0.24 |
| 4 | 53727450 | rs881743 | -2.459 | 1.951 | 1.263 | 1.99 | 0.04 | 0.092 | 0.092 | 0 | 0.23 | 0.47 | 0.21 | 0.34 | 0.28 | 0.35 |
| 4 | 53727487 | rs881744 | -1.703 | 1.951 | 1.2 | 1.972 | 0.005 | -0.015 | -0.015 | 0 | 0.17 | 0.25 | 0.14 | 0.08 | 0.2 | 0.24 |
| 4 | 53732445 | rs2279334 | -0.785 | 1.792 | 1.359 | 2.099 | 0.068 | 0.267 | 0.267 | 0.028 | 0.06 | 0.23 | 0.07 | 0.27 | 0.08 | 0.1 |
| 4 | 53732586 | rs8287 | -2.407 | 1.819 | 1.363 | 2.051 | 0.027 | 0.102 | 0.102 | 0 | 0.22 | 0.5 | 0.21 | 0.34 | 0.26 | 0.32 |
| 4 | 53732611 | rs1043975 | -0.766 | 1.813 | 1.365 | 2.049 | 0.073 | 0.271 | 0.271 | 0.028 | 0.06 | 0.23 | 0.07 | 0.27 | 0.09 | 0.1 |
| 4 | 53735661 | rs7699960 | -1.548 | 1.645 | 1.07 | 1.966 | 0.001 | -0.023 | -0.023 | 0 | 0.16 | 0.27 | 0.13 | 0.07 | 0.17 | 0.16 |
| 4 | 53750002 | rs6852148 | -2.038 | 1.639 | 1.256 | 1.485 | 0.006 | 0.033 | 0.033 | 0 | 0.22 | 0.35 | 0.18 | 0.28 | 0.25 | 0.27 |
| 4 | 53751932 | rs2306600 | -0.437 | 1.736 | 0.9 | 1.593 | 0.028 | 0.115 | 0.115 | 0.214 | 0.07 | 0.13 | 0.15 | 0.23 | 0.1 | 0.11 |
| 4 | 53757000 | rs2898681 | -1.523 | 1.492 | 0.32 | 0.982 | 0.011 | 0.035 | 0.035 | 0 | 0.17 | 0.2 | 0.16 | 0.28 | 0.22 | 0.21 |
| 4 | 53774933 | rs10866413 | 1.624 | 1.353 | -0.123 | 0.498 | -0.01 | 0.013 | 0.013 | 0.815 | 0.64 | 0.36 | 0.7 | 0.71 | 0.54 | 0.64 |
| 4 | 53776206 | rs10014061 | -1.145 | 1.138 | 0.452 | 1.035 | -0.008 | -0.022 | -0.022 | 0 | 0.07 | 0.06 | 0.03 | 0.01 | 0.04 | 0.15 |
| 4 | 53798146 | rs301119 | -2.077 | 1.182 | 0.299 | 1.082 | 0 | 0.103 | 0.103 | 1.886 | 0.14 | 0.53 | 0.13 | 0.28 | 0.12 | 0.23 |
| 4 | 53801092 | rs301113 | -1.366 | 0.834 | 0.353 | 0.426 | 0.009 | -0.03 | -0.03 | 0 | 0.1 | 0.42 | 0.07 | 0.01 | 0.04 | 0.14 |
| 4 | 53820155 | rs13104971 | -0.459 | 1.477 | -0.618 | 1.038 | -0.035 | -0.023 | -0.023 | 0 | 0.07 | 0.02 | 0.12 | 0.29 | 0.12 | 0.28 |
| 4 | 53821849 | rs301092 | -0.719 | 1.071 | -0.726 | 0.624 | 0 | 0.007 | 0.007 | 0.052 | 0.22 | 0.26 | 0.25 | 0.36 | 0.21 | 0.42 |
| 4 | 53828606 | rs17644026 | 0.499 | 1.023 | -0.587 | 0.6 | 0.004 | 0.006 | 0.006 | 1.417 | 0.13 | 0.08 | 0.12 | 0.07 | 0.09 | 0.14 |
| 4 | 53828649 | rs17082225 | 0.087 | 1.07 | -0.76 | 0.441 | 0.006 | 0.029 | 0.029 | 0 | 0.24 | 0.33 | 0.2 | 0.4 | 0.31 | 0.29 |
| 4 | 53830757 | rs301088 | 0.483 | 0.859 | -1.078 | 0.073 | -0.003 | -0.004 | -0.004 | 0.162 | 0.59 | 0.42 | 0.68 | 0.52 | 0.51 | 0.68 |
| 4 | 53844275 | rs2411289 | 0.017 | 0.71 | -1.293 | 0.075 | 0.014 | -0.016 | -0.016 | 0 | 0.11 | 0.09 | 0.03 | 0.18 | 0.04 | 0.13 |
| 4 | 53848506 | rs11133239 | 0.964 | 1.203 | -1.326 | -0.167 | 0.011 | -0.047 | -0.047 | 0.162 | 0.11 | 0.17 | 0.12 | 0.03 | 0.16 | 0.03 |
| 4 | 53850860 | rs1531836 | -1.405 | 1.159 | -1.246 | -0.08 | 0.077 | 0.238 | 0.238 | 0 | 0.95 | 0.81 | 0.88 | 0.66 | 0.9 | 0.8 |
| 4 | 53859627 | rs6849838 | -0.185 | 1.065 | -1.357 | -0.208 | 0.011 | 0.009 | 0.009 | 0 | 0.37 | 0.47 | 0.32 | 0.43 | 0.48 | 0.31 |
| 4 | 53863777 | rs6838924 | -0.597 | 0.892 | -1.478 | -0.404 | 0.016 | 0.019 | 0.019 | 0 | 0.51 | 0.67 | 0.41 | 0.44 | 0.58 | 0.49 |
| 4 | 53867132 | rs11929934 | 0.504 | 1.163 | -1.258 | -0.142 | 0.003 | 0.008 | 0.008 | 0 | 0.29 | 0.33 | 0.23 | 0.4 | 0.35 | 0.37 |
| 4 | 53869083 | rs11133240 | -1.114 | 1.198 | -1.675 | 0.391 | 0.075 | 0.227 | 0.227 | 0 | 0.95 | 0.83 | 0.88 | 0.65 | 0.9 | 0.81 |
| 4 | 53871388 | rs12641446 | 0.132 | 0.861 | -1.985 | 0.114 | 0.007 | 0 | 0 | 0 | 0.4 | 0.48 | 0.31 | 0.41 | 0.48 | 0.41 |
| 4 | 53872674 | rs17701605 | -0.281 | 0.473 | -2.5 | -0.281 | 0.034 | 0.093 | 0.093 | 0.001 | 0.33 | 0.13 | 0.27 | 0.16 | 0.25 | 0.2 |
| 4 | 53879085 | rs4864709 | -0.391 | 1.099 | -2.713 | -0.267 | -0.121 | 0.069 | 0.069 | 0 | 0.32 | 0.03 | 0.34 | 0.15 | 0.39 | 0.14 |
| 4 | 53879152 | rs10012324 | 0.194 | 0.486 | -2.288 | 0.019 | -0.021 | 0 | 0 | 0 | 0.4 | 0.46 | 0.34 | 0.37 | 0.3 | 0.49 |
| 4 | 53888072 | rs10024057 | -0.375 | -0.754 | -1.169 | -0.211 | -0.07 | -0.038 | -0.038 | 0.206 | 0.19 | 0.56 | 0.11 | 0.12 | 0.08 | 0.28 |
| 4 | 53893156 | rs17082301 | -0.027 | -0.836 | -0.425 | -0.244 | -0.07 | -0.006 | -0.006 | 0 | 0.15 | 0.47 | 0.09 | 0.12 | 0.05 | 0.26 |

| | | | | | | | | | | | | | | | | |
|---|---|---|---|---|---|---|---|---|---|---|---|---|---|---|---|---|
| 4 | 53907444 | rs4864715 | -0.98 | -1.03 | -0.547 | -0.407 | -0.072 | -0.057 | -0.057 | 0 | 0.23 | 0.59 | 0.11 | 0.12 | 0.1 | 0.29 |
| 4 | 53910138 | rs13143381 | 1.209 | -0.837 | -0.836 | 0.178 | 0.005 | 0.003 | 0.003 | 0.08 | 0.15 | 0.12 | 0.22 | 0.35 | 0.29 | 0.22 |
| 4 | 53915624 | rs10517278 | 0.287 | -1.185 | -0.604 | 0.216 | -0.092 | 0.017 | 0.017 | 0.067 | 0.77 | 0.53 | 0.88 | 0.78 | 0.9 | 0.7 |
| 4 | 53922626 | rs2046306 | 1.065 | -0.613 | -1.385 | 0.196 | -0.081 | -0.045 | -0.045 | 0.83 | 0.34 | 0.07 | 0.41 | 0.35 | 0.44 | 0.36 |
| 4 | 53939510 | rs2411290 | -0.339 | -0.725 | -1.315 | 0.136 | 0.017 | 0.002 | 0.002 | 0.02 | 0.46 | 0.39 | 0.4 | 0.38 | 0.29 | 0.43 |
| 4 | 53959141 | rs13141532 | 1.651 | -0.632 | -1.496 | 0.138 | -0.04 | -0.029 | -0.029 | 0 | 0.13 | 0.09 | 0.15 | 0.26 | 0.25 | 0.2 |
| 4 | 53960196 | rs11934112 | 0.715 | -0.74 | -1.461 | 0.018 | -0.007 | 0.006 | 0.006 | 0.962 | 0.41 | 0.31 | 0.44 | 0.35 | 0.47 | 0.37 |
| 4 | 53980677 | rs4864446 | 1.147 | -1.065 | -1.467 | -0.026 | -0.031 | -0.008 | -0.008 | 0 | 0.23 | 0.05 | 0.3 | 0.17 | 0.19 | 0.16 |
| 4 | 53986846 | rs4864723 | 0.172 | -1.036 | -0.881 | -0.066 | 0.055 | 0.056 | 0.056 | 0 | 0.55 | 0.28 | 0.6 | 0.71 | 0.72 | 0.58 |
| 4 | 53990594 | rs10517279 | -0.65 | -1.883 | -1.178 | -0.148 | 0.076 | 0.059 | 0.059 | 0.078 | 0.34 | 0.18 | 0.19 | 0.2 | 0.15 | 0.35 |
| 4 | 53991510 | rs10016091 | 0.757 | -1.359 | -0.637 | -0.033 | 0.008 | -0.01 | -0.01 | 0 | 0.33 | 0.33 | 0.34 | 0.23 | 0.45 | 0.23 |
| 4 | 54000949 | rs10028181 | -0.058 | -1.641 | -0.402 | 0.351 | -0.013 | -0.016 | -0.016 | 0 | 0.44 | 0.32 | 0.45 | 0.47 | 0.43 | 0.55 |
| 4 | 54008839 | rs7696426 | -1.643 | -1.737 | -0.236 | 0.335 | -0.001 | -0.001 | -0.001 | 0.083 | 0.87 | 0.87 | 0.82 | 0.77 | 0.92 | 0.81 |
| 4 | 54011424 | rs4864727 | -0.012 | -1.753 | -0.41 | 0.334 | -0.013 | -0.014 | -0.014 | 0 | 0.44 | 0.32 | 0.45 | 0.47 | 0.43 | 0.55 |
| 4 | 54011526 | rs7675987 | 1.326 | -1.053 | -0.708 | 0.27 | -0.02 | 0.02 | 0.02 | 0 | 0.28 | 0.19 | 0.28 | 0.21 | 0.39 | 0.22 |
| 4 | 54011603 | rs17082453 | 1.351 | -1.18 | -0.649 | 0.393 | 0.015 | -0.06 | -0.06 | 0.605 | 0.13 | 0.01 | 0.17 | 0.23 | 0.08 | 0.19 |
| 4 | 54016241 | rs6818457 | 0.953 | -0.998 | -0.555 | 0.305 | 0.001 | -0.02 | -0.02 | 0 | 0.09 | 0.05 | 0.07 | 0.09 | 0.08 | 0.04 |
| 4 | 54017971 | rs4864728 | 0.745 | -0.915 | -0.596 | 0.182 | 0 | 0.068 | 0.068 | 0 | 0.41 | 0.28 | 0.35 | 0.33 | 0.6 | 0.39 |
| 4 | 54021591 | rs1962230 | 0.519 | -1.629 | -0.23 | 0.23 | -0.059 | -0.019 | -0.019 | 0 | 0.18 | 0.2 | 0.06 | 0.1 | 0.05 | 0.25 |
| 4 | 54023351 | rs7694213 | 0.41 | -1.639 | -0.18 | 0.313 | 0.036 | -0.003 | -0.003 | 0 | 0.53 | 0.38 | 0.5 | 0.5 | 0.69 | 0.43 |
| 4 | 54026736 | rs9312637 | 0.778 | -1.557 | -0.205 | 0.269 | -0.059 | -0.017 | -0.017 | 0 | 0.16 | 0.2 | 0.06 | 0.1 | 0.05 | 0.25 |
| 4 | 54027595 | rs10517281 | 1.577 | -1.45 | -0.15 | 0.412 | 0 | 0 | 0 | 0.005 | 0.17 | 0.18 | 0.27 | 0.18 | 0.18 | 0.18 |
| 4 | 54033543 | rs17082492 | 1.321 | -1.582 | -0.166 | 0.379 | -0.062 | -0.016 | -0.016 | 0 | 0.13 | 0.2 | 0.04 | 0.09 | 0.03 | 0.17 |
| 4 | 54043211 | rs13111251 | 2.348 | -1.419 | -0.289 | 0.263 | -0.015 | 0.02 | 0.02 | 0 | 0.25 | 0.15 | 0.26 | 0.14 | 0.34 | 0.19 |
| 4 | 54058102 | rs11133248 | 1.963 | -1.683 | 0.234 | 0.454 | -0.021 | 0.028 | 0.028 | 0.17 | 0.41 | 0.16 | 0.51 | 0.31 | 0.52 | 0.32 |
| 4 | 54061989 | rs12507517 | 0.44 | -1.448 | 0.202 | 0.662 | -0.051 | -0.045 | -0.045 | 0 | 0.22 | 0.12 | 0.21 | 0.31 | 0.32 | 0.23 |
| 4 | 54074642 | rs4588514 | 0.831 | -1.463 | 0.23 | 0.645 | -0.001 | 0.002 | 0.002 | 0.924 | 0.58 | 0.46 | 0.59 | 0.45 | 0.58 | 0.58 |
| 4 | 54087096 | rs6818181 | -0.603 | -1.405 | 0.172 | 0.704 | -0.003 | -0.003 | -0.003 | 0 | 0.77 | 0.75 | 0.78 | 0.69 | 0.68 | 0.77 |
| 4 | 54089331 | rs11939189 | 1.602 | -1.521 | 0.227 | 0.737 | 0.009 | -0.07 | -0.07 | 0.389 | 0.12 | 0.01 | 0.18 | 0.24 | 0.09 | 0.19 |
| 4 | 54107382 | rs4472196 | -0.566 | -1.238 | 0.327 | 0.746 | 0.002 | 0.002 | 0.002 | 1.843 | 0.78 | 0.73 | 0.79 | 0.7 | 0.71 | 0.78 |
| 4 | 54140416 | rs17082557 | 0.901 | -1.384 | 0.391 | 0.736 | -0.053 | -0.001 | -0.001 | 0 | 0.14 | 0.2 | 0.06 | 0.17 | 0.06 | 0.25 |
| 4 | 54147428 | rs9884671 | 1.094 | -1.359 | 0.324 | 0.749 | -0.026 | -0.001 | -0.001 | 0 | 0.13 | 0.14 | 0.05 | 0.17 | 0.06 | 0.25 |
| 4 | 54181789 | rs4864747 | 0.482 | -0.84 | -0.063 | 0.872 | 0.006 | -0.004 | -0.004 | 0 | 0.18 | 0.18 | 0.18 | 0.22 | 0.08 | 0.2 |
| 4 | 54181814 | rs12512496 | 1.681 | -0.694 | -0.22 | 0.903 | -0.018 | -0.04 | -0.04 | 0 | 0.08 | 0 | 0.07 | 0.09 | 0.08 | 0.04 |
| 4 | 54203701 | rs11725172 | 0.585 | -0.863 | -0.105 | 0.742 | -0.001 | -0.01 | -0.01 | 0 | 0.61 | 0.59 | 0.6 | 0.48 | 0.61 | 0.58 |
| 4 | 54214528 | rs10517292 | 0.475 | -0.972 | -0.182 | 0.759 | 0.021 | -0.011 | -0.011 | 0 | 0.17 | 0.14 | 0.18 | 0.22 | 0.08 | 0.2 |
| 4 | 54221952 | rs1600795 | 0.199 | -0.461 | 0.02 | 0.883 | 0.007 | 0.03 | 0.03 | 0 | 0.73 | 0.6 | 0.69 | 0.65 | 0.72 | 0.72 |
| 4 | 54222481 | rs2128103 | -0.792 | -0.347 | -0.008 | 0.874 | -0.005 | 0.009 | 0.009 | 0 | 0.82 | 0.75 | 0.8 | 0.78 | 0.85 | 0.77 |
| 4 | 54232110 | rs1351514 | 1.406 | -0.352 | -0.143 | 0.871 | -0.04 | -0.068 | -0.068 | 0 | 0.13 | 0.01 | 0.18 | 0.22 | 0.14 | 0.23 |
| 4 | 54232658 | rs6554098 | 1.631 | -0.782 | -0.332 | 0.696 | -0.016 | 0.047 | 0.047 | 0.017 | 0.44 | 0.14 | 0.54 | 0.31 | 0.56 | 0.24 |
| 4 | 54235363 | rs6828559 | 1.281 | -0.866 | -0.242 | 0.714 | 0.067 | -0.009 | -0.009 | 0 | 0.55 | 0.21 | 0.63 | 0.47 | 0.68 | 0.39 |
| 4 | 54238009 | rs1481823 | 1.402 | -0.722 | -0.289 | 0.75 | -0.007 | 0.001 | 0.001 | 4.844 | 0.12 | 0.07 | 0.1 | 0.16 | 0.13 | 0.15 |
| 4 | 54243425 | rs11133267 | -0.882 | -0.827 | 0.063 | 0.849 | -0.005 | 0.011 | 0.011 | 0 | 0.82 | 0.75 | 0.8 | 0.78 | 0.85 | 0.77 |
| 4 | 54252882 | rs10517295 | 0.513 | -0.967 | 0.232 | 0.827 | -0.109 | 0.002 | 0.002 | 0 | 0.17 | 0.38 | 0.06 | 0.17 | 0.03 | 0.33 |
| 4 | 54254146 | rs1481819 | 0.104 | -0.879 | 0.204 | 0.833 | 0.004 | 0.016 | 0.016 | 0 | 0.73 | 0.66 | 0.69 | 0.65 | 0.71 | 0.72 |
| 4 | 54264543 | rs10025091 | 0.486 | -1.023 | 0.347 | 0.723 | -0.098 | -0.001 | -0.001 | 0.017 | 0.17 | 0.34 | 0.06 | 0.17 | 0.03 | 0.33 |
| 4 | 54299380 | rs11133274 | 0.76 | -1.002 | 0.488 | 0.796 | -0.021 | -0.002 | -0.002 | 0 | 0.66 | 0.4 | 0.75 | 0.68 | 0.8 | 0.45 |
| 4 | 54314418 | rs17326196 | 0.746 | -1.012 | 0.266 | 0.682 | -0.069 | 0 | 0 | 0.155 | 0.16 | 0.21 | 0.05 | 0.17 | 0.04 | 0.34 |
| 4 | 54324379 | rs2412487 | 0.508 | -0.984 | 0.086 | 0.796 | -0.11 | 0.003 | 0.003 | 0 | 0.17 | 0.38 | 0.06 | 0.17 | 0.04 | 0.34 |
| 4 | 54325990 | rs1014056 | -0.529 | -0.264 | -0.189 | 0.894 | -0.005 | 0.009 | 0.009 | 0.007 | 0.82 | 0.75 | 0.8 | 0.78 | 0.85 | 0.77 |
| 4 | 54339478 | rs12505040 | 1.043 | 0.247 | -0.261 | 0.942 | -0.011 | -0.012 | -0.012 | 0.245 | 0.08 | 0.08 | 0.11 | 0.13 | 0.14 | 0.05 |
| 4 | 54346483 | rs6835629 | 0.053 | -0.448 | 0.114 | 0.821 | -0.02 | 0.01 | 0.01 | 0 | 0.29 | 0.34 | 0.24 | 0.39 | 0.18 | 0.57 |
| 4 | 54355824 | rs6838260 | -0.73 | -0.364 | 0.107 | 0.757 | -0.009 | 0.006 | 0.006 | 0 | 0.41 | 0.53 | 0.37 | 0.52 | 0.34 | 0.64 |
| 4 | 54361640 | rs12642403 | -1.003 | -0.186 | -0.322 | 0.864 | 0.017 | -0.006 | -0.006 | 0.318 | 0.45 | 0.56 | 0.38 | 0.48 | 0.35 | 0.63 |
| 4 | 54362150 | rs6836800 | 1.439 | -0.204 | -0.571 | 0.731 | -0.051 | 0.012 | 0.012 | 0.108 | 0.41 | 0.17 | 0.5 | 0.34 | 0.51 | 0.22 |
| 4 | 54367136 | rs4864767 | -0.637 | -0.171 | -0.547 | 0.756 | 0.008 | -0.005 | -0.005 | 0 | 0.44 | 0.53 | 0.38 | 0.48 | 0.35 | 0.62 |
| 4 | 54373464 | rs10021772 | -0.485 | 0.201 | -0.456 | 0.752 | 0.001 | 0 | 0 | 0 | 0.47 | 0.47 | 0.36 | 0.52 | 0.31 | 0.72 |
| 4 | 54377540 | rs2164265 | -0.885 | 0.519 | -0.674 | 0.806 | 0.049 | 0.057 | 0.057 | 0 | 0.9 | 0.69 | 0.88 | 0.86 | 0.86 | 0.95 |
| 4 | 54377597 | rs11935144 | 1.521 | 0.612 | -1.046 | 0.517 | -0.049 | 0.028 | 0.028 | 0 | 0.22 | 0.06 | 0.23 | 0.14 | 0.33 | 0.12 |
| 4 | 54380052 | rs11723567 | 0.145 | 0.498 | -1.234 | 0.846 | -0.071 | -0.065 | -0.065 | 0.2 | 0.26 | 0.09 | 0.37 | 0.29 | 0.31 | 0.14 |
| 4 | 54381345 | rs9999406 | -0.697 | 0.011 | -0.747 | 0.527 | -0.001 | 0.001 | 0.001 | 0.013 | 0.56 | 0.46 | 0.54 | 0.68 | 0.51 | 0.55 |
| 4 | 54382119 | rs1541967 | -0.617 | -0.627 | -0.468 | 0.807 | -0.045 | 0.056 | 0.056 | 3.539 | 0.31 | 0.52 | 0.17 | 0.51 | 0.22 | 0.36 |
| 4 | 54382225 | rs4864777 | -0.664 | -0.618 | -0.862 | 0.438 | 0.006 | 0.003 | 0.003 | 1.591 | 0.51 | 0.54 | 0.4 | 0.54 | 0.42 | 0.42 |
| 4 | 54393115 | rs17082951 | -0.363 | -0.511 | -0.91 | 0.348 | 0.006 | -0.007 | -0.007 | 0 | 0.33 | 0.21 | 0.32 | 0.33 | 0.29 | 0.23 |
| 4 | 54394093 | rs17082953 | 0.595 | -0.856 | -0.45 | 0.292 | -0.004 | 0.006 | 0.006 | 0 | 0.23 | 0.28 | 0.12 | 0.36 | 0.2 | 0.23 |
| 4 | 54394410 | rs4864785 | 0.408 | -0.814 | -0.789 | 0.259 | -0.002 | -0.019 | -0.019 | 0 | 0.25 | 0.11 | 0.26 | 0.25 | 0.23 | 0.19 |
| 4 | 54395740 | rs10517300 | 1.488 | -0.687 | -0.34 | 0.423 | 0 | -0.007 | -0.007 | 0 | 0.1 | 0.15 | 0.08 | 0.08 | 0.1 | 0.06 |
| 4 | 54396077 | rs6812114 | -1.415 | -0.867 | 0.076 | 0.523 | 0.001 | 0.077 | 0.077 | 0 | 0.83 | 0.58 | 0.88 | 0.73 | 0.84 | 0.84 |
| 4 | 54398776 | rs1435221 | -0.345 | -0.817 | -0.319 | 0.557 | -0.028 | 0.025 | 0.025 | 0 | 0.74 | 0.38 | 0.85 | 0.67 | 0.8 | 0.77 |

| | | | | | | | | | | | | | | | | |
|---|---|---|---|---|---|---|---|---|---|---|---|---|---|---|---|---|
| 4 | 54401849 | rs1435219 | 0.636 | -0.289 | -1.043 | 0.333 | -0.04 | 0.04 | 0.04 | 5.872 | 0.47 | 0.12 | 0.5 | 0.34 | 0.55 | 0.37 |
| 4 | 54403898 | rs17082977 | 1.616 | -0.181 | -0.843 | 0.575 | -0.001 | 0 | 0 | 0 | 0.1 | 0.1 | 0.09 | 0.12 | 0.1 | 0.06 |
| 4 | 54409561 | rs12331079 | 0.655 | -0.12 | -1.007 | 0.451 | -0.017 | 0.015 | 0.015 | 2.592 | 0.41 | 0.19 | 0.48 | 0.31 | 0.49 | 0.34 |
| 4 | 54410224 | rs1040084 | -1.029 | -0.204 | -0.453 | 0.669 | 0 | 0.01 | 0.01 | 0 | 0.78 | 0.74 | 0.86 | 0.67 | 0.78 | 0.8 |
| 4 | 54410324 | rs1913484 | -0.004 | -0.459 | -0.44 | 0.596 | -0.024 | -0.002 | -0.002 | 0.456 | 0.67 | 0.53 | 0.84 | 0.63 | 0.74 | 0.73 |
| 4 | 54410671 | rs17660061 | -0.956 | -0.235 | -0.716 | 0.444 | -0.001 | 0 | 0 | 0 | 0.48 | 0.49 | 0.37 | 0.62 | 0.39 | 0.46 |
| 4 | 54411534 | rs1435215 | 1.017 | -0.137 | -1.203 | 0.509 | -0.008 | -0.056 | -0.056 | 0.193 | 0.22 | 0.08 | 0.28 | 0.23 | 0.17 | 0.38 |
| 4 | 54413069 | rs7697542 | 1.329 | -0.057 | -0.654 | 0.117 | 0.01 | -0.064 | -0.064 | 0.181 | 0.26 | 0.39 | 0.28 | 0.12 | 0.32 | 0.14 |
| 4 | 54414696 | rs6820391 | -0.104 | -0.239 | -0.729 | 0.32 | -0.006 | 0 | 0 | 0.573 | 0.27 | 0.15 | 0.3 | 0.27 | 0.29 | 0.22 |
| 4 | 54416496 | rs6815418 | 0.523 | -0.271 | -0.417 | 0.479 | 0.002 | -0.005 | -0.005 | 0 | 0.25 | 0.26 | 0.13 | 0.38 | 0.22 | 0.26 |
| 4 | 54417674 | rs2117600 | 0.473 | -0.358 | -0.501 | 0.431 | 0 | -0.001 | -0.001 | 0.004 | 0.43 | 0.42 | 0.5 | 0.35 | 0.49 | 0.37 |
| 4 | 54418635 | rs6554112 | -0.584 | -0.212 | -0.39 | 0.458 | -0.002 | 0.001 | 0.001 | 0 | 0.67 | 0.66 | 0.62 | 0.71 | 0.71 | 0.63 |
| 4 | 54425184 | rs4864795 | 0.594 | -0.32 | 0.024 | 0.427 | 0.015 | -0.03 | -0.03 | 0 | 0.13 | 0.37 | 0.11 | 0.06 | 0.11 | 0.04 |
| 4 | 54427925 | rs6850277 | -0.244 | -0.496 | -0.227 | 0.451 | 0.001 | -0.016 | -0.016 | 0 | 0.61 | 0.56 | 0.58 | 0.74 | 0.63 | 0.62 |
| 4 | 54439033 | rs4864800 | -0.429 | -0.432 | -0.199 | 0.413 | 0 | 0.001 | 0.001 | 0 | 0.59 | 0.58 | 0.6 | 0.48 | 0.63 | 0.48 |
| 4 | 54442182 | rs4864801 | -0.63 | -0.446 | 0.087 | 0.377 | -0.007 | -0.016 | -0.016 | 0 | 0.28 | 0.33 | 0.35 | 0.25 | 0.26 | 0.19 |
| 4 | 54446860 | rs1369355 | -0.586 | -0.293 | -0.108 | 0.352 | -0.005 | -0.011 | -0.011 | 1.496 | 0.28 | 0.34 | 0.35 | 0.24 | 0.26 | 0.2 |
| 4 | 54451175 | rs932039 | -1.597 | -0.008 | -0.298 | 0.55 | 0.002 | 0.01 | 0.01 | 0 | 0.89 | 0.89 | 0.93 | 0.8 | 0.89 | 0.88 |
| 4 | 54452487 | rs1347406 | 0.013 | 0.056 | -0.139 | 0.259 | -0.004 | -0.009 | -0.009 | 0.009 | 0.7 | 0.67 | 0.83 | 0.66 | 0.71 | 0.79 |
| 4 | 54452878 | rs979006 | -1.62 | -0.015 | -0.01 | 0.213 | 0 | 0.013 | 0.013 | 0 | 0.85 | 0.85 | 0.93 | 0.77 | 0.86 | 0.89 |
| 4 | 54453084 | rs11731437 | 1.783 | 0.168 | -0.209 | 0.078 | -0.003 | -0.003 | -0.003 | 0 | 0.14 | 0.12 | 0.15 | 0.25 | 0.22 | 0.11 |
| 4 | 54453676 | rs17083112 | 0.572 | 0.147 | -0.132 | -0.163 | -0.002 | -0.024 | -0.024 | 0.194 | 0.15 | 0.19 | 0.1 | 0.11 | 0.15 | 0.1 |
| 4 | 54456051 | rs9312642 | -0.56 | 0.261 | -0.363 | -0.057 | 0 | 0.007 | 0.007 | 0 | 0.75 | 0.77 | 0.66 | 0.53 | 0.75 | 0.77 |
| 4 | 54460531 | rs1028000 | 1.874 | 0.104 | -1.159 | -0.314 | -0.061 | -0.085 | -0.085 | 0.493 | 0.26 | 0.02 | 0.28 | 0.26 | 0.26 | 0.41 |
| 4 | 54460824 | rs1464028 | -0.264 | 0.317 | -0.615 | 0.037 | 0 | 0.001 | 0.001 | 0 | 0.24 | 0.24 | 0.37 | 0.33 | 0.27 | 0.25 |
| 4 | 54461548 | rs1015801 | 1.185 | 0.607 | -0.65 | 0.494 | -0.015 | -0.029 | -0.029 | 0 | 0.09 | 0.05 | 0.27 | 0.23 | 0.11 | 0.11 |
| 4 | 54462952 | rs1369351 | -1.78 | 0.373 | -0.28 | 0.341 | 0.003 | -0.001 | -0.001 | 0 | 0.86 | 0.84 | 0.84 | 0.83 | 0.77 | 0.93 |
| 4 | 54463384 | rs7668052 | 1.131 | 0.238 | -0.091 | 0.504 | -0.002 | 0.026 | 0.026 | 0 | 0.14 | 0.17 | 0.07 | 0.23 | 0.12 | 0.11 |
| 4 | 54471672 | rs2668568 | -0.453 | 0.417 | -0.204 | 0.04 | 0 | 0.016 | 0.016 | 0 | 0.51 | 0.43 | 0.52 | 0.64 | 0.51 | 0.4 |
| 4 | 54474244 | rs2616404 | -0.744 | 0.454 | -0.305 | 0.016 | -0.001 | 0 | 0 | 1.16 | 0.4 | 0.41 | 0.49 | 0.35 | 0.43 | 0.39 |
| 4 | 54483569 | rs959094 | -0.464 | 0.459 | -0.371 | 0.014 | -0.022 | -0.001 | -0.001 | 0 | 0.37 | 0.25 | 0.48 | 0.35 | 0.43 | 0.37 |
| 4 | 54485327 | rs2590816 | -1.403 | 0.49 | -0.304 | -0.315 | 0.009 | -0.041 | -0.041 | 0.108 | 0.79 | 0.7 | 0.8 | 0.9 | 0.74 | 0.92 |
| 4 | 54488165 | rs2616417 | -0.161 | 0.611 | -0.884 | -0.387 | -0.033 | 0.081 | 0.081 | 0 | 0.63 | 0.8 | 0.69 | 0.82 | 0.52 | 0.75 |
| 4 | 54489838 | rs10031866 | 0.936 | 0.482 | -0.713 | -0.295 | -0.012 | -0.001 | -0.001 | 0.411 | 0.41 | 0.5 | 0.31 | 0.59 | 0.3 | 0.55 |
| 4 | 54490116 | rs2572297 | -0.27 | 0.547 | -0.794 | -0.384 | -0.001 | -0.001 | -0.001 | 0 | 0.32 | 0.3 | 0.46 | 0.32 | 0.38 | 0.33 |
| 4 | 54493031 | rs2572290 | 0.243 | 0.504 | -0.922 | -0.405 | -0.022 | -0.014 | -0.014 | 1.213 | 0.28 | 0.22 | 0.45 | 0.31 | 0.37 | 0.32 |
| 4 | 54493383 | rs2572291 | -0.687 | 0.553 | -1.063 | -0.428 | -0.006 | 0.038 | 0.038 | 0 | 0.73 | 0.8 | 0.77 | 0.9 | 0.68 | 0.87 |
| 4 | 54499251 | rs1546526 | -1.454 | 0.692 | -0.964 | -0.319 | 0.002 | -0.029 | -0.029 | 0.045 | 0.78 | 0.76 | 0.81 | 0.91 | 0.73 | 0.92 |
| 4 | 54505850 | rs2668524 | -1.491 | 0.691 | -0.973 | -0.32 | -0.001 | -0.003 | -0.003 | 0 | 0.78 | 0.82 | 0.81 | 0.91 | 0.73 | 0.92 |
| 4 | 54511913 | rs4864809 | 0.242 | 0.669 | -0.995 | -0.182 | 0.011 | 0.009 | 0.009 | 0 | 0.48 | 0.59 | 0.35 | 0.59 | 0.36 | 0.59 |
| 4 | 54517341 | rs1077966 | 0.734 | 0.711 | -1.138 | -0.17 | 0.011 | -0.008 | -0.008 | 0 | 0.12 | 0.23 | 0.1 | 0.09 | 0.16 | 0.12 |
| 4 | 54517724 | rs17083248 | 1.485 | 0.719 | -1.953 | -0.007 | -0.038 | -0.103 | -0.103 | 0.172 | 0.22 | 0.01 | 0.21 | 0.27 | 0.17 | 0.43 |
| 4 | 54524939 | rs17083255 | 1.16 | 0.969 | -1.394 | -0.349 | -0.004 | 0 | 0 | 0 | 0.1 | 0.09 | 0.08 | 0.09 | 0.15 | 0.12 |
| 4 | 54542666 | rs2590809 | -1.386 | 0.967 | -1.413 | -0.338 | -0.061 | -0.072 | -0.072 | 0 | 0.86 | 0.99 | 0.67 | 0.79 | 0.81 | 0.88 |
| 4 | 54547031 | rs2199574 | -0.377 | 0.568 | -1.21 | -0.218 | 0.003 | -0.009 | -0.009 | 0 | 0.57 | 0.55 | 0.42 | 0.69 | 0.48 | 0.71 |
| 4 | 54547406 | rs2199572 | -0.047 | 0.625 | -1.181 | -0.048 | -0.002 | -0.007 | -0.007 | 0 | 0.48 | 0.51 | 0.37 | 0.69 | 0.41 | 0.66 |
| 4 | 54556087 | rs2668532 | -0.867 | 0.687 | -2.011 | -0.658 | -0.143 | -0.14 | -0.14 | 0 | 0.6 | 0.97 | 0.53 | 0.57 | 0.52 | 0.39 |
| 4 | 54558299 | rs991670 | -0.384 | 0.331 | -1.205 | -0.885 | -0.001 | -0.076 | -0.076 | 0 | 0.29 | 0.46 | 0.25 | 0.13 | 0.32 | 0.16 |
| 4 | 54559738 | rs2590783 | 0.425 | 0.205 | -0.967 | -0.768 | 0.018 | -0.032 | -0.032 | 0 | 0.15 | 0.35 | 0.15 | 0.09 | 0.16 | 0.06 |
| 4 | 54561980 | rs6827185 | 0.226 | -0.104 | -0.95 | -0.602 | -0.009 | 0.008 | 0.008 | 2.296 | 0.28 | 0.33 | 0.17 | 0.43 | 0.18 | 0.19 |
| 4 | 54564016 | rs1871029 | -0.917 | 0.417 | -1.154 | -0.739 | -0.037 | -0.094 | -0.094 | 0 | 0.67 | 0.89 | 0.55 | 0.58 | 0.61 | 0.48 |
| 4 | 54572307 | rs715155 | -0.937 | 0.375 | -0.289 | -0.907 | 0.014 | 0.005 | 0.005 | 0 | 0.5 | 0.62 | 0.46 | 0.48 | 0.51 | 0.41 |
| 4 | 54574146 | rs2668540 | -1.424 | 0.237 | -0.414 | -0.966 | -0.044 | -0.077 | -0.077 | 0 | 0.59 | 0.93 | 0.61 | 0.59 | 0.59 | 0.46 |
| 4 | 54574680 | rs728587 | -0.358 | 0.215 | -0.142 | -0.992 | -0.009 | 0.007 | 0.007 | 0 | 0.08 | 0.22 | 0.14 | 0.11 | 0.07 | 0.05 |
| 4 | 54575432 | rs2590784 | -0.231 | 0.35 | 0.014 | -1.144 | -0.001 | -0.001 | -0.001 | 0 | 0.38 | 0.57 | 0.38 | 0.42 | 0.35 | 0.28 |
| 4 | 54582132 | rs9684467 | 0.247 | 0.381 | -0.049 | -1.145 | -0.009 | 0.012 | 0.012 | 0 | 0.14 | 0.12 | 0.09 | 0.07 | 0.17 | 0.12 |
| 4 | 54589655 | rs2590828 | 0.53 | -0.284 | 0.429 | -0.964 | 0.034 | -0.023 | -0.023 | 0 | 0.13 | 0.22 | 0.15 | 0.05 | 0.21 | 0.06 |
| 4 | 54596466 | rs2245157 | -0.878 | -0.447 | 0.406 | -1.026 | 0.01 | -0.022 | -0.022 | 0 | 0.85 | 0.8 | 0.85 | 0.95 | 0.8 | 0.88 |
| 4 | 54596896 | rs2616411 | -1.99 | -0.509 | 0.264 | -1.046 | -0.145 | -0.092 | -0.092 | 0 | 0.59 | 0.95 | 0.57 | 0.59 | 0.53 | 0.5 |
| 4 | 54598787 | rs1382084 | -0.718 | -0.608 | 0.603 | -0.944 | -0.021 | 0.014 | 0.014 | 0 | 0.34 | 0.57 | 0.35 | 0.48 | 0.21 | 0.29 |
| 4 | 54608737 | rs3844504 | -1.491 | -0.55 | 0.553 | -0.935 | 0.075 | -0.043 | -0.043 | 0 | 0.44 | 0.81 | 0.42 | 0.54 | 0.33 | 0.37 |
| 4 | 54608763 | rs4082297 | -0.246 | -0.538 | 0.63 | -0.885 | -0.053 | 0.077 | 0.077 | 0 | 0.28 | 0.49 | 0.31 | 0.48 | 0.17 | 0.27 |
| 4 | 54609243 | rs3853200 | 0.484 | -0.544 | 0.724 | -0.879 | -0.012 | 0.024 | 0.024 | 0.01 | 0.22 | 0.26 | 0.21 | 0.37 | 0.14 | 0.23 |
| 4 | 54612049 | rs7664385 | -0.543 | -0.406 | 0.558 | -0.857 | -0.003 | 0.056 | 0.056 | 1.409 | 0.05 | 0.2 | 0.1 | 0.11 | 0.02 | 0.04 |
| 4 | 54612302 | rs3866377 | -0.554 | -0.464 | 0.425 | -0.869 | -0.022 | 0.019 | 0.019 | 0 | 0.35 | 0.47 | 0.27 | 0.43 | 0.25 | 0.34 |
| 4 | 54615216 | rs2590765 | -0.643 | -0.55 | 0.407 | -0.931 | -0.026 | 0.02 | 0.02 | 0.123 | 0.34 | 0.55 | 0.27 | 0.42 | 0.24 | 0.29 |
| 4 | 54616649 | rs2616434 | -2.311 | -0.489 | 0.079 | -1.04 | -0.175 | -0.146 | -0.146 | 0.377 | 0.63 | 0.97 | 0.56 | 0.56 | 0.53 | 0.5 |
| 4 | 54618574 | rs2590767 | 0.238 | -0.482 | 0.242 | -1.07 | 0 | -0.007 | -0.007 | 0 | 0.12 | 0.12 | 0.06 | 0.01 | 0.11 | 0.07 |
| 4 | 54629640 | rs4530695 | -0.6 | -0.528 | 0.2 | -1.108 | 0 | 0.033 | 0.033 | 0 | 0.23 | 0.17 | 0.18 | 0.06 | 0.25 | 0.17 |

| | | | | | | | | | | | | | | | | |
|---|---|---|---|---|---|---|---|---|---|---|---|---|---|---|---|---|
| 4 | 54632173 | rs6842213 | -1.554 | -0.645 | 0.078 | -0.996 | 0.038 | -0.004 | -0.004 | 0.367 | 0.44 | 0.65 | 0.38 | 0.5 | 0.3 | 0.36 |
| 4 | 54633159 | rs2412494 | 0.202 | -0.656 | -0.192 | -1.056 | -0.01 | -0.035 | -0.035 | 0 | 0.08 | 0.27 | 0.04 | 0 | 0.04 | 0.01 |
| 4 | 54638662 | rs4305579 | 0.197 | -0.596 | -0.143 | -1.002 | -0.001 | 0 | 0 | 0 | 0.23 | 0.21 | 0.22 | 0.38 | 0.16 | 0.2 |
| 4 | 54638699 | rs2412493 | 0.16 | -0.576 | -0.118 | -1.005 | -0.002 | -0.027 | -0.027 | 0.017 | 0.12 | 0.17 | 0.06 | 0.02 | 0.11 | 0.07 |
| 4 | 54639903 | rs11726943 | 0.253 | -0.326 | -0.017 | -0.745 | 0.013 | -0.011 | -0.011 | 0 | 0.3 | 0.21 | 0.25 | 0.38 | 0.23 | 0.23 |
| 4 | 54650286 | rs4501282 | 0.287 | 0.423 | -0.063 | -0.492 | -0.028 | 0.024 | 0.024 | 0.336 | 0.36 | 0.49 | 0.34 | 0.49 | 0.26 | 0.35 |
| 4 | 54653803 | rs2088582 | -0.896 | 0.854 | -1.033 | -0.33 | -0.135 | -0.14 | -0.14 | 0 | 0.58 | 0.96 | 0.53 | 0.55 | 0.49 | 0.48 |
| 4 | 54656460 | rs6811489 | -0.506 | 0.814 | -0.468 | -0.457 | 0.023 | -0.054 | -0.054 | 0 | 0.18 | 0.32 | 0.17 | 0.05 | 0.24 | 0.12 |
| 4 | 54657093 | rs6844333 | -0.563 | 0.443 | -0.835 | 0.11 | 0.038 | -0.021 | -0.021 | 0 | 0.44 | 0.73 | 0.42 | 0.58 | 0.32 | 0.44 |
| 4 | 54661158 | rs2412501 | 0.679 | 0.535 | 0.213 | -0.342 | 0.047 | -0.006 | -0.006 | 0 | 0.93 | 0.7 | 0.94 | 1 | 0.92 | 0.99 |
| 4 | 54666951 | rs1498821 | 1.077 | 0.669 | -0.938 | -0.885 | -0.068 | 0.153 | 0.153 | 0 | 0.58 | 0.05 | 0.7 | 0.85 | 0.61 | 0.73 |
| 4 | 54676030 | rs994965 | 0.904 | 0.719 | -0.861 | -1.005 | 0.028 | -0.029 | -0.029 | 0 | 0.76 | 0.66 | 0.81 | 0.89 | 0.7 | 0.84 |
| 4 | 54681059 | rs4864829 | 1.418 | 0.575 | -0.333 | -1.076 | -0.008 | -0.04 | -0.04 | 0 | 0.87 | 0.83 | 0.91 | 1 | 0.88 | 0.94 |
| 4 | 54698028 | rs13108970 | -0.551 | 0.735 | -0.194 | -1.055 | -0.02 | 0.002 | 0.002 | 0.932 | 0.14 | 0.08 | 0.12 | 0.11 | 0.2 | 0.11 |
| 4 | 54701833 | rs13125746 | -1.357 | 0.7 | 0.264 | -1.074 | -0.003 | 0.002 | 0.002 | 0 | 0.19 | 0.16 | 0.14 | 0.11 | 0.23 | 0.14 |
| 4 | 54706281 | rs17083558 | 1.781 | 0.618 | 1.159 | -1.122 | 0.012 | -0.034 | -0.034 | 0 | 0.69 | 0.38 | 0.73 | 0.87 | 0.63 | 0.81 |
| 4 | 54707699 | rs11133292 | 1.394 | 0.536 | 1.468 | -1.265 | 0.005 | -0.076 | -0.076 | 2.623 | 0.86 | 0.62 | 0.88 | 0.97 | 0.84 | 0.93 |
| 4 | 54713554 | rs6852162 | -1.906 | 0.692 | 1.517 | -1.334 | 0.049 | -0.078 | -0.078 | 0 | 0.25 | 0.55 | 0.2 | 0.1 | 0.26 | 0.19 |
| 4 | 54714868 | rs17083590 | -0.29 | 0.66 | 1.398 | -1.336 | -0.02 | 0.017 | 0.017 | 0 | 0.13 | 0.07 | 0.13 | 0.04 | 0.21 | 0.1 |
| 4 | 54718275 | rs17083601 | -0.253 | 0.47 | 1.465 | -1.28 | -0.017 | -0.003 | -0.003 | 0.086 | 0.05 | 0.08 | 0.01 | 0.03 | 0.01 | 0.07 |
| 4 | 54723742 | rs7662508 | -1.479 | 0.478 | 0.531 | -1.987 | 0.002 | -0.031 | -0.031 | 0 | 0.41 | 0.55 | 0.28 | 0.17 | 0.45 | 0.32 |
| 4 | 54724066 | rs13106140 | -0.358 | 0.606 | -0.576 | -1.942 | -0.171 | 0.088 | 0.088 | 0 | 0.27 | 0.02 | 0.2 | 0.11 | 0.38 | 0.24 |
| 4 | 54728147 | rs12500026 | 1.604 | 0.478 | -0.498 | -1.882 | -0.015 | 0.229 | 0.229 | 0.105 | 0.46 | 0.04 | 0.62 | 0.82 | 0.44 | 0.62 |
| 4 | 54734511 | rs1391758 | 0.727 | 0.35 | 0.413 | -1.729 | -0.018 | -0.067 | -0.067 | 0.058 | 0.87 | 0.61 | 0.93 | 0.97 | 0.92 | 0.9 |
| 4 | 54736988 | rs1498828 | -0.519 | 0.533 | 0.101 | -1.839 | -0.001 | 0 | 0 | 0 | 0.2 | 0.17 | 0.13 | 0.14 | 0.23 | 0.19 |
| 4 | 54738622 | rs6554139 | 1.602 | 0.093 | -0.186 | -2.2 | -0.015 | 0.106 | 0.106 | 0 | 0.56 | 0.27 | 0.71 | 0.8 | 0.61 | 0.66 |
| 4 | 54739506 | rs2898990 | -0.379 | 0.107 | -0.788 | -2.083 | -0.088 | 0.013 | 0.013 | 0.186 | 0.21 | 0.02 | 0.13 | 0.16 | 0.23 | 0.19 |
| 4 | 54740384 | rs4558921 | 1.4 | -0.21 | -1.583 | -2.687 | -0.082 | 0.128 | 0.128 | 0 | 0.26 | 0.02 | 0.16 | 0.05 | 0.31 | 0.19 |
| 4 | 54741949 | rs965126 | 0.763 | 0.143 | -0.752 | -2.092 | -0.011 | -0.067 | -0.067 | 0.785 | 0.72 | 0.35 | 0.81 | 0.94 | 0.8 | 0.85 |
| 4 | 54750921 | rs7659192 | 0.823 | 0.184 | -1.174 | -2.072 | 0.022 | 0.034 | 0.034 | 0.732 | 0.69 | 0.23 | 0.8 | 0.94 | 0.79 | 0.85 |
| 4 | 54756219 | rs1498827 | 1.217 | 0.287 | -1.511 | -1.897 | -0.034 | 0.188 | 0.188 | 0.042 | 0.58 | 0.1 | 0.73 | 0.81 | 0.65 | 0.69 |
| 4 | 54757388 | rs13133779 | -0.027 | 0.382 | -1.437 | -1.83 | -0.063 | 0 | 0 | 3.134 | 0.18 | 0.03 | 0.1 | 0.16 | 0.2 | 0.2 |
| 4 | 54757995 | rs13135139 | -0.098 | 0.385 | -1.481 | -1.907 | -0.075 | 0.025 | 0.025 | 2.804 | 0.18 | 0.02 | 0.1 | 0.13 | 0.2 | 0.19 |
| 4 | 54761404 | rs2102012 | -0.272 | 0.376 | -1.294 | -1.911 | -0.018 | 0.023 | 0.023 | 2.999 | 0.19 | 0.12 | 0.1 | 0.13 | 0.2 | 0.19 |
| 4 | 54762082 | rs1498820 | -0.08 | 0.324 | -1.052 | -1.957 | -0.023 | -0.131 | -0.131 | 0.005 | 0.8 | 0.58 | 0.88 | 0.97 | 0.86 | 0.9 |
| 4 | 54770332 | rs11133294 | 0.797 | 0.415 | -1.791 | -1.75 | -0.035 | 0.219 | 0.219 | 0.01 | 0.57 | 0.1 | 0.74 | 0.83 | 0.65 | 0.71 |
| 4 | 54775225 | rs10866415 | 0.832 | 0.389 | -1.808 | -1.764 | -0.035 | 0.232 | 0.232 | 0.049 | 0.56 | 0.1 | 0.73 | 0.83 | 0.64 | 0.7 |
| 4 | 54782044 | rs1109789 | 0.27 | 1.096 | -1.216 | -0.24 | 0.012 | 0.017 | 0.017 | 0 | 0.63 | 0.54 | 0.5 | 0.56 | 0.59 | 0.79 |
| 4 | 54784205 | rs1357133 | 0.15 | 1.029 | -1.15 | -0.272 | 0.013 | 0.017 | 0.017 | 0 | 0.65 | 0.59 | 0.5 | 0.56 | 0.59 | 0.79 |
| 4 | 54786223 | rs11723268 | 0.559 | 1.029 | -0.896 | -0.348 | 0.001 | 0.002 | 0.002 | 0 | 0.61 | 0.39 | 0.47 | 0.56 | 0.57 | 0.79 |
| 4 | 54791781 | rs11946709 | 0.736 | 1.29 | -1.251 | -0.195 | -0.042 | -0.047 | -0.047 | 0 | 0.13 | 0.01 | 0.33 | 0.16 | 0.14 | 0.06 |
| 4 | 54793060 | rs6811240 | 0.036 | 0.991 | -0.8 | -0.056 | 0.011 | -0.002 | -0.002 | 0.322 | 0.65 | 0.61 | 0.51 | 0.66 | 0.59 | 0.8 |
| 4 | 54796220 | rs11725926 | -0.192 | 0.033 | -1.095 | -0.071 | 0.001 | 0.003 | 0.003 | 1.184 | 0.23 | 0.19 | 0.21 | 0.2 | 0.19 | 0.35 |
| 4 | 54801549 | rs17083718 | -0.522 | 0.063 | -0.995 | 0.007 | -0.008 | -0.001 | -0.001 | 0 | 0.19 | 0.19 | 0.13 | 0.2 | 0.12 | 0.33 |
| 4 | 54801934 | rs11133301 | -1.029 | 0.058 | -0.955 | 0.109 | 0.002 | -0.002 | -0.002 | 0 | 0.6 | 0.59 | 0.43 | 0.62 | 0.52 | 0.78 |
| 4 | 54802080 | rs11737196 | -0.3 | 0.115 | -0.999 | -0.956 | -0.008 | -0.001 | -0.001 | 0 | 0.17 | 0.16 | 0.12 | 0.19 | 0.1 | 0.33 |
| 4 | 54804163 | rs17083734 | -0.018 | 0.13 | -0.88 | -0.843 | -0.028 | 0.009 | 0.009 | 0 | 0.13 | 0.2 | 0.11 | 0.19 | 0.06 | 0.27 |
| 4 | 54814760 | rs13133473 | -0.985 | 0.133 | -1.067 | -0.586 | -0.003 | 0.002 | 0.002 | 0 | 0.58 | 0.62 | 0.38 | 0.29 | 0.49 | 0.63 |
| 4 | 54819631 | rs7686432 | -0.919 | 0.134 | -1.071 | -0.652 | -0.003 | 0.003 | 0.003 | 0 | 0.58 | 0.61 | 0.38 | 0.28 | 0.48 | 0.62 |
| 4 | 54825255 | rs11133302 | -0.109 | 0.372 | -1.367 | -0.721 | -0.023 | 0.127 | 0.127 | 0 | 0.42 | 0.15 | 0.27 | 0.1 | 0.45 | 0.34 |
| 4 | 54829996 | rs13152774 | 0 | 0.293 | -1.488 | -0.652 | -0.014 | 0.135 | 0.135 | 0 | 0.39 | 0.1 | 0.2 | 0.11 | 0.38 | 0.33 |
| 4 | 54847381 | rs6828100 | 0.369 | -0.061 | -1.483 | -0.655 | -0.029 | -0.028 | -0.028 | 0.727 | 0.11 | 0.01 | 0.07 | 0.06 | 0.13 | 0.07 |
| 4 | 54850662 | rs13105833 | -0.467 | -0.089 | -1.651 | -0.846 | -0.012 | 0.133 | 0.133 | 0 | 0.38 | 0.07 | 0.19 | 0.1 | 0.36 | 0.33 |
| 4 | 54854377 | rs13134590 | 0.473 | -0.078 | -1.673 | -0.848 | -0.029 | -0.028 | -0.028 | 0 | 0.11 | 0.01 | 0.07 | 0.06 | 0.13 | 0.07 |
| 4 | 54866124 | rs6834659 | -0.996 | -0.362 | -0.901 | -0.985 | 0.018 | 0.039 | 0.039 | 0 | 0.44 | 0.65 | 0.25 | 0.1 | 0.37 | 0.36 |
| 4 | 54901801 | rs13145700 | -0.453 | -0.253 | -1.103 | -0.931 | -0.001 | 0.131 | 0.131 | 0 | 0.38 | 0.1 | 0.2 | 0.1 | 0.36 | 0.33 |
| 4 | 54914865 | rs2049170 | -0.699 | -0.209 | -0.502 | -0.897 | 0 | 0.003 | 0.003 | 0 | 0.85 | 0.84 | 0.84 | 0.8 | 0.88 | 0.76 |
| 4 | 54917484 | rs13120133 | -0.902 | -0.308 | -0.707 | -0.831 | 0.001 | -0.006 | -0.006 | 0 | 0.42 | 0.43 | 0.23 | 0.1 | 0.37 | 0.33 |
| 4 | 54917937 | rs13121173 | 0.213 | 0.01 | -1.123 | -0.581 | -0.034 | -0.028 | -0.028 | 0 | 0.11 | 0.01 | 0.07 | 0.06 | 0.13 | 0.07 |
| 4 | 54938543 | rs2412522 | -0.633 | -0.052 | -1.091 | -0.741 | 0.014 | 0.118 | 0.118 | 0 | 0.38 | 0.28 | 0.18 | 0.08 | 0.33 | 0.34 |
| 4 | 54943432 | rs13117345 | 0.154 | 0.362 | -1.193 | -0.57 | -0.038 | -0.004 | -0.004 | 0 | 0.12 | 0.03 | 0.09 | 0.06 | 0.14 | 0.07 |
| 4 | 54954323 | rs714860 | -0.206 | 0.051 | -0.891 | -0.696 | -0.025 | -0.064 | -0.064 | 2.979 | 0.29 | 0.44 | 0.16 | 0.14 | 0.23 | 0.3 |
| 4 | 54956686 | rs2412535 | -0.933 | 0.106 | -0.888 | -0.631 | -0.007 | -0.036 | -0.036 | 0 | 0.42 | 0.52 | 0.25 | 0.2 | 0.37 | 0.38 |
| 4 | 54957891 | rs6812385 | -0.177 | 0.244 | -1.416 | -0.171 | 0.011 | 0.013 | 0.013 | 0 | 0.24 | 0.16 | 0.15 | 0.18 | 0.22 | 0.28 |
| 4 | 54958646 | rs6554147 | 0.019 | 0.798 | -1.635 | -0.141 | -0.062 | 0.061 | 0.061 | 0.154 | 0.6 | 0.21 | 0.46 | 0.73 | 0.55 | 0.72 |
| 4 | 54967214 | rs6554148 | -0.363 | 0.693 | -0.73 | -0.296 | -0.007 | -0.035 | -0.035 | 0 | 0.77 | 0.69 | 0.85 | 0.91 | 0.76 | 0.72 |
| 4 | 54967709 | rs2278142 | 0.817 | 0.793 | -0.578 | -0.61 | 0.001 | 0.02 | 0.02 | 0 | 0.6 | 0.29 | 0.72 | 0.84 | 0.61 | 0.67 |
| 4 | 54970684 | rs11133304 | 0.35 | 0.713 | -0.638 | -0.675 | -0.059 | -0.004 | -0.004 | 0 | 0.11 | 0.01 | 0.09 | 0.07 | 0.15 | 0.05 |
| 4 | 54973200 | rs6554150 | 0.782 | 0.687 | -0.566 | -0.635 | 0.002 | 0.021 | 0.021 | 0 | 0.6 | 0.28 | 0.72 | 0.85 | 0.61 | 0.68 |

| | | | | | | | | | | | | | | | | |
|---|---|---|---|---|---|---|---|---|---|---|---|---|---|---|---|---|
| 4 | 54986885 | rs4864841 | 0.165 | 0.768 | -0.841 | -0.225 | 0.011 | 0.043 | 0.043 | 0 | 0.28 | 0.37 | 0.43 | 0.34 | 0.3 | 0.31 |
| 4 | 54986912 | rs6850083 | 0.163 | 0.508 | -0.016 | -0.645 | -0.032 | -0.085 | -0.085 | 0 | 0.26 | 0.42 | 0.17 | 0.07 | 0.21 | 0.28 |
| 4 | 55011123 | rs1017280 | 0.734 | 0.56 | -0.283 | -0.762 | 0.008 | 0.09 | 0.09 | 0.816 | 0.22 | 0.05 | 0.15 | 0.05 | 0.22 | 0.24 |
| 4 | 55016097 | rs6554152 | 0.725 | 0.614 | -0.204 | -0.672 | 0.008 | 0.014 | 0.014 | 0 | 0.61 | 0.28 | 0.73 | 0.84 | 0.65 | 0.67 |
| 4 | 55030375 | rs6840522 | 0.163 | 1.084 | -0.153 | -0.394 | 0.001 | -0.024 | -0.024 | 0.386 | 0.85 | 0.66 | 0.89 | 0.92 | 0.87 | 0.94 |
| 4 | 55045716 | rs10027676 | 0.563 | 0.694 | -0.901 | -0.496 | 0.001 | 0.099 | 0.099 | 0.186 | 0.25 | 0.03 | 0.16 | 0.06 | 0.22 | 0.24 |
| 4 | 55049751 | rs7659596 | -0.936 | 0.918 | -0.398 | -0.187 | -0.001 | -0.004 | -0.004 | 0.898 | 0.59 | 0.52 | 0.63 | 0.37 | 0.6 | 0.61 |
| 4 | 55061398 | rs2162136 | -0.419 | 1.304 | -0.368 | 0.021 | -0.005 | -0.017 | -0.017 | 0.044 | 0.19 | 0.2 | 0.37 | 0.24 | 0.22 | 0.27 |
| 4 | 55073270 | rs12233727 | -0.663 | 1.24 | -0.127 | -0.199 | 0 | 0 | 0 | 0 | 0.48 | 0.47 | 0.57 | 0.31 | 0.51 | 0.52 |
| 4 | 55073673 | rs1559715 | 0.85 | 1.206 | -0.232 | -0.252 | -0.053 | 0.113 | 0.113 | 0.299 | 0.28 | 0.03 | 0.17 | 0.06 | 0.28 | 0.26 |
| 4 | 55074368 | rs17084040 | 0.486 | 1.304 | -0.164 | -0.315 | -0.002 | 0.005 | 0.005 | 0.059 | 0.11 | 0.09 | 0.06 | 0.08 | 0.1 | 0.05 |
| 4 | 55076237 | rs6832597 | -0.237 | 1.526 | 0.076 | -0.018 | 0.003 | 0.018 | 0.018 | 0 | 0.19 | 0.26 | 0.39 | 0.25 | 0.23 | 0.26 |
| 4 | 55081757 | rs11133313 | -0.805 | 1.473 | 0.143 | -0.134 | -0.002 | 0.014 | 0.014 | 0 | 0.48 | 0.44 | 0.56 | 0.3 | 0.49 | 0.52 |
| 4 | 55087581 | rs17084051 | -0.748 | 1.321 | 0.823 | 0.208 | 0.009 | 0.03 | 0.03 | 0 | 0.19 | 0.36 | 0.39 | 0.22 | 0.21 | 0.26 |
| 4 | 55092626 | rs2114039 | -1.638 | 1.256 | 0.974 | 0.161 | -0.001 | -0.004 | -0.004 | 0 | 0.28 | 0.73 | 0.47 | 0.29 | 0.28 | 0.27 |
| 4 | 55093955 | rs6554162 | -1.563 | 1.307 | 0.672 | 0.186 | 0.002 | 0.013 | 0.013 | 1.465 | 0.27 | 0.67 | 0.46 | 0.29 | 0.28 | 0.27 |
| 4 | 55102425 | rs7678144 | -0.903 | 0.98 | 0.143 | 0.142 | 0.014 | 0.043 | 0.043 | 0 | 0.19 | 0.38 | 0.39 | 0.22 | 0.21 | 0.25 |
| 4 | 55124460 | rs7677751 | -0.777 | 1.196 | 0.232 | 0.348 | 0.027 | 0.079 | 0.079 | 0.605 | 0.14 | 0.32 | 0.28 | 0.22 | 0.14 | 0.25 |
| 4 | 55130154 | rs2307049 | -0.583 | 1.024 | -0.068 | 0.317 | 0.012 | 0.075 | 0.075 | 0.125 | 0.11 | 0.28 | 0.35 | 0.2 | 0.12 | 0.25 |
| 4 | 55133726 | rs4358459 | -0.388 | 1.112 | -0.026 | 0.36 | 0.021 | 0.088 | 0.088 | 0.1 | 0.11 | 0.27 | 0.35 | 0.2 | 0.12 | 0.25 |
| 4 | 55141843 | rs7656613 | 0.523 | 1.026 | -1.115 | 0.241 | 0.01 | -0.181 | -0.181 | 0 | 0.35 | 0.02 | 0.13 | 0.48 | 0.26 | 0.41 |
| 4 | 55146389 | rs1547904 | -0.11 | 1.428 | 0.084 | 0.242 | 0.072 | 0.063 | 0.063 | 0.253 | 0.12 | 0.34 | 0.28 | 0.18 | 0.15 | 0.24 |
| 4 | 55152040 | rs2228230 | -0.11 | 1.394 | 0.133 | 0.208 | 0.069 | 0.059 | 0.059 | 0.253 | 0.12 | 0.34 | 0.28 | 0.17 | 0.15 | 0.24 |
| 4 | 55161254 | rs3733540 | 0.124 | 1.43 | -0.74 | 0.239 | 0.046 | 0.037 | 0.037 | 1.504 | 0.86 | 0.44 | 0.71 | 0.83 | 0.85 | 0.75 |
| 4 | 55164866 | rs17739921 | -0.355 | 1.101 | -1.417 | 0.086 | -0.024 | 0.038 | 0.038 | 0 | 0.41 | 0.25 | 0.41 | 0.27 | 0.45 | 0.31 |
| 4 | 55193347 | rs4583821 | -0.395 | 0.551 | -1.293 | -0.105 | 0.019 | 0.025 | 0.025 | 0 | 0.51 | 0.38 | 0.46 | 0.38 | 0.44 | 0.54 |
| 4 | 55196620 | rs17084148 | -0.359 | 0.393 | -1.286 | -0.126 | -0.024 | -0.035 | -0.035 | 0.589 | 0.38 | 0.1 | 0.4 | 0.38 | 0.4 | 0.46 |
| 4 | 55198205 | rs11729535 | -0.45 | 0.531 | -1.209 | -0.017 | -0.042 | -0.041 | -0.041 | 0.054 | 0.34 | 0.12 | 0.42 | 0.38 | 0.46 | 0.45 |
| 4 | 55202124 | rs11736538 | -0.441 | 0.446 | -1.114 | 0.104 | -0.03 | -0.077 | -0.077 | 0 | 0.29 | 0.1 | 0.42 | 0.41 | 0.41 | 0.43 |
| 4 | 55202157 | rs17084178 | -0.614 | 0.483 | -0.794 | 0.121 | 0.002 | 0.041 | 0.041 | 0 | 0.33 | 0.51 | 0.46 | 0.42 | 0.41 | 0.43 |
| 4 | 55221467 | rs1492767 | -0.475 | 0.283 | -1.791 | -0.25 | -0.083 | 0.025 | 0.025 | 0 | 0.41 | 0.04 | 0.45 | 0.25 | 0.49 | 0.32 |
| 4 | 55225121 | rs2412563 | 0.558 | 0.021 | -0.688 | -0.356 | 0.011 | 0.03 | 0.03 | 0 | 0.53 | 0.45 | 0.5 | 0.75 | 0.51 | 0.68 |
| 4 | 55233465 | rs1444043 | -0.088 | 0.08 | -1.155 | -0.418 | -0.029 | 0.02 | 0.02 | 0 | 0.19 | 0.01 | 0.13 | 0.1 | 0.2 | 0.21 |
| 4 | 55239947 | rs2412564 | 0.86 | 0.023 | -0.776 | -0.555 | 0.022 | 0.074 | 0.074 | 0 | 0.49 | 0.31 | 0.51 | 0.73 | 0.53 | 0.64 |
| 4 | 55241063 | rs17747127 | 0.248 | 0.005 | -0.45 | -0.685 | 0.015 | 0.024 | 0.024 | 0.01 | 0.17 | 0.08 | 0.08 | 0.01 | 0.17 | 0.12 |
| 4 | 55242625 | rs1004564 | 1.117 | -0.346 | -0.45 | -0.783 | -0.002 | 0.149 | 0.149 | 0 | 0.48 | 0.1 | 0.49 | 0.76 | 0.49 | 0.65 |
| 4 | 55242960 | rs2166021 | -0.603 | -0.455 | 0.401 | -0.818 | 0.005 | 0.04 | 0.04 | 0.124 | 0.08 | 0.32 | 0.09 | 0.08 | 0.04 | 0.09 |
| 4 | 55245741 | rs4572929 | 0.993 | -0.51 | 0.177 | -0.846 | 0.051 | 0.074 | 0.074 | 0.588 | 0.72 | 0.17 | 0.8 | 0.89 | 0.78 | 0.79 |
| 4 | 55245897 | rs1353005 | 0.669 | -0.453 | 0.186 | -0.825 | -0.05 | 0.025 | 0.025 | 0 | 0.23 | 0.07 | 0.3 | 0.13 | 0.29 | 0.14 |
| 4 | 55251785 | rs1562058 | 1.18 | -0.514 | 0.397 | -1.059 | -0.007 | 0.098 | 0.098 | 0 | 0.46 | 0.29 | 0.49 | 0.79 | 0.45 | 0.64 |
| 4 | 55251828 | rs17691182 | 0.668 | -0.523 | 0.105 | -1.089 | -0.097 | 0.061 | 0.061 | 0 | 0.26 | 0.04 | 0.34 | 0.1 | 0.35 | 0.17 |
| 4 | 55256927 | rs11133321 | 1.457 | -0.553 | 0.137 | -1.05 | -0.07 | 0.124 | 0.124 | 0 | 0.45 | 0.08 | 0.47 | 0.77 | 0.46 | 0.65 |
| 4 | 55257611 | rs1492769 | -0.029 | -0.589 | 0.408 | -1.01 | 0.007 | -0.019 | -0.019 | 0 | 0.77 | 0.25 | 0.88 | 0.97 | 0.86 | 0.91 |
| 4 | 55259352 | rs1156181 | 0.814 | -0.49 | 0.428 | -0.954 | 0.021 | 0.083 | 0.083 | 0 | 0.5 | 0.4 | 0.51 | 0.77 | 0.49 | 0.65 |
| 4 | 55270873 | rs17691810 | 0.715 | -0.373 | -0.055 | -0.881 | -0.06 | 0.05 | 0.05 | 0 | 0.24 | 0.07 | 0.31 | 0.11 | 0.31 | 0.17 |
| 4 | 55286861 | rs11721618 | 0.053 | -0.481 | 0.546 | -0.838 | 0 | -0.02 | -0.02 | 0 | 0.67 | 0.42 | 0.56 | 0.78 | 0.63 | 0.76 |
| 4 | 55289826 | rs1396066 | 1.554 | -0.401 | -0.007 | -0.309 | -0.064 | -0.028 | -0.028 | 0.035 | 0.35 | 0.09 | 0.18 | 0.36 | 0.34 | 0.39 |
| 4 | 55295481 | rs13129356 | 1.14 | -0.226 | 0.042 | -0.247 | -0.013 | -0.043 | -0.043 | 1.454 | 0.13 | 0.06 | 0.21 | 0.4 | 0.13 | 0.28 |
| 4 | 55301708 | rs10517334 | -1.09 | -0.228 | 0.097 | -0.39 | -0.008 | 0.009 | 0.009 | 0 | 0.38 | 0.47 | 0.48 | 0.53 | 0.34 | 0.45 |
| 4 | 55311965 | rs17749557 | 1.128 | -0.282 | -0.28 | -0.54 | -0.023 | -0.014 | -0.014 | 0 | 0.32 | 0.2 | 0.18 | 0.37 | 0.33 | 0.38 |
| 4 | 55314638 | rs1507966 | 0.298 | 0.024 | -0.188 | -0.49 | 0.006 | -0.006 | -0.006 | 0 | 0.18 | 0.12 | 0.17 | 0.02 | 0.17 | 0.11 |
| 4 | 55342137 | rs6857523 | -0.49 | 0.064 | -0.044 | -0.445 | -0.004 | -0.043 | -0.043 | 0 | 0.81 | 0.81 | 0.9 | 0.99 | 0.84 | 0.87 |
| 4 | 55345209 | rs17693577 | 0.367 | 0.107 | 0.031 | -0.186 | -0.026 | -0.03 | -0.03 | 1.088 | 0.36 | 0.28 | 0.38 | 0.46 | 0.41 | 0.45 |
| 4 | 55351107 | rs12509670 | -0.487 | -0.158 | 0.348 | 0.039 | -0.006 | 0.017 | 0.017 | 0 | 0.24 | 0.37 | 0.22 | 0.29 | 0.2 | 0.31 |
| 4 | 55354275 | rs1507955 | -0.651 | -0.189 | 0.442 | 0.024 | -0.01 | 0.009 | 0.009 | 0 | 0.25 | 0.53 | 0.23 | 0.29 | 0.2 | 0.31 |
| 4 | 55357990 | rs7676607 | 0.944 | -0.091 | 0.076 | -0.025 | -0.061 | -0.004 | -0.004 | 0 | 0.38 | 0.02 | 0.38 | 0.26 | 0.39 | 0.26 |
| 4 | 55358779 | rs13151127 | -0.368 | -0.074 | 0.134 | -0.013 | -0.004 | 0.019 | 0.019 | 0 | 0.23 | 0.36 | 0.22 | 0.29 | 0.2 | 0.31 |
| 4 | 55361102 | rs1507952 | -0.554 | -0.12 | 0.458 | 0.15 | -0.001 | -0.001 | -0.001 | 0 | 0.64 | 0.57 | 0.61 | 0.55 | 0.59 | 0.58 |
| 4 | 55368702 | rs6838519 | -0.461 | -0.04 | 0.52 | 0.118 | 0.013 | 0.011 | 0.011 | 0 | 0.64 | 0.48 | 0.61 | 0.57 | 0.61 | 0.54 |
| 4 | 55371241 | rs6554190 | -0.047 | -0.086 | 0.674 | 0.014 | -0.005 | 0.002 | 0.002 | 0 | 0.41 | 0.53 | 0.34 | 0.38 | 0.43 | 0.43 |
| 4 | 55374968 | rs6554191 | -0.388 | -0.061 | 0.52 | 0.053 | -0.028 | 0.034 | 0.034 | 0 | 0.2 | 0.29 | 0.18 | 0.28 | 0.16 | 0.28 |
| 4 | 55380471 | rs11133333 | 0.972 | -0.033 | 0.376 | -0.047 | -0.06 | 0.05 | 0.05 | 0 | 0.42 | 0.07 | 0.42 | 0.28 | 0.45 | 0.26 |
| 4 | 55385637 | rs17084420 | 0.516 | -0.179 | 0.795 | 0.035 | -0.014 | -0.079 | -0.079 | 0.633 | 0.13 | 0.26 | 0.06 | 0 | 0.1 | 0.07 |
| 4 | 55390494 | rs170118 | -0.96 | 0.025 | 0.852 | 0.135 | 0.028 | 0.021 | 0.021 | 0 | 0.68 | 0.48 | 0.7 | 0.63 | 0.64 | 0.63 |
| 4 | 55392598 | rs10050226 | -0.071 | -0.047 | 0.438 | 0.125 | -0.03 | -0.115 | -0.115 | 0 | 0.15 | 0.47 | 0.09 | 0.01 | 0.1 | 0.07 |
| 4 | 55393419 | rs4864905 | 0.113 | -0.098 | 0.082 | 0.173 | -0.002 | 0.004 | 0.004 | 0.061 | 0.57 | 0.61 | 0.5 | 0.28 | 0.51 | 0.34 |
| 4 | 55394172 | rs218237 | -0.719 | -0.44 | 0.287 | 0.484 | -0.015 | 0.065 | 0.065 | 0.124 | 0.14 | 0.26 | 0.23 | 0.35 | 0.11 | 0.29 |
| 4 | 55402962 | rs6824783 | 0.319 | -0.097 | -0.173 | 0.255 | 0.01 | 0.001 | 0.001 | 1.05 | 0.11 | 0.03 | 0.04 | 0.01 | 0.08 | 0.07 |
| 4 | 55411087 | rs218268 | -0.548 | -0.106 | -0.273 | 0.456 | 0.003 | 0.004 | 0.004 | 3.201 | 0.33 | 0.71 | 0.26 | 0.18 | 0.2 | 0.29 |

| | | | | | | | | | | | | | | | | |
|---|---|---|---|---|---|---|---|---|---|---|---|---|---|---|---|---|
| 4 | 55415172 | rs218273 | -0.741 | 0.717 | -0.857 | 0.621 | -0.012 | 0.013 | 0.013 | 0 | 0.56 | 0.76 | 0.59 | 0.59 | 0.57 | 0.65 |
| 4 | 55418294 | rs2110152 | 0.663 | 0.748 | -1.468 | 0.493 | -0.092 | -0.055 | -0.055 | 0 | 0.25 | 0.03 | 0.31 | 0.29 | 0.36 | 0.36 |
| 4 | 55424289 | rs218280 | -0.429 | 0.648 | -1.027 | 0.22 | -0.022 | -0.021 | -0.021 | 0 | 0.33 | 0.62 | 0.27 | 0.19 | 0.24 | 0.29 |
| 4 | 55426561 | rs2537872 | -0.719 | 1.221 | -1.673 | 0.143 | -0.007 | 0.001 | 0.001 | 0 | 0.65 | 0.86 | 0.71 | 0.66 | 0.68 | 0.71 |
| 4 | 55426618 | rs2703489 | -0.32 | 1.032 | -1.534 | 0.259 | -0.019 | 0.078 | 0.078 | 0 | 0.54 | 0.81 | 0.65 | 0.68 | 0.54 | 0.66 |
| 4 | 55430051 | rs7657643 | -0.516 | 1.222 | -0.572 | -0.339 | 0.002 | -0.041 | -0.041 | 0 | 0.44 | 0.54 | 0.32 | 0.14 | 0.53 | 0.4 |
| 4 | 55434828 | rs2703487 | -1.224 | 1.198 | 0.044 | -0.154 | -0.003 | -0.004 | -0.004 | 0.386 | 0.26 | 0.27 | 0.17 | 0.16 | 0.23 | 0.39 |
| 4 | 55439893 | rs2703481 | -1.43 | 1.661 | 1.156 | -0.173 | 0.024 | -0.016 | -0.016 | 0.007 | 0.16 | 0.56 | 0.12 | 0.13 | 0.16 | 0.15 |
| 4 | 55448581 | rs17826456 | -0.883 | 1.769 | 1.178 | -0.275 | -0.041 | 0.035 | 0.035 | 0 | 0.21 | 0.03 | 0.12 | 0.11 | 0.22 | 0.39 |
| 4 | 55449616 | rs4864910 | -1.93 | 1.838 | 1.743 | -0.347 | 0.009 | -0.043 | -0.043 | 0.376 | 0.32 | 0.39 | 0.19 | 0.14 | 0.34 | 0.53 |
| 4 | 55457545 | rs10488850 | -0.664 | 1.869 | 1.678 | -0.337 | 0.022 | -0.027 | -0.027 | 0 | 0.08 | 0.13 | 0.05 | 0.03 | 0.12 | 0.13 |
| 4 | 55461034 | rs11733102 | -1.98 | 1.82 | 1.608 | -0.292 | 0.003 | -0.021 | -0.021 | 0 | 0.33 | 0.34 | 0.19 | 0.15 | 0.34 | 0.53 |
| 4 | 55471848 | rs2703476 | -1.284 | 1.528 | 1.648 | -0.249 | -0.011 | -0.033 | -0.033 | 0.252 | 0.19 | 0.31 | 0.1 | 0.09 | 0.14 | 0.28 |
| 4 | 55475409 | rs2703480 | -1.57 | 1.475 | 1.733 | -0.274 | -0.02 | -0.051 | -0.051 | 0 | 0.2 | 0.36 | 0.1 | 0.1 | 0.14 | 0.28 |
| 4 | 55477883 | rs17827110 | -2.233 | 2.459 | 1.838 | 0.042 | 0.09 | -0.027 | -0.027 | 0 | 0.22 | 0.42 | 0.28 | 0.13 | 0.35 | 0.31 |
| 4 | 55478047 | rs2537883 | -3.037 | 2.413 | 1.858 | 0.007 | -0.002 | 0.001 | 0.001 | 0 | 0.3 | 0.71 | 0.3 | 0.22 | 0.38 | 0.49 |
| 4 | 55478686 | rs17827152 | -1.303 | 2.902 | 1.641 | 0.024 | -0.065 | -0.004 | -0.004 | 0.141 | 0.12 | 0.02 | 0.2 | 0.07 | 0.24 | 0.2 |
| 4 | 55480024 | rs17230634 | -2.268 | 3.209 | 1.671 | 0.141 | 0.118 | -0.015 | -0.015 | 0 | 0.24 | 0.45 | 0.31 | 0.17 | 0.4 | 0.33 |
| 4 | 55484356 | rs2703457 | -2.238 | 3.078 | 1.39 | 0.221 | 0.043 | 0.009 | 0.009 | 0 | 0.1 | 0.27 | 0.25 | 0.11 | 0.14 | 0.19 |
| 4 | 55486791 | rs12512899 | -1.441 | 2.949 | 1.066 | -0.162 | -0.03 | 0.023 | 0.023 | 0.369 | 0.23 | 0.19 | 0.45 | 0.11 | 0.39 | 0.24 |
| 4 | 55490810 | rs17084534 | -1.558 | 2.951 | 1.383 | 0.523 | 0.019 | 0.078 | 0.078 | 0.198 | 0.11 | 0.18 | 0.11 | 0.32 | 0.14 | 0.32 |
| 4 | 55491904 | rs759082 | -1.219 | 2.928 | 1.402 | 0.516 | -0.08 | 0.054 | 0.054 | 0.291 | 0.23 | 0.11 | 0.45 | 0.11 | 0.39 | 0.24 |
| 4 | 55492212 | rs7655814 | -1.578 | 2.739 | 1.412 | 0.562 | 0.019 | 0.078 | 0.078 | 0.198 | 0.11 | 0.19 | 0.11 | 0.32 | 0.14 | 0.32 |
| 4 | 55499810 | rs6554194 | -2.828 | 2.657 | 1.629 | 0.519 | 0.007 | -0.001 | -0.001 | 0 | 0.4 | 0.45 | 0.56 | 0.43 | 0.53 | 0.56 |
| 4 | 55500226 | rs6831020 | -0.45 | 2.914 | 1.373 | 0.45 | -0.093 | 0.018 | 0.018 | 0 | 0.16 | 0.02 | 0.29 | 0.08 | 0.29 | 0.16 |
| 4 | 55512066 | rs7695300 | -3.026 | 2.228 | 1.857 | 0.403 | 0.001 | -0.003 | -0.003 | 0 | 0.41 | 0.54 | 0.56 | 0.42 | 0.53 | 0.53 |
| 4 | 55515359 | rs6825657 | -0.669 | 2.277 | 1.805 | 0.37 | -0.1 | 0.042 | 0.042 | 0.431 | 0.2 | 0.08 | 0.42 | 0.11 | 0.37 | 0.24 |
| 4 | 55517005 | rs4241987 | -1.568 | 2.268 | 1.993 | 0.418 | 0.036 | -0.017 | -0.017 | 0.179 | 0.25 | 0.34 | 0.48 | 0.16 | 0.44 | 0.34 |
| 4 | 55519748 | rs13147869 | -1.623 | 2.265 | 1.983 | 0.579 | 0.042 | -0.009 | -0.009 | 0 | 0.25 | 0.35 | 0.48 | 0.19 | 0.44 | 0.33 |
| 4 | 55521017 | rs2159935 | -2.516 | 2.234 | 2.151 | 0.609 | 0.02 | -0.002 | -0.002 | 0.041 | 0.37 | 0.42 | 0.52 | 0.4 | 0.47 | 0.43 |
| 4 | 55522160 | rs6554198 | -3.547 | 2.17 | 1.985 | 0.521 | 0.013 | 0.009 | 0.009 | 0.033 | 0.46 | 0.76 | 0.6 | 0.46 | 0.57 | 0.55 |
| 4 | 55526251 | rs2237035 | -0.725 | 2.22 | 1.936 | 0.659 | -0.105 | 0.009 | 0.009 | 0.102 | 0.19 | 0.07 | 0.42 | 0.14 | 0.37 | 0.23 |
| 4 | 55528653 | rs2237034 | -1.762 | 2.136 | 2.023 | 0.67 | -0.013 | 0.01 | 0.01 | 0 | 0.16 | 0.18 | 0.09 | 0.26 | 0.1 | 0.2 |
| 4 | 55531875 | rs2237032 | -2.369 | 2.143 | 2.032 | 0.663 | -0.021 | 0.041 | 0.041 | 0 | 0.21 | 0.38 | 0.14 | 0.31 | 0.17 | 0.3 |
| 4 | 55536375 | rs2237028 | -2.01 | 2.246 | 1.636 | 0.33 | 0.048 | -0.039 | -0.039 | 0 | 0.31 | 0.48 | 0.49 | 0.23 | 0.42 | 0.27 |
| 4 | 55541408 | rs6820303 | -2.048 | 2.055 | 1.553 | 0.402 | 0.036 | -0.037 | -0.037 | 0 | 0.33 | 0.49 | 0.49 | 0.22 | 0.42 | 0.28 |
| 4 | 55542869 | rs10488845 | -1.032 | 1.893 | 1.024 | 0.453 | -0.005 | 0.016 | 0.016 | 0.064 | 0.09 | 0.24 | 0.05 | 0.1 | 0.05 | 0.05 |
| 4 | 55552330 | rs13135792 | -0.869 | 1.99 | 0.419 | 0.383 | -0.126 | 0.053 | 0.053 | 0 | 0.24 | 0.03 | 0.41 | 0.12 | 0.34 | 0.24 |
| 4 | 55553801 | rs11946333 | -0.76 | 1.991 | 0.728 | 0.476 | 0.035 | 0.054 | 0.054 | 0 | 0.09 | 0.34 | 0.06 | 0.13 | 0.06 | 0.12 |
| 4 | 55554155 | rs2855777 | -1.698 | 1.785 | 0.51 | 0.262 | -0.003 | 0.004 | 0.004 | 0.002 | 0.42 | 0.6 | 0.54 | 0.24 | 0.49 | 0.41 |
| 4 | 55557984 | rs2237023 | -2.196 | 1.687 | 0.418 | 0.639 | -0.055 | 0.013 | 0.013 | 0.115 | 0.51 | 0.94 | 0.61 | 0.4 | 0.54 | 0.53 |
| 4 | 55565445 | rs17084644 | -0.245 | 1.664 | 0.441 | 0.495 | -0.015 | -0.025 | -0.025 | 0 | 0.08 | 0.02 | 0.07 | 0.09 | 0.1 | 0.05 |
| 4 | 55569146 | rs3020825 | 1.514 | 1.435 | 0.545 | 0.992 | -0.044 | -0.094 | -0.094 | 0 | 0.49 | 0.06 | 0.28 | 0.51 | 0.45 | 0.46 |
| 4 | 55574182 | rs3020826 | 0.924 | 1.411 | 0.675 | 0.887 | 0.003 | -0.009 | -0.009 | 0.058 | 0.58 | 0.43 | 0.45 | 0.74 | 0.5 | 0.58 |
| 4 | 55575144 | rs2237018 | -1.432 | 1.293 | 1.709 | 1.288 | 0.023 | 0.191 | 0.191 | 0 | 0.1 | 0.34 | 0.18 | 0.25 | 0.06 | 0.12 |
| 4 | 55588536 | rs2298976 | -1.224 | 1.671 | 1.509 | 0.498 | 0.023 | -0.014 | -0.014 | 0 | 0.08 | 0.17 | 0.06 | 0.05 | 0.08 | 0.12 |
| 4 | 55588719 | rs11735550 | -0.221 | 1.659 | 1.27 | 0.216 | -0.009 | 0.054 | 0.054 | 0 | 0.12 | 0.03 | 0.22 | 0.01 | 0.13 | 0.08 |
| 4 | 55590621 | rs4864920 | -2.598 | 1.708 | 1.995 | 0.345 | 0.021 | 0.001 | 0.001 | 2.9 | 0.18 | 0.62 | 0.17 | 0.15 | 0.19 | 0.35 |
| 4 | 55593464 | rs3822214 | -0.429 | 1.675 | 2.029 | 0.337 | -0.002 | 0.001 | 0.001 | 0.017 | 0.07 | 0.05 | 0.06 | 0.05 | 0.08 | 0.09 |
| 4 | 55599998 | rs2237012 | -1.53 | 1.71 | 2.395 | 0.355 | 0.065 | 0.007 | 0.007 | 0.433 | 0.1 | 0.47 | 0.08 | 0.06 | 0.08 | 0.12 |
| 4 | 55601327 | rs2213180 | -1.513 | 2.015 | 2.17 | 0.272 | 0.009 | 0.002 | 0.002 | 0 | 0.09 | 0.1 | 0.11 | 0.09 | 0.18 | 0.23 |
| 4 | 55604940 | rs17084733 | -0.729 | 1.09 | 2.218 | 0.345 | 0.018 | -0.014 | -0.014 | 0 | 0.08 | 0.15 | 0.06 | 0.05 | 0.08 | 0.11 |
| 4 | 55606427 | rs8022 | -1.158 | 1.164 | 2.107 | 0.333 | 0.074 | -0.005 | -0.005 | 0.559 | 0.09 | 0.32 | 0.08 | 0.05 | 0.1 | 0.12 |
| 4 | 55610192 | rs17084754 | -0.851 | 1.086 | 1.806 | 0.343 | 0.029 | 0.04 | 0.04 | 0.05 | 0.08 | 0.19 | 0.08 | 0.1 | 0.11 | 0.21 |
| 4 | 55610660 | rs4406072 | -1.705 | 0.98 | 1.484 | 0.459 | 0.006 | -0.021 | -0.021 | 0 | 0.24 | 0.39 | 0.32 | 0.21 | 0.23 | 0.24 |
| 4 | 55617208 | rs4864522 | -1.801 | 1.153 | 0.491 | 0.472 | 0.026 | 0.006 | 0.006 | 0 | 0.28 | 0.44 | 0.36 | 0.31 | 0.34 | 0.45 |
| 4 | 55624059 | rs10213491 | 0.92 | 0.526 | -0.594 | 0.465 | 0.003 | -0.02 | -0.02 | 0 | 0.67 | 0.24 | 0.78 | 0.61 | 0.72 | 0.59 |
| 4 | 55639547 | rs3109915 | -1.096 | 0.111 | -0.689 | 0.038 | 0.033 | 0.058 | 0.058 | 0 | 0.23 | 0.11 | 0.09 | 0.04 | 0.16 | 0.08 |
| 4 | 55643116 | rs11732323 | -1.054 | 0.623 | 0.082 | 0.933 | 0.023 | 0.08 | 0.08 | 0 | 0.09 | 0.15 | 0.11 | 0.3 | 0.16 | 0.27 |
| 4 | 55650767 | rs7686567 | -1.895 | 0.15 | 1.063 | 0.83 | 0.037 | 0.237 | 0.237 | 0.073 | 0.12 | 0.47 | 0.12 | 0.3 | 0.12 | 0.25 |
| 4 | 55656500 | rs2173229 | -0.551 | 0.028 | 0.695 | 0.659 | 0.017 | 0.018 | 0.018 | 0.159 | 0.08 | 0.03 | 0.03 | 0.04 | 0.05 | 0.06 |
| 4 | 55658124 | rs2646336 | -1.492 | 0.024 | 1.133 | 0.657 | -0.054 | -0.062 | -0.062 | 1.312 | 0.14 | 0.38 | 0.06 | 0.04 | 0.05 | 0.08 |
| 4 | 55668740 | rs2646325 | -1.559 | 0.026 | 1.044 | 0.658 | -0.059 | -0.068 | -0.068 | 0 | 0.15 | 0.43 | 0.07 | 0.04 | 0.05 | 0.08 |
| 4 | 55669590 | rs1982975 | -2.249 | 0.101 | 0.933 | 0.828 | -0.014 | 0.033 | 0.033 | 1.242 | 0.22 | 0.65 | 0.16 | 0.33 | 0.18 | 0.3 |
| 4 | 55673133 | rs6821963 | -0.574 | 0.355 | 0.239 | 0.889 | 0.035 | 0.147 | 0.147 | 0 | 0.07 | 0.15 | 0.09 | 0.31 | 0.12 | 0.19 |
| 4 | 55680855 | rs4864926 | -1.28 | 0.004 | -0.464 | 0.686 | 0.005 | -0.005 | -0.005 | 0 | 0.24 | 0.74 | 0.17 | 0.32 | 0.17 | 0.28 |
| 4 | 55682619 | rs6821967 | -1.132 | 0.695 | -1.448 | 0.517 | -0.069 | -0.063 | -0.063 | 0 | 0.47 | 0.92 | 0.44 | 0.53 | 0.5 | 0.63 |
| 4 | 55683735 | rs2412611 | -1.159 | 0.191 | -0.622 | 0.337 | -0.014 | -0.019 | -0.019 | 0.061 | 0.34 | 0.58 | 0.25 | 0.23 | 0.26 | 0.29 |
| 4 | 55683815 | rs904140 | -1.129 | -0.252 | 0.192 | 0.014 | -0.074 | -0.094 | -0.094 | 0.692 | 0.19 | 0.53 | 0.1 | 0.05 | 0.08 | 0.14 |

| | | | | | | | | | | | | | | | | |
|---|---|---|---|---|---|---|---|---|---|---|---|---|---|---|---|---|
| 4 | 55683987 | rs2726646 | -1.226 | 0.062 | 0.192 | -0.131 | -0.005 | -0.033 | -0.033 | 0 | 0.22 | 0.31 | 0.23 | 0.19 | 0.18 | 0.24 |
| 4 | 55684426 | rs2646356 | -1.156 | 0.284 | 0.23 | -0.331 | -0.002 | -0.049 | -0.049 | 0 | 0.24 | 0.38 | 0.16 | 0.12 | 0.24 | 0.28 |
| 4 | 55684500 | rs17084858 | 0.727 | 0.448 | -0.748 | -0.116 | 0.034 | -0.031 | -0.031 | 0 | 0.38 | 0.22 | 0.41 | 0.6 | 0.37 | 0.4 |
| 4 | 55684576 | rs2726647 | -1.449 | 0.845 | -0.933 | -0.408 | -0.038 | 0.062 | 0.062 | 0 | 0.78 | 0.95 | 0.89 | 0.92 | 0.81 | 0.81 |
| 4 | 55685764 | rs2646357 | 0.247 | 0.955 | -0.374 | -0.256 | -0.001 | 0.005 | 0.005 | 1.72 | 0.53 | 0.41 | 0.72 | 0.65 | 0.55 | 0.66 |
| 4 | 55689495 | rs1472411 | -0.43 | 0.873 | -0.081 | -0.005 | 0.002 | 0.012 | 0.012 | 0 | 0.28 | 0.5 | 0.2 | 0.33 | 0.27 | 0.32 |
| 4 | 55701312 | rs2646351 | -0.753 | 0.582 | -0.606 | 0.303 | 0.016 | -0.001 | -0.001 | 0 | 0.52 | 0.71 | 0.31 | 0.54 | 0.44 | 0.56 |
| 4 | 55703218 | rs2726626 | -0.724 | 0.666 | -0.391 | 0.321 | 0.011 | 0.003 | 0.003 | 1.328 | 0.52 | 0.68 | 0.31 | 0.54 | 0.44 | 0.52 |
| 4 | 55703257 | rs17763012 | 1.02 | 0.617 | -0.741 | 0.342 | 0.01 | 0 | 0 | 1.519 | 0.22 | 0.2 | 0.12 | 0.2 | 0.18 | 0.22 |
| 4 | 55706827 | rs1948607 | -0.427 | 1.153 | -0.898 | 0.453 | -0.019 | -0.016 | -0.016 | 0 | 0.54 | 0.77 | 0.44 | 0.54 | 0.46 | 0.55 |
| 4 | 55715633 | rs2270674 | -0.261 | 1.174 | -1.112 | 0.404 | 0.052 | 0.028 | 0.028 | 3.202 | 0.49 | 0.76 | 0.3 | 0.6 | 0.4 | 0.6 |
| 4 | 55716601 | rs1874653 | -0.254 | 1.121 | -1.084 | 0.361 | 0.059 | -0.007 | -0.007 | 0 | 0.49 | 0.74 | 0.29 | 0.54 | 0.37 | 0.58 |
| 4 | 55717962 | rs1497499 | -1.28 | 1.184 | -1.268 | 0.098 | -0.011 | 0.067 | 0.067 | 0 | 0.76 | 0.88 | 0.87 | 0.91 | 0.77 | 0.88 |
| 4 | 55721469 | rs1874647 | -1.333 | 1.221 | -1.172 | 0.116 | -0.016 | 0.069 | 0.069 | 0 | 0.76 | 0.93 | 0.88 | 0.9 | 0.77 | 0.88 |
| 4 | 55725382 | rs2173231 | -0.009 | 0.972 | -0.987 | 0.263 | 0.06 | -0.005 | -0.005 | 0 | 0.51 | 0.8 | 0.3 | 0.52 | 0.38 | 0.57 |
| 4 | 55733422 | rs11727341 | 0.015 | 0.983 | -1.309 | 0.284 | 0.015 | -0.027 | -0.027 | 0.029 | 0.48 | 0.13 | 0.57 | 0.49 | 0.58 | 0.38 |
| 4 | 55743651 | rs1532460 | -0.152 | 0.884 | -1.298 | 0.311 | 0.074 | 0.009 | 0.009 | 0.464 | 0.49 | 0.73 | 0.28 | 0.5 | 0.36 | 0.57 |
| 4 | 55746122 | rs6554207 | -0.126 | 0.837 | -1.317 | 0.292 | 0.051 | 0.003 | 0.003 | 0.567 | 0.48 | 0.73 | 0.41 | 0.5 | 0.39 | 0.59 |
| 4 | 55748445 | rs878120 | 0.071 | 0.496 | -1.138 | 0.279 | 0.012 | -0.012 | -0.012 | 0.03 | 0.4 | 0.62 | 0.26 | 0.54 | 0.32 | 0.56 |
| 4 | 55751539 | rs924223 | -0.857 | 0.737 | -1.348 | -0.084 | -0.02 | 0.093 | 0.093 | 0 | 0.75 | 0.89 | 0.86 | 0.94 | 0.77 | 0.89 |
| 4 | 55752730 | rs10010776 | -0.135 | 0.139 | -0.526 | 0.412 | -0.026 | 0.038 | 0.038 | 1.955 | 0.34 | 0.44 | 0.24 | 0.43 | 0.28 | 0.54 |
| 4 | 55782793 | rs6851808 | -0.415 | 0.361 | 1.709 | 0.06 | 0.11 | 0.038 | 0.038 | 0.028 | 0.05 | 0.53 | 0.16 | 0 | 0.05 | 0.09 |
| 4 | 55786946 | rs12642211 | 0.11 | 0.509 | 0.798 | 0.003 | -0.07 | -0.003 | -0.003 | 0 | 0.42 | 0.19 | 0.33 | 0.59 | 0.5 | 0.65 |
| 4 | 55787043 | rs17085113 | -0.013 | 0.549 | 0.835 | 0.019 | -0.02 | 0.025 | 0.025 | 0.647 | 0.46 | 0.34 | 0.35 | 0.59 | 0.5 | 0.65 |
| 4 | 55790393 | rs969340 | -0.439 | 0.541 | 0.722 | 0.017 | 0.009 | 0.015 | 0.015 | 2.33 | 0.49 | 0.61 | 0.49 | 0.59 | 0.55 | 0.74 |
| 4 | 55797131 | rs1497482 | -0.123 | 0.59 | -0.14 | 0.086 | -0.03 | 0.02 | 0.02 | 0.999 | 0.45 | 0.31 | 0.35 | 0.59 | 0.5 | 0.64 |
| 4 | 55800687 | rs2088402 | 0.083 | 0.802 | -0.501 | -0.212 | 0.001 | -0.001 | -0.001 | 0.013 | 0.18 | 0.17 | 0.38 | 0.14 | 0.23 | 0.13 |
| 4 | 55805555 | rs2067951 | 0.149 | 0.404 | -1.364 | -0.35 | -0.098 | 0.065 | 0.065 | 0 | 0.45 | 0.12 | 0.52 | 0.36 | 0.48 | 0.33 |
| 4 | 55814774 | rs11722442 | 0.246 | 0.683 | -0.117 | -0.431 | -0.003 | -0.015 | -0.015 | 0 | 0.63 | 0.46 | 0.54 | 0.71 | 0.61 | 0.42 |
| 4 | 55815650 | rs4426846 | 0.125 | 0.629 | 0.325 | -0.456 | 0 | -0.009 | -0.009 | 0 | 0.64 | 0.63 | 0.55 | 0.72 | 0.61 | 0.42 |
| 4 | 55818909 | rs4345237 | 0.199 | 0.744 | 0.352 | -0.402 | 0.006 | -0.017 | -0.017 | 0 | 0.64 | 0.49 | 0.61 | 0.7 | 0.59 | 0.45 |
| 4 | 55822867 | rs11728462 | 0.116 | 0.602 | -0.015 | -0.376 | 0 | 0.004 | 0.004 | 0 | 0.66 | 0.75 | 0.66 | 0.72 | 0.66 | 0.46 |
| 4 | 55823638 | rs7684211 | 0.604 | 0.734 | 0.055 | -0.497 | 0.014 | 0.03 | 0.03 | 0 | 0.49 | 0.28 | 0.27 | 0.6 | 0.48 | 0.28 |
| 4 | 55831233 | rs6554217 | 0.795 | 0.852 | 0.581 | -0.312 | 0.001 | -0.004 | -0.004 | 0 | 0.83 | 0.74 | 0.7 | 0.86 | 0.84 | 0.85 |
| 4 | 55831694 | rs6554218 | 0.189 | 0.847 | 0.524 | -0.313 | -0.002 | -0.005 | -0.005 | 0 | 0.87 | 0.95 | 0.83 | 0.88 | 0.89 | 0.88 |
| 4 | 55839848 | rs6835585 | 1.323 | 0.576 | 0.586 | -0.55 | -0.003 | -0.002 | -0.002 | 0 | 0.44 | 0.32 | 0.26 | 0.58 | 0.49 | 0.26 |
| 4 | 55862338 | rs11133355 | -0.271 | 0.576 | 0.589 | -0.544 | 0.002 | 0.01 | 0.01 | 0 | 0.39 | 0.33 | 0.43 | 0.29 | 0.35 | 0.6 |
| 4 | 55867442 | rs12502309 | -0.306 | 0.574 | 0.497 | -0.543 | -0.001 | -0.003 | -0.003 | 0 | 0.39 | 0.43 | 0.43 | 0.29 | 0.35 | 0.59 |
| 4 | 55873926 | rs6842830 | 0.479 | 0.775 | 0.489 | -0.132 | -0.004 | 0.01 | 0.01 | 0 | 0.51 | 0.67 | 0.62 | 0.57 | 0.54 | 0.31 |
| 4 | 55876463 | rs4262039 | -0.06 | 0.607 | 0.681 | 0.117 | 0.03 | 0.102 | 0.102 | 0 | 0.94 | 0.77 | 0.84 | 0.88 | 0.96 | 0.96 |
| 4 | 55877829 | rs13113419 | -0.955 | 0.741 | -0.143 | -0.134 | 0.029 | -0.011 | -0.011 | 0 | 0.09 | 0.21 | 0.37 | 0.06 | 0.16 | 0.12 |
| 4 | 55881411 | rs6841800 | -0.143 | -0.227 | -0.582 | -0.551 | 0.014 | 0.009 | 0.009 | 0.429 | 0.41 | 0.34 | 0.15 | 0.1 | 0.27 | 0.21 |
| 4 | 55881855 | rs4453992 | 0.167 | -0.639 | -0.373 | 0.37 | -0.022 | 0.074 | 0.074 | 0 | 0.11 | 0.16 | 0.06 | 0.37 | 0.04 | 0.2 |
| 4 | 55885310 | rs7658751 | -0.456 | -0.173 | -0.559 | -0.139 | -0.003 | -0.004 | -0.004 | 0 | 0.66 | 0.64 | 0.62 | 0.66 | 0.59 | 0.76 |
| 4 | 55891531 | rs4241988 | 0.097 | -0.497 | -0.695 | -0.351 | 0.004 | 0.011 | 0.011 | 0 | 0.37 | 0.42 | 0.43 | 0.59 | 0.36 | 0.48 |
| 4 | 55894238 | rs13105577 | 1.222 | -0.339 | -0.903 | -0.546 | -0.022 | 0.01 | 0.01 | 0 | 0.13 | 0.07 | 0.1 | 0 | 0.19 | 0.11 |
| 4 | 55899286 | rs13142103 | -0.599 | -0.416 | -0.957 | -0.484 | -0.007 | 0.034 | 0.034 | 0.34 | 0.54 | 0.85 | 0.55 | 0.67 | 0.41 | 0.59 |
| 4 | 55900647 | rs4864949 | -0.473 | -0.389 | -0.958 | -0.471 | 0.008 | 0.044 | 0.044 | 0.013 | 0.53 | 0.74 | 0.54 | 0.67 | 0.41 | 0.59 |
| 4 | 55913934 | rs7441352 | 0.592 | -0.29 | -1.056 | -0.387 | -0.028 | 0.066 | 0.066 | 0.116 | 0.23 | 0.32 | 0.33 | 0.58 | 0.17 | 0.37 |
| 4 | 55921535 | rs10461355 | 0.292 | 0.352 | -1.31 | 0.085 | 0.01 | 0.026 | 0.026 | 0.001 | 0.53 | 0.67 | 0.53 | 0.66 | 0.41 | 0.59 |
| 4 | 55931246 | rs4241991 | 0.519 | 0.298 | -1.09 | 0.111 | -0.002 | 0 | 0 | 0.001 | 0.78 | 0.79 | 0.61 | 0.83 | 0.76 | 0.76 |
| 4 | 55936768 | rs4536994 | 0.817 | -0.383 | -1.116 | 0.504 | 0.017 | -0.019 | -0.019 | 0.198 | 0.81 | 0.84 | 0.82 | 0.74 | 0.89 | 0.78 |
| 4 | 55943001 | rs7691507 | -0.694 | 0.068 | -0.786 | -0.049 | -0.021 | -0.035 | -0.035 | 0 | 0.33 | 0.58 | 0.12 | 0.15 | 0.18 | 0.2 |
| 4 | 55953483 | rs2125489 | -0.578 | 0.852 | -0.975 | 0.211 | -0.038 | 0.026 | 0.026 | 0 | 0.07 | 0 | 0.05 | 0.02 | 0.12 | 0.09 |
| 4 | 55955232 | rs1531289 | -0.759 | 0.611 | -0.931 | -0.27 | 0.021 | 0.027 | 0.027 | 0.212 | 0.54 | 0.63 | 0.72 | 0.8 | 0.71 | 0.56 |
| 4 | 55957723 | rs17709898 | -0.905 | 1.482 | -0.863 | 0.584 | 0.002 | 0.002 | 0.002 | 0.35 | 0.26 | 0.25 | 0.21 | 0.31 | 0.36 | 0.27 |
| 4 | 55961159 | rs2219471 | -0.233 | 1.255 | -0.674 | 0.991 | -0.004 | 0.012 | 0.012 | 0 | 0.17 | 0.14 | 0.15 | 0.46 | 0.24 | 0.12 |
| 4 | 55964101 | rs6838752 | -0.266 | 1.237 | -0.604 | 0.981 | -0.01 | -0.006 | -0.006 | 0 | 0.17 | 0.1 | 0.14 | 0.45 | 0.24 | 0.15 |
| 4 | 55966801 | rs6828477 | -1.083 | 1.118 | -0.123 | 0.888 | -0.002 | -0.018 | -0.018 | 0 | 0.45 | 0.52 | 0.66 | 0.78 | 0.6 | 0.45 |
| 4 | 55967483 | rs2168945 | 0.039 | 0.991 | -0.616 | 0.811 | -0.05 | -0.04 | -0.04 | 0.008 | 0.26 | 0.11 | 0.5 | 0.31 | 0.36 | 0.3 |
| 4 | 55970101 | rs11732292 | -0.135 | 1.016 | -0.572 | 0.823 | -0.001 | 0 | 0 | 0 | 0.28 | 0.29 | 0.52 | 0.31 | 0.36 | 0.3 |
| 4 | 55978210 | rs11941492 | -0.524 | 0.133 | -0.926 | 0.859 | 0.003 | -0.003 | -0.003 | 2.943 | 0.27 | 0.26 | 0.26 | 0.28 | 0.21 | 0.49 |
| 4 | 55979558 | rs2305948 | -0.268 | 0.422 | -0.696 | 0.926 | 0.015 | 0.049 | 0.049 | 0 | 0.1 | 0.24 | 0.1 | 0.17 | 0.09 | 0.12 |
| 4 | 55980239 | rs7692791 | -0.141 | 0.747 | -1.036 | 0.728 | 0.004 | 0 | 0 | 0 | 0.64 | 0.39 | 0.46 | 0.63 | 0.54 | 0.72 |
| 4 | 55985815 | rs6837735 | -0.477 | -0.085 | -1.368 | 0.914 | 0.022 | -0.037 | -0.037 | 0 | 0.25 | 0.19 | 0.22 | 0.41 | 0.19 | 0.41 |
| 4 | 55991042 | rs12502008 | -0.187 | 0.398 | -1.567 | 0.875 | -0.005 | -0.016 | -0.016 | 0.354 | 0.34 | 0.22 | 0.53 | 0.57 | 0.37 | 0.39 |
| 4 | 55992366 | rs2071559 | 0.328 | 0.285 | -1.736 | 0.382 | -0.002 | 0.043 | 0.043 | 0 | 0.51 | 0.67 | 0.36 | 0.35 | 0.49 | 0.55 |
| 4 | 55997340 | rs10013228 | 0.732 | 0.21 | -1.972 | -0.078 | -0.013 | 0.05 | 0.05 | 0 | 0.29 | 0.12 | 0.18 | 0.19 | 0.27 | 0.26 |
| 4 | 56002858 | rs10517342 | 1.107 | 0.144 | -1.708 | -0.13 | -0.011 | -0.021 | -0.021 | 0.698 | 0.34 | 0.2 | 0.54 | 0.58 | 0.37 | 0.34 |

| | | | | | | | | | | | | | | | |
|---|---|---|---|---|---|---|---|---|---|---|---|---|---|---|---|
| 4 | 56006102 | rs12505096 | 0.872 | 0.102 | -1.967 | -0.356 | -0.038 | 0.023 | 0.023 | 0 | 0.25 | 0.08 | 0.19 | 0.19 | 0.28 | 0.26 |
| 4 | 56011889 | rs12331507 | 0.226 | 0.09 | -1.943 | -0.069 | -0.037 | 0.024 | 0.024 | 0 | 0.32 | 0.11 | 0.21 | 0.25 | 0.32 | 0.31 |
| 4 | 56012149 | rs12331869 | -0.763 | 0.021 | -1.163 | -0.1 | -0.014 | -0.009 | -0.009 | 0 | 0.8 | 0.59 | 0.86 | 0.84 | 0.82 | 0.7 |
| 4 | 56014425 | rs2219469 | 0.377 | 0.128 | -1.157 | -0.042 | 0 | 0.018 | 0.018 | 0 | 0.49 | 0.38 | 0.65 | 0.6 | 0.51 | 0.44 |
| 4 | 56018295 | rs13126206 | 0.812 | 0.187 | -1.338 | 0.172 | -0.044 | -0.028 | -0.028 | 0 | 0.13 | 0.01 | 0.11 | 0.11 | 0.17 | 0.29 |
| 4 | 56025616 | rs4356965 | 1.218 | 0.162 | -1.365 | 0.601 | -0.013 | -0.024 | -0.024 | 0 | 0.33 | 0.19 | 0.48 | 0.38 | 0.37 | 0.33 |
| 4 | 56027518 | rs6816309 | 0.614 | 0.331 | -1.178 | 0.637 | -0.001 | -0.001 | -0.001 | 0.458 | 0.64 | 0.63 | 0.73 | 0.53 | 0.64 | 0.48 |
| 4 | 56027573 | rs2412627 | 0.015 | 0.312 | -1.453 | 0.896 | -0.02 | -0.045 | -0.045 | 0.092 | 0.17 | 0.05 | 0.12 | 0.27 | 0.19 | 0.23 |
| 4 | 56029474 | rs3986292 | 0.206 | 0.167 | -1.54 | 0.811 | -0.033 | -0.042 | -0.042 | 0 | 0.4 | 0.17 | 0.52 | 0.53 | 0.46 | 0.49 |
| 4 | 56029834 | rs11727885 | 0.249 | -0.769 | -0.516 | 1.006 | -0.036 | -0.004 | -0.004 | 0 | 0.72 | 0.69 | 0.77 | 0.79 | 0.81 | 0.93 |
| 4 | 56030064 | rs6554238 | -0.306 | -0.401 | 0.188 | 1.419 | 0.011 | 0.038 | 0.038 | 0.076 | 0.79 | 0.68 | 0.77 | 0.65 | 0.75 | 0.72 |
| 4 | 56030182 | rs7659869 | -0.945 | -0.671 | 0.218 | 1.187 | 0.018 | -0.038 | -0.038 | 0 | 0.39 | 0.19 | 0.28 | 0.41 | 0.35 | 0.35 |
| 4 | 56038620 | rs1993583 | -0.038 | -0.702 | 1.423 | 0.737 | -0.049 | -0.111 | -0.111 | 0 | 0.83 | 0.67 | 0.93 | 1 | 0.92 | 0.98 |
| 4 | 56038746 | rs1993585 | 0.618 | -0.732 | 1.431 | 0.703 | -0.038 | -0.094 | -0.094 | 0 | 0.8 | 0.31 | 0.89 | 1 | 0.89 | 0.96 |
| 4 | 56044260 | rs2085621 | 1.356 | -0.768 | 1.235 | 0.78 | 0.03 | 0.018 | 0.018 | 3.025 | 0.76 | 0.16 | 0.86 | 0.84 | 0.88 | 0.93 |
| 4 | 56046659 | rs12500524 | 0.541 | -0.472 | 1.071 | 0.902 | -0.09 | -0.136 | -0.136 | 0.63 | 0.17 | 0.02 | 0.25 | 0.45 | 0.23 | 0.28 |
| 4 | 56049425 | rs13139198 | 1.431 | -1.087 | 2.084 | 0.731 | -0.021 | -0.008 | -0.008 | 0.16 | 0.77 | 0.24 | 0.86 | 0.84 | 0.89 | 0.93 |
| 4 | 56089259 | rs10003420 | -2.584 | -0.321 | 1.516 | 1.235 | -0.032 | 0.115 | 0.115 | 3.762 | 0.09 | 0.66 | 0.08 | 0.19 | 0.04 | 0.05 |
| 4 | 56096263 | rs6554248 | -0.967 | -0.163 | 2.039 | 0.501 | 0.084 | 0.052 | 0.052 | 0.086 | 0.05 | 0.34 | 0.04 | 0 | 0.02 | 0.01 |
| 4 | 56118414 | rs6554254 | -2.178 | -0.176 | 2.072 | 0.492 | 0.054 | 0.003 | 0.003 | 1.018 | 0.08 | 0.48 | 0.06 | 0 | 0.04 | 0.03 |
| 4 | 56125042 | rs4241994 | -0.128 | -0.449 | 1.679 | 0.767 | 0.003 | 0.015 | 0.015 | 0 | 0.08 | 0.07 | 0.14 | 0.29 | 0.07 | 0.12 |
| 4 | 56131739 | rs4301169 | -1.315 | -0.467 | 2.099 | 0.755 | -0.026 | 0.156 | 0.156 | 0 | 0.13 | 0.36 | 0.18 | 0.29 | 0.09 | 0.13 |
| 4 | 56133629 | rs6837397 | -2.072 | -0.514 | 2.252 | 0.705 | -0.047 | 0.077 | 0.077 | 0 | 0.16 | 0.59 | 0.2 | 0.29 | 0.11 | 0.15 |
| 4 | 56134791 | rs6843086 | -1.592 | -0.427 | 1.889 | 0.754 | -0.037 | 0.149 | 0.149 | 0 | 0.14 | 0.41 | 0.19 | 0.29 | 0.1 | 0.13 |
| 4 | 56153732 | rs4864976 | -2.04 | -0.476 | 1.565 | 0.58 | -0.038 | 0.063 | 0.063 | 0 | 0.2 | 0.46 | 0.22 | 0.3 | 0.13 | 0.14 |
| 4 | 56155385 | rs7661296 | -1.075 | -0.476 | 1.669 | 0.572 | 0.028 | -0.049 | -0.049 | 0 | 0.1 | 0.31 | 0.07 | 0 | 0.07 | 0.01 |
| 4 | 56157142 | rs4077725 | -1.765 | -0.472 | 1.05 | 0.581 | -0.025 | 0.017 | 0.017 | 0 | 0.22 | 0.67 | 0.24 | 0.3 | 0.15 | 0.16 |
| 4 | 56176481 | rs4864539 | -0.644 | -0.219 | 0.302 | -0.151 | 0.019 | 0.024 | 0.024 | 5.946 | 0.42 | 0.7 | 0.33 | 0.32 | 0.34 | 0.19 |
| 4 | 56179883 | rs1399953 | -0.23 | -0.237 | 0.268 | -0.2 | 0.008 | 0.009 | 0.009 | 3.205 | 0.37 | 0.68 | 0.2 | 0.29 | 0.31 | 0.2 |
| 4 | 56180919 | rs6829598 | -1.076 | -0.221 | 1.023 | -0.236 | 0.026 | -0.057 | -0.057 | 0 | 0.1 | 0.32 | 0.08 | 0 | 0.09 | 0.04 |
| 4 | 56187375 | rs17776421 | -0.456 | 0.146 | 0.392 | -0.62 | 0.036 | 0.04 | 0.04 | 4.709 | 0.46 | 0.67 | 0.44 | 0.34 | 0.38 | 0.22 |
| 4 | 56188287 | rs11728305 | -0.35 | 0.121 | -0.212 | -1.159 | -0.011 | 0.029 | 0.029 | 0.11 | 0.78 | 0.87 | 0.74 | 0.98 | 0.74 | 0.77 |
| 4 | 56195626 | rs6815518 | 0.202 | 0.008 | 0.157 | -0.679 | -0.007 | -0.001 | -0.001 | 0 | 0.35 | 0.58 | 0.17 | 0.32 | 0.23 | 0.18 |
| 4 | 56214596 | rs9790808 | 0.329 | 0.364 | -0.486 | -1.129 | -0.041 | 0.207 | 0.207 | 2.179 | 0.55 | 0.18 | 0.54 | 0.86 | 0.49 | 0.53 |
| 4 | 56222599 | rs7664100 | 0.228 | 0.313 | -0.307 | -1.061 | -0.038 | 0.19 | 0.19 | 2.168 | 0.54 | 0.16 | 0.52 | 0.84 | 0.48 | 0.51 |
| 4 | 56231119 | rs7663650 | -0.242 | 0.474 | 0.217 | -0.91 | -0.001 | 0 | 0 | 0 | 0.35 | 0.34 | 0.46 | 0.55 | 0.33 | 0.38 |
| 4 | 56237137 | rs576732 | 0.331 | 0.57 | 0.431 | -1.134 | 0.001 | 0.027 | 0.027 | 0.09 | 0.29 | 0.24 | 0.24 | 0.11 | 0.32 | 0.44 |
| 4 | 56238263 | rs819269 | 0.327 | 0.573 | 0.432 | -1.132 | 0.001 | 0.028 | 0.028 | 0.112 | 0.29 | 0.24 | 0.24 | 0.11 | 0.32 | 0.44 |
| 4 | 56238847 | rs819271 | -1.525 | 0.623 | 0.447 | -1.143 | -0.012 | -0.021 | -0.021 | 0.583 | 0.67 | 0.86 | 0.72 | 0.65 | 0.66 | 0.82 |
| 4 | 56243522 | rs9993675 | -1.383 | 0.629 | 0.507 | -1.14 | -0.001 | -0.007 | -0.007 | 3.746 | 0.66 | 0.77 | 0.71 | 0.65 | 0.66 | 0.82 |
| 4 | 56246273 | rs552792 | -0.01 | 0.612 | 0.607 | -1.243 | -0.014 | -0.108 | -0.108 | 0 | 0.33 | 0.52 | 0.26 | 0.1 | 0.33 | 0.44 |
| 4 | 56260662 | rs693367 | -0.029 | 0.742 | 0.595 | -1.264 | -0.015 | -0.118 | -0.118 | 0 | 0.35 | 0.51 | 0.28 | 0.1 | 0.35 | 0.45 |
| 4 | 56267473 | rs12641881 | 0.55 | 0.78 | 0.228 | -1.238 | -0.089 | -0.126 | -0.126 | 0.063 | 0.26 | 0.02 | 0.4 | 0.57 | 0.25 | 0.38 |
| 4 | 56274132 | rs2048564 | 0.196 | 0.739 | 0.406 | -1.188 | -0.019 | -0.026 | -0.026 | 1.689 | 0.3 | 0.17 | 0.42 | 0.57 | 0.3 | 0.41 |
| 4 | 56276334 | rs11943456 | 0.341 | 0.69 | 0.37 | -1.289 | -0.016 | 0.162 | 0.162 | 0.459 | 0.54 | 0.21 | 0.51 | 0.88 | 0.46 | 0.55 |
| 4 | 56285305 | rs504836 | -0.415 | 0.645 | 0.551 | -1.306 | 0.06 | -0.072 | -0.072 | 0 | 0.85 | 0.48 | 0.78 | 0.98 | 0.81 | 0.96 |
| 4 | 56290220 | rs534654 | -1.645 | 0.647 | 0.574 | -1.305 | 0.024 | -0.025 | -0.025 | 0 | 0.89 | 0.8 | 0.8 | 0.98 | 0.81 | 0.96 |
| 4 | 56296763 | rs1056547 | -0.213 | 0.644 | 0.88 | -0.971 | -0.033 | -0.023 | -0.023 | 3.748 | 0.32 | 0.17 | 0.45 | 0.57 | 0.34 | 0.44 |
| 4 | 56301369 | rs1801260 | 0.091 | 0.632 | 0.945 | -0.984 | 0.013 | 0.075 | 0.075 | 0.153 | 0.3 | 0.15 | 0.24 | 0.1 | 0.31 | 0.38 |
| 4 | 56338793 | rs11931061 | -1.285 | 0.635 | 0.5 | -0.971 | 0.005 | 0.008 | 0.008 | 4.743 | 0.61 | 0.32 | 0.69 | 0.67 | 0.65 | 0.82 |
| 4 | 56346411 | rs17722979 | 0.095 | 0.639 | 0.46 | -0.976 | 0.013 | 0.074 | 0.074 | 0.132 | 0.3 | 0.15 | 0.24 | 0.1 | 0.3 | 0.38 |
| 4 | 56375981 | rs3817444 | -1.864 | 0.662 | 0.414 | -0.956 | -0.003 | 0.004 | 0.004 | 3.213 | 0.66 | 0.77 | 0.72 | 0.67 | 0.65 | 0.82 |
| 4 | 56380484 | rs12649507 | -0.158 | 0.72 | 0.265 | -0.897 | -0.019 | -0.025 | -0.025 | 0 | 0.31 | 0.17 | 0.42 | 0.57 | 0.3 | 0.41 |
| 4 | 56392813 | rs13132420 | -0.235 | 0.554 | 0.273 | -0.89 | -0.033 | -0.025 | -0.025 | 0 | 0.31 | 0.17 | 0.44 | 0.57 | 0.34 | 0.42 |
| 4 | 56413803 | rs4864548 | -0.235 | 0.554 | 0.273 | -0.89 | -0.033 | -0.025 | -0.025 | 0 | 0.31 | 0.17 | 0.45 | 0.57 | 0.34 | 0.44 |
| 4 | 56418755 | rs1979604 | -0.263 | 0.513 | 0.212 | -0.945 | 0.013 | 0.074 | 0.074 | 0.132 | 0.7 | 0.85 | 0.76 | 0.9 | 0.69 | 0.62 |
| 4 | 56439245 | rs7665846 | -0.305 | 0.514 | 0.187 | -0.945 | 0.012 | 0.084 | 0.084 | 0.103 | 0.7 | 0.86 | 0.76 | 0.9 | 0.69 | 0.62 |
| 4 | 56443809 | rs13133077 | -0.264 | 0.508 | 0.184 | -0.948 | 0.013 | 0.084 | 0.084 | 0.115 | 0.7 | 0.86 | 0.76 | 0.9 | 0.69 | 0.61 |
| 4 | 56444300 | rs1873092 | 1.472 | 0.556 | 0.21 | -0.887 | -0.013 | -0.002 | -0.002 | 0.079 | 0.34 | 0.15 | 0.28 | 0.33 | 0.35 | 0.18 |
| 4 | 56460085 | rs13132085 | -0.343 | 0.991 | 0.206 | -0.692 | -0.01 | -0.032 | -0.032 | 0.233 | 0.28 | 0.15 | 0.41 | 0.58 | 0.27 | 0.33 |
| 4 | 56461365 | rs2087319 | 0.135 | 1.104 | -0.218 | -0.699 | -0.048 | -0.127 | -0.127 | 0 | 0.23 | 0.02 | 0.39 | 0.58 | 0.22 | 0.3 |
| 4 | 56471433 | rs1873091 | 0.806 | 0.982 | -0.296 | -0.954 | -0.011 | 0.042 | 0.042 | 0 | 0.17 | 0.05 | 0.15 | 0.1 | 0.2 | 0.26 |
| 4 | 56475338 | rs3805383 | 0.385 | 1.084 | -0.279 | -0.963 | -0.023 | 0.032 | 0.032 | 0 | 0.2 | 0.14 | 0.18 | 0.1 | 0.28 | 0.3 |
| 4 | 56492499 | rs12499623 | -0.997 | 1.441 | -0.47 | -0.712 | -0.005 | 0.011 | 0.011 | 0.051 | 0.43 | 0.41 | 0.59 | 0.68 | 0.48 | 0.58 |
| 4 | 56494039 | rs12500837 | 0.007 | 1.309 | -0.547 | -0.652 | -0.047 | -0.132 | -0.132 | 0.086 | 0.23 | 0.02 | 0.39 | 0.57 | 0.22 | 0.3 |
| 4 | 56498249 | rs9999653 | -0.894 | 1.325 | -0.322 | 0.146 | 0.004 | 0.011 | 0.011 | 0 | 0.44 | 0.41 | 0.43 | 0.34 | 0.55 | 0.52 |
| 4 | 56499826 | rs9312664 | 1.459 | 1.376 | -1.224 | 0.247 | 0.002 | 0.045 | 0.045 | 3.639 | 0.42 | 0.14 | 0.34 | 0.32 | 0.39 | 0.18 |
| 4 | 56502178 | rs3792703 | -0.507 | 1.388 | -1.132 | 0.227 | -0.003 | -0.003 | -0.003 | 0 | 0.38 | 0.55 | 0.34 | 0.34 | 0.37 | 0.45 |
| 4 | 56512574 | rs1488938 | 0.615 | 1.758 | -1.222 | 0.183 | 0.046 | -0.026 | -0.026 | 1.256 | 0.95 | 0.8 | 0.95 | 1 | 0.92 | 0.99 |

| | | | | | | | | | | | | | | | | |
|---|---|---|---|---|---|---|---|---|---|---|---|---|---|---|---|---|
| 4 | 56519892 | rs10002268 | 0.635 | 1.76 | -1.926 | 0.335 | -0.029 | 0.074 | 0.074 | 3.669 | 0.46 | 0.2 | 0.37 | 0.32 | 0.49 | 0.24 |
| 4 | 56530796 | rs2634464 | -0.505 | 1.502 | -2.075 | 0.605 | -0.061 | -0.129 | -0.129 | 0 | 0.2 | 0.05 | 0.41 | 0.45 | 0.25 | 0.37 |
| 4 | 56530970 | rs13132077 | 0.564 | 1.474 | -1.927 | 0.444 | 0.003 | 0.007 | 0.007 | 2.846 | 0.44 | 0.14 | 0.34 | 0.45 | 0.42 | 0.18 |
| 4 | 56539449 | rs2634466 | -0.733 | 1.728 | -1.221 | 0.512 | -0.025 | -0.043 | -0.043 | 0.278 | 0.14 | 0.06 | 0.19 | 0.28 | 0.18 | 0.24 |
| 4 | 56547412 | rs4864552 | 0.642 | 1.639 | -0.734 | -0.611 | 0.008 | -0.076 | -0.076 | 0 | 0.79 | 0.76 | 0.85 | 1 | 0.76 | 0.91 |
| 4 | 56551185 | rs4288064 | -0.585 | 1.361 | -1.051 | -0.989 | 0.002 | 0.024 | 0.024 | 0 | 0.2 | 0.15 | 0.23 | 0.09 | 0.21 | 0.09 |
| 4 | 56551916 | rs6856478 | 0.626 | 1.299 | -1.114 | -1.657 | 0.004 | 0.01 | 0.01 | 0 | 0.76 | 0.85 | 0.86 | 1 | 0.76 | 0.87 |
| 4 | 56568521 | rs6828640 | 0.657 | 2.077 | -0.231 | -1.855 | 0.115 | -0.121 | -0.121 | 0.311 | 0.84 | 0.5 | 0.86 | 1 | 0.8 | 0.94 |
| 4 | 56581678 | rs7665149 | 1.423 | 1.554 | -0.633 | -2.304 | 0.01 | 0.114 | 0.114 | 0 | 0.52 | 0.29 | 0.54 | 0.91 | 0.44 | 0.74 |
| 4 | 56583590 | rs12648836 | -0.294 | 1.429 | -0.169 | -2.333 | -0.001 | -0.007 | -0.007 | 0 | 0.25 | 0.24 | 0.25 | 0.1 | 0.23 | 0.13 |
| 4 | 56584126 | rs7658424 | 1.639 | 1.322 | -0.154 | -2.38 | 0.018 | 0.164 | 0.164 | 0 | 0.5 | 0.27 | 0.53 | 0.9 | 0.44 | 0.74 |
| 4 | 56596137 | rs10014020 | -0.144 | 1.373 | 0.697 | -2.34 | 0.001 | -0.063 | -0.063 | 1.206 | 0.11 | 0.26 | 0.09 | 0 | 0.12 | 0.08 |
| 4 | 56596987 | rs4333263 | -0.143 | 1.415 | 0.613 | -2.354 | -0.049 | 0.046 | 0.046 | 0 | 0.88 | 0.97 | 0.89 | 1 | 0.79 | 0.91 |
| 4 | 56599963 | rs3923477 | 1.086 | 1.191 | 0.566 | -2.433 | 0.009 | -0.066 | -0.066 | 2.796 | 0.6 | 0.52 | 0.62 | 0.91 | 0.56 | 0.78 |
| 4 | 56609540 | rs4272068 | -0.101 | 1.383 | 0.344 | -2.324 | -0.01 | 0.034 | 0.034 | 0 | 0.88 | 0.96 | 0.92 | 1 | 0.88 | 0.98 |
| 4 | 56609872 | rs10031105 | -1.131 | 1.269 | 0.804 | -2.252 | 0.073 | -0.1 | -0.1 | 0.322 | 0.15 | 0.41 | 0.14 | 0 | 0.22 | 0.15 |
| 4 | 56615644 | rs13142719 | -1.222 | 0.968 | -0.057 | -1.769 | 0.007 | 0.031 | 0.031 | 0.058 | 0.59 | 0.8 | 0.72 | 0.62 | 0.61 | 0.5 |
| 4 | 56617884 | rs4519853 | 1.683 | 1.163 | -0.094 | -1.733 | 0.004 | -0.005 | -0.005 | 0 | 0.32 | 0.19 | 0.22 | 0.38 | 0.3 | 0.44 |
| 4 | 56623447 | rs7667111 | -0.19 | 1.384 | -0.017 | -1.779 | -0.002 | 0.007 | 0.007 | 0 | 0.31 | 0.32 | 0.41 | 0.52 | 0.28 | 0.34 |
| 4 | 56625468 | rs11725155 | -0.297 | 1.48 | -0.04 | -1.707 | -0.001 | 0 | 0 | 0 | 0.29 | 0.25 | 0.44 | 0.52 | 0.35 | 0.41 |
| 4 | 56628525 | rs12641893 | 0.066 | 1.446 | 0.005 | -1.648 | 0.045 | -0.066 | -0.066 | 2.021 | 0.87 | 0.77 | 0.88 | 1 | 0.8 | 0.9 |
| 4 | 56629191 | rs6816150 | 0.417 | 1.086 | -0.127 | -1.71 | 0.009 | -0.049 | -0.049 | 0 | 0.63 | 0.57 | 0.65 | 0.9 | 0.58 | 0.79 |
| 4 | 56629771 | rs4865034 | -0.746 | 0.785 | -0.492 | -1.272 | -0.007 | 0.024 | 0.024 | 0.255 | 0.92 | 0.97 | 0.94 | 1 | 0.91 | 0.97 |
| 4 | 56632288 | rs7691043 | 0.807 | 0.562 | -0.295 | -1.17 | 0.011 | -0.001 | -0.001 | 0 | 0.34 | 0.29 | 0.22 | 0.38 | 0.21 | 0.36 |
| 4 | 56641023 | rs10015409 | 0.538 | 0.512 | -0.631 | -1.425 | 0.022 | 0.045 | 0.045 | 0 | 0.43 | 0.22 | 0.36 | 0.63 | 0.32 | 0.57 |
| 4 | 56641680 | rs13113466 | 0.666 | 0.915 | -0.924 | -1.553 | 0 | 0.004 | 0.004 | 0.039 | 0.11 | 0.1 | 0.15 | 0.09 | 0.13 | 0.05 |
| 4 | 56641791 | rs10018255 | -0.635 | 1.315 | -0.993 | -1.285 | -0.007 | 0.005 | 0.005 | 0 | 0.86 | 0.93 | 0.9 | 1 | 0.86 | 0.97 |
| 4 | 56654246 | rs1955161 | 0.391 | 0.85 | -0.467 | -1.158 | -0.004 | -0.047 | -0.047 | 0 | 0.67 | 0.45 | 0.79 | 0.84 | 0.63 | 0.84 |
| 4 | 56657690 | rs895615 | -0.163 | 0.877 | -0.315 | -1.022 | -0.003 | -0.02 | -0.02 | 0 | 0.81 | 0.63 | 0.88 | 0.89 | 0.81 | 0.91 |
| 4 | 56664054 | rs2726736 | -0.866 | 0.646 | -1.03 | -1.039 | 0.01 | 0.015 | 0.015 | 1.371 | 0.8 | 0.89 | 0.91 | 0.89 | 0.82 | 0.87 |
| 4 | 56667225 | rs6554324 | -0.365 | 0.509 | -1.117 | -1.315 | 0.033 | 0.105 | 0.105 | 0 | 0.61 | 0.81 | 0.81 | 0.89 | 0.62 | 0.69 |
| 4 | 56668431 | rs753129 | 0.26 | 0.957 | -1.681 | -0.777 | -0.016 | -0.043 | -0.043 | 0 | 0.26 | 0.17 | 0.27 | 0.53 | 0.24 | 0.32 |
| 4 | 56671211 | rs2899048 | 0.726 | 1.079 | -1.59 | -0.747 | -0.034 | 0.003 | 0.003 | 0 | 0.23 | 0.04 | 0.15 | 0.16 | 0.25 | 0.11 |
| 4 | 56677652 | rs11736419 | -0.136 | 1.229 | -1.435 | -0.551 | -0.034 | -0.032 | -0.032 | 0 | 0.26 | 0.13 | 0.28 | 0.67 | 0.24 | 0.28 |
| 4 | 56680363 | rs13126824 | -0.116 | 1.226 | -1.452 | -0.553 | -0.038 | -0.036 | -0.036 | 0 | 0.26 | 0.12 | 0.28 | 0.67 | 0.24 | 0.28 |
| 4 | 56683846 | rs7679852 | -1.176 | 1.07 | -0.855 | -0.944 | 0 | 0.03 | 0.03 | 0.15 | 0.43 | 0.7 | 0.68 | 0.87 | 0.36 | 0.51 |
| 4 | 56700160 | rs12504347 | 1.147 | 1.533 | -0.329 | -0.915 | 0 | -0.012 | -0.012 | 2.599 | 0.08 | 0.08 | 0.06 | 0 | 0.07 | 0.02 |
| 4 | 56702995 | rs1447050 | -0.7 | 1.421 | -0.406 | -1.143 | -0.008 | -0.013 | -0.013 | 4.418 | 0.58 | 0.74 | 0.64 | 0.46 | 0.57 | 0.56 |
| 4 | 56706623 | rs6824915 | -1.709 | 1.358 | -0.418 | -1.195 | 0 | 0.026 | 0.026 | 0.687 | 0.74 | 0.85 | 0.83 | 0.98 | 0.7 | 0.76 |
| 4 | 56718312 | rs2412692 | -1.636 | 1.36 | -0.343 | -1.177 | -0.001 | 0.023 | 0.023 | 0.579 | 0.74 | 0.84 | 0.83 | 0.98 | 0.7 | 0.75 |
| 4 | 56734701 | rs3213948 | -0.351 | 1.176 | 0.319 | -1.155 | -0.003 | -0.011 | -0.011 | 0.027 | 0.57 | 0.47 | 0.54 | 0.68 | 0.58 | 0.47 |
| 4 | 56742205 | rs4865041 | -0.645 | 1.174 | 0.522 | -1.169 | 0 | 0.007 | 0.007 | 0 | 0.6 | 0.66 | 0.74 | 0.83 | 0.6 | 0.65 |
| 4 | 56756531 | rs2290016 | 0.064 | 1.316 | -0.009 | -1.139 | 0.008 | 0.022 | 0.022 | 0 | 0.25 | 0.26 | 0.28 | 0.53 | 0.28 | 0.28 |
| 4 | 56761548 | rs1447045 | 1.191 | 1.367 | 0.082 | -1.056 | 0.017 | 0.043 | 0.043 | 0.032 | 0.28 | 0.19 | 0.13 | 0.1 | 0.26 | 0.15 |
| 4 | 56771089 | rs11658 | 0.934 | 1.465 | -0.391 | -0.56 | -0.011 | -0.028 | -0.028 | 1.83 | 0.15 | 0.11 | 0.19 | 0.52 | 0.13 | 0.19 |
| 4 | 56776848 | rs2593079 | -0.283 | -0.111 | -0.183 | -0.934 | -0.005 | -0.005 | -0.005 | 0.967 | 0.46 | 0.47 | 0.57 | 0.4 | 0.41 | 0.45 |
| 4 | 56777124 | rs17086157 | 1.042 | -0.026 | -0.401 | -1.131 | 0.016 | 0.049 | 0.049 | 0.844 | 0.24 | 0.12 | 0.14 | 0.02 | 0.23 | 0.24 |
| 4 | 56780461 | rs12509278 | 0.363 | 0.212 | -0.815 | -0.722 | -0.007 | -0.049 | -0.049 | 0 | 0.16 | 0.04 | 0.27 | 0.24 | 0.11 | 0.25 |
| 4 | 56786589 | rs2593075 | -0.507 | 0.313 | -0.703 | -0.836 | -0.051 | -0.033 | -0.033 | 0 | 0.71 | 0.78 | 0.7 | 0.41 | 0.62 | 0.69 |
| 4 | 56786831 | rs10517365 | 1.512 | -0.148 | -0.529 | -0.785 | 0.01 | 0.032 | 0.032 | 0 | 0.18 | 0.07 | 0.08 | 0.09 | 0.17 | 0.12 |
| 4 | 56789617 | rs2593074 | -0.646 | -0.406 | -0.337 | -0.792 | 0.027 | 0.028 | 0.028 | 0 | 0.46 | 0.57 | 0.57 | 0.35 | 0.4 | 0.42 |
| 4 | 56813481 | rs2702322 | -0.17 | -0.352 | -0.014 | -0.602 | 0.009 | -0.009 | -0.009 | 0.601 | 0.2 | 0.18 | 0.35 | 0.29 | 0.12 | 0.28 |
| 4 | 56821806 | rs4865047 | 0.304 | -0.4 | 0.356 | -0.613 | -0.01 | 0.005 | 0.005 | 0.061 | 0.16 | 0.15 | 0.27 | 0.23 | 0.08 | 0.22 |
| 4 | 56829482 | rs2593068 | -1.119 | -0.476 | 0.236 | -0.788 | -0.011 | 0.012 | 0.012 | 0 | 0.57 | 0.69 | 0.64 | 0.39 | 0.51 | 0.47 |
| 4 | 56830934 | rs3806749 | 1.421 | -0.462 | 0.509 | -0.672 | 0 | 0.038 | 0.038 | 0 | 0.23 | 0.12 | 0.12 | 0.08 | 0.26 | 0.13 |
| 4 | 56839866 | rs6838426 | -0.078 | -0.466 | 0.501 | -0.682 | 0.032 | 0.021 | 0.021 | 0 | 0.39 | 0.17 | 0.38 | 0.31 | 0.35 | 0.37 |
| 4 | 56851023 | rs6815354 | 1.375 | -0.43 | 0.386 | -0.661 | 0.001 | 0.025 | 0.025 | 0 | 0.24 | 0.13 | 0.12 | 0.09 | 0.26 | 0.13 |
| 4 | 56855667 | rs4526032 | -0.847 | -0.444 | 0.273 | -0.676 | 0.028 | 0.029 | 0.029 | 0 | 0.48 | 0.62 | 0.59 | 0.39 | 0.42 | 0.43 |
| 4 | 56864032 | rs11133419 | 1.106 | -0.528 | -0.013 | -0.612 | 0 | -0.006 | -0.006 | 0.163 | 0.24 | 0.21 | 0.13 | 0.09 | 0.27 | 0.14 |
| 4 | 56868342 | rs13138432 | 0.389 | -0.508 | -0.706 | -0.469 | 0.036 | -0.06 | -0.06 | 2.283 | 0.16 | 0.04 | 0.26 | 0.26 | 0.08 | 0.24 |
| 4 | 56869217 | rs6856844 | 0.267 | -0.469 | -0.702 | -0.451 | -0.01 | 0.006 | 0.006 | 0.074 | 0.17 | 0.16 | 0.28 | 0.26 | 0.08 | 0.22 |
| 4 | 56874517 | rs3214045 | 0.938 | 0.128 | -1.075 | -0.547 | -0.008 | 0.049 | 0.049 | 0 | 0.23 | 0.09 | 0.12 | 0.09 | 0.27 | 0.14 |
| 4 | 56903251 | rs7660872 | 0.332 | -0.335 | -0.578 | -0.374 | -0.001 | -0.013 | -0.013 | 0 | 0.26 | 0.29 | 0.16 | 0.08 | 0.27 | 0.3 |
| 4 | 56904059 | rs12649879 | -0.226 | -0.515 | -0.965 | -0.106 | 0.008 | 0 | 0 | 0 | 0.15 | 0.13 | 0.25 | 0.19 | 0.1 | 0.21 |
| 4 | 56905444 | rs2951806 | -0.976 | -0.506 | -1.335 | 0.212 | -0.096 | -0.101 | -0.101 | 0 | 0.61 | 0.93 | 0.7 | 0.51 | 0.51 | 0.65 |
| 4 | 56908194 | rs2702340 | -1.195 | -1.163 | 0.001 | 0.468 | -0.03 | 0.009 | 0.009 | 0 | 0.26 | 0.35 | 0.4 | 0.34 | 0.17 | 0.35 |
| 4 | 56911058 | rs2702334 | -1.153 | -1.043 | 0.661 | 0.236 | -0.066 | -0.009 | -0.009 | 0.01 | 0.22 | 0.54 | 0.28 | 0.22 | 0.14 | 0.23 |
| 4 | 56924685 | rs6839870 | -0.905 | -0.926 | 0.427 | 0.005 | 0.001 | -0.005 | -0.005 | 0 | 0.31 | 0.65 | 0.26 | 0.14 | 0.3 | 0.29 |
| 4 | 56945325 | rs2412717 | -0.889 | -0.88 | 0.481 | 0.074 | -0.014 | -0.045 | -0.045 | 0.059 | 0.32 | 0.52 | 0.36 | 0.21 | 0.32 | 0.36 |

| | | | | | | | | | | | | | | | | |
|---|---|---|---|---|---|---|---|---|---|---|---|---|---|---|---|---|
| 4 | 56948718 | rs2412719 | -1.005 | -0.868 | -0.116 | 0.08 | 0.006 | 0.013 | 0.013 | 0.204 | 0.33 | 0.69 | 0.37 | 0.21 | 0.32 | 0.37 |
| 4 | 56951767 | rs2412723 | -0.887 | -1.342 | -0.006 | 0.193 | -0.015 | -0.012 | -0.012 | 0.51 | 0.28 | 0.65 | 0.32 | 0.21 | 0.2 | 0.29 |
| 4 | 56952163 | rs10019724 | -0.326 | -0.985 | -0.147 | 0.381 | -0.011 | 0.003 | 0.003 | 1.597 | 0.11 | 0.15 | 0.16 | 0.12 | 0.09 | 0.08 |
| 4 | 56953577 | rs10015524 | 0.333 | -1.003 | -0.105 | 0.42 | -0.018 | 0.016 | 0.016 | 2.352 | 0.08 | 0.13 | 0.08 | 0.12 | 0.04 | 0.06 |
| 4 | 56955556 | rs12331742 | 0.067 | -1.215 | -0.344 | -0.216 | 0.007 | 0.012 | 0.012 | 0 | 0.29 | 0.3 | 0.23 | 0.13 | 0.21 | 0.4 |
| 4 | 56965175 | rs4865050 | 0.701 | -0.711 | -0.465 | 0.774 | -0.001 | 0.017 | 0.017 | 0 | 0.06 | 0.05 | 0.05 | 0.19 | 0.05 | 0.07 |
| 4 | 56968671 | rs17799799 | -0.759 | -0.836 | -0.697 | 0.746 | 0.013 | -0.009 | -0.009 | 0 | 0.18 | 0.1 | 0.16 | 0.26 | 0.13 | 0.16 |
| 4 | 56986929 | rs11728328 | 0.208 | -0.26 | -0.925 | 0.609 | -0.017 | -0.003 | -0.003 | 1.605 | 0.57 | 0.72 | 0.55 | 0.42 | 0.45 | 0.65 |
| 4 | 56997350 | rs998064 | -0.424 | -0.522 | -0.652 | 0.877 | -0.007 | 0.011 | 0.011 | 0 | 0.32 | 0.34 | 0.33 | 0.43 | 0.26 | 0.33 |
| 4 | 57001112 | rs11133423 | -0.105 | -0.659 | -0.88 | 0.862 | 0.004 | 0.088 | 0.088 | 0.327 | 0.22 | 0.31 | 0.4 | 0.5 | 0.22 | 0.32 |
| 4 | 57004959 | rs10866432 | -0.908 | -0.74 | -1.058 | 0.723 | 0.015 | -0.019 | -0.019 | 0 | 0.34 | 0.22 | 0.32 | 0.43 | 0.26 | 0.33 |
| 4 | 57008387 | rs13143526 | -1.14 | -0.474 | -0.912 | 0.452 | -0.001 | -0.002 | -0.002 | 0 | 0.38 | 0.44 | 0.43 | 0.36 | 0.38 | 0.35 |
| 4 | 57008939 | rs13144710 | -0.855 | -0.316 | -1.189 | 0.169 | 0 | 0.01 | 0.01 | 0 | 0.34 | 0.28 | 0.22 | 0.29 | 0.33 | 0.25 |
| 4 | 57009127 | rs13148796 | -0.04 | 0.197 | -1.127 | 0.466 | -0.009 | -0.004 | -0.004 | 0 | 0.21 | 0.15 | 0.17 | 0.22 | 0.26 | 0.2 |
| 4 | 57010328 | rs11133424 | -0.522 | 0.537 | -1.296 | -0.132 | -0.04 | 0.105 | 0.105 | 0.729 | 0.4 | 0.27 | 0.64 | 0.81 | 0.48 | 0.56 |
| 4 | 57016020 | rs13105877 | 0.645 | 0.384 | -0.532 | 0.072 | 0.011 | 0.024 | 0.024 | 0.33 | 0.46 | 0.62 | 0.36 | 0.28 | 0.38 | 0.28 |
| 4 | 57017493 | rs1479008 | 0.081 | 0.602 | -0.741 | 0.336 | -0.041 | -0.013 | -0.013 | 0.102 | 0.22 | 0.15 | 0.35 | 0.19 | 0.32 | 0.32 |
| 4 | 57026928 | rs11736698 | 0.697 | 0.607 | -0.087 | 0.494 | 0.01 | 0.026 | 0.026 | 0.072 | 0.44 | 0.39 | 0.33 | 0.14 | 0.38 | 0.28 |
| 4 | 57036306 | rs2276901 | 0.389 | 0.701 | -0.462 | 0.893 | 0 | 0.074 | 0.074 | 0 | 0.07 | 0.07 | 0.13 | 0.42 | 0.05 | 0.18 |
| 4 | 57038461 | rs11133427 | 0.589 | 0.765 | -0.447 | -0.132 | 0.015 | 0.046 | 0.046 | 0.048 | 0.44 | 0.33 | 0.39 | 0.11 | 0.38 | 0.29 |
| 4 | 57039609 | rs10213659 | -0.318 | 0.786 | -0.565 | -0.108 | 0 | 0 | 0 | 0 | 0.7 | 0.71 | 0.72 | 0.3 | 0.69 | 0.61 |
| 4 | 57039646 | rs10213095 | -0.498 | 1.056 | -0.495 | 0.473 | 0.007 | -0.002 | -0.002 | 0.772 | 0.26 | 0.3 | 0.33 | 0.19 | 0.31 | 0.32 |
| 4 | 57051461 | rs11133431 | -0.451 | 0.532 | -0.834 | 0.332 | 0.033 | 0.063 | 0.063 | 0.129 | 0.47 | 0.75 | 0.56 | 0.72 | 0.38 | 0.66 |
| 4 | 57053541 | rs6856875 | 0.12 | 0.076 | -0.201 | 0.325 | -0.006 | -0.041 | -0.041 | 0 | 0.35 | 0.42 | 0.36 | 0.12 | 0.33 | 0.36 |
| 4 | 57061470 | rs11936569 | -0.33 | 0.343 | -0.238 | 0.397 | -0.024 | -0.019 | -0.019 | 0 | 0.3 | 0.25 | 0.42 | 0.73 | 0.39 | 0.5 |
| 4 | 57061920 | rs17744943 | 0.17 | -0.005 | -0.504 | 0.372 | -0.089 | 0.029 | 0.029 | 0.276 | 0.25 | 0.02 | 0.22 | 0.14 | 0.35 | 0.21 |
| 4 | 57067425 | rs2045984 | -0.11 | -0.015 | -0.133 | 0.434 | 0.002 | 0.07 | 0.07 | 0 | 0.32 | 0.18 | 0.25 | 0.13 | 0.34 | 0.22 |
| 4 | 57071639 | rs1993916 | -0.462 | 0.185 | -0.077 | 0.195 | -0.001 | -0.001 | -0.001 | 0 | 0.4 | 0.58 | 0.41 | 0.73 | 0.41 | 0.51 |
| 4 | 57087608 | rs7692217 | -0.17 | -0.039 | -0.03 | -0.331 | -0.027 | -0.059 | -0.059 | 0 | 0.31 | 0.62 | 0.18 | 0.07 | 0.25 | 0.23 |
| 4 | 57090811 | rs962488 | -1.18 | 0.633 | -0.477 | -0.824 | -0.015 | 0.026 | 0.026 | 0 | 0.77 | 0.86 | 0.73 | 0.94 | 0.74 | 0.92 |
| 4 | 57092292 | rs5012040 | 0.174 | 0.741 | 0.288 | -0.335 | -0.006 | -0.018 | -0.018 | 0.758 | 0.39 | 0.48 | 0.29 | 0.24 | 0.39 | 0.38 |
| 4 | 57093826 | rs4865070 | -0.115 | 0.754 | 0.317 | 0.243 | 0.005 | 0.105 | 0.105 | 0 | 0.14 | 0.26 | 0.31 | 0.58 | 0.1 | 0.25 |
| 4 | 57095679 | rs10020065 | 0.933 | 0.748 | -0.137 | -0.045 | 0.014 | 0.019 | 0.019 | 0 | 0.27 | 0.16 | 0.2 | 0.18 | 0.29 | 0.23 |
| 4 | 57095806 | rs7685126 | -0.619 | 0.923 | -0.08 | 0.273 | -0.034 | -0.042 | -0.042 | 0.038 | 0.65 | 0.79 | 0.46 | 0.45 | 0.63 | 0.67 |
| 4 | 57096491 | rs140645 | 0.279 | 0.828 | -0.237 | 0.273 | 0.019 | 0.016 | 0.016 | 0.089 | 0.53 | 0.31 | 0.37 | 0.38 | 0.54 | 0.46 |
| 4 | 57096602 | rs140643 | 0.279 | 0.828 | -0.241 | 0.273 | 0.019 | 0.016 | 0.016 | 0.089 | 0.53 | 0.3 | 0.37 | 0.38 | 0.54 | 0.46 |
| 4 | 57097742 | rs7698073 | -0.518 | 0.936 | -0.136 | -0.072 | 0.001 | -0.019 | -0.019 | 0 | 0.65 | 0.39 | 0.64 | 0.87 | 0.64 | 0.69 |
| 4 | 57099609 | rs1600268 | 0.424 | 0.856 | 0.15 | 0.24 | 0.015 | 0.019 | 0.019 | 0 | 0.27 | 0.16 | 0.2 | 0.18 | 0.29 | 0.23 |
| 4 | 57102386 | rs1479006 | -0.105 | 1.065 | 0.503 | 0.233 | 0.071 | -0.021 | -0.021 | 15.224 | 0.87 | 0.5 | 0.91 | 0.91 | 0.9 | 0.77 |
| 4 | 57103047 | rs10023405 | 0.284 | 0.851 | 0.017 | 0.242 | 0.013 | 0.019 | 0.019 | 0 | 0.27 | 0.16 | 0.2 | 0.18 | 0.3 | 0.23 |
| 4 | 57108271 | rs1038115 | -0.493 | 0.862 | -0.25 | 0.086 | -0.042 | 0.071 | 0.071 | 0 | 0.27 | 0.15 | 0.27 | 0.11 | 0.36 | 0.25 |
| 4 | 57110592 | rs1993917 | -0.542 | 0.683 | 0.67 | 0.654 | 0.036 | 0 | 0 | 0.057 | 0.77 | 0.64 | 0.79 | 0.79 | 0.73 | 0.74 |
| 4 | 57112435 | rs11734492 | 1.118 | 0.706 | 0.556 | 0.495 | 0.005 | -0.006 | -0.006 | 0 | 0.61 | 0.43 | 0.67 | 0.65 | 0.6 | 0.5 |
| 4 | 57113130 | rs4865075 | -0.638 | 1.019 | 0.687 | 0.647 | 0.006 | -0.004 | -0.004 | 0 | 0.78 | 0.75 | 0.8 | 0.81 | 0.75 | 0.75 |
| 4 | 57116323 | rs7690043 | -0.039 | 1.213 | 0.382 | 1.061 | 0.004 | 0.02 | 0.02 | 0 | 0.2 | 0.23 | 0.46 | 0.62 | 0.26 | 0.24 |
| 4 | 57118192 | rs11942391 | -0.663 | 0.742 | 0.645 | 1.383 | 0.017 | 0.095 | 0.095 | 0.678 | 0.14 | 0.29 | 0.35 | 0.7 | 0.13 | 0.2 |
| 4 | 57126378 | rs598210 | -0.255 | 0.959 | 0.25 | 0.963 | -0.027 | 0 | 0 | 0 | 0.78 | 0.87 | 0.82 | 0.76 | 0.71 | 0.7 |
| 4 | 57127117 | rs534208 | -0.142 | 0.644 | 0.36 | 0.509 | 0.005 | -0.049 | -0.049 | 4.308 | 0.21 | 0.32 | 0.25 | 0.03 | 0.26 | 0.19 |
| 4 | 57127848 | rs6554340 | 1.103 | 0.715 | 0.273 | 0.636 | -0.003 | -0.004 | -0.004 | 4.367 | 0.57 | 0.47 | 0.57 | 0.65 | 0.43 | 0.51 |
| 4 | 57129103 | rs572969 | -0.263 | 0.943 | -0.325 | 0.192 | 0 | -0.011 | -0.011 | 0 | 0.19 | 0.21 | 0.22 | 0.03 | 0.26 | 0.19 |
| 4 | 57134800 | rs536002 | -0.396 | 1.178 | -0.424 | 0.723 | 0.039 | 0.059 | 0.059 | 0 | 0.79 | 0.68 | 0.71 | 0.57 | 0.7 | 0.72 |
| 4 | 57139331 | rs551289 | -0.579 | 0.79 | -0.96 | 0.533 | -0.008 | -0.052 | -0.052 | 0.023 | 0.84 | 0.94 | 0.9 | 0.76 | 0.89 | 0.75 |
| 4 | 57142047 | rs562803 | 0.59 | 0.407 | -0.915 | 0.55 | 0.061 | 0.048 | 0.048 | 0 | 0.51 | 0.66 | 0.33 | 0.39 | 0.42 | 0.34 |
| 4 | 57143037 | rs629242 | 1.013 | -0.167 | -0.836 | 0.099 | -0.017 | 0.024 | 0.024 | 0 | 0.24 | 0.32 | 0.17 | 0.36 | 0.21 | 0.22 |
| 4 | 57149650 | rs533178 | 0.156 | 0.565 | -1.343 | -0.14 | -0.057 | -0.013 | -0.013 | 0 | 0.71 | 0.93 | 0.63 | 0.76 | 0.7 | 0.6 |
| 4 | 57154517 | rs493228 | 0.698 | 0.848 | -0.294 | 0.388 | 0.044 | -0.047 | -0.047 | 0.014 | 0.65 | 0.52 | 0.61 | 0.19 | 0.58 | 0.53 |
| 4 | 57154790 | rs490591 | 0.735 | 0.802 | -0.053 | 0.536 | 0.017 | 0.129 | 0.129 | 1.565 | 0.9 | 0.66 | 0.92 | 0.76 | 0.94 | 0.9 |
| 4 | 57156379 | rs510411 | 0.502 | 0.877 | -0.446 | 0.449 | 0.052 | 0.048 | 0.048 | 0 | 0.71 | 0.63 | 0.65 | 0.41 | 0.61 | 0.62 |
| 4 | 57162619 | rs7663149 | -0.011 | 0.092 | -0.825 | -0.079 | -0.016 | -0.008 | -0.008 | 0 | 0.27 | 0.34 | 0.38 | 0.73 | 0.28 | 0.34 |
| 4 | 57165323 | rs4337783 | -0.758 | 0.698 | -1.928 | -0.55 | -0.135 | 0.145 | 0.145 | 0 | 0.69 | 0.91 | 0.62 | 0.89 | 0.56 | 0.69 |
| 4 | 57167280 | rs13124016 | -0.031 | 0.401 | -0.758 | 0.23 | -0.003 | -0.039 | -0.039 | 0 | 0.13 | 0.24 | 0.08 | 0.03 | 0.09 | 0.08 |
| 4 | 57172978 | rs1401910 | -0.82 | 0.553 | -1.521 | 0.953 | 0.007 | -0.052 | -0.052 | 0.307 | 0.29 | 0.15 | 0.18 | 0.42 | 0.29 | 0.17 |
| 4 | 57173047 | rs7696217 | -0.793 | 0.662 | -1.24 | 1.083 | -0.001 | -0.012 | -0.012 | 0.544 | 0.13 | 0.11 | 0.12 | 0.28 | 0.16 | 0.11 |
| 4 | 57181995 | rs3796547 | -0.09 | 0.684 | -1.031 | 0.753 | 0.004 | 0.056 | 0.056 | 0.01 | 0.37 | 0.2 | 0.23 | 0.25 | 0.34 | 0.19 |
| 4 | 57182647 | rs3822003 | 0.137 | 0.579 | -1.186 | 0.689 | -0.086 | 0.004 | 0.004 | 0.07 | 0.33 | 0.02 | 0.21 | 0.25 | 0.34 | 0.19 |
| 4 | 57187665 | rs1401904 | 0.332 | 0.809 | -0.075 | 1.099 | -0.001 | -0.004 | -0.004 | 0 | 0.52 | 0.5 | 0.67 | 0.28 | 0.55 | 0.56 |
| 4 | 57188538 | rs6854916 | -0.745 | 0.689 | 1.087 | 1.227 | 0.052 | 0.355 | 0.355 | 0.556 | 0.11 | 0.42 | 0.13 | 0.44 | 0.13 | 0.24 |
| 4 | 57190242 | rs2292983 | -0.426 | 0.73 | 1.103 | 1.244 | 0 | 0.203 | 0.203 | 0.164 | 0.09 | 0.23 | 0.1 | 0.43 | 0.1 | 0.19 |
| 4 | 57190356 | rs7695701 | 0.85 | 0.82 | 0.847 | 0.97 | -0.09 | 0.208 | 0.208 | 0 | 0.39 | 0.1 | 0.27 | 0.07 | 0.41 | 0.34 |

| | | | | | | | | | | | | | | | | |
|---|---|---|---|---|---|---|---|---|---|---|---|---|---|---|---|---|
| 4 | 57193762 | rs6826795 | -0.816 | 0.794 | 1.017 | 0.99 | 0.006 | 0.017 | 0.017 | 0.312 | 0.62 | 0.7 | 0.77 | 0.72 | 0.67 | 0.81 |
| 4 | 57196155 | rs939293 | -0.061 | 0.792 | 1.078 | 0.984 | -0.001 | -0.009 | -0.009 | 0 | 0.53 | 0.49 | 0.67 | 0.29 | 0.55 | 0.57 |
| 4 | 57204777 | rs8340 | -0.043 | 0.724 | 1.048 | 0.972 | -0.007 | 0.037 | 0.037 | 0 | 0.13 | 0.22 | 0.38 | 0.22 | 0.13 | 0.23 |
| 4 | 57204960 | rs3733324 | 0.644 | 0.705 | 1.168 | 1.363 | -0.001 | 0.033 | 0.033 | 0 | 0.06 | 0.05 | 0.05 | 0.24 | 0.09 | 0.1 |
| 4 | 57214366 | rs13124696 | 0.358 | 0.702 | 1.237 | 1.362 | -0.004 | 0.014 | 0.014 | 0 | 0.46 | 0.38 | 0.35 | 0.31 | 0.5 | 0.43 |
| 4 | 57215677 | rs3796544 | 0.738 | 0.703 | 1.227 | 1.359 | -0.015 | 0.055 | 0.055 | 0 | 0.4 | 0.28 | 0.29 | 0.07 | 0.42 | 0.33 |
| 4 | 57219613 | rs17086696 | -0.015 | 0.7 | 1.235 | 1.381 | -0.006 | 0.04 | 0.04 | 0 | 0.13 | 0.22 | 0.38 | 0.22 | 0.13 | 0.23 |
| 4 | 57224519 | rs13102012 | 0.234 | 0.73 | 1.167 | 1.243 | -0.001 | 0 | 0 | 0 | 0.46 | 0.46 | 0.36 | 0.31 | 0.5 | 0.43 |
| 4 | 57237683 | rs6554354 | 0.648 | 0.743 | 0.984 | 1.082 | -0.015 | 0.055 | 0.055 | 0 | 0.4 | 0.28 | 0.29 | 0.07 | 0.42 | 0.33 |
| 4 | 57246019 | rs13150967 | 0.648 | 0.743 | 0.984 | 1.082 | -0.015 | 0.055 | 0.055 | 0 | 0.4 | 0.28 | 0.29 | 0.07 | 0.42 | 0.33 |
| 4 | 57256201 | rs7671868 | -0.287 | 0.7 | 1.027 | 1.192 | 0.004 | 0 | 0 | 1.295 | 0.54 | 0.6 | 0.71 | 0.48 | 0.57 | 0.65 |
| 4 | 57258537 | rs4493603 | -0.52 | 0.716 | 1.119 | 1.166 | 0.001 | 0.077 | 0.077 | 0 | 0.15 | 0.24 | 0.42 | 0.41 | 0.17 | 0.32 |
| 4 | 57259465 | rs1060472 | -0.09 | 0.709 | 1.193 | 1.218 | 0.006 | 0.027 | 0.027 | 0 | 0.13 | 0.2 | 0.39 | 0.22 | 0.16 | 0.24 |
| 4 | 57285163 | rs1401908 | 0.713 | 0.684 | 1.317 | 1.177 | -0.019 | 0.053 | 0.053 | 0 | 0.4 | 0.25 | 0.3 | 0.26 | 0.42 | 0.41 |
| 4 | 57297996 | rs13147824 | -0.709 | 0.726 | 1.339 | 1.289 | 0.008 | 0.015 | 0.015 | 0.155 | 0.61 | 0.68 | 0.77 | 0.72 | 0.67 | 0.8 |
| 4 | 57300557 | rs1520026 | -1.264 | 1.518 | 1.994 | 1.829 | 0 | 0.074 | 0.074 | 0 | 0.15 | 0.22 | 0.41 | 0.41 | 0.17 | 0.37 |
| 4 | 57305202 | rs1356787 | -1.867 | 1.49 | 2.143 | 1.863 | 0.011 | 0.176 | 0.176 | 0.129 | 0.22 | 0.4 | 0.46 | 0.48 | 0.25 | 0.4 |
| 4 | 57305699 | rs1967559 | -0.689 | 1.486 | 2.316 | 1.86 | 0.002 | 0.011 | 0.011 | 0 | 0.13 | 0.1 | 0.38 | 0.23 | 0.16 | 0.29 |
| 4 | 57327405 | rs6584 | -1.889 | 1.486 | 2.312 | 1.857 | 0.011 | 0.176 | 0.176 | 0.129 | 0.22 | 0.4 | 0.46 | 0.48 | 0.25 | 0.4 |
| 4 | 57341262 | rs4270628 | -1.601 | 1.938 | 2.264 | 1.628 | -0.002 | 0.001 | 0.001 | 1.918 | 0.41 | 0.54 | 0.61 | 0.69 | 0.53 | 0.5 |
| 4 | 57345195 | rs17086832 | -1.639 | 1.818 | 2.599 | 2.213 | 0.026 | 0.018 | 0.018 | 0.378 | 0.2 | 0.33 | 0.43 | 0.23 | 0.22 | 0.3 |
| 4 | 57353767 | rs12508008 | -1.354 | 1.826 | 2.558 | 2.21 | -0.004 | -0.002 | -0.002 | 0 | 0.18 | 0.12 | 0.41 | 0.23 | 0.22 | 0.25 |
| 4 | 57361627 | rs11934479 | -0.948 | 1.819 | 2.515 | 2.212 | 0.016 | 0.164 | 0.164 | 0 | 0.09 | 0.26 | 0.07 | 0.25 | 0.1 | 0.12 |
| 4 | 57368413 | rs4260596 | -0.3 | 1.911 | 1.701 | 2.041 | -0.005 | 0.004 | 0.004 | 0 | 0.1 | 0.05 | 0.14 | 0.07 | 0.09 | 0.08 |
| 4 | 57368759 | rs14211 | -1.383 | 1.917 | 1.669 | 2.048 | -0.004 | -0.002 | -0.002 | 0.119 | 0.18 | 0.12 | 0.41 | 0.23 | 0.22 | 0.25 |
| 4 | 57385455 | rs4131086 | -2.323 | 1.506 | 1.084 | 1.983 | -0.004 | 0.022 | 0.022 | 0.716 | 0.31 | 0.49 | 0.52 | 0.64 | 0.32 | 0.48 |
| 4 | 57387706 | rs4391105 | -1.213 | 0.956 | 1.031 | 2.203 | -0.008 | 0.031 | 0.031 | 0 | 0.14 | 0.26 | 0.18 | 0.25 | 0.09 | 0.19 |
| 4 | 57390650 | rs17086939 | -0.224 | 1.049 | 0.83 | 2.123 | -0.004 | 0.008 | 0.008 | 0 | 0.1 | 0.08 | 0.15 | 0.06 | 0.07 | 0.06 |
| 4 | 57397910 | rs4865115 | -1.485 | 1.097 | 0.831 | 2.123 | -0.003 | 0.051 | 0.051 | 0 | 0.2 | 0.26 | 0.25 | 0.49 | 0.19 | 0.25 |
| 4 | 57398752 | rs4280802 | -0.548 | 1.173 | 0.451 | 1.653 | 0.02 | 0.014 | 0.014 | 0 | 0.13 | 0.29 | 0.35 | 0.16 | 0.16 | 0.21 |
| 4 | 57410548 | rs4473708 | -0.648 | 0.716 | -0.104 | 1.569 | -0.001 | 0.026 | 0.026 | 0 | 0.1 | 0.14 | 0.08 | 0.18 | 0.13 | 0.13 |
| 4 | 57418400 | rs7691920 | 0.771 | 0.712 | -0.301 | 1.422 | -0.004 | 0.022 | 0.022 | 0.583 | 0.68 | 0.59 | 0.68 | 0.58 | 0.69 | 0.66 |
| 4 | 57426688 | rs7686452 | 0.884 | 0.447 | -0.526 | 1.54 | -0.002 | 0.016 | 0.016 | 0 | 0.95 | 0.93 | 0.96 | 0.83 | 0.96 | 0.93 |
| 4 | 57434245 | rs10213489 | -0.889 | -0.155 | -0.26 | 1.824 | -0.022 | 0.045 | 0.045 | 0 | 0.25 | 0.33 | 0.18 | 0.44 | 0.18 | 0.19 |
| 4 | 57442004 | rs4600976 | -1.289 | -0.463 | 0.704 | 1.792 | -0.022 | 0.03 | 0.03 | 0 | 0.29 | 0.67 | 0.21 | 0.48 | 0.18 | 0.3 |
| 4 | 57444413 | rs9996055 | -0.841 | -0.58 | 1.037 | 1.961 | -0.03 | 0.023 | 0.023 | 0 | 0.17 | 0.37 | 0.12 | 0.2 | 0.1 | 0.09 |
| 4 | 57450848 | rs4345238 | -0.777 | -0.491 | 0.659 | 2.027 | -0.011 | 0.046 | 0.046 | 0 | 0.11 | 0.15 | 0.06 | 0.24 | 0.08 | 0.18 |
| 4 | 57457048 | rs10006458 | -0.024 | 0.133 | -0.068 | 2.413 | -0.04 | -0.044 | -0.044 | 0 | 0.92 | 0.99 | 0.92 | 0.73 | 0.87 | 0.96 |
| 4 | 57457918 | rs7670824 | 0.346 | -0.415 | -0.666 | 1.662 | -0.005 | -0.077 | -0.077 | 0 | 0.72 | 0.93 | 0.75 | 0.61 | 0.7 | 0.7 |
| 4 | 57459490 | rs4864589 | -0.043 | -0.075 | -0.987 | 1.348 | -0.03 | -0.026 | -0.026 | 0 | 0.4 | 0.76 | 0.57 | 0.48 | 0.49 | 0.59 |
| 4 | 57477794 | rs7699631 | -0.836 | -0.263 | -0.837 | 1.193 | 0.019 | 0.004 | 0.004 | 0 | 0.44 | 0.63 | 0.44 | 0.63 | 0.38 | 0.38 |
| 4 | 57498446 | rs4865131 | -0.864 | -0.309 | -0.952 | 1.223 | 0.041 | -0.04 | -0.04 | 1.99 | 0.4 | 0.78 | 0.42 | 0.56 | 0.35 | 0.42 |
| 4 | 57503077 | rs7667406 | -0.543 | 0.091 | 0.172 | 1.327 | 0.024 | 0.131 | 0.131 | 0.005 | 0.12 | 0.39 | 0.12 | 0.21 | 0.12 | 0.09 |
| 4 | 57513912 | rs12649154 | -0.564 | 0.861 | 0.181 | 1.299 | 0.004 | 0.01 | 0.01 | 0 | 0.08 | 0.1 | 0.09 | 0.13 | 0.11 | 0.05 |
| 4 | 57516896 | rs4371677 | -0.549 | 0.572 | 0.057 | 0.953 | 0.015 | 0.017 | 0.017 | 2.156 | 0.51 | 0.67 | 0.36 | 0.47 | 0.48 | 0.35 |
| 4 | 57521325 | rs4406078 | -0.805 | 0.496 | 0.201 | 0.873 | 0.008 | 0.017 | 0.017 | 2.373 | 0.53 | 0.77 | 0.37 | 0.47 | 0.5 | 0.35 |
| 4 | 57525831 | rs4864593 | -0.32 | 0.056 | 0.87 | 0.704 | 0.003 | 0.145 | 0.145 | 0 | 0.11 | 0.4 | 0.09 | 0.26 | 0.08 | 0.05 |
| 4 | 57526349 | rs6817875 | -0.225 | 0.058 | 0.549 | 0.583 | 0.011 | 0.067 | 0.067 | 0.526 | 0.4 | 0.3 | 0.25 | 0.15 | 0.41 | 0.18 |
| 4 | 57526500 | rs6823211 | 0.754 | -0.089 | 0.24 | 0.412 | -0.001 | 0.167 | 0.167 | 0 | 0.3 | 0.06 | 0.14 | 0.02 | 0.3 | 0.14 |
| 4 | 57527444 | rs6824757 | -0.702 | -0.118 | 0.804 | 0.54 | -0.01 | 0.129 | 0.129 | 0.055 | 0.13 | 0.35 | 0.12 | 0.33 | 0.1 | 0.17 |
| 4 | 57527586 | rs6830271 | 0.408 | -0.556 | -0.378 | -0.017 | 0.023 | 0.118 | 0.118 | 0 | 0.29 | 0.09 | 0.11 | 0.01 | 0.27 | 0.15 |
| 4 | 57528443 | rs6836932 | -0.104 | -0.207 | -0.921 | 0.423 | -0.088 | -0.063 | -0.063 | 0 | 0.58 | 0.87 | 0.39 | 0.46 | 0.52 | 0.39 |
| 4 | 57529014 | rs4864594 | -1.068 | -0.462 | -1.271 | 0.668 | -0.043 | -0.088 | -0.088 | 0 | 0.76 | 0.98 | 0.74 | 0.77 | 0.76 | 0.63 |
| 4 | 57549528 | rs12642085 | 1.088 | -0.455 | -0.724 | 0.73 | -0.018 | -0.028 | -0.028 | 9.357 | 0.41 | 0.26 | 0.37 | 0.41 | 0.48 | 0.28 |
| 4 | 57549583 | rs4865142 | 0.199 | -0.627 | -0.296 | 0.73 | 0.027 | 0.007 | 0.007 | 0 | 0.56 | 0.77 | 0.61 | 0.57 | 0.59 | 0.41 |
| 4 | 57549603 | rs6829867 | 0.276 | -0.833 | -0.615 | 0.795 | 0.004 | 0.003 | 0.003 | 0 | 0.57 | 0.53 | 0.44 | 0.68 | 0.58 | 0.61 |
| 4 | 57552810 | rs899628 | 1.349 | -0.657 | -0.942 | 0.886 | -0.021 | -0.028 | -0.028 | 0 | 0.41 | 0.23 | 0.36 | 0.42 | 0.48 | 0.28 |
| 4 | 57555103 | rs6828831 | 1.041 | -1.098 | -0.727 | 0.924 | 0.025 | -0.005 | -0.005 | 1.222 | 0.44 | 0.31 | 0.39 | 0.57 | 0.57 | 0.42 |
| 4 | 57558426 | rs9968381 | -0.683 | -0.887 | -0.372 | 0.944 | -0.025 | 0.081 | 0.081 | 0 | 0.15 | 0.44 | 0.24 | 0.23 | 0.09 | 0.15 |
| 4 | 57564095 | rs6845520 | -0.63 | -1.057 | -0.459 | 0.87 | 0.026 | 0.049 | 0.049 | 0 | 0.67 | 0.87 | 0.67 | 0.9 | 0.7 | 0.73 |
| 4 | 57564293 | rs11729059 | 0.461 | -0.733 | -0.737 | 0.915 | 0.051 | 0.042 | 0.042 | 0.197 | 0.52 | 0.38 | 0.43 | 0.67 | 0.61 | 0.58 |
| 4 | 57573446 | rs12644752 | -0.58 | -0.033 | -0.91 | 0.977 | -0.04 | -0.039 | -0.039 | 0 | 0.09 | 0.01 | 0.19 | 0.21 | 0.12 | 0.21 |
| 4 | 57575299 | rs6816355 | -0.043 | -0.396 | -0.397 | 0.638 | 0.043 | 0.061 | 0.061 | 0 | 0.68 | 0.8 | 0.85 | 0.92 | 0.79 | 0.72 |
| 4 | 57586154 | rs11725915 | 0.621 | -0.467 | -0.302 | 1.001 | 0.016 | -0.009 | -0.009 | 3.385 | 0.59 | 0.66 | 0.67 | 0.41 | 0.71 | 0.43 |
| 4 | 57587795 | rs973031 | -0.267 | -0.316 | -0.317 | 1.091 | 0.022 | 0.267 | 0.267 | 0.107 | 0.08 | 0.22 | 0.18 | 0.51 | 0.08 | 0.27 |
| 4 | 57590071 | rs1383028 | -0.319 | -0.379 | -0.69 | 0.752 | 0.046 | 0.143 | 0.143 | 0 | 0.7 | 0.92 | 0.88 | 0.96 | 0.79 | 0.74 |
| 4 | 57597210 | rs4864599 | 0.166 | -0.268 | -0.357 | 0.98 | -0.029 | 0.011 | 0.011 | 0 | 0.59 | 0.52 | 0.64 | 0.4 | 0.69 | 0.45 |
| 4 | 57606444 | rs4865148 | -0.703 | 0.329 | -0.158 | 0.937 | 0.027 | 0.029 | 0.029 | 0 | 0.23 | 0.36 | 0.4 | 0.31 | 0.34 | 0.24 |
| 4 | 57608969 | rs1842295 | -0.831 | -0.035 | -0.775 | 0.376 | -0.027 | 0.015 | 0.015 | 0 | 0.69 | 0.78 | 0.67 | 0.75 | 0.61 | 0.73 |

| | | | | | | | | | | | | | | | | |
|---|---|---|---|---|---|---|---|---|---|---|---|---|---|---|---|---|
| 4 | 57614160 | rs10222715 | -1.226 | -0.483 | -0.305 | 0.404 | 0.007 | 0 | 0 | 0 | 0.42 | 0.39 | 0.48 | 0.36 | 0.4 | 0.39 |
| 4 | 57621843 | rs6830854 | -0.973 | -0.473 | -0.023 | 0.483 | 0 | 0.077 | 0.077 | 0 | 0.16 | 0.36 | 0.21 | 0.22 | 0.16 | 0.11 |
| 4 | 57624037 | rs6818605 | -1.387 | -0.55 | -0.274 | 0.371 | 0.021 | 0.011 | 0.011 | 0 | 0.35 | 0.71 | 0.31 | 0.23 | 0.25 | 0.32 |
| 4 | 57629710 | rs6811987 | -1.051 | -0.012 | -0.36 | 0.738 | -0.025 | 0.022 | 0.022 | 0 | 0.17 | 0.32 | 0.12 | 0.16 | 0.15 | 0.19 |
| 4 | 57637951 | rs10033814 | -0.438 | 0.546 | -1.282 | 0.608 | -0.009 | -0.001 | -0.001 | 0 | 0.32 | 0.2 | 0.34 | 0.26 | 0.34 | 0.47 |
| 4 | 57648046 | rs9992616 | -0.815 | 0.751 | -2.427 | -0.149 | -0.117 | 0.148 | 0.148 | 0.019 | 0.69 | 0.98 | 0.61 | 0.94 | 0.61 | 0.88 |
| 4 | 57649159 | rs2412757 | -0.125 | 0.185 | -2.222 | 0.19 | 0.036 | -0.008 | -0.008 | 1.887 | 0.41 | 0.81 | 0.36 | 0.38 | 0.33 | 0.4 |
| 4 | 57652380 | rs2008139 | -1.073 | -0.266 | -1.412 | 0.514 | -0.035 | 0.082 | 0.082 | 0 | 0.17 | 0.33 | 0.09 | 0.36 | 0.1 | 0.39 |
| 4 | 57674397 | rs13146442 | 0.841 | -0.305 | -1.799 | 0.165 | -0.005 | 0.016 | 0.016 | 0.057 | 0.1 | 0.04 | 0.04 | 0.03 | 0.09 | 0.09 |
| 4 | 57678832 | rs17087176 | -0.278 | -0.16 | -1.82 | 0.153 | 0.003 | 0.003 | 0.003 | 0 | 0.09 | 0.06 | 0.04 | 0.26 | 0.06 | 0.37 |
| 4 | 57680805 | rs13102386 | -0.99 | 0.037 | -1.485 | -0.188 | 0.006 | 0.056 | 0.056 | 0.062 | 0.58 | 0.79 | 0.71 | 0.72 | 0.57 | 0.51 |
| 4 | 57686657 | rs812991 | -0.234 | -0.011 | -0.633 | -0.12 | -0.009 | -0.052 | -0.052 | 0 | 0.09 | 0.21 | 0.05 | 0 | 0.05 | 0.02 |
| 4 | 57687304 | rs781542 | -1.394 | -0.297 | -1.195 | -0.271 | 0.053 | 0.039 | 0.039 | 0.106 | 0.5 | 0.76 | 0.5 | 0.39 | 0.36 | 0.57 |
| 4 | 57690713 | rs1708768 | -0.735 | -0.263 | -0.45 | -0.074 | -0.013 | 0.017 | 0.017 | 1.185 | 0.13 | 0.55 | 0.24 | 0.09 | 0.06 | 0.04 |
| 4 | 57694352 | rs1708772 | -1.215 | -0.395 | -1.272 | -0.234 | 0.064 | -0.011 | -0.011 | 1.034 | 0.43 | 0.75 | 0.41 | 0.41 | 0.28 | 0.55 |
| 4 | 57695126 | rs1399002 | -0.008 | 0.103 | -1.179 | 0.015 | 0.006 | 0.004 | 0.004 | 0 | 0.51 | 0.61 | 0.39 | 0.49 | 0.42 | 0.42 |
| 4 | 57696918 | rs2127495 | 1.333 | 0.519 | -1.439 | -0.132 | -0.009 | 0.025 | 0.025 | 0 | 0.17 | 0.14 | 0.19 | 0.08 | 0.2 | 0.07 |
| 4 | 57700528 | rs7668497 | 0.304 | 0.707 | -1.364 | 0.172 | 0.04 | 0.042 | 0.042 | 0 | 0.53 | 0.68 | 0.32 | 0.4 | 0.42 | 0.57 |
| 4 | 57705970 | rs781571 | -0.233 | 1.16 | -1.225 | 0.454 | -0.025 | -0.015 | -0.015 | 0 | 0.66 | 0.69 | 0.38 | 0.42 | 0.54 | 0.61 |
| 4 | 57705999 | rs781570 | -1.083 | 1.728 | -1.354 | 1.016 | -0.027 | -0.036 | -0.036 | 0 | 0.92 | 0.98 | 0.82 | 0.79 | 0.86 | 0.84 |
| 4 | 57707469 | rs781565 | -0.099 | 1.732 | -1.316 | 1.028 | -0.03 | -0.041 | -0.041 | 0.017 | 0.67 | 0.75 | 0.61 | 0.52 | 0.58 | 0.63 |
| 4 | 57708245 | rs781564 | -0.234 | 1.672 | -0.433 | 0.791 | 0.01 | 0.013 | 0.013 | 0 | 0.32 | 0.51 | 0.3 | 0.35 | 0.35 | 0.49 |
| 4 | 57713324 | rs751356 | 1.436 | 1.601 | 0.007 | 0.247 | 0.013 | -0.07 | -0.07 | 0 | 0.19 | 0.29 | 0.14 | 0.01 | 0.21 | 0.17 |
| 4 | 57726274 | rs1708774 | 0.849 | 1.377 | 0.101 | 0.583 | 0.008 | 0.028 | 0.028 | 0 | 0.15 | 0.24 | 0.33 | 0.24 | 0.18 | 0.09 |
| 4 | 57727311 | rs7684253 | 0.169 | 1.041 | -0.063 | 0.288 | 0.005 | -0.004 | -0.004 | 5.08 | 0.6 | 0.52 | 0.55 | 0.32 | 0.55 | 0.31 |
| 4 | 57730683 | rs796681 | 0.95 | 1.032 | -0.097 | 0.188 | 0.016 | -0.003 | -0.003 | 0 | 0.09 | 0.12 | 0.06 | 0.06 | 0.1 | 0.04 |
| 4 | 57731446 | rs9637714 | -0.694 | 0.603 | -0.267 | -0.15 | -0.006 | 0.012 | 0.012 | 5.182 | 0.29 | 0.3 | 0.31 | 0.66 | 0.27 | 0.57 |
| 4 | 57733604 | rs12650282 | -0.634 | 0.614 | -0.422 | -0.144 | 0.011 | -0.034 | -0.034 | 0 | 0.29 | 0.2 | 0.31 | 0.66 | 0.27 | 0.57 |
| 4 | 57738469 | rs11730321 | 1.988 | 0.625 | -0.248 | -0.085 | -0.009 | -0.002 | -0.002 | 0 | 0.3 | 0.32 | 0.38 | 0.22 | 0.25 | 0.15 |
| 4 | 57741415 | rs4560473 | -0.156 | 0.635 | -0.645 | 0.137 | 0 | -0.036 | -0.036 | 0.335 | 0.28 | 0.19 | 0.43 | 0.54 | 0.29 | 0.43 |
| 4 | 57746825 | rs2170073 | 0.382 | 0.797 | -0.514 | 0.5 | -0.007 | 0.011 | 0.011 | 0 | 0.57 | 0.62 | 0.57 | 0.55 | 0.47 | 0.54 |
| 4 | 57773330 | rs3806746 | -0.859 | 0.909 | -0.75 | 0.462 | -0.037 | -0.021 | -0.021 | 2.917 | 0.71 | 0.8 | 0.74 | 0.67 | 0.65 | 0.53 |
| 4 | 57786450 | rs1713985 | -1.082 | 1 | -0.863 | 0.435 | -0.037 | -0.067 | -0.067 | 0 | 0.9 | 0.99 | 0.89 | 0.69 | 0.91 | 0.73 |
| 4 | 57791864 | rs6847086 | 1.156 | 1.129 | -0.196 | 0.555 | 0.046 | 0.024 | 0.024 | 0 | 0.24 | 0.45 | 0.45 | 0.24 | 0.36 | 0.26 |
| 4 | 57797414 | rs3796529 | -0.205 | 0.738 | -0.257 | 0.324 | -0.002 | 0.048 | 0.048 | 1.001 | 0.18 | 0.27 | 0.22 | 0.38 | 0.18 | 0.23 |
| 4 | 57821309 | rs708547 | 1.986 | 0.971 | -0.437 | 0.261 | -0.017 | 0.02 | 0.02 | 0 | 0.18 | 0.16 | 0.16 | 0.02 | 0.25 | 0.21 |
| 4 | 57825190 | rs7668814 | -0.134 | 0.692 | -0.456 | 0.251 | -0.013 | -0.001 | -0.001 | 0 | 0.59 | 0.58 | 0.44 | 0.47 | 0.56 | 0.44 |
| 4 | 57827206 | rs1718839 | 0.921 | 0.755 | -0.434 | 0.259 | 0.002 | -0.003 | -0.003 | 1.039 | 0.31 | 0.41 | 0.46 | 0.22 | 0.35 | 0.28 |
| 4 | 57833203 | rs11932673 | 1.709 | 0.594 | -0.29 | 0.383 | -0.005 | 0.027 | 0.027 | 0 | 0.12 | 0.2 | 0.3 | 0.2 | 0.1 | 0.08 |
| 4 | 57838583 | rs17087335 | 0.147 | 0.515 | -0.202 | 0.31 | 0.002 | 0.032 | 0.032 | 0.366 | 0.16 | 0.21 | 0.22 | 0.38 | 0.19 | 0.21 |
| 4 | 57838690 | rs1107674 | 1.368 | 0.352 | -0.274 | 0.199 | 0.012 | 0.005 | 0.005 | 0 | 0.43 | 0.3 | 0.25 | 0.38 | 0.37 | 0.48 |
| 4 | 57838707 | rs1107675 | 1.705 | 0.545 | -0.415 | 0.271 | -0.003 | 0.028 | 0.028 | 0 | 0.13 | 0.21 | 0.3 | 0.2 | 0.1 | 0.08 |
| 4 | 57844902 | rs1713998 | 1.422 | 0.589 | -0.749 | -0.185 | -0.032 | 0.061 | 0.061 | 0.051 | 0.27 | 0.19 | 0.2 | 0.02 | 0.34 | 0.27 |
| 4 | 57845269 | rs13133792 | 2.278 | 0.613 | -0.711 | -0.126 | -0.001 | 0.054 | 0.054 | 0 | 0.11 | 0.04 | 0.05 | 0 | 0.07 | 0.05 |
| 4 | 57845740 | rs1625691 | 1.983 | 0.693 | -0.571 | 0.091 | 0.001 | 0.001 | 0.001 | 0 | 0.2 | 0.26 | 0.41 | 0.22 | 0.3 | 0.22 |
| 4 | 57856092 | rs1277281 | -1.46 | 0.864 | -0.635 | -0.105 | 0.006 | -0.009 | -0.009 | 0 | 0.92 | 0.91 | 0.95 | 0.98 | 0.9 | 0.96 |
| 4 | 57871203 | rs1718837 | 1.751 | 0.788 | -0.481 | 0.035 | -0.002 | -0.002 | -0.002 | 0 | 0.23 | 0.25 | 0.23 | 0.3 | 0.28 | 0.47 |
| 4 | 57872421 | rs7661194 | 1.696 | 0.588 | -0.371 | -0.125 | 0.001 | 0.001 | 0.001 | 0 | 0.37 | 0.29 | 0.4 | 0.23 | 0.29 | 0.22 |
| 4 | 57876955 | rs1718878 | 0.888 | 0.576 | -0.301 | -0.174 | -0.001 | -0.006 | -0.006 | 0 | 0.36 | 0.44 | 0.3 | 0.3 | 0.35 | 0.52 |
| 4 | 57881715 | rs1713982 | 0.888 | 0.576 | -0.301 | -0.174 | -0.001 | -0.006 | -0.006 | 0 | 0.36 | 0.43 | 0.3 | 0.3 | 0.35 | 0.52 |
| 4 | 57889288 | rs7440152 | 1.969 | 0.577 | -0.298 | -0.158 | -0.009 | -0.051 | -0.051 | 0 | 0.13 | 0.18 | 0.06 | 0 | 0.07 | 0.04 |
| 4 | 57889677 | rs1056364 | -0.488 | 0.609 | -0.377 | -0.052 | -0.001 | -0.013 | -0.013 | 0 | 0.56 | 0.46 | 0.64 | 0.68 | 0.58 | 0.44 |
| 4 | 57892643 | rs4865171 | -0.408 | 0.615 | -0.414 | -0.029 | -0.004 | 0.001 | 0.001 | 0 | 0.88 | 0.91 | 0.92 | 0.98 | 0.87 | 0.95 |
| 4 | 57897864 | rs1277308 | 1.195 | 0.59 | -0.359 | -0.019 | -0.001 | 0.006 | 0.006 | 0.019 | 0.35 | 0.38 | 0.3 | 0.3 | 0.35 | 0.52 |
| 4 | 57898230 | rs11573128 | 3.255 | -0.408 | -0.779 | -0.396 | -0.027 | 0.115 | 0.115 | 1.347 | 0.19 | 0.01 | 0.1 | 0 | 0.17 | 0.08 |
| 4 | 57899602 | rs2271808 | 1.298 | -0.045 | -0.327 | 0.015 | -0.003 | -0.003 | -0.003 | 0 | 0.4 | 0.39 | 0.44 | 0.3 | 0.38 | 0.24 |
| 4 | 57902276 | rs12642622 | 0.317 | 0.037 | -0.592 | -0.182 | -0.013 | -0.066 | -0.066 | 0.464 | 0.12 | 0.05 | 0.13 | 0.36 | 0.15 | 0.21 |
| 4 | 57902761 | rs1277311 | 0.746 | -0.038 | -0.305 | -0.22 | 0.006 | 0.004 | 0.004 | 0 | 0.4 | 0.57 | 0.61 | 0.52 | 0.43 | 0.54 |
| 4 | 57904852 | rs11573119 | 0.307 | -0.044 | -0.281 | 0.328 | -0.012 | 0.016 | 0.016 | 0 | 0.07 | 0.1 | 0.04 | 0.15 | 0.04 | 0.11 |
| 4 | 57905453 | rs1718885 | -0.573 | 0.192 | -0.394 | 0.104 | 0.001 | -0.008 | -0.008 | 0 | 0.92 | 0.91 | 0.96 | 0.97 | 0.93 | 0.97 |
| 4 | 57906335 | rs7687211 | -0.043 | 0.153 | -0.426 | 0.074 | 0 | 0.002 | 0.002 | 0 | 0.88 | 0.91 | 0.93 | 0.97 | 0.9 | 0.96 |
| 4 | 57909291 | rs1277273 | 0.93 | -0.115 | -0.473 | 0.227 | 0.004 | 0.008 | 0.008 | 0 | 0.68 | 0.24 | 0.72 | 0.68 | 0.67 | 0.78 |
| 4 | 57910214 | rs6852762 | -0.543 | 0.061 | 0.131 | 0.554 | 0.043 | 0.114 | 0.114 | 0 | 0.12 | 0.36 | 0.16 | 0.28 | 0.19 | 0.13 |
| 4 | 57915267 | rs3821996 | -0.722 | -0.167 | -0.196 | 0.547 | 0.001 | -0.001 | -0.001 | 4.964 | 0.32 | 0.35 | 0.27 | 0.48 | 0.32 | 0.33 |
| 4 | 57917346 | rs881382 | 0.695 | -0.24 | 0.016 | 0.347 | 0.001 | -0.004 | -0.004 | 0 | 0.27 | 0.35 | 0.38 | 0.21 | 0.29 | 0.18 |
| 4 | 57918245 | rs1713973 | 0.199 | -0.254 | 0.016 | 0.072 | -0.015 | 0.015 | 0.015 | 0 | 0.89 | 0.82 | 0.92 | 0.83 | 0.92 | 0.94 |
| 4 | 57918904 | rs1718834 | 1.334 | -0.346 | 0.312 | 0.061 | 0.111 | 0.076 | 0.076 | 0 | 0.94 | 0.69 | 0.9 | 0.94 | 0.86 | 0.92 |
| 4 | 57919919 | rs1401189 | -0.12 | -0.497 | 0.26 | 0.043 | -0.021 | 0.02 | 0.02 | 0 | 0.23 | 0.33 | 0.16 | 0.31 | 0.19 | 0.19 |
| 4 | 57921706 | rs1713963 | 0.563 | -0.591 | 0.373 | 0.02 | -0.005 | 0.029 | 0.029 | 0 | 0.71 | 0.39 | 0.74 | 0.62 | 0.67 | 0.73 |

| | | | | | | | | | | | | | | | | |
|---|---|---|---|---|---|---|---|---|---|---|---|---|---|---|---|---|
| 4 | 57922249 | rs11573086 | 0.268 | -0.425 | 0.329 | -0.386 | 0.015 | 0.103 | 0.103 | 0.376 | 0.32 | 0.13 | 0.16 | 0.06 | 0.25 | 0.17 |
| 4 | 57922990 | rs1713959 | 0.159 | -0.432 | 0.357 | -0.358 | 0.001 | -0.046 | -0.046 | 0 | 0.62 | 0.56 | 0.75 | 0.91 | 0.61 | 0.75 |
| 4 | 57926736 | rs7656865 | 1.055 | -0.436 | -0.069 | -0.151 | 0.004 | 0.055 | 0.055 | 0 | 0.2 | 0.08 | 0.1 | 0.02 | 0.16 | 0.11 |
| 4 | 57931341 | rs1277293 | 1.147 | -0.946 | 1.128 | 0.739 | 0.005 | -0.002 | -0.002 | 0.065 | 0.61 | 0.59 | 0.71 | 0.61 | 0.7 | 0.66 |
| 4 | 57933375 | rs7356193 | 0.072 | -0.956 | 1.007 | 0.693 | 0 | 0.001 | 0.001 | 0 | 0.06 | 0.04 | 0.03 | 0.05 | 0.06 | 0.05 |
| 4 | 57934091 | rs17761305 | -0.202 | -1.098 | 1.104 | 0.74 | 0.022 | 0 | 0 | 0 | 0.19 | 0.12 | 0.17 | 0.16 | 0.13 | 0.12 |
| 4 | 57934971 | rs4865174 | -0.193 | -0.972 | 1.139 | 0.786 | 0.033 | 0.07 | 0.07 | 0 | 0.23 | 0.11 | 0.18 | 0.07 | 0.16 | 0.14 |
| 4 | 57936735 | rs10516161 | -0.75 | -1.009 | 1.392 | 0.782 | -0.003 | -0.015 | -0.015 | 0 | 0.08 | 0.2 | 0.04 | 0.05 | 0.06 | 0.05 |
| 4 | 57937134 | rs1714020 | 0.916 | -1.036 | 1.432 | 0.796 | -0.011 | 0.021 | 0.021 | 0 | 0.77 | 0.6 | 0.85 | 0.72 | 0.8 | 0.76 |
| 4 | 57937511 | rs10516163 | 2.156 | -0.802 | 0.751 | 0.502 | -0.062 | -0.046 | -0.046 | 0 | 0.42 | 0.14 | 0.56 | 0.55 | 0.47 | 0.53 |
| 4 | 57937862 | rs13141016 | 2.042 | -1.222 | 0.84 | 0.622 | -0.027 | -0.048 | -0.048 | 0.049 | 0.43 | 0.14 | 0.58 | 0.54 | 0.52 | 0.54 |
| 4 | 57941026 | rs11934877 | -0.079 | -0.916 | 0.834 | 0.412 | 0.024 | 0.047 | 0.047 | 0 | 0.22 | 0.12 | 0.17 | 0.07 | 0.14 | 0.13 |
| 4 | 57941160 | rs1714014 | 0.927 | -0.899 | 1.238 | 0.383 | 0.044 | -0.061 | -0.061 | 0.187 | 0.85 | 0.71 | 0.87 | 0.98 | 0.77 | 0.85 |
| 4 | 57942581 | rs1718848 | 0.584 | -0.867 | 1.233 | 0.378 | 0.023 | -0.062 | -0.062 | 0 | 0.87 | 0.72 | 0.9 | 0.99 | 0.84 | 0.86 |
| 4 | 57942649 | rs1714010 | 1.099 | -0.836 | 1.195 | 0.421 | 0 | -0.039 | -0.039 | 0.016 | 0.63 | 0.59 | 0.7 | 0.88 | 0.63 | 0.72 |
| 4 | 57942687 | rs11573056 | -0.184 | -0.912 | 1.256 | 0.339 | -0.002 | -0.002 | -0.002 | 0 | 0.07 | 0.07 | 0.04 | 0.05 | 0.06 | 0.07 |
| 4 | 57943153 | rs1718845 | 1.058 | -0.966 | 1.326 | 0.491 | -0.002 | -0.038 | -0.038 | 0 | 0.65 | 0.64 | 0.74 | 0.91 | 0.7 | 0.74 |
| 4 | 57943382 | rs10009533 | 1.187 | -0.968 | 1.276 | 0.491 | 0 | -0.052 | -0.052 | 0 | 0.63 | 0.59 | 0.7 | 0.9 | 0.63 | 0.72 |
| 4 | 57944503 | rs1713976 | 1.205 | -0.957 | 1.295 | 0.577 | -0.003 | -0.032 | -0.032 | 0 | 0.63 | 0.58 | 0.7 | 0.84 | 0.65 | 0.72 |
| 4 | 57945439 | rs3806744 | -0.351 | -0.934 | 0.666 | 0.604 | 0.028 | 0.037 | 0.037 | 0 | 0.24 | 0.16 | 0.18 | 0.13 | 0.13 | 0.13 |
| 4 | 57948187 | rs1718858 | 0.563 | -0.711 | -0.046 | 0.689 | 0.01 | -0.008 | -0.008 | 0 | 0.63 | 0.34 | 0.69 | 0.54 | 0.75 | 0.73 |
| 4 | 57949699 | rs1718861 | 1.019 | -0.62 | -0.161 | 0.776 | 0.028 | -0.006 | -0.006 | 0 | 0.53 | 0.28 | 0.61 | 0.53 | 0.6 | 0.59 |
| 4 | 57955234 | rs7695536 | 1.152 | -0.56 | -0.515 | 0.785 | 0.029 | -0.021 | -0.021 | 0 | 0.48 | 0.12 | 0.58 | 0.48 | 0.56 | 0.53 |
| 4 | 57963469 | rs4865181 | -0.646 | -0.809 | 0.271 | 1.07 | -0.073 | 0.057 | 0.057 | 0.153 | 0.31 | 0.49 | 0.4 | 0.54 | 0.15 | 0.25 |
| 4 | 57963611 | rs4865182 | -0.711 | -0.637 | 0.249 | 1.072 | -0.062 | 0.054 | 0.054 | 0.215 | 0.31 | 0.52 | 0.41 | 0.54 | 0.16 | 0.25 |
| 4 | 57968444 | rs10004910 | 1.026 | -0.157 | -0.039 | 0.706 | 0.022 | 0.083 | 0.083 | 0 | 0.49 | 0.21 | 0.43 | 0.34 | 0.56 | 0.57 |
| 4 | 57981557 | rs13129156 | 1.064 | -0.319 | -0.327 | 0.766 | 0.03 | 0.099 | 0.099 | 0 | 0.48 | 0.15 | 0.43 | 0.34 | 0.57 | 0.58 |
| 4 | 57987433 | rs6828580 | -0.252 | -0.408 | -0.103 | 0.969 | -0.041 | 0.087 | 0.087 | 0 | 0.8 | 0.55 | 0.66 | 0.71 | 0.87 | 0.81 |
| 4 | 57999268 | rs4327548 | -0.39 | -0.344 | -0.285 | 0.906 | 0.001 | -0.003 | -0.003 | 0 | 0.31 | 0.32 | 0.21 | 0.39 | 0.29 | 0.24 |
| 4 | 58005459 | rs4444899 | 0.414 | 0.13 | 0.088 | 0.828 | -0.047 | 0.082 | 0.082 | 0 | 0.79 | 0.51 | 0.65 | 0.71 | 0.87 | 0.82 |
| 4 | 58012501 | rs6847938 | -0.806 | 0.813 | -0.301 | 0.365 | 0.004 | -0.03 | -0.03 | 0 | 0.1 | 0.11 | 0.06 | 0 | 0.12 | 0.05 |
| 4 | 58013351 | rs9996990 | -0.555 | 0.635 | 0.31 | 0.588 | 0.029 | 0.064 | 0.064 | 0 | 0.07 | 0.25 | 0.05 | 0.16 | 0.08 | 0.19 |
| 4 | 58015591 | rs7663355 | -0.358 | 0.613 | -0.594 | 0.35 | -0.002 | 0 | 0 | 0 | 0.57 | 0.62 | 0.58 | 0.59 | 0.63 | 0.54 |
| 4 | 58015899 | rs4579186 | 0.511 | 0.351 | -0.623 | 0.414 | -0.002 | 0.013 | 0.013 | 0 | 0.11 | 0.12 | 0.09 | 0.19 | 0.08 | 0.13 |
| 4 | 58020126 | rs6817385 | -0.749 | 0.508 | -0.95 | -0.293 | 0 | -0.006 | -0.006 | 0 | 0.7 | 0.75 | 0.69 | 0.78 | 0.73 | 0.69 |
| 4 | 58020760 | rs7677987 | 0.278 | 1.022 | -1.488 | -0.654 | -0.043 | 0.082 | 0.082 | 0 | 0.22 | 0.08 | 0.14 | 0 | 0.27 | 0.06 |
| 4 | 58027081 | rs10033422 | -0.721 | 1.134 | -1.652 | -0.279 | -0.015 | 0.103 | 0.103 | 0 | 0.54 | 0.81 | 0.67 | 0.76 | 0.53 | 0.56 |
| 4 | 58028209 | rs4516785 | -0.508 | 1.125 | -1.537 | -0.285 | -0.011 | 0.109 | 0.109 | 0 | 0.52 | 0.77 | 0.65 | 0.72 | 0.52 | 0.55 |
| 4 | 58031795 | rs12641603 | -0.264 | 1.186 | -1.442 | -0.153 | -0.01 | 0.032 | 0.032 | 0 | 0.21 | 0.11 | 0.29 | 0.11 | 0.22 | 0.13 |
| 4 | 58032848 | rs7654584 | -0.264 | 1.186 | -1.442 | -0.153 | -0.01 | 0.032 | 0.032 | 0 | 0.21 | 0.11 | 0.29 | 0.11 | 0.22 | 0.13 |
| 4 | 58036217 | rs11133488 | 0.516 | 0.651 | -1.422 | 0.409 | 0 | -0.002 | -0.002 | 0 | 0.87 | 0.87 | 0.87 | 0.77 | 0.93 | 0.93 |
| 4 | 58041751 | rs4412045 | 0.911 | 0.549 | -1.617 | 0.23 | 0.001 | -0.016 | -0.016 | 0 | 0.71 | 0.89 | 0.61 | 0.67 | 0.75 | 0.86 |
| 4 | 58053743 | rs6857279 | 1.271 | -0.196 | -2.61 | 0.418 | -0.104 | -0.032 | -0.032 | 0 | 0.37 | 0.07 | 0.39 | 0.37 | 0.51 | 0.45 |
| 4 | 58055260 | rs13145188 | 0.877 | -0.884 | -1.709 | 0.031 | -0.005 | -0.005 | -0.005 | 0 | 0.58 | 0.52 | 0.55 | 0.67 | 0.7 | 0.8 |
| 4 | 58056503 | rs17497594 | 1.031 | -0.66 | -1.705 | 0.014 | -0.011 | 0.025 | 0.025 | 0.001 | 0.17 | 0.07 | 0.26 | 0.1 | 0.18 | 0.08 |
| 4 | 58064464 | rs10014421 | -1.177 | -1.026 | -2.395 | 0.337 | 0.014 | -0.021 | -0.021 | 0 | 0.51 | 0.95 | 0.48 | 0.49 | 0.37 | 0.53 |
| 4 | 58066486 | rs10488926 | 0.76 | -0.983 | -1.981 | 0.353 | -0.005 | 0.022 | 0.022 | 4.039 | 0.09 | 0.15 | 0.04 | 0.17 | 0.07 | 0.18 |
| 4 | 58066825 | rs2215473 | -1.535 | -1.03 | -2.367 | 0.166 | -0.036 | 0.015 | 0.015 | 1.4 | 0.48 | 0.94 | 0.49 | 0.45 | 0.39 | 0.5 |
| 4 | 58072736 | rs2098159 | 0.758 | -0.969 | -2.132 | 0.48 | -0.03 | -0.048 | -0.048 | 0 | 0.08 | 0 | 0.06 | 0.2 | 0.09 | 0.15 |
| 4 | 58072932 | rs2079916 | -0.679 | -1.269 | -1.341 | 0.199 | 0.002 | -0.006 | -0.006 | 0.01 | 0.39 | 0.36 | 0.43 | 0.51 | 0.34 | 0.44 |
| 4 | 58075747 | rs6853264 | 0.351 | -1.446 | -1.071 | 0.052 | -0.051 | -0.079 | -0.079 | 1.782 | 0.12 | 0.39 | 0.06 | 0 | 0.04 | 0.05 |
| 4 | 58077421 | rs917674 | -0.806 | -0.652 | -1.397 | 0.382 | 0.003 | -0.015 | -0.015 | 0 | 0.41 | 0.63 | 0.49 | 0.51 | 0.38 | 0.45 |
| 4 | 58081173 | rs17425871 | 0.322 | -0.196 | -1.591 | -0.392 | 0.004 | 0.067 | 0.067 | 0.001 | 0.18 | 0.04 | 0.09 | 0.04 | 0.14 | 0.12 |
| 4 | 58081623 | rs1860622 | -0.435 | 0.181 | -1.812 | -0.478 | -0.019 | -0.011 | -0.011 | 0.21 | 0.73 | 0.83 | 0.8 | 0.68 | 0.7 | 0.75 |
| 4 | 58085224 | rs6812462 | 0.842 | 0.34 | -0.715 | -0.405 | 0 | 0.001 | 0.001 | 0 | 0.5 | 0.44 | 0.64 | 0.51 | 0.49 | 0.54 |
| 4 | 58087036 | rs1860621 | 0.998 | 0.084 | -1.148 | -0.942 | -0.04 | -0.024 | -0.024 | 0 | 0.29 | 0.17 | 0.6 | 0.62 | 0.43 | 0.43 |
| 4 | 58088414 | rs7680799 | -0.895 | 0.127 | -1.035 | -0.692 | -0.004 | -0.008 | -0.008 | 0 | 0.74 | 0.77 | 0.83 | 0.69 | 0.74 | 0.74 |
| 4 | 58094886 | rs2109589 | -1.397 | 0.057 | -1.095 | -1.122 | -0.007 | 0.042 | 0.042 | 0 | 0.78 | 0.83 | 0.87 | 0.93 | 0.76 | 0.82 |
| 4 | 58098249 | rs9995833 | 0.804 | -0.192 | -0.994 | -1.272 | -0.015 | -0.026 | -0.026 | 0 | 0.28 | 0.31 | 0.2 | 0.18 | 0.23 | 0.27 |
| 4 | 58098494 | rs6822293 | -1.449 | 0.215 | -1.339 | -1.253 | -0.038 | 0.099 | 0.099 | 0 | 0.77 | 0.92 | 0.88 | 0.93 | 0.76 | 0.82 |
| 4 | 58098554 | rs12500068 | 0.224 | 0.303 | -0.839 | -0.928 | -0.008 | 0.006 | 0.006 | 0 | 0.3 | 0.25 | 0.57 | 0.74 | 0.36 | 0.42 |
| 4 | 58100571 | rs17425986 | 0.565 | 0.369 | -0.804 | -1.017 | -0.011 | 0.012 | 0.012 | 0 | 0.05 | 0.04 | 0.04 | 0.02 | 0.1 | 0.16 |
| 4 | 58102272 | rs6856352 | -0.847 | 0.258 | -0.383 | -1.47 | 0 | -0.054 | -0.054 | 0.157 | 0.55 | 0.48 | 0.67 | 0.89 | 0.58 | 0.66 |
| 4 | 58102553 | rs9999844 | 0.419 | 0.057 | -0.41 | -1.619 | -0.013 | -0.066 | -0.066 | 0 | 0.31 | 0.3 | 0.2 | 0.07 | 0.28 | 0.29 |
| 4 | 58103527 | rs4865204 | -0.791 | 0.09 | -0.435 | -1.156 | 0.004 | 0.02 | 0.02 | 0 | 0.5 | 0.48 | 0.61 | 0.64 | 0.5 | 0.57 |
| 4 | 58104161 | rs2191567 | -0.741 | 0.108 | -0.435 | -1.162 | -0.001 | -0.002 | -0.002 | 0 | 0.49 | 0.44 | 0.61 | 0.64 | 0.53 | 0.59 |
| 4 | 58104571 | rs1981687 | -0.963 | 0.105 | -0.458 | -1.191 | -0.001 | -0.001 | -0.001 | 0 | 0.54 | 0.48 | 0.64 | 0.67 | 0.58 | 0.69 |
| 4 | 58105295 | rs11934184 | 0.344 | 0.152 | -0.661 | -1.518 | 0.004 | -0.04 | -0.04 | 1.455 | 0.29 | 0.22 | 0.17 | 0 | 0.24 | 0.22 |
| 4 | 58105688 | rs7694552 | -1.585 | 0.103 | -0.726 | -1.446 | -0.016 | 0.045 | 0.045 | 0 | 0.66 | 0.76 | 0.74 | 0.81 | 0.67 | 0.79 |

| | | | | | | | | | | | | | | | | |
|---|---|---|---|---|---|---|---|---|---|---|---|---|---|---|---|---|
| 4 | 58106366 | rs4865205 | -1.616 | 0.091 | -0.737 | -1.479 | -0.016 | 0.047 | 0.047 | 0 | 0.66 | 0.76 | 0.73 | 0.84 | 0.67 | 0.78 |
| 4 | 58107030 | rs7679409 | -1.694 | 0.18 | -0.681 | -1.457 | -0.016 | 0.044 | 0.044 | 0 | 0.66 | 0.75 | 0.74 | 0.85 | 0.67 | 0.78 |
| 4 | 58108131 | rs13147707 | -0.68 | -0.175 | 0.124 | -0.727 | 0 | 0.02 | 0.02 | 0 | 0.14 | 0.3 | 0.17 | 0.19 | 0.15 | 0.28 |
| 4 | 58111647 | rs10001448 | -0.077 | -0.259 | -0.174 | -0.971 | -0.002 | -0.053 | -0.053 | 0 | 0.27 | 0.23 | 0.18 | 0 | 0.26 | 0.19 |
| 4 | 58115768 | rs1990404 | -0.084 | -0.524 | -0.053 | -0.952 | 0.001 | -0.032 | -0.032 | 0 | 0.24 | 0.2 | 0.11 | 0.01 | 0.16 | 0.15 |
| 4 | 58116921 | rs11731338 | -1.813 | -0.39 | -0.413 | -0.783 | 0.002 | 0.047 | 0.047 | 0 | 0.54 | 0.88 | 0.44 | 0.43 | 0.5 | 0.59 |
| 4 | 58120471 | rs6815861 | -0.585 | -0.442 | 0.113 | -0.778 | -0.029 | -0.051 | -0.051 | 0 | 0.35 | 0.41 | 0.23 | 0.13 | 0.32 | 0.22 |
| 4 | 58120699 | rs6833012 | -1.768 | -0.438 | 0.223 | -0.764 | 0.066 | 0.112 | 0.112 | 0 | 0.49 | 0.73 | 0.4 | 0.29 | 0.44 | 0.52 |
| 4 | 58121404 | rs13132470 | 0.505 | -0.407 | 0.285 | -0.74 | -0.008 | -0.002 | -0.002 | 0 | 0.09 | 0.22 | 0.14 | 0.1 | 0.07 | 0.2 |
| 4 | 58121589 | rs10016362 | -1.406 | -0.422 | 0.335 | -0.634 | 0.031 | 0.046 | 0.046 | 0 | 0.45 | 0.62 | 0.3 | 0.34 | 0.42 | 0.38 |
| 4 | 58121668 | rs17087830 | -1.006 | -0.422 | 0.406 | -0.667 | 0.01 | 0.012 | 0.012 | 0 | 0.41 | 0.55 | 0.25 | 0.22 | 0.35 | 0.34 |
| 4 | 58122141 | rs917677 | -0.522 | -0.37 | -0.04 | -0.705 | -0.034 | -0.077 | -0.077 | 0 | 0.36 | 0.47 | 0.22 | 0.14 | 0.29 | 0.24 |
| 4 | 58122195 | rs13115696 | -1.439 | 0 | 0.255 | -0.38 | 0.007 | 0.039 | 0.039 | 0.866 | 0.18 | 0.39 | 0.22 | 0.28 | 0.21 | 0.35 |
| 4 | 58123111 | rs6825426 | -0.542 | 0.014 | -0.396 | -0.273 | -0.001 | 0.027 | 0.027 | 0.527 | 0.12 | 0.23 | 0.19 | 0.23 | 0.15 | 0.25 |
| 4 | 58136368 | rs17498104 | -0.686 | -0.183 | -1.014 | -0.659 | 0.047 | 0.11 | 0.11 | 0 | 0.29 | 0.09 | 0.18 | 0.07 | 0.27 | 0.19 |
| 4 | 58136698 | rs2191576 | -1.678 | 0.43 | -1.354 | -0.357 | -0.085 | 0.103 | 0.103 | 0.662 | 0.57 | 0.92 | 0.63 | 0.79 | 0.55 | 0.68 |
| 4 | 58138412 | rs10018606 | -0.052 | 0.135 | -0.692 | 0.184 | -0.002 | 0.035 | 0.035 | 0 | 0.22 | 0.28 | 0.4 | 0.46 | 0.28 | 0.28 |
| 4 | 58139970 | rs1476844 | -2.363 | 0.571 | -1.123 | -0.305 | -0.038 | 0.017 | 0.017 | 0.027 | 0.7 | 0.86 | 0.79 | 0.8 | 0.71 | 0.85 |
| 4 | 58140768 | rs17426309 | 0.569 | 0.125 | -1.25 | -0.48 | -0.008 | 0.111 | 0.111 | 0 | 0.21 | 0.04 | 0.17 | 0 | 0.2 | 0.17 |
| 4 | 58141367 | rs2191574 | -0.932 | -0.074 | -0.805 | -0.279 | 0.013 | -0.012 | -0.012 | 0 | 0.36 | 0.72 | 0.51 | 0.6 | 0.34 | 0.4 |
| 4 | 58145935 | rs2041437 | -1.904 | 0.254 | -0.014 | -0.124 | 0.001 | -0.012 | -0.012 | 0 | 0.63 | 0.54 | 0.73 | 0.81 | 0.67 | 0.81 |
| 4 | 58146502 | rs4865211 | -0.81 | 0.214 | -0.119 | 0.038 | 0.003 | -0.009 | -0.009 | 0 | 0.44 | 0.22 | 0.57 | 0.44 | 0.48 | 0.44 |
| 4 | 58146929 | rs7686067 | 1.046 | 0.25 | -0.16 | -0.004 | 0.001 | 0.05 | 0.05 | 0 | 0.13 | 0.17 | 0.38 | 0.44 | 0.15 | 0.27 |
| 4 | 58148600 | rs7675913 | -1.642 | 0.513 | 0.326 | 0.22 | -0.002 | 0.014 | 0.014 | 0.121 | 0.62 | 0.35 | 0.72 | 0.81 | 0.67 | 0.8 |
| 4 | 58148873 | rs6838965 | 1.071 | 0.337 | 0.252 | 0.212 | -0.02 | 0.04 | 0.04 | 0 | 0.11 | 0.03 | 0.07 | 0 | 0.14 | 0.11 |
| 4 | 58150716 | rs17498431 | -0.6 | 0.377 | 0.824 | 0.673 | 0.005 | 0.009 | 0.009 | 0 | 0.51 | 0.55 | 0.64 | 0.59 | 0.51 | 0.53 |
| 4 | 58170360 | rs6840839 | -1.031 | -0.081 | 1.279 | 0.762 | 0.009 | 0.095 | 0.095 | 0 | 0.06 | 0.25 | 0.06 | 0.15 | 0.02 | 0.09 |
| 4 | 58171793 | rs2691402 | -1.781 | -0.097 | 1.155 | 0.754 | 0.014 | 0.122 | 0.122 | 1.285 | 0.09 | 0.5 | 0.07 | 0.15 | 0.02 | 0.09 |
| 4 | 58178243 | rs2535523 | -0.858 | -0.07 | 0.943 | 0.697 | -0.003 | 0.016 | 0.016 | 0 | 0.53 | 0.69 | 0.65 | 0.59 | 0.54 | 0.55 |
| 4 | 58178960 | rs991920 | -1.545 | -0.097 | 1.01 | 0.731 | 0.035 | 0.164 | 0.164 | 0.574 | 0.08 | 0.43 | 0.07 | 0.15 | 0.03 | 0.09 |
| 4 | 58179993 | rs1524686 | -0.277 | -0.053 | 0.942 | 0.683 | 0.005 | 0 | 0 | 0 | 0.45 | 0.26 | 0.57 | 0.44 | 0.51 | 0.46 |
| 4 | 58184590 | rs2535516 | -0.606 | -0.021 | 0.873 | 0.695 | 0.013 | 0.007 | 0.007 | 0 | 0.48 | 0.58 | 0.6 | 0.45 | 0.5 | 0.46 |
| 4 | 58189742 | rs2535511 | 0.024 | -0.11 | 0.922 | 0.531 | -0.01 | 0.022 | 0.022 | 0.317 | 0.12 | 0.27 | 0.09 | 0.14 | 0.08 | 0.2 |
| 4 | 58191253 | rs715208 | -0.298 | -0.098 | 0.764 | 0.872 | 0.007 | 0.052 | 0.052 | 0 | 0.47 | 0.31 | 0.38 | 0.33 | 0.53 | 0.45 |
| 4 | 58193150 | rs2245144 | 0.065 | -0.235 | 0.774 | 0.505 | -0.005 | -0.019 | -0.019 | 1.078 | 0.14 | 0.19 | 0.16 | 0.11 | 0.17 | 0.11 |
| 4 | 58193776 | rs1524671 | -1.056 | -0.368 | 0.858 | 0.732 | -0.013 | 0.007 | 0.007 | 0 | 0.25 | 0.3 | 0.21 | 0.32 | 0.22 | 0.28 |
| 4 | 58194612 | rs1917335 | -1.004 | -0.415 | 1.129 | 0.703 | -0.035 | -0.003 | -0.003 | 0.047 | 0.24 | 0.43 | 0.17 | 0.25 | 0.21 | 0.22 |
| 4 | 58195063 | rs1829982 | 0.954 | -0.377 | 1.025 | 0.658 | -0.001 | 0.002 | 0.002 | 0 | 0.11 | 0.06 | 0.1 | 0 | 0.15 | 0.03 |
| 4 | 58195196 | rs2535503 | -1.607 | -0.362 | 0.843 | 0.747 | -0.013 | 0.016 | 0.016 | 0 | 0.64 | 0.68 | 0.7 | 0.79 | 0.61 | 0.74 |
| 4 | 58196071 | rs2535498 | -2.163 | -0.526 | 0.32 | 0.602 | -0.125 | 0.009 | 0.009 | 0.132 | 0.71 | 0.93 | 0.74 | 0.79 | 0.65 | 0.79 |
| 4 | 58196685 | rs1404370 | 0.423 | -0.535 | 0.376 | 0.644 | -0.002 | -0.052 | -0.052 | 0 | 0.37 | 0.2 | 0.49 | 0.43 | 0.39 | 0.44 |
| 4 | 58197731 | rs2177642 | -1.144 | -0.516 | 0.641 | 0.628 | -0.005 | -0.001 | -0.001 | 0 | 0.59 | 0.62 | 0.68 | 0.59 | 0.59 | 0.61 |
| 4 | 58198454 | rs940572 | -1.144 | -0.515 | 0.646 | 0.654 | -0.004 | -0.001 | -0.001 | 0.351 | 0.59 | 0.62 | 0.68 | 0.59 | 0.59 | 0.61 |
| 4 | 58199809 | rs1524672 | 0.591 | -0.501 | 0.778 | 0.649 | -0.004 | -0.055 | -0.055 | 0 | 0.13 | 0.23 | 0.11 | 0.05 | 0.15 | 0.04 |
| 4 | 58202128 | rs10021529 | -0.093 | -0.495 | 0.617 | 0.784 | -0.005 | 0.001 | 0.001 | 0 | 0.16 | 0.17 | 0.08 | 0.2 | 0.09 | 0.23 |
| 4 | 58207586 | rs17087996 | -0.913 | -0.593 | 0.737 | 0.787 | -0.063 | 0.079 | 0.079 | 1.5 | 0.21 | 0.48 | 0.17 | 0.33 | 0.13 | 0.31 |
| 4 | 58216884 | rs2222560 | -1.799 | -0.312 | 0.611 | 0.564 | -0.053 | 0.028 | 0.028 | 0.028 | 0.7 | 0.82 | 0.73 | 0.8 | 0.65 | 0.79 |
| 4 | 58219609 | rs6811739 | -0.853 | -0.375 | 0.379 | 0.668 | -0.001 | 0.004 | 0.004 | 0 | 0.54 | 0.45 | 0.57 | 0.64 | 0.49 | 0.67 |
| 4 | 58239085 | rs6853593 | -0.827 | -0.316 | -0.384 | 0.542 | -0.003 | 0.027 | 0.027 | 0.014 | 0.55 | 0.4 | 0.64 | 0.8 | 0.52 | 0.76 |
| 4 | 58248207 | rs10024454 | -0.142 | -0.514 | -1.532 | 0.89 | -0.04 | -0.058 | -0.058 | 0.641 | 0.3 | 0.07 | 0.45 | 0.33 | 0.33 | 0.41 |
| 4 | 58261728 | rs13129010 | -0.005 | -0.573 | -1.481 | 0.99 | -0.002 | -0.053 | -0.053 | 0 | 0.15 | 0.02 | 0.11 | 0.24 | 0.13 | 0.24 |
| 4 | 58263866 | rs10009611 | -0.185 | -0.424 | -1.358 | 0.882 | -0.037 | -0.053 | -0.053 | 0.095 | 0.29 | 0.09 | 0.45 | 0.35 | 0.33 | 0.39 |
| 4 | 58264562 | rs2204370 | -0.776 | -0.291 | -0.983 | 0.545 | 0.001 | 0.005 | 0.005 | 0 | 0.88 | 0.95 | 0.92 | 1 | 0.87 | 0.96 |
| 4 | 58271686 | rs10517394 | -0.115 | -0.205 | -0.994 | 0.949 | -0.011 | -0.065 | -0.065 | 0 | 0.13 | 0.01 | 0.06 | 0.21 | 0.11 | 0.2 |
| 4 | 58272001 | rs1012189 | -1.244 | -0.278 | -1.429 | 1.023 | 0.055 | -0.054 | -0.054 | 0 | 0.43 | 0.91 | 0.5 | 0.47 | 0.35 | 0.49 |
| 4 | 58272082 | rs1012188 | -2.002 | -0.272 | -1.398 | 0.931 | -0.01 | 0.05 | 0.05 | 0.353 | 0.56 | 0.92 | 0.56 | 0.68 | 0.46 | 0.69 |
| 4 | 58272528 | rs10517393 | -0.1 | -0.461 | -1.05 | 0.907 | 0.005 | 0.001 | 0.001 | 0 | 0.24 | 0.19 | 0.19 | 0.26 | 0.24 | 0.31 |
| 4 | 58275617 | rs6847118 | -1.087 | -0.235 | 0.031 | 0.61 | -0.003 | 0.009 | 0.009 | 0 | 0.39 | 0.43 | 0.37 | 0.48 | 0.38 | 0.52 |
| 4 | 58277976 | rs10155292 | 0.687 | -0.272 | 0.213 | 0.568 | -0.009 | -0.041 | -0.041 | 0 | 0.08 | 0.15 | 0.06 | 0 | 0.07 | 0.1 |
| 4 | 58285434 | rs9917898 | -0.62 | -0.207 | 0.142 | 0.749 | -0.035 | 0.024 | 0.024 | 2.082 | 0.1 | 0.29 | 0.09 | 0.13 | 0.04 | 0.1 |
| 4 | 58289649 | rs839021 | 0.77 | 0.002 | -0.367 | 0.426 | -0.044 | -0.077 | -0.077 | 0 | 0.62 | 0.87 | 0.63 | 0.53 | 0.56 | 0.48 |
| 4 | 58296141 | rs940580 | -1.109 | 0.333 | 0.449 | -0.273 | 0.017 | -0.074 | -0.074 | 0 | 0.23 | 0.53 | 0.19 | 0.11 | 0.26 | 0.15 |
| 4 | 58296826 | rs6554464 | 1.792 | 0.354 | -0.069 | -0.378 | -0.021 | 0.173 | 0.173 | 0 | 0.54 | 0.16 | 0.58 | 0.8 | 0.51 | 0.5 |
| 4 | 58298691 | rs1372088 | -0.183 | 0.081 | 0.377 | -0.263 | -0.007 | -0.035 | -0.035 | 0 | 0.86 | 0.77 | 0.89 | 0.93 | 0.89 | 0.75 |
| 4 | 58298856 | rs940578 | 2.036 | 0.282 | 0.052 | -0.196 | -0.102 | -0.115 | -0.115 | 0 | 0.37 | 0.05 | 0.36 | 0.51 | 0.41 | 0.34 |
| 4 | 58299763 | rs7668422 | 0.048 | 0.467 | 0.686 | -0.339 | -0.001 | -0.017 | -0.017 | 0 | 0.85 | 0.83 | 0.88 | 0.93 | 0.86 | 0.74 |
| 4 | 58301224 | rs7661860 | -0.719 | 0.357 | 1.291 | -0.254 | 0.051 | -0.049 | -0.049 | 0.051 | 0.11 | 0.45 | 0.13 | 0 | 0.12 | 0.06 |
| 4 | 58303358 | rs13101487 | -0.036 | 0.377 | 0.794 | -0.151 | -0.006 | -0.033 | -0.033 | 0 | 0.86 | 0.79 | 0.9 | 0.93 | 0.89 | 0.75 |
| 4 | 58308959 | rs6821078 | -0.528 | 0.533 | 1.035 | -0.108 | 0.066 | -0.037 | -0.037 | 3.458 | 0.09 | 0.33 | 0.13 | 0 | 0.13 | 0.06 |

| | | | | | | | | | | | | | | | | |
|---|---|---|---|---|---|---|---|---|---|---|---|---|---|---|---|---|
| 4 | 58309910 | rs6827688 | 0.277 | 0.294 | 0.974 | -0.144 | -0.015 | -0.031 | -0.031 | 0.482 | 0.83 | 0.57 | 0.86 | 0.88 | 0.85 | 0.86 |
| 4 | 58313925 | rs1561660 | -0.002 | 0.463 | 1.324 | 0.004 | 0.022 | 0.019 | 0.019 | 0 | 0.9 | 0.72 | 0.91 | 0.89 | 0.91 | 0.87 |
| 4 | 58314264 | rs11937087 | -0.822 | 0.443 | 0.981 | 0.157 | 0.009 | -0.034 | -0.034 | 0 | 0.37 | 0.27 | 0.4 | 0.45 | 0.32 | 0.49 |
| 4 | 58320842 | rs6855484 | 1.526 | 0.414 | 1.226 | 0.177 | 0.023 | -0.001 | -0.001 | 1.944 | 0.51 | 0.27 | 0.5 | 0.46 | 0.59 | 0.38 |
| 4 | 58324740 | rs11133493 | -0.753 | 0.445 | 0.777 | 0.2 | 0.007 | -0.029 | -0.029 | 0 | 0.37 | 0.29 | 0.4 | 0.45 | 0.32 | 0.49 |
| 4 | 58326265 | rs4864624 | -0.843 | 0.451 | 0.774 | 0.088 | 0.01 | -0.023 | -0.023 | 1.434 | 0.37 | 0.29 | 0.4 | 0.43 | 0.32 | 0.49 |
| 4 | 58332294 | rs7669775 | 0.184 | 0.415 | 0.464 | 0.081 | 0.002 | 0.006 | 0.006 | 0 | 0.13 | 0.13 | 0.1 | 0.07 | 0.1 | 0.25 |
| 4 | 58332757 | rs10020991 | -0.267 | 0.467 | 0.361 | 0.147 | 0.018 | 0.015 | 0.015 | 0 | 0.9 | 0.74 | 0.91 | 0.89 | 0.91 | 0.87 |
| 4 | 58340913 | rs12650171 | 1.649 | 0.561 | -0.459 | 0.116 | -0.137 | -0.132 | -0.132 | 0 | 0.45 | 0.02 | 0.38 | 0.46 | 0.47 | 0.33 |
| 4 | 58350599 | rs7654918 | -0.517 | 0.476 | 0.506 | 0.233 | -0.008 | 0.035 | 0.035 | 0.011 | 0.16 | 0.5 | 0.13 | 0.15 | 0.11 | 0.29 |
| 4 | 58350801 | rs13102887 | 0.21 | 0.448 | 0.292 | 0.242 | -0.041 | -0.076 | -0.076 | 0.023 | 0.24 | 0.03 | 0.28 | 0.3 | 0.22 | 0.21 |
| 4 | 58350816 | rs7696759 | -0.416 | 0.441 | 0.607 | 0.241 | -0.009 | 0.032 | 0.032 | 0.062 | 0.16 | 0.49 | 0.14 | 0.15 | 0.11 | 0.29 |
| 4 | 58352102 | rs12646772 | 0.754 | 0.414 | 0.446 | 0.24 | -0.014 | 0.007 | 0.007 | 0 | 0.09 | 0.03 | 0.08 | 0.07 | 0.11 | 0.21 |
| 4 | 58352237 | rs1345040 | -0.664 | 0.409 | 0.418 | 0.239 | -0.031 | -0.036 | -0.036 | 0 | 0.33 | 0.1 | 0.36 | 0.39 | 0.33 | 0.42 |
| 4 | 58352345 | rs17832655 | 1.316 | 0.495 | 0.266 | 0.153 | -0.014 | 0.019 | 0.019 | 0 | 0.05 | 0.02 | 0.1 | 0 | 0.11 | 0.04 |
| 4 | 58355015 | rs4865234 | -1.386 | 0.388 | 0.361 | 0.225 | -0.008 | 0.013 | 0.013 | 0.04 | 0.39 | 0.51 | 0.41 | 0.47 | 0.33 | 0.51 |
| 4 | 58355701 | rs11735670 | -0.141 | 0.709 | -0.1 | 0.235 | -0.041 | -0.073 | -0.073 | 0.995 | 0.24 | 0.03 | 0.28 | 0.32 | 0.22 | 0.21 |
| 4 | 58363063 | rs4865236 | 0.441 | 1.088 | 0.423 | 0.132 | -0.003 | 0.001 | 0.001 | 0 | 0.9 | 0.86 | 0.92 | 0.9 | 0.89 | 0.87 |
| 4 | 58381197 | rs1372094 | 0.768 | 1.02 | 0.89 | -0.247 | 0.028 | -0.103 | -0.103 | 0 | 0.79 | 0.7 | 0.79 | 1 | 0.72 | 0.87 |
| 4 | 58393319 | rs6829715 | -1.345 | 0.796 | 1.047 | 0.107 | 0.037 | 0.077 | 0.077 | 0 | 0.17 | 0.38 | 0.28 | 0.3 | 0.18 | 0.15 |
| 4 | 58396399 | rs1441073 | -0.146 | 0.824 | 1.139 | 0.083 | 0.001 | -0.004 | -0.004 | 0 | 0.9 | 0.92 | 0.88 | 1 | 0.89 | 0.94 |
| 4 | 58408330 | rs2028435 | 2.154 | 0.77 | 1.142 | 0.058 | -0.005 | 0.01 | 0.01 | 0.23 | 0.59 | 0.22 | 0.5 | 0.59 | 0.59 | 0.64 |
| 4 | 58412269 | rs17088360 | 0.415 | 0.826 | 1.407 | 0.055 | 0.005 | -0.02 | -0.02 | 0 | 0.1 | 0.06 | 0.08 | 0 | 0.16 | 0.07 |
| 4 | 58413456 | rs17088371 | -0.205 | 0.583 | 1.526 | 0.149 | 0 | 0 | 0 | 0 | 0.09 | 0.11 | 0.09 | 0.11 | 0.11 | 0.11 |
| 4 | 58445149 | rs1583881 | 0.087 | 0.504 | 1.565 | -0.05 | -0.035 | -0.072 | -0.072 | 0.108 | 0.85 | 0.68 | 0.84 | 0.99 | 0.88 | 0.87 |
| 4 | 58450181 | rs2899092 | -0.099 | 0.556 | 1.013 | 0 | -0.002 | -0.001 | -0.001 | 0 | 0.91 | 0.91 | 0.92 | 0.9 | 0.89 | 0.9 |
| 4 | 58457282 | rs13117119 | 2.087 | 1.007 | 0.723 | -0.189 | -0.029 | 0.039 | 0.039 | 0 | 0.55 | 0.24 | 0.45 | 0.59 | 0.53 | 0.61 |
| 4 | 58464353 | rs7693389 | -0.252 | 0.973 | 0.658 | -0.148 | -0.001 | 0 | 0 | 0.037 | 0.9 | 0.91 | 0.91 | 0.88 | 0.89 | 0.89 |
| 4 | 58470680 | rs4608868 | 1.735 | 1.045 | 0.448 | -0.147 | -0.04 | 0.039 | 0.039 | 0 | 0.54 | 0.19 | 0.45 | 0.56 | 0.53 | 0.61 |
| 4 | 58474857 | rs13129234 | 2.15 | 1.01 | 0.107 | -0.189 | -0.003 | -0.057 | -0.057 | 1.273 | 0.43 | 0.17 | 0.38 | 0.43 | 0.41 | 0.41 |
| 4 | 58477126 | rs7697011 | 1.282 | 0.847 | 0.694 | -0.189 | -0.026 | 0.015 | 0.015 | 0 | 0.71 | 0.42 | 0.59 | 0.59 | 0.75 | 0.69 |
| 4 | 58477962 | rs6840178 | 0.11 | 0.803 | 0.73 | -0.213 | -0.014 | -0.043 | -0.043 | 0 | 0.14 | 0.2 | 0.14 | 0 | 0.12 | 0.12 |
| 4 | 58479112 | rs12643621 | 0.796 | 0.955 | 0.503 | -0.115 | -0.002 | -0.01 | -0.01 | 0 | 0.11 | 0.06 | 0.07 | 0.15 | 0.11 | 0.2 |
| 4 | 58480567 | rs4599476 | 2.137 | 0.922 | 0.386 | -0.234 | 0.01 | 0.093 | 0.093 | 0 | 0.46 | 0.2 | 0.55 | 0.71 | 0.43 | 0.48 |
| 4 | 58481417 | rs4865252 | 0.619 | 0.919 | 0.824 | -0.28 | -0.009 | -0.01 | -0.01 | 3.736 | 0.73 | 0.69 | 0.78 | 0.85 | 0.76 | 0.67 |
| 4 | 58483725 | rs4438813 | 0.177 | 1.033 | 1.044 | -0.462 | -0.01 | -0.004 | -0.004 | 0 | 0.09 | 0.04 | 0.07 | 0.12 | 0.11 | 0.11 |
| 4 | 58484281 | rs4438814 | 0.177 | 1.033 | 1.044 | -0.462 | -0.01 | -0.004 | -0.004 | 0 | 0.09 | 0.04 | 0.07 | 0.12 | 0.11 | 0.11 |
| 4 | 58484885 | rs11731283 | 0.065 | 1.104 | 1.24 | -0.473 | -0.003 | -0.015 | -0.015 | 0 | 0.14 | 0.08 | 0.12 | 0 | 0.21 | 0.08 |
| 4 | 58491683 | rs1469235 | 1.426 | 1.053 | 1.32 | -0.451 | -0.001 | 0.006 | 0.006 | 0 | 0.65 | 0.38 | 0.52 | 0.7 | 0.64 | 0.72 |
| 4 | 58501865 | rs4865256 | -0.203 | 0.965 | 1.269 | -0.51 | -0.02 | -0.054 | -0.054 | 0 | 0.85 | 0.78 | 0.85 | 1 | 0.87 | 0.87 |
| 4 | 58513545 | rs17088477 | -0.113 | 0.98 | 1.373 | -0.512 | 0.026 | 0.013 | 0.013 | 0 | 0.1 | 0.21 | 0.08 | 0.12 | 0.11 | 0.11 |
| 4 | 58515319 | rs1425316 | -0.185 | 0.993 | 1.307 | -0.513 | -0.019 | -0.053 | -0.053 | 0 | 0.85 | 0.79 | 0.85 | 1 | 0.87 | 0.87 |
| 4 | 58522592 | rs1425320 | 1.571 | 1.219 | 0.591 | -0.533 | -0.032 | 0.096 | 0.096 | 0.331 | 0.5 | 0.17 | 0.36 | 0.57 | 0.5 | 0.53 |
| 4 | 58526596 | rs4865257 | 0.658 | 1.092 | 0.664 | -0.555 | 0.01 | 0.003 | 0.003 | 0 | 0.64 | 0.69 | 0.53 | 0.58 | 0.72 | 0.6 |
| 4 | 58528741 | rs17088485 | 1.332 | 0.995 | 0.547 | -0.496 | -0.011 | 0.074 | 0.074 | 0.962 | 0.52 | 0.36 | 0.38 | 0.6 | 0.52 | 0.53 |
| 4 | 58546539 | rs4276345 | -0.975 | 1.137 | 0.985 | -0.53 | -0.011 | 0.026 | 0.026 | 0.007 | 0.47 | 0.3 | 0.59 | 0.48 | 0.51 | 0.61 |
| 4 | 58560611 | rs343503 | -0.967 | 1.168 | 1.038 | -0.528 | -0.013 | 0.026 | 0.026 | 0.022 | 0.47 | 0.27 | 0.59 | 0.48 | 0.5 | 0.62 |
| 4 | 58566621 | rs343501 | -0.951 | 1.113 | 1.035 | -0.53 | -0.012 | 0.023 | 0.023 | 0.017 | 0.47 | 0.27 | 0.59 | 0.48 | 0.51 | 0.61 |
| 4 | 58573210 | rs11736736 | 0.849 | 1.102 | 1.079 | -0.435 | -0.004 | 0.07 | 0.07 | 0 | 0.49 | 0.29 | 0.54 | 0.62 | 0.49 | 0.38 |
| 4 | 58586920 | rs482084 | 1.634 | 1.316 | 1.998 | -0.29 | 0.106 | 0.072 | 0.072 | 1.831 | 0.9 | 0.56 | 0.74 | 0.9 | 0.89 | 0.96 |
| 4 | 58589128 | rs12647976 | -0.259 | 1.328 | 1.56 | -0.277 | -0.005 | -0.037 | -0.037 | 0 | 0.07 | 0 | 0.03 | 0.17 | 0.03 | 0.15 |
| 4 | 58592926 | rs17088636 | -0.704 | 1.345 | 1.652 | -0.307 | 0.076 | 0.052 | 0.052 | 0 | 0.07 | 0.19 | 0.24 | 0.1 | 0.11 | 0.04 |
| 4 | 58597247 | rs17088644 | -0.233 | 1.376 | 1.375 | -0.523 | 0.04 | -0.023 | -0.023 | 0 | 0.06 | 0.1 | 0.06 | 0 | 0.1 | 0.03 |
| 4 | 58598522 | rs9790425 | 1.186 | 0.79 | 1.609 | -0.389 | 0.008 | 0.043 | 0.043 | 0.465 | 0.91 | 0.7 | 0.94 | 0.87 | 0.97 | 0.88 |
| 4 | 58614509 | rs343549 | -1.362 | 0.567 | 1.861 | -0.847 | -0.008 | -0.024 | -0.024 | 0 | 0.26 | 0.54 | 0.42 | 0.19 | 0.26 | 0.28 |
| 4 | 58616162 | rs343554 | -0.839 | 0.539 | 1.812 | -0.839 | -0.011 | 0.034 | 0.034 | 0 | 0.2 | 0.12 | 0.27 | 0.1 | 0.26 | 0.15 |
| 4 | 58631640 | rs343592 | -1.571 | 0.499 | 1.95 | -0.883 | -0.008 | -0.02 | -0.02 | 4.407 | 0.27 | 0.52 | 0.42 | 0.19 | 0.26 | 0.28 |
| 4 | 58642656 | rs343566 | -0.414 | 0.478 | 1.358 | -0.893 | -0.068 | 0.029 | 0.029 | 0 | 0.17 | 0.01 | 0.25 | 0.1 | 0.25 | 0.15 |
| 4 | 58646143 | rs189353 | -1.708 | 0.491 | 1.446 | -0.884 | 0.009 | 0.019 | 0.019 | 0.157 | 0.28 | 0.68 | 0.42 | 0.19 | 0.26 | 0.28 |
| 4 | 58655901 | rs1509528 | 0.251 | 0.634 | 1.392 | -0.866 | 0.01 | -0.041 | -0.041 | 0.239 | 0.82 | 0.69 | 0.61 | 0.91 | 0.77 | 0.9 |
| 4 | 58662041 | rs10034684 | 0.796 | 0.729 | 0.708 | -1.163 | 0.006 | 0.015 | 0.015 | 0 | 0.54 | 0.54 | 0.4 | 0.75 | 0.41 | 0.55 |
| 4 | 58662227 | rs6814709 | -0.177 | 0.735 | 0.161 | -1.183 | -0.045 | 0.042 | 0.042 | 0 | 0.3 | 0.22 | 0.22 | 0.17 | 0.38 | 0.36 |
| 4 | 58662408 | rs6831719 | -0.177 | 0.735 | 0.161 | -1.183 | -0.045 | 0.042 | 0.042 | 0 | 0.3 | 0.22 | 0.22 | 0.17 | 0.38 | 0.36 |
| 4 | 58662671 | rs13126470 | 1.614 | 0.373 | -0.384 | -1.198 | 0.088 | -0.025 | -0.025 | 0.514 | 0.38 | 0.14 | 0.28 | 0.61 | 0.23 | 0.42 |
| 4 | 58664089 | rs13134774 | 0.076 | 0.476 | -0.122 | -1.128 | 0.004 | -0.005 | -0.005 | 1.742 | 0.13 | 0.13 | 0.1 | 0.14 | 0.16 | 0.12 |
| 4 | 58669378 | rs11933998 | 1.487 | -0.147 | -0.671 | -0.987 | 0.084 | -0.045 | -0.045 | 0 | 0.35 | 0.13 | 0.27 | 0.62 | 0.22 | 0.4 |
| 4 | 58670132 | rs2899102 | 0.235 | 0.135 | -0.487 | -1.149 | 0 | 0.011 | 0.011 | 0 | 0.79 | 0.82 | 0.72 | 0.9 | 0.76 | 0.77 |
| 4 | 58670994 | rs951232 | 0.045 | 0.008 | -1.017 | -1.172 | 0.004 | -0.014 | -0.014 | 0 | 0.39 | 0.32 | 0.29 | 0.2 | 0.47 | 0.4 |
| 4 | 58671636 | rs979119 | -0.347 | -0.234 | -1.934 | -0.683 | -0.027 | 0.003 | 0.003 | 0.235 | 0.11 | 0.01 | 0.05 | 0.11 | 0.1 | 0.17 |

| | | | | | | | | | | | | | | | | |
|---|---|---|---|---|---|---|---|---|---|---|---|---|---|---|---|---|
| 4 | 58674284 | rs4865283 | 0.803 | -0.354 | -1.551 | -0.373 | 0.064 | -0.03 | -0.03 | 0 | 0.32 | 0.16 | 0.38 | 0.66 | 0.21 | 0.34 |
| 4 | 58684651 | rs1509531 | -0.838 | 0.023 | -0.927 | -0.175 | 0.011 | 0.005 | 0.005 | 0.103 | 0.29 | 0.76 | 0.34 | 0.28 | 0.22 | 0.29 |
| 4 | 58689383 | rs6848712 | -1.175 | 0.298 | -1.002 | -0.391 | 0.01 | 0.026 | 0.026 | 0.211 | 0.38 | 0.7 | 0.55 | 0.29 | 0.38 | 0.39 |
| 4 | 58692467 | rs1509539 | 0.038 | 0.698 | -1.14 | -0.274 | -0.028 | 0.043 | 0.043 | 0 | 0.1 | 0.03 | 0.21 | 0.01 | 0.15 | 0.07 |
| 4 | 58696072 | rs7679186 | -0.347 | 0.323 | -0.803 | -0.214 | -0.003 | -0.038 | -0.038 | 0 | 0.25 | 0.56 | 0.27 | 0.15 | 0.29 | 0.26 |
| 4 | 58696511 | rs12505795 | -0.32 | -0.153 | -1.176 | 0.084 | 0.008 | -0.007 | -0.007 | 0 | 0.1 | 0.07 | 0.16 | 0.16 | 0.05 | 0.14 |
| 4 | 58698487 | rs10011661 | 0.68 | 0.27 | -1.885 | 0.197 | -0.011 | -0.119 | -0.119 | 0 | 0.28 | 0.07 | 0.12 | 0.51 | 0.2 | 0.23 |
| 4 | 58698638 | rs10019628 | 0.68 | 0.27 | -1.885 | 0.221 | -0.011 | -0.121 | -0.121 | 0 | 0.28 | 0.07 | 0.12 | 0.51 | 0.2 | 0.23 |
| 4 | 58700119 | rs9312691 | -0.212 | 0.753 | -1.191 | -0.458 | 0.003 | -0.056 | -0.056 | 0 | 0.64 | 0.42 | 0.55 | 0.9 | 0.57 | 0.72 |
| 4 | 58700901 | rs3843894 | 0.896 | 0.447 | -0.535 | 0.264 | -0.004 | 0.007 | 0.007 | 0 | 0.32 | 0.4 | 0.21 | 0.47 | 0.22 | 0.38 |
| 4 | 58702130 | rs13134527 | 0.838 | 0.757 | -0.284 | 0.084 | 0.004 | 0.009 | 0.009 | 0 | 0.27 | 0.24 | 0.24 | 0.14 | 0.25 | 0.27 |
| 4 | 58702467 | rs1904778 | 0.312 | 0.711 | -0.033 | 0.092 | 0.012 | -0.003 | -0.003 | 0 | 0.59 | 0.56 | 0.44 | 0.62 | 0.47 | 0.66 |
| 4 | 58703510 | rs983355 | -1.101 | 0.648 | -0.053 | 0.03 | -0.003 | 0.001 | 0.001 | 0 | 0.88 | 0.93 | 0.65 | 0.93 | 0.76 | 0.92 |
| 4 | 58704841 | rs1480321 | 0.079 | 0.527 | 0.102 | 0.088 | -0.001 | -0.001 | -0.001 | 0.826 | 0.28 | 0.22 | 0.2 | 0.31 | 0.29 | 0.25 |
| 4 | 58704924 | rs17088782 | 1.095 | 0.711 | -0.225 | -0.024 | -0.031 | 0.04 | 0.04 | 0 | 0.13 | 0.05 | 0.13 | 0.04 | 0.17 | 0.07 |
| 4 | 58706462 | rs10020207 | -0.897 | 0.661 | -0.103 | -0.058 | 0 | 0 | 0 | 0 | 0.55 | 0.44 | 0.44 | 0.45 | 0.53 | 0.53 |
| 4 | 58707016 | rs1480319 | -0.582 | 0.909 | -0.224 | 0.249 | 0.003 | 0.003 | 0.003 | 0 | 0.28 | 0.24 | 0.2 | 0.31 | 0.29 | 0.25 |
| 4 | 58714407 | rs922605 | -0.923 | 1.099 | -0.483 | -0.152 | -0.011 | 0.019 | 0.019 | 0 | 0.37 | 0.74 | 0.57 | 0.29 | 0.47 | 0.3 |
| 4 | 58727290 | rs1480301 | -1.14 | 0.983 | -0.47 | 0.143 | -0.038 | -0.019 | -0.019 | 0.719 | 0.38 | 0.83 | 0.6 | 0.46 | 0.47 | 0.32 |
| 4 | 58728612 | rs17088834 | -0.255 | 1.021 | 0.412 | 0.055 | 0.016 | -0.003 | -0.003 | 0 | 0.12 | 0.37 | 0.08 | 0.05 | 0.09 | 0.1 |
| 4 | 58738371 | rs17088838 | 0.133 | 1.371 | 0.138 | 0.049 | 0.002 | -0.001 | -0.001 | 0 | 0.09 | 0.08 | 0.07 | 0.04 | 0.11 | 0.07 |
| 4 | 58739725 | rs2412857 | 0.574 | 1.51 | 0.186 | 0.019 | -0.011 | -0.011 | -0.011 | 0 | 0.11 | 0.1 | 0.22 | 0.23 | 0.19 | 0.16 |
| 4 | 58750042 | rs10517417 | -0.825 | 1.459 | 0.231 | -0.509 | 0.029 | 0.052 | 0.052 | 0 | 0.22 | 0.47 | 0.48 | 0.43 | 0.34 | 0.21 |
| 4 | 58758713 | rs17088873 | 0.563 | 1.439 | 0.115 | -0.508 | 0.01 | 0.008 | 0.008 | 0 | 0.07 | 0.11 | 0.06 | 0.07 | 0.08 | 0.08 |
| 4 | 58760786 | rs17794379 | 2.704 | 1.915 | -0.154 | -0.618 | -0.034 | 0.028 | 0.028 | 0.012 | 0.06 | 0 | 0.04 | 0 | 0.11 | 0.03 |
| 4 | 58768454 | rs10517418 | -1.365 | 1.258 | 0.453 | -0.408 | 0.023 | 0.036 | 0.036 | 0 | 0.28 | 0.61 | 0.54 | 0.5 | 0.42 | 0.29 |
| 4 | 58768658 | rs6822760 | -0.658 | 1.21 | 1.088 | -0.469 | 0.033 | 0.019 | 0.019 | 0.762 | 0.11 | 0.44 | 0.12 | 0.07 | 0.1 | 0.09 |
| 4 | 58770663 | rs9884270 | -1.115 | 1.192 | 1.127 | -0.422 | 0.002 | 0.093 | 0.093 | 2.442 | 0.13 | 0.62 | 0.18 | 0.27 | 0.1 | 0.1 |
| 4 | 58772260 | rs9992403 | -0.852 | 1.176 | 1.245 | -0.401 | 0.026 | 0.002 | 0.002 | 0 | 0.2 | 0.27 | 0.31 | 0.23 | 0.34 | 0.2 |
| 4 | 58782909 | rs1480332 | -0.978 | 1.006 | 0.952 | -0.407 | -0.002 | 0.016 | 0.016 | 0 | 0.88 | 0.96 | 0.91 | 0.95 | 0.88 | 0.86 |
| 4 | 58796884 | rs1480308 | 0.235 | 1.401 | 1.013 | -0.156 | 0.015 | -0.018 | -0.018 | 0 | 0.11 | 0.18 | 0.2 | 0.07 | 0.19 | 0.11 |
| 4 | 58799916 | rs17088952 | -0.282 | 0.863 | 0.852 | 0.067 | 0.021 | 0.049 | 0.049 | 0 | 0.1 | 0.25 | 0.12 | 0.16 | 0.15 | 0.09 |
| 4 | 58807069 | rs6554506 | -1.075 | 0.495 | 0.379 | -0.005 | -0.003 | 0.004 | 0.004 | 0.13 | 0.24 | 0.73 | 0.25 | 0.39 | 0.23 | 0.2 |
| 4 | 58809621 | rs11133536 | -1.36 | 0.764 | 0.221 | -0.049 | 0.001 | -0.032 | -0.032 | 10.483 | 0.31 | 0.8 | 0.45 | 0.49 | 0.39 | 0.28 |
| 4 | 58810453 | rs17088964 | 0.85 | 0.728 | 0.628 | -0.087 | -0.021 | -0.007 | -0.007 | 0 | 0.11 | 0.21 | 0.06 | 0.06 | 0.08 | 0.1 |
| 4 | 58810492 | rs10001131 | -0.45 | 0.771 | 0.877 | -0.052 | 0.008 | 0.004 | 0.004 | 0 | 0.16 | 0.18 | 0.31 | 0.27 | 0.3 | 0.13 |
| 4 | 58815257 | rs17088981 | 0.826 | 0.683 | 0.901 | -0.131 | 0 | 0 | 0 | 0 | 0.11 | 0.16 | 0.08 | 0.06 | 0.09 | 0.14 |
| 4 | 58827259 | rs10015020 | -0.156 | 0.789 | 1.187 | -0.103 | 0.045 | 0.057 | 0.057 | 0 | 0.09 | 0.3 | 0.13 | 0.18 | 0.14 | 0.05 |
| 4 | 58832860 | rs11936360 | 0.798 | 0.793 | 1.059 | -0.118 | 0.039 | 0.009 | 0.009 | 0 | 0.08 | 0.17 | 0.2 | 0.1 | 0.17 | 0.09 |
| 4 | 58837610 | rs10008425 | 0.515 | 0.789 | 1.008 | -0.104 | -0.008 | -0.01 | -0.01 | 0.292 | 0.08 | 0.07 | 0.12 | 0.18 | 0.14 | 0.05 |
| 4 | 58840510 | rs17237378 | 1.518 | 0.774 | 0.589 | -0.423 | -0.047 | -0.041 | -0.041 | 0 | 0.52 | 0.09 | 0.43 | 0.48 | 0.43 | 0.51 |
| 4 | 58842939 | rs1383233 | 0.838 | 0.898 | 0.908 | -0.263 | 0.041 | 0.01 | 0.01 | 0 | 0.08 | 0.17 | 0.2 | 0.1 | 0.17 | 0.09 |
| 4 | 58851934 | rs11946236 | 0.008 | 0.855 | 1.093 | -0.229 | 0.025 | 0.04 | 0.04 | 0 | 0.09 | 0.23 | 0.13 | 0.18 | 0.14 | 0.08 |
| 4 | 58854092 | rs3912236 | 0.608 | 0.629 | 0.83 | -0.391 | -0.026 | -0.019 | -0.019 | 0.332 | 0.16 | 0.28 | 0.07 | 0.06 | 0.1 | 0.16 |
| 4 | 58857424 | rs13134586 | 0.833 | 0.689 | 0.514 | -0.385 | -0.002 | -0.001 | -0.001 | 0.375 | 0.15 | 0.22 | 0.07 | 0.06 | 0.1 | 0.16 |
| 4 | 58860052 | rs3913924 | -0.214 | 0.741 | 0.522 | -0.439 | -0.001 | -0.008 | -0.008 | 0 | 0.23 | 0.33 | 0.29 | 0.16 | 0.29 | 0.26 |
| 4 | 58861963 | rs270464 | -0.93 | 0.695 | 0.227 | -0.383 | 0.003 | -0.042 | -0.042 | 0 | 0.36 | 0.83 | 0.46 | 0.49 | 0.44 | 0.38 |
| 4 | 58884182 | rs1395880 | 1.129 | 0.782 | 0.163 | -0.311 | -0.003 | 0 | 0 | 0 | 0.08 | 0.06 | 0.19 | 0.1 | 0.18 | 0.09 |
| 4 | 58900069 | rs10013416 | 0.682 | 0.8 | 0.065 | -0.28 | 0.001 | -0.006 | -0.006 | 0 | 0.19 | 0.26 | 0.18 | 0.24 | 0.22 | 0.25 |
| 4 | 58904987 | rs4865293 | 1.245 | 0.769 | -0.321 | -0.569 | -0.062 | -0.032 | -0.032 | 1.267 | 0.47 | 0.08 | 0.41 | 0.5 | 0.39 | 0.46 |
| 4 | 58907784 | rs1507578 | -0.111 | 1.021 | -0.37 | -0.416 | -0.004 | 0.024 | 0.024 | 0 | 0.42 | 0.43 | 0.47 | 0.85 | 0.43 | 0.51 |
| 4 | 58908653 | rs1948974 | 1.189 | 1.057 | -0.637 | -0.437 | -0.016 | -0.054 | -0.054 | 0 | 0.21 | 0.09 | 0.27 | 0.48 | 0.2 | 0.27 |
| 4 | 58912736 | rs10028399 | 0.616 | 0.649 | -0.863 | -0.358 | 0 | -0.006 | -0.006 | 0 | 0.19 | 0.21 | 0.17 | 0.22 | 0.22 | 0.21 |
| 4 | 58913500 | rs1380196 | 0.553 | 0.691 | -0.874 | -0.165 | 0.018 | 0.016 | 0.016 | 0 | 0.32 | 0.38 | 0.47 | 0.58 | 0.38 | 0.4 |
| 4 | 58923439 | rs2174809 | -0.653 | 0.485 | -1.582 | -0.515 | -0.031 | 0.045 | 0.045 | 0 | 0.89 | 0.99 | 0.9 | 0.98 | 0.89 | 0.91 |
| 4 | 58933127 | rs270233 | -0.269 | 0.519 | -1.3 | -0.215 | 0.018 | 0.081 | 0.081 | 0 | 0.42 | 0.29 | 0.28 | 0.11 | 0.34 | 0.33 |
| 4 | 58935390 | rs930937 | -0.173 | 0.249 | -1.233 | 0.889 | 0.019 | -0.035 | -0.035 | 0 | 0.17 | 0.07 | 0.19 | 0.36 | 0.09 | 0.31 |
| 4 | 58935943 | rs2660943 | -0.438 | 0.823 | -1.275 | 0.107 | -0.018 | 0.035 | 0.035 | 0 | 0.7 | 0.82 | 0.66 | 0.91 | 0.62 | 0.71 |
| 4 | 58940551 | rs270245 | -0.128 | 1.108 | -0.638 | 0.301 | 0.024 | -0.012 | -0.012 | 4.502 | 0.23 | 0.39 | 0.21 | 0.2 | 0.27 | 0.15 |
| 4 | 58940617 | rs270244 | -0.067 | 0.506 | -0.653 | 0.508 | -0.013 | -0.014 | -0.014 | 0 | 0.42 | 0.37 | 0.55 | 0.55 | 0.51 | 0.46 |
| 4 | 58942148 | rs781588 | -0.323 | -0.421 | -0.245 | 0.272 | -0.072 | -0.019 | -0.019 | 0 | 0.73 | 0.57 | 0.88 | 0.79 | 0.89 | 0.77 |
| 4 | 58955664 | rs1507583 | 0.432 | -0.445 | -0.758 | 0.136 | 0.056 | 0.045 | 0.045 | 0 | 0.32 | 0.12 | 0.12 | 0.19 | 0.17 | 0.25 |
| 4 | 58960261 | rs1531305 | 0.589 | -0.412 | -0.758 | 0.122 | 0.046 | 0.038 | 0.038 | 0 | 0.3 | 0.12 | 0.12 | 0.19 | 0.17 | 0.25 |
| 4 | 58960389 | rs17290180 | 0.518 | -0.138 | -0.665 | 0.483 | -0.002 | 0.009 | 0.009 | 0 | 0.22 | 0.2 | 0.23 | 0.32 | 0.26 | 0.28 |
| 4 | 58960829 | rs17290187 | 0.612 | 0.93 | -0.605 | 0.545 | -0.011 | -0.018 | -0.018 | 0.579 | 0.14 | 0.16 | 0.13 | 0.33 | 0.14 | 0.18 |
| 4 | 58961150 | rs2610166 | -0.697 | 0.621 | -0.167 | 0.52 | 0.006 | -0.003 | -0.003 | 0 | 0.46 | 0.47 | 0.62 | 0.49 | 0.56 | 0.46 |
| 4 | 58967480 | rs1911605 | 0.132 | 0.898 | -0.448 | 0.382 | -0.084 | -0.063 | -0.063 | 1.072 | 0.31 | 0.18 | 0.43 | 0.42 | 0.38 | 0.35 |
| 4 | 58971838 | rs12504646 | 0.397 | 0.963 | -0.267 | 0.396 | 0.024 | -0.017 | -0.017 | 2.735 | 0.15 | 0.28 | 0.19 | 0.07 | 0.18 | 0.11 |
| 4 | 58974923 | rs17089110 | 0.379 | 0.964 | -0.267 | 0.396 | 0.023 | -0.017 | -0.017 | 0 | 0.15 | 0.29 | 0.19 | 0.07 | 0.18 | 0.11 |

| | | | | | | | | | | | | | | | | |
|---|---|---|---|---|---|---|---|---|---|---|---|---|---|---|---|---|
| 4 | 58984954 | rs1433418 | 0.73 | 0.772 | -0.622 | 0.412 | 0.036 | -0.003 | -0.003 | 0 | 0.51 | 0.27 | 0.35 | 0.51 | 0.44 | 0.53 |
| 4 | 58991684 | rs4864644 | 1.073 | 1.089 | -0.974 | 0.287 | -0.079 | 0.011 | 0.011 | 0 | 0.14 | 0.01 | 0.29 | 0.09 | 0.24 | 0.17 |
| 4 | 58994093 | rs17237553 | 1.014 | 1.061 | -0.966 | 0.276 | -0.048 | -0.004 | -0.004 | 0 | 0.12 | 0.02 | 0.18 | 0.07 | 0.19 | 0.11 |
| 4 | 58995218 | rs13115208 | 0.854 | 0.67 | -1.007 | 0.294 | 0.026 | 0.049 | 0.049 | 0 | 0.42 | 0.13 | 0.31 | 0.28 | 0.36 | 0.36 |
| 4 | 59003803 | rs12506735 | 0.963 | 0.881 | -1.057 | 0.187 | -0.054 | 0.002 | 0.002 | 0 | 0.13 | 0.03 | 0.2 | 0.06 | 0.22 | 0.12 |
| 4 | 59006847 | rs1433409 | 0.536 | 0.889 | -0.873 | 0.346 | -0.028 | -0.018 | -0.018 | 0.01 | 0.61 | 0.24 | 0.42 | 0.39 | 0.51 | 0.6 |
| 4 | 59006971 | rs1433408 | -1.088 | 0.878 | -0.438 | 0.392 | 0.021 | 0.029 | 0.029 | 0 | 0.86 | 0.81 | 0.87 | 0.74 | 0.87 | 0.86 |
| 4 | 59007188 | rs1563772 | 1.148 | 0.701 | -0.618 | 0.286 | 0.008 | 0.071 | 0.071 | 0 | 0.17 | 0.02 | 0.1 | 0.05 | 0.15 | 0.18 |
| 4 | 59008412 | rs10517437 | 0.882 | 1.39 | -0.045 | 0.739 | -0.018 | -0.012 | -0.012 | 0 | 0.61 | 0.28 | 0.43 | 0.39 | 0.51 | 0.61 |
| 4 | 59008961 | rs2117065 | 0.323 | 1.315 | 0.319 | 0.704 | 0.038 | -0.047 | -0.047 | 0.26 | 0.18 | 0.41 | 0.31 | 0.09 | 0.23 | 0.17 |
| 4 | 59026944 | rs1080057 | 0.392 | 1.152 | 0.101 | 0.877 | 0.049 | 0.031 | 0.031 | 0.476 | 0.68 | 0.42 | 0.43 | 0.61 | 0.46 | 0.61 |
| 4 | 59028886 | rs3943387 | -0.815 | 1.484 | 0.272 | 0.888 | 0.054 | 0.046 | 0.046 | 1.49 | 0.23 | 0.5 | 0.3 | 0.3 | 0.34 | 0.24 |
| 4 | 59031089 | rs892307 | -1.041 | 1.042 | 0.074 | 0.886 | 0.007 | 0.029 | 0.029 | 0.017 | 0.83 | 0.76 | 0.85 | 0.71 | 0.87 | 0.92 |
| 4 | 59035958 | rs12646025 | -0.978 | 0.115 | -0.073 | 0.367 | 0.007 | 0.005 | 0.005 | 0 | 0.64 | 0.68 | 0.89 | 0.87 | 0.83 | 0.6 |
| 4 | 59036816 | rs17827154 | 0.603 | 0.779 | -0.429 | 0.5 | -0.108 | 0 | 0 | 0 | 0.25 | 0.07 | 0.28 | 0.23 | 0.36 | 0.23 |
| 4 | 59042209 | rs10005074 | 0.12 | 0.65 | -0.052 | 0.452 | -0.019 | -0.01 | -0.01 | 0 | 0.34 | 0.19 | 0.56 | 0.64 | 0.46 | 0.37 |
| 4 | 59050934 | rs1865182 | -1.172 | 0.285 | 0.088 | 0.337 | -0.01 | -0.014 | -0.014 | 0 | 0.63 | 0.62 | 0.87 | 0.88 | 0.83 | 0.6 |
| 4 | 59071485 | rs2570100 | -0.223 | 0.7 | -0.052 | 0.891 | -0.029 | -0.029 | -0.029 | 0 | 0.37 | 0.24 | 0.57 | 0.53 | 0.53 | 0.38 |
| 4 | 59072672 | rs2611162 | -0.84 | 0.635 | -0.459 | 0.852 | -0.072 | -0.071 | -0.071 | 0.404 | 0.44 | 0.85 | 0.62 | 0.53 | 0.53 | 0.39 |
| 4 | 59108872 | rs2412882 | 0.381 | 0.674 | 0.133 | 0.87 | -0.017 | -0.017 | -0.017 | 0 | 0.58 | 0.33 | 0.37 | 0.48 | 0.46 | 0.6 |
| 4 | 59113239 | rs10007387 | 0.42 | 0.671 | 0.128 | 0.794 | -0.023 | -0.023 | -0.023 | 0 | 0.58 | 0.28 | 0.38 | 0.48 | 0.46 | 0.6 |
| 4 | 59117179 | rs2570085 | 1.27 | 0.434 | -0.209 | 0.741 | 0.09 | 0.012 | 0.012 | 0 | 0.34 | 0.13 | 0.12 | 0.26 | 0.17 | 0.42 |
| 4 | 59133280 | rs13105776 | 0.847 | 0.691 | -0.416 | 0.598 | -0.047 | 0.011 | 0.011 | 0 | 0.12 | 0.02 | 0.14 | 0.08 | 0.2 | 0.11 |
| 4 | 59139089 | rs2553297 | -0.101 | 0.856 | -0.393 | 0.524 | -0.016 | -0.086 | -0.086 | 0 | 0.72 | 0.87 | 0.72 | 0.57 | 0.71 | 0.79 |
| 4 | 59141562 | rs2553296 | 0.93 | 0.915 | -0.461 | 0.555 | -0.02 | -0.013 | -0.013 | 0 | 0.58 | 0.29 | 0.38 | 0.47 | 0.47 | 0.6 |
| 4 | 59156538 | rs17237658 | 1.618 | 1.011 | -0.876 | 0.455 | -0.055 | -0.017 | -0.017 | 0.266 | 0.09 | 0.02 | 0.29 | 0.1 | 0.24 | 0.19 |
| 4 | 59156851 | rs2570069 | -2.319 | 0.911 | -0.925 | 0.447 | -0.066 | -0.036 | -0.036 | 0.903 | 0.86 | 0.99 | 0.85 | 0.82 | 0.84 | 0.92 |
| 4 | 59177075 | rs1915903 | 1.541 | -0.073 | -0.783 | 0.293 | 0.066 | 0.072 | 0.072 | 0 | 0.28 | 0.1 | 0.06 | 0.11 | 0.15 | 0.29 |
| 4 | 59211129 | rs12648067 | -0.805 | 0.559 | 0.056 | 0.834 | -0.015 | 0.056 | 0.056 | 0 | 0.31 | 0.47 | 0.49 | 0.44 | 0.29 | 0.21 |
| 4 | 59217565 | rs17089256 | 0.36 | 1.131 | 0.756 | 0.95 | 0.047 | -0.035 | -0.035 | 0 | 0.12 | 0.25 | 0.12 | 0.03 | 0.24 | 0.16 |
| 4 | 59217612 | rs1522093 | -0.152 | 1.091 | 0.215 | 1.141 | -0.015 | -0.044 | -0.044 | 0 | 0.15 | 0.06 | 0.33 | 0.25 | 0.17 | 0.12 |
| 4 | 59221416 | rs1522094 | -1.29 | 1.011 | 0.265 | 1.109 | 0.007 | 0.004 | 0.004 | 0.609 | 0.31 | 0.66 | 0.49 | 0.36 | 0.42 | 0.31 |
| 4 | 59252047 | rs10015715 | -1.067 | 1.127 | -0.271 | 0.942 | 0.005 | 0.008 | 0.008 | 0.189 | 0.28 | 0.36 | 0.46 | 0.45 | 0.42 | 0.35 |
| 4 | 59273187 | rs12642751 | -0.598 | 0.504 | -0.684 | 1.029 | 0 | -0.003 | -0.003 | 0.03 | 0.17 | 0.14 | 0.34 | 0.35 | 0.17 | 0.16 |
| 4 | 59275372 | rs1222269 | -1.107 | -0.017 | -1.179 | 0.833 | -0.001 | -0.003 | -0.003 | 0 | 0.81 | 0.86 | 0.79 | 0.76 | 0.83 | 0.86 |
| 4 | 59279104 | rs2140069 | -0.575 | -0.375 | -0.708 | 0.995 | 0.02 | -0.036 | -0.036 | 0 | 0.18 | 0.58 | 0.31 | 0.11 | 0.24 | 0.22 |
| 4 | 59284649 | rs935999 | -0.204 | -0.378 | -0.618 | 0.993 | 0.061 | -0.071 | -0.071 | 0.19 | 0.16 | 0.45 | 0.3 | 0.04 | 0.23 | 0.21 |
| 4 | 59285092 | rs11937415 | -0.209 | -0.56 | -0.833 | 1.062 | 0.012 | -0.086 | -0.086 | 0 | 0.15 | 0.04 | 0.14 | 0.33 | 0.09 | 0.15 |
| 4 | 59304973 | rs1222271 | 0.51 | -1.064 | -1.083 | 0.878 | 0.007 | 0.005 | 0.005 | 0.019 | 0.19 | 0.19 | 0.07 | 0.11 | 0.1 | 0.26 |
| 4 | 59319223 | rs17218168 | 0.049 | -0.046 | -0.763 | 1.241 | 0.013 | -0.035 | -0.035 | 0 | 0.11 | 0.05 | 0.11 | 0.33 | 0.07 | 0.11 |
| 4 | 59321537 | rs2037458 | -1.335 | 0.049 | -0.904 | 1.283 | -0.006 | 0.028 | 0.028 | 1.412 | 0.3 | 0.6 | 0.42 | 0.44 | 0.31 | 0.34 |
| 4 | 59338580 | rs12647734 | -1.208 | 0.142 | -1.09 | 1.094 | -0.008 | 0.022 | 0.022 | 1.299 | 0.31 | 0.6 | 0.43 | 0.44 | 0.32 | 0.35 |
| 4 | 59345914 | rs11133566 | 0.035 | 1.368 | -1.713 | 0.559 | -0.019 | -0.043 | -0.043 | 0 | 0.11 | 0 | 0.09 | 0.16 | 0.08 | 0.09 |
| 4 | 59348360 | rs1463323 | -0.45 | 1.437 | -1.108 | 0.35 | -0.022 | 0.02 | 0.02 | 0 | 0.12 | 0.05 | 0.26 | 0.04 | 0.23 | 0.2 |
| 4 | 59360441 | rs13121688 | -0.511 | 0.887 | -0.896 | 0.374 | -0.002 | -0.007 | -0.007 | 0 | 0.45 | 0.49 | 0.49 | 0.31 | 0.45 | 0.55 |
| 4 | 59362260 | rs4864655 | -0.768 | -0.329 | -0.679 | 0.504 | 0.023 | 0.004 | 0.004 | 0.793 | 0.5 | 0.42 | 0.54 | 0.5 | 0.34 | 0.56 |
| 4 | 59369235 | rs10007576 | -0.708 | -0.615 | -0.332 | 0.971 | 0.066 | 0.074 | 0.074 | 0.778 | 0.48 | 0.71 | 0.61 | 0.67 | 0.34 | 0.56 |
| 4 | 59375202 | rs11937554 | -0.259 | 0.145 | -0.081 | 0.487 | 0.008 | -0.009 | -0.009 | 0.374 | 0.84 | 0.81 | 0.69 | 0.94 | 0.74 | 0.76 |
| 4 | 59375444 | rs10007616 | 0.059 | -0.493 | 0.316 | 0.826 | -0.04 | 0.087 | 0.087 | 0 | 0.22 | 0.33 | 0.22 | 0.51 | 0.15 | 0.24 |
| 4 | 59406749 | rs9999484 | -0.001 | -0.729 | -0.109 | 0.121 | 0.001 | 0 | 0 | 0 | 0.43 | 0.56 | 0.41 | 0.42 | 0.35 | 0.41 |
| 4 | 59408250 | rs6818886 | -0.92 | -0.624 | 0.026 | 0.049 | 0.014 | -0.02 | -0.02 | 0 | 0.84 | 0.77 | 0.7 | 0.95 | 0.76 | 0.74 |
| 4 | 59413750 | rs4242018 | 0.506 | -0.784 | 0.069 | -0.032 | 0.001 | 0.009 | 0.009 | 0 | 0.72 | 0.65 | 0.55 | 0.67 | 0.7 | 0.59 |
| 4 | 59413971 | rs13109943 | 0.61 | -0.839 | -0.011 | 0.002 | -0.016 | -0.033 | -0.033 | 0 | 0.41 | 0.23 | 0.29 | 0.53 | 0.41 | 0.33 |
| 4 | 59419273 | rs10517441 | 0.053 | -0.889 | -0.02 | 0.12 | -0.009 | 0.017 | 0.017 | 0 | 0.14 | 0.16 | 0.16 | 0.28 | 0.05 | 0.2 |
| 4 | 59423332 | rs13114819 | 0.966 | -0.951 | -0.049 | 0.088 | -0.007 | 0.035 | 0.035 | 0 | 0.15 | 0.15 | 0.12 | 0.49 | 0.11 | 0.18 |
| 4 | 59436076 | rs9996466 | 1.37 | -0.87 | -0.13 | -0.185 | -0.035 | 0.049 | 0.049 | 1.181 | 0.26 | 0.16 | 0.16 | 0.04 | 0.3 | 0.14 |
| 4 | 59444611 | rs282722 | -1.764 | -1.04 | 0.09 | -0.289 | -0.001 | -0.003 | -0.003 | 0 | 0.74 | 0.73 | 0.76 | 0.86 | 0.74 | 0.74 |
| 4 | 59445288 | rs282720 | -1.12 | -1.011 | 0.091 | -0.192 | -0.006 | -0.028 | -0.028 | 0.232 | 0.44 | 0.47 | 0.58 | 0.82 | 0.4 | 0.59 |
| 4 | 59452393 | rs282745 | 1.004 | -0.896 | -0.093 | -0.177 | -0.034 | 0.023 | 0.023 | 0.02 | 0.13 | 0.06 | 0.25 | 0.05 | 0.23 | 0.2 |
| 4 | 59481416 | rs12647295 | 0.952 | -1.001 | 0.079 | 0.366 | -0.022 | 0.051 | 0.051 | 0 | 0.15 | 0.18 | 0.13 | 0.42 | 0.11 | 0.19 |
| 4 | 59484288 | rs17218287 | -0.078 | -0.753 | 0.038 | 0.409 | -0.005 | 0.012 | 0.012 | 0 | 0.14 | 0.16 | 0.2 | 0.28 | 0.07 | 0.2 |
| 4 | 59489084 | rs1605774 | -0.135 | -0.732 | 0.061 | 0.405 | -0.014 | 0.031 | 0.031 | 0 | 0.15 | 0.2 | 0.21 | 0.28 | 0.07 | 0.2 |
| 4 | 59490196 | rs4640727 | -0.112 | -0.706 | -0.068 | 0.408 | -0.003 | 0.009 | 0.009 | 0 | 0.15 | 0.15 | 0.2 | 0.28 | 0.07 | 0.2 |
| 4 | 59491899 | rs2662841 | -1.026 | -0.657 | -0.079 | 0.325 | 0.008 | 0.064 | 0.064 | 0 | 0.71 | 0.84 | 0.76 | 0.87 | 0.71 | 0.73 |
| 4 | 59497894 | rs10031559 | -0.074 | -0.717 | 0.103 | 0.43 | -0.004 | 0.073 | 0.073 | 0 | 0.1 | 0.17 | 0.19 | 0.28 | 0.06 | 0.15 |
| 4 | 59502687 | rs282705 | 0.047 | -0.652 | 0.356 | 0.125 | 0.033 | -0.007 | -0.007 | 0.361 | 0.15 | 0.24 | 0.26 | 0.11 | 0.27 | 0.27 |
| 4 | 59502974 | rs282706 | -0.283 | -0.973 | 0.371 | -0.058 | 0.003 | -0.019 | -0.019 | 0 | 0.44 | 0.51 | 0.5 | 0.23 | 0.57 | 0.47 |
| 4 | 59507265 | rs282710 | 0.609 | -0.562 | 0.2 | 0.028 | 0.019 | -0.008 | -0.008 | 0 | 0.12 | 0.16 | 0.3 | 0.07 | 0.26 | 0.17 |
| 4 | 59515454 | rs1565822 | 0.54 | -0.967 | 0.36 | -0.108 | -0.008 | -0.04 | -0.04 | 0 | 0.32 | 0.42 | 0.2 | 0.12 | 0.31 | 0.2 |

| | | | | | | | | | | | | | | | | |
|---|---|---|---|---|---|---|---|---|---|---|---|---|---|---|---|---|
| 4 | 59520026 | rs1396579 | -0.078 | -1.113 | 0.425 | 0.246 | -0.017 | 0.002 | 0.002 | 0.033 | 0.39 | 0.55 | 0.31 | 0.37 | 0.32 | 0.41 |
| 4 | 59535533 | rs9994974 | 0.272 | -1.496 | 0.215 | 0.365 | 0.008 | -0.011 | -0.011 | 0.037 | 0.38 | 0.25 | 0.56 | 0.47 | 0.58 | 0.31 |
| 4 | 59541966 | rs1961726 | -0.481 | -1.811 | 0.223 | 0.104 | 0.043 | 0.02 | 0.02 | 0 | 0.73 | 0.84 | 0.9 | 0.83 | 0.88 | 0.67 |
| 4 | 59550505 | rs1828204 | 2.046 | -1.761 | 0.42 | 0.114 | 0 | 0.003 | 0.003 | 0.02 | 0.23 | 0.23 | 0.2 | 0.12 | 0.26 | 0.19 |
| 4 | 59559715 | rs10004285 | -0.141 | -1.632 | 0.516 | 0.232 | -0.008 | -0.007 | -0.007 | 0.1 | 0.44 | 0.5 | 0.67 | 0.68 | 0.58 | 0.39 |
| 4 | 59564300 | rs4865352 | 0.672 | -1.541 | 0.583 | 0.332 | -0.011 | -0.023 | -0.023 | 0 | 0.36 | 0.24 | 0.53 | 0.49 | 0.54 | 0.29 |
| 4 | 59565589 | rs4865353 | 0.011 | -1.45 | 0.616 | 0.348 | -0.004 | -0.009 | -0.009 | 0 | 0.42 | 0.49 | 0.64 | 0.68 | 0.54 | 0.38 |
| 4 | 59567739 | rs1396576 | 0.305 | -1.365 | 0.602 | 0.649 | 0.065 | -0.095 | -0.095 | 0 | 0.26 | 0.11 | 0.17 | 0.51 | 0.16 | 0.37 |
| 4 | 59572909 | rs13115335 | 1.432 | -1.328 | 0.582 | 0.493 | -0.001 | 0.024 | 0.024 | 0.056 | 0.31 | 0.3 | 0.22 | 0.12 | 0.31 | 0.24 |
| 4 | 59573186 | rs17089553 | 1.192 | -1.17 | 0.634 | 0.4 | 0.018 | 0.014 | 0.014 | 0 | 0.26 | 0.31 | 0.27 | 0.4 | 0.29 | 0.17 |
| 4 | 59576664 | rs6850334 | 1.003 | -1.05 | 0.424 | 0.356 | -0.049 | -0.008 | -0.008 | 0 | 0.16 | 0.04 | 0.31 | 0.14 | 0.26 | 0.25 |
| 4 | 59577435 | rs4864659 | 0.914 | -1.047 | 0.451 | 0.358 | -0.015 | 0.003 | 0.003 | 0 | 0.16 | 0.11 | 0.32 | 0.14 | 0.26 | 0.25 |
| 4 | 59582759 | rs1472926 | -0.355 | -1.067 | 0.505 | 0.368 | -0.038 | -0.018 | -0.018 | 0.052 | 0.74 | 0.71 | 0.89 | 0.85 | 0.86 | 0.71 |
| 4 | 59595781 | rs1027867 | -0.829 | -0.728 | 0.247 | 0.666 | -0.034 | -0.003 | -0.003 | 0 | 0.82 | 0.75 | 0.92 | 0.86 | 0.89 | 0.74 |
| 4 | 59604951 | rs10005244 | 0.912 | -0.559 | -0.451 | 0.564 | 0.013 | 0.092 | 0.092 | 0.319 | 0.37 | 0.2 | 0.22 | 0.12 | 0.33 | 0.28 |
| 4 | 59605202 | rs9993637 | 0.904 | -0.562 | -0.453 | 0.563 | 0.013 | 0.092 | 0.092 | 0.312 | 0.37 | 0.2 | 0.22 | 0.12 | 0.33 | 0.28 |
| 4 | 59621853 | rs1868416 | 0.358 | -0.476 | -0.203 | 0.873 | -0.009 | -0.003 | -0.003 | 0 | 0.64 | 0.47 | 0.58 | 0.65 | 0.63 | 0.51 |
| 4 | 59627178 | rs10517447 | -0.565 | 0.139 | 0.066 | 0.91 | -0.023 | -0.038 | -0.038 | 2.786 | 0.75 | 0.62 | 0.87 | 0.86 | 0.79 | 0.74 |
| 4 | 59629835 | rs6843830 | 0.512 | 0.822 | 0.215 | 1.576 | 0.015 | 0.028 | 0.028 | 0 | 0.2 | 0.28 | 0.24 | 0.34 | 0.21 | 0.15 |
| 4 | 59633011 | rs7666280 | 0.355 | 0.824 | -0.029 | 1.583 | 0.007 | 0.041 | 0.041 | 0 | 0.21 | 0.28 | 0.32 | 0.54 | 0.21 | 0.21 |
| 4 | 59636452 | rs10011039 | 0.698 | 0.96 | -0.213 | 1.045 | -0.015 | -0.02 | -0.02 | 0 | 0.61 | 0.69 | 0.44 | 0.46 | 0.5 | 0.58 |
| 4 | 59637473 | rs6829904 | 0.132 | 1.037 | -0.427 | 0.806 | -0.022 | 0 | 0 | 0 | 0.16 | 0.1 | 0.32 | 0.18 | 0.26 | 0.24 |
| 4 | 59637674 | rs6813611 | -0.278 | 1.064 | -0.541 | 0.802 | -0.058 | -0.003 | -0.003 | 0 | 0.19 | 0.05 | 0.32 | 0.18 | 0.3 | 0.26 |
| 4 | 59645426 | rs7691858 | -0.122 | 0.99 | -0.048 | 0.783 | 0.008 | -0.005 | -0.005 | 0 | 0.18 | 0.24 | 0.33 | 0.18 | 0.26 | 0.25 |
| 4 | 59667025 | rs10517449 | -0.396 | 0.989 | -0.04 | 0.774 | 0.005 | -0.004 | -0.004 | 0.003 | 0.21 | 0.24 | 0.34 | 0.19 | 0.3 | 0.26 |
| 4 | 59673037 | rs1992941 | -0.972 | 0.901 | -0.021 | 0.977 | -0.011 | -0.002 | -0.002 | 0.273 | 0.41 | 0.38 | 0.57 | 0.46 | 0.5 | 0.45 |
| 4 | 59673650 | rs1449031 | 0.903 | 0.411 | 0.104 | 0.564 | 0.014 | 0.02 | 0.02 | 0 | 0.38 | 0.31 | 0.18 | 0.11 | 0.28 | 0.31 |
| 4 | 59677511 | rs10517450 | 1.125 | 0.385 | -0.012 | 0.674 | -0.01 | 0.009 | 0.009 | 0 | 0.19 | 0.16 | 0.18 | 0.06 | 0.2 | 0.11 |
| 4 | 59686485 | rs13147033 | -0.379 | 0.328 | 0.043 | 0.899 | -0.016 | 0.046 | 0.046 | 0 | 0.42 | 0.34 | 0.51 | 0.25 | 0.51 | 0.38 |
| 4 | 59689961 | rs2122155 | -0.595 | 0.332 | 0.19 | 0.889 | 0.001 | -0.002 | -0.002 | 0 | 0.44 | 0.45 | 0.58 | 0.46 | 0.52 | 0.47 |
| 4 | 59697508 | rs17237994 | 0.933 | 0.516 | -0.187 | 0.606 | -0.075 | 0.124 | 0.124 | 0.376 | 0.23 | 0.04 | 0.14 | 0.04 | 0.2 | 0.16 |
| 4 | 59710106 | rs12502978 | -1.153 | 0.548 | 0.259 | 1.024 | 0.012 | 0.068 | 0.068 | 0.724 | 0.79 | 0.72 | 0.74 | 0.6 | 0.76 | 0.78 |
| 4 | 59713040 | rs1868419 | 1.241 | 0.736 | 0.082 | 1.077 | -0.006 | -0.005 | -0.005 | 0 | 0.12 | 0.14 | 0.35 | 0.24 | 0.26 | 0.23 |
| 4 | 59715417 | rs9998639 | 0.877 | 0.65 | 0.028 | 0.904 | -0.021 | 0.087 | 0.087 | 1.341 | 0.27 | 0.12 | 0.15 | 0.04 | 0.24 | 0.18 |
| 4 | 59720431 | rs7685041 | -1.365 | 0.553 | 0.107 | 1.124 | -0.002 | -0.001 | -0.001 | 0 | 0.8 | 0.82 | 0.74 | 0.6 | 0.76 | 0.78 |
| 4 | 59727134 | rs1900867 | -1.425 | 0.56 | -0.032 | 1.127 | -0.011 | -0.033 | -0.033 | 0.056 | 0.8 | 0.86 | 0.74 | 0.6 | 0.76 | 0.78 |
| 4 | 59799872 | rs9997522 | 0.676 | 0.359 | 0.212 | 0.933 | 0.014 | 0.017 | 0.017 | 0 | 0.37 | 0.34 | 0.17 | 0.12 | 0.26 | 0.26 |
| 4 | 59805404 | rs4074334 | 0.702 | 0.782 | 0.405 | 1.648 | -0.011 | -0.029 | -0.029 | 0 | 0.2 | 0.15 | 0.26 | 0.39 | 0.25 | 0.23 |
| 4 | 59806257 | rs7436625 | -0.406 | 0.719 | 0.515 | 1.629 | 0.012 | 0.116 | 0.116 | 0.215 | 0.77 | 0.62 | 0.67 | 0.4 | 0.75 | 0.67 |
| 4 | 59813594 | rs9685766 | -0.384 | 0.412 | 0.039 | 1.388 | -0.002 | 0.011 | 0.011 | 3.242 | 0.27 | 0.67 | 0.42 | 0.3 | 0.29 | 0.33 |
| 4 | 59819545 | rs6551609 | -0.873 | 0.612 | -0.068 | 1.544 | -0.003 | -0.016 | -0.016 | 0.12 | 0.79 | 0.84 | 0.74 | 0.66 | 0.78 | 0.77 |
| 4 | 59832091 | rs7659089 | -0.542 | 0.578 | 0.469 | 1.55 | -0.001 | 0.002 | 0.002 | 0.819 | 0.77 | 0.62 | 0.74 | 0.72 | 0.78 | 0.79 |
| 4 | 59835266 | rs6819968 | -0.407 | 0.564 | 0.523 | 1.28 | -0.01 | -0.003 | -0.003 | 0 | 0.76 | 0.55 | 0.74 | 0.72 | 0.78 | 0.79 |
| 4 | 59842999 | rs11932542 | 0.137 | 0.767 | 0.468 | 1.227 | 0.013 | -0.001 | -0.001 | 0 | 0.65 | 0.58 | 0.38 | 0.37 | 0.5 | 0.51 |
| 4 | 59843855 | rs9942216 | 0.064 | 0.082 | 0.536 | 0.994 | -0.066 | -0.034 | -0.034 | 0 | 0.16 | 0.42 | 0.13 | 0.11 | 0.07 | 0.16 |
| 4 | 59847627 | rs11734556 | -0.906 | 0.361 | 0.187 | 1.312 | -0.017 | -0.052 | -0.052 | 0 | 0.78 | 0.87 | 0.75 | 0.63 | 0.75 | 0.76 |
| 4 | 59866889 | rs13112623 | 0.243 | -0.026 | 0.546 | 1.271 | -0.031 | -0.022 | -0.022 | 0.543 | 0.15 | 0.25 | 0.11 | 0.08 | 0.07 | 0.16 |
| 4 | 59867437 | rs4860200 | -0.615 | 0.485 | 0.094 | 1.115 | -0.024 | -0.049 | -0.049 | 0 | 0.72 | 0.83 | 0.73 | 0.64 | 0.72 | 0.76 |
| 4 | 59869101 | rs11733784 | 0.158 | 0.771 | 0.385 | 1.141 | -0.003 | 0.006 | 0.006 | 0.006 | 0.59 | 0.59 | 0.36 | 0.35 | 0.43 | 0.48 |
| 4 | 59869371 | rs1495722 | 1.431 | 0.923 | 0.366 | 1.182 | -0.007 | -0.001 | -0.001 | 0 | 0.1 | 0.09 | 0.36 | 0.27 | 0.27 | 0.28 |
| 4 | 59878013 | rs12649958 | 1.391 | 0.922 | 0.352 | 1.295 | -0.01 | 0.002 | 0.002 | 0.957 | 0.11 | 0.09 | 0.36 | 0.32 | 0.28 | 0.3 |
| 4 | 59889898 | rs6849738 | -0.643 | 0.676 | 0.265 | 1.051 | -0.019 | -0.018 | -0.018 | 0 | 0.72 | 0.83 | 0.73 | 0.7 | 0.73 | 0.78 |
| 4 | 59899352 | rs10002245 | 0.696 | 0.55 | 0.666 | 0.994 | 0.012 | 0.031 | 0.031 | 0 | 0.52 | 0.39 | 0.28 | 0.26 | 0.39 | 0.39 |
| 4 | 59900914 | rs7667333 | 0.599 | 0.559 | 0.641 | 0.992 | -0.003 | -0.014 | -0.014 | 0 | 0.53 | 0.49 | 0.28 | 0.26 | 0.39 | 0.39 |
| 4 | 59906337 | rs13111434 | 0.539 | -0.063 | -0.121 | 0.565 | -0.032 | -0.063 | -0.063 | 0.018 | 0.31 | 0.13 | 0.5 | 0.51 | 0.4 | 0.49 |
| 4 | 59906632 | rs10452251 | 1.396 | -0.079 | -0.194 | 0.383 | -0.006 | 0.008 | 0.008 | 0 | 0.25 | 0.2 | 0.18 | 0.17 | 0.28 | 0.21 |
| 4 | 59908346 | rs11131211 | 1.858 | -0.166 | -0.431 | 0.437 | 0.046 | 0.008 | 0.008 | 0 | 0.17 | 0.03 | 0.06 | 0.1 | 0.08 | 0.11 |
| 4 | 59916674 | rs2036098 | 0.752 | 0.04 | -0.404 | 0.123 | -0.01 | 0.024 | 0.024 | 0.349 | 0.24 | 0.15 | 0.18 | 0.03 | 0.22 | 0.1 |
| 4 | 59930697 | rs10034214 | 0.003 | -0.162 | -0.461 | 0.227 | 0.004 | 0.018 | 0.018 | 0.59 | 0.38 | 0.23 | 0.29 | 0.32 | 0.31 | 0.26 |
| 4 | 59950731 | rs2333727 | 0.231 | -0.115 | -0.508 | 0.002 | -0.006 | -0.052 | -0.052 | 0.487 | 0.57 | 0.46 | 0.39 | 0.2 | 0.52 | 0.37 |
| 4 | 59965367 | rs11729950 | -0.08 | -0.1 | -0.768 | 0.277 | 0.024 | 0.017 | 0.017 | 0 | 0.53 | 0.14 | 0.67 | 0.55 | 0.61 | 0.6 |
| 4 | 59970766 | rs1155335 | -0.192 | 0.776 | -0.199 | 0.55 | 0.095 | 0.076 | 0.076 | 0.25 | 0.21 | 0.41 | 0.44 | 0.3 | 0.36 | 0.31 |
| 4 | 59980062 | rs13151697 | 0.735 | 0.286 | -0.745 | 0.253 | -0.018 | 0.032 | 0.032 | 0 | 0.31 | 0.15 | 0.25 | 0.27 | 0.31 | 0.35 |
| 4 | 59988095 | rs357833 | 1.105 | 0.428 | -0.551 | 0.685 | -0.009 | 0.001 | 0.001 | 0.059 | 0.12 | 0.1 | 0.37 | 0.33 | 0.28 | 0.25 |
| 4 | 59998696 | rs13134467 | 0.143 | -0.2 | -0.297 | 0.569 | -0.002 | 0.011 | 0.011 | 0.017 | 0.64 | 0.38 | 0.49 | 0.52 | 0.6 | 0.6 |
| 4 | 60005301 | rs3114023 | 0.826 | -0.356 | -0.665 | 0.08 | -0.044 | -0.054 | -0.054 | 0.057 | 0.21 | 0.11 | 0.42 | 0.35 | 0.34 | 0.3 |
| 4 | 60010116 | rs2055486 | 0.482 | -0.928 | -0.694 | -0.269 | 0.022 | 0.011 | 0.011 | 0.104 | 0.5 | 0.26 | 0.43 | 0.49 | 0.54 | 0.57 |
| 4 | 60010750 | rs439122 | -2.841 | -1.033 | -0.51 | -0.326 | -0.022 | -0.034 | -0.034 | 0 | 0.93 | 0.82 | 0.98 | 1 | 0.97 | 0.99 |
| 4 | 60013350 | rs10021662 | 2.812 | -1.036 | -0.467 | -0.328 | -0.007 | -0.016 | -0.016 | 0 | 0.06 | 0.1 | 0.02 | 0 | 0.03 | 0.01 |

| | | | | | | | | | | | | | | | | |
|---|---|---|---|---|---|---|---|---|---|---|---|---|---|---|---|---|
| 4 | 60013919 | rs13347305 | 1.968 | -1.076 | -0.124 | -0.35 | -0.022 | -0.009 | -0.009 | 0.058 | 0.16 | 0.19 | 0.1 | 0.16 | 0.08 | 0.14 |
| 4 | 60016213 | rs12498186 | 1.363 | -1.238 | -0.664 | -0.515 | -0.017 | -0.014 | -0.014 | 0 | 0.24 | 0.11 | 0.21 | 0.28 | 0.28 | 0.34 |
| 4 | 60016378 | rs357809 | 0.543 | -0.825 | -0.299 | -0.398 | -0.024 | -0.036 | -0.036 | 0 | 0.28 | 0.21 | 0.44 | 0.36 | 0.37 | 0.3 |
| 4 | 60048376 | rs2134445 | -1.202 | -0.483 | -0.63 | -0.224 | 0.005 | 0.01 | 0.01 | 0 | 0.78 | 0.8 | 0.57 | 0.64 | 0.66 | 0.71 |
| 4 | 60054426 | rs11722985 | -1.193 | -0.482 | -0.629 | -0.223 | 0.004 | 0.01 | 0.01 | 0 | 0.78 | 0.8 | 0.57 | 0.64 | 0.66 | 0.71 |
| 4 | 60059161 | rs4489000 | -1.193 | -0.482 | -0.629 | -0.223 | 0.004 | 0.01 | 0.01 | 0 | 0.78 | 0.8 | 0.57 | 0.64 | 0.66 | 0.71 |
| 4 | 60060761 | rs7664607 | 0.927 | -0.245 | -0.886 | -0.284 | -0.012 | 0.009 | 0.009 | 0 | 0.26 | 0.11 | 0.21 | 0.22 | 0.27 | 0.23 |
| 4 | 60079490 | rs7681389 | -0.17 | 0.025 | -0.401 | -0.084 | 0.002 | 0.007 | 0.007 | 0 | 0.61 | 0.49 | 0.46 | 0.48 | 0.58 | 0.56 |
| 4 | 60082188 | rs1354254 | -0.193 | 0.035 | -0.53 | -0.079 | 0.007 | 0.019 | 0.019 | 0 | 0.62 | 0.55 | 0.47 | 0.48 | 0.58 | 0.56 |
| 4 | 60088172 | rs964384 | 0.471 | 0.406 | -0.688 | -0.228 | -0.05 | -0.085 | -0.085 | 0.217 | 0.27 | 0.14 | 0.33 | 0.54 | 0.34 | 0.33 |
| 4 | 60096626 | rs12646524 | 0.212 | 0.132 | -0.305 | -0.643 | -0.007 | -0.037 | -0.037 | 0 | 0.31 | 0.36 | 0.23 | 0.18 | 0.32 | 0.37 |
| 4 | 60099156 | rs6855237 | -1.277 | 0.178 | -0.206 | -0.688 | -0.007 | -0.013 | -0.013 | 0 | 0.85 | 0.86 | 0.7 | 0.76 | 0.77 | 0.86 |
| 4 | 60104294 | rs995678 | 0.605 | 0.153 | -0.326 | -0.772 | -0.026 | -0.056 | -0.056 | 0.209 | 0.12 | 0.16 | 0.08 | 0.01 | 0.07 | 0.11 |
| 4 | 60106143 | rs990147 | -0.842 | 0.054 | -0.101 | -0.822 | 0.005 | 0.041 | 0.041 | 0 | 0.81 | 0.63 | 0.67 | 0.76 | 0.77 | 0.86 |
| 4 | 60111876 | rs17827688 | 0.688 | 0.08 | -0.482 | -0.806 | 0.015 | 0.103 | 0.103 | 0 | 0.44 | 0.26 | 0.27 | 0.22 | 0.38 | 0.42 |
| 4 | 60112029 | rs1502784 | 1.17 | 0.139 | -0.51 | -0.748 | -0.022 | -0.036 | -0.036 | 0 | 0.25 | 0.19 | 0.32 | 0.53 | 0.32 | 0.33 |
| 4 | 60127335 | rs17089537 | 1.382 | 0.171 | -0.952 | -0.827 | -0.008 | -0.021 | -0.021 | 0 | 0.08 | 0.06 | 0.07 | 0.01 | 0.05 | 0.08 |
| 4 | 60129168 | rs7663467 | 0.178 | 0.516 | -0.955 | -0.621 | 0.012 | 0.081 | 0.081 | 0 | 0.44 | 0.28 | 0.27 | 0.22 | 0.38 | 0.42 |
| 4 | 60129838 | rs10002811 | 1.2 | 0.529 | -1.029 | -0.255 | 0.021 | 0.087 | 0.087 | 0 | 0.22 | 0.08 | 0.11 | 0.05 | 0.11 | 0.15 |
| 4 | 60143686 | rs7693572 | -0.097 | 0.76 | -0.647 | 0.092 | 0 | 0.003 | 0.003 | 0 | 0.5 | 0.53 | 0.45 | 0.57 | 0.44 | 0.5 |
| 4 | 60149217 | rs13134179 | -0.198 | 0.931 | -0.515 | 0.094 | -0.001 | -0.001 | -0.001 | 0 | 0.63 | 0.64 | 0.5 | 0.68 | 0.65 | 0.67 |
| 4 | 60149698 | rs13135277 | -0.198 | 0.931 | -0.515 | 0.094 | -0.001 | -0.001 | -0.001 | 0 | 0.63 | 0.64 | 0.5 | 0.68 | 0.65 | 0.67 |
| 4 | 60155189 | rs7440788 | -0.736 | 0.563 | -0.232 | -0.026 | -0.027 | -0.079 | -0.079 | 0.094 | 0.71 | 0.64 | 0.84 | 0.92 | 0.86 | 0.8 |
| 4 | 60164589 | rs9312092 | -0.436 | 0.439 | -0.212 | 0.015 | -0.01 | -0.042 | -0.042 | 0.026 | 0.34 | 0.43 | 0.3 | 0.22 | 0.32 | 0.41 |
| 4 | 60176254 | rs17089560 | 0.276 | 0.374 | -0.136 | -0.109 | -0.006 | -0.03 | -0.03 | 0.118 | 0.06 | 0.11 | 0.03 | 0 | 0.04 | 0.05 |
| 4 | 60178068 | rs17089569 | 0.423 | 0.552 | -0.272 | -0.142 | 0 | 0.001 | 0.001 | 0 | 0.14 | 0.1 | 0.09 | 0.12 | 0.13 | 0.21 |
| 4 | 60179526 | rs10014071 | -0.777 | 0.333 | -0.288 | 0.048 | -0.009 | -0.022 | -0.022 | 0.074 | 0.37 | 0.54 | 0.35 | 0.28 | 0.32 | 0.43 |
| 4 | 60183194 | rs966998 | -0.777 | 0.333 | -0.288 | 0.048 | -0.009 | -0.022 | -0.022 | 0 | 0.37 | 0.54 | 0.35 | 0.28 | 0.32 | 0.43 |
| 4 | 60192397 | rs7683276 | 0.562 | 0.376 | -0.059 | 0.134 | 0 | -0.007 | -0.007 | 0 | 0.19 | 0.22 | 0.23 | 0.11 | 0.19 | 0.2 |
| 4 | 60192809 | rs17290640 | -1.321 | 0.058 | 0.14 | 0.33 | 0.038 | -0.063 | -0.063 | 0.459 | 0.39 | 0.82 | 0.33 | 0.51 | 0.3 | 0.5 |
| 4 | 60193870 | rs10008176 | -0.352 | 0.102 | 0.391 | -0.008 | -0.02 | -0.044 | -0.044 | 0 | 0.3 | 0.48 | 0.27 | 0.22 | 0.25 | 0.39 |
| 4 | 60200132 | rs17089590 | 0.197 | 0.129 | 0.306 | -0.008 | -0.001 | -0.011 | -0.011 | 0 | 0.24 | 0.25 | 0.24 | 0.11 | 0.2 | 0.22 |
| 4 | 60217601 | rs10022990 | -0.556 | 0.203 | 0.091 | 0.05 | -0.018 | -0.049 | -0.049 | 0 | 0.34 | 0.44 | 0.29 | 0.22 | 0.3 | 0.44 |
| 4 | 60228517 | rs1119950 | -0.507 | 0.474 | 0.056 | 0.297 | -0.002 | 0.001 | 0.001 | 1.022 | 0.68 | 0.75 | 0.78 | 0.37 | 0.64 | 0.74 |
| 4 | 60239646 | rs10866134 | -0.288 | 0.484 | -0.042 | 0.233 | -0.001 | 0.002 | 0.002 | 0.234 | 0.66 | 0.67 | 0.77 | 0.31 | 0.64 | 0.72 |
| 4 | 60249365 | rs1983239 | -0.474 | 0.685 | -0.251 | 0.779 | -0.001 | 0.04 | 0.04 | 0 | 0.91 | 0.89 | 0.91 | 0.65 | 0.88 | 0.94 |
| 4 | 60257120 | rs4860530 | 0.895 | 0.663 | -0.389 | -0.067 | 0.011 | 0.186 | 0.186 | 0 | 0.52 | 0.21 | 0.68 | 0.19 | 0.51 | 0.47 |
| 4 | 60257400 | rs4860532 | 0.117 | 0.724 | 0.188 | 0.195 | 0.001 | -0.015 | -0.015 | 0 | 0.65 | 0.42 | 0.73 | 0.31 | 0.64 | 0.7 |
| 4 | 60257454 | rs7681982 | -0.267 | 0.714 | 0.368 | 0.19 | -0.008 | 0.009 | 0.009 | 0.026 | 0.69 | 0.74 | 0.75 | 0.31 | 0.64 | 0.7 |
| 4 | 60268638 | rs1606137 | -0.659 | 0.658 | 0.303 | 0.767 | -0.011 | -0.045 | -0.045 | 0.107 | 0.74 | 0.83 | 0.84 | 0.6 | 0.7 | 0.75 |
| 4 | 60275069 | rs1079463 | 0.192 | 0.496 | 0.136 | 0.353 | 0.008 | 0.022 | 0.022 | 1.243 | 0.58 | 0.35 | 0.69 | 0.36 | 0.63 | 0.67 |
| 4 | 60283466 | rs1511404 | -0.123 | 0.572 | -0.081 | 0.389 | 0.001 | 0.002 | 0.002 | 1.834 | 0.62 | 0.63 | 0.75 | 0.41 | 0.64 | 0.7 |
| 4 | 60325010 | rs4860580 | -0.99 | 1.123 | -0.558 | 0.391 | -0.032 | -0.031 | -0.031 | 6.011 | 0.85 | 0.95 | 0.83 | 0.75 | 0.71 | 0.8 |
| 4 | 60325495 | rs13143351 | 0.4 | 0.658 | -0.777 | -0.138 | 0.015 | 0 | 0 | 0 | 0.47 | 0.4 | 0.29 | 0.49 | 0.38 | 0.34 |
| 4 | 60340339 | rs1858317 | 1.587 | 0.609 | -0.653 | -0.177 | -0.006 | 0.024 | 0.024 | 0 | 0.2 | 0.17 | 0.39 | 0.13 | 0.2 | 0.22 |
| 4 | 60344490 | rs1511408 | 0.011 | 0.026 | -0.946 | -0.438 | 0.018 | -0.023 | -0.023 | 0 | 0.37 | 0.22 | 0.22 | 0.45 | 0.33 | 0.38 |
| 4 | 60356966 | rs17238233 | 2.121 | 0.161 | -1.013 | -0.45 | 0.008 | 0 | 0 | 0.755 | 0.18 | 0.15 | 0.1 | 0.02 | 0.11 | 0.05 |
| 4 | 60367869 | rs6811282 | -0.645 | 0.421 | -0.681 | -0.189 | -0.003 | 0 | 0 | 0 | 0.42 | 0.42 | 0.28 | 0.58 | 0.39 | 0.54 |
| 4 | 60391854 | rs10007020 | 0.023 | 0.459 | -0.716 | -0.249 | -0.012 | -0.016 | -0.016 | 0 | 0.63 | 0.69 | 0.65 | 0.61 | 0.55 | 0.52 |
| 4 | 60393064 | rs11131575 | -0.823 | 0.453 | -0.816 | -0.233 | -0.067 | -0.089 | -0.089 | 0 | 0.74 | 0.96 | 0.77 | 0.69 | 0.65 | 0.69 |
| 4 | 60399836 | rs17218618 | 1.041 | 0.437 | -0.708 | -0.209 | -0.002 | -0.002 | -0.002 | 0 | 0.08 | 0.07 | 0.07 | 0.11 | 0.09 | 0.14 |
| 4 | 60401751 | rs11131579 | 0.416 | 0.205 | -0.981 | -0.486 | 0.006 | 0 | 0 | 0 | 0.4 | 0.57 | 0.24 | 0.44 | 0.31 | 0.29 |
| 4 | 60403334 | rs17089687 | 0.27 | 0.196 | -0.979 | -0.553 | 0.001 | 0 | 0 | 7.061 | 0.11 | 0.22 | 0.12 | 0.11 | 0.1 | 0.19 |
| 4 | 60405703 | rs1913311 | 0.525 | -0.079 | -0.699 | -0.642 | -0.006 | 0.035 | 0.035 | 0.228 | 0.53 | 0.42 | 0.58 | 0.26 | 0.46 | 0.42 |
| 4 | 60418595 | rs17089693 | 2.124 | -0.085 | -0.62 | -0.835 | 0.018 | 0.056 | 0.056 | 1.094 | 0.2 | 0.08 | 0.11 | 0.03 | 0.12 | 0.05 |
| 4 | 60428730 | rs1513536 | -0.389 | 0.288 | 0.05 | -0.215 | -0.002 | 0.118 | 0.118 | 0.679 | 0.21 | 0.58 | 0.2 | 0.49 | 0.22 | 0.33 |
| 4 | 60442490 | rs17290779 | 0.153 | 0.339 | 0.243 | -0.19 | 0.012 | 0.107 | 0.107 | 0 | 0.18 | 0.29 | 0.14 | 0.48 | 0.22 | 0.28 |
| 4 | 60458820 | rs970313 | -0.649 | 0.317 | -0.265 | -0.35 | -0.004 | 0.001 | 0.001 | 0 | 0.47 | 0.66 | 0.61 | 0.62 | 0.48 | 0.56 |
| 4 | 60465326 | rs1600954 | -0.167 | 0.373 | -0.194 | -0.234 | -0.007 | -0.015 | -0.015 | 1.449 | 0.57 | 0.66 | 0.41 | 0.56 | 0.47 | 0.49 |
| 4 | 60468539 | rs7439432 | 0.166 | 0.102 | 0.383 | 0.038 | 0.003 | -0.034 | -0.034 | 0 | 0.09 | 0.28 | 0.06 | 0.01 | 0.05 | 0.07 |
| 4 | 60475482 | rs1482701 | -1.214 | 0.247 | -0.001 | 0.089 | -0.048 | -0.077 | -0.077 | 1.38 | 0.85 | 0.97 | 0.83 | 0.69 | 0.76 | 0.69 |
| 4 | 60480763 | rs988744 | 0.444 | 0.206 | 0.162 | 0.041 | -0.003 | -0.036 | -0.036 | 0.871 | 0.09 | 0.23 | 0.06 | 0.01 | 0.05 | 0.07 |
| 4 | 60489510 | rs6842739 | 0.74 | 0.072 | 0.349 | -0.008 | 0.007 | 0.052 | 0.052 | 0.375 | 0.72 | 0.44 | 0.74 | 0.67 | 0.69 | 0.59 |
| 4 | 60493470 | rs1825238 | -1.278 | 0.216 | 0.112 | 0.064 | -0.01 | -0.006 | -0.006 | 1.085 | 0.85 | 0.93 | 0.83 | 0.69 | 0.75 | 0.69 |
| 4 | 60495394 | rs7683528 | -0.1 | -0.151 | -0.078 | 0.007 | -0.028 | -0.015 | -0.015 | 0 | 0.43 | 0.23 | 0.58 | 0.64 | 0.48 | 0.47 |
| 4 | 60496226 | rs17290786 | 0.526 | 0.768 | -0.109 | 0.196 | -0.064 | 0.009 | 0.009 | 0.182 | 0.16 | 0.03 | 0.4 | 0.14 | 0.25 | 0.21 |
| 4 | 60500708 | rs1482699 | -0.035 | -0.008 | 0.601 | 0.548 | 0 | -0.004 | -0.004 | 0 | 0.08 | 0.1 | 0.05 | 0.01 | 0.05 | 0.07 |
| 4 | 60512343 | rs10013542 | -0.052 | -0.058 | 0.656 | 0.581 | -0.004 | -0.021 | -0.021 | 0 | 0.08 | 0.15 | 0.05 | 0.03 | 0.05 | 0.07 |
| 4 | 60521038 | rs7670505 | -0.887 | 0.044 | 0.438 | 1.145 | -0.024 | -0.055 | -0.055 | 3.702 | 0.88 | 0.98 | 0.87 | 0.7 | 0.79 | 0.77 |

| | | | | | | | | | | | | | | | | |
|---|---|---|---|---|---|---|---|---|---|---|---|---|---|---|---|---|
| 4 | 60521175 | rs7655641 | 0.604 | -0.128 | 1.015 | 1.213 | -0.001 | 0.03 | 0.03 | 0 | 0.76 | 0.66 | 0.73 | 0.37 | 0.69 | 0.64 |
| 4 | 60529334 | rs1482704 | 1.482 | -0.172 | 1.234 | 1.174 | 0.001 | -0.001 | -0.001 | 0 | 0.66 | 0.31 | 0.7 | 0.35 | 0.64 | 0.57 |
| 4 | 60532584 | rs1513532 | 1.634 | -0.26 | 1.029 | 1.12 | 0.024 | 0.012 | 0.012 | 0.242 | 0.62 | 0.21 | 0.65 | 0.34 | 0.62 | 0.51 |
| 4 | 60536829 | rs35712489 | 0.233 | 0.141 | 0.339 | 1.579 | -0.004 | 0.039 | 0.039 | 0 | 0.1 | 0.16 | 0.13 | 0.34 | 0.1 | 0.13 |
| 4 | 60538549 | rs1032030 | -0.295 | 0.282 | 0.145 | 1.471 | -0.004 | -0.009 | -0.009 | 0 | 0.87 | 0.91 | 0.87 | 0.71 | 0.79 | 0.78 |
| 4 | 60549283 | rs1395652 | 0.087 | 0.228 | 0.216 | 1.441 | 0.002 | 0.015 | 0.015 | 0.067 | 0.83 | 0.73 | 0.85 | 0.79 | 0.78 | 0.72 |
| 4 | 60560780 | rs17218730 | 1.309 | 0.1 | -0.211 | 1.139 | -0.017 | -0.053 | -0.053 | 0.342 | 0.11 | 0 | 0.06 | 0.21 | 0.09 | 0.08 |
| 4 | 60568412 | rs9991718 | 0.291 | -0.098 | -0.057 | 0.692 | 0 | 0 | 0 | 0 | 0.46 | 0.51 | 0.55 | 0.4 | 0.47 | 0.5 |
| 4 | 60573507 | rs11936764 | -0.371 | 0.061 | 0.201 | 1.225 | -0.001 | 0 | 0 | 0 | 0.15 | 0.17 | 0.15 | 0.26 | 0.13 | 0.16 |
| 4 | 60578593 | rs6552029 | 0.881 | 0.104 | 0.205 | 1.27 | -0.004 | -0.034 | -0.034 | 0 | 0.69 | 0.73 | 0.7 | 0.46 | 0.63 | 0.6 |
| 4 | 60581437 | rs2880170 | -0.853 | 0.122 | 0.61 | 0.698 | -0.003 | 0.001 | 0.001 | 0 | 0.17 | 0.22 | 0.16 | 0.26 | 0.16 | 0.24 |
| 4 | 60616622 | rs17089786 | -0.283 | 0.349 | 0.137 | 0.484 | -0.012 | 0.012 | 0.012 | 0.034 | 0.09 | 0.18 | 0.08 | 0.09 | 0.08 | 0.1 |
| 4 | 60619937 | rs1507132 | -0.532 | 0.222 | -0.29 | 0.137 | 0.009 | 0.003 | 0.003 | 0.377 | 0.39 | 0.27 | 0.31 | 0.65 | 0.38 | 0.61 |
| 4 | 60621100 | rs7699730 | -1.066 | 0.061 | 0.112 | 0.057 | 0.02 | 0.043 | 0.043 | 3.628 | 0.5 | 0.64 | 0.4 | 0.74 | 0.46 | 0.72 |
| 4 | 60627772 | rs4600947 | -0.127 | 0.081 | 0.15 | 0.096 | -0.018 | 0.02 | 0.02 | 0.329 | 0.1 | 0.34 | 0.09 | 0.09 | 0.08 | 0.11 |
| 4 | 60639488 | rs10002873 | -0.071 | 0.082 | 0.178 | 0.138 | -0.014 | 0.044 | 0.044 | 0.349 | 0.1 | 0.36 | 0.13 | 0.11 | 0.08 | 0.11 |
| 4 | 60639822 | rs10005469 | -0.986 | 0.056 | -0.405 | -0.031 | 0.013 | 0.022 | 0.022 | 3.229 | 0.45 | 0.64 | 0.39 | 0.74 | 0.43 | 0.69 |
| 4 | 60673647 | rs7697784 | 0.787 | -0.044 | -0.739 | 0.438 | -0.002 | -0.019 | -0.019 | 0 | 0.18 | 0.1 | 0.14 | 0.27 | 0.16 | 0.23 |
| 4 | 60674843 | rs9312186 | -0.742 | 0.158 | -0.429 | -0.056 | 0.011 | 0.028 | 0.028 | 1.548 | 0.52 | 0.62 | 0.4 | 0.74 | 0.46 | 0.75 |
| 4 | 60678416 | rs1863308 | -0.858 | 0.577 | -0.785 | -0.508 | -0.034 | 0.023 | 0.023 | 0 | 0.81 | 0.9 | 0.56 | 0.91 | 0.66 | 0.83 |
| 4 | 60680656 | rs17089834 | 0.024 | 0.257 | -0.632 | -0.418 | -0.013 | 0.021 | 0.021 | 0.239 | 0.11 | 0.37 | 0.09 | 0.09 | 0.08 | 0.12 |
| 4 | 60687024 | rs1010849 | 0.271 | 0.344 | -1.162 | -0.366 | 0.018 | 0.009 | 0.009 | 0 | 0.4 | 0.2 | 0.3 | 0.65 | 0.38 | 0.61 |
| 4 | 60707711 | rs12499997 | 0.566 | 1.061 | -0.905 | -0.019 | -0.009 | -0.001 | -0.001 | 0 | 0.13 | 0.08 | 0.47 | 0.14 | 0.26 | 0.15 |
| 4 | 60715255 | rs4401481 | -0.757 | 0.95 | -0.979 | -0.151 | 0.004 | 0.008 | 0.008 | 0.311 | 0.58 | 0.74 | 0.72 | 0.37 | 0.54 | 0.42 |
| 4 | 60729234 | rs1304686 | 0.459 | 1.046 | -1.646 | -0.257 | 0.019 | 0.052 | 0.052 | 3.869 | 0.35 | 0.24 | 0.18 | 0.15 | 0.26 | 0.17 |
| 4 | 60730605 | rs6815012 | 0.011 | 0.389 | -1.402 | 0.056 | 0.015 | 0.062 | 0.062 | 1.294 | 0.5 | 0.32 | 0.65 | 0.29 | 0.52 | 0.32 |
| 4 | 60741765 | rs17089860 | -0.456 | -0.475 | -1.808 | 0.396 | 0.04 | 0.009 | 0.009 | 1.41 | 0.38 | 0.1 | 0.31 | 0.68 | 0.29 | 0.47 |
| 4 | 60751925 | rs13131266 | -0.437 | -0.651 | -1.009 | 1.113 | 0.02 | -0.026 | -0.026 | 0 | 0.37 | 0.13 | 0.25 | 0.34 | 0.29 | 0.35 |
| 4 | 60757025 | rs7655523 | -1.25 | -1.365 | 0.103 | 0.742 | -0.007 | 0.077 | 0.077 | 0 | 0.59 | 0.71 | 0.36 | 0.83 | 0.39 | 0.73 |
| 4 | 60774654 | rs1559703 | -0.07 | -1.212 | 0.417 | 1.635 | -0.013 | -0.036 | -0.036 | 0 | 0.06 | 0 | 0.03 | 0.14 | 0.06 | 0.08 |
| 4 | 60776281 | rs1834384 | -1.417 | -1.236 | 0.699 | 1.636 | -0.02 | 0.077 | 0.077 | 0.117 | 0.16 | 0.24 | 0.09 | 0.42 | 0.09 | 0.38 |
| 4 | 60793034 | rs1903298 | -1.389 | -1.311 | 0.517 | 0.661 | 0.027 | 0.132 | 0.132 | 0.004 | 0.51 | 0.7 | 0.31 | 0.84 | 0.39 | 0.72 |
| 4 | 60825149 | rs6833033 | -1.252 | -1.107 | 1.266 | 1.743 | -0.003 | 0.272 | 0.272 | 0.038 | 0.08 | 0.21 | 0.12 | 0.39 | 0.05 | 0.14 |
| 4 | 60826699 | rs13112895 | -0.559 | -1.075 | 0.905 | 1.759 | 0 | 0.097 | 0.097 | 0.043 | 0.06 | 0.05 | 0.11 | 0.39 | 0.05 | 0.14 |
| 4 | 60829098 | rs349247 | -1.571 | -1.499 | 0.526 | 0.399 | 0.058 | 0.191 | 0.191 | 0 | 0.46 | 0.84 | 0.31 | 0.84 | 0.32 | 0.63 |
| 4 | 60833758 | rs349235 | 1.189 | -1.291 | 1.225 | 0.449 | -0.01 | -0.041 | -0.041 | 0 | 0.59 | 0.5 | 0.67 | 0.15 | 0.64 | 0.3 |
| 4 | 60843933 | rs349215 | -0.617 | -1.251 | 1.229 | 0.451 | 0.049 | -0.008 | -0.008 | 0 | 0.07 | 0.28 | 0.03 | 0.01 | 0.04 | 0.02 |
| 4 | 60844380 | rs349211 | 1.247 | -1.329 | 1.236 | 0.404 | 0.001 | -0.008 | -0.008 | 0 | 0.58 | 0.54 | 0.67 | 0.16 | 0.64 | 0.3 |
| 4 | 60853613 | rs10461286 | -0.665 | -1.38 | 0.847 | 0.796 | 0.013 | 0.011 | 0.011 | 0 | 0.33 | 0.17 | 0.29 | 0.69 | 0.32 | 0.59 |
| 4 | 60854084 | rs11946874 | -0.943 | -1.319 | 0.888 | 0.629 | 0.016 | 0.046 | 0.046 | 0 | 0.38 | 0.31 | 0.32 | 0.78 | 0.36 | 0.67 |
| 4 | 60855477 | rs1993672 | 1.071 | -1.098 | 0.135 | 0.663 | 0.052 | 0.257 | 0.257 | 0.952 | 0.5 | 0.11 | 0.63 | 0.16 | 0.6 | 0.28 |
| 4 | 60873322 | rs9312045 | -1.29 | -0.792 | 1.055 | 1.507 | -0.01 | -0.013 | -0.013 | 3.28 | 0.08 | 0.12 | 0.03 | 0.06 | 0.04 | 0.1 |
| 4 | 60874182 | rs2880475 | -2.761 | -1.116 | 1.307 | 2.031 | -0.031 | 0.047 | 0.047 | 0 | 0.22 | 0.35 | 0.2 | 0.67 | 0.13 | 0.49 |
| 4 | 60879952 | rs12645408 | -1.743 | -1.026 | 1.494 | 2.825 | -0.031 | 0.289 | 0.289 | 0 | 0.11 | 0.29 | 0.15 | 0.52 | 0.04 | 0.35 |
| 4 | 60891956 | rs17291044 | -2.519 | -1.255 | 1.438 | 2.704 | -0.02 | 0.015 | 0.015 | 0 | 0.2 | 0.26 | 0.14 | 0.29 | 0.1 | 0.33 |
| 4 | 60892141 | rs11947191 | -2.82 | -0.66 | 1.851 | 2.567 | -0.021 | 0.036 | 0.036 | 0 | 0.25 | 0.45 | 0.2 | 0.3 | 0.24 | 0.36 |
| 4 | 60906671 | rs10000971 | -3.32 | -1.251 | 2.497 | 2.753 | -0.027 | 0.036 | 0.036 | 0 | 0.27 | 0.66 | 0.18 | 0.49 | 0.14 | 0.44 |
| 4 | 60908435 | rs2085777 | -2.772 | -1.229 | 2.693 | 2.847 | -0.062 | 0.151 | 0.151 | 0.019 | 0.18 | 0.48 | 0.14 | 0.35 | 0.09 | 0.33 |
| 4 | 60930593 | rs7656343 | -1.889 | -1.189 | 2.587 | 3.022 | 0.014 | 0.137 | 0.137 | 0.755 | 0.08 | 0.38 | 0.06 | 0.15 | 0.06 | 0.09 |
| 4 | 60931418 | rs531673 | 0.819 | -1.185 | 2.527 | 2.969 | -0.008 | -0.007 | -0.007 | 0 | 0.92 | 0.89 | 0.98 | 0.91 | 0.96 | 0.91 |
| 4 | 60931440 | rs531622 | 1.025 | -1.185 | 2.534 | 2.969 | -0.01 | -0.012 | -0.012 | 0 | 0.92 | 0.85 | 0.98 | 0.91 | 0.96 | 0.91 |
| 4 | 60941032 | rs2119551 | 0.989 | -1.184 | 2.531 | 2.97 | -0.01 | -0.011 | -0.011 | 0 | 0.92 | 0.87 | 0.98 | 0.91 | 0.96 | 0.91 |
| 4 | 60946148 | rs1109023 | 1.817 | -1.198 | 2.518 | 2.973 | -0.017 | -0.018 | -0.018 | 0 | 0.89 | 0.84 | 0.97 | 0.91 | 0.96 | 0.89 |
| 4 | 60946433 | rs6832310 | -2.881 | -1.089 | 2.465 | 3.088 | -0.029 | 0.213 | 0.213 | 0 | 0.15 | 0.39 | 0.14 | 0.41 | 0.1 | 0.33 |
| 4 | 60965262 | rs17291120 | -0.472 | -1.031 | 2.042 | 3.041 | 0.008 | -0.012 | -0.012 | 0 | 0.07 | 0.02 | 0.09 | 0.26 | 0.04 | 0.25 |
| 4 | 60969122 | rs7695675 | -2.666 | -1.011 | 2.396 | 3.025 | -0.028 | 0.155 | 0.155 | 0 | 0.17 | 0.47 | 0.14 | 0.37 | 0.08 | 0.33 |
| 4 | 60973349 | rs4530671 | 1.11 | -0.871 | 2.324 | 3.01 | -0.003 | 0.017 | 0.017 | 0 | 0.92 | 0.9 | 0.94 | 0.86 | 0.91 | 0.93 |
| 4 | 60984101 | rs6551519 | 3.26 | -0.932 | 2.13 | 3.027 | -0.026 | 0.027 | 0.027 | 0.372 | 0.72 | 0.38 | 0.75 | 0.38 | 0.81 | 0.51 |
| 4 | 61007315 | rs12511090 | 1.941 | -0.645 | 1.705 | 3.066 | 0.002 | -0.003 | -0.003 | 0.063 | 0.9 | 0.92 | 0.89 | 0.77 | 0.9 | 0.83 |
| 4 | 61008776 | rs10049506 | -2.103 | -0.595 | 1.718 | 2.517 | -0.001 | -0.014 | -0.014 | 0.466 | 0.1 | 0.06 | 0.13 | 0.28 | 0.11 | 0.31 |
| 4 | 61029597 | rs4146901 | 1.032 | -0.632 | 1.72 | 2.248 | 0.003 | 0.008 | 0.008 | 0 | 0.93 | 0.96 | 0.94 | 1 | 0.93 | 0.91 |
| 4 | 61074614 | rs7681003 | -2.404 | -0.522 | 2.668 | 2.11 | 0.066 | 0.156 | 0.156 | 0.367 | 0.07 | 0.36 | 0.07 | 0.22 | 0.07 | 0.04 |
| 4 | 61080364 | rs6820884 | -3.181 | -0.57 | 2.621 | 2.287 | 0 | 0.203 | 0.203 | 0.392 | 0.17 | 0.45 | 0.19 | 0.48 | 0.15 | 0.34 |
| 4 | 61080750 | rs11131228 | 3.422 | -0.68 | 2.537 | 2.31 | -0.02 | 0.096 | 0.096 | 1.563 | 0.74 | 0.42 | 0.79 | 0.49 | 0.79 | 0.53 |
| 4 | 61107712 | rs4277806 | -2.094 | -0.698 | 2.913 | 2.482 | -0.008 | 0.056 | 0.056 | 0.214 | 0.11 | 0.15 | 0.13 | 0.34 | 0.09 | 0.32 |
| 4 | 61110086 | rs3924779 | -3.838 | -0.712 | 2.997 | 2.485 | -0.029 | 0.207 | 0.207 | 1.363 | 0.21 | 0.49 | 0.2 | 0.51 | 0.18 | 0.42 |
| 4 | 61126692 | rs6834427 | -2.032 | -0.694 | 3.06 | 2.506 | 0.079 | 0.07 | 0.07 | 0 | 0.06 | 0.26 | 0.07 | 0.1 | 0.07 | 0.04 |
| 4 | 61133934 | rs2334124 | 1.409 | -0.694 | 3.037 | 2.507 | -0.01 | 0.043 | 0.043 | 0 | 0.95 | 0.86 | 0.98 | 0.86 | 0.97 | 0.92 |
| 4 | 61141856 | rs13434729 | -1.729 | -0.69 | 3.092 | 2.491 | 0.091 | 0.087 | 0.087 | 0 | 0.05 | 0.27 | 0.07 | 0.1 | 0.07 | 0.05 |

| | | | | | | | | | | | | | | | | |
|---|---|---|---|---|---|---|---|---|---|---|---|---|---|---|---|---|
| 4 | 61143085 | rs2333895 | 3.322 | -0.715 | 3.008 | 2.481 | -0.028 | 0.156 | 0.156 | 1.552 | 0.79 | 0.43 | 0.8 | 0.49 | 0.84 | 0.59 |
| 4 | 61154338 | rs10517502 | 3.272 | -0.704 | 3.014 | 2.487 | -0.026 | 0.162 | 0.162 | 1.786 | 0.79 | 0.43 | 0.8 | 0.49 | 0.84 | 0.58 |
| 4 | 61163718 | rs717252 | -3.116 | -0.696 | 3.041 | 2.479 | -0.003 | 0.169 | 0.169 | 1.108 | 0.16 | 0.4 | 0.18 | 0.37 | 0.14 | 0.34 |
| 4 | 61173570 | rs7672373 | -1.947 | -0.7 | 2.903 | 2.456 | -0.009 | 0.041 | 0.041 | 0.407 | 0.1 | 0.13 | 0.12 | 0.27 | 0.07 | 0.31 |
| 4 | 61177831 | rs6827408 | 2.675 | -0.697 | 2.94 | 2.453 | -0.026 | 0.148 | 0.148 | 1.819 | 0.79 | 0.43 | 0.8 | 0.51 | 0.83 | 0.57 |
| 4 | 61178793 | rs4860334 | 2.675 | -0.697 | 2.94 | 2.453 | -0.026 | 0.148 | 0.148 | 1.819 | 0.79 | 0.43 | 0.8 | 0.51 | 0.83 | 0.57 |
| 4 | 61181195 | rs6848703 | -3.192 | -0.699 | 2.986 | 2.453 | -0.021 | 0.206 | 0.206 | 1.306 | 0.2 | 0.49 | 0.19 | 0.49 | 0.17 | 0.43 |
| 4 | 61187284 | rs7673099 | 2.683 | -0.698 | 2.954 | 2.453 | -0.026 | 0.148 | 0.148 | 1.819 | 0.79 | 0.43 | 0.8 | 0.51 | 0.83 | 0.57 |
| 4 | 61194055 | rs1455731 | 2.813 | -0.707 | 2.95 | 2.449 | -0.027 | 0.144 | 0.144 | 1.3 | 0.79 | 0.43 | 0.81 | 0.51 | 0.83 | 0.57 |
| 4 | 61207237 | rs10517499 | 1.625 | -0.825 | 2.713 | 1.873 | 0.001 | -0.002 | -0.002 | 1.328 | 0.79 | 0.79 | 0.83 | 0.69 | 0.83 | 0.6 |
| 4 | 61210632 | rs12501945 | 1.942 | -0.828 | 2.876 | 1.858 | -0.039 | 0.038 | 0.038 | 0 | 0.77 | 0.56 | 0.82 | 0.69 | 0.83 | 0.58 |
| 4 | 61211180 | rs10866099 | 1.715 | -0.818 | 2.964 | 1.863 | -0.015 | 0.017 | 0.017 | 0 | 0.78 | 0.69 | 0.83 | 0.69 | 0.83 | 0.58 |
| 4 | 61216054 | rs1947493 | 2.296 | -0.824 | 2.959 | 1.853 | -0.022 | 0.028 | 0.028 | 0 | 0.74 | 0.53 | 0.78 | 0.66 | 0.77 | 0.55 |
| 4 | 61220878 | rs6551541 | 2.78 | -0.892 | 2.97 | 1.676 | 0.023 | -0.017 | -0.017 | 0 | 0.66 | 0.23 | 0.75 | 0.54 | 0.72 | 0.49 |
| 4 | 61226507 | rs6833291 | 1.837 | -0.855 | 2.977 | 1.548 | 0.04 | 0.035 | 0.035 | 0 | 0.89 | 0.61 | 0.93 | 0.86 | 0.9 | 0.92 |
| 4 | 61228770 | rs10019682 | 2.287 | -0.877 | 2.964 | 1.532 | -0.019 | -0.058 | -0.058 | 0 | 0.8 | 0.49 | 0.89 | 0.86 | 0.83 | 0.78 |
| 4 | 61231636 | rs17081849 | -1.163 | -0.926 | 2.78 | 1.249 | -0.007 | -0.032 | -0.032 | 1.089 | 0.1 | 0.11 | 0.04 | 0 | 0.07 | 0.14 |
| 4 | 61233219 | rs1463351 | 1.542 | -0.93 | 2.812 | 1.33 | -0.002 | -0.028 | -0.028 | 0 | 0.81 | 0.63 | 0.83 | 0.82 | 0.82 | 0.76 |
| 4 | 61238161 | rs11946495 | 2.836 | -0.885 | 2.664 | 1.296 | 0.034 | -0.009 | -0.009 | 0 | 0.63 | 0.13 | 0.71 | 0.59 | 0.71 | 0.48 |
| 4 | 61240227 | rs17090036 | -1.131 | -0.967 | 2.83 | 1.233 | -0.002 | -0.031 | -0.031 | 0.175 | 0.07 | 0.27 | 0.03 | 0 | 0.03 | 0.05 |
| 4 | 61244872 | rs13143552 | -0.64 | -0.948 | 2.779 | 1.233 | 0 | -0.017 | -0.017 | 0 | 0.08 | 0.07 | 0.05 | 0 | 0.09 | 0.12 |
| 4 | 61247315 | rs4860340 | 2.494 | -1.009 | 2.913 | 1.278 | 0.026 | 0.015 | 0.015 | 0 | 0.68 | 0.2 | 0.77 | 0.72 | 0.78 | 0.53 |
| 4 | 61249246 | rs4318679 | 1.494 | -0.972 | 3.021 | 1.302 | -0.048 | 0.001 | 0.001 | 0 | 0.79 | 0.59 | 0.84 | 0.77 | 0.88 | 0.67 |
| 4 | 61251815 | rs6852865 | -0.881 | -0.921 | 2.993 | 1.3 | -0.028 | 0.023 | 0.023 | 0 | 0.14 | 0.31 | 0.12 | 0.18 | 0.08 | 0.13 |
| 4 | 61258122 | rs6858288 | 2.849 | -1.045 | 2.657 | 1.201 | 0.034 | -0.019 | -0.019 | 0 | 0.61 | 0.1 | 0.71 | 0.59 | 0.71 | 0.48 |
| 4 | 61262697 | rs10517496 | -1.308 | -0.959 | 2.697 | 1.196 | 0 | -0.007 | -0.007 | 0 | 0.14 | 0.08 | 0.08 | 0 | 0.13 | 0.18 |
| 4 | 61264831 | rs1378566 | 1.362 | -0.813 | 2.771 | 1.34 | 0.036 | 0.056 | 0.056 | 0.463 | 0.95 | 0.83 | 0.94 | 0.88 | 0.94 | 0.95 |
| 4 | 61265932 | rs6551546 | 1.362 | -0.867 | 2.771 | 1.331 | 0.032 | 0.093 | 0.093 | 0.463 | 0.95 | 0.83 | 0.94 | 0.83 | 0.94 | 0.95 |
| 4 | 61266555 | rs4499722 | 2.477 | -0.872 | 2.885 | 1.328 | 0.016 | 0.117 | 0.117 | 1.08 | 0.91 | 0.71 | 0.91 | 0.76 | 0.93 | 0.82 |
| 4 | 61276747 | rs7690448 | 1.862 | -1.171 | 2.543 | 1.083 | -0.018 | 0.047 | 0.047 | 0.299 | 0.86 | 0.69 | 0.9 | 0.77 | 0.91 | 0.81 |
| 4 | 61278746 | rs9990505 | 1.829 | -1.156 | 2.538 | 1.091 | -0.017 | 0.046 | 0.046 | 0.32 | 0.86 | 0.69 | 0.9 | 0.77 | 0.91 | 0.8 |
| 4 | 61290298 | rs6853921 | -2.104 | -1.225 | 2.939 | 1.049 | -0.07 | 0.033 | 0.033 | 0 | 0.25 | 0.59 | 0.18 | 0.33 | 0.12 | 0.32 |
| 4 | 61296108 | rs980051 | -2.377 | -1.279 | 2.848 | 1.016 | -0.02 | 0.002 | 0.002 | 0 | 0.29 | 0.68 | 0.23 | 0.36 | 0.17 | 0.35 |
| 4 | 61305996 | rs3911391 | -1.688 | -1.251 | 2.869 | 1.027 | -0.023 | 0.041 | 0.041 | 0 | 0.2 | 0.54 | 0.18 | 0.28 | 0.13 | 0.2 |
| 4 | 61310371 | rs1350085 | -1.688 | -1.251 | 2.869 | 1.027 | -0.023 | 0.041 | 0.041 | 0 | 0.2 | 0.54 | 0.18 | 0.28 | 0.13 | 0.2 |
| 4 | 61316400 | rs17219303 | -1.339 | -1.136 | 2.991 | 0.776 | 0.043 | 0.109 | 0.109 | 0.652 | 0.05 | 0.19 | 0.05 | 0.16 | 0.06 | 0.05 |
| 4 | 61317176 | rs925395 | 1.82 | -1.236 | 3.008 | 0.667 | -0.028 | -0.075 | -0.075 | 2.173 | 0.77 | 0.64 | 0.87 | 0.92 | 0.84 | 0.69 |
| 4 | 61319052 | rs1553863 | 1.969 | -1.24 | 3.145 | 0.665 | -0.04 | -0.101 | -0.101 | 0.333 | 0.76 | 0.51 | 0.86 | 0.92 | 0.84 | 0.7 |
| 4 | 61321240 | rs6551553 | 1.943 | -1.197 | 2.987 | 0.652 | 0.02 | -0.022 | -0.022 | 0.128 | 0.81 | 0.6 | 0.87 | 0.84 | 0.81 | 0.79 |
| 4 | 61339224 | rs994342 | -1.205 | -1.154 | 2.805 | 0.659 | 0 | -0.062 | -0.062 | 0 | 0.14 | 0.2 | 0.08 | 0.01 | 0.13 | 0.16 |
| 4 | 61345972 | rs10517505 | -0.776 | -1.182 | 2.607 | 0.671 | -0.029 | -0.012 | -0.012 | 0 | 0.09 | 0.16 | 0.05 | 0.08 | 0.03 | 0.14 |
| 4 | 61352529 | rs10021608 | -1.944 | -1.187 | 2.357 | 0.696 | 0.008 | -0.021 | -0.021 | 0 | 0.2 | 0.39 | 0.14 | 0.16 | 0.2 | 0.22 |
| 4 | 61353754 | rs1397649 | 1.914 | -1.066 | 1.753 | 0.444 | -0.015 | -0.043 | -0.043 | 0 | 0.73 | 0.4 | 0.82 | 0.82 | 0.78 | 0.67 |
| 4 | 61356693 | rs997107 | -1.995 | -1.634 | 1.143 | 0.174 | 0.009 | 0.011 | 0.011 | 0 | 0.37 | 0.36 | 0.19 | 0.21 | 0.26 | 0.36 |
| 4 | 61360284 | rs4602520 | 0.257 | -1.732 | 1.002 | 0.754 | -0.015 | 0.101 | 0.101 | 0 | 0.13 | 0.24 | 0.12 | 0.37 | 0.09 | 0.18 |
| 4 | 61360679 | rs11728006 | 2.091 | -1.64 | 0.817 | 0.662 | 0.06 | -0.006 | -0.006 | 0 | 0.48 | 0.16 | 0.68 | 0.4 | 0.65 | 0.45 |
| 4 | 61363382 | rs9884208 | -1.368 | -1.284 | 1.025 | 0.726 | 0 | -0.003 | -0.003 | 0.021 | 0.16 | 0.16 | 0.12 | 0.09 | 0.19 | 0.17 |
| 4 | 61365718 | rs11131258 | 1.566 | -0.947 | -0.24 | 0.356 | -0.071 | -0.051 | -0.051 | 0 | 0.32 | 0.15 | 0.61 | 0.4 | 0.49 | 0.4 |
| 4 | 61367078 | rs6551565 | -0.42 | -0.699 | -0.344 | 0.461 | -0.001 | 0 | 0 | 0.461 | 0.45 | 0.4 | 0.3 | 0.52 | 0.37 | 0.42 |
| 4 | 61370159 | rs6852374 | 0.562 | -0.212 | -0.662 | 0.42 | 0 | -0.001 | -0.001 | 0 | 0.4 | 0.43 | 0.62 | 0.34 | 0.5 | 0.4 |
| 4 | 61371578 | rs1588514 | 0.262 | -0.244 | -1.269 | -0.036 | 0.026 | 0.093 | 0.093 | 0.135 | 0.67 | 0.89 | 0.83 | 0.88 | 0.75 | 0.66 |
| 4 | 61395456 | rs1553862 | -1.474 | -0.346 | -0.073 | 0.827 | -0.013 | 0.08 | 0.08 | 2.993 | 0.15 | 0.59 | 0.09 | 0.24 | 0.11 | 0.19 |
| 4 | 61402680 | rs9990993 | -0.73 | -0.363 | 0.309 | 0.893 | 0 | -0.001 | -0.001 | 0 | 0.08 | 0.09 | 0.04 | 0.12 | 0.08 | 0.14 |
| 4 | 61404788 | rs1912682 | -0.391 | -0.671 | 0.281 | 1.176 | 0.005 | -0.014 | -0.014 | 0 | 0.16 | 0.15 | 0.32 | 0.44 | 0.15 | 0.17 |
| 4 | 61422632 | rs12645705 | -1.265 | -0.658 | 0.31 | 1.213 | -0.021 | 0.186 | 0.186 | 0.183 | 0.18 | 0.53 | 0.34 | 0.56 | 0.15 | 0.18 |
| 4 | 61424207 | rs1439310 | 0.502 | -0.533 | 0.04 | 1.194 | -0.013 | -0.026 | -0.026 | 0 | 0.08 | 0.01 | 0.28 | 0.44 | 0.1 | 0.11 |
| 4 | 61443447 | rs13125772 | -0.903 | -0.466 | 0.164 | 0.736 | 0.012 | -0.001 | -0.001 | 0 | 0.1 | 0.22 | 0.05 | 0.11 | 0.08 | 0.14 |
| 4 | 61446054 | rs13146560 | 0.785 | 0.324 | -1.202 | -0.319 | -0.105 | 0.138 | 0.138 | 0 | 0.25 | 0.04 | 0.21 | 0 | 0.4 | 0.17 |
| 4 | 61484551 | rs11131268 | 1.041 | -0.387 | -1.034 | -0.249 | 0.042 | 0.143 | 0.143 | 1.113 | 0.5 | 0.26 | 0.42 | 0.22 | 0.61 | 0.49 |
| 4 | 61497064 | rs6843734 | 1.038 | -0.408 | -1.045 | -0.26 | 0.043 | 0.144 | 0.144 | 1.207 | 0.5 | 0.26 | 0.42 | 0.22 | 0.61 | 0.48 |
| 4 | 61504139 | rs2028315 | 0.789 | -0.506 | -1.128 | -0.053 | 0.019 | 0.049 | 0.049 | 0.762 | 0.58 | 0.27 | 0.78 | 0.72 | 0.69 | 0.57 |
| 4 | 61507007 | rs975734 | 0.153 | -0.71 | -0.959 | -0.274 | -0.005 | -0.005 | -0.005 | 0 | 0.7 | 0.4 | 0.82 | 0.77 | 0.75 | 0.69 |
| 4 | 61511878 | rs9996373 | -0.023 | -0.628 | -0.935 | -0.196 | -0.002 | -0.014 | -0.014 | 0 | 0.12 | 0.13 | 0.04 | 0.05 | 0.06 | 0.11 |
| 4 | 61513458 | rs17090230 | -0.46 | -0.654 | -0.401 | -0.194 | 0.021 | 0.126 | 0.126 | 1.082 | 0.11 | 0.36 | 0.07 | 0.2 | 0.1 | 0.17 |
| 4 | 61519303 | rs1898848 | -0.734 | -0.686 | 0.244 | -0.317 | -0.003 | -0.043 | -0.043 | 0 | 0.81 | 0.79 | 0.89 | 0.95 | 0.85 | 0.86 |
| 4 | 61528903 | rs1439317 | 1.372 | -0.218 | 0.267 | -0.242 | -0.001 | -0.005 | -0.005 | 0.573 | 0.55 | 0.46 | 0.43 | 0.26 | 0.6 | 0.5 |
| 4 | 61529037 | rs10024649 | -0.296 | -0.139 | 0.072 | -0.067 | 0.033 | 0.19 | 0.19 | 0 | 0.05 | 0.2 | 0.02 | 0.13 | 0.04 | 0.07 |
| 4 | 61533457 | rs1439299 | 1.015 | -0.297 | -0.232 | -0.136 | 0.017 | -0.03 | -0.03 | 0.012 | 0.09 | 0.02 | 0.36 | 0.55 | 0.08 | 0.18 |
| 4 | 61539305 | rs7662547 | 0.571 | -0.194 | -0.067 | -0.223 | -0.004 | 0.022 | 0.022 | 0 | 0.68 | 0.73 | 0.81 | 0.78 | 0.71 | 0.63 |

| | | | | | | | | | | | | | | | | |
|---|---|---|---|---|---|---|---|---|---|---|---|---|---|---|---|---|
| 4 | 61541738 | rs1439294 | -0.862 | -0.211 | 0.053 | -0.21 | 0.003 | 0.026 | 0.026 | 0 | 0.82 | 0.88 | 0.89 | 0.93 | 0.85 | 0.79 |
| 4 | 61552142 | rs11932579 | -0.594 | -0.113 | 0.115 | -0.162 | 0 | -0.002 | -0.002 | 0 | 0.14 | 0.15 | 0.08 | 0.15 | 0.15 | 0.16 |
| 4 | 61552933 | rs4535361 | -0.945 | -0.154 | 0.056 | -0.317 | 0.002 | 0.009 | 0.009 | 0.036 | 0.82 | 0.83 | 0.89 | 0.95 | 0.85 | 0.87 |
| 4 | 61555937 | rs9884903 | -1.44 | -0.25 | 0.269 | -0.126 | -0.01 | 0.031 | 0.031 | 1.139 | 0.3 | 0.44 | 0.47 | 0.67 | 0.26 | 0.36 |
| 4 | 61588040 | rs4860361 | 0.964 | -0.595 | 0.486 | -0.574 | -0.008 | -0.029 | -0.029 | 0 | 0.61 | 0.44 | 0.77 | 0.81 | 0.7 | 0.69 |
| 4 | 61590894 | rs4309865 | 0.485 | -0.682 | 0.822 | -0.66 | -0.003 | -0.034 | -0.034 | 0 | 0.19 | 0.17 | 0.1 | 0.03 | 0.16 | 0.1 |
| 4 | 61592332 | rs10012618 | 0.391 | -0.742 | 0.998 | -0.614 | -0.008 | -0.005 | -0.005 | 0 | 0.84 | 0.77 | 0.86 | 0.82 | 0.86 | 0.81 |
| 4 | 61602645 | rs10050170 | 1.182 | -0.644 | 1.023 | -0.725 | -0.011 | -0.039 | -0.039 | 0.555 | 0.61 | 0.47 | 0.74 | 0.79 | 0.71 | 0.69 |
| 4 | 61603501 | rs7679019 | 0.431 | -0.652 | 1.144 | -0.708 | -0.015 | -0.007 | -0.007 | 0.277 | 0.84 | 0.69 | 0.87 | 0.82 | 0.88 | 0.82 |
| 4 | 61603973 | rs4510484 | 0.245 | -0.554 | 0.941 | -0.591 | -0.011 | -0.062 | -0.062 | 0.174 | 0.22 | 0.28 | 0.12 | 0.03 | 0.16 | 0.12 |
| 4 | 61612126 | rs4527515 | 1.127 | -0.552 | 0.793 | -0.557 | -0.006 | -0.021 | -0.021 | 0 | 0.6 | 0.43 | 0.74 | 0.79 | 0.7 | 0.7 |
| 4 | 61621387 | rs4394027 | 0.654 | -0.483 | 1.035 | -0.467 | 0.012 | 0.02 | 0.02 | 0.332 | 0.89 | 0.67 | 0.89 | 0.87 | 0.89 | 0.86 |
| 4 | 61626653 | rs12504712 | 0.964 | -0.599 | 0.27 | 0.158 | -0.012 | 0.001 | 0.001 | 0 | 0.07 | 0.01 | 0.39 | 0.5 | 0.11 | 0.09 |
| 4 | 61628548 | rs6849233 | 0.778 | -0.627 | 0.404 | 0.108 | 0 | 0 | 0 | 0 | 0.07 | 0.09 | 0.07 | 0.14 | 0.08 | 0.1 |
| 4 | 61629652 | rs13109616 | 2.014 | -1.096 | -0.316 | -0.562 | -0.147 | 0.234 | 0.234 | 0.517 | 0.36 | 0.07 | 0.27 | 0 | 0.49 | 0.23 |
| 4 | 61641564 | rs4496609 | -0.986 | -0.864 | 0.312 | 0.162 | -0.002 | 0.105 | 0.105 | 0 | 0.92 | 0.89 | 0.59 | 0.49 | 0.89 | 0.89 |
| 4 | 61654288 | rs4072938 | 0.644 | -0.951 | 0.525 | -0.314 | -0.009 | -0.056 | -0.056 | 0 | 0.2 | 0.27 | 0.11 | 0.02 | 0.15 | 0.09 |
| 4 | 61666777 | rs12648062 | -0.502 | -0.838 | 0.57 | -0.065 | -0.005 | -0.016 | -0.016 | 0 | 0.91 | 0.8 | 0.91 | 0.96 | 0.91 | 0.88 |
| 4 | 61675083 | rs17081884 | -1.464 | -0.938 | 0.637 | 0.05 | -0.024 | 0.001 | 0.001 | 0 | 0.28 | 0.5 | 0.56 | 0.78 | 0.25 | 0.33 |
| 4 | 61685448 | rs17090281 | 0.581 | -0.821 | 0.937 | -0.036 | -0.008 | -0.06 | -0.06 | 0 | 0.19 | 0.25 | 0.1 | 0.02 | 0.15 | 0.09 |
| 4 | 61688725 | rs1378360 | 0.069 | -0.886 | 1.081 | 0.281 | -0.012 | 0.101 | 0.101 | 0.069 | 0.22 | 0.51 | 0.48 | 0.55 | 0.19 | 0.19 |
| 4 | 61695508 | rs17090284 | 0.614 | -0.897 | 0.69 | -0.101 | -0.007 | -0.045 | -0.045 | 0.103 | 0.19 | 0.2 | 0.09 | 0.03 | 0.15 | 0.11 |
| 4 | 61696791 | rs4611956 | 1.251 | -0.511 | -0.227 | -0.434 | 0 | 0.046 | 0.046 | 0.259 | 0.13 | 0.03 | 0.08 | 0.02 | 0.13 | 0.06 |
| 4 | 61709761 | rs17291739 | -0.331 | -0.486 | 0.083 | -0.162 | -0.002 | 0.007 | 0.007 | 0.16 | 0.34 | 0.29 | 0.54 | 0.48 | 0.31 | 0.23 |
| 4 | 61724114 | rs11725567 | 1.879 | -0.785 | -0.334 | -0.382 | -0.012 | 0.052 | 0.052 | 0.918 | 0.1 | 0.01 | 0.06 | 0.01 | 0.1 | 0.05 |
| 4 | 61725276 | rs10019200 | 1.179 | -0.767 | -0.269 | -0.392 | 0.006 | 0.011 | 0.011 | 0.248 | 0.13 | 0.06 | 0.07 | 0.01 | 0.11 | 0.06 |
| 4 | 61729184 | rs10012645 | -0.767 | -1.011 | -0.28 | -0.72 | -0.017 | -0.071 | -0.071 | 0 | 0.3 | 0.35 | 0.18 | 0.07 | 0.2 | 0.25 |
| 4 | 61738094 | rs1455268 | -0.39 | -1.014 | -0.433 | -0.649 | -0.004 | 0.015 | 0.015 | 0 | 0.32 | 0.57 | 0.55 | 0.49 | 0.28 | 0.31 |
| 4 | 61738470 | rs17090295 | 0.683 | -0.849 | -1.105 | -0.534 | -0.014 | 0.024 | 0.024 | 0 | 0.08 | 0.04 | 0.07 | 0.03 | 0.1 | 0.11 |
| 4 | 61739960 | rs6826243 | -1.22 | -0.87 | -1.156 | -0.548 | 0.011 | -0.059 | -0.059 | 6.011 | 0.53 | 0.83 | 0.68 | 0.53 | 0.4 | 0.44 |
| 4 | 61753646 | rs1455286 | 0.558 | -0.705 | -1.072 | -0.682 | -0.028 | 0.038 | 0.038 | 0 | 0.09 | 0.01 | 0.07 | 0.03 | 0.11 | 0.11 |
| 4 | 61762647 | rs7670391 | 0.942 | -0.972 | -0.844 | -0.739 | 0.002 | -0.006 | -0.006 | 0 | 0.2 | 0.12 | 0.46 | 0.45 | 0.17 | 0.19 |
| 4 | 61767946 | rs716167 | 1.186 | -0.266 | -1.376 | -1.464 | -0.001 | 0.014 | 0.014 | 1.039 | 0.15 | 0.11 | 0.43 | 0.42 | 0.23 | 0.24 |
| 4 | 61771431 | rs1455280 | 1.156 | -0.469 | -0.871 | -1.349 | 0.01 | 0.01 | 0.01 | 0 | 0.2 | 0.12 | 0.08 | 0.01 | 0.12 | 0.13 |
| 4 | 61773762 | rs10000438 | -1.252 | -0.598 | -0.956 | -1.463 | 0.079 | 0.202 | 0.202 | 0 | 0.49 | 0.77 | 0.29 | 0.1 | 0.33 | 0.34 |
| 4 | 61773790 | rs9312059 | 0.876 | -0.557 | -0.608 | -1.387 | -0.056 | -0.109 | -0.109 | 0 | 0.19 | 0.46 | 0.13 | 0.06 | 0.1 | 0.11 |
| 4 | 61781801 | rs6823258 | 0.639 | -0.466 | -1.061 | -1.724 | -0.033 | 0.101 | 0.101 | 0 | 0.37 | 0.18 | 0.26 | 0.01 | 0.38 | 0.2 |
| 4 | 61788908 | rs10022351 | -0.991 | -0.498 | -0.98 | -1.758 | 0 | -0.053 | -0.053 | 0.52 | 0.63 | 0.66 | 0.46 | 0.09 | 0.59 | 0.42 |
| 4 | 61795416 | rs10517518 | 0.881 | -1.417 | -0.315 | -0.964 | 0.014 | 0.252 | 0.252 | 0.095 | 0.05 | 0.11 | 0.39 | 0.44 | 0.06 | 0.1 |
| 4 | 61799685 | rs6851977 | 0.914 | -1.555 | -0.475 | -1.759 | 0.008 | -0.003 | -0.003 | 0 | 0.21 | 0.15 | 0.08 | 0.01 | 0.12 | 0.13 |
| 4 | 61800185 | rs2045199 | 1.046 | -1.563 | -0.476 | -1.762 | -0.018 | 0.039 | 0.039 | 0 | 0.08 | 0.03 | 0.07 | 0.01 | 0.12 | 0.11 |
| 4 | 61803301 | rs2604632 | -0.028 | -1.478 | -0.427 | -1.69 | -0.03 | -0.124 | -0.124 | 0.009 | 0.73 | 0.52 | 0.8 | 0.93 | 0.79 | 0.79 |
| 4 | 61809210 | rs2631054 | 0.302 | -0.843 | -0.417 | -1.234 | 0.023 | 0.007 | 0.007 | 0 | 0.53 | 0.37 | 0.71 | 0.92 | 0.66 | 0.66 |
| 4 | 61830190 | rs1486492 | -0.139 | -1.339 | -0.282 | -0.542 | 0.052 | 0.04 | 0.04 | 0.494 | 0.59 | 0.33 | 0.77 | 0.7 | 0.78 | 0.73 |
| 4 | 61832236 | rs901388 | 0.119 | -1.333 | -0.428 | -0.814 | 0.049 | 0.076 | 0.076 | 1.077 | 0.48 | 0.33 | 0.7 | 0.91 | 0.67 | 0.54 |
| 4 | 61836190 | rs1486493 | -0.679 | -1.34 | -0.464 | -0.812 | 0.02 | 0.008 | 0.008 | 0.288 | 0.63 | 0.36 | 0.79 | 0.93 | 0.8 | 0.81 |
| 4 | 61837133 | rs1906073 | 1.222 | -1.228 | -0.643 | -0.425 | -0.071 | 0.045 | 0.045 | 1.219 | 0.36 | 0.1 | 0.31 | 0.15 | 0.59 | 0.33 |
| 4 | 61846956 | rs8180200 | -1.076 | -1.213 | -0.213 | -0.496 | 0.019 | 0.015 | 0.015 | 1 | 0.68 | 0.75 | 0.82 | 0.93 | 0.8 | 0.81 |
| 4 | 61848154 | rs13129872 | 0.705 | -0.597 | -1.593 | -0.169 | -0.025 | 0.089 | 0.089 | 0 | 0.14 | 0.03 | 0.09 | 0.02 | 0.13 | 0.27 |
| 4 | 61860937 | rs12641314 | -0.087 | -0.604 | -1.501 | -0.229 | 0.008 | 0.013 | 0.013 | 1.944 | 0.47 | 0.47 | 0.69 | 0.85 | 0.64 | 0.55 |
| 4 | 61866264 | rs6831772 | -0.628 | -0.368 | -1.033 | 0.155 | 0 | 0.104 | 0.104 | 0.067 | 0.07 | 0.23 | 0.05 | 0.23 | 0.04 | 0.07 |
| 4 | 61871415 | rs13119110 | -0.422 | -0.425 | -1.36 | -0.674 | -0.009 | -0.016 | -0.016 | 0 | 0.59 | 0.49 | 0.78 | 0.87 | 0.76 | 0.79 |
| 4 | 61872079 | rs1486498 | 0.108 | 0.279 | -2.041 | -0.826 | -0.122 | 0.15 | 0.15 | 1.017 | 0.35 | 0.07 | 0.29 | 0.07 | 0.56 | 0.36 |
| 4 | 61894078 | rs10006348 | -0.134 | -0.13 | -1.8 | -0.781 | -0.034 | 0.187 | 0.187 | 0.008 | 0.42 | 0.11 | 0.31 | 0.12 | 0.6 | 0.46 |
| 4 | 61900003 | rs4860078 | 0.028 | -0.076 | -1.899 | -0.855 | -0.139 | 0.138 | 0.138 | 0.397 | 0.38 | 0.03 | 0.25 | 0.11 | 0.55 | 0.38 |
| 4 | 61903366 | rs12501471 | -0.235 | -0.542 | -0.823 | -1.042 | 0.024 | 0.007 | 0.007 | 0 | 0.95 | 0.75 | 0.97 | 1 | 0.97 | 1 |
| 4 | 61908999 | rs6813806 | 0.516 | -0.717 | -0.399 | -0.684 | -0.01 | -0.01 | -0.01 | 2.621 | 0.45 | 0.48 | 0.59 | 0.66 | 0.29 | 0.41 |
| 4 | 61911862 | rs725761 | -1.229 | -0.596 | 0.122 | -0.492 | 0 | 0.013 | 0.013 | 2.245 | 0.2 | 0.68 | 0.26 | 0.28 | 0.19 | 0.29 |
| 4 | 61917658 | rs966084 | -0.655 | -1.38 | 0.448 | -0.175 | 0.047 | 0.005 | 0.005 | 0.061 | 0.58 | 0.74 | 0.52 | 0.45 | 0.71 | 0.63 |
| 4 | 61921234 | rs17090351 | -0.609 | -0.841 | 0.733 | 0.106 | 0.018 | 0.164 | 0.164 | 0 | 0.06 | 0.24 | 0.03 | 0.22 | 0.03 | 0.13 |
| 4 | 61946476 | rs9884795 | -0.525 | -0.833 | 0.028 | 0.054 | 0.033 | -0.007 | -0.007 | 0 | 0.57 | 0.74 | 0.51 | 0.5 | 0.66 | 0.67 |
| 4 | 61948488 | rs7699691 | -0.02 | -0.846 | 0.204 | 0.078 | 0.038 | 0.093 | 0.093 | 0 | 0.48 | 0.27 | 0.47 | 0.28 | 0.63 | 0.53 |
| 4 | 61950476 | rs10030728 | -0.26 | -0.264 | 0.348 | 0.277 | 0.038 | 0.011 | 0.011 | 0.611 | 0.48 | 0.72 | 0.45 | 0.5 | 0.58 | 0.65 |
| 4 | 61966273 | rs4599440 | 0.359 | -0.226 | -0.168 | 0.596 | -0.037 | 0.002 | 0.002 | 0 | 0.63 | 0.48 | 0.83 | 0.43 | 0.78 | 0.51 |
| 4 | 61967025 | rs7439943 | 0.279 | -0.147 | -0.666 | 0.625 | 0.033 | -0.013 | -0.013 | 0 | 0.65 | 0.74 | 0.85 | 0.43 | 0.78 | 0.52 |
| 4 | 61975533 | rs1449618 | -0.731 | 0.457 | -0.927 | 0.438 | -0.011 | 0.017 | 0.017 | 0 | 0.24 | 0.17 | 0.34 | 0.16 | 0.26 | 0.25 |
| 4 | 61978764 | rs2053864 | 0.988 | 1.306 | -0.346 | 1.334 | 0.018 | 0.009 | 0.009 | 0 | 0.63 | 0.53 | 0.47 | 0.44 | 0.52 | 0.54 |
| 4 | 61982096 | rs10517524 | -0.21 | 1.293 | -0.463 | 1.349 | -0.045 | -0.048 | -0.048 | 0 | 0.11 | 0.03 | 0.19 | 0.4 | 0.22 | 0.2 |
| 4 | 61984568 | rs2034448 | -0.342 | 1.243 | -0.315 | 1.092 | -0.003 | 0.006 | 0.006 | 0 | 0.24 | 0.21 | 0.34 | 0.16 | 0.26 | 0.27 |

| | | | | | | | | | | | | | | | | |
|---|---|---|---|---|---|---|---|---|---|---|---|---|---|---|---|---|
| 4 | 61987337 | rs12506236 | -0.806 | 1.217 | -0.411 | 1.601 | -0.03 | -0.039 | -0.039 | 1.006 | 0.13 | 0.07 | 0.24 | 0.52 | 0.22 | 0.19 |
| 4 | 61992249 | rs1900957 | -1.312 | 0.877 | -0.449 | 1.455 | -0.002 | 0.012 | 0.012 | 0 | 0.16 | 0.17 | 0.24 | 0.53 | 0.22 | 0.2 |
| 4 | 61998057 | rs10866114 | -0.52 | 0.301 | -1.067 | 0.421 | -0.011 | 0.069 | 0.069 | 0 | 0.42 | 0.28 | 0.38 | 0.78 | 0.46 | 0.5 |
| 4 | 62002328 | rs10006264 | 0.892 | 0.303 | -1.061 | 0.431 | 0.002 | -0.011 | -0.011 | 0.282 | 0.26 | 0.13 | 0.15 | 0.25 | 0.23 | 0.3 |
| 4 | 62004333 | rs930668 | -0.591 | -0.494 | -0.312 | 0.338 | -0.029 | 0.022 | 0.022 | 0 | 0.91 | 0.68 | 0.87 | 0.94 | 0.97 | 0.96 |
| 4 | 62007979 | rs1470645 | -0.404 | -0.638 | -0.455 | 0.164 | 0.039 | 0.078 | 0.078 | 0.428 | 0.58 | 0.29 | 0.66 | 0.82 | 0.67 | 0.66 |
| 4 | 62015576 | rs12186207 | 0.808 | -0.845 | -0.786 | -0.128 | -0.077 | 0.009 | 0.009 | 0 | 0.37 | 0.14 | 0.39 | 0.29 | 0.39 | 0.45 |
| 4 | 62017472 | rs1375680 | 1.171 | -1.21 | -0.725 | -0.145 | -0.008 | -0.068 | -0.068 | 0 | 0.32 | 0.36 | 0.22 | 0.11 | 0.31 | 0.29 |
| 4 | 62044559 | rs1449613 | 1.148 | -1.281 | -0.761 | -0.18 | -0.008 | -0.066 | -0.066 | 1.232 | 0.32 | 0.37 | 0.22 | 0.11 | 0.3 | 0.29 |
| 4 | 62049284 | rs2218741 | 0.701 | -0.833 | -1.189 | -0.087 | -0.087 | 0.019 | 0.019 | 0 | 0.37 | 0.16 | 0.42 | 0.29 | 0.46 | 0.47 |
| 4 | 62052467 | rs998666 | 0.762 | -0.975 | -1.202 | 0.142 | -0.086 | 0.093 | 0.093 | 0 | 0.36 | 0.15 | 0.41 | 0.16 | 0.43 | 0.44 |
| 4 | 62061051 | rs6841219 | 0.877 | -0.848 | -0.769 | 0.185 | -0.007 | 0.034 | 0.034 | 0.069 | 0.31 | 0.15 | 0.2 | 0.24 | 0.33 | 0.34 |
| 4 | 62074758 | rs1505659 | -0.49 | -0.765 | -1.034 | 0.49 | -0.001 | -0.001 | -0.001 | 0 | 0.6 | 0.47 | 0.65 | 0.68 | 0.65 | 0.56 |
| 4 | 62076443 | rs6854914 | 0.7 | -0.961 | -1.127 | 0.472 | -0.011 | 0.036 | 0.036 | 0.069 | 0.42 | 0.35 | 0.45 | 0.19 | 0.46 | 0.4 |
| 4 | 62080215 | rs6845545 | -0.63 | -0.93 | -1.218 | 0.553 | -0.002 | -0.058 | -0.058 | 0 | 0.18 | 0.08 | 0.22 | 0.51 | 0.2 | 0.16 |
| 4 | 62084538 | rs6825047 | -0.323 | -0.873 | -1.159 | 0.672 | -0.007 | -0.007 | -0.007 | 0 | 0.6 | 0.44 | 0.67 | 0.72 | 0.67 | 0.57 |
| 4 | 62088597 | rs12499198 | -0.319 | -0.437 | -1.206 | 1.094 | -0.025 | -0.03 | -0.03 | 0 | 0.09 | 0.01 | 0.15 | 0.5 | 0.1 | 0.09 |
| 4 | 62098937 | rs17090416 | 0.031 | -0.178 | -0.976 | 0.516 | -0.03 | -0.024 | -0.024 | 0 | 0.33 | 0.44 | 0.31 | 0.28 | 0.27 | 0.39 |
| 4 | 62100860 | rs924940 | -0.948 | 0.244 | -1.354 | 1.147 | -0.008 | -0.088 | -0.088 | 0.287 | 0.15 | 0.02 | 0.2 | 0.41 | 0.18 | 0.14 |
| 4 | 62104112 | rs1827770 | -0.438 | 0.476 | -0.948 | -0.025 | 0.001 | -0.029 | -0.029 | 0 | 0.17 | 0.18 | 0.27 | 0.04 | 0.18 | 0.18 |
| 4 | 62106989 | rs11131314 | 0.497 | 0.536 | -0.882 | -0.2 | 0.003 | -0.004 | -0.004 | 0 | 0.41 | 0.43 | 0.41 | 0.61 | 0.48 | 0.42 |
| 4 | 62111856 | rs1505682 | 0.815 | 0.797 | -0.881 | 0.151 | -0.04 | -0.051 | -0.051 | 0 | 0.23 | 0.12 | 0.25 | 0.37 | 0.33 | 0.14 |
| 4 | 62113587 | rs1979111 | 0.459 | 0.464 | -0.738 | -0.279 | -0.002 | -0.001 | -0.001 | 0.62 | 0.26 | 0.25 | 0.2 | 0.29 | 0.23 | 0.29 |
| 4 | 62121348 | rs974541 | -0.881 | -0.115 | -0.434 | -0.209 | 0.003 | -0.006 | -0.006 | 0 | 0.75 | 0.77 | 0.68 | 0.74 | 0.77 | 0.6 |
| 4 | 62124053 | rs2342456 | 0.091 | -0.185 | -0.369 | -0.326 | 0.014 | 0.031 | 0.031 | 0 | 0.56 | 0.67 | 0.57 | 0.69 | 0.64 | 0.46 |
| 4 | 62130085 | rs10517529 | 1.195 | -0.327 | -0.442 | -0.322 | -0.001 | 0.049 | 0.049 | 0 | 0.18 | 0.09 | 0.1 | 0.05 | 0.13 | 0.13 |
| 4 | 62132090 | rs1505663 | 0.607 | -0.516 | -0.427 | -0.363 | -0.014 | 0.004 | 0.004 | 0 | 0.24 | 0.3 | 0.25 | 0.31 | 0.2 | 0.31 |
| 4 | 62132345 | rs1505664 | 0.883 | -0.274 | -0.446 | -0.342 | -0.027 | -0.013 | -0.013 | 0 | 0.31 | 0.24 | 0.33 | 0.37 | 0.47 | 0.16 |
| 4 | 62136915 | rs4860406 | -0.079 | -0.409 | -0.375 | -0.327 | -0.01 | -0.002 | -0.002 | 0 | 0.45 | 0.53 | 0.43 | 0.57 | 0.36 | 0.69 |
| 4 | 62140177 | rs12505990 | 0.581 | 0.198 | -0.472 | 0.054 | -0.038 | -0.033 | -0.033 | 0 | 0.28 | 0.17 | 0.31 | 0.41 | 0.41 | 0.14 |
| 4 | 62143023 | rs9312075 | -0.922 | -0.06 | -0.355 | 0.027 | -0.046 | 0.024 | 0.024 | 0 | 0.79 | 0.64 | 0.75 | 0.71 | 0.83 | 0.69 |
| 4 | 62150059 | rs7435420 | 0.291 | 0.048 | -0.517 | 0.045 | 0.001 | 0.018 | 0.018 | 0 | 0.48 | 0.4 | 0.43 | 0.31 | 0.39 | 0.55 |
| 4 | 62159915 | rs9998310 | 0.265 | 0.727 | -1.012 | 0.371 | -0.121 | -0.104 | -0.104 | 1.088 | 0.27 | 0.06 | 0.38 | 0.4 | 0.42 | 0.14 |
| 4 | 62160473 | rs12508911 | 0.124 | 0.764 | -0.99 | 0.389 | -0.04 | -0.033 | -0.033 | 0 | 0.28 | 0.17 | 0.4 | 0.4 | 0.42 | 0.14 |
| 4 | 62162828 | rs17828306 | -0.264 | -0.214 | -0.728 | 0.146 | 0.028 | 0.005 | 0.005 | 0 | 0.37 | 0.31 | 0.22 | 0.29 | 0.26 | 0.48 |
| 4 | 62163043 | rs17090424 | -0.219 | -0.46 | -0.151 | 0.077 | -0.022 | -0.002 | -0.002 | 0 | 0.34 | 0.56 | 0.24 | 0.29 | 0.25 | 0.44 |
| 4 | 62163509 | rs7695405 | 0.554 | -0.06 | 0.334 | 0.33 | 0.021 | 0.059 | 0.059 | 0 | 0.88 | 0.7 | 0.8 | 0.84 | 0.86 | 0.94 |
| 4 | 62167906 | rs2880363 | 0.813 | -0.034 | -0.877 | -0.131 | -0.014 | 0.115 | 0.115 | 0.106 | 0.49 | 0.17 | 0.53 | 0.69 | 0.57 | 0.32 |
| 4 | 62172534 | rs13151711 | -0.381 | -0.101 | -0.459 | -0.521 | 0.031 | -0.055 | -0.055 | 0 | 0.84 | 0.75 | 0.75 | 0.97 | 0.8 | 0.86 |
| 4 | 62188118 | rs2135242 | 1.102 | -0.256 | -0.466 | -0.737 | 0.004 | -0.08 | -0.08 | 0 | 0.71 | 0.65 | 0.72 | 0.97 | 0.74 | 0.74 |
| 4 | 62198201 | rs1505679 | -0.209 | -0.216 | -0.43 | -0.703 | 0.024 | -0.071 | -0.071 | 0 | 0.83 | 0.7 | 0.76 | 0.98 | 0.82 | 0.89 |
| 4 | 62208968 | rs10032294 | 2.297 | -0.562 | -1.225 | -0.616 | -0.001 | 0.056 | 0.056 | 0 | 0.46 | 0.17 | 0.52 | 0.66 | 0.59 | 0.29 |
| 4 | 62210266 | rs11729528 | -0.249 | -0.369 | -1.287 | -0.653 | 0.002 | -0.038 | -0.038 | 0 | 0.16 | 0.02 | 0.12 | 0.19 | 0.14 | 0.2 |
| 4 | 62213988 | rs1456863 | -0.191 | -0.525 | -0.919 | -0.479 | 0 | -0.018 | -0.018 | 0 | 0.1 | 0.13 | 0.03 | 0.01 | 0.08 | 0.16 |
| 4 | 62225605 | rs17828324 | -1.424 | -0.571 | -1.061 | -0.491 | -0.004 | 0.104 | 0.104 | 0.02 | 0.38 | 0.14 | 0.33 | 0.21 | 0.32 | 0.5 |
| 4 | 62229733 | rs4860411 | 1.355 | 0.023 | -0.919 | 0.052 | 0.003 | -0.028 | -0.028 | 0 | 0.6 | 0.27 | 0.58 | 0.51 | 0.66 | 0.42 |
| 4 | 62240526 | rs6858223 | -0.291 | 0.216 | -0.895 | 0.263 | -0.043 | -0.011 | -0.011 | 0 | 0.21 | 0.08 | 0.3 | 0.2 | 0.27 | 0.28 |
| 4 | 62242432 | rs10866117 | 0.859 | 0.13 | -0.376 | -0.018 | 0.003 | 0.007 | 0.007 | 0 | 0.69 | 0.7 | 0.61 | 0.51 | 0.66 | 0.42 |
| 4 | 62244228 | rs17219773 | 0.377 | 0.557 | -0.721 | 0.107 | -0.043 | -0.025 | -0.025 | 2.999 | 0.12 | 0.02 | 0.09 | 0.13 | 0.15 | 0.15 |
| 4 | 62251012 | rs9684510 | -0.474 | 0.265 | -0.486 | -0.048 | -0.001 | -0.025 | -0.025 | 0 | 0.52 | 0.47 | 0.28 | 0.18 | 0.41 | 0.46 |
| 4 | 62251077 | rs9312076 | 1.378 | 0.409 | -0.876 | 0.087 | -0.032 | -0.03 | -0.03 | 1.084 | 0.33 | 0.2 | 0.4 | 0.38 | 0.42 | 0.21 |
| 4 | 62255222 | rs17090446 | 0.332 | 0.275 | -0.589 | 0.243 | 0 | 0.006 | 0.006 | 0 | 0.08 | 0.04 | 0.02 | 0.04 | 0.05 | 0.2 |
| 4 | 62256029 | rs2124793 | -0.222 | 0.351 | -0.488 | 0.496 | -0.004 | 0.095 | 0.095 | 0 | 0.84 | 0.75 | 0.69 | 0.55 | 0.84 | 0.67 |
| 4 | 62256068 | rs17238968 | 0.491 | 0.818 | -0.832 | 0.615 | -0.066 | -0.038 | -0.038 | 0 | 0.36 | 0.19 | 0.45 | 0.41 | 0.5 | 0.22 |
| 4 | 62258836 | rs2061266 | 0.246 | 0.291 | 0.172 | 0.887 | 0.018 | 0.228 | 0.228 | 0.009 | 0.92 | 0.65 | 0.95 | 0.76 | 0.97 | 0.82 |
| 4 | 62268153 | rs17238975 | 0.972 | 0.751 | -0.805 | 0.858 | -0.068 | -0.027 | -0.027 | 0 | 0.27 | 0.06 | 0.48 | 0.25 | 0.34 | 0.17 |
| 4 | 62269277 | rs13127162 | 0.3 | 0.359 | -0.685 | 0.351 | 0.007 | 0.036 | 0.036 | 0 | 0.33 | 0.21 | 0.18 | 0.24 | 0.32 | 0.43 |
| 4 | 62276356 | rs1350246 | 0.084 | 0.276 | 0.024 | 0.389 | 0.006 | -0.003 | -0.003 | 0.297 | 0.59 | 0.39 | 0.77 | 0.51 | 0.65 | 0.4 |
| 4 | 62284310 | rs17238982 | 0.481 | 0.982 | -0.441 | -0.268 | -0.038 | 0.037 | 0.037 | 0 | 0.11 | 0.04 | 0.09 | 0.02 | 0.16 | 0.05 |
| 4 | 62290252 | rs6824906 | -0.563 | 0.571 | -0.145 | -0.211 | 0.006 | 0.035 | 0.035 | 0 | 0.52 | 0.4 | 0.38 | 0.36 | 0.52 | 0.62 |
| 4 | 62290634 | rs2124794 | -0.608 | 0.6 | -0.305 | -0.274 | 0.004 | 0.061 | 0.061 | 0 | 0.53 | 0.27 | 0.44 | 0.37 | 0.54 | 0.62 |
| 4 | 62293449 | rs6821028 | -0.729 | 0.36 | -0.255 | -1.018 | -0.001 | 0.136 | 0.136 | 0 | 0.46 | 0.26 | 0.41 | 0.16 | 0.45 | 0.51 |
| 4 | 62303163 | rs1026419 | 1.231 | 0.755 | 0.237 | -0.801 | -0.013 | 0.075 | 0.075 | 0 | 0.58 | 0.33 | 0.58 | 0.8 | 0.57 | 0.45 |
| 4 | 62303629 | rs7661450 | -0.166 | 0.9 | 0.104 | -0.89 | -0.013 | 0.029 | 0.029 | 0 | 0.21 | 0.1 | 0.23 | 0.14 | 0.18 | 0.2 |
| 4 | 62316572 | rs2643036 | 1.362 | 0.801 | 0.811 | -0.724 | -0.001 | -0.051 | -0.051 | 0.259 | 0.76 | 0.39 | 0.82 | 0.91 | 0.75 | 0.65 |
| 4 | 62317170 | rs10009335 | -1.011 | 0.828 | 0.626 | -0.711 | 0.002 | -0.013 | -0.013 | 0 | 0.18 | 0.23 | 0.15 | 0.09 | 0.25 | 0.35 |
| 4 | 62333348 | rs335336 | 0.98 | 0.408 | 0.755 | -0.288 | -0.002 | 0.02 | 0.02 | 1.424 | 0.9 | 0.64 | 0.95 | 0.95 | 0.95 | 0.95 |
| 4 | 62349873 | rs335324 | 0.734 | 0.617 | 0.683 | -0.667 | -0.02 | -0.116 | -0.116 | 0 | 0.78 | 0.55 | 0.88 | 0.96 | 0.85 | 0.91 |
| 4 | 62359319 | rs10016690 | -0.033 | 0.953 | 0.27 | -0.729 | 0 | -0.011 | -0.011 | 0 | 0.09 | 0.08 | 0.1 | 0.01 | 0.1 | 0.15 |

| | | | | | | | | | | | | | | | | |
|---|---|---|---|---|---|---|---|---|---|---|---|---|---|---|---|---|
| 4 | 62361459 | rs335271 | 0.258 | 0.909 | 0.397 | -0.721 | 0.001 | -0.072 | -0.072 | 0 | 0.82 | 0.69 | 0.89 | 0.96 | 0.85 | 0.91 |
| 4 | 62366666 | rs10008278 | -0.072 | 1.164 | 0.07 | -0.539 | 0.002 | -0.018 | -0.018 | 0 | 0.07 | 0.06 | 0.05 | 0.01 | 0.09 | 0.14 |
| 4 | 62368762 | rs335286 | -0.304 | 1.231 | -0.271 | -0.86 | 0.009 | -0.029 | -0.029 | 0 | 0.68 | 0.61 | 0.52 | 0.82 | 0.61 | 0.78 |
| 4 | 62373539 | rs17090492 | -0.173 | 0.848 | -0.224 | -0.554 | -0.005 | 0.008 | 0.008 | 0 | 0.08 | 0.16 | 0.04 | 0.08 | 0.09 | 0.1 |
| 4 | 62380911 | rs10029192 | 0.134 | 0.773 | -0.715 | -0.639 | -0.002 | 0.058 | 0.058 | 0.496 | 0.6 | 0.25 | 0.48 | 0.71 | 0.61 | 0.74 |
| 4 | 62381539 | rs335306 | -0.652 | 0.864 | -0.633 | -0.812 | -0.014 | 0.06 | 0.06 | 0 | 0.75 | 0.85 | 0.62 | 0.94 | 0.71 | 0.85 |
| 4 | 62387140 | rs335317 | -0.702 | 0.853 | -0.638 | -0.829 | -0.014 | 0.06 | 0.06 | 0 | 0.75 | 0.85 | 0.62 | 0.94 | 0.71 | 0.85 |
| 4 | 62393760 | rs335305 | 0.397 | 0.564 | -0.786 | -0.699 | -0.012 | -0.007 | -0.007 | 0 | 0.42 | 0.3 | 0.37 | 0.48 | 0.46 | 0.45 |
| 4 | 62398938 | rs6813884 | 0.521 | 0.576 | -0.609 | -0.889 | -0.002 | -0.004 | -0.004 | 0 | 0.34 | 0.32 | 0.23 | 0.29 | 0.28 | 0.44 |
| 4 | 62399244 | rs17226207 | 2.084 | 0.558 | -0.763 | -0.885 | 0 | -0.028 | -0.028 | 0 | 0.19 | 0.06 | 0.11 | 0.22 | 0.15 | 0.33 |
| 4 | 62402002 | rs11734047 | 0.859 | 0.65 | -0.657 | -0.889 | -0.003 | 0.018 | 0.018 | 0 | 0.22 | 0.17 | 0.35 | 0.07 | 0.25 | 0.11 |
| 4 | 62406445 | rs904243 | 1.485 | 0.717 | -1.095 | -0.75 | -0.022 | 0.082 | 0.082 | 0.374 | 0.24 | 0.04 | 0.16 | 0.06 | 0.23 | 0.17 |
| 4 | 62407327 | rs1565901 | 0.002 | 0.471 | -0.776 | -0.592 | -0.008 | 0.031 | 0.031 | 0 | 0.19 | 0.32 | 0.2 | 0.28 | 0.15 | 0.29 |
| 4 | 62410156 | rs17090504 | 0.142 | 0.634 | -0.952 | -0.992 | -0.001 | -0.045 | -0.045 | 0 | 0.4 | 0.54 | 0.46 | 0.24 | 0.4 | 0.28 |
| 4 | 62411676 | rs7656189 | -1.055 | 0.122 | -0.493 | -0.731 | 0.027 | 0.108 | 0.108 | 0.025 | 0.45 | 0.78 | 0.39 | 0.75 | 0.35 | 0.56 |
| 4 | 62412292 | rs9312078 | -1.27 | 0.165 | -0.58 | -0.785 | -0.001 | -0.029 | -0.029 | 0 | 0.58 | 0.53 | 0.69 | 0.93 | 0.6 | 0.69 |
| 4 | 62416499 | rs2132074 | -0.447 | 0.158 | -0.556 | -0.736 | -0.001 | -0.015 | -0.015 | 0.063 | 0.39 | 0.37 | 0.37 | 0.74 | 0.35 | 0.56 |
| 4 | 62416593 | rs1497917 | 0.543 | 0.146 | -0.47 | -0.679 | 0.053 | 0.182 | 0.182 | 0 | 0.05 | 0.41 | 0.06 | 0.14 | 0.01 | 0 |
| 4 | 62418658 | rs10011995 | 0.912 | 0.252 | -1.062 | -0.869 | -0.032 | 0.049 | 0.049 | 0 | 0.18 | 0.08 | 0.28 | 0.04 | 0.26 | 0.11 |
| 4 | 62419426 | rs2343249 | -1.63 | 0.246 | -1.057 | -0.761 | -0.033 | 0.156 | 0.156 | 0.032 | 0.64 | 0.92 | 0.71 | 0.92 | 0.61 | 0.69 |
| 4 | 62434848 | rs958862 | -1.508 | 0.114 | -0.793 | -0.755 | -0.061 | 0.276 | 0.276 | 0 | 0.58 | 0.94 | 0.68 | 0.96 | 0.54 | 0.68 |
| 4 | 62434977 | rs723120 | 0.68 | 0.418 | -0.367 | -0.614 | 0.005 | -0.026 | -0.026 | 0 | 0.16 | 0.22 | 0.13 | 0.04 | 0.23 | 0.09 |
| 4 | 62455462 | rs17239080 | -1.505 | 0.096 | -0.226 | -0.031 | -0.013 | 0.096 | 0.096 | 0.851 | 0.5 | 0.9 | 0.65 | 0.66 | 0.51 | 0.47 |
| 4 | 62457454 | rs11131334 | -1.02 | 0.181 | -0.164 | 0.012 | -0.003 | 0.025 | 0.025 | 0 | 0.45 | 0.66 | 0.64 | 0.66 | 0.51 | 0.47 |
| 4 | 62462234 | rs35550109 | -0.138 | 0.328 | -0.213 | 0.126 | 0.005 | 0.068 | 0.068 | 0 | 0.28 | 0.39 | 0.46 | 0.49 | 0.28 | 0.38 |
| 4 | 62468670 | rs9996810 | 0.054 | 0.635 | -0.026 | 0.028 | 0.02 | -0.072 | -0.072 | 1.236 | 0.19 | 0.33 | 0.14 | 0.04 | 0.23 | 0.09 |
| 4 | 62495735 | rs17239101 | 0.091 | 0.156 | -0.259 | 0.573 | 0.014 | 0.016 | 0.016 | 0.086 | 0.14 | 0.04 | 0.18 | 0.11 | 0.1 | 0.12 |
| 4 | 62503837 | rs12642037 | -0.897 | 0.104 | -0.316 | 0.513 | 0.007 | -0.007 | -0.007 | 0 | 0.36 | 0.72 | 0.5 | 0.54 | 0.31 | 0.5 |
| 4 | 62526281 | rs1497921 | 1.348 | 0.141 | -0.681 | 0.22 | -0.016 | 0.247 | 0.247 | 0 | 0.45 | 0.15 | 0.33 | 0.07 | 0.46 | 0.38 |
| 4 | 62542957 | rs4860425 | 0.207 | 0.225 | -0.229 | 0.341 | 0 | 0.009 | 0.009 | 0 | 0.18 | 0.13 | 0.25 | 0.12 | 0.17 | 0.14 |
| 4 | 62545435 | rs12233859 | 0.075 | 0.431 | -0.328 | 0.069 | -0.003 | 0.031 | 0.031 | 0 | 0.33 | 0.44 | 0.49 | 0.49 | 0.31 | 0.43 |
| 4 | 62563504 | rs12510774 | -0.157 | 0.389 | -0.58 | 0.008 | -0.003 | 0 | 0 | 0 | 0.65 | 0.42 | 0.49 | 0.42 | 0.69 | 0.5 |
| 4 | 62568688 | rs6843311 | 0.628 | 0.384 | -0.924 | -0.747 | -0.002 | 0.206 | 0.206 | 0 | 0.47 | 0.2 | 0.33 | 0.13 | 0.46 | 0.46 |
| 4 | 62585789 | rs6848408 | 1.581 | 0.369 | -1.121 | -0.958 | 0.001 | 0.04 | 0.04 | 0 | 0.11 | 0.04 | 0.09 | 0 | 0.08 | 0.03 |
| 4 | 62586313 | rs9998713 | -1.21 | 0.423 | -1.125 | -0.919 | -0.059 | 0.135 | 0.135 | 0 | 0.69 | 0.89 | 0.77 | 0.95 | 0.62 | 0.68 |
| 4 | 62599289 | rs9312082 | -1.144 | 0.429 | -1.142 | -0.927 | -0.069 | 0.136 | 0.136 | 0 | 0.7 | 0.89 | 0.77 | 0.95 | 0.62 | 0.68 |
| 4 | 62606930 | rs4640696 | -0.343 | -0.179 | -0.318 | -0.283 | -0.002 | 0.095 | 0.095 | 0.054 | 0.06 | 0.25 | 0.03 | 0.11 | 0.02 | 0.02 |
| 4 | 62616180 | rs12646895 | -0.783 | -0.167 | -0.718 | -0.392 | 0.001 | 0.235 | 0.235 | 0 | 0.53 | 0.89 | 0.68 | 0.81 | 0.44 | 0.49 |
| 4 | 62626565 | rs2343577 | -0.429 | 0.203 | -0.436 | -0.185 | -0.005 | -0.007 | -0.007 | 0 | 0.47 | 0.49 | 0.6 | 0.61 | 0.41 | 0.49 |
| 4 | 62637158 | rs2345049 | -0.408 | 0.138 | -0.409 | -0.213 | 0.009 | 0.048 | 0.048 | 0 | 0.51 | 0.69 | 0.64 | 0.67 | 0.44 | 0.51 |
| 4 | 62645032 | rs12509742 | -0.392 | 0.023 | -0.216 | -0.161 | -0.001 | -0.002 | -0.002 | 0 | 0.46 | 0.49 | 0.56 | 0.59 | 0.41 | 0.49 |
| 4 | 62651583 | rs6815347 | 1.252 | -0.101 | -0.448 | -0.335 | 0.012 | 0.038 | 0.038 | 0 | 0.12 | 0.06 | 0.09 | 0 | 0.08 | 0.03 |
| 4 | 62660731 | rs1901223 | -0.751 | -0.234 | -0.355 | -0.317 | -0.001 | -0.001 | -0.001 | 0 | 0.47 | 0.54 | 0.56 | 0.58 | 0.41 | 0.49 |
| 4 | 62665721 | rs4493573 | -0.961 | -0.371 | -0.331 | -0.454 | 0.022 | 0.109 | 0.109 | 0 | 0.52 | 0.83 | 0.63 | 0.72 | 0.42 | 0.5 |
| 4 | 62676117 | rs7683391 | 0.494 | -0.272 | -0.258 | -0.362 | -0.016 | -0.012 | -0.012 | 0 | 0.15 | 0.28 | 0.16 | 0.13 | 0.1 | 0.04 |
| 4 | 62686490 | rs11724057 | 0.968 | -0.272 | -0.257 | -0.391 | 0.001 | -0.02 | -0.02 | 0 | 0.13 | 0.13 | 0.1 | 0 | 0.09 | 0.03 |
| 4 | 62693692 | rs11734607 | -0.11 | -0.298 | -0.14 | -0.298 | -0.006 | 0.011 | 0.011 | 0 | 0.35 | 0.46 | 0.47 | 0.58 | 0.32 | 0.43 |
| 4 | 62698357 | rs2345041 | -0.256 | -0.533 | -0.239 | -0.287 | -0.013 | 0.004 | 0.004 | 0 | 0.34 | 0.59 | 0.41 | 0.59 | 0.22 | 0.4 |
| 4 | 62707476 | rs10021525 | -0.359 | -0.531 | -0.424 | -0.316 | 0.003 | 0.003 | 0.003 | 0.086 | 0.35 | 0.67 | 0.46 | 0.74 | 0.23 | 0.41 |
| 4 | 62707653 | rs10021694 | -0.337 | -0.534 | -0.433 | -0.318 | 0.01 | 0.007 | 0.007 | 0.295 | 0.35 | 0.7 | 0.46 | 0.74 | 0.23 | 0.41 |
| 4 | 62716210 | rs6551661 | 0.407 | -0.429 | -0.415 | -0.205 | -0.012 | 0.034 | 0.034 | 0 | 0.12 | 0.38 | 0.13 | 0.16 | 0.08 | 0.04 |
| 4 | 62726874 | rs10446786 | 0.482 | -0.586 | -0.657 | -0.225 | 0.059 | 0.063 | 0.063 | 0 | 0.58 | 0.17 | 0.47 | 0.27 | 0.7 | 0.54 |
| 4 | 62728124 | rs34855831 | -0.47 | 0.109 | -0.783 | -0.157 | -0.002 | 0.001 | 0.001 | 0 | 0.44 | 0.58 | 0.39 | 0.41 | 0.46 | 0.39 |
| 4 | 62739541 | rs6551665 | 0.301 | 0.265 | -0.763 | 0.051 | 0.003 | 0.002 | 0.002 | 0.025 | 0.59 | 0.43 | 0.69 | 0.67 | 0.65 | 0.66 |
| 4 | 62739674 | rs6846033 | 0.308 | 0.255 | -0.306 | 0.113 | -0.008 | -0.01 | -0.01 | 0 | 0.22 | 0.27 | 0.13 | 0.17 | 0.16 | 0.14 |
| 4 | 62744545 | rs1947275 | -0.662 | 0.661 | -0.473 | 0.16 | 0.008 | 0.019 | 0.019 | 0 | 0.84 | 0.7 | 0.82 | 0.84 | 0.81 | 0.81 |
| 4 | 62748384 | rs9683662 | -0.133 | 0.93 | -1.384 | 0.248 | -0.041 | 0.017 | 0.017 | 0 | 0.22 | 0.08 | 0.18 | 0.19 | 0.26 | 0.22 |
| 4 | 62754320 | rs6858066 | -0.074 | 0.677 | -1.083 | 0.466 | 0.005 | 0.011 | 0.011 | 0 | 0.54 | 0.37 | 0.35 | 0.63 | 0.53 | 0.6 |
| 4 | 62759270 | rs11131347 | -0.167 | 0.886 | -0.129 | 0.579 | -0.002 | 0.002 | 0.002 | 0 | 0.43 | 0.56 | 0.5 | 0.53 | 0.45 | 0.44 |
| 4 | 62766445 | rs1349408 | -0.35 | 1.04 | -0.364 | 0.138 | -0.014 | 0 | 0 | 0.265 | 0.79 | 0.82 | 0.87 | 0.99 | 0.72 | 0.74 |
| 4 | 62773995 | rs4860442 | 0.73 | 0.278 | -0.092 | 0.454 | 0.004 | -0.011 | -0.011 | 0 | 0.32 | 0.28 | 0.35 | 0.47 | 0.26 | 0.27 |
| 4 | 62779822 | rs1355368 | -0.41 | 0.19 | 0.218 | 0.45 | -0.011 | 0.002 | 0.002 | 0 | 0.62 | 0.51 | 0.54 | 0.38 | 0.68 | 0.5 |
| 4 | 62788111 | rs12648453 | 0.452 | -0.243 | -0.087 | 0.148 | -0.024 | 0.155 | 0.155 | 0.408 | 0.42 | 0.14 | 0.34 | 0.09 | 0.57 | 0.37 |
| 4 | 62788474 | rs10023425 | -0.764 | 0.002 | 0.187 | 0.337 | 0.007 | -0.005 | -0.005 | 0 | 0.64 | 0.72 | 0.56 | 0.38 | 0.69 | 0.52 |
| 4 | 62800554 | rs2305339 | -1.033 | -0.018 | -0.02 | 0.293 | 0.01 | -0.045 | -0.045 | 0 | 0.67 | 0.8 | 0.61 | 0.54 | 0.73 | 0.61 |
| 4 | 62800728 | rs734644 | -1.111 | -0.017 | -0.019 | 0.548 | 0.008 | -0.031 | -0.031 | 0 | 0.68 | 0.8 | 0.62 | 0.59 | 0.73 | 0.61 |
| 4 | 62830533 | rs1510925 | 1.148 | -0.224 | -0.109 | 0.373 | 0.019 | -0.121 | -0.121 | 0.241 | 0.28 | 0.08 | 0.35 | 0.45 | 0.2 | 0.3 |
| 4 | 62834741 | rs1510923 | 0.373 | -0.229 | 0.466 | 0.39 | -0.017 | 0.009 | 0.009 | 0 | 0.36 | 0.43 | 0.43 | 0.61 | 0.26 | 0.5 |
| 4 | 62835607 | rs17239219 | 1.082 | -0.197 | 0.117 | 0.318 | 0.003 | -0.002 | -0.002 | 0 | 0.12 | 0.1 | 0.1 | 0.13 | 0.1 | 0.1 |

| | | | | | | | | | | | | | | | | |
|---|---|---|---|---|---|---|---|---|---|---|---|---|---|---|---|---|
| 4 | 62838634 | rs7672853 | 1 | -0.129 | 0.527 | 0.418 | 0.036 | 0.109 | 0.109 | 0 | 0.9 | 0.63 | 0.92 | 0.84 | 0.94 | 0.81 |
| 4 | 62845490 | rs1397548 | -0.518 | -0.201 | -0.215 | 0.235 | 0.007 | -0.003 | -0.003 | 0 | 0.63 | 0.35 | 0.57 | 0.39 | 0.74 | 0.51 |
| 4 | 62850522 | rs4860106 | -0.274 | -0.354 | -0.803 | -0.119 | 0.006 | 0.03 | 0.03 | 0 | 0.55 | 0.37 | 0.39 | 0.79 | 0.41 | 0.6 |
| 4 | 62851157 | rs17292128 | 1.179 | -0.491 | -1.412 | 0.085 | -0.035 | -0.048 | -0.048 | 0 | 0.13 | 0.03 | 0.16 | 0.23 | 0.15 | 0.15 |
| 4 | 62856659 | rs10017760 | 1.588 | -0.78 | -1.575 | 0.01 | 0.03 | -0.128 | -0.128 | 0 | 0.27 | 0.07 | 0.17 | 0.45 | 0.19 | 0.38 |
| 4 | 62859511 | rs954645 | -0.683 | -0.53 | -1.221 | 0.002 | 0.008 | -0.006 | -0.006 | 0 | 0.26 | 0.21 | 0.22 | 0.33 | 0.22 | 0.22 |
| 4 | 62862615 | rs6824673 | -0.159 | -0.285 | -0.939 | -0.121 | -0.01 | -0.006 | -0.006 | 0.514 | 0.14 | 0.18 | 0.07 | 0.12 | 0.08 | 0.07 |
| 4 | 62863411 | rs17828558 | 0.924 | 0.703 | -1.61 | -0.274 | -0.084 | 0.033 | 0.033 | 0 | 0.21 | 0.01 | 0.16 | 0.06 | 0.26 | 0.21 |
| 4 | 62883431 | rs1510920 | -0.509 | 0.302 | -0.084 | -0.403 | -0.007 | 0.067 | 0.067 | 0.34 | 0.9 | 0.72 | 0.94 | 0.84 | 0.94 | 0.95 |
| 4 | 62895592 | rs1510921 | 0.94 | 0.261 | -0.884 | -0.529 | 0.023 | -0.075 | -0.075 | 0 | 0.31 | 0.09 | 0.22 | 0.39 | 0.24 | 0.44 |
| 4 | 62911349 | rs2058764 | -0.633 | 0.214 | -0.504 | -0.55 | 0.029 | 0.122 | 0.122 | 0.546 | 0.92 | 0.68 | 0.93 | 0.86 | 0.95 | 0.96 |
| 4 | 62915024 | rs1397546 | -0.047 | 0.373 | -1.174 | -1.16 | -0.084 | 0.075 | 0.075 | 0 | 0.28 | 0.09 | 0.44 | 0.16 | 0.37 | 0.21 |
| 4 | 62923797 | rs11736888 | 1.297 | 0.028 | -0.725 | -0.96 | 0.008 | 0.026 | 0.026 | 0 | 0.22 | 0.14 | 0.13 | 0.03 | 0.22 | 0.13 |
| 4 | 62944634 | rs10032372 | 0.322 | 0.358 | -0.628 | -0.788 | -0.009 | 0.008 | 0.008 | 0 | 0.63 | 0.43 | 0.65 | 0.42 | 0.69 | 0.7 |
| 4 | 62945462 | rs3922274 | 0.775 | 0.552 | -0.702 | -0.845 | -0.002 | 0 | 0 | 0 | 0.57 | 0.51 | 0.61 | 0.42 | 0.61 | 0.64 |
| 4 | 62990100 | rs7697748 | -0.462 | 0.599 | -0.653 | -0.5 | -0.012 | 0.015 | 0.015 | 1.082 | 0.75 | 0.62 | 0.81 | 0.74 | 0.79 | 0.91 |
| 4 | 62994770 | rs12715708 | 0.379 | 0.622 | -0.59 | -0.503 | -0.012 | 0.015 | 0.015 | 0 | 0.63 | 0.44 | 0.65 | 0.42 | 0.69 | 0.7 |
| 4 | 63000769 | rs7676979 | 0.903 | 0.603 | -0.936 | -0.473 | 0.008 | -0.021 | -0.021 | 0.273 | 0.13 | 0.09 | 0.16 | 0.33 | 0.1 | 0.19 |
| 4 | 63012413 | rs6857895 | 0.187 | 0.57 | -0.935 | -1.1 | -0.008 | 0.025 | 0.025 | 0.302 | 0.27 | 0.67 | 0.28 | 0.41 | 0.21 | 0.14 |
| 4 | 63037324 | rs1124974 | 0.51 | 0.536 | -0.876 | -1.141 | -0.009 | 0.093 | 0.093 | 0.025 | 0.25 | 0.53 | 0.27 | 0.4 | 0.21 | 0.14 |
| 4 | 63038015 | rs7440496 | 0.438 | 0.541 | -0.907 | -1.139 | -0.011 | 0.045 | 0.045 | 0.233 | 0.26 | 0.63 | 0.27 | 0.4 | 0.21 | 0.14 |
| 4 | 63042975 | rs2193120 | -0.936 | 0.686 | -0.945 | -1.13 | -0.002 | -0.014 | -0.014 | 0 | 0.63 | 0.71 | 0.66 | 0.58 | 0.59 | 0.44 |
| 4 | 63059629 | rs17292254 | 1.863 | 0.599 | -0.853 | -1.034 | 0 | 0 | 0 | 0.828 | 0.1 | 0.08 | 0.09 | 0.18 | 0.12 | 0.13 |
| 4 | 63060580 | rs4446347 | -0.521 | 0.75 | -0.651 | -1.069 | -0.004 | -0.015 | -0.015 | 0.344 | 0.63 | 0.71 | 0.66 | 0.58 | 0.59 | 0.43 |
| 4 | 63067995 | rs1563796 | -0.412 | 0.827 | -0.341 | -0.781 | -0.002 | 0.005 | 0.005 | 0 | 0.64 | 0.42 | 0.64 | 0.39 | 0.67 | 0.71 |
| 4 | 63084429 | rs6823272 | 0.268 | 1.4 | -0.351 | -1.303 | -0.003 | -0.002 | -0.002 | 0 | 0.25 | 0.25 | 0.19 | 0.03 | 0.29 | 0.2 |
| 4 | 63096403 | rs7694238 | 0.259 | 1.411 | -0.229 | -1.283 | 0.002 | -0.059 | -0.059 | 0 | 0.25 | 0.3 | 0.19 | 0.03 | 0.28 | 0.19 |
| 4 | 63195550 | rs1478528 | 0.901 | 1.471 | -0.987 | -1.164 | -0.001 | -0.005 | -0.005 | 0.023 | 0.07 | 0.07 | 0.04 | 0 | 0.08 | 0 |
| 4 | 63214913 | rs9994927 | 0.199 | 1.612 | -0.927 | -1.049 | 0.004 | -0.056 | -0.056 | 0 | 0.25 | 0.34 | 0.16 | 0.03 | 0.29 | 0.32 |
| 4 | 63229194 | rs13137382 | -0.252 | 1.576 | -0.415 | -0.435 | 0.001 | -0.036 | -0.036 | 0.002 | 0.25 | 0.3 | 0.16 | 0.03 | 0.29 | 0.32 |
| 4 | 63229994 | rs10001214 | -0.246 | 1.702 | -0.639 | -0.375 | -0.005 | 0.001 | 0.001 | 0 | 0.24 | 0.23 | 0.15 | 0.03 | 0.29 | 0.32 |
| 4 | 63239289 | rs7692732 | -0.659 | 1.85 | -0.339 | 0.877 | 0.025 | 0.177 | 0.177 | 0 | 0.1 | 0.16 | 0.17 | 0.49 | 0.15 | 0.33 |
| 4 | 63244085 | rs778940 | 0.335 | 2.25 | -0.451 | 0.445 | -0.008 | -0.013 | -0.013 | 0 | 0.9 | 0.94 | 0.9 | 0.81 | 0.87 | 0.93 |
| 4 | 63246560 | rs10033123 | 0.481 | 2.328 | -0.929 | -0.218 | -0.008 | 0.073 | 0.073 | 0 | 0.21 | 0.1 | 0.13 | 0.02 | 0.22 | 0.27 |
| 4 | 63247984 | rs2169814 | 1.008 | 2.204 | -0.147 | -0.596 | 0.022 | 0.001 | 0.001 | 0 | 0.78 | 0.48 | 0.8 | 0.8 | 0.76 | 0.85 |
| 4 | 63258627 | rs10517821 | 0.102 | 2.186 | 0.202 | -0.619 | 0.008 | -0.021 | -0.021 | 0 | 0.09 | 0.12 | 0.08 | 0.01 | 0.11 | 0.09 |
| 4 | 63258843 | rs801454 | 0.427 | 2.086 | 0.418 | -0.495 | 0.05 | 0.091 | 0.091 | 0.013 | 0.87 | 0.64 | 0.89 | 0.81 | 0.87 | 0.93 |
| 4 | 63263936 | rs778950 | 0.03 | 2.022 | 0.215 | -0.348 | 0.002 | 0.004 | 0.004 | 0 | 0.25 | 0.29 | 0.28 | 0.73 | 0.29 | 0.48 |
| 4 | 63281335 | rs1983497 | 0.055 | 1.798 | -0.105 | -1.002 | 0.001 | 0.01 | 0.01 | 0 | 0.2 | 0.31 | 0.32 | 0.79 | 0.18 | 0.35 |
| 4 | 63288507 | rs1830310 | 0.42 | 1.811 | -0.093 | -0.871 | -0.002 | -0.007 | -0.007 | 0 | 0.17 | 0.21 | 0.2 | 0.12 | 0.15 | 0.18 |
| 4 | 63293938 | rs1590068 | 0.382 | 2.445 | -0.324 | -1.344 | 0.002 | -0.001 | -0.001 | 0 | 0.82 | 0.83 | 0.66 | 0.8 | 0.69 | 0.81 |
| 4 | 63296955 | rs1528375 | -0.775 | 2.531 | -0.349 | -1.333 | -0.02 | -0.026 | -0.026 | 0 | 0.91 | 0.97 | 0.9 | 0.81 | 0.86 | 0.94 |
| 4 | 63309789 | rs778572 | 0.023 | 2.565 | -0.321 | -1.344 | -0.001 | 0.003 | 0.003 | 0 | 0.22 | 0.31 | 0.32 | 0.79 | 0.2 | 0.37 |
| 4 | 63312810 | rs11946748 | 1.245 | 2.198 | -0.863 | -1.781 | -0.049 | 0.191 | 0.191 | 0.255 | 0.28 | 0.02 | 0.18 | 0.01 | 0.27 | 0.3 |
| 4 | 63314242 | rs17828750 | 0.217 | 2.29 | -0.799 | -1.763 | 0.025 | -0.026 | -0.026 | 0.901 | 0.1 | 0.14 | 0.24 | 0.01 | 0.18 | 0.13 |
| 4 | 63315877 | rs2697201 | 0.061 | 1.712 | -1.099 | -1.764 | 0.009 | 0.031 | 0.031 | 0 | 0.47 | 0.4 | 0.45 | 0.78 | 0.4 | 0.49 |
| 4 | 63319526 | rs17239472 | 0.982 | 1.638 | -0.848 | -1.484 | 0.021 | 0.062 | 0.062 | 0 | 0.27 | 0.11 | 0.13 | 0 | 0.22 | 0.16 |
| 4 | 63322306 | rs778554 | -0.053 | 1.612 | -0.86 | -1.104 | 0.005 | 0.089 | 0.089 | 0.028 | 0.48 | 0.34 | 0.5 | 0.8 | 0.45 | 0.64 |
| 4 | 63328348 | rs1378353 | -0.158 | 1.457 | -0.927 | -0.773 | 0.006 | 0.01 | 0.01 | 0 | 0.49 | 0.52 | 0.55 | 0.3 | 0.39 | 0.44 |
| 4 | 63338415 | rs12640895 | -0.438 | 1.716 | -1.448 | -0.919 | -0.009 | 0.144 | 0.144 | 0.494 | 0.48 | 0.22 | 0.6 | 0.79 | 0.5 | 0.62 |
| 4 | 63341773 | rs13140211 | 0.36 | 1.883 | -1.719 | -0.933 | -0.043 | 0.165 | 0.165 | 0.263 | 0.28 | 0.07 | 0.21 | 0.02 | 0.28 | 0.33 |
| 4 | 63350062 | rs13107417 | 0.172 | 1.572 | -1.755 | -0.929 | -0.192 | 0.198 | 0.198 | 0.136 | 0.35 | 0.02 | 0.31 | 0.03 | 0.49 | 0.24 |
| 4 | 63354303 | rs6843214 | -0.346 | 1.425 | -0.686 | -0.654 | -0.004 | -0.021 | -0.021 | 0 | 0.44 | 0.51 | 0.35 | 0.27 | 0.56 | 0.33 |
| 4 | 63361327 | rs17082777 | -0.993 | 1.533 | -0.152 | -0.498 | 0.026 | -0.033 | -0.033 | 0 | 0.09 | 0.16 | 0.05 | 0.01 | 0.13 | 0.05 |
| 4 | 63366728 | rs10027570 | -1.148 | 1.188 | -0.218 | -0.044 | 0 | 0 | 0 | 0.152 | 0.16 | 0.17 | 0.16 | 0.22 | 0.18 | 0.22 |
| 4 | 63370926 | rs1000226 | -1.806 | 1.013 | 0.639 | -0.305 | 0.019 | -0.001 | -0.001 | 0 | 0.24 | 0.48 | 0.21 | 0.24 | 0.24 | 0.39 |
| 4 | 63372524 | rs12108508 | -1.806 | 1 | 0.775 | -0.307 | 0.019 | -0.001 | -0.001 | 0 | 0.24 | 0.48 | 0.22 | 0.24 | 0.24 | 0.39 |
| 4 | 63378266 | rs13126803 | 0.702 | 0.575 | -0.23 | -0.881 | 0.018 | 0.046 | 0.046 | 0 | 0.25 | 0.16 | 0.08 | 0.01 | 0.17 | 0.15 |
| 4 | 63379063 | rs6830305 | -1.081 | 0.882 | 0.143 | -0.109 | 0.001 | 0.002 | 0.002 | 0 | 0.16 | 0.19 | 0.16 | 0.2 | 0.18 | 0.23 |
| 4 | 63391971 | rs6817682 | -1.305 | 0.623 | -0.019 | -0.334 | 0 | 0.022 | 0.022 | 0 | 0.18 | 0.16 | 0.15 | 0.03 | 0.18 | 0.17 |
| 4 | 63397548 | rs6848319 | 0.39 | 0.61 | 0.312 | -0.338 | 0.005 | -0.026 | -0.026 | 0 | 0.85 | 0.72 | 0.84 | 0.91 | 0.81 | 0.87 |
| 4 | 63403138 | rs969607 | 0.33 | 0.357 | 0.36 | -0.375 | -0.005 | -0.019 | -0.019 | 0 | 0.84 | 0.78 | 0.85 | 0.91 | 0.83 | 0.87 |
| 4 | 63407474 | rs7699771 | 0.045 | 0.342 | 0.36 | -0.383 | 0 | -0.012 | -0.012 | 0 | 0.12 | 0.14 | 0.08 | 0.06 | 0.16 | 0.07 |
| 4 | 63407579 | rs2168267 | -0.58 | 0.346 | 0.495 | -0.103 | -0.009 | -0.015 | -0.015 | 0 | 0.28 | 0.2 | 0.38 | 0.69 | 0.32 | 0.39 |
| 4 | 63416030 | rs7670184 | 0.138 | 0.35 | 1.121 | -0.129 | -0.004 | -0.045 | -0.045 | 0 | 0.84 | 0.6 | 0.84 | 0.91 | 0.82 | 0.87 |
| 4 | 63424089 | rs11944965 | -0.796 | 0.284 | 0.664 | -0.062 | 0 | -0.001 | -0.001 | 0.224 | 0.3 | 0.33 | 0.36 | 0.7 | 0.32 | 0.39 |
| 4 | 63448853 | rs6854419 | 0.209 | 0.421 | 0.645 | 0.042 | -0.018 | -0.012 | -0.012 | 0.251 | 0.94 | 0.83 | 0.93 | 0.97 | 0.98 | 0.94 |
| 4 | 63458896 | rs4379088 | -1.359 | 0.549 | 0.306 | 0.66 | -0.001 | 0.043 | 0.043 | 0 | 0.29 | 0.51 | 0.25 | 0.39 | 0.28 | 0.27 |
| 4 | 63467034 | rs4348128 | 0.274 | 0.992 | -0.338 | 1.004 | 0.004 | 0.03 | 0.03 | 0 | 0.11 | 0.15 | 0.1 | 0.29 | 0.11 | 0.14 |

| | | | | | | | | | | | | | | | | |
|---|---|---|---|---|---|---|---|---|---|---|---|---|---|---|---|---|
| 4 | 63482692 | rs4355419 | 0.254 | 0.931 | -0.73 | 0 | -0.005 | -0.006 | -0.006 | 0 | 0.33 | 0.31 | 0.26 | 0.1 | 0.39 | 0.28 |
| 4 | 63487723 | rs7690075 | 0.273 | 0.093 | -1.446 | 0.906 | 0.032 | -0.087 | -0.087 | 0.026 | 0.17 | 0.05 | 0.21 | 0.38 | 0.14 | 0.21 |
| 4 | 63493963 | rs11941453 | 0.618 | 0.216 | -1.623 | 1.119 | -0.012 | -0.029 | -0.029 | 0.438 | 0.14 | 0.07 | 0.12 | 0.25 | 0.17 | 0.1 |
| 4 | 63509409 | rs12500171 | -0.94 | 0.23 | -1.402 | 0.412 | 0.003 | -0.033 | -0.033 | 0 | 0.57 | 0.52 | 0.69 | 0.88 | 0.51 | 0.6 |
| 4 | 63525643 | rs3887103 | -0.014 | 0.173 | -1.421 | 0.681 | -0.043 | -0.157 | -0.157 | 0.115 | 0.34 | 0.04 | 0.36 | 0.49 | 0.31 | 0.34 |
| 4 | 63533684 | rs13137221 | 1.663 | 0.174 | -1.442 | 0.646 | -0.008 | -0.018 | -0.018 | 0.002 | 0.1 | 0.08 | 0.08 | 0.22 | 0.14 | 0.09 |
| 4 | 63535629 | rs11933934 | -0.663 | 0.098 | -1.375 | 0.572 | -0.027 | 0.059 | 0.059 | 0 | 0.45 | 0.14 | 0.44 | 0.71 | 0.45 | 0.44 |
| 4 | 63539931 | rs4280764 | -1.628 | 0.113 | -1.448 | 0.324 | -0.004 | -0.016 | -0.016 | 0 | 0.8 | 0.78 | 0.87 | 0.94 | 0.82 | 0.89 |
| 4 | 63540378 | rs10013579 | 1.653 | 0.136 | -1.397 | 0.619 | -0.014 | 0.019 | 0.019 | 0 | 0.27 | 0.23 | 0.21 | 0.1 | 0.33 | 0.29 |
| 4 | 63541157 | rs12509725 | 0.812 | -0.003 | -0.92 | 0.884 | -0.001 | 0.026 | 0.026 | 0 | 0.06 | 0.07 | 0.19 | 0.16 | 0.04 | 0.16 |
| 4 | 63558975 | rs6551714 | -1.052 | 0.03 | -0.863 | 0.756 | -0.001 | -0.002 | -0.002 | 0.051 | 0.79 | 0.79 | 0.72 | 0.87 | 0.81 | 0.8 |
| 4 | 63562828 | rs11131395 | -0.148 | 0.031 | -1.09 | 0.669 | 0.024 | -0.039 | -0.039 | 0 | 0.37 | 0.18 | 0.37 | 0.55 | 0.3 | 0.44 |
| 4 | 63566758 | rs4860481 | 0.761 | 0.735 | -1.078 | 0.777 | -0.023 | 0.061 | 0.061 | 0.143 | 0.39 | 0.25 | 0.3 | 0.15 | 0.41 | 0.33 |
| 4 | 63579914 | rs6815661 | -0.658 | 1.013 | -0.827 | 1.643 | 0 | 0.002 | 0.002 | 0 | 0.88 | 0.86 | 0.87 | 0.77 | 0.86 | 0.95 |
| 4 | 63614842 | rs3922569 | -0.663 | 0.876 | -0.941 | 1.641 | -0.003 | -0.028 | -0.028 | 0.2 | 0.15 | 0.11 | 0.16 | 0.49 | 0.15 | 0.29 |
| 4 | 63648085 | rs4473678 | -1.127 | 0.941 | -0.847 | 1.378 | -0.002 | 0.015 | 0.015 | 0 | 0.9 | 0.94 | 0.9 | 1 | 0.88 | 0.99 |
| 4 | 63667799 | rs11131413 | -0.927 | 1.298 | -0.5 | 2.067 | -0.002 | -0.006 | -0.006 | 0 | 0.21 | 0.24 | 0.22 | 0.72 | 0.25 | 0.43 |
| 4 | 63679630 | rs11131415 | -0.352 | 1.057 | -0.686 | 2.157 | -0.008 | -0.043 | -0.043 | 0 | 0.17 | 0.09 | 0.17 | 0.39 | 0.21 | 0.31 |
| 4 | 63689854 | rs10027721 | 0.025 | 1.109 | -0.951 | 1.219 | -0.04 | 0.046 | 0.046 | 0 | 0.16 | 0.02 | 0.09 | 0.05 | 0.14 | 0.07 |
| 4 | 63701231 | rs11727577 | -0.919 | 1.012 | 0.303 | 2.079 | -0.001 | 0.009 | 0.009 | 2.02 | 0.22 | 0.19 | 0.19 | 0.5 | 0.24 | 0.4 |
| 4 | 63718434 | rs12649725 | -1.386 | 0.599 | 0.909 | 1.971 | 0.016 | 0.201 | 0.201 | 1.444 | 0.08 | 0.39 | 0.14 | 0.29 | 0.05 | 0.1 |
| 4 | 63730296 | rs6551725 | 0.678 | 0.885 | 0.689 | 2.175 | -0.005 | 0.065 | 0.065 | 1.082 | 0.8 | 0.78 | 0.69 | 0.5 | 0.8 | 0.64 |
| 4 | 63731916 | rs6551726 | 1.015 | 0.777 | 0.778 | 1.949 | -0.002 | 0.004 | 0.004 | 1.422 | 0.78 | 0.74 | 0.66 | 0.25 | 0.78 | 0.57 |
| 4 | 63734689 | rs906345 | 0.562 | 0.754 | 0.703 | 1.975 | -0.007 | 0.016 | 0.016 | 0.19 | 0.82 | 0.79 | 0.63 | 0.74 | 0.84 | 0.75 |
| 4 | 63735874 | rs17658230 | 0.796 | 0.85 | -0.369 | 0.897 | -0.066 | 0.035 | 0.035 | 0 | 0.23 | 0.02 | 0.16 | 0.1 | 0.26 | 0.2 |
| 4 | 63739803 | rs1395738 | 1.086 | 0.764 | -0.041 | 0.712 | -0.017 | 0.244 | 0.244 | 0.194 | 0.5 | 0.28 | 0.31 | 0.1 | 0.49 | 0.35 |
| 4 | 63747908 | rs4470662 | 0.229 | 0.587 | 0.067 | 0.182 | -0.003 | -0.024 | -0.024 | 0.336 | 0.56 | 0.51 | 0.4 | 0.18 | 0.61 | 0.46 |
| 4 | 63748261 | rs4269203 | 0.785 | 0.58 | 0.115 | 0.577 | 0.011 | -0.027 | -0.027 | 0 | 0.33 | 0.44 | 0.27 | 0.27 | 0.36 | 0.31 |
| 4 | 63750632 | rs10213657 | 1.272 | 0.651 | -0.222 | 0.479 | -0.066 | 0.035 | 0.035 | 0 | 0.23 | 0.02 | 0.16 | 0.11 | 0.26 | 0.19 |
| 4 | 63763762 | rs1507317 | -0.952 | 0.833 | 0.127 | 0.385 | 0.033 | -0.013 | -0.013 | 0.306 | 0.08 | 0.33 | 0.04 | 0.03 | 0.04 | 0.11 |
| 4 | 63768826 | rs13107261 | 0.179 | 0.865 | 0.065 | 0.373 | 0 | -0.009 | -0.009 | 0 | 0.35 | 0.55 | 0.27 | 0.26 | 0.36 | 0.32 |
| 4 | 63769930 | rs4565107 | 1.051 | 0.869 | 0.02 | 0.312 | -0.002 | -0.032 | -0.032 | 0 | 0.28 | 0.25 | 0.2 | 0.04 | 0.31 | 0.25 |
| 4 | 63779288 | rs12374362 | 0.351 | 0.928 | -0.814 | 0.706 | -0.05 | -0.104 | -0.104 | 0 | 0.14 | 0.01 | 0.17 | 0.37 | 0.18 | 0.17 |
| 4 | 63785436 | rs11932508 | -0.395 | 0.691 | -0.501 | 0.031 | -0.011 | 0.062 | 0.062 | 0.629 | 0.55 | 0.67 | 0.35 | 0.29 | 0.5 | 0.43 |
| 4 | 63788331 | rs7668525 | 0.64 | 0.682 | -0.524 | 0.118 | -0.037 | -0.009 | -0.009 | 0.395 | 0.25 | 0.53 | 0.13 | 0.24 | 0.15 | 0.15 |
| 4 | 63824453 | rs1507315 | 0.502 | 0.76 | -2.278 | -0.722 | -0.062 | 0.157 | 0.157 | 0 | 0.29 | 0.11 | 0.21 | 0.06 | 0.35 | 0.3 |
| 4 | 63835544 | rs17083151 | 0.5 | 0.617 | -1.772 | -0.112 | 0.003 | 0.075 | 0.075 | 0 | 0.24 | 0.12 | 0.12 | 0 | 0.22 | 0.12 |
| 4 | 63838796 | rs10025992 | 0.2 | 0.732 | -1.522 | 0.328 | -0.003 | 0.119 | 0.119 | 0 | 0.36 | 0.14 | 0.17 | 0.15 | 0.34 | 0.31 |
| 4 | 63842062 | rs2660598 | -0.5 | 0.844 | -0.674 | 1.006 | -0.002 | -0.001 | -0.001 | 0.171 | 0.32 | 0.22 | 0.28 | 0.31 | 0.32 | 0.37 |
| 4 | 63856523 | rs1395733 | -0.558 | 0.779 | 0.637 | 0.772 | -0.021 | 0.038 | 0.038 | 0 | 0.12 | 0.04 | 0.08 | 0.03 | 0.13 | 0.07 |
| 4 | 63859056 | rs1507308 | -0.924 | 1.402 | 0.501 | 1.401 | -0.012 | -0.017 | -0.017 | 0 | 0.15 | 0.1 | 0.11 | 0.28 | 0.17 | 0.22 |
| 4 | 63867349 | rs2683405 | 1.42 | 0.82 | 0.841 | 1.916 | -0.019 | 0.111 | 0.111 | 0.022 | 0.41 | 0.16 | 0.4 | 0.15 | 0.44 | 0.29 |
| 4 | 63872125 | rs9312100 | -0.411 | 0.791 | 0.894 | 1.868 | -0.007 | -0.046 | -0.046 | 0 | 0.39 | 0.49 | 0.36 | 0.2 | 0.37 | 0.28 |
| 4 | 63878877 | rs10029207 | -1.599 | 0.525 | 0.884 | 1.857 | 0.068 | 0.035 | 0.035 | 2.321 | 0.24 | 0.41 | 0.2 | 0.32 | 0.3 | 0.29 |
| 4 | 63879707 | rs9996786 | -1.091 | 0.576 | 0.856 | 1.883 | 0.024 | 0.015 | 0.015 | 0 | 0.51 | 0.59 | 0.4 | 0.46 | 0.42 | 0.43 |
| 4 | 63883185 | rs10213271 | 0.399 | 0.561 | 0.825 | 1.676 | 0.036 | 0.075 | 0.075 | 0.466 | 0.3 | 0.06 | 0.27 | 0.16 | 0.24 | 0.21 |
| 4 | 63887305 | rs17083254 | -0.044 | 0.661 | 0.832 | 1.798 | 0.003 | -0.006 | -0.006 | 0.09 | 0.12 | 0.06 | 0.07 | 0.14 | 0.09 | 0.16 |
| 4 | 63895857 | rs2346648 | -1.55 | 0.724 | 0.892 | 1.81 | -0.007 | 0.022 | 0.022 | 0 | 0.54 | 0.61 | 0.47 | 0.35 | 0.52 | 0.45 |
| 4 | 63901724 | rs2346649 | -1.334 | 0.802 | 0.948 | 1.772 | -0.037 | 0.009 | 0.009 | 1.305 | 0.2 | 0.12 | 0.18 | 0.17 | 0.28 | 0.24 |
| 4 | 63975710 | rs4860122 | 0.025 | 0.78 | 1.138 | 1.762 | -0.011 | -0.002 | -0.002 | 0 | 0.64 | 0.57 | 0.61 | 0.33 | 0.72 | 0.53 |
| 4 | 63992411 | rs2171748 | 0.361 | 0.781 | 0.876 | 1.783 | 0.042 | 0.019 | 0.019 | 0.171 | 0.31 | 0.06 | 0.26 | 0.23 | 0.24 | 0.21 |
| 4 | 64001013 | rs12502749 | 0.614 | 0.891 | 0.922 | 1.726 | 0.004 | 0.013 | 0.013 | 0 | 0.57 | 0.34 | 0.53 | 0.28 | 0.6 | 0.46 |
| 4 | 64040427 | rs7679758 | 0.157 | 0.852 | 0.764 | 1.424 | 0.037 | 0.038 | 0.038 | 0.007 | 0.32 | 0.22 | 0.27 | 0.19 | 0.24 | 0.22 |
| 4 | 64076557 | rs13127900 | 1.361 | 0.958 | 0.305 | 1.454 | -0.04 | 0.05 | 0.05 | 0.04 | 0.43 | 0.13 | 0.43 | 0.3 | 0.46 | 0.3 |
| 4 | 64076807 | rs976921 | -1.534 | 1.436 | 0.872 | 1.212 | 0.021 | -0.01 | -0.01 | 0.149 | 0.16 | 0.34 | 0.13 | 0.13 | 0.15 | 0.15 |
| 4 | 64103715 | rs13144816 | 0.758 | 1.26 | 1.019 | 1.124 | 0.184 | 0.09 | 0.09 | 0 | 0.94 | 0.71 | 0.92 | 0.95 | 0.89 | 0.93 |
| 4 | 64118963 | rs6837749 | -1.06 | 0.497 | 0.376 | 1.015 | 0.016 | -0.031 | -0.031 | 0 | 0.06 | 0.13 | 0.05 | 0 | 0.06 | 0.01 |
| 4 | 64119829 | rs9996258 | -1.179 | -0.022 | 0.106 | 1.33 | -0.016 | 0.011 | 0.011 | 0 | 0.21 | 0.27 | 0.25 | 0.25 | 0.17 | 0.34 |
| 4 | 64120486 | rs9999371 | -0.781 | 0.116 | -0.33 | 1.073 | 0.013 | 0.007 | 0.007 | 0 | 0.1 | 0.03 | 0.11 | 0.07 | 0.09 | 0.17 |
| 4 | 64122349 | rs1381737 | 0.168 | 0.319 | -0.919 | 1.11 | -0.074 | -0.086 | -0.086 | 0 | 0.4 | 0.1 | 0.32 | 0.48 | 0.48 | 0.37 |
| 4 | 64125181 | rs2346937 | -0.409 | 0.429 | -0.222 | 0.745 | 0 | -0.018 | -0.018 | 0 | 0.59 | 0.6 | 0.59 | 0.8 | 0.65 | 0.66 |
| 4 | 64171187 | rs2346939 | 0.205 | -0.199 | -0.809 | 1.298 | 0.068 | -0.051 | -0.051 | 0.073 | 0.62 | 0.15 | 0.67 | 0.49 | 0.74 | 0.68 |
| 4 | 64174876 | rs17671382 | -0.479 | 0.315 | -0.39 | 0.795 | 0.007 | -0.024 | -0.024 | 0.008 | 0.2 | 0.3 | 0.32 | 0.1 | 0.24 | 0.18 |
| 4 | 64176939 | rs17083621 | 0.148 | -0.512 | -0.69 | 1.027 | 0.024 | -0.029 | -0.029 | 3.271 | 0.16 | 0.09 | 0.06 | 0.28 | 0.1 | 0.05 |
| 4 | 64182495 | rs10005499 | -0.005 | -0.915 | -0.307 | 1.702 | -0.029 | 0.119 | 0.119 | 0 | 0.09 | 0.17 | 0.21 | 0.25 | 0.04 | 0.18 |
| 4 | 64184644 | rs1982616 | 0.831 | -1.029 | 0.488 | 1.519 | 0.002 | -0.005 | -0.005 | 0.159 | 0.69 | 0.35 | 0.67 | 0.48 | 0.83 | 0.77 |
| 4 | 64214915 | rs7662456 | 0.808 | -0.558 | 0.107 | 0.582 | 0.05 | 0.233 | 0.233 | 0 | 0.5 | 0.26 | 0.52 | 0.15 | 0.63 | 0.4 |
| 4 | 64217228 | rs10050307 | 1.143 | -0.463 | 0.202 | 0.714 | 0.001 | 0.024 | 0.024 | 0 | 0.08 | 0.09 | 0.21 | 0.25 | 0.04 | 0.18 |
| 4 | 64229023 | rs12503849 | 0.515 | 0.218 | 0.195 | 0.461 | 0.016 | 0.04 | 0.04 | 0 | 0.21 | 0.33 | 0.18 | 0.33 | 0.2 | 0.36 |

| | | | | | | | | | | | | | | | | |
|---|---|---|---|---|---|---|---|---|---|---|---|---|---|---|---|---|
| 4 | 64238559 | rs1462622 | 0.496 | 0.223 | 0.251 | 0.411 | 0.021 | 0.058 | 0.058 | 0 | 0.2 | 0.33 | 0.17 | 0.33 | 0.19 | 0.35 |
| 4 | 64243373 | rs17672372 | 1.942 | 0.509 | -0.601 | -0.145 | -0.04 | 0.109 | 0.109 | 0 | 0.2 | 0.05 | 0.13 | 0 | 0.23 | 0.07 |
| 4 | 64253631 | rs13353631 | 0.498 | 0.427 | -0.589 | -0.108 | -0.043 | 0.075 | 0.075 | 0 | 0.32 | 0.21 | 0.4 | 0.15 | 0.41 | 0.35 |
| 4 | 64259569 | rs10007651 | 1.035 | 0.332 | -0.144 | 0.055 | -0.036 | 0.165 | 0.165 | 0 | 0.11 | 0.27 | 0.22 | 0.25 | 0.04 | 0.19 |
| 4 | 64275466 | rs549815 | -0.194 | 0.373 | -0.176 | 0.004 | -0.001 | 0.048 | 0.048 | 0 | 0.49 | 0.31 | 0.48 | 0.42 | 0.54 | 0.4 |
| 4 | 64282069 | rs12501961 | 1.132 | 0.341 | -0.031 | 0.07 | -0.035 | 0.147 | 0.147 | 0 | 0.1 | 0.22 | 0.22 | 0.25 | 0.04 | 0.19 |
| 4 | 64295154 | rs13105340 | 1.572 | 0.391 | -0.288 | -0.124 | -0.073 | -0.001 | -0.001 | 0.265 | 0.19 | 0.02 | 0.32 | 0.11 | 0.26 | 0.22 |
| 4 | 64299915 | rs12647475 | 0.64 | 0.49 | -0.186 | -0.121 | 0.005 | -0.013 | -0.013 | 0 | 0.4 | 0.43 | 0.29 | 0.34 | 0.41 | 0.41 |
| 4 | 64305801 | rs2347407 | -2.036 | 0.358 | -0.22 | -0.068 | 0.008 | -0.013 | -0.013 | 0 | 0.83 | 0.88 | 0.92 | 0.73 | 0.87 | 0.94 |
| 4 | 64308401 | rs17627822 | 1.279 | -0.038 | -0.35 | -0.346 | -0.018 | 0.006 | 0.006 | 0 | 0.15 | 0.14 | 0.3 | 0.11 | 0.22 | 0.21 |
| 4 | 64309441 | rs1514950 | 1.059 | -0.111 | -0.395 | -0.383 | -0.007 | 0.005 | 0.005 | 0 | 0.16 | 0.19 | 0.3 | 0.11 | 0.22 | 0.21 |
| 4 | 64313453 | rs6551774 | -2.284 | -0.14 | -0.418 | -0.369 | 0.008 | -0.018 | -0.018 | 0 | 0.83 | 0.9 | 0.92 | 0.73 | 0.87 | 0.94 |
| 4 | 64315104 | rs7689785 | 0.711 | 0.528 | -0.512 | -0.545 | -0.004 | -0.014 | -0.014 | 2.301 | 0.45 | 0.48 | 0.29 | 0.32 | 0.43 | 0.48 |
| 4 | 64315661 | rs1399531 | -0.567 | 0.629 | -0.542 | -0.522 | 0.002 | 0.002 | 0.002 | 0 | 0.37 | 0.4 | 0.63 | 0.42 | 0.44 | 0.46 |
| 4 | 64316145 | rs17628227 | 1.756 | 0.613 | -0.512 | -0.451 | -0.007 | 0.023 | 0.023 | 0 | 0.08 | 0.04 | 0.03 | 0 | 0.08 | 0.09 |
| 4 | 64329169 | rs972651 | 1.592 | 0.542 | -0.355 | -0.487 | -0.001 | 0.003 | 0.003 | 0 | 0.17 | 0.12 | 0.12 | 0.31 | 0.13 | 0.33 |
| 4 | 64330224 | rs2102701 | 0.237 | 0.399 | -0.574 | -0.264 | -0.004 | 0.069 | 0.069 | 0 | 0.26 | 0.33 | 0.33 | 0.38 | 0.23 | 0.32 |
| 4 | 64332005 | rs13133029 | 2.43 | 0.418 | -0.658 | -0.401 | -0.029 | 0.104 | 0.104 | 0 | 0.19 | 0.05 | 0.13 | 0 | 0.22 | 0.07 |
| 4 | 64333810 | rs10021833 | -1.037 | 0.563 | -0.519 | -0.273 | -0.007 | -0.011 | -0.011 | 0 | 0.68 | 0.73 | 0.62 | 0.63 | 0.66 | 0.74 |
| 4 | 64337600 | rs2090189 | -0.618 | 0.466 | -0.393 | 0.148 | -0.002 | 0.005 | 0.005 | 0 | 0.38 | 0.35 | 0.36 | 0.59 | 0.34 | 0.56 |
| 4 | 64342824 | rs11131461 | 0.137 | 0.3 | -0.558 | 0.374 | 0.001 | -0.001 | -0.001 | 0 | 0.52 | 0.52 | 0.41 | 0.55 | 0.36 | 0.63 |
| 4 | 64354567 | rs9996032 | 0.691 | 0.168 | -0.483 | 0.528 | 0.014 | 0.008 | 0.008 | 0 | 0.39 | 0.34 | 0.27 | 0.36 | 0.25 | 0.37 |
| 4 | 64359342 | rs1399529 | 0.576 | 0.086 | -0.22 | 0.498 | -0.029 | -0.017 | -0.017 | 0 | 0.42 | 0.5 | 0.28 | 0.36 | 0.26 | 0.39 |
| 4 | 64359382 | rs1399530 | 0.716 | 0.162 | -0.479 | 0.536 | 0.014 | 0.008 | 0.008 | 0 | 0.39 | 0.34 | 0.27 | 0.36 | 0.26 | 0.37 |
| 4 | 64360700 | rs11131464 | 0.871 | 0.543 | -0.33 | 0.963 | 0 | 0.023 | 0.023 | 0 | 0.07 | 0.07 | 0.16 | 0.24 | 0.1 | 0.21 |
| 4 | 64390049 | rs13118127 | 0.359 | 1.164 | -0.906 | 0.506 | -0.091 | -0.022 | -0.022 | 0 | 0.25 | 0.04 | 0.29 | 0.21 | 0.32 | 0.26 |
| 4 | 64404377 | rs6855459 | 0.074 | 1.114 | -0.574 | 0.362 | -0.001 | -0.002 | -0.002 | 1.393 | 0.66 | 0.66 | 0.53 | 0.58 | 0.61 | 0.68 |
| 4 | 64442439 | rs7663493 | 0.654 | 0.221 | -0.45 | 0.335 | 0.029 | -0.003 | -0.003 | 0 | 0.42 | 0.67 | 0.34 | 0.61 | 0.27 | 0.5 |
| 4 | 64461794 | rs1020711 | 1.958 | -0.025 | -0.417 | 0.433 | -0.005 | 0.001 | 0.001 | 0.164 | 0.26 | 0.29 | 0.22 | 0.35 | 0.16 | 0.25 |
| 4 | 64468573 | rs12507316 | 0.553 | 0.085 | -0.455 | 0.503 | 0.002 | -0.008 | -0.008 | 0 | 0.14 | 0.11 | 0.1 | 0.25 | 0.12 | 0.25 |
| 4 | 64474350 | rs12648875 | 0.382 | 0.252 | -0.368 | 0.206 | -0.022 | 0.032 | 0.032 | 0 | 0.33 | 0.22 | 0.49 | 0.13 | 0.39 | 0.35 |
| 4 | 64475210 | rs12500270 | 1.608 | -0.374 | 0.331 | 0.677 | -0.033 | 0.005 | 0.005 | 0.108 | 0.27 | 0.39 | 0.22 | 0.34 | 0.16 | 0.22 |
| 4 | 64476938 | rs4413438 | 0.601 | -0.137 | -0.135 | 0.736 | 0.002 | -0.003 | -0.003 | 0 | 0.13 | 0.1 | 0.1 | 0.26 | 0.09 | 0.25 |
| 4 | 64483900 | rs17083921 | 1.635 | -0.304 | 0.069 | 0.687 | -0.044 | 0.009 | 0.009 | 0.005 | 0.27 | 0.39 | 0.24 | 0.37 | 0.16 | 0.24 |
| 4 | 64484803 | rs10452252 | -0.404 | -0.206 | -0.253 | 0.744 | -0.008 | 0.069 | 0.069 | 1.99 | 0.85 | 0.68 | 0.89 | 0.76 | 0.89 | 0.75 |
| 4 | 64484975 | rs12507969 | 0.659 | -0.127 | -1.256 | 0.783 | 0.002 | -0.024 | -0.024 | 2.95 | 0.13 | 0.1 | 0.1 | 0.24 | 0.11 | 0.25 |
| 4 | 64485837 | rs1456848 | -0.822 | 0.204 | -1.225 | 0.258 | 0.087 | -0.025 | -0.025 | 0.061 | 0.57 | 0.2 | 0.65 | 0.39 | 0.73 | 0.52 |
| 4 | 64489926 | rs1157023 | -0.978 | 0.256 | -1.201 | 0.279 | 0.078 | -0.032 | -0.032 | 0 | 0.58 | 0.2 | 0.65 | 0.39 | 0.73 | 0.52 |
| 4 | 64500994 | rs7692181 | -0.605 | 0.898 | -0.949 | -0.026 | -0.058 | 0.027 | 0.027 | 0 | 0.35 | 0.13 | 0.22 | 0.25 | 0.41 | 0.28 |
| 4 | 64514978 | rs1531202 | 0.509 | 0.521 | -0.128 | -0.597 | -0.01 | 0.037 | 0.037 | 0.617 | 0.64 | 0.29 | 0.58 | 0.78 | 0.61 | 0.83 |
| 4 | 64535584 | rs2881716 | 0.644 | 0.549 | -0.069 | -0.6 | -0.01 | 0.03 | 0.03 | 0.582 | 0.63 | 0.33 | 0.58 | 0.75 | 0.6 | 0.8 |
| 4 | 64611539 | rs10025060 | 2.076 | -0.392 | -0.346 | -1.068 | 0.067 | -0.065 | -0.065 | 2.61 | 0.26 | 0.06 | 0.25 | 0.77 | 0.18 | 0.36 |
| 4 | 64612436 | rs7659497 | 0.415 | -0.213 | 0.198 | -1.086 | -0.053 | -0.078 | -0.078 | 0 | 0.87 | 0.4 | 0.94 | 1 | 0.97 | 0.96 |
| 4 | 64625606 | rs13146719 | 0.615 | 0.001 | 0.157 | -1.146 | -0.005 | 0.018 | 0.018 | 0.753 | 0.65 | 0.35 | 0.69 | 0.9 | 0.62 | 0.83 |
| 4 | 64662510 | rs4860553 | 0.215 | 0.095 | 0.236 | -1.229 | -0.055 | 0.058 | 0.058 | 0 | 0.23 | 0.08 | 0.18 | 0.07 | 0.34 | 0.13 |
| 4 | 64662777 | rs17636187 | 0.172 | 0.093 | 0.333 | -1.229 | -0.039 | 0.049 | 0.049 | 0 | 0.23 | 0.12 | 0.18 | 0.07 | 0.34 | 0.13 |
| 4 | 64663792 | rs2347824 | 1.239 | 0.219 | -0.011 | -1.466 | 0.054 | 0.071 | 0.071 | 0.237 | 0.46 | 0.34 | 0.36 | 0.81 | 0.37 | 0.59 |
| 4 | 64668177 | rs17636490 | 0.307 | 0.427 | 0.007 | -1.305 | -0.072 | 0.062 | 0.062 | 0 | 0.23 | 0.05 | 0.18 | 0.07 | 0.34 | 0.13 |
| 4 | 64686848 | rs2347833 | -0.09 | 0.358 | 0.395 | -1.284 | 0.035 | -0.076 | -0.076 | 0 | 0.73 | 0.52 | 0.76 | 0.92 | 0.62 | 0.88 |
| 4 | 64693032 | rs12715741 | -0.101 | 0.359 | 0.395 | -1.284 | 0.035 | -0.076 | -0.076 | 0 | 0.73 | 0.52 | 0.76 | 0.92 | 0.62 | 0.88 |
| 4 | 64727794 | rs11131484 | 0.784 | 0.429 | -0.077 | -1.289 | -0.099 | -0.014 | -0.014 | 0 | 0.19 | 0.03 | 0.39 | 0.11 | 0.25 | 0.24 |
| 4 | 64759742 | rs1375469 | 2.214 | 0.043 | -0.168 | -1.329 | 0.072 | -0.043 | -0.043 | 0.288 | 0.27 | 0.04 | 0.25 | 0.79 | 0.19 | 0.36 |
| 4 | 64769162 | rs13116801 | 0.621 | 0.589 | 1.29 | -0.969 | 0.001 | -0.025 | -0.025 | 0 | 0.67 | 0.41 | 0.76 | 0.92 | 0.61 | 0.83 |
| 4 | 64772647 | rs1349182 | 0.046 | 0.705 | 1.355 | -0.846 | -0.022 | 0.03 | 0.03 | 0 | 0.75 | 0.82 | 0.79 | 0.92 | 0.62 | 0.86 |
| 4 | 64775396 | rs2678065 | 0.112 | 0.716 | 1.5 | -0.841 | 0.012 | -0.031 | -0.031 | 0.316 | 0.75 | 0.71 | 0.79 | 0.92 | 0.62 | 0.86 |
| 4 | 64791788 | rs1878564 | 0.228 | 0.73 | 1.535 | -0.842 | 0.032 | -0.059 | -0.059 | 0 | 0.74 | 0.62 | 0.79 | 0.92 | 0.62 | 0.86 |
| 4 | 64853008 | rs2122206 | 0.086 | 0.514 | 1.555 | -0.91 | -0.025 | 0.026 | 0.026 | 0 | 0.29 | 0.17 | 0.43 | 0.12 | 0.31 | 0.35 |
| 4 | 64881166 | rs1449191 | 0.081 | 0.518 | 1.576 | -0.887 | -0.025 | 0.023 | 0.023 | 0 | 0.29 | 0.18 | 0.43 | 0.16 | 0.31 | 0.36 |
| 4 | 64888674 | rs12647624 | 1.028 | 0.483 | 1.658 | -0.906 | -0.018 | -0.015 | -0.015 | 0 | 0.45 | 0.52 | 0.36 | 0.76 | 0.33 | 0.46 |
| 4 | 64892027 | rs11932534 | 1.226 | 0.473 | 1.726 | -0.911 | 0.017 | 0.015 | 0.015 | 0 | 0.43 | 0.37 | 0.35 | 0.76 | 0.33 | 0.5 |
| 4 | 64900987 | rs2881806 | -0.949 | -0.134 | 2.032 | -0.65 | 0.008 | -0.036 | -0.036 | 0 | 0.38 | 0.46 | 0.47 | 0.19 | 0.34 | 0.49 |
| 4 | 64933680 | rs1449187 | 0.059 | -0.104 | 1.167 | -0.599 | -0.038 | 0.025 | 0.025 | 0.494 | 0.27 | 0.13 | 0.42 | 0.17 | 0.3 | 0.39 |
| 4 | 64948753 | rs17697056 | -0.226 | -0.054 | 1.013 | -0.519 | -0.022 | -0.001 | -0.001 | 0.086 | 0.2 | 0.04 | 0.14 | 0.17 | 0.2 | 0.12 |
| 4 | 64970948 | rs1402142 | 1.705 | 0.2 | 1.385 | -0.286 | 0.014 | 0.033 | 0.033 | 0 | 0.46 | 0.28 | 0.39 | 0.66 | 0.5 | 0.39 |
| 4 | 64976578 | rs10002861 | -0.181 | 0.551 | 1.2 | -0.171 | -0.032 | 0.027 | 0.027 | 0.611 | 0.27 | 0.13 | 0.43 | 0.16 | 0.3 | 0.42 |
| 4 | 65085452 | rs9312127 | -0.413 | 0.185 | 1.349 | 0.053 | 0.01 | -0.023 | -0.023 | 0 | 0.31 | 0.35 | 0.47 | 0.18 | 0.32 | 0.45 |
| 4 | 65085505 | rs12649001 | 0.039 | 0.13 | 0.862 | 0.059 | 0 | 0.004 | 0.004 | 0.24 | 0.14 | 0.21 | 0.08 | 0.16 | 0.13 | 0.06 |
| 4 | 65097893 | rs1483716 | 1.763 | -0.048 | 0.716 | 0.012 | -0.076 | -0.104 | -0.104 | 0 | 0.45 | 0.05 | 0.3 | 0.54 | 0.51 | 0.37 |
| 4 | 65102656 | rs1531864 | -0.111 | 0.038 | 0.875 | -0.139 | 0.003 | 0.016 | 0.016 | 0.727 | 0.85 | 0.69 | 0.91 | 0.83 | 0.87 | 0.94 |

| | | | | | | | | | | | | | | | |
|---|---|---|---|---|---|---|---|---|---|---|---|---|---|---|---|
| 4 | 65108929 | rs2128887 | 1.592 | 0.038 | 0.355 | -0.138 | -0.009 | 0.051 | 0.051 | 0.009 | 0.47 | 0.15 | 0.43 | 0.66 | 0.52 | 0.4 |
| 4 | 65114074 | rs2200231 | -0.259 | 0.387 | 0.173 | 0.116 | -0.049 | 0.031 | 0.031 | 1.224 | 0.28 | 0.1 | 0.44 | 0.17 | 0.32 | 0.44 |
| 4 | 65117251 | rs7691568 | -0.343 | 0.082 | 0.438 | 0.391 | -0.001 | -0.001 | -0.001 | 0.013 | 0.14 | 0.16 | 0.08 | 0.17 | 0.13 | 0.06 |
| 4 | 65134786 | rs972537 | -0.218 | 0.149 | 0.501 | 0.184 | -0.004 | 0.006 | 0.006 | 0.606 | 0.26 | 0.16 | 0.39 | 0.18 | 0.25 | 0.4 |
| 4 | 65134819 | rs972536 | 0.729 | -0.03 | 1.248 | 0.79 | -0.011 | 0.021 | 0.021 | 0 | 0.76 | 0.55 | 0.74 | 0.73 | 0.78 | 0.76 |
| 4 | 65142710 | rs1483713 | 0.708 | -0.018 | 1.278 | 0.787 | -0.007 | 0.012 | 0.012 | 0 | 0.76 | 0.63 | 0.72 | 0.71 | 0.77 | 0.74 |
| 4 | 65164696 | rs11930994 | 1.38 | -0.076 | 1.264 | 0.739 | -0.006 | -0.033 | -0.033 | 0 | 0.93 | 0.81 | 0.94 | 1 | 0.96 | 0.96 |
| 4 | 65170415 | rs7656244 | -0.775 | -0.149 | 1.193 | 0.714 | 0.01 | -0.015 | -0.015 | 0 | 0.25 | 0.23 | 0.25 | 0.29 | 0.22 | 0.25 |
| 4 | 65189839 | rs10011095 | 0.559 | -0.036 | 0.929 | 0.676 | -0.038 | -0.061 | -0.061 | 0 | 0.12 | 0.01 | 0.15 | 0.13 | 0.1 | 0.22 |
| 4 | 65200217 | rs17084662 | -1.046 | -0.215 | 1.782 | 0.518 | -0.005 | 0.006 | 0.006 | 0 | 0.25 | 0.32 | 0.18 | 0.32 | 0.22 | 0.25 |
| 4 | 65215475 | rs1384725 | 1.583 | -0.201 | 1.788 | 0.509 | -0.029 | 0.004 | 0.004 | 0 | 0.67 | 0.44 | 0.67 | 0.67 | 0.74 | 0.71 |
| 4 | 65217836 | rs1552211 | 1.775 | -0.061 | 1.71 | 0.368 | 0.009 | -0.032 | -0.032 | 3.301 | 0.93 | 0.74 | 0.93 | 1 | 0.96 | 0.97 |
| 4 | 65225307 | rs17084731 | -1.231 | -0.103 | 1.18 | 0.511 | 0.008 | -0.005 | -0.005 | 0.653 | 0.27 | 0.27 | 0.27 | 0.29 | 0.22 | 0.28 |
| 4 | 65252628 | rs10049939 | 1.64 | -0.174 | 1.698 | 0.438 | 0.037 | 0.001 | 0.001 | 0.007 | 0.66 | 0.16 | 0.64 | 0.71 | 0.73 | 0.68 |
| 4 | 65258550 | rs17084784 | -1.184 | -0.099 | 1.872 | 0.435 | -0.002 | 0 | 0 | 0 | 0.27 | 0.3 | 0.33 | 0.29 | 0.23 | 0.29 |
| 4 | 65264183 | rs13151311 | 1.617 | -0.099 | 1.988 | 0.43 | 0.015 | 0.004 | 0.004 | 0 | 0.66 | 0.24 | 0.59 | 0.71 | 0.72 | 0.68 |
| 4 | 65265753 | rs11727042 | 1.444 | -0.073 | 1.861 | 0.443 | -0.01 | -0.002 | -0.002 | 0 | 0.68 | 0.39 | 0.6 | 0.71 | 0.72 | 0.68 |
| 4 | 65265927 | rs7690664 | 1.121 | -0.083 | 1.541 | 0.452 | 0.093 | 0.025 | 0.025 | 0.114 | 0.95 | 0.7 | 0.93 | 1 | 0.95 | 0.96 |
| 4 | 65270357 | rs1483288 | 1.381 | -0.136 | 0.577 | 0.534 | 0.003 | 0 | 0 | 0 | 0.67 | 0.3 | 0.59 | 0.68 | 0.72 | 0.68 |
| 4 | 65273961 | rs17084842 | -0.489 | -0.322 | 0.348 | 0.49 | 0.01 | -0.046 | -0.046 | 0.71 | 0.08 | 0 | 0.04 | 0.15 | 0.06 | 0.08 |
| 4 | 65275649 | rs1384720 | 1.516 | -0.22 | 0.731 | 0.314 | -0.043 | -0.066 | -0.066 | 0 | 0.89 | 0.29 | 0.86 | 0.99 | 0.96 | 0.9 |
| 4 | 65304222 | rs6828446 | -0.839 | -0.054 | 0.645 | 0.753 | -0.009 | 0.096 | 0.096 | 0 | 0.07 | 0.21 | 0.06 | 0.18 | 0.05 | 0.1 |
| 4 | 65305086 | rs4510490 | 0.163 | 0.325 | 0.058 | 0.375 | -0.019 | 0.024 | 0.024 | 0 | 0.08 | 0.04 | 0.07 | 0 | 0.14 | 0.02 |
| 4 | 65309678 | rs1842955 | 0.73 | 0.115 | -0.247 | -0.1 | -0.004 | 0.057 | 0.057 | 0 | 0.43 | 0.25 | 0.39 | 0.13 | 0.39 | 0.39 |
| 4 | 65309971 | rs12501821 | -0.095 | 0.195 | -0.923 | -0.089 | -0.135 | -0.029 | -0.029 | 0.361 | 0.39 | 0.07 | 0.46 | 0.69 | 0.5 | 0.39 |
| 4 | 65313065 | rs10002960 | 0.695 | 0.1 | 0.246 | 0.599 | 0.015 | 0.206 | 0.206 | 0.396 | 0.91 | 0.67 | 0.91 | 0.76 | 0.94 | 0.89 |
| 4 | 65313497 | rs10517889 | 0.172 | -0.013 | -0.683 | 0.31 | -0.082 | -0.096 | -0.096 | 0 | 0.4 | 0.08 | 0.43 | 0.61 | 0.49 | 0.43 |
| 4 | 65313753 | rs13434443 | 0.004 | -0.055 | -0.117 | 0.451 | -0.021 | 0.078 | 0.078 | 0 | 0.1 | 0.19 | 0.04 | 0.25 | 0.06 | 0.06 |
| 4 | 65324353 | rs2086735 | -0.67 | 0.126 | 0.285 | 0.359 | -0.006 | 0.041 | 0.041 | 0 | 0.54 | 0.43 | 0.54 | 0.83 | 0.57 | 0.51 |
| 4 | 65343461 | rs11725219 | -0.515 | -0.142 | 0.255 | 0.363 | -0.008 | 0.027 | 0.027 | 0 | 0.15 | 0.2 | 0.1 | 0.25 | 0.11 | 0.1 |
| 4 | 65346967 | rs2086731 | 0.505 | -0.465 | -0.314 | -0.325 | -0.014 | -0.01 | -0.01 | 0 | 0.5 | 0.13 | 0.45 | 0.6 | 0.56 | 0.54 |
| 4 | 65351771 | rs6851961 | 0.448 | -0.134 | -0.386 | -0.288 | -0.041 | -0.048 | -0.048 | 0 | 0.51 | 0.12 | 0.44 | 0.56 | 0.54 | 0.51 |
| 4 | 65361943 | rs10471121 | -0.039 | -0.141 | -0.344 | -0.216 | -0.005 | 0.008 | 0.008 | 0 | 0.16 | 0.14 | 0.1 | 0.23 | 0.12 | 0.1 |
| 4 | 65382034 | rs13105217 | 1.371 | -0.188 | 0.255 | -0.268 | -0.021 | 0.007 | 0.007 | 0 | 0.23 | 0.14 | 0.39 | 0.14 | 0.25 | 0.3 |
| 4 | 65384957 | rs1425395 | 0.702 | -0.704 | 0.003 | -0.304 | -0.001 | -0.001 | -0.001 | 0 | 0.76 | 0.78 | 0.85 | 0.73 | 0.78 | 0.81 |
| 4 | 65392212 | rs1425392 | 0.777 | -1.017 | -0.694 | 0.027 | 0.028 | 0.003 | 0.003 | 0.026 | 0.33 | 0.07 | 0.31 | 0.31 | 0.28 | 0.36 |
| 4 | 65403406 | rs10032135 | -0.199 | -0.958 | -0.092 | 0.069 | 0.015 | -0.019 | -0.019 | 0 | 0.17 | 0.11 | 0.09 | 0.26 | 0.11 | 0.1 |
| 4 | 65408723 | rs1035709 | 0.436 | -0.913 | 0.239 | 0.093 | -0.04 | 0.019 | 0.019 | 0.761 | 0.75 | 0.61 | 0.87 | 0.68 | 0.83 | 0.83 |
| 4 | 65412694 | rs17663093 | 1.059 | -0.385 | -0.314 | 0.02 | -0.018 | 0.002 | 0.002 | 0 | 0.21 | 0.13 | 0.38 | 0.14 | 0.24 | 0.31 |
| 4 | 65423635 | rs9918005 | 0.19 | -0.578 | -0.002 | -0.032 | 0.002 | -0.042 | -0.042 | 0.402 | 0.31 | 0.54 | 0.42 | 0.14 | 0.27 | 0.34 |
| 4 | 65436643 | rs1425377 | -0.252 | -0.844 | -0.191 | 0.156 | -0.007 | 0.028 | 0.028 | 0 | 0.83 | 0.74 | 0.89 | 0.71 | 0.87 | 0.86 |
| 4 | 65437395 | rs1425376 | 0.149 | -0.758 | -0.203 | -0.328 | 0.001 | -0.016 | -0.016 | 0 | 0.35 | 0.59 | 0.45 | 0.17 | 0.34 | 0.4 |
| 4 | 65439260 | rs7688529 | 0.253 | -0.685 | -0.13 | -0.074 | -0.003 | 0.041 | 0.041 | 0 | 0.49 | 0.25 | 0.36 | 0.45 | 0.54 | 0.48 |
| 4 | 65451606 | rs6551838 | 0.454 | -0.876 | -0.299 | -0.155 | -0.055 | -0.007 | -0.007 | 0 | 0.42 | 0.17 | 0.38 | 0.46 | 0.5 | 0.44 |
| 4 | 65453210 | rs10024736 | -0.439 | -1.218 | 0.271 | -0.238 | -0.014 | 0.006 | 0.006 | 0 | 0.32 | 0.38 | 0.28 | 0.42 | 0.23 | 0.23 |
| 4 | 65459781 | rs7664375 | -0.477 | -1.222 | 0.388 | -0.241 | -0.043 | 0.022 | 0.022 | 0 | 0.32 | 0.49 | 0.29 | 0.42 | 0.23 | 0.23 |
| 4 | 65464297 | rs10021202 | -0.035 | -1.273 | 0.492 | -0.27 | -0.002 | 0.002 | 0.002 | 0 | 0.25 | 0.28 | 0.26 | 0.42 | 0.18 | 0.22 |
| 4 | 65475782 | rs10212780 | -1.236 | -0.851 | 0.533 | -0.068 | 0.006 | -0.006 | -0.006 | 0.738 | 0.76 | 0.75 | 0.67 | 0.88 | 0.73 | 0.67 |
| 4 | 65480701 | rs10007849 | -0.138 | -0.867 | 0.547 | -0.02 | -0.055 | 0.019 | 0.019 | 0 | 0.33 | 0.55 | 0.29 | 0.41 | 0.23 | 0.23 |
| 4 | 65493709 | rs1991222 | -0.22 | -0.322 | 0.271 | -0.115 | -0.03 | -0.006 | -0.006 | 0.32 | 0.43 | 0.22 | 0.37 | 0.47 | 0.5 | 0.44 |
| 4 | 65501422 | rs11943670 | -0.279 | -0.323 | 0.294 | -0.116 | -0.015 | -0.001 | -0.001 | 0.397 | 0.43 | 0.28 | 0.38 | 0.47 | 0.5 | 0.44 |
| 4 | 65501761 | rs1425390 | -0.263 | -0.301 | -0.017 | -0.116 | -0.038 | -0.009 | -0.009 | 0 | 0.43 | 0.19 | 0.38 | 0.47 | 0.51 | 0.44 |
| 4 | 65508643 | rs11131555 | 0.173 | 0.134 | -0.305 | -0.107 | -0.127 | -0.107 | -0.107 | 0 | 0.25 | 0.07 | 0.35 | 0.46 | 0.45 | 0.35 |
| 4 | 65510169 | rs7340852 | 0.086 | 0.108 | -0.243 | -0.107 | -0.089 | -0.077 | -0.077 | 0 | 0.26 | 0.1 | 0.35 | 0.46 | 0.45 | 0.35 |
| 4 | 65510635 | rs4403082 | 0.039 | 0.141 | -0.13 | -0.101 | -0.042 | -0.037 | -0.037 | 0 | 0.26 | 0.19 | 0.36 | 0.47 | 0.44 | 0.36 |
| 4 | 65510729 | rs4403083 | 0.039 | 0.141 | -0.13 | -0.101 | -0.042 | -0.037 | -0.037 | 0 | 0.26 | 0.19 | 0.36 | 0.47 | 0.44 | 0.36 |
| 4 | 65512619 | rs766818 | -0.026 | 0.237 | -0.153 | -0.122 | -0.056 | -0.042 | -0.042 | 0.533 | 0.31 | 0.19 | 0.38 | 0.47 | 0.47 | 0.36 |
| 4 | 65513391 | rs11937285 | -0.162 | 0.077 | 0.003 | -0.261 | -0.012 | -0.047 | -0.047 | 0 | 0.09 | 0.15 | 0.08 | 0 | 0.05 | 0.08 |
| 4 | 65517439 | rs10028130 | -0.663 | 0.135 | -0.242 | -0.09 | -0.01 | -0.004 | -0.004 | 5.491 | 0.4 | 0.34 | 0.46 | 0.47 | 0.52 | 0.44 |
| 4 | 65519108 | rs11131559 | 0.227 | 0.202 | -0.729 | -0.061 | -0.073 | -0.028 | -0.028 | 0.136 | 0.61 | 0.8 | 0.58 | 0.53 | 0.5 | 0.54 |
| 4 | 65519866 | rs7678201 | 0.261 | 0.193 | -0.614 | -0.067 | -0.042 | -0.014 | -0.014 | 0 | 0.61 | 0.73 | 0.58 | 0.53 | 0.5 | 0.53 |
| 4 | 65521501 | rs4604083 | 0.638 | 0.7 | -0.867 | -0.019 | -0.066 | -0.041 | -0.041 | 0.446 | 0.17 | 0.01 | 0.17 | 0.19 | 0.23 | 0.15 |
| 4 | 65535470 | rs6831411 | 0.728 | -0.064 | -0.209 | -0.358 | -0.002 | -0.003 | -0.003 | 0 | 0.47 | 0.47 | 0.55 | 0.53 | 0.44 | 0.47 |
| 4 | 65549349 | rs6811404 | -1.139 | -0.065 | -0.468 | -0.364 | -0.001 | -0.001 | -0.001 | 1.199 | 0.55 | 0.5 | 0.46 | 0.47 | 0.56 | 0.53 |
| 4 | 65556925 | rs6828768 | -0.523 | 0.433 | -0.925 | 0.001 | -0.043 | -0.014 | -0.014 | 0 | 0.39 | 0.26 | 0.42 | 0.47 | 0.5 | 0.47 |
| 4 | 65557183 | rs4355422 | -0.496 | 0.477 | -1.109 | -0.003 | -0.062 | -0.025 | -0.025 | 0.058 | 0.38 | 0.2 | 0.42 | 0.47 | 0.5 | 0.47 |
| 4 | 65563678 | rs4075197 | -0.555 | 0.383 | -0.735 | -0.138 | 0 | -0.003 | -0.003 | 0 | 0.46 | 0.48 | 0.36 | 0.46 | 0.46 | 0.43 |
| 4 | 65572162 | rs9995377 | -0.025 | -0.234 | -1.184 | -0.179 | 0.033 | -0.044 | -0.044 | 0.236 | 0.28 | 0.16 | 0.32 | 0.38 | 0.2 | 0.37 |
| 4 | 65574910 | rs6819000 | 0.556 | -0.37 | -0.627 | -0.527 | 0.037 | 0.01 | 0.01 | 0.072 | 0.49 | 0.36 | 0.43 | 0.58 | 0.58 | 0.49 |

| | | | | | | | | | | | | | | | | |
|---|---|---|---|---|---|---|---|---|---|---|---|---|---|---|---|---|
| 4 | 65576097 | rs7685391 | 0.226 | -0.225 | -0.191 | -0.494 | 0.009 | -0.055 | -0.055 | 0 | 0.9 | 0.59 | 0.94 | 1 | 0.95 | 0.98 |
| 4 | 65586360 | rs7695636 | 0.108 | -0.096 | -0.546 | -0.638 | 0.007 | 0.013 | 0.013 | 0 | 0.46 | 0.65 | 0.51 | 0.59 | 0.5 | 0.45 |
| 4 | 65592228 | rs12639640 | 0.367 | 0.012 | -0.402 | -0.517 | -0.011 | -0.009 | -0.009 | 0 | 0.37 | 0.36 | 0.44 | 0.46 | 0.43 | 0.36 |
| 4 | 65603201 | rs17085254 | 1.722 | 0.029 | -0.444 | -0.632 | 0.002 | -0.019 | -0.019 | 0 | 0.07 | 0.03 | 0.05 | 0.12 | 0.07 | 0.09 |
| 4 | 65613269 | rs6847583 | -0.187 | -0.04 | -0.254 | -0.417 | -0.001 | 0 | 0 | 0.75 | 0.46 | 0.5 | 0.47 | 0.54 | 0.5 | 0.45 |
| 4 | 65616248 | rs1367342 | 1.625 | 0.634 | -0.569 | -0.326 | -0.07 | -0.063 | -0.063 | 0.506 | 0.12 | 0.01 | 0.27 | 0.17 | 0.18 | 0.18 |
| 4 | 65620087 | rs1074431 | 0.786 | -0.214 | -0.555 | -0.668 | -0.03 | 0.07 | 0.07 | 0 | 0.13 | 0.02 | 0.1 | 0.02 | 0.17 | 0.12 |
| 4 | 65624721 | rs1430519 | -0.113 | -0.094 | -0.283 | -0.452 | 0.003 | -0.003 | -0.003 | 0 | 0.45 | 0.42 | 0.47 | 0.54 | 0.5 | 0.45 |
| 4 | 65640223 | rs17085296 | -0.449 | -0.391 | -0.28 | -0.224 | -0.002 | -0.002 | -0.002 | 0 | 0.15 | 0.15 | 0.09 | 0.12 | 0.14 | 0.2 |
| 4 | 65643064 | rs6551871 | 0.024 | -0.695 | -0.49 | -0.469 | 0.001 | -0.013 | -0.013 | 0.108 | 0.36 | 0.29 | 0.45 | 0.45 | 0.34 | 0.43 |
| 4 | 65651730 | rs1346841 | -0.838 | -0.934 | -0.455 | -0.574 | 0.014 | 0.009 | 0.009 | 0 | 0.51 | 0.59 | 0.52 | 0.55 | 0.42 | 0.58 |
| 4 | 65654988 | rs11731709 | 0.986 | -0.854 | -0.483 | -0.399 | -0.014 | -0.094 | -0.094 | 0.084 | 0.25 | 0.09 | 0.38 | 0.44 | 0.24 | 0.32 |
| 4 | 65662699 | rs1430512 | -0.252 | -0.839 | -0.29 | -0.366 | -0.003 | 0.007 | 0.007 | 0.023 | 0.38 | 0.45 | 0.45 | 0.45 | 0.34 | 0.43 |
| 4 | 65675052 | rs1346839 | -0.942 | -0.809 | -0.369 | -0.38 | 0.002 | 0.001 | 0.001 | 0 | 0.48 | 0.47 | 0.51 | 0.55 | 0.44 | 0.57 |
| 4 | 65682693 | rs978067 | -1.147 | -0.881 | -0.308 | -0.678 | -0.038 | -0.062 | -0.062 | 0.185 | 0.27 | 0.54 | 0.14 | 0.11 | 0.19 | 0.24 |
| 4 | 65697696 | rs17085407 | 0.284 | -0.706 | -0.515 | -0.504 | 0.023 | 0.011 | 0.011 | 0 | 0.15 | 0.07 | 0.06 | 0.11 | 0.1 | 0.14 |
| 4 | 65711998 | rs7654975 | -0.87 | -0.754 | -0.447 | -0.421 | 0.002 | 0.001 | 0.001 | 0 | 0.47 | 0.55 | 0.52 | 0.63 | 0.43 | 0.56 |
| 4 | 65712811 | rs6818012 | -0.827 | -0.763 | -0.239 | -0.425 | 0.008 | 0.006 | 0.006 | 0 | 0.46 | 0.47 | 0.52 | 0.63 | 0.43 | 0.56 |
| 4 | 65716998 | rs986971 | 1.35 | -0.795 | -1.031 | -0.825 | -0.076 | -0.045 | -0.045 | 0 | 0.23 | 0.01 | 0.1 | 0.24 | 0.25 | 0.14 |
| 4 | 65736609 | rs17085449 | 0.164 | -1.671 | -0.397 | -0.663 | 0.025 | 0.012 | 0.012 | 0 | 0.17 | 0.09 | 0.07 | 0.13 | 0.11 | 0.14 |
| 4 | 65737833 | rs1367337 | 0.838 | -1.667 | -0.304 | -0.7 | 0 | -0.018 | -0.018 | 0.396 | 0.08 | 0.13 | 0.05 | 0.01 | 0.09 | 0.09 |
| 4 | 65740387 | rs17085463 | -0.556 | -1.622 | -0.227 | -0.631 | -0.015 | 0.007 | 0.007 | 0 | 0.39 | 0.44 | 0.46 | 0.62 | 0.33 | 0.46 |
| 4 | 65750033 | rs7674285 | 1.13 | -1.883 | -0.149 | -0.688 | 0.014 | 0.019 | 0.019 | 0 | 0.46 | 0.34 | 0.45 | 0.36 | 0.57 | 0.42 |
| 4 | 65750373 | rs10517902 | -0.338 | -2.038 | -0.168 | -0.756 | 0.017 | 0.037 | 0.037 | 0 | 0.25 | 0.12 | 0.12 | 0.14 | 0.2 | 0.22 |
| 4 | 65751266 | rs1864522 | 0.876 | -2.073 | -0.011 | -0.775 | -0.027 | -0.058 | -0.058 | 1.442 | 0.67 | 0.49 | 0.84 | 0.86 | 0.77 | 0.74 |
| 4 | 65755777 | rs10023055 | 0.269 | -2.11 | -0.026 | -0.604 | 0.016 | -0.074 | -0.074 | 1.228 | 0.26 | 0.15 | 0.4 | 0.5 | 0.23 | 0.32 |
| 4 | 65774664 | rs12502234 | -0.358 | -2.035 | 0.242 | -0.614 | 0.027 | -0.016 | -0.016 | 0.887 | 0.93 | 0.68 | 0.97 | 1 | 0.98 | 0.96 |
| 4 | 65774726 | rs12504921 | 0.285 | -2.058 | -0.192 | -0.542 | 0.02 | -0.071 | -0.071 | 0 | 0.25 | 0.13 | 0.39 | 0.5 | 0.22 | 0.33 |
| 4 | 65776107 | rs10008353 | 1.029 | -2.043 | -0.143 | -0.534 | -0.009 | -0.113 | -0.113 | 0.777 | 0.2 | 0.06 | 0.38 | 0.5 | 0.21 | 0.32 |
| 4 | 65778213 | rs10005825 | 0.236 | -2.004 | -0.036 | -0.516 | 0.002 | -0.004 | -0.004 | 0 | 0.26 | 0.25 | 0.4 | 0.5 | 0.22 | 0.33 |
| 4 | 65784245 | rs6857709 | 0.215 | -1.828 | -0.101 | -0.411 | 0.03 | -0.053 | -0.053 | 0 | 0.31 | 0.17 | 0.41 | 0.5 | 0.25 | 0.33 |
| 4 | 65791276 | rs1372029 | 0.336 | -1.705 | 0.245 | -0.222 | 0.017 | -0.021 | -0.021 | 0 | 0.34 | 0.25 | 0.43 | 0.46 | 0.27 | 0.28 |
| 4 | 65798746 | rs1595974 | 0.591 | -1.857 | 0.427 | -0.334 | 0.011 | 0.005 | 0.005 | 0 | 0.45 | 0.38 | 0.45 | 0.41 | 0.56 | 0.45 |
| 4 | 65801134 | rs968827 | -0.261 | -1.931 | 0.624 | -0.364 | -0.014 | 0.006 | 0.006 | 0.39 | 0.6 | 0.54 | 0.55 | 0.54 | 0.7 | 0.67 |
| 4 | 65807695 | rs2033944 | 0.123 | -1.719 | 0.553 | -0.19 | -0.003 | -0.002 | -0.002 | 0.072 | 0.16 | 0.13 | 0.09 | 0.13 | 0.15 | 0.22 |
| 4 | 65817923 | rs1372020 | -0.063 | -1.725 | 0.561 | -0.192 | -0.017 | -0.012 | -0.012 | 0.003 | 0.17 | 0.21 | 0.1 | 0.13 | 0.15 | 0.22 |
| 4 | 65822976 | rs2119436 | -0.563 | -1.848 | 0.56 | -0.237 | 0.04 | 0.018 | 0.018 | 0 | 0.49 | 0.33 | 0.51 | 0.59 | 0.42 | 0.49 |
| 4 | 65824824 | rs6837932 | -0.528 | -2.013 | 0.77 | -0.365 | -0.048 | -0.042 | -0.042 | 0 | 0.21 | 0.43 | 0.13 | 0.13 | 0.16 | 0.23 |
| 4 | 65825862 | rs920508 | 0.099 | -1.987 | 0.489 | -0.338 | 0.012 | 0.007 | 0.007 | 0.205 | 0.17 | 0.11 | 0.1 | 0.13 | 0.15 | 0.22 |
| 4 | 65831184 | rs3943667 | -0.275 | -2.072 | 0.625 | -0.326 | 0.041 | -0.02 | -0.02 | 0 | 0.61 | 0.3 | 0.54 | 0.55 | 0.7 | 0.71 |
| 4 | 65840580 | rs1440923 | -0.739 | -2.115 | 0.84 | -0.362 | 0 | -0.002 | -0.002 | 0 | 0.65 | 0.68 | 0.59 | 0.54 | 0.72 | 0.7 |
| 4 | 65846651 | rs13106164 | 0.303 | -2.196 | 0.843 | -0.42 | 0.024 | 0.01 | 0.01 | 0 | 0.48 | 0.41 | 0.47 | 0.42 | 0.58 | 0.49 |
| 4 | 65863772 | rs10155085 | 0.344 | -1.996 | 0.893 | -0.33 | -0.005 | -0.003 | -0.003 | 0.083 | 0.16 | 0.18 | 0.1 | 0.13 | 0.14 | 0.21 |
| 4 | 65871723 | rs10011331 | -0.984 | -2.064 | 0.9 | -0.301 | -0.015 | -0.01 | -0.01 | 0 | 0.24 | 0.68 | 0.15 | 0.14 | 0.16 | 0.23 |
| 4 | 65873197 | rs17085702 | 0.21 | -2.008 | 0.75 | -0.271 | -0.022 | -0.005 | -0.005 | 0.238 | 0.17 | 0.24 | 0.1 | 0.14 | 0.14 | 0.21 |
| 4 | 65881161 | rs6811034 | -0.582 | -2.1 | 0.563 | -0.352 | -0.021 | 0.014 | 0.014 | 0.22 | 0.63 | 0.48 | 0.55 | 0.55 | 0.71 | 0.72 |
| 4 | 65883560 | rs1440949 | 0.538 | -2.044 | -0.032 | -0.361 | 0.005 | -0.047 | -0.047 | 0.177 | 0.44 | 0.12 | 0.44 | 0.42 | 0.56 | 0.51 |
| 4 | 65886260 | rs12186163 | -1.066 | -1.85 | 0.211 | -0.262 | 0.018 | -0.041 | -0.041 | 0 | 0.68 | 0.79 | 0.59 | 0.52 | 0.72 | 0.73 |
| 4 | 65890760 | rs9917886 | -0.579 | -1.827 | 0.988 | -0.201 | 0.003 | 0.002 | 0.002 | 0 | 0.5 | 0.45 | 0.51 | 0.61 | 0.42 | 0.48 |
| 4 | 65897053 | rs6819203 | -0.268 | -1.586 | 0.551 | -0.189 | -0.023 | -0.011 | -0.011 | 1.985 | 0.18 | 0.23 | 0.1 | 0.13 | 0.14 | 0.22 |
| 4 | 65902264 | rs1440931 | -0.927 | -2.032 | 0.267 | -0.843 | -0.022 | -0.092 | -0.092 | 0 | 0.29 | 0.62 | 0.14 | 0.01 | 0.18 | 0.25 |
| 4 | 65903597 | rs13152245 | 0.908 | -1.818 | -0.409 | -0.433 | -0.085 | -0.106 | -0.106 | 0.081 | 0.36 | 0.1 | 0.43 | 0.52 | 0.51 | 0.38 |
| 4 | 65905019 | rs17085830 | 1.468 | -1.528 | -0.932 | -0.499 | 0.024 | -0.077 | -0.077 | 0 | 0.21 | 0.02 | 0.1 | 0.35 | 0.16 | 0.16 |
| 4 | 65906658 | rs7693800 | 0.19 | -1.578 | -0.032 | -0.675 | -0.012 | -0.07 | -0.07 | 0 | 0.2 | 0.22 | 0.11 | 0.01 | 0.16 | 0.22 |
| 4 | 65907274 | rs9312133 | 0.742 | -1.353 | -0.319 | 0.229 | 0.02 | 0.01 | 0.01 | 0 | 0.32 | 0.22 | 0.34 | 0.27 | 0.26 | 0.27 |
| 4 | 65934317 | rs1348098 | -0.008 | -1.561 | 0.304 | 0.232 | -0.05 | -0.029 | -0.029 | 0.388 | 0.72 | 0.56 | 0.82 | 0.84 | 0.83 | 0.73 |
| 4 | 65940796 | rs10009571 | 0.723 | -1.741 | 0.202 | 0.249 | 0.076 | 0.049 | 0.049 | 0 | 0.63 | 0.18 | 0.67 | 0.76 | 0.76 | 0.64 |
| 4 | 65945051 | rs7668959 | 0.098 | -1.791 | 0.679 | 0.258 | -0.02 | -0.017 | -0.017 | 0 | 0.69 | 0.62 | 0.77 | 0.82 | 0.77 | 0.66 |
| 4 | 65957836 | rs2083551 | 0.745 | -1.844 | 0.498 | 0.258 | 0.079 | 0.044 | 0.044 | 0 | 0.63 | 0.19 | 0.68 | 0.76 | 0.77 | 0.65 |
| 4 | 65966308 | rs1877116 | -0.502 | -1.758 | 0.675 | 0.26 | -0.003 | -0.001 | -0.001 | 0 | 0.75 | 0.72 | 0.84 | 0.85 | 0.83 | 0.72 |
| 4 | 65979199 | rs329461 | -0.421 | -1.781 | 0.665 | 0.248 | -0.016 | -0.009 | -0.009 | 0 | 0.74 | 0.7 | 0.84 | 0.84 | 0.83 | 0.72 |
| 4 | 65996048 | rs682680 | 0.227 | -1.835 | 0.707 | 0.198 | -0.045 | -0.043 | -0.043 | 0 | 0.66 | 0.49 | 0.75 | 0.82 | 0.75 | 0.67 |
| 4 | 65996512 | rs10013406 | 1.311 | -0.762 | -0.059 | 0.236 | -0.058 | 0.01 | 0.01 | 0 | 0.29 | 0.09 | 0.22 | 0.22 | 0.33 | 0.23 |
| 4 | 66024351 | rs11721592 | 1.323 | -0.294 | -0.49 | 0.458 | -0.054 | -0.051 | -0.051 | 0 | 0.12 | 0.03 | 0.32 | 0.19 | 0.23 | 0.24 |
| 4 | 66058606 | rs371925 | -0.827 | -0.67 | -0.145 | 0.196 | -0.015 | -0.011 | -0.011 | 0 | 0.72 | 0.67 | 0.83 | 0.84 | 0.8 | 0.73 |
| 4 | 66079826 | rs1514267 | -0.329 | -0.655 | -0.237 | 0.199 | -0.044 | -0.042 | -0.042 | 0 | 0.66 | 0.51 | 0.75 | 0.82 | 0.75 | 0.69 |
| 4 | 66083947 | rs2349420 | 1.228 | -0.486 | -0.663 | 0.287 | -0.13 | -0.035 | -0.035 | 0 | 0.28 | 0.02 | 0.22 | 0.21 | 0.33 | 0.23 |
| 4 | 66111370 | rs13141982 | 0.591 | -0.353 | -0.403 | 0.31 | -0.003 | -0.012 | -0.012 | 0 | 0.05 | 0.08 | 0.06 | 0.02 | 0.05 | 0.05 |
| 4 | 66117152 | rs12504837 | 0.472 | -0.66 | -0.721 | 0.329 | -0.085 | -0.051 | -0.051 | 0.474 | 0.38 | 0.1 | 0.52 | 0.62 | 0.46 | 0.51 |

| | | | | | | | | | | | | | | | | |
|---|---|---|---|---|---|---|---|---|---|---|---|---|---|---|---|---|
| 4 | 66119496 | rs54808 | 0.86 | -0.732 | -0.318 | 0.343 | 0.002 | -0.001 | -0.001 | 0.028 | 0.32 | 0.34 | 0.29 | 0.28 | 0.34 | 0.24 |
| 4 | 66126518 | rs12499393 | -0.043 | -0.388 | -0.643 | 0.516 | -0.058 | 0.005 | 0.005 | 0.372 | 0.42 | 0.15 | 0.58 | 0.64 | 0.5 | 0.56 |
| 4 | 66131067 | rs1510896 | 0.319 | -0.038 | -0.542 | 0.521 | -0.053 | -0.065 | -0.065 | 0 | 0.33 | 0.16 | 0.42 | 0.49 | 0.42 | 0.47 |
| 4 | 66136223 | rs10008833 | 1.23 | 0.281 | -0.82 | 0.318 | -0.039 | 0.026 | 0.026 | 0 | 0.22 | 0.01 | 0.15 | 0.13 | 0.21 | 0.24 |
| 4 | 66142596 | rs7692514 | -0.469 | 0.519 | 0.058 | 0.497 | 0.026 | 0.015 | 0.015 | 0 | 0.05 | 0.25 | 0.18 | 0.03 | 0.06 | 0.04 |
| 4 | 66147444 | rs4860646 | 0.544 | 0.526 | -0.537 | 0.542 | -0.095 | -0.104 | -0.104 | 0 | 0.33 | 0.07 | 0.42 | 0.48 | 0.41 | 0.49 |
| 4 | 66150055 | rs7688176 | 0.035 | 0.552 | 0.112 | 0.526 | -0.009 | -0.01 | -0.01 | 0 | 0.37 | 0.3 | 0.48 | 0.49 | 0.43 | 0.53 |
| 4 | 66167253 | rs1376409 | 1.287 | 0.28 | 0 | 0.389 | 0.006 | 0.016 | 0.016 | 0 | 0.23 | 0.13 | 0.24 | 0.15 | 0.21 | 0.22 |
| 4 | 66168773 | rs11737511 | 0.193 | -0.016 | -0.049 | 0.186 | -0.055 | -0.015 | -0.015 | 0 | 0.44 | 0.13 | 0.38 | 0.47 | 0.47 | 0.51 |
| 4 | 66173126 | rs1160158 | -1.222 | -0.083 | 0.323 | 0.207 | -0.007 | 0.046 | 0.046 | 0 | 0.94 | 0.78 | 0.93 | 0.97 | 1 | 0.99 |
| 4 | 66174910 | rs2882226 | -0.953 | -0.122 | 0.276 | 0.142 | -0.039 | -0.055 | -0.055 | 0 | 0.81 | 0.68 | 0.85 | 0.97 | 0.9 | 0.95 |
| 4 | 66181030 | rs1451160 | 0.382 | -0.096 | 0.17 | 0.165 | 0.023 | 0.024 | 0.024 | 0 | 0.54 | 0.18 | 0.54 | 0.62 | 0.59 | 0.7 |
| 4 | 66187627 | rs4860650 | -0.535 | -0.009 | 0.434 | 0.296 | 0.004 | 0.067 | 0.067 | 0 | 0.93 | 0.67 | 0.93 | 0.97 | 1 | 0.99 |
| 4 | 66188295 | rs4860651 | 0.655 | 0.035 | 0.214 | 0.228 | 0.008 | 0.098 | 0.098 | 0 | 0.36 | 0.14 | 0.32 | 0.15 | 0.32 | 0.25 |
| 4 | 66195601 | rs2122722 | 0.369 | 0.341 | 0.409 | 0.302 | 0.017 | 0.027 | 0.027 | 0 | 0.27 | 0.46 | 0.31 | 0.36 | 0.31 | 0.26 |
| 4 | 66197804 | rs7349683 | 0.859 | -0.058 | 0.063 | 0.363 | -0.079 | -0.091 | -0.091 | 0.036 | 0.31 | 0.09 | 0.31 | 0.46 | 0.37 | 0.47 |
| 4 | 66202345 | rs17086092 | 1.138 | 0.017 | -0.28 | 0.477 | -0.002 | 0.001 | 0.001 | 0 | 0.2 | 0.21 | 0.09 | 0.33 | 0.16 | 0.21 |
| 4 | 66204540 | rs12645455 | 1.189 | 0.291 | -0.513 | 0.575 | -0.058 | -0.096 | -0.096 | 1.583 | 0.15 | 0.02 | 0.19 | 0.32 | 0.19 | 0.24 |
| 4 | 66206492 | rs986208 | 0.239 | -0.16 | -0.128 | 0.106 | 0.006 | 0.066 | 0.066 | 0 | 0.52 | 0.35 | 0.4 | 0.79 | 0.54 | 0.68 |
| 4 | 66208717 | rs2349723 | -0.234 | 0.002 | 0.241 | 0.278 | 0.004 | -0.012 | -0.012 | 0.4 | 0.65 | 0.43 | 0.49 | 0.79 | 0.63 | 0.71 |
| 4 | 66219433 | rs1451180 | -0.522 | -0.073 | 0.095 | 0.314 | -0.001 | -0.018 | -0.018 | 0 | 0.69 | 0.44 | 0.66 | 0.82 | 0.71 | 0.74 |
| 4 | 66228727 | rs13135753 | -0.36 | -0.119 | 0.21 | 0.253 | 0.001 | -0.01 | -0.01 | 0.214 | 0.65 | 0.66 | 0.6 | 0.82 | 0.65 | 0.7 |
| 4 | 66234071 | rs11737618 | 1.184 | -0.463 | 0.061 | 0.203 | 0.015 | 0.023 | 0.023 | 0 | 0.13 | 0.08 | 0.09 | 0.03 | 0.1 | 0.04 |
| 4 | 66238145 | rs17086137 | 1.052 | -0.887 | -0.002 | 0.434 | -0.024 | 0.012 | 0.012 | 0 | 0.17 | 0.2 | 0.07 | 0.28 | 0.11 | 0.26 |
| 4 | 66244369 | rs2034537 | 0.593 | -0.651 | 0.287 | 0.133 | 0.018 | 0.077 | 0.077 | 0.001 | 0.55 | 0.24 | 0.51 | 0.8 | 0.6 | 0.69 |
| 4 | 66244695 | rs11726788 | -0.466 | -0.205 | 0.608 | 0.413 | 0.002 | -0.013 | -0.013 | 0 | 0.06 | 0.2 | 0.03 | 0 | 0.03 | 0.06 |
| 4 | 66246438 | rs1451173 | -0.717 | -0.02 | 0.412 | 0.493 | 0.017 | -0.002 | -0.002 | 0.741 | 0.77 | 0.69 | 0.55 | 0.82 | 0.75 | 0.78 |
| 4 | 66254728 | rs4860656 | -0.279 | 0.138 | 0.247 | 0.323 | 0.004 | 0.054 | 0.054 | 0 | 0.51 | 0.59 | 0.65 | 0.79 | 0.45 | 0.65 |
| 4 | 66256159 | rs10002887 | -0.341 | 0.223 | 0.225 | 0.822 | 0.024 | 0.075 | 0.075 | 0 | 0.15 | 0.27 | 0.18 | 0.35 | 0.22 | 0.32 |
| 4 | 66258273 | rs1376416 | -0.764 | 0.212 | 0.145 | 0.665 | -0.05 | 0.189 | 0.189 | 0 | 0.56 | 0.85 | 0.63 | 0.83 | 0.5 | 0.73 |
| 4 | 66260257 | rs10033393 | 0.915 | 0.028 | 0.34 | 1.08 | -0.068 | 0.013 | 0.013 | 0 | 0.27 | 0.5 | 0.18 | 0.24 | 0.13 | 0.18 |
| 4 | 66260526 | rs6828375 | 1.183 | 0.293 | -0.05 | 1.15 | -0.026 | -0.029 | -0.029 | 0 | 0.09 | 0.05 | 0.09 | 0.24 | 0.15 | 0.24 |
| 4 | 66262049 | rs2084682 | -0.124 | 0.25 | 0.167 | 1.134 | -0.011 | -0.014 | -0.014 | 1.533 | 0.25 | 0.23 | 0.41 | 0.5 | 0.33 | 0.48 |
| 4 | 66264302 | rs1597611 | 1 | 0.263 | 0.161 | 0.851 | -0.003 | -0.001 | -0.001 | 0 | 0.16 | 0.15 | 0.1 | 0.16 | 0.13 | 0.06 |
| 4 | 66272179 | rs2198103 | -0.757 | 0.224 | 0.394 | 0.797 | -0.025 | 0.004 | 0.004 | 0 | 0.81 | 0.7 | 0.79 | 0.81 | 0.84 | 0.94 |
| 4 | 66272411 | rs13120093 | 1.392 | 0.091 | 0.394 | 0.744 | 0.021 | 0.018 | 0.018 | 0 | 0.24 | 0.18 | 0.17 | 0.1 | 0.13 | 0.15 |
| 4 | 66273350 | rs17086199 | 1.488 | 0.303 | 0.254 | 0.822 | -0.011 | -0.013 | -0.013 | 0 | 0.13 | 0.06 | 0.3 | 0.28 | 0.17 | 0.22 |
| 4 | 66277278 | rs17086207 | 0.978 | 0.304 | 0.306 | 0.816 | -0.002 | -0.003 | -0.003 | 0.017 | 0.1 | 0.09 | 0.1 | 0.23 | 0.15 | 0.25 |
| 4 | 66278087 | rs12509018 | 0.292 | -0.124 | 0.57 | 1.004 | 0.094 | -0.008 | -0.008 | 1.246 | 0.52 | 0.24 | 0.57 | 0.63 | 0.67 | 0.71 |
| 4 | 66280171 | rs2305351 | 1.266 | 0.149 | 0.385 | 1.163 | -0.063 | -0.081 | -0.081 | 2.607 | 0.12 | 0.02 | 0.3 | 0.29 | 0.17 | 0.22 |
| 4 | 66283093 | rs10027864 | -0.219 | -0.359 | 0.855 | 0.9 | 0.039 | 0.161 | 0.161 | 0 | 0.05 | 0.3 | 0.06 | 0.09 | 0.05 | 0.07 |
| 4 | 66294649 | rs7672488 | -0.916 | -0.419 | -0.141 | 0.148 | -0.051 | -0.054 | -0.054 | 1.421 | 0.74 | 0.63 | 0.81 | 0.91 | 0.87 | 0.85 |
| 4 | 66300425 | rs6831652 | 0.049 | 0.538 | -0.942 | 0.271 | -0.053 | 0.081 | 0.081 | 0 | 0.39 | 0.13 | 0.29 | 0.34 | 0.54 | 0.49 |
| 4 | 66304962 | rs10013782 | -0.306 | 0.379 | -0.475 | 0.37 | 0.031 | 0.01 | 0.01 | 0 | 0.51 | 0.44 | 0.45 | 0.52 | 0.63 | 0.54 |
| 4 | 66305150 | rs10003638 | -0.544 | 0.797 | -0.507 | -0.212 | -0.035 | 0.056 | 0.056 | 0 | 0.65 | 0.75 | 0.76 | 0.92 | 0.6 | 0.73 |
| 4 | 66306421 | rs10007066 | 0.424 | 0.748 | 0.245 | 0.552 | 0.007 | 0.101 | 0.101 | 0 | 0.89 | 0.77 | 0.81 | 0.77 | 0.92 | 0.87 |
| 4 | 66322437 | rs4860665 | 0.999 | 1.019 | -0.367 | 0.154 | 0.017 | 0.065 | 0.065 | 0.018 | 0.34 | 0.15 | 0.3 | 0.12 | 0.29 | 0.2 |
| 4 | 66327854 | rs9991256 | 1.429 | 1.452 | -0.703 | 0.458 | -0.047 | -0.066 | -0.066 | 0.051 | 0.11 | 0.02 | 0.25 | 0.38 | 0.16 | 0.22 |
| 4 | 66335098 | rs4267782 | 1.422 | 1.485 | -0.736 | 0.313 | -0.015 | 0.107 | 0.107 | 0 | 0.32 | 0.03 | 0.2 | 0.09 | 0.26 | 0.21 |
| 4 | 66340967 | rs4425430 | -0.356 | 1.372 | -0.35 | 0.284 | -0.005 | 0.071 | 0.071 | 0 | 0.13 | 0.25 | 0.2 | 0.24 | 0.11 | 0.15 |
| 4 | 66343670 | rs4637457 | -0.143 | 1.582 | -0.232 | 0.116 | -0.006 | -0.009 | -0.009 | 0.577 | 0.44 | 0.33 | 0.45 | 0.45 | 0.47 | 0.45 |
| 4 | 66344954 | rs11936348 | 0.955 | 1.477 | -0.831 | -0.135 | -0.033 | 0.178 | 0.178 | 0 | 0.3 | 0.06 | 0.18 | 0.08 | 0.31 | 0.22 |
| 4 | 66348548 | rs7694306 | 1.405 | 1.417 | -0.517 | 0.154 | -0.009 | 0.105 | 0.105 | 0 | 0.32 | 0.04 | 0.2 | 0.09 | 0.26 | 0.21 |
| 4 | 66353220 | rs4516775 | 0.767 | 1.986 | 0.079 | 0.434 | 0.017 | 0.014 | 0.014 | 0 | 0.09 | 0.14 | 0.11 | 0.22 | 0.15 | 0.14 |
| 4 | 66355104 | rs11724363 | 0.006 | 1.71 | -0.28 | 0.159 | -0.004 | -0.006 | -0.006 | 0.089 | 0.43 | 0.22 | 0.48 | 0.51 | 0.4 | 0.44 |
| 4 | 66359577 | rs4337775 | -0.406 | 1.645 | 0.035 | 0.556 | -0.042 | 0.006 | 0.006 | 0 | 0.6 | 0.16 | 0.54 | 0.62 | 0.56 | 0.6 |
| 4 | 66362487 | rs6815080 | 1.102 | 1.29 | 0.503 | 0.518 | -0.011 | -0.048 | -0.048 | 0.078 | 0.28 | 0.47 | 0.23 | 0.13 | 0.27 | 0.24 |
| 4 | 66378192 | rs4541571 | -1.036 | 1.335 | 0.128 | 0.562 | 0.006 | 0.023 | 0.023 | 0 | 0.68 | 0.27 | 0.65 | 0.84 | 0.7 | 0.75 |
| 4 | 66378789 | rs13109328 | 0.292 | 1.319 | -0.223 | 0.571 | -0.016 | 0.122 | 0.122 | 0.068 | 0.46 | 0.08 | 0.35 | 0.28 | 0.39 | 0.34 |
| 4 | 66381581 | rs4132313 | 0.658 | 1.583 | 0.335 | 0.686 | 0.002 | 0.094 | 0.094 | 0 | 0.16 | 0.28 | 0.28 | 0.34 | 0.17 | 0.27 |
| 4 | 66382111 | rs7673091 | -1.555 | 1.577 | 0.336 | 0.649 | -0.011 | -0.022 | -0.022 | 0 | 0.71 | 0.66 | 0.79 | 0.87 | 0.74 | 0.77 |
| 4 | 66382160 | rs13149946 | 0.429 | 1.583 | 0.032 | 0.669 | -0.008 | 0.117 | 0.117 | 0.178 | 0.42 | 0.08 | 0.32 | 0.25 | 0.35 | 0.31 |
| 4 | 66383283 | rs7679950 | -0.122 | 1.66 | 0.452 | 0.637 | 0.07 | 0.062 | 0.062 | 0 | 0.12 | 0.26 | 0.17 | 0.28 | 0.21 | 0.19 |
| 4 | 66383430 | rs11131599 | 0.724 | 1.637 | 0.11 | 0.632 | 0.034 | 0.017 | 0.017 | 0 | 0.09 | 0.14 | 0.12 | 0.21 | 0.17 | 0.15 |
| 4 | 66389701 | rs7684022 | -0.982 | 0.959 | -0.072 | 0.446 | -0.011 | -0.015 | -0.015 | 0 | 0.68 | 0.63 | 0.74 | 0.8 | 0.71 | 0.74 |
| 4 | 66394160 | rs11947260 | -0.025 | 0.942 | -0.213 | 0.397 | 0.004 | 0.008 | 0.008 | 0 | 0.31 | 0.31 | 0.29 | 0.22 | 0.25 | 0.23 |
| 4 | 66397114 | rs10032173 | -0.781 | 1.059 | 0.114 | 0.556 | 0.008 | 0.006 | 0.006 | 0 | 0.89 | 0.83 | 0.9 | 0.88 | 0.88 | 0.87 |
| 4 | 66398855 | rs10016747 | 0.653 | 1.125 | -0.101 | 1.055 | -0.004 | -0.048 | -0.048 | 0.214 | 0.13 | 0.06 | 0.25 | 0.28 | 0.13 | 0.22 |
| 4 | 66401110 | rs4860679 | -0.96 | 1.142 | -0.024 | 1.037 | 0.008 | 0.006 | 0.006 | 0 | 0.89 | 0.83 | 0.9 | 0.88 | 0.88 | 0.87 |

| | | | | | | | | | | | | | | | | |
|---|---|---|---|---|---|---|---|---|---|---|---|---|---|---|---|---|
| 4 | 66403567 | rs6551933 | 1.171 | 1.087 | -0.001 | 0.968 | 0.002 | -0.007 | -0.007 | 0 | 0.45 | 0.48 | 0.36 | 0.38 | 0.51 | 0.42 |
| 4 | 66421666 | rs4518304 | 0.403 | 1.032 | 0.041 | 1.008 | -0.001 | 0.043 | 0.043 | 0.094 | 0.75 | 0.66 | 0.64 | 0.6 | 0.75 | 0.65 |
| 4 | 66431909 | rs11931601 | 0.025 | 1.24 | -0.143 | 0.516 | 0.006 | -0.053 | -0.053 | 0 | 0.38 | 0.18 | 0.47 | 0.45 | 0.34 | 0.36 |
| 4 | 66438114 | rs9997603 | -0.546 | 1.619 | 0.793 | 0.839 | 0.002 | 0.058 | 0.058 | 0 | 0.89 | 0.77 | 0.9 | 0.8 | 0.9 | 0.89 |
| 4 | 66443874 | rs13127631 | 0.533 | 1.441 | 0.37 | 0.647 | -0.037 | -0.101 | -0.101 | 0.603 | 0.28 | 0.14 | 0.44 | 0.44 | 0.28 | 0.24 |
| 4 | 66454374 | rs13144830 | -0.77 | 1.883 | 0.646 | 0.485 | 0.083 | 0.117 | 0.117 | 0.449 | 0.21 | 0.51 | 0.26 | 0.35 | 0.32 | 0.34 |
| 4 | 66454906 | rs7440592 | -1.253 | 1.834 | 0.649 | 0.453 | -0.002 | 0.01 | 0.01 | 0 | 0.9 | 0.89 | 0.91 | 0.81 | 0.9 | 0.86 |
| 4 | 66456092 | rs6855605 | 0.649 | 1.811 | 0.585 | 0.373 | -0.018 | -0.093 | -0.093 | 0 | 0.29 | 0.12 | 0.44 | 0.46 | 0.27 | 0.26 |
| 4 | 66468381 | rs4860188 | 0.918 | 1.839 | 0.905 | -0.108 | 0.023 | 0.029 | 0.029 | 1.107 | 0.69 | 0.39 | 0.64 | 0.59 | 0.58 | 0.53 |
| 4 | 66470077 | rs11947738 | 1.117 | 1.849 | 0.844 | -0.103 | -0.036 | -0.044 | -0.044 | 0 | 0.67 | 0.2 | 0.63 | 0.59 | 0.58 | 0.53 |
| 4 | 66483593 | rs4320186 | -0.822 | 1.353 | 0.871 | -0.199 | 0.009 | 0.054 | 0.054 | 0 | 0.89 | 0.76 | 0.9 | 0.81 | 0.9 | 0.86 |
| 4 | 66504490 | rs4505878 | 0.502 | 1.471 | -0.097 | -0.182 | -0.005 | -0.075 | -0.075 | 0 | 0.3 | 0.13 | 0.44 | 0.59 | 0.27 | 0.27 |
| 4 | 66526101 | rs4565127 | -0.296 | 0.603 | 0.163 | -1.099 | -0.006 | 0.031 | 0.031 | 0 | 0.87 | 0.76 | 0.84 | 0.81 | 0.9 | 0.85 |
| 4 | 66527955 | rs7655988 | -0.624 | 0.846 | 0.061 | -1.117 | 0.045 | 0.072 | 0.072 | 0 | 0.52 | 0.64 | 0.66 | 0.8 | 0.64 | 0.6 |
| 4 | 66550619 | rs4371664 | -0.14 | 1.001 | 0.365 | -0.564 | 0.004 | 0.051 | 0.051 | 0 | 0.87 | 0.75 | 0.82 | 0.78 | 0.87 | 0.8 |
| 4 | 66553425 | rs11131613 | 0.032 | 1.448 | -0.105 | -0.885 | -0.023 | 0.01 | 0.01 | 0 | 0.37 | 0.26 | 0.48 | 0.7 | 0.36 | 0.43 |
| 4 | 66554209 | rs4498177 | 0.661 | 1.408 | -0.475 | -0.954 | 0.003 | 0.042 | 0.042 | 0.021 | 0.46 | 0.3 | 0.32 | 0.04 | 0.49 | 0.36 |
| 4 | 66562536 | rs7699335 | -0.354 | 1.96 | -1.028 | -0.387 | -0.09 | -0.013 | -0.013 | 0.022 | 0.37 | 0.1 | 0.49 | 0.68 | 0.4 | 0.43 |
| 4 | 66563086 | rs9997431 | -0.923 | 1.738 | -0.762 | -0.612 | -0.015 | 0.024 | 0.024 | 0 | 0.08 | 0.04 | 0.06 | 0.02 | 0.09 | 0.11 |
| 4 | 66565043 | rs9784450 | 0.68 | 1.785 | -0.588 | -0.715 | 0.002 | -0.025 | -0.025 | 0.057 | 0.85 | 0.87 | 0.88 | 0.98 | 0.75 | 0.83 |
| 4 | 66567179 | rs13127866 | -0.014 | 1.981 | -0.548 | -0.854 | -0.138 | -0.001 | -0.001 | 0 | 0.32 | 0.07 | 0.47 | 0.76 | 0.4 | 0.44 |
| 4 | 66571857 | rs11727254 | 0.034 | 2.159 | -0.352 | -0.735 | 0.008 | -0.008 | -0.008 | 0 | 0.06 | 0.09 | 0.07 | 0.01 | 0.12 | 0.1 |
| 4 | 66571998 | rs4529111 | -0.545 | 2.106 | -0.338 | -0.733 | -0.005 | -0.043 | -0.043 | 0 | 0.91 | 0.88 | 0.94 | 1 | 0.87 | 0.93 |
| 4 | 66576310 | rs4860189 | 0.414 | 2.369 | -0.412 | -0.706 | 0.001 | 0.003 | 0.003 | 0 | 0.26 | 0.26 | 0.36 | 0.66 | 0.28 | 0.33 |
| 4 | 66578235 | rs13144348 | 0.111 | 2.36 | -0.418 | -0.702 | 0.008 | -0.008 | -0.008 | 0.936 | 0.06 | 0.09 | 0.07 | 0.01 | 0.12 | 0.1 |
| 4 | 66583502 | rs4860701 | 1.448 | 2.216 | -0.57 | -0.778 | 0.076 | 0.132 | 0.132 | 0.301 | 0.46 | 0.12 | 0.34 | 0.22 | 0.35 | 0.38 |
| 4 | 66583906 | rs4629506 | 0.603 | 2.243 | -0.535 | -0.782 | 0.007 | -0.049 | -0.049 | 0 | 0.77 | 0.39 | 0.77 | 0.89 | 0.75 | 0.8 |
| 4 | 66584922 | rs4860703 | -0.106 | 2.299 | -0.605 | -0.712 | -0.013 | -0.006 | -0.006 | 0 | 0.31 | 0.26 | 0.42 | 0.68 | 0.4 | 0.42 |
| 4 | 66589229 | rs6551952 | 0.048 | 2.155 | -0.059 | -0.537 | -0.004 | -0.043 | -0.043 | 0 | 0.91 | 0.88 | 0.94 | 1 | 0.87 | 0.93 |
| 4 | 66596857 | rs11131615 | 0.404 | 1.709 | 0.09 | -0.479 | 0.045 | 0.014 | 0.014 | 0.564 | 0.87 | 0.58 | 0.82 | 0.9 | 0.88 | 0.87 |
| 4 | 66597539 | rs11728136 | -0.998 | 1.448 | -0.748 | -0.311 | 0.001 | -0.025 | -0.025 | 0 | 0.14 | 0.13 | 0.09 | 0.01 | 0.18 | 0.16 |
| 4 | 66601246 | rs6848595 | -0.04 | 1.241 | -0.972 | -0.35 | 0 | 0.015 | 0.015 | 0 | 0.8 | 0.85 | 0.73 | 0.94 | 0.81 | 0.81 |
| 4 | 66610307 | rs4241669 | 0.024 | 0.518 | -1.064 | -0.767 | 0 | -0.001 | -0.001 | 1.18 | 0.3 | 0.29 | 0.2 | 0.21 | 0.22 | 0.36 |
| 4 | 66611856 | rs4588513 | 0.063 | 1.157 | -0.718 | 0.324 | -0.006 | -0.022 | -0.022 | 0 | 0.7 | 0.79 | 0.62 | 0.48 | 0.7 | 0.68 |
| 4 | 66613120 | rs10009221 | 0.364 | 1.554 | -0.817 | 0.122 | 0.11 | 0.045 | 0.045 | 0 | 0.17 | 0.43 | 0.29 | 0.21 | 0.3 | 0.19 |
| 4 | 66615653 | rs6813706 | 0.511 | 1.508 | -0.782 | 0.117 | 0.014 | 0.026 | 0.026 | 0 | 0.55 | 0.39 | 0.33 | 0.27 | 0.41 | 0.49 |
| 4 | 66616643 | rs4440279 | 0.054 | 1.101 | -0.953 | 0.411 | -0.004 | -0.015 | -0.015 | 0 | 0.71 | 0.76 | 0.63 | 0.48 | 0.71 | 0.68 |
| 4 | 66622386 | rs10031256 | -0.295 | 0.615 | -0.388 | 0.374 | -0.013 | 0.017 | 0.017 | 0 | 0.1 | 0.22 | 0.04 | 0.09 | 0.04 | 0.06 |
| 4 | 66622515 | rs2350336 | -0.569 | 0.73 | -1.051 | 0.492 | -0.015 | 0.001 | 0.001 | 2.433 | 0.63 | 0.86 | 0.58 | 0.66 | 0.6 | 0.72 |
| 4 | 66626260 | rs4368661 | 0.379 | 1.37 | -1.191 | 0.267 | -0.012 | 0.007 | 0.007 | 0.458 | 0.14 | 0.1 | 0.2 | 0.11 | 0.21 | 0.19 |
| 4 | 66627407 | rs17086349 | 0.506 | 0.987 | -1.256 | 0.384 | 0 | -0.033 | -0.033 | 1.103 | 0.22 | 0.1 | 0.19 | 0.31 | 0.21 | 0.26 |
| 4 | 66628221 | rs4382087 | 0.527 | 0.671 | -0.183 | 0.147 | 0 | -0.002 | -0.002 | 1.779 | 0.62 | 0.41 | 0.62 | 0.71 | 0.59 | 0.52 |
| 4 | 66628728 | rs4373205 | -0.348 | 0.564 | 0.218 | 0.312 | -0.016 | 0.038 | 0.038 | 0 | 0.86 | 0.7 | 0.87 | 0.82 | 0.9 | 0.78 |
| 4 | 66628975 | rs2350339 | 0.143 | 0.544 | -0.369 | 0.278 | -0.007 | 0.012 | 0.012 | 0.028 | 0.15 | 0.11 | 0.09 | 0.1 | 0.16 | 0.2 |
| 4 | 66629765 | rs2882388 | -0.741 | 0.504 | -0.092 | 0.419 | -0.028 | -0.021 | -0.021 | 0 | 0.31 | 0.24 | 0.42 | 0.4 | 0.37 | 0.46 |
| 4 | 66638816 | rs4860707 | 0.437 | 0.493 | 0.001 | 0.304 | 0.01 | -0.001 | -0.001 | 0 | 0.18 | 0.13 | 0.17 | 0.18 | 0.14 | 0.24 |
| 4 | 66645823 | rs2882317 | -1.144 | 0.599 | 0.611 | 0.294 | 0.057 | -0.002 | -0.002 | 0 | 0.9 | 0.68 | 0.94 | 0.96 | 0.9 | 0.94 |
| 4 | 66655689 | rs6845738 | -2.169 | 0.028 | 0.43 | 0.086 | 0.001 | 0.006 | 0.006 | 4.588 | 0.64 | 0.53 | 0.6 | 0.62 | 0.67 | 0.79 |
| 4 | 66655790 | rs17086372 | 1.047 | 0.114 | 0.471 | -0.086 | 0 | -0.034 | -0.034 | 0 | 0.13 | 0.17 | 0.09 | 0 | 0.14 | 0.18 |
| 4 | 66657060 | rs1074532 | 2.353 | 0.104 | 0.544 | -0.09 | -0.007 | -0.011 | -0.011 | 0 | 0.34 | 0.22 | 0.38 | 0.38 | 0.33 | 0.21 |
| 4 | 66658538 | rs11131618 | 0.912 | 0.123 | 0.226 | -0.011 | -0.015 | -0.026 | -0.026 | 0 | 0.13 | 0.05 | 0.22 | 0.22 | 0.15 | 0.21 |
| 4 | 66663257 | rs17086389 | 1.034 | 0.065 | 0.281 | 0 | 0 | -0.034 | -0.034 | 0 | 0.13 | 0.17 | 0.09 | 0 | 0.14 | 0.17 |
| 4 | 66664021 | rs4146479 | -1.388 | 0.185 | 0.125 | 0.247 | -0.003 | 0.036 | 0.036 | 0.112 | 0.51 | 0.34 | 0.51 | 0.62 | 0.53 | 0.61 |
| 4 | 66681736 | rs4484355 | -1.041 | 0.171 | 0.128 | 0.301 | 0.001 | 0.002 | 0.002 | 0 | 0.46 | 0.33 | 0.49 | 0.46 | 0.49 | 0.55 |
| 4 | 66683829 | rs11735605 | 0.084 | -0.079 | -0.01 | 0.292 | 0.037 | 0 | 0 | 0 | 0.23 | 0.12 | 0.19 | 0.19 | 0.18 | 0.28 |
| 4 | 66688425 | rs2882314 | 0.426 | -0.119 | 0.328 | 0.323 | -0.042 | 0.04 | 0.04 | 0.436 | 0.68 | 0.48 | 0.74 | 0.59 | 0.76 | 0.65 |
| 4 | 66689367 | rs7660878 | 1.12 | -0.157 | -0.191 | -0.499 | 0.017 | 0.019 | 0.019 | 0 | 0.13 | 0.1 | 0.07 | 0.03 | 0.12 | 0.08 |
| 4 | 66694545 | rs10517941 | 2.09 | 0.468 | -0.618 | -0.809 | -0.046 | 0.052 | 0.052 | 0 | 0.12 | 0.02 | 0.22 | 0 | 0.21 | 0.08 |
| 4 | 66697780 | rs17082137 | 0.643 | 0.26 | 0.101 | -0.324 | -0.006 | 0.075 | 0.075 | 0 | 0.07 | 0.13 | 0.03 | 0.2 | 0.04 | 0.07 |
| 4 | 66709201 | rs2350064 | -0.632 | 0.314 | -0.201 | -0.624 | 0 | 0.022 | 0.022 | 0 | 0.31 | 0.24 | 0.4 | 0.15 | 0.31 | 0.44 |
| 4 | 66712131 | rs10866165 | 1.126 | 0.49 | -0.005 | -0.675 | 0 | 0.007 | 0.007 | 0.054 | 0.53 | 0.53 | 0.4 | 0.69 | 0.45 | 0.46 |
| 4 | 66712685 | rs17755896 | 1.464 | 0.555 | -0.216 | -0.725 | -0.002 | -0.016 | -0.016 | 0 | 0.1 | 0.08 | 0.09 | 0 | 0.16 | 0.07 |
| 4 | 66716472 | rs2350070 | 1.234 | 0.55 | -0.202 | -0.66 | -0.021 | -0.007 | -0.007 | 0.07 | 0.14 | 0.11 | 0.19 | 0.16 | 0.24 | 0.1 |
| 4 | 66716573 | rs4146476 | 1.504 | 0.413 | 0.011 | -0.658 | 0.001 | -0.001 | -0.001 | 0.039 | 0.47 | 0.47 | 0.37 | 0.5 | 0.4 | 0.39 |
| 4 | 66729586 | rs10517944 | -0.026 | 0.611 | -0.237 | -0.879 | -0.007 | 0.037 | 0.037 | 0 | 0.22 | 0.13 | 0.27 | 0.07 | 0.25 | 0.27 |
| 4 | 66734932 | rs17086487 | 1.44 | 0.439 | -0.386 | -0.59 | -0.001 | -0.023 | -0.023 | 0.152 | 0.07 | 0 | 0.04 | 0.28 | 0.04 | 0.07 |
| 4 | 66738581 | rs6836494 | 0.559 | 0.319 | -0.212 | -0.979 | 0.002 | -0.004 | -0.004 | 0 | 0.59 | 0.62 | 0.63 | 0.52 | 0.61 | 0.62 |
| 4 | 66745310 | rs2350343 | 0.888 | 0.367 | -0.223 | -1.013 | 0.005 | 0.007 | 0.007 | 0 | 0.5 | 0.57 | 0.51 | 0.45 | 0.52 | 0.45 |
| 4 | 66750577 | rs6551968 | -0.165 | 0.373 | -0.192 | -1.009 | -0.002 | -0.015 | -0.015 | 0 | 0.34 | 0.34 | 0.32 | 0.2 | 0.35 | 0.31 |

| | | | | | | | | | | | | | | | | |
|---|---|---|---|---|---|---|---|---|---|---|---|---|---|---|---|---|
| 4 | 66751139 | rs17086506 | 1.035 | 0.363 | -0.185 | -1.023 | 0.007 | -0.038 | -0.038 | 0 | 0.21 | 0.3 | 0.14 | 0.12 | 0.24 | 0.1 |
| 4 | 66751426 | rs17086507 | 1.216 | 0.358 | -0.226 | -1.029 | -0.023 | 0.034 | 0.034 | 0 | 0.19 | 0.09 | 0.13 | 0.12 | 0.24 | 0.1 |
| 4 | 66759828 | rs10517949 | 0.962 | 0.736 | -0.25 | -1.051 | 0.006 | 0.01 | 0.01 | 0 | 0.49 | 0.59 | 0.49 | 0.45 | 0.45 | 0.44 |
| 4 | 66760539 | rs2576511 | -1.803 | 0.094 | -0.21 | -1.035 | -0.001 | 0 | 0 | 0 | 0.9 | 0.94 | 0.87 | 0.94 | 0.91 | 0.83 |
| 4 | 66763683 | rs11930963 | 1.042 | 0.092 | -0.199 | -1.042 | 0.004 | -0.026 | -0.026 | 0 | 0.21 | 0.27 | 0.14 | 0.12 | 0.24 | 0.1 |
| 4 | 66763742 | rs7690714 | -1.796 | 0.106 | -0.202 | -1.024 | -0.003 | -0.001 | -0.001 | 0.074 | 0.9 | 0.95 | 0.87 | 0.94 | 0.91 | 0.83 |
| 4 | 66771088 | rs12513262 | 0.706 | 0.091 | -0.217 | -1.028 | 0.002 | 0.017 | 0.017 | 0 | 0.56 | 0.61 | 0.55 | 0.73 | 0.57 | 0.51 |
| 4 | 66774464 | rs1897694 | 1.863 | -0.088 | -0.481 | -1.265 | -0.017 | 0.07 | 0.07 | 0 | 0.14 | 0.03 | 0.07 | 0 | 0.13 | 0.16 |
| 4 | 66777240 | rs7660970 | 2.737 | -0.137 | -0.156 | -1.257 | -0.006 | -0.015 | -0.015 | 0 | 0.24 | 0.18 | 0.34 | 0.43 | 0.27 | 0.22 |
| 4 | 66777683 | rs7679437 | 0.839 | -0.137 | -0.271 | -1.058 | -0.011 | -0.009 | -0.009 | 0 | 0.15 | 0.17 | 0.18 | 0.12 | 0.1 | 0.22 |
| 4 | 66779689 | rs12505828 | -1.621 | 0.022 | -0.165 | -0.953 | 0 | 0 | 0 | 0.052 | 0.9 | 0.91 | 0.87 | 0.94 | 0.91 | 0.83 |
| 4 | 66783098 | rs10022875 | -1.591 | 0.019 | -0.166 | -0.955 | 0 | 0 | 0 | 0.218 | 0.9 | 0.91 | 0.87 | 0.94 | 0.91 | 0.83 |
| 4 | 66783283 | rs10023064 | 0.429 | 0.014 | -0.147 | -0.985 | 0 | -0.002 | -0.002 | 0 | 0.63 | 0.63 | 0.69 | 0.55 | 0.63 | 0.65 |
| 4 | 66786261 | rs6825624 | 1.004 | 0.319 | 0.117 | -1.882 | -0.007 | 0.017 | 0.017 | 0 | 0.19 | 0.13 | 0.15 | 0.12 | 0.25 | 0.11 |
| 4 | 66791167 | rs17086628 | 0.926 | 0.311 | 0.004 | -1.885 | 0.001 | -0.014 | -0.014 | 0 | 0.2 | 0.21 | 0.15 | 0.12 | 0.25 | 0.11 |
| 4 | 66796556 | rs1897691 | -1.453 | 0.313 | -0.008 | -1.86 | 0 | -0.006 | -0.006 | 0.094 | 0.9 | 0.87 | 0.86 | 0.87 | 0.91 | 0.82 |
| 4 | 66797440 | rs6815570 | -0.147 | 0.386 | -0.11 | -1.812 | 0 | -0.007 | -0.007 | 0 | 0.69 | 0.65 | 0.7 | 0.78 | 0.66 | 0.72 |
| 4 | 66803361 | rs11131630 | -0.177 | 0.419 | -0.251 | -1.808 | 0 | -0.017 | -0.017 | 0.054 | 0.32 | 0.37 | 0.29 | 0.22 | 0.34 | 0.3 |
| 4 | 66805499 | rs11733341 | -0.232 | 0.417 | -0.252 | -1.802 | -0.001 | -0.011 | -0.011 | 0 | 0.32 | 0.4 | 0.29 | 0.25 | 0.34 | 0.3 |
| 4 | 66811323 | rs10517951 | -0.378 | 0.596 | -0.273 | -1.655 | -0.003 | 0.019 | 0.019 | 0 | 0.22 | 0.2 | 0.23 | 0.12 | 0.23 | 0.25 |
| 4 | 66812771 | rs11945164 | -2.356 | 0.535 | -0.32 | -1.806 | 0 | 0.001 | 0.001 | 0.375 | 0.66 | 0.67 | 0.57 | 0.35 | 0.65 | 0.68 |
| 4 | 66822729 | rs17708692 | 0.728 | 0.786 | -0.662 | -1.587 | -0.006 | 0.019 | 0.019 | 0 | 0.12 | 0.05 | 0.16 | 0.09 | 0.11 | 0.18 |
| 4 | 66827845 | rs17708914 | 1.212 | 0.855 | -1.283 | -1.59 | -0.036 | 0.016 | 0.016 | 0 | 0.16 | 0.06 | 0.12 | 0.12 | 0.21 | 0.06 |
| 4 | 66881057 | rs7683217 | -0.332 | 0.952 | -1.293 | -1.497 | -0.008 | 0.03 | 0.03 | 0 | 0.21 | 0.17 | 0.24 | 0.09 | 0.23 | 0.24 |
| 4 | 66882613 | rs17709448 | -0.157 | 0.992 | -1.392 | -1.477 | -0.019 | 0.047 | 0.047 | 0 | 0.2 | 0.11 | 0.23 | 0.09 | 0.23 | 0.24 |
| 4 | 66899906 | rs13102215 | 1.635 | 1.058 | -1.393 | -1.329 | -0.047 | -0.022 | -0.022 | 0 | 0.1 | 0.01 | 0.08 | 0.12 | 0.14 | 0.05 |
| 4 | 66911964 | rs1979342 | 0.843 | 0.751 | -1.361 | -1.45 | -0.012 | 0.09 | 0.09 | 0.055 | 0.16 | 0.03 | 0.08 | 0 | 0.13 | 0.17 |
| 4 | 66917610 | rs6855976 | -0.43 | 0.753 | -1.306 | -1.492 | -0.003 | 0.105 | 0.105 | 0.075 | 0.28 | 0.12 | 0.25 | 0.09 | 0.24 | 0.35 |
| 4 | 66923059 | rs2588665 | -0.474 | 0.741 | -1.303 | -1.498 | -0.001 | 0.105 | 0.105 | 0.147 | 0.29 | 0.14 | 0.25 | 0.09 | 0.24 | 0.35 |
| 4 | 66949475 | rs2588681 | 0.003 | 0.741 | -1.057 | -1.603 | 0.009 | -0.036 | -0.036 | 1.72 | 0.63 | 0.51 | 0.67 | 0.78 | 0.59 | 0.67 |
| 4 | 66965155 | rs6850299 | -0.464 | 0.848 | -0.729 | -1.267 | 0.003 | 0.082 | 0.082 | 0.022 | 0.28 | 0.16 | 0.25 | 0.09 | 0.24 | 0.34 |
| 4 | 66984331 | rs13149202 | 0.942 | 0.975 | -0.557 | -1.219 | 0.002 | 0.001 | 0.001 | 0.001 | 0.21 | 0.2 | 0.21 | 0.21 | 0.22 | 0.24 |
| 4 | 66991420 | rs13110147 | 0.512 | 0.863 | -0.449 | -1.141 | 0.011 | -0.007 | -0.007 | 0 | 0.26 | 0.37 | 0.24 | 0.22 | 0.26 | 0.26 |
| 4 | 67005499 | rs11942020 | -0.198 | 0.447 | -0.446 | -1.177 | 0.002 | 0.001 | 0.001 | 0 | 0.25 | 0.22 | 0.24 | 0.09 | 0.21 | 0.34 |
| 4 | 67009474 | rs1461827 | -0.148 | 0.582 | -0.465 | -1.219 | 0.016 | -0.02 | -0.02 | 0.097 | 0.66 | 0.57 | 0.69 | 0.78 | 0.61 | 0.68 |
| 4 | 67012685 | rs6551997 | 1.43 | 0.621 | -0.675 | -1.122 | 0.007 | 0.004 | 0.004 | 1.155 | 0.4 | 0.3 | 0.45 | 0.66 | 0.4 | 0.33 |
| 4 | 67022718 | rs960865 | -0.591 | 0.58 | -0.94 | -0.88 | -0.004 | -0.026 | -0.026 | 0 | 0.26 | 0.26 | 0.24 | 0.08 | 0.2 | 0.34 |
| 4 | 67033158 | rs1459417 | -0.011 | 0.977 | -1.275 | -0.581 | -0.005 | 0.001 | 0.001 | 0 | 0.25 | 0.2 | 0.21 | 0.22 | 0.27 | 0.27 |
| 4 | 67033505 | rs10517955 | -0.25 | 0.554 | -1.137 | -0.351 | 0.001 | -0.008 | -0.008 | 0 | 0.11 | 0.09 | 0.07 | 0 | 0.08 | 0.14 |
| 4 | 67036815 | rs2588666 | -0.864 | 0.269 | -1.401 | -0.237 | 0.023 | 0.035 | 0.035 | 0 | 0.17 | 0.06 | 0.2 | 0.15 | 0.13 | 0.24 |
| 4 | 67041359 | rs2611059 | 0.479 | 0.395 | -1.988 | -0.106 | 0.013 | -0.025 | -0.025 | 3.008 | 0.47 | 0.16 | 0.54 | 0.45 | 0.54 | 0.38 |
| 4 | 67053769 | rs11725618 | 1.193 | 1.013 | -2.397 | -0.092 | 0.002 | 0.062 | 0.062 | 0 | 0.32 | 0.05 | 0.42 | 0.17 | 0.31 | 0.17 |
| 4 | 67055090 | rs13102033 | -0.04 | 0.861 | -2.011 | -0.215 | 0.03 | 0.008 | 0.008 | 0 | 0.49 | 0.25 | 0.54 | 0.44 | 0.54 | 0.38 |
| 4 | 67067806 | rs3919827 | -0.183 | 0.651 | -1.005 | -0.352 | 0.007 | -0.055 | -0.055 | 6.573 | 0.23 | 0.36 | 0.29 | 0.16 | 0.28 | 0.22 |
| 4 | 67069001 | rs2061459 | 0.911 | 0.642 | -1.182 | -0.308 | -0.022 | -0.059 | -0.059 | 0 | 0.45 | 0.06 | 0.53 | 0.45 | 0.54 | 0.37 |
| 4 | 67069599 | rs1993644 | -1.175 | 0.088 | -0.473 | -0.298 | -0.002 | -0.003 | -0.003 | 0.202 | 0.61 | 0.61 | 0.54 | 0.79 | 0.68 | 0.83 |
| 4 | 67075938 | rs2350804 | -1.216 | 0.115 | -0.412 | -0.313 | -0.014 | -0.019 | -0.019 | 1.222 | 0.62 | 0.56 | 0.54 | 0.79 | 0.68 | 0.83 |
| 4 | 67082181 | rs7680820 | 1.524 | 0.136 | -0.847 | -0.28 | 0.011 | 0.03 | 0.03 | 0 | 0.33 | 0.06 | 0.43 | 0.22 | 0.32 | 0.17 |
| 4 | 67087489 | rs7654157 | 1.183 | 0.405 | -0.673 | -0.148 | 0.012 | 0.079 | 0.079 | 0 | 0.39 | 0.06 | 0.46 | 0.21 | 0.41 | 0.21 |
| 4 | 67093452 | rs10011993 | -0.587 | 1.458 | 0.443 | 0.222 | -0.075 | -0.047 | -0.047 | 0.022 | 0.25 | 0.1 | 0.32 | 0.39 | 0.33 | 0.35 |
| 4 | 67104099 | rs6834031 | 1.675 | 0.827 | 1.563 | 0.127 | 0.061 | -0.005 | -0.005 | 0.985 | 0.92 | 0.55 | 0.95 | 1 | 0.93 | 0.98 |
| 4 | 67118667 | rs400524 | -1.71 | 1.162 | 1.453 | 0.187 | 0.13 | -0.028 | -0.028 | 0 | 0.1 | 0.42 | 0.08 | 0.01 | 0.1 | 0.07 |
| 4 | 67124222 | rs450788 | -1.25 | 1.006 | 1.595 | 0.164 | 0.033 | -0.022 | -0.022 | 0.398 | 0.07 | 0.29 | 0.04 | 0 | 0.05 | 0.04 |
| 4 | 67124441 | rs17086929 | -0.932 | 0.334 | 1.554 | 0.662 | -0.064 | 0.117 | 0.117 | 0.042 | 0.17 | 0.36 | 0.09 | 0.34 | 0.1 | 0.2 |
| 4 | 67125477 | rs535324 | -1.483 | 0.338 | 1.622 | 0.583 | 0.021 | -0.036 | -0.036 | 0 | 0.09 | 0.42 | 0.05 | 0 | 0.05 | 0.04 |
| 4 | 67126645 | rs434623 | -1.867 | 0.555 | 0.88 | 0.657 | 0.017 | -0.029 | -0.029 | 1.456 | 0.28 | 0.8 | 0.17 | 0.36 | 0.21 | 0.23 |
| 4 | 67139606 | rs17086968 | 0.468 | 0.699 | 1.118 | 0.951 | -0.017 | -0.061 | -0.061 | 0.102 | 0.1 | 0.01 | 0.13 | 0.18 | 0.08 | 0.11 |
| 4 | 67140834 | rs1397755 | -1.559 | 0.688 | 2.005 | 0.729 | -0.005 | 0.02 | 0.02 | 0.15 | 0.28 | 0.63 | 0.19 | 0.37 | 0.25 | 0.26 |
| 4 | 67147791 | rs12645637 | -0.872 | 0.66 | 2.26 | 0.726 | -0.01 | 0.017 | 0.017 | 0 | 0.2 | 0.31 | 0.13 | 0.35 | 0.15 | 0.19 |
| 4 | 67169493 | rs12642661 | -2.013 | 0.565 | 2.417 | 0.695 | 0.001 | -0.003 | -0.003 | 0.242 | 0.3 | 0.74 | 0.2 | 0.36 | 0.25 | 0.26 |
| 4 | 67183218 | rs12649988 | -1.66 | 0.505 | 2.437 | 0.708 | -0.025 | 0.016 | 0.016 | 1.039 | 0.27 | 0.67 | 0.18 | 0.35 | 0.2 | 0.24 |
| 4 | 67188285 | rs17087006 | -0.602 | 0.507 | 2.64 | 0.699 | 0.019 | -0.015 | -0.015 | 0 | 0.06 | 0.21 | 0.04 | 0 | 0.04 | 0.05 |
| 4 | 67188696 | rs17087008 | -0.818 | 0.534 | 2.302 | 0.751 | -0.037 | 0.069 | 0.069 | 0 | 0.2 | 0.44 | 0.14 | 0.35 | 0.15 | 0.19 |
| 4 | 67203187 | rs10009967 | -0.759 | 0.604 | 1.397 | 0.475 | 0.004 | -0.009 | -0.009 | 0 | 0.41 | 0.65 | 0.32 | 0.55 | 0.37 | 0.44 |
| 4 | 67210563 | rs2942544 | -2.128 | 0.997 | 0.22 | 0.094 | -0.061 | 0.061 | 0.061 | 0.388 | 0.78 | 0.98 | 0.61 | 0.94 | 0.73 | 0.88 |
| 4 | 67218350 | rs10034030 | -0.265 | 0.789 | 0.685 | 0.251 | 0.002 | -0.018 | -0.018 | 0 | 0.39 | 0.32 | 0.3 | 0.55 | 0.37 | 0.42 |
| 4 | 67227162 | rs13127224 | -0.045 | 0.8 | 0.623 | 0.41 | 0.022 | 0.011 | 0.011 | 0 | 0.37 | 0.24 | 0.27 | 0.4 | 0.34 | 0.39 |
| 4 | 67230762 | rs11729723 | 1.095 | 0.944 | 0.405 | -0.28 | -0.022 | -0.042 | -0.042 | 0 | 0.08 | 0.01 | 0.03 | 0.14 | 0.08 | 0.14 |
| 4 | 67232011 | rs1033095 | -0.602 | 0.881 | 1.113 | -0.461 | 0.126 | 0.005 | 0.005 | 0.526 | 0.08 | 0.45 | 0.06 | 0.01 | 0.06 | 0.04 |

| | | | | | | | | | | | | | | | | |
|---|---|---|---|---|---|---|---|---|---|---|---|---|---|---|---|---|
| 4 | 67234581 | rs1511279 | -0.045 | 1.041 | 0.715 | -0.398 | 0.028 | 0.004 | 0.004 | 0 | 0.38 | 0.22 | 0.4 | 0.65 | 0.35 | 0.39 |
| 4 | 67236121 | rs11945575 | -1.54 | 1.117 | 1.226 | -0.353 | 0.071 | 0.038 | 0.038 | 0.941 | 0.06 | 0.43 | 0.04 | 0 | 0.01 | 0.02 |
| 4 | 67237446 | rs2942543 | -2.041 | 1.191 | 0.653 | -0.437 | -0.059 | 0.1 | 0.1 | 1.395 | 0.78 | 0.98 | 0.63 | 0.97 | 0.73 | 0.85 |
| 4 | 67253757 | rs6552006 | -0.112 | 1.053 | 1.692 | 0.064 | 0.144 | 0.04 | 0.04 | 0.263 | 0.06 | 0.4 | 0.05 | 0.01 | 0.05 | 0.03 |
| 4 | 67257460 | rs10517980 | 0.181 | 0.88 | 1.643 | -0.081 | 0.002 | -0.002 | -0.002 | 0 | 0.12 | 0.12 | 0.09 | 0.01 | 0.11 | 0.1 |
| 4 | 67262225 | rs6823164 | -1.01 | 0.887 | 1.503 | 0.076 | 0.002 | 0.011 | 0.011 | 0 | 0.44 | 0.63 | 0.47 | 0.66 | 0.42 | 0.44 |
| 4 | 67287728 | rs6838213 | -0.538 | 0.918 | 0.537 | 0.078 | 0.097 | 0.004 | 0.004 | 2.054 | 0.07 | 0.58 | 0.06 | 0.01 | 0.05 | 0.02 |
| 4 | 67292325 | rs2175431 | -0.42 | 1.08 | 0.169 | 0.047 | 0.005 | 0.008 | 0.008 | 0 | 0.4 | 0.24 | 0.43 | 0.63 | 0.41 | 0.44 |
| 4 | 67293088 | rs4860743 | -1.093 | 1.101 | -0.321 | 0.132 | -0.05 | 0.057 | 0.057 | 0.001 | 0.48 | 0.9 | 0.5 | 0.65 | 0.46 | 0.46 |
| 4 | 67300802 | rs2009314 | -0.383 | 0.902 | 0.411 | -0.067 | 0.002 | 0.004 | 0.004 | 0.166 | 0.24 | 0.63 | 0.29 | 0.28 | 0.21 | 0.21 |
| 4 | 67301889 | rs1877515 | -0.806 | 0.45 | 0.066 | -0.468 | -0.002 | 0.005 | 0.005 | 0 | 0.38 | 0.33 | 0.43 | 0.35 | 0.37 | 0.44 |
| 4 | 67311736 | rs1996076 | 0.163 | 0.465 | -0.384 | -0.496 | 0.005 | 0.035 | 0.035 | 0 | 0.2 | 0.13 | 0.18 | 0.07 | 0.2 | 0.21 |
| 4 | 67314393 | rs1996074 | 0.188 | 0.43 | -0.178 | -0.455 | 0.011 | 0.041 | 0.041 | 0 | 0.21 | 0.41 | 0.27 | 0.28 | 0.17 | 0.22 |
| 4 | 67315227 | rs9647495 | 0.905 | 0.381 | -0.67 | -0.388 | 0.018 | -0.057 | -0.057 | 0.126 | 0.17 | 0.06 | 0.23 | 0.29 | 0.12 | 0.2 |
| 4 | 67334039 | rs1512965 | -0.546 | 0.589 | -0.544 | -0.278 | -0.002 | -0.005 | -0.005 | 0 | 0.42 | 0.55 | 0.45 | 0.35 | 0.37 | 0.44 |
| 4 | 67337614 | rs7663168 | -0.498 | 0.435 | -1.12 | -0.781 | 0.012 | 0.045 | 0.045 | 2.201 | 0.25 | 0.86 | 0.14 | 0.03 | 0.17 | 0.1 |
| 4 | 67345098 | rs11731608 | 0.539 | 0.962 | -1.425 | -1.16 | -0.026 | 0.112 | 0.112 | 0.94 | 0.6 | 0.2 | 0.53 | 0.93 | 0.53 | 0.73 |
| 4 | 67345899 | rs7669583 | -0.192 | 1.186 | -0.319 | -1.272 | -0.005 | -0.007 | -0.007 | 0 | 0.67 | 0.67 | 0.58 | 0.93 | 0.6 | 0.76 |
| 4 | 67347879 | rs17087196 | 0.454 | 1.188 | 0.192 | -1.223 | 0.2 | 0.032 | 0.032 | 0.076 | 0.06 | 0.33 | 0.04 | 0 | 0.07 | 0.02 |
| 4 | 67350474 | rs4860756 | 1.706 | 0.618 | -0.605 | -0.956 | -0.013 | -0.099 | -0.099 | 0 | 0.16 | 0.01 | 0.23 | 0.32 | 0.13 | 0.2 |
| 4 | 67358218 | rs1398548 | 1.141 | 0.456 | -0.69 | -1.108 | 0.018 | -0.054 | -0.054 | 0.022 | 0.21 | 0.09 | 0.26 | 0.32 | 0.16 | 0.24 |
| 4 | 67358778 | rs1398547 | 0.384 | 0.441 | -0.552 | -1.092 | -0.001 | -0.001 | -0.001 | 0 | 0.46 | 0.46 | 0.32 | 0.58 | 0.43 | 0.51 |
| 4 | 67362872 | rs1512945 | 1.008 | 0.489 | -0.382 | -0.907 | 0.016 | -0.031 | -0.031 | 0 | 0.21 | 0.17 | 0.26 | 0.33 | 0.16 | 0.24 |
| 4 | 67377865 | rs9312156 | 1.524 | 0.695 | -0.526 | -0.969 | -0.093 | 0.015 | 0.015 | 0 | 0.14 | 0.02 | 0.33 | 0.05 | 0.29 | 0.18 |
| 4 | 67382811 | rs1022126 | -0.159 | 0.119 | 0.028 | -0.774 | 0.201 | 0.02 | 0.02 | 1.655 | 0.93 | 0.54 | 0.95 | 1 | 0.93 | 0.97 |
| 4 | 67386470 | rs7676173 | 1.551 | -0.459 | -0.584 | -0.947 | 0.028 | -0.078 | -0.078 | 0 | 0.2 | 0.07 | 0.12 | 0.51 | 0.16 | 0.27 |
| 4 | 67389529 | rs8180319 | -0.19 | -0.37 | -0.293 | -0.781 | 0 | -0.009 | -0.009 | 0.039 | 0.7 | 0.21 | 0.67 | 0.67 | 0.77 | 0.73 |
| 4 | 67401602 | rs1510740 | -1.006 | -0.465 | 0.455 | -0.77 | 0.004 | -0.011 | -0.011 | 0.022 | 0.78 | 0.79 | 0.73 | 0.63 | 0.83 | 0.76 |
| 4 | 67423095 | rs17731697 | 1.181 | -0.359 | 0.491 | -0.834 | 0.025 | -0.022 | -0.022 | 2.316 | 0.05 | 0.1 | 0.02 | 0 | 0.07 | 0.02 |
| 4 | 67424395 | rs2203018 | -0.556 | -0.636 | 0.661 | -0.974 | 0.007 | -0.002 | -0.002 | 4.216 | 0.34 | 0.7 | 0.36 | 0.37 | 0.24 | 0.28 |
| 4 | 67432445 | rs1510742 | -0.588 | -0.654 | 0.499 | -0.943 | 0.013 | 0.004 | 0.004 | 0.205 | 0.35 | 0.76 | 0.32 | 0.33 | 0.25 | 0.28 |
| 4 | 67436220 | rs1510743 | -1.248 | -0.893 | 0.275 | -1.048 | 0.017 | -0.008 | -0.008 | 0.062 | 0.44 | 0.82 | 0.44 | 0.43 | 0.35 | 0.42 |
| 4 | 67438833 | rs2882598 | 0.765 | -0.917 | 0.513 | -1.069 | -0.014 | 0.015 | 0.015 | 0 | 0.24 | 0.29 | 0.23 | 0.32 | 0.17 | 0.25 |
| 4 | 67439729 | rs10013019 | -0.771 | -0.888 | 0.393 | -1.067 | 0.019 | -0.053 | -0.053 | 0.362 | 0.2 | 0.47 | 0.2 | 0.1 | 0.18 | 0.18 |
| 4 | 67448037 | rs10021655 | 1.104 | -1.148 | 0.124 | -1.378 | -0.13 | 0.183 | 0.183 | 0.064 | 0.35 | 0.1 | 0.44 | 0.09 | 0.48 | 0.29 |
| 4 | 67451328 | rs17087338 | 1.281 | -0.892 | -0.67 | -1.196 | 0.006 | -0.019 | -0.019 | 0 | 0.22 | 0.16 | 0.22 | 0.32 | 0.17 | 0.24 |
| 4 | 67453634 | rs17732645 | 0.424 | -0.557 | -0.98 | -0.928 | -0.058 | -0.018 | -0.018 | 0 | 0.14 | 0.02 | 0.13 | 0.13 | 0.17 | 0.17 |
| 4 | 67459961 | rs1397470 | -1.615 | -0.722 | -0.728 | -0.755 | 0.02 | -0.058 | -0.058 | 1.421 | 0.8 | 0.91 | 0.87 | 0.53 | 0.83 | 0.71 |
| 4 | 67463707 | rs4032303 | -0.379 | -0.653 | -0.322 | -0.706 | 0.037 | 0.005 | 0.005 | 0 | 0.34 | 0.71 | 0.31 | 0.33 | 0.2 | 0.26 |
| 4 | 67468459 | rs1510745 | -1.334 | -0.274 | 0.276 | -0.85 | -0.01 | -0.113 | -0.113 | 0.017 | 0.26 | 0.47 | 0.2 | 0.06 | 0.26 | 0.13 |
| 4 | 67470362 | rs11934386 | -0.386 | -0.309 | 0.521 | -0.851 | -0.017 | -0.063 | -0.063 | 0 | 0.15 | 0.19 | 0.09 | 0 | 0.12 | 0.04 |
| 4 | 67471664 | rs4860765 | 0.2 | -0.44 | 0.407 | -0.781 | 0 | 0.009 | 0.009 | 0 | 0.09 | 0.1 | 0.15 | 0.26 | 0.06 | 0.11 |
| 4 | 67471682 | rs4860766 | -1.818 | -0.374 | 0.618 | -0.801 | -0.008 | -0.013 | -0.013 | 0 | 0.35 | 0.53 | 0.35 | 0.31 | 0.32 | 0.24 |
| 4 | 67482764 | rs2136822 | 0.086 | -0.045 | 0.793 | -0.794 | -0.016 | 0.015 | 0.015 | 0 | 0.08 | 0.04 | 0.1 | 0.03 | 0.15 | 0.09 |
| 4 | 67501423 | rs1397453 | -1.675 | -0.098 | 1.003 | -0.783 | 0.018 | -0.01 | -0.01 | 0 | 0.25 | 0.55 | 0.33 | 0.27 | 0.27 | 0.21 |
| 4 | 67507665 | rs2203017 | -0.36 | -0.133 | 1.197 | -0.728 | -0.004 | 0.052 | 0.052 | 0 | 0.15 | 0.35 | 0.21 | 0.24 | 0.12 | 0.12 |
| 4 | 67523957 | rs4624710 | -0.549 | -0.121 | 1.199 | -0.444 | -0.007 | 0.126 | 0.126 | 0 | 0.16 | 0.49 | 0.22 | 0.29 | 0.12 | 0.12 |
| 4 | 67525255 | rs7695566 | 0.024 | -0.107 | 1.05 | -0.434 | 0.001 | -0.002 | -0.002 | 0 | 0.13 | 0.12 | 0.19 | 0.29 | 0.12 | 0.12 |
| 4 | 67567104 | rs4860771 | -1.499 | 0.259 | -0.295 | -0.232 | -0.062 | -0.183 | -0.183 | 0 | 0.73 | 0.98 | 0.66 | 0.51 | 0.68 | 0.67 |
| 4 | 67572774 | rs11733560 | -1.267 | 0.25 | 0.757 | 0.07 | 0.016 | 0.028 | 0.028 | 0 | 0.24 | 0.47 | 0.29 | 0.32 | 0.26 | 0.21 |
| 4 | 67589776 | rs17790467 | -0.121 | 0.543 | 0.032 | 0.105 | -0.002 | 0 | 0 | 0.002 | 0.06 | 0.04 | 0.13 | 0.07 | 0.11 | 0.05 |
| 4 | 67594125 | rs713435 | 1.15 | 0.643 | -0.192 | 0.006 | -0.039 | 0.015 | 0.015 | 0.13 | 0.49 | 0.11 | 0.5 | 0.58 | 0.54 | 0.54 |
| 4 | 67596504 | rs1530759 | 0.028 | 0.127 | 0.25 | -0.125 | -0.022 | -0.078 | -0.078 | 0 | 0.25 | 0.42 | 0.14 | 0.05 | 0.18 | 0.26 |
| 4 | 67599908 | rs10518014 | 0.202 | 0.275 | -0.297 | 0.086 | 0.005 | 0.019 | 0.019 | 0 | 0.75 | 0.69 | 0.7 | 0.67 | 0.77 | 0.8 |
| 4 | 67600200 | rs10518013 | -0.239 | -0.119 | -0.748 | -0.34 | 0 | -0.01 | -0.01 | 0 | 0.24 | 0.2 | 0.14 | 0.05 | 0.22 | 0.25 |
| 4 | 67606895 | rs10866173 | 0.971 | -0.519 | -0.779 | 0.331 | 0.05 | 0.01 | 0.01 | 0.48 | 0.59 | 0.23 | 0.61 | 0.63 | 0.7 | 0.65 |
| 4 | 67607007 | rs7655881 | 0.389 | -0.03 | -0.897 | 0.858 | 0.021 | -0.046 | -0.046 | 0.01 | 0.78 | 0.9 | 0.71 | 0.64 | 0.81 | 0.74 |
| 4 | 67613272 | rs13144587 | 0.816 | 0.199 | 0.193 | 1.151 | -0.006 | 0.073 | 0.073 | 0.192 | 0.78 | 0.35 | 0.72 | 0.58 | 0.79 | 0.65 |
| 4 | 67624510 | rs9996690 | -0.575 | -0.141 | 0.517 | 0.235 | -0.018 | -0.105 | -0.105 | 0 | 0.22 | 0.51 | 0.18 | 0.04 | 0.22 | 0.13 |
| 4 | 67630679 | rs1375890 | 1.148 | -1.225 | 0.234 | 0.595 | 0.057 | -0.011 | -0.011 | 0 | 0.62 | 0.21 | 0.7 | 0.58 | 0.74 | 0.69 |
| 4 | 67635469 | rs1375885 | 0.191 | -1.268 | 0.909 | 0.591 | -0.091 | 0.19 | 0.19 | 0 | 0.85 | 0.45 | 0.85 | 0.61 | 0.96 | 0.85 |
| 4 | 67641168 | rs6834814 | 0.111 | -1.278 | 1.397 | 0.668 | -0.082 | 0.147 | 0.147 | 0 | 0.85 | 0.59 | 0.87 | 0.66 | 0.96 | 0.85 |
| 4 | 67642371 | rs6842084 | -0.434 | -0.75 | 1.552 | 0.833 | 0.005 | -0.005 | -0.005 | 0 | 0.23 | 0.24 | 0.15 | 0.03 | 0.22 | 0.16 |
| 4 | 67644546 | rs1545429 | 1.299 | -0.42 | 1.793 | 1.219 | 0.014 | 0.409 | 0.409 | 0 | 0.94 | 0.68 | 0.89 | 0.61 | 0.98 | 0.87 |
| 4 | 67645785 | rs11131652 | 1.67 | -0.424 | 1.343 | 1.155 | 0.018 | 0.419 | 0.419 | 0.134 | 0.92 | 0.46 | 0.87 | 0.66 | 0.98 | 0.87 |
| 4 | 67654478 | rs1450038 | -1.301 | 1.128 | 0.993 | 0.637 | 0.035 | -0.04 | -0.04 | 0 | 0.1 | 0.2 | 0.13 | 0 | 0.22 | 0.1 |
| 4 | 67657239 | rs1450036 | 1.269 | 1.265 | 0.958 | 0.861 | 0.058 | 0.069 | 0.069 | 0.445 | 0.77 | 0.4 | 0.69 | 0.61 | 0.65 | 0.64 |
| 4 | 67661122 | rs7697495 | 1.525 | 1.312 | 1.293 | 0.497 | 0.272 | 0.015 | 0.015 | 0 | 0.93 | 0.5 | 0.89 | 1 | 0.84 | 0.91 |
| 4 | 67668580 | rs997185 | -0.722 | 1.244 | 1.18 | 0.648 | 0 | 0 | 0 | 0.429 | 0.16 | 0.11 | 0.19 | 0.26 | 0.18 | 0.26 |

| | | | | | | | | | | | | | | | | |
|---|---|---|---|---|---|---|---|---|---|---|---|---|---|---|---|---|
| 4 | 67673566 | rs1495103 | 1.445 | 1.417 | 0.935 | 0.635 | 0.043 | 0.001 | 0.001 | 0.586 | 0.76 | 0.36 | 0.69 | 0.72 | 0.62 | 0.59 |
| 4 | 67677853 | rs1353320 | 0.934 | 0.606 | 0.893 | 0.714 | -0.002 | 0.029 | 0.029 | 0.016 | 0.8 | 0.43 | 0.77 | 0.72 | 0.78 | 0.67 |
| 4 | 67686485 | rs17736022 | -0.674 | 0.554 | -0.109 | 0.376 | -0.034 | 0.035 | 0.035 | 0 | 0.26 | 0.06 | 0.17 | 0.18 | 0.27 | 0.37 |
| 4 | 67708427 | rs12506243 | 0.738 | 0.065 | -0.84 | 0.155 | -0.178 | -0.033 | -0.033 | 0 | 0.27 | 0.02 | 0.47 | 0.18 | 0.46 | 0.19 |
| 4 | 67716110 | rs1450024 | 0.686 | 0.118 | 0.268 | 0.166 | 0 | -0.002 | -0.002 | 0 | 0.67 | 0.34 | 0.69 | 0.66 | 0.7 | 0.59 |
| 4 | 67716733 | rs12507395 | 1.069 | -0.109 | 0.81 | 0.308 | 0.02 | 0.187 | 0.187 | 1.704 | 0.93 | 0.43 | 0.87 | 0.84 | 0.98 | 0.95 |
| 4 | 67722262 | rs12512195 | -0.006 | -0.054 | 0.495 | -0.424 | -0.01 | -0.047 | -0.047 | 0.003 | 0.8 | 0.68 | 0.84 | 0.88 | 0.87 | 0.66 |
| 4 | 67733165 | rs11726617 | -0.128 | -0.047 | 0.529 | -0.421 | -0.007 | -0.033 | -0.033 | 0 | 0.81 | 0.75 | 0.85 | 0.88 | 0.86 | 0.66 |
| 4 | 67733857 | rs1155865 | 0.139 | -0.214 | 1.235 | -0.502 | -0.019 | -0.066 | -0.066 | 0.008 | 0.79 | 0.53 | 0.84 | 0.88 | 0.86 | 0.66 |
| 4 | 67734753 | rs1868599 | -0.038 | -0.211 | 1.248 | -0.501 | 0.001 | 0.008 | 0.008 | 0.825 | 0.18 | 0.14 | 0.14 | 0.12 | 0.14 | 0.34 |
| 4 | 67741837 | rs17799404 | 0.864 | -0.031 | 0.272 | -1.059 | -0.105 | 0.113 | 0.113 | 0 | 0.28 | 0.09 | 0.48 | 0.04 | 0.5 | 0.17 |
| 4 | 67758372 | rs9999249 | -0.274 | -0.079 | 0.739 | -1.002 | -0.018 | -0.07 | -0.07 | 0 | 0.79 | 0.55 | 0.84 | 0.88 | 0.86 | 0.66 |
| 4 | 67759254 | rs1481253 | -0.346 | -0.078 | 0.776 | -0.976 | -0.015 | -0.055 | -0.055 | 0 | 0.79 | 0.58 | 0.84 | 0.88 | 0.86 | 0.66 |
| 4 | 67762025 | rs7671625 | 0.409 | -0.656 | 0.218 | -1.363 | 0.004 | 0.009 | 0.009 | 0 | 0.3 | 0.25 | 0.18 | 0.05 | 0.27 | 0.19 |
| 4 | 67762959 | rs6818708 | 0.443 | -0.657 | 0.219 | -1.343 | -0.001 | -0.034 | -0.034 | 4.849 | 0.31 | 0.32 | 0.19 | 0.06 | 0.27 | 0.19 |
| 4 | 67763820 | rs1481245 | -0.946 | -0.615 | -0.142 | -1.323 | -0.015 | 0.165 | 0.165 | 0 | 0.55 | 0.84 | 0.37 | 0.17 | 0.44 | 0.55 |
| 4 | 67774097 | rs1585959 | -0.237 | -0.846 | 0.127 | -0.87 | -0.024 | -0.061 | -0.061 | 0.214 | 0.77 | 0.49 | 0.82 | 0.88 | 0.85 | 0.65 |
| 4 | 67802992 | rs923994 | -0.67 | -0.066 | -0.501 | -0.999 | -0.014 | 0.011 | 0.011 | 0 | 0.87 | 0.92 | 0.81 | 0.94 | 0.79 | 0.84 |
| 4 | 67807854 | rs11947612 | 0.901 | -0.608 | -0.666 | -0.409 | -0.026 | 0.003 | 0.003 | 0 | 0.6 | 0.13 | 0.63 | 0.62 | 0.63 | 0.59 |
| 4 | 67808949 | rs1390067 | 0.116 | -0.625 | -0.424 | -0.402 | -0.005 | 0.085 | 0.085 | 0.028 | 0.06 | 0.11 | 0.08 | 0.28 | 0.05 | 0.05 |
| 4 | 67809075 | rs1390066 | -0.379 | -0.633 | -0.424 | -0.408 | 0.015 | -0.007 | -0.007 | 0 | 0.27 | 0.79 | 0.18 | 0.32 | 0.16 | 0.25 |
| 4 | 67811092 | rs9999351 | 0.645 | -0.626 | -0.076 | -0.415 | -0.036 | -0.083 | -0.083 | 0 | 0.2 | 0.39 | 0.09 | 0.04 | 0.11 | 0.2 |
| 4 | 67813017 | rs2164300 | 1.779 | -1.043 | -0.647 | -0.761 | -0.18 | 0.111 | 0.111 | 0 | 0.26 | 0.02 | 0.39 | 0.04 | 0.49 | 0.18 |
| 4 | 67813997 | rs9998536 | 0.877 | -1.038 | -0.484 | -0.705 | -0.036 | -0.083 | -0.083 | 0.146 | 0.2 | 0.39 | 0.09 | 0.04 | 0.11 | 0.2 |
| 4 | 67814133 | rs10031872 | 0.304 | -0.991 | -0.463 | -0.667 | -0.005 | 0.085 | 0.085 | 0.028 | 0.06 | 0.11 | 0.08 | 0.28 | 0.05 | 0.05 |
| 4 | 67815068 | rs11938473 | 0.054 | -1.285 | -0.719 | -0.814 | -0.021 | -0.052 | -0.052 | 0 | 0.37 | 0.41 | 0.14 | 0.08 | 0.2 | 0.35 |
| 4 | 67820174 | rs2035366 | -0.248 | -1.368 | -0.736 | -0.746 | -0.013 | -0.002 | -0.002 | 0.993 | 0.39 | 0.41 | 0.19 | 0.36 | 0.22 | 0.4 |
| 4 | 67826009 | rs2019834 | -0.241 | -1.371 | -0.728 | -0.743 | -0.008 | 0 | 0 | 0 | 0.39 | 0.39 | 0.2 | 0.39 | 0.22 | 0.4 |
| 4 | 67828945 | rs10518015 | 1.057 | -1.509 | -0.737 | -0.784 | -0.039 | -0.061 | -0.061 | 0.014 | 0.19 | 0.37 | 0.09 | 0.06 | 0.09 | 0.2 |
| 4 | 67839637 | rs11736089 | 1.327 | -1.487 | -0.907 | -0.788 | 0.027 | 0.054 | 0.054 | 0 | 0.17 | 0.01 | 0.05 | 0.06 | 0.1 | 0.16 |
| 4 | 67843376 | rs12500685 | 0.633 | -1.238 | -0.674 | -0.763 | 0 | 0.002 | 0.002 | 0 | 0.47 | 0.48 | 0.61 | 0.58 | 0.56 | 0.44 |
| 4 | 67872428 | rs987811 | -2.594 | -1.023 | -0.811 | -0.729 | 0.001 | -0.02 | -0.02 | 1.255 | 0.93 | 0.95 | 0.94 | 1 | 0.9 | 0.99 |
| 4 | 67878373 | rs17746145 | 0.945 | -0.967 | -0.863 | -0.703 | -0.028 | 0.011 | 0.011 | 0 | 0.06 | 0.01 | 0.13 | 0.04 | 0.12 | 0.15 |
| 4 | 67884593 | rs1435423 | -0.454 | -1.159 | -0.72 | -0.736 | 0.001 | 0 | 0 | 0 | 0.45 | 0.42 | 0.33 | 0.42 | 0.35 | 0.55 |
| 4 | 67891641 | rs13141210 | 1.507 | -1.042 | -0.955 | -0.107 | -0.123 | 0.02 | 0.02 | 0 | 0.26 | 0.09 | 0.41 | 0.16 | 0.5 | 0.19 |
| 4 | 67899235 | rs12506221 | 1.12 | -0.416 | -0.424 | -0.174 | -0.019 | -0.001 | -0.001 | 1.86 | 0.43 | 0.29 | 0.59 | 0.56 | 0.54 | 0.43 |
| 4 | 67904595 | rs787361 | -1.119 | -0.33 | -0.488 | -0.122 | -0.001 | 0.008 | 0.008 | 0 | 0.54 | 0.59 | 0.4 | 0.44 | 0.44 | 0.57 |
| 4 | 67906156 | rs787363 | 0.168 | 0.387 | -0.591 | 0.346 | 0.046 | -0.007 | -0.007 | 0 | 0.06 | 0.08 | 0.13 | 0.06 | 0.12 | 0.14 |
| 4 | 67910084 | rs11940944 | -0.051 | -0.141 | -0.887 | 0.543 | 0 | -0.111 | -0.111 | 2.678 | 0.2 | 0.02 | 0.11 | 0.35 | 0.14 | 0.21 |
| 4 | 67911404 | rs787370 | 1.146 | 0.267 | -0.103 | 0.158 | 0.004 | -0.023 | -0.023 | 2.457 | 0.92 | 0.72 | 0.87 | 0.92 | 0.92 | 0.78 |
| 4 | 67921630 | rs12645870 | -0.674 | 0.53 | -0.294 | 0.187 | 0.002 | 0 | 0 | 0 | 0.09 | 0.1 | 0.05 | 0.12 | 0.12 | 0.13 |
| 4 | 67925197 | rs12186186 | -1.152 | 0.143 | -0.523 | -0.092 | 0.013 | 0.027 | 0.027 | 0 | 0.38 | 0.26 | 0.23 | 0.33 | 0.32 | 0.44 |
| 4 | 67939089 | rs1425664 | -1.421 | 0.239 | -0.776 | -0.358 | 0.008 | 0.083 | 0.083 | 0 | 0.47 | 0.26 | 0.31 | 0.33 | 0.43 | 0.45 |
| 4 | 67954517 | rs1425653 | -0.477 | 0.455 | -0.439 | 0.122 | 0.019 | 0.09 | 0.09 | 0 | 0.95 | 0.83 | 0.88 | 0.82 | 0.95 | 0.89 |
| 4 | 67958264 | rs10010944 | 0.03 | 0.366 | -0.607 | -0.006 | -0.041 | 0.032 | 0.032 | 0 | 0.18 | 0.06 | 0.12 | 0.12 | 0.24 | 0.14 |
| 4 | 67961296 | rs10018066 | -0.623 | 0.15 | -0.639 | -0.122 | -0.007 | 0.012 | 0.012 | 0 | 0.23 | 0.19 | 0.17 | 0.19 | 0.27 | 0.22 |
| 4 | 67972601 | rs17088003 | 0.192 | 0.105 | -0.774 | -0.083 | 0.037 | 0.051 | 0.051 | 0 | 0.26 | 0.08 | 0.15 | 0.14 | 0.17 | 0.23 |
| 4 | 67994618 | rs10866177 | -0.463 | 0.244 | -0.713 | -0.061 | -0.019 | 0.028 | 0.028 | 0 | 0.3 | 0.1 | 0.18 | 0.25 | 0.28 | 0.36 |
| 4 | 68006895 | rs6818650 | 0.082 | 0.212 | -0.583 | -0.169 | 0.001 | 0.007 | 0.007 | 0 | 0.17 | 0.14 | 0.12 | 0.08 | 0.17 | 0.11 |
| 4 | 68007388 | rs10518020 | 0.757 | 0.067 | -0.721 | -0.264 | 0.009 | 0.082 | 0.082 | 0 | 0.18 | 0.04 | 0.06 | 0.03 | 0.12 | 0.14 |
| 4 | 68007509 | rs11131679 | -0.021 | 0.114 | -0.356 | -0.124 | -0.001 | -0.017 | -0.017 | 0 | 0.18 | 0.21 | 0.13 | 0.08 | 0.17 | 0.11 |
| 4 | 68019294 | rs933830 | 0.481 | 0.166 | -0.192 | 0.041 | -0.016 | 0.001 | 0.001 | 0 | 0.74 | 0.56 | 0.74 | 0.73 | 0.78 | 0.79 |
| 4 | 68024153 | rs730934 | 1.813 | 0.088 | -0.356 | 0.072 | -0.021 | -0.021 | -0.021 | 0.428 | 0.44 | 0.25 | 0.53 | 0.45 | 0.5 | 0.41 |
| 4 | 68038559 | rs7669686 | -1.061 | 0.14 | -0.409 | -0.115 | -0.019 | 0.059 | 0.059 | 0.15 | 0.4 | 0.23 | 0.28 | 0.27 | 0.42 | 0.4 |
| 4 | 68039482 | rs28534578 | 0.706 | 0.023 | 0.059 | 0.086 | 0.001 | -0.032 | -0.032 | 0 | 0.13 | 0.15 | 0.1 | 0.02 | 0.14 | 0.03 |
| 4 | 68056613 | rs7690543 | 0.565 | -0.183 | 0.155 | 0.45 | -0.04 | 0.101 | 0.101 | 0 | 0.84 | 0.47 | 0.82 | 0.75 | 0.92 | 0.81 |
| 4 | 68058793 | rs1425660 | -0.525 | 1.082 | -0.546 | 0.048 | -0.013 | 0.028 | 0.028 | 0 | 0.39 | 0.26 | 0.35 | 0.25 | 0.51 | 0.4 |
| 4 | 68059850 | rs6854985 | -0.586 | 1.196 | -0.867 | -0.324 | -0.016 | 0.045 | 0.045 | 0 | 0.17 | 0.08 | 0.14 | 0.05 | 0.21 | 0.14 |
| 4 | 68064440 | rs10518025 | -0.088 | 1.052 | -0.287 | -0.307 | 0.005 | 0.007 | 0.007 | 0 | 0.09 | 0.17 | 0.15 | 0.11 | 0.12 | 0.19 |
| 4 | 68064531 | rs1425658 | -1.632 | 1.069 | -1.463 | -1.279 | -0.024 | 0.096 | 0.096 | 1.744 | 0.74 | 0.92 | 0.84 | 0.98 | 0.69 | 0.91 |
| 4 | 68065150 | rs9312170 | -0.831 | 0.889 | -1.866 | -1.233 | -0.063 | 0.107 | 0.107 | 0 | 0.34 | 0.15 | 0.38 | 0.08 | 0.45 | 0.31 |
| 4 | 68067034 | rs11942221 | 1.018 | 0.856 | -1.99 | -0.15 | -0.027 | 0.039 | 0.039 | 0 | 0.17 | 0.08 | 0.25 | 0.02 | 0.23 | 0.17 |
| 4 | 68069022 | rs12639670 | 0.215 | -0.12 | -1.974 | 0.351 | 0.079 | -0.087 | -0.087 | 0 | 0.28 | 0.07 | 0.28 | 0.46 | 0.14 | 0.33 |
| 4 | 68070195 | rs17088117 | 0.67 | 0.864 | -1.821 | 0.362 | -0.04 | -0.019 | -0.019 | 0.388 | 0.07 | 0 | 0.06 | 0.08 | 0.11 | 0.08 |
| 4 | 68071781 | rs13151209 | -0.096 | 0.845 | -1.875 | 0.39 | -0.01 | -0.002 | -0.002 | 0 | 0.7 | 0.75 | 0.62 | 0.69 | 0.65 | 0.66 |
| 4 | 68074583 | rs1549448 | 1.07 | -0.271 | -2.201 | 0.312 | 0.076 | 0.024 | 0.024 | 0.368 | 0.37 | 0.11 | 0.3 | 0.27 | 0.24 | 0.3 |
| 4 | 68075795 | rs6814470 | 0.45 | -0.1 | -1.793 | 0.475 | 0.021 | 0.01 | 0.01 | 0 | 0.48 | 0.58 | 0.36 | 0.37 | 0.29 | 0.36 |
| 4 | 68085262 | rs1425655 | -0.799 | 0.118 | -1.572 | 0.189 | 0.01 | 0.009 | 0.009 | 0.021 | 0.49 | 0.41 | 0.65 | 0.42 | 0.51 | 0.61 |
| 4 | 68088235 | rs6819869 | 0.741 | 0.231 | -1.733 | 0.276 | 0 | 0 | 0 | 0.015 | 0.46 | 0.58 | 0.3 | 0.39 | 0.41 | 0.29 |

| | | | | | | | | | | | | | | | | |
|---|---|---|---|---|---|---|---|---|---|---|---|---|---|---|---|---|
| 4 | 68093400 | rs4860238 | 0.655 | 0.348 | -2.093 | 0.274 | 0.017 | -0.037 | -0.037 | 1.65 | 0.23 | 0.04 | 0.25 | 0.22 | 0.16 | 0.31 |
| 4 | 68095870 | rs4639131 | 0.008 | 0.644 | -1.713 | 0.386 | 0.003 | 0.001 | 0.001 | 0 | 0.7 | 0.7 | 0.55 | 0.61 | 0.58 | 0.6 |
| 4 | 68101830 | rs1866784 | 0.375 | 0.565 | -1.783 | 0.222 | 0.001 | -0.002 | -0.002 | 0 | 0.25 | 0.21 | 0.39 | 0.12 | 0.33 | 0.25 |
| 4 | 68102182 | rs17766472 | 1.453 | 0.175 | -2.015 | -0.087 | -0.028 | 0.199 | 0.199 | 0.393 | 0.33 | 0.06 | 0.19 | 0.01 | 0.35 | 0.1 |
| 4 | 68104490 | rs4133270 | 0.641 | 0.254 | -1.886 | 0.263 | -0.016 | -0.021 | -0.021 | 0.212 | 0.07 | 0.02 | 0.06 | 0.28 | 0.1 | 0.16 |
| 4 | 68106106 | rs11131687 | 0.263 | 0.369 | -1.789 | -0.197 | 0 | -0.002 | -0.002 | 0.128 | 0.63 | 0.36 | 0.63 | 0.24 | 0.57 | 0.49 |
| 4 | 68113850 | rs10005602 | -0.19 | 0.373 | -1.539 | 0.287 | 0.009 | 0.003 | 0.003 | 0 | 0.3 | 0.23 | 0.43 | 0.27 | 0.22 | 0.4 |
| 4 | 68116539 | rs1158408 | -0.347 | 0.617 | -0.976 | 0.552 | 0.078 | 0.19 | 0.19 | 0.479 | 0.83 | 0.53 | 0.74 | 0.63 | 0.78 | 0.74 |
| 4 | 68117200 | rs4860822 | 1.137 | 0.636 | -1.231 | 0.571 | -0.07 | -0.068 | -0.068 | 16.546 | 0.15 | 0.01 | 0.23 | 0.22 | 0.2 | 0.24 |
| 4 | 68123521 | rs2218192 | 1.306 | 0.677 | -1.071 | 0.598 | -0.014 | -0.002 | -0.002 | 0 | 0.09 | 0.06 | 0.17 | 0.1 | 0.19 | 0.11 |
| 4 | 68123616 | rs1440556 | -1.205 | 0.582 | -1.018 | 0.618 | 0.032 | 0.324 | 0.324 | 0.54 | 0.93 | 0.6 | 0.91 | 0.73 | 0.97 | 0.85 |
| 4 | 68127826 | rs1371864 | -0.258 | 0.61 | -1.548 | 0.38 | 0.047 | 0.136 | 0.136 | 1.382 | 0.82 | 0.44 | 0.73 | 0.62 | 0.77 | 0.74 |
| 4 | 68132509 | rs17767450 | 1.281 | 0.834 | -2.345 | -0.448 | -0.031 | 0.008 | 0.008 | 0.377 | 0.1 | 0.05 | 0.17 | 0.07 | 0.2 | 0.11 |
| 4 | 68154530 | rs1946918 | 1.154 | 0.807 | -2.395 | -0.414 | -0.041 | -0.049 | -0.049 | 0 | 0.15 | 0.05 | 0.21 | 0.26 | 0.2 | 0.24 |
| 4 | 68155364 | rs17818614 | 1.019 | 0.532 | -1.907 | -0.188 | -0.022 | 0.007 | 0.007 | 0.306 | 0.1 | 0.07 | 0.18 | 0.07 | 0.2 | 0.11 |
| 4 | 68159587 | rs11724259 | 0.846 | 0.322 | -2.219 | -0.71 | 0.02 | 0.017 | 0.017 | 3.579 | 0.24 | 0.17 | 0.16 | 0.01 | 0.18 | 0.06 |
| 4 | 68159785 | rs6852744 | -0.443 | 0.763 | -2.366 | -0.098 | -0.026 | 0.048 | 0.048 | 0.056 | 0.47 | 0.08 | 0.4 | 0.34 | 0.58 | 0.45 |
| 4 | 68167336 | rs28373882 | 0.016 | 0.969 | -1.62 | -0.18 | -0.037 | 0.049 | 0.049 | 0 | 0.09 | 0.01 | 0.08 | 0 | 0.15 | 0.02 |
| 4 | 68174701 | rs1440559 | 0.03 | 0.405 | -1.474 | -0.615 | 0.009 | 0.012 | 0.012 | 0 | 0.29 | 0.25 | 0.24 | 0.08 | 0.24 | 0.2 |
| 4 | 68176106 | rs9985687 | -0.684 | 0.24 | -1.273 | -1.163 | 0 | -0.055 | -0.055 | 0 | 0.54 | 0.51 | 0.41 | 0.09 | 0.5 | 0.28 |
| 4 | 68183082 | rs12504640 | -0.178 | 0.107 | -1.099 | -0.092 | -0.004 | 0.005 | 0.005 | 1.421 | 0.28 | 0.17 | 0.31 | 0.25 | 0.29 | 0.24 |
| 4 | 68186030 | rs1899130 | 0.36 | 0.101 | -1.29 | 0.052 | 0.014 | 0.042 | 0.042 | 0 | 0.4 | 0.27 | 0.29 | 0.27 | 0.35 | 0.44 |
| 4 | 68192816 | rs1371863 | -1.731 | 0.011 | -0.603 | -0.127 | -0.012 | 0.107 | 0.107 | 0 | 0.89 | 0.81 | 0.85 | 0.65 | 0.93 | 0.86 |
| 4 | 68193125 | rs1348058 | 1.01 | 0.016 | -0.384 | -0.398 | 0.001 | 0.004 | 0.004 | 0.44 | 0.2 | 0.2 | 0.36 | 0.35 | 0.28 | 0.32 |
| 4 | 68198578 | rs746877 | 0.47 | -0.051 | -0.427 | -0.399 | 0.007 | 0.005 | 0.005 | 0.393 | 0.56 | 0.42 | 0.5 | 0.4 | 0.6 | 0.6 |
| 4 | 68199483 | rs17088281 | 0.901 | 0.078 | -0.419 | -0.72 | 0.006 | -0.022 | -0.022 | 2.445 | 0.06 | 0.18 | 0.04 | 0.01 | 0.04 | 0.02 |
| 4 | 68207449 | rs1946920 | 0.271 | 0.079 | -0.353 | -0.717 | 0 | -0.038 | -0.038 | 0 | 0.28 | 0.21 | 0.19 | 0.02 | 0.3 | 0.1 |
| 4 | 68215003 | rs1497431 | 0.692 | 0.228 | -0.323 | -0.317 | 0.019 | 0.036 | 0.036 | 1.795 | 0.2 | 0.31 | 0.34 | 0.35 | 0.28 | 0.33 |
| 4 | 68222094 | rs356310 | -0.351 | 0.158 | -0.335 | -1.171 | 0.016 | -0.011 | -0.011 | 0 | 0.67 | 0.61 | 0.56 | 0.31 | 0.64 | 0.51 |
| 4 | 68222294 | rs10518036 | 0.603 | -0.28 | -0.743 | -1.778 | 0.006 | 0.027 | 0.027 | 0 | 0.33 | 0.09 | 0.34 | 0.25 | 0.31 | 0.36 |
| 4 | 68224591 | rs356306 | 2.085 | -0.322 | -0.585 | -1.807 | 0.004 | -0.017 | -0.017 | 0.343 | 0.09 | 0.23 | 0.06 | 0.06 | 0.07 | 0.04 |
| 4 | 68228744 | rs356335 | -1.6 | -0.289 | -0.5 | -1.801 | -0.01 | 0.077 | 0.077 | 0 | 0.84 | 0.74 | 0.73 | 0.42 | 0.89 | 0.68 |
| 4 | 68237962 | rs10008670 | 0.188 | -0.308 | -0.506 | -1.902 | -0.01 | 0.076 | 0.076 | 0.323 | 0.53 | 0.31 | 0.46 | 0.25 | 0.51 | 0.43 |
| 4 | 68241071 | rs6824881 | 0.569 | -0.33 | -0.461 | -1.916 | -0.002 | -0.024 | -0.024 | 0.446 | 0.37 | 0.31 | 0.3 | 0.11 | 0.47 | 0.26 |
| 4 | 68241081 | rs920482 | -0.894 | -0.343 | -0.383 | -1.816 | -0.01 | 0.038 | 0.038 | 0 | 0.8 | 0.71 | 0.7 | 0.42 | 0.85 | 0.65 |
| 4 | 68250856 | rs7669658 | 0.206 | -0.599 | -0.484 | -2.025 | -0.005 | -0.001 | -0.001 | 0.486 | 0.44 | 0.34 | 0.49 | 0.13 | 0.52 | 0.3 |
| 4 | 68252674 | rs13103397 | 0.913 | -0.578 | -0.539 | -2.01 | -0.008 | 0.016 | 0.016 | 0 | 0.26 | 0.16 | 0.16 | 0.23 | 0.26 | 0.3 |
| 4 | 68253127 | rs356343 | 2.258 | -0.553 | -0.568 | -1.944 | 0.002 | -0.01 | -0.01 | 0.068 | 0.08 | 0.19 | 0.05 | 0.06 | 0.08 | 0.04 |
| 4 | 68262262 | rs1497427 | 0.904 | -0.556 | -0.646 | -1.99 | 0.003 | 0.039 | 0.039 | 2.06 | 0.25 | 0.13 | 0.32 | 0.02 | 0.26 | 0.12 |
| 4 | 68263831 | rs3932174 | -1.611 | -0.428 | -0.617 | -1.898 | -0.002 | -0.021 | -0.021 | 0.038 | 0.88 | 0.77 | 0.91 | 0.93 | 0.89 | 0.94 |
| 4 | 68267437 | rs17428139 | 1.973 | -0.417 | -0.639 | -1.895 | -0.008 | 0.007 | 0.007 | 1.414 | 0.17 | 0.11 | 0.17 | 0.11 | 0.25 | 0.18 |
| 4 | 68269400 | rs3935662 | 0.752 | -0.511 | -0.661 | -1.863 | -0.007 | 0.011 | 0.011 | 0 | 0.27 | 0.17 | 0.15 | 0.24 | 0.26 | 0.3 |
| 4 | 68289606 | rs4860843 | -0.82 | -0.32 | -0.612 | -1.848 | 0.014 | -0.043 | -0.043 | 0 | 0.71 | 0.62 | 0.76 | 0.83 | 0.65 | 0.76 |
| 4 | 68293285 | rs7696609 | 0.881 | -0.47 | -0.711 | -1.952 | 0.003 | 0.094 | 0.094 | 4.509 | 0.27 | 0.09 | 0.34 | 0.02 | 0.28 | 0.13 |
| 4 | 68337435 | rs1874633 | -0.767 | -0.164 | -0.694 | -1.887 | 0.007 | -0.031 | -0.031 | 0 | 0.69 | 0.62 | 0.76 | 0.83 | 0.63 | 0.72 |
| 4 | 68369968 | rs17088445 | 2.646 | -0.139 | -0.729 | -1.914 | -0.023 | -0.01 | -0.01 | 0 | 0.07 | 0 | 0.02 | 0.07 | 0.07 | 0.03 |
| 4 | 68377472 | rs355466 | -0.746 | -0.166 | -0.708 | -1.903 | 0.008 | -0.036 | -0.036 | 0 | 0.69 | 0.62 | 0.75 | 0.83 | 0.63 | 0.72 |
| 4 | 68379299 | rs355462 | -1.559 | -0.171 | -0.668 | -1.831 | -0.006 | -0.022 | -0.022 | 0 | 0.88 | 0.77 | 0.93 | 0.93 | 0.9 | 0.94 |
| 4 | 68381367 | rs173317 | 1.376 | -0.311 | -0.63 | -1.69 | -0.009 | 0.074 | 0.074 | 0 | 0.16 | 0.25 | 0.26 | 0.57 | 0.1 | 0.33 |
| 4 | 68385372 | rs13104130 | 1.766 | -0.258 | -0.811 | -1.932 | -0.024 | 0.086 | 0.086 | 2.166 | 0.2 | 0.04 | 0.15 | 0.01 | 0.2 | 0.07 |
| 4 | 68410009 | rs17625547 | 1.269 | 0.01 | -0.693 | -1.739 | -0.005 | 0.003 | 0.003 | 2.397 | 0.19 | 0.14 | 0.18 | 0.1 | 0.28 | 0.22 |
| 4 | 68412148 | rs957582 | 0.339 | -0.143 | -0.562 | -1.734 | -0.007 | -0.056 | -0.056 | 0.142 | 0.16 | 0.37 | 0.09 | 0.07 | 0.12 | 0.1 |
| 4 | 68415286 | rs13119483 | 0.602 | -0.295 | -0.562 | -1.737 | -0.002 | 0.214 | 0.214 | 0 | 0.48 | 0.17 | 0.52 | 0.12 | 0.56 | 0.37 |
| 4 | 68419485 | rs7700004 | -0.281 | -0.377 | -0.476 | -1.617 | -0.024 | 0.028 | 0.028 | 0 | 0.72 | 0.47 | 0.67 | 0.35 | 0.8 | 0.6 |
| 4 | 68428733 | rs7681780 | -0.897 | -0.378 | -0.451 | -1.627 | -0.026 | 0.127 | 0.127 | 0 | 0.79 | 0.49 | 0.7 | 0.41 | 0.86 | 0.63 |
| 4 | 68428980 | rs7682206 | 0.671 | -0.294 | -0.532 | -1.658 | -0.023 | -0.013 | -0.013 | 0 | 0.22 | 0.08 | 0.14 | 0.23 | 0.24 | 0.25 |
| 4 | 68436883 | rs13111555 | -1.743 | -0.323 | -0.477 | -1.586 | -0.009 | 0.099 | 0.099 | 0 | 0.84 | 0.73 | 0.74 | 0.43 | 0.89 | 0.66 |
| 4 | 68441091 | rs2306810 | 2.117 | -0.102 | -1.027 | -1.588 | -0.054 | 0.026 | 0.026 | 1.513 | 0.16 | 0.02 | 0.17 | 0.1 | 0.24 | 0.15 |
| 4 | 68443308 | rs17088473 | 0.007 | -0.004 | -1.062 | -1.601 | -0.008 | -0.004 | -0.004 | 0.133 | 0.27 | 0.17 | 0.33 | 0.26 | 0.31 | 0.34 |
| 4 | 68444102 | rs13131164 | 1.474 | 0.047 | -1.108 | -1.621 | -0.065 | -0.038 | -0.038 | 0.083 | 0.13 | 0.01 | 0.06 | 0.13 | 0.19 | 0.15 |
| 4 | 68444180 | rs3775866 | 0.235 | 0.178 | -0.838 | -1.535 | -0.006 | 0.002 | 0.002 | 0 | 0.29 | 0.17 | 0.33 | 0.26 | 0.31 | 0.36 |
| 4 | 68444192 | rs11725922 | 1.379 | 0.404 | -0.822 | -1.399 | -0.005 | 0.044 | 0.044 | 1.798 | 0.26 | 0.12 | 0.16 | 0.01 | 0.25 | 0.12 |
| 4 | 68447200 | rs2242331 | 0.723 | 0.408 | -0.782 | -1.392 | 0.01 | -0.026 | -0.026 | 0 | 0.25 | 0.33 | 0.21 | 0.17 | 0.3 | 0.19 |
| 4 | 68447249 | rs2242330 | 0.502 | 0.05 | -1.431 | -0.918 | -0.018 | -0.018 | -0.018 | 0.033 | 0.23 | 0.11 | 0.14 | 0.25 | 0.25 | 0.3 |
| 4 | 68450302 | rs2087176 | 0.236 | 0.571 | -1.309 | -0.926 | -0.023 | 0.088 | 0.088 | 0 | 0.42 | 0.21 | 0.33 | 0.1 | 0.48 | 0.27 |
| 4 | 68453189 | rs17632290 | 0.102 | 0.646 | -1.202 | -0.669 | 0.017 | 0.118 | 0.118 | 0 | 0.49 | 0.26 | 0.36 | 0.17 | 0.55 | 0.3 |
| 4 | 68457695 | rs10000695 | 0.028 | 0.908 | -0.909 | -0.034 | -0.002 | 0.009 | 0.009 | 7.319 | 0.51 | 0.63 | 0.37 | 0.45 | 0.53 | 0.58 |
| 4 | 68458808 | rs7681638 | -0.251 | 0.229 | 0.05 | -0.524 | -0.032 | -0.051 | -0.051 | 0 | 0.11 | 0.36 | 0.1 | 0.01 | 0.05 | 0.04 |
| 4 | 68460255 | rs6830199 | -0.575 | 0.25 | 0.091 | -0.321 | -0.032 | 0.012 | 0.012 | 0 | 0.12 | 0.42 | 0.18 | 0.1 | 0.05 | 0.04 |

| | | | | | | | | | | | | | | | | |
|---|---|---|---|---|---|---|---|---|---|---|---|---|---|---|---|---|
| 4 | 68470455 | rs900791 | 0.32 | 0.895 | -0.149 | 0.256 | -0.006 | 0.01 | 0.01 | 0.582 | 0.14 | 0.13 | 0.11 | 0.52 | 0.19 | 0.34 |
| 4 | 68470522 | rs354883 | 0.495 | 0.799 | -0.643 | -0.251 | -0.014 | 0.015 | 0.015 | 0.268 | 0.37 | 0.34 | 0.33 | 0.25 | 0.46 | 0.32 |
| 4 | 68472428 | rs1054593 | 0.272 | 0.96 | -0.444 | 0.173 | -0.007 | 0.005 | 0.005 | 1.05 | 0.15 | 0.16 | 0.12 | 0.53 | 0.2 | 0.36 |
| 4 | 68472604 | rs354885 | -0.347 | 0.411 | 0.184 | -0.439 | -0.033 | -0.048 | -0.048 | 0 | 0.9 | 0.72 | 0.91 | 0.99 | 0.95 | 0.96 |
| 4 | 68480525 | rs4860262 | 0.753 | 0.691 | -0.197 | -0.866 | 0.014 | 0.058 | 0.058 | 0 | 0.47 | 0.36 | 0.37 | 0.28 | 0.41 | 0.3 |
| 4 | 68483472 | rs1048447 | 1.703 | 0.471 | -0.318 | -1.087 | 0.028 | 0.033 | 0.033 | 0 | 0.24 | 0.15 | 0.12 | 0.11 | 0.16 | 0.22 |
| 4 | 68493076 | rs3796715 | 1.703 | 0.471 | -0.238 | -1.087 | 0.023 | 0.026 | 0.026 | 0 | 0.24 | 0.16 | 0.12 | 0.11 | 0.17 | 0.22 |
| 4 | 68515155 | rs354872 | -0.567 | 0.277 | 0.102 | -1.288 | -0.017 | -0.044 | -0.044 | 2.317 | 0.88 | 0.81 | 0.9 | 0.99 | 0.92 | 0.97 |
| 4 | 68524526 | rs10006851 | 0.527 | 0.376 | -0.006 | -1.322 | 0.014 | 0.066 | 0.066 | 0.031 | 0.48 | 0.36 | 0.34 | 0.28 | 0.42 | 0.31 |
| 4 | 68534392 | rs10010188 | 1.823 | 0.322 | -0.041 | -1.339 | 0.03 | 0.034 | 0.034 | 0 | 0.25 | 0.15 | 0.12 | 0.11 | 0.17 | 0.22 |
| 4 | 68536679 | rs1017878 | 0.877 | 0.38 | 0.012 | -1.322 | -0.015 | 0.026 | 0.026 | 2.409 | 0.24 | 0.16 | 0.23 | 0.17 | 0.26 | 0.08 |
| 4 | 68536966 | rs2170872 | -0.655 | 0.711 | 0.322 | -1.504 | -0.005 | 0.005 | 0.005 | 0.557 | 0.67 | 0.72 | 0.51 | 0.29 | 0.59 | 0.36 |
| 4 | 68562365 | rs17572160 | 0.848 | 0.208 | 0.42 | -1.242 | -0.01 | 0.018 | 0.018 | 2.409 | 0.24 | 0.19 | 0.23 | 0.17 | 0.26 | 0.08 |
| 4 | 68591252 | rs2630489 | -0.304 | 0.176 | 0.475 | -1.259 | -0.008 | 0.009 | 0.009 | 0.671 | 0.61 | 0.67 | 0.45 | 0.29 | 0.5 | 0.35 |
| 4 | 68598694 | rs2711144 | 0.215 | 0.159 | 0.55 | -1.266 | -0.03 | -0.071 | -0.071 | 0.834 | 0.12 | 0.23 | 0.1 | 0.01 | 0.08 | 0.04 |
| 4 | 68598998 | rs1843593 | 2.047 | 0.227 | 0.181 | -1.218 | 0.033 | 0.046 | 0.046 | 0 | 0.23 | 0.05 | 0.11 | 0.11 | 0.17 | 0.22 |
| 4 | 68603702 | rs1038427 | 0.418 | 0.184 | 0.188 | -1.176 | 0.015 | 0.069 | 0.069 | 0.01 | 0.48 | 0.36 | 0.35 | 0.28 | 0.42 | 0.31 |
| 4 | 68613969 | rs17573883 | 0.57 | 0.479 | 0.007 | -1.287 | -0.016 | 0.043 | 0.043 | 2.42 | 0.24 | 0.15 | 0.18 | 0.08 | 0.28 | 0.08 |
| 4 | 68615500 | rs974483 | -0.311 | 0.568 | 0.067 | -1.263 | -0.006 | 0.009 | 0.009 | 1.374 | 0.35 | 0.33 | 0.29 | 0.09 | 0.37 | 0.11 |
| 4 | 68617365 | rs972072 | -0.371 | 0.455 | -0.01 | -0.727 | -0.018 | -0.008 | -0.008 | 0 | 0.63 | 0.73 | 0.5 | 0.47 | 0.53 | 0.36 |
| 4 | 68618451 | rs6828922 | 0.646 | -0.141 | 0.314 | -0.872 | -0.026 | -0.081 | -0.081 | 0 | 0.85 | 0.31 | 0.85 | 0.99 | 0.91 | 0.91 |
| 4 | 68619601 | rs4986942 | 0.286 | -0.103 | 0.397 | -0.853 | 0 | -0.021 | -0.021 | 0.219 | 0.08 | 0.09 | 0.09 | 0.01 | 0.08 | 0.07 |
| 4 | 68620957 | rs6552113 | 0.11 | -0.198 | 0.247 | -0.927 | -0.067 | -0.008 | -0.008 | 0 | 0.38 | 0.19 | 0.57 | 0.61 | 0.5 | 0.59 |
| 4 | 68624442 | rs10005857 | 1.469 | -0.33 | 0.21 | -0.831 | 0.049 | -0.008 | -0.008 | 0.097 | 0.26 | 0.06 | 0.12 | 0.22 | 0.17 | 0.26 |
| 4 | 68631416 | rs13102739 | 0.591 | -0.054 | 0.686 | -0.498 | 0.04 | 0.093 | 0.093 | 0 | 0.47 | 0.22 | 0.32 | 0.27 | 0.41 | 0.33 |
| 4 | 68654739 | rs2582358 | -1.149 | -0.27 | 0.868 | -0.452 | -0.024 | -0.021 | -0.021 | 0.446 | 0.24 | 0.69 | 0.14 | 0.15 | 0.11 | 0.14 |
| 4 | 68672416 | rs6552116 | -0.062 | 0.395 | 0.976 | -0.343 | -0.006 | -0.084 | -0.084 | 0 | 0.19 | 0.27 | 0.1 | 0.01 | 0.17 | 0.08 |
| 4 | 68673583 | rs4504308 | 1.586 | 0.675 | 0.603 | 0.015 | -0.028 | -0.028 | -0.028 | 0 | 0.52 | 0.15 | 0.6 | 0.51 | 0.52 | 0.63 |
| 4 | 68686507 | rs6552123 | 0.446 | 0.54 | 1.402 | 0.133 | 0.023 | 0.226 | 0.226 | 0 | 0.89 | 0.65 | 0.86 | 0.65 | 0.87 | 0.8 |
| 4 | 68697199 | rs13142201 | 0.976 | 0.553 | 0.563 | 0.109 | 0.026 | 0.256 | 0.256 | 0 | 0.86 | 0.51 | 0.84 | 0.65 | 0.86 | 0.8 |
| 4 | 68701771 | rs1371928 | 1.414 | 0.621 | -0.469 | 0.08 | 0.01 | -0.012 | -0.012 | 0 | 0.61 | 0.15 | 0.7 | 0.64 | 0.64 | 0.69 |
| 4 | 68706863 | rs10022223 | 1.394 | 0.833 | 0.352 | 0.907 | -0.005 | 0.036 | 0.036 | 0.764 | 0.82 | 0.28 | 0.84 | 0.67 | 0.82 | 0.78 |
| 4 | 68715140 | rs1371927 | 0.992 | 0.996 | 0.417 | 1.177 | 0.035 | 0.239 | 0.239 | 2.129 | 0.89 | 0.43 | 0.87 | 0.67 | 0.87 | 0.87 |
| 4 | 68716465 | rs718591 | -1.113 | 1.269 | -0.326 | 0.422 | 0.001 | -0.005 | -0.005 | 0.606 | 0.14 | 0.18 | 0.14 | 0.1 | 0.16 | 0.1 |
| 4 | 68718486 | rs1872581 | 0.194 | 1.229 | -1.06 | 0.532 | -0.022 | -0.069 | -0.069 | 0.007 | 0.28 | 0.13 | 0.52 | 0.54 | 0.33 | 0.4 |
| 4 | 68718552 | rs1348079 | -0.311 | 1.195 | -0.943 | -0.323 | 0.011 | -0.006 | -0.006 | 0 | 0.52 | 0.54 | 0.74 | 0.89 | 0.59 | 0.72 |
| 4 | 68725776 | rs1542664 | -0.128 | 1.057 | -0.944 | -0.424 | 0.014 | 0.046 | 0.046 | 0 | 0.75 | 0.42 | 0.59 | 0.64 | 0.75 | 0.58 |
| 4 | 68727562 | rs11131729 | -0.129 | 1.062 | -0.942 | -0.399 | 0.014 | 0.032 | 0.032 | 0 | 0.75 | 0.42 | 0.59 | 0.66 | 0.75 | 0.58 |
| 4 | 68737593 | rs17577246 | 0.228 | 0.231 | -1.508 | -1.216 | -0.033 | 0.134 | 0.134 | 0.488 | 0.21 | 0.03 | 0.13 | 0 | 0.21 | 0.09 |
| 4 | 68738544 | rs6811522 | 0.205 | 0.418 | -1.135 | -0.665 | -0.008 | 0.049 | 0.049 | 1.569 | 0.14 | 0.04 | 0.08 | 0 | 0.16 | 0.07 |
| 4 | 68750409 | rs960994 | -0.423 | 1.024 | -0.859 | -0.924 | -0.009 | 0.01 | 0.01 | 0 | 0.25 | 0.22 | 0.26 | 0.12 | 0.32 | 0.3 |
| 4 | 68751635 | rs1440743 | -0.688 | 0.505 | -0.618 | -1.434 | -0.001 | -0.024 | -0.024 | 4.253 | 0.58 | 0.62 | 0.51 | 0.82 | 0.58 | 0.67 |
| 4 | 68763794 | rs959636 | -0.953 | 0.4 | -0.528 | -1.437 | -0.015 | 0.122 | 0.122 | 0 | 0.62 | 0.89 | 0.58 | 0.82 | 0.63 | 0.74 |
| 4 | 68773898 | rs2165365 | -0.612 | 0.427 | -0.111 | -0.627 | 0.007 | 0.03 | 0.03 | 0 | 0.54 | 0.64 | 0.52 | 0.66 | 0.58 | 0.59 |
| 4 | 68775707 | rs1440740 | -0.509 | 0.433 | -0.152 | -0.614 | 0.01 | 0.056 | 0.056 | 0 | 0.49 | 0.33 | 0.43 | 0.66 | 0.53 | 0.52 |
| 4 | 68780041 | rs353172 | 0.069 | 0.472 | -0.793 | -0.684 | -0.055 | 0.012 | 0.012 | 0 | 0.28 | 0.14 | 0.29 | 0.28 | 0.37 | 0.31 |
| 4 | 68789778 | rs6839279 | 0.628 | -0.051 | -0.385 | 0.516 | 0.008 | -0.018 | -0.018 | 0 | 0.19 | 0.05 | 0.15 | 0.19 | 0.16 | 0.2 |
| 4 | 68791400 | rs353158 | 0.57 | 0.292 | 0.117 | 0.376 | 0.022 | 0.077 | 0.077 | 0 | 0.06 | 0.17 | 0.06 | 0.16 | 0.05 | 0.08 |
| 4 | 68806909 | rs6849403 | 0.231 | 0.551 | -0.695 | -0.423 | -0.058 | 0.072 | 0.072 | 0.561 | 0.34 | 0.23 | 0.37 | 0.17 | 0.48 | 0.36 |
| 4 | 68808967 | rs168482 | -1.118 | -0.329 | -0.448 | -0.649 | -0.004 | -0.018 | -0.018 | 0 | 0.69 | 0.7 | 0.59 | 0.81 | 0.73 | 0.75 |
| 4 | 68810246 | rs1370840 | 0.335 | -0.994 | 0.058 | -1.104 | -0.023 | -0.07 | -0.07 | 0.243 | 0.26 | 0.55 | 0.38 | 0.19 | 0.22 | 0.24 |
| 4 | 68811137 | rs12642405 | 0.816 | -0.853 | -0.584 | -0.367 | -0.008 | -0.17 | -0.17 | 0 | 0.29 | 0.03 | 0.18 | 0.39 | 0.24 | 0.36 |
| 4 | 68812283 | rs11930532 | -0.604 | -0.888 | -0.231 | -0.425 | 0.009 | 0.018 | 0.018 | 0 | 0.73 | 0.25 | 0.6 | 0.81 | 0.78 | 0.78 |
| 4 | 68814218 | rs2319691 | -0.592 | -0.888 | -0.236 | -0.425 | 0.01 | 0.021 | 0.021 | 0 | 0.73 | 0.25 | 0.6 | 0.81 | 0.79 | 0.76 |
| 4 | 68818382 | rs12643884 | 0.925 | -0.974 | -0.406 | -0.347 | 0.021 | -0.104 | -0.104 | 4.995 | 0.26 | 0.07 | 0.16 | 0.43 | 0.21 | 0.33 |
| 4 | 68821282 | rs1561385 | 1.019 | -0.888 | -0.432 | -0.265 | -0.006 | -0.157 | -0.157 | 0 | 0.26 | 0.03 | 0.14 | 0.42 | 0.22 | 0.32 |
| 4 | 68822218 | rs727534 | -1.529 | -0.938 | -0.002 | -0.396 | 0 | -0.01 | -0.01 | 0 | 0.8 | 0.78 | 0.64 | 0.81 | 0.79 | 0.78 |
| 4 | 68829207 | rs6552134 | 0.177 | -1.125 | -0.087 | -0.532 | 0.045 | 0.096 | 0.096 | 0 | 0.62 | 0.21 | 0.54 | 0.82 | 0.72 | 0.71 |
| 4 | 68829223 | rs11947613 | -0.058 | -1.149 | 0.143 | -0.567 | 0.068 | 0.057 | 0.057 | 2.401 | 0.06 | 0.45 | 0.03 | 0 | 0 | 0.01 |
| 4 | 68829603 | rs17088850 | 1.345 | -0.921 | -0.016 | -0.516 | -0.023 | -0.058 | -0.058 | 0 | 0.12 | 0.21 | 0.08 | 0 | 0.06 | 0.03 |
| 4 | 68832165 | rs1866325 | 1.37 | -0.985 | -0.09 | -0.514 | -0.011 | -0.034 | -0.034 | 0 | 0.12 | 0.16 | 0.08 | 0 | 0.06 | 0.03 |
| 4 | 68873083 | rs4362874 | 0.766 | -0.775 | -0.106 | -0.384 | 0.007 | 0.112 | 0.112 | 0 | 0.29 | 0.09 | 0.4 | 0.13 | 0.27 | 0.26 |
| 4 | 68874618 | rs6826271 | -0.504 | -0.438 | 0.323 | -0.489 | 0.027 | 0.004 | 0.004 | 0 | 0.63 | 0.46 | 0.6 | 0.59 | 0.55 | 0.61 |
| 4 | 68877522 | rs3860694 | 0.848 | -0.413 | 0.104 | -0.453 | -0.079 | -0.04 | -0.04 | 0 | 0.32 | 0.16 | 0.37 | 0.41 | 0.45 | 0.36 |
| 4 | 68882510 | rs7661514 | 0.228 | -0.468 | 0.241 | -0.534 | 0.025 | -0.04 | -0.04 | 0 | 0.36 | 0.23 | 0.19 | 0.46 | 0.29 | 0.37 |
| 4 | 68890221 | rs3910547 | -0.502 | -0.437 | 0.351 | -0.489 | 0.014 | 0.01 | 0.01 | 0 | 0.64 | 0.37 | 0.56 | 0.87 | 0.73 | 0.74 |
| 4 | 68899971 | rs4324597 | 2.285 | -0.779 | 0.035 | -0.124 | -0.003 | -0.058 | -0.058 | 0 | 0.12 | 0 | 0.07 | 0.17 | 0.1 | 0.13 |
| 4 | 68911041 | rs2164937 | -0.64 | -0.71 | 0.22 | -0.459 | 0.002 | -0.022 | -0.022 | 0 | 0.64 | 0.4 | 0.54 | 0.87 | 0.73 | 0.74 |
| 4 | 68913256 | rs6817194 | -0.66 | -0.71 | 0.224 | -0.454 | -0.001 | -0.034 | -0.034 | 0.099 | 0.64 | 0.42 | 0.55 | 0.87 | 0.73 | 0.74 |

| | | | | | | | | | | | | | | | | |
|---|---|---|---|---|---|---|---|---|---|---|---|---|---|---|---|---|
| 4 | 68914769 | rs6851972 | -0.705 | -0.719 | 0.339 | -0.457 | -0.009 | -0.053 | -0.053 | 0.196 | 0.65 | 0.43 | 0.55 | 0.87 | 0.73 | 0.75 |
| 4 | 68915798 | rs13112221 | 1.167 | -0.765 | 0.003 | -0.464 | -0.038 | -0.149 | -0.149 | 0 | 0.19 | 0.01 | 0.14 | 0.45 | 0.22 | 0.23 |
| 4 | 68919617 | rs10018943 | -0.612 | -0.532 | 0.305 | -0.339 | -0.009 | -0.053 | -0.053 | 0 | 0.65 | 0.43 | 0.55 | 0.87 | 0.73 | 0.75 |
| 4 | 68920638 | rs1438445 | -0.752 | -0.547 | 0.509 | -0.346 | -0.03 | -0.12 | -0.12 | 0 | 0.66 | 0.55 | 0.56 | 0.87 | 0.73 | 0.73 |
| 4 | 68921021 | rs2164935 | 0.923 | -0.586 | 0.729 | -0.366 | -0.005 | -0.043 | -0.043 | 0 | 0.31 | 0.27 | 0.42 | 0.13 | 0.27 | 0.27 |
| 4 | 68925617 | rs1370828 | -0.472 | -0.599 | 0.659 | -0.373 | 0.014 | 0.008 | 0.008 | 0 | 0.64 | 0.36 | 0.54 | 0.87 | 0.73 | 0.73 |
| 4 | 68933046 | rs1898644 | -0.912 | -0.605 | 0.793 | -0.363 | -0.026 | -0.113 | -0.113 | 3.114 | 0.68 | 0.56 | 0.57 | 0.86 | 0.73 | 0.73 |
| 4 | 68934306 | rs10004522 | -0.468 | -0.605 | 0.784 | -0.371 | 0.014 | 0.008 | 0.008 | 0 | 0.64 | 0.36 | 0.55 | 0.86 | 0.73 | 0.73 |
| 4 | 68940208 | rs1370830 | -0.525 | -0.621 | 0.776 | -0.379 | 0.012 | 0.005 | 0.005 | 0 | 0.65 | 0.36 | 0.55 | 0.87 | 0.73 | 0.74 |
| 4 | 68942150 | rs6552145 | -0.845 | -0.614 | 0.751 | -0.375 | -0.021 | -0.095 | -0.095 | 0.062 | 0.68 | 0.53 | 0.57 | 0.87 | 0.73 | 0.74 |
| 4 | 68949916 | rs4860900 | 0.774 | -0.649 | 0.496 | -0.39 | -0.178 | -0.099 | -0.099 | 0 | 0.33 | 0.04 | 0.34 | 0.41 | 0.45 | 0.39 |
| 4 | 68956040 | rs1370834 | 0.192 | -0.654 | 0.628 | -0.389 | 0.01 | -0.022 | -0.022 | 0 | 0.31 | 0.3 | 0.2 | 0.46 | 0.28 | 0.34 |
| 4 | 68956856 | rs17089042 | 1.103 | -0.667 | 0.652 | -0.404 | 0.01 | 0.041 | 0.041 | 0 | 0.29 | 0.14 | 0.41 | 0.13 | 0.27 | 0.25 |
| 4 | 68958290 | rs7665380 | 0.381 | -0.619 | 0.346 | -0.377 | -0.033 | -0.005 | -0.005 | 0.096 | 0.37 | 0.26 | 0.36 | 0.41 | 0.45 | 0.39 |
| 4 | 68962223 | rs966827 | -0.801 | -0.528 | -0.094 | -0.337 | -0.013 | -0.063 | -0.063 | 0.046 | 0.66 | 0.47 | 0.55 | 0.87 | 0.73 | 0.75 |
| 4 | 68963631 | rs7676484 | 0.565 | -0.474 | -0.279 | -0.3 | -0.1 | -0.04 | -0.04 | 1.776 | 0.34 | 0.1 | 0.34 | 0.41 | 0.45 | 0.39 |
| 4 | 68982038 | rs10019959 | 0.109 | -0.466 | -0.189 | -0.269 | 0.007 | -0.015 | -0.015 | 0.069 | 0.32 | 0.33 | 0.21 | 0.46 | 0.28 | 0.34 |
| 4 | 68990767 | rs10031258 | 0.634 | -0.507 | -0.265 | -0.324 | -0.146 | -0.069 | -0.069 | 0.016 | 0.34 | 0.06 | 0.34 | 0.41 | 0.46 | 0.4 |
| 4 | 68995529 | rs10030708 | 0.139 | -0.458 | -0.244 | -0.285 | 0.006 | -0.015 | -0.015 | 0 | 0.31 | 0.32 | 0.21 | 0.46 | 0.28 | 0.34 |
| 4 | 69013001 | rs6819568 | 0.365 | -0.296 | 0.16 | -0.427 | -0.021 | -0.088 | -0.088 | 0.013 | 0.33 | 0.39 | 0.44 | 0.13 | 0.27 | 0.24 |
| 4 | 69050703 | rs17590869 | -0.263 | 0.071 | -0.264 | -0.193 | 0.001 | 0.006 | 0.006 | 0 | 0.11 | 0.06 | 0.11 | 0.1 | 0.11 | 0.16 |
| 4 | 69052317 | rs6845487 | -0.551 | 0.315 | -0.079 | -0.677 | -0.005 | -0.128 | -0.128 | 0 | 0.72 | 0.47 | 0.6 | 0.99 | 0.73 | 0.9 |
| 4 | 69083563 | rs10008636 | -0.551 | 0.315 | -0.079 | -0.677 | -0.005 | -0.128 | -0.128 | 0.842 | 0.72 | 0.47 | 0.6 | 0.99 | 0.73 | 0.9 |
| 4 | 69087267 | rs7434495 | 0.541 | 0.28 | 0.114 | -0.695 | 0.001 | -0.169 | -0.169 | 0 | 0.26 | 0.4 | 0.38 | 0.01 | 0.27 | 0.1 |
| 4 | 69091168 | rs4860919 | -0.439 | 0.286 | -0.073 | -0.692 | -0.004 | -0.092 | -0.092 | 0.185 | 0.71 | 0.37 | 0.6 | 0.99 | 0.73 | 0.9 |
| 4 | 69095197 | rs12331141 | -0.495 | 0.287 | -0.071 | -0.692 | -0.006 | -0.117 | -0.117 | 0 | 0.72 | 0.43 | 0.6 | 0.99 | 0.73 | 0.9 |
| 4 | 69098006 | rs6552175 | 0.441 | 0.3 | -0.143 | -0.693 | -0.003 | -0.124 | -0.124 | 0.3 | 0.27 | 0.49 | 0.39 | 0.01 | 0.27 | 0.1 |
| 4 | 69126345 | rs6840820 | 0.206 | 1.058 | -1.023 | 0.426 | -0.061 | -0.075 | -0.075 | 0 | 0.2 | 0.08 | 0.13 | 0.39 | 0.29 | 0.25 |
| 4 | 69132978 | rs11933242 | 0.152 | 1.339 | -1.04 | 0.454 | -0.056 | -0.044 | -0.044 | 0 | 0.22 | 0.08 | 0.21 | 0.32 | 0.3 | 0.22 |
| 4 | 69138386 | rs7691074 | 0.045 | 0.441 | -0.318 | -0.096 | -0.048 | -0.06 | -0.06 | 0 | 0.1 | 0.38 | 0.06 | 0 | 0.03 | 0.01 |
| 4 | 69138643 | rs7670560 | -0.027 | 0.327 | -0.339 | -0.721 | -0.009 | -0.087 | -0.087 | 1.319 | 0.33 | 0.63 | 0.43 | 0.01 | 0.26 | 0.12 |
| 4 | 69141870 | rs1996598 | 0.721 | 0.877 | -0.691 | -0.341 | -0.016 | -0.043 | -0.043 | 0 | 0.19 | 0.08 | 0.17 | 0.32 | 0.22 | 0.23 |
| 4 | 69149571 | rs10866204 | 1.018 | 0.789 | -0.447 | -0.491 | -0.016 | -0.083 | -0.083 | 0 | 0.24 | 0.32 | 0.37 | 0.01 | 0.17 | 0.1 |
| 4 | 69151046 | rs7678448 | 0.053 | 0.632 | -0.307 | -0.571 | -0.038 | -0.125 | -0.125 | 0 | 0.34 | 0.56 | 0.43 | 0.01 | 0.2 | 0.12 |
| 4 | 69155649 | rs7659406 | 0.781 | 0.887 | -0.381 | -0.423 | -0.03 | -0.05 | -0.05 | 0 | 0.2 | 0.08 | 0.18 | 0.34 | 0.25 | 0.24 |
| 4 | 69160724 | rs1962172 | 0.496 | 0.837 | -0.16 | -0.968 | -0.048 | -0.128 | -0.128 | 0 | 0.28 | 0.48 | 0.39 | 0.04 | 0.17 | 0.1 |
| 4 | 69174057 | rs1730875 | -1.661 | 1.074 | -0.387 | -0.643 | -0.007 | -0.039 | -0.039 | 0 | 0.8 | 0.77 | 0.64 | 0.96 | 0.84 | 0.9 |
| 4 | 69176798 | rs1715081 | 0.323 | 1.23 | -0.483 | -0.45 | 0.003 | 0.01 | 0.01 | 0 | 0.48 | 0.4 | 0.4 | 0.39 | 0.55 | 0.65 |
| 4 | 69177138 | rs17592288 | 0.779 | 1.339 | -0.568 | -0.695 | -0.001 | 0 | 0 | 0 | 0.11 | 0.11 | 0.14 | 0.07 | 0.18 | 0.22 |
| 4 | 69178920 | rs2293595 | 0.464 | 1.127 | -0.69 | -0.479 | 0.003 | 0.013 | 0.013 | 0 | 0.47 | 0.39 | 0.4 | 0.38 | 0.56 | 0.65 |
| 4 | 69215034 | rs7666993 | 0.477 | 0.819 | -0.659 | -0.586 | 0.001 | 0.009 | 0.009 | 0 | 0.47 | 0.4 | 0.4 | 0.39 | 0.55 | 0.65 |
| 4 | 69219049 | rs4860928 | -0.366 | 0.273 | -0.342 | -1.063 | -0.014 | -0.054 | -0.054 | 0 | 0.73 | 0.71 | 0.6 | 0.96 | 0.8 | 0.87 |
| 4 | 69244539 | rs34527218 | -0.78 | 0.784 | -0.712 | -0.73 | -0.028 | -0.096 | -0.096 | 2.024 | 0.77 | 0.6 | 0.62 | 0.96 | 0.84 | 0.91 |
| 4 | 69252261 | rs35377534 | -0.024 | 1.276 | -1.116 | -0.604 | 0.012 | 0.003 | 0.003 | 0 | 0.64 | 0.51 | 0.47 | 0.36 | 0.6 | 0.71 |
| 4 | 69280975 | rs4860941 | -0.648 | 1.233 | -1.924 | -1.496 | -0.081 | 0.205 | 0.205 | 1.89 | 0.36 | 0.07 | 0.46 | 0.06 | 0.39 | 0.33 |
| 4 | 69282119 | rs11722296 | -0.105 | 1.308 | -2.046 | -1.604 | -0.096 | 0.158 | 0.158 | 0 | 0.24 | 0.03 | 0.39 | 0 | 0.3 | 0.11 |
| 4 | 69283448 | rs4860296 | -0.608 | 1.258 | -1.717 | -1.42 | -0.046 | 0.169 | 0.169 | 1.786 | 0.37 | 0.12 | 0.47 | 0.06 | 0.39 | 0.32 |
| 4 | 69341037 | rs2708699 | -0.108 | 0.889 | -0.86 | -0.762 | 0.009 | 0.061 | 0.061 | 0 | 0.49 | 0.6 | 0.45 | 0.77 | 0.44 | 0.61 |
| 4 | 69341579 | rs1454247 | -0.108 | 0.889 | -0.86 | -0.762 | 0.009 | 0.061 | 0.061 | 0 | 0.49 | 0.6 | 0.45 | 0.77 | 0.44 | 0.61 |
| 4 | 69349034 | rs2708697 | -0.557 | 0.867 | -0.902 | -0.793 | 0.001 | 0.011 | 0.011 | 0 | 0.61 | 0.64 | 0.76 | 0.85 | 0.63 | 0.79 |
| 4 | 69367074 | rs2603152 | 0.821 | 0.411 | -0.375 | -0.032 | -0.003 | -0.036 | -0.036 | 0.064 | 0.75 | 0.5 | 0.64 | 0.85 | 0.72 | 0.65 |
| 4 | 69500664 | rs36069550 | -0.559 | 0.263 | -0.66 | 0.895 | -0.001 | 0.005 | 0.005 | 0.005 | 0.35 | 0.4 | 0.34 | 0.51 | 0.31 | 0.4 |
| 4 | 69512637 | rs4148271 | -0.542 | 0.141 | -0.793 | 1.593 | 0.03 | -0.028 | -0.028 | 0.228 | 0.07 | 0.01 | 0.05 | 0.27 | 0.01 | 0.09 |
| 4 | 69590353 | rs293433 | -1.352 | 0.244 | -0.109 | 1.344 | -0.023 | 0.047 | 0.047 | 1.109 | 0.27 | 0.45 | 0.47 | 0.61 | 0.22 | 0.3 |
| 4 | 69590669 | rs293431 | -1.367 | 0.485 | -0.073 | 1.333 | -0.012 | 0.051 | 0.051 | 1.36 | 0.27 | 0.47 | 0.47 | 0.61 | 0.24 | 0.31 |
| 4 | 69591612 | rs293429 | -1.67 | 0.607 | -0.181 | 1.224 | -0.007 | 0.031 | 0.031 | 1.095 | 0.3 | 0.49 | 0.48 | 0.61 | 0.28 | 0.37 |
| 4 | 69595144 | rs376360 | -0.115 | 0.274 | 0.049 | 1.351 | -0.013 | 0.096 | 0.096 | 0 | 0.14 | 0.22 | 0.16 | 0.56 | 0.1 | 0.2 |
| 4 | 69598506 | rs4694270 | -0.87 | 0.386 | 0.316 | 1.314 | -0.007 | -0.046 | -0.046 | 0 | 0.14 | 0.37 | 0.32 | 0.06 | 0.13 | 0.12 |
| 4 | 69601124 | rs293408 | -0.086 | 0.822 | -0.648 | 1.192 | -0.103 | -0.081 | -0.081 | 0 | 0.81 | 0.99 | 0.78 | 0.73 | 0.74 | 0.83 |
| 4 | 69606943 | rs407182 | -1.049 | 0.217 | 0.347 | 1.545 | -0.004 | -0.004 | -0.004 | 0.647 | 0.25 | 0.66 | 0.52 | 0.24 | 0.24 | 0.15 |
| 4 | 69611505 | rs397678 | -1.049 | 0.217 | 0.366 | 1.545 | -0.005 | -0.004 | -0.004 | 0 | 0.25 | 0.66 | 0.52 | 0.24 | 0.24 | 0.15 |
| 4 | 69613030 | rs403552 | -1.049 | 0.217 | 0.361 | 1.545 | -0.004 | -0.004 | -0.004 | 0 | 0.25 | 0.66 | 0.52 | 0.24 | 0.24 | 0.15 |
| 4 | 69627526 | rs4235087 | -0.583 | 0.151 | 0.586 | 2.158 | -0.004 | -0.01 | -0.01 | 0 | 0.2 | 0.16 | 0.42 | 0.3 | 0.24 | 0.13 |
| 4 | 69628948 | rs1119049 | -1.343 | 0.118 | 0.56 | 2.139 | 0 | -0.001 | -0.001 | 0 | 0.27 | 0.74 | 0.53 | 0.43 | 0.25 | 0.18 |
| 4 | 69630828 | rs1513557 | -1.728 | -0.6 | 0.336 | 1.75 | 0.055 | -0.032 | -0.032 | 0 | 0.36 | 0.85 | 0.55 | 0.42 | 0.28 | 0.19 |
| 4 | 69632909 | rs9993883 | -1.192 | -0.512 | 0.416 | 1.788 | -0.004 | 0.014 | 0.014 | 0 | 0.27 | 0.7 | 0.53 | 0.43 | 0.25 | 0.18 |
| 4 | 69646009 | rs11726899 | -0.326 | -0.422 | 0.347 | 1.829 | 0.011 | 0.041 | 0.041 | 0 | 0.05 | 0.09 | 0.08 | 0.19 | 0.1 | 0.06 |
| 4 | 69665970 | rs294755 | 0.511 | -0.26 | 0.094 | 2.07 | -0.041 | -0.04 | -0.04 | 0 | 0.08 | 0 | 0.14 | 0.31 | 0.13 | 0.16 |
| 4 | 69676142 | rs4694340 | 0.364 | -0.518 | 0.886 | 1.913 | -0.025 | -0.013 | -0.013 | 0.549 | 0.15 | 0.01 | 0.33 | 0.12 | 0.15 | 0.07 |

| | | | | | | | | | | | | | | | | |
|---|---|---|---|---|---|---|---|---|---|---|---|---|---|---|---|---|
| 4 | 69686221 | rs835317 | 1.313 | -0.478 | 1.085 | 1.872 | 0.051 | 0.042 | 0.042 | 1.813 | 0.91 | 0.38 | 0.84 | 0.91 | 0.91 | 0.94 |
| 4 | 69701067 | rs11737566 | -0.962 | -0.7 | 1.37 | 1.969 | -0.013 | 0.015 | 0.015 | 0 | 0.15 | 0.26 | 0.1 | 0.18 | 0.12 | 0.13 |
| 4 | 69703527 | rs1379809 | 1.671 | -0.824 | 1.343 | 1.921 | 0.06 | -0.055 | -0.055 | 0 | 0.63 | 0.15 | 0.46 | 0.44 | 0.73 | 0.71 |
| 4 | 69706273 | rs6843220 | -0.878 | -0.819 | 1.368 | 1.927 | -0.011 | 0.008 | 0.008 | 0 | 0.14 | 0.24 | 0.1 | 0.18 | 0.12 | 0.13 |
| 4 | 69708718 | rs1841043 | 1.295 | -0.803 | 1.346 | 1.934 | -0.004 | 0.003 | 0.003 | 0.884 | 0.67 | 0.41 | 0.53 | 0.54 | 0.73 | 0.75 |
| 4 | 69717920 | rs1841046 | 0.426 | -0.701 | 1.159 | 1.925 | -0.025 | -0.006 | -0.006 | 0.162 | 0.15 | 0 | 0.33 | 0.11 | 0.14 | 0.07 |
| 4 | 69725006 | rs6600855 | -0.431 | -0.679 | 1.195 | 1.942 | 0.015 | 0.046 | 0.046 | 0 | 0.06 | 0.14 | 0.08 | 0.18 | 0.09 | 0.06 |
| 4 | 69727682 | rs7674919 | -0.431 | -0.679 | 1.195 | 1.942 | 0.015 | 0.046 | 0.046 | 0 | 0.06 | 0.14 | 0.08 | 0.18 | 0.09 | 0.06 |
| 4 | 69736745 | rs12500933 | 0.299 | -0.724 | 1.145 | 1.911 | 0.005 | 0.014 | 0.014 | 0.641 | 0.16 | 0.04 | 0.33 | 0.11 | 0.14 | 0.07 |
| 4 | 69749311 | rs1841037 | 0.312 | -0.611 | 0.779 | 1.624 | 0.005 | 0.013 | 0.013 | 0.827 | 0.16 | 0.05 | 0.33 | 0.11 | 0.14 | 0.08 |
| 4 | 69752468 | rs11724856 | -1.303 | -0.653 | 1.133 | 1.65 | -0.014 | -0.006 | -0.006 | 0 | 0.32 | 0.36 | 0.44 | 0.29 | 0.26 | 0.22 |
| 4 | 69762953 | rs7678798 | -0.815 | -0.532 | 1.16 | 1.599 | -0.021 | 0.015 | 0.015 | 0 | 0.16 | 0.31 | 0.1 | 0.18 | 0.12 | 0.15 |
| 4 | 69771935 | rs12651295 | -0.86 | -0.44 | 1.121 | 1.599 | -0.014 | 0.01 | 0.01 | 0 | 0.16 | 0.3 | 0.11 | 0.17 | 0.12 | 0.14 |
| 4 | 69786431 | rs10019843 | 0.41 | -0.922 | 1.674 | 1.722 | -0.031 | 0.061 | 0.061 | 0 | 0.78 | 0.49 | 0.6 | 0.69 | 0.84 | 0.81 |
| 4 | 69787683 | rs12648504 | 0.396 | -0.921 | 1.675 | 1.722 | -0.031 | 0.062 | 0.062 | 0 | 0.78 | 0.49 | 0.6 | 0.69 | 0.84 | 0.81 |
| 4 | 69818151 | rs17605102 | 0.339 | -0.898 | 1.154 | 1.588 | -0.024 | 0.061 | 0.061 | 0 | 0.15 | 0.02 | 0.29 | 0.04 | 0.15 | 0.09 |
| 4 | 69822928 | rs1824264 | 0.531 | -1.025 | 1.332 | 1.779 | -0.094 | 0.059 | 0.059 | 3.072 | 0.82 | 0.4 | 0.88 | 0.75 | 0.96 | 0.89 |
| 4 | 69830542 | rs3966085 | -0.122 | -0.885 | 1.12 | 1.885 | 0.03 | 0.232 | 0.232 | 0 | 0.13 | 0.27 | 0.19 | 0.53 | 0.17 | 0.29 |
| 4 | 69830871 | rs1458241 | 0.433 | -0.924 | 1.141 | 1.875 | -0.099 | 0.058 | 0.058 | 3.036 | 0.82 | 0.42 | 0.88 | 0.75 | 0.96 | 0.89 |
| 4 | 69833007 | rs3886642 | 1.339 | -0.916 | 1.138 | 1.969 | 0.024 | -0.023 | -0.023 | 0 | 0.71 | 0.25 | 0.69 | 0.4 | 0.79 | 0.71 |
| 4 | 69850452 | rs6818594 | 0.417 | -1.017 | 1.147 | 1.544 | -0.105 | 0.002 | 0.002 | 1.663 | 0.79 | 0.44 | 0.88 | 0.79 | 0.95 | 0.87 |
| 4 | 69853849 | rs4590113 | 0.46 | -0.969 | 1.137 | 1.631 | 0.025 | 0.122 | 0.122 | 0 | 0.11 | 0.17 | 0.19 | 0.35 | 0.17 | 0.16 |
| 4 | 69859702 | rs4694490 | 1.093 | -0.992 | 1.113 | 1.626 | 0.032 | -0.033 | -0.033 | 0 | 0.69 | 0.26 | 0.69 | 0.44 | 0.78 | 0.73 |
| 4 | 69860688 | rs7678207 | 1.093 | -0.992 | 1.113 | 1.626 | 0.032 | -0.033 | -0.033 | 0.687 | 0.69 | 0.26 | 0.69 | 0.44 | 0.78 | 0.73 |
| 4 | 69862775 | rs1841050 | 0.276 | -0.795 | 1.101 | 1.618 | 0.032 | 0.146 | 0.146 | 0 | 0.11 | 0.21 | 0.2 | 0.35 | 0.17 | 0.16 |
| 4 | 69862918 | rs2018024 | 0.227 | -0.795 | 1.095 | 1.618 | 0.037 | 0.164 | 0.164 | 0 | 0.11 | 0.24 | 0.2 | 0.35 | 0.17 | 0.16 |
| 4 | 69883503 | rs4576082 | 1.362 | -1.161 | 0.408 | 1.143 | 0.096 | 0.086 | 0.086 | 0.505 | 0.52 | 0.14 | 0.38 | 0.31 | 0.64 | 0.57 |
| 4 | 69893854 | rs4235108 | 0.932 | -1.249 | 1.003 | 1.248 | -0.009 | 0.004 | 0.004 | 0.485 | 0.71 | 0.31 | 0.68 | 0.39 | 0.79 | 0.71 |
| 4 | 69901663 | rs13147648 | 0.945 | -1.249 | 1.003 | 1.248 | -0.009 | 0.004 | 0.004 | 0.469 | 0.71 | 0.31 | 0.68 | 0.39 | 0.79 | 0.71 |
| 4 | 69907597 | rs4299666 | 0.911 | -1.248 | 0.995 | 1.249 | -0.014 | 0.008 | 0.008 | 0.475 | 0.71 | 0.32 | 0.68 | 0.39 | 0.79 | 0.71 |
| 4 | 69916384 | rs7438808 | -0.767 | -1.044 | 0.107 | 0.423 | 0.027 | 0.02 | 0.02 | 0 | 0.46 | 0.73 | 0.41 | 0.61 | 0.4 | 0.43 |
| 4 | 69923096 | rs6833862 | 0.826 | -0.986 | 0.351 | 0.514 | 0.009 | 0.112 | 0.112 | 0 | 0.12 | 0.21 | 0.23 | 0.44 | 0.16 | 0.21 |
| 4 | 69927974 | rs7662632 | -0.085 | -0.988 | -0.113 | 0.269 | -0.003 | -0.009 | -0.009 | 0.034 | 0.35 | 0.71 | 0.38 | 0.6 | 0.36 | 0.42 |
| 4 | 69932587 | rs9991142 | -0.071 | -1.012 | -0.199 | 0.052 | 0.026 | 0.032 | 0.032 | 1.792 | 0.28 | 0.17 | 0.35 | 0.04 | 0.19 | 0.13 |
| 4 | 69954151 | rs4694169 | -1.04 | -0.944 | -0.701 | 0.112 | -0.091 | 0.002 | 0.002 | 0 | 0.65 | 0.88 | 0.72 | 0.73 | 0.55 | 0.61 |
| 4 | 69956930 | rs6600876 | -0.439 | -1.026 | -0.47 | 0.065 | -0.013 | 0.082 | 0.082 | 0 | 0.53 | 0.77 | 0.68 | 0.72 | 0.51 | 0.6 |
| 4 | 69961127 | rs11940316 | -0.439 | -1.026 | -0.47 | 0.065 | -0.013 | 0.082 | 0.082 | 0 | 0.53 | 0.77 | 0.68 | 0.73 | 0.51 | 0.6 |
| 4 | 69961339 | rs7438135 | -0.439 | -0.996 | -0.47 | 0.065 | -0.016 | 0.082 | 0.082 | 0 | 0.53 | 0.77 | 0.68 | 0.73 | 0.51 | 0.6 |
| 4 | 69961912 | rs7662029 | -0.439 | -1.008 | -0.47 | 0.065 | -0.013 | 0.082 | 0.082 | 0 | 0.53 | 0.77 | 0.68 | 0.72 | 0.51 | 0.6 |
| 4 | 69962078 | rs7668258 | -0.439 | -1.008 | -0.47 | 0.065 | -0.013 | 0.082 | 0.082 | 0 | 0.53 | 0.77 | 0.68 | 0.73 | 0.51 | 0.6 |
| 4 | 69964554 | rs7441774 | -0.432 | -1.063 | -0.402 | 0.029 | -0.013 | 0.083 | 0.083 | 0 | 0.52 | 0.77 | 0.68 | 0.73 | 0.52 | 0.6 |
| 4 | 69965502 | rs4293848 | -0.432 | -1.065 | -0.361 | 0.029 | -0.012 | 0.08 | 0.08 | 0 | 0.52 | 0.77 | 0.68 | 0.72 | 0.51 | 0.6 |
| 4 | 69966024 | rs12506962 | -0.44 | -1.062 | -0.379 | 0.03 | -0.013 | 0.082 | 0.082 | 0 | 0.53 | 0.77 | 0.68 | 0.73 | 0.51 | 0.6 |
| 4 | 69969006 | rs7439152 | -0.494 | -1.07 | -0.381 | 0.026 | -0.037 | 0.089 | 0.089 | 0 | 0.53 | 0.83 | 0.69 | 0.73 | 0.51 | 0.6 |
| 4 | 69969543 | rs6858558 | -0.494 | -1.07 | -0.381 | 0.026 | -0.037 | 0.089 | 0.089 | 0 | 0.53 | 0.83 | 0.69 | 0.73 | 0.51 | 0.6 |
| 4 | 69970937 | rs10028494 | 1.495 | -0.7 | -0.925 | -0.47 | -0.015 | -0.028 | -0.028 | 0 | 0.17 | 0.08 | 0.14 | 0.21 | 0.19 | 0.15 |
| 4 | 69971335 | rs7435335 | 1.318 | -1.201 | -0.111 | -0.388 | -0.004 | -0.055 | -0.055 | 0.823 | 0.16 | 0.26 | 0.33 | 0.04 | 0.16 | 0.12 |
| 4 | 69971596 | rs6600891 | -0.289 | -1.189 | -0.281 | -0.382 | -0.034 | 0.095 | 0.095 | 0 | 0.52 | 0.83 | 0.69 | 0.72 | 0.51 | 0.55 |
| 4 | 69972086 | rs12513195 | -0.289 | -1.189 | -0.281 | -0.382 | -0.034 | 0.095 | 0.095 | 0 | 0.52 | 0.83 | 0.69 | 0.72 | 0.51 | 0.55 |
| 4 | 69972463 | rs4356975 | 0.454 | -1.209 | -0.196 | -0.393 | 0.005 | -0.001 | -0.001 | 0.241 | 0.35 | 0.47 | 0.35 | 0.68 | 0.36 | 0.43 |
| 4 | 69973525 | rs4521414 | -0.158 | -1.069 | -0.868 | -0.286 | -0.034 | 0.095 | 0.095 | 0 | 0.52 | 0.83 | 0.69 | 0.72 | 0.51 | 0.55 |
| 4 | 69987249 | rs6600897 | -0.189 | -1.072 | -0.808 | -0.289 | -0.025 | 0.087 | 0.087 | 0 | 0.53 | 0.8 | 0.69 | 0.73 | 0.52 | 0.55 |
| 4 | 69995509 | rs4318715 | -0.821 | -1.091 | -0.847 | -0.496 | -0.057 | 0.032 | 0.032 | 0 | 0.66 | 0.82 | 0.72 | 0.78 | 0.56 | 0.59 |
| 4 | 69997633 | rs11249529 | 1.256 | -1.131 | -0.251 | -0.574 | -0.003 | -0.037 | -0.037 | 0 | 0.16 | 0.19 | 0.33 | 0.04 | 0.15 | 0.12 |
| 4 | 70003963 | rs6822259 | 1.61 | -1.197 | -0.35 | -0.435 | -0.013 | 0.001 | 0.001 | 0 | 0.18 | 0.1 | 0.1 | 0.18 | 0.2 | 0.15 |
| 4 | 70004010 | rs4694181 | 1.265 | -1.176 | -0.013 | -0.49 | -0.015 | -0.081 | -0.081 | 1.967 | 0.17 | 0.34 | 0.33 | 0.05 | 0.14 | 0.12 |
| 4 | 70005443 | rs4415024 | -0.561 | -1.414 | -0.514 | -0.617 | -0.037 | 0.069 | 0.069 | 0 | 0.6 | 0.79 | 0.7 | 0.77 | 0.53 | 0.62 |
| 4 | 70013537 | rs6600900 | -0.276 | -1.528 | -0.3 | -0.563 | 0.011 | 0.02 | 0.02 | 2.269 | 0.53 | 0.51 | 0.49 | 0.3 | 0.38 | 0.39 |
| 4 | 70019933 | rs4273535 | 0.715 | -1.328 | -0.23 | -0.388 | 0.001 | 0.002 | 0.002 | 0 | 0.25 | 0.17 | 0.13 | 0.25 | 0.21 | 0.26 |
| 4 | 70031620 | rs4593203 | 0.624 | -1.032 | 0.009 | -0.096 | 0.001 | 0.053 | 0.053 | 0 | 0.17 | 0.24 | 0.23 | 0.47 | 0.18 | 0.24 |
| 4 | 70057930 | rs6825376 | 1.203 | -1.14 | 0.132 | -0.459 | -0.026 | -0.066 | -0.066 | 0.018 | 0.15 | 0.38 | 0.19 | 0.05 | 0.11 | 0.16 |
| 4 | 70058114 | rs4342251 | 1.261 | -1.138 | 0.172 | -0.458 | -0.025 | -0.062 | -0.062 | 0 | 0.14 | 0.37 | 0.19 | 0.05 | 0.11 | 0.16 |
| 4 | 70058335 | rs4326078 | 0.071 | -1.213 | -0.1 | -0.387 | -0.003 | 0.041 | 0.041 | 0 | 0.24 | 0.35 | 0.28 | 0.47 | 0.24 | 0.37 |
| 4 | 70059980 | rs4694199 | -1.467 | -1.257 | -0.176 | -0.814 | -0.013 | 0.061 | 0.061 | 0.204 | 0.77 | 0.91 | 0.9 | 1 | 0.81 | 0.89 |
| 4 | 70088601 | rs6817250 | 1.149 | -1.2 | -0.733 | -0.616 | -0.073 | 0 | 0 | 0 | 0.31 | 0.16 | 0.38 | 0.29 | 0.42 | 0.31 |
| 4 | 70112313 | rs10032551 | 0.848 | -1.239 | -0.759 | -0.518 | -0.046 | 0 | 0 | 0 | 0.34 | 0.19 | 0.38 | 0.33 | 0.42 | 0.31 |
| 4 | 70115470 | rs11939111 | 1.112 | -1.207 | -0.431 | -0.614 | -0.032 | -0.042 | -0.042 | 0 | 0.18 | 0.34 | 0.21 | 0.11 | 0.16 | 0.26 |
| 4 | 70120699 | rs6830865 | 0.269 | -1.227 | -0.556 | -0.61 | 0.003 | 0.025 | 0.025 | 1.259 | 0.3 | 0.18 | 0.16 | 0.18 | 0.24 | 0.16 |
| 4 | 70234419 | rs7693378 | 0.707 | -0.691 | -0.322 | -0.383 | -0.031 | -0.04 | -0.04 | 0 | 0.18 | 0.33 | 0.21 | 0.11 | 0.16 | 0.26 |

| | | | | | | | | | | | | | | | | |
|---|---|---|---|---|---|---|---|---|---|---|---|---|---|---|---|---|
| 4 | 70247163 | rs12501393 | 0.714 | -0.69 | -0.354 | -0.381 | -0.027 | -0.035 | -0.035 | 0.057 | 0.18 | 0.29 | 0.2 | 0.11 | 0.16 | 0.26 |
| 4 | 70255961 | rs12505338 | 1.302 | -0.706 | -0.353 | -0.387 | -0.026 | -0.049 | -0.049 | 0 | 0.14 | 0.27 | 0.19 | 0.06 | 0.11 | 0.24 |
| 4 | 70261746 | rs12506196 | 0.903 | -0.657 | -0.274 | -0.359 | -0.027 | -0.037 | -0.037 | 0 | 0.18 | 0.29 | 0.2 | 0.11 | 0.16 | 0.26 |
| 4 | 70274557 | rs12509072 | 0.893 | -0.535 | -0.405 | -0.444 | -0.028 | -0.037 | -0.037 | 0 | 0.18 | 0.29 | 0.2 | 0.11 | 0.16 | 0.26 |
| 4 | 70287303 | rs2736509 | -0.772 | -0.06 | -0.975 | -0.529 | 0.004 | -0.025 | -0.025 | 0.714 | 0.7 | 0.75 | 0.72 | 0.61 | 0.73 | 0.78 |
| 4 | 70288522 | rs2254718 | 0.028 | -0.031 | -1.037 | -0.497 | 0.004 | -0.001 | -0.001 | 0 | 0.52 | 0.46 | 0.51 | 0.49 | 0.57 | 0.52 |
| 4 | 70297879 | rs2736488 | 1.979 | 0.883 | -1.455 | -0.703 | -0.045 | -0.014 | -0.014 | 0 | 0.1 | 0.01 | 0.06 | 0.12 | 0.17 | 0.14 |
| 4 | 70299686 | rs2736486 | -0.035 | 0.533 | -1.071 | -0.638 | -0.001 | 0.01 | 0.01 | 0 | 0.61 | 0.42 | 0.57 | 0.77 | 0.62 | 0.58 |
| 4 | 70311222 | rs7670842 | 0.622 | 1.268 | -1.287 | -0.442 | -0.047 | 0.051 | 0.051 | 0 | 0.18 | 0.07 | 0.24 | 0 | 0.24 | 0.13 |
| 4 | 70314776 | rs1845545 | 0.524 | 0.517 | -1.208 | -0.554 | 0.048 | 0.062 | 0.062 | 1.479 | 0.43 | 0.28 | 0.34 | 0.26 | 0.33 | 0.37 |
| 4 | 70317551 | rs1845549 | -0.757 | 0.546 | -1.206 | -0.556 | 0.048 | 0.064 | 0.064 | 1.42 | 0.57 | 0.71 | 0.66 | 0.74 | 0.67 | 0.63 |
| 4 | 70321293 | rs12505081 | 1.104 | 1.173 | -0.785 | -0.952 | 0.008 | -0.055 | -0.055 | 0 | 0.21 | 0.2 | 0.24 | 0 | 0.26 | 0.12 |
| 4 | 70324048 | rs4694743 | 1.194 | 1.18 | -0.799 | -0.948 | -0.023 | 0.004 | 0.004 | 0 | 0.2 | 0.13 | 0.24 | 0 | 0.26 | 0.12 |
| 4 | 70325709 | rs2736467 | -0.869 | 1.264 | -0.28 | -0.437 | 0.013 | 0.007 | 0.007 | 0.798 | 0.51 | 0.45 | 0.59 | 0.57 | 0.61 | 0.6 |
| 4 | 70328040 | rs1353280 | -1.289 | 1.253 | -0.141 | -0.436 | 0 | -0.001 | -0.001 | 0.175 | 0.55 | 0.56 | 0.65 | 0.74 | 0.67 | 0.63 |
| 4 | 70330250 | rs10518060 | 1.273 | 1.345 | -0.491 | -0.582 | -0.036 | 0.027 | 0.027 | 0 | 0.2 | 0.09 | 0.24 | 0 | 0.26 | 0.12 |
| 4 | 70336172 | rs2736520 | 2.872 | 1.524 | -0.441 | -0.166 | -0.02 | -0.015 | -0.015 | 0 | 0.07 | 0.03 | 0.06 | 0.13 | 0.14 | 0.1 |
| 4 | 70336501 | rs903446 | -0.748 | 1.375 | -0.098 | -0.3 | -0.001 | 0.001 | 0.001 | 0.037 | 0.34 | 0.35 | 0.4 | 0.74 | 0.41 | 0.51 |
| 4 | 70341392 | rs1569344 | 1.72 | 1.732 | -0.318 | -0.505 | -0.017 | 0.003 | 0.003 | 0 | 0.15 | 0.09 | 0.22 | 0 | 0.22 | 0.1 |
| 4 | 70341463 | rs1569343 | -1.355 | 1.113 | -0.467 | -0.722 | 0.025 | 0.114 | 0.114 | 0 | 0.53 | 0.72 | 0.62 | 0.76 | 0.55 | 0.64 |
| 4 | 70341550 | rs1569342 | 1.887 | 1.258 | -0.457 | -0.562 | -0.02 | 0.007 | 0.007 | 0 | 0.16 | 0.09 | 0.22 | 0 | 0.23 | 0.1 |
| 4 | 70345669 | rs2736446 | -0.888 | 0.684 | -0.167 | -0.938 | 0.004 | -0.042 | -0.042 | 0 | 0.68 | 0.64 | 0.68 | 0.9 | 0.7 | 0.87 |
| 4 | 70345904 | rs1131878 | -0.888 | 0.684 | -0.156 | -0.938 | 0.004 | -0.04 | -0.04 | 0 | 0.68 | 0.64 | 0.68 | 0.9 | 0.7 | 0.87 |
| 4 | 70346127 | rs1966151 | 0.575 | 0.291 | -0.018 | -0.94 | 0 | 0.044 | 0.044 | 0 | 0.2 | 0.32 | 0.17 | 0.42 | 0.25 | 0.24 |
| 4 | 70350006 | rs4557343 | 0.561 | 0.37 | -0.039 | -0.923 | 0.006 | 0.006 | 0.006 | 0 | 0.5 | 0.4 | 0.53 | 0.47 | 0.45 | 0.63 |
| 4 | 70350357 | rs11249442 | 0.139 | 0.348 | 0.001 | -0.976 | -0.02 | 0.04 | 0.04 | 0 | 0.67 | 0.5 | 0.75 | 0.47 | 0.67 | 0.73 |
| 4 | 70350444 | rs1121124 | 0.544 | 0.322 | 0.003 | -0.975 | 0.004 | 0.004 | 0.004 | 0 | 0.51 | 0.41 | 0.53 | 0.47 | 0.45 | 0.63 |
| 4 | 70350467 | rs12645716 | 0.596 | 0.326 | -0.015 | -0.972 | 0.006 | 0.006 | 0.006 | 0 | 0.5 | 0.4 | 0.53 | 0.47 | 0.45 | 0.63 |
| 4 | 70354575 | rs2013562 | 0.779 | 0.454 | 0.006 | -1.067 | 0 | 0 | 0 | 0 | 0.48 | 0.25 | 0.52 | 0.47 | 0.45 | 0.63 |
| 4 | 70354794 | rs1845557 | 0.234 | 0.453 | 0.041 | -1.071 | -0.01 | 0.013 | 0.013 | 0 | 0.65 | 0.38 | 0.74 | 0.47 | 0.67 | 0.73 |
| 4 | 70354811 | rs13111134 | 1.989 | 0.681 | -0.194 | -1.262 | -0.022 | 0.012 | 0.012 | 0 | 0.17 | 0.09 | 0.23 | 0 | 0.25 | 0.1 |
| 4 | 70356195 | rs3822179 | 1.217 | 0.686 | -0.051 | -1.336 | -0.002 | 0.113 | 0.113 | 0 | 0.09 | 0.23 | 0.1 | 0.33 | 0.08 | 0.14 |
| 4 | 70357876 | rs13139888 | 2.17 | 1.023 | -0.796 | -1.591 | -0.017 | 0 | 0 | 0 | 0.16 | 0.09 | 0.22 | 0 | 0.23 | 0.1 |
| 4 | 70358845 | rs17671289 | 2.069 | 0.871 | -1.032 | -1.85 | 0.006 | 0.121 | 0.121 | 0 | 0.28 | 0.07 | 0.25 | 0.02 | 0.26 | 0.29 |
| 4 | 70359068 | rs13145834 | 2.096 | 1.044 | -0.687 | -1.425 | -0.017 | 0.001 | 0.001 | 0 | 0.16 | 0.09 | 0.22 | 0 | 0.23 | 0.1 |
| 4 | 70359694 | rs7441743 | 0.78 | 0.94 | -0.889 | -0.804 | -0.008 | 0.055 | 0.055 | 0 | 0.42 | 0.18 | 0.43 | 0.27 | 0.39 | 0.46 |
| 4 | 70360229 | rs17614939 | 1.747 | 0.888 | -0.735 | -1.628 | -0.061 | 0.069 | 0.069 | 0 | 0.13 | 0.01 | 0.09 | 0 | 0.19 | 0.09 |
| 4 | 70371555 | rs7657880 | -0.183 | 0.654 | -0.207 | -1.612 | -0.001 | -0.001 | -0.001 | 0 | 0.24 | 0.26 | 0.18 | 0.22 | 0.19 | 0.2 |
| 4 | 70394123 | rs10461367 | 0.901 | 0.868 | -0.335 | -1.807 | 0.018 | 0.114 | 0.114 | 0 | 0.46 | 0.21 | 0.43 | 0.75 | 0.43 | 0.52 |
| 4 | 70395452 | rs13151633 | 1.438 | 0.908 | -0.237 | -1.827 | -0.001 | -0.032 | -0.032 | 0.43 | 0.18 | 0.17 | 0.21 | 0 | 0.2 | 0.17 |
| 4 | 70395945 | rs1603117 | 0.752 | 1.104 | 0.099 | -1.429 | 0 | -0.006 | -0.006 | 0.072 | 0.64 | 0.4 | 0.64 | 0.75 | 0.63 | 0.69 |
| 4 | 70405837 | rs6825148 | 0.116 | 0.688 | -0.052 | -1.329 | 0 | -0.001 | -0.001 | 0.018 | 0.76 | 0.74 | 0.82 | 0.79 | 0.81 | 0.81 |
| 4 | 70438197 | rs12502677 | -0.085 | 0.744 | -0.14 | -1.159 | -0.007 | 0.02 | 0.02 | 0 | 0.44 | 0.32 | 0.52 | 0.58 | 0.44 | 0.47 |
| 4 | 70438763 | rs17616972 | 1.675 | 0.984 | -0.006 | -1.052 | 0.021 | -0.027 | -0.027 | 1.239 | 0.08 | 0.16 | 0.15 | 0.03 | 0.17 | 0.13 |
| 4 | 70443083 | rs4463132 | -0.107 | 0.808 | 0.018 | -1.202 | 0 | 0.009 | 0.009 | 0 | 0.27 | 0.2 | 0.19 | 0.18 | 0.22 | 0.27 |
| 4 | 70448535 | rs11939113 | 0.138 | 0.792 | -0.073 | -1.21 | -0.003 | 0.011 | 0.011 | 0 | 0.26 | 0.18 | 0.19 | 0.18 | 0.22 | 0.26 |
| 4 | 70449652 | rs7687676 | 0.107 | 0.818 | -0.051 | -1.197 | -0.002 | 0.014 | 0.014 | 0 | 0.26 | 0.18 | 0.19 | 0.18 | 0.22 | 0.26 |
| 4 | 70453516 | rs11929908 | -1.094 | 0.768 | 0.075 | -1.279 | 0.001 | -0.026 | -0.026 | 0.07 | 0.66 | 0.56 | 0.68 | 0.75 | 0.58 | 0.72 |
| 4 | 70453638 | rs11929943 | -0.151 | 0.75 | 0.098 | -1.265 | -0.001 | -0.006 | -0.006 | 0.799 | 0.28 | 0.26 | 0.2 | 0.18 | 0.23 | 0.27 |
| 4 | 70455478 | rs2288741 | -1.736 | 0.762 | 0.087 | -1.307 | -0.015 | -0.017 | -0.017 | 0 | 0.74 | 0.69 | 0.83 | 0.78 | 0.75 | 0.85 |
| 4 | 70455650 | rs1438537 | 0.441 | 0.772 | 0.088 | -1.32 | -0.001 | 0.002 | 0.002 | 0 | 0.38 | 0.3 | 0.47 | 0.57 | 0.36 | 0.46 |
| 4 | 70455884 | rs4148309 | 0.061 | 0.776 | 0.09 | -1.323 | -0.001 | -0.005 | -0.005 | 0 | 0.27 | 0.24 | 0.2 | 0.18 | 0.22 | 0.27 |
| 4 | 70456186 | rs4148308 | -1.068 | 0.777 | 0.089 | -1.321 | 0.001 | -0.025 | -0.025 | 0 | 0.66 | 0.57 | 0.68 | 0.75 | 0.58 | 0.72 |
| 4 | 70457877 | rs1432327 | -1.068 | 0.777 | 0.061 | -1.321 | 0.001 | -0.027 | -0.027 | 0 | 0.66 | 0.57 | 0.68 | 0.75 | 0.58 | 0.72 |
| 4 | 70458182 | rs1432324 | 0.413 | 0.84 | -0.569 | -1.289 | -0.001 | 0.002 | 0.002 | 0 | 0.38 | 0.3 | 0.47 | 0.57 | 0.36 | 0.46 |
| 4 | 70460328 | rs4148304 | 1.959 | 0.843 | -0.571 | -1.287 | -0.003 | -0.003 | -0.003 | 0 | 0.08 | 0.13 | 0.15 | 0.03 | 0.17 | 0.12 |
| 4 | 70462621 | rs1158439 | -1.909 | 0.821 | -0.505 | -1.298 | 0.016 | 0.018 | 0.018 | 0.108 | 0.76 | 0.88 | 0.85 | 0.78 | 0.76 | 0.85 |
| 4 | 70462669 | rs1821185 | 0.422 | 0.834 | -0.535 | -1.288 | -0.001 | 0.002 | 0.002 | 0 | 0.38 | 0.3 | 0.47 | 0.57 | 0.36 | 0.46 |
| 4 | 70462803 | rs2010207 | -1.185 | 0.834 | -0.488 | -1.288 | -0.004 | 0.02 | 0.02 | 0 | 0.67 | 0.69 | 0.69 | 0.75 | 0.59 | 0.72 |
| 4 | 70462893 | rs1319811 | -1.185 | 0.834 | -0.488 | -1.288 | -0.004 | 0.02 | 0.02 | 0 | 0.67 | 0.69 | 0.69 | 0.75 | 0.59 | 0.72 |
| 4 | 70463600 | rs1432316 | 0.818 | 0.829 | -0.566 | -1.416 | 0 | 0.02 | 0.02 | 0.282 | 0.15 | 0.17 | 0.36 | 0.57 | 0.16 | 0.27 |
| 4 | 70463662 | rs1432315 | 0.818 | 0.829 | -0.566 | -1.416 | 0 | 0.02 | 0.02 | 0.282 | 0.15 | 0.17 | 0.36 | 0.57 | 0.16 | 0.27 |
| 4 | 70463854 | rs3775783 | 0.073 | 0.827 | -0.563 | -1.413 | -0.004 | -0.033 | -0.033 | 0 | 0.28 | 0.36 | 0.21 | 0.18 | 0.22 | 0.27 |
| 4 | 70464701 | rs4148298 | 0.073 | 0.827 | -0.563 | -1.413 | -0.004 | -0.033 | -0.033 | 0 | 0.28 | 0.36 | 0.21 | 0.18 | 0.23 | 0.27 |
| 4 | 70465493 | rs4148296 | 0.218 | 0.828 | -0.57 | -1.414 | -0.001 | 0.002 | 0.002 | 0 | 0.38 | 0.3 | 0.47 | 0.57 | 0.36 | 0.46 |
| 4 | 70465584 | rs10023782 | 0.818 | 0.838 | -0.572 | -1.419 | 0.001 | 0.02 | 0.02 | 0.282 | 0.15 | 0.17 | 0.37 | 0.57 | 0.16 | 0.27 |
| 4 | 70465626 | rs10033801 | -1.32 | 0.811 | -0.565 | -1.425 | -0.003 | 0.018 | 0.018 | 0 | 0.67 | 0.69 | 0.69 | 0.75 | 0.59 | 0.72 |
| 4 | 70465737 | rs10033879 | -0.048 | 0.806 | -0.601 | -1.427 | 0 | 0 | 0 | 0.425 | 0.4 | 0.37 | 0.48 | 0.57 | 0.36 | 0.46 |
| 4 | 70466810 | rs1432313 | -0.048 | 0.806 | -0.602 | -1.417 | -0.007 | -0.036 | -0.036 | 0 | 0.29 | 0.38 | 0.22 | 0.18 | 0.23 | 0.27 |

| | | | | | | | | | | | | | | | | |
|---|---|---|---|---|---|---|---|---|---|---|---|---|---|---|---|---|
| 4 | 70467109 | rs1036160 | 0.8 | 0.813 | -0.623 | -1.39 | 0 | 0.02 | 0.02 | 0.282 | 0.15 | 0.17 | 0.36 | 0.57 | 0.16 | 0.27 |
| 4 | 70474310 | rs6832167 | -1.912 | 0.817 | -0.764 | -1.385 | 0.01 | 0.01 | 0.01 | 0.05 | 0.75 | 0.81 | 0.84 | 0.78 | 0.76 | 0.85 |
| 4 | 70475866 | rs4148295 | -0.421 | 0.854 | -0.755 | -1.377 | 0.008 | -0.021 | -0.021 | 0 | 0.36 | 0.42 | 0.36 | 0.21 | 0.4 | 0.39 |
| 4 | 70476595 | rs1432336 | 1.721 | 0.902 | -0.9 | -1.414 | -0.005 | -0.003 | -0.003 | 0 | 0.09 | 0.11 | 0.16 | 0.03 | 0.17 | 0.12 |
| 4 | 70477328 | rs11728546 | -0.296 | 0.738 | -1.13 | -1.307 | 0.015 | -0.05 | -0.05 | 0.691 | 0.37 | 0.51 | 0.37 | 0.21 | 0.39 | 0.39 |
| 4 | 70477984 | rs6600793 | -0.296 | 0.738 | -1.13 | -1.307 | 0.015 | -0.05 | -0.05 | 0 | 0.37 | 0.51 | 0.37 | 0.21 | 0.4 | 0.39 |
| 4 | 70478169 | rs6600794 | -0.296 | 0.738 | -1.13 | -1.307 | 0.015 | -0.05 | -0.05 | 0.715 | 0.37 | 0.51 | 0.37 | 0.21 | 0.4 | 0.39 |
| 4 | 70478971 | rs6848997 | 1.411 | 0.698 | -1.208 | -1.313 | -0.007 | 0.078 | 0.078 | 0 | 0.22 | 0.09 | 0.1 | 0 | 0.2 | 0.19 |
| 4 | 70480327 | rs10011912 | -1.63 | 0.608 | -1.145 | -1.182 | 0.01 | 0.012 | 0.012 | 0.05 | 0.75 | 0.81 | 0.84 | 0.79 | 0.76 | 0.85 |
| 4 | 70480926 | rs4694222 | -0.614 | 1.15 | -0.887 | -1.044 | 0.016 | -0.045 | -0.045 | 0 | 0.37 | 0.51 | 0.37 | 0.22 | 0.4 | 0.39 |
| 4 | 70481880 | rs7670819 | -0.403 | 0.943 | -0.714 | -0.88 | -0.005 | -0.024 | -0.024 | 0 | 0.55 | 0.52 | 0.45 | 0.25 | 0.56 | 0.51 |
| 4 | 70482097 | rs7676249 | -0.036 | 0.604 | -0.647 | -0.907 | -0.004 | -0.037 | -0.037 | 0.619 | 0.39 | 0.51 | 0.24 | 0.22 | 0.32 | 0.4 |
| 4 | 70483323 | rs4148291 | -0.003 | 0.599 | -0.593 | -0.91 | -0.005 | -0.036 | -0.036 | 0.785 | 0.4 | 0.47 | 0.25 | 0.22 | 0.33 | 0.38 |
| 4 | 70484250 | rs4148288 | 1.292 | 0.6 | -0.506 | -0.931 | -0.001 | 0.001 | 0.001 | 0 | 0.07 | 0.07 | 0.03 | 0.07 | 0.03 | 0.07 |
| 4 | 70485627 | rs6824830 | -1.66 | 0.471 | -0.518 | -0.97 | 0.013 | 0.019 | 0.019 | 0.038 | 0.73 | 0.81 | 0.83 | 0.81 | 0.76 | 0.86 |
| 4 | 70485663 | rs6855271 | 0.998 | 0.455 | -0.336 | -0.977 | -0.001 | 0.021 | 0.021 | 0 | 0.31 | 0.24 | 0.2 | 0.14 | 0.3 | 0.31 |
| 4 | 70488786 | rs11249453 | 1.002 | 0.48 | -0.317 | -0.965 | -0.002 | 0.023 | 0.023 | 0 | 0.31 | 0.24 | 0.2 | 0.14 | 0.3 | 0.31 |
| 4 | 70488849 | rs10903210 | 0.076 | 0.481 | -0.366 | -0.962 | -0.004 | -0.039 | -0.039 | 0.107 | 0.39 | 0.47 | 0.24 | 0.22 | 0.33 | 0.37 |
| 4 | 70491891 | rs1835829 | 0.139 | 0.396 | -0.326 | -1.023 | -0.017 | -0.008 | -0.008 | 0 | 0.3 | 0.23 | 0.56 | 0.6 | 0.38 | 0.4 |
| 4 | 70491970 | rs10001991 | 0.06 | 0.633 | -0.3 | -0.899 | -0.004 | -0.037 | -0.037 | 0.049 | 0.39 | 0.47 | 0.25 | 0.22 | 0.33 | 0.37 |
| 4 | 70492055 | rs10023877 | 1.514 | 0.645 | -0.3 | -0.897 | -0.001 | 0.001 | 0.001 | 0 | 0.06 | 0.07 | 0.02 | 0.07 | 0.03 | 0.06 |
| 4 | 70494631 | rs10022954 | 1.514 | 0.626 | -0.291 | -0.897 | -0.002 | 0.002 | 0.002 | 0 | 0.06 | 0.07 | 0.03 | 0.07 | 0.03 | 0.06 |
| 4 | 70494901 | rs6854350 | 0.721 | 0.626 | -0.294 | -0.899 | -0.002 | -0.045 | -0.045 | 0.001 | 0.33 | 0.4 | 0.22 | 0.14 | 0.3 | 0.32 |
| 4 | 70494933 | rs17674565 | 1.605 | 0.621 | -0.281 | -0.824 | 0.022 | 0.041 | 0.041 | 0.331 | 0.25 | 0.09 | 0.15 | 0.19 | 0.24 | 0.14 |
| 4 | 70494984 | rs10011630 | 1.44 | 0.646 | -0.262 | -0.825 | -0.001 | 0.001 | 0.001 | 0 | 0.06 | 0.07 | 0.03 | 0.07 | 0.03 | 0.06 |
| 4 | 70495961 | rs17618178 | 1.278 | 0.683 | -0.328 | -0.862 | -0.003 | -0.114 | -0.114 | 0 | 0.26 | 0.38 | 0.18 | 0.01 | 0.25 | 0.17 |
| 4 | 70496262 | rs17674648 | 1.278 | 0.683 | -0.328 | -0.862 | -0.003 | -0.114 | -0.114 | 0.206 | 0.26 | 0.39 | 0.18 | 0.01 | 0.25 | 0.17 |
| 4 | 70497601 | rs7665571 | 1.496 | 0.683 | -0.326 | -0.862 | -0.001 | -0.028 | -0.028 | 0.531 | 0.24 | 0.22 | 0.17 | 0.01 | 0.25 | 0.16 |
| 4 | 70498165 | rs17147521 | 1.082 | 0.679 | -0.319 | -0.867 | -0.002 | 0.002 | 0.002 | 0 | 0.12 | 0.09 | 0.06 | 0.11 | 0.1 | 0.23 |
| 4 | 70499234 | rs11249454 | 1.398 | 0.751 | -0.35 | -0.413 | -0.017 | -0.051 | -0.051 | 3.093 | 0.13 | 0.05 | 0.31 | 0.24 | 0.14 | 0.11 |
| 4 | 70500830 | rs6824954 | 0.969 | 0.135 | -0.041 | -1.32 | -0.001 | -0.153 | -0.153 | 0.013 | 0.26 | 0.41 | 0.18 | 0.01 | 0.25 | 0.17 |
| 4 | 70501980 | rs7662309 | -1.577 | 0.127 | -0.085 | -1.327 | 0.023 | 0.042 | 0.042 | 0.266 | 0.75 | 0.9 | 0.85 | 0.79 | 0.76 | 0.86 |
| 4 | 70502386 | rs7692074 | 1.15 | 0.206 | -0.071 | -1.29 | 0.002 | -0.077 | -0.077 | 1.246 | 0.24 | 0.28 | 0.17 | 0.01 | 0.25 | 0.16 |
| 4 | 70502779 | rs17618520 | 0.005 | 0.19 | -0.075 | -1.291 | 0 | 0.01 | 0.01 | 0 | 0.36 | 0.33 | 0.22 | 0.14 | 0.35 | 0.39 |
| 4 | 70503978 | rs10518063 | 1.163 | 0.207 | -0.072 | -1.285 | 0 | -0.028 | -0.028 | 0 | 0.24 | 0.22 | 0.16 | 0.01 | 0.25 | 0.16 |
| 4 | 70504313 | rs9992698 | 0.854 | 0.195 | -0.059 | -1.289 | -0.009 | -0.004 | -0.004 | 0.888 | 0.3 | 0.28 | 0.57 | 0.6 | 0.38 | 0.41 |
| 4 | 70505904 | rs17147532 | 0.591 | 0.229 | -0.042 | -1.276 | 0 | -0.001 | -0.001 | 0.056 | 0.12 | 0.13 | 0.06 | 0.13 | 0.12 | 0.24 |
| 4 | 70506120 | rs11936310 | 1.149 | 0.263 | -0.025 | -1.263 | -0.001 | -0.028 | -0.028 | 0 | 0.24 | 0.22 | 0.16 | 0.01 | 0.25 | 0.16 |
| 4 | 70506518 | rs988672 | -0.05 | 0.262 | -0.022 | -1.263 | 0 | -0.058 | -0.058 | 0 | 0.37 | 0.42 | 0.23 | 0.14 | 0.37 | 0.4 |
| 4 | 70506828 | rs988671 | 1.126 | 0.273 | -0.041 | -1.255 | 0.001 | -0.075 | -0.075 | 4.913 | 0.24 | 0.27 | 0.17 | 0.01 | 0.25 | 0.16 |
| 4 | 70507156 | rs17147542 | 0.034 | 0.259 | -0.253 | -1.249 | 0 | -0.048 | -0.048 | 0 | 0.36 | 0.4 | 0.22 | 0.14 | 0.35 | 0.4 |
| 4 | 70507794 | rs2331685 | -0.059 | 0.437 | -1.342 | -1.17 | -0.003 | -0.076 | -0.076 | 0 | 0.38 | 0.46 | 0.23 | 0.12 | 0.34 | 0.4 |
| 4 | 70509530 | rs4148284 | 1.025 | 0.584 | -1.306 | -1.097 | 0 | 0 | 0 | 0 | 0.07 | 0.06 | 0.03 | 0.08 | 0.03 | 0.06 |
| 4 | 70511200 | rs4148282 | 1.205 | 0.62 | -1.288 | -1.079 | 0 | -0.002 | -0.002 | 0 | 0.06 | 0.04 | 0.03 | 0.07 | 0.03 | 0.05 |
| 4 | 70513710 | rs1432329 | -0.849 | 0.615 | -1.308 | -1.084 | 0.018 | 0.022 | 0.022 | 0 | 0.67 | 0.77 | 0.81 | 0.72 | 0.72 | 0.78 |
| 4 | 70514358 | rs4148279 | 0.267 | 0.938 | -0.98 | -0.59 | -0.008 | -0.003 | -0.003 | 0 | 0.29 | 0.27 | 0.57 | 0.61 | 0.38 | 0.4 |
| 4 | 70515025 | rs4148275 | 0.131 | 0.909 | -0.96 | -0.591 | -0.008 | -0.096 | -0.096 | 0 | 0.32 | 0.46 | 0.21 | 0.09 | 0.28 | 0.21 |
| 4 | 70516671 | rs10000435 | -0.624 | 1.564 | -1.023 | 0.611 | -0.007 | -0.004 | -0.004 | 0 | 0.3 | 0.27 | 0.56 | 0.61 | 0.37 | 0.39 |
| 4 | 70517952 | rs7668703 | -0.659 | 1.329 | -1.166 | 0.475 | -0.033 | 0.031 | 0.031 | 0 | 0.38 | 0.28 | 0.58 | 0.66 | 0.48 | 0.49 |
| 4 | 70519417 | rs4395582 | 0.709 | 0.914 | -0.659 | 0.569 | -0.003 | -0.005 | -0.005 | 1.156 | 0.73 | 0.35 | 0.78 | 0.76 | 0.78 | 0.75 |
| 4 | 70519904 | rs10033854 | -0.633 | 0.982 | -1.135 | 0.67 | 0.004 | -0.009 | -0.009 | 0 | 0.43 | 0.49 | 0.44 | 0.63 | 0.47 | 0.45 |
| 4 | 70520597 | rs10518065 | 0.706 | 1.827 | -1.309 | 0.484 | -0.013 | 0.018 | 0.018 | 0 | 0.05 | 0.02 | 0.09 | 0.01 | 0.15 | 0.12 |
| 4 | 70520830 | rs13134357 | -0.072 | 1.589 | -1.163 | 0.522 | -0.004 | -0.006 | -0.006 | 0 | 0.33 | 0.3 | 0.34 | 0.4 | 0.4 | 0.3 |
| 4 | 70521755 | rs1347047 | -0.54 | 1.472 | -0.405 | 0.392 | 0.001 | -0.001 | -0.001 | 0 | 0.37 | 0.41 | 0.4 | 0.59 | 0.42 | 0.4 |
| 4 | 70540370 | rs1948828 | 0.71 | 1.042 | -0.034 | 1.018 | -0.004 | 0.023 | 0.023 | 0 | 0.58 | 0.29 | 0.57 | 0.34 | 0.56 | 0.57 |
| 4 | 70540563 | rs11249456 | 0.71 | 1.042 | -0.034 | 1.018 | -0.004 | 0.023 | 0.023 | 0 | 0.58 | 0.29 | 0.58 | 0.34 | 0.56 | 0.57 |
| 4 | 70567104 | rs1587765 | 1.679 | 1.183 | 0.018 | 0.758 | 0.018 | 0.06 | 0.06 | 0 | 0.37 | 0.14 | 0.44 | 0.22 | 0.33 | 0.41 |
| 4 | 70570637 | rs17681469 | 0.219 | 1.204 | 0.005 | 0.664 | -0.011 | 0.019 | 0.019 | 0 | 0.16 | 0.1 | 0.08 | 0 | 0.16 | 0.04 |
| 4 | 70584038 | rs4694242 | 0.407 | 1.041 | 0.363 | 0.798 | -0.004 | -0.03 | -0.03 | 0.61 | 0.63 | 0.55 | 0.64 | 0.72 | 0.62 | 0.63 |
| 4 | 70586610 | rs7696239 | 0.854 | 1.049 | 0.71 | 0.868 | 0.001 | 0.023 | 0.023 | 0 | 0.58 | 0.29 | 0.57 | 0.6 | 0.56 | 0.56 |
| 4 | 70587338 | rs17620185 | 0.489 | 1.251 | 0.67 | 0.828 | -0.002 | 0.016 | 0.016 | 0 | 0.16 | 0.1 | 0.06 | 0 | 0.14 | 0.04 |
| 4 | 70588309 | rs2292090 | 0.558 | 1.123 | 0.461 | 0.682 | -0.069 | 0.016 | 0.016 | 0 | 0.23 | 0.04 | 0.19 | 0.14 | 0.27 | 0.22 |
| 4 | 70588741 | rs9685617 | 0.718 | 1.148 | 0.529 | 1.016 | 0.007 | 0.142 | 0.142 | 0 | 0.05 | 0.12 | 0.06 | 0.37 | 0.07 | 0.07 |
| 4 | 70592242 | rs4149419 | 2.312 | 1.142 | 0.513 | 1.006 | -0.078 | 0.01 | 0.01 | 0 | 0.37 | 0.02 | 0.44 | 0.23 | 0.35 | 0.46 |
| 4 | 70592790 | rs2292092 | 1.905 | 1.126 | 0.554 | 1.005 | -0.019 | 0.031 | 0.031 | 0 | 0.42 | 0.2 | 0.51 | 0.6 | 0.42 | 0.52 |
| 4 | 70597081 | rs6817798 | 0.684 | 1.122 | 1.084 | 1.135 | 0.006 | 0.139 | 0.139 | 0 | 0.05 | 0.11 | 0.06 | 0.37 | 0.07 | 0.07 |
| 4 | 70597380 | rs3775780 | 2.294 | 1.121 | 1.073 | 1.126 | -0.015 | 0.051 | 0.051 | 0 | 0.37 | 0.08 | 0.45 | 0.24 | 0.35 | 0.46 |
| 4 | 70599073 | rs1847366 | 1.834 | 1.12 | 1.175 | 1.124 | -0.007 | 0.028 | 0.028 | 0 | 0.43 | 0.25 | 0.52 | 0.6 | 0.42 | 0.52 |
| 4 | 70599386 | rs1394038 | 2.3 | 1.111 | 1.107 | 1.084 | -0.017 | 0.05 | 0.05 | 0 | 0.37 | 0.08 | 0.45 | 0.24 | 0.35 | 0.46 |

| | | | | | | | | | | | | | | | | |
|---|---|---|---|---|---|---|---|---|---|---|---|---|---|---|---|---|
| 4 | 70600256 | rs17626012 | 0.859 | 1.108 | 1.116 | 1.052 | -0.038 | 0.079 | 0.079 | 0 | 0.14 | 0 | 0.06 | 0 | 0.15 | 0.04 |
| 4 | 70601326 | rs1847364 | 1.688 | 1.042 | 1.467 | 1.051 | 0.002 | 0.015 | 0.015 | 0 | 0.46 | 0.43 | 0.53 | 0.6 | 0.42 | 0.53 |
| 4 | 70602840 | rs11731028 | 2.262 | 1.046 | 1.348 | 1.06 | -0.013 | 0.058 | 0.058 | 0 | 0.37 | 0.09 | 0.45 | 0.24 | 0.34 | 0.46 |
| 4 | 70604618 | rs2089075 | 2.262 | 1.046 | 1.348 | 1.06 | -0.013 | 0.058 | 0.058 | 0 | 0.37 | 0.09 | 0.45 | 0.24 | 0.34 | 0.46 |
| 4 | 70609040 | rs1847361 | 2.272 | 1.046 | 1.348 | 1.06 | -0.013 | 0.057 | 0.057 | 0 | 0.37 | 0.08 | 0.45 | 0.24 | 0.34 | 0.46 |
| 4 | 70612712 | rs11727948 | 2.272 | 1.046 | 1.347 | 1.06 | -0.015 | 0.057 | 0.057 | 0 | 0.37 | 0.08 | 0.46 | 0.24 | 0.34 | 0.46 |
| 4 | 70614259 | rs1604747 | 1.018 | 1.08 | 1.411 | 1.053 | 0.001 | -0.006 | -0.006 | 0.014 | 0.59 | 0.49 | 0.6 | 0.35 | 0.55 | 0.56 |
| 4 | 70616322 | rs1354360 | 2.303 | 1.238 | 1.385 | 1.062 | -0.008 | 0.057 | 0.057 | 0 | 0.37 | 0.08 | 0.45 | 0.24 | 0.33 | 0.46 |
| 4 | 70621074 | rs11569731 | 0.982 | 1.166 | 1.766 | 0.975 | 0.003 | -0.011 | -0.011 | 0.023 | 0.06 | 0.11 | 0.05 | 0.03 | 0.05 | 0.05 |
| 4 | 70629413 | rs1847359 | 0.544 | 1.192 | 1.428 | 0.999 | -0.008 | -0.034 | -0.034 | 0 | 0.63 | 0.51 | 0.65 | 0.72 | 0.62 | 0.63 |
| 4 | 70667521 | rs12499134 | 2.227 | 1.394 | 0.727 | 0.96 | -0.031 | 0.036 | 0.036 | 0 | 0.36 | 0.04 | 0.35 | 0.21 | 0.3 | 0.43 |
| 4 | 70667744 | rs1813685 | 0.402 | 1.47 | 0.626 | 0.989 | -0.064 | 0.019 | 0.019 | 0 | 0.25 | 0.04 | 0.2 | 0.16 | 0.3 | 0.22 |
| 4 | 70671743 | rs2014777 | 1.222 | 1.474 | 0.841 | 1.112 | -0.004 | -0.069 | -0.069 | 0.212 | 0.67 | 0.15 | 0.68 | 0.49 | 0.7 | 0.72 |
| 4 | 70677967 | rs1220709 | 0.265 | 1.474 | 1.219 | 1.059 | 0.005 | -0.1 | -0.1 | 0 | 0.17 | 0.33 | 0.1 | 0 | 0.15 | 0.04 |
| 4 | 70679571 | rs1503721 | 2.391 | 1.472 | 1.154 | 1.092 | -0.03 | 0.025 | 0.025 | 0 | 0.36 | 0.05 | 0.35 | 0.23 | 0.3 | 0.44 |
| 4 | 70684350 | rs1581680 | 0.775 | 1.473 | 1.414 | 1.122 | 0.025 | 0.268 | 0.268 | 0 | 0.85 | 0.57 | 0.82 | 0.52 | 0.85 | 0.79 |
| 4 | 70694646 | rs1220718 | 0.672 | 1.474 | 1.306 | 1.063 | -0.017 | 0.055 | 0.055 | 0 | 0.15 | 0.06 | 0.07 | 0 | 0.15 | 0.04 |
| 4 | 70702006 | rs1529039 | 0.643 | 1.454 | 1.382 | 1.072 | -0.009 | 0.024 | 0.024 | 0 | 0.15 | 0.08 | 0.07 | 0 | 0.15 | 0.04 |
| 4 | 70704206 | rs1220726 | 0.622 | 1.448 | 1.474 | 1.068 | -0.003 | -0.011 | -0.011 | 0.607 | 0.15 | 0.11 | 0.08 | 0 | 0.15 | 0.04 |
| 4 | 70704542 | rs1910465 | 2.633 | 1.423 | 1.348 | 1.056 | -0.05 | 0.02 | 0.02 | 0 | 0.36 | 0.02 | 0.35 | 0.22 | 0.3 | 0.44 |
| 4 | 70704877 | rs13102618 | 0.438 | 1.46 | 1.345 | 1.084 | -0.069 | 0.011 | 0.011 | 0 | 0.25 | 0.03 | 0.2 | 0.15 | 0.29 | 0.21 |
| 4 | 70706366 | rs1220725 | 0.628 | 1.455 | 1.467 | 1.099 | 0.015 | 0.245 | 0.245 | 0 | 0.84 | 0.58 | 0.82 | 0.52 | 0.85 | 0.79 |
| 4 | 70707945 | rs1220724 | 0.644 | 1.439 | 1.457 | 1.356 | 0.014 | 0.22 | 0.22 | 0 | 0.84 | 0.58 | 0.82 | 0.56 | 0.85 | 0.79 |
| 4 | 70708461 | rs2331821 | 2.535 | 1.425 | 1.416 | 1.347 | 0.003 | 0.058 | 0.058 | 0 | 0.37 | 0.13 | 0.36 | 0.22 | 0.3 | 0.44 |
| 4 | 70709432 | rs3775778 | 2.526 | 1.425 | 1.438 | 1.407 | 0 | 0.058 | 0.058 | 0 | 0.37 | 0.12 | 0.36 | 0.22 | 0.3 | 0.44 |
| 4 | 70709510 | rs3775777 | 0.357 | 1.428 | 1.431 | 1.403 | -0.069 | 0.011 | 0.011 | 0 | 0.25 | 0.03 | 0.2 | 0.15 | 0.29 | 0.21 |
| 4 | 70714171 | rs4149535 | 0.797 | 1.688 | 1.362 | 1.338 | -0.037 | -0.036 | -0.036 | 0 | 0.07 | 0.02 | 0.14 | 0.14 | 0.18 | 0.11 |
| 4 | 70716627 | rs1220715 | 0.621 | 1.571 | 1.341 | 1.661 | -0.008 | 0.019 | 0.019 | 0 | 0.15 | 0.09 | 0.07 | 0 | 0.15 | 0.04 |
| 4 | 70718005 | rs4149534 | 0.321 | 1.546 | 1.254 | 1.662 | -0.07 | 0.01 | 0.01 | 0 | 0.25 | 0.03 | 0.2 | 0.15 | 0.29 | 0.21 |
| 4 | 70718036 | rs4149533 | 0.321 | 1.546 | 1.254 | 1.662 | -0.07 | 0.01 | 0.01 | 0 | 0.25 | 0.03 | 0.2 | 0.15 | 0.29 | 0.21 |
| 4 | 70718282 | rs3775775 | -0.136 | 1.553 | 1.299 | 1.656 | -0.003 | -0.03 | -0.03 | 0 | 0.11 | 0.27 | 0.1 | 0.07 | 0.09 | 0.1 |
| 4 | 70719238 | rs3775774 | 2.394 | 1.39 | 0.212 | 1.572 | -0.001 | 0.054 | 0.054 | 0 | 0.36 | 0.13 | 0.36 | 0.22 | 0.3 | 0.44 |
| 4 | 70719533 | rs10019305 | 0.237 | 1.4 | 0.184 | 1.575 | -0.033 | 0.03 | 0.03 | 0 | 0.25 | 0.07 | 0.21 | 0.15 | 0.29 | 0.21 |
| 4 | 70722297 | rs4694036 | 2.303 | 1.379 | 0.178 | 1.592 | 0 | 0.073 | 0.073 | 0 | 0.37 | 0.12 | 0.36 | 0.2 | 0.3 | 0.44 |
| 4 | 70724270 | rs3775770 | -0.018 | 1.273 | 0.227 | 1.015 | -0.031 | 0.017 | 0.017 | 0 | 0.25 | 0.07 | 0.2 | 0.17 | 0.28 | 0.21 |
| 4 | 70724996 | rs4149527 | -0.018 | 1.325 | -0.05 | 1.042 | -0.066 | -0.006 | -0.006 | 0 | 0.25 | 0.03 | 0.2 | 0.17 | 0.27 | 0.21 |
| 4 | 70725112 | rs3775768 | -0.019 | 1.349 | -0.02 | 1.043 | -0.031 | 0.017 | 0.017 | 0 | 0.25 | 0.07 | 0.2 | 0.17 | 0.28 | 0.21 |
| 4 | 70725698 | rs1534672 | 0.471 | 1.575 | 0.071 | 1.021 | -0.034 | 0.099 | 0.099 | 0 | 0.58 | 0.79 | 0.61 | 0.8 | 0.51 | 0.71 |
| 4 | 70726399 | rs4149525 | -0.855 | 1.067 | 1.055 | 1.104 | 0.014 | 0.163 | 0.163 | 0 | 0.11 | 0.28 | 0.18 | 0.25 | 0.14 | 0.2 |
| 4 | 70734059 | rs1590128 | 0.725 | 0.875 | -0.488 | -0.205 | -0.023 | 0.151 | 0.151 | 0.343 | 0.37 | 0.07 | 0.39 | 0.2 | 0.26 | 0.45 |
| 4 | 70753524 | rs13122069 | 0.565 | 1.162 | 0.026 | -0.051 | -0.006 | 0.022 | 0.022 | 0 | 0.11 | 0.08 | 0.19 | 0.43 | 0.31 | 0.12 |
| 4 | 70756086 | rs12507764 | -0.211 | 0.965 | 0.341 | -0.165 | 0.037 | 0.044 | 0.044 | 0 | 0.48 | 0.6 | 0.4 | 0.32 | 0.43 | 0.36 |
| 4 | 70759142 | rs17147830 | 1.054 | 0.359 | -0.02 | -0.727 | 0.004 | 0.01 | 0.01 | 0 | 0.1 | 0.01 | 0.14 | 0.1 | 0.05 | 0.07 |
| 4 | 70760559 | rs17628741 | 0.748 | 0.541 | 0.058 | -0.66 | -0.022 | 0.148 | 0.148 | 0.318 | 0.37 | 0.07 | 0.39 | 0.2 | 0.26 | 0.45 |
| 4 | 70765416 | rs2172159 | 0.266 | 0.339 | 0.78 | -0.594 | 0.002 | 0 | 0 | 0 | 0.48 | 0.48 | 0.49 | 0.29 | 0.37 | 0.52 |
| 4 | 70768646 | rs10518069 | -0.552 | 0.315 | 0.877 | -0.606 | 0.007 | 0.019 | 0.019 | 1.057 | 0.49 | 0.39 | 0.49 | 0.71 | 0.63 | 0.44 |
| 4 | 70774400 | rs7437229 | 1.871 | 0.169 | 0.821 | -0.725 | -0.027 | 0.068 | 0.068 | 4.613 | 0.24 | 0.12 | 0.28 | 0.1 | 0.23 | 0.34 |
| 4 | 70778543 | rs17147844 | 0.945 | 0.174 | 0.807 | -0.713 | 0.012 | 0.057 | 0.057 | 0 | 0.19 | 0.01 | 0.14 | 0.11 | 0.11 | 0.12 |
| 4 | 70783348 | rs12650947 | 0.885 | 0.174 | 0.996 | -0.712 | 0 | -0.005 | -0.005 | 0 | 0.2 | 0.16 | 0.16 | 0.11 | 0.11 | 0.12 |
| 4 | 70784305 | rs1220823 | -0.611 | 0.175 | 0.995 | -0.712 | -0.032 | -0.052 | -0.052 | 0 | 0.77 | 0.67 | 0.82 | 0.89 | 0.89 | 0.79 |
| 4 | 70789170 | rs4694257 | 1.552 | 0.279 | 0.771 | -0.598 | -0.006 | 0.013 | 0.013 | 5.089 | 0.26 | 0.22 | 0.32 | 0.14 | 0.26 | 0.35 |
| 4 | 70796390 | rs17696835 | 1.67 | 0.325 | 0.461 | -0.603 | -0.032 | 0.071 | 0.071 | 3.268 | 0.26 | 0.1 | 0.29 | 0.14 | 0.27 | 0.35 |
| 4 | 70807771 | rs10030475 | 0.273 | 0.368 | 0.619 | -0.563 | 0.005 | 0.116 | 0.116 | 0.471 | 0.48 | 0.3 | 0.46 | 0.29 | 0.41 | 0.48 |
| 4 | 70811806 | rs1237375 | -0.352 | 0.305 | 0.952 | -0.451 | 0.004 | 0.062 | 0.062 | 0.634 | 0.47 | 0.3 | 0.48 | 0.64 | 0.59 | 0.47 |
| 4 | 70813971 | rs1002082 | -0.394 | 0.314 | 1.015 | -0.447 | 0.002 | 0.021 | 0.021 | 0.499 | 0.48 | 0.39 | 0.49 | 0.64 | 0.59 | 0.47 |
| 4 | 70818296 | rs1388354 | -0.391 | 0.319 | 1.018 | -0.445 | 0.002 | 0.021 | 0.021 | 0.485 | 0.48 | 0.39 | 0.49 | 0.64 | 0.59 | 0.47 |
| 4 | 70833723 | rs9994584 | 0.798 | 0.317 | 0.897 | -0.421 | 0.027 | 0.027 | 0.027 | 0 | 0.2 | 0.08 | 0.17 | 0.17 | 0.12 | 0.11 |
| 4 | 70836155 | rs776840 | -1.224 | 0.798 | 1.009 | -0.093 | 0 | 0.031 | 0.031 | 0 | 0.9 | 0.91 | 0.84 | 0.65 | 0.77 | 0.85 |
| 4 | 70846936 | rs1356118 | 1.205 | 0.825 | 0.942 | -0.084 | -0.01 | 0.022 | 0.022 | 0 | 0.09 | 0.06 | 0.15 | 0.35 | 0.23 | 0.15 |
| 4 | 70863901 | rs17641990 | 1.626 | 0.315 | 0.62 | -0.579 | -0.029 | 0.043 | 0.043 | 0 | 0.08 | 0.01 | 0.06 | 0 | 0.09 | 0.03 |
| 4 | 70871257 | rs12648068 | 0.722 | 0.126 | 0.578 | -0.506 | 0.006 | -0.025 | -0.025 | 0 | 0.22 | 0.02 | 0.17 | 0.24 | 0.14 | 0.13 |
| 4 | 70892669 | rs776847 | -2.085 | 0.18 | 0.624 | -0.581 | -0.032 | 0.063 | 0.063 | 4.714 | 0.76 | 0.94 | 0.71 | 0.87 | 0.73 | 0.66 |
| 4 | 70910928 | rs2331856 | -2.072 | -0.061 | 0.69 | -0.564 | -0.032 | 0.06 | 0.06 | 4.587 | 0.75 | 0.94 | 0.71 | 0.87 | 0.73 | 0.66 |
| 4 | 70911720 | rs2138269 | -0.464 | -0.167 | 0.951 | -0.604 | 0.005 | 0.034 | 0.034 | 1.166 | 0.48 | 0.38 | 0.48 | 0.63 | 0.59 | 0.47 |
| 4 | 70917429 | rs6854359 | -1.947 | -0.149 | 0.781 | -0.631 | -0.017 | 0.032 | 0.032 | 3.802 | 0.74 | 0.86 | 0.69 | 0.83 | 0.7 | 0.62 |
| 4 | 70920812 | rs2009025 | 1.5 | -1.773 | 1.141 | -0.53 | 0 | 0.003 | 0.003 | 0 | 0.12 | 0.11 | 0.2 | 0.23 | 0.16 | 0.1 |
| 4 | 70932305 | rs17644018 | 0.138 | -1.307 | 1.207 | -0.396 | 0.018 | 0.026 | 0.026 | 0.04 | 0.37 | 0.21 | 0.31 | 0.27 | 0.33 | 0.25 |
| 4 | 70934328 | rs11249476 | 0.711 | -1.892 | 1.773 | -0.086 | -0.047 | -0.038 | -0.038 | 0 | 0.71 | 0.44 | 0.75 | 0.74 | 0.83 | 0.8 |
| 4 | 70934473 | rs11249478 | 0.795 | -1.906 | 1.781 | -0.088 | -0.017 | -0.012 | -0.012 | 0.155 | 0.7 | 0.34 | 0.74 | 0.74 | 0.83 | 0.81 |

| | | | | | | | | | | | | | | | |
|---|---|---|---|---|---|---|---|---|---|---|---|---|---|---|---|
| 4 | 70935067 | rs2063818 | 0.017 | -1.869 | 1.472 | -0.07 | 0.028 | 0.015 | 0.015 | 0 | 0.23 | 0.07 | 0.16 | 0.21 | 0.14 | 0.16 |
| 4 | 70957090 | rs9998140 | -0.392 | -1.855 | 1.569 | -0.063 | 0.014 | 0.012 | 0.012 | 0 | 0.26 | 0.24 | 0.18 | 0.21 | 0.14 | 0.2 |
| 4 | 70974315 | rs1717600 | -0.07 | -1.651 | 0.959 | 0.042 | -0.043 | -0.003 | -0.003 | 0 | 0.31 | 0.51 | 0.32 | 0.3 | 0.25 | 0.24 |
| 4 | 70974741 | rs17148162 | 0.466 | -1.732 | 0.281 | 0.002 | 0.027 | 0.02 | 0.02 | 0 | 0.19 | 0.06 | 0.15 | 0.16 | 0.11 | 0.12 |
| 4 | 70979385 | rs6855744 | -0.085 | -1.745 | 0.335 | 0 | -0.02 | -0.001 | -0.001 | 0 | 0.67 | 0.41 | 0.63 | 0.66 | 0.72 | 0.76 |
| 4 | 70980547 | rs13107309 | 1.208 | -1.73 | 0.192 | 0.006 | 0.032 | -0.012 | -0.012 | 0 | 0.46 | 0.3 | 0.47 | 0.46 | 0.58 | 0.6 |
| 4 | 70982011 | rs17148188 | 0.377 | -1.623 | -0.266 | 0.058 | 0.011 | -0.006 | -0.006 | 0 | 0.27 | 0.18 | 0.29 | 0.28 | 0.25 | 0.2 |
| 4 | 70985025 | rs7685261 | 0.197 | -1.661 | -0.871 | 0.139 | 0.02 | 0.021 | 0.021 | 0 | 0.2 | 0.04 | 0.14 | 0.16 | 0.13 | 0.12 |
| 4 | 70990224 | rs1607989 | 0.148 | -1.643 | -0.897 | 0.198 | 0.025 | 0.011 | 0.011 | 0 | 0.2 | 0.1 | 0.15 | 0.19 | 0.11 | 0.12 |
| 4 | 70999619 | rs1399252 | -0.222 | -1.436 | -0.92 | 0.262 | -0.046 | 0.009 | 0.009 | 0 | 0.68 | 0.53 | 0.67 | 0.65 | 0.75 | 0.78 |
| 4 | 71008940 | rs3775759 | -0.365 | -0.686 | -1.465 | 0.161 | -0.012 | -0.013 | -0.013 | 0 | 0.2 | 0.01 | 0.18 | 0.21 | 0.16 | 0.12 |
| 4 | 71018758 | rs708235 | 0.184 | -1.053 | -0.971 | -0.089 | -0.028 | -0.033 | -0.033 | 0 | 0.92 | 0.87 | 0.99 | 1 | 0.99 | 1 |
| 4 | 71021940 | rs17148265 | -0.049 | -1.601 | -0.758 | 0.366 | -0.009 | 0.06 | 0.06 | 0.026 | 0.18 | 0.17 | 0.3 | 0.45 | 0.15 | 0.12 |
| 4 | 71024240 | rs1613461 | 0.516 | -1.991 | -0.999 | 0.116 | 0.007 | 0.005 | 0.005 | 0 | 0.05 | 0.06 | 0.02 | 0 | 0.01 | 0.11 |
| 4 | 71025548 | rs11735430 | 1.083 | -2.446 | -1.332 | -0.594 | -0.03 | -0.035 | -0.035 | 0 | 0.24 | 0.17 | 0.43 | 0.43 | 0.51 | 0.34 |
| 4 | 71026763 | rs11942432 | 0.562 | -2.677 | -1.362 | -0.885 | 0.059 | 0.235 | 0.235 | 2.857 | 0.4 | 0.05 | 0.23 | 0.13 | 0.31 | 0.37 |
| 4 | 71032811 | rs6600807 | 0.238 | -3.164 | -0.772 | -1.213 | -0.076 | -0.163 | -0.163 | 0 | 0.71 | 0.53 | 0.88 | 1 | 0.88 | 0.75 |
| 4 | 71041750 | rs17647132 | 1.004 | -3.265 | -1.164 | -1.284 | 0.006 | 0.107 | 0.107 | 0.145 | 0.22 | 0.02 | 0.13 | 0.1 | 0.21 | 0.22 |
| 4 | 71043494 | rs13105632 | 0.236 | -2.539 | -0.978 | -0.961 | 0.063 | -0.02 | -0.02 | 0 | 0.55 | 0.23 | 0.63 | 0.49 | 0.68 | 0.57 |
| 4 | 71045118 | rs13151337 | 0.734 | -1.877 | -0.608 | -0.633 | -0.061 | -0.044 | -0.044 | 0.561 | 0.33 | 0.21 | 0.5 | 0.4 | 0.48 | 0.32 |
| 4 | 71046288 | rs10049496 | 0.578 | -1.267 | 0.061 | -0.562 | -0.011 | -0.054 | -0.054 | 0 | 0.16 | 0.21 | 0.09 | 0.01 | 0.09 | 0.13 |
| 4 | 71046529 | rs1709276 | -0.566 | -1.263 | 0.025 | -0.393 | -0.005 | -0.033 | -0.033 | 0.05 | 0.23 | 0.4 | 0.17 | 0.18 | 0.17 | 0.3 |
| 4 | 71048803 | rs777390 | -0.414 | -0.605 | -0.387 | 0.143 | 0.016 | 0.031 | 0.031 | 0 | 0.07 | 0.11 | 0.09 | 0.18 | 0.08 | 0.17 |
| 4 | 71051031 | rs17652789 | -0.272 | -0.365 | -0.671 | -0.087 | 0.008 | 0.071 | 0.071 | 0.149 | 0.53 | 0.33 | 0.3 | 0.33 | 0.43 | 0.56 |
| 4 | 71052152 | rs869894 | -0.748 | -0.21 | -0.127 | 0.274 | -0.008 | -0.01 | -0.01 | 0 | 0.71 | 0.74 | 0.55 | 0.55 | 0.59 | 0.71 |
| 4 | 71058996 | rs17148330 | 0.991 | -1.343 | -0.711 | -0.234 | 0.059 | -0.014 | -0.014 | 0.011 | 0.2 | 0.05 | 0.09 | 0.29 | 0.14 | 0.24 |
| 4 | 71061842 | rs1383436 | 0.595 | -0.541 | -0.786 | -0.433 | 0.002 | -0.007 | -0.007 | 0 | 0.06 | 0.06 | 0.22 | 0 | 0.13 | 0.05 |
| 4 | 71062426 | rs17704351 | -0.058 | -0.466 | -0.836 | -0.62 | 0.147 | 0.143 | 0.143 | 0 | 0.48 | 0.16 | 0.2 | 0.3 | 0.28 | 0.52 |
| 4 | 71063359 | rs1514393 | -0.009 | -0.505 | -0.856 | -0.639 | 0.142 | 0.139 | 0.139 | 0.008 | 0.47 | 0.16 | 0.2 | 0.3 | 0.28 | 0.52 |
| 4 | 71063374 | rs1514394 | 0.992 | -0.508 | -0.849 | -0.649 | 0.043 | 0.104 | 0.104 | 0 | 0.26 | 0.09 | 0.11 | 0.01 | 0.14 | 0.26 |
| 4 | 71067317 | rs4694273 | -1.642 | -0.353 | -0.574 | -0.567 | 0.005 | 0.014 | 0.014 | 0 | 0.73 | 0.72 | 0.71 | 0.55 | 0.63 | 0.75 |
| 4 | 71068489 | rs3196714 | 0.038 | -0.406 | -0.755 | -0.554 | 0.142 | 0.138 | 0.138 | 0.005 | 0.47 | 0.16 | 0.2 | 0.3 | 0.28 | 0.5 |
| 4 | 71069113 | rs3775756 | 2.118 | -0.493 | 0.069 | -0.539 | -0.002 | 0.003 | 0.003 | 0 | 0.13 | 0.1 | 0.17 | 0.22 | 0.11 | 0.14 |
| 4 | 71079192 | rs12504890 | 2.118 | -0.493 | 0.069 | -0.557 | -0.002 | 0.003 | 0.003 | 0 | 0.13 | 0.1 | 0.17 | 0.22 | 0.11 | 0.14 |
| 4 | 71081038 | rs1828826 | -0.002 | -0.523 | 0.04 | -0.59 | 0.146 | 0.142 | 0.142 | 0 | 0.48 | 0.16 | 0.2 | 0.3 | 0.28 | 0.52 |
| 4 | 71088563 | rs28563113 | 1.904 | -0.189 | 0.061 | -0.307 | -0.002 | 0.003 | 0.003 | 0 | 0.13 | 0.1 | 0.16 | 0.22 | 0.11 | 0.14 |
| 4 | 71090658 | rs10518072 | 0.299 | 0.104 | 0.012 | -0.489 | 0.038 | -0.03 | -0.03 | 0 | 0.06 | 0.11 | 0.23 | 0 | 0.13 | 0.05 |
| 4 | 71091123 | rs1514399 | 0.261 | -0.002 | -0.29 | -0.521 | 0.006 | -0.014 | -0.014 | 0 | 0.26 | 0.14 | 0.17 | 0.32 | 0.26 | 0.3 |
| 4 | 71106017 | rs2249813 | 0.3 | 0.02 | -0.128 | -0.549 | 0.004 | -0.008 | -0.008 | 0.242 | 0.66 | 0.58 | 0.59 | 0.68 | 0.61 | 0.64 |
| 4 | 71109108 | rs3775753 | 1.639 | -0.288 | -0.857 | -0.483 | -0.027 | -0.001 | -0.001 | 0 | 0.16 | 0 | 0.11 | 0.13 | 0.12 | 0.06 |
| 4 | 71113124 | rs17654723 | 0.427 | -0.055 | -0.424 | -0.274 | -0.027 | 0.025 | 0.025 | 1.017 | 0.06 | 0.02 | 0.22 | 0 | 0.13 | 0.05 |
| 4 | 71115060 | rs3775739 | 0.947 | -0.273 | -0.471 | -0.264 | 0.068 | -0.052 | -0.052 | 0 | 0.2 | 0.01 | 0.09 | 0.29 | 0.14 | 0.25 |
| 4 | 71117202 | rs1399253 | -0.501 | -0.156 | -0.268 | -0.309 | -0.016 | 0.013 | 0.013 | 0.691 | 0.32 | 0.26 | 0.4 | 0.32 | 0.39 | 0.36 |
| 4 | 71121681 | rs2046106 | -0.678 | -0.272 | -0.018 | -0.367 | 0.016 | 0.01 | 0.01 | 2.037 | 0.5 | 0.48 | 0.68 | 0.65 | 0.74 | 0.52 |
| 4 | 71126287 | rs970681 | -1.285 | -0.282 | 0.135 | -0.51 | -0.004 | -0.022 | -0.022 | 0.756 | 0.95 | 0.93 | 0.98 | 1 | 0.96 | 0.97 |
| 4 | 71127403 | rs17148451 | -0.639 | -0.135 | 0.042 | -0.49 | 0.021 | -0.042 | -0.042 | 1.198 | 0.3 | 0.41 | 0.41 | 0.22 | 0.37 | 0.3 |
| 4 | 71137121 | rs17148491 | 0.109 | 0.07 | 0.209 | -0.389 | 0.14 | -0.003 | -0.003 | 0 | 0.07 | 0.27 | 0.09 | 0.02 | 0.12 | 0.03 |
| 4 | 71142199 | rs13127641 | 1.585 | -0.143 | -0.221 | -0.249 | -0.123 | -0.147 | -0.147 | 1.071 | 0.2 | 0.03 | 0.27 | 0.44 | 0.37 | 0.21 |
| 4 | 71147013 | rs12510290 | 1.479 | -0.837 | 0.488 | -0.347 | -0.003 | 0 | 0 | 1.961 | 0.16 | 0.11 | 0.13 | 0.16 | 0.11 | 0.11 |
| 4 | 71150652 | rs1842469 | -0.556 | -0.741 | 0.603 | -0.455 | -0.021 | -0.037 | -0.037 | 0 | 0.25 | 0.38 | 0.34 | 0.19 | 0.24 | 0.28 |
| 4 | 71151940 | rs17148526 | 0.481 | -0.75 | 0.268 | -0.466 | 0.042 | -0.005 | -0.005 | 0 | 0.17 | 0.04 | 0.11 | 0.19 | 0.14 | 0.21 |
| 4 | 71154114 | rs1812922 | -0.659 | -0.49 | 0.486 | -0.28 | -0.004 | -0.02 | -0.02 | 0 | 0.95 | 0.93 | 0.98 | 1 | 0.96 | 0.97 |
| 4 | 71156301 | rs1976001 | 0.318 | -0.343 | 0.291 | -0.092 | -0.027 | 0.027 | 0.027 | 0 | 0.05 | 0.01 | 0.2 | 0 | 0.1 | 0.05 |
| 4 | 71165706 | rs729314 | 0.058 | -0.174 | -0.313 | -0.006 | 0.044 | -0.033 | -0.033 | 0.312 | 0.06 | 0.17 | 0.22 | 0 | 0.1 | 0.06 |
| 4 | 71166102 | rs17146493 | 0.598 | -0.93 | -0.842 | 0.22 | 0.04 | -0.035 | -0.035 | 0.065 | 0.13 | 0.01 | 0.06 | 0.19 | 0.08 | 0.19 |
| 4 | 71171720 | rs10007509 | 0.686 | -1.021 | -0.516 | 0.264 | -0.007 | 0.001 | 0.001 | 1.797 | 0.18 | 0.15 | 0.15 | 0.17 | 0.11 | 0.1 |
| 4 | 71181917 | rs10024824 | 0.655 | -0.904 | -0.516 | 0.249 | -0.003 | 0 | 0 | 1.622 | 0.19 | 0.16 | 0.15 | 0.18 | 0.12 | 0.16 |
| 4 | 71184850 | rs4421051 | 1.392 | -0.626 | -0.927 | 0.302 | -0.042 | -0.02 | -0.02 | 0 | 0.23 | 0.12 | 0.38 | 0.22 | 0.35 | 0.18 |
| 4 | 71185226 | rs9329032 | -0.424 | -0.936 | -0.455 | -0.077 | 0.016 | -0.013 | -0.013 | 0 | 0.62 | 0.44 | 0.5 | 0.78 | 0.49 | 0.75 |
| 4 | 71185783 | rs1904851 | -0.947 | -0.959 | 0.004 | -0.108 | 0.014 | -0.021 | -0.021 | 0 | 0.94 | 0.78 | 0.96 | 1 | 0.96 | 0.98 |
| 4 | 71190111 | rs2013685 | 1.188 | -1.381 | -0.213 | -0.245 | -0.017 | -0.021 | -0.021 | 0.19 | 0.26 | 0.18 | 0.27 | 0.4 | 0.27 | 0.38 |
| 4 | 71190301 | rs717360 | 0.222 | -1.437 | -0.094 | -0.249 | 0.035 | -0.004 | -0.004 | 0.655 | 0.36 | 0.25 | 0.23 | 0.38 | 0.22 | 0.37 |
| 4 | 71201388 | rs1351419 | -1.073 | -1.389 | -0.077 | -0.238 | 0.014 | -0.021 | -0.021 | 0 | 0.94 | 0.78 | 0.96 | 1 | 0.96 | 0.98 |
| 4 | 71201648 | rs2291182 | 0.935 | -1.387 | -0.491 | -0.208 | 0.041 | -0.028 | -0.028 | 0.026 | 0.17 | 0.01 | 0.07 | 0.2 | 0.1 | 0.21 |
| 4 | 71201943 | rs13039 | 0.629 | -1.234 | -0.256 | -0.129 | 0.002 | 0.001 | 0.001 | 0.14 | 0.24 | 0.22 | 0.19 | 0.17 | 0.19 | 0.2 |
| 4 | 71206687 | rs9992346 | -0.145 | -1.202 | -0.248 | -0.149 | 0.002 | 0.009 | 0.009 | 0 | 0.49 | 0.4 | 0.45 | 0.58 | 0.46 | 0.57 |
| 4 | 71207398 | rs4694051 | -0.251 | -1.258 | -0.115 | -0.196 | 0.002 | 0.002 | 0.002 | 0 | 0.5 | 0.53 | 0.55 | 0.42 | 0.57 | 0.45 |
| 4 | 71214585 | rs17148650 | 0.722 | -1.274 | 0.083 | -0.2 | 0.027 | -0.012 | -0.012 | 0 | 0.06 | 0.26 | 0.05 | 0 | 0.04 | 0.02 |
| 4 | 71217628 | rs1480757 | -0.216 | -1.036 | -0.038 | -0.228 | -0.001 | 0.001 | 0.001 | 0 | 0.49 | 0.53 | 0.55 | 0.42 | 0.55 | 0.44 |

| | | | | | | | | | | | | | | | | |
|---|---|---|---|---|---|---|---|---|---|---|---|---|---|---|---|---|
| 4 | 71218054 | rs6600813 | -0.521 | -1.099 | 0.054 | -0.268 | -0.033 | -0.04 | -0.04 | 0.207 | 0.27 | 0.41 | 0.17 | 0.2 | 0.22 | 0.26 |
| 4 | 71218510 | rs12647928 | 1.422 | -1.184 | 0.018 | -0.272 | -0.002 | 0 | 0 | 0.964 | 0.2 | 0.18 | 0.15 | 0.17 | 0.15 | 0.16 |
| 4 | 71219358 | rs17660273 | 0.846 | -1.032 | -0.005 | -0.264 | 0.003 | -0.014 | -0.014 | 0 | 0.05 | 0.07 | 0.06 | 0 | 0.11 | 0.03 |
| 4 | 71221837 | rs17148759 | 1.196 | -1.019 | -0.064 | -0.257 | 0.023 | -0.02 | -0.02 | 0 | 0.14 | 0.04 | 0.08 | 0.2 | 0.11 | 0.23 |
| 4 | 71226962 | rs17148794 | -0.596 | -0.779 | 0.167 | -0.243 | -0.001 | -0.082 | -0.082 | 0.148 | 0.13 | 0.35 | 0.08 | 0 | 0.12 | 0.03 |
| 4 | 71231202 | rs13137718 | 0.028 | -0.086 | -0.192 | 0.144 | 0.063 | 0.05 | 0.05 | 0 | 0.06 | 0.15 | 0.11 | 0.2 | 0.11 | 0.03 |
| 4 | 71231241 | rs13435339 | 0.235 | -0.299 | -0.667 | 0.093 | -0.025 | 0.021 | 0.021 | 0.211 | 0.6 | 0.24 | 0.62 | 0.63 | 0.58 | 0.81 |
| 4 | 71233986 | rs1813261 | -0.838 | -0.706 | -0.926 | 0.024 | -0.013 | -0.011 | -0.011 | 4.721 | 0.72 | 0.77 | 0.7 | 0.59 | 0.69 | 0.52 |
| 4 | 71236847 | rs17661300 | 1.898 | -0.626 | -0.991 | 0.098 | -0.063 | -0.045 | -0.045 | 0.021 | 0.22 | 0.07 | 0.38 | 0.23 | 0.33 | 0.16 |
| 4 | 71237421 | rs11249497 | 0.339 | -0.295 | -0.695 | 0.242 | 0.014 | 0.004 | 0.004 | 0.417 | 0.18 | 0.07 | 0.12 | 0.2 | 0.17 | 0.23 |
| 4 | 71252241 | rs6600824 | -0.294 | -0.273 | -0.448 | 0.299 | -0.015 | -0.01 | -0.01 | 0 | 0.7 | 0.77 | 0.7 | 0.59 | 0.68 | 0.61 |
| 4 | 71260539 | rs1490482 | 0.109 | -0.516 | -0.052 | 0.49 | -0.014 | 0.006 | 0.006 | 0 | 0.63 | 0.42 | 0.72 | 0.59 | 0.7 | 0.63 |
| 4 | 71261913 | rs1387964 | 0.533 | -0.907 | -0.305 | 0.326 | 0.079 | 0.002 | 0.002 | 0 | 0.54 | 0.25 | 0.69 | 0.59 | 0.68 | 0.45 |
| 4 | 71263790 | rs4426856 | 0.42 | -1.21 | -0.192 | 0.36 | 0.076 | 0.02 | 0.02 | 0 | 0.53 | 0.67 | 0.75 | 0.57 | 0.73 | 0.45 |
| 4 | 71264191 | rs11938536 | 1.353 | -1.192 | -0.178 | 0.366 | -0.018 | -0.009 | -0.009 | 0 | 0.3 | 0.23 | 0.54 | 0.38 | 0.53 | 0.3 |
| 4 | 71265172 | rs1844635 | -0.652 | -1.205 | -0.159 | 0.348 | 0 | -0.006 | -0.006 | 0.051 | 0.6 | 0.54 | 0.42 | 0.61 | 0.46 | 0.52 |
| 4 | 71265840 | rs7682761 | 1.007 | -1.108 | -0.096 | 0.345 | -0.022 | 0.014 | 0.014 | 0.032 | 0.23 | 0.44 | 0.2 | 0.19 | 0.19 | 0.14 |
| 4 | 71272547 | rs6827846 | -0.431 | -0.686 | -0.14 | 0.445 | 0.026 | 0.016 | 0.016 | 0 | 0.89 | 0.81 | 0.87 | 0.8 | 0.87 | 0.79 |
| 4 | 71290636 | rs17662455 | 0.719 | -0.78 | -0.304 | 0.419 | -0.03 | -0.001 | -0.001 | 0 | 0.4 | 0.27 | 0.51 | 0.4 | 0.51 | 0.49 |
| 4 | 71296033 | rs10018283 | 1.202 | -1.021 | -0.5 | 0.268 | 0.013 | -0.039 | -0.039 | 1.438 | 0.17 | 0.06 | 0.14 | 0.18 | 0.13 | 0.16 |
| 4 | 71296050 | rs17148915 | 1.271 | -1.112 | -0.195 | 0.237 | 0.039 | 0.018 | 0.018 | 0 | 0.25 | 0.12 | 0.16 | 0.21 | 0.16 | 0.15 |
| 4 | 71309233 | rs962922 | 0.703 | -0.853 | 0.302 | 0.414 | 0.011 | 0.028 | 0.028 | 0 | 0.71 | 0.57 | 0.76 | 0.58 | 0.7 | 0.58 |
| 4 | 71310245 | rs12645253 | 1.568 | -0.88 | 0.013 | 0.469 | -0.004 | -0.057 | -0.057 | 0.795 | 0.1 | 0.01 | 0.06 | 0.21 | 0.09 | 0.15 |
| 4 | 71311646 | rs2172006 | 0.011 | -0.852 | 0.358 | 0.406 | 0.083 | 0.065 | 0.065 | 0 | 0.88 | 0.65 | 0.86 | 0.79 | 0.87 | 0.75 |
| 4 | 71329490 | rs1109501 | 0.549 | -0.995 | 0.115 | 0.386 | 0.045 | -0.093 | -0.093 | 0 | 0.4 | 0.06 | 0.26 | 0.39 | 0.29 | 0.3 |
| 4 | 71346701 | rs6826961 | -0.209 | -0.842 | 0.335 | 0.435 | 0.013 | 0.038 | 0.038 | 0.153 | 0.21 | 0.3 | 0.2 | 0.41 | 0.22 | 0.4 |
| 4 | 71347854 | rs3733491 | -1.188 | -0.907 | 0.396 | 0.386 | 0.02 | -0.032 | -0.032 | 0 | 0.63 | 0.55 | 0.5 | 0.8 | 0.51 | 0.73 |
| 4 | 71351436 | rs4694317 | 1.852 | -0.941 | 0.463 | 0.35 | -0.034 | 0.016 | 0.016 | 0 | 0.28 | 0.17 | 0.44 | 0.2 | 0.41 | 0.24 |
| 4 | 71353586 | rs6837080 | 1.406 | -0.975 | 0.497 | 0.378 | -0.011 | 0.026 | 0.026 | 0.213 | 0.11 | 0.14 | 0.07 | 0.2 | 0.09 | 0.2 |
| 4 | 71357077 | rs17838972 | -2.472 | -1.048 | 0.297 | 0.369 | 0.006 | -0.07 | -0.07 | 0 | 0.87 | 0.97 | 0.87 | 0.8 | 0.92 | 0.88 |
| 4 | 71362025 | rs17148999 | 0.742 | -1.002 | 0.376 | 0.341 | 0.008 | -0.052 | -0.052 | 0 | 0.09 | 0.28 | 0.06 | 0 | 0.08 | 0.02 |
| 4 | 71362151 | rs17732035 | 0.16 | -0.96 | 0.374 | 0.339 | 0.001 | 0.004 | 0.004 | 0 | 0.47 | 0.54 | 0.57 | 0.4 | 0.59 | 0.46 |
| 4 | 71363289 | rs17675439 | 0.954 | -0.946 | 0.332 | 0.354 | -0.023 | -0.007 | -0.007 | 0 | 0.38 | 0.26 | 0.51 | 0.4 | 0.51 | 0.44 |
| 4 | 71368508 | rs3864157 | 1.158 | -0.898 | 0.432 | 0.343 | 0.008 | -0.044 | -0.044 | 0 | 0.08 | 0.21 | 0.05 | 0 | 0.08 | 0.02 |
| 4 | 71372099 | rs3852091 | -0.943 | -0.875 | 0.453 | 0.364 | 0.024 | 0.036 | 0.036 | 0 | 0.28 | 0.44 | 0.24 | 0.41 | 0.3 | 0.44 |
| 4 | 71372596 | rs17149002 | -1.238 | -0.932 | 0.442 | 0.348 | -0.014 | -0.01 | -0.01 | 0 | 0.57 | 0.68 | 0.42 | 0.6 | 0.51 | 0.64 |
| 4 | 71376022 | rs17732605 | 0.98 | -0.934 | 0.287 | 0.336 | 0.016 | 0.017 | 0.017 | 0 | 0.3 | 0.26 | 0.18 | 0.19 | 0.21 | 0.2 |
| 4 | 71376829 | rs3852093 | 0.076 | -0.933 | 0.303 | 0.332 | -0.024 | -0.044 | -0.044 | 0.165 | 0.38 | 0.46 | 0.23 | 0.19 | 0.29 | 0.22 |
| 4 | 71385167 | rs28420645 | 0.982 | -1.065 | 0.255 | 0.386 | -0.014 | 0.02 | 0.02 | 0 | 0.38 | 0.32 | 0.51 | 0.2 | 0.52 | 0.28 |
| 4 | 71388551 | rs7660807 | 1.623 | -0.966 | 0.106 | 0.474 | -0.009 | -0.056 | -0.056 | 0.251 | 0.1 | 0.01 | 0.06 | 0.2 | 0.1 | 0.17 |
| 4 | 71403159 | rs4024031 | 0.11 | -0.913 | 0.309 | 0.373 | 0.069 | 0.049 | 0.049 | 0 | 0.1 | 0.28 | 0.14 | 0.21 | 0.13 | 0.23 |
| 4 | 71407154 | rs28408923 | -0.321 | -1.151 | -0.061 | 0.344 | 0.105 | -0.017 | -0.017 | 3.656 | 0.57 | 0.79 | 0.75 | 0.58 | 0.75 | 0.7 |
| 4 | 71408454 | rs28770195 | 0.863 | -0.984 | -0.105 | 0.186 | 0.09 | 0.025 | 0.025 | 0 | 0.3 | 0.03 | 0.11 | 0.2 | 0.16 | 0.2 |
| 4 | 71419827 | rs28719939 | 0.537 | -0.893 | -0.014 | 0.235 | 0.056 | 0.03 | 0.03 | 0 | 0.3 | 0.17 | 0.13 | 0.2 | 0.16 | 0.2 |
| 4 | 71422796 | rs3852105 | 0.238 | -0.346 | -0.011 | 0.506 | 0 | 0 | 0 | 0 | 0.39 | 0.37 | 0.19 | 0.2 | 0.24 | 0.26 |
| 4 | 71439354 | rs1490470 | -0.381 | 0.586 | 0.218 | 0.909 | -0.002 | 0.018 | 0.018 | 0 | 0.63 | 0.49 | 0.4 | 0.53 | 0.44 | 0.58 |
| 4 | 71447093 | rs17733915 | -1.016 | 0.824 | -0.776 | 0.832 | -0.054 | 0 | 0 | 0 | 0.7 | 0.82 | 0.5 | 0.74 | 0.55 | 0.76 |
| 4 | 71453028 | rs28661403 | 0.123 | 0.163 | -0.724 | 0.912 | 0.015 | 0.009 | 0.009 | 0.545 | 0.5 | 0.52 | 0.33 | 0.38 | 0.34 | 0.32 |
| 4 | 71455685 | rs28660083 | -0.838 | 0.422 | -0.903 | 1.047 | -0.045 | -0.037 | -0.037 | 0.088 | 0.7 | 0.84 | 0.46 | 0.56 | 0.52 | 0.66 |
| 4 | 71455961 | rs28432331 | -0.624 | 0.432 | -0.894 | 1.052 | -0.043 | -0.036 | -0.036 | 0.847 | 0.7 | 0.84 | 0.46 | 0.56 | 0.52 | 0.7 |
| 4 | 71466451 | rs28509200 | -0.027 | -0.709 | -0.229 | 0.767 | -0.057 | -0.029 | -0.029 | 0 | 0.28 | 0.39 | 0.15 | 0.19 | 0.13 | 0.17 |
| 4 | 71466914 | rs4694075 | -0.431 | -0.657 | -0.359 | 0.722 | -0.003 | 0.004 | 0.004 | 0 | 0.48 | 0.67 | 0.42 | 0.54 | 0.47 | 0.43 |
| 4 | 71469604 | rs7439186 | 0.201 | -1.406 | 0.405 | 0.898 | -0.026 | 0.025 | 0.025 | 0.774 | 0.12 | 0.15 | 0.07 | 0.15 | 0.06 | 0.14 |
| 4 | 71472426 | rs7680880 | -1.095 | -1.397 | 0.982 | 0.724 | -0.052 | 0.011 | 0.011 | 0.958 | 0.14 | 0.68 | 0.13 | 0.15 | 0.08 | 0.17 |
| 4 | 71506412 | rs12640848 | 0.65 | -1.441 | -0.43 | 0.249 | 0.146 | 0.121 | 0.121 | 0 | 0.49 | 0.03 | 0.34 | 0.23 | 0.67 | 0.47 |
| 4 | 71509086 | rs7671281 | -1.066 | -0.908 | 1.069 | 0.217 | -0.071 | -0.095 | -0.095 | 0.105 | 0.15 | 0.53 | 0.11 | 0.02 | 0.05 | 0.06 |
| 4 | 71509431 | rs3796704 | -0.78 | -0.967 | 1.091 | -0.013 | -0.05 | -0.068 | -0.068 | 0 | 0.13 | 0.38 | 0.1 | 0.02 | 0.05 | 0.05 |
| 4 | 71510914 | rs7679569 | -0.746 | -0.949 | 1.077 | -0.004 | -0.05 | -0.068 | -0.068 | 0 | 0.13 | 0.38 | 0.1 | 0.02 | 0.05 | 0.05 |
| 4 | 71520178 | rs13146209 | 0.296 | 0.408 | -1.277 | 0.679 | -0.058 | -0.019 | -0.019 | 0.242 | 0.42 | 0.14 | 0.45 | 0.35 | 0.49 | 0.42 |
| 4 | 71520630 | rs16845299 | -0.638 | -0.016 | -1.092 | 0.595 | 0.007 | -0.014 | -0.014 | 0 | 0.35 | 0.21 | 0.19 | 0.35 | 0.31 | 0.5 |
| 4 | 71540886 | rs16845359 | -1.56 | -0.725 | 1.619 | -0.178 | -0.029 | -0.071 | -0.071 | 0 | 0.16 | 0.7 | 0.08 | 0 | 0.05 | 0.06 |
| 4 | 71552398 | rs16845383 | -0.832 | -0.461 | 2.406 | -0.072 | -0.012 | -0.036 | -0.036 | 0.151 | 0.07 | 0.18 | 0.03 | 0 | 0.03 | 0.01 |
| 4 | 71554275 | rs3733488 | -1.092 | -0.549 | 1.901 | -0.263 | -0.011 | 0.031 | 0.031 | 0 | 0.08 | 0.01 | 0.05 | 0.03 | 0.09 | 0.08 |
| 4 | 71564925 | rs7697206 | -1.53 | -0.517 | 2.18 | -0.207 | 0.001 | -0.037 | -0.037 | 0 | 0.09 | 0.34 | 0.04 | 0 | 0.03 | 0.02 |
| 4 | 71583087 | rs7678779 | -2.712 | -0.874 | 2.457 | 0.342 | -0.038 | 0.012 | 0.012 | 1.3 | 0.18 | 0.66 | 0.13 | 0.21 | 0.05 | 0.15 |
| 4 | 71587040 | rs28475626 | -1.616 | -0.886 | 2.873 | 0.308 | -0.006 | -0.065 | -0.065 | 0 | 0.11 | 0.31 | 0.05 | 0 | 0.04 | 0.05 |
| 4 | 71597069 | rs10009233 | -2.848 | -0.901 | 2.692 | 0.33 | -0.037 | 0.016 | 0.016 | 1.662 | 0.17 | 0.66 | 0.13 | 0.21 | 0.05 | 0.14 |
| 4 | 71625511 | rs16845433 | -0.112 | -0.869 | 2.7 | 0.329 | -0.007 | -0.02 | -0.02 | 0 | 0.05 | 0.07 | 0.03 | 0 | 0.02 | 0.01 |
| 4 | 71626572 | rs3775730 | -2.023 | -0.889 | 2.797 | 0.34 | -0.018 | 0.017 | 0.017 | 0 | 0.11 | 0.13 | 0.09 | 0.21 | 0.03 | 0.11 |

| | | | | | | | | | | | | | | | | |
|---|---|---|---|---|---|---|---|---|---|---|---|---|---|---|---|---|
| 4 | 71630161 | rs2278134 | -1.625 | -0.889 | 3.125 | 0.339 | -0.016 | 0.038 | 0.038 | 0 | 0.06 | 0.08 | 0.06 | 0.21 | 0.01 | 0.1 |
| 4 | 71658684 | rs10012829 | -1.066 | -0.879 | 3.253 | 0.332 | 0.035 | -0.001 | -0.001 | 0 | 0.06 | 0.24 | 0.02 | 0 | 0.01 | 0.04 |
| 4 | 71672653 | rs1043903 | -2.919 | -0.733 | 3.15 | 0.329 | 0.049 | 0.014 | 0.014 | 0.581 | 0.48 | 0.78 | 0.41 | 0.4 | 0.38 | 0.47 |
| 4 | 71672811 | rs1043906 | -0.809 | -0.633 | 3.241 | 0.539 | 0.035 | -0.001 | -0.001 | 0.287 | 0.06 | 0.24 | 0.02 | 0 | 0.01 | 0.04 |
| 4 | 71673137 | rs7681549 | -1.602 | -0.64 | 3.142 | 0.54 | -0.027 | 0.037 | 0.037 | 0 | 0.1 | 0.16 | 0.09 | 0.21 | 0.03 | 0.1 |
| 4 | 71680954 | rs11737063 | 1.212 | -0.602 | 3.215 | 0.426 | 0.05 | 0.007 | 0.007 | 2.478 | 0.93 | 0.62 | 0.97 | 1 | 0.99 | 0.96 |
| 4 | 71685092 | rs10938024 | 1.244 | -0.602 | 3.227 | 0.426 | 0.049 | 0.006 | 0.006 | 2.53 | 0.93 | 0.62 | 0.97 | 1 | 0.99 | 0.96 |
| 4 | 71686450 | rs7686378 | 0.141 | -0.599 | 3.116 | 0.428 | -0.007 | -0.02 | -0.02 | 0.026 | 0.05 | 0.07 | 0.02 | 0 | 0.02 | 0.01 |
| 4 | 71700247 | rs13112870 | 2.242 | -0.621 | 3.198 | 0.481 | -0.001 | 0.011 | 0.011 | 1.383 | 0.87 | 0.65 | 0.92 | 0.9 | 0.91 | 0.89 |
| 4 | 71702684 | rs3775729 | 2.555 | -0.613 | 3.198 | 0.533 | -0.044 | 0.01 | 0.01 | 1.009 | 0.82 | 0.36 | 0.87 | 0.8 | 0.96 | 0.86 |
| 4 | 71719138 | rs13137120 | 1.901 | -0.602 | 3.198 | 0.579 | -0.012 | 0.107 | 0.107 | 0.749 | 0.9 | 0.66 | 0.93 | 0.84 | 0.98 | 0.9 |
| 4 | 71724706 | rs6446909 | 2.256 | -0.599 | 2.909 | 0.575 | -0.062 | 0.034 | 0.034 | 0.828 | 0.83 | 0.47 | 0.87 | 0.8 | 0.96 | 0.86 |
| 4 | 71748245 | rs12649012 | 2.423 | -0.509 | 3.076 | 0.582 | -0.035 | 0.082 | 0.082 | 0.099 | 0.86 | 0.58 | 0.87 | 0.8 | 0.96 | 0.89 |
| 4 | 71761726 | rs10433674 | 2.33 | -0.542 | 3.383 | 0.586 | -0.048 | 0.045 | 0.045 | 0.597 | 0.84 | 0.41 | 0.86 | 0.79 | 0.95 | 0.88 |
| 4 | 71776935 | rs12512502 | 2.422 | -0.506 | 3.214 | 0.436 | 0.02 | 0.022 | 0.022 | 1.562 | 0.52 | 0.15 | 0.57 | 0.62 | 0.59 | 0.51 |
| 4 | 71791502 | rs6446945 | -1.595 | -0.477 | 3.452 | 0.404 | -0.024 | 0.084 | 0.084 | 0.055 | 0.07 | 0.18 | 0.07 | 0.16 | 0.01 | 0.07 |
| 4 | 71794649 | rs6826930 | 0.302 | -0.477 | 3.45 | 0.395 | -0.007 | -0.027 | -0.027 | 0.228 | 0.05 | 0.13 | 0.03 | 0 | 0.02 | 0.01 |
| 4 | 71806349 | rs10003046 | -0.19 | -0.466 | 3.408 | 0.393 | -0.009 | -0.031 | -0.031 | 0.955 | 0.06 | 0.13 | 0.04 | 0 | 0.03 | 0.01 |
| 4 | 71817710 | rs28673341 | -2.429 | -0.475 | 3.447 | 0.399 | -0.048 | 0.065 | 0.065 | 0.013 | 0.13 | 0.42 | 0.12 | 0.16 | 0.04 | 0.08 |
| 4 | 71849888 | rs6842838 | 2.323 | -0.476 | 3.349 | 0.399 | -0.048 | 0.044 | 0.044 | 0.331 | 0.84 | 0.39 | 0.86 | 0.8 | 0.95 | 0.88 |
| 4 | 71850271 | rs3733399 | -0.531 | -0.546 | 3.203 | 0.324 | -0.057 | 0.043 | 0.043 | 0 | 0.15 | 0.01 | 0.08 | 0.05 | 0.21 | 0.19 |
| 4 | 71852410 | rs1385985 | 2.293 | -0.543 | 3.145 | 0.381 | -0.048 | 0.044 | 0.044 | 0.331 | 0.84 | 0.4 | 0.86 | 0.8 | 0.95 | 0.88 |
| 4 | 71858122 | rs7689093 | 0.261 | -0.369 | 2.956 | -0.062 | -0.007 | -0.027 | -0.027 | 0 | 0.05 | 0.14 | 0.03 | 0 | 0.02 | 0.01 |
| 4 | 71874331 | rs10805074 | 2.333 | -0.405 | 2.487 | -0.082 | -0.03 | -0.048 | -0.048 | 0.085 | 0.87 | 0.33 | 0.9 | 0.96 | 0.96 | 0.96 |
| 4 | 71875237 | rs7684954 | 2.333 | -0.405 | 2.487 | -0.082 | -0.03 | -0.048 | -0.048 | 0 | 0.87 | 0.33 | 0.9 | 0.96 | 0.96 | 0.96 |
| 4 | 71893864 | rs4694362 | 2.461 | -0.139 | 2.004 | -0.005 | -0.013 | 0.112 | 0.112 | 2.066 | 0.55 | 0.14 | 0.61 | 0.77 | 0.59 | 0.57 |
| 4 | 71913507 | rs1843614 | 1.949 | -0.535 | 2.347 | -0.088 | -0.005 | -0.023 | -0.023 | 0.862 | 0.68 | 0.44 | 0.79 | 0.85 | 0.71 | 0.71 |
| 4 | 71917988 | rs10518081 | 2.041 | -0.464 | 2.424 | -0.034 | -0.004 | -0.019 | -0.019 | 0 | 0.72 | 0.42 | 0.81 | 0.85 | 0.74 | 0.71 |
| 4 | 71926314 | rs10805077 | 1.527 | -0.516 | 2.892 | -0.037 | 0 | -0.018 | -0.018 | 0.634 | 0.75 | 0.7 | 0.84 | 0.85 | 0.74 | 0.73 |
| 4 | 71937445 | rs11941079 | -1.262 | -0.408 | 1.733 | 0.17 | 0 | 0.003 | 0.003 | 0.251 | 0.07 | 0.1 | 0.06 | 0.1 | 0.08 | 0.09 |
| 4 | 71948304 | rs10518083 | -0.331 | -0.405 | 1.758 | 0.133 | -0.006 | -0.008 | -0.008 | 0 | 0.1 | 0.12 | 0.14 | 0.04 | 0.06 | 0.12 |
| 4 | 71958094 | rs6816128 | -1.949 | -0.278 | 1.891 | 0.155 | -0.001 | 0.001 | 0.001 | 0.445 | 0.24 | 0.22 | 0.15 | 0.16 | 0.24 | 0.24 |
| 4 | 71975779 | rs4694373 | 2.363 | -0.396 | 1.635 | 0.573 | 0.039 | 0.008 | 0.008 | 0.323 | 0.64 | 0.25 | 0.73 | 0.69 | 0.79 | 0.57 |
| 4 | 71978043 | rs6831852 | -1.749 | -0.418 | 1.917 | 0.58 | -0.028 | 0.03 | 0.03 | 0.409 | 0.2 | 0.38 | 0.1 | 0.26 | 0.13 | 0.29 |
| 4 | 71979558 | rs12330929 | 0.203 | -0.384 | 1.643 | 0.57 | -0.003 | -0.016 | -0.016 | 0 | 0.06 | 0.08 | 0.04 | 0 | 0.03 | 0.03 |
| 4 | 71982160 | rs16845785 | -0.443 | -0.323 | 1.261 | 0.631 | -0.004 | -0.002 | -0.002 | 0 | 0.09 | 0.11 | 0.12 | 0.05 | 0.05 | 0.11 |
| 4 | 71997259 | rs17688758 | 0.412 | 0.324 | 0.688 | 0.818 | -0.034 | -0.032 | -0.032 | 1.716 | 0.06 | 0 | 0.03 | 0.15 | 0.08 | 0.11 |
| 4 | 71999712 | rs2123430 | 1.872 | -0.421 | 0.936 | 0.504 | -0.002 | -0.002 | -0.002 | 0 | 0.68 | 0.37 | 0.8 | 0.74 | 0.78 | 0.64 |
| 4 | 72006446 | rs7694411 | 1.847 | -0.084 | 1.003 | 0.851 | -0.014 | 0.007 | 0.007 | 0.401 | 0.75 | 0.46 | 0.84 | 0.74 | 0.81 | 0.64 |
| 4 | 72016176 | rs6845432 | -1.328 | -0.097 | 0.41 | 0.989 | -0.015 | -0.039 | -0.039 | 0.053 | 0.1 | 0.02 | 0.04 | 0.14 | 0.07 | 0.23 |
| 4 | 72017058 | rs16845929 | -0.872 | -0.191 | 0.624 | 0.626 | 0 | -0.007 | -0.007 | 0 | 0.09 | 0.08 | 0.29 | 0.02 | 0.05 | 0.21 |
| 4 | 72026724 | rs16845941 | -2.242 | -0.301 | 1.291 | 0.866 | -0.008 | -0.035 | -0.035 | 0 | 0.17 | 0.26 | 0.33 | 0.12 | 0.12 | 0.39 |
| 4 | 72027476 | rs1979537 | 2.74 | -0.484 | 1.319 | 0.872 | 0.029 | 0.085 | 0.085 | 0.652 | 0.69 | 0.12 | 0.59 | 0.78 | 0.84 | 0.43 |
| 4 | 72032466 | rs873954 | -1.433 | -0.142 | 1.98 | 1.222 | 0.001 | -0.005 | -0.005 | 0 | 0.1 | 0.23 | 0.05 | 0.1 | 0.04 | 0.19 |
| 4 | 72040651 | rs10938136 | 2.156 | -0.141 | 1.92 | 1.222 | -0.017 | -0.024 | -0.024 | 0 | 0.87 | 0.38 | 0.93 | 0.9 | 0.96 | 0.81 |
| 4 | 72063044 | rs2198341 | -1.451 | -0.178 | 1.539 | 1.64 | -0.004 | 0.1 | 0.1 | 0.418 | 0.1 | 0.27 | 0.09 | 0.21 | 0.05 | 0.26 |
| 4 | 72069713 | rs2579302 | -1.929 | -0.189 | 0.885 | 1.636 | -0.018 | 0.01 | 0.01 | 0 | 0.22 | 0.65 | 0.41 | 0.24 | 0.1 | 0.45 |
| 4 | 72084653 | rs4521314 | -1.75 | -0.492 | 0.981 | 2.19 | -0.001 | 0.004 | 0.004 | 0 | 0.16 | 0.17 | 0.12 | 0.33 | 0.12 | 0.38 |
| 4 | 72105088 | rs2579362 | 1.626 | -0.808 | 2.888 | 1.954 | 0.032 | 0.034 | 0.034 | 0 | 0.95 | 0.73 | 0.96 | 0.99 | 0.99 | 0.91 |
| 4 | 72108593 | rs12506660 | 2.896 | -1.106 | 2.822 | 2.511 | 0.02 | -0.024 | -0.024 | 0 | 0.73 | 0.26 | 0.71 | 0.55 | 0.87 | 0.54 |
| 4 | 72115097 | rs9790757 | -2.14 | -1.113 | 2.835 | 2.595 | -0.015 | 0.028 | 0.028 | 0 | 0.18 | 0.23 | 0.23 | 0.38 | 0.12 | 0.34 |
| 4 | 72117691 | rs2060903 | 2.867 | -1.119 | 2.362 | 2.601 | 0.014 | -0.018 | -0.018 | 1.104 | 0.72 | 0.29 | 0.71 | 0.54 | 0.87 | 0.54 |
| 4 | 72135455 | rs2602098 | -2.712 | -0.738 | 2.299 | 2.508 | -0.01 | 0.019 | 0.019 | 0 | 0.24 | 0.57 | 0.26 | 0.33 | 0.21 | 0.42 |
| 4 | 72136575 | rs1377287 | -1.764 | -0.657 | 1.859 | 2.691 | 0 | -0.001 | -0.001 | 0.227 | 0.11 | 0.09 | 0.16 | 0.32 | 0.12 | 0.3 |
| 4 | 72138216 | rs2363719 | -1.808 | -0.988 | 1.942 | 2.411 | 0.002 | -0.003 | -0.003 | 0 | 0.12 | 0.11 | 0.06 | 0.16 | 0.09 | 0.21 |
| 4 | 72147070 | rs2602049 | -2.228 | -1.696 | 2.273 | 2.181 | -0.035 | 0.004 | 0.004 | 1.052 | 0.3 | 0.56 | 0.37 | 0.31 | 0.16 | 0.51 |
| 4 | 72147944 | rs1031420 | -2.762 | -1.834 | 1.978 | 2.316 | 0.085 | -0.045 | -0.045 | 0 | 0.39 | 0.73 | 0.47 | 0.54 | 0.23 | 0.67 |
| 4 | 72149030 | rs1901709 | -1.477 | -1.896 | 2.352 | 2.294 | -0.039 | 0.082 | 0.082 | 0 | 0.13 | 0.28 | 0.28 | 0.23 | 0.05 | 0.22 |
| 4 | 72163134 | rs930754 | -1.48 | -1.902 | 2.35 | 2.305 | -0.039 | 0.082 | 0.082 | 0 | 0.13 | 0.28 | 0.28 | 0.23 | 0.05 | 0.22 |
| 4 | 72183646 | rs1377286 | -1.043 | -1.871 | 1.876 | 2.312 | 0.027 | -0.046 | -0.046 | 0 | 0.11 | 0.03 | 0.27 | 0.23 | 0.06 | 0.2 |
| 4 | 72199788 | rs11727507 | 2.223 | -1.98 | 2.084 | 2.158 | 0.053 | -0.032 | -0.032 | 1.357 | 0.63 | 0.32 | 0.54 | 0.46 | 0.8 | 0.35 |
| 4 | 72237431 | rs16846305 | -0.378 | -1.82 | 2.078 | 1.76 | -0.001 | 0.007 | 0.007 | 0 | 0.05 | 0.06 | 0.04 | 0.15 | 0.01 | 0.15 |
| 4 | 72239375 | rs16846318 | -0.621 | -1.824 | 2.344 | 1.758 | -0.015 | 0.07 | 0.07 | 0 | 0.05 | 0.14 | 0.04 | 0.15 | 0.01 | 0.15 |
| 4 | 72257715 | rs6852522 | 1.199 | -2.006 | 2.474 | 1.717 | -0.048 | -0.002 | -0.002 | 0 | 0.84 | 0.68 | 0.91 | 0.83 | 0.93 | 0.79 |
| 4 | 72278430 | rs4314260 | -1.161 | -2.021 | 2.169 | 1.427 | -0.002 | -0.001 | -0.001 | 0 | 0.23 | 0.2 | 0.36 | 0.23 | 0.16 | 0.45 |
| 4 | 72282753 | rs6854303 | 2.204 | -2.06 | 2.392 | 1.413 | 0.046 | 0.01 | 0.01 | 0 | 0.6 | 0.29 | 0.53 | 0.6 | 0.77 | 0.35 |
| 4 | 72304002 | rs13124079 | -1.117 | -1.79 | 2.127 | 1.46 | -0.003 | -0.001 | -0.001 | 0.088 | 0.22 | 0.2 | 0.36 | 0.23 | 0.16 | 0.44 |
| 4 | 72322773 | rs4130912 | -0.441 | -1.728 | 1.797 | 1.065 | -0.003 | 0.045 | 0.045 | 0.002 | 0.17 | 0.06 | 0.29 | 0.06 | 0.15 | 0.3 |
| 4 | 72335972 | rs6840014 | -1.429 | -2.355 | 1.84 | 1.074 | -0.074 | 0.053 | 0.053 | 0 | 0.28 | 0.58 | 0.21 | 0.58 | 0.11 | 0.24 |

| | | | | | | | | | | | | | | | | |
|---|---|---|---|---|---|---|---|---|---|---|---|---|---|---|---|---|
| 4 | 72337490 | rs4600889 | 0.179 | -2.406 | 2.055 | 1.098 | -0.046 | 0.142 | 0.142 | 0 | 0.12 | 0.2 | 0.12 | 0.44 | 0.03 | 0.07 |
| 4 | 72340242 | rs4639040 | -1.533 | -2.385 | 1.826 | 1.097 | -0.074 | 0.053 | 0.053 | 0 | 0.28 | 0.58 | 0.21 | 0.58 | 0.11 | 0.24 |
| 4 | 72347491 | rs6846301 | -0.662 | -2.093 | 1.762 | 1.193 | -0.035 | 0.001 | 0.001 | 0 | 0.13 | 0.27 | 0.08 | 0.14 | 0.08 | 0.17 |
| 4 | 72360478 | rs4469035 | 0.813 | -2.269 | 1.763 | 1.176 | -0.045 | 0.067 | 0.067 | 0 | 0.71 | 0.56 | 0.59 | 0.53 | 0.83 | 0.62 |
| 4 | 72361951 | rs6858376 | 1.591 | -2.163 | 0.641 | 1.072 | 0.127 | -0.014 | -0.014 | 0 | 0.57 | 0.16 | 0.49 | 0.37 | 0.77 | 0.42 |
| 4 | 72382471 | rs10031807 | 0.146 | -2.064 | 0.59 | 0.826 | -0.009 | 0.043 | 0.043 | 0.155 | 0.16 | 0.06 | 0.29 | 0.07 | 0.14 | 0.19 |
| 4 | 72386350 | rs6842334 | 1.453 | -2.124 | 0.631 | 0.867 | 0.055 | 0 | 0 | 0.22 | 0.58 | 0.29 | 0.5 | 0.37 | 0.77 | 0.44 |
| 4 | 72387348 | rs1377527 | -0.76 | -2.137 | 0.71 | 0.854 | -0.103 | 0.116 | 0.116 | 0 | 0.25 | 0.47 | 0.2 | 0.56 | 0.09 | 0.36 |
| 4 | 72387926 | rs2886107 | 1.356 | -2.024 | 0.336 | 0.86 | 0.102 | 0.01 | 0.01 | 0.525 | 0.54 | 0.25 | 0.49 | 0.37 | 0.76 | 0.39 |
| 4 | 72391695 | rs1453445 | 1.156 | -1.939 | 0.247 | 0.902 | 0.025 | -0.003 | -0.003 | 0 | 0.58 | 0.33 | 0.5 | 0.38 | 0.77 | 0.46 |
| 4 | 72396727 | rs7683903 | -0.027 | -1.625 | -0.366 | 0.583 | 0.023 | 0.038 | 0.038 | 0.419 | 0.2 | 0.05 | 0.12 | 0.12 | 0.17 | 0.11 |
| 4 | 72397710 | rs3775162 | 1.172 | -1.987 | -0.512 | 0.202 | 0.186 | 0.262 | 0.262 | 1.85 | 0.55 | 0.08 | 0.64 | 0.1 | 0.77 | 0.44 |
| 4 | 72405132 | rs7693843 | 0.126 | -1.412 | -0.177 | 0.332 | -0.006 | 0.038 | 0.038 | 0.591 | 0.73 | 0.65 | 0.77 | 0.59 | 0.73 | 0.71 |
| 4 | 72406365 | rs1869377 | -0.133 | -1.361 | -0.509 | 0.358 | -0.002 | -0.052 | -0.052 | 0 | 0.24 | 0.14 | 0.2 | 0.41 | 0.27 | 0.26 |
| 4 | 72412029 | rs3733362 | -0.142 | -1.054 | -0.295 | 0.508 | -0.085 | 0.199 | 0.199 | 0 | 0.18 | 0.43 | 0.16 | 0.48 | 0.05 | 0.3 |
| 4 | 72415556 | rs11937245 | 0.567 | 0.144 | -1.219 | -0.407 | -0.143 | 0.249 | 0.249 | 1.202 | 0.37 | 0.04 | 0.31 | 0.04 | 0.53 | 0.22 |
| 4 | 72421709 | rs16846575 | -1.082 | 1.048 | -0.272 | 0.749 | -0.002 | 0 | 0 | 0.14 | 0.16 | 0.16 | 0.36 | 0.13 | 0.19 | 0.2 |
| 4 | 72423624 | rs1453474 | 0.543 | -0.08 | 0.503 | 1.146 | -0.066 | 0.101 | 0.101 | 0.07 | 0.85 | 0.62 | 0.89 | 0.74 | 0.96 | 0.8 |
| 4 | 72425863 | rs1453458 | 0.543 | -0.08 | 0.53 | 1.146 | -0.066 | 0.103 | 0.103 | 0.07 | 0.85 | 0.61 | 0.86 | 0.75 | 0.96 | 0.8 |
| 4 | 72427450 | rs1031453 | 0.844 | 0.22 | -0.143 | 0.484 | -0.023 | -0.04 | -0.04 | 0 | 0.67 | 0.53 | 0.66 | 0.19 | 0.73 | 0.47 |
| 4 | 72433300 | rs3816468 | -1.142 | 0.011 | -0.28 | 0.406 | -0.006 | -0.029 | -0.029 | 0 | 0.26 | 0.19 | 0.44 | 0.34 | 0.22 | 0.34 |
| 4 | 72433545 | rs1062677 | -0.037 | 0.092 | -0.191 | 0.429 | -0.009 | 0 | 0 | 0 | 0.14 | 0.1 | 0.35 | 0.12 | 0.18 | 0.19 |
| 4 | 72436235 | rs1043218 | 0.223 | 0.031 | 0.423 | 0.486 | 0.003 | 0.111 | 0.111 | 0 | 0.77 | 0.59 | 0.77 | 0.43 | 0.76 | 0.61 |
| 4 | 72438028 | rs16846687 | -0.641 | -0.053 | -0.028 | 0.176 | -0.021 | 0.02 | 0.02 | 0 | 0.21 | 0.11 | 0.38 | 0.12 | 0.24 | 0.25 |
| 4 | 72447421 | rs1377523 | -1.311 | -0.155 | 0.84 | 0.757 | -0.01 | 0.096 | 0.096 | 0.01 | 0.2 | 0.54 | 0.22 | 0.31 | 0.14 | 0.21 |
| 4 | 72469183 | rs1597978 | 0.635 | -0.134 | 0.423 | 0.136 | 0.019 | -0.006 | -0.006 | 0.049 | 0.66 | 0.77 | 0.53 | 0.34 | 0.67 | 0.52 |
| 4 | 72476761 | rs17763915 | 0.036 | -0.191 | 1.024 | 0.196 | 0.001 | -0.014 | -0.014 | 0 | 0.15 | 0.2 | 0.11 | 0.1 | 0.14 | 0.09 |
| 4 | 72481753 | rs10212827 | 1.609 | -0.203 | 1.146 | 0.178 | 0.012 | 0.039 | 0.039 | 0 | 0.5 | 0.42 | 0.41 | 0.24 | 0.54 | 0.43 |
| 4 | 72513178 | rs1526691 | -0.523 | -0.271 | 0.981 | 0.252 | -0.005 | -0.009 | -0.009 | 1.571 | 0.84 | 0.82 | 0.68 | 0.92 | 0.86 | 0.82 |
| 4 | 72515260 | rs2219070 | 1.896 | -0.573 | 0.777 | 0.051 | -0.006 | 0.118 | 0.118 | 0 | 0.44 | 0.25 | 0.32 | 0.02 | 0.53 | 0.27 |
| 4 | 72528017 | rs9654062 | 1.912 | -0.569 | 0.834 | 0.053 | -0.006 | 0.107 | 0.107 | 0 | 0.44 | 0.26 | 0.32 | 0.02 | 0.53 | 0.27 |
| 4 | 72531761 | rs2177921 | 0.202 | -0.586 | 1.043 | 0.214 | -0.003 | 0.119 | 0.119 | 0 | 0.13 | 0.28 | 0.18 | 0.29 | 0.1 | 0.21 |
| 4 | 72531863 | rs13150933 | -0.12 | -0.626 | 0.678 | 0.221 | -0.012 | -0.135 | -0.135 | 0.395 | 0.2 | 0.02 | 0.15 | 0.59 | 0.19 | 0.28 |
| 4 | 72541092 | rs12501894 | -0.239 | -0.356 | 1.162 | 0.358 | -0.005 | 0.178 | 0.178 | 0 | 0.14 | 0.39 | 0.19 | 0.34 | 0.1 | 0.22 |
| 4 | 72543907 | rs6446756 | 0.538 | -0.376 | 0.727 | 0.986 | 0.013 | -0.016 | -0.016 | 0 | 0.6 | 0.68 | 0.51 | 0.48 | 0.63 | 0.48 |
| 4 | 72545947 | rs4694421 | 1.389 | 0.26 | 0.176 | 0.543 | -0.031 | 0.056 | 0.056 | 0 | 0.42 | 0.27 | 0.36 | 0.25 | 0.47 | 0.36 |
| 4 | 72569065 | rs377687 | -1.301 | 0.541 | 0.176 | 0.336 | 0 | -0.01 | -0.01 | 0 | 0.63 | 0.62 | 0.6 | 0.8 | 0.65 | 0.75 |
| 4 | 72569693 | rs6845026 | 0.304 | 0.443 | -0.382 | 0.127 | 0.017 | 0.069 | 0.069 | 0 | 0.22 | 0.01 | 0.17 | 0.1 | 0.17 | 0.27 |
| 4 | 72578724 | rs1526692 | -0.04 | 0.641 | 0.005 | 0.726 | 0.038 | 0.023 | 0.023 | 0 | 0.26 | 0.48 | 0.26 | 0.27 | 0.39 | 0.31 |
| 4 | 72578956 | rs842990 | -0.513 | 0.639 | -0.34 | 0.764 | 0.031 | -0.078 | -0.078 | 0 | 0.36 | 0.12 | 0.33 | 0.54 | 0.27 | 0.44 |
| 4 | 72592214 | rs4694105 | 0.198 | 1.172 | 0.11 | 0.966 | -0.013 | -0.027 | -0.027 | 0.021 | 0.2 | 0.15 | 0.22 | 0.26 | 0.3 | 0.29 |
| 4 | 72596821 | rs6837549 | 0.861 | 0.856 | 0.366 | 0.55 | -0.009 | 0.014 | 0.014 | 0 | 0.6 | 0.39 | 0.57 | 0.3 | 0.52 | 0.52 |
| 4 | 72601331 | rs12512631 | 1.272 | 0.575 | 0.374 | 0.541 | 0.006 | 0.026 | 0.026 | 0.009 | 0.4 | 0.35 | 0.43 | 0.23 | 0.35 | 0.26 |
| 4 | 72608115 | rs705117 | 1.129 | 0.407 | 0.962 | 1.287 | 0.009 | -0.036 | -0.036 | 0.238 | 0.76 | 0.17 | 0.74 | 0.51 | 0.83 | 0.83 |
| 4 | 72608383 | rs2282679 | 0.308 | 0.528 | 0.732 | 1.233 | -0.036 | -0.06 | -0.06 | 0.112 | 0.21 | 0.05 | 0.21 | 0.26 | 0.25 | 0.3 |
| 4 | 72610208 | rs222047 | -1.886 | 0.535 | 0.751 | 1.185 | -0.019 | 0.154 | 0.154 | 0.721 | 0.45 | 0.88 | 0.47 | 0.75 | 0.41 | 0.47 |
| 4 | 72613036 | rs705119 | 1.644 | 0.656 | 1.363 | 1.473 | -0.023 | 0.095 | 0.095 | 0.852 | 0.56 | 0.09 | 0.54 | 0.3 | 0.6 | 0.55 |
| 4 | 72613566 | rs1491709 | 0.829 | 0.648 | 1.53 | 1.789 | 0.003 | -0.008 | -0.008 | 0.036 | 0.13 | 0.08 | 0.06 | 0.17 | 0.08 | 0.12 |
| 4 | 72614140 | rs705120 | 1.034 | 0.649 | 1.558 | 1.837 | 0.012 | 0.025 | 0.025 | 0.04 | 0.63 | 0.53 | 0.69 | 0.56 | 0.61 | 0.57 |
| 4 | 72618323 | rs4588 | 0.274 | 0.637 | 1.191 | 1.601 | -0.046 | -0.068 | -0.068 | 0 | 0.2 | 0.07 | 0.21 | 0.26 | 0.25 | 0.3 |
| 4 | 72618334 | rs7041 | 1.479 | 0.649 | 1.247 | 1.252 | -0.034 | 0.098 | 0.098 | 1.135 | 0.56 | 0.09 | 0.54 | 0.3 | 0.58 | 0.54 |
| 4 | 72623347 | rs4364228 | -1.271 | 0.618 | 1.949 | 1.083 | 0.082 | 0.066 | 0.066 | 1.307 | 0.08 | 0.46 | 0.1 | 0.09 | 0.09 | 0.01 |
| 4 | 72632643 | rs705124 | 1.61 | 0.361 | 1.848 | 1.121 | 0.011 | -0.013 | -0.013 | 0 | 0.9 | 0.72 | 0.88 | 0.92 | 0.88 | 0.96 |
| 4 | 72634975 | rs222016 | 1.292 | 0.235 | 1.776 | 1.326 | -0.018 | 0.088 | 0.088 | 0 | 0.79 | 0.37 | 0.79 | 0.58 | 0.84 | 0.87 |
| 4 | 72636272 | rs222020 | 1.292 | 0.235 | 1.776 | 1.326 | -0.018 | 0.088 | 0.088 | 0 | 0.79 | 0.37 | 0.79 | 0.58 | 0.84 | 0.87 |
| 4 | 72637080 | rs222023 | -0.016 | 0.233 | 1.799 | 1.333 | -0.044 | 0.214 | 0.214 | 0 | 0.85 | 0.45 | 0.83 | 0.59 | 0.95 | 0.9 |
| 4 | 72641279 | rs1491718 | -0.26 | 0.403 | 1.349 | 1.085 | -0.002 | -0.007 | -0.007 | 0 | 0.06 | 0.09 | 0.04 | 0 | 0.12 | 0.03 |
| 4 | 72647749 | rs1352844 | 0.592 | -0.019 | 1.105 | 0.732 | -0.001 | -0.024 | -0.024 | 0 | 0.12 | 0.15 | 0.08 | 0 | 0.13 | 0.04 |
| 4 | 72647762 | rs1352845 | -1.095 | -0.142 | 1.523 | 0.895 | -0.025 | 0.035 | 0.035 | 0 | 0.27 | 0.43 | 0.22 | 0.4 | 0.2 | 0.16 |
| 4 | 72649774 | rs3733359 | 0.173 | -0.124 | 1.389 | 0.932 | -0.027 | 0.116 | 0.116 | 0 | 0.14 | 0.27 | 0.15 | 0.4 | 0.05 | 0.11 |
| 4 | 72672158 | rs11939173 | 1.61 | -0.008 | 0.815 | 0.656 | -0.016 | 0.221 | 0.221 | 0.087 | 0.48 | 0.1 | 0.48 | 0.16 | 0.56 | 0.53 |
| 4 | 72686078 | rs842889 | -0.68 | -0.101 | 1.286 | 0.648 | -0.034 | -0.016 | -0.016 | 0 | 0.27 | 0.53 | 0.22 | 0.25 | 0.2 | 0.12 |
| 4 | 72689926 | rs11737702 | 1.415 | 0.226 | 0.481 | 0.231 | -0.056 | 0.013 | 0.013 | 0 | 0.17 | 0.01 | 0.14 | 0.12 | 0.23 | 0.21 |
| 4 | 72695059 | rs842872 | -1.465 | 0.289 | 0.366 | 0.36 | -0.03 | 0.217 | 0.217 | 0.094 | 0.52 | 0.9 | 0.52 | 0.84 | 0.47 | 0.47 |
| 4 | 72695479 | rs16847105 | 0.832 | 0.106 | 0.748 | 0.706 | -0.006 | 0.017 | 0.017 | 0 | 0.12 | 0.13 | 0.13 | 0.25 | 0.06 | 0.08 |
| 4 | 72699705 | rs9291163 | 0.787 | 0.23 | 0.477 | 0.225 | 0.02 | 0.058 | 0.058 | 0 | 0.49 | 0.37 | 0.58 | 0.25 | 0.59 | 0.51 |
| 4 | 72699778 | rs1520498 | 1.34 | 0.49 | 0.32 | 0.136 | -0.051 | 0.016 | 0.016 | 0 | 0.16 | 0.05 | 0.17 | 0.12 | 0.22 | 0.2 |
| 4 | 72700747 | rs9992215 | 1.538 | 0.443 | 0.494 | 0.151 | -0.004 | 0.015 | 0.015 | 0 | 0.34 | 0.29 | 0.4 | 0.13 | 0.37 | 0.31 |
| 4 | 72704949 | rs6837292 | 0.611 | 0.227 | 0.619 | 0.667 | -0.005 | 0.012 | 0.012 | 0 | 0.13 | 0.14 | 0.14 | 0.26 | 0.06 | 0.08 |

| | | | | | | | | | | | | | | | | |
|---|---|---|---|---|---|---|---|---|---|---|---|---|---|---|---|---|
| 4 | 72707118 | rs6851050 | 0.016 | 0.256 | 0.838 | 0.566 | -0.002 | -0.001 | -0.001 | 0.407 | 0.25 | 0.27 | 0.21 | 0.26 | 0.21 | 0.12 |
| 4 | 72707400 | rs962226 | 1.632 | 0.339 | 0.84 | 0.472 | -0.065 | 0.21 | 0.21 | 0 | 0.45 | 0.11 | 0.51 | 0.16 | 0.52 | 0.48 |
| 4 | 72707517 | rs962227 | 0.665 | 0.298 | 1.425 | 0.852 | -0.002 | 0.015 | 0.015 | 0.806 | 0.71 | 0.38 | 0.72 | 0.42 | 0.73 | 0.61 |
| 4 | 72707821 | rs1851024 | -0.318 | 0.266 | 1.345 | 0.871 | 0.01 | 0.121 | 0.121 | 0 | 0.95 | 0.8 | 0.95 | 0.85 | 0.97 | 0.92 |
| 4 | 72709580 | rs7681566 | 0.298 | 0.296 | 1.079 | 0.866 | 0.003 | -0.037 | -0.037 | 0 | 0.2 | 0.14 | 0.16 | 0.34 | 0.17 | 0.29 |
| 4 | 72712872 | rs12639968 | 0.298 | 0.296 | 1.079 | 0.866 | 0.003 | -0.037 | -0.037 | 0.072 | 0.2 | 0.15 | 0.16 | 0.34 | 0.17 | 0.29 |
| 4 | 72723948 | rs7683624 | 0.772 | 0.095 | 1.539 | 0.858 | 0.07 | 0.182 | 0.182 | 0 | 0.91 | 0.67 | 0.88 | 0.75 | 0.9 | 0.89 |
| 4 | 72730996 | rs6845801 | 0.358 | 0.089 | 1.696 | 0.838 | 0.134 | 0.328 | 0.328 | 0.546 | 0.95 | 0.64 | 0.88 | 0.76 | 0.92 | 0.93 |
| 4 | 72735132 | rs6810725 | -0.896 | -0.006 | 1.459 | 0.494 | -0.009 | -0.017 | -0.017 | 0 | 0.62 | 0.5 | 0.5 | 0.71 | 0.66 | 0.63 |
| 4 | 72739098 | rs13113067 | 0.454 | 0.013 | 1.524 | 0.587 | 0.005 | 0.014 | 0.014 | 0.973 | 0.68 | 0.43 | 0.67 | 0.46 | 0.66 | 0.6 |
| 4 | 72743474 | rs10033936 | 0.355 | 0.028 | 1.452 | 0.652 | -0.008 | -0.023 | -0.023 | 0 | 0.22 | 0.14 | 0.14 | 0.29 | 0.24 | 0.3 |
| 4 | 72743549 | rs10022720 | 0.09 | 0.006 | 1.684 | 0.614 | 0 | 0 | 0 | 0 | 0.56 | 0.53 | 0.53 | 0.43 | 0.55 | 0.59 |
| 4 | 72744626 | rs2365520 | -0.514 | 0.013 | 1.603 | 0.615 | 0.006 | 0.124 | 0.124 | 0 | 0.95 | 0.78 | 0.88 | 0.85 | 0.97 | 0.92 |
| 4 | 72751686 | rs16847200 | 0.146 | 0.087 | 1.574 | 0.6 | -0.029 | -0.004 | -0.004 | 0 | 0.38 | 0.2 | 0.35 | 0.41 | 0.42 | 0.33 |
| 4 | 72756782 | rs4694440 | 0.241 | 0.071 | 1.067 | 0.648 | -0.012 | -0.031 | -0.031 | 0 | 0.22 | 0.12 | 0.14 | 0.27 | 0.24 | 0.3 |
| 4 | 72757529 | rs4694441 | -1.643 | 0.024 | 0.853 | 0.606 | 0.023 | 0.022 | 0.022 | 0 | 0.67 | 0.77 | 0.57 | 0.78 | 0.74 | 0.66 |
| 4 | 72769012 | rs1474198 | -0.289 | 0.106 | 0.98 | 0.689 | -0.01 | -0.001 | -0.001 | 0.023 | 0.43 | 0.38 | 0.41 | 0.51 | 0.49 | 0.37 |
| 4 | 72784005 | rs12504562 | -1.099 | -0.046 | 0.471 | 0.579 | -0.015 | 0.059 | 0.059 | 0 | 0.74 | 0.88 | 0.74 | 0.89 | 0.71 | 0.77 |
| 4 | 72791849 | rs1520499 | -0.401 | -0.069 | 0.67 | 0.597 | 0.131 | 0.365 | 0.365 | 1.038 | 0.07 | 0.42 | 0.13 | 0.27 | 0.07 | 0.07 |
| 4 | 72792667 | rs10035034 | 1.713 | -0.188 | 0.109 | 0.42 | 0.032 | 0.123 | 0.123 | 0.113 | 0.3 | 0.12 | 0.33 | 0.05 | 0.24 | 0.27 |
| 4 | 72797923 | rs1973750 | -0.768 | 0.312 | 0.888 | 1.06 | -0.012 | -0.014 | -0.014 | 0.138 | 0.37 | 0.27 | 0.28 | 0.57 | 0.4 | 0.43 |
| 4 | 72800493 | rs1520497 | -1.355 | 0.318 | 0.275 | 0.657 | -0.022 | 0.089 | 0.089 | 0.51 | 0.44 | 0.73 | 0.41 | 0.84 | 0.48 | 0.52 |
| 4 | 72801285 | rs17774208 | -0.448 | 0.695 | -0.009 | 1.086 | -0.12 | -0.082 | -0.082 | 1.173 | 0.25 | 0.02 | 0.17 | 0.24 | 0.35 | 0.35 |
| 4 | 72806951 | rs2139639 | -0.128 | 0.225 | 0.55 | 1.316 | -0.008 | -0.008 | -0.008 | 0 | 0.64 | 0.22 | 0.57 | 0.65 | 0.62 | 0.64 |
| 4 | 72809000 | rs10937987 | -0.67 | 0.227 | 1.034 | 1.308 | -0.004 | 0.007 | 0.007 | 0 | 0.69 | 0.7 | 0.64 | 0.82 | 0.62 | 0.67 |
| 4 | 72811523 | rs6446772 | -0.601 | 0.218 | 0.876 | 1.303 | -0.021 | 0.03 | 0.03 | 0 | 0.7 | 0.82 | 0.66 | 0.82 | 0.62 | 0.67 |
| 4 | 72822599 | rs11726886 | -0.355 | 0.244 | 0.54 | 1.353 | -0.076 | -0.064 | -0.064 | 0 | 0.24 | 0.05 | 0.16 | 0.28 | 0.3 | 0.26 |
| 4 | 72834313 | rs11733593 | -0.101 | 0.408 | 0.935 | 1.396 | -0.024 | -0.026 | -0.026 | 0.011 | 0.64 | 0.19 | 0.6 | 0.65 | 0.63 | 0.64 |
| 4 | 72841348 | rs4583729 | 1.473 | 0.189 | 1.469 | 1.449 | 0.011 | 0.213 | 0.213 | 4.894 | 0.94 | 0.39 | 0.91 | 0.84 | 1 | 0.97 |
| 4 | 72845549 | rs7683913 | 1.337 | 0.208 | 1.657 | 1.458 | 0.055 | 0.296 | 0.296 | 2.915 | 0.95 | 0.46 | 0.92 | 0.84 | 1 | 0.97 |
| 4 | 72875597 | rs11723796 | -0.055 | 0.237 | 1.837 | 1.277 | 0 | -0.013 | -0.013 | 0.356 | 0.64 | 0.19 | 0.6 | 0.65 | 0.67 | 0.64 |
| 4 | 72885899 | rs7699080 | 0.266 | 0.047 | 1.789 | 0.999 | 0.007 | 0.01 | 0.01 | 0.005 | 0.59 | 0.3 | 0.54 | 0.33 | 0.64 | 0.54 |
| 4 | 72891467 | rs10028416 | -0.319 | 0.134 | 1.949 | 1.019 | -0.01 | -0.012 | -0.012 | 0.682 | 0.4 | 0.56 | 0.43 | 0.68 | 0.33 | 0.45 |
| 4 | 72896212 | rs6446795 | -0.327 | 0.129 | 1.695 | 1.015 | 0 | 0 | 0 | 0.046 | 0.4 | 0.61 | 0.43 | 0.67 | 0.33 | 0.45 |
| 4 | 72903792 | rs12503480 | -0.339 | 0.165 | 1.55 | 0.97 | 0.009 | 0.011 | 0.011 | 0 | 0.39 | 0.31 | 0.42 | 0.67 | 0.34 | 0.46 |
| 4 | 72905089 | rs12512773 | -0.349 | 0.172 | 1.553 | 0.964 | 0.009 | 0.01 | 0.01 | 0 | 0.4 | 0.31 | 0.42 | 0.67 | 0.34 | 0.46 |
| 4 | 72921718 | rs7675387 | 1.454 | 0.195 | 1.258 | 0.737 | 0.011 | 0.161 | 0.161 | 0.683 | 0.35 | 0.14 | 0.19 | 0.02 | 0.33 | 0.2 |
| 4 | 72927226 | rs7678974 | 0.817 | 0.103 | 1.691 | 1.085 | -0.031 | 0.36 | 0.36 | 0.04 | 0.91 | 0.72 | 0.87 | 0.51 | 0.98 | 0.88 |
| 4 | 72938462 | rs4296652 | 0.689 | 0.121 | 1.627 | 1.088 | -0.031 | 0.329 | 0.329 | 0 | 0.92 | 0.75 | 0.88 | 0.52 | 0.98 | 0.88 |
| 4 | 72947764 | rs7674293 | 0.579 | 0.313 | 1.674 | 0.983 | -0.013 | 0.045 | 0.045 | 0 | 0.31 | 0.16 | 0.31 | 0.18 | 0.33 | 0.33 |
| 4 | 73000885 | rs9790741 | -0.915 | 0.686 | 2.157 | 2.152 | -0.086 | -0.085 | -0.085 | 0 | 0.21 | 0.04 | 0.35 | 0.3 | 0.31 | 0.32 |
| 4 | 73061027 | rs1513895 | -1.665 | 0.302 | 2.817 | 2.628 | -0.019 | 0.427 | 0.427 | 0.007 | 0.08 | 0.33 | 0.3 | 0.49 | 0.01 | 0.13 |
| 4 | 73073061 | rs10805048 | -1.233 | 0.371 | 2.199 | 2.557 | -0.032 | -0.031 | -0.031 | 0.594 | 0.21 | 0.15 | 0.17 | 0.31 | 0.32 | 0.35 |
| 4 | 73091934 | rs10029245 | -1.524 | 0.374 | 2.275 | 2.558 | 0.008 | 0.007 | 0.007 | 0.174 | 0.23 | 0.28 | 0.19 | 0.31 | 0.32 | 0.35 |
| 4 | 73096392 | rs10938012 | -1.57 | 0.372 | 2.257 | 2.557 | 0.031 | 0.029 | 0.029 | 0.18 | 0.23 | 0.34 | 0.2 | 0.31 | 0.32 | 0.35 |
| 4 | 73106712 | rs16847697 | -1.348 | -0.265 | 2.164 | 2.716 | -0.03 | 0.198 | 0.198 | 0 | 0.07 | 0.17 | 0.29 | 0.46 | 0.01 | 0.13 |
| 4 | 73117353 | rs1554017 | -1.534 | -0.026 | 1.363 | 2.409 | 0.016 | 0.034 | 0.034 | 0 | 0.2 | 0.27 | 0.13 | 0.33 | 0.24 | 0.32 |
| 4 | 73140229 | rs768029 | 1.861 | -0.309 | 1.153 | 2.092 | -0.039 | 0.167 | 0.167 | 0.338 | 0.87 | 0.51 | 0.89 | 0.71 | 0.97 | 0.89 |
| 4 | 73144182 | rs1369095 | 1.271 | -0.226 | 1.076 | 2.051 | 0.001 | 0.203 | 0.203 | 0.651 | 0.9 | 0.6 | 0.88 | 0.69 | 0.96 | 0.87 |
| 4 | 73146274 | rs9994557 | -1.388 | 0.129 | 0.246 | 2.005 | -0.018 | 0.066 | 0.066 | 0 | 0.25 | 0.41 | 0.33 | 0.47 | 0.19 | 0.41 |
| 4 | 73195704 | rs1897814 | 0.459 | 0.142 | -0.943 | 1.957 | 0.016 | -0.08 | -0.08 | 0 | 0.73 | 0.91 | 0.74 | 0.56 | 0.79 | 0.6 |
| 4 | 73245191 | rs1369093 | -1.439 | 0.221 | 0.83 | 1.729 | 0.017 | 0.197 | 0.197 | 1.322 | 0.09 | 0.38 | 0.33 | 0.24 | 0.03 | 0.14 |
| 4 | 73257944 | rs17722955 | -1.227 | 0.609 | 0.483 | 1.139 | 0.009 | -0.025 | -0.025 | 1.477 | 0.06 | 0.14 | 0.08 | 0 | 0.11 | 0.02 |
| 4 | 73264368 | rs10019624 | -0.204 | 0.812 | -0.253 | 1.027 | -0.064 | 0.023 | 0.023 | 0.61 | 0.42 | 0.14 | 0.55 | 0.36 | 0.49 | 0.54 |
| 4 | 73273605 | rs13112404 | -0.356 | 0.617 | 0.821 | 1.664 | 0.016 | 0.08 | 0.08 | 0 | 0.71 | 0.48 | 0.79 | 0.57 | 0.65 | 0.8 |
| 4 | 73284282 | rs922739 | -0.417 | 0.617 | 1.034 | 1.678 | 0 | 0.101 | 0.101 | 0 | 0.72 | 0.5 | 0.81 | 0.57 | 0.69 | 0.81 |
| 4 | 73293130 | rs1481667 | -0.732 | 0.619 | 1.287 | 1.679 | 0.003 | 0.045 | 0.045 | 0 | 0.74 | 0.68 | 0.83 | 0.57 | 0.69 | 0.81 |
| 4 | 73295961 | rs11727604 | -0.732 | 0.619 | 1.287 | 1.679 | 0.003 | 0.045 | 0.045 | 0 | 0.74 | 0.68 | 0.83 | 0.57 | 0.69 | 0.81 |
| 4 | 73328204 | rs10024063 | -0.567 | 0.616 | 1.438 | 1.676 | 0.007 | 0.031 | 0.031 | 0 | 0.73 | 0.63 | 0.8 | 0.57 | 0.65 | 0.81 |
| 4 | 73333387 | rs17725271 | 0.61 | 0.595 | 1.464 | 1.706 | -0.03 | -0.013 | -0.013 | 0 | 0.24 | 0.1 | 0.15 | 0.28 | 0.31 | 0.12 |
| 4 | 73334051 | rs1383930 | -0.262 | 0.592 | 1.482 | 1.65 | 0.017 | 0.076 | 0.076 | 0 | 0.71 | 0.45 | 0.79 | 0.57 | 0.65 | 0.8 |
| 4 | 73335098 | rs6823875 | 0.545 | 0.591 | 1.431 | 1.666 | -0.013 | -0.067 | -0.067 | 0.271 | 0.25 | 0.16 | 0.16 | 0.43 | 0.31 | 0.2 |
| 4 | 73353894 | rs788919 | -0.233 | 0.541 | 1.412 | 1.612 | -0.001 | -0.013 | -0.013 | 0 | 0.26 | 0.31 | 0.3 | 0.22 | 0.26 | 0.2 |
| 4 | 73363929 | rs11730719 | -0.177 | 0.525 | 1.402 | 1.702 | 0 | 0.006 | 0.006 | 0.029 | 0.25 | 0.21 | 0.29 | 0.22 | 0.26 | 0.2 |
| 4 | 73379012 | rs788935 | -0.555 | 0.657 | 0.332 | 1.261 | -0.006 | 0.043 | 0.043 | 0.085 | 0.57 | 0.77 | 0.49 | 0.65 | 0.6 | 0.42 |
| 4 | 73404916 | rs6446834 | 0.415 | 0.569 | 0.055 | 1.263 | 0.004 | 0.045 | 0.045 | 0.483 | 0.42 | 0.15 | 0.49 | 0.35 | 0.36 | 0.57 |
| 4 | 73409668 | rs700513 | -0.611 | 0.62 | 0.091 | 1.263 | 0.017 | 0.048 | 0.048 | 0.509 | 0.57 | 0.84 | 0.51 | 0.67 | 0.64 | 0.45 |
| 4 | 73412231 | rs788912 | -0.631 | 0.619 | 0.091 | 1.263 | 0.015 | 0.047 | 0.047 | 0.536 | 0.58 | 0.86 | 0.51 | 0.66 | 0.64 | 0.45 |
| 4 | 73414286 | rs788908 | -0.612 | 0.623 | 0.09 | 1.265 | 0.017 | 0.048 | 0.048 | 0.509 | 0.57 | 0.84 | 0.51 | 0.67 | 0.64 | 0.45 |

| | | | | | | | | | | | | | | | | |
|---|---|---|---|---|---|---|---|---|---|---|---|---|---|---|---|---|
| 4 | 73431464 | rs12645174 | 0.133 | 0.435 | 0.324 | 1.266 | 0.002 | -0.005 | -0.005 | 4.87 | 0.48 | 0.3 | 0.35 | 0.61 | 0.41 | 0.65 |
| 4 | 73433875 | rs1518492 | -0.381 | 0.257 | 1.05 | 1.195 | 0.002 | 0.001 | 0.001 | 0 | 0.48 | 0.28 | 0.61 | 0.4 | 0.59 | 0.35 |
| 4 | 73435404 | rs7670571 | 0.549 | 0.303 | 0.765 | 0.9 | -0.07 | -0.132 | -0.132 | 0 | 0.28 | 0.04 | 0.44 | 0.4 | 0.36 | 0.28 |
| 4 | 73449917 | rs753944 | 0.018 | 0.071 | 0.983 | 1.02 | -0.05 | -0.04 | -0.04 | 0.485 | 0.42 | 0.14 | 0.57 | 0.41 | 0.56 | 0.31 |
| 4 | 73454114 | rs719382 | 0.645 | 0.152 | 1.087 | 1.209 | 0.019 | 0.114 | 0.114 | 0 | 0.45 | 0.14 | 0.24 | 0.32 | 0.35 | 0.59 |
| 4 | 73454510 | rs11933831 | 0.722 | 0.137 | 1.094 | 1.175 | 0 | -0.01 | -0.01 | 0 | 0.07 | 0.11 | 0.03 | 0 | 0.03 | 0.03 |
| 4 | 73455382 | rs13104467 | 0.536 | 0.151 | 1.132 | 1.193 | 0.001 | -0.002 | -0.002 | 0 | 0.47 | 0.23 | 0.34 | 0.59 | 0.41 | 0.65 |
| 4 | 73470972 | rs1518485 | -0.479 | 0.147 | 2.157 | 1.191 | 0.006 | 0.005 | 0.005 | 0 | 0.5 | 0.48 | 0.63 | 0.41 | 0.59 | 0.35 |
| 4 | 73506817 | rs1401144 | 0.566 | 0.082 | 1.751 | 1.115 | -0.009 | -0.01 | -0.01 | 0 | 0.44 | 0.2 | 0.33 | 0.58 | 0.39 | 0.6 |
| 4 | 73508425 | rs11939290 | 0.239 | 0.121 | 1.548 | 1.1 | -0.002 | -0.018 | -0.018 | 0 | 0.09 | 0.14 | 0.04 | 0 | 0.04 | 0.06 |
| 4 | 73509826 | rs11728435 | 0.368 | 0.152 | 1.657 | 1.141 | 0.001 | 0.001 | 0.001 | 0 | 0.46 | 0.52 | 0.35 | 0.58 | 0.39 | 0.6 |
| 4 | 73518909 | rs12644269 | 0.603 | 0.17 | 1.417 | 1.33 | 0.01 | -0.043 | -0.043 | 0 | 0.43 | 0.22 | 0.25 | 0.46 | 0.37 | 0.55 |
| 4 | 73527617 | rs9784530 | 0.366 | 0.166 | 1.435 | 1.285 | 0 | -0.008 | -0.008 | 0.117 | 0.46 | 0.48 | 0.36 | 0.72 | 0.43 | 0.61 |
| 4 | 73528682 | rs13103561 | 0.93 | 0.245 | 0.974 | 0.984 | 0 | -0.084 | -0.084 | 0 | 0.34 | 0.12 | 0.19 | 0.42 | 0.27 | 0.46 |
| 4 | 73530700 | rs16848482 | 0.28 | -0.061 | 0.058 | 0.594 | -0.004 | -0.022 | -0.022 | 1.074 | 0.09 | 0.14 | 0.04 | 0 | 0.05 | 0.05 |
| 4 | 73543031 | rs2086471 | 0.127 | 0.907 | -0.144 | 0.804 | -0.045 | 0.047 | 0.047 | 0 | 0.41 | 0.14 | 0.58 | 0.26 | 0.51 | 0.32 |
| 4 | 73572187 | rs16848533 | -0.782 | 0.792 | 0.4 | 1.305 | -0.012 | 0.049 | 0.049 | 0 | 0.06 | 0.1 | 0.1 | 0.18 | 0.04 | 0.14 |
| 4 | 73575548 | rs10518108 | -0.474 | 0.619 | 1.345 | 1.039 | 0.004 | -0.067 | -0.067 | 0 | 0.11 | 0.43 | 0.06 | 0 | 0.06 | 0.05 |
| 4 | 73578725 | rs986397 | -1.389 | 0.919 | 0.58 | 0.926 | 0.014 | 0.014 | 0.014 | 0 | 0.51 | 0.67 | 0.54 | 0.6 | 0.51 | 0.52 |
| 4 | 73586055 | rs9291178 | -0.822 | 0.903 | 0.653 | 0.891 | -0.051 | -0.006 | -0.006 | 0 | 0.42 | 0.23 | 0.39 | 0.41 | 0.46 | 0.37 |
| 4 | 73614025 | rs316431 | 1.292 | 0.897 | 1.105 | 0.875 | 0.029 | 0.023 | 0.023 | 0 | 0.49 | 0.2 | 0.45 | 0.4 | 0.47 | 0.48 |
| 4 | 73617478 | rs4694517 | -0.098 | 0.89 | 0.997 | 0.881 | -0.154 | -0.1 | -0.1 | 0.019 | 0.33 | 0.06 | 0.37 | 0.42 | 0.43 | 0.32 |
| 4 | 73618256 | rs4694519 | -1.104 | 0.905 | 1.174 | 0.895 | -0.037 | 0.011 | 0.011 | 1.285 | 0.17 | 0.61 | 0.18 | 0.18 | 0.1 | 0.19 |
| 4 | 73618440 | rs4694523 | -0.667 | 0.88 | 1.005 | 0.913 | -0.021 | 0.005 | 0.005 | 0 | 0.15 | 0.28 | 0.14 | 0.18 | 0.09 | 0.19 |
| 4 | 73633046 | rs2366447 | -1.422 | 0.84 | 0.861 | 0.89 | 0.014 | 0.012 | 0.012 | 0.003 | 0.52 | 0.71 | 0.56 | 0.6 | 0.54 | 0.52 |
| 4 | 73637255 | rs1006667 | 1.144 | 1.095 | 0.784 | 1.006 | 0.001 | 0.01 | 0.01 | 0 | 0.53 | 0.25 | 0.48 | 0.42 | 0.51 | 0.53 |
| 4 | 73637861 | rs1383626 | 0.204 | 1.532 | 0.492 | 0.918 | -0.03 | -0.007 | -0.007 | 3.986 | 0.18 | 0.1 | 0.23 | 0.19 | 0.25 | 0.13 |
| 4 | 73640679 | rs316438 | 1.442 | 1.508 | 0.528 | 0.884 | 0.001 | 0.011 | 0.011 | 0.04 | 0.52 | 0.22 | 0.48 | 0.42 | 0.51 | 0.54 |
| 4 | 73640771 | rs316440 | -0.256 | 1.738 | 0.651 | 1.26 | 0.042 | 0.073 | 0.073 | 0 | 0.85 | 0.68 | 0.83 | 0.72 | 0.83 | 0.84 |
| 4 | 73651309 | rs316424 | 0.23 | 1.673 | 0.643 | 1.317 | 0.002 | 0.011 | 0.011 | 0 | 0.76 | 0.63 | 0.67 | 0.68 | 0.74 | 0.77 |
| 4 | 73657570 | rs316452 | 1.119 | 1.745 | 0.753 | 1.332 | 0.014 | 0.008 | 0.008 | 0 | 0.62 | 0.5 | 0.51 | 0.41 | 0.57 | 0.58 |
| 4 | 73673070 | rs1381252 | 0.249 | 1.649 | 0.71 | 1.724 | 0.002 | 0.049 | 0.049 | 0 | 0.76 | 0.61 | 0.68 | 0.59 | 0.74 | 0.77 |
| 4 | 73674204 | rs1842553 | 0.249 | 1.649 | 0.71 | 1.724 | 0.002 | 0.049 | 0.049 | 0 | 0.76 | 0.62 | 0.68 | 0.59 | 0.74 | 0.77 |
| 4 | 73674780 | rs1841587 | 0.953 | 1.522 | 0.612 | 1.479 | 0.022 | -0.023 | -0.023 | 0.132 | 0.62 | 0.44 | 0.51 | 0.31 | 0.57 | 0.58 |
| 4 | 73676145 | rs1383623 | -1.311 | 0.904 | 0.55 | 1.438 | -0.002 | 0 | 0 | 0 | 0.45 | 0.6 | 0.58 | 0.47 | 0.47 | 0.34 |
| 4 | 73680605 | rs4146484 | 0.15 | 0.574 | 0.313 | 0.327 | -0.001 | 0.001 | 0.001 | 0 | 0.37 | 0.32 | 0.26 | 0.31 | 0.39 | 0.4 |
| 4 | 73681946 | rs4694548 | 0.845 | 0.528 | 0.357 | 0.359 | -0.003 | -0.006 | -0.006 | 0 | 0.25 | 0.21 | 0.22 | 0.33 | 0.28 | 0.44 |
| 4 | 73683824 | rs4694549 | -0.193 | 0.399 | 0.582 | 0.218 | -0.003 | 0.012 | 0.012 | 0 | 0.53 | 0.53 | 0.43 | 0.64 | 0.49 | 0.49 |
| 4 | 73686750 | rs221593 | -1.704 | 0.359 | 0.653 | 0.237 | 0.006 | -0.012 | -0.012 | 0.164 | 0.85 | 0.87 | 0.83 | 0.66 | 0.9 | 0.87 |
| 4 | 73689660 | rs6446885 | 0.709 | 0.35 | 0.747 | 0.228 | -0.002 | 0.004 | 0.004 | 0 | 0.37 | 0.31 | 0.26 | 0.3 | 0.39 | 0.36 |
| 4 | 73693542 | rs315335 | -1.757 | 0.349 | 0.738 | 0.228 | 0.005 | -0.011 | -0.011 | 0 | 0.85 | 0.87 | 0.83 | 0.66 | 0.9 | 0.88 |
| 4 | 73694092 | rs221594 | -0.053 | 0.351 | 0.749 | 0.227 | -0.003 | 0.012 | 0.012 | 0 | 0.47 | 0.47 | 0.57 | 0.36 | 0.51 | 0.51 |
| 4 | 73694511 | rs315337 | 1.368 | 0.366 | 0.687 | 0.211 | -0.001 | -0.035 | -0.035 | 0 | 0.22 | 0.26 | 0.35 | 0.03 | 0.23 | 0.03 |
| 4 | 73711378 | rs2126207 | -0.141 | 0.387 | 0.439 | 0.251 | -0.001 | 0 | 0 | 0 | 0.48 | 0.55 | 0.57 | 0.36 | 0.51 | 0.51 |
| 4 | 73720317 | rs315359 | -1.723 | 0.404 | 0.345 | 0.261 | 0.005 | -0.011 | -0.011 | 0 | 0.85 | 0.87 | 0.83 | 0.66 | 0.9 | 0.88 |
| 4 | 73722977 | rs6822170 | 0.844 | 0.336 | 0.284 | 0.197 | -0.014 | 0.036 | 0.036 | 0 | 0.34 | 0.16 | 0.24 | 0.23 | 0.37 | 0.3 |
| 4 | 73740674 | rs186403 | -0.916 | 0.362 | -0.02 | -0.076 | -0.008 | -0.06 | -0.06 | 0.136 | 0.77 | 0.86 | 0.77 | 0.66 | 0.81 | 0.84 |
| 4 | 73742111 | rs315383 | -0.63 | 0.36 | 0.112 | -0.171 | -0.001 | 0.007 | 0.007 | 0 | 0.75 | 0.72 | 0.76 | 0.59 | 0.79 | 0.8 |
| 4 | 73744442 | rs315390 | -0.63 | 0.36 | 0.112 | -0.171 | -0.001 | 0.007 | 0.007 | 0 | 0.75 | 0.72 | 0.75 | 0.59 | 0.79 | 0.8 |
| 4 | 73757679 | rs9291180 | -0.642 | 0.364 | 0.124 | -0.174 | -0.001 | 0.008 | 0.008 | 0 | 0.75 | 0.72 | 0.76 | 0.59 | 0.79 | 0.8 |
| 4 | 73759117 | rs7698530 | 0.715 | 0.302 | 0.092 | -0.246 | -0.001 | -0.001 | -0.001 | 0 | 0.28 | 0.22 | 0.19 | 0.23 | 0.29 | 0.3 |
| 4 | 73761638 | rs17779822 | 1.676 | 0.375 | 0.017 | -0.289 | -0.001 | -0.013 | -0.013 | 0.039 | 0.21 | 0.2 | 0.17 | 0 | 0.23 | 0.03 |
| 4 | 73762643 | rs7677865 | -0.5 | 0.147 | -0.121 | 0.106 | -0.002 | -0.025 | -0.025 | 0 | 0.75 | 0.77 | 0.76 | 0.59 | 0.79 | 0.81 |
| 4 | 73768487 | rs17779898 | 0.523 | 0.17 | -0.553 | 0.045 | -0.016 | 0.01 | 0.01 | 0 | 0.13 | 0.06 | 0.11 | 0.11 | 0.22 | 0.16 |
| 4 | 73769708 | rs11930953 | 0.523 | 0.194 | -0.449 | 0.078 | 0.002 | 0.01 | 0.01 | 0.06 | 0.27 | 0.17 | 0.18 | 0.21 | 0.28 | 0.28 |
| 4 | 73770378 | rs919047 | 0.079 | 0.35 | -0.268 | 0.355 | 0.002 | -0.015 | -0.015 | 0 | 0.72 | 0.74 | 0.75 | 0.61 | 0.78 | 0.81 |
| 4 | 73774717 | rs1346133 | -0.483 | -0.059 | -0.159 | 0.586 | -0.018 | 0.071 | 0.071 | 0 | 0.8 | 0.7 | 0.8 | 0.61 | 0.87 | 0.84 |
| 4 | 73781518 | rs6814202 | -0.151 | 0.052 | -0.997 | 0.163 | 0.04 | 0.032 | 0.032 | 2.307 | 0.62 | 0.82 | 0.64 | 0.79 | 0.69 | 0.75 |
| 4 | 73784340 | rs11941162 | 0.599 | 0.347 | -0.768 | -0.049 | -0.004 | 0.022 | 0.022 | 0 | 0.37 | 0.27 | 0.31 | 0.26 | 0.4 | 0.34 |
| 4 | 73799039 | rs1895855 | -1.512 | 0.549 | -0.865 | -0.107 | 0.005 | 0.002 | 0.002 | 0 | 0.81 | 0.85 | 0.85 | 1 | 0.82 | 0.97 |
| 4 | 73803268 | rs7688718 | 0.47 | 0.934 | -0.39 | 0.468 | 0.029 | 0.099 | 0.099 | 0.342 | 0.21 | 0.36 | 0.12 | 0.5 | 0.26 | 0.4 |
| 4 | 73812806 | rs1381015 | 0.803 | 0.845 | -0.689 | 0.38 | -0.019 | 0.013 | 0.013 | 0.349 | 0.12 | 0.06 | 0.14 | 0.09 | 0.24 | 0.12 |
| 4 | 73821904 | rs13102676 | 0.404 | 0.204 | -0.239 | 0.186 | -0.009 | -0.008 | -0.008 | 0 | 0.39 | 0.36 | 0.26 | 0.5 | 0.48 | 0.38 |
| 4 | 73824798 | rs2059569 | -0.767 | -0.195 | -0.489 | 0.159 | -0.028 | 0.04 | 0.04 | 0 | 0.66 | 0.86 | 0.74 | 0.74 | 0.62 | 0.7 |
| 4 | 73832518 | rs2366711 | 0.145 | -0.077 | -0.182 | 0.283 | -0.012 | -0.006 | -0.006 | 0 | 0.44 | 0.65 | 0.47 | 0.51 | 0.49 | 0.39 |
| 4 | 73833081 | rs6446910 | 0.588 | 0.01 | -0.358 | 0.345 | -0.008 | -0.005 | -0.005 | 0 | 0.39 | 0.35 | 0.26 | 0.48 | 0.47 | 0.38 |
| 4 | 73843550 | rs2045680 | 0.366 | -0.051 | -1.268 | 0.252 | -0.02 | 0.037 | 0.037 | 0 | 0.36 | 0.15 | 0.27 | 0.25 | 0.38 | 0.28 |
| 4 | 73851964 | rs16849052 | 0.844 | -0.099 | -1.373 | 0.426 | 0.012 | -0.063 | -0.063 | 0.306 | 0.19 | 0.04 | 0.25 | 0.24 | 0.13 | 0.31 |
| 4 | 73854103 | rs2366715 | 1.027 | 0.163 | -1.546 | 0.685 | -0.128 | -0.021 | -0.021 | 0 | 0.29 | 0.02 | 0.2 | 0.28 | 0.42 | 0.32 |

| | | | | | | | | | | | | | | | | |
|---|---|---|---|---|---|---|---|---|---|---|---|---|---|---|---|---|
| 4 | 73854191 | rs961873 | -0.496 | 0.013 | -1.078 | 0.313 | -0.007 | 0.007 | 0.007 | 0 | 0.66 | 0.69 | 0.74 | 0.75 | 0.63 | 0.7 |
| 4 | 73854746 | rs13137265 | 0.508 | -0.233 | -0.966 | 0.713 | 0.035 | -0.038 | -0.038 | 1.134 | 0.25 | 0.12 | 0.26 | 0.29 | 0.14 | 0.31 |
| 4 | 73855253 | rs10009409 | -0.098 | 0.091 | -0.447 | 0.569 | -0.004 | 0.001 | 0.001 | 0 | 0.38 | 0.41 | 0.55 | 0.52 | 0.33 | 0.37 |
| 4 | 73855515 | rs16849080 | 0.179 | -0.226 | -0.563 | 0.28 | -0.016 | -0.015 | -0.015 | 0.842 | 0.27 | 0.32 | 0.3 | 0.28 | 0.2 | 0.17 |
| 4 | 73856466 | rs1381016 | 0.348 | -0.257 | -0.943 | 0.281 | 0.007 | -0.003 | -0.003 | 0 | 0.19 | 0.07 | 0.11 | 0.14 | 0.1 | 0.24 |
| 4 | 73857746 | rs13141048 | 0.336 | 0.062 | -0.535 | 0.171 | -0.012 | 0.01 | 0.01 | 0 | 0.39 | 0.46 | 0.5 | 0.47 | 0.31 | 0.33 |
| 4 | 73858464 | rs7666763 | -0.601 | 0.097 | -0.361 | 0.163 | -0.004 | 0.009 | 0.009 | 0 | 0.59 | 0.62 | 0.62 | 0.6 | 0.41 | 0.58 |
| 4 | 73862030 | rs1381014 | 0.3 | 0.038 | -0.405 | 0.091 | -0.019 | 0.018 | 0.018 | 0.06 | 0.39 | 0.45 | 0.49 | 0.47 | 0.31 | 0.32 |
| 4 | 73862194 | rs7675007 | 0.09 | -0.11 | -0.76 | 0.037 | 0.022 | 0.033 | 0.033 | 0 | 0.22 | 0.1 | 0.12 | 0.13 | 0.11 | 0.23 |
| 4 | 73891050 | rs6446913 | 0.596 | 0.341 | -0.724 | 0.247 | -0.009 | -0.005 | -0.005 | 0.922 | 0.16 | 0.08 | 0.29 | 0.19 | 0.26 | 0.08 |
| 4 | 73920622 | rs1052080 | 0.948 | 0.09 | -0.775 | 0.098 | -0.057 | -0.013 | -0.013 | 0 | 0.34 | 0.15 | 0.29 | 0.39 | 0.45 | 0.4 |
| 4 | 73931170 | rs2306057 | 0.025 | -0.323 | -0.443 | -0.096 | -0.051 | -0.058 | -0.058 | 0 | 0.24 | 0.45 | 0.14 | 0.13 | 0.1 | 0.19 |
| 4 | 73937533 | rs6845845 | 0.029 | -0.244 | -0.278 | -0.068 | -0.023 | -0.031 | -0.031 | 0 | 0.73 | 0.41 | 0.82 | 0.87 | 0.88 | 0.78 |
| 4 | 73940743 | rs10092 | 2.081 | -0.263 | -0.405 | 0.012 | -0.037 | -0.107 | -0.107 | 3.415 | 0.2 | 0.01 | 0.22 | 0.29 | 0.16 | 0.26 |
| 4 | 73942678 | rs2306058 | 2.16 | -0.246 | -0.403 | 0.014 | -0.041 | -0.107 | -0.107 | 0 | 0.2 | 0.01 | 0.22 | 0.29 | 0.17 | 0.26 |
| 4 | 73956736 | rs6855349 | 0.666 | -0.038 | -0.121 | -0.037 | -0.033 | 0.002 | 0.002 | 0 | 0.37 | 0.24 | 0.3 | 0.38 | 0.45 | 0.41 |
| 4 | 73988492 | rs10050235 | 0.334 | -0.032 | -0.081 | -0.069 | -0.002 | 0.007 | 0.007 | 0 | 0.39 | 0.5 | 0.55 | 0.48 | 0.42 | 0.35 |
| 4 | 74010834 | rs11939939 | 0.735 | -0.246 | -0.239 | -0.034 | -0.045 | -0.006 | -0.006 | 0.158 | 0.36 | 0.19 | 0.29 | 0.39 | 0.45 | 0.4 |
| 4 | 74036166 | rs5020219 | -1.059 | -0.189 | -0.375 | 0.029 | -0.049 | -0.112 | -0.112 | 0.455 | 0.64 | 0.92 | 0.48 | 0.52 | 0.58 | 0.65 |
| 4 | 74039463 | rs10035001 | -1.021 | -0.174 | -0.337 | 0.012 | -0.031 | -0.085 | -0.085 | 0.013 | 0.64 | 0.89 | 0.48 | 0.52 | 0.58 | 0.65 |
| 4 | 74058406 | rs10028486 | -0.45 | -0.388 | -0.004 | -0.073 | -0.04 | -0.064 | -0.064 | 1.54 | 0.26 | 0.47 | 0.17 | 0.13 | 0.13 | 0.22 |
| 4 | 74141448 | rs6820167 | -0.463 | -0.307 | -0.09 | 0.01 | 0.001 | 0.003 | 0.003 | 0 | 0.59 | 0.36 | 0.58 | 0.45 | 0.45 | 0.55 |
| 4 | 74151780 | rs6853164 | 0.411 | -0.35 | 0.084 | -0.035 | -0.004 | -0.006 | -0.006 | 0 | 0.41 | 0.4 | 0.4 | 0.55 | 0.55 | 0.43 |
| 4 | 74208199 | rs714825 | -0.215 | -0.4 | 0.109 | -0.08 | 0.005 | -0.004 | -0.004 | 0 | 0.54 | 0.34 | 0.56 | 0.45 | 0.42 | 0.5 |
| 4 | 74236264 | rs13125875 | 0.094 | -0.259 | 0.084 | -0.085 | -0.002 | -0.005 | -0.005 | 0.908 | 0.44 | 0.43 | 0.41 | 0.55 | 0.55 | 0.47 |
| 4 | 74254678 | rs4364229 | 1.021 | -0.456 | -0.03 | -0.142 | 0.006 | -0.001 | -0.001 | 0 | 0.32 | 0.23 | 0.21 | 0.29 | 0.24 | 0.25 |
| 4 | 74268658 | rs7688274 | 0.202 | -0.427 | -0.049 | -0.141 | 0 | -0.003 | -0.003 | 0.001 | 0.45 | 0.46 | 0.42 | 0.56 | 0.58 | 0.53 |
| 4 | 74275515 | rs3775486 | -0.383 | -0.419 | -0.06 | -0.138 | 0 | -0.001 | -0.001 | 1.467 | 0.54 | 0.42 | 0.57 | 0.44 | 0.42 | 0.46 |
| 4 | 74278809 | rs10022167 | 0.196 | -0.405 | -0.012 | -0.14 | 0 | 0.001 | 0.001 | 0 | 0.45 | 0.48 | 0.42 | 0.56 | 0.55 | 0.47 |
| 4 | 74279549 | rs2051512 | -0.394 | -0.405 | -0.056 | -0.137 | 0 | -0.001 | -0.001 | 0 | 0.54 | 0.42 | 0.57 | 0.44 | 0.42 | 0.47 |
| 4 | 74284428 | rs7678298 | 0.668 | -0.441 | -0.251 | -0.136 | -0.001 | 0.028 | 0.028 | 0 | 0.19 | 0.08 | 0.16 | 0.11 | 0.17 | 0.2 |
| 4 | 74284686 | rs3926327 | 1.161 | 0.046 | -0.289 | -0.036 | -0.016 | -0.014 | -0.014 | 0 | 0.07 | 0.06 | 0.1 | 0.13 | 0.17 | 0.09 |
| 4 | 74292407 | rs16849364 | 1.218 | -0.19 | -0.301 | -0.118 | -0.038 | -0.085 | -0.085 | 0 | 0.29 | 0.01 | 0.19 | 0.29 | 0.23 | 0.22 |
| 4 | 74296480 | rs6829532 | 0.918 | 0.009 | -0.166 | -0.174 | 0.007 | 0.002 | 0.002 | 0 | 0.33 | 0.22 | 0.22 | 0.29 | 0.27 | 0.22 |
| 4 | 74300880 | rs12651581 | 0.661 | -0.009 | 0.007 | -0.055 | -0.003 | 0.035 | 0.035 | 0 | 0.19 | 0.08 | 0.16 | 0.11 | 0.16 | 0.19 |
| 4 | 74301313 | rs4018 | 0.405 | -0.147 | 0.009 | -0.063 | -0.001 | -0.002 | -0.002 | 0 | 0.46 | 0.47 | 0.42 | 0.58 | 0.54 | 0.48 |
| 4 | 74302870 | rs16849384 | -0.295 | -0.126 | -0.162 | -0.045 | 0.009 | 0.001 | 0.001 | 0 | 0.51 | 0.34 | 0.55 | 0.42 | 0.42 | 0.45 |
| 4 | 74303200 | rs4640638 | 0.93 | 0.121 | -0.218 | -0.06 | 0.002 | -0.003 | -0.003 | 0 | 0.29 | 0.22 | 0.37 | 0.28 | 0.24 | 0.2 |
| 4 | 74310844 | rs2298839 | 0.045 | 0.219 | -0.22 | 0.119 | 0 | 0.002 | 0.002 | 0 | 0.46 | 0.46 | 0.42 | 0.67 | 0.55 | 0.46 |
| 4 | 74322512 | rs9784480 | -0.595 | 0.181 | -0.38 | 0.329 | 0.003 | 0.001 | 0.001 | 0 | 0.6 | 0.61 | 0.72 | 0.57 | 0.6 | 0.58 |
| 4 | 74344535 | rs16849461 | 0.197 | 0.499 | -0.382 | 0.367 | 0.026 | -0.003 | -0.003 | 0.582 | 0.37 | 0.24 | 0.43 | 0.34 | 0.3 | 0.36 |
| 4 | 74357851 | rs11733262 | 0.049 | 0.449 | -0.256 | 0.323 | 0.009 | 0 | 0 | 0 | 0.57 | 0.38 | 0.49 | 0.64 | 0.69 | 0.6 |
| 4 | 74358277 | rs12510870 | 1.17 | 0.359 | -0.592 | 0.459 | -0.049 | -0.013 | -0.013 | 0.003 | 0.27 | 0.01 | 0.34 | 0.2 | 0.24 | 0.19 |
| 4 | 74373060 | rs4694170 | 0.504 | 1.029 | 0.091 | 1.213 | -0.043 | -0.027 | -0.027 | 0 | 0.28 | 0.03 | 0.43 | 0.23 | 0.27 | 0.23 |
| 4 | 74402790 | rs4574378 | 0.528 | 0.871 | 0.325 | 1.272 | 0.001 | -0.001 | -0.001 | 0 | 0.59 | 0.39 | 0.48 | 0.64 | 0.68 | 0.6 |
| 4 | 74413589 | rs3756073 | -0.253 | 0.907 | 0.435 | 1.167 | 0.028 | 0.032 | 0.032 | 0 | 0.67 | 0.88 | 0.54 | 0.76 | 0.71 | 0.71 |
| 4 | 74413975 | rs9996759 | -1.069 | 1.104 | 0.717 | 1.076 | 0.052 | 0.026 | 0.026 | 0.457 | 0.06 | 0.39 | 0.05 | 0 | 0.03 | 0.03 |
| 4 | 74426173 | rs10018058 | 0.424 | 1.375 | 0.727 | 1.183 | -0.03 | -0.021 | -0.021 | 0 | 0.09 | 0.04 | 0.1 | 0.13 | 0.2 | 0.1 |
| 4 | 74428908 | rs2367154 | -1.178 | 1.241 | 0.849 | 1.037 | 0.071 | 0.043 | 0.043 | 0.483 | 0.06 | 0.44 | 0.05 | 0 | 0.03 | 0.03 |
| 4 | 74440552 | rs17838981 | -1.762 | 1.443 | 0.837 | 1.3 | 0.179 | 0.32 | 0.32 | 0 | 0.05 | 0.31 | 0.05 | 0.18 | 0.07 | 0.12 |
| 4 | 74440784 | rs10003721 | -1.762 | 1.46 | 0.837 | 1.3 | 0.186 | 0.32 | 0.32 | 0 | 0.05 | 0.31 | 0.05 | 0.18 | 0.07 | 0.12 |
| 4 | 74441786 | rs3756068 | -1.685 | 1.452 | 0.853 | 1.311 | 0.163 | 0.296 | 0.296 | 0 | 0.05 | 0.3 | 0.05 | 0.18 | 0.07 | 0.12 |
| 4 | 74447104 | rs1745300 | 0.271 | 1.406 | 0.646 | 1.27 | -0.001 | -0.001 | -0.001 | 4.213 | 0.63 | 0.58 | 0.49 | 0.7 | 0.65 | 0.66 |
| 4 | 74447452 | rs1247670 | -0.5 | 1.374 | 0.649 | 1.245 | 0 | -0.006 | -0.006 | 0 | 0.72 | 0.72 | 0.55 | 0.9 | 0.74 | 0.83 |
| 4 | 74451073 | rs12507775 | -0.289 | 1.373 | 0.582 | 1.244 | 0.015 | 0.032 | 0.032 | 0 | 0.33 | 0.28 | 0.47 | 0.11 | 0.28 | 0.23 |
| 4 | 74452469 | rs1528916 | -0.289 | 1.373 | 0.574 | 1.244 | 0.016 | 0.034 | 0.034 | 0 | 0.33 | 0.27 | 0.47 | 0.11 | 0.28 | 0.23 |
| 4 | 74453782 | rs10009048 | 0.584 | 1.385 | 0.352 | 1.266 | 0.001 | -0.004 | -0.004 | 0 | 0.61 | 0.46 | 0.47 | 0.7 | 0.62 | 0.63 |
| 4 | 74453857 | rs10011536 | 0.63 | 1.381 | 0.337 | 1.247 | 0 | -0.003 | -0.003 | 2.033 | 0.6 | 0.43 | 0.46 | 0.7 | 0.62 | 0.63 |
| 4 | 74455381 | rs7658159 | -0.466 | 1.406 | 0.29 | 1.32 | -0.008 | -0.002 | -0.002 | 0 | 0.08 | 0.08 | 0.06 | 0.2 | 0.09 | 0.18 |
| 4 | 74458223 | rs6446943 | 0.916 | 0.858 | 0.321 | 0.793 | -0.01 | -0.017 | -0.017 | 0.574 | 0.06 | 0.07 | 0.03 | 0.01 | 0.02 | 0.06 |
| 4 | 74466409 | rs6827519 | 0.612 | 1.032 | 0.359 | 0.891 | 0.005 | -0.012 | -0.012 | 0 | 0.59 | 0.42 | 0.4 | 0.68 | 0.51 | 0.62 |
| 4 | 74489271 | rs1247583 | -0.878 | 1.221 | 0.191 | 0.765 | -0.005 | 0.007 | 0.007 | 0 | 0.41 | 0.4 | 0.61 | 0.29 | 0.55 | 0.36 |
| 4 | 74494008 | rs4694626 | 0.528 | 0.886 | -0.727 | 0.884 | -0.061 | 0.08 | 0.08 | 0 | 0.25 | 0.02 | 0.43 | 0.08 | 0.26 | 0.18 |
| 4 | 74504614 | rs7655717 | -0.279 | 0.98 | -0.041 | 1.287 | 0 | 0.001 | 0.001 | 0 | 0.1 | 0.07 | 0.12 | 0.21 | 0.2 | 0.13 |
| 4 | 74521045 | rs1247574 | 0.2 | 0.522 | 0.098 | 1.483 | -0.029 | 0.11 | 0.11 | 0 | 0.92 | 0.78 | 0.96 | 0.71 | 0.97 | 0.83 |
| 4 | 74525356 | rs4694627 | 1.05 | 0.477 | -0.083 | 1.055 | -0.009 | 0.036 | 0.036 | 0 | 0.46 | 0.25 | 0.48 | 0.41 | 0.39 | 0.59 |
| 4 | 74527971 | rs1247683 | 0.578 | 0.449 | 0.633 | 0.674 | 0.043 | 0.011 | 0.011 | 0.012 | 0.93 | 0.62 | 0.94 | 1 | 0.96 | 0.96 |
| 4 | 74528627 | rs6855668 | 0.309 | 0.633 | 0.17 | 0.685 | -0.003 | 0.019 | 0.019 | 0 | 0.28 | 0.21 | 0.24 | 0.06 | 0.31 | 0.14 |
| 4 | 74534437 | rs2091588 | -0.526 | 0.425 | 0.721 | 1.491 | -0.004 | -0.002 | -0.002 | 0 | 0.11 | 0.07 | 0.19 | 0.25 | 0.22 | 0.13 |

| | | | | | | | | | | | | | | | | |
|---|---|---|---|---|---|---|---|---|---|---|---|---|---|---|---|---|
| 4 | 74555477 | rs1358594 | 1.572 | -0.55 | 1.564 | 1.545 | -0.075 | 0.156 | 0.156 | 0.494 | 0.83 | 0.44 | 0.89 | 0.67 | 0.94 | 0.74 |
| 4 | 74561560 | rs1247620 | -1.137 | -0.542 | 1.399 | 1.548 | 0.023 | -0.028 | -0.028 | 0.805 | 0.12 | 0.08 | 0.06 | 0.32 | 0.06 | 0.17 |
| 4 | 74572039 | rs1951703 | 2.022 | -0.604 | 1.539 | 1.516 | -0.043 | 0.043 | 0.043 | 0.658 | 0.81 | 0.31 | 0.88 | 0.67 | 0.93 | 0.72 |
| 4 | 74589784 | rs7657356 | -1.175 | -0.263 | -0.06 | 1.691 | 0.029 | -0.039 | -0.039 | 0 | 0.12 | 0.07 | 0.05 | 0.32 | 0.06 | 0.17 |
| 4 | 74592161 | rs12506479 | -1.307 | 0.191 | -0.19 | 1.446 | -0.005 | -0.044 | -0.044 | 1.962 | 0.25 | 0.18 | 0.43 | 0.43 | 0.25 | 0.31 |
| 4 | 74594043 | rs10031141 | -0.682 | -0.108 | -0.446 | 0.705 | -0.071 | -0.061 | -0.061 | 1.534 | 0.32 | 0.07 | 0.26 | 0.33 | 0.37 | 0.23 |
| 4 | 74607910 | rs2227543 | -1.107 | -0.77 | 0.505 | 0.411 | -0.049 | -0.051 | -0.051 | 0 | 0.37 | 0.1 | 0.27 | 0.38 | 0.39 | 0.36 |
| 4 | 74609045 | rs1126647 | -1.06 | -0.736 | 0.507 | 0.413 | -0.054 | -0.053 | -0.053 | 0 | 0.37 | 0.1 | 0.27 | 0.38 | 0.39 | 0.35 |
| 4 | 74617282 | rs1951242 | 1.456 | -0.904 | 1.235 | 0.358 | 0.015 | 0.013 | 0.013 | 0.121 | 0.53 | 0.16 | 0.62 | 0.59 | 0.58 | 0.61 |
| 4 | 74626161 | rs10938095 | -0.697 | -0.94 | 1.071 | 0.454 | -0.036 | -0.038 | -0.038 | 0 | 0.35 | 0.09 | 0.26 | 0.39 | 0.37 | 0.31 |
| 4 | 74626945 | rs2886920 | -0.913 | -0.958 | 1.206 | 0.445 | -0.017 | -0.028 | -0.028 | 0.661 | 0.37 | 0.16 | 0.3 | 0.42 | 0.39 | 0.32 |
| 4 | 74628455 | rs9999446 | -1.646 | -0.977 | 1.201 | 0.436 | 0.029 | 0.016 | 0.016 | 0.01 | 0.46 | 0.77 | 0.36 | 0.42 | 0.39 | 0.35 |
| 4 | 74657255 | rs9998340 | -1.696 | -1.254 | 1.059 | 0.275 | 0.039 | 0.034 | 0.034 | 0.173 | 0.51 | 0.8 | 0.39 | 0.41 | 0.42 | 0.39 |
| 4 | 74667727 | rs1951235 | -1.589 | -1.235 | 1.232 | 0.277 | 0.009 | 0.009 | 0.009 | 0 | 0.46 | 0.63 | 0.36 | 0.41 | 0.42 | 0.37 |
| 4 | 74679286 | rs6851997 | -0.294 | -1.298 | 1.113 | 0.234 | -0.066 | -0.05 | -0.05 | 0 | 0.32 | 0.08 | 0.26 | 0.33 | 0.37 | 0.22 |
| 4 | 74688555 | rs6834586 | -1.886 | -1.228 | 1.605 | 0.327 | -0.08 | -0.061 | -0.061 | 0 | 0.15 | 0.59 | 0.1 | 0.07 | 0.05 | 0.15 |
| 4 | 74699333 | rs10805069 | 1.854 | -1.306 | 1.253 | 0.357 | -0.093 | -0.054 | -0.054 | 3.402 | 0.82 | 0.44 | 0.9 | 0.92 | 0.94 | 0.83 |
| 4 | 74703999 | rs16850073 | -0.792 | -1.309 | 0.515 | 0.356 | -0.035 | -0.028 | -0.028 | 0.294 | 0.3 | 0.12 | 0.27 | 0.32 | 0.35 | 0.21 |
| 4 | 74714003 | rs6815457 | -1.684 | -2.206 | 0.781 | 0.305 | 0.056 | 0.022 | 0.022 | 1.071 | 0.44 | 0.72 | 0.27 | 0.34 | 0.24 | 0.27 |
| 4 | 74714084 | rs16850078 | -1.655 | -2.267 | 0.741 | 0.198 | -0.053 | -0.113 | -0.113 | 4.282 | 0.21 | 0.61 | 0.11 | 0.03 | 0.09 | 0.16 |
| 4 | 74718101 | rs872914 | 1.963 | -2.172 | -0.092 | 0.172 | 0.148 | -0.014 | -0.014 | 0 | 0.48 | 0.06 | 0.68 | 0.6 | 0.71 | 0.63 |
| 4 | 74718941 | rs941758 | 1.97 | -2.146 | -0.19 | 0.193 | 0.134 | -0.045 | -0.045 | 0.331 | 0.48 | 0.04 | 0.68 | 0.6 | 0.71 | 0.63 |
| 4 | 74719925 | rs11723868 | 1.449 | -1.575 | 0.692 | 0.459 | -0.026 | -0.037 | -0.037 | 0.168 | 0.92 | 0.77 | 0.97 | 0.99 | 0.99 | 0.87 |
| 4 | 74724385 | rs7695276 | 0.337 | -1.397 | 0.52 | 0.74 | -0.014 | 0.009 | 0.009 | 0.034 | 0.76 | 0.73 | 0.82 | 0.67 | 0.85 | 0.9 |
| 4 | 74727832 | rs960821 | 1.065 | -0.895 | 0.416 | 0.471 | -0.001 | 0.007 | 0.007 | 0 | 0.93 | 0.88 | 0.78 | 0.89 | 0.93 | 0.89 |
| 4 | 74728327 | rs7438101 | -0.34 | -1.166 | 0.323 | 0.741 | 0.009 | -0.014 | -0.014 | 0 | 0.24 | 0.14 | 0.26 | 0.27 | 0.2 | 0.18 |
| 4 | 74728508 | rs2145614 | -1.63 | -1.353 | 1.174 | 0.679 | 0.007 | 0.044 | 0.044 | 2.844 | 0.1 | 0.43 | 0.23 | 0.06 | 0.05 | 0.05 |
| 4 | 74730921 | rs3111693 | -1.63 | -1.353 | 1.183 | 0.679 | 0.008 | 0.045 | 0.045 | 2.844 | 0.1 | 0.43 | 0.23 | 0.06 | 0.05 | 0.05 |
| 4 | 74731278 | rs3117598 | -1.925 | -1.529 | 0.853 | 0.479 | 0.083 | -0.017 | -0.017 | 0.428 | 0.43 | 0.87 | 0.53 | 0.38 | 0.28 | 0.4 |
| 4 | 74734668 | rs3117604 | -2.013 | -1.582 | 0.803 | 0.618 | 0.089 | -0.058 | -0.058 | 0.081 | 0.44 | 0.87 | 0.53 | 0.43 | 0.29 | 0.4 |
| 4 | 74736144 | rs4074 | 2.249 | -1.824 | 0.439 | 0.185 | 0 | -0.088 | -0.088 | 0 | 0.45 | 0.03 | 0.43 | 0.58 | 0.65 | 0.45 |
| 4 | 74737930 | rs1429638 | 0.248 | -1.89 | 0.655 | 0.274 | 0.032 | -0.02 | -0.02 | 0 | 0.22 | 0.09 | 0.37 | 0.26 | 0.16 | 0.16 |
| 4 | 74748288 | rs3097412 | -1.884 | -1.895 | 0.69 | 0.258 | 0.122 | -0.031 | -0.031 | 3.034 | 0.42 | 0.86 | 0.5 | 0.41 | 0.23 | 0.39 |
| 4 | 74748325 | rs6816849 | -0.892 | -1.95 | 1.014 | 0.249 | -0.026 | 0 | 0 | 0 | 0.06 | 0.15 | 0.02 | 0.05 | 0.01 | 0.14 |
| 4 | 74750051 | rs10518117 | -0.618 | -1.969 | 1.485 | 0.246 | -0.006 | 0.041 | 0.041 | 0 | 0.07 | 0.17 | 0.06 | 0.1 | 0.05 | 0.06 |
| 4 | 74750147 | rs516224 | -1.825 | -1.953 | 1.451 | 0.259 | 0.111 | -0.028 | -0.028 | 2.06 | 0.42 | 0.82 | 0.49 | 0.42 | 0.23 | 0.38 |
| 4 | 74760458 | rs4694649 | 0.117 | -1.562 | 1.543 | 0.259 | 0.012 | -0.008 | -0.008 | 1.34 | 0.23 | 0.18 | 0.38 | 0.26 | 0.18 | 0.16 |
| 4 | 74762971 | rs482239 | -1.503 | -1.529 | 1.181 | 0.209 | 0.082 | -0.053 | -0.053 | 0.861 | 0.39 | 0.85 | 0.49 | 0.41 | 0.23 | 0.27 |
| 4 | 74776218 | rs629315 | -1.834 | -1.06 | 2.132 | 0.329 | 0.005 | 0.112 | 0.112 | 1.789 | 0.1 | 0.36 | 0.13 | 0.14 | 0.05 | 0.06 |
| 4 | 74785503 | rs13122604 | 2.099 | -1.108 | 2.019 | 0.228 | -0.012 | -0.013 | -0.013 | 0 | 0.77 | 0.32 | 0.78 | 0.84 | 0.84 | 0.73 |
| 4 | 74795131 | rs487834 | 0.552 | -1.092 | 2.215 | 0.222 | -0.002 | -0.054 | -0.054 | 0.028 | 0.87 | 0.8 | 0.92 | 0.99 | 0.89 | 0.84 |
| 4 | 74797139 | rs352024 | 0.352 | -1.093 | 2.208 | 0.222 | 0 | -0.013 | -0.013 | 0 | 0.88 | 0.89 | 0.93 | 0.99 | 0.89 | 0.84 |
| 4 | 74805322 | rs352018 | 0.352 | -1.093 | 2.208 | 0.222 | 0 | -0.013 | -0.013 | 0 | 0.88 | 0.89 | 0.93 | 0.99 | 0.89 | 0.84 |
| 4 | 74806082 | rs4694655 | 2.437 | -1.199 | 2.171 | 0.16 | 0.052 | 0.079 | 0.079 | 3.697 | 0.69 | 0.11 | 0.75 | 0.83 | 0.82 | 0.59 |
| 4 | 74811199 | rs4694656 | 0.904 | -1.196 | 2.277 | 0.141 | -0.016 | -0.109 | -0.109 | 0 | 0.83 | 0.44 | 0.89 | 0.99 | 0.89 | 0.81 |
| 4 | 74826087 | rs10938105 | 2.031 | -1.162 | 1.173 | 0.276 | 0.048 | 0.039 | 0.039 | 3.48 | 0.69 | 0.16 | 0.7 | 0.8 | 0.82 | 0.58 |
| 4 | 74834656 | rs1595009 | 2.071 | -1.127 | 0.453 | 0.396 | 0.055 | 0.006 | 0.006 | 3.523 | 0.69 | 0.1 | 0.48 | 0.74 | 0.82 | 0.58 |
| 4 | 74837402 | rs4694180 | 2.062 | -1.12 | 0.456 | 0.306 | 0.055 | 0.039 | 0.039 | 3.546 | 0.69 | 0.11 | 0.69 | 0.79 | 0.82 | 0.58 |
| 4 | 74855231 | rs352010 | 0.454 | -0.829 | 1.273 | 0.236 | 0.021 | -0.082 | -0.082 | 1.122 | 0.85 | 0.55 | 0.9 | 0.96 | 0.88 | 0.91 |
| 4 | 74857708 | rs2472649 | 2.034 | -0.897 | 0.883 | 0.175 | 0.034 | 0.059 | 0.059 | 4.819 | 0.72 | 0.1 | 0.73 | 0.82 | 0.82 | 0.75 |
| 4 | 74860238 | rs2437285 | 0.991 | -0.87 | 1.324 | -0.114 | -0.028 | -0.113 | -0.113 | 0.367 | 0.8 | 0.54 | 0.86 | 0.96 | 0.88 | 0.88 |
| 4 | 74862852 | rs3775488 | 0.517 | -0.713 | -0.452 | -0.685 | -0.04 | 0.018 | 0.018 | 0 | 0.35 | 0.04 | 0.48 | 0.23 | 0.33 | 0.38 |
| 4 | 74863997 | rs425535 | 0.467 | -0.431 | 0.347 | -0.577 | -0.009 | -0.113 | -0.113 | 0 | 0.82 | 0.53 | 0.88 | 0.96 | 0.89 | 0.92 |
| 4 | 74877895 | rs6446974 | 0.125 | -0.377 | -0.573 | -0.345 | -0.039 | 0.107 | 0.107 | 0 | 0.45 | 0.2 | 0.53 | 0.24 | 0.44 | 0.45 |
| 4 | 74881968 | rs422140 | 0.416 | -0.368 | -0.327 | -0.323 | -0.009 | 0.035 | 0.035 | 0.466 | 0.42 | 0.31 | 0.36 | 0.73 | 0.5 | 0.45 |
| 4 | 74887033 | rs6840660 | -0.181 | -0.143 | -0.351 | -0.254 | -0.017 | 0.069 | 0.069 | 0 | 0.45 | 0.35 | 0.54 | 0.22 | 0.47 | 0.41 |
| 4 | 74894373 | rs2115694 | 0.11 | -0.396 | -0.831 | -0.097 | -0.003 | 0.044 | 0.044 | 0 | 0.57 | 0.16 | 0.67 | 0.33 | 0.66 | 0.58 |
| 4 | 74903134 | rs370655 | 0.17 | -0.503 | 0.265 | 0.555 | -0.001 | -0.004 | -0.004 | 0 | 0.41 | 0.6 | 0.34 | 0.5 | 0.39 | 0.3 |
| 4 | 74915427 | rs10019219 | 0.063 | -0.541 | 0.153 | 0.544 | -0.017 | 0.066 | 0.066 | 0.988 | 0.54 | 0.14 | 0.61 | 0.34 | 0.61 | 0.64 |
| 4 | 74918698 | rs352038 | 0.764 | -0.55 | 0.746 | 0.511 | -0.012 | 0.118 | 0.118 | 0 | 0.95 | 0.84 | 0.96 | 0.83 | 0.99 | 0.94 |
| 4 | 74959093 | rs1837559 | 0.658 | -0.632 | 0.02 | 0.225 | -0.061 | -0.16 | -0.16 | 1.149 | 0.36 | 0.05 | 0.22 | 0.52 | 0.36 | 0.29 |
| 4 | 74963049 | rs9131 | 0.302 | -0.762 | 0.036 | 0.205 | 0.068 | 0.09 | 0.09 | 1.205 | 0.5 | 0.18 | 0.65 | 0.34 | 0.62 | 0.55 |
| 4 | 74971196 | rs549280 | -0.353 | -0.61 | 0.231 | 0.254 | 0.003 | -0.005 | -0.005 | 0.045 | 0.59 | 0.36 | 0.73 | 0.48 | 0.64 | 0.73 |
| 4 | 74973129 | rs1893319 | 0.693 | -0.395 | -0.835 | 0.365 | -0.083 | -0.184 | -0.184 | 0 | 0.34 | 0.03 | 0.22 | 0.52 | 0.35 | 0.27 |
| 4 | 74999784 | rs1072389 | -0.301 | 0.53 | -0.066 | 0.544 | -0.13 | -0.035 | -0.035 | 0 | 0.25 | 0.04 | 0.51 | 0.21 | 0.38 | 0.41 |
| 4 | 75013441 | rs538775 | -2.035 | 0.016 | 1.56 | 0.944 | -0.032 | 0.069 | 0.069 | 0 | 0.13 | 0.58 | 0.12 | 0.16 | 0.04 | 0.19 |
| 4 | 75037867 | rs977598 | 2.482 | -0.083 | 1.929 | 0.879 | -0.035 | 0.013 | 0.013 | 0.412 | 0.85 | 0.29 | 0.89 | 0.84 | 0.96 | 0.81 |
| 4 | 75040696 | rs16850531 | -1.598 | -0.093 | 2.729 | 0.896 | 0.009 | 0.083 | 0.083 | 0 | 0.09 | 0.32 | 0.05 | 0.12 | 0.04 | 0.08 |
| 4 | 75054234 | rs564793 | -2.206 | -0.052 | 2.64 | 0.867 | -0.018 | 0.1 | 0.1 | 0 | 0.12 | 0.49 | 0.08 | 0.15 | 0.04 | 0.18 |

| | | | | | | | | | | | | | | | | |
|---|---|---|---|---|---|---|---|---|---|---|---|---|---|---|---|---|
| 4 | 75060567 | rs477975 | -2.585 | -0.128 | 2.698 | 0.76 | -0.003 | 0.056 | 0.056 | 0.057 | 0.09 | 0.4 | 0.07 | 0.04 | 0.01 | 0.13 |
| 4 | 75067427 | rs480090 | 1.406 | -0.159 | 2.688 | 0.865 | 0.015 | 0.086 | 0.086 | 0.002 | 0.91 | 0.62 | 0.78 | 0.87 | 0.96 | 0.93 |
| 4 | 75067842 | rs4694666 | 2.531 | -0.158 | 1.981 | 0.863 | 0.025 | -0.004 | -0.004 | 0 | 0.82 | 0.16 | 0.71 | 0.83 | 0.96 | 0.79 |
| 4 | 75095061 | rs1265327 | -1.489 | 0.653 | 2.46 | 0.657 | 0.046 | -0.022 | -0.022 | 0 | 0.2 | 0.42 | 0.35 | 0.2 | 0.24 | 0.21 |
| 4 | 75102993 | rs1460016 | 1.576 | 0.637 | 2.461 | 0.803 | 0.015 | 0.005 | 0.005 | 0 | 0.76 | 0.39 | 0.6 | 0.74 | 0.74 | 0.7 |
| 4 | 75106485 | rs10518120 | 0.579 | 0.521 | 1.943 | 0.857 | 0.001 | -0.027 | -0.027 | 0 | 0.19 | 0.09 | 0.25 | 0.23 | 0.19 | 0.19 |
| 4 | 75110540 | rs1246591 | -1.813 | 0.792 | 2.697 | 0.67 | 0.038 | -0.007 | -0.007 | 0 | 0.22 | 0.43 | 0.35 | 0.21 | 0.25 | 0.21 |
| 4 | 75132798 | rs7665105 | -2.264 | 0.787 | 2.561 | 0.612 | -0.004 | 0.002 | 0.002 | 0.676 | 0.26 | 0.76 | 0.42 | 0.26 | 0.27 | 0.3 |
| 4 | 75152436 | rs9684145 | -0.716 | 1.014 | 2.98 | 1.052 | 0.008 | -0.025 | -0.025 | 0 | 0.13 | 0.2 | 0.11 | 0.03 | 0.21 | 0.11 |
| 4 | 75153710 | rs10023519 | -1.521 | 0.91 | 3.007 | 1.099 | 0.036 | -0.052 | -0.052 | 0 | 0.19 | 0.4 | 0.14 | 0.12 | 0.25 | 0.16 |
| 4 | 75168571 | rs10011730 | -1.753 | 0.905 | 2.907 | 1.1 | 0.071 | 0.161 | 0.161 | 3.663 | 0.06 | 0.42 | 0.08 | 0.06 | 0.01 | 0.09 |
| 4 | 75172809 | rs16850892 | -1.631 | 0.917 | 2.104 | 1.106 | 0.056 | 0.141 | 0.141 | 3.788 | 0.06 | 0.37 | 0.07 | 0.06 | 0.01 | 0.09 |
| 4 | 75186254 | rs715844 | -1.435 | 0.952 | 0.973 | 1.14 | 0.012 | -0.013 | -0.013 | 0.721 | 0.25 | 0.66 | 0.23 | 0.19 | 0.28 | 0.26 |
| 4 | 75199609 | rs16850959 | -1.767 | 0.323 | 0.937 | 1.44 | -0.015 | 0.057 | 0.057 | 0.631 | 0.15 | 0.38 | 0.15 | 0.21 | 0.09 | 0.25 |
| 4 | 75202277 | rs17282228 | -1.597 | 0.291 | 1.154 | 1.499 | -0.007 | 0.063 | 0.063 | 0.008 | 0.08 | 0.24 | 0.12 | 0.12 | 0.05 | 0.19 |
| 4 | 75205133 | rs12512256 | -0.705 | 0.278 | 1.595 | 1.995 | 0 | 0.041 | 0.041 | 0 | 0.22 | 0.25 | 0.37 | 0.36 | 0.21 | 0.35 |
| 4 | 75206039 | rs10938119 | -2.101 | 0.301 | 1.65 | 1.987 | -0.003 | 0 | 0 | 0 | 0.36 | 0.59 | 0.53 | 0.57 | 0.3 | 0.55 |
| 4 | 75208519 | rs2125398 | -1.596 | 0.307 | 1.639 | 1.97 | -0.007 | 0.061 | 0.061 | 0 | 0.29 | 0.48 | 0.49 | 0.48 | 0.26 | 0.49 |
| 4 | 75210003 | rs7679352 | -1.363 | 0.292 | 1.796 | 1.933 | 0.001 | 0.086 | 0.086 | 0.141 | 0.08 | 0.27 | 0.13 | 0.12 | 0.05 | 0.19 |
| 4 | 75216238 | rs16851013 | -2.071 | 0.243 | 2.036 | 1.863 | 0.011 | 0.002 | 0.002 | 0.223 | 0.37 | 0.72 | 0.56 | 0.57 | 0.3 | 0.55 |
| 4 | 75216365 | rs16851014 | -1.265 | 0.287 | 2.143 | 2.104 | -0.004 | 0.069 | 0.069 | 0 | 0.23 | 0.38 | 0.26 | 0.4 | 0.22 | 0.46 |
| 4 | 75216675 | rs16851016 | -0.986 | 0.283 | 2.261 | 2.176 | -0.003 | 0.084 | 0.084 | 0 | 0.21 | 0.37 | 0.2 | 0.36 | 0.19 | 0.45 |
| 4 | 75224590 | rs1350666 | 1.796 | 0.178 | 2.705 | 2.191 | -0.013 | 0.052 | 0.052 | 0.04 | 0.71 | 0.48 | 0.76 | 0.52 | 0.77 | 0.55 |
| 4 | 75227649 | rs968061 | -0.11 | 0.19 | 2.531 | 2.223 | -0.007 | -0.049 | -0.049 | 0 | 0.84 | 0.9 | 0.87 | 0.71 | 0.83 | 0.72 |
| 4 | 75230302 | rs3806794 | -0.687 | 0.176 | 2.579 | 2.244 | 0.001 | -0.001 | -0.001 | 0 | 0.06 | 0.04 | 0.03 | 0.08 | 0.04 | 0.08 |
| 4 | 75243702 | rs4694683 | 0.009 | 0.203 | 2.136 | 2.151 | -0.006 | -0.047 | -0.047 | 0.374 | 0.83 | 0.89 | 0.87 | 0.7 | 0.82 | 0.73 |
| 4 | 75248434 | rs2367707 | 0.344 | 0.312 | 2.251 | 2.131 | 0 | 0.013 | 0.013 | 0 | 0.81 | 0.79 | 0.83 | 0.7 | 0.81 | 0.72 |
| 4 | 75248544 | rs2367708 | 0.198 | 0.336 | 1.973 | 2.135 | -0.007 | -0.043 | -0.043 | 0 | 0.82 | 0.88 | 0.85 | 0.7 | 0.81 | 0.72 |
| 4 | 75252350 | rs1017733 | 0.129 | 0.341 | 1.988 | 2.151 | -0.002 | -0.009 | -0.009 | 0 | 0.83 | 0.83 | 0.83 | 0.7 | 0.81 | 0.72 |
| 4 | 75254258 | rs1460007 | 0.078 | 0.34 | 1.988 | 2.225 | -0.011 | -0.041 | -0.041 | 0 | 0.83 | 0.89 | 0.85 | 0.73 | 0.81 | 0.72 |
| 4 | 75254282 | rs1460008 | 0.137 | 0.343 | 1.95 | 2.227 | -0.002 | -0.009 | -0.009 | 0 | 0.83 | 0.83 | 0.84 | 0.73 | 0.81 | 0.72 |
| 4 | 75258429 | rs1460013 | 0.137 | 0.343 | 1.925 | 2.227 | -0.002 | -0.013 | -0.013 | 0.401 | 0.83 | 0.83 | 0.84 | 0.73 | 0.81 | 0.72 |
| 4 | 75258678 | rs6816797 | -1.431 | -0.326 | 1.645 | 1.788 | -0.035 | 0.092 | 0.092 | 0.013 | 0.22 | 0.46 | 0.35 | 0.31 | 0.12 | 0.25 |
| 4 | 75259405 | rs6845297 | 1.185 | -0.383 | 0.135 | 1.236 | 0.054 | 0.083 | 0.083 | 0.089 | 0.49 | 0.15 | 0.61 | 0.38 | 0.62 | 0.32 |
| 4 | 75261407 | rs10002688 | -0.037 | -0.159 | 0.562 | 0.223 | -0.001 | -0.053 | -0.053 | 0 | 0.68 | 0.69 | 0.69 | 0.97 | 0.69 | 0.74 |
| 4 | 75266446 | rs16851150 | 0.107 | -0.193 | -0.166 | 0.107 | -0.027 | -0.059 | -0.059 | 1.357 | 0.18 | 0.1 | 0.4 | 0.33 | 0.23 | 0.24 |
| 4 | 75268730 | rs2047193 | 1.096 | -0.402 | 0.306 | -0.128 | 0.004 | 0.005 | 0.005 | 0 | 0.4 | 0.28 | 0.24 | 0.63 | 0.39 | 0.39 |
| 4 | 75272863 | rs7693713 | 0.04 | -0.381 | 0.112 | 0.03 | -0.026 | -0.061 | -0.061 | 5.191 | 0.18 | 0.09 | 0.41 | 0.32 | 0.23 | 0.25 |
| 4 | 75273080 | rs13139174 | 0.309 | -0.621 | 0.492 | -0.132 | 0.009 | 0.031 | 0.031 | 0 | 0.5 | 0.42 | 0.27 | 0.66 | 0.44 | 0.48 |
| 4 | 75276524 | rs718607 | 0.182 | -0.736 | 0.315 | -0.192 | 0.006 | 0.013 | 0.013 | 0 | 0.49 | 0.56 | 0.28 | 0.64 | 0.43 | 0.5 |
| 4 | 75282847 | rs1494878 | 0.196 | -0.734 | 0.443 | -0.191 | 0.008 | 0.017 | 0.017 | 0 | 0.49 | 0.56 | 0.28 | 0.64 | 0.43 | 0.5 |
| 4 | 75324290 | rs1691274 | 0.947 | -1.088 | 0.14 | -0.185 | -0.007 | -0.017 | -0.017 | 0 | 0.38 | 0.33 | 0.38 | 0.47 | 0.42 | 0.41 |
| 4 | 75328108 | rs11725706 | 0.086 | -1.164 | 0.222 | -0.404 | -0.001 | -0.001 | -0.001 | 4.569 | 0.18 | 0.18 | 0.37 | 0.19 | 0.18 | 0.23 |
| 4 | 75329122 | rs1353295 | -0.675 | -1.189 | 0.243 | -0.762 | -0.004 | 0.026 | 0.026 | 0 | 0.63 | 0.75 | 0.78 | 0.98 | 0.65 | 0.82 |
| 4 | 75336464 | rs1246415 | -0.76 | -0.973 | 0.62 | -0.804 | -0.003 | -0.136 | -0.136 | 0 | 0.65 | 0.54 | 0.78 | 0.99 | 0.68 | 0.82 |
| 4 | 75341240 | rs2609204 | 0.556 | -1.037 | 0.488 | -0.787 | -0.018 | -0.006 | -0.006 | 0 | 0.1 | 0.13 | 0.12 | 0.09 | 0.04 | 0.16 |
| 4 | 75344134 | rs7668817 | 1.497 | -1.136 | 0.606 | -0.686 | -0.028 | -0.043 | -0.043 | 0 | 0.33 | 0.2 | 0.46 | 0.42 | 0.39 | 0.32 |
| 4 | 75356565 | rs1494893 | -1.146 | -1.078 | 0.624 | -0.815 | -0.015 | 0.095 | 0.095 | 0 | 0.68 | 0.86 | 0.8 | 0.99 | 0.68 | 0.82 |
| 4 | 75362636 | rs1603131 | 0.463 | -0.96 | 0.281 | -0.695 | -0.006 | -0.002 | -0.002 | 0 | 0.1 | 0.09 | 0.12 | 0.09 | 0.05 | 0.17 |
| 4 | 75377358 | rs6447000 | -0.143 | -0.885 | 0.224 | -0.688 | -0.002 | -0.039 | -0.039 | 0.199 | 0.57 | 0.41 | 0.66 | 0.88 | 0.64 | 0.66 |
| 4 | 75377395 | rs6828372 | 0.855 | -0.83 | -0.249 | -0.505 | 0 | -0.004 | -0.004 | 0.009 | 0.15 | 0.14 | 0.15 | 0.21 | 0.18 | 0.18 |
| 4 | 75387973 | rs7656798 | 1.096 | -0.624 | -0.358 | -0.44 | -0.046 | -0.076 | -0.076 | 0 | 0.34 | 0.17 | 0.48 | 0.43 | 0.4 | 0.33 |
| 4 | 75395312 | rs13104811 | 0.548 | -0.452 | 0.264 | -0.685 | 0.008 | 0.053 | 0.053 | 0 | 0.52 | 0.38 | 0.64 | 0.63 | 0.58 | 0.55 |
| 4 | 75400315 | rs1994941 | 0.18 | -0.493 | 0.399 | -0.652 | -0.02 | -0.003 | -0.003 | 0 | 0.1 | 0.16 | 0.12 | 0.1 | 0.04 | 0.09 |
| 4 | 75404702 | rs4694196 | -0.882 | -0.516 | -0.022 | -0.846 | -0.035 | 0.116 | 0.116 | 0 | 0.72 | 0.88 | 0.82 | 0.99 | 0.69 | 0.79 |
| 4 | 75406050 | rs10001945 | 0.729 | -0.544 | 0.185 | -0.795 | 0.01 | 0.059 | 0.059 | 0 | 0.52 | 0.36 | 0.64 | 0.63 | 0.58 | 0.55 |
| 4 | 75409327 | rs6447006 | -0.404 | -0.145 | -0.368 | -0.597 | -0.022 | 0.033 | 0.033 | 0 | 0.68 | 0.82 | 0.81 | 0.74 | 0.65 | 0.66 |
| 4 | 75416565 | rs7435073 | -0.559 | -0.251 | -0.528 | -1.047 | -0.047 | 0.13 | 0.13 | 0 | 0.72 | 0.89 | 0.82 | 0.99 | 0.68 | 0.78 |
| 4 | 75510963 | rs9307637 | 0.151 | -0.33 | 0.036 | -1.141 | 0.002 | -0.053 | -0.053 | 0 | 0.45 | 0.59 | 0.28 | 0.01 | 0.38 | 0.23 |
| 4 | 75515889 | rs10029098 | 0.474 | -0.298 | -0.055 | -0.872 | -0.002 | 0.062 | 0.062 | 0 | 0.41 | 0.25 | 0.25 | 0.18 | 0.39 | 0.31 |
| 4 | 75525096 | rs10050072 | -0.652 | -0.6 | 0.037 | -0.95 | -0.024 | -0.02 | -0.02 | 0.03 | 0.86 | 0.79 | 0.87 | 0.89 | 0.93 | 0.83 |
| 4 | 75526108 | rs7435213 | 0.558 | -0.49 | -0.245 | -0.782 | -0.012 | -0.011 | -0.011 | 0 | 0.24 | 0.49 | 0.25 | 0.22 | 0.22 | 0.26 |
| 4 | 75528603 | rs11097585 | -0.989 | -0.195 | -0.718 | -0.583 | -0.025 | -0.02 | -0.02 | 0.233 | 0.87 | 0.82 | 0.88 | 0.9 | 0.94 | 0.84 |
| 4 | 75529098 | rs7437891 | 0.318 | -0.146 | -0.716 | -0.573 | -0.007 | -0.001 | -0.001 | 0 | 0.65 | 0.74 | 0.5 | 0.4 | 0.61 | 0.53 |
| 4 | 75534357 | rs11097821 | 0.518 | 0.135 | -1.258 | -0.402 | -0.07 | -0.105 | -0.105 | 0.012 | 0.2 | 0.06 | 0.37 | 0.49 | 0.32 | 0.27 |
| 4 | 75546594 | rs4596322 | -0.029 | -0.404 | -0.731 | -0.442 | -0.012 | -0.014 | -0.014 | 0.538 | 0.15 | 0.19 | 0.15 | 0.11 | 0.1 | 0.17 |
| 4 | 75547013 | rs7659874 | -0.149 | -0.463 | -0.498 | -0.323 | -0.017 | 0.053 | 0.053 | 0.251 | 0.26 | 0.57 | 0.25 | 0.41 | 0.23 | 0.37 |
| 4 | 75547732 | rs6534012 | -0.086 | -0.873 | -1.168 | -1.16 | -0.005 | 0.121 | 0.121 | 0 | 0.61 | 0.81 | 0.7 | 0.88 | 0.63 | 0.73 |
| 4 | 75547845 | rs13124944 | -0.332 | -0.616 | -0.99 | -1.148 | -0.009 | -0.104 | -0.104 | 0 | 0.36 | 0.5 | 0.23 | 0.1 | 0.33 | 0.38 |

| | | | | | | | | | | | | | | | | |
|---|---|---|---|---|---|---|---|---|---|---|---|---|---|---|---|---|
| 4 | 75555721 | rs4333264 | -0.036 | -0.249 | -2.122 | -0.607 | -0.166 | -0.045 | -0.045 | 0 | 0.32 | 0.05 | 0.31 | 0.33 | 0.47 | 0.38 |
| 4 | 75564110 | rs7664990 | -0.187 | -1.139 | -1.819 | -1.183 | 0.004 | 0.001 | 0.001 | 0 | 0.32 | 0.69 | 0.18 | 0.11 | 0.14 | 0.23 |
| 4 | 75575898 | rs11099492 | 0.224 | -1.542 | -1.561 | -1.583 | 0 | 0.006 | 0.006 | 0 | 0.33 | 0.3 | 0.21 | 0.05 | 0.31 | 0.15 |
| 4 | 75579453 | rs3923243 | -0.617 | -1.517 | -1.405 | -1.558 | 0.013 | 0.125 | 0.125 | 0 | 0.53 | 0.72 | 0.33 | 0.12 | 0.4 | 0.38 |
| 4 | 75589514 | rs6836453 | -0.573 | -1.543 | -1.37 | -1.57 | 0.012 | 0.083 | 0.083 | 0 | 0.52 | 0.67 | 0.33 | 0.12 | 0.4 | 0.38 |
| 4 | 75599687 | rs7663639 | 1.142 | -1.437 | -1.312 | -1.564 | -0.003 | -0.032 | -0.032 | 0 | 0.21 | 0.19 | 0.16 | 0 | 0.26 | 0.15 |
| 4 | 75605512 | rs11930214 | 0.461 | -1.533 | -1.354 | -1.599 | -0.049 | -0.084 | -0.084 | 0.238 | 0.21 | 0.41 | 0.12 | 0.07 | 0.09 | 0.22 |
| 4 | 75608390 | rs11097027 | 0.405 | -1.555 | -2.272 | -1.241 | -0.098 | -0.053 | -0.053 | 0 | 0.35 | 0.13 | 0.36 | 0.41 | 0.48 | 0.39 |
| 4 | 75619533 | rs6848228 | -0.921 | -1.136 | -1.669 | -0.653 | 0.023 | -0.045 | -0.045 | 0 | 0.79 | 0.85 | 0.65 | 0.54 | 0.83 | 0.76 |
| 4 | 75622361 | rs6849245 | 0.217 | -1.119 | -1.782 | -0.735 | -0.043 | 0.003 | 0.003 | 0.172 | 0.42 | 0.18 | 0.4 | 0.42 | 0.52 | 0.57 |
| 4 | 75624475 | rs7691380 | 0.166 | -0.531 | -1.161 | -0.075 | -0.006 | 0.014 | 0.014 | 0.639 | 0.73 | 0.72 | 0.62 | 0.53 | 0.78 | 0.74 |
| 4 | 75629315 | rs10025144 | 0.052 | -0.409 | -1.69 | -0.964 | 0.048 | 0.168 | 0.168 | 0 | 0.56 | 0.81 | 0.29 | 0.03 | 0.34 | 0.34 |
| 4 | 75632076 | rs4412061 | -0.346 | -0.672 | -1.193 | -0.3 | -0.001 | 0 | 0 | 0 | 0.53 | 0.5 | 0.51 | 0.45 | 0.66 | 0.52 |
| 4 | 75639740 | rs4859540 | -0.194 | -0.772 | -0.924 | -0.18 | 0.077 | 0.033 | 0.033 | 0 | 0.51 | 0.29 | 0.48 | 0.38 | 0.67 | 0.51 |
| 4 | 75654169 | rs7686112 | -0.359 | -0.295 | -0.796 | -0.758 | -0.012 | -0.062 | -0.062 | 0 | 0.27 | 0.42 | 0.23 | 0.12 | 0.27 | 0.24 |
| 4 | 75654722 | rs7676863 | 0.223 | 0.034 | -1.071 | -0.817 | -0.013 | 0.023 | 0.023 | 0 | 0.14 | 0.07 | 0.14 | 0.04 | 0.2 | 0.17 |
| 4 | 75658650 | rs10022199 | 0.528 | -0.117 | -0.938 | -0.6 | -0.03 | -0.034 | -0.034 | 0 | 0.31 | 0.23 | 0.21 | 0.42 | 0.42 | 0.28 |
| 4 | 75661832 | rs11736577 | 0.558 | -0.107 | -0.98 | -0.595 | -0.038 | -0.043 | -0.043 | 0.301 | 0.3 | 0.21 | 0.2 | 0.42 | 0.42 | 0.28 |
| 4 | 75664504 | rs12503079 | -0.119 | -0.467 | -0.752 | -0.938 | 0 | -0.053 | -0.053 | 0 | 0.26 | 0.25 | 0.2 | 0.04 | 0.25 | 0.25 |
| 4 | 75667398 | rs4302527 | 1.304 | -0.895 | -1.036 | -0.888 | -0.014 | -0.071 | -0.071 | 0 | 0.17 | 0.03 | 0.22 | 0.35 | 0.18 | 0.24 |
| 4 | 75668334 | rs7667014 | -0.643 | -0.678 | -0.33 | -0.905 | 0.024 | -0.017 | -0.017 | 0 | 0.59 | 0.69 | 0.44 | 0.46 | 0.68 | 0.53 |
| 4 | 75670619 | rs11727548 | -0.593 | -0.677 | -0.312 | -0.905 | 0.017 | -0.011 | -0.011 | 6.387 | 0.58 | 0.65 | 0.44 | 0.46 | 0.68 | 0.53 |
| 4 | 75673363 | rs4585380 | -0.263 | -0.358 | -0.224 | -1.874 | 0.001 | -0.054 | -0.054 | 0 | 0.26 | 0.25 | 0.2 | 0.04 | 0.26 | 0.25 |
| 4 | 75681217 | rs11734944 | 0.723 | -0.234 | -0.644 | -1.596 | -0.002 | 0 | 0 | 0 | 0.32 | 0.31 | 0.48 | 0.63 | 0.42 | 0.37 |
| 4 | 75682484 | rs9307118 | 0.591 | -0.231 | -0.712 | -1.59 | 0.016 | -0.004 | -0.004 | 0 | 0.33 | 0.42 | 0.49 | 0.63 | 0.42 | 0.37 |
| 4 | 75683594 | rs4352548 | -0.455 | -0.636 | -1.043 | -1.625 | -0.023 | 0.109 | 0.109 | 0.011 | 0.22 | 0.05 | 0.15 | 0.04 | 0.21 | 0.16 |
| 4 | 75694208 | rs10009801 | -0.591 | -1.494 | -0.284 | -1.618 | -0.041 | -0.088 | -0.088 | 0 | 0.32 | 0.61 | 0.2 | 0.05 | 0.12 | 0.25 |
| 4 | 75712104 | rs1377032 | -0.065 | -1.515 | 0.195 | -1.584 | -0.076 | -0.111 | -0.111 | 0.577 | 0.24 | 0.45 | 0.08 | 0.04 | 0.07 | 0.18 |
| 4 | 75717601 | rs977266 | -0.861 | -1.279 | -0.527 | -1.694 | -0.012 | 0.027 | 0.027 | 0.508 | 0.63 | 0.72 | 0.47 | 0.31 | 0.46 | 0.57 |
| 4 | 75718366 | rs1869172 | 0.254 | -1.281 | -0.204 | -1.69 | -0.077 | -0.113 | -0.113 | 0.038 | 0.24 | 0.43 | 0.08 | 0.05 | 0.07 | 0.19 |
| 4 | 75718547 | rs1349634 | -0.402 | -0.43 | 0.132 | -1.131 | 0.007 | 0.027 | 0.027 | 0 | 0.4 | 0.33 | 0.39 | 0.28 | 0.39 | 0.39 |
| 4 | 75730241 | rs16999874 | -0.305 | -0.479 | -0.219 | -1.004 | 0 | -0.011 | -0.011 | 0.107 | 0.07 | 0.07 | 0.11 | 0.01 | 0.06 | 0.07 |
| 4 | 75731348 | rs7667066 | 0.355 | -0.142 | -0.484 | -1.073 | -0.067 | 0.081 | 0.081 | 0 | 0.35 | 0.1 | 0.29 | 0.29 | 0.42 | 0.35 |
| 4 | 75738792 | rs1377028 | -0.176 | 0.029 | -0.365 | -0.83 | 0 | -0.014 | -0.014 | 0 | 0.46 | 0.43 | 0.43 | 0.3 | 0.49 | 0.41 |
| 4 | 75745515 | rs6827871 | -0.336 | 0.366 | -0.552 | -0.572 | -0.003 | 0.014 | 0.014 | 0.035 | 0.55 | 0.42 | 0.47 | 0.36 | 0.55 | 0.53 |
| 4 | 75746665 | rs6854845 | -1.295 | 0.328 | -0.17 | -0.141 | 0.007 | -0.027 | -0.027 | 2.934 | 0.09 | 0.21 | 0.14 | 0.04 | 0.1 | 0.09 |
| 4 | 75748121 | rs896577 | 0.335 | -0.179 | -0.641 | 0.185 | 0.023 | 0.005 | 0.005 | 1.072 | 0.57 | 0.38 | 0.53 | 0.4 | 0.67 | 0.62 |
| 4 | 75761496 | rs4859467 | 0.204 | 0.029 | -0.377 | 0.467 | 0 | -0.001 | -0.001 | 0.031 | 0.53 | 0.49 | 0.54 | 0.44 | 0.5 | 0.63 |
| 4 | 75771799 | rs6837250 | 0.028 | -0.498 | -0.199 | 0.516 | -0.003 | -0.003 | -0.003 | 0.79 | 0.56 | 0.49 | 0.55 | 0.39 | 0.58 | 0.65 |
| 4 | 75797003 | rs16999967 | -0.21 | -0.32 | 0.253 | 0.573 | 0.024 | -0.02 | -0.02 | 0.154 | 0.08 | 0.31 | 0.06 | 0 | 0.08 | 0.01 |
| 4 | 75805790 | rs6833056 | 0.205 | -0.609 | -0.149 | 0.484 | 0 | 0 | 0 | 0 | 0.52 | 0.48 | 0.56 | 0.43 | 0.57 | 0.71 |
| 4 | 75808541 | rs17248137 | 0.147 | -0.449 | -0.619 | 0.097 | -0.001 | -0.01 | -0.01 | 0 | 0.06 | 0.11 | 0.05 | 0 | 0.08 | 0.01 |
| 4 | 75809063 | rs2902802 | 0.736 | -0.462 | -1.039 | 0.009 | -0.169 | 0.189 | 0.189 | 0.366 | 0.37 | 0.02 | 0.32 | 0.1 | 0.48 | 0.53 |
| 4 | 75826870 | rs4859853 | -0.019 | -0.767 | -0.659 | 0.16 | -0.01 | 0.007 | 0.007 | 0 | 0.09 | 0.02 | 0.18 | 0.06 | 0.08 | 0.08 |
| 4 | 75844039 | rs16999999 | -0.288 | -0.789 | 0.148 | 0.137 | -0.004 | -0.018 | -0.018 | 0 | 0.11 | 0.18 | 0.2 | 0.06 | 0.09 | 0.08 |
| 4 | 75845127 | rs3886161 | 0.398 | -0.967 | 0.306 | 0.4 | 0.065 | -0.003 | -0.003 | 0 | 0.4 | 0.19 | 0.35 | 0.41 | 0.28 | 0.28 |
| 4 | 75845559 | rs1452303 | -0.15 | -0.883 | 0.602 | 0.379 | 0.02 | 0.009 | 0.009 | 0.001 | 0.52 | 0.45 | 0.57 | 0.58 | 0.63 | 0.71 |
| 4 | 75847533 | rs1377027 | -0.356 | -0.804 | 0.542 | 0.391 | 0.028 | -0.019 | -0.019 | 0.193 | 0.94 | 0.74 | 0.94 | 1 | 0.91 | 0.99 |
| 4 | 75847586 | rs17248704 | 0.324 | -0.783 | 0.424 | 0.418 | 0.003 | -0.017 | -0.017 | 0 | 0.06 | 0.13 | 0.06 | 0 | 0.09 | 0.01 |
| 4 | 75851372 | rs2060849 | 0.683 | -0.816 | 0.332 | 0.511 | -0.028 | -0.055 | -0.055 | 0 | 0.38 | 0.15 | 0.35 | 0.46 | 0.54 | 0.59 |
| 4 | 75857836 | rs4859880 | -0.283 | -0.849 | 0.587 | 0.419 | 0.004 | 0.002 | 0.002 | 0.046 | 0.54 | 0.47 | 0.61 | 0.58 | 0.72 | 0.73 |
| 4 | 75860438 | rs17248859 | -0.185 | -0.203 | 0.505 | 0.26 | -0.013 | 0.023 | 0.023 | 0 | 0.06 | 0.03 | 0.05 | 0 | 0.09 | 0.01 |
| 4 | 75898443 | rs7656977 | 0.85 | -0.422 | 0.291 | 0.237 | -0.002 | -0.01 | -0.01 | 0.002 | 0.05 | 0.06 | 0.01 | 0.01 | 0.02 | 0.07 |
| 4 | 75899133 | rs1007044 | -1.111 | -1.45 | 0.443 | 0.133 | 0.002 | 0.002 | 0.002 | 1.423 | 0.36 | 0.7 | 0.35 | 0.37 | 0.19 | 0.25 |
| 4 | 75911362 | rs6838268 | 0.451 | -1.441 | 0.47 | 0.196 | 0.003 | 0.003 | 0.003 | 0 | 0.06 | 0.04 | 0.05 | 0 | 0.05 | 0.03 |
| 4 | 75911758 | rs1816181 | 0.215 | -1.504 | 0.77 | 0.152 | 0.034 | -0.028 | -0.028 | 0 | 0.21 | 0.13 | 0.23 | 0.36 | 0.11 | 0.15 |
| 4 | 75929358 | rs11098369 | -0.848 | -1.454 | 0.584 | 0.179 | -0.036 | -0.001 | -0.001 | 0 | 0.31 | 0.64 | 0.32 | 0.36 | 0.17 | 0.18 |
| 4 | 75937971 | rs3822109 | 0.804 | -0.392 | 0.311 | -0.55 | -0.001 | 0 | 0 | 0.002 | 0.31 | 0.29 | 0.28 | 0.16 | 0.46 | 0.47 |
| 4 | 75938179 | rs3775527 | 0.881 | -0.284 | 0.379 | -0.544 | -0.011 | 0.007 | 0.007 | 0.693 | 0.31 | 0.25 | 0.26 | 0.16 | 0.45 | 0.47 |
| 4 | 75938236 | rs1062293 | -0.047 | -0.556 | 0.405 | -0.571 | 0 | -0.004 | -0.004 | 0 | 0.41 | 0.42 | 0.35 | 0.21 | 0.53 | 0.53 |
| 4 | 75942134 | rs6824037 | -0.206 | -0.51 | 0.447 | -0.461 | -0.018 | -0.015 | -0.015 | 3.213 | 0.1 | 0.18 | 0.07 | 0.05 | 0.07 | 0.05 |
| 4 | 75945366 | rs10049553 | 0.966 | -0.582 | 0.123 | -0.603 | -0.145 | 0.138 | 0.138 | 1.555 | 0.33 | 0.05 | 0.23 | 0.11 | 0.47 | 0.47 |
| 4 | 75952499 | rs4859933 | -0.169 | -0.636 | 0.208 | -0.623 | -0.001 | 0.003 | 0.003 | 0 | 0.46 | 0.59 | 0.37 | 0.23 | 0.55 | 0.52 |
| 4 | 75967200 | rs10028207 | -0.41 | -0.697 | -0.179 | -0.655 | -0.005 | -0.006 | -0.006 | 0 | 0.41 | 0.58 | 0.48 | 0.72 | 0.34 | 0.37 |
| 4 | 75971764 | rs3796683 | 0.021 | -0.721 | -0.291 | -0.614 | -0.01 | 0.001 | 0.001 | 0.619 | 0.11 | 0.45 | 0.09 | 0.05 | 0.09 | 0.05 |
| 4 | 75971997 | rs10012438 | 0.021 | -0.721 | -0.291 | -0.614 | -0.01 | 0.001 | 0.001 | 0.619 | 0.11 | 0.45 | 0.09 | 0.05 | 0.09 | 0.05 |
| 4 | 75972413 | rs9090 | 0.059 | -0.763 | -0.277 | -0.609 | -0.013 | -0.002 | -0.002 | 0 | 0.11 | 0.44 | 0.09 | 0.05 | 0.09 | 0.05 |
| 4 | 75972812 | rs10015446 | -0.46 | -0.677 | -0.255 | -0.618 | -0.007 | -0.009 | -0.009 | 0 | 0.4 | 0.56 | 0.48 | 0.73 | 0.36 | 0.37 |
| 4 | 75973830 | rs1043633 | 0.286 | -0.593 | -0.359 | -0.581 | -0.02 | -0.011 | -0.011 | 0 | 0.1 | 0.33 | 0.08 | 0.05 | 0.09 | 0.05 |

| | | | | | | | | | | | | | | | | |
|---|---|---|---|---|---|---|---|---|---|---|---|---|---|---|---|---|
| 4 | 75974353 | rs1061570 | 0.493 | -0.134 | -1.254 | -0.327 | -0.006 | -0.052 | -0.052 | 0 | 0.28 | 0.09 | 0.39 | 0.67 | 0.27 | 0.32 |
| 4 | 75980925 | rs11944489 | 0.099 | 0.684 | -0.819 | -0.158 | -0.054 | 0.051 | 0.051 | 0.467 | 0.34 | 0.17 | 0.29 | 0.18 | 0.47 | 0.44 |
| 4 | 75992533 | rs17000192 | -0.472 | 0.36 | 0.074 | -0.093 | 0.01 | -0.058 | -0.058 | 0 | 0.1 | 0.28 | 0.08 | 0.01 | 0.08 | 0.13 |
| 4 | 75996025 | rs1426064 | -0.593 | -0.247 | -0.278 | 0.177 | -0.036 | 0.034 | 0.034 | 0 | 0.28 | 0.46 | 0.54 | 0.7 | 0.26 | 0.26 |
| 4 | 75998962 | rs17000203 | -0.336 | 0.02 | -0.633 | 0.855 | -0.022 | -0.04 | -0.04 | 0 | 0.09 | 0.02 | 0.2 | 0.37 | 0.12 | 0.16 |
| 4 | 76010215 | rs6837969 | -1.269 | -0.144 | 0.231 | 0.545 | 0.05 | -0.081 | -0.081 | 0 | 0.13 | 0.52 | 0.13 | 0.01 | 0.15 | 0.1 |
| 4 | 76010255 | rs10000988 | 0.764 | -0.172 | 0.223 | 0.776 | 0.018 | -0.047 | -0.047 | 0 | 0.13 | 0.05 | 0.2 | 0.34 | 0.08 | 0.14 |
| 4 | 76012856 | rs4464630 | -0.695 | -0.166 | 0.178 | 0.778 | 0.049 | -0.021 | -0.021 | 0 | 0.09 | 0.48 | 0.08 | 0.01 | 0.06 | 0.06 |
| 4 | 76014897 | rs4859943 | -1.45 | -0.302 | 0.207 | 0.724 | 0.016 | -0.013 | -0.013 | 0 | 0.28 | 0.79 | 0.35 | 0.35 | 0.23 | 0.24 |
| 4 | 76031576 | rs6534607 | -2.679 | -0.146 | 0.751 | 1.061 | 0.005 | -0.01 | -0.01 | 3.546 | 0.19 | 0.8 | 0.22 | 0.32 | 0.17 | 0.18 |
| 4 | 76031652 | rs6534608 | -1.678 | 0.187 | -0.07 | 0.586 | -0.065 | 0.302 | 0.302 | 0.101 | 0.51 | 0.93 | 0.55 | 0.89 | 0.52 | 0.5 |
| 4 | 76034046 | rs17000262 | -0.447 | 0.477 | 0.192 | 0.819 | 0.008 | -0.02 | -0.02 | 0 | 0.05 | 0.11 | 0.06 | 0 | 0.11 | 0.03 |
| 4 | 76051855 | rs6853754 | -1.659 | 0.483 | 0.202 | 0.875 | 0.037 | 0.156 | 0.156 | 0 | 0.1 | 0.29 | 0.13 | 0.31 | 0.14 | 0.11 |
| 4 | 76072424 | rs769325 | 0.296 | 0.494 | -0.49 | 0.311 | -0.047 | -0.076 | -0.076 | 0.859 | 0.23 | 0.05 | 0.13 | 0.22 | 0.23 | 0.17 |
| 4 | 76075268 | rs7699626 | 0.614 | 0.563 | -0.422 | -0.258 | -0.041 | 0.024 | 0.024 | 0 | 0.11 | 0.03 | 0.12 | 0.05 | 0.21 | 0.18 |
| 4 | 76075792 | rs6534849 | -0.74 | 0.468 | -0.285 | -0.384 | -0.028 | 0.026 | 0.026 | 0 | 0.18 | 0.14 | 0.19 | 0.08 | 0.32 | 0.23 |
| 4 | 76078680 | rs7349670 | 1.784 | -0.2 | -0.613 | -0.685 | -0.079 | 0.102 | 0.102 | 0 | 0.43 | 0.07 | 0.42 | 0.19 | 0.47 | 0.49 |
| 4 | 76081120 | rs980761 | -1.27 | -0.288 | 0.226 | -0.678 | -0.006 | -0.015 | -0.015 | 0.045 | 0.36 | 0.56 | 0.45 | 0.81 | 0.36 | 0.46 |
| 4 | 76081553 | rs1391010 | -0.719 | -0.441 | 0.354 | -0.61 | -0.029 | 0.021 | 0.021 | 0.28 | 0.24 | 0.38 | 0.33 | 0.76 | 0.18 | 0.28 |
| 4 | 76082809 | rs1497209 | 1.468 | 0.095 | -0.174 | -0.59 | -0.051 | 0.078 | 0.078 | 0.022 | 0.4 | 0.07 | 0.4 | 0.19 | 0.39 | 0.47 |
| 4 | 76086206 | rs1391013 | 0.276 | 0.074 | 0.609 | -0.006 | -0.007 | 0.006 | 0.006 | 0 | 0.12 | 0.11 | 0.12 | 0.06 | 0.19 | 0.18 |
| 4 | 76088845 | rs13113687 | 1.495 | -0.251 | 1.086 | -0.035 | 0.048 | 0.018 | 0.018 | 0 | 0.61 | 0.14 | 0.62 | 0.25 | 0.7 | 0.72 |
| 4 | 76093188 | rs12642121 | -0.978 | -0.283 | 1.53 | -0.047 | 0.015 | 0.098 | 0.098 | 0 | 0.1 | 0.19 | 0.18 | 0.73 | 0.13 | 0.17 |
| 4 | 76106090 | rs7685095 | -0.823 | -0.353 | 1.815 | -0.056 | -0.012 | 0.016 | 0.016 | 0 | 0.23 | 0.3 | 0.21 | 0.7 | 0.16 | 0.17 |
| 4 | 76119783 | rs4859946 | 1.167 | -0.348 | 1.957 | -0.058 | 0.011 | -0.096 | -0.096 | 0 | 0.86 | 0.48 | 0.84 | 0.98 | 0.87 | 0.89 |
| 4 | 76130475 | rs1497214 | -0.887 | -0.269 | 1.86 | -0.043 | 0.005 | 0.052 | 0.052 | 0 | 0.1 | 0.14 | 0.19 | 0.69 | 0.13 | 0.18 |
| 4 | 76137551 | rs6855222 | 0.731 | -0.288 | 1.891 | -0.07 | -0.055 | -0.143 | -0.143 | 0.408 | 0.74 | 0.6 | 0.83 | 0.98 | 0.84 | 0.89 |
| 4 | 76138373 | rs2055215 | 0.731 | -0.288 | 1.88 | -0.07 | -0.055 | -0.143 | -0.143 | 0.151 | 0.74 | 0.61 | 0.83 | 0.98 | 0.84 | 0.89 |
| 4 | 76139891 | rs7677444 | 0.967 | -0.207 | 1.796 | -0.033 | 0.008 | -0.08 | -0.08 | 0.058 | 0.87 | 0.67 | 0.85 | 0.98 | 0.87 | 0.89 |
| 4 | 76161222 | rs7681167 | -0.757 | -0.208 | 1.62 | -0.01 | -0.027 | 0.047 | 0.047 | 0 | 0.24 | 0.44 | 0.23 | 0.7 | 0.16 | 0.17 |
| 4 | 76164287 | rs7691721 | 1.444 | -0.174 | 0.85 | 0.025 | 0.04 | 0.004 | 0.004 | 0.01 | 0.62 | 0.15 | 0.62 | 0.29 | 0.7 | 0.72 |
| 4 | 76180442 | rs10021538 | -0.5 | -0.341 | 0.91 | 0.336 | 0.02 | -0.022 | -0.022 | 0 | 0.23 | 0.17 | 0.18 | 0.7 | 0.12 | 0.2 |
| 4 | 76181972 | rs11099470 | -1.12 | -0.403 | 1.503 | 0.353 | -0.047 | 0.034 | 0.034 | 0.003 | 0.3 | 0.47 | 0.26 | 0.69 | 0.19 | 0.25 |
| 4 | 76191643 | rs11734596 | 1.37 | -0.238 | 1.51 | 0.207 | 0.006 | 0.03 | 0.03 | 0 | 0.58 | 0.32 | 0.59 | 0.22 | 0.61 | 0.61 |
| 4 | 76192871 | rs1826876 | 1.37 | -0.238 | 1.51 | 0.207 | 0.006 | 0.03 | 0.03 | 0 | 0.58 | 0.32 | 0.59 | 0.22 | 0.61 | 0.61 |
| 4 | 76193549 | rs13147022 | -0.634 | -0.244 | 1.519 | 0.171 | -0.006 | -0.043 | -0.043 | 0 | 0.09 | 0.18 | 0.09 | 0.02 | 0.07 | 0.09 |
| 4 | 76207445 | rs4859952 | 2.055 | -0.263 | 1.238 | 0.162 | -0.061 | 0.109 | 0.109 | 0 | 0.48 | 0.06 | 0.5 | 0.21 | 0.53 | 0.53 |
| 4 | 76209438 | rs7655689 | 0.227 | 0.052 | 1.778 | 0.371 | -0.01 | 0.005 | 0.005 | 0 | 0.76 | 0.71 | 0.78 | 0.28 | 0.84 | 0.79 |
| 4 | 76209646 | rs10518137 | 0.327 | 0.18 | 1.511 | 0.459 | -0.005 | 0.003 | 0.003 | 0 | 0.12 | 0.11 | 0.14 | 0.09 | 0.2 | 0.14 |
| 4 | 76210182 | rs1497216 | 1.159 | 0.157 | 1.412 | 0.435 | 0 | 0.002 | 0.002 | 0 | 0.58 | 0.41 | 0.59 | 0.19 | 0.6 | 0.62 |
| 4 | 76212995 | rs6535270 | -1.815 | 0.08 | 0.587 | 0.583 | -0.027 | -0.008 | -0.008 | 0.04 | 0.53 | 0.89 | 0.49 | 0.63 | 0.46 | 0.51 |
| 4 | 76231406 | rs9993892 | -0.416 | 0.009 | 0.743 | 0.876 | 0.007 | -0.036 | -0.036 | 0 | 0.11 | 0.06 | 0.1 | 0.55 | 0.1 | 0.2 |
| 4 | 76232965 | rs13138899 | -0.051 | -0.081 | 1.131 | 0.696 | -0.022 | -0.089 | -0.089 | 1.024 | 0.2 | 0.31 | 0.14 | 0.01 | 0.13 | 0.1 |
| 4 | 76235455 | rs2131688 | -1.328 | -0.013 | 0.728 | 0.784 | 0.031 | -0.022 | -0.022 | 0 | 0.37 | 0.72 | 0.29 | 0.59 | 0.26 | 0.36 |
| 4 | 76248474 | rs2220799 | 1.492 | 0.661 | 1.453 | 1.053 | 0.025 | 0.064 | 0.064 | 0 | 0.93 | 0.65 | 0.95 | 0.95 | 0.97 | 0.95 |
| 4 | 76249812 | rs4859957 | 1.753 | 1.04 | 1.509 | 1.241 | 0.004 | 0.048 | 0.048 | 0.441 | 0.71 | 0.42 | 0.74 | 0.4 | 0.66 | 0.6 |
| 4 | 76250849 | rs1389441 | 1.858 | 1.021 | 1.828 | 1.2 | -0.007 | -0.032 | -0.032 | 0 | 0.67 | 0.23 | 0.72 | 0.37 | 0.63 | 0.55 |
| 4 | 76251731 | rs10019688 | -0.419 | 1.037 | 1.873 | 1.228 | 0.012 | 0 | 0 | 0.038 | 0.1 | 0.09 | 0.1 | 0.49 | 0.1 | 0.16 |
| 4 | 76252397 | rs2008051 | 0.882 | 1.029 | 1.949 | 1.212 | 0.061 | -0.059 | -0.059 | 0 | 0.81 | 0.56 | 0.84 | 0.89 | 0.76 | 0.77 |
| 4 | 76263405 | rs6813112 | -1.526 | 0.943 | 1.86 | 1.239 | -0.014 | 0.228 | 0.228 | 0.032 | 0.15 | 0.37 | 0.12 | 0.53 | 0.13 | 0.21 |
| 4 | 76270417 | rs12501796 | -1.341 | 1.238 | 1.826 | 1.167 | 0.068 | -0.059 | -0.059 | 0 | 0.2 | 0.38 | 0.22 | 0.11 | 0.31 | 0.25 |
| 4 | 76279001 | rs7680886 | 2.049 | 1.215 | 1.301 | 1.213 | -0.02 | -0.017 | -0.017 | 0 | 0.64 | 0.26 | 0.66 | 0.36 | 0.55 | 0.54 |
| 4 | 76281282 | rs7667024 | -2.26 | 1.379 | 1.088 | 1.289 | 0.011 | 0.022 | 0.022 | 0.166 | 0.31 | 0.61 | 0.32 | 0.61 | 0.4 | 0.39 |
| 4 | 76281630 | rs10026677 | -0.657 | 1.234 | 0.079 | 1.574 | 0.011 | -0.003 | -0.003 | 0 | 0.1 | 0.08 | 0.11 | 0.51 | 0.1 | 0.16 |
| 4 | 76283179 | rs4269201 | -1.468 | 1.279 | 0.127 | 1.47 | 0.003 | 0.026 | 0.026 | 0.565 | 0.22 | 0.28 | 0.21 | 0.58 | 0.3 | 0.34 |
| 4 | 76284744 | rs1493705 | -0.591 | 1.52 | 0.152 | 1.506 | 0.014 | 0.022 | 0.022 | 0 | 0.07 | 0.1 | 0.08 | 0.15 | 0.1 | 0.16 |
| 4 | 76285970 | rs1857913 | -1.021 | -0.039 | -0.728 | 1.214 | 0.133 | 0.091 | 0.091 | 0.841 | 0.49 | 0.86 | 0.37 | 0.67 | 0.37 | 0.38 |
| 4 | 76291076 | rs3853187 | -0.518 | 1.181 | -0.447 | 0.626 | -0.014 | 0 | 0 | 1.339 | 0.57 | 0.68 | 0.43 | 0.59 | 0.51 | 0.36 |
| 4 | 76295629 | rs17000522 | 1.157 | 0.381 | -0.058 | 1.038 | 0.058 | 0.026 | 0.026 | 0 | 0.26 | 0.14 | 0.21 | 0.14 | 0.16 | 0.09 |
| 4 | 76296421 | rs2201331 | -0.108 | 0.409 | 0.106 | 0.98 | -0.03 | 0.004 | 0.004 | 1.538 | 0.41 | 0.55 | 0.27 | 0.52 | 0.27 | 0.21 |
| 4 | 76299595 | rs7674576 | -0.827 | 0.414 | 0.195 | 0.958 | 0.087 | 0.063 | 0.063 | 0.545 | 0.49 | 0.73 | 0.35 | 0.67 | 0.37 | 0.38 |
| 4 | 76318507 | rs6841628 | -0.178 | 0.054 | 1.036 | 0.864 | -0.022 | 0.072 | 0.072 | 13.888 | 0.18 | 0.32 | 0.12 | 0.46 | 0.15 | 0.13 |
| 4 | 76321147 | rs1026694 | -0.358 | 0.055 | 1.087 | 0.865 | -0.048 | 0.2 | 0.2 | 12.792 | 0.19 | 0.55 | 0.13 | 0.46 | 0.15 | 0.13 |
| 4 | 76323522 | rs7683530 | 0.488 | 0.083 | 0.935 | 0.874 | 0.022 | 0.013 | 0.013 | 0.189 | 0.3 | 0.19 | 0.22 | 0.21 | 0.22 | 0.24 |
| 4 | 76332975 | rs2903698 | 0.934 | 0.283 | 0.691 | 0.865 | -0.037 | 0.031 | 0.031 | 0 | 0.16 | 0.07 | 0.15 | 0.07 | 0.25 | 0.15 |
| 4 | 76350536 | rs6535429 | -0.724 | 0.18 | 0.789 | 0.912 | 0.007 | 0.013 | 0.013 | 7.702 | 0.33 | 0.43 | 0.35 | 0.53 | 0.39 | 0.28 |
| 4 | 76352987 | rs7677609 | -1.706 | 0.185 | 0.364 | 0.884 | -0.114 | -0.023 | -0.023 | 0.851 | 0.65 | 0.97 | 0.61 | 0.74 | 0.61 | 0.52 |
| 4 | 76379793 | rs11099595 | -1.674 | 0.185 | 0.398 | 0.884 | -0.074 | 0.006 | 0.006 | 0.877 | 0.65 | 0.92 | 0.6 | 0.74 | 0.61 | 0.52 |
| 4 | 76382176 | rs6814729 | -1.695 | 0.186 | 0.364 | 1.21 | -0.088 | -0.072 | -0.072 | 0 | 0.65 | 0.94 | 0.6 | 0.65 | 0.61 | 0.52 |

| | | | | | | | | | | | | | | | | |
|---|---|---|---|---|---|---|---|---|---|---|---|---|---|---|---|---|
| 4 | 76393189 | rs1478174 | -1.725 | 0.159 | 0.301 | 1.309 | -0.117 | -0.07 | -0.07 | 0 | 0.66 | 0.97 | 0.6 | 0.63 | 0.61 | 0.52 |
| 4 | 76406846 | rs2126852 | 0.124 | 0.499 | 0.515 | 1.232 | -0.016 | -0.008 | -0.008 | 0 | 0.23 | 0.19 | 0.22 | 0.26 | 0.3 | 0.18 |
| 4 | 76415985 | rs6833496 | -1.972 | 0.353 | 0.53 | 1.247 | -0.087 | -0.051 | -0.051 | 0 | 0.66 | 0.91 | 0.6 | 0.62 | 0.61 | 0.52 |
| 4 | 76417401 | rs10518139 | -0.244 | 0.345 | 0.554 | 1.219 | -0.041 | -0.008 | -0.008 | 0 | 0.27 | 0.17 | 0.28 | 0.29 | 0.35 | 0.26 |
| 4 | 76419565 | rs17000594 | 0.387 | 0.272 | 0.857 | 1.215 | -0.007 | 0.059 | 0.059 | 0.797 | 0.06 | 0.19 | 0.07 | 0.1 | 0.04 | 0.01 |
| 4 | 76421029 | rs11731909 | 1.498 | 0.137 | 0.791 | 1.108 | 0.069 | 0.059 | 0.059 | 0.187 | 0.21 | 0.07 | 0.14 | 0.06 | 0.12 | 0.08 |
| 4 | 76431119 | rs10020976 | -0.227 | 0.236 | 0.798 | 1.156 | -0.024 | -0.004 | -0.004 | 0 | 0.28 | 0.21 | 0.28 | 0.29 | 0.35 | 0.26 |
| 4 | 76436884 | rs2903701 | 0.359 | 0.097 | 1.127 | 1.154 | -0.019 | -0.008 | -0.008 | 0 | 0.32 | 0.37 | 0.24 | 0.22 | 0.22 | 0.25 |
| 4 | 76443568 | rs17000603 | 0.421 | 0.082 | 1.126 | 1.164 | -0.006 | 0.117 | 0.117 | 0.278 | 0.06 | 0.19 | 0.13 | 0.15 | 0.04 | 0.01 |
| 4 | 76453149 | rs2126854 | 0.487 | 0.108 | 1.001 | 0.937 | 0.031 | 0.032 | 0.032 | 0.38 | 0.31 | 0.2 | 0.16 | 0.17 | 0.22 | 0.24 |
| 4 | 76461012 | rs11721925 | 0.315 | 0.111 | 0.907 | 0.947 | -0.026 | -0.025 | -0.025 | 0 | 0.32 | 0.38 | 0.18 | 0.17 | 0.22 | 0.24 |
| 4 | 76462087 | rs7681108 | -0.601 | 0.296 | 0.738 | 0.928 | 0.008 | 0.053 | 0.053 | 0 | 0.5 | 0.33 | 0.36 | 0.3 | 0.47 | 0.34 |
| 4 | 76464584 | rs10857225 | -0.391 | 0.644 | 0.47 | 1.1 | 0.051 | 0.025 | 0.025 | 3.185 | 0.44 | 0.66 | 0.24 | 0.35 | 0.32 | 0.27 |
| 4 | 76464991 | rs17000620 | 0.367 | 0.751 | 0.032 | 1.191 | -0.018 | 0.05 | 0.05 | 0.801 | 0.06 | 0.15 | 0.06 | 0.12 | 0.03 | 0.01 |
| 4 | 76475750 | rs9998529 | -1.527 | 0.916 | -0.419 | 1.257 | -0.163 | -0.093 | -0.093 | 0 | 0.67 | 0.96 | 0.5 | 0.6 | 0.56 | 0.51 |
| 4 | 76480880 | rs2061742 | 0.588 | 0.518 | -1.172 | 0.082 | -0.095 | -0.023 | -0.023 | 0.318 | 0.4 | 0.13 | 0.59 | 0.62 | 0.45 | 0.59 |
| 4 | 76482371 | rs6811713 | 0.805 | 0.754 | -0.821 | 0.038 | 0.03 | 0.071 | 0.071 | 0.245 | 0.31 | 0.16 | 0.16 | 0.13 | 0.24 | 0.17 |
| 4 | 76483340 | rs12646094 | 1.464 | 0.838 | -1.045 | 0.126 | -0.015 | -0.057 | -0.057 | 0 | 0.1 | 0.01 | 0.06 | 0.2 | 0.09 | 0.22 |
| 4 | 76483511 | rs11099642 | 1.026 | 0.835 | -0.98 | 0.136 | 0.002 | -0.008 | -0.008 | 0 | 0.12 | 0.1 | 0.06 | 0.2 | 0.1 | 0.22 |
| 4 | 76487438 | rs17000658 | -1.318 | 0.986 | -1.291 | 0 | -0.115 | -0.11 | -0.11 | 0 | 0.64 | 0.97 | 0.46 | 0.58 | 0.58 | 0.61 |
| 4 | 76489345 | rs2306175 | 0.004 | 1.061 | -0.36 | 0.068 | 0.048 | 0.058 | 0.058 | 0.012 | 0.16 | 0.32 | 0.17 | 0.24 | 0.23 | 0.2 |
| 4 | 76489582 | rs2306174 | 1.11 | 1.009 | -0.241 | 0.041 | -0.004 | 0.013 | 0.013 | 0.177 | 0.12 | 0.17 | 0.07 | 0.19 | 0.09 | 0.22 |
| 4 | 76491568 | rs6857229 | 0.47 | 1.194 | 0.38 | -0.473 | -0.003 | -0.013 | -0.013 | 0 | 0.32 | 0.3 | 0.17 | 0.14 | 0.25 | 0.16 |
| 4 | 76493256 | rs6851864 | -0.537 | 1.101 | -0.074 | -0.44 | -0.002 | 0.029 | 0.029 | 0 | 0.49 | 0.65 | 0.35 | 0.4 | 0.48 | 0.37 |
| 4 | 76502748 | rs1871114 | -0.516 | 1.416 | 0.295 | -1.813 | 0.001 | -0.044 | -0.044 | 0 | 0.38 | 0.53 | 0.28 | 0.07 | 0.4 | 0.12 |
| 4 | 76503197 | rs6829402 | -0.9 | 1.613 | 0.242 | -1.9 | 0.002 | -0.004 | -0.004 | 0 | 0.9 | 0.86 | 0.87 | 0.92 | 0.84 | 0.89 |
| 4 | 76509634 | rs2867918 | 1.045 | 1.695 | 0.187 | -2.004 | -0.026 | 0.016 | 0.016 | 0 | 0.1 | 0.03 | 0.07 | 0.04 | 0.17 | 0.05 |
| 4 | 76514964 | rs4610362 | 0.653 | 1.62 | 0.068 | -2.155 | 0 | 0.013 | 0.013 | 0 | 0.17 | 0.11 | 0.12 | 0.1 | 0.15 | 0.41 |
| 4 | 76519096 | rs4859966 | 1.963 | 1.591 | 0.1 | -2.205 | 0.004 | 0.052 | 0.052 | 0 | 0.33 | 0.13 | 0.46 | 0.76 | 0.29 | 0.37 |
| 4 | 76535086 | rs2055997 | 0.28 | 1.543 | 0.14 | -2.215 | -0.002 | 0.007 | 0.007 | 0 | 0.73 | 0.75 | 0.75 | 0.82 | 0.7 | 0.48 |
| 4 | 76535466 | rs13123545 | 0.302 | 1.569 | 0.397 | -2.335 | 0.014 | 0.121 | 0.121 | 0 | 0.32 | 0.12 | 0.17 | 0.05 | 0.3 | 0.09 |
| 4 | 76554604 | rs1553985 | -0.389 | 1.548 | 0.468 | -2.316 | 0.002 | -0.054 | -0.054 | 0 | 0.38 | 0.53 | 0.28 | 0.07 | 0.4 | 0.12 |
| 4 | 76558024 | rs12498415 | 0.515 | 1.479 | 0.488 | -2.324 | 0 | -0.01 | -0.01 | 3.318 | 0.18 | 0.18 | 0.13 | 0.1 | 0.15 | 0.39 |
| 4 | 76563470 | rs10856853 | 0.706 | 1.482 | 0.491 | -2.322 | 0 | 0.015 | 0.015 | 0 | 0.17 | 0.11 | 0.12 | 0.1 | 0.15 | 0.39 |
| 4 | 76565970 | rs11097041 | 0.501 | 1.482 | 0.493 | -2.322 | 0 | -0.01 | -0.01 | 0 | 0.18 | 0.18 | 0.13 | 0.1 | 0.15 | 0.39 |
| 4 | 76566975 | rs2037012 | 0.262 | 1.496 | 0.936 | -2.328 | 0.176 | 0.033 | 0.033 | 0 | 0.06 | 0.33 | 0.11 | 0.02 | 0.1 | 0.02 |
| 4 | 76569553 | rs11556146 | 0.707 | 1.499 | 0.755 | -2.324 | 0 | 0.015 | 0.015 | 0 | 0.17 | 0.11 | 0.12 | 0.1 | 0.15 | 0.4 |
| 4 | 76572191 | rs2175766 | -1.254 | 1.234 | 0.521 | -2.235 | -0.001 | 0.116 | 0.116 | 0 | 0.56 | 0.7 | 0.4 | 0.17 | 0.53 | 0.52 |
| 4 | 76576116 | rs9306994 | 2.02 | 1.226 | 0.281 | -2.226 | -0.031 | 0.047 | 0.047 | 0.13 | 0.33 | 0.07 | 0.47 | 0.75 | 0.31 | 0.37 |
| 4 | 76585823 | rs13141160 | 1.301 | 1.264 | 0.325 | -2.145 | -0.044 | 0.004 | 0.004 | 0 | 0.1 | 0.01 | 0.07 | 0.05 | 0.15 | 0.05 |
| 4 | 76585975 | rs13141390 | 0.786 | 1.245 | 0.714 | -1.811 | 0.016 | 0.062 | 0.062 | 0 | 0.31 | 0.17 | 0.17 | 0.05 | 0.28 | 0.1 |
| 4 | 76622077 | rs11097070 | -0.631 | 1.171 | 0.137 | -1.571 | 0 | -0.005 | -0.005 | 0 | 0.89 | 0.86 | 0.87 | 0.92 | 0.84 | 0.9 |
| 4 | 76629797 | rs6827809 | -0.659 | 1.176 | 0.124 | -1.569 | 0.001 | -0.009 | -0.009 | 0 | 0.89 | 0.85 | 0.87 | 0.92 | 0.84 | 0.9 |
| 4 | 76633325 | rs7665811 | 1.213 | 1.163 | -0.517 | -1.574 | -0.013 | 0.057 | 0.057 | 0.022 | 0.33 | 0.13 | 0.47 | 0.76 | 0.32 | 0.44 |
| 4 | 76650443 | rs7662940 | -0.628 | 1.245 | -0.081 | -1.309 | -0.004 | 0.009 | 0.009 | 0 | 0.9 | 0.93 | 0.88 | 0.91 | 0.84 | 0.9 |
| 4 | 76660500 | rs4859406 | 1.039 | 1.266 | 0.025 | -1.521 | -0.021 | 0.082 | 0.082 | 0.134 | 0.63 | 0.23 | 0.64 | 0.8 | 0.6 | 0.54 |
| 4 | 76692898 | rs226416 | -0.814 | 1.282 | -0.065 | -1.514 | -0.031 | 0.009 | 0.009 | 0 | 0.89 | 0.99 | 0.88 | 0.91 | 0.84 | 0.9 |
| 4 | 76695302 | rs324709 | -0.494 | 1.263 | 0.12 | -1.272 | 0.003 | -0.004 | -0.004 | 0.008 | 0.26 | 0.66 | 0.23 | 0.23 | 0.24 | 0.37 |
| 4 | 76697211 | rs2089253 | 0.409 | 1.335 | 0.519 | -1.705 | -0.012 | 0 | 0 | 0 | 0.11 | 0.29 | 0.1 | 0.1 | 0.06 | 0.23 |
| 4 | 76698424 | rs324706 | -0.925 | 1.368 | 0.264 | -1.681 | 0.012 | -0.068 | -0.068 | 0 | 0.86 | 0.76 | 0.91 | 0.99 | 0.85 | 0.87 |
| 4 | 76712173 | rs10212995 | 1.08 | 1.055 | 0.083 | -1.636 | -0.013 | 0.053 | 0.053 | 0.107 | 0.63 | 0.27 | 0.65 | 0.8 | 0.6 | 0.54 |
| 4 | 76720885 | rs13119659 | 0.396 | 1.095 | 0.147 | -1.624 | 0.015 | -0.044 | -0.044 | 0 | 0.75 | 0.7 | 0.76 | 0.9 | 0.65 | 0.77 |
| 4 | 76722353 | rs324726 | 0.498 | 1.088 | 0.211 | -1.674 | 0.003 | -0.031 | -0.031 | 0 | 0.75 | 0.34 | 0.77 | 0.89 | 0.76 | 0.64 |
| 4 | 76722579 | rs324725 | 0.097 | 1.32 | -0.305 | -1.575 | -0.008 | 0.018 | 0.018 | 0 | 0.12 | 0.07 | 0.12 | 0.09 | 0.16 | 0.1 |
| 4 | 76730195 | rs324689 | 0.321 | 1.386 | -0.65 | -1.542 | -0.031 | 0.01 | 0.01 | 0 | 0.11 | 0.01 | 0.12 | 0.09 | 0.16 | 0.1 |
| 4 | 76730908 | rs324694 | -0.578 | 1.464 | -0.545 | -1.456 | 0.059 | -0.09 | -0.09 | 0.673 | 0.85 | 0.69 | 0.88 | 0.99 | 0.82 | 0.87 |
| 4 | 76738361 | rs186562 | -0.023 | 0.825 | -0.976 | -0.939 | -0.004 | -0.006 | -0.006 | 0 | 0.1 | 0.04 | 0.09 | 0.14 | 0.15 | 0.09 |
| 4 | 76741116 | rs6531940 | -0.011 | 0.87 | -1.147 | -1.268 | -0.017 | 0.111 | 0.111 | 0 | 0.29 | 0.13 | 0.16 | 0.05 | 0.3 | 0.09 |
| 4 | 76750356 | rs6847598 | -0.56 | 0.827 | -0.717 | -1.044 | -0.009 | 0.048 | 0.048 | 0 | 0.53 | 0.67 | 0.41 | 0.29 | 0.51 | 0.46 |
| 4 | 76756059 | rs10856860 | 0.498 | 0.054 | 0.062 | -1.543 | -0.057 | -0.076 | -0.076 | 0 | 0.78 | 0.67 | 0.9 | 0.88 | 0.9 | 0.73 |
| 4 | 76761164 | rs7691847 | -0.097 | 0.627 | -0.671 | -2.236 | -0.102 | 0.131 | 0.131 | 0 | 0.31 | 0.11 | 0.31 | 0.05 | 0.49 | 0.27 |
| 4 | 76769442 | rs924937 | 0.046 | 0.295 | -0.016 | -2.213 | -0.057 | -0.044 | -0.044 | 0 | 0.84 | 0.4 | 0.9 | 0.88 | 0.96 | 0.75 |
| 4 | 76780087 | rs6811965 | -0.337 | 0.251 | 0.552 | -2.506 | -0.004 | -0.043 | -0.043 | 0 | 0.36 | 0.29 | 0.34 | 0.02 | 0.53 | 0.26 |
| 4 | 76780943 | rs1013588 | 0.982 | 0.162 | 0.407 | -2.529 | 0.047 | 0.173 | 0.173 | 0 | 0.5 | 0.29 | 0.63 | 0.87 | 0.42 | 0.49 |
| 4 | 76788268 | rs3817099 | 0.432 | 0.8 | 0.982 | -2.357 | -0.027 | 0.02 | 0.02 | 0 | 0.65 | 0.26 | 0.71 | 0.97 | 0.53 | 0.75 |
| 4 | 76809074 | rs13120456 | -0.138 | 1.519 | 1.523 | -2.498 | -0.02 | 0.008 | 0.008 | 0 | 0.22 | 0.16 | 0.16 | 0.01 | 0.29 | 0.13 |
| 4 | 76817221 | rs10007472 | -0.234 | 1.633 | 1.568 | -2.52 | 0.043 | -0.087 | -0.087 | 0 | 0.14 | 0.53 | 0.15 | 0.02 | 0.16 | 0.1 |
| 4 | 76818367 | rs13147538 | -0.222 | 1.807 | 1.324 | -2.502 | 0.112 | -0.08 | -0.08 | 0 | 0.87 | 0.49 | 0.85 | 0.98 | 0.81 | 0.9 |
| 4 | 76824078 | rs9784511 | 0.515 | 1.498 | 0.586 | -2.532 | -0.017 | -0.01 | -0.01 | 0.267 | 0.63 | 0.25 | 0.69 | 0.97 | 0.51 | 0.76 |

| | | | | | | | | | | | | | | | | |
|---|---|---|---|---|---|---|---|---|---|---|---|---|---|---|---|---|
| 4 | 76834209 | rs2047985 | 0.506 | 1.038 | 0.772 | -1.585 | 0.02 | 0.087 | 0.087 | 0 | 0.9 | 0.73 | 0.87 | 0.79 | 0.92 | 0.85 |
| 4 | 76834545 | rs2047987 | -0.243 | 0.597 | -0.837 | -1.083 | -0.02 | -0.166 | -0.166 | 0 | 0.63 | 0.97 | 0.7 | 0.54 | 0.71 | 0.7 |
| 4 | 76836137 | rs6823734 | -0.263 | 0.599 | -0.836 | -1.077 | -0.022 | -0.163 | -0.163 | 0.067 | 0.63 | 0.98 | 0.7 | 0.54 | 0.71 | 0.71 |
| 4 | 76839108 | rs6819442 | 0.842 | 0.922 | -0.559 | -0.962 | 0.047 | -0.134 | -0.134 | 0.857 | 0.24 | 0.41 | 0.18 | 0.02 | 0.34 | 0.24 |
| 4 | 76839729 | rs2869456 | 1.033 | 0.258 | -1.245 | -0.455 | -0.016 | -0.114 | -0.114 | 0 | 0.18 | 0.01 | 0.16 | 0.3 | 0.16 | 0.09 |
| 4 | 76846755 | rs937788 | 0.088 | 0.366 | -0.722 | -0.541 | 0.01 | -0.007 | -0.007 | 0 | 0.58 | 0.36 | 0.5 | 0.5 | 0.69 | 0.68 |
| 4 | 76848886 | rs7670521 | 0.04 | 0.408 | -0.986 | -0.153 | 0.007 | -0.049 | -0.049 | 0 | 0.31 | 0.19 | 0.34 | 0.42 | 0.31 | 0.43 |
| 4 | 76857309 | rs4859571 | 0.433 | 0.613 | -1.016 | 0.165 | -0.023 | -0.051 | -0.051 | 0 | 0.15 | 0.08 | 0.3 | 0.36 | 0.23 | 0.19 |
| 4 | 76857388 | rs4859572 | 0.433 | 0.613 | -1.016 | 0.165 | -0.023 | -0.051 | -0.051 | 0 | 0.15 | 0.08 | 0.29 | 0.36 | 0.23 | 0.18 |
| 4 | 76874996 | rs17001275 | 0.269 | 0.648 | 0.294 | -0.079 | 0.061 | 0.015 | 0.015 | 0 | 0.09 | 0.37 | 0.19 | 0.06 | 0.07 | 0.06 |
| 4 | 76877402 | rs9790644 | -0.794 | 0.214 | -0.247 | -0.771 | 0.004 | 0.193 | 0.193 | 0 | 0.49 | 0.22 | 0.58 | 0.87 | 0.5 | 0.49 |
| 4 | 76878688 | rs2242470 | 0.974 | 0.138 | -0.301 | -0.738 | -0.013 | -0.042 | -0.042 | 1.909 | 0.17 | 0.1 | 0.31 | 0.36 | 0.25 | 0.19 |
| 4 | 76878956 | rs2242473 | 1.485 | -0.103 | -0.102 | -0.839 | -0.017 | 0.047 | 0.047 | 0 | 0.37 | 0.24 | 0.21 | 0.07 | 0.42 | 0.45 |
| 4 | 76881979 | rs1857821 | 1.111 | -0.133 | -0.147 | -0.808 | -0.015 | -0.042 | -0.042 | 0.534 | 0.17 | 0.1 | 0.31 | 0.36 | 0.25 | 0.19 |
| 4 | 76885423 | rs2271527 | 0.546 | -0.26 | -0.305 | -0.896 | 0.012 | -0.075 | -0.075 | 0.017 | 0.29 | 0.1 | 0.27 | 0.51 | 0.24 | 0.3 |
| 4 | 76902739 | rs3733238 | 1.24 | -0.226 | -0.338 | -0.871 | -0.055 | -0.118 | -0.118 | 2.164 | 0.16 | 0.02 | 0.3 | 0.37 | 0.25 | 0.19 |
| 4 | 76903036 | rs10008757 | -1.757 | -0.239 | -0.255 | -0.911 | -0.017 | 0.047 | 0.047 | 0 | 0.64 | 0.76 | 0.79 | 0.93 | 0.58 | 0.55 |
| 4 | 76921814 | rs2869460 | 0.637 | -0.309 | -0.278 | -0.952 | 0.032 | -0.047 | -0.047 | 0 | 0.25 | 0.14 | 0.25 | 0.51 | 0.18 | 0.28 |
| 4 | 76922988 | rs10336 | -1.886 | -0.303 | -0.25 | -1.006 | -0.021 | 0.085 | 0.085 | 0 | 0.58 | 0.76 | 0.76 | 0.94 | 0.51 | 0.53 |
| 4 | 76923998 | rs3733236 | 1.038 | -0.325 | 0.136 | -1.02 | 0.053 | 0.012 | 0.012 | 0 | 0.08 | 0.3 | 0.19 | 0.05 | 0.08 | 0.06 |
| 4 | 76924933 | rs10031452 | -1.898 | -0.328 | 0.044 | -1.049 | -0.021 | 0.085 | 0.085 | 0 | 0.58 | 0.75 | 0.76 | 0.94 | 0.51 | 0.53 |
| 4 | 76933445 | rs10518143 | 0.986 | -0.337 | -0.001 | -1.001 | -0.013 | -0.048 | -0.048 | 0 | 0.16 | 0.08 | 0.3 | 0.37 | 0.24 | 0.19 |
| 4 | 76934975 | rs868007 | 1.88 | -1.261 | 0.067 | -0.755 | 0.03 | -0.005 | -0.005 | 0 | 0.1 | 0.03 | 0.08 | 0.12 | 0.03 | 0.15 |
| 4 | 76935263 | rs884304 | 0.041 | -1.259 | 0.08 | -0.745 | 0.006 | -0.019 | -0.019 | 0 | 0.27 | 0.21 | 0.39 | 0.49 | 0.27 | 0.34 |
| 4 | 76942300 | rs8878 | -2.078 | -1.279 | 0.097 | -0.755 | -0.015 | 0.075 | 0.075 | 0 | 0.58 | 0.73 | 0.75 | 0.94 | 0.51 | 0.53 |
| 4 | 76945522 | rs4256246 | 1.171 | -1.187 | 0.077 | -0.772 | -0.01 | -0.083 | -0.083 | 0 | 0.15 | 0.03 | 0.15 | 0.39 | 0.14 | 0.13 |
| 4 | 76946097 | rs4508917 | -0.378 | -1.029 | 0.245 | -0.622 | 0.013 | -0.021 | -0.021 | 0 | 0.31 | 0.21 | 0.4 | 0.49 | 0.28 | 0.34 |
| 4 | 76947304 | rs12504339 | -0.362 | -1.029 | 0.245 | -0.622 | -0.015 | 0.057 | 0.057 | 0.167 | 0.28 | 0.52 | 0.35 | 0.45 | 0.23 | 0.19 |
| 4 | 76952879 | rs11097212 | -1.961 | -0.976 | 0.251 | -0.624 | -0.015 | 0.075 | 0.075 | 0 | 0.58 | 0.73 | 0.75 | 0.94 | 0.51 | 0.53 |
| 4 | 76955229 | rs7436646 | -1.961 | -0.976 | 0.251 | -0.624 | -0.015 | 0.075 | 0.075 | 0 | 0.58 | 0.73 | 0.75 | 0.94 | 0.51 | 0.53 |
| 4 | 76955248 | rs9994667 | 0.96 | -0.986 | 0.246 | -0.618 | -0.04 | -0.108 | -0.108 | 5.534 | 0.17 | 0.04 | 0.3 | 0.39 | 0.24 | 0.19 |
| 4 | 76955406 | rs10017431 | -1.898 | -1.019 | 0.288 | -0.733 | -0.015 | 0.075 | 0.075 | 0 | 0.58 | 0.73 | 0.75 | 0.94 | 0.51 | 0.53 |
| 4 | 76955914 | rs6532111 | -1.898 | -1.019 | 0.288 | -0.733 | -0.015 | 0.075 | 0.075 | 0 | 0.58 | 0.73 | 0.75 | 0.94 | 0.51 | 0.53 |
| 4 | 76957974 | rs6532114 | -1.898 | -1.019 | 0.327 | -0.733 | -0.01 | 0.036 | 0.036 | 0 | 0.58 | 0.71 | 0.75 | 0.94 | 0.51 | 0.53 |
| 4 | 76963187 | rs7679689 | 1.377 | -1.144 | 0.449 | -0.792 | -0.006 | -0.024 | -0.024 | 0.591 | 0.13 | 0.2 | 0.19 | 0.05 | 0.08 | 0.06 |
| 4 | 76987496 | rs11097223 | -1.778 | -0.997 | 0.112 | -0.716 | -0.004 | -0.005 | -0.005 | 0 | 0.58 | 0.64 | 0.74 | 0.94 | 0.51 | 0.53 |
| 4 | 76991897 | rs7693606 | 0.338 | -0.497 | 0.295 | -0.384 | -0.03 | -0.084 | -0.084 | 0.12 | 0.18 | 0.55 | 0.22 | 0.05 | 0.1 | 0.07 |
| 4 | 76994500 | rs7666094 | 0.338 | -0.497 | 0.295 | -0.384 | -0.03 | -0.084 | -0.084 | 1.846 | 0.18 | 0.55 | 0.22 | 0.05 | 0.1 | 0.07 |
| 4 | 76995049 | rs6532170 | 0.338 | -0.497 | 0.295 | -0.384 | -0.03 | -0.084 | -0.084 | 0.013 | 0.18 | 0.55 | 0.22 | 0.05 | 0.1 | 0.07 |
| 4 | 76999197 | rs4859609 | 0.552 | -0.538 | -0.269 | -0.13 | -0.04 | -0.077 | -0.077 | 0 | 0.16 | 0.05 | 0.3 | 0.42 | 0.26 | 0.17 |
| 4 | 76999841 | rs4859418 | -0.978 | -0.656 | -0.182 | 0.077 | -0.001 | 0.037 | 0.037 | 0 | 0.9 | 0.91 | 0.85 | 0.59 | 0.87 | 0.88 |
| 4 | 77007912 | rs17001357 | -0.638 | -0.674 | 0.029 | -0.06 | 0.002 | 0.009 | 0.009 | 0 | 0.32 | 0.37 | 0.4 | 0.42 | 0.37 | 0.28 |
| 4 | 77019581 | rs6829592 | -0.921 | -1.749 | 0.265 | 0.078 | -0.01 | 0.003 | 0.003 | 0.166 | 0.4 | 0.41 | 0.48 | 0.57 | 0.28 | 0.38 |
| 4 | 77023238 | rs10022137 | -1.623 | -1.789 | 0.437 | -0.148 | 0.006 | -0.034 | -0.034 | 0 | 0.52 | 0.52 | 0.66 | 0.98 | 0.44 | 0.55 |
| 4 | 77024534 | rs12510869 | -0.269 | -1.954 | 0.48 | -0.078 | -0.034 | -0.019 | -0.019 | 0 | 0.29 | 0.35 | 0.35 | 0.22 | 0.13 | 0.28 |
| 4 | 77025136 | rs11097233 | 1.102 | -2.013 | 0.693 | -0.105 | -0.015 | -0.032 | -0.032 | 0 | 0.15 | 0.19 | 0.17 | 0.03 | 0.08 | 0.05 |
| 4 | 77025273 | rs7682766 | -1.89 | -1.981 | 0.555 | -0.118 | 0.009 | 0.011 | 0.011 | 0.018 | 0.5 | 0.41 | 0.64 | 0.92 | 0.45 | 0.55 |
| 4 | 77025368 | rs7697711 | -2.279 | -1.777 | 0.442 | 0.011 | -0.139 | 0.191 | 0.191 | 0 | 0.68 | 0.93 | 0.75 | 0.98 | 0.54 | 0.57 |
| 4 | 77027110 | rs10006380 | 0.903 | -1.694 | 0.483 | 0.175 | -0.049 | -0.033 | -0.033 | 0 | 0.09 | 0.01 | 0.16 | 0.37 | 0.18 | 0.15 |
| 4 | 77030147 | rs6836703 | -0.014 | -1.71 | 0.54 | 0.133 | -0.011 | 0.102 | 0.102 | 0 | 0.15 | 0.22 | 0.15 | 0.41 | 0.13 | 0.13 |
| 4 | 77030971 | rs4241583 | -1.207 | -1.678 | 0.381 | 0.176 | 0.021 | 0.032 | 0.032 | 0 | 0.56 | 0.89 | 0.63 | 0.64 | 0.41 | 0.44 |
| 4 | 77037045 | rs4859426 | 0.339 | -1.559 | 0.722 | 0.266 | -0.062 | 0.018 | 0.018 | 0.319 | 0.17 | 0.42 | 0.21 | 0.19 | 0.06 | 0.23 |
| 4 | 77038807 | rs7657846 | 0.66 | -1.433 | 0.374 | 0.393 | -0.001 | -0.021 | -0.021 | 2.996 | 0.12 | 0.06 | 0.13 | 0.35 | 0.13 | 0.13 |
| 4 | 77044956 | rs4538520 | -0.561 | -1.444 | 0.492 | 0.375 | 0.001 | 0.001 | 0.001 | 0 | 0.2 | 0.18 | 0.3 | 0.71 | 0.31 | 0.28 |
| 4 | 77064939 | rs11736405 | 0.626 | -2.126 | 0.368 | 0.448 | -0.001 | -0.021 | -0.021 | 0.249 | 0.12 | 0.06 | 0.13 | 0.35 | 0.13 | 0.13 |
| 4 | 77072597 | rs10008834 | 1.043 | -1.957 | 0.34 | 0.489 | -0.039 | -0.017 | -0.017 | 1.986 | 0.08 | 0 | 0.15 | 0.36 | 0.15 | 0.14 |
| 4 | 77076451 | rs1465920 | 0.268 | -2.075 | 0.672 | 0.483 | 0 | 0.124 | 0.124 | 0 | 0.13 | 0.25 | 0.14 | 0.35 | 0.13 | 0.18 |
| 4 | 77087704 | rs7664889 | -0.387 | -2.327 | 0.528 | 0.249 | 0.008 | -0.007 | -0.007 | 0 | 0.2 | 0.76 | 0.13 | 0.26 | 0.07 | 0.17 |
| 4 | 77094831 | rs2869851 | -0.916 | -2.332 | 0.648 | 0.376 | -0.034 | 0.035 | 0.035 | 1.316 | 0.3 | 0.57 | 0.24 | 0.57 | 0.19 | 0.37 |
| 4 | 77096118 | rs17001561 | 0.694 | -2.525 | 0.339 | 0.186 | 0.059 | 0.087 | 0.087 | 1.433 | 0.28 | 0.17 | 0.22 | 0.03 | 0.18 | 0.07 |
| 4 | 77096220 | rs3796496 | 0.334 | -2.426 | 0.586 | 0.282 | -0.008 | 0.212 | 0.212 | 0 | 0.15 | 0.44 | 0.17 | 0.36 | 0.13 | 0.2 |
| 4 | 77099384 | rs17001573 | 1.113 | -2.13 | -0.324 | 0.088 | -0.037 | -0.02 | -0.02 | 0.174 | 0.07 | 0.01 | 0.26 | 0.33 | 0.14 | 0.16 |
| 4 | 77105700 | rs11946079 | -0.555 | -1.949 | -0.07 | 0.202 | 0.054 | -0.021 | -0.021 | 0 | 0.32 | 0.76 | 0.26 | 0.62 | 0.19 | 0.37 |
| 4 | 77128026 | rs12508223 | -1.129 | -0.85 | -1.548 | -0.23 | -0.076 | 0.159 | 0.159 | 0 | 0.67 | 0.89 | 0.74 | 0.99 | 0.5 | 0.62 |
| 4 | 77129840 | rs894251 | -0.677 | -0.819 | -0.707 | 1.108 | -0.027 | 0.072 | 0.072 | 0.009 | 0.17 | 0.22 | 0.15 | 0.4 | 0.12 | 0.22 |
| 4 | 77130285 | rs9991821 | -0.201 | -0.452 | -1.196 | 0.88 | -0.05 | -0.045 | -0.045 | 0 | 0.17 | 0.03 | 0.29 | 0.21 | 0.23 | 0.15 |
| 4 | 77131069 | rs17001659 | -0.359 | -1.123 | -0.261 | 0.812 | -0.09 | 0.122 | 0.122 | 0 | 0.18 | 0.4 | 0.18 | 0.34 | 0.06 | 0.19 |
| 4 | 77139510 | rs7685696 | -0.97 | -0.87 | -0.999 | -0.34 | 0.029 | 0.144 | 0.144 | 1.379 | 0.43 | 0.2 | 0.58 | 0.94 | 0.39 | 0.67 |
| 4 | 77144682 | rs2034004 | -0.837 | -1.487 | -0.695 | -0.423 | 0.045 | 0.066 | 0.066 | 0 | 0.5 | 0.4 | 0.49 | 0.77 | 0.39 | 0.57 |

| | | | | | | | | | | | | | | | | |
|---|---|---|---|---|---|---|---|---|---|---|---|---|---|---|---|---|
| 4 | 77149119 | rs12505005 | 0.737 | -1.896 | -0.848 | -0.531 | -0.023 | 0.042 | 0.042 | 0 | 0.08 | 0 | 0.02 | 0.01 | 0.07 | 0.06 |
| 4 | 77151474 | rs1596116 | 0.747 | -2.251 | -0.74 | 0.078 | 0.02 | -0.057 | -0.057 | 0 | 0.17 | 0.1 | 0.26 | 0.35 | 0.14 | 0.23 |
| 4 | 77151880 | rs1441904 | -1.532 | -2.483 | -0.446 | -0.051 | 0.055 | 0.06 | 0.06 | 0 | 0.41 | 0.29 | 0.41 | 0.78 | 0.28 | 0.53 |
| 4 | 77151938 | rs6846983 | -0.489 | -2.402 | -0.525 | -0.082 | 0.023 | -0.06 | -0.06 | 0 | 0.27 | 0.15 | 0.28 | 0.36 | 0.21 | 0.3 |
| 4 | 77154865 | rs11097297 | -0.832 | -2.618 | -0.022 | -0.423 | -0.023 | -0.025 | -0.025 | 0.659 | 0.38 | 0.43 | 0.44 | 0.81 | 0.27 | 0.48 |
| 4 | 77159700 | rs1441922 | 0.68 | -2.386 | 0.193 | -0.321 | 0.006 | -0.019 | -0.019 | 0 | 0.79 | 0.52 | 0.77 | 0.83 | 0.8 | 0.8 |
| 4 | 77167242 | rs1348206 | -0.591 | -2.28 | 0.144 | -0.234 | 0.04 | 0.081 | 0.081 | 0 | 0.89 | 0.7 | 0.81 | 0.84 | 0.87 | 0.85 |
| 4 | 77173914 | rs13117238 | -0.25 | -2.579 | -0.14 | -0.238 | 0.029 | -0.038 | -0.038 | 0 | 0.26 | 0.13 | 0.28 | 0.36 | 0.17 | 0.28 |
| 4 | 77174469 | rs10856883 | 0.174 | -2.588 | 0.186 | -0.227 | -0.036 | 0.081 | 0.081 | 1.639 | 0.73 | 0.49 | 0.55 | 0.5 | 0.76 | 0.63 |
| 4 | 77175743 | rs717239 | 1.761 | -2.796 | -0.055 | -0.474 | 0.011 | 0.154 | 0.154 | 0.524 | 0.42 | 0.23 | 0.33 | 0.02 | 0.53 | 0.31 |
| 4 | 77176552 | rs7671407 | 0.3 | -2.868 | 0.28 | -0.348 | 0.006 | -0.019 | -0.019 | 0.493 | 0.75 | 0.56 | 0.72 | 0.78 | 0.72 | 0.8 |
| 4 | 77176768 | rs7655536 | -0.607 | -2.754 | 0.214 | -0.243 | -0.023 | -0.004 | -0.004 | 0.263 | 0.31 | 0.34 | 0.38 | 0.75 | 0.19 | 0.5 |
| 4 | 77177027 | rs17001726 | 0.633 | -2.699 | 0.115 | -0.102 | -0.007 | -0.037 | -0.037 | 0 | 0.08 | 0 | 0.11 | 0.37 | 0.08 | 0.23 |
| 4 | 77179830 | rs6851219 | 0.762 | -2.791 | 0.165 | -0.078 | 0.016 | -0.021 | -0.021 | 0 | 0.18 | 0.1 | 0.23 | 0.33 | 0.11 | 0.27 |
| 4 | 77182033 | rs2904221 | 0.519 | -2.951 | 0.337 | -0.074 | 0.004 | -0.025 | -0.025 | 0.438 | 0.74 | 0.5 | 0.72 | 0.79 | 0.72 | 0.8 |
| 4 | 77186415 | rs17308336 | 0.991 | -2.591 | -0.311 | -0.521 | -0.022 | 0.005 | 0.005 | 0.627 | 0.42 | 0.18 | 0.43 | 0.32 | 0.41 | 0.35 |
| 4 | 77192838 | rs3733251 | 0.526 | -2.661 | 0.046 | -0.544 | -0.004 | -0.036 | -0.036 | 0 | 0.17 | 0.28 | 0.09 | 0.05 | 0.16 | 0.08 |
| 4 | 77192868 | rs3733250 | 1.072 | -2.7 | -0.023 | -0.57 | -0.024 | 0.004 | 0.004 | 0.634 | 0.41 | 0.17 | 0.43 | 0.32 | 0.41 | 0.35 |
| 4 | 77196476 | rs17236661 | 1.798 | -2.721 | -0.028 | -0.61 | -0.017 | 0.056 | 0.056 | 0 | 0.2 | 0.13 | 0.14 | 0.03 | 0.23 | 0.23 |
| 4 | 77198756 | rs6532316 | -1.099 | -2.732 | 0.017 | -0.572 | 0.053 | 0.078 | 0.078 | 0 | 0.91 | 0.73 | 0.81 | 0.84 | 0.87 | 0.87 |
| 4 | 77198986 | rs6812193 | 0.024 | -2.749 | 0.052 | -0.649 | -0.001 | -0.031 | -0.031 | 0 | 0.39 | 0.47 | 0.25 | 0.08 | 0.39 | 0.31 |
| 4 | 77199414 | rs964051 | 1.11 | -2.739 | -0.066 | -0.61 | -0.035 | -0.005 | -0.005 | 0.223 | 0.41 | 0.14 | 0.43 | 0.35 | 0.41 | 0.34 |
| 4 | 77201487 | rs1036788 | 0.237 | -2.571 | -0.069 | -0.406 | -0.003 | 0.142 | 0.142 | 0.919 | 0.82 | 0.7 | 0.68 | 0.43 | 0.8 | 0.66 |
| 4 | 77202861 | rs1530297 | 0.151 | -2.574 | -0.062 | -0.439 | 0 | 0.008 | 0.008 | 0 | 0.6 | 0.55 | 0.54 | 0.4 | 0.57 | 0.42 |
| 4 | 77204500 | rs1542096 | -0.321 | -2.369 | 0.046 | -0.671 | 0.003 | -0.049 | -0.049 | 0 | 0.66 | 0.49 | 0.66 | 0.77 | 0.64 | 0.64 |
| 4 | 77208412 | rs6532327 | -0.966 | -2.058 | 0.081 | -0.905 | -0.003 | -0.082 | -0.082 | 0.101 | 0.74 | 0.67 | 0.85 | 0.99 | 0.75 | 0.75 |
| 4 | 77218240 | rs1508335 | 1.339 | -2.038 | -0.566 | -0.784 | -0.188 | 0.042 | 0.042 | 0.019 | 0.27 | 0.02 | 0.34 | 0.08 | 0.38 | 0.11 |
| 4 | 77219402 | rs6830162 | 0.483 | -2.188 | 0.015 | -0.58 | 0.004 | 0.025 | 0.025 | 0 | 0.29 | 0.24 | 0.18 | 0.01 | 0.28 | 0.26 |
| 4 | 77220837 | rs12506745 | 1.184 | -1.84 | -0.069 | -0.347 | -0.125 | 0.076 | 0.076 | 0 | 0.3 | 0.09 | 0.36 | 0.13 | 0.44 | 0.16 |
| 4 | 77222933 | rs10019831 | -0.517 | -2.045 | 0.443 | 0.133 | -0.002 | 0.053 | 0.053 | 0 | 0.87 | 0.69 | 0.92 | 0.84 | 0.89 | 0.96 |
| 4 | 77225313 | rs13104573 | -0.032 | -2.115 | 0.433 | 0.121 | 0.024 | -0.008 | -0.008 | 0 | 0.31 | 0.25 | 0.4 | 0.69 | 0.19 | 0.57 |
| 4 | 77227485 | rs3733249 | -0.399 | -1.802 | 0.675 | 0.252 | -0.01 | 0.013 | 0.013 | 0 | 0.85 | 0.8 | 0.92 | 0.84 | 0.86 | 0.94 |
| 4 | 77228724 | rs3796491 | -0.189 | -1.793 | 0.766 | 0.065 | -0.004 | -0.019 | -0.019 | 0 | 0.59 | 0.54 | 0.76 | 0.77 | 0.61 | 0.7 |
| 4 | 77231294 | rs3733248 | -0.559 | -1.874 | 0.76 | 0.063 | -0.011 | 0.016 | 0.016 | 0 | 0.86 | 0.81 | 0.92 | 0.84 | 0.87 | 0.95 |
| 4 | 77252196 | rs7658585 | 0.272 | -1.893 | 0.679 | 0.071 | 0.004 | -0.005 | -0.005 | 0 | 0.28 | 0.25 | 0.38 | 0.7 | 0.18 | 0.55 |
| 4 | 77262103 | rs1911737 | -0.575 | -1.855 | 0.725 | 0.067 | -0.016 | 0.019 | 0.019 | 0 | 0.85 | 0.76 | 0.91 | 0.84 | 0.87 | 0.95 |
| 4 | 77269262 | rs10002448 | -0.003 | -1.811 | 0.638 | 0.092 | 0.006 | 0.017 | 0.017 | 0.017 | 0.58 | 0.38 | 0.75 | 0.77 | 0.61 | 0.7 |
| 4 | 77289952 | rs11097338 | -0.374 | -1.766 | 0.416 | 0.115 | -0.008 | -0.032 | -0.032 | 0 | 0.62 | 0.54 | 0.77 | 0.8 | 0.64 | 0.72 |
| 4 | 77292054 | rs13116030 | 0.435 | -1.669 | 0.179 | 0.269 | -0.008 | 0.009 | 0.009 | 1.184 | 0.25 | 0.25 | 0.35 | 0.47 | 0.16 | 0.39 |
| 4 | 77305346 | rs17001890 | 0.828 | -1.047 | -0.296 | -0.576 | -0.001 | -0.066 | -0.066 | 0 | 0.27 | 0.34 | 0.16 | 0.07 | 0.26 | 0.25 |
| 4 | 77317124 | rs6857452 | 0.758 | -0.216 | -0.453 | 0.035 | -0.106 | 0.077 | 0.077 | 0 | 0.3 | 0.1 | 0.36 | 0.08 | 0.43 | 0.15 |
| 4 | 77330682 | rs4272041 | 0.758 | -0.216 | -0.453 | 0.035 | -0.106 | 0.077 | 0.077 | 0 | 0.3 | 0.1 | 0.36 | 0.08 | 0.43 | 0.15 |
| 4 | 77356246 | rs10000169 | 0.028 | -1.397 | 0.048 | 0.504 | -0.034 | 0 | 0 | 2.237 | 0.65 | 0.58 | 0.58 | 0.39 | 0.75 | 0.59 |
| 4 | 77357747 | rs10006043 | -0.143 | -1.376 | -0.487 | -0.167 | 0.007 | 0.015 | 0.015 | 0 | 0.47 | 0.39 | 0.42 | 0.37 | 0.53 | 0.38 |
| 4 | 77359677 | rs11097356 | -0.327 | -1.234 | -0.498 | -0.109 | -0.008 | -0.001 | -0.001 | 0 | 0.17 | 0.2 | 0.08 | 0.21 | 0.14 | 0.24 |
| 4 | 77359789 | rs11737407 | -0.286 | -1.234 | -0.468 | -0.087 | 0 | -0.001 | -0.001 | 0 | 0.17 | 0.14 | 0.07 | 0.21 | 0.14 | 0.24 |
| 4 | 77362699 | rs13102317 | 0.012 | -1.308 | -0.006 | 0.164 | 0.006 | 0.025 | 0.025 | 0.763 | 0.11 | 0.24 | 0.17 | 0.22 | 0.11 | 0.23 |
| 4 | 77364126 | rs7677847 | -0.82 | -1.417 | -0.151 | 0.749 | 0.008 | -0.009 | -0.009 | 0 | 0.35 | 0.29 | 0.41 | 0.7 | 0.26 | 0.61 |
| 4 | 77364799 | rs4859673 | -1.504 | -1.5 | -0.14 | 0.706 | -0.007 | 0.001 | 0.001 | 0 | 0.74 | 0.6 | 0.8 | 0.8 | 0.73 | 0.76 |
| 4 | 77367688 | rs1398016 | 1.437 | -1.658 | -0.401 | 0.521 | -0.089 | 0.056 | 0.056 | 0.252 | 0.28 | 0.1 | 0.34 | 0.09 | 0.41 | 0.13 |
| 4 | 77370038 | rs17001970 | 1.172 | -1.632 | -0.262 | 0.537 | -0.013 | -0.031 | -0.031 | 0 | 0.11 | 0.14 | 0.04 | 0.06 | 0.06 | 0.02 |
| 4 | 77386372 | rs4859678 | 0.778 | -1.778 | -0.229 | 0.867 | 0 | 0 | 0 | 0.849 | 0.2 | 0.18 | 0.19 | 0.2 | 0.1 | 0.16 |
| 4 | 77386678 | rs1511817 | -0.616 | -1.888 | -0.228 | 0.907 | -0.004 | -0.004 | -0.004 | 0.008 | 0.62 | 0.37 | 0.76 | 0.74 | 0.68 | 0.74 |
| 4 | 77392092 | rs6843293 | -1.594 | -1.868 | -0.233 | 0.954 | -0.002 | 0.025 | 0.025 | 0 | 0.74 | 0.6 | 0.79 | 0.68 | 0.73 | 0.77 |
| 4 | 77397585 | rs4129337 | 1.307 | -1.933 | -0.43 | 0.558 | 0.025 | -0.048 | -0.048 | 0 | 0.16 | 0 | 0.16 | 0.17 | 0.08 | 0.15 |
| 4 | 77397830 | rs10213666 | 1.635 | -1.814 | -0.322 | 0.519 | -0.008 | -0.027 | -0.027 | 0 | 0.1 | 0.08 | 0.03 | 0 | 0.05 | 0.01 |
| 4 | 77398015 | rs10032549 | 0.689 | -1.798 | -0.441 | 0.843 | -0.029 | 0.003 | 0.003 | 0 | 0.37 | 0.22 | 0.48 | 0.27 | 0.52 | 0.34 |
| 4 | 77408781 | rs6532418 | 1.025 | -1.792 | -0.436 | 0.83 | 0.011 | 0 | 0 | 0 | 0.07 | 0.13 | 0.14 | 0.07 | 0.08 | 0.2 |
| 4 | 77409795 | rs7654754 | 0.635 | -1.802 | -0.452 | 0.832 | -0.02 | 0.005 | 0.005 | 0 | 0.37 | 0.25 | 0.49 | 0.27 | 0.52 | 0.34 |
| 4 | 77410318 | rs4859682 | 1.277 | -1.756 | -0.571 | 0.86 | -0.111 | -0.019 | -0.019 | 2.617 | 0.3 | 0.08 | 0.35 | 0.2 | 0.43 | 0.14 |
| 4 | 77411996 | rs12651108 | 1.099 | -1.597 | -0.402 | 0.999 | -0.024 | -0.042 | -0.042 | 0.087 | 0.07 | 0 | 0.1 | 0.26 | 0.08 | 0.26 |
| 4 | 77417060 | rs2068064 | 0.533 | -1.809 | -0.137 | 0.432 | -0.001 | 0 | 0 | 0.064 | 0.09 | 0.1 | 0.15 | 0.07 | 0.09 | 0.22 |
| 4 | 77417756 | rs13152154 | -1.421 | -1.959 | 0.103 | 0.405 | 0 | -0.022 | -0.022 | 0 | 0.74 | 0.65 | 0.81 | 0.83 | 0.73 | 0.76 |
| 4 | 77418681 | rs13106227 | -0.262 | -1.985 | -0.002 | 0.466 | 0.013 | -0.003 | -0.003 | 0 | 0.56 | 0.38 | 0.64 | 0.51 | 0.65 | 0.61 |
| 4 | 77420784 | rs1986734 | 1.096 | -2.012 | -0.084 | 0.403 | -0.053 | -0.031 | -0.031 | 0 | 0.37 | 0.18 | 0.46 | 0.55 | 0.51 | 0.4 |
| 4 | 77424167 | rs7696372 | -0.697 | -1.827 | 0.31 | 0.602 | -0.029 | -0.014 | -0.014 | 0.029 | 0.2 | 0.28 | 0.19 | 0.17 | 0.14 | 0.24 |
| 4 | 77430136 | rs4452451 | -0.893 | -1.553 | 0.071 | 0.558 | -0.034 | -0.003 | -0.003 | 1.982 | 0.37 | 0.47 | 0.35 | 0.37 | 0.21 | 0.37 |
| 4 | 77430773 | rs17254217 | -0.463 | -0.149 | 0.085 | 0.529 | -0.001 | 0 | 0 | 0 | 0.09 | 0.1 | 0.15 | 0.07 | 0.09 | 0.22 |
| 4 | 77432584 | rs17002031 | -0.57 | -0.135 | 0.01 | 0.536 | -0.002 | -0.017 | -0.017 | 0 | 0.11 | 0.22 | 0.16 | 0.07 | 0.09 | 0.22 |

| | | | | | | | | | | | | | | | | |
|---|---|---|---|---|---|---|---|---|---|---|---|---|---|---|---|---|
| 4 | 77439979 | rs4859688 | -0.111 | -0.006 | -0.246 | 0.505 | -0.01 | -0.034 | -0.034 | 0 | 0.75 | 0.58 | 0.79 | 0.84 | 0.74 | 0.76 |
| 4 | 77440267 | rs4371638 | 0.241 | 1.052 | -0.994 | 0.916 | -0.061 | -0.051 | -0.051 | 0.001 | 0.76 | 0.88 | 0.55 | 0.59 | 0.55 | 0.66 |
| 4 | 77441772 | rs9999448 | 0.258 | 1.632 | -1.096 | 0.216 | -0.014 | 0.032 | 0.032 | 0 | 0.71 | 0.8 | 0.76 | 0.91 | 0.63 | 0.82 |
| 4 | 77442504 | rs10002311 | -0.567 | 2.653 | -0.549 | 0.859 | 0.05 | 0.065 | 0.065 | 0 | 0.1 | 0.17 | 0.12 | 0.32 | 0.16 | 0.21 |
| 4 | 77446681 | rs6848817 | 0.597 | 2.42 | -0.148 | 0.835 | -0.018 | 0.003 | 0.003 | 0 | 0.64 | 0.73 | 0.67 | 0.59 | 0.53 | 0.65 |
| 4 | 77448199 | rs13151434 | 0.795 | 1.834 | -0.63 | 1.211 | 0.01 | -0.038 | -0.038 | 0 | 0.11 | 0.03 | 0.07 | 0.19 | 0.08 | 0.08 |
| 4 | 77452151 | rs6810716 | 1.027 | 1.422 | -0.583 | 1.056 | 0.027 | 0.04 | 0.04 | 0 | 0.49 | 0.79 | 0.6 | 0.38 | 0.43 | 0.57 |
| 4 | 77454497 | rs10023611 | -0.064 | 1.214 | 0.118 | 1.209 | -0.011 | 0.137 | 0.137 | 0 | 0.11 | 0.3 | 0.09 | 0.2 | 0.07 | 0.08 |
| 4 | 77455181 | rs10024176 | -1.608 | 1.366 | 0.238 | 1.183 | 0.017 | 0.154 | 0.154 | 0 | 0.19 | 0.33 | 0.16 | 0.53 | 0.18 | 0.29 |
| 4 | 77456093 | rs9999005 | -0.395 | 1.406 | 0.612 | 0.604 | -0.002 | -0.019 | -0.019 | 5.394 | 0.69 | 0.67 | 0.81 | 0.93 | 0.62 | 0.82 |
| 4 | 77459600 | rs11097403 | 0.789 | 1.285 | 0.737 | 0.906 | 0 | -0.011 | -0.011 | 6.139 | 0.56 | 0.51 | 0.67 | 0.8 | 0.52 | 0.75 |
| 4 | 77461011 | rs6817593 | 1.501 | 1.202 | 0.991 | 0.974 | 0.025 | -0.013 | -0.013 | 0 | 0.47 | 0.28 | 0.62 | 0.49 | 0.42 | 0.58 |
| 4 | 77464016 | rs17002074 | 0.334 | 1.159 | 1.083 | 1.075 | -0.015 | 0.019 | 0.019 | 0.033 | 0.12 | 0.14 | 0.12 | 0.14 | 0.1 | 0.07 |
| 4 | 77465074 | rs10027450 | -0.793 | 1.228 | 0.978 | 1.101 | 0.06 | 0.142 | 0.142 | 0.23 | 0.1 | 0.29 | 0.06 | 0.31 | 0.1 | 0.18 |
| 4 | 77467624 | rs9996716 | -1.258 | 1.182 | 0.961 | 1.078 | 0.003 | 0.083 | 0.083 | 0 | 0.2 | 0.31 | 0.17 | 0.45 | 0.2 | 0.24 |
| 4 | 77468196 | rs11932470 | -0.194 | 1.347 | 0.584 | 1.023 | -0.065 | 0.044 | 0.044 | 0 | 0.35 | 0.08 | 0.28 | 0.21 | 0.45 | 0.23 |
| 4 | 77468594 | rs3923380 | 0.512 | 1.423 | 1.205 | 0.9 | 0.003 | 0 | 0 | 0 | 0.61 | 0.43 | 0.74 | 0.56 | 0.52 | 0.66 |
| 4 | 77469221 | rs6532473 | -1.507 | 1.257 | 1.201 | 1.16 | -0.001 | 0.031 | 0.031 | 0 | 0.22 | 0.26 | 0.17 | 0.46 | 0.2 | 0.28 |
| 4 | 77470300 | rs10005964 | 0.082 | 1.609 | 1.008 | 1.026 | -0.022 | -0.026 | -0.026 | 0.288 | 0.3 | 0.22 | 0.24 | 0.44 | 0.41 | 0.34 |
| 4 | 77471116 | rs13120833 | -1.074 | 1.469 | 1.035 | 0.766 | -0.001 | 0 | 0 | 0 | 0.52 | 0.55 | 0.39 | 0.54 | 0.57 | 0.45 |
| 4 | 77484202 | rs4380545 | -0.712 | 1.76 | 1.264 | 1.076 | 0.002 | 0.018 | 0.018 | 0 | 0.12 | 0.14 | 0.17 | 0.31 | 0.15 | 0.21 |
| 4 | 77487977 | rs7687284 | 1.886 | 1.77 | 1.427 | 1.12 | 0.015 | 0.042 | 0.042 | 0 | 0.71 | 0.57 | 0.63 | 0.44 | 0.66 | 0.57 |
| 4 | 77492155 | rs6824297 | -1.386 | 1.648 | 1.268 | 0.986 | 0.003 | 0.008 | 0.008 | 0 | 0.59 | 0.67 | 0.56 | 0.68 | 0.65 | 0.57 |
| 4 | 77497756 | rs4380546 | 0.203 | 1.657 | 1.347 | 0.797 | 0 | -0.011 | -0.011 | 0 | 0.35 | 0.47 | 0.29 | 0.26 | 0.4 | 0.26 |
| 4 | 77500947 | rs17002091 | -0.376 | 1.618 | 1.064 | 0.758 | 0.009 | 0.037 | 0.037 | 0 | 0.11 | 0.16 | 0.17 | 0.28 | 0.14 | 0.21 |
| 4 | 77503120 | rs12503111 | 0.105 | 1.523 | 0.953 | 0.55 | -0.031 | 0.061 | 0.061 | 0 | 0.44 | 0.19 | 0.45 | 0.36 | 0.51 | 0.41 |
| 4 | 77504400 | rs13148163 | -0.892 | 1.568 | 1.087 | 0.58 | -0.001 | -0.001 | -0.001 | 1.381 | 0.59 | 0.44 | 0.55 | 0.43 | 0.62 | 0.51 |
| 4 | 77507863 | rs13140663 | -0.035 | 1.504 | 0.847 | 0.44 | -0.003 | 0.013 | 0.013 | 0 | 0.13 | 0.01 | 0.09 | 0.07 | 0.11 | 0.1 |
| 4 | 77515370 | rs4859453 | -1.227 | 1.591 | 1.246 | 0.678 | 0.05 | 0.22 | 0.22 | 0 | 0.16 | 0.45 | 0.31 | 0.45 | 0.21 | 0.32 |
| 4 | 77517128 | rs6846198 | -0.032 | 1.515 | 1.368 | 0.532 | 0.014 | 0.111 | 0.111 | 0 | 0.51 | 0.69 | 0.62 | 0.86 | 0.52 | 0.7 |
| 4 | 77521826 | rs12501546 | -0.169 | 1.467 | 1.299 | 0.603 | 0.021 | 0.093 | 0.093 | 0 | 0.11 | 0.22 | 0.18 | 0.29 | 0.13 | 0.22 |
| 4 | 77523129 | rs17322390 | 1.066 | 1.477 | 0.905 | 0.49 | -0.079 | 0.194 | 0.194 | 0 | 0.34 | 0.02 | 0.28 | 0.07 | 0.39 | 0.2 |
| 4 | 77531337 | rs12640320 | 1.21 | 1.502 | 1.194 | 0.647 | 0 | -0.013 | -0.013 | 0 | 0.32 | 0.25 | 0.3 | 0.4 | 0.29 | 0.37 |
| 4 | 77540746 | rs11734813 | 0.454 | 1.373 | 1.202 | 0.464 | 0.001 | 0.011 | 0.011 | 0 | 0.11 | 0.04 | 0.08 | 0.05 | 0.09 | 0.09 |
| 4 | 77542398 | rs7669852 | 0.471 | 1.471 | 1.337 | 0.475 | -0.009 | -0.006 | -0.006 | 0.025 | 0.4 | 0.35 | 0.44 | 0.56 | 0.4 | 0.5 |
| 4 | 77543863 | rs13104324 | -0.979 | 1.477 | 1.187 | 0.481 | -0.003 | -0.009 | -0.009 | 0.665 | 0.18 | 0.12 | 0.2 | 0.19 | 0.17 | 0.2 |
| 4 | 77545018 | rs4330370 | -0.987 | 1.554 | 1.39 | 0.4 | 0.01 | 0.108 | 0.108 | 0 | 0.6 | 0.75 | 0.7 | 0.93 | 0.59 | 0.78 |
| 4 | 77547939 | rs10223020 | -1.484 | 1.55 | 1.519 | 0.447 | -0.004 | -0.014 | -0.014 | 0.18 | 0.24 | 0.22 | 0.36 | 0.52 | 0.27 | 0.4 |
| 4 | 77551310 | rs7695340 | -0.99 | 1.569 | 1.511 | 0.425 | 0.012 | 0.107 | 0.107 | 0 | 0.6 | 0.74 | 0.7 | 0.93 | 0.59 | 0.78 |
| 4 | 77551856 | rs11722153 | -0.76 | 1.558 | 1.448 | 0.467 | 0 | -0.021 | -0.021 | 0 | 0.19 | 0.11 | 0.23 | 0.37 | 0.18 | 0.29 |
| 4 | 77552782 | rs4582178 | 1.041 | 1.562 | 1.483 | 0.469 | -0.005 | -0.012 | -0.012 | 0 | 0.33 | 0.26 | 0.32 | 0.42 | 0.32 | 0.39 |
| 4 | 77555857 | rs4859699 | -1.097 | 1.598 | 1.455 | 0.515 | -0.003 | 0.088 | 0.088 | 0 | 0.21 | 0.37 | 0.27 | 0.37 | 0.19 | 0.29 |
| 4 | 77560811 | rs17322529 | 0.325 | 1.543 | 1.507 | 0.461 | 0.001 | 0.034 | 0.034 | 0 | 0.46 | 0.39 | 0.43 | 0.21 | 0.5 | 0.32 |
| 4 | 77562958 | rs4859456 | -0.974 | 1.503 | 1.506 | 0.462 | 0 | 0.012 | 0.012 | 0.024 | 0.64 | 0.49 | 0.65 | 0.59 | 0.68 | 0.61 |
| 4 | 77576402 | rs17002139 | 0.09 | 1.345 | 1.456 | 0.701 | 0.016 | 0.127 | 0.127 | 0 | 0.08 | 0.19 | 0.07 | 0.31 | 0.1 | 0.25 |
| 4 | 77576975 | rs11727631 | -1.448 | 1.373 | 1.036 | 0.589 | -0.001 | 0.111 | 0.111 | 0 | 0.27 | 0.47 | 0.37 | 0.52 | 0.26 | 0.45 |
| 4 | 77594917 | rs13130505 | -0.448 | 1.366 | 0.39 | 0.524 | -0.002 | -0.024 | -0.024 | 0 | 0.17 | 0.11 | 0.22 | 0.36 | 0.17 | 0.32 |
| 4 | 77601715 | rs1493363 | 0.019 | 1.464 | 0.284 | 0.358 | 0.019 | 0.003 | 0.003 | 0.328 | 0.44 | 0.78 | 0.47 | 0.58 | 0.39 | 0.44 |
| 4 | 77612903 | rs2172505 | 0.183 | 1.326 | 0.792 | 0.519 | 0.014 | 0.169 | 0.169 | 0 | 0.12 | 0.23 | 0.21 | 0.36 | 0.09 | 0.26 |
| 4 | 77619296 | rs344117 | 0.786 | 1.508 | 0.416 | 0.263 | -0.028 | 0.142 | 0.142 | 0 | 0.37 | 0.18 | 0.3 | 0.06 | 0.43 | 0.25 |
| 4 | 77621705 | rs2035945 | 0.902 | 1.466 | 0.672 | 0.453 | 0 | 0.003 | 0.003 | 0 | 0.32 | 0.35 | 0.31 | 0.41 | 0.29 | 0.34 |
| 4 | 77626887 | rs13121538 | -0.898 | 1.601 | 0.738 | 0.333 | 0.021 | 0.025 | 0.025 | 0 | 0.15 | 0.28 | 0.19 | 0.22 | 0.17 | 0.16 |
| 4 | 77634028 | rs6849066 | -0.916 | 1.557 | 0.989 | 0.311 | 0.026 | 0.031 | 0.031 | 0 | 0.15 | 0.32 | 0.19 | 0.22 | 0.17 | 0.16 |
| 4 | 77636643 | rs10013334 | -0.189 | 1.495 | 1.153 | 0.397 | -0.018 | 0.078 | 0.078 | 0 | 0.16 | 0.25 | 0.21 | 0.33 | 0.11 | 0.3 |
| 4 | 77637502 | rs344122 | -1.532 | 1.454 | 0.877 | 0.209 | 0 | -0.009 | -0.009 | 0 | 0.69 | 0.72 | 0.7 | 0.58 | 0.71 | 0.7 |
| 4 | 77640617 | rs11734394 | -1.521 | 1.479 | 0.906 | 0.232 | -0.002 | 0.013 | 0.013 | 0 | 0.31 | 0.59 | 0.4 | 0.55 | 0.28 | 0.46 |
| 4 | 77647590 | rs344130 | -0.239 | 1.446 | 0.809 | 0.186 | 0.009 | 0.033 | 0.033 | 0 | 0.53 | 0.44 | 0.49 | 0.27 | 0.6 | 0.4 |
| 4 | 77649374 | rs11735775 | 1.297 | 1.487 | 0.768 | 0.18 | -0.021 | 0.031 | 0.031 | 0 | 0.07 | 0.03 | 0.04 | 0 | 0.09 | 0.03 |
| 4 | 77650897 | rs724096 | 0.045 | 1.471 | 0.803 | 0.185 | 0.043 | 0.159 | 0.159 | 0 | 0.08 | 0.29 | 0.15 | 0.23 | 0.08 | 0.13 |
| 4 | 77652219 | rs344135 | 0.895 | 1.458 | 0.604 | 0.141 | -0.059 | 0.232 | 0.232 | 0 | 0.39 | 0.13 | 0.31 | 0.04 | 0.43 | 0.23 |
| 4 | 77660162 | rs344140 | -1.354 | 1.369 | 0.729 | 0.324 | -0.002 | 0.01 | 0.01 | 0 | 0.69 | 0.61 | 0.7 | 0.6 | 0.71 | 0.73 |
| 4 | 77665279 | rs4859711 | -0.156 | 1.937 | 0.97 | 0.768 | 0.028 | 0.092 | 0.092 | 0 | 0.16 | 0.28 | 0.25 | 0.34 | 0.18 | 0.31 |
| 4 | 77666254 | rs10518155 | -0.404 | 1.934 | 0.983 | 0.958 | 0.038 | 0.093 | 0.093 | 0 | 0.08 | 0.21 | 0.14 | 0.23 | 0.12 | 0.18 |
| 4 | 77669728 | rs10022778 | -1.152 | 1.988 | 0.508 | 0.599 | 0.002 | -0.007 | -0.007 | 0 | 0.45 | 0.7 | 0.52 | 0.57 | 0.57 | 0.56 |
| 4 | 77684099 | rs13113476 | -1.766 | 1.981 | 0.674 | 1.416 | 0.048 | 0.078 | 0.078 | 0 | 0.14 | 0.37 | 0.18 | 0.24 | 0.19 | 0.21 |
| 4 | 77688152 | rs13129187 | 0.542 | 1.898 | 0.154 | 1.319 | 0.001 | -0.019 | -0.019 | 0 | 0.06 | 0.1 | 0.02 | 0 | 0.06 | 0.01 |
| 4 | 77701909 | rs7661695 | -1.237 | 1.479 | 0.086 | 1.193 | 0.012 | 0.029 | 0.029 | 0 | 0.12 | 0.18 | 0.18 | 0.22 | 0.15 | 0.24 |
| 4 | 77717083 | rs716525 | -0.706 | 1.613 | -0.246 | 1.307 | 0 | 0.004 | 0.004 | 0.005 | 0.09 | 0.09 | 0.17 | 0.22 | 0.14 | 0.21 |
| 4 | 77718624 | rs344108 | 0.249 | 1.304 | -0.399 | 0.913 | 0 | 0 | 0 | 0 | 0.6 | 0.45 | 0.53 | 0.44 | 0.6 | 0.55 |

| | | | | | | | | | | | | | | | | |
|---|---|---|---|---|---|---|---|---|---|---|---|---|---|---|---|---|
| 4 | 77719779 | rs11097519 | -0.832 | 1.143 | -0.433 | 1.311 | 0 | 0.005 | 0.005 | 0 | 0.29 | 0.26 | 0.29 | 0.48 | 0.31 | 0.48 |
| 4 | 77719812 | rs344107 | 0.611 | 1.058 | 0.36 | 0.842 | 0.016 | 0.035 | 0.035 | 0 | 0.68 | 0.48 | 0.64 | 0.55 | 0.66 | 0.57 |
| 4 | 77719935 | rs17324961 | 0.056 | 0.455 | -0.428 | -0.979 | 0.019 | 0.016 | 0.016 | 2.931 | 0.38 | 0.28 | 0.18 | 0 | 0.34 | 0.14 |
| 4 | 77720135 | rs12646332 | -0.135 | 0.284 | -0.097 | -0.643 | -0.001 | 0.075 | 0.075 | 0 | 0.29 | 0.5 | 0.28 | 0.53 | 0.29 | 0.35 |
| 4 | 77721997 | rs9998681 | 0.134 | 0.455 | -0.674 | -0.976 | -0.008 | -0.01 | -0.01 | 0.667 | 0.13 | 0.12 | 0.18 | 0.01 | 0.17 | 0.1 |
| 4 | 77724484 | rs12054540 | 1.173 | -0.048 | -1.167 | -1.253 | -0.064 | -0.089 | -0.089 | 0 | 0.39 | 0.11 | 0.46 | 0.46 | 0.39 | 0.45 |
| 4 | 77725026 | rs6821442 | 0.416 | -0.838 | -0.153 | -1.318 | -0.008 | -0.092 | -0.092 | 0 | 0.23 | 0.27 | 0.1 | 0.02 | 0.19 | 0.09 |
| 4 | 77725091 | rs17002218 | 0.457 | -0.93 | 0.099 | -1.311 | -0.02 | 0.11 | 0.11 | 0 | 0.19 | 0.35 | 0.2 | 0.49 | 0.18 | 0.29 |
| 4 | 77725242 | rs6853653 | -1.225 | -0.601 | 0.422 | -2.053 | -0.018 | -0.126 | -0.126 | 0 | 0.35 | 0.38 | 0.17 | 0.02 | 0.25 | 0.14 |
| 4 | 77726159 | rs6840749 | -1.062 | -0.409 | 0.46 | -1.849 | -0.033 | -0.155 | -0.155 | 0.445 | 0.31 | 0.39 | 0.13 | 0.02 | 0.21 | 0.14 |
| 4 | 77727505 | rs13124337 | -0.492 | 0.064 | 0.367 | -1.367 | -0.005 | -0.053 | -0.053 | 0 | 0.2 | 0.23 | 0.09 | 0.02 | 0.15 | 0.09 |
| 4 | 77730260 | rs7663030 | -0.409 | 0.084 | 0.227 | -1.338 | -0.016 | -0.105 | -0.105 | 0 | 0.2 | 0.32 | 0.11 | 0.02 | 0.15 | 0.09 |
| 4 | 77734939 | rs2172496 | 0.675 | 0.37 | -0.122 | -1.096 | 0.015 | 0.145 | 0.145 | 0 | 0.39 | 0.13 | 0.25 | 0.11 | 0.32 | 0.23 |
| 4 | 77735406 | rs1586933 | 0.051 | 0.254 | 0.5 | -0.604 | 0.027 | 0.002 | 0.002 | 0 | 0.54 | 0.69 | 0.41 | 0.52 | 0.37 | 0.48 |
| 4 | 77735709 | rs2865147 | 0.063 | 0.125 | 0.976 | -0.091 | -0.014 | 0.135 | 0.135 | 0 | 0.09 | 0.14 | 0.13 | 0.32 | 0.05 | 0.23 |
| 4 | 77739602 | rs11938295 | 0.277 | 0.489 | 0.719 | -0.321 | -0.027 | -0.072 | -0.072 | 0.04 | 0.24 | 0.12 | 0.35 | 0.45 | 0.32 | 0.12 |
| 4 | 77742277 | rs17002240 | -0.543 | 1.112 | 1.503 | 0.055 | -0.015 | 0.185 | 0.185 | 0 | 0.09 | 0.19 | 0.13 | 0.34 | 0.05 | 0.23 |
| 4 | 77747798 | rs1493305 | 0.986 | 1.829 | 1.387 | -1.08 | 0.039 | -0.098 | -0.098 | 0 | 0.78 | 0.67 | 0.75 | 0.99 | 0.71 | 0.75 |
| 4 | 77751647 | rs9307199 | 1.035 | 1.819 | 1.369 | -1.081 | 0.063 | -0.13 | -0.13 | 0 | 0.78 | 0.59 | 0.75 | 0.99 | 0.71 | 0.72 |
| 4 | 77754140 | rs11097539 | 1.618 | 0.968 | 1.327 | -0.647 | 0.038 | -0.066 | -0.066 | 0.454 | 0.82 | 0.66 | 0.86 | 0.92 | 0.72 | 0.88 |
| 4 | 77754619 | rs12500486 | 0.824 | 0.657 | 1.303 | -0.156 | 0.039 | -0.032 | -0.032 | 3.771 | 0.94 | 0.88 | 0.92 | 1 | 0.84 | 0.97 |
| 4 | 77754699 | rs12500522 | 0.943 | 0.662 | 1.286 | -0.146 | 0.035 | -0.025 | -0.025 | 0 | 0.93 | 0.86 | 0.92 | 0.98 | 0.84 | 0.97 |
| 4 | 77763831 | rs4859466 | 1.408 | 0.673 | 0.888 | -0.214 | 0.101 | 0.003 | 0.003 | 0.698 | 0.9 | 0.47 | 0.89 | 0.92 | 0.84 | 0.94 |
| 4 | 77767645 | rs1994854 | -0.674 | -0.309 | -0.221 | -0.445 | 0.015 | 0.014 | 0.014 | 0 | 0.29 | 0.21 | 0.09 | 0.11 | 0.16 | 0.26 |
| 4 | 77768463 | rs1108255 | -1.363 | 0.241 | 0.348 | 0.239 | -0.006 | 0.041 | 0.041 | 0.128 | 0.11 | 0.28 | 0.07 | 0.1 | 0.07 | 0.14 |
| 4 | 77770388 | rs17260538 | -1.37 | 0.273 | 0.504 | -0.171 | 0.01 | -0.061 | -0.061 | 0 | 0.13 | 0.22 | 0.14 | 0 | 0.12 | 0.19 |
| 4 | 77771345 | rs17002253 | -1.105 | 0.078 | 1.112 | -0.099 | -0.001 | 0.071 | 0.071 | 0 | 0.06 | 0.23 | 0.03 | 0.07 | 0.03 | 0.12 |
| 4 | 77772872 | rs10518160 | -1.105 | 0.078 | 1.129 | -0.102 | -0.01 | 0.045 | 0.045 | 0 | 0.06 | 0.18 | 0.03 | 0.07 | 0.03 | 0.12 |
| 4 | 77773437 | rs4859723 | 0.866 | -0.55 | 0.486 | 0.017 | -0.004 | 0 | 0 | 0 | 0.61 | 0.43 | 0.61 | 0.6 | 0.72 | 0.59 |
| 4 | 77773610 | rs6829573 | -0.087 | -0.394 | -0.292 | -0.214 | -0.009 | -0.023 | -0.023 | 1.051 | 0.76 | 0.96 | 0.83 | 0.81 | 0.84 | 0.79 |
| 4 | 77778487 | rs344837 | -1.146 | -0.152 | 0.031 | -0.11 | -0.003 | 0.008 | 0.008 | 0.559 | 0.1 | 0.05 | 0.08 | 0.14 | 0.1 | 0.14 |
| 4 | 77780095 | rs6835485 | -0.65 | 1.404 | 0.399 | -0.002 | 0.133 | 0.024 | 0.024 | 0 | 0.06 | 0.19 | 0.1 | 0.04 | 0.17 | 0.04 |
| 4 | 77783243 | rs17261150 | -1.292 | 1.394 | 0.281 | -0.012 | 0.042 | -0.003 | -0.003 | 0 | 0.08 | 0.15 | 0.12 | 0.04 | 0.19 | 0.07 |
| 4 | 77783901 | rs17261171 | -1.381 | 1.387 | 0.295 | -0.015 | 0.041 | -0.004 | -0.004 | 0 | 0.08 | 0.17 | 0.12 | 0.04 | 0.19 | 0.07 |
| 4 | 77784395 | rs7693322 | -1.381 | 1.387 | 0.292 | -0.015 | 0.041 | -0.004 | -0.004 | 0.43 | 0.08 | 0.17 | 0.12 | 0.04 | 0.19 | 0.07 |
| 4 | 77784736 | rs401382 | -0.205 | 1.357 | 0.272 | -0.024 | -0.002 | -0.002 | -0.002 | 0.222 | 0.05 | 0.05 | 0.04 | 0 | 0.04 | 0.08 |
| 4 | 77785183 | rs6821290 | -1.408 | 1.382 | 0.292 | 0.032 | 0.038 | -0.003 | -0.003 | 0 | 0.08 | 0.17 | 0.12 | 0.04 | 0.19 | 0.07 |
| 4 | 77792007 | rs6810774 | 0.445 | 0.666 | 0.219 | 0.185 | 0.011 | 0.086 | 0.086 | 0 | 0.27 | 0.46 | 0.28 | 0.43 | 0.3 | 0.34 |
| 4 | 77793015 | rs237616 | -0.667 | 0.452 | 0.006 | 0.004 | 0.003 | -0.011 | -0.011 | 0 | 0.42 | 0.67 | 0.38 | 0.57 | 0.42 | 0.47 |
| 4 | 77793375 | rs1472201 | 0.612 | 0.357 | 0.202 | -0.052 | 0.015 | 0.08 | 0.08 | 0 | 0.27 | 0.37 | 0.28 | 0.43 | 0.31 | 0.34 |
| 4 | 77793829 | rs6532676 | -1.318 | 0.555 | 0.042 | -0.067 | -0.025 | 0.065 | 0.065 | 0 | 0.49 | 0.86 | 0.43 | 0.66 | 0.49 | 0.55 |
| 4 | 77799849 | rs6828780 | -0.216 | 0.459 | 0.123 | -0.153 | -0.002 | 0.015 | 0.015 | 0 | 0.07 | 0.03 | 0.1 | 0 | 0.06 | 0.12 |
| 4 | 77800625 | rs2132766 | -0.546 | 0.353 | 0.171 | -0.147 | -0.002 | 0 | 0 | 0 | 0.1 | 0.07 | 0.05 | 0.14 | 0.08 | 0.06 |
| 4 | 77804009 | rs6532686 | -1.692 | 0.387 | 0.219 | -0.111 | -0.009 | -0.002 | -0.002 | 0.209 | 0.17 | 0.28 | 0.1 | 0.22 | 0.15 | 0.14 |
| 4 | 77806097 | rs7667074 | -1.4 | 0.401 | 0.508 | -0.121 | -0.02 | -0.043 | -0.043 | 0 | 0.18 | 0.26 | 0.15 | 0.14 | 0.14 | 0.18 |
| 4 | 77812990 | rs6853383 | -2.244 | 0.124 | 0.359 | -0.275 | -0.029 | -0.044 | -0.044 | 0 | 0.27 | 0.53 | 0.22 | 0.22 | 0.23 | 0.3 |
| 4 | 77814209 | rs11947481 | 0.814 | 0.326 | -0.887 | -0.762 | -0.036 | -0.012 | -0.012 | 0 | 0.22 | 0.02 | 0.3 | 0.18 | 0.2 | 0.14 |
| 4 | 77815545 | rs4859725 | -1.067 | 0.404 | -0.891 | -0.767 | -0.05 | -0.012 | -0.012 | 0.725 | 0.78 | 0.99 | 0.69 | 0.82 | 0.79 | 0.87 |
| 4 | 77817874 | rs2703130 | -0.601 | 0.992 | -0.421 | -0.573 | -0.002 | 0.002 | 0.002 | 0 | 0.11 | 0.14 | 0.07 | 0.09 | 0.15 | 0.07 |
| 4 | 77820005 | rs2645683 | -0.8 | 1.111 | -0.32 | -0.591 | -0.003 | -0.017 | -0.017 | 0 | 0.17 | 0.26 | 0.16 | 0.14 | 0.18 | 0.23 |
| 4 | 77824442 | rs2703115 | -1.626 | 1.139 | -0.354 | -0.559 | 0.001 | -0.012 | -0.012 | 0 | 0.27 | 0.39 | 0.23 | 0.23 | 0.34 | 0.3 |
| 4 | 77831307 | rs2703147 | -1.456 | 0.995 | -0.376 | -0.562 | -0.002 | -0.015 | -0.015 | 0 | 0.25 | 0.36 | 0.19 | 0.2 | 0.3 | 0.3 |
| 4 | 77836665 | rs12331206 | -0.522 | 0.816 | -0.565 | -0.402 | -0.009 | -0.011 | -0.011 | 0.951 | 0.38 | 0.6 | 0.3 | 0.31 | 0.34 | 0.29 |
| 4 | 77851537 | rs6847460 | 1.155 | 1.043 | -0.786 | -0.075 | 0.003 | -0.008 | -0.008 | 0 | 0.62 | 0.42 | 0.53 | 0.67 | 0.56 | 0.48 |
| 4 | 77854979 | rs2703137 | -0.401 | 1.491 | -1.087 | 0.176 | -0.022 | -0.019 | -0.019 | 0.011 | 0.92 | 0.96 | 0.86 | 0.82 | 0.82 | 0.93 |
| 4 | 77857753 | rs1874565 | 0.176 | 0.982 | -1.118 | 0.852 | -0.004 | -0.018 | -0.018 | 0.648 | 0.66 | 0.74 | 0.62 | 0.34 | 0.64 | 0.63 |
| 4 | 77863211 | rs11722689 | 0.868 | 0.857 | -1.223 | 0.734 | 0.002 | 0.039 | 0.039 | 0 | 0.11 | 0.04 | 0.12 | 0 | 0.09 | 0.18 |
| 4 | 77864488 | rs2645651 | 0.275 | 1.014 | -0.945 | 1.27 | -0.005 | -0.023 | -0.023 | 0 | 0.65 | 0.73 | 0.61 | 0.37 | 0.62 | 0.62 |
| 4 | 77866648 | rs4859727 | 0.193 | 0.546 | -0.762 | 1.13 | 0.017 | -0.022 | -0.022 | 0 | 0.26 | 0.22 | 0.24 | 0.48 | 0.17 | 0.3 |
| 4 | 77868088 | rs2645656 | -0.981 | 1.815 | -0.652 | 0.409 | 0.016 | -0.025 | -0.025 | 0 | 0.24 | 0.32 | 0.21 | 0.08 | 0.41 | 0.2 |
| 4 | 77868340 | rs10009464 | -0.871 | 0.378 | -0.413 | 0.694 | 0.006 | 0.018 | 0.018 | 0 | 0.51 | 0.37 | 0.57 | 0.6 | 0.47 | 0.58 |
| 4 | 77871016 | rs873090 | 0.142 | 0.131 | -0.593 | 0.338 | -0.008 | -0.016 | -0.016 | 0 | 0.65 | 0.71 | 0.7 | 0.48 | 0.63 | 0.65 |
| 4 | 77878392 | rs10213389 | 0.507 | -0.078 | -0.307 | 0.029 | 0.003 | -0.001 | -0.001 | 0 | 0.6 | 0.39 | 0.53 | 0.34 | 0.61 | 0.59 |
| 4 | 77878521 | rs12650893 | 2.053 | -0.217 | -0.394 | -0.049 | 0.002 | 0.003 | 0.003 | 0 | 0.28 | 0.26 | 0.26 | 0.27 | 0.34 | 0.2 |
| 4 | 77882431 | rs17002297 | 0.733 | 0.022 | -0.092 | 0.432 | 0.001 | -0.023 | -0.023 | 0.062 | 0.22 | 0.19 | 0.24 | 0.34 | 0.19 | 0.2 |
| 4 | 77894220 | rs2132768 | 0.361 | 0.084 | -0.139 | 0.399 | -0.018 | -0.011 | -0.011 | 0 | 0.65 | 0.69 | 0.66 | 0.59 | 0.65 | 0.61 |
| 4 | 77894399 | rs2703109 | -1.073 | 0.154 | -0.43 | 0.291 | -0.059 | 0.025 | 0.025 | 0 | 0.9 | 1 | 0.92 | 0.95 | 0.84 | 0.88 |
| 4 | 77915665 | rs883698 | 1.338 | 0.141 | -0.373 | 0.229 | 0.001 | 0.015 | 0.015 | 0 | 0.43 | 0.29 | 0.37 | 0.28 | 0.45 | 0.39 |
| 4 | 77916705 | rs6836567 | 0.488 | 0.158 | -0.378 | 0.233 | 0 | 0 | 0 | 0 | 0.62 | 0.35 | 0.54 | 0.34 | 0.62 | 0.6 |

| | | | | | | | | | | | | | | | | |
|---|---|---|---|---|---|---|---|---|---|---|---|---|---|---|---|---|
| 4 | 77918966 | rs7692644 | 0.273 | 0.298 | -0.545 | 0.037 | 0.008 | 0.021 | 0.021 | 0.02 | 0.64 | 0.39 | 0.55 | 0.46 | 0.64 | 0.61 |
| 4 | 77921005 | rs878729 | -0.31 | -0.111 | -0.831 | 0.124 | -0.003 | 0.068 | 0.068 | 0 | 0.19 | 0.07 | 0.16 | 0.06 | 0.18 | 0.22 |
| 4 | 77927538 | rs6826681 | 0.313 | -0.315 | -0.209 | 0.779 | 0 | -0.001 | -0.001 | 0.564 | 0.18 | 0.21 | 0.19 | 0.27 | 0.17 | 0.08 |
| 4 | 77943171 | rs4859731 | -0.21 | -0.293 | 0.058 | 0.556 | -0.009 | 0.01 | 0.01 | 0 | 0.09 | 0.19 | 0.15 | 0.14 | 0.07 | 0.18 |
| 4 | 77958765 | rs1127595 | -0.5 | -0.87 | 0.413 | 0.429 | 0.027 | 0.022 | 0.022 | 0.064 | 0.47 | 0.68 | 0.54 | 0.64 | 0.38 | 0.44 |
| 4 | 77961377 | rs10000606 | -0.5 | -0.87 | 0.413 | 0.429 | 0.027 | 0.022 | 0.022 | 0.064 | 0.47 | 0.68 | 0.54 | 0.65 | 0.38 | 0.45 |
| 4 | 77974545 | rs4241599 | -0.424 | -1.024 | 0.565 | 0.42 | 0.029 | 0.023 | 0.023 | 0.043 | 0.46 | 0.71 | 0.53 | 0.64 | 0.37 | 0.43 |
| 4 | 77974776 | rs4252913 | -0.685 | -1.022 | 0.958 | 0.505 | 0.018 | 0.13 | 0.13 | 0 | 0.06 | 0.32 | 0.03 | 0.09 | 0.02 | 0.02 |
| 4 | 77984155 | rs4252845 | -1.135 | -1.054 | 0.824 | 0.489 | -0.044 | 0.052 | 0.052 | 0.226 | 0.15 | 0.59 | 0.2 | 0.22 | 0.09 | 0.21 |
| 4 | 77986728 | rs4252828 | 0.304 | -0.958 | 0.647 | 0.503 | -0.016 | 0.026 | 0.026 | 0 | 0.09 | 0.23 | 0.15 | 0.14 | 0.05 | 0.17 |
| 4 | 77987351 | rs4241602 | -1.691 | -0.739 | 0.406 | 0.368 | -0.029 | 0.034 | 0.034 | 0 | 0.94 | 1 | 0.92 | 1 | 0.92 | 0.95 |
| 4 | 77994734 | rs4252782 | 0.722 | -1.195 | -0.793 | 0.289 | 0.017 | 0.062 | 0.062 | 0 | 0.13 | 0.04 | 0.05 | 0 | 0.08 | 0.16 |
| 4 | 78004668 | rs4502705 | 0.109 | -1.259 | -1.019 | 0.414 | 0.065 | 0.045 | 0.045 | 2.063 | 0.5 | 0.2 | 0.45 | 0.37 | 0.63 | 0.57 |
| 4 | 78016353 | rs11097670 | -0.009 | -0.942 | -1.008 | 0.562 | 0.015 | -0.001 | -0.001 | 0.692 | 0.56 | 0.35 | 0.47 | 0.41 | 0.63 | 0.67 |
| 4 | 78019799 | rs4518273 | 1.001 | -0.816 | -1.811 | 0.658 | -0.058 | -0.077 | -0.077 | 0.254 | 0.31 | 0.06 | 0.33 | 0.39 | 0.29 | 0.2 |
| 4 | 78022904 | rs4461545 | 0.178 | -0.163 | -1.306 | 0.912 | -0.016 | 0.003 | 0.003 | 0.043 | 0.4 | 0.26 | 0.41 | 0.37 | 0.54 | 0.41 |
| 4 | 78045685 | rs11097684 | -0.238 | 0.378 | -2.202 | 0.784 | -0.029 | -0.055 | -0.055 | 0 | 0.54 | 0.91 | 0.59 | 0.46 | 0.6 | 0.6 |
| 4 | 78053975 | rs4272042 | 0.262 | 0.461 | -1.774 | 0.733 | 0.001 | -0.021 | -0.021 | 0 | 0.36 | 0.23 | 0.37 | 0.5 | 0.33 | 0.29 |
| 4 | 78063308 | rs7659891 | -0.47 | 0.235 | -1.137 | 0.492 | 0.011 | -0.012 | -0.012 | 0.714 | 0.08 | 0.27 | 0.06 | 0.04 | 0.06 | 0.09 |
| 4 | 78064766 | rs4382065 | -0.404 | 0.2 | -1.668 | 0.585 | -0.064 | -0.067 | -0.067 | 0 | 0.67 | 0.88 | 0.68 | 0.6 | 0.58 | 0.45 |
| 4 | 78069066 | rs4518274 | -0.45 | 0.029 | -1.747 | 0.442 | -0.029 | -0.039 | -0.039 | 0 | 0.72 | 0.75 | 0.7 | 0.59 | 0.61 | 0.46 |
| 4 | 78073655 | rs6849534 | 0.5 | -0.277 | -0.915 | 0.216 | 0.01 | 0.005 | 0.005 | 0 | 0.59 | 0.46 | 0.61 | 0.53 | 0.53 | 0.39 |
| 4 | 78075622 | rs10010725 | 0.601 | -0.247 | -0.856 | 0.23 | -0.021 | -0.017 | -0.017 | 0.182 | 0.13 | 0.3 | 0.09 | 0.06 | 0.08 | 0.07 |
| 4 | 78079772 | rs4150052 | 0.197 | -0.214 | -0.803 | 0.205 | -0.009 | -0.023 | -0.023 | 0 | 0.09 | 0.12 | 0.04 | 0.04 | 0.07 | 0.06 |
| 4 | 78083125 | rs4150074 | 1.679 | 0.244 | -1.039 | 0.345 | -0.062 | -0.079 | -0.079 | 0 | 0.13 | 0.02 | 0.15 | 0.28 | 0.22 | 0.28 |
| 4 | 78084904 | rs4150084 | 0.27 | 0.051 | -0.912 | 0.243 | -0.009 | -0.023 | -0.023 | 0.051 | 0.09 | 0.12 | 0.04 | 0.04 | 0.07 | 0.06 |
| 4 | 78098806 | rs881997 | 2.102 | -0.367 | -1.462 | 0.174 | -0.091 | -0.019 | -0.019 | 0 | 0.26 | 0.01 | 0.27 | 0.24 | 0.29 | 0.21 |
| 4 | 78103458 | rs6848898 | 0.12 | -0.414 | -0.587 | 0.327 | -0.055 | -0.062 | -0.062 | 0 | 0.2 | 0.57 | 0.14 | 0.06 | 0.11 | 0.14 |
| 4 | 78106123 | rs28503908 | 1.725 | -0.277 | -0.953 | 0.413 | -0.052 | -0.037 | -0.037 | 0.107 | 0.44 | 0.08 | 0.52 | 0.48 | 0.43 | 0.28 |
| 4 | 78106441 | rs6822800 | -0.963 | -0.38 | -0.748 | -0.007 | 0.017 | 0.018 | 0.018 | 3.059 | 0.29 | 0.72 | 0.19 | 0.1 | 0.18 | 0.2 |
| 4 | 78115334 | rs11097730 | 1.607 | -0.188 | -0.712 | 0.291 | -0.044 | -0.044 | -0.044 | 1.23 | 0.44 | 0.1 | 0.43 | 0.5 | 0.44 | 0.29 |
| 4 | 78120747 | rs10025838 | -0.498 | -0.282 | -0.696 | 0.246 | -0.073 | -0.082 | -0.082 | 0.005 | 0.75 | 0.88 | 0.64 | 0.57 | 0.61 | 0.51 |
| 4 | 78123574 | rs11943130 | 0.818 | -0.454 | 0.401 | 0.162 | -0.032 | -0.027 | -0.027 | 0.502 | 0.15 | 0.27 | 0.1 | 0.06 | 0.09 | 0.08 |
| 4 | 78124348 | rs4859756 | -0.16 | -0.326 | 0.324 | 0.118 | 0.002 | 0.003 | 0.003 | 2.27 | 0.72 | 0.71 | 0.62 | 0.57 | 0.59 | 0.52 |
| 4 | 78129387 | rs6840915 | -1.361 | -0.196 | 0.136 | 0.242 | -0.005 | 0 | 0 | 0.081 | 0.47 | 0.64 | 0.48 | 0.45 | 0.51 | 0.64 |
| 4 | 78136933 | rs11937061 | -1.026 | 0.134 | -0.228 | 0.324 | 0 | 0 | 0 | 0 | 0.37 | 0.37 | 0.28 | 0.29 | 0.37 | 0.41 |
| 4 | 78137036 | rs11937096 | 1.571 | 0.126 | -0.619 | 0.406 | -0.008 | 0.003 | 0.003 | 0 | 0.08 | 0.1 | 0.04 | 0.06 | 0.05 | 0.08 |
| 4 | 78137234 | rs1477314 | -1.676 | 0.3 | -0.707 | 0.535 | -0.04 | -0.074 | -0.074 | 0 | 0.57 | 0.92 | 0.55 | 0.5 | 0.57 | 0.72 |
| 4 | 78139160 | rs1009141 | 0.71 | 0.489 | -0.731 | 0.585 | -0.004 | -0.003 | -0.003 | 0 | 0.17 | 0.13 | 0.25 | 0.19 | 0.2 | 0.26 |
| 4 | 78145293 | rs716024 | 1.63 | 1.028 | -1.12 | 0.61 | -0.056 | -0.079 | -0.079 | 0.121 | 0.18 | 0.02 | 0.18 | 0.25 | 0.19 | 0.1 |
| 4 | 78153905 | rs11097744 | -1.434 | 1.399 | 0.398 | 0.764 | -0.001 | 0.002 | 0.002 | 0 | 0.35 | 0.34 | 0.27 | 0.26 | 0.36 | 0.41 |
| 4 | 78154524 | rs28735825 | -0.88 | 0.783 | 0.374 | 0.574 | -0.036 | -0.054 | -0.054 | 0 | 0.22 | 0.37 | 0.12 | 0.07 | 0.13 | 0.16 |
| 4 | 78161220 | rs2009091 | -1.344 | 1.635 | 0.288 | 1.172 | -0.029 | -0.023 | -0.023 | 0.568 | 0.31 | 0.22 | 0.34 | 0.37 | 0.43 | 0.52 |
| 4 | 78161257 | rs7661051 | -0.572 | 1.616 | 0.407 | 0.665 | 0.005 | -0.007 | -0.007 | 0 | 0.12 | 0.24 | 0.1 | 0.1 | 0.1 | 0.27 |
| 4 | 78161745 | rs13144211 | -0.441 | 2.155 | -0.312 | 0.616 | -0.074 | -0.029 | -0.029 | 0 | 0.13 | 0.03 | 0.2 | 0.16 | 0.28 | 0.17 |
| 4 | 78161843 | rs2865830 | -0.317 | 1.1 | -0.113 | 0 | 0.002 | 0.087 | 0.087 | 0 | 0.49 | 0.58 | 0.39 | 0.1 | 0.48 | 0.5 |
| 4 | 78162739 | rs10491462 | -0.599 | 1.615 | 0.543 | 0.72 | 0.006 | 0.039 | 0.039 | 0 | 0.13 | 0.15 | 0.3 | 0.42 | 0.19 | 0.29 |
| 4 | 78168932 | rs6835713 | -0.537 | 1.323 | 0.486 | 0.489 | -0.002 | 0 | 0 | 0 | 0.47 | 0.57 | 0.59 | 0.48 | 0.55 | 0.48 |
| 4 | 78181653 | rs736601 | -0.573 | 1.809 | 0.795 | 0.521 | 0.026 | 0.014 | 0.014 | 0 | 0.09 | 0.26 | 0.09 | 0.08 | 0.07 | 0.07 |
| 4 | 78194110 | rs17643945 | 0.91 | 1.509 | 0.614 | 0.324 | -0.056 | 0.194 | 0.194 | 0.629 | 0.3 | 0.05 | 0.24 | 0.01 | 0.32 | 0.17 |
| 4 | 78203795 | rs11732260 | -0.925 | 1.613 | 0.953 | 0.512 | 0.082 | 0.176 | 0.176 | 1.809 | 0.21 | 0.53 | 0.3 | 0.49 | 0.26 | 0.34 |
| 4 | 78210296 | rs17396361 | -0.514 | 1.674 | 0.909 | 0.58 | 0.037 | 0.015 | 0.015 | 0.542 | 0.17 | 0.37 | 0.25 | 0.24 | 0.21 | 0.21 |
| 4 | 78211866 | rs4859484 | -0.201 | 1.671 | 0.938 | 0.579 | -0.018 | -0.01 | -0.01 | 0 | 0.16 | 0.14 | 0.23 | 0.24 | 0.21 | 0.21 |
| 4 | 78231196 | rs4859782 | -0.209 | 1.376 | 0.196 | 0.728 | 0.005 | 0.001 | 0.001 | 0.895 | 0.16 | 0.19 | 0.24 | 0.24 | 0.21 | 0.21 |
| 4 | 78237955 | rs4859487 | -0.039 | 0.531 | -0.042 | 0.748 | 0.039 | -0.009 | -0.009 | 0 | 0.52 | 0.71 | 0.55 | 0.5 | 0.58 | 0.51 |
| 4 | 78268226 | rs17002564 | -0.952 | 0.571 | 0.008 | 0.59 | -0.009 | -0.015 | -0.015 | 0 | 0.17 | 0.16 | 0.11 | 0.1 | 0.1 | 0.17 |
| 4 | 78268641 | rs11726925 | -1.649 | 1.523 | -0.21 | 1.22 | -0.044 | -0.046 | -0.046 | 0 | 0.09 | 0.02 | 0.1 | 0.17 | 0.15 | 0.23 |
| 4 | 78270126 | rs11936421 | -0.931 | 1.298 | -0.161 | 1.366 | -0.015 | -0.009 | -0.009 | 0 | 0.14 | 0.09 | 0.16 | 0.21 | 0.17 | 0.17 |
| 4 | 78271822 | rs10023142 | -1.392 | 1.157 | 0.169 | 1.523 | -0.004 | 0.042 | 0.042 | 0 | 0.07 | 0.08 | 0.07 | 0.19 | 0.06 | 0.17 |
| 4 | 78276420 | rs1559121 | -1.545 | 1.132 | 0.519 | 1.585 | -0.001 | 0.105 | 0.105 | 0 | 0.07 | 0.14 | 0.07 | 0.22 | 0.06 | 0.16 |
| 4 | 78280329 | rs10034168 | -1.445 | 1.138 | 0.46 | 1.592 | -0.004 | 0.048 | 0.048 | 0 | 0.07 | 0.09 | 0.07 | 0.22 | 0.06 | 0.17 |
| 4 | 78291269 | rs7375525 | 2.09 | 0.922 | 0.842 | 1.57 | 0.001 | -0.034 | -0.034 | 0 | 0.73 | 0.29 | 0.73 | 0.54 | 0.76 | 0.64 |
| 4 | 78292049 | rs2866281 | 1.411 | 0.894 | 2.555 | 1.576 | -0.003 | -0.01 | -0.01 | 0.496 | 0.79 | 0.79 | 0.77 | 0.61 | 0.77 | 0.66 |
| 4 | 78292203 | rs12500776 | -0.563 | 0.922 | 2.541 | 1.58 | -0.032 | -0.029 | -0.029 | 0 | 0.13 | 0.02 | 0.14 | 0.2 | 0.13 | 0.17 |
| 4 | 78293878 | rs4859824 | 1.937 | 0.936 | 2.922 | 1.586 | 0.008 | 0.066 | 0.066 | 0 | 0.77 | 0.59 | 0.76 | 0.61 | 0.77 | 0.66 |
| 4 | 78307791 | rs1806786 | -1.543 | 1.047 | 3.053 | 1.561 | 0.035 | 0.029 | 0.029 | 0 | 0.18 | 0.48 | 0.18 | 0.23 | 0.15 | 0.19 |
| 4 | 78308601 | rs12502190 | 1.397 | 1.053 | 2.753 | 1.582 | 0.007 | 0.094 | 0.094 | 0 | 0.92 | 0.82 | 0.92 | 0.78 | 0.91 | 0.83 |
| 4 | 78328101 | rs1372784 | -2.348 | 1.045 | 2.644 | 1.546 | 0.012 | 0.051 | 0.051 | 0 | 0.25 | 0.5 | 0.25 | 0.43 | 0.24 | 0.36 |
| 4 | 78329877 | rs6533029 | -2.156 | 1.045 | 2.36 | 1.546 | 0.027 | 0.088 | 0.088 | 0 | 0.23 | 0.35 | 0.23 | 0.43 | 0.24 | 0.36 |

| | | | | | | | | | | | | | | | | |
|---|---|---|---|---|---|---|---|---|---|---|---|---|---|---|---|---|
| 4 | 78330433 | rs1372786 | 2.111 | 1.064 | 1.338 | 1.539 | -0.007 | -0.046 | -0.046 | 0 | 0.73 | 0.27 | 0.73 | 0.54 | 0.75 | 0.65 |
| 4 | 78333798 | rs17002672 | -1.351 | 0.925 | 1.182 | 1.463 | 0.003 | 0.045 | 0.045 | 0 | 0.07 | 0.09 | 0.07 | 0.17 | 0.09 | 0.16 |
| 4 | 78334386 | rs2197167 | -1.513 | 0.481 | 1.187 | 1.599 | -0.035 | 0.033 | 0.033 | 0 | 0.27 | 0.42 | 0.25 | 0.36 | 0.22 | 0.22 |
| 4 | 78334882 | rs6841674 | -0.73 | 1.305 | 0.335 | 0.909 | -0.031 | -0.021 | -0.021 | 0 | 0.28 | 0.09 | 0.33 | 0.26 | 0.35 | 0.18 |
| 4 | 78335569 | rs11931577 | -0.24 | 1.244 | 0.333 | 1.027 | 0.001 | 0.003 | 0.003 | 0.489 | 0.11 | 0.12 | 0.16 | 0.17 | 0.12 | 0.21 |
| 4 | 78336408 | rs13138737 | -0.657 | 2.107 | 0.517 | 0.772 | -0.031 | 0.002 | 0.002 | 0 | 0.33 | 0.2 | 0.39 | 0.27 | 0.47 | 0.24 |
| 4 | 78339282 | rs11721695 | -0.288 | 1.099 | 0.311 | 0.243 | -0.013 | -0.016 | -0.016 | 0.926 | 0.36 | 0.46 | 0.3 | 0.34 | 0.29 | 0.34 |
| 4 | 78339441 | rs11726795 | -0.373 | 1.096 | 0.284 | 0.254 | -0.012 | -0.014 | -0.014 | 3.103 | 0.36 | 0.46 | 0.3 | 0.35 | 0.3 | 0.34 |
| 4 | 78341476 | rs6851181 | -1.709 | 1.024 | 0.247 | 0.343 | -0.001 | 0 | 0 | 0 | 0.68 | 0.66 | 0.69 | 0.61 | 0.76 | 0.58 |
| 4 | 78343820 | rs12649862 | 0.062 | 1.624 | 1.113 | 0.768 | 0.018 | 0.077 | 0.077 | 0 | 0.08 | 0.13 | 0.18 | 0.26 | 0.13 | 0.15 |
| 4 | 78344132 | rs34107162 | -0.46 | 1.572 | 1.332 | 0.732 | 0 | -0.016 | -0.016 | 0.184 | 0.31 | 0.27 | 0.29 | 0.44 | 0.3 | 0.3 |
| 4 | 78346794 | rs17002676 | -0.433 | 1.594 | 1.336 | 0.611 | 0.026 | 0.015 | 0.015 | 0.071 | 0.16 | 0.31 | 0.24 | 0.18 | 0.21 | 0.21 |
| 4 | 78349135 | rs10020156 | -1.275 | 1.572 | 1.25 | 0.601 | 0.009 | 0.016 | 0.016 | 0 | 0.42 | 0.61 | 0.37 | 0.36 | 0.38 | 0.35 |
| 4 | 78351064 | rs1442696 | 0.128 | 1.339 | 1.237 | 0.671 | 0.022 | 0.031 | 0.031 | 0 | 0.24 | 0.16 | 0.12 | 0.17 | 0.17 | 0.15 |
| 4 | 78351923 | rs1073284 | -0.092 | 1.032 | 1.432 | 0.649 | -0.026 | -0.019 | -0.019 | 0 | 0.24 | 0.32 | 0.12 | 0.18 | 0.14 | 0.13 |
| 4 | 78352799 | rs4280762 | 0.044 | 1.002 | 1.636 | 0.828 | -0.038 | -0.001 | -0.001 | 0 | 0.17 | 0.35 | 0.09 | 0.18 | 0.09 | 0.12 |
| 4 | 78355165 | rs12510042 | -0.583 | 0.919 | 1.183 | 0.983 | 0 | -0.025 | -0.025 | 0 | 0.22 | 0.18 | 0.24 | 0.44 | 0.21 | 0.26 |
| 4 | 78356862 | rs11939449 | -0.54 | 0.923 | 1.258 | 0.894 | 0.038 | 0.044 | 0.044 | 0.101 | 0.14 | 0.36 | 0.16 | 0.18 | 0.18 | 0.18 |
| 4 | 78366298 | rs7685130 | -0.538 | 0.861 | 0.427 | 0.87 | 0.002 | 0.001 | 0.001 | 0 | 0.84 | 0.83 | 0.93 | 0.83 | 0.91 | 0.88 |
| 4 | 78373045 | rs1348299 | 1.672 | 0.891 | 0.328 | 0.867 | -0.009 | -0.018 | -0.018 | 2.466 | 0.58 | 0.11 | 0.56 | 0.38 | 0.6 | 0.55 |
| 4 | 78389293 | rs6831175 | 1.786 | 0.877 | 0.401 | 0.848 | 0.013 | 0.003 | 0.003 | 0.02 | 0.57 | 0.17 | 0.57 | 0.38 | 0.59 | 0.55 |
| 4 | 78391533 | rs12501028 | 0.096 | 0.971 | 0.364 | 0.957 | 0 | 0.036 | 0.036 | 0 | 0.07 | 0.1 | 0.17 | 0.28 | 0.13 | 0.15 |
| 4 | 78396544 | rs1596230 | 1.334 | 0.913 | 0.59 | 0.89 | 0.03 | 0.082 | 0.082 | 0.561 | 0.76 | 0.44 | 0.65 | 0.57 | 0.69 | 0.65 |
| 4 | 78404249 | rs11735457 | 0.012 | 0.905 | 0.052 | 0.882 | -0.012 | -0.002 | -0.002 | 0 | 0.19 | 0.22 | 0.08 | 0.2 | 0.1 | 0.11 |
| 4 | 78413360 | rs12510689 | 2.003 | 0.918 | -0.358 | 0.865 | -0.011 | -0.007 | -0.007 | 0.31 | 0.52 | 0.09 | 0.56 | 0.38 | 0.58 | 0.52 |
| 4 | 78417537 | rs17002700 | -0.204 | 1.11 | -0.368 | 0.987 | -0.013 | -0.007 | -0.007 | 0.057 | 0.14 | 0.07 | 0.14 | 0.17 | 0.18 | 0.21 |
| 4 | 78419683 | rs10518178 | -0.316 | 0.69 | -0.213 | 0.872 | 0.022 | 0.013 | 0.013 | 0 | 0.2 | 0.13 | 0.08 | 0.18 | 0.1 | 0.11 |
| 4 | 78420568 | rs10518177 | -0.326 | 1.301 | -0.18 | 1.107 | -0.036 | -0.028 | -0.028 | 2.249 | 0.07 | 0.01 | 0.16 | 0.25 | 0.13 | 0.15 |
| 4 | 78424381 | rs1442691 | 0.971 | 0.793 | 0.458 | 0.382 | 0.053 | 0.014 | 0.014 | 1.789 | 0.94 | 0.67 | 0.96 | 1 | 0.97 | 0.94 |
| 4 | 78426029 | rs2053526 | -1.697 | 0.74 | -0.739 | 0.371 | -0.016 | -0.037 | -0.037 | 0 | 0.46 | 0.89 | 0.3 | 0.49 | 0.39 | 0.52 |
| 4 | 78456502 | rs17406477 | -0.093 | 0.438 | -0.511 | 0.089 | -0.013 | 0.023 | 0.023 | 0 | 0.12 | 0.01 | 0.1 | 0.05 | 0.11 | 0.12 |
| 4 | 78459361 | rs1077234 | -0.301 | 0.398 | -0.61 | 0.138 | -0.036 | -0.058 | -0.058 | 0 | 0.21 | 0.07 | 0.13 | 0.3 | 0.22 | 0.32 |
| 4 | 78464698 | rs1596231 | -2.089 | 0.342 | -0.573 | 0.124 | 0.006 | 0 | 0 | 1.029 | 0.43 | 0.88 | 0.29 | 0.38 | 0.36 | 0.48 |
| 4 | 78478506 | rs355679 | -0.438 | 0.366 | 0.348 | 0.443 | 0.001 | 0.047 | 0.047 | 0.338 | 0.2 | 0.27 | 0.17 | 0.37 | 0.18 | 0.3 |
| 4 | 78500085 | rs189587 | -1.816 | 0.565 | 0.489 | 0.233 | 0.001 | -0.005 | -0.005 | 0 | 0.43 | 0.62 | 0.35 | 0.44 | 0.43 | 0.48 |
| 4 | 78505372 | rs355686 | -0.888 | 0.046 | 0.52 | 0.48 | 0.001 | -0.003 | -0.003 | 0 | 0.24 | 0.22 | 0.19 | 0.33 | 0.2 | 0.35 |
| 4 | 78505578 | rs355687 | -1.06 | 0.072 | 0.548 | 0.5 | -0.009 | 0.011 | 0.011 | 0.249 | 0.27 | 0.31 | 0.21 | 0.33 | 0.24 | 0.38 |
| 4 | 78511203 | rs920666 | -0.855 | -0.802 | 0.877 | -0.151 | -0.015 | -0.025 | -0.025 | 0 | 0.36 | 0.49 | 0.24 | 0.17 | 0.25 | 0.28 |
| 4 | 78513987 | rs17002731 | -0.025 | -0.778 | 1.002 | -0.347 | 0.007 | -0.038 | -0.038 | 0 | 0.11 | 0.19 | 0.07 | 0.02 | 0.1 | 0.02 |
| 4 | 78521525 | rs171388 | -1.697 | -0.958 | 1.219 | -0.373 | -0.019 | 0.007 | 0.007 | 0 | 0.23 | 0.63 | 0.21 | 0.21 | 0.15 | 0.2 |
| 4 | 78523092 | rs355661 | -0.271 | -1.005 | 1.04 | -0.469 | -0.012 | 0.013 | 0.013 | 0 | 0.1 | 0.16 | 0.12 | 0.15 | 0.05 | 0.18 |
| 4 | 78532232 | rs1052563 | 0.18 | -0.875 | -0.049 | -0.515 | -0.02 | 0.049 | 0.049 | 0 | 0.1 | 0.03 | 0.06 | 0 | 0.1 | 0.04 |
| 4 | 78533252 | rs10022693 | -1.363 | -1.012 | -0.109 | -0.576 | 0.079 | 0.102 | 0.102 | 0 | 0.43 | 0.76 | 0.27 | 0.17 | 0.27 | 0.35 |
| 4 | 78542597 | rs1566485 | 0.505 | -0.745 | -0.733 | -0.809 | 0.023 | 0.089 | 0.089 | 0 | 0.45 | 0.24 | 0.62 | 0.72 | 0.59 | 0.55 |
| 4 | 78544132 | rs9993375 | 0.082 | -0.976 | -0.794 | -1.161 | 0 | -0.035 | -0.035 | 0.298 | 0.18 | 0.17 | 0.09 | 0 | 0.15 | 0.09 |
| 4 | 78544487 | rs17416271 | 0.553 | -0.815 | -0.997 | -0.938 | -0.005 | 0.021 | 0.021 | 0 | 0.1 | 0.06 | 0.05 | 0.03 | 0.09 | 0.11 |
| 4 | 78549559 | rs2173755 | 0.206 | -1.183 | -0.462 | -0.9 | -0.004 | -0.001 | -0.001 | 0 | 0.75 | 0.76 | 0.77 | 0.77 | 0.83 | 0.75 |
| 4 | 78555443 | rs2866441 | 0.005 | -1.263 | -0.8 | -0.527 | 0.012 | -0.032 | -0.032 | 0.562 | 0.85 | 0.95 | 0.91 | 0.77 | 0.85 | 0.88 |
| 4 | 78558585 | rs2866442 | 1.328 | -1.152 | -1.042 | -1.112 | -0.02 | 0.03 | 0.03 | 0 | 0.42 | 0.22 | 0.63 | 0.65 | 0.52 | 0.52 |
| 4 | 78562040 | rs972755 | -0.073 | -1.466 | -1.021 | -1.378 | 0.015 | 0.088 | 0.088 | 0 | 0.23 | 0.1 | 0.12 | 0.03 | 0.21 | 0.14 |
| 4 | 78562573 | rs4859868 | 0.515 | -1.224 | -0.306 | -1.041 | -0.014 | -0.034 | -0.034 | 0 | 0.68 | 0.45 | 0.8 | 0.74 | 0.72 | 0.8 |
| 4 | 78597444 | rs17416515 | -0.236 | -1.166 | -1.06 | -0.766 | 0.015 | 0.06 | 0.06 | 0 | 0.12 | 0.03 | 0.05 | 0.03 | 0.09 | 0.1 |
| 4 | 78598313 | rs10009981 | 0.866 | -1.384 | -1.084 | -0.751 | 0.092 | 0.001 | 0.001 | 0.747 | 0.63 | 0.16 | 0.62 | 0.67 | 0.75 | 0.65 |
| 4 | 78605369 | rs958617 | 1.023 | -1.59 | -1.595 | -0.767 | 0.124 | -0.014 | -0.014 | 0 | 0.6 | 0.12 | 0.61 | 0.63 | 0.73 | 0.62 |
| 4 | 78624604 | rs17002781 | -0.707 | 0.506 | -0.734 | -0.876 | 0.051 | -0.022 | -0.022 | 0 | 0.1 | 0.2 | 0.11 | 0.06 | 0.21 | 0.05 |
| 4 | 78638491 | rs3205281 | 0.536 | -0.167 | -0.344 | -0.933 | -0.031 | -0.03 | -0.03 | 0 | 0.75 | 0.59 | 0.87 | 0.81 | 0.87 | 0.74 |
| 4 | 78638730 | rs2134048 | 0.536 | -0.167 | -0.344 | -0.933 | -0.031 | -0.03 | -0.03 | 0 | 0.75 | 0.59 | 0.87 | 0.81 | 0.87 | 0.74 |
| 4 | 78643791 | rs13104947 | 0.942 | 0.565 | -1.151 | -0.611 | -0.069 | -0.109 | -0.109 | 0.005 | 0.27 | 0.07 | 0.28 | 0.47 | 0.33 | 0.3 |
| 4 | 78655778 | rs17448771 | 1.485 | 0.372 | -0.995 | -0.751 | -0.003 | -0.025 | -0.025 | 0.892 | 0.07 | 0.14 | 0.03 | 0 | 0.05 | 0.04 |
| 4 | 78682349 | rs10005741 | -0.707 | 0.333 | -0.926 | -0.766 | -0.017 | -0.005 | -0.005 | 0 | 0.82 | 0.78 | 0.9 | 0.82 | 0.92 | 0.78 |
| 4 | 78682581 | rs10005935 | 0.652 | 0.492 | -0.954 | -0.78 | 0.003 | 0.014 | 0.014 | 0 | 0.5 | 0.52 | 0.61 | 0.35 | 0.58 | 0.45 |
| 4 | 78708696 | rs17002804 | 0.788 | 0.971 | -0.528 | -1.012 | -0.001 | 0.003 | 0.003 | 0 | 0.47 | 0.42 | 0.6 | 0.35 | 0.53 | 0.44 |
| 4 | 78713254 | rs4859878 | -1.095 | 0.885 | -0.571 | -0.984 | -0.017 | -0.005 | -0.005 | 0.01 | 0.82 | 0.78 | 0.9 | 0.82 | 0.92 | 0.78 |
| 4 | 78722315 | rs4859879 | 1.639 | 0.958 | -0.602 | -0.987 | -0.091 | -0.154 | -0.154 | 0.081 | 0.24 | 0.03 | 0.26 | 0.47 | 0.29 | 0.29 |
| 4 | 78732149 | rs10222901 | 0.097 | 1.15 | -0.574 | -1.01 | -0.031 | -0.03 | -0.03 | 0 | 0.75 | 0.59 | 0.87 | 0.82 | 0.87 | 0.74 |
| 4 | 78734563 | rs1587563 | 0.122 | 1.346 | -0.661 | -1.1 | 0.003 | 0.004 | 0.004 | 0 | 0.54 | 0.52 | 0.66 | 0.35 | 0.63 | 0.45 |
| 4 | 78738089 | rs1814033 | -0.891 | 0.791 | -0.462 | -0.809 | -0.018 | 0.001 | 0.001 | 0 | 0.82 | 0.78 | 0.9 | 0.81 | 0.92 | 0.78 |
| 4 | 78747771 | rs4365752 | -0.944 | 0.757 | -0.398 | -0.897 | -0.01 | -0.003 | -0.003 | 0.624 | 0.83 | 0.8 | 0.91 | 0.81 | 0.92 | 0.78 |
| 4 | 78748977 | rs1996740 | 0.679 | 0.7 | 0.202 | -0.891 | 0 | 0.045 | 0.045 | 0 | 0.38 | 0.18 | 0.51 | 0.22 | 0.42 | 0.4 |

| | | | | | | | | | | | | | | | | |
|---|---|---|---|---|---|---|---|---|---|---|---|---|---|---|---|---|
| 4 | 78749042 | rs2866688 | 0.079 | 0.45 | 0.255 | -0.935 | -0.031 | -0.03 | -0.03 | 0 | 0.75 | 0.59 | 0.87 | 0.81 | 0.87 | 0.74 |
| 4 | 78757015 | rs10518182 | 1.321 | 0.314 | -0.101 | -1.274 | 0.024 | 0.111 | 0.111 | 0.392 | 0.28 | 0.1 | 0.21 | 0.03 | 0.23 | 0.15 |
| 4 | 78776883 | rs6817305 | -0.238 | 0.51 | 0.12 | -0.964 | -0.026 | -0.033 | -0.033 | 0 | 0.25 | 0.39 | 0.13 | 0.19 | 0.14 | 0.26 |
| 4 | 78794541 | rs10026176 | -0.289 | 0.609 | 0.143 | -0.934 | 0.001 | 0.006 | 0.006 | 0 | 0.49 | 0.43 | 0.61 | 0.32 | 0.57 | 0.46 |
| 4 | 78806253 | rs6849145 | -0.16 | 0.606 | 0.214 | -0.935 | 0.003 | 0.044 | 0.044 | 0 | 0.48 | 0.28 | 0.6 | 0.32 | 0.57 | 0.46 |
| 4 | 78817124 | rs6816113 | 1.273 | 0.563 | 0.354 | -0.845 | -0.007 | -0.036 | -0.036 | 0 | 0.08 | 0.18 | 0.03 | 0 | 0.05 | 0.04 |
| 4 | 78820513 | rs10005531 | -0.277 | 0.398 | 0.293 | -0.881 | -0.001 | 0.001 | 0.001 | 0 | 0.47 | 0.43 | 0.6 | 0.32 | 0.57 | 0.45 |
| 4 | 78834863 | rs7671941 | -0.446 | 0.343 | 0.281 | -0.902 | -0.036 | -0.033 | -0.033 | 0.558 | 0.25 | 0.42 | 0.13 | 0.21 | 0.14 | 0.26 |
| 4 | 78840447 | rs4859512 | -0.222 | 0.386 | 0.176 | -0.843 | 0.001 | 0.04 | 0.04 | 0 | 0.48 | 0.29 | 0.59 | 0.33 | 0.56 | 0.4 |
| 4 | 78841872 | rs7439347 | -0.393 | 0.469 | -0.218 | -0.797 | -0.027 | -0.025 | -0.025 | 0 | 0.23 | 0.4 | 0.13 | 0.2 | 0.14 | 0.32 |
| 4 | 78849088 | rs924684 | -0.225 | 0.469 | -0.222 | -0.805 | 0.006 | 0.03 | 0.03 | 0 | 0.5 | 0.55 | 0.61 | 0.33 | 0.56 | 0.39 |
| 4 | 78854942 | rs7687500 | 1.571 | 0.403 | -0.304 | -0.78 | -0.007 | -0.031 | -0.031 | 2.346 | 0.07 | 0.15 | 0.03 | 0 | 0.05 | 0.04 |
| 4 | 78858515 | rs4577601 | -2.289 | 0.436 | -0.412 | -0.705 | -0.059 | -0.135 | -0.135 | 0 | 0.73 | 0.94 | 0.73 | 0.54 | 0.71 | 0.71 |
| 4 | 78868343 | rs17450279 | 1.608 | 0.325 | -0.625 | -0.901 | -0.052 | 0.076 | 0.076 | 0 | 0.13 | 0.01 | 0.09 | 0.01 | 0.18 | 0.12 |
| 4 | 78875974 | rs13101418 | 1.783 | 0.275 | -0.089 | -0.41 | -0.01 | -0.049 | -0.049 | 0 | 0.28 | 0.17 | 0.29 | 0.47 | 0.3 | 0.29 |
| 4 | 78879461 | rs7699718 | 1.285 | 0.271 | 0.073 | -0.348 | -0.007 | -0.039 | -0.039 | 0 | 0.07 | 0.2 | 0.03 | 0 | 0.05 | 0.04 |
| 4 | 78889803 | rs12504644 | 1.139 | 0.323 | 0.061 | -0.376 | -0.016 | -0.016 | -0.016 | 2.001 | 0.11 | 0.08 | 0.31 | 0.21 | 0.23 | 0.22 |
| 4 | 78889868 | rs7683766 | 1.012 | -0.317 | 0.515 | -0.122 | -0.007 | -0.039 | -0.039 | 0 | 0.07 | 0.2 | 0.03 | 0 | 0.05 | 0.04 |
| 4 | 78909717 | rs17450634 | 1.207 | -0.292 | 0.424 | -0.109 | -0.005 | -0.026 | -0.026 | 0.009 | 0.07 | 0.14 | 0.03 | 0 | 0.05 | 0.04 |
| 4 | 78915841 | rs6533374 | -0.426 | -0.313 | 0.442 | -0.117 | -0.013 | -0.002 | -0.002 | 1.903 | 0.84 | 0.79 | 0.91 | 0.8 | 0.92 | 0.73 |
| 4 | 78918906 | rs4859894 | 0.603 | -0.305 | 0.438 | -0.151 | -0.026 | -0.027 | -0.027 | 0.02 | 0.77 | 0.64 | 0.87 | 0.81 | 0.87 | 0.69 |
| 4 | 78924968 | rs12641050 | 0.559 | -0.136 | 0.183 | -0.024 | -0.001 | -0.006 | -0.006 | 0 | 0.35 | 0.37 | 0.31 | 0.47 | 0.35 | 0.37 |
| 4 | 78937771 | rs11940556 | 0.267 | -0.116 | 0.149 | -0.058 | 0 | -0.008 | -0.008 | 0 | 0.4 | 0.56 | 0.42 | 0.71 | 0.38 | 0.56 |
| 4 | 78947327 | rs7667122 | 1.121 | -0.09 | 0.248 | 0.063 | -0.004 | -0.015 | -0.015 | 0 | 0.29 | 0.26 | 0.28 | 0.5 | 0.31 | 0.34 |
| 4 | 78949372 | rs1546318 | -0.286 | -0.354 | -0.092 | -0.361 | -0.001 | -0.03 | -0.03 | 0 | 0.85 | 0.9 | 0.93 | 1 | 0.9 | 0.89 |
| 4 | 78967046 | rs1875649 | 0.176 | -0.224 | 0.036 | -0.089 | 0.006 | 0.023 | 0.023 | 0.166 | 0.4 | 0.7 | 0.41 | 0.72 | 0.37 | 0.56 |
| 4 | 78968798 | rs1502877 | 0.204 | -0.209 | 0.094 | -0.048 | -0.001 | -0.004 | -0.004 | 1.733 | 0.38 | 0.57 | 0.4 | 0.72 | 0.37 | 0.54 |
| 4 | 78970022 | rs7696815 | -0.167 | -0.808 | 0.512 | 0.031 | 0.009 | 0.016 | 0.016 | 0 | 0.16 | 0.13 | 0.14 | 0.03 | 0.17 | 0.07 |
| 4 | 78970864 | rs7656678 | 0.879 | -1.057 | 1.211 | 0.424 | -0.039 | -0.014 | -0.014 | 0 | 0.73 | 0.56 | 0.78 | 0.75 | 0.77 | 0.79 |
| 4 | 78974336 | rs2036460 | 0.46 | -1.107 | 0.904 | 1.039 | 0.014 | 0.063 | 0.063 | 0 | 0.13 | 0.05 | 0.11 | 0 | 0.15 | 0.04 |
| 4 | 78974867 | rs7677984 | -0.137 | -0.87 | 1.003 | 0.81 | 0.001 | -0.004 | -0.004 | 0.678 | 0.3 | 0.24 | 0.3 | 0.27 | 0.29 | 0.37 |
| 4 | 78975137 | rs1910301 | 0.759 | -0.891 | 1.516 | 0.73 | -0.016 | -0.007 | -0.007 | 0 | 0.71 | 0.36 | 0.78 | 0.75 | 0.77 | 0.79 |
| 4 | 78983383 | rs1502887 | -0.227 | -0.509 | 1.964 | 0.81 | -0.005 | -0.03 | -0.03 | 0 | 0.16 | 0.17 | 0.14 | 0 | 0.12 | 0.06 |
| 4 | 78984082 | rs10518184 | -2.313 | -0.414 | 1.848 | 0.938 | 0.038 | 0.015 | 0.015 | 0 | 0.49 | 0.75 | 0.47 | 0.54 | 0.42 | 0.49 |
| 4 | 78987157 | rs4859905 | -2.328 | -0.389 | 1.846 | 0.951 | 0.038 | 0.016 | 0.016 | 0 | 0.49 | 0.75 | 0.47 | 0.54 | 0.42 | 0.49 |
| 4 | 78989939 | rs873455 | -1.606 | -0.517 | 1.463 | 1.114 | -0.008 | 0.008 | 0.008 | 0.512 | 0.28 | 0.34 | 0.23 | 0.39 | 0.21 | 0.24 |
| 4 | 78991030 | rs4591635 | -1.4 | -0.402 | 1.151 | 0.36 | -0.009 | -0.003 | -0.003 | 0.343 | 0.25 | 0.31 | 0.21 | 0.22 | 0.19 | 0.25 |
| 4 | 78992568 | rs9284617 | -0.39 | -0.265 | 0.855 | 0.233 | 0.015 | 0.007 | 0.007 | 0.684 | 0.17 | 0.13 | 0.13 | 0 | 0.12 | 0.06 |
| 4 | 78996420 | rs17002919 | -0.608 | -0.311 | 1.049 | 0.34 | 0.012 | 0.009 | 0.009 | 0.313 | 0.18 | 0.15 | 0.14 | 0.11 | 0.12 | 0.18 |
| 4 | 78997515 | rs4241622 | 0.35 | 0.687 | 1.687 | 0.609 | 0.016 | -0.06 | -0.06 | 0 | 0.82 | 0.76 | 0.78 | 0.96 | 0.79 | 0.88 |
| 4 | 79002597 | rs6843142 | 1.598 | 0.705 | 1.579 | 0.592 | 0.003 | -0.066 | -0.066 | 4.628 | 0.7 | 0.48 | 0.68 | 0.87 | 0.7 | 0.75 |
| 4 | 79023544 | rs7699864 | -0.031 | 0.756 | 1.535 | 1.203 | -0.003 | 0.145 | 0.145 | 0 | 0.92 | 0.87 | 0.96 | 0.66 | 0.94 | 0.87 |
| 4 | 79025681 | rs10518187 | -0.97 | 0.629 | 1.779 | 1.154 | -0.014 | 0.034 | 0.034 | 0.016 | 0.09 | 0.26 | 0.11 | 0.07 | 0.05 | 0.13 |
| 4 | 79033115 | rs13130161 | -1.629 | 0.757 | 1.82 | 1.084 | 0.004 | -0.041 | -0.041 | 0 | 0.23 | 0.39 | 0.31 | 0.12 | 0.26 | 0.22 |
| 4 | 79033428 | rs6847174 | -1.629 | 0.757 | 1.82 | 1.084 | 0.004 | -0.041 | -0.041 | 0.03 | 0.23 | 0.39 | 0.31 | 0.12 | 0.26 | 0.22 |
| 4 | 79048490 | rs6845871 | 2.346 | 0.747 | 1.621 | 1.025 | 0.016 | -0.032 | -0.032 | 0.182 | 0.59 | 0.25 | 0.6 | 0.51 | 0.62 | 0.55 |
| 4 | 79052851 | rs2078481 | -0.696 | 1.433 | 1.484 | 0.845 | -0.001 | -0.003 | -0.003 | 0 | 0.15 | 0.13 | 0.2 | 0.05 | 0.2 | 0.09 |
| 4 | 79056713 | rs1317018 | -0.691 | 1.465 | 1.393 | 0.862 | -0.005 | 0.006 | 0.006 | 0 | 0.15 | 0.12 | 0.2 | 0.05 | 0.2 | 0.09 |
| 4 | 79061464 | rs17002971 | -0.599 | 1.629 | 1.564 | 1.014 | 0.007 | -0.015 | -0.015 | 0 | 0.07 | 0.19 | 0.08 | 0 | 0.06 | 0.01 |
| 4 | 79081466 | rs9307352 | -1.663 | 1.683 | 1.465 | 1.005 | 0.027 | -0.071 | -0.071 | 0 | 0.19 | 0.32 | 0.22 | 0.06 | 0.25 | 0.1 |
| 4 | 79083490 | rs10518194 | -0.89 | 1.868 | 0.744 | 1.244 | 0.008 | -0.019 | -0.019 | 0 | 0.14 | 0.15 | 0.15 | 0.05 | 0.2 | 0.09 |
| 4 | 79083751 | rs9284618 | 0.679 | 1.542 | 0.444 | 1.703 | -0.003 | 0.13 | 0.13 | 0 | 0.92 | 0.88 | 0.96 | 0.66 | 0.94 | 0.87 |
| 4 | 79093399 | rs10518195 | -1.231 | 2.4 | 0.531 | 0.776 | 0.158 | -0.043 | -0.043 | 0 | 0.1 | 0.38 | 0.14 | 0.01 | 0.19 | 0.05 |
| 4 | 79095407 | rs2903455 | -1.095 | 1.979 | -0.334 | 1.475 | -0.024 | -0.073 | -0.073 | 0.084 | 0.17 | 0.08 | 0.11 | 0.36 | 0.19 | 0.16 |
| 4 | 79095777 | rs4309859 | -0.679 | 1.937 | -0.166 | 1.054 | 0.001 | 0.003 | 0.003 | 0.279 | 0.53 | 0.45 | 0.46 | 0.66 | 0.55 | 0.47 |
| 4 | 79099678 | rs7688981 | -0.887 | 2.363 | -0.473 | 1.334 | -0.083 | -0.081 | -0.081 | 3.052 | 0.17 | 0.07 | 0.17 | 0.37 | 0.35 | 0.14 |
| 4 | 79103687 | rs17003025 | -0.186 | 1.919 | 0.096 | 1.086 | -0.007 | -0.025 | -0.025 | 0 | 0.05 | 0.18 | 0.02 | 0 | 0.02 | 0.01 |
| 4 | 79104112 | rs17003028 | 0.676 | 1.735 | 0.504 | 0.894 | 0.009 | -0.038 | -0.038 | 0 | 0.3 | 0.37 | 0.3 | 0.15 | 0.33 | 0.33 |
| 4 | 79108589 | rs11098085 | 0.025 | 1.482 | 0.54 | 0.86 | -0.006 | 0.029 | 0.029 | 0 | 0.6 | 0.17 | 0.59 | 0.68 | 0.62 | 0.63 |
| 4 | 79118533 | rs1996389 | -1.308 | 1.437 | 0.613 | 0.956 | 0.051 | 0.044 | 0.044 | 0.08 | 0.55 | 0.68 | 0.53 | 0.68 | 0.7 | 0.48 |
| 4 | 79120256 | rs7695549 | -0.469 | 1.71 | 0.675 | 1.056 | -0.072 | -0.082 | -0.082 | 0 | 0.17 | 0.09 | 0.18 | 0.36 | 0.33 | 0.14 |
| 4 | 79122800 | rs17003043 | 0.623 | 1.821 | 0.84 | 1.074 | 0.011 | 0.003 | 0.003 | 0.019 | 0.3 | 0.35 | 0.34 | 0.31 | 0.33 | 0.34 |
| 4 | 79126005 | rs10518188 | -1.008 | 1.733 | 0.862 | 1.136 | 0.065 | 0.064 | 0.064 | 0.219 | 0.51 | 0.7 | 0.54 | 0.65 | 0.67 | 0.48 |
| 4 | 79136032 | rs2866993 | 0.722 | 1.749 | 0.57 | 1.333 | 0.101 | 0.08 | 0.08 | 0 | 0.78 | 0.52 | 0.79 | 0.69 | 0.65 | 0.87 |
| 4 | 79138857 | rs7686101 | 0.529 | 1.655 | 0.437 | 1.194 | 0.01 | 0.007 | 0.007 | 0.065 | 0.3 | 0.35 | 0.34 | 0.34 | 0.34 | 0.34 |
| 4 | 79148953 | rs11941469 | -0.723 | 1.299 | 0.429 | 0.935 | 0.047 | 0.039 | 0.039 | 0 | 0.49 | 0.36 | 0.64 | 0.64 | 0.65 | 0.45 |
| 4 | 79158128 | rs11725082 | -0.868 | 1.767 | 0.797 | 1.175 | 0.018 | 0.006 | 0.006 | 0 | 0.2 | 0.2 | 0.25 | 0.28 | 0.35 | 0.14 |
| 4 | 79163162 | rs1869646 | 1.006 | 1.341 | 1.111 | 1.066 | 0.09 | 0.063 | 0.063 | 0 | 0.44 | 0.15 | 0.28 | 0.32 | 0.31 | 0.51 |
| 4 | 79173335 | rs13117388 | 0.763 | 1.519 | 1.22 | 0.958 | -0.06 | -0.061 | -0.061 | 0 | 0.31 | 0.1 | 0.41 | 0.38 | 0.35 | 0.34 |

| | | | | | | | | | | | | | | | | |
|---|---|---|---|---|---|---|---|---|---|---|---|---|---|---|---|---|
| 4 | 79176094 | rs10010118 | -0.462 | 1.473 | 2.239 | 0.952 | 0.002 | 0.011 | 0.011 | 0 | 0.64 | 0.49 | 0.54 | 0.59 | 0.65 | 0.65 |
| 4 | 79177408 | rs12510741 | 0.734 | 1.398 | 2.387 | 0.847 | 0.059 | 0.016 | 0.016 | 0 | 0.47 | 0.28 | 0.32 | 0.59 | 0.33 | 0.59 |
| 4 | 79185399 | rs1840392 | 0.746 | 1.389 | 2.349 | 0.86 | -0.033 | -0.035 | -0.035 | 0 | 0.32 | 0.16 | 0.42 | 0.38 | 0.35 | 0.34 |
| 4 | 79198744 | rs13108102 | 0.746 | 1.389 | 2.349 | 0.86 | -0.033 | -0.035 | -0.035 | 0 | 0.32 | 0.16 | 0.42 | 0.38 | 0.35 | 0.34 |
| 4 | 79205747 | rs6815945 | 0.414 | 1.425 | 2.416 | 0.863 | 0.01 | 0.006 | 0.006 | 0 | 0.34 | 0.39 | 0.44 | 0.38 | 0.37 | 0.34 |
| 4 | 79209463 | rs13136648 | 0.837 | 1.468 | 1.978 | 0.86 | -0.11 | -0.091 | -0.091 | 1.334 | 0.31 | 0.07 | 0.41 | 0.38 | 0.37 | 0.34 |
| 4 | 79210546 | rs17456396 | -0.089 | 1.497 | 1.945 | 0.813 | -0.09 | 0.085 | 0.085 | 0 | 0.16 | 0.04 | 0.21 | 0 | 0.32 | 0.06 |
| 4 | 79212910 | rs1484344 | 0.712 | 1.381 | 2.022 | 0.901 | -0.032 | -0.024 | -0.024 | 0.006 | 0.32 | 0.17 | 0.42 | 0.38 | 0.37 | 0.34 |
| 4 | 79222347 | rs12511889 | 0.835 | 1.464 | 1.793 | 0.939 | -0.049 | -0.033 | -0.033 | 0 | 0.31 | 0.1 | 0.4 | 0.34 | 0.34 | 0.32 |
| 4 | 79229302 | rs345528 | -0.787 | 1.341 | 2.261 | 0.905 | 0.039 | 0.088 | 0.088 | 0.1 | 0.06 | 0.44 | 0.04 | 0.04 | 0 | 0.03 |
| 4 | 79229609 | rs12505887 | 0.641 | 1.416 | 2.212 | 0.905 | -0.012 | -0.024 | -0.024 | 0.172 | 0.33 | 0.22 | 0.4 | 0.38 | 0.34 | 0.35 |
| 4 | 79231933 | rs1385133 | -1.08 | 1.298 | 2.268 | 1.15 | 0.038 | 0.279 | 0.279 | 0.02 | 0.05 | 0.22 | 0.05 | 0.29 | 0.03 | 0.08 |
| 4 | 79236212 | rs345518 | -1.024 | 1.383 | 1.837 | 0.994 | 0.094 | 0.026 | 0.026 | 0 | 0.55 | 0.8 | 0.67 | 0.38 | 0.68 | 0.43 |
| 4 | 79238448 | rs345515 | -0.52 | 1.025 | 1.483 | 0.668 | -0.014 | 0.075 | 0.075 | 0 | 0.49 | 0.89 | 0.51 | 0.67 | 0.42 | 0.51 |
| 4 | 79240063 | rs345513 | 0.235 | 1.099 | 1.762 | 0.647 | -0.004 | -0.002 | -0.002 | 0 | 0.42 | 0.51 | 0.45 | 0.4 | 0.39 | 0.39 |
| 4 | 79245028 | rs11724700 | -0.789 | 1.637 | 1.572 | 0.618 | 0.091 | -0.101 | -0.101 | 0 | 0.17 | 0.29 | 0.24 | 0 | 0.32 | 0.06 |
| 4 | 79248325 | rs391275 | -0.949 | 0.569 | 0.609 | 0.955 | 0.048 | -0.084 | -0.084 | 0 | 0.64 | 0.92 | 0.72 | 0.56 | 0.71 | 0.49 |
| 4 | 79254058 | rs168282 | -1.108 | 0.303 | 1.115 | 0.557 | 0.011 | -0.012 | -0.012 | 0 | 0.64 | 0.69 | 0.68 | 0.47 | 0.73 | 0.51 |
| 4 | 79261323 | rs429977 | -1.602 | 1.157 | 1.842 | 0.297 | 0.075 | -0.012 | -0.012 | 0 | 0.11 | 0.41 | 0.09 | 0.06 | 0.11 | 0.06 |
| 4 | 79262696 | rs366463 | -1.602 | 1.158 | 1.844 | 0.296 | 0.075 | -0.012 | -0.012 | 0 | 0.11 | 0.41 | 0.09 | 0.06 | 0.11 | 0.06 |
| 4 | 79265536 | rs10015470 | -1.606 | 1.157 | 1.877 | 0.31 | 0.059 | -0.014 | -0.014 | 1.605 | 0.11 | 0.39 | 0.09 | 0.06 | 0.11 | 0.06 |
| 4 | 79268802 | rs368313 | -1.444 | 1.177 | 1.739 | 0.325 | 0.091 | 0.015 | 0.015 | 0 | 0.09 | 0.39 | 0.09 | 0.06 | 0.11 | 0.06 |
| 4 | 79276588 | rs2008485 | 0.265 | 1.069 | 1.711 | 0.236 | -0.037 | 0.081 | 0.081 | 0 | 0.4 | 0.22 | 0.56 | 0.2 | 0.55 | 0.37 |
| 4 | 79283643 | rs345545 | -0.454 | 1.032 | 1.658 | 0.243 | 0.004 | 0.003 | 0.003 | 0 | 0.47 | 0.59 | 0.65 | 0.26 | 0.66 | 0.43 |
| 4 | 79284428 | rs6833805 | 0.171 | 0.942 | 1.54 | 0.241 | -0.022 | 0.059 | 0.059 | 0 | 0.39 | 0.28 | 0.57 | 0.2 | 0.56 | 0.37 |
| 4 | 79284694 | rs6835769 | 0.297 | 0.94 | 1.55 | 0.24 | -0.08 | 0.108 | 0.108 | 0 | 0.38 | 0.13 | 0.56 | 0.2 | 0.56 | 0.37 |
| 4 | 79285958 | rs6837091 | -0.724 | 0.668 | 1.533 | 0.358 | -0.024 | 0.041 | 0.041 | 0 | 0.21 | 0.31 | 0.12 | 0.34 | 0.11 | 0.13 |
| 4 | 79287631 | rs11932185 | -0.962 | 0.755 | 1.458 | 0.398 | -0.001 | 0 | 0 | 0.037 | 0.59 | 0.43 | 0.67 | 0.54 | 0.66 | 0.5 |
| 4 | 79288238 | rs345542 | -2.003 | 0.792 | 1.074 | 0.416 | 0.038 | -0.125 | -0.125 | 0 | 0.69 | 0.97 | 0.77 | 0.6 | 0.77 | 0.56 |
| 4 | 79291535 | rs11098160 | -0.97 | 0.79 | 1.11 | 0.387 | -0.01 | 0.009 | 0.009 | 0 | 0.6 | 0.57 | 0.68 | 0.54 | 0.66 | 0.5 |
| 4 | 79295156 | rs7687731 | -1.2 | 0.896 | 1.342 | 0.706 | -0.029 | 0.089 | 0.089 | 0.082 | 0.21 | 0.54 | 0.14 | 0.34 | 0.13 | 0.13 |
| 4 | 79296723 | rs17003169 | 1.247 | 1.185 | 0.615 | 0.618 | 0.012 | -0.042 | -0.042 | 0 | 0.2 | 0.09 | 0.3 | 0.25 | 0.2 | 0.19 |
| 4 | 79300173 | rs923019 | -1.081 | 1.12 | 1.242 | 0.428 | 0.01 | -0.006 | -0.006 | 0 | 0.56 | 0.66 | 0.67 | 0.54 | 0.63 | 0.39 |
| 4 | 79300280 | rs923017 | -1.077 | 1.133 | 1.249 | 0.434 | 0.011 | -0.006 | -0.006 | 1.641 | 0.56 | 0.66 | 0.67 | 0.54 | 0.63 | 0.39 |
| 4 | 79300993 | rs12512164 | 1.287 | 1.536 | 1.408 | 0.421 | -0.012 | 0.006 | 0.006 | 0 | 0.23 | 0.12 | 0.35 | 0.22 | 0.27 | 0.18 |
| 4 | 79302857 | rs7679527 | -1.138 | 1.542 | 1.426 | 0.425 | 0.01 | -0.006 | -0.006 | 0 | 0.56 | 0.66 | 0.67 | 0.54 | 0.63 | 0.39 |
| 4 | 79303200 | rs1471748 | -0.898 | 1.541 | 1.591 | 0.423 | 0.012 | -0.003 | -0.003 | 0.478 | 0.33 | 0.44 | 0.31 | 0.32 | 0.36 | 0.2 |
| 4 | 79304226 | rs10033307 | -1.95 | 1.552 | 1.379 | 0.415 | 0.035 | -0.109 | -0.109 | 0.551 | 0.65 | 0.95 | 0.73 | 0.6 | 0.72 | 0.52 |
| 4 | 79304598 | rs17003174 | 1.108 | 1.663 | 1.664 | -0.225 | -0.016 | 0.005 | 0.005 | 0 | 0.23 | 0.11 | 0.35 | 0.22 | 0.27 | 0.18 |
| 4 | 79306297 | rs12644161 | 0.228 | 1.575 | 1.796 | -0.214 | 0.016 | -0.017 | -0.017 | 0 | 0.31 | 0.4 | 0.41 | 0.27 | 0.36 | 0.31 |
| 4 | 79307872 | rs869349 | -0.267 | 1.511 | 1.842 | -0.243 | -0.008 | 0.026 | 0.026 | 0 | 0.35 | 0.74 | 0.44 | 0.27 | 0.36 | 0.31 |
| 4 | 79319428 | rs12651417 | -0.079 | 1.524 | 1.86 | -0.236 | -0.008 | 0.016 | 0.016 | 0 | 0.33 | 0.7 | 0.43 | 0.27 | 0.36 | 0.31 |
| 4 | 79335148 | rs1484334 | 0.542 | 1.544 | 1.788 | -0.207 | -0.031 | -0.004 | -0.004 | 0 | 0.28 | 0.14 | 0.39 | 0.31 | 0.36 | 0.31 |
| 4 | 79339332 | rs1484332 | -0.24 | 1.698 | 1.963 | -0.345 | 0.071 | -0.084 | -0.084 | 0.111 | 0.14 | 0.32 | 0.18 | 0 | 0.23 | 0.09 |
| 4 | 79346759 | rs6825981 | -0.64 | 1.422 | 1.792 | -0.27 | -0.001 | -0.005 | -0.005 | 0.429 | 0.43 | 0.55 | 0.57 | 0.31 | 0.59 | 0.39 |
| 4 | 79350115 | rs2054722 | -1.327 | 1.224 | 1.734 | -0.153 | -0.035 | 0.015 | 0.015 | 0 | 0.62 | 0.6 | 0.71 | 0.51 | 0.75 | 0.51 |
| 4 | 79350769 | rs17427362 | 1.144 | 1.67 | 1.83 | -0.124 | -0.01 | 0.004 | 0.004 | 0 | 0.08 | 0.06 | 0.12 | 0 | 0.2 | 0.06 |
| 4 | 79359885 | rs2170899 | 0.78 | 1.579 | 1.749 | -0.113 | -0.076 | -0.029 | -0.029 | 0.349 | 0.27 | 0.1 | 0.44 | 0.31 | 0.36 | 0.32 |
| 4 | 79362209 | rs2035510 | 0.615 | 1.581 | 1.958 | -0.116 | -0.023 | -0.001 | -0.001 | 0 | 0.28 | 0.21 | 0.45 | 0.31 | 0.36 | 0.31 |
| 4 | 79372856 | rs10026662 | -1.454 | 1.551 | 2.058 | -0.095 | -0.003 | -0.001 | -0.001 | 0.917 | 0.59 | 0.62 | 0.71 | 0.53 | 0.73 | 0.48 |
| 4 | 79372930 | rs753752 | -1.306 | 1.558 | 2.009 | -0.091 | -0.023 | 0.005 | 0.005 | 0 | 0.58 | 0.51 | 0.7 | 0.53 | 0.73 | 0.48 |
| 4 | 79378809 | rs2627654 | -1.596 | 1.544 | 1.871 | -0.098 | 0.063 | -0.05 | -0.05 | 0.487 | 0.62 | 0.8 | 0.72 | 0.53 | 0.73 | 0.48 |
| 4 | 79385719 | rs6851427 | 0.505 | 1.506 | 1.914 | -0.085 | -0.036 | -0.007 | -0.007 | 0 | 0.28 | 0.14 | 0.44 | 0.32 | 0.36 | 0.32 |
| 4 | 79391988 | rs13114980 | 0.602 | 1.554 | 1.903 | -0.372 | -0.004 | 0.005 | 0.005 | 0 | 0.68 | 0.33 | 0.71 | 0.79 | 0.63 | 0.84 |
| 4 | 79398481 | rs1995060 | 0.511 | 1.567 | 1.861 | -0.366 | 0.026 | -0.039 | -0.039 | 0.307 | 0.69 | 0.48 | 0.72 | 0.79 | 0.63 | 0.84 |
| 4 | 79401434 | rs12509281 | 0.165 | 1.523 | 1.801 | -0.353 | 0.006 | -0.006 | -0.006 | 0 | 0.19 | 0.12 | 0.12 | 0.24 | 0.16 | 0.1 |
| 4 | 79402688 | rs17409828 | 0.263 | 1.597 | 1.62 | -0.369 | -0.015 | -0.033 | -0.033 | 0 | 0.19 | 0.04 | 0.12 | 0.21 | 0.17 | 0.1 |
| 4 | 79403747 | rs924149 | 1.341 | 1.467 | 1.594 | -0.332 | 0.07 | -0.054 | -0.054 | 0 | 0.38 | 0.12 | 0.28 | 0.52 | 0.27 | 0.52 |
| 4 | 79409610 | rs7686157 | -0.104 | 1.664 | 1.779 | -0.317 | 0.144 | -0.079 | -0.079 | 0 | 0.88 | 0.53 | 0.85 | 1 | 0.81 | 0.94 |
| 4 | 79409899 | rs1496604 | -0.055 | 1.665 | 1.654 | -0.317 | 0.123 | -0.08 | -0.08 | 0.129 | 0.88 | 0.48 | 0.85 | 1 | 0.81 | 0.94 |
| 4 | 79416629 | rs6825278 | 0.517 | 1.666 | 1.527 | -0.316 | 0.07 | -0.1 | -0.1 | 5.309 | 0.86 | 0.44 | 0.84 | 1 | 0.8 | 0.94 |
| 4 | 79420616 | rs4975132 | -0.138 | 1.776 | 1.075 | -0.188 | -0.008 | 0.004 | 0.004 | 0 | 0.65 | 0.63 | 0.45 | 0.68 | 0.48 | 0.64 |
| 4 | 79421011 | rs11933630 | 1.093 | 1.911 | 1.023 | -0.171 | 0.001 | -0.008 | -0.008 | 0 | 0.08 | 0.07 | 0.11 | 0 | 0.18 | 0.05 |
| 4 | 79421698 | rs1496585 | 1.2 | 1.734 | 1.294 | -0.217 | -0.03 | 0.018 | 0.018 | 0 | 0.39 | 0.45 | 0.29 | 0.47 | 0.27 | 0.52 |
| 4 | 79422054 | rs4975135 | -0.042 | 1.736 | 1.32 | -0.23 | -0.001 | -0.001 | -0.001 | 0 | 0.23 | 0.17 | 0.14 | 0.22 | 0.2 | 0.12 |
| 4 | 79422073 | rs2867447 | 1.229 | 1.734 | 1.32 | -0.23 | -0.03 | 0.019 | 0.019 | 0 | 0.39 | 0.45 | 0.29 | 0.46 | 0.27 | 0.52 |
| 4 | 79431352 | rs11940606 | -0.681 | 1.736 | 1.324 | -0.23 | -0.011 | 0.007 | 0.007 | 0 | 0.3 | 0.25 | 0.25 | 0.23 | 0.39 | 0.17 |
| 4 | 79434343 | rs3792608 | 1.216 | 1.627 | 1.405 | -0.248 | -0.001 | -0.001 | -0.001 | 0 | 0.39 | 0.38 | 0.29 | 0.46 | 0.27 | 0.52 |
| 4 | 79439935 | rs6844368 | 0.697 | 1.649 | 1.16 | -0.237 | 0 | -0.001 | -0.001 | 2.193 | 0.47 | 0.51 | 0.4 | 0.46 | 0.45 | 0.57 |

| | | | | | | | | | | | | | | | | |
|---|---|---|---|---|---|---|---|---|---|---|---|---|---|---|---|---|
| 4 | 79442003 | rs7669587 | 0.404 | 1.644 | 1.222 | -0.24 | 0.015 | -0.002 | -0.002 | 0 | 0.5 | 0.61 | 0.4 | 0.46 | 0.45 | 0.57 |
| 4 | 79443837 | rs931605 | 0.81 | 2.337 | 1.675 | 0.442 | 0.001 | -0.008 | -0.008 | 0.003 | 0.08 | 0.07 | 0.1 | 0 | 0.18 | 0.05 |
| 4 | 79443850 | rs931606 | 0.694 | 2.294 | 1.623 | 0.419 | 0.013 | 0.013 | 0.013 | 0 | 0.58 | 0.5 | 0.45 | 0.46 | 0.55 | 0.59 |
| 4 | 79445470 | rs10028215 | 1.23 | 2.207 | 1.604 | 0.423 | 0.017 | -0.001 | -0.001 | 0.23 | 0.51 | 0.43 | 0.35 | 0.45 | 0.36 | 0.53 |
| 4 | 79450373 | rs4975137 | -0.293 | 2.376 | 1.414 | 0.352 | -0.002 | 0 | 0 | 0 | 0.72 | 0.69 | 0.54 | 0.68 | 0.66 | 0.69 |
| 4 | 79454315 | rs1496594 | 0.532 | 1.898 | 1.208 | 0.23 | -0.004 | 0.01 | 0.01 | 0 | 0.59 | 0.35 | 0.45 | 0.78 | 0.46 | 0.63 |
| 4 | 79455733 | rs4975139 | 0.537 | 1.901 | 1.079 | 0.232 | -0.008 | 0.032 | 0.032 | 0 | 0.58 | 0.31 | 0.44 | 0.78 | 0.46 | 0.63 |
| 4 | 79456303 | rs12639791 | -0.545 | 2.078 | 1.267 | 0.303 | 0.038 | 0.096 | 0.096 | 0 | 0.11 | 0.22 | 0.1 | 0.33 | 0.12 | 0.1 |
| 4 | 79458537 | rs6533756 | 0.079 | 2.153 | 1.305 | 0.174 | 0.015 | -0.029 | -0.029 | 0 | 0.65 | 0.59 | 0.51 | 0.83 | 0.5 | 0.74 |
| 4 | 79458598 | rs7689740 | 1.106 | 1.547 | 0.037 | 0.331 | 0.118 | -0.052 | -0.052 | 0 | 0.49 | 0.08 | 0.34 | 0.45 | 0.36 | 0.53 |
| 4 | 79464092 | rs17003321 | -0.695 | 0.648 | 0.749 | 0.874 | 0.025 | 0.181 | 0.181 | 0 | 0.11 | 0.3 | 0.06 | 0.35 | 0.07 | 0.08 |
| 4 | 79464432 | rs3811747 | 0.31 | 0.754 | 0.165 | 0.387 | -0.001 | -0.023 | -0.023 | 3.277 | 0.7 | 0.56 | 0.56 | 0.81 | 0.69 | 0.69 |
| 4 | 79465559 | rs3749482 | 0.305 | 0.719 | 0.063 | 0.378 | -0.005 | -0.021 | -0.021 | 0 | 0.66 | 0.56 | 0.55 | 0.81 | 0.66 | 0.62 |
| 4 | 79485876 | rs931602 | 0.9 | -0.098 | -0.513 | 0.723 | 0.093 | -0.028 | -0.028 | 0 | 0.56 | 0.16 | 0.62 | 0.48 | 0.71 | 0.62 |
| 4 | 79487060 | rs6533780 | 0.337 | -0.191 | 0.295 | 0.33 | -0.043 | -0.081 | -0.081 | 0 | 0.71 | 0.52 | 0.69 | 0.87 | 0.8 | 0.75 |
| 4 | 79488270 | rs4975147 | 0.241 | -0.181 | 0.325 | 0.312 | -0.033 | -0.075 | -0.075 | 0 | 0.73 | 0.53 | 0.68 | 0.86 | 0.8 | 0.71 |
| 4 | 79494430 | rs5945 | -0.844 | -0.223 | 0.654 | 0.886 | -0.011 | 0.073 | 0.073 | 0 | 0.12 | 0.16 | 0.07 | 0.3 | 0.08 | 0.16 |
| 4 | 79505884 | rs7656476 | 1.164 | -0.127 | 0.309 | 0.238 | -0.002 | -0.042 | -0.042 | 0 | 0.7 | 0.31 | 0.53 | 0.87 | 0.72 | 0.69 |
| 4 | 79506864 | rs7679218 | 0.716 | -0.211 | 0.767 | 0.196 | -0.017 | 0.049 | 0.049 | 0 | 0.76 | 0.82 | 0.57 | 0.87 | 0.73 | 0.7 |
| 4 | 79513215 | rs2867461 | 1.414 | -0.103 | 1.304 | 0.267 | 0.003 | 0.011 | 0.011 | 0 | 0.65 | 0.47 | 0.5 | 0.57 | 0.67 | 0.57 |
| 4 | 79522598 | rs6844356 | -1.216 | -0.071 | 2.449 | 0.151 | -0.012 | -0.042 | -0.042 | 0 | 0.08 | 0.19 | 0.04 | 0 | 0.05 | 0.04 |
| 4 | 79524420 | rs6533821 | 0.944 | -0.452 | 2.74 | -0.086 | -0.003 | 0.001 | 0.001 | 0 | 0.78 | 0.78 | 0.59 | 0.88 | 0.74 | 0.72 |
| 4 | 79525404 | rs5942 | -1.441 | -0.627 | 3.013 | 0.341 | -0.017 | 0.191 | 0.191 | 0 | 0.09 | 0.26 | 0.1 | 0.3 | 0.04 | 0.16 |
| 4 | 79528957 | rs7661370 | -1.432 | -0.656 | 2.859 | 0.33 | 0.009 | -0.016 | -0.016 | 0 | 0.13 | 0.11 | 0.12 | 0.3 | 0.09 | 0.17 |
| 4 | 79529206 | rs10029214 | 2.595 | -0.769 | 3.14 | 0.295 | 0.013 | -0.011 | -0.011 | 1.28 | 0.6 | 0.22 | 0.44 | 0.59 | 0.65 | 0.55 |
| 4 | 79531202 | rs5950 | -1.766 | -0.777 | 3.155 | 0.276 | 0.011 | -0.019 | -0.019 | 0.072 | 0.16 | 0.15 | 0.13 | 0.29 | 0.11 | 0.19 |
| 4 | 79531308 | rs4612 | 2.807 | -0.799 | 3.109 | 0.278 | 0.017 | 0.004 | 0.004 | 0 | 0.6 | 0.21 | 0.44 | 0.63 | 0.65 | 0.55 |
| 4 | 79537360 | rs6825328 | 2.66 | -0.795 | 3.151 | 0.28 | 0.005 | 0 | 0 | 0 | 0.62 | 0.3 | 0.45 | 0.63 | 0.66 | 0.55 |
| 4 | 79546144 | rs12643488 | 1.084 | -0.736 | 3.671 | 0.302 | 0.014 | -0.037 | -0.037 | 0 | 0.81 | 0.7 | 0.6 | 0.91 | 0.77 | 0.74 |
| 4 | 79552176 | rs7680766 | 1.334 | -0.737 | 3.699 | 0.301 | 0.03 | -0.079 | -0.079 | 0 | 0.79 | 0.52 | 0.59 | 0.91 | 0.77 | 0.74 |
| 4 | 79572289 | rs11736602 | 1.383 | -0.779 | 3.675 | 0.281 | 0.032 | -0.084 | -0.084 | 0.886 | 0.79 | 0.47 | 0.59 | 0.91 | 0.77 | 0.74 |
| 4 | 79584433 | rs11942573 | 1.392 | -0.838 | 3.596 | 0.172 | 0.027 | -0.089 | -0.089 | 0 | 0.77 | 0.48 | 0.58 | 0.91 | 0.76 | 0.73 |
| 4 | 79589045 | rs10007186 | 1.698 | -0.853 | 3.58 | 1.081 | 0.009 | 0.002 | 0.002 | 0.064 | 0.75 | 0.4 | 0.55 | 0.75 | 0.76 | 0.73 |
| 4 | 79602718 | rs1134124 | 2.478 | -0.844 | 3.292 | 1.154 | 0.012 | -0.023 | -0.023 | 0 | 0.59 | 0.21 | 0.41 | 0.46 | 0.64 | 0.54 |
| 4 | 79612009 | rs4619914 | 2.23 | -0.769 | 3.398 | 1.229 | -0.002 | 0 | 0 | 0 | 0.63 | 0.4 | 0.47 | 0.62 | 0.66 | 0.58 |
| 4 | 79616556 | rs1834519 | 1.99 | -0.82 | 3.134 | 1.181 | 0 | -0.002 | -0.002 | 2.108 | 0.64 | 0.39 | 0.45 | 0.46 | 0.66 | 0.58 |
| 4 | 79625240 | rs1346145 | 2.663 | -1.038 | 2.54 | 0.837 | 0.011 | -0.071 | -0.071 | 0 | 0.58 | 0.08 | 0.44 | 0.59 | 0.66 | 0.54 |
| 4 | 79629350 | rs7694971 | 2.372 | -1.311 | 2.773 | 0.626 | -0.011 | -0.007 | -0.007 | 0 | 0.58 | 0.48 | 0.58 | 0.65 | 0.66 | 0.58 |
| 4 | 79634522 | rs1966537 | 2.633 | -1.297 | 2.236 | 0.632 | 0.056 | 0.024 | 0.024 | 0 | 0.54 | 0.08 | 0.55 | 0.65 | 0.66 | 0.57 |
| 4 | 79645394 | rs6811782 | 1.627 | -1.081 | 2.398 | 0.46 | -0.001 | -0.03 | -0.03 | 0 | 0.65 | 0.38 | 0.6 | 0.9 | 0.69 | 0.66 |
| 4 | 79646371 | rs11724233 | 2.704 | -1.107 | 2.356 | 0.528 | 0.044 | 0.048 | 0.048 | 0 | 0.53 | 0.24 | 0.55 | 0.65 | 0.64 | 0.53 |
| 4 | 79655286 | rs12648219 | -1.579 | -1.505 | 2.541 | 0.583 | -0.075 | 0.126 | 0.126 | 0.015 | 0.15 | 0.45 | 0.07 | 0.25 | 0.05 | 0.13 |
| 4 | 79685513 | rs12644278 | -0.143 | -1.467 | 2.253 | 0.618 | -0.014 | 0.085 | 0.085 | 0 | 0.06 | 0.09 | 0.05 | 0.25 | 0.05 | 0.12 |
| 4 | 79689207 | rs13138117 | -1.914 | -1.495 | 2.429 | 0.52 | 0.013 | 0.015 | 0.015 | 0 | 0.35 | 0.36 | 0.32 | 0.1 | 0.2 | 0.29 |
| 4 | 79700596 | rs1816184 | -3.431 | -1.431 | 2.443 | 0.583 | 0.068 | 0.063 | 0.063 | 4.59 | 0.05 | 0.47 | 0.03 | 0 | 0 | 0.01 |
| 4 | 79701528 | rs1426138 | -0.947 | -1.463 | 1.882 | 0.564 | -0.021 | 0.037 | 0.037 | 0 | 0.2 | 0.03 | 0.15 | 0.09 | 0.15 | 0.28 |
| 4 | 79720870 | rs3775518 | 0.144 | -1.358 | 2.016 | 0.664 | -0.003 | 0.041 | 0.041 | 0 | 0.06 | 0.04 | 0.04 | 0.24 | 0.04 | 0.11 |
| 4 | 79800080 | rs10518205 | -0.157 | -1.356 | 2.025 | 0.681 | -0.018 | 0.103 | 0.103 | 0 | 0.06 | 0.11 | 0.05 | 0.23 | 0.04 | 0.11 |
| 4 | 79813622 | rs6830661 | -1.355 | -1.361 | 2.124 | 0.678 | -0.004 | 0.247 | 0.247 | 0 | 0.09 | 0.38 | 0.06 | 0.23 | 0.04 | 0.11 |
| 4 | 79824233 | rs17003488 | -0.353 | -1.318 | 2.051 | 0.7 | -0.02 | 0.103 | 0.103 | 0 | 0.07 | 0.12 | 0.05 | 0.24 | 0.04 | 0.11 |
| 4 | 79832509 | rs12649750 | -0.312 | -1.325 | 2.011 | 0.711 | -0.023 | 0.097 | 0.097 | 0 | 0.07 | 0.11 | 0.05 | 0.24 | 0.04 | 0.11 |
| 4 | 79834326 | rs17008951 | -0.278 | -1.304 | 0.951 | 0.721 | -0.023 | 0.096 | 0.096 | 0 | 0.07 | 0.11 | 0.05 | 0.22 | 0.04 | 0.11 |
| 4 | 79840498 | rs3811766 | -0.16 | -1.304 | 0.952 | 0.717 | -0.02 | 0.079 | 0.079 | 0 | 0.06 | 0.09 | 0.04 | 0.24 | 0.04 | 0.11 |
| 4 | 79840956 | rs1564458 | -0.258 | -1.303 | 0.938 | 0.717 | -0.023 | 0.097 | 0.097 | 0 | 0.07 | 0.11 | 0.04 | 0.24 | 0.04 | 0.11 |
| 4 | 79845007 | rs13108247 | 1.624 | -1.342 | 0.833 | 0.662 | -0.002 | 0 | 0 | 0 | 0.72 | 0.73 | 0.7 | 0.66 | 0.83 | 0.63 |
| 4 | 79862591 | rs7664339 | 0.638 | -1.284 | 0.131 | 0.767 | 0.011 | 0.017 | 0.017 | 0 | 0.8 | 0.86 | 0.75 | 0.88 | 0.89 | 0.74 |
| 4 | 79875172 | rs9993426 | 1.66 | -1.107 | -0.257 | 0.77 | 0.11 | 0.008 | 0.008 | 0 | 0.56 | 0.15 | 0.62 | 0.62 | 0.77 | 0.59 |
| 4 | 79883086 | rs10015978 | 0.587 | -1.257 | -0.324 | 0.583 | 0.019 | -0.019 | -0.019 | 0 | 0.72 | 0.8 | 0.67 | 0.64 | 0.83 | 0.61 |
| 4 | 79887043 | rs13150370 | 0.854 | -1.373 | -1.99 | -0.366 | -0.102 | -0.133 | -0.133 | 0 | 0.39 | 0.03 | 0.46 | 0.56 | 0.56 | 0.37 |
| 4 | 79892811 | rs1480713 | 0.342 | -1.449 | -1.457 | -0.596 | -0.006 | -0.036 | -0.036 | 0 | 0.07 | 0.16 | 0.04 | 0 | 0.04 | 0.05 |
| 4 | 79904876 | rs4975154 | 1.128 | -1.844 | -1.725 | -0.908 | 0.093 | 0.151 | 0.151 | 1.697 | 0.67 | 0.1 | 0.83 | 0.96 | 0.8 | 0.9 |
| 4 | 79907023 | rs2046105 | 1.215 | -1.828 | -1.533 | -0.971 | 0.067 | 0.112 | 0.112 | 2.055 | 0.66 | 0.14 | 0.82 | 0.96 | 0.78 | 0.89 |
| 4 | 79913463 | rs11728698 | 1.691 | -1.344 | -1.687 | -0.952 | -0.161 | -0.146 | -0.146 | 0 | 0.38 | 0.03 | 0.43 | 0.56 | 0.5 | 0.38 |
| 4 | 79913491 | rs17003548 | 0.701 | -1.485 | -1.2 | -1.027 | -0.017 | -0.036 | -0.036 | 0.66 | 0.09 | 0.17 | 0.03 | 0 | 0.06 | 0.01 |
| 4 | 79914407 | rs1545805 | 1.812 | -1.592 | -1.113 | -1.093 | 0.021 | 0.018 | 0.018 | 2.013 | 0.75 | 0.18 | 0.84 | 0.96 | 0.85 | 0.91 |
| 4 | 79924121 | rs4306981 | 0.42 | -1.442 | -1.119 | -0.967 | -0.028 | -0.085 | -0.085 | 0 | 0.28 | 0.07 | 0.38 | 0.4 | 0.28 | 0.51 |
| 4 | 79927368 | rs1585920 | 0.575 | -1.667 | 0.197 | -1.523 | 0.007 | 0.004 | 0.004 | 0 | 0.92 | 0.71 | 0.93 | 0.96 | 0.95 | 0.96 |
| 4 | 79939309 | rs1480720 | -0.674 | -1.526 | -0.18 | -1.674 | -0.028 | -0.126 | -0.126 | 0 | 0.17 | 0.44 | 0.08 | 0 | 0.11 | 0.06 |
| 4 | 79947170 | rs286479 | -0.572 | -1.371 | -0.344 | -1.583 | -0.005 | -0.081 | -0.081 | 0 | 0.15 | 0.24 | 0.08 | 0 | 0.12 | 0.07 |

| | | | | | | | | | | | | | | | | |
|---|---|---|---|---|---|---|---|---|---|---|---|---|---|---|---|---|
| 4 | 79949599 | rs17003582 | -0.023 | -1.39 | -0.193 | -1.426 | -0.01 | -0.071 | -0.071 | 0 | 0.12 | 0.24 | 0.04 | 0 | 0.08 | 0.04 |
| 4 | 79951750 | rs7687803 | 1.034 | -0.537 | -0.925 | -1.259 | -0.054 | 0.037 | 0.037 | 0 | 0.38 | 0.13 | 0.39 | 0.28 | 0.39 | 0.32 |
| 4 | 79957848 | rs6852257 | 0.598 | -0.906 | -0.561 | -1.719 | -0.02 | -0.011 | -0.011 | 0 | 0.2 | 0.07 | 0.11 | 0.13 | 0.17 | 0.1 |
| 4 | 79967383 | rs286451 | 1.153 | -1.228 | -0.276 | -1.968 | 0.087 | 0.177 | 0.177 | 2.169 | 0.54 | 0.15 | 0.77 | 0.88 | 0.67 | 0.83 |
| 4 | 79970208 | rs10032909 | -1.184 | -1.142 | 0.173 | -2.518 | 0.051 | 0.047 | 0.047 | 7.718 | 0.05 | 0.51 | 0.04 | 0 | 0 | 0 |
| 4 | 79981840 | rs2622351 | 1.34 | -1.232 | -0.11 | -2.593 | 0.074 | 0.298 | 0.298 | 2.729 | 0.54 | 0.06 | 0.72 | 0.93 | 0.67 | 0.84 |
| 4 | 79983945 | rs2604828 | 1.132 | -1.187 | 0.094 | -2.506 | 0.029 | 0.009 | 0.009 | 2.369 | 0.75 | 0.22 | 0.87 | 1 | 0.83 | 0.93 |
| 4 | 79985443 | rs2127782 | 0.286 | -1.211 | 0.26 | -2.509 | -0.023 | -0.08 | -0.08 | 0.424 | 0.8 | 0.72 | 0.91 | 1 | 0.84 | 0.93 |
| 4 | 79991221 | rs1455297 | -1.461 | -1.198 | 0.065 | -2.502 | 0.03 | 0.009 | 0.009 | 2.369 | 0.25 | 0.78 | 0.13 | 0 | 0.17 | 0.07 |
| 4 | 80008023 | rs10518213 | -0.875 | -1.12 | 0.219 | -2.465 | -0.044 | -0.12 | -0.12 | 0.499 | 0.21 | 0.38 | 0.1 | 0 | 0.17 | 0.07 |
| 4 | 80009984 | rs1154761 | -1.488 | -0.849 | -0.51 | -2.596 | -0.015 | 0.338 | 0.338 | 1.99 | 0.51 | 0.94 | 0.38 | 0.06 | 0.44 | 0.2 |
| 4 | 80010684 | rs1378376 | -1.29 | -0.825 | 0.269 | -2.492 | -0.007 | -0.15 | -0.15 | 0 | 0.2 | 0.57 | 0.21 | 0 | 0.21 | 0.09 |
| 4 | 80012823 | rs6844930 | -0.837 | -0.713 | 0.342 | -2.399 | 0.035 | -0.088 | -0.088 | 0 | 0.13 | 0.6 | 0.17 | 0 | 0.13 | 0.06 |
| 4 | 80033504 | rs4588482 | 1.059 | -0.72 | -0.206 | -2.416 | 0 | -0.022 | -0.022 | 1.709 | 0.85 | 0.17 | 0.81 | 1 | 0.86 | 0.94 |
| 4 | 80034047 | rs1378378 | 0.686 | -0.815 | -0.451 | -2.551 | -0.007 | 0.055 | 0.055 | 0 | 0.2 | 0.07 | 0.14 | 0.06 | 0.17 | 0.07 |
| 4 | 80049788 | rs10001729 | 0.039 | -0.408 | -0.06 | -2.265 | 0.022 | -0.051 | -0.051 | 0 | 0.91 | 0.81 | 0.86 | 1 | 0.86 | 0.94 |
| 4 | 80057589 | rs10020375 | 0.866 | -0.529 | -0.748 | -2.134 | -0.019 | 0.279 | 0.279 | 1.858 | 0.46 | 0.07 | 0.6 | 0.94 | 0.54 | 0.78 |
| 4 | 80059123 | rs12648730 | 0.685 | -0.574 | -0.622 | -2.048 | 0.033 | 0.262 | 0.262 | 1.504 | 0.48 | 0.1 | 0.6 | 0.94 | 0.54 | 0.81 |
| 4 | 80084425 | rs1078234 | -0.75 | -0.583 | -0.04 | -1.391 | -0.003 | -0.042 | -0.042 | 0 | 0.18 | 0.17 | 0.16 | 0 | 0.13 | 0.03 |
| 4 | 80085942 | rs10518214 | 1.501 | -1.33 | 0.113 | -1.979 | 0.046 | 0.031 | 0.031 | 5.67 | 0.66 | 0.27 | 0.77 | 1 | 0.83 | 0.88 |
| 4 | 80091507 | rs1455308 | 0.48 | -1.358 | 0.494 | -1.951 | -0.061 | -0.111 | -0.111 | 0 | 0.84 | 0.46 | 0.91 | 1 | 0.93 | 0.92 |
| 4 | 80108181 | rs1026394 | -1.411 | -1.532 | 0.366 | -1.814 | -0.002 | -0.009 | -0.009 | 0 | 0.31 | 0.29 | 0.22 | 0.11 | 0.16 | 0.14 |
| 4 | 80135060 | rs7667438 | 0.399 | -1.506 | 0.726 | -1.776 | -0.053 | -0.032 | -0.032 | 4.625 | 0.83 | 0.6 | 0.88 | 0.89 | 0.93 | 0.91 |
| 4 | 80146690 | rs4975172 | 1.27 | -1.537 | 0.366 | -1.83 | -0.019 | -0.039 | -0.039 | 1.483 | 0.64 | 0.41 | 0.75 | 0.89 | 0.83 | 0.86 |
| 4 | 80160465 | rs4571360 | 1.264 | -1.539 | 0.365 | -1.831 | -0.019 | -0.039 | -0.039 | 0 | 0.65 | 0.41 | 0.75 | 0.88 | 0.83 | 0.85 |
| 4 | 80179372 | rs11098609 | 1.324 | -1.037 | 0.394 | -1.854 | 0.008 | 0.003 | 0.003 | 3.299 | 0.64 | 0.37 | 0.73 | 0.88 | 0.8 | 0.85 |
| 4 | 80189105 | rs17003697 | -0.533 | -0.794 | 0.556 | -1.505 | -0.016 | 0.007 | 0.007 | 0 | 0.07 | 0.1 | 0.05 | 0.07 | 0.05 | 0.06 |
| 4 | 80202960 | rs6534293 | 0.229 | -0.124 | -0.251 | -1.836 | -0.072 | 0.025 | 0.025 | 0.896 | 0.7 | 0.89 | 0.59 | 0.26 | 0.54 | 0.54 |
| 4 | 80208634 | rs968256 | 0.204 | -0.355 | -0.355 | -2.075 | -0.059 | 0.062 | 0.062 | 0.863 | 0.65 | 0.85 | 0.56 | 0.25 | 0.52 | 0.53 |
| 4 | 80217155 | rs11930784 | 0.066 | -0.254 | -0.105 | -1.795 | -0.005 | 0.002 | 0.002 | 0 | 0.07 | 0.05 | 0.06 | 0.07 | 0.05 | 0.06 |
| 4 | 80226156 | rs4401473 | -0.379 | 0.119 | -0.107 | -2.039 | -0.047 | 0.05 | 0.05 | 0.243 | 0.34 | 0.18 | 0.48 | 0.76 | 0.48 | 0.53 |
| 4 | 80234648 | rs1484140 | -0.978 | 0.358 | 0.282 | -2.196 | -0.004 | 0.053 | 0.053 | 0 | 0.45 | 0.61 | 0.55 | 0.83 | 0.55 | 0.6 |
| 4 | 80238693 | rs1385056 | 0.526 | 0.3 | 0.371 | -2.171 | -0.008 | 0.025 | 0.025 | 0 | 0.07 | 0.17 | 0.05 | 0.07 | 0.04 | 0.06 |
| 4 | 80239092 | rs13106792 | -1.089 | 0.413 | -0.119 | -2.353 | 0.025 | 0.112 | 0.112 | 0.037 | 0.55 | 0.69 | 0.59 | 0.83 | 0.63 | 0.64 |
| 4 | 80243645 | rs6820325 | -0.881 | 0.166 | -0.498 | -2.013 | -0.004 | 0.027 | 0.027 | 0 | 0.44 | 0.33 | 0.49 | 0.79 | 0.49 | 0.53 |
| 4 | 80255768 | rs1545886 | 0.236 | -0.148 | -0.871 | -0.606 | 0.047 | -0.047 | -0.047 | 0 | 0.67 | 0.82 | 0.6 | 0.43 | 0.71 | 0.53 |
| 4 | 80262521 | rs11737437 | -1.38 | 0.046 | -0.595 | -1.169 | 0.004 | -0.056 | -0.056 | 0.049 | 0.2 | 0.65 | 0.27 | 0.03 | 0.21 | 0.12 |
| 4 | 80272792 | rs10004440 | -1.074 | 0.043 | 0.155 | -1.17 | 0.058 | -0.075 | -0.075 | 0 | 0.16 | 0.5 | 0.25 | 0.03 | 0.2 | 0.11 |
| 4 | 80309825 | rs10017376 | -0.246 | -0.531 | -0.494 | -1.106 | -0.01 | 0.008 | 0.008 | 0 | 0.13 | 0.16 | 0.08 | 0.19 | 0.12 | 0.1 |
| 4 | 80312214 | rs6855734 | 0.321 | -0.643 | -1.073 | -0.964 | 0.012 | 0.023 | 0.023 | 0.38 | 0.1 | 0.02 | 0.04 | 0 | 0.06 | 0.04 |
| 4 | 80313510 | rs17003742 | -0.46 | -0.935 | -1.1 | -1.083 | -0.004 | 0.002 | 0.002 | 0.323 | 0.2 | 0.19 | 0.24 | 0.04 | 0.25 | 0.19 |
| 4 | 80316079 | rs6534361 | -2.06 | -1.144 | -1.291 | -1.149 | 0.042 | 0.171 | 0.171 | 0 | 0.51 | 0.87 | 0.41 | 0.22 | 0.44 | 0.36 |
| 4 | 80317772 | rs17003749 | -0.646 | -0.758 | -0.839 | -0.978 | -0.011 | 0.026 | 0.026 | 0.173 | 0.2 | 0.16 | 0.24 | 0.04 | 0.25 | 0.18 |
| 4 | 80322959 | rs17003752 | -0.758 | -1.346 | -0.744 | -1.201 | 0.009 | -0.014 | -0.014 | 0.388 | 0.18 | 0.16 | 0.09 | 0 | 0.13 | 0.05 |
| 4 | 80326400 | rs10003904 | 0.149 | -1.275 | -0.771 | -1.158 | -0.003 | -0.02 | -0.02 | 0 | 0.08 | 0.12 | 0.05 | 0 | 0.06 | 0.01 |
| 4 | 80331240 | rs1922285 | -1.691 | -1.824 | 0.284 | -1.478 | -0.058 | -0.133 | -0.133 | 0 | 0.27 | 0.58 | 0.14 | 0 | 0.16 | 0.11 |
| 4 | 80332039 | rs10049840 | -1.992 | -1.526 | 0.161 | -1.373 | 0.048 | 0.062 | 0.062 | 0 | 0.4 | 0.72 | 0.22 | 0.16 | 0.29 | 0.21 |
| 4 | 80339467 | rs9993682 | 0.158 | -1.607 | 0.008 | -0.467 | -0.004 | -0.021 | -0.021 | 0 | 0.09 | 0.14 | 0.04 | 0 | 0.06 | 0.01 |
| 4 | 80352632 | rs3924829 | -1.355 | -2.314 | -0.693 | -0.308 | -0.021 | -0.013 | -0.013 | 0.059 | 0.65 | 0.78 | 0.42 | 0.6 | 0.45 | 0.75 |
| 4 | 80353066 | rs12645140 | 1.726 | -2.493 | -0.841 | -0.299 | -0.001 | -0.07 | -0.07 | 0.014 | 0.19 | 0.08 | 0.17 | 0.41 | 0.14 | 0.48 |
| 4 | 80365765 | rs6838184 | 0.666 | -2.679 | -0.671 | -0.406 | 0.013 | -0.021 | -0.021 | 0.165 | 0.27 | 0.2 | 0.19 | 0.41 | 0.18 | 0.55 |
| 4 | 80375367 | rs3920463 | -1.509 | -2.6 | -0.762 | -0.701 | -0.016 | -0.011 | -0.011 | 3.607 | 0.66 | 0.74 | 0.39 | 0.59 | 0.45 | 0.76 |
| 4 | 80381867 | rs12644205 | 1.821 | -2.102 | -1.281 | -0.736 | -0.02 | -0.014 | -0.014 | 0 | 0.09 | 0 | 0.23 | 0.07 | 0.1 | 0.1 |
| 4 | 80386263 | rs7655458 | 1.738 | -2.928 | -0.935 | -0.751 | 0.001 | -0.054 | -0.054 | 0.001 | 0.2 | 0.1 | 0.14 | 0.46 | 0.15 | 0.5 |
| 4 | 80387141 | rs10518219 | 0.415 | -2.56 | -0.592 | -0.63 | -0.01 | 0.002 | 0.002 | 0 | 0.41 | 0.38 | 0.23 | 0.46 | 0.25 | 0.6 |
| 4 | 80390999 | rs7687998 | -1.039 | -2.059 | -1.014 | -0.353 | -0.013 | -0.02 | -0.02 | 0 | 0.66 | 0.71 | 0.38 | 0.55 | 0.45 | 0.76 |
| 4 | 80393032 | rs10033347 | 0.042 | -1.365 | -0.877 | -0.358 | 0.012 | 0.014 | 0.014 | 0.394 | 0.1 | 0.11 | 0.09 | 0.03 | 0.07 | 0.07 |
| 4 | 80394763 | rs17443595 | 0.242 | -0.858 | -0.757 | -0.386 | 0.006 | -0.02 | -0.02 | 0.552 | 0.1 | 0.16 | 0.07 | 0.06 | 0.12 | 0.08 |
| 4 | 80394862 | rs7685006 | -0.514 | -1.388 | -1.293 | -0.311 | 0.026 | -0.017 | -0.017 | 0.027 | 0.53 | 0.71 | 0.31 | 0.51 | 0.4 | 0.72 |
| 4 | 80398708 | rs10027646 | 0.948 | -0.577 | -1.472 | -0.228 | -0.189 | -0.175 | -0.175 | 0 | 0.33 | 0.02 | 0.52 | 0.48 | 0.44 | 0.21 |
| 4 | 80412577 | rs1115518 | 0.364 | -0.48 | -0.533 | 0.105 | -0.012 | 0.027 | 0.027 | 0 | 0.69 | 0.56 | 0.76 | 0.57 | 0.77 | 0.41 |
| 4 | 80416491 | rs7691918 | -0.223 | -0.455 | -0.641 | 0.099 | 0.008 | -0.035 | -0.035 | 0 | 0.76 | 0.81 | 0.82 | 0.57 | 0.82 | 0.43 |
| 4 | 80420140 | rs1561639 | 0.079 | -0.273 | -0.736 | 0.21 | 0.001 | -0.017 | -0.017 | 0 | 0.74 | 0.78 | 0.77 | 0.57 | 0.77 | 0.41 |
| 4 | 80425028 | rs11943366 | -0.379 | -0.165 | -0.412 | 0.101 | 0.019 | -0.027 | -0.027 | 0.489 | 0.07 | 0.28 | 0.06 | 0 | 0.05 | 0.02 |
| 4 | 80429770 | rs998437 | 0.529 | -0.413 | -0.593 | 0.137 | -0.005 | 0.016 | 0.016 | 0 | 0.7 | 0.63 | 0.76 | 0.57 | 0.77 | 0.41 |
| 4 | 80442759 | rs9994690 | 1.084 | -0.581 | -1.479 | -0.3 | -0.189 | -0.175 | -0.175 | 0.047 | 0.33 | 0.02 | 0.52 | 0.48 | 0.44 | 0.21 |
| 4 | 80450322 | rs2119421 | 0.568 | -0.766 | 0.166 | 0.398 | -0.015 | 0.041 | 0.041 | 0 | 0.69 | 0.55 | 0.75 | 0.57 | 0.76 | 0.41 |
| 4 | 80461045 | rs12507692 | 0.735 | -0.749 | 0.133 | 0.403 | -0.006 | 0.015 | 0.015 | 0 | 0.7 | 0.64 | 0.76 | 0.57 | 0.76 | 0.41 |
| 4 | 80482400 | rs13112577 | 0.111 | -0.553 | 0.237 | 0.466 | -0.007 | 0.046 | 0.046 | 0.007 | 0.75 | 0.67 | 0.81 | 0.56 | 0.81 | 0.4 |

| | | | | | | | | | | | | | | | | |
|---|---|---|---|---|---|---|---|---|---|---|---|---|---|---|---|---|
| 4 | 80501938 | rs6850999 | -0.531 | -0.527 | 0.024 | 0.411 | -0.009 | -0.057 | -0.057 | 0.199 | 0.8 | 0.9 | 0.82 | 0.56 | 0.81 | 0.4 |
| 4 | 80506647 | rs1440853 | -0.561 | -0.527 | 0.024 | 0.411 | -0.009 | -0.056 | -0.056 | 0 | 0.8 | 0.91 | 0.82 | 0.56 | 0.81 | 0.4 |
| 4 | 80513004 | rs12501981 | 0.479 | -0.547 | 0.198 | 0.374 | -0.015 | 0.073 | 0.073 | 0 | 0.69 | 0.5 | 0.75 | 0.52 | 0.75 | 0.38 |
| 4 | 80539001 | rs11098778 | 0.461 | -0.517 | -0.167 | 0.194 | -0.003 | -0.002 | -0.002 | 0 | 0.64 | 0.57 | 0.62 | 0.63 | 0.69 | 0.35 |
| 4 | 80543277 | rs1808870 | 1.072 | -0.395 | -0.453 | -0.369 | -0.001 | 0.006 | 0.006 | 0 | 0.45 | 0.58 | 0.29 | 0.11 | 0.5 | 0.21 |
| 4 | 80544474 | rs13120966 | 1.146 | -0.436 | 0.171 | -0.393 | -0.009 | 0.065 | 0.065 | 0 | 0.44 | 0.3 | 0.27 | 0.1 | 0.5 | 0.21 |
| 4 | 80545715 | rs1371990 | 0.841 | -0.41 | -0.228 | -0.378 | 0.005 | 0.076 | 0.076 | 0 | 0.49 | 0.65 | 0.3 | 0.1 | 0.5 | 0.21 |
| 4 | 80550505 | rs11098789 | 0.039 | -0.36 | -0.308 | -0.321 | -0.002 | -0.021 | -0.021 | 0 | 0.71 | 0.84 | 0.7 | 0.63 | 0.73 | 0.38 |
| 4 | 80552199 | rs3109166 | -1.007 | -0.46 | -0.535 | -0.378 | -0.031 | -0.067 | -0.067 | 0 | 0.81 | 0.96 | 0.77 | 0.66 | 0.79 | 0.41 |
| 4 | 80554217 | rs3111788 | 1.149 | -0.803 | -0.559 | -0.443 | -0.003 | 0.017 | 0.017 | 0 | 0.39 | 0.33 | 0.24 | 0.29 | 0.42 | 0.15 |
| 4 | 80558107 | rs6819600 | -0.294 | -0.877 | -0.201 | -0.521 | -0.014 | -0.108 | -0.108 | 0.006 | 0.15 | 0.39 | 0.14 | 0 | 0.12 | 0.06 |
| 4 | 80558341 | rs17004018 | 0.544 | -0.744 | -0.192 | -0.4 | 0.005 | -0.028 | -0.028 | 0 | 0.05 | 0.16 | 0.06 | 0 | 0.05 | 0.03 |
| 4 | 80561055 | rs3109164 | -0.551 | -0.735 | -0.667 | -0.349 | -0.017 | 0.075 | 0.075 | 0 | 0.45 | 0.28 | 0.62 | 0.71 | 0.47 | 0.79 |
| 4 | 80561681 | rs4490487 | 0.793 | -0.504 | -1.086 | -0.217 | -0.045 | -0.133 | -0.133 | 0.037 | 0.21 | 0.02 | 0.35 | 0.37 | 0.19 | 0.14 |
| 4 | 80564868 | rs10518222 | 0.739 | -0.419 | -1.01 | -0.243 | -0.022 | -0.063 | -0.063 | 0 | 0.23 | 0.1 | 0.36 | 0.37 | 0.22 | 0.14 |
| 4 | 80566060 | rs7696440 | -1.088 | -0.6 | -1.053 | -0.759 | 0.005 | 0.018 | 0.018 | 0 | 0.49 | 0.68 | 0.58 | 0.36 | 0.48 | 0.21 |
| 4 | 80570500 | rs10518221 | 1.381 | -0.476 | -0.59 | -0.755 | -0.011 | -0.054 | -0.054 | 0 | 0.13 | 0.16 | 0.09 | 0 | 0.14 | 0.01 |
| 4 | 80574242 | rs12503047 | -0.292 | -0.18 | -0.753 | -0.178 | -0.004 | -0.031 | -0.031 | 0 | 0.23 | 0.08 | 0.27 | 0.37 | 0.25 | 0.58 |
| 4 | 80575387 | rs7340867 | -0.491 | -0.013 | -0.563 | -0.226 | 0.01 | 0.003 | 0.003 | 0.231 | 0.95 | 0.68 | 0.98 | 0.99 | 1 | 0.96 |
| 4 | 80579291 | rs4975062 | -0.162 | 0.005 | -0.678 | -0.193 | -0.009 | -0.028 | -0.028 | 0 | 0.22 | 0.08 | 0.28 | 0.36 | 0.26 | 0.57 |
| 4 | 80588739 | rs4975105 | 0.068 | 0.107 | -0.711 | -0.131 | -0.003 | -0.01 | -0.01 | 0 | 0.67 | 0.58 | 0.62 | 0.79 | 0.7 | 0.74 |
| 4 | 80598713 | rs4975107 | -0.604 | 0.127 | -0.646 | -0.062 | 0.003 | 0.015 | 0.015 | 0.009 | 0.95 | 0.73 | 0.98 | 0.94 | 1 | 0.96 |
| 4 | 80598796 | rs13123790 | 1.389 | 0.117 | -0.838 | -0.047 | 0.001 | 0 | 0 | 0 | 0.45 | 0.5 | 0.34 | 0.43 | 0.44 | 0.18 |
| 4 | 80601457 | rs4286542 | -0.186 | 0.257 | -0.771 | 0.225 | 0.005 | 0.011 | 0.011 | 0 | 0.76 | 0.63 | 0.68 | 0.74 | 0.8 | 0.79 |
| 4 | 80602627 | rs17004052 | 0.567 | 0.571 | -1.651 | 0.362 | 0.005 | -0.029 | -0.029 | 0 | 0.15 | 0.06 | 0.23 | 0.31 | 0.17 | 0.52 |
| 4 | 80602865 | rs17447715 | 0.675 | 0.655 | -1.614 | 0.399 | -0.062 | -0.07 | -0.07 | 0 | 0.18 | 0.03 | 0.3 | 0.21 | 0.19 | 0.16 |
| 4 | 80605580 | rs17004060 | -0.411 | 0.497 | -1.037 | 0.421 | -0.015 | -0.029 | -0.029 | 0 | 0.2 | 0.07 | 0.26 | 0.31 | 0.24 | 0.53 |
| 4 | 80606999 | rs4975109 | 0.364 | 0.213 | -0.489 | 0.397 | 0 | -0.001 | -0.001 | 0.077 | 0.47 | 0.55 | 0.44 | 0.44 | 0.45 | 0.15 |
| 4 | 80607964 | rs10029533 | 0.901 | 0.094 | -0.578 | 0.206 | 0.004 | -0.001 | -0.001 | 0.73 | 0.11 | 0.07 | 0.08 | 0 | 0.11 | 0.06 |
| 4 | 80608250 | rs4580675 | -0.207 | 0.076 | -0.41 | 0.209 | -0.014 | -0.014 | -0.014 | 0 | 0.59 | 0.74 | 0.52 | 0.44 | 0.56 | 0.21 |
| 4 | 80609149 | rs4547818 | -0.944 | 0.029 | -0.498 | 0.333 | -0.007 | 0.006 | 0.006 | 0 | 0.74 | 0.79 | 0.81 | 0.81 | 0.67 | 0.67 |
| 4 | 80618658 | rs13127915 | 0.972 | -0.126 | -0.048 | 0.284 | 0.001 | 0 | 0 | 0.001 | 0.45 | 0.43 | 0.41 | 0.44 | 0.42 | 0.14 |
| 4 | 80620235 | rs6534511 | -0.149 | -0.139 | -0.336 | 0.256 | -0.014 | -0.059 | -0.059 | 0 | 0.84 | 0.66 | 0.89 | 0.93 | 0.85 | 0.86 |
| 4 | 80620555 | rs4464597 | -0.185 | 0.017 | -0.797 | 0.428 | -0.065 | -0.099 | -0.099 | 0 | 0.37 | 0.12 | 0.47 | 0.49 | 0.43 | 0.72 |
| 4 | 80621795 | rs11944230 | 0.88 | 0.236 | -0.653 | 0.259 | -0.068 | 0.048 | 0.048 | 0 | 0.22 | 0.06 | 0.18 | 0.12 | 0.31 | 0.26 |
| 4 | 80634958 | rs9991949 | 0.876 | 0.011 | -0.385 | 0.257 | 0.004 | 0.035 | 0.035 | 0 | 0.1 | 0.03 | 0.08 | 0 | 0.11 | 0.06 |
| 4 | 80635570 | rs4356930 | -0.039 | -0.091 | 0.123 | 0.595 | -0.018 | 0.08 | 0.08 | 0 | 0.8 | 0.72 | 0.68 | 0.56 | 0.84 | 0.46 |
| 4 | 80645990 | rs7678939 | -0.011 | -0.092 | 0.148 | 0.585 | -0.019 | 0.083 | 0.083 | 0 | 0.8 | 0.72 | 0.67 | 0.56 | 0.84 | 0.46 |
| 4 | 80649730 | rs7658299 | -0.011 | -0.092 | 0.148 | 0.585 | -0.019 | 0.083 | 0.083 | 0 | 0.8 | 0.72 | 0.68 | 0.56 | 0.84 | 0.46 |
| 4 | 80652842 | rs7657079 | 0.245 | -0.052 | 0.034 | 0.402 | -0.036 | 0.083 | 0.083 | 0 | 0.34 | 0.18 | 0.26 | 0.12 | 0.42 | 0.31 |
| 4 | 80656305 | rs7694021 | 0.752 | -0.172 | 0.021 | 0.447 | -0.01 | -0.103 | -0.103 | 1.767 | 0.14 | 0.01 | 0.29 | 0.37 | 0.12 | 0.46 |
| 4 | 80661930 | rs12510706 | -0.027 | -0.341 | 0.486 | 0.352 | 0.005 | -0.033 | -0.033 | 0 | 0.31 | 0.24 | 0.45 | 0.51 | 0.3 | 0.49 |
| 4 | 80663590 | rs3935850 | 1.2 | -0.27 | 0.564 | 0.33 | -0.001 | 0 | 0 | 0 | 0.3 | 0.33 | 0.25 | 0.29 | 0.24 | 0.1 |
| 4 | 80666077 | rs10000154 | -0.766 | -0.27 | 0.544 | 0.328 | -0.004 | 0 | 0 | 0 | 0.64 | 0.41 | 0.71 | 0.63 | 0.72 | 0.81 |
| 4 | 80667570 | rs4270608 | 1.136 | -0.254 | 0.457 | 0.307 | -0.004 | 0 | 0 | 0 | 0.31 | 0.37 | 0.25 | 0.29 | 0.24 | 0.11 |
| 4 | 80672520 | rs4447889 | 1.477 | -0.176 | 0.358 | 0.09 | -0.005 | 0.003 | 0.003 | 0 | 0.15 | 0.11 | 0.15 | 0.14 | 0.18 | 0.03 |
| 4 | 80675751 | rs13134009 | 0.55 | -0.095 | 0.389 | 0.005 | -0.074 | 0.116 | 0.116 | 0 | 0.33 | 0.07 | 0.26 | 0.12 | 0.42 | 0.32 |
| 4 | 80681293 | rs7665764 | 0.2 | -0.546 | 0.326 | -0.061 | -0.019 | 0.158 | 0.158 | 0 | 0.55 | 0.18 | 0.66 | 0.81 | 0.55 | 0.6 |
| 4 | 80688131 | rs4359935 | 0.209 | -0.594 | 0.688 | -0.082 | 0.001 | -0.022 | -0.022 | 0 | 0.37 | 0.41 | 0.3 | 0.11 | 0.43 | 0.31 |
| 4 | 80688257 | rs4285108 | 0.626 | -0.625 | 0.478 | -0.098 | -0.081 | 0.124 | 0.124 | 0 | 0.34 | 0.08 | 0.27 | 0.11 | 0.43 | 0.31 |
| 4 | 80688396 | rs6534560 | 0.066 | -0.551 | 0.556 | -0.054 | -0.008 | 0.085 | 0.085 | 0 | 0.57 | 0.28 | 0.67 | 0.81 | 0.55 | 0.62 |
| 4 | 80706259 | rs17004380 | 0.916 | -0.989 | 0.457 | 0.038 | -0.016 | 0.031 | 0.031 | 0.041 | 0.23 | 0.28 | 0.17 | 0.29 | 0.14 | 0.07 |
| 4 | 80719386 | rs6813693 | 0.577 | -0.963 | 0.259 | 0.117 | -0.03 | -0.03 | -0.03 | 0 | 0.18 | 0.05 | 0.14 | 0.28 | 0.18 | 0.54 |
| 4 | 80722214 | rs7670590 | -0.686 | -1.046 | 0.426 | 0.079 | -0.004 | -0.02 | -0.02 | 0.791 | 0.76 | 0.27 | 0.76 | 0.89 | 0.77 | 0.93 |
| 4 | 80732878 | rs4381994 | -0.787 | -1.045 | 0.53 | 0.08 | -0.009 | -0.059 | -0.059 | 0 | 0.77 | 0.39 | 0.77 | 0.89 | 0.77 | 0.93 |
| 4 | 80735714 | rs4632591 | -0.572 | -1.057 | 0.49 | 0.073 | -0.013 | -0.051 | -0.051 | 0.274 | 0.75 | 0.35 | 0.74 | 0.89 | 0.76 | 0.92 |
| 4 | 80740009 | rs10011959 | 0.815 | -1.081 | 0.102 | 0.09 | -0.097 | -0.039 | -0.039 | 0 | 0.35 | 0.05 | 0.45 | 0.31 | 0.45 | 0.34 |
| 4 | 80742791 | rs4690145 | 0.534 | -1.119 | 0.066 | 0.108 | -0.026 | 0.062 | 0.062 | 0 | 0.55 | 0.08 | 0.61 | 0.6 | 0.59 | 0.4 |
| 4 | 80743976 | rs11943513 | 1.894 | -1.17 | -0.1 | -0.018 | -0.099 | -0.094 | -0.094 | 0 | 0.22 | 0.02 | 0.36 | 0.31 | 0.32 | 0.27 |
| 4 | 80745103 | rs11098907 | 0.792 | -1.122 | 0.055 | 0.031 | -0.026 | -0.026 | -0.026 | 0.588 | 0.18 | 0.08 | 0.14 | 0.29 | 0.18 | 0.52 |
| 4 | 80749973 | rs4546189 | -1.099 | -1.081 | 0.418 | 0.009 | -0.004 | -0.056 | -0.056 | 2.578 | 0.78 | 0.64 | 0.79 | 0.89 | 0.76 | 0.92 |
| 4 | 80750928 | rs4441708 | 0.636 | -1.042 | 0.102 | 0.056 | -0.059 | -0.009 | -0.009 | 2.285 | 0.36 | 0.09 | 0.45 | 0.31 | 0.45 | 0.34 |
| 4 | 80751289 | rs13141201 | 0.652 | -1.062 | 0.096 | 0.05 | -0.06 | -0.01 | -0.01 | 0 | 0.35 | 0.09 | 0.45 | 0.31 | 0.45 | 0.34 |
| 4 | 80753792 | rs10031299 | 1.115 | -1.153 | 0.503 | 0.078 | -0.02 | 0.034 | 0.034 | 0.045 | 0.23 | 0.38 | 0.19 | 0.29 | 0.14 | 0.06 |
| 4 | 80758914 | rs7680943 | 0.884 | -1.155 | 0.576 | 0.076 | -0.037 | 0.054 | 0.054 | 0.201 | 0.25 | 0.46 | 0.19 | 0.29 | 0.14 | 0.06 |
| 4 | 80769739 | rs6855167 | -1.088 | -1.122 | 0.449 | -0.02 | 0 | -0.067 | -0.067 | 0 | 0.78 | 0.58 | 0.79 | 0.89 | 0.76 | 0.92 |
| 4 | 80770939 | rs10014970 | 0.696 | -1.031 | 0.098 | -0.01 | -0.08 | -0.023 | -0.023 | 0.984 | 0.35 | 0.08 | 0.45 | 0.31 | 0.45 | 0.33 |
| 4 | 80773461 | rs6857026 | -0.108 | -0.88 | 0.314 | 0.11 | -0.003 | -0.018 | -0.018 | 0.797 | 0.32 | 0.37 | 0.25 | 0.28 | 0.3 | 0.59 |
| 4 | 80778464 | rs4271959 | -0.013 | -0.869 | 0.473 | 0.158 | -0.006 | 0.008 | 0.008 | 0.205 | 0.6 | 0.51 | 0.61 | 0.6 | 0.63 | 0.87 |

| | | | | | | | | | | | | | | | | |
|---|---|---|---|---|---|---|---|---|---|---|---|---|---|---|---|---|
| 4 | 80788782 | rs4690123 | -1.088 | -0.854 | 0.225 | 0.201 | 0.005 | -0.011 | -0.011 | 0 | 0.78 | 0.79 | 0.82 | 0.7 | 0.86 | 0.94 |
| 4 | 80791037 | rs6534620 | 1.059 | -0.83 | 0.24 | -0.089 | -0.005 | -0.033 | -0.033 | 0 | 0.06 | 0.16 | 0.05 | 0 | 0.06 | 0.02 |
| 4 | 80793114 | rs11735728 | 0.042 | -0.902 | -0.291 | 0.105 | 0.013 | -0.022 | -0.022 | 0 | 0.7 | 0.25 | 0.71 | 0.65 | 0.7 | 0.9 |
| 4 | 80796202 | rs11733071 | 0.239 | -0.839 | -0.489 | -0.695 | -0.01 | -0.011 | -0.011 | 0 | 0.56 | 0.49 | 0.56 | 0.31 | 0.69 | 0.42 |
| 4 | 80799867 | rs10011562 | 0.312 | -0.846 | -0.493 | -0.698 | -0.006 | -0.007 | -0.007 | 0 | 0.55 | 0.48 | 0.56 | 0.31 | 0.69 | 0.4 |
| 4 | 80800327 | rs11727192 | -0.408 | -0.781 | -0.543 | -0.746 | -0.001 | 0.006 | 0.006 | 0 | 0.76 | 0.75 | 0.65 | 0.38 | 0.81 | 0.47 |
| 4 | 80801911 | rs10010217 | 0.038 | -0.761 | -0.609 | -0.865 | -0.002 | -0.057 | -0.057 | 0 | 0.26 | 0.61 | 0.22 | 0.05 | 0.28 | 0.08 |
| 4 | 80805430 | rs10020809 | -0.403 | -0.668 | -0.47 | -0.475 | -0.007 | 0.024 | 0.024 | 0 | 0.79 | 0.75 | 0.67 | 0.38 | 0.86 | 0.47 |
| 4 | 80809997 | rs6534625 | -0.639 | -0.236 | -0.38 | -1.015 | 0 | -0.08 | -0.08 | 0 | 0.43 | 0.49 | 0.29 | 0.06 | 0.43 | 0.15 |
| 4 | 80812960 | rs6847975 | -0.212 | -0.083 | -0.467 | -0.933 | 0 | -0.028 | -0.028 | 0 | 0.35 | 0.31 | 0.24 | 0.06 | 0.37 | 0.12 |
| 4 | 80816632 | rs6534636 | -0.821 | -0.24 | -0.243 | -0.932 | 0.011 | -0.028 | -0.028 | 0.208 | 0.13 | 0.38 | 0.09 | 0.06 | 0.09 | 0.05 |
| 4 | 80818417 | rs11939401 | -0.762 | -0.49 | -0.159 | -1.44 | 0.006 | -0.078 | -0.078 | 1.314 | 0.82 | 0.76 | 0.85 | 1 | 0.76 | 0.9 |
| 4 | 80821366 | rs4264776 | 0.51 | -1.13 | -0.632 | -1.432 | 0.006 | 0.011 | 0.011 | 0.456 | 0.52 | 0.56 | 0.48 | 0.32 | 0.57 | 0.42 |
| 4 | 80827062 | rs7747 | -1.11 | -0.846 | -0.516 | -1.519 | 0.008 | -0.075 | -0.075 | 0.051 | 0.82 | 0.77 | 0.86 | 1 | 0.76 | 0.9 |
| 4 | 80850805 | rs4234848 | -0.195 | -1.076 | -1.087 | -1.939 | -0.002 | -0.107 | -0.107 | 0.12 | 0.63 | 0.63 | 0.8 | 1 | 0.64 | 0.79 |
| 4 | 80857151 | rs6824433 | -0.8 | -1.1 | -0.979 | -1.828 | 0.012 | -0.065 | -0.065 | 0.519 | 0.76 | 0.71 | 0.82 | 0.96 | 0.7 | 0.84 |
| 4 | 80871932 | rs4554013 | 1.179 | -1.138 | -1.138 | -1.845 | 0.03 | 0.026 | 0.026 | 0.329 | 0.19 | 0.1 | 0.06 | 0 | 0.1 | 0.11 |
| 4 | 80872086 | rs4511947 | 1.179 | -1.138 | -1.138 | -1.845 | 0.03 | 0.026 | 0.026 | 0.329 | 0.19 | 0.1 | 0.06 | 0 | 0.1 | 0.11 |
| 4 | 80887556 | rs10033596 | 1.912 | -1.264 | -1.288 | -1.572 | -0.144 | -0.079 | -0.079 | 0 | 0.33 | 0.03 | 0.51 | 0.29 | 0.45 | 0.31 |
| 4 | 80887969 | rs11098964 | 1.043 | -1.268 | -1.152 | -1.608 | -0.036 | 0.212 | 0.212 | 0.024 | 0.44 | 0.16 | 0.67 | 0.95 | 0.5 | 0.68 |
| 4 | 80906131 | rs4444771 | -0.233 | -1.273 | -1.13 | -1.622 | -0.001 | -0.019 | -0.019 | 0 | 0.7 | 0.31 | 0.78 | 0.96 | 0.7 | 0.84 |
| 4 | 80927001 | rs12504282 | 1.022 | -1.227 | -1.172 | -1.584 | -0.036 | 0.206 | 0.206 | 0.024 | 0.44 | 0.17 | 0.67 | 0.93 | 0.5 | 0.68 |
| 4 | 80931982 | rs4690135 | 0.428 | -1.237 | -1.173 | -1.581 | 0.04 | 0.229 | 0.229 | 0.249 | 0.52 | 0.21 | 0.72 | 0.94 | 0.59 | 0.74 |
| 4 | 80933225 | rs7682733 | 1.064 | -1.208 | -1.222 | -1.518 | 0.009 | 0.015 | 0.015 | 0 | 0.1 | 0.07 | 0.15 | 0.64 | 0.05 | 0.37 |
| 4 | 80943688 | rs10000471 | 0.154 | -1.244 | -1.074 | -1.605 | -0.004 | -0.081 | -0.081 | 0 | 0.56 | 0.55 | 0.75 | 0.95 | 0.59 | 0.74 |
| 4 | 80946541 | rs4640621 | 0.34 | -0.913 | -0.944 | -1.417 | -0.011 | 0.007 | 0.007 | 0 | 0.63 | 0.36 | 0.74 | 0.94 | 0.6 | 0.78 |
| 4 | 80948670 | rs12645809 | 0.393 | -0.924 | -0.95 | -1.404 | 0.032 | -0.022 | -0.022 | 0 | 0.29 | 0.22 | 0.21 | 0.65 | 0.16 | 0.47 |
| 4 | 80949829 | rs4333130 | 0.789 | -0.092 | -0.332 | -1.434 | -0.011 | 0.007 | 0.007 | 0 | 0.63 | 0.35 | 0.74 | 0.94 | 0.6 | 0.78 |
| 4 | 80959926 | rs7667228 | -0.967 | 0.048 | -0.115 | -0.657 | 0.048 | -0.066 | -0.066 | 0 | 0.12 | 0.37 | 0.09 | 0.01 | 0.09 | 0.06 |
| 4 | 80968006 | rs13132967 | 0.106 | 0.299 | -1.015 | -0.76 | -0.04 | 0.034 | 0.034 | 0.212 | 0.07 | 0.01 | 0.05 | 0.01 | 0.09 | 0.06 |
| 4 | 80970213 | rs7668399 | -0.361 | 0.291 | -0.917 | -1.057 | -0.002 | 0 | 0 | 0 | 0.38 | 0.41 | 0.52 | 0.24 | 0.51 | 0.29 |
| 4 | 80978467 | rs4308327 | 0.121 | 0.52 | -0.993 | -0.922 | 0.016 | -0.006 | -0.006 | 1.187 | 0.3 | 0.35 | 0.52 | 0.24 | 0.48 | 0.26 |
| 4 | 80984670 | rs7668376 | -0.176 | 0.254 | -1.718 | -0.774 | -0.072 | 0.081 | 0.081 | 0.588 | 0.63 | 0.88 | 0.39 | 0.27 | 0.57 | 0.48 |
| 4 | 80989462 | rs17004812 | 0.103 | -0.031 | -1.058 | 0.098 | 0.005 | 0.058 | 0.058 | 0 | 0.05 | 0.11 | 0.06 | 0.22 | 0.06 | 0.18 |
| 4 | 80989630 | rs17509725 | -0.319 | -0.635 | -1.287 | -0.188 | 0.085 | -0.011 | -0.011 | 0 | 0.45 | 0.81 | 0.38 | 0.54 | 0.29 | 0.31 |
| 4 | 80993796 | rs13140055 | -0.175 | -0.226 | -0.474 | -0.554 | 0.002 | 0.001 | 0.001 | 0 | 0.54 | 0.43 | 0.25 | 0.21 | 0.37 | 0.58 |
| 4 | 81001940 | rs2867698 | -1.569 | -0.368 | -0.056 | -0.195 | -0.005 | -0.003 | -0.003 | 0 | 0.13 | 0.15 | 0.05 | 0.08 | 0.05 | 0.21 |
| 4 | 81008844 | rs7660651 | -2.014 | -0.127 | 0.642 | -0.304 | -0.005 | -0.074 | -0.074 | 0 | 0.15 | 0.48 | 0.09 | 0.04 | 0.1 | 0.11 |
| 4 | 81055721 | rs9307622 | -1.887 | -0.033 | 0.556 | -0.337 | 0.007 | -0.089 | -0.089 | 1.431 | 0.14 | 0.5 | 0.08 | 0.02 | 0.1 | 0.1 |
| 4 | 81057927 | rs17464115 | -1.173 | 0.224 | -0.557 | -0.142 | 0 | -0.012 | -0.012 | 0 | 0.09 | 0.09 | 0.04 | 0.02 | 0.09 | 0.1 |
| 4 | 81061233 | rs2867701 | -0.687 | -0.121 | -1.145 | -0.486 | 0.06 | 0.142 | 0.142 | 0.035 | 0.45 | 0.81 | 0.61 | 0.83 | 0.34 | 0.56 |
| 4 | 81061422 | rs12513014 | -0.137 | -0.758 | -1.221 | 0.205 | 0.014 | -0.051 | -0.051 | 0.67 | 0.16 | 0.1 | 0.41 | 0.62 | 0.14 | 0.29 |
| 4 | 81068101 | rs13129263 | -1.319 | -1.282 | -0.343 | 0.38 | -0.031 | 0.018 | 0.018 | 0 | 0.25 | 0.3 | 0.3 | 0.68 | 0.13 | 0.49 |
| 4 | 81069624 | rs13119154 | -1.533 | -1.305 | -0.332 | 0.386 | -0.023 | 0.008 | 0.008 | 0 | 0.28 | 0.32 | 0.31 | 0.68 | 0.15 | 0.5 |
| 4 | 81073276 | rs2867696 | -0.983 | -0.734 | -0.394 | -0.343 | -0.096 | 0.028 | 0.028 | 0.222 | 0.84 | 0.99 | 0.87 | 0.95 | 0.76 | 0.87 |
| 4 | 81073672 | rs13102102 | -1.69 | -0.962 | -0.125 | -0.303 | 0.084 | 0.076 | 0.076 | 0 | 0.43 | 0.82 | 0.44 | 0.74 | 0.28 | 0.53 |
| 4 | 81073695 | rs13102278 | 0.384 | -0.934 | -0.047 | -0.075 | 0.005 | 0.013 | 0.013 | 0 | 0.14 | 0.12 | 0.25 | 0.57 | 0.09 | 0.27 |
| 4 | 81080290 | rs12646498 | -0.482 | -1.066 | 0.125 | -0.23 | 0.006 | 0.008 | 0.008 | 0 | 0.17 | 0.11 | 0.08 | 0.11 | 0.12 | 0.07 |
| 4 | 81087073 | rs13138779 | 0.329 | -0.53 | 0.5 | 0.235 | -0.001 | 0.035 | 0.035 | 0 | 0.13 | 0.14 | 0.25 | 0.57 | 0.09 | 0.28 |
| 4 | 81103562 | rs3755912 | -1.582 | -0.547 | 0.904 | 0.125 | -0.011 | 0.009 | 0.009 | 0 | 0.18 | 0.39 | 0.13 | 0.17 | 0.13 | 0.22 |
| 4 | 81104548 | rs3822010 | -1.539 | -0.545 | 0.908 | 0.126 | -0.009 | 0.011 | 0.011 | 0 | 0.18 | 0.39 | 0.13 | 0.17 | 0.13 | 0.22 |
| 4 | 81105762 | rs3755913 | -0.601 | -0.498 | 0.927 | 0.13 | 0.021 | 0.011 | 0.011 | 0 | 0.12 | 0.26 | 0.08 | 0.1 | 0.11 | 0.05 |
| 4 | 81106182 | rs1996455 | 1.034 | -0.831 | 1.057 | -0.109 | -0.008 | 0.012 | 0.012 | 0 | 0.67 | 0.38 | 0.73 | 0.83 | 0.65 | 0.84 |
| 4 | 81107049 | rs17004832 | -0.711 | -0.987 | 0.864 | -0.141 | 0.014 | -0.003 | -0.003 | 0.003 | 0.14 | 0.29 | 0.08 | 0.1 | 0.11 | 0.05 |
| 4 | 81108264 | rs6834136 | -0.493 | -0.729 | 0.685 | -0.037 | -0.034 | 0.027 | 0.027 | 0 | 0.83 | 0.93 | 0.85 | 0.93 | 0.77 | 0.88 |
| 4 | 81109088 | rs17530504 | -0.241 | -0.763 | 0.662 | -0.007 | -0.006 | 0.008 | 0.008 | 0 | 0.18 | 0.13 | 0.11 | 0.13 | 0.19 | 0.12 |
| 4 | 81113195 | rs1458043 | -2.225 | -0.98 | 0.414 | -0.069 | 0.016 | 0.122 | 0.122 | 0 | 0.49 | 0.87 | 0.5 | 0.7 | 0.4 | 0.59 |
| 4 | 81126083 | rs2168813 | -1.061 | -0.895 | 0.895 | 0.119 | -0.01 | 0.006 | 0.006 | 0 | 0.32 | 0.36 | 0.36 | 0.63 | 0.27 | 0.41 |
| 4 | 81130714 | rs34319350 | 0.141 | -0.889 | 0.48 | 0.315 | 0.027 | -0.037 | -0.037 | 0.132 | 0.13 | 0.05 | 0.24 | 0.56 | 0.08 | 0.31 |
| 4 | 81134893 | rs12650744 | -0.219 | -0.8 | 0.363 | 1.044 | 0.006 | -0.045 | -0.045 | 0.028 | 0.11 | 0.01 | 0.08 | 0.23 | 0.08 | 0.13 |
| 4 | 81135131 | rs6854711 | -2.255 | -0.737 | 0.77 | 1.077 | 0.008 | -0.005 | -0.005 | 0 | 0.32 | 0.67 | 0.4 | 0.66 | 0.25 | 0.5 |
| 4 | 81140615 | rs1458027 | -0.375 | -0.577 | 0.698 | 1.316 | 0.001 | 0.076 | 0.076 | 0 | 0.05 | 0.06 | 0.19 | 0.33 | 0.03 | 0.16 |
| 4 | 81149169 | rs6824301 | -1.732 | -0.715 | 0.754 | 0.961 | 0.013 | -0.003 | -0.003 | 0 | 0.42 | 0.72 | 0.35 | 0.41 | 0.36 | 0.3 |
| 4 | 81149690 | rs17004839 | 0.249 | -0.765 | 1.114 | 0.97 | 0.012 | -0.007 | -0.007 | 0 | 0.07 | 0.02 | 0.13 | 0.3 | 0.04 | 0.14 |
| 4 | 81150109 | rs11934116 | -0.998 | -0.699 | 1.313 | 0.871 | -0.016 | -0.01 | -0.01 | 0 | 0.33 | 0.41 | 0.26 | 0.29 | 0.27 | 0.28 |
| 4 | 81151514 | rs11727345 | -1.473 | -0.651 | 0.718 | 0.85 | 0.016 | 0.029 | 0.029 | 0 | 0.42 | 0.72 | 0.35 | 0.34 | 0.36 | 0.3 |
| 4 | 81152247 | rs10518234 | -1.687 | -0.538 | 0.645 | 1.128 | 0.011 | -0.034 | -0.034 | 0.005 | 0.44 | 0.81 | 0.39 | 0.51 | 0.37 | 0.55 |
| 4 | 81152804 | rs17004844 | -0.568 | -0.608 | 0.201 | 1.136 | 0.01 | -0.01 | -0.01 | 0.202 | 0.1 | 0.05 | 0.16 | 0.29 | 0.06 | 0.15 |
| 4 | 81154975 | rs7682543 | -1.768 | -0.528 | 0.284 | 0.492 | 0.005 | 0.108 | 0.108 | 0 | 0.52 | 0.69 | 0.53 | 0.81 | 0.44 | 0.71 |

| | | | | | | | | | | | | | | | | |
|---|---|---|---|---|---|---|---|---|---|---|---|---|---|---|---|---|
| 4 | 81158221 | rs17004846 | -1.474 | -0.162 | 0.181 | 0.998 | -0.007 | -0.04 | -0.04 | 0 | 0.35 | 0.79 | 0.38 | 0.54 | 0.35 | 0.31 |
| 4 | 81158545 | rs13149993 | -0.617 | -0.195 | 0.862 | 1.463 | -0.011 | -0.021 | -0.021 | 0 | 0.24 | 0.22 | 0.31 | 0.38 | 0.3 | 0.23 |
| 4 | 81160847 | rs7664429 | -0.662 | -1.017 | 0.594 | 1.516 | -0.005 | 0.011 | 0.011 | 0 | 0.12 | 0.18 | 0.15 | 0.26 | 0.06 | 0.37 |
| 4 | 81164723 | rs1458038 | -0.347 | -1.033 | 0.517 | 1.547 | -0.022 | -0.101 | -0.101 | 0 | 0.24 | 0.04 | 0.27 | 0.39 | 0.27 | 0.25 |
| 4 | 81174237 | rs6534834 | -1.783 | -0.963 | 1.411 | 1.188 | 0.022 | 0.051 | 0.051 | 0.023 | 0.15 | 0.49 | 0.11 | 0.16 | 0.12 | 0.2 |
| 4 | 81179173 | rs10009490 | -0.672 | -0.953 | 1.377 | 1.142 | 0.025 | -0.062 | -0.062 | 0 | 0.11 | 0.25 | 0.08 | 0 | 0.11 | 0.16 |
| 4 | 81181388 | rs12054583 | -2.199 | -1.018 | 0.949 | 1.15 | 0.032 | 0.266 | 0.266 | 0.023 | 0.51 | 0.9 | 0.57 | 0.8 | 0.42 | 0.65 |
| 4 | 81183900 | rs1379706 | -1.497 | -0.645 | 1.471 | 1.422 | -0.012 | 0.035 | 0.035 | 0 | 0.16 | 0.47 | 0.11 | 0.15 | 0.11 | 0.05 |
| 4 | 81185974 | rs17004857 | 0.57 | -0.602 | 1.359 | 1.012 | -0.001 | -0.009 | -0.009 | 0 | 0.06 | 0.05 | 0.07 | 0 | 0.1 | 0.01 |
| 4 | 81187895 | rs1047245 | -1.224 | -0.575 | 1.305 | 1.05 | 0.002 | -0.045 | -0.045 | 0 | 0.15 | 0.44 | 0.11 | 0.07 | 0.11 | 0.05 |
| 4 | 81190135 | rs716678 | -1.174 | -0.594 | 1.057 | 1.07 | -0.002 | -0.043 | -0.043 | 0.93 | 0.15 | 0.39 | 0.11 | 0.07 | 0.11 | 0.05 |
| 4 | 81192363 | rs3796605 | 0.083 | -0.404 | -0.149 | 1.351 | 0.002 | 0.044 | 0.044 | 0 | 0.08 | 0.06 | 0.09 | 0.34 | 0.04 | 0.16 |
| 4 | 81200645 | rs6827834 | -2.15 | -0.428 | -0.581 | 1.07 | -0.005 | 0.197 | 0.197 | 0 | 0.49 | 0.95 | 0.52 | 0.77 | 0.4 | 0.58 |
| 4 | 81202382 | rs7683390 | -1.246 | 0.461 | 0.558 | 1.37 | 0.015 | 0.138 | 0.138 | 0 | 0.17 | 0.54 | 0.18 | 0.35 | 0.15 | 0.29 |
| 4 | 81207963 | rs3733336 | -0.922 | 0.488 | -0.035 | 0.824 | 0.009 | 0.003 | 0.003 | 0 | 0.26 | 0.3 | 0.35 | 0.25 | 0.35 | 0.21 |
| 4 | 81209842 | rs10006753 | -1.829 | -0.869 | 1.22 | 0.34 | 0.022 | 0.005 | 0.005 | 0 | 0.08 | 0.43 | 0.05 | 0.01 | 0.01 | 0.03 |
| 4 | 81223154 | rs12650677 | -0.039 | -2.223 | 0.939 | 0.011 | 0.021 | 0.005 | 0.005 | 0.775 | 0.57 | 0.61 | 0.7 | 0.65 | 0.73 | 0.62 |
| 4 | 81225455 | rs6837827 | 0.357 | -2.345 | 1.349 | -0.073 | -0.003 | 0.001 | 0.001 | 0.111 | 0.49 | 0.48 | 0.66 | 0.65 | 0.7 | 0.49 |
| 4 | 81227042 | rs747125 | -0.917 | -3.152 | 0.842 | -0.338 | -0.004 | -0.063 | -0.063 | 2.11 | 0.8 | 0.96 | 0.77 | 0.76 | 0.82 | 0.83 |
| 4 | 81227964 | rs7664124 | 1.079 | -3.163 | 0.866 | -0.344 | 0.007 | 0.005 | 0.005 | 0.494 | 0.52 | 0.55 | 0.67 | 0.65 | 0.7 | 0.49 |
| 4 | 81233244 | rs7698854 | -0.77 | -2.878 | 0.751 | -0.242 | 0.004 | -0.04 | -0.04 | 0.273 | 0.8 | 0.9 | 0.77 | 0.76 | 0.81 | 0.83 |
| 4 | 81253275 | rs10009812 | -0.679 | -3.063 | 1.41 | -0.381 | -0.08 | -0.054 | -0.054 | 0 | 0.27 | 0.57 | 0.12 | 0.12 | 0.1 | 0.33 |
| 4 | 81253388 | rs10033567 | -0.151 | -3.023 | 1.671 | -0.333 | -0.087 | -0.049 | -0.049 | 0.414 | 0.23 | 0.44 | 0.17 | 0.13 | 0.1 | 0.32 |
| 4 | 81254610 | rs1458045 | -0.756 | -2.932 | 1.415 | -0.286 | 0 | -0.067 | -0.067 | 0.208 | 0.79 | 0.97 | 0.85 | 0.77 | 0.82 | 0.85 |
| 4 | 81283916 | rs17004886 | -1.223 | -3.044 | 1.767 | -0.388 | 0.036 | 0.018 | 0.018 | 0 | 0.05 | 0.29 | 0.03 | 0 | 0.01 | 0 |
| 4 | 81307025 | rs17004889 | 0.399 | -3.14 | 1.24 | -0.379 | 0.005 | -0.062 | -0.062 | 0 | 0.22 | 0.03 | 0.15 | 0.24 | 0.18 | 0.16 |
| 4 | 81307865 | rs1874337 | 2.274 | -3.195 | 2.142 | -0.403 | 0.048 | 0.013 | 0.013 | 0.72 | 0.94 | 0.68 | 0.96 | 1 | 0.98 | 0.93 |
| 4 | 81310385 | rs10031438 | 1.358 | -3.381 | 1.501 | -0.413 | 0.086 | 0.033 | 0.033 | 0.233 | 0.61 | 0.19 | 0.69 | 0.73 | 0.77 | 0.64 |
| 4 | 81346212 | rs6821676 | 1.199 | -3.407 | 1.623 | -0.431 | 0.057 | 0.025 | 0.025 | 0.253 | 0.62 | 0.27 | 0.7 | 0.74 | 0.77 | 0.64 |
| 4 | 81385386 | rs7672761 | 0.5 | -3.504 | 1.854 | -0.476 | -0.011 | -0.003 | -0.003 | 0 | 0.18 | 0.21 | 0.06 | 0.12 | 0.08 | 0.19 |
| 4 | 81397872 | rs11721676 | -0.145 | -3.52 | 1.951 | -0.522 | 0.01 | -0.004 | -0.004 | 0 | 0.27 | 0.18 | 0.19 | 0.23 | 0.21 | 0.3 |
| 4 | 81402774 | rs1039053 | -1.859 | -3.47 | 1.601 | -0.529 | 0.101 | 0.033 | 0.033 | 0.011 | 0.5 | 0.7 | 0.36 | 0.37 | 0.31 | 0.53 |
| 4 | 81411451 | rs2132264 | 1.224 | -3.334 | 1.565 | -0.462 | -0.001 | -0.005 | -0.005 | 1.208 | 0.08 | 0.08 | 0.06 | 0 | 0.09 | 0.01 |
| 4 | 81413618 | rs11937407 | -2.326 | -3.55 | 1.733 | -0.557 | 0.035 | 0.007 | 0.007 | 0 | 0.45 | 0.65 | 0.24 | 0.36 | 0.25 | 0.51 |
| 4 | 81430314 | rs10857170 | -0.867 | -3.533 | 1.811 | -0.552 | -0.038 | -0.051 | -0.051 | 0.895 | 0.29 | 0.44 | 0.14 | 0.12 | 0.18 | 0.21 |
| 4 | 81441082 | rs7689698 | -0.105 | -3.502 | 1.772 | -0.512 | -0.01 | 0.054 | 0.054 | 0 | 0.1 | 0.13 | 0.07 | 0.23 | 0.04 | 0.2 |
| 4 | 81441407 | rs12645025 | 0.016 | -3.556 | 1.884 | -0.548 | 0.02 | 0.016 | 0.016 | 0 | 0.21 | 0.17 | 0.07 | 0.12 | 0.09 | 0.19 |
| 4 | 81446625 | rs11728439 | 1.118 | -3.548 | 1.983 | -0.556 | 0.023 | -0.046 | -0.046 | 3.853 | 0.08 | 0.27 | 0.07 | 0 | 0.09 | 0.01 |
| 4 | 81447179 | rs11933716 | -0.937 | -3.561 | 2.002 | -0.54 | -0.009 | 0.067 | 0.067 | 0.299 | 0.18 | 0.4 | 0.14 | 0.23 | 0.14 | 0.2 |
| 4 | 81448749 | rs989123 | -0.653 | -3.598 | 2.117 | -0.563 | -0.012 | -0.071 | -0.071 | 0 | 0.14 | 0.22 | 0.1 | 0 | 0.11 | 0.1 |
| 4 | 81449030 | rs13111397 | 0.616 | -3.592 | 2.048 | -0.556 | 0.005 | 0.005 | 0.005 | 0 | 0.18 | 0.17 | 0.06 | 0.12 | 0.08 | 0.19 |
| 4 | 81450152 | rs17004930 | -0.038 | -3.592 | 2.02 | -0.553 | -0.01 | 0.054 | 0.054 | 0 | 0.1 | 0.13 | 0.07 | 0.23 | 0.04 | 0.19 |
| 4 | 81453383 | rs12650756 | 0.586 | -3.485 | 1.927 | -0.502 | 0.028 | 0.014 | 0.014 | 0 | 0.18 | 0.12 | 0.06 | 0.12 | 0.08 | 0.19 |
| 4 | 81460931 | rs11940032 | -2.454 | -3.582 | 1.918 | -0.545 | 0.08 | 0.011 | 0.011 | 0 | 0.43 | 0.74 | 0.25 | 0.36 | 0.24 | 0.48 |
| 4 | 81461715 | rs11945984 | -1.117 | -3.617 | 1.931 | -0.539 | -0.014 | -0.036 | -0.036 | 0 | 0.07 | 0.16 | 0.04 | 0 | 0.02 | 0.09 |
| 4 | 81462163 | rs1531959 | 0.534 | -3.596 | 1.729 | -0.525 | -0.001 | 0.002 | 0.002 | 0 | 0.17 | 0.17 | 0.06 | 0.13 | 0.08 | 0.19 |
| 4 | 81488421 | rs7666589 | -1.827 | -3.608 | 1.711 | -0.493 | 0.016 | -0.004 | -0.004 | 1.699 | 0.37 | 0.68 | 0.19 | 0.37 | 0.16 | 0.47 |
| 4 | 81494123 | rs1601513 | -1.69 | -3.5 | 1.331 | -0.489 | 0 | -0.015 | -0.015 | 0 | 0.93 | 0.92 | 0.94 | 1 | 0.92 | 0.99 |
| 4 | 81529088 | rs13141796 | -0.476 | -3.614 | 1.284 | -0.479 | 0.025 | 0.02 | 0.02 | 0 | 0.23 | 0.16 | 0.09 | 0.12 | 0.11 | 0.24 |
| 4 | 81535393 | rs1385890 | 2.172 | -3.721 | 1.354 | -0.484 | 0.133 | 0.007 | 0.007 | 0 | 0.55 | 0.18 | 0.76 | 0.63 | 0.77 | 0.52 |
| 4 | 81543568 | rs1486025 | -0.447 | -3.678 | 1.283 | -0.491 | 0.025 | 0.02 | 0.02 | 0 | 0.23 | 0.16 | 0.09 | 0.12 | 0.11 | 0.24 |
| 4 | 81549972 | rs4693532 | 1.275 | -4.021 | 0.139 | -0.533 | 0.024 | 0.03 | 0.03 | 0 | 0.22 | 0.01 | 0.18 | 0.16 | 0.18 | 0.21 |
| 4 | 81552878 | rs13136169 | 2.141 | -3.304 | 1.099 | 0.036 | 0.151 | 0.021 | 0.021 | 0 | 0.52 | 0.18 | 0.75 | 0.62 | 0.75 | 0.49 |
| 4 | 81554785 | rs16998198 | -0.722 | -2.669 | 1.608 | 0.305 | -0.008 | 0.06 | 0.06 | 0 | 0.09 | 0.15 | 0.07 | 0.2 | 0.04 | 0.2 |
| 4 | 81573178 | rs3925214 | 2.141 | -2.778 | 2.066 | 0.244 | -0.016 | 0.006 | 0.006 | 0.684 | 0.76 | 0.32 | 0.75 | 0.77 | 0.85 | 0.71 |
| 4 | 81624639 | rs7668590 | -1.665 | -2.618 | 2.062 | 0.288 | -0.016 | 0.031 | 0.031 | 0.104 | 0.94 | 0.99 | 0.95 | 1 | 0.92 | 0.99 |
| 4 | 81633586 | rs432324 | 1.365 | -2.648 | 2.256 | 0.297 | -0.015 | -0.09 | -0.09 | 0 | 0.85 | 0.47 | 0.82 | 0.97 | 0.89 | 0.89 |
| 4 | 81648384 | rs11734172 | -2.498 | -2.944 | 2.213 | 0.435 | 0.037 | 0.002 | 0.002 | 0.015 | 0.42 | 0.68 | 0.3 | 0.35 | 0.25 | 0.46 |
| 4 | 81653826 | rs11936472 | -1.609 | -2.948 | 2.144 | 0.448 | -0.021 | 0.008 | 0.008 | 0 | 0.3 | 0.33 | 0.22 | 0.35 | 0.15 | 0.38 |
| 4 | 81667695 | rs1493177 | -2.778 | -2.955 | 2.006 | 0.431 | 0.136 | 0.015 | 0.015 | 0 | 0.45 | 0.83 | 0.32 | 0.36 | 0.25 | 0.47 |
| 4 | 81703379 | rs2903736 | -0.1 | -2.736 | 1.978 | 0.585 | -0.016 | -0.008 | -0.008 | 0 | 0.17 | 0.15 | 0.07 | 0.1 | 0.1 | 0.22 |
| 4 | 81708094 | rs17004990 | -0.819 | -2.705 | 1.844 | 0.626 | 0.004 | -0.014 | -0.014 | 0.393 | 0.1 | 0.05 | 0.06 | 0.2 | 0.04 | 0.13 |
| 4 | 81735968 | rs2047039 | -2.699 | -2.706 | 2.02 | 0.6 | 0.013 | 0.001 | 0.001 | 0 | 0.43 | 0.61 | 0.31 | 0.36 | 0.24 | 0.46 |
| 4 | 81759688 | rs10031019 | -0.029 | -2.629 | 1.656 | 0.46 | 0.006 | -0.044 | -0.044 | 0.008 | 0.07 | 0 | 0.02 | 0.13 | 0.03 | 0.1 |
| 4 | 81764113 | rs6849668 | -1.331 | -2.629 | 2.077 | 0.4 | -0.047 | -0.038 | -0.038 | 0 | 0.27 | 0.37 | 0.22 | 0.16 | 0.19 | 0.26 |
| 4 | 81773914 | rs10028057 | 2.678 | -2.238 | 2.124 | 1.067 | 0.083 | -0.069 | -0.069 | 0 | 0.59 | 0.04 | 0.72 | 0.65 | 0.81 | 0.58 |
| 4 | 81773932 | rs13131042 | -2.675 | -2.097 | 2.535 | 0.76 | 0.003 | 0.002 | 0.002 | 0 | 0.39 | 0.62 | 0.26 | 0.28 | 0.19 | 0.42 |
| 4 | 81776186 | rs13107344 | -1.385 | -2.102 | 2.699 | 0.752 | -0.024 | -0.029 | -0.029 | 0 | 0.24 | 0.25 | 0.11 | 0.1 | 0.12 | 0.2 |
| 4 | 81789570 | rs1493182 | -2.951 | -2.1 | 2.727 | 0.761 | 0.081 | 0.034 | 0.034 | 0 | 0.39 | 0.83 | 0.27 | 0.28 | 0.19 | 0.42 |

| | | | | | | | | | | | | | | | | |
|---|---|---|---|---|---|---|---|---|---|---|---|---|---|---|---|---|
| 4 | 81790211 | rs1027295 | -2.139 | -2.632 | 2.662 | 0.776 | -0.055 | -0.038 | -0.038 | 0 | 0.29 | 0.56 | 0.21 | 0.16 | 0.14 | 0.24 |
| 4 | 81790266 | rs1027294 | -1.013 | -2.471 | 2.246 | 0.828 | -0.006 | -0.008 | -0.008 | 0 | 0.2 | 0.16 | 0.1 | 0.1 | 0.12 | 0.19 |
| 4 | 81794590 | rs34620960 | -1.013 | -2.471 | 2.246 | 0.828 | -0.006 | -0.008 | -0.008 | 0 | 0.2 | 0.16 | 0.1 | 0.1 | 0.12 | 0.19 |
| 4 | 81801291 | rs10026644 | -2.846 | -2.547 | 2.513 | 1.081 | -0.068 | 0.079 | 0.079 | 0 | 0.21 | 0.51 | 0.19 | 0.28 | 0.09 | 0.24 |
| 4 | 81802673 | rs960168 | -2.456 | -2.464 | 2.572 | 1.13 | -0.053 | 0.152 | 0.152 | 0.073 | 0.16 | 0.51 | 0.16 | 0.28 | 0.06 | 0.22 |
| 4 | 81829660 | rs10007113 | -1.742 | -2.463 | 2.387 | 0.584 | -0.043 | -0.039 | -0.039 | 0.185 | 0.13 | 0.33 | 0.06 | 0.05 | 0.03 | 0.15 |
| 4 | 81842668 | rs17005004 | -0.492 | -2.489 | 1.403 | 0.599 | 0.045 | 0.041 | 0.041 | 0 | 0.17 | 0.04 | 0.07 | 0.11 | 0.1 | 0.15 |
| 4 | 81844526 | rs2055177 | -2.124 | -2.329 | 1.534 | 0.625 | 0.07 | 0.033 | 0.033 | 0 | 0.37 | 0.79 | 0.25 | 0.31 | 0.16 | 0.36 |
| 4 | 81849352 | rs1353388 | -2.038 | -2.313 | 1.518 | 0.613 | 0.022 | 0.013 | 0.013 | 0 | 0.36 | 0.7 | 0.25 | 0.31 | 0.16 | 0.36 |
| 4 | 81883741 | rs11099454 | 1.48 | -1.131 | 1.757 | 0.547 | 0.017 | -0.04 | -0.04 | 0.343 | 0.82 | 0.69 | 0.85 | 0.94 | 0.84 | 0.85 |
| 4 | 81884722 | rs1052325 | 1.547 | -1.141 | 1.76 | 0.542 | 0.018 | -0.046 | -0.046 | 0.165 | 0.81 | 0.64 | 0.85 | 0.94 | 0.84 | 0.85 |
| 4 | 81899545 | rs10011602 | -1.407 | -1.749 | 1.794 | 0.638 | -0.008 | -0.015 | -0.015 | 0 | 0.21 | 0.19 | 0.09 | 0.06 | 0.1 | 0.12 |
| 4 | 81906024 | rs7667446 | 2.282 | -2.301 | 1.9 | 0.918 | 0.108 | 0.008 | 0.008 | 0 | 0.59 | 0.16 | 0.68 | 0.67 | 0.8 | 0.54 |
| 4 | 81913993 | rs11099457 | 1.542 | -2.19 | 2.712 | 0.953 | -0.055 | 0.006 | 0.006 | 0.372 | 0.75 | 0.48 | 0.84 | 0.79 | 0.89 | 0.84 |
| 4 | 81927206 | rs4458448 | 0.994 | -2.113 | 2.655 | 0.93 | -0.034 | -0.063 | -0.063 | 0 | 0.79 | 0.59 | 0.88 | 0.94 | 0.88 | 0.87 |
| 4 | 81941456 | rs6814223 | -0.963 | -2.171 | 2.422 | 0.89 | 0.015 | 0.033 | 0.033 | 1.919 | 0.18 | 0.08 | 0.1 | 0.06 | 0.13 | 0.12 |
| 4 | 81943313 | rs17005022 | -1.472 | -2.177 | 2.662 | 0.945 | -0.017 | 0.002 | 0.002 | 2.683 | 0.21 | 0.28 | 0.16 | 0.2 | 0.14 | 0.17 |
| 4 | 81947080 | rs1353386 | 1.936 | -2.461 | 2.588 | 0.848 | -0.013 | -0.003 | -0.003 | 0 | 0.72 | 0.33 | 0.79 | 0.8 | 0.82 | 0.82 |
| 4 | 81947402 | rs6845445 | -0.849 | -2.423 | 2.627 | 0.876 | 0.001 | -0.002 | -0.002 | 0 | 0.18 | 0.14 | 0.11 | 0.06 | 0.12 | 0.12 |
| 4 | 81950655 | rs13138361 | 0.111 | -2.377 | 1.965 | 0.875 | 0.013 | -0.008 | -0.008 | 0.225 | 0.12 | 0.07 | 0.12 | 0.12 | 0.08 | 0.29 |
| 4 | 81954086 | rs10516738 | 0.221 | -2.343 | 1.958 | 0.889 | 0.01 | -0.011 | -0.011 | 0 | 0.11 | 0.07 | 0.12 | 0.13 | 0.07 | 0.28 |
| 4 | 81961697 | rs10002189 | 2.193 | -2.239 | 1.208 | 0.979 | -0.137 | -0.088 | -0.088 | 2.753 | 0.38 | 0.03 | 0.33 | 0.45 | 0.42 | 0.31 |
| 4 | 81963418 | rs1495634 | 0.265 | -1.726 | 2.108 | 0.874 | -0.034 | 0.029 | 0.029 | 0 | 0.86 | 0.63 | 0.85 | 0.87 | 0.92 | 0.72 |
| 4 | 81963960 | rs10019046 | 0.357 | -1.729 | 2.165 | 0.872 | -0.038 | 0.034 | 0.034 | 0 | 0.86 | 0.56 | 0.84 | 0.87 | 0.92 | 0.72 |
| 4 | 81975674 | rs1495637 | 1.482 | -1.585 | 1.049 | 0.894 | 0.112 | 0.086 | 0.086 | 0 | 0.62 | 0.11 | 0.7 | 0.82 | 0.78 | 0.59 |
| 4 | 81977690 | rs10007784 | 1.482 | -1.527 | 0.033 | 0.925 | 0.111 | 0.086 | 0.086 | 0 | 0.62 | 0.1 | 0.71 | 0.82 | 0.78 | 0.59 |
| 4 | 81981434 | rs4693810 | -0.995 | -1.037 | 0.655 | 1.079 | -0.018 | 0.013 | 0.013 | 0 | 0.34 | 0.25 | 0.34 | 0.26 | 0.44 | 0.34 |
| 4 | 81981592 | rs2685471 | 1.397 | -1.233 | 1.447 | 1.46 | 0.003 | 0.044 | 0.044 | 0 | 0.52 | 0.31 | 0.45 | 0.3 | 0.47 | 0.4 |
| 4 | 81984196 | rs2643925 | 2.019 | -2.111 | 2.978 | 1.742 | -0.009 | -0.014 | -0.014 | 0 | 0.73 | 0.37 | 0.75 | 0.86 | 0.8 | 0.72 |
| 4 | 81986784 | rs2643921 | 2.255 | -2.125 | 2.904 | 1.735 | 0.017 | 0.024 | 0.024 | 0 | 0.71 | 0.24 | 0.74 | 0.86 | 0.8 | 0.71 |
| 4 | 81996925 | rs2685462 | 2.187 | -2.151 | 2.943 | 1.969 | -0.026 | 0.018 | 0.018 | 0 | 0.67 | 0.43 | 0.66 | 0.55 | 0.8 | 0.53 |
| 4 | 82009707 | rs3796804 | -2.618 | -2.339 | 2.913 | 2.12 | -0.019 | 0.016 | 0.016 | 0.367 | 0.19 | 0.76 | 0.23 | 0.39 | 0.08 | 0.29 |
| 4 | 82012206 | rs2868132 | -2.745 | -2.33 | 3.617 | 2.177 | 0.066 | 0.451 | 0.451 | 0.287 | 0.06 | 0.41 | 0.1 | 0.33 | 0.01 | 0.16 |
| 4 | 82023516 | rs11723025 | -1.452 | -2.287 | 3.374 | 1.814 | -0.033 | -0.012 | -0.012 | 0.001 | 0.12 | 0.25 | 0.11 | 0.06 | 0.07 | 0.11 |
| 4 | 82029505 | rs17005056 | -1.794 | -2.328 | 3.254 | 1.649 | -0.031 | -0.073 | -0.073 | 0.127 | 0.18 | 0.33 | 0.13 | 0.05 | 0.09 | 0.12 |
| 4 | 82030410 | rs6821258 | -2.856 | -2.425 | 3.391 | 1.921 | -0.051 | 0.032 | 0.032 | 0 | 0.27 | 0.56 | 0.26 | 0.43 | 0.12 | 0.33 |
| 4 | 82035067 | rs7675843 | -2.812 | -2.421 | 3.28 | 1.965 | -0.05 | 0.041 | 0.041 | 0 | 0.27 | 0.48 | 0.26 | 0.44 | 0.12 | 0.33 |
| 4 | 82035993 | rs17005061 | -1.705 | -2.314 | 2.927 | 1.483 | -0.001 | -0.007 | -0.007 | 1.498 | 0.18 | 0.21 | 0.12 | 0.06 | 0.09 | 0.12 |
| 4 | 82062931 | rs6837293 | -2.259 | -2.724 | 1.68 | 1.926 | 0.156 | -0.107 | -0.107 | 0.051 | 0.4 | 0.94 | 0.32 | 0.57 | 0.18 | 0.39 |
| 4 | 82069449 | rs7688672 | -2.219 | -2.709 | 1.729 | 1.893 | 0.154 | -0.1 | -0.1 | 0 | 0.4 | 0.91 | 0.32 | 0.56 | 0.18 | 0.39 |
| 4 | 82083445 | rs11722928 | -2.03 | -2.74 | 2.316 | 1.873 | 0.001 | -0.004 | -0.004 | 0 | 0.37 | 0.59 | 0.29 | 0.56 | 0.18 | 0.41 |
| 4 | 82092005 | rs17005078 | -0.739 | -2.8 | 2.239 | 1.88 | 0.043 | 0.053 | 0.053 | 0 | 0.21 | 0.08 | 0.11 | 0.07 | 0.12 | 0.17 |
| 4 | 82092967 | rs9992933 | -2.402 | -2.83 | 2.332 | 1.972 | 0.095 | -0.031 | -0.031 | 0.978 | 0.39 | 0.76 | 0.3 | 0.56 | 0.19 | 0.4 |
| 4 | 82094069 | rs1972127 | -2.043 | -2.812 | 2.324 | 1.964 | -0.037 | 0.077 | 0.077 | 0 | 0.18 | 0.69 | 0.19 | 0.48 | 0.07 | 0.23 |
| 4 | 82097836 | rs10005912 | -1.961 | -2.815 | 2.495 | 1.96 | -0.055 | 0.148 | 0.148 | 0 | 0.18 | 0.62 | 0.19 | 0.48 | 0.07 | 0.23 |
| 4 | 82104348 | rs12330930 | -1.799 | -2.806 | 2.416 | 1.965 | -0.056 | 0.166 | 0.166 | 0 | 0.17 | 0.61 | 0.19 | 0.48 | 0.07 | 0.23 |
| 4 | 82114503 | rs3733553 | -2.503 | -2.537 | 2.142 | 1.847 | 0.117 | -0.125 | -0.125 | 0 | 0.41 | 0.92 | 0.34 | 0.48 | 0.24 | 0.4 |
| 4 | 82118352 | rs4149 | -0.538 | -2.37 | 2.272 | 1.736 | -0.017 | 0.061 | 0.061 | 0 | 0.09 | 0.13 | 0.1 | 0.27 | 0.05 | 0.15 |
| 4 | 82118996 | rs6849299 | -1.313 | -1.931 | 2.063 | 0.872 | -0.03 | -0.043 | -0.043 | 0.347 | 0.23 | 0.27 | 0.19 | 0.07 | 0.13 | 0.18 |
| 4 | 82120029 | rs7656323 | -1.108 | -1.925 | 1.504 | 0.892 | 0.008 | 0.01 | 0.01 | 0 | 0.22 | 0.17 | 0.19 | 0.07 | 0.18 | 0.18 |
| 4 | 82123118 | rs6850526 | -1.37 | -1.754 | 1.325 | 0.965 | -0.041 | -0.05 | -0.05 | 0 | 0.28 | 0.51 | 0.27 | 0.16 | 0.18 | 0.2 |
| 4 | 82133264 | rs989445 | -2.231 | -1.476 | 1.785 | 1.753 | -0.022 | 0.243 | 0.243 | 0 | 0.09 | 0.26 | 0.26 | 0.34 | 0.04 | 0.18 |
| 4 | 82134787 | rs710839 | -1.076 | -1.666 | 0.88 | 0.813 | -0.002 | -0.018 | -0.018 | 0 | 0.29 | 0.68 | 0.28 | 0.18 | 0.24 | 0.21 |
| 4 | 82135230 | rs710840 | -0.792 | -1.708 | 1.615 | 0.794 | 0.002 | -0.028 | -0.028 | 0 | 0.26 | 0.37 | 0.26 | 0.17 | 0.24 | 0.21 |
| 4 | 82149016 | rs17005096 | -1.206 | -2.165 | 1.307 | 0.509 | -0.005 | 0.002 | 0.002 | 0.143 | 0.19 | 0.16 | 0.32 | 0.3 | 0.12 | 0.19 |
| 4 | 82154710 | rs1712374 | -0.981 | -1.972 | 1.6 | 0.396 | 0 | -0.004 | -0.004 | 0.524 | 0.3 | 0.71 | 0.3 | 0.18 | 0.32 | 0.22 |
| 4 | 82166904 | rs10516652 | -1.266 | -2.038 | 1.402 | 0.427 | 0.062 | -0.041 | -0.041 | 0 | 0.21 | 0.1 | 0.31 | 0.3 | 0.09 | 0.2 |
| 4 | 82184234 | rs1878528 | -1.216 | -2.02 | 1.737 | 0.235 | -0.001 | 0 | 0 | 0 | 0.31 | 0.62 | 0.31 | 0.18 | 0.32 | 0.2 |
| 4 | 82184539 | rs7687899 | -0.981 | -2.093 | 1.718 | 0.188 | -0.001 | -0.016 | -0.016 | 0 | 0.06 | 0.2 | 0.03 | 0 | 0.02 | 0.01 |
| 4 | 82187030 | rs1901568 | -1.727 | -2.069 | 0.886 | 0.202 | -0.017 | 0.115 | 0.115 | 0.003 | 0.33 | 0.86 | 0.32 | 0.18 | 0.33 | 0.2 |
| 4 | 82192994 | rs6535239 | -1.523 | -2.054 | 0.918 | 0.205 | 0.006 | 0.085 | 0.085 | 0.013 | 0.32 | 0.78 | 0.31 | 0.18 | 0.27 | 0.21 |
| 4 | 82226349 | rs2034629 | -1.296 | -2.043 | 1.15 | 0.204 | 0.007 | -0.043 | -0.043 | 0 | 0.29 | 0.52 | 0.29 | 0.17 | 0.33 | 0.21 |
| 4 | 82241612 | rs7681976 | -2.799 | -2.145 | 0.976 | 0.266 | 0.053 | 0.015 | 0.015 | 0.213 | 0.5 | 0.88 | 0.62 | 0.47 | 0.41 | 0.39 |
| 4 | 82274298 | rs4277772 | -0.828 | -2.134 | 0.853 | 0.074 | 0.018 | -0.01 | -0.01 | 0.861 | 0.16 | 0.11 | 0.12 | 0.21 | 0.07 | 0.17 |
| 4 | 82276379 | rs10488933 | -0.791 | -2.131 | 0.855 | 0.075 | 0.017 | -0.01 | -0.01 | 0 | 0.16 | 0.11 | 0.12 | 0.21 | 0.07 | 0.17 |
| 4 | 82284055 | rs1452373 | -1.198 | -2.137 | 0.859 | 0.07 | -0.008 | -0.001 | -0.001 | 0 | 0.18 | 0.22 | 0.13 | 0.21 | 0.07 | 0.18 |
| 4 | 82287652 | rs13435795 | -1.459 | -2.126 | 0.856 | 0.076 | -0.057 | 0.003 | 0.003 | 0 | 0.2 | 0.34 | 0.14 | 0.21 | 0.07 | 0.18 |
| 4 | 82287755 | rs11723619 | -1.279 | -2.162 | 0.799 | 0.04 | -0.001 | -0.025 | -0.025 | 0 | 0.3 | 0.56 | 0.3 | 0.17 | 0.27 | 0.2 |
| 4 | 82290018 | rs6535244 | -2.251 | -2.098 | 0.387 | 0.042 | 0.05 | 0.052 | 0.052 | 0.424 | 0.5 | 0.97 | 0.45 | 0.39 | 0.35 | 0.39 |

| | | | | | | | | | | | | | | | | |
|---|---|---|---|---|---|---|---|---|---|---|---|---|---|---|---|---|
| 4 | 82315470 | rs17005123 | -0.645 | -2.124 | 0.645 | -0.099 | 0.001 | -0.025 | -0.025 | 0.785 | 0.09 | 0.33 | 0.04 | 0 | 0.02 | 0.01 |
| 4 | 82318524 | rs10028610 | -2.259 | -2.055 | -0.81 | -0.008 | 0.049 | 0.04 | 0.04 | 0.77 | 0.51 | 0.93 | 0.45 | 0.37 | 0.35 | 0.38 |
| 4 | 82324437 | rs17500100 | -2.3 | -2.065 | -0.825 | 0.009 | -0.004 | 0.02 | 0.02 | 0.519 | 0.51 | 0.97 | 0.46 | 0.38 | 0.41 | 0.39 |
| 4 | 82324877 | rs17005129 | -1.671 | -2.113 | -0.647 | -0.433 | 0.022 | 0.024 | 0.024 | 1.187 | 0.24 | 0.71 | 0.16 | 0.1 | 0.06 | 0.03 |
| 4 | 82325636 | rs17500149 | -0.538 | -1.46 | -0.601 | -0.216 | -0.028 | 0.016 | 0.016 | 0 | 0.21 | 0.11 | 0.25 | 0.19 | 0.25 | 0.19 |
| 4 | 82329840 | rs10488931 | -0.636 | -1.486 | -0.474 | -0.247 | -0.013 | -0.008 | -0.008 | 0 | 0.13 | 0.13 | 0.11 | 0.06 | 0.04 | 0.02 |
| 4 | 82330924 | rs10488930 | -0.396 | -0.831 | -0.794 | 0.043 | -0.061 | -0.053 | -0.053 | 0.084 | 0.15 | 0.04 | 0.27 | 0.26 | 0.27 | 0.18 |
| 4 | 82332999 | rs10488928 | -1.157 | -1.912 | -0.665 | -0.53 | 0.009 | 0.016 | 0.016 | 1.589 | 0.26 | 0.78 | 0.15 | 0.09 | 0.07 | 0.17 |
| 4 | 82338731 | rs4692955 | 0.623 | -0.718 | -1.169 | -1.01 | 0.018 | 0.181 | 0.181 | 0 | 0.51 | 0.21 | 0.65 | 0.84 | 0.58 | 0.65 |
| 4 | 82339079 | rs6836031 | -1.561 | -1.086 | -0.279 | -1.22 | -0.04 | -0.084 | -0.084 | 0 | 0.2 | 0.33 | 0.1 | 0.03 | 0.09 | 0.13 |
| 4 | 82339516 | rs6823416 | -1.559 | -0.878 | -0.536 | -1.206 | -0.001 | -0.023 | -0.023 | 0 | 0.13 | 0.12 | 0.08 | 0 | 0.1 | 0.11 |
| 4 | 82340952 | rs17005133 | -1.086 | -0.882 | -0.488 | -1.089 | -0.003 | -0.023 | -0.023 | 0 | 0.06 | 0.11 | 0.03 | 0 | 0.06 | 0.04 |
| 4 | 82347502 | rs1037405 | -1.041 | -0.962 | -0.928 | -0.976 | -0.016 | -0.029 | -0.029 | 0 | 0.17 | 0.06 | 0.1 | 0.2 | 0.16 | 0.31 |
| 4 | 82355593 | rs2159657 | -1.98 | -0.985 | -0.194 | -1.402 | 0 | -0.011 | -0.011 | 0 | 0.32 | 0.63 | 0.2 | 0.22 | 0.29 | 0.41 |
| 4 | 82364962 | rs10010385 | -1.091 | -0.939 | 0.51 | -1.331 | -0.001 | -0.079 | -0.079 | 1.005 | 0.15 | 0.57 | 0.1 | 0.02 | 0.12 | 0.1 |
| 4 | 82365496 | rs6839120 | -2.606 | -1.586 | 0.112 | -1.653 | 0.02 | 0.08 | 0.08 | 0 | 0.4 | 0.87 | 0.23 | 0.23 | 0.31 | 0.41 |
| 4 | 82375936 | rs17558056 | 0.202 | -1.68 | 0.899 | -1.73 | -0.005 | -0.013 | -0.013 | 0 | 0.07 | 0.05 | 0.03 | 0 | 0.03 | 0.05 |
| 4 | 82376621 | rs1444891 | -0.874 | -1.595 | 0.935 | -1.76 | 0.009 | -0.078 | -0.078 | 0.102 | 0.15 | 0.57 | 0.11 | 0.03 | 0.14 | 0.1 |
| 4 | 82377329 | rs2868215 | -0.881 | -1.558 | 0.948 | -1.765 | 0.017 | -0.079 | -0.079 | 0 | 0.15 | 0.56 | 0.11 | 0.03 | 0.15 | 0.1 |
| 4 | 82381313 | rs895086 | -1.436 | -1.627 | 0.842 | -1.79 | 0.005 | 0.008 | 0.008 | 0 | 0.23 | 0.2 | 0.12 | 0.18 | 0.19 | 0.3 |
| 4 | 82381627 | rs748436 | -2.432 | -1.597 | 0.581 | -1.699 | 0.014 | 0.075 | 0.075 | 0 | 0.39 | 0.79 | 0.24 | 0.22 | 0.34 | 0.4 |
| 4 | 82382526 | rs7670580 | -1.905 | -1.433 | 0.764 | -1.781 | 0.012 | 0.007 | 0.007 | 0 | 0.26 | 0.75 | 0.16 | 0.03 | 0.19 | 0.16 |
| 4 | 82393862 | rs13353696 | -0.749 | -1.483 | 1.239 | -1.722 | -0.032 | -0.052 | -0.052 | 3.629 | 0.1 | 0.15 | 0.06 | 0 | 0.03 | 0.05 |
| 4 | 82401332 | rs10015146 | -1.596 | -1.426 | 1.302 | -1.951 | -0.054 | -0.112 | -0.112 | 1.066 | 0.24 | 0.46 | 0.26 | 0.07 | 0.16 | 0.18 |
| 4 | 82402588 | rs17005142 | -2.026 | -1.281 | 1.376 | -2.213 | -0.003 | -0.066 | -0.066 | 0 | 0.37 | 0.57 | 0.35 | 0.07 | 0.36 | 0.32 |
| 4 | 82403942 | rs6819741 | -1.295 | -1.296 | 1.379 | -2.176 | -0.045 | -0.096 | -0.096 | 0 | 0.24 | 0.41 | 0.25 | 0.07 | 0.16 | 0.18 |
| 4 | 82404274 | rs17005144 | 0.184 | -1.162 | 0.752 | -2.106 | -0.003 | -0.023 | -0.023 | 0 | 0.13 | 0.2 | 0.19 | 0.07 | 0.12 | 0.12 |
| 4 | 82417143 | rs11939325 | -0.058 | -1.003 | 0.355 | -2.018 | 0.029 | -0.066 | -0.066 | 0 | 0.13 | 0.23 | 0.09 | 0.01 | 0.21 | 0.12 |
| 4 | 82422327 | rs2166181 | 1.729 | -1.25 | -0.455 | -1.965 | -0.029 | 0.112 | 0.112 | 0.136 | 0.45 | 0.25 | 0.55 | 0.74 | 0.48 | 0.45 |
| 4 | 82422433 | rs10018581 | -0.102 | -0.599 | -0.441 | -1.225 | -0.04 | -0.051 | -0.051 | 1.354 | 0.88 | 0.99 | 0.92 | 0.8 | 0.87 | 0.76 |
| 4 | 82426814 | rs11930241 | 0.042 | -0.911 | -0.083 | -1.314 | -0.009 | -0.033 | -0.033 | 0 | 0.18 | 0.28 | 0.34 | 0.08 | 0.17 | 0.15 |
| 4 | 82427419 | rs6840449 | -0.463 | -1.134 | -0.041 | -0.997 | -0.002 | -0.023 | -0.023 | 0 | 0.24 | 0.34 | 0.2 | 0.2 | 0.27 | 0.2 |
| 4 | 82429283 | rs13147840 | -0.039 | -1.122 | 0.128 | -1.002 | -0.014 | -0.054 | -0.054 | 0 | 0.2 | 0.46 | 0.35 | 0.08 | 0.18 | 0.16 |
| 4 | 82430252 | rs882043 | -1.569 | -1.139 | 0.079 | -1.011 | 0.005 | 0.088 | 0.088 | 0 | 0.43 | 0.85 | 0.54 | 0.27 | 0.43 | 0.35 |
| 4 | 82432524 | rs17005153 | -0.655 | -1.065 | 0.157 | -0.967 | -0.001 | -0.029 | -0.029 | 9.607 | 0.25 | 0.41 | 0.19 | 0.2 | 0.26 | 0.2 |
| 4 | 82440730 | rs1011631 | -2.331 | -1.289 | -0.665 | -0.63 | -0.101 | -0.072 | -0.072 | 0.097 | 0.61 | 0.89 | 0.62 | 0.5 | 0.56 | 0.6 |
| 4 | 82441160 | rs7672307 | -1.682 | -1.29 | -0.498 | -0.587 | 0.041 | 0.036 | 0.036 | 0 | 0.44 | 0.75 | 0.29 | 0.42 | 0.39 | 0.45 |
| 4 | 82441375 | rs35142427 | -0.741 | -1.503 | -0.356 | -0.223 | 0.021 | 0.007 | 0.007 | 0 | 0.2 | 0.17 | 0.1 | 0.22 | 0.14 | 0.26 |
| 4 | 82448309 | rs6841069 | -1.049 | -1.388 | -0.259 | -0.253 | 0.024 | -0.008 | -0.008 | 0 | 0.25 | 0.78 | 0.19 | 0.31 | 0.19 | 0.3 |
| 4 | 82452878 | rs1348601 | 0.626 | -1.291 | -0.602 | -0.305 | -0.028 | 0.024 | 0.024 | 0 | 0.17 | 0.03 | 0.39 | 0.09 | 0.2 | 0.19 |
| 4 | 82452933 | rs10019003 | -0.192 | -0.915 | 0.022 | -0.044 | -0.013 | 0.023 | 0.023 | 0 | 0.19 | 0.23 | 0.15 | 0.31 | 0.18 | 0.29 |
| 4 | 82459940 | rs17005157 | -0.143 | -0.922 | 0.05 | -0.038 | -0.027 | 0.034 | 0.034 | 0 | 0.21 | 0.54 | 0.16 | 0.31 | 0.18 | 0.29 |
| 4 | 82473932 | rs1373637 | -0.856 | -0.295 | -1.88 | 0.219 | 0.013 | -0.042 | -0.042 | 1.887 | 0.53 | 0.9 | 0.62 | 0.51 | 0.46 | 0.57 |
| 4 | 82476030 | rs2018726 | 0.426 | 0.042 | -1.116 | 0.704 | -0.001 | -0.004 | -0.004 | 0 | 0.92 | 0.96 | 0.93 | 0.73 | 0.91 | 0.87 |
| 4 | 82482584 | rs9307785 | 0.201 | 0.239 | -1.291 | 0.285 | -0.025 | -0.018 | -0.018 | 0 | 0.32 | 0.22 | 0.23 | 0.37 | 0.39 | 0.34 |
| 4 | 82484170 | rs10022264 | -0.346 | -0.194 | -0.589 | -0.602 | -0.001 | -0.001 | -0.001 | 0 | 0.56 | 0.61 | 0.56 | 0.44 | 0.56 | 0.65 |
| 4 | 82492363 | rs10033009 | 0.201 | -0.425 | -0.006 | -0.708 | -0.009 | 0.033 | 0.033 | 0 | 0.45 | 0.31 | 0.32 | 0.37 | 0.51 | 0.48 |
| 4 | 82493067 | rs10033652 | -0.443 | -0.625 | -0.055 | -0.751 | -0.057 | 0.021 | 0.021 | 0 | 0.57 | 0.85 | 0.41 | 0.39 | 0.57 | 0.56 |
| 4 | 82506162 | rs4693298 | 0.435 | -0.64 | -0.047 | -0.905 | -0.04 | 0.006 | 0.006 | 0.015 | 0.38 | 0.16 | 0.58 | 0.65 | 0.41 | 0.43 |
| 4 | 82518513 | rs6812919 | 0.863 | -0.564 | 0.436 | -0.63 | -0.063 | -0.122 | -0.122 | 0 | 0.81 | 0.51 | 0.89 | 0.98 | 0.9 | 0.92 |
| 4 | 82519643 | rs6535261 | 0.144 | -0.602 | 0.495 | -0.587 | 0 | -0.002 | -0.002 | 0.344 | 0.55 | 0.56 | 0.35 | 0.37 | 0.55 | 0.56 |
| 4 | 82547364 | rs11933681 | 0.983 | -0.467 | 1.172 | -0.43 | -0.028 | 0.04 | 0.04 | 0.169 | 0.88 | 0.6 | 0.96 | 0.91 | 0.96 | 0.9 |
| 4 | 82549474 | rs7662480 | 1.338 | -0.494 | 1.131 | -0.444 | -0.045 | 0.026 | 0.026 | 0.237 | 0.87 | 0.49 | 0.95 | 0.91 | 0.95 | 0.9 |
| 4 | 82556945 | rs7657174 | 0.773 | -0.528 | 0.668 | -0.349 | -0.039 | 0.022 | 0.022 | 0 | 0.9 | 0.71 | 0.97 | 0.91 | 0.97 | 0.91 |
| 4 | 82564162 | rs4396981 | -0.079 | -0.119 | 0.465 | -0.295 | 0.018 | -0.02 | -0.02 | 0 | 0.93 | 0.73 | 0.92 | 1 | 0.93 | 0.95 |
| 4 | 82569389 | rs10434096 | 0.659 | -0.083 | -0.371 | -0.337 | -0.073 | -0.143 | -0.143 | 0 | 0.33 | 0.03 | 0.43 | 0.56 | 0.33 | 0.34 |
| 4 | 82572441 | rs4073341 | 0.77 | 0.049 | -0.069 | -0.368 | -0.041 | -0.07 | -0.07 | 0.219 | 0.34 | 0.1 | 0.52 | 0.57 | 0.35 | 0.39 |
| 4 | 82584423 | rs6853493 | -0.55 | -0.51 | 0.245 | -0.301 | -0.022 | 0.023 | 0.023 | 0 | 0.12 | 0.27 | 0.09 | 0.12 | 0.11 | 0.2 |
| 4 | 82587050 | rs11099493 | 0.973 | -0.348 | -0.065 | -0.582 | -0.081 | 0 | 0 | 0 | 0.27 | 0.09 | 0.21 | 0.22 | 0.35 | 0.26 |
| 4 | 82595257 | rs4437242 | -0.769 | -0.054 | 0.21 | 0.117 | -0.023 | -0.086 | -0.086 | 0 | 0.64 | 0.91 | 0.5 | 0.44 | 0.69 | 0.62 |
| 4 | 82597792 | rs4074490 | -1.983 | 0.123 | 0.609 | 0.323 | -0.032 | 0.026 | 0.026 | 0 | 0.21 | 0.52 | 0.17 | 0.22 | 0.18 | 0.34 |
| 4 | 82600430 | rs9307786 | 0.051 | 0.199 | 0.815 | 0.54 | -0.023 | 0.052 | 0.052 | 0 | 0.07 | 0.18 | 0.03 | 0.12 | 0.04 | 0.15 |
| 4 | 82602057 | rs9985504 | -0.467 | 0.206 | 0.655 | 0.523 | 0.001 | -0.005 | -0.005 | 0 | 0.61 | 0.63 | 0.44 | 0.44 | 0.64 | 0.57 |
| 4 | 82616045 | rs6535273 | -0.682 | 0.288 | -0.155 | 0.608 | -0.029 | -0.07 | -0.07 | 0 | 0.63 | 0.87 | 0.46 | 0.44 | 0.63 | 0.57 |
| 4 | 82619170 | rs9992825 | -0.003 | 0.363 | -0.339 | 0.652 | -0.022 | 0.037 | 0.037 | 0 | 0.07 | 0.18 | 0.03 | 0.11 | 0.04 | 0.16 |
| 4 | 82620131 | rs4401444 | -0.026 | 0.422 | -0.457 | 0.61 | 0.005 | 0.014 | 0.014 | 0 | 0.55 | 0.58 | 0.42 | 0.33 | 0.59 | 0.42 |
| 4 | 82628567 | rs7683827 | -0.964 | 0.504 | -0.731 | 1.026 | -0.001 | -0.001 | -0.001 | 0 | 0.07 | 0.05 | 0.05 | 0.11 | 0.06 | 0.14 |
| 4 | 82634810 | rs6834681 | -1.042 | 0.661 | -0.939 | 1.075 | 0.004 | -0.011 | -0.011 | 0 | 0.12 | 0.15 | 0.08 | 0.23 | 0.1 | 0.3 |
| 4 | 82643252 | rs7697168 | -0.289 | 0.829 | -0.815 | 0.553 | 0 | 0.002 | 0.002 | 0 | 0.48 | 0.58 | 0.35 | 0.35 | 0.53 | 0.41 |

| | | | | | | | | | | | | | | | | |
|---|---|---|---|---|---|---|---|---|---|---|---|---|---|---|---|---|
| 4 | 82644654 | rs11099497 | -0.256 | 0.936 | -0.779 | -0.476 | -0.001 | 0.006 | 0.006 | 0.35 | 0.42 | 0.42 | 0.71 | 0.85 | 0.59 | 0.58 |
| 4 | 82654242 | rs6822657 | -1.034 | 0.668 | 0.035 | -0.793 | 0.016 | -0.089 | -0.089 | 0.247 | 0.14 | 0.41 | 0.11 | 0.01 | 0.11 | 0.03 |
| 4 | 82659400 | rs1353663 | -0.092 | 0.479 | 0.171 | -1.711 | -0.002 | -0.076 | -0.076 | 0 | 0.8 | 0.69 | 0.89 | 0.98 | 0.83 | 0.69 |
| 4 | 82659919 | rs1497784 | -0.842 | 0.187 | 0.045 | -0.655 | 0 | -0.007 | -0.007 | 0.377 | 0.2 | 0.16 | 0.17 | 0.41 | 0.18 | 0.11 |
| 4 | 82664674 | rs7676794 | -0.419 | -0.127 | -0.118 | -1.19 | 0.016 | -0.072 | -0.072 | 0.264 | 0.88 | 0.7 | 0.91 | 0.99 | 0.88 | 0.98 |
| 4 | 82666782 | rs7680648 | -0.62 | -0.133 | -0.059 | -1.202 | 0 | -0.067 | -0.067 | 0.376 | 0.3 | 0.36 | 0.19 | 0.03 | 0.29 | 0.34 |
| 4 | 82667620 | rs6819863 | -1.495 | -1.033 | 0.022 | -1.543 | 0.011 | 0 | 0 | 0 | 0.5 | 0.61 | 0.31 | 0.42 | 0.38 | 0.44 |
| 4 | 82671938 | rs7672440 | 0.954 | -1.206 | -0.056 | -1.554 | 0.03 | -0.003 | -0.003 | 0 | 0.49 | 0.32 | 0.69 | 0.57 | 0.62 | 0.54 |
| 4 | 82677938 | rs6535279 | 0.266 | -1.089 | -0.06 | -1.631 | -0.018 | -0.09 | -0.09 | 0 | 0.82 | 0.71 | 0.91 | 0.99 | 0.88 | 0.96 |
| 4 | 82687275 | rs7696872 | -0.666 | -1.045 | -0.328 | -1.498 | 0.005 | 0.003 | 0.003 | 0 | 0.29 | 0.28 | 0.14 | 0.06 | 0.23 | 0.35 |
| 4 | 82687523 | rs4693005 | 1.108 | -0.399 | -0.606 | -0.972 | -0.013 | -0.001 | -0.001 | 0 | 0.36 | 0.32 | 0.62 | 0.45 | 0.52 | 0.44 |
| 4 | 82688257 | rs17568540 | 0.426 | -0.103 | -0.262 | -0.813 | -0.006 | 0.029 | 0.029 | 0.241 | 0.21 | 0.13 | 0.13 | 0.06 | 0.21 | 0.33 |
| 4 | 82690494 | rs9307787 | -0.051 | 0.403 | -0.04 | -0.245 | 0.012 | -0.006 | -0.006 | 0 | 0.17 | 0.25 | 0.31 | 0.19 | 0.26 | 0.13 |
| 4 | 82698629 | rs2055263 | -0.443 | 0.475 | 0.025 | 0.307 | 0.007 | 0.023 | 0.023 | 0.218 | 0.16 | 0.24 | 0.17 | 0.26 | 0.23 | 0.13 |
| 4 | 82720331 | rs523329 | 1.285 | -0.008 | 0.678 | 0.042 | -0.016 | 0.015 | 0.015 | 0 | 0.77 | 0.55 | 0.8 | 0.72 | 0.76 | 0.82 |
| 4 | 82723964 | rs751104 | 1.435 | -0.05 | -0.397 | 0.238 | -0.146 | 0.015 | 0.015 | 0 | 0.36 | 0.07 | 0.47 | 0.27 | 0.46 | 0.4 |
| 4 | 82730697 | rs515108 | 0.252 | -0.639 | -0.21 | 0.046 | -0.039 | 0.176 | 0.176 | 0 | 0.19 | 0.41 | 0.22 | 0.4 | 0.13 | 0.15 |
| 4 | 82734906 | rs1497770 | 0.362 | -0.694 | -0.097 | -0.09 | -0.005 | 0.002 | 0.002 | 0.314 | 0.69 | 0.51 | 0.62 | 0.33 | 0.64 | 0.72 |
| 4 | 82738099 | rs6535280 | -0.23 | -0.841 | -0.088 | -0.261 | -0.028 | -0.085 | -0.085 | 0 | 0.32 | 0.42 | 0.15 | 0.07 | 0.17 | 0.31 |
| 4 | 82744243 | rs17570404 | 0.925 | -0.873 | -0.322 | -0.254 | -0.007 | 0.08 | 0.08 | 0 | 0.2 | 0.06 | 0.1 | 0.07 | 0.15 | 0.28 |
| 4 | 82744299 | rs4566646 | 1.64 | -0.712 | -0.332 | 0.028 | -0.077 | 0.015 | 0.015 | 0 | 0.39 | 0.11 | 0.49 | 0.31 | 0.48 | 0.44 |
| 4 | 82752721 | rs685815 | -1.302 | -1.349 | -0.067 | 0.076 | 0.073 | 0.017 | 0.017 | 0 | 0.48 | 0.71 | 0.35 | 0.42 | 0.29 | 0.44 |
| 4 | 82766046 | rs583138 | 0.354 | -1.329 | 0.135 | 0.132 | -0.069 | 0.147 | 0.147 | 0 | 0.2 | 0.51 | 0.21 | 0.34 | 0.11 | 0.15 |
| 4 | 82767175 | rs7653998 | 0.694 | -1.358 | 0.695 | 0.116 | -0.022 | 0.041 | 0.041 | 0 | 0.18 | 0.28 | 0.2 | 0.34 | 0.11 | 0.15 |
| 4 | 82767300 | rs4693387 | 0.216 | -1.389 | 0.706 | 0.1 | -0.053 | 0.07 | 0.07 | 0 | 0.21 | 0.42 | 0.21 | 0.34 | 0.12 | 0.15 |
| 4 | 82773619 | rs666149 | 0.584 | -1.395 | 0.641 | 0.105 | -0.049 | 0.112 | 0.112 | 0 | 0.19 | 0.37 | 0.2 | 0.35 | 0.11 | 0.16 |
| 4 | 82779892 | rs667443 | 0.42 | -0.803 | 0.409 | 0.038 | -0.003 | 0.001 | 0.001 | 0 | 0.61 | 0.64 | 0.66 | 0.6 | 0.52 | 0.59 |
| 4 | 82783136 | rs1824657 | -1.364 | -0.647 | 0.079 | 0.103 | -0.077 | -0.076 | -0.076 | 0 | 0.86 | 0.97 | 0.8 | 0.66 | 0.68 | 0.86 |
| 4 | 82786911 | rs10002393 | 1.901 | -0.733 | 0.075 | 0.06 | -0.062 | 0.056 | 0.056 | 0 | 0.36 | 0.09 | 0.43 | 0.24 | 0.39 | 0.41 |
| 4 | 82805077 | rs17571905 | 1.945 | -0.716 | 0.042 | 0.063 | -0.076 | 0.052 | 0.052 | 0 | 0.36 | 0.07 | 0.43 | 0.24 | 0.39 | 0.4 |
| 4 | 82812093 | rs7663795 | -0.861 | -0.542 | 0.281 | 0.257 | 0.006 | 0.007 | 0.007 | 0.441 | 0.85 | 0.88 | 0.8 | 0.68 | 0.69 | 0.85 |
| 4 | 82816806 | rs1462694 | 0.633 | -0.751 | 0.384 | -0.042 | -0.018 | 0.029 | 0.029 | 0 | 0.18 | 0.26 | 0.2 | 0.35 | 0.1 | 0.15 |
| 4 | 82820008 | rs2199487 | -0.176 | -0.633 | 0.44 | -0.058 | 0.023 | 0.008 | 0.008 | 0 | 0.81 | 0.82 | 0.77 | 0.7 | 0.66 | 0.83 |
| 4 | 82820406 | rs12500182 | 0.228 | -0.705 | 0.61 | -0.072 | -0.083 | 0.131 | 0.131 | 0 | 0.22 | 0.44 | 0.22 | 0.35 | 0.1 | 0.15 |
| 4 | 82825171 | rs7673236 | 1.371 | -0.703 | 0.072 | -0.105 | -0.008 | 0.041 | 0.041 | 0 | 0.55 | 0.12 | 0.53 | 0.35 | 0.55 | 0.68 |
| 4 | 82835295 | rs1982618 | -0.681 | -0.383 | 0.527 | -0.106 | 0.018 | 0.009 | 0.009 | 0 | 0.85 | 0.83 | 0.79 | 0.73 | 0.69 | 0.86 |
| 4 | 82843061 | rs4693011 | 0.279 | -0.557 | 0.694 | -0.274 | 0.018 | -0.008 | -0.008 | 0 | 0.65 | 0.64 | 0.63 | 0.64 | 0.52 | 0.6 |
| 4 | 82843970 | rs10516668 | -0.045 | -0.578 | 0.626 | -0.301 | -0.025 | -0.102 | -0.102 | 0.147 | 0.24 | 0.42 | 0.16 | 0.09 | 0.19 | 0.29 |
| 4 | 82865191 | rs2120368 | 0.808 | -0.602 | 0.481 | -0.229 | -0.019 | 0.025 | 0.025 | 0 | 0.59 | 0.38 | 0.59 | 0.64 | 0.48 | 0.57 |
| 4 | 82875045 | rs320754 | 0.273 | -0.602 | 0.471 | -0.312 | -0.008 | -0.068 | -0.068 | 0.101 | 0.23 | 0.25 | 0.19 | 0.09 | 0.21 | 0.27 |
| 4 | 82889237 | rs320770 | 0.253 | -0.607 | 0.451 | -0.329 | -0.006 | -0.067 | -0.067 | 0 | 0.23 | 0.26 | 0.2 | 0.09 | 0.22 | 0.27 |
| 4 | 82895808 | rs11099512 | 1.918 | -0.893 | -0.078 | -0.314 | -0.096 | -0.004 | -0.004 | 0 | 0.33 | 0.06 | 0.39 | 0.26 | 0.36 | 0.4 |
| 4 | 82896567 | rs6838368 | 0.668 | -1.052 | -0.085 | -0.447 | 0 | 0.044 | 0.044 | 0 | 0.21 | 0.13 | 0.17 | 0.09 | 0.18 | 0.27 |
| 4 | 82896862 | rs11099514 | 0.669 | -1.068 | 0.124 | -0.413 | 0.007 | 0.02 | 0.02 | 0 | 0.58 | 0.3 | 0.56 | 0.38 | 0.55 | 0.67 |
| 4 | 82900372 | rs11730183 | 1.029 | -1.107 | 0.02 | -0.453 | 0.01 | 0.049 | 0.049 | 0 | 0.56 | 0.13 | 0.55 | 0.35 | 0.54 | 0.67 |
| 4 | 82905378 | rs17515274 | 1.987 | -1.022 | 0.039 | -0.419 | -0.046 | 0.019 | 0.019 | 0 | 0.34 | 0.17 | 0.39 | 0.26 | 0.36 | 0.4 |
| 4 | 82910126 | rs1443785 | 0.211 | -1.134 | 0.339 | -0.346 | 0.05 | 0.026 | 0.026 | 0.315 | 0.19 | 0.34 | 0.23 | 0.27 | 0.32 | 0.16 |
| 4 | 82913135 | rs12650942 | 2.143 | -1.218 | 0.14 | -0.4 | -0.103 | -0.005 | -0.005 | 0.008 | 0.33 | 0.06 | 0.39 | 0.26 | 0.38 | 0.41 |
| 4 | 82914118 | rs12711232 | 0.153 | -1.221 | 0.269 | -0.344 | 0.04 | 0.018 | 0.018 | 0.444 | 0.21 | 0.38 | 0.2 | 0.3 | 0.31 | 0.16 |
| 4 | 82915079 | rs13109155 | 0.56 | -1.284 | 0.166 | -0.357 | 0 | -0.001 | -0.001 | 0.752 | 0.57 | 0.63 | 0.61 | 0.53 | 0.69 | 0.57 |
| 4 | 82915702 | rs4693013 | 0.443 | -1.349 | 0.263 | -0.406 | 0.043 | -0.056 | -0.056 | 0.538 | 0.21 | 0.15 | 0.23 | 0.38 | 0.13 | 0.17 |
| 4 | 82917515 | rs7670373 | -0.113 | -1.337 | 0.242 | -0.4 | -0.06 | 0.055 | 0.055 | 0.062 | 0.75 | 0.67 | 0.76 | 0.63 | 0.86 | 0.83 |
| 4 | 82922668 | rs2868278 | 0.272 | -1.249 | 0.124 | -0.357 | 0.027 | -0.034 | -0.034 | 0 | 0.22 | 0.2 | 0.23 | 0.38 | 0.13 | 0.16 |
| 4 | 82928877 | rs7678127 | -0.04 | -1.202 | 0.242 | -0.382 | 0.049 | 0.02 | 0.02 | 0.254 | 0.22 | 0.48 | 0.2 | 0.3 | 0.31 | 0.16 |
| 4 | 82928896 | rs11939555 | 1.025 | -1.123 | 0.081 | -0.28 | 0.009 | 0.008 | 0.008 | 0 | 0.15 | 0.15 | 0.15 | 0.3 | 0.27 | 0.14 |
| 4 | 82932359 | rs6814999 | -0.071 | -1.229 | -0.081 | -0.147 | 0.03 | -0.036 | -0.036 | 0 | 0.22 | 0.2 | 0.23 | 0.37 | 0.13 | 0.16 |
| 4 | 82949189 | rs1443769 | 0.039 | -0.581 | 0.021 | -0.274 | -0.054 | 0.087 | 0.087 | 0 | 0.19 | 0.01 | 0.17 | 0.03 | 0.21 | 0.24 |
| 4 | 82955013 | rs1470338 | 0.768 | -1.067 | 0.169 | -0.218 | -0.009 | 0.087 | 0.087 | 0 | 0.48 | 0.22 | 0.49 | 0.28 | 0.46 | 0.48 |
| 4 | 82956948 | rs4693416 | 0.12 | -1.293 | 0.156 | -0.047 | -0.001 | -0.019 | -0.019 | 0 | 0.18 | 0.12 | 0.28 | 0.4 | 0.18 | 0.28 |
| 4 | 82959120 | rs1443772 | 0.555 | -1.353 | 0.517 | -0.2 | -0.026 | -0.027 | -0.027 | 0.089 | 0.05 | 0.16 | 0.02 | 0 | 0 | 0 |
| 4 | 82962095 | rs1562030 | 0.079 | -1.4 | 0.425 | -0.268 | -0.001 | -0.003 | -0.003 | 0.53 | 0.05 | 0.05 | 0.03 | 0 | 0.04 | 0.06 |
| 4 | 82964509 | rs7671690 | -0.835 | -1.374 | 0.537 | -0.294 | -0.011 | 0.02 | 0.02 | 0 | 0.44 | 0.41 | 0.48 | 0.74 | 0.52 | 0.49 |
| 4 | 82974329 | rs7654293 | 1.26 | -1.086 | 0.384 | -0.17 | -0.022 | 0.035 | 0.035 | 0 | 0.4 | 0.13 | 0.43 | 0.26 | 0.41 | 0.47 |
| 4 | 82975345 | rs971825 | 1.451 | -1.145 | 0.222 | -0.1 | -0.101 | 0.008 | 0.008 | 0 | 0.37 | 0.03 | 0.26 | 0.22 | 0.4 | 0.42 |
| 4 | 82977205 | rs7664786 | 0.424 | -1.229 | 0.756 | -0.088 | 0.018 | 0.007 | 0.007 | 0 | 0.25 | 0.43 | 0.22 | 0.34 | 0.34 | 0.21 |
| 4 | 82978391 | rs12508988 | 0.336 | -1.273 | 0.59 | -0.103 | 0.011 | -0.069 | -0.069 | 0.009 | 0.25 | 0.07 | 0.31 | 0.38 | 0.18 | 0.29 |
| 4 | 82980833 | rs2131361 | -1.266 | -1.28 | 0.827 | -0.116 | -0.001 | -0.001 | -0.001 | 0.05 | 0.57 | 0.62 | 0.56 | 0.72 | 0.57 | 0.53 |
| 4 | 82981181 | rs4693418 | -0.092 | -1.214 | 1.169 | 0.017 | 0.015 | -0.058 | -0.058 | 4.918 | 0.25 | 0.08 | 0.31 | 0.38 | 0.18 | 0.29 |
| 4 | 82981940 | rs1373238 | -0.091 | -1.2 | 1.381 | 0.024 | 0.015 | -0.058 | -0.058 | 0 | 0.25 | 0.08 | 0.31 | 0.38 | 0.19 | 0.29 |

| | | | | | | | | | | | | | | | | |
|---|---|---|---|---|---|---|---|---|---|---|---|---|---|---|---|---|
| 4 | 82990379 | rs10010220 | 1.424 | -0.586 | 2.551 | -0.111 | 0.071 | 0.005 | 0.005 | 0 | 0.92 | 0.57 | 0.95 | 1 | 0.96 | 0.97 |
| 4 | 83007475 | rs2035912 | 1.175 | -0.383 | 0.544 | -0.001 | -0.023 | -0.04 | -0.04 | 0 | 0.84 | 0.31 | 0.75 | 0.94 | 0.93 | 0.92 |
| 4 | 83019396 | rs4693421 | 0.411 | -0.149 | 0.005 | 0.135 | -0.004 | 0.004 | 0.004 | 0 | 0.58 | 0.61 | 0.59 | 0.3 | 0.52 | 0.69 |
| 4 | 83026416 | rs6535307 | -1.06 | -0.52 | 1.066 | -0.197 | 0.009 | -0.048 | -0.048 | 0 | 0.1 | 0.29 | 0.05 | 0 | 0.04 | 0.03 |
| 4 | 83028150 | rs2011786 | 0.773 | -0.06 | 0.694 | 0.172 | -0.022 | -0.017 | -0.017 | 0.373 | 0.18 | 0.12 | 0.18 | 0.3 | 0.35 | 0.22 |
| 4 | 83029942 | rs10009173 | 0.415 | -1.651 | 0.853 | -0.31 | 0.023 | 0.068 | 0.068 | 0 | 0.39 | 0.28 | 0.43 | 0.23 | 0.36 | 0.39 |
| 4 | 83044475 | rs7677948 | 0.737 | -1.64 | 1.254 | -0.317 | 0.013 | 0.031 | 0.031 | 0 | 0.38 | 0.31 | 0.26 | 0.21 | 0.34 | 0.36 |
| 4 | 83045736 | rs11943170 | 0.119 | -1.563 | 1.208 | -0.268 | -0.007 | -0.014 | -0.014 | 0 | 0.46 | 0.58 | 0.46 | 0.25 | 0.37 | 0.4 |
| 4 | 83058150 | rs962753 | 1.04 | -1.509 | 0.765 | -0.541 | -0.044 | -0.084 | -0.084 | 0 | 0.81 | 0.44 | 0.74 | 0.96 | 0.92 | 0.93 |
| 4 | 83059714 | rs13109978 | 0.136 | -1.435 | 0.214 | -0.452 | 0.005 | 0.096 | 0.096 | 0.124 | 0.43 | 0.2 | 0.48 | 0.75 | 0.6 | 0.58 |
| 4 | 83062902 | rs6812383 | 0.578 | -1.141 | 0.666 | 0.065 | -0.015 | -0.018 | -0.018 | 0 | 0.91 | 0.71 | 0.88 | 0.98 | 0.97 | 0.96 |
| 4 | 83067671 | rs17005475 | -0.553 | -0.838 | -0.268 | 0.081 | -0.018 | -0.047 | -0.047 | 0 | 0.09 | 0.17 | 0.03 | 0 | 0.03 | 0.03 |
| 4 | 83068042 | rs1492848 | -0.526 | -0.465 | -0.816 | 0.652 | 0.015 | -0.095 | -0.095 | 0 | 0.13 | 0.01 | 0.27 | 0.47 | 0.08 | 0.1 |
| 4 | 83085123 | rs6825436 | 0.025 | -0.235 | -0.424 | -0.501 | 0.009 | 0.016 | 0.016 | 0.012 | 0.47 | 0.38 | 0.36 | 0.22 | 0.36 | 0.38 |
| 4 | 83087694 | rs2868298 | 0.238 | -0.351 | -0.081 | -0.398 | -0.005 | -0.009 | -0.009 | 0.06 | 0.94 | 0.84 | 0.96 | 0.98 | 0.96 | 0.96 |
| 4 | 83094251 | rs13116695 | 0.102 | 0.068 | -0.152 | -0.963 | 0.002 | -0.016 | -0.016 | 0 | 0.27 | 0.3 | 0.2 | 0.12 | 0.35 | 0.31 |
| 4 | 83096524 | rs2101680 | 0.372 | -0.698 | -0.291 | -1.576 | -0.054 | -0.113 | -0.113 | 0.147 | 0.77 | 0.47 | 0.89 | 0.98 | 0.9 | 0.88 |
| 4 | 83102836 | rs10019349 | 0.815 | -0.692 | -0.996 | -1.393 | -0.043 | -0.133 | -0.133 | 0 | 0.18 | 0.02 | 0.29 | 0.56 | 0.17 | 0.14 |
| 4 | 83112989 | rs12507749 | 0.329 | -0.871 | -0.261 | -1.617 | 0.001 | 0.012 | 0.012 | 0 | 0.53 | 0.4 | 0.71 | 0.89 | 0.55 | 0.72 |
| 4 | 83114530 | rs1389044 | -0.649 | -0.695 | -0.226 | -1.474 | 0.003 | 0.069 | 0.069 | 0 | 0.75 | 0.53 | 0.68 | 0.41 | 0.79 | 0.83 |
| 4 | 83115166 | rs2035910 | 0.757 | -0.885 | -0.268 | -1.596 | -0.033 | 0.052 | 0.052 | 0 | 0.29 | 0.21 | 0.21 | 0.09 | 0.39 | 0.2 |
| 4 | 83115740 | rs12510027 | -0.73 | -0.888 | -0.257 | -1.59 | 0.003 | 0.069 | 0.069 | 0 | 0.75 | 0.53 | 0.68 | 0.41 | 0.79 | 0.83 |
| 4 | 83115904 | rs4146415 | 0.12 | -0.886 | -0.937 | -1.589 | 0.016 | -0.003 | -0.003 | 0 | 0.64 | 0.34 | 0.65 | 0.41 | 0.77 | 0.79 |
| 4 | 83116329 | rs4693432 | 0.975 | -0.641 | -0.877 | -1.444 | -0.01 | 0.015 | 0.015 | 0 | 0.35 | 0.13 | 0.44 | 0.32 | 0.37 | 0.59 |
| 4 | 83130688 | rs11941179 | 0.422 | 0.164 | -0.804 | -0.918 | 0.005 | 0.013 | 0.013 | 0 | 0.52 | 0.51 | 0.51 | 0.32 | 0.45 | 0.68 |
| 4 | 83132921 | rs4693023 | -0.305 | 0.519 | -1.098 | -1.112 | -0.024 | 0.033 | 0.033 | 0 | 0.75 | 0.8 | 0.81 | 0.91 | 0.65 | 0.85 |
| 4 | 83149478 | rs4693026 | -0.407 | -0.182 | -1.192 | -1.19 | 0.023 | 0.03 | 0.03 | 1.979 | 0.31 | 0.18 | 0.39 | 0.22 | 0.25 | 0.51 |
| 4 | 83153179 | rs2583636 | -0.944 | 0.319 | -2.025 | -1.288 | -0.079 | 0.129 | 0.129 | 0 | 0.48 | 0.91 | 0.31 | 0.75 | 0.48 | 0.3 |
| 4 | 83160983 | rs2556543 | -1.188 | 1.014 | -1.305 | -1.368 | -0.038 | 0.024 | 0.024 | 0 | 0.88 | 1 | 0.7 | 0.97 | 0.88 | 0.74 |
| 4 | 83164184 | rs7660665 | -1.422 | 1.197 | -0.398 | -0.964 | 0.009 | 0.001 | 0.001 | 0 | 0.33 | 0.6 | 0.26 | 0.66 | 0.4 | 0.22 |
| 4 | 83170172 | rs2114693 | -2.191 | 1.093 | -0.529 | -1.044 | -0.024 | 0.17 | 0.17 | 0 | 0.5 | 0.94 | 0.34 | 0.79 | 0.53 | 0.3 |
| 4 | 83171331 | rs7678274 | -0.519 | 1.294 | 0.366 | -0.786 | 0.029 | 0.064 | 0.064 | 0 | 0.19 | 0.33 | 0.17 | 0.61 | 0.28 | 0.19 |
| 4 | 83177773 | rs2303571 | 0.242 | 1.316 | 0.335 | -0.789 | 0.003 | 0.021 | 0.021 | 0 | 0.13 | 0.09 | 0.13 | 0.61 | 0.26 | 0.15 |
| 4 | 83189631 | rs6815242 | -0.825 | 1.136 | 0.14 | -0.69 | 0.036 | 0.042 | 0.042 | 0 | 0.26 | 0.42 | 0.21 | 0.61 | 0.35 | 0.21 |
| 4 | 83196536 | rs13119500 | -0.145 | 1.01 | -0.215 | -0.599 | -0.007 | -0.033 | -0.033 | 0 | 0.17 | 0.3 | 0.08 | 0.13 | 0.12 | 0.07 |
| 4 | 83209493 | rs7692570 | -1.618 | 1.13 | -1.271 | -0.74 | -0.012 | 0.21 | 0.21 | 0 | 0.49 | 0.9 | 0.35 | 0.79 | 0.54 | 0.29 |
| 4 | 83210000 | rs17005574 | 0.2 | 1.548 | -1.321 | -0.578 | -0.048 | -0.054 | -0.054 | 0.148 | 0.12 | 0.03 | 0.13 | 0.57 | 0.26 | 0.15 |
| 4 | 83218230 | rs6822283 | -0.877 | 1.476 | -0.736 | -0.681 | 0.006 | 0.025 | 0.025 | 0 | 0.32 | 0.73 | 0.2 | 0.16 | 0.23 | 0.13 |
| 4 | 83237661 | rs1116664 | -0.246 | 1.433 | 0.056 | -0.707 | 0.075 | 0.182 | 0.182 | 0.343 | 0.06 | 0.41 | 0.05 | 0.12 | 0.02 | 0.02 |
| 4 | 83266333 | rs1961547 | -0.699 | 1.437 | -0.555 | -0.73 | -0.029 | 0.091 | 0.091 | 0 | 0.23 | 0.05 | 0.12 | 0.04 | 0.22 | 0.11 |
| 4 | 83273140 | rs34471118 | -0.526 | 1.44 | -0.554 | -0.728 | -0.03 | 0.086 | 0.086 | 0 | 0.22 | 0.05 | 0.12 | 0.04 | 0.22 | 0.11 |
| 4 | 83274634 | rs2288338 | 0.006 | 1.438 | -0.571 | -0.647 | -0.058 | -0.074 | -0.074 | 0.033 | 0.13 | 0.03 | 0.13 | 0.57 | 0.26 | 0.15 |
| 4 | 83278778 | rs1820577 | -0.632 | 1.338 | -0.471 | -0.857 | -0.03 | 0.086 | 0.086 | 0 | 0.22 | 0.05 | 0.12 | 0.04 | 0.22 | 0.11 |
| 4 | 83305278 | rs3762885 | -1.457 | 1.45 | 0.251 | -0.323 | 0.058 | 0.144 | 0.144 | 0.076 | 0.17 | 0.46 | 0.13 | 0.63 | 0.22 | 0.12 |
| 4 | 83309685 | rs6843699 | -1.294 | 0.997 | 0.04 | -0.719 | -0.014 | -0.05 | -0.05 | 0 | 0.31 | 0.4 | 0.25 | 0.15 | 0.28 | 0.17 |
| 4 | 83312849 | rs6816230 | -1.63 | 0.395 | 0.229 | -0.341 | -0.084 | -0.079 | -0.079 | 0 | 0.3 | 0.5 | 0.23 | 0.12 | 0.11 | 0.12 |
| 4 | 83317753 | rs13105860 | -0.806 | 1.315 | -0.294 | -0.668 | -0.103 | 0.054 | 0.054 | 0 | 0.28 | 0.08 | 0.31 | 0.79 | 0.39 | 0.27 |
| 4 | 83319611 | rs2004987 | -0.58 | 1.314 | -0.301 | -0.674 | -0.105 | 0.043 | 0.043 | 0 | 0.26 | 0.07 | 0.3 | 0.79 | 0.39 | 0.27 |
| 4 | 83321808 | rs17005633 | -1.048 | 1.003 | 0.147 | -0.916 | 0.012 | -0.038 | -0.038 | 0.297 | 0.14 | 0.34 | 0.13 | 0.07 | 0.16 | 0.07 |
| 4 | 83323062 | rs7695741 | -0.528 | 1.04 | 0.243 | -0.898 | 0.003 | -0.016 | -0.016 | 0 | 0.11 | 0.15 | 0.12 | 0.07 | 0.16 | 0.07 |
| 4 | 83325574 | rs13125359 | -0.04 | 0.877 | 0.27 | -0.967 | -0.048 | -0.079 | -0.079 | 0 | 0.87 | 0.64 | 0.95 | 1 | 0.97 | 0.93 |
| 4 | 83340605 | rs1559949 | 1.551 | 0.943 | 0.278 | -0.886 | -0.034 | -0.048 | -0.048 | 0.043 | 0.73 | 0.36 | 0.79 | 0.91 | 0.87 | 0.86 |
| 4 | 83343745 | rs11737188 | 0.635 | 1.135 | 0.043 | -0.496 | -0.085 | -0.094 | -0.094 | 0 | 0.17 | 0.04 | 0.2 | 0.61 | 0.31 | 0.21 |
| 4 | 83345146 | rs997720 | 0.241 | 0.906 | 0.048 | -0.49 | 0.004 | -0.001 | -0.001 | 0 | 0.11 | 0.03 | 0.11 | 0.08 | 0.08 | 0.06 |
| 4 | 83345292 | rs997721 | 0.267 | 0.908 | 0.049 | -0.489 | 0.004 | -0.001 | -0.001 | 0 | 0.11 | 0.03 | 0.11 | 0.08 | 0.08 | 0.06 |
| 4 | 83348978 | rs6826022 | 2.252 | 0.851 | 0.022 | -0.528 | -0.023 | 0.167 | 0.167 | 0 | 0.45 | 0.09 | 0.49 | 0.22 | 0.43 | 0.6 |
| 4 | 83351821 | rs8732 | 0.037 | 0.778 | 0.123 | -0.484 | -0.009 | -0.005 | -0.005 | 0 | 0.12 | 0.2 | 0.12 | 0.08 | 0.08 | 0.06 |
| 4 | 83358762 | rs10516674 | -0.28 | 0.799 | -0.114 | -0.39 | -0.05 | -0.081 | -0.081 | 0 | 0.13 | 0.36 | 0.05 | 0 | 0.03 | 0.07 |
| 4 | 83362192 | rs1878494 | 1.641 | 0.899 | -0.545 | -0.307 | 0.009 | 0.141 | 0.141 | 0 | 0.46 | 0.21 | 0.52 | 0.23 | 0.45 | 0.61 |
| 4 | 83368980 | rs1113576 | -0.572 | 0.9 | -0.071 | 0.37 | -0.04 | -0.069 | -0.069 | 0.054 | 0.12 | 0.31 | 0.05 | 0 | 0.03 | 0.07 |
| 4 | 83369287 | rs4285076 | -0.123 | 1.767 | -0.838 | 0.824 | -0.06 | -0.093 | -0.093 | 0 | 0.84 | 0.97 | 0.82 | 0.39 | 0.75 | 0.79 |
| 4 | 83378872 | rs1980187 | -0.323 | 1.173 | -0.493 | 0.712 | 0.001 | 0.001 | 0.001 | 0.74 | 0.53 | 0.49 | 0.61 | 0.52 | 0.53 | 0.69 |
| 4 | 83379826 | rs11929894 | -0.409 | 1.123 | -0.307 | 0.701 | -0.003 | -0.003 | -0.003 | 0 | 0.54 | 0.57 | 0.71 | 0.53 | 0.53 | 0.7 |
| 4 | 83381564 | rs3796470 | -0.442 | 0.861 | -0.429 | 0.842 | 0.013 | -0.017 | -0.017 | 0 | 0.07 | 0.02 | 0.11 | 0.14 | 0.03 | 0.04 |
| 4 | 83403563 | rs13114300 | -0.311 | 0.89 | -0.188 | 0.51 | 0.008 | 0.005 | 0.005 | 1.238 | 0.41 | 0.49 | 0.65 | 0.46 | 0.45 | 0.57 |
| 4 | 83405383 | rs6818677 | -0.491 | 1.068 | -0.019 | 0.482 | -0.001 | -0.002 | -0.002 | 0 | 0.45 | 0.67 | 0.69 | 0.47 | 0.53 | 0.62 |
| 4 | 83418930 | rs4693448 | -0.991 | 0.707 | 1.025 | 0.514 | 0 | 0.031 | 0.031 | 0 | 0.09 | 0.37 | 0.32 | 0.07 | 0.03 | 0.21 |
| 4 | 83421440 | rs10003787 | -0.953 | 1.409 | 0.212 | 0.512 | -0.03 | 0.001 | 0.001 | 0 | 0.1 | 0.04 | 0.14 | 0.1 | 0.2 | 0.09 |
| 4 | 83427464 | rs4693450 | -0.919 | 0.118 | 0.501 | 0.862 | -0.055 | 0.048 | 0.048 | 0.109 | 0.27 | 0.41 | 0.46 | 0.43 | 0.16 | 0.31 |

| | | | | | | | | | | | | | | | | |
|---|---|---|---|---|---|---|---|---|---|---|---|---|---|---|---|---|
| 4 | 83431048 | rs10016201 | -0.852 | 0.748 | 1.11 | 0.959 | 0.015 | 0.111 | 0.111 | 0 | 0.21 | 0.34 | 0.48 | 0.43 | 0.23 | 0.3 |
| 4 | 83433406 | rs953186 | -0.834 | 0.756 | 1.103 | 0.955 | 0.011 | 0.091 | 0.091 | 0 | 0.21 | 0.27 | 0.47 | 0.43 | 0.23 | 0.3 |
| 4 | 83436006 | rs12506540 | -1.281 | 0.314 | 1.297 | 0.501 | -0.001 | 0.016 | 0.016 | 0.347 | 0.08 | 0.2 | 0.07 | 0.1 | 0.05 | 0.05 |
| 4 | 83439160 | rs6535350 | -1.711 | 0.396 | 1.457 | 0.552 | 0.04 | 0.071 | 0.071 | 2.112 | 0.15 | 0.4 | 0.15 | 0.23 | 0.17 | 0.1 |
| 4 | 83441867 | rs7671872 | 0.162 | -0.549 | 0.92 | -0.429 | -0.013 | -0.028 | -0.028 | 0 | 0.72 | 0.77 | 0.89 | 0.93 | 0.82 | 0.87 |
| 4 | 83443054 | rs6836241 | -1.424 | -0.561 | 1.94 | -0.556 | 0.006 | -0.015 | -0.015 | 0 | 0.11 | 0.29 | 0.05 | 0.04 | 0.07 | 0.07 |
| 4 | 83446505 | rs17005759 | -1.882 | -1.356 | 1.454 | -0.765 | 0.072 | 0.079 | 0.079 | 0.022 | 0.37 | 0.8 | 0.19 | 0.14 | 0.22 | 0.21 |
| 4 | 83447641 | rs10014422 | -1.801 | -1.398 | 1.747 | -0.848 | 0.015 | -0.066 | -0.066 | 0 | 0.15 | 0.59 | 0.1 | 0.02 | 0.11 | 0.08 |
| 4 | 83448507 | rs4693453 | -0.817 | -1.418 | 1.882 | -0.794 | -0.05 | -0.032 | -0.032 | 0 | 0.25 | 0.57 | 0.21 | 0.14 | 0.16 | 0.14 |
| 4 | 83448764 | rs4693454 | -1.438 | -1.394 | 1.976 | -0.783 | -0.044 | -0.05 | -0.05 | 3.182 | 0.33 | 0.45 | 0.22 | 0.15 | 0.22 | 0.21 |
| 4 | 83458667 | rs11946794 | -1.787 | -1.25 | 0.478 | -0.728 | 0.032 | 0.04 | 0.04 | 0.472 | 0.35 | 0.72 | 0.25 | 0.13 | 0.22 | 0.19 |
| 4 | 83462935 | rs10516675 | -0.207 | -1.093 | 0.081 | -0.444 | 0.045 | 0.013 | 0.013 | 0.024 | 0.18 | 0.07 | 0.15 | 0.13 | 0.1 | 0.12 |
| 4 | 83485723 | rs4693458 | -0.174 | -0.191 | -0.495 | -0.796 | 0 | 0.01 | 0.01 | 0 | 0.59 | 0.64 | 0.56 | 0.8 | 0.64 | 0.5 |
| 4 | 83486981 | rs2176401 | 0.546 | -0.049 | -1.269 | -0.893 | -0.004 | 0.013 | 0.013 | 0 | 0.2 | 0.02 | 0.26 | 0.12 | 0.15 | 0.19 |
| 4 | 83491535 | rs11728793 | -0.127 | 0.456 | -0.922 | 0.213 | 0.013 | 0.119 | 0.119 | 0 | 0.17 | 0.27 | 0.27 | 0.43 | 0.21 | 0.22 |
| 4 | 83503020 | rs4693460 | 0.621 | 0.456 | -1.88 | -0.22 | -0.061 | -0.002 | -0.002 | 0 | 0.35 | 0.1 | 0.24 | 0.35 | 0.44 | 0.23 |
| 4 | 83507794 | rs11723799 | -0.168 | 0.556 | -1.381 | 0.411 | -0.038 | -0.122 | -0.122 | 0 | 0.15 | 0.02 | 0.22 | 0.42 | 0.16 | 0.23 |
| 4 | 83521262 | rs4693465 | 0.466 | 0.591 | -1.612 | 0.28 | -0.069 | -0.13 | -0.13 | 0 | 0.19 | 0.02 | 0.39 | 0.39 | 0.21 | 0.25 |
| 4 | 83527257 | rs12645956 | 0.518 | 0.312 | -1.922 | 0.126 | -0.054 | -0.138 | -0.138 | 0 | 0.2 | 0.01 | 0.15 | 0.38 | 0.24 | 0.19 |
| 4 | 83527733 | rs17005836 | -0.799 | 0.072 | -1.125 | -0.187 | -0.001 | -0.001 | -0.001 | 0.003 | 0.36 | 0.64 | 0.27 | 0.38 | 0.37 | 0.38 |
| 4 | 83534854 | rs6845080 | -0.636 | -0.093 | -0.81 | -0.272 | 0.002 | -0.006 | -0.006 | 0 | 0.32 | 0.31 | 0.24 | 0.38 | 0.33 | 0.23 |
| 4 | 83536279 | rs11734355 | -0.469 | -0.012 | -0.365 | -0.426 | 0 | -0.009 | -0.009 | 0 | 0.09 | 0.08 | 0.05 | 0 | 0.06 | 0.06 |
| 4 | 83538430 | rs6853729 | -0.332 | -0.034 | -0.046 | -0.439 | 0.073 | 0.013 | 0.013 | 0 | 0.06 | 0.31 | 0.06 | 0 | 0.05 | 0.03 |
| 4 | 83539606 | rs6842536 | -1.44 | -0.079 | -0.02 | -0.463 | 0.007 | -0.066 | -0.066 | 0 | 0.12 | 0.45 | 0.08 | 0 | 0.06 | 0.06 |
| 4 | 83547584 | rs4693468 | 0.73 | -0.102 | -0.339 | -0.405 | 0 | 0.021 | 0.021 | 0.132 | 0.68 | 0.56 | 0.73 | 0.64 | 0.65 | 0.61 |
| 4 | 83551007 | rs6840 | 0.734 | -0.106 | -0.367 | -0.407 | 0 | 0.02 | 0.02 | 0.137 | 0.68 | 0.56 | 0.73 | 0.64 | 0.65 | 0.61 |
| 4 | 83551119 | rs1065403 | 0.655 | -0.107 | -0.368 | -0.407 | 0 | 0.021 | 0.021 | 0.19 | 0.68 | 0.56 | 0.73 | 0.64 | 0.65 | 0.61 |
| 4 | 83554880 | rs7672297 | 0.242 | -0.134 | -0.322 | -0.419 | 0.092 | 0.294 | 0.294 | 0.154 | 0.13 | 0.47 | 0.18 | 0.36 | 0.21 | 0.12 |
| 4 | 83559578 | rs1395370 | -0.718 | -0.133 | -0.404 | -0.569 | -0.007 | 0.024 | 0.024 | 0 | 0.9 | 0.94 | 0.93 | 0.99 | 0.88 | 0.79 |
| 4 | 83560949 | rs9991687 | 0.371 | -0.206 | -0.424 | -0.611 | 0.022 | 0.041 | 0.041 | 0 | 0.79 | 0.88 | 0.9 | 0.99 | 0.86 | 0.72 |
| 4 | 83561503 | rs6535360 | 0.224 | -0.17 | -0.355 | -0.475 | 0.006 | 0.021 | 0.021 | 0.263 | 0.79 | 0.75 | 0.77 | 0.62 | 0.68 | 0.68 |
| 4 | 83564114 | rs1506606 | 1.629 | -0.38 | -0.528 | -0.643 | 0.002 | -0.001 | -0.001 | 1.7 | 0.59 | 0.39 | 0.71 | 0.62 | 0.6 | 0.58 |
| 4 | 83564947 | rs996465 | 0.549 | -0.314 | -0.59 | -0.596 | 0.001 | 0.017 | 0.017 | 0.876 | 0.74 | 0.7 | 0.77 | 0.63 | 0.71 | 0.77 |
| 4 | 83567535 | rs1506604 | -0.25 | -0.349 | -0.755 | -0.697 | 0.019 | 0.04 | 0.04 | 0 | 0.86 | 0.95 | 0.94 | 1 | 0.92 | 0.89 |
| 4 | 83569708 | rs1317366 | -0.226 | -0.349 | -0.732 | -0.697 | 0.007 | -0.004 | -0.004 | 0.49 | 0.85 | 0.89 | 0.93 | 1 | 0.92 | 0.89 |
| 4 | 83573730 | rs11935974 | 0.152 | -0.334 | -0.67 | -0.674 | -0.027 | -0.073 | -0.073 | 0 | 0.15 | 0.28 | 0.07 | 0 | 0.11 | 0.19 |
| 4 | 83576973 | rs12500897 | 1.254 | -0.171 | -1.316 | -0.802 | -0.087 | -0.114 | -0.114 | 4.931 | 0.23 | 0.03 | 0.46 | 0.37 | 0.27 | 0.3 |
| 4 | 83582064 | rs3733228 | 1.174 | -0.239 | -1.224 | -1.058 | -0.067 | -0.13 | -0.13 | 2.827 | 0.25 | 0.08 | 0.49 | 0.48 | 0.29 | 0.31 |
| 4 | 83586671 | rs2135507 | -0.944 | -0.348 | -0.537 | -0.709 | -0.007 | 0.09 | 0.09 | 0 | 0.27 | 0.49 | 0.28 | 0.41 | 0.31 | 0.31 |
| 4 | 83590099 | rs6811012 | 0.856 | -0.495 | -0.616 | -0.704 | -0.013 | 0.061 | 0.061 | 1.135 | 0.73 | 0.58 | 0.73 | 0.59 | 0.78 | 0.76 |
| 4 | 83596745 | rs10030631 | -0.137 | -0.467 | -0.625 | -0.688 | -0.003 | 0.118 | 0.118 | 0 | 0.18 | 0.33 | 0.22 | 0.41 | 0.18 | 0.29 |
| 4 | 83597815 | rs7689308 | -0.379 | -0.468 | -0.582 | -0.698 | 0.008 | 0.059 | 0.059 | 1.278 | 0.86 | 0.81 | 0.77 | 0.59 | 0.8 | 0.84 |
| 4 | 83598784 | rs7695607 | -0.232 | -0.508 | -0.608 | -0.731 | -0.017 | -0.047 | -0.047 | 1.76 | 0.17 | 0.21 | 0.07 | 0 | 0.08 | 0.26 |
| 4 | 83602150 | rs2276883 | 1.579 | -0.559 | -0.482 | -0.768 | 0.007 | -0.005 | -0.005 | 0 | 0.6 | 0.38 | 0.68 | 0.59 | 0.67 | 0.57 |
| 4 | 83602396 | rs11938637 | 0.68 | -0.348 | -1.38 | -0.634 | -0.015 | -0.045 | -0.045 | 6.636 | 0.09 | 0.2 | 0.05 | 0 | 0.06 | 0.18 |
| 4 | 83609415 | rs6834939 | 1.155 | -0.185 | -1.776 | -0.535 | 0.004 | 0.094 | 0.094 | 1.04 | 0.09 | 0.13 | 0.18 | 0.41 | 0.13 | 0.11 |
| 4 | 83613357 | rs17353688 | 1.433 | -0.311 | -2.479 | -0.845 | -0.018 | 0.086 | 0.086 | 0.487 | 0.3 | 0.04 | 0.16 | 0.17 | 0.26 | 0.25 |
| 4 | 83617849 | rs953534 | 0.445 | 0.27 | -1.96 | -0.404 | 0.028 | 0.04 | 0.04 | 0.002 | 0.52 | 0.53 | 0.24 | 0.16 | 0.34 | 0.37 |
| 4 | 83623978 | rs6852540 | -0.055 | 0.403 | -2.715 | -0.186 | -0.071 | -0.055 | -0.055 | 0 | 0.62 | 0.84 | 0.43 | 0.57 | 0.47 | 0.49 |
| 4 | 83626382 | rs1876218 | 0.369 | 0.87 | -2.707 | 0.135 | -0.12 | -0.124 | -0.124 | 0 | 0.27 | 0.07 | 0.48 | 0.43 | 0.37 | 0.35 |
| 4 | 83627583 | rs4693491 | -0.429 | 0.66 | -1.522 | 0.746 | 0.004 | -0.009 | -0.009 | 0 | 0.17 | 0.14 | 0.2 | 0.4 | 0.15 | 0.18 |
| 4 | 83637368 | rs4693496 | -0.097 | 0.813 | -1.621 | 0.192 | -0.052 | -0.003 | -0.003 | 0.358 | 0.27 | 0.06 | 0.28 | 0.2 | 0.32 | 0.19 |
| 4 | 83638620 | rs4693497 | -1.168 | 0.426 | -2.003 | 0.191 | -0.05 | -0.046 | -0.046 | 0 | 0.54 | 0.87 | 0.68 | 0.44 | 0.52 | 0.43 |
| 4 | 83639461 | rs10025210 | -0.231 | 0.323 | -1.458 | 0.104 | 0 | 0 | 0 | 0.574 | 0.14 | 0.18 | 0.18 | 0.1 | 0.11 | 0.2 |
| 4 | 83644676 | rs10516678 | 0.025 | 0.51 | -1.035 | 0.344 | -0.001 | -0.001 | -0.001 | 0.512 | 0.06 | 0.07 | 0.14 | 0.1 | 0.07 | 0.12 |
| 4 | 83648552 | rs1358055 | -0.248 | 0.386 | -0.537 | 0.358 | -0.004 | -0.001 | -0.001 | 0 | 0.36 | 0.34 | 0.43 | 0.31 | 0.39 | 0.37 |
| 4 | 83649829 | rs10009661 | -1.161 | 0.324 | 0.377 | 0.45 | -0.009 | -0.001 | -0.001 | 0 | 0.47 | 0.71 | 0.63 | 0.49 | 0.48 | 0.43 |
| 4 | 83650157 | rs4693501 | -1.341 | 0.259 | -0.144 | 0.311 | -0.12 | -0.108 | -0.108 | 0 | 0.54 | 0.97 | 0.69 | 0.49 | 0.55 | 0.44 |
| 4 | 83652094 | rs17006191 | -0.087 | -0.336 | 0.665 | 0.274 | -0.005 | 0.035 | 0.035 | 0 | 0.1 | 0.17 | 0.19 | 0.2 | 0.07 | 0.1 |
| 4 | 83653968 | rs6822001 | -1.157 | -0.197 | 1.17 | 0.262 | 0.012 | 0.014 | 0.014 | 0.165 | 0.51 | 0.74 | 0.66 | 0.51 | 0.49 | 0.43 |
| 4 | 83654463 | rs4693052 | -0.306 | -0.192 | 1.417 | 0.26 | 0.003 | 0.007 | 0.007 | 0.016 | 0.4 | 0.36 | 0.46 | 0.31 | 0.42 | 0.32 |
| 4 | 83656703 | rs13124051 | 1.031 | -0.29 | 1.032 | 0.167 | -0.034 | -0.02 | -0.02 | 0.042 | 0.45 | 0.15 | 0.32 | 0.5 | 0.49 | 0.55 |
| 4 | 83659028 | rs951500 | -0.332 | -0.661 | 1.329 | 0.303 | 0.015 | 0.049 | 0.049 | 0 | 0.06 | 0.15 | 0.03 | 0.11 | 0.05 | 0.11 |
| 4 | 83659628 | rs11733801 | 1.282 | -0.613 | 0.988 | 0.238 | -0.053 | -0.096 | -0.096 | 0 | 0.44 | 0.04 | 0.44 | 0.49 | 0.53 | 0.55 |
| 4 | 83659974 | rs4626189 | -1.345 | -0.701 | 1.403 | 0.285 | -0.024 | 0.064 | 0.064 | 0 | 0.14 | 0.39 | 0.09 | 0.16 | 0.07 | 0.19 |
| 4 | 83660842 | rs17354029 | 0.586 | -0.927 | 1.298 | 0.102 | 0.005 | 0.041 | 0.041 | 6.982 | 0.12 | 0.06 | 0.08 | 0 | 0.14 | 0.02 |
| 4 | 83661968 | rs12650548 | -1.26 | -0.86 | 1.285 | 0.155 | -0.025 | 0.068 | 0.068 | 0 | 0.14 | 0.4 | 0.09 | 0.16 | 0.07 | 0.18 |
| 4 | 83662588 | rs10019834 | 0.549 | -0.856 | 1.4 | 0.099 | -0.007 | -0.005 | -0.005 | 0 | 0.59 | 0.51 | 0.53 | 0.64 | 0.6 | 0.74 |
| 4 | 83668072 | rs13148235 | 0.254 | -0.585 | 1.642 | 0.528 | -0.002 | 0 | 0 | 1.934 | 0.63 | 0.39 | 0.57 | 0.59 | 0.72 | 0.7 |

| | | | | | | | | | | | | | | | | |
|---|---|---|---|---|---|---|---|---|---|---|---|---|---|---|---|---|
| 4 | 83668410 | rs11933135 | -1.789 | -0.507 | 1.444 | 0.563 | 0.002 | 0.03 | 0.03 | 3.625 | 0.14 | 0.76 | 0.19 | 0.25 | 0.13 | 0.15 |
| 4 | 83673572 | rs13139428 | 0.07 | -1.737 | 0.224 | 0.195 | -0.017 | -0.001 | -0.001 | 1.155 | 0.67 | 0.64 | 0.72 | 0.74 | 0.79 | 0.76 |
| 4 | 83680552 | rs11732247 | -1.25 | -1.137 | 0.642 | 0.074 | -0.027 | -0.044 | -0.044 | 0.051 | 0.21 | 0.38 | 0.1 | 0.1 | 0.14 | 0.17 |
| 4 | 83682200 | rs11729909 | -0.8 | -1.053 | 0.709 | 0.138 | -0.005 | 0.001 | 0.001 | 0 | 0.07 | 0.21 | 0.05 | 0.03 | 0.03 | 0.04 |
| 4 | 83687425 | rs11726264 | -1.774 | -1.229 | 0.574 | -0.08 | -0.026 | -0.05 | -0.05 | 8.495 | 0.32 | 0.5 | 0.13 | 0.17 | 0.21 | 0.21 |
| 4 | 83693278 | rs17351947 | 0.169 | -1.227 | 0.414 | -0.107 | -0.017 | 0.046 | 0.046 | 0 | 0.08 | 0.03 | 0.03 | 0 | 0.06 | 0.03 |
| 4 | 83694283 | rs1980360 | 1.712 | -1.048 | -0.187 | 0.168 | -0.08 | 0.131 | 0.131 | 0.102 | 0.51 | 0.05 | 0.72 | 0.25 | 0.61 | 0.5 |
| 4 | 83697854 | rs11099549 | -1.735 | -1.21 | 0.345 | -0.013 | -0.017 | -0.033 | -0.033 | 0 | 0.32 | 0.47 | 0.14 | 0.18 | 0.21 | 0.2 |
| 4 | 83700755 | rs11099551 | -1.06 | -0.854 | 0.467 | 0.192 | -0.017 | -0.06 | -0.06 | 0 | 0.16 | 0.35 | 0.09 | 0.04 | 0.09 | 0.08 |
| 4 | 83710092 | rs3857139 | -0.234 | -0.827 | 0.566 | 0.613 | 0.015 | 0.027 | 0.027 | 0 | 0.09 | 0.15 | 0.04 | 0.16 | 0.11 | 0.11 |
| 4 | 83719944 | rs6841081 | -1.8 | -0.744 | 0.996 | 0.45 | -0.011 | -0.025 | -0.025 | 0 | 0.05 | 0.19 | 0.02 | 0 | 0.02 | 0.01 |
| 4 | 83725273 | rs17354301 | 0.193 | -1.694 | 0.457 | 0.301 | 0.071 | 0.003 | 0.003 | 0 | 0.33 | 0.08 | 0.13 | 0.26 | 0.22 | 0.31 |
| 4 | 83733766 | rs4235052 | 0.355 | -1.485 | 0.683 | 0.563 | -0.001 | 0.001 | 0.001 | 0 | 0.74 | 0.27 | 0.81 | 0.51 | 0.79 | 0.7 |
| 4 | 83740718 | rs12331100 | 1.019 | -1.61 | 0.627 | 0.479 | -0.101 | 0.021 | 0.021 | 0 | 0.37 | 0.11 | 0.67 | 0.25 | 0.56 | 0.34 |
| 4 | 83758827 | rs1136882 | 0.432 | -1.578 | 0.672 | 0.528 | 0.062 | 0.025 | 0.025 | 0 | 0.35 | 0.17 | 0.14 | 0.26 | 0.23 | 0.36 |
| 4 | 83778839 | rs7658703 | 0.206 | -1.555 | 0.909 | 0.535 | 0.046 | 0.138 | 0.138 | 0.021 | 0.21 | 0.51 | 0.17 | 0.49 | 0.19 | 0.29 |
| 4 | 83791959 | rs3762901 | -0.294 | -1.562 | 0.937 | 0.527 | 0.011 | 0.008 | 0.008 | 0 | 0.61 | 0.64 | 0.84 | 0.74 | 0.77 | 0.64 |
| 4 | 83795806 | rs10025654 | 0.837 | -1.465 | 0.877 | 0.654 | -0.092 | 0.011 | 0.011 | 0 | 0.39 | 0.1 | 0.68 | 0.26 | 0.56 | 0.34 |
| 4 | 83798446 | rs6830527 | 0.771 | -1.528 | 1.284 | 0.509 | -0.026 | -0.04 | -0.04 | 0.674 | 0.16 | 0.1 | 0.13 | 0.49 | 0.19 | 0.29 |
| 4 | 83800428 | rs7656058 | -0.63 | -1.475 | 0.868 | 0.547 | -0.003 | 0.041 | 0.041 | 0 | 0.55 | 0.64 | 0.31 | 0.75 | 0.42 | 0.65 |
| 4 | 83807801 | rs7682147 | -0.38 | -1.413 | 1.29 | 0.581 | -0.047 | -0.024 | -0.024 | 0.012 | 0.62 | 0.53 | 0.83 | 0.74 | 0.77 | 0.64 |
| 4 | 83814018 | rs1020748 | 0.165 | -1.439 | 1.335 | 0.703 | 0.005 | 0.049 | 0.049 | 0 | 0.75 | 0.35 | 0.83 | 0.51 | 0.8 | 0.7 |
| 4 | 83817693 | rs1031640 | 0.25 | -1.448 | 1.298 | 0.695 | 0.004 | 0.036 | 0.036 | 0 | 0.74 | 0.33 | 0.83 | 0.51 | 0.8 | 0.7 |
| 4 | 83819464 | rs1350386 | 0.261 | -1.454 | 1.301 | 0.752 | 0.004 | 0.031 | 0.031 | 0 | 0.74 | 0.34 | 0.83 | 0.53 | 0.8 | 0.7 |
| 4 | 83820170 | rs4302451 | -0.151 | -1.37 | 1.135 | 0.776 | 0 | 0.013 | 0.013 | 0 | 0.69 | 0.36 | 0.79 | 0.52 | 0.76 | 0.69 |
| 4 | 83833473 | rs10857215 | -0.933 | -1.432 | 0.925 | 0.365 | -0.055 | 0.066 | 0.066 | 0 | 0.63 | 0.81 | 0.35 | 0.8 | 0.43 | 0.65 |
| 4 | 83838262 | rs897945 | 0.964 | -1.407 | 0.929 | 0.361 | -0.125 | 0.047 | 0.047 | 0 | 0.34 | 0.07 | 0.65 | 0.2 | 0.55 | 0.34 |
| 4 | 83840906 | rs1047564 | -0.932 | -1.405 | 0.942 | 0.362 | -0.056 | 0.067 | 0.067 | 0 | 0.63 | 0.82 | 0.35 | 0.8 | 0.43 | 0.65 |
| 4 | 83848388 | rs13132580 | 0.702 | -1.358 | 1.011 | 0.398 | 0.048 | -0.001 | -0.001 | 0 | 0.31 | 0.16 | 0.14 | 0.27 | 0.18 | 0.33 |
| 4 | 83862479 | rs1473394 | 0.702 | -1.358 | 0.979 | 0.398 | 0.05 | -0.002 | -0.002 | 0 | 0.31 | 0.16 | 0.14 | 0.27 | 0.19 | 0.33 |
| 4 | 83873373 | rs11938691 | 0.463 | -1.362 | 0.98 | 0.396 | 0.035 | 0.004 | 0.004 | 0 | 0.33 | 0.22 | 0.15 | 0.27 | 0.2 | 0.34 |
| 4 | 83899764 | rs10000180 | -0.02 | -1.432 | 1.022 | 0.417 | 0.003 | 0.038 | 0.038 | 0.015 | 0.31 | 0.55 | 0.2 | 0.51 | 0.24 | 0.32 |
| 4 | 83903849 | rs1020750 | 0.71 | -1.424 | 0.725 | 0.421 | 0.051 | -0.002 | -0.002 | 0 | 0.31 | 0.16 | 0.14 | 0.27 | 0.18 | 0.34 |
| 4 | 83907105 | rs11099563 | 0.134 | -1.259 | 0.895 | 0.498 | 0.005 | 0.024 | 0.024 | 0 | 0.28 | 0.35 | 0.18 | 0.51 | 0.24 | 0.32 |
| 4 | 83923869 | rs9307811 | -0.196 | -1.288 | 0.944 | 0.501 | 0.013 | 0.055 | 0.055 | 0 | 0.7 | 0.53 | 0.81 | 0.49 | 0.74 | 0.67 |
| 4 | 83929394 | rs7691430 | 0.869 | -1.292 | 0.555 | 0.501 | -0.126 | 0.024 | 0.024 | 0 | 0.34 | 0.07 | 0.65 | 0.22 | 0.55 | 0.33 |
| 4 | 83939171 | rs10033399 | -0.783 | -1.267 | 0.398 | 0.464 | 0.037 | 0.005 | 0.005 | 0 | 0.67 | 0.77 | 0.85 | 0.73 | 0.81 | 0.66 |
| 4 | 83954383 | rs6821110 | 0.117 | 0.085 | -0.45 | 0.299 | -0.172 | -0.015 | -0.015 | 2.497 | 0.33 | 0.02 | 0.65 | 0.23 | 0.55 | 0.34 |
| 4 | 83997158 | rs11947785 | -1.477 | 0.049 | 0.676 | 0.536 | 0.034 | 0.026 | 0.026 | 2.008 | 0.05 | 0.3 | 0.03 | 0 | 0 | 0 |
| 4 | 84005377 | rs11732191 | 0.625 | 0.118 | 0.186 | 0.814 | -0.006 | 0.009 | 0.009 | 0.066 | 0.75 | 0.64 | 0.89 | 0.68 | 0.83 | 0.77 |
| 4 | 84011131 | rs11722733 | -1.511 | -0.611 | 1.075 | 0.284 | -0.045 | -0.065 | -0.065 | 0.109 | 0.12 | 0.31 | 0.05 | 0.01 | 0.05 | 0.15 |
| 4 | 84013292 | rs11099568 | 0.234 | -0.252 | -0.157 | 0.264 | -0.072 | -0.05 | -0.05 | 0.005 | 0.4 | 0.14 | 0.64 | 0.42 | 0.54 | 0.45 |
| 4 | 84013509 | rs11099569 | 0.058 | -0.349 | 0.011 | 0.227 | -0.017 | -0.007 | -0.007 | 0 | 0.42 | 0.27 | 0.65 | 0.42 | 0.54 | 0.45 |
| 4 | 84017945 | rs7692926 | 0.826 | -0.614 | 0.324 | -0.334 | 0.001 | 0.023 | 0.023 | 0 | 0.32 | 0.33 | 0.22 | 0.58 | 0.3 | 0.28 |
| 4 | 84018272 | rs6846520 | -0.05 | -0.773 | 0.211 | -0.561 | 0.006 | 0.002 | 0.002 | 0 | 0.85 | 0.88 | 0.92 | 1 | 0.9 | 0.88 |
| 4 | 84026991 | rs7688441 | -0.234 | -0.283 | 0.715 | -0.191 | 0.007 | 0.004 | 0.004 | 0 | 0.46 | 0.42 | 0.27 | 0.39 | 0.39 | 0.31 |
| 4 | 84034675 | rs2868763 | 0.652 | 0.011 | 0.244 | -0.467 | -0.044 | -0.021 | -0.021 | 0 | 0.4 | 0.19 | 0.51 | 0.59 | 0.49 | 0.57 |
| 4 | 84045915 | rs2868750 | 0.289 | -0.187 | 0.619 | -0.645 | 0.013 | -0.052 | -0.052 | 1.34 | 0.89 | 0.8 | 0.81 | 1 | 0.88 | 0.94 |
| 4 | 84053945 | rs12510343 | 0.4 | -0.615 | 0.688 | -0.645 | -0.001 | -0.001 | -0.001 | 0 | 0.46 | 0.48 | 0.56 | 0.62 | 0.58 | 0.56 |
| 4 | 84058227 | rs2868752 | -0.041 | -1.043 | 0.033 | -0.978 | 0.01 | 0.001 | 0.001 | 0 | 0.62 | 0.69 | 0.77 | 0.63 | 0.72 | 0.68 |
| 4 | 84059844 | rs11940883 | -0.779 | -0.76 | 0.736 | -1.079 | 0.024 | -0.144 | -0.144 | 0 | 0.18 | 0.46 | 0.25 | 0.01 | 0.17 | 0.11 |
| 4 | 84065038 | rs6852740 | 1.505 | -0.873 | 0.926 | -0.796 | 0.036 | 0.026 | 0.026 | 0.244 | 0.94 | 0.59 | 0.96 | 0.99 | 0.99 | 0.99 |
| 4 | 84072132 | rs7657984 | -0.071 | -0.886 | 0.366 | -0.763 | -0.016 | -0.004 | -0.004 | 0 | 0.27 | 0.35 | 0.19 | 0.25 | 0.25 | 0.21 |
| 4 | 84072737 | rs9999963 | -0.698 | -1.632 | 0.202 | -1.057 | 0.004 | 0.021 | 0.021 | 0 | 0.05 | 0.01 | 0.04 | 0 | 0.03 | 0.05 |
| 4 | 84074826 | rs10516683 | 1.681 | -2.014 | 0.069 | -1.534 | 0.045 | 0.093 | 0.093 | 0.161 | 0.61 | 0.2 | 0.68 | 0.99 | 0.76 | 0.79 |
| 4 | 84091444 | rs7673451 | 1.462 | -1.726 | 0.757 | -1.11 | -0.063 | -0.14 | -0.14 | 1.713 | 0.8 | 0.45 | 0.89 | 1 | 0.92 | 0.92 |
| 4 | 84100592 | rs2868760 | 0.749 | -1.158 | 0.168 | -0.005 | -0.024 | 0.029 | 0.029 | 0 | 0.06 | 0.01 | 0.12 | 0.34 | 0.07 | 0.12 |
| 4 | 84120705 | rs7679817 | -0.036 | -1.686 | 0.951 | -1.097 | -0.024 | 0.025 | 0.025 | 0.586 | 0.12 | 0.41 | 0.07 | 0.1 | 0.05 | 0.08 |
| 4 | 84123698 | rs2903913 | 1.053 | -1.714 | 0.879 | -1.242 | 0.044 | -0.056 | -0.056 | 1.782 | 0.9 | 0.57 | 0.9 | 1 | 0.92 | 0.99 |
| 4 | 84130153 | rs12508182 | 1.264 | -2.367 | -0.189 | -1.746 | 0.041 | 0.164 | 0.164 | 0 | 0.45 | 0.05 | 0.7 | 0.77 | 0.64 | 0.67 |
| 4 | 84131775 | rs10033388 | 0.568 | -1.478 | 0.873 | -1.202 | -0.014 | 0.021 | 0.021 | 0.652 | 0.1 | 0.2 | 0.06 | 0.1 | 0.05 | 0.08 |
| 4 | 84133315 | rs11731396 | 1.013 | -1.451 | 0.35 | -1.204 | -0.019 | -0.038 | -0.038 | 0.021 | 0.8 | 0.42 | 0.86 | 0.9 | 0.87 | 0.91 |
| 4 | 84140400 | rs6854272 | 0.801 | -1.386 | 0.082 | -1.217 | 0.005 | 0.006 | 0.006 | 1.244 | 0.78 | 0.26 | 0.72 | 0.82 | 0.84 | 0.88 |
| 4 | 84143589 | rs6856697 | -1.226 | -1.961 | -0.433 | -1.664 | 0.083 | 0.186 | 0.186 | 0 | 0.48 | 0.91 | 0.41 | 0.28 | 0.32 | 0.34 |
| 4 | 84145617 | rs11942687 | -0.288 | -2.034 | 0.235 | -1.659 | -0.021 | -0.023 | -0.023 | 0 | 0.39 | 0.46 | 0.34 | 0.28 | 0.26 | 0.29 |
| 4 | 84151064 | rs17352615 | 1.358 | -1.769 | -0.11 | -1.664 | -0.042 | -0.005 | -0.005 | 0 | 0.09 | 0.01 | 0.05 | 0.08 | 0.1 | 0.04 |
| 4 | 84152973 | rs13122305 | 1.175 | -2.391 | 0.128 | -1.825 | 0.074 | 0.139 | 0.139 | 0 | 0.53 | 0.07 | 0.59 | 0.7 | 0.69 | 0.68 |
| 4 | 84153610 | rs6840620 | -0.738 | -2.413 | 0.445 | -1.814 | 0.044 | -0.023 | -0.023 | 0.169 | 0.06 | 0.28 | 0.05 | 0 | 0.06 | 0.05 |
| 4 | 84156465 | rs6819374 | 0.016 | -2.386 | 0.381 | -1.801 | 0.027 | -0.027 | -0.027 | 2.426 | 0.05 | 0.16 | 0.04 | 0 | 0.06 | 0.05 |

| | | | | | | | | | | | | | | | | |
|---|---|---|---|---|---|---|---|---|---|---|---|---|---|---|---|---|
| 4 | 84168653 | rs4396979 | -0.058 | -2.391 | 0.734 | -1.838 | -0.03 | -0.094 | -0.094 | 0 | 0.33 | 0.57 | 0.16 | 0.12 | 0.21 | 0.25 |
| 4 | 84175235 | rs4346637 | 0.019 | -2.405 | 0.673 | -1.783 | -0.03 | 0.022 | 0.022 | 0 | 0.16 | 0.39 | 0.24 | 0.18 | 0.11 | 0.08 |
| 4 | 84177179 | rs10002954 | 0.327 | -2.443 | 0.64 | -1.818 | -0.008 | -0.037 | -0.037 | 0 | 0.3 | 0.42 | 0.15 | 0.12 | 0.21 | 0.25 |
| 4 | 84179480 | rs6535450 | 0.298 | -2.46 | 0.62 | -1.826 | -0.007 | -0.035 | -0.035 | 0.043 | 0.29 | 0.41 | 0.15 | 0.12 | 0.2 | 0.25 |
| 4 | 84182602 | rs10021552 | -0.922 | -2.319 | 0.234 | -1.681 | 0.073 | 0.114 | 0.114 | 0 | 0.46 | 0.81 | 0.39 | 0.31 | 0.32 | 0.33 |
| 4 | 84189869 | rs4693596 | 0.995 | -2.113 | 0.66 | -2.345 | 0.047 | 0.102 | 0.102 | 0 | 0.5 | 0.31 | 0.55 | 0.87 | 0.64 | 0.67 |
| 4 | 84201479 | rs11099588 | 0.736 | -1.81 | 1.226 | -1.76 | 0.021 | 0.004 | 0.004 | 0.135 | 0.93 | 0.72 | 0.96 | 0.99 | 0.97 | 0.95 |
| 4 | 84202683 | rs6849677 | 1.487 | -1.787 | 1.117 | -1.76 | 0.018 | -0.001 | -0.001 | 0 | 0.92 | 0.63 | 0.96 | 0.99 | 0.97 | 0.95 |
| 4 | 84216941 | rs10025120 | 0.661 | -1.82 | 0.456 | -1.785 | 0.011 | 0.009 | 0.009 | 0 | 0.58 | 0.35 | 0.65 | 0.88 | 0.73 | 0.7 |
| 4 | 84220612 | rs9307824 | 1.274 | -1.012 | 1.005 | -0.95 | 0.021 | 0.004 | 0.004 | 0.061 | 0.94 | 0.77 | 0.97 | 0.99 | 0.97 | 0.95 |
| 4 | 84226063 | rs11726245 | -1.252 | -0.919 | 0.673 | -0.961 | -0.007 | -0.034 | -0.034 | 0 | 0.11 | 0.25 | 0.21 | 0.06 | 0.1 | 0.11 |
| 4 | 84230619 | rs11099592 | 0.897 | -0.818 | 0.344 | -1.099 | -0.003 | 0.026 | 0.026 | 0.099 | 0.8 | 0.88 | 0.68 | 0.87 | 0.75 | 0.83 |
| 4 | 84233449 | rs6535458 | 0.917 | -0.811 | 0.348 | -1.095 | -0.004 | 0.023 | 0.023 | 0.157 | 0.8 | 0.88 | 0.68 | 0.87 | 0.75 | 0.83 |
| 4 | 84241357 | rs4693608 | 0.335 | -0.685 | 0.304 | -1.263 | -0.035 | 0.122 | 0.122 | 0 | 0.56 | 0.82 | 0.54 | 0.8 | 0.5 | 0.68 |
| 4 | 84243549 | rs4328905 | -0.264 | -0.704 | 1.673 | -0.432 | -0.039 | 0.143 | 0.143 | 0 | 0.18 | 0.3 | 0.19 | 0.42 | 0.12 | 0.29 |
| 4 | 84246090 | rs7670729 | 1.261 | -0.712 | 1.034 | -0.499 | 0.044 | -0.06 | -0.06 | 0 | 0.67 | 0.23 | 0.58 | 0.58 | 0.74 | 0.59 |
| 4 | 84247056 | rs11722753 | -1.512 | 0.083 | 1.506 | -1.202 | -0.002 | -0.056 | -0.056 | 0 | 0.13 | 0.2 | 0.21 | 0.01 | 0.15 | 0.12 |
| 4 | 84253150 | rs4693612 | 1.287 | -0.026 | 1.293 | -1.058 | -0.002 | -0.073 | -0.073 | 0.234 | 0.87 | 0.65 | 0.89 | 0.99 | 0.87 | 0.88 |
| 4 | 84254494 | rs12507864 | -1.024 | 0.099 | 0.366 | -0.97 | -0.006 | -0.06 | -0.06 | 0 | 0.13 | 0.29 | 0.1 | 0.01 | 0.12 | 0.13 |
| 4 | 84260521 | rs4693614 | -0.89 | 0.325 | 0.187 | -0.722 | -0.001 | -0.001 | -0.001 | 0.026 | 0.25 | 0.22 | 0.31 | 0.1 | 0.24 | 0.16 |
| 4 | 84282828 | rs6831788 | -0.805 | -0.4 | 0.541 | -0.94 | -0.04 | -0.128 | -0.128 | 0 | 0.29 | 0.43 | 0.29 | 0.02 | 0.24 | 0.22 |
| 4 | 84288014 | rs7436836 | 0.087 | -1.455 | -0.05 | -1.499 | 0.05 | 0.058 | 0.058 | 0.024 | 0.58 | 0.34 | 0.75 | 0.83 | 0.75 | 0.7 |
| 4 | 84288613 | rs6854611 | -0.217 | 0.382 | -0.737 | -0.499 | -0.081 | -0.107 | -0.107 | 0 | 0.19 | 0.08 | 0.28 | 0.51 | 0.32 | 0.32 |
| 4 | 84289142 | rs6535469 | 0.048 | -0.229 | -0.632 | -1.039 | -0.001 | -0.038 | -0.038 | 0 | 0.72 | 0.45 | 0.77 | 0.84 | 0.77 | 0.72 |
| 4 | 84290641 | rs4311309 | 0.514 | -0.146 | -0.837 | -0.864 | -0.093 | -0.118 | -0.118 | 0 | 0.19 | 0.06 | 0.28 | 0.51 | 0.32 | 0.33 |
| 4 | 84295261 | rs12650866 | 0.207 | -0.818 | -0.273 | -1.238 | 0.027 | 0.077 | 0.077 | 0.842 | 0.68 | 0.16 | 0.76 | 0.84 | 0.79 | 0.72 |
| 4 | 84298192 | rs13128996 | -0.028 | -0.755 | 0.999 | -1.293 | 0.098 | 0.014 | 0.014 | 0.044 | 0.09 | 0.34 | 0.06 | 0.06 | 0.07 | 0.07 |
| 4 | 84304734 | rs6831180 | -1.53 | -0.757 | 0.757 | -1.425 | 0.018 | 0.052 | 0.052 | 0.208 | 0.29 | 0.82 | 0.23 | 0.15 | 0.17 | 0.25 |
| 4 | 84310711 | rs10516684 | -0.569 | -0.965 | 1.316 | -1.525 | -0.009 | -0.049 | -0.049 | 1.216 | 0.18 | 0.22 | 0.09 | 0 | 0.1 | 0.11 |
| 4 | 84312759 | rs10004897 | -1.756 | -0.876 | 1.264 | -1.538 | 0.022 | 0.047 | 0.047 | 0.475 | 0.29 | 0.8 | 0.23 | 0.1 | 0.16 | 0.25 |
| 4 | 84317759 | rs7681313 | -1.766 | -0.952 | 1.063 | -1.77 | 0.027 | 0.086 | 0.086 | 0 | 0.3 | 0.85 | 0.16 | 0.01 | 0.17 | 0.18 |
| 4 | 84325901 | rs17006780 | -0.559 | -0.96 | 2.025 | -1.771 | 0.062 | -0.044 | -0.044 | 0.311 | 0.1 | 0.4 | 0.05 | 0.01 | 0.06 | 0.06 |
| 4 | 84330059 | rs17006802 | -0.374 | -0.988 | 2.125 | -1.76 | 0.018 | 0.056 | 0.056 | 0 | 0.17 | 0.1 | 0.08 | 0 | 0.11 | 0.12 |
| 4 | 84331768 | rs4693623 | -1.571 | -0.163 | 0.958 | -1.562 | -0.006 | 0.047 | 0.047 | 1.187 | 0.45 | 0.79 | 0.66 | 0.69 | 0.42 | 0.58 |
| 4 | 84364751 | rs7665103 | -0.576 | -0.173 | 1.146 | -1.428 | 0.042 | -0.006 | -0.006 | 0 | 0.31 | 0.45 | 0.54 | 0.67 | 0.38 | 0.52 |
| 4 | 84370091 | rs13141136 | -0.142 | -0.219 | 1.207 | -1.443 | -0.072 | -0.018 | -0.018 | 0 | 0.27 | 0.11 | 0.52 | 0.68 | 0.38 | 0.52 |
| 4 | 84372998 | rs4693626 | -2.253 | -0.707 | 0.413 | -1.693 | -0.019 | 0.04 | 0.04 | 0.283 | 0.86 | 0.98 | 0.86 | 0.91 | 0.82 | 0.86 |
| 4 | 84374480 | rs1494961 | -1.684 | -0.55 | 0.931 | -1.497 | 0.008 | 0.119 | 0.119 | 0.176 | 0.51 | 0.83 | 0.64 | 0.68 | 0.49 | 0.63 |
| 4 | 84374692 | rs17006837 | -1.321 | -0.546 | 1.087 | -1.488 | 0.065 | 0.061 | 0.061 | 1.699 | 0.05 | 0.49 | 0.03 | 0 | 0 | 0 |
| 4 | 84380955 | rs12505955 | 0.352 | -0.548 | 0.764 | -1.497 | -0.057 | -0.013 | -0.013 | 0 | 0.27 | 0.12 | 0.53 | 0.68 | 0.38 | 0.52 |
| 4 | 84382763 | rs11099601 | -1.7 | -0.557 | 0.772 | -1.505 | 0.01 | 0.117 | 0.117 | 0.15 | 0.51 | 0.82 | 0.64 | 0.68 | 0.49 | 0.63 |
| 4 | 84383810 | rs12642536 | 0.942 | -0.5 | 0.765 | -1.508 | -0.057 | -0.013 | -0.013 | 0.12 | 0.27 | 0.12 | 0.53 | 0.68 | 0.38 | 0.52 |
| 4 | 84384995 | rs10516689 | 0.942 | -0.5 | 0.765 | -1.508 | -0.057 | -0.013 | -0.013 | 0 | 0.27 | 0.12 | 0.52 | 0.68 | 0.38 | 0.52 |
| 4 | 84387157 | rs12499395 | 0.942 | -0.5 | 0.765 | -1.508 | -0.057 | -0.013 | -0.013 | 0 | 0.27 | 0.12 | 0.53 | 0.68 | 0.38 | 0.52 |
| 4 | 84388915 | rs17355027 | 1.015 | -0.473 | 0.856 | -1.451 | -0.006 | -0.045 | -0.045 | 0 | 0.17 | 0.17 | 0.08 | 0 | 0.11 | 0.11 |
| 4 | 84400120 | rs17006851 | -1.478 | -0.46 | 0.755 | -1.369 | 0.048 | 0.044 | 0.044 | 2.548 | 0.06 | 0.58 | 0.04 | 0 | 0 | 0 |
| 4 | 84414282 | rs28488925 | 1.21 | -0.411 | 0.21 | -1.413 | -0.006 | 0.128 | 0.128 | 0.068 | 0.5 | 0.13 | 0.35 | 0.32 | 0.51 | 0.38 |
| 4 | 84434011 | rs17352915 | 0.967 | -0.298 | 0.379 | -1.344 | -0.003 | -0.042 | -0.042 | 0 | 0.17 | 0.14 | 0.09 | 0 | 0.11 | 0.08 |
| 4 | 84444097 | rs9999564 | -0.92 | -0.141 | -0.186 | -1.019 | -0.054 | -0.012 | -0.012 | 1.046 | 0.71 | 0.85 | 0.48 | 0.33 | 0.62 | 0.45 |
| 4 | 84456409 | rs9992543 | 0.877 | 0.055 | -0.463 | -0.837 | 0.006 | 0.165 | 0.165 | 1.297 | 0.5 | 0.14 | 0.36 | 0.26 | 0.5 | 0.36 |
| 4 | 84465613 | rs10516690 | 0.686 | -0.31 | -0.465 | -0.825 | 0.061 | 0.14 | 0.14 | 0.364 | 0.47 | 0.15 | 0.33 | 0.26 | 0.42 | 0.33 |
| 4 | 84515775 | rs1474250 | 0.838 | -0.134 | 0.33 | -0.534 | 0.005 | 0.007 | 0.007 | 0.24 | 0.36 | 0.27 | 0.28 | 0.28 | 0.34 | 0.28 |
| 4 | 84518315 | rs3733328 | -0.445 | -0.017 | -0.046 | -0.051 | 0.022 | 0.021 | 0.021 | 0.018 | 0.72 | 0.33 | 0.57 | 0.59 | 0.63 | 0.63 |
| 4 | 84518558 | rs3733329 | 0.396 | 0.002 | -0.191 | -0.042 | -0.043 | -0.058 | -0.058 | 1.774 | 0.37 | 0.03 | 0.29 | 0.31 | 0.29 | 0.35 |
| 4 | 84519076 | rs1379974 | -0.684 | 0.087 | -0.004 | 0.005 | 0.076 | 0.073 | 0.073 | 0 | 0.75 | 0.48 | 0.58 | 0.59 | 0.63 | 0.63 |
| 4 | 84526275 | rs1521531 | -1.115 | 0.094 | -0.165 | 0.009 | 0.045 | 0.045 | 0.045 | 0 | 0.76 | 0.63 | 0.59 | 0.59 | 0.64 | 0.63 |
| 4 | 84526536 | rs1521532 | 0.218 | 0.117 | -1.177 | 0.038 | -0.019 | -0.033 | -0.033 | 0 | 0.37 | 0.08 | 0.29 | 0.31 | 0.29 | 0.35 |
| 4 | 84550759 | rs1379977 | 0.394 | -0.214 | -0.561 | 0.011 | 0.004 | 0.005 | 0.005 | 0.773 | 0.36 | 0.32 | 0.28 | 0.28 | 0.34 | 0.28 |
| 4 | 84552920 | rs6818782 | -0.103 | -0.121 | -1.003 | -0.313 | -0.016 | 0.049 | 0.049 | 0 | 0.53 | 0.66 | 0.5 | 0.59 | 0.46 | 0.52 |
| 4 | 84554429 | rs17359809 | 0.401 | -0.346 | -0.971 | -0.422 | -0.046 | 0.014 | 0.014 | 0.021 | 0.32 | 0.15 | 0.25 | 0.27 | 0.36 | 0.36 |
| 4 | 84558656 | rs781443 | -0.235 | -0.738 | -0.402 | -0.796 | -0.004 | -0.006 | -0.006 | 0 | 0.54 | 0.45 | 0.55 | 0.34 | 0.55 | 0.49 |
| 4 | 84560219 | rs11099608 | 0.092 | -0.579 | -0.115 | -1.115 | 0.012 | -0.009 | -0.009 | 0.146 | 0.38 | 0.43 | 0.46 | 0.34 | 0.41 | 0.4 |
| 4 | 84562362 | rs1458642 | 0.303 | -0.83 | -0.295 | -0.848 | 0 | 0.002 | 0.002 | 0 | 0.37 | 0.38 | 0.32 | 0.29 | 0.33 | 0.32 |
| 4 | 84566027 | rs1458650 | -0.85 | -0.973 | -0.605 | -1.115 | -0.032 | -0.067 | -0.067 | 0 | 0.64 | 0.79 | 0.54 | 0.48 | 0.54 | 0.53 |
| 4 | 84566072 | rs2061408 | 1.688 | -0.879 | -0.905 | -1.077 | -0.007 | 0.08 | 0.08 | 0 | 0.15 | 0.03 | 0.08 | 0 | 0.11 | 0.08 |
| 4 | 84569279 | rs13151073 | 0.25 | -0.845 | -0.656 | -0.979 | 0.004 | -0.004 | -0.004 | 0 | 0.28 | 0.24 | 0.17 | 0.28 | 0.21 | 0.24 |
| 4 | 84571718 | rs1674338 | 1.774 | -0.725 | -1.117 | -0.961 | -0.006 | -0.004 | -0.004 | 0 | 0.21 | 0.13 | 0.23 | 0.19 | 0.22 | 0.23 |
| 4 | 84578678 | rs17007085 | 0.326 | -0.524 | -0.992 | -0.653 | -0.011 | -0.043 | -0.043 | 0.146 | 0.25 | 0.1 | 0.17 | 0.3 | 0.22 | 0.24 |
| 4 | 84580763 | rs7694734 | 0.969 | -0.733 | -0.778 | -0.875 | 0.002 | 0.004 | 0.004 | 0.001 | 0.3 | 0.27 | 0.37 | 0.23 | 0.34 | 0.28 |

| | | | | | | | | | | | | | | | | |
|---|---|---|---|---|---|---|---|---|---|---|---|---|---|---|---|---|
| 4 | 84584157 | rs13118676 | 0.136 | -0.806 | -0.371 | -0.863 | -0.011 | 0.018 | 0.018 | 0 | 0.28 | 0.44 | 0.18 | 0.3 | 0.22 | 0.29 |
| 4 | 84584220 | rs4693096 | 0.238 | -0.88 | -0.542 | -0.954 | -0.003 | 0 | 0 | 0 | 0.62 | 0.4 | 0.66 | 0.62 | 0.7 | 0.61 |
| 4 | 84587179 | rs1547507 | 0.182 | -0.885 | -0.43 | -0.971 | -0.009 | 0.001 | 0.001 | 0.018 | 0.63 | 0.44 | 0.66 | 0.62 | 0.7 | 0.63 |
| 4 | 84588508 | rs17360129 | 0.142 | -0.815 | -0.949 | -1.028 | -0.02 | 0.072 | 0.072 | 0.198 | 0.45 | 0.18 | 0.36 | 0.68 | 0.49 | 0.6 |
| 4 | 84589904 | rs11099613 | 1.57 | -0.688 | -1.156 | -0.942 | -0.068 | -0.087 | -0.087 | 0.594 | 0.2 | 0.04 | 0.19 | 0.39 | 0.27 | 0.32 |
| 4 | 84597304 | rs7694527 | 0.214 | -0.772 | -1.098 | -1.03 | -0.007 | 0.003 | 0.003 | 0 | 0.56 | 0.36 | 0.53 | 0.53 | 0.56 | 0.53 |
| 4 | 84598254 | rs17007149 | 1.092 | -0.395 | -0.776 | -0.874 | 0.006 | -0.027 | -0.027 | 0 | 0.12 | 0.19 | 0.14 | 0.05 | 0.14 | 0.08 |
| 4 | 84598282 | rs17007150 | -0.007 | -0.371 | -0.502 | -0.686 | -0.014 | -0.112 | -0.112 | 0 | 0.15 | 0.43 | 0.11 | 0 | 0.09 | 0.05 |
| 4 | 84602082 | rs7693390 | 0.66 | -0.1 | -0.615 | -0.512 | 0 | 0.001 | 0.001 | 0 | 0.31 | 0.28 | 0.38 | 0.22 | 0.34 | 0.24 |
| 4 | 84603816 | rs13117154 | 0.483 | 0.058 | -0.49 | -0.309 | 0.106 | 0.384 | 0.384 | 0 | 0.07 | 0.31 | 0.05 | 0.3 | 0.06 | 0.13 |
| 4 | 84603852 | rs17356247 | 1.109 | 0.046 | -1.039 | -0.392 | -0.051 | -0.071 | -0.071 | 0 | 0.29 | 0.05 | 0.24 | 0.4 | 0.32 | 0.43 |
| 4 | 84604639 | rs7340961 | -0.591 | 0.36 | -0.054 | 0.239 | 0.112 | 0.435 | 0.435 | 0.26 | 0.08 | 0.48 | 0.06 | 0.3 | 0.06 | 0.13 |
| 4 | 84606737 | rs6839928 | -0.98 | 0.277 | -0.685 | -0.541 | 0 | 0 | 0 | 1.087 | 0.39 | 0.37 | 0.49 | 0.22 | 0.4 | 0.29 |
| 4 | 84606822 | rs17356270 | -0.27 | 0.047 | -0.752 | -0.498 | -0.001 | -0.003 | -0.003 | 0.006 | 0.08 | 0.05 | 0.08 | 0 | 0.07 | 0.05 |
| 4 | 84624474 | rs7677300 | -0.549 | -0.023 | -0.585 | -0.444 | 0 | -0.014 | -0.014 | 0 | 0.08 | 0.07 | 0.04 | 0 | 0.05 | 0.02 |
| 4 | 84626069 | rs6849805 | -1.3 | -0.656 | -0.684 | -1.025 | 0.033 | 0.062 | 0.062 | 2.575 | 0.34 | 0.73 | 0.16 | 0.08 | 0.21 | 0.12 |
| 4 | 84633401 | rs10516696 | -1.284 | -0.728 | -0.498 | -1.066 | 0.023 | 0.037 | 0.037 | 1.502 | 0.33 | 0.68 | 0.16 | 0.08 | 0.21 | 0.12 |
| 4 | 84636196 | rs17007216 | 0.194 | -0.749 | 0.305 | -1.049 | -0.005 | -0.025 | -0.025 | 0 | 0.17 | 0.13 | 0.06 | 0 | 0.08 | 0.07 |
| 4 | 84636731 | rs6814874 | 0.129 | -0.482 | 1.062 | -0.884 | 0.047 | 0.021 | 0.021 | 0 | 0.06 | 0.13 | 0.04 | 0.08 | 0.08 | 0.03 |
| 4 | 84639697 | rs7660262 | -0.773 | -0.483 | 1.495 | -0.885 | 0.107 | 0.06 | 0.06 | 0.258 | 0.08 | 0.35 | 0.06 | 0.08 | 0.08 | 0.03 |
| 4 | 84648524 | rs4082474 | 0.267 | -0.272 | 1.003 | -0.459 | 0.045 | 0.019 | 0.019 | 0.005 | 0.23 | 0.25 | 0.42 | 0.29 | 0.38 | 0.41 |
| 4 | 84652935 | rs6821747 | 0.111 | -0.312 | 1.151 | -0.474 | 0.066 | 0.029 | 0.029 | 0.014 | 0.24 | 0.29 | 0.43 | 0.29 | 0.39 | 0.41 |
| 4 | 84654205 | rs1352436 | -1.217 | -1.132 | 0.66 | -0.965 | 0.005 | -0.023 | -0.023 | 0 | 0.63 | 0.78 | 0.62 | 0.39 | 0.65 | 0.54 |
| 4 | 84658591 | rs1458649 | -1.271 | -1.401 | 0.805 | -1.179 | 0.011 | -0.007 | -0.007 | 0 | 0.82 | 0.88 | 0.94 | 1 | 0.9 | 0.93 |
| 4 | 84658805 | rs2125426 | 1.022 | -0.935 | 0.809 | -1.042 | -0.04 | -0.081 | -0.081 | 0 | 0.37 | 0.17 | 0.39 | 0.52 | 0.42 | 0.45 |
| 4 | 84659390 | rs9307828 | -0.158 | -0.944 | 0.945 | -1.078 | 0.036 | 0.013 | 0.013 | 0 | 0.63 | 0.73 | 0.82 | 1 | 0.72 | 0.88 |
| 4 | 84664343 | rs10004130 | 1.016 | -0.938 | 0.982 | -1.068 | -0.024 | -0.056 | -0.056 | 0 | 0.37 | 0.24 | 0.4 | 0.52 | 0.42 | 0.46 |
| 4 | 84671227 | rs1458647 | 0.722 | -1.101 | 1.165 | -0.952 | -0.006 | -0.006 | -0.006 | 0 | 0.23 | 0.17 | 0.4 | 0.38 | 0.3 | 0.38 |
| 4 | 84673934 | rs1488652 | 0.489 | -1.089 | 0.679 | -0.939 | 0.068 | 0.097 | 0.097 | 0 | 0.25 | 0.4 | 0.41 | 0.39 | 0.31 | 0.38 |
| 4 | 84676270 | rs4693641 | 0.634 | -2.105 | 0.802 | -1.528 | -0.025 | -0.089 | -0.089 | 0.002 | 0.57 | 0.43 | 0.78 | 0.93 | 0.7 | 0.84 |
| 4 | 84684230 | rs1993570 | 0.685 | -1.995 | 0.444 | -1.486 | 0.024 | 0.017 | 0.017 | 0.98 | 0.56 | 0.58 | 0.78 | 0.89 | 0.7 | 0.85 |
| 4 | 84684277 | rs17356652 | 0.129 | -1.429 | 0.088 | -1.219 | 0.013 | 0.032 | 0.032 | 0 | 0.23 | 0.15 | 0.13 | 0 | 0.2 | 0.06 |
| 4 | 84700945 | rs4629440 | -0.848 | -0.892 | 0.468 | -0.736 | 0.071 | 0.114 | 0.114 | 7.508 | 0.1 | 0.47 | 0.08 | 0.14 | 0.09 | 0.07 |
| 4 | 84706464 | rs7679666 | 0.578 | -1.288 | -0.986 | -0.828 | -0.111 | -0.048 | -0.048 | 0.084 | 0.33 | 0.05 | 0.49 | 0.66 | 0.44 | 0.58 |
| 4 | 84712638 | rs11099616 | 1.042 | -2.003 | -0.867 | -1.056 | 0.012 | -0.038 | -0.038 | 0.022 | 0.16 | 0.02 | 0.21 | 0.17 | 0.16 | 0.21 |
| 4 | 84721814 | rs4693646 | 0.2 | -1.455 | -0.623 | -0.949 | -0.016 | 0.075 | 0.075 | 0.841 | 0.38 | 0.07 | 0.55 | 0.75 | 0.53 | 0.66 |
| 4 | 84737580 | rs13120080 | 0.145 | -1.309 | -0.495 | -1.14 | -0.002 | -0.016 | -0.016 | 0.37 | 0.41 | 0.46 | 0.58 | 0.82 | 0.53 | 0.66 |
| 4 | 84750283 | rs10018145 | 0.168 | -1.007 | -0.295 | -0.65 | 0.033 | 0.025 | 0.025 | 0 | 0.84 | 0.94 | 0.93 | 1 | 0.91 | 0.93 |
| 4 | 84755196 | rs1352437 | -0.024 | -0.177 | 0.204 | -0.118 | -0.001 | 0.001 | 0.001 | 0.017 | 0.52 | 0.55 | 0.66 | 0.6 | 0.51 | 0.65 |
| 4 | 84755605 | rs13140644 | -0.543 | 0.238 | 0.265 | -0.36 | 0.02 | -0.033 | -0.033 | 0 | 0.13 | 0.2 | 0.17 | 0.03 | 0.25 | 0.09 |
| 4 | 84759300 | rs7689018 | -0.3 | -0.185 | 0.04 | -0.24 | -0.011 | 0.026 | 0.026 | 0.111 | 0.18 | 0.23 | 0.18 | 0.3 | 0.1 | 0.2 |
| 4 | 84762151 | rs11099621 | 0.259 | -0.183 | 0.065 | -0.327 | 0 | 0.082 | 0.082 | 0.735 | 0.43 | 0.17 | 0.56 | 0.33 | 0.59 | 0.46 |
| 4 | 84762853 | rs9999718 | 0.212 | -0.216 | 0.027 | -0.216 | 0.011 | -0.043 | -0.043 | 0.688 | 0.15 | 0.09 | 0.17 | 0.27 | 0.1 | 0.2 |
| 4 | 84764167 | rs4557253 | 1.26 | -0.214 | -0.036 | -0.23 | -0.054 | 0.014 | 0.014 | 0.184 | 0.3 | 0.1 | 0.39 | 0.3 | 0.34 | 0.35 |
| 4 | 84764738 | rs10005962 | -0.616 | -0.421 | 0.215 | -0.138 | 0.008 | 0.001 | 0.001 | 0.082 | 0.59 | 0.41 | 0.74 | 0.63 | 0.69 | 0.66 |
| 4 | 84777648 | rs11939071 | -0.823 | -0.216 | 0.596 | -0.093 | 0.008 | 0.045 | 0.045 | 4.162 | 0.06 | 0.62 | 0.04 | 0.03 | 0 | 0 |
| 4 | 84778595 | rs17357139 | 0.059 | -0.534 | 0.104 | -0.271 | 0.021 | 0.068 | 0.068 | 0 | 0.5 | 0.72 | 0.52 | 0.65 | 0.43 | 0.53 |
| 4 | 84779091 | rs10003040 | 0.365 | -0.597 | 0.006 | -0.376 | -0.019 | 0.015 | 0.015 | 0.06 | 0.14 | 0.1 | 0.16 | 0.03 | 0.26 | 0.13 |
| 4 | 84779231 | rs10516704 | 1.549 | -0.552 | -0.137 | -0.28 | -0.056 | -0.008 | -0.008 | 0 | 0.31 | 0.1 | 0.4 | 0.33 | 0.36 | 0.34 |
| 4 | 84779373 | rs9307830 | -0.658 | -0.606 | 0.471 | -0.374 | -0.005 | -0.003 | -0.003 | 0 | 0.61 | 0.58 | 0.66 | 0.68 | 0.69 | 0.66 |
| 4 | 84787315 | rs2017394 | -0.448 | -0.575 | 0.092 | -0.21 | -0.053 | 0.151 | 0.151 | 0 | 0.18 | 0.49 | 0.11 | 0.37 | 0.08 | 0.2 |
| 4 | 84788998 | rs1994511 | -0.415 | -0.566 | 0.062 | -0.205 | -0.055 | 0.178 | 0.178 | 0 | 0.18 | 0.46 | 0.11 | 0.37 | 0.08 | 0.2 |
| 4 | 84790267 | rs10021381 | 0.186 | -0.5 | 0.037 | -0.256 | 0.017 | -0.032 | -0.032 | 0 | 0.15 | 0.2 | 0.17 | 0.03 | 0.26 | 0.13 |
| 4 | 84791189 | rs7696864 | -1.365 | -0.5 | -0.167 | -0.248 | -0.001 | -0.004 | -0.004 | 0 | 0.33 | 0.67 | 0.28 | 0.4 | 0.34 | 0.33 |
| 4 | 84793018 | rs721461 | 0.261 | -0.433 | -0.316 | -0.434 | -0.06 | 0.008 | 0.008 | 4.038 | 0.34 | 0.45 | 0.18 | 0.37 | 0.2 | 0.25 |
| 4 | 84793814 | rs1906578 | 0.275 | -0.386 | -0.279 | -0.447 | 0.033 | -0.047 | -0.047 | 0.016 | 0.14 | 0.24 | 0.18 | 0.03 | 0.26 | 0.13 |
| 4 | 84793918 | rs1906579 | -0.723 | -0.341 | -0.386 | -0.412 | 0.023 | 0.037 | 0.037 | 0.259 | 0.48 | 0.68 | 0.36 | 0.4 | 0.46 | 0.38 |
| 4 | 84797545 | rs1488656 | 0.028 | -0.278 | 0.05 | -0.307 | 0.056 | 0.116 | 0.116 | 0 | 0.51 | 0.82 | 0.61 | 0.33 | 0.62 | 0.45 |
| 4 | 84802420 | rs1988722 | 0.042 | 0.091 | 0.222 | -0.026 | 0 | 0.004 | 0.004 | 0 | 0.45 | 0.59 | 0.68 | 0.33 | 0.55 | 0.41 |
| 4 | 84808625 | rs6822630 | 0.097 | 0.117 | 0.771 | -0.087 | 0 | 0 | 0 | 0 | 0.46 | 0.56 | 0.67 | 0.32 | 0.55 | 0.4 |
| 4 | 84820870 | rs13117766 | 0.054 | 0.083 | 1.09 | -0.106 | -0.001 | -0.004 | -0.004 | 0 | 0.46 | 0.55 | 0.67 | 0.33 | 0.55 | 0.4 |
| 4 | 84823287 | rs11735575 | -0.125 | -0.537 | 0.915 | -0.482 | 0.079 | 0.137 | 0.137 | 0.01 | 0.53 | 0.25 | 0.56 | 0.17 | 0.67 | 0.51 |
| 4 | 84823736 | rs2048005 | 0.36 | -0.061 | 1.147 | -0.281 | 0 | -0.025 | -0.025 | 0 | 0.45 | 0.51 | 0.46 | 0.23 | 0.52 | 0.41 |
| 4 | 84825877 | rs4106571 | 0.513 | -0.116 | 0.758 | -0.313 | 0.001 | -0.019 | -0.019 | 1.679 | 0.41 | 0.45 | 0.45 | 0.23 | 0.5 | 0.36 |
| 4 | 84831724 | rs6535510 | -0.592 | -0.636 | 0.008 | 0.007 | 0.046 | -0.053 | -0.053 | 0 | 0.67 | 0.84 | 0.81 | 0.46 | 0.76 | 0.64 |
| 4 | 84837046 | rs1505773 | 1.234 | -0.534 | -0.426 | -0.454 | -0.188 | 0.139 | 0.139 | 0.001 | 0.34 | 0.02 | 0.42 | 0.15 | 0.46 | 0.26 |
| 4 | 84839094 | rs1995597 | -0.428 | -0.465 | 0.309 | -0.431 | 0.001 | 0 | 0 | 0 | 0.61 | 0.62 | 0.75 | 0.36 | 0.71 | 0.52 |
| 4 | 84841250 | rs17007739 | -0.783 | -0.412 | 0.552 | -0.412 | 0 | 0 | 0 | 0.263 | 0.27 | 0.74 | 0.34 | 0.21 | 0.26 | 0.26 |
| 4 | 84853269 | rs7656326 | -0.381 | -0.658 | -0.335 | -0.559 | 0.05 | 0.074 | 0.074 | 0 | 0.62 | 0.91 | 0.62 | 0.25 | 0.7 | 0.5 |

| | | | | | | | | | | | | | | | | |
|---|---|---|---|---|---|---|---|---|---|---|---|---|---|---|---|---|
| 4 | 84854633 | rs6847383 | 0.876 | -0.449 | -0.267 | -0.783 | -0.049 | 0.118 | 0.118 | 0 | 0.38 | 0.16 | 0.35 | 0.14 | 0.45 | 0.27 |
| 4 | 84854953 | rs1827814 | -0.54 | -1.001 | -0.564 | -1.021 | 0.033 | 0.068 | 0.068 | 1.368 | 0.74 | 0.95 | 0.86 | 0.89 | 0.74 | 0.8 |
| 4 | 84858835 | rs12640485 | -1.621 | -0.811 | -0.196 | -0.834 | 0.026 | 0.214 | 0.214 | 1.669 | 0.5 | 0.8 | 0.62 | 0.86 | 0.38 | 0.71 |
| 4 | 84870572 | rs9992996 | -0.021 | -0.928 | 0.422 | -0.943 | 0.014 | -0.091 | -0.091 | 0 | 0.17 | 0.3 | 0.19 | 0 | 0.14 | 0.14 |
| 4 | 84879860 | rs12503076 | 0.671 | -1.017 | 0.14 | -0.018 | -0.014 | -0.042 | -0.042 | 0 | 0.09 | 0.01 | 0.19 | 0.34 | 0.05 | 0.22 |
| 4 | 84880846 | rs11099628 | 0.599 | -1.012 | 0.229 | 0.176 | -0.003 | -0.027 | -0.027 | 0.01 | 0.1 | 0.04 | 0.18 | 0.42 | 0.11 | 0.23 |
| 4 | 84881829 | rs17007783 | 0.468 | -0.993 | 0.298 | 0.185 | 0.003 | -0.015 | -0.015 | 0 | 0.12 | 0.07 | 0.18 | 0.42 | 0.12 | 0.23 |
| 4 | 84883128 | rs1857836 | 0.357 | -0.991 | 0.318 | 0.187 | 0.001 | 0.019 | 0.019 | 0 | 0.12 | 0.12 | 0.18 | 0.42 | 0.12 | 0.23 |
| 4 | 84884893 | rs12650954 | -0.629 | -0.901 | 0.206 | 0.251 | 0.003 | 0.039 | 0.039 | 1.763 | 0.28 | 0.39 | 0.38 | 0.42 | 0.26 | 0.37 |
| 4 | 84893974 | rs905991 | 0.376 | -0.694 | 0.471 | 0.581 | -0.019 | -0.016 | -0.016 | 0.097 | 0.84 | 0.7 | 0.9 | 0.9 | 0.87 | 0.86 |
| 4 | 84894991 | rs1847819 | -0.38 | -0.291 | -0.193 | 0.977 | -0.008 | 0.112 | 0.112 | 0.1 | 0.1 | 0.21 | 0.2 | 0.34 | 0.05 | 0.22 |
| 4 | 84901327 | rs1876117 | 0.425 | -0.34 | -0.336 | 0.852 | 0.001 | 0.013 | 0.013 | 4.862 | 0.73 | 0.73 | 0.6 | 0.65 | 0.74 | 0.64 |
| 4 | 84902964 | rs9999681 | -0.692 | -1.033 | -0.6 | 0.491 | -0.015 | -0.011 | -0.011 | 0.244 | 0.18 | 0.24 | 0.11 | 0.1 | 0.16 | 0.1 |
| 4 | 84906728 | rs17362048 | 0.237 | -1.55 | -1.431 | 0.61 | 0.133 | -0.016 | -0.016 | 0 | 0.53 | 0.2 | 0.47 | 0.4 | 0.74 | 0.44 |
| 4 | 84907924 | rs17007811 | 0.359 | -2.244 | -1.602 | 0.816 | 0.036 | -0.017 | -0.017 | 0 | 0.24 | 0.11 | 0.09 | 0.26 | 0.14 | 0.23 |
| 4 | 84908159 | rs2135284 | 0.861 | -2.665 | -2.016 | 0.643 | 0.014 | -0.011 | -0.011 | 0.096 | 0.16 | 0.03 | 0.07 | 0.13 | 0.11 | 0.07 |
| 4 | 84909466 | rs4693665 | 0.892 | -2.967 | -0.95 | 0.45 | -0.012 | -0.017 | -0.017 | 0.732 | 0.43 | 0.5 | 0.42 | 0.27 | 0.62 | 0.41 |
| 4 | 84930230 | rs4308354 | -0.37 | -2.837 | -0.105 | 0.728 | -0.022 | 0.046 | 0.046 | 0 | 0.84 | 0.76 | 0.6 | 0.66 | 0.9 | 0.72 |
| 4 | 84939404 | rs13149123 | 1.132 | -3.261 | -0.539 | 0.588 | 0.131 | -0.069 | -0.069 | 2.217 | 0.58 | 0.17 | 0.48 | 0.51 | 0.78 | 0.6 |
| 4 | 84947233 | rs17007862 | -0.527 | -3.26 | -0.458 | 0.589 | 0.062 | -0.089 | -0.089 | 0 | 0.29 | 0.11 | 0.4 | 0.47 | 0.18 | 0.35 |
| 4 | 84953207 | rs17007881 | -0.446 | -3.304 | 0.281 | 0.429 | -0.042 | -0.023 | -0.023 | 0.287 | 0.26 | 0.3 | 0.11 | 0.16 | 0.13 | 0.12 |
| 4 | 84957198 | rs1921880 | 0.46 | -3.282 | 0.07 | 0.426 | 0.041 | 0.024 | 0.024 | 0.013 | 0.19 | 0.05 | 0.09 | 0.12 | 0.11 | 0.08 |
| 4 | 84961080 | rs2091745 | 1.288 | -3.587 | 0.231 | 0.484 | 0.162 | -0.068 | -0.068 | 0 | 0.55 | 0.18 | 0.48 | 0.51 | 0.77 | 0.57 |
| 4 | 84964281 | rs1921873 | 0.108 | -3.142 | 0.726 | 0.655 | -0.043 | 0.147 | 0.147 | 0.02 | 0.83 | 0.49 | 0.59 | 0.64 | 0.9 | 0.71 |
| 4 | 84969104 | rs17007921 | 1.128 | -3.387 | -0.477 | 0.336 | 0.002 | -0.014 | -0.014 | 0 | 0.25 | 0.01 | 0.12 | 0.23 | 0.19 | 0.29 |
| 4 | 84970969 | rs1921871 | 0.03 | -2.763 | 0.602 | 0.69 | -0.029 | 0.123 | 0.123 | 0 | 0.84 | 0.7 | 0.6 | 0.63 | 0.9 | 0.71 |
| 4 | 84974648 | rs1343881 | -1.105 | -1.654 | 0.175 | 0.391 | 0.018 | 0.001 | 0.001 | 0 | 0.48 | 0.74 | 0.61 | 0.58 | 0.43 | 0.43 |
| 4 | 84975437 | rs2178602 | -0.881 | -1.853 | 0.773 | 0.281 | 0.002 | 0.008 | 0.008 | 0 | 0.35 | 0.59 | 0.52 | 0.45 | 0.35 | 0.41 |
| 4 | 84976857 | rs12499536 | 0.362 | -1.749 | 0.854 | 0.245 | -0.039 | -0.017 | -0.017 | 0 | 0.17 | 0.26 | 0.09 | 0.14 | 0.09 | 0.07 |
| 4 | 84978944 | rs10000976 | -0.336 | -1.676 | 0.701 | 0.264 | 0.024 | -0.066 | -0.066 | 0 | 0.29 | 0.15 | 0.47 | 0.45 | 0.23 | 0.43 |
| 4 | 84984952 | rs17007945 | -0.265 | -1.645 | 1.196 | 0.204 | -0.002 | -0.036 | -0.036 | 0 | 0.21 | 0.57 | 0.16 | 0.13 | 0.2 | 0.07 |
| 4 | 84991230 | rs10022529 | -1.563 | -1.88 | 0.952 | 0.145 | -0.002 | -0.026 | -0.026 | 0 | 0.55 | 0.85 | 0.63 | 0.57 | 0.42 | 0.54 |
| 4 | 85003358 | rs7696640 | 0.01 | -1.794 | 1.311 | 0.243 | 0.031 | -0.025 | -0.025 | 0 | 0.19 | 0.46 | 0.15 | 0.14 | 0.2 | 0.09 |
| 4 | 85005873 | rs12508400 | 0.324 | -1.792 | 1.304 | 0.244 | 0.016 | -0.014 | -0.014 | 0 | 0.18 | 0.29 | 0.14 | 0.14 | 0.2 | 0.09 |
| 4 | 85006975 | rs1343762 | -1.573 | -1.809 | 0.544 | 0.268 | 0.011 | -0.017 | -0.017 | 0 | 0.54 | 0.8 | 0.63 | 0.56 | 0.42 | 0.54 |
| 4 | 85024046 | rs13107199 | 0.776 | -1.778 | 0.703 | 0.304 | -0.009 | 0.003 | 0.003 | 0 | 0.16 | 0.12 | 0.12 | 0.14 | 0.2 | 0.09 |
| 4 | 85029546 | rs7683351 | -1.06 | -1.687 | 0.142 | 0.344 | 0.027 | 0.011 | 0.011 | 0 | 0.5 | 0.62 | 0.62 | 0.56 | 0.4 | 0.52 |
| 4 | 85029894 | rs1950128 | -1.077 | -1.661 | 0.155 | 0.356 | 0.025 | 0.009 | 0.009 | 0 | 0.51 | 0.62 | 0.62 | 0.56 | 0.4 | 0.52 |
| 4 | 85036481 | rs1921848 | -1.267 | -1.551 | -0.018 | 0.285 | 0.03 | 0.03 | 0.03 | 0 | 0.52 | 0.79 | 0.61 | 0.61 | 0.41 | 0.52 |
| 4 | 85042254 | rs1921847 | -1.299 | -1.554 | -0.022 | 0.284 | 0.027 | 0.028 | 0.028 | 0 | 0.53 | 0.79 | 0.62 | 0.61 | 0.41 | 0.52 |
| 4 | 85054547 | rs6820732 | 0.026 | -1.51 | 0.185 | 0.299 | 0.026 | -0.028 | -0.028 | 0.158 | 0.2 | 0.45 | 0.15 | 0.14 | 0.2 | 0.09 |
| 4 | 85056551 | rs1921867 | -0.543 | -1.524 | 0.189 | 0.271 | -0.001 | -0.007 | -0.007 | 0.01 | 0.25 | 0.69 | 0.17 | 0.15 | 0.21 | 0.13 |
| 4 | 85068653 | rs1950130 | -0.478 | -1.614 | 0.039 | 0.307 | -0.003 | 0.001 | 0.001 | 0 | 0.33 | 0.32 | 0.46 | 0.47 | 0.21 | 0.43 |
| 4 | 85080471 | rs6815073 | -0.764 | -1.058 | -0.337 | 0.403 | 0.02 | 0.013 | 0.013 | 0 | 0.48 | 0.59 | 0.6 | 0.6 | 0.41 | 0.49 |
| 4 | 85081980 | rs10023423 | -1.031 | -0.275 | -0.104 | 0.456 | -0.023 | -0.039 | -0.039 | 0.007 | 0.29 | 0.51 | 0.18 | 0.18 | 0.22 | 0.16 |
| 4 | 85082238 | rs6824813 | 0.268 | -0.113 | -0.509 | 0.541 | 0.007 | 0.031 | 0.031 | 0.554 | 0.35 | 0.28 | 0.33 | 0.24 | 0.33 | 0.36 |
| 4 | 85082294 | rs4693676 | -0.266 | -0.141 | -0.856 | 0.676 | 0.007 | -0.003 | -0.003 | 0 | 0.54 | 0.64 | 0.58 | 0.52 | 0.45 | 0.66 |
| 4 | 85090749 | rs2178603 | -1.145 | -0.507 | -1.267 | 0.828 | 0.066 | -0.144 | -0.144 | 0 | 0.36 | 0.95 | 0.36 | 0.47 | 0.23 | 0.29 |
| 4 | 85118112 | rs6825533 | -1.141 | -0.389 | 0.588 | 0.596 | 0.006 | -0.008 | -0.008 | 0.394 | 0.19 | 0.19 | 0.17 | 0.25 | 0.15 | 0.21 |
| 4 | 85136552 | rs13122998 | -0.669 | -0.005 | 0.816 | 0.261 | -0.001 | -0.004 | -0.004 | 0 | 0.14 | 0.21 | 0.11 | 0.13 | 0.14 | 0.21 |
| 4 | 85138894 | rs1319771 | -1.437 | -0.101 | 1.614 | 0.24 | -0.014 | -0.024 | -0.024 | 0 | 0.18 | 0.62 | 0.13 | 0.14 | 0.16 | 0.21 |
| 4 | 85149128 | rs6827060 | -1.492 | 0.47 | 1.132 | -0.496 | 0.002 | -0.141 | -0.141 | 0 | 0.29 | 0.53 | 0.19 | 0.01 | 0.31 | 0.25 |
| 4 | 85150623 | rs4380514 | 1.378 | 0.311 | 1.496 | -0.169 | -0.003 | -0.023 | -0.023 | 0 | 0.71 | 0.39 | 0.79 | 0.81 | 0.72 | 0.76 |
| 4 | 85159891 | rs1532358 | 0.57 | 0.828 | 1.281 | -0.512 | 0.032 | -0.019 | -0.019 | 0 | 0.72 | 0.4 | 0.67 | 0.77 | 0.59 | 0.83 |
| 4 | 85160070 | rs1532357 | 0.388 | 0.848 | 1.396 | -0.516 | 0.054 | -0.025 | -0.025 | 0 | 0.73 | 0.45 | 0.67 | 0.78 | 0.59 | 0.83 |
| 4 | 85164682 | rs903521 | 1.381 | 0.79 | 1.336 | -0.584 | -0.01 | 0.015 | 0.015 | 0 | 0.61 | 0.32 | 0.6 | 0.77 | 0.51 | 0.66 |
| 4 | 85165381 | rs348701 | 1.344 | 0.323 | 1.912 | -0.444 | -0.039 | -0.035 | -0.035 | 0.273 | 0.85 | 0.6 | 0.89 | 0.93 | 0.91 | 0.83 |
| 4 | 85169677 | rs17008412 | 0.944 | 0.899 | 1.508 | -0.765 | -0.003 | 0.011 | 0.011 | 0 | 0.61 | 0.32 | 0.5 | 0.78 | 0.43 | 0.68 |
| 4 | 85174676 | rs17008416 | -0.164 | 0.918 | 1.525 | -0.785 | 0.009 | 0.009 | 0.009 | 0 | 0.1 | 0.07 | 0.07 | 0.01 | 0.07 | 0.15 |
| 4 | 85178558 | rs10020342 | -0.094 | 1.056 | 1.846 | -0.818 | 0.046 | -0.021 | -0.021 | 0 | 0.73 | 0.64 | 0.58 | 0.81 | 0.52 | 0.83 |
| 4 | 85189813 | rs10030773 | 1.126 | 0.97 | 1.842 | -0.839 | -0.009 | 0.062 | 0.062 | 0 | 0.59 | 0.22 | 0.49 | 0.78 | 0.43 | 0.68 |
| 4 | 85199286 | rs6816852 | -0.514 | 0.834 | 1.872 | -0.77 | 0.087 | -0.044 | -0.044 | 0 | 0.77 | 0.65 | 0.65 | 0.88 | 0.55 | 0.86 |
| 4 | 85201560 | rs7662217 | 0.172 | -0.111 | 1.131 | -1.295 | 0 | -0.004 | -0.004 | 0.122 | 0.2 | 0.16 | 0.14 | 0.1 | 0.14 | 0.19 |
| 4 | 85205214 | rs1495151 | 1.194 | -0.312 | 0.87 | -1.393 | 0.062 | 0.131 | 0.131 | 0 | 0.45 | 0.07 | 0.43 | 0.71 | 0.33 | 0.54 |
| 4 | 85206535 | rs11932874 | 0.017 | -0.312 | 1.088 | -1.365 | -0.018 | -0.042 | -0.042 | 0 | 0.11 | 0.37 | 0.08 | 0.03 | 0.08 | 0.13 |
| 4 | 85215142 | rs1495145 | -1.194 | -0.295 | 1.018 | -1.359 | -0.022 | -0.057 | -0.057 | 0 | 0.32 | 0.59 | 0.22 | 0.13 | 0.22 | 0.32 |
| 4 | 85244068 | rs6850349 | -1.165 | -0.645 | 1.046 | -1.357 | -0.037 | -0.062 | -0.062 | 0 | 0.29 | 0.59 | 0.2 | 0.13 | 0.19 | 0.28 |
| 4 | 85260011 | rs13109014 | 0.329 | -0.626 | 1.072 | -1.343 | 0.002 | 0.001 | 0.001 | 0 | 0.16 | 0.13 | 0.11 | 0.1 | 0.11 | 0.13 |
| 4 | 85260621 | rs882109 | 1.19 | -1.622 | 0.129 | -0.925 | 0.006 | -0.113 | -0.113 | 0 | 0.19 | 0 | 0.23 | 0.31 | 0.12 | 0.23 |

| | | | | | | | | | | | | | | | | |
|---|---|---|---|---|---|---|---|---|---|---|---|---|---|---|---|---|
| 4 | 85268860 | rs1899392 | -0.108 | -1.87 | 0.786 | -1.144 | -0.021 | -0.047 | -0.047 | 0.87 | 0.12 | 0.4 | 0.08 | 0.03 | 0.08 | 0.15 |
| 4 | 85272572 | rs4264821 | 0.187 | -1.856 | 0.744 | -1.12 | -0.003 | -0.011 | -0.011 | 1.828 | 0.13 | 0.15 | 0.12 | 0.04 | 0.1 | 0.18 |
| 4 | 85274557 | rs1442163 | -1.21 | -1.883 | 0.759 | -1.136 | -0.029 | -0.046 | -0.046 | 0 | 0.33 | 0.59 | 0.26 | 0.15 | 0.22 | 0.31 |
| 4 | 85274948 | rs1442164 | -1.068 | -1.901 | 0.811 | -1.153 | 0.035 | -0.018 | -0.018 | 0 | 0.75 | 0.69 | 0.65 | 0.83 | 0.56 | 0.81 |
| 4 | 85277790 | rs6817184 | -1.81 | -2.489 | 2.037 | -0.625 | -0.029 | -0.046 | -0.046 | 0 | 0.33 | 0.59 | 0.26 | 0.15 | 0.23 | 0.31 |
| 4 | 85283718 | rs11729834 | -1.15 | -1.968 | 1.67 | -0.339 | -0.038 | -0.06 | -0.06 | 1.057 | 0.16 | 0.5 | 0.15 | 0.05 | 0.12 | 0.18 |
| 4 | 85294388 | rs1442173 | -1.089 | -2.213 | 0.914 | -0.479 | 0.009 | 0.008 | 0.008 | 0.04 | 0.31 | 0.71 | 0.26 | 0.13 | 0.2 | 0.23 |
| 4 | 85309078 | rs385194 | 1.384 | -2.503 | -0.012 | -0.146 | 0.046 | -0.062 | -0.062 | 0.19 | 0.48 | 0.04 | 0.4 | 0.43 | 0.69 | 0.37 |
| 4 | 85315581 | rs6849720 | -0.678 | -2.541 | 0.67 | -0.266 | 0.021 | 0.053 | 0.053 | 0 | 0.22 | 0.13 | 0.14 | 0.08 | 0.13 | 0.22 |
| 4 | 85316658 | rs446951 | 0.795 | -2.22 | 1.419 | 0.212 | 0.003 | 0.087 | 0.087 | 1.082 | 0.93 | 0.68 | 0.87 | 0.88 | 0.98 | 0.94 |
| 4 | 85324586 | rs12650179 | -1.225 | -2.253 | 1.642 | 0.201 | -0.014 | 0.069 | 0.069 | 1.194 | 0.08 | 0.36 | 0.12 | 0.13 | 0.02 | 0.07 |
| 4 | 85328262 | rs4101237 | 0.597 | -2.65 | 1.246 | 0.245 | 0.019 | -0.012 | -0.012 | 0 | 0.66 | 0.71 | 0.66 | 0.6 | 0.78 | 0.56 |
| 4 | 85339322 | rs7686486 | -1.259 | -2.691 | 1.615 | 0.201 | -0.019 | 0.052 | 0.052 | 2.816 | 0.1 | 0.49 | 0.11 | 0.11 | 0.02 | 0.1 |
| 4 | 85341562 | rs17008894 | 1.516 | -2.73 | 1.52 | 0.217 | 0.079 | -0.025 | -0.025 | 0 | 0.57 | 0.23 | 0.56 | 0.49 | 0.76 | 0.47 |
| 4 | 85342849 | rs13434947 | -1.28 | -2.447 | 1.614 | 0.183 | -0.009 | 0.066 | 0.066 | 2.376 | 0.1 | 0.45 | 0.1 | 0.11 | 0.02 | 0.1 |
| 4 | 85359985 | rs1562147 | 0.55 | -2.37 | 0.92 | 0.137 | -0.026 | -0.004 | -0.004 | 0 | 0.1 | 0.01 | 0.14 | 0.1 | 0.08 | 0.06 |
| 4 | 85360873 | rs12233901 | 1.089 | -2.31 | 1.754 | 0.16 | -0.025 | 0.024 | 0.024 | 3.289 | 0.91 | 0.44 | 0.95 | 0.94 | 0.98 | 0.95 |
| 4 | 85363446 | rs1444976 | 1.156 | -2.477 | 1.562 | 0.156 | 0.05 | -0.004 | -0.004 | 0 | 0.55 | 0.27 | 0.55 | 0.5 | 0.74 | 0.46 |
| 4 | 85368880 | rs752402 | 1.171 | -2.395 | 1.497 | 0.179 | 0.091 | -0.056 | -0.056 | 0.087 | 0.55 | 0.13 | 0.55 | 0.46 | 0.74 | 0.48 |
| 4 | 85377040 | rs1444957 | 0.984 | -2.389 | 1.667 | 0.181 | 0.031 | -0.011 | -0.011 | 0 | 0.57 | 0.33 | 0.55 | 0.46 | 0.75 | 0.48 |
| 4 | 85387480 | rs1822836 | -0.507 | -2.332 | 1.156 | 0.206 | 0.005 | -0.009 | -0.009 | 0 | 0.34 | 0.34 | 0.41 | 0.48 | 0.24 | 0.47 |
| 4 | 85401169 | rs7668923 | 1.016 | -2.388 | 0.107 | 0.111 | 0.082 | -0.072 | -0.072 | 0.066 | 0.48 | 0.05 | 0.52 | 0.43 | 0.72 | 0.48 |
| 4 | 85407784 | rs11721853 | 0.612 | -2.052 | 0.367 | 0.255 | 0.071 | -0.025 | -0.025 | 0 | 0.47 | 0.18 | 0.52 | 0.45 | 0.68 | 0.52 |
| 4 | 85416797 | rs2278671 | -0.518 | -1.813 | -0.388 | 0.347 | 0.011 | 0.007 | 0.007 | 0 | 0.45 | 0.32 | 0.39 | 0.38 | 0.32 | 0.36 |
| 4 | 85419973 | rs3806750 | -0.065 | -1.686 | -1.246 | 0.381 | 0.017 | -0.002 | -0.002 | 0.408 | 0.37 | 0.15 | 0.35 | 0.36 | 0.29 | 0.31 |
| 4 | 85428178 | rs1545330 | -0.546 | -1.138 | -1.735 | 0.476 | 0.055 | -0.053 | -0.053 | 0 | 0.53 | 0.87 | 0.49 | 0.51 | 0.34 | 0.5 |
| 4 | 85434987 | rs1017560 | -0.486 | -1.139 | -1.789 | 0.338 | 0.081 | 0.05 | 0.05 | 0 | 0.43 | 0.79 | 0.4 | 0.29 | 0.31 | 0.35 |
| 4 | 85447672 | rs6812841 | -0.76 | -1.811 | -1.308 | -0.138 | 0.065 | 0.039 | 0.039 | 0 | 0.41 | 0.74 | 0.28 | 0.25 | 0.27 | 0.43 |
| 4 | 85447778 | rs6835003 | -0.692 | -1.735 | -1.277 | -0.413 | 0.018 | 0.019 | 0.019 | 0.005 | 0.35 | 0.74 | 0.29 | 0.22 | 0.27 | 0.36 |
| 4 | 85449323 | rs6849318 | -0.452 | -1.866 | -0.341 | -0.585 | -0.017 | -0.041 | -0.041 | 1.185 | 0.27 | 0.38 | 0.15 | 0.17 | 0.23 | 0.31 |
| 4 | 85457113 | rs1017557 | 0.288 | -0.801 | -1.016 | -0.188 | -0.027 | -0.03 | -0.03 | 0 | 0.28 | 0.25 | 0.47 | 0.4 | 0.37 | 0.35 |
| 4 | 85469315 | rs974130 | -1.394 | -0.927 | -1.013 | -0.216 | 0.009 | 0.002 | 0.002 | 0 | 0.6 | 0.7 | 0.63 | 0.57 | 0.61 | 0.69 |
| 4 | 85475077 | rs12642236 | 0.003 | -0.722 | -0.922 | -0.097 | -0.033 | -0.037 | -0.037 | 0 | 0.29 | 0.23 | 0.47 | 0.4 | 0.37 | 0.35 |
| 4 | 85483258 | rs7695015 | -0.871 | -1.469 | -1.347 | -0.57 | 0.033 | 0.051 | 0.051 | 1.22 | 0.38 | 0.72 | 0.2 | 0.21 | 0.26 | 0.36 |
| 4 | 85483427 | rs7699871 | -1.201 | -1.547 | -1.256 | -0.333 | 0.001 | 0.001 | 0.001 | 0 | 0.59 | 0.61 | 0.62 | 0.65 | 0.61 | 0.7 |
| 4 | 85483950 | rs11099650 | -0.053 | -1.311 | -0.585 | -0.356 | -0.015 | -0.02 | -0.02 | 0 | 0.33 | 0.28 | 0.47 | 0.45 | 0.4 | 0.36 |
| 4 | 85485329 | rs4449405 | -1.512 | -1.171 | -0.673 | -0.386 | -0.004 | -0.003 | -0.003 | 0 | 0.63 | 0.72 | 0.61 | 0.64 | 0.62 | 0.74 |
| 4 | 85491947 | rs17009161 | 0.561 | -1.2 | -0.61 | -0.404 | -0.002 | -0.001 | -0.001 | 0 | 0.25 | 0.2 | 0.11 | 0.19 | 0.18 | 0.3 |
| 4 | 85496761 | rs2868914 | -1.18 | -1.222 | -0.249 | -0.426 | -0.001 | -0.001 | -0.001 | 0 | 0.6 | 0.56 | 0.59 | 0.64 | 0.62 | 0.74 |
| 4 | 85505474 | rs11097023 | -1.192 | -1.237 | -0.216 | -0.293 | -0.001 | -0.005 | -0.005 | 0 | 0.61 | 0.58 | 0.6 | 0.66 | 0.62 | 0.74 |
| 4 | 85510643 | rs17367226 | 0.051 | -1.323 | -0.48 | -0.271 | 0.028 | 0.084 | 0.084 | 0 | 0.29 | 0.07 | 0.15 | 0.14 | 0.24 | 0.37 |
| 4 | 85520643 | rs2868919 | -1.254 | -1.193 | -0.148 | -0.136 | 0 | -0.001 | -0.001 | 0 | 0.6 | 0.61 | 0.59 | 0.67 | 0.6 | 0.73 |
| 4 | 85532915 | rs2165786 | -1.219 | -1.006 | -0.008 | -0.125 | 0 | 0 | 0 | 2.328 | 0.59 | 0.58 | 0.59 | 0.63 | 0.57 | 0.72 |
| 4 | 85532930 | rs1020278 | -1.219 | -1.004 | -0.008 | -0.125 | 0 | 0 | 0 | 0 | 0.59 | 0.58 | 0.6 | 0.63 | 0.6 | 0.72 |
| 4 | 85545644 | rs4490450 | -2.262 | -1.473 | -0.481 | -0.312 | -0.12 | -0.025 | -0.025 | 0 | 0.81 | 0.99 | 0.7 | 0.83 | 0.7 | 0.82 |
| 4 | 85558840 | rs6829806 | -0.467 | -1.562 | 0.037 | -0.361 | -0.002 | 0 | 0 | 0 | 0.58 | 0.42 | 0.57 | 0.63 | 0.57 | 0.74 |
| 4 | 85566809 | rs1946998 | -2.268 | -1.498 | -0.398 | -0.356 | -0.12 | -0.026 | -0.026 | 0 | 0.81 | 0.99 | 0.7 | 0.83 | 0.7 | 0.82 |
| 4 | 85570364 | rs1838038 | 0.946 | -2.009 | -0.252 | -0.623 | -0.03 | -0.071 | -0.071 | 2.619 | 0.25 | 0.15 | 0.42 | 0.48 | 0.34 | 0.3 |
| 4 | 85570767 | rs6827228 | 0.454 | -2.008 | -0.296 | -0.599 | 0.02 | 0.025 | 0.025 | 0 | 0.3 | 0.19 | 0.13 | 0.15 | 0.2 | 0.36 |
| 4 | 85571339 | rs1372971 | -1.136 | -2.008 | -0.25 | -0.606 | -0.027 | -0.013 | -0.013 | 0 | 0.62 | 0.75 | 0.6 | 0.63 | 0.57 | 0.73 |
| 4 | 85572374 | rs1120 | 0.419 | -1.862 | -0.312 | -0.615 | 0.016 | 0.022 | 0.022 | 0 | 0.28 | 0.18 | 0.13 | 0.15 | 0.2 | 0.36 |
| 4 | 85583567 | rs17367858 | 0.884 | -1.779 | -0.612 | -0.617 | 0.035 | 0.071 | 0.071 | 0 | 0.25 | 0.07 | 0.09 | 0.12 | 0.17 | 0.27 |
| 4 | 85612737 | rs3747680 | 0.815 | -1.665 | -0.346 | -0.64 | -0.006 | 0.007 | 0.007 | 0.106 | 0.34 | 0.38 | 0.28 | 0.43 | 0.33 | 0.26 |
| 4 | 85620879 | rs35617206 | 2.878 | -1.541 | -0.745 | -0.776 | -0.02 | 0.076 | 0.076 | 0 | 0.13 | 0.01 | 0.1 | 0 | 0.13 | 0.02 |
| 4 | 85621772 | rs6816534 | 1.6 | -1.562 | -0.541 | -0.658 | -0.024 | -0.057 | -0.057 | 0 | 0.24 | 0.14 | 0.27 | 0.42 | 0.31 | 0.24 |
| 4 | 85625701 | rs1838039 | -2.321 | -1.525 | -0.809 | -0.667 | -0.162 | -0.106 | -0.106 | 0 | 0.77 | 0.99 | 0.64 | 0.75 | 0.61 | 0.78 |
| 4 | 85634488 | rs13108722 | 1.28 | -1.51 | -0.725 | -0.64 | -0.006 | -0.072 | -0.072 | 0.016 | 0.33 | 0.1 | 0.27 | 0.41 | 0.32 | 0.27 |
| 4 | 85640591 | rs10516727 | 1.872 | -1.518 | -0.598 | -0.703 | 0.036 | 0.002 | 0.002 | 0 | 0.25 | 0.08 | 0.1 | 0.26 | 0.17 | 0.28 |
| 4 | 85730752 | rs7681742 | -0.114 | -2.007 | -0.442 | -0.236 | 0.014 | 0.011 | 0.011 | 0.166 | 0.09 | 0.3 | 0.19 | 0.05 | 0.07 | 0.12 |
| 4 | 85730914 | rs6531775 | -0.496 | -2.206 | -0.597 | -0.256 | -0.002 | 0 | 0 | 1.495 | 0.45 | 0.58 | 0.48 | 0.48 | 0.4 | 0.38 |
| 4 | 85734049 | rs1484662 | -0.339 | -1.438 | -0.092 | 0.217 | 0.012 | 0.006 | 0.006 | 0.135 | 0.09 | 0.28 | 0.19 | 0.05 | 0.07 | 0.12 |
| 4 | 85769856 | rs17009317 | 0.76 | -1.463 | -0.171 | 0.214 | -0.002 | -0.006 | -0.006 | 0 | 0.26 | 0.26 | 0.3 | 0.43 | 0.31 | 0.25 |
| 4 | 85779738 | rs11936391 | 1.335 | -1.466 | -0.792 | 0.208 | -0.003 | 0.007 | 0.007 | 0 | 0.05 | 0.13 | 0.17 | 0.05 | 0.04 | 0.05 |
| 4 | 85782946 | rs11726635 | -0.791 | -1.479 | -0.773 | 0.197 | 0.011 | 0.004 | 0.004 | 0.409 | 0.1 | 0.32 | 0.19 | 0.05 | 0.07 | 0.12 |
| 4 | 85786828 | rs12501525 | 0.529 | -1.458 | -0.864 | 0.172 | -0.01 | -0.024 | -0.024 | 0 | 0.28 | 0.22 | 0.31 | 0.43 | 0.34 | 0.28 |
| 4 | 85803607 | rs11097026 | -0.153 | -1.791 | -0.785 | -0.212 | 0 | -0.022 | -0.022 | 0 | 0.33 | 0.47 | 0.47 | 0.25 | 0.35 | 0.35 |
| 4 | 85824232 | rs10029496 | -0.662 | -1.788 | -1.228 | -0.389 | -0.004 | -0.038 | -0.038 | 0 | 0.36 | 0.87 | 0.49 | 0.37 | 0.35 | 0.35 |
| 4 | 85827192 | rs12501332 | -1.98 | -1.769 | -1.904 | -0.473 | -0.185 | -0.193 | -0.193 | 0 | 0.66 | 0.99 | 0.62 | 0.5 | 0.57 | 0.75 |
| 4 | 85844157 | rs10516733 | 0.937 | -1.581 | -1.698 | -0.491 | -0.004 | 0.008 | 0.008 | 0.013 | 0.06 | 0.15 | 0.17 | 0.05 | 0.04 | 0.05 |

| | | | | | | | | | | | | | | | | |
|---|---|---|---|---|---|---|---|---|---|---|---|---|---|---|---|---|
| 4 | 85858329 | rs6848127 | -2.236 | -1.557 | -1.911 | -0.471 | -0.177 | -0.144 | -0.144 | 0 | 0.7 | 0.99 | 0.64 | 0.63 | 0.59 | 0.76 |
| 4 | 85872414 | rs4627854 | 0.8 | -1.066 | -0.96 | -0.661 | 0.016 | 0.069 | 0.069 | 0 | 0.17 | 0.01 | 0.05 | 0.06 | 0.11 | 0.19 |
| 4 | 85898741 | rs4590017 | -0.707 | -0.156 | 0.128 | 0.047 | -0.003 | 0.001 | 0.001 | 0 | 0.27 | 0.23 | 0.29 | 0.27 | 0.32 | 0.25 |
| 4 | 85915570 | rs4533765 | -1.371 | 0.06 | -0.027 | 0.115 | -0.005 | -0.005 | -0.005 | 0 | 0.35 | 0.48 | 0.47 | 0.32 | 0.36 | 0.39 |
| 4 | 85974024 | rs4287999 | -1.362 | 0.038 | 0.161 | 0.724 | -0.001 | 0.029 | 0.029 | 0.012 | 0.24 | 0.39 | 0.36 | 0.32 | 0.24 | 0.39 |
| 4 | 85981276 | rs10516730 | -0.59 | -0.016 | 0.273 | 0.689 | 0.006 | 0.043 | 0.043 | 0.94 | 0.15 | 0.29 | 0.17 | 0.27 | 0.16 | 0.24 |
| 4 | 85993738 | rs4129430 | -1.614 | -0.392 | 0.534 | 0.313 | 0.003 | 0.003 | 0.003 | 4.567 | 0.26 | 0.71 | 0.28 | 0.24 | 0.24 | 0.4 |
| 4 | 85998783 | rs10516731 | -0.911 | -0.484 | 1.345 | 0.261 | -0.018 | -0.052 | -0.052 | 0.049 | 0.1 | 0.41 | 0.07 | 0 | 0.04 | 0.12 |
| 4 | 86000455 | rs11097033 | -0.518 | -0.449 | 1.316 | 0.283 | 0.008 | 0.015 | 0.015 | 0.023 | 0.15 | 0.25 | 0.21 | 0.24 | 0.2 | 0.28 |
| 4 | 86005206 | rs4235071 | 1.933 | -0.839 | 1.103 | 0.083 | 0.02 | 0.017 | 0.017 | 2.994 | 0.64 | 0.29 | 0.6 | 0.67 | 0.69 | 0.43 |
| 4 | 86009525 | rs6835154 | -1.343 | -0.841 | 1.011 | 0.086 | -0.007 | -0.018 | -0.018 | 0.749 | 0.24 | 0.16 | 0.32 | 0.33 | 0.26 | 0.43 |
| 4 | 86021602 | rs4558866 | 1.369 | -0.868 | 1.308 | 0.046 | -0.047 | -0.066 | -0.066 | 0 | 0.88 | 0.34 | 0.92 | 1 | 0.95 | 0.87 |
| 4 | 86024239 | rs4507353 | 0.615 | -0.856 | 1.433 | 0.05 | -0.013 | -0.045 | -0.045 | 0 | 0.9 | 0.53 | 0.93 | 1 | 0.96 | 0.89 |
| 4 | 86033441 | rs11941410 | -1.973 | -0.926 | 1.662 | 0.163 | -0.007 | -0.002 | -0.002 | 0 | 0.25 | 0.68 | 0.26 | 0.26 | 0.22 | 0.4 |
| 4 | 86034744 | rs10034520 | 1.988 | -1.011 | 0.427 | -0.21 | -0.17 | -0.193 | -0.193 | 0 | 0.3 | 0.02 | 0.45 | 0.43 | 0.42 | 0.19 |
| 4 | 86036723 | rs17009476 | -0.616 | -0.87 | 1.015 | -0.124 | -0.01 | -0.043 | -0.043 | 0 | 0.1 | 0.47 | 0.07 | 0 | 0.04 | 0.11 |
| 4 | 86042277 | rs17009493 | 0.13 | -0.936 | 0.771 | -0.173 | -0.006 | 0.004 | 0.004 | 0 | 0.1 | 0 | 0.12 | 0.09 | 0.07 | 0.15 |
| 4 | 86044561 | rs6820757 | -0.147 | -1.109 | 0.973 | -0.525 | -0.006 | -0.108 | -0.108 | 2.242 | 0.18 | 0.37 | 0.12 | 0 | 0.14 | 0.11 |
| 4 | 86049136 | rs6835768 | -2.205 | -1.087 | 0.753 | -0.479 | 0.027 | 0.071 | 0.071 | 0.467 | 0.44 | 0.81 | 0.45 | 0.32 | 0.38 | 0.53 |
| 4 | 86057003 | rs13145163 | 1.018 | -0.925 | 0.761 | -0.42 | 0.01 | -0.063 | -0.063 | 0.751 | 0.12 | 0.25 | 0.07 | 0 | 0.1 | 0.03 |
| 4 | 86064719 | rs2869268 | -0.38 | -0.964 | 0.648 | -0.418 | -0.001 | -0.029 | -0.029 | 2.864 | 0.21 | 0.22 | 0.17 | 0.05 | 0.17 | 0.17 |
| 4 | 86078683 | rs11728848 | 0.801 | -0.966 | 0.468 | -0.315 | -0.039 | -0.067 | -0.067 | 0.901 | 0.11 | 0.01 | 0.19 | 0.27 | 0.17 | 0.26 |
| 4 | 86080861 | rs2869291 | 0.024 | -0.988 | 0.521 | -0.343 | 0.009 | 0.02 | 0.02 | 0 | 0.11 | 0.03 | 0.11 | 0.04 | 0.07 | 0.15 |
| 4 | 86088655 | rs17009514 | 1.336 | -0.979 | 0.739 | -0.361 | 0.025 | -0.064 | -0.064 | 1.378 | 0.11 | 0.29 | 0.07 | 0 | 0.1 | 0.03 |
| 4 | 86094786 | rs7349600 | 0.594 | -0.985 | 0.869 | -0.348 | -0.001 | -0.001 | -0.001 | 0 | 0.12 | 0.14 | 0.19 | 0.27 | 0.17 | 0.26 |
| 4 | 86098946 | rs12507778 | -1.096 | -1.088 | 0.968 | -0.429 | -0.002 | -0.007 | -0.007 | 0.496 | 0.31 | 0.66 | 0.32 | 0.27 | 0.31 | 0.37 |
| 4 | 86101484 | rs12510879 | -0.843 | -1.079 | 0.931 | -0.416 | -0.009 | 0.02 | 0.02 | 0 | 0.2 | 0.37 | 0.25 | 0.27 | 0.21 | 0.34 |
| 4 | 86101771 | rs6822829 | 2.253 | -1.05 | 0.523 | -0.461 | -0.151 | -0.183 | -0.183 | 0 | 0.35 | 0.02 | 0.44 | 0.48 | 0.41 | 0.29 |
| 4 | 86107271 | rs17009586 | -0.734 | -0.788 | 1.2 | -0.311 | -0.009 | -0.079 | -0.079 | 2.015 | 0.23 | 0.36 | 0.19 | 0.05 | 0.17 | 0.17 |
| 4 | 86109620 | rs2869302 | 0.468 | -0.741 | 1.055 | -0.262 | 0.004 | 0.028 | 0.028 | 0 | 0.12 | 0.22 | 0.2 | 0.27 | 0.17 | 0.26 |
| 4 | 86129563 | rs7666306 | -1.947 | -0.761 | 1.014 | -0.273 | 0.012 | 0.031 | 0.031 | 0.121 | 0.42 | 0.69 | 0.43 | 0.31 | 0.38 | 0.52 |
| 4 | 86153942 | rs6818991 | -0.413 | -0.689 | 1.095 | -0.251 | 0.006 | 0.001 | 0.001 | 0 | 0.23 | 0.34 | 0.26 | 0.27 | 0.27 | 0.29 |
| 4 | 86167397 | rs6841256 | -1.108 | -0.818 | 1.099 | -0.289 | -0.035 | -0.074 | -0.074 | 0 | 0.19 | 0.35 | 0.18 | 0.05 | 0.12 | 0.24 |
| 4 | 86167708 | rs11730221 | -0.499 | -0.798 | 1.119 | -0.267 | 0.005 | 0.003 | 0.003 | 0 | 0.24 | 0.39 | 0.25 | 0.27 | 0.26 | 0.28 |
| 4 | 86167996 | rs2869329 | -1.312 | -0.693 | 0.989 | 0.076 | -0.037 | -0.075 | -0.075 | 0.31 | 0.19 | 0.37 | 0.18 | 0.05 | 0.12 | 0.24 |
| 4 | 86178448 | rs12504718 | -1.83 | -0.527 | 1.067 | 0.162 | -0.014 | 0.006 | 0.006 | 0.456 | 0.28 | 0.51 | 0.37 | 0.31 | 0.28 | 0.46 |
| 4 | 86178701 | rs11097048 | 0.014 | -0.408 | 0.906 | 0.303 | 0 | 0.01 | 0.01 | 0 | 0.13 | 0.18 | 0.19 | 0.27 | 0.17 | 0.26 |
| 4 | 86191083 | rs7673714 | -1.631 | -0.354 | 0.78 | 0.496 | -0.006 | 0.082 | 0.082 | 0.019 | 0.24 | 0.51 | 0.36 | 0.37 | 0.25 | 0.41 |
| 4 | 86210281 | rs11732437 | -1.218 | -0.353 | 0.745 | 0.519 | -0.007 | 0.063 | 0.063 | 0.041 | 0.2 | 0.38 | 0.26 | 0.29 | 0.21 | 0.35 |
| 4 | 86222977 | rs7678568 | -1.213 | -0.334 | 0.903 | 0.296 | -0.008 | -0.007 | -0.007 | 0 | 0.12 | 0.28 | 0.16 | 0.09 | 0.08 | 0.16 |
| 4 | 86253489 | rs434193 | -0.671 | -0.433 | -0.074 | 0.043 | 0.016 | 0.036 | 0.036 | 0.148 | 0.4 | 0.67 | 0.24 | 0.19 | 0.33 | 0.3 |
| 4 | 86264971 | rs1955237 | -0.589 | -0.503 | -0.115 | 0.012 | 0.028 | 0.067 | 0.067 | 0.295 | 0.4 | 0.73 | 0.25 | 0.19 | 0.33 | 0.3 |
| 4 | 86266527 | rs342944 | -0.176 | -0.499 | 0.025 | 0.01 | -0.009 | -0.023 | -0.023 | 0.215 | 0.35 | 0.58 | 0.2 | 0.19 | 0.29 | 0.22 |
| 4 | 86276070 | rs1817149 | -0.771 | -0.279 | 0.332 | 0.177 | 0.001 | 0.011 | 0.011 | 0.131 | 0.37 | 0.68 | 0.35 | 0.27 | 0.37 | 0.37 |
| 4 | 86288423 | rs11938951 | -0.178 | -0.301 | 0.277 | 0.158 | -0.002 | -0.043 | -0.043 | 0 | 0.32 | 0.46 | 0.22 | 0.2 | 0.32 | 0.32 |
| 4 | 86289106 | rs391925 | -1.096 | -0.287 | 0.133 | 0.401 | 0.036 | 0.17 | 0.17 | 0.461 | 0.18 | 0.49 | 0.33 | 0.35 | 0.22 | 0.3 |
| 4 | 86289320 | rs403870 | -0.753 | -0.268 | 0.175 | 0.411 | 0.047 | 0.175 | 0.175 | 0 | 0.16 | 0.43 | 0.32 | 0.35 | 0.22 | 0.3 |
| 4 | 86289637 | rs412856 | -0.444 | -0.193 | -0.122 | 0.301 | -0.001 | 0.051 | 0.051 | 0.012 | 0.35 | 0.81 | 0.26 | 0.22 | 0.34 | 0.33 |
| 4 | 86292692 | rs10222882 | -1.105 | 0.095 | -0.06 | 0.252 | -0.027 | -0.029 | -0.029 | 0 | 0.45 | 0.82 | 0.43 | 0.48 | 0.5 | 0.58 |
| 4 | 86292752 | rs6842048 | -1.494 | 0.081 | -0.108 | 0.242 | -0.02 | 0.002 | 0.002 | 0.128 | 0.49 | 0.87 | 0.54 | 0.53 | 0.53 | 0.63 |
| 4 | 86294733 | rs1377362 | -0.332 | 0.066 | 0.05 | 0.687 | -0.001 | -0.001 | -0.001 | 0 | 0.39 | 0.41 | 0.24 | 0.43 | 0.35 | 0.38 |
| 4 | 86297661 | rs9306997 | 0.063 | 0.208 | -0.29 | 0.772 | -0.009 | -0.011 | -0.011 | 0.578 | 0.13 | 0.13 | 0.2 | 0.27 | 0.21 | 0.29 |
| 4 | 86309824 | rs381032 | -1.401 | 0.161 | -0.748 | 0.563 | -0.037 | 0.058 | 0.058 | 1.542 | 0.58 | 0.92 | 0.58 | 0.74 | 0.59 | 0.73 |
| 4 | 86321876 | rs384878 | -1.348 | 0.26 | -0.053 | 0.807 | 0.026 | 0.1 | 0.1 | 0 | 0.28 | 0.52 | 0.39 | 0.53 | 0.32 | 0.4 |
| 4 | 86324046 | rs405074 | 0.121 | 0.55 | -0.383 | 0.958 | -0.012 | -0.007 | -0.007 | 0 | 0.11 | 0.07 | 0.2 | 0.29 | 0.19 | 0.3 |
| 4 | 86324520 | rs1453071 | -0.981 | 0.547 | -0.252 | 0.915 | 0.026 | 0.086 | 0.086 | 0.04 | 0.16 | 0.33 | 0.32 | 0.33 | 0.23 | 0.35 |
| 4 | 86335426 | rs340185 | 0.415 | 0.725 | -1.04 | 0.243 | 0.005 | -0.023 | -0.023 | 0 | 0.38 | 0.49 | 0.42 | 0.25 | 0.38 | 0.16 |
| 4 | 86337028 | rs744597 | 0.059 | 0.804 | -1.471 | 0.315 | -0.002 | 0.002 | 0.002 | 0.667 | 0.41 | 0.4 | 0.5 | 0.31 | 0.42 | 0.23 |
| 4 | 86339453 | rs4693115 | 0.255 | 1.044 | -2.533 | 0.233 | -0.017 | -0.055 | -0.055 | 0 | 0.09 | 0.01 | 0.18 | 0.23 | 0.09 | 0.07 |
| 4 | 86339956 | rs340192 | 0.699 | 0.792 | -2.273 | -0.095 | -0.047 | 0.029 | 0.029 | 2.419 | 0.35 | 0.12 | 0.38 | 0.25 | 0.39 | 0.16 |
| 4 | 86340696 | rs168153 | 0.343 | 0.507 | -1.235 | 0.294 | -0.084 | -0.115 | -0.115 | 0 | 0.82 | 0.98 | 0.8 | 0.7 | 0.72 | 0.57 |
| 4 | 86350683 | rs340199 | -1.025 | -0.74 | -0.391 | -0.975 | -0.005 | -0.036 | -0.036 | 0 | 0.41 | 0.52 | 0.27 | 0.08 | 0.38 | 0.31 |
| 4 | 86351156 | rs340200 | -1.028 | -0.817 | -0.15 | -1.013 | -0.017 | -0.083 | -0.083 | 0.112 | 0.39 | 0.47 | 0.26 | 0.08 | 0.37 | 0.29 |
| 4 | 86353091 | rs975967 | -0.688 | -1.037 | 0.309 | -0.881 | 0.025 | 0.125 | 0.125 | 2.156 | 0.12 | 0.48 | 0.06 | 0.18 | 0.06 | 0.07 |
| 4 | 86357821 | rs13118367 | -0.487 | -1.342 | 0.122 | -1.124 | -0.01 | 0.044 | 0.044 | 1.073 | 0.23 | 0.13 | 0.15 | 0.08 | 0.28 | 0.22 |
| 4 | 86358621 | rs1481775 | -2.226 | -1.426 | 0.374 | -1.151 | 0.045 | 0.12 | 0.12 | 0.464 | 0.5 | 0.82 | 0.32 | 0.26 | 0.43 | 0.38 |
| 4 | 86366812 | rs10516748 | -1.446 | -1.625 | 0.579 | -1.169 | -0.009 | 0.003 | 0.003 | 0.76 | 0.27 | 0.64 | 0.13 | 0.24 | 0.18 | 0.19 |
| 4 | 86369449 | rs17010244 | -0.465 | -1.63 | 0.556 | -1.206 | -0.008 | -0.013 | -0.013 | 0 | 0.17 | 0.18 | 0.06 | 0.06 | 0.1 | 0.12 |
| 4 | 86381967 | rs9995572 | 0.571 | -1.443 | 0.536 | -1.216 | -0.003 | -0.043 | -0.043 | 0.012 | 0.79 | 0.79 | 0.8 | 0.98 | 0.75 | 0.83 |

| | | | | | | | | | | | | | | | | |
|---|---|---|---|---|---|---|---|---|---|---|---|---|---|---|---|---|
| 4 | 86384329 | rs1383972 | 1.428 | -1.351 | 0.283 | -1.167 | 0.007 | 0.05 | 0.05 | 4.882 | 0.66 | 0.19 | 0.71 | 0.81 | 0.65 | 0.73 |
| 4 | 86397877 | rs10034597 | -0.72 | -2.027 | 0.61 | -1.626 | -0.002 | -0.006 | -0.006 | 0.5 | 0.28 | 0.28 | 0.34 | 0.09 | 0.19 | 0.15 |
| 4 | 86414621 | rs11097055 | 1.198 | -1.78 | 0.982 | -1.656 | -0.061 | -0.124 | -0.124 | 0.236 | 0.72 | 0.46 | 0.84 | 0.97 | 0.8 | 0.83 |
| 4 | 86426882 | rs10015995 | 1.436 | -1.411 | 0.47 | -1.413 | -0.006 | -0.026 | -0.026 | 0 | 0.62 | 0.42 | 0.81 | 0.86 | 0.74 | 0.8 |
| 4 | 86432381 | rs4611924 | -0.94 | -0.966 | 0.458 | -1.331 | -0.028 | -0.077 | -0.077 | 0 | 0.21 | 0.34 | 0.1 | 0.07 | 0.12 | 0.08 |
| 4 | 86434204 | rs1383976 | 1.031 | -0.674 | -0.242 | -1.222 | 0.009 | -0.002 | -0.002 | 0 | 0.62 | 0.35 | 0.81 | 0.9 | 0.73 | 0.81 |
| 4 | 86434268 | rs9307000 | -0.776 | -0.584 | -0.225 | -1.097 | -0.032 | -0.114 | -0.114 | 0.024 | 0.29 | 0.51 | 0.14 | 0.1 | 0.22 | 0.17 |
| 4 | 86436599 | rs9985756 | 0.703 | -0.444 | -0.589 | -1.108 | 0.008 | -0.002 | -0.002 | 0 | 0.62 | 0.35 | 0.81 | 0.9 | 0.73 | 0.81 |
| 4 | 86437529 | rs4693701 | 0.105 | -0.127 | -1.295 | -1.379 | 0.004 | 0.059 | 0.059 | 0 | 0.67 | 0.77 | 0.78 | 0.79 | 0.7 | 0.66 |
| 4 | 86437794 | rs2078049 | 0.327 | 0.563 | -2.096 | -0.73 | -0.081 | -0.158 | -0.158 | 0 | 0.26 | 0.03 | 0.28 | 0.55 | 0.28 | 0.31 |
| 4 | 86440758 | rs17387199 | -0.121 | 0.194 | -2.039 | -1.24 | -0.01 | 0.014 | 0.014 | 0.753 | 0.27 | 0.05 | 0.46 | 0.8 | 0.28 | 0.48 |
| 4 | 86445949 | rs10020489 | -0.148 | -0.055 | -1.264 | -1.34 | -0.006 | 0.014 | 0.014 | 0.687 | 0.27 | 0.07 | 0.45 | 0.79 | 0.28 | 0.48 |
| 4 | 86455224 | rs6845803 | -0.388 | -0.834 | -0.684 | -1.777 | -0.035 | -0.037 | -0.037 | 0 | 0.13 | 0.24 | 0.06 | 0.03 | 0.04 | 0.05 |
| 4 | 86461983 | rs11942163 | -0.74 | -0.921 | -0.231 | -1.839 | 0.013 | 0.009 | 0.009 | 0.777 | 0.07 | 0.47 | 0.06 | 0.02 | 0.02 | 0.03 |
| 4 | 86463980 | rs11097059 | 1.124 | -0.417 | 0.231 | -1.874 | 0.017 | 0.054 | 0.054 | 0.003 | 0.56 | 0.27 | 0.43 | 0.15 | 0.63 | 0.43 |
| 4 | 86464117 | rs1545823 | 0.147 | -0.177 | 0.273 | -1.648 | -0.045 | 0.01 | 0.01 | 0.82 | 0.27 | 0.06 | 0.46 | 0.79 | 0.31 | 0.49 |
| 4 | 86467056 | rs6829120 | -0.883 | -0.161 | 0.37 | -1.579 | 0.014 | 0.011 | 0.011 | 0.684 | 0.07 | 0.43 | 0.06 | 0.02 | 0.02 | 0.03 |
| 4 | 86475051 | rs2869356 | 0.731 | -0.121 | 0.205 | -1.72 | -0.028 | -0.088 | -0.088 | 0 | 0.86 | 0.48 | 0.92 | 0.98 | 0.94 | 0.97 |
| 4 | 86478104 | rs12510597 | 0.279 | -0.07 | 0.078 | -1.76 | -0.022 | 0.009 | 0.009 | 0.442 | 0.26 | 0.08 | 0.44 | 0.79 | 0.27 | 0.45 |
| 4 | 86482468 | rs964837 | 0.439 | -0.079 | 0.074 | -1.693 | -0.006 | -0.044 | -0.044 | 0 | 0.62 | 0.49 | 0.49 | 0.19 | 0.67 | 0.52 |
| 4 | 86487089 | rs2128101 | 0.129 | -0.466 | 0.816 | -1.231 | -0.017 | 0.005 | 0.005 | 0.3 | 0.22 | 0.28 | 0.42 | 0.77 | 0.18 | 0.43 |
| 4 | 86498937 | rs11930449 | 0.699 | -0.745 | 0.729 | -1.335 | 0.016 | 0.023 | 0.023 | 0.052 | 0.62 | 0.32 | 0.49 | 0.16 | 0.73 | 0.5 |
| 4 | 86501891 | rs1384137 | -0.905 | -0.75 | 0.796 | -1.338 | -0.027 | -0.059 | -0.059 | 0 | 0.37 | 0.56 | 0.5 | 0.84 | 0.27 | 0.49 |
| 4 | 86508815 | rs1161262 | -0.798 | -0.961 | 0.812 | -1.562 | -0.035 | -0.068 | -0.068 | 0.008 | 0.84 | 0.7 | 0.9 | 0.97 | 0.92 | 0.87 |
| 4 | 86510607 | rs2623770 | -1.142 | -0.972 | 0.765 | -1.567 | 0.007 | 0.006 | 0.006 | 0 | 0.87 | 0.91 | 0.91 | 0.97 | 0.91 | 0.87 |
| 4 | 86520349 | rs13149991 | 1.409 | -0.702 | 0.422 | -1.853 | -0.074 | 0.112 | 0.112 | 0 | 0.35 | 0.2 | 0.31 | 0.12 | 0.46 | 0.35 |
| 4 | 86523580 | rs1482090 | -0.708 | -1.138 | 0.339 | -1.337 | -0.038 | -0.074 | -0.074 | 0 | 0.84 | 0.67 | 0.9 | 0.97 | 0.91 | 0.87 |
| 4 | 86528170 | rs6849280 | 1.068 | -1.185 | 0.248 | -1.416 | 0.022 | 0.04 | 0.04 | 0 | 0.48 | 0.41 | 0.41 | 0.13 | 0.62 | 0.39 |
| 4 | 86533932 | rs13103706 | 0.834 | -1.19 | 0.301 | -1.418 | 0.019 | 0.023 | 0.023 | 0 | 0.5 | 0.53 | 0.42 | 0.13 | 0.62 | 0.39 |
| 4 | 86549594 | rs12504294 | -0.707 | -0.73 | 0.528 | -0.853 | 0.015 | 0.038 | 0.038 | 0.013 | 0.36 | 0.27 | 0.49 | 0.84 | 0.3 | 0.49 |
| 4 | 86559173 | rs1482088 | 0.62 | -0.64 | -0.323 | -0.241 | -0.034 | -0.162 | -0.162 | 0 | 0.21 | 0.01 | 0.33 | 0.57 | 0.19 | 0.42 |
| 4 | 86565198 | rs1384135 | -0.244 | -0.358 | -0.041 | -1.217 | 0.026 | 0.079 | 0.079 | 0.257 | 0.36 | 0.17 | 0.48 | 0.84 | 0.3 | 0.49 |
| 4 | 86581005 | rs4693122 | -0.562 | -0.354 | -0.135 | -1.215 | 0.01 | 0.015 | 0.015 | 0 | 0.38 | 0.32 | 0.5 | 0.84 | 0.3 | 0.49 |
| 4 | 86594072 | rs1482097 | -1.041 | -0.06 | 0.717 | -0.457 | 0.042 | 0.038 | 0.038 | 1.364 | 0.06 | 0.41 | 0.04 | 0 | 0 | 0 |
| 4 | 86601669 | rs1026873 | -0.263 | 0.393 | -0.248 | -0.817 | -0.008 | 0.01 | 0.01 | 0.096 | 0.41 | 0.64 | 0.37 | 0.13 | 0.47 | 0.37 |
| 4 | 86604729 | rs7698203 | -1.146 | -0.926 | 0.201 | -1.014 | -0.068 | -0.074 | -0.074 | 0.018 | 0.25 | 0.61 | 0.15 | 0.09 | 0.15 | 0.2 |
| 4 | 86614203 | rs6823441 | 0.479 | -1.261 | 0.631 | -1.267 | -0.052 | -0.087 | -0.087 | 1.333 | 0.8 | 0.66 | 0.91 | 0.97 | 0.87 | 0.86 |
| 4 | 86615391 | rs10017047 | 1.456 | -1.356 | 1.137 | -0.425 | -0.018 | 0.001 | 0.001 | 0 | 0.69 | 0.34 | 0.65 | 0.66 | 0.74 | 0.63 |
| 4 | 86632771 | rs4413396 | 1.564 | -1.369 | 1.148 | -0.432 | 0.017 | -0.003 | -0.003 | 0 | 0.68 | 0.29 | 0.64 | 0.66 | 0.74 | 0.63 |
| 4 | 86652248 | rs2062098 | -1.707 | -2.223 | 1.211 | -1.004 | -0.029 | -0.096 | -0.096 | 3.345 | 0.39 | 0.48 | 0.57 | 0.91 | 0.33 | 0.61 |
| 4 | 86653361 | rs1482091 | 0.053 | -2.196 | 0.969 | -0.749 | 0.005 | 0.133 | 0.133 | 0.02 | 0.07 | 0.11 | 0.25 | 0.33 | 0.12 | 0.21 |
| 4 | 86669762 | rs13106553 | 0.252 | -2.199 | 0.702 | -0.734 | 0.014 | -0.045 | -0.045 | 0 | 0.17 | 0.11 | 0.26 | 0.58 | 0.11 | 0.33 |
| 4 | 86675725 | rs6848047 | 1.298 | -1.665 | -0.214 | -1.91 | -0.017 | 0.05 | 0.05 | 0 | 0.24 | 0.13 | 0.13 | 0.05 | 0.24 | 0.16 |
| 4 | 86689557 | rs13150113 | -0.001 | -1.605 | 0.514 | -1.456 | 0.04 | 0.036 | 0.036 | 3.25 | 0.94 | 0.59 | 0.96 | 1 | 1 | 1 |
| 4 | 86691595 | rs11945319 | 0.288 | -1.429 | 0.232 | -1.243 | 0.016 | -0.032 | -0.032 | 0.416 | 0.17 | 0.17 | 0.28 | 0.68 | 0.12 | 0.34 |
| 4 | 86692665 | rs6831420 | -0.135 | -1.117 | 0.213 | -1.081 | -0.022 | 0.018 | 0.018 | 0 | 0.2 | 0.27 | 0.31 | 0.23 | 0.22 | 0.28 |
| 4 | 86703976 | rs10516756 | 0.529 | -0.884 | -0.127 | -0.228 | -0.001 | 0.043 | 0.043 | 0 | 0.07 | 0.06 | 0.23 | 0.64 | 0.07 | 0.28 |
| 4 | 86704339 | rs7670874 | -1.018 | -1.105 | -0.517 | -0.607 | -0.011 | -0.092 | -0.092 | 0.243 | 0.65 | 0.43 | 0.76 | 0.9 | 0.69 | 0.78 |
| 4 | 86704797 | rs10516757 | 0.994 | -0.419 | -0.687 | -0.78 | -0.01 | -0.003 | -0.003 | 0.013 | 0.05 | 0.01 | 0.18 | 0.16 | 0.11 | 0.22 |
| 4 | 86705664 | rs1452681 | 2.24 | -0.199 | -1.011 | -1.1 | -0.131 | 0.142 | 0.142 | 0.258 | 0.21 | 0.01 | 0.16 | 0 | 0.28 | 0.16 |
| 4 | 86706193 | rs9307010 | 1.791 | -0.337 | -0.538 | -1.013 | 0 | -0.024 | -0.024 | 0 | 0.28 | 0.3 | 0.17 | 0.03 | 0.28 | 0.15 |
| 4 | 86709899 | rs7662174 | -1.206 | -0.699 | -0.419 | -0.472 | -0.011 | -0.01 | -0.01 | 0 | 0.37 | 0.36 | 0.37 | 0.26 | 0.35 | 0.34 |
| 4 | 86713124 | rs17010925 | -0.278 | -0.761 | -0.294 | -0.583 | -0.049 | -0.072 | -0.072 | 3.475 | 0.26 | 0.37 | 0.16 | 0.09 | 0.19 | 0.11 |
| 4 | 86719165 | rs17010957 | -0.629 | -0.429 | -0.54 | -0.064 | 0 | -0.009 | -0.009 | 0 | 0.12 | 0.1 | 0.22 | 0.17 | 0.14 | 0.22 |
| 4 | 86720955 | rs12504839 | 0.808 | -0.592 | -0.323 | 0.814 | -0.006 | 0.103 | 0.103 | 0 | 0.07 | 0.08 | 0.15 | 0.32 | 0.04 | 0.25 |
| 4 | 86723173 | rs7689056 | -0.352 | -1.458 | 0.114 | 0.372 | -0.07 | 0.178 | 0.178 | 0 | 0.21 | 0.47 | 0.22 | 0.5 | 0.16 | 0.19 |
| 4 | 86725502 | rs1452676 | -2.224 | -1.327 | -0.323 | -0.232 | -0.001 | -0.023 | -0.023 | 0 | 0.72 | 0.74 | 0.83 | 0.98 | 0.73 | 0.86 |
| 4 | 86726240 | rs17395916 | 1.484 | -1.28 | -0.34 | -0.218 | -0.138 | 0.234 | 0.234 | 0.991 | 0.34 | 0.04 | 0.23 | 0 | 0.39 | 0.2 |
| 4 | 86730514 | rs6815441 | 0.313 | -1.405 | -0.056 | 0.255 | -0.04 | 0.013 | 0.013 | 0 | 0.16 | 0.47 | 0.12 | 0.13 | 0.11 | 0.07 |
| 4 | 86730714 | rs4693128 | -1.012 | -0.801 | 0.062 | 0.645 | -0.001 | 0.046 | 0.046 | 0 | 0.18 | 0.19 | 0.38 | 0.47 | 0.17 | 0.47 |
| 4 | 86732728 | rs12512001 | 1.407 | -0.646 | 0.076 | 0.743 | -0.003 | 0.127 | 0.127 | 0 | 0.06 | 0.08 | 0.15 | 0.32 | 0.04 | 0.26 |
| 4 | 86739805 | rs12498568 | 0.126 | -0.563 | 0.223 | 0.482 | 0.021 | 0.079 | 0.079 | 0.713 | 0.07 | 0.19 | 0.15 | 0.16 | 0.08 | 0.2 |
| 4 | 86744000 | rs1823762 | 0.691 | -0.551 | 0.037 | 0.495 | 0.001 | 0.013 | 0.013 | 0.046 | 0.06 | 0.09 | 0.14 | 0.16 | 0.09 | 0.21 |
| 4 | 86750699 | rs345363 | 0.751 | -0.647 | -0.027 | 0.147 | -0.099 | 0.156 | 0.156 | 0.344 | 0.35 | 0.12 | 0.26 | 0.05 | 0.45 | 0.18 |
| 4 | 86751384 | rs17011024 | 0.233 | -0.71 | -0.176 | 0.035 | -0.012 | -0.013 | -0.013 | 0 | 0.18 | 0.08 | 0.21 | 0.16 | 0.22 | 0.25 |
| 4 | 86751718 | rs3796625 | -0.293 | -0.969 | 0.255 | -0.183 | -0.008 | 0.067 | 0.067 | 0 | 0.17 | 0.21 | 0.29 | 0.5 | 0.18 | 0.33 |
| 4 | 86752397 | rs6844781 | 1.073 | -1.011 | 0.483 | -0.114 | 0.02 | 0.063 | 0.063 | 0 | 0.38 | 0.26 | 0.23 | 0.12 | 0.31 | 0.21 |
| 4 | 86756230 | rs345370 | -0.43 | -1.198 | 0.768 | 0.002 | 0.01 | 0.38 | 0.38 | 0.039 | 0.9 | 0.54 | 0.78 | 0.62 | 0.95 | 0.72 |
| 4 | 86763097 | rs10008894 | 0.585 | -0.977 | 0.451 | -0.18 | -0.001 | 0 | 0 | 0 | 0.46 | 0.47 | 0.44 | 0.5 | 0.36 | 0.49 |

| | | | | | | | | | | | | | | | | |
|---|---|---|---|---|---|---|---|---|---|---|---|---|---|---|---|---|
| 4 | 86763901 | rs345328 | 1.215 | -0.617 | -0.104 | -0.831 | -0.164 | 0.152 | 0.152 | 0 | 0.23 | 0.01 | 0.18 | 0 | 0.34 | 0.14 |
| 4 | 86768945 | rs6834349 | 0.152 | -0.581 | 0.441 | -0.704 | -0.005 | -0.005 | -0.005 | 0 | 0.57 | 0.63 | 0.62 | 0.86 | 0.51 | 0.7 |
| 4 | 86776488 | rs7672187 | 0.67 | -0.509 | -0.078 | -0.641 | 0 | -0.011 | -0.011 | 0.037 | 0.13 | 0.1 | 0.09 | 0.01 | 0.15 | 0.05 |
| 4 | 86795362 | rs2601864 | -0.106 | -0.182 | 0.469 | -0.426 | 0.02 | -0.102 | -0.102 | 0 | 0.35 | 0.46 | 0.32 | 0.09 | 0.39 | 0.26 |
| 4 | 86808963 | rs10032626 | 0.254 | -0.174 | 0.367 | 0.106 | -0.014 | 0.029 | 0.029 | 0 | 0.57 | 0.35 | 0.51 | 0.68 | 0.56 | 0.48 |
| 4 | 86810287 | rs6817562 | 0.444 | -0.355 | 1.088 | 0.554 | -0.015 | 0.114 | 0.114 | 0.101 | 0.86 | 0.65 | 0.71 | 0.67 | 0.89 | 0.64 |
| 4 | 86812285 | rs10009525 | -0.923 | -0.379 | 1.173 | 0.205 | -0.004 | 0.013 | 0.013 | 0.961 | 0.1 | 0.18 | 0.09 | 0.12 | 0.11 | 0.14 |
| 4 | 86819377 | rs1016510 | -1.304 | -0.619 | 1.276 | 0.123 | -0.015 | -0.017 | -0.017 | 0 | 0.27 | 0.44 | 0.35 | 0.26 | 0.27 | 0.45 |
| 4 | 86821509 | rs10023650 | -1.578 | -0.846 | 0.875 | -0.222 | -0.102 | 0.168 | 0.168 | 0 | 0.67 | 0.93 | 0.76 | 0.99 | 0.56 | 0.81 |
| 4 | 86829529 | rs7696763 | -0.171 | -0.831 | 0.918 | -0.177 | -0.001 | -0.011 | -0.011 | 0 | 0.08 | 0.13 | 0.09 | 0.08 | 0.11 | 0.14 |
| 4 | 86832756 | rs723570 | 0.326 | -0.852 | 0.92 | -0.175 | 0.001 | -0.004 | -0.004 | 0 | 0.17 | 0.11 | 0.25 | 0.18 | 0.16 | 0.31 |
| 4 | 86836381 | rs346510 | 0.615 | -0.84 | 1.29 | -0.147 | 0.008 | 0.004 | 0.004 | 0 | 0.37 | 0.36 | 0.39 | 0.63 | 0.29 | 0.28 |
| 4 | 86837535 | rs10006487 | -0.366 | -0.863 | 1.383 | -0.105 | 0.007 | -0.009 | -0.009 | 1.005 | 0.09 | 0.2 | 0.09 | 0.08 | 0.11 | 0.17 |
| 4 | 86837843 | rs9307013 | -0.561 | -0.738 | 1.316 | -0.039 | -0.009 | 0.009 | 0.009 | 0 | 0.27 | 0.35 | 0.36 | 0.36 | 0.27 | 0.53 |
| 4 | 86847640 | rs346521 | 0.531 | -0.901 | 0.934 | 0.213 | 0.008 | -0.018 | -0.018 | 0 | 0.34 | 0.28 | 0.36 | 0.53 | 0.28 | 0.27 |
| 4 | 86851560 | rs346498 | 0.032 | -0.784 | 0.938 | 0.317 | 0.016 | -0.004 | -0.004 | 0 | 0.42 | 0.31 | 0.46 | 0.59 | 0.35 | 0.46 |
| 4 | 86853558 | rs346500 | -1.197 | -0.498 | 0.932 | 0.269 | -0.062 | 0.098 | 0.098 | 0 | 0.64 | 0.88 | 0.72 | 0.9 | 0.51 | 0.65 |
| 4 | 86864518 | rs2665869 | -1.345 | -0.311 | 0.31 | 0.409 | -0.05 | 0.061 | 0.061 | 0 | 0.67 | 0.88 | 0.74 | 0.82 | 0.54 | 0.7 |
| 4 | 86871282 | rs13110329 | 0.565 | -0.053 | -0.262 | 0.596 | 0 | -0.029 | -0.029 | 1.353 | 0.22 | 0.1 | 0.29 | 0.31 | 0.22 | 0.41 |
| 4 | 86880673 | rs346480 | -0.553 | 0.008 | -0.313 | 0.807 | -0.003 | 0.002 | 0.002 | 0.069 | 0.4 | 0.4 | 0.43 | 0.51 | 0.32 | 0.29 |
| 4 | 86891199 | rs12498346 | 0.213 | 0.202 | -1.114 | 0.485 | 0.002 | 0.025 | 0.025 | 0 | 0.16 | 0.17 | 0.29 | 0.41 | 0.19 | 0.19 |
| 4 | 86912951 | rs11724681 | -0.337 | 0.492 | -0.886 | 0.714 | 0.01 | -0.017 | -0.017 | 1.266 | 0.62 | 0.73 | 0.59 | 0.57 | 0.68 | 0.71 |
| 4 | 86930207 | rs2575676 | -0.241 | 1.068 | -0.803 | 0.358 | -0.026 | 0.048 | 0.048 | 0 | 0.37 | 0.21 | 0.25 | 0.16 | 0.47 | 0.27 |
| 4 | 86930609 | rs7438872 | 0.872 | 0.639 | 0.18 | 1.201 | -0.027 | 0.135 | 0.135 | 0.224 | 0.81 | 0.48 | 0.87 | 0.7 | 0.85 | 0.92 |
| 4 | 86931091 | rs7439032 | 1.07 | 1.195 | 0.658 | 1.396 | 0.027 | 0.143 | 0.143 | 0.829 | 0.82 | 0.38 | 0.68 | 0.64 | 0.81 | 0.81 |
| 4 | 86931482 | rs2589525 | 1.247 | 1.377 | -0.474 | 1.062 | -0.063 | 0.038 | 0.038 | 0 | 0.53 | 0.07 | 0.49 | 0.41 | 0.46 | 0.61 |
| 4 | 86932486 | rs4640664 | -1.645 | 1.315 | -0.381 | 0.753 | -0.041 | 0.005 | 0.005 | 0.277 | 0.44 | 0.89 | 0.33 | 0.35 | 0.46 | 0.33 |
| 4 | 86933060 | rs17011297 | -0.193 | 1.26 | -0.348 | 0.817 | 0.003 | -0.033 | -0.033 | 2.985 | 0.08 | 0.02 | 0.06 | 0.19 | 0.07 | 0.1 |
| 4 | 86933202 | rs2589523 | -1.369 | 1.383 | 0.331 | 0.653 | -0.007 | 0.002 | 0.002 | 0 | 0.19 | 0.26 | 0.09 | 0.19 | 0.13 | 0.1 |
| 4 | 86935890 | rs2575675 | -1.57 | 1.745 | 0.635 | 0.48 | 0.001 | 0.002 | 0.002 | 0 | 0.25 | 0.23 | 0.17 | 0.19 | 0.24 | 0.12 |
| 4 | 86936468 | rs1201 | -1.194 | 1.499 | 1.24 | 0.53 | -0.002 | 0.007 | 0.007 | 1.507 | 0.18 | 0.19 | 0.13 | 0.25 | 0.13 | 0.27 |
| 4 | 86937874 | rs958 | -1.3 | 1.59 | 1.239 | 0.529 | 0.001 | 0.003 | 0.003 | 1.085 | 0.19 | 0.19 | 0.17 | 0.25 | 0.19 | 0.28 |
| 4 | 86940532 | rs2589514 | -2.794 | 1.623 | 1.601 | 0.494 | -0.024 | 0.004 | 0.004 | 0.295 | 0.42 | 0.78 | 0.32 | 0.35 | 0.43 | 0.34 |
| 4 | 86946141 | rs3775167 | -1.225 | 1.807 | 1.513 | 0.429 | -0.001 | 0 | 0 | 0 | 0.2 | 0.19 | 0.18 | 0.25 | 0.24 | 0.3 |
| 4 | 86948942 | rs2043649 | -0.939 | 1.76 | 1.563 | 0.364 | 0.06 | 0.001 | 0.001 | 0 | 0.15 | 0.51 | 0.1 | 0.1 | 0.15 | 0.05 |
| 4 | 86954867 | rs3775173 | -0.138 | 1.715 | 1.619 | 0.358 | -0.01 | 0.007 | 0.007 | 0.075 | 0.12 | 0.11 | 0.13 | 0.06 | 0.16 | 0.17 |
| 4 | 86956026 | rs10856856 | -2.202 | 1.591 | 2.049 | 0.299 | -0.009 | 0.008 | 0.008 | 0.963 | 0.34 | 0.49 | 0.25 | 0.33 | 0.33 | 0.32 |
| 4 | 86966327 | rs6815306 | 2.86 | 1.545 | 1.579 | 0.337 | -0.047 | -0.07 | -0.07 | 0 | 0.48 | 0.07 | 0.49 | 0.52 | 0.49 | 0.58 |
| 4 | 86979583 | rs3775182 | -0.152 | 1.493 | 1.657 | 0.259 | -0.02 | -0.05 | -0.05 | 0 | 0.09 | 0.01 | 0.05 | 0.18 | 0.08 | 0.11 |
| 4 | 86983232 | rs3775187 | -2.131 | 1.339 | 1.858 | 0.177 | -0.011 | -0.003 | -0.003 | 0 | 0.36 | 0.69 | 0.28 | 0.34 | 0.37 | 0.34 |
| 4 | 87021505 | rs10009993 | -1.912 | 1.262 | 1.922 | 0.439 | 0.015 | 0.023 | 0.023 | 0.769 | 0.29 | 0.58 | 0.26 | 0.34 | 0.32 | 0.34 |
| 4 | 87021938 | rs12644947 | -2.154 | 1.267 | 1.872 | 0.426 | 0.009 | 0.013 | 0.013 | 1.065 | 0.3 | 0.64 | 0.26 | 0.33 | 0.33 | 0.33 |
| 4 | 87023394 | rs12640395 | 1.244 | 0.444 | 2.196 | -0.136 | -0.024 | -0.042 | -0.042 | 2.362 | 0.78 | 0.71 | 0.86 | 0.89 | 0.83 | 0.83 |
| 4 | 87028072 | rs12505595 | -1.552 | 0.495 | 2.396 | -0.058 | 0.049 | 0.054 | 0.054 | 0 | 0.09 | 0.26 | 0.19 | 0.14 | 0.14 | 0.07 |
| 4 | 87037379 | rs5006575 | 0.971 | 0.41 | 2.349 | -0.125 | -0.025 | -0.044 | -0.044 | 0.914 | 0.81 | 0.72 | 0.87 | 0.89 | 0.85 | 0.88 |
| 4 | 87038481 | rs4403040 | 2.623 | 0.348 | 2.489 | -0.153 | 0.024 | 0.046 | 0.046 | 0.047 | 0.63 | 0.2 | 0.65 | 0.74 | 0.68 | 0.75 |
| 4 | 87054813 | rs12647506 | -0.927 | 0.495 | 2.437 | -0.047 | 0.01 | 0.013 | 0.013 | 0 | 0.07 | 0.1 | 0.18 | 0.14 | 0.13 | 0.06 |
| 4 | 87062363 | rs4352468 | 2.355 | 0.468 | 2.303 | -0.039 | 0.008 | 0.016 | 0.016 | 0 | 0.67 | 0.25 | 0.66 | 0.74 | 0.68 | 0.76 |
| 4 | 87074032 | rs10001480 | -2.042 | 0.619 | 2.113 | -0.073 | 0.068 | 0.071 | 0.071 | 0 | 0.11 | 0.41 | 0.2 | 0.14 | 0.15 | 0.07 |
| 4 | 87076360 | rs6822478 | 1.105 | -0.102 | 1.537 | -0.087 | -0.031 | -0.062 | -0.062 | 0 | 0.79 | 0.62 | 0.87 | 0.88 | 0.85 | 0.85 |
| 4 | 87084039 | rs7440491 | 2.163 | -0.247 | 1.395 | -0.362 | 0.005 | -0.001 | -0.001 | 0 | 0.72 | 0.3 | 0.83 | 0.92 | 0.79 | 0.81 |
| 4 | 87087320 | rs7678480 | -1.095 | -0.314 | 2.026 | -0.403 | -0.017 | -0.03 | -0.03 | 1.248 | 0.16 | 0.23 | 0.1 | 0.07 | 0.14 | 0.09 |
| 4 | 87095840 | rs1903360 | -2.592 | -0.31 | 2.245 | -0.402 | -0.029 | -0.055 | -0.055 | 0 | 0.27 | 0.61 | 0.16 | 0.08 | 0.2 | 0.19 |
| 4 | 87096662 | rs1460761 | -2.437 | -0.315 | 2.312 | -0.405 | -0.044 | -0.072 | -0.072 | 5.864 | 0.24 | 0.59 | 0.14 | 0.07 | 0.17 | 0.15 |
| 4 | 87098956 | rs1460763 | -2.893 | -0.311 | 2.232 | -0.403 | 0.005 | -0.001 | -0.001 | 0 | 0.28 | 0.7 | 0.17 | 0.08 | 0.21 | 0.19 |
| 4 | 87108742 | rs11097100 | -0.481 | -0.306 | 2.419 | -0.399 | -0.014 | -0.034 | -0.034 | 0.011 | 0.08 | 0.26 | 0.03 | 0 | 0.02 | 0.06 |
| 4 | 87122922 | rs10034428 | -2.869 | -0.252 | 2.324 | -0.4 | 0.003 | -0.004 | -0.004 | 0 | 0.28 | 0.69 | 0.17 | 0.08 | 0.21 | 0.19 |
| 4 | 87132771 | rs2904090 | -1.033 | -0.306 | 2.372 | -0.417 | 0.003 | 0.003 | 0.003 | 0 | 0.16 | 0.12 | 0.1 | 0.07 | 0.14 | 0.1 |
| 4 | 87135124 | rs1599313 | -3.007 | -0.31 | 2.422 | -0.423 | 0.007 | 0.001 | 0.001 | 1.232 | 0.29 | 0.69 | 0.18 | 0.08 | 0.21 | 0.2 |
| 4 | 87138738 | rs12650052 | -1.111 | -0.322 | 2.472 | -0.429 | -0.008 | -0.018 | -0.018 | 0 | 0.16 | 0.16 | 0.11 | 0.07 | 0.14 | 0.1 |
| 4 | 87139714 | rs10010011 | -2.542 | -0.316 | 2.495 | -0.411 | -0.022 | -0.047 | -0.047 | 0 | 0.26 | 0.62 | 0.15 | 0.07 | 0.18 | 0.17 |
| 4 | 87151567 | rs17011547 | -1.122 | -0.301 | 2.507 | -0.397 | -0.014 | -0.027 | -0.027 | 0 | 0.16 | 0.19 | 0.11 | 0.07 | 0.14 | 0.1 |
| 4 | 87172098 | rs10516770 | -1.529 | -0.247 | 1.972 | -0.382 | -0.009 | -0.048 | -0.048 | 0 | 0.11 | 0.42 | 0.06 | 0.01 | 0.06 | 0.1 |
| 4 | 87180112 | rs11725943 | -0.589 | -0.646 | 0.586 | -0.829 | -0.007 | -0.019 | -0.019 | 0.043 | 0.24 | 0.27 | 0.2 | 0.07 | 0.17 | 0.2 |
| 4 | 87182910 | rs1460756 | -1.836 | -0.521 | -0.221 | -0.743 | 0.066 | 0.124 | 0.124 | 0.634 | 0.46 | 0.72 | 0.35 | 0.14 | 0.35 | 0.34 |
| 4 | 87185006 | rs2035058 | -0.215 | -0.409 | -0.169 | -0.491 | 0.002 | 0.006 | 0.006 | 0 | 0.11 | 0.1 | 0.09 | 0.07 | 0.12 | 0.08 |
| 4 | 87185848 | rs17011645 | 0.707 | -0.502 | -0.356 | -0.541 | -0.017 | -0.02 | -0.02 | 0 | 0.08 | 0 | 0.14 | 0.07 | 0.08 | 0.14 |
| 4 | 87190830 | rs6855169 | -2.177 | -0.914 | -0.214 | -0.703 | 0.033 | 0.042 | 0.042 | 1.925 | 0.35 | 0.73 | 0.28 | 0.15 | 0.25 | 0.29 |
| 4 | 87197433 | rs17011655 | -0.293 | -0.346 | -0.314 | -0.444 | 0 | -0.001 | -0.001 | 0 | 0.11 | 0.12 | 0.09 | 0.07 | 0.12 | 0.08 |

| | | | | | | | | | | | | | | | | |
|---|---|---|---|---|---|---|---|---|---|---|---|---|---|---|---|---|
| 4 | 87219895 | rs1460751 | 0.854 | -0.572 | -0.351 | 0.024 | 0.002 | -0.003 | -0.003 | 0.043 | 0.59 | 0.46 | 0.61 | 0.54 | 0.64 | 0.75 |
| 4 | 87231768 | rs7655217 | 0.704 | -0.608 | -0.166 | -0.005 | -0.001 | 0 | 0 | 0 | 0.62 | 0.47 | 0.61 | 0.54 | 0.65 | 0.75 |
| 4 | 87231849 | rs4693763 | 1.002 | -0.405 | -1.168 | -0.312 | -0.023 | 0.005 | 0.005 | 0 | 0.28 | 0.15 | 0.34 | 0.27 | 0.34 | 0.31 |
| 4 | 87232968 | rs10213444 | 1.591 | -0.634 | -1.231 | -0.433 | -0.077 | 0.001 | 0.001 | 0.625 | 0.27 | 0.08 | 0.24 | 0.26 | 0.3 | 0.41 |
| 4 | 87235780 | rs4299551 | 0.994 | -0.53 | -1.13 | -0.422 | -0.023 | 0.005 | 0.005 | 0 | 0.28 | 0.15 | 0.34 | 0.27 | 0.34 | 0.31 |
| 4 | 87240020 | rs6531915 | -0.239 | -0.902 | -0.73 | -0.278 | -0.017 | -0.003 | -0.003 | 0 | 0.73 | 0.66 | 0.8 | 0.78 | 0.75 | 0.88 |
| 4 | 87250736 | rs2290882 | 1.757 | -0.851 | -1.025 | -0.027 | -0.05 | -0.031 | -0.031 | 0 | 0.25 | 0.1 | 0.23 | 0.34 | 0.28 | 0.43 |
| 4 | 87252259 | rs4693142 | -0.175 | -1.018 | -0.815 | -0.18 | -0.022 | -0.021 | -0.021 | 0 | 0.73 | 0.54 | 0.77 | 0.8 | 0.77 | 0.91 |
| 4 | 87254477 | rs7689808 | -0.069 | -1.035 | -0.74 | -0.188 | -0.024 | -0.024 | -0.024 | 0.825 | 0.72 | 0.53 | 0.77 | 0.8 | 0.77 | 0.91 |
| 4 | 87255851 | rs13132206 | 2.021 | -1.033 | -0.799 | -0.188 | -0.079 | -0.054 | -0.054 | 0.727 | 0.25 | 0.07 | 0.23 | 0.34 | 0.28 | 0.43 |
| 4 | 87259268 | rs4608765 | -0.052 | -1.019 | -0.702 | -0.18 | 0.003 | -0.007 | -0.007 | 0 | 0.45 | 0.35 | 0.53 | 0.46 | 0.5 | 0.48 |
| 4 | 87259805 | rs7688651 | 1.508 | -1.129 | -0.761 | -0.181 | -0.028 | 0.005 | 0.005 | 0 | 0.27 | 0.14 | 0.34 | 0.24 | 0.33 | 0.29 |
| 4 | 87263901 | rs6531925 | 0.02 | -1.071 | -0.417 | -0.12 | -0.004 | 0.014 | 0.014 | 0.064 | 0.17 | 0.17 | 0.19 | 0.22 | 0.17 | 0.2 |
| 4 | 87269343 | rs11097109 | -0.06 | -1.052 | -0.429 | -0.117 | -0.019 | -0.019 | -0.019 | 0 | 0.73 | 0.59 | 0.77 | 0.8 | 0.77 | 0.92 |
| 4 | 87282216 | rs10008238 | 1.351 | -1.125 | -0.624 | -0.15 | 0.035 | -0.027 | -0.027 | 0.179 | 0.55 | 0.24 | 0.57 | 0.57 | 0.61 | 0.72 |
| 4 | 87287934 | rs12644920 | -0.314 | -1.072 | -0.564 | -0.122 | -0.01 | 0.016 | 0.016 | 0.078 | 0.19 | 0.24 | 0.2 | 0.22 | 0.17 | 0.2 |
| 4 | 87296065 | rs13115030 | 0.192 | -0.986 | -0.551 | -0.069 | -0.019 | -0.019 | -0.019 | 0.183 | 0.72 | 0.44 | 0.77 | 0.8 | 0.77 | 0.91 |
| 4 | 87321260 | rs2904095 | -0.754 | -0.545 | -0.65 | -0.034 | -0.006 | 0.009 | 0.009 | 0.352 | 0.21 | 0.25 | 0.21 | 0.22 | 0.2 | 0.29 |
| 4 | 87322336 | rs4235080 | -0.634 | -0.48 | -0.788 | -0.002 | 0 | 0 | 0 | 0.466 | 0.19 | 0.15 | 0.21 | 0.22 | 0.2 | 0.29 |
| 4 | 87336776 | rs17418062 | 1.914 | -0.186 | -1.06 | 0.065 | -0.017 | -0.015 | -0.015 | 0 | 0.05 | 0.02 | 0.04 | 0.06 | 0.09 | 0.02 |
| 4 | 87351740 | rs12507758 | -0.503 | -0.435 | -1.072 | -0.128 | -0.002 | -0.002 | -0.002 | 0 | 0.42 | 0.44 | 0.53 | 0.52 | 0.47 | 0.37 |
| 4 | 87362834 | rs4488910 | 0.404 | -0.095 | -1.061 | 0.062 | 0.006 | 0.028 | 0.028 | 0.505 | 0.12 | 0.15 | 0.17 | 0.21 | 0.15 | 0.17 |
| 4 | 87369063 | rs12503051 | -0.051 | -0.115 | -1.075 | 0.05 | 0.016 | 0.06 | 0.06 | 0.679 | 0.14 | 0.27 | 0.18 | 0.22 | 0.15 | 0.18 |
| 4 | 87389117 | rs7658485 | 1.161 | -0.431 | -1.625 | -0.113 | -0.11 | -0.076 | -0.076 | 0.172 | 0.32 | 0.06 | 0.27 | 0.36 | 0.37 | 0.49 |
| 4 | 87404608 | rs4693147 | 0.61 | -0.369 | -1.516 | -0.144 | -0.086 | -0.067 | -0.067 | 0 | 0.23 | 0.02 | 0.33 | 0.3 | 0.28 | 0.16 |
| 4 | 87407518 | rs11947775 | -0.258 | -0.426 | -1.126 | -0.218 | 0.001 | -0.001 | -0.001 | 0 | 0.3 | 0.33 | 0.38 | 0.31 | 0.34 | 0.19 |
| 4 | 87457636 | rs4607180 | 0.64 | -0.408 | -1.24 | -0.202 | -0.045 | -0.035 | -0.035 | 0.019 | 0.23 | 0.06 | 0.32 | 0.3 | 0.27 | 0.16 |
| 4 | 87463813 | rs4529013 | -0.124 | -0.409 | -1.182 | -0.204 | -0.01 | -0.001 | -0.001 | 0 | 0.28 | 0.26 | 0.36 | 0.31 | 0.32 | 0.19 |
| 4 | 87465741 | rs6847139 | -0.124 | -0.409 | -1.182 | -0.204 | -0.01 | -0.001 | -0.001 | 0 | 0.28 | 0.26 | 0.36 | 0.31 | 0.32 | 0.19 |
| 4 | 87473776 | rs4478147 | 0.733 | -0.542 | -1.381 | -0.316 | -0.03 | 0.002 | 0.002 | 5.744 | 0.48 | 0.1 | 0.44 | 0.57 | 0.54 | 0.69 |
| 4 | 87500298 | rs4277747 | 0.821 | -0.367 | -1.331 | -0.211 | -0.045 | -0.032 | -0.032 | 0.264 | 0.21 | 0.06 | 0.32 | 0.3 | 0.27 | 0.16 |
| 4 | 87517609 | rs7660964 | -0.94 | -0.966 | -0.438 | -0.051 | -0.052 | -0.029 | -0.029 | 0 | 0.79 | 0.7 | 0.83 | 0.88 | 0.86 | 0.89 |
| 4 | 87541252 | rs6531932 | -0.072 | -0.271 | -0.727 | 0.06 | -0.003 | 0.01 | 0.01 | 0 | 0.21 | 0.3 | 0.33 | 0.31 | 0.22 | 0.17 |
| 4 | 87542149 | rs10023787 | 0.138 | -0.216 | -0.903 | 0.088 | 0 | 0 | 0 | 0 | 0.2 | 0.2 | 0.32 | 0.31 | 0.22 | 0.17 |
| 4 | 87543847 | rs9307020 | 0.432 | -0.178 | -0.996 | 0.106 | -0.001 | -0.012 | -0.012 | 0 | 0.18 | 0.12 | 0.31 | 0.3 | 0.21 | 0.17 |
| 4 | 87552903 | rs10029671 | 0.475 | -0.167 | -0.991 | 0.11 | -0.002 | -0.011 | -0.011 | 0 | 0.18 | 0.12 | 0.31 | 0.3 | 0.21 | 0.16 |
| 4 | 87580297 | rs1021101 | 0.994 | -0.181 | -0.958 | 0.114 | 0.004 | -0.02 | -0.02 | 0 | 0.16 | 0.1 | 0.28 | 0.3 | 0.17 | 0.15 |
| 4 | 87626028 | rs1212166 | 1.023 | 0.149 | -1.807 | 0.023 | -0.141 | -0.036 | -0.036 | 0 | 0.35 | 0.03 | 0.28 | 0.33 | 0.36 | 0.5 |
| 4 | 87632672 | rs1420621 | 0.848 | 0.11 | -1.629 | 0.248 | -0.041 | -0.06 | -0.06 | 0 | 0.16 | 0.04 | 0.26 | 0.27 | 0.21 | 0.17 |
| 4 | 87641219 | rs4693780 | 1.237 | 0.063 | -1.578 | 0.25 | -0.023 | -0.056 | -0.056 | 0 | 0.14 | 0.05 | 0.23 | 0.27 | 0.17 | 0.16 |
| 4 | 87644842 | rs12508339 | 0.461 | 0.216 | -1.674 | 0.161 | -0.011 | 0.011 | 0.011 | 0 | 0.06 | 0.03 | 0.07 | 0.01 | 0.11 | 0.04 |
| 4 | 87656992 | rs10024860 | -0.526 | 0.123 | -1.272 | 0.114 | 0.036 | -0.034 | -0.034 | 1.645 | 0.08 | 0.23 | 0.08 | 0.01 | 0.11 | 0.04 |
| 4 | 87659465 | rs10033679 | -1.003 | 0.062 | -1.062 | 0.084 | 0.052 | -0.036 | -0.036 | 0 | 0.09 | 0.29 | 0.08 | 0.01 | 0.11 | 0.04 |
| 4 | 87664854 | rs7664860 | -1.003 | 0.062 | -1.062 | 0.084 | 0.052 | -0.036 | -0.036 | 0 | 0.09 | 0.29 | 0.08 | 0.01 | 0.11 | 0.04 |
| 4 | 87667171 | rs13112139 | 0.652 | 0.067 | -1.079 | 0.086 | -0.027 | 0.023 | 0.023 | 0 | 0.05 | 0.01 | 0.07 | 0.01 | 0.11 | 0.04 |
| 4 | 87670795 | rs10029565 | 0.401 | 0.068 | -1.077 | 0.087 | -0.01 | 0.009 | 0.009 | 0 | 0.06 | 0.03 | 0.07 | 0.01 | 0.11 | 0.04 |
| 4 | 87673221 | rs2054593 | 0.222 | 0.073 | -1.087 | 0.087 | -0.056 | -0.057 | -0.057 | 0 | 0.19 | 0.04 | 0.3 | 0.28 | 0.27 | 0.19 |
| 4 | 87679642 | rs1420624 | -1.049 | -0.136 | -0.977 | -0.006 | -0.046 | 0 | 0 | 0.429 | 0.6 | 0.88 | 0.7 | 0.63 | 0.61 | 0.49 |
| 4 | 87684031 | rs710832 | -1.389 | -0.127 | -0.831 | 0.021 | -0.008 | 0.003 | 0.003 | 0 | 0.66 | 0.84 | 0.75 | 0.69 | 0.72 | 0.52 |
| 4 | 87689268 | rs11097118 | -1.086 | -0.575 | -0.074 | -0.115 | 0.055 | -0.035 | -0.035 | 0 | 0.09 | 0.3 | 0.08 | 0.01 | 0.11 | 0.04 |
| 4 | 87690998 | rs2230600 | 1.14 | -0.592 | -0.375 | -0.024 | -0.042 | -0.084 | -0.084 | 0.709 | 0.12 | 0.02 | 0.23 | 0.32 | 0.16 | 0.14 |
| 4 | 87696476 | rs12499471 | -0.287 | -0.567 | -0.229 | -0.201 | 0.021 | -0.027 | -0.027 | 0 | 0.07 | 0.11 | 0.07 | 0.01 | 0.11 | 0.04 |
| 4 | 87706506 | rs989902 | -1.278 | -1.033 | -0.358 | -0.312 | 0.035 | 0.003 | 0.003 | 1.108 | 0.44 | 0.83 | 0.46 | 0.45 | 0.4 | 0.27 |
| 4 | 87728676 | rs17694112 | 0.523 | -0.606 | -0.255 | -0.008 | -0.033 | -0.087 | -0.087 | 0 | 0.15 | 0.02 | 0.22 | 0.31 | 0.15 | 0.13 |
| 4 | 87737110 | rs10016456 | -0.565 | -0.607 | -0.177 | -0.035 | 0.009 | -0.024 | -0.024 | 0 | 0.07 | 0.1 | 0.07 | 0.01 | 0.11 | 0.04 |
| 4 | 87738967 | rs10000226 | -1.104 | -0.597 | -0.118 | -0.03 | 0.039 | -0.043 | -0.043 | 0.654 | 0.1 | 0.31 | 0.08 | 0.01 | 0.11 | 0.04 |
| 4 | 87743956 | rs7688011 | 0.839 | -0.898 | -0.352 | -0.125 | 0.049 | 0.05 | 0.05 | 0.212 | 0.09 | 0 | 0.01 | 0 | 0.01 | 0.02 |
| 4 | 87743984 | rs7688019 | -0.838 | -0.041 | 0.491 | 0.323 | 0.005 | 0.018 | 0.018 | 0 | 0.25 | 0.43 | 0.3 | 0.31 | 0.26 | 0.17 |
| 4 | 87754713 | rs7678571 | -0.855 | -0.121 | 0.326 | 0.392 | -0.019 | 0.076 | 0.076 | 0.849 | 0.24 | 0.61 | 0.31 | 0.38 | 0.21 | 0.25 |
| 4 | 87760103 | rs6531935 | -1.5 | -0.457 | -0.267 | -0.266 | 0.014 | -0.005 | -0.005 | 0 | 0.45 | 0.89 | 0.46 | 0.38 | 0.42 | 0.28 |
| 4 | 87769929 | rs13106574 | 0.264 | -0.386 | 0.15 | -0.322 | 0.001 | -0.028 | -0.028 | 3.506 | 0.18 | 0.12 | 0.13 | 0 | 0.18 | 0.07 |
| 4 | 87773193 | rs8180256 | 0.768 | -0.388 | 0.309 | -0.348 | 0.006 | 0.007 | 0.007 | 0.034 | 0.67 | 0.32 | 0.75 | 0.89 | 0.7 | 0.76 |
| 4 | 87793546 | rs727143 | 0.111 | -0.269 | 0.76 | -0.279 | -0.001 | -0.054 | -0.054 | 0 | 0.19 | 0.2 | 0.12 | 0.01 | 0.18 | 0.1 |
| 4 | 87798196 | rs1374934 | 0.082 | -0.26 | 1.866 | -0.272 | 0.005 | 0.091 | 0.091 | 4.629 | 0.94 | 0.71 | 0.95 | 0.91 | 1 | 0.92 |
| 4 | 87799216 | rs6811350 | 0.311 | -0.312 | 1.941 | -0.305 | 0 | -0.018 | -0.018 | 0.313 | 0.19 | 0.12 | 0.11 | 0.01 | 0.19 | 0.11 |
| 4 | 87800577 | rs2121871 | -0.832 | -0.319 | 1.835 | -0.153 | -0.017 | 0.007 | 0.007 | 0 | 0.36 | 0.21 | 0.39 | 0.33 | 0.45 | 0.24 |
| 4 | 87809387 | rs1550931 | 0.086 | -0.315 | 1.618 | -0.15 | 0.003 | 0.089 | 0.089 | 4.873 | 0.94 | 0.71 | 0.95 | 0.91 | 1 | 0.92 |
| 4 | 87815387 | rs1868244 | 1.193 | -0.266 | 0.81 | -0.178 | -0.002 | -0.001 | -0.001 | 0.086 | 0.57 | 0.2 | 0.54 | 0.58 | 0.55 | 0.68 |
| 4 | 87834607 | rs17751557 | 0.638 | -0.018 | 0.931 | 0.132 | -0.001 | -0.003 | -0.003 | 0 | 0.14 | 0.13 | 0.23 | 0.3 | 0.16 | 0.09 |

| | | | | | | | | | | | | | | | |
|---|---|---|---|---|---|---|---|---|---|---|---|---|---|---|---|
| 4 | 87852108 | rs7657530 | 0.482 | -0.149 | 1.253 | 0.322 | 0.005 | 0.102 | 0.102 | 3.559 | 0.77 | 0.43 | 0.68 | 0.53 | 0.74 | 0.79 |
| 4 | 87863770 | rs6836128 | -0.404 | -0.325 | 0.711 | 0.266 | -0.008 | -0.004 | -0.004 | 0 | 0.21 | 0.41 | 0.24 | 0.18 | 0.2 | 0.22 |
| 4 | 87873164 | rs1992876 | -0.536 | -0.456 | 0.479 | 0.086 | 0.001 | -0.002 | -0.002 | 0 | 0.19 | 0.09 | 0.21 | 0.18 | 0.19 | 0.21 |
| 4 | 87877291 | rs725030 | 1.229 | -0.401 | 0.304 | 0.059 | -0.036 | -0.044 | -0.044 | 0.858 | 0.08 | 0.01 | 0.17 | 0.18 | 0.13 | 0.17 |
| 4 | 87883837 | rs4274814 | 1.89 | -0.546 | 0.149 | -0.198 | -0.133 | -0.017 | -0.017 | 0.67 | 0.4 | 0.05 | 0.39 | 0.33 | 0.43 | 0.51 |
| 4 | 87884766 | rs12651081 | 1.459 | -0.518 | 0.19 | -0.159 | -0.01 | 0 | 0 | 0 | 0.58 | 0.06 | 0.49 | 0.33 | 0.62 | 0.61 |
| 4 | 87895960 | rs6531941 | -1.12 | -0.518 | 0.829 | -0.004 | 0.002 | 0.223 | 0.223 | 5.015 | 0.84 | 0.64 | 0.75 | 0.52 | 0.84 | 0.83 |
| 4 | 87928204 | rs340638 | 0.225 | -0.334 | 0.928 | -0.038 | -0.002 | 0.013 | 0.013 | 3.297 | 0.75 | 0.26 | 0.69 | 0.51 | 0.76 | 0.81 |
| 4 | 87930760 | rs340636 | 1.115 | -0.434 | 0.613 | -0.152 | -0.026 | -0.085 | -0.085 | 0 | 0.48 | 0.06 | 0.55 | 0.5 | 0.54 | 0.67 |
| 4 | 87951393 | rs340641 | 1.115 | -0.434 | 0.613 | -0.145 | -0.026 | -0.081 | -0.081 | 0.128 | 0.48 | 0.06 | 0.55 | 0.51 | 0.54 | 0.67 |
| 4 | 87955141 | rs10032200 | 0.298 | -0.441 | 0.384 | -0.131 | -0.018 | -0.035 | -0.035 | 0.029 | 0.07 | 0.14 | 0.03 | 0 | 0.04 | 0.04 |
| 4 | 87956886 | rs340628 | -0.656 | -0.213 | -0.114 | 0.055 | -0.01 | 0.091 | 0.091 | 0 | 0.95 | 0.85 | 0.97 | 0.84 | 0.97 | 0.91 |
| 4 | 87972465 | rs375795 | -0.918 | 0.014 | -0.803 | 0.182 | -0.017 | -0.061 | -0.061 | 0 | 0.51 | 0.92 | 0.44 | 0.49 | 0.47 | 0.34 |
| 4 | 87976178 | rs6819155 | 0.739 | 0.663 | 0.045 | 0.86 | 0.015 | 0.057 | 0.057 | 4.329 | 0.13 | 0.21 | 0.24 | 0.33 | 0.19 | 0.09 |
| 4 | 87978626 | rs3775229 | 0.262 | 0.887 | -0.53 | 0.529 | -0.044 | -0.049 | -0.049 | 0.357 | 0.11 | 0.01 | 0.21 | 0.17 | 0.16 | 0.19 |
| 4 | 87979212 | rs4693802 | 0.15 | 0.895 | 0.168 | 0.849 | 0.047 | 0.162 | 0.162 | 4.439 | 0.17 | 0.47 | 0.26 | 0.38 | 0.22 | 0.1 |
| 4 | 87982790 | rs384059 | -0.843 | 0.547 | -0.772 | 0.189 | -0.003 | 0.037 | 0.037 | 0 | 0.68 | 0.82 | 0.71 | 0.79 | 0.75 | 0.67 |
| 4 | 87983880 | rs10006308 | 0.18 | 0.415 | -0.453 | 0.093 | -0.009 | -0.03 | -0.03 | 0 | 0.06 | 0.17 | 0.02 | 0 | 0.03 | 0.04 |
| 4 | 87994787 | rs10489045 | 0.045 | 0.382 | -0.153 | 0.077 | 0.007 | -0.017 | -0.017 | 0.849 | 0.06 | 0.21 | 0.02 | 0 | 0.03 | 0.04 |
| 4 | 87996745 | rs17605615 | 0.288 | 0.132 | -0.185 | -0.067 | 0.004 | -0.016 | -0.016 | 0 | 0.34 | 0.25 | 0.35 | 0.38 | 0.38 | 0.19 |
| 4 | 88005203 | rs236996 | -0.268 | 0.044 | -0.033 | -0.083 | 0.006 | 0.003 | 0.003 | 0 | 0.36 | 0.51 | 0.33 | 0.41 | 0.4 | 0.55 |
| 4 | 88010601 | rs236675 | -1.791 | 0.051 | -0.19 | -0.061 | -0.019 | 0.007 | 0.007 | 0 | 0.73 | 0.9 | 0.69 | 0.79 | 0.78 | 0.74 |
| 4 | 88024738 | rs3755985 | 1.506 | 0.019 | -0.407 | -0.186 | -0.041 | -0.115 | -0.115 | 0 | 0.17 | 0.01 | 0.23 | 0.38 | 0.2 | 0.1 |
| 4 | 88030486 | rs3822041 | 0.226 | -0.233 | -0.517 | -0.397 | -0.168 | -0.168 | -0.168 | 0 | 0.31 | 0.02 | 0.46 | 0.56 | 0.44 | 0.36 |
| 4 | 88033352 | rs447990 | 0.58 | -0.308 | -0.033 | -0.314 | 0.002 | 0.008 | 0.008 | 0 | 0.48 | 0.47 | 0.44 | 0.21 | 0.4 | 0.35 |
| 4 | 88041256 | rs19026 | 2.022 | -0.306 | -0.1 | -0.329 | -0.028 | 0.009 | 0.009 | 0 | 0.28 | 0.09 | 0.3 | 0.2 | 0.22 | 0.26 |
| 4 | 88052219 | rs342467 | 0.085 | -0.323 | 0.27 | -0.296 | 0.009 | 0.004 | 0.004 | 0 | 0.65 | 0.54 | 0.67 | 0.6 | 0.61 | 0.46 |
| 4 | 88057353 | rs1408 | 0.222 | -0.25 | 0.334 | -0.341 | -0.007 | -0.003 | -0.003 | 6.645 | 0.63 | 0.77 | 0.66 | 0.6 | 0.58 | 0.44 |
| 4 | 88057481 | rs342462 | 1.991 | -0.194 | -0.187 | -0.334 | -0.008 | 0.019 | 0.019 | 0 | 0.28 | 0.14 | 0.3 | 0.2 | 0.22 | 0.27 |
| 4 | 88058472 | rs342461 | 0.638 | -0.159 | -0.173 | -0.347 | 0.009 | 0.08 | 0.08 | 0 | 0.48 | 0.35 | 0.43 | 0.21 | 0.4 | 0.37 |
| 4 | 88059415 | rs342459 | 0.518 | -0.177 | -0.191 | -0.344 | 0.002 | 0.01 | 0.01 | 0 | 0.48 | 0.47 | 0.44 | 0.21 | 0.4 | 0.36 |
| 4 | 88060025 | rs342458 | 0.584 | -0.181 | -0.075 | -0.346 | 0.006 | 0.038 | 0.038 | 0 | 0.48 | 0.41 | 0.44 | 0.21 | 0.4 | 0.36 |
| 4 | 88062611 | rs2926511 | 0.884 | -0.084 | -0.53 | -0.167 | -0.014 | 0.038 | 0.038 | 0.101 | 0.46 | 0.22 | 0.54 | 0.6 | 0.42 | 0.36 |
| 4 | 88067504 | rs17012512 | 1.36 | 0.064 | -0.544 | -0.003 | -0.013 | -0.073 | -0.073 | 0 | 0.17 | 0.05 | 0.23 | 0.38 | 0.21 | 0.1 |
| 4 | 88068531 | rs4693803 | -0.282 | 0.025 | -0.977 | -0.601 | -0.025 | 0.106 | 0.106 | 0.369 | 0.35 | 0.17 | 0.33 | 0.13 | 0.39 | 0.36 |
| 4 | 88068599 | rs1466364 | 0.847 | -0.016 | -1.153 | -0.36 | 0.006 | 0.112 | 0.112 | 0.219 | 0.48 | 0.15 | 0.39 | 0.26 | 0.4 | 0.38 |
| 4 | 88069143 | rs17012516 | 0.668 | 0.234 | -1.079 | 0.058 | -0.006 | -0.115 | -0.115 | 0 | 0.24 | 0.05 | 0.26 | 0.46 | 0.26 | 0.15 |
| 4 | 88073178 | rs13145898 | 0.637 | 0.278 | -1.006 | -0.078 | -0.009 | 0.004 | 0.004 | 0 | 0.05 | 0.02 | 0.08 | 0 | 0.11 | 0.04 |
| 4 | 88082526 | rs6840470 | 1.534 | 0.191 | -1.066 | -0.187 | -0.021 | 0.135 | 0.135 | 0.088 | 0.41 | 0.18 | 0.27 | 0.14 | 0.38 | 0.38 |
| 4 | 88083107 | rs3193672 | 1.113 | 0.147 | -1.019 | -0.013 | -0.001 | -0.121 | -0.121 | 2.185 | 0.23 | 0.04 | 0.25 | 0.45 | 0.22 | 0.14 |
| 4 | 88087832 | rs17754933 | 1.312 | 0.194 | -0.857 | 0.019 | -0.028 | -0.014 | -0.014 | 0 | 0.12 | 0.07 | 0.3 | 0.18 | 0.19 | 0.25 |
| 4 | 88088507 | rs964739 | -0.199 | 0.111 | -0.001 | 0.164 | -0.028 | 0.164 | 0.164 | 0 | 0.11 | 0.49 | 0.08 | 0.22 | 0.04 | 0.15 |
| 4 | 88117272 | rs6531956 | 0.772 | 0.268 | -0.411 | 0.033 | -0.006 | 0.193 | 0.193 | 0 | 0.48 | 0.2 | 0.3 | 0.14 | 0.43 | 0.43 |
| 4 | 88148812 | rs6834470 | -0.593 | 0.395 | -0.097 | 0.212 | 0.008 | 0.006 | 0.006 | 0 | 0.35 | 0.4 | 0.47 | 0.42 | 0.4 | 0.48 |
| 4 | 88148895 | rs6531964 | -0.105 | 0.137 | 0.222 | 0.298 | -0.019 | 0.142 | 0.142 | 0 | 0.09 | 0.31 | 0.07 | 0.22 | 0.04 | 0.15 |
| 4 | 88156939 | rs11097130 | -0.897 | 0.507 | -0.531 | 0.056 | 0 | 0 | 0 | 0.347 | 0.69 | 0.69 | 0.8 | 0.76 | 0.73 | 0.74 |
| 4 | 88159429 | rs13137873 | 1.23 | 0.883 | -1.086 | 0.137 | -0.051 | -0.111 | -0.111 | 0 | 0.19 | 0.04 | 0.13 | 0.42 | 0.22 | 0.17 |
| 4 | 88159940 | rs7668684 | 1.668 | 1.097 | -0.803 | -0.102 | 0.025 | -0.04 | -0.04 | 0 | 0.1 | 0.18 | 0.13 | 0.01 | 0.14 | 0.02 |
| 4 | 88160018 | rs10006766 | 1.175 | 1.012 | -0.482 | -0.12 | 0.006 | 0.047 | 0.047 | 0.827 | 0.2 | 0.34 | 0.15 | 0.42 | 0.22 | 0.17 |
| 4 | 88162693 | rs4345139 | -1.518 | 0.83 | -0.734 | -0.319 | 0.004 | -0.004 | -0.004 | 0.048 | 0.71 | 0.79 | 0.7 | 0.72 | 0.76 | 0.7 |
| 4 | 88162837 | rs4134363 | 2.055 | 0.81 | -0.979 | -0.473 | -0.037 | 0.005 | 0.005 | 0 | 0.19 | 0.04 | 0.31 | 0.13 | 0.21 | 0.28 |
| 4 | 88180631 | rs6847980 | 0.416 | 0.843 | -0.664 | -0.28 | 0.029 | 0.018 | 0.018 | 0.02 | 0.29 | 0.46 | 0.28 | 0.36 | 0.35 | 0.18 |
| 4 | 88189651 | rs3923441 | -1.901 | 0.614 | -0.815 | -0.277 | 0.012 | -0.008 | -0.008 | 0.804 | 0.74 | 0.79 | 0.73 | 0.71 | 0.82 | 0.71 |
| 4 | 88190169 | rs7668929 | -0.51 | 0.597 | -0.808 | -0.254 | 0.047 | -0.03 | -0.03 | 0 | 0.57 | 0.77 | 0.66 | 0.48 | 0.65 | 0.5 |
| 4 | 88191100 | rs7699556 | -0.819 | 0.595 | -0.809 | -0.408 | 0.03 | 0.033 | 0.033 | 0 | 0.61 | 0.78 | 0.67 | 0.71 | 0.68 | 0.6 |
| 4 | 88198088 | rs7436142 | 2.062 | 0.956 | -0.286 | -0.072 | 0.024 | -0.029 | -0.029 | 0 | 0.09 | 0.15 | 0.14 | 0.01 | 0.16 | 0.03 |
| 4 | 88199484 | rs6531968 | -0.075 | 0.829 | -0.366 | -0.218 | 0.001 | 0.001 | 0.001 | 0 | 0.47 | 0.48 | 0.43 | 0.37 | 0.44 | 0.52 |
| 4 | 88200095 | rs4693811 | -0.635 | 0.829 | -0.385 | -0.215 | -0.004 | 0.013 | 0.013 | 0 | 0.57 | 0.64 | 0.57 | 0.37 | 0.59 | 0.55 |
| 4 | 88211056 | rs10084835 | -0.766 | 1.03 | -0.602 | -0.332 | -0.034 | 0.005 | 0.005 | 0.814 | 0.38 | 0.75 | 0.36 | 0.35 | 0.44 | 0.25 |
| 4 | 88214144 | rs4607179 | 1.176 | 1.066 | -0.69 | -0.309 | -0.065 | -0.09 | -0.09 | 1.058 | 0.2 | 0.05 | 0.16 | 0.34 | 0.25 | 0.17 |
| 4 | 88217223 | rs13133311 | 0.101 | 1.054 | -0.639 | -0.31 | -0.026 | -0.01 | -0.01 | 0 | 0.3 | 0.26 | 0.3 | 0.34 | 0.41 | 0.2 |
| 4 | 88220112 | rs13102451 | 1.147 | 1.043 | -0.688 | -0.283 | -0.065 | -0.092 | -0.092 | 3.07 | 0.2 | 0.05 | 0.16 | 0.34 | 0.25 | 0.17 |
| 4 | 88224998 | rs11945520 | 0.539 | 0.815 | -0.528 | -0.286 | 0.019 | -0.007 | -0.007 | 1.572 | 0.08 | 0.49 | 0.06 | 0 | 0.04 | 0.06 |
| 4 | 88226231 | rs11735092 | 0.296 | 0.894 | -0.738 | -0.297 | -0.076 | -0.041 | -0.041 | 0 | 0.29 | 0.15 | 0.29 | 0.34 | 0.41 | 0.2 |
| 4 | 88226480 | rs6850131 | -0.59 | 0.961 | -0.916 | -0.237 | -0.021 | 0.003 | 0.003 | 0.69 | 0.38 | 0.71 | 0.36 | 0.35 | 0.45 | 0.26 |
| 4 | 88228228 | rs10022237 | -0.59 | 0.961 | -0.916 | -0.237 | -0.02 | 0.003 | 0.003 | 0.69 | 0.38 | 0.7 | 0.36 | 0.34 | 0.45 | 0.26 |
| 4 | 88230737 | rs6531972 | -0.879 | 0.976 | -1.39 | -0.106 | -0.046 | -0.096 | -0.096 | 0 | 0.67 | 0.91 | 0.51 | 0.44 | 0.65 | 0.62 |
| 4 | 88232510 | rs9992651 | 1.055 | 1.07 | -1.321 | -0.109 | -0.059 | -0.089 | -0.089 | 0 | 0.19 | 0.05 | 0.16 | 0.31 | 0.23 | 0.15 |
| 4 | 88246162 | rs6531977 | -2.046 | 0.388 | -0.467 | 0.009 | -0.015 | -0.055 | -0.055 | 0 | 0.81 | 0.96 | 0.72 | 0.74 | 0.84 | 0.77 |

| | | | | | | | | | | | | | | | | |
|---|---|---|---|---|---|---|---|---|---|---|---|---|---|---|---|---|
| 4 | 88264614 | rs7698689 | -0.949 | 0.559 | -0.365 | -0.055 | -0.003 | -0.067 | -0.067 | 1.038 | 0.65 | 0.88 | 0.64 | 0.43 | 0.71 | 0.64 |
| 4 | 88266006 | rs11097131 | 1.159 | 0.836 | -0.136 | -0.515 | 0.034 | -0.064 | -0.064 | 0 | 0.13 | 0.24 | 0.15 | 0.01 | 0.18 | 0.06 |
| 4 | 88274740 | rs11734330 | 1.098 | 0.834 | -0.135 | -0.516 | 0.039 | -0.067 | -0.067 | 0 | 0.13 | 0.27 | 0.2 | 0.01 | 0.18 | 0.06 |
| 4 | 88277927 | rs6531978 | 0.283 | 0.7 | -0.367 | -0.684 | 0 | 0.008 | 0.008 | 0 | 0.42 | 0.38 | 0.41 | 0.09 | 0.43 | 0.39 |
| 4 | 88280502 | rs6845304 | -0.123 | 0.78 | -0.406 | -0.51 | -0.003 | -0.003 | -0.003 | 0 | 0.56 | 0.42 | 0.5 | 0.46 | 0.55 | 0.5 |
| 4 | 88290265 | rs10028441 | 0.114 | 0.885 | -0.152 | -0.506 | 0.003 | 0.004 | 0.004 | 1.403 | 0.3 | 0.58 | 0.3 | 0.31 | 0.3 | 0.18 |
| 4 | 88291439 | rs11737402 | 1.063 | 1.208 | 0.114 | -0.901 | 0.023 | -0.051 | -0.051 | 5.123 | 0.12 | 0.18 | 0.19 | 0 | 0.18 | 0.06 |
| 4 | 88293887 | rs6531985 | -0.003 | 1.133 | -0.004 | -1.097 | 0.005 | -0.133 | -0.133 | 0 | 0.28 | 0.58 | 0.29 | 0.01 | 0.33 | 0.14 |
| 4 | 88295008 | rs10222732 | 0.921 | 1.168 | -0.336 | -1.151 | 0 | -0.005 | -0.005 | 0 | 0.23 | 0.22 | 0.32 | 0.1 | 0.25 | 0.27 |
| 4 | 88304358 | rs7682516 | -0.272 | 0.941 | 0.024 | -0.948 | 0.007 | -0.17 | -0.17 | 0 | 0.28 | 0.49 | 0.29 | 0.01 | 0.33 | 0.14 |
| 4 | 88308415 | rs11097148 | -0.647 | 1.003 | 0.003 | -0.666 | 0.052 | -0.125 | -0.125 | 0 | 0.8 | 0.6 | 0.77 | 1 | 0.75 | 0.86 |
| 4 | 88312504 | rs13138404 | 0.316 | 0.771 | -0.717 | -0.342 | -0.115 | 0.095 | 0.095 | 0 | 0.31 | 0.02 | 0.36 | 0.11 | 0.36 | 0.28 |
| 4 | 88313179 | rs12644152 | -0.193 | 0.622 | -0.274 | 0.216 | -0.012 | 0.062 | 0.062 | 0 | 0.1 | 0.19 | 0.06 | 0.28 | 0.1 | 0.17 |
| 4 | 88326380 | rs11097151 | -1.412 | 0.497 | -0.479 | 0.461 | -0.023 | -0.015 | -0.015 | 0.268 | 0.63 | 0.85 | 0.55 | 0.63 | 0.61 | 0.48 |
| 4 | 88330987 | rs6827817 | 0.044 | 0.367 | -0.45 | 0.337 | -0.096 | -0.005 | -0.005 | 0.4 | 0.41 | 0.14 | 0.45 | 0.35 | 0.48 | 0.31 |
| 4 | 88347368 | rs10017282 | -1.36 | 0.449 | -0.132 | 0.519 | -0.023 | -0.004 | -0.004 | 0 | 0.62 | 0.85 | 0.55 | 0.64 | 0.61 | 0.49 |
| 4 | 88349022 | rs7673763 | 0.719 | 0.483 | -0.051 | 0.558 | 0 | 0.008 | 0.008 | 3.247 | 0.16 | 0.19 | 0.11 | 0.24 | 0.15 | 0.07 |
| 4 | 88349200 | rs11097155 | 0.745 | 0.45 | -0.158 | 0.557 | 0.01 | -0.016 | -0.016 | 0 | 0.11 | 0.06 | 0.05 | 0.3 | 0.1 | 0.15 |
| 4 | 88356799 | rs7690728 | 0.566 | 0.6 | -0.078 | 0.418 | 0.001 | 0.022 | 0.022 | 4.051 | 0.16 | 0.27 | 0.12 | 0.24 | 0.15 | 0.07 |
| 4 | 88365083 | rs13119333 | -0.054 | 0.613 | -0.108 | 0.547 | 0.012 | -0.019 | -0.019 | 0 | 0.36 | 0.25 | 0.39 | 0.47 | 0.37 | 0.35 |
| 4 | 88381080 | rs9968511 | -0.976 | 0.597 | -0.492 | 0.324 | -0.001 | -0.018 | -0.018 | 0 | 0.59 | 0.56 | 0.54 | 0.71 | 0.61 | 0.48 |
| 4 | 88382475 | rs9307036 | -1.069 | 0.725 | -1.542 | 0.387 | -0.016 | 0.028 | 0.028 | 0.056 | 0.62 | 0.83 | 0.55 | 0.71 | 0.61 | 0.48 |
| 4 | 88386769 | rs4405983 | -0.195 | 0.853 | -1.63 | 0.566 | -0.048 | -0.092 | -0.092 | 0 | 0.3 | 0.03 | 0.36 | 0.4 | 0.34 | 0.29 |
| 4 | 88389081 | rs10024099 | 0.043 | 0.912 | -1.439 | 0.739 | -0.06 | -0.115 | -0.115 | 0.958 | 0.2 | 0.01 | 0.3 | 0.38 | 0.23 | 0.25 |
| 4 | 88392755 | rs7683311 | -0.331 | 0.829 | -0.991 | 0.684 | -0.002 | -0.01 | -0.01 | 1.077 | 0.19 | 0.18 | 0.09 | 0.31 | 0.21 | 0.19 |
| 4 | 88393780 | rs9994839 | -0.822 | 0.864 | -0.326 | 0.689 | 0.004 | 0.02 | 0.02 | 0.37 | 0.33 | 0.63 | 0.21 | 0.5 | 0.33 | 0.28 |
| 4 | 88398749 | rs4693164 | 0.562 | 0.397 | -0.096 | 0.545 | -0.033 | -0.006 | -0.006 | 0 | 0.39 | 0.44 | 0.37 | 0.28 | 0.24 | 0.48 |
| 4 | 88400110 | rs4610302 | -0.862 | 0.877 | -0.164 | 0.547 | -0.056 | -0.05 | -0.05 | 0 | 0.69 | 0.9 | 0.79 | 0.63 | 0.6 | 0.78 |
| 4 | 88404126 | rs10461161 | 0.742 | 1.067 | 0.203 | 0.352 | 0.023 | -0.017 | -0.017 | 0 | 0.16 | 0.51 | 0.13 | 0.15 | 0.12 | 0.06 |
| 4 | 88415884 | rs3763392 | 1.097 | 1.194 | -0.217 | 0.367 | -0.079 | -0.048 | -0.048 | 0 | 0.13 | 0.02 | 0.29 | 0.19 | 0.22 | 0.21 |
| 4 | 88424737 | rs9993980 | 0.335 | 0.605 | 0.54 | 0.284 | -0.003 | -0.012 | -0.012 | 0.188 | 0.49 | 0.58 | 0.49 | 0.78 | 0.37 | 0.58 |
| 4 | 88429759 | rs10003497 | 0.237 | 0.524 | 1.072 | 0.232 | 0.01 | 0.014 | 0.014 | 0 | 0.52 | 0.46 | 0.49 | 0.78 | 0.39 | 0.64 |
| 4 | 88433880 | rs2869678 | 0.109 | 0.566 | 1.054 | 0.243 | 0.009 | -0.012 | -0.012 | 0 | 0.19 | 0.22 | 0.16 | 0.17 | 0.21 | 0.11 |
| 4 | 88435789 | rs1870881 | 1.301 | 0.499 | 1.006 | 0.254 | 0.069 | 0 | 0 | 0.85 | 0.34 | 0.17 | 0.36 | 0.62 | 0.26 | 0.52 |
| 4 | 88437710 | rs10001608 | -0.367 | 0.46 | 1.084 | 0.216 | -0.051 | 0.142 | 0.142 | 0 | 0.57 | 0.91 | 0.56 | 0.78 | 0.47 | 0.65 |
| 4 | 88438011 | rs4693829 | 1.224 | 0.497 | 1.359 | 0.252 | 0.029 | 0.006 | 0.006 | 0 | 0.35 | 0.28 | 0.37 | 0.62 | 0.26 | 0.52 |
| 4 | 88438520 | rs12643464 | 1.224 | 0.497 | 1.359 | 0.252 | 0.029 | 0.006 | 0.006 | 0 | 0.35 | 0.28 | 0.37 | 0.62 | 0.26 | 0.52 |
| 4 | 88439106 | rs12640408 | 0.964 | 0.738 | 1.088 | 0.293 | -0.033 | 0.013 | 0.013 | 4.817 | 0.15 | 0.12 | 0.15 | 0.16 | 0.19 | 0.06 |
| 4 | 88443685 | rs17012782 | 1.129 | 0.119 | 1.016 | 0.412 | 0.003 | -0.002 | -0.002 | 0 | 0.29 | 0.23 | 0.34 | 0.26 | 0.22 | 0.35 |
| 4 | 88448026 | rs2011326 | 0.641 | 0.194 | 0.971 | 0.558 | -0.059 | 0.019 | 0.019 | 0 | 0.34 | 0.48 | 0.36 | 0.56 | 0.24 | 0.49 |
| 4 | 88450348 | rs1049539 | 0.695 | 0.194 | 0.975 | 0.624 | -0.058 | 0.013 | 0.013 | 0 | 0.33 | 0.47 | 0.35 | 0.28 | 0.23 | 0.43 |
| 4 | 88451096 | rs17012830 | 1.389 | 0.462 | 0.766 | 0.557 | -0.003 | -0.004 | -0.004 | 0 | 0.12 | 0.08 | 0.26 | 0.22 | 0.14 | 0.23 |
| 4 | 88451182 | rs1031794 | -0.465 | 0.728 | 0.943 | 0.738 | 0.01 | -0.002 | -0.002 | 0 | 0.49 | 0.63 | 0.51 | 0.46 | 0.42 | 0.5 |
| 4 | 88451579 | rs1031796 | -0.337 | 0.726 | 0.77 | 0.543 | 0.026 | 0.091 | 0.091 | 0 | 0.5 | 0.75 | 0.53 | 0.74 | 0.44 | 0.56 |
| 4 | 88456441 | rs10027979 | 0.866 | 0.883 | 0.418 | 0.517 | -0.035 | 0.006 | 0.006 | 3.953 | 0.15 | 0.12 | 0.16 | 0.18 | 0.2 | 0.07 |
| 4 | 88466135 | rs4693841 | -0.315 | 0.787 | 0.411 | 0.582 | 0.001 | 0.012 | 0.012 | 0.053 | 0.47 | 0.48 | 0.47 | 0.77 | 0.45 | 0.56 |
| 4 | 88468386 | rs7685612 | 0.943 | 0.612 | -0.092 | 0.662 | 0.029 | -0.058 | -0.058 | 0 | 0.29 | 0.18 | 0.3 | 0.54 | 0.25 | 0.48 |
| 4 | 88468852 | rs1381633 | -0.997 | 0.975 | -0.327 | 0.634 | -0.037 | -0.001 | -0.001 | 0 | 0.83 | 0.89 | 0.83 | 0.77 | 0.77 | 0.91 |
| 4 | 88471575 | rs16996086 | 0.115 | 0.943 | -0.362 | 0.741 | 0.02 | 0.067 | 0.067 | 3.15 | 0.77 | 0.34 | 0.79 | 0.49 | 0.74 | 0.81 |
| 4 | 88474171 | rs13144424 | 1.098 | 1.015 | -1.101 | 0.525 | -0.064 | 0.054 | 0.054 | 0 | 0.15 | 0.01 | 0.13 | 0.02 | 0.22 | 0.06 |
| 4 | 88477532 | rs2904166 | 0.461 | 0.479 | -0.61 | 0.995 | 0.032 | -0.019 | -0.019 | 0 | 0.27 | 0.13 | 0.27 | 0.26 | 0.2 | 0.44 |
| 4 | 88478750 | rs1462372 | -0.218 | 0.673 | -0.615 | 0.696 | 0.002 | -0.001 | -0.001 | 0 | 0.4 | 0.63 | 0.37 | 0.53 | 0.34 | 0.52 |
| 4 | 88489929 | rs7435692 | 0.042 | 1.132 | -1.035 | 0.446 | -0.032 | 0.084 | 0.084 | 1.168 | 0.35 | 0.2 | 0.25 | 0.08 | 0.41 | 0.17 |
| 4 | 88491716 | rs2199458 | 0.677 | 1.186 | -0.278 | 1.946 | 0.04 | 0.294 | 0.294 | 0 | 0.92 | 0.78 | 0.85 | 0.5 | 0.9 | 0.89 |
| 4 | 88502776 | rs2869680 | 0.103 | 0.638 | -1.057 | 1.046 | 0.006 | 0.001 | 0.001 | 0.117 | 0.71 | 0.89 | 0.78 | 0.69 | 0.76 | 0.51 |
| 4 | 88505945 | rs13435625 | 0.806 | 0.871 | -0.757 | 1.114 | 0.001 | 0.028 | 0.028 | 1.354 | 0.55 | 0.57 | 0.6 | 0.29 | 0.58 | 0.24 |
| 4 | 88509540 | rs13137502 | -0.654 | 1.001 | -1.113 | 0.912 | 0.025 | 0.003 | 0.003 | 0.118 | 0.07 | 0.16 | 0.09 | 0.09 | 0.13 | 0.14 |
| 4 | 88509940 | rs4693847 | -0.02 | 0.606 | -1.297 | 0.951 | 0.005 | -0.004 | -0.004 | 1.047 | 0.66 | 0.95 | 0.77 | 0.67 | 0.74 | 0.46 |
| 4 | 88522478 | rs2615485 | 0.363 | -0.151 | -0.676 | 0.508 | -0.046 | -0.056 | -0.056 | 0.443 | 0.66 | 0.48 | 0.87 | 0.88 | 0.8 | 0.65 |
| 4 | 88530463 | rs2615487 | 0.557 | 0.058 | -1.508 | 0.572 | -0.061 | -0.065 | -0.065 | 0 | 0.23 | 0.09 | 0.27 | 0.6 | 0.32 | 0.4 |
| 4 | 88533441 | rs13131929 | 0.747 | 0.135 | -1.788 | 0.334 | 0.047 | 0.106 | 0.106 | 2.921 | 0.51 | 0.17 | 0.5 | 0.71 | 0.57 | 0.52 |
| 4 | 88534235 | rs2736982 | 0.058 | 0.358 | -1.531 | 0.028 | -0.002 | 0.003 | 0.003 | 0.036 | 0.67 | 0.76 | 0.58 | 0.71 | 0.65 | 0.61 |
| 4 | 88537737 | rs2615489 | -0.618 | -0.569 | -1.924 | -0.523 | 0.038 | 0.044 | 0.044 | 0 | 0.78 | 0.9 | 0.93 | 0.89 | 0.86 | 0.7 |
| 4 | 88539776 | rs10016823 | 0.559 | 0.072 | -1.74 | -0.131 | 0.008 | 0.115 | 0.115 | 4.636 | 0.3 | 0.05 | 0.25 | 0.1 | 0.27 | 0.15 |
| 4 | 88539948 | rs10028688 | 0.425 | 0.098 | -1.727 | -0.118 | 0.006 | 0.113 | 0.113 | 4.785 | 0.3 | 0.05 | 0.25 | 0.1 | 0.27 | 0.15 |
| 4 | 88552784 | rs1381626 | 0.114 | -0.143 | -0.791 | -0.055 | -0.039 | -0.048 | -0.048 | 0 | 0.75 | 0.56 | 0.9 | 0.88 | 0.84 | 0.67 |
| 4 | 88553414 | rs6532013 | 0.109 | 0.121 | -1.273 | -0.256 | -0.005 | 0.064 | 0.064 | 0 | 0.49 | 0.69 | 0.57 | 0.67 | 0.52 | 0.3 |
| 4 | 88553549 | rs2045743 | -1.029 | 0.086 | -1.408 | -0.195 | 0.025 | 0.025 | 0.025 | 0 | 0.83 | 0.95 | 0.94 | 0.89 | 0.88 | 0.69 |
| 4 | 88554115 | rs2627657 | 0.607 | 0.082 | -1.883 | 0.324 | -0.057 | -0.049 | -0.049 | 0 | 0.54 | 0.15 | 0.5 | 0.57 | 0.51 | 0.44 |

| | | | | | | | | | | | | | | | | |
|---|---|---|---|---|---|---|---|---|---|---|---|---|---|---|---|---|
| 4 | 88564721 | rs2627669 | -0.141 | -0.103 | -1.16 | 0.314 | -0.018 | -0.011 | -0.011 | 0.246 | 0.37 | 0.47 | 0.21 | 0.27 | 0.24 | 0.36 |
| 4 | 88567295 | rs2627696 | -1.053 | 0.421 | -1.171 | 0.451 | -0.066 | 0.107 | 0.107 | 0.631 | 0.54 | 0.89 | 0.56 | 0.69 | 0.49 | 0.61 |
| 4 | 88569583 | rs2904169 | 0.258 | 0.427 | -0.685 | 0.456 | 0.077 | 0.18 | 0.18 | 0 | 0.15 | 0.26 | 0.33 | 0.42 | 0.23 | 0.21 |
| 4 | 88570528 | rs1462367 | 1.317 | 0.389 | -0.916 | 0.334 | -0.088 | 0.017 | 0.017 | 0.011 | 0.19 | 0.01 | 0.16 | 0.1 | 0.28 | 0.27 |
| 4 | 88578487 | rs1545735 | -0.975 | 0.423 | -0.819 | 0.373 | -0.062 | 0.105 | 0.105 | 0.558 | 0.54 | 0.88 | 0.56 | 0.69 | 0.49 | 0.61 |
| 4 | 88583135 | rs10019009 | 0.252 | 0.64 | -0.641 | 0.499 | 0.075 | 0.184 | 0.184 | 0 | 0.15 | 0.26 | 0.33 | 0.43 | 0.23 | 0.21 |
| 4 | 88601464 | rs10034250 | 0.245 | 0.621 | -0.618 | 0.49 | 0.108 | 0.252 | 0.252 | 0 | 0.15 | 0.36 | 0.34 | 0.42 | 0.23 | 0.21 |
| 4 | 88611344 | rs11097166 | -0.52 | 0.275 | -0.654 | 0.295 | -0.037 | 0.04 | 0.04 | 0 | 0.43 | 0.74 | 0.54 | 0.66 | 0.46 | 0.55 |
| 4 | 88615772 | rs17012948 | 1.009 | 0.281 | -0.595 | 0.424 | -0.003 | 0.001 | 0.001 | 0 | 0.25 | 0.36 | 0.17 | 0.26 | 0.21 | 0.28 |
| 4 | 88629174 | rs2869704 | -0.535 | 0.288 | -0.939 | 0.411 | -0.036 | 0.057 | 0.057 | 0 | 0.43 | 0.74 | 0.54 | 0.7 | 0.46 | 0.55 |
| 4 | 88638522 | rs7680291 | -1.766 | 0.256 | -1.024 | 0.357 | -0.048 | 0.029 | 0.029 | 0 | 0.82 | 0.96 | 0.85 | 0.9 | 0.73 | 0.74 |
| 4 | 88640064 | rs10016081 | -0.519 | 0.28 | -0.851 | 0.413 | -0.032 | 0.052 | 0.052 | 0 | 0.43 | 0.73 | 0.54 | 0.7 | 0.46 | 0.54 |
| 4 | 88652577 | rs4416440 | 0.966 | 0.206 | -0.874 | 0.436 | -0.02 | -0.045 | -0.045 | 0.086 | 0.24 | 0.07 | 0.15 | 0.3 | 0.21 | 0.28 |
| 4 | 88655984 | rs2732171 | -1.482 | 0.285 | -0.657 | 0.328 | -0.007 | 0.007 | 0.007 | 0 | 0.81 | 0.84 | 0.84 | 0.9 | 0.73 | 0.73 |
| 4 | 88665977 | rs7661141 | -0.139 | -0.023 | -0.612 | 0.243 | -0.026 | -0.021 | -0.021 | 0 | 0.11 | 0.15 | 0.03 | 0.02 | 0.03 | 0.08 |
| 4 | 88665994 | rs7685567 | -0.139 | -0.023 | -0.612 | 0.25 | -0.026 | -0.019 | -0.019 | 0.021 | 0.11 | 0.15 | 0.03 | 0.02 | 0.03 | 0.09 |
| 4 | 88677931 | rs2627704 | -1.107 | 0.127 | -0.004 | 0.362 | 0.008 | 0.074 | 0.074 | 0.266 | 0.28 | 0.49 | 0.39 | 0.41 | 0.28 | 0.28 |
| 4 | 88678096 | rs4693868 | 1.813 | 0.007 | -0.148 | 0.311 | 0.021 | 0.048 | 0.048 | 3.478 | 0.28 | 0.07 | 0.27 | 0.19 | 0.23 | 0.12 |
| 4 | 88681801 | rs2627723 | -1.377 | 0.073 | -0.029 | 0.353 | 0 | 0.001 | 0.001 | 1.142 | 0.3 | 0.7 | 0.46 | 0.48 | 0.29 | 0.33 |
| 4 | 88687693 | rs2627714 | 1.422 | 0.074 | -0.142 | 0.309 | -0.042 | 0.026 | 0.026 | 0 | 0.18 | 0.07 | 0.15 | 0.11 | 0.27 | 0.28 |
| 4 | 88689765 | rs4693869 | -1.015 | 0.053 | -0.13 | 0.333 | 0.011 | 0.118 | 0.118 | 0.151 | 0.27 | 0.49 | 0.43 | 0.48 | 0.27 | 0.28 |
| 4 | 88696086 | rs7661149 | -1.152 | 0.087 | -0.323 | 0.337 | -0.031 | 0.02 | 0.02 | 1.642 | 0.47 | 0.86 | 0.62 | 0.59 | 0.55 | 0.56 |
| 4 | 88698345 | rs2100889 | -0.314 | -0.541 | -0.46 | 0.471 | 0.014 | 0.08 | 0.08 | 0 | 0.24 | 0.43 | 0.44 | 0.43 | 0.23 | 0.25 |
| 4 | 88701522 | rs6857696 | 1.496 | -0.548 | -0.606 | 0.403 | -0.047 | 0.004 | 0.004 | 0 | 0.19 | 0.05 | 0.15 | 0.13 | 0.27 | 0.28 |
| 4 | 88703641 | rs2045839 | -0.214 | -0.946 | -0.759 | 0.191 | -0.006 | 0.003 | 0.003 | 0.405 | 0.69 | 0.6 | 0.81 | 0.69 | 0.74 | 0.61 |
| 4 | 88705132 | rs12650760 | 1.295 | -0.962 | -1.04 | 0.22 | -0.04 | 0.098 | 0.098 | 1.938 | 0.44 | 0.13 | 0.37 | 0.25 | 0.5 | 0.36 |
| 4 | 88707081 | rs13136331 | -0.07 | -0.901 | -0.745 | 0.233 | 0.015 | 0.021 | 0.021 | 0 | 0.4 | 0.35 | 0.22 | 0.17 | 0.33 | 0.42 |
| 4 | 88709783 | rs17761748 | 1.884 | -0.821 | -0.982 | 0.335 | 0.013 | 0.044 | 0.044 | 2.716 | 0.26 | 0.12 | 0.23 | 0.12 | 0.23 | 0.08 |
| 4 | 88710784 | rs2732166 | -0.902 | -0.172 | -0.987 | 0.588 | -0.029 | -0.026 | -0.026 | 0 | 0.8 | 0.9 | 0.86 | 0.71 | 0.77 | 0.71 |
| 4 | 88715324 | rs13137552 | 0.574 | -0.297 | -1.496 | 0.028 | -0.05 | 0.179 | 0.179 | 0 | 0.46 | 0.15 | 0.37 | 0.14 | 0.55 | 0.39 |
| 4 | 88723210 | rs2616262 | 0.404 | 0.006 | -0.991 | 0.339 | 0.018 | 0.015 | 0.015 | 0.013 | 0.31 | 0.36 | 0.55 | 0.58 | 0.47 | 0.51 |
| 4 | 88724600 | rs1870964 | 0.597 | 0.079 | -0.702 | 0.84 | 0.087 | 0.105 | 0.105 | 0 | 0.07 | 0.16 | 0.2 | 0.21 | 0.13 | 0.1 |
| 4 | 88724967 | rs3805374 | 1.198 | -0.198 | -0.692 | 0.683 | -0.09 | 0.136 | 0.136 | 0.336 | 0.4 | 0.11 | 0.36 | 0.14 | 0.52 | 0.39 |
| 4 | 88726959 | rs17013175 | 0.516 | -0.016 | -0.699 | 0.836 | 0.086 | 0.123 | 0.123 | 0 | 0.07 | 0.17 | 0.2 | 0.24 | 0.13 | 0.1 |
| 4 | 88728904 | rs1381965 | -0.85 | -0.028 | -0.694 | 0.835 | 0.078 | 0.316 | 0.316 | 0 | 0.11 | 0.34 | 0.28 | 0.47 | 0.15 | 0.17 |
| 4 | 88732692 | rs1054627 | 1.878 | 0.002 | -0.885 | 0.788 | -0.1 | 0.004 | 0.004 | 0.026 | 0.21 | 0.02 | 0.28 | 0.11 | 0.33 | 0.31 |
| 4 | 88732746 | rs13144371 | -1.071 | -0.158 | -0.613 | 0.808 | -0.01 | -0.061 | -0.061 | 0 | 0.7 | 0.91 | 0.73 | 0.59 | 0.76 | 0.65 |
| 4 | 88732763 | rs17013181 | -0.83 | 0.087 | 0.2 | 1.118 | 0.023 | 0.005 | 0.005 | 0 | 0.2 | 0.4 | 0.27 | 0.23 | 0.19 | 0.17 |
| 4 | 88732911 | rs1054628 | -0.053 | 0.505 | -0.17 | 1.46 | 0.122 | 0.24 | 0.24 | 0 | 0.07 | 0.27 | 0.25 | 0.29 | 0.13 | 0.11 |
| 4 | 88742318 | rs7681895 | -0.768 | -0.37 | -1.533 | 0.736 | 0.011 | -0.041 | -0.041 | 0 | 0.66 | 0.94 | 0.8 | 0.64 | 0.77 | 0.74 |
| 4 | 88746734 | rs13117929 | 1.615 | 0.147 | -1.072 | 0.456 | 0.008 | -0.017 | -0.017 | 0 | 0.23 | 0.28 | 0.35 | 0.1 | 0.32 | 0.33 |
| 4 | 88749701 | rs10007111 | -1.353 | 0.392 | -1.064 | 0.629 | 0.011 | 0.002 | 0.002 | 0 | 0.74 | 0.67 | 0.62 | 0.72 | 0.62 | 0.62 |
| 4 | 88752878 | rs11097172 | 1.5 | 0.339 | -0.812 | 0.58 | 0.023 | -0.059 | -0.059 | 0 | 0.24 | 0.39 | 0.35 | 0.08 | 0.32 | 0.36 |
| 4 | 88753045 | rs1585616 | -0.585 | 0.038 | -0.845 | 0.382 | 0.002 | -0.014 | -0.014 | 0 | 0.86 | 0.88 | 0.93 | 0.81 | 0.89 | 0.89 |
| 4 | 88754060 | rs17013212 | -0.069 | 0.061 | -0.53 | 0.158 | 0.004 | 0.124 | 0.124 | 0 | 0.14 | 0.27 | 0.25 | 0.35 | 0.16 | 0.17 |
| 4 | 88754681 | rs1463114 | 0.668 | 0.146 | -0.488 | -0.18 | 0.001 | -0.045 | -0.045 | 0 | 0.34 | 0.42 | 0.4 | 0.08 | 0.37 | 0.39 |
| 4 | 88755967 | rs11097173 | 2.103 | 0.111 | -0.472 | -0.271 | -0.005 | -0.002 | -0.002 | 2.345 | 0.15 | 0.13 | 0.14 | 0.02 | 0.23 | 0.1 |
| 4 | 88756245 | rs2045836 | 0.013 | 0.217 | -0.569 | -0.134 | 0.004 | -0.013 | -0.013 | 0 | 0.5 | 0.56 | 0.54 | 0.1 | 0.6 | 0.52 |
| 4 | 88756675 | rs2045837 | 1.841 | 0.351 | -1.044 | 0.057 | -0.101 | 0.064 | 0.064 | 0 | 0.22 | 0.03 | 0.33 | 0.08 | 0.32 | 0.32 |
| 4 | 88756978 | rs1381966 | -0.531 | -0.255 | -0.958 | 0.18 | 0.016 | -0.013 | -0.013 | 0.067 | 0.66 | 0.81 | 0.71 | 0.38 | 0.73 | 0.67 |
| 4 | 88757008 | rs2045838 | -0.282 | -0.248 | -0.91 | 0.051 | -0.001 | 0.019 | 0.019 | 0 | 0.64 | 0.8 | 0.67 | 0.33 | 0.68 | 0.67 |
| 4 | 88757703 | rs17712558 | -1.803 | 0.433 | -0.025 | 0.103 | 0.026 | 0.009 | 0.009 | 0 | 0.78 | 0.68 | 0.68 | 0.74 | 0.64 | 0.66 |
| 4 | 88765516 | rs13127257 | 1.378 | 0.214 | -0.526 | -0.291 | -0.042 | 0.08 | 0.08 | 1.653 | 0.17 | 0.05 | 0.11 | 0.01 | 0.21 | 0.11 |
| 4 | 88767008 | rs17013285 | 1.002 | -0.217 | -0.801 | -0.193 | 0.067 | -0.033 | -0.033 | 0.06 | 0.31 | 0.05 | 0.18 | 0.28 | 0.16 | 0.23 |
| 4 | 88770890 | rs10018094 | 1.816 | 0.417 | -0.282 | -0.067 | -0.002 | -0.01 | -0.01 | 0 | 0.23 | 0.28 | 0.29 | 0.05 | 0.28 | 0.3 |
| 4 | 88774073 | rs1471399 | 0.052 | 0.389 | -0.034 | 0.132 | -0.002 | -0.001 | -0.001 | 0.663 | 0.66 | 0.67 | 0.75 | 0.67 | 0.64 | 0.67 |
| 4 | 88774247 | rs1471400 | 0.232 | 0.34 | 0.015 | 0.118 | -0.006 | -0.008 | -0.008 | 0.249 | 0.42 | 0.39 | 0.47 | 0.62 | 0.37 | 0.37 |
| 4 | 88777743 | rs7655902 | -0.323 | 0.556 | 0.128 | 0.236 | -0.004 | -0.007 | -0.007 | 0 | 0.58 | 0.46 | 0.59 | 0.63 | 0.58 | 0.5 |
| 4 | 88778446 | rs931032 | 1.086 | 0.239 | 0.106 | 0.324 | -0.003 | 0.008 | 0.008 | 0 | 0.09 | 0.07 | 0.18 | 0.27 | 0.15 | 0.12 |
| 4 | 88783177 | rs7667813 | 0.821 | 0.158 | 0.073 | 0.323 | 0.008 | 0.003 | 0.003 | 0 | 0.13 | 0.1 | 0.1 | 0.11 | 0.07 | 0.15 |
| 4 | 88788036 | rs1463091 | 0.155 | 0.224 | 0.237 | 0.338 | 0.006 | 0.053 | 0.053 | 0 | 0.49 | 0.28 | 0.34 | 0.3 | 0.45 | 0.37 |
| 4 | 88789140 | rs1026663 | -0.346 | 0.079 | 0.336 | 0.418 | -0.004 | -0.001 | -0.001 | 0 | 0.58 | 0.5 | 0.53 | 0.57 | 0.59 | 0.51 |
| 4 | 88792438 | rs11728139 | 0.753 | 0.062 | 0.315 | 0.426 | 0.008 | 0.003 | 0.003 | 0 | 0.13 | 0.1 | 0.1 | 0.11 | 0.07 | 0.15 |
| 4 | 88793888 | rs17013421 | 1.06 | -0.002 | 0.276 | 0.307 | 0.015 | -0.001 | -0.001 | 0 | 0.33 | 0.23 | 0.22 | 0.3 | 0.21 | 0.24 |
| 4 | 88795989 | rs6815121 | -0.446 | 0.049 | 0.349 | 0.329 | -0.01 | 0.061 | 0.061 | 0 | 0.83 | 0.77 | 0.8 | 0.62 | 0.87 | 0.81 |
| 4 | 88797100 | rs12510860 | 1.738 | 0.067 | 0.321 | 0.282 | 0 | 0.004 | 0.004 | 0 | 0.24 | 0.26 | 0.28 | 0.05 | 0.27 | 0.3 |
| 4 | 88798284 | rs7672749 | 0.604 | 0.131 | -0.005 | 0.382 | 0.009 | -0.002 | -0.002 | 0 | 0.14 | 0.11 | 0.17 | 0.16 | 0.08 | 0.16 |
| 4 | 88799710 | rs1463104 | 0.239 | 0.18 | 0.021 | 0.373 | -0.002 | 0 | 0 | 0 | 0.27 | 0.27 | 0.32 | 0.27 | 0.39 | 0.26 |
| 4 | 88804039 | rs10222708 | 1.983 | -0.006 | -0.144 | 0.181 | -0.022 | 0.086 | 0.086 | 0 | 0.22 | 0.07 | 0.26 | 0.05 | 0.27 | 0.3 |

| | | | | | | | | | | | | | | | | |
|---|---|---|---|---|---|---|---|---|---|---|---|---|---|---|---|---|
| 4 | 88805208 | rs12644436 | -0.569 | 0.09 | -0.008 | 0.374 | -0.001 | -0.001 | -0.001 | 0 | 0.6 | 0.63 | 0.55 | 0.57 | 0.59 | 0.5 |
| 4 | 88807246 | rs10004908 | -0.588 | -0.094 | -0.266 | 0.548 | -0.026 | 0.025 | 0.025 | 0 | 0.65 | 0.48 | 0.76 | 0.53 | 0.78 | 0.76 |
| 4 | 88808272 | rs17013516 | 0.671 | 0.649 | -0.602 | 0.74 | -0.003 | 0.003 | 0.003 | 0 | 0.12 | 0.12 | 0.2 | 0.31 | 0.17 | 0.14 |
| 4 | 88817116 | rs4693901 | 0.884 | -0.266 | -0.556 | 0.68 | -0.062 | 0.004 | 0.004 | 0 | 0.35 | 0.18 | 0.48 | 0.36 | 0.47 | 0.49 |
| 4 | 88821873 | rs12651696 | 0.551 | -0.669 | -0.371 | 0.531 | 0.028 | -0.009 | -0.009 | 0.126 | 0.29 | 0.16 | 0.2 | 0.32 | 0.18 | 0.2 |
| 4 | 88823410 | rs713538 | 1.199 | -0.63 | -0.552 | 0.497 | -0.088 | -0.003 | -0.003 | 0 | 0.36 | 0.11 | 0.47 | 0.36 | 0.47 | 0.5 |
| 4 | 88823643 | rs17013544 | 0.078 | -0.573 | -0.04 | 0.599 | -0.054 | 0.036 | 0.036 | 0 | 0.33 | 0.44 | 0.27 | 0.43 | 0.19 | 0.28 |
| 4 | 88834454 | rs7683960 | -0.654 | -0.018 | -0.199 | 0.698 | -0.007 | -0.002 | -0.002 | 0.805 | 0.8 | 0.83 | 0.78 | 0.79 | 0.69 | 0.84 |
| 4 | 88836002 | rs716202 | 0.495 | -0.529 | -0.269 | 0.612 | -0.006 | 0.004 | 0.004 | 0 | 0.23 | 0.21 | 0.18 | 0.32 | 0.15 | 0.17 |
| 4 | 88840386 | rs6846509 | -0.538 | -0.062 | -0.293 | 0.677 | -0.002 | -0.001 | -0.001 | 0.049 | 0.8 | 0.81 | 0.78 | 0.79 | 0.7 | 0.84 |
| 4 | 88848691 | rs16996103 | 0.649 | -0.382 | -0.303 | 0.73 | -0.04 | 0.059 | 0.059 | 0 | 0.17 | 0.29 | 0.09 | 0.26 | 0.09 | 0.12 |
| 4 | 88849669 | rs1477602 | -0.779 | -0.34 | -0.332 | 0.679 | -0.018 | -0.004 | -0.004 | 0 | 0.8 | 0.85 | 0.79 | 0.79 | 0.7 | 0.84 |
| 4 | 88852643 | rs10005067 | 1.541 | -0.647 | -1.008 | 0.517 | -0.078 | 0.027 | 0.027 | 0 | 0.44 | 0.08 | 0.5 | 0.38 | 0.5 | 0.54 |
| 4 | 88856229 | rs1477600 | 0.94 | -0.511 | -0.792 | 0.333 | 0.001 | -0.001 | -0.001 | 0 | 0.62 | 0.37 | 0.59 | 0.64 | 0.6 | 0.66 |
| 4 | 88859131 | rs11723364 | 0.894 | 0.094 | -1.476 | 0.362 | -0.081 | 0.089 | 0.089 | 0.859 | 0.14 | 0.01 | 0.12 | 0.01 | 0.23 | 0.11 |
| 4 | 88864775 | rs4693920 | 1.04 | 0.008 | -1.166 | 0.783 | -0.01 | -0.016 | -0.016 | 0.013 | 0.06 | 0.01 | 0.1 | 0.21 | 0.07 | 0.05 |
| 4 | 88867961 | rs6532035 | 0.587 | -0.235 | -0.987 | 0.665 | -0.001 | -0.003 | -0.003 | 0 | 0.52 | 0.37 | 0.58 | 0.5 | 0.53 | 0.67 |
| 4 | 88868506 | rs7675985 | 0.754 | -0.26 | -0.977 | 0.709 | 0 | 0.037 | 0.037 | 0 | 0.5 | 0.36 | 0.52 | 0.4 | 0.52 | 0.58 |
| 4 | 88868955 | rs6532037 | 0.596 | -0.215 | -0.995 | 0.531 | -0.001 | -0.001 | -0.001 | 0 | 0.52 | 0.37 | 0.58 | 0.5 | 0.53 | 0.66 |
| 4 | 88878504 | rs17013600 | 0.612 | -0.422 | -1.716 | 0.145 | 0.009 | -0.012 | -0.012 | 0 | 0.48 | 0.12 | 0.55 | 0.44 | 0.57 | 0.65 |
| 4 | 88888113 | rs4693923 | -1.408 | -0.65 | -1.662 | 0.153 | -0.014 | -0.046 | -0.046 | 0.508 | 0.88 | 0.99 | 0.87 | 0.8 | 0.89 | 0.92 |
| 4 | 88889605 | rs6833161 | 0.791 | -0.807 | -1.704 | 0.102 | 0.025 | 0.006 | 0.006 | 0 | 0.5 | 0.17 | 0.56 | 0.43 | 0.56 | 0.65 |
| 4 | 88894235 | rs6813526 | -0.172 | -0.903 | -0.618 | -0.061 | 0.005 | 0.016 | 0.016 | 0 | 0.3 | 0.42 | 0.22 | 0.37 | 0.33 | 0.27 |
| 4 | 88896248 | rs2853744 | 0.499 | -1.412 | -0.414 | -0.102 | -0.047 | 0.052 | 0.052 | 0 | 0.18 | 0.29 | 0.09 | 0.37 | 0.09 | 0.17 |
| 4 | 88896421 | rs11730582 | 0.985 | -1.241 | -0.772 | -0.159 | -0.099 | 0.032 | 0.032 | 0 | 0.45 | 0.06 | 0.46 | 0.35 | 0.46 | 0.49 |
| 4 | 88902405 | rs6811536 | 0.87 | -1.07 | -0.667 | -0.561 | 0.023 | -0.113 | -0.113 | 0.249 | 0.27 | 0.42 | 0.18 | 0.04 | 0.28 | 0.36 |
| 4 | 88902692 | rs4754 | 0.693 | -1.264 | -0.744 | -0.438 | 0.043 | -0.012 | -0.012 | 0 | 0.32 | 0.15 | 0.38 | 0.71 | 0.28 | 0.36 |
| 4 | 88904186 | rs1126772 | 1.41 | -1.235 | -0.791 | -0.471 | -0.01 | -0.068 | -0.068 | 0.208 | 0.25 | 0.02 | 0.24 | 0.3 | 0.21 | 0.15 |
| 4 | 88905795 | rs10012150 | 0.93 | -1.336 | -0.413 | -0.518 | -0.01 | -0.002 | -0.002 | 0 | 0.29 | 0.3 | 0.25 | 0.3 | 0.21 | 0.14 |
| 4 | 88906458 | rs7685225 | 0.505 | -1.33 | -0.63 | -0.86 | 0.031 | -0.152 | -0.152 | 0.49 | 0.31 | 0.44 | 0.21 | 0 | 0.37 | 0.24 |
| 4 | 88908998 | rs7675246 | 0.957 | -1.209 | -0.57 | -0.705 | 0.007 | -0.066 | -0.066 | 0 | 0.26 | 0.33 | 0.17 | 0.04 | 0.27 | 0.35 |
| 4 | 88919106 | rs2725236 | -0.767 | -1.116 | -0.778 | -0.361 | 0.001 | -0.008 | -0.008 | 0 | 0.47 | 0.56 | 0.58 | 0.65 | 0.5 | 0.45 |
| 4 | 88926020 | rs12511728 | 0.053 | -0.872 | -1.563 | -0.172 | -0.011 | 0.083 | 0.083 | 0 | 0.11 | 0.18 | 0.15 | 0.4 | 0.11 | 0.21 |
| 4 | 88927707 | rs6841030 | 0.864 | -0.905 | -1.542 | -0.565 | 0.005 | -0.092 | -0.092 | 0 | 0.23 | 0.5 | 0.16 | 0.04 | 0.22 | 0.33 |
| 4 | 88928399 | rs13110188 | 0.874 | -0.709 | -1.803 | -0.556 | -0.107 | 0.126 | 0.126 | 0 | 0.23 | 0.07 | 0.18 | 0 | 0.34 | 0.17 |
| 4 | 88933584 | rs2725233 | -1.308 | -0.928 | -1.74 | -0.36 | 0.018 | 0.001 | 0.001 | 0 | 0.7 | 0.84 | 0.74 | 0.7 | 0.75 | 0.84 |
| 4 | 88935231 | rs2725230 | -1.328 | -0.954 | -1.676 | -0.369 | 0.016 | -0.01 | -0.01 | 0 | 0.7 | 0.91 | 0.75 | 0.7 | 0.76 | 0.84 |
| 4 | 88939000 | rs4484262 | 0.403 | -1.085 | -1.446 | 0.013 | -0.009 | 0.059 | 0.059 | 0 | 0.09 | 0.11 | 0.14 | 0.39 | 0.08 | 0.21 |
| 4 | 88941549 | rs10516801 | 0.039 | -1.022 | -1.598 | -0.147 | -0.004 | 0.005 | 0.005 | 1.243 | 0.32 | 0.59 | 0.3 | 0.44 | 0.31 | 0.54 |
| 4 | 88944511 | rs2725227 | -0.736 | -1.061 | -1.671 | -0.207 | 0.017 | 0.032 | 0.032 | 0 | 0.44 | 0.61 | 0.54 | 0.7 | 0.38 | 0.61 |
| 4 | 88947869 | rs4312723 | 0.072 | -1.062 | -1.67 | -0.198 | -0.002 | 0.002 | 0.002 | 1.283 | 0.32 | 0.59 | 0.3 | 0.44 | 0.31 | 0.54 |
| 4 | 88949175 | rs2467052 | -1.026 | -1.054 | -1.657 | -0.244 | 0.044 | 0.102 | 0.102 | 0.15 | 0.47 | 0.78 | 0.57 | 0.7 | 0.41 | 0.67 |
| 4 | 88952076 | rs2725225 | -0.745 | -1.01 | -1.717 | -0.208 | 0.02 | 0.037 | 0.037 | 0 | 0.44 | 0.62 | 0.54 | 0.7 | 0.38 | 0.61 |
| 4 | 88956521 | rs6532045 | -0.025 | -0.824 | -1.617 | -0.199 | -0.002 | 0.003 | 0.003 | 1.279 | 0.32 | 0.59 | 0.3 | 0.44 | 0.31 | 0.54 |
| 4 | 88957562 | rs17786456 | 0.367 | -0.821 | -1.956 | -0.35 | -0.079 | 0.1 | 0.1 | 0 | 0.26 | 0.1 | 0.17 | 0 | 0.34 | 0.13 |
| 4 | 88959381 | rs2725221 | -1.263 | -0.729 | -2.103 | 0.067 | -0.062 | -0.033 | -0.033 | 0.315 | 0.6 | 0.95 | 0.64 | 0.66 | 0.57 | 0.56 |
| 4 | 88959745 | rs17013735 | 0.175 | -1.074 | -1.862 | 0.166 | 0.006 | -0.011 | -0.011 | 0.846 | 0.16 | 0.13 | 0.19 | 0.42 | 0.08 | 0.19 |
| 4 | 88959998 | rs7696304 | -0.613 | -0.179 | -1.488 | -0.087 | -0.02 | 0.092 | 0.092 | 0 | 0.41 | 0.74 | 0.26 | 0.01 | 0.41 | 0.29 |
| 4 | 88961736 | rs2728108 | -0.931 | -0.138 | -1.585 | 0.161 | -0.019 | 0.085 | 0.085 | 0 | 0.46 | 0.82 | 0.39 | 0.26 | 0.47 | 0.37 |
| 4 | 88966000 | rs4555590 | 0.723 | -0.328 | -1.781 | 0.108 | 0.023 | 0.031 | 0.031 | 0 | 0.13 | 0 | 0.02 | 0.03 | 0.06 | 0.1 |
| 4 | 88970375 | rs2725211 | 0.401 | 0.229 | -1.335 | 0.617 | -0.026 | -0.008 | -0.008 | 1.722 | 0.05 | 0.02 | 0.14 | 0.23 | 0.07 | 0.07 |
| 4 | 88972067 | rs2728105 | -1.571 | 0.04 | -1.362 | 0.8 | -0.066 | -0.047 | -0.047 | 0.006 | 0.4 | 0.85 | 0.42 | 0.48 | 0.47 | 0.42 |
| 4 | 88973006 | rs2728104 | 0.224 | -0.111 | -1.207 | 0.965 | -0.023 | -0.024 | -0.024 | 0 | 0.07 | 0.01 | 0.14 | 0.26 | 0.1 | 0.08 |
| 4 | 88979529 | rs2725207 | -1.862 | -0.236 | -0.823 | 0.85 | -0.019 | 0.039 | 0.039 | 0.971 | 0.21 | 0.72 | 0.24 | 0.47 | 0.14 | 0.29 |
| 4 | 88981690 | rs2728133 | -1.984 | -0.253 | -0.826 | 0.84 | -0.014 | 0.027 | 0.027 | 1.103 | 0.22 | 0.74 | 0.24 | 0.46 | 0.14 | 0.29 |
| 4 | 88985711 | rs2725205 | -1.753 | -0.275 | -0.731 | 0.839 | -0.03 | 0.077 | 0.077 | 1.103 | 0.2 | 0.67 | 0.24 | 0.46 | 0.14 | 0.28 |
| 4 | 88987916 | rs4336187 | -1.917 | -0.273 | -0.746 | 0.837 | -0.019 | 0.039 | 0.039 | 1.575 | 0.21 | 0.72 | 0.24 | 0.46 | 0.14 | 0.3 |
| 4 | 88995795 | rs2725203 | -2.079 | -0.272 | -0.762 | 0.816 | -0.011 | 0.019 | 0.019 | 1.201 | 0.22 | 0.74 | 0.24 | 0.46 | 0.14 | 0.29 |
| 4 | 88997102 | rs2728121 | -0.86 | -0.213 | -0.612 | 0.9 | -0.001 | 0.063 | 0.063 | 0.36 | 0.09 | 0.13 | 0.17 | 0.43 | 0.09 | 0.16 |
| 4 | 88999306 | rs2725201 | -1.485 | 0.049 | -0.867 | 0.736 | -0.064 | -0.035 | -0.035 | 0 | 0.4 | 0.85 | 0.43 | 0.46 | 0.49 | 0.42 |
| 4 | 89004957 | rs17013808 | -0.403 | -0.47 | -0.38 | 0.844 | -0.004 | 0.041 | 0.041 | 0.493 | 0.08 | 0.51 | 0.05 | 0.03 | 0.03 | 0.01 |
| 4 | 89007921 | rs2725270 | -0.772 | 0.117 | -0.91 | 0.789 | -0.146 | -0.096 | -0.096 | 0 | 0.33 | 0.08 | 0.36 | 0.44 | 0.48 | 0.34 |
| 4 | 89016175 | rs2725267 | 1.494 | -0.221 | -0.398 | 1.275 | -0.04 | 0.018 | 0.018 | 3.647 | 0.86 | 0.25 | 0.73 | 0.77 | 0.95 | 0.85 |
| 4 | 89016782 | rs2231162 | 0.621 | -0.21 | 0.319 | 1.287 | -0.048 | 0.193 | 0.193 | 0.828 | 0.89 | 0.5 | 0.77 | 0.78 | 0.95 | 0.87 |
| 4 | 89022362 | rs2231153 | 1.458 | -0.177 | -0.519 | 1.377 | -0.04 | 0.018 | 0.018 | 3.582 | 0.86 | 0.25 | 0.73 | 0.77 | 0.95 | 0.83 |
| 4 | 89023224 | rs4693924 | 0.603 | -0.021 | -0.764 | 1.371 | -0.026 | 0 | 0 | 0 | 0.05 | 0.03 | 0.13 | 0.25 | 0.07 | 0.07 |
| 4 | 89026109 | rs2725264 | 1.36 | -0.176 | -0.816 | 1.17 | 0.041 | -0.023 | -0.023 | 0.389 | 0.82 | 0.15 | 0.7 | 0.78 | 0.91 | 0.81 |
| 4 | 89026428 | rs2725263 | 1.378 | -0.06 | -1.2 | 0.955 | -0.044 | -0.021 | -0.021 | 0.075 | 0.58 | 0.14 | 0.48 | 0.56 | 0.51 | 0.51 |
| 4 | 89027542 | rs2622610 | 1.018 | -0.102 | -1.132 | 0.977 | 0.026 | -0.017 | -0.017 | 0 | 0.83 | 0.17 | 0.7 | 0.78 | 0.91 | 0.81 |

| | | | | | | | | | | | | | | | | |
|---|---|---|---|---|---|---|---|---|---|---|---|---|---|---|---|---|
| 4 | 89028478 | rs2231148 | -0.075 | 0.068 | -1.236 | 0.996 | -0.13 | 0.008 | 0.008 | 0 | 0.25 | 0.03 | 0.22 | 0.21 | 0.4 | 0.29 |
| 4 | 89029252 | rs2622628 | 0.51 | -0.182 | -0.261 | 1.015 | -0.05 | 0.181 | 0.181 | 0.406 | 0.89 | 0.52 | 0.77 | 0.78 | 0.95 | 0.87 |
| 4 | 89030920 | rs2622621 | 0.592 | -0.027 | -1.382 | 0.476 | -0.012 | -0.05 | -0.05 | 0 | 0.35 | 0.12 | 0.36 | 0.58 | 0.3 | 0.39 |
| 4 | 89039629 | rs1871744 | 1.099 | 0.072 | -0.903 | 0.178 | -0.03 | -0.05 | -0.05 | 0 | 0.22 | 0.07 | 0.14 | 0.31 | 0.22 | 0.14 |
| 4 | 89050998 | rs2725256 | 1.221 | -0.137 | -0.57 | 0.283 | -0.001 | -0.008 | -0.008 | 0 | 0.37 | 0.34 | 0.32 | 0.29 | 0.33 | 0.39 |
| 4 | 89053718 | rs2231138 | 0.635 | -0.249 | -0.356 | 0.34 | 0.01 | -0.009 | -0.009 | 0 | 0.08 | 0.04 | 0.2 | 0.12 | 0.04 | 0.1 |
| 4 | 89054486 | rs17013859 | 0.423 | -0.313 | -0.315 | 0.291 | -0.034 | -0.047 | -0.047 | 0.001 | 0.1 | 0.38 | 0.04 | 0 | 0.02 | 0.16 |
| 4 | 89059819 | rs17731631 | 1.723 | -0.298 | -0.915 | 0.261 | -0.035 | 0.053 | 0.053 | 0.088 | 0.19 | 0.03 | 0.1 | 0.09 | 0.2 | 0.09 |
| 4 | 89059917 | rs12641369 | -1.034 | -0.347 | -0.709 | 0.269 | -0.093 | 0.073 | 0.073 | 0.087 | 0.21 | 0.52 | 0.29 | 0.33 | 0.08 | 0.29 |
| 4 | 89060909 | rs4148152 | 0.24 | -0.263 | -0.715 | 0.319 | 0.019 | -0.018 | -0.018 | 0 | 0.1 | 0.05 | 0.24 | 0.33 | 0.06 | 0.16 |
| 4 | 89061114 | rs2231137 | 0.254 | -0.272 | -0.719 | 0.314 | 0.016 | -0.014 | -0.014 | 0.278 | 0.1 | 0.06 | 0.24 | 0.33 | 0.06 | 0.15 |
| 4 | 89061265 | rs1564481 | 1.812 | -0.446 | -0.781 | 0.175 | -0.005 | 0.003 | 0.003 | 0 | 0.33 | 0.21 | 0.3 | 0.28 | 0.33 | 0.39 |
| 4 | 89061910 | rs2725252 | 1.641 | -0.372 | -0.433 | 0.454 | 0.015 | 0.021 | 0.021 | 0.234 | 0.52 | 0.28 | 0.41 | 0.41 | 0.53 | 0.48 |
| 4 | 89062189 | rs4148150 | -0.165 | -0.338 | -0.139 | -0.089 | 0.016 | -0.015 | -0.015 | 0.391 | 0.1 | 0.06 | 0.24 | 0.3 | 0.06 | 0.15 |
| 4 | 89062285 | rs4148149 | -1.434 | -0.313 | -0.149 | -0.128 | -0.006 | -0.008 | -0.008 | 0.002 | 0.37 | 0.58 | 0.45 | 0.3 | 0.37 | 0.39 |
| 4 | 89062513 | rs13137622 | -0.662 | -0.026 | -0.084 | -0.349 | 0.011 | -0.17 | -0.17 | 0 | 0.26 | 0.48 | 0.2 | 0 | 0.32 | 0.24 |
| 4 | 89063363 | rs2046134 | -0.25 | -0.169 | 0.143 | -0.144 | -0.032 | -0.048 | -0.048 | 0 | 0.1 | 0.46 | 0.05 | 0.01 | 0.02 | 0.14 |
| 4 | 89063985 | rs2725250 | 0.35 | -0.094 | -0.509 | 0.037 | -0.009 | -0.01 | -0.01 | 3.551 | 0.73 | 0.78 | 0.61 | 0.68 | 0.64 | 0.75 |
| 4 | 89064602 | rs3109823 | 0.778 | 0.146 | -0.102 | -0.251 | 0.018 | -0.015 | -0.015 | 0 | 0.79 | 0.39 | 0.74 | 0.81 | 0.74 | 0.76 |
| 4 | 89065076 | rs17013881 | 1.043 | 0.088 | -0.453 | -0.349 | 0.005 | 0.047 | 0.047 | 0 | 0.09 | 0.03 | 0.04 | 0 | 0.07 | 0.08 |
| 4 | 89065353 | rs2622627 | 0.616 | 0.233 | -0.785 | -0.67 | -0.001 | 0.051 | 0.051 | 0 | 0.51 | 0.3 | 0.54 | 0.65 | 0.47 | 0.55 |
| 4 | 89066075 | rs6857600 | 0.179 | 0.207 | -0.701 | -0.736 | -0.032 | -0.054 | -0.054 | 0 | 0.34 | 0.51 | 0.22 | 0.16 | 0.25 | 0.31 |
| 4 | 89066715 | rs2622626 | 0.692 | 0.244 | -0.78 | -0.72 | -0.001 | 0.053 | 0.053 | 0 | 0.51 | 0.3 | 0.54 | 0.65 | 0.47 | 0.55 |
| 4 | 89067526 | rs6532049 | 0.112 | 0.206 | -0.764 | -0.713 | -0.013 | -0.025 | -0.025 | 0.366 | 0.35 | 0.61 | 0.23 | 0.16 | 0.25 | 0.3 |
| 4 | 89068498 | rs2725246 | 1.126 | 0.312 | -0.775 | -0.658 | -0.007 | 0.03 | 0.03 | 0 | 0.42 | 0.22 | 0.5 | 0.65 | 0.39 | 0.47 |
| 4 | 89069406 | rs2622624 | 1.135 | 0.313 | -0.774 | -0.658 | -0.007 | 0.029 | 0.029 | 0 | 0.42 | 0.22 | 0.5 | 0.64 | 0.39 | 0.47 |
| 4 | 89078924 | rs2622604 | -0.045 | 0.492 | -0.802 | -0.474 | -0.016 | -0.011 | -0.011 | 0 | 0.87 | 0.89 | 0.8 | 0.8 | 0.76 | 0.88 |
| 4 | 89079386 | rs2622605 | 1.099 | 0.38 | -0.903 | -0.663 | -0.01 | 0.014 | 0.014 | 0 | 0.39 | 0.21 | 0.51 | 0.63 | 0.4 | 0.46 |
| 4 | 89081441 | rs3114019 | -0.364 | 0.717 | -0.895 | -0.64 | -0.025 | -0.018 | -0.018 | 0.056 | 0.88 | 0.91 | 0.8 | 0.8 | 0.76 | 0.89 |
| 4 | 89083666 | rs3114020 | 0.886 | 0.412 | -0.779 | -0.564 | -0.009 | 0.017 | 0.017 | 0 | 0.39 | 0.21 | 0.51 | 0.63 | 0.4 | 0.47 |
| 4 | 89090877 | rs10011796 | -1.003 | 0.367 | -0.915 | -0.576 | -0.002 | 0.018 | 0.018 | 0 | 0.57 | 0.73 | 0.48 | 0.37 | 0.61 | 0.52 |
| 4 | 89094064 | rs2622629 | 1.323 | -0.01 | -1.504 | -0.451 | -0.107 | -0.187 | -0.187 | 0 | 0.31 | 0.02 | 0.4 | 0.49 | 0.3 | 0.4 |
| 4 | 89096641 | rs7658584 | 0.316 | -0.525 | -0.814 | -1.126 | -0.036 | -0.09 | -0.09 | 0.305 | 0.25 | 0.5 | 0.14 | 0.07 | 0.19 | 0.22 |
| 4 | 89111468 | rs2127863 | 0.19 | 0.362 | -0.732 | -1.026 | -0.008 | -0.006 | -0.006 | 0.86 | 0.88 | 0.88 | 0.89 | 1 | 0.79 | 0.93 |
| 4 | 89119659 | rs10856870 | -0.424 | -0.094 | -0.668 | -1.534 | 0.002 | -0.063 | -0.063 | 0.611 | 0.48 | 0.44 | 0.72 | 0.99 | 0.53 | 0.7 |
| 4 | 89129484 | rs11723264 | -0.781 | -0.271 | -0.792 | -1.462 | 0.074 | 0.103 | 0.103 | 0.943 | 0.56 | 0.3 | 0.8 | 1 | 0.74 | 0.74 |
| 4 | 89139002 | rs4693933 | 0.657 | -0.153 | -0.494 | -1.287 | -0.016 | -0.065 | -0.065 | 0.112 | 0.46 | 0.48 | 0.27 | 0.09 | 0.3 | 0.25 |
| 4 | 89139275 | rs4693935 | 0.586 | 0.122 | -0.582 | -0.724 | 0.003 | 0.014 | 0.014 | 0 | 0.76 | 0.67 | 0.76 | 0.67 | 0.72 | 0.81 |
| 4 | 89139832 | rs2904185 | 0.514 | 0.317 | -0.686 | -0.609 | -0.022 | -0.025 | -0.025 | 0.243 | 0.29 | 0.18 | 0.48 | 0.57 | 0.41 | 0.55 |
| 4 | 89149570 | rs9307048 | 0.075 | 0.232 | -1.159 | -1.037 | -0.031 | 0.034 | 0.034 | 0 | 0.19 | 0.11 | 0.16 | 0 | 0.3 | 0.06 |
| 4 | 89159819 | rs1114568 | -0.481 | -0.094 | -0.761 | -0.702 | 0.003 | -0.053 | -0.053 | 0.007 | 0.42 | 0.48 | 0.62 | 0.89 | 0.45 | 0.7 |
| 4 | 89160677 | rs13120819 | 0.363 | 0.082 | -0.81 | -0.508 | 0.002 | 0.003 | 0.003 | 5.143 | 0.31 | 0.33 | 0.52 | 0.55 | 0.41 | 0.56 |
| 4 | 89163923 | rs6532059 | 1.472 | 0.046 | -1.074 | -0.475 | -0.009 | 0.045 | 0.045 | 0.491 | 0.1 | 0.01 | 0.06 | 0 | 0.09 | 0.02 |
| 4 | 89165613 | rs6532061 | 0.162 | 0.24 | -0.909 | -0.36 | 0.022 | 0.028 | 0.028 | 4.4 | 0.33 | 0.43 | 0.53 | 0.54 | 0.42 | 0.56 |
| 4 | 89165737 | rs11932727 | 0.617 | 0.151 | -1.004 | -0.856 | 0.013 | 0.08 | 0.08 | 0.522 | 0.42 | 0.24 | 0.25 | 0.07 | 0.4 | 0.26 |
| 4 | 89167407 | rs1351441 | -0.056 | 0.833 | -0.902 | -0.164 | -0.004 | 0.008 | 0.008 | 0.895 | 0.88 | 0.89 | 0.91 | 0.96 | 0.88 | 0.9 |
| 4 | 89171059 | rs11729997 | 0.058 | 0.731 | -0.765 | 0.354 | -0.001 | -0.001 | -0.001 | 0.961 | 0.32 | 0.3 | 0.54 | 0.56 | 0.42 | 0.54 |
| 4 | 89174193 | rs11733577 | 0.035 | -0.027 | -1.217 | 0.209 | 0.1 | -0.047 | -0.047 | 0 | 0.46 | 0.12 | 0.66 | 0.55 | 0.69 | 0.61 |
| 4 | 89191346 | rs7682161 | 0.437 | -0.562 | -0.641 | -0.234 | -0.022 | -0.056 | -0.056 | 0 | 0.1 | 0.24 | 0.05 | 0 | 0.06 | 0.01 |
| 4 | 89196821 | rs7656367 | -0.66 | -0.164 | -0.373 | -0.153 | -0.013 | 0.012 | 0.012 | 0 | 0.6 | 0.71 | 0.38 | 0.3 | 0.52 | 0.39 |
| 4 | 89201007 | rs7661312 | -0.605 | -0.162 | -0.252 | -0.152 | -0.04 | 0.03 | 0.03 | 0 | 0.61 | 0.8 | 0.38 | 0.3 | 0.52 | 0.39 |
| 4 | 89203197 | rs17013978 | 0.617 | -0.15 | -0.318 | -0.192 | -0.024 | -0.057 | -0.057 | 1.782 | 0.1 | 0.24 | 0.05 | 0 | 0.06 | 0.01 |
| 4 | 89204093 | rs17732955 | 1.548 | -0.441 | -0.962 | -0.434 | -0.113 | 0.167 | 0.167 | 0 | 0.24 | 0.01 | 0.15 | 0 | 0.31 | 0.1 |
| 4 | 89209681 | rs958325 | -0.526 | -0.308 | -0.576 | -0.086 | -0.015 | 0.017 | 0.017 | 0 | 0.59 | 0.72 | 0.37 | 0.3 | 0.51 | 0.39 |
| 4 | 89222164 | rs12641788 | -0.642 | 0.002 | -0.399 | 0.461 | 0.002 | 0.003 | 0.003 | 0 | 0.08 | 0.08 | 0.12 | 0.33 | 0.09 | 0.12 |
| 4 | 89226422 | rs1440581 | -0.723 | 0.044 | -0.221 | 0.446 | 0.003 | -0.009 | -0.009 | 0 | 0.53 | 0.73 | 0.48 | 0.55 | 0.53 | 0.48 |
| 4 | 89227094 | rs17013995 | 0.198 | 0.004 | -0.419 | 0.397 | -0.028 | 0.01 | 0.01 | 0 | 0.12 | 0.24 | 0.15 | 0.18 | 0.05 | 0.18 |
| 4 | 89227121 | rs17789621 | 1.232 | 0.082 | -0.953 | 0.306 | -0.068 | 0.118 | 0.118 | 0 | 0.28 | 0.13 | 0.18 | 0.01 | 0.36 | 0.15 |
| 4 | 89228928 | rs6841731 | -0.114 | 0.3 | -0.821 | 0.859 | 0.004 | -0.002 | -0.002 | 0 | 0.52 | 0.68 | 0.47 | 0.55 | 0.5 | 0.48 |
| 4 | 89236735 | rs893970 | 0.118 | 0.504 | -0.846 | 0.302 | 0.004 | 0.034 | 0.034 | 0 | 0.52 | 0.62 | 0.41 | 0.3 | 0.49 | 0.44 |
| 4 | 89239492 | rs1545207 | -0.667 | 0.787 | -0.834 | 0.74 | 0.026 | 0.074 | 0.074 | 0 | 0.21 | 0.31 | 0.27 | 0.55 | 0.27 | 0.4 |
| 4 | 89243818 | rs10022462 | 1.019 | 0.104 | -1.144 | -0.403 | 0.002 | -0.143 | -0.143 | 2.297 | 0.4 | 0.43 | 0.27 | 0.02 | 0.43 | 0.24 |
| 4 | 89244007 | rs2869929 | 0.136 | -1.045 | -1.447 | 0.569 | 0.091 | -0.015 | -0.015 | 0.394 | 0.77 | 0.93 | 0.82 | 0.79 | 0.88 | 0.79 |
| 4 | 89249061 | rs4693211 | -0.44 | -0.414 | -0.971 | 0.913 | -0.002 | 0.049 | 0.049 | 0 | 0.07 | 0.13 | 0.11 | 0.22 | 0.05 | 0.15 |
| 4 | 89249372 | rs10001661 | -0.208 | -0.053 | -1.287 | 1.056 | -0.053 | -0.082 | -0.082 | 2.848 | 0.16 | 0.04 | 0.23 | 0.46 | 0.25 | 0.32 |
| 4 | 89252068 | rs12512051 | -0.199 | -0.153 | -1.245 | 1.116 | -0.003 | 0.046 | 0.046 | 0 | 0.07 | 0.13 | 0.11 | 0.22 | 0.05 | 0.15 |
| 4 | 89252659 | rs17014031 | -0.293 | -0.814 | -1.55 | 0.784 | 0.065 | -0.013 | -0.013 | 0 | 0.19 | 0.08 | 0.16 | 0.23 | 0.08 | 0.11 |
| 4 | 89253079 | rs7657298 | -1.278 | -0.84 | -1.359 | 0.737 | 0.008 | -0.01 | -0.01 | 0 | 0.28 | 0.31 | 0.29 | 0.45 | 0.14 | 0.26 |
| 4 | 89258623 | rs17014050 | -0.504 | -0.865 | -1.469 | 0.204 | 0.041 | 0.002 | 0.002 | 1.405 | 0.21 | 0.14 | 0.19 | 0.22 | 0.11 | 0.12 |

| | | | | | | | | | | | | | | | | |
|---|---|---|---|---|---|---|---|---|---|---|---|---|---|---|---|---|
| 4 | 89261908 | rs2869928 | -1.697 | -0.91 | -0.882 | 0.176 | 0.013 | 0.049 | 0.049 | 0 | 0.46 | 0.69 | 0.54 | 0.91 | 0.39 | 0.59 |
| 4 | 89270121 | rs4693953 | -1.222 | -0.822 | -1.032 | 0.162 | -0.001 | -0.008 | -0.008 | 0.949 | 0.41 | 0.59 | 0.42 | 0.69 | 0.37 | 0.48 |
| 4 | 89277494 | rs4693954 | -1.7 | -0.643 | -1.163 | -0.105 | -0.001 | -0.04 | -0.04 | 0.116 | 0.49 | 0.54 | 0.54 | 0.92 | 0.44 | 0.66 |
| 4 | 89283496 | rs4693212 | -0.62 | -0.101 | -1.292 | 0.637 | -0.044 | -0.055 | -0.055 | 0 | 0.23 | 0.12 | 0.34 | 0.63 | 0.28 | 0.37 |
| 4 | 89287096 | rs4693957 | -0.438 | -0.934 | -1.128 | 0.37 | -0.062 | 0.024 | 0.024 | 0 | 0.26 | 0.4 | 0.28 | 0.39 | 0.13 | 0.3 |
| 4 | 89287709 | rs12651141 | -0.765 | -0.955 | -1.182 | -0.363 | -0.061 | -0.014 | -0.014 | 0.227 | 0.32 | 0.5 | 0.4 | 0.78 | 0.19 | 0.53 |
| 4 | 89293265 | rs10016511 | -0.887 | -0.222 | -1.492 | -0.305 | 0.002 | -0.003 | -0.003 | 0 | 0.3 | 0.33 | 0.46 | 0.83 | 0.3 | 0.61 |
| 4 | 89293297 | rs10029067 | -1.174 | 0.215 | -2.102 | -1.363 | -0.011 | 0.12 | 0.12 | 0 | 0.64 | 0.85 | 0.75 | 0.99 | 0.58 | 0.87 |
| 4 | 89299174 | rs28694742 | -0.208 | 0.332 | -1.264 | -0.159 | -0.025 | 0.028 | 0.028 | 0 | 0.14 | 0.09 | 0.13 | 0.03 | 0.18 | 0.09 |
| 4 | 89306659 | rs7677237 | -0.431 | -0.379 | -0.617 | 0.261 | -0.024 | -0.063 | -0.063 | 0 | 0.15 | 0.23 | 0.07 | 0 | 0.08 | 0.02 |
| 4 | 89311962 | rs12510688 | -0.446 | 0.041 | -0.996 | 0.721 | -0.006 | -0.009 | -0.009 | 0 | 0.14 | 0 | 0.18 | 0.12 | 0.09 | 0.07 |
| 4 | 89314310 | rs17014114 | -0.664 | 0.028 | -0.387 | 1.602 | 0 | 0 | 0 | 0.02 | 0.12 | 0.11 | 0.09 | 0.23 | 0.07 | 0.15 |
| 4 | 89316229 | rs17014115 | -0.91 | 0.286 | 0.548 | 1.651 | 0.037 | 0.053 | 0.053 | 0.429 | 0.14 | 0.41 | 0.15 | 0.2 | 0.14 | 0.08 |
| 4 | 89320406 | rs7662660 | -0.899 | 0.296 | 0.565 | 1.914 | -0.009 | 0.028 | 0.028 | 0 | 0.12 | 0.14 | 0.13 | 0.24 | 0.13 | 0.15 |
| 4 | 89383662 | rs10516808 | -0.919 | 1.173 | 1.098 | 1.861 | -0.039 | -0.032 | -0.032 | 0 | 0.08 | 0.01 | 0.18 | 0.14 | 0.13 | 0.07 |
| 4 | 89417500 | rs11722508 | -0.861 | 1.061 | 0.892 | 1.875 | -0.036 | -0.031 | -0.031 | 0 | 0.07 | 0.01 | 0.18 | 0.14 | 0.12 | 0.06 |
| 4 | 89433700 | rs7665582 | -1.859 | 0.72 | 1.469 | 2.33 | 0.009 | 0.205 | 0.205 | 1.258 | 0.19 | 0.5 | 0.34 | 0.54 | 0.16 | 0.3 |
| 4 | 89441779 | rs2717533 | 1.202 | 0.985 | 1.055 | 1.956 | 0.031 | 0.045 | 0.045 | 0.256 | 0.95 | 0.88 | 0.9 | 0.85 | 0.92 | 0.9 |
| 4 | 89445906 | rs4693959 | -1.851 | 0.486 | 1.201 | 2.028 | -0.022 | 0.087 | 0.087 | 0 | 0.11 | 0.31 | 0.12 | 0.22 | 0.06 | 0.18 |
| 4 | 89448232 | rs6823968 | -1.028 | 0.053 | 1.244 | 1.441 | -0.014 | -0.008 | -0.008 | 0 | 0.26 | 0.32 | 0.37 | 0.82 | 0.23 | 0.51 |
| 4 | 89451509 | rs3846281 | -1.079 | 0.036 | 1.23 | 1.756 | -0.015 | 0.008 | 0.008 | 0.108 | 0.26 | 0.33 | 0.37 | 0.76 | 0.23 | 0.51 |
| 4 | 89452479 | rs10031689 | -0.829 | 0.192 | 1.32 | 1.353 | 0.018 | 0.027 | 0.027 | 0.983 | 0.15 | 0.3 | 0.16 | 0.14 | 0.16 | 0.12 |
| 4 | 89453522 | rs2869665 | -1.241 | -0.107 | 1.325 | 1.524 | -0.031 | 0.002 | 0.002 | 0.649 | 0.29 | 0.46 | 0.39 | 0.78 | 0.23 | 0.54 |
| 4 | 89456219 | rs7435410 | -2.056 | 0.277 | 2.153 | 2.234 | 0.01 | 0.136 | 0.136 | 0.001 | 0.19 | 0.36 | 0.3 | 0.51 | 0.19 | 0.42 |
| 4 | 89457478 | rs17014179 | -0.75 | 0.259 | 1.942 | 2.13 | -0.001 | -0.005 | -0.005 | 0.02 | 0.08 | 0.11 | 0.04 | 0.01 | 0.06 | 0.28 |
| 4 | 89458051 | rs11736472 | -1.548 | 0.312 | 1.832 | 2.314 | 0.014 | 0.113 | 0.113 | 0 | 0.11 | 0.16 | 0.26 | 0.47 | 0.15 | 0.15 |
| 4 | 89477899 | rs17014220 | -1.536 | 0.379 | 1.779 | 2.51 | -0.011 | 0.074 | 0.074 | 0.397 | 0.08 | 0.14 | 0.04 | 0.25 | 0.05 | 0.06 |
| 4 | 89482021 | rs6850944 | -2.041 | -0.251 | 1.585 | 1.91 | 0.001 | 0.019 | 0.019 | 9.449 | 0.31 | 0.75 | 0.37 | 0.26 | 0.28 | 0.48 |
| 4 | 89492405 | rs16996122 | -0.642 | -0.295 | 1.53 | 2.013 | -0.026 | 0.06 | 0.06 | 0.517 | 0.06 | 0.08 | 0.01 | 0.24 | 0 | 0.04 |
| 4 | 89494828 | rs7671998 | -2.561 | -0.305 | 1.774 | 2.022 | 0.027 | -0.052 | -0.052 | 0 | 0.36 | 0.83 | 0.29 | 0.5 | 0.27 | 0.5 |
| 4 | 89512853 | rs10084787 | -2.561 | -0.305 | 1.774 | 2.022 | 0.027 | -0.052 | -0.052 | 0.131 | 0.36 | 0.83 | 0.3 | 0.5 | 0.27 | 0.5 |
| 4 | 89514572 | rs6857847 | -0.963 | -0.288 | 1.983 | 2.119 | 0.048 | 0.095 | 0.095 | 0 | 0.05 | 0.1 | 0.16 | 0.29 | 0.17 | 0.15 |
| 4 | 89522699 | rs3822084 | -2.227 | -0.325 | 1.979 | 1.926 | 0.026 | 0.004 | 0.004 | 11.805 | 0.31 | 0.73 | 0.13 | 0.29 | 0.1 | 0.36 |
| 4 | 89525053 | rs17014252 | -2.101 | -0.381 | 2.786 | 1.896 | -0.07 | 0.094 | 0.094 | 1.064 | 0.17 | 0.53 | 0.09 | 0.26 | 0.06 | 0.3 |
| 4 | 89526143 | rs17014254 | -0.611 | -0.358 | 2.524 | 1.865 | 0.001 | -0.013 | -0.013 | 0 | 0.14 | 0.13 | 0.04 | 0.03 | 0.04 | 0.06 |
| 4 | 89527323 | rs17799056 | -0.99 | -0.264 | 2.513 | 1.967 | 0.042 | 0.041 | 0.041 | 0 | 0.05 | 0.08 | 0.16 | 0.22 | 0.17 | 0.14 |
| 4 | 89532682 | rs2924942 | -2.811 | -0.345 | 3.09 | 1.895 | -0.006 | 0.011 | 0.011 | 5.237 | 0.32 | 0.62 | 0.25 | 0.49 | 0.24 | 0.48 |
| 4 | 89539581 | rs11935902 | -0.605 | -0.285 | 3.22 | 1.88 | -0.004 | -0.025 | -0.025 | 0 | 0.08 | 0.2 | 0.03 | 0.02 | 0.06 | 0.28 |
| 4 | 89540209 | rs10028621 | -0.268 | -0.286 | 3.235 | 1.879 | -0.007 | -0.02 | -0.02 | 0 | 0.07 | 0.14 | 0.03 | 0.02 | 0.06 | 0.28 |
| 4 | 89542281 | rs2869663 | -1.683 | -0.278 | 3.059 | 1.903 | 0.076 | 0.047 | 0.047 | 0 | 0.14 | 0.35 | 0.2 | 0.21 | 0.21 | 0.39 |
| 4 | 89555107 | rs2972023 | -0.609 | -0.28 | 2.389 | 1.934 | -0.03 | 0.131 | 0.131 | 0.401 | 0.06 | 0.15 | 0.02 | 0.24 | 0 | 0.04 |
| 4 | 89562091 | rs2869664 | -3.161 | -0.701 | 2.021 | 1.835 | 0.043 | -0.102 | -0.102 | 0 | 0.39 | 0.89 | 0.29 | 0.52 | 0.26 | 0.6 |
| 4 | 89567767 | rs3017925 | -3.166 | -0.708 | 1.955 | 1.823 | 0.039 | -0.108 | -0.108 | 0 | 0.39 | 0.9 | 0.29 | 0.52 | 0.26 | 0.59 |
| 4 | 89584883 | rs2924939 | -0.5 | -0.41 | 1.97 | 1.934 | -0.029 | 0.099 | 0.099 | 0.637 | 0.06 | 0.1 | 0.02 | 0.24 | 0.01 | 0.04 |
| 4 | 89585781 | rs17014374 | -0.328 | -0.316 | 1.666 | 1.695 | 0.012 | 0.009 | 0.009 | 5.659 | 0.11 | 0.08 | 0.02 | 0.03 | 0.03 | 0.06 |
| 4 | 89612444 | rs1020593 | -2.248 | -0.383 | 2.03 | 1.853 | -0.021 | 0.074 | 0.074 | 4.572 | 0.29 | 0.43 | 0.24 | 0.51 | 0.21 | 0.44 |
| 4 | 89615729 | rs2924358 | -1.463 | -0.694 | 1.719 | 1.781 | 0.017 | -0.021 | -0.021 | 3.202 | 0.14 | 0.11 | 0.03 | 0.28 | 0.04 | 0.37 |
| 4 | 89622904 | rs2924924 | -2.933 | -0.621 | 2.497 | 1.761 | -0.002 | 0.009 | 0.009 | 1.91 | 0.33 | 0.58 | 0.26 | 0.52 | 0.27 | 0.56 |
| 4 | 89629000 | rs1063143 | -2.868 | -0.685 | 2.805 | 1.288 | -0.033 | 0.02 | 0.02 | 0.059 | 0.29 | 0.57 | 0.15 | 0.31 | 0.23 | 0.53 |
| 4 | 89639032 | rs2860408 | -1.616 | -0.861 | 3.002 | 1.347 | -0.029 | 0.064 | 0.064 | 0.007 | 0.16 | 0.31 | 0.05 | 0.28 | 0.1 | 0.43 |
| 4 | 89641514 | rs2924936 | -1.539 | -0.865 | 3.01 | 1.352 | -0.027 | 0.065 | 0.065 | 0 | 0.16 | 0.31 | 0.05 | 0.28 | 0.1 | 0.43 |
| 4 | 89644197 | rs2924941 | -2.73 | -0.859 | 2.998 | 1.349 | -0.007 | 0.03 | 0.03 | 1.406 | 0.32 | 0.53 | 0.25 | 0.52 | 0.26 | 0.56 |
| 4 | 89647070 | rs2860500 | -3.293 | -0.903 | 2.884 | 1.325 | 0.04 | -0.061 | -0.061 | 0.823 | 0.4 | 0.84 | 0.31 | 0.54 | 0.3 | 0.65 |
| 4 | 89648845 | rs13130 | -2.113 | -0.764 | 3.167 | 1.394 | -0.047 | 0.077 | 0.077 | 0.624 | 0.18 | 0.41 | 0.06 | 0.28 | 0.1 | 0.43 |
| 4 | 89649491 | rs3017895 | -1.902 | -0.778 | 3.169 | 1.434 | -0.041 | 0.093 | 0.093 | 0.583 | 0.17 | 0.38 | 0.06 | 0.29 | 0.1 | 0.37 |
| 4 | 89649659 | rs9224 | -2.7 | -1.539 | 2.545 | 1.208 | 0.056 | 0.172 | 0.172 | 0.088 | 0.47 | 0.77 | 0.51 | 0.83 | 0.34 | 0.64 |
| 4 | 89653257 | rs1801915 | -0.979 | -0.866 | 2.712 | 1.339 | 0.006 | 0.005 | 0.005 | 0 | 0.16 | 0.2 | 0.19 | 0.23 | 0.16 | 0.2 |
| 4 | 89665193 | rs3775391 | -1.938 | -0.805 | 2.457 | 1.41 | -0.012 | 0.133 | 0.133 | 0.031 | 0.14 | 0.34 | 0.06 | 0.3 | 0.1 | 0.39 |
| 4 | 89667591 | rs17014483 | -1.726 | -0.803 | 2.54 | 1.411 | -0.011 | 0.091 | 0.091 | 0 | 0.13 | 0.24 | 0.05 | 0.3 | 0.1 | 0.39 |
| 4 | 89668859 | rs7657817 | -1.36 | -0.825 | 2.359 | 1.431 | 0.031 | 0.048 | 0.048 | 0.364 | 0.18 | 0.47 | 0.23 | 0.25 | 0.18 | 0.2 |
| 4 | 89669852 | rs11944668 | -2.567 | -0.826 | 2.423 | 1.428 | 0.001 | -0.018 | -0.018 | 0 | 0.32 | 0.73 | 0.28 | 0.54 | 0.28 | 0.59 |
| 4 | 89672805 | rs7662156 | 0.886 | -0.823 | 2.106 | 1.5 | 0.044 | 0.058 | 0.058 | 0.405 | 0.82 | 0.53 | 0.68 | 0.74 | 0.82 | 0.81 |
| 4 | 89677537 | rs1965869 | 2.096 | -0.555 | 1.709 | 1.315 | -0.002 | 0.041 | 0.041 | 3.677 | 0.67 | 0.51 | 0.62 | 0.44 | 0.72 | 0.43 |
| 4 | 89684640 | rs17014513 | -1.055 | -0.521 | 1.674 | 1.367 | -0.028 | 0.079 | 0.079 | 0.325 | 0.11 | 0.18 | 0.04 | 0.3 | 0.04 | 0.32 |
| 4 | 89693098 | rs3816453 | 1.038 | -0.092 | 1.273 | 1.25 | 0.025 | 0.062 | 0.062 | 0.331 | 0.82 | 0.59 | 0.68 | 0.69 | 0.82 | 0.78 |
| 4 | 89703883 | rs10017416 | -0.978 | -0.081 | 1.154 | 1.132 | -0.001 | -0.01 | -0.01 | 0.188 | 0.08 | 0.25 | 0.1 | 0.02 | 0.07 | 0.11 |
| 4 | 89713121 | rs9991328 | -0.488 | -0.073 | 0.4 | 0.172 | -0.002 | 0.032 | 0.032 | 0 | 0.56 | 0.63 | 0.55 | 0.7 | 0.49 | 0.65 |
| 4 | 89717990 | rs7680994 | -1.201 | -0.268 | 0.143 | 0.745 | -0.027 | 0.072 | 0.072 | 0 | 0.15 | 0.25 | 0.14 | 0.32 | 0.09 | 0.32 |
| 4 | 89724236 | rs17014580 | -0.171 | -0.122 | -0.185 | 0.775 | 0.007 | 0.009 | 0.009 | 0 | 0.09 | 0.07 | 0.09 | 0.28 | 0.06 | 0.31 |

| | | | | | | | | | | | | | | | | |
|---|---|---|---|---|---|---|---|---|---|---|---|---|---|---|---|---|
| 4 | 89739808 | rs3775380 | -0.294 | 0.077 | 0.037 | 0.221 | 0 | -0.001 | -0.001 | 0.373 | 0.55 | 0.54 | 0.55 | 0.7 | 0.49 | 0.67 |
| 4 | 89742764 | rs2869950 | -0.298 | -0.149 | -0.082 | 0.201 | 0.002 | 0.003 | 0.003 | 0 | 0.5 | 0.45 | 0.44 | 0.66 | 0.42 | 0.55 |
| 4 | 89743821 | rs3775373 | -0.649 | -0.462 | -0.014 | 0.382 | -0.048 | 0.022 | 0.022 | 0.292 | 0.34 | 0.44 | 0.27 | 0.52 | 0.21 | 0.26 |
| 4 | 89744242 | rs2305934 | -0.11 | 0.757 | 0.261 | 0.575 | 0.045 | 0.215 | 0.215 | 0 | 0.11 | 0.37 | 0.33 | 0.39 | 0.12 | 0.12 |
| 4 | 89751858 | rs7660000 | -0.65 | 0.934 | -0.292 | 0.034 | 0.031 | -0.004 | -0.004 | 0 | 0.39 | 0.27 | 0.28 | 0.53 | 0.3 | 0.58 |
| 4 | 89770583 | rs10461165 | -0.287 | 0.626 | -0.008 | 0.217 | -0.011 | 0.009 | 0.009 | 0 | 0.33 | 0.38 | 0.24 | 0.39 | 0.25 | 0.54 |
| 4 | 89789478 | rs1979290 | 0.428 | 1.171 | -0.065 | 0.241 | -0.008 | -0.008 | -0.008 | 0 | 0.07 | 0.05 | 0.09 | 0.13 | 0.14 | 0.11 |
| 4 | 89797638 | rs10516823 | -0.047 | 1.334 | -0.062 | 0.245 | 0 | 0.003 | 0.003 | 0 | 0.07 | 0.11 | 0.07 | 0.13 | 0.11 | 0.1 |
| 4 | 89798954 | rs17816002 | 0.827 | 0.837 | -0.772 | -0.562 | 0.001 | 0.08 | 0.08 | 0 | 0.32 | 0.15 | 0.26 | 0.16 | 0.35 | 0.16 |
| 4 | 89799575 | rs13131811 | 0.81 | 0.876 | -0.779 | -0.569 | -0.001 | 0.082 | 0.082 | 0 | 0.33 | 0.15 | 0.26 | 0.16 | 0.35 | 0.16 |
| 4 | 89811195 | rs2609255 | -0.833 | 0.51 | -0.451 | -0.638 | -0.011 | 0.007 | 0.007 | 0 | 0.66 | 0.7 | 0.7 | 0.58 | 0.76 | 0.52 |
| 4 | 89828080 | rs2609264 | -0.863 | 0.499 | -0.415 | -0.677 | 0.014 | -0.015 | -0.015 | 0 | 0.66 | 0.72 | 0.7 | 0.58 | 0.76 | 0.55 |
| 4 | 89831335 | rs13109149 | 2.059 | 0.553 | -0.543 | -0.793 | 0.046 | 0.129 | 0.129 | 0 | 0.25 | 0.06 | 0.16 | 0 | 0.18 | 0.05 |
| 4 | 89832532 | rs10019681 | 0.761 | 1.435 | -0.19 | -0.763 | 0.098 | 0.114 | 0.114 | 0 | 0.14 | 0.33 | 0.3 | 0.38 | 0.22 | 0.26 |
| 4 | 89835017 | rs1458555 | 0.703 | 1.406 | -0.227 | -0.813 | -0.009 | 0.013 | 0.013 | 0.024 | 0.19 | 0.16 | 0.14 | 0.05 | 0.19 | 0.11 |
| 4 | 89839840 | rs4693976 | 0.732 | 1.422 | -0.22 | -0.79 | 0.115 | 0.132 | 0.132 | 0 | 0.14 | 0.4 | 0.3 | 0.38 | 0.22 | 0.26 |
| 4 | 89849772 | rs2704588 | 0.427 | 1.109 | -0.444 | -0.824 | 0.007 | -0.028 | -0.028 | 0.14 | 0.2 | 0.22 | 0.14 | 0.05 | 0.19 | 0.11 |
| 4 | 89853598 | rs1458562 | 0.168 | 1.171 | -0.5 | -0.659 | -0.001 | -0.001 | -0.001 | 0.003 | 0.46 | 0.53 | 0.56 | 0.53 | 0.58 | 0.45 |
| 4 | 89879663 | rs2464526 | 0.966 | 1.109 | -0.783 | -0.821 | 0.02 | 0.116 | 0.116 | 0 | 0.35 | 0.14 | 0.25 | 0.05 | 0.27 | 0.1 |
| 4 | 89887620 | rs1458558 | 0.03 | 1.178 | -0.268 | -0.679 | -0.003 | 0.053 | 0.053 | 0.015 | 0.52 | 0.65 | 0.41 | 0.24 | 0.52 | 0.51 |
| 4 | 89895944 | rs2085601 | -0.464 | 1.668 | 0.059 | -0.354 | 0.037 | 0.028 | 0.028 | 0 | 0.8 | 0.7 | 0.71 | 0.61 | 0.73 | 0.68 |
| 4 | 89896036 | rs6838424 | 1.713 | 1.551 | 0.436 | -0.518 | 0.031 | 0.007 | 0.007 | 0 | 0.54 | 0.38 | 0.45 | 0.37 | 0.35 | 0.22 |
| 4 | 89897373 | rs17817631 | 0.054 | 1.715 | 0.534 | -0.551 | 0.041 | -0.003 | -0.003 | 0 | 0.14 | 0.21 | 0.11 | 0.07 | 0.15 | 0.29 |
| 4 | 89897522 | rs16996144 | 0.977 | 1.725 | 0.454 | -0.533 | 0.015 | 0.01 | 0.01 | 0 | 0.65 | 0.48 | 0.57 | 0.54 | 0.54 | 0.37 |
| 4 | 89898476 | rs11737182 | -0.484 | 1.746 | 0.126 | -0.465 | -0.01 | -0.008 | -0.008 | 0 | 0.8 | 0.83 | 0.69 | 0.61 | 0.69 | 0.66 |
| 4 | 89917135 | rs10007590 | -0.381 | 1.679 | 0.114 | -0.532 | 0.006 | -0.009 | -0.009 | 0 | 0.18 | 0.19 | 0.14 | 0.07 | 0.2 | 0.3 |
| 4 | 89918629 | rs9307061 | -1.424 | 1.912 | -0.03 | -0.43 | -0.006 | 0 | 0 | 0 | 0.89 | 0.89 | 0.75 | 0.61 | 0.8 | 0.8 |
| 4 | 89920082 | rs10516825 | 1.613 | 1.918 | -0.105 | -0.49 | -0.037 | 0.047 | 0.047 | 0.011 | 0.08 | 0 | 0.05 | 0 | 0.11 | 0.14 |
| 4 | 89922143 | rs1921684 | -1.152 | 1.896 | 0.132 | -0.526 | 0.035 | 0.057 | 0.057 | 0 | 0.88 | 0.81 | 0.74 | 0.61 | 0.8 | 0.8 |
| 4 | 89933630 | rs1795734 | -1.219 | 1.389 | 0.187 | -1.017 | 0.006 | -0.015 | -0.015 | 0.88 | 0.61 | 0.48 | 0.54 | 0.61 | 0.6 | 0.74 |
| 4 | 89938564 | rs1708661 | -1.795 | 1.644 | 0.277 | -1.049 | 0.021 | -0.011 | -0.011 | 2.212 | 0.34 | 0.61 | 0.3 | 0.24 | 0.44 | 0.57 |
| 4 | 89940992 | rs6822256 | 0.524 | 1.684 | 0.151 | -0.978 | -0.043 | -0.038 | -0.038 | 0 | 0.1 | 0.02 | 0.11 | 0.17 | 0.2 | 0.16 |
| 4 | 89945459 | rs9993181 | 0.808 | 1.048 | -0.168 | -1.371 | 0 | 0.042 | 0.042 | 0 | 0.54 | 0.47 | 0.38 | 0.18 | 0.53 | 0.53 |
| 4 | 89947462 | rs1343921 | 0.808 | 1.048 | -0.168 | -1.371 | 0 | 0.042 | 0.042 | 0 | 0.54 | 0.47 | 0.38 | 0.19 | 0.53 | 0.53 |
| 4 | 89954027 | rs13140085 | 1.533 | 0.999 | -0.176 | -1.388 | 0.028 | 0.014 | 0.014 | 0 | 0.44 | 0.45 | 0.27 | 0.02 | 0.33 | 0.37 |
| 4 | 89964456 | rs1533291 | 0.262 | 1.059 | -0.144 | -1.37 | 0.003 | -0.014 | -0.014 | 0.072 | 0.62 | 0.65 | 0.47 | 0.24 | 0.62 | 0.55 |
| 4 | 89975033 | rs6826407 | 1.43 | 1.083 | -0.113 | -1.367 | 0.047 | 0.06 | 0.06 | 0 | 0.45 | 0.41 | 0.27 | 0.02 | 0.34 | 0.37 |
| 4 | 89976882 | rs1795731 | -0.108 | 1.086 | -0.119 | -1.352 | 0.032 | 0.014 | 0.014 | 0.026 | 0.18 | 0.24 | 0.2 | 0.22 | 0.28 | 0.18 |
| 4 | 89978281 | rs11097210 | 1.388 | 1.007 | -0.133 | -1.463 | 0.041 | 0.048 | 0.048 | 0 | 0.45 | 0.42 | 0.27 | 0.02 | 0.34 | 0.36 |
| 4 | 89992347 | rs9790655 | 1.382 | 0.96 | -0.081 | -1.306 | -0.033 | 0.057 | 0.057 | 0 | 0.18 | 0.23 | 0.23 | 0.37 | 0.09 | 0.13 |
| 4 | 90002785 | rs13148714 | -0.02 | 1.263 | -0.263 | -1.268 | 0.039 | 0.018 | 0.018 | 0.087 | 0.81 | 0.71 | 0.79 | 0.78 | 0.72 | 0.82 |
| 4 | 90010941 | rs12649385 | 1.461 | 1.343 | -0.311 | -1.276 | -0.007 | 0 | 0 | 0 | 0.11 | 0.06 | 0.25 | 0.39 | 0.2 | 0.19 |
| 4 | 90011642 | rs6825998 | 1.209 | 1.103 | -0.298 | -1.322 | -0.005 | 0.011 | 0.011 | 0 | 0.62 | 0.59 | 0.49 | 0.39 | 0.43 | 0.5 |
| 4 | 90024739 | rs4627822 | 0.084 | 0.816 | -0.139 | -1.106 | -0.047 | 0.079 | 0.079 | 0 | 0.24 | 0.57 | 0.29 | 0.39 | 0.15 | 0.39 |
| 4 | 90034030 | rs17015025 | 0.245 | 0.983 | -0.179 | -1.031 | 0.022 | 0.02 | 0.02 | 0 | 0.11 | 0.14 | 0.11 | 0.16 | 0.19 | 0.17 |
| 4 | 90035549 | rs2280099 | 0.621 | 1.028 | -0.52 | -1.009 | -0.05 | -0.047 | -0.047 | 0.015 | 0.1 | 0.01 | 0.11 | 0.16 | 0.19 | 0.17 |
| 4 | 90044292 | rs4367140 | -0.228 | 0.725 | -0.457 | -1.071 | -0.034 | 0.04 | 0.04 | 0.314 | 0.31 | 0.48 | 0.33 | 0.39 | 0.24 | 0.52 |
| 4 | 90046848 | rs13138280 | -0.067 | 1.089 | -0.931 | -1.096 | -0.053 | -0.047 | -0.047 | 0.012 | 0.23 | 0.11 | 0.32 | 0.41 | 0.35 | 0.58 |
| 4 | 90049495 | rs17821540 | 1.502 | 0.817 | -0.675 | -1.018 | 0.016 | 0.063 | 0.063 | 0.094 | 0.11 | 0 | 0.03 | 0 | 0.07 | 0.04 |
| 4 | 90049968 | rs7687207 | -1.323 | 0.918 | -0.485 | -1.057 | 0 | 0.127 | 0.127 | 0 | 0.52 | 0.71 | 0.67 | 0.93 | 0.56 | 0.55 |
| 4 | 90057814 | rs1398944 | 0.262 | 1.591 | 0.209 | -0.536 | 0.082 | 0.131 | 0.131 | 0.427 | 0.13 | 0.34 | 0.27 | 0.4 | 0.2 | 0.17 |
| 4 | 90059434 | rs6828137 | -0.764 | 0.9 | -0.164 | -0.506 | -0.037 | -0.043 | -0.043 | 0.035 | 0.31 | 0.2 | 0.39 | 0.57 | 0.46 | 0.39 |
| 4 | 90062409 | rs7690336 | 0.533 | 0.877 | -0.064 | -0.751 | 0.061 | 0.029 | 0.029 | 0 | 0.73 | 0.61 | 0.56 | 0.55 | 0.57 | 0.74 |
| 4 | 90067301 | rs4552429 | -0.776 | 1.101 | -0.026 | -0.7 | 0.002 | -0.012 | -0.012 | 0 | 0.88 | 0.85 | 0.9 | 0.96 | 0.9 | 0.92 |
| 4 | 90069964 | rs756227 | -0.153 | 1.13 | -0.145 | -0.642 | 0.005 | 0.001 | 0.001 | 0.084 | 0.19 | 0.21 | 0.09 | 0.17 | 0.14 | 0.19 |
| 4 | 90073214 | rs756345 | 0.152 | 1.798 | 0.095 | -0.329 | 0.076 | 0.062 | 0.062 | 0 | 0.13 | 0.22 | 0.35 | 0.42 | 0.34 | 0.19 |
| 4 | 90091547 | rs6532123 | -0.173 | 0.726 | -0.154 | -1.51 | 0.009 | -0.017 | -0.017 | 0.242 | 0.63 | 0.6 | 0.35 | 0.18 | 0.51 | 0.66 |
| 4 | 90099188 | rs17771380 | 0.187 | 0.202 | -0.486 | -1.854 | -0.032 | 0.105 | 0.105 | 0.116 | 0.25 | 0.1 | 0.13 | 0.03 | 0.29 | 0.41 |
| 4 | 90107063 | rs9990512 | 0.48 | -0.083 | -0.365 | -1.95 | -0.006 | 0.038 | 0.038 | 0 | 0.12 | 0.34 | 0.13 | 0.14 | 0.1 | 0.15 |
| 4 | 90115197 | rs2116325 | 0.378 | 0.017 | -0.773 | -1.947 | -0.031 | 0.106 | 0.106 | 0.153 | 0.25 | 0.11 | 0.12 | 0.03 | 0.29 | 0.41 |
| 4 | 90115680 | rs524907 | 0.816 | -0.348 | -0.925 | -2.042 | -0.007 | 0.034 | 0.034 | 0 | 0.12 | 0.05 | 0.1 | 0.03 | 0.11 | 0.08 |
| 4 | 90116188 | rs11734085 | 1.101 | -0.335 | -0.987 | -2.089 | -0.009 | 0.12 | 0.12 | 0 | 0.19 | 0 | 0.09 | 0 | 0.16 | 0.12 |
| 4 | 90116432 | rs13353765 | 1.02 | -0.414 | -0.851 | -2.122 | 0.028 | 0.034 | 0.034 | 0.169 | 0.33 | 0.22 | 0.2 | 0.14 | 0.2 | 0.2 |
| 4 | 90123667 | rs6821473 | -0.304 | -0.387 | -0.453 | -2.243 | 0.002 | 0.147 | 0.147 | 1.33 | 0.31 | 0.11 | 0.19 | 0.03 | 0.27 | 0.2 |
| 4 | 90124654 | rs695053 | -0.162 | -0.629 | -0.127 | -1.897 | -0.01 | -0.044 | -0.044 | 0 | 0.41 | 0.45 | 0.22 | 0.11 | 0.22 | 0.19 |
| 4 | 90125419 | rs7688911 | 0.889 | -0.376 | -0.545 | -1.755 | -0.012 | -0.062 | -0.062 | 0 | 0.26 | 0.22 | 0.11 | 0 | 0.1 | 0.05 |
| 4 | 90126051 | rs678701 | -0.275 | 0.815 | -0.6 | -0.957 | -0.006 | -0.002 | -0.002 | 0 | 0.68 | 0.68 | 0.39 | 0.06 | 0.53 | 0.67 |
| 4 | 90126331 | rs679672 | -0.632 | 0.426 | -0.428 | -0.021 | -0.003 | 0.004 | 0.004 | 0.033 | 0.09 | 0.08 | 0.09 | 0.03 | 0.1 | 0.06 |
| 4 | 90126898 | rs7674810 | -0.992 | -0.158 | -0.014 | 0.641 | -0.053 | 0.089 | 0.089 | 0.104 | 0.17 | 0.34 | 0.35 | 0.36 | 0.09 | 0.29 |

| | | | | | | | | | | | | | | | | |
|---|---|---|---|---|---|---|---|---|---|---|---|---|---|---|---|---|
| 4 | 90126909 | rs7696471 | -1.089 | -0.155 | -0.2 | 0.567 | -0.061 | 0.067 | 0.067 | 0.697 | 0.25 | 0.47 | 0.38 | 0.39 | 0.12 | 0.33 |
| 4 | 90133795 | rs499411 | -0.35 | 0.553 | -0.826 | -0.35 | -0.035 | 0.04 | 0.04 | 0 | 0.66 | 0.75 | 0.72 | 0.93 | 0.56 | 0.81 |
| 4 | 90137336 | rs518156 | -0.057 | -0.3 | -0.811 | 0.305 | 0.009 | -0.022 | -0.022 | 0 | 0.29 | 0.22 | 0.14 | 0.42 | 0.26 | 0.33 |
| 4 | 90137884 | rs656063 | -1.079 | -0.251 | -0.88 | 0.128 | -0.046 | 0.065 | 0.065 | 0 | 0.66 | 0.78 | 0.72 | 0.93 | 0.56 | 0.8 |
| 4 | 90138063 | rs17015190 | 0.408 | -0.698 | -0.916 | 0.384 | 0.01 | -0.018 | -0.018 | 0 | 0.14 | 0.07 | 0.1 | 0.24 | 0.12 | 0.3 |
| 4 | 90146649 | rs9307064 | -1.656 | -0.154 | -1.05 | 0.07 | -0.027 | 0.042 | 0.042 | 0 | 0.69 | 0.79 | 0.75 | 0.93 | 0.64 | 0.9 |
| 4 | 90149589 | rs17773361 | 0.529 | -0.499 | -1.139 | -0.056 | -0.003 | 0.088 | 0.088 | 0 | 0.21 | 0.05 | 0.11 | 0 | 0.19 | 0.15 |
| 4 | 90152565 | rs1978673 | -0.037 | -0.565 | -0.565 | 0.062 | 0 | -0.002 | -0.002 | 0 | 0.18 | 0.19 | 0.13 | 0.24 | 0.2 | 0.41 |
| 4 | 90155694 | rs6847228 | -0.154 | -1.007 | -0.48 | 0.955 | -0.011 | 0.005 | 0.005 | 0.201 | 0.13 | 0.1 | 0.03 | 0.28 | 0.06 | 0.14 |
| 4 | 90161863 | rs7660707 | -0.724 | -1.079 | 0.396 | 1.221 | 0.004 | 0.019 | 0.019 | 0.01 | 0.33 | 0.43 | 0.25 | 0.4 | 0.38 | 0.57 |
| 4 | 90166159 | rs891674 | -0.831 | -1.213 | 0.316 | 1.232 | 0.001 | 0.035 | 0.035 | 0 | 0.35 | 0.48 | 0.25 | 0.45 | 0.38 | 0.59 |
| 4 | 90166778 | rs6532146 | -1.848 | -1.243 | 0.003 | 1.066 | -0.003 | 0.035 | 0.035 | 0 | 0.53 | 0.64 | 0.33 | 0.68 | 0.51 | 0.8 |
| 4 | 90166921 | rs6532148 | -0.686 | -1.215 | 0.02 | 1.083 | -0.001 | 0.032 | 0.032 | 0 | 0.36 | 0.54 | 0.26 | 0.45 | 0.38 | 0.59 |
| 4 | 90167445 | rs10516830 | -0.547 | -0.973 | -0.04 | 1.046 | 0.006 | 0 | 0 | 0.983 | 0.17 | 0.1 | 0.08 | 0.23 | 0.13 | 0.18 |
| 4 | 90168572 | rs1431548 | -1.682 | -1.218 | -0.092 | 1.107 | 0.001 | -0.004 | -0.004 | 0 | 0.48 | 0.61 | 0.33 | 0.5 | 0.49 | 0.64 |
| 4 | 90168825 | rs7672015 | -1.31 | -1.359 | 0.126 | 1.064 | 0.002 | -0.011 | -0.011 | 0 | 0.41 | 0.58 | 0.28 | 0.34 | 0.46 | 0.49 |
| 4 | 90169925 | rs7653897 | -1.759 | -0.979 | -0.083 | 0.684 | -0.011 | 0.029 | 0.029 | 1.693 | 0.52 | 0.76 | 0.35 | 0.55 | 0.57 | 0.75 |
| 4 | 90171503 | rs2298756 | -0.091 | -0.828 | -0.334 | 0.492 | -0.018 | 0.056 | 0.056 | 1.081 | 0.22 | 0.09 | 0.14 | 0.05 | 0.26 | 0.2 |
| 4 | 90180437 | rs6532149 | -1.489 | -0.875 | 0.149 | 0.73 | 0 | 0.012 | 0.012 | 0.287 | 0.5 | 0.48 | 0.35 | 0.55 | 0.56 | 0.71 |
| 4 | 90188353 | rs6532151 | -1.558 | -0.925 | 0.853 | 0.854 | 0.023 | 0.12 | 0.12 | 0 | 0.15 | 0.4 | 0.16 | 0.26 | 0.17 | 0.34 |
| 4 | 90193727 | rs6830724 | -0.646 | -1.435 | 0.575 | 0.204 | -0.017 | -0.011 | -0.011 | 0 | 0.24 | 0.37 | 0.15 | 0.23 | 0.16 | 0.22 |
| 4 | 90194997 | rs4399946 | -0.845 | -1.573 | 0.138 | -0.475 | -0.037 | -0.068 | -0.068 | 0.116 | 0.12 | 0.24 | 0.06 | 0 | 0.07 | 0.12 |
| 4 | 90198463 | rs1560489 | -1.571 | -1.004 | -0.331 | -0.232 | 0.095 | -0.02 | -0.02 | 0.178 | 0.43 | 0.91 | 0.46 | 0.37 | 0.27 | 0.47 |
| 4 | 90202027 | rs4557230 | -1.917 | -0.746 | 0.689 | -0.26 | 0.016 | -0.026 | -0.026 | 0 | 0.08 | 0.61 | 0.04 | 0 | 0.04 | 0.12 |
| 4 | 90205253 | rs12640416 | 1.42 | -0.492 | -0.675 | -0.282 | 0.054 | 0.015 | 0.015 | 0.416 | 0.49 | 0.08 | 0.47 | 0.59 | 0.62 | 0.48 |
| 4 | 90207602 | rs17015332 | 0.399 | -0.297 | -0.136 | -0.3 | 0.009 | -0.083 | -0.083 | 1.802 | 0.15 | 0.25 | 0.12 | 0 | 0.18 | 0.04 |
| 4 | 90209723 | rs937719 | 1.182 | -0.295 | -0.068 | -0.265 | -0.002 | -0.001 | -0.001 | 0.062 | 0.57 | 0.4 | 0.52 | 0.59 | 0.65 | 0.61 |
| 4 | 90211918 | rs1079070 | 1.345 | -0.312 | -0.395 | -0.266 | 0.04 | 0.012 | 0.012 | 0.403 | 0.55 | 0.16 | 0.5 | 0.59 | 0.65 | 0.6 |
| 4 | 90214108 | rs1019875 | 0.384 | -0.296 | -0.843 | -0.228 | 0.002 | -0.003 | -0.003 | 0.028 | 0.71 | 0.65 | 0.64 | 0.6 | 0.83 | 0.64 |
| 4 | 90216747 | rs7687888 | 0.877 | -0.001 | -1.285 | -0.396 | 0 | -0.009 | -0.009 | 0 | 0.59 | 0.59 | 0.47 | 0.51 | 0.63 | 0.46 |
| 4 | 90225835 | rs1560488 | -0.103 | 0.298 | -1.638 | -0.6 | 0.001 | -0.051 | -0.051 | 0.71 | 0.81 | 0.78 | 0.87 | 1 | 0.78 | 0.9 |
| 4 | 90227469 | rs7653939 | 1.23 | 0.438 | -2.205 | -0.056 | -0.095 | -0.123 | -0.123 | 0 | 0.4 | 0.03 | 0.32 | 0.54 | 0.44 | 0.43 |
| 4 | 90228238 | rs17015404 | -0.672 | 0.408 | -0.985 | 0.228 | 0.004 | 0.005 | 0.005 | 0 | 0.35 | 0.34 | 0.42 | 0.31 | 0.3 | 0.27 |
| 4 | 90233839 | rs4693992 | 0.638 | 0.098 | -1.305 | -0.111 | 0.016 | 0.103 | 0.103 | 0 | 0.77 | 0.95 | 0.88 | 0.97 | 0.84 | 0.93 |
| 4 | 90235488 | rs1431559 | -0.554 | 0.313 | -0.922 | 0.653 | 0.001 | 0.02 | 0.02 | 0.042 | 0.18 | 0.21 | 0.23 | 0.46 | 0.22 | 0.25 |
| 4 | 90238073 | rs10015406 | -1.142 | -0.232 | 0.017 | 0.615 | -0.029 | -0.008 | -0.008 | 0 | 0.28 | 0.37 | 0.33 | 0.24 | 0.2 | 0.21 |
| 4 | 90241029 | rs13104039 | -1.036 | -0.24 | 0.72 | 0.627 | 0.06 | 0.019 | 0.019 | 0 | 0.05 | 0.18 | 0.05 | 0.03 | 0.07 | 0.04 |
| 4 | 90242059 | rs10027938 | -1.29 | -0.355 | 0.794 | 0.599 | -0.05 | 0.004 | 0.004 | 0.016 | 0.25 | 0.46 | 0.3 | 0.23 | 0.15 | 0.18 |
| 4 | 90244476 | rs4106153 | -0.134 | -0.429 | 0.709 | 0.541 | -0.004 | -0.068 | -0.068 | 0 | 0.8 | 0.91 | 0.73 | 0.5 | 0.78 | 0.71 |
| 4 | 90255167 | rs1463908 | 0.999 | -0.423 | 0.882 | 0.657 | 0.027 | 0.006 | 0.006 | 0.454 | 0.6 | 0.71 | 0.48 | 0.34 | 0.7 | 0.57 |
| 4 | 90256211 | rs1875863 | 1.603 | -0.521 | 1.096 | 0.521 | 0.059 | 0.1 | 0.1 | 0.054 | 0.5 | 0.23 | 0.4 | 0.3 | 0.6 | 0.53 |
| 4 | 90259075 | rs17015445 | 0.076 | -0.378 | 1.119 | 0.56 | 0.035 | 0.005 | 0.005 | 0 | 0.05 | 0.27 | 0.04 | 0.02 | 0.04 | 0.01 |
| 4 | 90271894 | rs17015458 | 0.383 | -0.481 | 0.469 | 0.583 | -0.005 | -0.048 | -0.048 | 1.172 | 0.08 | 0.01 | 0.23 | 0.2 | 0.07 | 0.14 |
| 4 | 90272120 | rs10516833 | 0.756 | -0.299 | 0.535 | 0.443 | -0.023 | -0.007 | -0.007 | 1.067 | 0.59 | 0.51 | 0.47 | 0.34 | 0.71 | 0.58 |
| 4 | 90277638 | rs12511565 | 1.106 | -0.193 | -0.451 | 0.444 | 0.047 | 0.104 | 0.104 | 0 | 0.51 | 0.12 | 0.4 | 0.29 | 0.6 | 0.53 |
| 4 | 90280688 | rs6846715 | 0.52 | 0.235 | 0.378 | 0.821 | -0.001 | 0.007 | 0.007 | 0 | 0.76 | 0.74 | 0.78 | 0.55 | 0.78 | 0.75 |
| 4 | 90285544 | rs6821767 | -1.646 | 0.579 | 0.62 | 0.582 | 0.008 | -0.052 | -0.052 | 0 | 0.34 | 0.78 | 0.44 | 0.52 | 0.32 | 0.27 |
| 4 | 90295681 | rs1504484 | -0.806 | 0.413 | 0.804 | 0.63 | -0.007 | 0.03 | 0.03 | 0 | 0.25 | 0.3 | 0.23 | 0.52 | 0.21 | 0.23 |
| 4 | 90306233 | rs6827974 | -1.383 | 0.439 | 1.091 | 0.632 | -0.009 | 0.013 | 0.013 | 1.069 | 0.31 | 0.64 | 0.39 | 0.52 | 0.23 | 0.27 |
| 4 | 90309312 | rs12650758 | -0.777 | 0.431 | 1.16 | 0.638 | -0.005 | 0.019 | 0.019 | 0 | 0.26 | 0.31 | 0.24 | 0.52 | 0.22 | 0.23 |
| 4 | 90310346 | rs1105137 | -1.411 | 0.431 | 1.106 | 0.637 | -0.009 | 0.013 | 0.013 | 1.007 | 0.32 | 0.64 | 0.39 | 0.52 | 0.23 | 0.27 |
| 4 | 90310693 | rs1910679 | -0.858 | 0.657 | 0.316 | 0.814 | -0.003 | 0.011 | 0.011 | 0 | 0.26 | 0.29 | 0.24 | 0.52 | 0.22 | 0.23 |
| 4 | 90371779 | rs12643268 | -0.309 | 0.982 | 0.548 | 1.44 | -0.014 | 0.014 | 0.014 | 0 | 0.07 | 0.01 | 0.22 | 0.47 | 0.08 | 0.18 |
| 4 | 90371934 | rs17778445 | -0.291 | 1.518 | 0.119 | 1.284 | -0.037 | -0.016 | -0.016 | 0 | 0.08 | 0.01 | 0.08 | 0.09 | 0.11 | 0.1 |
| 4 | 90372566 | rs772492 | -0.986 | 1.451 | 0.006 | 0.084 | 0.007 | -0.089 | -0.089 | 0 | 0.73 | 0.71 | 0.8 | 1 | 0.7 | 0.89 |
| 4 | 90373071 | rs772494 | 0.03 | 1.401 | 0.92 | 0.026 | 0.003 | 0.003 | 0.003 | 0 | 0.62 | 0.44 | 0.53 | 0.53 | 0.54 | 0.7 |
| 4 | 90373435 | rs772496 | -0.36 | 1.3 | 0.664 | 0.051 | -0.003 | -0.004 | -0.004 | 1.066 | 0.66 | 0.7 | 0.58 | 0.53 | 0.62 | 0.71 |
| 4 | 90375951 | rs772501 | -0.227 | 1.306 | 0.655 | 0.084 | 0.006 | 0.017 | 0.017 | 0 | 0.65 | 0.61 | 0.58 | 0.53 | 0.62 | 0.71 |
| 4 | 90384672 | rs7691267 | 0.73 | 1.219 | 0.363 | -0.156 | -0.059 | 0.098 | 0.098 | 0 | 0.27 | 0.1 | 0.2 | 0 | 0.33 | 0.11 |
| 4 | 90385574 | rs2172895 | -0.477 | 0.589 | 0.742 | -0.152 | -0.001 | 0.002 | 0.002 | 0 | 0.45 | 0.45 | 0.48 | 0.49 | 0.49 | 0.27 |
| 4 | 90388994 | rs976595 | 0.834 | 1.21 | 0.823 | -0.332 | -0.038 | 0.055 | 0.055 | 0 | 0.28 | 0.17 | 0.21 | 0 | 0.35 | 0.11 |
| 4 | 90389965 | rs1495553 | -0.356 | 0.988 | 1.366 | 0.031 | 0.003 | 0.008 | 0.008 | 0 | 0.41 | 0.48 | 0.35 | 0.46 | 0.49 | 0.28 |
| 4 | 90395266 | rs9998962 | -0.431 | 1.293 | 1.45 | 0.017 | 0.002 | 0.007 | 0.007 | 0 | 0.41 | 0.54 | 0.35 | 0.45 | 0.49 | 0.31 |
| 4 | 90395499 | rs12644111 | -0.638 | 1.307 | 1.581 | 0.317 | 0.005 | 0.201 | 0.201 | 0.082 | 0.11 | 0.27 | 0.12 | 0.44 | 0.08 | 0.18 |
| 4 | 90404263 | rs2132281 | -1.373 | 1.207 | 1.566 | 0.232 | 0.047 | 0.248 | 0.248 | 0.842 | 0.16 | 0.4 | 0.16 | 0.45 | 0.15 | 0.16 |
| 4 | 90404465 | rs2172894 | -1.51 | 1.192 | 1.617 | 0.225 | 0.051 | 0.278 | 0.278 | 1.677 | 0.16 | 0.54 | 0.17 | 0.45 | 0.15 | 0.16 |
| 4 | 90410551 | rs12648284 | -0.024 | 1.214 | 0.674 | 0.311 | -0.006 | 0.034 | 0.034 | 0.144 | 0.09 | 0.05 | 0.1 | 0.44 | 0.08 | 0.15 |
| 4 | 90418192 | rs17174961 | 0.783 | 1.183 | 0.17 | -0.321 | -0.086 | 0.184 | 0.184 | 0 | 0.29 | 0.03 | 0.17 | 0 | 0.35 | 0.13 |
| 4 | 90421700 | rs1908556 | -1.598 | 0.285 | -0.487 | -0.643 | 0.04 | 0.032 | 0.032 | 0 | 0.82 | 0.96 | 0.81 | 0.87 | 0.87 | 0.66 |

| | | | | | | | | | | | | | | | | |
|---|---|---|---|---|---|---|---|---|---|---|---|---|---|---|---|---|
| 4 | 90422209 | rs10008041 | 0.634 | 0.808 | -0.106 | -0.46 | 0.04 | 0.003 | 0.003 | 0 | 0.06 | 0.11 | 0.12 | 0.07 | 0.12 | 0.15 |
| 4 | 90426646 | rs17015705 | 0.822 | 0.928 | -0.011 | -0.051 | -0.01 | 0.036 | 0.036 | 0.012 | 0.08 | 0.04 | 0.1 | 0.42 | 0.08 | 0.09 |
| 4 | 90429040 | rs1495552 | 0.481 | 1.559 | -0.396 | -0.643 | -0.007 | 0.002 | 0.002 | 0 | 0.36 | 0.32 | 0.33 | 0.14 | 0.47 | 0.29 |
| 4 | 90430695 | rs903600 | -0.818 | 1.215 | -0.693 | -0.785 | -0.003 | -0.005 | -0.005 | 0 | 0.74 | 0.75 | 0.64 | 0.7 | 0.69 | 0.85 |
| 4 | 90431418 | rs13146919 | 1.138 | 1.256 | -1.192 | -0.885 | -0.04 | 0.127 | 0.127 | 0.224 | 0.32 | 0.12 | 0.22 | 0.07 | 0.33 | 0.24 |
| 4 | 90432746 | rs6840873 | -0.45 | 1.465 | -0.863 | -0.543 | -0.015 | -0.005 | -0.005 | 0.838 | 0.73 | 0.76 | 0.62 | 0.73 | 0.62 | 0.84 |
| 4 | 90434672 | rs6854472 | 1.385 | 1.558 | -1.12 | -0.633 | -0.083 | 0.086 | 0.086 | 0 | 0.26 | 0.08 | 0.23 | 0.07 | 0.38 | 0.24 |
| 4 | 90435011 | rs729686 | -0.115 | 1.608 | -0.801 | -0.198 | -0.004 | -0.001 | -0.001 | 0 | 0.68 | 0.65 | 0.59 | 0.71 | 0.56 | 0.84 |
| 4 | 90439694 | rs17015738 | 0.937 | 1.328 | -0.75 | -0.01 | -0.005 | -0.052 | -0.052 | 0.015 | 0.31 | 0.13 | 0.28 | 0.51 | 0.29 | 0.35 |
| 4 | 90441242 | rs990695 | -0.935 | 0.945 | -0.499 | 0.479 | -0.022 | 0.055 | 0.055 | 0.52 | 0.8 | 0.98 | 0.87 | 0.93 | 0.81 | 0.92 |
| 4 | 90441824 | rs11097225 | 0.519 | 0.842 | -0.44 | 0.503 | -0.001 | -0.001 | -0.001 | 0 | 0.71 | 0.66 | 0.61 | 0.71 | 0.65 | 0.86 |
| 4 | 90450579 | rs4694010 | 0.965 | -0.121 | -0.329 | 0.713 | 0.002 | 0.02 | 0.02 | 0 | 0.05 | 0.06 | 0.16 | 0.14 | 0.08 | 0.04 |
| 4 | 90458283 | rs1495556 | 0.615 | -0.963 | -0.329 | 0.882 | 0.009 | 0.027 | 0.027 | 0 | 0.37 | 0.16 | 0.29 | 0.29 | 0.36 | 0.49 |
| 4 | 90461313 | rs4147250 | -0.032 | -1.006 | -0.007 | -0.076 | 0.002 | -0.019 | -0.019 | 0 | 0.4 | 0.48 | 0.33 | 0.25 | 0.39 | 0.32 |
| 4 | 90478149 | rs17780881 | 1.095 | -1.145 | -0.542 | -0.128 | 0.021 | -0.021 | -0.021 | 0 | 0.09 | 0.04 | 0.04 | 0.14 | 0.05 | 0.08 |
| 4 | 90484613 | rs1826739 | -1.019 | -0.754 | -0.178 | -0.517 | -0.008 | 0.037 | 0.037 | 0 | 0.83 | 0.93 | 0.85 | 0.91 | 0.81 | 0.89 |
| 4 | 90492766 | rs1156115 | -0.702 | -0.739 | -0.203 | -0.509 | 0.001 | 0.009 | 0.009 | 0 | 0.81 | 0.85 | 0.84 | 0.91 | 0.8 | 0.88 |
| 4 | 90494272 | rs1495557 | 0.516 | -0.774 | -0.095 | -0.594 | 0.013 | -0.059 | -0.059 | 0.213 | 0.32 | 0.56 | 0.3 | 0.11 | 0.34 | 0.24 |
| 4 | 90495946 | rs2870006 | -0.092 | -1.186 | -0.215 | -0.121 | 0.019 | 0.138 | 0.138 | 0 | 0.49 | 0.28 | 0.53 | 0.8 | 0.47 | 0.65 |
| 4 | 90522480 | rs6852411 | -1.646 | -2.176 | 0.436 | 0.547 | 0.067 | -0.024 | -0.024 | 0.5 | 0.34 | 0.77 | 0.36 | 0.54 | 0.16 | 0.37 |
| 4 | 90527344 | rs6850389 | -0.392 | -2.13 | 0.417 | 0.573 | 0.079 | -0.08 | -0.08 | 0.159 | 0.22 | 0.08 | 0.2 | 0.53 | 0.08 | 0.24 |
| 4 | 90538108 | rs10000875 | -0.425 | -1.621 | 1.068 | 0.662 | 0.034 | 0.003 | 0.003 | 2.631 | 0.06 | 0.26 | 0.04 | 0 | 0.03 | 0.03 |
| 4 | 90552920 | rs1430961 | -0.288 | -2.034 | 0.587 | 0.909 | 0.006 | 0 | 0 | 0.419 | 0.2 | 0.17 | 0.14 | 0.15 | 0.08 | 0.21 |
| 4 | 90555520 | rs12513009 | -0.658 | -2.258 | 0.83 | 0.762 | -0.023 | 0.208 | 0.208 | 0 | 0.13 | 0.31 | 0.19 | 0.38 | 0.07 | 0.12 |
| 4 | 90565218 | rs11932021 | 0.054 | -2.244 | 0.03 | 0.735 | 0.049 | -0.008 | -0.008 | 0 | 0.19 | 0.1 | 0.14 | 0.16 | 0.08 | 0.21 |
| 4 | 90572322 | rs3733449 | -1.797 | -2.368 | 0.018 | 0.672 | 0.109 | -0.016 | -0.016 | 0 | 0.39 | 0.77 | 0.38 | 0.54 | 0.18 | 0.37 |
| 4 | 90600620 | rs10213062 | -1.084 | -2.113 | 0.439 | 0.91 | -0.042 | 0.214 | 0.214 | 0.465 | 0.17 | 0.51 | 0.1 | 0.38 | 0.1 | 0.15 |
| 4 | 90602795 | rs13108925 | 0.179 | -1.569 | -0.294 | 0.848 | -0.065 | -0.041 | -0.041 | 0.074 | 0.23 | 0.05 | 0.22 | 0.27 | 0.26 | 0.28 |
| 4 | 90603419 | rs12644119 | -0.632 | -2.101 | 0.624 | 0.978 | -0.042 | 0.081 | 0.081 | 0 | 0.22 | 0.31 | 0.17 | 0.48 | 0.11 | 0.26 |
| 4 | 90606518 | rs3857047 | -1.001 | -1.708 | 0.447 | -0.515 | -0.067 | -0.147 | -0.147 | 1.126 | 0.2 | 0.45 | 0.13 | 0 | 0.15 | 0.06 |
| 4 | 90614899 | rs17016018 | -0.765 | -1.382 | 0.815 | -0.23 | 0.004 | -0.035 | -0.035 | 0.283 | 0.09 | 0.43 | 0.05 | 0 | 0.04 | 0.05 |
| 4 | 90618264 | rs17286290 | -0.509 | -2.024 | 0.282 | -0.483 | -0.006 | -0.001 | -0.001 | 0 | 0.19 | 0.15 | 0.06 | 0.07 | 0.08 | 0.05 |
| 4 | 90618526 | rs3906628 | -1.272 | -1.924 | -0.215 | -0.429 | 0.032 | 0.032 | 0.032 | 0.069 | 0.3 | 0.71 | 0.12 | 0.07 | 0.12 | 0.1 |
| 4 | 90625419 | rs3857050 | -1.156 | -1.944 | -0.443 | -0.438 | 0.072 | 0.068 | 0.068 | 0.047 | 0.3 | 0.77 | 0.13 | 0.07 | 0.12 | 0.14 |
| 4 | 90630901 | rs356174 | -0.125 | -0.967 | -0.579 | -1.143 | -0.012 | 0.041 | 0.041 | 0.867 | 0.67 | 0.77 | 0.81 | 0.93 | 0.64 | 0.84 |
| 4 | 90641340 | rs356220 | 0.406 | -1.308 | -0.256 | -0.219 | -0.017 | -0.028 | -0.028 | 0.027 | 0.61 | 0.29 | 0.55 | 0.48 | 0.61 | 0.58 |
| 4 | 90644454 | rs8180209 | -0.802 | -1.529 | -0.025 | -0.294 | -0.055 | 0.099 | 0.099 | 7.978 | 0.21 | 0.59 | 0.29 | 0.52 | 0.07 | 0.32 |
| 4 | 90674431 | rs356168 | 1.296 | -1.636 | -0.213 | -0.442 | -0.043 | 0.001 | 0.001 | 0 | 0.48 | 0.18 | 0.49 | 0.41 | 0.52 | 0.48 |
| 4 | 90678541 | rs2736990 | 1.296 | -1.636 | -0.213 | -0.442 | -0.043 | 0.001 | 0.001 | 0 | 0.48 | 0.18 | 0.49 | 0.41 | 0.52 | 0.48 |
| 4 | 90687507 | rs3796661 | 0.28 | -1.415 | 0.513 | -0.026 | -0.015 | 0.202 | 0.202 | 0.724 | 0.07 | 0.13 | 0.25 | 0.54 | 0.02 | 0.28 |
| 4 | 90709741 | rs3775439 | -1.367 | -1.507 | 1.117 | -0.104 | -0.07 | 0.134 | 0.134 | 1.316 | 0.25 | 0.52 | 0.31 | 0.54 | 0.12 | 0.37 |
| 4 | 90712629 | rs10014396 | -0.933 | -1.51 | 1.257 | 0.464 | -0.014 | 0.042 | 0.042 | 0.019 | 0.14 | 0.34 | 0.19 | 0.18 | 0.1 | 0.26 |
| 4 | 90713064 | rs12502363 | 1.061 | -2.108 | 0.377 | -0.021 | -0.055 | -0.022 | -0.022 | 0 | 0.3 | 0.12 | 0.26 | 0.29 | 0.37 | 0.25 |
| 4 | 90715231 | rs3775442 | 1.636 | -2.077 | 0.415 | 0.025 | -0.029 | 0.067 | 0.067 | 5.19 | 0.1 | 0.15 | 0.13 | 0.36 | 0.02 | 0.11 |
| 4 | 90716852 | rs3889917 | 0.153 | -2.132 | 0.498 | 0.002 | -0.018 | 0.017 | 0.017 | 0 | 0.12 | 0.19 | 0.16 | 0.18 | 0.07 | 0.23 |
| 4 | 90734535 | rs894278 | 0.824 | -2.102 | 0.377 | 0.022 | -0.028 | 0.171 | 0.171 | 2.272 | 0.13 | 0.35 | 0.15 | 0.36 | 0.06 | 0.14 |
| 4 | 90736113 | rs1442144 | -1.184 | -2.054 | 0.289 | -0.153 | 0.025 | 0.022 | 0.022 | 0 | 0.72 | 0.8 | 0.73 | 0.84 | 0.8 | 0.71 |
| 4 | 90736517 | rs3822095 | 1.005 | -1.904 | 0.365 | -0.058 | -0.029 | -0.009 | -0.009 | 0 | 0.31 | 0.2 | 0.27 | 0.29 | 0.37 | 0.25 |
| 4 | 90739539 | rs1812923 | -0.044 | -1.9 | 0.325 | -0.072 | -0.005 | 0.009 | 0.009 | 0.046 | 0.44 | 0.57 | 0.42 | 0.65 | 0.43 | 0.38 |
| 4 | 90743331 | rs10005233 | -0.682 | -1.863 | 0.202 | -0.167 | -0.016 | 0.124 | 0.124 | 1.095 | 0.56 | 0.76 | 0.58 | 0.83 | 0.5 | 0.61 |
| 4 | 90746709 | rs11097238 | 0.974 | -1.818 | 0.165 | -0.118 | -0.04 | -0.013 | -0.013 | 0 | 0.31 | 0.15 | 0.26 | 0.29 | 0.37 | 0.25 |
| 4 | 90758945 | rs2301134 | -0.692 | -1.777 | 0.168 | -0.255 | -0.012 | 0.118 | 0.118 | 0.322 | 0.53 | 0.69 | 0.59 | 0.84 | 0.49 | 0.6 |
| 4 | 90761412 | rs1023777 | 0.913 | -1.762 | 0.192 | -0.197 | -0.025 | -0.015 | -0.015 | 0 | 0.28 | 0.21 | 0.27 | 0.29 | 0.37 | 0.22 |
| 4 | 90770255 | rs12331318 | -0.487 | -1.822 | 0.312 | -0.213 | -0.001 | 0.099 | 0.099 | 0.305 | 0.5 | 0.63 | 0.5 | 0.84 | 0.48 | 0.55 |
| 4 | 90770408 | rs17016279 | 0.961 | -1.761 | 0.276 | -0.035 | -0.002 | 0.011 | 0.011 | 0.316 | 0.09 | 0.1 | 0.19 | 0.23 | 0.06 | 0.18 |
| 4 | 90775212 | rs10516848 | 0.853 | -1.677 | 0.187 | -0.074 | -0.025 | -0.015 | -0.015 | 0 | 0.28 | 0.21 | 0.27 | 0.29 | 0.37 | 0.22 |
| 4 | 90779823 | rs2737026 | -0.847 | -1.833 | 0.152 | -0.215 | 0.016 | 0.016 | 0.016 | 0 | 0.67 | 0.71 | 0.65 | 0.85 | 0.78 | 0.64 |
| 4 | 90780902 | rs6532194 | -1.24 | -1.597 | 0.853 | 0.273 | -0.09 | 0.115 | 0.115 | 0.429 | 0.25 | 0.51 | 0.32 | 0.55 | 0.09 | 0.32 |
| 4 | 90781261 | rs6532196 | -0.061 | -1.524 | 1.087 | 0.284 | -0.069 | 0.072 | 0.072 | 1.657 | 0.15 | 0.3 | 0.05 | 0.31 | 0.03 | 0.09 |
| 4 | 90789539 | rs10516849 | -0.365 | -1.495 | 1.143 | 0.279 | -0.041 | 0.05 | 0.05 | 1.27 | 0.16 | 0.28 | 0.06 | 0.31 | 0.07 | 0.13 |
| 4 | 90795804 | rs11097239 | -0.082 | -1.44 | 1.05 | 0.372 | -0.008 | -0.019 | -0.019 | 0.866 | 0.34 | 0.23 | 0.4 | 0.45 | 0.37 | 0.42 |
| 4 | 90797301 | rs6532197 | -0.287 | -1.722 | 1.164 | 0.446 | -0.052 | 0.084 | 0.084 | 1.274 | 0.16 | 0.37 | 0.08 | 0.32 | 0.07 | 0.13 |
| 4 | 90802998 | rs1899389 | -1.285 | -1.628 | 1.095 | 0.314 | -0.004 | 0.059 | 0.059 | 0 | 0.53 | 0.68 | 0.6 | 0.77 | 0.51 | 0.57 |
| 4 | 90816023 | rs1442140 | -1.428 | -1.633 | 1.114 | 0.312 | -0.014 | 0.082 | 0.082 | 0 | 0.54 | 0.75 | 0.61 | 0.77 | 0.5 | 0.56 |
| 4 | 90816294 | rs1442138 | -0.632 | -1.479 | 1.154 | 0.444 | 0.014 | -0.024 | -0.024 | 1.399 | 0.05 | 0.17 | 0.05 | 0 | 0.06 | 0.05 |
| 4 | 90818857 | rs1442137 | 0.512 | -1.593 | 0.746 | 0.512 | 0.009 | -0.01 | -0.01 | 0 | 0.12 | 0.09 | 0.2 | 0.23 | 0.06 | 0.23 |
| 4 | 90826760 | rs3775464 | 0.525 | -1.586 | 0.824 | 0.488 | -0.045 | 0.072 | 0.072 | 0.888 | 0.13 | 0.36 | 0.04 | 0.29 | 0.03 | 0.08 |
| 4 | 90827466 | rs3775465 | 0.544 | -1.513 | 0.739 | 0.49 | -0.016 | 0.044 | 0.044 | 1.035 | 0.13 | 0.24 | 0.04 | 0.29 | 0.07 | 0.11 |
| 4 | 90833372 | rs9307077 | -1.638 | -1.281 | 0.452 | 0.258 | 0.006 | 0.046 | 0.046 | 0 | 0.55 | 0.7 | 0.62 | 0.76 | 0.55 | 0.58 |

| | | | | | | | | | | | | | | | | |
|---|---|---|---|---|---|---|---|---|---|---|---|---|---|---|---|---|
| 4 | 90833421 | rs10011790 | 0.674 | -0.675 | -0.024 | 0.426 | -0.037 | -0.006 | -0.006 | 4.466 | 0.22 | 0.1 | 0.28 | 0.21 | 0.35 | 0.27 |
| 4 | 90837127 | rs10516850 | -0.637 | -1.09 | -0.041 | 0.403 | -0.03 | 0.027 | 0.027 | 0.98 | 0.18 | 0.29 | 0.06 | 0.29 | 0.08 | 0.12 |
| 4 | 90842840 | rs3775478 | -0.636 | -1.067 | -0.167 | 0.415 | -0.037 | 0.037 | 0.037 | 1.188 | 0.18 | 0.31 | 0.07 | 0.29 | 0.08 | 0.12 |
| 4 | 90844848 | rs2241884 | 0.695 | -0.834 | -0.611 | 0.612 | -0.016 | 0.022 | 0.022 | 0.544 | 0.12 | 0.15 | 0.03 | 0.29 | 0.03 | 0.08 |
| 4 | 90862372 | rs10030931 | 0.337 | -0.184 | -1.06 | 0.022 | -0.11 | -0.034 | -0.034 | 0 | 0.23 | 0.06 | 0.28 | 0.23 | 0.38 | 0.31 |
| 4 | 90870359 | rs3775483 | -1.033 | -0.473 | -0.763 | -0.367 | -0.001 | 0.011 | 0.011 | 0 | 0.61 | 0.65 | 0.64 | 0.89 | 0.58 | 0.74 |
| 4 | 90871662 | rs1479429 | 0.507 | -0.32 | -0.938 | -0.345 | -0.047 | 0.001 | 0.001 | 0 | 0.26 | 0.16 | 0.26 | 0.24 | 0.34 | 0.3 |
| 4 | 90875910 | rs13109927 | -2.019 | -0.55 | -0.838 | -0.375 | -0.037 | 0.1 | 0.1 | 0 | 0.68 | 0.87 | 0.67 | 0.88 | 0.6 | 0.73 |
| 4 | 90879406 | rs7667606 | -0.109 | -0.751 | -0.788 | -0.478 | -0.015 | 0.025 | 0.025 | 0 | 0.37 | 0.26 | 0.32 | 0.25 | 0.4 | 0.34 |
| 4 | 90880403 | rs12233759 | -0.426 | -0.784 | -0.744 | -0.478 | -0.029 | 0.021 | 0.021 | 0 | 0.3 | 0.55 | 0.34 | 0.64 | 0.2 | 0.39 |
| 4 | 90882154 | rs11939062 | 0.46 | -0.737 | -1.018 | -0.455 | -0.072 | 0.016 | 0.016 | 0 | 0.32 | 0.1 | 0.3 | 0.25 | 0.39 | 0.33 |
| 4 | 90883047 | rs11931138 | -1.657 | -0.413 | -0.838 | -0.469 | -0.025 | 0.089 | 0.089 | 0 | 0.66 | 0.82 | 0.67 | 0.88 | 0.6 | 0.73 |
| 4 | 90895017 | rs13107488 | -1.767 | -0.401 | -0.874 | -0.46 | -0.035 | 0.092 | 0.092 | 0 | 0.68 | 0.85 | 0.67 | 0.89 | 0.6 | 0.74 |
| 4 | 90910930 | rs6532210 | -1.003 | -0.433 | -0.031 | -0.405 | -0.001 | 0.007 | 0.007 | 0 | 0.59 | 0.4 | 0.53 | 0.86 | 0.57 | 0.65 |
| 4 | 90916437 | rs10516854 | -1.715 | -0.539 | -0.074 | -0.402 | -0.011 | 0.057 | 0.057 | 0 | 0.66 | 0.79 | 0.67 | 0.88 | 0.61 | 0.74 |
| 4 | 90928959 | rs13113259 | -1.308 | -0.726 | 0.049 | -0.632 | 0.013 | 0.053 | 0.053 | 0 | 0.59 | 0.71 | 0.75 | 0.88 | 0.61 | 0.72 |
| 4 | 90938575 | rs11728457 | -1.527 | -0.907 | -0.166 | -0.481 | 0.034 | 0.004 | 0.004 | 0 | 0.39 | 0.7 | 0.49 | 0.63 | 0.24 | 0.39 |
| 4 | 90942824 | rs6812321 | -1.488 | -0.817 | -0.132 | -0.505 | 0.015 | 0.038 | 0.038 | 0 | 0.57 | 0.66 | 0.66 | 0.86 | 0.62 | 0.65 |
| 4 | 90947793 | rs17808648 | 0.88 | -0.799 | -0.2 | -0.499 | -0.138 | -0.052 | -0.052 | 0.012 | 0.26 | 0.03 | 0.28 | 0.25 | 0.38 | 0.33 |
| 4 | 90958046 | rs17016459 | -0.487 | -0.795 | 0.085 | -0.475 | -0.023 | 0.046 | 0.046 | 0 | 0.28 | 0.4 | 0.27 | 0.61 | 0.23 | 0.31 |
| 4 | 90963446 | rs724454 | -0.625 | -0.801 | 0.028 | -0.466 | -0.028 | 0.05 | 0.05 | 0 | 0.29 | 0.43 | 0.27 | 0.61 | 0.23 | 0.31 |
| 4 | 90968156 | rs1443797 | -2.203 | -0.824 | 0.006 | -0.529 | 0 | 0.018 | 0.018 | 0 | 0.66 | 0.74 | 0.77 | 0.88 | 0.63 | 0.72 |
| 4 | 90969565 | rs11929845 | -0.56 | -0.843 | 0.191 | -0.168 | -0.027 | 0.074 | 0.074 | 0 | 0.29 | 0.41 | 0.27 | 0.58 | 0.23 | 0.32 |
| 4 | 90971745 | rs722937 | -2.206 | -0.773 | 0.002 | -0.139 | -0.003 | 0.041 | 0.041 | 0 | 0.66 | 0.8 | 0.77 | 0.84 | 0.63 | 0.72 |
| 4 | 90971800 | rs722936 | 0.638 | -0.786 | 0.15 | 0.597 | -0.008 | -0.023 | -0.023 | 0 | 0.14 | 0.09 | 0.06 | 0.33 | 0.16 | 0.19 |
| 4 | 90976413 | rs13114320 | 0.361 | -0.611 | -0.108 | 0.354 | -0.04 | 0.005 | 0.005 | 0 | 0.28 | 0.21 | 0.39 | 0.24 | 0.39 | 0.33 |
| 4 | 90993172 | rs894803 | -2.034 | -0.846 | -0.301 | 0.285 | -0.01 | 0.065 | 0.065 | 0 | 0.63 | 0.83 | 0.57 | 0.84 | 0.6 | 0.66 |
| 4 | 91000334 | rs2197291 | -1.711 | -0.949 | -0.288 | 0.253 | -0.002 | 0.077 | 0.077 | 0 | 0.58 | 0.73 | 0.52 | 0.84 | 0.55 | 0.63 |
| 4 | 91025850 | rs34001515 | 0.725 | -0.74 | -0.358 | 0.365 | -0.086 | -0.002 | -0.002 | 0.188 | 0.25 | 0.1 | 0.27 | 0.23 | 0.38 | 0.28 |
| 4 | 91033300 | rs7674520 | -2.091 | -0.806 | -0.365 | 0.34 | -0.013 | 0.085 | 0.085 | 0 | 0.62 | 0.82 | 0.57 | 0.85 | 0.59 | 0.65 |
| 4 | 91046719 | rs12644131 | -1.691 | -1.331 | 0.205 | 0.818 | 0.027 | -0.009 | -0.009 | 5.248 | 0.37 | 0.7 | 0.3 | 0.62 | 0.21 | 0.37 |
| 4 | 91069888 | rs6837278 | -1.163 | -1.121 | 0.436 | 1.018 | -0.038 | 0.036 | 0.036 | 0 | 0.29 | 0.53 | 0.23 | 0.63 | 0.17 | 0.31 |
| 4 | 91123665 | rs10461168 | -1.208 | -0.166 | 0.049 | 0.999 | -0.019 | 0.014 | 0.014 | 0 | 0.58 | 0.75 | 0.5 | 0.59 | 0.47 | 0.64 |
| 4 | 91123874 | rs13109951 | 0.383 | -0.251 | -0.115 | 1.232 | -0.026 | -0.048 | -0.048 | 0 | 0.22 | 0.07 | 0.29 | 0.22 | 0.2 | 0.31 |
| 4 | 91156917 | rs4356886 | 0.09 | -0.144 | 0.677 | 1.265 | 0.049 | 0.227 | 0.227 | 0 | 0.08 | 0.34 | 0.07 | 0.31 | 0.1 | 0.12 |
| 4 | 91194806 | rs1350855 | -1.081 | -0.349 | 0.774 | 1.189 | 0.022 | 0.027 | 0.027 | 0 | 0.48 | 0.38 | 0.41 | 0.29 | 0.35 | 0.48 |
| 4 | 91210048 | rs13147804 | -1.179 | -0.357 | 0.645 | 1.226 | 0.029 | 0.035 | 0.035 | 0.062 | 0.49 | 0.39 | 0.41 | 0.29 | 0.35 | 0.47 |
| 4 | 91212092 | rs10516867 | 0.322 | -0.523 | 0.496 | 1.084 | 0.034 | 0.024 | 0.024 | 0 | 0.19 | 0.04 | 0.08 | 0.1 | 0.13 | 0.12 |
| 4 | 91227681 | rs1037926 | -0.88 | 0.067 | 0.761 | 1.304 | 0.008 | 0.002 | 0.002 | 0 | 0.51 | 0.45 | 0.51 | 0.51 | 0.38 | 0.56 |
| 4 | 91230579 | rs12647859 | 0.506 | 0.178 | 1.004 | 1.386 | 0.028 | 0.148 | 0.148 | 0 | 0.08 | 0.24 | 0.06 | 0.3 | 0.11 | 0.11 |
| 4 | 91234498 | rs2870246 | -1.563 | 0.213 | 0.621 | 0.982 | -0.012 | 0.028 | 0.028 | 0 | 0.61 | 0.73 | 0.59 | 0.82 | 0.52 | 0.71 |
| 4 | 91247973 | rs11723677 | 0.84 | -0.012 | 0.29 | 1.331 | -0.058 | -0.066 | -0.066 | 0.254 | 0.17 | 0.01 | 0.23 | 0.2 | 0.19 | 0.26 |
| 4 | 91253355 | rs10015182 | -0.01 | 0.176 | 0.739 | 1.524 | -0.028 | 0.077 | 0.077 | 3.288 | 0.27 | 0.49 | 0.33 | 0.4 | 0.2 | 0.37 |
| 4 | 91254945 | rs7693233 | -1.694 | 0.306 | 0.545 | 1.323 | -0.052 | 0.072 | 0.072 | 0 | 0.64 | 0.82 | 0.5 | 0.82 | 0.49 | 0.75 |
| 4 | 91269591 | rs12642077 | -1.953 | 0.325 | 0.515 | 1.386 | -0.096 | 0.099 | 0.099 | 0 | 0.66 | 0.87 | 0.54 | 0.82 | 0.53 | 0.79 |
| 4 | 91271176 | rs8180269 | -0.596 | 0.252 | 0.6 | 1.414 | 0.013 | -0.011 | -0.011 | 0 | 0.37 | 0.29 | 0.17 | 0.43 | 0.29 | 0.38 |
| 4 | 91280571 | rs4693236 | -1.538 | 0.338 | 0.52 | 1.403 | -0.058 | 0.011 | 0.011 | 0 | 0.63 | 0.8 | 0.44 | 0.65 | 0.49 | 0.69 |
| 4 | 91287263 | rs17016912 | -0.992 | 0.317 | 0.49 | 0.972 | 0.004 | 0.007 | 0.007 | 0 | 0.54 | 0.55 | 0.37 | 0.33 | 0.41 | 0.61 |
| 4 | 91289294 | rs6842950 | -0.037 | 0.534 | 1.161 | 1.639 | -0.043 | 0.002 | 0.002 | 4.559 | 0.25 | 0.51 | 0.26 | 0.22 | 0.16 | 0.3 |
| 4 | 91296454 | rs1903577 | -0.23 | 0.996 | 1.561 | 1.617 | -0.016 | -0.004 | -0.004 | 0.402 | 0.4 | 0.51 | 0.32 | 0.33 | 0.28 | 0.45 |
| 4 | 91301321 | rs7688033 | -0.347 | 1.015 | 1.275 | 1.646 | 0.017 | 0.229 | 0.229 | 0 | 0.12 | 0.3 | 0.17 | 0.49 | 0.12 | 0.18 |
| 4 | 91307525 | rs2169398 | -0.624 | 0.881 | 1.202 | 1.501 | -0.012 | -0.003 | -0.003 | 0.135 | 0.4 | 0.45 | 0.32 | 0.33 | 0.28 | 0.45 |
| 4 | 91319038 | rs6834254 | -2.382 | 1.588 | 2.024 | 1.848 | 0.004 | 0.072 | 0.072 | 1.917 | 0.14 | 0.68 | 0.17 | 0.34 | 0.13 | 0.17 |
| 4 | 91340811 | rs12645541 | -1.365 | 1.513 | 2.571 | 1.89 | 0.008 | 0.207 | 0.207 | 0 | 0.07 | 0.2 | 0.12 | 0.35 | 0.06 | 0.13 |
| 4 | 91367607 | rs10002613 | -1.799 | 1.775 | 3.057 | 1.823 | 0.1 | 0.279 | 0.279 | 0.007 | 0.1 | 0.41 | 0.16 | 0.36 | 0.13 | 0.16 |
| 4 | 91371416 | rs6843585 | -2.036 | 1.768 | 3.104 | 1.82 | 0.082 | 0.249 | 0.249 | 0.479 | 0.12 | 0.52 | 0.17 | 0.36 | 0.13 | 0.18 |
| 4 | 91428124 | rs11097253 | -2.255 | 1.737 | 2.318 | 1.792 | 0.038 | 0.158 | 0.158 | 1.119 | 0.14 | 0.61 | 0.17 | 0.35 | 0.14 | 0.19 |
| 4 | 91460581 | rs2036153 | -1.973 | 1.637 | 0.52 | 1.205 | -0.065 | -0.114 | -0.114 | 0.434 | 0.35 | 0.94 | 0.31 | 0.47 | 0.34 | 0.4 |
| 4 | 91464162 | rs17017104 | -0.987 | 1.832 | 1.536 | 1.131 | 0.025 | -0.118 | -0.118 | 0 | 0.17 | 0.41 | 0.14 | 0.01 | 0.18 | 0.16 |
| 4 | 91465389 | rs6532233 | -2.35 | 1.839 | 1.979 | 1.133 | -0.007 | -0.015 | -0.015 | 0.221 | 0.32 | 0.77 | 0.29 | 0.47 | 0.34 | 0.39 |
| 4 | 91465511 | rs6849123 | -1.269 | 1.809 | 1.756 | 1.541 | 0.031 | 0.233 | 0.233 | 0 | 0.08 | 0.25 | 0.12 | 0.42 | 0.12 | 0.1 |
| 4 | 91469221 | rs6822624 | -0.754 | 1.801 | 1.39 | 1.505 | -0.002 | 0.001 | 0.001 | 0 | 0.07 | 0.06 | 0.03 | 0.04 | 0.04 | 0.13 |
| 4 | 91475787 | rs10010423 | -1.723 | 1.787 | 1.552 | 1.433 | 0.039 | 0.224 | 0.224 | 0.008 | 0.15 | 0.43 | 0.14 | 0.45 | 0.16 | 0.21 |
| 4 | 91481271 | rs6845935 | -1.29 | 1.826 | 1.26 | 0.714 | 0.002 | -0.043 | -0.043 | 0 | 0.21 | 0.25 | 0.15 | 0.02 | 0.22 | 0.2 |
| 4 | 91500284 | rs2008600 | -2.025 | 2.13 | 2.598 | 1.152 | 0.046 | 0.105 | 0.105 | 0 | 0.28 | 0.47 | 0.29 | 0.5 | 0.33 | 0.27 |
| 4 | 91501614 | rs1510795 | -2.206 | 2.097 | 2.739 | 1.139 | -0.006 | -0.013 | -0.013 | 1.156 | 0.31 | 0.7 | 0.31 | 0.51 | 0.34 | 0.38 |
| 4 | 91514884 | rs2048398 | -2.435 | 1.597 | 2.356 | 1.371 | -0.023 | -0.082 | -0.082 | 0.948 | 0.35 | 0.88 | 0.33 | 0.56 | 0.31 | 0.4 |
| 4 | 91516040 | rs10031325 | -1.35 | 1.716 | 2.899 | 1.459 | 0.043 | 0.132 | 0.132 | 1.392 | 0.18 | 0.3 | 0.26 | 0.53 | 0.26 | 0.24 |
| 4 | 91516422 | rs17017247 | -1.709 | 1.715 | 2.98 | 1.458 | 0.053 | 0.174 | 0.174 | 1.062 | 0.19 | 0.45 | 0.28 | 0.53 | 0.26 | 0.24 |

| | | | | | | | | | | | | | | | | |
|---|---|---|---|---|---|---|---|---|---|---|---|---|---|---|---|---|
| 4 | 91526531 | rs7687903 | -0.684 | 1.718 | 2.735 | 1.976 | -0.012 | -0.016 | -0.016 | 0 | 0.12 | 0.09 | 0.16 | 0.34 | 0.18 | 0.11 |
| 4 | 91532461 | rs9884326 | -2.661 | 1.465 | 2.318 | 1.279 | -0.015 | -0.061 | -0.061 | 0.089 | 0.43 | 0.87 | 0.32 | 0.55 | 0.37 | 0.45 |
| 4 | 91533617 | rs17240838 | -1.148 | 1.381 | 2.557 | 1.022 | -0.001 | -0.044 | -0.044 | 0 | 0.19 | 0.23 | 0.05 | 0.01 | 0.11 | 0.11 |
| 4 | 91546380 | rs9790562 | -1.637 | 1.436 | 2.323 | 1.181 | 0 | 0 | 0 | 0.63 | 0.32 | 0.41 | 0.22 | 0.36 | 0.29 | 0.21 |
| 4 | 91566875 | rs7672803 | -0.919 | 1.881 | 1.213 | 1.072 | -0.01 | 0.004 | 0.004 | 0 | 0.2 | 0.12 | 0.2 | 0.2 | 0.25 | 0.17 |
| 4 | 91567112 | rs950127 | -1.524 | 2.707 | 1.233 | 2.037 | -0.007 | 0.006 | 0.006 | 0 | 0.09 | 0.07 | 0.29 | 0.32 | 0.16 | 0.18 |
| 4 | 91567979 | rs2132834 | -0.405 | 2.601 | 1.353 | 1.561 | 0.005 | 0.104 | 0.104 | 0 | 0.21 | 0.46 | 0.17 | 0.36 | 0.22 | 0.37 |
| 4 | 91568501 | rs10014992 | 0.332 | 1.93 | 0.116 | 0.63 | 0.002 | -0.002 | -0.002 | 0 | 0.56 | 0.39 | 0.63 | 0.55 | 0.6 | 0.49 |
| 4 | 91573596 | rs17017347 | 0.707 | 1.916 | 0.151 | 1.099 | 0.024 | 0.012 | 0.012 | 0.073 | 0.62 | 0.49 | 0.43 | 0.42 | 0.55 | 0.58 |
| 4 | 91574795 | rs2088386 | 0.707 | 1.916 | 0.151 | 1.075 | 0.024 | 0.014 | 0.014 | 0.073 | 0.62 | 0.49 | 0.43 | 0.44 | 0.55 | 0.58 |
| 4 | 91574914 | rs10022603 | 0.107 | 1.977 | -0.674 | 0.677 | -0.033 | 0.023 | 0.023 | 0 | 0.05 | 0 | 0.04 | 0.02 | 0.1 | 0.14 |
| 4 | 91576994 | rs1399409 | -0.888 | 1.557 | -0.468 | 0.711 | 0.006 | 0.022 | 0.022 | 0 | 0.79 | 0.84 | 0.82 | 0.91 | 0.83 | 0.86 |
| 4 | 91591467 | rs1567929 | 0.801 | 1.704 | -0.197 | 0.713 | 0.003 | -0.011 | -0.011 | 0 | 0.57 | 0.51 | 0.37 | 0.24 | 0.51 | 0.49 |
| 4 | 91606389 | rs13113267 | -0.988 | 1.575 | -0.585 | 0.587 | -0.001 | -0.019 | -0.019 | 0 | 0.87 | 0.87 | 0.9 | 0.99 | 0.88 | 0.9 |
| 4 | 91607363 | rs6856591 | 0.137 | 1.704 | -0.576 | 1.018 | -0.005 | -0.006 | -0.006 | 0.279 | 0.8 | 0.82 | 0.63 | 0.69 | 0.76 | 0.73 |
| 4 | 91630043 | rs17017439 | -1.157 | 1.307 | -0.327 | 0.953 | 0 | 0.004 | 0.004 | 0 | 0.3 | 0.61 | 0.33 | 0.38 | 0.29 | 0.29 |
| 4 | 91632410 | rs13150060 | 0.066 | 1.764 | -0.419 | 1.205 | 0 | 0.022 | 0.022 | 0.013 | 0.08 | 0.09 | 0.24 | 0.28 | 0.15 | 0.2 |
| 4 | 91633300 | rs7660162 | -0.563 | 1.629 | 0.157 | 1.146 | -0.072 | 0.013 | 0.013 | 0 | 0.21 | 0.45 | 0.22 | 0.27 | 0.12 | 0.19 |
| 4 | 91636530 | rs17017443 | -0.319 | 1.548 | 0.523 | 1.164 | -0.037 | 0.024 | 0.024 | 0 | 0.13 | 0.34 | 0.08 | 0.15 | 0.07 | 0.15 |
| 4 | 91638789 | rs17183896 | 1.144 | 1.065 | -0.403 | 0.629 | -0.007 | -0.02 | -0.02 | 0 | 0.23 | 0.11 | 0.17 | 0.26 | 0.24 | 0.28 |
| 4 | 91653148 | rs17017488 | -1.058 | 1.083 | -0.348 | 0.634 | 0.03 | -0.008 | -0.008 | 0 | 0.45 | 0.66 | 0.4 | 0.53 | 0.35 | 0.46 |
| 4 | 91661745 | rs1355993 | 2.651 | 0.788 | -0.561 | 0.385 | 0.015 | 0.133 | 0.133 | 0.244 | 0.25 | 0.01 | 0.17 | 0.02 | 0.19 | 0.14 |
| 4 | 91664577 | rs1399419 | -0.462 | 0.993 | -0.31 | 0.706 | 0.058 | 0.078 | 0.078 | 0 | 0.17 | 0.6 | 0.19 | 0.3 | 0.23 | 0.21 |
| 4 | 91667562 | rs1114878 | -0.486 | 0.958 | -0.154 | 0.686 | 0.06 | 0.084 | 0.084 | 0 | 0.17 | 0.58 | 0.19 | 0.29 | 0.24 | 0.21 |
| 4 | 91671650 | rs17244178 | 1.595 | 0.915 | -0.416 | 0.702 | -0.049 | -0.035 | -0.035 | 0.153 | 0.21 | 0.02 | 0.13 | 0.22 | 0.21 | 0.24 |
| 4 | 91674460 | rs6532239 | -1.197 | 0.924 | -0.304 | 0.657 | -0.006 | 0.026 | 0.026 | 0 | 0.47 | 0.69 | 0.48 | 0.67 | 0.52 | 0.52 |
| 4 | 91676401 | rs6833239 | 0.825 | 0.923 | -0.448 | 0.682 | -0.002 | -0.044 | -0.044 | 0 | 0.3 | 0.12 | 0.3 | 0.37 | 0.28 | 0.31 |
| 4 | 91680245 | rs1514742 | -1.601 | 1.054 | -0.507 | 0.654 | 0.044 | 0.029 | 0.029 | 0 | 0.56 | 0.78 | 0.56 | 0.69 | 0.65 | 0.68 |
| 4 | 91687322 | rs7667003 | 1.052 | 1.062 | -0.264 | 0.71 | 0.012 | 0.003 | 0.003 | 0 | 0.07 | 0.07 | 0.09 | 0.14 | 0.16 | 0.1 |
| 4 | 91691313 | rs7693519 | 1.716 | 0.991 | -0.281 | 0.603 | -0.02 | 0.051 | 0.051 | 0.083 | 0.33 | 0.2 | 0.27 | 0.16 | 0.35 | 0.24 |
| 4 | 91695730 | rs7685883 | -1.003 | 1.397 | 0.36 | 1.225 | 0.057 | 0.081 | 0.081 | 0 | 0.15 | 0.62 | 0.19 | 0.29 | 0.22 | 0.19 |
| 4 | 91700548 | rs11727861 | 0.95 | 1.453 | 0.245 | 1.189 | -0.033 | -0.033 | -0.033 | 0.819 | 0.06 | 0.02 | 0.09 | 0.14 | 0.16 | 0.1 |
| 4 | 91709484 | rs1869539 | -1.058 | 1.298 | 0.245 | 1.105 | -0.019 | -0.016 | -0.016 | 0 | 0.39 | 0.7 | 0.35 | 0.56 | 0.45 | 0.46 |
| 4 | 91711665 | rs11097264 | -0.946 | 1.238 | 0.174 | 1.058 | -0.029 | -0.026 | -0.026 | 0 | 0.37 | 0.73 | 0.37 | 0.56 | 0.45 | 0.46 |
| 4 | 91715008 | rs11932857 | -0.666 | 1.367 | 0.784 | 1.192 | 0.096 | 0.149 | 0.149 | 0 | 0.19 | 0.47 | 0.31 | 0.4 | 0.27 | 0.21 |
| 4 | 91722598 | rs17246676 | 1.032 | 1.344 | 0.482 | 1.187 | -0.051 | -0.052 | -0.052 | 0 | 0.23 | 0.03 | 0.17 | 0.26 | 0.25 | 0.3 |
| 4 | 91725803 | rs10856879 | -1.674 | 1.305 | 0.668 | 1.21 | 0.001 | 0.032 | 0.032 | 0 | 0.47 | 0.64 | 0.5 | 0.68 | 0.54 | 0.54 |
| 4 | 91732390 | rs17187375 | 0.452 | 1.503 | 0.933 | 1.349 | -0.005 | -0.014 | -0.014 | 0 | 0.06 | 0.02 | 0.08 | 0.14 | 0.17 | 0.1 |
| 4 | 91737500 | rs17017652 | -0.749 | 1.445 | 1.113 | 1.299 | 0.082 | 0.136 | 0.136 | 0 | 0.18 | 0.4 | 0.3 | 0.42 | 0.27 | 0.21 |
| 4 | 91740550 | rs10516879 | -0.554 | 1.581 | 1.185 | 1.337 | 0.165 | 0.21 | 0.21 | 0 | 0.09 | 0.29 | 0.16 | 0.3 | 0.2 | 0.17 |
| 4 | 91748644 | rs17017704 | 0.135 | 0.895 | 0.944 | 0.809 | -0.035 | -0.037 | -0.037 | 0 | 0.07 | 0.18 | 0.03 | 0 | 0.02 | 0.01 |
| 4 | 91762984 | rs10021524 | -1.162 | 0.751 | 0.801 | 0.806 | 0.009 | 0.001 | 0.001 | 0.185 | 0.47 | 0.44 | 0.52 | 0.47 | 0.64 | 0.53 |
| 4 | 91764098 | rs11725878 | -1.383 | 0.682 | 0.94 | 0.786 | 0.007 | 0.001 | 0.001 | 0.457 | 0.49 | 0.45 | 0.52 | 0.49 | 0.65 | 0.53 |
| 4 | 91764932 | rs6822939 | 1.231 | 1.098 | 0.753 | 0.783 | -0.014 | 0.002 | 0.002 | 0 | 0.21 | 0.09 | 0.17 | 0.2 | 0.24 | 0.28 |
| 4 | 91767861 | rs9998288 | -0.055 | 1.365 | 0.995 | 0.758 | 0.023 | 0.006 | 0.006 | 0 | 0.17 | 0.18 | 0.27 | 0.25 | 0.3 | 0.14 |
| 4 | 91768115 | rs10010346 | 0.153 | 1.477 | 0.996 | 0.781 | 0.134 | 0.053 | 0.053 | 0 | 0.08 | 0.16 | 0.13 | 0.14 | 0.22 | 0.1 |
| 4 | 91772546 | rs17195440 | 0.355 | 1.442 | 0.99 | 0.768 | -0.001 | 0.008 | 0.008 | 0 | 0.31 | 0.18 | 0.32 | 0.3 | 0.31 | 0.32 |
| 4 | 91775333 | rs11936158 | -1.414 | 1.43 | 0.698 | 0.76 | 0.013 | 0.009 | 0.009 | 0 | 0.46 | 0.62 | 0.48 | 0.46 | 0.56 | 0.44 |
| 4 | 91775468 | rs17248761 | 0.041 | 1.635 | 0.461 | 0.888 | 0.114 | 0.043 | 0.043 | 0 | 0.08 | 0.16 | 0.13 | 0.14 | 0.22 | 0.1 |
| 4 | 91787375 | rs17196007 | 1.389 | 1.065 | -0.29 | 0.328 | -0.004 | -0.008 | -0.008 | 0 | 0.1 | 0 | 0.14 | 0.1 | 0.08 | 0.04 |
| 4 | 91788611 | rs13143215 | 0.073 | 1.109 | -0.181 | 0.265 | -0.005 | 0.002 | 0.002 | 0 | 0.1 | 0.11 | 0.08 | 0.02 | 0.13 | 0.1 |
| 4 | 91788661 | rs4693247 | -1.798 | 0.828 | -0.265 | 0.263 | 0.077 | 0.043 | 0.043 | 0.01 | 0.58 | 0.73 | 0.7 | 0.7 | 0.71 | 0.61 |
| 4 | 91790439 | rs10004104 | -1.695 | 1.43 | -0.018 | -0.094 | 0.017 | -0.046 | -0.046 | 0.096 | 0.31 | 0.59 | 0.36 | 0.11 | 0.41 | 0.25 |
| 4 | 91806782 | rs9994175 | -1.177 | 1.517 | 0.029 | 0.088 | 0.051 | -0.07 | -0.07 | 0.106 | 0.22 | 0.55 | 0.29 | 0.1 | 0.29 | 0.17 |
| 4 | 91816783 | rs11946311 | -0.755 | 0.085 | -0.783 | -0.202 | -0.028 | 0.061 | 0.061 | 0 | 0.38 | 0.25 | 0.36 | 0.16 | 0.45 | 0.26 |
| 4 | 91819988 | rs6828263 | -0.248 | 0.2 | -0.757 | -0.141 | -0.037 | 0.053 | 0.053 | 0 | 0.32 | 0.18 | 0.22 | 0.1 | 0.4 | 0.24 |
| 4 | 91823910 | rs1377917 | 0.951 | 0.168 | -1.423 | 0.569 | -0.116 | -0.117 | -0.117 | 0.095 | 0.21 | 0.02 | 0.21 | 0.39 | 0.31 | 0.25 |
| 4 | 91825652 | rs1037910 | -0.456 | 0.085 | -0.693 | 0.594 | -0.003 | 0 | 0 | 0 | 0.36 | 0.34 | 0.38 | 0.3 | 0.46 | 0.28 |
| 4 | 91827528 | rs17017809 | 0.34 | 0.179 | -0.214 | 0.31 | 0.009 | -0.078 | -0.078 | 0.044 | 0.27 | 0.27 | 0.18 | 0.03 | 0.32 | 0.15 |
| 4 | 91828373 | rs11097273 | -0.526 | 0.013 | -0.205 | 0.457 | -0.022 | -0.012 | -0.012 | 0 | 0.34 | 0.74 | 0.38 | 0.28 | 0.39 | 0.24 |
| 4 | 91829443 | rs7676836 | 2.478 | -0.224 | -0.378 | 0.454 | 0.02 | -0.028 | -0.028 | 0 | 0.06 | 0 | 0.17 | 0.11 | 0.04 | 0.01 |
| 4 | 91830496 | rs7683238 | -0.907 | -0.258 | -0.318 | 0.442 | -0.031 | -0.055 | -0.055 | 0.033 | 0.41 | 0.82 | 0.57 | 0.4 | 0.43 | 0.25 |
| 4 | 91831513 | rs9994801 | -1.716 | -0.214 | -0.276 | 0.398 | -0.033 | -0.028 | -0.028 | 0 | 0.49 | 0.84 | 0.62 | 0.42 | 0.51 | 0.32 |
| 4 | 91847796 | rs10012191 | -0.245 | -0.285 | -0.102 | 0.323 | 0.016 | -0.004 | -0.004 | 0.47 | 0.33 | 0.55 | 0.34 | 0.24 | 0.4 | 0.23 |
| 4 | 91860898 | rs11097280 | -1.801 | -0.569 | -0.922 | 0.328 | 0.061 | 0.083 | 0.083 | 0 | 0.68 | 0.84 | 0.8 | 0.88 | 0.81 | 0.56 |
| 4 | 91862940 | rs2870278 | -1.385 | -0.527 | -1.012 | 0.26 | 0.064 | -0.035 | -0.035 | 0.323 | 0.64 | 0.88 | 0.67 | 0.67 | 0.76 | 0.46 |
| 4 | 91866171 | rs10433957 | 0.275 | 0.146 | -0.656 | 0.596 | -0.069 | -0.074 | -0.074 | 0.012 | 0.26 | 0.11 | 0.23 | 0.47 | 0.36 | 0.3 |
| 4 | 91869258 | rs34473363 | -0.263 | -0.637 | 0.008 | 0.654 | 0 | 0 | 0 | 0 | 0.09 | 0.08 | 0.07 | 0.03 | 0.09 | 0.09 |
| 4 | 91871784 | rs11097282 | 0.644 | -1.419 | 0.325 | 0.424 | -0.011 | -0.005 | -0.005 | 0 | 0.57 | 0.59 | 0.55 | 0.67 | 0.72 | 0.54 |

| | | | | | | | | | | | | | | | | |
|---|---|---|---|---|---|---|---|---|---|---|---|---|---|---|---|---|
| 4 | 91872431 | rs1947442 | 0.578 | -1.668 | 0.363 | 0.346 | -0.027 | -0.02 | -0.02 | 0 | 0.82 | 0.68 | 0.83 | 0.85 | 0.85 | 0.73 |
| 4 | 91877301 | rs11097283 | 2.089 | -1.586 | 0.057 | 0.249 | -0.14 | -0.073 | -0.073 | 0 | 0.27 | 0.05 | 0.24 | 0.38 | 0.4 | 0.28 |
| 4 | 91881730 | rs1869538 | 0.337 | -1.859 | 0.045 | 0.03 | -0.044 | -0.029 | -0.029 | 0 | 0.67 | 0.63 | 0.75 | 0.82 | 0.82 | 0.57 |
| 4 | 91884244 | rs7672960 | 1.473 | -1.685 | 0.177 | -0.17 | -0.017 | 0.058 | 0.058 | 0 | 0.21 | 0.09 | 0.13 | 0.06 | 0.25 | 0.13 |
| 4 | 91889469 | rs7676210 | 1.357 | -1.628 | 0.352 | -0.099 | -0.013 | 0.048 | 0.048 | 0 | 0.23 | 0.13 | 0.14 | 0.06 | 0.29 | 0.14 |
| 4 | 91894603 | rs10016048 | 0.234 | -1.796 | 0.777 | 0.018 | -0.007 | -0.001 | -0.001 | 0 | 0.66 | 0.71 | 0.6 | 0.71 | 0.82 | 0.61 |
| 4 | 91900701 | rs1903496 | 1.461 | -1.674 | 0.384 | 0.038 | 0.03 | 0.025 | 0.025 | 0.648 | 0.16 | 0.06 | 0.09 | 0.03 | 0.03 | 0.18 |
| 4 | 91902620 | rs2219630 | -0.628 | -1.701 | 0.475 | 0.037 | 0.026 | 0.025 | 0.025 | 0.098 | 0.34 | 0.23 | 0.37 | 0.19 | 0.2 | 0.31 |
| 4 | 91910883 | rs6849036 | -1.155 | -1.407 | 0.684 | 0.271 | -0.011 | 0.091 | 0.091 | 0 | 0.26 | 0.46 | 0.45 | 0.39 | 0.21 | 0.24 |
| 4 | 91911417 | rs11727994 | -0.207 | -1.34 | 0.554 | 0.215 | -0.004 | -0.012 | -0.012 | 0 | 0.53 | 0.53 | 0.67 | 0.78 | 0.6 | 0.52 |
| 4 | 91917416 | rs2035116 | 1.835 | -1.168 | 0.41 | 0.317 | -0.13 | -0.084 | -0.084 | 0.262 | 0.27 | 0.05 | 0.22 | 0.39 | 0.39 | 0.27 |
| 4 | 91919207 | rs17017924 | -0.578 | -1.202 | 0.756 | 0.282 | -0.02 | 0.012 | 0.012 | 0.584 | 0.11 | 0.3 | 0.07 | 0.07 | 0.07 | 0.11 |
| 4 | 91925480 | rs2035115 | 0.84 | -1.147 | 0.289 | 0.281 | -0.076 | -0.019 | -0.019 | 0 | 0.35 | 0.15 | 0.27 | 0.42 | 0.45 | 0.37 |
| 4 | 91927412 | rs9992741 | 0.265 | -0.809 | 0.537 | 0.194 | -0.003 | -0.01 | -0.01 | 0 | 0.07 | 0.09 | 0.04 | 0.03 | 0.06 | 0.1 |
| 4 | 91931508 | rs1599455 | -0.895 | -0.781 | 0.601 | 0.24 | -0.014 | 0.055 | 0.055 | 0.16 | 0.93 | 0.82 | 0.92 | 0.87 | 0.96 | 0.84 |
| 4 | 91941868 | rs6830306 | 1.107 | -0.961 | 0.026 | 0.059 | -0.069 | -0.052 | -0.052 | 0 | 0.42 | 0.13 | 0.36 | 0.57 | 0.45 | 0.41 |
| 4 | 91941942 | rs10031177 | -0.171 | -1.067 | 0.768 | -0.092 | -0.019 | -0.018 | -0.018 | 0 | 0.13 | 0.36 | 0.1 | 0.07 | 0.07 | 0.21 |
| 4 | 91942194 | rs6830608 | -0.788 | -0.951 | 0.378 | -0.156 | -0.011 | 0.04 | 0.04 | 0 | 0.69 | 0.84 | 0.74 | 0.81 | 0.66 | 0.68 |
| 4 | 91952313 | rs6819554 | 0.191 | -0.943 | 0.392 | -0.099 | -0.019 | 0.053 | 0.053 | 0.02 | 0.56 | 0.75 | 0.48 | 0.69 | 0.51 | 0.6 |
| 4 | 91954105 | rs10856881 | -0.542 | -0.925 | 0.665 | 0 | -0.003 | 0.005 | 0.005 | 0 | 0.86 | 0.87 | 0.85 | 0.84 | 0.9 | 0.68 |
| 4 | 91954768 | rs1531403 | -0.094 | -0.984 | 1.158 | -0.512 | -0.02 | -0.024 | -0.024 | 0.271 | 0.84 | 0.6 | 0.87 | 0.9 | 0.89 | 0.68 |
| 4 | 91957343 | rs10002981 | 0.248 | -0.983 | 0.693 | -0.571 | 0.008 | 0.023 | 0.023 | 0.227 | 0.07 | 0.04 | 0.03 | 0.04 | 0.04 | 0.03 |
| 4 | 91959750 | rs2126218 | -0.874 | -1 | 0.735 | -0.571 | 0.003 | 0.003 | 0.003 | 0 | 0.87 | 0.92 | 0.89 | 0.9 | 0.9 | 0.7 |
| 4 | 91962013 | rs1461605 | 1.184 | -1.067 | 0.754 | -0.696 | -0.015 | 0.017 | 0.017 | 0 | 0.42 | 0.32 | 0.41 | 0.64 | 0.46 | 0.42 |
| 4 | 91962477 | rs10516887 | 1.054 | -1.236 | 0.361 | -0.741 | -0.012 | 0.055 | 0.055 | 0 | 0.25 | 0.14 | 0.18 | 0.05 | 0.29 | 0.16 |
| 4 | 91966364 | rs13133514 | 1.003 | -0.938 | 0.142 | -0.451 | -0.079 | 0.008 | 0.008 | 0 | 0.4 | 0.14 | 0.4 | 0.63 | 0.46 | 0.37 |
| 4 | 91973424 | rs1381294 | -0.84 | -0.76 | 0.584 | -0.506 | -0.001 | -0.001 | -0.001 | 0 | 0.19 | 0.21 | 0.27 | 0.18 | 0.15 | 0.09 |
| 4 | 91974658 | rs6849961 | -0.05 | -0.639 | 0.426 | -0.449 | 0.002 | 0.012 | 0.012 | 0.138 | 0.07 | 0.04 | 0.03 | 0.04 | 0.05 | 0.03 |
| 4 | 91989547 | rs10009574 | -1.059 | -0.674 | 0.81 | -0.488 | -0.004 | -0.005 | -0.005 | 0 | 0.19 | 0.36 | 0.32 | 0.2 | 0.16 | 0.11 |
| 4 | 91991096 | rs7660020 | 0.285 | -0.871 | 0.857 | -0.867 | 0 | -0.041 | -0.041 | 0 | 0.37 | 0.3 | 0.21 | 0.06 | 0.37 | 0.41 |
| 4 | 91996010 | rs12513240 | -0.518 | -1.131 | 0.502 | -0.663 | 0.005 | 0.014 | 0.014 | 0 | 0.57 | 0.6 | 0.42 | 0.33 | 0.44 | 0.66 |
| 4 | 92014627 | rs12646841 | 0.831 | -1.009 | 0.32 | -0.478 | 0.017 | -0.018 | -0.018 | 0 | 0.18 | 0.14 | 0.2 | 0.26 | 0.07 | 0.24 |
| 4 | 92016267 | rs11097298 | 0.973 | -1.014 | 0.321 | -0.481 | 0.043 | -0.05 | -0.05 | 0.383 | 0.17 | 0.07 | 0.2 | 0.26 | 0.07 | 0.24 |
| 4 | 92017500 | rs12508842 | 0.546 | -0.972 | 0.39 | -0.517 | -0.037 | 0.077 | 0.077 | 0 | 0.42 | 0.25 | 0.5 | 0.73 | 0.48 | 0.48 |
| 4 | 92025683 | rs13114934 | 1.064 | -0.955 | 0.301 | -0.495 | 0.034 | -0.037 | -0.037 | 0.004 | 0.17 | 0.09 | 0.19 | 0.26 | 0.07 | 0.24 |
| 4 | 92028168 | rs6817115 | 0.285 | -0.423 | 0.371 | -0.254 | 0 | 0 | 0 | 0 | 0.06 | 0.09 | 0.07 | 0.07 | 0.06 | 0.03 |
| 4 | 92028969 | rs3944131 | 1.431 | -0.367 | 0.333 | -0.263 | -0.022 | -0.023 | -0.023 | 0.979 | 0.25 | 0.23 | 0.31 | 0.47 | 0.41 | 0.25 |
| 4 | 92036269 | rs17243008 | 1.455 | -0.292 | 0.217 | -0.261 | -0.058 | -0.062 | -0.062 | 0.25 | 0.25 | 0.14 | 0.31 | 0.47 | 0.41 | 0.24 |
| 4 | 92038546 | rs17018037 | -0.734 | -0.463 | 0.272 | -0.287 | 0.004 | 0.02 | 0.02 | 0 | 0.13 | 0.25 | 0.25 | 0.2 | 0.12 | 0.1 |
| 4 | 92042065 | rs11936964 | 0.045 | -0.297 | 0.409 | -0.39 | 0.006 | -0.009 | -0.009 | 0 | 0.4 | 0.56 | 0.55 | 0.67 | 0.53 | 0.35 |
| 4 | 92053582 | rs7667572 | -0.452 | -0.477 | 0.322 | 0.054 | 0.005 | 0.05 | 0.05 | 0 | 0.11 | 0.2 | 0.26 | 0.25 | 0.09 | 0.08 |
| 4 | 92054563 | rs17018078 | 0.354 | -1.065 | -0.154 | -0.515 | 0.065 | -0.082 | -0.082 | 0.393 | 0.22 | 0.03 | 0.21 | 0.26 | 0.09 | 0.21 |
| 4 | 92066776 | rs13104402 | 0.649 | -1.325 | 0.225 | -0.741 | -0.022 | -0.003 | -0.003 | 0.056 | 0.62 | 0.42 | 0.67 | 0.69 | 0.8 | 0.59 |
| 4 | 92076288 | rs10016028 | -1.27 | -1.283 | -0.065 | -0.723 | 0.017 | 0.016 | 0.016 | 0.006 | 0.9 | 0.97 | 0.95 | 0.95 | 0.93 | 0.83 |
| 4 | 92079569 | rs1541365 | 0.29 | -1.051 | 0.139 | -0.635 | -0.03 | -0.003 | -0.003 | 0 | 0.64 | 0.6 | 0.68 | 0.69 | 0.8 | 0.6 |
| 4 | 92114903 | rs17268740 | 0.678 | -0.327 | -0.339 | -0.736 | 0.001 | -0.047 | -0.047 | 0 | 0.16 | 0.15 | 0.12 | 0.01 | 0.2 | 0.14 |
| 4 | 92141455 | rs872669 | -0.199 | -0.443 | 0.267 | -0.94 | 0.01 | 0.042 | 0.042 | 0 | 0.58 | 0.71 | 0.6 | 0.94 | 0.6 | 0.68 |
| 4 | 92142940 | rs13146287 | 0.455 | -0.003 | 0.117 | -1.126 | -0.003 | 0.004 | 0.004 | 0 | 0.4 | 0.41 | 0.48 | 0.86 | 0.41 | 0.49 |
| 4 | 92142966 | rs13122462 | 0.169 | -0.003 | 0.115 | -1.126 | 0.002 | -0.048 | -0.048 | 0.201 | 0.44 | 0.54 | 0.49 | 0.87 | 0.43 | 0.55 |
| 4 | 92151187 | rs4693271 | 1.193 | -0.207 | 0.034 | -0.458 | 0.02 | -0.025 | -0.025 | 0.712 | 0.12 | 0.07 | 0.15 | 0.3 | 0.03 | 0.12 |
| 4 | 92154401 | rs7671793 | -1.054 | 0.256 | 0.361 | -1.097 | 0.015 | -0.015 | -0.015 | 0 | 0.09 | 0.41 | 0.06 | 0.01 | 0.06 | 0.09 |
| 4 | 92159830 | rs1544017 | -0.108 | 0.67 | -0.789 | -0.772 | -0.019 | 0.026 | 0.026 | 0.065 | 0.14 | 0.05 | 0.09 | 0.07 | 0.17 | 0.14 |
| 4 | 92163756 | rs2204703 | -0.176 | 0.235 | -0.454 | -0.988 | -0.005 | -0.048 | -0.048 | 0 | 0.58 | 0.58 | 0.73 | 0.93 | 0.62 | 0.64 |
| 4 | 92172833 | rs7675354 | 0.174 | 0.769 | -0.005 | -0.396 | 0.033 | 0.011 | 0.011 | 0.131 | 0.07 | 0.14 | 0.06 | 0.09 | 0.11 | 0.15 |
| 4 | 92178955 | rs13151372 | -0.064 | 0.757 | 0.373 | 0.061 | 0.005 | 0.072 | 0.072 | 0 | 0.2 | 0.38 | 0.19 | 0.33 | 0.16 | 0.14 |
| 4 | 92179229 | rs6815212 | 0.745 | 0.433 | 0.139 | 0.191 | -0.017 | -0.007 | -0.007 | 0 | 0.12 | 0.13 | 0.05 | 0.11 | 0.06 | 0.06 |
| 4 | 92184080 | rs10028174 | -0.008 | 0.168 | -0.883 | -0.211 | 0 | 0.086 | 0.086 | 0 | 0.25 | 0.03 | 0.23 | 0.08 | 0.17 | 0.26 |
| 4 | 92185156 | rs9986071 | -1.563 | 0.088 | -0.584 | -0.134 | -0.028 | -0.039 | -0.039 | 0 | 0.59 | 0.79 | 0.42 | 0.49 | 0.46 | 0.57 |
| 4 | 92193156 | rs4331741 | 0.509 | 0.445 | -0.76 | 0.203 | -0.033 | -0.013 | -0.013 | 0 | 0.14 | 0.25 | 0.08 | 0.11 | 0.07 | 0.06 |
| 4 | 92200866 | rs10003471 | 0.255 | 1.188 | -0.958 | 0.425 | -0.028 | 0 | 0 | 0 | 0.12 | 0.02 | 0.07 | 0.07 | 0.11 | 0.19 |
| 4 | 92201568 | rs10006439 | -0.49 | 1.273 | -0.535 | 0.463 | 0.022 | 0.012 | 0.012 | 0 | 0.27 | 0.37 | 0.24 | 0.31 | 0.26 | 0.29 |
| 4 | 92218323 | rs10033018 | -0.377 | 1.115 | -1.038 | 0.607 | -0.008 | 0.01 | 0.01 | 0 | 0.15 | 0.12 | 0.23 | 0.1 | 0.18 | 0.18 |
| 4 | 92225277 | rs1969531 | -0.188 | 1.185 | -0.973 | 0.592 | -0.006 | 0.002 | 0.002 | 1.72 | 0.36 | 0.31 | 0.31 | 0.31 | 0.38 | 0.28 |
| 4 | 92231314 | rs11725743 | -0.584 | 0.462 | -0.867 | 0.976 | 0.007 | -0.011 | -0.011 | 0.015 | 0.77 | 0.79 | 0.81 | 0.67 | 0.79 | 0.7 |
| 4 | 92244300 | rs17018345 | 0.674 | 0.798 | -0.544 | 0.936 | 0.009 | 0.018 | 0.018 | 0 | 0.22 | 0.26 | 0.28 | 0.34 | 0.27 | 0.25 |
| 4 | 92251839 | rs6532299 | -0.239 | 0.584 | -0.742 | -0.584 | 0 | -0.008 | -0.008 | 0 | 0.67 | 0.61 | 0.67 | 0.79 | 0.69 | 0.64 |
| 4 | 92267559 | rs2870313 | 0.212 | 0.35 | -1.293 | -0.921 | 0.008 | 0.139 | 0.139 | 0 | 0.36 | 0.12 | 0.25 | 0.16 | 0.36 | 0.33 |
| 4 | 92280933 | rs12507382 | 0.18 | 0.079 | -0.854 | -1.06 | 0.006 | -0.017 | -0.017 | 1.026 | 0.36 | 0.33 | 0.22 | 0.07 | 0.31 | 0.19 |
| 4 | 92281057 | rs12507411 | 0.22 | 0.066 | -0.861 | -1.086 | 0.005 | -0.021 | -0.021 | 0 | 0.36 | 0.32 | 0.22 | 0.07 | 0.3 | 0.19 |

| | | | | | | | | | | | | | | | | |
|---|---|---|---|---|---|---|---|---|---|---|---|---|---|---|---|---|
| 4 | 92283771 | rs4128705 | 0.169 | 0.102 | -0.922 | -1.077 | 0.009 | -0.01 | -0.01 | 2.332 | 0.37 | 0.33 | 0.24 | 0.08 | 0.31 | 0.2 |
| 4 | 92286605 | rs1919225 | -0.849 | 0.249 | -1.443 | -0.768 | -0.031 | 0.013 | 0.013 | 0.013 | 0.81 | 0.98 | 0.77 | 0.91 | 0.82 | 0.86 |
| 4 | 92304224 | rs6837355 | 0.051 | 0.088 | -1.337 | -1.021 | 0 | 0.112 | 0.112 | 0 | 0.65 | 0.85 | 0.73 | 0.86 | 0.73 | 0.76 |
| 4 | 92309543 | rs10856885 | 0.197 | -0.048 | -0.384 | -1.062 | -0.006 | -0.034 | -0.034 | 2.209 | 0.63 | 0.55 | 0.7 | 0.86 | 0.73 | 0.76 |
| 4 | 92316484 | rs4411947 | 0.638 | -0.12 | -0.744 | -1.465 | 0.034 | 0.174 | 0.174 | 0 | 0.35 | 0.11 | 0.22 | 0.07 | 0.31 | 0.21 |
| 4 | 92349263 | rs13123650 | -0.285 | -0.262 | -0.316 | -1.225 | -0.006 | -0.024 | -0.024 | 0 | 0.75 | 0.67 | 0.76 | 0.88 | 0.81 | 0.88 |
| 4 | 92350342 | rs6845679 | 0.566 | -0.446 | -0.47 | -1.526 | 0.005 | 0.019 | 0.019 | 0.291 | 0.49 | 0.42 | 0.56 | 0.82 | 0.55 | 0.71 |
| 4 | 92352498 | rs10022644 | -0.814 | -0.426 | -0.776 | -1.48 | -0.004 | 0.057 | 0.057 | 0.453 | 0.77 | 0.93 | 0.78 | 0.88 | 0.8 | 0.88 |
| 4 | 92354457 | rs6824640 | 0.349 | -0.432 | -0.161 | -1.606 | -0.012 | -0.069 | -0.069 | 0.11 | 0.56 | 0.53 | 0.66 | 0.9 | 0.64 | 0.78 |
| 4 | 92357499 | rs1922217 | -0.477 | -0.411 | -0.1 | -1.45 | 0.012 | 0.063 | 0.063 | 0 | 0.75 | 0.89 | 0.73 | 0.92 | 0.78 | 0.84 |
| 4 | 92358383 | rs7659358 | 0.87 | -0.333 | 0.158 | -1.454 | 0.104 | 0.011 | 0.011 | 0 | 0.07 | 0.29 | 0.1 | 0.03 | 0.11 | 0.07 |
| 4 | 92366263 | rs1922216 | -0.456 | -0.382 | -0.418 | -1.437 | 0.018 | 0.07 | 0.07 | 0 | 0.75 | 0.89 | 0.74 | 0.92 | 0.79 | 0.85 |
| 4 | 92368346 | rs1922233 | -0.46 | -0.386 | -0.467 | -1.443 | 0.017 | 0.08 | 0.08 | 0 | 0.75 | 0.9 | 0.76 | 0.93 | 0.79 | 0.85 |
| 4 | 92380924 | rs12646638 | 0.834 | -0.265 | -0.273 | -1.399 | 0.039 | -0.014 | -0.014 | 0 | 0.07 | 0.14 | 0.07 | 0.03 | 0.11 | 0.07 |
| 4 | 92381080 | rs12649871 | -0.526 | -0.37 | -0.267 | -1.491 | 0.016 | 0.078 | 0.078 | 0.119 | 0.76 | 0.85 | 0.76 | 0.94 | 0.79 | 0.84 |
| 4 | 92381094 | rs12646710 | 1.199 | -0.286 | -0.23 | -1.411 | 0.033 | -0.012 | -0.012 | 0 | 0.06 | 0.12 | 0.06 | 0.03 | 0.11 | 0.07 |
| 4 | 92399884 | rs7675583 | 0.6 | -0.533 | -0.393 | -1.636 | 0.014 | 0.121 | 0.121 | 0.361 | 0.51 | 0.65 | 0.6 | 0.88 | 0.53 | 0.7 |
| 4 | 92420171 | rs6838229 | -0.191 | -0.2 | 0.077 | -0.682 | -0.012 | 0.005 | 0.005 | 0 | 0.7 | 0.42 | 0.55 | 0.67 | 0.72 | 0.63 |
| 4 | 92421296 | rs11731642 | -0.286 | -0.182 | 0.055 | -0.677 | -0.004 | 0.011 | 0.011 | 0.242 | 0.73 | 0.61 | 0.56 | 0.67 | 0.73 | 0.64 |
| 4 | 92431729 | rs7677659 | 0.495 | -0.485 | -0.113 | -0.201 | -0.002 | -0.002 | -0.002 | 0 | 0.46 | 0.5 | 0.43 | 0.58 | 0.51 | 0.39 |
| 4 | 92434672 | rs1607075 | -0.238 | -0.514 | -0.23 | -0.237 | -0.003 | 0.002 | 0.002 | 0.546 | 0.71 | 0.68 | 0.66 | 0.73 | 0.72 | 0.7 |
| 4 | 92439678 | rs12641499 | -0.757 | -0.277 | -0.188 | -0.112 | -0.001 | -0.001 | -0.001 | 0 | 0.07 | 0.08 | 0.07 | 0.1 | 0.1 | 0.23 |
| 4 | 92451147 | rs1516700 | 1.409 | -0.143 | -0.195 | -0.162 | -0.002 | 0.01 | 0.01 | 0 | 0.16 | 0.15 | 0.31 | 0.44 | 0.23 | 0.13 |
| 4 | 92452205 | rs6852056 | -0.831 | -0.791 | -0.515 | -0.552 | -0.02 | 0.029 | 0.029 | 0 | 0.62 | 0.86 | 0.63 | 0.74 | 0.63 | 0.59 |
| 4 | 92454931 | rs17283537 | 2.206 | -0.935 | -0.731 | -0.65 | -0.023 | -0.051 | -0.051 | 0.988 | 0.1 | 0.01 | 0.06 | 0.16 | 0.13 | 0.13 |
| 4 | 92457618 | rs7697141 | 0.599 | -0.907 | -0.647 | -0.654 | -0.001 | -0.006 | -0.006 | 0 | 0.05 | 0.09 | 0.03 | 0.03 | 0.05 | 0.02 |
| 4 | 92466958 | rs6532317 | -0.225 | -0.937 | -0.318 | -0.613 | -0.003 | 0.014 | 0.014 | 0 | 0.53 | 0.61 | 0.53 | 0.64 | 0.53 | 0.35 |
| 4 | 92472139 | rs12152593 | 0.712 | -0.866 | 0.149 | -0.556 | 0.014 | 0.02 | 0.02 | 0 | 0.27 | 0.34 | 0.41 | 0.61 | 0.37 | 0.27 |
| 4 | 92477566 | rs17018613 | 1.705 | -0.785 | -0.111 | -0.522 | -0.001 | 0.012 | 0.012 | 0 | 0.16 | 0.15 | 0.32 | 0.44 | 0.23 | 0.14 |
| 4 | 92483806 | rs17284306 | 1.083 | -1.06 | -0.286 | -0.947 | 0.042 | 0.091 | 0.091 | 0.194 | 0.23 | 0.08 | 0.11 | 0.02 | 0.15 | 0.08 |
| 4 | 92496463 | rs7675138 | -0.715 | -0.543 | 0.227 | -0.578 | 0.004 | 0.001 | 0.001 | 0 | 0.08 | 0.14 | 0.08 | 0.09 | 0.1 | 0.24 |
| 4 | 92502367 | rs1516708 | 0.444 | -0.681 | 0.114 | -0.749 | 0.005 | 0.021 | 0.021 | 0 | 0.48 | 0.61 | 0.49 | 0.66 | 0.48 | 0.36 |
| 4 | 92520419 | rs9991015 | -0.559 | -0.618 | 0.346 | -0.718 | 0.002 | -0.062 | -0.062 | 0 | 0.15 | 0.28 | 0.11 | 0.03 | 0.13 | 0.04 |
| 4 | 92539867 | rs13113814 | 1.274 | -0.739 | -0.035 | -0.989 | 0.004 | 0.038 | 0.038 | 0.92 | 0.17 | 0.1 | 0.06 | 0 | 0.1 | 0.11 |
| 4 | 92544404 | rs11947273 | 1.027 | -0.641 | -0.016 | -0.755 | 0 | 0.005 | 0.005 | 0 | 0.45 | 0.39 | 0.46 | 0.66 | 0.48 | 0.34 |
| 4 | 92566775 | rs11946091 | 0.79 | -0.463 | -0.626 | -0.654 | 0.009 | 0.049 | 0.049 | 0 | 0.51 | 0.71 | 0.5 | 0.66 | 0.49 | 0.36 |
| 4 | 92572499 | rs12649563 | -0.283 | 0.102 | -0.115 | -0.644 | 0.003 | -0.005 | -0.005 | 0.638 | 0.12 | 0.12 | 0.12 | 0.09 | 0.15 | 0.24 |
| 4 | 92591560 | rs12647856 | 0.007 | 0.083 | -0.061 | -0.679 | 0.006 | -0.024 | -0.024 | 0 | 0.1 | 0.12 | 0.09 | 0.03 | 0.11 | 0.03 |
| 4 | 92615579 | rs4508848 | -0.989 | 0.074 | -0.372 | -0.78 | -0.001 | -0.033 | -0.033 | 0 | 0.41 | 0.52 | 0.38 | 0.2 | 0.38 | 0.37 |
| 4 | 92634711 | rs11731487 | -0.353 | 0.555 | -0.017 | -0.597 | 0.052 | -0.05 | -0.05 | 0 | 0.17 | 0.33 | 0.17 | 0.09 | 0.18 | 0.12 |
| 4 | 92651110 | rs6532345 | 0.329 | 0.952 | -0.51 | -0.451 | -0.001 | -0.001 | -0.001 | 0.037 | 0.74 | 0.75 | 0.5 | 0.23 | 0.68 | 0.58 |
| 4 | 92661382 | rs13141601 | 0.748 | -0.203 | -0.479 | -1.24 | 0.045 | 0.112 | 0.112 | 0.03 | 0.53 | 0.26 | 0.41 | 0.23 | 0.6 | 0.51 |
| 4 | 92662425 | rs4348059 | -0.445 | -0.071 | -0.611 | -0.162 | -0.002 | 0.007 | 0.007 | 1.339 | 0.79 | 0.79 | 0.76 | 0.66 | 0.8 | 0.69 |
| 4 | 92666216 | rs6820916 | -0.352 | -0.083 | -0.617 | -0.165 | -0.002 | 0.006 | 0.006 | 1.481 | 0.78 | 0.79 | 0.76 | 0.64 | 0.8 | 0.69 |
| 4 | 92667871 | rs12505028 | -0.133 | 0.029 | -0.757 | -1.143 | -0.003 | -0.045 | -0.045 | 0 | 0.35 | 0.65 | 0.23 | 0.07 | 0.26 | 0.22 |
| 4 | 92668180 | rs4314258 | 1.093 | -0.47 | -0.197 | -1.058 | -0.056 | -0.092 | -0.092 | 0.626 | 0.18 | 0.33 | 0.08 | 0 | 0.08 | 0.07 |
| 4 | 92670313 | rs4128277 | -0.578 | -0.081 | -0.339 | -0.851 | -0.001 | 0.001 | 0.001 | 0 | 0.25 | 0.26 | 0.46 | 0.71 | 0.23 | 0.44 |
| 4 | 92672743 | rs17221444 | 1.701 | 0.305 | -0.865 | -1.253 | -0.041 | 0.065 | 0.065 | 0 | 0.14 | 0.03 | 0.1 | 0 | 0.17 | 0.05 |
| 4 | 92673841 | rs4455379 | -0.491 | 0.258 | -0.757 | -1.329 | -0.015 | 0.039 | 0.039 | 0 | 0.71 | 0.79 | 0.81 | 0.89 | 0.67 | 0.83 |
| 4 | 92678634 | rs17287631 | 1.351 | 0.148 | -0.362 | -1.39 | -0.016 | 0.027 | 0.027 | 0 | 0.29 | 0.2 | 0.28 | 0.19 | 0.35 | 0.32 |
| 4 | 92684140 | rs6811317 | 0.586 | 0.113 | -0.584 | -0.951 | 0.005 | -0.008 | -0.008 | 0 | 0.41 | 0.48 | 0.48 | 0.36 | 0.46 | 0.49 |
| 4 | 92685286 | rs13106681 | -0.21 | 0.518 | -0.476 | -0.27 | -0.004 | -0.011 | -0.011 | 0 | 0.79 | 0.83 | 0.69 | 0.36 | 0.77 | 0.67 |
| 4 | 92688140 | rs13114360 | 1.081 | 0.558 | -0.304 | -0.096 | -0.001 | 0.013 | 0.013 | 0 | 0.12 | 0.17 | 0.19 | 0.16 | 0.1 | 0.17 |
| 4 | 92688358 | rs9307104 | 0.928 | 0.477 | -0.374 | -0.4 | -0.003 | -0.033 | -0.033 | 0 | 0.44 | 0.49 | 0.34 | 0.19 | 0.42 | 0.38 |
| 4 | 92690410 | rs4615142 | 0.711 | 0.389 | -0.268 | 0.014 | 0.005 | 0.035 | 0.035 | 0 | 0.12 | 0.26 | 0.21 | 0.18 | 0.1 | 0.17 |
| 4 | 92691639 | rs11097328 | -0.618 | 0.128 | -0.534 | -0.095 | 0.046 | 0.041 | 0.041 | 0.193 | 0.4 | 0.76 | 0.56 | 0.71 | 0.29 | 0.45 |
| 4 | 92692108 | rs11937169 | 0.043 | 0.723 | -0.599 | 0.023 | 0.008 | 0 | 0 | 0 | 0.16 | 0.17 | 0.23 | 0.19 | 0.23 | 0.2 |
| 4 | 92696668 | rs4273429 | -0.748 | 0.071 | -0.273 | -0.03 | 0.065 | 0.061 | 0.061 | 6.14 | 0.05 | 0.52 | 0.04 | 0 | 0.01 | 0 |
| 4 | 92706551 | rs1564958 | -0.832 | -0.02 | -1.291 | -0.024 | 0.046 | 0.078 | 0.078 | 0 | 0.49 | 0.76 | 0.42 | 0.37 | 0.39 | 0.45 |
| 4 | 92718180 | rs4622965 | -1.947 | 0.187 | -1.555 | 0.168 | -0.032 | -0.037 | -0.037 | 0 | 0.93 | 1 | 0.96 | 0.88 | 0.9 | 0.95 |
| 4 | 92720480 | rs1003076 | 1.314 | -0.02 | -1.793 | -0.196 | -0.104 | 0.042 | 0.042 | 0 | 0.3 | 0.03 | 0.25 | 0.19 | 0.37 | 0.3 |
| 4 | 92734290 | rs1485991 | 0.071 | -0.932 | -1.005 | -0.581 | -0.064 | -0.01 | -0.01 | 0.021 | 0.19 | 0.56 | 0.2 | 0.18 | 0.08 | 0.15 |
| 4 | 92750394 | rs970694 | -0.525 | -0.202 | -1.596 | -0.651 | -0.147 | 0.007 | 0.007 | 0.445 | 0.79 | 0.98 | 0.8 | 0.85 | 0.64 | 0.77 |
| 4 | 92758836 | rs1485989 | 1.128 | -0.783 | -1.729 | -1.045 | -0.052 | -0.017 | -0.017 | 0 | 0.28 | 0.14 | 0.5 | 0.67 | 0.42 | 0.47 |
| 4 | 92761582 | rs1485988 | 0.39 | -0.72 | -1.314 | -1.114 | -0.021 | 0.079 | 0.079 | 0 | 0.59 | 0.71 | 0.73 | 0.85 | 0.55 | 0.68 |
| 4 | 92770887 | rs11941387 | -0.339 | -0.422 | -0.984 | -0.779 | -0.083 | 0.034 | 0.034 | 0.356 | 0.78 | 0.93 | 0.79 | 0.85 | 0.63 | 0.77 |
| 4 | 92790543 | rs17018839 | 0.536 | -0.894 | -0.599 | -0.443 | -0.027 | -0.058 | -0.058 | 4.817 | 0.12 | 0.21 | 0.04 | 0 | 0.04 | 0.07 |
| 4 | 92803775 | rs10028425 | -0.437 | -0.449 | -0.343 | -0.026 | 0.014 | 0.023 | 0.023 | 0 | 0.41 | 0.69 | 0.3 | 0.37 | 0.34 | 0.33 |
| 4 | 92816815 | rs991345 | 0.773 | -0.585 | 0.388 | -0.417 | -0.052 | -0.041 | -0.041 | 0 | 0.24 | 0.3 | 0.09 | 0.15 | 0.08 | 0.1 |

| | | | | | | | | | | | | | | | | |
|---|---|---|---|---|---|---|---|---|---|---|---|---|---|---|---|---|
| 4 | 92817301 | rs1471822 | 0.882 | -0.564 | 0.386 | -0.418 | -0.03 | -0.018 | -0.018 | 0 | 0.08 | 0.05 | 0.12 | 0.13 | 0.22 | 0.14 |
| 4 | 92820451 | rs1120646 | 0.758 | -0.543 | 0.425 | -0.448 | -0.051 | -0.042 | -0.042 | 0 | 0.24 | 0.3 | 0.09 | 0.15 | 0.08 | 0.1 |
| 4 | 92837584 | rs17303960 | 0.817 | -0.567 | 0.54 | -0.427 | 0.025 | 0.002 | 0.002 | 0 | 0.08 | 0.11 | 0.13 | 0.13 | 0.22 | 0.15 |
| 4 | 92838346 | rs1564957 | 0.095 | -0.71 | 0.662 | -0.484 | -0.06 | -0.052 | -0.052 | 0 | 0.31 | 0.36 | 0.13 | 0.15 | 0.11 | 0.18 |
| 4 | 92839908 | rs11944581 | 0.325 | -0.7 | 0.461 | -0.471 | 0.007 | 0.004 | 0.004 | 0 | 0.07 | 0.07 | 0.05 | 0.01 | 0.03 | 0.09 |
| 4 | 92845437 | rs321723 | 0.759 | -0.655 | 0.321 | -0.114 | 0.024 | 0.008 | 0.008 | 0 | 0.54 | 0.28 | 0.65 | 0.52 | 0.59 | 0.58 |
| 4 | 92867432 | rs1844346 | 0.599 | -0.579 | 0.524 | -0.655 | 0.012 | 0.032 | 0.032 | 0 | 0.58 | 0.36 | 0.72 | 0.72 | 0.66 | 0.68 |
| 4 | 92873306 | rs17291988 | 0.018 | -0.677 | 0.752 | -0.695 | -0.076 | -0.067 | -0.067 | 1.665 | 0.32 | 0.38 | 0.14 | 0.15 | 0.11 | 0.18 |
| 4 | 92883857 | rs1489557 | 0.053 | -0.682 | 0.872 | -0.697 | -0.058 | -0.05 | -0.05 | 0 | 0.32 | 0.33 | 0.14 | 0.15 | 0.11 | 0.18 |
| 4 | 92889426 | rs6532349 | -0.802 | -0.622 | 0.83 | -0.672 | -0.002 | -0.003 | -0.003 | 0 | 0.91 | 0.93 | 0.87 | 0.88 | 0.77 | 0.86 |
| 4 | 92890930 | rs6817844 | -0.077 | -0.69 | 0.944 | -0.765 | -0.071 | -0.056 | -0.056 | 0 | 0.32 | 0.38 | 0.15 | 0.15 | 0.11 | 0.18 |
| 4 | 92897624 | rs7672071 | 1.396 | -0.778 | 0.571 | -1.243 | 0.001 | -0.021 | -0.021 | 0 | 0.2 | 0.17 | 0.08 | 0 | 0.14 | 0.1 |
| 4 | 92906792 | rs966867 | 1.045 | -0.407 | 1.102 | -0.371 | 0.014 | 0.039 | 0.039 | 0 | 0.57 | 0.36 | 0.7 | 0.74 | 0.64 | 0.67 |
| 4 | 92919069 | rs3864150 | -0.533 | -0.276 | 0.955 | -0.301 | -0.027 | -0.008 | -0.008 | 0.526 | 0.91 | 0.94 | 0.85 | 0.89 | 0.75 | 0.85 |
| 4 | 92930924 | rs4693287 | 0.518 | -0.59 | 1.014 | -0.35 | 0.063 | 0.034 | 0.034 | 1.91 | 0.23 | 0.05 | 0.08 | 0.15 | 0.08 | 0.09 |
| 4 | 93009187 | rs10032505 | -0.451 | -0.603 | 1.258 | -0.388 | -0.087 | -0.083 | -0.083 | 0 | 0.32 | 0.47 | 0.15 | 0.16 | 0.11 | 0.18 |
| 4 | 93016862 | rs730293 | 1.993 | 0.264 | 1.803 | 0.17 | 0.001 | -0.021 | -0.021 | 0 | 0.75 | 0.35 | 0.75 | 0.85 | 0.7 | 0.74 |
| 4 | 93030390 | rs10023188 | -0.424 | 0.385 | 1.328 | 0.22 | -0.027 | -0.032 | -0.032 | 0 | 0.08 | 0.12 | 0.05 | 0.01 | 0.04 | 0.09 |
| 4 | 93035922 | rs6837448 | -0.033 | 0.46 | 1.041 | 0.248 | -0.015 | -0.004 | -0.004 | 0 | 0.9 | 0.94 | 0.86 | 0.89 | 0.76 | 0.85 |
| 4 | 93044488 | rs6532359 | -0.504 | 0.299 | 1.014 | 0.258 | -0.033 | -0.039 | -0.039 | 0.11 | 0.08 | 0.15 | 0.05 | 0.01 | 0.04 | 0.09 |
| 4 | 93062161 | rs2169202 | -0.332 | 0.433 | 0.764 | 0.316 | 0.009 | -0.001 | -0.001 | 0 | 0.91 | 0.91 | 0.84 | 0.88 | 0.76 | 0.83 |
| 4 | 93064187 | rs17019139 | -1.2 | -0.14 | 0.76 | 0.465 | -0.026 | 0.002 | 0.002 | 0 | 0.14 | 0.19 | 0.11 | 0.17 | 0.05 | 0.15 |
| 4 | 93088664 | rs1585428 | -1.33 | -0.15 | 0.803 | 0.459 | -0.065 | 0.013 | 0.013 | 0.373 | 0.15 | 0.32 | 0.13 | 0.17 | 0.06 | 0.14 |
| 4 | 93107648 | rs6848613 | 0.134 | 0.016 | 0.548 | 0.537 | -0.019 | 0.036 | 0.036 | 0 | 0.05 | 0.12 | 0.03 | 0.15 | 0.01 | 0.06 |
| 4 | 93113274 | rs17019186 | -1.776 | -0.103 | 1.258 | 0.406 | -0.087 | 0.024 | 0.024 | 0.908 | 0.17 | 0.5 | 0.13 | 0.15 | 0.07 | 0.15 |
| 4 | 93118886 | rs1227423 | 1.047 | -0.222 | 0.768 | -0.246 | 0.008 | 0.026 | 0.026 | 0.029 | 0.31 | 0.2 | 0.18 | 0.05 | 0.27 | 0.16 |
| 4 | 93136397 | rs7683443 | -1.634 | 0.147 | 1.182 | 0.778 | -0.051 | 0.029 | 0.029 | 0.086 | 0.16 | 0.32 | 0.12 | 0.23 | 0.07 | 0.15 |
| 4 | 93136797 | rs2199291 | 2.118 | 0.436 | 0.93 | 0.854 | -0.002 | 0.018 | 0.018 | 1.3 | 0.73 | 0.35 | 0.71 | 0.65 | 0.7 | 0.67 |
| 4 | 93146144 | rs10049958 | -2.042 | -0.278 | 0.76 | 0.751 | -0.094 | 0.075 | 0.075 | 0.821 | 0.17 | 0.51 | 0.13 | 0.23 | 0.07 | 0.15 |
| 4 | 93152870 | rs10516905 | -0.67 | -0.197 | 0.534 | 0.792 | -0.016 | 0.177 | 0.177 | 0 | 0.07 | 0.29 | 0.05 | 0.22 | 0.01 | 0.07 |
| 4 | 93161851 | rs17019255 | -0.139 | -0.181 | 0.275 | 0.77 | -0.015 | 0.042 | 0.042 | 0 | 0.06 | 0.08 | 0.03 | 0.22 | 0.01 | 0.07 |
| 4 | 93184838 | rs1913301 | -1.029 | -0.153 | 0.353 | 0.597 | -0.035 | -0.06 | -0.06 | 0 | 0.11 | 0.41 | 0.1 | 0.01 | 0.06 | 0.09 |
| 4 | 93190198 | rs17019283 | 0.283 | 0.077 | -0.326 | 0.904 | -0.05 | -0.068 | -0.068 | 0 | 0.2 | 0.02 | 0.11 | 0.3 | 0.24 | 0.32 |
| 4 | 93198845 | rs8180202 | -1.606 | -0.11 | 0.517 | 0.727 | -0.065 | 0.058 | 0.058 | 0.655 | 0.17 | 0.5 | 0.13 | 0.22 | 0.09 | 0.16 |
| 4 | 93242743 | rs17238193 | -0.99 | 0.027 | -0.095 | 0.292 | 0.019 | 0.033 | 0.033 | 0 | 0.08 | 0.02 | 0.07 | 0.01 | 0.09 | 0.07 |
| 4 | 93250536 | rs6824860 | -2.224 | -0.251 | 0.501 | 0.524 | -0.075 | 0.036 | 0.036 | 0 | 0.16 | 0.42 | 0.13 | 0.16 | 0.1 | 0.14 |
| 4 | 93255959 | rs17019318 | -0.962 | -0.25 | 0.589 | 0.529 | -0.01 | 0.146 | 0.146 | 0 | 0.08 | 0.33 | 0.05 | 0.15 | 0.01 | 0.07 |
| 4 | 93264193 | rs10009241 | -2.009 | -0.241 | 0.603 | 0.528 | -0.068 | 0.035 | 0.035 | 0 | 0.16 | 0.37 | 0.13 | 0.16 | 0.1 | 0.14 |
| 4 | 93285927 | rs17019342 | -0.351 | -0.107 | 0.459 | 0.606 | -0.012 | 0.012 | 0.012 | 0 | 0.05 | 0.06 | 0.03 | 0.11 | 0.01 | 0.04 |
| 4 | 93288641 | rs7685835 | -0.599 | 0.646 | 0.328 | 0.334 | 0.021 | -0.01 | -0.01 | 0 | 0.25 | 0.42 | 0.18 | 0.25 | 0.31 | 0.35 |
| 4 | 93300188 | rs2089990 | 0.051 | 0.865 | -0.319 | 0.261 | -0.026 | -0.054 | -0.054 | 0 | 0.15 | 0.07 | 0.47 | 0.61 | 0.23 | 0.29 |
| 4 | 93309526 | rs1513460 | -1.174 | 0.557 | -0.178 | 0.068 | 0 | -0.016 | -0.016 | 0.028 | 0.25 | 0.19 | 0.54 | 0.67 | 0.33 | 0.4 |
| 4 | 93318804 | rs13122471 | -0.733 | 0.244 | -0.097 | -0.073 | -0.001 | 0.001 | 0.001 | 0 | 0.1 | 0.1 | 0.07 | 0.05 | 0.1 | 0.12 |
| 4 | 93322977 | rs4133324 | -0.524 | 0.227 | 0.099 | -0.276 | 0.007 | 0.005 | 0.005 | 0 | 0.28 | 0.18 | 0.15 | 0.23 | 0.23 | 0.38 |
| 4 | 93344512 | rs4374594 | -1.787 | 0.241 | -0.041 | -0.881 | -0.007 | 0.073 | 0.073 | 3.912 | 0.63 | 0.79 | 0.82 | 0.9 | 0.74 | 0.8 |
| 4 | 93346743 | rs5021051 | -1.837 | 0.319 | -0.068 | -0.908 | 0.009 | 0.066 | 0.066 | 3.708 | 0.64 | 0.79 | 0.85 | 0.9 | 0.78 | 0.8 |
| 4 | 93354048 | rs6844474 | -1.143 | -0.457 | 0.691 | -0.645 | -0.04 | -0.006 | -0.006 | 0 | 0.16 | 0.35 | 0.1 | 0.13 | 0.1 | 0.16 |
| 4 | 93368860 | rs7668752 | -1.207 | -0.547 | 0.569 | -0.67 | -0.02 | -0.018 | -0.018 | 0 | 0.34 | 0.58 | 0.22 | 0.24 | 0.29 | 0.39 |
| 4 | 93372969 | rs4629410 | 0.844 | -0.485 | 0.416 | -0.542 | -0.018 | 0.004 | 0.004 | 0 | 0.17 | 0.02 | 0.11 | 0.13 | 0.2 | 0.24 |
| 4 | 93391877 | rs12645728 | 0.502 | -0.44 | 0.467 | -0.664 | -0.001 | 0.051 | 0.051 | 0 | 0.2 | 0.06 | 0.14 | 0.12 | 0.23 | 0.24 |
| 4 | 93398759 | rs12643954 | -1.264 | -0.457 | 0.656 | -0.662 | -0.033 | -0.041 | -0.041 | 0.046 | 0.38 | 0.54 | 0.26 | 0.24 | 0.32 | 0.4 |
| 4 | 93416752 | rs7684294 | -0.506 | -0.393 | 0.21 | -0.572 | -0.007 | 0.009 | 0.009 | 0 | 0.07 | 0.11 | 0.07 | 0.08 | 0.05 | 0.06 |
| 4 | 93428335 | rs7659159 | -1.394 | -0.289 | -0.018 | -0.31 | 0.015 | 0.012 | 0.012 | 0.147 | 0.42 | 0.66 | 0.44 | 0.36 | 0.38 | 0.44 |
| 4 | 93428415 | rs7659338 | -0.009 | 0.132 | -0.625 | -0.192 | -0.018 | 0.001 | 0.001 | 0 | 0.24 | 0.1 | 0.33 | 0.23 | 0.28 | 0.29 |
| 4 | 93449941 | rs4692972 | -0.415 | -0.016 | -0.459 | -0.229 | 0 | -0.009 | -0.009 | 0 | 0.05 | 0.05 | 0.08 | 0 | 0.08 | 0.05 |
| 4 | 93450349 | rs4692973 | -1.396 | -0.731 | -0.762 | -0.61 | 0.049 | 0.051 | 0.051 | 0 | 0.33 | 0.26 | 0.2 | 0.18 | 0.25 | 0.24 |
| 4 | 93451160 | rs11944800 | -1.275 | -0.749 | -0.339 | -0.628 | -0.011 | -0.021 | -0.021 | 0 | 0.43 | 0.52 | 0.49 | 0.31 | 0.42 | 0.44 |
| 4 | 93451321 | rs13140156 | -0.301 | -0.789 | -0.623 | -0.683 | -0.014 | 0.03 | 0.03 | 0 | 0.29 | 0.08 | 0.36 | 0.25 | 0.32 | 0.34 |
| 4 | 93455880 | rs7661958 | -1.968 | -1.031 | -0.393 | -0.576 | 0.012 | 0.004 | 0.004 | 0 | 0.48 | 0.58 | 0.43 | 0.42 | 0.41 | 0.49 |
| 4 | 93458820 | rs17319190 | 0.073 | -1.173 | -0.524 | -0.671 | 0.025 | 0.061 | 0.061 | 0 | 0.13 | 0.03 | 0.07 | 0 | 0.09 | 0.03 |
| 4 | 93467486 | rs1993030 | 0.228 | -1.232 | -0.386 | -0.655 | -0.009 | -0.01 | -0.01 | 0 | 0.22 | 0.09 | 0.29 | 0.26 | 0.26 | 0.3 |
| 4 | 93467552 | rs17019538 | -1.671 | -1.246 | -0.39 | -0.674 | -0.062 | -0.021 | -0.021 | 0 | 0.26 | 0.44 | 0.14 | 0.16 | 0.15 | 0.18 |
| 4 | 93480717 | rs7663643 | -1.791 | -1.131 | -0.221 | -0.444 | -0.013 | 0.004 | 0.004 | 0.017 | 0.4 | 0.51 | 0.42 | 0.43 | 0.36 | 0.44 |
| 4 | 93485733 | rs2060676 | 0.117 | -1.091 | -0.354 | -0.44 | -0.023 | -0.021 | -0.021 | 1.173 | 0.22 | 0.08 | 0.3 | 0.26 | 0.26 | 0.3 |
| 4 | 93489287 | rs7682842 | -0.512 | -1.074 | -0.106 | -0.363 | -0.002 | 0.012 | 0.012 | 0 | 0.28 | 0.17 | 0.34 | 0.27 | 0.31 | 0.32 |
| 4 | 93504230 | rs9685970 | -1.612 | -1.084 | 0.054 | -0.399 | 0.01 | -0.008 | -0.008 | 0 | 0.39 | 0.39 | 0.41 | 0.43 | 0.36 | 0.44 |
| 4 | 93508181 | rs13101814 | 1.848 | -0.96 | 0.019 | -1.039 | 0.006 | -0.015 | -0.015 | 0.901 | 0.31 | 0.21 | 0.12 | 0 | 0.19 | 0.16 |
| 4 | 93508896 | rs9307120 | -0.607 | -0.937 | 0.239 | -0.147 | 0 | 0.013 | 0.013 | 0 | 0.28 | 0.18 | 0.34 | 0.26 | 0.31 | 0.32 |
| 4 | 93539532 | rs7668740 | -1.309 | -0.905 | 0.619 | 0.11 | -0.013 | 0.005 | 0.005 | 1.092 | 0.35 | 0.59 | 0.37 | 0.42 | 0.32 | 0.44 |

| | | | | | | | | | | | | | | | | |
|---|---|---|---|---|---|---|---|---|---|---|---|---|---|---|---|---|
| 4 | 93542768 | rs1349350 | -1.287 | -0.897 | 0.685 | 0.122 | -0.022 | 0.013 | 0.013 | 0.797 | 0.35 | 0.55 | 0.37 | 0.42 | 0.32 | 0.46 |
| 4 | 93553252 | rs921355 | -0.412 | -0.919 | 0.443 | 0.13 | 0.001 | 0.003 | 0.003 | 0.021 | 0.25 | 0.24 | 0.3 | 0.27 | 0.27 | 0.34 |
| 4 | 93559571 | rs2169945 | -1.329 | -0.914 | 0.447 | 0.136 | 0 | -0.001 | -0.001 | 0 | 0.34 | 0.4 | 0.36 | 0.43 | 0.32 | 0.46 |
| 4 | 93579550 | rs12648061 | -0.186 | -0.762 | 0.206 | 0.194 | 0.013 | -0.023 | -0.023 | 1.908 | 0.08 | 0.04 | 0.05 | 0.16 | 0.05 | 0.15 |
| 4 | 93592657 | rs34905965 | -1.235 | -0.826 | 0.292 | 0.249 | -0.008 | 0.007 | 0.007 | 0.054 | 0.34 | 0.4 | 0.36 | 0.46 | 0.32 | 0.43 |
| 4 | 93602855 | rs9994093 | -0.775 | -1.051 | 0.276 | 0.109 | -0.035 | -0.035 | -0.035 | 0 | 0.29 | 0.51 | 0.33 | 0.28 | 0.23 | 0.26 |
| 4 | 93612903 | rs17019763 | -0.057 | -1.032 | -0.195 | 0.345 | 0.017 | -0.023 | -0.023 | 0.18 | 0.2 | 0.13 | 0.23 | 0.35 | 0.15 | 0.32 |
| 4 | 93619350 | rs17257464 | 0.456 | -1.382 | -0.31 | 0.038 | 0.008 | 0.009 | 0.009 | 0.293 | 0.1 | 0.05 | 0.07 | 0.01 | 0.06 | 0.03 |
| 4 | 93621420 | rs11734837 | -0.22 | -1.128 | -0.296 | 0.273 | 0 | -0.002 | -0.002 | 0.011 | 0.09 | 0.06 | 0.06 | 0.09 | 0.07 | 0.07 |
| 4 | 93621793 | rs2061889 | -0.851 | -1.122 | -0.288 | 0.255 | 0.012 | -0.019 | -0.019 | 0 | 0.29 | 0.22 | 0.31 | 0.45 | 0.22 | 0.39 |
| 4 | 93627019 | rs12643444 | -1.022 | -1.142 | -0.12 | 0.245 | -0.013 | 0.012 | 0.012 | 0 | 0.3 | 0.37 | 0.3 | 0.44 | 0.22 | 0.39 |
| 4 | 93635505 | rs1836439 | -0.623 | -1.278 | -0.423 | 0.406 | 0.008 | -0.011 | -0.011 | 0.37 | 0.27 | 0.23 | 0.27 | 0.45 | 0.18 | 0.39 |
| 4 | 93638479 | rs17019814 | 0.104 | -1.261 | -0.429 | 0.447 | -0.007 | 0.01 | 0.01 | 0 | 0.2 | 0.22 | 0.25 | 0.37 | 0.15 | 0.33 |
| 4 | 93641094 | rs10032050 | -0.854 | -1.301 | -0.488 | 0.499 | 0.026 | -0.018 | -0.018 | 0.22 | 0.32 | 0.24 | 0.33 | 0.44 | 0.24 | 0.4 |
| 4 | 93653338 | rs6532381 | -0.81 | -1.349 | -0.6 | 1.022 | 0.048 | -0.074 | -0.074 | 0.001 | 0.32 | 0.16 | 0.36 | 0.56 | 0.25 | 0.4 |
| 4 | 93659471 | rs1369166 | -0.718 | -1.497 | -0.986 | 0.79 | 0.059 | 0.022 | 0.022 | 0.812 | 0.44 | 0.15 | 0.38 | 0.42 | 0.31 | 0.38 |
| 4 | 93660667 | rs10516914 | -1.134 | -1.533 | -0.358 | 0.862 | 0.044 | 0.013 | 0.013 | 0 | 0.5 | 0.39 | 0.44 | 0.59 | 0.32 | 0.5 |
| 4 | 93660958 | rs10516916 | 0.563 | -1.516 | -0.512 | 0.703 | 0.002 | 0.03 | 0.03 | 0 | 0.07 | 0.02 | 0.06 | 0 | 0.07 | 0.03 |
| 4 | 93665778 | rs954467 | -1.576 | -1.263 | -0.086 | 0.347 | -0.016 | -0.015 | -0.015 | 0 | 0.75 | 0.79 | 0.63 | 0.58 | 0.5 | 0.61 |
| 4 | 93673153 | rs17019871 | 1.328 | -1.408 | 0.292 | 0.137 | -0.018 | -0.052 | -0.052 | 0 | 0.22 | 0.29 | 0.1 | 0.05 | 0.12 | 0.17 |
| 4 | 93674890 | rs17019874 | -0.222 | -1.426 | 0.369 | 0.135 | -0.01 | 0.019 | 0.019 | 0.208 | 0.08 | 0.15 | 0.06 | 0.12 | 0.06 | 0.06 |
| 4 | 93675391 | rs13141798 | 1.284 | -1.419 | 0.479 | 0.141 | -0.027 | -0.067 | -0.067 | 0 | 0.22 | 0.31 | 0.09 | 0.06 | 0.12 | 0.17 |
| 4 | 93678655 | rs12507379 | 0.306 | -1.402 | 0.257 | 0.238 | 0.029 | 0.076 | 0.076 | 0.455 | 0.42 | 0.15 | 0.45 | 0.33 | 0.33 | 0.39 |
| 4 | 93691318 | rs6852643 | -0.176 | -1.314 | 0.375 | 0.288 | -0.011 | 0.026 | 0.026 | 0 | 0.08 | 0.15 | 0.07 | 0.12 | 0.06 | 0.06 |
| 4 | 93707044 | rs17259489 | 0.204 | -0.848 | 0.162 | 0.242 | 0 | 0.092 | 0.092 | 0.721 | 0.44 | 0.14 | 0.48 | 0.32 | 0.38 | 0.41 |
| 4 | 93707288 | rs7671095 | -0.224 | -1.287 | 0.347 | 0.163 | -0.029 | -0.048 | -0.048 | 0.023 | 0.35 | 0.49 | 0.18 | 0.18 | 0.21 | 0.18 |
| 4 | 93710269 | rs28409430 | 0.766 | -1.229 | 0.114 | -0.083 | -0.001 | -0.058 | -0.058 | 0.177 | 0.28 | 0.27 | 0.14 | 0 | 0.16 | 0.12 |
| 4 | 93723466 | rs10856898 | 0.042 | -1.128 | -0.219 | 0.161 | 0.035 | 0.095 | 0.095 | 0.982 | 0.42 | 0.17 | 0.45 | 0.3 | 0.33 | 0.39 |
| 4 | 93786775 | rs6532382 | -0.334 | -1.099 | -0.047 | 0.485 | -0.005 | -0.002 | -0.002 | 0.022 | 0.45 | 0.44 | 0.57 | 0.45 | 0.33 | 0.44 |
| 4 | 93803563 | rs1456359 | -0.11 | -1.122 | -0.446 | 0.498 | 0.023 | 0.096 | 0.096 | 1.304 | 0.48 | 0.18 | 0.58 | 0.34 | 0.37 | 0.39 |
| 4 | 93833561 | rs1456362 | -1.287 | -1.208 | -0.208 | 0.866 | -0.064 | -0.043 | -0.043 | 0.054 | 0.72 | 0.8 | 0.73 | 0.57 | 0.47 | 0.53 |
| 4 | 93847904 | rs2168489 | 0.299 | -1.473 | 0.176 | 0.782 | 0.013 | 0.038 | 0.038 | 2.03 | 0.45 | 0.3 | 0.56 | 0.29 | 0.35 | 0.38 |
| 4 | 93849612 | rs3911386 | -1.729 | -1.349 | -0.049 | 0.856 | -0.055 | -0.051 | -0.051 | 1.554 | 0.75 | 0.87 | 0.75 | 0.56 | 0.51 | 0.53 |
| 4 | 93872906 | rs17327937 | 0.953 | -1.549 | 0.052 | 0.544 | 0.031 | 0.043 | 0.043 | 0 | 0.19 | 0.08 | 0.09 | 0 | 0.09 | 0.09 |
| 4 | 93892487 | rs17020045 | -1.766 | -1.333 | 0.07 | 0.82 | -0.093 | -0.067 | -0.067 | 0.161 | 0.75 | 0.89 | 0.75 | 0.61 | 0.51 | 0.59 |
| 4 | 93902882 | rs2045316 | -0.115 | -1.243 | 0.756 | 0.753 | 0.003 | 0.011 | 0.011 | 0.327 | 0.5 | 0.4 | 0.61 | 0.34 | 0.4 | 0.39 |
| 4 | 93918343 | rs1039938 | 0.063 | -0.835 | 0.851 | 0.751 | 0.012 | 0.023 | 0.023 | 0.498 | 0.46 | 0.34 | 0.53 | 0.33 | 0.36 | 0.38 |
| 4 | 93969165 | rs17263644 | 0.464 | -1.073 | 0.2 | 0.451 | -0.032 | 0.077 | 0.077 | 0.39 | 0.19 | 0.06 | 0.21 | 0.06 | 0.2 | 0.12 |
| 4 | 93975772 | rs13135367 | -0.228 | -1.318 | 0.237 | 0.707 | -0.002 | -0.007 | -0.007 | 0 | 0.43 | 0.45 | 0.58 | 0.37 | 0.38 | 0.36 |
| 4 | 93985925 | rs17329945 | -1.201 | -1.505 | 0.256 | 0.81 | -0.035 | 0.003 | 0.003 | 0.549 | 0.59 | 0.84 | 0.7 | 0.6 | 0.41 | 0.5 |
| 4 | 93987251 | rs13126969 | -0.484 | -1.51 | 0.503 | 0.8 | 0.006 | 0.005 | 0.005 | 1.656 | 0.5 | 0.43 | 0.57 | 0.41 | 0.36 | 0.39 |
| 4 | 93989892 | rs1912718 | -0.413 | -1.355 | 0.556 | 0.817 | 0.001 | 0.004 | 0.004 | 0.056 | 0.47 | 0.49 | 0.61 | 0.35 | 0.39 | 0.4 |
| 4 | 93993208 | rs10032153 | -0.257 | -1.357 | 0.527 | 0.704 | -0.001 | -0.001 | -0.001 | 0.798 | 0.44 | 0.42 | 0.62 | 0.42 | 0.37 | 0.43 |
| 4 | 93994938 | rs1511294 | -0.342 | -1.477 | 0.519 | 0.711 | -0.001 | 0 | 0 | 0.675 | 0.46 | 0.43 | 0.67 | 0.48 | 0.4 | 0.41 |
| 4 | 93998463 | rs6857940 | -0.424 | -1.384 | 0.488 | 0.754 | 0.011 | -0.005 | -0.005 | 0.233 | 0.46 | 0.57 | 0.64 | 0.49 | 0.37 | 0.41 |
| 4 | 94011311 | rs4502650 | -0.581 | -1.382 | 0.504 | 0.821 | 0.015 | 0.006 | 0.006 | 0 | 0.5 | 0.57 | 0.68 | 0.53 | 0.41 | 0.42 |
| 4 | 94014156 | rs12506519 | -0.225 | -1.365 | 0.452 | 0.813 | 0.004 | 0.032 | 0.032 | 0 | 0.44 | 0.3 | 0.63 | 0.34 | 0.39 | 0.37 |
| 4 | 94034683 | rs3843409 | -0.622 | -1.445 | 0.362 | 0.837 | 0.041 | 0.06 | 0.06 | 0 | 0.51 | 0.77 | 0.69 | 0.61 | 0.39 | 0.48 |
| 4 | 94037375 | rs6849575 | 0.298 | -1.334 | 0.442 | 0.541 | 0.01 | 0.041 | 0.041 | 0 | 0.05 | 0.22 | 0.04 | 0.08 | 0.04 | 0.06 |
| 4 | 94052135 | rs17264265 | -0.081 | -1.45 | 0.423 | 0.336 | 0.011 | 0.055 | 0.055 | 0 | 0.44 | 0.21 | 0.62 | 0.33 | 0.38 | 0.38 |
| 4 | 94064059 | rs11097358 | -1.214 | -1.565 | 0.169 | 0.31 | -0.032 | 0.017 | 0.017 | 0 | 0.63 | 0.77 | 0.74 | 0.7 | 0.45 | 0.52 |
| 4 | 94075263 | rs1354296 | -0.71 | -1.53 | 0.222 | 0.352 | -0.008 | 0.023 | 0.023 | 0 | 0.55 | 0.76 | 0.72 | 0.6 | 0.44 | 0.48 |
| 4 | 94092083 | rs995906 | 0.183 | -1.625 | 0.003 | 0.279 | -0.034 | 0.103 | 0.103 | 0 | 0.22 | 0.06 | 0.26 | 0.06 | 0.23 | 0.12 |
| 4 | 94100410 | rs6821061 | 0.525 | -1.548 | 0.597 | 0.894 | -0.004 | 0.042 | 0.042 | 0 | 0.07 | 0.11 | 0.07 | 0.18 | 0.05 | 0.04 |
| 4 | 94106172 | rs12645274 | -0.789 | -1.566 | 0.792 | 0.9 | -0.003 | 0.21 | 0.21 | 0.392 | 0.09 | 0.37 | 0.1 | 0.26 | 0.05 | 0.1 |
| 4 | 94115801 | rs1503223 | 0.132 | -1.538 | 0.732 | 0.839 | 0.002 | 0.13 | 0.13 | 0 | 0.08 | 0.19 | 0.07 | 0.23 | 0.06 | 0.07 |
| 4 | 94116539 | rs989927 | -0.935 | -1.321 | 0.176 | 0.594 | -0.018 | -0.001 | -0.001 | 0.011 | 0.67 | 0.76 | 0.79 | 0.66 | 0.54 | 0.55 |
| 4 | 94117721 | rs6857551 | -1.199 | -1.446 | 0.543 | 0.681 | -0.002 | 0.305 | 0.305 | 1.734 | 0.11 | 0.5 | 0.1 | 0.31 | 0.06 | 0.13 |
| 4 | 94120374 | rs1155646 | 0.196 | -1.257 | -0.069 | 0.485 | 0.026 | 0.046 | 0.046 | 0 | 0.5 | 0.37 | 0.73 | 0.65 | 0.56 | 0.5 |
| 4 | 94123529 | rs7667127 | -1.447 | -0.142 | 0.423 | 0.816 | 0.008 | 0.007 | 0.007 | 0 | 0.83 | 0.83 | 0.85 | 0.64 | 0.63 | 0.67 |
| 4 | 94125424 | rs12507496 | 0.739 | -0.255 | 0.379 | 0.785 | -0.011 | 0.014 | 0.014 | 0 | 0.6 | 0.24 | 0.72 | 0.35 | 0.54 | 0.5 |
| 4 | 94146093 | rs3796675 | -0.828 | -0.878 | 0.455 | 0.746 | -0.062 | 0.035 | 0.035 | 0 | 0.22 | 0.39 | 0.12 | 0.28 | 0.08 | 0.18 |
| 4 | 94159658 | rs17020286 | -0.099 | 0.353 | -0.605 | 1.047 | -0.001 | 0.059 | 0.059 | 0 | 0.07 | 0.08 | 0.06 | 0.26 | 0.05 | 0.07 |
| 4 | 94162154 | rs7686280 | 0.352 | 0.179 | -0.775 | 0.724 | 0.081 | 0.074 | 0.074 | 1.419 | 0.47 | 0.64 | 0.27 | 0.27 | 0.26 | 0.3 |
| 4 | 94162560 | rs1433654 | 0.745 | 0.742 | 0.269 | 1.722 | -0.002 | -0.002 | -0.002 | 0 | 0.72 | 0.66 | 0.8 | 0.52 | 0.63 | 0.51 |
| 4 | 94172097 | rs1821342 | -0.183 | 0.913 | 0.241 | 1.695 | -0.014 | -0.015 | -0.015 | 0 | 0.79 | 0.81 | 0.84 | 0.6 | 0.65 | 0.64 |
| 4 | 94185705 | rs7691723 | -0.176 | 0.911 | 0.218 | 1.682 | -0.05 | -0.056 | -0.056 | 0 | 0.79 | 0.87 | 0.84 | 0.6 | 0.65 | 0.63 |
| 4 | 94219390 | rs7684707 | -0.165 | 0.946 | -0.755 | 1.694 | -0.042 | -0.072 | -0.072 | 0 | 0.8 | 0.89 | 0.87 | 0.6 | 0.71 | 0.65 |
| 4 | 94227970 | rs6813949 | -1.196 | 0.243 | -0.214 | 1.274 | -0.013 | 0.083 | 0.083 | 0.186 | 0.11 | 0.31 | 0.09 | 0.28 | 0.08 | 0.19 |

| | | | | | | | | | | | | | | | | |
|---|---|---|---|---|---|---|---|---|---|---|---|---|---|---|---|---|
| 4 | 94233967 | rs6854968 | -0.511 | 0.023 | -0.572 | 0.252 | -0.005 | 0.005 | 0.005 | 0.105 | 0.37 | 0.61 | 0.38 | 0.58 | 0.39 | 0.5 |
| 4 | 94239926 | rs1433676 | -0.3 | -0.835 | -0.94 | -0.805 | 0.054 | 0.061 | 0.061 | 0 | 0.45 | 0.6 | 0.22 | 0.13 | 0.27 | 0.37 |
| 4 | 94242858 | rs7698899 | 0.756 | -1.289 | -1.023 | -1.028 | 0.009 | 0.02 | 0.02 | 0.392 | 0.13 | 0.06 | 0.08 | 0 | 0.11 | 0.1 |
| 4 | 94251270 | rs733942 | 0.172 | -1.281 | -1.184 | -1.103 | -0.007 | 0 | 0 | 0 | 0.38 | 0.43 | 0.38 | 0.61 | 0.31 | 0.42 |
| 4 | 94252360 | rs1433666 | 0.137 | -0.748 | -0.866 | -1.25 | -0.005 | 0.04 | 0.04 | 0 | 0.55 | 0.7 | 0.52 | 0.72 | 0.59 | 0.61 |
| 4 | 94252683 | rs10002068 | 0.213 | -0.643 | -0.129 | -1.113 | 0.042 | 0.038 | 0.038 | 0.54 | 0.08 | 0.39 | 0.06 | 0.05 | 0.07 | 0.14 |
| 4 | 94252722 | rs11943480 | -0.748 | -0.789 | -0.609 | -1.345 | -0.088 | -0.06 | -0.06 | 1.449 | 0.73 | 0.93 | 0.59 | 0.73 | 0.68 | 0.68 |
| 4 | 94255091 | rs7696884 | -0.945 | -1.479 | -0.182 | -1.311 | -0.047 | -0.072 | -0.072 | 1.352 | 0.21 | 0.6 | 0.15 | 0.05 | 0.13 | 0.26 |
| 4 | 94258848 | rs919866 | -0.935 | -1.488 | -0.206 | -1.31 | -0.043 | -0.064 | -0.064 | 0 | 0.21 | 0.62 | 0.15 | 0.05 | 0.13 | 0.26 |
| 4 | 94262808 | rs10028824 | -0.655 | -1.452 | -0.093 | -1.278 | -0.042 | -0.041 | -0.041 | 0 | 0.13 | 0.57 | 0.07 | 0.05 | 0.08 | 0.17 |
| 4 | 94283482 | rs4693315 | 1.644 | -1.961 | -0.779 | -1.523 | -0.101 | 0.011 | 0.011 | 0 | 0.18 | 0.01 | 0.19 | 0.15 | 0.31 | 0.17 |
| 4 | 94290979 | rs6834311 | 0.112 | -2.209 | -0.776 | -1.799 | -0.004 | 0.039 | 0.039 | 0.579 | 0.54 | 0.22 | 0.44 | 0.68 | 0.49 | 0.48 |
| 4 | 94294991 | rs1368716 | -0.068 | -2.159 | -0.533 | -1.78 | -0.011 | -0.05 | -0.05 | 0.826 | 0.14 | 0.49 | 0.13 | 0.05 | 0.12 | 0.22 |
| 4 | 94297664 | rs1905708 | -0.368 | -2.314 | -0.562 | -1.833 | -0.031 | -0.038 | -0.038 | 0 | 0.11 | 0.63 | 0.09 | 0.04 | 0.08 | 0.21 |
| 4 | 94300038 | rs11737556 | -1.66 | -2.345 | -0.59 | -1.943 | -0.076 | -0.036 | -0.036 | 1.317 | 0.75 | 0.89 | 0.59 | 0.73 | 0.63 | 0.75 |
| 4 | 94301291 | rs2082553 | 1.182 | -2.342 | -0.195 | -1.932 | 0.019 | -0.017 | -0.017 | 0 | 0.05 | 0.19 | 0.05 | 0.01 | 0.06 | 0.09 |
| 4 | 94312343 | rs1948016 | 0.282 | -2.676 | -0.264 | -1.927 | 0.002 | 0.005 | 0.005 | 0 | 0.32 | 0.37 | 0.42 | 0.64 | 0.39 | 0.42 |
| 4 | 94314479 | rs1994253 | -0.733 | -2.425 | -0.259 | -1.885 | -0.045 | -0.09 | -0.09 | 0 | 0.17 | 0.54 | 0.15 | 0.04 | 0.13 | 0.22 |
| 4 | 94316763 | rs1385405 | -1.807 | -2.438 | -0.307 | -1.894 | -0.153 | -0.088 | -0.088 | 3.915 | 0.77 | 0.96 | 0.6 | 0.73 | 0.63 | 0.75 |
| 4 | 94319668 | rs9631724 | -0.873 | -2.347 | -0.235 | -1.883 | -0.041 | -0.096 | -0.096 | 0 | 0.18 | 0.54 | 0.16 | 0.04 | 0.15 | 0.21 |
| 4 | 94324475 | rs11727464 | 0.71 | -2.557 | -0.507 | -2.026 | -0.027 | -0.021 | -0.021 | 0.403 | 0.36 | 0.16 | 0.38 | 0.66 | 0.41 | 0.41 |
| 4 | 94329254 | rs7662289 | -1.032 | -2.247 | -0.136 | -1.821 | -0.036 | -0.077 | -0.077 | 0 | 0.17 | 0.62 | 0.16 | 0.04 | 0.15 | 0.22 |
| 4 | 94331320 | rs11938668 | -0.543 | -2.003 | 0.025 | -1.633 | -0.011 | -0.034 | -0.034 | 0.039 | 0.11 | 0.54 | 0.1 | 0.04 | 0.1 | 0.21 |
| 4 | 94344042 | rs10034345 | 0.331 | -1.987 | 0.067 | -1.627 | -0.004 | -0.038 | -0.038 | 0 | 0.07 | 0.18 | 0.06 | 0.02 | 0.08 | 0.16 |
| 4 | 94358023 | rs10022753 | -1.309 | -1.928 | -0.242 | -1.625 | -0.01 | -0.006 | -0.006 | 0 | 0.75 | 0.79 | 0.59 | 0.72 | 0.63 | 0.73 |
| 4 | 94359448 | rs10006372 | -1.395 | -1.912 | -0.26 | -1.616 | -0.028 | -0.015 | -0.015 | 0.03 | 0.76 | 0.85 | 0.59 | 0.72 | 0.64 | 0.73 |
| 4 | 94361687 | rs10012811 | -0.282 | -1.973 | -0.027 | -1.552 | -0.042 | -0.059 | -0.059 | 0.587 | 0.13 | 0.54 | 0.09 | 0.04 | 0.09 | 0.18 |
| 4 | 94367364 | rs11097364 | 0.327 | -2.142 | -0.627 | -1.685 | -0.027 | 0.011 | 0.011 | 0.482 | 0.42 | 0.17 | 0.44 | 0.67 | 0.47 | 0.39 |
| 4 | 94372827 | rs6532407 | -0.608 | -1.468 | 0.17 | -1.243 | -0.043 | -0.069 | -0.069 | 0 | 0.13 | 0.48 | 0.09 | 0.04 | 0.08 | 0.19 |
| 4 | 94380515 | rs17020631 | 0.311 | -1.428 | 0.063 | -1.216 | -0.025 | -0.01 | -0.01 | 0 | 0.08 | 0.28 | 0.03 | 0.03 | 0.03 | 0.08 |
| 4 | 94382396 | rs1905724 | -0.323 | -1.613 | -1.177 | -1.11 | 0.039 | -0.003 | -0.003 | 0 | 0.56 | 0.7 | 0.41 | 0.53 | 0.3 | 0.51 |
| 4 | 94390998 | rs2200375 | 0.381 | -1.157 | -1.906 | -0.895 | 0.084 | -0.054 | -0.054 | 0 | 0.38 | 0.17 | 0.33 | 0.53 | 0.24 | 0.3 |
| 4 | 94402365 | rs11097368 | 0.466 | -0.098 | -1.745 | -0.578 | -0.077 | 0.051 | 0.051 | 0 | 0.22 | 0.04 | 0.16 | 0.14 | 0.32 | 0.18 |
| 4 | 94406148 | rs10030500 | -0.054 | -0.65 | -1.475 | -0.527 | 0.082 | 0.009 | 0.009 | 0 | 0.55 | 0.81 | 0.51 | 0.61 | 0.34 | 0.58 |
| 4 | 94413681 | rs7676645 | 0.021 | 0.305 | -0.785 | 0.246 | 0.027 | 0.045 | 0.045 | 0 | 0.81 | 0.63 | 0.68 | 0.75 | 0.78 | 0.73 |
| 4 | 94417245 | rs1905719 | -0.591 | 0.267 | -0.418 | 0.499 | 0.008 | 0.023 | 0.023 | 0 | 0.84 | 0.82 | 0.71 | 0.69 | 0.8 | 0.77 |
| 4 | 94423150 | rs7655363 | 0.848 | 0.631 | -1.141 | 0.322 | -0.083 | -0.001 | -0.001 | 0.047 | 0.19 | 0.02 | 0.14 | 0.12 | 0.3 | 0.14 |
| 4 | 94426942 | rs6851153 | 0.52 | 0.543 | -0.887 | 0.583 | -0.027 | -0.026 | -0.026 | 0.69 | 0.22 | 0.13 | 0.16 | 0.27 | 0.32 | 0.18 |
| 4 | 94430437 | rs4557232 | 0.594 | -0.115 | -1.245 | 0.052 | -0.011 | 0.033 | 0.033 | 0 | 0.26 | 0.37 | 0.34 | 0.43 | 0.22 | 0.35 |
| 4 | 94430964 | rs13144042 | 1.531 | 0.002 | -1.86 | -0.284 | -0.069 | 0.179 | 0.179 | 0 | 0.26 | 0.01 | 0.13 | 0.01 | 0.23 | 0.16 |
| 4 | 94440026 | rs4286488 | -1.136 | 0.179 | -1.824 | -0.179 | -0.084 | -0.107 | -0.107 | 0 | 0.8 | 0.98 | 0.68 | 0.71 | 0.75 | 0.74 |
| 4 | 94446458 | rs13131107 | 0.952 | 0.031 | -1.299 | -0.625 | -0.017 | 0.101 | 0.101 | 0 | 0.28 | 0.14 | 0.16 | 0.02 | 0.25 | 0.21 |
| 4 | 94458391 | rs1435473 | -0.26 | 0.139 | -0.967 | -0.551 | 0.003 | 0.034 | 0.034 | 0 | 0.54 | 0.6 | 0.36 | 0.27 | 0.58 | 0.45 |
| 4 | 94461705 | rs6836589 | 1.392 | 0.158 | -1.022 | -0.711 | -0.044 | 0.126 | 0.126 | 0 | 0.24 | 0.05 | 0.14 | 0.01 | 0.23 | 0.15 |
| 4 | 94494455 | rs11942069 | 0.858 | 0.096 | -0.547 | -0.738 | 0 | -0.08 | -0.08 | 0 | 0.29 | 0.28 | 0.17 | 0.02 | 0.27 | 0.21 |
| 4 | 94505733 | rs6824100 | -0.326 | -0.159 | -0.387 | -0.887 | 0 | 0.002 | 0.002 | 0 | 0.61 | 0.59 | 0.4 | 0.22 | 0.63 | 0.41 |
| 4 | 94516290 | rs13121519 | 1.019 | -0.073 | -0.288 | -0.757 | -0.035 | 0.153 | 0.153 | 0 | 0.28 | 0.08 | 0.15 | 0.02 | 0.25 | 0.21 |
| 4 | 94518801 | rs13117931 | 0.553 | -0.044 | -0.099 | -0.718 | 0 | -0.027 | -0.027 | 0 | 0.32 | 0.27 | 0.19 | 0.1 | 0.28 | 0.27 |
| 4 | 94528666 | rs1435474 | 0.593 | 0 | -0.43 | -0.67 | -0.001 | -0.025 | -0.025 | 0 | 0.32 | 0.27 | 0.19 | 0.1 | 0.28 | 0.27 |
| 4 | 94543233 | rs12503522 | -0.205 | 0.174 | -0.792 | 0.207 | 0 | -0.032 | -0.032 | 0 | 0.25 | 0.19 | 0.3 | 0.57 | 0.26 | 0.35 |
| 4 | 94543882 | rs11728147 | 1.556 | 0.164 | -0.883 | 0.571 | -0.027 | -0.036 | -0.036 | 0 | 0.13 | 0.03 | 0.09 | 0.17 | 0.17 | 0.09 |
| 4 | 94552306 | rs10516928 | 0.031 | 0.124 | -0.945 | 0.206 | 0.001 | 0.049 | 0.049 | 0 | 0.5 | 0.31 | 0.44 | 0.64 | 0.5 | 0.51 |
| 4 | 94566830 | rs6856229 | 0.36 | -0.366 | -1.231 | -0.23 | 0.027 | 0.132 | 0.132 | 0.788 | 0.51 | 0.71 | 0.29 | 0.14 | 0.4 | 0.32 |
| 4 | 94569114 | rs9993464 | -0.033 | -0.379 | -1.169 | -0.243 | -0.034 | -0.066 | -0.066 | 0 | 0.27 | 0.59 | 0.15 | 0.08 | 0.17 | 0.17 |
| 4 | 94570408 | rs10516933 | 2.355 | -0.063 | -1.527 | -0.122 | -0.036 | 0.091 | 0.091 | 0 | 0.23 | 0.09 | 0.13 | 0.06 | 0.23 | 0.15 |
| 4 | 94572516 | rs10516936 | 1.517 | 0.053 | -1.129 | -0.019 | -0.001 | -0.007 | -0.007 | 0 | 0.13 | 0.12 | 0.09 | 0.07 | 0.13 | 0.1 |
| 4 | 94572864 | rs6848888 | -0.928 | 0.568 | -1.928 | 0.259 | -0.05 | -0.038 | -0.038 | 0 | 0.77 | 0.9 | 0.6 | 0.73 | 0.67 | 0.69 |
| 4 | 94579992 | rs12505284 | 1.134 | 0.231 | -1.263 | 0.914 | -0.026 | -0.052 | -0.052 | 0.238 | 0.13 | 0.03 | 0.09 | 0.22 | 0.17 | 0.09 |
| 4 | 94582661 | rs6855368 | -1.282 | -0.203 | -1.492 | 0.513 | 0.06 | 0.057 | 0.057 | 0.294 | 0.51 | 0.81 | 0.46 | 0.66 | 0.42 | 0.53 |
| 4 | 94583396 | rs12509478 | 1.436 | 0.008 | -1.458 | 0.635 | -0.031 | -0.054 | -0.054 | 0.173 | 0.12 | 0.01 | 0.07 | 0.18 | 0.14 | 0.1 |
| 4 | 94585087 | rs6532424 | 0.031 | 0.041 | -1.131 | 0.811 | -0.001 | 0.004 | 0.004 | 0 | 0.17 | 0.17 | 0.27 | 0.42 | 0.18 | 0.26 |
| 4 | 94591898 | rs17020960 | -0.263 | -0.288 | -0.351 | 0.38 | -0.031 | -0.096 | -0.096 | 0.054 | 0.16 | 0.35 | 0.09 | 0.01 | 0.1 | 0.11 |
| 4 | 94594199 | rs10516940 | -0.105 | -0.231 | -0.474 | 0.456 | -0.001 | -0.007 | -0.007 | 0.544 | 0.29 | 0.28 | 0.33 | 0.6 | 0.31 | 0.34 |
| 4 | 94594845 | rs2859980 | -1.559 | -0.117 | -0.786 | 0.481 | 0.024 | 0.053 | 0.053 | 0 | 0.52 | 0.8 | 0.46 | 0.67 | 0.43 | 0.49 |
| 4 | 94605867 | rs6821896 | -0.368 | 0.335 | -0.537 | 0.732 | 0 | -0.001 | -0.001 | 0 | 0.27 | 0.68 | 0.37 | 0.29 | 0.25 | 0.3 |
| 4 | 94605968 | rs13135407 | -0.145 | 0.652 | -0.582 | 0.547 | -0.019 | -0.016 | -0.016 | 1.431 | 0.26 | 0.21 | 0.4 | 0.72 | 0.33 | 0.47 |
| 4 | 94616814 | rs4692987 | -0.93 | 0.47 | -0.333 | 0.236 | -0.008 | 0.013 | 0.013 | 0 | 0.41 | 0.38 | 0.51 | 0.74 | 0.47 | 0.54 |
| 4 | 94617165 | rs1036640 | -0.589 | 0.481 | -0.412 | 0.242 | -0.015 | 0.009 | 0.009 | 0 | 0.36 | 0.33 | 0.47 | 0.73 | 0.44 | 0.53 |
| 4 | 94618455 | rs4693334 | 1.965 | 0.619 | -1.148 | -0.322 | -0.003 | 0.041 | 0.041 | 0 | 0.24 | 0.16 | 0.16 | 0.06 | 0.26 | 0.12 |

| | | | | | | | | | | | | | | | | |
|---|---|---|---|---|---|---|---|---|---|---|---|---|---|---|---|---|
| 4 | 94623649 | rs1371357 | -0.933 | 0.68 | -0.467 | -0.066 | -0.001 | -0.03 | -0.03 | 0.703 | 0.86 | 0.89 | 0.92 | 0.97 | 0.88 | 0.86 |
| 4 | 94629367 | rs7659517 | -0.842 | 0.605 | -0.366 | -0.049 | 0.001 | -0.011 | -0.011 | 0.08 | 0.42 | 0.6 | 0.53 | 0.75 | 0.47 | 0.54 |
| 4 | 94634302 | rs6850113 | 1.65 | 0.631 | -0.741 | 0.038 | 0.016 | 0.083 | 0.083 | 0.206 | 0.42 | 0.18 | 0.37 | 0.19 | 0.38 | 0.29 |
| 4 | 94650978 | rs1898905 | 0.594 | 0.93 | -0.525 | 0.667 | -0.009 | 0.008 | 0.008 | 0 | 0.09 | 0.08 | 0.1 | 0.41 | 0.12 | 0.21 |
| 4 | 94676316 | rs12642295 | -0.712 | 0.737 | -0.362 | 0.084 | -0.001 | -0.008 | -0.008 | 2.734 | 0.4 | 0.59 | 0.48 | 0.75 | 0.4 | 0.57 |
| 4 | 94680540 | rs13101891 | -0.284 | 0.703 | -0.673 | -0.109 | 0 | -0.011 | -0.011 | 0.005 | 0.79 | 0.84 | 0.86 | 0.95 | 0.81 | 0.82 |
| 4 | 94680811 | rs4693337 | -0.935 | 0.795 | -0.652 | 0.049 | 0.003 | -0.016 | -0.016 | 2.4 | 0.88 | 0.89 | 0.92 | 0.95 | 0.88 | 0.87 |
| 4 | 94699009 | rs4619859 | -0.576 | 0.753 | -0.879 | 0.008 | -0.002 | 0.018 | 0.018 | 0 | 0.38 | 0.34 | 0.46 | 0.78 | 0.4 | 0.57 |
| 4 | 94701354 | rs732020 | -0.554 | 0.773 | -0.877 | 0.009 | -0.002 | 0.017 | 0.017 | 0 | 0.37 | 0.34 | 0.46 | 0.77 | 0.4 | 0.57 |
| 4 | 94702001 | rs10084947 | -0.55 | 0.772 | -1.121 | 0.009 | -0.007 | 0.04 | 0.04 | 0 | 0.37 | 0.25 | 0.45 | 0.77 | 0.4 | 0.57 |
| 4 | 94702064 | rs17021105 | 1.134 | 0.808 | -1.856 | 0.566 | -0.041 | -0.071 | -0.071 | 0 | 0.12 | 0.01 | 0.29 | 0.24 | 0.16 | 0.14 |
| 4 | 94704547 | rs4693340 | 1.161 | 0.418 | -1.575 | 0.17 | -0.009 | 0.042 | 0.042 | 0 | 0.37 | 0.21 | 0.35 | 0.2 | 0.38 | 0.25 |
| 4 | 94706706 | rs735256 | 0.607 | 0.522 | -1.422 | -0.038 | 0.006 | 0.06 | 0.06 | 0 | 0.48 | 0.34 | 0.44 | 0.2 | 0.49 | 0.3 |
| 4 | 94714996 | rs12644979 | 0.221 | 0.251 | -1.623 | 0.625 | -0.012 | -0.107 | -0.107 | 3.168 | 0.14 | 0.01 | 0.1 | 0.47 | 0.12 | 0.23 |
| 4 | 94717163 | rs11097387 | 0.42 | 0.64 | -1.411 | -0.08 | -0.008 | 0.091 | 0.091 | 0 | 0.42 | 0.23 | 0.49 | 0.19 | 0.41 | 0.24 |
| 4 | 94718878 | rs2218120 | -0.46 | 0.221 | -0.839 | 0.321 | 0.011 | 0.016 | 0.016 | 0 | 0.34 | 0.51 | 0.39 | 0.38 | 0.36 | 0.39 |
| 4 | 94719712 | rs10009431 | 0.623 | 0.124 | -0.942 | 0.09 | -0.009 | 0.034 | 0.034 | 0 | 0.56 | 0.35 | 0.48 | 0.35 | 0.55 | 0.48 |
| 4 | 94720590 | rs10024051 | 0.742 | 0.004 | -1.05 | 0.025 | -0.029 | 0.076 | 0.076 | 0 | 0.53 | 0.18 | 0.46 | 0.35 | 0.54 | 0.35 |
| 4 | 94721315 | rs1371356 | 1.505 | -0.068 | -1.09 | 0.061 | -0.026 | -0.053 | -0.053 | 0 | 0.08 | 0.01 | 0.14 | 0.19 | 0.1 | 0.11 |
| 4 | 94721747 | rs6844311 | 0.618 | -0.07 | -1.019 | 0.023 | -0.022 | 0.049 | 0.049 | 0 | 0.56 | 0.27 | 0.48 | 0.35 | 0.55 | 0.49 |
| 4 | 94725853 | rs10033683 | 1.634 | -0.082 | -1.429 | -0.122 | -0.07 | 0.029 | 0.029 | 0.001 | 0.23 | 0.07 | 0.26 | 0.19 | 0.3 | 0.21 |
| 4 | 94727622 | rs726899 | 0.481 | 0.35 | -0.266 | 0.438 | 0 | -0.002 | -0.002 | 0 | 0.57 | 0.47 | 0.49 | 0.34 | 0.55 | 0.49 |
| 4 | 94729848 | rs10024195 | 1.133 | 0.491 | -0.013 | 0.372 | -0.027 | 0.095 | 0.095 | 0 | 0.36 | 0.15 | 0.36 | 0.21 | 0.4 | 0.27 |
| 4 | 94730209 | rs17021163 | 0.96 | 0.498 | -0.043 | 0.392 | -0.027 | -0.055 | -0.055 | 0 | 0.09 | 0.01 | 0.14 | 0.18 | 0.11 | 0.11 |
| 4 | 94731641 | rs993775 | 0.055 | 0.518 | 0.231 | 0.4 | -0.003 | 0.041 | 0.041 | 0 | 0.51 | 0.73 | 0.55 | 0.39 | 0.52 | 0.52 |
| 4 | 94733470 | rs9991641 | 1.007 | 0.523 | 0.289 | 0.4 | -0.005 | 0.057 | 0.057 | 0 | 0.37 | 0.25 | 0.37 | 0.21 | 0.4 | 0.28 |
| 4 | 94734922 | rs9284593 | -0.833 | 0.609 | 0.032 | 0.547 | 0.011 | -0.029 | -0.029 | 0 | 0.6 | 0.77 | 0.72 | 0.54 | 0.66 | 0.62 |
| 4 | 94736376 | rs9307131 | 1.021 | 0.612 | 0.093 | 0.544 | 0 | -0.015 | -0.015 | 0 | 0.37 | 0.38 | 0.38 | 0.21 | 0.4 | 0.28 |
| 4 | 94739787 | rs12499415 | 0.701 | 0.614 | -0.156 | 0.545 | -0.001 | -0.016 | -0.016 | 0 | 0.4 | 0.57 | 0.41 | 0.21 | 0.41 | 0.3 |
| 4 | 94741522 | rs10022714 | -0.693 | 0.646 | -0.833 | 0.567 | 0.011 | -0.03 | -0.03 | 0 | 0.6 | 0.77 | 0.72 | 0.54 | 0.66 | 0.63 |
| 4 | 94745528 | rs11731687 | 0.842 | 0.759 | -0.877 | 0.631 | -0.048 | -0.042 | -0.042 | 0 | 0.08 | 0 | 0.17 | 0.15 | 0.13 | 0.1 |
| 4 | 94746245 | rs9996938 | -0.487 | 0.228 | -0.424 | 0.278 | -0.005 | 0.04 | 0.04 | 0 | 0.52 | 0.73 | 0.56 | 0.39 | 0.52 | 0.52 |
| 4 | 94746370 | rs10021588 | -0.32 | 0.38 | -1.117 | -0.304 | 0.001 | 0.011 | 0.011 | 0 | 0.4 | 0.39 | 0.4 | 0.21 | 0.41 | 0.41 |
| 4 | 94748289 | rs10015253 | -0.018 | 0.75 | -2.156 | -0.094 | -0.025 | 0.093 | 0.093 | 0 | 0.36 | 0.15 | 0.36 | 0.21 | 0.4 | 0.27 |
| 4 | 94759826 | rs7435060 | -0.392 | 0.375 | -1.769 | -0.037 | -0.011 | -0.016 | -0.016 | 0.694 | 0.16 | 0.02 | 0.11 | 0.13 | 0.13 | 0.13 |
| 4 | 94760262 | rs4264798 | 0.026 | 0.075 | -1.296 | -1.022 | 0.001 | -0.055 | -0.055 | 1.617 | 0.61 | 0.47 | 0.73 | 0.91 | 0.63 | 0.74 |
| 4 | 94760857 | rs7664530 | -0.583 | -0.611 | -0.964 | -0.194 | 0.002 | -0.003 | -0.003 | 0.346 | 0.16 | 0.14 | 0.28 | 0.25 | 0.1 | 0.15 |
| 4 | 94761097 | rs7689734 | -0.18 | -1.005 | -0.626 | -0.256 | -0.014 | 0.036 | 0.036 | 0 | 0.23 | 0.26 | 0.2 | 0.57 | 0.18 | 0.43 |
| 4 | 94761237 | rs4485792 | -0.361 | -1.09 | -0.229 | -0.296 | -0.016 | 0.043 | 0.043 | 0 | 0.17 | 0.35 | 0.32 | 0.23 | 0.13 | 0.16 |
| 4 | 94761551 | rs4558831 | -1.545 | -1.179 | -0.946 | -0.758 | -0.073 | 0.211 | 0.211 | 0.187 | 0.58 | 0.97 | 0.64 | 0.87 | 0.44 | 0.66 |
| 4 | 94766621 | rs7693338 | -1.102 | -1.188 | -0.36 | -0.493 | 0.002 | -0.002 | -0.002 | 0.243 | 0.33 | 0.74 | 0.44 | 0.35 | 0.31 | 0.22 |
| 4 | 94769055 | rs3924764 | -0.009 | -0.919 | -0.198 | -0.65 | 0.045 | -0.012 | -0.012 | 0 | 0.12 | 0.3 | 0.1 | 0.09 | 0.17 | 0.06 |
| 4 | 94769686 | rs7688836 | -0.265 | -1.002 | -0.279 | -0.688 | -0.018 | -0.001 | -0.001 | 2.907 | 0.23 | 0.32 | 0.36 | 0.22 | 0.19 | 0.17 |
| 4 | 94775266 | rs4568209 | -1.044 | -0.176 | -0.652 | -1.047 | -0.142 | 0.187 | 0.187 | 0.195 | 0.6 | 0.97 | 0.67 | 0.85 | 0.54 | 0.66 |
| 4 | 94777861 | rs4476567 | -1.92 | 0.269 | 0.831 | -0.168 | -0.001 | -0.031 | -0.031 | 0.189 | 0.18 | 0.65 | 0.14 | 0.1 | 0.19 | 0.09 |
| 4 | 94778298 | rs12643691 | -0.9 | 0.735 | 0.146 | 0.124 | 0.062 | -0.009 | -0.009 | 0.027 | 0.11 | 0.3 | 0.11 | 0.09 | 0.17 | 0.07 |
| 4 | 94793323 | rs4693349 | -0.698 | 0.434 | -0.534 | 0.383 | 0.006 | 0.005 | 0.005 | 0 | 0.35 | 0.34 | 0.39 | 0.66 | 0.29 | 0.51 |
| 4 | 94799694 | rs7666492 | -0.977 | 0.552 | 0.166 | 0.217 | -0.001 | -0.007 | -0.007 | 3.021 | 0.4 | 0.42 | 0.46 | 0.71 | 0.38 | 0.53 |
| 4 | 94802707 | rs6818437 | -0.907 | 0.394 | 1.35 | 0.23 | -0.002 | -0.01 | -0.01 | 0 | 0.37 | 0.43 | 0.35 | 0.65 | 0.37 | 0.5 |
| 4 | 94803402 | rs7667683 | -0.64 | 0.374 | 1.174 | 0.84 | -0.023 | -0.024 | -0.024 | 0 | 0.1 | 0.04 | 0.16 | 0.14 | 0.16 | 0.09 |
| 4 | 94804559 | rs7674525 | -0.27 | 0.209 | 1.175 | 0.742 | -0.005 | -0.097 | -0.097 | 0 | 0.22 | 0.09 | 0.15 | 0.51 | 0.2 | 0.38 |
| 4 | 94807875 | rs7698518 | -1.889 | 0.22 | 1.238 | 0.717 | -0.003 | -0.011 | -0.011 | 0 | 0.38 | 0.38 | 0.36 | 0.66 | 0.38 | 0.51 |
| 4 | 94814676 | rs7673218 | -1.952 | -0.031 | 1.237 | 0.679 | -0.013 | 0.057 | 0.057 | 0.062 | 0.12 | 0.35 | 0.16 | 0.13 | 0.1 | 0.11 |
| 4 | 94816144 | rs7692683 | -2.388 | -1.124 | 0.927 | -0.014 | 0.068 | 0.089 | 0.089 | 0.87 | 0.47 | 0.73 | 0.49 | 0.71 | 0.35 | 0.54 |
| 4 | 94816722 | rs9998896 | -1.797 | -1.101 | 1.001 | 0.061 | -0.046 | -0.02 | -0.02 | 0 | 0.24 | 0.64 | 0.33 | 0.17 | 0.15 | 0.15 |
| 4 | 94830161 | rs6813248 | -1.24 | -0.967 | 1.359 | -0.334 | -0.007 | -0.012 | -0.012 | 0 | 0.4 | 0.45 | 0.36 | 0.68 | 0.36 | 0.53 |
| 4 | 94830314 | rs2578165 | -1.24 | -0.967 | 1.359 | -0.334 | -0.007 | -0.012 | -0.012 | 0.013 | 0.4 | 0.45 | 0.36 | 0.68 | 0.36 | 0.53 |
| 4 | 94835289 | rs2578160 | -1.222 | -0.975 | 1.355 | -0.337 | -0.007 | -0.012 | -0.012 | 1.1 | 0.4 | 0.45 | 0.36 | 0.68 | 0.36 | 0.53 |
| 4 | 94839483 | rs1898638 | -0.725 | -0.98 | 0.877 | -1.053 | 0.004 | -0.018 | -0.018 | 0 | 0.57 | 0.48 | 0.59 | 0.69 | 0.56 | 0.71 |
| 4 | 94842776 | rs13105254 | 0.662 | -1.071 | 1.221 | -1.104 | -0.005 | 0.012 | 0.012 | 0 | 0.41 | 0.35 | 0.31 | 0.31 | 0.43 | 0.27 |
| 4 | 94854453 | rs1595502 | -0.601 | -1.1 | 1.277 | -0.628 | 0 | -0.007 | -0.007 | 0 | 0.57 | 0.45 | 0.59 | 0.68 | 0.56 | 0.71 |
| 4 | 94862267 | rs1583661 | 0.924 | -1.202 | 1.305 | -0.677 | -0.039 | 0.025 | 0.025 | 0 | 0.37 | 0.17 | 0.3 | 0.32 | 0.43 | 0.27 |
| 4 | 94866291 | rs1438374 | 0.563 | -1.186 | 1.294 | -0.669 | -0.003 | 0.007 | 0.007 | 0 | 0.41 | 0.38 | 0.31 | 0.32 | 0.43 | 0.28 |
| 4 | 94877244 | rs2618766 | -0.118 | -1.397 | 1.087 | -0.805 | -0.011 | -0.004 | -0.004 | 0 | 0.46 | 0.28 | 0.35 | 0.47 | 0.48 | 0.52 |
| 4 | 94881319 | rs2578125 | 0.884 | -1.455 | 1.177 | -0.811 | -0.029 | 0.035 | 0.035 | 0 | 0.86 | 0.71 | 0.67 | 0.79 | 0.92 | 0.81 |
| 4 | 94882341 | rs10017390 | 1.508 | -1.622 | 0.84 | -1.05 | -0.108 | -0.016 | -0.016 | 0 | 0.31 | 0.09 | 0.3 | 0.32 | 0.46 | 0.28 |
| 4 | 94883373 | rs1898636 | 0.098 | -1.561 | 0.959 | -1.006 | -0.001 | -0.003 | -0.003 | 0 | 0.44 | 0.36 | 0.31 | 0.47 | 0.4 | 0.51 |
| 4 | 94887031 | rs13130787 | 1.069 | -1.522 | 0.69 | -0.988 | -0.046 | 0.026 | 0.026 | 0.01 | 0.37 | 0.16 | 0.3 | 0.32 | 0.43 | 0.29 |
| 4 | 94889283 | rs2578179 | 1.374 | -1.521 | 0.274 | -0.497 | -0.071 | -0.023 | -0.023 | 0 | 0.25 | 0.01 | 0.13 | 0.22 | 0.23 | 0.33 |

| | | | | | | | | | | | | | | | | |
|---|---|---|---|---|---|---|---|---|---|---|---|---|---|---|---|---|
| 4 | 94901470 | rs1370823 | 0.084 | -1.158 | 0.825 | -0.462 | 0.009 | 0.009 | 0.009 | 0 | 0.51 | 0.29 | 0.49 | 0.46 | 0.55 | 0.54 |
| 4 | 94907486 | rs1438383 | 0.084 | -1.158 | 0.825 | -0.468 | 0.009 | 0.01 | 0.01 | 0 | 0.51 | 0.29 | 0.49 | 0.46 | 0.55 | 0.54 |
| 4 | 94914066 | rs2618758 | -0.19 | -1.121 | 0.88 | -0.464 | -0.001 | -0.001 | -0.001 | 0 | 0.48 | 0.51 | 0.49 | 0.54 | 0.45 | 0.46 |
| 4 | 94927323 | rs11944674 | 0.428 | -1.142 | 0.09 | -0.468 | -0.008 | -0.011 | -0.011 | 0.21 | 0.43 | 0.28 | 0.47 | 0.54 | 0.43 | 0.46 |
| 4 | 94949180 | rs280067 | -0.508 | -1.028 | -0.156 | -0.461 | -0.03 | -0.003 | -0.003 | 0.027 | 0.53 | 0.85 | 0.55 | 0.54 | 0.51 | 0.43 |
| 4 | 94953631 | rs2618764 | 0.041 | -1.206 | 0.253 | -0.32 | 0.005 | 0.165 | 0.165 | 0 | 0.88 | 0.49 | 0.73 | 0.77 | 0.89 | 0.86 |
| 4 | 94954308 | rs9685344 | 0.264 | -1.138 | 0.137 | -0.323 | -0.006 | 0.009 | 0.009 | 0 | 0.48 | 0.17 | 0.43 | 0.44 | 0.46 | 0.56 |
| 4 | 94959526 | rs13121813 | 1.105 | -1.141 | -0.381 | -0.307 | -0.061 | 0.006 | 0.006 | 0 | 0.36 | 0.14 | 0.29 | 0.32 | 0.42 | 0.31 |
| 4 | 94962497 | rs12641765 | -0.3 | -1.241 | -0.251 | -0.287 | -0.018 | 0.183 | 0.183 | 0 | 0.12 | 0.51 | 0.25 | 0.22 | 0.09 | 0.14 |
| 4 | 94970758 | rs280117 | -1.243 | -1.13 | -0.785 | -0.281 | -0.052 | 0.008 | 0.008 | 0 | 0.64 | 0.83 | 0.69 | 0.68 | 0.57 | 0.7 |
| 4 | 94976716 | rs11732124 | 0.104 | -0.901 | -1.154 | -0.285 | -0.03 | 0.004 | 0.004 | 0 | 0.48 | 0.12 | 0.43 | 0.45 | 0.46 | 0.56 |
| 4 | 94993624 | rs2048299 | 0.278 | -0.836 | -0.892 | -0.179 | 0.039 | 0.043 | 0.043 | 0 | 0.45 | 0.73 | 0.56 | 0.65 | 0.33 | 0.6 |
| 4 | 94994695 | rs17366966 | 0.404 | -0.74 | -0.967 | -0.294 | -0.001 | -0.012 | -0.012 | 0 | 0.06 | 0.07 | 0.06 | 0 | 0.09 | 0.02 |
| 4 | 94995494 | rs1509951 | -0.058 | -0.748 | -0.994 | -0.273 | 0.03 | 0.083 | 0.083 | 0.012 | 0.51 | 0.8 | 0.62 | 0.65 | 0.43 | 0.62 |
| 4 | 95012684 | rs1904096 | -0.401 | -0.323 | -1.205 | -0.367 | -0.044 | 0.113 | 0.113 | 0.899 | 0.57 | 0.91 | 0.66 | 0.75 | 0.45 | 0.62 |
| 4 | 95019816 | rs899135 | -0.395 | -0.304 | -1.201 | -0.363 | -0.044 | 0.114 | 0.114 | 1.001 | 0.57 | 0.91 | 0.66 | 0.75 | 0.45 | 0.62 |
| 4 | 95035175 | rs1841925 | -0.273 | -0.546 | -0.871 | -0.35 | -0.008 | 0.083 | 0.083 | 0.544 | 0.56 | 0.74 | 0.64 | 0.75 | 0.45 | 0.62 |
| 4 | 95047893 | rs17021364 | 0.74 | -0.538 | -0.707 | -0.286 | 0.01 | 0.01 | 0.01 | 0 | 0.41 | 0.6 | 0.52 | 0.68 | 0.29 | 0.57 |
| 4 | 95049123 | rs17301787 | 0.741 | -0.313 | -0.673 | -0.186 | 0.011 | 0.035 | 0.035 | 0.34 | 0.16 | 0.08 | 0.12 | 0 | 0.15 | 0.04 |
| 4 | 95055821 | rs1509944 | 0.777 | -0.268 | -0.7 | -0.185 | -0.002 | 0.056 | 0.056 | 0.248 | 0.15 | 0.06 | 0.11 | 0 | 0.16 | 0.04 |
| 4 | 95056300 | rs10011158 | 0.261 | -0.33 | -0.458 | -0.168 | -0.001 | -0.007 | -0.007 | 4.308 | 0.18 | 0.23 | 0.13 | 0.07 | 0.17 | 0.05 |
| 4 | 95058293 | rs9992640 | 0.107 | -0.404 | -0.064 | 0.329 | -0.005 | 0.007 | 0.007 | 0.012 | 0.19 | 0.21 | 0.31 | 0.2 | 0.18 | 0.15 |
| 4 | 95063631 | rs1605923 | 1.1 | -0.783 | -0.377 | 0.108 | 0.002 | -0.092 | -0.092 | 0 | 0.35 | 0.09 | 0.3 | 0.48 | 0.27 | 0.41 |
| 4 | 95081378 | rs4693373 | -0.007 | -0.78 | -0.114 | 0.104 | -0.017 | 0.036 | 0.036 | 0.279 | 0.21 | 0.36 | 0.32 | 0.28 | 0.18 | 0.16 |
| 4 | 95086164 | rs4693374 | -0.265 | -0.783 | -0.094 | 0.037 | -0.003 | 0.022 | 0.022 | 0.865 | 0.57 | 0.44 | 0.62 | 0.76 | 0.46 | 0.57 |
| 4 | 95099194 | rs7676797 | -0.395 | -0.801 | 0.29 | 0.023 | -0.003 | 0.009 | 0.009 | 0.863 | 0.58 | 0.66 | 0.63 | 0.78 | 0.47 | 0.57 |
| 4 | 95103545 | rs2171384 | -0.022 | -1.034 | 0.135 | -0.083 | 0.011 | 0.059 | 0.059 | 0 | 0.5 | 0.42 | 0.61 | 0.71 | 0.45 | 0.56 |
| 4 | 95115302 | rs4693002 | -0.375 | -1.015 | 0.166 | -0.072 | 0 | -0.012 | -0.012 | 0 | 0.57 | 0.51 | 0.62 | 0.79 | 0.47 | 0.57 |
| 4 | 95120289 | rs6848742 | 1.142 | -1.034 | 0.136 | -0.096 | 0.001 | 0.03 | 0.03 | 0.419 | 0.17 | 0.11 | 0.11 | 0 | 0.17 | 0.04 |
| 4 | 95136164 | rs7654919 | -0.332 | -1.037 | 0.153 | -0.089 | -0.001 | 0.002 | 0.002 | 1.076 | 0.57 | 0.5 | 0.62 | 0.71 | 0.46 | 0.56 |
| 4 | 95155505 | rs17309887 | 1.024 | -0.859 | 0.157 | -0.047 | -0.145 | 0.021 | 0.021 | 2.872 | 0.26 | 0.01 | 0.25 | 0.18 | 0.37 | 0.23 |
| 4 | 95155719 | rs10007463 | 0.857 | -0.755 | 0.355 | 0.104 | -0.025 | 0.082 | 0.082 | 0.093 | 0.15 | 0.02 | 0.11 | 0 | 0.17 | 0.04 |
| 4 | 95165662 | rs11097408 | -0.316 | -0.76 | 0.369 | 0.075 | 0.004 | -0.006 | -0.006 | 0.143 | 0.63 | 0.65 | 0.68 | 0.71 | 0.56 | 0.63 |
| 4 | 95170839 | rs11722476 | 1.743 | -0.79 | 0.344 | 0.057 | -0.01 | 0.042 | 0.042 | 0 | 0.38 | 0.2 | 0.54 | 0.7 | 0.38 | 0.59 |
| 4 | 95173779 | rs7439869 | -0.533 | -0.812 | 0.381 | 0.046 | -0.122 | 0.017 | 0.017 | 0.629 | 0.65 | 0.96 | 0.71 | 0.71 | 0.57 | 0.64 |
| 4 | 95185854 | rs10029551 | 0.61 | -0.762 | 0.406 | 0.077 | 0.001 | 0.029 | 0.029 | 0.413 | 0.17 | 0.11 | 0.12 | 0 | 0.17 | 0.04 |
| 4 | 95186055 | rs2306802 | 2.164 | -0.773 | 0.404 | 0.136 | -0.068 | -0.133 | -0.133 | 0 | 0.34 | 0.07 | 0.35 | 0.5 | 0.37 | 0.48 |
| 4 | 95197520 | rs6823404 | 0.452 | -0.485 | 0.216 | -0.573 | -0.004 | 0.058 | 0.058 | 0 | 0.55 | 0.34 | 0.66 | 0.7 | 0.56 | 0.63 |
| 4 | 95207438 | rs10027578 | 0.7 | -0.484 | 0.199 | -0.574 | -0.008 | 0.059 | 0.059 | 0.249 | 0.16 | 0.07 | 0.11 | 0 | 0.17 | 0.04 |
| 4 | 95211610 | rs8336 | -0.159 | -0.457 | 0.204 | -0.56 | 0.008 | -0.013 | -0.013 | 0 | 0.62 | 0.5 | 0.68 | 0.71 | 0.56 | 0.63 |
| 4 | 95212185 | rs8026 | 1.863 | -0.444 | -0.168 | -0.553 | -0.01 | 0.041 | 0.041 | 0 | 0.38 | 0.2 | 0.54 | 0.7 | 0.38 | 0.59 |
| 4 | 95223698 | rs10028613 | -0.419 | -0.319 | -0.209 | -0.48 | -0.126 | 0.085 | 0.085 | 0.685 | 0.64 | 0.97 | 0.71 | 0.79 | 0.57 | 0.64 |
| 4 | 95239462 | rs10033662 | -1.318 | -0.35 | -0.084 | -0.425 | 0.028 | 0.05 | 0.05 | 0 | 0.57 | 0.72 | 0.46 | 0.31 | 0.65 | 0.44 |
| 4 | 95254575 | rs10084984 | 0.488 | -0.393 | -0.059 | -0.472 | 0.003 | 0.025 | 0.025 | 0.413 | 0.17 | 0.12 | 0.12 | 0 | 0.18 | 0.04 |
| 4 | 95255212 | rs2059605 | -1.241 | -0.377 | 0.067 | -0.407 | 0.013 | 0.022 | 0.022 | 0 | 0.57 | 0.62 | 0.45 | 0.31 | 0.65 | 0.44 |
| 4 | 95255637 | rs2289186 | -2.146 | -0.41 | -0.143 | -0.411 | -0.002 | 0.022 | 0.022 | 0 | 0.65 | 0.8 | 0.47 | 0.31 | 0.66 | 0.44 |
| 4 | 95262665 | rs11727030 | -1.081 | -0.14 | -0.267 | -0.252 | 0.03 | 0.041 | 0.041 | 0 | 0.55 | 0.28 | 0.43 | 0.34 | 0.64 | 0.44 |
| 4 | 95268272 | rs1991316 | -1.285 | -0.131 | -0.249 | -0.242 | 0.011 | 0.015 | 0.015 | 0 | 0.56 | 0.43 | 0.44 | 0.34 | 0.64 | 0.44 |
| 4 | 95277099 | rs6847989 | -0.194 | 0.126 | -0.301 | -0.227 | -0.123 | 0.006 | 0.006 | 0.593 | 0.38 | 0.07 | 0.31 | 0.33 | 0.47 | 0.4 |
| 4 | 95277311 | rs6532481 | -1.437 | 0.11 | -0.223 | -0.236 | 0.004 | 0.004 | 0.004 | 0 | 0.57 | 0.48 | 0.46 | 0.33 | 0.65 | 0.45 |
| 4 | 95285174 | rs7696494 | 0.06 | 0.903 | -0.022 | 0.294 | -0.002 | -0.06 | -0.06 | 2.222 | 0.17 | 0.28 | 0.14 | 0 | 0.18 | 0.04 |
| 4 | 95298847 | rs12108275 | 0.098 | 0.949 | -0.192 | 0.319 | -0.002 | -0.062 | -0.062 | 0 | 0.17 | 0.23 | 0.14 | 0 | 0.18 | 0.04 |
| 4 | 95301887 | rs1609424 | -0.173 | 1.274 | -0.813 | 0.578 | -0.122 | 0.064 | 0.064 | 0.487 | 0.32 | 0.06 | 0.26 | 0.16 | 0.41 | 0.35 |
| 4 | 95313228 | rs17311887 | -0.502 | 1.288 | -0.763 | 0.611 | -0.096 | 0.033 | 0.033 | 0.918 | 0.35 | 0.09 | 0.29 | 0.26 | 0.45 | 0.4 |
| 4 | 95328925 | rs7685534 | -1.352 | 1.465 | 0.643 | 1.294 | -0.043 | -0.008 | -0.008 | 0.401 | 0.41 | 0.8 | 0.62 | 0.42 | 0.5 | 0.37 |
| 4 | 95329157 | rs7664215 | -0.061 | 1.592 | 1.409 | 1.488 | 0 | 0 | 0 | 0.361 | 0.26 | 0.25 | 0.33 | 0.23 | 0.37 | 0.29 |
| 4 | 95339333 | rs13435066 | -1.412 | 1.374 | 1.742 | 1.501 | 0.026 | 0.024 | 0.024 | 0 | 0.52 | 0.79 | 0.69 | 0.42 | 0.55 | 0.38 |
| 4 | 95342789 | rs4699852 | -1.309 | 1.338 | 1.902 | 1.285 | 0.02 | 0.022 | 0.022 | 0 | 0.51 | 0.67 | 0.67 | 0.41 | 0.55 | 0.38 |
| 4 | 95356328 | rs10008257 | -1.463 | 1.344 | 1.537 | 1.254 | 0.016 | 0.046 | 0.046 | 0.071 | 0.53 | 0.84 | 0.69 | 0.39 | 0.55 | 0.38 |
| 4 | 95360346 | rs7697295 | -1.528 | 1.212 | 1.627 | 1.203 | -0.04 | 0.166 | 0.166 | 0.725 | 0.52 | 0.87 | 0.38 | 0.26 | 0.51 | 0.25 |
| 4 | 95370805 | rs2433320 | -0.83 | 1.401 | 2.063 | 1.165 | -0.006 | 0.091 | 0.091 | 0.071 | 0.41 | 0.25 | 0.29 | 0.19 | 0.45 | 0.2 |
| 4 | 95389193 | rs6843496 | -0.844 | 1.146 | 2.685 | 1.166 | 0.035 | 0.021 | 0.021 | 0.033 | 0.08 | 0.39 | 0.04 | 0.08 | 0.05 | 0.06 |
| 4 | 95407821 | rs2438140 | -1.687 | 1.383 | 2.088 | 1.325 | -0.037 | 0.088 | 0.088 | 0.256 | 0.53 | 0.87 | 0.39 | 0.39 | 0.52 | 0.25 |
| 4 | 95409802 | rs2452554 | 0.146 | 1.595 | 1.898 | 1.282 | -0.025 | -0.012 | -0.012 | 0 | 0.19 | 0.13 | 0.15 | 0.26 | 0.25 | 0.11 |
| 4 | 95410168 | rs2510802 | 0.312 | 1.6 | 1.899 | 1.307 | -0.061 | -0.032 | -0.032 | 0 | 0.2 | 0.03 | 0.15 | 0.26 | 0.27 | 0.16 |
| 4 | 95410364 | rs774982 | 0.864 | 1.601 | 2.05 | 1.301 | -0.008 | 0.05 | 0.05 | 0.004 | 0.58 | 0.7 | 0.71 | 0.68 | 0.54 | 0.8 |
| 4 | 95412692 | rs3805301 | -0.111 | 1.244 | 2.084 | 1.313 | 0 | -0.009 | -0.009 | 0 | 0.22 | 0.23 | 0.14 | 0.06 | 0.19 | 0.08 |
| 4 | 95424226 | rs2452563 | -1.795 | 1.43 | 1.758 | 1.42 | -0.079 | 0.088 | 0.088 | 0.047 | 0.52 | 0.89 | 0.38 | 0.41 | 0.51 | 0.27 |
| 4 | 95428800 | rs2433317 | 0.038 | 1.921 | 2.036 | 1.525 | -0.021 | 0 | 0 | 0 | 0.21 | 0.09 | 0.18 | 0.27 | 0.27 | 0.12 |

| | | | | | | | | | | | | | | | | |
|---|---|---|---|---|---|---|---|---|---|---|---|---|---|---|---|---|
| 4 | 95438406 | rs2510788 | 0.028 | 1.955 | 2.052 | 1.509 | -0.015 | -0.003 | -0.003 | 0 | 0.22 | 0.09 | 0.18 | 0.28 | 0.27 | 0.12 |
| 4 | 95439154 | rs2639795 | -0.53 | 1.886 | 2.242 | 1.624 | -0.001 | -0.009 | -0.009 | 0.019 | 0.83 | 0.79 | 0.88 | 0.92 | 0.82 | 0.91 |
| 4 | 95439956 | rs2433324 | -0.47 | 1.825 | 2.198 | 1.584 | 0.001 | -0.002 | -0.002 | 0 | 0.26 | 0.23 | 0.22 | 0.28 | 0.31 | 0.13 |
| 4 | 95447392 | rs2162346 | -1.153 | 1.978 | 2.316 | 1.76 | -0.001 | -0.004 | -0.004 | 0.083 | 0.31 | 0.62 | 0.21 | 0.44 | 0.31 | 0.24 |
| 4 | 95450770 | rs3805288 | -0.216 | 1.837 | 2.353 | 1.813 | -0.003 | -0.018 | -0.018 | 0 | 0.19 | 0.26 | 0.16 | 0.08 | 0.2 | 0.07 |
| 4 | 95458029 | rs3805284 | 0.417 | 1.787 | 2.119 | 2.008 | -0.029 | 0.011 | 0.011 | 0 | 0.06 | 0.01 | 0.23 | 0.26 | 0.1 | 0.15 |
| 4 | 95468433 | rs10213504 | -0.335 | 1.666 | 2.674 | 1.832 | -0.012 | -0.03 | -0.03 | 0.282 | 0.19 | 0.29 | 0.14 | 0.08 | 0.16 | 0.09 |
| 4 | 95470815 | rs2510772 | -1.594 | 1.758 | 3.082 | 1.839 | -0.013 | -0.02 | -0.02 | 0 | 0.4 | 0.71 | 0.34 | 0.5 | 0.47 | 0.22 |
| 4 | 95472630 | rs1493112 | -0.474 | 1.46 | 3.169 | 1.868 | 0.024 | 0.054 | 0.054 | 0.009 | 0.22 | 0.29 | 0.19 | 0.42 | 0.31 | 0.15 |
| 4 | 95487714 | rs2452593 | 0.439 | 1.64 | 3.105 | 1.703 | 0.004 | -0.054 | -0.054 | 0 | 0.78 | 0.36 | 0.79 | 0.92 | 0.79 | 0.89 |
| 4 | 95491516 | rs17021818 | -0.392 | 1.544 | 2.912 | 1.719 | -0.008 | -0.014 | -0.014 | 0 | 0.08 | 0.13 | 0.03 | 0.05 | 0.04 | 0.08 |
| 4 | 95496882 | rs2452600 | -0.258 | 1.568 | 2.868 | 1.829 | -0.036 | -0.054 | -0.054 | 0 | 0.2 | 0.11 | 0.24 | 0.38 | 0.31 | 0.13 |
| 4 | 95500628 | rs3828518 | -0.387 | 1.569 | 2.942 | 1.811 | 0.002 | -0.017 | -0.017 | 0 | 0.2 | 0.27 | 0.17 | 0.11 | 0.2 | 0.09 |
| 4 | 95505975 | rs12639887 | 1.77 | 1.474 | 2.055 | 1.818 | -0.035 | -0.058 | -0.058 | 0 | 0.5 | 0.06 | 0.53 | 0.46 | 0.44 | 0.67 |
| 4 | 95506827 | rs11722955 | -1.055 | 1.375 | 2.188 | 1.739 | -0.007 | -0.02 | -0.02 | 0 | 0.1 | 0.13 | 0.04 | 0.05 | 0.07 | 0.11 |
| 4 | 95525186 | rs12639623 | 1.189 | 1.398 | 2.36 | 1.906 | 0.001 | 0.002 | 0.002 | 0 | 0.6 | 0.39 | 0.57 | 0.5 | 0.48 | 0.73 |
| 4 | 95529444 | rs9307146 | 1.028 | 1.429 | 2.194 | 1.822 | -0.004 | -0.018 | -0.018 | 0 | 0.91 | 0.76 | 0.96 | 0.96 | 0.95 | 0.9 |
| 4 | 95544162 | rs4699312 | -0.876 | 1.623 | 1.621 | 1.925 | -0.007 | -0.016 | -0.016 | 0 | 0.23 | 0.18 | 0.23 | 0.43 | 0.28 | 0.17 |
| 4 | 95545821 | rs6824181 | 0.984 | 1.694 | 1.741 | 1.682 | 0.013 | 0.002 | 0.002 | 0 | 0.7 | 0.39 | 0.62 | 0.7 | 0.68 | 0.71 |
| 4 | 95555833 | rs11732687 | 1.749 | 1.749 | 1.324 | 1.803 | 0.026 | 0.009 | 0.009 | 0 | 0.52 | 0.26 | 0.56 | 0.54 | 0.44 | 0.67 |
| 4 | 95561459 | rs7690296 | 1.779 | 1.724 | 1.261 | 1.836 | 0.025 | 0.047 | 0.047 | 0 | 0.52 | 0.15 | 0.43 | 0.4 | 0.44 | 0.62 |
| 4 | 95563972 | rs1509620 | 0.199 | 1.635 | 1.356 | 1.849 | -0.002 | 0.001 | 0.001 | 0 | 0.17 | 0.14 | 0.2 | 0.3 | 0.24 | 0.1 |
| 4 | 95575601 | rs2280003 | -0.925 | 1.767 | 1.407 | 1.648 | -0.01 | -0.006 | -0.006 | 0 | 0.3 | 0.22 | 0.26 | 0.33 | 0.35 | 0.18 |
| 4 | 95577290 | rs13121500 | -0.941 | 1.767 | 1.4 | 1.648 | -0.011 | -0.006 | -0.006 | 0.046 | 0.3 | 0.22 | 0.26 | 0.33 | 0.35 | 0.18 |
| 4 | 95579800 | rs11097433 | -1.833 | 1.794 | 1.473 | 1.637 | 0.001 | 0.002 | 0.002 | 0.025 | 0.44 | 0.59 | 0.57 | 0.59 | 0.56 | 0.34 |
| 4 | 95586224 | rs14082 | -1.887 | 1.771 | 1.441 | 1.635 | 0.006 | 0.007 | 0.007 | 0 | 0.47 | 0.59 | 0.57 | 0.59 | 0.57 | 0.37 |
| 4 | 95587675 | rs1043848 | -0.938 | 1.786 | 1.341 | 1.65 | -0.009 | -0.002 | -0.002 | 0 | 0.32 | 0.26 | 0.26 | 0.33 | 0.35 | 0.2 |
| 4 | 95588274 | rs1043853 | 1.518 | 1.81 | 0.938 | 1.002 | 0.032 | 0.042 | 0.042 | 0.119 | 0.5 | 0.14 | 0.41 | 0.41 | 0.43 | 0.59 |
| 4 | 95589337 | rs7285 | 0.078 | 1.867 | 0.894 | 0.679 | 0.043 | -0.024 | -0.024 | 0 | 0.14 | 0.31 | 0.18 | 0.11 | 0.2 | 0.11 |
| 4 | 95594616 | rs13133845 | -0.462 | 1.194 | -1.23 | 0.691 | -0.001 | 0.004 | 0.004 | 0.039 | 0.13 | 0.08 | 0.06 | 0.05 | 0.1 | 0.09 |
| 4 | 95597814 | rs6831015 | 0.368 | 0.965 | -1.327 | 0.986 | -0.008 | 0.011 | 0.011 | 0.192 | 0.5 | 0.23 | 0.41 | 0.44 | 0.46 | 0.59 |
| 4 | 95602550 | rs17022003 | -1.476 | 0.728 | -0.289 | 0.921 | -0.002 | 0.005 | 0.005 | 0 | 0.09 | 0.25 | 0.09 | 0.05 | 0.03 | 0.01 |
| 4 | 95606136 | rs17022008 | -1.056 | 0.669 | -0.714 | 1.76 | -0.008 | -0.015 | -0.015 | 0 | 0.07 | 0.04 | 0.16 | 0.29 | 0.07 | 0.11 |
| 4 | 95606958 | rs17399013 | -0.469 | 0.698 | -0.656 | 1.53 | 0.001 | -0.01 | -0.01 | 0.662 | 0.13 | 0.21 | 0.13 | 0.1 | 0.14 | 0.31 |
| 4 | 95607190 | rs4699578 | -0.151 | 0.693 | -0.665 | 0.992 | 0 | -0.026 | -0.026 | 0 | 0.09 | 0.09 | 0.07 | 0 | 0.1 | 0.02 |
| 4 | 95608023 | rs6532503 | -1.447 | 0.642 | -0.709 | 0.004 | -0.006 | 0.012 | 0.012 | 1.753 | 0.57 | 0.67 | 0.57 | 0.4 | 0.62 | 0.52 |
| 4 | 95612497 | rs7680214 | -1 | 0.347 | -0.831 | -0.374 | 0.026 | 0.054 | 0.054 | 0 | 0.47 | 0.36 | 0.47 | 0.32 | 0.44 | 0.22 |
| 4 | 95620254 | rs6850621 | 0.127 | 0.717 | -0.765 | 0.519 | 0.034 | 0.103 | 0.103 | 1.149 | 0.21 | 0.5 | 0.25 | 0.35 | 0.23 | 0.42 |
| 4 | 95622305 | rs11931151 | -0.458 | 0.548 | -0.959 | 0.237 | -0.001 | -0.002 | -0.002 | 0 | 0.37 | 0.61 | 0.36 | 0.35 | 0.39 | 0.47 |
| 4 | 95626444 | rs6812065 | 0.648 | 0.509 | -0.913 | 0.209 | -0.002 | -0.012 | -0.012 | 0 | 0.19 | 0.22 | 0.2 | 0.05 | 0.19 | 0.06 |
| 4 | 95628481 | rs11735755 | -0.267 | 0.786 | -0.837 | 0.707 | 0.001 | 0.022 | 0.022 | 0.014 | 0.12 | 0.1 | 0.2 | 0.28 | 0.13 | 0.13 |
| 4 | 95631307 | rs6846079 | -0.835 | 0.743 | -0.807 | 0.44 | 0.006 | 0.011 | 0.011 | 2.365 | 0.33 | 0.64 | 0.42 | 0.6 | 0.36 | 0.55 |
| 4 | 95642908 | rs9631727 | -0.365 | 0.641 | -0.664 | 0.381 | -0.001 | 0.004 | 0.004 | 1.72 | 0.39 | 0.36 | 0.24 | 0.34 | 0.37 | 0.19 |
| 4 | 95644727 | rs1553745 | -0.601 | 0.254 | 0.07 | 0.617 | 0.005 | 0.004 | 0.004 | 2.59 | 0.46 | 0.61 | 0.32 | 0.5 | 0.39 | 0.28 |
| 4 | 95648755 | rs4699340 | -0.664 | 0.24 | 0.089 | 0.635 | 0.012 | 0.01 | 0.01 | 3.042 | 0.46 | 0.7 | 0.34 | 0.49 | 0.39 | 0.29 |
| 4 | 95649261 | rs7663879 | -0.63 | 0.254 | 0.078 | 0.438 | 0.007 | 0.03 | 0.03 | 4.232 | 0.45 | 0.67 | 0.28 | 0.38 | 0.38 | 0.21 |
| 4 | 95653664 | rs978429 | 0.045 | -0.129 | -0.232 | 0.111 | 0.018 | 0.022 | 0.022 | 0.567 | 0.14 | 0.08 | 0.06 | 0 | 0.09 | 0.02 |
| 4 | 95678443 | rs4348103 | -0.012 | 0.604 | -0.2 | 0.607 | 0.001 | -0.002 | -0.002 | 0 | 0.7 | 0.7 | 0.71 | 0.69 | 0.67 | 0.62 |
| 4 | 95684380 | rs10049681 | 0.492 | 0.283 | -0.049 | 0.618 | -0.024 | -0.008 | -0.008 | 0.411 | 0.42 | 0.55 | 0.46 | 0.41 | 0.28 | 0.34 |
| 4 | 95691583 | rs4401458 | 0.3 | 1.636 | 0.163 | 0.318 | 0 | 0.006 | 0.006 | 0 | 0.51 | 0.41 | 0.61 | 0.77 | 0.51 | 0.36 |
| 4 | 95692315 | rs7694754 | -1.447 | 1.597 | -0.331 | 0.182 | -0.019 | -0.07 | -0.07 | 4.531 | 0.83 | 0.99 | 0.82 | 0.8 | 0.83 | 0.69 |
| 4 | 95703047 | rs11733843 | 0.327 | 1.633 | 0.067 | 0.493 | 0 | 0 | 0 | 0 | 0.55 | 0.52 | 0.64 | 0.59 | 0.55 | 0.36 |
| 4 | 95706285 | rs12501965 | 0.437 | 1.631 | 0.179 | 0.458 | -0.019 | 0.062 | 0.062 | 0 | 0.27 | 0.22 | 0.16 | 0.02 | 0.28 | 0.31 |
| 4 | 95708561 | rs4599420 | 0.317 | 1.718 | 0.109 | 0.58 | 0.001 | -0.008 | -0.008 | 0 | 0.55 | 0.53 | 0.64 | 0.7 | 0.55 | 0.37 |
| 4 | 95709925 | rs17022227 | 1.259 | 1.812 | 0.002 | 0.725 | -0.054 | -0.043 | -0.043 | 0 | 0.08 | 0 | 0.15 | 0.29 | 0.2 | 0.05 |
| 4 | 95713679 | rs2510543 | 1.929 | 1.478 | 0.072 | 0.426 | 0.065 | 0.049 | 0.049 | 0.032 | 0.35 | 0.13 | 0.37 | 0.28 | 0.24 | 0.2 |
| 4 | 95718070 | rs7677261 | -0.083 | 1.437 | 0.31 | 0.277 | 0.009 | -0.029 | -0.029 | 0.925 | 0.3 | 0.46 | 0.18 | 0.19 | 0.28 | 0.33 |
| 4 | 95723924 | rs11945770 | 0.962 | 1.76 | -0.02 | 0.089 | -0.023 | 0.022 | 0.022 | 0 | 0.15 | 0.11 | 0.09 | 0.01 | 0.19 | 0.17 |
| 4 | 95724988 | rs4456963 | -0.485 | 1.612 | 0.161 | 0.459 | 0.001 | 0.001 | 0.001 | 0 | 0.79 | 0.78 | 0.76 | 0.79 | 0.77 | 0.69 |
| 4 | 95732301 | rs11723423 | 0.43 | 1.505 | 0.452 | 0.446 | 0.159 | 0.252 | 0.252 | 0 | 0.06 | 0.33 | 0.05 | 0.16 | 0.06 | 0.04 |
| 4 | 95733906 | rs10000023 | 0.642 | 1.599 | 0.234 | 0.247 | 0.001 | 0.001 | 0.001 | 0 | 0.61 | 0.45 | 0.64 | 0.6 | 0.61 | 0.56 |
| 4 | 95738226 | rs7692640 | -0.812 | 1.472 | 0.251 | 0.242 | -0.001 | 0.003 | 0.003 | 0 | 0.37 | 0.56 | 0.33 | 0.39 | 0.32 | 0.45 |
| 4 | 95748634 | rs17022328 | 0.296 | 1.371 | 0.329 | 0.295 | 0.016 | 0.141 | 0.141 | 0 | 0.22 | 0.43 | 0.23 | 0.39 | 0.2 | 0.33 |
| 4 | 95750839 | rs6831418 | -1.886 | 1.289 | -0.239 | -0.066 | -0.031 | -0.084 | -0.084 | 0 | 0.57 | 0.89 | 0.45 | 0.49 | 0.56 | 0.71 |
| 4 | 95750866 | rs17428494 | 0.64 | 1.312 | -0.186 | -0.225 | -0.007 | 0.015 | 0.015 | 0 | 0.15 | 0.12 | 0.1 | 0 | 0.13 | 0.12 |
| 4 | 95769791 | rs13127472 | 0.182 | 1.302 | -0.083 | -0.234 | -0.001 | -0.027 | -0.027 | 1.585 | 0.19 | 0.24 | 0.11 | 0.09 | 0.23 | 0.25 |
| 4 | 95770843 | rs10005096 | -1.861 | 1.291 | -0.429 | -0.15 | -0.022 | -0.076 | -0.076 | 0 | 0.57 | 0.85 | 0.45 | 0.48 | 0.56 | 0.7 |
| 4 | 95775101 | rs1544392 | -0.743 | 1.292 | -0.368 | -0.161 | -0.004 | -0.027 | -0.027 | 0 | 0.41 | 0.66 | 0.35 | 0.48 | 0.43 | 0.58 |
| 4 | 95775606 | rs4254781 | -1.643 | 1.276 | -0.51 | -0.002 | -0.013 | -0.063 | -0.063 | 0 | 0.57 | 0.81 | 0.45 | 0.48 | 0.56 | 0.7 |

| | | | | | | | | | | | | | | | | |
|---|---|---|---|---|---|---|---|---|---|---|---|---|---|---|---|---|
| 4 | 95793041 | rs17428994 | 0.37 | 1.28 | -0.306 | -0.606 | -0.004 | 0.029 | 0.029 | 0.018 | 0.16 | 0.1 | 0.09 | 0.1 | 0.2 | 0.26 |
| 4 | 95794060 | rs2719175 | 1.068 | 1.599 | -0.577 | -0.486 | -0.051 | -0.052 | -0.052 | 0 | 0.09 | 0.01 | 0.27 | 0.21 | 0.2 | 0.11 |
| 4 | 95796952 | rs17500892 | 0.778 | 1.544 | -0.569 | -0.714 | -0.013 | 0.025 | 0.025 | 0 | 0.13 | 0.07 | 0.1 | 0 | 0.13 | 0.12 |
| 4 | 95803113 | rs7678784 | -1.398 | 1.374 | -0.045 | -0.742 | -0.018 | 0.075 | 0.075 | 0 | 0.53 | 0.9 | 0.46 | 0.34 | 0.54 | 0.71 |
| 4 | 95810540 | rs1011961 | 0.226 | 1.262 | 0.464 | -0.679 | 0.034 | -0.072 | -0.072 | 1.384 | 0.17 | 0.52 | 0.13 | 0.09 | 0.22 | 0.24 |
| 4 | 95811997 | rs13116658 | 0.931 | 1.243 | 0.505 | -0.686 | -0.001 | 0.003 | 0.003 | 0 | 0.21 | 0.2 | 0.22 | 0.23 | 0.18 | 0.34 |
| 4 | 95817017 | rs17429196 | 1.479 | 1.178 | 0.479 | -0.704 | -0.007 | -0.005 | -0.005 | 0.755 | 0.11 | 0.11 | 0.08 | 0.01 | 0.18 | 0.21 |
| 4 | 95821404 | rs3775009 | 0.811 | 1.176 | 0.543 | -0.703 | 0.004 | -0.015 | -0.015 | 0 | 0.14 | 0.18 | 0.09 | 0.09 | 0.2 | 0.24 |
| 4 | 95834034 | rs1859156 | 0.095 | 1.192 | 0.259 | -0.691 | 0.038 | -0.07 | -0.07 | 0 | 0.17 | 0.47 | 0.13 | 0.1 | 0.22 | 0.24 |
| 4 | 95841123 | rs1295450 | 1.994 | 1.12 | -0.108 | -0.674 | 0.042 | -0.07 | -0.07 | 0 | 0.35 | 0.08 | 0.25 | 0.43 | 0.26 | 0.15 |
| 4 | 95841960 | rs7669822 | 0.325 | 1.156 | 0.231 | -0.626 | 0.024 | -0.051 | -0.051 | 0.029 | 0.15 | 0.42 | 0.11 | 0.11 | 0.2 | 0.24 |
| 4 | 95843673 | rs3775017 | 1.32 | 0.734 | 0.085 | -0.448 | -0.007 | -0.066 | -0.066 | 1.88 | 0.16 | 0.01 | 0.19 | 0.2 | 0.13 | 0.32 |
| 4 | 95845911 | rs1444930 | -0.478 | 0.812 | 0.181 | -0.525 | 0.001 | -0.023 | -0.023 | 0 | 0.34 | 0.39 | 0.3 | 0.23 | 0.29 | 0.48 |
| 4 | 95849555 | rs11939949 | 1.316 | 0.839 | -0.003 | -0.562 | -0.018 | 0.022 | 0.022 | 0 | 0.09 | 0.04 | 0.06 | 0 | 0.1 | 0.08 |
| 4 | 95856681 | rs10021303 | -0.727 | 0.819 | 0.122 | -0.537 | 0 | 0.001 | 0.001 | 0 | 0.37 | 0.65 | 0.32 | 0.34 | 0.32 | 0.51 |
| 4 | 95859913 | rs7671870 | 0.979 | 1.179 | 0.354 | -0.655 | 0.004 | 0.005 | 0.005 | 0.436 | 0.26 | 0.18 | 0.25 | 0.2 | 0.24 | 0.34 |
| 4 | 95861615 | rs6842585 | 1.674 | 1.109 | 0.183 | -0.782 | -0.002 | -0.052 | -0.052 | 1.457 | 0.28 | 0.12 | 0.22 | 0.42 | 0.24 | 0.15 |
| 4 | 95867338 | rs3796413 | 0.708 | 0.396 | 0.575 | -1.4 | 0 | 0 | 0 | 0.522 | 0.27 | 0.22 | 0.25 | 0.2 | 0.24 | 0.36 |
| 4 | 95868525 | rs1373651 | -0.793 | 0.629 | 0.813 | -1.108 | -0.027 | 0.057 | 0.057 | 0 | 0.57 | 0.73 | 0.43 | 0.24 | 0.51 | 0.64 |
| 4 | 95876278 | rs1444928 | -0.733 | 0.591 | 0.782 | -1.157 | -0.021 | 0.05 | 0.05 | 0 | 0.57 | 0.68 | 0.43 | 0.23 | 0.51 | 0.64 |
| 4 | 95876313 | rs1444927 | -0.508 | 0.796 | 0.169 | -1.118 | -0.001 | 0 | 0 | 0 | 0.52 | 0.4 | 0.4 | 0.21 | 0.51 | 0.59 |
| 4 | 95889271 | rs17402559 | 0.311 | 0.959 | -0.779 | -1.047 | -0.003 | 0.046 | 0.046 | 0 | 0.15 | 0.04 | 0.09 | 0 | 0.15 | 0.09 |
| 4 | 95892711 | rs1373649 | -0.014 | 0.882 | -0.901 | -0.316 | -0.001 | 0.003 | 0.003 | 0 | 0.53 | 0.54 | 0.4 | 0.21 | 0.51 | 0.59 |
| 4 | 95901481 | rs17343504 | -0.867 | 0.554 | -1.246 | -0.266 | 0.003 | 0.074 | 0.074 | 0.042 | 0.14 | 0.03 | 0.08 | 0.01 | 0.12 | 0.06 |
| 4 | 95905449 | rs3775031 | 0.649 | 0.582 | -1.122 | -0.592 | -0.019 | 0.077 | 0.077 | 0 | 0.45 | 0.23 | 0.55 | 0.7 | 0.45 | 0.36 |
| 4 | 95914424 | rs1545326 | 0.945 | 0.169 | -0.458 | -0.92 | -0.034 | -0.056 | -0.056 | 0.015 | 0.72 | 0.45 | 0.82 | 0.88 | 0.79 | 0.81 |
| 4 | 95928570 | rs7662504 | -0.882 | 0.153 | -0.986 | -1.513 | 0.006 | 0.014 | 0.014 | 1.834 | 0.48 | 0.51 | 0.57 | 0.71 | 0.4 | 0.25 |
| 4 | 95933337 | rs12500473 | 0.571 | 0.091 | -0.898 | -1.581 | -0.024 | 0.067 | 0.067 | 0 | 0.45 | 0.27 | 0.55 | 0.7 | 0.46 | 0.37 |
| 4 | 95938386 | rs1347345 | -1.2 | 0.057 | -0.88 | -1.602 | -0.026 | 0 | 0 | 0 | 0.72 | 0.82 | 0.71 | 0.72 | 0.64 | 0.47 |
| 4 | 95948640 | rs17022885 | -0.691 | 0.039 | -0.796 | -1.598 | -0.048 | -0.141 | -0.141 | 0 | 0.27 | 0.54 | 0.16 | 0.02 | 0.18 | 0.11 |
| 4 | 95954962 | rs1434546 | 0.814 | 0.436 | -0.975 | -0.334 | -0.011 | -0.021 | -0.021 | 0 | 0.7 | 0.37 | 0.82 | 0.88 | 0.79 | 0.82 |
| 4 | 95955095 | rs17022918 | 1.791 | 0.413 | -1.078 | -0.373 | -0.014 | -0.093 | -0.093 | 0 | 0.18 | 0.02 | 0.27 | 0.29 | 0.13 | 0.06 |
| 4 | 95961926 | rs13103406 | 1.08 | 0.458 | -0.621 | -0.001 | -0.068 | 0.037 | 0.037 | 0.305 | 0.26 | 0.11 | 0.27 | 0.18 | 0.33 | 0.44 |
| 4 | 95969759 | rs1434545 | 0.996 | 0.491 | -0.405 | 0.054 | -0.043 | 0.03 | 0.03 | 0.458 | 0.27 | 0.15 | 0.27 | 0.2 | 0.33 | 0.39 |
| 4 | 95984379 | rs2114534 | 0.288 | 0.354 | 0.006 | 0.008 | -0.005 | 0.038 | 0.038 | 0 | 0.55 | 0.35 | 0.6 | 0.68 | 0.51 | 0.47 |
| 4 | 95991417 | rs1863652 | 1.432 | 0.311 | 0.048 | 0.027 | -0.002 | 0.037 | 0.037 | 0.005 | 0.39 | 0.15 | 0.38 | 0.26 | 0.36 | 0.33 |
| 4 | 96004341 | rs1158793 | 1.196 | 0.286 | 0.979 | 0.257 | -0.01 | 0.048 | 0.048 | 0 | 0.68 | 0.61 | 0.68 | 0.47 | 0.7 | 0.75 |
| 4 | 96011233 | rs6816834 | 1.207 | 0.286 | 0.979 | 0.257 | -0.011 | 0.047 | 0.047 | 0 | 0.68 | 0.6 | 0.68 | 0.47 | 0.7 | 0.74 |
| 4 | 96015307 | rs723367 | 1.254 | 0.317 | 1.113 | 0.292 | -0.016 | 0.065 | 0.065 | 0.011 | 0.68 | 0.5 | 0.66 | 0.46 | 0.7 | 0.73 |
| 4 | 96020228 | rs1560909 | -0.064 | 0.368 | 0.735 | 0.195 | -0.012 | -0.016 | -0.016 | 0.061 | 0.83 | 0.78 | 0.89 | 0.88 | 0.84 | 0.87 |
| 4 | 96030402 | rs3796432 | 1.106 | 0.322 | 0.254 | 0.031 | -0.005 | 0.065 | 0.065 | 0 | 0.42 | 0.14 | 0.4 | 0.25 | 0.38 | 0.31 |
| 4 | 96031455 | rs3796433 | -1.073 | 0.363 | 1.254 | 0.683 | 0.021 | 0.111 | 0.111 | 0 | 0.12 | 0.35 | 0.19 | 0.24 | 0.09 | 0.07 |
| 4 | 96031577 | rs3796434 | 0.923 | 0.302 | 1.05 | 0.605 | -0.014 | 0.152 | 0.152 | 0 | 0.51 | 0.16 | 0.47 | 0.24 | 0.5 | 0.35 |
| 4 | 96033978 | rs1369087 | 0.425 | 0.259 | 1.402 | 0.706 | 0.005 | 0.204 | 0.204 | 0 | 0.91 | 0.76 | 0.95 | 0.65 | 0.93 | 0.81 |
| 4 | 96047368 | rs6826261 | -0.01 | 0.115 | 1.617 | 0.63 | 0.01 | 0.026 | 0.026 | 0.007 | 0.07 | 0.33 | 0.05 | 0.03 | 0.02 | 0.02 |
| 4 | 96052186 | rs17023083 | -1.01 | 0.064 | 1.629 | 0.592 | -0.015 | -0.02 | -0.02 | 0 | 0.12 | 0.52 | 0.09 | 0.04 | 0.05 | 0.03 |
| 4 | 96053225 | rs3796441 | -1.202 | 0.062 | 1.627 | 0.571 | -0.026 | -0.035 | -0.035 | 0 | 0.12 | 0.53 | 0.1 | 0.04 | 0.05 | 0.04 |
| 4 | 96054027 | rs3796442 | -0.011 | 0.083 | 1.447 | 0.599 | -0.007 | -0.021 | -0.021 | 0 | 0.08 | 0.03 | 0.14 | 0.2 | 0.08 | 0.08 |
| 4 | 96061509 | rs3796443 | 0.709 | 0.149 | 1.623 | 0.553 | -0.032 | -0.018 | -0.018 | 0 | 0.56 | 0.2 | 0.59 | 0.45 | 0.56 | 0.44 |
| 4 | 96062784 | rs3755876 | 0.69 | 0.148 | 1.609 | 0.555 | -0.038 | -0.024 | -0.024 | 0 | 0.56 | 0.18 | 0.59 | 0.45 | 0.56 | 0.45 |
| 4 | 96066672 | rs1434538 | 0.606 | 0.164 | 1.693 | 0.667 | -0.014 | 0.012 | 0.012 | 0 | 0.57 | 0.29 | 0.6 | 0.39 | 0.56 | 0.43 |
| 4 | 96068624 | rs3775042 | 0.692 | 0.141 | 1.642 | 0.635 | -0.063 | 0.005 | 0.005 | 0 | 0.55 | 0.09 | 0.59 | 0.39 | 0.56 | 0.45 |
| 4 | 96070525 | rs1970801 | -0.675 | 0.124 | 1.765 | 0.644 | -0.008 | -0.008 | -0.008 | 1.082 | 0.69 | 0.71 | 0.68 | 0.42 | 0.6 | 0.47 |
| 4 | 96075965 | rs1434536 | 0.62 | 0.181 | 1.351 | 0.687 | -0.028 | 0.014 | 0.014 | 0 | 0.57 | 0.18 | 0.6 | 0.38 | 0.56 | 0.43 |
| 4 | 96087973 | rs1813074 | 0.561 | 0.255 | 1.419 | 0.724 | -0.028 | 0.007 | 0.007 | 0 | 0.57 | 0.21 | 0.6 | 0.37 | 0.56 | 0.3 |
| 4 | 96088849 | rs10516959 | -0.435 | 0.637 | 1.915 | 0.912 | 0.003 | 0.017 | 0.017 | 0 | 0.05 | 0.2 | 0.04 | 0.03 | 0.02 | 0.02 |
| 4 | 96117230 | rs3775059 | -0.06 | 1.175 | 1.304 | 0.99 | 0.032 | 0.081 | 0.081 | 0 | 0.87 | 0.76 | 0.87 | 0.67 | 0.81 | 0.66 |
| 4 | 96118696 | rs1991946 | 1.63 | 1.157 | 1.257 | 0.876 | 0.006 | 0.024 | 0.024 | 1.158 | 0.74 | 0.34 | 0.77 | 0.57 | 0.71 | 0.55 |
| 4 | 96122615 | rs11737434 | 1.281 | 1.194 | 0.74 | 0.443 | -0.023 | -0.019 | -0.019 | 0 | 0.72 | 0.72 | 0.49 | 0.45 | 0.57 | 0.54 |
| 4 | 96126231 | rs3775050 | 0.466 | 1.152 | 1.102 | 0.518 | 0.029 | 0.07 | 0.07 | 0.157 | 0.82 | 0.77 | 0.69 | 0.46 | 0.75 | 0.61 |
| 4 | 96130145 | rs2162448 | -1.881 | 1.348 | 0.966 | 0.367 | 0.022 | -0.029 | -0.029 | 0 | 0.06 | 0.17 | 0.03 | 0 | 0.05 | 0.08 |
| 4 | 96160117 | rs4699412 | -1.758 | 0.768 | 0.517 | -0.069 | 0.003 | 0.003 | 0.003 | 0 | 0.36 | 0.4 | 0.4 | 0.6 | 0.35 | 0.61 |
| 4 | 96161342 | rs7434638 | -1.8 | 0.737 | 0.624 | -0.086 | 0.01 | 0.008 | 0.008 | 0 | 0.37 | 0.44 | 0.41 | 0.6 | 0.37 | 0.61 |
| 4 | 96163244 | rs11730446 | -2.08 | 0.673 | 0.664 | -0.147 | 0.003 | 0.001 | 0.001 | 0 | 0.4 | 0.46 | 0.42 | 0.59 | 0.37 | 0.59 |
| 4 | 96165142 | rs12642020 | -0.36 | 0.737 | 0.715 | -0.052 | 0.048 | 0.086 | 0.086 | 0.018 | 0.18 | 0.25 | 0.34 | 0.55 | 0.28 | 0.42 |
| 4 | 96170439 | rs4699837 | -0.957 | 0.776 | 0.367 | -0.039 | 0.028 | 0.034 | 0.034 | 0 | 0.32 | 0.47 | 0.41 | 0.6 | 0.35 | 0.61 |
| 4 | 96171211 | rs3775003 | -1.926 | 0.804 | -0.75 | -0.413 | -0.075 | 0.149 | 0.149 | 0.771 | 0.63 | 0.98 | 0.59 | 0.87 | 0.62 | 0.7 |
| 4 | 96179750 | rs6838035 | -0.362 | 1.642 | 0.417 | 0.276 | 0.024 | -0.028 | -0.028 | 0 | 0.05 | 0.14 | 0.05 | 0 | 0.08 | 0.01 |
| 4 | 96180560 | rs17352477 | -0.73 | 1.201 | -0.001 | 0.355 | 0 | -0.009 | -0.009 | 0.403 | 0.09 | 0.1 | 0.03 | 0 | 0.06 | 0.03 |

| | | | | | | | | | | | | | | | | |
|---|---|---|---|---|---|---|---|---|---|---|---|---|---|---|---|---|
| 4 | 96181408 | rs6532537 | -1.122 | 1.05 | -0.032 | 0.653 | -0.009 | 0.001 | 0.001 | 0 | 0.42 | 0.65 | 0.6 | 0.57 | 0.43 | 0.59 |
| 4 | 96200216 | rs10010112 | 0.819 | 0.927 | -0.013 | 0.424 | 0.005 | -0.027 | -0.027 | 0 | 0.66 | 0.57 | 0.53 | 0.84 | 0.62 | 0.54 |
| 4 | 96201243 | rs2865421 | -0.136 | 0.754 | -0.47 | 0.875 | 0.013 | 0.234 | 0.234 | 0 | 0.12 | 0.38 | 0.24 | 0.36 | 0.08 | 0.15 |
| 4 | 96204556 | rs11097465 | 1.38 | 0.925 | -1.904 | 0.878 | -0.069 | -0.034 | -0.034 | 0 | 0.52 | 0.11 | 0.28 | 0.47 | 0.51 | 0.36 |
| 4 | 96206860 | rs10032274 | 0.727 | 1.38 | -1.537 | 0.375 | -0.052 | 0.037 | 0.037 | 2.994 | 0.7 | 0.87 | 0.55 | 0.81 | 0.61 | 0.63 |
| 4 | 96209804 | rs4504262 | 0.436 | 1.637 | -1.702 | 0.422 | -0.101 | -0.016 | -0.016 | 0 | 0.82 | 0.99 | 0.68 | 0.88 | 0.71 | 0.84 |
| 4 | 96213821 | rs17023470 | -0.966 | 1.345 | -1.666 | 0.269 | -0.033 | 0.029 | 0.029 | 0 | 0.12 | 0.02 | 0.09 | 0.06 | 0.16 | 0.07 |
| 4 | 96213883 | rs6857589 | 1.214 | 1.278 | -1.555 | 0.151 | -0.02 | 0.027 | 0.027 | 0 | 0.62 | 0.73 | 0.6 | 0.75 | 0.57 | 0.71 |
| 4 | 96213928 | rs6831808 | 0.987 | 1.451 | -1.426 | 0.223 | -0.005 | 0.003 | 0.003 | 0 | 0.72 | 0.75 | 0.67 | 0.8 | 0.65 | 0.76 |
| 4 | 96220207 | rs4699847 | 0.825 | 1.57 | -1.9 | 0.042 | -0.101 | 0.005 | 0.005 | 0.021 | 0.78 | 0.96 | 0.75 | 0.85 | 0.67 | 0.69 |
| 4 | 96226609 | rs17433092 | 0.248 | 1.334 | -2.492 | 0.45 | -0.024 | -0.06 | -0.06 | 0.034 | 0.12 | 0.01 | 0.23 | 0.19 | 0.11 | 0.24 |
| 4 | 96236932 | rs10856912 | 0.229 | 1.432 | -1.874 | 0.849 | 0.02 | 0.007 | 0.007 | 0.267 | 0.87 | 0.85 | 0.79 | 0.85 | 0.79 | 0.8 |
| 4 | 96238873 | rs6532549 | 0.185 | 1.424 | -1.866 | 0.845 | 0.016 | 0.005 | 0.005 | 0.2 | 0.87 | 0.88 | 0.8 | 0.85 | 0.79 | 0.8 |
| 4 | 96240018 | rs17023493 | -0.584 | 1.262 | -1.089 | 0.693 | 0 | -0.045 | -0.045 | 0 | 0.12 | 0.18 | 0.07 | 0.01 | 0.12 | 0.11 |
| 4 | 96247498 | rs2621462 | -0.269 | 0.367 | -0.325 | -0.109 | -0.037 | -0.13 | -0.13 | 2.699 | 0.74 | 0.64 | 0.86 | 1 | 0.81 | 0.91 |
| 4 | 96255088 | rs2626053 | 0.378 | 0.442 | -0.441 | -0.008 | -0.008 | -0.01 | -0.01 | 0 | 0.1 | 0.06 | 0.2 | 0.16 | 0.19 | 0.18 |
| 4 | 96255814 | rs10516965 | -0.725 | 0.4 | -0.559 | -0.188 | -0.001 | -0.039 | -0.039 | 0 | 0.17 | 0.19 | 0.11 | 0.01 | 0.17 | 0.17 |
| 4 | 96259062 | rs4130507 | 0.412 | 0.386 | -0.55 | -0.196 | 0.009 | -0.042 | -0.042 | 0 | 0.64 | 0.51 | 0.66 | 0.84 | 0.62 | 0.73 |
| 4 | 96260842 | rs1552225 | 1.505 | 0.43 | -0.646 | 0.528 | 0.006 | 0.059 | 0.059 | 0.034 | 0.47 | 0.29 | 0.55 | 0.7 | 0.46 | 0.55 |
| 4 | 96262520 | rs2626055 | 1.04 | 0.413 | -0.695 | -0.004 | 0.06 | 0.127 | 0.127 | 0 | 0.57 | 0.22 | 0.74 | 0.99 | 0.65 | 0.74 |
| 4 | 96286793 | rs2626035 | -1.614 | 0.18 | -1.042 | 0.195 | -0.028 | 0.071 | 0.071 | 0 | 0.54 | 0.89 | 0.72 | 0.29 | 0.53 | 0.44 |
| 4 | 96292381 | rs1542616 | -0.182 | 0.041 | -0.382 | -0.334 | -0.048 | -0.058 | -0.058 | 0 | 0.28 | 0.39 | 0.4 | 0.13 | 0.18 | 0.09 |
| 4 | 96295569 | rs2626040 | -0.767 | 0.253 | -0.346 | -0.347 | -0.003 | 0.019 | 0.019 | 1.356 | 0.39 | 0.65 | 0.58 | 0.79 | 0.4 | 0.37 |
| 4 | 96298575 | rs2621456 | -2.012 | 0.156 | -0.665 | -0.578 | -0.05 | 0.211 | 0.211 | 0 | 0.63 | 0.92 | 0.76 | 0.96 | 0.62 | 0.73 |
| 4 | 96299024 | rs1994175 | 0.734 | 0.122 | -0.592 | -0.466 | 0.019 | -0.009 | -0.009 | 0 | 0.14 | 0.03 | 0.1 | 0.12 | 0.09 | 0.09 |
| 4 | 96299591 | rs12644448 | -0.739 | 0.24 | -0.344 | -0.376 | 0 | -0.003 | -0.003 | 0.8 | 0.39 | 0.63 | 0.59 | 0.79 | 0.4 | 0.37 |
| 4 | 96300827 | rs10222858 | -0.125 | -0.193 | -0.073 | -0.976 | -0.01 | -0.117 | -0.117 | 0.017 | 0.45 | 0.48 | 0.25 | 0.06 | 0.38 | 0.46 |
| 4 | 96303313 | rs4353913 | -0.722 | -0.166 | 0.193 | -0.621 | -0.007 | 0.023 | 0.023 | 0 | 0.57 | 0.7 | 0.42 | 0.72 | 0.59 | 0.65 |
| 4 | 96304008 | rs10461127 | -1.93 | -0.262 | 0.029 | -0.764 | -0.054 | 0.083 | 0.083 | 0 | 0.74 | 0.91 | 0.8 | 0.96 | 0.68 | 0.83 |
| 4 | 96304133 | rs10461172 | 0.63 | -0.32 | 0.341 | -0.798 | 0.021 | 0.051 | 0.051 | 0 | 0.14 | 0.21 | 0.19 | 0.66 | 0.2 | 0.25 |
| 4 | 96306407 | rs9998018 | 0.457 | -0.392 | 0.385 | -0.863 | 0.01 | -0.023 | -0.023 | 0 | 0.41 | 0.35 | 0.23 | 0.06 | 0.38 | 0.4 |
| 4 | 96307840 | rs6532555 | -1.949 | -0.079 | 0.253 | -0.852 | -0.048 | 0.09 | 0.09 | 0 | 0.73 | 0.91 | 0.79 | 0.94 | 0.7 | 0.82 |
| 4 | 96309211 | rs1384862 | 0.297 | 0.053 | 0.268 | -0.657 | 0.01 | -0.021 | -0.021 | 0.03 | 0.38 | 0.33 | 0.21 | 0.06 | 0.32 | 0.36 |
| 4 | 96310163 | rs10516969 | -0.043 | 0.638 | 0.259 | 0.296 | 0.034 | 0.079 | 0.079 | 0 | 0.14 | 0.26 | 0.19 | 0.66 | 0.2 | 0.25 |
| 4 | 96311797 | rs1351770 | -0.208 | 0.142 | -0.118 | 0.129 | -0.001 | 0.002 | 0.002 | 0 | 0.06 | 0.07 | 0.03 | 0.05 | 0.04 | 0.19 |
| 4 | 96314856 | rs10012134 | -0.455 | -0.039 | 0.364 | 0.653 | -0.039 | 0.067 | 0.067 | 0.171 | 0.18 | 0.32 | 0.33 | 0.74 | 0.13 | 0.24 |
| 4 | 96314874 | rs7679033 | -0.798 | 0.387 | -0.283 | 0.037 | 0.003 | -0.007 | -0.007 | 0 | 0.38 | 0.4 | 0.48 | 0.76 | 0.4 | 0.42 |
| 4 | 96315629 | rs6854948 | -0.979 | 0.159 | 0.14 | 0.768 | 0.01 | -0.002 | -0.002 | 0 | 0.4 | 0.47 | 0.52 | 0.45 | 0.47 | 0.44 |
| 4 | 96315706 | rs6832475 | 0.031 | -0.043 | 0.423 | 0.753 | 0.014 | 0.068 | 0.068 | 0 | 0.06 | 0.25 | 0.06 | 0.08 | 0.05 | 0.09 |
| 4 | 96316032 | rs9998674 | 0.618 | 0.1 | 0.285 | 0.777 | 0.011 | 0.019 | 0.019 | 0 | 0.53 | 0.4 | 0.47 | 0.66 | 0.45 | 0.47 |
| 4 | 96316619 | rs17023753 | -1.295 | -0.066 | 0.671 | 1.14 | -0.002 | 0.087 | 0.087 | 0 | 0.13 | 0.41 | 0.26 | 0.17 | 0.11 | 0.25 |
| 4 | 96322804 | rs4406027 | -1.283 | 0.003 | 0.654 | 1.042 | 0.01 | 0.023 | 0.023 | 0 | 0.13 | 0.4 | 0.1 | 0.12 | 0.12 | 0.21 |
| 4 | 96330417 | rs7691901 | -1.029 | -0.712 | 0.314 | 1.135 | -0.03 | -0.002 | -0.002 | 0 | 0.29 | 0.66 | 0.21 | 0.3 | 0.18 | 0.35 |
| 4 | 96330516 | rs9307158 | -0.172 | -0.7 | 0.357 | 1.128 | -0.061 | -0.014 | -0.014 | 0 | 0.21 | 0.46 | 0.14 | 0.21 | 0.11 | 0.21 |
| 4 | 96333037 | rs17023779 | -0.286 | -0.601 | -0.347 | 1.167 | 0.047 | -0.019 | -0.019 | 0 | 0.23 | 0.11 | 0.18 | 0.29 | 0.14 | 0.29 |
| 4 | 96341046 | rs10516971 | -1.257 | -0.372 | -0.557 | 1.096 | 0.007 | 0.011 | 0.011 | 0 | 0.37 | 0.34 | 0.31 | 0.32 | 0.32 | 0.35 |
| 4 | 96341886 | rs9307159 | -0.16 | -0.343 | -0.522 | 1.12 | -0.002 | -0.007 | -0.007 | 0 | 0.24 | 0.33 | 0.19 | 0.22 | 0.2 | 0.23 |
| 4 | 96342750 | rs10017332 | 0.656 | -0.344 | -0.846 | 1.196 | 0 | -0.002 | -0.002 | 0 | 0.18 | 0.08 | 0.17 | 0.19 | 0.15 | 0.14 |
| 4 | 96344267 | rs900617 | 0.816 | 0.219 | -0.752 | 1.464 | 0.007 | -0.016 | -0.016 | 0 | 0.09 | 0.02 | 0.09 | 0.09 | 0.07 | 0.12 |
| 4 | 96347978 | rs10004779 | -0.466 | 0.653 | -0.433 | 2.009 | 0.009 | 0.068 | 0.068 | 0.029 | 0.11 | 0.16 | 0.18 | 0.32 | 0.14 | 0.16 |
| 4 | 96348686 | rs4277782 | -0.863 | 1.067 | -0.674 | 1.2 | -0.032 | 0.015 | 0.015 | 0 | 0.82 | 0.92 | 0.86 | 0.9 | 0.71 | 0.83 |
| 4 | 96349407 | rs4380535 | 0.46 | 0.918 | -0.643 | 1.245 | -0.012 | -0.009 | -0.009 | 0 | 0.7 | 0.76 | 0.67 | 0.47 | 0.57 | 0.67 |
| 4 | 96349420 | rs1483749 | 0.77 | 0.653 | -0.918 | 0.422 | 0.006 | 0.02 | 0.02 | 0 | 0.3 | 0.22 | 0.24 | 0.17 | 0.26 | 0.25 |
| 4 | 96353173 | rs1483748 | 0.63 | 0.669 | -1.501 | 0.467 | -0.084 | 0.062 | 0.062 | 0.003 | 0.29 | 0.02 | 0.24 | 0.17 | 0.29 | 0.25 |
| 4 | 96356534 | rs1026926 | 0.089 | 0.348 | -0.778 | 0.821 | 0.021 | 0.067 | 0.067 | 0 | 0.48 | 0.29 | 0.4 | 0.23 | 0.55 | 0.39 |
| 4 | 96357927 | rs13116794 | 0.247 | 0.254 | -0.619 | 1.05 | 0.036 | 0.011 | 0.011 | 0.359 | 0.51 | 0.37 | 0.56 | 0.56 | 0.65 | 0.42 |
| 4 | 96358203 | rs6854070 | -0.468 | 0.29 | -0.848 | 1.016 | -0.017 | 0.016 | 0.016 | 0 | 0.9 | 0.83 | 0.95 | 0.88 | 0.92 | 0.85 |
| 4 | 96360642 | rs3846457 | 0.399 | 0.563 | -1.245 | 0.75 | 0.092 | 0.102 | 0.102 | 0.976 | 0.32 | 0.06 | 0.31 | 0.15 | 0.2 | 0.3 |
| 4 | 96362183 | rs17023881 | -0.891 | 0.824 | -0.66 | 0.793 | 0.058 | 0.127 | 0.127 | 0 | 0.05 | 0.24 | 0.05 | 0.16 | 0.07 | 0.14 |
| 4 | 96363909 | rs4699266 | -0.077 | 0.725 | -1.38 | 0.376 | 0.005 | 0.003 | 0.003 | 0.003 | 0.41 | 0.14 | 0.47 | 0.37 | 0.35 | 0.46 |
| 4 | 96368463 | rs7662972 | -0.881 | 1.256 | -0.311 | -0.09 | 0.025 | -0.057 | -0.057 | 0 | 0.23 | 0.37 | 0.23 | 0.08 | 0.31 | 0.11 |
| 4 | 96369736 | rs9995935 | -0.902 | 1.315 | -0.259 | -0.362 | -0.005 | -0.001 | -0.001 | 0.048 | 0.27 | 0.29 | 0.22 | 0.11 | 0.4 | 0.19 |
| 4 | 96370093 | rs17381344 | -0.778 | 0.962 | -0.527 | -0.145 | -0.004 | -0.02 | -0.02 | 0.069 | 0.71 | 0.76 | 0.5 | 0.55 | 0.71 | 0.6 |
| 4 | 96372611 | rs265052 | -0.906 | 0.675 | 0.018 | -0.256 | 0.02 | -0.029 | -0.029 | 0.718 | 0.3 | 0.53 | 0.27 | 0.22 | 0.35 | 0.23 |
| 4 | 96373680 | rs265047 | 0.551 | 0.713 | -0.06 | -0.193 | 0.006 | 0.065 | 0.065 | 0 | 0.1 | 0.28 | 0.1 | 0.16 | 0.08 | 0.14 |
| 4 | 96374120 | rs265044 | 0.049 | 0.825 | -0.035 | -0.246 | 0.007 | 0.042 | 0.042 | 0 | 0.11 | 0.28 | 0.09 | 0.16 | 0.11 | 0.09 |
| 4 | 96374272 | rs13119662 | 0.316 | 0.842 | -0.293 | -0.279 | -0.036 | 0.044 | 0.044 | 0 | 0.16 | 0.07 | 0.15 | 0.05 | 0.26 | 0.09 |
| 4 | 96375542 | rs17023932 | 0.735 | 0.358 | -0.01 | -0.032 | -0.01 | -0.003 | -0.003 | 0 | 0.36 | 0.37 | 0.18 | 0.31 | 0.23 | 0.27 |
| 4 | 96375818 | rs265042 | -0.678 | 0.409 | -0.301 | 0.036 | -0.021 | -0.023 | -0.023 | 0.896 | 0.58 | 0.76 | 0.38 | 0.52 | 0.48 | 0.52 |

| | | | | | | | | | | | | | | | | |
|---|---|---|---|---|---|---|---|---|---|---|---|---|---|---|---|---|
| 4 | 96376995 | rs17023936 | -0.347 | 0.142 | -0.18 | -0.107 | -0.014 | -0.007 | -0.007 | 0.006 | 0.43 | 0.49 | 0.23 | 0.34 | 0.33 | 0.35 |
| 4 | 96377584 | rs11730759 | -0.111 | 0.836 | -0.33 | -0.008 | -0.002 | 0.01 | 0.01 | 0 | 0.1 | 0.07 | 0.08 | 0.06 | 0.14 | 0.11 |
| 4 | 96378484 | rs7669439 | -0.9 | 0.63 | 0.157 | -0.157 | 0.009 | -0.111 | -0.111 | 0.001 | 0.21 | 0.53 | 0.15 | 0.02 | 0.22 | 0.09 |
| 4 | 96379021 | rs11936864 | -1.481 | 0.476 | 0.261 | 0.631 | -0.003 | 0.053 | 0.053 | 0 | 0.07 | 0.14 | 0.03 | 0.13 | 0.05 | 0.09 |
| 4 | 96379903 | rs3912476 | 0.262 | 0.073 | 0.214 | 0.396 | -0.002 | 0.003 | 0.003 | 0.052 | 0.47 | 0.43 | 0.66 | 0.57 | 0.46 | 0.49 |
| 4 | 96382027 | rs265033 | -0.156 | -0.14 | 0.958 | 0.463 | -0.001 | -0.009 | -0.009 | 0 | 0.54 | 0.47 | 0.62 | 0.77 | 0.57 | 0.67 |
| 4 | 96383256 | rs3846452 | 1.012 | 0.007 | 0.466 | 1.06 | -0.083 | -0.041 | -0.041 | 0.206 | 0.33 | 0.1 | 0.47 | 0.37 | 0.4 | 0.39 |
| 4 | 96385457 | rs265030 | -0.876 | -0.207 | 0.969 | 1.125 | 0.003 | 0.028 | 0.028 | 0 | 0.21 | 0.28 | 0.12 | 0.35 | 0.17 | 0.29 |
| 4 | 96385593 | rs265029 | -0.504 | -0.501 | 0.817 | 0.862 | 0.005 | 0.017 | 0.017 | 0 | 0.6 | 0.63 | 0.6 | 0.76 | 0.58 | 0.73 |
| 4 | 96388651 | rs4444836 | -1.113 | -0.437 | 1.027 | 1.076 | -0.006 | 0.054 | 0.054 | 0.356 | 0.22 | 0.35 | 0.13 | 0.36 | 0.17 | 0.3 |
| 4 | 96390834 | rs10856917 | 0.623 | -0.404 | 0.57 | 0.994 | 0.011 | -0.002 | -0.002 | 2.875 | 0.32 | 0.35 | 0.38 | 0.26 | 0.4 | 0.34 |
| 4 | 96393102 | rs265061 | -0.885 | -1.36 | 0.613 | 0.769 | -0.053 | -0.004 | -0.004 | 0 | 0.29 | 0.48 | 0.23 | 0.28 | 0.13 | 0.17 |
| 4 | 96394332 | rs17383133 | 1.341 | -1.16 | 0.397 | 0.927 | -0.033 | 0.01 | 0.01 | 0 | 0.26 | 0.13 | 0.34 | 0.21 | 0.36 | 0.28 |
| 4 | 96395955 | rs10005557 | 1.07 | -1.325 | 0.587 | 0.89 | -0.022 | -0.026 | -0.026 | 0.943 | 0.05 | 0.11 | 0.01 | 0 | 0.01 | 0.01 |
| 4 | 96397988 | rs265057 | -0.968 | -1.587 | 0.842 | 0.945 | -0.064 | 0.034 | 0.034 | 0.206 | 0.27 | 0.54 | 0.24 | 0.37 | 0.14 | 0.2 |
| 4 | 96406950 | rs4505826 | -1.045 | -1.599 | 0.374 | 0.726 | -0.062 | -0.051 | -0.051 | 3.737 | 0.63 | 0.85 | 0.64 | 0.58 | 0.56 | 0.5 |
| 4 | 96407772 | rs6834411 | 1.18 | -1.687 | 0.773 | 0.545 | -0.019 | 0.01 | 0.01 | 0 | 0.29 | 0.25 | 0.4 | 0.21 | 0.44 | 0.29 |
| 4 | 96407962 | rs2626047 | -0.195 | -1.776 | 0.993 | 0.595 | -0.005 | 0.01 | 0.01 | 0 | 0.06 | 0.13 | 0.15 | 0.05 | 0.02 | 0.08 |
| 4 | 96408615 | rs17383596 | -0.255 | -1.81 | 1.001 | 0.56 | 0.002 | -0.01 | -0.01 | 0.092 | 0.45 | 0.55 | 0.56 | 0.28 | 0.49 | 0.4 |
| 4 | 96409200 | rs2629833 | -0.303 | -1.807 | 1.119 | 0.572 | -0.012 | 0.04 | 0.04 | 0 | 0.46 | 0.71 | 0.58 | 0.33 | 0.49 | 0.41 |
| 4 | 96409981 | rs2626048 | -0.926 | -1.77 | 1.083 | 0.582 | -0.024 | 0.004 | 0.004 | 0 | 0.11 | 0.28 | 0.17 | 0.07 | 0.04 | 0.12 |
| 4 | 96410355 | rs17383903 | 0.944 | -1.541 | 0.991 | 0.717 | 0.005 | -0.003 | -0.003 | 0 | 0.3 | 0.33 | 0.4 | 0.25 | 0.43 | 0.29 |
| 4 | 96416472 | rs6532561 | 0.101 | -1.443 | 0.552 | 0.468 | -0.001 | -0.001 | -0.001 | 0 | 0.8 | 0.75 | 0.89 | 0.74 | 0.81 | 0.88 |
| 4 | 96424923 | rs10003330 | -0.303 | -1.448 | 0.705 | 0.443 | -0.008 | 0.013 | 0.013 | 0 | 0.07 | 0.31 | 0.03 | 0.04 | 0.02 | 0.02 |
| 4 | 96427086 | rs13107936 | 1.662 | -1.404 | -0.371 | 0.525 | -0.049 | 0.02 | 0.02 | 0 | 0.24 | 0.07 | 0.3 | 0.17 | 0.32 | 0.27 |
| 4 | 96428992 | rs7699716 | 0.09 | -1.295 | -0.634 | 0.649 | 0.041 | 0.007 | 0.007 | 0 | 0.79 | 0.88 | 0.94 | 0.77 | 0.91 | 0.9 |
| 4 | 96434992 | rs1905850 | 0.483 | -0.548 | -0.559 | 0.99 | -0.082 | -0.081 | -0.081 | 1.856 | 0.4 | 0.1 | 0.4 | 0.51 | 0.45 | 0.5 |
| 4 | 96441556 | rs11934067 | -0.884 | -0.217 | 0.274 | 0.965 | 0.003 | 0.019 | 0.019 | 0 | 0.05 | 0.22 | 0.02 | 0.04 | 0.01 | 0.01 |
| 4 | 96448687 | rs1564896 | 0.351 | 0.011 | -0.83 | 1.08 | -0.017 | -0.051 | -0.051 | 0.333 | 0.41 | 0.2 | 0.33 | 0.49 | 0.43 | 0.35 |
| 4 | 96450093 | rs10021966 | 0.251 | 0.312 | -0.925 | 1.002 | 0 | 0.023 | 0.023 | 0 | 0.18 | 0.12 | 0.26 | 0.05 | 0.15 | 0.16 |
| 4 | 96450925 | rs4699429 | 0.381 | 0.334 | -1.166 | 1.498 | -0.068 | -0.085 | -0.085 | 2.214 | 0.47 | 0.05 | 0.32 | 0.53 | 0.47 | 0.45 |
| 4 | 96454775 | rs1545919 | -0.346 | 0.322 | -0.298 | 1.378 | -0.024 | 0.073 | 0.073 | 0 | 0.27 | 0.48 | 0.35 | 0.33 | 0.2 | 0.32 |
| 4 | 96456602 | rs17386118 | 0.634 | 0.664 | -1.302 | 1.094 | -0.076 | 0.097 | 0.097 | 0 | 0.44 | 0.05 | 0.27 | 0.28 | 0.43 | 0.32 |
| 4 | 96458433 | rs10032931 | -0.57 | 1.34 | -0.04 | 1.225 | 0.01 | 0 | 0 | 1.04 | 0.78 | 0.78 | 0.67 | 0.75 | 0.68 | 0.7 |
| 4 | 96460537 | rs970689 | -0.517 | 1.261 | -0.07 | 1.006 | -0.002 | 0.001 | 0.001 | 0 | 0.77 | 0.83 | 0.68 | 0.82 | 0.65 | 0.7 |
| 4 | 96461841 | rs10008641 | 0.133 | 1.33 | -0.727 | 0.848 | -0.019 | -0.029 | -0.029 | 0.355 | 0.66 | 0.28 | 0.6 | 0.59 | 0.62 | 0.62 |
| 4 | 96467503 | rs11097470 | -1.334 | 1.665 | -0.146 | 0.542 | 0.065 | 0.003 | 0.003 | 0 | 0.07 | 0.19 | 0.08 | 0.06 | 0.14 | 0.03 |
| 4 | 96468160 | rs13149449 | -1.225 | 1.497 | -0.129 | 0.443 | 0.007 | -0.02 | -0.02 | 0 | 0.05 | 0.05 | 0.05 | 0 | 0.08 | 0.01 |
| 4 | 96475314 | rs1351999 | 0.264 | 1.005 | -0.305 | -0.484 | 0 | -0.021 | -0.021 | 0 | 0.63 | 0.64 | 0.76 | 0.94 | 0.67 | 0.77 |
| 4 | 96475689 | rs17024180 | -0.878 | 1.033 | 0.293 | -0.416 | 0.004 | -0.007 | -0.007 | 0.451 | 0.1 | 0.28 | 0.08 | 0.06 | 0.07 | 0.06 |
| 4 | 96480555 | rs13152256 | 1.054 | 1.257 | 0.213 | -0.046 | 0.008 | 0.004 | 0.004 | 0 | 0.76 | 0.69 | 0.74 | 0.76 | 0.72 | 0.77 |
| 4 | 96484360 | rs9307161 | 1.389 | 0.45 | -0.286 | -0.833 | 0.007 | 0.023 | 0.023 | 0 | 0.51 | 0.39 | 0.64 | 0.77 | 0.57 | 0.71 |
| 4 | 96485224 | rs6818365 | 0.048 | 0.407 | -0.579 | -0.624 | 0.036 | -0.007 | -0.007 | 0.623 | 0.84 | 0.95 | 0.87 | 0.85 | 0.9 | 0.88 |
| 4 | 96494451 | rs1385674 | 0.111 | 1.076 | -0.584 | 0.107 | -0.009 | -0.014 | -0.014 | 7.287 | 0.2 | 0.11 | 0.24 | 0.25 | 0.21 | 0.09 |
| 4 | 96494806 | rs11931784 | -1.08 | 0.905 | 0.095 | -0.03 | 0.091 | 0.003 | 0.003 | 0 | 0.14 | 0.59 | 0.16 | 0.1 | 0.22 | 0.14 |
| 4 | 96506439 | rs17453026 | 0.271 | 0.293 | -0.318 | 0.163 | 0.005 | 0.009 | 0.009 | 0.006 | 0.2 | 0.1 | 0.16 | 0.2 | 0.18 | 0.1 |
| 4 | 96506611 | rs4699434 | 1.533 | -0.145 | 0.26 | 0.875 | 0.006 | 0.017 | 0.017 | 0 | 0.48 | 0.41 | 0.6 | 0.39 | 0.55 | 0.63 |
| 4 | 96508701 | rs1385677 | 0.929 | -0.718 | -0.132 | 0.332 | 0.034 | 0.008 | 0.008 | 1.669 | 0.23 | 0.14 | 0.21 | 0.25 | 0.11 | 0.36 |
| 4 | 96509012 | rs7658734 | 0.184 | -0.759 | 0.289 | 0.293 | 0.005 | -0.006 | -0.006 | 0 | 0.16 | 0.15 | 0.11 | 0.08 | 0.2 | 0.11 |
| 4 | 96510283 | rs7665819 | 0.002 | -1.098 | 0.291 | 0.27 | -0.02 | 0.01 | 0.01 | 0 | 0.62 | 0.49 | 0.61 | 0.44 | 0.74 | 0.41 |
| 4 | 96512246 | rs11097473 | 2.18 | -1.463 | -0.225 | 0.03 | -0.141 | 0.003 | 0.003 | 0 | 0.27 | 0.02 | 0.4 | 0.18 | 0.42 | 0.23 |
| 4 | 96516310 | rs1485611 | 1.278 | -1.379 | 0.295 | 0.102 | -0.018 | 0.025 | 0.025 | 0 | 0.43 | 0.33 | 0.51 | 0.37 | 0.55 | 0.31 |
| 4 | 96517899 | rs10024436 | 1.316 | -1.484 | 0.266 | 0.062 | 0.035 | 0.004 | 0.004 | 0.699 | 0.22 | 0.13 | 0.21 | 0.25 | 0.11 | 0.34 |
| 4 | 96518564 | rs1564893 | -0.349 | -1.43 | 0.336 | 0.157 | -0.007 | 0.068 | 0.068 | 0.043 | 0.84 | 0.74 | 0.86 | 0.73 | 0.86 | 0.78 |
| 4 | 96521263 | rs11097474 | -0.33 | -1.43 | 0.343 | 0.157 | -0.007 | 0.071 | 0.071 | 0.057 | 0.84 | 0.74 | 0.86 | 0.73 | 0.86 | 0.78 |
| 4 | 96522796 | rs17387847 | -0.233 | -1.431 | 0.276 | 0.137 | 0.004 | -0.016 | -0.016 | 0 | 0.21 | 0.28 | 0.14 | 0.11 | 0.21 | 0.12 |
| 4 | 96524095 | rs17024229 | 1.628 | -1.418 | 0.208 | 0.117 | 0.02 | -0.031 | -0.031 | 0 | 0.06 | 0.11 | 0.03 | 0 | 0.07 | 0.05 |
| 4 | 96527272 | rs264298 | -1.005 | -1.413 | 0.135 | 0.147 | -0.001 | 0.019 | 0.019 | 0 | 0.9 | 0.91 | 0.9 | 0.73 | 0.93 | 0.81 |
| 4 | 96527801 | rs12499437 | 1.379 | -1.341 | 0.341 | 0.151 | -0.061 | 0.028 | 0.028 | 0 | 0.42 | 0.16 | 0.5 | 0.37 | 0.54 | 0.31 |
| 4 | 96531822 | rs264305 | 0.79 | -1.312 | 0.44 | -0.016 | 0 | 0 | 0 | 0 | 0.24 | 0.23 | 0.25 | 0.26 | 0.18 | 0.39 |
| 4 | 96532925 | rs6833223 | 0.804 | -1.279 | 0.425 | 0.025 | 0 | 0 | 0 | 0.018 | 0.46 | 0.5 | 0.55 | 0.46 | 0.55 | 0.39 |
| 4 | 96537724 | rs11937354 | 0.426 | -1.273 | 0.4 | 0.039 | 0 | 0 | 0 | 0.2 | 0.49 | 0.5 | 0.57 | 0.47 | 0.59 | 0.42 |
| 4 | 96541556 | rs10516981 | 0.861 | -1.259 | -0.181 | 0.078 | -0.049 | -0.049 | -0.049 | 0 | 0.44 | 0.15 | 0.53 | 0.47 | 0.56 | 0.41 |
| 4 | 96543990 | rs2043656 | -0.655 | -1.485 | 0.18 | 0.244 | -0.059 | -0.007 | -0.007 | 0 | 0.72 | 0.53 | 0.69 | 0.67 | 0.86 | 0.61 |
| 4 | 96545816 | rs1369985 | -0.89 | -1.475 | 0.29 | 0.226 | 0.011 | -0.002 | -0.002 | 0.44 | 0.33 | 0.56 | 0.22 | 0.25 | 0.35 | 0.23 |
| 4 | 96546107 | rs977737 | 1.047 | -1.562 | 0.294 | 0.196 | -0.015 | -0.019 | -0.019 | 2.554 | 0.45 | 0.29 | 0.59 | 0.51 | 0.55 | 0.44 |
| 4 | 96546227 | rs977738 | 1.047 | -1.562 | 0.294 | 0.196 | -0.015 | -0.019 | -0.019 | 0 | 0.45 | 0.3 | 0.59 | 0.51 | 0.55 | 0.44 |
| 4 | 96546552 | rs11941922 | -1.438 | -1.536 | 0.178 | 0.24 | 0.034 | -0.003 | -0.003 | 0.093 | 0.78 | 0.89 | 0.79 | 0.73 | 0.89 | 0.65 |
| 4 | 96547093 | rs987382 | -0.204 | -1.387 | 0.182 | 0.266 | 0.019 | 0.001 | 0.001 | 0.279 | 0.28 | 0.38 | 0.16 | 0.22 | 0.3 | 0.19 |

| | | | | | | | | | | | | | | | | |
|---|---|---|---|---|---|---|---|---|---|---|---|---|---|---|---|---|
| 4 | 96557708 | rs1436591 | 0.944 | -1.43 | -0.269 | -0.013 | -0.027 | -0.006 | -0.006 | 0 | 0.44 | 0.2 | 0.51 | 0.44 | 0.56 | 0.39 |
| 4 | 96560136 | rs992852 | -1.592 | -1.732 | -0.313 | 0.076 | 0.019 | -0.017 | -0.017 | 0.471 | 0.78 | 0.84 | 0.79 | 0.64 | 0.9 | 0.65 |
| 4 | 96560457 | rs13106700 | 1.435 | -1.625 | -0.469 | 0.026 | -0.065 | -0.045 | -0.045 | 0 | 0.36 | 0.15 | 0.49 | 0.43 | 0.54 | 0.36 |
| 4 | 96562035 | rs12644144 | 1.042 | -1.614 | -0.237 | 0.035 | -0.017 | -0.021 | -0.021 | 0 | 0.39 | 0.28 | 0.52 | 0.46 | 0.55 | 0.39 |
| 4 | 96565112 | rs10016927 | 1.366 | -1.629 | -0.357 | 0.041 | -0.067 | -0.048 | -0.048 | 0.016 | 0.37 | 0.14 | 0.49 | 0.43 | 0.54 | 0.37 |
| 4 | 96567263 | rs11727175 | -0.942 | -1.739 | -0.116 | -0.012 | -0.002 | -0.021 | -0.021 | 0 | 0.36 | 0.59 | 0.22 | 0.17 | 0.34 | 0.21 |
| 4 | 96568238 | rs7698569 | -0.904 | -1.58 | 0.054 | 0.174 | -0.028 | 0.013 | 0.013 | 0 | 0.73 | 0.7 | 0.71 | 0.61 | 0.88 | 0.57 |
| 4 | 96577171 | rs2865709 | -1.09 | -1.574 | -0.164 | 0.208 | -0.002 | -0.003 | -0.003 | 0 | 0.74 | 0.78 | 0.71 | 0.6 | 0.89 | 0.59 |
| 4 | 96577810 | rs7658020 | 0.9 | -1.452 | 0.009 | 0.058 | 0.003 | 0.001 | 0.001 | 0 | 0.14 | 0.11 | 0.05 | 0.12 | 0.1 | 0.11 |
| 4 | 96581831 | rs6841996 | -0.483 | -1.472 | 0.162 | 0.039 | 0.015 | -0.032 | -0.032 | 0.085 | 0.26 | 0.39 | 0.16 | 0.14 | 0.29 | 0.18 |
| 4 | 96600219 | rs1436576 | -1.019 | -1.622 | -0.224 | 0.26 | -0.05 | 0.057 | 0.057 | 0 | 0.78 | 0.68 | 0.72 | 0.63 | 0.89 | 0.62 |
| 4 | 96610670 | rs13112339 | -0.597 | -1.625 | -0.047 | 0.229 | -0.045 | 0.046 | 0.046 | 0 | 0.75 | 0.41 | 0.69 | 0.63 | 0.86 | 0.61 |
| 4 | 96614776 | rs17390627 | 1.78 | -1.751 | -0.272 | 0.189 | 0.008 | -0.001 | -0.001 | 0 | 0.06 | 0.02 | 0.09 | 0.05 | 0.03 | 0.05 |
| 4 | 96614904 | rs11097480 | 0.148 | -1.804 | -0.323 | -0.108 | 0.035 | -0.027 | -0.027 | 0.051 | 0.7 | 0.23 | 0.64 | 0.6 | 0.85 | 0.57 |
| 4 | 96625942 | rs1865848 | 0.63 | -1.893 | -0.697 | -0.388 | -0.095 | 0.094 | 0.094 | 0 | 0.18 | 0.02 | 0.11 | 0.02 | 0.23 | 0.13 |
| 4 | 96636651 | rs1579627 | -0.737 | -1.809 | -0.205 | -0.08 | -0.054 | 0.058 | 0.058 | 0 | 0.77 | 0.66 | 0.72 | 0.63 | 0.89 | 0.59 |
| 4 | 96640243 | rs7685363 | 2.397 | -1.698 | -1.085 | -0.112 | -0.143 | -0.02 | -0.02 | 1.049 | 0.23 | 0.01 | 0.35 | 0.17 | 0.41 | 0.21 |
| 4 | 96641816 | rs12508603 | 0.309 | -2.304 | -0.03 | 0.42 | 0.051 | -0.038 | -0.038 | 0.027 | 0.7 | 0.21 | 0.64 | 0.6 | 0.87 | 0.57 |
| 4 | 96642580 | rs1985147 | 0.728 | -2.124 | -0.113 | 0.455 | 0.167 | -0.046 | -0.046 | 5.596 | 0.58 | 0.17 | 0.64 | 0.49 | 0.84 | 0.46 |
| 4 | 96663991 | rs1946742 | 0.301 | -1.081 | 0.393 | 0.82 | -0.037 | 0.035 | 0.035 | 0 | 0.72 | 0.64 | 0.69 | 0.64 | 0.82 | 0.61 |
| 4 | 96664225 | rs1946741 | 0.876 | -1.163 | 0.084 | 0.772 | -0.007 | 0.002 | 0.002 | 0 | 0.64 | 0.39 | 0.61 | 0.39 | 0.76 | 0.55 |
| 4 | 96664321 | rs1946740 | 0.371 | -0.974 | -0.04 | 0.7 | -0.031 | 0.064 | 0.064 | 0 | 0.75 | 0.65 | 0.68 | 0.6 | 0.81 | 0.59 |
| 4 | 96672472 | rs7681417 | 0.61 | -1.176 | -0.633 | -0.217 | 0.003 | 0.004 | 0.004 | 0 | 0.19 | 0.17 | 0.08 | 0.07 | 0.12 | 0.15 |
| 4 | 96685664 | rs2196515 | -0.471 | -1.44 | -1.217 | -0.262 | 0.02 | -0.096 | -0.096 | 0.218 | 0.78 | 0.95 | 0.65 | 0.4 | 0.85 | 0.6 |
| 4 | 96686886 | rs1595201 | -0.457 | -1.361 | -1.219 | -0.263 | 0.014 | -0.096 | -0.096 | 0.304 | 0.78 | 0.95 | 0.65 | 0.4 | 0.85 | 0.6 |
| 4 | 96688897 | rs6851495 | 0.002 | -1.498 | -0.832 | -0.543 | 0.005 | 0.006 | 0.006 | 0 | 0.26 | 0.73 | 0.11 | 0.09 | 0.14 | 0.17 |
| 4 | 96691264 | rs4293789 | 1.129 | -1.47 | -0.866 | -0.525 | 0.013 | 0.013 | 0.013 | 0 | 0.18 | 0.13 | 0.07 | 0.06 | 0.12 | 0.15 |
| 4 | 96691621 | rs4699458 | 1.213 | -0.582 | -1.067 | -0.95 | -0.04 | 0.056 | 0.056 | 0.018 | 0.29 | 0.16 | 0.28 | 0.16 | 0.38 | 0.2 |
| 4 | 96704454 | rs7669620 | -0.168 | -0.575 | -1.311 | -0.957 | 0.045 | 0.188 | 0.188 | 0 | 0.48 | 0.8 | 0.41 | 0.2 | 0.58 | 0.32 |
| 4 | 96724242 | rs13116310 | 1.136 | -0.615 | -1.017 | -0.829 | -0.012 | 0.008 | 0.008 | 0.359 | 0.3 | 0.29 | 0.28 | 0.13 | 0.43 | 0.2 |
| 4 | 96724483 | rs4463073 | -0.219 | -0.919 | -1.172 | -0.708 | -0.01 | -0.045 | -0.045 | 0 | 0.31 | 0.17 | 0.32 | 0.6 | 0.29 | 0.48 |
| 4 | 96725043 | rs11097485 | 1.207 | -1.236 | -1.043 | -0.706 | 0.016 | -0.01 | -0.01 | 0 | 0.28 | 0.32 | 0.42 | 0.21 | 0.44 | 0.24 |
| 4 | 96727544 | rs13132449 | -0.5 | -1.516 | -1.048 | -0.997 | -0.003 | -0.052 | -0.052 | 0 | 0.67 | 0.48 | 0.74 | 0.84 | 0.68 | 0.84 |
| 4 | 96728233 | rs5012933 | -0.164 | -1.505 | -1.037 | -0.722 | -0.004 | -0.016 | -0.016 | 0.196 | 0.6 | 0.47 | 0.72 | 0.76 | 0.64 | 0.73 |
| 4 | 96728284 | rs5012929 | -0.164 | -1.505 | -1.037 | -0.722 | -0.004 | -0.016 | -0.016 | 0.07 | 0.6 | 0.47 | 0.72 | 0.76 | 0.64 | 0.73 |
| 4 | 96729750 | rs1583544 | 0.349 | -1.284 | -1.152 | -0.481 | -0.006 | 0.075 | 0.075 | 0.021 | 0.16 | 0.24 | 0.28 | 0.56 | 0.13 | 0.33 |
| 4 | 96730586 | rs17024592 | -0.104 | -1.168 | -1.09 | -0.471 | 0.017 | -0.007 | -0.007 | 1.198 | 0.34 | 0.28 | 0.35 | 0.58 | 0.24 | 0.51 |
| 4 | 96732967 | rs4352481 | 0.636 | -1.373 | -1.031 | -0.61 | 0.001 | 0.076 | 0.076 | 0.096 | 0.55 | 0.18 | 0.55 | 0.65 | 0.59 | 0.65 |
| 4 | 96733861 | rs11938262 | -0.734 | -1.932 | -0.511 | -0.875 | -0.009 | -0.044 | -0.044 | 0 | 0.32 | 0.61 | 0.25 | 0.13 | 0.25 | 0.29 |
| 4 | 96734419 | rs11939292 | -0.673 | -1.944 | -0.284 | -0.881 | -0.014 | -0.072 | -0.072 | 0 | 0.31 | 0.56 | 0.25 | 0.13 | 0.25 | 0.29 |
| 4 | 96736162 | rs1436554 | 0.825 | -1.79 | -0.204 | -0.845 | 0.008 | -0.027 | -0.027 | 0 | 0.1 | 0.1 | 0.08 | 0 | 0.15 | 0.07 |
| 4 | 96742723 | rs1436548 | -0.579 | -2.315 | 0.223 | -1.091 | -0.057 | -0.054 | -0.054 | 2.913 | 0.79 | 0.68 | 0.85 | 0.89 | 0.92 | 0.8 |
| 4 | 96743842 | rs6819351 | -1.816 | -2.272 | 0.217 | -1.071 | 0.007 | 0.008 | 0.008 | 0.589 | 0.87 | 0.9 | 0.88 | 0.93 | 0.93 | 0.94 |
| 4 | 96743973 | rs6819078 | -0.814 | -2.254 | 0.396 | -1.06 | 0.041 | 0.047 | 0.047 | 0 | 0.66 | 0.73 | 0.78 | 0.87 | 0.79 | 0.75 |
| 4 | 96747103 | rs1836899 | -0.78 | -2.223 | 0.363 | -1.044 | 0.033 | 0.036 | 0.036 | 0 | 0.66 | 0.71 | 0.78 | 0.87 | 0.79 | 0.76 |
| 4 | 96759448 | rs16996519 | 0.894 | -2.2 | 0.375 | -1.024 | -0.007 | -0.011 | -0.011 | 0.619 | 0.11 | 0.15 | 0.06 | 0.05 | 0.08 | 0.06 |
| 4 | 96760489 | rs11733585 | 0.742 | -2.306 | 0.557 | -1.081 | -0.038 | -0.052 | -0.052 | 0 | 0.09 | 0.21 | 0.03 | 0.01 | 0.03 | 0.15 |
| 4 | 96760945 | rs1076860 | -0.314 | -2.173 | 0.581 | -0.62 | -0.036 | -0.052 | -0.052 | 0.019 | 0.58 | 0.53 | 0.77 | 0.86 | 0.78 | 0.62 |
| 4 | 96761233 | rs10516983 | 0.656 | -1.814 | 0.627 | -0.218 | -0.007 | -0.011 | -0.011 | 2.554 | 0.11 | 0.14 | 0.06 | 0.05 | 0.08 | 0.06 |
| 4 | 96764785 | rs1436536 | 1.108 | -1.692 | 0.295 | -0.234 | -0.078 | -0.039 | -0.039 | 0 | 0.3 | 0.12 | 0.48 | 0.58 | 0.49 | 0.43 |
| 4 | 96765029 | rs7684431 | 0.964 | -1.824 | 0.319 | -0.715 | -0.038 | -0.043 | -0.043 | 0 | 0.08 | 0.18 | 0.01 | 0.01 | 0.01 | 0.14 |
| 4 | 96783302 | rs2054159 | -0.402 | -1.341 | -0.009 | -0.486 | 0.006 | -0.012 | -0.012 | 0.141 | 0.49 | 0.45 | 0.68 | 0.86 | 0.63 | 0.57 |
| 4 | 96794027 | rs2014345 | 0.186 | -0.678 | -0.58 | -0.05 | -0.028 | 0.055 | 0.055 | 0 | 0.36 | 0.18 | 0.61 | 0.77 | 0.56 | 0.43 |
| 4 | 96798214 | rs10007534 | 0.066 | -0.586 | -0.59 | -0.107 | -0.014 | 0.035 | 0.035 | 0 | 0.37 | 0.31 | 0.62 | 0.8 | 0.56 | 0.51 |
| 4 | 96814281 | rs13131407 | -0.324 | -1.079 | -0.46 | 0.923 | -0.006 | 0.003 | 0.003 | 0 | 0.36 | 0.33 | 0.27 | 0.55 | 0.27 | 0.26 |
| 4 | 96816411 | rs6848902 | -0.449 | -1.164 | -0.461 | 0.847 | -0.022 | -0.008 | -0.008 | 0.074 | 0.41 | 0.48 | 0.22 | 0.34 | 0.27 | 0.38 |
| 4 | 96828109 | rs6532586 | -1.062 | -0.859 | -1.333 | 0.998 | -0.007 | -0.106 | -0.106 | 0.042 | 0.84 | 0.98 | 0.76 | 0.44 | 0.85 | 0.73 |
| 4 | 96830011 | rs10516985 | 0.292 | -0.753 | -1.492 | 0.926 | -0.026 | -0.034 | -0.034 | 0 | 0.07 | 0 | 0.09 | 0.24 | 0.1 | 0.07 |
| 4 | 96831739 | rs10516987 | 0.073 | -1.003 | -1.099 | -0.073 | 0.031 | 0.038 | 0.038 | 0 | 0.35 | 0.25 | 0.16 | 0.05 | 0.2 | 0.34 |
| 4 | 96860000 | rs9997683 | 0.512 | -0.33 | -1.177 | 0.179 | -0.032 | 0.069 | 0.069 | 0 | 0.37 | 0.22 | 0.47 | 0.14 | 0.54 | 0.3 |
| 4 | 96875436 | rs6532589 | -0.637 | -0.339 | -0.852 | 0.645 | 0.034 | -0.005 | -0.005 | 1.179 | 0.45 | 0.63 | 0.3 | 0.55 | 0.28 | 0.44 |
| 4 | 96877723 | rs17024928 | 0.512 | -0.974 | -1.16 | 0.591 | -0.016 | 0.022 | 0.022 | 0 | 0.19 | 0.03 | 0.1 | 0.13 | 0.16 | 0.13 |
| 4 | 96879552 | rs10027211 | -0.621 | -0.987 | -0.975 | 0.663 | -0.052 | 0.048 | 0.048 | 0 | 0.26 | 0.6 | 0.2 | 0.42 | 0.12 | 0.31 |
| 4 | 96880695 | rs17024939 | -0.004 | -1.043 | -0.908 | 0.651 | 0.009 | -0.011 | -0.011 | 0.005 | 0.16 | 0.14 | 0.12 | 0.4 | 0.09 | 0.24 |
| 4 | 96893798 | rs4699479 | -1.373 | -0.214 | -1.203 | 0.336 | -0.041 | 0.085 | 0.085 | 0.065 | 0.59 | 0.8 | 0.54 | 0.82 | 0.49 | 0.75 |
| 4 | 96897523 | rs6823094 | -0.678 | -0.237 | -0.619 | 0.348 | 0.016 | -0.004 | -0.004 | 0.22 | 0.44 | 0.34 | 0.31 | 0.47 | 0.34 | 0.43 |
| 4 | 96906648 | rs17024965 | -0.303 | -0.162 | -0.643 | -0.519 | -0.004 | -0.04 | -0.04 | 0 | 0.35 | 0.41 | 0.38 | 0.12 | 0.28 | 0.25 |
| 4 | 96908427 | rs4699481 | 0.611 | 0.45 | -1.024 | -0.816 | -0.024 | 0.165 | 0.165 | 0 | 0.46 | 0.2 | 0.3 | 0.12 | 0.53 | 0.27 |
| 4 | 96918295 | rs6855600 | 0.045 | 0.592 | -0.489 | -0.992 | -0.001 | -0.063 | -0.063 | 0 | 0.29 | 0.3 | 0.37 | 0.01 | 0.26 | 0.21 |

| | | | | | | | | | | | | | | | | |
|---|---|---|---|---|---|---|---|---|---|---|---|---|---|---|---|---|
| 4 | 96923483 | rs1514974 | -0.454 | 0.365 | -0.504 | -0.688 | 0 | -0.001 | -0.001 | 0 | 0.68 | 0.68 | 0.6 | 0.31 | 0.66 | 0.52 |
| 4 | 96933043 | rs12503593 | 0.341 | 0.343 | -0.497 | -0.392 | 0.012 | -0.051 | -0.051 | 0 | 0.11 | 0.26 | 0.25 | 0.02 | 0.1 | 0.11 |
| 4 | 96939471 | rs17425319 | 1.416 | 0.213 | -0.916 | -0.491 | -0.098 | -0.016 | -0.016 | 0 | 0.31 | 0.04 | 0.16 | 0.22 | 0.36 | 0.25 |
| 4 | 96944577 | rs971524 | 0.032 | 0.285 | -0.612 | -0.403 | -0.001 | 0.013 | 0.013 | 0.808 | 0.48 | 0.58 | 0.45 | 0.28 | 0.47 | 0.37 |
| 4 | 96946680 | rs7689120 | -0.469 | 0.348 | -0.584 | -0.193 | -0.002 | -0.001 | -0.001 | 0 | 0.67 | 0.64 | 0.59 | 0.28 | 0.65 | 0.51 |
| 4 | 96952180 | rs3737480 | 0.81 | 0.491 | -1.262 | -0.35 | -0.079 | -0.074 | -0.074 | 0 | 0.14 | 0.01 | 0.13 | 0.23 | 0.22 | 0.16 |
| 4 | 96961891 | rs1567957 | -1.627 | 0.365 | -1.256 | -0.074 | -0.015 | -0.013 | -0.013 | 0 | 0.94 | 0.99 | 0.79 | 0.73 | 0.91 | 0.72 |
| 4 | 96963418 | rs997076 | -1.603 | 0.371 | -1.254 | -0.067 | -0.015 | -0.013 | -0.013 | 0 | 0.94 | 0.99 | 0.79 | 0.72 | 0.91 | 0.71 |
| 4 | 96965849 | rs1514976 | -0.736 | 0.175 | -0.632 | -1.28 | -0.001 | -0.003 | -0.003 | 0.063 | 0.64 | 0.64 | 0.49 | 0.26 | 0.6 | 0.43 |
| 4 | 96967431 | rs2203442 | -0.436 | 0.44 | -1.01 | -1.2 | -0.01 | 0.01 | 0.01 | 0 | 0.37 | 0.33 | 0.29 | 0.24 | 0.44 | 0.3 |
| 4 | 96977876 | rs7695763 | 0.995 | 0.568 | -0.939 | -1.366 | -0.001 | -0.009 | -0.009 | 0 | 0.2 | 0.1 | 0.14 | 0 | 0.2 | 0.14 |
| 4 | 96979391 | rs10026084 | -0.396 | 0.569 | -0.81 | -1.321 | 0.005 | -0.007 | -0.007 | 0 | 0.36 | 0.32 | 0.29 | 0.22 | 0.43 | 0.3 |
| 4 | 96982735 | rs17025056 | -0.103 | 0.92 | -0.753 | -0.953 | 0.028 | 0.022 | 0.022 | 0 | 0.15 | 0.15 | 0.14 | 0.22 | 0.22 | 0.16 |
| 4 | 97008338 | rs6825966 | -0.488 | 0.614 | -0.598 | -1.145 | 0.001 | 0.002 | 0.002 | 0 | 0.48 | 0.41 | 0.41 | 0.26 | 0.54 | 0.36 |
| 4 | 97014317 | rs7686416 | -0.093 | 0.838 | -0.731 | -1.28 | 0.008 | -0.03 | -0.03 | 0 | 0.43 | 0.31 | 0.31 | 0.02 | 0.36 | 0.24 |
| 4 | 97018808 | rs4318657 | 1.638 | 0.789 | -0.85 | -1.391 | 0.015 | 0.009 | 0.009 | 0.016 | 0.18 | 0.1 | 0.06 | 0.01 | 0.12 | 0.08 |
| 4 | 97024525 | rs6851001 | -0.231 | 0.809 | -0.929 | -1.36 | -0.001 | -0.085 | -0.085 | 2.509 | 0.43 | 0.35 | 0.31 | 0.02 | 0.36 | 0.24 |
| 4 | 97037142 | rs6814006 | 1.03 | 1.057 | -0.933 | -0.968 | -0.027 | 0.082 | 0.082 | 0.148 | 0.2 | 0.04 | 0.14 | 0 | 0.2 | 0.15 |
| 4 | 97059594 | rs6848939 | -0.213 | 0.736 | -0.603 | -1.354 | -0.011 | -0.141 | -0.141 | 0.031 | 0.31 | 0.4 | 0.31 | 0.01 | 0.28 | 0.16 |
| 4 | 97062525 | rs10516990 | 0.742 | 0.85 | -0.91 | -1.041 | -0.008 | 0.031 | 0.031 | 0 | 0.06 | 0.01 | 0.04 | 0 | 0.07 | 0.02 |
| 4 | 97074573 | rs6844234 | 0.018 | 0.855 | -1.04 | -0.899 | -0.075 | 0.007 | 0.007 | 0 | 0.33 | 0.09 | 0.28 | 0.29 | 0.42 | 0.28 |
| 4 | 97075010 | rs6819348 | 0.107 | 1.06 | -1.073 | -0.879 | -0.021 | -0.02 | -0.02 | 0 | 0.12 | 0.05 | 0.14 | 0.29 | 0.25 | 0.16 |
| 4 | 97082410 | rs7679128 | -0.827 | 0.443 | -0.823 | -1.442 | 0.005 | -0.008 | -0.008 | 0 | 0.55 | 0.57 | 0.7 | 0.88 | 0.62 | 0.69 |
| 4 | 97083012 | rs1840042 | -1.751 | 0.151 | -1.292 | -1.702 | -0.067 | 0.095 | 0.095 | 0 | 0.82 | 0.99 | 0.87 | 0.98 | 0.78 | 0.89 |
| 4 | 97086618 | rs7671563 | 0.451 | -0.003 | -0.712 | -1.398 | -0.022 | 0.05 | 0.05 | 0 | 0.18 | 0.29 | 0.3 | 0.61 | 0.17 | 0.4 |
| 4 | 97094011 | rs2042624 | 1.172 | 0.303 | -0.524 | -1.282 | 0.003 | 0.063 | 0.063 | 0.363 | 0.23 | 0.14 | 0.24 | 0.01 | 0.19 | 0.14 |
| 4 | 97104881 | rs6818950 | 0.692 | 0.086 | -0.46 | -1.522 | 0.003 | -0.056 | -0.056 | 0 | 0.35 | 0.31 | 0.16 | 0.02 | 0.24 | 0.23 |
| 4 | 97125086 | rs7692294 | -0.895 | 0.231 | -0.492 | -1.458 | -0.078 | -0.017 | -0.017 | 0 | 0.71 | 0.86 | 0.67 | 0.65 | 0.55 | 0.71 |
| 4 | 97135777 | rs6852164 | 1.105 | 0.326 | -0.255 | -1.205 | 0.003 | 0.008 | 0.008 | 0.256 | 0.23 | 0.22 | 0.25 | 0.01 | 0.19 | 0.14 |
| 4 | 97161412 | rs17025209 | 0.974 | 0.333 | -0.183 | -0.966 | 0.001 | 0.259 | 0.259 | 0.082 | 0.08 | 0.24 | 0.22 | 0.56 | 0.06 | 0.31 |
| 4 | 97185259 | rs4699496 | 0.126 | 0.316 | -0.182 | -1.354 | -0.017 | -0.001 | -0.001 | 0 | 0.33 | 0.49 | 0.47 | 0.63 | 0.26 | 0.45 |
| 4 | 97216464 | rs7682601 | 1.031 | 0.335 | -0.286 | -1.399 | 0.003 | 0 | 0 | 0 | 0.24 | 0.24 | 0.25 | 0.01 | 0.19 | 0.14 |
| 4 | 97245414 | rs11728676 | 1.676 | 0.319 | -0.393 | -1.414 | -0.021 | 0.1 | 0.1 | 0.05 | 0.2 | 0.03 | 0.14 | 0 | 0.18 | 0.13 |
| 4 | 97278349 | rs11097505 | 0.051 | 0.234 | -0.253 | -1.387 | -0.019 | -0.006 | -0.006 | 0 | 0.33 | 0.53 | 0.47 | 0.63 | 0.26 | 0.45 |
| 4 | 97293102 | rs6817554 | 1.176 | 0.504 | -0.448 | -1.335 | -0.001 | 0.067 | 0.067 | 0.363 | 0.23 | 0.14 | 0.24 | 0.02 | 0.2 | 0.14 |
| 4 | 97308914 | rs4699509 | -0.141 | 0.522 | -0.186 | -1.339 | 0.009 | 0.007 | 0.007 | 0 | 0.36 | 0.68 | 0.52 | 0.63 | 0.31 | 0.48 |
| 4 | 97395575 | rs9884206 | 0.97 | 0.609 | -0.977 | -1.504 | 0 | 0.064 | 0.064 | 0.963 | 0.24 | 0.15 | 0.24 | 0.01 | 0.2 | 0.14 |
| 4 | 97419088 | rs7693049 | 0.284 | 0.452 | -0.776 | -1.575 | -0.012 | -0.079 | -0.079 | 0 | 0.38 | 0.4 | 0.26 | 0.03 | 0.25 | 0.23 |
| 4 | 97430897 | rs1607062 | 0.094 | 0.48 | -0.78 | -1.58 | -0.005 | -0.068 | -0.068 | 1.39 | 0.39 | 0.4 | 0.29 | 0.03 | 0.3 | 0.26 |
| 4 | 97435210 | rs4699523 | -0.942 | 0.551 | -0.736 | -1.487 | -0.052 | 0.003 | 0.003 | 0.255 | 0.7 | 0.84 | 0.67 | 0.67 | 0.55 | 0.7 |
| 4 | 97435859 | rs12646509 | 0.451 | 0.436 | -0.661 | -1.252 | -0.005 | 0.001 | 0.001 | 0 | 0.3 | 0.38 | 0.37 | 0.64 | 0.26 | 0.44 |
| 4 | 97438464 | rs2090376 | 0.379 | 0.441 | -0.683 | -1.249 | -0.011 | -0.001 | -0.001 | 0 | 0.31 | 0.44 | 0.38 | 0.64 | 0.26 | 0.44 |
| 4 | 97460916 | rs1516662 | 0.436 | 0.442 | -0.659 | -1.237 | -0.003 | -0.001 | -0.001 | 0 | 0.3 | 0.35 | 0.37 | 0.63 | 0.26 | 0.43 |
| 4 | 97472001 | rs17025630 | 0.996 | 0.389 | -0.671 | -0.71 | -0.012 | 0.205 | 0.205 | 0 | 0.08 | 0.19 | 0.23 | 0.52 | 0.06 | 0.18 |
| 4 | 97475233 | rs13124250 | 0.237 | 0.705 | -0.614 | -0.998 | -0.002 | -0.004 | -0.004 | 0.179 | 0.56 | 0.41 | 0.33 | 0.14 | 0.48 | 0.47 |
| 4 | 97476439 | rs10033523 | 0.026 | 0.705 | -0.604 | -0.998 | 0.004 | -0.024 | -0.024 | 0.702 | 0.58 | 0.54 | 0.33 | 0.14 | 0.48 | 0.47 |
| 4 | 97489174 | rs6834178 | -0.286 | 0.745 | -0.657 | -0.998 | -0.019 | 0.06 | 0.06 | 0 | 0.59 | 0.71 | 0.39 | 0.14 | 0.53 | 0.49 |
| 4 | 97490815 | rs954241 | 0.806 | 0.649 | -0.829 | -1.078 | -0.003 | -0.014 | -0.014 | 0.712 | 0.32 | 0.33 | 0.18 | 0.13 | 0.27 | 0.33 |
| 4 | 97496521 | rs1516666 | -1.471 | 1.008 | -1.312 | -1.003 | -0.008 | 0.077 | 0.077 | 0 | 0.73 | 0.91 | 0.78 | 0.89 | 0.71 | 0.82 |
| 4 | 97503973 | rs17514862 | 1.705 | 0.482 | -1.077 | -0.932 | -0.035 | 0.083 | 0.083 | 0 | 0.19 | 0.04 | 0.14 | 0 | 0.2 | 0.12 |
| 4 | 97505284 | rs1400338 | -0.438 | 0.406 | -0.982 | -0.884 | -0.01 | 0.031 | 0.031 | 0 | 0.59 | 0.66 | 0.38 | 0.14 | 0.53 | 0.49 |
| 4 | 97507670 | rs11732439 | 1.462 | 0.445 | -1.131 | -0.808 | -0.015 | 0.025 | 0.025 | 0 | 0.21 | 0.1 | 0.18 | 0 | 0.25 | 0.15 |
| 4 | 97511566 | rs4699293 | -1.161 | 0.576 | -1.26 | -0.673 | -0.062 | -0.007 | -0.007 | 0.907 | 0.68 | 0.9 | 0.62 | 0.66 | 0.61 | 0.67 |
| 4 | 97523338 | rs7674376 | -0.353 | 0.561 | -0.497 | -0.029 | -0.004 | 0.047 | 0.047 | 0 | 0.34 | 0.52 | 0.45 | 0.51 | 0.34 | 0.32 |
| 4 | 97524510 | rs6838767 | 0.397 | 0.473 | -0.624 | -0.159 | -0.01 | -0.008 | -0.008 | 0 | 0.19 | 0.36 | 0.13 | 0.14 | 0.16 | 0.27 |
| 4 | 97528878 | rs7673132 | 0.78 | 0.523 | -1.045 | -0.173 | -0.002 | -0.033 | -0.033 | 0 | 0.23 | 0.2 | 0.19 | 0 | 0.25 | 0.15 |
| 4 | 97529769 | rs13151319 | 0.882 | 0.545 | -1.136 | -0.162 | -0.003 | -0.023 | -0.023 | 0 | 0.23 | 0.19 | 0.19 | 0 | 0.25 | 0.15 |
| 4 | 97534146 | rs6816701 | -0.625 | 0.69 | -1.529 | 0.324 | -0.059 | 0.072 | 0.072 | 0.005 | 0.53 | 0.89 | 0.59 | 0.64 | 0.5 | 0.59 |
| 4 | 97534436 | rs7349632 | -0.407 | 0.443 | -0.435 | 1.462 | -0.006 | 0.199 | 0.199 | 0 | 0.1 | 0.26 | 0.26 | 0.51 | 0.09 | 0.17 |
| 4 | 97540873 | rs7670682 | 0.224 | 0.718 | -0.948 | 1.273 | -0.017 | -0.024 | -0.024 | 0 | 0.68 | 0.74 | 0.58 | 0.68 | 0.61 | 0.57 |
| 4 | 97551175 | rs7680300 | -0.678 | -0.065 | -0.387 | 1.36 | -0.025 | 0.03 | 0.03 | 0 | 0.22 | 0.34 | 0.34 | 0.36 | 0.19 | 0.27 |
| 4 | 97554148 | rs10010351 | 1.161 | -0.344 | -0.525 | 1.395 | -0.034 | -0.026 | -0.026 | 0.055 | 0.4 | 0.22 | 0.36 | 0.44 | 0.48 | 0.35 |
| 4 | 97560105 | rs7679806 | -0.38 | -0.389 | -0.279 | 1.202 | -0.002 | 0.002 | 0.002 | 0 | 0.69 | 0.71 | 0.77 | 0.81 | 0.74 | 0.72 |
| 4 | 97560757 | rs7685841 | 1.605 | -0.39 | -0.315 | 1.214 | -0.032 | -0.029 | -0.029 | 0 | 0.38 | 0.22 | 0.34 | 0.44 | 0.46 | 0.35 |
| 4 | 97564929 | rs4699530 | 1.081 | -0.403 | -0.287 | 1.225 | -0.003 | 0.004 | 0.004 | 0.114 | 0.46 | 0.33 | 0.42 | 0.44 | 0.55 | 0.46 |
| 4 | 97567998 | rs6855625 | 1.575 | -0.269 | -0.313 | 1.156 | -0.045 | 0.123 | 0.123 | 2.947 | 0.42 | 0.15 | 0.33 | 0.24 | 0.46 | 0.28 |
| 4 | 97570607 | rs11735922 | 1.759 | -0.299 | -0.301 | 1.141 | -0.069 | 0.132 | 0.132 | 3.136 | 0.41 | 0.11 | 0.32 | 0.24 | 0.46 | 0.28 |
| 4 | 97591085 | rs7686119 | 0.442 | -0.238 | -0.243 | 1.247 | -0.006 | 0.005 | 0.005 | 0 | 0.73 | 0.64 | 0.59 | 0.66 | 0.73 | 0.66 |
| 4 | 97593469 | rs4476677 | 1.139 | -0.296 | -0.373 | 0.89 | 0.016 | -0.001 | -0.001 | 0 | 0.52 | 0.28 | 0.45 | 0.47 | 0.58 | 0.46 |

| | | | | | | | | | | | | | | | | |
|---|---|---|---|---|---|---|---|---|---|---|---|---|---|---|---|---|
| 4 | 97599993 | rs2178763 | 0.459 | -0.333 | -0.136 | 0.875 | 0.005 | 0.011 | 0.011 | 0 | 0.64 | 0.64 | 0.61 | 0.48 | 0.64 | 0.56 |
| 4 | 97611764 | rs17025751 | 1.092 | -0.458 | -0.18 | 0.815 | -0.002 | -0.013 | -0.013 | 0.556 | 0.14 | 0.12 | 0.13 | 0.02 | 0.15 | 0.12 |
| 4 | 97617981 | rs6532644 | 1.07 | -0.463 | -0.183 | 0.812 | -0.002 | -0.012 | -0.012 | 0 | 0.14 | 0.12 | 0.15 | 0.02 | 0.15 | 0.12 |
| 4 | 97621871 | rs10024940 | 1.962 | -0.621 | -0.294 | 0.795 | -0.091 | 0.123 | 0.123 | 1.898 | 0.41 | 0.06 | 0.32 | 0.25 | 0.42 | 0.26 |
| 4 | 97623215 | rs4308434 | 1.193 | -0.643 | -0.113 | 0.564 | -0.001 | -0.026 | -0.026 | 0 | 0.14 | 0.1 | 0.08 | 0 | 0.11 | 0.1 |
| 4 | 97627858 | rs10004820 | 1.003 | -0.649 | -0.212 | 0.719 | -0.008 | -0.024 | -0.024 | 0 | 0.14 | 0.17 | 0.13 | 0.02 | 0.15 | 0.12 |
| 4 | 97637435 | rs10856922 | 0.309 | -0.601 | -0.486 | 0.808 | 0.004 | 0.006 | 0.006 | 0 | 0.64 | 0.64 | 0.6 | 0.45 | 0.63 | 0.56 |
| 4 | 97644876 | rs10009024 | 1.315 | -0.265 | -1.317 | 0.498 | -0.148 | 0.126 | 0.126 | 2.313 | 0.4 | 0.02 | 0.31 | 0.22 | 0.43 | 0.25 |
| 4 | 97652057 | rs1922857 | -1.244 | -0.264 | -0.886 | 0.804 | 0.009 | -0.031 | -0.031 | 1.668 | 0.46 | 0.77 | 0.59 | 0.5 | 0.4 | 0.53 |
| 4 | 97653130 | rs1950166 | -1.504 | -0.346 | -0.15 | 0.374 | -0.003 | -0.003 | -0.003 | 0.205 | 0.56 | 0.53 | 0.66 | 0.58 | 0.6 | 0.72 |
| 4 | 97656445 | rs17025844 | 0.848 | -0.299 | -0.172 | 0.326 | -0.001 | -0.008 | -0.008 | 0.126 | 0.16 | 0.2 | 0.18 | 0.06 | 0.19 | 0.15 |
| 4 | 97662437 | rs17025848 | 1.003 | -0.29 | -0.176 | 0.112 | -0.002 | 0.007 | 0.007 | 0 | 0.16 | 0.13 | 0.17 | 0.02 | 0.19 | 0.15 |
| 4 | 97662454 | rs17456988 | -0.827 | -0.283 | -0.201 | 0.201 | -0.005 | 0 | 0 | 0 | 0.4 | 0.34 | 0.48 | 0.59 | 0.41 | 0.57 |
| 4 | 97680655 | rs1922848 | 0.466 | -0.201 | -0.053 | 0.021 | -0.002 | -0.001 | -0.001 | 0 | 0.07 | 0.04 | 0.05 | 0.06 | 0.09 | 0.17 |
| 4 | 97680928 | rs1922847 | -1.436 | -0.385 | -0.152 | -0.152 | 0.024 | 0.119 | 0.119 | 0.876 | 0.44 | 0.73 | 0.45 | 0.13 | 0.39 | 0.46 |
| 4 | 97685296 | rs6532649 | -0.923 | -0.437 | -0.081 | -0.123 | -0.015 | -0.044 | -0.044 | 0.555 | 0.27 | 0.57 | 0.27 | 0.11 | 0.2 | 0.29 |
| 4 | 97689917 | rs10516415 | 0.033 | -0.444 | 0.553 | -0.164 | -0.016 | -0.049 | -0.049 | 0.015 | 0.18 | 0.26 | 0.2 | 0.04 | 0.11 | 0.12 |
| 4 | 97695401 | rs12509878 | -1.439 | -0.393 | 0.418 | -0.17 | 0.024 | 0.119 | 0.119 | 1.013 | 0.45 | 0.74 | 0.45 | 0.13 | 0.39 | 0.46 |
| 4 | 97698070 | rs7695101 | -0.812 | -0.35 | 0.462 | -0.113 | -0.016 | -0.053 | -0.053 | 1.191 | 0.27 | 0.57 | 0.27 | 0.1 | 0.2 | 0.29 |
| 4 | 97705034 | rs1922850 | -1.018 | -0.387 | 0.452 | -0.132 | -0.019 | -0.054 | -0.054 | 4.683 | 0.29 | 0.57 | 0.27 | 0.1 | 0.2 | 0.29 |
| 4 | 97737582 | rs4579126 | -1.265 | -0.252 | 0.458 | 0.066 | -0.004 | -0.017 | -0.017 | 0 | 0.42 | 0.69 | 0.57 | 0.5 | 0.45 | 0.51 |
| 4 | 97742963 | rs4550934 | 0.567 | -0.273 | 0.287 | -0.036 | 0.001 | 0.013 | 0.013 | 0 | 0.13 | 0.06 | 0.06 | 0 | 0.09 | 0.05 |
| 4 | 97748436 | rs1922854 | -0.429 | -0.304 | 0.657 | -0.028 | 0.014 | 0.054 | 0.054 | 0.116 | 0.3 | 0.42 | 0.38 | 0.46 | 0.35 | 0.39 |
| 4 | 97751638 | rs12509492 | 0.007 | -0.289 | 0.548 | -0.28 | -0.012 | -0.034 | -0.034 | 0 | 0.12 | 0.22 | 0.18 | 0.05 | 0.09 | 0.13 |
| 4 | 97759914 | rs17025930 | -0.499 | -0.229 | 0.259 | -0.409 | -0.004 | -0.105 | -0.105 | 0.04 | 0.3 | 0.42 | 0.25 | 0.02 | 0.28 | 0.21 |
| 4 | 97765720 | rs17025935 | 0.747 | -0.222 | 0.031 | -0.245 | -0.001 | 0.005 | 0.005 | 0.47 | 0.16 | 0.14 | 0.18 | 0.02 | 0.2 | 0.16 |
| 4 | 97774817 | rs17025949 | -1.046 | -0.416 | -0.469 | -0.298 | -0.002 | -0.002 | -0.002 | 0.006 | 0.47 | 0.53 | 0.48 | 0.45 | 0.47 | 0.5 |
| 4 | 97776070 | rs6532654 | -1.844 | -0.256 | -1.41 | -0.242 | -0.131 | -0.144 | -0.144 | 0.015 | 0.64 | 0.96 | 0.69 | 0.5 | 0.61 | 0.64 |
| 4 | 97776909 | rs9997488 | 0.451 | -0.117 | -1.212 | -0.397 | -0.004 | 0.024 | 0.024 | 0 | 0.13 | 0.11 | 0.23 | 0.43 | 0.18 | 0.27 |
| 4 | 97779789 | rs11936397 | -0.181 | -0.182 | -1.093 | -0.418 | -0.002 | -0.001 | -0.001 | 0 | 0.27 | 0.23 | 0.29 | 0.45 | 0.27 | 0.33 |
| 4 | 97781855 | rs11725440 | -0.406 | -0.481 | -0.777 | -0.748 | 0.007 | -0.016 | -0.016 | 0 | 0.33 | 0.73 | 0.37 | 0.04 | 0.29 | 0.28 |
| 4 | 97782987 | rs7688717 | -0.779 | 0.265 | -1.093 | -0.279 | -0.021 | -0.006 | -0.006 | 0.053 | 0.63 | 0.84 | 0.77 | 0.61 | 0.63 | 0.64 |
| 4 | 97799998 | rs929290 | -0.469 | -0.265 | -0.149 | -0.526 | 0.014 | -0.002 | -0.002 | 0.791 | 0.46 | 0.58 | 0.37 | 0.47 | 0.43 | 0.33 |
| 4 | 97805511 | rs12651219 | -0.137 | -0.345 | 0.246 | -0.559 | 0.009 | -0.011 | -0.011 | 0.196 | 0.41 | 0.31 | 0.32 | 0.47 | 0.39 | 0.28 |
| 4 | 97807484 | rs9994928 | -0.3 | -0.413 | 0.036 | -0.529 | -0.003 | 0.003 | 0.003 | 0.07 | 0.42 | 0.43 | 0.34 | 0.47 | 0.43 | 0.29 |
| 4 | 97853451 | rs10488826 | -1.496 | -0.741 | 0.349 | -1.104 | 0.005 | -0.001 | -0.001 | 0 | 0.32 | 0.67 | 0.2 | 0.12 | 0.26 | 0.27 |
| 4 | 97860526 | rs2107405 | -1.196 | -0.762 | 0.744 | -1.121 | -0.008 | -0.027 | -0.027 | 0.098 | 0.27 | 0.3 | 0.15 | 0.12 | 0.23 | 0.25 |
| 4 | 97861141 | rs10488827 | -0.574 | -0.76 | 0.734 | -1.009 | -0.014 | -0.007 | -0.007 | 0.673 | 0.13 | 0.18 | 0.07 | 0.12 | 0.08 | 0.19 |
| 4 | 97865889 | rs10011515 | -1.426 | -0.634 | 0.937 | -0.984 | -0.025 | -0.079 | -0.079 | 0 | 0.25 | 0.47 | 0.16 | 0.12 | 0.23 | 0.2 |
| 4 | 97874682 | rs6825085 | -0.118 | -0.306 | 0.677 | -0.822 | 0.002 | -0.093 | -0.093 | 0 | 0.13 | 0.43 | 0.09 | 0 | 0.12 | 0.05 |
| 4 | 97874724 | rs6848042 | -0.14 | -0.255 | 0.674 | -0.825 | 0.009 | -0.094 | -0.094 | 1.361 | 0.14 | 0.43 | 0.1 | 0 | 0.13 | 0.05 |
| 4 | 97880409 | rs7694207 | -1.08 | -1.103 | -0.188 | -1.268 | -0.043 | -0.077 | -0.077 | 0 | 0.37 | 0.47 | 0.2 | 0.12 | 0.23 | 0.34 |
| 4 | 97889973 | rs6830362 | -0.333 | -1.224 | -0.448 | -1.348 | -0.031 | -0.044 | -0.044 | 0 | 0.23 | 0.32 | 0.16 | 0.11 | 0.19 | 0.25 |
| 4 | 97891970 | rs981391 | -1.467 | -0.551 | -1.357 | -0.916 | -0.081 | -0.087 | -0.087 | 0 | 0.74 | 0.9 | 0.75 | 0.57 | 0.67 | 0.84 |
| 4 | 97906295 | rs6834049 | -1.251 | -1.321 | -0.663 | -1.018 | 0.108 | 0.154 | 0.154 | 0 | 0.41 | 0.79 | 0.23 | 0.12 | 0.24 | 0.36 |
| 4 | 97907354 | rs13109296 | -0.538 | -0.362 | -0.319 | -0.375 | 0.005 | -0.016 | -0.016 | 0 | 0.14 | 0.3 | 0.14 | 0.11 | 0.18 | 0.21 |
| 4 | 97908904 | rs2525795 | 0.043 | 0.016 | -1.081 | -0.1 | -0.017 | -0.017 | -0.017 | 2.773 | 0.42 | 0.22 | 0.41 | 0.5 | 0.54 | 0.35 |
| 4 | 97909469 | rs2189572 | -0.612 | 0.067 | -0.667 | -0.048 | 0.008 | 0.008 | 0.008 | 0 | 0.18 | 0.13 | 0.15 | 0.11 | 0.14 | 0.24 |
| 4 | 97915370 | rs7659471 | -0.541 | 0.412 | -0.585 | 0.009 | -0.016 | -0.001 | -0.001 | 0 | 0.37 | 0.32 | 0.28 | 0.41 | 0.49 | 0.32 |
| 4 | 97917772 | rs7673377 | -0.347 | 0.994 | -0.845 | 0.106 | -0.007 | 0.01 | 0.01 | 0 | 0.21 | 0.22 | 0.16 | 0.11 | 0.27 | 0.23 |
| 4 | 97918476 | rs733323 | -1.465 | 0.48 | -1.058 | -0.158 | 0.007 | -0.018 | -0.018 | 0.517 | 0.55 | 0.85 | 0.37 | 0.48 | 0.57 | 0.52 |
| 4 | 97921027 | rs2903086 | 1.19 | 0.686 | -1.029 | 0.026 | -0.087 | -0.093 | -0.093 | 0.006 | 0.16 | 0.01 | 0.12 | 0.32 | 0.22 | 0.1 |
| 4 | 97934806 | rs17247004 | 0.279 | 0.504 | -0.984 | -0.286 | -0.042 | 0.083 | 0.083 | 0 | 0.14 | 0.03 | 0.1 | 0 | 0.17 | 0.1 |
| 4 | 97935617 | rs4141303 | -1.358 | 0.118 | -0.38 | -0.367 | 0.015 | 0.074 | 0.074 | 0 | 0.39 | 0.78 | 0.25 | 0.19 | 0.35 | 0.4 |
| 4 | 97935662 | rs4141304 | -0.482 | 0.218 | 0.409 | -0.344 | 0.032 | -0.058 | -0.058 | 4.459 | 0.22 | 0.44 | 0.18 | 0.11 | 0.27 | 0.21 |
| 4 | 97949728 | rs17026302 | 1.063 | -0.248 | -0.257 | 0.27 | -0.016 | -0.04 | -0.04 | 0.242 | 0.12 | 0.07 | 0.2 | 0.31 | 0.15 | 0.16 |
| 4 | 97962970 | rs4627872 | -1.153 | -0.35 | -0.313 | 0.072 | 0.045 | 0.101 | 0.101 | 0 | 0.38 | 0.86 | 0.22 | 0.19 | 0.28 | 0.4 |
| 4 | 97963926 | rs2865775 | 0.253 | -0.378 | -0.094 | 0.05 | -0.006 | 0.013 | 0.013 | 0 | 0.18 | 0.18 | 0.12 | 0.11 | 0.19 | 0.2 |
| 4 | 97968711 | rs7695114 | 1.267 | -0.351 | 0.455 | 0.077 | 0.001 | 0.023 | 0.023 | 0 | 0.11 | 0.33 | 0.07 | 0.07 | 0.08 | 0.16 |
| 4 | 97974722 | rs13109193 | -1.105 | -0.335 | 0.267 | 0.078 | 0.041 | 0.087 | 0.087 | 0 | 0.37 | 0.82 | 0.21 | 0.19 | 0.28 | 0.4 |
| 4 | 97984954 | rs11097537 | -1.38 | -0.167 | -0.179 | 0.163 | 0.009 | -0.057 | -0.057 | 0 | 0.55 | 0.9 | 0.39 | 0.5 | 0.6 | 0.5 |
| 4 | 97985065 | rs11942031 | 0.033 | 0.016 | -0.101 | 0.147 | -0.001 | -0.045 | -0.045 | 0.004 | 0.22 | 0.38 | 0.14 | 0.11 | 0.2 | 0.2 |
| 4 | 97989968 | rs10516420 | 1.113 | 0.263 | -0.864 | 0.31 | -0.116 | -0.086 | -0.086 | 0 | 0.17 | 0.02 | 0.17 | 0.31 | 0.32 | 0.11 |
| 4 | 98017954 | rs9997830 | -1.205 | -0.2 | -0.342 | 0.369 | -0.013 | -0.078 | -0.078 | 3.924 | 0.5 | 0.94 | 0.43 | 0.49 | 0.42 | 0.6 |
| 4 | 98045861 | rs2865763 | -1.298 | 0.004 | -0.403 | 0.399 | 0.015 | -0.027 | -0.027 | 2.085 | 0.47 | 0.84 | 0.42 | 0.49 | 0.42 | 0.55 |
| 4 | 98050429 | rs4131679 | -0.241 | 0.354 | -0.396 | 0.765 | 0.001 | 0.009 | 0.009 | 0 | 0.18 | 0.24 | 0.27 | 0.38 | 0.24 | 0.26 |
| 4 | 98053277 | rs4078016 | 0.871 | -0.115 | -0.561 | 0.599 | 0.003 | -0.019 | -0.019 | 0.903 | 0.09 | 0.03 | 0.05 | 0.11 | 0.08 | 0.16 |
| 4 | 98055658 | rs4074142 | -0.091 | -0.436 | -0.571 | 0.441 | 0.037 | 0.013 | 0.013 | 0 | 0.15 | 0.07 | 0.07 | 0.1 | 0.09 | 0.27 |
| 4 | 98065231 | rs11942998 | 0.576 | -0.275 | -0.664 | 0.711 | -0.025 | -0.037 | -0.037 | 0.179 | 0.1 | 0.04 | 0.13 | 0.28 | 0.12 | 0.14 |

| | | | | | | | | | | | | | | | | |
|---|---|---|---|---|---|---|---|---|---|---|---|---|---|---|---|---|
| 4 | 98069011 | rs7693495 | -1.962 | -0.929 | -0.973 | 0.214 | 0.08 | -0.008 | -0.008 | 1.691 | 0.47 | 0.92 | 0.32 | 0.38 | 0.31 | 0.5 |
| 4 | 98098473 | rs9715346 | -0.078 | -1.002 | 0.745 | 0.073 | -0.009 | -0.05 | -0.05 | 0 | 0.16 | 0.17 | 0.07 | 0 | 0.1 | 0.09 |
| 4 | 98100051 | rs11942542 | 0.438 | -1.022 | 0.697 | 0.066 | 0.028 | 0.006 | 0.006 | 0.112 | 0.15 | 0.08 | 0.07 | 0.13 | 0.09 | 0.23 |
| 4 | 98119717 | rs11721581 | -0.853 | -1.019 | 0.704 | 0.188 | -0.012 | -0.005 | -0.005 | 0.021 | 0.28 | 0.39 | 0.2 | 0.24 | 0.23 | 0.24 |
| 4 | 98121286 | rs13111598 | -0.476 | -1.002 | 0.561 | 0.196 | 0.001 | -0.001 | -0.001 | 0.063 | 0.25 | 0.2 | 0.19 | 0.24 | 0.22 | 0.24 |
| 4 | 98127149 | rs4642255 | -0.798 | -1.006 | 1.15 | 0.179 | -0.013 | -0.009 | -0.009 | 0 | 0.27 | 0.49 | 0.21 | 0.24 | 0.22 | 0.24 |
| 4 | 98160778 | rs10009327 | -1.839 | -0.964 | 0.818 | 0.215 | 0.081 | 0.037 | 0.037 | 0 | 0.47 | 0.84 | 0.3 | 0.36 | 0.32 | 0.47 |
| 4 | 98170598 | rs13126649 | -0.539 | -0.86 | 0.88 | 0.246 | 0.001 | 0 | 0 | 0 | 0.25 | 0.2 | 0.19 | 0.23 | 0.22 | 0.24 |
| 4 | 98182517 | rs11938519 | -1.488 | -1.022 | 1.39 | 0.357 | 0 | 0.001 | 0.001 | 0 | 0.44 | 0.55 | 0.28 | 0.29 | 0.31 | 0.49 |
| 4 | 98198773 | rs7690692 | -0.344 | -1.054 | 1.319 | 0.223 | 0.005 | 0.007 | 0.007 | 0 | 0.23 | 0.19 | 0.14 | 0.13 | 0.18 | 0.35 |
| 4 | 98200637 | rs1462758 | -1.227 | -1.065 | 1.107 | 0.416 | -0.021 | -0.02 | -0.02 | 0 | 0.34 | 0.4 | 0.34 | 0.21 | 0.28 | 0.42 |
| 4 | 98201071 | rs1982619 | -1.894 | -1.163 | 0.913 | 0.444 | 0.059 | -0.037 | -0.037 | 0.6 | 0.51 | 0.86 | 0.48 | 0.44 | 0.4 | 0.57 |
| 4 | 98206435 | rs11097557 | 2.073 | -0.521 | -0.031 | 0.155 | -0.015 | 0.144 | 0.144 | 0 | 0.35 | 0.04 | 0.31 | 0.15 | 0.31 | 0.3 |
| 4 | 98208206 | rs9307203 | -1.881 | -0.061 | 0.118 | 0.605 | -0.073 | -0.07 | -0.07 | 1.564 | 0.89 | 0.98 | 0.86 | 0.67 | 0.73 | 0.92 |
| 4 | 98209654 | rs9997401 | -0.228 | -0.761 | 0.84 | 0.4 | 0.002 | 0.014 | 0.014 | 0 | 0.22 | 0.33 | 0.2 | 0.23 | 0.21 | 0.22 |
| 4 | 98210045 | rs17026605 | 0.82 | -0.628 | 0.469 | 0.421 | -0.016 | -0.006 | -0.006 | 0 | 0.15 | 0.19 | 0.08 | 0.13 | 0.09 | 0.27 |
| 4 | 98219875 | rs10213670 | -0.681 | -0.523 | 0.705 | 0.507 | 0.001 | 0.017 | 0.017 | 0 | 0.26 | 0.32 | 0.37 | 0.31 | 0.28 | 0.27 |
| 4 | 98227375 | rs13119820 | -1.355 | -0.633 | 0.782 | 0.433 | 0.002 | -0.002 | -0.002 | 0 | 0.44 | 0.54 | 0.44 | 0.44 | 0.38 | 0.52 |
| 4 | 98228716 | rs11939157 | -1.811 | -0.757 | 0.869 | 0.378 | 0.04 | -0.006 | -0.006 | 0 | 0.48 | 0.67 | 0.46 | 0.44 | 0.38 | 0.56 |
| 4 | 98230356 | rs11936871 | 0.855 | -0.752 | 0.846 | 0.391 | -0.017 | -0.006 | -0.006 | 0 | 0.16 | 0.18 | 0.07 | 0.13 | 0.09 | 0.25 |
| 4 | 98237090 | rs1462778 | -0.657 | -0.836 | 0.986 | 0.404 | -0.059 | -0.01 | -0.01 | 0.746 | 0.27 | 0.5 | 0.28 | 0.21 | 0.18 | 0.34 |
| 4 | 98238605 | rs1462777 | -0.441 | -0.746 | 1.092 | 0.48 | 0 | 0.053 | 0.053 | 0 | 0.1 | 0.34 | 0.2 | 0.08 | 0.08 | 0.05 |
| 4 | 98240651 | rs11737720 | -0.44 | -0.704 | 0.856 | 0.567 | 0 | -0.007 | -0.007 | 0 | 0.23 | 0.19 | 0.2 | 0.34 | 0.21 | 0.28 |
| 4 | 98241354 | rs7686247 | -0.163 | -0.685 | 0.731 | 0.577 | -0.002 | -0.017 | -0.017 | 0 | 0.21 | 0.17 | 0.2 | 0.34 | 0.2 | 0.26 |
| 4 | 98243379 | rs1381822 | 0.83 | -0.723 | 0.727 | 0.527 | -0.009 | 0.004 | 0.004 | 0 | 0.07 | 0.08 | 0.05 | 0.12 | 0.03 | 0.08 |
| 4 | 98258036 | rs11728761 | -0.153 | -0.644 | 0.708 | 0.511 | -0.003 | -0.017 | -0.017 | 0 | 0.21 | 0.17 | 0.21 | 0.34 | 0.22 | 0.26 |
| 4 | 98264960 | rs10516422 | 0.324 | -0.487 | 0.536 | 0.697 | -0.004 | 0.03 | 0.03 | 0 | 0.07 | 0.15 | 0.19 | 0.1 | 0.08 | 0.06 |
| 4 | 98266404 | rs12512414 | -0.174 | -0.458 | 0.511 | 0.693 | -0.01 | 0.026 | 0.026 | 0 | 0.09 | 0.18 | 0.19 | 0.1 | 0.09 | 0.07 |
| 4 | 98270800 | rs1462767 | -1.254 | -0.368 | -0.038 | 0.577 | 0.043 | -0.003 | -0.003 | 0 | 0.49 | 0.69 | 0.46 | 0.45 | 0.39 | 0.54 |
| 4 | 98279501 | rs13151057 | -0.129 | -0.143 | -0.645 | 0.337 | 0.002 | -0.003 | -0.003 | 0.016 | 0.56 | 0.37 | 0.63 | 0.44 | 0.58 | 0.75 |
| 4 | 98289559 | rs2055957 | 0.694 | -0.154 | -1.374 | 0.193 | -0.034 | 0.023 | 0.023 | 0 | 0.24 | 0.03 | 0.21 | 0.17 | 0.22 | 0.34 |
| 4 | 98291379 | rs1350971 | 0.088 | 0.421 | -1.506 | 0.066 | -0.059 | -0.048 | -0.048 | 0 | 0.6 | 0.13 | 0.63 | 0.51 | 0.55 | 0.61 |
| 4 | 98292053 | rs10011676 | -1.226 | 0.532 | -0.275 | 0.412 | 0.169 | 0.265 | 0.265 | 0.352 | 0.88 | 0.58 | 0.76 | 0.58 | 0.74 | 0.85 |
| 4 | 98299136 | rs4699559 | 0.442 | 0.375 | -0.944 | 0.339 | -0.006 | -0.036 | -0.036 | 0 | 0.74 | 0.18 | 0.7 | 0.51 | 0.69 | 0.8 |
| 4 | 98330337 | rs6825634 | 1.164 | 0.026 | -1.231 | 0.279 | -0.029 | -0.036 | -0.036 | 0.077 | 0.06 | 0.01 | 0.13 | 0.12 | 0.1 | 0.11 |
| 4 | 98333918 | rs17026725 | -0.702 | -0.326 | -1.003 | 0.283 | -0.019 | -0.003 | -0.003 | 0 | 0.18 | 0.77 | 0.11 | 0.1 | 0.06 | 0.1 |
| 4 | 98370425 | rs9999251 | 0.85 | -0.341 | -0.676 | 0.273 | -0.026 | -0.009 | -0.009 | 3.305 | 0.09 | 0.17 | 0.05 | 0.06 | 0.05 | 0.03 |
| 4 | 98411916 | rs35632782 | -0.044 | -0.281 | -0.662 | 0.29 | -0.03 | 0.001 | 0.001 | 5.8 | 0.08 | 0.59 | 0.06 | 0.04 | 0.01 | 0.07 |
| 4 | 98414540 | rs11097562 | -1.359 | 0.297 | -1.839 | 0.532 | -0.074 | -0.094 | -0.094 | 0 | 0.77 | 0.95 | 0.78 | 0.58 | 0.7 | 0.71 |
| 4 | 98425582 | rs2865798 | 0.737 | 0.411 | -1.933 | 0.366 | -0.011 | 0.046 | 0.046 | 0.026 | 0.52 | 0.1 | 0.47 | 0.37 | 0.56 | 0.54 |
| 4 | 98435890 | rs6822100 | 1.533 | 0.612 | -1.833 | 0.147 | -0.053 | 0.061 | 0.061 | 0.045 | 0.26 | 0.07 | 0.21 | 0.16 | 0.32 | 0.32 |
| 4 | 98444547 | rs9307206 | 1.253 | 0.285 | -1.157 | -0.096 | -0.014 | 0.035 | 0.035 | 0 | 0.25 | 0.06 | 0.18 | 0.17 | 0.24 | 0.12 |
| 4 | 98466559 | rs960450 | 0.625 | 0.382 | -0.842 | 0.013 | 0.012 | -0.01 | -0.01 | 0 | 0.55 | 0.28 | 0.57 | 0.47 | 0.55 | 0.69 |
| 4 | 98471647 | rs2136080 | 2.801 | 0.511 | -1.047 | 0.058 | -0.05 | -0.011 | -0.011 | 0 | 0.15 | 0.01 | 0.05 | 0.11 | 0.13 | 0.21 |
| 4 | 98513928 | rs12649979 | -0.496 | 0.469 | -0.778 | -0.086 | -0.038 | -0.066 | -0.066 | 1.503 | 0.67 | 0.88 | 0.65 | 0.51 | 0.61 | 0.74 |
| 4 | 98518164 | rs13130146 | 1.943 | 0.395 | -0.829 | -0.069 | -0.016 | 0.007 | 0.007 | 0 | 0.2 | 0.05 | 0.15 | 0.17 | 0.2 | 0.11 |
| 4 | 98567966 | rs6832426 | -0.155 | 0.438 | -0.559 | 0.08 | -0.034 | -0.024 | -0.024 | 0 | 0.62 | 0.76 | 0.62 | 0.51 | 0.55 | 0.71 |
| 4 | 98575319 | rs1508470 | -0.155 | 0.438 | -0.518 | 0.08 | -0.028 | -0.02 | -0.02 | 0.004 | 0.62 | 0.75 | 0.62 | 0.51 | 0.55 | 0.71 |
| 4 | 98596032 | rs10856927 | -0.009 | 0.891 | -0.515 | 0.049 | -0.014 | -0.013 | -0.013 | 0 | 0.61 | 0.7 | 0.59 | 0.51 | 0.47 | 0.7 |
| 4 | 98597993 | rs11097568 | 0.761 | 0.04 | -0.367 | 0.096 | -0.029 | -0.035 | -0.035 | 0 | 0.54 | 0.18 | 0.56 | 0.41 | 0.54 | 0.66 |
| 4 | 98625497 | rs877376 | -0.162 | 0.091 | -0.419 | 0.126 | -0.022 | -0.014 | -0.014 | 0.063 | 0.62 | 0.7 | 0.61 | 0.51 | 0.54 | 0.72 |
| 4 | 98635524 | rs11097573 | 0.286 | 0.14 | -0.288 | -0.019 | -0.018 | -0.026 | -0.026 | 0 | 0.58 | 0.27 | 0.57 | 0.41 | 0.54 | 0.68 |
| 4 | 98652689 | rs17545594 | 1.667 | 0.025 | -0.687 | -0.115 | -0.109 | 0.05 | 0.05 | 0 | 0.34 | 0.08 | 0.35 | 0.18 | 0.44 | 0.38 |
| 4 | 98657190 | rs11729826 | 0.557 | -0.041 | -0.371 | -0.098 | -0.008 | 0.055 | 0.055 | 0 | 0.45 | 0.3 | 0.53 | 0.24 | 0.48 | 0.46 |
| 4 | 98743020 | rs7657354 | 2.21 | -0.251 | -0.757 | -0.343 | -0.095 | 0.033 | 0.033 | 0 | 0.19 | 0.04 | 0.18 | 0.09 | 0.28 | 0.28 |
| 4 | 98761900 | rs11735193 | 0.189 | -0.803 | -0.11 | -0.169 | -0.002 | -0.002 | -0.002 | 0.352 | 0.46 | 0.43 | 0.29 | 0.37 | 0.4 | 0.41 |
| 4 | 98819843 | rs722024 | 2.56 | -0.857 | 0.075 | -0.12 | -0.032 | 0.078 | 0.078 | 0 | 0.14 | 0.28 | 0.06 | 0.21 | 0.06 | 0.22 |
| 4 | 98837863 | rs968854 | -0.752 | -0.633 | 0.102 | -0.13 | -0.011 | 0.054 | 0.054 | 0 | 0.56 | 0.73 | 0.47 | 0.76 | 0.52 | 0.53 |
| 4 | 98838339 | rs921521 | 0.144 | -0.62 | -0.017 | -0.131 | 0.006 | 0.005 | 0.005 | 0 | 0.47 | 0.37 | 0.29 | 0.37 | 0.4 | 0.43 |
| 4 | 98848093 | rs1228208 | -1.558 | -0.496 | -0.039 | -0.019 | 0.037 | 0.137 | 0.137 | 0 | 0.93 | 0.82 | 0.83 | 0.62 | 0.88 | 0.9 |
| 4 | 98863009 | rs1215054 | 0.541 | -0.587 | -0.193 | -0.143 | -0.005 | 0.036 | 0.036 | 0 | 0.45 | 0.37 | 0.54 | 0.24 | 0.48 | 0.47 |
| 4 | 98885670 | rs783961 | 2.948 | -0.664 | -0.326 | -0.29 | 0.02 | 0.068 | 0.068 | 0 | 0.17 | 0.03 | 0.09 | 0 | 0.13 | 0.05 |
| 4 | 98893437 | rs783960 | 0.918 | -0.691 | 0.322 | 0.337 | -0.005 | 0.036 | 0.036 | 0 | 0.45 | 0.37 | 0.54 | 0.24 | 0.47 | 0.47 |
| 4 | 98919415 | rs13133355 | -0.081 | -0.69 | 0.373 | 0.267 | -0.005 | -0.004 | -0.004 | 0 | 0.45 | 0.45 | 0.29 | 0.37 | 0.4 | 0.4 |
| 4 | 98927532 | rs11097592 | 0.721 | -0.864 | 0.361 | -0.18 | -0.002 | 0.006 | 0.006 | 0 | 0.34 | 0.17 | 0.23 | 0.28 | 0.32 | 0.35 |
| 4 | 98965696 | rs7677523 | 2.974 | -0.94 | 0.361 | -0.199 | 0.004 | -0.005 | -0.005 | 0 | 0.12 | 0.07 | 0.04 | 0.16 | 0.06 | 0.18 |
| 4 | 98966141 | rs1967977 | -0.523 | -0.94 | -0.081 | -0.199 | -0.005 | -0.011 | -0.011 | 0.034 | 0.47 | 0.43 | 0.29 | 0.28 | 0.4 | 0.42 |
| 4 | 98967484 | rs4699323 | 1.235 | -0.907 | -0.21 | -0.196 | -0.017 | 0.032 | 0.032 | 0 | 0.22 | 0.1 | 0.19 | 0.12 | 0.26 | 0.17 |
| 4 | 99033352 | rs2865983 | 0.633 | -0.769 | -0.245 | -0.107 | -0.002 | -0.006 | -0.006 | 0.282 | 0.35 | 0.33 | 0.24 | 0.27 | 0.32 | 0.35 |

| | | | | | | | | | | | | | | | | |
|---|---|---|---|---|---|---|---|---|---|---|---|---|---|---|---|---|
| 4 | 99057457 | rs2903153 | -2.114 | -0.664 | -0.945 | 0.073 | -0.11 | -0.054 | -0.054 | 0.972 | 0.67 | 0.92 | 0.65 | 0.68 | 0.57 | 0.61 |
| 4 | 99065480 | rs6854802 | -0.724 | -0.274 | -0.755 | 0.33 | -0.005 | 0.019 | 0.019 | 0 | 0.53 | 0.71 | 0.45 | 0.63 | 0.52 | 0.5 |
| 4 | 99078105 | rs17027299 | 0.979 | -1.265 | -0.617 | -0.428 | -0.035 | -0.014 | -0.014 | 0.889 | 0.1 | 0.18 | 0.19 | 0.06 | 0.04 | 0.08 |
| 4 | 99094892 | rs13150465 | 1.205 | -1.141 | -0.865 | -0.473 | -0.018 | 0.021 | 0.021 | 0 | 0.18 | 0.07 | 0.15 | 0.12 | 0.18 | 0.15 |
| 4 | 99097492 | rs4699611 | -0.982 | -0.875 | -0.648 | -0.247 | 0.006 | 0.014 | 0.014 | 0 | 0.58 | 0.68 | 0.47 | 0.34 | 0.41 | 0.52 |
| 4 | 99101605 | rs9994641 | -0.397 | -0.006 | -0.691 | 0.018 | -0.003 | -0.007 | -0.007 | 1.435 | 0.44 | 0.5 | 0.29 | 0.28 | 0.37 | 0.41 |
| 4 | 99104879 | rs13134014 | 0.358 | -0.199 | -1.18 | -0.092 | 0.025 | -0.009 | -0.009 | 0 | 0.26 | 0.09 | 0.15 | 0.23 | 0.17 | 0.3 |
| 4 | 99105055 | rs6820691 | -0.275 | -0.83 | -0.544 | -0.313 | 0.014 | 0.012 | 0.012 | 0 | 0.4 | 0.59 | 0.35 | 0.23 | 0.22 | 0.37 |
| 4 | 99125659 | rs6820913 | -0.468 | -0.859 | 0.219 | -0.505 | -0.045 | -0.062 | -0.062 | 1.537 | 0.15 | 0.52 | 0.2 | 0.04 | 0.08 | 0.1 |
| 4 | 99147896 | rs2866006 | -0.887 | -0.858 | 0.285 | -0.574 | 0.057 | 0.07 | 0.07 | 0 | 0.5 | 0.69 | 0.41 | 0.28 | 0.33 | 0.47 |
| 4 | 99152631 | rs13150538 | 1.602 | -0.87 | 0.052 | -0.655 | -0.054 | 0.047 | 0.047 | 0 | 0.16 | 0 | 0.13 | 0.08 | 0.15 | 0.1 |
| 4 | 99172175 | rs3974886 | -0.851 | -0.851 | 0.776 | -0.303 | -0.066 | -0.086 | -0.086 | 1.924 | 0.18 | 0.47 | 0.22 | 0.04 | 0.08 | 0.1 |
| 4 | 99187874 | rs2034677 | -1.422 | -0.745 | 0.721 | -0.21 | 0.113 | 0.026 | 0.026 | 3.023 | 0.46 | 0.83 | 0.42 | 0.33 | 0.29 | 0.45 |
| 4 | 99208535 | rs10028275 | -0.111 | -0.856 | 0.853 | -0.286 | -0.039 | -0.054 | -0.054 | 0 | 0.15 | 0.34 | 0.21 | 0.04 | 0.07 | 0.08 |
| 4 | 99208603 | rs10005811 | -0.111 | -0.856 | 0.853 | -0.286 | -0.039 | -0.054 | -0.054 | 1.27 | 0.15 | 0.34 | 0.21 | 0.04 | 0.07 | 0.08 |
| 4 | 99214851 | rs603215 | 0.674 | -0.499 | 0.515 | 0.141 | -0.028 | 0.008 | 0.008 | 0 | 0.06 | 0.01 | 0.13 | 0.39 | 0.08 | 0.07 |
| 4 | 99235747 | rs17027456 | 1.343 | -0.627 | 0.567 | -0.574 | 0.038 | 0.017 | 0.017 | 0 | 0.16 | 0.05 | 0.05 | 0.08 | 0.07 | 0.24 |
| 4 | 99237439 | rs10516428 | -1.471 | -0.861 | 0.58 | -0.573 | 0.123 | 0.059 | 0.059 | 2.465 | 0.46 | 0.82 | 0.42 | 0.31 | 0.27 | 0.42 |
| 4 | 99259807 | rs6846417 | 0.023 | -0.822 | 0.77 | -0.554 | -0.04 | -0.061 | -0.061 | 0 | 0.15 | 0.41 | 0.21 | 0.04 | 0.08 | 0.07 |
| 4 | 99261097 | rs6854095 | 1.215 | -0.839 | 0.631 | -0.538 | 0.001 | -0.015 | -0.015 | 2.582 | 0.09 | 0.05 | 0.13 | 0.19 | 0.12 | 0.08 |
| 4 | 99265074 | rs6532732 | -0.059 | -0.907 | 0.67 | -0.598 | -0.047 | -0.06 | -0.06 | 0 | 0.16 | 0.33 | 0.2 | 0.04 | 0.07 | 0.09 |
| 4 | 99298166 | rs10002767 | -0.723 | -0.429 | 1.033 | -0.276 | -0.07 | -0.093 | -0.093 | 0.194 | 0.18 | 0.49 | 0.21 | 0.04 | 0.07 | 0.09 |
| 4 | 99304544 | rs3775540 | 1.107 | -0.42 | 1.006 | -0.263 | -0.004 | -0.011 | -0.011 | 0 | 0.09 | 0.05 | 0.14 | 0.2 | 0.13 | 0.08 |
| 4 | 99309404 | rs3756088 | -2.233 | -0.19 | 0.821 | -0.239 | -0.13 | 0.214 | 0.214 | 0.822 | 0.6 | 0.97 | 0.73 | 0.9 | 0.58 | 0.59 |
| 4 | 99317251 | rs13103626 | 0.49 | -0.741 | 1.753 | 0.142 | -0.008 | -0.009 | -0.009 | 0 | 0.12 | 0.07 | 0.17 | 0.19 | 0.2 | 0.12 |
| 4 | 99322274 | rs17027564 | -0.682 | -0.642 | 1.842 | 0.111 | -0.046 | -0.061 | -0.061 | 0.002 | 0.16 | 0.35 | 0.2 | 0.04 | 0.07 | 0.1 |
| 4 | 99322767 | rs2063745 | -0.682 | -0.642 | 1.836 | 0.111 | -0.046 | -0.06 | -0.06 | 0 | 0.16 | 0.35 | 0.2 | 0.04 | 0.08 | 0.1 |
| 4 | 99328306 | rs1399243 | 0.277 | -0.638 | 1.641 | 0.114 | -0.005 | -0.006 | -0.006 | 1.326 | 0.1 | 0.09 | 0.17 | 0.04 | 0.07 | 0.06 |
| 4 | 99330630 | rs7671104 | -1.786 | -0.646 | 1.622 | 0.123 | 0.073 | 0.069 | 0.069 | 1.873 | 0.47 | 0.76 | 0.44 | 0.3 | 0.35 | 0.43 |
| 4 | 99337881 | rs10516430 | -0.107 | -0.463 | 1.252 | 0.202 | -0.002 | -0.015 | -0.015 | 2.455 | 0.26 | 0.12 | 0.22 | 0.25 | 0.28 | 0.33 |
| 4 | 99364621 | rs4699634 | -0.161 | -0.235 | 1.459 | 0.177 | -0.016 | -0.017 | -0.017 | 0 | 0.11 | 0.15 | 0.17 | 0.04 | 0.07 | 0.06 |
| 4 | 99366583 | rs13119703 | -0.062 | -0.232 | 1.29 | 0.179 | -0.004 | -0.004 | -0.004 | 0 | 0.1 | 0.11 | 0.17 | 0.04 | 0.07 | 0.06 |
| 4 | 99369061 | rs12503056 | -1.537 | -0.194 | 1.111 | 0.187 | -0.046 | -0.075 | -0.075 | 0 | 0.19 | 0.52 | 0.23 | 0.06 | 0.09 | 0.1 |
| 4 | 99381148 | rs2004316 | -1.551 | -0.186 | 0.941 | 0.192 | -0.046 | -0.072 | -0.072 | 0.129 | 0.19 | 0.53 | 0.23 | 0.06 | 0.09 | 0.1 |
| 4 | 99387827 | rs2090118 | -1.044 | -0.175 | 0.476 | 0.196 | -0.044 | -0.046 | -0.046 | 1.551 | 0.16 | 0.32 | 0.21 | 0.05 | 0.08 | 0.09 |
| 4 | 99392101 | rs17027607 | -0.715 | 0.137 | 0.057 | 0.282 | 0.01 | -0.037 | -0.037 | 2.175 | 0.09 | 0.37 | 0.04 | 0 | 0.02 | 0.04 |
| 4 | 99399748 | rs4303985 | -0.708 | 0.172 | -1.212 | 0.593 | 0.053 | 0.023 | 0.023 | 1.432 | 0.46 | 0.77 | 0.44 | 0.33 | 0.36 | 0.38 |
| 4 | 99407200 | rs6854915 | 0.618 | 0.243 | -1.365 | 0.837 | 0.035 | -0.005 | -0.005 | 0.39 | 0.51 | 0.23 | 0.52 | 0.45 | 0.62 | 0.59 |
| 4 | 99411237 | rs7677026 | 0.046 | 0.65 | -0.943 | 1.081 | -0.007 | -0.011 | -0.011 | 0.759 | 0.78 | 0.82 | 0.84 | 0.59 | 0.74 | 0.87 |
| 4 | 99421741 | rs4527483 | -0.82 | 0.012 | -0.663 | 0.78 | -0.001 | -0.001 | -0.001 | 0.007 | 0.38 | 0.59 | 0.48 | 0.38 | 0.3 | 0.4 |
| 4 | 99422081 | rs4699638 | -0.669 | -0.099 | -0.427 | 0.806 | -0.029 | 0.008 | 0.008 | 0 | 0.35 | 0.43 | 0.46 | 0.37 | 0.26 | 0.35 |
| 4 | 99423280 | rs10009693 | -0.485 | -0.072 | -0.572 | 0.746 | -0.006 | 0.001 | 0.001 | 0 | 0.12 | 0.15 | 0.24 | 0.13 | 0.08 | 0.17 |
| 4 | 99425353 | rs10023791 | -0.375 | -0.321 | -1.574 | 0.521 | 0.002 | -0.045 | -0.045 | 13.108 | 0.54 | 0.86 | 0.58 | 0.53 | 0.45 | 0.62 |
| 4 | 99428163 | rs955931 | -0.33 | -1.346 | -0.872 | 0.279 | -0.036 | 0.063 | 0.063 | 0.116 | 0.28 | 0.53 | 0.35 | 0.43 | 0.19 | 0.34 |
| 4 | 99441349 | rs17499015 | 0.291 | -1.434 | -0.784 | 0.005 | 0.004 | 0.007 | 0.007 | 0 | 0.21 | 0.19 | 0.13 | 0.09 | 0.19 | 0.26 |
| 4 | 99443585 | rs7695093 | 0.081 | -1.292 | -0.716 | -0.342 | -0.006 | 0.001 | 0.001 | 0 | 0.47 | 0.34 | 0.37 | 0.44 | 0.45 | 0.54 |
| 4 | 99447433 | rs10004753 | -0.386 | -1.537 | -0.731 | -0.231 | -0.027 | 0.014 | 0.014 | 0 | 0.57 | 0.75 | 0.58 | 0.56 | 0.49 | 0.61 |
| 4 | 99458977 | rs6835899 | 1.502 | -1.831 | -0.187 | 0.272 | 0 | 0.068 | 0.068 | 0 | 0.08 | 0.2 | 0.05 | 0.17 | 0.06 | 0.03 |
| 4 | 99459528 | rs2037832 | -1.652 | -1.601 | -0.083 | 0.38 | -0.086 | -0.093 | -0.093 | 0 | 0.8 | 0.97 | 0.72 | 0.65 | 0.71 | 0.9 |
| 4 | 99463989 | rs1919211 | -0.581 | -1.788 | 0.072 | 0.299 | -0.021 | -0.053 | -0.053 | 0 | 0.7 | 0.83 | 0.52 | 0.59 | 0.67 | 0.85 |
| 4 | 99468567 | rs6532748 | -0.641 | -1.801 | 0.132 | 0.282 | -0.013 | -0.033 | -0.033 | 0.065 | 0.7 | 0.79 | 0.52 | 0.59 | 0.67 | 0.85 |
| 4 | 99474774 | rs1554849 | 0.584 | -1.578 | 0.387 | 0.317 | -0.016 | -0.042 | -0.042 | 0 | 0.17 | 0.26 | 0.1 | 0.09 | 0.15 | 0.27 |
| 4 | 99478756 | rs13141997 | 0.057 | -1.389 | 0.276 | 0.676 | 0.003 | 0.002 | 0.002 | 0 | 0.67 | 0.42 | 0.44 | 0.35 | 0.65 | 0.79 |
| 4 | 99483238 | rs10021296 | -0.655 | -1.281 | 0.093 | 0.65 | 0 | 0.001 | 0.001 | 0 | 0.75 | 0.75 | 0.52 | 0.62 | 0.67 | 0.87 |
| 4 | 99488312 | rs1534555 | -0.759 | -1.064 | -0.315 | 0.689 | -0.023 | -0.023 | -0.023 | 0.008 | 0.76 | 0.82 | 0.7 | 0.65 | 0.69 | 0.89 |
| 4 | 99490578 | rs17027758 | -0.337 | -1.377 | -0.15 | 0.841 | -0.046 | 0.023 | 0.023 | 0 | 0.1 | 0.22 | 0.03 | 0.09 | 0.04 | 0.05 |
| 4 | 99491187 | rs7691465 | 0.35 | -1.325 | -0.354 | 0.769 | 0.004 | -0.009 | -0.009 | 0 | 0.62 | 0.65 | 0.65 | 0.57 | 0.64 | 0.8 |
| 4 | 99493776 | rs11944636 | -1.053 | -1.464 | -0.428 | 0.909 | -0.009 | -0.001 | -0.001 | 0 | 0.22 | 0.18 | 0.11 | 0.2 | 0.13 | 0.35 |
| 4 | 99493955 | rs1547382 | 0.305 | -1.45 | -0.433 | 0.733 | 0.006 | 0 | 0 | 0 | 0.12 | 0.09 | 0.08 | 0.11 | 0.09 | 0.29 |
| 4 | 99495460 | rs1406311 | -1.095 | -1.462 | -0.448 | 0.84 | -0.007 | 0.002 | 0.002 | 0.028 | 0.22 | 0.18 | 0.19 | 0.39 | 0.13 | 0.38 |
| 4 | 99498899 | rs10026446 | -0.503 | -0.962 | -0.384 | 0.438 | -0.023 | -0.01 | -0.01 | 0 | 0.14 | 0.31 | 0.11 | 0.11 | 0.12 | 0.25 |
| 4 | 99504635 | rs1358548 | 0.342 | -0.842 | -0.79 | 0.379 | 0 | 0 | 0 | 0 | 0.6 | 0.61 | 0.64 | 0.55 | 0.59 | 0.63 |
| 4 | 99509133 | rs4699353 | 0.599 | -0.481 | -0.545 | 0.309 | 0.001 | -0.002 | -0.002 | 0.294 | 0.58 | 0.56 | 0.62 | 0.57 | 0.55 | 0.73 |
| 4 | 99509229 | rs17027806 | 0.418 | -0.443 | -0.873 | 0.33 | 0.009 | 0.017 | 0.017 | 0 | 0.09 | 0.02 | 0.08 | 0.07 | 0.09 | 0.11 |
| 4 | 99510279 | rs1893714 | 0.257 | -0.307 | -0.699 | 0.423 | -0.003 | 0.004 | 0.004 | 0 | 0.58 | 0.65 | 0.62 | 0.58 | 0.55 | 0.72 |
| 4 | 99527572 | rs17027845 | -0.674 | -0.254 | -0.7 | 0.417 | -0.025 | -0.015 | -0.015 | 0.947 | 0.07 | 0.16 | 0.03 | 0.06 | 0.03 | 0.06 |
| 4 | 99528449 | rs846015 | -0.095 | 0.387 | -0.838 | 0.457 | 0.012 | -0.007 | -0.007 | 0 | 0.66 | 0.64 | 0.61 | 0.74 | 0.59 | 0.75 |
| 4 | 99528486 | rs2000814 | -0.412 | 0.191 | -1.192 | 0.599 | -0.011 | -0.041 | -0.041 | 0.336 | 0.11 | 0.03 | 0.15 | 0.22 | 0.14 | 0.17 |
| 4 | 99534928 | rs4699661 | -0.74 | -0.054 | -1.501 | -0.141 | 0.012 | 0.051 | 0.051 | 0 | 0.33 | 0.11 | 0.39 | 0.17 | 0.31 | 0.47 |

| | | | | | | | | | | | | | | | | |
|---|---|---|---|---|---|---|---|---|---|---|---|---|---|---|---|---|
| 4 | 99539290 | rs7659885 | -0.645 | -0.362 | -0.47 | -0.236 | -0.042 | -0.075 | -0.075 | 0.016 | 0.13 | 0.28 | 0.07 | 0 | 0.08 | 0.23 |
| 4 | 99539498 | rs2866015 | -1.045 | -0.348 | -0.444 | -0.835 | -0.021 | 0.064 | 0.064 | 0 | 0.6 | 0.76 | 0.52 | 0.18 | 0.46 | 0.53 |
| 4 | 99542125 | rs9684705 | -1.116 | -0.475 | -0.425 | -0.883 | -0.06 | 0.043 | 0.043 | 0 | 0.63 | 0.82 | 0.56 | 0.21 | 0.5 | 0.6 |
| 4 | 99546654 | rs12504979 | -0.026 | -0.703 | -0.074 | -0.99 | 0 | -0.02 | -0.02 | 2.195 | 0.44 | 0.43 | 0.48 | 0.18 | 0.41 | 0.33 |
| 4 | 99546928 | rs12509170 | -0.066 | -0.694 | -0.182 | -0.986 | 0 | -0.019 | -0.019 | 0 | 0.44 | 0.46 | 0.48 | 0.18 | 0.41 | 0.33 |
| 4 | 99548182 | rs3941 | -0.998 | -0.218 | -0.559 | -0.673 | -0.059 | 0.021 | 0.021 | 0 | 0.63 | 0.82 | 0.56 | 0.23 | 0.5 | 0.61 |
| 4 | 99551907 | rs10026666 | -0.812 | -0.385 | -0.211 | -0.684 | -0.008 | 0.011 | 0.011 | 0 | 0.61 | 0.69 | 0.55 | 0.22 | 0.49 | 0.63 |
| 4 | 99555740 | rs12509052 | -0.71 | -0.156 | -0.164 | -0.995 | 0.014 | 0.025 | 0.025 | 0 | 0.13 | 0.05 | 0.19 | 0.06 | 0.13 | 0.35 |
| 4 | 99556496 | rs10489126 | -1.081 | 0.486 | -0.256 | -1.108 | -0.014 | 0.012 | 0.012 | 0 | 0.07 | 0.05 | 0.13 | 0.03 | 0.11 | 0.09 |
| 4 | 99561045 | rs1528545 | -1.454 | -0.129 | 0.269 | -1.125 | 0.005 | 0.053 | 0.053 | 0.984 | 0.5 | 0.61 | 0.54 | 0.31 | 0.51 | 0.57 |
| 4 | 99561995 | rs871473 | -1.751 | -0.347 | 0.557 | -1.207 | -0.003 | 0.009 | 0.009 | 0.304 | 0.22 | 0.28 | 0.35 | 0.31 | 0.24 | 0.47 |
| 4 | 99563676 | rs883509 | -0.1 | -0.301 | 0.714 | -1.255 | -0.013 | 0.04 | 0.04 | 0 | 0.33 | 0.21 | 0.31 | 0.03 | 0.36 | 0.2 |
| 4 | 99565852 | rs10856938 | -0.709 | -0.377 | 0.487 | -1.053 | 0.008 | 0.015 | 0.015 | 0 | 0.14 | 0.11 | 0.19 | 0.06 | 0.14 | 0.37 |
| 4 | 99571085 | rs6829297 | -0.593 | -0.027 | 0.743 | -0.877 | 0.02 | -0.028 | -0.028 | 0.119 | 0.07 | 0.17 | 0.14 | 0.03 | 0.09 | 0.1 |
| 4 | 99573350 | rs12507328 | -1.278 | -0.103 | 0.295 | -0.999 | 0 | 0.11 | 0.11 | 0.085 | 0.46 | 0.81 | 0.54 | 0.28 | 0.44 | 0.45 |
| 4 | 99573992 | rs17027889 | -0.359 | -0.196 | 0.696 | -0.714 | -0.002 | -0.008 | -0.008 | 0 | 0.09 | 0.11 | 0.18 | 0.04 | 0.08 | 0.25 |
| 4 | 99583513 | rs12512460 | -0.692 | -0.302 | 0.904 | -0.765 | -0.009 | -0.02 | -0.02 | 0 | 0.1 | 0.29 | 0.19 | 0.04 | 0.07 | 0.25 |
| 4 | 99583639 | rs7675896 | -0.994 | -0.348 | 0.855 | -0.684 | -0.024 | 0.052 | 0.052 | 0 | 0.12 | 0.27 | 0.21 | 0.22 | 0.09 | 0.29 |
| 4 | 99595604 | rs6819703 | 0.054 | -0.635 | 0.543 | -1.405 | -0.014 | -0.088 | -0.088 | 0.014 | 0.36 | 0.39 | 0.19 | 0.03 | 0.28 | 0.21 |
| 4 | 99598549 | rs4699665 | -0.297 | -0.68 | 0.258 | -1.412 | -0.021 | -0.017 | -0.017 | 0 | 0.11 | 0.25 | 0.18 | 0.07 | 0.06 | 0.25 |
| 4 | 99599578 | rs1528009 | -0.869 | -0.425 | -0.345 | -1.052 | 0.101 | 0.169 | 0.169 | 0 | 0.5 | 0.8 | 0.4 | 0.23 | 0.35 | 0.47 |
| 4 | 99608363 | rs7680513 | -1.048 | 0.082 | -1.253 | -0.878 | -0.069 | -0.111 | -0.111 | 0 | 0.73 | 0.99 | 0.75 | 0.32 | 0.7 | 0.83 |
| 4 | 99613485 | rs4621437 | 0.27 | -0.3 | -0.35 | -1.393 | -0.003 | -0.067 | -0.067 | 0.176 | 0.37 | 0.37 | 0.2 | 0.01 | 0.28 | 0.35 |
| 4 | 99617267 | rs1003174 | -0.106 | -0.016 | 0.355 | -0.998 | 0.044 | -0.042 | -0.042 | 0 | 0.33 | 0.52 | 0.52 | 0.19 | 0.4 | 0.47 |
| 4 | 99619286 | rs12507086 | -0.457 | 0.049 | 0.717 | -1.034 | 0.057 | 0.059 | 0.059 | 0 | 0.06 | 0.34 | 0.17 | 0.06 | 0.06 | 0.1 |
| 4 | 99620154 | rs2178125 | 0.562 | 0.087 | 0.72 | -1.026 | -0.032 | 0.029 | 0.029 | 0.005 | 0.27 | 0.17 | 0.35 | 0.12 | 0.34 | 0.37 |
| 4 | 99628067 | rs1528007 | 0.102 | 0.423 | 1.232 | -0.967 | 0.061 | 0.001 | 0.001 | 0 | 0.07 | 0.23 | 0.14 | 0.03 | 0.09 | 0.1 |
| 4 | 99632473 | rs4146004 | -1.702 | 0.433 | 1.383 | -0.949 | 0.01 | 0.033 | 0.033 | 0.546 | 0.15 | 0.66 | 0.36 | 0.22 | 0.18 | 0.2 |
| 4 | 99636215 | rs17027976 | -0.246 | 0.296 | 1.3 | -1.062 | -0.005 | -0.013 | -0.013 | 0 | 0.13 | 0.19 | 0.19 | 0.06 | 0.11 | 0.31 |
| 4 | 99636952 | rs7668409 | -0.493 | 0.286 | 1.397 | -1.066 | 0.07 | 0.134 | 0.134 | 0.093 | 0.09 | 0.38 | 0.17 | 0.15 | 0.09 | 0.1 |
| 4 | 99640223 | rs10016647 | -1.65 | 0.23 | 1.31 | -1.115 | -0.003 | -0.001 | -0.001 | 0 | 0.22 | 0.69 | 0.37 | 0.22 | 0.2 | 0.41 |
| 4 | 99643472 | rs17566763 | 0.876 | 0.149 | 1 | -1.165 | -0.085 | 0.048 | 0.048 | 0.202 | 0.23 | 0.04 | 0.21 | 0.11 | 0.26 | 0.25 |
| 4 | 99651480 | rs10213420 | -0.208 | 0.156 | 1.319 | -1.086 | 0.085 | 0.009 | 0.009 | 0.009 | 0.09 | 0.36 | 0.15 | 0.02 | 0.09 | 0.1 |
| 4 | 99657397 | rs13134776 | -0.787 | 0.136 | 0.966 | -0.991 | 0.002 | -0.007 | -0.007 | 0.022 | 0.39 | 0.62 | 0.47 | 0.3 | 0.38 | 0.56 |
| 4 | 99661738 | rs4699667 | -0.395 | 0.262 | 0.428 | -0.925 | -0.008 | 0.003 | 0.003 | 0 | 0.35 | 0.34 | 0.43 | 0.3 | 0.36 | 0.56 |
| 4 | 99665433 | rs4129895 | 0.513 | 0.504 | -0.098 | -0.963 | -0.083 | 0.053 | 0.053 | 0.164 | 0.24 | 0.04 | 0.21 | 0.11 | 0.26 | 0.25 |
| 4 | 99668311 | rs4235430 | -1.067 | 0.671 | -0.036 | -1.135 | -0.002 | -0.003 | -0.003 | 0 | 0.73 | 0.72 | 0.72 | 0.85 | 0.65 | 0.81 |
| 4 | 99672392 | rs4699670 | 0.038 | 0.293 | 0.14 | -0.918 | 0.021 | 0.02 | 0.02 | 0.004 | 0.39 | 0.27 | 0.45 | 0.23 | 0.34 | 0.52 |
| 4 | 99674593 | rs4493540 | -0.878 | 0.601 | 0.368 | -0.877 | 0.019 | -0.057 | -0.057 | 0 | 0.71 | 0.58 | 0.72 | 0.84 | 0.65 | 0.81 |
| 4 | 99676081 | rs971169 | -0.215 | 0.291 | 0.409 | -0.9 | -0.005 | -0.034 | -0.034 | 1.04 | 0.59 | 0.46 | 0.48 | 0.77 | 0.43 | 0.66 |
| 4 | 99687168 | rs11097632 | -0.989 | 0.325 | 0.392 | -0.926 | -0.024 | 0.031 | 0.031 | 0 | 0.66 | 0.72 | 0.65 | 0.85 | 0.51 | 0.78 |
| 4 | 99698254 | rs10010115 | 0.727 | 0.208 | -0.166 | -0.895 | 0.019 | 0.032 | 0.032 | 0 | 0.05 | 0.14 | 0.16 | 0.07 | 0.08 | 0.12 |
| 4 | 99699141 | rs4616760 | -0.073 | 0.225 | -0.092 | -0.934 | 0.003 | 0.006 | 0.006 | 0 | 0.57 | 0.36 | 0.46 | 0.79 | 0.4 | 0.68 |
| 4 | 99700351 | rs17028075 | 0.597 | 0.512 | -0.353 | -0.72 | 0.019 | 0.03 | 0.03 | 0 | 0.05 | 0.14 | 0.16 | 0.07 | 0.08 | 0.12 |
| 4 | 99701724 | rs17028081 | 0.389 | 0.513 | -0.434 | -0.719 | 0.027 | 0.04 | 0.04 | 0 | 0.06 | 0.21 | 0.17 | 0.07 | 0.08 | 0.12 |
| 4 | 99702148 | rs7438406 | 0.758 | 0.287 | -0.795 | -0.769 | 0.018 | -0.033 | -0.033 | 0 | 0.24 | 0.14 | 0.16 | 0.63 | 0.15 | 0.24 |
| 4 | 99703142 | rs6532765 | -1.152 | 0.545 | -0.172 | -0.844 | 0.038 | 0.144 | 0.144 | 0.407 | 0.48 | 0.77 | 0.52 | 0.14 | 0.42 | 0.57 |
| 4 | 99707614 | rs894810 | -0.537 | 0.725 | 0.454 | -0.384 | -0.02 | -0.12 | -0.12 | 1.38 | 0.19 | 0.42 | 0.12 | 0.01 | 0.14 | 0.09 |
| 4 | 99708057 | rs2083865 | -0.852 | 0.735 | -0.66 | -0.306 | -0.001 | -0.007 | -0.007 | 0 | 0.25 | 0.7 | 0.29 | 0.08 | 0.22 | 0.21 |
| 4 | 99714727 | rs2903158 | -0.025 | 1.355 | -0.794 | 0.442 | -0.011 | -0.011 | -0.011 | 0.082 | 0.89 | 0.91 | 0.86 | 0.72 | 0.82 | 0.91 |
| 4 | 99719961 | rs1867343 | -0.19 | 0.209 | 0.285 | 0.858 | -0.054 | 0.372 | 0.372 | 0 | 0.13 | 0.44 | 0.1 | 0.46 | 0.04 | 0.08 |
| 4 | 99723669 | rs7696219 | -0.975 | -0.012 | 0.055 | 0.062 | -0.018 | 0.046 | 0.046 | 0.033 | 0.34 | 0.47 | 0.26 | 0.46 | 0.29 | 0.15 |
| 4 | 99728276 | rs12649579 | 1.137 | 0.004 | 0.23 | 0.111 | 0.01 | 0.028 | 0.028 | 0 | 0.1 | 0.06 | 0.07 | 0.46 | 0.04 | 0.09 |
| 4 | 99730246 | rs10011147 | 0.741 | -0.352 | -0.838 | -0.766 | -0.05 | 0.171 | 0.171 | 0.232 | 0.39 | 0.03 | 0.42 | 0.2 | 0.41 | 0.33 |
| 4 | 99730691 | rs4699675 | -0.391 | -0.336 | -0.125 | -0.297 | 0.058 | -0.081 | -0.081 | 0.217 | 0.12 | 0.39 | 0.11 | 0 | 0.13 | 0.03 |
| 4 | 99740952 | rs13114194 | 0.568 | -0.347 | -0.095 | -0.289 | -0.019 | 0.169 | 0.169 | 0 | 0.77 | 0.53 | 0.82 | 0.54 | 0.83 | 0.88 |
| 4 | 99743545 | rs12331010 | -1.716 | -0.386 | -0.248 | -0.257 | 0.007 | 0.038 | 0.038 | 0 | 0.9 | 0.93 | 0.93 | 0.56 | 0.96 | 0.92 |
| 4 | 99745705 | rs6822279 | 1.6 | -0.58 | -0.726 | -0.508 | 0.008 | 0.183 | 0.183 | 0.713 | 0.5 | 0.06 | 0.42 | 0.24 | 0.56 | 0.58 |
| 4 | 99748988 | rs12649469 | -0.011 | -0.506 | -0.561 | -0.471 | 0.01 | -0.022 | -0.022 | 0 | 0.25 | 0.11 | 0.38 | 0.32 | 0.27 | 0.3 |
| 4 | 99772547 | rs10446681 | 0.991 | -0.388 | 0.62 | -0.323 | -0.031 | 0.049 | 0.049 | 0 | 0.73 | 0.52 | 0.65 | 0.35 | 0.79 | 0.83 |
| 4 | 99774183 | rs4401453 | 0.686 | -0.328 | 0.524 | -0.292 | 0 | -0.001 | -0.001 | 0 | 0.75 | 0.77 | 0.66 | 0.35 | 0.78 | 0.83 |
| 4 | 99774466 | rs6851217 | 1.25 | -0.313 | 0.481 | -0.284 | 0.01 | -0.011 | -0.011 | 0 | 0.71 | 0.29 | 0.62 | 0.35 | 0.78 | 0.83 |
| 4 | 99781443 | rs11097638 | -0.724 | 0.15 | 0.594 | -0.161 | 0.004 | -0.005 | -0.005 | 0.567 | 0.15 | 0.2 | 0.21 | 0.21 | 0.16 | 0.09 |
| 4 | 99783669 | rs6532770 | -1.071 | 0.045 | 0.266 | -0.353 | -0.004 | -0.01 | -0.01 | 0 | 0.28 | 0.69 | 0.3 | 0.21 | 0.27 | 0.21 |
| 4 | 99783733 | rs2165944 | 0.669 | 0.154 | 0.064 | -0.414 | -0.026 | 0.092 | 0.092 | 0 | 0.15 | 0.01 | 0.1 | 0 | 0.19 | 0.12 |
| 4 | 99788605 | rs7669736 | -1.5 | -0.124 | 0.099 | -0.285 | 0.009 | 0.008 | 0.008 | 0 | 0.4 | 0.7 | 0.43 | 0.66 | 0.39 | 0.35 |
| 4 | 99789979 | rs10032966 | -1.335 | -0.052 | 0.168 | -0.495 | 0.004 | 0.031 | 0.031 | 0 | 0.41 | 0.64 | 0.4 | 0.24 | 0.42 | 0.28 |
| 4 | 99800310 | rs10433891 | -0.51 | -0.093 | 0.423 | -0.477 | -0.011 | 0.032 | 0.032 | 0.017 | 0.16 | 0.33 | 0.23 | 0.24 | 0.15 | 0.13 |
| 4 | 99808872 | rs994636 | -0.213 | -0.205 | 0.448 | -0.457 | -0.022 | 0.023 | 0.023 | 0 | 0.15 | 0.22 | 0.19 | 0.24 | 0.11 | 0.09 |

| | | | | | | | | | | | | | | | | |
|---|---|---|---|---|---|---|---|---|---|---|---|---|---|---|---|---|
| 4 | 99822905 | rs17570252 | 0.355 | -0.149 | -0.022 | -0.567 | -0.071 | 0.14 | 0.14 | 0 | 0.21 | 0.02 | 0.15 | 0 | 0.27 | 0.16 |
| 4 | 99850126 | rs11723037 | -1.901 | -0.211 | -0.415 | -0.464 | -0.004 | 0.08 | 0.08 | 0 | 0.46 | 0.9 | 0.48 | 0.68 | 0.46 | 0.42 |
| 4 | 99852395 | rs13107892 | 0.25 | -0.107 | -0.344 | -0.509 | -0.042 | 0.092 | 0.092 | 0 | 0.22 | 0.06 | 0.16 | 0 | 0.29 | 0.16 |
| 4 | 99863585 | rs9999702 | -0.8 | -0.174 | -0.049 | -0.412 | -0.035 | 0.017 | 0.017 | 0.252 | 0.19 | 0.58 | 0.23 | 0.24 | 0.14 | 0.13 |
| 4 | 99867860 | rs4699693 | -1.61 | -0.15 | -0.451 | -0.282 | -0.001 | 0.08 | 0.08 | 0 | 0.47 | 0.89 | 0.48 | 0.68 | 0.46 | 0.42 |
| 4 | 99877445 | rs6810489 | -1.154 | -0.028 | -0.384 | -0.51 | 0.008 | 0.082 | 0.082 | 0 | 0.42 | 0.74 | 0.38 | 0.1 | 0.43 | 0.29 |
| 4 | 99896306 | rs1230194 | 0.516 | 0.575 | -0.111 | -0.142 | -0.005 | 0.066 | 0.066 | 0 | 0.08 | 0.12 | 0.23 | 0.25 | 0.06 | 0.04 |
| 4 | 99909462 | rs1230173 | 0.573 | 0.608 | -0.089 | -0.175 | -0.001 | 0.01 | 0.01 | 0 | 0.08 | 0.06 | 0.23 | 0.28 | 0.07 | 0.04 |
| 4 | 99918661 | rs10035037 | 0.485 | 0.554 | -0.305 | -0.811 | -0.032 | 0.075 | 0.075 | 0 | 0.66 | 0.78 | 0.46 | 0.18 | 0.6 | 0.67 |
| 4 | 99938721 | rs1230182 | 0.086 | 0.547 | -0.309 | -0.795 | -0.002 | -0.015 | -0.015 | 0.082 | 0.11 | 0.11 | 0.33 | 0.81 | 0.12 | 0.16 |
| 4 | 99944097 | rs6841819 | 1.645 | 0.367 | -1.218 | -1.618 | -0.121 | 0.165 | 0.165 | 0 | 0.29 | 0.02 | 0.22 | 0.05 | 0.37 | 0.27 |
| 4 | 99952607 | rs1037476 | -1.049 | 0.341 | -1.2 | -1.621 | -0.042 | 0.099 | 0.099 | 0 | 0.77 | 0.92 | 0.79 | 0.99 | 0.72 | 0.83 |
| 4 | 99952710 | rs1037475 | 0.77 | 0.258 | -0.645 | -1.69 | -0.013 | 0.085 | 0.085 | 0.048 | 0.6 | 0.28 | 0.41 | 0.18 | 0.57 | 0.67 |
| 4 | 99959932 | rs17216080 | 1.766 | 0.473 | -1.369 | -1.681 | -0.086 | 0.149 | 0.149 | 0 | 0.27 | 0.05 | 0.21 | 0.05 | 0.34 | 0.24 |
| 4 | 99962758 | rs1230185 | -0.861 | 0.537 | -0.974 | -1.197 | -0.043 | 0.096 | 0.096 | 0 | 0.77 | 0.92 | 0.79 | 0.99 | 0.72 | 0.83 |
| 4 | 99963407 | rs1230186 | 0.408 | 0.643 | -1.048 | -1.177 | -0.001 | -0.002 | -0.002 | 0 | 0.11 | 0.15 | 0.33 | 0.81 | 0.13 | 0.17 |
| 4 | 99967691 | rs1230210 | -0.633 | 0.581 | -1.156 | -1.271 | -0.04 | 0.094 | 0.094 | 0 | 0.77 | 0.94 | 0.8 | 0.99 | 0.72 | 0.82 |
| 4 | 99986373 | rs1230165 | -1.009 | 0.472 | -1.233 | -1.274 | -0.004 | 0.108 | 0.108 | 0 | 0.81 | 0.96 | 0.85 | 1 | 0.81 | 0.9 |
| 4 | 99988659 | rs1230154 | 0.026 | 0.449 | -0.847 | -1.573 | -0.011 | 0.091 | 0.091 | 0 | 0.72 | 0.85 | 0.78 | 1 | 0.71 | 0.88 |
| 4 | 99988713 | rs17595186 | 1.898 | 0.63 | -1.525 | -2.154 | -0.115 | 0.117 | 0.117 | 0 | 0.19 | 0.01 | 0.15 | 0.03 | 0.28 | 0.12 |
| 4 | 99989259 | rs1230155 | -0.077 | 0.538 | -0.979 | -1.672 | -0.001 | -0.046 | -0.046 | 0.17 | 0.7 | 0.69 | 0.76 | 1 | 0.69 | 0.87 |
| 4 | 99989808 | rs1230157 | -1.482 | 0.523 | -0.988 | -1.678 | 0.001 | 0.101 | 0.101 | 0 | 0.81 | 0.94 | 0.85 | 1 | 0.81 | 0.91 |
| 4 | 99991642 | rs7662987 | 0.808 | 0.534 | -0.993 | -1.671 | -0.005 | -0.005 | -0.005 | 0.852 | 0.09 | 0.07 | 0.07 | 0 | 0.1 | 0.03 |
| 4 | 99992602 | rs7669660 | -0.286 | 0.505 | -0.999 | -1.687 | 0.028 | -0.075 | -0.075 | 1.516 | 0.13 | 0.29 | 0.09 | 0 | 0.12 | 0.04 |
| 4 | 99992793 | rs11547772 | 0.583 | 0.505 | -0.999 | -1.687 | -0.001 | -0.017 | -0.017 | 0.775 | 0.1 | 0.12 | 0.07 | 0 | 0.1 | 0.03 |
| 4 | 99993151 | rs1803037 | 0.872 | 0.515 | -0.999 | -1.687 | -0.009 | 0.003 | 0.003 | 0 | 0.09 | 0.06 | 0.07 | 0 | 0.1 | 0.03 |
| 4 | 99993869 | rs17595424 | 0.807 | 0.512 | -1.063 | -1.684 | -0.005 | -0.005 | -0.005 | 0.492 | 0.09 | 0.07 | 0.07 | 0 | 0.1 | 0.03 |
| 4 | 99995138 | rs7683802 | 0.7 | 0.512 | -1.063 | -1.684 | -0.003 | -0.01 | -0.01 | 0 | 0.09 | 0.08 | 0.07 | 0 | 0.1 | 0.03 |
| 4 | 99999033 | rs1154415 | 1.097 | 0.077 | -1.29 | -1.867 | -0.103 | 0.244 | 0.244 | 0 | 0.43 | 0.09 | 0.32 | 0.13 | 0.45 | 0.54 |
| 4 | 99999130 | rs13119035 | 0.939 | 0.132 | -1.223 | -1.832 | -0.005 | -0.005 | -0.005 | 0 | 0.09 | 0.07 | 0.07 | 0 | 0.1 | 0.03 |
| 4 | 99999160 | rs13146416 | 0.865 | 0.132 | -1.223 | -1.832 | -0.003 | -0.01 | -0.01 | 0.031 | 0.09 | 0.08 | 0.07 | 0 | 0.1 | 0.03 |
| 4 | 99999848 | rs13125919 | 1.004 | 0.131 | -1.226 | -1.833 | -0.009 | 0.004 | 0.004 | 0 | 0.09 | 0.06 | 0.07 | 0 | 0.1 | 0.03 |
| 4 | 100001932 | rs6852848 | 0.902 | 0.173 | -1.16 | -1.77 | -0.005 | -0.005 | -0.005 | 0 | 0.09 | 0.07 | 0.07 | 0 | 0.1 | 0.03 |
| 4 | 100004226 | rs7683704 | 0.03 | 0.162 | -1.134 | -1.775 | 0.01 | -0.07 | -0.07 | 0 | 0.12 | 0.28 | 0.08 | 0 | 0.1 | 0.03 |
| 4 | 100005538 | rs3018048 | -0.96 | 0.207 | -1.056 | -1.685 | -0.024 | 0.11 | 0.11 | 0 | 0.74 | 0.91 | 0.83 | 1 | 0.72 | 0.9 |
| 4 | 100006645 | rs1154410 | -0.98 | 0.256 | -1.047 | -1.688 | -0.028 | 0.108 | 0.108 | 0 | 0.74 | 0.92 | 0.83 | 1 | 0.72 | 0.9 |
| 4 | 100008750 | rs1154405 | -0.968 | 0.213 | -0.979 | -1.682 | -0.024 | 0.109 | 0.109 | 0 | 0.74 | 0.91 | 0.83 | 1 | 0.72 | 0.9 |
| 4 | 100010010 | rs1154400 | -0.4 | 0.237 | -0.817 | -1.692 | 0 | -0.067 | -0.067 | 0 | 0.7 | 0.69 | 0.81 | 1 | 0.7 | 0.89 |
| 4 | 100010127 | rs2851301 | 0.931 | 0.2 | -0.936 | -1.712 | -0.107 | 0.242 | 0.242 | 0 | 0.43 | 0.09 | 0.31 | 0.13 | 0.45 | 0.53 |
| 4 | 100012261 | rs1311615 | -1.036 | 0.133 | -0.936 | -1.707 | -0.031 | 0.122 | 0.122 | 0 | 0.74 | 0.93 | 0.83 | 1 | 0.72 | 0.9 |
| 4 | 100012303 | rs1311616 | -1.022 | 0.133 | -0.936 | -1.707 | -0.025 | 0.108 | 0.108 | 0 | 0.74 | 0.91 | 0.83 | 1 | 0.72 | 0.9 |
| 4 | 100012357 | rs1312200 | 1.998 | 0.032 | -1.03 | -1.408 | 0.001 | -0.093 | -0.093 | 0 | 0.14 | 0.01 | 0.09 | 0.73 | 0.13 | 0.31 |
| 4 | 100012425 | rs1311617 | -1.026 | 0.108 | -0.932 | -1.362 | -0.025 | 0.108 | 0.108 | 0 | 0.74 | 0.91 | 0.83 | 1 | 0.73 | 0.9 |
| 4 | 100014805 | rs2602836 | 0.649 | 0.084 | -0.542 | -1.363 | -0.002 | 0.06 | 0.06 | 0 | 0.59 | 0.31 | 0.41 | 0.86 | 0.58 | 0.84 |
| 4 | 100022910 | rs2851277 | -0.948 | 0.117 | -0.535 | -1.345 | -0.025 | 0.108 | 0.108 | 0 | 0.74 | 0.91 | 0.83 | 1 | 0.72 | 0.9 |
| 4 | 100034841 | rs2602865 | -0.948 | 0.141 | -0.647 | -1.333 | -0.025 | 0.108 | 0.108 | 0 | 0.74 | 0.91 | 0.83 | 1 | 0.72 | 0.9 |
| 4 | 100039953 | rs2602878 | -0.961 | 0.142 | -0.66 | -1.332 | -0.031 | 0.122 | 0.122 | 0 | 0.74 | 0.93 | 0.83 | 1 | 0.72 | 0.9 |
| 4 | 100042221 | rs6532795 | -0.795 | 0.026 | -0.675 | -1.331 | -0.012 | 0.091 | 0.091 | 0 | 0.73 | 0.86 | 0.77 | 1 | 0.7 | 0.88 |
| 4 | 100042242 | rs6532796 | -0.795 | 0.026 | -0.675 | -1.331 | -0.012 | 0.091 | 0.091 | 0 | 0.73 | 0.86 | 0.77 | 1 | 0.7 | 0.88 |
| 4 | 100042262 | rs3828541 | 0.806 | -0.1 | -0.6 | -1.453 | -0.018 | 0.101 | 0.101 | 0 | 0.44 | 0.29 | 0.32 | 0.12 | 0.45 | 0.52 |
| 4 | 100043284 | rs2602891 | -1.119 | -0.105 | -0.893 | -1.452 | -0.036 | 0.14 | 0.14 | 0 | 0.74 | 0.93 | 0.83 | 1 | 0.72 | 0.9 |
| 4 | 100043505 | rs2851253 | -1.119 | -0.105 | -0.893 | -1.452 | -0.036 | 0.14 | 0.14 | 0 | 0.74 | 0.93 | 0.83 | 1 | 0.72 | 0.9 |
| 4 | 100044640 | rs2924583 | -1.093 | -0.11 | -0.896 | -1.455 | -0.035 | 0.142 | 0.142 | 0 | 0.74 | 0.93 | 0.83 | 1 | 0.72 | 0.9 |
| 4 | 100045574 | rs1042364 | -1.118 | -0.104 | -0.963 | -1.452 | -0.036 | 0.14 | 0.14 | 0 | 0.74 | 0.93 | 0.83 | 1 | 0.72 | 0.9 |
| 4 | 100045616 | rs1126673 | -0.888 | -0.101 | -0.96 | -1.452 | -0.011 | 0.092 | 0.092 | 0 | 0.73 | 0.86 | 0.77 | 1 | 0.7 | 0.88 |
| 4 | 100045840 | rs2602895 | -1.118 | -0.122 | -0.96 | -1.452 | -0.036 | 0.14 | 0.14 | 0 | 0.74 | 0.93 | 0.83 | 1 | 0.72 | 0.9 |
| 4 | 100045974 | rs2851248 | -1.118 | -0.122 | -0.96 | -1.452 | -0.036 | 0.14 | 0.14 | 0 | 0.74 | 0.93 | 0.83 | 1 | 0.72 | 0.9 |
| 4 | 100046091 | rs2851247 | -1.118 | -0.122 | -0.96 | -1.452 | -0.036 | 0.14 | 0.14 | 0 | 0.74 | 0.93 | 0.83 | 1 | 0.72 | 0.9 |
| 4 | 100046104 | rs2851246 | -1.118 | -0.122 | -0.96 | -1.452 | -0.036 | 0.14 | 0.14 | 0 | 0.74 | 0.93 | 0.83 | 1 | 0.72 | 0.9 |
| 4 | 100046207 | rs2602896 | -1.118 | -0.122 | -0.96 | -1.452 | -0.036 | 0.14 | 0.14 | 0 | 0.74 | 0.93 | 0.83 | 1 | 0.72 | 0.9 |
| 4 | 100047812 | rs1126672 | 0.896 | -0.122 | -0.96 | -1.452 | -0.036 | 0.14 | 0.14 | 0 | 0.26 | 0.07 | 0.17 | 0 | 0.28 | 0.1 |
| 4 | 100048414 | rs1126671 | -0.888 | -0.122 | -0.96 | -1.452 | -0.014 | 0.092 | 0.092 | 0 | 0.73 | 0.86 | 0.77 | 1 | 0.7 | 0.88 |
| 4 | 100049145 | rs6837685 | -0.888 | -0.122 | -0.96 | -1.452 | -0.014 | 0.092 | 0.092 | 0 | 0.73 | 0.86 | 0.77 | 1 | 0.7 | 0.88 |
| 4 | 100051819 | rs4699710 | -0.899 | -0.123 | -0.961 | -1.453 | -0.014 | 0.092 | 0.092 | 0 | 0.73 | 0.86 | 0.77 | 1 | 0.7 | 0.88 |
| 4 | 100054192 | rs4148888 | 1.979 | -0.146 | -0.989 | -1.415 | 0.002 | -0.094 | -0.094 | 0.85 | 0.14 | 0.01 | 0.09 | 0.73 | 0.13 | 0.31 |
| 4 | 100054312 | rs17217949 | 0.924 | -0.146 | -0.989 | -1.416 | -0.036 | 0.14 | 0.14 | 0 | 0.26 | 0.07 | 0.17 | 0 | 0.28 | 0.1 |
| 4 | 100054827 | rs6532798 | -0.916 | -0.146 | -0.989 | -1.416 | -0.014 | 0.092 | 0.092 | 0 | 0.73 | 0.86 | 0.77 | 1 | 0.7 | 0.88 |
| 4 | 100054970 | rs13143133 | 0.916 | -0.135 | -1.154 | -1.405 | -0.036 | 0.14 | 0.14 | 0 | 0.26 | 0.07 | 0.17 | 0 | 0.28 | 0.1 |
| 4 | 100055739 | rs17218073 | 0.916 | -0.089 | -1.681 | -1.382 | -0.036 | 0.14 | 0.14 | 0 | 0.26 | 0.07 | 0.17 | 0 | 0.28 | 0.1 |

| | | | | | | | | | | | | | | | | |
|---|---|---|---|---|---|---|---|---|---|---|---|---|---|---|---|---|
| 4 | 100055825 | rs13112176 | 0.916 | -0.089 | -1.681 | -1.382 | -0.036 | 0.14 | 0.14 | 0 | 0.26 | 0.07 | 0.17 | 0 | 0.28 | 0.1 |
| 4 | 100055862 | rs4148887 | 1.971 | -0.089 | -1.682 | -1.382 | 0.002 | -0.094 | -0.094 | 2.979 | 0.14 | 0.01 | 0.09 | 0.73 | 0.13 | 0.31 |
| 4 | 100055908 | rs17218141 | 0.859 | -0.014 | -1.593 | -1.351 | -0.036 | 0.14 | 0.14 | 0 | 0.26 | 0.07 | 0.17 | 0 | 0.28 | 0.1 |
| 4 | 100056060 | rs17218162 | 0.859 | -0.014 | -1.593 | -1.351 | -0.036 | 0.14 | 0.14 | 0 | 0.26 | 0.07 | 0.17 | 0 | 0.28 | 0.1 |
| 4 | 100057020 | rs17218239 | 0.859 | -0.006 | -1.589 | -1.633 | -0.036 | 0.14 | 0.14 | 0 | 0.26 | 0.07 | 0.17 | 0 | 0.28 | 0.1 |
| 4 | 100058727 | rs13133647 | 0.859 | -0.006 | -1.589 | -1.633 | -0.036 | 0.14 | 0.14 | 0 | 0.26 | 0.07 | 0.17 | 0 | 0.28 | 0.1 |
| 4 | 100059312 | rs7669636 | 0.859 | -0.006 | -1.589 | -1.633 | -0.036 | 0.14 | 0.14 | 0 | 0.26 | 0.07 | 0.17 | 0 | 0.28 | 0.1 |
| 4 | 100059568 | rs7670060 | 0.872 | -0.007 | -1.589 | -1.633 | -0.042 | 0.15 | 0.15 | 0 | 0.25 | 0.05 | 0.17 | 0 | 0.28 | 0.1 |
| 4 | 100059619 | rs7670241 | 0.861 | -0.004 | -1.589 | -1.631 | -0.036 | 0.14 | 0.14 | 0 | 0.26 | 0.07 | 0.17 | 0 | 0.28 | 0.1 |
| 4 | 100060513 | rs4699713 | 0.861 | -0.004 | -1.589 | -1.631 | -0.036 | 0.14 | 0.14 | 0 | 0.26 | 0.07 | 0.17 | 0 | 0.28 | 0.1 |
| 4 | 100060538 | rs4699714 | 0.861 | -0.004 | -1.589 | -1.631 | -0.036 | 0.14 | 0.14 | 0 | 0.26 | 0.07 | 0.17 | 0 | 0.28 | 0.1 |
| 4 | 100060623 | rs17817868 | 0.861 | -0.004 | -1.589 | -1.631 | -0.036 | 0.14 | 0.14 | 0 | 0.26 | 0.07 | 0.17 | 0 | 0.28 | 0.1 |
| 4 | 100060794 | rs17218560 | 0.861 | -0.004 | -1.589 | -1.631 | -0.036 | 0.14 | 0.14 | 0 | 0.26 | 0.07 | 0.17 | 0 | 0.28 | 0.1 |
| 4 | 100060836 | rs13148577 | 0.924 | -0.048 | -1.621 | -1.669 | -0.1 | 0.247 | 0.247 | 0 | 0.43 | 0.09 | 0.31 | 0.12 | 0.45 | 0.52 |
| 4 | 100062430 | rs13110176 | 0.645 | 0.149 | -1.298 | -1.415 | -0.036 | 0.14 | 0.14 | 0 | 0.26 | 0.07 | 0.17 | 0 | 0.28 | 0.1 |
| 4 | 100062466 | rs13110764 | 0.645 | 0.149 | -1.298 | -1.415 | -0.036 | 0.14 | 0.14 | 0 | 0.26 | 0.07 | 0.17 | 0 | 0.28 | 0.1 |
| 4 | 100065509 | rs1800759 | 0.573 | 0.091 | -1.228 | -1.34 | 0.013 | 0.114 | 0.114 | 0 | 0.59 | 0.18 | 0.41 | 0.86 | 0.6 | 0.84 |
| 4 | 100066084 | rs3762894 | 1.329 | 0.033 | -0.966 | -1.422 | -0.009 | 0.03 | 0.03 | 0 | 0.17 | 0.23 | 0.11 | 0.74 | 0.16 | 0.32 |
| 4 | 100066125 | rs4699716 | 0.898 | -0.05 | -1.012 | -1.572 | -0.025 | 0.126 | 0.126 | 0 | 0.26 | 0.07 | 0.17 | 0 | 0.27 | 0.1 |
| 4 | 100066137 | rs4699717 | 0.898 | -0.05 | -1.012 | -1.572 | -0.025 | 0.126 | 0.126 | 0 | 0.26 | 0.07 | 0.17 | 0 | 0.27 | 0.1 |
| 4 | 100066287 | rs4148884 | 1.253 | -0.091 | -0.395 | -1.542 | 0.03 | 0.144 | 0.144 | 0.016 | 0.1 | 0.3 | 0.34 | 0.14 | 0.09 | 0.04 |
| 4 | 100066332 | rs4148883 | 0.106 | -0.077 | -0.466 | -1.54 | 0.007 | 0.025 | 0.025 | 0 | 0.56 | 0.62 | 0.66 | 0.27 | 0.54 | 0.56 |
| 4 | 100067337 | rs2226896 | 1.658 | -0.027 | -0.119 | -2.367 | -0.003 | -0.006 | -0.006 | 0 | 0.08 | 0.05 | 0.31 | 0.14 | 0.09 | 0.04 |
| 4 | 100073261 | rs4699720 | -0.873 | -0.155 | -0.292 | -2.852 | -0.016 | -0.106 | -0.106 | 2.16 | 0.77 | 0.63 | 0.81 | 1 | 0.77 | 0.9 |
| 4 | 100077749 | rs16996598 | 0.524 | -0.186 | -0.309 | -2.844 | 0.015 | 0.047 | 0.047 | 0 | 0.27 | 0.12 | 0.44 | 0.1 | 0.27 | 0.1 |
| 4 | 100082154 | rs1540053 | -0.877 | -0.188 | -0.328 | -2.839 | -0.016 | -0.107 | -0.107 | 0 | 0.77 | 0.63 | 0.81 | 1 | 0.78 | 0.91 |
| 4 | 100122819 | rs6814114 | 1.284 | -0.281 | -0.491 | -2.864 | 0.011 | 0.112 | 0.112 | 0 | 0.19 | 0.05 | 0.12 | 0 | 0.17 | 0.06 |
| 4 | 100123186 | rs2051428 | 1.054 | -0.123 | -0.327 | -2.981 | 0.002 | 0.004 | 0.004 | 0 | 0.22 | 0.25 | 0.17 | 0.78 | 0.24 | 0.46 |
| 4 | 100123776 | rs2000864 | -0.697 | -0.158 | -0.462 | -3.084 | 0.007 | 0.074 | 0.074 | 0 | 0.52 | 0.65 | 0.63 | 0.88 | 0.51 | 0.56 |
| 4 | 100126896 | rs17219626 | 1.502 | -0.175 | -0.78 | -3.067 | 0.008 | 0.107 | 0.107 | 0 | 0.18 | 0.05 | 0.12 | 0 | 0.17 | 0.06 |
| 4 | 100127311 | rs4147547 | 0.854 | -0.005 | -0.68 | -2.979 | -0.106 | 0.258 | 0.258 | 0 | 0.43 | 0.08 | 0.3 | 0.12 | 0.43 | 0.41 |
| 4 | 100134103 | rs2187483 | -0.934 | 0.056 | -0.689 | -2.969 | -0.077 | 0.255 | 0.255 | 0 | 0.55 | 0.88 | 0.65 | 0.88 | 0.55 | 0.57 |
| 4 | 100137534 | rs4699733 | -0.479 | 0.479 | -0.655 | -3.379 | -0.003 | -0.025 | -0.025 | 0 | 0.66 | 0.62 | 0.74 | 0.91 | 0.73 | 0.67 |
| 4 | 100138538 | rs9995799 | 1.06 | 0.368 | -0.667 | -3.488 | 0.002 | 0.002 | 0.002 | 0 | 0.22 | 0.25 | 0.18 | 0.78 | 0.24 | 0.5 |
| 4 | 100142302 | rs10008281 | 1.06 | 0.374 | -0.667 | -3.488 | 0.003 | 0.002 | 0.002 | 0 | 0.22 | 0.25 | 0.18 | 0.78 | 0.24 | 0.5 |
| 4 | 100143184 | rs6830685 | -0.82 | 0.305 | -0.87 | -3.556 | 0.033 | 0.118 | 0.118 | 0 | 0.7 | 0.89 | 0.77 | 0.91 | 0.76 | 0.68 |
| 4 | 100147101 | rs12510722 | 0.748 | 0.347 | -0.828 | -3.523 | -0.072 | 0.146 | 0.146 | 0 | 0.4 | 0.15 | 0.56 | 0.13 | 0.49 | 0.17 |
| 4 | 100171379 | rs2173199 | 0.746 | 0.359 | -0.829 | -3.52 | -0.069 | 0.146 | 0.146 | 0 | 0.4 | 0.15 | 0.56 | 0.13 | 0.49 | 0.17 |
| 4 | 100178151 | rs13142106 | 1.385 | 0.371 | -0.942 | -3.551 | -0.141 | 0.231 | 0.231 | 0 | 0.32 | 0.06 | 0.24 | 0.02 | 0.4 | 0.13 |
| 4 | 100185431 | rs12506882 | 1.572 | 0.352 | -0.673 | -3.558 | 0.01 | 0.042 | 0.042 | 0.021 | 0.08 | 0.17 | 0.33 | 0.1 | 0.09 | 0.04 |
| 4 | 100192460 | rs1566141 | 1.163 | 0.296 | -0.638 | -3.712 | 0.005 | -0.003 | -0.003 | 2.171 | 0.22 | 0.25 | 0.18 | 0.8 | 0.25 | 0.5 |
| 4 | 100194517 | rs13123099 | 1.434 | 0.295 | -0.642 | -3.713 | -0.141 | 0.232 | 0.232 | 0 | 0.32 | 0.06 | 0.24 | 0.02 | 0.4 | 0.13 |
| 4 | 100194977 | rs6811453 | 1.434 | 0.295 | -0.642 | -3.713 | -0.141 | 0.232 | 0.232 | 0 | 0.32 | 0.06 | 0.24 | 0.02 | 0.4 | 0.13 |
| 4 | 100195815 | rs12512110 | 1.9 | 0.306 | -0.616 | -3.701 | -0.001 | -0.002 | -0.002 | 0 | 0.08 | 0.06 | 0.32 | 0.09 | 0.09 | 0.04 |
| 4 | 100200509 | rs3819197 | 1.155 | 0.296 | -0.631 | -3.694 | 0.005 | -0.003 | -0.003 | 0 | 0.22 | 0.25 | 0.18 | 0.8 | 0.25 | 0.5 |
| 4 | 100202078 | rs1229976 | -0.976 | 0.285 | -1.225 | -3.65 | 0.041 | 0.122 | 0.122 | 0.001 | 0.71 | 0.93 | 0.83 | 0.91 | 0.81 | 0.71 |
| 4 | 100203447 | rs2276332 | 1.573 | -0.102 | -0.977 | -3.429 | -0.001 | -0.001 | -0.001 | 0 | 0.08 | 0.06 | 0.32 | 0.09 | 0.09 | 0.04 |
| 4 | 100211786 | rs4147532 | 0.462 | -0.06 | -0.992 | -3.452 | -0.037 | 0.11 | 0.11 | 0 | 0.41 | 0.22 | 0.57 | 0.11 | 0.5 | 0.17 |
| 4 | 100212197 | rs4147531 | 0.462 | -0.08 | -0.992 | -3.452 | -0.035 | 0.11 | 0.11 | 0 | 0.41 | 0.22 | 0.57 | 0.11 | 0.5 | 0.17 |
| 4 | 100214460 | rs35743364 | 1.014 | -0.098 | -1.085 | -3.477 | -0.097 | 0.186 | 0.186 | 0 | 0.32 | 0.11 | 0.24 | 0.03 | 0.4 | 0.13 |
| 4 | 100217743 | rs1826909 | 1.014 | -0.098 | -1.085 | -3.477 | -0.097 | 0.186 | 0.186 | 0 | 0.32 | 0.11 | 0.24 | 0.03 | 0.4 | 0.13 |
| 4 | 100220240 | rs13103321 | 1.086 | -0.106 | -1.12 | -3.482 | -0.132 | 0.225 | 0.225 | 0 | 0.32 | 0.06 | 0.24 | 0.03 | 0.4 | 0.12 |
| 4 | 100238998 | rs2075633 | 0.951 | -0.356 | -1.53 | -3.671 | -0.062 | -0.018 | -0.018 | 0 | 0.23 | 0.07 | 0.17 | 0.8 | 0.3 | 0.54 |
| 4 | 100239112 | rs4147536 | -0.646 | -0.393 | -1.333 | -3.677 | -0.055 | -0.109 | -0.109 | 0 | 0.68 | 0.55 | 0.77 | 0.91 | 0.82 | 0.71 |
| 4 | 100241575 | rs1353621 | 1.034 | -0.069 | -2.346 | -3.452 | -0.135 | 0.228 | 0.228 | 0 | 0.32 | 0.06 | 0.24 | 0.03 | 0.4 | 0.13 |
| 4 | 100243009 | rs1159918 | 0.531 | -0.368 | -2.17 | -3.311 | 0.065 | 0.15 | 0.15 | 0 | 0.56 | 0.24 | 0.4 | 0.83 | 0.67 | 0.66 |
| 4 | 100243932 | rs1229982 | -0.234 | -0.986 | -1.79 | -3.575 | -0.032 | -0.104 | -0.104 | 0 | 0.69 | 0.51 | 0.81 | 0.94 | 0.81 | 0.84 |
| 4 | 100244221 | rs3811802 | 0.04 | -0.425 | -2.649 | -3.345 | 0.003 | -0.046 | -0.046 | 0 | 0.38 | 0.51 | 0.34 | 0.09 | 0.47 | 0.27 |
| 4 | 100245489 | rs1693439 | -1.486 | -0.021 | -2.516 | -2.897 | 0.001 | -0.014 | -0.014 | 0 | 0.88 | 0.91 | 0.96 | 1 | 0.95 | 0.96 |
| 4 | 100247351 | rs2213041 | 0.104 | 0.128 | -2.594 | -2.946 | -0.048 | 0.025 | 0.025 | 0 | 0.11 | 0.01 | 0.15 | 0.05 | 0.15 | 0.17 |
| 4 | 100250419 | rs1789891 | 0.104 | 0.128 | -2.588 | -2.946 | -0.043 | 0.027 | 0.027 | 0 | 0.11 | 0.01 | 0.15 | 0.05 | 0.15 | 0.17 |
| 4 | 100254935 | rs3846448 | 1.567 | 0.119 | -2.798 | -3.15 | -0.052 | 0.083 | 0.083 | 0 | 0.22 | 0.09 | 0.13 | 0.02 | 0.26 | 0.11 |
| 4 | 100255953 | rs1229979 | 1.47 | -0.146 | -2.627 | -2.99 | -0.072 | 0.101 | 0.101 | 0 | 0.24 | 0.09 | 0.17 | 0.02 | 0.32 | 0.17 |
| 4 | 100256199 | rs1229978 | 0.684 | -0.098 | -2.605 | -2.969 | -0.114 | 0.163 | 0.163 | 0.59 | 0.32 | 0.1 | 0.27 | 0.08 | 0.41 | 0.28 |
| 4 | 100256984 | rs1789896 | -0.014 | -0.131 | -2.614 | -2.993 | -0.021 | 0.181 | 0.181 | 0.001 | 0.56 | 0.8 | 0.69 | 0.92 | 0.54 | 0.69 |
| 4 | 100257432 | rs3114046 | -1.607 | -0.121 | -2.609 | -2.988 | 0.001 | -0.014 | -0.014 | 0 | 0.88 | 0.91 | 0.96 | 1 | 0.95 | 0.96 |
| 4 | 100258381 | rs1662033 | 1.38 | -0.224 | -2.497 | -2.877 | -0.072 | 0.105 | 0.105 | 0.059 | 0.25 | 0.09 | 0.17 | 0.02 | 0.32 | 0.17 |
| 4 | 100263383 | rs904095 | 0.612 | -0.226 | -2.496 | -2.876 | -0.114 | 0.163 | 0.163 | 0.59 | 0.32 | 0.1 | 0.27 | 0.08 | 0.4 | 0.28 |
| 4 | 100263584 | rs904096 | 0.612 | -0.226 | -2.495 | -2.876 | -0.11 | 0.159 | 0.159 | 0.59 | 0.32 | 0.1 | 0.27 | 0.08 | 0.41 | 0.28 |

| | | | | | | | | | | | | | | | | |
|---|---|---|---|---|---|---|---|---|---|---|---|---|---|---|---|---|
| 4 | 100263778 | rs1789911 | 0.612 | -0.226 | -2.495 | -2.876 | -0.11 | 0.159 | 0.159 | 0.59 | 0.32 | 0.1 | 0.27 | 0.08 | 0.4 | 0.28 |
| 4 | 100269202 | rs1789921 | 1.377 | -0.205 | -2.482 | -2.865 | -0.069 | 0.1 | 0.1 | 0.059 | 0.25 | 0.09 | 0.17 | 0.02 | 0.32 | 0.17 |
| 4 | 100270607 | rs283415 | -0.386 | 0.101 | -2.332 | -2.714 | -0.046 | 0.154 | 0.154 | 0 | 0.58 | 0.81 | 0.7 | 0.92 | 0.58 | 0.7 |
| 4 | 100274286 | rs1789924 | 0.303 | 0.649 | -2.63 | -2.45 | -0.107 | 0.179 | 0.179 | 0.102 | 0.35 | 0.1 | 0.27 | 0.08 | 0.4 | 0.28 |
| 4 | 100274749 | rs11499823 | 0.598 | 0.649 | -2.63 | -2.45 | -0.029 | 0.008 | 0.008 | 0 | 0.08 | 0.01 | 0.1 | 0.05 | 0.09 | 0.1 |
| 4 | 100278697 | rs4699741 | 1.544 | 0.578 | -2.697 | -2.251 | 0.025 | -0.034 | -0.034 | 0 | 0.14 | 0.06 | 0.36 | 0.16 | 0.08 | 0.04 |
| 4 | 100278819 | rs1662037 | 1.041 | 0.784 | -2.48 | -2.002 | -0.067 | 0.122 | 0.122 | 0 | 0.27 | 0.09 | 0.17 | 0.03 | 0.32 | 0.18 |
| 4 | 100279332 | rs1614377 | 1.041 | 0.763 | -2.48 | -2.002 | -0.065 | 0.122 | 0.122 | 0 | 0.27 | 0.09 | 0.17 | 0.03 | 0.31 | 0.18 |
| 4 | 100279824 | rs2851300 | 0.568 | 0.551 | -2.212 | -1.742 | -0.107 | 0.179 | 0.179 | 0.102 | 0.35 | 0.1 | 0.27 | 0.08 | 0.4 | 0.28 |
| 4 | 100282765 | rs166892 | 0.585 | 0.814 | -1.33 | -0.636 | -0.001 | -0.011 | -0.011 | 0 | 0.06 | 0.07 | 0.05 | 0 | 0.08 | 0.01 |
| 4 | 100284199 | rs7661978 | -0.592 | 1.234 | -1.27 | -0.326 | 0.008 | 0.011 | 0.011 | 0 | 0.26 | 0.24 | 0.46 | 0.16 | 0.23 | 0.07 |
| 4 | 100304545 | rs1372680 | -0.628 | 1.625 | -0.993 | 0.128 | 0.003 | 0.002 | 0.002 | 0 | 0.06 | 0.06 | 0.14 | 0.17 | 0.1 | 0.09 |
| 4 | 100305393 | rs10516440 | -1.224 | 1.646 | -0.857 | 0.04 | 0.011 | 0.004 | 0.004 | 0 | 0.08 | 0.11 | 0.16 | 0.17 | 0.12 | 0.1 |
| 4 | 100307378 | rs2851293 | -0.473 | 2.05 | -1.308 | 0 | -0.043 | -0.036 | -0.036 | 0 | 0.86 | 0.99 | 0.83 | 0.86 | 0.88 | 0.95 |
| 4 | 100310907 | rs284782 | 1.03 | 1.752 | -1.036 | -0.927 | 0.005 | 0.027 | 0.027 | 0 | 0.5 | 0.38 | 0.48 | 0.04 | 0.46 | 0.33 |
| 4 | 100318099 | rs7667212 | -0.676 | 1.998 | -0.986 | -0.666 | 0.013 | 0.005 | 0.005 | 0 | 0.08 | 0.11 | 0.15 | 0.17 | 0.12 | 0.1 |
| 4 | 100321443 | rs4699748 | -0.464 | 2.03 | -0.971 | -0.65 | 0.017 | 0.007 | 0.007 | 0 | 0.08 | 0.11 | 0.16 | 0.17 | 0.12 | 0.09 |
| 4 | 100321573 | rs2584461 | -0.759 | 1.724 | -1.034 | -0.602 | -0.013 | -0.026 | -0.026 | 0 | 0.86 | 0.93 | 0.81 | 0.83 | 0.88 | 0.95 |
| 4 | 100322445 | rs991316 | 0.369 | 1.49 | -0.744 | -1.003 | 0.004 | -0.064 | -0.064 | 3.712 | 0.58 | 0.54 | 0.48 | 0.04 | 0.55 | 0.34 |
| 4 | 100324071 | rs17587689 | -0.141 | 2.104 | -1.187 | -0.851 | -0.054 | 0.055 | 0.055 | 0 | 0.1 | 0.02 | 0.1 | 0.01 | 0.19 | 0.06 |
| 4 | 100325630 | rs1348276 | 0.028 | 1.373 | -1.367 | -1.125 | 0.008 | 0.006 | 0.006 | 0 | 0.4 | 0.69 | 0.34 | 0.56 | 0.38 | 0.5 |
| 4 | 100326403 | rs1442490 | 0.425 | 0.946 | -1.265 | -1.273 | -0.027 | -0.078 | -0.078 | 0 | 0.29 | 0.15 | 0.23 | 0.5 | 0.31 | 0.38 |
| 4 | 100327803 | rs994771 | -0.187 | 0.974 | -0.972 | -1.302 | -0.004 | -0.009 | -0.009 | 0.788 | 0.36 | 0.22 | 0.3 | 0.56 | 0.38 | 0.5 |
| 4 | 100330309 | rs284792 | -0.735 | 1.287 | -0.964 | -1.291 | 0.165 | 0.029 | 0.029 | 2.358 | 0.07 | 0.5 | 0.09 | 0.02 | 0.1 | 0.05 |
| 4 | 100333267 | rs729147 | -1.038 | 0.842 | -1.414 | -1.702 | -0.005 | -0.025 | -0.025 | 0 | 0.76 | 0.85 | 0.57 | 0.59 | 0.81 | 0.68 |
| 4 | 100333556 | rs284787 | 0.613 | 0.509 | -1.008 | -1.2 | 0.026 | -0.105 | -0.105 | 0.185 | 0.26 | 0.11 | 0.18 | 0.48 | 0.22 | 0.33 |
| 4 | 100333932 | rs3805331 | 0.333 | 0.534 | -0.951 | -1.178 | 0.004 | -0.001 | -0.001 | 0 | 0.07 | 0.05 | 0.07 | 0.06 | 0.06 | 0.13 |
| 4 | 100334943 | rs4147553 | -0.398 | 0.648 | -0.707 | -1.129 | 0.062 | 0.094 | 0.094 | 0.07 | 0.07 | 0.41 | 0.08 | 0.06 | 0.06 | 0.12 |
| 4 | 100335586 | rs284785 | -0.254 | 0.623 | -0.772 | -1.159 | -0.02 | 0.007 | 0.007 | 0 | 0.34 | 0.52 | 0.27 | 0.54 | 0.28 | 0.45 |
| 4 | 100336102 | rs1827567 | 0.433 | 0.74 | -1.234 | -1.02 | 0.027 | -0.075 | -0.075 | 0.174 | 0.26 | 0.13 | 0.19 | 0.48 | 0.22 | 0.34 |
| 4 | 100338342 | rs1154454 | -1.266 | 0.706 | -0.259 | -1.342 | 0.035 | -0.101 | -0.101 | 0 | 0.14 | 0.46 | 0.09 | 0 | 0.15 | 0.1 |
| 4 | 100339596 | rs1154456 | 0.133 | 1.257 | -0.986 | -1.074 | -0.041 | 0.001 | 0.001 | 0 | 0.23 | 0.12 | 0.37 | 0.24 | 0.32 | 0.28 |
| 4 | 100339793 | rs1154457 | 0.146 | 1.259 | -1.015 | -1.073 | -0.063 | -0.008 | -0.008 | 0.384 | 0.23 | 0.08 | 0.37 | 0.23 | 0.32 | 0.28 |
| 4 | 100340586 | rs17529377 | 0.656 | 0.626 | -1.943 | -1.282 | 0.006 | 0.049 | 0.049 | 0 | 0.16 | 0.04 | 0.07 | 0.13 | 0.15 | 0.14 |
| 4 | 100341103 | rs1154459 | 0.613 | 0.701 | -1.888 | -1.22 | -0.06 | -0.007 | -0.007 | 1.136 | 0.23 | 0.08 | 0.37 | 0.24 | 0.32 | 0.28 |
| 4 | 100341643 | rs1154460 | -0.148 | 0.274 | -1.391 | -1.087 | 0.004 | -0.004 | -0.004 | 0 | 0.39 | 0.46 | 0.5 | 0.36 | 0.44 | 0.36 |
| 4 | 100341861 | rs971074 | 1.439 | 0.217 | -1.782 | -0.968 | 0.005 | -0.001 | -0.001 | 0 | 0.15 | 0.14 | 0.13 | 0.12 | 0.12 | 0.09 |
| 4 | 100356179 | rs1154469 | 1.249 | 0.379 | -2.097 | -0.815 | -0.051 | 0.003 | 0.003 | 0 | 0.23 | 0.11 | 0.37 | 0.24 | 0.33 | 0.28 |
| 4 | 100356466 | rs17537595 | 1.346 | 0.258 | -2.268 | -0.936 | -0.005 | 0.047 | 0.047 | 0 | 0.16 | 0.03 | 0.07 | 0.13 | 0.15 | 0.15 |
| 4 | 100357018 | rs2851028 | -1.557 | 0.313 | -2.651 | -0.744 | -0.048 | -0.138 | -0.138 | 0 | 0.82 | 1 | 0.77 | 0.51 | 0.84 | 0.71 |
| 4 | 100360117 | rs1154474 | 1.676 | 0.208 | -2.366 | -0.808 | -0.106 | -0.024 | -0.024 | 0 | 0.23 | 0.03 | 0.37 | 0.24 | 0.32 | 0.27 |
| 4 | 100360727 | rs1154476 | 1.109 | 0.117 | -2.303 | -0.769 | -0.069 | 0.023 | 0.023 | 0 | 0.27 | 0.11 | 0.4 | 0.24 | 0.38 | 0.29 |
| 4 | 100361769 | rs1154477 | 1.052 | 0.116 | -2.299 | -0.763 | -0.066 | 0.022 | 0.022 | 0 | 0.27 | 0.14 | 0.4 | 0.24 | 0.38 | 0.29 |
| 4 | 100363569 | rs1154480 | 1.052 | 0.116 | -2.285 | -0.763 | -0.061 | 0.022 | 0.022 | 0 | 0.27 | 0.14 | 0.4 | 0.24 | 0.38 | 0.29 |
| 4 | 100380148 | rs2119888 | 0.472 | -0.261 | -2.373 | -0.835 | -0.091 | 0.115 | 0.115 | 0 | 0.4 | 0.12 | 0.48 | 0.24 | 0.49 | 0.32 |
| 4 | 100391104 | rs2851024 | 1.379 | -0.727 | -2.251 | -0.751 | -0.09 | 0.006 | 0.006 | 0.059 | 0.26 | 0.15 | 0.42 | 0.26 | 0.46 | 0.3 |
| 4 | 100392060 | rs12505135 | 0.577 | -1.415 | -1.605 | -0.529 | -0.016 | 0.063 | 0.063 | 0 | 0.08 | 0.17 | 0.06 | 0.13 | 0.03 | 0.07 |
| 4 | 100394210 | rs1121886 | 0.143 | -1.869 | -1.574 | -0.678 | 0.001 | -0.007 | -0.007 | 0.446 | 0.5 | 0.52 | 0.57 | 0.72 | 0.62 | 0.39 |
| 4 | 100399736 | rs1442482 | 1.037 | -1.866 | -1.61 | -0.683 | -0.047 | 0.095 | 0.095 | 0.389 | 0.38 | 0.09 | 0.46 | 0.24 | 0.59 | 0.32 |
| 4 | 100423715 | rs1032827 | -2.106 | -1.945 | -1.823 | -0.877 | -0.025 | -0.044 | -0.044 | 0 | 0.86 | 1 | 0.77 | 0.86 | 0.87 | 0.77 |
| 4 | 100424345 | rs2654838 | -1.146 | -2.328 | -1.509 | -1.106 | -0.025 | 0.048 | 0.048 | 0 | 0.72 | 0.86 | 0.63 | 0.86 | 0.67 | 0.71 |
| 4 | 100430924 | rs7669241 | 0.798 | -2.265 | -1.315 | -0.992 | -0.092 | 0.211 | 0.211 | 0.163 | 0.17 | 0.45 | 0.12 | 0.5 | 0.06 | 0.07 |
| 4 | 100437553 | rs2851025 | 1.861 | -2.267 | -1.559 | -1.059 | -0.122 | 0.049 | 0.049 | 0.751 | 0.33 | 0.05 | 0.45 | 0.23 | 0.56 | 0.31 |
| 4 | 100441886 | rs1354368 | 0.852 | -2.625 | -1.317 | -1.079 | -0.011 | -0.035 | -0.035 | 0 | 0.43 | 0.46 | 0.71 | 0.75 | 0.65 | 0.4 |
| 4 | 100443720 | rs13143848 | 0.235 | -2.906 | -1.221 | -1.221 | 0.015 | 0.023 | 0.023 | 1.082 | 0.42 | 0.34 | 0.22 | 0.24 | 0.3 | 0.44 |
| 4 | 100443782 | rs13119384 | -1.512 | -2.811 | -1.461 | -1.075 | -0.033 | -0.056 | -0.056 | 0 | 0.61 | 0.93 | 0.35 | 0.4 | 0.36 | 0.65 |
| 4 | 100443800 | rs17029087 | 0.982 | -2.73 | -1.358 | -1.358 | 0 | 0.045 | 0.045 | 0.141 | 0.28 | 0.18 | 0.19 | 0.12 | 0.27 | 0.27 |
| 4 | 100444178 | rs10021082 | -0.08 | -2.656 | -1.18 | -1.172 | 0.008 | -0.009 | -0.009 | 0 | 0.27 | 0.71 | 0.12 | 0.63 | 0.07 | 0.38 |
| 4 | 100444684 | rs13126513 | -1.994 | -1.863 | -1.368 | -1.082 | -0.149 | 0.056 | 0.056 | 1.931 | 0.71 | 0.97 | 0.39 | 0.79 | 0.47 | 0.71 |
| 4 | 100450712 | rs884275 | -1.199 | -2.302 | -0.871 | -1.017 | 0.042 | 0.012 | 0.012 | 2.433 | 0.46 | 0.69 | 0.29 | 0.35 | 0.32 | 0.47 |
| 4 | 100457841 | rs10031512 | -0.664 | -2.371 | -0.606 | -1.071 | -0.025 | -0.001 | -0.001 | 0.128 | 0.41 | 0.49 | 0.25 | 0.35 | 0.28 | 0.43 |
| 4 | 100467879 | rs1491235 | -0.612 | -2.381 | -0.612 | -1.076 | -0.026 | 0 | 0 | 0.042 | 0.41 | 0.49 | 0.25 | 0.35 | 0.28 | 0.43 |
| 4 | 100472636 | rs13135826 | 1.161 | -2.161 | -0.528 | -1.067 | 0.011 | -0.022 | -0.022 | 0.192 | 0.05 | 0.19 | 0.04 | 0 | 0.04 | 0.04 |
| 4 | 100477462 | rs11935423 | 0.065 | -2.327 | -0.631 | -1.072 | 0.009 | 0.018 | 0.018 | 0 | 0.25 | 0.24 | 0.12 | 0.13 | 0.18 | 0.34 |
| 4 | 100484543 | rs10026790 | -0.59 | -2.351 | -0.634 | -1.085 | 0.039 | 0.067 | 0.067 | 0 | 0.35 | 0.26 | 0.18 | 0.14 | 0.26 | 0.39 |
| 4 | 100486295 | rs11097669 | 1.576 | -2.415 | -0.603 | -0.864 | 0.002 | 0.001 | 0.001 | 0 | 0.2 | 0.15 | 0.1 | 0.45 | 0.12 | 0.19 |
| 4 | 100487712 | rs12647527 | 1.616 | -2.464 | -0.704 | -0.863 | -0.026 | 0.093 | 0.093 | 0 | 0.2 | 0.28 | 0.1 | 0.45 | 0.12 | 0.19 |
| 4 | 100490438 | rs1503777 | -0.026 | -2.196 | -1.019 | -0.951 | 0.009 | 0.017 | 0.017 | 0 | 0.25 | 0.24 | 0.12 | 0.13 | 0.18 | 0.34 |
| 4 | 100490688 | rs1503778 | -0.627 | -2.135 | -0.983 | -0.918 | 0.039 | 0.067 | 0.067 | 0 | 0.35 | 0.26 | 0.18 | 0.14 | 0.26 | 0.39 |

| | | | | | | | | | | | | | | | | |
|---|---|---|---|---|---|---|---|---|---|---|---|---|---|---|---|---|
| 4 | 100491772 | rs11933337 | 1.028 | -2.171 | -1.005 | -0.944 | 0.002 | 0.004 | 0.004 | 0 | 0.19 | 0.2 | 0.11 | 0.13 | 0.16 | 0.33 |
| 4 | 100495817 | rs1800804 | -0.519 | -2.039 | -0.93 | -0.858 | 0.038 | 0.067 | 0.067 | 0 | 0.35 | 0.26 | 0.18 | 0.14 | 0.26 | 0.39 |
| 4 | 100504664 | rs3816873 | -0.519 | -2.039 | -0.93 | -0.858 | 0.038 | 0.067 | 0.067 | 0 | 0.35 | 0.26 | 0.18 | 0.14 | 0.26 | 0.39 |
| 4 | 100509321 | rs17029173 | -0.028 | -2.177 | -0.59 | -0.952 | -0.016 | -0.062 | -0.062 | 0 | 0.21 | 0.27 | 0.11 | 0 | 0.15 | 0.1 |
| 4 | 100510859 | rs991811 | -1.163 | -2.614 | -0.943 | -1.037 | 0.157 | 0.141 | 0.141 | 0.008 | 0.5 | 0.82 | 0.31 | 0.67 | 0.31 | 0.44 |
| 4 | 100512412 | rs982424 | 1.005 | -2.554 | -0.421 | -0.717 | -0.013 | 0.153 | 0.153 | 0.53 | 0.09 | 0.25 | 0.1 | 0.22 | 0.05 | 0.15 |
| 4 | 100518247 | rs17029213 | 1.234 | -2.508 | -0.457 | -0.756 | 0.007 | -0.022 | -0.022 | 0 | 0.05 | 0.19 | 0.04 | 0 | 0.04 | 0.04 |
| 4 | 100521805 | rs17029215 | 1.234 | -2.508 | -0.457 | -0.756 | 0.007 | -0.022 | -0.022 | 0 | 0.05 | 0.19 | 0.04 | 0 | 0.04 | 0.04 |
| 4 | 100532637 | rs745075 | 1.163 | -2.584 | -0.574 | -0.809 | 0.04 | 0.048 | 0.048 | 0 | 0.16 | 0.07 | 0.07 | 0 | 0.1 | 0.06 |
| 4 | 100539896 | rs1032355 | -0.525 | -2.573 | -0.494 | -0.722 | -0.074 | 0.003 | 0.003 | 1.493 | 0.27 | 0.47 | 0.17 | 0.22 | 0.16 | 0.14 |
| 4 | 100543304 | rs881980 | 1.701 | -2.597 | -0.374 | -0.71 | -0.004 | 0.017 | 0.017 | 0 | 0.2 | 0.18 | 0.1 | 0.45 | 0.12 | 0.19 |
| 4 | 100543745 | rs2255119 | 1.554 | -2.353 | -0.612 | -0.62 | 0.006 | 0.008 | 0.008 | 0 | 0.17 | 0.14 | 0.08 | 0.13 | 0.13 | 0.23 |
| 4 | 100553223 | rs958868 | 2.131 | -2.132 | -0.271 | -0.288 | -0.019 | 0.179 | 0.179 | 0.823 | 0.06 | 0.2 | 0.07 | 0.21 | 0.02 | 0.04 |
| 4 | 100554402 | rs7659550 | -1.136 | -1.929 | -0.775 | -0.619 | 0.078 | 0.085 | 0.085 | 0 | 0.51 | 0.7 | 0.32 | 0.67 | 0.35 | 0.45 |
| 4 | 100558594 | rs12641768 | -1.129 | -1.936 | -0.766 | -0.621 | 0.076 | 0.082 | 0.082 | 0 | 0.51 | 0.69 | 0.32 | 0.67 | 0.35 | 0.45 |
| 4 | 100563999 | rs17600719 | 1.087 | -1.325 | -0.794 | -0.936 | -0.006 | 0.064 | 0.064 | 0.307 | 0.15 | 0.05 | 0.12 | 0.01 | 0.16 | 0.15 |
| 4 | 100571001 | rs1491242 | 1.537 | -1.49 | -0.823 | -0.877 | -0.019 | 0.078 | 0.078 | 5.578 | 0.13 | 0.01 | 0.11 | 0.01 | 0.14 | 0.14 |
| 4 | 100588306 | rs7696714 | 0.051 | -1.484 | 0.035 | -0.895 | 0.004 | -0.031 | -0.031 | 0.585 | 0.07 | 0.22 | 0.05 | 0 | 0.07 | 0.05 |
| 4 | 100613859 | rs1027223 | 0.904 | -1.692 | 0.257 | -0.63 | -0.034 | 0.103 | 0.103 | 2.005 | 0.08 | 0.17 | 0.15 | 0.22 | 0.01 | 0.04 |
| 4 | 100626549 | rs10516451 | -1.768 | -1.22 | -0.42 | -1.014 | -0.05 | 0.021 | 0.021 | 1.141 | 0.76 | 0.86 | 0.55 | 0.8 | 0.58 | 0.7 |
| 4 | 100637756 | rs1586757 | -0.934 | -1.888 | -0.797 | -0.986 | -0.021 | 0.004 | 0.004 | 2.269 | 0.24 | 0.66 | 0.27 | 0.23 | 0.11 | 0.1 |
| 4 | 100638882 | rs12648843 | -1.161 | -2.043 | -1.047 | -1.116 | 0.118 | 0.049 | 0.049 | 1.683 | 0.41 | 0.83 | 0.39 | 0.24 | 0.28 | 0.27 |
| 4 | 100642113 | rs13123292 | 0.807 | -1.479 | -0.715 | -0.809 | 0.002 | -0.008 | -0.008 | 0 | 0.18 | 0.16 | 0.14 | 0.01 | 0.2 | 0.18 |
| 4 | 100643484 | rs13112253 | -1.253 | -1.425 | -0.748 | -0.833 | 0.065 | 0.008 | 0.008 | 0.713 | 0.36 | 0.84 | 0.38 | 0.24 | 0.28 | 0.27 |
| 4 | 100646108 | rs7670607 | -1.033 | -1.414 | -0.719 | -0.811 | -0.025 | 0.021 | 0.021 | 1.923 | 0.18 | 0.67 | 0.24 | 0.23 | 0.08 | 0.09 |
| 4 | 100651270 | rs13150420 | -1.156 | -1.269 | -1.308 | -0.786 | 0.058 | 0.017 | 0.017 | 0.258 | 0.38 | 0.84 | 0.4 | 0.24 | 0.31 | 0.28 |
| 4 | 100658372 | rs13116934 | 0.071 | -0.902 | -0.49 | -0.149 | -0.023 | 0.185 | 0.185 | 3.529 | 0.1 | 0.31 | 0.18 | 0.21 | 0.03 | 0.04 |
| 4 | 100663166 | rs1108961 | -0.27 | -0.696 | -1.014 | -0.555 | -0.006 | 0.001 | 0.001 | 0 | 0.43 | 0.45 | 0.19 | 0.53 | 0.34 | 0.49 |
| 4 | 100666364 | rs7666347 | 0.767 | 0.063 | -0.845 | -0.35 | 0.004 | 0.1 | 0.1 | 0.109 | 0.08 | 0.3 | 0.04 | 0.14 | 0.03 | 0.02 |
| 4 | 100668696 | rs11729480 | 0.691 | 0.487 | -2.298 | -0.427 | -0.072 | 0.112 | 0.112 | 0 | 0.2 | 0.04 | 0.11 | 0.01 | 0.28 | 0.19 |
| 4 | 100671057 | rs4699760 | 0.242 | 0.859 | -1.939 | 0.327 | 0 | -0.095 | -0.095 | 0.277 | 0.3 | 0.08 | 0.2 | 0.47 | 0.29 | 0.44 |
| 4 | 100671124 | rs1426730 | -0.6 | 0.542 | -1.914 | 0.295 | -0.023 | -0.04 | -0.04 | 0 | 0.75 | 0.86 | 0.43 | 0.62 | 0.68 | 0.72 |
| 4 | 100673830 | rs12649867 | -0.454 | 0.029 | -1.437 | 0.699 | 0.035 | -0.098 | -0.098 | 0.189 | 0.29 | 0.08 | 0.19 | 0.47 | 0.25 | 0.43 |
| 4 | 100678905 | rs1426736 | -0.695 | 0.556 | 0.055 | 0.493 | 0.075 | 0.042 | 0.042 | 1.2 | 0.06 | 0.39 | 0.02 | 0.01 | 0.01 | 0 |
| 4 | 100682937 | rs6849369 | -0.457 | 0.514 | -0.049 | 1.294 | 0.001 | 0.057 | 0.057 | 0 | 0.09 | 0.12 | 0.14 | 0.43 | 0.09 | 0.32 |
| 4 | 100686270 | rs6834229 | 0.14 | 0.848 | -0.235 | 1.168 | -0.007 | -0.003 | -0.003 | 0 | 0.85 | 0.89 | 0.82 | 0.84 | 0.82 | 0.8 |
| 4 | 100705272 | rs6831238 | 1.095 | 0.787 | -0.031 | 1.156 | 0 | 0.006 | 0.006 | 0 | 0.76 | 0.71 | 0.67 | 0.42 | 0.73 | 0.48 |
| 4 | 100705602 | rs2555482 | -0.062 | 0.731 | -0.11 | 1.259 | -0.001 | 0.057 | 0.057 | 0 | 0.91 | 0.88 | 0.85 | 0.58 | 0.9 | 0.68 |
| 4 | 100712375 | rs12152555 | 0.79 | 0.867 | -0.665 | 0.397 | 0.021 | 0.032 | 0.032 | 0 | 0.49 | 0.27 | 0.47 | 0.42 | 0.4 | 0.26 |
| 4 | 100728698 | rs1426738 | -0.398 | 0.877 | -0.516 | 0.432 | -0.001 | -0.002 | -0.002 | 0 | 0.15 | 0.13 | 0.18 | 0.16 | 0.17 | 0.2 |
| 4 | 100734264 | rs1365495 | -0.674 | 0.843 | -0.668 | 0.335 | 0.006 | 0.006 | 0.006 | 1.906 | 0.44 | 0.66 | 0.41 | 0.42 | 0.45 | 0.29 |
| 4 | 100740709 | rs4699764 | -0.695 | 0.857 | -1.141 | -0.308 | -0.015 | 0.017 | 0.017 | 1.477 | 0.54 | 0.75 | 0.46 | 0.43 | 0.52 | 0.37 |
| 4 | 100743389 | rs9994042 | -0.196 | 0.519 | -0.566 | -0.597 | -0.017 | -0.059 | -0.059 | 0.094 | 0.13 | 0.17 | 0.04 | 0 | 0.08 | 0.09 |
| 4 | 100743513 | rs3775500 | 0.385 | 0.49 | -0.478 | -0.618 | -0.017 | 0.057 | 0.057 | 1.056 | 0.28 | 0.51 | 0.33 | 0.43 | 0.28 | 0.2 |
| 4 | 100747164 | rs9307241 | -0.115 | 0.523 | -0.483 | -0.65 | -0.008 | -0.037 | -0.037 | 0 | 0.13 | 0.11 | 0.04 | 0 | 0.08 | 0.09 |
| 4 | 100753527 | rs1465518 | -0.67 | 0.424 | -0.257 | -0.378 | 0.001 | 0.001 | 0.001 | 0.387 | 0.38 | 0.6 | 0.31 | 0.18 | 0.33 | 0.24 |
| 4 | 100755107 | rs6813099 | 0.536 | 0.459 | -0.256 | 0.254 | -0.008 | 0.043 | 0.043 | 0.629 | 0.26 | 0.43 | 0.27 | 0.26 | 0.26 | 0.15 |
| 4 | 100770454 | rs6848666 | 0.655 | 0.36 | 0.173 | 0.113 | -0.001 | -0.004 | -0.004 | 2.4 | 0.23 | 0.31 | 0.22 | 0.15 | 0.25 | 0.14 |
| 4 | 100770696 | rs6849599 | -1.343 | 0.194 | 0.267 | -0.028 | 0.009 | 0.032 | 0.032 | 0.16 | 0.49 | 0.69 | 0.39 | 0.41 | 0.45 | 0.32 |
| 4 | 100784980 | rs3737484 | 0.564 | 0.229 | 0.365 | -0.045 | -0.006 | -0.015 | -0.015 | 0.364 | 0.25 | 0.47 | 0.24 | 0.15 | 0.26 | 0.14 |
| 4 | 100790307 | rs15924 | 0.866 | 0.202 | 0.361 | -0.07 | 0 | -0.022 | -0.022 | 0 | 0.13 | 0.1 | 0.04 | 0 | 0.06 | 0.09 |
| 4 | 100790328 | rs6532834 | 0.62 | 0.212 | 0.358 | -0.064 | -0.005 | -0.014 | -0.014 | 3.187 | 0.25 | 0.46 | 0.24 | 0.15 | 0.26 | 0.14 |
| 4 | 100799961 | rs11944405 | 0.572 | -0.033 | 0.362 | -0.095 | -0.003 | -0.012 | -0.012 | 0 | 0.25 | 0.44 | 0.24 | 0.15 | 0.26 | 0.14 |
| 4 | 100801273 | rs11723286 | -0.158 | -0.006 | 0.36 | -0.099 | -0.012 | -0.034 | -0.034 | 0 | 0.3 | 0.52 | 0.25 | 0.15 | 0.28 | 0.17 |
| 4 | 100805207 | rs6532835 | -1.492 | -0.553 | 0.048 | -0.381 | -0.01 | -0.002 | -0.002 | 0 | 0.57 | 0.82 | 0.45 | 0.42 | 0.46 | 0.42 |
| 4 | 100807221 | rs10016741 | 0.897 | -0.551 | 0.041 | -0.373 | 0.002 | -0.021 | -0.021 | 1.018 | 0.14 | 0.04 | 0.13 | 0.26 | 0.12 | 0.17 |
| 4 | 100808909 | rs11937985 | 0.779 | -0.544 | 0.055 | -0.38 | -0.004 | -0.013 | -0.013 | 0 | 0.25 | 0.46 | 0.24 | 0.15 | 0.26 | 0.14 |
| 4 | 100810611 | rs11945303 | 0.842 | -0.557 | 0.048 | -0.386 | -0.002 | -0.012 | -0.012 | 0 | 0.24 | 0.44 | 0.24 | 0.15 | 0.26 | 0.14 |
| 4 | 100823058 | rs11931053 | 0.072 | -0.557 | 0.036 | -0.381 | -0.012 | -0.036 | -0.036 | 0 | 0.3 | 0.52 | 0.25 | 0.15 | 0.28 | 0.17 |
| 4 | 100829746 | rs7676564 | -0.918 | -0.503 | 0.005 | -0.349 | 0.019 | 0.015 | 0.015 | 0.001 | 0.46 | 0.68 | 0.39 | 0.41 | 0.4 | 0.33 |
| 4 | 100832060 | rs2114442 | -1.499 | -0.528 | -0.008 | -0.366 | -0.013 | 0.001 | 0.001 | 0.363 | 0.57 | 0.82 | 0.45 | 0.41 | 0.46 | 0.42 |
| 4 | 100847105 | rs17029662 | 0.126 | -0.489 | 0 | -0.323 | -0.014 | -0.036 | -0.036 | 0 | 0.31 | 0.56 | 0.25 | 0.15 | 0.28 | 0.17 |
| 4 | 100848810 | rs10516458 | 1.464 | -0.59 | -0.24 | -0.373 | -0.065 | -0.046 | -0.046 | 0.235 | 0.19 | 0.02 | 0.21 | 0.15 | 0.26 | 0.14 |
| 4 | 100852921 | rs11097681 | 0.82 | -0.449 | -0.139 | -0.295 | -0.001 | -0.005 | -0.005 | 1.491 | 0.24 | 0.32 | 0.23 | 0.15 | 0.26 | 0.14 |
| 4 | 100872365 | rs17029673 | 0.652 | -0.318 | -0.4 | -0.265 | -0.002 | -0.016 | -0.016 | 0 | 0.25 | 0.47 | 0.24 | 0.16 | 0.26 | 0.14 |
| 4 | 100873826 | rs2070738 | -0.866 | -0.343 | -0.375 | -0.277 | 0.05 | 0.112 | 0.112 | 0 | 0.46 | 0.74 | 0.3 | 0.16 | 0.34 | 0.29 |
| 4 | 100876648 | rs11736218 | 0.52 | -0.32 | -0.363 | -0.261 | -0.004 | -0.016 | -0.016 | 0 | 0.25 | 0.47 | 0.24 | 0.16 | 0.26 | 0.14 |
| 4 | 100945695 | rs2866180 | -0.737 | -0.004 | -0.996 | -0.031 | 0.054 | 0.043 | 0.043 | 0 | 0.5 | 0.78 | 0.39 | 0.42 | 0.4 | 0.37 |
| 4 | 100971095 | rs13136235 | -0.715 | 0.731 | -1.216 | 0.414 | -0.069 | -0.063 | -0.063 | 0 | 0.9 | 0.99 | 0.71 | 0.8 | 0.79 | 0.87 |

| | | | | | | | | | | | | | | | | |
|---|---|---|---|---|---|---|---|---|---|---|---|---|---|---|---|---|
| 4 | 100979170 | rs4699769 | -1.312 | -0.17 | -1.239 | 0.038 | 0.005 | -0.067 | -0.067 | 0.053 | 0.75 | 0.93 | 0.79 | 0.62 | 0.76 | 0.62 |
| 4 | 101006705 | rs7340886 | -0.108 | -0.094 | -0.196 | -0.196 | -0.004 | -0.018 | -0.018 | 0 | 0.36 | 0.63 | 0.29 | 0.15 | 0.32 | 0.24 |
| 4 | 101016395 | rs4699773 | -0.563 | -0.098 | 0.427 | -0.105 | 0.006 | 0.005 | 0.005 | 0.439 | 0.5 | 0.51 | 0.67 | 0.67 | 0.57 | 0.48 |
| 4 | 101026248 | rs11735694 | 1.144 | -0.044 | 0.438 | -0.376 | 0.004 | -0.002 | -0.002 | 0.339 | 0.2 | 0.26 | 0.16 | 0.15 | 0.25 | 0.14 |
| 4 | 101046193 | rs4602501 | 1.293 | 0.017 | 0.399 | -0.382 | 0.006 | -0.002 | -0.002 | 0.433 | 0.19 | 0.28 | 0.14 | 0.15 | 0.25 | 0.15 |
| 4 | 101055012 | rs6855061 | 0.712 | 0.181 | 0.112 | -0.145 | -0.075 | -0.096 | -0.096 | 0 | 0.13 | 0.01 | 0.23 | 0.53 | 0.22 | 0.19 |
| 4 | 101057346 | rs11097688 | -0.909 | 0.094 | 0.3 | -0.247 | 0.001 | 0 | 0 | 0.586 | 0.49 | 0.51 | 0.64 | 0.68 | 0.58 | 0.48 |
| 4 | 101072734 | rs2866185 | 0.63 | 0.288 | -0.203 | -0.242 | -0.075 | -0.103 | -0.103 | 0 | 0.13 | 0.02 | 0.36 | 0.58 | 0.22 | 0.19 |
| 4 | 101082077 | rs7654712 | -0.586 | -0.095 | 0.55 | -0.783 | 0 | 0.002 | 0.002 | 0.153 | 0.48 | 0.55 | 0.57 | 0.72 | 0.53 | 0.44 |
| 4 | 101082943 | rs6817883 | -1.373 | -0.403 | -0.161 | -1.115 | -0.004 | 0.002 | 0.002 | 0.608 | 0.67 | 0.85 | 0.73 | 0.72 | 0.68 | 0.57 |
| 4 | 101085258 | rs6854962 | -0.132 | -0.209 | -0.062 | -0.92 | -0.005 | 0.059 | 0.059 | 0.123 | 0.45 | 0.3 | 0.54 | 0.72 | 0.53 | 0.44 |
| 4 | 101094284 | rs6532846 | -1.019 | -0.003 | -0.06 | -0.785 | -0.001 | 0 | 0 | 1.147 | 0.73 | 0.85 | 0.79 | 0.75 | 0.74 | 0.66 |
| 4 | 101100658 | rs2866194 | -0.71 | 0.104 | -0.092 | -0.715 | 0.002 | 0.004 | 0.004 | 0.304 | 0.67 | 0.7 | 0.76 | 0.75 | 0.7 | 0.64 |
| 4 | 101106316 | rs2866197 | 1.026 | 0.273 | 0.028 | -0.706 | 0.003 | -0.043 | -0.043 | 0.103 | 0.18 | 0.24 | 0.12 | 0.03 | 0.23 | 0.1 |
| 4 | 101107276 | rs1063821 | -0.54 | 0.1 | 0.187 | -0.932 | -0.001 | -0.017 | -0.017 | 0 | 0.45 | 0.46 | 0.61 | 0.74 | 0.43 | 0.54 |
| 4 | 101107476 | rs3819183 | 0.699 | 0.186 | 0.139 | -0.981 | -0.003 | -0.017 | -0.017 | 0.068 | 0.23 | 0.24 | 0.14 | 0.01 | 0.25 | 0.16 |
| 4 | 101108008 | rs11734827 | 1.276 | 0.198 | 0 | -0.951 | -0.008 | 0.009 | 0.009 | 0 | 0.19 | 0.16 | 0.13 | 0.01 | 0.23 | 0.09 |
| 4 | 101108057 | rs11726613 | -0.374 | 0.047 | 0.511 | -0.649 | -0.003 | 0.009 | 0.009 | 0 | 0.39 | 0.59 | 0.54 | 0.53 | 0.39 | 0.3 |
| 4 | 101108069 | rs11734833 | -0.565 | 0.01 | 0.877 | -0.672 | -0.002 | 0.002 | 0.002 | 0 | 0.43 | 0.59 | 0.55 | 0.53 | 0.41 | 0.36 |
| 4 | 101108132 | rs11726694 | -0.365 | 0.02 | 0.818 | -0.661 | -0.001 | 0.001 | 0.001 | 0 | 0.39 | 0.5 | 0.54 | 0.53 | 0.39 | 0.3 |
| 4 | 101108191 | rs11726650 | -0.365 | 0.02 | 0.818 | -0.65 | -0.001 | 0.001 | 0.001 | 0 | 0.39 | 0.5 | 0.54 | 0.53 | 0.39 | 0.3 |
| 4 | 101108332 | rs10007201 | -0.387 | 0.425 | 0.826 | -0.473 | -0.006 | -0.041 | -0.041 | 0 | 0.31 | 0.44 | 0.23 | 0.19 | 0.29 | 0.32 |
| 4 | 101114549 | rs17029785 | 1.068 | 0.458 | 0.721 | -0.584 | -0.006 | -0.076 | -0.076 | 0.106 | 0.13 | 0.24 | 0.08 | 0 | 0.09 | 0.11 |
| 4 | 101114999 | rs3737483 | 0.201 | 0.447 | 0.76 | -0.596 | 0 | -0.001 | -0.001 | 0 | 0.18 | 0.14 | 0.14 | 0.18 | 0.19 | 0.21 |
| 4 | 101118465 | rs10033894 | -0.638 | 0.467 | 0.723 | -0.541 | -0.006 | -0.047 | -0.047 | 0 | 0.31 | 0.41 | 0.22 | 0.2 | 0.28 | 0.32 |
| 4 | 101126405 | rs10030715 | 0.089 | 0.531 | 0.429 | -0.646 | -0.001 | -0.152 | -0.152 | 0 | 0.35 | 0.45 | 0.22 | 0.01 | 0.36 | 0.22 |
| 4 | 101128211 | rs868911 | -0.164 | 0.579 | 0.146 | -0.157 | 0 | -0.002 | -0.002 | 0 | 0.4 | 0.56 | 0.46 | 0.46 | 0.4 | 0.41 |
| 4 | 101131209 | rs7699773 | -0.667 | 0.511 | 0.054 | -0.427 | 0.001 | 0.012 | 0.012 | 0.236 | 0.55 | 0.6 | 0.58 | 0.63 | 0.58 | 0.56 |
| 4 | 101131793 | rs2866209 | -0.827 | 0.975 | -0.123 | 0.026 | 0.016 | 0.008 | 0.008 | 0 | 0.1 | 0.15 | 0.17 | 0.11 | 0.13 | 0.14 |
| 4 | 101132253 | rs12506292 | -0.36 | 0.831 | -0.442 | 0.009 | -0.008 | -0.001 | -0.001 | 0 | 0.39 | 0.34 | 0.49 | 0.63 | 0.47 | 0.39 |
| 4 | 101133509 | rs4352484 | -0.364 | 1.05 | -0.469 | -0.815 | 0.017 | -0.101 | -0.101 | 2.383 | 0.39 | 0.5 | 0.36 | 0.07 | 0.53 | 0.33 |
| 4 | 101135216 | rs4396998 | -0.375 | 0.187 | -0.884 | -0.651 | 0.023 | 0.008 | 0.008 | 0.225 | 0.64 | 0.76 | 0.6 | 0.44 | 0.72 | 0.55 |
| 4 | 101135989 | rs11097701 | 0.448 | 1.322 | -1.367 | -0.662 | -0.041 | -0.001 | -0.001 | 0 | 0.16 | 0.04 | 0.25 | 0.16 | 0.24 | 0.18 |
| 4 | 101137084 | rs10001464 | 0.411 | 1.335 | -1.11 | -0.649 | -0.012 | 0.002 | 0.002 | 0 | 0.16 | 0.12 | 0.26 | 0.17 | 0.24 | 0.18 |
| 4 | 101137500 | rs6820091 | 1.026 | 1.681 | -1.364 | -1.05 | -0.024 | 0.021 | 0.021 | 0 | 0.23 | 0.06 | 0.14 | 0.23 | 0.28 | 0.15 |
| 4 | 101140986 | rs13103925 | 0.9 | 1.729 | -1.338 | -1.124 | -0.025 | 0.025 | 0.025 | 0 | 0.1 | 0.08 | 0.11 | 0.05 | 0.13 | 0.17 |
| 4 | 101153391 | rs4440226 | 0.543 | 1.635 | -1.164 | -1.176 | -0.007 | 0.004 | 0.004 | 0 | 0.11 | 0.13 | 0.11 | 0.05 | 0.13 | 0.17 |
| 4 | 101168332 | rs6825670 | 0.798 | 1.669 | -0.648 | -1.037 | 0 | -0.015 | -0.015 | 0.041 | 0.26 | 0.29 | 0.15 | 0.24 | 0.29 | 0.15 |
| 4 | 101170196 | rs4557267 | 0.473 | 1.808 | -0.89 | -0.287 | -0.011 | -0.004 | -0.004 | 0 | 0.16 | 0.12 | 0.27 | 0.21 | 0.25 | 0.19 |
| 4 | 101176062 | rs11734053 | -0.64 | 1.082 | -1.03 | 0.354 | -0.004 | 0.07 | 0.07 | 0 | 0.88 | 0.86 | 0.69 | 0.58 | 0.91 | 0.72 |
| 4 | 101179982 | rs4699395 | 0.041 | 0.965 | -1.218 | 0.141 | -0.002 | -0.009 | -0.009 | 0 | 0.8 | 0.83 | 0.65 | 0.5 | 0.8 | 0.66 |
| 4 | 101191677 | rs4235448 | -0.664 | 0.548 | -1.572 | 0.424 | -0.01 | -0.056 | -0.056 | 0 | 0.9 | 0.96 | 0.78 | 0.83 | 0.91 | 0.84 |
| 4 | 101193982 | rs4699786 | 0.384 | 0.805 | -1.523 | -0.042 | -0.04 | 0.043 | 0.043 | 0 | 0.27 | 0.17 | 0.28 | 0.1 | 0.35 | 0.22 |
| 4 | 101197518 | rs10026337 | -0.226 | 0.378 | -1.208 | 0.856 | 0.005 | 0.099 | 0.099 | 0.014 | 0.14 | 0.28 | 0.25 | 0.35 | 0.11 | 0.25 |
| 4 | 101210715 | rs4235450 | 0.029 | 0.007 | -2.129 | 0.108 | 0.015 | 0.033 | 0.033 | 0 | 0.46 | 0.19 | 0.47 | 0.36 | 0.57 | 0.38 |
| 4 | 101213802 | rs6836741 | -1.361 | -0.052 | -2.389 | 0.115 | -0.044 | 0.016 | 0.016 | 3.286 | 0.81 | 0.99 | 0.91 | 0.92 | 0.83 | 0.74 |
| 4 | 101214573 | rs4128643 | 0.446 | 0.134 | -2.175 | 0.211 | -0.075 | -0.002 | -0.002 | 0 | 0.38 | 0.11 | 0.43 | 0.3 | 0.53 | 0.36 |
| 4 | 101216854 | rs4130108 | -0.019 | 0.1 | -1.341 | -0.046 | 0 | 0 | 0 | 0 | 0.62 | 0.37 | 0.69 | 0.77 | 0.72 | 0.57 |
| 4 | 101219932 | rs7659715 | 0.513 | 0.546 | -1.053 | -0.005 | -0.003 | -0.004 | -0.004 | 0.324 | 0.43 | 0.44 | 0.42 | 0.33 | 0.39 | 0.5 |
| 4 | 101222442 | rs10000959 | 0.515 | 1.676 | -1.35 | 0.659 | -0.006 | -0.003 | -0.003 | 0 | 0.09 | 0.06 | 0.08 | 0.1 | 0.1 | 0.1 |
| 4 | 101228493 | rs4306962 | 0.235 | 1.32 | -1.258 | 0.617 | -0.016 | -0.026 | -0.026 | 0 | 0.7 | 0.76 | 0.63 | 0.44 | 0.65 | 0.66 |
| 4 | 101250176 | rs11934689 | 0.141 | 0.638 | -1.32 | -0.11 | 0.053 | 0.052 | 0.052 | 0 | 0.41 | 0.23 | 0.57 | 0.73 | 0.34 | 0.53 |
| 4 | 101251778 | rs11097709 | 0.064 | 0.753 | -0.228 | -0.416 | 0.029 | 0.001 | 0.001 | 0 | 0.89 | 0.65 | 0.88 | 0.94 | 0.9 | 0.95 |
| 4 | 101262840 | rs4478196 | 0.419 | 0.607 | 0.366 | -0.489 | 0.027 | -0.001 | -0.001 | 0 | 0.88 | 0.49 | 0.87 | 0.94 | 0.91 | 0.95 |
| 4 | 101283279 | rs4458459 | 0.464 | 0.544 | 0.548 | -0.562 | 0.053 | 0.059 | 0.059 | 0 | 0.42 | 0.34 | 0.58 | 0.78 | 0.34 | 0.53 |
| 4 | 101299450 | rs12647890 | -0.029 | 0.854 | -0.302 | -1.464 | -0.062 | 0.112 | 0.112 | 0 | 0.36 | 0.21 | 0.6 | 0.92 | 0.5 | 0.58 |
| 4 | 101306586 | rs1982086 | 0.013 | 0.823 | -0.338 | -1.621 | -0.002 | 0.003 | 0.003 | 3.454 | 0.25 | 0.23 | 0.48 | 0.81 | 0.24 | 0.47 |
| 4 | 101306906 | rs13122464 | 1.288 | 0.845 | -0.454 | -1.604 | -0.011 | 0.036 | 0.036 | 3.18 | 0.15 | 0.05 | 0.07 | 0.02 | 0.13 | 0.15 |
| 4 | 101310665 | rs1019947 | 0.889 | 0.853 | -0.468 | -1.598 | -0.13 | 0.038 | 0.038 | 0 | 0.25 | 0.03 | 0.19 | 0.13 | 0.39 | 0.25 |
| 4 | 101319165 | rs4071 | -0.186 | 0.705 | 0.189 | -1.663 | -0.005 | 0.022 | 0.022 | 0.3 | 0.75 | 0.84 | 0.82 | 0.92 | 0.77 | 0.8 |
| 4 | 101319332 | rs17552409 | 1.32 | 0.702 | 0.201 | -1.671 | -0.015 | 0.046 | 0.046 | 3.285 | 0.15 | 0.04 | 0.07 | 0.02 | 0.13 | 0.15 |
| 4 | 101320412 | rs2116992 | -0.616 | 0.723 | 0.206 | -1.662 | 0 | -0.002 | -0.002 | 0.564 | 0.9 | 0.9 | 0.89 | 0.94 | 0.9 | 0.95 |
| 4 | 101321764 | rs1056659 | -0.192 | 0.659 | 0.18 | -1.685 | -0.005 | 0.025 | 0.025 | 0.302 | 0.75 | 0.85 | 0.82 | 0.92 | 0.77 | 0.8 |
| 4 | 101330587 | rs17552895 | 1.799 | 0.381 | 0.164 | -1.698 | 0.064 | 0.137 | 0.137 | 0.087 | 0.31 | 0.09 | 0.17 | 0 | 0.23 | 0.22 |
| 4 | 101341678 | rs1347197 | 0.311 | 0.458 | 0.305 | -1.609 | -0.028 | -0.011 | -0.011 | 0.046 | 0.88 | 0.71 | 0.87 | 0.94 | 0.92 | 0.92 |
| 4 | 101342922 | rs1392887 | -0.364 | 0.6 | -0.037 | -1.616 | -0.143 | 0.039 | 0.039 | 0 | 0.67 | 0.92 | 0.71 | 0.81 | 0.51 | 0.7 |
| 4 | 101348412 | rs7674306 | 0.31 | 0.568 | 0.139 | -1.62 | 0 | -0.077 | -0.077 | 0 | 0.21 | 0.24 | 0.1 | 0.02 | 0.17 | 0.15 |
| 4 | 101353122 | rs357652 | -0.43 | 0.809 | -0.259 | -1.656 | 0.019 | 0.194 | 0.194 | 0 | 0.44 | 0.09 | 0.27 | 0.15 | 0.56 | 0.4 |
| 4 | 101368154 | rs1501075 | 0.904 | 1.047 | -0.211 | -1.51 | -0.157 | 0.018 | 0.018 | 0 | 0.23 | 0.02 | 0.18 | 0.13 | 0.38 | 0.24 |

| | | | | | | | | | | | | | | | | |
|---|---|---|---|---|---|---|---|---|---|---|---|---|---|---|---|---|
| 4 | 101383092 | rs242596 | 0.044 | 0.732 | 0.39 | -1.435 | 0.024 | 0.04 | 0.04 | 0 | 0.9 | 0.56 | 0.87 | 0.94 | 0.91 | 0.95 |
| 4 | 101388148 | rs11097720 | 0.804 | 0.674 | 0.369 | -1.591 | 0.019 | 0.026 | 0.026 | 0 | 0.43 | 0.37 | 0.59 | 0.8 | 0.35 | 0.55 |
| 4 | 101398205 | rs924531 | 1.313 | 0.476 | 0.421 | -1.757 | -0.021 | -0.117 | -0.117 | 0 | 0.35 | 0.41 | 0.2 | 0.01 | 0.24 | 0.23 |
| 4 | 101402501 | rs992834 | -0.524 | 1.366 | 0.367 | -1.714 | -0.008 | 0.002 | 0.002 | 0 | 0.69 | 0.74 | 0.74 | 0.75 | 0.62 | 0.72 |
| 4 | 101423326 | rs1587549 | 0.647 | 1.102 | 0.533 | -1.247 | -0.008 | -0.005 | -0.005 | 1.418 | 0.45 | 0.57 | 0.47 | 0.69 | 0.34 | 0.55 |
| 4 | 101432085 | rs3775367 | 0.614 | 1.028 | 0.502 | -1.248 | -0.004 | 0.077 | 0.077 | 0 | 0.11 | 0.19 | 0.27 | 0.69 | 0.1 | 0.31 |
| 4 | 101432849 | rs1501096 | 0.688 | 0.996 | 0.513 | -1.276 | -0.007 | -0.004 | -0.004 | 0.049 | 0.43 | 0.48 | 0.45 | 0.69 | 0.34 | 0.55 |
| 4 | 101436201 | rs1318710 | 0.653 | 1.024 | 0.391 | -1.185 | -0.003 | 0.058 | 0.058 | 0 | 0.11 | 0.16 | 0.26 | 0.71 | 0.1 | 0.31 |
| 4 | 101442540 | rs1501089 | 0.842 | 1.194 | 0.386 | -1.306 | 0.002 | -0.008 | -0.008 | 0 | 0.18 | 0.16 | 0.24 | 0.07 | 0.18 | 0.14 |
| 4 | 101443803 | rs2036357 | -0.166 | 1.277 | 0.259 | -1.279 | -0.007 | 0.006 | 0.006 | 0.075 | 0.64 | 0.69 | 0.71 | 0.82 | 0.54 | 0.69 |
| 4 | 101452208 | rs357670 | -0.377 | 1.174 | 0.127 | -1.115 | -0.001 | -0.015 | -0.015 | 0.048 | 0.16 | 0.16 | 0.32 | 0.73 | 0.18 | 0.37 |
| 4 | 101454173 | rs1846870 | 0.217 | 1.274 | 0.215 | -1.098 | 0 | 0.007 | 0.007 | 0 | 0.55 | 0.43 | 0.56 | 0.73 | 0.51 | 0.61 |
| 4 | 101466820 | rs1392874 | 0.288 | 1.344 | -0.036 | -0.86 | -0.007 | -0.024 | -0.024 | 0.407 | 0.38 | 0.52 | 0.32 | 0.25 | 0.37 | 0.33 |
| 4 | 101470231 | rs977668 | 0.336 | 0.701 | 0.016 | -0.832 | -0.026 | -0.019 | -0.019 | 0 | 0.37 | 0.53 | 0.28 | 0.25 | 0.35 | 0.27 |
| 4 | 101471756 | rs2567398 | -1.084 | 0.697 | 0.217 | -0.831 | 0 | 0.108 | 0.108 | 0 | 0.14 | 0.33 | 0.13 | 0.24 | 0.14 | 0.15 |
| 4 | 101475765 | rs10021695 | -0.984 | 0.721 | 0.333 | -0.858 | 0.045 | -0.023 | -0.023 | 0 | 0.09 | 0.45 | 0.05 | 0 | 0.06 | 0.04 |
| 4 | 101479155 | rs2013390 | 0.288 | 0.914 | -0.427 | -0.783 | -0.027 | -0.089 | -0.089 | 0.02 | 0.32 | 0.12 | 0.55 | 0.53 | 0.33 | 0.42 |
| 4 | 101486889 | rs2651581 | -0.153 | 0.898 | -0.343 | -0.883 | 0.054 | 0.071 | 0.071 | 0.876 | 0.46 | 0.75 | 0.32 | 0.26 | 0.42 | 0.38 |
| 4 | 101488171 | rs17030274 | -0.341 | 0.774 | 0.232 | -0.868 | 0.044 | 0.02 | 0.02 | 0 | 0.06 | 0.31 | 0.02 | 0 | 0.03 | 0.03 |
| 4 | 101494460 | rs11936939 | 0.25 | 0.66 | 0.313 | -0.957 | 0.016 | 0.015 | 0.015 | 0.184 | 0.4 | 0.37 | 0.29 | 0.28 | 0.38 | 0.32 |
| 4 | 101494857 | rs1827303 | -0.569 | 0.78 | 0.214 | -0.856 | 0.027 | -0.032 | -0.032 | 0 | 0.72 | 0.61 | 0.86 | 0.82 | 0.71 | 0.72 |
| 4 | 101501761 | rs2567388 | -0.747 | 0.467 | 0.347 | -0.966 | 0.004 | 0.127 | 0.127 | 0 | 0.15 | 0.38 | 0.13 | 0.28 | 0.13 | 0.13 |
| 4 | 101503277 | rs6848407 | 1.414 | 0.377 | 0.133 | -1.11 | 0.033 | 0.036 | 0.036 | 2.395 | 0.27 | 0.19 | 0.15 | 0.02 | 0.2 | 0.2 |
| 4 | 101503451 | rs2651555 | -1.474 | 0.489 | 0.06 | -1.132 | -0.039 | 0.006 | 0.006 | 0 | 0.78 | 0.92 | 0.89 | 0.82 | 0.75 | 0.76 |
| 4 | 101505168 | rs17030283 | 1.307 | 0.294 | 0.178 | -1.144 | -0.011 | -0.064 | -0.064 | 0 | 0.27 | 0.29 | 0.16 | 0 | 0.2 | 0.19 |
| 4 | 101506795 | rs2651562 | 0.396 | 0.486 | 0.187 | -0.956 | 0.004 | 0.067 | 0.067 | 0 | 0.09 | 0.09 | 0.1 | 0.28 | 0.11 | 0.11 |
| 4 | 101507299 | rs13148793 | 0.53 | 0.696 | 0.075 | -0.986 | -0.02 | -0.058 | -0.058 | 0 | 0.32 | 0.19 | 0.57 | 0.55 | 0.32 | 0.42 |
| 4 | 101508622 | rs2651569 | -0.373 | 0.415 | 0.056 | -1.007 | 0.001 | -0.001 | -0.001 | 0 | 0.6 | 0.57 | 0.52 | 0.34 | 0.55 | 0.41 |
| 4 | 101510054 | rs1501104 | -0.456 | -0.042 | 0.061 | -0.812 | 0.059 | 0.053 | 0.053 | 0.013 | 0.45 | 0.79 | 0.32 | 0.31 | 0.42 | 0.37 |
| 4 | 101510537 | rs6830402 | 0.234 | -0.059 | 0.193 | -0.862 | 0 | -0.044 | -0.044 | 0 | 0.36 | 0.67 | 0.21 | 0.04 | 0.3 | 0.26 |
| 4 | 101511108 | rs2567373 | 0.697 | 0.015 | 0.09 | -0.671 | -0.002 | 0.014 | 0.014 | 0 | 0.08 | 0.06 | 0.1 | 0.24 | 0.11 | 0.1 |
| 4 | 101512875 | rs17030327 | 0.234 | -0.12 | 0.116 | -0.795 | 0.003 | -0.025 | -0.025 | 0 | 0.36 | 0.67 | 0.21 | 0.07 | 0.3 | 0.27 |
| 4 | 101514587 | rs6857393 | 1.785 | -0.313 | 0.344 | -0.909 | -0.005 | -0.046 | -0.046 | 0.891 | 0.22 | 0.24 | 0.12 | 0 | 0.18 | 0.16 |
| 4 | 101515453 | rs7671093 | 0.023 | -0.295 | 0.466 | -0.881 | 0.004 | 0.004 | 0.004 | 0 | 0.59 | 0.37 | 0.71 | 0.72 | 0.62 | 0.66 |
| 4 | 101523312 | rs2866217 | 0.672 | -0.337 | 0.08 | -0.896 | -0.013 | -0.04 | -0.04 | 0 | 0.32 | 0.19 | 0.56 | 0.53 | 0.31 | 0.44 |
| 4 | 101525948 | rs2866223 | 0.747 | -0.756 | 0.118 | -0.837 | -0.002 | 0.03 | 0.03 | 0.38 | 0.82 | 0.76 | 0.85 | 0.72 | 0.83 | 0.81 |
| 4 | 101528207 | rs2866224 | 0.05 | -0.774 | 0.109 | -0.854 | -0.019 | -0.013 | -0.013 | 0.328 | 0.61 | 0.52 | 0.73 | 0.72 | 0.66 | 0.67 |
| 4 | 101534643 | rs12641140 | 0.978 | -0.22 | -0.109 | -0.427 | -0.002 | 0.03 | 0.03 | 0.38 | 0.82 | 0.74 | 0.85 | 0.72 | 0.83 | 0.81 |
| 4 | 101551216 | rs7658792 | -0.742 | 0.039 | -0.454 | -0.314 | -0.003 | -0.006 | -0.006 | 0 | 0.91 | 0.94 | 0.91 | 0.83 | 0.89 | 0.9 |
| 4 | 101568956 | rs10025419 | -0.174 | 0.039 | -0.468 | -0.335 | -0.008 | -0.013 | -0.013 | 0.745 | 0.89 | 0.92 | 0.9 | 0.83 | 0.85 | 0.9 |
| 4 | 101573688 | rs9759759 | -0.299 | 0.014 | -0.329 | -0.426 | 0.002 | 0.015 | 0.015 | 0 | 0.89 | 0.83 | 0.9 | 0.84 | 0.88 | 0.9 |
| 4 | 101579340 | rs13118526 | 1.369 | 0.21 | -0.829 | -0.615 | -0.068 | 0.017 | 0.017 | 0 | 0.23 | 0.06 | 0.13 | 0.18 | 0.3 | 0.18 |
| 4 | 101579972 | rs4586953 | 0.509 | -0.047 | -0.711 | -0.615 | -0.007 | -0.046 | -0.046 | 0 | 0.32 | 0.18 | 0.56 | 0.51 | 0.3 | 0.44 |
| 4 | 101586719 | rs13131732 | -0.669 | 0.104 | -0.739 | -0.549 | -0.001 | 0 | 0 | 0 | 0.9 | 0.92 | 0.9 | 0.84 | 0.88 | 0.9 |
| 4 | 101595019 | rs7693246 | -0.765 | 0.123 | -0.736 | -0.54 | -0.001 | 0 | 0 | 0.01 | 0.91 | 0.92 | 0.9 | 0.85 | 0.88 | 0.9 |
| 4 | 101599854 | rs6848941 | 1.282 | 0.144 | -0.806 | -0.657 | -0.042 | 0.022 | 0.022 | 0 | 0.24 | 0.1 | 0.26 | 0.18 | 0.3 | 0.19 |
| 4 | 101599908 | rs10516468 | 0.201 | 0.085 | -0.609 | -0.473 | -0.006 | -0.045 | -0.045 | 0.051 | 0.33 | 0.19 | 0.43 | 0.53 | 0.3 | 0.41 |
| 4 | 101616281 | rs10030146 | 0.859 | 0.083 | -0.714 | -0.664 | -0.027 | 0.04 | 0.04 | 0 | 0.29 | 0.19 | 0.18 | 0.15 | 0.33 | 0.19 |
| 4 | 101619817 | rs925169 | -0.546 | -0.006 | -0.702 | -0.725 | -0.007 | 0.005 | 0.005 | 0.571 | 0.77 | 0.78 | 0.81 | 0.83 | 0.76 | 0.82 |
| 4 | 101627782 | rs7691456 | 0.093 | -0.095 | -0.73 | -0.762 | 0.038 | 0.044 | 0.044 | 0 | 0.5 | 0.61 | 0.64 | 0.68 | 0.44 | 0.65 |
| 4 | 101627867 | rs7667413 | 0.852 | -0.251 | -0.469 | -0.82 | -0.004 | 0.039 | 0.039 | 0 | 0.22 | 0.24 | 0.48 | 0.53 | 0.2 | 0.38 |
| 4 | 101634693 | rs1396287 | -0.289 | 0.493 | -0.37 | -0.908 | 0.036 | 0.059 | 0.059 | 0 | 0.92 | 0.73 | 0.89 | 0.87 | 0.89 | 0.92 |
| 4 | 101640802 | rs10008030 | 0.834 | 0.435 | -0.468 | -1.003 | -0.046 | 0.072 | 0.072 | 0 | 0.28 | 0.14 | 0.18 | 0.11 | 0.32 | 0.18 |
| 4 | 101644204 | rs9995730 | -0.469 | 0.308 | -0.41 | -1.027 | 0.001 | 0.009 | 0.009 | 0 | 0.46 | 0.42 | 0.29 | 0.16 | 0.46 | 0.28 |
| 4 | 101651113 | rs981272 | -0.63 | 0.333 | -0.592 | -1.027 | 0.009 | 0.045 | 0.045 | 0 | 0.48 | 0.41 | 0.3 | 0.15 | 0.5 | 0.29 |
| 4 | 101651209 | rs17572230 | 1.286 | 0.833 | -1.011 | -0.822 | -0.06 | 0.027 | 0.027 | 0.16 | 0.14 | 0.01 | 0.09 | 0.04 | 0.14 | 0.09 |
| 4 | 101651417 | rs981271 | 0.601 | 0.605 | -0.763 | -0.866 | -0.041 | 0.068 | 0.068 | 0 | 0.28 | 0.13 | 0.17 | 0.11 | 0.32 | 0.18 |
| 4 | 101683999 | rs1508421 | 0.952 | 0.948 | 0.029 | -0.363 | -0.001 | 0 | 0 | 0 | 0.72 | 0.74 | 0.81 | 0.84 | 0.74 | 0.69 |
| 4 | 101698883 | rs6839885 | -0.346 | 1.028 | -0.397 | -0.353 | -0.074 | 0.017 | 0.017 | 1.385 | 0.82 | 0.98 | 0.86 | 0.92 | 0.8 | 0.85 |
| 4 | 101699382 | rs2089540 | 0.819 | 0.994 | -0.186 | -0.267 | 0.006 | 0.006 | 0.006 | 1.716 | 0.7 | 0.58 | 0.77 | 0.71 | 0.74 | 0.66 |
| 4 | 101712104 | rs10010623 | 1.107 | 0.949 | -0.433 | -0.974 | 0.001 | 0.08 | 0.08 | 0 | 0.55 | 0.3 | 0.6 | 0.91 | 0.56 | 0.61 |
| 4 | 101713756 | rs12507027 | 0.296 | 1.424 | -0.055 | -0.547 | 0.005 | -0.01 | -0.01 | 0 | 0.84 | 0.8 | 0.79 | 0.91 | 0.79 | 0.89 |
| 4 | 101714921 | rs6853889 | 1.056 | 1.501 | 0.189 | -0.413 | 0.044 | -0.001 | -0.001 | 0 | 0.73 | 0.41 | 0.67 | 0.73 | 0.63 | 0.77 |
| 4 | 101722485 | rs4699824 | 1.057 | 0.765 | 0.35 | -0.779 | -0.001 | -0.001 | -0.001 | 0.355 | 0.66 | 0.29 | 0.69 | 0.81 | 0.69 | 0.79 |
| 4 | 101725061 | rs1567282 | 1.122 | 1.067 | 0.25 | -0.645 | -0.037 | 0.004 | 0.004 | 0 | 0.88 | 0.94 | 0.82 | 0.91 | 0.8 | 0.9 |
| 4 | 101728821 | rs10008442 | -0.53 | 0.619 | 0.72 | -1.043 | -0.014 | -0.132 | -0.132 | 0 | 0.34 | 0.47 | 0.26 | 0.05 | 0.28 | 0.17 |
| 4 | 101742816 | rs13130138 | -0.692 | 0.614 | 0.808 | -1.061 | -0.009 | -0.113 | -0.113 | 0 | 0.36 | 0.55 | 0.29 | 0.04 | 0.3 | 0.18 |
| 4 | 101748186 | rs6419156 | -0.158 | 0.562 | 0.863 | -1.06 | -0.036 | -0.137 | -0.137 | 1.959 | 0.29 | 0.45 | 0.18 | 0.04 | 0.19 | 0.14 |
| 4 | 101766650 | rs17030500 | -1.509 | 0.394 | 0.962 | -0.952 | 0.032 | 0.051 | 0.051 | 0.147 | 0.07 | 0.41 | 0.06 | 0.04 | 0.01 | 0.05 |

| | | | | | | | | | | | | | | | | |
|---|---|---|---|---|---|---|---|---|---|---|---|---|---|---|---|---|
| 4 | 101774201 | rs6847657 | -0.687 | 0.425 | 0.707 | -0.977 | -0.007 | -0.071 | -0.071 | 1.391 | 0.4 | 0.52 | 0.31 | 0.1 | 0.36 | 0.28 |
| 4 | 101798797 | rs11725047 | 0.754 | 0.797 | 1.073 | -0.649 | -0.002 | -0.004 | -0.004 | 0 | 0.05 | 0.04 | 0.11 | 0.11 | 0.11 | 0.04 |
| 4 | 101799187 | rs4699162 | -0.029 | 0.466 | 0.86 | -0.919 | 0.013 | 0.009 | 0.009 | 0 | 0.16 | 0.07 | 0.12 | 0.1 | 0.13 | 0.15 |
| 4 | 101800766 | rs978155 | -1.36 | 0.523 | 0.877 | -0.929 | -0.011 | -0.061 | -0.061 | 0.621 | 0.24 | 0.48 | 0.23 | 0.1 | 0.22 | 0.22 |
| 4 | 101804019 | rs12646623 | -0.049 | 0.055 | 0.925 | -0.785 | 0.013 | 0.008 | 0.008 | 0 | 0.15 | 0.07 | 0.12 | 0.1 | 0.13 | 0.15 |
| 4 | 101851136 | rs13121097 | 0.011 | 0.035 | 0.925 | -0.788 | 0.01 | 0.011 | 0.011 | 0 | 0.15 | 0.08 | 0.1 | 0.07 | 0.13 | 0.15 |
| 4 | 101851511 | rs10001585 | -0.981 | 0.018 | 0.876 | -0.795 | -0.062 | -0.052 | -0.052 | 0 | 0.21 | 0.5 | 0.16 | 0.1 | 0.13 | 0.19 |
| 4 | 101855402 | rs17826594 | 1.004 | 0.069 | 0.341 | -0.903 | -0.03 | 0.093 | 0.093 | 0.038 | 0.26 | 0.08 | 0.19 | 0 | 0.27 | 0.11 |
| 4 | 101858763 | rs4698838 | -0.315 | 0.058 | 0.56 | -0.403 | -0.024 | -0.011 | -0.011 | 0 | 0.16 | 0.27 | 0.1 | 0.09 | 0.12 | 0.14 |
| 4 | 101867277 | rs4698981 | -0.05 | 0.06 | -0.23 | -0.402 | 0.019 | 0.003 | 0.003 | 0 | 0.15 | 0.04 | 0.09 | 0.09 | 0.12 | 0.14 |
| 4 | 101868169 | rs17030661 | -0.63 | 0.133 | -0.333 | -0.368 | -0.055 | -0.02 | -0.02 | 0 | 0.2 | 0.54 | 0.15 | 0.12 | 0.12 | 0.15 |
| 4 | 101868917 | rs4626209 | -0.348 | 0.743 | -0.724 | 0.638 | -0.018 | -0.02 | -0.02 | 0 | 0.65 | 0.85 | 0.76 | 0.59 | 0.63 | 0.47 |
| 4 | 101881356 | rs979665 | -0.145 | 0.697 | -0.691 | 0.25 | 0.001 | -0.003 | -0.003 | 0.786 | 0.67 | 0.67 | 0.48 | 0.25 | 0.6 | 0.46 |
| 4 | 101886921 | rs2645370 | -0.484 | 0.649 | -1.197 | 0.386 | 0.006 | 0.018 | 0.018 | 0 | 0.22 | 0.09 | 0.12 | 0.17 | 0.18 | 0.12 |
| 4 | 101896676 | rs12649554 | -0.053 | 0.538 | -0.524 | 0.894 | -0.001 | 0.037 | 0.037 | 0 | 0.7 | 0.6 | 0.48 | 0.41 | 0.66 | 0.56 |
| 4 | 101904475 | rs2645373 | 1.187 | 0.974 | -0.046 | 1.435 | 0.034 | 0.129 | 0.129 | 0 | 0.87 | 0.77 | 0.63 | 0.54 | 0.82 | 0.79 |
| 4 | 101906493 | rs4699014 | -0.37 | 0.207 | -0.823 | 1.325 | 0.033 | -0.069 | -0.069 | 0 | 0.14 | 0.02 | 0.1 | 0.26 | 0.09 | 0.29 |
| 4 | 101924060 | rs2851048 | 1.297 | -0.286 | -0.674 | 0.137 | 0 | 0.091 | 0.091 | 0.272 | 0.48 | 0.25 | 0.38 | 0.27 | 0.53 | 0.21 |
| 4 | 101933417 | rs2851057 | 0.998 | -0.348 | 0.184 | -0.06 | 0 | 0 | 0 | 0.052 | 0.6 | 0.43 | 0.47 | 0.33 | 0.66 | 0.36 |
| 4 | 101942483 | rs10000538 | -0.877 | -0.203 | 0.551 | -0.794 | -0.004 | 0 | 0 | 0.056 | 0.28 | 0.28 | 0.21 | 0.03 | 0.29 | 0.38 |
| 4 | 101945139 | rs6826912 | -1.557 | -0.355 | 1.265 | -0.829 | 0.034 | -0.019 | -0.019 | 0 | 0.08 | 0.33 | 0.06 | 0.01 | 0.05 | 0.07 |
| 4 | 101946948 | rs1042094 | -1.816 | -0.246 | 1.569 | -0.622 | 0.006 | 0.031 | 0.031 | 0.113 | 0.12 | 0.36 | 0.06 | 0.11 | 0.08 | 0.23 |
| 4 | 101953182 | rs1880942 | 2.083 | -0.798 | 0.961 | -1.281 | -0.027 | 0.078 | 0.078 | 0.433 | 0.39 | 0.19 | 0.29 | 0 | 0.46 | 0.17 |
| 4 | 101953322 | rs2851060 | 0.294 | -0.717 | 0.783 | -1.213 | -0.004 | 0.035 | 0.035 | 0 | 0.75 | 0.91 | 0.54 | 0.13 | 0.75 | 0.7 |
| 4 | 101958617 | rs2037773 | 0.248 | -0.829 | 0.758 | -1.252 | -0.008 | 0.048 | 0.048 | 0 | 0.19 | 0.12 | 0.15 | 0.02 | 0.21 | 0.27 |
| 4 | 101965295 | rs10025768 | -0.566 | -0.749 | 0.89 | -1.135 | 0.005 | -0.021 | -0.021 | 1.257 | 0.06 | 0.11 | 0.04 | 0.01 | 0.05 | 0.07 |
| 4 | 101970512 | rs2178006 | -1.955 | -0.909 | 0.886 | -1.543 | 0.003 | -0.059 | -0.059 | 0.041 | 0.58 | 0.49 | 0.7 | 0.9 | 0.55 | 0.66 |
| 4 | 101971451 | rs1358312 | -0.039 | -0.698 | 1.282 | -1.063 | 0.005 | 0.028 | 0.028 | 0.058 | 0.24 | 0.16 | 0.17 | 0.02 | 0.23 | 0.28 |
| 4 | 101983607 | rs6851231 | -1.191 | -0.692 | 1.577 | -1.065 | -0.001 | -0.054 | -0.054 | 0 | 0.35 | 0.65 | 0.26 | 0.03 | 0.3 | 0.39 |
| 4 | 101989580 | rs10020845 | -0.887 | -0.485 | 1.426 | -0.68 | 0.013 | -0.024 | -0.024 | 0 | 0.07 | 0.14 | 0.04 | 0.01 | 0.05 | 0.07 |
| 4 | 101996812 | rs17030739 | -1.635 | -0.458 | 1.178 | -0.8 | -0.015 | -0.129 | -0.129 | 0 | 0.2 | 0.54 | 0.13 | 0.01 | 0.14 | 0.11 |
| 4 | 102002123 | rs3804357 | -0.482 | -0.303 | 0.882 | -0.561 | -0.002 | 0.003 | 0.003 | 0 | 0.34 | 0.27 | 0.22 | 0.18 | 0.33 | 0.56 |
| 4 | 102010038 | rs17030741 | -1.738 | -0.018 | 1.307 | -0.155 | -0.003 | -0.087 | -0.087 | 1.111 | 0.18 | 0.36 | 0.11 | 0.01 | 0.14 | 0.1 |
| 4 | 102016486 | rs17030746 | -1.209 | 0.051 | 1.412 | -0.018 | 0.085 | 0.009 | 0.009 | 0 | 0.06 | 0.22 | 0.05 | 0.01 | 0.07 | 0.05 |
| 4 | 102021290 | rs17030753 | -1.472 | 0.021 | 1.191 | -0.039 | 0.001 | -0.079 | -0.079 | 0.048 | 0.16 | 0.36 | 0.09 | 0.01 | 0.11 | 0.1 |
| 4 | 102026137 | rs17030758 | -1.172 | -0.401 | 1.307 | 0.034 | -0.041 | -0.093 | -0.093 | 0 | 0.13 | 0.42 | 0.09 | 0 | 0.05 | 0.05 |
| 4 | 102031775 | rs2732510 | -1.691 | -0.612 | 1.191 | 0.647 | -0.061 | 0.015 | 0.015 | 0 | 0.22 | 0.4 | 0.17 | 0.27 | 0.11 | 0.13 |
| 4 | 102033997 | rs2732512 | -1.265 | -0.578 | 1.293 | 0.691 | -0.003 | 0.154 | 0.154 | 0 | 0.11 | 0.31 | 0.12 | 0.27 | 0.06 | 0.08 |
| 4 | 102040091 | rs12639641 | -0.818 | -0.426 | 0.956 | 1.593 | -0.015 | 0.115 | 0.115 | 0 | 0.08 | 0.2 | 0.31 | 0.31 | 0.03 | 0.22 |
| 4 | 102041236 | rs6820738 | -2.941 | -0.129 | 0.394 | 1.392 | 0.022 | 0.008 | 0.008 | 0 | 0.41 | 0.84 | 0.56 | 0.62 | 0.3 | 0.6 |
| 4 | 102048076 | rs17030795 | -0.695 | 0.352 | 0.264 | 1.414 | -0.01 | -0.039 | -0.039 | 0 | 0.16 | 0.14 | 0.44 | 0.31 | 0.2 | 0.29 |
| 4 | 102061730 | rs2732518 | -0.879 | 0.423 | 0.733 | 1.265 | -0.002 | 0.165 | 0.165 | 0 | 0.11 | 0.42 | 0.12 | 0.25 | 0.04 | 0.06 |
| 4 | 102065589 | rs10516472 | -0.365 | 0.762 | -0.052 | 0.997 | -0.028 | -0.011 | -0.011 | 0.015 | 0.08 | 0.02 | 0.04 | 0.06 | 0.12 | 0.17 |
| 4 | 102074134 | rs1405688 | 0.765 | 0.596 | 0.065 | 0.922 | -0.04 | 0.048 | 0.048 | 0 | 0.09 | 0.01 | 0.1 | 0 | 0.15 | 0.03 |
| 4 | 102084331 | rs1527351 | -2.069 | 0.648 | 0.563 | 1.704 | -0.002 | 0 | 0 | 0 | 0.35 | 0.63 | 0.35 | 0.4 | 0.34 | 0.39 |
| 4 | 102089849 | rs13142024 | 2.332 | 0.69 | 0.844 | 2.055 | -0.012 | 0.028 | 0.028 | 0 | 0.59 | 0.28 | 0.38 | 0.27 | 0.59 | 0.47 |
| 4 | 102094453 | rs4699110 | 0.072 | 0.678 | 0.767 | 2.256 | -0.01 | -0.007 | -0.007 | 0 | 0.06 | 0.03 | 0.12 | 0.18 | 0.09 | 0.19 |
| 4 | 102094764 | rs2850979 | 1.128 | 0.698 | 1.018 | 2.302 | -0.006 | -0.041 | -0.041 | 0.241 | 0.71 | 0.52 | 0.78 | 0.74 | 0.76 | 0.8 |
| 4 | 102095497 | rs2659504 | -0.969 | 0.814 | 0.825 | 2.282 | 0.02 | 0.174 | 0.174 | 0.013 | 0.07 | 0.2 | 0.28 | 0.29 | 0.08 | 0.14 |
| 4 | 102098066 | rs17030887 | -0.325 | 0.812 | 0.681 | 2.044 | -0.005 | 0.003 | 0.003 | 0 | 0.06 | 0.04 | 0.12 | 0.19 | 0.09 | 0.19 |
| 4 | 102115186 | rs1395475 | -2.191 | 0.892 | 0.899 | 2.016 | -0.001 | 0.127 | 0.127 | 1.249 | 0.12 | 0.61 | 0.33 | 0.25 | 0.1 | 0.15 |
| 4 | 102122755 | rs2850992 | 0.984 | 0.71 | 0.359 | 2.157 | -0.123 | -0.063 | -0.063 | 0 | 0.74 | 0.98 | 0.73 | 0.63 | 0.67 | 0.62 |
| 4 | 102158450 | rs2850950 | 1.676 | 0.864 | 1.284 | 2.699 | -0.057 | -0.078 | -0.078 | 2.045 | 0.86 | 0.98 | 0.83 | 0.61 | 0.81 | 0.65 |
| 4 | 102165917 | rs17030994 | -1.494 | 0.471 | 1.814 | 2.386 | 0.046 | 0.042 | 0.042 | 2.742 | 0.05 | 0.4 | 0.03 | 0.01 | 0 | 0.02 |
| 4 | 102168234 | rs2850373 | -0.553 | 0.585 | 1.78 | 2.514 | 0.024 | 0.211 | 0.211 | 0 | 0.05 | 0.12 | 0.3 | 0.29 | 0.07 | 0.13 |
| 4 | 102176098 | rs2850962 | -1.167 | 0.267 | 1.341 | 1.718 | -0.021 | -0.018 | -0.018 | 0.642 | 0.1 | 0.01 | 0.06 | 0.17 | 0.1 | 0.15 |
| 4 | 102261268 | rs17031191 | -1.064 | 0.125 | 1.566 | 1.83 | -0.02 | -0.031 | -0.031 | 0 | 0.08 | 0 | 0.07 | 0.16 | 0.06 | 0.14 |
| 4 | 102301819 | rs13135111 | 0.581 | 0.009 | 1.748 | 1.925 | -0.005 | -0.006 | -0.006 | 0 | 0.75 | 0.81 | 0.56 | 0.49 | 0.72 | 0.57 |
| 4 | 102323959 | rs10015209 | -0.824 | -0.038 | 2.204 | 1.921 | -0.009 | 0.017 | 0.017 | 5.789 | 0.06 | 0.3 | 0.03 | 0.03 | 0.01 | 0 |
| 4 | 102325626 | rs17031254 | -0.709 | -0.038 | 2.201 | 1.921 | -0.007 | 0.019 | 0.019 | 5.937 | 0.06 | 0.3 | 0.03 | 0.03 | 0.01 | 0 |
| 4 | 102333865 | rs2044044 | 0.44 | -0.051 | 2.278 | 1.938 | 0.007 | 0.02 | 0.02 | 0 | 0.76 | 0.77 | 0.55 | 0.49 | 0.72 | 0.58 |
| 4 | 102343571 | rs4699215 | 1.061 | -0.081 | 2.143 | 1.916 | 0.008 | 0.034 | 0.034 | 0.689 | 0.69 | 0.41 | 0.51 | 0.45 | 0.71 | 0.62 |
| 4 | 102353934 | rs990960 | 1.071 | -0.08 | 1.866 | 1.917 | -0.002 | -0.007 | -0.007 | 0 | 0.69 | 0.32 | 0.51 | 0.45 | 0.71 | 0.62 |
| 4 | 102382498 | rs4398526 | -0.409 | -0.205 | 1.705 | 1.626 | -0.027 | 0.002 | 0.002 | 6.151 | 0.09 | 0.45 | 0.05 | 0.03 | 0.01 | 0.02 |
| 4 | 102387446 | rs6855746 | -0.699 | -0.219 | 1.753 | 1.619 | -0.043 | -0.013 | -0.013 | 0.484 | 0.1 | 0.53 | 0.05 | 0.03 | 0.01 | 0.02 |
| 4 | 102395086 | rs17031304 | -0.685 | -0.217 | 1.754 | 1.62 | -0.047 | -0.016 | -0.016 | 0 | 0.1 | 0.53 | 0.05 | 0.03 | 0.01 | 0.02 |
| 4 | 102396292 | rs9307257 | -1.666 | -0.326 | 1.314 | 1.433 | 0.004 | 0.178 | 0.178 | 0.098 | 0.53 | 0.87 | 0.7 | 0.76 | 0.54 | 0.56 |
| 4 | 102399209 | rs6532939 | -1.495 | -0.163 | 1.66 | 1.632 | -0.002 | -0.004 | -0.004 | 0.102 | 0.36 | 0.67 | 0.54 | 0.6 | 0.34 | 0.48 |
| 4 | 102407109 | rs1863575 | -1.586 | -0.127 | 1.503 | 1.638 | -0.007 | -0.013 | -0.013 | 0.118 | 0.37 | 0.74 | 0.54 | 0.6 | 0.34 | 0.49 |

| | | | | | | | | | | | | | | | | |
|---|---|---|---|---|---|---|---|---|---|---|---|---|---|---|---|---|
| 4 | 102427214 | rs6825984 | -0.734 | -0.278 | 1.392 | 1.565 | -0.055 | 0.023 | 0.023 | 6.127 | 0.11 | 0.56 | 0.06 | 0.1 | 0.02 | 0.03 |
| 4 | 102432702 | rs890448 | -0.717 | 0.056 | 0.62 | 1.593 | -0.011 | -0.015 | -0.015 | 0.01 | 0.26 | 0.2 | 0.51 | 0.59 | 0.33 | 0.48 |
| 4 | 102449905 | rs1426538 | -1.001 | -0.128 | 0.268 | 1.31 | -0.025 | -0.039 | -0.039 | 0.891 | 0.37 | 0.71 | 0.57 | 0.55 | 0.4 | 0.53 |
| 4 | 102451022 | rs13434873 | -1.19 | -0.868 | 0.663 | 0.836 | -0.015 | 0 | 0 | 0 | 0.63 | 0.78 | 0.45 | 0.66 | 0.59 | 0.66 |
| 4 | 102470700 | rs10022442 | 0.418 | -0.748 | 0.736 | 0.931 | -0.048 | -0.027 | -0.027 | 0 | 0.35 | 0.18 | 0.28 | 0.36 | 0.44 | 0.35 |
| 4 | 102484972 | rs17031441 | -0.34 | -0.642 | 0.919 | 0.921 | -0.008 | 0.028 | 0.028 | 0 | 0.17 | 0.28 | 0.17 | 0.28 | 0.12 | 0.23 |
| 4 | 102492986 | rs2723054 | 0.807 | -0.506 | 0.05 | 0.945 | -0.023 | 0.01 | 0.01 | 0 | 0.39 | 0.13 | 0.52 | 0.32 | 0.42 | 0.37 |
| 4 | 102508948 | rs2723058 | 0.295 | -0.147 | -0.529 | 1.12 | -0.018 | -0.002 | -0.002 | 0.104 | 0.74 | 0.32 | 0.79 | 0.69 | 0.86 | 0.75 |
| 4 | 102515133 | rs2615973 | 0.094 | 1.022 | -0.769 | 1.179 | -0.028 | 0.007 | 0.007 | 0 | 0.38 | 0.14 | 0.52 | 0.32 | 0.42 | 0.37 |
| 4 | 102520862 | rs2717833 | 1.159 | 1.152 | 0.315 | 1.478 | 0.041 | 0.086 | 0.086 | 0.05 | 0.94 | 0.89 | 0.9 | 0.77 | 0.9 | 0.89 |
| 4 | 102525045 | rs7661775 | -1.53 | 0.726 | 0.25 | 1.242 | 0.001 | 0.036 | 0.036 | 0 | 0.06 | 0.13 | 0.05 | 0.2 | 0.07 | 0.12 |
| 4 | 102526313 | rs11930552 | -1.068 | 0.551 | 0.42 | 0.913 | 0.002 | -0.038 | -0.038 | 0.378 | 0.08 | 0.23 | 0.07 | 0.01 | 0.04 | 0.04 |
| 4 | 102545130 | rs17031472 | -1.398 | 0.414 | 0.299 | 1.506 | -0.016 | 0.079 | 0.079 | 0 | 0.07 | 0.12 | 0.04 | 0.29 | 0.06 | 0.18 |
| 4 | 102548422 | rs12643950 | -1.493 | 0.439 | 0.498 | 1.583 | -0.012 | 0.079 | 0.079 | 0.016 | 0.07 | 0.15 | 0.04 | 0.24 | 0.06 | 0.2 |
| 4 | 102551731 | rs17031503 | -0.869 | -0.288 | 1.128 | 1.756 | -0.035 | 0.024 | 0.024 | 0.493 | 0.17 | 0.35 | 0.14 | 0.17 | 0.1 | 0.1 |
| 4 | 102560877 | rs7666463 | -0.752 | -0.174 | 0.865 | 1.075 | -0.002 | -0.044 | -0.044 | 0 | 0.14 | 0.14 | 0.11 | 0.01 | 0.13 | 0.05 |
| 4 | 102563794 | rs4699246 | -0.649 | 0.316 | 0.714 | 1.018 | 0.003 | 0.026 | 0.026 | 0 | 0.28 | 0.4 | 0.21 | 0.43 | 0.31 | 0.54 |
| 4 | 102565480 | rs7663310 | -0.695 | 0.238 | 0.766 | 0.964 | 0.003 | -0.006 | -0.006 | 0 | 0.28 | 0.42 | 0.2 | 0.32 | 0.31 | 0.56 |
| 4 | 102566810 | rs2016504 | 0.05 | 0.204 | 0.525 | 0.886 | -0.002 | 0.028 | 0.028 | 0 | 0.08 | 0.11 | 0.11 | 0.18 | 0.1 | 0.03 |
| 4 | 102569368 | rs2866627 | -0.663 | 0.267 | -0.397 | 0.551 | 0.015 | 0.107 | 0.107 | 0 | 0.56 | 0.74 | 0.53 | 0.77 | 0.64 | 0.65 |
| 4 | 102569411 | rs1009257 | -0.226 | 0.021 | -0.427 | 0.736 | -0.038 | -0.038 | -0.038 | 0 | 0.43 | 0.24 | 0.33 | 0.49 | 0.5 | 0.58 |
| 4 | 102570449 | rs10009721 | -0.659 | 0.659 | -1.135 | 0.392 | -0.071 | 0.067 | 0.067 | 0 | 0.72 | 0.93 | 0.75 | 0.83 | 0.69 | 0.5 |
| 4 | 102570753 | rs17250305 | 1.457 | 0.049 | -1.809 | -0.225 | -0.003 | 0.094 | 0.094 | 0.017 | 0.22 | 0.07 | 0.15 | 0.06 | 0.19 | 0.19 |
| 4 | 102571890 | rs7681723 | -1.109 | 0.212 | -1.645 | -0.06 | -0.065 | 0.046 | 0.046 | 0 | 0.72 | 0.93 | 0.58 | 0.81 | 0.69 | 0.52 |
| 4 | 102572251 | rs7682523 | -0.612 | 0.221 | -1.791 | -0.078 | 0.012 | 0.092 | 0.092 | 0 | 0.44 | 0.8 | 0.37 | 0.71 | 0.43 | 0.31 |
| 4 | 102574367 | rs7676236 | -0.937 | 0.551 | -1.581 | 0.244 | -0.068 | 0.015 | 0.015 | 0 | 0.72 | 0.93 | 0.58 | 0.77 | 0.69 | 0.5 |
| 4 | 102603328 | rs2866422 | -0.187 | 0.4 | -0.329 | 0.86 | 0 | -0.004 | -0.004 | 0 | 0.25 | 0.26 | 0.14 | 0.11 | 0.27 | 0.12 |
| 4 | 102606843 | rs7669395 | -0.834 | 0.754 | -0.352 | 1.345 | -0.001 | -0.001 | -0.001 | 0 | 0.25 | 0.27 | 0.24 | 0.43 | 0.26 | 0.23 |
| 4 | 102608288 | rs17251102 | 0.244 | 0.745 | -0.242 | 0.605 | 0.003 | 0.044 | 0.044 | 0.639 | 0.49 | 0.38 | 0.33 | 0.19 | 0.5 | 0.4 |
| 4 | 102613885 | rs12644085 | 0.868 | 0.695 | -0.067 | 1.066 | -0.01 | 0.067 | 0.067 | 0 | 0.07 | 0.08 | 0.12 | 0.31 | 0.04 | 0.09 |
| 4 | 102614169 | rs7669524 | 0.228 | 0.688 | -0.08 | 0.709 | 0.006 | -0.016 | -0.016 | 0.023 | 0.19 | 0.25 | 0.12 | 0.09 | 0.23 | 0.09 |
| 4 | 102616166 | rs11723683 | 0.15 | 0.566 | 0.071 | 0.705 | 0 | -0.009 | -0.009 | 0 | 0.48 | 0.51 | 0.34 | 0.2 | 0.49 | 0.39 |
| 4 | 102624280 | rs13149930 | -1.484 | 0.644 | -0.72 | 0.879 | -0.022 | -0.053 | -0.053 | 0 | 0.69 | 0.9 | 0.61 | 0.64 | 0.69 | 0.58 |
| 4 | 102634710 | rs17252142 | 1.666 | 0.724 | -0.813 | 0.479 | -0.082 | 0.119 | 0.119 | 0.117 | 0.29 | 0.01 | 0.21 | 0.11 | 0.27 | 0.31 |
| 4 | 102635952 | rs768290 | 0.216 | 0.919 | -0.268 | 0.784 | -0.006 | 0.01 | 0.01 | 0 | 0.18 | 0.13 | 0.11 | 0.09 | 0.22 | 0.1 |
| 4 | 102638137 | rs12512546 | 1.057 | 0.9 | -0.501 | 0.931 | -0.024 | -0.088 | -0.088 | 1.097 | 0.27 | 0.08 | 0.31 | 0.38 | 0.23 | 0.4 |
| 4 | 102638201 | rs2866430 | -0.477 | 1.011 | 0.048 | 0.969 | 0.003 | 0.142 | 0.142 | 0 | 0.16 | 0.34 | 0.21 | 0.4 | 0.19 | 0.18 |
| 4 | 102641428 | rs10011961 | -0.952 | 1.291 | -0.453 | 0.92 | -0.028 | 0.01 | 0.01 | 0.249 | 0.84 | 0.93 | 0.85 | 0.91 | 0.77 | 0.91 |
| 4 | 102642012 | rs12501586 | 0.705 | 1.337 | -0.451 | 0.912 | 0 | -0.004 | -0.004 | 0 | 0.35 | 0.34 | 0.39 | 0.37 | 0.31 | 0.4 |
| 4 | 102642526 | rs12506517 | 1.573 | 1.535 | 0.666 | 1.586 | 0.021 | 0.059 | 0.059 | 0 | 0.67 | 0.53 | 0.62 | 0.5 | 0.58 | 0.73 |
| 4 | 102656452 | rs13103794 | 0.886 | 1.64 | 0.642 | 1.504 | -0.019 | 0.02 | 0.02 | 0 | 0.07 | 0.02 | 0.06 | 0 | 0.11 | 0.04 |
| 4 | 102665975 | rs6818863 | 1.498 | 1.595 | 0.184 | 1.776 | 0.018 | 0.041 | 0.041 | 0 | 0.67 | 0.51 | 0.65 | 0.59 | 0.59 | 0.73 |
| 4 | 102675023 | rs7675129 | 1.45 | 1.561 | 0.164 | 1.566 | 0.016 | 0.048 | 0.048 | 0 | 0.67 | 0.58 | 0.63 | 0.5 | 0.58 | 0.73 |
| 4 | 102677450 | rs6815355 | -0.289 | 1.642 | -0.02 | 1.577 | -0.035 | 0.012 | 0.012 | 0.599 | 0.84 | 0.94 | 0.85 | 0.91 | 0.76 | 0.91 |
| 4 | 102679337 | rs871061 | 0.735 | 1.627 | -0.146 | 1.483 | 0.019 | 0.056 | 0.056 | 0 | 0.67 | 0.53 | 0.62 | 0.5 | 0.59 | 0.73 |
| 4 | 102696219 | rs10003889 | -0.871 | 1.523 | -0.378 | 1.052 | 0 | -0.004 | -0.004 | 0 | 0.07 | 0.1 | 0.05 | 0.04 | 0.07 | 0.04 |
| 4 | 102701232 | rs7695738 | -1.051 | 1.673 | -0.131 | 1.226 | 0.032 | 0.024 | 0.024 | 0 | 0.08 | 0.2 | 0.13 | 0.17 | 0.11 | 0.12 |
| 4 | 102701262 | rs7695470 | -1.105 | 1.447 | -0.129 | 1.201 | 0.001 | -0.001 | -0.001 | 0 | 0.32 | 0.68 | 0.31 | 0.35 | 0.31 | 0.27 |
| 4 | 102708235 | rs4371620 | 0.063 | 1.376 | -0.511 | 1.568 | 0 | -0.006 | -0.006 | 0 | 0.84 | 0.87 | 0.9 | 0.76 | 0.81 | 0.9 |
| 4 | 102709119 | rs7375516 | 0.36 | 1.381 | -0.734 | 2.223 | -0.004 | -0.007 | -0.007 | 0 | 0.93 | 0.97 | 0.9 | 0.76 | 0.94 | 0.8 |
| 4 | 102714886 | rs10031210 | 1.268 | 1.788 | -1.009 | 2.065 | -0.032 | -0.037 | -0.037 | 0.091 | 0.66 | 0.79 | 0.51 | 0.6 | 0.57 | 0.73 |
| 4 | 102718795 | rs11944613 | -1.069 | 1.853 | -0.838 | 2.214 | -0.008 | -0.005 | -0.005 | 0 | 0.24 | 0.21 | 0.21 | 0.28 | 0.31 | 0.14 |
| 4 | 102723640 | rs13146194 | -0.972 | 1.85 | -0.906 | 2.095 | -0.025 | 0.018 | 0.018 | 0 | 0.23 | 0.11 | 0.17 | 0.17 | 0.31 | 0.13 |
| 4 | 102726005 | rs4699262 | -0.985 | 1.826 | -0.906 | 2.099 | -0.023 | 0.004 | 0.004 | 0 | 0.23 | 0.11 | 0.17 | 0.19 | 0.31 | 0.13 |
| 4 | 102726073 | rs6833249 | 0.63 | 1.902 | -0.79 | 2.188 | -0.038 | -0.053 | -0.053 | 0 | 0.91 | 0.99 | 0.71 | 0.84 | 0.87 | 0.87 |
| 4 | 102736456 | rs13136297 | -1.969 | 1.742 | 0.223 | 2.121 | 0.021 | 0.043 | 0.043 | 0 | 0.28 | 0.4 | 0.46 | 0.39 | 0.38 | 0.28 |
| 4 | 102737250 | rs10028805 | -2.166 | 1.627 | 0.96 | 2.064 | 0.002 | 0.005 | 0.005 | 0 | 0.29 | 0.64 | 0.47 | 0.39 | 0.37 | 0.28 |
| 4 | 102739536 | rs1421627 | -1.336 | 1.615 | 1.157 | 1.909 | 0.038 | 0.02 | 0.02 | 0 | 0.22 | 0.43 | 0.23 | 0.26 | 0.3 | 0.19 |
| 4 | 102753408 | rs4637409 | -1.224 | 1.614 | 1.034 | 1.799 | 0.006 | -0.007 | -0.007 | 0 | 0.21 | 0.22 | 0.19 | 0.19 | 0.3 | 0.19 |
| 4 | 102764358 | rs17031708 | -1.197 | 1.455 | 0.758 | 1.96 | -0.004 | 0.013 | 0.013 | 0 | 0.15 | 0.2 | 0.08 | 0.23 | 0.13 | 0.12 |
| 4 | 102774964 | rs1421624 | -0.202 | 1.35 | 0.422 | 1.984 | -0.004 | -0.042 | -0.042 | 0 | 0.08 | 0.02 | 0.23 | 0.17 | 0.08 | 0.1 |
| 4 | 102776083 | rs6849308 | 0.014 | 0.947 | 0.857 | 2.083 | 0.001 | 0.055 | 0.055 | 0 | 0.06 | 0.2 | 0.04 | 0.11 | 0.03 | 0.17 |
| 4 | 102783351 | rs4572884 | -1.991 | 0.907 | 1.23 | 2.092 | -0.001 | 0 | 0 | 0.224 | 0.32 | 0.63 | 0.27 | 0.35 | 0.38 | 0.35 |
| 4 | 102786685 | rs13101528 | -1.41 | 0.902 | 1.223 | 2.077 | 0.028 | -0.015 | -0.015 | 0 | 0.25 | 0.42 | 0.23 | 0.24 | 0.34 | 0.18 |
| 4 | 102790613 | rs12331846 | -0.455 | 0.881 | 1.284 | 2.069 | -0.011 | 0.043 | 0.043 | 0 | 0.08 | 0.24 | 0.04 | 0.11 | 0.04 | 0.17 |
| 4 | 102790818 | rs12331849 | -2.717 | 0.875 | 1.24 | 2.243 | -0.002 | -0.001 | -0.001 | 0 | 0.41 | 0.78 | 0.52 | 0.59 | 0.46 | 0.45 |
| 4 | 102792085 | rs10516484 | 0.131 | 0.789 | 1.141 | 2.236 | -0.004 | -0.041 | -0.041 | 0 | 0.08 | 0.01 | 0.23 | 0.17 | 0.08 | 0.1 |
| 4 | 102792354 | rs17031755 | -0.491 | 0.789 | 1.153 | 2.232 | -0.009 | 0.086 | 0.086 | 0 | 0.09 | 0.17 | 0.26 | 0.24 | 0.08 | 0.1 |
| 4 | 102792983 | rs17031758 | -0.505 | 0.755 | 1.067 | 2.215 | -0.008 | 0.063 | 0.063 | 0 | 0.09 | 0.14 | 0.26 | 0.24 | 0.08 | 0.1 |

| | | | | | | | | | | | | | | | | |
|---|---|---|---|---|---|---|---|---|---|---|---|---|---|---|---|---|
| 4 | 102821381 | rs12505747 | 1.242 | 0.744 | 1.1 | 2.17 | -0.042 | 0.114 | 0.114 | 0 | 0.83 | 0.62 | 0.71 | 0.72 | 0.88 | 0.73 |
| 4 | 102834387 | rs6532979 | 1.032 | 0.705 | 0.922 | 2.131 | -0.027 | 0.123 | 0.123 | 0 | 0.84 | 0.58 | 0.72 | 0.72 | 0.88 | 0.73 |
| 4 | 102839008 | rs6812874 | 0.233 | 0.707 | 0.943 | 2.106 | -0.008 | 0.041 | 0.041 | 0 | 0.92 | 0.69 | 0.94 | 0.9 | 0.95 | 0.83 |
| 4 | 102839287 | rs3733197 | -1.16 | 0.981 | 0.989 | 2.094 | -0.022 | 0.001 | 0.001 | 0.019 | 0.23 | 0.15 | 0.22 | 0.25 | 0.35 | 0.18 |
| 4 | 102841605 | rs13119516 | 0.774 | 1.265 | 0.784 | 1.883 | -0.039 | 0.043 | 0.043 | 0 | 0.08 | 0.01 | 0.09 | 0 | 0.18 | 0.01 |
| 4 | 102849920 | rs2631255 | 1.337 | 0.808 | 0.888 | 2.009 | -0.036 | 0.161 | 0.161 | 0 | 0.82 | 0.57 | 0.69 | 0.65 | 0.87 | 0.73 |
| 4 | 102854390 | rs17208914 | 2.097 | 1.166 | 0.782 | 1.897 | 0.004 | 0.002 | 0.002 | 0 | 0.58 | 0.44 | 0.45 | 0.47 | 0.46 | 0.54 |
| 4 | 102883425 | rs2112139 | 0.691 | 1.141 | 0.755 | 1.916 | -0.022 | 0.137 | 0.137 | 0 | 0.84 | 0.66 | 0.7 | 0.65 | 0.88 | 0.71 |
| 4 | 102893323 | rs2850374 | 0.82 | 1.151 | 0.777 | 1.818 | -0.024 | 0.084 | 0.084 | 0 | 0.84 | 0.67 | 0.71 | 0.7 | 0.87 | 0.71 |
| 4 | 102899504 | rs2631273 | -0.517 | 1.171 | 0.388 | 1.812 | -0.015 | -0.046 | -0.046 | 0 | 0.92 | 1 | 0.78 | 0.83 | 0.92 | 0.89 |
| 4 | 102904490 | rs1027013 | -0.905 | 1.239 | 0.411 | 1.68 | -0.048 | 0.008 | 0.008 | 0.002 | 0.25 | 0.13 | 0.26 | 0.25 | 0.42 | 0.19 |
| 4 | 102909994 | rs2851320 | -0.161 | 1.237 | 0.261 | 1.687 | -0.111 | -0.044 | -0.044 | 0 | 0.18 | 0.02 | 0.23 | 0.23 | 0.38 | 0.09 |
| 4 | 102912159 | rs2850377 | -0.797 | 1.278 | 0.414 | 1.72 | -0.004 | 0.001 | 0.001 | 0.022 | 0.26 | 0.23 | 0.26 | 0.25 | 0.41 | 0.19 |
| 4 | 102914396 | rs2631271 | 0.713 | 0.871 | 0.786 | 1.562 | -0.026 | 0.095 | 0.095 | 0 | 0.84 | 0.4 | 0.7 | 0.72 | 0.88 | 0.71 |
| 4 | 102936592 | rs6844449 | 1.336 | 1.042 | -0.758 | 0.797 | 0.005 | 0.192 | 0.192 | 0 | 0.47 | 0.1 | 0.3 | 0.25 | 0.36 | 0.46 |
| 4 | 102943996 | rs2850390 | 0.303 | 1.171 | -0.823 | 1.385 | -0.078 | -0.088 | -0.088 | 0 | 0.7 | 0.89 | 0.73 | 0.59 | 0.59 | 0.64 |
| 4 | 102948472 | rs2631267 | 0.672 | 1.173 | -0.868 | 1.5 | -0.042 | -0.06 | -0.06 | 0.601 | 0.09 | 0.01 | 0.25 | 0.19 | 0.13 | 0.12 |
| 4 | 102948730 | rs2631268 | 0.668 | 0.262 | -0.956 | 0.359 | 0.003 | -0.006 | -0.006 | 1.187 | 0.36 | 0.32 | 0.43 | 0.35 | 0.29 | 0.29 |
| 4 | 102948991 | rs7685012 | 1.877 | 0.116 | -1.359 | 0.189 | 0.019 | 0.153 | 0.153 | 0 | 0.23 | 0.01 | 0.09 | 0 | 0.12 | 0.12 |
| 4 | 102953681 | rs1872701 | 0.128 | 0.549 | -0.619 | 0.765 | -0.045 | -0.037 | -0.037 | 3.105 | 0.68 | 0.83 | 0.71 | 0.61 | 0.58 | 0.64 |
| 4 | 102968082 | rs2850395 | -0.031 | 0.587 | -0.594 | 0.751 | -0.047 | -0.038 | -0.038 | 0 | 0.69 | 0.84 | 0.71 | 0.61 | 0.56 | 0.64 |
| 4 | 102970332 | rs2658534 | 0.328 | 0.578 | -0.026 | 0.775 | 0 | 0 | 0 | 0 | 0.66 | 0.63 | 0.68 | 0.59 | 0.56 | 0.64 |
| 4 | 102977140 | rs17211702 | 1.464 | 0.593 | -0.54 | 0.747 | -0.067 | 0.08 | 0.08 | 0 | 0.46 | 0.03 | 0.3 | 0.28 | 0.38 | 0.43 |
| 4 | 102979904 | rs17031983 | 0.015 | 0.54 | 0.364 | 0.862 | 0.036 | 0.138 | 0.138 | 0 | 0.16 | 0.42 | 0.34 | 0.27 | 0.19 | 0.18 |
| 4 | 102983720 | rs4550926 | 1.553 | 0.509 | 0.189 | 0.801 | -0.068 | 0.079 | 0.079 | 0 | 0.46 | 0.03 | 0.3 | 0.28 | 0.38 | 0.43 |
| 4 | 102988188 | rs4698987 | 0.31 | 0.481 | 0.357 | 0.866 | -0.002 | -0.001 | -0.001 | 0 | 0.63 | 0.66 | 0.64 | 0.56 | 0.54 | 0.61 |
| 4 | 102995263 | rs6842661 | -0.24 | 0.497 | 0.302 | 0.848 | 0.019 | 0.107 | 0.107 | 0 | 0.17 | 0.59 | 0.35 | 0.27 | 0.18 | 0.18 |
| 4 | 102998867 | rs7675363 | 0.296 | 0.528 | 0.264 | 0.848 | -0.002 | 0.001 | 0.001 | 0 | 0.64 | 0.67 | 0.66 | 0.61 | 0.55 | 0.61 |
| 4 | 103008712 | rs6856488 | 1.249 | 0.519 | 0.305 | 0.836 | 0.021 | 0.038 | 0.038 | 0 | 0.48 | 0.34 | 0.34 | 0.36 | 0.36 | 0.43 |
| 4 | 103013903 | rs6532990 | 1.293 | 0.49 | 0.219 | 0.804 | 0.024 | 0.048 | 0.048 | 0 | 0.48 | 0.32 | 0.34 | 0.34 | 0.36 | 0.43 |
| 4 | 103021348 | rs238487 | -0.639 | 0.439 | 0.085 | 0.739 | 0.004 | 0.002 | 0.002 | 0 | 0.36 | 0.44 | 0.36 | 0.44 | 0.45 | 0.4 |
| 4 | 103023963 | rs238490 | 0.493 | 0.458 | -0.514 | 0.675 | -0.111 | -0.034 | -0.034 | 0 | 0.21 | 0.01 | 0.22 | 0.22 | 0.37 | 0.18 |
| 4 | 103058514 | rs236755 | -0.636 | 0.458 | -0.051 | 0.564 | 0.003 | -0.005 | -0.005 | 0 | 0.38 | 0.58 | 0.35 | 0.37 | 0.45 | 0.4 |
| 4 | 103075330 | rs151476 | -0.28 | 0.603 | -0.809 | 0.617 | -0.022 | 0.005 | 0.005 | 0 | 0.34 | 0.26 | 0.32 | 0.36 | 0.45 | 0.4 |
| 4 | 103084840 | rs236780 | 0.666 | 0.49 | -0.653 | 0.715 | -0.003 | 0.022 | 0.022 | 0 | 0.79 | 0.33 | 0.61 | 0.65 | 0.82 | 0.84 |
| 4 | 103088491 | rs17032126 | 0.538 | 0.847 | -1.184 | 0.628 | -0.04 | -0.064 | -0.064 | 0 | 0.1 | 0.01 | 0.25 | 0.22 | 0.13 | 0.1 |
| 4 | 103091829 | rs13131758 | 0.669 | 0.683 | -1.44 | -0.851 | 0.01 | -0.004 | -0.004 | 0 | 0.41 | 0.36 | 0.23 | 0 | 0.33 | 0.21 |
| 4 | 103098004 | rs11727065 | 0.268 | 0.051 | -1.483 | -0.739 | 0.001 | -0.045 | -0.045 | 0 | 0.27 | 0.28 | 0.12 | 0 | 0.16 | 0.18 |
| 4 | 103105685 | rs190223 | -0.436 | 0.705 | -1.45 | 0.417 | -0.037 | -0.079 | -0.079 | 0 | 0.85 | 0.96 | 0.7 | 0.64 | 0.82 | 0.78 |
| 4 | 103106466 | rs1573494 | -0.198 | 0.75 | -1.623 | 0.702 | 0.002 | -0.055 | -0.055 | 0.361 | 0.17 | 0.09 | 0.2 | 0.3 | 0.16 | 0.27 |
| 4 | 103107786 | rs439538 | 0.376 | 0.607 | -0.97 | 0.253 | 0 | -0.001 | -0.001 | 0 | 0.67 | 0.62 | 0.47 | 0.33 | 0.66 | 0.51 |
| 4 | 103112813 | rs238449 | 0.064 | 0.752 | -0.687 | -0.134 | 0.019 | -0.021 | -0.021 | 0 | 0.71 | 0.64 | 0.62 | 0.77 | 0.67 | 0.82 |
| 4 | 103113866 | rs451461 | 0.275 | 0.614 | -1.411 | -0.307 | 0.001 | 0.087 | 0.087 | 0 | 0.31 | 0.09 | 0.15 | 0.16 | 0.24 | 0.43 |
| 4 | 103115022 | rs7665350 | 0.054 | 0.453 | -1.015 | -0.115 | -0.015 | 0.042 | 0.042 | 0 | 0.17 | 0.45 | 0.17 | 0.18 | 0.16 | 0.22 |
| 4 | 103117289 | rs6419158 | 1.415 | 0.489 | -1.625 | -0.23 | 0.002 | -0.022 | -0.022 | 0 | 0.26 | 0.19 | 0.28 | 0.4 | 0.25 | 0.16 |
| 4 | 103117826 | rs151382 | 0.357 | 0.799 | -1.416 | -0.118 | -0.017 | -0.002 | -0.002 | 0.303 | 0.58 | 0.34 | 0.44 | 0.56 | 0.49 | 0.59 |
| 4 | 103121043 | rs151378 | -0.43 | 1.064 | -1.943 | -0.358 | -0.111 | 0.154 | 0.154 | 0.276 | 0.43 | 0.08 | 0.31 | 0.24 | 0.48 | 0.63 |
| 4 | 103121726 | rs151377 | -1.16 | 1.003 | -1.663 | -0.012 | -0.004 | -0.041 | -0.041 | 0 | 0.84 | 0.94 | 0.72 | 0.6 | 0.85 | 0.82 |
| 4 | 103122348 | rs151375 | -1.212 | 1.073 | -1.825 | -0.059 | -0.052 | -0.108 | -0.108 | 0 | 0.84 | 0.99 | 0.71 | 0.61 | 0.83 | 0.84 |
| 4 | 103125285 | rs1440235 | 0.573 | 0.963 | -1.71 | -0.315 | 0.046 | -0.02 | -0.02 | 0 | 0.4 | 0.78 | 0.39 | 0.38 | 0.33 | 0.17 |
| 4 | 103125383 | rs11735227 | 1.341 | 0.688 | -1.24 | -0.496 | 0.011 | -0.028 | -0.028 | 0 | 0.3 | 0.17 | 0.3 | 0.38 | 0.28 | 0.16 |
| 4 | 103133044 | rs7667281 | -0.507 | 0.859 | -1.734 | -0.757 | -0.092 | -0.099 | -0.099 | 0 | 0.69 | 0.91 | 0.58 | 0.55 | 0.61 | 0.61 |
| 4 | 103135867 | rs151417 | 0.343 | 0.941 | -1.706 | -0.696 | -0.08 | -0.057 | -0.057 | 0 | 0.57 | 0.9 | 0.53 | 0.44 | 0.55 | 0.42 |
| 4 | 103137577 | rs17032168 | 0.65 | 0.864 | -1.752 | -0.831 | 0.008 | -0.001 | -0.001 | 12.419 | 0.09 | 0.07 | 0.11 | 0.09 | 0.09 | 0.22 |
| 4 | 103137977 | rs1540052 | -0.138 | 0.891 | -1.719 | -0.793 | -0.018 | 0.062 | 0.062 | 0 | 0.27 | 0.09 | 0.19 | 0.17 | 0.27 | 0.41 |
| 4 | 103144754 | rs142493 | 0.267 | 0.757 | -1.526 | -0.92 | -0.062 | -0.055 | -0.055 | 0 | 0.57 | 0.88 | 0.52 | 0.43 | 0.55 | 0.42 |
| 4 | 103173749 | rs13126885 | 0.466 | 0.608 | -0.33 | -0.993 | 0 | 0 | 0 | 0 | 0.54 | 0.56 | 0.5 | 0.44 | 0.56 | 0.42 |
| 4 | 103176770 | rs151369 | -1.517 | 0.741 | -0.463 | -1.085 | -0.056 | -0.122 | -0.122 | 1.831 | 0.81 | 0.95 | 0.64 | 0.55 | 0.78 | 0.79 |
| 4 | 103177071 | rs2298752 | -0.077 | 0.72 | -0.415 | -1.106 | 0.006 | 0.084 | 0.084 | 0 | 0.24 | 0.11 | 0.12 | 0.11 | 0.22 | 0.37 |
| 4 | 103180149 | rs170871 | -1.136 | 0.739 | -0.467 | -1.267 | -0.002 | -0.059 | -0.059 | 1.735 | 0.45 | 0.52 | 0.28 | 0.15 | 0.43 | 0.59 |
| 4 | 103183218 | rs9331 | -0.912 | 0.911 | -0.11 | -1.104 | 0.028 | 0.173 | 0.173 | 0.054 | 0.78 | 0.56 | 0.61 | 0.54 | 0.78 | 0.79 |
| 4 | 103183510 | rs151390 | 0.262 | 0.934 | -0.147 | -1.165 | 0.006 | -0.065 | -0.065 | 0 | 0.23 | 0.33 | 0.17 | 0.04 | 0.27 | 0.25 |
| 4 | 103183531 | rs9705 | -0.835 | 0.962 | -0.384 | -0.838 | 0.015 | 0.105 | 0.105 | 0.015 | 0.78 | 0.66 | 0.62 | 0.54 | 0.78 | 0.79 |
| 4 | 103183935 | rs151392 | -0.835 | 0.962 | -0.384 | -0.838 | 0.015 | 0.105 | 0.105 | 0.015 | 0.78 | 0.66 | 0.62 | 0.54 | 0.78 | 0.79 |
| 4 | 103190128 | rs151401 | 0.974 | 0.887 | -0.69 | -1.024 | -0.017 | 0.037 | 0.037 | 0.244 | 0.17 | 0.11 | 0.13 | 0.05 | 0.21 | 0.21 |
| 4 | 103190229 | rs17032400 | 0.952 | 0.76 | -0.83 | -1.097 | -0.021 | -0.039 | -0.039 | 0.38 | 0.47 | 0.12 | 0.4 | 0.47 | 0.47 | 0.33 |
| 4 | 103190486 | rs151402 | -0.302 | 0.725 | -0.478 | -1.037 | 0.006 | 0.064 | 0.064 | 0 | 0.68 | 0.49 | 0.55 | 0.51 | 0.68 | 0.54 |
| 4 | 103200577 | rs10014145 | -0.375 | 0.695 | -0.402 | -1.106 | 0 | -0.031 | -0.031 | 0.821 | 0.32 | 0.34 | 0.19 | 0.12 | 0.33 | 0.35 |
| 4 | 103203500 | rs2165265 | 1.706 | 0.711 | -1.375 | -1.027 | -0.003 | -0.057 | -0.057 | 0 | 0.35 | 0.12 | 0.35 | 0.39 | 0.36 | 0.19 |

| | | | | | | | | | | | | | | | | |
|---|---|---|---|---|---|---|---|---|---|---|---|---|---|---|---|---|
| 4 | 103204358 | rs233821 | -0.645 | 0.691 | -1.401 | -1.186 | -0.004 | 0.072 | 0.072 | 0 | 0.36 | 0.24 | 0.24 | 0.15 | 0.38 | 0.51 |
| 4 | 103204734 | rs233818 | -1.249 | 0.729 | -1.513 | -1.055 | -0.022 | -0.064 | -0.064 | 0 | 0.8 | 0.89 | 0.63 | 0.5 | 0.75 | 0.75 |
| 4 | 103205126 | rs448195 | 0.094 | 1.004 | -1.821 | -1.063 | -0.049 | 0.087 | 0.087 | 0.077 | 0.23 | 0.07 | 0.17 | 0.14 | 0.29 | 0.25 |
| 4 | 103205668 | rs17226148 | 0.719 | 1.045 | -1.771 | -1.01 | -0.022 | 0.045 | 0.045 | 0 | 0.13 | 0.03 | 0.04 | 0.08 | 0.11 | 0.13 |
| 4 | 103206152 | rs17824524 | 0.621 | 1.022 | -1.397 | -0.787 | -0.001 | -0.008 | -0.008 | 0 | 0.09 | 0.1 | 0.06 | 0.03 | 0.09 | 0.21 |
| 4 | 103212864 | rs233805 | -0.217 | 1.08 | -1.524 | -0.807 | -0.011 | 0.04 | 0.04 | 0 | 0.26 | 0.21 | 0.2 | 0.09 | 0.29 | 0.38 |
| 4 | 103217980 | rs4698844 | 0.41 | 1.011 | -1.529 | -0.384 | 0.014 | -0.046 | -0.046 | 0 | 0.37 | 0.17 | 0.36 | 0.39 | 0.36 | 0.21 |
| 4 | 103218446 | rs7699390 | 0.448 | 1.058 | -1.618 | -0.362 | 0.011 | -0.055 | -0.055 | 0 | 0.37 | 0.15 | 0.35 | 0.39 | 0.36 | 0.21 |
| 4 | 103220401 | rs170869 | 0.043 | 0.939 | -1.21 | -1.083 | -0.033 | 0.052 | 0.052 | 0.677 | 0.15 | 0.07 | 0.13 | 0.03 | 0.21 | 0.18 |
| 4 | 103224534 | rs971752 | -0.054 | 0.461 | -0.802 | -1.003 | 0.005 | 0.004 | 0.004 | 0 | 0.19 | 0.16 | 0.18 | 0.12 | 0.16 | 0.21 |
| 4 | 103232168 | rs13103835 | -1.111 | 0.614 | -0.241 | -1.497 | 0.004 | -0.047 | -0.047 | 3.164 | 0.33 | 0.4 | 0.33 | 0.14 | 0.36 | 0.25 |
| 4 | 103243101 | rs17278473 | -0.341 | -0.013 | -0.812 | -2.273 | -0.022 | 0.096 | 0.096 | 2.392 | 0.37 | 0.17 | 0.26 | 0.04 | 0.39 | 0.15 |
| 4 | 103244676 | rs998719 | 1.159 | -0.049 | -0.968 | -2.338 | 0.029 | 0 | 0 | 0 | 0.17 | 0.09 | 0.12 | 0.11 | 0.14 | 0.19 |
| 4 | 103255692 | rs7664683 | 1.091 | -0.298 | -0.566 | -2.506 | 0.006 | 0.007 | 0.007 | 0 | 0.21 | 0.19 | 0.07 | 0.07 | 0.13 | 0.3 |
| 4 | 103271888 | rs11931217 | 0.524 | 0.309 | 0.12 | -2.151 | 0.023 | 0.033 | 0.033 | 0 | 0.42 | 0.38 | 0.21 | 0.18 | 0.3 | 0.5 |
| 4 | 103275840 | rs13152048 | 0.49 | 0.469 | -0.119 | -2.209 | -0.044 | 0.155 | 0.155 | 0.857 | 0.27 | 0.05 | 0.18 | 0.02 | 0.25 | 0.12 |
| 4 | 103285705 | rs10516494 | -0.489 | 0.487 | 0.09 | -2.228 | -0.018 | 0.078 | 0.078 | 2.078 | 0.38 | 0.19 | 0.25 | 0.04 | 0.39 | 0.15 |
| 4 | 103306670 | rs4394001 | 0.503 | 0.49 | 0.034 | -2.222 | -0.044 | 0.155 | 0.155 | 0.857 | 0.27 | 0.05 | 0.18 | 0.02 | 0.25 | 0.12 |
| 4 | 103308071 | rs6843751 | -1.591 | 0.357 | 0.266 | -1.913 | -0.008 | -0.009 | -0.009 | 0 | 0.8 | 0.82 | 0.55 | 0.28 | 0.69 | 0.65 |
| 4 | 103327149 | rs12508086 | 0.933 | 0.282 | 0.145 | -1.947 | 0.038 | 0.022 | 0.022 | 0.021 | 0.37 | 0.15 | 0.24 | 0.24 | 0.3 | 0.49 |
| 4 | 103327536 | rs10856967 | -0.28 | 0.351 | 0.084 | -2.022 | -0.027 | 0.068 | 0.068 | 2.424 | 0.34 | 0.16 | 0.23 | 0.04 | 0.38 | 0.12 |
| 4 | 103337685 | rs12510124 | -1.048 | 0.487 | -0.032 | -1.735 | 0.005 | 0.013 | 0.013 | 0 | 0.75 | 0.73 | 0.54 | 0.3 | 0.7 | 0.64 |
| 4 | 103340510 | rs12513242 | 0.141 | 0.454 | -0.108 | -1.802 | -0.068 | 0.093 | 0.093 | 0.112 | 0.24 | 0.09 | 0.21 | 0.02 | 0.35 | 0.09 |
| 4 | 103343133 | rs9996947 | -0.107 | 0.305 | 0.339 | -1.403 | 0.036 | 0.035 | 0.035 | 0 | 0.51 | 0.64 | 0.33 | 0.28 | 0.35 | 0.55 |
| 4 | 103349049 | rs964504 | 0.406 | 0.11 | 0.274 | -1.198 | 0.032 | -0.005 | -0.005 | 0 | 0.42 | 0.3 | 0.24 | 0.42 | 0.29 | 0.5 |
| 4 | 103349740 | rs6533014 | -0.826 | 0.262 | 0.297 | -1.285 | 0.002 | 0.002 | 0.002 | 0 | 0.67 | 0.67 | 0.44 | 0.44 | 0.5 | 0.62 |
| 4 | 103359744 | rs6846971 | -0.036 | 0.289 | 0.173 | -1.618 | 0 | -0.065 | -0.065 | 0.012 | 0.23 | 0.28 | 0.19 | 0.02 | 0.2 | 0.11 |
| 4 | 103364762 | rs926163 | 1.051 | 0.356 | 0.113 | -1.625 | 0.016 | 0.03 | 0.03 | 1.145 | 0.17 | 0.1 | 0.09 | 0.03 | 0.1 | 0.06 |
| 4 | 103365532 | rs13115343 | -0.715 | 0.353 | 0.436 | -1.73 | 0.004 | -0.114 | -0.114 | 0 | 0.3 | 0.59 | 0.23 | 0.03 | 0.31 | 0.11 |
| 4 | 103369480 | rs10031415 | -1.165 | 0.571 | 0.416 | -1.601 | 0.125 | -0.096 | -0.096 | 1.921 | 0.13 | 0.48 | 0.17 | 0 | 0.21 | 0.05 |
| 4 | 103373286 | rs13116385 | 0.284 | 0.669 | 0.103 | -1.421 | 0.078 | -0.021 | -0.021 | 0 | 0.05 | 0.19 | 0.1 | 0 | 0.09 | 0.04 |
| 4 | 103374154 | rs10000030 | 0.606 | 0.35 | -0.369 | -1.501 | -0.005 | -0.003 | -0.003 | 0 | 0.23 | 0.23 | 0.11 | 0.22 | 0.14 | 0.09 |
| 4 | 103375303 | rs1313924 | 0.891 | 1.139 | -0.457 | -1.028 | 0.026 | 0.019 | 0.019 | 0.901 | 0.49 | 0.41 | 0.33 | 0.44 | 0.35 | 0.44 |
| 4 | 103376474 | rs1585607 | -0.76 | 1.654 | -0.27 | -0.42 | 0.074 | -0.08 | -0.08 | 0.269 | 0.13 | 0.36 | 0.15 | 0 | 0.21 | 0.05 |
| 4 | 103377576 | rs1313925 | -0.278 | 1.686 | -0.489 | -0.449 | -0.038 | -0.035 | -0.035 | 0 | 0.73 | 0.86 | 0.52 | 0.45 | 0.61 | 0.58 |
| 4 | 103378866 | rs1314336 | 0.828 | 1.644 | -0.485 | -0.42 | 0.003 | -0.007 | -0.007 | 0 | 0.6 | 0.48 | 0.37 | 0.44 | 0.41 | 0.53 |
| 4 | 103387419 | rs230490 | -0.295 | 1.671 | -1.455 | 0.121 | -0.038 | -0.036 | -0.036 | 0 | 0.27 | 0.14 | 0.48 | 0.55 | 0.39 | 0.42 |
| 4 | 103388441 | rs230489 | -1.205 | 1.292 | -1.135 | -0.088 | 0 | -0.021 | -0.021 | 0 | 0.1 | 0.12 | 0.09 | 0 | 0.11 | 0.04 |
| 4 | 103389409 | rs230487 | 0.165 | 0.983 | -1.697 | 0.5 | 0.002 | -0.036 | -0.036 | 0 | 0.65 | 0.84 | 0.68 | 0.39 | 0.72 | 0.51 |
| 4 | 103396333 | rs2903281 | 0.426 | 1.173 | -2.399 | 0.382 | -0.056 | -0.075 | -0.075 | 1.194 | 0.21 | 0.02 | 0.31 | 0.27 | 0.24 | 0.15 |
| 4 | 103397014 | rs2085548 | -0.324 | 1.262 | -1.098 | 0.104 | 0.013 | 0.014 | 0.014 | 0 | 0.28 | 0.49 | 0.37 | 0.32 | 0.3 | 0.21 |
| 4 | 103427476 | rs3774934 | -0.804 | 1.075 | -1.505 | 0.921 | -0.021 | -0.049 | -0.049 | 0.014 | 0.91 | 0.99 | 0.91 | 0.56 | 0.9 | 0.71 |
| 4 | 103432974 | rs17032705 | -0.34 | 1.183 | -1.278 | 0.61 | -0.015 | -0.011 | -0.011 | 1.96 | 0.34 | 0.31 | 0.47 | 0.38 | 0.4 | 0.27 |
| 4 | 103443569 | rs1599961 | -0.536 | 0.834 | -0.773 | 0.439 | 0 | 0 | 0 | 0.299 | 0.37 | 0.43 | 0.48 | 0.38 | 0.4 | 0.28 |
| 4 | 103446115 | rs1598856 | -0.911 | 0.409 | -1.05 | -0.342 | -0.028 | -0.042 | -0.042 | 0 | 0.65 | 0.81 | 0.59 | 0.43 | 0.54 | 0.48 |
| 4 | 103448582 | rs230535 | -1.065 | 0.494 | -0.954 | -0.131 | -0.012 | -0.034 | -0.034 | 0.11 | 0.69 | 0.83 | 0.57 | 0.56 | 0.66 | 0.76 |
| 4 | 103463007 | rs118882 | -1.132 | 0.437 | -0.755 | -0.151 | -0.036 | -0.068 | -0.068 | 0 | 0.69 | 0.89 | 0.57 | 0.56 | 0.66 | 0.76 |
| 4 | 103478703 | rs13117745 | 0.685 | 0.281 | -0.182 | -0.266 | -0.018 | -0.059 | -0.059 | 0.185 | 0.27 | 0.3 | 0.11 | 0.05 | 0.14 | 0.2 |
| 4 | 103510511 | rs1020759 | 0.217 | 0.234 | -0.19 | -0.288 | 0.001 | 0.001 | 0.001 | 0 | 0.37 | 0.42 | 0.49 | 0.5 | 0.41 | 0.3 |
| 4 | 103527068 | rs4648090 | 1.108 | 0.007 | -0.209 | -0.508 | -0.003 | -0.036 | -0.036 | 0 | 0.18 | 0.17 | 0.09 | 0 | 0.12 | 0.11 |
| 4 | 103536261 | rs230547 | 1.11 | 0.063 | -0.306 | 0.044 | 0 | 0.04 | 0.04 | 0 | 0.1 | 0.14 | 0.1 | 0.27 | 0.1 | 0.2 |
| 4 | 103537442 | rs1609798 | 1.156 | -0.084 | -0.441 | -0.103 | -0.041 | -0.064 | -0.064 | 0.022 | 0.28 | 0.07 | 0.41 | 0.41 | 0.31 | 0.28 |
| 4 | 103542005 | rs997476 | 1.427 | -0.116 | -0.447 | -0.088 | -0.033 | -0.028 | -0.028 | 12.676 | 0.07 | 0 | 0.03 | 0.08 | 0.06 | 0.06 |
| 4 | 103542201 | rs10489113 | -0.178 | -0.34 | -0.28 | -0.209 | -0.031 | -0.088 | -0.088 | 0 | 0.28 | 0.48 | 0.16 | 0.08 | 0.18 | 0.17 |
| 4 | 103545253 | rs11733293 | 0.642 | -0.075 | -0.516 | -0.033 | 0.016 | -0.005 | -0.005 | 0 | 0.34 | 0.26 | 0.5 | 0.41 | 0.31 | 0.27 |
| 4 | 103549149 | rs10213251 | -0.241 | -0.272 | -0.243 | -0.212 | -0.036 | -0.081 | -0.081 | 0 | 0.29 | 0.5 | 0.24 | 0.08 | 0.17 | 0.17 |
| 4 | 103552709 | rs1054037 | -0.66 | -0.119 | -0.093 | -0.107 | -0.013 | -0.01 | -0.01 | 0 | 0.61 | 0.66 | 0.65 | 0.52 | 0.49 | 0.41 |
| 4 | 103552813 | rs4013 | -0.66 | -0.119 | -0.093 | -0.107 | -0.013 | -0.01 | -0.01 | 1.202 | 0.61 | 0.66 | 0.65 | 0.52 | 0.49 | 0.41 |
| 4 | 103553053 | rs1054029 | -0.66 | -0.229 | -0.098 | -0.084 | -0.013 | -0.01 | -0.01 | 0 | 0.61 | 0.66 | 0.65 | 0.52 | 0.49 | 0.41 |
| 4 | 103555992 | rs2272697 | -0.645 | -0.225 | -0.086 | -0.083 | -0.013 | -0.01 | -0.01 | 0.025 | 0.61 | 0.66 | 0.65 | 0.52 | 0.49 | 0.41 |
| 4 | 103557077 | rs2866413 | -0.645 | -0.225 | -0.086 | -0.083 | -0.013 | -0.01 | -0.01 | 0 | 0.61 | 0.66 | 0.65 | 0.52 | 0.49 | 0.41 |
| 4 | 103558731 | rs6833962 | 0.698 | -0.427 | -0.109 | -0.192 | -0.068 | -0.114 | -0.114 | 0 | 0.2 | 0.37 | 0.1 | 0 | 0.08 | 0.11 |
| 4 | 103578637 | rs228614 | 0.235 | 0.056 | -0.093 | -0.037 | -0.003 | -0.002 | -0.002 | 0 | 0.4 | 0.57 | 0.34 | 0.5 | 0.49 | 0.57 |
| 4 | 103607635 | rs227284 | -0.706 | -0.329 | -0.124 | -0.317 | -0.005 | -0.013 | -0.013 | 0.113 | 0.63 | 0.39 | 0.61 | 0.79 | 0.67 | 0.8 |
| 4 | 103611845 | rs227368 | 0.832 | -0.147 | -0.288 | -0.02 | -0.097 | -0.075 | -0.075 | 0 | 0.36 | 0.12 | 0.35 | 0.47 | 0.5 | 0.52 |
| 4 | 103618023 | rs227375 | 0.754 | -0.4 | -0.285 | -0.013 | -0.079 | -0.071 | -0.071 | 1.11 | 0.37 | 0.12 | 0.38 | 0.47 | 0.53 | 0.52 |
| 4 | 103640758 | rs223502 | 0.902 | -0.494 | -0.198 | -0.235 | -0.01 | -0.016 | -0.016 | 0 | 0.27 | 0.4 | 0.37 | 0.2 | 0.27 | 0.13 |
| 4 | 103645082 | rs6857760 | -0.501 | -0.414 | -0.015 | -0.343 | 0.086 | 0.043 | 0.043 | 0 | 0.05 | 0.38 | 0.02 | 0 | 0.02 | 0.01 |
| 4 | 103647047 | rs223500 | 0.61 | -0.548 | -0.38 | -0.211 | 0.01 | 0.017 | 0.017 | 0.918 | 0.3 | 0.77 | 0.36 | 0.19 | 0.26 | 0.13 |

| | | | | | | | | | | | | | | | |
|---|---|---|---|---|---|---|---|---|---|---|---|---|---|---|---|
| 4 | 103651962 | rs223498 | -1.024 | -0.287 | -0.726 | -0.222 | -0.088 | -0.037 | -0.037 | 0 | 0.61 | 0.89 | 0.62 | 0.5 | 0.48 | 0.47 |
| 4 | 103679117 | rs223490 | -0.742 | -0.547 | -0.748 | -0.443 | 0.089 | 0.088 | 0.088 | 0 | 0.49 | 0.81 | 0.48 | 0.38 | 0.38 | 0.39 |
| 4 | 103680984 | rs223489 | 0.159 | -0.631 | -0.398 | -0.494 | -0.003 | -0.003 | -0.003 | 0 | 0.43 | 0.47 | 0.47 | 0.38 | 0.36 | 0.38 |
| 4 | 103687208 | rs223482 | -1.061 | -0.681 | -0.37 | -0.233 | -0.018 | -0.005 | -0.005 | 0.067 | 0.59 | 0.71 | 0.53 | 0.5 | 0.49 | 0.49 |
| 4 | 103730493 | rs223418 | -1.157 | -0.707 | -0.345 | -0.242 | -0.061 | -0.04 | -0.04 | 0 | 0.61 | 0.84 | 0.55 | 0.5 | 0.49 | 0.49 |
| 4 | 103733600 | rs223412 | -1.091 | -0.7 | -0.343 | -0.24 | -0.042 | -0.021 | -0.021 | 0 | 0.59 | 0.8 | 0.55 | 0.5 | 0.49 | 0.49 |
| 4 | 103734219 | rs10005010 | 0.965 | -0.7 | -0.316 | -0.27 | 0 | -0.001 | -0.001 | 0 | 0.16 | 0.14 | 0.07 | 0.12 | 0.16 | 0.1 |
| 4 | 103746753 | rs3774997 | 1.349 | -0.702 | -0.348 | -0.239 | -0.006 | 0.002 | 0.002 | 0 | 0.14 | 0.08 | 0.07 | 0.13 | 0.12 | 0.09 |
| 4 | 103748224 | rs223391 | 2.633 | -1.01 | -0.52 | -0.365 | -0.029 | -0.041 | -0.041 | 0 | 0.17 | 0.02 | 0.27 | 0.19 | 0.2 | 0.12 |
| 4 | 103758301 | rs223374 | 0.285 | -0.895 | -0.321 | -0.311 | -0.006 | -0.003 | -0.003 | 0 | 0.41 | 0.47 | 0.47 | 0.38 | 0.33 | 0.4 |
| 4 | 103765156 | rs223366 | 0.416 | -0.587 | -0.748 | -0.788 | 0.003 | 0.014 | 0.014 | 0.016 | 0.53 | 0.48 | 0.51 | 0.37 | 0.62 | 0.42 |
| 4 | 103775704 | rs223353 | -1.297 | -0.806 | -0.417 | -0.408 | -0.061 | -0.04 | -0.04 | 0 | 0.61 | 0.84 | 0.55 | 0.5 | 0.49 | 0.49 |
| 4 | 103785051 | rs223340 | -2.384 | -1.016 | -0.545 | -0.028 | -0.167 | -0.011 | -0.011 | 0 | 0.73 | 0.98 | 0.75 | 0.86 | 0.59 | 0.74 |
| 4 | 103788211 | rs223334 | -1.637 | -0.6 | 0.123 | 0.198 | -0.034 | -0.021 | -0.021 | 0 | 0.61 | 0.77 | 0.54 | 0.5 | 0.49 | 0.49 |
| 4 | 103789782 | rs2198199 | 1.061 | -0.593 | 0.147 | 0.199 | -0.009 | 0.004 | 0.004 | 0 | 0.16 | 0.1 | 0.18 | 0.13 | 0.15 | 0.1 |
| 4 | 103805443 | rs223315 | -1.538 | -0.523 | 0.129 | 0.209 | -0.022 | -0.009 | -0.009 | 0.038 | 0.59 | 0.72 | 0.54 | 0.5 | 0.49 | 0.49 |
| 4 | 103807720 | rs223313 | -1.619 | -0.517 | 0.087 | 0.212 | -0.053 | -0.034 | -0.034 | 0 | 0.61 | 0.82 | 0.55 | 0.5 | 0.49 | 0.49 |
| 4 | 103919065 | rs9994654 | 1.973 | -0.667 | 0.486 | 0.233 | -0.001 | 0 | 0 | 0 | 0.19 | 0.22 | 0.17 | 0.19 | 0.2 | 0.14 |
| 4 | 103936001 | rs6843738 | -0.122 | -0.617 | 0.651 | 0.279 | 0.042 | 0.027 | 0.027 | 0 | 0.4 | 0.31 | 0.35 | 0.38 | 0.33 | 0.39 |
| 4 | 103937595 | rs28445579 | -1.527 | -0.633 | 0.803 | 0.281 | -0.019 | -0.01 | -0.01 | 0 | 0.58 | 0.71 | 0.55 | 0.51 | 0.49 | 0.48 |
| 4 | 103937933 | rs7676765 | -1.896 | -0.661 | 0.816 | 0.282 | -0.059 | -0.046 | -0.046 | 0.049 | 0.61 | 0.84 | 0.57 | 0.51 | 0.49 | 0.48 |
| 4 | 103937974 | rs7698608 | -1.55 | -0.655 | 0.677 | 0.288 | -0.02 | -0.011 | -0.011 | 0 | 0.58 | 0.71 | 0.56 | 0.51 | 0.49 | 0.48 |
| 4 | 103942165 | rs10516496 | -0.283 | -0.628 | 0.808 | 0.278 | -0.018 | -0.014 | -0.014 | 0 | 0.42 | 0.47 | 0.36 | 0.37 | 0.34 | 0.39 |
| 4 | 103942498 | rs13150953 | -0.354 | -0.628 | 0.744 | 0.278 | -0.022 | -0.018 | -0.018 | 0.175 | 0.42 | 0.52 | 0.36 | 0.37 | 0.34 | 0.39 |
| 4 | 103942714 | rs10516497 | 2.083 | -0.663 | 0.627 | 0.274 | -0.001 | 0 | 0 | 0.138 | 0.19 | 0.22 | 0.17 | 0.19 | 0.2 | 0.13 |
| 4 | 103944742 | rs11946020 | -0.22 | -0.491 | 0.719 | 0.372 | -0.004 | -0.003 | -0.003 | 0.26 | 0.42 | 0.47 | 0.36 | 0.37 | 0.34 | 0.39 |
| 4 | 103950425 | rs1080081 | -1.769 | -0.455 | 0.074 | -0.358 | -0.048 | -0.035 | -0.035 | 0 | 0.61 | 0.82 | 0.55 | 0.51 | 0.49 | 0.48 |
| 4 | 103951624 | rs4699049 | -1.761 | -0.433 | 0.085 | -0.347 | -0.048 | -0.034 | -0.034 | 0 | 0.61 | 0.82 | 0.55 | 0.51 | 0.49 | 0.48 |
| 4 | 103954851 | rs34475639 | -0.126 | -0.443 | 0.145 | -0.353 | 0.022 | 0.018 | 0.018 | 0 | 0.41 | 0.37 | 0.35 | 0.37 | 0.34 | 0.39 |
| 4 | 103956956 | rs13130741 | -1.724 | -0.371 | -0.335 | -0.324 | -0.049 | -0.035 | -0.035 | 2.029 | 0.61 | 0.82 | 0.56 | 0.51 | 0.49 | 0.48 |
| 4 | 103960768 | rs7660298 | -1.887 | -0.301 | -0.461 | -0.342 | -0.053 | -0.04 | -0.04 | 0 | 0.62 | 0.82 | 0.57 | 0.51 | 0.5 | 0.48 |
| 4 | 103962532 | rs7694724 | -1.848 | -0.306 | -0.46 | -0.341 | -0.05 | -0.039 | -0.039 | 0 | 0.62 | 0.82 | 0.56 | 0.51 | 0.5 | 0.48 |
| 4 | 103964529 | rs2276976 | 0.279 | -0.659 | 0.168 | -0.251 | 0.03 | 0.019 | 0.019 | 0 | 0.18 | 0.06 | 0.16 | 0.18 | 0.11 | 0.24 |
| 4 | 103974122 | rs13113099 | -1.091 | -0.409 | 0.354 | -0.333 | 0.08 | 0.098 | 0.098 | 0.019 | 0.5 | 0.85 | 0.41 | 0.4 | 0.39 | 0.41 |
| 4 | 103977700 | rs13149311 | -0.497 | -0.64 | -0.223 | -0.294 | 0.009 | 0.012 | 0.012 | 0 | 0.43 | 0.39 | 0.36 | 0.4 | 0.39 | 0.4 |
| 4 | 103987899 | rs7691873 | -0.944 | -0.69 | -1.316 | -0.312 | 0.009 | 0.071 | 0.071 | 0.014 | 0.52 | 0.93 | 0.42 | 0.4 | 0.42 | 0.42 |
| 4 | 104000498 | rs7666067 | -0.278 | -1.033 | -0.723 | -0.706 | 0.011 | 0.058 | 0.058 | 0 | 0.23 | 0.06 | 0.18 | 0.16 | 0.17 | 0.28 |
| 4 | 104000922 | rs12508269 | 0.316 | -1.066 | -0.786 | -0.743 | -0.015 | -0.003 | -0.003 | 1.043 | 0.15 | 0.01 | 0.14 | 0.16 | 0.11 | 0.23 |
| 4 | 104001975 | rs3796983 | -1.025 | -1.145 | 0.08 | -0.786 | -0.019 | -0.048 | -0.048 | 0 | 0.31 | 0.37 | 0.23 | 0.16 | 0.21 | 0.33 |
| 4 | 104003148 | rs2306478 | -0.445 | -1.11 | 0.03 | -0.769 | 0.015 | 0.052 | 0.052 | 0 | 0.25 | 0.13 | 0.2 | 0.16 | 0.2 | 0.32 |
| 4 | 104004362 | rs3775972 | -0.647 | -1.107 | 0.017 | -0.766 | 0.021 | 0.071 | 0.071 | 0 | 0.27 | 0.12 | 0.2 | 0.16 | 0.2 | 0.32 |
| 4 | 104007105 | rs6825519 | -0.934 | -1.114 | 0.158 | -0.764 | -0.021 | -0.051 | -0.051 | 0 | 0.3 | 0.39 | 0.23 | 0.16 | 0.2 | 0.33 |
| 4 | 104007996 | rs7675057 | -0.724 | -1.105 | 0.211 | -0.755 | -0.021 | -0.046 | -0.046 | 1.294 | 0.27 | 0.37 | 0.23 | 0.16 | 0.18 | 0.32 |
| 4 | 104010027 | rs10516498 | -0.165 | -1.2 | 0.31 | -0.727 | -0.001 | -0.005 | -0.005 | 0 | 0.24 | 0.27 | 0.19 | 0.15 | 0.15 | 0.3 |
| 4 | 104011067 | rs746829 | -0.61 | -1.216 | 0.489 | -0.74 | -0.025 | -0.047 | -0.047 | 0 | 0.29 | 0.41 | 0.22 | 0.15 | 0.18 | 0.32 |
| 4 | 104014114 | rs7690059 | -0.238 | -1.165 | 0.373 | -0.715 | -0.002 | -0.009 | -0.009 | 0.005 | 0.27 | 0.31 | 0.22 | 0.15 | 0.18 | 0.32 |
| 4 | 104026547 | rs2720447 | 1.032 | -1.164 | -0.198 | -0.673 | -0.017 | 0.005 | 0.005 | 0.211 | 0.18 | 0.06 | 0.12 | 0.14 | 0.22 | 0.08 |
| 4 | 104030792 | rs2623064 | -0.922 | -1.375 | -0.068 | -0.806 | 0.043 | 0.083 | 0.083 | 0 | 0.46 | 0.29 | 0.33 | 0.3 | 0.4 | 0.42 |
| 4 | 104031440 | rs2711896 | 1.111 | -1.364 | -0.071 | -0.8 | -0.017 | 0.005 | 0.005 | 0.211 | 0.18 | 0.06 | 0.12 | 0.14 | 0.22 | 0.08 |
| 4 | 104054686 | rs2720460 | -0.865 | -1.386 | -0.046 | -0.811 | 0.04 | 0.079 | 0.079 | 0.03 | 0.46 | 0.3 | 0.33 | 0.3 | 0.4 | 0.41 |
| 4 | 104059542 | rs2243682 | 1.245 | -1.375 | -0.168 | -0.801 | -0.043 | -0.018 | -0.018 | 0 | 0.17 | 0.02 | 0.12 | 0.14 | 0.22 | 0.08 |
| 4 | 104066461 | rs2615542 | 1.068 | -1.377 | -0.165 | -0.791 | -0.017 | 0.005 | 0.005 | 0.211 | 0.18 | 0.06 | 0.12 | 0.14 | 0.22 | 0.08 |
| 4 | 104098605 | rs11724309 | -0.117 | -1.406 | -0.139 | -0.808 | 0.028 | 0.042 | 0.042 | 0 | 0.28 | 0.17 | 0.22 | 0.15 | 0.18 | 0.33 |
| 4 | 104117191 | rs2290943 | 1.106 | -1.378 | -0.188 | -0.773 | -0.023 | -0.003 | -0.003 | 0 | 0.18 | 0.06 | 0.12 | 0.14 | 0.22 | 0.08 |
| 4 | 104120041 | rs6850461 | -1.193 | -1.604 | 0.545 | -0.882 | 0.033 | 0.057 | 0.057 | 0 | 0.51 | 0.68 | 0.37 | 0.3 | 0.41 | 0.42 |
| 4 | 104120119 | rs17217592 | -0.062 | -1.531 | 0.662 | -0.87 | 0.032 | 0.058 | 0.058 | 0.012 | 0.28 | 0.15 | 0.23 | 0.15 | 0.19 | 0.33 |
| 4 | 104127786 | rs6815702 | -0.746 | -1.514 | 0.571 | -0.843 | -0.023 | -0.04 | -0.04 | 0 | 0.33 | 0.61 | 0.25 | 0.16 | 0.19 | 0.33 |
| 4 | 104128452 | rs6821646 | -0.664 | -1.515 | 0.602 | -0.843 | -0.051 | -0.079 | -0.079 | 0 | 0.33 | 0.5 | 0.24 | 0.16 | 0.19 | 0.33 |
| 4 | 104133376 | rs6533059 | -0.88 | -1.565 | 0.207 | -0.701 | 0.013 | 0.016 | 0.016 | 0 | 0.5 | 0.58 | 0.36 | 0.37 | 0.39 | 0.44 |
| 4 | 104134846 | rs35199266 | 0.374 | -1.618 | 0.135 | -0.745 | -0.001 | 0.058 | 0.058 | 0.149 | 0.25 | 0.03 | 0.16 | 0.16 | 0.17 | 0.33 |
| 4 | 104137790 | rs4699052 | -1.022 | -1.565 | 0.189 | -0.716 | 0.01 | 0.013 | 0.013 | 0 | 0.49 | 0.58 | 0.36 | 0.37 | 0.39 | 0.44 |
| 4 | 104140848 | rs1580278 | -1.882 | -1.37 | -0.637 | -0.59 | -0.175 | -0.137 | -0.137 | 0.611 | 0.68 | 0.97 | 0.58 | 0.55 | 0.55 | 0.56 |
| 4 | 104165717 | rs7676474 | 1.232 | -0.693 | -0.034 | -1.125 | -0.032 | -0.048 | -0.048 | 0 | 0.17 | 0.05 | 0.14 | 0.2 | 0.21 | 0.11 |
| 4 | 104179569 | rs17033496 | 0.479 | -0.821 | 0.095 | -1.252 | 0.02 | 0.056 | 0.056 | 0 | 0.23 | 0.09 | 0.16 | 0.14 | 0.16 | 0.31 |
| 4 | 104185435 | rs11723084 | -0.635 | -0.959 | 0.5 | -1.32 | -0.008 | -0.027 | -0.027 | 0 | 0.33 | 0.65 | 0.22 | 0.15 | 0.17 | 0.31 |
| 4 | 104194329 | rs7656832 | 0.536 | -0.946 | 0.473 | -1.314 | 0.02 | 0.06 | 0.06 | 0 | 0.23 | 0.09 | 0.16 | 0.14 | 0.16 | 0.3 |
| 4 | 104204409 | rs7687989 | -0.545 | -0.899 | 0.36 | -1.291 | -0.018 | -0.042 | -0.042 | 0.886 | 0.32 | 0.63 | 0.22 | 0.15 | 0.17 | 0.31 |
| 4 | 104214616 | rs10516502 | -0.881 | -0.791 | 0.364 | -1.226 | 0.006 | 0.008 | 0.008 | 0 | 0.47 | 0.46 | 0.35 | 0.35 | 0.38 | 0.42 |
| 4 | 104216086 | rs6820979 | 1.136 | -0.678 | -0.133 | -1.155 | -0.018 | -0.035 | -0.035 | 1.039 | 0.18 | 0.05 | 0.14 | 0.2 | 0.22 | 0.11 |

| | | | | | | | | | | | | | | | | |
|---|---|---|---|---|---|---|---|---|---|---|---|---|---|---|---|---|
| 4 | 104225413 | rs1607094 | -0.502 | -0.491 | 0.101 | -1.489 | 0.012 | 0.035 | 0.035 | 0.003 | 0.45 | 0.3 | 0.34 | 0.38 | 0.42 | 0.42 |
| 4 | 104235380 | rs17219677 | 1.201 | -0.476 | -0.008 | -1.471 | -0.005 | 0.008 | 0.008 | 0.747 | 0.19 | 0.09 | 0.13 | 0.15 | 0.21 | 0.09 |
| 4 | 104241267 | rs6533083 | -0.841 | -1.018 | 0.521 | -1.399 | 0.016 | 0.037 | 0.037 | 0.105 | 0.46 | 0.32 | 0.34 | 0.38 | 0.42 | 0.4 |
| 4 | 104242990 | rs6836157 | -1.221 | -1.072 | 0.742 | -1.435 | 0.026 | 0.071 | 0.071 | 0 | 0.52 | 0.79 | 0.42 | 0.38 | 0.44 | 0.42 |
| 4 | 104245131 | rs7682050 | 0.069 | -0.992 | 0.796 | -1.349 | 0.006 | 0.013 | 0.013 | 0 | 0.26 | 0.19 | 0.2 | 0.23 | 0.21 | 0.31 |
| 4 | 104250324 | rs1996346 | -1.084 | -0.827 | 0.355 | -1.166 | 0.029 | 0.045 | 0.045 | 0 | 0.52 | 0.73 | 0.37 | 0.4 | 0.43 | 0.44 |
| 4 | 104257925 | rs4699068 | -1.479 | -0.865 | -0.1 | -1.205 | -0.015 | 0.012 | 0.012 | 0 | 0.57 | 0.73 | 0.52 | 0.43 | 0.5 | 0.48 |
| 4 | 104258396 | rs7662107 | -0.01 | -1.293 | 0.334 | -1.042 | 0.005 | 0.015 | 0.015 | 0 | 0.28 | 0.25 | 0.16 | 0.17 | 0.28 | 0.2 |
| 4 | 104270805 | rs10856971 | 0.627 | -1.356 | 0.215 | -1.009 | -0.017 | -0.036 | -0.036 | 0 | 0.17 | 0.03 | 0.15 | 0.22 | 0.14 | 0.23 |
| 4 | 104271269 | rs12649701 | 0.65 | -1.352 | 0.217 | -1.007 | -0.017 | -0.037 | -0.037 | 0 | 0.17 | 0.03 | 0.15 | 0.22 | 0.14 | 0.23 |
| 4 | 104275452 | rs17286262 | 0.638 | -1.374 | 0.214 | -1.014 | -0.003 | 0.005 | 0.005 | 0.275 | 0.22 | 0.08 | 0.13 | 0.16 | 0.23 | 0.18 |
| 4 | 104282146 | rs13101586 | 0.663 | -1.359 | 0.094 | -1.005 | -0.017 | -0.037 | -0.037 | 0 | 0.17 | 0.03 | 0.15 | 0.22 | 0.14 | 0.23 |
| 4 | 104288620 | rs11734572 | -1.273 | -1.354 | 0.224 | -1.017 | -0.005 | -0.006 | -0.006 | 0 | 0.47 | 0.5 | 0.33 | 0.39 | 0.42 | 0.44 |
| 4 | 104293598 | rs10009242 | -1.525 | -1.349 | 0.167 | -1.046 | 0 | 0 | 0 | 0 | 0.49 | 0.49 | 0.39 | 0.39 | 0.46 | 0.51 |
| 4 | 104294789 | rs6533086 | 0.418 | -1.22 | -0.59 | -0.992 | 0.001 | -0.037 | -0.037 | 0 | 0.07 | 0.16 | 0.04 | 0 | 0.05 | 0.02 |
| 4 | 104308097 | rs13111985 | 0.538 | -1.144 | -1.294 | -0.918 | -0.02 | -0.04 | -0.04 | 0.841 | 0.17 | 0.02 | 0.15 | 0.22 | 0.14 | 0.26 |
| 4 | 104335667 | rs6828829 | -1.223 | -0.962 | -1.103 | -0.769 | 0.033 | 0.043 | 0.043 | 0 | 0.52 | 0.73 | 0.39 | 0.39 | 0.44 | 0.48 |
| 4 | 104372029 | rs17221044 | -0.276 | -0.724 | -0.589 | -0.363 | -0.012 | -0.007 | -0.007 | 0 | 0.22 | 0.07 | 0.22 | 0.17 | 0.24 | 0.19 |
| 4 | 104374596 | rs10026360 | -0.736 | -0.543 | -0.182 | -0.241 | -0.027 | -0.055 | -0.055 | 0.732 | 0.34 | 0.49 | 0.31 | 0.17 | 0.32 | 0.23 |
| 4 | 104382243 | rs340479 | -1.184 | -0.528 | -0.237 | 0.464 | 0.003 | -0.024 | -0.024 | 0 | 0.17 | 0.09 | 0.12 | 0.26 | 0.18 | 0.22 |
| 4 | 104385495 | rs17033669 | -0.772 | -1.34 | 0.453 | -0.725 | -0.041 | -0.101 | -0.101 | 0 | 0.27 | 0.46 | 0.22 | 0.08 | 0.18 | 0.24 |
| 4 | 104396376 | rs10011202 | -0.261 | -1.535 | 0.876 | -0.707 | -0.034 | 0.012 | 0.012 | 0 | 0.12 | 0.34 | 0.16 | 0.1 | 0.06 | 0.14 |
| 4 | 104401567 | rs437156 | -0.304 | -1.621 | 1.271 | -0.807 | 0.002 | -0.062 | -0.062 | 0.251 | 0.12 | 0.18 | 0.09 | 0 | 0.12 | 0.03 |
| 4 | 104402247 | rs7694802 | -2.305 | -1.91 | 1.633 | -0.958 | 0.04 | 0.099 | 0.099 | 0 | 0.39 | 0.72 | 0.33 | 0.1 | 0.3 | 0.31 |
| 4 | 104403357 | rs17033694 | -0.116 | -1.913 | 1.485 | -0.958 | -0.001 | -0.032 | -0.032 | 0 | 0.11 | 0.11 | 0.09 | 0 | 0.12 | 0.03 |
| 4 | 104413137 | rs17221890 | -0.361 | -1.924 | 1.608 | -0.948 | -0.005 | -0.044 | -0.044 | 0.026 | 0.14 | 0.14 | 0.07 | 0.02 | 0.12 | 0.14 |
| 4 | 104414511 | rs420934 | 1.829 | -1.902 | 2.032 | -0.919 | 0.024 | 0.042 | 0.042 | 0.098 | 0.61 | 0.35 | 0.67 | 0.88 | 0.7 | 0.68 |
| 4 | 104414948 | rs402681 | 1.086 | -1.834 | 2.004 | -0.901 | -0.017 | -0.163 | -0.163 | 3.288 | 0.73 | 0.56 | 0.83 | 0.98 | 0.76 | 0.83 |
| 4 | 104418239 | rs340468 | 1.749 | -2.007 | 0.82 | -0.755 | 0.074 | 0.182 | 0.182 | 0 | 0.61 | 0.18 | 0.77 | 0.93 | 0.7 | 0.71 |
| 4 | 104418307 | rs894996 | -1.014 | -1.525 | 1.079 | -0.313 | -0.006 | -0.003 | -0.003 | 0 | 0.09 | 0.17 | 0.06 | 0.06 | 0.07 | 0.13 |
| 4 | 104421641 | rs13115641 | -1.085 | -1.476 | 0.789 | -0.295 | -0.049 | -0.019 | -0.019 | 0 | 0.22 | 0.42 | 0.22 | 0.17 | 0.15 | 0.27 |
| 4 | 104424598 | rs7691671 | -1.03 | -1.236 | 0.883 | -0.541 | 0 | -0.133 | -0.133 | 1.105 | 0.17 | 0.44 | 0.13 | 0 | 0.16 | 0.03 |
| 4 | 104424934 | rs340451 | 0.813 | -2.09 | 0.262 | -0.468 | 0.079 | 0.002 | 0.002 | 0 | 0.47 | 0.21 | 0.6 | 0.48 | 0.64 | 0.37 |
| 4 | 104447091 | rs445761 | 0.109 | -2.59 | 0.698 | -0.656 | 0.001 | -0.002 | -0.002 | 0 | 0.69 | 0.34 | 0.7 | 0.51 | 0.79 | 0.53 |
| 4 | 104455234 | rs448991 | 0.096 | -2.582 | 0.702 | -0.652 | 0.001 | -0.002 | -0.002 | 0 | 0.7 | 0.34 | 0.7 | 0.51 | 0.79 | 0.53 |
| 4 | 104457398 | rs451907 | 0.096 | -2.582 | 0.701 | -0.652 | 0.003 | -0.004 | -0.004 | 0.095 | 0.7 | 0.33 | 0.7 | 0.51 | 0.79 | 0.53 |
| 4 | 104461487 | rs543330 | 0.096 | -2.582 | 0.701 | -0.652 | 0.003 | -0.004 | -0.004 | 0 | 0.7 | 0.33 | 0.7 | 0.51 | 0.79 | 0.53 |
| 4 | 104467267 | rs10002719 | 0.31 | -2.582 | 1.009 | -0.652 | -0.035 | 0.046 | 0.046 | 0 | 0.7 | 0.47 | 0.71 | 0.51 | 0.79 | 0.53 |
| 4 | 104472071 | rs405717 | -0.52 | -2.83 | 1.195 | -0.793 | 0.023 | -0.073 | -0.073 | 0.008 | 0.88 | 0.73 | 0.86 | 0.98 | 0.85 | 0.93 |
| 4 | 104474749 | rs419391 | 1.212 | -2.906 | 1.073 | -0.835 | 0.073 | -0.035 | -0.035 | 0.284 | 0.61 | 0.19 | 0.68 | 0.51 | 0.74 | 0.52 |
| 4 | 104480799 | rs449075 | 0.611 | -2.727 | 1.307 | -0.716 | -0.008 | 0.007 | 0.007 | 0.045 | 0.69 | 0.34 | 0.7 | 0.51 | 0.79 | 0.54 |
| 4 | 104483419 | rs434269 | 0.656 | -2.741 | 1.278 | -0.755 | 0.001 | -0.003 | -0.003 | 0 | 0.68 | 0.32 | 0.7 | 0.51 | 0.79 | 0.54 |
| 4 | 104490633 | rs2588901 | 0.597 | -2.734 | 1.296 | -0.739 | -0.009 | 0.009 | 0.009 | 0.045 | 0.69 | 0.34 | 0.7 | 0.51 | 0.79 | 0.54 |
| 4 | 104493370 | rs2588907 | 0.597 | -2.734 | 1.237 | -0.739 | 0.023 | -0.026 | -0.026 | 0 | 0.69 | 0.26 | 0.69 | 0.51 | 0.79 | 0.54 |
| 4 | 104495075 | rs2588910 | 0.604 | -2.737 | 1.232 | -0.741 | 0.023 | -0.026 | -0.026 | 0.005 | 0.69 | 0.25 | 0.69 | 0.51 | 0.79 | 0.54 |
| 4 | 104495431 | rs11932144 | 0.149 | -2.72 | 1.194 | -0.871 | -0.032 | -0.086 | -0.086 | 0 | 0.17 | 0.27 | 0.1 | 0 | 0.11 | 0.16 |
| 4 | 104500455 | rs13124966 | 0.535 | -2.672 | 0.731 | -0.835 | 0.028 | 0.043 | 0.043 | 0 | 0.15 | 0.06 | 0.08 | 0 | 0.1 | 0.15 |
| 4 | 104510766 | rs2765 | 0.906 | -2.596 | 0.892 | -0.542 | 0.01 | -0.001 | -0.001 | 0 | 0.55 | 0.58 | 0.62 | 0.49 | 0.69 | 0.39 |
| 4 | 104528309 | rs7654507 | 0.929 | -2.596 | 0.941 | -0.548 | -0.004 | 0 | 0 | 0 | 0.55 | 0.45 | 0.62 | 0.5 | 0.68 | 0.41 |
| 4 | 104532648 | rs10013026 | -0.177 | -2.719 | 0.817 | -0.586 | -0.019 | 0.038 | 0.038 | 0 | 0.18 | 0.22 | 0.16 | 0.48 | 0.06 | 0.36 |
| 4 | 104532950 | rs11722288 | 0.46 | -2.454 | 1.134 | -0.783 | -0.017 | -0.116 | -0.116 | 0.563 | 0.72 | 0.63 | 0.77 | 0.98 | 0.75 | 0.78 |
| 4 | 104542360 | rs11935705 | -0.655 | -2.643 | 0.715 | -0.886 | -0.028 | -0.109 | -0.109 | 0.532 | 0.31 | 0.36 | 0.2 | 0.04 | 0.21 | 0.25 |
| 4 | 104543891 | rs17303520 | 0.913 | -1.881 | -0.163 | -0.803 | -0.005 | -0.01 | -0.01 | 0 | 0.27 | 0.25 | 0.18 | 0.31 | 0.33 | 0.23 |
| 4 | 104545046 | rs17033923 | 0.528 | -1.969 | -0.518 | -0.87 | 0 | -0.034 | -0.034 | 0 | 0.22 | 0.21 | 0.12 | 0.04 | 0.12 | 0.17 |
| 4 | 104545224 | rs3796974 | -0.96 | -2.183 | -0.668 | -0.983 | -0.064 | 0.009 | 0.009 | 0.316 | 0.64 | 0.86 | 0.4 | 0.35 | 0.54 | 0.49 |
| 4 | 104546313 | rs1564613 | 0.565 | -2.022 | -0.735 | -0.894 | -0.025 | -0.071 | -0.071 | 0 | 0.22 | 0.28 | 0.11 | 0.04 | 0.12 | 0.17 |
| 4 | 104552687 | rs12641703 | -0.354 | -1.119 | -1.196 | -0.348 | -0.079 | 0.021 | 0.021 | 0 | 0.6 | 0.91 | 0.4 | 0.35 | 0.53 | 0.49 |
| 4 | 104553287 | rs10516503 | 0.307 | -0.622 | -0.567 | -0.132 | -0.002 | -0.038 | -0.038 | 0 | 0.09 | 0.12 | 0.08 | 0.01 | 0.09 | 0.08 |
| 4 | 104555791 | rs3796972 | 0.167 | 0.068 | -1.259 | 0.55 | -0.066 | -0.089 | -0.089 | 0 | 0.27 | 0.09 | 0.36 | 0.45 | 0.3 | 0.37 |
| 4 | 104556666 | rs3796969 | -0.965 | 0.421 | -1.441 | 0.327 | -0.011 | 0.04 | 0.04 | 0 | 0.35 | 0.83 | 0.32 | 0.24 | 0.33 | 0.33 |
| 4 | 104574265 | rs3796962 | -0.885 | 0.518 | -1.17 | 0.336 | -0.004 | 0.027 | 0.027 | 0 | 0.33 | 0.74 | 0.31 | 0.23 | 0.33 | 0.32 |
| 4 | 104583111 | rs11733295 | -0.171 | 1.059 | -1.046 | 0.477 | -0.013 | 0.001 | 0.001 | 0 | 0.08 | 0.07 | 0.1 | 0.06 | 0.17 | 0.09 |
| 4 | 104595253 | rs10516506 | -0.214 | 0.448 | -0.317 | 1.082 | -0.013 | 0.072 | 0.072 | 0 | 0.15 | 0.25 | 0.17 | 0.41 | 0.11 | 0.31 |
| 4 | 104615281 | rs10019555 | -1.717 | 0.188 | -0.473 | 0.923 | 0.03 | -0.05 | -0.05 | 0.742 | 0.26 | 0.83 | 0.23 | 0.42 | 0.16 | 0.36 |
| 4 | 104640935 | rs3733632 | -1.944 | 0.223 | 0.213 | 0.906 | 0.039 | -0.007 | -0.007 | 0.149 | 0.29 | 0.82 | 0.23 | 0.34 | 0.17 | 0.39 |
| 4 | 104650673 | rs17034020 | -0.256 | 0.162 | 0.802 | 0.845 | 0.048 | -0.001 | -0.001 | 0 | 0.05 | 0.24 | 0.04 | 0 | 0.04 | 0.02 |
| 4 | 104657070 | rs233992 | -1.426 | 0.358 | 0.099 | 0.992 | -0.067 | 0.108 | 0.108 | 0 | 0.51 | 0.92 | 0.63 | 0.72 | 0.46 | 0.6 |
| 4 | 104665738 | rs233991 | 0.341 | 0.498 | 0.24 | 1.103 | -0.005 | -0.005 | -0.005 | 0 | 0.28 | 0.23 | 0.44 | 0.38 | 0.34 | 0.23 |
| 4 | 104672280 | rs4327490 | -0.268 | 0.154 | 0.478 | 1.014 | -0.009 | 0.017 | 0.017 | 0 | 0.17 | 0.23 | 0.15 | 0.33 | 0.11 | 0.31 |

| | | | | | | | | | | | | | | | | |
|---|---|---|---|---|---|---|---|---|---|---|---|---|---|---|---|---|
| 4 | 104674280 | rs6833446 | -0.075 | 0.186 | 0.264 | 1.03 | 0.014 | -0.03 | -0.03 | 4.427 | 0.16 | 0.11 | 0.15 | 0.33 | 0.11 | 0.31 |
| 4 | 104695124 | rs2189223 | -0.134 | 0.19 | 0.261 | 0.849 | 0.006 | 0.012 | 0.012 | 0 | 0.14 | 0.1 | 0.05 | 0.14 | 0.1 | 0.26 |
| 4 | 104712494 | rs12502493 | -0.178 | 1.122 | 0.048 | 0.87 | -0.006 | -0.007 | -0.007 | 0.018 | 0.28 | 0.22 | 0.48 | 0.44 | 0.37 | 0.25 |
| 4 | 104719501 | rs4698905 | -1.973 | 0.916 | -0.213 | 0.513 | 0.005 | 0.137 | 0.137 | 0.152 | 0.6 | 0.84 | 0.74 | 0.85 | 0.6 | 0.7 |
| 4 | 104721398 | rs12640158 | -0.767 | 0.851 | 0.273 | 0.504 | -0.004 | 0.004 | 0.004 | 0 | 0.3 | 0.37 | 0.24 | 0.4 | 0.22 | 0.41 |
| 4 | 104722062 | rs6533109 | -2.053 | 0.827 | 0.053 | 0.42 | -0.032 | 0.151 | 0.151 | 0.126 | 0.6 | 0.93 | 0.74 | 0.85 | 0.6 | 0.7 |
| 4 | 104725088 | rs2189164 | 0.018 | 0.772 | 0.1 | 0.471 | 0.019 | 0.023 | 0.023 | 0 | 0.19 | 0.12 | 0.08 | 0.16 | 0.11 | 0.27 |
| 4 | 104737913 | rs6837683 | -2.266 | 0.81 | 0.087 | 0.434 | -0.007 | 0.151 | 0.151 | 0.114 | 0.6 | 0.89 | 0.74 | 0.85 | 0.59 | 0.67 |
| 4 | 104757193 | rs36096554 | -0.181 | 0.928 | 0.292 | 0.269 | 0.011 | 0.014 | 0.014 | 0 | 0.19 | 0.2 | 0.09 | 0.15 | 0.11 | 0.28 |
| 4 | 104763949 | rs10488855 | -1.906 | 0.955 | 0.202 | 0.355 | -0.029 | -0.064 | -0.064 | 0 | 0.54 | 0.93 | 0.63 | 0.59 | 0.54 | 0.57 |
| 4 | 104766986 | rs6831167 | -1.294 | 0.894 | 0.491 | 0.322 | -0.013 | -0.039 | -0.039 | 0 | 0.27 | 0.69 | 0.14 | 0.15 | 0.16 | 0.32 |
| 4 | 104776075 | rs11933069 | 0.053 | 1.082 | 0.505 | 0.364 | 0.053 | 0.067 | 0.067 | 0.059 | 0.29 | 0.37 | 0.51 | 0.45 | 0.4 | 0.26 |
| 4 | 104778948 | rs17034148 | -0.424 | 0.632 | 0.218 | 0.176 | 0.012 | 0.014 | 0.014 | 0 | 0.19 | 0.19 | 0.09 | 0.15 | 0.11 | 0.27 |
| 4 | 104789707 | rs2106987 | -0.624 | 0.891 | -0.133 | 0.561 | 0.018 | 0.003 | 0.003 | 0.016 | 0.3 | 0.51 | 0.36 | 0.25 | 0.29 | 0.24 |
| 4 | 104792271 | rs11937825 | -0.946 | 0.607 | 0.644 | 0.109 | 0.106 | 0.003 | 0.003 | 1.332 | 0.08 | 0.47 | 0.05 | 0 | 0.05 | 0.04 |
| 4 | 104792456 | rs11937871 | -1.169 | 0.603 | -0.063 | 0.152 | -0.015 | -0.026 | -0.026 | 0 | 0.2 | 0.69 | 0.18 | 0.16 | 0.11 | 0.24 |
| 4 | 104830724 | rs3857207 | -0.797 | 0.922 | -1.596 | -0.228 | -0.098 | 0.111 | 0.111 | 0 | 0.68 | 0.95 | 0.74 | 0.87 | 0.58 | 0.62 |
| 4 | 104831771 | rs12511381 | -0.183 | 0.536 | -1.077 | -0.013 | -0.004 | -0.009 | -0.009 | 0 | 0.36 | 0.24 | 0.44 | 0.59 | 0.35 | 0.3 |
| 4 | 104834423 | rs6845362 | -0.743 | 0.298 | -0.544 | -0.069 | 0.018 | 0.033 | 0.033 | 0 | 0.43 | 0.71 | 0.54 | 0.66 | 0.36 | 0.38 |
| 4 | 104838047 | rs17266258 | -0.113 | 0.308 | -0.536 | -0.04 | -0.005 | -0.014 | -0.014 | 0 | 0.35 | 0.24 | 0.42 | 0.6 | 0.34 | 0.28 |
| 4 | 104838774 | rs12513169 | -0.039 | 0.279 | -0.803 | -0.055 | -0.031 | -0.051 | -0.051 | 0 | 0.34 | 0.09 | 0.41 | 0.6 | 0.34 | 0.28 |
| 4 | 104850169 | rs6533118 | -0.392 | 0.201 | -0.73 | -0.052 | 0.015 | 0.003 | 0.003 | 0 | 0.38 | 0.63 | 0.48 | 0.66 | 0.3 | 0.34 |
| 4 | 104882439 | rs6533126 | 0.366 | 0.354 | -0.418 | -0.188 | -0.002 | -0.028 | -0.028 | 0.017 | 0.59 | 0.54 | 0.66 | 0.8 | 0.62 | 0.56 |
| 4 | 104890866 | rs4272013 | 0.296 | 0.31 | -0.608 | -0.384 | 0.003 | 0.037 | 0.037 | 0 | 0.6 | 0.67 | 0.69 | 0.84 | 0.62 | 0.76 |
| 4 | 104903537 | rs2866672 | -0.018 | 0.445 | -0.595 | -0.153 | 0.003 | -0.026 | -0.026 | 0 | 0.68 | 0.6 | 0.73 | 0.84 | 0.68 | 0.79 |
| 4 | 104903879 | rs2866673 | 0.2 | 0.412 | -1.591 | -0.174 | -0.004 | 0.078 | 0.078 | 0.579 | 0.63 | 0.18 | 0.7 | 0.84 | 0.66 | 0.77 |
| 4 | 104919109 | rs2951111 | -0.444 | 0.252 | -2.154 | 0.713 | -0.02 | -0.185 | -0.185 | 0 | 0.25 | 0.01 | 0.31 | 0.52 | 0.2 | 0.27 |
| 4 | 104944893 | rs1406639 | -0.104 | 0.175 | -2.193 | -0.419 | 0.039 | 0.078 | 0.078 | 0 | 0.58 | 0.82 | 0.57 | 0.74 | 0.44 | 0.47 |
| 4 | 104949606 | rs11097839 | -0.406 | 1.687 | -2.712 | -0.833 | -0.132 | 0.033 | 0.033 | 0 | 0.81 | 0.99 | 0.83 | 0.94 | 0.7 | 0.8 |
| 4 | 104955958 | rs12651382 | 0.686 | -0.195 | -3.049 | -1.692 | 0.053 | -0.007 | -0.007 | 0.262 | 0.27 | 0.14 | 0.4 | 0.74 | 0.23 | 0.3 |
| 4 | 104963745 | rs12645446 | 0.693 | -0.236 | -2.527 | -1.712 | 0.04 | 0.001 | 0.001 | 0 | 0.27 | 0.16 | 0.4 | 0.74 | 0.23 | 0.3 |
| 4 | 104965585 | rs11725491 | 0.775 | -0.304 | -2.424 | -1.768 | -0.006 | -0.068 | -0.068 | 0 | 0.3 | 0.64 | 0.15 | 0 | 0.19 | 0.15 |
| 4 | 104972057 | rs2134026 | 0.761 | -0.473 | -1.349 | -1.851 | -0.044 | -0.152 | -0.152 | 0.121 | 0.28 | 0.46 | 0.14 | 0 | 0.19 | 0.15 |
| 4 | 104978031 | rs950882 | 0.376 | 0.025 | -1.522 | -1.945 | 0.012 | -0.022 | -0.022 | 0 | 0.33 | 0.66 | 0.17 | 0 | 0.24 | 0.15 |
| 4 | 104988768 | rs9996114 | 0.675 | -0.384 | -1.451 | -1.893 | -0.008 | -0.072 | -0.072 | 0 | 0.3 | 0.66 | 0.15 | 0 | 0.19 | 0.15 |
| 4 | 104997276 | rs10019375 | -0.349 | -0.442 | -1.508 | -1.918 | 0.046 | 0.076 | 0.076 | 0 | 0.58 | 0.81 | 0.55 | 0.74 | 0.42 | 0.46 |
| 4 | 105013863 | rs1304013 | 0.196 | -0.328 | -1.516 | -1.949 | 0.032 | 0.045 | 0.045 | 0 | 0.35 | 0.75 | 0.19 | 0.07 | 0.26 | 0.26 |
| 4 | 105039193 | rs17306764 | -0.535 | -0.377 | -1.684 | -2.104 | -0.028 | 0.106 | 0.106 | 0 | 0.63 | 0.9 | 0.66 | 0.8 | 0.47 | 0.57 |
| 4 | 105053161 | rs10470905 | 0.563 | -0.353 | -1.19 | -2.02 | -0.039 | -0.161 | -0.161 | 0 | 0.3 | 0.47 | 0.17 | 0 | 0.24 | 0.16 |
| 4 | 105054988 | rs13140777 | -0.08 | -0.381 | -0.718 | -2.239 | -0.014 | -0.02 | -0.02 | 1.735 | 0.55 | 0.46 | 0.44 | 0.33 | 0.63 | 0.44 |
| 4 | 105059801 | rs882920 | 0.638 | -0.417 | -0.739 | -2.446 | -0.069 | 0.118 | 0.118 | 0 | 0.38 | 0.09 | 0.36 | 0.2 | 0.42 | 0.35 |
| 4 | 105061306 | rs13108980 | -0.471 | -0.492 | -0.376 | -2.368 | 0.027 | 0.046 | 0.046 | 0 | 0.58 | 0.67 | 0.5 | 0.33 | 0.61 | 0.44 |
| 4 | 105078507 | rs795121 | 0.381 | -0.652 | -0.465 | -2.399 | 0.035 | 0.045 | 0.045 | 0 | 0.37 | 0.2 | 0.48 | 0.69 | 0.37 | 0.52 |
| 4 | 105108352 | rs9307291 | -1.13 | -0.706 | 0.555 | -2.473 | 0.033 | 0.122 | 0.122 | 5.505 | 0.06 | 0.41 | 0.05 | 0.11 | 0.01 | 0.01 |
| 4 | 105142122 | rs17034369 | 0.599 | -0.624 | 0.398 | -2.584 | 0.002 | -0.038 | -0.038 | 0 | 0.13 | 0.16 | 0.09 | 0.01 | 0.17 | 0.1 |
| 4 | 105151560 | rs4698914 | -0.726 | -0.522 | 0.214 | -2.692 | -0.002 | 0.052 | 0.052 | 0 | 0.57 | 0.7 | 0.64 | 0.92 | 0.56 | 0.63 |
| 4 | 105157058 | rs886329 | 0.212 | -0.44 | 0.257 | -2.544 | 0.006 | -0.02 | -0.02 | 0.011 | 0.16 | 0.41 | 0.11 | 0.12 | 0.17 | 0.09 |
| 4 | 105183237 | rs1528377 | -0.399 | -0.74 | 0.408 | -2.272 | -0.003 | -0.015 | -0.015 | 0.542 | 0.43 | 0.54 | 0.54 | 0.81 | 0.39 | 0.56 |
| 4 | 105184685 | rs4698917 | -0.507 | -0.733 | 0.384 | -2.269 | -0.017 | -0.042 | -0.042 | 0 | 0.44 | 0.59 | 0.55 | 0.81 | 0.39 | 0.57 |
| 4 | 105231571 | rs10002598 | -0.702 | -0.329 | -0.191 | -2.269 | 0.002 | -0.068 | -0.068 | 0 | 0.56 | 0.59 | 0.63 | 0.89 | 0.5 | 0.65 |
| 4 | 105238884 | rs11097857 | 0.143 | -0.448 | -0.45 | -2.347 | -0.053 | 0.105 | 0.105 | 0 | 0.39 | 0.08 | 0.54 | 0.8 | 0.41 | 0.51 |
| 4 | 105239875 | rs17268078 | 0.011 | 0.117 | 0.161 | -1.876 | 0 | 0.177 | 0.177 | 0 | 0.52 | 0.23 | 0.6 | 0.8 | 0.49 | 0.61 |
| 4 | 105259892 | rs11933178 | -1.967 | 0.64 | 1.107 | -0.557 | 0.031 | 0.132 | 0.132 | 5.885 | 0.05 | 0.33 | 0.03 | 0.1 | 0 | 0.04 |
| 4 | 105311957 | rs11731936 | 0.208 | 1.286 | -0.002 | 0.636 | -0.028 | -0.044 | -0.044 | 0 | 0.84 | 0.9 | 0.64 | 0.31 | 0.79 | 0.67 |
| 4 | 105333354 | rs10007721 | 0.288 | 1.247 | 0.093 | 0.659 | -0.028 | -0.04 | -0.04 | 0 | 0.75 | 0.84 | 0.61 | 0.33 | 0.7 | 0.62 |
| 4 | 105375423 | rs7657799 | -1.772 | 0.581 | 0.625 | 0.764 | -0.022 | 0.031 | 0.031 | 8.553 | 0.1 | 0.79 | 0.06 | 0.16 | 0.02 | 0.07 |
| 4 | 105391999 | rs2298730 | -1.865 | 0.593 | 0.974 | 0.759 | -0.021 | 0.02 | 0.02 | 7.211 | 0.1 | 0.8 | 0.06 | 0.14 | 0.02 | 0.07 |
| 4 | 105400344 | rs10013053 | -0.283 | 0.958 | 0.864 | 0.528 | -0.134 | -0.098 | -0.098 | 1.879 | 0.7 | 0.97 | 0.56 | 0.35 | 0.58 | 0.65 |
| 4 | 105401714 | rs11940318 | -1.59 | 0.82 | 1.02 | 0.512 | -0.038 | 0.028 | 0.028 | 6.177 | 0.09 | 0.77 | 0.05 | 0.13 | 0.01 | 0.03 |
| 4 | 105405240 | rs9307295 | -0.039 | 0.995 | 0.806 | 0.566 | -0.039 | -0.018 | -0.018 | 0 | 0.68 | 0.8 | 0.55 | 0.34 | 0.57 | 0.64 |
| 4 | 105409654 | rs4129288 | -0.208 | 1.06 | 0.494 | 0.578 | -0.105 | -0.061 | -0.061 | 0 | 0.7 | 0.93 | 0.55 | 0.33 | 0.57 | 0.62 |
| 4 | 105409696 | rs3897938 | -1.037 | 0.745 | 1.327 | 0.806 | 0.079 | 0.279 | 0.279 | 0.516 | 0.06 | 0.38 | 0.03 | 0.15 | 0.01 | 0.03 |
| 4 | 105422865 | rs4699128 | -1.087 | 0.837 | 1.023 | 0.841 | 0.058 | 0.184 | 0.184 | 0 | 0.05 | 0.27 | 0.05 | 0.13 | 0.03 | 0.05 |
| 4 | 105425576 | rs17237942 | 0.837 | 0.11 | -0.389 | -0.115 | 0.037 | 0.217 | 0.217 | 0.003 | 0.3 | 0.02 | 0.22 | 0 | 0.21 | 0.2 |
| 4 | 105426879 | rs11097864 | -0.741 | 0.763 | -0.536 | 0.441 | -0.073 | -0.057 | -0.057 | 0.347 | 0.67 | 0.85 | 0.59 | 0.44 | 0.58 | 0.63 |
| 4 | 105428417 | rs7682418 | 0.446 | 0.716 | -0.158 | 0.358 | -0.005 | 0.004 | 0.004 | 0 | 0.25 | 0.17 | 0.17 | 0.06 | 0.31 | 0.28 |
| 4 | 105432249 | rs4414954 | -1.094 | 1.022 | -0.369 | 0.647 | -0.111 | -0.137 | -0.137 | 0.033 | 0.73 | 0.96 | 0.67 | 0.57 | 0.63 | 0.74 |
| 4 | 105434230 | rs7675016 | -1.028 | 0.996 | 0.501 | 0.429 | 0.039 | 0.097 | 0.097 | 0 | 0.11 | 0.4 | 0.11 | 0.15 | 0.11 | 0.16 |
| 4 | 105437362 | rs10516510 | 0.178 | 0.79 | 0.258 | 0.467 | -0.002 | 0 | 0 | 0 | 0.23 | 0.19 | 0.24 | 0.27 | 0.26 | 0.28 |

| | | | | | | | | | | | | | | | | |
|---|---|---|---|---|---|---|---|---|---|---|---|---|---|---|---|---|
| 4 | 105444337 | rs10488865 | -1.128 | 0.785 | 0.442 | 0.275 | 0.035 | 0.087 | 0.087 | 0 | 0.09 | 0.3 | 0.09 | 0.14 | 0.09 | 0.16 |
| 4 | 105450085 | rs2905627 | 0.209 | 1.047 | -0.076 | 0.353 | 0.001 | 0.001 | 0.001 | 0 | 0.32 | 0.58 | 0.27 | 0.29 | 0.32 | 0.38 |
| 4 | 105450529 | rs2051788 | -0.512 | 0.808 | -0.526 | 0.005 | -0.01 | 0.004 | 0.004 | 0 | 0.37 | 0.35 | 0.51 | 0.7 | 0.46 | 0.45 |
| 4 | 105452721 | rs2860529 | -1.697 | 0.772 | -0.933 | 0.137 | 0.045 | 0.226 | 0.226 | 0 | 0.69 | 0.98 | 0.78 | 1 | 0.78 | 0.83 |
| 4 | 105454573 | rs1859157 | -0.157 | 1.378 | -0.501 | 0.558 | -0.04 | -0.055 | -0.055 | 0 | 0.17 | 0.11 | 0.38 | 0.65 | 0.24 | 0.26 |
| 4 | 105473684 | rs3017684 | 0.849 | 1.372 | -0.555 | 0.236 | -0.076 | 0.016 | 0.016 | 0 | 0.26 | 0.11 | 0.26 | 0.2 | 0.41 | 0.18 |
| 4 | 105474268 | rs3017685 | -0.145 | 1.389 | 0.151 | 0.609 | -0.005 | -0.001 | -0.001 | 0 | 0.38 | 0.37 | 0.46 | 0.58 | 0.43 | 0.35 |
| 4 | 105475331 | rs3017687 | -1.289 | 1.186 | 0.205 | 0.404 | 0.029 | 0.037 | 0.037 | 0.056 | 0.63 | 0.75 | 0.62 | 0.74 | 0.69 | 0.71 |
| 4 | 105475940 | rs3017689 | -1.461 | 1.079 | 0.116 | 0.414 | 0.059 | 0.055 | 0.055 | 0.221 | 0.64 | 0.91 | 0.63 | 0.74 | 0.71 | 0.72 |
| 4 | 105475979 | rs6840786 | -0.313 | 1.166 | 0.399 | 0.433 | 0.026 | -0.045 | -0.045 | 0 | 0.26 | 0.49 | 0.17 | 0.16 | 0.28 | 0.36 |
| 4 | 105476932 | rs17269632 | 0.417 | 1.111 | 0.487 | 0.411 | -0.004 | 0.002 | 0.002 | 0 | 0.19 | 0.15 | 0.1 | 0.05 | 0.21 | 0.24 |
| 4 | 105492066 | rs17034632 | 0.386 | 1.19 | 0.173 | 0.666 | -0.044 | -0.077 | -0.077 | 0 | 0.14 | 0.03 | 0.24 | 0.38 | 0.15 | 0.18 |
| 4 | 105492822 | rs1468231 | -1.198 | 1.224 | 0.063 | 0.466 | 0.022 | 0.065 | 0.065 | 0.19 | 0.63 | 0.91 | 0.6 | 0.74 | 0.67 | 0.71 |
| 4 | 105493321 | rs3017707 | -1.027 | 1.074 | 0.201 | 0.487 | 0.052 | 0.062 | 0.062 | 0.183 | 0.62 | 0.8 | 0.62 | 0.74 | 0.67 | 0.64 |
| 4 | 105493835 | rs7668924 | -1.02 | 1.11 | 0.415 | 0.594 | -0.008 | -0.006 | -0.006 | 0.461 | 0.39 | 0.62 | 0.39 | 0.54 | 0.4 | 0.47 |
| 4 | 105497463 | rs11934708 | -0.981 | 1.149 | 0.061 | 0.613 | 0 | 0 | 0 | 0 | 0.39 | 0.61 | 0.39 | 0.54 | 0.39 | 0.47 |
| 4 | 105507674 | rs3017722 | -0.619 | 0.554 | -0.585 | 0.126 | -0.011 | 0.002 | 0.002 | 0 | 0.61 | 0.81 | 0.42 | 0.37 | 0.62 | 0.61 |
| 4 | 105509600 | rs494660 | 1.466 | 0.405 | -0.721 | -0.076 | -0.026 | -0.003 | -0.003 | 0 | 0.23 | 0.09 | 0.2 | 0.21 | 0.27 | 0.18 |
| 4 | 105511810 | rs17034684 | -0.05 | 0.235 | -0.194 | 0.243 | -0.001 | 0.002 | 0.002 | 0 | 0.29 | 0.28 | 0.17 | 0.26 | 0.3 | 0.33 |
| 4 | 105512358 | rs607012 | -0.931 | 0.218 | -0.885 | 0.239 | -0.012 | -0.045 | -0.045 | 0 | 0.66 | 0.81 | 0.52 | 0.49 | 0.69 | 0.66 |
| 4 | 105515047 | rs2074395 | 0.993 | 0.55 | -1.059 | 0.346 | -0.051 | -0.04 | -0.04 | 0.576 | 0.08 | 0.01 | 0.07 | 0.15 | 0.18 | 0.16 |
| 4 | 105515423 | rs488076 | 0.574 | -0.026 | -0.65 | 0.063 | -0.016 | 0.003 | 0.003 | 0 | 0.26 | 0.16 | 0.23 | 0.24 | 0.33 | 0.21 |
| 4 | 105517973 | rs9307296 | 0.221 | 0.066 | -0.427 | 0.128 | 0.003 | 0.003 | 0.003 | 0 | 0.41 | 0.41 | 0.37 | 0.33 | 0.38 | 0.39 |
| 4 | 105518122 | rs555039 | -0.658 | 0.357 | -0.388 | 0.006 | 0.013 | 0.015 | 0.015 | 0.135 | 0.9 | 0.84 | 0.83 | 0.84 | 0.88 | 0.82 |
| 4 | 105519735 | rs639114 | -0.63 | 0.285 | -0.755 | -0.065 | -0.073 | -0.047 | -0.047 | 0 | 0.88 | 0.99 | 0.76 | 0.87 | 0.82 | 0.75 |
| 4 | 105528713 | rs680550 | 0.529 | -0.749 | -0.729 | -0.796 | 0.034 | 0.089 | 0.089 | 0 | 0.53 | 0.66 | 0.62 | 0.82 | 0.61 | 0.48 |
| 4 | 105528877 | rs477451 | 1.675 | -0.608 | -0.702 | -0.775 | -0.019 | -0.007 | -0.007 | 0.572 | 0.2 | 0.12 | 0.29 | 0.27 | 0.28 | 0.16 |
| 4 | 105543843 | rs578265 | 1.623 | -0.644 | -0.618 | -0.794 | -0.013 | -0.003 | -0.003 | 0.305 | 0.21 | 0.14 | 0.29 | 0.27 | 0.28 | 0.16 |
| 4 | 105565218 | rs728123 | 0.764 | -0.93 | -0.712 | -1.002 | 0.065 | 0.078 | 0.078 | 0 | 0.29 | 0.16 | 0.13 | 0.05 | 0.17 | 0.17 |
| 4 | 105566441 | rs624909 | 0.502 | -0.462 | -0.374 | -0.778 | 0.029 | 0.071 | 0.071 | 0 | 0.49 | 0.32 | 0.41 | 0.33 | 0.45 | 0.31 |
| 4 | 105581394 | rs4698923 | 0.606 | -0.454 | 0.015 | -0.763 | -0.025 | -0.049 | -0.049 | 6.563 | 0.29 | 0.13 | 0.31 | 0.55 | 0.34 | 0.28 |
| 4 | 105598022 | rs506932 | -1.465 | -0.236 | 0.164 | -0.742 | -0.063 | -0.033 | -0.033 | 0 | 0.89 | 0.99 | 0.8 | 0.88 | 0.84 | 0.75 |
| 4 | 105619927 | rs2522515 | 1.574 | -0.545 | -0.12 | -0.893 | -0.035 | -0.036 | -0.036 | 0.386 | 0.24 | 0.07 | 0.3 | 0.31 | 0.29 | 0.16 |
| 4 | 105624313 | rs2651339 | 0.179 | -0.562 | 0.061 | -0.897 | 0.006 | 0.001 | 0.001 | 0 | 0.4 | 0.42 | 0.54 | 0.44 | 0.46 | 0.5 |
| 4 | 105653263 | rs2522502 | 0.366 | -0.577 | -0.323 | -0.933 | 0.006 | 0.015 | 0.015 | 0 | 0.56 | 0.42 | 0.63 | 0.82 | 0.65 | 0.45 |
| 4 | 105665680 | rs2522507 | -0.711 | -0.626 | -0.646 | -1.203 | -0.063 | -0.124 | -0.124 | 0 | 0.69 | 0.51 | 0.85 | 0.95 | 0.82 | 0.71 |
| 4 | 105691743 | rs4698925 | 0.771 | -0.163 | -1.593 | -1.005 | -0.085 | -0.144 | -0.144 | 1.028 | 0.29 | 0.02 | 0.3 | 0.54 | 0.33 | 0.28 |
| 4 | 105691934 | rs9996250 | 0.676 | 0.404 | -1.046 | -0.666 | 0.015 | -0.01 | -0.01 | 0 | 0.12 | 0.15 | 0.22 | 0.09 | 0.16 | 0.25 |
| 4 | 105696569 | rs2522517 | 0.321 | 0.415 | -1.367 | -0.652 | 0.006 | -0.002 | -0.002 | 0 | 0.14 | 0.15 | 0.22 | 0.14 | 0.17 | 0.27 |
| 4 | 105700542 | rs2725762 | 0.909 | 0.067 | -1.65 | -0.726 | -0.091 | -0.087 | -0.087 | 0.441 | 0.24 | 0.01 | 0.29 | 0.3 | 0.28 | 0.16 |
| 4 | 105704161 | rs1981542 | -0.118 | -0.395 | -0.78 | -0.847 | -0.024 | -0.094 | -0.094 | 0.963 | 0.75 | 0.68 | 0.86 | 1 | 0.83 | 0.84 |
| 4 | 105706327 | rs2725759 | 0.534 | 0.182 | -1.778 | -0.719 | -0.146 | -0.018 | -0.018 | 0.661 | 0.35 | 0.03 | 0.34 | 0.33 | 0.38 | 0.29 |
| 4 | 105708053 | rs2522524 | 0.159 | -0.75 | -0.685 | -1.188 | -0.036 | -0.073 | -0.073 | 0.071 | 0.67 | 0.56 | 0.83 | 0.91 | 0.78 | 0.74 |
| 4 | 105715931 | rs2107003 | -0.572 | -0.911 | -1.218 | -1.267 | 0.055 | 0.101 | 0.101 | 0 | 0.78 | 0.94 | 0.89 | 1 | 0.84 | 0.87 |
| 4 | 105716540 | rs2051780 | 0.429 | -0.904 | -0.519 | -1.248 | -0.024 | -0.051 | -0.051 | 3.536 | 0.65 | 0.46 | 0.83 | 0.9 | 0.78 | 0.68 |
| 4 | 105747315 | rs2725750 | -0.481 | -0.602 | -0.576 | -0.76 | -0.021 | -0.007 | -0.007 | 0 | 0.44 | 0.51 | 0.46 | 0.64 | 0.32 | 0.44 |
| 4 | 105758739 | rs2725771 | -0.522 | -0.703 | -0.335 | -0.81 | 0.072 | 0.028 | 0.028 | 0 | 0.45 | 0.67 | 0.47 | 0.63 | 0.32 | 0.46 |
| 4 | 105763065 | rs2079112 | -0.75 | -0.093 | -0.785 | -1.035 | -0.005 | 0.021 | 0.021 | 0.096 | 0.57 | 0.8 | 0.55 | 0.64 | 0.43 | 0.55 |
| 4 | 105768938 | rs2522482 | -0.882 | 0.401 | -0.018 | -0.617 | -0.005 | 0.001 | 0.001 | 0.447 | 0.38 | 0.44 | 0.49 | 0.62 | 0.35 | 0.43 |
| 4 | 105772094 | rs3886330 | -1.037 | 0.399 | -0.357 | -0.364 | 0.016 | -0.03 | -0.03 | 0 | 0.38 | 0.71 | 0.46 | 0.45 | 0.31 | 0.4 |
| 4 | 105781935 | rs4698926 | -1.472 | 0.607 | -0.581 | -0.431 | 0.009 | -0.036 | -0.036 | 0 | 0.46 | 0.8 | 0.54 | 0.46 | 0.41 | 0.38 |
| 4 | 105792576 | rs6850952 | 1.528 | 1.434 | -0.782 | -0.563 | 0.043 | -0.002 | -0.002 | 0 | 0.45 | 0.19 | 0.39 | 0.45 | 0.39 | 0.55 |
| 4 | 105794260 | rs6533163 | 1.51 | 1.4 | -0.721 | -0.455 | 0.012 | 0.003 | 0.003 | 0.035 | 0.47 | 0.16 | 0.41 | 0.46 | 0.43 | 0.55 |
| 4 | 105803313 | rs4585303 | -0.56 | 1.545 | -0.288 | -0.368 | 0.049 | 0 | 0 | 0 | 0.9 | 0.8 | 0.91 | 0.93 | 0.81 | 0.92 |
| 4 | 105816898 | rs9998015 | -1.887 | 1.263 | 0.189 | -0.521 | 0.037 | -0.017 | -0.017 | 0 | 0.48 | 0.73 | 0.57 | 0.49 | 0.41 | 0.36 |
| 4 | 105821431 | rs10516511 | -1.506 | 1.026 | 0.179 | -0.504 | 0.037 | -0.02 | -0.02 | 0 | 0.4 | 0.68 | 0.51 | 0.47 | 0.3 | 0.31 |
| 4 | 105835961 | rs4264829 | 0.27 | 1.646 | -0.256 | -0.189 | 0 | 0.003 | 0.003 | 0 | 0.06 | 0.03 | 0.04 | 0 | 0.05 | 0.01 |
| 4 | 105838702 | rs7695587 | -1.093 | 1.624 | -0.205 | -0.199 | -0.009 | -0.039 | -0.039 | 0.04 | 0.11 | 0.2 | 0.11 | 0.01 | 0.07 | 0.04 |
| 4 | 105844273 | rs10005603 | -1.307 | 1.6 | -0.205 | -0.192 | -0.007 | -0.055 | -0.055 | 2.224 | 0.12 | 0.3 | 0.12 | 0.02 | 0.07 | 0.04 |
| 4 | 105845011 | rs9637619 | -0.433 | 1.743 | -1.208 | -0.084 | 0.002 | -0.144 | -0.144 | 0.816 | 0.19 | 0.02 | 0.33 | 0.45 | 0.16 | 0.25 |
| 4 | 105849384 | rs11935593 | 0.09 | 1.927 | -0.882 | -0.123 | -0.001 | -0.022 | -0.022 | 0 | 0.07 | 0.05 | 0.04 | 0 | 0.05 | 0.01 |
| 4 | 105860001 | rs10032763 | -0.642 | 2.216 | -1.009 | 0.38 | 0.002 | -0.002 | -0.002 | 0 | 0.09 | 0.09 | 0.18 | 0.09 | 0.05 | 0.05 |
| 4 | 105866724 | rs17239519 | 0.588 | 2.675 | -0.9 | 0.435 | -0.001 | -0.003 | -0.003 | 0.453 | 0.07 | 0.05 | 0.09 | 0 | 0.15 | 0.06 |
| 4 | 105869983 | rs7660636 | -0.256 | 0.834 | -0.713 | 0.892 | 0 | 0.016 | 0.016 | 0 | 0.05 | 0.09 | 0.17 | 0.09 | 0.04 | 0.04 |
| 4 | 105872720 | rs4698929 | -0.559 | 0.881 | -1.004 | 0.945 | -0.005 | -0.04 | -0.04 | 0 | 0.16 | 0.07 | 0.22 | 0.39 | 0.16 | 0.3 |
| 4 | 105890017 | rs1490586 | -1.039 | 0.539 | -0.739 | 0.705 | 0.004 | -0.009 | -0.009 | 0 | 0.32 | 0.31 | 0.44 | 0.52 | 0.24 | 0.37 |
| 4 | 105890515 | rs2291507 | -0.815 | 0.456 | -0.779 | 0.712 | 0.011 | -0.1 | -0.1 | 0.018 | 0.25 | 0.09 | 0.4 | 0.48 | 0.19 | 0.35 |
| 4 | 105892565 | rs17035170 | 0.51 | 1.33 | -0.026 | -0.009 | 0.064 | -0.039 | -0.039 | 0.321 | 0.08 | 0.19 | 0.11 | 0 | 0.17 | 0.04 |
| 4 | 105897524 | rs7688673 | 0.815 | 0.392 | -0.911 | -0.699 | 0.017 | 0.021 | 0.021 | 0 | 0.19 | 0.14 | 0.16 | 0.05 | 0.12 | 0.32 |

| | | | | | | | | | | | | | | | | |
|---|---|---|---|---|---|---|---|---|---|---|---|---|---|---|---|---|
| 4 | 105897585 | rs1490579 | -0.5 | -0.17 | 0.152 | -1.066 | -0.015 | -0.004 | -0.004 | 0 | 0.1 | 0.41 | 0.09 | 0.04 | 0.04 | 0.03 |
| 4 | 105911182 | rs1490583 | -0.158 | -0.075 | -0.179 | -1.939 | -0.002 | 0.037 | 0.037 | 0 | 0.42 | 0.33 | 0.35 | 0.06 | 0.45 | 0.45 |
| 4 | 105921314 | rs12500787 | 0.038 | -0.018 | -0.16 | -1.944 | -0.013 | 0.1 | 0.1 | 0 | 0.4 | 0.23 | 0.35 | 0.06 | 0.45 | 0.44 |
| 4 | 105923047 | rs902444 | -0.762 | 0.015 | -0.063 | -1.912 | 0.006 | 0.158 | 0.158 | 0 | 0.51 | 0.71 | 0.44 | 0.11 | 0.49 | 0.47 |
| 4 | 105925099 | rs1873361 | 0.146 | -0.212 | -0.019 | -1.954 | -0.009 | -0.047 | -0.047 | 0 | 0.35 | 0.35 | 0.29 | 0.07 | 0.27 | 0.36 |
| 4 | 105933321 | rs4699145 | 0.719 | 0.223 | -1.005 | -1.599 | -0.071 | -0.116 | -0.116 | 0 | 0.23 | 0.03 | 0.31 | 0.65 | 0.24 | 0.43 |
| 4 | 105936370 | rs13151168 | 1.44 | 0.19 | -1.031 | -1.905 | -0.01 | 0.05 | 0.05 | 0 | 0.17 | 0.01 | 0.2 | 0.06 | 0.15 | 0.25 |
| 4 | 105937236 | rs10027139 | -1.306 | 0.617 | -1.049 | -2.078 | -0.011 | 0.011 | 0.011 | 0 | 0.91 | 0.93 | 0.92 | 0.98 | 0.85 | 0.98 |
| 4 | 105945776 | rs1490589 | -0.67 | 0.582 | -1.064 | -2.09 | -0.037 | 0.093 | 0.093 | 0 | 0.74 | 0.92 | 0.71 | 0.91 | 0.71 | 0.73 |
| 4 | 105986597 | rs195149 | -0.952 | 0.525 | -0.836 | -2.187 | 0.013 | 0.127 | 0.127 | 0 | 0.47 | 0.67 | 0.4 | 0.09 | 0.44 | 0.39 |
| 4 | 105987213 | rs6811026 | 0.557 | 0.651 | -0.594 | -1.955 | -0.03 | -0.122 | -0.122 | 0 | 0.17 | 0.38 | 0.1 | 0 | 0.12 | 0.11 |
| 4 | 105988057 | rs195148 | 0.476 | 0.991 | -1.162 | -1.561 | -0.01 | 0.009 | 0.009 | 0.081 | 0.1 | 0.09 | 0.09 | 0.02 | 0.17 | 0.02 |
| 4 | 106005985 | rs2079135 | -0.609 | 0.788 | -1.272 | -1.917 | 0.009 | 0.036 | 0.036 | 0 | 0.55 | 0.57 | 0.47 | 0.1 | 0.59 | 0.43 |
| 4 | 106006036 | rs2079134 | 0.352 | 0.921 | -1.397 | -2.215 | -0.048 | 0.068 | 0.068 | 0 | 0.18 | 0.07 | 0.14 | 0 | 0.27 | 0.13 |
| 4 | 106009763 | rs2301718 | 0.628 | 0.551 | -1.622 | -2.118 | -0.035 | -0.08 | -0.08 | 1.909 | 0.24 | 0.01 | 0.39 | 0.72 | 0.23 | 0.44 |
| 4 | 106010433 | rs12509636 | 0.083 | 1.022 | -1.409 | -1.976 | 0.005 | -0.004 | -0.004 | 2.613 | 0.39 | 0.41 | 0.54 | 0.92 | 0.34 | 0.51 |
| 4 | 106016627 | rs764275 | -1.103 | 1.081 | -1.664 | -2.155 | 0 | -0.04 | -0.04 | 0.045 | 0.66 | 0.67 | 0.75 | 0.95 | 0.73 | 0.62 |
| 4 | 106022992 | rs4141270 | 2.06 | 1.277 | -1.725 | -1.501 | -0.014 | -0.039 | -0.039 | 0 | 0.09 | 0 | 0.3 | 0.6 | 0.09 | 0.21 |
| 4 | 106028073 | rs17429675 | 0.423 | 1.422 | -1.361 | -2.458 | 0.004 | -0.011 | -0.011 | 1.12 | 0.4 | 0.41 | 0.55 | 0.92 | 0.34 | 0.51 |
| 4 | 106032945 | rs2214409 | -0.049 | 1.539 | -1.41 | -2.502 | -0.005 | 0.049 | 0.049 | 0 | 0.48 | 0.42 | 0.64 | 0.92 | 0.49 | 0.55 |
| 4 | 106034478 | rs4698931 | -0.59 | 0.729 | -1.105 | -2.302 | -0.001 | -0.047 | -0.047 | 0.292 | 0.54 | 0.49 | 0.66 | 0.94 | 0.57 | 0.59 |
| 4 | 106036545 | rs17035270 | -0.018 | 0.796 | -0.814 | -1.115 | 0.018 | -0.055 | -0.055 | 0 | 0.16 | 0.23 | 0.11 | 0.03 | 0.22 | 0.06 |
| 4 | 106048291 | rs17035289 | -0.228 | -0.331 | -0.726 | -1.11 | -0.04 | -0.048 | -0.048 | 0 | 0.3 | 0.39 | 0.21 | 0.16 | 0.19 | 0.25 |
| 4 | 106061534 | rs7679673 | -0.813 | 0.036 | -0.957 | -1.329 | 0.008 | 0.025 | 0.025 | 0 | 0.56 | 0.65 | 0.52 | 0.81 | 0.39 | 0.55 |
| 4 | 106064683 | rs17617028 | 1.124 | -0.162 | -1.054 | -1.389 | -0.01 | -0.057 | -0.057 | 0 | 0.22 | 0.05 | 0.3 | 0.64 | 0.21 | 0.28 |
| 4 | 106066293 | rs17035311 | 0.318 | -0.239 | -0.973 | -1.402 | -0.047 | -0.044 | -0.044 | 0 | 0.3 | 0.41 | 0.2 | 0.18 | 0.16 | 0.25 |
| 4 | 106071064 | rs974801 | 0.014 | -0.137 | -1.164 | -1.546 | -0.014 | 0.006 | 0.006 | 0 | 0.34 | 0.27 | 0.4 | 0.64 | 0.41 | 0.34 |
| 4 | 106075498 | rs2189234 | -1.573 | -0.745 | -1.284 | -2.059 | 0.003 | -0.014 | -0.014 | 0 | 0.64 | 0.58 | 0.61 | 0.63 | 0.63 | 0.53 |
| 4 | 106084778 | rs9790517 | 1.627 | -0.58 | -1.419 | -1.882 | -0.017 | -0.061 | -0.061 | 0 | 0.22 | 0.05 | 0.29 | 0.63 | 0.21 | 0.28 |
| 4 | 106091978 | rs17035323 | 1.424 | -0.484 | -1.201 | -1.884 | -0.022 | 0.02 | 0.02 | 0.007 | 0.11 | 0.06 | 0.1 | 0 | 0.2 | 0.06 |
| 4 | 106108902 | rs17508261 | 1.309 | -0.481 | -0.664 | -1.541 | 0.002 | 0.008 | 0.008 | 0 | 0.16 | 0.11 | 0.09 | 0.06 | 0.14 | 0.09 |
| 4 | 106115450 | rs1015521 | 0.291 | -0.559 | -0.725 | -1.238 | 0.003 | -0.018 | -0.018 | 0.639 | 0.38 | 0.15 | 0.38 | 0.54 | 0.34 | 0.39 |
| 4 | 106118756 | rs7698522 | 0.21 | -0.661 | -0.437 | -1.085 | -0.009 | -0.021 | -0.021 | 0 | 0.11 | 0.2 | 0.11 | 0.06 | 0.08 | 0.09 |
| 4 | 106125715 | rs9637601 | 2.189 | -0.828 | -0.722 | -1.002 | 0.038 | -0.039 | -0.039 | 0 | 0.08 | 0.01 | 0.09 | 0.2 | 0.02 | 0.12 |
| 4 | 106137033 | rs2047409 | 0.778 | -0.966 | -0.466 | -1.338 | 0.031 | 0.103 | 0.103 | 0 | 0.44 | 0.28 | 0.47 | 0.19 | 0.63 | 0.44 |
| 4 | 106144735 | rs6839705 | 0.132 | -1.042 | -0.213 | -1.365 | 0.03 | 0.052 | 0.052 | 0 | 0.52 | 0.66 | 0.52 | 0.26 | 0.64 | 0.48 |
| 4 | 106152984 | rs7683416 | 1.052 | -1.037 | -0.053 | -1.28 | -0.004 | 0.028 | 0.028 | 0 | 0.39 | 0.35 | 0.4 | 0.2 | 0.57 | 0.38 |
| 4 | 106155751 | rs6843141 | -0.539 | -1.06 | -0.099 | -1.293 | -0.01 | 0.024 | 0.024 | 1.564 | 0.07 | 0.29 | 0.03 | 0.06 | 0.02 | 0.03 |
| 4 | 106158216 | rs3796927 | -0.038 | -1.143 | -0.085 | -1.315 | 0.016 | 0.069 | 0.069 | 4.852 | 0.05 | 0.28 | 0.02 | 0.06 | 0.01 | 0 |
| 4 | 106160365 | rs7670522 | 1.243 | -1.105 | -0.247 | -1.349 | -0.013 | 0.052 | 0.052 | 0 | 0.38 | 0.3 | 0.4 | 0.2 | 0.56 | 0.38 |
| 4 | 106177882 | rs4464576 | 1.002 | -1.14 | -0.304 | -1.417 | -0.006 | 0.062 | 0.062 | 0 | 0.21 | 0.09 | 0.17 | 0.05 | 0.2 | 0.16 |
| 4 | 106192457 | rs1498126 | 1.011 | -1.146 | -0.31 | -1.42 | -0.006 | 0.059 | 0.059 | 0 | 0.21 | 0.09 | 0.18 | 0.05 | 0.21 | 0.16 |
| 4 | 106195541 | rs2726492 | -1.98 | -1.23 | -0.376 | -1.269 | -0.006 | -0.104 | -0.104 | 0 | 0.8 | 0.96 | 0.72 | 0.51 | 0.82 | 0.75 |
| 4 | 106196951 | rs2454206 | 2.33 | -1.183 | -0.402 | -1.431 | -0.102 | 0.053 | 0.053 | 0 | 0.29 | 0.07 | 0.28 | 0.19 | 0.37 | 0.31 |
| 4 | 106201556 | rs2647256 | -1.943 | -1.228 | -0.328 | -1.341 | -0.003 | -0.107 | -0.107 | 0 | 0.79 | 0.95 | 0.72 | 0.51 | 0.82 | 0.69 |
| 4 | 106202633 | rs904274 | -0.835 | -0.986 | 0.269 | -0.989 | -0.093 | 0.039 | 0.039 | 0 | 0.19 | 0.57 | 0.14 | 0.27 | 0.05 | 0.16 |
| 4 | 106213874 | rs6533185 | 0.098 | -0.811 | 0.015 | -0.879 | -0.087 | 0.071 | 0.071 | 0.138 | 0.14 | 0.32 | 0.11 | 0.27 | 0.03 | 0.19 |
| 4 | 106216205 | rs12639764 | 1.123 | -0.374 | -0.603 | -1.155 | -0.058 | 0.06 | 0.06 | 0 | 0.3 | 0.15 | 0.3 | 0.19 | 0.38 | 0.32 |
| 4 | 106239038 | rs6827660 | -0.708 | -0.227 | -0.208 | -0.795 | -0.002 | 0 | 0 | 0 | 0.2 | 0.31 | 0.22 | 0.22 | 0.21 | 0.22 |
| 4 | 106253209 | rs17035461 | 0.419 | -0.112 | -0.191 | -0.786 | 0.025 | -0.028 | -0.028 | 0 | 0.09 | 0.12 | 0.1 | 0.01 | 0.18 | 0.07 |
| 4 | 106256521 | rs2647259 | 0.522 | -0.299 | -0.39 | -0.912 | -0.028 | 0.068 | 0.068 | 0 | 0.36 | 0.24 | 0.31 | 0.17 | 0.41 | 0.36 |
| 4 | 106263450 | rs2726485 | 0.433 | -0.252 | -0.395 | -0.867 | -0.023 | 0.017 | 0.017 | 0 | 0.41 | 0.12 | 0.39 | 0.38 | 0.41 | 0.48 |
| 4 | 106263681 | rs6823995 | 1.483 | 0.254 | -0.22 | -0.686 | 0.075 | -0.02 | -0.02 | 0 | 0.05 | 0.1 | 0.09 | 0.01 | 0.16 | 0.06 |
| 4 | 106264681 | rs2726486 | -2.239 | 0.008 | -0.72 | -0.791 | 0.008 | -0.095 | -0.095 | 0.545 | 0.87 | 0.99 | 0.78 | 0.48 | 0.92 | 0.77 |
| 4 | 106270291 | rs2647266 | 1.094 | 0.049 | -0.566 | -0.695 | -0.061 | 0.073 | 0.073 | 0 | 0.33 | 0.11 | 0.32 | 0.23 | 0.39 | 0.4 |
| 4 | 106282165 | rs924283 | -1.227 | 0.11 | -0.337 | -0.553 | 0.002 | 0.03 | 0.03 | 0 | 0.83 | 0.85 | 0.71 | 0.56 | 0.77 | 0.8 |
| 4 | 106285090 | rs2647254 | 0.3 | 0.056 | -0.042 | -0.675 | 0.016 | 0.027 | 0.027 | 0 | 0.41 | 0.37 | 0.35 | 0.3 | 0.35 | 0.33 |
| 4 | 106288827 | rs2726496 | 0.482 | 0.113 | 0.038 | -0.662 | 0.03 | 0.169 | 0.169 | 0 | 0.21 | 0.39 | 0.32 | 0.44 | 0.25 | 0.18 |
| 4 | 106293125 | rs13108489 | -0.091 | 0.09 | -0.182 | -0.644 | 0.018 | 0.132 | 0.132 | 0 | 0.24 | 0.45 | 0.33 | 0.44 | 0.28 | 0.2 |
| 4 | 106294756 | rs2636693 | 1.477 | 0.076 | -0.316 | -0.625 | 0.014 | -0.03 | -0.03 | 0.642 | 0.14 | 0.1 | 0.22 | 0.43 | 0.1 | 0.12 |
| 4 | 106297376 | rs2713833 | -0.233 | 0.226 | -0.11 | -0.693 | -0.001 | -0.003 | -0.003 | 0 | 0.56 | 0.5 | 0.62 | 0.68 | 0.64 | 0.65 |
| 4 | 106298394 | rs2726503 | 0.361 | 0.162 | -1.028 | -0.51 | -0.03 | 0.087 | 0.087 | 0 | 0.46 | 0.18 | 0.52 | 0.68 | 0.46 | 0.57 |
| 4 | 106316427 | rs2726528 | 1.038 | 0.266 | -1.117 | -0.514 | -0.065 | 0.052 | 0.052 | 0 | 0.31 | 0.08 | 0.3 | 0.25 | 0.36 | 0.41 |
| 4 | 106318456 | rs959086 | 1.4 | -0.062 | -1.106 | -1.007 | 0.002 | -0.02 | -0.02 | 0 | 0.13 | 0.13 | 0.06 | 0 | 0.1 | 0.07 |
| 4 | 106321423 | rs17035553 | -0.049 | 0.034 | -0.24 | -1.157 | 0.092 | -0.065 | -0.065 | 0.075 | 0.11 | 0.41 | 0.12 | 0.01 | 0.18 | 0.07 |
| 4 | 106325106 | rs2713871 | 0.57 | -0.187 | 0.206 | -1.134 | 0 | 0.015 | 0.015 | 0 | 0.5 | 0.44 | 0.55 | 0.71 | 0.49 | 0.57 |
| 4 | 106335490 | rs2726471 | 1.088 | 0.803 | -0.162 | -1.129 | 0.038 | 0.049 | 0.049 | 0 | 0.44 | 0.21 | 0.45 | 0.67 | 0.34 | 0.54 |
| 4 | 106338434 | rs2636740 | -2.404 | 0.27 | -0.319 | -1.069 | 0.005 | -0.05 | -0.05 | 0 | 0.92 | 1 | 0.9 | 0.77 | 0.97 | 0.84 |
| 4 | 106338490 | rs11721982 | 2.561 | 0.424 | -0.288 | -1.097 | -0.011 | 0 | 0 | 0 | 0.06 | 0 | 0.03 | 0.06 | 0.04 | 0.02 |

| | | | | | | | | | | | | | | | | |
|---|---|---|---|---|---|---|---|---|---|---|---|---|---|---|---|---|
| 4 | 106346206 | rs2726516 | 0.126 | 0.666 | -0.1 | -1.118 | 0 | 0.001 | 0.001 | 0 | 0.53 | 0.52 | 0.6 | 0.67 | 0.59 | 0.62 |
| 4 | 106353784 | rs753972 | -0.153 | 0.638 | -0.021 | -1.106 | 0.001 | 0.002 | 0.002 | 0 | 0.58 | 0.57 | 0.63 | 0.67 | 0.64 | 0.63 |
| 4 | 106356395 | rs3898404 | 0.398 | 0.688 | 0.12 | -1.135 | 0.081 | -0.056 | -0.056 | 1.183 | 0.1 | 0.31 | 0.11 | 0.01 | 0.18 | 0.07 |
| 4 | 106356581 | rs2636743 | -0.016 | 0.648 | 0.117 | -1.121 | -0.005 | -0.007 | -0.007 | 0 | 0.56 | 0.45 | 0.63 | 0.67 | 0.64 | 0.63 |
| 4 | 106364129 | rs923587 | 0.844 | 0.585 | -0.471 | -1.077 | -0.042 | 0.074 | 0.074 | 0 | 0.46 | 0.14 | 0.52 | 0.67 | 0.46 | 0.56 |
| 4 | 106366209 | rs2636751 | 1.14 | 0.188 | -0.465 | -0.812 | -0.031 | 0.037 | 0.037 | 0 | 0.42 | 0.09 | 0.48 | 0.67 | 0.42 | 0.55 |
| 4 | 106366687 | rs7671891 | -0.128 | -0.333 | -0.077 | -0.584 | 0.003 | -0.026 | -0.026 | 0 | 0.07 | 0.28 | 0.03 | 0 | 0.03 | 0.02 |
| 4 | 106382268 | rs2029513 | -2.841 | -0.342 | -0.733 | -0.011 | 0.013 | -0.075 | -0.075 | 0 | 0.89 | 1 | 0.89 | 0.66 | 0.94 | 0.91 |
| 4 | 106386760 | rs12643660 | -0.08 | -0.261 | -0.63 | -0.294 | 0.008 | 0.052 | 0.052 | 0 | 0.42 | 0.28 | 0.33 | 0.29 | 0.36 | 0.36 |
| 4 | 106388702 | rs4699176 | -1.894 | -1.115 | -0.245 | 0.337 | 0.01 | -0.068 | -0.068 | 0 | 0.87 | 0.95 | 0.8 | 0.58 | 0.89 | 0.89 |
| 4 | 106390734 | rs2298733 | -0.091 | -1.434 | -0.085 | 0.136 | -0.022 | 0 | 0 | 0 | 0.27 | 0.57 | 0.2 | 0.26 | 0.14 | 0.23 |
| 4 | 106391093 | rs2298732 | -1.1 | -1.47 | -0.111 | 0.097 | 0.013 | -0.041 | -0.041 | 0 | 0.82 | 0.84 | 0.77 | 0.58 | 0.85 | 0.88 |
| 4 | 106394380 | rs2074396 | 1.039 | -1.485 | -0.108 | 0.079 | 0.103 | 0.112 | 0.112 | 0 | 0.55 | 0.23 | 0.56 | 0.32 | 0.71 | 0.63 |
| 4 | 106399576 | rs12507480 | 0.018 | -1.447 | -0.041 | 0.135 | -0.021 | 0 | 0 | 0.07 | 0.26 | 0.58 | 0.2 | 0.26 | 0.14 | 0.23 |
| 4 | 106408926 | rs11097891 | -0.712 | -1.162 | -0.063 | 0.299 | -0.005 | 0.029 | 0.029 | 0 | 0.81 | 0.74 | 0.76 | 0.58 | 0.85 | 0.88 |
| 4 | 106412529 | rs757176 | -1.798 | -1.084 | -0.181 | 0.354 | 0.004 | 0.011 | 0.011 | 0 | 0.86 | 0.87 | 0.79 | 0.58 | 0.89 | 0.89 |
| 4 | 106429568 | rs17617523 | 1.039 | -1.161 | -0.257 | -0.011 | 0.005 | -0.007 | -0.007 | 0 | 0.11 | 0.11 | 0.13 | 0.22 | 0.06 | 0.17 |
| 4 | 106442518 | rs9307305 | -2.321 | -1.079 | -0.359 | 0.368 | 0.006 | -0.042 | -0.042 | 0 | 0.88 | 0.95 | 0.77 | 0.62 | 0.9 | 0.9 |
| 4 | 106447752 | rs3864227 | -0.649 | -1.665 | -0.267 | -0.046 | -0.025 | -0.014 | -0.014 | 0.252 | 0.34 | 0.58 | 0.24 | 0.2 | 0.17 | 0.24 |
| 4 | 106448900 | rs4082504 | 0.539 | -1.631 | -0.287 | -0.115 | 0.03 | 0.009 | 0.009 | 0 | 0.55 | 0.4 | 0.54 | 0.42 | 0.74 | 0.67 |
| 4 | 106459205 | rs1388040 | -2.546 | -1.615 | -0.788 | -0.02 | 0.009 | -0.036 | -0.036 | 3.506 | 0.88 | 0.93 | 0.77 | 0.6 | 0.89 | 0.9 |
| 4 | 106463713 | rs11730243 | 0.819 | -1.752 | -0.78 | -0.121 | 0.116 | 0.044 | 0.044 | 0 | 0.52 | 0.26 | 0.52 | 0.41 | 0.71 | 0.64 |
| 4 | 106467126 | rs6858204 | 0.875 | -1.37 | -0.193 | -0.209 | -0.07 | -0.115 | -0.115 | 0 | 0.15 | 0.47 | 0.07 | 0 | 0.06 | 0.05 |
| 4 | 106467767 | rs17617811 | 1.652 | -1.174 | -0.455 | -0.384 | -0.032 | -0.049 | -0.049 | 0.488 | 0.19 | 0.06 | 0.2 | 0.33 | 0.24 | 0.44 |
| 4 | 106472259 | rs3756258 | -0.315 | -1.575 | 0.307 | -0.364 | -0.007 | -0.001 | -0.001 | 0.34 | 0.18 | 0.21 | 0.16 | 0.19 | 0.09 | 0.19 |
| 4 | 106474078 | rs10032241 | 0.031 | -0.776 | 0.124 | 0.176 | -0.068 | -0.114 | -0.114 | 0.631 | 0.15 | 0.45 | 0.07 | 0 | 0.06 | 0.05 |
| 4 | 106474576 | rs3796915 | 0.77 | -0.517 | -1.157 | 0.322 | 0.008 | -0.029 | -0.029 | 0 | 0.08 | 0.03 | 0.13 | 0.19 | 0.04 | 0.16 |
| 4 | 106477417 | rs6816893 | 0.25 | -0.086 | -0.837 | -0.257 | -0.037 | -0.065 | -0.065 | 1.445 | 0.12 | 0.21 | 0.06 | 0 | 0.06 | 0.05 |
| 4 | 106482235 | rs4699189 | -0.212 | 0.766 | -1.71 | 0.096 | -0.058 | -0.032 | -0.032 | 0.573 | 0.39 | 0.09 | 0.45 | 0.39 | 0.58 | 0.61 |
| 4 | 106482715 | rs9996656 | -0.909 | 0.896 | -0.295 | -0.497 | 0.046 | -0.048 | -0.048 | 0.368 | 0.12 | 0.41 | 0.1 | 0.02 | 0.12 | 0.03 |
| 4 | 106482887 | rs6533209 | 0.295 | 0.754 | -0.533 | -0.677 | -0.011 | 0.033 | 0.033 | 0 | 0.56 | 0.33 | 0.47 | 0.31 | 0.46 | 0.45 |
| 4 | 106483327 | rs6533210 | -0.503 | 0.162 | 0.115 | -1.03 | -0.023 | -0.027 | -0.027 | 0 | 0.82 | 0.71 | 0.86 | 0.9 | 0.91 | 0.96 |
| 4 | 106483898 | rs17584487 | 0.499 | 0.212 | -0.395 | -0.997 | 0.011 | 0.012 | 0.012 | 0 | 0.17 | 0.12 | 0.13 | 0.11 | 0.09 | 0.04 |
| 4 | 106485194 | rs10433994 | -0.066 | 0.286 | -0.65 | -1.364 | -0.011 | 0.019 | 0.019 | 0.195 | 0.79 | 0.86 | 0.88 | 1 | 0.77 | 0.94 |
| 4 | 106485474 | rs11097894 | 0.688 | 0.472 | -0.755 | -1.207 | 0.015 | 0.045 | 0.045 | 0.586 | 0.57 | 0.32 | 0.71 | 0.89 | 0.68 | 0.86 |
| 4 | 106485745 | rs6813330 | 1.247 | 0.512 | -1.133 | -1.002 | -0.008 | -0.024 | -0.024 | 0 | 0.34 | 0.16 | 0.36 | 0.63 | 0.38 | 0.67 |
| 4 | 106485836 | rs6533212 | 0.087 | 0.005 | -0.303 | -1.458 | 0.014 | 0.042 | 0.042 | 0 | 0.76 | 0.85 | 0.84 | 0.89 | 0.85 | 0.95 |
| 4 | 106504943 | rs199330 | 1.102 | 0.027 | -0.336 | -1.314 | 0.028 | 0.079 | 0.079 | 0.14 | 0.58 | 0.27 | 0.71 | 0.89 | 0.69 | 0.87 |
| 4 | 106506244 | rs11935777 | -1.063 | 0.519 | 0.195 | -0.694 | 0.003 | -0.149 | -0.149 | 1.04 | 0.19 | 0.45 | 0.11 | 0 | 0.17 | 0.09 |
| 4 | 106510470 | rs2276970 | -1.329 | 0.619 | -0.421 | -0.032 | -0.001 | -0.006 | -0.006 | 0.215 | 0.09 | 0.09 | 0.08 | 0.11 | 0.11 | 0.05 |
| 4 | 106516028 | rs715706 | -1.906 | 0.718 | 0.326 | -0.619 | 0.018 | -0.077 | -0.077 | 0 | 0.22 | 0.4 | 0.18 | 0.06 | 0.27 | 0.09 |
| 4 | 106520742 | rs17035917 | -1.083 | 0.639 | 0.625 | -0.71 | -0.001 | -0.038 | -0.038 | 0.593 | 0.08 | 0.12 | 0.06 | 0 | 0.07 | 0.02 |
| 4 | 106521514 | rs11721732 | -2.296 | 0.602 | 0.719 | -0.75 | 0 | 0.007 | 0.007 | 0 | 0.15 | 0.1 | 0.12 | 0.03 | 0.14 | 0.07 |
| 4 | 106523977 | rs1374530 | -3.106 | 0.728 | 1.169 | -0.788 | 0.012 | -0.089 | -0.089 | 0.043 | 0.22 | 0.41 | 0.17 | 0.06 | 0.26 | 0.11 |
| 4 | 106527106 | rs7687176 | -1.079 | 0.529 | 1.123 | -0.769 | 0 | 0.004 | 0.004 | 0 | 0.08 | 0.06 | 0.06 | 0.03 | 0.07 | 0.05 |
| 4 | 106531758 | rs13138544 | -2.661 | -0.469 | 1.514 | -1.138 | 0.025 | 0.032 | 0.032 | 0 | 0.32 | 0.78 | 0.25 | 0.11 | 0.23 | 0.15 |
| 4 | 106534110 | rs10516520 | -2.25 | -0.405 | 1.534 | -1.132 | 0.001 | -0.006 | -0.006 | 0 | 0.17 | 0.19 | 0.13 | 0.03 | 0.16 | 0.07 |
| 4 | 106549096 | rs10007278 | -2.242 | -0.392 | 1.462 | -1.153 | -0.002 | -0.016 | -0.016 | 0 | 0.26 | 0.72 | 0.19 | 0.07 | 0.17 | 0.08 |
| 4 | 106552414 | rs2044536 | -2.656 | -0.483 | 1.914 | -1.157 | 0.004 | -0.001 | -0.001 | 0 | 0.31 | 0.71 | 0.24 | 0.1 | 0.22 | 0.12 |
| 4 | 106553247 | rs10516521 | -1.39 | -0.477 | 1.972 | -1.151 | -0.001 | -0.001 | -0.001 | 0 | 0.07 | 0.09 | 0.07 | 0.03 | 0.08 | 0.05 |
| 4 | 106559378 | rs1447096 | -2.684 | -0.41 | 1.941 | -1.156 | 0.025 | 0.04 | 0.04 | 0 | 0.32 | 0.78 | 0.25 | 0.1 | 0.23 | 0.12 |
| 4 | 106572456 | rs1374534 | 0.367 | -0.826 | 0.906 | -2.09 | 0.007 | 0.027 | 0.027 | 0 | 0.19 | 0.07 | 0.1 | 0 | 0.18 | 0.13 |
| 4 | 106575269 | rs17036076 | -0.178 | -0.777 | 1.054 | -1.902 | -0.002 | 0.014 | 0.014 | 0 | 0.07 | 0.02 | 0.06 | 0 | 0.07 | 0.02 |
| 4 | 106588509 | rs10019217 | -1.398 | -1.151 | 1.331 | -2.324 | -0.037 | -0.06 | -0.06 | 1.991 | 0.22 | 0.63 | 0.18 | 0.08 | 0.14 | 0.08 |
| 4 | 106593850 | rs4235415 | -1.7 | -1.16 | 1.237 | -2.307 | -0.001 | -0.007 | -0.007 | 0 | 0.26 | 0.78 | 0.2 | 0.08 | 0.14 | 0.09 |
| 4 | 106594625 | rs11930965 | -0.07 | -1.15 | 1.624 | -2.254 | -0.017 | 0.02 | 0.02 | 0 | 0.1 | 0.35 | 0.09 | 0.07 | 0.06 | 0.05 |
| 4 | 106595055 | rs17036093 | -0.606 | -1.171 | 2.321 | -2.27 | -0.026 | 0.01 | 0.01 | 0 | 0.12 | 0.43 | 0.14 | 0.07 | 0.07 | 0.07 |
| 4 | 106601989 | rs4699193 | -0.318 | -1.15 | 2.47 | -2.285 | -0.019 | 0.007 | 0.007 | 0 | 0.11 | 0.33 | 0.12 | 0.06 | 0.07 | 0.08 |
| 4 | 106603098 | rs17036102 | -1.742 | -1.258 | 2.522 | -2.371 | -0.041 | -0.072 | -0.072 | 0 | 0.21 | 0.5 | 0.21 | 0.07 | 0.14 | 0.1 |
| 4 | 106613821 | rs6820671 | -1.742 | -1.258 | 2.522 | -2.371 | -0.041 | -0.072 | -0.072 | 0.111 | 0.21 | 0.5 | 0.21 | 0.07 | 0.14 | 0.1 |
| 4 | 106614806 | rs17262443 | -0.413 | -1.321 | 2.443 | -2.456 | -0.004 | 0.002 | 0.002 | 0 | 0.07 | 0.05 | 0.06 | 0.03 | 0.1 | 0.06 |
| 4 | 106618362 | rs17262457 | -0.413 | -1.321 | 2.443 | -2.463 | -0.004 | 0.003 | 0.003 | 0 | 0.07 | 0.05 | 0.06 | 0.03 | 0.1 | 0.06 |
| 4 | 106621643 | rs7659526 | -1.715 | -1.328 | 2.452 | -2.466 | -0.042 | -0.072 | -0.072 | 2.352 | 0.21 | 0.52 | 0.21 | 0.07 | 0.14 | 0.08 |
| 4 | 106622190 | rs17036123 | -0.234 | -1.329 | 2.449 | -2.467 | 0.001 | 0.022 | 0.022 | 0 | 0.08 | 0.03 | 0.06 | 0 | 0.06 | 0.02 |
| 4 | 106625263 | rs10516523 | -2.778 | -1.331 | 2.48 | -2.51 | 0.047 | 0.106 | 0.106 | 0 | 0.33 | 0.91 | 0.29 | 0.1 | 0.24 | 0.15 |
| 4 | 106629009 | rs17262764 | -2.778 | -1.331 | 2.468 | -2.51 | 0.047 | 0.106 | 0.106 | 0 | 0.33 | 0.91 | 0.29 | 0.1 | 0.24 | 0.15 |
| 4 | 106630949 | rs4699196 | -2.781 | -1.333 | 2.542 | -2.511 | 0.043 | 0.084 | 0.084 | 0.006 | 0.33 | 0.86 | 0.29 | 0.1 | 0.24 | 0.15 |
| 4 | 106632436 | rs10516524 | -0.386 | -1.33 | 2.539 | -2.509 | -0.005 | 0.003 | 0.003 | 0 | 0.07 | 0.05 | 0.06 | 0.03 | 0.1 | 0.06 |
| 4 | 106632657 | rs17036139 | -0.159 | -1.329 | 2.539 | -2.508 | 0.001 | 0.022 | 0.022 | 0 | 0.08 | 0.03 | 0.06 | 0 | 0.06 | 0.02 |

| | | | | | | | | | | | | | | | | |
|---|---|---|---|---|---|---|---|---|---|---|---|---|---|---|---|---|
| 4 | 106638256 | rs11724839 | -0.159 | -1.329 | 2.539 | -2.508 | 0.001 | 0.022 | 0.022 | 0 | 0.08 | 0.03 | 0.06 | 0 | 0.06 | 0.02 |
| 4 | 106641923 | rs4699197 | -0.07 | -1.33 | 2.559 | -2.508 | -0.021 | 0.002 | 0.002 | 0 | 0.11 | 0.29 | 0.12 | 0.06 | 0.07 | 0.06 |
| 4 | 106643445 | rs6820200 | -2.747 | -1.34 | 2.448 | -2.513 | 0.044 | 0.085 | 0.085 | 0.003 | 0.33 | 0.86 | 0.29 | 0.1 | 0.24 | 0.15 |
| 4 | 106657151 | rs1294671 | 0.193 | -1.329 | 2.433 | -2.507 | -0.004 | 0.003 | 0.003 | 0 | 0.93 | 0.95 | 0.94 | 0.97 | 0.9 | 0.94 |
| 4 | 106662118 | rs12511885 | -1.629 | -1.332 | 2.448 | -2.506 | -0.046 | -0.08 | -0.08 | 0 | 0.21 | 0.48 | 0.21 | 0.07 | 0.14 | 0.08 |
| 4 | 106663541 | rs11934139 | 2.363 | -1.338 | 2.391 | -2.512 | 0.045 | 0.086 | 0.086 | 0 | 0.67 | 0.13 | 0.71 | 0.9 | 0.76 | 0.85 |
| 4 | 106666359 | rs11097896 | 0.199 | -1.323 | 2.393 | -2.496 | -0.005 | 0.003 | 0.003 | 0 | 0.93 | 0.95 | 0.94 | 0.97 | 0.9 | 0.94 |
| 4 | 106666773 | rs11722731 | 1.298 | -1.33 | 2.464 | -2.501 | -0.022 | -0.027 | -0.027 | 0 | 0.81 | 0.52 | 0.79 | 0.9 | 0.83 | 0.88 |
| 4 | 106668358 | rs7676226 | 0.204 | -1.305 | 2.349 | -2.486 | -0.005 | 0.003 | 0.003 | 0 | 0.93 | 0.95 | 0.94 | 0.97 | 0.9 | 0.94 |
| 4 | 106669594 | rs9991722 | -1.056 | -1.34 | 2.364 | -2.499 | 0.037 | 0.024 | 0.024 | 0.17 | 0.05 | 0.35 | 0.02 | 0 | 0.01 | 0.01 |
| 4 | 106672198 | rs4698951 | -1.161 | -1.365 | 2.603 | -2.512 | -0.049 | -0.034 | -0.034 | 0 | 0.15 | 0.65 | 0.11 | 0.06 | 0.08 | 0.06 |
| 4 | 106680764 | rs2193852 | -0.116 | -1.424 | 2.586 | -2.541 | -0.023 | -0.001 | -0.001 | 0 | 0.11 | 0.3 | 0.09 | 0.06 | 0.07 | 0.05 |
| 4 | 106681251 | rs2866781 | 1.183 | -1.415 | 2.594 | -2.542 | 0.021 | -0.007 | -0.007 | 2.391 | 0.91 | 0.77 | 0.88 | 0.97 | 0.9 | 0.93 |
| 4 | 106687733 | rs2544402 | 1.577 | -1.456 | 2.57 | -2.57 | -0.004 | -0.051 | -0.051 | 0 | 0.83 | 0.75 | 0.82 | 0.97 | 0.83 | 0.91 |
| 4 | 106688976 | rs2544401 | 1.422 | -1.465 | 2.565 | -2.575 | 0.009 | -0.016 | -0.016 | 0 | 0.9 | 0.78 | 0.88 | 0.97 | 0.89 | 0.93 |
| 4 | 106696476 | rs2553446 | 1.752 | -1.466 | 2.565 | -2.575 | -0.006 | -0.054 | -0.054 | 0 | 0.82 | 0.73 | 0.82 | 0.97 | 0.83 | 0.91 |
| 4 | 106698892 | rs2553449 | 2.094 | -1.766 | 2.227 | -2.891 | 0.005 | 0.266 | 0.266 | 0.341 | 0.48 | 0.03 | 0.62 | 0.9 | 0.57 | 0.72 |
| 4 | 106704166 | rs6825379 | 0.062 | -1.493 | 2.534 | -2.545 | -0.015 | 0.008 | 0.008 | 0 | 0.1 | 0.3 | 0.09 | 0.06 | 0.06 | 0.05 |
| 4 | 106705339 | rs11733287 | -0.356 | -1.481 | 2.188 | -2.542 | 0.001 | 0.023 | 0.023 | 0 | 0.08 | 0.03 | 0.06 | 0 | 0.06 | 0.02 |
| 4 | 106705863 | rs2544395 | 2.511 | -1.481 | 2.184 | -2.542 | 0.053 | 0.116 | 0.116 | 0 | 0.67 | 0.08 | 0.71 | 0.9 | 0.76 | 0.85 |
| 4 | 106709583 | rs2553471 | 0.554 | -1.465 | 2.203 | -2.532 | 0 | -0.004 | -0.004 | 6.544 | 0.93 | 0.92 | 0.91 | 0.97 | 0.9 | 0.93 |
| 4 | 106713673 | rs2553470 | 1.167 | -1.466 | 2.221 | -2.532 | 0.013 | -0.012 | -0.012 | 0 | 0.91 | 0.79 | 0.88 | 0.97 | 0.89 | 0.93 |
| 4 | 106717797 | rs2544426 | 1.251 | -1.391 | 2.202 | -2.532 | 0.019 | -0.012 | -0.012 | 3.016 | 0.91 | 0.78 | 0.88 | 0.97 | 0.89 | 0.93 |
| 4 | 106718316 | rs2544424 | 1.263 | -1.419 | 2.202 | -2.532 | 0.014 | -0.013 | -0.013 | 0 | 0.91 | 0.78 | 0.88 | 0.97 | 0.89 | 0.93 |
| 4 | 106722298 | rs2553468 | 2.455 | -1.426 | 2.152 | -2.538 | 0.05 | 0.113 | 0.113 | 0 | 0.67 | 0.08 | 0.71 | 0.9 | 0.76 | 0.85 |
| 4 | 106724328 | rs2553467 | 0.555 | -1.392 | 2.155 | -2.475 | 0.001 | -0.003 | -0.003 | 0.116 | 0.93 | 0.92 | 0.9 | 0.96 | 0.89 | 0.93 |
| 4 | 106727328 | rs2193857 | 0.63 | -1.437 | 2.147 | -2.5 | 0 | -0.004 | -0.004 | 0 | 0.92 | 0.92 | 0.91 | 0.97 | 0.9 | 0.93 |
| 4 | 106731410 | rs2553461 | 1.729 | -1.441 | 2.17 | -2.504 | -0.004 | -0.051 | -0.051 | 0 | 0.83 | 0.75 | 0.82 | 0.97 | 0.83 | 0.91 |
| 4 | 106732183 | rs2553460 | 1.729 | -1.441 | 2.17 | -2.504 | -0.004 | -0.051 | -0.051 | 0 | 0.83 | 0.75 | 0.82 | 0.97 | 0.83 | 0.91 |
| 4 | 106736355 | rs17036254 | -1.111 | -1.465 | 2.239 | -2.445 | -0.046 | -0.031 | -0.031 | 0 | 0.15 | 0.65 | 0.11 | 0.06 | 0.07 | 0.06 |
| 4 | 106737796 | rs2553458 | 2.508 | -1.479 | 2.197 | -2.454 | 0.05 | 0.113 | 0.113 | 0 | 0.67 | 0.08 | 0.71 | 0.91 | 0.76 | 0.85 |
| 4 | 106741974 | rs2544418 | 1.479 | -1.465 | 2.25 | -2.438 | 0.011 | -0.016 | -0.016 | 0.152 | 0.9 | 0.77 | 0.88 | 0.97 | 0.89 | 0.93 |
| 4 | 106742958 | rs2193856 | 1.887 | -1.48 | 2.252 | -2.446 | -0.006 | -0.058 | -0.058 | 0 | 0.82 | 0.74 | 0.82 | 0.97 | 0.83 | 0.91 |
| 4 | 106745405 | rs2544417 | 1.855 | -1.469 | 2.245 | -2.441 | -0.006 | -0.057 | -0.057 | 0 | 0.82 | 0.74 | 0.82 | 0.97 | 0.83 | 0.91 |
| 4 | 106745751 | rs11726674 | -0.503 | -1.468 | 2.229 | -2.44 | 0.002 | 0.019 | 0.019 | 0 | 0.08 | 0.03 | 0.06 | 0 | 0.06 | 0.02 |
| 4 | 106746012 | rs11731417 | -0.459 | -1.468 | 2.229 | -2.44 | 0.002 | 0.019 | 0.019 | 0 | 0.08 | 0.03 | 0.06 | 0 | 0.06 | 0.02 |
| 4 | 106750214 | rs2216515 | 1.915 | -1.469 | 2.258 | -2.441 | -0.009 | -0.072 | -0.072 | 0 | 0.81 | 0.67 | 0.82 | 0.97 | 0.83 | 0.91 |
| 4 | 106751028 | rs2553456 | 1.935 | -1.47 | 2.258 | -2.441 | -0.01 | -0.072 | -0.072 | 0 | 0.81 | 0.67 | 0.82 | 0.97 | 0.83 | 0.91 |
| 4 | 106751231 | rs2112051 | 2.267 | -1.443 | 2.069 | -2.422 | 0.05 | 0.117 | 0.117 | 0 | 0.67 | 0.08 | 0.71 | 0.91 | 0.75 | 0.85 |
| 4 | 106753845 | rs6533218 | 2.267 | -1.443 | 2.069 | -2.422 | 0.05 | 0.117 | 0.117 | 0 | 0.67 | 0.08 | 0.71 | 0.91 | 0.76 | 0.85 |
| 4 | 106761837 | rs2544412 | 2.267 | -1.442 | 2.047 | -2.422 | 0.05 | 0.117 | 0.117 | 0 | 0.67 | 0.08 | 0.71 | 0.91 | 0.75 | 0.85 |
| 4 | 106765915 | rs10050159 | -1.134 | -1.259 | 1.923 | -2.324 | -0.008 | -0.062 | -0.062 | 0 | 0.11 | 0.23 | 0.09 | 0 | 0.07 | 0.02 |
| 4 | 106808172 | rs2553453 | 1.117 | -1.124 | 1.567 | -2.188 | -0.02 | -0.03 | -0.03 | 0 | 0.81 | 0.63 | 0.84 | 0.9 | 0.83 | 0.8 |
| 4 | 106820344 | rs11731982 | -1.87 | -0.967 | 1.097 | -2.033 | 0.037 | -0.018 | -0.018 | 2.208 | 0.07 | 0.24 | 0.05 | 0 | 0.05 | 0.02 |
| 4 | 106829645 | rs4600917 | -1.401 | -1.054 | -0.076 | -2.496 | 0.029 | 0.166 | 0.166 | 0 | 0.52 | 0.9 | 0.42 | 0.18 | 0.39 | 0.31 |
| 4 | 106830232 | rs6811135 | -1.257 | -1.04 | -0.045 | -2.489 | 0.048 | 0.178 | 0.178 | 0.013 | 0.5 | 0.88 | 0.4 | 0.18 | 0.39 | 0.29 |
| 4 | 106857657 | rs4507363 | 1.35 | -0.18 | 0.099 | -2.075 | 0.064 | -0.027 | -0.027 | 0 | 0.92 | 0.72 | 0.92 | 1 | 0.92 | 0.96 |
| 4 | 106861730 | rs4340795 | 1.35 | -0.18 | 0.074 | -2.075 | 0.062 | -0.028 | -0.028 | 0 | 0.92 | 0.72 | 0.92 | 1 | 0.92 | 0.96 |
| 4 | 106872861 | rs7677557 | -0.071 | -0.288 | -0.273 | -2.161 | -0.012 | 0.002 | 0.002 | 0 | 0.15 | 0.03 | 0.07 | 0.11 | 0.14 | 0.07 |
| 4 | 106887163 | rs7680832 | 0.719 | -0.078 | -1.507 | -1.752 | 0.012 | 0.095 | 0.095 | 1.241 | 0.54 | 0.1 | 0.46 | 0.79 | 0.52 | 0.62 |
| 4 | 106888263 | rs4597836 | -0.526 | 0.105 | -0.979 | -1.942 | -0.01 | -0.008 | -0.008 | 0 | 0.37 | 0.52 | 0.42 | 0.21 | 0.39 | 0.39 |
| 4 | 106891531 | rs6817700 | 0.631 | 0.095 | -1.059 | -1.928 | 0.008 | 0.022 | 0.022 | 0.24 | 0.58 | 0.37 | 0.52 | 0.79 | 0.53 | 0.61 |
| 4 | 106915834 | rs4345186 | -0.149 | 0.319 | -1.377 | 0.678 | -0.014 | 0.013 | 0.013 | 0 | 0.13 | 0.22 | 0.21 | 0.24 | 0.08 | 0.25 |
| 4 | 106921594 | rs4367177 | 0.369 | 0.59 | -1.235 | 0.682 | 0.001 | -0.017 | -0.017 | 0 | 0.94 | 0.92 | 0.94 | 1 | 0.9 | 0.98 |
| 4 | 106923626 | rs994370 | 0.19 | 0.508 | -1.804 | 0.922 | -0.001 | 0.087 | 0.087 | 1.466 | 0.52 | 0.29 | 0.67 | 0.76 | 0.49 | 0.6 |
| 4 | 106926748 | rs17199327 | 0.804 | 0.563 | -2.382 | 0.906 | -0.15 | -0.046 | -0.046 | 0 | 0.39 | 0.03 | 0.25 | 0.4 | 0.37 | 0.36 |
| 4 | 106936604 | rs10001483 | -0.772 | 0.253 | -1.404 | 0.783 | -0.008 | 0.003 | 0.003 | 0 | 0.15 | 0.16 | 0.28 | 0.12 | 0.09 | 0.06 |
| 4 | 106938490 | rs4611932 | 0.761 | 0.874 | -1.356 | 0.571 | 0.002 | -0.013 | -0.013 | 0 | 0.94 | 0.95 | 0.94 | 1 | 0.9 | 0.98 |
| 4 | 106947487 | rs6853554 | -0.366 | 0.795 | -1.437 | 0.727 | -0.004 | -0.003 | -0.003 | 0 | 0.28 | 0.25 | 0.33 | 0.44 | 0.29 | 0.48 |
| 4 | 106958170 | rs6842919 | -0.824 | 0.165 | -1.437 | 0.292 | -0.001 | -0.006 | -0.006 | 0 | 0.39 | 0.36 | 0.26 | 0.15 | 0.36 | 0.33 |
| 4 | 106964945 | rs17036535 | 0.954 | 0.161 | -1.429 | 0.291 | -0.008 | -0.033 | -0.033 | 0.168 | 0.12 | 0.12 | 0.06 | 0 | 0.07 | 0.03 |
| 4 | 106970272 | rs7683106 | -0.302 | 0.194 | -1.423 | 0.264 | -0.009 | -0.064 | -0.064 | 0 | 0.17 | 0.2 | 0.1 | 0 | 0.15 | 0.06 |
| 4 | 106973602 | rs10023586 | 0.33 | 0.204 | -1.455 | 0.254 | -0.001 | -0.008 | -0.008 | 0.596 | 0.05 | 0.04 | 0.04 | 0 | 0.07 | 0.02 |
| 4 | 106982793 | rs13101737 | 1.008 | 0 | -2.151 | -0.153 | -0.083 | 0.019 | 0.019 | 0.314 | 0.25 | 0.02 | 0.16 | 0.15 | 0.29 | 0.16 |
| 4 | 107022503 | rs17272618 | 0.085 | 0.305 | -1.87 | -0.016 | -0.012 | -0.009 | -0.009 | 6.582 | 0.28 | 0.23 | 0.33 | 0.43 | 0.29 | 0.49 |
| 4 | 107128360 | rs13101612 | 0.361 | 0.381 | -1.484 | 0.016 | -0.004 | -0.013 | -0.013 | 0 | 0.59 | 0.52 | 0.6 | 0.56 | 0.63 | 0.48 |
| 4 | 107171131 | rs13106209 | 1.081 | 0.038 | -1.61 | 0.058 | -0.08 | -0.157 | -0.157 | 4.97 | 0.25 | 0.02 | 0.36 | 0.36 | 0.29 | 0.24 |
| 4 | 107172182 | rs12641015 | 1.563 | 0.068 | -1.605 | 0.044 | -0.002 | 0.001 | 0.001 | 0 | 0.19 | 0.23 | 0.13 | 0.15 | 0.18 | 0.28 |
| 4 | 107174698 | rs13129572 | 1.63 | -0.066 | -1.779 | 0.206 | -0.093 | -0.131 | -0.131 | 0 | 0.21 | 0.01 | 0.35 | 0.31 | 0.28 | 0.19 |

| | | | | | | | | | | | | | | | | |
|---|---|---|---|---|---|---|---|---|---|---|---|---|---|---|---|---|
| 4 | 107178254 | rs12504721 | 1.905 | -0.016 | -1.584 | 0.211 | -0.069 | -0.009 | -0.009 | 0 | 0.17 | 0.01 | 0.12 | 0.15 | 0.18 | 0.27 |
| 4 | 107179126 | rs727618 | 0.895 | -0.006 | -1.574 | 0.216 | -0.002 | -0.042 | -0.042 | 0 | 0.17 | 0.17 | 0.1 | 0 | 0.15 | 0.05 |
| 4 | 107185855 | rs7662395 | 1.499 | 0.201 | -1.605 | 0.463 | -0.014 | 0.015 | 0.015 | 0 | 0.18 | 0.12 | 0.13 | 0.15 | 0.18 | 0.28 |
| 4 | 107236648 | rs3762947 | 1.443 | 0.227 | -1.592 | 0.477 | 0 | -0.002 | -0.002 | 0.041 | 0.18 | 0.18 | 0.13 | 0.15 | 0.18 | 0.28 |
| 4 | 107237395 | rs6814166 | 1.515 | 0.226 | -1.591 | 0.476 | -0.008 | 0.009 | 0.009 | 0 | 0.18 | 0.1 | 0.12 | 0.15 | 0.18 | 0.28 |
| 4 | 107246136 | rs3737498 | 1.451 | 0.232 | -1.623 | 0.479 | -0.001 | 0 | 0 | 0 | 0.18 | 0.17 | 0.13 | 0.15 | 0.18 | 0.28 |
| 4 | 107266883 | rs12642454 | 1.512 | 0.253 | -1.711 | 0.49 | -0.011 | 0.012 | 0.012 | 0 | 0.18 | 0.09 | 0.12 | 0.15 | 0.18 | 0.28 |
| 4 | 107268949 | rs3805410 | 1.512 | 0.253 | -1.711 | 0.494 | -0.011 | 0.011 | 0.011 | 0 | 0.18 | 0.09 | 0.12 | 0.15 | 0.18 | 0.28 |
| 4 | 107279450 | rs7654865 | 1.495 | 0.25 | -1.712 | 0.486 | -0.01 | 0.012 | 0.012 | 0 | 0.18 | 0.1 | 0.12 | 0.15 | 0.18 | 0.28 |
| 4 | 107281920 | rs3018058 | 0.871 | 0.053 | -1.421 | 0.431 | -0.002 | -0.008 | -0.008 | 0 | 0.34 | 0.32 | 0.22 | 0.2 | 0.35 | 0.24 |
| 4 | 107282566 | rs3018059 | 0.475 | 0.208 | -2.216 | 0.044 | -0.017 | -0.007 | -0.007 | 0 | 0.22 | 0.39 | 0.28 | 0.29 | 0.19 | 0.24 |
| 4 | 107288449 | rs2949646 | 0.425 | 0.41 | -2.48 | 0.064 | -0.007 | 0.002 | 0.002 | 1.66 | 0.24 | 0.3 | 0.27 | 0.32 | 0.19 | 0.29 |
| 4 | 107289526 | rs3018066 | -0.376 | 0.619 | -2.496 | 0.073 | 0.003 | 0.005 | 0.005 | 0.127 | 0.43 | 0.39 | 0.4 | 0.48 | 0.37 | 0.57 |
| 4 | 107295179 | rs3108151 | -0.857 | 1.394 | -2.696 | 0.416 | -0.088 | -0.052 | -0.052 | 0 | 0.8 | 0.99 | 0.64 | 0.86 | 0.72 | 0.81 |
| 4 | 107296620 | rs7697673 | 0.228 | 1.12 | -1.697 | 0.277 | -0.033 | 0.016 | 0.016 | 0 | 0.18 | 0.09 | 0.13 | 0.16 | 0.18 | 0.29 |
| 4 | 107302260 | rs2949619 | -0.323 | 0.893 | -1.959 | 0.135 | -0.059 | 0.048 | 0.048 | 0 | 0.28 | 0.05 | 0.18 | 0.24 | 0.33 | 0.27 |
| 4 | 107303131 | rs3108146 | -0.821 | -0.297 | -2.101 | -0.053 | 0.084 | 0.063 | 0.063 | 1.565 | 0.4 | 0.85 | 0.26 | 0.31 | 0.33 | 0.38 |
| 4 | 107304010 | rs2949623 | -0.687 | -0.388 | -1.41 | -0.088 | 0.038 | 0.035 | 0.035 | 1.053 | 0.39 | 0.7 | 0.25 | 0.3 | 0.31 | 0.36 |
| 4 | 107308215 | rs3113243 | -0.854 | -0.372 | -1.325 | -0.099 | 0.078 | 0.062 | 0.062 | 1.039 | 0.4 | 0.84 | 0.25 | 0.31 | 0.32 | 0.38 |
| 4 | 107310178 | rs2949640 | -0.107 | -0.457 | -0.247 | -0.137 | -0.023 | -0.06 | -0.06 | 0 | 0.3 | 0.45 | 0.21 | 0.24 | 0.3 | 0.27 |
| 4 | 107310444 | rs2949641 | -0.768 | -0.44 | -0.432 | -0.13 | 0.028 | 0.027 | 0.027 | 1.038 | 0.38 | 0.69 | 0.25 | 0.3 | 0.31 | 0.36 |
| 4 | 107318580 | rs4340862 | 0.997 | -0.555 | -0.307 | -0.119 | -0.032 | 0.014 | 0.014 | 0 | 0.06 | 0.15 | 0.03 | 0.05 | 0.01 | 0.02 |
| 4 | 107329974 | rs3018056 | 0.312 | -0.431 | -0.559 | -0.127 | 0.008 | 0.045 | 0.045 | 0 | 0.27 | 0.05 | 0.17 | 0.22 | 0.27 | 0.27 |
| 4 | 107334044 | rs3108140 | -0.959 | -0.516 | -0.544 | -0.182 | 0.12 | 0.081 | 0.081 | 1.847 | 0.41 | 0.87 | 0.25 | 0.28 | 0.3 | 0.36 |
| 4 | 107334361 | rs17276911 | 0.978 | 0.013 | -0.995 | -0.297 | -0.064 | -0.018 | -0.018 | 0 | 0.12 | 0.01 | 0.12 | 0.15 | 0.18 | 0.21 |
| 4 | 107353688 | rs940617 | 0.052 | -0.187 | -0.488 | -0.039 | 0.02 | 0.034 | 0.034 | 0.178 | 0.54 | 0.78 | 0.67 | 0.56 | 0.52 | 0.5 |
| 4 | 107370685 | rs2903421 | 1.089 | 0.13 | -0.001 | 0.586 | -0.084 | -0.074 | -0.074 | 0 | 0.4 | 0.08 | 0.38 | 0.44 | 0.44 | 0.33 |
| 4 | 107394976 | rs10516535 | 0.244 | 0.044 | 0.159 | 0.545 | 0.029 | 0.039 | 0.039 | 0.294 | 0.49 | 0.3 | 0.65 | 0.56 | 0.52 | 0.43 |
| 4 | 107419151 | rs4495119 | 0.318 | 0.209 | -0.068 | 0.446 | -0.009 | 0.042 | 0.042 | 0.828 | 0.47 | 0.24 | 0.62 | 0.56 | 0.5 | 0.44 |
| 4 | 107423164 | rs13148020 | -0.185 | 0.207 | 0.026 | 0.34 | -0.002 | 0.002 | 0.002 | 0 | 0.57 | 0.44 | 0.66 | 0.64 | 0.5 | 0.53 |
| 4 | 107427351 | rs1522221 | 0.241 | 0.24 | -0.102 | 0.466 | -0.001 | -0.004 | -0.004 | 0.018 | 0.4 | 0.45 | 0.5 | 0.37 | 0.38 | 0.41 |
| 4 | 107431789 | rs1522218 | -0.031 | 0.386 | 0.176 | 0.598 | 0.036 | 0.167 | 0.167 | 0 | 0.1 | 0.3 | 0.2 | 0.25 | 0.11 | 0.1 |
| 4 | 107448046 | rs7690115 | -1.399 | 0.231 | 0.044 | 0.424 | -0.046 | 0.053 | 0.053 | 0 | 0.66 | 0.87 | 0.75 | 0.65 | 0.56 | 0.56 |
| 4 | 107470497 | rs7672970 | 0.993 | 0.508 | -0.032 | 0.532 | -0.01 | -0.062 | -0.062 | 0 | 0.31 | 0.17 | 0.43 | 0.36 | 0.32 | 0.39 |
| 4 | 107471140 | rs2686308 | -0.075 | 0.517 | 0.003 | 0.125 | 0.035 | 0.164 | 0.164 | 0 | 0.1 | 0.3 | 0.2 | 0.26 | 0.11 | 0.1 |
| 4 | 107477804 | rs2648048 | -0.302 | 0.521 | -0.043 | 0.127 | 0.037 | 0.172 | 0.172 | 0 | 0.9 | 0.69 | 0.79 | 0.74 | 0.89 | 0.9 |
| 4 | 107486817 | rs769169 | 0.692 | 0.411 | -0.779 | -0.418 | -0.013 | 0.004 | 0.004 | 0.027 | 0.28 | 0.11 | 0.21 | 0.22 | 0.29 | 0.27 |
| 4 | 107498301 | rs10024795 | -0.489 | 0.222 | 0.001 | -0.452 | -0.025 | -0.02 | -0.02 | 0.45 | 0.11 | 0.15 | 0.03 | 0.07 | 0.04 | 0.03 |
| 4 | 107499928 | rs10008290 | -0.438 | 0.234 | -0.035 | -0.544 | -0.024 | -0.033 | -0.033 | 0 | 0.11 | 0.15 | 0.03 | 0.03 | 0.04 | 0.02 |
| 4 | 107501341 | rs10034263 | -1.377 | 0.696 | 0.118 | -0.548 | 0.016 | -0.077 | -0.077 | 0 | 0.17 | 0.32 | 0.14 | 0.06 | 0.17 | 0.16 |
| 4 | 107504175 | rs764538 | 0.402 | 0.589 | -0.781 | -1.257 | 0.008 | 0.008 | 0.008 | 0 | 0.72 | 0.79 | 0.83 | 0.99 | 0.8 | 0.8 |
| 4 | 107509092 | rs2648031 | 1.014 | 0.602 | -0.687 | -1.319 | 0.002 | -0.023 | -0.023 | 1.527 | 0.62 | 0.64 | 0.75 | 0.99 | 0.67 | 0.77 |
| 4 | 107520859 | rs2648037 | 1.048 | 0.857 | -1.378 | -0.687 | 0.002 | 0.015 | 0.015 | 0 | 0.58 | 0.62 | 0.5 | 0.61 | 0.6 | 0.63 |
| 4 | 107526541 | rs2686311 | 0.4 | 0.845 | -1.505 | -0.702 | 0.01 | 0.029 | 0.029 | 0 | 0.69 | 0.71 | 0.7 | 0.89 | 0.75 | 0.74 |
| 4 | 107527205 | rs2553205 | 0.859 | 0.846 | -1.504 | -0.708 | 0.016 | 0.049 | 0.049 | 0 | 0.63 | 0.66 | 0.68 | 0.89 | 0.72 | 0.73 |
| 4 | 107533620 | rs2686305 | 1.496 | 0.89 | -1.557 | -0.76 | 0.02 | 0.107 | 0.107 | 0.259 | 0.56 | 0.66 | 0.62 | 0.89 | 0.64 | 0.7 |
| 4 | 107536850 | rs4956261 | 0.684 | 0.92 | -1.57 | -0.711 | -0.032 | -0.015 | -0.015 | 0 | 0.05 | 0.01 | 0.18 | 0.28 | 0.12 | 0.1 |
| 4 | 107537390 | rs1023792 | 1.827 | 0.921 | -1.568 | -0.714 | -0.007 | 0.06 | 0.06 | 0.205 | 0.51 | 0.65 | 0.44 | 0.61 | 0.52 | 0.6 |
| 4 | 107547518 | rs4340861 | 0.557 | 0.785 | -1.424 | -0.813 | 0.005 | 0.099 | 0.099 | 0 | 0.67 | 0.79 | 0.79 | 0.99 | 0.72 | 0.79 |
| 4 | 107549400 | rs2686297 | 0.553 | 0.903 | -1.274 | -0.535 | -0.012 | 0.004 | 0.004 | 0 | 0.72 | 0.77 | 0.65 | 0.71 | 0.68 | 0.72 |
| 4 | 107583181 | rs1425261 | -0.845 | 0.796 | -1.041 | -0.633 | -0.001 | -0.024 | -0.024 | 0 | 0.1 | 0.12 | 0.1 | 0 | 0.09 | 0.04 |
| 4 | 107587752 | rs2587136 | -0.108 | 0.871 | -0.92 | -0.78 | 0 | -0.025 | -0.025 | 0.019 | 0.11 | 0.16 | 0.08 | 0 | 0.1 | 0.05 |
| 4 | 107588257 | rs13141555 | 1.485 | 0.975 | -0.9 | -0.769 | 0 | -0.027 | -0.027 | 0.799 | 0.77 | 0.68 | 0.8 | 0.92 | 0.78 | 0.85 |
| 4 | 107591650 | rs4956263 | 0.085 | 1.164 | -1.074 | -0.67 | -0.029 | -0.038 | -0.038 | 0 | 0.07 | 0.01 | 0.06 | 0.16 | 0.1 | 0.11 |
| 4 | 107601518 | rs10010360 | 0.818 | 1.011 | -0.931 | -0.797 | 0.005 | -0.04 | -0.04 | 0 | 0.84 | 0.71 | 0.87 | 0.93 | 0.84 | 0.89 |
| 4 | 107608787 | rs1425258 | 0.794 | 1.021 | -0.991 | -0.821 | 0.001 | -0.005 | -0.005 | 0 | 0.88 | 0.92 | 0.89 | 1 | 0.88 | 0.95 |
| 4 | 107609695 | rs2553207 | 0.519 | 1.003 | -0.965 | -0.796 | 0.001 | -0.009 | -0.009 | 0 | 0.89 | 0.93 | 0.9 | 1 | 0.89 | 0.95 |
| 4 | 107613144 | rs12646341 | 1.472 | 1.016 | -0.961 | -0.792 | -0.001 | -0.011 | -0.011 | 0 | 0.76 | 0.73 | 0.8 | 0.93 | 0.78 | 0.86 |
| 4 | 107625430 | rs9784417 | 0.9 | 1.065 | -0.834 | -0.617 | 0.004 | -0.028 | -0.028 | 0 | 0.85 | 0.76 | 0.88 | 0.93 | 0.85 | 0.91 |
| 4 | 107637162 | rs1443893 | 0.429 | 1.109 | -1.063 | -0.592 | -0.008 | 0.003 | 0.003 | 0.647 | 0.88 | 0.92 | 0.89 | 1 | 0.87 | 0.95 |
| 4 | 107654106 | rs11725068 | 0.029 | 1.111 | -1.065 | -0.591 | -0.002 | 0.01 | 0.01 | 0.175 | 0.1 | 0.04 | 0.07 | 0 | 0.09 | 0.03 |
| 4 | 107654943 | rs7695009 | 0.104 | 1.11 | -1.066 | -0.592 | -0.006 | 0.022 | 0.022 | 0.36 | 0.1 | 0.03 | 0.07 | 0 | 0.09 | 0.03 |
| 4 | 107663696 | rs11097937 | 0.392 | 1.117 | -1.062 | -0.591 | -0.002 | -0.002 | -0.002 | 0.448 | 0.88 | 0.92 | 0.9 | 1 | 0.88 | 0.96 |
| 4 | 107666927 | rs1425249 | 1.512 | 1.321 | -0.701 | -0.511 | 0.003 | -0.016 | -0.016 | 0.07 | 0.68 | 0.63 | 0.71 | 0.82 | 0.66 | 0.83 |
| 4 | 107685776 | rs12508455 | 1.332 | 1.191 | -0.692 | -0.475 | 0 | -0.006 | -0.006 | 0.009 | 0.69 | 0.71 | 0.72 | 0.81 | 0.69 | 0.83 |
| 4 | 107708035 | rs817147 | 1.374 | 1.239 | -1.054 | -1.001 | 0.002 | 0.007 | 0.007 | 0 | 0.5 | 0.52 | 0.56 | 0.71 | 0.46 | 0.65 |
| 4 | 107745306 | rs12510474 | 0.818 | 1.02 | -1.841 | -0.247 | -0.034 | -0.126 | -0.126 | 0.472 | 0.16 | 0.01 | 0.32 | 0.45 | 0.14 | 0.18 |
| 4 | 107751089 | rs11733406 | -0.276 | 0.893 | -1.28 | -0.592 | 0.011 | 0.043 | 0.043 | 0.716 | 0.14 | 0.03 | 0.07 | 0.01 | 0.09 | 0.03 |
| 4 | 107769255 | rs11943338 | -0.276 | 1.166 | -1.051 | -0.399 | -0.001 | -0.001 | -0.001 | 0.067 | 0.19 | 0.14 | 0.16 | 0.23 | 0.18 | 0.16 |

| | | | | | | | | | | | | | | | | |
|---|---|---|---|---|---|---|---|---|---|---|---|---|---|---|---|---|
| 4 | 107773179 | rs10014676 | -0.385 | 1.407 | -0.881 | -0.259 | -0.006 | -0.002 | -0.002 | 0 | 0.13 | 0.08 | 0.1 | 0 | 0.13 | 0.05 |
| 4 | 107826705 | rs2324 | 0.033 | 1.534 | -0.494 | 0.298 | 0.024 | -0.013 | -0.013 | 1.545 | 0.26 | 0.45 | 0.19 | 0.25 | 0.33 | 0.24 |
| 4 | 107830261 | rs450533 | -0.052 | 2.004 | -0.62 | 0.177 | 0.018 | -0.015 | -0.015 | 0.046 | 0.27 | 0.46 | 0.22 | 0.25 | 0.35 | 0.23 |
| 4 | 107841693 | rs422661 | 0.545 | 2.032 | -0.496 | 0.321 | 0.011 | 0.008 | 0.008 | 0 | 0.22 | 0.38 | 0.14 | 0.27 | 0.27 | 0.21 |
| 4 | 107842959 | rs402586 | 0.52 | 2.039 | -0.442 | 0.344 | 0.009 | 0.007 | 0.007 | 0 | 0.22 | 0.38 | 0.14 | 0.27 | 0.27 | 0.21 |
| 4 | 107843493 | rs10488898 | 0.637 | 1.345 | -0.096 | -0.089 | -0.018 | -0.042 | -0.042 | 0 | 0.11 | 0.16 | 0.04 | 0.06 | 0.07 | 0.11 |
| 4 | 107844092 | rs419558 | 0.841 | 1.347 | -0.144 | -0.088 | 0.008 | 0.005 | 0.005 | 0 | 0.22 | 0.38 | 0.14 | 0.27 | 0.27 | 0.21 |
| 4 | 107844157 | rs419764 | 0.429 | 1.379 | -0.165 | -0.09 | 0.021 | -0.004 | -0.004 | 0.159 | 0.26 | 0.44 | 0.19 | 0.27 | 0.34 | 0.24 |
| 4 | 107845794 | rs17037102 | 1.233 | 1.825 | -0.982 | 0.603 | -0.042 | -0.04 | -0.04 | 0 | 0.08 | 0.01 | 0.31 | 0.39 | 0.11 | 0.11 |
| 4 | 107851275 | rs433201 | -0.329 | 1.651 | -0.725 | 0.157 | -0.003 | 0.008 | 0.008 | 0 | 0.37 | 0.67 | 0.23 | 0.32 | 0.41 | 0.35 |
| 4 | 107888737 | rs17431749 | -0.153 | 0.896 | -0.655 | -0.229 | -0.011 | -0.041 | -0.041 | 0 | 0.12 | 0.2 | 0.06 | 0.06 | 0.1 | 0.11 |
| 4 | 107889798 | rs396196 | -0.505 | 1.145 | -0.882 | -0.271 | 0.003 | -0.048 | -0.048 | 0.039 | 0.44 | 0.58 | 0.62 | 0.88 | 0.49 | 0.54 |
| 4 | 107903545 | rs2051756 | -1.107 | 1.266 | -0.831 | 1.197 | -0.003 | 0.009 | 0.009 | 2.923 | 0.16 | 0.14 | 0.31 | 0.41 | 0.17 | 0.27 |
| 4 | 107905508 | rs2866908 | -0.875 | 1.289 | -0.836 | 1.195 | 0 | 0.039 | 0.039 | 4.826 | 0.12 | 0.12 | 0.29 | 0.41 | 0.14 | 0.26 |
| 4 | 107939938 | rs4956274 | -0.607 | 1.314 | -0.75 | 1.413 | 0.005 | 0.031 | 0.031 | 3.596 | 0.08 | 0.09 | 0.18 | 0.24 | 0.09 | 0.17 |
| 4 | 107946161 | rs4956275 | -1.681 | 1.437 | -0.978 | 1.77 | -0.033 | -0.058 | -0.058 | 0.493 | 0.14 | 0.05 | 0.32 | 0.52 | 0.2 | 0.29 |
| 4 | 107947618 | rs2212966 | -0.401 | 1.261 | -0.886 | 1.458 | -0.021 | -0.007 | -0.007 | 0.367 | 0.07 | 0.01 | 0.14 | 0.23 | 0.11 | 0.09 |
| 4 | 107951360 | rs10021344 | -0.269 | 1.137 | -1.036 | -0.093 | -0.052 | 0.066 | 0.066 | 0 | 0.34 | 0.22 | 0.27 | 0.02 | 0.47 | 0.17 |
| 4 | 107953879 | rs956137 | -0.018 | 0.366 | -0.831 | -0.269 | -0.005 | -0.03 | -0.03 | 0 | 0.13 | 0.11 | 0.07 | 0 | 0.08 | 0.11 |
| 4 | 107956905 | rs3733635 | -0.358 | 0.35 | -1.03 | 0.388 | 0.003 | -0.012 | -0.012 | 1.266 | 0.07 | 0.03 | 0.12 | 0.26 | 0.1 | 0.16 |
| 4 | 107958376 | rs2704335 | -0.421 | -0.292 | -0.777 | 0.116 | -0.004 | 0.003 | 0.003 | 0.244 | 0.65 | 0.65 | 0.67 | 0.55 | 0.65 | 0.45 |
| 4 | 107960572 | rs13147822 | 1.034 | -0.458 | -0.891 | 0.014 | -0.063 | 0.098 | 0.098 | 0 | 0.33 | 0.19 | 0.28 | 0.05 | 0.44 | 0.19 |
| 4 | 107961767 | rs383567 | -1.171 | -0.41 | -0.757 | 0.037 | -0.001 | -0.003 | -0.003 | 0 | 0.71 | 0.74 | 0.78 | 0.8 | 0.74 | 0.61 |
| 4 | 107970647 | rs7685856 | -0.01 | -0.39 | -0.812 | 0.101 | -0.002 | 0.026 | 0.026 | 0.111 | 0.08 | 0.1 | 0.12 | 0.25 | 0.1 | 0.16 |
| 4 | 107974736 | rs17584885 | 0.813 | -0.538 | -0.915 | 0.604 | -0.065 | -0.028 | -0.028 | 0 | 0.36 | 0.18 | 0.3 | 0.39 | 0.47 | 0.28 |
| 4 | 107976020 | rs419783 | 0.682 | -0.773 | -0.315 | 0.476 | -0.001 | 0.012 | 0.012 | 0 | 0.16 | 0.32 | 0.3 | 0.15 | 0.11 | 0.05 |
| 4 | 107980550 | rs398093 | 0.008 | -0.941 | -0.316 | 0.225 | -0.001 | -0.039 | -0.039 | 0.002 | 0.19 | 0.18 | 0.1 | 0.01 | 0.14 | 0.13 |
| 4 | 107984723 | rs2704324 | -0.74 | -0.618 | -0.364 | 0.372 | -0.007 | -0.002 | -0.002 | 0 | 0.63 | 0.53 | 0.51 | 0.62 | 0.67 | 0.66 |
| 4 | 107988210 | rs441202 | 0.464 | -0.583 | -0.777 | 0.461 | 0.007 | 0.02 | 0.02 | 0 | 0.45 | 0.26 | 0.33 | 0.37 | 0.52 | 0.4 |
| 4 | 107993687 | rs386499 | -1.346 | -0.569 | -0.647 | 0.532 | -0.014 | 0.031 | 0.031 | 0.002 | 0.68 | 0.83 | 0.74 | 0.77 | 0.64 | 0.65 |
| 4 | 108000247 | rs378405 | -0.492 | -0.585 | -0.628 | 0.542 | -0.008 | 0.116 | 0.116 | 0.286 | 0.23 | 0.54 | 0.43 | 0.41 | 0.21 | 0.21 |
| 4 | 108007908 | rs6849760 | 0.48 | -0.65 | -0.625 | 0.52 | 0.001 | 0.011 | 0.011 | 0 | 0.08 | 0.05 | 0.12 | 0.25 | 0.1 | 0.16 |
| 4 | 108015733 | rs403608 | -1.309 | -0.599 | -0.531 | 0.565 | -0.005 | 0.01 | 0.01 | 0 | 0.67 | 0.79 | 0.59 | 0.74 | 0.65 | 0.65 |
| 4 | 108016162 | rs425273 | -0.78 | -0.32 | -0.394 | 0.819 | -0.015 | 0.083 | 0.083 | 0.239 | 0.24 | 0.54 | 0.29 | 0.38 | 0.21 | 0.21 |
| 4 | 108017272 | rs81299 | 0.058 | -0.246 | -0.369 | 0.533 | 0.007 | 0.005 | 0.005 | 0.059 | 0.17 | 0.48 | 0.16 | 0.13 | 0.11 | 0.05 |
| 4 | 108018123 | rs382525 | 0.127 | -0.232 | -0.395 | 0.524 | 0.001 | -0.003 | -0.003 | 0 | 0.16 | 0.41 | 0.16 | 0.13 | 0.11 | 0.05 |
| 4 | 108020599 | rs435887 | -0.426 | -0.061 | -0.855 | 0.215 | 0.003 | 0.032 | 0.032 | 0 | 0.58 | 0.73 | 0.55 | 0.74 | 0.6 | 0.55 |
| 4 | 108040129 | rs373586 | -0.188 | -0.201 | -0.822 | 0.336 | 0.007 | 0.008 | 0.008 | 0 | 0.61 | 0.38 | 0.57 | 0.31 | 0.62 | 0.63 |
| 4 | 108040751 | rs404662 | -0.022 | -0.292 | -1.234 | 0.456 | 0.021 | -0.007 | -0.007 | 0 | 0.46 | 0.71 | 0.37 | 0.51 | 0.38 | 0.26 |
| 4 | 108041804 | rs412908 | -0.159 | -0.267 | -0.9 | 0.913 | -0.003 | 0.023 | 0.023 | 0 | 0.89 | 0.88 | 0.82 | 0.75 | 0.88 | 0.79 |
| 4 | 108048311 | rs17510449 | 0.329 | -0.366 | -1.25 | 0.659 | -0.069 | -0.028 | -0.028 | 0.041 | 0.33 | 0.07 | 0.34 | 0.34 | 0.33 | 0.39 |
| 4 | 108048977 | rs2158182 | -0.003 | -0.316 | -1.423 | 0.485 | -0.021 | 0.009 | 0.009 | 0.26 | 0.77 | 0.9 | 0.75 | 0.8 | 0.76 | 0.69 |
| 4 | 108051500 | rs718820 | 0.515 | -0.035 | -0.824 | 0.48 | 0 | -0.001 | -0.001 | 0.029 | 0.41 | 0.62 | 0.37 | 0.4 | 0.41 | 0.27 |
| 4 | 108052407 | rs2522457 | 0.215 | 0.04 | -1.246 | 0.376 | 0.021 | -0.027 | -0.027 | 0 | 0.44 | 0.76 | 0.4 | 0.46 | 0.42 | 0.3 |
| 4 | 108071458 | rs28513708 | 0.301 | -0.11 | -0.478 | 0.626 | -0.04 | 0.008 | 0.008 | 0 | 0.31 | 0.16 | 0.39 | 0.28 | 0.43 | 0.29 |
| 4 | 108083471 | rs1807847 | 0.336 | -0.29 | -0.008 | 0.75 | -0.022 | 0.005 | 0.005 | 0 | 0.3 | 0.19 | 0.39 | 0.28 | 0.41 | 0.28 |
| 4 | 108084614 | rs7694364 | -0.146 | -0.314 | 0.191 | 0.755 | 0.015 | 0.006 | 0.006 | 0 | 0.34 | 0.41 | 0.43 | 0.35 | 0.41 | 0.34 |
| 4 | 108086688 | rs12512114 | 0.285 | -0.304 | 0.013 | 0.749 | -0.078 | -0.004 | -0.004 | 0 | 0.3 | 0.1 | 0.38 | 0.28 | 0.41 | 0.28 |
| 4 | 108094650 | rs17037473 | 0.19 | -0.363 | 0.442 | 0.714 | 0.076 | 0.198 | 0.198 | 0.644 | 0.06 | 0.41 | 0.05 | 0.07 | 0.01 | 0.05 |
| 4 | 108105844 | rs12511906 | 0.34 | -0.361 | 0.067 | 0.752 | -0.079 | -0.014 | -0.014 | 0.197 | 0.3 | 0.1 | 0.38 | 0.29 | 0.41 | 0.28 |
| 4 | 108109894 | rs950252 | -0.528 | -0.547 | -0.825 | 0.745 | -0.015 | 0.008 | 0.008 | 0 | 0.72 | 0.64 | 0.8 | 0.66 | 0.79 | 0.81 |
| 4 | 108115022 | rs17483781 | 0.401 | -0.365 | -1.523 | 0.591 | -0.07 | 0.018 | 0.018 | 0.229 | 0.36 | 0.09 | 0.36 | 0.3 | 0.37 | 0.56 |
| 4 | 108117467 | rs1027253 | -0.655 | -0.087 | -0.927 | 0.651 | 0.018 | -0.02 | -0.02 | 0 | 0.79 | 0.82 | 0.82 | 0.7 | 0.85 | 0.84 |
| 4 | 108118301 | rs17549820 | 0.033 | 0.193 | -1.193 | 0.452 | -0.035 | 0.055 | 0.055 | 0 | 0.34 | 0.16 | 0.38 | 0.18 | 0.42 | 0.19 |
| 4 | 108119422 | rs2029635 | 0.269 | 0.179 | -1.29 | 0.345 | -0.058 | 0.049 | 0.049 | 0 | 0.31 | 0.15 | 0.35 | 0.18 | 0.43 | 0.19 |
| 4 | 108120442 | rs4403120 | -0.691 | 0.013 | -1.165 | 0.503 | 0 | -0.001 | -0.001 | 0.424 | 0.56 | 0.43 | 0.57 | 0.49 | 0.62 | 0.36 |
| 4 | 108120955 | rs17037525 | -0.37 | -0.357 | -1.485 | 0.685 | 0.032 | -0.069 | -0.069 | 0 | 0.27 | 0.06 | 0.27 | 0.3 | 0.24 | 0.19 |
| 4 | 108127486 | rs17037547 | -0.315 | -0.341 | -0.711 | 0.631 | -0.034 | 0.013 | 0.013 | 0.088 | 0.11 | 0.25 | 0.07 | 0.14 | 0.08 | 0.11 |
| 4 | 108132331 | rs4956005 | 0.547 | -0.461 | 0.175 | 0.75 | -0.039 | 0.092 | 0.092 | 0.177 | 0.88 | 0.56 | 0.9 | 0.79 | 0.91 | 0.89 |
| 4 | 108133070 | rs7691374 | 0.084 | -0.403 | -1.152 | 0.189 | 0.001 | 0.005 | 0.005 | 0 | 0.48 | 0.53 | 0.24 | 0.2 | 0.44 | 0.26 |
| 4 | 108135377 | rs6827724 | -0.628 | -0.707 | -0.652 | -0.071 | -0.005 | 0.054 | 0.054 | 0 | 0.56 | 0.72 | 0.77 | 0.91 | 0.55 | 0.8 |
| 4 | 108135681 | rs6828013 | 0.103 | -0.697 | -0.322 | 0.037 | -0.044 | 0.057 | 0.057 | 0 | 0.12 | 0.37 | 0.09 | 0.22 | 0.09 | 0.12 |
| 4 | 108145936 | rs1492473 | -0.606 | -0.652 | -0.835 | -0.07 | -0.005 | 0.054 | 0.054 | 0 | 0.57 | 0.72 | 0.78 | 0.91 | 0.55 | 0.8 |
| 4 | 108146502 | rs12646290 | 0.34 | -0.23 | -1.211 | -0.294 | 0.002 | -0.013 | -0.013 | 0 | 0.18 | 0.27 | 0.1 | 0.1 | 0.15 | 0.15 |
| 4 | 108148408 | rs959482 | 0.518 | -0.309 | -1.325 | -0.007 | -0.165 | -0.057 | -0.057 | 0 | 0.38 | 0.03 | 0.66 | 0.61 | 0.46 | 0.63 |
| 4 | 108156306 | rs1388845 | -1.202 | -0.279 | -1.392 | -0.266 | 0 | -0.029 | -0.029 | 0.886 | 0.87 | 0.76 | 0.93 | 0.92 | 0.86 | 0.95 |
| 4 | 108158105 | rs1021420 | -1.01 | -0.3 | -1.669 | 0.205 | 0 | -0.008 | -0.008 | 0 | 0.7 | 0.69 | 0.75 | 0.77 | 0.67 | 0.88 |
| 4 | 108164882 | rs4956081 | 0.407 | 0.112 | -2.133 | 0.487 | -0.155 | -0.144 | -0.144 | 0 | 0.39 | 0.03 | 0.57 | 0.49 | 0.46 | 0.59 |
| 4 | 108167905 | rs11722765 | 0.237 | 0.237 | -0.801 | 0.6 | -0.044 | 0.033 | 0.033 | 0.188 | 0.88 | 0.68 | 0.92 | 0.87 | 0.92 | 0.88 |

| | | | | | | | | | | | | | | | | |
|---|---|---|---|---|---|---|---|---|---|---|---|---|---|---|---|---|
| 4 | 108174195 | rs1873710 | -0.02 | 1.275 | -1.899 | 0.626 | -0.065 | 0.002 | 0.002 | 0 | 0.18 | 0.06 | 0.21 | 0.13 | 0.25 | 0.29 |
| 4 | 108174391 | rs931481 | 0.196 | 0.629 | -1.736 | 0.697 | 0.011 | -0.016 | -0.016 | 2.409 | 0.82 | 0.84 | 0.87 | 0.81 | 0.82 | 0.8 |
| 4 | 108177310 | rs17037583 | 0.26 | 0.657 | -2.021 | 0.944 | -0.037 | -0.095 | -0.095 | 0.168 | 0.14 | 0.01 | 0.27 | 0.37 | 0.14 | 0.24 |
| 4 | 108181819 | rs10019755 | -0.279 | 0.576 | -2.687 | 0.568 | -0.008 | -0.028 | -0.028 | 2.244 | 0.46 | 0.9 | 0.39 | 0.37 | 0.41 | 0.28 |
| 4 | 108189246 | rs1038854 | 0.04 | 0.558 | -1.878 | 0.505 | 0.013 | -0.03 | -0.03 | 1.108 | 0.38 | 0.25 | 0.51 | 0.55 | 0.34 | 0.36 |
| 4 | 108197243 | rs1393587 | 0.07 | 0.465 | -1.852 | 0.253 | 0.013 | -0.036 | -0.036 | 3.053 | 0.37 | 0.22 | 0.51 | 0.55 | 0.34 | 0.36 |
| 4 | 108197862 | rs1388850 | 0.104 | 0.451 | -1.828 | 0.246 | 0.014 | -0.048 | -0.048 | 1.008 | 0.37 | 0.17 | 0.51 | 0.55 | 0.34 | 0.36 |
| 4 | 108222771 | rs1056079 | -0.093 | 0.268 | -0.786 | 0.362 | 0.001 | -0.013 | -0.013 | 0.221 | 0.42 | 0.3 | 0.56 | 0.52 | 0.43 | 0.35 |
| 4 | 108231167 | rs6533295 | -0.421 | 0.176 | -0.785 | 0.318 | 0.015 | 0.007 | 0.007 | 0 | 0.47 | 0.72 | 0.59 | 0.61 | 0.43 | 0.36 |
| 4 | 108233063 | rs717435 | -1.117 | 0.135 | -0.875 | 0.283 | -0.128 | -0.005 | -0.005 | 0.201 | 0.61 | 0.92 | 0.69 | 0.7 | 0.55 | 0.52 |
| 4 | 108239396 | rs1023514 | 0.096 | 0.292 | -1.076 | 0.025 | 0.004 | -0.001 | -0.001 | 0 | 0.44 | 0.61 | 0.59 | 0.61 | 0.43 | 0.36 |
| 4 | 108266027 | rs1502773 | 0.427 | 0.286 | -1.092 | 0.02 | -0.008 | -0.028 | -0.028 | 0.007 | 0.41 | 0.2 | 0.55 | 0.52 | 0.43 | 0.35 |
| 4 | 108284661 | rs17218497 | 0.358 | 0.387 | -0.913 | 0.29 | -0.011 | -0.005 | -0.005 | 0 | 0.15 | 0.2 | 0.11 | 0.09 | 0.12 | 0.16 |
| 4 | 108296579 | rs203721 | -1.577 | 0.626 | -1.188 | 0.254 | -0.073 | -0.054 | -0.054 | 0 | 0.77 | 0.96 | 0.79 | 0.74 | 0.7 | 0.62 |
| 4 | 108304505 | rs7680755 | 1.34 | 0.56 | -0.769 | 0.356 | -0.104 | -0.055 | -0.055 | 1.69 | 0.39 | 0.06 | 0.25 | 0.33 | 0.4 | 0.44 |
| 4 | 108312043 | rs6847720 | 0.376 | 0.471 | -0.355 | 0.37 | -0.004 | -0.004 | -0.004 | 0 | 0.62 | 0.6 | 0.53 | 0.62 | 0.64 | 0.59 |
| 4 | 108313081 | rs10488872 | 1.438 | 0.34 | -0.421 | 0.295 | 0.004 | 0.009 | 0.009 | 0 | 0.07 | 0.01 | 0.07 | 0.05 | 0.06 | 0.05 |
| 4 | 108320885 | rs203217 | 1.409 | 0.397 | -0.371 | 0.28 | -0.038 | 0.073 | 0.073 | 0 | 0.13 | 0.01 | 0.08 | 0.02 | 0.14 | 0.09 |
| 4 | 108324658 | rs203221 | -0.114 | 0.381 | -0.185 | 0.276 | -0.002 | 0 | 0 | 0 | 0.79 | 0.79 | 0.75 | 0.8 | 0.77 | 0.86 |
| 4 | 108326530 | rs203223 | 0.001 | 0.381 | -0.167 | 0.276 | 0.02 | -0.002 | -0.002 | 0 | 0.79 | 0.53 | 0.74 | 0.8 | 0.77 | 0.86 |
| 4 | 108327835 | rs203227 | 1.353 | 0.529 | -0.189 | 0.381 | -0.038 | 0.073 | 0.073 | 0 | 0.13 | 0.01 | 0.08 | 0.02 | 0.14 | 0.09 |
| 4 | 108347993 | rs10007314 | -0.178 | 0.367 | 0.028 | 0.554 | -0.019 | 0.04 | 0.04 | 0 | 0.68 | 0.78 | 0.67 | 0.77 | 0.65 | 0.77 |
| 4 | 108383641 | rs10023658 | 1.19 | 0.369 | -0.232 | 0.301 | -0.021 | -0.061 | -0.061 | 0 | 0.11 | 0.03 | 0.27 | 0.28 | 0.12 | 0.2 |
| 4 | 108387300 | rs2214389 | 1.157 | 0.369 | -0.033 | 0.257 | -0.001 | -0.004 | -0.004 | 1.005 | 0.42 | 0.37 | 0.28 | 0.42 | 0.42 | 0.43 |
| 4 | 108390499 | rs7692560 | -0.445 | 0.378 | -0.095 | 0.198 | -0.024 | 0.024 | 0.024 | 0 | 0.73 | 0.87 | 0.69 | 0.77 | 0.69 | 0.81 |
| 4 | 108392974 | rs10005025 | 0.007 | 0.428 | 0.005 | 0.22 | -0.003 | 0.016 | 0.016 | 0 | 0.66 | 0.69 | 0.67 | 0.79 | 0.66 | 0.76 |
| 4 | 108396913 | rs17501731 | 0.035 | 0.428 | 0.014 | 0.237 | -0.004 | 0.019 | 0.019 | 0 | 0.25 | 0.33 | 0.39 | 0.34 | 0.24 | 0.33 |
| 4 | 108414125 | rs2189201 | 0.065 | 0.492 | -0.53 | 0.16 | -0.003 | 0.017 | 0.017 | 0 | 0.66 | 0.69 | 0.67 | 0.8 | 0.66 | 0.76 |
| 4 | 108419815 | rs1399403 | -0.255 | 0.628 | -0.515 | 0.233 | -0.008 | 0.015 | 0.015 | 0 | 0.72 | 0.8 | 0.68 | 0.81 | 0.69 | 0.83 |
| 4 | 108458314 | rs11942323 | -0.499 | 0.427 | -0.05 | 0.358 | -0.001 | -0.005 | -0.005 | 0.417 | 0.24 | 0.22 | 0.38 | 0.38 | 0.24 | 0.34 |
| 4 | 108462490 | rs2672469 | -0.769 | 0.495 | -0.635 | 0.43 | 0 | -0.001 | -0.001 | 0 | 0.31 | 0.71 | 0.43 | 0.38 | 0.27 | 0.39 |
| 4 | 108474586 | rs1514728 | 0.072 | 0.621 | -0.538 | 0.497 | 0.001 | -0.004 | -0.004 | 0.803 | 0.66 | 0.65 | 0.64 | 0.5 | 0.67 | 0.45 |
| 4 | 108493334 | rs2726692 | 0.521 | 0.617 | -0.592 | -0.19 | 0 | -0.001 | -0.001 | 0.594 | 0.49 | 0.41 | 0.51 | 0.5 | 0.5 | 0.28 |
| 4 | 108493705 | rs10516545 | 0.174 | 0.726 | -0.606 | -0.165 | -0.016 | 0.035 | 0.035 | 0 | 0.14 | 0.05 | 0.11 | 0 | 0.16 | 0.12 |
| 4 | 108506584 | rs932100 | -0.117 | 0.669 | -0.477 | 0.071 | 0.008 | -0.005 | -0.005 | 0 | 0.54 | 0.63 | 0.56 | 0.38 | 0.58 | 0.42 |
| 4 | 108517790 | rs12648912 | -0.231 | 0.589 | -0.338 | 0.002 | -0.03 | -0.042 | -0.042 | 0 | 0.59 | 0.81 | 0.57 | 0.38 | 0.57 | 0.43 |
| 4 | 108530885 | rs1514730 | -0.775 | 0.977 | 0.292 | -0.184 | 0.042 | 0.106 | 0.106 | 0 | 0.26 | 0.49 | 0.32 | 0.48 | 0.32 | 0.2 |
| 4 | 108534358 | rs2726180 | -0.787 | 0.918 | 0.431 | -0.297 | 0.001 | 0.026 | 0.026 | 0 | 0.65 | 0.55 | 0.52 | 0.49 | 0.65 | 0.44 |
| 4 | 108535335 | rs9569 | 0.73 | 0.996 | 0.394 | -0.218 | -0.052 | -0.074 | -0.074 | 0.742 | 0.14 | 0.04 | 0.26 | 0.35 | 0.23 | 0.15 |
| 4 | 108537766 | rs2672487 | -0.845 | 0.936 | 0.456 | -0.241 | 0.003 | 0.026 | 0.026 | 0 | 0.66 | 0.59 | 0.52 | 0.49 | 0.65 | 0.45 |
| 4 | 108541282 | rs2726673 | 0.07 | 0.995 | 0.843 | -0.202 | 0.008 | 0.037 | 0.037 | 0 | 0.09 | 0.28 | 0.05 | 0.11 | 0.06 | 0.02 |
| 4 | 108547865 | rs4956111 | 0.483 | 1.085 | 0.342 | -0.189 | -0.012 | -0.011 | -0.011 | 0.612 | 0.15 | 0.11 | 0.27 | 0.35 | 0.23 | 0.15 |
| 4 | 108557559 | rs2672458 | -0.205 | 1.135 | 0.568 | -0.241 | 0.047 | 0.052 | 0.052 | 0 | 0.1 | 0.42 | 0.06 | 0.11 | 0.07 | 0.03 |
| 4 | 108562555 | rs10461141 | 0.447 | 1.164 | 0.61 | -0.162 | 0.029 | 0.095 | 0.095 | 0.045 | 0.15 | 0.25 | 0.26 | 0.35 | 0.23 | 0.15 |
| 4 | 108565215 | rs2726659 | -0.197 | 1.158 | 0.612 | -0.165 | 0.047 | 0.052 | 0.052 | 0 | 0.1 | 0.42 | 0.06 | 0.11 | 0.07 | 0.03 |
| 4 | 108572567 | rs2726655 | -0.197 | 1.158 | 0.612 | -0.165 | 0.047 | 0.052 | 0.052 | 0 | 0.1 | 0.42 | 0.06 | 0.11 | 0.07 | 0.03 |
| 4 | 108575989 | rs3733633 | 0.305 | 1.261 | 0.436 | 0.061 | 0.029 | 0.095 | 0.095 | 0.045 | 0.15 | 0.25 | 0.26 | 0.35 | 0.23 | 0.16 |
| 4 | 108584007 | rs3805342 | 0.416 | 1.298 | -0.232 | 0.08 | -0.017 | -0.022 | -0.022 | 0.043 | 0.14 | 0.09 | 0.25 | 0.35 | 0.23 | 0.16 |
| 4 | 108592665 | rs4956024 | 0.436 | 1.284 | -0.474 | 0.082 | -0.044 | -0.076 | -0.076 | 0 | 0.14 | 0.05 | 0.25 | 0.35 | 0.22 | 0.15 |
| 4 | 108593062 | rs1476086 | 0.353 | 1.187 | -0.564 | -0.042 | -0.015 | -0.05 | -0.05 | 0.178 | 0.36 | 0.23 | 0.48 | 0.53 | 0.38 | 0.58 |
| 4 | 108602062 | rs3805347 | -0.093 | 1.207 | -0.25 | 0.635 | -0.044 | -0.076 | -0.076 | 0.341 | 0.14 | 0.05 | 0.25 | 0.35 | 0.22 | 0.16 |
| 4 | 108605212 | rs12502059 | 0.148 | 1.258 | -0.117 | 0.726 | -0.021 | -0.026 | -0.026 | 0 | 0.12 | 0.07 | 0.23 | 0.36 | 0.2 | 0.14 |
| 4 | 108609628 | rs2522420 | -0.251 | 1.069 | -0.236 | 0.771 | -0.057 | -0.109 | -0.109 | 4.504 | 0.23 | 0.05 | 0.38 | 0.47 | 0.32 | 0.3 |
| 4 | 108611941 | rs2522422 | 0.335 | 0.773 | 0.582 | 0.643 | 0.003 | -0.005 | -0.005 | 0 | 0.89 | 0.41 | 0.88 | 0.87 | 0.93 | 0.96 |
| 4 | 108618232 | rs2522425 | 0.269 | 0.745 | 1.168 | 0.629 | 0.025 | 0.015 | 0.015 | 0 | 0.89 | 0.49 | 0.88 | 0.87 | 0.93 | 0.96 |
| 4 | 108621483 | rs2522426 | 0.54 | 0.746 | 1.223 | 0.629 | -0.004 | -0.012 | -0.012 | 1.781 | 0.89 | 0.4 | 0.88 | 0.87 | 0.93 | 0.96 |
| 4 | 108622475 | rs13110240 | 0.93 | 0.713 | 1.263 | 0.618 | -0.015 | -0.034 | -0.034 | 0 | 0.64 | 0.22 | 0.49 | 0.4 | 0.6 | 0.67 |
| 4 | 108633506 | rs2726176 | 0.194 | 0.724 | 1.322 | 0.585 | 0.024 | 0.014 | 0.014 | 0 | 0.89 | 0.53 | 0.89 | 0.88 | 0.93 | 0.96 |
| 4 | 108639491 | rs2726207 | -0.232 | 0.786 | 1.017 | 0.72 | -0.017 | -0.053 | -0.053 | 0.447 | 0.25 | 0.14 | 0.38 | 0.48 | 0.29 | 0.29 |
| 4 | 108641300 | rs973126 | 0.046 | 0.983 | 1.016 | 0.774 | 0.019 | -0.002 | -0.002 | 0.613 | 0.92 | 0.8 | 0.9 | 0.88 | 0.9 | 0.96 |
| 4 | 108655834 | rs2189158 | 0.899 | 0.715 | 0.566 | 0.555 | -0.048 | 0.026 | 0.026 | 0 | 0.41 | 0.18 | 0.4 | 0.31 | 0.47 | 0.31 |
| 4 | 108671723 | rs11732603 | -0.526 | 1.508 | 1.785 | 0.989 | 0.028 | 0.103 | 0.103 | 0 | 0.24 | 0.4 | 0.38 | 0.47 | 0.31 | 0.27 |
| 4 | 108681819 | rs4585383 | 2.169 | 1.026 | 2.334 | 0.282 | 0.029 | 0.082 | 0.082 | 1.094 | 0.86 | 0.45 | 0.8 | 0.78 | 0.84 | 0.91 |
| 4 | 108691131 | rs10027815 | -1.763 | 1.025 | 2.321 | 0.282 | 0.032 | 0.081 | 0.081 | 0.116 | 0.12 | 0.39 | 0.19 | 0.22 | 0.16 | 0.09 |
| 4 | 108717195 | rs1826712 | -2.008 | 1.102 | 2.043 | 0.309 | 0.032 | 0.088 | 0.088 | 0 | 0.14 | 0.54 | 0.21 | 0.22 | 0.16 | 0.1 |
| 4 | 108731771 | rs7665836 | -0.555 | 1.201 | 1.316 | 0.047 | 0.032 | -0.039 | -0.039 | 0 | 0.08 | 0.2 | 0.05 | 0 | 0.08 | 0.04 |
| 4 | 108732508 | rs10008418 | -1.107 | 1.151 | 1.034 | 0.076 | 0.066 | 0.1 | 0.1 | 0 | 0.1 | 0.38 | 0.13 | 0.11 | 0.12 | 0.09 |
| 4 | 108758579 | rs1389167 | 0.028 | 1.053 | -0.021 | -0.784 | -0.011 | 0.093 | 0.093 | 0 | 0.54 | 0.67 | 0.34 | 0.05 | 0.49 | 0.29 |
| 4 | 108784514 | rs10516546 | 0.029 | 1.083 | 0.569 | -0.667 | 0 | 0.061 | 0.061 | 0 | 0.57 | 0.7 | 0.44 | 0.16 | 0.52 | 0.24 |

| | | | | | | | | | | | | | | | | |
|---|---|---|---|---|---|---|---|---|---|---|---|---|---|---|---|---|
| 4 | 108788680 | rs11940819 | 0.804 | 0.887 | 0.38 | -1.248 | -0.021 | 0.023 | 0.023 | 0 | 0.33 | 0.21 | 0.32 | 0.16 | 0.36 | 0.14 |
| 4 | 108792735 | rs7671347 | 0.452 | 0.723 | 0.153 | -0.65 | 0.003 | -0.004 | -0.004 | 0.104 | 0.46 | 0.39 | 0.48 | 0.43 | 0.51 | 0.56 |
| 4 | 108794544 | rs975219 | -0.689 | 0.793 | -0.783 | -0.555 | -0.011 | -0.138 | -0.138 | 4.447 | 0.7 | 0.91 | 0.6 | 0.43 | 0.67 | 0.65 |
| 4 | 108794813 | rs17038153 | 0.193 | 0.75 | -0.267 | -0.878 | 0.003 | 0.025 | 0.025 | 0 | 0.13 | 0.06 | 0.07 | 0 | 0.11 | 0.04 |
| 4 | 108802411 | rs6811703 | -0.32 | 0.863 | -0.047 | -0.82 | -0.006 | -0.031 | -0.031 | 0 | 0.3 | 0.58 | 0.27 | 0.11 | 0.27 | 0.14 |
| 4 | 108802762 | rs12648528 | -0.511 | 0.78 | -0.159 | -0.247 | -0.001 | 0 | 0 | 0 | 0.28 | 0.28 | 0.38 | 0.75 | 0.28 | 0.56 |
| 4 | 108818757 | rs1160677 | 0.735 | 1.022 | 0.523 | -0.385 | 0.055 | 0.011 | 0.011 | 0 | 0.9 | 0.56 | 0.95 | 0.98 | 0.93 | 0.94 |
| 4 | 108823320 | rs6826111 | -0.958 | 0.909 | 0.095 | -0.439 | 0.001 | -0.017 | -0.017 | 0 | 0.07 | 0.12 | 0.04 | 0 | 0.06 | 0.02 |
| 4 | 108829343 | rs13144835 | -1.086 | 0.809 | 0.592 | -1.346 | -0.008 | -0.141 | -0.141 | 0.422 | 0.32 | 0.5 | 0.19 | 0.02 | 0.28 | 0.09 |
| 4 | 108832286 | rs11945554 | -1.095 | 0.774 | 0.587 | -1.351 | -0.008 | -0.141 | -0.141 | 0 | 0.33 | 0.5 | 0.19 | 0.02 | 0.28 | 0.09 |
| 4 | 108834529 | rs13103176 | -0.014 | 0.722 | 0.908 | -1.361 | -0.009 | -0.035 | -0.035 | 0 | 0.07 | 0.17 | 0.02 | 0 | 0.02 | 0.03 |
| 4 | 108848793 | rs4956140 | -1.462 | 0.437 | 0.913 | -1.558 | -0.001 | -0.049 | -0.049 | 0 | 0.34 | 0.66 | 0.2 | 0.02 | 0.28 | 0.08 |
| 4 | 108851574 | rs4956029 | -0.254 | 0.525 | 1.143 | -1.518 | 0.005 | -0.044 | -0.044 | 0.008 | 0.19 | 0.23 | 0.12 | 0.02 | 0.22 | 0.04 |
| 4 | 108855776 | rs2131463 | 0.35 | 0.695 | -0.042 | -1.275 | -0.007 | -0.002 | -0.002 | 0.509 | 0.3 | 0.44 | 0.45 | 0.66 | 0.33 | 0.47 |
| 4 | 108857140 | rs4956031 | 0.822 | 0.746 | -0.013 | -1.285 | -0.006 | 0.002 | 0.002 | 0 | 0.17 | 0.14 | 0.12 | 0.02 | 0.22 | 0.04 |
| 4 | 108860906 | rs1493131 | 0.86 | 0.746 | -0.015 | -1.285 | -0.012 | 0.02 | 0.02 | 0 | 0.17 | 0.11 | 0.12 | 0.02 | 0.22 | 0.04 |
| 4 | 108861140 | rs6533327 | -1.042 | 0.723 | -0.033 | -1.292 | -0.001 | 0.042 | 0.042 | 0 | 0.5 | 0.59 | 0.59 | 0.69 | 0.57 | 0.52 |
| 4 | 108873285 | rs8727 | 0.597 | 0.777 | -0.077 | -1.268 | -0.004 | -0.007 | -0.007 | 0 | 0.19 | 0.16 | 0.13 | 0.02 | 0.23 | 0.05 |
| 4 | 108874039 | rs3756271 | -0.455 | 0.618 | -0.019 | -0.858 | -0.007 | -0.003 | -0.003 | 0.202 | 0.7 | 0.6 | 0.55 | 0.34 | 0.67 | 0.53 |
| 4 | 108921367 | rs221330 | 0.983 | 1.34 | -1.01 | -1.627 | -0.022 | -0.001 | -0.001 | 2.919 | 0.3 | 0.1 | 0.29 | 0.36 | 0.3 | 0.42 |
| 4 | 108931039 | rs4956145 | -0.092 | 1.021 | 0.058 | -1.639 | 0.077 | 0.046 | 0.046 | 0.17 | 0.92 | 0.67 | 0.95 | 0.98 | 0.92 | 0.92 |
| 4 | 108941283 | rs7691506 | -0.092 | 1.021 | 0.058 | -1.639 | 0.077 | 0.046 | 0.046 | 0.17 | 0.92 | 0.67 | 0.95 | 0.98 | 0.92 | 0.92 |
| 4 | 108944101 | rs7664031 | 1.347 | 0.802 | 0.266 | -1.779 | -0.007 | -0.061 | -0.061 | 0 | 0.82 | 0.43 | 0.88 | 0.97 | 0.84 | 0.89 |
| 4 | 108946378 | rs4245924 | -0.198 | 0.777 | 0.269 | -1.765 | 0.077 | 0.046 | 0.046 | 0.17 | 0.92 | 0.67 | 0.95 | 0.98 | 0.92 | 0.92 |
| 4 | 108956423 | rs6831530 | -0.6 | 0.766 | 0.423 | -1.772 | 0.062 | 0.021 | 0.021 | 0.371 | 0.1 | 0.41 | 0.05 | 0.02 | 0.08 | 0.08 |
| 4 | 108959282 | rs6851442 | 0.168 | 0.766 | 0.478 | -1.771 | 0.063 | 0.035 | 0.035 | 0.044 | 0.07 | 0.28 | 0.05 | 0.02 | 0.08 | 0.08 |
| 4 | 108960709 | rs4956035 | 1.327 | 0.768 | 0.329 | -1.752 | -0.009 | -0.056 | -0.056 | 0 | 0.82 | 0.43 | 0.88 | 0.97 | 0.84 | 0.89 |
| 4 | 108967846 | rs4245926 | 1.318 | 0.767 | 0.328 | -1.753 | -0.007 | -0.055 | -0.055 | 0 | 0.83 | 0.42 | 0.89 | 0.96 | 0.85 | 0.89 |
| 4 | 108969590 | rs4245927 | 1.314 | 0.772 | 0.331 | -1.751 | -0.007 | -0.055 | -0.055 | 0.983 | 0.83 | 0.42 | 0.88 | 0.96 | 0.85 | 0.89 |
| 4 | 108978450 | rs10034874 | 1.379 | 0.966 | 0.362 | -1.736 | 0.001 | -0.043 | -0.043 | 0 | 0.82 | 0.33 | 0.89 | 0.97 | 0.84 | 0.89 |
| 4 | 108980772 | rs6826687 | -0.271 | 0.987 | 0.299 | -1.732 | 0.055 | 0.042 | 0.042 | 0 | 0.93 | 0.73 | 0.96 | 0.97 | 0.93 | 0.93 |
| 4 | 108985388 | rs4613638 | -0.427 | 0.982 | 0.312 | -1.73 | 0.069 | 0.039 | 0.039 | 0.05 | 0.93 | 0.76 | 0.95 | 0.97 | 0.91 | 0.92 |
| 4 | 108994012 | rs1291490 | 1.15 | 0.806 | -0.151 | -2.139 | -0.018 | 0.096 | 0.096 | 0 | 0.17 | 0.02 | 0.08 | 0 | 0.15 | 0.04 |
| 4 | 108999366 | rs4956157 | 1.414 | 0.984 | 0.331 | -1.842 | 0 | -0.045 | -0.045 | 0 | 0.83 | 0.34 | 0.89 | 0.97 | 0.84 | 0.9 |
| 4 | 109004975 | rs4403119 | 0.263 | 0.98 | 0.141 | -1.763 | -0.009 | -0.028 | -0.028 | 0 | 0.25 | 0.16 | 0.46 | 0.63 | 0.32 | 0.39 |
| 4 | 109009087 | rs4547871 | 0.717 | 0.976 | 0.345 | -1.761 | 0.045 | 0.002 | 0.002 | 0.876 | 0.89 | 0.6 | 0.93 | 0.97 | 0.9 | 0.92 |
| 4 | 109016824 | rs898518 | 0.121 | 0.992 | 0.339 | -1.735 | 0.013 | -0.008 | -0.008 | 0.407 | 0.68 | 0.6 | 0.49 | 0.35 | 0.59 | 0.54 |
| 4 | 109033979 | rs10022956 | -0.884 | 1.023 | 0.262 | -1.747 | -0.002 | -0.022 | -0.022 | 2.263 | 0.78 | 0.85 | 0.55 | 0.37 | 0.73 | 0.63 |
| 4 | 109055567 | rs6533354 | -0.758 | 1.072 | 0.436 | -1.733 | 0.09 | 0.056 | 0.056 | 0.028 | 0.93 | 0.72 | 0.95 | 0.96 | 0.91 | 0.92 |
| 4 | 109056791 | rs17038630 | -1.31 | 1.164 | 0.455 | -1.766 | 0.015 | -0.05 | -0.05 | 0.027 | 0.09 | 0.37 | 0.06 | 0 | 0.08 | 0.03 |
| 4 | 109060364 | rs922167 | 0.512 | 0.903 | 0.255 | -1.708 | 0.001 | -0.028 | -0.028 | 0.01 | 0.23 | 0.15 | 0.46 | 0.63 | 0.28 | 0.39 |
| 4 | 109062510 | rs12503104 | 0.738 | 1.092 | -0.206 | -2.048 | -0.002 | 0.058 | 0.058 | 0 | 0.54 | 0.33 | 0.31 | 0.28 | 0.44 | 0.47 |
| 4 | 109066366 | rs922163 | 0.221 | 0.97 | 0.209 | -1.493 | 0.011 | -0.004 | -0.004 | 0 | 0.7 | 0.56 | 0.49 | 0.33 | 0.63 | 0.53 |
| 4 | 109104682 | rs2343113 | -0.401 | 1.165 | -1.138 | -1.134 | -0.001 | 0.001 | 0.001 | 0 | 0.73 | 0.71 | 0.52 | 0.38 | 0.66 | 0.6 |
| 4 | 109106451 | rs10000124 | -0.281 | 1.25 | -1.126 | -0.338 | 0.041 | 0.027 | 0.027 | 0 | 0.07 | 0.2 | 0.04 | 0.05 | 0.08 | 0.08 |
| 4 | 109111753 | rs2221145 | -0.163 | 0.956 | -2.085 | -0.035 | -0.038 | -0.055 | -0.055 | 0.935 | 0.22 | 0.02 | 0.16 | 0.28 | 0.19 | 0.39 |
| 4 | 109113914 | rs4956041 | 0.023 | 0.726 | -1.496 | 0.096 | 0.024 | 0.002 | 0.002 | 0.021 | 0.51 | 0.32 | 0.67 | 0.55 | 0.57 | 0.48 |
| 4 | 109119628 | rs17567325 | -0.791 | 0.523 | -1.699 | -0.754 | 0.012 | 0.066 | 0.066 | 0 | 0.27 | 0.1 | 0.15 | 0.01 | 0.26 | 0.17 |
| 4 | 109142299 | rs4956042 | -1.055 | 0.798 | -2.033 | -1.404 | -0.023 | 0.106 | 0.106 | 0 | 0.35 | 0.16 | 0.22 | 0.01 | 0.4 | 0.21 |
| 4 | 109143439 | rs12501237 | 1.164 | 0.068 | -1.465 | -1.944 | -0.012 | 0.034 | 0.034 | 0 | 0.43 | 0.37 | 0.65 | 0.87 | 0.47 | 0.63 |
| 4 | 109151467 | rs7671754 | 0.444 | -0.323 | -1.568 | -2.012 | -0.006 | 0.038 | 0.038 | 0 | 0.11 | 0.02 | 0.09 | 0.06 | 0.11 | 0.13 |
| 4 | 109162596 | rs10026586 | 0.384 | -0.377 | -1.388 | -2.037 | -0.041 | -0.049 | -0.049 | 0 | 0.8 | 0.53 | 0.88 | 0.88 | 0.87 | 0.84 |
| 4 | 109163035 | rs11097974 | 1.395 | -0.361 | -1.275 | -1.752 | -0.017 | 0.067 | 0.067 | 0 | 0.43 | 0.3 | 0.67 | 0.83 | 0.49 | 0.63 |
| 4 | 109163364 | rs13140104 | -0.642 | -0.267 | -0.394 | -1.898 | -0.003 | -0.106 | -0.106 | 0.087 | 0.41 | 0.41 | 0.22 | 0.01 | 0.39 | 0.2 |
| 4 | 109172913 | rs931771 | -0.639 | -0.532 | -0.128 | -2.15 | 0.034 | 0.085 | 0.085 | 0 | 0.33 | 0.16 | 0.2 | 0.07 | 0.28 | 0.22 |
| 4 | 109180551 | rs7665502 | 1.614 | -0.602 | 0.231 | -1.511 | 0.006 | 0.072 | 0.072 | 0 | 0.47 | 0.33 | 0.69 | 0.78 | 0.53 | 0.66 |
| 4 | 109182392 | rs11934029 | 0.311 | -0.564 | 0.242 | -1.494 | 0.025 | 0.011 | 0.011 | 0 | 0.8 | 0.6 | 0.88 | 0.86 | 0.79 | 0.87 |
| 4 | 109192876 | rs3901937 | -1.761 | -0.638 | -0.028 | -1.528 | 0.026 | 0.08 | 0.08 | 0 | 0.52 | 0.68 | 0.31 | 0.22 | 0.44 | 0.34 |
| 4 | 109193333 | rs13139506 | -0.688 | -0.419 | -0.364 | -1.527 | 0.028 | 0.072 | 0.072 | 0 | 0.31 | 0.16 | 0.19 | 0.07 | 0.26 | 0.21 |
| 4 | 109196994 | rs11725852 | -0.716 | -0.415 | -0.348 | -1.525 | 0.026 | 0.058 | 0.058 | 0 | 0.31 | 0.16 | 0.19 | 0.07 | 0.26 | 0.21 |
| 4 | 109211978 | rs12507470 | 0.724 | 0.458 | -0.927 | -0.541 | -0.062 | -0.131 | -0.131 | 0.002 | 0.2 | 0.04 | 0.27 | 0.42 | 0.25 | 0.46 |
| 4 | 109235238 | rs220634 | 0.04 | 0.249 | -0.399 | -1.103 | 0.006 | 0.01 | 0.01 | 0 | 0.82 | 0.85 | 0.9 | 0.93 | 0.86 | 0.88 |
| 4 | 109235352 | rs220636 | 0.054 | 0.241 | -0.403 | -1.107 | 0.006 | 0.01 | 0.01 | 0 | 0.82 | 0.85 | 0.9 | 0.93 | 0.86 | 0.88 |
| 4 | 109236454 | rs852269 | 0.052 | 0.295 | -0.402 | -1.106 | 0.005 | 0.01 | 0.01 | 0 | 0.82 | 0.85 | 0.9 | 0.93 | 0.86 | 0.88 |
| 4 | 109238322 | rs220612 | 0.245 | 0.258 | -0.318 | -1.047 | -0.001 | -0.011 | -0.011 | 0 | 0.88 | 0.84 | 0.93 | 0.93 | 0.9 | 0.91 |
| 4 | 109242752 | rs220620 | -0.013 | 0.251 | -0.261 | -1.049 | -0.001 | -0.013 | -0.013 | 0 | 0.89 | 0.87 | 0.93 | 0.93 | 0.91 | 0.91 |
| 4 | 109243057 | rs220623 | 0.858 | 0.137 | 0.052 | -1.338 | 0.005 | -0.085 | -0.085 | 4.451 | 0.76 | 0.62 | 0.84 | 0.93 | 0.74 | 0.83 |
| 4 | 109251708 | rs17608469 | 1.19 | 0.067 | -0.473 | -1.813 | 0.003 | 0.048 | 0.048 | 0.003 | 0.37 | 0.19 | 0.2 | 0.04 | 0.34 | 0.21 |
| 4 | 109254423 | rs17038822 | 0.495 | 0.341 | -0.482 | -1.532 | 0.004 | -0.056 | -0.056 | 0 | 0.12 | 0.21 | 0.09 | 0 | 0.16 | 0.07 |

| | | | | | | | | | | | | | | | | |
|---|---|---|---|---|---|---|---|---|---|---|---|---|---|---|---|---|
| 4 | 109263671 | rs13138626 | 0.331 | 0.398 | -0.855 | -1.063 | -0.076 | -0.042 | -0.042 | 0.02 | 0.37 | 0.06 | 0.56 | 0.64 | 0.43 | 0.64 |
| 4 | 109264055 | rs12108639 | 0.415 | 0.597 | -0.104 | -0.791 | 0.022 | -0.063 | -0.063 | 0 | 0.11 | 0.28 | 0.08 | 0 | 0.11 | 0.02 |
| 4 | 109272140 | rs11940782 | -0.535 | 0.556 | 0.005 | -0.816 | 0.02 | -0.115 | -0.115 | 1.059 | 0.15 | 0.48 | 0.1 | 0 | 0.13 | 0.04 |
| 4 | 109275454 | rs917413 | -0.112 | 0.782 | -0.479 | -0.69 | 0.035 | -0.087 | -0.087 | 0.659 | 0.84 | 0.66 | 0.89 | 1 | 0.83 | 0.95 |
| 4 | 109276830 | rs1558172 | 0.692 | 0.841 | -1.062 | -0.133 | -0.061 | -0.105 | -0.105 | 0 | 0.25 | 0.06 | 0.47 | 0.45 | 0.34 | 0.54 |
| 4 | 109282683 | rs10489025 | 1.607 | 0.473 | -0.867 | -0.629 | -0.001 | -0.092 | -0.092 | 1.278 | 0.31 | 0.32 | 0.19 | 0.04 | 0.31 | 0.21 |
| 4 | 109286645 | rs7667905 | 1.572 | 0.421 | -0.848 | -0.496 | 0 | -0.11 | -0.11 | 2.146 | 0.29 | 0.35 | 0.19 | 0.04 | 0.3 | 0.21 |
| 4 | 109288854 | rs12374269 | 0.629 | 0.491 | -1.369 | -0.133 | -0.044 | -0.078 | -0.078 | 0.172 | 0.3 | 0.08 | 0.47 | 0.44 | 0.35 | 0.52 |
| 4 | 109289367 | rs6844729 | 0.917 | 0.212 | -1.246 | -0.199 | -0.028 | -0.025 | -0.025 | 0 | 0.06 | 0.16 | 0.03 | 0.01 | 0.02 | 0.03 |
| 4 | 109290374 | rs17038931 | 1.128 | 0.227 | -1.461 | -0.045 | 0.005 | -0.057 | -0.057 | 0 | 0.14 | 0.01 | 0.1 | 0.18 | 0.1 | 0.07 |
| 4 | 109292368 | rs219457 | -0.006 | 0.495 | -1.246 | -0.331 | 0.007 | -0.018 | -0.018 | 0 | 0.36 | 0.29 | 0.5 | 0.45 | 0.36 | 0.56 |
| 4 | 109295549 | rs11097983 | -0.207 | 0.832 | -1.645 | -0.992 | -0.039 | 0.223 | 0.223 | 0 | 0.43 | 0.09 | 0.63 | 0.82 | 0.43 | 0.61 |
| 4 | 109295714 | rs188720 | -0.207 | 0.832 | -1.645 | -0.992 | -0.039 | 0.223 | 0.223 | 0 | 0.43 | 0.09 | 0.63 | 0.82 | 0.43 | 0.61 |
| 4 | 109296917 | rs188721 | 0.141 | 0.968 | -1.269 | -1.065 | 0.026 | -0.034 | -0.034 | 0 | 0.82 | 0.73 | 0.9 | 0.9 | 0.77 | 0.86 |
| 4 | 109298935 | rs737605 | 1.093 | 0.92 | -1.294 | -1.202 | -0.004 | -0.062 | -0.062 | 0 | 0.38 | 0.57 | 0.24 | 0.08 | 0.33 | 0.25 |
| 4 | 109299202 | rs740053 | -0.139 | 0.84 | -0.955 | -0.563 | 0.001 | -0.004 | -0.004 | 0 | 0.12 | 0.16 | 0.06 | 0.1 | 0.17 | 0.11 |
| 4 | 109301049 | rs13132148 | 0.039 | 0.823 | -1.215 | -0.356 | -0.003 | -0.103 | -0.103 | 0.454 | 0.18 | 0.03 | 0.17 | 0.29 | 0.16 | 0.18 |
| 4 | 109301111 | rs219467 | 1.099 | 0.797 | -0.776 | -0.313 | 0.005 | 0.004 | 0.004 | 0 | 0.62 | 0.52 | 0.71 | 0.6 | 0.61 | 0.68 |
| 4 | 109305397 | rs1452397 | -0.912 | 0.499 | -0.656 | -0.233 | 0.002 | 0.018 | 0.018 | 0 | 0.29 | 0.47 | 0.25 | 0.29 | 0.28 | 0.2 |
| 4 | 109311641 | rs219472 | -0.35 | 0.215 | -0.792 | -0.79 | -0.015 | 0.035 | 0.035 | 0 | 0.42 | 0.37 | 0.65 | 0.73 | 0.45 | 0.5 |
| 4 | 109312549 | rs17039002 | 0.873 | 0.359 | -0.475 | -0.984 | 0.035 | -0.042 | -0.042 | 0 | 0.08 | 0.16 | 0.08 | 0 | 0.14 | 0.02 |
| 4 | 109316327 | rs121967 | -0.3 | -0.007 | -0.783 | -1.169 | -0.001 | -0.01 | -0.01 | 0 | 0.95 | 0.96 | 0.96 | 1 | 0.94 | 0.97 |
| 4 | 109320729 | rs219456 | -0.189 | -0.041 | -0.855 | -1.229 | -0.008 | 0.034 | 0.034 | 0 | 0.43 | 0.4 | 0.64 | 0.74 | 0.43 | 0.46 |
| 4 | 109322782 | rs219451 | 0.003 | -0.153 | -0.95 | -1.279 | -0.018 | 0.035 | 0.035 | 0 | 0.38 | 0.35 | 0.62 | 0.78 | 0.43 | 0.49 |
| 4 | 109338471 | rs172666 | -0.766 | -0.155 | -0.732 | -1.286 | 0.011 | 0.016 | 0.016 | 0 | 0.48 | 0.6 | 0.7 | 0.78 | 0.6 | 0.51 |
| 4 | 109340406 | rs6811130 | 0.642 | -0.083 | -0.716 | -1.278 | 0.023 | -0.08 | -0.08 | 0 | 0.14 | 0.25 | 0.1 | 0 | 0.18 | 0.02 |
| 4 | 109340680 | rs219481 | 1.373 | 0.222 | -1.122 | -0.766 | -0.1 | -0.144 | -0.144 | 0 | 0.18 | 0.05 | 0.49 | 0.53 | 0.3 | 0.35 |
| 4 | 109346722 | rs17039028 | 1.092 | 0.396 | -1.005 | -0.634 | -0.035 | -0.059 | -0.059 | 0 | 0.1 | 0.01 | 0.1 | 0.24 | 0.11 | 0.14 |
| 4 | 109349464 | rs1377065 | 0.477 | 0.249 | -0.928 | -0.605 | 0.037 | 0.05 | 0.05 | 0 | 0.5 | 0.37 | 0.26 | 0.26 | 0.37 | 0.48 |
| 4 | 109350880 | rs219493 | -0.775 | 0.427 | -1.556 | -0.301 | -0.025 | -0.045 | -0.045 | 0 | 0.86 | 0.94 | 0.86 | 0.76 | 0.85 | 0.85 |
| 4 | 109362923 | rs219500 | -0.539 | -0.325 | -1.245 | -0.945 | 0.067 | 0.115 | 0.115 | 0.1 | 0.6 | 0.86 | 0.78 | 0.77 | 0.68 | 0.57 |
| 4 | 109369135 | rs11936528 | -0.879 | 0.227 | -0.54 | 0.033 | -0.011 | -0.029 | -0.029 | 0 | 0.29 | 0.51 | 0.16 | 0.22 | 0.28 | 0.22 |
| 4 | 109372293 | rs7667423 | -0.467 | -0.152 | -0.894 | 0.329 | 0.014 | 0.004 | 0.004 | 0 | 0.34 | 0.64 | 0.27 | 0.36 | 0.25 | 0.34 |
| 4 | 109374717 | rs1452401 | 0.329 | 0.251 | -0.661 | 0.318 | -0.001 | 0 | 0 | 0 | 0.6 | 0.39 | 0.59 | 0.61 | 0.59 | 0.54 |
| 4 | 109396732 | rs719786 | 0.119 | 0.394 | -0.561 | 0.556 | -0.007 | 0.032 | 0.032 | 0 | 0.16 | 0.19 | 0.2 | 0.33 | 0.13 | 0.35 |
| 4 | 109407499 | rs7654972 | 0.061 | 0.622 | -0.378 | 0.051 | 0.015 | 0.02 | 0.02 | 0 | 0.31 | 0.39 | 0.32 | 0.38 | 0.36 | 0.46 |
| 4 | 109408608 | rs1562975 | 0.703 | 0.444 | -0.523 | -0.261 | -0.008 | 0.024 | 0.024 | 0.024 | 0.28 | 0.19 | 0.18 | 0.04 | 0.3 | 0.16 |
| 4 | 109411079 | rs6838036 | -0.438 | 0.047 | -0.599 | -1.144 | -0.004 | -0.023 | -0.023 | 0 | 0.72 | 0.75 | 0.77 | 1 | 0.7 | 0.91 |
| 4 | 109413413 | rs12508024 | 1.554 | -0.389 | -0.335 | -1.19 | 0.005 | 0.085 | 0.085 | 0 | 0.1 | 0.15 | 0.18 | 0.35 | 0.13 | 0.34 |
| 4 | 109417821 | rs1115882 | 1.153 | -0.436 | -0.492 | -1.334 | -0.01 | -0.059 | -0.059 | 0 | 0.13 | 0.18 | 0.09 | 0 | 0.1 | 0.07 |
| 4 | 109439995 | rs4996943 | -0.503 | 0.333 | -0.146 | -0.931 | 0.001 | 0.003 | 0.003 | 0.059 | 0.39 | 0.52 | 0.43 | 0.4 | 0.41 | 0.51 |
| 4 | 109441230 | rs11737760 | 0.929 | 0.364 | -0.099 | -0.801 | 0.036 | 0.235 | 0.235 | 0 | 0.11 | 0.32 | 0.25 | 0.36 | 0.13 | 0.35 |
| 4 | 109461985 | rs17583662 | 0.202 | 0.168 | -0.574 | -1.098 | -0.032 | 0.107 | 0.107 | 0.005 | 0.34 | 0.15 | 0.2 | 0.08 | 0.38 | 0.25 |
| 4 | 109465726 | rs7688462 | -0.667 | -0.05 | -0.462 | -1.024 | 0.002 | 0 | 0 | 0.257 | 0.46 | 0.49 | 0.46 | 0.47 | 0.5 | 0.59 |
| 4 | 109474255 | rs1017432 | 0.521 | -0.231 | -0.341 | -0.73 | 0.011 | 0.199 | 0.199 | 0 | 0.12 | 0.25 | 0.26 | 0.43 | 0.12 | 0.34 |
| 4 | 109477779 | rs7686569 | -0.572 | -0.859 | -0.526 | -1.245 | 0.002 | 0.021 | 0.021 | 0 | 0.5 | 0.64 | 0.59 | 0.57 | 0.52 | 0.59 |
| 4 | 109486382 | rs11734670 | 1.77 | -0.849 | -0.507 | -1.303 | 0.022 | 0.071 | 0.071 | 0 | 0.12 | 0.14 | 0.37 | 0.53 | 0.2 | 0.38 |
| 4 | 109488632 | rs7676871 | 0.136 | -0.949 | -0.475 | -1.486 | 0.029 | 0.049 | 0.049 | 0 | 0.33 | 0.2 | 0.18 | 0.04 | 0.28 | 0.18 |
| 4 | 109493253 | rs17800095 | -0.004 | -0.947 | -0.496 | -1.485 | -0.008 | -0.053 | -0.053 | 1.816 | 0.34 | 0.39 | 0.19 | 0.04 | 0.28 | 0.19 |
| 4 | 109517946 | rs12646462 | 0.966 | -1.051 | -1.31 | -1.018 | 0.039 | 0.063 | 0.063 | 0 | 0.24 | 0.34 | 0.45 | 0.55 | 0.34 | 0.45 |
| 4 | 109529238 | rs17584310 | 1.362 | -1.264 | -1.636 | -1.102 | -0.043 | -0.007 | -0.007 | 0.182 | 0.27 | 0.01 | 0.18 | 0.18 | 0.22 | 0.2 |
| 4 | 109532430 | rs1865845 | 1.5 | -0.802 | -1.332 | -0.803 | -0.006 | -0.002 | -0.002 | 0 | 0.19 | 0.16 | 0.42 | 0.56 | 0.31 | 0.44 |
| 4 | 109536616 | rs7691236 | 1.484 | -0.804 | -1.333 | -0.804 | -0.007 | -0.003 | -0.003 | 0.278 | 0.19 | 0.16 | 0.42 | 0.56 | 0.31 | 0.44 |
| 4 | 109552881 | rs11097999 | 1.588 | -0.801 | -1.389 | -0.804 | -0.024 | -0.029 | -0.029 | 1.729 | 0.19 | 0.12 | 0.41 | 0.56 | 0.31 | 0.43 |
| 4 | 109553331 | rs2880787 | -0.438 | -1.069 | -0.414 | -0.937 | 0.04 | 0.076 | 0.076 | 0 | 0.23 | 0.12 | 0.11 | 0.04 | 0.16 | 0.13 |
| 4 | 109583785 | rs1369962 | -0.557 | 0.165 | 0.016 | -0.146 | -0.003 | -0.017 | -0.017 | 0 | 0.82 | 0.83 | 0.85 | 1 | 0.78 | 0.96 |
| 4 | 109586178 | rs2575632 | -0.557 | 0.165 | -0.127 | -0.141 | -0.007 | 0.009 | 0.009 | 0 | 0.82 | 0.85 | 0.85 | 1 | 0.78 | 0.96 |
| 4 | 109613948 | rs6852328 | -0.901 | 0.182 | -0.385 | -0.104 | -0.024 | -0.001 | -0.001 | 0.087 | 0.29 | 0.39 | 0.23 | 0.33 | 0.24 | 0.29 |
| 4 | 109621884 | rs2077246 | 1.352 | 0.131 | -0.766 | -0.095 | 0.022 | 0.099 | 0.099 | 0 | 0.52 | 0.21 | 0.61 | 0.66 | 0.54 | 0.65 |
| 4 | 109628025 | rs10029851 | -0.19 | 0.131 | -0.937 | 0.54 | 0 | -0.009 | -0.009 | 0 | 0.18 | 0.18 | 0.11 | 0.03 | 0.17 | 0.15 |
| 4 | 109630951 | rs1436506 | -0.542 | 0.525 | -1.415 | 0.696 | -0.031 | 0.005 | 0.005 | 0 | 0.29 | 0.46 | 0.23 | 0.35 | 0.24 | 0.29 |
| 4 | 109636348 | rs2575625 | 0.033 | 0.98 | -2.432 | 1.114 | -0.055 | 0.009 | 0.009 | 0 | 0.45 | 0.12 | 0.55 | 0.63 | 0.48 | 0.58 |
| 4 | 109642764 | rs1812429 | -0.702 | 0.983 | -1.718 | 1.8 | -0.058 | -0.146 | -0.146 | 0.066 | 0.2 | 0.02 | 0.3 | 0.36 | 0.22 | 0.35 |
| 4 | 109657931 | rs2575657 | -1.134 | 0.914 | -1.822 | 1.695 | -0.056 | -0.181 | -0.181 | 0 | 0.25 | 0.02 | 0.34 | 0.5 | 0.26 | 0.38 |
| 4 | 109658910 | rs12710996 | 1.419 | 1.161 | -0.439 | 2.285 | 0.094 | 0.271 | 0.271 | 0 | 0.94 | 0.73 | 0.91 | 0.71 | 0.92 | 0.96 |
| 4 | 109660715 | rs12649328 | -0.412 | 1.079 | -1.11 | 1.695 | -0.028 | 0.029 | 0.029 | 0 | 0.35 | 0.19 | 0.36 | 0.24 | 0.38 | 0.4 |
| 4 | 109663371 | rs4956208 | -1 | 1.098 | -0.5 | 2.438 | 0.006 | 0.194 | 0.194 | 0 | 0.06 | 0.09 | 0.11 | 0.5 | 0.05 | 0.18 |
| 4 | 109664667 | rs7663958 | -1.628 | 1.272 | -0.285 | 2.493 | 0.013 | 0.174 | 0.174 | 0 | 0.06 | 0.13 | 0.13 | 0.48 | 0.08 | 0.2 |
| 4 | 109665016 | rs10035050 | -0.351 | 0.383 | 0.653 | 2.024 | -0.056 | -0.073 | -0.073 | 0 | 0.12 | 0.35 | 0.05 | 0 | 0.03 | 0.07 |

| | | | | | | | | | | | | | | | | |
|---|---|---|---|---|---|---|---|---|---|---|---|---|---|---|---|---|
| 4 | 109670312 | rs1078132 | -0.921 | 0.466 | 0.437 | 2.123 | 0.005 | 0.182 | 0.182 | 0 | 0.06 | 0.1 | 0.11 | 0.5 | 0.05 | 0.18 |
| 4 | 109674116 | rs1377210 | -1.084 | 0.392 | 1.165 | 2.076 | -0.028 | 0.195 | 0.195 | 0 | 0.16 | 0.3 | 0.15 | 0.5 | 0.08 | 0.25 |
| 4 | 109684830 | rs12108563 | -1.244 | 0.562 | 1.177 | 1.674 | 0.001 | 0.017 | 0.017 | 0 | 0.17 | 0.4 | 0.09 | 0.16 | 0.12 | 0.23 |
| 4 | 109704907 | rs6814067 | 0.623 | 0.466 | 0.336 | 1.221 | -0.002 | -0.002 | -0.002 | 0 | 0.53 | 0.3 | 0.5 | 0.57 | 0.53 | 0.54 |
| 4 | 109707619 | rs6852039 | 1.18 | 0.107 | -0.468 | 0.933 | -0.072 | 0.006 | 0.006 | 0 | 0.26 | 0.03 | 0.28 | 0.25 | 0.29 | 0.39 |
| 4 | 109707993 | rs11732844 | 1.011 | 0.344 | -0.382 | 1.084 | -0.044 | 0.004 | 0.004 | 0 | 0.16 | 0.07 | 0.14 | 0.14 | 0.27 | 0.14 |
| 4 | 109708479 | rs2880902 | -0.751 | 0.042 | 0.331 | 1.11 | -0.025 | -0.004 | -0.004 | 0 | 0.84 | 0.82 | 0.8 | 0.93 | 0.93 | 0.76 |
| 4 | 109711994 | rs1840063 | 0.396 | 0.091 | 0.73 | 1.084 | 0.019 | 0.006 | 0.006 | 0 | 0.7 | 0.61 | 0.71 | 0.63 | 0.6 | 0.77 |
| 4 | 109714131 | rs17039317 | 1.368 | -0.208 | 0.916 | 0.636 | 0.002 | 0 | 0 | 0 | 0.45 | 0.38 | 0.35 | 0.27 | 0.39 | 0.44 |
| 4 | 109714155 | rs7692671 | -0.198 | -0.182 | 1.077 | 0.684 | -0.004 | 0.006 | 0.006 | 0 | 0.32 | 0.31 | 0.36 | 0.2 | 0.37 | 0.38 |
| 4 | 109715558 | rs9994524 | 0.347 | -0.122 | 1.191 | 0.845 | -0.004 | 0.032 | 0.032 | 0 | 0.12 | 0.19 | 0.12 | 0.24 | 0.1 | 0.08 |
| 4 | 109722566 | rs9631731 | 0.487 | -0.077 | 1.142 | 0.784 | 0.019 | 0.066 | 0.066 | 0.475 | 0.76 | 0.63 | 0.69 | 0.47 | 0.73 | 0.82 |
| 4 | 109730810 | rs1812590 | 1.287 | -0.034 | 0.909 | 0.595 | 0.002 | -0.021 | -0.021 | 0.053 | 0.56 | 0.26 | 0.51 | 0.52 | 0.58 | 0.58 |
| 4 | 109734959 | rs2305437 | 0.91 | 0.22 | 0.414 | 0.625 | -0.041 | -0.001 | -0.001 | 0 | 0.14 | 0.05 | 0.11 | 0.11 | 0.22 | 0.16 |
| 4 | 109750774 | rs1452689 | -0.721 | -0.127 | 0.487 | 0.532 | 0.002 | -0.001 | -0.001 | 0.222 | 0.18 | 0.16 | 0.14 | 0.24 | 0.12 | 0.08 |
| 4 | 109754899 | rs6829773 | 0.946 | 0.087 | 0.627 | 0.624 | -0.001 | 0.008 | 0.008 | 0 | 0.61 | 0.44 | 0.55 | 0.56 | 0.62 | 0.63 |
| 4 | 109759469 | rs4956063 | 1.472 | 0.112 | 0.356 | 0.193 | 0.009 | 0.064 | 0.064 | 0 | 0.53 | 0.22 | 0.62 | 0.63 | 0.54 | 0.72 |
| 4 | 109762297 | rs1349677 | 0.822 | 0.216 | 0.817 | -0.466 | -0.004 | -0.127 | -0.127 | 0.186 | 0.67 | 0.49 | 0.73 | 0.94 | 0.67 | 0.82 |
| 4 | 109766290 | rs11098004 | 1.021 | 0.55 | 1.296 | 0.285 | 0.029 | 0.028 | 0.028 | 0 | 0.89 | 0.76 | 0.87 | 0.85 | 0.85 | 0.91 |
| 4 | 109767869 | rs1563005 | 1.475 | 0.526 | 1.134 | 0.254 | 0.026 | 0.022 | 0.022 | 0 | 0.87 | 0.63 | 0.84 | 0.85 | 0.84 | 0.9 |
| 4 | 109772715 | rs12642704 | 1.582 | 0.393 | 0.283 | 0.428 | -0.003 | 0.008 | 0.008 | 0 | 0.71 | 0.35 | 0.67 | 0.63 | 0.71 | 0.76 |
| 4 | 109774351 | rs2198329 | 1.582 | 0.401 | 0.119 | 0.416 | -0.001 | 0 | 0 | 0 | 0.71 | 0.32 | 0.67 | 0.63 | 0.71 | 0.76 |
| 4 | 109781788 | rs965935 | 1.376 | 0.863 | -0.402 | -0.506 | -0.025 | 0.076 | 0.076 | 0.017 | 0.64 | 0.22 | 0.64 | 0.94 | 0.61 | 0.8 |
| 4 | 109803577 | rs7686077 | 1.516 | 1.648 | 0.533 | -0.819 | 0.01 | -0.097 | -0.097 | 1.521 | 0.65 | 0.41 | 0.67 | 1 | 0.57 | 0.82 |
| 4 | 109805696 | rs884049 | 1.134 | 1.892 | 1.232 | -0.945 | 0.003 | -0.08 | -0.08 | 0 | 0.76 | 0.68 | 0.84 | 1 | 0.77 | 0.84 |
| 4 | 109817962 | rs10856995 | 2.032 | 1.863 | 1.413 | -0.94 | 0.023 | -0.148 | -0.148 | 0 | 0.64 | 0.56 | 0.68 | 1 | 0.55 | 0.8 |
| 4 | 109821503 | rs7655700 | 2.4 | 1.474 | 0.989 | -1.201 | -0.017 | 0.12 | 0.12 | 0 | 0.57 | 0.28 | 0.62 | 0.95 | 0.52 | 0.77 |
| 4 | 109822990 | rs12643975 | 2.434 | 1.474 | 0.953 | -1.202 | -0.03 | 0.186 | 0.186 | 0.009 | 0.56 | 0.2 | 0.61 | 0.95 | 0.52 | 0.77 |
| 4 | 109826319 | rs17596124 | -0.477 | 1.542 | 1.14 | -1.155 | 0.048 | -0.055 | -0.055 | 1.255 | 0.12 | 0.14 | 0.16 | 0 | 0.21 | 0.04 |
| 4 | 109834239 | rs1526131 | 1.385 | 1.495 | 1.153 | -1.201 | -0.003 | -0.178 | -0.178 | 0 | 0.72 | 0.47 | 0.83 | 1 | 0.75 | 0.84 |
| 4 | 109841334 | rs12504037 | -0.514 | 1.543 | 1.108 | -1.22 | 0.077 | -0.069 | -0.069 | 0.352 | 0.12 | 0.19 | 0.17 | 0 | 0.21 | 0.04 |
| 4 | 109841743 | rs17596705 | -0.514 | 1.543 | 1.122 | -1.22 | 0.082 | -0.07 | -0.07 | 0 | 0.12 | 0.19 | 0.17 | 0 | 0.21 | 0.04 |
| 4 | 109843603 | rs13104889 | 2.116 | 1.451 | 0.776 | -1.175 | -0.064 | 0.194 | 0.194 | 0.276 | 0.59 | 0.16 | 0.63 | 0.95 | 0.54 | 0.81 |
| 4 | 109863159 | rs7677050 | 0.785 | 0.743 | 1.052 | -1.1 | 0.003 | -0.066 | -0.066 | 1.857 | 0.88 | 0.75 | 0.91 | 1 | 0.89 | 0.95 |
| 4 | 109870169 | rs2881015 | 2.529 | 0.503 | 0.637 | -0.79 | -0.037 | 0.067 | 0.067 | 0 | 0.57 | 0.21 | 0.53 | 0.76 | 0.52 | 0.7 |
| 4 | 109880276 | rs6853519 | 2.873 | -0.131 | 0.16 | -0.52 | -0.028 | 0.021 | 0.021 | 0 | 0.39 | 0.17 | 0.21 | 0.38 | 0.37 | 0.48 |
| 4 | 109884269 | rs10009816 | -0.076 | -0.147 | 0.206 | -0.37 | 0.001 | -0.001 | -0.001 | 0 | 0.23 | 0.2 | 0.36 | 0.39 | 0.17 | 0.26 |
| 4 | 109888212 | rs17039593 | 0.376 | -0.005 | 0.203 | -0.577 | 0.024 | -0.112 | -0.112 | 1.739 | 0.79 | 0.67 | 0.82 | 1 | 0.75 | 0.86 |
| 4 | 109889251 | rs2194861 | -1.429 | -0.297 | 0.38 | -0.333 | -0.015 | -0.015 | -0.015 | 0.43 | 0.4 | 0.47 | 0.61 | 0.62 | 0.38 | 0.38 |
| 4 | 109898846 | rs3863122 | 0.904 | -0.074 | 0.215 | -0.478 | -0.024 | -0.168 | -0.168 | 0.355 | 0.74 | 0.53 | 0.82 | 1 | 0.79 | 0.85 |
| 4 | 109898887 | rs3863123 | -0.112 | 0.075 | 0.219 | -0.299 | -0.013 | -0.09 | -0.09 | 0 | 0.83 | 0.73 | 0.9 | 1 | 0.88 | 0.9 |
| 4 | 109898914 | rs3863124 | 0.223 | 0.144 | -0.322 | 0.193 | -0.017 | -0.084 | -0.084 | 3.203 | 0.12 | 0 | 0.34 | 0.4 | 0.15 | 0.18 |
| 4 | 109903445 | rs11098020 | 0.141 | 0.169 | -0.288 | 0.05 | -0.003 | -0.043 | -0.043 | 0 | 0.12 | 0.06 | 0.37 | 0.45 | 0.15 | 0.19 |
| 4 | 109922064 | rs10516554 | -0.412 | 0.053 | 0.002 | -0.328 | 0.001 | -0.003 | -0.003 | 0 | 0.2 | 0.2 | 0.24 | 0.16 | 0.23 | 0.16 |
| 4 | 109926806 | rs4547872 | 0.57 | -0.504 | 0.515 | -0.355 | -0.023 | -0.128 | -0.128 | 0.086 | 0.8 | 0.6 | 0.87 | 1 | 0.86 | 0.89 |
| 4 | 109928033 | rs1648012 | -1.989 | -1.339 | 0.361 | -0.236 | 0.066 | 0.132 | 0.132 | 0 | 0.48 | 0.8 | 0.62 | 0.72 | 0.42 | 0.47 |
| 4 | 109931951 | rs1405160 | -2.012 | -1.6 | 0.459 | -0.118 | 0.084 | 0.136 | 0.136 | 0 | 0.47 | 0.84 | 0.61 | 0.69 | 0.38 | 0.46 |
| 4 | 109938078 | rs2081466 | -0.658 | -1.672 | 0.942 | 0.322 | 0.008 | -0.016 | -0.016 | 0 | 0.28 | 0.26 | 0.43 | 0.45 | 0.23 | 0.33 |
| 4 | 109945016 | rs12502197 | -2.2 | -1.756 | 0.361 | 0.324 | 0.09 | 0.11 | 0.11 | 0.007 | 0.48 | 0.84 | 0.59 | 0.67 | 0.38 | 0.46 |
| 4 | 109948511 | rs2194860 | -1.873 | -2.04 | 0.604 | 0.301 | 0.052 | 0.025 | 0.025 | 0.842 | 0.4 | 0.69 | 0.49 | 0.65 | 0.17 | 0.41 |
| 4 | 109948640 | rs1648045 | -2.151 | -1.708 | 0.16 | 0.342 | 0.087 | 0.078 | 0.078 | 0.196 | 0.53 | 0.82 | 0.55 | 0.66 | 0.37 | 0.49 |
| 4 | 109952249 | rs752673 | -0.423 | -1.531 | 0.381 | 0.227 | -0.004 | 0.024 | 0.024 | 0 | 0.26 | 0.28 | 0.44 | 0.54 | 0.25 | 0.39 |
| 4 | 109952756 | rs794139 | -1.032 | -1.434 | -0.008 | -0.005 | -0.026 | -0.122 | -0.122 | 0.433 | 0.19 | 0.38 | 0.13 | 0 | 0.13 | 0.06 |
| 4 | 109953266 | rs794144 | -1.62 | -1.194 | -1.005 | 0.151 | 0.068 | 0.114 | 0.114 | 0 | 0.5 | 0.76 | 0.58 | 0.65 | 0.39 | 0.47 |
| 4 | 109954423 | rs7672923 | 0.162 | -1.078 | -1 | 0.17 | 0.006 | -0.047 | -0.047 | 0.285 | 0.09 | 0.19 | 0.05 | 0 | 0.08 | 0.04 |
| 4 | 109957769 | rs12648799 | 0.111 | -0.785 | -1.891 | 0.411 | -0.018 | -0.152 | -0.152 | 2.927 | 0.2 | 0.01 | 0.4 | 0.45 | 0.18 | 0.31 |
| 4 | 109970465 | rs3108923 | -0.361 | -0.03 | -1.182 | -0.115 | 0.061 | 0.1 | 0.1 | 0.131 | 0.51 | 0.72 | 0.58 | 0.66 | 0.39 | 0.46 |
| 4 | 109975633 | rs1863293 | -0.5 | 0.086 | -1.181 | -0.08 | 0.045 | 0.086 | 0.086 | 0.555 | 0.53 | 0.75 | 0.58 | 0.66 | 0.4 | 0.46 |
| 4 | 109981685 | rs1425353 | -1.272 | 0.101 | -0.306 | -0.341 | -0.017 | -0.112 | -0.112 | 0 | 0.19 | 0.55 | 0.13 | 0 | 0.19 | 0.03 |
| 4 | 109987288 | rs1895742 | -0.576 | 0.004 | -0.797 | -0.609 | -0.02 | 0.031 | 0.031 | 0 | 0.15 | 0.08 | 0.18 | 0.03 | 0.2 | 0.06 |
| 4 | 109987381 | rs3113687 | -1.909 | -0.391 | -1.224 | -0.173 | 0.026 | 0.036 | 0.036 | 0 | 0.48 | 0.79 | 0.68 | 0.64 | 0.45 | 0.45 |
| 4 | 109989467 | rs876365 | -1.205 | -0.532 | 0.048 | -0.064 | 0.006 | -0.051 | -0.051 | 0 | 0.28 | 0.42 | 0.24 | 0.14 | 0.3 | 0.17 |
| 4 | 110000675 | rs1991219 | -1.306 | -0.818 | 1.19 | 0.834 | -0.002 | 0.216 | 0.216 | 0.787 | 0.16 | 0.47 | 0.16 | 0.4 | 0.16 | 0.27 |
| 4 | 110006432 | rs17039766 | -0.283 | -0.87 | 0.932 | 1.255 | 0.003 | -0.014 | -0.014 | 0.217 | 0.08 | 0.05 | 0.32 | 0.17 | 0.07 | 0.06 |
| 4 | 110017206 | rs4632736 | -0.132 | -0.88 | 1.255 | 1.219 | 0.018 | 0.205 | 0.205 | 0.009 | 0.06 | 0.16 | 0.11 | 0.29 | 0.06 | 0.16 |
| 4 | 110030410 | rs4368681 | 1.614 | -1.33 | -0.014 | 0.255 | 0.097 | 0.028 | 0.028 | 0 | 0.5 | 0.1 | 0.41 | 0.37 | 0.63 | 0.47 |
| 4 | 110034004 | rs7679372 | 1.388 | -1.332 | 0.308 | 0.204 | 0.062 | -0.002 | -0.002 | 0 | 0.53 | 0.27 | 0.44 | 0.43 | 0.65 | 0.52 |
| 4 | 110035050 | rs1317984 | -0.717 | -0.978 | 0.456 | -0.042 | -0.004 | -0.044 | -0.044 | 0 | 0.83 | 0.81 | 0.94 | 1 | 0.88 | 0.88 |
| 4 | 110037163 | rs3113727 | -0.776 | -0.723 | 0.693 | -0.137 | -0.016 | -0.091 | -0.091 | 0.359 | 0.28 | 0.42 | 0.18 | 0.06 | 0.2 | 0.27 |

| | | | | | | | | | | | | | | | | |
|---|---|---|---|---|---|---|---|---|---|---|---|---|---|---|---|---|
| 4 | 110048492 | rs17039896 | -0.467 | -0.787 | 0.683 | -0.039 | -0.01 | 0.022 | 0.022 | 0 | 0.05 | 0.16 | 0.02 | 0.12 | 0.03 | 0.04 |
| 4 | 110048604 | rs11947505 | -1.271 | -1.106 | 0.722 | 0.396 | -0.026 | 0.078 | 0.078 | 0 | 0.18 | 0.38 | 0.15 | 0.38 | 0.11 | 0.19 |
| 4 | 110052853 | rs17039902 | -0.383 | -1.085 | 0.797 | 0.025 | -0.007 | -0.156 | -0.156 | 0.152 | 0.27 | 0.48 | 0.21 | 0 | 0.22 | 0.17 |
| 4 | 110057072 | rs17039914 | -0.715 | -1.013 | 0.667 | 0.244 | -0.006 | 0.04 | 0.04 | 0 | 0.06 | 0.22 | 0.02 | 0.12 | 0.03 | 0.04 |
| 4 | 110069083 | rs17288525 | 1.225 | -1.276 | 0.346 | 0.323 | 0.036 | 0.04 | 0.04 | 0 | 0.58 | 0.23 | 0.53 | 0.65 | 0.65 | 0.64 |
| 4 | 110077762 | rs4956247 | 0.652 | -1.116 | 0.067 | 0.747 | -0.021 | -0.042 | -0.042 | 0 | 0.07 | 0.01 | 0.19 | 0.23 | 0.12 | 0.13 |
| 4 | 110097265 | rs17223850 | 2.33 | -1.281 | -0.233 | 0.657 | -0.015 | 0.007 | 0.007 | 0 | 0.44 | 0.13 | 0.32 | 0.37 | 0.52 | 0.39 |
| 4 | 110097567 | rs2526434 | -0.981 | -1.385 | -0.152 | 0.569 | -0.018 | 0.055 | 0.055 | 0 | 0.81 | 0.95 | 0.84 | 1 | 0.8 | 0.87 |
| 4 | 110099651 | rs13149803 | 1.852 | -1.34 | -0.191 | 0.764 | 0.08 | 0.106 | 0.106 | 0.014 | 0.51 | 0.14 | 0.53 | 0.64 | 0.64 | 0.52 |
| 4 | 110103485 | rs2704102 | 0.424 | -1.353 | -0.048 | 0.59 | -0.01 | -0.024 | -0.024 | 0 | 0.65 | 0.59 | 0.56 | 0.76 | 0.68 | 0.69 |
| 4 | 110122500 | rs2189708 | 2.451 | -0.896 | -0.599 | 0.502 | -0.181 | -0.105 | -0.105 | 0 | 0.38 | 0.02 | 0.28 | 0.39 | 0.45 | 0.36 |
| 4 | 110131702 | rs987695 | -2.09 | -1.153 | -0.382 | 0.644 | 0.066 | 0.019 | 0.019 | 0 | 0.52 | 0.86 | 0.65 | 0.6 | 0.42 | 0.58 |
| 4 | 110132858 | rs10049987 | -1.879 | -1.16 | -0.335 | 0.596 | 0.099 | 0.038 | 0.038 | 0.039 | 0.45 | 0.85 | 0.46 | 0.38 | 0.31 | 0.45 |
| 4 | 110136612 | rs2007788 | -0.563 | -0.954 | -0.037 | 0.63 | 0.005 | 0.019 | 0.019 | 0.094 | 0.06 | 0.28 | 0.03 | 0.07 | 0.04 | 0.04 |
| 4 | 110153951 | rs4956253 | -1.624 | -1.104 | 0.118 | 0.635 | -0.002 | 0 | 0 | 0 | 0.42 | 0.6 | 0.61 | 0.54 | 0.42 | 0.44 |
| 4 | 110155038 | rs17304953 | 1.044 | -1.136 | 0.083 | 0.339 | -0.002 | 0.029 | 0.029 | 0 | 0.15 | 0.05 | 0.13 | 0 | 0.15 | 0.1 |
| 4 | 110155157 | rs12640304 | -0.684 | -0.971 | 0.274 | 0.586 | -0.002 | 0.054 | 0.054 | 0.021 | 0.13 | 0.27 | 0.22 | 0.3 | 0.15 | 0.17 |
| 4 | 110158446 | rs17040162 | -0.906 | -0.893 | 0.284 | 0.962 | -0.001 | 0.132 | 0.132 | 0.01 | 0.05 | 0.22 | 0.03 | 0.1 | 0.01 | 0.14 |
| 4 | 110162634 | rs17226518 | -1.195 | -0.383 | -0.37 | 0.731 | -0.027 | 0.028 | 0.028 | 0 | 0.95 | 1 | 0.96 | 1 | 0.93 | 0.96 |
| 4 | 110178174 | rs7675657 | -1.902 | -1.125 | -0.07 | 1.015 | 0.109 | 0.024 | 0.024 | 0 | 0.47 | 0.83 | 0.57 | 0.61 | 0.33 | 0.55 |
| 4 | 110179436 | rs7682615 | -1.301 | -0.985 | 0.227 | 1.05 | -0.019 | 0.157 | 0.157 | 0.56 | 0.12 | 0.43 | 0.22 | 0.24 | 0.09 | 0.16 |
| 4 | 110181439 | rs7695378 | -0.197 | -0.935 | 0.065 | 1.08 | 0 | 0.116 | 0.116 | 0 | 0.06 | 0.15 | 0.18 | 0.24 | 0.07 | 0.14 |
| 4 | 110189698 | rs4512071 | -0.306 | -0.539 | -0.155 | 0.712 | -0.002 | -0.001 | -0.001 | 0 | 0.26 | 0.17 | 0.33 | 0.24 | 0.26 | 0.25 |
| 4 | 110192405 | rs1297941 | -1.029 | -0.38 | -0.32 | 0.514 | -0.006 | -0.039 | -0.039 | 0 | 0.1 | 0.45 | 0.05 | 0 | 0.05 | 0.05 |
| 4 | 110198260 | rs1297940 | -1.473 | 0 | -1.931 | 0.673 | -0.14 | -0.107 | -0.107 | 0 | 0.57 | 0.98 | 0.68 | 0.58 | 0.51 | 0.62 |
| 4 | 110200344 | rs917022 | 0.217 | -0.329 | -1.498 | 0.728 | 0.002 | -0.028 | -0.028 | 5.352 | 0.11 | 0.05 | 0.19 | 0.22 | 0.1 | 0.12 |
| 4 | 110207927 | rs2346029 | -1.44 | -0.804 | -1.201 | 0.588 | -0.028 | -0.001 | -0.001 | 0 | 0.27 | 0.65 | 0.25 | 0.29 | 0.12 | 0.3 |
| 4 | 110209365 | rs2214375 | -1.716 | -0.442 | -1.706 | 0.636 | 0.089 | -0.041 | -0.041 | 0 | 0.47 | 0.87 | 0.55 | 0.51 | 0.34 | 0.53 |
| 4 | 110211727 | rs6820019 | 0.114 | 0.113 | -1.126 | 0.651 | -0.038 | -0.061 | -0.061 | 0 | 0.11 | 0.01 | 0.19 | 0.23 | 0.12 | 0.15 |
| 4 | 110216053 | rs961695 | -1.152 | -0.318 | -0.244 | 0.29 | -0.061 | -0.025 | -0.025 | 0 | 0.12 | 0.61 | 0.06 | 0.08 | 0.02 | 0.15 |
| 4 | 110220384 | rs6822076 | -0.097 | 0.475 | -0.773 | 0.469 | 0.029 | 0.044 | 0.044 | 0 | 0.19 | 0.24 | 0.37 | 0.36 | 0.31 | 0.22 |
| 4 | 110223320 | rs1859143 | -1.405 | 0.423 | -1.255 | 0.104 | -0.092 | -0.03 | -0.03 | 0 | 0.44 | 0.91 | 0.45 | 0.42 | 0.46 | 0.46 |
| 4 | 110226406 | rs714586 | -0.661 | -0.057 | -0.02 | -0.121 | -0.026 | -0.129 | -0.129 | 0 | 0.19 | 0.53 | 0.1 | 0 | 0.11 | 0.1 |
| 4 | 110228717 | rs13128445 | -0.706 | -0.071 | 0.08 | -0.128 | -0.025 | -0.112 | -0.112 | 0.945 | 0.19 | 0.57 | 0.11 | 0 | 0.11 | 0.1 |
| 4 | 110250481 | rs10005272 | -0.491 | -0.181 | -0.044 | -0.186 | -0.004 | -0.003 | -0.003 | 0 | 0.07 | 0.13 | 0.03 | 0.05 | 0.05 | 0.01 |
| 4 | 110254092 | rs11935942 | -1.829 | -0.226 | -0.01 | -0.179 | 0.007 | 0.007 | 0.007 | 0.991 | 0.28 | 0.72 | 0.15 | 0.1 | 0.17 | 0.21 |
| 4 | 110255140 | rs13104799 | -1.531 | -0.242 | 0.344 | -0.197 | -0.017 | -0.059 | -0.059 | 0 | 0.25 | 0.65 | 0.13 | 0.05 | 0.16 | 0.11 |
| 4 | 110278217 | rs10031399 | -0.57 | -0.225 | 0.399 | -0.196 | -0.005 | -0.006 | -0.006 | 0.68 | 0.07 | 0.16 | 0.03 | 0.04 | 0.04 | 0.01 |
| 4 | 110278483 | rs7673867 | -1.896 | -0.247 | 0.405 | -0.172 | 0.006 | 0.005 | 0.005 | 1.048 | 0.28 | 0.71 | 0.15 | 0.11 | 0.17 | 0.21 |
| 4 | 110278816 | rs1557804 | -1.545 | -0.25 | 0.409 | -0.203 | -0.024 | -0.078 | -0.078 | 0.77 | 0.26 | 0.62 | 0.14 | 0.04 | 0.16 | 0.11 |
| 4 | 110283708 | rs1859144 | -1.597 | -0.24 | 0.359 | -0.198 | -0.028 | -0.096 | -0.096 | 0 | 0.25 | 0.54 | 0.14 | 0.04 | 0.16 | 0.11 |
| 4 | 110299443 | rs6854151 | -0.542 | -0.281 | 0.536 | -0.184 | -0.029 | -0.139 | -0.139 | 0.574 | 0.18 | 0.45 | 0.1 | 0 | 0.11 | 0.1 |
| 4 | 110324528 | rs11098033 | -1.203 | -0.307 | 0.559 | -0.099 | -0.013 | 0.012 | 0.012 | 0.471 | 0.1 | 0.26 | 0.03 | 0.12 | 0.04 | 0.1 |
| 4 | 110350409 | rs11098035 | -0.685 | -0.316 | 0.585 | -0.102 | -0.012 | 0.007 | 0.007 | 0 | 0.08 | 0.15 | 0.03 | 0.13 | 0.04 | 0.1 |
| 4 | 110365426 | rs17040436 | -1.384 | -0.339 | 0.548 | -0.237 | -0.012 | -0.036 | -0.036 | 0 | 0.22 | 0.32 | 0.09 | 0.13 | 0.14 | 0.18 |
| 4 | 110369533 | rs6852062 | -0.825 | -0.485 | 0.745 | -0.31 | 0.086 | 0.082 | 0.082 | 3.405 | 0.05 | 0.47 | 0.03 | 0 | 0 | 0 |
| 4 | 110373682 | rs10009463 | -0.989 | -0.496 | 0.642 | -0.316 | 0.06 | 0.056 | 0.056 | 4.501 | 0.05 | 0.49 | 0.04 | 0 | 0 | 0 |
| 4 | 110378688 | rs1396709 | 1.541 | -0.23 | -0.044 | -0.278 | -0.066 | -0.041 | -0.041 | 4.416 | 0.5 | 0.06 | 0.54 | 0.52 | 0.53 | 0.6 |
| 4 | 110388034 | rs9985632 | -1.742 | -0.223 | -0.049 | -0.274 | 0.038 | 0.067 | 0.067 | 0.435 | 0.28 | 0.81 | 0.12 | 0.13 | 0.14 | 0.18 |
| 4 | 110395146 | rs10516556 | 0.733 | -0.236 | -0.069 | -0.28 | -0.069 | -0.088 | -0.088 | 0 | 0.21 | 0.08 | 0.34 | 0.35 | 0.34 | 0.21 |
| 4 | 110395406 | rs10516557 | 0.095 | -0.244 | 0.177 | -0.511 | 0.004 | 0.005 | 0.005 | 0 | 0.07 | 0.07 | 0.02 | 0.05 | 0.04 | 0.01 |
| 4 | 110409978 | rs6856291 | -0.951 | -0.303 | 0.222 | -0.537 | -0.005 | -0.015 | -0.015 | 0.262 | 0.22 | 0.27 | 0.09 | 0.13 | 0.13 | 0.18 |
| 4 | 110446454 | rs6812906 | -1.547 | -0.389 | -0.202 | -0.485 | -0.113 | -0.023 | -0.023 | 0.213 | 0.63 | 0.98 | 0.62 | 0.7 | 0.6 | 0.58 |
| 4 | 110454306 | rs10003981 | -1.128 | -0.436 | 0.982 | -0.378 | 0.067 | 0.062 | 0.062 | 4.259 | 0.05 | 0.48 | 0.03 | 0 | 0 | 0 |
| 4 | 110466413 | rs3775962 | -1.805 | -0.356 | 0.529 | -0.404 | 0.026 | 0.051 | 0.051 | 0.543 | 0.28 | 0.79 | 0.14 | 0.13 | 0.16 | 0.15 |
| 4 | 110466758 | rs17040519 | -0.721 | -0.446 | 0.637 | -0.336 | -0.01 | -0.013 | -0.013 | 0 | 0.08 | 0.17 | 0.03 | 0.05 | 0.04 | 0.01 |
| 4 | 110474321 | rs6533434 | -1.119 | -0.538 | 0.612 | -0.394 | -0.034 | -0.096 | -0.096 | 0 | 0.24 | 0.63 | 0.1 | 0 | 0.08 | 0.08 |
| 4 | 110486492 | rs7687245 | -1.482 | -0.515 | 0.411 | -0.305 | 0.026 | 0.033 | 0.033 | 0.106 | 0.28 | 0.77 | 0.1 | 0.13 | 0.14 | 0.15 |
| 4 | 110491665 | rs6831048 | -0.857 | -0.264 | 0.741 | -0.023 | -0.02 | -0.032 | -0.032 | 0 | 0.08 | 0.32 | 0.02 | 0 | 0.01 | 0 |
| 4 | 110498602 | rs197203 | -0.115 | -0.275 | 0.219 | 0.102 | -0.003 | -0.001 | -0.001 | 0 | 0.53 | 0.49 | 0.59 | 0.54 | 0.58 | 0.68 |
| 4 | 110510080 | rs7694402 | -0.782 | 0.228 | 0.251 | -0.028 | 0.059 | -0.012 | -0.012 | 0 | 0.13 | 0.38 | 0.14 | 0.09 | 0.13 | 0.03 |
| 4 | 110515471 | rs149800 | 1.179 | 0.228 | -0.392 | 0.045 | -0.063 | -0.027 | -0.027 | 0 | 0.26 | 0.03 | 0.29 | 0.24 | 0.28 | 0.34 |
| 4 | 110515484 | rs197230 | -0.319 | 0.138 | 0.142 | 0.11 | -0.005 | 0.003 | 0.003 | 0 | 0.42 | 0.61 | 0.44 | 0.41 | 0.42 | 0.47 |
| 4 | 110523044 | rs197212 | -0.532 | -0.297 | 0.166 | -0.142 | 0.002 | 0.004 | 0.004 | 0 | 0.61 | 0.44 | 0.52 | 0.46 | 0.55 | 0.72 |
| 4 | 110537567 | rs17440077 | 1.169 | -0.426 | -0.431 | -0.197 | -0.074 | -0.016 | -0.016 | 0 | 0.3 | 0.02 | 0.3 | 0.24 | 0.31 | 0.38 |
| 4 | 110538429 | rs6533438 | -1.574 | -0.426 | -0.124 | -0.191 | -0.005 | -0.036 | -0.036 | 0 | 0.72 | 0.76 | 0.61 | 0.57 | 0.69 | 0.77 |
| 4 | 110539442 | rs7687878 | -0.144 | -0.383 | 0.235 | -0.292 | -0.001 | -0.004 | -0.004 | 0.783 | 0.28 | 0.27 | 0.16 | 0.15 | 0.26 | 0.29 |
| 4 | 110544696 | rs6832174 | -0.165 | -0.392 | 0.494 | -0.288 | -0.004 | 0.016 | 0.016 | 0 | 0.55 | 0.3 | 0.46 | 0.39 | 0.54 | 0.64 |
| 4 | 110546567 | rs7677585 | 0.751 | -0.454 | 0.443 | -0.382 | -0.006 | -0.003 | -0.003 | 0.783 | 0.14 | 0.1 | 0.12 | 0.15 | 0.16 | 0.24 |

| | | | | | | | | | | | | | | | | |
|---|---|---|---|---|---|---|---|---|---|---|---|---|---|---|---|---|
| 4 | 110548394 | rs10017480 | 1.31 | -0.531 | 0.443 | -0.372 | 0.003 | -0.011 | -0.011 | 0.04 | 0.13 | 0.12 | 0.04 | 0 | 0.1 | 0.05 |
| 4 | 110555678 | rs11938222 | -0.252 | -0.544 | 0.931 | -0.302 | 0 | -0.001 | -0.001 | 0 | 0.52 | 0.54 | 0.64 | 0.71 | 0.58 | 0.51 |
| 4 | 110562936 | rs4698773 | -0.711 | -0.594 | 0.776 | -0.357 | 0.002 | -0.006 | -0.006 | 0.485 | 0.27 | 0.34 | 0.24 | 0.23 | 0.26 | 0.29 |
| 4 | 110567473 | rs10032323 | 0.461 | -0.788 | 0.78 | -0.394 | 0.002 | -0.005 | -0.005 | 0 | 0.42 | 0.6 | 0.46 | 0.54 | 0.42 | 0.36 |
| 4 | 110568156 | rs13128388 | -1.548 | -0.699 | 0.685 | -0.402 | -0.048 | 0.018 | 0.018 | 0 | 0.69 | 0.94 | 0.7 | 0.77 | 0.69 | 0.64 |
| 4 | 110568711 | rs9990765 | -1.6 | -0.767 | 0.738 | -0.431 | -0.047 | -0.107 | -0.107 | 0.642 | 0.7 | 0.89 | 0.6 | 0.52 | 0.66 | 0.75 |
| 4 | 110571181 | rs1029452 | 1.15 | -0.765 | 0.965 | -0.428 | -0.023 | -0.026 | -0.026 | 0 | 0.14 | 0.25 | 0.06 | 0.06 | 0.1 | 0.13 |
| 4 | 110577363 | rs13103963 | 1.15 | -0.765 | 0.965 | -0.428 | -0.023 | -0.026 | -0.026 | 0 | 0.14 | 0.25 | 0.06 | 0.06 | 0.1 | 0.13 |
| 4 | 110582100 | rs17440280 | -1.212 | -0.797 | 0.935 | -0.442 | -0.007 | -0.033 | -0.033 | 0.109 | 0.4 | 0.48 | 0.29 | 0.24 | 0.37 | 0.34 |
| 4 | 110584287 | rs3796911 | -1.451 | -0.801 | 1.069 | -0.267 | -0.05 | -0.11 | -0.11 | 0 | 0.7 | 0.89 | 0.61 | 0.52 | 0.66 | 0.75 |
| 4 | 110590821 | rs4698776 | 1.039 | -0.813 | 1.113 | -0.272 | 0.012 | 0.07 | 0.07 | 0 | 0.29 | 0.49 | 0.41 | 0.54 | 0.32 | 0.25 |
| 4 | 110593460 | rs13103167 | 1.497 | -0.807 | 1.123 | -0.268 | 0.002 | 0.013 | 0.013 | 0 | 0.26 | 0.23 | 0.39 | 0.48 | 0.32 | 0.23 |
| 4 | 110602430 | rs4698780 | -1.27 | -0.891 | 1.06 | -0.24 | -0.05 | -0.111 | -0.111 | 1.208 | 0.7 | 0.89 | 0.61 | 0.52 | 0.66 | 0.75 |
| 4 | 110603784 | rs3733611 | -1.447 | -0.894 | 1.229 | -0.207 | -0.029 | -0.076 | -0.076 | 0 | 0.7 | 0.86 | 0.62 | 0.52 | 0.66 | 0.75 |
| 4 | 110606713 | rs2285712 | -0.899 | -0.898 | 1.253 | -0.208 | 0.001 | 0.035 | 0.035 | 0 | 0.66 | 0.51 | 0.59 | 0.46 | 0.65 | 0.69 |
| 4 | 110608967 | rs5030606 | -0.606 | -0.786 | 1.081 | -0.101 | -0.001 | -0.017 | -0.017 | 1.906 | 0.57 | 0.72 | 0.57 | 0.52 | 0.55 | 0.69 |
| 4 | 110609481 | rs3212153 | 0.8 | -0.571 | 0.837 | -0.037 | -0.013 | -0.004 | -0.004 | 0 | 0.06 | 0.04 | 0.12 | 0.09 | 0.14 | 0.04 |
| 4 | 110613658 | rs1800627 | 0.049 | -1.71 | 1 | 0.056 | -0.004 | 0.007 | 0.007 | 0.012 | 0.46 | 0.5 | 0.47 | 0.34 | 0.47 | 0.47 |
| 4 | 110628894 | rs1541373 | -1.178 | -1.479 | 0.576 | 0.402 | -0.034 | -0.032 | -0.032 | 0 | 0.67 | 0.82 | 0.57 | 0.55 | 0.6 | 0.75 |
| 4 | 110631621 | rs3087494 | 0.042 | -1.619 | 0.459 | 0.299 | 0.007 | 0.001 | 0.001 | 0 | 0.51 | 0.33 | 0.51 | 0.46 | 0.53 | 0.64 |
| 4 | 110638810 | rs2285714 | 1.531 | -1.524 | -0.851 | 0.147 | -0.139 | -0.031 | -0.031 | 0.001 | 0.29 | 0.02 | 0.37 | 0.2 | 0.39 | 0.39 |
| 4 | 110648124 | rs10004377 | -1.147 | -1.213 | -0.891 | 0.149 | -0.067 | -0.052 | -0.052 | 0 | 0.68 | 0.88 | 0.58 | 0.57 | 0.59 | 0.75 |
| 4 | 110654640 | rs2346841 | 0.808 | -0.867 | -0.578 | -0.145 | 0.01 | 0.001 | 0.001 | 0 | 0.13 | 0.08 | 0.04 | 0 | 0.07 | 0.06 |
| 4 | 110656730 | rs7439493 | 0.015 | -0.73 | -0.872 | 0.276 | -0.042 | -0.061 | -0.061 | 0.098 | 0.39 | 0.14 | 0.49 | 0.55 | 0.45 | 0.43 |
| 4 | 110659067 | rs10033900 | -0.084 | -0.057 | -0.105 | 0.279 | 0.005 | 0.024 | 0.024 | 0 | 0.51 | 0.66 | 0.53 | 0.39 | 0.54 | 0.31 |
| 4 | 110664630 | rs11726949 | 0.833 | -0.522 | 0.555 | 0.592 | -0.032 | 0.008 | 0.008 | 0.44 | 0.83 | 0.43 | 0.94 | 0.82 | 0.94 | 0.83 |
| 4 | 110665481 | rs13104777 | 0.202 | -0.443 | -0.762 | 0.156 | -0.064 | 0.085 | 0.085 | 0 | 0.36 | 0.13 | 0.48 | 0.19 | 0.42 | 0.23 |
| 4 | 110666292 | rs6822976 | 0.78 | -0.601 | -0.759 | 0.23 | -0.055 | 0.024 | 0.024 | 0.2 | 0.44 | 0.18 | 0.45 | 0.6 | 0.51 | 0.53 |
| 4 | 110667345 | rs9998151 | -1.578 | -1.444 | 0.651 | 0.465 | -0.023 | -0.006 | -0.006 | 0 | 0.11 | 0.29 | 0.04 | 0.09 | 0.03 | 0.09 |
| 4 | 110668651 | rs7438961 | 1.296 | -1.568 | 0.401 | 0.656 | 0.081 | -0.03 | -0.03 | 0 | 0.56 | 0.22 | 0.51 | 0.51 | 0.7 | 0.58 |
| 4 | 110681505 | rs2298749 | -0.639 | -1.657 | 1.23 | 0.659 | -0.017 | -0.006 | -0.006 | 0 | 0.33 | 0.36 | 0.44 | 0.31 | 0.25 | 0.23 |
| 4 | 110682487 | rs4541508 | 1.748 | -1.922 | 0.794 | 0.553 | 0.093 | -0.044 | -0.044 | 0 | 0.48 | 0.11 | 0.38 | 0.5 | 0.67 | 0.58 |
| 4 | 110682953 | rs17610314 | 0.536 | -1.921 | 0.885 | 0.581 | 0.008 | 0.047 | 0.047 | 0 | 0.22 | 0.06 | 0.38 | 0.13 | 0.23 | 0.16 |
| 4 | 110683550 | rs3822272 | 0.73 | -1.899 | 1.369 | 0.583 | 0.007 | 0.004 | 0.004 | 0.583 | 0.63 | 0.42 | 0.55 | 0.69 | 0.75 | 0.77 |
| 4 | 110684744 | rs7439356 | 1.037 | -1.924 | 1.339 | 0.561 | 0.053 | -0.009 | -0.009 | 0.044 | 0.61 | 0.33 | 0.54 | 0.55 | 0.75 | 0.71 |
| 4 | 110708517 | rs4626205 | 0.239 | -1.642 | 1.092 | 0.486 | -0.016 | 0.06 | 0.06 | 0 | 0.24 | 0.06 | 0.42 | 0.13 | 0.28 | 0.17 |
| 4 | 110713964 | rs10029485 | -2.196 | -1.684 | 1.623 | 0.59 | -0.068 | 0.111 | 0.111 | 0.401 | 0.18 | 0.48 | 0.19 | 0.34 | 0.05 | 0.2 |
| 4 | 110719534 | rs4698788 | -2.159 | -1.683 | 1.626 | 0.591 | -0.069 | 0.121 | 0.121 | 0.28 | 0.18 | 0.47 | 0.19 | 0.34 | 0.05 | 0.2 |
| 4 | 110726322 | rs884711 | -0.789 | -1.555 | 1.753 | 0.679 | -0.011 | 0.178 | 0.178 | 0 | 0.07 | 0.18 | 0.04 | 0.32 | 0.02 | 0.2 |
| 4 | 110727560 | rs4698790 | 0.999 | -1.679 | 1.878 | 0.594 | 0.002 | -0.001 | -0.001 | 0.189 | 0.61 | 0.45 | 0.51 | 0.57 | 0.7 | 0.61 |
| 4 | 110729304 | rs4698792 | 0.037 | -1.509 | 1.63 | 0.498 | -0.006 | -0.04 | -0.04 | 0.11 | 0.72 | 0.71 | 0.56 | 0.89 | 0.72 | 0.82 |
| 4 | 110734498 | rs11930711 | -0.907 | -1.167 | 1.653 | 1.075 | 0.009 | 0.091 | 0.091 | 0 | 0.1 | 0.18 | 0.06 | 0.25 | 0.07 | 0.2 |
| 4 | 110745938 | rs4353905 | -1.508 | -1.339 | 1.604 | 1.032 | -0.002 | -0.003 | -0.003 | 0 | 0.15 | 0.16 | 0.38 | 0.28 | 0.15 | 0.21 |
| 4 | 110754499 | rs12513108 | -1.971 | -1.442 | 1.861 | 1.142 | 0 | 0.038 | 0.038 | 0 | 0.17 | 0.26 | 0.39 | 0.43 | 0.15 | 0.32 |
| 4 | 110758769 | rs4698795 | -0.284 | -1.425 | 1.763 | 1.128 | -0.017 | 0.001 | 0.001 | 0 | 0.09 | 0.04 | 0.17 | 0.08 | 0.13 | 0.11 |
| 4 | 110763413 | rs7438763 | 2.154 | -2.013 | 1.403 | 0.813 | 0.065 | -0.039 | -0.039 | 4.669 | 0.49 | 0.08 | 0.36 | 0.51 | 0.64 | 0.43 |
| 4 | 110765372 | rs7657133 | 1.706 | -1.992 | 1.708 | 0.829 | -0.011 | 0.054 | 0.054 | 0.291 | 0.78 | 0.64 | 0.61 | 0.59 | 0.85 | 0.68 |
| 4 | 110765496 | rs6839680 | -0.051 | -2.007 | 1.637 | 0.756 | 0.004 | 0.07 | 0.07 | 0 | 0.21 | 0.04 | 0.2 | 0.08 | 0.21 | 0.25 |
| 4 | 110773067 | rs4698797 | 2.308 | -2.057 | 1.982 | 0.8 | 0.004 | -0.003 | -0.003 | 0 | 0.55 | 0.4 | 0.39 | 0.49 | 0.64 | 0.43 |
| 4 | 110773433 | rs10857002 | -0.309 | -2.069 | 1.962 | 0.827 | 0.001 | 0.032 | 0.032 | 0 | 0.81 | 0.95 | 0.81 | 0.91 | 0.8 | 0.75 |
| 4 | 110779516 | rs2078040 | -1.665 | -2.076 | 2.087 | 0.858 | 0.003 | 0.03 | 0.03 | 0.387 | 0.17 | 0.31 | 0.38 | 0.27 | 0.15 | 0.21 |
| 4 | 110784379 | rs7654660 | 2.126 | -2.089 | 2.152 | 0.855 | 0.008 | -0.004 | -0.004 | 0 | 0.55 | 0.39 | 0.4 | 0.49 | 0.64 | 0.43 |
| 4 | 110785384 | rs17040920 | -0.635 | -2.012 | 2.15 | 0.856 | -0.012 | 0.051 | 0.051 | 0 | 0.07 | 0.18 | 0.21 | 0.2 | 0.02 | 0.1 |
| 4 | 110786290 | rs7679048 | 1.879 | -2.021 | 2.181 | 0.805 | -0.008 | 0.004 | 0.004 | 0.056 | 0.63 | 0.53 | 0.45 | 0.63 | 0.68 | 0.58 |
| 4 | 110791413 | rs2347132 | -1.41 | -1.839 | 1.899 | 0.877 | 0.002 | 0.031 | 0.031 | 0.052 | 0.17 | 0.27 | 0.38 | 0.32 | 0.15 | 0.21 |
| 4 | 110791453 | rs9994891 | -1.306 | -1.795 | 2.028 | 0.702 | -0.015 | -0.026 | -0.026 | 0 | 0.09 | 0.42 | 0.03 | 0 | 0 | 0 |
| 4 | 110795962 | rs11722475 | 1.351 | -1.61 | 1.171 | 1.641 | -0.011 | 0.002 | 0.002 | 0 | 0.63 | 0.63 | 0.51 | 0.39 | 0.71 | 0.48 |
| 4 | 110800020 | rs6847467 | 1.026 | -1.586 | 1.184 | 1.884 | -0.045 | 0.011 | 0.011 | 0 | 0.69 | 0.62 | 0.6 | 0.58 | 0.8 | 0.56 |
| 4 | 110803816 | rs9685587 | 0.255 | -1.475 | 0.778 | 1.795 | 0.001 | 0.018 | 0.018 | 0 | 0.06 | 0.08 | 0.09 | 0.18 | 0.06 | 0.08 |
| 4 | 110808173 | rs11937583 | 1.142 | -1.541 | 0.997 | 1.749 | -0.043 | -0.023 | -0.023 | 0 | 0.71 | 0.63 | 0.6 | 0.74 | 0.8 | 0.68 |
| 4 | 110808321 | rs11736783 | 0.296 | -1.217 | 0.162 | 1.919 | 0.019 | -0.005 | -0.005 | 0.34 | 0.78 | 0.91 | 0.63 | 0.74 | 0.81 | 0.68 |
| 4 | 110810499 | rs7689017 | -1.626 | -0.69 | 0.304 | 1.954 | -0.015 | -0.03 | -0.03 | 0.372 | 0.16 | 0.12 | 0.25 | 0.31 | 0.2 | 0.19 |
| 4 | 110812598 | rs2024111 | -0.729 | -0.549 | 0.305 | 1.447 | -0.016 | -0.016 | -0.016 | 0 | 0.1 | 0.04 | 0.17 | 0.13 | 0.14 | 0.11 |
| 4 | 110824115 | rs718768 | 0.92 | -0.699 | 0.296 | 0.979 | -0.001 | 0.015 | 0.015 | 0 | 0.74 | 0.84 | 0.75 | 0.93 | 0.71 | 0.78 |
| 4 | 110825634 | rs17316633 | -0.63 | -0.246 | 0.455 | 1.129 | -0.001 | -0.002 | -0.002 | 0 | 0.19 | 0.16 | 0.24 | 0.1 | 0.23 | 0.19 |
| 4 | 110826275 | rs17238095 | -0.737 | -0.264 | 0.653 | 1.12 | 0.005 | -0.016 | -0.016 | 0.012 | 0.19 | 0.2 | 0.25 | 0.1 | 0.23 | 0.19 |
| 4 | 110834110 | rs4444903 | -2.295 | -0.868 | 0.865 | 1.903 | 0.008 | 0.048 | 0.048 | 0.1 | 0.43 | 0.76 | 0.58 | 0.69 | 0.39 | 0.54 |
| 4 | 110837667 | rs2298978 | -2.202 | -0.889 | 0.858 | 2.124 | 0.004 | -0.012 | -0.012 | 0.011 | 0.42 | 0.75 | 0.58 | 0.46 | 0.39 | 0.38 |
| 4 | 110846796 | rs2298982 | -1.094 | -0.913 | 1.542 | 2.178 | -0.033 | 0.127 | 0.127 | 0.18 | 0.15 | 0.36 | 0.28 | 0.37 | 0.06 | 0.18 |

| | | | | | | | | | | | | | | | | |
|---|---|---|---|---|---|---|---|---|---|---|---|---|---|---|---|---|
| 4 | 110865271 | rs10470911 | -1.113 | -1.02 | 2.675 | 2.054 | 0.003 | -0.007 | -0.007 | 0 | 0.27 | 0.32 | 0.46 | 0.27 | 0.33 | 0.27 |
| 4 | 110866442 | rs4698756 | -1.432 | -1.044 | 2.563 | 2.232 | 0.006 | 0.079 | 0.079 | 0 | 0.29 | 0.45 | 0.47 | 0.51 | 0.33 | 0.42 |
| 4 | 110866508 | rs4698800 | -1.162 | -1.028 | 2.542 | 2.11 | 0.008 | -0.016 | -0.016 | 0 | 0.28 | 0.38 | 0.47 | 0.27 | 0.33 | 0.27 |
| 4 | 110870765 | rs3796947 | -1.143 | -1.043 | 2.645 | 2.138 | -0.023 | 0.073 | 0.073 | 0.317 | 0.15 | 0.44 | 0.12 | 0.19 | 0.06 | 0.1 |
| 4 | 110882590 | rs9992755 | -1.38 | -1.059 | 2.467 | 2.077 | 0.01 | -0.026 | -0.026 | 0.305 | 0.29 | 0.44 | 0.47 | 0.27 | 0.33 | 0.27 |
| 4 | 110883121 | rs11568943 | -0.893 | -1.038 | 2.395 | 2.214 | -0.023 | 0.046 | 0.046 | 0.016 | 0.13 | 0.3 | 0.12 | 0.19 | 0.06 | 0.1 |
| 4 | 110889909 | rs11568978 | -0.989 | -0.933 | 2.033 | 2.354 | -0.011 | 0.177 | 0.177 | 0 | 0.11 | 0.22 | 0.11 | 0.42 | 0.06 | 0.25 |
| 4 | 110891543 | rs2255355 | -1.523 | -0.979 | 2.014 | 2.355 | -0.017 | 0.096 | 0.096 | 0 | 0.13 | 0.22 | 0.27 | 0.61 | 0.06 | 0.34 |
| 4 | 110894301 | rs9991367 | -0.801 | -0.836 | 2.069 | 2.266 | -0.005 | 0.037 | 0.037 | 0 | 0.1 | 0.19 | 0.1 | 0.2 | 0.06 | 0.08 |
| 4 | 110897535 | rs11568994 | -0.964 | -0.898 | 2.118 | 1.888 | -0.005 | 0.014 | 0.014 | 0 | 0.26 | 0.24 | 0.29 | 0.09 | 0.33 | 0.2 |
| 4 | 110911907 | rs2298999 | -2.436 | -0.98 | 1.54 | 1.659 | 0.024 | 0.187 | 0.187 | 0 | 0.5 | 0.88 | 0.55 | 0.76 | 0.38 | 0.67 |
| 4 | 110913193 | rs11569061 | -1.85 | -0.677 | 2.403 | 1.99 | -0.024 | -0.034 | -0.034 | 0 | 0.07 | 0.28 | 0.01 | 0 | 0 | 0.01 |
| 4 | 110929128 | rs11569121 | -1.599 | -0.658 | 2.311 | 2.207 | -0.019 | 0.05 | 0.05 | 0.273 | 0.11 | 0.32 | 0.09 | 0.16 | 0.05 | 0.07 |
| 4 | 110929653 | rs11569126 | 0.437 | -0.597 | 1.723 | 2.245 | -0.011 | -0.028 | -0.028 | 0 | 0.05 | 0.01 | 0.07 | 0.16 | 0.05 | 0.07 |
| 4 | 110933004 | rs3733625 | -0.273 | -0.609 | 1.735 | 2.227 | -0.001 | -0.02 | -0.02 | 0 | 0.07 | 0.04 | 0.08 | 0.16 | 0.06 | 0.09 |
| 4 | 110940045 | rs4698804 | -2.224 | -0.725 | 1.862 | 2.189 | -0.054 | 0.035 | 0.035 | 7.93 | 0.12 | 0.67 | 0.2 | 0.15 | 0.01 | 0.07 |
| 4 | 110945256 | rs17041245 | -1.362 | -0.367 | 1.535 | 2.181 | 0.012 | 0.024 | 0.024 | 0.122 | 0.24 | 0.27 | 0.28 | 0.33 | 0.3 | 0.4 |
| 4 | 110945966 | rs720419 | -1.362 | -0.367 | 1.535 | 2.181 | 0.012 | 0.024 | 0.024 | 0.122 | 0.24 | 0.27 | 0.28 | 0.33 | 0.3 | 0.4 |
| 4 | 110946249 | rs7440841 | -0.794 | -0.315 | 1.092 | 1.901 | -0.028 | 0.037 | 0.037 | 0.245 | 0.2 | 0.11 | 0.26 | 0.09 | 0.3 | 0.21 |
| 4 | 110953909 | rs6854522 | -1.942 | -0.519 | 1.381 | 2.035 | -0.009 | 0.02 | 0.02 | 0.051 | 0.35 | 0.5 | 0.43 | 0.45 | 0.36 | 0.34 |
| 4 | 110958164 | rs6845765 | -0.948 | -0.479 | 0.982 | 1.814 | -0.007 | 0.037 | 0.037 | 0.833 | 0.23 | 0.11 | 0.18 | 0.11 | 0.31 | 0.2 |
| 4 | 110964272 | rs9990430 | -2.673 | -0.551 | 1.117 | 2.493 | 0.026 | -0.033 | -0.033 | 0 | 0.4 | 0.82 | 0.44 | 0.53 | 0.34 | 0.44 |
| 4 | 110964362 | rs10012953 | -0.885 | -0.434 | 1.151 | 2.366 | 0 | -0.002 | -0.002 | 0 | 0.18 | 0.16 | 0.16 | 0.13 | 0.26 | 0.15 |
| 4 | 110966120 | rs11098064 | -1.304 | -0.564 | -0.012 | 1.892 | -0.068 | -0.091 | -0.091 | 0.783 | 0.82 | 0.98 | 0.66 | 0.68 | 0.77 | 0.74 |
| 4 | 110967310 | rs9030 | -1.826 | -0.916 | 1.227 | 2.35 | 0 | -0.002 | -0.002 | 0 | 0.31 | 0.7 | 0.39 | 0.44 | 0.3 | 0.42 |
| 4 | 110967788 | rs3813826 | 0.198 | -0.765 | 1.092 | 1.926 | -0.009 | 0.018 | 0.018 | 0 | 0.08 | 0.11 | 0.06 | 0.16 | 0.05 | 0.09 |
| 4 | 110968599 | rs12108641 | 0.354 | -0.606 | 1.168 | 1.606 | 0.025 | -0.041 | -0.041 | 2.808 | 0.08 | 0.13 | 0.06 | 0 | 0.12 | 0.02 |
| 4 | 110975433 | rs7662161 | -1.37 | -0.814 | 1.47 | 1.894 | -0.028 | 0.016 | 0.016 | 0.491 | 0.17 | 0.35 | 0.25 | 0.23 | 0.14 | 0.15 |
| 4 | 110976538 | rs7690709 | -1.236 | -0.691 | 1.388 | 2.163 | -0.001 | 0 | 0 | 0 | 0.16 | 0.17 | 0.27 | 0.35 | 0.19 | 0.33 |
| 4 | 110977053 | rs9997849 | 0.655 | -0.601 | 1.374 | 2.017 | 0.011 | -0.031 | -0.031 | 0 | 0.07 | 0.09 | 0.05 | 0 | 0.1 | 0.02 |
| 4 | 110979707 | rs17041309 | -0.141 | -0.675 | 1.117 | 2.262 | -0.026 | -0.034 | -0.034 | 0 | 0.06 | 0 | 0.08 | 0.11 | 0.07 | 0.14 |
| 4 | 110982227 | rs17041315 | -0.103 | -0.943 | 1.25 | 2.429 | 0.004 | -0.008 | -0.008 | 0 | 0.09 | 0.05 | 0.19 | 0.27 | 0.1 | 0.15 |
| 4 | 110982248 | rs12498468 | -0.467 | -0.947 | 1.275 | 2.475 | 0.002 | 0.019 | 0.019 | 0 | 0.1 | 0.08 | 0.36 | 0.41 | 0.1 | 0.25 |
| 4 | 110984555 | rs17041323 | -1.686 | -0.776 | 1.139 | 2.18 | -0.017 | 0.095 | 0.095 | 0.075 | 0.17 | 0.36 | 0.24 | 0.39 | 0.15 | 0.27 |
| 4 | 110989719 | rs16997129 | -1.699 | -1.254 | 0.553 | 1.369 | -0.029 | -0.005 | -0.005 | 0 | 0.33 | 0.62 | 0.44 | 0.7 | 0.2 | 0.35 |
| 4 | 110994243 | rs11737840 | -0.475 | -0.845 | 0.102 | 0.573 | 0.01 | -0.031 | -0.031 | 0 | 0.23 | 0.24 | 0.16 | 0.06 | 0.29 | 0.28 |
| 4 | 110994792 | rs2040710 | -2.224 | -0.876 | 0.011 | 0.767 | -0.023 | 0.066 | 0.066 | 0.821 | 0.54 | 0.84 | 0.44 | 0.64 | 0.49 | 0.6 |
| 4 | 110995658 | rs6858635 | -2.25 | -0.868 | -0.303 | 0.772 | -0.036 | 0.063 | 0.063 | 1.258 | 0.55 | 0.86 | 0.43 | 0.64 | 0.49 | 0.6 |
| 4 | 110995710 | rs10033691 | 0.622 | -0.92 | -0.28 | 1.016 | -0.007 | 0.007 | 0.007 | 0 | 0.11 | 0.14 | 0.06 | 0.16 | 0.07 | 0.12 |
| 4 | 110997775 | rs2005701 | -1.915 | -0.82 | -0.745 | 0.607 | 0.004 | 0.089 | 0.089 | 1.899 | 0.51 | 0.77 | 0.43 | 0.64 | 0.48 | 0.6 |
| 4 | 111004545 | rs11726880 | 0.506 | -0.473 | -0.949 | 0.568 | 0.024 | -0.057 | -0.057 | 0 | 0.11 | 0.16 | 0.08 | 0 | 0.15 | 0.1 |
| 4 | 111007345 | rs34415611 | -0.526 | -0.608 | -0.923 | 0.866 | -0.009 | 0.002 | 0.002 | 0.117 | 0.26 | 0.17 | 0.17 | 0.19 | 0.33 | 0.33 |
| 4 | 111016014 | rs10005776 | -1.579 | -0.274 | -1.398 | 0.534 | -0.046 | 0.055 | 0.055 | 1.127 | 0.55 | 0.9 | 0.43 | 0.64 | 0.48 | 0.6 |
| 4 | 111026520 | rs3733623 | -0.565 | -0.646 | -1.279 | 0.171 | -0.039 | 0.022 | 0.022 | 0.006 | 0.27 | 0.41 | 0.3 | 0.69 | 0.21 | 0.4 |
| 4 | 111026927 | rs10011926 | -0.569 | -0.644 | -1.286 | 0.172 | -0.04 | 0.023 | 0.023 | 0.016 | 0.27 | 0.41 | 0.3 | 0.69 | 0.21 | 0.39 |
| 4 | 111030787 | rs6533491 | -1.048 | 0.191 | -2.182 | 0.081 | -0.163 | 0.102 | 0.102 | 0.018 | 0.61 | 0.95 | 0.65 | 0.8 | 0.53 | 0.69 |
| 4 | 111032048 | rs1005595 | -0.452 | 0.331 | -1.102 | 0.528 | 0.003 | 0.002 | 0.002 | 0 | 0.38 | 0.42 | 0.6 | 0.47 | 0.44 | 0.31 |
| 4 | 111032489 | rs1005594 | -0.997 | -0.049 | -0.975 | 0.022 | 0.005 | 0.055 | 0.055 | 0 | 0.63 | 0.76 | 0.77 | 0.82 | 0.64 | 0.58 |
| 4 | 111032602 | rs1001857 | -0.484 | -0.109 | -0.401 | 0.496 | 0.016 | -0.006 | -0.006 | 1.433 | 0.37 | 0.5 | 0.46 | 0.62 | 0.51 | 0.39 |
| 4 | 111033180 | rs11098070 | -0.206 | 0.159 | -0.508 | 0.592 | -0.001 | -0.001 | -0.001 | 0.306 | 0.33 | 0.32 | 0.5 | 0.58 | 0.5 | 0.38 |
| 4 | 111036402 | rs1125819 | 0.787 | 1.043 | -0.271 | 1.027 | 0.071 | 0.154 | 0.154 | 0 | 0.1 | 0.2 | 0.33 | 0.45 | 0.19 | 0.16 |
| 4 | 111036781 | rs1026883 | -0.693 | 0.561 | -0.704 | 0.833 | -0.028 | 0.044 | 0.044 | 0.474 | 0.77 | 0.88 | 0.82 | 0.92 | 0.68 | 0.75 |
| 4 | 111043210 | rs2170553 | -0.554 | 0.983 | -0.5 | 1.154 | -0.003 | 0.023 | 0.023 | 0 | 0.66 | 0.88 | 0.79 | 0.77 | 0.65 | 0.65 |
| 4 | 111051486 | rs10010167 | -0.871 | 1.878 | 0.654 | 1.782 | 0.107 | 0.112 | 0.112 | 0 | 0.23 | 0.44 | 0.39 | 0.46 | 0.33 | 0.27 |
| 4 | 111055176 | rs6533493 | -0.312 | 0.862 | 0.271 | 1.475 | 0.068 | -0.03 | -0.03 | 0 | 0.08 | 0.34 | 0.04 | 0 | 0.09 | 0.02 |
| 4 | 111055558 | rs11936079 | -1.155 | 0.608 | -0.849 | 2.023 | -0.011 | 0.071 | 0.071 | 0.228 | 0.1 | 0.2 | 0.19 | 0.29 | 0.06 | 0.21 |
| 4 | 111064092 | rs6835929 | -0.686 | 0.908 | -1.382 | 1.799 | -0.04 | -0.083 | -0.083 | 3.954 | 0.12 | 0.01 | 0.25 | 0.36 | 0.15 | 0.18 |
| 4 | 111065534 | rs4698812 | -0.299 | 0.599 | -1.593 | 0.842 | 0.039 | 0.053 | 0.053 | 0 | 0.46 | 0.28 | 0.56 | 0.66 | 0.6 | 0.42 |
| 4 | 111067650 | rs4146696 | -0.673 | 0.002 | -1.919 | 0.393 | 0.002 | 0.048 | 0.048 | 0 | 0.8 | 0.94 | 0.88 | 0.9 | 0.81 | 0.69 |
| 4 | 111072985 | rs6813255 | 0.547 | 0.082 | -1.196 | -0.065 | -0.009 | -0.011 | -0.011 | 0 | 0.58 | 0.56 | 0.64 | 0.66 | 0.65 | 0.48 |
| 4 | 111073867 | rs11733718 | 1.096 | -0.097 | -1.396 | -0.315 | -0.033 | 0.041 | 0.041 | 0 | 0.43 | 0.23 | 0.53 | 0.66 | 0.51 | 0.39 |
| 4 | 111079337 | rs2200106 | -0.207 | -0.183 | -0.644 | -0.1 | 0.005 | 0.009 | 0.009 | 0 | 0.71 | 0.76 | 0.72 | 0.81 | 0.77 | 0.64 |
| 4 | 111084981 | rs2035415 | 1.203 | -0.242 | -0.654 | -0.099 | -0.056 | 0.009 | 0.009 | 0 | 0.41 | 0.22 | 0.51 | 0.63 | 0.51 | 0.39 |
| 4 | 111092664 | rs10028161 | -1.015 | -0.166 | 0.599 | 0.038 | 0.038 | -0.022 | -0.022 | 0 | 0.19 | 0.4 | 0.27 | 0.13 | 0.21 | 0.26 |
| 4 | 111093616 | rs10008897 | 0.313 | -0.114 | 0.358 | -0.059 | 0.007 | -0.048 | -0.048 | 0.461 | 0.11 | 0.15 | 0.1 | 0 | 0.14 | 0.1 |
| 4 | 111105442 | rs1384331 | -1.062 | -0.354 | 0.244 | 0.052 | 0.011 | -0.005 | -0.005 | 0 | 0.35 | 0.49 | 0.38 | 0.31 | 0.36 | 0.51 |
| 4 | 111109020 | rs13145689 | -0.933 | -0.436 | 0.216 | 0.055 | -0.006 | -0.005 | -0.005 | 0 | 0.35 | 0.41 | 0.37 | 0.31 | 0.31 | 0.49 |
| 4 | 111112788 | rs6533498 | 0.612 | -0.3 | -0.896 | -0.063 | 0.027 | 0.017 | 0.017 | 0.393 | 0.39 | 0.28 | 0.32 | 0.3 | 0.33 | 0.28 |
| 4 | 111117247 | rs11098071 | -0.405 | -0.221 | -1.136 | -0.077 | -0.006 | 0.011 | 0.011 | 0 | 0.52 | 0.61 | 0.53 | 0.6 | 0.47 | 0.46 |

| | | | | | | | | | | | | | | | | |
|---|---|---|---|---|---|---|---|---|---|---|---|---|---|---|---|---|
| 4 | 111120713 | rs6824447 | 0.521 | 0.017 | -1.612 | 0.313 | -0.012 | 0.056 | 0.056 | 0 | 0.55 | 0.76 | 0.4 | 0.33 | 0.47 | 0.45 |
| 4 | 111121841 | rs6831073 | -0.112 | 0.199 | -1.14 | 0.569 | -0.011 | -0.002 | -0.002 | 0 | 0.84 | 0.89 | 0.82 | 0.87 | 0.79 | 0.69 |
| 4 | 111125483 | rs4698814 | -0.933 | 0.416 | -0.921 | 0.886 | -0.002 | -0.002 | -0.002 | 0.051 | 0.37 | 0.38 | 0.57 | 0.64 | 0.35 | 0.45 |
| 4 | 111126874 | rs17041421 | 0.25 | 0.737 | -1.117 | 1.382 | -0.057 | -0.086 | -0.086 | 0 | 0.12 | 0.03 | 0.25 | 0.45 | 0.17 | 0.18 |
| 4 | 111128587 | rs5022521 | 0.101 | 0.602 | -0.718 | 1.589 | 0.006 | 0.036 | 0.036 | 0.037 | 0.86 | 0.75 | 0.74 | 0.79 | 0.85 | 0.82 |
| 4 | 111129756 | rs12499874 | -0.718 | -0.14 | -0.656 | 1.05 | -0.01 | 0.028 | 0.028 | 0.183 | 0.3 | 0.43 | 0.32 | 0.45 | 0.24 | 0.22 |
| 4 | 111132642 | rs12510756 | -0.959 | 0.07 | -1.085 | 1.072 | 0.002 | -0.001 | -0.001 | 0.741 | 0.34 | 0.63 | 0.35 | 0.35 | 0.28 | 0.46 |
| 4 | 111132948 | rs17041448 | -0.447 | 0.486 | -1.411 | 1.319 | -0.044 | -0.095 | -0.095 | 0.009 | 0.19 | 0.06 | 0.3 | 0.41 | 0.23 | 0.22 |
| 4 | 111143870 | rs4698761 | -0.203 | 0.205 | -1.606 | 1.191 | -0.065 | -0.14 | -0.14 | 0 | 0.23 | 0.04 | 0.33 | 0.41 | 0.26 | 0.22 |
| 4 | 111144049 | rs4698816 | -0.623 | 0.088 | -1.921 | 0.61 | 0.009 | 0.048 | 0.048 | 0 | 0.62 | 0.76 | 0.77 | 0.79 | 0.63 | 0.67 |
| 4 | 111144505 | rs7686686 | 0.099 | 0.1 | -0.852 | 0.494 | 0.013 | -0.007 | -0.007 | 0 | 0.33 | 0.49 | 0.41 | 0.32 | 0.36 | 0.3 |
| 4 | 111146079 | rs4557340 | -0.187 | -0.751 | -0.961 | -0.288 | 0.02 | 0.034 | 0.034 | 0 | 0.74 | 0.83 | 0.85 | 0.9 | 0.79 | 0.76 |
| 4 | 111151421 | rs7697518 | -0.329 | -0.784 | -0.966 | -0.303 | 0.019 | 0.04 | 0.04 | 0 | 0.76 | 0.84 | 0.85 | 0.91 | 0.8 | 0.87 |
| 4 | 111151810 | rs11098072 | 0.445 | -1.597 | -0.077 | -0.643 | -0.053 | -0.078 | -0.078 | 0 | 0.63 | 0.47 | 0.81 | 0.88 | 0.78 | 0.65 |
| 4 | 111158481 | rs13105344 | 1.102 | -1.447 | 0.231 | -0.58 | -0.034 | -0.021 | -0.021 | 0 | 0.07 | 0.18 | 0.03 | 0.02 | 0.02 | 0.01 |
| 4 | 111164592 | rs17041514 | 0.491 | -1.487 | 0.196 | -0.586 | 0 | -0.001 | -0.001 | 0 | 0.17 | 0.13 | 0.14 | 0.19 | 0.21 | 0.2 |
| 4 | 111166848 | rs6533505 | -0.382 | -1.655 | 0.343 | -0.708 | -0.011 | -0.026 | -0.026 | 0 | 0.69 | 0.64 | 0.82 | 0.91 | 0.78 | 0.87 |
| 4 | 111166999 | rs7661205 | 1.263 | -1.578 | 0.003 | -0.691 | -0.045 | -0.029 | -0.029 | 0.013 | 0.3 | 0.19 | 0.56 | 0.61 | 0.43 | 0.46 |
| 4 | 111168797 | rs11735814 | 1.062 | -1.462 | -0.062 | -0.612 | -0.007 | -0.018 | -0.018 | 0 | 0.16 | 0.09 | 0.14 | 0.21 | 0.21 | 0.17 |
| 4 | 111171406 | rs6823387 | 1.203 | -1.428 | 0.123 | -0.677 | 0.016 | 0.027 | 0.027 | 0 | 0.19 | 0.15 | 0.1 | 0.09 | 0.13 | 0.12 |
| 4 | 111173378 | rs1482470 | 0.679 | -1.369 | 0.184 | -0.496 | -0.029 | -0.023 | -0.023 | 0 | 0.32 | 0.23 | 0.57 | 0.54 | 0.44 | 0.58 |
| 4 | 111181565 | rs10009070 | 0.608 | -1.414 | 0.331 | -0.587 | -0.018 | -0.029 | -0.029 | 0 | 0.25 | 0.29 | 0.13 | 0.09 | 0.15 | 0.12 |
| 4 | 111187683 | rs7656124 | -0.192 | -1.343 | 0.278 | -0.59 | -0.002 | -0.002 | -0.002 | 0 | 0.57 | 0.57 | 0.69 | 0.62 | 0.56 | 0.56 |
| 4 | 111191861 | rs10516561 | 0.5 | -1.091 | 0.118 | -0.585 | -0.023 | -0.042 | -0.042 | 0.44 | 0.25 | 0.32 | 0.14 | 0.09 | 0.17 | 0.13 |
| 4 | 111195127 | rs7694909 | -0.154 | -0.988 | -0.092 | -0.562 | 0.006 | 0 | 0 | 0 | 0.62 | 0.65 | 0.73 | 0.61 | 0.63 | 0.57 |
| 4 | 111197373 | rs17542839 | 1.074 | -1.012 | -1.096 | -0.654 | -0.033 | 0.032 | 0.032 | 0 | 0.19 | 0.05 | 0.18 | 0.09 | 0.22 | 0.11 |
| 4 | 111198328 | rs2713949 | 1.265 | -0.782 | -0.806 | -0.398 | -0.015 | -0.058 | -0.058 | 2.579 | 0.13 | 0.02 | 0.07 | 0.27 | 0.14 | 0.18 |
| 4 | 111211272 | rs417693 | 0.345 | -0.98 | -0.766 | -0.598 | 0.01 | 0.12 | 0.12 | 0 | 0.39 | 0.15 | 0.29 | 0.16 | 0.38 | 0.28 |
| 4 | 111214289 | rs7659944 | 1.122 | -0.452 | -0.927 | -0.177 | -0.027 | -0.018 | -0.018 | 0 | 0.16 | 0.06 | 0.33 | 0.23 | 0.25 | 0.15 |
| 4 | 111214726 | rs12510061 | -0.026 | -0.448 | -0.96 | -0.157 | -0.051 | -0.065 | -0.065 | 0 | 0.29 | 0.14 | 0.4 | 0.49 | 0.39 | 0.33 |
| 4 | 111216329 | rs244053 | -0.912 | -0.01 | -0.527 | 0.052 | 0 | -0.008 | -0.008 | 0 | 0.74 | 0.75 | 0.73 | 0.67 | 0.78 | 0.74 |
| 4 | 111218256 | rs763298 | 0.792 | 0.276 | -1.146 | 0.15 | -0.028 | -0.021 | -0.021 | 0 | 0.16 | 0.06 | 0.33 | 0.24 | 0.25 | 0.15 |
| 4 | 111227771 | rs244043 | -1.528 | -0.254 | -0.748 | 0.09 | -0.067 | -0.094 | -0.094 | 0.629 | 0.83 | 0.98 | 0.85 | 0.71 | 0.8 | 0.76 |
| 4 | 111227913 | rs9990445 | 0.208 | -0.862 | 0.017 | -0.368 | -0.036 | -0.023 | -0.023 | 1.154 | 0.1 | 0.44 | 0.04 | 0.02 | 0.01 | 0.01 |
| 4 | 111231836 | rs244036 | -0.394 | -0.283 | -1.103 | -0.217 | 0.01 | 0.02 | 0.02 | 0 | 0.24 | 0.18 | 0.19 | 0.06 | 0.22 | 0.11 |
| 4 | 111239014 | rs244055 | 0.561 | 0.205 | -1.32 | 0.274 | 0.009 | 0.021 | 0.021 | 1.401 | 0.34 | 0.13 | 0.5 | 0.32 | 0.35 | 0.36 |
| 4 | 111241892 | rs13113943 | -0.148 | 0.338 | -1.094 | 0.54 | 0.016 | -0.001 | -0.001 | 0 | 0.4 | 0.46 | 0.31 | 0.39 | 0.4 | 0.22 |
| 4 | 111252829 | rs10011752 | 0.469 | 0.734 | -1.535 | 0.629 | -0.003 | -0.011 | -0.011 | 0 | 0.16 | 0.24 | 0.11 | 0.13 | 0.11 | 0.06 |
| 4 | 111254360 | rs244019 | 0.146 | 0.661 | -2.186 | 0.762 | 0.012 | 0.025 | 0.025 | 1.518 | 0.34 | 0.13 | 0.51 | 0.33 | 0.37 | 0.38 |
| 4 | 111255917 | rs244017 | -0.048 | 0.676 | -1.483 | 1.142 | -0.014 | -0.057 | -0.057 | 0 | 0.8 | 0.9 | 0.84 | 0.71 | 0.79 | 0.77 |
| 4 | 111257452 | rs17586996 | -0.409 | 0.46 | -0.989 | 1.06 | -0.002 | 0.011 | 0.011 | 0 | 0.13 | 0.2 | 0.1 | 0.32 | 0.11 | 0.39 |
| 4 | 111257748 | rs244016 | -0.543 | 0.178 | -1.866 | 0.275 | -0.033 | 0.098 | 0.098 | 0 | 0.77 | 0.99 | 0.9 | 0.95 | 0.79 | 0.85 |
| 4 | 111262618 | rs244005 | 0.589 | 0.667 | -1.622 | 0.717 | -0.031 | -0.019 | -0.019 | 0 | 0.27 | 0.14 | 0.23 | 0.23 | 0.32 | 0.17 |
| 4 | 111272563 | rs409267 | 0.896 | 0.594 | -0.69 | 0.64 | -0.082 | -0.076 | -0.076 | 0 | 0.24 | 0.03 | 0.2 | 0.23 | 0.28 | 0.17 |
| 4 | 111275822 | rs6836372 | 0.593 | 0.566 | -0.6 | 0.831 | -0.012 | -0.029 | -0.029 | 0.038 | 0.16 | 0.08 | 0.13 | 0.3 | 0.17 | 0.33 |
| 4 | 111278172 | rs400966 | -0.367 | 0.569 | -0.557 | 0.867 | -0.001 | -0.001 | -0.001 | 0 | 0.45 | 0.43 | 0.37 | 0.53 | 0.47 | 0.55 |
| 4 | 111280757 | rs243981 | 0.033 | 0.707 | -0.439 | 0.704 | 0.023 | -0.022 | -0.022 | 0 | 0.38 | 0.43 | 0.32 | 0.21 | 0.46 | 0.22 |
| 4 | 111281932 | rs243978 | -0.661 | 0.819 | -0.607 | 0.745 | 0.005 | -0.003 | -0.003 | 0 | 0.55 | 0.54 | 0.47 | 0.51 | 0.66 | 0.53 |
| 4 | 111286032 | rs243973 | -0.369 | 0.866 | -0.625 | 0.93 | -0.001 | -0.001 | -0.001 | 0 | 0.48 | 0.41 | 0.39 | 0.51 | 0.51 | 0.5 |
| 4 | 111288232 | rs243970 | -0.449 | 0.872 | -0.728 | 0.903 | -0.001 | -0.001 | -0.001 | 0.059 | 0.48 | 0.51 | 0.4 | 0.55 | 0.51 | 0.6 |
| 4 | 111288254 | rs17625855 | 0.562 | 0.856 | -0.741 | 0.916 | -0.006 | -0.019 | -0.019 | 0.459 | 0.18 | 0.09 | 0.14 | 0.29 | 0.2 | 0.32 |
| 4 | 111295923 | rs243957 | -1.145 | 0.805 | -0.635 | 0.977 | 0.033 | 0.036 | 0.036 | 0 | 0.6 | 0.69 | 0.63 | 0.78 | 0.69 | 0.72 |
| 4 | 111297688 | rs243952 | -1.108 | 0.799 | -0.616 | 0.974 | 0.025 | 0.027 | 0.027 | 0 | 0.6 | 0.66 | 0.63 | 0.78 | 0.69 | 0.72 |
| 4 | 111306263 | rs10516562 | 0.748 | 1.23 | -0.117 | 1.027 | 0.015 | -0.031 | -0.031 | 0 | 0.08 | 0.11 | 0.08 | 0 | 0.16 | 0.04 |
| 4 | 111313114 | rs17041746 | 0.257 | 1.219 | -0.174 | 1.063 | 0.02 | 0.026 | 0.026 | 0 | 0.12 | 0.17 | 0.22 | 0.23 | 0.18 | 0.12 |
| 4 | 111313570 | rs2348205 | -0.575 | 1.231 | -0.306 | 1.044 | 0.009 | -0.007 | -0.007 | 0 | 0.7 | 0.61 | 0.74 | 0.78 | 0.69 | 0.81 |
| 4 | 111319763 | rs1027045 | -0.374 | 1.399 | -0.818 | 1.235 | 0.034 | 0.049 | 0.049 | 0 | 0.12 | 0.22 | 0.23 | 0.27 | 0.18 | 0.21 |
| 4 | 111324679 | rs10017096 | -0.366 | 0.826 | -1.815 | 0.944 | -0.096 | -0.064 | -0.064 | 0.65 | 0.74 | 0.95 | 0.77 | 0.75 | 0.7 | 0.85 |
| 4 | 111324997 | rs12506286 | -0.432 | -0.221 | -1.277 | 0.472 | -0.003 | -0.006 | -0.006 | 0 | 0.39 | 0.39 | 0.39 | 0.28 | 0.31 | 0.26 |
| 4 | 111335604 | rs1879056 | 0.282 | -1.214 | -0.974 | 0.51 | -0.041 | 0.052 | 0.052 | 0.647 | 0.67 | 0.42 | 0.73 | 0.48 | 0.79 | 0.57 |
| 4 | 111335722 | rs1879057 | 0.282 | -1.214 | -0.961 | 0.51 | -0.042 | 0.054 | 0.054 | 0.647 | 0.67 | 0.43 | 0.73 | 0.48 | 0.79 | 0.57 |
| 4 | 111356532 | rs9942210 | 0.164 | -1.301 | -1.258 | 0.217 | -0.004 | 0.033 | 0.033 | 0 | 0.16 | 0.05 | 0.19 | 0.08 | 0.16 | 0.13 |
| 4 | 111363108 | rs17008091 | 0.728 | -1.392 | -0.725 | 0.373 | -0.016 | -0.003 | -0.003 | 0 | 0.13 | 0.13 | 0.04 | 0.08 | 0.04 | 0.04 |
| 4 | 111364167 | rs2017368 | -1.089 | -1.177 | -0.763 | 0.5 | -0.03 | -0.042 | -0.042 | 0.405 | 0.91 | 0.97 | 0.93 | 0.76 | 0.87 | 0.79 |
| 4 | 111388265 | rs6814283 | -0.069 | -1.209 | -0.507 | 0.488 | 0.007 | 0.038 | 0.038 | 0.013 | 0.87 | 0.69 | 0.91 | 0.71 | 0.87 | 0.73 |
| 4 | 111392963 | rs17041821 | -0.428 | -1.458 | -0.121 | 0.278 | -0.043 | 0.043 | 0.043 | 0 | 0.19 | 0.32 | 0.16 | 0.23 | 0.09 | 0.17 |
| 4 | 111393691 | rs6823980 | 0.389 | -1.488 | -0.158 | -0.015 | -0.018 | -0.007 | -0.007 | 0 | 0.14 | 0.16 | 0.05 | 0.07 | 0.05 | 0.07 |
| 4 | 111396948 | rs1879053 | 0.098 | -1.478 | -0.488 | -0.441 | 0 | -0.007 | -0.007 | 0 | 0.61 | 0.61 | 0.74 | 0.74 | 0.55 | 0.55 |
| 4 | 111398208 | rs10004516 | -0.704 | -1.099 | -0.333 | -0.181 | -0.002 | -0.002 | -0.002 | 0 | 0.87 | 0.88 | 0.93 | 0.79 | 0.86 | 0.79 |

| | | | | | | | | | | | | | | | | |
|---|---|---|---|---|---|---|---|---|---|---|---|---|---|---|---|---|
| 4 | 111416282 | rs2881913 | -0.024 | -1.177 | -0.351 | -0.346 | 0.003 | 0.015 | 0.015 | 0 | 0.6 | 0.64 | 0.55 | 0.66 | 0.54 | 0.56 |
| 4 | 111418814 | rs17551860 | -0.229 | -1.237 | -0.343 | -0.387 | 0.018 | 0.015 | 0.015 | 0 | 0.25 | 0.16 | 0.08 | 0.17 | 0.15 | 0.25 |
| 4 | 111419299 | rs17041828 | -0.34 | -1.212 | -0.425 | -0.381 | -0.031 | -0.026 | -0.026 | 0 | 0.28 | 0.34 | 0.1 | 0.18 | 0.15 | 0.31 |
| 4 | 111420577 | rs939213 | -1.006 | -0.849 | -0.676 | -0.185 | -0.012 | -0.013 | -0.013 | 0 | 0.89 | 0.95 | 0.82 | 0.79 | 0.87 | 0.81 |
| 4 | 111426341 | rs11731078 | 0.372 | -1.014 | -0.561 | -0.32 | 0.001 | 0.004 | 0.004 | 0 | 0.29 | 0.27 | 0.27 | 0.13 | 0.32 | 0.25 |
| 4 | 111437127 | rs2348433 | 0.153 | -1.007 | -0.691 | -0.303 | 0.002 | 0.015 | 0.015 | 0 | 0.6 | 0.67 | 0.55 | 0.65 | 0.55 | 0.56 |
| 4 | 111454631 | rs670014 | -1.251 | -0.615 | -1.101 | -0.127 | -0.004 | -0.004 | -0.004 | 0.349 | 0.68 | 0.68 | 0.55 | 0.52 | 0.61 | 0.75 |
| 4 | 111462203 | rs557785 | -0.051 | -1.112 | -1.195 | -0.331 | -0.017 | -0.026 | -0.026 | 0 | 0.86 | 0.82 | 0.96 | 0.92 | 0.92 | 0.85 |
| 4 | 111463277 | rs478454 | 0.141 | -0.541 | -1.591 | 0.011 | 0.034 | 0.049 | 0.049 | 0 | 0.44 | 0.24 | 0.62 | 0.69 | 0.55 | 0.48 |
| 4 | 111473366 | rs9997349 | 0.583 | -0.041 | -1.492 | -0.111 | 0.017 | 0.07 | 0.07 | 0 | 0.79 | 0.93 | 0.92 | 0.92 | 0.82 | 0.88 |
| 4 | 111473485 | rs17041900 | -0.32 | 0.551 | -1.567 | 0.239 | -0.004 | -0.005 | -0.005 | 0 | 0.13 | 0.13 | 0.19 | 0.26 | 0.16 | 0.22 |
| 4 | 111476701 | rs3796879 | 0.435 | 0.169 | -1.999 | 0.267 | 0.017 | -0.014 | -0.014 | 0 | 0.12 | 0.06 | 0.06 | 0.09 | 0.08 | 0.14 |
| 4 | 111477949 | rs2029073 | 1.596 | -0.125 | -1.003 | 0.115 | 0.015 | 0.015 | 0.015 | 0 | 0.5 | 0.39 | 0.66 | 0.57 | 0.58 | 0.53 |
| 4 | 111484143 | rs2290105 | 1.645 | -0.165 | -0.869 | 0.159 | 0 | 0 | 0 | 0 | 0.5 | 0.48 | 0.67 | 0.56 | 0.59 | 0.53 |
| 4 | 111485171 | rs7682249 | -0.251 | -0.031 | -1.525 | 0.212 | -0.013 | -0.004 | -0.004 | 0 | 0.17 | 0.06 | 0.1 | 0.09 | 0.14 | 0.19 |
| 4 | 111492369 | rs1562640 | 0.009 | 0.033 | -1.089 | 0.345 | 0.028 | 0.033 | 0.033 | 0.02 | 0.85 | 0.94 | 0.95 | 0.92 | 0.88 | 0.91 |
| 4 | 111496844 | rs1466097 | -0.328 | 0.41 | -1.076 | 0.398 | 0.003 | 0.03 | 0.03 | 0 | 0.15 | 0.06 | 0.05 | 0.1 | 0.15 | 0.1 |
| 4 | 111501269 | rs11721423 | 0.681 | 0.544 | -0.722 | 0.6 | 0.002 | 0.028 | 0.028 | 0.143 | 0.87 | 0.72 | 0.84 | 0.75 | 0.89 | 0.83 |
| 4 | 111509740 | rs1900829 | 0.705 | 0.791 | -0.97 | 0.771 | 0.001 | 0.007 | 0.007 | 0 | 0.94 | 0.9 | 0.89 | 0.84 | 0.9 | 0.88 |
| 4 | 111510571 | rs1470618 | -0.722 | 0.682 | -1.195 | 1.104 | -0.001 | -0.029 | -0.029 | 0 | 0.18 | 0.06 | 0.27 | 0.23 | 0.2 | 0.12 |
| 4 | 111520926 | rs17553596 | -0.917 | 0.598 | -1.383 | 1.193 | -0.023 | -0.05 | -0.05 | 0 | 0.16 | 0.04 | 0.22 | 0.29 | 0.23 | 0.14 |
| 4 | 111530168 | rs9790621 | 0.652 | 0.243 | -0.526 | 0.841 | 0.001 | -0.001 | -0.001 | 0 | 0.73 | 0.34 | 0.82 | 0.7 | 0.82 | 0.75 |
| 4 | 111538121 | rs10024173 | -0.641 | 0.11 | -0.096 | 0.821 | 0.004 | 0.006 | 0.006 | 0 | 0.2 | 0.14 | 0.13 | 0.17 | 0.16 | 0.17 |
| 4 | 111540468 | rs3796902 | -1.161 | 0.092 | 0.461 | 0.822 | -0.03 | -0.001 | -0.001 | 0 | 0.24 | 0.39 | 0.16 | 0.27 | 0.16 | 0.23 |
| 4 | 111545323 | rs2595110 | 1.287 | 0.352 | -0.833 | -0.198 | -0.051 | 0.174 | 0.174 | 0.158 | 0.28 | 0.08 | 0.14 | 0 | 0.31 | 0.24 |
| 4 | 111548596 | rs7668322 | -0.577 | 0.112 | -0.138 | 0.421 | -0.003 | 0 | 0 | 4.251 | 0.18 | 0.2 | 0.13 | 0.24 | 0.13 | 0.23 |
| 4 | 111570223 | rs1448818 | -0.896 | -0.142 | 0.529 | 1.042 | 0.014 | 0.085 | 0.085 | 0 | 0.2 | 0.38 | 0.26 | 0.31 | 0.24 | 0.15 |
| 4 | 111571042 | rs1982360 | 1.402 | -0.188 | 0.479 | 1.139 | -0.007 | 0.046 | 0.046 | 0 | 0.7 | 0.5 | 0.67 | 0.48 | 0.7 | 0.68 |
| 4 | 111584419 | rs12498374 | -0.034 | -0.073 | -0.032 | 1.105 | -0.025 | -0.021 | -0.021 | 0 | 0.1 | 0.03 | 0.18 | 0.3 | 0.19 | 0.14 |
| 4 | 111604483 | rs2723296 | 1.076 | -0.153 | 0.281 | 0.98 | -0.004 | 0.016 | 0.016 | 0.041 | 0.69 | 0.42 | 0.64 | 0.44 | 0.69 | 0.63 |
| 4 | 111608485 | rs2723298 | 0.586 | -0.163 | 0.06 | 1.153 | -0.002 | 0.001 | 0.001 | 0 | 0.6 | 0.61 | 0.61 | 0.43 | 0.67 | 0.6 |
| 4 | 111622134 | rs7667461 | -0.256 | -0.169 | -0.11 | 1.403 | -0.006 | -0.021 | -0.021 | 0 | 0.15 | 0.06 | 0.19 | 0.26 | 0.16 | 0.2 |
| 4 | 111625925 | rs969642 | -1.106 | -0.323 | -0.083 | 1.424 | 0.014 | -0.056 | -0.056 | 0 | 0.38 | 0.16 | 0.4 | 0.52 | 0.34 | 0.35 |
| 4 | 111627356 | rs2595093 | -0.336 | -0.342 | 0.104 | 1.4 | -0.008 | 0.012 | 0.012 | 0 | 0.75 | 0.72 | 0.79 | 0.68 | 0.81 | 0.78 |
| 4 | 111627489 | rs2723313 | 0.962 | -0.331 | 0.071 | 1.339 | 0.007 | -0.008 | -0.008 | 0 | 0.61 | 0.65 | 0.59 | 0.42 | 0.65 | 0.58 |
| 4 | 111629039 | rs16997168 | 0.14 | -0.33 | 0.062 | 1.336 | 0.012 | -0.008 | -0.008 | 0 | 0.24 | 0.17 | 0.21 | 0.26 | 0.18 | 0.16 |
| 4 | 111634287 | rs2723316 | 0.586 | -0.169 | 0.063 | 1.13 | 0.014 | -0.021 | -0.021 | 0 | 0.61 | 0.73 | 0.6 | 0.43 | 0.65 | 0.59 |
| 4 | 111634826 | rs10222783 | -0.077 | -0.406 | 0.308 | 1.511 | 0.07 | 0.408 | 0.408 | 0 | 0.07 | 0.36 | 0.25 | 0.34 | 0.05 | 0.18 |
| 4 | 111641053 | rs1448817 | -0.901 | -0.65 | -0.128 | 1.197 | -0.03 | -0.012 | -0.012 | 0.146 | 0.41 | 0.46 | 0.47 | 0.72 | 0.32 | 0.39 |
| 4 | 111649869 | rs10019689 | 0.098 | -0.848 | -0.021 | 1.623 | -0.025 | 0.026 | 0.026 | 0 | 0.25 | 0.3 | 0.29 | 0.41 | 0.16 | 0.21 |
| 4 | 111665783 | rs2634073 | 0.48 | -0.888 | -0.018 | 1.498 | -0.056 | 0 | 0 | 0 | 0.66 | 0.56 | 0.57 | 0.28 | 0.78 | 0.64 |
| 4 | 111677722 | rs10516563 | 0.454 | -0.973 | 0.01 | 1.548 | 0.024 | -0.033 | -0.033 | 1.742 | 0.22 | 0.16 | 0.25 | 0.39 | 0.15 | 0.12 |
| 4 | 111685081 | rs4605724 | 0.557 | -0.971 | -0.033 | 1.563 | 0.038 | -0.059 | -0.059 | 0 | 0.21 | 0.11 | 0.25 | 0.38 | 0.15 | 0.12 |
| 4 | 111687514 | rs6533527 | 0.53 | -0.972 | 0 | 1.58 | 0.025 | -0.045 | -0.045 | 0 | 0.22 | 0.17 | 0.25 | 0.45 | 0.15 | 0.18 |
| 4 | 111688752 | rs2723334 | 0.56 | -0.858 | 0 | 1.511 | -0.038 | -0.002 | -0.002 | 0.016 | 0.64 | 0.49 | 0.56 | 0.29 | 0.76 | 0.62 |
| 4 | 111704143 | rs2129982 | -0.663 | -0.743 | -0.53 | 1.24 | -0.007 | 0.001 | 0.001 | 0 | 0.33 | 0.3 | 0.41 | 0.71 | 0.24 | 0.32 |
| 4 | 111705768 | rs6817105 | 0.222 | -0.76 | -0.546 | 1.278 | -0.02 | 0.031 | 0.031 | 0 | 0.23 | 0.25 | 0.28 | 0.45 | 0.16 | 0.14 |
| 4 | 111714419 | rs13143308 | 0.499 | -0.76 | -0.666 | 1.271 | 0 | -0.001 | -0.001 | 0 | 0.66 | 0.69 | 0.57 | 0.29 | 0.75 | 0.68 |
| 4 | 111714889 | rs2220427 | 0.571 | -0.699 | -0.743 | 1.301 | 0.034 | -0.064 | -0.064 | 0.328 | 0.21 | 0.12 | 0.27 | 0.45 | 0.15 | 0.13 |
| 4 | 111719685 | rs11931959 | 0.058 | -0.332 | -1.111 | 0.512 | 0.001 | 0.001 | 0.001 | 0 | 0.42 | 0.57 | 0.49 | 0.64 | 0.3 | 0.4 |
| 4 | 111720761 | rs10033464 | -0.123 | -0.278 | -1.08 | 0.534 | -0.012 | 0.035 | 0.035 | 0 | 0.89 | 0.81 | 0.85 | 0.75 | 0.9 | 0.8 |
| 4 | 111724501 | rs723363 | 0.444 | -0.29 | -1.045 | 0.51 | 0.059 | 0.187 | 0.187 | 0 | 0.48 | 0.24 | 0.38 | 0.1 | 0.6 | 0.4 |
| 4 | 111728614 | rs13141190 | 0.459 | -0.243 | -1.19 | 0.533 | 0.057 | 0.181 | 0.181 | 0 | 0.48 | 0.25 | 0.38 | 0.1 | 0.6 | 0.4 |
| 4 | 111729740 | rs3853441 | 0.797 | 0.431 | -0.858 | 1.298 | -0.012 | 0.034 | 0.034 | 0 | 0.89 | 0.81 | 0.85 | 0.73 | 0.9 | 0.8 |
| 4 | 111730243 | rs4124159 | 0.797 | 0.431 | -0.858 | 1.298 | -0.012 | 0.034 | 0.034 | 0 | 0.89 | 0.81 | 0.85 | 0.73 | 0.9 | 0.8 |
| 4 | 111734136 | rs3853444 | -0.708 | 1.434 | -1.777 | 1.345 | -0.118 | -0.138 | -0.138 | 0.088 | 0.27 | 0.05 | 0.44 | 0.62 | 0.37 | 0.42 |
| 4 | 111735539 | rs880309 | -0.233 | 0.385 | -1.435 | 0.795 | -0.03 | 0.011 | 0.011 | 0 | 0.29 | 0.53 | 0.17 | 0.29 | 0.21 | 0.34 |
| 4 | 111739156 | rs13130446 | -0.083 | 0.912 | -1.998 | 0.514 | -0.051 | -0.07 | -0.07 | 0.173 | 0.39 | 0.16 | 0.51 | 0.47 | 0.49 | 0.46 |
| 4 | 111739292 | rs10516564 | 0.581 | 1.184 | -1.668 | 0.484 | -0.028 | 0.03 | 0.03 | 0.274 | 0.29 | 0.12 | 0.31 | 0.25 | 0.3 | 0.2 |
| 4 | 111745640 | rs3866836 | 0.658 | 1.16 | -0.613 | 0.807 | 0.012 | 0.019 | 0.019 | 0 | 0.84 | 0.63 | 0.8 | 0.85 | 0.83 | 0.87 |
| 4 | 111753815 | rs3866838 | 0.723 | 1.126 | -0.596 | 0.802 | 0.023 | 0.03 | 0.03 | 0 | 0.85 | 0.66 | 0.82 | 0.85 | 0.84 | 0.86 |
| 4 | 111761487 | rs3853445 | -0.061 | 1.217 | -1.201 | 0.867 | -0.024 | -0.055 | -0.055 | 0 | 0.25 | 0.08 | 0.32 | 0.31 | 0.24 | 0.32 |
| 4 | 111765495 | rs6838973 | -0.625 | 1.121 | -1.063 | 0.248 | -0.026 | 0.009 | 0.009 | 0.77 | 0.4 | 0.19 | 0.44 | 0.58 | 0.4 | 0.6 |
| 4 | 111773867 | rs4631108 | -1.075 | 0.73 | -0.617 | 0.013 | 0.006 | 0.041 | 0.041 | 0.07 | 0.57 | 0.62 | 0.64 | 0.74 | 0.58 | 0.72 |
| 4 | 111780482 | rs10213323 | -0.065 | 0.864 | 0.041 | 0.229 | 0.006 | -0.001 | -0.001 | 0 | 0.17 | 0.17 | 0.19 | 0.16 | 0.18 | 0.11 |
| 4 | 111783306 | rs4033104 | 0.692 | 0.696 | 0.598 | 0.029 | 0.007 | -0.04 | -0.04 | 0.666 | 0.91 | 0.66 | 0.94 | 0.99 | 0.94 | 0.93 |
| 4 | 111787380 | rs695183 | 0.111 | 0.703 | 0.619 | 0.031 | 0.009 | -0.032 | -0.032 | 1.545 | 0.93 | 0.74 | 0.95 | 0.99 | 0.94 | 0.98 |
| 4 | 111787455 | rs4833470 | 0.214 | 0.662 | 0.328 | 0.347 | 0.002 | -0.005 | -0.005 | 0 | 0.69 | 0.65 | 0.67 | 0.62 | 0.73 | 0.55 |
| 4 | 111798201 | rs11098092 | 1.37 | 0.62 | 0.474 | 0.302 | 0.01 | 0.011 | 0.011 | 0 | 0.51 | 0.34 | 0.46 | 0.47 | 0.55 | 0.43 |

| | | | | | | | | | | | | | | | | |
|---|---|---|---|---|---|---|---|---|---|---|---|---|---|---|---|---|
| 4 | 111798258 | rs542331 | 0.464 | 0.598 | 0.46 | 0.299 | -0.017 | -0.098 | -0.098 | 0.163 | 0.23 | 0.07 | 0.29 | 0.37 | 0.21 | 0.41 |
| 4 | 111799417 | rs6843250 | -0.184 | 0.572 | 0.87 | 0.115 | 0.01 | -0.002 | -0.002 | 5.851 | 0.18 | 0.32 | 0.2 | 0.15 | 0.18 | 0.11 |
| 4 | 111804131 | rs577781 | 0.863 | 0.598 | 0.49 | 0.152 | -0.006 | -0.024 | -0.024 | 0 | 0.74 | 0.4 | 0.75 | 0.83 | 0.76 | 0.84 |
| 4 | 111811620 | rs537772 | 0.603 | 0.742 | 0.705 | 0.177 | 0.044 | -0.051 | -0.051 | 0 | 0.91 | 0.75 | 0.94 | 0.99 | 0.9 | 0.93 |
| 4 | 111812288 | rs10488914 | -0.277 | 0.739 | 0.753 | 0.175 | 0.003 | -0.02 | -0.02 | 0 | 0.07 | 0.07 | 0.05 | 0 | 0.1 | 0.06 |
| 4 | 111812327 | rs7679158 | 0.505 | 0.565 | 0.985 | 0.366 | -0.001 | 0.008 | 0.008 | 0 | 0.69 | 0.45 | 0.64 | 0.66 | 0.71 | 0.53 |
| 4 | 111812462 | rs536099 | 0.213 | 0.55 | 0.821 | 0.443 | -0.002 | 0.012 | 0.012 | 0 | 0.21 | 0.23 | 0.3 | 0.34 | 0.18 | 0.4 |
| 4 | 111812531 | rs11947581 | 0.151 | 0.772 | 0.744 | 0.481 | -0.006 | 0.011 | 0.011 | 0 | 0.17 | 0.07 | 0.19 | 0.12 | 0.2 | 0.11 |
| 4 | 111825573 | rs17042473 | -0.502 | 0.762 | 0.993 | 0.485 | -0.001 | -0.012 | -0.012 | 0 | 0.08 | 0.06 | 0.04 | 0 | 0.11 | 0.07 |
| 4 | 111845147 | rs17587726 | -0.046 | 0.726 | 1.192 | 0.858 | -0.01 | 0.078 | 0.078 | 0 | 0.09 | 0.17 | 0.07 | 0.27 | 0.06 | 0.13 |
| 4 | 111846955 | rs17587740 | -0.086 | 0.747 | 1.23 | 0.857 | -0.008 | 0.09 | 0.09 | 0 | 0.09 | 0.2 | 0.07 | 0.27 | 0.06 | 0.13 |
| 4 | 111849422 | rs374582 | -1.221 | 0.858 | 0.744 | 0.692 | 0.007 | 0.058 | 0.058 | 0 | 0.57 | 0.88 | 0.64 | 0.66 | 0.6 | 0.68 |
| 4 | 111858116 | rs886840 | -1.723 | 0.886 | 1.916 | 1.017 | 0 | 0.072 | 0.072 | 2.514 | 0.19 | 0.54 | 0.15 | 0.28 | 0.16 | 0.18 |
| 4 | 111866337 | rs10004479 | -1.94 | 0.884 | 1.677 | 1.016 | -0.008 | 0.032 | 0.032 | 1.605 | 0.2 | 0.64 | 0.16 | 0.28 | 0.16 | 0.18 |
| 4 | 111872144 | rs11943026 | -1.959 | 0.883 | 1.676 | 1.016 | -0.009 | 0.033 | 0.033 | 1.454 | 0.2 | 0.64 | 0.15 | 0.28 | 0.16 | 0.18 |
| 4 | 111927642 | rs10002518 | -2.317 | 0.815 | 1.357 | 1.256 | 0.029 | 0.131 | 0.131 | 0.094 | 0.15 | 0.52 | 0.12 | 0.29 | 0.15 | 0.14 |
| 4 | 111928603 | rs11726961 | -0.89 | 1.029 | 0.59 | 1.212 | 0.036 | -0.038 | -0.038 | 0 | 0.07 | 0.15 | 0.08 | 0 | 0.12 | 0.04 |
| 4 | 111932510 | rs6850589 | -1.254 | 0.715 | 0.233 | 1.578 | -0.004 | 0.196 | 0.196 | 0 | 0.06 | 0.17 | 0.03 | 0.29 | 0.03 | 0.09 |
| 4 | 111934020 | rs1010816 | -0.703 | 0.641 | -0.831 | 1.294 | -0.016 | -0.059 | -0.059 | 0 | 0.24 | 0.11 | 0.35 | 0.26 | 0.29 | 0.06 |
| 4 | 111937018 | rs542046 | -0.018 | 1.051 | -1.348 | 0.532 | -0.173 | 0.092 | 0.092 | 0.613 | 0.33 | 0.05 | 0.28 | 0.08 | 0.46 | 0.34 |
| 4 | 111953270 | rs1386389 | -0.964 | 0.893 | 0.811 | 1.154 | 0.028 | 0.096 | 0.096 | 0 | 0.23 | 0.44 | 0.34 | 0.46 | 0.26 | 0.27 |
| 4 | 111960686 | rs159471 | -1.927 | 0.713 | 0.82 | 1.092 | -0.013 | 0 | 0 | 0 | 0.2 | 0.71 | 0.15 | 0.21 | 0.1 | 0.11 |
| 4 | 111964082 | rs977013 | -0.568 | 0.91 | 0.69 | 1.263 | 0.024 | 0.01 | 0.01 | 0 | 0.18 | 0.31 | 0.27 | 0.23 | 0.24 | 0.23 |
| 4 | 111966090 | rs6840447 | -1.674 | 0.886 | 0.507 | 1.231 | 0.013 | 0.051 | 0.051 | 0 | 0.31 | 0.46 | 0.39 | 0.47 | 0.33 | 0.29 |
| 4 | 111980281 | rs159463 | -1.618 | 0.842 | 0.382 | 1.224 | -0.033 | 0.05 | 0.05 | 0.4 | 0.17 | 0.51 | 0.15 | 0.21 | 0.12 | 0.07 |
| 4 | 111984518 | rs159439 | -1.367 | 0.813 | 0.335 | 1.243 | -0.027 | 0.058 | 0.058 | 0.025 | 0.16 | 0.46 | 0.15 | 0.21 | 0.11 | 0.07 |
| 4 | 111985054 | rs159438 | -0.577 | 0.831 | 0.347 | 1.233 | 0.043 | 0.039 | 0.039 | 2.304 | 0.05 | 0.39 | 0.04 | 0 | 0.01 | 0 |
| 4 | 111986708 | rs212282 | -0.605 | 0.952 | 0.293 | 1.269 | 0.016 | 0.017 | 0.017 | 0 | 0.25 | 0.44 | 0.31 | 0.28 | 0.27 | 0.3 |
| 4 | 111986946 | rs190042 | -0.585 | 0.972 | 0.302 | 1.118 | 0.017 | 0.008 | 0.008 | 0 | 0.25 | 0.44 | 0.31 | 0.26 | 0.27 | 0.3 |
| 4 | 112004589 | rs297012 | 0.524 | 0.999 | -0.145 | 0.886 | 0.004 | 0.023 | 0.023 | 0 | 0.64 | 0.46 | 0.58 | 0.54 | 0.63 | 0.64 |
| 4 | 112024192 | rs439192 | -0.742 | 0.808 | -0.09 | 0.366 | 0.03 | -0.042 | -0.042 | 0 | 0.34 | 0.46 | 0.34 | 0.23 | 0.48 | 0.36 |
| 4 | 112024388 | rs11932853 | 0.737 | -0.337 | -0.484 | 0.319 | 0.011 | -0.011 | -0.011 | 1.741 | 0.38 | 0.22 | 0.52 | 0.55 | 0.33 | 0.39 |
| 4 | 112050006 | rs159228 | 0.448 | -0.462 | -0.081 | 0.9 | -0.022 | 0.077 | 0.077 | 0 | 0.85 | 0.83 | 0.76 | 0.58 | 0.9 | 0.82 |
| 4 | 112051379 | rs294941 | -0.455 | -0.573 | 0.203 | 0.706 | 0.007 | 0.001 | 0.001 | 0.271 | 0.27 | 0.25 | 0.35 | 0.42 | 0.46 | 0.46 |
| 4 | 112051824 | rs294942 | 1.15 | 0.066 | 0.174 | 0.646 | 0.002 | -0.002 | -0.002 | 0.001 | 0.07 | 0.07 | 0.06 | 0.04 | 0.16 | 0.13 |
| 4 | 112055872 | rs10446719 | 1.119 | -0.638 | 0.017 | 0.535 | 0.013 | 0.021 | 0.021 | 0 | 0.13 | 0.07 | 0.06 | 0.02 | 0.11 | 0.1 |
| 4 | 112060015 | rs9884929 | 0.407 | -0.262 | 0.188 | 0.576 | 0.015 | -0.001 | -0.001 | 0 | 0.18 | 0.18 | 0.3 | 0.23 | 0.4 | 0.35 |
| 4 | 112061855 | rs17042789 | 0.644 | -0.304 | 0.16 | 0.591 | 0.001 | -0.001 | -0.001 | 0 | 0.09 | 0.06 | 0.05 | 0.19 | 0.05 | 0.1 |
| 4 | 112077613 | rs12508528 | 0.243 | -0.604 | 0.53 | 0.434 | 0.002 | 0.002 | 0.002 | 0 | 0.63 | 0.73 | 0.75 | 0.77 | 0.76 | 0.75 |
| 4 | 112082915 | rs1601667 | -1.111 | -0.59 | 0.55 | 0.393 | 0.002 | -0.015 | -0.015 | 0 | 0.86 | 0.91 | 0.94 | 1 | 0.92 | 0.98 |
| 4 | 112093354 | rs11098102 | -0.837 | -0.687 | 0.571 | 0.389 | -0.026 | -0.066 | -0.066 | 0 | 0.86 | 0.83 | 0.94 | 1 | 0.92 | 0.99 |
| 4 | 112100154 | rs7660826 | 1.904 | -0.667 | 0.597 | 0.397 | 0.003 | 0 | 0 | 0 | 0.35 | 0.38 | 0.39 | 0.35 | 0.3 | 0.3 |
| 4 | 112126757 | rs1546053 | -0.768 | -0.439 | 0.449 | 0.137 | -0.016 | -0.05 | -0.05 | 0 | 0.86 | 0.82 | 0.94 | 1 | 0.92 | 0.99 |
| 4 | 112129311 | rs1552445 | 1.401 | -0.425 | 0.476 | 0.18 | 0.001 | -0.002 | -0.002 | 1.638 | 0.44 | 0.46 | 0.45 | 0.55 | 0.36 | 0.39 |
| 4 | 112129705 | rs17042928 | 0.716 | -0.351 | -0.095 | 0.215 | -0.002 | -0.024 | -0.024 | 0.251 | 0.13 | 0.13 | 0.06 | 0 | 0.08 | 0.02 |
| 4 | 112131789 | rs17042944 | -1.199 | 0.273 | -0.284 | 0.689 | -0.023 | -0.022 | -0.022 | 0.946 | 0.4 | 0.27 | 0.49 | 0.45 | 0.56 | 0.59 |
| 4 | 112145870 | rs17042997 | 0.012 | 0.345 | -0.843 | 0.782 | 0.044 | -0.038 | -0.038 | 0.02 | 0.22 | 0.08 | 0.19 | 0.22 | 0.17 | 0.24 |
| 4 | 112157798 | rs1490901 | -0.634 | 1.261 | -0.368 | 0.584 | 0.045 | -0.002 | -0.002 | 0 | 0.19 | 0.25 | 0.3 | 0.23 | 0.4 | 0.35 |
| 4 | 112172938 | rs2220490 | 0.301 | 0.048 | -0.817 | 0.535 | 0.025 | 0.007 | 0.007 | 0 | 0.78 | 0.88 | 0.88 | 0.79 | 0.87 | 0.85 |
| 4 | 112173532 | rs11098109 | 1.071 | -0.004 | -0.892 | 0.437 | 0.007 | 0.037 | 0.037 | 0 | 0.37 | 0.22 | 0.4 | 0.29 | 0.35 | 0.24 |
| 4 | 112176909 | rs7669813 | 1.029 | -0.014 | -0.774 | 0.431 | 0.009 | 0.034 | 0.034 | 0 | 0.38 | 0.27 | 0.4 | 0.29 | 0.35 | 0.24 |
| 4 | 112179424 | rs1388152 | 0.223 | 0.619 | -0.491 | 0.613 | 0.017 | 0.035 | 0.035 | 0 | 0.34 | 0.53 | 0.41 | 0.4 | 0.36 | 0.47 |
| 4 | 112192062 | rs17043025 | -0.238 | 0.379 | -0.386 | 0.482 | 0.007 | 0.005 | 0.005 | 0 | 0.41 | 0.49 | 0.47 | 0.4 | 0.44 | 0.48 |
| 4 | 112193351 | rs7694111 | -0.457 | 0.579 | -0.789 | 0.745 | -0.004 | -0.004 | -0.004 | 0.248 | 0.39 | 0.62 | 0.48 | 0.49 | 0.52 | 0.6 |
| 4 | 112198777 | rs17043035 | -0.083 | 0.57 | -0.81 | 0.904 | 0.028 | -0.019 | -0.019 | 0 | 0.13 | 0.05 | 0.06 | 0.21 | 0.05 | 0.14 |
| 4 | 112201846 | rs17576345 | 0.587 | 1.213 | -0.588 | 0.782 | -0.032 | -0.027 | -0.027 | 0.025 | 0.05 | 0 | 0.06 | 0.09 | 0.16 | 0.13 |
| 4 | 112207954 | rs11098115 | -0.625 | 0.734 | -0.453 | 0.621 | 0.003 | 0.015 | 0.015 | 0 | 0.45 | 0.37 | 0.46 | 0.57 | 0.43 | 0.45 |
| 4 | 112208694 | rs6826534 | 0.279 | 0.442 | -0.4 | 1.245 | -0.018 | 0.161 | 0.161 | 0 | 0.06 | 0.15 | 0.19 | 0.27 | 0.03 | 0.08 |
| 4 | 112209616 | rs6833281 | -0.425 | 0.406 | -0.412 | 1.206 | -0.017 | 0.041 | 0.041 | 0 | 0.39 | 0.22 | 0.23 | 0.3 | 0.4 | 0.38 |
| 4 | 112217523 | rs17576773 | 1.005 | 0.286 | -0.705 | 0.845 | 0.011 | 0.06 | 0.06 | 0 | 0.15 | 0.05 | 0.05 | 0 | 0.12 | 0.11 |
| 4 | 112217944 | rs4833619 | -1.789 | 0.137 | -0.616 | 0.251 | 0.013 | -0.105 | -0.105 | 0.106 | 0.69 | 0.53 | 0.73 | 0.88 | 0.66 | 0.8 |
| 4 | 112220233 | rs17043075 | -0.356 | 0.041 | -0.108 | 0.275 | -0.023 | 0.176 | 0.176 | 0.155 | 0.1 | 0.25 | 0.23 | 0.29 | 0.05 | 0.09 |
| 4 | 112220899 | rs4834781 | 0.132 | 0.079 | -0.043 | 0.293 | -0.033 | 0.025 | 0.025 | 0.015 | 0.26 | 0.41 | 0.29 | 0.29 | 0.17 | 0.2 |
| 4 | 112226088 | rs7692766 | -0.914 | -0.004 | 0.404 | 0.29 | 0.002 | 0.006 | 0.006 | 0 | 0.49 | 0.54 | 0.5 | 0.59 | 0.45 | 0.46 |
| 4 | 112229861 | rs7667177 | -0.785 | -0.082 | 0.184 | 0.324 | 0.007 | 0.04 | 0.04 | 0 | 0.51 | 0.74 | 0.47 | 0.58 | 0.45 | 0.45 |
| 4 | 112233572 | rs9307366 | 0.751 | -0.013 | -0.012 | 0.556 | 0 | -0.002 | -0.002 | 0 | 0.27 | 0.29 | 0.27 | 0.23 | 0.27 | 0.35 |
| 4 | 112244790 | rs7684254 | -1.215 | -0.319 | -0.35 | 0.791 | -0.029 | -0.077 | -0.077 | 3.912 | 0.73 | 0.91 | 0.72 | 0.55 | 0.7 | 0.72 |
| 4 | 112245519 | rs12642421 | -0.447 | -0.054 | 0.195 | 0.489 | 0 | 0 | 0 | 0 | 0.47 | 0.52 | 0.43 | 0.58 | 0.45 | 0.54 |
| 4 | 112252039 | rs2130816 | -0.387 | -0.055 | 0.175 | 0.454 | 0.017 | -0.049 | -0.049 | 6.064 | 0.64 | 0.53 | 0.68 | 0.84 | 0.58 | 0.69 |

| | | | | | | | | | | | | | | | | |
|---|---|---|---|---|---|---|---|---|---|---|---|---|---|---|---|---|
| 4 | 112254270 | rs11727195 | -0.21 | 0.275 | 0.202 | 0.37 | 0.044 | -0.011 | -0.011 | 0 | 0.1 | 0.22 | 0.07 | 0.05 | 0.14 | 0.11 |
| 4 | 112264996 | rs1352734 | -0.361 | 0.258 | 0.191 | 0.36 | 0.001 | 0 | 0 | 0 | 0.11 | 0.11 | 0.08 | 0.06 | 0.15 | 0.12 |
| 4 | 112265165 | rs6827570 | -0.263 | 0.254 | 0.181 | 0.385 | 0.018 | -0.031 | -0.031 | 0.251 | 0.64 | 0.53 | 0.68 | 0.81 | 0.58 | 0.69 |
| 4 | 112265437 | rs1388156 | -1.888 | 0.271 | 0.081 | 0.41 | -0.001 | 0.001 | 0.001 | 0 | 0.78 | 0.79 | 0.78 | 0.87 | 0.73 | 0.81 |
| 4 | 112272793 | rs1388159 | -1.854 | 0.198 | 0.074 | 0.403 | -0.001 | 0.002 | 0.002 | 0 | 0.77 | 0.79 | 0.78 | 0.87 | 0.73 | 0.82 |
| 4 | 112277270 | rs2661762 | -1.158 | 0.244 | 0.155 | 0.423 | -0.002 | -0.014 | -0.014 | 0 | 0.73 | 0.69 | 0.76 | 0.85 | 0.73 | 0.81 |
| 4 | 112279167 | rs1388158 | -1.33 | 0.225 | 0.14 | 0.423 | -0.002 | 0.011 | 0.011 | 0 | 0.74 | 0.79 | 0.76 | 0.85 | 0.72 | 0.81 |
| 4 | 112288658 | rs1847354 | 0.577 | 0.379 | 0.064 | 0.507 | 0.017 | -0.001 | -0.001 | 0 | 0.07 | 0.1 | 0.06 | 0.06 | 0.13 | 0.11 |
| 4 | 112289444 | rs1910452 | -0.528 | 0.306 | 0.011 | 0.478 | 0.02 | -0.021 | -0.021 | 0 | 0.66 | 0.58 | 0.69 | 0.78 | 0.58 | 0.69 |
| 4 | 112293391 | rs12499979 | 0.239 | 0.617 | 0.201 | 0.73 | -0.005 | 0.011 | 0.011 | 0 | 0.34 | 0.39 | 0.35 | 0.49 | 0.29 | 0.42 |
| 4 | 112293846 | rs2714997 | -0.315 | 0.824 | 0.139 | 0.793 | 0.03 | -0.001 | -0.001 | 0 | 0.08 | 0.12 | 0.06 | 0.06 | 0.13 | 0.07 |
| 4 | 112306505 | rs9685906 | -0.439 | 0.712 | 0.087 | 0.765 | 0.018 | -0.006 | -0.006 | 0 | 0.08 | 0.1 | 0.06 | 0.05 | 0.11 | 0.08 |
| 4 | 112307357 | rs1394001 | 0.445 | 0.818 | 0.112 | 0.616 | 0.001 | -0.001 | -0.001 | 0 | 0.25 | 0.24 | 0.39 | 0.24 | 0.37 | 0.21 |
| 4 | 112312860 | rs7669293 | -0.803 | 0.142 | 0.27 | 0.794 | -0.01 | 0.023 | 0.023 | 0 | 0.31 | 0.46 | 0.29 | 0.37 | 0.23 | 0.37 |
| 4 | 112314841 | rs1503666 | -1.372 | -0.041 | 0.24 | 1.004 | 0.006 | 0.056 | 0.056 | 0 | 0.6 | 0.73 | 0.7 | 0.82 | 0.6 | 0.67 |
| 4 | 112321704 | rs4833646 | 1.196 | -0.181 | 0.272 | 1.006 | 0.012 | -0.024 | -0.024 | 0 | 0.15 | 0.08 | 0.16 | 0.25 | 0.09 | 0.26 |
| 4 | 112323147 | rs1110906 | 0.106 | -0.171 | 0.394 | 1.011 | 0.017 | -0.019 | -0.019 | 0 | 0.22 | 0.16 | 0.2 | 0.29 | 0.13 | 0.3 |
| 4 | 112336798 | rs6833355 | 0.446 | -0.122 | 0.477 | 0.999 | 0.038 | 0.009 | 0.009 | 0.629 | 0.08 | 0.15 | 0.08 | 0.08 | 0.11 | 0.07 |
| 4 | 112345629 | rs1910455 | 0.052 | -0.126 | 0.446 | 1.07 | -0.006 | -0.031 | -0.031 | 0 | 0.35 | 0.23 | 0.42 | 0.5 | 0.37 | 0.33 |
| 4 | 112357791 | rs3109360 | -1.358 | -0.045 | 0.552 | 1.115 | 0.005 | -0.011 | -0.011 | 0 | 0.84 | 0.87 | 0.82 | 0.75 | 0.89 | 0.74 |
| 4 | 112371737 | rs3105288 | -1.897 | -0.101 | 0.297 | 1.111 | 0.001 | 0.011 | 0.011 | 0 | 0.87 | 0.98 | 0.94 | 0.9 | 0.91 | 0.81 |
| 4 | 112390143 | rs2661781 | 0.635 | -0.007 | 0.386 | 1.008 | -0.012 | -0.022 | -0.022 | 0 | 0.29 | 0.18 | 0.39 | 0.45 | 0.35 | 0.29 |
| 4 | 112398613 | rs2654735 | 1.208 | 0.202 | 0.455 | 0.875 | 0.005 | 0.07 | 0.07 | 0 | 0.41 | 0.28 | 0.32 | 0.18 | 0.43 | 0.34 |
| 4 | 112405007 | rs2661773 | 1.418 | 0.301 | 0.125 | 1.048 | -0.015 | -0.01 | -0.01 | 0.471 | 0.23 | 0.15 | 0.35 | 0.3 | 0.33 | 0.2 |
| 4 | 112413651 | rs2714992 | 2.301 | 0.182 | -0.315 | 0.852 | -0.046 | 0.045 | 0.045 | 0 | 0.2 | 0.06 | 0.18 | 0.09 | 0.24 | 0.1 |
| 4 | 112419559 | rs924758 | -1.041 | 0.649 | 0.426 | 1.398 | -0.009 | -0.005 | -0.005 | 0 | 0.43 | 0.6 | 0.51 | 0.56 | 0.46 | 0.43 |
| 4 | 112424765 | rs10027375 | 0.489 | 0.671 | 0.46 | 1.408 | 0.005 | -0.003 | -0.003 | 0 | 0.12 | 0.12 | 0.08 | 0.08 | 0.11 | 0.11 |
| 4 | 112428657 | rs1394002 | -0.897 | 0.681 | 0.452 | 1.426 | -0.008 | -0.005 | -0.005 | 0 | 0.43 | 0.6 | 0.5 | 0.55 | 0.46 | 0.4 |
| 4 | 112462458 | rs12498306 | 0.914 | 0.676 | 0.352 | 1.415 | 0.006 | 0.003 | 0.003 | 0 | 0.24 | 0.29 | 0.36 | 0.3 | 0.33 | 0.22 |
| 4 | 112466269 | rs17043348 | 0.076 | 0.709 | 0.272 | 1.368 | 0.029 | -0.01 | -0.01 | 0 | 0.14 | 0.3 | 0.1 | 0.08 | 0.11 | 0.11 |
| 4 | 112467251 | rs1806506 | -1.764 | 0.739 | -0.336 | 1.397 | 0.005 | 0.04 | 0.04 | 0 | 0.6 | 0.68 | 0.68 | 0.82 | 0.57 | 0.69 |
| 4 | 112476534 | rs10001625 | 0.376 | 0.859 | -0.79 | 1.466 | 0.009 | -0.005 | -0.005 | 1.071 | 0.12 | 0.14 | 0.09 | 0.08 | 0.12 | 0.11 |
| 4 | 112497952 | rs9992389 | 0.616 | 0.821 | -0.933 | 1.48 | 0 | -0.003 | -0.003 | 0 | 0.12 | 0.13 | 0.07 | 0.08 | 0.09 | 0.11 |
| 4 | 112506540 | rs4324606 | 0.073 | 1.093 | -1.498 | 1.485 | -0.022 | -0.038 | -0.038 | 0 | 0.28 | 0.17 | 0.39 | 0.43 | 0.35 | 0.32 |
| 4 | 112510973 | rs12648416 | 0.467 | 1.135 | -1.29 | 1 | -0.022 | -0.01 | -0.01 | 1.095 | 0.24 | 0.17 | 0.36 | 0.29 | 0.34 | 0.22 |
| 4 | 112514407 | rs1450782 | -0.224 | 0.689 | -1.467 | 0.592 | 0.006 | 0.058 | 0.058 | 0 | 0.61 | 0.72 | 0.69 | 0.82 | 0.58 | 0.69 |
| 4 | 112519126 | rs6830890 | -0.674 | 0.93 | -1.259 | 1.267 | -0.012 | -0.013 | -0.013 | 0 | 0.08 | 0.03 | 0.19 | 0.2 | 0.13 | 0.19 |
| 4 | 112521730 | rs2717196 | -0.593 | 0.476 | -1.259 | 1.006 | -0.001 | 0.001 | 0.001 | 0 | 0.22 | 0.24 | 0.29 | 0.26 | 0.25 | 0.2 |
| 4 | 112524605 | rs1470716 | -0.108 | 0.506 | -1.486 | 1.009 | -0.052 | -0.056 | -0.056 | 0 | 0.13 | 0.01 | 0.22 | 0.25 | 0.2 | 0.15 |
| 4 | 112524778 | rs10015214 | 0.75 | 0.073 | -1.374 | 0.541 | 0.002 | 0.015 | 0.015 | 0.32 | 0.49 | 0.46 | 0.4 | 0.28 | 0.51 | 0.57 |
| 4 | 112530606 | rs2597863 | 0.027 | -0.035 | -1.157 | 0.514 | -0.003 | 0.006 | 0.006 | 0 | 0.89 | 0.88 | 0.9 | 0.81 | 0.93 | 0.95 |
| 4 | 112536566 | rs2597875 | -0.252 | 0.033 | -1.232 | 0.489 | 0.001 | -0.008 | -0.008 | 0 | 0.9 | 0.95 | 0.92 | 0.82 | 0.93 | 0.95 |
| 4 | 112538240 | rs980387 | 0.68 | -0.243 | -1.365 | 0.115 | -0.004 | -0.028 | -0.028 | 0 | 0.45 | 0.5 | 0.38 | 0.2 | 0.5 | 0.48 |
| 4 | 112542699 | rs6845692 | 0.484 | -0.286 | -1.032 | 0.443 | -0.01 | -0.022 | -0.022 | 2.592 | 0.54 | 0.51 | 0.44 | 0.2 | 0.6 | 0.52 |
| 4 | 112550848 | rs2717207 | 0.554 | 0.114 | -0.927 | 0.807 | -0.005 | 0.005 | 0.005 | 0 | 0.88 | 0.87 | 0.9 | 0.82 | 0.92 | 0.95 |
| 4 | 112557798 | rs983966 | -0.445 | -0.249 | -1.509 | 0.345 | 0.035 | 0.097 | 0.097 | 0.018 | 0.44 | 0.12 | 0.35 | 0.25 | 0.43 | 0.38 |
| 4 | 112562714 | rs2443057 | 0.041 | -0.759 | -1.875 | 0.021 | -0.063 | 0.031 | 0.031 | 0 | 0.31 | 0.02 | 0.33 | 0.18 | 0.35 | 0.35 |
| 4 | 112568464 | rs10025530 | 0.496 | -1.475 | -1.339 | -0.418 | -0.031 | -0.04 | -0.04 | 0 | 0.08 | 0.22 | 0.05 | 0 | 0.02 | 0.02 |
| 4 | 112576171 | rs4833869 | 0.564 | -2.125 | -0.262 | -0.083 | 0.059 | 0.046 | 0.046 | 0 | 0.5 | 0.32 | 0.52 | 0.65 | 0.6 | 0.62 |
| 4 | 112584467 | rs10014697 | 0.118 | -2.11 | 0.172 | -0.086 | -0.038 | -0.05 | -0.05 | 0 | 0.09 | 0.33 | 0.06 | 0 | 0.02 | 0.02 |
| 4 | 112588951 | rs1450781 | -0.352 | -2.36 | -0.04 | -0.198 | 0.021 | 0.087 | 0.087 | 0.125 | 0.42 | 0.13 | 0.36 | 0.25 | 0.43 | 0.38 |
| 4 | 112591777 | rs6533579 | 0.76 | -2.137 | 0.141 | -0.044 | 0.051 | 0.043 | 0.043 | 0 | 0.51 | 0.35 | 0.52 | 0.66 | 0.61 | 0.63 |
| 4 | 112628325 | rs2597899 | -0.491 | -1.915 | 0.697 | 0.09 | -0.02 | -0.035 | -0.035 | 0 | 0.93 | 0.73 | 0.96 | 1 | 0.97 | 0.99 |
| 4 | 112669571 | rs326896 | -0.381 | -0.71 | 0.68 | 0.472 | 0.005 | 0.002 | 0.002 | 0 | 0.49 | 0.85 | 0.49 | 0.52 | 0.39 | 0.51 |
| 4 | 112673539 | rs6823743 | 0.821 | -0.675 | 0.865 | 0.428 | -0.022 | -0.034 | -0.034 | 0 | 0.06 | 0.19 | 0.04 | 0 | 0.03 | 0.01 |
| 4 | 112676788 | rs326902 | -0.486 | -0.029 | 0.858 | 0.38 | 0.018 | -0.032 | -0.032 | 0 | 0.94 | 0.85 | 0.96 | 1 | 0.94 | 0.98 |
| 4 | 112696927 | rs12508492 | 1.175 | -0.143 | 0.501 | 1.094 | -0.002 | 0.007 | 0.007 | 0 | 0.07 | 0.04 | 0.19 | 0.33 | 0.07 | 0.17 |
| 4 | 112712361 | rs6815054 | 0.479 | -0.261 | 0.756 | 0.902 | -0.005 | -0.016 | -0.016 | 0 | 0.34 | 0.25 | 0.33 | 0.42 | 0.31 | 0.48 |
| 4 | 112722748 | rs966055 | -0.124 | -0.294 | 0.901 | 0.94 | 0 | 0 | 0 | 0.083 | 0.39 | 0.37 | 0.35 | 0.42 | 0.34 | 0.48 |
| 4 | 112729781 | rs326880 | 0.248 | -0.4 | 0.809 | 0.807 | 0.027 | 0.002 | 0.002 | 0 | 0.56 | 0.23 | 0.54 | 0.46 | 0.67 | 0.53 |
| 4 | 112733434 | rs326872 | -0.079 | -0.355 | 1.074 | 0.792 | 0.001 | -0.003 | -0.003 | 0 | 0.6 | 0.4 | 0.56 | 0.46 | 0.66 | 0.51 |
| 4 | 112739250 | rs326866 | 0.035 | -0.4 | 1.268 | 0.816 | 0 | 0 | 0 | 0 | 0.58 | 0.39 | 0.56 | 0.46 | 0.63 | 0.5 |
| 4 | 112744390 | rs7671351 | 0.779 | -0.249 | 1.14 | 0.738 | -0.002 | -0.022 | -0.022 | 3.896 | 0.06 | 0.22 | 0.04 | 0 | 0.03 | 0.01 |
| 4 | 112756367 | rs13128359 | 0.696 | -0.062 | 0.698 | 0.766 | -0.001 | -0.001 | -0.001 | 0 | 0.17 | 0.15 | 0.23 | 0.34 | 0.17 | 0.22 |
| 4 | 112762156 | rs2199342 | 0.391 | -0.056 | 0.531 | 0.826 | -0.003 | -0.001 | -0.001 | 0 | 0.55 | 0.38 | 0.6 | 0.58 | 0.6 | 0.5 |
| 4 | 112771435 | rs400245 | 0.652 | 0.155 | 0.399 | 0.518 | -0.005 | -0.005 | -0.005 | 0 | 0.72 | 0.44 | 0.71 | 0.73 | 0.74 | 0.87 |
| 4 | 112777194 | rs10516580 | -0.374 | 0.106 | 0.771 | 0.518 | 0.003 | -0.024 | -0.024 | 0 | 0.23 | 0.34 | 0.21 | 0.16 | 0.24 | 0.11 |
| 4 | 112793563 | rs13435138 | -0.366 | 0.121 | 0.717 | 0.614 | 0.005 | -0.029 | -0.029 | 0 | 0.23 | 0.35 | 0.21 | 0.16 | 0.24 | 0.11 |
| 4 | 112799771 | rs6843121 | 0.585 | 0.415 | 0.463 | 0.46 | 0.025 | -0.007 | -0.007 | 0 | 0.71 | 0.64 | 0.68 | 0.73 | 0.64 | 0.82 |

| | | | | | | | | | | | | | | | | |
|---|---|---|---|---|---|---|---|---|---|---|---|---|---|---|---|---|
| 4 | 112805191 | rs17627147 | 1.304 | 0.12 | 0.047 | 0.223 | 0.001 | 0.028 | 0.028 | 0 | 0.08 | 0 | 0.02 | 0.04 | 0.04 | 0.13 |
| 4 | 112809145 | rs3850475 | -0.01 | 0.144 | 0.218 | 0.263 | 0.001 | -0.006 | -0.006 | 0 | 0.75 | 0.76 | 0.76 | 0.88 | 0.65 | 0.83 |
| 4 | 112813419 | rs6845600 | 0.373 | 0.126 | 0.224 | 0.303 | 0.011 | -0.002 | -0.002 | 0.072 | 0.72 | 0.69 | 0.69 | 0.73 | 0.64 | 0.82 |
| 4 | 112824982 | rs3913813 | 1.461 | 0.092 | 0.069 | 0.201 | 0.005 | 0.019 | 0.019 | 0 | 0.52 | 0.37 | 0.62 | 0.63 | 0.57 | 0.68 |
| 4 | 112827769 | rs4256285 | 1.103 | -0.04 | -1.071 | -0.018 | 0.045 | 0.081 | 0.081 | 0 | 0.37 | 0.17 | 0.3 | 0.22 | 0.34 | 0.45 |
| 4 | 112830162 | rs227906 | -1.84 | -0.875 | -1.236 | -0.041 | -0.056 | 0.088 | 0.088 | 10.447 | 0.65 | 0.91 | 0.69 | 0.79 | 0.57 | 0.51 |
| 4 | 112845980 | rs722052 | -0.122 | -1.104 | -1.135 | 0.044 | -0.002 | -0.023 | -0.023 | 0.009 | 0.35 | 0.78 | 0.4 | 0.58 | 0.31 | 0.31 |
| 4 | 112847457 | rs7678991 | -0.873 | -1.109 | -1.182 | 0.009 | -0.018 | 0.06 | 0.06 | 3.103 | 0.42 | 0.87 | 0.51 | 0.71 | 0.37 | 0.32 |
| 4 | 112874121 | rs17527615 | 0.656 | -1.213 | -1.19 | -0.016 | 0.002 | 0.047 | 0.047 | 0 | 0.22 | 0.06 | 0.19 | 0.08 | 0.21 | 0.2 |
| 4 | 112880414 | rs2016740 | 0.485 | -1.259 | -0.688 | 0.052 | 0.021 | 0.08 | 0.08 | 0 | 0.3 | 0.54 | 0.38 | 0.49 | 0.29 | 0.3 |
| 4 | 112885796 | rs1981567 | -1.658 | -1.215 | -0.807 | 0.041 | -0.057 | 0.123 | 0.123 | 3.017 | 0.59 | 0.87 | 0.66 | 0.77 | 0.53 | 0.5 |
| 4 | 112900775 | rs4834106 | -1.741 | -1.675 | -0.786 | 0.034 | -0.06 | 0.023 | 0.023 | 8.225 | 0.61 | 0.88 | 0.6 | 0.65 | 0.52 | 0.49 |
| 4 | 112902209 | rs6812046 | -0.556 | -1.58 | -0.605 | 0.113 | 0.006 | 0.008 | 0.008 | 2.346 | 0.32 | 0.28 | 0.23 | 0.16 | 0.25 | 0.2 |
| 4 | 112905223 | rs7654718 | -0.535 | -1.549 | -0.572 | 0.171 | -0.008 | 0 | 0 | 0.662 | 0.29 | 0.3 | 0.27 | 0.28 | 0.22 | 0.2 |
| 4 | 112909820 | rs7660014 | -0.1 | -1.504 | -0.64 | 0.152 | 0.008 | 0.012 | 0.012 | 0.24 | 0.26 | 0.2 | 0.2 | 0.16 | 0.23 | 0.2 |
| 4 | 112914966 | rs3850482 | 0.173 | -2.435 | -0.252 | 0.057 | -0.001 | 0.037 | 0.037 | 0.252 | 0.31 | 0.53 | 0.41 | 0.56 | 0.28 | 0.32 |
| 4 | 112922517 | rs6820458 | 0.103 | -2.639 | -0.528 | -0.047 | -0.003 | -0.009 | -0.009 | 0.601 | 0.15 | 0.14 | 0.09 | 0.06 | 0.11 | 0.15 |
| 4 | 112927422 | rs3850485 | -2.575 | -2.879 | -0.906 | -0.186 | -0.139 | 0.088 | 0.088 | 2.862 | 0.66 | 0.94 | 0.63 | 0.81 | 0.47 | 0.56 |
| 4 | 112929419 | rs10488956 | 2.293 | -2.769 | -0.649 | -0.023 | -0.003 | 0.002 | 0.002 | 0 | 0.08 | 0.07 | 0.04 | 0.03 | 0.11 | 0.02 |
| 4 | 112929836 | rs12500823 | -2.137 | -2.734 | -0.779 | 0.011 | -0.045 | 0.136 | 0.136 | 3.514 | 0.58 | 0.9 | 0.59 | 0.78 | 0.42 | 0.47 |
| 4 | 112971781 | rs4404614 | 1.809 | -2.64 | -0.606 | 0.126 | 0.002 | 0.01 | 0.01 | 1.01 | 0.06 | 0.04 | 0.03 | 0 | 0.04 | 0.01 |
| 4 | 112979568 | rs2018700 | -1.935 | -2.683 | -0.606 | 0.121 | 0.003 | 0.119 | 0.119 | 2.793 | 0.55 | 0.84 | 0.59 | 0.71 | 0.42 | 0.46 |
| 4 | 112987361 | rs12506001 | -0.569 | -2.764 | -0.492 | 0.248 | 0.002 | -0.008 | -0.008 | 0 | 0.36 | 0.66 | 0.44 | 0.49 | 0.32 | 0.31 |
| 4 | 112995439 | rs4240282 | 0.047 | -2.757 | -0.423 | 0.255 | -0.002 | 0.05 | 0.05 | 0 | 0.3 | 0.54 | 0.4 | 0.49 | 0.27 | 0.3 |
| 4 | 113003174 | rs6842261 | -0.082 | -2.762 | -0.422 | 0.267 | -0.027 | 0.085 | 0.085 | 0.632 | 0.06 | 0.14 | 0.09 | 0.22 | 0.01 | 0.01 |
| 4 | 113014647 | rs9307378 | -1.217 | -2.767 | -0.451 | 0.233 | 0.031 | 0.069 | 0.069 | 0.031 | 0.42 | 0.77 | 0.52 | 0.72 | 0.33 | 0.31 |
| 4 | 113019167 | rs10004030 | -0.623 | -2.692 | -0.336 | 0.43 | 0.003 | 0.002 | 0.002 | 0 | 0.34 | 0.65 | 0.41 | 0.68 | 0.27 | 0.3 |
| 4 | 113025264 | rs10516583 | 0.187 | -2.682 | -0.317 | 0.439 | -0.003 | -0.001 | -0.001 | 0.891 | 0.15 | 0.14 | 0.14 | 0.1 | 0.09 | 0.15 |
| 4 | 113032743 | rs12644031 | -1.179 | -2.649 | -0.409 | 0.427 | 0.026 | 0.047 | 0.047 | 0.159 | 0.43 | 0.72 | 0.47 | 0.68 | 0.34 | 0.31 |
| 4 | 113033220 | rs6533596 | 0.47 | -2.651 | -0.372 | 0.443 | -0.003 | 0 | 0 | 0.055 | 0.13 | 0.12 | 0.14 | 0.16 | 0.09 | 0.02 |
| 4 | 113034504 | rs1968489 | 0.213 | -2.626 | -0.329 | 0.456 | 0.008 | 0.115 | 0.115 | 0 | 0.27 | 0.47 | 0.32 | 0.46 | 0.25 | 0.29 |
| 4 | 113035523 | rs7699106 | 0.02 | -2.669 | -0.237 | 0.124 | -0.013 | 0.002 | 0.002 | 1.971 | 0.16 | 0.19 | 0.15 | 0.2 | 0.09 | 0.02 |
| 4 | 113036309 | rs6815972 | -0.185 | -2.669 | -0.167 | 0.125 | -0.031 | 0.008 | 0.008 | 2.353 | 0.16 | 0.26 | 0.16 | 0.2 | 0.09 | 0.02 |
| 4 | 113039827 | rs6836984 | -1.207 | -2.668 | -0.224 | 0.109 | 0.02 | 0.066 | 0.066 | 0.916 | 0.44 | 0.7 | 0.49 | 0.71 | 0.37 | 0.33 |
| 4 | 113042939 | rs7682303 | -1.157 | -2.668 | -0.255 | 0.109 | 0.011 | 0.033 | 0.033 | 0.789 | 0.44 | 0.63 | 0.49 | 0.71 | 0.37 | 0.33 |
| 4 | 113050302 | rs1521477 | -2.194 | -2.722 | -0.406 | 0.062 | -0.066 | 0.14 | 0.14 | 5.814 | 0.61 | 0.89 | 0.63 | 0.8 | 0.46 | 0.48 |
| 4 | 113055291 | rs9995910 | -0.028 | -2.428 | -0.082 | 0.436 | 0.01 | 0.06 | 0.06 | 0 | 0.28 | 0.42 | 0.34 | 0.52 | 0.28 | 0.31 |
| 4 | 113071518 | rs2117917 | -1.308 | -2.505 | -0.142 | 0.418 | 0.016 | 0.053 | 0.053 | 1.332 | 0.44 | 0.69 | 0.5 | 0.71 | 0.38 | 0.35 |
| 4 | 113079391 | rs1040278 | -1.897 | -2.59 | -0.15 | 0.419 | 0.006 | 0.063 | 0.063 | 5.748 | 0.55 | 0.77 | 0.54 | 0.64 | 0.44 | 0.48 |
| 4 | 113081283 | rs11098145 | -2.153 | -2.588 | -0.163 | 0.415 | -0.02 | 0.149 | 0.149 | 5.806 | 0.58 | 0.82 | 0.62 | 0.8 | 0.45 | 0.48 |
| 4 | 113092380 | rs2351236 | 0.158 | -2.269 | 0.156 | 0.615 | -0.014 | -0.003 | -0.003 | 0.181 | 0.13 | 0.14 | 0.12 | 0.09 | 0.06 | 0.13 |
| 4 | 113106286 | rs10212874 | -1.233 | -2.288 | 0.055 | 0.604 | 0.01 | 0.012 | 0.012 | 0.274 | 0.42 | 0.61 | 0.47 | 0.6 | 0.36 | 0.45 |
| 4 | 113120268 | rs4834241 | -0.494 | -1.658 | -0.163 | 0.762 | 0.011 | 0.052 | 0.052 | 0 | 0.29 | 0.43 | 0.34 | 0.51 | 0.3 | 0.32 |
| 4 | 113123832 | rs9994166 | -1.481 | -1.719 | -0.808 | 0.705 | -0.028 | 0.012 | 0.012 | 2.137 | 0.6 | 0.74 | 0.54 | 0.58 | 0.49 | 0.39 |
| 4 | 113125894 | rs7437464 | 1.785 | -1.75 | -0.968 | 0.686 | -0.092 | 0.058 | 0.058 | 1.821 | 0.33 | 0.13 | 0.35 | 0.24 | 0.44 | 0.58 |
| 4 | 113126582 | rs1554467 | -0.113 | -2.18 | -0.921 | 0.747 | 0.035 | 0.026 | 0.026 | 0 | 0.15 | 0.07 | 0.12 | 0.06 | 0.08 | 0.15 |
| 4 | 113150559 | rs4834244 | -1.994 | -2.106 | -1.063 | 0.75 | -0.079 | 0.074 | 0.074 | 2.221 | 0.64 | 0.83 | 0.63 | 0.76 | 0.51 | 0.4 |
| 4 | 113151732 | rs7699435 | -1.977 | -2.112 | -1.066 | 0.875 | -0.079 | 0.049 | 0.049 | 2.139 | 0.64 | 0.83 | 0.59 | 0.73 | 0.5 | 0.4 |
| 4 | 113152490 | rs2280499 | -2.085 | -2.1 | -1.17 | 0.881 | -0.135 | 0.034 | 0.034 | 2.185 | 0.65 | 0.91 | 0.59 | 0.73 | 0.51 | 0.4 |
| 4 | 113169092 | rs17044338 | 0.454 | -1.984 | -1.04 | 0.696 | 0.03 | 0.001 | 0.001 | 0.307 | 0.11 | 0.05 | 0.09 | 0.09 | 0.02 | 0.12 |
| 4 | 113174700 | rs10026870 | 0.274 | -1.816 | -0.904 | 0.764 | 0.01 | 0.095 | 0.095 | 1.036 | 0.24 | 0.38 | 0.29 | 0.4 | 0.26 | 0.12 |
| 4 | 113185053 | rs3775969 | -2.015 | -1.746 | -0.964 | 0.765 | -0.13 | 0.075 | 0.075 | 3.453 | 0.65 | 0.91 | 0.63 | 0.75 | 0.51 | 0.38 |
| 4 | 113195610 | rs17604169 | 0.887 | -1.764 | -0.984 | 0.737 | -0.019 | 0.025 | 0.025 | 0 | 0.08 | 0 | 0.05 | 0.04 | 0.07 | 0.07 |
| 4 | 113196545 | rs3822289 | -1.733 | -1.348 | -0.723 | 0.938 | -0.114 | 0.075 | 0.075 | 3.485 | 0.64 | 0.89 | 0.63 | 0.75 | 0.51 | 0.38 |
| 4 | 113196906 | rs3202899 | -1.733 | -1.348 | -0.715 | 0.924 | -0.104 | 0.072 | 0.072 | 3.485 | 0.64 | 0.88 | 0.63 | 0.75 | 0.51 | 0.38 |
| 4 | 113199075 | rs701759 | -1.733 | -1.324 | -0.756 | 0.924 | -0.12 | 0.071 | 0.071 | 3.485 | 0.64 | 0.89 | 0.63 | 0.75 | 0.51 | 0.38 |
| 4 | 113200382 | rs12510189 | 0.252 | -1.398 | -0.73 | 0.939 | 0.009 | 0.115 | 0.115 | 1.563 | 0.23 | 0.38 | 0.29 | 0.39 | 0.24 | 0.1 |
| 4 | 113204153 | rs10003654 | -0.997 | -0.453 | -1.442 | 0.836 | -0.115 | 0.07 | 0.07 | 3.359 | 0.64 | 0.89 | 0.63 | 0.75 | 0.51 | 0.38 |
| 4 | 113207277 | rs17604670 | -0.327 | -0.242 | -0.233 | 1.198 | 0.008 | 0 | 0 | 0 | 0.08 | 0.12 | 0.07 | 0.06 | 0.08 | 0.04 |
| 4 | 113232651 | rs17627730 | 0.039 | -0.625 | -0.672 | 1.218 | 0.018 | -0.02 | -0.02 | 0.027 | 0.06 | 0.02 | 0.07 | 0.17 | 0.01 | 0 |
| 4 | 113240529 | rs6839324 | -0.783 | -1.039 | -0.716 | 1.172 | 0.002 | -0.002 | -0.002 | 4.206 | 0.09 | 0.09 | 0.2 | 0.11 | 0.05 | 0.11 |
| 4 | 113243265 | rs6810537 | -1.372 | -1.129 | -0.906 | 1.218 | 0.041 | -0.022 | -0.022 | 0 | 0.4 | 0.75 | 0.61 | 0.61 | 0.31 | 0.29 |
| 4 | 113243456 | rs12648308 | -0.523 | -1.076 | -0.914 | 1.243 | 0.003 | -0.007 | -0.007 | 0 | 0.31 | 0.65 | 0.4 | 0.51 | 0.27 | 0.17 |
| 4 | 113244313 | rs1544382 | -1.057 | -1.04 | -0.884 | 1.026 | 0.031 | -0.014 | -0.014 | 0 | 0.39 | 0.7 | 0.6 | 0.62 | 0.31 | 0.29 |
| 4 | 113244979 | rs4834259 | -0.429 | -0.358 | -0.528 | 0.676 | -0.011 | 0.006 | 0.006 | 0.264 | 0.32 | 0.47 | 0.51 | 0.38 | 0.33 | 0.2 |
| 4 | 113246438 | rs17044418 | -0.61 | -0.485 | -0.842 | 0.293 | 0.003 | 0.004 | 0.004 | 0 | 0.16 | 0.13 | 0.05 | 0.09 | 0.1 | 0.11 |
| 4 | 113246972 | rs2351451 | 0.613 | 0.41 | -0.567 | -0.07 | 0.01 | -0.019 | -0.019 | 0.849 | 0.75 | 0.7 | 0.79 | 0.81 | 0.67 | 0.76 |
| 4 | 113247435 | rs10018567 | -0.997 | 0.021 | 0.053 | -0.313 | -0.026 | -0.049 | -0.049 | 0 | 0.1 | 0.23 | 0.13 | 0 | 0.05 | 0.11 |
| 4 | 113248536 | rs2051776 | 0.439 | -0.233 | 0.191 | -0.784 | 0.01 | -0.043 | -0.043 | 0.159 | 0.75 | 0.64 | 0.77 | 0.9 | 0.7 | 0.8 |

| | | | | | | | | | | | | | | | | |
|---|---|---|---|---|---|---|---|---|---|---|---|---|---|---|---|---|
| 4 | 113251247 | rs4834262 | 0.505 | -0.325 | 0.576 | -0.83 | 0.016 | -0.071 | -0.071 | 0 | 0.74 | 0.53 | 0.77 | 0.9 | 0.7 | 0.81 |
| 4 | 113251677 | rs6829831 | 0.633 | -0.46 | 0.721 | -0.917 | 0 | -0.04 | -0.04 | 0 | 0.73 | 0.38 | 0.75 | 0.9 | 0.7 | 0.81 |
| 4 | 113260465 | rs2351454 | -0.191 | -0.299 | -0.472 | -0.875 | 0.003 | -0.016 | -0.016 | 0 | 0.32 | 0.27 | 0.42 | 0.61 | 0.28 | 0.48 |
| 4 | 113262777 | rs17528404 | 0.997 | -0.022 | -1.21 | -1.574 | -0.027 | 0.079 | 0.079 | 0 | 0.27 | 0.06 | 0.14 | 0.14 | 0.24 | 0.16 |
| 4 | 113264524 | rs2240033 | 0.056 | -0.105 | -1.079 | -1.627 | 0.004 | -0.027 | -0.027 | 0 | 0.57 | 0.53 | 0.66 | 0.78 | 0.5 | 0.73 |
| 4 | 113269731 | rs11098152 | -0.29 | -0.052 | -0.665 | -1.307 | 0.019 | -0.071 | -0.071 | 0.341 | 0.74 | 0.58 | 0.79 | 0.92 | 0.71 | 0.81 |
| 4 | 113278511 | rs4834267 | -0.048 | -0.439 | -0.691 | -1.105 | -0.009 | 0.009 | 0.009 | 0.935 | 0.32 | 0.36 | 0.49 | 0.63 | 0.31 | 0.55 |
| 4 | 113280747 | rs1009475 | 0.581 | 0.093 | -0.039 | -0.658 | 0.013 | 0.075 | 0.075 | 0.415 | 0.79 | 0.55 | 0.76 | 0.69 | 0.74 | 0.69 |
| 4 | 113282802 | rs2097745 | -0.063 | 0.101 | 0.012 | -0.625 | 0 | 0 | 0 | 0.161 | 0.87 | 0.86 | 0.85 | 0.88 | 0.82 | 0.81 |
| 4 | 113290045 | rs1859140 | -0.11 | 0.095 | 0.225 | -0.614 | 0.005 | 0.099 | 0.099 | 0 | 0.08 | 0.24 | 0.08 | 0.19 | 0.08 | 0.13 |
| 4 | 113293314 | rs7694057 | 0.341 | 0.096 | -0.044 | -0.345 | 0 | 0.003 | 0.003 | 0.24 | 0.84 | 0.83 | 0.84 | 0.78 | 0.82 | 0.75 |
| 4 | 113298305 | rs2023824 | -0.699 | 0.086 | 0.195 | 0.039 | -0.008 | 0.059 | 0.059 | 0 | 0.05 | 0.18 | 0.02 | 0.09 | 0.01 | 0.06 |
| 4 | 113302357 | rs2023825 | -0.988 | 0.237 | -0.076 | -0.006 | 0 | 0.001 | 0.001 | 0 | 0.91 | 0.92 | 0.89 | 0.89 | 0.86 | 0.7 |
| 4 | 113302869 | rs7673053 | -0.732 | -0.363 | -0.154 | -0.485 | 0.048 | 0.108 | 0.108 | 2.557 | 0.47 | 0.71 | 0.6 | 0.83 | 0.41 | 0.58 |
| 4 | 113306879 | rs2051777 | 0.448 | -0.481 | -0.138 | -0.214 | -0.001 | -0.001 | -0.001 | 0 | 0.32 | 0.36 | 0.53 | 0.73 | 0.35 | 0.47 |
| 4 | 113310981 | rs965013 | -1.012 | -0.436 | -0.697 | -0.353 | 0.029 | 0.163 | 0.163 | 1.78 | 0.53 | 0.81 | 0.65 | 0.77 | 0.46 | 0.62 |
| 4 | 113311741 | rs7664678 | 0.967 | -0.141 | -0.387 | 0.03 | -0.001 | 0.009 | 0.009 | 0 | 0.28 | 0.28 | 0.48 | 0.54 | 0.31 | 0.42 |
| 4 | 113317475 | rs2882661 | -0.956 | 0.117 | -0.697 | -0.328 | -0.019 | 0.002 | 0.002 | 0.846 | 0.79 | 0.89 | 0.82 | 0.79 | 0.74 | 0.74 |
| 4 | 113317583 | rs2351458 | -0.589 | -0.127 | -0.441 | -0.324 | 0.016 | 0.046 | 0.046 | 0 | 0.51 | 0.64 | 0.33 | 0.27 | 0.42 | 0.29 |
| 4 | 113320553 | rs7663969 | -0.506 | -0.23 | -0.161 | -0.272 | -0.035 | 0.061 | 0.061 | 0 | 0.15 | 0.37 | 0.12 | 0.17 | 0.09 | 0.13 |
| 4 | 113323634 | rs13104183 | 0.224 | -0.176 | -0.593 | -0.521 | -0.002 | 0.052 | 0.052 | 0 | 0.34 | 0.28 | 0.24 | 0.1 | 0.34 | 0.17 |
| 4 | 113327945 | rs6838405 | -0.568 | -0.185 | -0.52 | 0.175 | -0.003 | -0.002 | -0.002 | 0.175 | 0.44 | 0.52 | 0.34 | 0.36 | 0.32 | 0.27 |
| 4 | 113329345 | rs7687819 | -0.648 | -0.289 | -0.586 | 0.07 | 0.002 | 0.003 | 0.003 | 0.159 | 0.25 | 0.18 | 0.21 | 0.28 | 0.25 | 0.35 |
| 4 | 113329465 | rs7688190 | -1.111 | -0.473 | -0.513 | -0.078 | -0.019 | -0.013 | -0.013 | 0 | 0.41 | 0.45 | 0.33 | 0.38 | 0.33 | 0.41 |
| 4 | 113330772 | rs17628268 | 1.671 | -0.239 | -0.79 | 0.07 | 0.001 | -0.023 | -0.023 | 0 | 0.07 | 0.03 | 0.1 | 0.21 | 0.08 | 0.23 |
| 4 | 113330940 | rs17044631 | 0.622 | -0.359 | -0.536 | -0.295 | -0.023 | -0.016 | -0.016 | 0 | 0.19 | 0.23 | 0.13 | 0.1 | 0.12 | 0.07 |
| 4 | 113331585 | rs6811550 | -0.233 | -0.032 | -0.671 | -0.383 | -0.007 | 0.03 | 0.03 | 0.216 | 0.15 | 0.06 | 0.1 | 0.07 | 0.17 | 0.11 |
| 4 | 113339161 | rs6829461 | -0.897 | 0.058 | -0.632 | -0.026 | -0.002 | -0.011 | -0.011 | 0 | 0.48 | 0.45 | 0.49 | 0.72 | 0.46 | 0.5 |
| 4 | 113341588 | rs7686496 | 0.71 | 0.085 | -0.866 | -0.121 | -0.066 | 0.052 | 0.052 | 0.059 | 0.15 | 0.03 | 0.11 | 0.09 | 0.23 | 0.15 |
| 4 | 113343224 | rs2106988 | -1.719 | 0.033 | -0.447 | -0.127 | 0 | -0.019 | -0.019 | 0 | 0.59 | 0.54 | 0.63 | 0.79 | 0.59 | 0.52 |
| 4 | 113344203 | rs11733852 | 0.503 | -0.001 | -0.679 | -0.018 | -0.001 | 0 | 0 | 0 | 0.13 | 0.1 | 0.15 | 0.08 | 0.16 | 0.03 |
| 4 | 113345147 | rs6533616 | 1.771 | 0.115 | -0.696 | -0.083 | -0.002 | -0.007 | -0.007 | 3.067 | 0.09 | 0.05 | 0.07 | 0 | 0.12 | 0.02 |
| 4 | 113346714 | rs916868 | -1.209 | 0.069 | -0.659 | -0.088 | 0 | -0.002 | -0.002 | 0 | 0.53 | 0.48 | 0.55 | 0.72 | 0.55 | 0.53 |
| 4 | 113346996 | rs10017209 | 0.52 | 0.339 | -0.777 | 0.047 | 0.009 | -0.014 | -0.014 | 0 | 0.16 | 0.15 | 0.12 | 0.09 | 0.23 | 0.15 |
| 4 | 113350665 | rs12649485 | -0.257 | 0.013 | -0.921 | 0.183 | 0.008 | -0.002 | -0.002 | 0 | 0.32 | 0.24 | 0.4 | 0.63 | 0.29 | 0.34 |
| 4 | 113352397 | rs2074388 | -0.879 | 0.475 | -1.256 | 0.286 | 0.029 | 0.021 | 0.021 | 0 | 0.56 | 0.72 | 0.56 | 0.64 | 0.62 | 0.55 |
| 4 | 113352628 | rs13148353 | -1.239 | 0.39 | -1.297 | 0.287 | 0.015 | 0.027 | 0.027 | 0 | 0.61 | 0.75 | 0.58 | 0.7 | 0.65 | 0.56 |
| 4 | 113352899 | rs2074379 | -0.826 | 0.39 | -1.297 | 0.309 | 0.029 | 0.02 | 0.02 | 0 | 0.56 | 0.72 | 0.56 | 0.64 | 0.62 | 0.55 |
| 4 | 113353285 | rs11726117 | -1.233 | 0.401 | -1.293 | 0.315 | 0.015 | 0.027 | 0.027 | 0 | 0.61 | 0.75 | 0.58 | 0.7 | 0.65 | 0.56 |
| 4 | 113354150 | rs13110304 | 0.883 | 0.33 | -1.238 | 0.712 | -0.044 | 0.011 | 0.011 | 0 | 0.26 | 0.56 | 0.21 | 0.2 | 0.16 | 0.2 |
| 4 | 113356927 | rs17628432 | -0.16 | 0.753 | -1.136 | 0.711 | -0.012 | -0.012 | -0.012 | 0.095 | 0.22 | 0.17 | 0.27 | 0.45 | 0.31 | 0.35 |
| 4 | 113356961 | rs2023828 | 0.112 | 0.731 | -1.147 | 0.642 | -0.008 | -0.012 | -0.012 | 0.011 | 0.34 | 0.54 | 0.29 | 0.2 | 0.31 | 0.22 |
| 4 | 113359703 | rs231247 | -1.491 | 0.898 | -1.225 | 0.648 | 0.016 | 0.024 | 0.024 | 0 | 0.61 | 0.75 | 0.57 | 0.7 | 0.66 | 0.57 |
| 4 | 113362463 | rs11730442 | 1.416 | 1.39 | -1.125 | 0.705 | -0.026 | 0.024 | 0.024 | 1.467 | 0.09 | 0.06 | 0.08 | 0 | 0.15 | 0.02 |
| 4 | 113370913 | rs231259 | 0.565 | 1.307 | -1.134 | 0.825 | -0.06 | 0.036 | 0.036 | 0.211 | 0.14 | 0.03 | 0.11 | 0.11 | 0.21 | 0.14 |
| 4 | 113372448 | rs231263 | -0.623 | 1.243 | -1.024 | 0.91 | 0.001 | 0.004 | 0.004 | 0.897 | 0.23 | 0.2 | 0.27 | 0.5 | 0.31 | 0.34 |
| 4 | 113373974 | rs17444829 | 0.963 | 1.18 | -1.118 | 0.653 | -0.007 | -0.003 | -0.003 | 0.729 | 0.1 | 0.07 | 0.08 | 0 | 0.15 | 0.02 |
| 4 | 113374323 | rs6815237 | 0.147 | 1.224 | -1.275 | 0.895 | 0 | 0.019 | 0.019 | 0 | 0.11 | 0.12 | 0.09 | 0.4 | 0.1 | 0.26 |
| 4 | 113375494 | rs17590516 | -0.607 | 1.399 | -1.334 | 0.854 | -0.014 | -0.023 | -0.023 | 0 | 0.17 | 0.13 | 0.16 | 0.41 | 0.23 | 0.27 |
| 4 | 113377758 | rs2074384 | 1.619 | 1.344 | -1.335 | 0.857 | -0.01 | -0.031 | -0.031 | 0.204 | 0.06 | 0.01 | 0.06 | 0.28 | 0.07 | 0.18 |
| 4 | 113379501 | rs879068 | -1.334 | 1.223 | -1.361 | 0.767 | -0.01 | -0.028 | -0.028 | 0.043 | 0.23 | 0.19 | 0.19 | 0.4 | 0.27 | 0.28 |
| 4 | 113380119 | rs7662734 | -1.477 | 1.19 | -1.325 | 0.75 | -0.005 | -0.018 | -0.018 | 0.438 | 0.24 | 0.23 | 0.21 | 0.41 | 0.27 | 0.28 |
| 4 | 113382048 | rs28806888 | -0.896 | 1.081 | -1.332 | 0.773 | 0.018 | -0.025 | -0.025 | 0.734 | 0.18 | 0.14 | 0.12 | 0.4 | 0.14 | 0.27 |
| 4 | 113393443 | rs1486860 | 1.065 | 0.881 | -1.225 | 0.641 | 0 | 0 | 0 | 0 | 0.45 | 0.4 | 0.51 | 0.35 | 0.4 | 0.46 |
| 4 | 113406671 | rs17044746 | 0.816 | 0.945 | -1.208 | 0.641 | 0.003 | 0.012 | 0.012 | 0 | 0.5 | 0.41 | 0.58 | 0.35 | 0.53 | 0.48 |
| 4 | 113409270 | rs11936846 | 0.224 | 0.944 | -1.242 | 0.593 | -0.002 | -0.003 | -0.003 | 0 | 0.58 | 0.49 | 0.66 | 0.64 | 0.61 | 0.66 |
| 4 | 113415402 | rs7699752 | 0.252 | 0.963 | -1.25 | 0.531 | -0.019 | -0.022 | -0.022 | 0 | 0.11 | 0.15 | 0.06 | 0.07 | 0.06 | 0.07 |
| 4 | 113430752 | rs12641188 | 1.201 | 0.908 | -1.339 | 0.453 | -0.003 | 0.004 | 0.004 | 0 | 0.39 | 0.28 | 0.5 | 0.35 | 0.41 | 0.48 |
| 4 | 113433294 | rs6848238 | 1.18 | 0.908 | -1.46 | 0.467 | -0.002 | 0.002 | 0.002 | 0 | 0.39 | 0.29 | 0.5 | 0.35 | 0.41 | 0.48 |
| 4 | 113443282 | rs1994376 | 0.619 | 0.802 | -1.516 | 0.573 | -0.007 | 0.006 | 0.006 | 0 | 0.45 | 0.3 | 0.57 | 0.58 | 0.48 | 0.68 |
| 4 | 113447137 | rs2129405 | 1.05 | 0.816 | -1.528 | 0.478 | -0.003 | -0.004 | -0.004 | 0.258 | 0.38 | 0.3 | 0.49 | 0.41 | 0.4 | 0.48 |
| 4 | 113459405 | rs11726579 | 1.683 | 0.766 | -1.538 | 0.632 | -0.017 | -0.04 | -0.04 | 0 | 0.07 | 0.01 | 0.07 | 0.22 | 0.08 | 0.2 |
| 4 | 113459416 | rs13106629 | 1.019 | 0.838 | -1.499 | 0.617 | -0.003 | -0.004 | -0.004 | 0 | 0.38 | 0.29 | 0.49 | 0.41 | 0.4 | 0.48 |
| 4 | 113465651 | rs1486865 | -1.102 | 0.705 | -1.363 | 0.549 | -0.002 | -0.038 | -0.038 | 0 | 0.63 | 0.61 | 0.74 | 0.94 | 0.56 | 0.8 |
| 4 | 113465971 | rs12054589 | -0.589 | 0.682 | -1.345 | 0.557 | 0.012 | -0.019 | -0.019 | 0.375 | 0.19 | 0.17 | 0.23 | 0.5 | 0.15 | 0.28 |
| 4 | 113472958 | rs13144672 | 1.053 | 0.693 | -1.279 | 0.609 | -0.004 | -0.004 | -0.004 | 0 | 0.39 | 0.29 | 0.5 | 0.41 | 0.42 | 0.51 |
| 4 | 113494607 | rs6533629 | 0.096 | 0.788 | -1.291 | 0.636 | 0.02 | 0.025 | 0.025 | 0 | 0.49 | 0.6 | 0.6 | 0.57 | 0.43 | 0.56 |
| 4 | 113519607 | rs11724428 | -0.388 | 0.699 | -1.372 | 0.694 | 0.015 | -0.02 | -0.02 | 0 | 0.18 | 0.15 | 0.21 | 0.39 | 0.13 | 0.26 |
| 4 | 113533796 | rs9993534 | -0.376 | 0.634 | -1.411 | 0.667 | 0.013 | -0.017 | -0.017 | 0.009 | 0.18 | 0.15 | 0.21 | 0.39 | 0.12 | 0.25 |

| | | | | | | | | | | | | | | | | |
|---|---|---|---|---|---|---|---|---|---|---|---|---|---|---|---|---|
| 4 | 113538280 | rs10002557 | -1.706 | 0.667 | -1.472 | 0.213 | -0.001 | -0.061 | -0.061 | 0.059 | 0.71 | 0.65 | 0.82 | 0.94 | 0.73 | 0.83 |
| 4 | 113539969 | rs7696816 | 0.894 | 0.565 | -1.203 | 0.398 | -0.005 | -0.003 | -0.003 | 0 | 0.39 | 0.3 | 0.51 | 0.41 | 0.43 | 0.51 |
| 4 | 113546107 | rs1471880 | 0.267 | 0.325 | -1.02 | 0.585 | -0.001 | 0.003 | 0.003 | 0 | 0.45 | 0.47 | 0.53 | 0.44 | 0.44 | 0.54 |
| 4 | 113549989 | rs12054518 | 0.89 | 0.444 | -0.961 | 0.552 | -0.005 | -0.004 | -0.004 | 0 | 0.39 | 0.3 | 0.51 | 0.41 | 0.43 | 0.51 |
| 4 | 113558602 | rs13146963 | -1.176 | 0.542 | -1.494 | 0.153 | -0.004 | -0.034 | -0.034 | 0 | 0.63 | 0.61 | 0.74 | 0.94 | 0.56 | 0.8 |
| 4 | 113570885 | rs7656819 | 0.645 | -0.181 | -1.257 | 0.537 | 0.056 | 0.12 | 0.12 | 0.545 | 0.06 | 0.36 | 0.04 | 0.06 | 0.01 | 0.01 |
| 4 | 113578266 | rs6533631 | -0.408 | 0.021 | -1.415 | 0.548 | 0.025 | -0.01 | -0.01 | 0 | 0.2 | 0.15 | 0.17 | 0.21 | 0.09 | 0.18 |
| 4 | 113578609 | rs1129065 | -0.675 | 0.066 | -1.394 | 0.536 | 0.036 | -0.051 | -0.051 | 0.195 | 0.22 | 0.16 | 0.22 | 0.5 | 0.13 | 0.29 |
| 4 | 113579162 | rs12644868 | -1.072 | 0.787 | -1.111 | 0.065 | 0.004 | -0.068 | -0.068 | 2.195 | 0.69 | 0.63 | 0.81 | 0.95 | 0.68 | 0.81 |
| 4 | 113609023 | rs7664015 | -0.18 | 1.08 | -1.573 | 1.003 | -0.005 | -0.022 | -0.022 | 0 | 0.14 | 0.06 | 0.16 | 0.21 | 0.13 | 0.22 |
| 4 | 113617694 | rs6533636 | 0.158 | 1.375 | -1.194 | 1.513 | 0.015 | 0.072 | 0.072 | 0 | 0.92 | 0.8 | 0.79 | 0.68 | 0.88 | 0.9 |
| 4 | 113622050 | rs11731671 | 0.432 | 1.365 | -2.025 | 1.151 | -0.09 | -0.082 | -0.082 | 0.414 | 0.31 | 0.08 | 0.37 | 0.35 | 0.39 | 0.34 |
| 4 | 113658555 | rs10021207 | -1.033 | 0.47 | -1.867 | 0.183 | -0.039 | 0.211 | 0.211 | 0 | 0.54 | 0.84 | 0.73 | 0.91 | 0.54 | 0.63 |
| 4 | 113674445 | rs6855396 | 1.058 | 0.2 | -1.019 | 0.154 | -0.028 | -0.013 | -0.013 | 0.98 | 0.09 | 0.16 | 0.04 | 0.03 | 0.03 | 0.09 |
| 4 | 113685804 | rs7654314 | 0.489 | 0.06 | -0.665 | 0.249 | -0.012 | -0.034 | -0.034 | 0 | 0.3 | 0.23 | 0.33 | 0.36 | 0.33 | 0.33 |
| 4 | 113698005 | rs17607420 | 0.796 | 0.051 | -1.041 | -0.186 | -0.019 | 0.048 | 0.048 | 0 | 0.13 | 0.02 | 0.07 | 0.05 | 0.12 | 0.12 |
| 4 | 113710173 | rs11098171 | -0.785 | -0.076 | -0.227 | -0.023 | -0.016 | 0.117 | 0.117 | 0 | 0.21 | 0.51 | 0.38 | 0.58 | 0.18 | 0.29 |
| 4 | 113719791 | rs6829775 | 0.272 | -0.201 | 0.393 | -0.09 | -0.008 | 0.011 | 0.011 | 0 | 0.35 | 0.56 | 0.38 | 0.41 | 0.34 | 0.35 |
| 4 | 113724018 | rs10434020 | -1.141 | -0.191 | 0.32 | -0.073 | -0.001 | 0.082 | 0.082 | 0 | 0.5 | 0.64 | 0.68 | 0.88 | 0.5 | 0.62 |
| 4 | 113737479 | rs11732297 | 0.439 | -0.299 | 0.131 | -0.075 | -0.053 | -0.039 | -0.039 | 0.065 | 0.13 | 0.31 | 0.05 | 0.03 | 0.05 | 0.09 |
| 4 | 113762103 | rs10516590 | 0.67 | -0.25 | -0.431 | -0.074 | -0.013 | -0.021 | -0.021 | 0 | 0.31 | 0.25 | 0.34 | 0.36 | 0.34 | 0.34 |
| 4 | 113765546 | rs10013796 | 0.464 | -0.388 | -0.476 | -0.447 | -0.04 | -0.03 | -0.03 | 0.476 | 0.12 | 0.22 | 0.05 | 0.03 | 0.05 | 0.09 |
| 4 | 113772458 | rs11098178 | -1.425 | -0.087 | -1.068 | -0.422 | -0.044 | 0.165 | 0.165 | 0 | 0.64 | 0.89 | 0.78 | 0.95 | 0.62 | 0.73 |
| 4 | 113775868 | rs17672142 | 0.037 | -0.5 | -1.153 | -0.288 | 0.005 | 0.044 | 0.044 | 0 | 0.28 | 0.16 | 0.25 | 0.15 | 0.28 | 0.11 |
| 4 | 113775901 | rs17672170 | -1.216 | -0.245 | -0.848 | -0.246 | -0.001 | 0.006 | 0.006 | 0 | 0.63 | 0.67 | 0.76 | 0.76 | 0.63 | 0.55 |
| 4 | 113779891 | rs1448221 | -1.889 | -1.104 | -0.598 | -0.391 | -0.01 | 0.159 | 0.159 | 0 | 0.49 | 0.74 | 0.67 | 0.85 | 0.51 | 0.53 |
| 4 | 113781667 | rs17045073 | 0.186 | -1.086 | -0.37 | -0.336 | -0.018 | -0.03 | -0.03 | 0 | 0.06 | 0.13 | 0.04 | 0 | 0.03 | 0.01 |
| 4 | 113782926 | rs895763 | -1.329 | -0.976 | -0.6 | -0.287 | 0.007 | 0.072 | 0.072 | 0 | 0.43 | 0.7 | 0.62 | 0.82 | 0.45 | 0.5 |
| 4 | 113784482 | rs7657978 | 0.502 | -0.565 | -1.434 | -0.113 | -0.001 | 0.009 | 0.009 | 0 | 0.19 | 0.15 | 0.2 | 0.14 | 0.24 | 0.08 |
| 4 | 113789510 | rs6835163 | -1.292 | -0.304 | -1.248 | 0.312 | -0.126 | 0.212 | 0.212 | 0 | 0.64 | 0.97 | 0.72 | 0.94 | 0.61 | 0.71 |
| 4 | 113808126 | rs4834308 | -0.588 | -0.086 | 0.212 | 2.004 | -0.008 | 0.049 | 0.049 | 0 | 0.06 | 0.05 | 0.27 | 0.35 | 0.16 | 0.21 |
| 4 | 113817332 | rs17045113 | -1.229 | -1.115 | 0.581 | 1.931 | -0.032 | 0.15 | 0.15 | 0 | 0.08 | 0.19 | 0.1 | 0.34 | 0.02 | 0.26 |
| 4 | 113817807 | rs12513140 | -0.693 | -1.073 | 0.579 | 1.951 | -0.024 | 0.152 | 0.152 | 0 | 0.06 | 0.16 | 0.09 | 0.34 | 0.02 | 0.25 |
| 4 | 113825869 | rs7675254 | -0.906 | -1.373 | 0.562 | 1.547 | -0.028 | -0.029 | -0.029 | 0 | 0.25 | 0.58 | 0.19 | 0.13 | 0.19 | 0.08 |
| 4 | 113830885 | rs10016365 | -0.893 | -1.372 | 0.573 | 1.547 | -0.03 | -0.03 | -0.03 | 0.728 | 0.25 | 0.58 | 0.15 | 0.12 | 0.19 | 0.08 |
| 4 | 113835185 | rs7680555 | -0.482 | -1.347 | 1.013 | 1.527 | 0.011 | -0.007 | -0.007 | 0 | 0.07 | 0.26 | 0.04 | 0 | 0.03 | 0 |
| 4 | 113858241 | rs4404544 | -1.514 | -1.431 | 0.106 | 1.801 | 0.035 | 0.005 | 0.005 | 0.229 | 0.35 | 0.67 | 0.3 | 0.64 | 0.24 | 0.41 |
| 4 | 113866368 | rs10015915 | -1.145 | -1.358 | 0.066 | 1.712 | 0 | -0.001 | -0.001 | 0 | 0.31 | 0.62 | 0.22 | 0.29 | 0.23 | 0.15 |
| 4 | 113866381 | rs12506747 | -0.334 | -1.247 | -0.574 | 1.295 | -0.002 | -0.002 | -0.002 | 0 | 0.21 | 0.22 | 0.16 | 0.13 | 0.18 | 0.08 |
| 4 | 113867271 | rs9996317 | -1.395 | -1.122 | -1.103 | 0.906 | 0.12 | 0.094 | 0.094 | 0.308 | 0.42 | 0.87 | 0.34 | 0.71 | 0.28 | 0.46 |
| 4 | 113868607 | rs7678728 | -0.828 | -0.404 | -0.374 | 1.134 | 0.003 | -0.008 | -0.008 | 0 | 0.32 | 0.69 | 0.44 | 0.47 | 0.29 | 0.36 |
| 4 | 113915647 | rs6820742 | 0.525 | -0.396 | -0.08 | 0.633 | -0.002 | -0.012 | -0.012 | 0 | 0.75 | 0.72 | 0.85 | 0.9 | 0.77 | 0.85 |
| 4 | 113916575 | rs12503954 | 0.711 | -0.379 | -0.021 | 1.072 | -0.021 | -0.003 | -0.003 | 0.182 | 0.63 | 0.39 | 0.77 | 0.66 | 0.67 | 0.74 |
| 4 | 113918544 | rs6818050 | -1.184 | 0.014 | 0.375 | 1.569 | 0.001 | 0.071 | 0.071 | 0.018 | 0.27 | 0.47 | 0.29 | 0.42 | 0.29 | 0.4 |
| 4 | 113918577 | rs1354680 | -1.612 | -0.49 | 1.217 | 1.878 | 0.001 | -0.003 | -0.003 | 0 | 0.33 | 0.68 | 0.32 | 0.54 | 0.3 | 0.51 |
| 4 | 113921105 | rs6812106 | -1.526 | -0.689 | 2.009 | 2.144 | -0.006 | 0.071 | 0.071 | 0.07 | 0.18 | 0.32 | 0.13 | 0.28 | 0.16 | 0.23 |
| 4 | 113921253 | rs7655703 | -2.329 | -0.788 | 1.954 | 2.109 | -0.025 | 0.034 | 0.034 | 0 | 0.32 | 0.57 | 0.28 | 0.4 | 0.27 | 0.4 |
| 4 | 113924858 | rs6833846 | 0.833 | -0.503 | 1.812 | 1.984 | 0.048 | 0.032 | 0.032 | 0.114 | 0.86 | 0.68 | 0.86 | 0.85 | 0.81 | 0.87 |
| 4 | 113927365 | rs2174558 | -1.288 | -0.533 | 1.584 | 2.169 | 0.011 | -0.016 | -0.016 | 2.924 | 0.16 | 0.15 | 0.15 | 0.37 | 0.13 | 0.39 |
| 4 | 113931773 | rs7687726 | 0.697 | -0.445 | 1.905 | 2.13 | 0.007 | 0.002 | 0.002 | 0.07 | 0.87 | 0.82 | 0.87 | 0.85 | 0.8 | 0.88 |
| 4 | 113935712 | rs1506064 | 1.98 | -0.568 | 2.14 | 1.975 | 0.017 | -0.033 | -0.033 | 0 | 0.66 | 0.24 | 0.69 | 0.47 | 0.68 | 0.48 |
| 4 | 113948790 | rs10014072 | 1.324 | -0.968 | 2.198 | 2.036 | -0.035 | 0.068 | 0.068 | 0 | 0.69 | 0.54 | 0.77 | 0.48 | 0.67 | 0.51 |
| 4 | 113948864 | rs13103796 | 0.863 | -1.43 | 1.339 | 1.224 | 0.023 | -0.02 | -0.02 | 0 | 0.61 | 0.24 | 0.57 | 0.49 | 0.73 | 0.68 |
| 4 | 113951572 | rs10155397 | 0.822 | -1.452 | 1.418 | 1.204 | 0.003 | -0.004 | -0.004 | 2.452 | 0.61 | 0.32 | 0.58 | 0.49 | 0.73 | 0.67 |
| 4 | 113951743 | rs17045384 | -0.433 | -1.466 | 1.414 | 1.186 | 0.005 | -0.053 | -0.053 | 1.294 | 0.2 | 0.09 | 0.16 | 0.4 | 0.25 | 0.36 |
| 4 | 113953383 | rs13115459 | 0.803 | -1.488 | 1.332 | 1.191 | -0.002 | 0 | 0 | 0 | 0.62 | 0.33 | 0.58 | 0.49 | 0.73 | 0.68 |
| 4 | 113953581 | rs1395114 | 0.121 | -1.525 | 1.253 | 1.085 | 0.012 | -0.079 | -0.079 | 0.151 | 0.24 | 0.1 | 0.32 | 0.4 | 0.17 | 0.22 |
| 4 | 113954301 | rs12504906 | 0.624 | -1.744 | 1.077 | 1.078 | 0.016 | 0.003 | 0.003 | 0.017 | 0.1 | 0.06 | 0.07 | 0.08 | 0.02 | 0.06 |
| 4 | 113954926 | rs1354681 | -0.056 | -1.749 | 1.232 | 1.054 | -0.009 | -0.069 | -0.069 | 0.284 | 0.88 | 0.68 | 0.92 | 0.98 | 0.9 | 0.88 |
| 4 | 113960961 | rs10015472 | 1.035 | -1.548 | 1.02 | 0.989 | -0.01 | 0.02 | 0.02 | 0.132 | 0.65 | 0.43 | 0.74 | 0.49 | 0.63 | 0.51 |
| 4 | 113961039 | rs10015551 | 1.617 | -1.465 | 0.92 | 0.713 | 0.015 | 0.007 | 0.007 | 0 | 0.55 | 0.39 | 0.68 | 0.4 | 0.61 | 0.46 |
| 4 | 113963746 | rs7654153 | 0.071 | -1.088 | 0.848 | 1.017 | -0.004 | -0.062 | -0.062 | 0 | 0.89 | 0.73 | 0.92 | 0.98 | 0.9 | 0.88 |
| 4 | 113964288 | rs716051 | -0.174 | -0.806 | 0.204 | 1.321 | 0 | 0.003 | 0.003 | 0 | 0.16 | 0.18 | 0.27 | 0.33 | 0.15 | 0.16 |
| 4 | 113967494 | rs313938 | -0.173 | -0.761 | 0.245 | 1.104 | 0 | -0.001 | -0.001 | 0.252 | 0.11 | 0.12 | 0.02 | 0.12 | 0.04 | 0.04 |
| 4 | 113967603 | rs313937 | 0.659 | -0.892 | 0.178 | 0.763 | 0.003 | -0.003 | -0.003 | 0 | 0.57 | 0.42 | 0.48 | 0.47 | 0.52 | 0.45 |
| 4 | 113968160 | rs6856101 | 1.183 | -0.681 | 0.709 | 0.05 | -0.005 | -0.07 | -0.07 | 0 | 0.74 | 0.45 | 0.8 | 0.93 | 0.74 | 0.7 |
| 4 | 113968893 | rs313936 | -1.387 | -0.837 | 0.823 | 0.572 | -0.036 | 0.146 | 0.146 | 0 | 0.15 | 0.4 | 0.07 | 0.27 | 0.09 | 0.12 |
| 4 | 113969404 | rs13128057 | 1.663 | -0.795 | 0.539 | -0.002 | 0.02 | -0.007 | -0.007 | 0 | 0.58 | 0.29 | 0.64 | 0.6 | 0.6 | 0.6 |
| 4 | 113969945 | rs7663732 | 0.543 | -0.352 | 0.394 | -0.122 | -0.017 | 0.003 | 0.003 | 0 | 0.82 | 0.91 | 0.8 | 0.86 | 0.77 | 0.7 |

| | | | | | | | | | | | | | | | | |
|---|---|---|---|---|---|---|---|---|---|---|---|---|---|---|---|---|
| 4 | 113971374 | rs1026975 | -1.558 | -0.327 | 0.965 | -0.055 | -0.019 | -0.023 | -0.023 | 0.115 | 0.23 | 0.36 | 0.14 | 0.17 | 0.19 | 0.16 |
| 4 | 113972856 | rs313942 | 1.743 | -0.591 | 0.895 | -0.234 | -0.01 | 0.011 | 0.011 | 0 | 0.64 | 0.54 | 0.71 | 0.61 | 0.66 | 0.7 |
| 4 | 113973952 | rs931838 | 2.232 | -0.449 | 0.544 | -0.224 | 0.024 | 0.004 | 0.004 | 0 | 0.54 | 0.2 | 0.61 | 0.59 | 0.57 | 0.59 |
| 4 | 113976831 | rs10007543 | -1.19 | -0.017 | 0.424 | -0.558 | -0.003 | -0.087 | -0.087 | 0.064 | 0.14 | 0.36 | 0.13 | 0.01 | 0.15 | 0.09 |
| 4 | 113978279 | rs313985 | 0.402 | 0.299 | -0.246 | -0.444 | -0.024 | -0.005 | -0.005 | 0 | 0.84 | 0.89 | 0.8 | 0.86 | 0.78 | 0.77 |
| 4 | 113981730 | rs313980 | 0.519 | 0.316 | -0.312 | -0.443 | -0.033 | 0.008 | 0.008 | 0 | 0.85 | 0.93 | 0.86 | 0.89 | 0.81 | 0.83 |
| 4 | 113986740 | rs678035 | 0.066 | 0.385 | -0.275 | -0.438 | -0.035 | 0.009 | 0.009 | 0 | 0.87 | 0.95 | 0.87 | 0.89 | 0.82 | 0.87 |
| 4 | 113988513 | rs632895 | -1.362 | 0.276 | -0.066 | -0.524 | -0.014 | -0.076 | -0.076 | 0 | 0.19 | 0.52 | 0.19 | 0.04 | 0.17 | 0.17 |
| 4 | 113989119 | rs689117 | 0.381 | 0.252 | -0.124 | -0.532 | -0.035 | 0.003 | 0.003 | 0 | 0.83 | 0.91 | 0.78 | 0.86 | 0.76 | 0.8 |
| 4 | 113989326 | rs582050 | -1.322 | 0.016 | 0.31 | -0.565 | -0.027 | -0.123 | -0.123 | 0.157 | 0.17 | 0.49 | 0.12 | 0.01 | 0.13 | 0.11 |
| 4 | 113991016 | rs2279891 | -0.534 | -0.076 | -0.518 | -0.638 | -0.021 | -0.037 | -0.037 | 0.012 | 0.36 | 0.38 | 0.21 | 0.2 | 0.26 | 0.35 |
| 4 | 113998638 | rs884555 | 0.415 | 0.8 | -1.875 | -0.743 | -0.163 | 0.12 | 0.12 | 0 | 0.39 | 0.04 | 0.49 | 0.25 | 0.46 | 0.29 |
| 4 | 114001887 | rs2135354 | -0.069 | 0.476 | -1.398 | -0.565 | 0.038 | 0.052 | 0.052 | 0 | 0.45 | 0.64 | 0.29 | 0.25 | 0.34 | 0.46 |
| 4 | 114008326 | rs12500552 | 0.011 | 1.258 | -1.883 | -0.813 | -0.05 | -0.027 | -0.027 | 0 | 0.3 | 0.22 | 0.36 | 0.58 | 0.44 | 0.35 |
| 4 | 114009586 | rs7656666 | 1.129 | 0.776 | -1.748 | -1.011 | -0.038 | -0.033 | -0.033 | 0.586 | 0.34 | 0.46 | 0.21 | 0.24 | 0.23 | 0.4 |
| 4 | 114009885 | rs649022 | -0.818 | 1.458 | -1.933 | -1.213 | -0.12 | 0.141 | 0.141 | 0 | 0.64 | 0.96 | 0.56 | 0.84 | 0.61 | 0.78 |
| 4 | 114010341 | rs6819998 | 0.981 | 1.114 | -1.829 | -1.102 | 0.003 | 0.023 | 0.023 | 0 | 0.15 | 0.17 | 0.24 | 0.6 | 0.18 | 0.24 |
| 4 | 114011088 | rs10029516 | 0.314 | 1.125 | -1.819 | -1.13 | 0 | -0.01 | -0.01 | 0 | 0.48 | 0.47 | 0.32 | 0.21 | 0.48 | 0.5 |
| 4 | 114013008 | rs17045537 | 1.344 | 1.091 | -1.824 | -1.137 | -0.038 | -0.072 | -0.072 | 0.426 | 0.33 | 0.45 | 0.18 | 0.14 | 0.23 | 0.39 |
| 4 | 114013286 | rs592670 | -0.058 | 1.251 | -2.032 | -1.081 | 0.023 | 0.076 | 0.076 | 0 | 0.63 | 0.72 | 0.57 | 0.82 | 0.67 | 0.75 |
| 4 | 114013698 | rs7683186 | 0.077 | 1.249 | -2.026 | -1.065 | 0.013 | 0.019 | 0.019 | 0 | 0.62 | 0.64 | 0.54 | 0.72 | 0.67 | 0.74 |
| 4 | 114014365 | rs17045555 | 1.039 | 1.234 | -2.024 | -1.079 | -0.002 | -0.007 | -0.007 | 0 | 0.11 | 0.15 | 0.07 | 0.02 | 0.1 | 0.07 |
| 4 | 114014463 | rs684932 | 0.176 | 1.249 | -2.03 | -1.071 | -0.025 | 0.003 | 0.003 | 0.076 | 0.14 | 0.33 | 0.11 | 0.12 | 0.1 | 0.08 |
| 4 | 114016236 | rs17476342 | 1.089 | 1.282 | -2.084 | -1.084 | -0.101 | 0.065 | 0.065 | 0 | 0.16 | 0.02 | 0.14 | 0.08 | 0.26 | 0.11 |
| 4 | 114016978 | rs626267 | -1.124 | 1.141 | -1.913 | -0.927 | -0.029 | 0.046 | 0.046 | 0 | 0.76 | 0.97 | 0.65 | 0.84 | 0.77 | 0.82 |
| 4 | 114017352 | rs17476377 | -0.03 | 1.151 | -1.901 | -0.92 | -0.027 | 0.045 | 0.045 | 0 | 0.28 | 0.27 | 0.23 | 0.1 | 0.35 | 0.18 |
| 4 | 114017964 | rs695002 | 0.527 | 1.213 | -1.998 | -0.877 | 0.034 | 0.104 | 0.104 | 0.011 | 0.48 | 0.74 | 0.41 | 0.64 | 0.41 | 0.63 |
| 4 | 114020087 | rs6851877 | -0.357 | 1.216 | -1.513 | -0.942 | -0.007 | 0.006 | 0.006 | 0 | 0.29 | 0.31 | 0.25 | 0.2 | 0.36 | 0.19 |
| 4 | 114023849 | rs313956 | -0.201 | 0.666 | -1.769 | -1.088 | -0.018 | 0.116 | 0.116 | 0 | 0.75 | 0.95 | 0.88 | 0.94 | 0.76 | 0.83 |
| 4 | 114024092 | rs313958 | -0.011 | 0.46 | -1.645 | -1.329 | -0.004 | 0.013 | 0.013 | 0 | 0.22 | 0.18 | 0.12 | 0.13 | 0.2 | 0.24 |
| 4 | 114025922 | rs9759975 | -0.225 | 0.061 | -1.203 | -1.271 | 0.031 | 0.024 | 0.024 | 0 | 0.56 | 0.68 | 0.74 | 0.66 | 0.68 | 0.69 |
| 4 | 114026519 | rs1385662 | 0.443 | 0.513 | -0.769 | -1.314 | 0.044 | -0.002 | -0.002 | 0.099 | 0.11 | 0.2 | 0.13 | 0.12 | 0.19 | 0.13 |
| 4 | 114027334 | rs4834315 | -0.423 | 0.517 | -0.799 | -1.32 | 0.031 | 0.023 | 0.023 | 0 | 0.57 | 0.68 | 0.75 | 0.66 | 0.68 | 0.69 |
| 4 | 114030580 | rs7679106 | 0.363 | 0.478 | -0.541 | -1.33 | -0.015 | 0.003 | 0.003 | 0 | 0.44 | 0.34 | 0.6 | 0.53 | 0.48 | 0.56 |
| 4 | 114032596 | rs11731561 | 0.339 | 0.574 | -0.544 | -1.269 | 0.067 | 0.004 | 0.004 | 0.007 | 0.12 | 0.23 | 0.13 | 0.12 | 0.19 | 0.13 |
| 4 | 114033802 | rs7699082 | -0.324 | 0.205 | -0.402 | -1.247 | 0.032 | 0.025 | 0.025 | 0 | 0.57 | 0.71 | 0.74 | 0.66 | 0.68 | 0.69 |
| 4 | 114035610 | rs17045612 | 0.774 | -0.161 | -0.638 | -1.476 | 0.011 | -0.006 | -0.006 | 0 | 0.29 | 0.22 | 0.2 | 0.36 | 0.22 | 0.27 |
| 4 | 114038963 | rs10022885 | -0.417 | 0.398 | -0.306 | -1.055 | 0.028 | 0.019 | 0.019 | 0 | 0.56 | 0.66 | 0.74 | 0.64 | 0.67 | 0.69 |
| 4 | 114044885 | rs12502247 | 0.473 | 0.416 | -0.336 | -1.31 | -0.005 | 0.066 | 0.066 | 0 | 0.16 | 0.27 | 0.4 | 0.49 | 0.13 | 0.35 |
| 4 | 114047406 | rs1506063 | 0.124 | 0.392 | -0.391 | -1.419 | -0.001 | -0.005 | -0.005 | 0 | 0.41 | 0.38 | 0.34 | 0.15 | 0.55 | 0.39 |
| 4 | 114051112 | rs13149379 | -0.777 | 0.486 | -0.527 | -1.34 | 0.041 | 0.03 | 0.03 | 0 | 0.57 | 0.72 | 0.76 | 0.66 | 0.68 | 0.69 |
| 4 | 114052710 | rs10003143 | 0.054 | 0.467 | -0.489 | -1.359 | 0 | -0.051 | -0.051 | 0 | 0.41 | 0.5 | 0.35 | 0.16 | 0.55 | 0.34 |
| 4 | 114058373 | rs7664927 | 0.595 | 0.221 | -0.52 | -0.952 | -0.009 | 0.099 | 0.099 | 0.792 | 0.1 | 0.19 | 0.38 | 0.51 | 0.08 | 0.32 |
| 4 | 114061536 | rs17676256 | 0.249 | -0.005 | -0.663 | -1.08 | 0.009 | -0.012 | -0.012 | 0 | 0.19 | 0.19 | 0.41 | 0.53 | 0.12 | 0.45 |
| 4 | 114062948 | rs10020548 | 0.5 | 0.63 | -0.265 | -0.715 | -0.011 | 0.171 | 0.171 | 1.086 | 0.1 | 0.23 | 0.36 | 0.49 | 0.08 | 0.28 |
| 4 | 114063451 | rs10021192 | -0.117 | 1.046 | -0.507 | -0.622 | 0 | 0.002 | 0.002 | 0 | 0.49 | 0.49 | 0.63 | 0.54 | 0.52 | 0.55 |
| 4 | 114066333 | rs17579878 | -0.053 | 1.342 | -0.666 | -0.544 | 0.052 | -0.011 | -0.011 | 0 | 0.12 | 0.26 | 0.12 | 0.11 | 0.17 | 0.12 |
| 4 | 114067139 | rs3112980 | 0.473 | 1.351 | -1.195 | -0.476 | -0.031 | 0.053 | 0.053 | 0 | 0.13 | 0.02 | 0.07 | 0.02 | 0.15 | 0.19 |
| 4 | 114067233 | rs3112981 | 0.024 | 1.189 | -0.73 | -0.295 | 0.006 | 0.015 | 0.015 | 0 | 0.36 | 0.48 | 0.55 | 0.53 | 0.37 | 0.3 |
| 4 | 114068831 | rs13146022 | -0.596 | 1.509 | -0.638 | -0.499 | 0.001 | -0.001 | -0.001 | 0.493 | 0.18 | 0.16 | 0.14 | 0.11 | 0.24 | 0.31 |
| 4 | 114078457 | rs17482885 | -0.368 | 1.478 | -0.01 | -0.537 | 0.017 | -0.023 | -0.023 | 0 | 0.1 | 0.16 | 0.09 | 0.05 | 0.11 | 0.14 |
| 4 | 114078812 | rs10516593 | -0.812 | 1.295 | -0.702 | -0.26 | 0.014 | 0.031 | 0.031 | 0 | 0.55 | 0.84 | 0.64 | 0.59 | 0.62 | 0.51 |
| 4 | 114078931 | rs13134375 | 0.72 | 1.197 | -0.552 | -0.494 | -0.038 | 0.084 | 0.084 | 0 | 0.31 | 0.17 | 0.2 | 0.04 | 0.38 | 0.21 |
| 4 | 114080240 | rs17590572 | 0.268 | 0.746 | -0.227 | -0.132 | -0.001 | 0.005 | 0.005 | 0.133 | 0.26 | 0.29 | 0.44 | 0.55 | 0.25 | 0.41 |
| 4 | 114082021 | rs17590593 | -0.088 | 0.556 | -0.322 | 0.007 | -0.006 | -0.03 | -0.03 | 0 | 0.34 | 0.22 | 0.39 | 0.46 | 0.35 | 0.45 |
| 4 | 114082323 | rs4627864 | -0.345 | 0.719 | 0.127 | -0.207 | 0.013 | -0.012 | -0.012 | 3.905 | 0.38 | 0.42 | 0.4 | 0.32 | 0.43 | 0.45 |
| 4 | 114086121 | rs7697503 | -1.585 | 0.069 | -0.127 | -0.247 | -0.003 | 0.013 | 0.013 | 0 | 0.35 | 0.66 | 0.23 | 0.1 | 0.36 | 0.28 |
| 4 | 114086242 | rs404950 | -1.591 | 0.043 | -0.211 | -0.224 | -0.01 | -0.109 | -0.109 | 0 | 0.63 | 0.91 | 0.57 | 0.46 | 0.67 | 0.65 |
| 4 | 114087355 | rs6814622 | -0.447 | 0.162 | 0.498 | -0.311 | 0.077 | -0.078 | -0.078 | 0 | 0.14 | 0.34 | 0.11 | 0.02 | 0.19 | 0.09 |
| 4 | 114089743 | rs4833422 | -0.558 | -0.125 | 0.009 | -0.139 | -0.023 | -0.04 | -0.04 | 3.318 | 0.19 | 0.24 | 0.09 | 0.04 | 0.13 | 0.17 |
| 4 | 114092317 | rs399754 | -1.046 | -0.089 | -0.32 | 0.118 | 0.026 | -0.061 | -0.061 | 0.545 | 0.62 | 0.84 | 0.57 | 0.46 | 0.68 | 0.65 |
| 4 | 114105776 | rs6858425 | 0.541 | 0.318 | 1.064 | 0.189 | 0.047 | -0.062 | -0.062 | 0 | 0.26 | 0.43 | 0.26 | 0.14 | 0.29 | 0.34 |
| 4 | 114108991 | rs4834321 | -1.294 | 0.206 | 1.127 | 0.539 | 0.001 | 0.032 | 0.032 | 0.001 | 0.47 | 0.7 | 0.47 | 0.43 | 0.5 | 0.52 |
| 4 | 114109593 | rs11933707 | -1.498 | -0.294 | 1.381 | 0.725 | -0.003 | 0.017 | 0.017 | 0 | 0.16 | 0.27 | 0.21 | 0.29 | 0.19 | 0.15 |
| 4 | 114117472 | rs2285710 | -1.181 | -0.47 | 1.349 | 0.746 | -0.006 | 0.004 | 0.004 | 0 | 0.14 | 0.2 | 0.17 | 0.23 | 0.14 | 0.14 |
| 4 | 114118650 | rs967099 | -0.937 | -0.565 | 1.642 | 0.948 | 0.036 | -0.003 | -0.003 | 0 | 0.51 | 0.69 | 0.53 | 0.53 | 0.59 | 0.59 |
| 4 | 114122971 | rs6533674 | 0.8 | -0.55 | 1.649 | 0.985 | -0.08 | 0.006 | 0.006 | 0 | 0.31 | 0.17 | 0.3 | 0.23 | 0.41 | 0.42 |
| 4 | 114127850 | rs11943237 | -0.641 | -0.634 | 2.027 | 1.177 | -0.001 | 0.029 | 0.029 | 0 | 0.11 | 0.2 | 0.15 | 0.25 | 0.12 | 0.14 |
| 4 | 114135827 | rs17629019 | -0.376 | -0.607 | 1.944 | 1.34 | 0.013 | 0.034 | 0.034 | 0.725 | 0.1 | 0.1 | 0.19 | 0.26 | 0.15 | 0.2 |

| | | | | | | | | | | | | | | | | |
|---|---|---|---|---|---|---|---|---|---|---|---|---|---|---|---|---|
| 4 | 114139428 | rs362492 | 0.074 | -0.689 | 1.885 | 1.341 | 0.001 | 0.009 | 0.009 | 0.43 | 0.09 | 0.06 | 0.15 | 0.19 | 0.12 | 0.19 |
| 4 | 114164585 | rs362503 | 0.626 | -0.899 | 1.731 | 1.126 | -0.088 | 0.03 | 0.03 | 0 | 0.38 | 0.12 | 0.32 | 0.24 | 0.45 | 0.42 |
| 4 | 114168845 | rs362496 | -1.065 | -0.911 | 2.11 | 1.151 | -0.014 | -0.012 | -0.012 | 0.174 | 0.61 | 0.58 | 0.77 | 0.78 | 0.72 | 0.79 |
| 4 | 114171605 | rs29436 | -1.067 | -0.873 | 2.074 | 1.16 | -0.031 | -0.028 | -0.028 | 0 | 0.61 | 0.5 | 0.75 | 0.71 | 0.71 | 0.78 |
| 4 | 114193571 | rs29401 | -2.254 | -0.867 | 2.176 | 1.314 | -0.007 | 0.09 | 0.09 | 0.212 | 0.26 | 0.54 | 0.43 | 0.48 | 0.26 | 0.36 |
| 4 | 114194411 | rs17629353 | -0.975 | -0.876 | 2.24 | 1.35 | -0.02 | 0.052 | 0.052 | 0 | 0.09 | 0.29 | 0.04 | 0.19 | 0.05 | 0.05 |
| 4 | 114197064 | rs29394 | -0.893 | -0.874 | 2.241 | 1.351 | -0.018 | 0.055 | 0.055 | 0 | 0.08 | 0.29 | 0.04 | 0.19 | 0.05 | 0.05 |
| 4 | 114201136 | rs29331 | 0.69 | -0.846 | 1.82 | 1.423 | -0.032 | -0.041 | -0.041 | 0 | 0.07 | 0 | 0.23 | 0.18 | 0.08 | 0.17 |
| 4 | 114201353 | rs29382 | -1.561 | -0.864 | 2.076 | 1.228 | 0.016 | 0.059 | 0.059 | 0 | 0.17 | 0.27 | 0.39 | 0.29 | 0.21 | 0.3 |
| 4 | 114207666 | rs29327 | -0.034 | -0.936 | 1.987 | 1.061 | 0.026 | -0.004 | -0.004 | 0 | 0.45 | 0.27 | 0.56 | 0.46 | 0.56 | 0.61 |
| 4 | 114208433 | rs29362 | -0.433 | -0.563 | 2.022 | 1.272 | 0.036 | 0.04 | 0.04 | 0.054 | 0.56 | 0.66 | 0.66 | 0.72 | 0.64 | 0.67 |
| 4 | 114211892 | rs29324 | -1.334 | -0.619 | 1.942 | 1.26 | 0.051 | -0.023 | -0.023 | 0 | 0.65 | 0.86 | 0.76 | 0.54 | 0.72 | 0.78 |
| 4 | 114216407 | rs3025713 | 0.187 | -0.503 | 1.976 | 1.329 | 0.011 | 0.073 | 0.073 | 0 | 0.08 | 0.13 | 0.25 | 0.24 | 0.1 | 0.17 |
| 4 | 114216903 | rs29306 | -1.443 | -0.53 | 1.736 | 1.317 | 0.061 | 0.053 | 0.053 | 0.271 | 0.66 | 0.86 | 0.8 | 0.79 | 0.75 | 0.8 |
| 4 | 114216924 | rs29305 | 0.094 | -0.496 | 1.629 | 1.313 | 0.001 | 0.005 | 0.005 | 0 | 0.47 | 0.47 | 0.36 | 0.23 | 0.51 | 0.49 |
| 4 | 114217997 | rs29319 | -0.613 | -0.81 | 0.981 | 0.658 | 0.002 | -0.005 | -0.005 | 0.178 | 0.08 | 0.13 | 0.14 | 0.07 | 0.11 | 0.13 |
| 4 | 114224592 | rs29302 | 0.071 | -0.828 | 0.957 | 0.587 | 0.023 | 0.009 | 0.009 | 0 | 0.5 | 0.39 | 0.6 | 0.47 | 0.58 | 0.63 |
| 4 | 114228154 | rs3025705 | -1.47 | -0.008 | 1.248 | 0.021 | 0.001 | -0.006 | -0.006 | 0 | 0.08 | 0.12 | 0.14 | 0.06 | 0.11 | 0.13 |
| 4 | 114231125 | rs6837523 | -1.967 | -0.189 | 1.443 | 0.129 | -0.015 | -0.009 | -0.009 | 0 | 0.12 | 0.25 | 0.15 | 0.09 | 0.11 | 0.09 |
| 4 | 114235039 | rs10020183 | -2.1 | -0.431 | 1.847 | 0.225 | -0.046 | 0.035 | 0.035 | 0.13 | 0.12 | 0.42 | 0.13 | 0.12 | 0.06 | 0.08 |
| 4 | 114236227 | rs10023495 | -2.368 | -0.285 | 1.834 | 0.192 | -0.019 | 0.008 | 0.008 | 0.01 | 0.14 | 0.38 | 0.14 | 0.12 | 0.1 | 0.09 |
| 4 | 114248543 | rs7676554 | -1.807 | -0.515 | 1.485 | 0.16 | -0.013 | 0.002 | 0.002 | 0 | 0.1 | 0.11 | 0.11 | 0.11 | 0.05 | 0.11 |
| 4 | 114249897 | rs11940206 | -2.543 | -0.413 | 1.96 | 0.045 | -0.031 | -0.02 | -0.02 | 1.12 | 0.18 | 0.57 | 0.2 | 0.13 | 0.12 | 0.13 |
| 4 | 114257201 | rs3736575 | -2.732 | -0.61 | 2.186 | -0.025 | -0.039 | -0.025 | -0.025 | 0.538 | 0.2 | 0.62 | 0.2 | 0.12 | 0.1 | 0.12 |
| 4 | 114260538 | rs2272230 | -2.53 | -0.642 | 2.325 | -0.041 | -0.042 | -0.018 | -0.018 | 1.695 | 0.18 | 0.51 | 0.2 | 0.12 | 0.1 | 0.12 |
| 4 | 114276880 | rs28377576 | -1.457 | -0.858 | 2.099 | -0.432 | -0.001 | -0.032 | -0.032 | 0.148 | 0.16 | 0.2 | 0.08 | 0 | 0.13 | 0.02 |
| 4 | 114278277 | rs3733617 | -1.664 | -0.891 | 2.151 | -0.441 | -0.034 | 0.048 | 0.048 | 1.081 | 0.12 | 0.42 | 0.16 | 0.11 | 0.04 | 0.12 |
| 4 | 114279422 | rs10013743 | -1.436 | -0.888 | 2.011 | -0.452 | -0.001 | -0.032 | -0.032 | 1.468 | 0.16 | 0.19 | 0.08 | 0 | 0.13 | 0.02 |
| 4 | 114285321 | rs11930964 | -2.638 | -0.862 | 1.738 | -0.438 | -0.017 | -0.027 | -0.027 | 0 | 0.29 | 0.66 | 0.24 | 0.11 | 0.17 | 0.14 |
| 4 | 114294308 | rs2293324 | -2.502 | -0.861 | 1.637 | -0.239 | -0.001 | 0 | 0 | 0 | 0.28 | 0.73 | 0.24 | 0.14 | 0.17 | 0.14 |
| 4 | 114295540 | rs11946150 | -1.42 | -0.645 | 1.889 | -0.091 | -0.009 | 0.001 | 0.001 | 0 | 0.09 | 0.1 | 0.13 | 0.09 | 0.04 | 0.12 |
| 4 | 114305184 | rs6826949 | 0.095 | -0.026 | 1.328 | -0.069 | 0.031 | 0.036 | 0.036 | 0 | 0.31 | 0.35 | 0.4 | 0.44 | 0.42 | 0.58 |
| 4 | 114307861 | rs1534093 | -1.866 | 0.075 | 1.873 | -0.179 | 0.027 | -0.048 | -0.048 | 4.001 | 0.09 | 0.34 | 0.07 | 0 | 0.08 | 0.01 |
| 4 | 114309577 | rs6533680 | -0.83 | -0.122 | 1.147 | -0.304 | 0.001 | -0.007 | -0.007 | 0 | 0.53 | 0.84 | 0.52 | 0.42 | 0.57 | 0.68 |
| 4 | 114311645 | rs6846326 | -0.297 | -0.172 | 1.198 | -0.34 | -0.015 | 0.003 | 0.003 | 0 | 0.45 | 0.24 | 0.46 | 0.41 | 0.51 | 0.68 |
| 4 | 114312389 | rs17580458 | -0.046 | -0.031 | 0.279 | -0.357 | -0.014 | 0.021 | 0.021 | 0.754 | 0.06 | 0.01 | 0.05 | 0.02 | 0.06 | 0.02 |
| 4 | 114314698 | rs2037673 | -0.836 | -0.07 | 0.363 | -0.27 | -0.044 | -0.061 | -0.061 | 0.605 | 0.54 | 0.92 | 0.53 | 0.42 | 0.57 | 0.69 |
| 4 | 114315944 | rs13130205 | -1.141 | 0.103 | 0.778 | -0.232 | 0.054 | -0.006 | -0.006 | 6.507 | 0.06 | 0.31 | 0.06 | 0 | 0.07 | 0.01 |
| 4 | 114318689 | rs11098192 | -0.532 | 0.147 | 0.355 | -0.118 | -0.003 | 0.01 | 0.01 | 0 | 0.48 | 0.56 | 0.51 | 0.41 | 0.49 | 0.62 |
| 4 | 114329115 | rs7664824 | -1.491 | 0.134 | 0.732 | -0.313 | -0.003 | -0.078 | -0.078 | 0.948 | 0.16 | 0.44 | 0.09 | 0.04 | 0.11 | 0.08 |
| 4 | 114332302 | rs10015062 | -1.101 | 0.094 | 0.704 | -0.746 | -0.014 | -0.098 | -0.098 | 0 | 0.17 | 0.59 | 0.12 | 0 | 0.13 | 0.06 |
| 4 | 114332705 | rs1525004 | -0.012 | 0.422 | -0.591 | -0.738 | -0.018 | -0.013 | -0.013 | 0 | 0.23 | 0.17 | 0.14 | 0.29 | 0.24 | 0.18 |
| 4 | 114334319 | rs17676757 | 1.015 | 0.191 | -0.898 | -0.203 | 0.013 | -0.033 | -0.033 | 0 | 0.06 | 0 | 0.13 | 0.13 | 0.03 | 0.01 |
| 4 | 114352881 | rs13129231 | 0.854 | 0.455 | -1.738 | -0.237 | -0.051 | -0.088 | -0.088 | 0 | 0.19 | 0.06 | 0.36 | 0.44 | 0.28 | 0.45 |
| 4 | 114354856 | rs17046034 | 1.056 | 0.303 | -1.606 | -0.289 | 0.002 | 0.014 | 0.014 | 0 | 0.1 | 0.05 | 0.06 | 0.07 | 0.09 | 0.06 |
| 4 | 114358532 | rs7658972 | 0.746 | 0.379 | -0.853 | -0.416 | -0.001 | -0.03 | -0.03 | 0 | 0.13 | 0.23 | 0.09 | 0.07 | 0.11 | 0.08 |
| 4 | 114360599 | rs17620306 | 1.718 | 0.346 | -0.982 | -0.326 | 0.014 | -0.035 | -0.035 | 0 | 0.06 | 0 | 0.13 | 0.13 | 0.04 | 0.01 |
| 4 | 114365275 | rs6533687 | -0.469 | -0.098 | -0.866 | -0.775 | -0.001 | 0.041 | 0.041 | 0 | 0.6 | 0.72 | 0.75 | 0.73 | 0.62 | 0.67 |
| 4 | 114369065 | rs13117519 | 2.133 | -0.026 | -1.138 | -0.835 | -0.032 | 0.009 | 0.009 | 0 | 0.16 | 0.04 | 0.13 | 0.08 | 0.18 | 0.12 |
| 4 | 114373068 | rs17676882 | 1.537 | -0.109 | -1.201 | -0.806 | -0.069 | -0.111 | -0.111 | 0.05 | 0.18 | 0.04 | 0.35 | 0.43 | 0.28 | 0.38 |
| 4 | 114373255 | rs3733619 | 0.869 | -0.291 | -0.779 | -0.866 | -0.007 | -0.034 | -0.034 | 0 | 0.15 | 0.25 | 0.09 | 0.07 | 0.12 | 0.12 |
| 4 | 114379379 | rs1880529 | -0.481 | -0.116 | -0.987 | -0.978 | -0.07 | 0.015 | 0.015 | 0 | 0.63 | 0.87 | 0.65 | 0.65 | 0.57 | 0.61 |
| 4 | 114379548 | rs1524998 | -0.504 | -0.122 | -1.049 | -0.981 | -0.092 | 0.002 | 0.002 | 0 | 0.63 | 0.91 | 0.65 | 0.65 | 0.58 | 0.61 |
| 4 | 114384869 | rs1525000 | 1.09 | 0.062 | -0.889 | -0.566 | -0.055 | -0.081 | -0.081 | 0 | 0.34 | 0.13 | 0.33 | 0.44 | 0.41 | 0.5 |
| 4 | 114391404 | rs6835747 | 1.521 | 0.236 | -1.061 | -0.522 | -0.016 | -0.062 | -0.062 | 0 | 0.14 | 0.04 | 0.16 | 0.35 | 0.13 | 0.37 |
| 4 | 114392393 | rs916874 | -1.121 | 0.106 | -1.309 | -0.698 | -0.113 | -0.101 | -0.101 | 0.252 | 0.78 | 0.98 | 0.86 | 0.68 | 0.7 | 0.79 |
| 4 | 114392629 | rs6842886 | 1.434 | 0.156 | -1.15 | -0.69 | -0.011 | -0.032 | -0.032 | 0 | 0.15 | 0.07 | 0.29 | 0.35 | 0.17 | 0.38 |
| 4 | 114416596 | rs2158196 | 0.114 | -0.067 | -0.876 | -0.926 | -0.004 | 0.003 | 0.003 | 0 | 0.55 | 0.58 | 0.53 | 0.12 | 0.53 | 0.4 |
| 4 | 114429407 | rs11098193 | 2.386 | -0.101 | -0.985 | -0.898 | -0.054 | 0.021 | 0.021 | 0 | 0.18 | 0.03 | 0.16 | 0.09 | 0.27 | 0.13 |
| 4 | 114432073 | rs17046126 | 0.952 | -0.172 | -0.797 | -0.894 | -0.001 | 0.006 | 0.006 | 0.002 | 0.28 | 0.33 | 0.37 | 0.31 | 0.33 | 0.15 |
| 4 | 114432254 | rs17630012 | 2.228 | -0.06 | -0.894 | -0.813 | 0 | -0.03 | -0.03 | 0 | 0.06 | 0 | 0.16 | 0.22 | 0.05 | 0.01 |
| 4 | 114435877 | rs3815072 | 1.629 | 0.124 | -0.953 | -0.774 | -0.07 | -0.077 | -0.077 | 5.557 | 0.15 | 0.01 | 0.05 | 0.24 | 0.16 | 0.13 |
| 4 | 114437456 | rs6533691 | -0.573 | -0.491 | -0.502 | -1.103 | -0.099 | -0.014 | -0.014 | 0 | 0.62 | 0.89 | 0.57 | 0.34 | 0.53 | 0.41 |
| 4 | 114439090 | rs17531026 | 0.151 | -0.501 | -0.517 | -1.166 | 0.018 | 0.002 | 0.002 | 0 | 0.36 | 0.27 | 0.34 | 0.04 | 0.25 | 0.29 |
| 4 | 114449719 | rs11938486 | 1.768 | 0.027 | -1.199 | -0.816 | 0.002 | -0.007 | -0.007 | 0 | 0.19 | 0.2 | 0.16 | 0.08 | 0.28 | 0.14 |
| 4 | 114450380 | rs13133676 | 1.093 | 0.467 | -1.428 | -0.189 | -0.002 | -0.029 | -0.029 | 0 | 0.14 | 0.12 | 0.22 | 0.03 | 0.13 | 0.16 |
| 4 | 114452724 | rs4240286 | 0.268 | 0.953 | -1.494 | 0.428 | -0.108 | -0.034 | -0.034 | 1.186 | 0.66 | 0.9 | 0.71 | 0.33 | 0.58 | 0.44 |
| 4 | 114477154 | rs10027475 | 0.312 | 0.884 | -1.107 | 1.11 | -0.005 | -0.049 | -0.049 | 0.844 | 0.75 | 0.85 | 0.85 | 0.6 | 0.74 | 0.74 |
| 4 | 114478653 | rs11728021 | 0.496 | 0.88 | -0.27 | 0.347 | 0.012 | -0.093 | -0.093 | 8.831 | 0.18 | 0.29 | 0.12 | 0.04 | 0.17 | 0.17 |

| | | | | | | | | | | | | | | | | |
|---|---|---|---|---|---|---|---|---|---|---|---|---|---|---|---|---|
| 4 | 114505498 | rs7665102 | 0.156 | 0.62 | 0.018 | 0.995 | 0.003 | 0.025 | 0.025 | 0 | 0.74 | 0.69 | 0.63 | 0.56 | 0.74 | 0.65 |
| 4 | 114506450 | rs17447193 | 0.713 | 0.449 | -0.297 | 0.754 | -0.014 | -0.041 | -0.041 | 0.385 | 0.1 | 0.02 | 0.18 | 0.2 | 0.11 | 0.12 |
| 4 | 114520105 | rs7681248 | -1.023 | 0.446 | 0.034 | 0.751 | 0.002 | 0.007 | 0.007 | 0.001 | 0.34 | 0.52 | 0.55 | 0.63 | 0.34 | 0.44 |
| 4 | 114524255 | rs10025791 | -1.91 | 0.207 | 0.258 | 0.448 | -0.001 | -0.02 | -0.02 | 0 | 0.55 | 0.63 | 0.66 | 0.96 | 0.51 | 0.65 |
| 4 | 114527882 | rs10033037 | -0.201 | 0.154 | 0.463 | 0.843 | -0.001 | -0.001 | -0.001 | 0 | 0.15 | 0.17 | 0.14 | 0.18 | 0.17 | 0.19 |
| 4 | 114533473 | rs9307393 | -0.152 | 0.149 | 0.444 | 1.077 | -0.002 | -0.002 | -0.002 | 0 | 0.14 | 0.16 | 0.09 | 0.28 | 0.17 | 0.19 |
| 4 | 114534723 | rs10488958 | -0.48 | 0.068 | 0.491 | 1.091 | 0 | 0.029 | 0.029 | 0 | 0.25 | 0.42 | 0.36 | 0.32 | 0.23 | 0.3 |
| 4 | 114543214 | rs6826859 | 0.172 | -0.205 | 0.356 | 1.145 | -0.043 | -0.043 | -0.043 | 0 | 0.13 | 0.02 | 0.07 | 0.18 | 0.17 | 0.19 |
| 4 | 114546689 | rs10000775 | -2.179 | -0.567 | 0.634 | 0.995 | -0.023 | 0.148 | 0.148 | 0.007 | 0.55 | 0.82 | 0.66 | 0.82 | 0.44 | 0.61 |
| 4 | 114546704 | rs10023113 | 0.163 | -0.597 | 0.846 | 1.085 | 0.015 | 0.034 | 0.034 | 0 | 0.13 | 0.33 | 0.1 | 0.14 | 0.13 | 0.09 |
| 4 | 114548301 | rs916913 | -2.063 | -0.769 | 0.849 | 1.162 | 0.064 | 0.17 | 0.17 | 0.427 | 0.46 | 0.86 | 0.6 | 0.78 | 0.32 | 0.47 |
| 4 | 114555661 | rs2189363 | -0.272 | -0.647 | 0.724 | 1.275 | -0.017 | -0.008 | -0.008 | 0 | 0.19 | 0.21 | 0.11 | 0.16 | 0.09 | 0.15 |
| 4 | 114559741 | rs12510043 | 0.311 | -0.573 | 0.664 | 1.288 | -0.008 | -0.046 | -0.046 | 0.277 | 0.18 | 0.09 | 0.32 | 0.32 | 0.19 | 0.29 |
| 4 | 114560543 | rs17630766 | -0.469 | -0.698 | 0.789 | 1.309 | -0.007 | -0.001 | -0.001 | 0 | 0.21 | 0.2 | 0.12 | 0.23 | 0.12 | 0.15 |
| 4 | 114561123 | rs12512019 | -0.249 | -0.731 | 0.47 | 1.407 | 0.012 | -0.05 | -0.05 | 0 | 0.23 | 0.08 | 0.31 | 0.26 | 0.2 | 0.32 |
| 4 | 114561733 | rs10488959 | -0.518 | -0.516 | 1.212 | 1.738 | 0.064 | 0.4 | 0.4 | 1.207 | 0.08 | 0.51 | 0.18 | 0.33 | 0.06 | 0.08 |
| 4 | 114572090 | rs12506881 | -0.901 | -0.535 | 0.645 | 1.686 | -0.003 | -0.023 | -0.023 | 3.57 | 0.35 | 0.1 | 0.39 | 0.29 | 0.32 | 0.48 |
| 4 | 114573947 | rs17046372 | 0.387 | -0.508 | 0.922 | 1.658 | 0.014 | -0.034 | -0.034 | 0 | 0.12 | 0.22 | 0.09 | 0.03 | 0.12 | 0.1 |
| 4 | 114581207 | rs6845568 | -0.616 | -0.501 | 0.194 | 1.69 | 0.005 | 0.091 | 0.091 | 0.078 | 0.19 | 0.62 | 0.25 | 0.38 | 0.17 | 0.16 |
| 4 | 114585800 | rs2107172 | -1.734 | -0.554 | 0.028 | 1.398 | 0.019 | 0.052 | 0.052 | 0 | 0.45 | 0.7 | 0.57 | 0.64 | 0.39 | 0.54 |
| 4 | 114588367 | rs12512765 | -0.085 | -0.519 | -0.197 | 1.259 | 0.02 | -0.121 | -0.121 | 0 | 0.24 | 0.05 | 0.31 | 0.36 | 0.19 | 0.38 |
| 4 | 114593466 | rs1011973 | -1.741 | -0.4 | -0.037 | 0.52 | -0.014 | 0.145 | 0.145 | 0 | 0.55 | 0.79 | 0.65 | 0.81 | 0.45 | 0.64 |
| 4 | 114594271 | rs6827138 | -0.882 | -0.082 | 0.679 | 0.91 | 0.047 | -0.02 | -0.02 | 0 | 0.18 | 0.44 | 0.13 | 0.11 | 0.17 | 0.11 |
| 4 | 114611948 | rs7660775 | -1.207 | -0.922 | -0.248 | 1.204 | 0.046 | -0.036 | -0.036 | 0 | 0.31 | 0.78 | 0.18 | 0.41 | 0.16 | 0.33 |
| 4 | 114616823 | rs4475149 | -0.651 | -0.8 | 0.562 | 1.102 | 0.021 | 0.002 | 0.002 | 0 | 0.08 | 0.26 | 0.05 | 0.03 | 0.05 | 0.11 |
| 4 | 114617815 | rs764830 | -0.864 | -0.634 | -0.446 | 0.852 | -0.048 | 0.03 | 0.03 | 0 | 0.72 | 0.9 | 0.72 | 0.74 | 0.62 | 0.65 |
| 4 | 114618602 | rs987694 | 1.697 | -0.965 | -0.528 | 0.68 | -0.048 | -0.059 | -0.059 | 0.171 | 0.32 | 0.13 | 0.52 | 0.38 | 0.41 | 0.31 |
| 4 | 114621830 | rs7687511 | -0.855 | -1.194 | -0.141 | 0.467 | -0.063 | -0.109 | -0.109 | 0.137 | 0.21 | 0.5 | 0.1 | 0.03 | 0.08 | 0.15 |
| 4 | 114626186 | rs6533711 | -1.239 | -1.206 | 0.008 | 0.466 | -0.021 | 0.023 | 0.023 | 0 | 0.71 | 0.84 | 0.72 | 0.75 | 0.62 | 0.64 |
| 4 | 114626620 | rs6842343 | -0.848 | -1.209 | -0.007 | 0.465 | -0.063 | -0.109 | -0.109 | 0 | 0.21 | 0.5 | 0.1 | 0.03 | 0.08 | 0.15 |
| 4 | 114628988 | rs1978139 | -1.282 | -1.213 | -0.096 | 0.463 | -0.059 | 0.037 | 0.037 | 0 | 0.71 | 0.92 | 0.72 | 0.75 | 0.62 | 0.64 |
| 4 | 114631347 | rs2040742 | -0.94 | -1.233 | 0.027 | 0.451 | -0.059 | -0.103 | -0.103 | 0.868 | 0.21 | 0.49 | 0.09 | 0.03 | 0.07 | 0.15 |
| 4 | 114635044 | rs2107170 | 1.897 | -1.157 | -0.603 | 0.573 | -0.076 | -0.081 | -0.081 | 0 | 0.31 | 0.09 | 0.51 | 0.36 | 0.41 | 0.31 |
| 4 | 114635433 | rs9993234 | -1.698 | -1.145 | -0.413 | 0.665 | 0.083 | 0.08 | 0.08 | 0.171 | 0.41 | 0.8 | 0.19 | 0.21 | 0.22 | 0.3 |
| 4 | 114635782 | rs11098200 | 0.335 | -1.106 | -0.457 | 0.718 | -0.002 | -0.001 | -0.001 | 0 | 0.2 | 0.22 | 0.08 | 0.18 | 0.14 | 0.15 |
| 4 | 114661681 | rs1859229 | -1.245 | -0.941 | -1.095 | 0.797 | 0.1 | 0.005 | 0.005 | 0.235 | 0.41 | 0.82 | 0.21 | 0.38 | 0.21 | 0.33 |
| 4 | 114664962 | rs4834354 | -1.187 | -0.885 | -1.232 | 0.277 | -0.055 | 0.078 | 0.078 | 0 | 0.74 | 0.89 | 0.71 | 0.92 | 0.61 | 0.71 |
| 4 | 114665832 | rs10488961 | 1.473 | -0.934 | -1.207 | 0.731 | -0.134 | -0.145 | -0.145 | 0.916 | 0.31 | 0.02 | 0.51 | 0.41 | 0.39 | 0.36 |
| 4 | 114671843 | rs6812422 | 1.424 | -0.879 | -1.118 | 0.627 | -0.07 | -0.095 | -0.095 | 0.927 | 0.31 | 0.09 | 0.51 | 0.43 | 0.38 | 0.38 |
| 4 | 114678642 | rs10006113 | -0.375 | -0.633 | -0.541 | 0.798 | -0.06 | 0.035 | 0.035 | 0.441 | 0.26 | 0.57 | 0.15 | 0.34 | 0.16 | 0.24 |
| 4 | 114678786 | rs10018022 | -0.781 | -0.315 | -1.433 | 0.496 | -0.069 | 0.046 | 0.046 | 0 | 0.69 | 0.88 | 0.7 | 0.77 | 0.59 | 0.66 |
| 4 | 114679682 | rs4141007 | -0.141 | -0.258 | -1.154 | 0.544 | -0.001 | 0 | 0 | 0 | 0.19 | 0.18 | 0.09 | 0.12 | 0.13 | 0.06 |
| 4 | 114682714 | rs11731775 | 0.189 | 0.1 | -1.562 | 0.772 | -0.028 | -0.045 | -0.045 | 0 | 0.4 | 0.14 | 0.53 | 0.43 | 0.41 | 0.4 |
| 4 | 114690439 | rs11935493 | -0.757 | 0.113 | -1.215 | 0.751 | 0.008 | -0.027 | -0.027 | 0 | 0.38 | 0.71 | 0.34 | 0.49 | 0.32 | 0.45 |
| 4 | 114692396 | rs7676980 | -0.757 | 0.113 | -1.221 | 0.762 | 0.009 | -0.027 | -0.027 | 0 | 0.37 | 0.71 | 0.34 | 0.49 | 0.32 | 0.45 |
| 4 | 114696373 | rs1557815 | -0.911 | 0.143 | -1.638 | 0.234 | -0.063 | -0.058 | -0.058 | 0 | 0.67 | 0.87 | 0.49 | 0.54 | 0.55 | 0.51 |
| 4 | 114698696 | rs1476569 | -0.047 | -0.216 | -0.297 | 0.232 | -0.016 | -0.024 | -0.024 | 0 | 0.43 | 0.46 | 0.31 | 0.24 | 0.31 | 0.2 |
| 4 | 114702819 | rs7662446 | 0.866 | -0.067 | -0.572 | 0.385 | -0.024 | 0.005 | 0.005 | 0 | 0.16 | 0.06 | 0.09 | 0.13 | 0.18 | 0.18 |
| 4 | 114703071 | rs11945240 | -0.023 | -0.077 | -0.59 | 0.384 | 0.005 | 0.007 | 0.007 | 2.762 | 0.44 | 0.53 | 0.31 | 0.24 | 0.31 | 0.21 |
| 4 | 114703773 | rs10461151 | -0.937 | 0.049 | -1.045 | 0.469 | -0.143 | -0.093 | -0.093 | 0 | 0.64 | 0.94 | 0.43 | 0.37 | 0.49 | 0.45 |
| 4 | 114703967 | rs17532826 | 0.847 | -0.213 | -0.76 | 0.178 | 0.034 | 0.1 | 0.1 | 3.057 | 0.29 | 0.1 | 0.13 | 0.05 | 0.21 | 0.05 |
| 4 | 114709273 | rs13134836 | 0.335 | 0.299 | -0.417 | 0.653 | 0.002 | -0.023 | -0.023 | 0.948 | 0.33 | 0.22 | 0.29 | 0.52 | 0.31 | 0.52 |
| 4 | 114710145 | rs11731601 | 0.429 | 0.321 | -0.286 | 0.769 | 0.001 | -0.003 | -0.003 | 1.014 | 0.14 | 0.13 | 0.18 | 0.2 | 0.13 | 0.17 |
| 4 | 114732529 | rs1081010 | 1.24 | 0.336 | -0.421 | 0.861 | -0.034 | -0.065 | -0.065 | 0 | 0.25 | 0.02 | 0.21 | 0.37 | 0.24 | 0.41 |
| 4 | 114741032 | rs10010935 | 1.24 | 0.323 | -0.121 | 0.795 | -0.034 | -0.065 | -0.065 | 0.802 | 0.25 | 0.02 | 0.21 | 0.37 | 0.24 | 0.41 |
| 4 | 114741276 | rs7699538 | -0.311 | 0.271 | 0.206 | 0.723 | -0.009 | 0.059 | 0.059 | 0.204 | 0.43 | 0.21 | 0.45 | 0.72 | 0.44 | 0.62 |
| 4 | 114742249 | rs7659107 | 1.077 | 0.375 | 0.402 | 0.595 | -0.005 | 0.003 | 0.003 | 0 | 0.17 | 0.18 | 0.25 | 0.13 | 0.2 | 0.13 |
| 4 | 114742637 | rs11730230 | 1.562 | 0.36 | 0.613 | 0.507 | 0.018 | 0.018 | 0.018 | 2.585 | 0.28 | 0.18 | 0.18 | 0.15 | 0.18 | 0.06 |
| 4 | 114748150 | rs4345183 | 1.18 | 0.766 | 1.131 | 0.65 | -0.007 | -0.009 | -0.009 | 2.157 | 0.71 | 0.74 | 0.66 | 0.67 | 0.6 | 0.6 |
| 4 | 114751650 | rs7684832 | 1.495 | 0.663 | 1.343 | 0.582 | 0.014 | 0 | 0 | 0 | 0.52 | 0.36 | 0.39 | 0.54 | 0.39 | 0.47 |
| 4 | 114752809 | rs9918007 | -1.159 | 0.943 | 0.973 | 0.353 | -0.026 | 0.022 | 0.022 | 0 | 0.92 | 0.97 | 0.9 | 1 | 0.81 | 0.88 |
| 4 | 114754302 | rs13126791 | 0.284 | 0.92 | 1.185 | 0.368 | 0.006 | -0.012 | -0.012 | 0.51 | 0.19 | 0.3 | 0.26 | 0.13 | 0.21 | 0.19 |
| 4 | 114759474 | rs4426777 | 0.256 | 0.895 | 1.209 | 0.315 | -0.012 | 0.011 | 0.011 | 0 | 0.73 | 0.76 | 0.64 | 0.87 | 0.59 | 0.65 |
| 4 | 114762343 | rs4834359 | 0.61 | 0.872 | 1.397 | 0.303 | 0.03 | -0.068 | -0.068 | 0.125 | 0.69 | 0.44 | 0.62 | 0.87 | 0.59 | 0.59 |
| 4 | 114768317 | rs6836841 | 0.679 | 1.036 | 0.909 | 0.455 | -0.027 | 0.006 | 0.006 | 0 | 0.71 | 0.75 | 0.58 | 0.74 | 0.51 | 0.64 |
| 4 | 114772555 | rs6816415 | -0.83 | 1.094 | 0.729 | 0.425 | -0.002 | -0.01 | -0.01 | 0 | 0.92 | 0.93 | 0.9 | 1 | 0.82 | 0.88 |
| 4 | 114797011 | rs13142831 | -1.034 | 0.664 | 0.963 | 1.036 | -0.002 | -0.013 | -0.013 | 0 | 0.32 | 0.75 | 0.42 | 0.45 | 0.32 | 0.39 |
| 4 | 114800468 | rs7437325 | 0.963 | 0.664 | 1.147 | 1.036 | -0.012 | -0.001 | -0.001 | 0 | 0.15 | 0.11 | 0.25 | 0.16 | 0.21 | 0.15 |
| 4 | 114803632 | rs13136741 | 0.689 | 0.682 | 1.165 | 1.046 | 0.01 | -0.002 | -0.002 | 0 | 0.17 | 0.22 | 0.26 | 0.16 | 0.21 | 0.21 |

| | | | | | | | | | | | | | | | | |
|---|---|---|---|---|---|---|---|---|---|---|---|---|---|---|---|---|
| 4 | 114816474 | rs7674158 | 0.58 | 0.709 | 0.803 | 1.198 | -0.039 | 0.103 | 0.103 | 2.478 | 0.08 | 0.2 | 0.1 | 0.19 | 0.02 | 0.07 |
| 4 | 114816990 | rs2374218 | 1.171 | 0.705 | 0.651 | 1.205 | -0.042 | -0.011 | -0.011 | 0 | 0.6 | 0.12 | 0.5 | 0.45 | 0.61 | 0.53 |
| 4 | 114821256 | rs6816611 | 1.531 | 0.707 | 0.654 | 1.118 | 0.005 | 0.01 | 0.01 | 4.273 | 0.06 | 0.04 | 0.09 | 0.19 | 0.02 | 0.05 |
| 4 | 114822485 | rs2374216 | 0.329 | 0.723 | 0.856 | 1.113 | -0.012 | 0.042 | 0.042 | 0 | 0.72 | 0.34 | 0.61 | 0.6 | 0.76 | 0.68 |
| 4 | 114833153 | rs6533728 | 0.231 | 0.436 | 0.63 | 1.144 | -0.013 | 0.07 | 0.07 | 0 | 0.72 | 0.41 | 0.62 | 0.6 | 0.76 | 0.67 |
| 4 | 114833290 | rs6533729 | 0.231 | 0.436 | 0.63 | 1.144 | -0.013 | 0.07 | 0.07 | 0 | 0.72 | 0.41 | 0.62 | 0.6 | 0.76 | 0.67 |
| 4 | 114842742 | rs12645879 | -0.726 | 0.439 | 0.169 | 0.819 | 0.003 | 0.024 | 0.024 | 0 | 0.51 | 0.53 | 0.56 | 0.78 | 0.57 | 0.63 |
| 4 | 114854312 | rs2374196 | -0.467 | -0.029 | -0.149 | 1.249 | -0.002 | -0.001 | -0.001 | 0 | 0.37 | 0.33 | 0.43 | 0.68 | 0.3 | 0.41 |
| 4 | 114855353 | rs7437812 | -0.073 | 0.577 | -0.134 | 1.077 | -0.002 | 0.002 | 0.002 | 0.391 | 0.31 | 0.28 | 0.34 | 0.49 | 0.28 | 0.36 |
| 4 | 114856627 | rs10012647 | -0.38 | 0.929 | 0.302 | 0.794 | 0.029 | -0.003 | -0.003 | 1.654 | 0.09 | 0.23 | 0.09 | 0.07 | 0.08 | 0.06 |
| 4 | 114865091 | rs4364328 | -0.039 | 0.811 | -0.283 | 0.942 | 0.002 | 0.001 | 0.001 | 0 | 0.8 | 0.75 | 0.72 | 0.78 | 0.78 | 0.79 |
| 4 | 114867878 | rs11098213 | -0.349 | 0.704 | -0.8 | 1.066 | -0.011 | -0.025 | -0.025 | 0 | 0.36 | 0.22 | 0.28 | 0.51 | 0.38 | 0.34 |
| 4 | 114884901 | rs728313 | 0.235 | 0.025 | -0.852 | 1.137 | 0.03 | -0.004 | -0.004 | 0 | 0.16 | 0.05 | 0.18 | 0.21 | 0.11 | 0.16 |
| 4 | 114885155 | rs2374219 | 0.386 | 0.289 | -0.778 | 1.279 | -0.001 | -0.004 | -0.004 | 2.168 | 0.31 | 0.31 | 0.45 | 0.51 | 0.34 | 0.37 |
| 4 | 114898691 | rs12505253 | 0.037 | 0.91 | -0.886 | 0.897 | -0.008 | 0.006 | 0.006 | 0 | 0.66 | 0.65 | 0.73 | 0.81 | 0.58 | 0.62 |
| 4 | 114914110 | rs17581675 | -0.124 | 0.322 | -1.398 | 1.13 | 0 | 0.035 | 0.035 | 0 | 0.36 | 0.13 | 0.21 | 0.25 | 0.33 | 0.38 |
| 4 | 114938478 | rs4833439 | -0.094 | -0.919 | -0.563 | 1.032 | -0.071 | -0.013 | -0.013 | 0.52 | 0.71 | 0.46 | 0.75 | 0.67 | 0.85 | 0.72 |
| 4 | 114942909 | rs2374223 | -0.052 | -1.114 | -0.457 | 1.022 | 0.03 | 0.007 | 0.007 | 0.108 | 0.68 | 0.23 | 0.73 | 0.64 | 0.85 | 0.69 |
| 4 | 114946868 | rs882308 | 0.315 | -1.399 | 0.266 | 0.954 | 0.01 | -0.004 | -0.004 | 0 | 0.24 | 0.17 | 0.22 | 0.34 | 0.15 | 0.25 |
| 4 | 114957205 | rs6533743 | 0.364 | -1.437 | 0.51 | 0.931 | 0.001 | 0.001 | 0.001 | 0.143 | 0.67 | 0.29 | 0.74 | 0.63 | 0.85 | 0.69 |
| 4 | 114957962 | rs7440732 | 0.749 | -1.483 | 0.979 | 0.906 | 0.005 | 0.036 | 0.036 | 5.901 | 0.92 | 0.55 | 0.96 | 0.97 | 1 | 0.96 |
| 4 | 114958543 | rs13112787 | 0.418 | -1.482 | 0.663 | 0.907 | 0.043 | -0.028 | -0.028 | 0 | 0.23 | 0.07 | 0.22 | 0.34 | 0.15 | 0.25 |
| 4 | 114985792 | rs10020382 | -0.675 | -1.439 | 0.694 | 0.937 | 0.003 | 0 | 0 | 0.108 | 0.33 | 0.72 | 0.27 | 0.37 | 0.15 | 0.31 |
| 4 | 115005026 | rs6835349 | -0.804 | -1.402 | 0.123 | 0.369 | 0.093 | 0.085 | 0.085 | 0.085 | 0.5 | 0.74 | 0.38 | 0.37 | 0.32 | 0.47 |
| 4 | 115013043 | rs17046808 | -0.228 | -1.485 | 0.903 | 0.233 | -0.02 | -0.102 | -0.102 | 0 | 0.25 | 0.43 | 0.17 | 0.02 | 0.18 | 0.22 |
| 4 | 115016497 | rs17621440 | 0.978 | -0.318 | 0.544 | 0.802 | -0.065 | -0.061 | -0.061 | 2.826 | 0.15 | 0.04 | 0.14 | 0.3 | 0.27 | 0.16 |
| 4 | 115023489 | rs11098226 | 0.677 | -0.584 | 0.968 | 0.622 | -0.001 | -0.001 | -0.001 | 0 | 0.4 | 0.54 | 0.53 | 0.35 | 0.41 | 0.36 |
| 4 | 115041483 | rs7340953 | 0.408 | -0.668 | 1.528 | 0.585 | 0.009 | 0.013 | 0.013 | 0 | 0.43 | 0.68 | 0.54 | 0.35 | 0.41 | 0.36 |
| 4 | 115044037 | rs4834387 | -0.783 | -0.691 | 1.432 | 0.653 | 0.046 | 0.027 | 0.027 | 0.286 | 0.61 | 0.84 | 0.68 | 0.65 | 0.7 | 0.68 |
| 4 | 115047815 | rs1497606 | 1.098 | -0.685 | 1.505 | 0.645 | -0.007 | -0.004 | -0.004 | 1.588 | 0.36 | 0.25 | 0.5 | 0.35 | 0.41 | 0.36 |
| 4 | 115059118 | rs1497591 | 0.713 | -0.656 | 1.478 | 0.659 | -0.002 | -0.005 | -0.005 | 0 | 0.4 | 0.47 | 0.52 | 0.35 | 0.41 | 0.36 |
| 4 | 115075724 | rs1112496 | -0.761 | -0.668 | 1.291 | 0.649 | 0.047 | 0.029 | 0.029 | 0.169 | 0.61 | 0.84 | 0.68 | 0.66 | 0.7 | 0.68 |
| 4 | 115078294 | rs1497590 | 0.935 | -0.663 | 1.327 | 0.618 | -0.009 | -0.003 | -0.003 | 0 | 0.37 | 0.24 | 0.5 | 0.35 | 0.41 | 0.36 |
| 4 | 115084166 | rs6832117 | 0.19 | -0.68 | 1.397 | 0.609 | 0.004 | 0.008 | 0.008 | 0 | 0.44 | 0.65 | 0.54 | 0.36 | 0.43 | 0.53 |
| 4 | 115091159 | rs1909020 | 0.895 | -0.604 | 1.169 | 0.649 | -0.009 | -0.002 | -0.002 | 0.046 | 0.37 | 0.25 | 0.5 | 0.35 | 0.41 | 0.36 |
| 4 | 115095176 | rs1497586 | 0.913 | -0.625 | 1.121 | 0.638 | -0.022 | -0.01 | -0.01 | 0 | 0.36 | 0.19 | 0.5 | 0.35 | 0.41 | 0.36 |
| 4 | 115101499 | rs6851287 | 0.594 | -0.615 | 1.162 | 0.644 | -0.001 | -0.004 | -0.004 | 0.149 | 0.4 | 0.54 | 0.53 | 0.35 | 0.41 | 0.36 |
| 4 | 115113997 | rs1002287 | 1.482 | -1.02 | 0.753 | 0.644 | 0.037 | -0.051 | -0.051 | 0 | 0.15 | 0.01 | 0.14 | 0.24 | 0.09 | 0.14 |
| 4 | 115134768 | rs13128386 | -0.058 | -1.016 | 0.994 | 0.606 | 0.011 | 0.009 | 0.009 | 0 | 0.56 | 0.59 | 0.68 | 0.66 | 0.69 | 0.53 |
| 4 | 115137619 | rs1497597 | -0.2 | -1.057 | 1.154 | 0.584 | -0.003 | -0.002 | -0.002 | 0.052 | 0.57 | 0.58 | 0.67 | 0.67 | 0.7 | 0.69 |
| 4 | 115146328 | rs1497595 | -0.079 | -1.023 | 1.204 | 0.578 | -0.002 | -0.001 | -0.001 | 0 | 0.56 | 0.46 | 0.67 | 0.67 | 0.7 | 0.69 |
| 4 | 115147720 | rs11732278 | 0.759 | -0.879 | 1.218 | 0.648 | 0.029 | 0.027 | 0.027 | 0.011 | 0.18 | 0.22 | 0.15 | 0.3 | 0.28 | 0.17 |
| 4 | 115149454 | rs11944255 | -1.981 | -0.869 | 1.408 | 0.773 | 0.006 | 0.033 | 0.033 | 1.676 | 0.08 | 0.35 | 0.06 | 0.05 | 0.02 | 0.16 |
| 4 | 115151872 | rs12513381 | 0.141 | -1.093 | 1.322 | 0.774 | 0.025 | 0.01 | 0.01 | 0 | 0.55 | 0.39 | 0.63 | 0.58 | 0.69 | 0.55 |
| 4 | 115161980 | rs1603622 | 0.83 | -1.078 | 1.366 | 0.763 | -0.001 | -0.015 | -0.015 | 0 | 0.39 | 0.46 | 0.49 | 0.3 | 0.42 | 0.37 |
| 4 | 115163958 | rs2646653 | -0.758 | -1.039 | 1.202 | 0.784 | 0.033 | 0.002 | 0.002 | 0.015 | 0.63 | 0.8 | 0.69 | 0.62 | 0.7 | 0.71 |
| 4 | 115167948 | rs2725520 | 0.439 | -0.978 | 1.04 | 0.832 | 0.028 | 0.019 | 0.019 | 0 | 0.21 | 0.3 | 0.16 | 0.27 | 0.28 | 0.19 |
| 4 | 115171607 | rs12499919 | 1.702 | -0.904 | 0.305 | 0.686 | -0.066 | -0.006 | -0.006 | 0 | 0.16 | 0.02 | 0.34 | 0.1 | 0.24 | 0.22 |
| 4 | 115172667 | rs9998695 | 0.349 | -0.907 | 0.508 | 0.759 | 0 | 0.004 | 0.004 | 0 | 0.42 | 0.42 | 0.52 | 0.33 | 0.43 | 0.52 |
| 4 | 115196903 | rs13134083 | 0.403 | -0.921 | 0.398 | 0.832 | -0.001 | 0.01 | 0.01 | 0 | 0.61 | 0.49 | 0.66 | 0.44 | 0.64 | 0.6 |
| 4 | 115199110 | rs11947200 | 1.139 | -0.997 | 0.744 | 0.722 | 0.001 | 0.003 | 0.003 | 0 | 0.23 | 0.21 | 0.18 | 0.13 | 0.21 | 0.22 |
| 4 | 115223284 | rs9790620 | 0.752 | -0.798 | 0.413 | 0.911 | -0.006 | -0.014 | -0.014 | 0.305 | 0.44 | 0.47 | 0.32 | 0.33 | 0.39 | 0.38 |
| 4 | 115225571 | rs4549468 | -0.925 | -0.132 | 0.065 | 1.003 | -0.009 | 0.018 | 0.018 | 0 | 0.06 | 0.28 | 0.05 | 0.05 | 0.01 | 0.11 |
| 4 | 115227215 | rs4597902 | -0.279 | 0.003 | -0.515 | 1.055 | -0.121 | -0.088 | -0.088 | 0.899 | 0.35 | 0.1 | 0.48 | 0.38 | 0.51 | 0.38 |
| 4 | 115239529 | rs12644990 | 0.283 | 0.231 | -0.662 | 1.011 | -0.056 | -0.045 | -0.045 | 1.073 | 0.17 | 0.09 | 0.14 | 0.26 | 0.27 | 0.15 |
| 4 | 115240188 | rs12498367 | 0.326 | 0.242 | -0.814 | 1.08 | -0.002 | -0.02 | -0.02 | 0 | 0.21 | 0.1 | 0.34 | 0.2 | 0.22 | 0.32 |
| 4 | 115242791 | rs11098233 | -0.362 | 0.129 | -0.525 | 0.971 | -0.07 | -0.069 | -0.069 | 0 | 0.35 | 0.18 | 0.48 | 0.46 | 0.49 | 0.38 |
| 4 | 115243741 | rs11098234 | -0.754 | 0.197 | -0.206 | 0.965 | 0.001 | 0.001 | 0.001 | 0.735 | 0.4 | 0.43 | 0.51 | 0.46 | 0.5 | 0.47 |
| 4 | 115245137 | rs10034750 | -0.68 | 0.5 | -0.3 | 0.656 | -0.011 | -0.013 | -0.013 | 1.155 | 0.6 | 0.48 | 0.62 | 0.59 | 0.61 | 0.58 |
| 4 | 115250531 | rs11098236 | 0.232 | 0.368 | -0.678 | 0.73 | -0.001 | -0.001 | -0.001 | 0.698 | 0.61 | 0.57 | 0.52 | 0.49 | 0.52 | 0.6 |
| 4 | 115251586 | rs17046991 | 0.805 | 0.306 | -0.751 | 0.664 | -0.038 | -0.062 | -0.062 | 0 | 0.24 | 0.49 | 0.19 | 0.1 | 0.14 | 0.26 |
| 4 | 115253765 | rs4444907 | 0.368 | 0.683 | -0.688 | 0.917 | 0.012 | 0 | 0 | 1.669 | 0.4 | 0.32 | 0.5 | 0.35 | 0.35 | 0.52 |
| 4 | 115255278 | rs10516601 | 0.119 | 0.912 | -1.391 | 0.754 | 0.006 | 0.008 | 0.008 | 0 | 0.38 | 0.28 | 0.34 | 0.39 | 0.38 | 0.33 |
| 4 | 115264835 | rs2353563 | -0.07 | 1.027 | -1.43 | 0.883 | 0.009 | 0.014 | 0.014 | 1.612 | 0.7 | 0.46 | 0.73 | 0.56 | 0.7 | 0.65 |
| 4 | 115264948 | rs4418044 | -0.008 | 1.449 | -1.116 | 1.425 | -0.018 | -0.017 | -0.017 | 3.591 | 0.18 | 0.17 | 0.37 | 0.28 | 0.31 | 0.37 |
| 4 | 115271601 | rs7667542 | 0.093 | -0.018 | -1.117 | 0.991 | 0.013 | 0.002 | 0.002 | 0.305 | 0.38 | 0.34 | 0.29 | 0.31 | 0.23 | 0.22 |
| 4 | 115283928 | rs4455493 | -0.595 | 1.351 | -1.115 | 1.294 | -0.051 | -0.065 | -0.065 | 0 | 0.14 | 0.03 | 0.07 | 0.23 | 0.16 | 0.19 |
| 4 | 115287331 | rs4458522 | 0.423 | 1.243 | -0.6 | 0.98 | 0.011 | -0.026 | -0.026 | 0 | 0.93 | 0.84 | 0.96 | 1 | 0.95 | 0.92 |
| 4 | 115289204 | rs4989308 | 1.053 | 1.239 | -0.912 | 1.19 | 0.01 | 0.024 | 0.024 | 0.619 | 0.63 | 0.49 | 0.61 | 0.55 | 0.67 | 0.45 |

| | | | | | | | | | | | | | | | | |
|---|---|---|---|---|---|---|---|---|---|---|---|---|---|---|---|---|
| 4 | 115292482 | rs12643524 | -0.734 | 1.053 | -1.475 | 1.136 | -0.03 | -0.053 | -0.053 | 0.591 | 0.13 | 0.01 | 0.26 | 0.2 | 0.12 | 0.25 |
| 4 | 115299073 | rs4834400 | 0.61 | 0.888 | -2.304 | 0.656 | -0.099 | 0.054 | 0.054 | 0 | 0.41 | 0.12 | 0.39 | 0.29 | 0.43 | 0.23 |
| 4 | 115308760 | rs12511321 | -0.385 | 0.474 | -2.038 | 0.721 | -0.01 | -0.037 | -0.037 | 0 | 0.09 | 0.02 | 0.16 | 0.13 | 0.09 | 0.16 |
| 4 | 115313167 | rs13122061 | -1.295 | 0.396 | -1.381 | 0.364 | 0 | -0.002 | -0.002 | 0 | 0.27 | 0.32 | 0.17 | 0.2 | 0.23 | 0.25 |
| 4 | 115313315 | rs1962495 | 1.649 | -0.272 | -1.575 | 0.434 | 0.05 | 0.097 | 0.097 | 0 | 0.52 | 0.15 | 0.56 | 0.69 | 0.61 | 0.58 |
| 4 | 115315128 | rs4834401 | 1.32 | 0.256 | -0.677 | 0.711 | 0.014 | 0.033 | 0.033 | 0 | 0.81 | 0.71 | 0.77 | 0.69 | 0.79 | 0.81 |
| 4 | 115316659 | rs6843850 | -1.595 | 0.061 | -0.075 | 0.533 | -0.002 | -0.017 | -0.017 | 1.194 | 0.29 | 0.25 | 0.3 | 0.13 | 0.3 | 0.19 |
| 4 | 115316768 | rs11935578 | 1.588 | 0.03 | -0.047 | 0.503 | -0.004 | -0.06 | -0.06 | 0 | 0.67 | 0.54 | 0.67 | 0.87 | 0.64 | 0.8 |
| 4 | 115318983 | rs4834403 | 2.604 | 0.006 | -0.065 | 0.513 | -0.003 | 0.005 | 0.005 | 0 | 0.52 | 0.45 | 0.48 | 0.55 | 0.48 | 0.63 |
| 4 | 115319437 | rs4440296 | -0.541 | -0.027 | -0.044 | 0.268 | -0.008 | 0.006 | 0.006 | 0 | 0.14 | 0.06 | 0.21 | 0.09 | 0.19 | 0.12 |
| 4 | 115321016 | rs4561982 | 2.394 | -0.253 | -0.287 | 0.888 | 0 | 0.11 | 0.11 | 0 | 0.64 | 0.18 | 0.63 | 0.86 | 0.63 | 0.72 |
| 4 | 115327659 | rs10857025 | 1.663 | -0.421 | 0.024 | 0.798 | -0.043 | -0.082 | -0.082 | 0.091 | 0.87 | 0.35 | 0.92 | 1 | 0.94 | 0.9 |
| 4 | 115329332 | rs4446396 | -1.599 | -0.306 | -0.255 | 0.884 | -0.019 | 0.025 | 0.025 | 0.798 | 0.33 | 0.22 | 0.44 | 0.22 | 0.41 | 0.29 |
| 4 | 115339308 | rs13120563 | -1.77 | -0.415 | 0.555 | 1.07 | -0.009 | -0.022 | -0.022 | 0 | 0.38 | 0.24 | 0.48 | 0.43 | 0.42 | 0.38 |
| 4 | 115359532 | rs1841432 | -0.829 | -1.177 | 0.678 | 0.793 | 0.005 | 0.022 | 0.022 | 0 | 0.15 | 0.09 | 0.11 | 0.03 | 0.18 | 0.08 |
| 4 | 115368816 | rs1460775 | -2.026 | -1.381 | 1.184 | 1.175 | 0.037 | -0.001 | -0.001 | 0.104 | 0.51 | 0.67 | 0.5 | 0.47 | 0.46 | 0.44 |
| 4 | 115370762 | rs13434798 | -0.26 | -1.429 | 1.323 | 1.138 | -0.018 | -0.021 | -0.021 | 0.199 | 0.1 | 0.14 | 0.03 | 0.03 | 0.05 | 0.07 |
| 4 | 115374527 | rs7660554 | 0.911 | -1.635 | 1.434 | 1.162 | -0.017 | 0.003 | 0.003 | 0 | 0.67 | 0.55 | 0.65 | 0.66 | 0.67 | 0.73 |
| 4 | 115381844 | rs4834411 | 1.897 | -1.538 | 0.876 | 1.059 | 0.074 | 0.001 | 0.001 | 0 | 0.51 | 0.09 | 0.5 | 0.48 | 0.58 | 0.57 |
| 4 | 115385540 | rs2388996 | 1.705 | -1.512 | 0.996 | 1.072 | 0.065 | 0.006 | 0.006 | 0 | 0.53 | 0.17 | 0.52 | 0.48 | 0.58 | 0.58 |
| 4 | 115392483 | rs6819276 | 0.759 | -1.746 | 1.093 | 0.339 | 0.001 | 0.012 | 0.012 | 0 | 0.06 | 0.04 | 0.03 | 0 | 0.07 | 0 |
| 4 | 115398200 | rs1870653 | 1.57 | -1.79 | 1.46 | 0.33 | -0.066 | -0.098 | -0.098 | 2.26 | 0.78 | 0.46 | 0.9 | 0.97 | 0.89 | 0.85 |
| 4 | 115399738 | rs10857027 | 0.839 | -1.654 | 1.386 | 0.33 | -0.034 | -0.095 | -0.095 | 0.903 | 0.84 | 0.65 | 0.92 | 1 | 0.9 | 0.95 |
| 4 | 115405345 | rs1460770 | -0.874 | -1.501 | 1.396 | 0.489 | 0 | 0.004 | 0.004 | 0.701 | 0.3 | 0.32 | 0.4 | 0.38 | 0.39 | 0.28 |
| 4 | 115411284 | rs11722041 | 2.934 | -1.774 | 1.111 | 0.438 | -0.189 | 0.064 | 0.064 | 0 | 0.33 | 0.02 | 0.25 | 0.18 | 0.41 | 0.24 |
| 4 | 115412047 | rs4833457 | -0.759 | -1.727 | 1.23 | 0.48 | 0 | 0.003 | 0.003 | 0.701 | 0.3 | 0.31 | 0.4 | 0.38 | 0.39 | 0.29 |
| 4 | 115412233 | rs9995562 | 0.062 | -1.758 | 1.245 | 0.468 | -0.003 | -0.027 | -0.027 | 0 | 0.12 | 0.11 | 0.06 | 0 | 0.1 | 0.04 |
| 4 | 115415910 | rs6839599 | 1.738 | -1.753 | 1.332 | 0.473 | 0.011 | 0.007 | 0.007 | 0.023 | 0.55 | 0.37 | 0.52 | 0.62 | 0.51 | 0.67 |
| 4 | 115423197 | rs6839694 | 0.865 | -1.751 | 1.382 | 0.475 | -0.004 | 0.013 | 0.013 | 0 | 0.67 | 0.41 | 0.58 | 0.62 | 0.61 | 0.72 |
| 4 | 115451608 | rs2100782 | -1.676 | -1.701 | 0.785 | 0.489 | 0.002 | 0 | 0 | 0 | 0.42 | 0.38 | 0.45 | 0.38 | 0.49 | 0.33 |
| 4 | 115458057 | rs11938586 | -1.098 | -1.708 | 0.786 | 0.497 | 0.003 | -0.002 | -0.002 | 0 | 0.2 | 0.28 | 0.25 | 0.19 | 0.27 | 0.19 |
| 4 | 115458168 | rs11938628 | -0.412 | -1.706 | 0.753 | 0.514 | -0.036 | 0.013 | 0.013 | 0.753 | 0.2 | 0.45 | 0.21 | 0.18 | 0.19 | 0.1 |
| 4 | 115462264 | rs1903364 | -0.538 | -1.23 | 0.938 | 0.851 | 0.003 | 0.004 | 0.004 | 0 | 0.14 | 0.18 | 0.22 | 0.2 | 0.2 | 0.19 |
| 4 | 115467459 | rs10516603 | 0.254 | -1.212 | 0.967 | 0.861 | -0.011 | 0.014 | 0.014 | 0 | 0.16 | 0.26 | 0.16 | 0.18 | 0.16 | 0.1 |
| 4 | 115484827 | rs10017302 | -0.158 | -1.058 | 1.102 | 0.951 | 0.033 | 0.005 | 0.005 | 0.021 | 0.94 | 0.78 | 0.96 | 1 | 0.98 | 1 |
| 4 | 115490751 | rs17678871 | -0.362 | -1.133 | 1.045 | 0.968 | -0.025 | -0.025 | -0.025 | 1.062 | 0.13 | 0.09 | 0.21 | 0.2 | 0.19 | 0.18 |
| 4 | 115490786 | rs17678877 | 0.584 | -1.179 | 0.974 | 0.986 | 0.016 | -0.038 | -0.038 | 0 | 0.13 | 0.05 | 0.22 | 0.44 | 0.07 | 0.33 |
| 4 | 115499117 | rs12504058 | -0.428 | -0.783 | 1.262 | 0.894 | -0.041 | -0.041 | -0.041 | 0 | 0.13 | 0.05 | 0.21 | 0.2 | 0.19 | 0.19 |
| 4 | 115499674 | rs12650546 | -1.803 | -0.801 | 1.252 | 0.871 | -0.003 | -0.001 | -0.001 | 0.84 | 0.41 | 0.52 | 0.43 | 0.38 | 0.45 | 0.33 |
| 4 | 115506203 | rs17047249 | 1.358 | -0.26 | 0.867 | 0.971 | 0.001 | 0.008 | 0.008 | 0 | 0.05 | 0.03 | 0.02 | 0.01 | 0.03 | 0.04 |
| 4 | 115514975 | rs7436074 | -0.598 | -0.048 | 0.962 | 1.177 | -0.057 | -0.052 | -0.052 | 0.188 | 0.12 | 0.03 | 0.21 | 0.19 | 0.19 | 0.19 |
| 4 | 115530635 | rs4130705 | 0.655 | -0.02 | 1.072 | 1.079 | 0.007 | 0.027 | 0.027 | 0 | 0.23 | 0.17 | 0.12 | 0.06 | 0.21 | 0.16 |
| 4 | 115534816 | rs4585328 | 1.001 | -0.066 | 1.111 | 1.133 | 0.069 | 0.086 | 0.086 | 0.045 | 0.59 | 0.19 | 0.74 | 0.82 | 0.69 | 0.78 |
| 4 | 115541961 | rs9685397 | -0.831 | -0.041 | 1.168 | 1.108 | -0.01 | -0.032 | -0.032 | 0.927 | 0.07 | 0.22 | 0.04 | 0 | 0.02 | 0 |
| 4 | 115554439 | rs7677294 | 0.813 | -0.011 | 1.045 | 0.977 | -0.008 | 0.083 | 0.083 | 0 | 0.22 | 0.03 | 0.1 | 0.06 | 0.2 | 0.16 |
| 4 | 115558241 | rs6533803 | 0.664 | -0.011 | 1.053 | 0.977 | 0.01 | 0.027 | 0.027 | 0 | 0.23 | 0.17 | 0.11 | 0.06 | 0.2 | 0.16 |
| 4 | 115565659 | rs13152031 | 0.923 | -0.027 | 1.055 | 0.987 | 0.068 | 0.09 | 0.09 | 0.034 | 0.59 | 0.2 | 0.74 | 0.82 | 0.69 | 0.78 |
| 4 | 115566754 | rs4277792 | 0.718 | -0.024 | 1.054 | 0.849 | -0.023 | -0.053 | -0.053 | 0 | 0.9 | 0.53 | 0.94 | 1 | 0.97 | 1 |
| 4 | 115570238 | rs4377607 | 0.656 | -0.024 | 1.054 | 0.849 | -0.016 | -0.048 | -0.048 | 0.337 | 0.9 | 0.57 | 0.95 | 1 | 0.97 | 1 |
| 4 | 115583723 | rs4834426 | 0.225 | 0.012 | 0.424 | 0.866 | 0.017 | -0.016 | -0.016 | 0.878 | 0.92 | 0.58 | 0.95 | 1 | 0.97 | 1 |
| 4 | 115586696 | rs4833463 | 0.851 | 0.009 | 0.436 | 0.865 | -0.025 | -0.056 | -0.056 | 0 | 0.9 | 0.54 | 0.94 | 1 | 0.97 | 1 |
| 4 | 115589159 | rs6831964 | 0.237 | 0.053 | 0.458 | 0.887 | 0.02 | -0.014 | -0.014 | 0 | 0.92 | 0.59 | 0.95 | 1 | 0.97 | 1 |
| 4 | 115595647 | rs6815181 | 0.081 | 0.091 | 0.108 | 0.905 | -0.014 | -0.047 | -0.047 | 0.186 | 0.9 | 0.58 | 0.95 | 1 | 0.97 | 1 |
| 4 | 115596092 | rs7686301 | 0.181 | 0.095 | -0.203 | 0.907 | -0.027 | -0.057 | -0.057 | 0.016 | 0.9 | 0.53 | 0.94 | 1 | 0.97 | 1 |
| 4 | 115605064 | rs4834428 | 0.605 | 0.34 | -1.047 | 1.06 | 0.068 | 0.1 | 0.1 | 0.009 | 0.59 | 0.21 | 0.74 | 0.83 | 0.69 | 0.78 |
| 4 | 115607637 | rs11098263 | 1.261 | 0.477 | -1.117 | 1.101 | -0.043 | -0.021 | -0.021 | 0 | 0.5 | 0.08 | 0.51 | 0.6 | 0.51 | 0.64 |
| 4 | 115638209 | rs4269197 | -1.452 | 1.027 | 0.338 | 2.227 | -0.003 | 0.001 | 0.001 | 2.932 | 0.19 | 0.2 | 0.43 | 0.62 | 0.24 | 0.52 |
| 4 | 115640581 | rs12510781 | -1.037 | 0.663 | 0.26 | 1.619 | -0.006 | 0.027 | 0.027 | 0 | 0.06 | 0.13 | 0.07 | 0.12 | 0.04 | 0.06 |
| 4 | 115656533 | rs7700052 | 1.19 | 0.692 | -0.173 | 1.344 | 0 | -0.002 | -0.002 | 0.061 | 0.75 | 0.26 | 0.63 | 0.66 | 0.74 | 0.8 |
| 4 | 115665320 | rs6533817 | -1.113 | 0.578 | 0.266 | 1.343 | -0.004 | 0.106 | 0.106 | 0.12 | 0.14 | 0.37 | 0.2 | 0.2 | 0.12 | 0.06 |
| 4 | 115667187 | rs4833467 | 0 | 0.693 | -0.789 | 0.506 | -0.041 | 0.101 | 0.101 | 0.07 | 0.61 | 0.81 | 0.64 | 0.82 | 0.54 | 0.75 |
| 4 | 115672966 | rs4286532 | 0.33 | 0.164 | -1.252 | 0.895 | -0.03 | 0.008 | 0.008 | 4.047 | 0.37 | 0.12 | 0.38 | 0.3 | 0.35 | 0.45 |
| 4 | 115678239 | rs4834440 | 0.451 | -0.203 | -1.744 | 0.525 | -0.088 | -0.094 | -0.094 | 0.482 | 0.18 | 0.03 | 0.31 | 0.27 | 0.22 | 0.18 |
| 4 | 115681827 | rs4834442 | -0.356 | -0.296 | -1.405 | 0.439 | -0.037 | -0.038 | -0.038 | 0 | 0.79 | 0.89 | 0.68 | 0.72 | 0.76 | 0.8 |
| 4 | 115688354 | rs11723161 | 0.755 | -0.676 | -0.938 | 0.89 | 0.014 | -0.001 | -0.001 | 0.132 | 0.53 | 0.27 | 0.5 | 0.54 | 0.58 | 0.52 |
| 4 | 115692870 | rs4508906 | -1.735 | -0.835 | 0.675 | 0.653 | 0.02 | 0.128 | 0.128 | 5.156 | 0.1 | 0.52 | 0.12 | 0.16 | 0.07 | 0.06 |
| 4 | 115720115 | rs4376167 | 1.126 | -1.116 | 0.945 | 0.557 | -0.09 | 0.157 | 0.157 | 0.647 | 0.44 | 0.12 | 0.39 | 0.19 | 0.52 | 0.27 |
| 4 | 115739428 | rs4574434 | -0.242 | -1.45 | 0.85 | 0.522 | -0.048 | -0.061 | -0.061 | 0 | 0.71 | 0.92 | 0.65 | 0.4 | 0.69 | 0.61 |
| 4 | 115749005 | rs6843860 | -0.88 | -1.518 | 1.554 | 0.533 | -0.052 | -0.037 | -0.037 | 0 | 0.23 | 0.55 | 0.25 | 0.16 | 0.17 | 0.32 |

| | | | | | | | | | | | | | | | | |
|---|---|---|---|---|---|---|---|---|---|---|---|---|---|---|---|---|
| 4 | 115755760 | rs11098270 | 0.161 | -1.587 | 1.47 | 0.465 | 0.006 | 0.004 | 0.004 | 1.898 | 0.16 | 0.1 | 0.14 | 0.06 | 0.11 | 0.09 |
| 4 | 115769330 | rs10022934 | -2.026 | -1.501 | 1.655 | 0.461 | 0.02 | 0.037 | 0.037 | 0 | 0.12 | 0.43 | 0.21 | 0.14 | 0.13 | 0.08 |
| 4 | 115786656 | rs13120543 | 1.6 | -1.144 | 1.058 | 0.255 | -0.099 | 0.17 | 0.17 | 0.235 | 0.41 | 0.09 | 0.37 | 0.13 | 0.48 | 0.24 |
| 4 | 115787479 | rs17486233 | 0.764 | -1.242 | 1.201 | 0.435 | 0.01 | 0.013 | 0.013 | 0.841 | 0.11 | 0.01 | 0.04 | 0.07 | 0.08 | 0.07 |
| 4 | 115788728 | rs7665509 | -0.667 | -1.066 | 1.577 | 0.547 | -0.01 | -0.011 | -0.011 | 1.295 | 0.06 | 0.1 | 0.1 | 0.06 | 0.05 | 0.06 |
| 4 | 115813013 | rs11938588 | -1.682 | -1.445 | 1.412 | 0.135 | -0.055 | -0.066 | -0.066 | 0 | 0.19 | 0.5 | 0.16 | 0.12 | 0.12 | 0.3 |
| 4 | 115823991 | rs11945934 | -0.828 | -1.402 | 1.483 | 0.21 | -0.009 | 0.008 | 0.008 | 0.019 | 0.11 | 0.29 | 0.14 | 0.12 | 0.1 | 0.06 |
| 4 | 115829615 | rs7694381 | -0.66 | -1.103 | -0.015 | -0.075 | -0.017 | -0.048 | -0.048 | 0 | 0.67 | 0.88 | 0.64 | 0.45 | 0.66 | 0.57 |
| 4 | 115829919 | rs7675371 | -2.186 | -0.797 | 0.714 | 0.212 | -0.026 | 0.031 | 0.031 | 0.28 | 0.22 | 0.55 | 0.24 | 0.32 | 0.18 | 0.31 |
| 4 | 115844385 | rs17486477 | 1.884 | -0.832 | -0.194 | -0.205 | -0.071 | 0.056 | 0.056 | 0.028 | 0.25 | 0.09 | 0.33 | 0.12 | 0.38 | 0.17 |
| 4 | 115844634 | rs11939954 | -2.399 | -0.848 | 0.701 | 0.51 | -0.023 | 0.034 | 0.034 | 0 | 0.23 | 0.63 | 0.24 | 0.43 | 0.18 | 0.33 |
| 4 | 115851183 | rs1036757 | -1.422 | -0.61 | 0.67 | 0.668 | -0.031 | -0.089 | -0.089 | 0 | 0.14 | 0.36 | 0.07 | 0 | 0.09 | 0.06 |
| 4 | 115858331 | rs1020219 | -1.522 | -0.642 | 0.535 | 0.198 | -0.035 | -0.102 | -0.102 | 0 | 0.15 | 0.4 | 0.07 | 0 | 0.09 | 0.06 |
| 4 | 115867213 | rs1441639 | 0.797 | -0.902 | -0.267 | 0.702 | 0.068 | 0.029 | 0.029 | 0 | 0.52 | 0.31 | 0.66 | 0.56 | 0.68 | 0.47 |
| 4 | 115876042 | rs2018486 | 0.4 | -1.072 | 0.108 | 0.524 | 0.021 | 0.001 | 0.001 | 0.003 | 0.59 | 0.73 | 0.69 | 0.57 | 0.65 | 0.49 |
| 4 | 115897055 | rs6820993 | 1.702 | -1.456 | 0.072 | 0.299 | -0.038 | 0.044 | 0.044 | 0 | 0.31 | 0.16 | 0.43 | 0.17 | 0.54 | 0.15 |
| 4 | 115910256 | rs2028481 | 0.031 | -1.511 | 0.188 | 0.347 | 0.025 | 0.002 | 0.002 | 0 | 0.43 | 0.33 | 0.32 | 0.43 | 0.34 | 0.5 |
| 4 | 115912013 | rs279505 | -1.981 | -1.532 | 0.195 | 0.37 | 0.019 | -0.051 | -0.051 | 0 | 0.83 | 0.92 | 0.79 | 0.6 | 0.9 | 0.68 |
| 4 | 115919123 | rs4082347 | 1.538 | -1.864 | 0.015 | 0.064 | -0.027 | -0.054 | -0.054 | 2.716 | 0.12 | 0.21 | 0.12 | 0.01 | 0.07 | 0.05 |
| 4 | 115920956 | rs2583515 | 1.483 | -1.775 | -0.265 | 0.254 | -0.035 | -0.079 | -0.079 | 0 | 0.21 | 0.05 | 0.14 | 0.32 | 0.18 | 0.39 |
| 4 | 115952955 | rs979513 | 1.016 | -1.635 | 0.2 | 0.169 | 0.011 | 0.026 | 0.026 | 0.211 | 0.45 | 0.36 | 0.55 | 0.18 | 0.63 | 0.26 |
| 4 | 115955380 | rs1552025 | 0.4 | -1.082 | 0.65 | 0.661 | -0.052 | 0.16 | 0.16 | 0.195 | 0.22 | 0.48 | 0.27 | 0.39 | 0.12 | 0.22 |
| 4 | 115964380 | rs2046050 | 1.327 | -0.887 | -0.026 | 0.778 | 0.004 | 0.001 | 0.001 | 0 | 0.16 | 0.11 | 0.22 | 0.39 | 0.11 | 0.22 |
| 4 | 115973568 | rs1383001 | 0.851 | -0.592 | -0.065 | 0.346 | -0.023 | 0.031 | 0.031 | 0.024 | 0.36 | 0.21 | 0.49 | 0.27 | 0.54 | 0.25 |
| 4 | 115995713 | rs541329 | 0.941 | -0.485 | 0.445 | 0.808 | -0.005 | 0.037 | 0.037 | 0 | 0.19 | 0.21 | 0.23 | 0.47 | 0.14 | 0.25 |
| 4 | 115996988 | rs509127 | 0.638 | -0.562 | 0.826 | 0.77 | -0.035 | 0.175 | 0.175 | 0 | 0.21 | 0.5 | 0.26 | 0.47 | 0.14 | 0.26 |
| 4 | 116007803 | rs13104877 | 0.795 | -0.119 | 0.613 | 0.483 | -0.063 | 0.05 | 0.05 | 0 | 0.3 | 0.14 | 0.42 | 0.22 | 0.46 | 0.19 |
| 4 | 116012782 | rs512349 | -0.034 | -0.299 | 0.86 | 0.503 | 0.001 | -0.016 | -0.016 | 0.065 | 0.43 | 0.53 | 0.51 | 0.32 | 0.54 | 0.24 |
| 4 | 116012863 | rs511497 | -0.957 | -0.435 | 0.29 | 0.401 | -0.046 | 0.006 | 0.006 | 1.296 | 0.65 | 0.95 | 0.76 | 0.77 | 0.7 | 0.52 |
| 4 | 116013647 | rs566198 | -0.815 | -0.349 | 0.216 | 0.6 | -0.006 | 0.001 | 0.001 | 1.085 | 0.64 | 0.89 | 0.73 | 0.67 | 0.67 | 0.52 |
| 4 | 116028808 | rs505277 | -0.775 | -0.559 | 0.378 | 0.299 | 0.027 | 0.022 | 0.022 | 0.644 | 0.64 | 0.8 | 0.73 | 0.75 | 0.72 | 0.52 |
| 4 | 116032339 | rs13105185 | -0.01 | -0.198 | 0.124 | 0.22 | -0.001 | 0.003 | 0.003 | 0 | 0.42 | 0.43 | 0.5 | 0.31 | 0.57 | 0.26 |
| 4 | 116074962 | rs1940336 | 0.006 | -0.104 | -0.296 | 0.131 | 0.021 | 0.002 | 0.002 | 0.005 | 0.57 | 0.68 | 0.67 | 0.63 | 0.63 | 0.49 |
| 4 | 116077415 | rs10018147 | -0.894 | -0.167 | -0.61 | 0.103 | -0.022 | -0.073 | -0.073 | 0.307 | 0.68 | 0.91 | 0.72 | 0.68 | 0.7 | 0.54 |
| 4 | 116081714 | rs2155516 | -0.894 | -0.167 | -0.61 | 0.115 | -0.022 | -0.076 | -0.076 | 0 | 0.68 | 0.91 | 0.72 | 0.68 | 0.7 | 0.54 |
| 4 | 116083584 | rs1940319 | -0.513 | -0.2 | -0.565 | 0.084 | 0.01 | -0.037 | -0.037 | 0 | 0.64 | 0.86 | 0.71 | 0.68 | 0.69 | 0.5 |
| 4 | 116084489 | rs10516609 | -0.798 | -0.197 | -0.561 | 0.1 | -0.021 | -0.073 | -0.073 | 0 | 0.67 | 0.91 | 0.72 | 0.68 | 0.7 | 0.53 |
| 4 | 116089855 | rs10027823 | -0.724 | -0.196 | -0.533 | 0.1 | 0.002 | -0.044 | -0.044 | 0 | 0.66 | 0.86 | 0.72 | 0.68 | 0.7 | 0.53 |
| 4 | 116091803 | rs7678790 | -0.456 | -0.222 | -0.345 | 0.1 | -0.009 | -0.019 | -0.019 | 0 | 0.09 | 0.16 | 0.04 | 0.05 | 0.06 | 0.04 |
| 4 | 116093537 | rs948186 | 1.386 | -0.289 | -0.484 | 0.029 | 0.005 | -0.058 | -0.058 | 0 | 0.18 | 0.07 | 0.19 | 0.43 | 0.17 | 0.26 |
| 4 | 116099668 | rs4834457 | 1.449 | -0.322 | -0.599 | 0.013 | -0.013 | -0.117 | -0.117 | 0 | 0.18 | 0.02 | 0.19 | 0.43 | 0.17 | 0.26 |
| 4 | 116105196 | rs17685799 | 1.496 | -0.298 | -0.586 | 0.063 | -0.009 | -0.105 | -0.105 | 0 | 0.18 | 0.03 | 0.19 | 0.43 | 0.17 | 0.26 |
| 4 | 116109043 | rs11723597 | -1.079 | -0.176 | -0.61 | -0.006 | -0.027 | -0.075 | -0.075 | 0.611 | 0.67 | 0.91 | 0.72 | 0.67 | 0.69 | 0.53 |
| 4 | 116122612 | rs6856972 | -1.07 | -0.176 | -0.61 | -0.006 | -0.027 | -0.074 | -0.074 | 0 | 0.67 | 0.91 | 0.72 | 0.67 | 0.69 | 0.53 |
| 4 | 116126250 | rs11940517 | 0.895 | -0.231 | -0.011 | -0.13 | -0.039 | 0.049 | 0.049 | 0 | 0.33 | 0.18 | 0.43 | 0.19 | 0.44 | 0.22 |
| 4 | 116130120 | rs1960049 | 0.861 | -0.244 | -0.014 | -0.116 | -0.032 | 0.039 | 0.039 | 0 | 0.33 | 0.19 | 0.43 | 0.19 | 0.44 | 0.22 |
| 4 | 116143319 | rs11946458 | -1.088 | -0.266 | 0.293 | -0.093 | 0.047 | 0.036 | 0.036 | 7.181 | 0.07 | 0.48 | 0.04 | 0 | 0.01 | 0 |
| 4 | 116151157 | rs10029472 | -0.572 | -0.343 | 0.142 | -0.087 | 0.006 | -0.01 | -0.01 | 0 | 0.33 | 0.69 | 0.29 | 0.48 | 0.25 | 0.31 |
| 4 | 116164027 | rs2048163 | -0.188 | -0.346 | 0.284 | -0.003 | 0.006 | 0.001 | 0.001 | 0 | 0.6 | 0.33 | 0.66 | 0.67 | 0.68 | 0.52 |
| 4 | 116181809 | rs13113622 | -0.319 | -0.314 | 0.447 | 0.013 | -0.013 | -0.003 | -0.003 | 0.312 | 0.61 | 0.5 | 0.68 | 0.67 | 0.68 | 0.52 |
| 4 | 116194139 | rs7694946 | -0.505 | -0.393 | 0.343 | -0.015 | -0.02 | -0.018 | -0.018 | 0 | 0.67 | 0.52 | 0.73 | 0.75 | 0.7 | 0.52 |
| 4 | 116211711 | rs13128882 | 0.88 | 0.391 | -0.222 | -0.278 | -0.1 | -0.013 | -0.013 | 0.216 | 0.3 | 0.05 | 0.31 | 0.28 | 0.36 | 0.19 |
| 4 | 116218720 | rs6853566 | -1.307 | 0.344 | -0.265 | -0.396 | -0.039 | -0.008 | -0.008 | 0.599 | 0.74 | 0.86 | 0.61 | 0.75 | 0.68 | 0.49 |
| 4 | 116230563 | rs1989243 | 1.2 | 0.161 | -0.345 | -0.466 | -0.004 | -0.137 | -0.137 | 0.03 | 0.25 | 0.03 | 0.16 | 0.41 | 0.23 | 0.25 |
| 4 | 116237539 | rs17686110 | -1.464 | 0.016 | -0.219 | -0.403 | -0.064 | -0.019 | -0.019 | 0 | 0.75 | 0.89 | 0.61 | 0.75 | 0.68 | 0.49 |
| 4 | 116243567 | rs11727958 | 0.043 | 0.18 | 0.469 | -0.403 | 0 | -0.01 | -0.01 | 1.104 | 0.61 | 0.41 | 0.51 | 0.69 | 0.61 | 0.46 |
| 4 | 116246035 | rs11725144 | 0.786 | 0.026 | 0.626 | -0.381 | -0.002 | 0.003 | 0.003 | 0 | 0.28 | 0.22 | 0.18 | 0.41 | 0.24 | 0.25 |
| 4 | 116261419 | rs13144926 | 1.088 | 0.05 | 0.585 | -0.378 | -0.025 | 0.02 | 0.02 | 0.061 | 0.31 | 0.17 | 0.33 | 0.28 | 0.36 | 0.19 |
| 4 | 116266582 | rs6829587 | -0.018 | 0.046 | 0.699 | -0.377 | 0.004 | -0.013 | -0.013 | 0.048 | 0.6 | 0.42 | 0.51 | 0.69 | 0.6 | 0.45 |
| 4 | 116276218 | rs7434749 | 0.976 | 0.048 | 0.644 | -0.379 | -0.022 | 0.022 | 0.022 | 0.273 | 0.32 | 0.17 | 0.33 | 0.28 | 0.36 | 0.2 |
| 4 | 116279442 | rs12651062 | -1.338 | 0.028 | 0.833 | -0.395 | -0.025 | -0.003 | -0.003 | 0.512 | 0.73 | 0.8 | 0.6 | 0.75 | 0.68 | 0.49 |
| 4 | 116283010 | rs10000358 | -0.608 | 0.069 | 0.721 | -0.355 | 0.016 | -0.019 | -0.019 | 0 | 0.66 | 0.5 | 0.55 | 0.75 | 0.63 | 0.49 |
| 4 | 116291436 | rs7437820 | 0.969 | 0.088 | 0.698 | -0.357 | -0.022 | 0.024 | 0.024 | 0.273 | 0.32 | 0.17 | 0.33 | 0.28 | 0.36 | 0.2 |
| 4 | 116316085 | rs4834476 | 0.759 | 0.089 | 0.718 | -0.354 | -0.002 | 0.003 | 0.003 | 0 | 0.28 | 0.23 | 0.18 | 0.41 | 0.24 | 0.25 |
| 4 | 116332722 | rs6836582 | -0.564 | 0.1 | 0.66 | -0.352 | 0.02 | -0.024 | -0.024 | 0.055 | 0.66 | 0.46 | 0.55 | 0.75 | 0.63 | 0.49 |
| 4 | 116336119 | rs6533873 | -0.568 | 0.092 | 0.638 | -0.355 | 0.017 | -0.021 | -0.021 | 0.432 | 0.66 | 0.49 | 0.55 | 0.75 | 0.63 | 0.49 |
| 4 | 116340193 | rs6854632 | -0.578 | 0.1 | 0.642 | -0.352 | 0.018 | -0.021 | -0.021 | 0.062 | 0.66 | 0.49 | 0.55 | 0.75 | 0.63 | 0.49 |
| 4 | 116349081 | rs6830739 | -0.273 | -0.585 | 0.57 | -0.171 | -0.008 | 0.007 | 0.007 | 0 | 0.35 | 0.32 | 0.22 | 0.46 | 0.28 | 0.3 |
| 4 | 116352641 | rs4833482 | 0.746 | -0.223 | 0.155 | 0.024 | 0.015 | -0.048 | -0.048 | 0.543 | 0.26 | 0.14 | 0.18 | 0.41 | 0.23 | 0.25 |

| | | | | | | | | | | | | | | | | |
|---|---|---|---|---|---|---|---|---|---|---|---|---|---|---|---|---|
| 4 | 116375611 | rs12508013 | 0.175 | -0.233 | 0.339 | -0.026 | 0.003 | -0.006 | -0.006 | 0 | 0.31 | 0.31 | 0.22 | 0.41 | 0.28 | 0.25 |
| 4 | 116377844 | rs13131555 | 0.789 | -0.183 | 0.344 | -0.228 | -0.038 | 0.038 | 0.038 | 0.279 | 0.36 | 0.19 | 0.36 | 0.27 | 0.41 | 0.24 |
| 4 | 116378717 | rs6850894 | 0.16 | -0.247 | 0.343 | 0.059 | 0.012 | -0.025 | -0.025 | 0.09 | 0.31 | 0.28 | 0.22 | 0.4 | 0.28 | 0.3 |
| 4 | 116387928 | rs4834488 | 0.17 | -0.292 | 0.288 | 0.146 | 0.028 | -0.074 | -0.074 | 0 | 0.29 | 0.17 | 0.19 | 0.46 | 0.25 | 0.29 |
| 4 | 116389374 | rs12509776 | 0.198 | -0.3 | 0.484 | 0.142 | 0.009 | -0.017 | -0.017 | 0 | 0.3 | 0.26 | 0.2 | 0.46 | 0.25 | 0.29 |
| 4 | 116395664 | rs10026917 | -0.388 | -0.417 | 0.53 | -0.172 | -0.003 | -0.002 | -0.002 | 0 | 0.62 | 0.55 | 0.57 | 0.66 | 0.68 | 0.49 |
| 4 | 116399649 | rs6533913 | -0.805 | -0.444 | 0.514 | -0.168 | -0.004 | -0.004 | -0.004 | 0.113 | 0.66 | 0.61 | 0.59 | 0.72 | 0.7 | 0.51 |
| 4 | 116406659 | rs7668500 | -0.687 | -0.486 | 0.731 | -0.196 | -0.007 | -0.007 | -0.007 | 0 | 0.65 | 0.57 | 0.58 | 0.72 | 0.7 | 0.51 |
| 4 | 116409580 | rs9884192 | -0.036 | -0.34 | 0.753 | -0.208 | 0.003 | -0.007 | -0.007 | 0 | 0.41 | 0.53 | 0.42 | 0.32 | 0.47 | 0.24 |
| 4 | 116411907 | rs13129771 | 0.891 | -0.242 | 0.615 | -0.224 | -0.028 | 0.015 | 0.015 | 0 | 0.29 | 0.19 | 0.34 | 0.26 | 0.37 | 0.22 |
| 4 | 116420387 | rs10012098 | -0.784 | -0.281 | 0.85 | -0.243 | 0.003 | -0.018 | -0.018 | 0 | 0.11 | 0.34 | 0.09 | 0.06 | 0.1 | 0.02 |
| 4 | 116421669 | rs6533922 | -1.01 | -0.444 | 0.843 | -0.303 | 0.005 | 0.005 | 0.005 | 0.21 | 0.67 | 0.7 | 0.6 | 0.72 | 0.7 | 0.51 |
| 4 | 116426581 | rs7435768 | -0.904 | -0.388 | 1.034 | -0.04 | 0.002 | -0.018 | -0.018 | 0 | 0.11 | 0.34 | 0.09 | 0.06 | 0.1 | 0.02 |
| 4 | 116432420 | rs7438394 | -0.358 | -0.416 | 0.843 | 0.159 | -0.011 | 0.013 | 0.013 | 0.035 | 0.36 | 0.54 | 0.26 | 0.43 | 0.33 | 0.3 |
| 4 | 116464839 | rs4522939 | -0.559 | -0.354 | 0.956 | 0.188 | -0.001 | 0.036 | 0.036 | 0.08 | 0.05 | 0.22 | 0.01 | 0.06 | 0.01 | 0.01 |
| 4 | 116490478 | rs4292399 | -0.351 | -0.296 | 0.396 | 0.157 | -0.002 | 0.001 | 0.001 | 0.565 | 0.36 | 0.6 | 0.26 | 0.43 | 0.33 | 0.29 |
| 4 | 116509605 | rs13105584 | 0.648 | -0.147 | -0.623 | 0.386 | -0.102 | -0.029 | -0.029 | 0 | 0.36 | 0.12 | 0.35 | 0.36 | 0.44 | 0.26 |
| 4 | 116527361 | rs7664444 | 0.59 | -0.035 | -0.597 | 0.335 | -0.073 | 0.028 | 0.028 | 0 | 0.37 | 0.18 | 0.36 | 0.28 | 0.42 | 0.25 |
| 4 | 116529926 | rs17048021 | -0.005 | -0.133 | -0.312 | 0.198 | 0 | -0.025 | -0.025 | 0 | 0.07 | 0.13 | 0.06 | 0 | 0.07 | 0.01 |
| 4 | 116529968 | rs11098314 | 0.304 | -0.588 | -0.199 | 0.269 | 0.003 | -0.004 | -0.004 | 0 | 0.28 | 0.28 | 0.27 | 0.35 | 0.26 | 0.46 |
| 4 | 116530435 | rs1545763 | 0.153 | -0.557 | -0.542 | 0.313 | 0.021 | 0.003 | 0.003 | 0 | 0.62 | 0.34 | 0.64 | 0.65 | 0.66 | 0.53 |
| 4 | 116541424 | rs1871306 | 0.361 | -0.64 | -0.982 | 0.176 | -0.016 | -0.069 | -0.069 | 0 | 0.27 | 0.07 | 0.3 | 0.35 | 0.26 | 0.31 |
| 4 | 116547100 | rs4146409 | 1.536 | -1.01 | -0.307 | -0.204 | -0.01 | -0.02 | -0.02 | 0.174 | 0.81 | 0.29 | 0.86 | 0.93 | 0.9 | 0.94 |
| 4 | 116548435 | rs1478926 | 1.009 | -1.185 | -0.602 | -0.11 | 0.054 | 0.076 | 0.076 | 0 | 0.59 | 0.22 | 0.64 | 0.68 | 0.67 | 0.51 |
| 4 | 116560690 | rs10002695 | -1.784 | -1.132 | -0.038 | -0.204 | -0.022 | -0.052 | -0.052 | 0 | 0.2 | 0.64 | 0.15 | 0.07 | 0.12 | 0.07 |
| 4 | 116598539 | rs7677317 | -1.406 | -1.166 | 0.743 | -0.271 | -0.026 | -0.091 | -0.091 | 0.316 | 0.14 | 0.46 | 0.11 | 0.02 | 0.09 | 0.03 |
| 4 | 116606013 | rs6844072 | 0.174 | -1.086 | 1.024 | -0.234 | 0.001 | -0.028 | -0.028 | 0 | 0.06 | 0.13 | 0.06 | 0 | 0.07 | 0.01 |
| 4 | 116612109 | rs11942806 | -1.276 | -1.129 | 1.509 | -0.255 | 0.015 | -0.055 | -0.055 | 0 | 0.1 | 0.42 | 0.11 | 0 | 0.08 | 0.01 |
| 4 | 116658760 | rs13112862 | -1.253 | -1.425 | 0.885 | -0.419 | 0.032 | 0.049 | 0.049 | 0 | 0.39 | 0.71 | 0.41 | 0.3 | 0.36 | 0.46 |
| 4 | 116658800 | rs10017879 | -0.721 | -1.331 | 0.963 | -0.247 | 0.013 | -0.046 | -0.046 | 0.526 | 0.09 | 0.28 | 0.09 | 0 | 0.08 | 0.01 |
| 4 | 116667561 | rs10029219 | -0.165 | -1.272 | 0.668 | -0.219 | 0 | -0.042 | -0.042 | 0 | 0.08 | 0.2 | 0.08 | 0 | 0.07 | 0.01 |
| 4 | 116684526 | rs7677410 | 0.175 | -1.268 | 0.344 | -0.057 | 0.009 | -0.012 | -0.012 | 0 | 0.23 | 0.08 | 0.26 | 0.29 | 0.24 | 0.41 |
| 4 | 116710985 | rs2389089 | 0.159 | -0.55 | 0.688 | -0.048 | 0.008 | -0.029 | -0.029 | 0.392 | 0.31 | 0.41 | 0.26 | 0.23 | 0.3 | 0.2 |
| 4 | 116718238 | rs10516621 | 0.288 | -0.524 | 0.216 | -0.035 | -0.038 | 0.046 | 0.046 | 0 | 0.29 | 0.17 | 0.25 | 0.23 | 0.3 | 0.2 |
| 4 | 116727973 | rs1013411 | 0.834 | -0.501 | -0.012 | 0.054 | -0.013 | 0.016 | 0.016 | 0 | 0.34 | 0.12 | 0.23 | 0.25 | 0.34 | 0.2 |
| 4 | 116730435 | rs7680117 | -1.281 | -0.475 | 0.348 | -0.05 | 0 | -0.005 | -0.005 | 0 | 0.84 | 0.84 | 0.76 | 0.87 | 0.81 | 0.72 |
| 4 | 116740301 | rs13106872 | 0.759 | -0.509 | 0.412 | -0.047 | -0.077 | 0.044 | 0.044 | 0.183 | 0.29 | 0.08 | 0.24 | 0.23 | 0.3 | 0.22 |
| 4 | 116766626 | rs1549805 | 0.155 | -0.748 | 0.523 | -0.141 | -0.002 | 0.057 | 0.057 | 0 | 0.37 | 0.2 | 0.26 | 0.23 | 0.31 | 0.25 |
| 4 | 116769746 | rs13151706 | 0.204 | -0.74 | 0.341 | -0.137 | -0.01 | 0.082 | 0.082 | 0 | 0.37 | 0.15 | 0.26 | 0.23 | 0.31 | 0.25 |
| 4 | 116786795 | rs9999955 | -1.461 | -0.668 | 0.031 | -0.113 | -0.001 | -0.006 | -0.006 | 0 | 0.84 | 0.85 | 0.77 | 0.87 | 0.81 | 0.72 |
| 4 | 116804645 | rs1430934 | 0.639 | -0.613 | 0.145 | -0.061 | -0.004 | 0.004 | 0.004 | 0 | 0.43 | 0.38 | 0.48 | 0.59 | 0.5 | 0.44 |
| 4 | 116834084 | rs1821011 | 0.56 | -0.103 | -0.397 | -0.237 | -0.067 | 0.05 | 0.05 | 0.096 | 0.29 | 0.1 | 0.25 | 0.23 | 0.3 | 0.22 |
| 4 | 116844501 | rs11098326 | -0.865 | 0.073 | -0.244 | -0.152 | 0.001 | -0.022 | -0.022 | 0 | 0.8 | 0.73 | 0.72 | 0.9 | 0.77 | 0.72 |
| 4 | 116848573 | rs1025397 | 0.701 | 0.106 | -0.649 | -0.127 | -0.031 | 0.031 | 0.031 | 0.02 | 0.28 | 0.11 | 0.21 | 0.27 | 0.26 | 0.24 |
| 4 | 116862170 | rs9307416 | -0.31 | 0.114 | -0.323 | 0.219 | -0.005 | 0.053 | 0.053 | 0 | 0.52 | 0.62 | 0.5 | 0.63 | 0.51 | 0.47 |
| 4 | 116880079 | rs6824665 | -0.55 | 0.509 | -0.094 | 0.314 | 0.015 | 0.014 | 0.014 | 0 | 0.69 | 0.6 | 0.68 | 0.57 | 0.67 | 0.67 |
| 4 | 116889740 | rs1456365 | 0.023 | 0.188 | -1.033 | 0.001 | -0.061 | -0.113 | -0.113 | 0 | 0.35 | 0.14 | 0.52 | 0.44 | 0.44 | 0.37 |
| 4 | 116893245 | rs17586724 | 0.97 | 0.132 | -1.036 | -0.081 | 0.035 | 0.069 | 0.069 | 0 | 0.27 | 0.07 | 0.09 | 0.13 | 0.18 | 0.26 |
| 4 | 116894599 | rs13151079 | -1.306 | 0.031 | -1.065 | -0.164 | -0.099 | -0.097 | -0.097 | 0.552 | 0.73 | 0.94 | 0.71 | 0.57 | 0.67 | 0.67 |
| 4 | 116895632 | rs17502284 | 0.94 | -0.329 | -0.797 | -0.23 | 0.03 | 0.052 | 0.052 | 0 | 0.28 | 0.1 | 0.1 | 0.13 | 0.18 | 0.26 |
| 4 | 116908711 | rs7669751 | -0.6 | 0.205 | 0.144 | 0.094 | -0.006 | -0.004 | -0.004 | 0.104 | 0.7 | 0.69 | 0.65 | 0.56 | 0.62 | 0.6 |
| 4 | 116916813 | rs11098331 | -0.152 | 0.139 | 0.796 | 0.028 | 0.022 | 0 | 0 | 0.379 | 0.63 | 0.49 | 0.68 | 0.6 | 0.56 | 0.57 |
| 4 | 116917134 | rs10776499 | -0.103 | 0.132 | 0.984 | 0.024 | 0.015 | -0.001 | -0.001 | 0 | 0.63 | 0.42 | 0.66 | 0.6 | 0.56 | 0.57 |
| 4 | 116920093 | rs2045320 | 0.309 | 0.113 | 0.686 | 0.144 | 0 | -0.004 | -0.004 | 2.068 | 0.56 | 0.5 | 0.43 | 0.23 | 0.43 | 0.48 |
| 4 | 116927517 | rs10033887 | -0.108 | 0.071 | 0.876 | 0.124 | -0.008 | 0.026 | 0.026 | 0.049 | 0.6 | 0.63 | 0.44 | 0.23 | 0.44 | 0.51 |
| 4 | 116927825 | rs10011809 | -1.501 | 0.079 | 1.06 | 0.203 | -0.01 | 0.189 | 0.189 | 1.121 | 0.17 | 0.55 | 0.32 | 0.37 | 0.19 | 0.29 |
| 4 | 116932834 | rs13135582 | 1.06 | -0.003 | 0.589 | -0.036 | -0.039 | 0.042 | 0.042 | 0.014 | 0.17 | 0.02 | 0.26 | 0.09 | 0.18 | 0.19 |
| 4 | 116942805 | rs6854280 | -0.538 | 0.299 | 1.284 | 0.46 | 0.015 | 0.151 | 0.151 | 0 | 0.11 | 0.25 | 0.29 | 0.37 | 0.18 | 0.26 |
| 4 | 116954591 | rs6851349 | -0.38 | 0.242 | 1.04 | 0.475 | -0.008 | 0.13 | 0.13 | 0 | 0.11 | 0.2 | 0.26 | 0.38 | 0.13 | 0.25 |
| 4 | 116973405 | rs12509225 | -0.334 | 0.287 | 0.616 | 0.377 | 0.002 | 0.011 | 0.011 | 0 | 0.13 | 0.09 | 0.27 | 0.36 | 0.16 | 0.25 |
| 4 | 116986729 | rs814382 | 0.84 | 0.306 | 0.435 | 0.245 | 0.003 | 0.082 | 0.082 | 0.688 | 0.54 | 0.3 | 0.42 | 0.21 | 0.42 | 0.48 |
| 4 | 116987639 | rs814383 | -0.755 | 0.75 | 0.232 | 0.427 | -0.068 | -0.066 | -0.066 | 0 | 0.75 | 0.89 | 0.75 | 0.57 | 0.64 | 0.77 |
| 4 | 116999572 | rs814397 | -0.752 | 0.885 | -0.362 | 0.495 | -0.112 | -0.109 | -0.109 | 0 | 0.75 | 0.94 | 0.75 | 0.57 | 0.63 | 0.77 |
| 4 | 117017017 | rs701779 | 0.391 | 0.385 | -0.124 | 0.644 | -0.03 | 0.067 | 0.067 | 0.108 | 0.81 | 0.71 | 0.71 | 0.69 | 0.84 | 0.74 |
| 4 | 117042009 | rs701793 | 0.14 | 1.122 | -0.885 | 0.637 | -0.069 | -0.059 | -0.059 | 0.09 | 0.24 | 0.11 | 0.28 | 0.42 | 0.4 | 0.23 |
| 4 | 117047226 | rs292911 | 0.488 | 0.231 | -0.466 | 0.441 | -0.03 | -0.008 | -0.008 | 0 | 0.77 | 0.28 | 0.71 | 0.79 | 0.86 | 0.7 |
| 4 | 117055445 | rs13108295 | 0.298 | 0.81 | -1.305 | 0.349 | -0.078 | -0.126 | -0.126 | 0.244 | 0.22 | 0.03 | 0.36 | 0.43 | 0.36 | 0.28 |
| 4 | 117073520 | rs6858305 | -0.434 | 0.218 | -0.538 | -0.779 | 0.012 | -0.025 | -0.025 | 0 | 0.31 | 0.43 | 0.33 | 0.21 | 0.31 | 0.38 |
| 4 | 117078327 | rs1506732 | 0.426 | 0.2 | -0.306 | -0.788 | 0 | -0.005 | -0.005 | 0 | 0.66 | 0.31 | 0.64 | 0.79 | 0.69 | 0.62 |

| | | | | | | | | | | | | | | | | |
|---|---|---|---|---|---|---|---|---|---|---|---|---|---|---|---|---|
| 4 | 117084740 | rs1588040 | 0.808 | -0.056 | -0.875 | -0.94 | -0.065 | 0.11 | 0.11 | 2.063 | 0.61 | 0.05 | 0.61 | 0.79 | 0.66 | 0.57 |
| 4 | 117099219 | rs17048339 | -0.792 | -1.91 | -0.274 | -1.161 | 0 | -0.002 | -0.002 | 0.017 | 0.28 | 0.71 | 0.23 | 0.15 | 0.2 | 0.14 |
| 4 | 117099769 | rs6533958 | 0.519 | -1.868 | 0.169 | -1.054 | -0.04 | -0.04 | -0.04 | 0 | 0.16 | 0.45 | 0.13 | 0.08 | 0.08 | 0.09 |
| 4 | 117135380 | rs1525760 | 0.36 | -1.984 | -0.013 | -1.116 | 0.076 | 0.084 | 0.084 | 7.674 | 0.73 | 0.13 | 0.79 | 0.91 | 0.89 | 0.85 |
| 4 | 117147006 | rs1588041 | 0.367 | -1.972 | -0.173 | -1.111 | 0.078 | 0.087 | 0.087 | 7.436 | 0.73 | 0.13 | 0.79 | 0.91 | 0.89 | 0.72 |
| 4 | 117148675 | rs1032940 | 1.266 | -1.911 | -0.282 | -0.989 | -0.02 | 0.005 | 0.005 | 0 | 0.13 | 0 | 0.08 | 0.06 | 0.11 | 0.1 |
| 4 | 117159811 | rs1965299 | -0.494 | -2.326 | 0.201 | -1.313 | -0.006 | -0.032 | -0.032 | 0 | 0.68 | 0.71 | 0.83 | 0.93 | 0.85 | 0.75 |
| 4 | 117163200 | rs1387218 | -0.561 | -2.329 | 0.15 | -1.309 | 0.026 | 0.018 | 0.018 | 0 | 0.71 | 0.79 | 0.88 | 0.94 | 0.87 | 0.77 |
| 4 | 117172342 | rs727235 | 1.464 | -2.353 | -0.22 | -1.476 | -0.047 | 0.05 | 0.05 | 0 | 0.17 | 0.01 | 0.12 | 0.04 | 0.15 | 0.1 |
| 4 | 117211030 | rs11938979 | -0.079 | -2.242 | 0.736 | -1.208 | 0.055 | 0.051 | 0.051 | 3.012 | 0.05 | 0.4 | 0.04 | 0 | 0.01 | 0.05 |
| 4 | 117219559 | rs7654510 | -0.648 | -2.291 | 0.349 | -1.244 | 0.025 | 0.015 | 0.015 | 0.137 | 0.69 | 0.77 | 0.88 | 0.94 | 0.87 | 0.74 |
| 4 | 117226309 | rs6821696 | -0.609 | -2.179 | 0.501 | -0.504 | 0.07 | 0 | 0 | 0.016 | 0.66 | 0.78 | 0.74 | 0.7 | 0.82 | 0.71 |
| 4 | 117227330 | rs10428494 | -1.099 | -2.028 | 0.781 | -0.654 | -0.001 | -0.012 | -0.012 | 0.079 | 0.07 | 0.31 | 0.08 | 0.01 | 0.03 | 0.02 |
| 4 | 117237644 | rs11725289 | 0.857 | -1.834 | 0.185 | -0.528 | -0.044 | 0.038 | 0.038 | 0.094 | 0.3 | 0.03 | 0.19 | 0.21 | 0.29 | 0.24 |
| 4 | 117245430 | rs1489001 | 0.008 | -2.13 | 1.354 | -0.269 | -0.073 | -0.018 | -0.018 | 0.203 | 0.68 | 0.55 | 0.81 | 0.77 | 0.86 | 0.67 |
| 4 | 117257732 | rs1906699 | 0.554 | -2.113 | 1.098 | -0.305 | -0.024 | -0.006 | -0.006 | 0 | 0.63 | 0.39 | 0.74 | 0.77 | 0.84 | 0.65 |
| 4 | 117262160 | rs1873103 | -0.704 | -1.932 | 0.254 | -0.216 | -0.013 | -0.002 | -0.002 | 0.016 | 0.34 | 0.38 | 0.23 | 0.23 | 0.16 | 0.35 |
| 4 | 117272840 | rs1489012 | -0.307 | -1.612 | -0.06 | -0.137 | -0.011 | -0.002 | -0.002 | 0.346 | 0.31 | 0.31 | 0.17 | 0.21 | 0.14 | 0.33 |
| 4 | 117298571 | rs6533970 | -1.624 | -1.07 | 0.796 | -0.066 | 0.032 | 0.028 | 0.028 | 2.839 | 0.07 | 0.47 | 0.06 | 0 | 0.01 | 0.02 |
| 4 | 117300599 | rs7675851 | -0.252 | -1.155 | 0.075 | -0.153 | 0.057 | 0.054 | 0.054 | 2.205 | 0.51 | 0.66 | 0.45 | 0.32 | 0.38 | 0.31 |
| 4 | 117374128 | rs6533974 | -0.341 | -1.215 | 0.602 | -0.244 | -0.004 | -0.005 | -0.005 | 2.933 | 0.48 | 0.5 | 0.38 | 0.29 | 0.37 | 0.25 |
| 4 | 117379825 | rs970943 | -0.498 | -1.386 | 0.584 | -0.392 | -0.031 | -0.055 | -0.055 | 3.236 | 0.19 | 0.3 | 0.17 | 0.07 | 0.12 | 0.12 |
| 4 | 117387380 | rs6826566 | 0.093 | -1.399 | 0.722 | -0.406 | -0.006 | -0.012 | -0.012 | 0 | 0.15 | 0.16 | 0.15 | 0.07 | 0.11 | 0.09 |
| 4 | 117406228 | rs6849811 | 0.981 | -1.017 | 0.298 | -0.619 | -0.067 | -0.032 | -0.032 | 0 | 0.3 | 0.05 | 0.23 | 0.25 | 0.34 | 0.12 |
| 4 | 117407113 | rs7697881 | -1.262 | -0.855 | 0.299 | -0.568 | -0.037 | -0.056 | -0.056 | 0.324 | 0.78 | 0.84 | 0.76 | 0.52 | 0.69 | 0.68 |
| 4 | 117412778 | rs7663989 | -0.486 | -0.832 | 0.703 | -0.628 | 0.004 | 0.051 | 0.051 | 0.041 | 0.53 | 0.63 | 0.4 | 0.27 | 0.47 | 0.25 |
| 4 | 117414663 | rs11942266 | -0.432 | -0.775 | 0.224 | -0.616 | -0.029 | -0.11 | -0.11 | 0 | 0.2 | 0.31 | 0.14 | 0.02 | 0.12 | 0.12 |
| 4 | 117426252 | rs10222978 | -0.282 | -0.77 | 0.194 | -0.594 | 0.01 | 0.041 | 0.041 | 0 | 0.5 | 0.37 | 0.38 | 0.27 | 0.46 | 0.25 |
| 4 | 117429799 | rs13102076 | 1.015 | -0.629 | -0.262 | -0.472 | -0.066 | -0.023 | -0.023 | 0.022 | 0.3 | 0.05 | 0.24 | 0.25 | 0.35 | 0.12 |
| 4 | 117438399 | rs13127996 | 0.204 | -0.73 | -0.007 | -0.415 | 0.014 | 0.102 | 0.102 | 0.993 | 0.43 | 0.2 | 0.34 | 0.22 | 0.39 | 0.21 |
| 4 | 117449878 | rs2042651 | 0.163 | -0.656 | 0.337 | -0.387 | 0.003 | 0.009 | 0.009 | 0 | 0.39 | 0.37 | 0.45 | 0.29 | 0.39 | 0.19 |
| 4 | 117463727 | rs7661211 | 0.822 | -0.709 | 0.197 | -0.406 | -0.002 | 0.001 | 0.001 | 0 | 0.32 | 0.21 | 0.25 | 0.29 | 0.35 | 0.13 |
| 4 | 117468423 | rs10024264 | -0.206 | -0.845 | 0.256 | -0.337 | -0.022 | -0.022 | -0.022 | 0 | 0.37 | 0.45 | 0.34 | 0.27 | 0.26 | 0.52 |
| 4 | 117484810 | rs12639860 | 1.054 | -0.604 | -0.199 | -0.247 | -0.065 | -0.061 | -0.061 | 0.16 | 0.31 | 0.05 | 0.23 | 0.29 | 0.35 | 0.13 |
| 4 | 117509065 | rs1582716 | -0.271 | -0.668 | 0.103 | -0.309 | 0.001 | 0.001 | 0.001 | 0 | 0.5 | 0.49 | 0.55 | 0.32 | 0.46 | 0.31 |
| 4 | 117512637 | rs1895794 | -0.699 | -1.053 | 0.11 | -0.287 | -0.025 | -0.124 | -0.124 | 0 | 0.2 | 0.51 | 0.33 | 0.03 | 0.11 | 0.18 |
| 4 | 117513027 | rs1425632 | 0.818 | -1.046 | 0.166 | -0.293 | -0.004 | -0.032 | -0.032 | 0 | 0.11 | 0.12 | 0.11 | 0.03 | 0.07 | 0.11 |
| 4 | 117516679 | rs17632683 | 1.187 | -0.928 | -0.22 | -0.289 | -0.042 | -0.075 | -0.075 | 0.059 | 0.3 | 0.03 | 0.2 | 0.29 | 0.31 | 0.13 |
| 4 | 117520731 | rs4833516 | -0.465 | -0.849 | -0.812 | -0.256 | -0.022 | -0.124 | -0.124 | 0 | 0.2 | 0.51 | 0.33 | 0.03 | 0.12 | 0.18 |
| 4 | 117525693 | rs4557348 | 0.005 | -0.862 | -0.814 | -0.261 | 0.013 | 0.025 | 0.025 | 0.004 | 0.5 | 0.45 | 0.48 | 0.68 | 0.58 | 0.68 |
| 4 | 117527892 | rs4398591 | 0.41 | -0.496 | -0.45 | -0.714 | -0.023 | -0.125 | -0.125 | 0.316 | 0.8 | 0.49 | 0.67 | 0.97 | 0.88 | 0.82 |
| 4 | 117540588 | rs13114676 | 0.113 | -0.632 | -1.167 | -0.366 | 0.015 | -0.039 | -0.039 | 0 | 0.49 | 0.1 | 0.39 | 0.43 | 0.45 | 0.52 |
| 4 | 117543972 | rs4560484 | -0.027 | -0.183 | -0.951 | -0.051 | 0.007 | 0.009 | 0.009 | 0 | 0.39 | 0.47 | 0.39 | 0.57 | 0.5 | 0.4 |
| 4 | 117552737 | rs10005909 | -0.107 | -0.226 | -1.727 | -0.484 | 0.037 | 0.155 | 0.155 | 0 | 0.35 | 0.1 | 0.19 | 0.1 | 0.28 | 0.15 |
| 4 | 117568631 | rs13124273 | 0.175 | 0.682 | -0.926 | 0.03 | 0.012 | 0.004 | 0.004 | 0 | 0.62 | 0.43 | 0.48 | 0.33 | 0.44 | 0.64 |
| 4 | 117569267 | rs4415029 | 0.098 | 1.173 | -0.584 | 0.133 | -0.009 | -0.009 | -0.009 | 0 | 0.23 | 0.22 | 0.39 | 0.61 | 0.39 | 0.31 |
| 4 | 117569522 | rs4640736 | -0.164 | 1.142 | -0.452 | 0.104 | 0.009 | -0.028 | -0.028 | 0 | 0.85 | 0.71 | 0.87 | 0.94 | 0.84 | 0.96 |
| 4 | 117585086 | rs13118630 | 0.967 | -0.059 | -0.784 | -0.618 | 0.036 | 0.064 | 0.064 | 0 | 0.36 | 0.2 | 0.2 | 0.07 | 0.24 | 0.15 |
| 4 | 117594889 | rs10012562 | -0.021 | -0.006 | -0.847 | -0.549 | -0.008 | -0.003 | -0.003 | 0 | 0.6 | 0.31 | 0.47 | 0.33 | 0.44 | 0.64 |
| 4 | 117597044 | rs12507901 | 0.155 | -0.022 | -0.695 | -0.216 | 0.001 | -0.011 | -0.011 | 0 | 0.22 | 0.17 | 0.21 | 0.26 | 0.19 | 0.46 |
| 4 | 117609491 | rs12648967 | -0.275 | -0.595 | -1.177 | -0.32 | 0.024 | 0.011 | 0.011 | 2.075 | 0.26 | 0.16 | 0.19 | 0.23 | 0.16 | 0.28 |
| 4 | 117612897 | rs4130340 | 1.043 | 0.5 | -1.054 | 0.284 | -0.008 | 0.044 | 0.044 | 0 | 0.08 | 0.06 | 0.22 | 0.37 | 0.18 | 0.3 |
| 4 | 117618126 | rs11939225 | -0.681 | -0.217 | -0.447 | -0.076 | -0.032 | -0.008 | -0.008 | 0 | 0.24 | 0.38 | 0.13 | 0.23 | 0.13 | 0.29 |
| 4 | 117623903 | rs4077692 | -1.482 | -0.354 | 0.145 | -0.187 | -0.002 | -0.002 | -0.002 | 0 | 0.35 | 0.66 | 0.39 | 0.59 | 0.32 | 0.6 |
| 4 | 117624139 | rs4274920 | -1.728 | 0.064 | -0.37 | -0.5 | -0.026 | -0.028 | -0.028 | 0 | 0.85 | 0.9 | 0.65 | 0.66 | 0.67 | 0.75 |
| 4 | 117626118 | rs7654840 | 0.192 | -0.166 | -0.116 | -0.375 | 0.002 | -0.003 | -0.003 | 0 | 0.19 | 0.18 | 0.11 | 0.23 | 0.12 | 0.27 |
| 4 | 117630240 | rs7659215 | -1.531 | -0.373 | -0.292 | -0.619 | 0.008 | 0.063 | 0.063 | 0 | 0.49 | 0.76 | 0.48 | 0.59 | 0.39 | 0.59 |
| 4 | 117630678 | rs7682047 | 1.119 | -0.677 | -0.154 | -0.742 | -0.017 | -0.019 | -0.019 | 0 | 0.05 | 0.18 | 0.02 | 0 | 0.01 | 0.02 |
| 4 | 117638802 | rs11942536 | -0.138 | -0.896 | -0.497 | -0.771 | 0.015 | 0.017 | 0.017 | 0.416 | 0.3 | 0.22 | 0.19 | 0.23 | 0.19 | 0.29 |
| 4 | 117653901 | rs1500818 | -0.221 | -0.925 | -0.545 | -0.811 | -0.042 | -0.015 | -0.015 | 0.347 | 0.29 | 0.43 | 0.33 | 0.29 | 0.18 | 0.29 |
| 4 | 117659634 | rs10021925 | -0.997 | -0.878 | -1.214 | -0.608 | -0.113 | -0.066 | -0.066 | 0.353 | 0.54 | 0.92 | 0.72 | 0.62 | 0.53 | 0.56 |
| 4 | 117665686 | rs6814762 | -0.649 | -0.952 | -0.941 | -0.638 | -0.009 | 0.021 | 0.021 | 0.433 | 0.5 | 0.77 | 0.71 | 0.62 | 0.53 | 0.56 |
| 4 | 117683039 | rs1604230 | -0.128 | -1.046 | -0.758 | -0.697 | 0.042 | 0.039 | 0.039 | 0 | 0.42 | 0.66 | 0.38 | 0.29 | 0.23 | 0.33 |
| 4 | 117684483 | rs12505108 | -0.033 | -1.068 | -0.783 | -0.708 | 0.047 | 0.044 | 0.044 | 0 | 0.42 | 0.68 | 0.38 | 0.29 | 0.23 | 0.33 |
| 4 | 117687055 | rs11726029 | 1.863 | -0.925 | -0.785 | -0.679 | -0.045 | -0.043 | -0.043 | 1.909 | 0.17 | 0.09 | 0.36 | 0.32 | 0.35 | 0.27 |
| 4 | 117689811 | rs7675219 | 1.15 | -1.03 | -0.844 | -0.847 | 0.008 | 0.101 | 0.101 | 0 | 0.28 | 0.1 | 0.15 | 0.09 | 0.25 | 0.11 |
| 4 | 117715353 | rs12649281 | 0.096 | -1.143 | -0.474 | -0.721 | -0.051 | -0.033 | -0.033 | 1.924 | 0.38 | 0.47 | 0.24 | 0.29 | 0.19 | 0.33 |
| 4 | 117716133 | rs13114307 | -0.503 | -1.068 | -0.654 | -0.701 | 0.109 | 0.098 | 0.098 | 0 | 0.45 | 0.74 | 0.39 | 0.3 | 0.23 | 0.35 |
| 4 | 117724087 | rs10009620 | 0.383 | -1.109 | -0.395 | -0.628 | -0.014 | -0.008 | -0.008 | 0 | 0.39 | 0.39 | 0.24 | 0.28 | 0.2 | 0.33 |

| | | | | | | | | | | | | | | | | |
|---|---|---|---|---|---|---|---|---|---|---|---|---|---|---|---|---|
| 4 | 117733111 | rs17048820 | -1.058 | -1.143 | -0.179 | -0.57 | 0.079 | 0.031 | 0.031 | 4.148 | 0.06 | 0.34 | 0.16 | 0.01 | 0.04 | 0.02 |
| 4 | 117743263 | rs4834589 | 1.582 | -0.942 | -0.467 | -0.482 | -0.064 | -0.056 | -0.056 | 0 | 0.18 | 0.08 | 0.37 | 0.31 | 0.35 | 0.27 |
| 4 | 117761449 | rs1392764 | -0.168 | -1.138 | -0.644 | -0.525 | 0.005 | 0.039 | 0.039 | 0 | 0.48 | 0.26 | 0.55 | 0.41 | 0.59 | 0.38 |
| 4 | 117763136 | rs6856485 | -1.45 | -0.809 | -0.387 | -0.324 | 0.027 | 0.074 | 0.074 | 0 | 0.91 | 0.88 | 0.93 | 0.7 | 0.83 | 0.73 |
| 4 | 117772331 | rs10033111 | -0.352 | -0.921 | -0.494 | -0.453 | -0.001 | -0.001 | -0.001 | 0.493 | 0.7 | 0.7 | 0.55 | 0.4 | 0.49 | 0.48 |
| 4 | 117773396 | rs6534004 | -0.23 | -0.954 | -0.198 | -0.475 | 0.04 | 0.007 | 0.007 | 0 | 0.68 | 0.58 | 0.54 | 0.39 | 0.48 | 0.48 |
| 4 | 117779937 | rs10008401 | -0.21 | -1.126 | -0.297 | -0.592 | -0.02 | -0.024 | -0.024 | 2.254 | 0.63 | 0.72 | 0.75 | 0.6 | 0.6 | 0.61 |
| 4 | 117780360 | rs1875303 | 1.034 | -1.095 | -0.419 | -0.579 | -0.028 | 0.067 | 0.067 | 0 | 0.24 | 0.06 | 0.17 | 0.11 | 0.22 | 0.12 |
| 4 | 117783592 | rs7668517 | -0.84 | -1.033 | -0.267 | -0.576 | -0.042 | -0.047 | -0.047 | 0.563 | 0.7 | 0.76 | 0.76 | 0.6 | 0.61 | 0.63 |
| 4 | 117785317 | rs9684450 | 0.332 | -1.138 | -0.239 | -0.647 | -0.017 | -0.017 | -0.017 | 0 | 0.46 | 0.45 | 0.37 | 0.28 | 0.26 | 0.38 |
| 4 | 117795282 | rs12505094 | -0.529 | -0.779 | 0.111 | -0.545 | 0.002 | -0.032 | -0.032 | 0 | 0.06 | 0.15 | 0.14 | 0 | 0.04 | 0.02 |
| 4 | 117795603 | rs10033291 | -0.462 | -0.52 | 0.204 | -0.334 | 0.078 | 0.189 | 0.189 | 0 | 0.87 | 0.72 | 0.9 | 0.65 | 0.8 | 0.74 |
| 4 | 117796865 | rs2047636 | 0.007 | -0.962 | -0.112 | -0.496 | 0.016 | 0.001 | 0.001 | 0 | 0.67 | 0.6 | 0.41 | 0.39 | 0.44 | 0.46 |
| 4 | 117799215 | rs12498144 | -0.317 | -0.868 | -0.032 | -0.431 | 0.005 | -0.03 | -0.03 | 0.481 | 0.06 | 0.17 | 0.14 | 0 | 0.04 | 0.02 |
| 4 | 117799718 | rs2389247 | -0.162 | -0.799 | -0.113 | -0.359 | 0.076 | 0.137 | 0.137 | 0 | 0.85 | 0.7 | 0.79 | 0.7 | 0.79 | 0.72 |
| 4 | 117802073 | rs17505738 | 1.344 | -0.851 | -0.627 | -0.422 | -0.041 | -0.031 | -0.031 | 0 | 0.18 | 0.1 | 0.36 | 0.29 | 0.32 | 0.26 |
| 4 | 117803013 | rs11098361 | 0.336 | -1.405 | -0.927 | -0.75 | 0.037 | 0.028 | 0.028 | 0.021 | 0.4 | 0.23 | 0.3 | 0.36 | 0.33 | 0.3 |
| 4 | 117803266 | rs11098362 | 0.669 | -1.495 | -0.967 | -0.843 | 0.018 | 0.062 | 0.062 | 0.775 | 0.31 | 0.22 | 0.22 | 0.12 | 0.26 | 0.13 |
| 4 | 117830724 | rs10516632 | 0.492 | -1.74 | -0.616 | -1.001 | 0 | 0 | 0 | 0 | 0.29 | 0.3 | 0.32 | 0.12 | 0.28 | 0.14 |
| 4 | 117832770 | rs11940805 | 0.563 | -1.751 | -0.628 | -1.006 | 0.001 | -0.014 | -0.014 | 0 | 0.45 | 0.27 | 0.52 | 0.55 | 0.46 | 0.55 |
| 4 | 117836020 | rs1440312 | -1.511 | -1.419 | -0.563 | -0.843 | 0.045 | 0.114 | 0.114 | 0 | 0.89 | 0.83 | 0.92 | 0.69 | 0.81 | 0.72 |
| 4 | 117844745 | rs17505791 | 1.574 | -1.426 | -0.988 | -0.827 | -0.052 | 0.014 | 0.014 | 0 | 0.19 | 0.05 | 0.16 | 0.11 | 0.22 | 0.11 |
| 4 | 117848472 | rs6534006 | 1.826 | -1.478 | -0.77 | -0.741 | 0.005 | -0.011 | -0.011 | 0.013 | 0.09 | 0.05 | 0.09 | 0.23 | 0.07 | 0.08 |
| 4 | 117850456 | rs979961 | 0.639 | -1.678 | -0.535 | -0.867 | 0.025 | 0.013 | 0.013 | 0 | 0.27 | 0.19 | 0.16 | 0.27 | 0.14 | 0.2 |
| 4 | 117851859 | rs4834594 | -0.504 | -0.799 | -0.154 | -0.414 | 0.036 | 0.088 | 0.088 | 0 | 0.83 | 0.71 | 0.77 | 0.67 | 0.75 | 0.6 |
| 4 | 117856960 | rs1822050 | -0.697 | -0.811 | -0.36 | -0.462 | -0.046 | -0.051 | -0.051 | 0 | 0.72 | 0.8 | 0.78 | 0.59 | 0.58 | 0.58 |
| 4 | 117863537 | rs4834599 | 1.182 | -1.186 | -0.363 | -0.582 | 0.038 | 0.013 | 0.013 | 0.076 | 0.24 | 0.15 | 0.16 | 0.23 | 0.15 | 0.11 |
| 4 | 117863553 | rs12508441 | 0.813 | -0.641 | 0.025 | -0.315 | -0.013 | -0.026 | -0.026 | 0 | 0.09 | 0.14 | 0.16 | 0.02 | 0.06 | 0.11 |
| 4 | 117867406 | rs10050215 | -1.918 | -0.15 | -0.05 | -0.082 | 0.039 | 0.104 | 0.104 | 0 | 0.92 | 0.88 | 0.93 | 0.69 | 0.81 | 0.69 |
| 4 | 117868349 | rs1822046 | 0.311 | -0.33 | -0.027 | -0.182 | 0.074 | 0.072 | 0.072 | 0 | 0.5 | 0.4 | 0.41 | 0.3 | 0.26 | 0.35 |
| 4 | 117868882 | rs717904 | 1.469 | -0.179 | -0.015 | -0.189 | -0.032 | -0.025 | -0.025 | 0 | 0.17 | 0.08 | 0.34 | 0.28 | 0.31 | 0.23 |
| 4 | 117879013 | rs12506385 | 0.033 | -0.566 | -0.019 | -0.144 | 0.026 | 0.026 | 0.026 | 0 | 0.52 | 0.47 | 0.41 | 0.3 | 0.26 | 0.36 |
| 4 | 117885030 | rs12331356 | 0.027 | -0.402 | -0.152 | -0.206 | 0.002 | -0.004 | -0.004 | 0.012 | 0.41 | 0.48 | 0.52 | 0.39 | 0.54 | 0.34 |
| 4 | 117888361 | rs1440304 | 0.036 | -0.378 | -0.274 | -0.194 | 0.002 | -0.003 | -0.003 | 0 | 0.41 | 0.42 | 0.52 | 0.39 | 0.54 | 0.34 |
| 4 | 117897658 | rs13109536 | 0.068 | 0.251 | -0.57 | 0.13 | -0.068 | -0.03 | -0.03 | 0.782 | 0.3 | 0.16 | 0.39 | 0.43 | 0.44 | 0.31 |
| 4 | 117897700 | rs10016819 | 0.788 | -0.188 | -0.386 | -0.049 | 0.028 | 0.043 | 0.043 | 0 | 0.36 | 0.25 | 0.3 | 0.18 | 0.28 | 0.15 |
| 4 | 117912395 | rs7697486 | -0.639 | -0.23 | -0.057 | -0.001 | 0 | 0.005 | 0.005 | 0 | 0.55 | 0.58 | 0.63 | 0.5 | 0.56 | 0.55 |
| 4 | 117927811 | rs17690630 | 1.07 | 0.084 | 0.273 | 0.07 | -0.009 | -0.009 | -0.009 | 0.01 | 0.2 | 0.15 | 0.31 | 0.37 | 0.33 | 0.23 |
| 4 | 117931447 | rs11734096 | 1.28 | -0.074 | 0.519 | -0.372 | -0.011 | 0.006 | 0.006 | 0 | 0.1 | 0.06 | 0.08 | 0.09 | 0.12 | 0.06 |
| 4 | 117946646 | rs12649613 | -0.436 | -0.138 | 0.471 | -0.288 | 0.001 | -0.001 | -0.001 | 0 | 0.45 | 0.55 | 0.63 | 0.48 | 0.45 | 0.49 |
| 4 | 117950600 | rs11098370 | 1.029 | -0.077 | 0.53 | -0.249 | 0.008 | 0.021 | 0.021 | 0 | 0.21 | 0.23 | 0.4 | 0.42 | 0.34 | 0.23 |
| 4 | 117953708 | rs1899071 | -1.973 | -0.046 | 0.424 | -0.217 | -0.038 | -0.015 | -0.015 | 0 | 0.92 | 0.98 | 0.94 | 0.7 | 0.84 | 0.66 |
| 4 | 117954643 | rs17048984 | 0.662 | -0.176 | 0.653 | -0.3 | 0.006 | -0.018 | -0.018 | 2.652 | 0.13 | 0.26 | 0.1 | 0.09 | 0.12 | 0.06 |
| 4 | 117961014 | rs298924 | -0.653 | -0.243 | 0.65 | -0.375 | -0.001 | -0.016 | -0.016 | 0.516 | 0.49 | 0.43 | 0.67 | 0.79 | 0.61 | 0.83 |
| 4 | 117962388 | rs298926 | 1.823 | -0.256 | 0.488 | -0.338 | -0.037 | -0.011 | -0.011 | 0 | 0.08 | 0.02 | 0.06 | 0.3 | 0.17 | 0.34 |
| 4 | 117963914 | rs13150008 | -0.455 | -0.295 | 0.514 | -0.344 | -0.02 | -0.021 | -0.021 | 0 | 0.38 | 0.65 | 0.64 | 0.52 | 0.51 | 0.4 |
| 4 | 117964138 | rs298928 | -1.45 | -0.274 | 0.428 | -0.335 | 0.049 | 0.093 | 0.093 | 0 | 0.63 | 0.75 | 0.78 | 0.86 | 0.73 | 0.88 |
| 4 | 117976279 | rs11098373 | 0.23 | -0.162 | 0.237 | -0.269 | 0.025 | 0.033 | 0.033 | 0 | 0.31 | 0.35 | 0.48 | 0.52 | 0.46 | 0.29 |
| 4 | 117981294 | rs298941 | -1.114 | -0.366 | 0.334 | -0.367 | 0.009 | -0.001 | -0.001 | 0.604 | 0.59 | 0.62 | 0.64 | 0.86 | 0.69 | 0.78 |
| 4 | 117987578 | rs298942 | -1.387 | -0.428 | 0.343 | -0.338 | 0.031 | 0.052 | 0.052 | 0 | 0.64 | 0.75 | 0.77 | 0.88 | 0.73 | 0.88 |
| 4 | 117993779 | rs2196715 | -0.91 | -0.169 | 0.638 | 0.067 | -0.035 | -0.094 | -0.094 | 0 | 0.35 | 0.54 | 0.31 | 0.15 | 0.23 | 0.31 |
| 4 | 118000844 | rs298944 | 1.128 | 0.013 | 0.372 | 0.211 | 0.002 | 0.045 | 0.045 | 0 | 0.09 | 0.05 | 0.07 | 0.3 | 0.17 | 0.35 |
| 4 | 118003302 | rs17049001 | -0.001 | -0.624 | 0.463 | -0.484 | 0.016 | 0.003 | 0.003 | 0 | 0.2 | 0.21 | 0.09 | 0.03 | 0.07 | 0.15 |
| 4 | 118007156 | rs7683967 | -0.444 | -0.628 | 0.009 | -0.397 | 0.011 | -0.05 | -0.05 | 0 | 0.18 | 0.48 | 0.24 | 0.11 | 0.16 | 0.16 |
| 4 | 118014813 | rs2043805 | 0.682 | -0.614 | -0.159 | -0.399 | 0.049 | 0.078 | 0.078 | 0.657 | 0.16 | 0.03 | 0.06 | 0.03 | 0.06 | 0.14 |
| 4 | 118020477 | rs3943702 | -0.835 | 0.22 | 0.524 | 0.225 | 0.013 | -0.033 | -0.033 | 1.88 | 0.18 | 0.48 | 0.24 | 0.13 | 0.17 | 0.16 |
| 4 | 118039852 | rs1438217 | -0.379 | 0.489 | 0.106 | 0.596 | 0.008 | 0.03 | 0.03 | 0 | 0.34 | 0.43 | 0.52 | 0.61 | 0.42 | 0.58 |
| 4 | 118043043 | rs4395584 | -0.09 | 0.167 | 0.353 | 0.85 | -0.019 | -0.012 | -0.012 | 0 | 0.2 | 0.27 | 0.14 | 0.23 | 0.15 | 0.22 |
| 4 | 118043203 | rs7676913 | -0.858 | -0.104 | 0.301 | 0.403 | 0.007 | 0.052 | 0.052 | 0 | 0.5 | 0.75 | 0.62 | 0.65 | 0.53 | 0.66 |
| 4 | 118043617 | rs11933950 | -0.917 | -0.069 | 0.37 | 0.421 | 0.001 | 0.021 | 0.021 | 0 | 0.47 | 0.64 | 0.61 | 0.65 | 0.53 | 0.66 |
| 4 | 118072289 | rs2196712 | -0.044 | -0.355 | 0.496 | 0.381 | -0.013 | 0.006 | 0.006 | 0 | 0.45 | 0.3 | 0.55 | 0.42 | 0.54 | 0.46 |
| 4 | 118078654 | rs12504828 | -0.045 | -0.39 | 0.595 | 0.376 | -0.002 | 0.004 | 0.004 | 0 | 0.46 | 0.42 | 0.56 | 0.42 | 0.54 | 0.46 |
| 4 | 118079907 | rs10516635 | 0.421 | -0.394 | 0.134 | 0.801 | 0.005 | -0.007 | -0.007 | 0 | 0.16 | 0.16 | 0.1 | 0.25 | 0.13 | 0.19 |
| 4 | 118080628 | rs13110184 | -0.125 | -0.394 | -0.079 | 0.264 | -0.024 | -0.064 | -0.064 | 0 | 0.27 | 0.18 | 0.35 | 0.53 | 0.33 | 0.44 |
| 4 | 118081229 | rs12644587 | 0.13 | -0.4 | -0.063 | 0.268 | -0.009 | -0.033 | -0.033 | 3.858 | 0.71 | 0.74 | 0.64 | 0.47 | 0.67 | 0.56 |
| 4 | 118093624 | rs11098380 | 0.334 | -0.433 | 0.146 | 0.096 | 0.001 | 0 | 0 | 0 | 0.43 | 0.36 | 0.71 | 0.43 | 0.54 | 0.46 |
| 4 | 118109122 | rs1438220 | -0.829 | 0.089 | -0.284 | 0.124 | -0.021 | -0.022 | -0.022 | 0 | 0.8 | 0.87 | 0.85 | 0.66 | 0.72 | 0.75 |
| 4 | 118122688 | rs10022128 | 1.036 | -0.51 | -0.014 | -0.363 | -0.019 | -0.082 | -0.082 | 0.948 | 0.24 | 0.25 | 0.07 | 0 | 0.12 | 0.09 |
| 4 | 118131899 | rs10025815 | -0.184 | -0.466 | -0.409 | -0.22 | -0.04 | -0.055 | -0.055 | 0 | 0.66 | 0.8 | 0.75 | 0.5 | 0.57 | 0.66 |

| | | | | | | | | | | | | | | | | |
|---|---|---|---|---|---|---|---|---|---|---|---|---|---|---|---|---|
| 4 | 118135356 | rs4498198 | 0.87 | -0.865 | -0.345 | -0.182 | -0.012 | -0.017 | -0.017 | 0.861 | 0.3 | 0.22 | 0.62 | 0.37 | 0.41 | 0.52 |
| 4 | 118145893 | rs4535398 | -0.089 | -0.972 | 0.03 | -0.18 | -0.004 | -0.004 | -0.004 | 0 | 0.64 | 0.61 | 0.74 | 0.49 | 0.59 | 0.65 |
| 4 | 118155066 | rs12639976 | 0.699 | -0.773 | -0.479 | -0.101 | -0.011 | -0.019 | -0.019 | 0.043 | 0.36 | 0.23 | 0.65 | 0.39 | 0.45 | 0.53 |
| 4 | 118157777 | rs9999009 | 0.62 | -0.74 | -0.446 | -0.105 | -0.007 | -0.012 | -0.012 | 0 | 0.37 | 0.25 | 0.65 | 0.39 | 0.45 | 0.53 |
| 4 | 118163167 | rs7690351 | 0.735 | -0.733 | -0.854 | -0.093 | -0.023 | -0.037 | -0.037 | 0 | 0.36 | 0.18 | 0.65 | 0.39 | 0.45 | 0.53 |
| 4 | 118164717 | rs13147087 | 1.055 | -0.58 | -1.681 | -0.465 | -0.116 | -0.061 | -0.061 | 0.911 | 0.27 | 0.03 | 0.47 | 0.25 | 0.4 | 0.48 |
| 4 | 118185220 | rs6534029 | -0.506 | -0.544 | -1.441 | -0.651 | 0.002 | 0.007 | 0.007 | 0 | 0.37 | 0.65 | 0.38 | 0.75 | 0.36 | 0.44 |
| 4 | 118190581 | rs12644393 | 0.358 | -0.768 | -1.65 | -0.849 | -0.009 | 0.023 | 0.023 | 0 | 0.43 | 0.35 | 0.57 | 0.25 | 0.51 | 0.47 |
| 4 | 118204404 | rs7681687 | -0.818 | -0.693 | -1.843 | -0.888 | -0.127 | 0.114 | 0.114 | 0.041 | 0.5 | 0.97 | 0.47 | 0.75 | 0.47 | 0.45 |
| 4 | 118208216 | rs11930731 | 0.147 | -0.549 | -1.193 | -1.187 | 0.009 | -0.018 | -0.018 | 0.001 | 0.15 | 0.12 | 0.1 | 0.27 | 0.13 | 0.13 |
| 4 | 118215853 | rs10033190 | -0.872 | -0.222 | -1.036 | -1.21 | 0.01 | 0.025 | 0.025 | 0 | 0.66 | 0.76 | 0.86 | 1 | 0.77 | 0.9 |
| 4 | 118218045 | rs6853512 | 0.132 | 0.168 | -0.438 | -0.075 | 0.002 | 0.01 | 0.01 | 0 | 0.11 | 0.17 | 0.09 | 0.18 | 0.12 | 0.08 |
| 4 | 118221551 | rs4361453 | 0.024 | -0.138 | -0.745 | -0.366 | -0.014 | -0.02 | -0.02 | 0 | 0.57 | 0.53 | 0.46 | 0.35 | 0.66 | 0.57 |
| 4 | 118235444 | rs13137355 | 0.413 | 0.392 | -0.479 | -0.433 | 0.002 | 0.049 | 0.049 | 0.03 | 0.14 | 0.14 | 0.41 | 0.54 | 0.19 | 0.34 |
| 4 | 118239302 | rs4833543 | 0.077 | 0.163 | -0.612 | -0.967 | 0.014 | 0.024 | 0.024 | 0 | 0.15 | 0.09 | 0.06 | 0.01 | 0.09 | 0.09 |
| 4 | 118244984 | rs9998109 | -0.98 | 0.408 | -0.023 | -1.368 | -0.021 | -0.131 | -0.131 | 0.48 | 0.29 | 0.5 | 0.16 | 0.03 | 0.2 | 0.15 |
| 4 | 118246863 | rs4074599 | -0.648 | 0.383 | 0.098 | -1.383 | -0.004 | -0.07 | -0.07 | 0.919 | 0.26 | 0.32 | 0.13 | 0.01 | 0.19 | 0.11 |
| 4 | 118254355 | rs4455494 | 1.139 | 0.733 | -1.576 | -1.885 | -0.029 | -0.071 | -0.071 | 0.47 | 0.3 | 0.14 | 0.29 | 0.42 | 0.36 | 0.31 |
| 4 | 118254457 | rs17861758 | 1.181 | 1.262 | -1.22 | -1.373 | -0.006 | 0.002 | 0.002 | 0 | 0.08 | 0.08 | 0.05 | 0 | 0.1 | 0.02 |
| 4 | 118254740 | rs12507136 | 0.927 | 1.164 | -1.111 | -1.053 | -0.003 | 0.016 | 0.016 | 0.048 | 0.14 | 0.09 | 0.41 | 0.55 | 0.18 | 0.39 |
| 4 | 118262620 | rs4416544 | -1.159 | 0.918 | -0.984 | -1.501 | -0.023 | 0.101 | 0.101 | 0 | 0.77 | 0.96 | 0.91 | 1 | 0.8 | 0.89 |
| 4 | 118265683 | rs17861768 | 0.599 | 0.91 | -0.657 | -1.748 | 0.002 | -0.029 | -0.029 | 0 | 0.12 | 0.14 | 0.11 | 0.01 | 0.15 | 0.08 |
| 4 | 118276666 | rs4409028 | -1.012 | 0.915 | -0.697 | -1.524 | -0.012 | -0.019 | -0.019 | 0 | 0.8 | 0.83 | 0.44 | 0.45 | 0.68 | 0.6 |
| 4 | 118281116 | rs13138792 | 1.064 | 0.345 | 0.527 | -1.728 | -0.024 | -0.011 | -0.011 | 1.094 | 0.07 | 0.14 | 0.02 | 0.02 | 0.02 | 0.07 |
| 4 | 118281370 | rs11723116 | 0.768 | 0.171 | 0.235 | -2.03 | 0.024 | -0.05 | -0.05 | 0 | 0.28 | 0.14 | 0.12 | 0.42 | 0.22 | 0.26 |
| 4 | 118292314 | rs3925016 | -0.405 | 0.264 | -0.029 | -1.986 | 0.022 | -0.011 | -0.011 | 0 | 0.4 | 0.64 | 0.19 | 0.44 | 0.25 | 0.33 |
| 4 | 118297556 | rs10034473 | 0.624 | 0.308 | -0.192 | -1.844 | -0.029 | -0.015 | -0.015 | 0 | 0.08 | 0.25 | 0.04 | 0.02 | 0.02 | 0.07 |
| 4 | 118297655 | rs4309912 | 1.317 | 0.695 | -0.505 | -1.778 | -0.027 | -0.061 | -0.061 | 0 | 0.17 | 0.07 | 0.5 | 0.54 | 0.26 | 0.38 |
| 4 | 118306228 | rs7695817 | -0.104 | 0.561 | -0.304 | -2.257 | -0.002 | 0.004 | 0.004 | 0 | 0.48 | 0.37 | 0.28 | 0.04 | 0.45 | 0.34 |
| 4 | 118309390 | rs13113229 | 1.194 | 0.892 | -1 | -2.04 | -0.042 | 0.146 | 0.146 | 0 | 0.26 | 0.06 | 0.12 | 0.01 | 0.26 | 0.16 |
| 4 | 118310450 | rs17861840 | 1.308 | 0.889 | -0.717 | -1.816 | -0.007 | 0.008 | 0.008 | 0 | 0.05 | 0.04 | 0.05 | 0 | 0.09 | 0.01 |
| 4 | 118313760 | rs4834614 | 0.222 | 0.882 | -0.578 | -2.257 | -0.026 | 0.053 | 0.053 | 0 | 0.41 | 0.3 | 0.3 | 0.04 | 0.5 | 0.31 |
| 4 | 118315560 | rs4339281 | 0.449 | 0.738 | -0.375 | -2.166 | 0.006 | -0.015 | -0.015 | 0 | 0.1 | 0.15 | 0.09 | 0.03 | 0.11 | 0.09 |
| 4 | 118319741 | rs17861858 | 1.143 | 0.074 | -0.472 | -2.157 | 0.028 | -0.034 | -0.034 | 0 | 0.21 | 0.13 | 0.23 | 0.32 | 0.11 | 0.22 |
| 4 | 118320277 | rs6839566 | 1.62 | 0.368 | -0.237 | -2.192 | -0.001 | -0.015 | -0.015 | 0 | 0.1 | 0.1 | 0.09 | 0.01 | 0.12 | 0.06 |
| 4 | 118325168 | rs17861866 | 0.701 | 0.399 | 0.364 | -2.296 | 0.004 | -0.132 | -0.132 | 0.445 | 0.4 | 0.47 | 0.26 | 0.01 | 0.42 | 0.26 |
| 4 | 118326937 | rs7692931 | -1.559 | 0.451 | 0.271 | -2.131 | -0.018 | -0.025 | -0.025 | 0.04 | 0.81 | 0.88 | 0.61 | 0.36 | 0.68 | 0.62 |
| 4 | 118327461 | rs6851671 | -0.706 | 0.417 | 0.308 | -2.115 | -0.007 | -0.012 | -0.012 | 1.089 | 0.74 | 0.82 | 0.59 | 0.36 | 0.66 | 0.58 |
| 4 | 118335241 | rs7664551 | -0.76 | 0.039 | 0.328 | -2.009 | -0.009 | -0.001 | -0.001 | 0.05 | 0.42 | 0.45 | 0.36 | 0.37 | 0.27 | 0.36 |
| 4 | 118339156 | rs4619965 | 0.765 | 0.263 | -0.21 | -2.384 | 0.004 | -0.042 | -0.042 | 0 | 0.27 | 0.13 | 0.28 | 0.35 | 0.21 | 0.28 |
| 4 | 118340170 | rs4385147 | -2.414 | 0.39 | -0.127 | -2.509 | 0.002 | 0.045 | 0.045 | 0 | 0.88 | 0.89 | 0.66 | 0.4 | 0.78 | 0.69 |
| 4 | 118340693 | rs17861884 | 2.199 | 0.395 | -0.477 | -2.669 | -0.061 | 0.145 | 0.145 | 0 | 0.26 | 0.04 | 0.13 | 0 | 0.29 | 0.2 |
| 4 | 118340708 | rs17861885 | 1.94 | 0.347 | -0.341 | -2.704 | -0.001 | -0.01 | -0.01 | 0 | 0.09 | 0.07 | 0.08 | 0.01 | 0.12 | 0.06 |
| 4 | 118340731 | rs17875070 | 2.237 | 0.345 | -0.353 | -2.68 | -0.061 | 0.148 | 0.148 | 0 | 0.26 | 0.04 | 0.14 | 0.01 | 0.29 | 0.2 |
| 4 | 118340767 | rs6534034 | -1.021 | 0.315 | -0.05 | -2.698 | 0.017 | 0.017 | 0.017 | 0 | 0.49 | 0.46 | 0.41 | 0.39 | 0.37 | 0.43 |
| 4 | 118340829 | rs17866961 | 1.846 | 0.212 | -0.038 | -2.737 | -0.017 | -0.024 | -0.024 | 0 | 0.07 | 0.11 | 0.03 | 0 | 0.02 | 0.04 |
| 4 | 118341781 | rs7658609 | -1.057 | 0.444 | -0.008 | -2.738 | 0.011 | 0.016 | 0.016 | 0 | 0.49 | 0.47 | 0.43 | 0.39 | 0.4 | 0.46 |
| 4 | 118344201 | rs6834348 | -0.184 | 0.493 | -0.343 | -2.59 | -0.002 | 0.005 | 0.005 | 0.002 | 0.69 | 0.39 | 0.52 | 0.38 | 0.64 | 0.6 |
| 4 | 118348459 | rs6825806 | 1.763 | 0.471 | -0.735 | -2.5 | -0.001 | -0.01 | -0.01 | 0 | 0.09 | 0.07 | 0.08 | 0.01 | 0.12 | 0.06 |
| 4 | 118351381 | rs4240307 | 0.169 | 0.159 | -0.847 | -2.468 | -0.011 | 0.033 | 0.033 | 0.703 | 0.57 | 0.19 | 0.42 | 0.34 | 0.51 | 0.49 |
| 4 | 118353315 | rs6812138 | 0.214 | 0.048 | -0.561 | -1.882 | -0.004 | -0.058 | -0.058 | 0 | 0.26 | 0.11 | 0.28 | 0.34 | 0.21 | 0.28 |
| 4 | 118358899 | rs4632732 | -0.755 | -0.993 | -0.108 | -2.504 | -0.015 | -0.058 | -0.058 | 0 | 0.31 | 0.39 | 0.15 | 0.12 | 0.21 | 0.3 |
| 4 | 118361718 | rs12644762 | 1.281 | -1.06 | -0.539 | -2.679 | -0.092 | -0.011 | -0.011 | 0 | 0.24 | 0.07 | 0.41 | 0.73 | 0.45 | 0.41 |
| 4 | 118362073 | rs17861918 | 1.305 | -1.032 | -0.547 | -2.681 | -0.093 | -0.015 | -0.015 | 0 | 0.23 | 0.07 | 0.4 | 0.73 | 0.43 | 0.41 |
| 4 | 118370511 | rs13119978 | 0.636 | -1.33 | -0.453 | -2.733 | -0.025 | 0.031 | 0.031 | 0 | 0.12 | 0.04 | 0.05 | 0.04 | 0.13 | 0.09 |
| 4 | 118376267 | rs4282258 | -1.73 | -1.539 | -0.399 | -2.868 | -0.001 | 0.047 | 0.047 | 0.125 | 0.75 | 0.83 | 0.83 | 0.91 | 0.75 | 0.85 |
| 4 | 118376553 | rs17861942 | -0.011 | -1.517 | -0.713 | -2.866 | 0 | -0.02 | -0.02 | 0 | 0.27 | 0.27 | 0.13 | 0.05 | 0.2 | 0.15 |
| 4 | 118379516 | rs7675550 | 1.281 | -0.764 | -1.03 | -2.336 | -0.034 | 0.096 | 0.096 | 0 | 0.23 | 0.06 | 0.16 | 0.09 | 0.25 | 0.15 |
| 4 | 118384890 | rs7679074 | 0.357 | -0.025 | -0.243 | -1.585 | -0.004 | 0 | 0 | 0.684 | 0.24 | 0.21 | 0.43 | 0.73 | 0.45 | 0.41 |
| 4 | 118390464 | rs6823688 | -0.274 | -0.688 | 0.076 | -1.585 | 0.006 | -0.006 | -0.006 | 0 | 0.52 | 0.39 | 0.55 | 0.88 | 0.59 | 0.73 |
| 4 | 118393611 | rs17861959 | 0.233 | -0.737 | 0.092 | -1.627 | 0.001 | 0.001 | 0.001 | 0 | 0.08 | 0.07 | 0.04 | 0.06 | 0.04 | 0.22 |
| 4 | 118393990 | rs12108495 | -0.558 | -1.109 | 0.519 | -1.782 | -0.032 | -0.007 | -0.007 | 0 | 0.09 | 0.27 | 0.17 | 0.05 | 0.03 | 0.07 |
| 4 | 118394656 | rs4645284 | 0.937 | -1.15 | 0.502 | -1.869 | -0.112 | 0.039 | 0.039 | 0.378 | 0.34 | 0.12 | 0.44 | 0.76 | 0.48 | 0.47 |
| 4 | 118395930 | rs6827477 | -0.79 | -1.147 | 0.961 | -1.882 | -0.011 | -0.035 | -0.035 | 0 | 0.13 | 0.41 | 0.08 | 0.06 | 0.07 | 0.04 |
| 4 | 118396765 | rs17861965 | 0.072 | -1.054 | 0.751 | -1.837 | 0.002 | -0.013 | -0.013 | 0.046 | 0.1 | 0.16 | 0.06 | 0.06 | 0.06 | 0.04 |
| 4 | 118396886 | rs12651122 | 0.003 | -0.769 | 0.455 | -1.616 | -0.017 | 0.114 | 0.114 | 0.044 | 0.43 | 0.24 | 0.49 | 0.82 | 0.52 | 0.69 |
| 4 | 118402189 | rs12510650 | -0.121 | -0.865 | 0.975 | -1.459 | 0 | -0.023 | -0.023 | 0.001 | 0.44 | 0.47 | 0.66 | 0.8 | 0.55 | 0.65 |
| 4 | 118404415 | rs11725039 | 0.256 | -1.093 | 1.041 | -1.686 | 0.003 | -0.001 | -0.001 | 0 | 0.46 | 0.43 | 0.25 | 0.2 | 0.35 | 0.34 |
| 4 | 118408245 | rs6839724 | 0.48 | -1.169 | 1.026 | -1.748 | 0 | -0.06 | -0.06 | 1.938 | 0.83 | 0.63 | 0.86 | 0.93 | 0.85 | 0.94 |

| | | | | | | | | | | | | | | | | |
|---|---|---|---|---|---|---|---|---|---|---|---|---|---|---|---|---|
| 4 | 118408638 | rs10014124 | -0.809 | -1.138 | 0.821 | -1.728 | -0.002 | -0.032 | -0.032 | 0 | 0.91 | 0.88 | 0.92 | 1 | 0.89 | 0.98 |
| 4 | 118413749 | rs6852609 | 0.67 | -1.126 | 0.958 | -1.715 | 0.026 | 0.07 | 0.07 | 0 | 0.41 | 0.28 | 0.23 | 0.13 | 0.31 | 0.3 |
| 4 | 118417254 | rs6842580 | -0.293 | -0.976 | 1.086 | -1.528 | 0.006 | -0.019 | -0.019 | 0.553 | 0.12 | 0.23 | 0.1 | 0.06 | 0.13 | 0.05 |
| 4 | 118420131 | rs7679549 | -0.136 | -0.991 | 0.874 | -1.548 | 0.001 | 0.009 | 0.009 | 0.233 | 0.55 | 0.64 | 0.34 | 0.16 | 0.44 | 0.35 |
| 4 | 118420302 | rs17865839 | -0.349 | -0.694 | 1.088 | -1.171 | 0.029 | 0.048 | 0.048 | 0 | 0.06 | 0.16 | 0.04 | 0.07 | 0.04 | 0.05 |
| 4 | 118435901 | rs10006777 | -1.029 | -0.895 | 1.093 | -1.197 | -0.003 | 0.021 | 0.021 | 0 | 0.08 | 0.18 | 0.03 | 0.06 | 0.03 | 0.05 |
| 4 | 118444804 | rs11562936 | -1.61 | -1.117 | 1.249 | -1.346 | -0.046 | -0.052 | -0.052 | 0 | 0.23 | 0.55 | 0.14 | 0.1 | 0.15 | 0.1 |
| 4 | 118445839 | rs6829877 | -1.649 | -1.123 | 1.287 | -1.363 | -0.036 | -0.043 | -0.043 | 0.007 | 0.24 | 0.58 | 0.14 | 0.09 | 0.16 | 0.1 |
| 4 | 118450681 | rs10034912 | -0.589 | -1.114 | 1.287 | -1.353 | -0.022 | -0.021 | -0.021 | 0 | 0.09 | 0.2 | 0.03 | 0.03 | 0.03 | 0.05 |
| 4 | 118450767 | rs17866755 | 0.286 | -1.111 | 1.336 | -1.367 | -0.009 | -0.068 | -0.068 | 0 | 0.11 | 0.29 | 0.09 | 0 | 0.1 | 0.01 |
| 4 | 118453775 | rs11933680 | -0.511 | -1.126 | 0.768 | -1.361 | -0.029 | -0.048 | -0.048 | 0 | 0.16 | 0.48 | 0.12 | 0.06 | 0.12 | 0.05 |
| 4 | 118454987 | rs1948245 | -0.058 | -1.093 | 0.061 | -1.611 | 0.01 | 0.034 | 0.034 | 0 | 0.53 | 0.37 | 0.38 | 0.17 | 0.37 | 0.38 |
| 4 | 118463264 | rs1017964 | 0.371 | -0.949 | -0.214 | -1.61 | 0.032 | 0.151 | 0.151 | 0 | 0.44 | 0.2 | 0.23 | 0.13 | 0.34 | 0.32 |
| 4 | 118479461 | rs2635348 | -0.048 | -0.723 | -0.292 | -1.279 | 0.004 | 0.01 | 0.01 | 0.094 | 0.44 | 0.39 | 0.69 | 0.89 | 0.59 | 0.69 |
| 4 | 118480441 | rs11562861 | -0.536 | -0.791 | -0.266 | -0.118 | -0.006 | 0.018 | 0.018 | 0.047 | 0.13 | 0.19 | 0.09 | 0.22 | 0.13 | 0.07 |
| 4 | 118481560 | rs17863856 | 0.907 | -1.16 | -0.514 | -0.903 | 0.009 | -0.012 | -0.012 | 0.167 | 0.22 | 0.15 | 0.1 | 0 | 0.15 | 0.1 |
| 4 | 118486543 | rs1948226 | 0.389 | -1.247 | -1.143 | -0.867 | -0.061 | -0.009 | -0.009 | 0.313 | 0.37 | 0.16 | 0.67 | 0.59 | 0.54 | 0.37 |
| 4 | 118488900 | rs6813241 | 1.103 | -1.21 | -0.985 | -0.915 | -0.002 | -0.039 | -0.039 | 0 | 0.09 | 0.17 | 0.08 | 0.01 | 0.1 | 0.01 |
| 4 | 118496919 | rs3811755 | -0.016 | -0.589 | -1.363 | -0.978 | -0.025 | -0.09 | -0.09 | 0.086 | 0.38 | 0.1 | 0.24 | 0.48 | 0.37 | 0.58 |
| 4 | 118497403 | rs2389380 | 1.898 | -0.728 | -1.06 | -0.961 | 0.006 | -0.026 | -0.026 | 0 | 0.23 | 0.16 | 0.1 | 0 | 0.15 | 0.1 |
| 4 | 118497425 | rs17865200 | 1.246 | -1.083 | -1.263 | -1.067 | 0.027 | 0.024 | 0.024 | 0 | 0.11 | 0.04 | 0.02 | 0 | 0.03 | 0.07 |
| 4 | 118497883 | rs2245104 | -0.016 | -0.911 | -0.622 | -0.972 | -0.017 | 0.04 | 0.04 | 0.014 | 0.66 | 0.42 | 0.78 | 0.51 | 0.69 | 0.49 |
| 4 | 118499387 | rs11562932 | 1.122 | -0.515 | -0.314 | -0.702 | -0.001 | -0.044 | -0.044 | 0 | 0.09 | 0.21 | 0.08 | 0.01 | 0.1 | 0.01 |
| 4 | 118505637 | rs13129382 | 0.701 | -0.692 | -0.108 | -0.64 | 0.018 | -0.008 | -0.008 | 0.042 | 0.44 | 0.33 | 0.21 | 0.48 | 0.36 | 0.56 |
| 4 | 118505700 | rs1352573 | -1.529 | -0.509 | 0.154 | -0.414 | 0.048 | -0.021 | -0.021 | 0 | 0.76 | 0.7 | 0.5 | 0.79 | 0.65 | 0.79 |
| 4 | 118510812 | rs6850876 | -1.5 | -0.514 | 0.068 | -0.419 | 0.01 | -0.005 | -0.005 | 0 | 0.76 | 0.76 | 0.5 | 0.79 | 0.65 | 0.79 |
| 4 | 118511981 | rs6814620 | 0.829 | -0.51 | -0.11 | 0.002 | 0.001 | 0.03 | 0.03 | 0.295 | 0.09 | 0.2 | 0.08 | 0.14 | 0.1 | 0.04 |
| 4 | 118512999 | rs17863870 | 0.097 | -0.56 | -0.141 | -0.03 | -0.008 | 0 | 0 | 0 | 0.16 | 0.18 | 0.09 | 0.17 | 0.06 | 0.16 |
| 4 | 118513265 | rs13135130 | -0.632 | -0.047 | -0.288 | -0.016 | -0.001 | -0.001 | -0.001 | 0.167 | 0.17 | 0.22 | 0.2 | 0.14 | 0.24 | 0.07 |
| 4 | 118519146 | rs2635191 | 0.876 | 0.063 | -0.568 | 0.047 | -0.015 | -0.024 | -0.024 | 0.002 | 0.06 | 0 | 0.05 | 0.06 | 0.07 | 0.04 |
| 4 | 118519506 | rs10019567 | -0.629 | -0.259 | -0.175 | -0.182 | -0.003 | -0.012 | -0.012 | 0.9 | 0.22 | 0.31 | 0.17 | 0.14 | 0.21 | 0.11 |
| 4 | 118531857 | rs12511367 | 0.845 | -0.164 | 0.119 | 0.265 | -0.003 | 0.084 | 0.084 | 0 | 0.05 | 0.11 | 0.23 | 0.22 | 0.03 | 0.09 |
| 4 | 118532328 | rs2635295 | -1.854 | -0.414 | 0.087 | -0.228 | 0.031 | 0.147 | 0.147 | 0 | 0.48 | 0.74 | 0.54 | 0.82 | 0.42 | 0.66 |
| 4 | 118538842 | rs17862054 | -0.159 | -0.316 | 0.297 | -0.198 | -0.013 | -0.006 | -0.006 | 0 | 0.18 | 0.32 | 0.1 | 0.13 | 0.13 | 0.09 |
| 4 | 118541595 | rs9999101 | 1.239 | -0.228 | -0.117 | -0.577 | 0 | -0.003 | -0.003 | 0 | 0.33 | 0.27 | 0.18 | 0.13 | 0.27 | 0.18 |
| 4 | 118542986 | rs2220497 | 0.897 | -0.242 | -0.434 | -0.535 | -0.018 | -0.005 | -0.005 | 1.118 | 0.1 | 0 | 0.05 | 0.06 | 0.1 | 0.05 |
| 4 | 118543044 | rs11562909 | 2.173 | 0.17 | -0.024 | -0.209 | -0.002 | -0.06 | -0.06 | 0 | 0.12 | 0.04 | 0.07 | 0.42 | 0.11 | 0.41 |
| 4 | 118543358 | rs2635299 | -1.111 | 0.214 | 0.212 | -0.171 | -0.005 | 0.002 | 0.002 | 0.073 | 0.7 | 0.71 | 0.64 | 0.83 | 0.59 | 0.77 |
| 4 | 118544344 | rs923441 | 0.382 | -0.073 | 0.529 | -0.09 | -0.004 | 0.119 | 0.119 | 0.304 | 0.06 | 0.18 | 0.24 | 0.22 | 0.03 | 0.1 |
| 4 | 118545387 | rs1948223 | 0.363 | -0.082 | 0.356 | -0.127 | 0.009 | 0.029 | 0.029 | 2.18 | 0.44 | 0.3 | 0.24 | 0.19 | 0.37 | 0.23 |
| 4 | 118547300 | rs1304736 | -0.177 | 0.046 | -0.092 | -0.074 | 0.023 | 0.006 | 0.006 | 0.011 | 0.52 | 0.61 | 0.49 | 0.41 | 0.39 | 0.33 |
| 4 | 118548943 | rs1948221 | 0.589 | 0.425 | 0.478 | -0.068 | 0.057 | 0.055 | 0.055 | 0.795 | 0.86 | 0.52 | 0.86 | 0.86 | 0.84 | 0.94 |
| 4 | 118551128 | rs1948250 | 0.979 | 0.514 | 0.626 | 0.198 | 0.024 | 0.114 | 0.114 | 1.176 | 0.83 | 0.38 | 0.75 | 0.62 | 0.81 | 0.82 |
| 4 | 118556306 | rs2247756 | -0.268 | 0.44 | 0.947 | 0.192 | 0.015 | 0.085 | 0.085 | 0 | 0.91 | 0.71 | 0.92 | 0.87 | 0.92 | 0.95 |
| 4 | 118557318 | rs17862059 | 0.239 | 0.715 | 0.262 | -0.046 | -0.07 | 0.067 | 0.067 | 0.196 | 0.29 | 0.12 | 0.35 | 0.14 | 0.37 | 0.19 |
| 4 | 118558034 | rs17865601 | 1.156 | 0.688 | 0.321 | 0.385 | -0.013 | -0.04 | -0.04 | 0.057 | 0.11 | 0.01 | 0.07 | 0.11 | 0.14 | 0.07 |
| 4 | 118560782 | rs2635304 | 1.974 | 0.646 | 0.214 | 0.274 | -0.014 | -0.036 | -0.036 | 0 | 0.16 | 0.1 | 0.25 | 0.38 | 0.21 | 0.4 |
| 4 | 118567841 | rs2635199 | 0.402 | 0.611 | 0.305 | 0.217 | -0.003 | -0.036 | -0.036 | 0.353 | 0.28 | 0.22 | 0.33 | 0.48 | 0.34 | 0.45 |
| 4 | 118587270 | rs2389381 | 0.286 | 0.491 | 0.585 | 0.084 | -0.023 | 0.041 | 0.041 | 0 | 0.79 | 0.64 | 0.84 | 0.76 | 0.79 | 0.88 |
| 4 | 118599194 | rs1459545 | 0.418 | 0.51 | 0.645 | 0.035 | 0.003 | 0.085 | 0.085 | 0 | 0.09 | 0.29 | 0.07 | 0.13 | 0.07 | 0.05 |
| 4 | 118608250 | rs4834632 | -1.082 | 0.539 | 0.363 | 0.047 | -0.014 | 0.036 | 0.036 | 0 | 0.26 | 0.6 | 0.27 | 0.49 | 0.25 | 0.24 |
| 4 | 118610383 | rs1459551 | -1.075 | 0.467 | 0.395 | 0.011 | -0.011 | 0.03 | 0.03 | 0.022 | 0.27 | 0.59 | 0.26 | 0.49 | 0.25 | 0.24 |
| 4 | 118612335 | rs17862984 | 2.084 | 0.638 | -0.118 | -0.14 | -0.011 | 0.028 | 0.028 | 0.18 | 0.07 | 0.02 | 0.05 | 0 | 0.07 | 0 |
| 4 | 118613005 | rs4551001 | -1.966 | 0.261 | 0.148 | 0.014 | -0.061 | -0.085 | -0.085 | 0 | 0.73 | 0.87 | 0.53 | 0.49 | 0.65 | 0.52 |
| 4 | 118617008 | rs10011738 | -1.557 | -0.983 | 0.923 | 0.619 | -0.018 | 0.057 | 0.057 | 0 | 0.26 | 0.57 | 0.27 | 0.49 | 0.25 | 0.24 |
| 4 | 118622573 | rs17862070 | -1.162 | -0.936 | 0.823 | 0.64 | -0.024 | 0.118 | 0.118 | 0 | 0.21 | 0.39 | 0.21 | 0.38 | 0.2 | 0.22 |
| 4 | 118627589 | rs1120641 | -1.432 | -0.951 | 0.855 | 0.6 | -0.024 | 0.061 | 0.061 | 0 | 0.23 | 0.56 | 0.22 | 0.38 | 0.2 | 0.22 |
| 4 | 118629807 | rs10022014 | 0.269 | -0.957 | 0.96 | 0.637 | -0.001 | 0 | 0 | 0 | 0.12 | 0.08 | 0.08 | 0.11 | 0.14 | 0.07 |
| 4 | 118638662 | rs6834486 | -0.331 | -0.84 | 0.371 | 0.494 | 0.016 | 0.039 | 0.039 | 0.016 | 0.12 | 0.33 | 0.11 | 0.14 | 0.11 | 0.03 |
| 4 | 118641047 | rs10009424 | -0.959 | -0.801 | 0.38 | 0.517 | -0.029 | 0.129 | 0.129 | 0 | 0.22 | 0.42 | 0.21 | 0.38 | 0.2 | 0.22 |
| 4 | 118646907 | rs11098403 | 0.194 | -0.662 | 0.216 | 0.581 | 0.024 | 0.046 | 0.046 | 0.643 | 0.37 | 0.25 | 0.21 | 0.14 | 0.27 | 0.19 |
| 4 | 118650042 | rs11562926 | 0.023 | -0.631 | 0.181 | 0.32 | 0.01 | 0.009 | 0.009 | 0.24 | 0.39 | 0.35 | 0.34 | 0.22 | 0.28 | 0.2 |
| 4 | 118653376 | rs11562852 | -1.204 | -0.455 | 0.243 | 0.434 | -0.031 | 0.111 | 0.111 | 0 | 0.23 | 0.46 | 0.22 | 0.37 | 0.2 | 0.23 |
| 4 | 118663416 | rs7678262 | -1.147 | -1.165 | -0.718 | 0.271 | -0.042 | 0.004 | 0.004 | 0.033 | 0.6 | 0.84 | 0.54 | 0.57 | 0.44 | 0.41 |
| 4 | 118667234 | rs11945758 | -0.346 | -1.336 | -0.359 | 0.263 | 0.037 | 0.002 | 0.002 | 0.385 | 0.47 | 0.57 | 0.45 | 0.47 | 0.3 | 0.33 |
| 4 | 118667520 | rs11562835 | 0.406 | -1.289 | -0.43 | 0.179 | -0.001 | -0.002 | -0.002 | 2.982 | 0.1 | 0.1 | 0.08 | 0.1 | 0.14 | 0.07 |
| 4 | 118689954 | rs1350618 | -1.153 | -1.316 | -0.5 | 0.194 | -0.038 | 0.017 | 0.017 | 0 | 0.6 | 0.87 | 0.55 | 0.58 | 0.45 | 0.39 |
| 4 | 118705004 | rs4585377 | -1.122 | -1.319 | -0.487 | 0.193 | -0.027 | 0.028 | 0.028 | 0 | 0.6 | 0.87 | 0.55 | 0.58 | 0.45 | 0.39 |
| 4 | 118705939 | rs17869797 | 0.345 | -1.463 | -0.606 | 0.076 | -0.013 | 0.048 | 0.048 | 0 | 0.09 | 0.01 | 0.05 | 0.01 | 0.06 | 0.04 |

| | | | | | | | | | | | | | | | | |
|---|---|---|---|---|---|---|---|---|---|---|---|---|---|---|---|---|
| 4 | 118707424 | rs2125709 | -1.02 | -1.108 | -0.381 | 0.335 | -0.029 | 0.022 | 0.022 | 0 | 0.6 | 0.87 | 0.55 | 0.58 | 0.45 | 0.39 |
| 4 | 118717350 | rs7698673 | 0.102 | -1.117 | -0.376 | 0.348 | 0 | -0.003 | -0.003 | 0 | 0.11 | 0.1 | 0.09 | 0.1 | 0.14 | 0.07 |
| 4 | 118722033 | rs4377657 | -0.301 | -1.096 | -0.289 | 0.372 | 0.108 | 0.019 | 0.019 | 0 | 0.51 | 0.71 | 0.48 | 0.48 | 0.32 | 0.33 |
| 4 | 118727351 | rs11562865 | -0.269 | -1.059 | -0.118 | 0.376 | -0.006 | 0.001 | 0.001 | 0 | 0.12 | 0.22 | 0.09 | 0.1 | 0.12 | 0.07 |
| 4 | 118731430 | rs994619 | -0.945 | -0.988 | -0.134 | 0.365 | 0.004 | 0.034 | 0.034 | 0 | 0.59 | 0.69 | 0.54 | 0.58 | 0.44 | 0.39 |
| 4 | 118756418 | rs6811518 | -0.307 | -0.79 | 0.251 | 0.5 | 0 | 0.018 | 0.018 | 0 | 0.11 | 0.26 | 0.1 | 0.13 | 0.12 | 0.07 |
| 4 | 118758795 | rs4597906 | -1.132 | -1.141 | -0.136 | 0.402 | -0.004 | 0.049 | 0.049 | 0 | 0.59 | 0.89 | 0.5 | 0.62 | 0.43 | 0.4 |
| 4 | 118760390 | rs6813598 | 0.137 | -1.09 | -0.106 | 0.428 | 0 | -0.002 | -0.002 | 0 | 0.1 | 0.11 | 0.08 | 0.13 | 0.12 | 0.07 |
| 4 | 118777687 | rs4580714 | -0.154 | -1.167 | 0.245 | 0.384 | -0.018 | -0.003 | -0.003 | 0 | 0.45 | 0.52 | 0.41 | 0.49 | 0.3 | 0.32 |
| 4 | 118784946 | rs12503813 | 1.022 | -0.626 | -0.161 | 0.337 | -0.081 | 0.043 | 0.043 | 0 | 0.36 | 0.13 | 0.44 | 0.34 | 0.48 | 0.59 |
| 4 | 118789715 | rs7696793 | 0.173 | -0.901 | 0.035 | 0.606 | 0.03 | 0.026 | 0.026 | 0 | 0.5 | 0.4 | 0.63 | 0.72 | 0.63 | 0.77 |
| 4 | 118789858 | rs7675389 | -0.03 | -0.849 | -0.019 | 0.629 | -0.002 | -0.002 | -0.002 | 0 | 0.52 | 0.57 | 0.65 | 0.72 | 0.63 | 0.77 |
| 4 | 118802784 | rs4131119 | -1.038 | -1.064 | -0.487 | 0.339 | 0.08 | 0.096 | 0.096 | 0 | 0.55 | 0.91 | 0.48 | 0.65 | 0.36 | 0.38 |
| 4 | 118803489 | rs6534063 | -0.569 | -0.951 | -0.14 | 0.417 | 0.086 | 0.003 | 0.003 | 0 | 0.43 | 0.65 | 0.37 | 0.52 | 0.23 | 0.3 |
| 4 | 118820791 | rs4543205 | -1.059 | -0.128 | 0.703 | 0.079 | -0.013 | -0.021 | -0.021 | 0.197 | 0.34 | 0.6 | 0.23 | 0.24 | 0.3 | 0.25 |
| 4 | 118851421 | rs6834344 | -0.861 | -0.142 | 0.6 | 0.094 | -0.009 | -0.015 | -0.015 | 0 | 0.33 | 0.46 | 0.21 | 0.23 | 0.3 | 0.22 |
| 4 | 118868678 | rs11946339 | 0.313 | -0.168 | 0.737 | 0.081 | 0.001 | 0.003 | 0.003 | 0 | 0.69 | 0.7 | 0.8 | 0.77 | 0.7 | 0.78 |
| 4 | 118870933 | rs6534066 | 0.313 | -0.168 | 0.737 | 0.081 | 0.001 | 0.003 | 0.003 | 0 | 0.69 | 0.7 | 0.8 | 0.77 | 0.7 | 0.78 |
| 4 | 118891359 | rs11098416 | 0.809 | -0.139 | 0.777 | 0.077 | -0.019 | -0.026 | -0.026 | 0 | 0.65 | 0.47 | 0.78 | 0.76 | 0.7 | 0.78 |
| 4 | 118895571 | rs2892782 | 0.322 | -0.146 | 0.77 | 0.074 | 0.003 | 0.008 | 0.008 | 0 | 0.7 | 0.8 | 0.81 | 0.76 | 0.7 | 0.78 |
| 4 | 118919497 | rs4833558 | 0.506 | -0.008 | 0.375 | 0.116 | -0.002 | -0.004 | -0.004 | 1.814 | 0.68 | 0.64 | 0.79 | 0.77 | 0.7 | 0.75 |
| 4 | 118932958 | rs9997637 | -0.55 | 0.321 | -0.843 | 0.387 | -0.005 | 0.003 | 0.003 | 0.203 | 0.3 | 0.13 | 0.21 | 0.24 | 0.32 | 0.33 |
| 4 | 118939953 | rs11098419 | 0.43 | 0.132 | -1.037 | 0.317 | 0.042 | 0.043 | 0.043 | 0 | 0.58 | 0.19 | 0.65 | 0.75 | 0.68 | 0.78 |
| 4 | 118940052 | rs2389487 | 0.198 | -0.14 | -0.034 | 0.028 | -0.013 | 0.016 | 0.016 | 0 | 0.72 | 0.47 | 0.7 | 0.74 | 0.76 | 0.89 |
| 4 | 118941387 | rs2389489 | 0.211 | -0.242 | 0.05 | -0.04 | -0.03 | 0.015 | 0.015 | 0.108 | 0.75 | 0.47 | 0.83 | 0.77 | 0.83 | 0.91 |
| 4 | 118941742 | rs6849073 | 0.725 | -0.154 | 0.136 | -0.134 | -0.016 | -0.009 | -0.009 | 0 | 0.68 | 0.41 | 0.79 | 0.77 | 0.75 | 0.82 |
| 4 | 118948220 | rs10000049 | 0.197 | -0.24 | 0.193 | -0.167 | -0.031 | 0.011 | 0.011 | 0.34 | 0.75 | 0.43 | 0.76 | 0.76 | 0.82 | 0.55 |
| 4 | 118953901 | rs6534078 | 0.187 | -0.242 | 0.188 | -0.171 | -0.03 | 0.012 | 0.012 | 0.296 | 0.75 | 0.46 | 0.83 | 0.77 | 0.82 | 0.91 |
| 4 | 118963085 | rs10857057 | 0.809 | -0.234 | 0.304 | -0.156 | 0.027 | 0.017 | 0.017 | 0 | 0.66 | 0.25 | 0.78 | 0.77 | 0.75 | 0.83 |
| 4 | 118985104 | rs2389508 | 0.463 | -0.549 | 0.678 | -0.049 | -0.03 | 0.146 | 0.146 | 4.013 | 0.88 | 0.56 | 0.94 | 0.81 | 0.94 | 0.97 |
| 4 | 119001513 | rs2389513 | 0.463 | -0.549 | 0.678 | -0.049 | -0.03 | 0.146 | 0.146 | 4.013 | 0.88 | 0.56 | 0.94 | 0.81 | 0.94 | 0.97 |
| 4 | 119020848 | rs4428355 | 1.476 | -0.669 | -0.208 | -0.727 | 0.052 | 0.064 | 0.064 | 1.161 | 0.56 | 0.07 | 0.72 | 0.73 | 0.65 | 0.66 |
| 4 | 119024014 | rs4240315 | 1.475 | -0.686 | -0.216 | -0.735 | 0.052 | 0.065 | 0.065 | 1.07 | 0.56 | 0.07 | 0.72 | 0.73 | 0.65 | 0.67 |
| 4 | 119047274 | rs4833561 | 0.403 | -0.639 | -0.05 | -0.746 | -0.02 | 0.059 | 0.059 | 3.492 | 0.84 | 0.41 | 0.9 | 0.8 | 0.89 | 0.92 |
| 4 | 119068637 | rs13114867 | -0.782 | -0.668 | 0.02 | -0.81 | -0.004 | -0.047 | -0.047 | 0 | 0.3 | 0.36 | 0.21 | 0.06 | 0.29 | 0.3 |
| 4 | 119072862 | rs6853879 | -0.852 | -0.678 | 0.036 | -0.815 | -0.017 | -0.107 | -0.107 | 0.003 | 0.31 | 0.4 | 0.21 | 0.06 | 0.29 | 0.3 |
| 4 | 119115366 | rs12505156 | 0.183 | -0.628 | 0.424 | -0.667 | 0.008 | 0.138 | 0.138 | 2.602 | 0.87 | 0.57 | 0.93 | 0.8 | 0.92 | 0.96 |
| 4 | 119134308 | rs2892800 | 0.292 | -0.312 | -0.929 | -0.688 | 0.001 | -0.022 | -0.022 | 0 | 0.25 | 0.18 | 0.38 | 0.53 | 0.27 | 0.33 |
| 4 | 119142770 | rs2204269 | 0.777 | -0.193 | -0.992 | -0.804 | -0.001 | -0.024 | -0.024 | 0 | 0.15 | 0.17 | 0.14 | 0.05 | 0.19 | 0.24 |
| 4 | 119155229 | rs788663 | 1.539 | -0.168 | -1.149 | -0.84 | -0.132 | 0.075 | 0.075 | 0 | 0.37 | 0.02 | 0.36 | 0.21 | 0.37 | 0.36 |
| 4 | 119161795 | rs617430 | 1.463 | -0.182 | -1.119 | -0.846 | -0.088 | 0.103 | 0.103 | 0 | 0.38 | 0.05 | 0.36 | 0.21 | 0.37 | 0.36 |
| 4 | 119170256 | rs687492 | 1.557 | -0.163 | -1.139 | -0.714 | -0.125 | 0.179 | 0.179 | 0 | 0.37 | 0.02 | 0.35 | 0.12 | 0.36 | 0.36 |
| 4 | 119175904 | rs631271 | 1.459 | -0.176 | -1.107 | -0.705 | -0.064 | 0.106 | 0.106 | 0 | 0.38 | 0.07 | 0.36 | 0.21 | 0.37 | 0.36 |
| 4 | 119199879 | rs1573955 | 0.288 | -0.244 | -0.503 | -0.677 | -0.001 | -0.001 | -0.001 | 0 | 0.27 | 0.26 | 0.4 | 0.53 | 0.29 | 0.33 |
| 4 | 119203708 | rs11098432 | -0.574 | 0.083 | -0.497 | -0.774 | 0 | -0.001 | -0.001 | 0.009 | 0.4 | 0.37 | 0.51 | 0.58 | 0.45 | 0.53 |
| 4 | 119204720 | rs4833571 | -1.077 | 0.062 | -0.469 | -0.775 | 0 | 0.003 | 0.003 | 0 | 0.51 | 0.51 | 0.58 | 0.67 | 0.56 | 0.6 |
| 4 | 119228883 | rs11730327 | 1.385 | 0.05 | -0.533 | -0.695 | -0.027 | 0.032 | 0.032 | 0 | 0.12 | 0.01 | 0.1 | 0.05 | 0.16 | 0.2 |
| 4 | 119240618 | rs4834678 | 0.09 | 0.127 | -0.558 | 0.021 | 0 | 0.008 | 0.008 | 0 | 0.24 | 0.25 | 0.39 | 0.4 | 0.27 | 0.22 |
| 4 | 119266964 | rs1523743 | 0.862 | 0.213 | -0.707 | 0.408 | -0.044 | -0.067 | -0.067 | 0.057 | 0.17 | 0.04 | 0.36 | 0.35 | 0.27 | 0.2 |
| 4 | 119272958 | rs7674055 | 1.122 | 0.525 | -0.612 | 0.342 | 0.012 | -0.023 | -0.023 | 0.426 | 0.59 | 0.21 | 0.74 | 0.47 | 0.64 | 0.57 |
| 4 | 119277490 | rs10016452 | 1.122 | 0.585 | -0.296 | 0.475 | -0.033 | 0.037 | 0.037 | 0.785 | 0.08 | 0.01 | 0.03 | 0 | 0.11 | 0.01 |
| 4 | 119279098 | rs1514657 | 0.085 | 0.526 | -0.045 | 0.579 | -0.003 | 0.162 | 0.162 | 2.771 | 0.88 | 0.48 | 0.93 | 0.8 | 0.94 | 0.94 |
| 4 | 119299078 | rs2892812 | -0.928 | 0.504 | 0.117 | 0.577 | -0.038 | 0.052 | 0.052 | 3.35 | 0.15 | 0.63 | 0.09 | 0.19 | 0.06 | 0.06 |
| 4 | 119302960 | rs11932636 | -1.15 | 0.473 | 0.203 | 0.562 | -0.026 | 0.013 | 0.013 | 4.189 | 0.16 | 0.72 | 0.09 | 0.19 | 0.06 | 0.06 |
| 4 | 119303024 | rs7696047 | -0.269 | 0.48 | -0.141 | 0.725 | -0.028 | -0.05 | -0.05 | 0 | 0.13 | 0.08 | 0.18 | 0.27 | 0.16 | 0.18 |
| 4 | 119311524 | rs12498329 | -0.898 | 0.472 | 0.045 | 0.721 | 0.011 | 0.05 | 0.05 | 0.298 | 0.17 | 0.27 | 0.19 | 0.27 | 0.16 | 0.18 |
| 4 | 119322004 | rs1818521 | -1.152 | 0.238 | 0.238 | 0.674 | 0.008 | 0.029 | 0.029 | 0.511 | 0.32 | 0.43 | 0.5 | 0.59 | 0.33 | 0.37 |
| 4 | 119331363 | rs10518293 | -0.411 | 0.301 | 0.119 | 0.87 | 0 | 0 | 0 | 0 | 0.2 | 0.17 | 0.34 | 0.44 | 0.22 | 0.36 |
| 4 | 119337263 | rs6811485 | -0.388 | 0.315 | 0.166 | 0.901 | -0.003 | 0.04 | 0.04 | 0 | 0.21 | 0.23 | 0.34 | 0.44 | 0.22 | 0.36 |
| 4 | 119371173 | rs2631108 | -0.702 | -0.42 | -0.174 | 1.697 | 0.005 | 0.02 | 0.02 | 0 | 0.26 | 0.32 | 0.31 | 0.41 | 0.32 | 0.31 |
| 4 | 119399111 | rs17693682 | -0.066 | -0.346 | -0.269 | 1.61 | -0.001 | -0.027 | -0.027 | 0.233 | 0.21 | 0.12 | 0.16 | 0.26 | 0.24 | 0.31 |
| 4 | 119407189 | rs10021843 | -1.06 | -0.01 | -0.705 | 1.128 | -0.001 | -0.001 | -0.001 | 0 | 0.44 | 0.44 | 0.48 | 0.42 | 0.58 | 0.51 |
| 4 | 119408351 | rs17642621 | 0.778 | -0.725 | -0.806 | 1.128 | 0.073 | -0.043 | -0.043 | 0 | 0.21 | 0.07 | 0.04 | 0.28 | 0.08 | 0.06 |
| 4 | 119437972 | rs2035600 | -0.9 | 0.179 | -0.679 | 0.883 | 0 | -0.007 | -0.007 | 0 | 0.45 | 0.45 | 0.35 | 0.64 | 0.48 | 0.39 |
| 4 | 119441680 | rs4001461 | -0.966 | 0.181 | -0.687 | 0.884 | 0 | -0.007 | -0.007 | 0 | 0.45 | 0.43 | 0.36 | 0.64 | 0.48 | 0.37 |
| 4 | 119455055 | rs17595265 | 0.927 | 0.165 | -0.44 | 1.073 | 0 | 0.006 | 0.006 | 0 | 0.21 | 0.12 | 0.18 | 0.17 | 0.23 | 0.18 |
| 4 | 119462685 | rs2200517 | -0.34 | -0.097 | -0.326 | 1.09 | 0.009 | -0.001 | -0.001 | 0 | 0.42 | 0.38 | 0.3 | 0.52 | 0.32 | 0.33 |
| 4 | 119483415 | rs11728902 | 0.15 | 0.493 | -0.313 | 1.243 | 0.02 | 0.011 | 0.011 | 0.179 | 0.24 | 0.3 | 0.35 | 0.27 | 0.39 | 0.21 |
| 4 | 119506139 | rs7689541 | -0.23 | 0.325 | -0.173 | 0.863 | 0.042 | -0.005 | -0.005 | 0.03 | 0.26 | 0.37 | 0.25 | 0.19 | 0.38 | 0.24 |

| | | | | | | | | | | | | | | | | |
|---|---|---|---|---|---|---|---|---|---|---|---|---|---|---|---|---|
| 4 | 119510402 | rs7669896 | 0.483 | 0.48 | -0.313 | 0.89 | -0.046 | -0.02 | -0.02 | 0 | 0.21 | 0.11 | 0.21 | 0.18 | 0.33 | 0.23 |
| 4 | 119595364 | rs299005 | -0.014 | -0.426 | -0.688 | 0.692 | 0.001 | 0.001 | 0.001 | 0.798 | 0.59 | 0.42 | 0.76 | 0.69 | 0.69 | 0.69 |
| 4 | 119605009 | rs298982 | -0.739 | -0.852 | -0.835 | 0.538 | 0.028 | -0.024 | -0.024 | 0 | 0.81 | 0.47 | 0.81 | 0.86 | 0.77 | 0.76 |
| 4 | 119607376 | rs298980 | 2.166 | -0.268 | -1.311 | 0.354 | -0.057 | -0.168 | -0.168 | 0.083 | 0.29 | 0.04 | 0.48 | 0.51 | 0.26 | 0.4 |
| 4 | 119613045 | rs6821350 | 0.803 | -0.054 | -0.866 | 0.378 | -0.065 | -0.002 | -0.002 | 0 | 0.25 | 0.11 | 0.23 | 0.17 | 0.37 | 0.29 |
| 4 | 119624641 | rs6813357 | 0.747 | -0.05 | -0.803 | 0.38 | -0.038 | 0.008 | 0.008 | 1.107 | 0.26 | 0.16 | 0.24 | 0.17 | 0.37 | 0.29 |
| 4 | 119630402 | rs441216 | -0.166 | -0.105 | -0.626 | 0.207 | -0.029 | -0.027 | -0.027 | 0 | 0.64 | 0.46 | 0.81 | 0.79 | 0.77 | 0.83 |
| 4 | 119643010 | rs12640223 | 0.908 | -0.062 | -0.787 | 0.249 | -0.05 | 0.004 | 0.004 | 1.288 | 0.25 | 0.12 | 0.23 | 0.17 | 0.37 | 0.29 |
| 4 | 119646000 | rs298991 | 0.229 | -0.189 | -0.769 | 0.434 | 0.013 | 0.005 | 0.005 | 0.355 | 0.6 | 0.39 | 0.72 | 0.6 | 0.69 | 0.69 |
| 4 | 119648399 | rs298988 | -0.863 | 0.132 | -0.713 | 0.517 | 0.037 | 0.015 | 0.015 | 0 | 0.82 | 0.52 | 0.78 | 0.77 | 0.77 | 0.76 |
| 4 | 119651565 | rs960738 | 1.592 | 0.376 | -1.454 | 0.626 | -0.063 | -0.085 | -0.085 | 1.228 | 0.32 | 0.08 | 0.48 | 0.41 | 0.32 | 0.4 |
| 4 | 119655888 | rs12509236 | 1.743 | 0.308 | -1.86 | 0.545 | -0.077 | -0.16 | -0.16 | 0 | 0.31 | 0.02 | 0.45 | 0.43 | 0.27 | 0.4 |
| 4 | 119663911 | rs3775835 | -1.337 | 0.442 | -1.524 | 0.493 | -0.007 | 0.006 | 0.006 | 1.778 | 0.19 | 0.75 | 0.2 | 0.26 | 0.16 | 0.22 |
| 4 | 119664228 | rs4834700 | -1.196 | 1.292 | -2.201 | 0.23 | -0.03 | -0.006 | -0.006 | 0.794 | 0.83 | 0.9 | 0.82 | 0.82 | 0.72 | 0.81 |
| 4 | 119675900 | rs6827740 | -0.041 | 1.028 | -2.286 | 0.098 | -0.04 | 0.032 | 0.032 | 0.743 | 0.26 | 0.12 | 0.23 | 0.15 | 0.37 | 0.29 |
| 4 | 119678580 | rs4340779 | 0.538 | 0.301 | -2.361 | -0.023 | 0.063 | 0.001 | 0.001 | 0.975 | 0.6 | 0.19 | 0.72 | 0.6 | 0.71 | 0.7 |
| 4 | 119691624 | rs4834703 | 1.789 | 0.385 | -2.253 | -0.099 | -0.003 | 0.029 | 0.029 | 0 | 0.09 | 0.04 | 0.07 | 0.06 | 0.11 | 0.09 |
| 4 | 119717634 | rs10518325 | 0.543 | 0.246 | -2.076 | 0.056 | 0.064 | 0.074 | 0.074 | 0 | 0.19 | 0.03 | 0.04 | 0.05 | 0.09 | 0.1 |
| 4 | 119733366 | rs4834706 | 0.811 | 1.082 | -1.954 | 0.408 | -0.012 | -0.101 | -0.101 | 0 | 0.38 | 0.09 | 0.55 | 0.55 | 0.38 | 0.47 |
| 4 | 119743682 | rs6843239 | -0.122 | 1.417 | -1.481 | 0.267 | 0.001 | 0.001 | 0.001 | 0.488 | 0.56 | 0.41 | 0.68 | 0.65 | 0.6 | 0.6 |
| 4 | 119752920 | rs13139244 | 0.873 | 1.415 | -1.515 | 0.317 | 0.014 | -0.057 | -0.057 | 1.031 | 0.39 | 0.16 | 0.56 | 0.55 | 0.38 | 0.47 |
| 4 | 119761678 | rs11730732 | -0.659 | 1.541 | -1.771 | 0.157 | -0.022 | 0.138 | 0.138 | 0 | 0.55 | 0.78 | 0.36 | 0.15 | 0.5 | 0.39 |
| 4 | 119762951 | rs2162400 | 0.768 | 1.753 | -1.739 | 0.241 | -0.012 | 0.004 | 0.004 | 0.253 | 0.14 | 0.1 | 0.09 | 0.1 | 0.17 | 0.12 |
| 4 | 119768598 | rs7676892 | -0.054 | 1.742 | -1.711 | 0.498 | 0.009 | 0.063 | 0.063 | 0 | 0.94 | 0.92 | 0.9 | 0.7 | 0.88 | 0.86 |
| 4 | 119770915 | rs10019751 | -0.633 | 1.675 | -1.832 | 0.201 | -0.038 | 0.169 | 0.169 | 0 | 0.57 | 0.81 | 0.33 | 0.14 | 0.45 | 0.38 |
| 4 | 119781338 | rs10857061 | 0.759 | 1.382 | -1.463 | 0.646 | 0.069 | 0.142 | 0.142 | 0 | 0.92 | 0.81 | 0.87 | 0.69 | 0.82 | 0.85 |
| 4 | 119785381 | rs2114461 | 0.791 | 1.185 | -1.842 | 0.455 | -0.08 | -0.119 | -0.119 | 0.507 | 0.4 | 0.08 | 0.61 | 0.56 | 0.48 | 0.48 |
| 4 | 119789448 | rs2389655 | 0.145 | 0.975 | -1.692 | 0.331 | 0.076 | 0.037 | 0.037 | 0.063 | 0.54 | 0.15 | 0.71 | 0.65 | 0.68 | 0.61 |
| 4 | 119790013 | rs7679385 | 0.198 | 0.929 | -1.53 | 0.249 | -0.022 | -0.073 | -0.073 | 0 | 0.37 | 0.56 | 0.18 | 0.06 | 0.2 | 0.26 |
| 4 | 119792297 | rs4240322 | 0.248 | 1.472 | -1.381 | 0.838 | 0.032 | 0.14 | 0.14 | 0 | 0.93 | 0.82 | 0.87 | 0.71 | 0.88 | 0.85 |
| 4 | 119793776 | rs4834711 | 0.742 | 1.423 | -2.007 | 0.856 | -0.075 | -0.058 | -0.058 | 0 | 0.35 | 0.06 | 0.46 | 0.4 | 0.41 | 0.46 |
| 4 | 119795642 | rs1426819 | 0.603 | 1.567 | -1.453 | 1.066 | -0.004 | -0.004 | -0.004 | 0 | 0.1 | 0.15 | 0.11 | 0.23 | 0.14 | 0.13 |
| 4 | 119800081 | rs6534110 | -1.151 | 0.923 | -1.345 | 0.694 | 0.001 | 0.001 | 0.001 | 0.133 | 0.83 | 0.9 | 0.9 | 0.9 | 0.84 | 0.78 |
| 4 | 119800779 | rs7690236 | -0.628 | 1.265 | -1.12 | 0.981 | -0.005 | -0.003 | -0.003 | 0 | 0.6 | 0.48 | 0.68 | 0.75 | 0.7 | 0.71 |
| 4 | 119801012 | rs7695141 | 0.961 | 2.009 | -1.066 | 0.859 | -0.052 | -0.041 | -0.041 | 1.074 | 0.1 | 0.01 | 0.08 | 0.12 | 0.13 | 0.1 |
| 4 | 119807193 | rs11940444 | -0.178 | 1.859 | -1.051 | 0.786 | -0.071 | 0.037 | 0.037 | 0 | 0.33 | 0.16 | 0.33 | 0.25 | 0.49 | 0.24 |
| 4 | 119811735 | rs13117019 | -0.814 | 1.814 | -0.75 | 0.976 | -0.005 | 0 | 0 | 0 | 0.42 | 0.44 | 0.42 | 0.64 | 0.47 | 0.42 |
| 4 | 119817622 | rs6831957 | 0.216 | 1.973 | -0.438 | 0.733 | -0.017 | 0.001 | 0.001 | 0 | 0.7 | 0.77 | 0.69 | 0.71 | 0.57 | 0.76 |
| 4 | 119825219 | rs4834713 | -0.129 | 2.17 | -0.165 | 0.606 | 0.047 | 0.011 | 0.011 | 0 | 0.87 | 0.78 | 0.85 | 0.88 | 0.78 | 0.9 |
| 4 | 119827183 | rs955607 | -0.582 | 1.732 | 0.069 | 0.567 | 0.033 | 0.01 | 0.01 | 0 | 0.89 | 0.77 | 0.91 | 0.93 | 0.87 | 0.9 |
| 4 | 119831193 | rs12506322 | -0.418 | 1.723 | 0.181 | 0.567 | 0.042 | 0.015 | 0.015 | 0 | 0.88 | 0.71 | 0.9 | 0.93 | 0.87 | 0.9 |
| 4 | 119833292 | rs736965 | 0.735 | 1.389 | 0.599 | 0.26 | -0.004 | 0.034 | 0.034 | 0.055 | 0.63 | 0.28 | 0.71 | 0.79 | 0.66 | 0.76 |
| 4 | 119835148 | rs7676506 | 0.464 | 1.377 | 0.968 | 0.252 | 0.002 | -0.027 | -0.027 | 0 | 0.2 | 0.28 | 0.17 | 0.12 | 0.19 | 0.13 |
| 4 | 119844851 | rs2162398 | -1.054 | 1.413 | 0.834 | 0.264 | 0.022 | 0.005 | 0.005 | 0 | 0.89 | 0.81 | 0.91 | 0.93 | 0.87 | 0.91 |
| 4 | 119846806 | rs754397 | 0.56 | 1.363 | 0.604 | 0.281 | 0 | -0.008 | -0.008 | 0 | 0.65 | 0.34 | 0.72 | 0.8 | 0.66 | 0.77 |
| 4 | 119853638 | rs13138512 | 0.49 | 1.64 | 0.035 | -0.005 | -0.003 | -0.001 | -0.001 | 0.85 | 0.63 | 0.37 | 0.71 | 0.81 | 0.6 | 0.78 |
| 4 | 119859344 | rs7672077 | 0.123 | 1.266 | 0.463 | 0.69 | 0.024 | 0.07 | 0.07 | 0 | 0.85 | 0.46 | 0.88 | 0.8 | 0.84 | 0.9 |
| 4 | 119860592 | rs1565475 | 0.122 | 1.269 | 0.463 | 0.69 | 0.023 | 0.069 | 0.069 | 0 | 0.85 | 0.46 | 0.88 | 0.8 | 0.84 | 0.9 |
| 4 | 119862144 | rs6826016 | 0.757 | 0.73 | -0.291 | 0.383 | 0 | -0.004 | -0.004 | 0 | 0.62 | 0.43 | 0.74 | 0.7 | 0.6 | 0.68 |
| 4 | 119864315 | rs17261234 | 0.025 | 0.8 | -0.719 | 0.351 | 0 | 0.049 | 0.049 | 0 | 0.23 | 0.12 | 0.14 | 0.1 | 0.24 | 0.22 |
| 4 | 119864599 | rs17327171 | -0.029 | 0.854 | -0.776 | 0.317 | -0.01 | 0.066 | 0.066 | 0 | 0.23 | 0.07 | 0.15 | 0.1 | 0.25 | 0.22 |
| 4 | 119865067 | rs4834729 | 0.58 | 0.746 | -0.866 | 0.928 | -0.03 | -0.078 | -0.078 | 0 | 0.19 | 0.02 | 0.14 | 0.24 | 0.21 | 0.21 |
| 4 | 119867342 | rs1021377 | 0.982 | 0.946 | -0.572 | 0.948 | 0.003 | -0.011 | -0.011 | 0 | 0.56 | 0.44 | 0.63 | 0.3 | 0.53 | 0.56 |
| 4 | 119873248 | rs13130144 | -0.717 | 0.966 | -0.488 | 1.124 | -0.003 | -0.008 | -0.008 | 0.04 | 0.35 | 0.34 | 0.29 | 0.58 | 0.36 | 0.38 |
| 4 | 119874955 | rs10025067 | 0.708 | 0.15 | -0.961 | 0.51 | 0.041 | 0.039 | 0.039 | 0.04 | 0.15 | 0.08 | 0.05 | 0 | 0.06 | 0.11 |
| 4 | 119875492 | rs1387981 | -1.22 | 0.223 | -0.835 | 0.562 | 0.046 | 0.065 | 0.065 | 0 | 0.89 | 0.65 | 0.9 | 0.89 | 0.88 | 0.94 |
| 4 | 119877794 | rs12508843 | 0.417 | 0.249 | -0.9 | 0.603 | -0.001 | -0.005 | -0.005 | 0 | 0.67 | 0.58 | 0.76 | 0.69 | 0.72 | 0.69 |
| 4 | 119881072 | rs6858546 | 1.016 | -0.004 | -0.812 | 0.396 | -0.004 | 0.004 | 0.004 | 0 | 0.45 | 0.41 | 0.61 | 0.43 | 0.51 | 0.53 |
| 4 | 119883564 | rs10518316 | 0.971 | 0.001 | -0.833 | 0.467 | 0.004 | -0.057 | -0.057 | 0 | 0.2 | 0.06 | 0.13 | 0.26 | 0.18 | 0.17 |
| 4 | 119884361 | rs7697974 | -0.364 | 0.09 | -0.72 | 0.498 | -0.008 | -0.013 | -0.013 | 0 | 0.81 | 0.74 | 0.89 | 0.87 | 0.83 | 0.9 |
| 4 | 119886179 | rs1490510 | -1.474 | 0.155 | -0.707 | 0.526 | 0.054 | 0.057 | 0.057 | 0 | 0.9 | 0.77 | 0.91 | 0.87 | 0.86 | 0.94 |
| 4 | 119890139 | rs17050209 | 1.274 | 0.199 | -0.789 | 0.551 | -0.002 | -0.001 | -0.001 | 0 | 0.08 | 0.08 | 0.1 | 0.19 | 0.1 | 0.12 |
| 4 | 119890772 | rs1038771 | 1.58 | 0.109 | -1.043 | 0.304 | -0.001 | -0.002 | -0.002 | 0 | 0.36 | 0.36 | 0.55 | 0.26 | 0.41 | 0.52 |
| 4 | 119890865 | rs17050208 | 1.134 | 0.135 | -1.161 | 0.352 | 0.012 | -0.049 | -0.049 | 0 | 0.21 | 0.07 | 0.13 | 0.26 | 0.18 | 0.17 |
| 4 | 119892879 | rs6855192 | 0.486 | 0.171 | -0.978 | 0.386 | 0.011 | 0.014 | 0.014 | 0.397 | 0.72 | 0.57 | 0.78 | 0.67 | 0.73 | 0.79 |
| 4 | 119896186 | rs11098467 | 1.062 | 0.099 | -1.012 | 0.301 | 0 | -0.032 | -0.032 | 0.169 | 0.38 | 0.49 | 0.57 | 0.26 | 0.41 | 0.53 |
| 4 | 119896331 | rs9307465 | -0.047 | 0.567 | -1.229 | 0.503 | 0.008 | -0.062 | -0.062 | 0.609 | 0.35 | 0.12 | 0.22 | 0.41 | 0.32 | 0.26 |
| 4 | 119896633 | rs17050196 | 0.872 | 0.966 | -1.012 | 0.761 | -0.002 | 0 | 0 | 0 | 0.07 | 0.08 | 0.1 | 0.2 | 0.1 | 0.12 |
| 4 | 119899664 | rs1387983 | 0.638 | 0.723 | -1.013 | 0.324 | -0.02 | -0.012 | -0.012 | 2.612 | 0.15 | 0.01 | 0.07 | 0.13 | 0.11 | 0.03 |

| | | | | | | | | | | | | | | | | |
|---|---|---|---|---|---|---|---|---|---|---|---|---|---|---|---|---|
| 4 | 119902065 | rs4833599 | -0.954 | 0.897 | -0.566 | 0.418 | 0.039 | 0.052 | 0.052 | 0 | 0.91 | 0.82 | 0.8 | 0.85 | 0.86 | 0.94 |
| 4 | 119903064 | rs1847 | -0.812 | 0.895 | -0.561 | 0.416 | 0.048 | 0.059 | 0.059 | 0 | 0.9 | 0.74 | 0.79 | 0.85 | 0.86 | 0.94 |
| 4 | 119907402 | rs4834733 | 0.238 | 0.408 | -0.684 | -0.076 | -0.001 | 0 | 0 | 0.045 | 0.42 | 0.37 | 0.46 | 0.26 | 0.45 | 0.6 |
| 4 | 119912286 | rs7437916 | -0.427 | 0.845 | -0.524 | 0.155 | -0.001 | 0.003 | 0.003 | 0 | 0.36 | 0.34 | 0.4 | 0.24 | 0.36 | 0.36 |
| 4 | 119913577 | rs17328610 | 0.355 | 0.902 | -0.646 | 0.112 | 0.001 | 0.041 | 0.041 | 0 | 0.24 | 0.16 | 0.3 | 0.12 | 0.22 | 0.33 |
| 4 | 119914670 | rs2061443 | -0.64 | 0.885 | -0.663 | 0.107 | 0 | -0.015 | -0.015 | 0 | 0.37 | 0.41 | 0.41 | 0.24 | 0.36 | 0.36 |
| 4 | 119922111 | rs1380163 | 0.046 | 0.887 | -0.332 | 0.481 | 0.02 | -0.009 | -0.009 | 0.161 | 0.15 | 0.34 | 0.12 | 0.14 | 0.14 | 0.03 |
| 4 | 119927268 | rs1994630 | -0.089 | 0.949 | -1.141 | 0.806 | -0.039 | -0.044 | -0.044 | 0.316 | 0.24 | 0.06 | 0.17 | 0.26 | 0.29 | 0.29 |
| 4 | 119929733 | rs983417 | -0.089 | 1.016 | -0.246 | 0.993 | 0.036 | 0.073 | 0.073 | 0 | 0.93 | 0.81 | 0.8 | 0.86 | 0.89 | 0.95 |
| 4 | 119932620 | rs1380158 | -0.201 | 0.647 | -0.794 | 0.662 | 0.024 | 0.041 | 0.041 | 0 | 0.48 | 0.23 | 0.43 | 0.38 | 0.42 | 0.6 |
| 4 | 119934384 | rs1380159 | 0.248 | -0.207 | 0.225 | 0.373 | -0.072 | -0.034 | -0.034 | 2.458 | 0.8 | 0.5 | 0.77 | 0.82 | 0.91 | 0.75 |
| 4 | 119941399 | rs1459056 | 0.438 | 0.104 | -0.196 | 0.396 | -0.012 | -0.002 | -0.002 | 0 | 0.67 | 0.67 | 0.66 | 0.67 | 0.71 | 0.59 |
| 4 | 119947135 | rs6828669 | -0.077 | 0.328 | -0.354 | 0.313 | -0.001 | -0.002 | -0.002 | 0 | 0.45 | 0.48 | 0.38 | 0.6 | 0.41 | 0.49 |
| 4 | 119947188 | rs7671665 | 0.598 | 0.346 | -0.216 | 0.834 | -0.001 | -0.003 | -0.003 | 0.01 | 0.3 | 0.33 | 0.4 | 0.41 | 0.3 | 0.3 |
| 4 | 119947362 | rs12511929 | -0.009 | 0.555 | -0.219 | 0.547 | -0.012 | -0.059 | -0.059 | 0 | 0.19 | 0.1 | 0.38 | 0.4 | 0.23 | 0.24 |
| 4 | 119948046 | rs17263971 | 0.606 | 0.697 | -0.232 | -0.005 | -0.006 | 0.015 | 0.015 | 0.38 | 0.24 | 0.21 | 0.22 | 0.16 | 0.26 | 0.26 |
| 4 | 119950186 | rs2045547 | 0.761 | 0.527 | -0.049 | 0.127 | -0.028 | -0.088 | -0.088 | 0.005 | 0.15 | 0.05 | 0.36 | 0.41 | 0.2 | 0.24 |
| 4 | 119951647 | rs7698598 | -1.47 | 0.248 | 0.349 | 0.194 | -0.001 | 0.009 | 0.009 | 0.975 | 0.87 | 0.81 | 0.9 | 0.87 | 0.87 | 0.8 |
| 4 | 119954533 | rs13837 | -0.876 | 0.175 | 0.39 | 0.274 | -0.011 | 0.115 | 0.115 | 1.237 | 0.83 | 0.59 | 0.87 | 0.74 | 0.86 | 0.77 |
| 4 | 119954710 | rs2279506 | 0.13 | 0.155 | -0.194 | 0.066 | -0.007 | -0.042 | -0.042 | 0 | 0.27 | 0.2 | 0.43 | 0.49 | 0.33 | 0.26 |
| 4 | 119955787 | rs3733511 | 0.837 | 0.343 | -0.57 | -0.51 | -0.015 | 0.03 | 0.03 | 0.054 | 0.25 | 0.18 | 0.24 | 0.16 | 0.29 | 0.27 |
| 4 | 119956089 | rs752097 | 0.563 | 0.363 | -0.699 | -0.189 | -0.001 | 0.121 | 0.121 | 4.693 | 0.3 | 0.05 | 0.19 | 0.08 | 0.24 | 0.24 |
| 4 | 119962741 | rs10518313 | -0.708 | 0.209 | -0.477 | -0.188 | 0.024 | 0.025 | 0.025 | 1.805 | 0.68 | 0.84 | 0.81 | 0.8 | 0.76 | 0.68 |
| 4 | 119963526 | rs17264415 | 1.044 | 0.4 | -0.8 | -0.29 | -0.035 | -0.001 | -0.001 | 0 | 0.09 | 0 | 0.06 | 0.08 | 0.12 | 0.03 |
| 4 | 119963646 | rs9784534 | 0.546 | 0.468 | -0.011 | -0.501 | 0.007 | -0.047 | -0.047 | 1.196 | 0.29 | 0.47 | 0.26 | 0.16 | 0.31 | 0.22 |
| 4 | 119966718 | rs1037763 | 0.555 | 0.429 | 0.165 | -0.495 | -0.016 | -0.035 | -0.035 | 0 | 0.28 | 0.14 | 0.46 | 0.57 | 0.34 | 0.44 |
| 4 | 119970203 | rs1380162 | -0.666 | 0.357 | 0.449 | -0.547 | 0.014 | 0.068 | 0.068 | 2.642 | 0.6 | 0.79 | 0.75 | 0.84 | 0.64 | 0.68 |
| 4 | 119971043 | rs13137064 | 0.356 | 0.37 | 0.521 | -0.353 | -0.005 | -0.014 | -0.014 | 0 | 0.28 | 0.23 | 0.47 | 0.57 | 0.33 | 0.44 |
| 4 | 119974540 | rs12506805 | 0.19 | 0.387 | 0.759 | -0.359 | 0.009 | 0.013 | 0.013 | 2.046 | 0.31 | 0.37 | 0.5 | 0.56 | 0.38 | 0.44 |
| 4 | 119980501 | rs17050125 | -0.312 | 0.052 | 0.959 | -0.659 | 0.039 | 0.052 | 0.052 | 0.934 | 0.49 | 0.63 | 0.3 | 0.23 | 0.34 | 0.25 |
| 4 | 119980640 | rs1058450 | 0.771 | -0.397 | 0.556 | -1.033 | -0.008 | 0.046 | 0.046 | 0.807 | 0.19 | 0.04 | 0.11 | 0.09 | 0.2 | 0.05 |
| 4 | 119981296 | rs17050116 | 0.463 | -0.377 | 0.809 | -0.836 | -0.036 | -0.02 | -0.02 | 0 | 0.28 | 0.4 | 0.17 | 0.14 | 0.13 | 0.2 |
| 4 | 119981828 | rs1047894 | 0.251 | -0.347 | 0.587 | -0.821 | -0.065 | -0.039 | -0.039 | 0 | 0.3 | 0.57 | 0.18 | 0.14 | 0.14 | 0.2 |
| 4 | 119992822 | rs4438744 | 0.687 | -0.002 | 0.013 | -0.657 | -0.028 | -0.033 | -0.033 | 0 | 0.27 | 0.19 | 0.43 | 0.56 | 0.37 | 0.4 |
| 4 | 119994554 | rs12509365 | 0.814 | -0.026 | -0.341 | -0.67 | -0.054 | -0.065 | -0.065 | 0 | 0.26 | 0.12 | 0.43 | 0.56 | 0.36 | 0.39 |
| 4 | 120005147 | rs11098479 | 0.129 | -0.648 | -0.313 | -1.069 | 0.048 | 0.048 | 0.048 | 0.215 | 0.55 | 0.31 | 0.74 | 0.76 | 0.72 | 0.76 |
| 4 | 120005814 | rs2219497 | 0.097 | -0.645 | -0.274 | -1.068 | 0.016 | 0.016 | 0.016 | 0.166 | 0.56 | 0.37 | 0.74 | 0.76 | 0.72 | 0.76 |
| 4 | 120008227 | rs7441142 | -0.292 | -0.369 | -0.255 | -0.898 | 0.001 | -0.017 | -0.017 | 0 | 0.59 | 0.51 | 0.63 | 0.65 | 0.57 | 0.6 |
| 4 | 120008569 | rs7670862 | 0.669 | -0.211 | -0.564 | -1.057 | -0.048 | -0.056 | -0.056 | 0.244 | 0.3 | 0.16 | 0.45 | 0.51 | 0.38 | 0.37 |
| 4 | 120009168 | rs10857064 | -0.237 | -0.021 | -0.343 | -0.72 | 0.003 | -0.018 | -0.018 | 0 | 0.6 | 0.58 | 0.54 | 0.62 | 0.57 | 0.64 |
| 4 | 120013157 | rs11729054 | -0.106 | 0.005 | -0.503 | -0.519 | -0.007 | 0.012 | 0.012 | 0 | 0.18 | 0.28 | 0.12 | 0.2 | 0.16 | 0.1 |
| 4 | 120016617 | rs11098483 | 0.648 | 0.918 | -0.836 | 0.382 | -0.07 | -0.084 | -0.084 | 1.391 | 0.12 | 0.01 | 0.21 | 0.36 | 0.21 | 0.17 |
| 4 | 120018082 | rs11098484 | 0.114 | 0.36 | -0.87 | -0.158 | -0.025 | -0.01 | -0.01 | 0 | 0.35 | 0.25 | 0.45 | 0.61 | 0.43 | 0.41 |
| 4 | 120025002 | rs4261975 | 0.066 | 0.257 | -0.383 | -0.442 | 0.011 | 0.017 | 0.017 | 0 | 0.39 | 0.37 | 0.25 | 0.16 | 0.31 | 0.27 |
| 4 | 120029841 | rs10029388 | 1.951 | 0.356 | 0.21 | -0.56 | 0.002 | -0.001 | -0.001 | 0 | 0.17 | 0.15 | 0.28 | 0.22 | 0.22 | 0.3 |
| 4 | 120043874 | rs2168987 | 0.031 | 0.362 | 0.242 | -0.494 | -0.03 | 0.009 | 0.009 | 0.383 | 0.38 | 0.27 | 0.46 | 0.62 | 0.48 | 0.43 |
| 4 | 120048884 | rs9307468 | -0.21 | 0.279 | 0.344 | -0.496 | 0.002 | -0.009 | -0.009 | 0.054 | 0.45 | 0.53 | 0.5 | 0.62 | 0.51 | 0.43 |
| 4 | 120053224 | rs4833607 | 0.99 | -0.299 | -0.229 | -0.712 | 0.022 | 0.014 | 0.014 | 0 | 0.15 | 0.07 | 0.05 | 0 | 0.06 | 0.12 |
| 4 | 120055799 | rs4834752 | 0.426 | -0.02 | -0.179 | -0.62 | -0.025 | 0.001 | 0.001 | 0 | 0.38 | 0.22 | 0.44 | 0.44 | 0.43 | 0.48 |
| 4 | 120056815 | rs1459055 | 0.288 | 0.165 | -0.153 | -0.336 | -0.022 | -0.02 | -0.02 | 0 | 0.23 | 0.18 | 0.32 | 0.39 | 0.43 | 0.22 |
| 4 | 120063715 | rs7694500 | 1.369 | 0.089 | -0.104 | -0.501 | -0.001 | 0 | 0 | 0.474 | 0.22 | 0.23 | 0.35 | 0.23 | 0.26 | 0.38 |
| 4 | 120067703 | rs6844238 | 0.978 | -0.489 | 0.027 | -0.715 | -0.003 | -0.004 | -0.004 | 0 | 0.35 | 0.33 | 0.25 | 0.21 | 0.27 | 0.31 |
| 4 | 120072585 | rs4452412 | -1.51 | -0.523 | -0.286 | -0.684 | 0.007 | -0.005 | -0.005 | 0.245 | 0.83 | 0.86 | 0.94 | 0.83 | 0.97 | 0.93 |
| 4 | 120077220 | rs7669431 | -0.19 | -0.189 | -0.395 | -0.595 | 0.073 | 0 | 0 | 0 | 0.46 | 0.25 | 0.67 | 0.63 | 0.7 | 0.62 |
| 4 | 120081606 | rs11098489 | 0.089 | 0.16 | -0.004 | -0.191 | -0.067 | -0.056 | -0.056 | 0.017 | 0.24 | 0.11 | 0.32 | 0.4 | 0.43 | 0.24 |
| 4 | 120085448 | rs7687613 | 1.054 | -0.089 | 0.29 | -0.243 | -0.026 | -0.024 | -0.024 | 0.181 | 0.07 | 0.2 | 0.04 | 0.04 | 0.02 | 0.07 |
| 4 | 120108603 | rs9995277 | 0.421 | 0.369 | 0.124 | -0.147 | -0.012 | -0.02 | -0.02 | 0 | 0.18 | 0.14 | 0.28 | 0.53 | 0.31 | 0.22 |
| 4 | 120112049 | rs17049978 | 1.336 | 0.059 | 0.272 | -0.445 | -0.01 | -0.022 | -0.022 | 0.035 | 0.27 | 0.31 | 0.21 | 0.13 | 0.21 | 0.24 |
| 4 | 120115228 | rs10022508 | 0.612 | -0.09 | 0.151 | -0.313 | -0.012 | -0.018 | -0.018 | 0 | 0.18 | 0.13 | 0.28 | 0.53 | 0.31 | 0.22 |
| 4 | 120127117 | rs11737086 | 0.542 | -0.052 | 0.147 | -0.446 | -0.004 | 0.019 | 0.019 | 0 | 0.36 | 0.31 | 0.25 | 0.17 | 0.34 | 0.33 |
| 4 | 120132690 | rs4834762 | 1.771 | -0.237 | -0.019 | -0.589 | -0.066 | 0.007 | 0.007 | 0.008 | 0.23 | 0.05 | 0.35 | 0.24 | 0.29 | 0.31 |
| 4 | 120133549 | rs12644283 | 1.726 | -0.24 | -0.006 | -0.59 | -0.062 | 0.011 | 0.011 | 0 | 0.23 | 0.07 | 0.35 | 0.23 | 0.29 | 0.31 |
| 4 | 120133892 | rs6833072 | 0.427 | -0.224 | 0.046 | -0.581 | -0.001 | 0.001 | 0.001 | 0 | 0.38 | 0.39 | 0.25 | 0.17 | 0.34 | 0.33 |
| 4 | 120147460 | rs4833611 | 0.98 | -0.316 | 0.206 | -0.625 | 0.002 | -0.014 | -0.014 | 0.307 | 0.3 | 0.42 | 0.43 | 0.29 | 0.33 | 0.4 |
| 4 | 120150467 | rs951570 | -0.239 | -0.497 | 0.115 | -0.374 | 0.001 | -0.029 | -0.029 | 5.655 | 0.69 | 0.81 | 0.68 | 0.46 | 0.67 | 0.73 |
| 4 | 120151452 | rs10031033 | -0.041 | -0.482 | 0.22 | -0.603 | 0.001 | -0.004 | -0.004 | 0 | 0.67 | 0.75 | 0.67 | 0.59 | 0.67 | 0.73 |
| 4 | 120151624 | rs6853317 | -0.778 | -0.676 | 0.482 | -0.657 | -0.063 | -0.048 | -0.048 | 0.08 | 0.82 | 0.67 | 0.94 | 0.94 | 0.97 | 0.94 |
| 4 | 120156040 | rs13113112 | 0.63 | -0.194 | 0.672 | -0.404 | 0.002 | 0.012 | 0.012 | 0 | 0.17 | 0.18 | 0.29 | 0.39 | 0.3 | 0.18 |
| 4 | 120159334 | rs11944880 | 0.452 | -0.199 | 0.703 | -0.407 | 0.022 | 0.062 | 0.062 | 0 | 0.18 | 0.26 | 0.3 | 0.39 | 0.3 | 0.18 |

| | | | | | | | | | | | | | | | | |
|---|---|---|---|---|---|---|---|---|---|---|---|---|---|---|---|---|
| 4 | 120167111 | rs7356185 | -1.443 | -0.278 | 0.589 | -0.603 | 0.003 | -0.005 | -0.005 | 0 | 0.7 | 0.75 | 0.79 | 0.76 | 0.77 | 0.71 |
| 4 | 120168588 | rs13139045 | 0.074 | -0.328 | 0.586 | -0.624 | 0.007 | 0.043 | 0.043 | 0 | 0.22 | 0.25 | 0.33 | 0.45 | 0.33 | 0.26 |
| 4 | 120177708 | rs7668423 | 0.272 | -0.335 | 0.598 | -0.684 | 0.004 | 0.004 | 0.004 | 2.967 | 0.41 | 0.43 | 0.43 | 0.33 | 0.33 | 0.44 |
| 4 | 120180790 | rs6823331 | -0.776 | -0.298 | 0.499 | -0.678 | -0.008 | 0.01 | 0.01 | 0.029 | 0.6 | 0.68 | 0.72 | 0.72 | 0.63 | 0.62 |
| 4 | 120187119 | rs11098498 | -0.531 | -0.107 | 0.343 | -0.484 | 0.036 | 0.029 | 0.029 | 0.361 | 0.3 | 0.38 | 0.37 | 0.43 | 0.44 | 0.27 |
| 4 | 120187611 | rs11098499 | 0.672 | -0.09 | 0.336 | -0.474 | 0.01 | 0.033 | 0.033 | 0 | 0.17 | 0.21 | 0.29 | 0.39 | 0.3 | 0.18 |
| 4 | 120214030 | rs3749591 | 0.726 | -0.091 | 0.306 | -0.47 | -0.015 | -0.024 | -0.024 | 0 | 0.17 | 0.12 | 0.28 | 0.39 | 0.3 | 0.18 |
| 4 | 120215314 | rs1052633 | -0.414 | -0.106 | 0.221 | -0.483 | 0.029 | 0.023 | 0.023 | 0.284 | 0.3 | 0.35 | 0.37 | 0.43 | 0.44 | 0.27 |
| 4 | 120222883 | rs13127517 | 1.415 | -0.032 | 0.17 | -0.402 | -0.05 | 0.045 | 0.045 | 0 | 0.1 | 0.02 | 0.06 | 0.04 | 0.14 | 0.09 |
| 4 | 120230446 | rs4834769 | 1.642 | -0.135 | 0.048 | -0.456 | -0.053 | -0.033 | -0.033 | 0 | 0.21 | 0.04 | 0.32 | 0.26 | 0.26 | 0.33 |
| 4 | 120231150 | rs1397607 | -0.441 | -0.205 | 0.308 | -0.427 | 0.012 | 0.01 | 0.01 | 0 | 0.5 | 0.56 | 0.69 | 0.69 | 0.68 | 0.59 |
| 4 | 120235125 | rs10518301 | 0.305 | -0.133 | 0.29 | -0.414 | -0.005 | 0.01 | 0.01 | 0 | 0.36 | 0.34 | 0.41 | 0.3 | 0.39 | 0.42 |
| 4 | 120237564 | rs878374 | -0.126 | -0.386 | 0.349 | -0.504 | -0.001 | 0.017 | 0.017 | 0.808 | 0.5 | 0.42 | 0.65 | 0.65 | 0.57 | 0.54 |
| 4 | 120238780 | rs878372 | -0.048 | -0.397 | 0.355 | -0.509 | -0.006 | 0.014 | 0.014 | 0 | 0.3 | 0.4 | 0.33 | 0.39 | 0.31 | 0.2 |
| 4 | 120239878 | rs11724758 | -0.792 | -0.397 | 0.424 | -0.537 | 0.001 | -0.001 | -0.001 | 0 | 0.41 | 0.56 | 0.41 | 0.44 | 0.45 | 0.28 |
| 4 | 120239944 | rs9999724 | -0.058 | -0.409 | 0.457 | -0.543 | -0.003 | 0.011 | 0.011 | 0 | 0.29 | 0.35 | 0.32 | 0.39 | 0.31 | 0.2 |
| 4 | 120240238 | rs1511025 | -0.026 | -0.395 | 0.427 | -0.536 | -0.006 | 0.014 | 0.014 | 0 | 0.3 | 0.4 | 0.33 | 0.39 | 0.31 | 0.2 |
| 4 | 120241849 | rs4834770 | -0.363 | -0.414 | 0.671 | -0.545 | -0.001 | -0.001 | -0.001 | 0 | 0.53 | 0.55 | 0.65 | 0.65 | 0.57 | 0.53 |
| 4 | 120241902 | rs1799883 | -1.697 | -0.512 | 0.602 | -0.534 | -0.004 | -0.005 | -0.005 | 0 | 0.71 | 0.78 | 0.77 | 0.75 | 0.73 | 0.69 |
| 4 | 120243352 | rs1397614 | 1.559 | -0.504 | 0.565 | -0.527 | 0.024 | -0.018 | -0.018 | 0.04 | 0.06 | 0 | 0.04 | 0.06 | 0.03 | 0.07 |
| 4 | 120244085 | rs10034661 | -1.717 | -0.444 | 0.609 | -0.551 | -0.001 | -0.001 | -0.001 | 3.834 | 0.7 | 0.72 | 0.77 | 0.75 | 0.73 | 0.69 |
| 4 | 120264996 | rs9995716 | -0.537 | -0.432 | 0.484 | -0.516 | -0.004 | 0.002 | 0.002 | 0 | 0.36 | 0.4 | 0.37 | 0.45 | 0.34 | 0.27 |
| 4 | 120277395 | rs4834773 | 1.503 | -0.33 | 0.22 | -0.477 | -0.009 | 0.003 | 0.003 | 0.171 | 0.23 | 0.13 | 0.32 | 0.26 | 0.26 | 0.33 |
| 4 | 120289042 | rs11722431 | 1.609 | -0.196 | 0.17 | -0.464 | -0.049 | 0.044 | 0.044 | 0 | 0.1 | 0.01 | 0.06 | 0.04 | 0.14 | 0.08 |
| 4 | 120309500 | rs4833618 | 0.206 | -0.405 | 0.377 | -0.524 | 0.003 | 0.045 | 0.045 | 0 | 0.52 | 0.31 | 0.35 | 0.29 | 0.44 | 0.42 |
| 4 | 120326323 | rs6846442 | 0.463 | -0.376 | 1.146 | -0.192 | 0.048 | 0.119 | 0.119 | 0.758 | 0.14 | 0.25 | 0.29 | 0.39 | 0.28 | 0.17 |
| 4 | 120371729 | rs7659551 | 0.463 | -0.376 | 1.146 | -0.192 | 0.048 | 0.119 | 0.119 | 0.758 | 0.14 | 0.25 | 0.29 | 0.39 | 0.28 | 0.17 |
| 4 | 120381341 | rs2892848 | 0.316 | -0.382 | 1.147 | -0.195 | 0.044 | 0.112 | 0.112 | 0.717 | 0.15 | 0.26 | 0.29 | 0.39 | 0.28 | 0.17 |
| 4 | 120388406 | rs7654587 | 0.316 | -0.382 | 1.147 | -0.195 | 0.044 | 0.112 | 0.112 | 0.717 | 0.15 | 0.26 | 0.29 | 0.39 | 0.28 | 0.17 |
| 4 | 120390032 | rs11098524 | 0.316 | -0.382 | 1.147 | -0.195 | 0.044 | 0.112 | 0.112 | 0.717 | 0.15 | 0.26 | 0.29 | 0.39 | 0.28 | 0.17 |
| 4 | 120391632 | rs10032158 | 0.316 | -0.382 | 1.147 | -0.195 | 0.044 | 0.112 | 0.112 | 0.717 | 0.15 | 0.26 | 0.29 | 0.39 | 0.29 | 0.17 |
| 4 | 120397829 | rs10034623 | -0.023 | -0.403 | 1.208 | -0.205 | 0.04 | 0.114 | 0.114 | 0.808 | 0.17 | 0.32 | 0.29 | 0.39 | 0.29 | 0.17 |
| 4 | 120399505 | rs988146 | -0.162 | -0.492 | 1.139 | -0.285 | -0.013 | 0.107 | 0.107 | 0.711 | 0.24 | 0.08 | 0.12 | 0.06 | 0.18 | 0.11 |
| 4 | 120401835 | rs10518329 | -1.183 | -0.539 | 1.251 | -0.273 | -0.003 | 0.048 | 0.048 | 0.049 | 0.25 | 0.58 | 0.35 | 0.44 | 0.32 | 0.24 |
| 4 | 120408931 | rs10006525 | 0.385 | -0.525 | 1.262 | -0.265 | 0.045 | 0.115 | 0.115 | 0.717 | 0.15 | 0.26 | 0.29 | 0.39 | 0.28 | 0.17 |
| 4 | 120416445 | rs13124532 | 1.295 | -0.544 | 1.226 | -0.27 | -0.058 | 0.024 | 0.024 | 0 | 0.1 | 0 | 0.07 | 0.07 | 0.15 | 0.09 |
| 4 | 120416788 | rs11729521 | 0.419 | -0.526 | 1.27 | -0.258 | 0.028 | 0.078 | 0.078 | 0.845 | 0.15 | 0.23 | 0.29 | 0.39 | 0.28 | 0.17 |
| 4 | 120416872 | rs11734241 | 0.419 | -0.526 | 1.27 | -0.258 | 0.028 | 0.078 | 0.078 | 0.845 | 0.15 | 0.23 | 0.28 | 0.39 | 0.28 | 0.17 |
| 4 | 120423729 | rs3733524 | 0.629 | -0.522 | 1.33 | -0.249 | -0.006 | 0.003 | 0.003 | 0.883 | 0.14 | 0.11 | 0.28 | 0.39 | 0.28 | 0.17 |
| 4 | 120424087 | rs6843229 | -2.265 | -0.514 | 1.297 | -0.25 | 0.007 | 0.003 | 0.003 | 0 | 0.49 | 0.67 | 0.47 | 0.51 | 0.49 | 0.36 |
| 4 | 120424580 | rs13107334 | -0.334 | -0.469 | 0.634 | -0.197 | -0.01 | 0.101 | 0.101 | 0.54 | 0.25 | 0.08 | 0.13 | 0.1 | 0.18 | 0.11 |
| 4 | 120425246 | rs4834779 | -0.439 | -0.426 | 0.805 | -0.06 | 0.001 | 0.041 | 0.041 | 0 | 0.15 | 0.19 | 0.21 | 0.38 | 0.2 | 0.17 |
| 4 | 120428404 | rs13121569 | -0.417 | -0.484 | 0.603 | -0.015 | 0.004 | 0.045 | 0.045 | 0.521 | 0.25 | 0.14 | 0.13 | 0.07 | 0.18 | 0.12 |
| 4 | 120445720 | rs3822192 | 0.582 | -0.419 | 0.524 | 0.03 | 0.003 | 0.021 | 0.021 | 0.503 | 0.14 | 0.14 | 0.28 | 0.39 | 0.29 | 0.17 |
| 4 | 120449810 | rs1155577 | 1.863 | -0.48 | 0.607 | -0.003 | 0.004 | 0.002 | 0.002 | 0 | 0.5 | 0.38 | 0.54 | 0.49 | 0.5 | 0.64 |
| 4 | 120450159 | rs1155576 | 0.048 | -0.448 | 0.626 | -0.01 | 0.031 | 0.091 | 0.091 | 0.467 | 0.17 | 0.31 | 0.29 | 0.39 | 0.29 | 0.17 |
| 4 | 120491714 | rs17358524 | 1.091 | -0.473 | 0.473 | -0.084 | -0.059 | 0.026 | 0.026 | 0 | 0.11 | 0.01 | 0.08 | 0.06 | 0.15 | 0.09 |
| 4 | 120502461 | rs12646525 | 1.09 | -0.488 | 0.532 | -0.086 | 0.001 | -0.013 | -0.013 | 0.532 | 0.66 | 0.65 | 0.83 | 0.89 | 0.79 | 0.81 |
| 4 | 120508298 | rs10012485 | 0.954 | -0.483 | 0.508 | -0.083 | 0.014 | 0.02 | 0.02 | 0.812 | 0.67 | 0.72 | 0.83 | 0.89 | 0.79 | 0.81 |
| 4 | 120522038 | rs2389870 | -0.585 | -0.478 | 0.479 | -0.087 | 0.009 | 0.04 | 0.04 | 0.428 | 0.27 | 0.18 | 0.12 | 0.07 | 0.18 | 0.12 |
| 4 | 120522172 | rs10518335 | -1.3 | -0.495 | 0.479 | -0.095 | 0.03 | 0.063 | 0.063 | 0.645 | 0.33 | 0.22 | 0.17 | 0.11 | 0.21 | 0.19 |
| 4 | 120522934 | rs10518336 | -1.27 | -0.402 | 0.387 | -0.034 | -0.012 | -0.004 | -0.004 | 0 | 0.4 | 0.33 | 0.42 | 0.46 | 0.47 | 0.29 |
| 4 | 120528064 | rs17051276 | 1.073 | -0.384 | 0.354 | 0.024 | 0.024 | -0.013 | -0.013 | 0 | 0.06 | 0 | 0.04 | 0.04 | 0.03 | 0.07 |
| 4 | 120534734 | rs6847164 | 0.34 | -0.375 | 0.427 | 0.004 | 0 | -0.017 | -0.017 | 0 | 0.8 | 0.77 | 0.86 | 0.9 | 0.81 | 0.84 |
| 4 | 120538823 | rs6534146 | 0.494 | -0.383 | 0.453 | 0 | -0.002 | -0.033 | -0.033 | 0 | 0.79 | 0.71 | 0.86 | 0.9 | 0.81 | 0.84 |
| 4 | 120546136 | rs6832740 | -0.805 | -0.734 | 0.797 | 0.445 | 0.001 | -0.005 | -0.005 | 0 | 0.31 | 0.3 | 0.33 | 0.4 | 0.32 | 0.2 |
| 4 | 120551085 | rs3806808 | -0.997 | -0.73 | 0.771 | 0.424 | -0.006 | 0.01 | 0.01 | 0 | 0.33 | 0.41 | 0.34 | 0.41 | 0.32 | 0.2 |
| 4 | 120558608 | rs10009566 | -0.984 | -0.727 | 0.773 | 0.425 | -0.006 | 0.01 | 0.01 | 0 | 0.32 | 0.41 | 0.34 | 0.41 | 0.32 | 0.2 |
| 4 | 120569563 | rs6844418 | -0.077 | -0.955 | 0.556 | 0.1 | -0.08 | -0.114 | -0.114 | 0 | 0.82 | 0.63 | 0.93 | 1 | 0.97 | 0.97 |
| 4 | 120576177 | rs2389891 | 0.751 | -0.955 | 0.347 | 0.127 | -0.125 | -0.117 | -0.117 | 2.324 | 0.75 | 0.5 | 0.88 | 0.95 | 0.93 | 0.89 |
| 4 | 120585497 | rs4833624 | -0.829 | -0.082 | 0.08 | 0.734 | -0.003 | 0.02 | 0.02 | 0 | 0.28 | 0.34 | 0.44 | 0.54 | 0.31 | 0.19 |
| 4 | 120587803 | rs9994651 | -1.112 | -0.299 | 0.048 | 0.694 | -0.005 | 0.008 | 0.008 | 0.314 | 0.32 | 0.35 | 0.46 | 0.56 | 0.31 | 0.24 |
| 4 | 120589750 | rs1986695 | 0.089 | -0.132 | 0.031 | 0.951 | -0.012 | -0.018 | -0.018 | 0 | 0.16 | 0.14 | 0.4 | 0.56 | 0.29 | 0.18 |
| 4 | 120595698 | rs10003106 | -0.391 | -0.247 | 0.111 | 0.868 | -0.028 | -0.014 | -0.014 | 0 | 0.16 | 0.2 | 0.08 | 0.06 | 0.06 | 0.08 |
| 4 | 120601731 | rs13122890 | 1.327 | -0.357 | 0.646 | 0.855 | -0.024 | -0.002 | -0.002 | 0.212 | 0.65 | 0.5 | 0.51 | 0.32 | 0.67 | 0.71 |
| 4 | 120606495 | rs2253404 | 0.008 | -0.348 | 0.658 | 0.843 | -0.021 | 0.029 | 0.029 | 0 | 0.94 | 0.86 | 0.97 | 0.89 | 0.98 | 0.92 |
| 4 | 120609868 | rs2715023 | 0.565 | -0.365 | 0.682 | 0.835 | -0.065 | -0.018 | -0.018 | 0.052 | 0.82 | 0.66 | 0.9 | 0.87 | 0.94 | 0.89 |
| 4 | 120612853 | rs7683474 | -1.46 | -0.342 | 0.635 | 0.856 | -0.004 | 0.001 | 0.001 | 0.004 | 0.33 | 0.38 | 0.48 | 0.62 | 0.33 | 0.27 |
| 4 | 120613018 | rs17295253 | -0.445 | -0.376 | 0.196 | 0.097 | 0.002 | -0.03 | -0.03 | 0 | 0.32 | 0.31 | 0.2 | 0.02 | 0.25 | 0.14 |

| | | | | | | | | | | | | | | | | |
|---|---|---|---|---|---|---|---|---|---|---|---|---|---|---|---|---|
| 4 | 120618359 | rs12510467 | -0.62 | -0.344 | 0.213 | 0.156 | -0.004 | 0.029 | 0.029 | 0 | 0.29 | 0.37 | 0.46 | 0.56 | 0.32 | 0.21 |
| 4 | 120619421 | rs4094 | -1.117 | -0.151 | 0.194 | 0.1 | -0.002 | 0.001 | 0.001 | 0 | 0.34 | 0.38 | 0.48 | 0.61 | 0.36 | 0.27 |
| 4 | 120627421 | rs1009077 | -0.889 | -0.14 | 0.147 | 0.21 | -0.003 | 0.02 | 0.02 | 0 | 0.79 | 0.88 | 0.87 | 0.91 | 0.79 | 0.88 |
| 4 | 120628627 | rs2714961 | 1.531 | -0.18 | 0.317 | 0.158 | -0.004 | 0.014 | 0.014 | 0 | 0.45 | 0.36 | 0.37 | 0.34 | 0.44 | 0.61 |
| 4 | 120643124 | rs11726524 | 0.168 | -0.121 | 0.712 | 0.662 | -0.041 | -0.047 | -0.047 | 0 | 0.13 | 0.31 | 0.07 | 0.02 | 0.04 | 0.03 |
| 4 | 120663649 | rs749526 | -0.834 | -0.057 | 0.651 | 0.751 | -0.001 | 0.013 | 0.013 | 0 | 0.27 | 0.29 | 0.44 | 0.51 | 0.32 | 0.21 |
| 4 | 120670520 | rs13112498 | 0.098 | 0.046 | 0.295 | 0.801 | -0.005 | 0.002 | 0.002 | 0 | 0.17 | 0.14 | 0.39 | 0.49 | 0.28 | 0.18 |
| 4 | 120680950 | rs4447843 | -0.523 | 0.019 | 0.439 | 0.765 | -0.003 | 0.03 | 0.03 | 0 | 0.28 | 0.36 | 0.47 | 0.49 | 0.32 | 0.21 |
| 4 | 120685312 | rs11736455 | -0.513 | -0.008 | 0.4 | 0.767 | -0.003 | 0.025 | 0.025 | 0 | 0.28 | 0.35 | 0.46 | 0.49 | 0.32 | 0.21 |
| 4 | 120690702 | rs17005814 | 0.629 | -0.004 | -0.053 | 0.828 | -0.001 | 0.005 | 0.005 | 0 | 0.05 | 0.04 | 0.03 | 0.15 | 0.04 | 0.08 |
| 4 | 120695322 | rs2389957 | 0.591 | 0.178 | -0.046 | 0.616 | -0.009 | 0.003 | 0.003 | 0 | 0.63 | 0.39 | 0.5 | 0.36 | 0.65 | 0.69 |
| 4 | 120704766 | rs6534152 | -0.114 | 1.173 | -0.266 | 0.669 | 0.003 | 0.001 | 0.001 | 0 | 0.17 | 0.21 | 0.2 | 0.2 | 0.29 | 0.15 |
| 4 | 120709561 | rs11729954 | 0.59 | 0.641 | -0.479 | 0.175 | 0.016 | 0.031 | 0.031 | 0 | 0.17 | 0.1 | 0.08 | 0.1 | 0.14 | 0.08 |
| 4 | 120714886 | rs17006113 | 1.309 | 0.09 | -0.427 | -0.181 | 0.004 | -0.008 | -0.008 | 0 | 0.26 | 0.3 | 0.22 | 0.23 | 0.26 | 0.35 |
| 4 | 120716967 | rs7699214 | 0.127 | 0.057 | -0.553 | -0.104 | -0.007 | 0.005 | 0.005 | 0 | 0.44 | 0.53 | 0.42 | 0.43 | 0.55 | 0.49 |
| 4 | 120721063 | rs2389954 | 0.602 | -0.053 | -0.66 | -0.15 | 0.019 | -0.067 | -0.067 | 0.831 | 0.33 | 0.11 | 0.47 | 0.43 | 0.29 | 0.37 |
| 4 | 120731697 | rs2389923 | 0.411 | 0.005 | -0.686 | -0.239 | -0.065 | -0.044 | -0.044 | 0 | 0.3 | 0.1 | 0.24 | 0.3 | 0.4 | 0.23 |
| 4 | 120736449 | rs10004197 | 0.569 | -0.028 | -0.285 | -0.248 | -0.001 | -0.013 | -0.013 | 2.927 | 0.34 | 0.59 | 0.27 | 0.27 | 0.3 | 0.4 |
| 4 | 120756608 | rs17363983 | 0.638 | 0.267 | -0.319 | -0.069 | 0.002 | -0.052 | -0.052 | 0 | 0.32 | 0.09 | 0.49 | 0.39 | 0.3 | 0.34 |
| 4 | 120757419 | rs1503538 | 0.992 | 0.048 | -0.41 | -0.226 | -0.004 | -0.047 | -0.047 | 0 | 0.14 | 0.16 | 0.06 | 0 | 0.13 | 0.06 |
| 4 | 120758258 | rs7690791 | 0.091 | 0.053 | -0.267 | -0.186 | 0.018 | 0.014 | 0.014 | 0 | 0.52 | 0.64 | 0.44 | 0.57 | 0.56 | 0.57 |
| 4 | 120760872 | rs7688177 | 1.358 | -0.038 | -0.641 | -0.174 | -0.009 | -0.006 | -0.006 | 1.056 | 0.13 | 0.08 | 0.4 | 0.37 | 0.21 | 0.14 |
| 4 | 120773133 | rs1124166 | -0.229 | -0.012 | -0.865 | -0.446 | -0.018 | 0.077 | 0.077 | 0 | 0.56 | 0.28 | 0.67 | 0.72 | 0.55 | 0.59 |
| 4 | 120775160 | rs17006835 | 0.884 | -0.095 | -0.424 | -0.511 | 0.013 | -0.016 | -0.016 | 0.974 | 0.06 | 0.26 | 0.04 | 0 | 0.03 | 0.01 |
| 4 | 120775448 | rs1123767 | -0.663 | -0.067 | -0.545 | -0.495 | 0.018 | -0.037 | -0.037 | 0 | 0.61 | 0.51 | 0.71 | 0.72 | 0.57 | 0.59 |
| 4 | 120776279 | rs10452195 | -2.105 | -0.193 | -0.785 | -0.549 | 0.005 | 0.009 | 0.009 | 0 | 0.88 | 0.9 | 0.94 | 1 | 0.89 | 0.94 |
| 4 | 120776752 | rs722766 | 1.365 | -0.12 | -0.885 | -0.425 | -0.008 | -0.052 | -0.052 | 0 | 0.2 | 0.09 | 0.17 | 0.3 | 0.22 | 0.2 |
| 4 | 120784320 | rs17364898 | 1.295 | -0.111 | -0.846 | -0.586 | 0.032 | 0.088 | 0.088 | 3.116 | 0.2 | 0.01 | 0.09 | 0.02 | 0.1 | 0.2 |
| 4 | 120788956 | rs17006982 | 1.39 | 0.482 | -0.529 | -0.482 | 0.005 | -0.016 | -0.016 | 0 | 0.23 | 0.29 | 0.18 | 0.22 | 0.22 | 0.19 |
| 4 | 120790884 | rs1818167 | 0.37 | 0.453 | -0.722 | -0.279 | 0.01 | -0.044 | -0.044 | 0 | 0.33 | 0.11 | 0.5 | 0.4 | 0.3 | 0.37 |
| 4 | 120791979 | rs1032818 | 0.962 | 0.05 | -0.295 | -0.291 | -0.013 | -0.022 | -0.022 | 0 | 0.33 | 0.27 | 0.57 | 0.56 | 0.42 | 0.35 |
| 4 | 120797098 | rs4834821 | 0.794 | 0.258 | 0.292 | 0.138 | -0.006 | 0.005 | 0.005 | 0.786 | 0.49 | 0.32 | 0.62 | 0.59 | 0.52 | 0.47 |
| 4 | 120812306 | rs1503540 | -0.682 | 0.184 | 0.291 | 0.328 | -0.005 | -0.03 | -0.03 | 0 | 0.78 | 0.89 | 0.81 | 0.73 | 0.76 | 0.8 |
| 4 | 120821994 | rs13132107 | 0.726 | -0.24 | 0.645 | 0.165 | -0.012 | -0.001 | -0.001 | 0.01 | 0.5 | 0.24 | 0.65 | 0.54 | 0.53 | 0.43 |
| 4 | 120842452 | rs17300152 | 0.986 | -0.326 | 0.523 | 0.069 | 0.04 | 0.058 | 0.058 | 0.984 | 0.15 | 0.05 | 0.05 | 0.02 | 0.06 | 0.14 |
| 4 | 120843895 | rs11098551 | -0.711 | -0.298 | 0.565 | 0.104 | -0.052 | -0.052 | -0.052 | 0 | 0.74 | 0.9 | 0.79 | 0.69 | 0.66 | 0.8 |
| 4 | 120846553 | rs10018121 | 0.708 | -0.262 | 0.472 | 0.125 | 0.003 | 0.007 | 0.007 | 0 | 0.53 | 0.34 | 0.68 | 0.6 | 0.58 | 0.61 |
| 4 | 120847421 | rs12508689 | -0.153 | -0.039 | -0.073 | 0.232 | 0.019 | 0.02 | 0.02 | 0.005 | 0.7 | 0.59 | 0.76 | 0.64 | 0.64 | 0.77 |
| 4 | 120880067 | rs2389925 | -0.532 | -0.343 | -0.3 | 0.024 | -0.026 | -0.026 | -0.026 | 0 | 0.24 | 0.34 | 0.1 | 0.14 | 0.14 | 0.38 |
| 4 | 120898509 | rs1503535 | -0.18 | 0.006 | -0.47 | 0.254 | 0 | 0 | 0 | 0 | 0.51 | 0.5 | 0.52 | 0.61 | 0.35 | 0.47 |
| 4 | 120900282 | rs13134800 | -0.353 | 0.598 | -0.853 | 0.155 | -0.028 | 0.016 | 0.016 | 0 | 0.72 | 0.82 | 0.82 | 0.84 | 0.69 | 0.8 |
| 4 | 120905242 | rs17366987 | 0.634 | -0.03 | -1.204 | -0.074 | 0.006 | 0.035 | 0.035 | 0 | 0.13 | 0.01 | 0.07 | 0.06 | 0.09 | 0.08 |
| 4 | 120909594 | rs7678743 | -1.191 | -0.509 | -0.606 | 0.101 | -0.012 | 0.014 | 0.014 | 0.161 | 0.76 | 0.74 | 0.72 | 0.69 | 0.8 | 0.71 |
| 4 | 120910200 | rs1604711 | 0.689 | -0.008 | -0.097 | 0.17 | -0.012 | -0.002 | -0.002 | 0 | 0.28 | 0.26 | 0.39 | 0.35 | 0.34 | 0.26 |
| 4 | 120912054 | rs2389926 | -0.116 | -0.289 | 0.258 | 0.217 | -0.009 | 0.001 | 0.001 | 0.324 | 0.57 | 0.48 | 0.61 | 0.56 | 0.67 | 0.46 |
| 4 | 120930676 | rs17008785 | -0.278 | -0.126 | 0.29 | 0.3 | 0.04 | 0.122 | 0.122 | 0 | 0.05 | 0.24 | 0.17 | 0.19 | 0.05 | 0.19 |
| 4 | 120939273 | rs1847333 | 0.375 | 0.115 | -0.112 | 0.136 | -0.018 | 0.035 | 0.035 | 0 | 0.32 | 0.18 | 0.34 | 0.17 | 0.31 | 0.23 |
| 4 | 120939685 | rs1503534 | -0.462 | 0.724 | -0.055 | 0.125 | 0.019 | -0.004 | -0.004 | 0 | 0.39 | 0.54 | 0.57 | 0.4 | 0.45 | 0.43 |
| 4 | 120942372 | rs6856643 | 0.319 | 0.75 | -1.013 | -0.031 | -0.027 | 0.042 | 0.042 | 0 | 0.31 | 0.16 | 0.35 | 0.19 | 0.31 | 0.23 |
| 4 | 120946927 | rs1967119 | 0.404 | 0.582 | -0.933 | -0.006 | -0.019 | 0.014 | 0.014 | 0 | 0.27 | 0.12 | 0.23 | 0.24 | 0.31 | 0.25 |
| 4 | 120952953 | rs6823111 | 0.209 | 0.782 | -0.4 | -0.24 | -0.017 | 0.007 | 0.007 | 0 | 0.27 | 0.24 | 0.37 | 0.18 | 0.36 | 0.21 |
| 4 | 120953490 | rs10006314 | -0.571 | 0.38 | -0.458 | -0.562 | 0.009 | 0.08 | 0.08 | 0 | 0.44 | 0.23 | 0.38 | 0.21 | 0.36 | 0.29 |
| 4 | 120962385 | rs1845344 | -0.228 | 0.507 | -0.066 | -0.535 | -0.009 | 0.051 | 0.051 | 0 | 0.34 | 0.28 | 0.22 | 0.13 | 0.33 | 0.21 |
| 4 | 120968070 | rs6828161 | 0.666 | 0.465 | -0.195 | -0.586 | -0.073 | 0.065 | 0.065 | 0 | 0.24 | 0.05 | 0.16 | 0.1 | 0.26 | 0.17 |
| 4 | 120969089 | rs7677531 | -0.981 | 0.238 | 0.074 | -0.649 | 0.007 | -0.019 | -0.019 | 0.503 | 0.43 | 0.52 | 0.41 | 0.31 | 0.44 | 0.29 |
| 4 | 120971860 | rs6827271 | -0.46 | 0.356 | 0.32 | -0.927 | 0.019 | -0.045 | -0.045 | 0 | 0.36 | 0.51 | 0.3 | 0.18 | 0.41 | 0.35 |
| 4 | 120972650 | rs17009453 | -1.141 | 0.451 | 0.221 | -0.886 | 0.066 | -0.032 | -0.032 | 1.661 | 0.11 | 0.35 | 0.11 | 0.06 | 0.14 | 0.19 |
| 4 | 120973238 | rs13126776 | -1.164 | 0.188 | 0.028 | -0.962 | -0.007 | 0.202 | 0.202 | 0.078 | 0.48 | 0.85 | 0.36 | 0.21 | 0.48 | 0.4 |
| 4 | 120983259 | rs2908989 | -0.753 | 0.507 | -0.079 | -0.948 | -0.038 | 0.042 | 0.042 | 0.598 | 0.77 | 0.87 | 0.81 | 0.87 | 0.72 | 0.7 |
| 4 | 120984369 | rs2908988 | -0.807 | 0.507 | -0.085 | -0.948 | -0.046 | 0.045 | 0.045 | 0.685 | 0.77 | 0.89 | 0.81 | 0.87 | 0.72 | 0.7 |
| 4 | 120986142 | rs2934379 | -0.187 | 0.4 | -0.041 | -0.781 | -0.013 | 0.021 | 0.021 | 0 | 0.72 | 0.79 | 0.79 | 0.82 | 0.7 | 0.66 |
| 4 | 120989853 | rs12711084 | -1.098 | 0.266 | 0.25 | -0.91 | -0.007 | 0.012 | 0.012 | 0 | 0.4 | 0.64 | 0.31 | 0.21 | 0.41 | 0.41 |
| 4 | 120990081 | rs13121806 | -0.081 | 0.296 | 0.609 | -0.926 | -0.001 | -0.001 | -0.001 | 0 | 0.29 | 0.29 | 0.21 | 0.18 | 0.29 | 0.34 |
| 4 | 120992642 | rs2892890 | -0.625 | 0.306 | 0.542 | -0.925 | 0.03 | -0.043 | -0.043 | 0 | 0.33 | 0.5 | 0.27 | 0.18 | 0.37 | 0.37 |
| 4 | 121005634 | rs6855113 | 0.436 | 0.323 | 0.521 | -0.921 | -0.012 | 0.017 | 0.017 | 0.865 | 0.24 | 0.22 | 0.2 | 0.11 | 0.27 | 0.3 |
| 4 | 121017399 | rs972311 | -1.428 | -0.283 | 0.221 | -1.551 | 0.005 | 0.095 | 0.095 | 0 | 0.58 | 0.76 | 0.43 | 0.2 | 0.53 | 0.49 |
| 4 | 121021737 | rs6831851 | 1.054 | -0.304 | 0.156 | -1.561 | -0.034 | 0.044 | 0.044 | 1.207 | 0.23 | 0.13 | 0.18 | 0.11 | 0.27 | 0.3 |
| 4 | 121033187 | rs1908169 | -0.972 | -0.496 | 0.302 | -1.565 | 0.007 | 0.029 | 0.029 | 0 | 0.5 | 0.54 | 0.39 | 0.16 | 0.48 | 0.45 |
| 4 | 121034480 | rs17010476 | 0.356 | -0.439 | -0.042 | -1.481 | -0.035 | -0.071 | -0.071 | 0 | 0.25 | 0.3 | 0.15 | 0.05 | 0.14 | 0.15 |

| | | | | | | | | | | | | | | | | |
|---|---|---|---|---|---|---|---|---|---|---|---|---|---|---|---|---|
| 4 | 121042212 | rs7695341 | -0.962 | -0.232 | -0.767 | -1.181 | 0.004 | 0.102 | 0.102 | 0 | 0.58 | 0.79 | 0.43 | 0.2 | 0.53 | 0.49 |
| 4 | 121056511 | rs9307484 | -0.84 | -0.043 | -1.352 | -0.913 | -0.044 | 0.013 | 0.013 | 0.422 | 0.7 | 0.94 | 0.58 | 0.77 | 0.64 | 0.69 |
| 4 | 121058095 | rs6842087 | -0.13 | -0.371 | -0.843 | -1.049 | -0.002 | 0.005 | 0.005 | 0 | 0.38 | 0.45 | 0.43 | 0.39 | 0.37 | 0.41 |
| 4 | 121063357 | rs13142346 | 1.056 | -0.428 | -1.063 | -1.071 | -0.024 | -0.047 | -0.047 | 0 | 0.26 | 0.1 | 0.37 | 0.31 | 0.3 | 0.33 |
| 4 | 121063683 | rs6854848 | -1.193 | -0.599 | -1.282 | -0.931 | -0.055 | 0.037 | 0.037 | 0 | 0.74 | 0.91 | 0.72 | 0.78 | 0.66 | 0.69 |
| 4 | 121078848 | rs10022765 | 1.3 | -0.617 | -0.671 | -0.972 | 0.015 | -0.006 | -0.006 | 0 | 0.33 | 0.24 | 0.23 | 0.39 | 0.24 | 0.28 |
| 4 | 121095379 | rs12644766 | 0.485 | -0.612 | -0.327 | -1 | -0.02 | -0.026 | -0.026 | 0 | 0.33 | 0.26 | 0.42 | 0.37 | 0.38 | 0.37 |
| 4 | 121100036 | rs13102180 | 0.6 | -0.746 | -0.282 | -0.977 | 0 | -0.026 | -0.026 | 0 | 0.33 | 0.26 | 0.4 | 0.37 | 0.32 | 0.39 |
| 4 | 121105615 | rs13116644 | 1.072 | -0.708 | -0.179 | -0.973 | -0.035 | -0.045 | -0.045 | 0 | 0.27 | 0.1 | 0.38 | 0.31 | 0.32 | 0.36 |
| 4 | 121107275 | rs1509002 | 0.926 | -0.631 | 0.04 | -1.422 | -0.019 | 0.01 | 0.01 | 0.082 | 0.31 | 0.39 | 0.25 | 0.39 | 0.23 | 0.25 |
| 4 | 121108446 | rs2662756 | -0.356 | -0.646 | -0.146 | -1.433 | 0.048 | 0.004 | 0.004 | 1.127 | 0.42 | 0.68 | 0.3 | 0.42 | 0.29 | 0.32 |
| 4 | 121110203 | rs1509001 | 1.142 | -0.621 | -0.212 | -1.417 | 0.022 | -0.025 | -0.025 | 0.925 | 0.3 | 0.16 | 0.22 | 0.38 | 0.23 | 0.25 |
| 4 | 121114622 | rs1876701 | 2.806 | -0.703 | -0.221 | -1.399 | 0.016 | -0.049 | -0.049 | 0.048 | 0.12 | 0.04 | 0.09 | 0.38 | 0.1 | 0.16 |
| 4 | 121117729 | rs6851191 | -1.066 | -0.709 | -0.117 | -1.825 | -0.03 | -0.039 | -0.039 | 0 | 0.12 | 0.42 | 0.06 | 0.04 | 0.05 | 0.07 |
| 4 | 121139889 | rs13127309 | 0.917 | -0.59 | -0.421 | -1.776 | 0 | 0 | 0 | 0 | 0.29 | 0.23 | 0.37 | 0.23 | 0.31 | 0.34 |
| 4 | 121140144 | rs2662692 | -0.121 | -0.673 | -0.371 | -1.837 | 0.003 | -0.002 | -0.002 | 2.895 | 0.44 | 0.43 | 0.31 | 0.47 | 0.31 | 0.37 |
| 4 | 121149877 | rs13134323 | 0.83 | -0.528 | -0.735 | -1.744 | -0.003 | -0.002 | -0.002 | 0 | 0.31 | 0.31 | 0.37 | 0.23 | 0.31 | 0.35 |
| 4 | 121164119 | rs9684604 | 1.277 | -0.16 | -1.165 | -1.836 | 0.001 | -0.009 | -0.009 | 0 | 0.14 | 0.1 | 0.1 | 0.01 | 0.11 | 0.02 |
| 4 | 121167953 | rs2662738 | -0.788 | -0.14 | -1.217 | -1.649 | -0.106 | -0.005 | -0.005 | 0 | 0.74 | 0.94 | 0.7 | 0.69 | 0.62 | 0.69 |
| 4 | 121168762 | rs2715990 | -1.803 | 0.049 | -1.095 | -1.636 | -0.056 | 0.052 | 0.052 | 0 | 0.87 | 0.98 | 0.89 | 0.96 | 0.75 | 0.82 |
| 4 | 121183954 | rs1355112 | -0.257 | -0.071 | -0.602 | -1.44 | 0.039 | 0.106 | 0.106 | 0.932 | 0.5 | 0.72 | 0.54 | 0.66 | 0.44 | 0.58 |
| 4 | 121206490 | rs2715970 | -0.979 | -0.127 | -0.238 | -1.481 | 0.014 | -0.01 | -0.01 | 0 | 0.7 | 0.6 | 0.66 | 0.7 | 0.61 | 0.68 |
| 4 | 121207648 | rs2662744 | -0.25 | -0.199 | -0.165 | -1.525 | 0.02 | -0.028 | -0.028 | 0.008 | 0.63 | 0.45 | 0.63 | 0.67 | 0.57 | 0.64 |
| 4 | 121229736 | rs2662715 | -1.539 | -0.129 | -0.278 | -1.515 | -0.086 | -0.03 | -0.03 | 0.634 | 0.76 | 0.96 | 0.74 | 0.7 | 0.68 | 0.7 |
| 4 | 121232742 | rs1543277 | 0.508 | -0.348 | -0.381 | -1.515 | 0.026 | -0.036 | -0.036 | 0 | 0.34 | 0.15 | 0.21 | 0.42 | 0.26 | 0.31 |
| 4 | 121239258 | rs2715952 | 1.343 | -0.239 | -0.707 | -1.519 | -0.132 | -0.026 | -0.026 | 0 | 0.25 | 0.05 | 0.3 | 0.3 | 0.39 | 0.3 |
| 4 | 121242474 | rs6850328 | 0.404 | -0.23 | -0.609 | -1.549 | -0.006 | -0.009 | -0.009 | 0 | 0.07 | 0.1 | 0.03 | 0.03 | 0.04 | 0.04 |
| 4 | 121248446 | rs1405241 | -0.766 | -0.26 | -0.527 | -1.507 | -0.001 | 0 | 0 | 0.977 | 0.7 | 0.72 | 0.61 | 0.76 | 0.7 | 0.67 |
| 4 | 121261806 | rs2715992 | 0.655 | -0.101 | -0.472 | -1.406 | -0.022 | 0.008 | 0.008 | 0 | 0.31 | 0.15 | 0.29 | 0.33 | 0.36 | 0.34 |
| 4 | 121263497 | rs1160185 | -0.807 | -0.096 | -0.349 | -1.411 | 0 | 0 | 0 | 7.797 | 0.68 | 0.71 | 0.61 | 0.76 | 0.7 | 0.66 |
| 4 | 121269728 | rs950658 | -0.047 | 0.055 | -0.318 | -1.306 | -0.009 | -0.006 | -0.006 | 0 | 0.6 | 0.47 | 0.57 | 0.73 | 0.68 | 0.6 |
| 4 | 121270939 | rs7683870 | 0.233 | 0.075 | -0.426 | -1.374 | 0.032 | -0.166 | -0.166 | 0.953 | 0.23 | 0.48 | 0.22 | 0 | 0.23 | 0.1 |
| 4 | 121292818 | rs6817848 | 0.251 | 0.016 | -1.082 | -1.351 | -0.019 | 0.063 | 0.063 | 0 | 0.44 | 0.23 | 0.39 | 0.75 | 0.49 | 0.55 |
| 4 | 121292945 | rs1374988 | -0.621 | -0.021 | -1.008 | -1.286 | -0.02 | -0.007 | -0.007 | 2.25 | 0.67 | 0.57 | 0.6 | 0.75 | 0.71 | 0.64 |
| 4 | 121298764 | rs1526326 | -0.602 | -0.034 | -1.045 | -1.285 | -0.02 | -0.007 | -0.007 | 0.528 | 0.66 | 0.53 | 0.6 | 0.75 | 0.71 | 0.64 |
| 4 | 121323123 | rs1448065 | -0.248 | 0.341 | -1.133 | -1.193 | 0.004 | 0.014 | 0.014 | 0 | 0.51 | 0.55 | 0.47 | 0.78 | 0.57 | 0.6 |
| 4 | 121323918 | rs2166838 | -1.346 | 0.19 | -1.381 | -1.274 | -0.052 | -0.055 | -0.055 | 0.087 | 0.78 | 0.97 | 0.65 | 0.8 | 0.75 | 0.75 |
| 4 | 121339394 | rs7694494 | 0.654 | 0.515 | -0.843 | -0.498 | -0.074 | -0.052 | -0.052 | 0 | 0.23 | 0.04 | 0.25 | 0.31 | 0.33 | 0.32 |
| 4 | 121342283 | rs1405240 | -0.815 | 0.448 | -0.74 | -0.457 | 0.004 | 0.018 | 0.018 | 0 | 0.51 | 0.56 | 0.47 | 0.77 | 0.57 | 0.59 |
| 4 | 121342367 | rs1808015 | 0.384 | 0.289 | -1.018 | 0.257 | -0.01 | -0.108 | -0.108 | 0 | 0.15 | 0.03 | 0.11 | 0.42 | 0.13 | 0.21 |
| 4 | 121347367 | rs705103 | 0.203 | 0.474 | -1.016 | -0.593 | -0.024 | 0.089 | 0.089 | 0 | 0.29 | 0.1 | 0.17 | 0.12 | 0.33 | 0.32 |
| 4 | 121347730 | rs697364 | 0.276 | 0.474 | -1.081 | -0.593 | -0.057 | 0.102 | 0.102 | 0 | 0.29 | 0.04 | 0.16 | 0.12 | 0.33 | 0.32 |
| 4 | 121351003 | rs1026083 | 0.782 | 0.66 | -1.068 | -0.32 | 0.003 | -0.067 | -0.067 | 0.645 | 0.14 | 0.06 | 0.2 | 0.6 | 0.12 | 0.23 |
| 4 | 121351589 | rs7667780 | -0.354 | 0.811 | -0.433 | -0.611 | 0.094 | -0.039 | -0.039 | 2.601 | 0.1 | 0.49 | 0.11 | 0.01 | 0.11 | 0.03 |
| 4 | 121353265 | rs11938648 | 0.465 | 0.295 | -0.46 | -0.518 | 0.023 | -0.009 | -0.009 | 0 | 0.07 | 0.33 | 0.05 | 0 | 0.03 | 0.02 |
| 4 | 121361500 | rs1448083 | -0.919 | 0.528 | -1.245 | -0.309 | -0.043 | 0.026 | 0.026 | 0 | 0.63 | 0.9 | 0.52 | 0.77 | 0.61 | 0.65 |
| 4 | 121368939 | rs843613 | 0.142 | 0.562 | -1.339 | -0.27 | -0.001 | -0.021 | -0.021 | 0.866 | 0.42 | 0.59 | 0.28 | 0.14 | 0.44 | 0.38 |
| 4 | 121371794 | rs1685593 | -0.093 | 0.904 | -0.851 | 0.003 | -0.001 | 0.004 | 0.004 | 0 | 0.61 | 0.65 | 0.5 | 0.77 | 0.61 | 0.65 |
| 4 | 121393821 | rs863620 | -0.069 | 0.956 | -1.422 | 0.165 | -0.019 | 0.06 | 0.06 | 0 | 0.55 | 0.8 | 0.53 | 0.68 | 0.55 | 0.58 |
| 4 | 121395169 | rs843596 | 0.075 | 0.946 | -1.179 | 0.385 | 0.001 | -0.005 | -0.005 | 0.014 | 0.57 | 0.46 | 0.63 | 0.47 | 0.62 | 0.5 |
| 4 | 121396721 | rs7697117 | -0.464 | 2.014 | -1.361 | -0.004 | -0.011 | 0.008 | 0.008 | 0.682 | 0.14 | 0.1 | 0.19 | 0.1 | 0.31 | 0.21 |
| 4 | 121397402 | rs12501789 | -0.464 | 2.014 | -1.361 | -0.004 | -0.011 | 0.008 | 0.008 | 0.682 | 0.14 | 0.1 | 0.19 | 0.1 | 0.31 | 0.21 |
| 4 | 121402735 | rs17339087 | -1.159 | 1.38 | -1.025 | -0.465 | 0.019 | -0.036 | -0.036 | 0 | 0.39 | 0.55 | 0.33 | 0.19 | 0.46 | 0.32 |
| 4 | 121403974 | rs4833161 | 0.52 | 0.417 | -1.498 | -0.089 | 0.022 | -0.03 | -0.03 | 0 | 0.14 | 0.05 | 0.27 | 0.26 | 0.14 | 0.13 |
| 4 | 121406055 | rs1994475 | 0.298 | 0.453 | -1.259 | -0.125 | 0.002 | -0.1 | -0.1 | 0 | 0.16 | 0.35 | 0.11 | 0 | 0.1 | 0.07 |
| 4 | 121417757 | rs17050662 | 0.283 | 0.454 | -1.258 | -0.124 | 0 | -0.1 | -0.1 | 0.652 | 0.16 | 0.35 | 0.11 | 0 | 0.1 | 0.07 |
| 4 | 121426884 | rs843605 | -0.77 | 0.263 | -1.727 | -0.009 | -0.055 | -0.155 | -0.155 | 0.095 | 0.76 | 0.98 | 0.81 | 0.55 | 0.76 | 0.68 |
| 4 | 121427700 | rs17050741 | 0.523 | 0.456 | -0.451 | -0.027 | 0.02 | -0.043 | -0.043 | 0 | 0.08 | 0.3 | 0.05 | 0 | 0.05 | 0.02 |
| 4 | 121429845 | rs17339852 | 0.243 | 0.449 | -0.649 | 0.219 | -0.001 | 0.034 | 0.034 | 0 | 0.07 | 0.17 | 0.05 | 0.1 | 0.08 | 0.03 |
| 4 | 121450846 | rs10024986 | -0.044 | 0.373 | -1.107 | 0.326 | 0.014 | 0 | 0 | 0 | 0.28 | 0.14 | 0.31 | 0.33 | 0.21 | 0.32 |
| 4 | 121457685 | rs2390022 | 0.048 | 0.48 | -0.396 | 0.714 | 0.007 | 0.111 | 0.111 | 0 | 0.86 | 0.78 | 0.81 | 0.57 | 0.82 | 0.77 |
| 4 | 121474572 | rs4833652 | 0.175 | 0.155 | -0.8 | 0.076 | 0.003 | -0.004 | -0.004 | 0 | 0.44 | 0.56 | 0.25 | 0.1 | 0.3 | 0.17 |
| 4 | 121474967 | rs1386814 | -0.111 | 0.49 | -1.062 | 0.895 | -0.001 | 0.055 | 0.055 | 0 | 0.89 | 0.87 | 0.81 | 0.56 | 0.89 | 0.74 |
| 4 | 121478712 | rs12511769 | 0.915 | 0.344 | -1.33 | 1.064 | 0.007 | -0.025 | -0.025 | 0 | 0.08 | 0.03 | 0.2 | 0.15 | 0.04 | 0.11 |
| 4 | 121479039 | rs6855878 | -0.804 | 0.265 | -1.952 | 0.779 | -0.071 | -0.143 | -0.143 | 0.285 | 0.67 | 0.94 | 0.7 | 0.57 | 0.72 | 0.69 |
| 4 | 121479398 | rs6856719 | 0.111 | -0.019 | -1.46 | 0.571 | -0.004 | 0.01 | 0.01 | 0 | 0.09 | 0.11 | 0.06 | 0.11 | 0.08 | 0.04 |
| 4 | 121481950 | rs4395533 | -0.213 | 0.101 | -1.062 | 0.307 | 0.006 | 0.019 | 0.019 | 0 | 0.5 | 0.45 | 0.45 | 0.4 | 0.62 | 0.53 |
| 4 | 121487559 | rs11098574 | -0.17 | 0.067 | -1.011 | 0.505 | -0.012 | -0.006 | -0.006 | 0 | 0.44 | 0.41 | 0.39 | 0.4 | 0.48 | 0.37 |
| 4 | 121489983 | rs1532055 | 0.687 | -0.012 | -0.879 | 0.591 | -0.003 | 0.002 | 0.002 | 0.036 | 0.41 | 0.53 | 0.4 | 0.45 | 0.48 | 0.49 |

| | | | | | | | | | | | | | | | | |
|---|---|---|---|---|---|---|---|---|---|---|---|---|---|---|---|---|
| 4 | 121491629 | rs6854696 | -1.098 | 0.03 | -0.91 | 0.582 | 0.014 | -0.008 | -0.008 | 0.08 | 0.86 | 0.94 | 0.8 | 0.85 | 0.95 | 0.86 |
| 4 | 121525149 | rs13121429 | 0.7 | 0.028 | -0.985 | 0.563 | -0.022 | -0.009 | -0.009 | 0.497 | 0.11 | 0 | 0.06 | 0.06 | 0.12 | 0.04 |
| 4 | 121526447 | rs10002802 | -0.778 | -0.1 | -0.888 | 0.511 | -0.005 | -0.005 | -0.005 | 0.038 | 0.65 | 0.68 | 0.66 | 0.66 | 0.63 | 0.66 |
| 4 | 121535293 | rs1353473 | -0.033 | 0.24 | -0.811 | 0.275 | 0.003 | 0.008 | 0.008 | 0 | 0.53 | 0.55 | 0.46 | 0.44 | 0.59 | 0.52 |
| 4 | 121536939 | rs10518365 | 0.8 | 0.181 | -0.846 | -0.089 | 0.009 | -0.021 | -0.021 | 0 | 0.4 | 0.43 | 0.39 | 0.38 | 0.48 | 0.48 |
| 4 | 121538689 | rs11098579 | -0.713 | 0.905 | -1.143 | 0.061 | -0.014 | -0.014 | -0.014 | 0 | 0.44 | 0.4 | 0.39 | 0.47 | 0.48 | 0.37 |
| 4 | 121539450 | rs1463782 | -0.048 | 0.883 | -1.031 | -0.764 | -0.009 | 0.059 | 0.059 | 0.186 | 0.28 | 0.24 | 0.2 | 0 | 0.27 | 0.1 |
| 4 | 121545138 | rs17051145 | -0.924 | 1.148 | 0.025 | -0.139 | 0.052 | -0.014 | -0.014 | 0 | 0.12 | 0.39 | 0.12 | 0.12 | 0.14 | 0.22 |
| 4 | 121553198 | rs17051152 | -1.269 | 0.192 | 0.596 | -0.54 | -0.066 | -0.079 | -0.079 | 1.399 | 0.13 | 0.56 | 0.06 | 0 | 0.02 | 0.03 |
| 4 | 121575053 | rs7667818 | -0.208 | 0.954 | 0.174 | -0.303 | -0.011 | -0.009 | -0.009 | 0 | 0.06 | 0.05 | 0.19 | 0.09 | 0.13 | 0.14 |
| 4 | 121581769 | rs6818888 | 0.462 | 0.703 | -0.474 | -1.187 | 0.02 | 0.083 | 0.083 | 0.203 | 0.4 | 0.22 | 0.33 | 0.16 | 0.35 | 0.29 |
| 4 | 121582628 | rs13138888 | 0.364 | 0.843 | -0.825 | -1.211 | -0.009 | 0.031 | 0.031 | 0 | 0.2 | 0.1 | 0.14 | 0.09 | 0.22 | 0.07 |
| 4 | 121587269 | rs1580743 | -0.731 | 1.046 | -0.168 | -1.154 | 0.044 | 0.048 | 0.048 | 0 | 0.93 | 0.78 | 0.82 | 0.9 | 0.91 | 0.86 |
| 4 | 121587550 | rs1580742 | 1.7 | 0.685 | -0.605 | -1.538 | -0.051 | 0.192 | 0.192 | 0.591 | 0.31 | 0.03 | 0.13 | 0.06 | 0.32 | 0.21 |
| 4 | 121588603 | rs1496270 | 0.502 | 0.52 | 0.137 | -1.128 | 0.004 | -0.011 | -0.011 | 0.153 | 0.72 | 0.63 | 0.69 | 0.82 | 0.69 | 0.8 |
| 4 | 121594207 | rs10027783 | 0.888 | 0.407 | 0.137 | -0.799 | -0.032 | 0.034 | 0.034 | 0 | 0.08 | 0.2 | 0.2 | 0.1 | 0.01 | 0.08 |
| 4 | 121623057 | rs1874442 | -0.445 | 0.705 | -0.033 | -0.76 | 0.045 | 0.046 | 0.046 | 0 | 0.93 | 0.78 | 0.82 | 0.91 | 0.91 | 0.87 |
| 4 | 121623516 | rs1496276 | 1.143 | 0.607 | -0.393 | -1.212 | 0.054 | 0.05 | 0.05 | 0 | 0.56 | 0.2 | 0.45 | 0.72 | 0.68 | 0.69 |
| 4 | 121626824 | rs1390563 | 0.552 | 0.168 | -0.433 | -1.711 | 0.002 | 0.049 | 0.049 | 0.187 | 0.52 | 0.36 | 0.66 | 0.75 | 0.55 | 0.55 |
| 4 | 121627113 | rs724818 | 2.291 | -0.483 | -0.847 | -1.953 | 0.052 | -0.031 | -0.031 | 0 | 0.11 | 0 | 0.17 | 0.11 | 0.04 | 0.07 |
| 4 | 121627842 | rs17345994 | 0.847 | -0.559 | -0.628 | -2.1 | -0.015 | 0.013 | 0.013 | 0.018 | 0.18 | 0.13 | 0.16 | 0.1 | 0.25 | 0.08 |
| 4 | 121630332 | rs11730092 | 0.962 | -0.536 | -0.619 | -2.092 | -0.015 | 0.011 | 0.011 | 0 | 0.18 | 0.13 | 0.16 | 0.1 | 0.25 | 0.08 |
| 4 | 121633580 | rs7657769 | 1.379 | -0.015 | -0.765 | -2.11 | -0.037 | -0.016 | -0.016 | 0 | 0.08 | 0.04 | 0.33 | 0.61 | 0.22 | 0.31 |
| 4 | 121637960 | rs10518375 | 1.081 | -0.503 | -0.106 | -2.178 | -0.018 | -0.055 | -0.055 | 0 | 0.24 | 0.29 | 0.11 | 0.09 | 0.13 | 0.23 |
| 4 | 121642180 | rs6838858 | 1.328 | -0.47 | -0.027 | -2.177 | 0.002 | -0.006 | -0.006 | 0 | 0.23 | 0.2 | 0.09 | 0.09 | 0.12 | 0.22 |
| 4 | 121645012 | rs843575 | -0.53 | -0.376 | -0.018 | -2.229 | -0.026 | -0.027 | -0.027 | 0.079 | 0.6 | 0.46 | 0.69 | 0.79 | 0.83 | 0.76 |
| 4 | 121648617 | rs10518377 | 0.753 | -0.096 | -0.557 | -2.152 | -0.008 | 0.005 | 0.005 | 0 | 0.4 | 0.37 | 0.48 | 0.66 | 0.51 | 0.49 |
| 4 | 121661208 | rs10518378 | 0.503 | -0.395 | -0.778 | -2.007 | 0.056 | 0.06 | 0.06 | 2.831 | 0.36 | 0.19 | 0.27 | 0.21 | 0.17 | 0.25 |
| 4 | 121669224 | rs7661193 | 2.481 | 0.316 | -1.07 | -1.893 | -0.072 | 0.092 | 0.092 | 0.02 | 0.15 | 0.01 | 0.08 | 0.02 | 0.22 | 0.09 |
| 4 | 121671965 | rs11725662 | 0.38 | 0.604 | -0.409 | -1.341 | 0.015 | 0.021 | 0.021 | 0 | 0.28 | 0.34 | 0.41 | 0.5 | 0.39 | 0.38 |
| 4 | 121683755 | rs10857072 | -0.057 | 0.066 | -0.556 | -1.691 | 0.025 | -0.011 | -0.011 | 0.138 | 0.34 | 0.18 | 0.41 | 0.32 | 0.35 | 0.17 |
| 4 | 121684267 | rs6813562 | 1.462 | -0.061 | -0.261 | -1.772 | -0.053 | -0.005 | -0.005 | 0.066 | 0.17 | 0.31 | 0.23 | 0.15 | 0.08 | 0.09 |
| 4 | 121685814 | rs13147610 | 2.619 | 0.141 | -0.241 | -1.759 | -0.013 | 0.007 | 0.007 | 0 | 0.18 | 0.09 | 0.1 | 0.02 | 0.24 | 0.1 |
| 4 | 121691899 | rs907298 | -0.458 | 0.073 | -0.209 | -1.751 | 0.009 | 0.01 | 0.01 | 0 | 0.52 | 0.47 | 0.63 | 0.68 | 0.68 | 0.48 |
| 4 | 121703782 | rs17051254 | 2.595 | -0.006 | -0.345 | -1.693 | 0.03 | -0.007 | -0.007 | 0.731 | 0.13 | 0.05 | 0.19 | 0.14 | 0.05 | 0.07 |
| 4 | 121705238 | rs2390053 | 1.909 | 0.107 | -0.255 | -1.66 | 0.006 | 0.1 | 0.1 | 0.138 | 0.09 | 0.15 | 0.28 | 0.47 | 0.14 | 0.26 |
| 4 | 121706201 | rs12499000 | 1.242 | 0.128 | -0.434 | -1.753 | -0.063 | -0.069 | -0.069 | 1.369 | 0.18 | 0.01 | 0.19 | 0.19 | 0.27 | 0.08 |
| 4 | 121709930 | rs13111620 | 2.415 | 0.221 | -0.255 | -1.777 | -0.003 | -0.024 | -0.024 | 0 | 0.19 | 0.16 | 0.12 | 0.02 | 0.24 | 0.1 |
| 4 | 121714045 | rs6827372 | 0.929 | -0.043 | -0.5 | -1.673 | 0.01 | 0.018 | 0.018 | 0 | 0.29 | 0.36 | 0.4 | 0.49 | 0.39 | 0.36 |
| 4 | 121719932 | rs3804174 | -0.071 | -0.109 | -0.439 | -1.551 | 0.016 | -0.009 | -0.009 | 0 | 0.33 | 0.25 | 0.4 | 0.35 | 0.32 | 0.15 |
| 4 | 121738049 | rs343192 | 0.095 | -0.116 | -0.408 | -1.537 | 0.024 | -0.019 | -0.019 | 0 | 0.33 | 0.21 | 0.39 | 0.36 | 0.32 | 0.15 |
| 4 | 121746462 | rs1511301 | -2.319 | 0.047 | -0.545 | -1.678 | 0.024 | 0.051 | 0.051 | 0 | 0.78 | 0.87 | 0.93 | 0.9 | 0.87 | 0.79 |
| 4 | 121750566 | rs1033097 | -2.313 | 0.068 | -0.555 | -1.687 | 0.016 | 0.054 | 0.054 | 0 | 0.78 | 0.88 | 0.92 | 0.9 | 0.86 | 0.78 |
| 4 | 121755299 | rs7673521 | 2.194 | 0.121 | -0.676 | -1.678 | -0.027 | 0.002 | 0.002 | 0 | 0.08 | 0.01 | 0.06 | 0.06 | 0.1 | 0.21 |
| 4 | 121759300 | rs343175 | -3.117 | 0.034 | -0.744 | -1.746 | 0.005 | 0.093 | 0.093 | 0 | 0.8 | 0.97 | 0.93 | 0.9 | 0.87 | 0.79 |
| 4 | 121759338 | rs343176 | -1.599 | 0.071 | -0.563 | -1.699 | 0.008 | 0.051 | 0.051 | 0.12 | 0.69 | 0.82 | 0.86 | 0.85 | 0.77 | 0.57 |
| 4 | 121759963 | rs12509870 | 0.822 | 0.182 | -0.385 | -1.483 | -0.011 | -0.016 | -0.016 | 0 | 0.18 | 0.09 | 0.21 | 0.19 | 0.27 | 0.08 |
| 4 | 121765527 | rs6534216 | 0.258 | 0.096 | -0.267 | -1.502 | 0.008 | 0.008 | 0.008 | 0 | 0.23 | 0.42 | 0.25 | 0.21 | 0.3 | 0.12 |
| 4 | 121766616 | rs1588523 | 0.1 | 0.239 | -0.301 | -1.401 | -0.005 | -0.004 | -0.004 | 0 | 0.37 | 0.36 | 0.47 | 0.55 | 0.5 | 0.58 |
| 4 | 121774515 | rs3115310 | 1.3 | -0.219 | -0.15 | -1.71 | 0.049 | -0.105 | -0.105 | 0.5 | 0.19 | 0.42 | 0.13 | 0.04 | 0.24 | 0.1 |
| 4 | 121775592 | rs10518380 | 1.529 | -0.52 | -0.507 | -1.719 | 0.064 | -0.021 | -0.021 | 0 | 0.16 | 0.02 | 0.2 | 0.14 | 0.07 | 0.08 |
| 4 | 121784013 | rs1450476 | 0.912 | -0.375 | -0.429 | -1.753 | 0 | -0.001 | -0.001 | 0 | 0.14 | 0.14 | 0.11 | 0.15 | 0.16 | 0.06 |
| 4 | 121786273 | rs2597553 | -1.3 | -0.433 | -0.429 | -1.928 | 0.008 | 0.056 | 0.056 | 0 | 0.66 | 0.78 | 0.85 | 0.85 | 0.76 | 0.56 |
| 4 | 121789730 | rs7675777 | 0.986 | -0.377 | -0.284 | -1.918 | 0.049 | 0.134 | 0.134 | 0 | 0.17 | 0.32 | 0.42 | 0.52 | 0.29 | 0.29 |
| 4 | 121792785 | rs17051273 | 1.68 | -0.556 | 0.199 | -1.773 | 0.008 | -0.036 | -0.036 | 0.741 | 0.18 | 0.2 | 0.11 | 0.04 | 0.24 | 0.1 |
| 4 | 121794535 | rs6844159 | 0.471 | -0.497 | 0.203 | -1.742 | 0.007 | 0.031 | 0.031 | 0 | 0.34 | 0.25 | 0.32 | 0.18 | 0.31 | 0.18 |
| 4 | 121795956 | rs343202 | -0.208 | -0.638 | 0.315 | -1.503 | 0.002 | 0.004 | 0.004 | 0.08 | 0.49 | 0.48 | 0.44 | 0.33 | 0.47 | 0.25 |
| 4 | 121801457 | rs17051275 | 0.854 | -1.727 | 0.758 | -0.84 | 0.058 | -0.002 | -0.002 | 0 | 0.17 | 0.04 | 0.21 | 0.15 | 0.07 | 0.08 |
| 4 | 121814232 | rs11723050 | 2.009 | -1.675 | 0.734 | -0.922 | -0.129 | -0.11 | -0.11 | 0 | 0.31 | 0.1 | 0.49 | 0.5 | 0.53 | 0.4 |
| 4 | 121816501 | rs343168 | -0.58 | -1.722 | 1.196 | -0.887 | 0.017 | 0.064 | 0.064 | 0 | 0.13 | 0.34 | 0.12 | 0.16 | 0.13 | 0.06 |
| 4 | 121830057 | rs2122512 | 0.615 | -1.654 | 0.669 | -0.913 | 0.008 | -0.013 | -0.013 | 0 | 0.11 | 0.17 | 0.08 | 0.08 | 0.1 | 0.21 |
| 4 | 121831385 | rs11939905 | 0.568 | -1.293 | 0.59 | -0.715 | -0.001 | 0.008 | 0.008 | 0 | 0.08 | 0.09 | 0.07 | 0.14 | 0.09 | 0.04 |
| 4 | 121831539 | rs6844456 | 0.8 | -1.375 | 0.491 | -0.803 | -0.012 | -0.052 | -0.052 | 0 | 0.27 | 0.33 | 0.1 | 0.11 | 0.15 | 0.2 |
| 4 | 121837791 | rs1450478 | -0.143 | -1.364 | 0.423 | -0.983 | 0.001 | -0.001 | -0.001 | 0.364 | 0.34 | 0.36 | 0.16 | 0.19 | 0.23 | 0.22 |
| 4 | 121842567 | rs1470696 | 0.587 | -1.316 | 0.462 | -0.874 | 0.007 | -0.001 | -0.001 | 0 | 0.58 | 0.43 | 0.59 | 0.61 | 0.69 | 0.61 |
| 4 | 121857957 | rs6848268 | 0.557 | -1.193 | 0.478 | -1.17 | 0.009 | 0.005 | 0.005 | 0.163 | 0.59 | 0.41 | 0.57 | 0.65 | 0.68 | 0.62 |
| 4 | 121859395 | rs7673618 | 1.713 | -1.19 | 0.24 | -1.246 | -0.136 | -0.064 | -0.064 | 0.299 | 0.32 | 0.08 | 0.48 | 0.58 | 0.53 | 0.43 |
| 4 | 121873063 | rs1450471 | -0.692 | -1.297 | 0.412 | -1.273 | 0.012 | 0.007 | 0.007 | 0.034 | 0.73 | 0.84 | 0.71 | 0.77 | 0.82 | 0.73 |
| 4 | 121875311 | rs13132472 | 0.184 | -1.002 | 0.42 | -1.117 | -0.009 | -0.004 | -0.004 | 0.727 | 0.6 | 0.51 | 0.57 | 0.64 | 0.68 | 0.62 |

| | | | | | | | | | | | | | | | | |
|---|---|---|---|---|---|---|---|---|---|---|---|---|---|---|---|---|
| 4 | 121875396 | rs13132115 | 1.11 | -0.211 | -0.005 | -0.635 | -0.127 | -0.064 | -0.064 | 0 | 0.33 | 0.08 | 0.48 | 0.58 | 0.54 | 0.44 |
| 4 | 121888708 | rs11098592 | -0.242 | -0.749 | 0.287 | -1.037 | 0.009 | 0.004 | 0.004 | 2.093 | 0.69 | 0.28 | 0.82 | 0.89 | 0.8 | 0.8 |
| 4 | 121889053 | rs6855125 | 0.666 | -0.745 | 0.646 | -1.02 | 0.04 | 0.066 | 0.066 | 1.277 | 0.94 | 0.69 | 0.95 | 0.93 | 0.95 | 0.96 |
| 4 | 121893179 | rs12648040 | 0.698 | -0.879 | 0.274 | -1.11 | 0.052 | 0.069 | 0.069 | 2.738 | 0.31 | 0.09 | 0.27 | 0.21 | 0.19 | 0.36 |
| 4 | 121893316 | rs12641971 | 0.07 | -0.632 | 0.674 | -1.204 | 0.011 | 0.014 | 0.014 | 2.205 | 0.7 | 0.26 | 0.82 | 0.89 | 0.81 | 0.78 |
| 4 | 121894207 | rs6841260 | 0.585 | -0.65 | 0.591 | -1.205 | -0.015 | -0.099 | -0.099 | 0 | 0.23 | 0.3 | 0.12 | 0.04 | 0.15 | 0.15 |
| 4 | 121898421 | rs6855437 | 0.593 | -0.589 | 0.475 | -1.178 | 0.015 | 0.049 | 0.049 | 1.986 | 0.65 | 0.25 | 0.75 | 0.84 | 0.71 | 0.77 |
| 4 | 121904225 | rs6824931 | 1.422 | -0.721 | 0.439 | -0.688 | 0.036 | 0.069 | 0.069 | 0.113 | 0.88 | 0.57 | 0.87 | 0.88 | 0.86 | 0.94 |
| 4 | 121938296 | rs1807071 | 0.185 | -1.424 | -0.646 | -0.66 | 0.047 | 0.05 | 0.05 | 0.736 | 0.34 | 0.11 | 0.34 | 0.24 | 0.2 | 0.33 |
| 4 | 121940998 | rs11931955 | 1.046 | -1.536 | -0.459 | -0.793 | -0.018 | -0.091 | -0.091 | 0.697 | 0.21 | 0.3 | 0.12 | 0.03 | 0.14 | 0.14 |
| 4 | 121944932 | rs12509928 | 0.104 | -1.528 | -0.748 | -0.786 | -0.011 | -0.006 | -0.006 | 0 | 0.59 | 0.46 | 0.6 | 0.67 | 0.75 | 0.59 |
| 4 | 121956610 | rs10461204 | 1.325 | -1.274 | -1.651 | -0.615 | -0.121 | -0.097 | -0.097 | 0.146 | 0.32 | 0.08 | 0.44 | 0.6 | 0.57 | 0.43 |
| 4 | 121957051 | rs1045080 | -0.603 | -1.293 | -1.655 | -0.625 | 0.032 | 0.053 | 0.053 | 4.513 | 0.71 | 0.18 | 0.81 | 0.88 | 0.82 | 0.77 |
| 4 | 121958091 | rs3733558 | 0.747 | -1.268 | -1.647 | -0.61 | -0.074 | -0.015 | -0.015 | 0 | 0.37 | 0.09 | 0.47 | 0.65 | 0.61 | 0.45 |
| 4 | 121958697 | rs3733560 | 1.279 | -1.211 | -1.608 | -0.574 | -0.121 | -0.097 | -0.097 | 0.812 | 0.32 | 0.08 | 0.44 | 0.6 | 0.57 | 0.43 |
| 4 | 121972302 | rs17051284 | -0.594 | -1.213 | -1.58 | -0.586 | 0.03 | 0.051 | 0.051 | 3.637 | 0.7 | 0.2 | 0.81 | 0.88 | 0.8 | 0.78 |
| 4 | 121972352 | rs11098600 | -0.091 | -0.334 | -1.417 | -0.076 | -0.008 | 0.008 | 0.008 | 1.919 | 0.38 | 0.3 | 0.49 | 0.65 | 0.6 | 0.45 |
| 4 | 121974645 | rs11931196 | -1.186 | -1.028 | -0.404 | -0.406 | 0.024 | 0.067 | 0.067 | 0 | 0.06 | 0.24 | 0.05 | 0.05 | 0.04 | 0.02 |
| 4 | 121976444 | rs17051286 | -0.667 | -1.746 | -1.629 | -0.979 | 0.075 | 0.094 | 0.094 | 2.612 | 0.25 | 0.02 | 0.06 | 0.11 | 0.09 | 0.17 |
| 4 | 121978877 | rs4833705 | 0.649 | -2.616 | -1.242 | -1.506 | 0.057 | 0.051 | 0.051 | 0 | 0.58 | 0.32 | 0.76 | 0.74 | 0.82 | 0.52 |
| 4 | 121982736 | rs882045 | -0.249 | -3.058 | -0.719 | -1.725 | -0.003 | -0.001 | -0.001 | 0.497 | 0.83 | 0.85 | 0.84 | 0.81 | 0.89 | 0.79 |
| 4 | 121984023 | rs6829190 | 1.312 | -3.157 | -0.688 | -1.738 | 0.074 | 0.064 | 0.064 | 0 | 0.57 | 0.3 | 0.76 | 0.73 | 0.81 | 0.51 |
| 4 | 121986137 | rs2390070 | -0.256 | -3.198 | -0.513 | -1.76 | -0.013 | -0.006 | -0.006 | 0.569 | 0.84 | 0.81 | 0.84 | 0.81 | 0.89 | 0.79 |
| 4 | 121988791 | rs2136947 | -0.411 | -3.211 | -0.298 | -1.809 | -0.087 | -0.105 | -0.105 | 1.149 | 0.23 | 0.46 | 0.07 | 0.04 | 0.07 | 0.27 |
| 4 | 121997348 | rs6823534 | 0.919 | -3.088 | -0.405 | -1.74 | 0.017 | 0.016 | 0.016 | 0 | 0.6 | 0.37 | 0.76 | 0.73 | 0.82 | 0.52 |
| 4 | 122010915 | rs1105274 | 0.851 | -3.021 | -1.026 | -2.37 | 0.02 | 0.093 | 0.093 | 2.316 | 0.28 | 0.12 | 0.17 | 0.05 | 0.2 | 0.15 |
| 4 | 122015591 | rs13128284 | 0.437 | -3.065 | -0.961 | -2.323 | 0.011 | 0.012 | 0.012 | 0 | 0.6 | 0.38 | 0.76 | 0.73 | 0.81 | 0.52 |
| 4 | 122024540 | rs13142482 | 0.61 | -2.619 | -1.071 | -2.419 | 0.003 | -0.004 | -0.004 | 0 | 0.3 | 0.24 | 0.21 | 0.04 | 0.24 | 0.15 |
| 4 | 122028784 | rs1397643 | 0.754 | -2.799 | -1.08 | -2.502 | -0.034 | 0.083 | 0.083 | 0 | 0.36 | 0.18 | 0.65 | 0.81 | 0.6 | 0.4 |
| 4 | 122030864 | rs6833755 | 0.401 | -2.769 | -0.446 | -2.671 | -0.028 | -0.135 | -0.135 | 0 | 0.29 | 0.41 | 0.2 | 0.03 | 0.21 | 0.15 |
| 4 | 122039475 | rs1027976 | -0.363 | -2.241 | -0.752 | -2.498 | -0.014 | 0.029 | 0.029 | 0 | 0.94 | 0.99 | 0.95 | 1 | 0.9 | 0.98 |
| 4 | 122052641 | rs11098602 | 0.36 | -2.49 | -0.315 | -2.735 | 0.003 | -0.037 | -0.037 | 0 | 0.33 | 0.65 | 0.21 | 0.05 | 0.21 | 0.16 |
| 4 | 122053223 | rs6846553 | -0.522 | -2.591 | -0.516 | -2.823 | 0.089 | 0.174 | 0.174 | 0.134 | 0.49 | 0.72 | 0.26 | 0.1 | 0.28 | 0.4 |
| 4 | 122055956 | rs1858291 | 0.488 | -2.565 | -0.491 | -2.807 | 0.009 | -0.021 | -0.021 | 0 | 0.33 | 0.68 | 0.21 | 0.05 | 0.22 | 0.16 |
| 4 | 122063735 | rs3817171 | 0.939 | -2.483 | -0.109 | -2.79 | 0.004 | -0.005 | -0.005 | 0 | 0.28 | 0.27 | 0.18 | 0.07 | 0.22 | 0.16 |
| 4 | 122074463 | rs959268 | -0.487 | -2.683 | -0.208 | -2.826 | 0.004 | 0.018 | 0.018 | 0 | 0.85 | 0.91 | 0.84 | 0.86 | 0.85 | 0.92 |
| 4 | 122075739 | rs17051298 | 0.555 | -2.699 | -0.227 | -2.821 | 0.021 | 0.022 | 0.022 | 0 | 0.18 | 0.09 | 0.04 | 0.07 | 0.07 | 0.37 |
| 4 | 122083428 | rs6821331 | 0.849 | -2.635 | -0.076 | -2.815 | -0.022 | -0.109 | -0.109 | 0 | 0.31 | 0.47 | 0.2 | 0.07 | 0.22 | 0.16 |
| 4 | 122088141 | rs7661427 | 1.821 | -2.698 | -0.809 | -2.799 | -0.173 | -0.021 | -0.021 | 0 | 0.26 | 0.05 | 0.53 | 0.7 | 0.5 | 0.35 |
| 4 | 122091233 | rs13129450 | -0.924 | -2.221 | -0.218 | -2.39 | 0.083 | 0.183 | 0.183 | 0.044 | 0.49 | 0.73 | 0.26 | 0.13 | 0.29 | 0.41 |
| 4 | 122096742 | rs6838000 | 0.484 | -1.444 | 0.007 | -1.929 | -0.001 | 0.109 | 0.109 | 0 | 0.24 | 0.07 | 0.15 | 0.08 | 0.2 | 0.14 |
| 4 | 122137121 | rs13104875 | -0.131 | -1.334 | -0.266 | -1.815 | -0.01 | -0.05 | -0.05 | 0 | 0.32 | 0.64 | 0.23 | 0.09 | 0.22 | 0.14 |
| 4 | 122142224 | rs10518385 | 0.642 | -1.08 | -0.461 | -1.366 | -0.01 | -0.005 | -0.005 | 0.002 | 0.78 | 0.51 | 0.64 | 0.8 | 0.77 | 0.91 |
| 4 | 122143664 | rs6835704 | 0.1 | -0.489 | -1.278 | -0.765 | 0.015 | 0.053 | 0.053 | 0 | 0.53 | 0.62 | 0.4 | 0.71 | 0.54 | 0.78 |
| 4 | 122149001 | rs12639692 | -0.95 | -0.592 | -1.363 | -0.194 | 0.028 | -0.038 | -0.038 | 0 | 0.11 | 0.01 | 0.23 | 0.12 | 0.07 | 0.04 |
| 4 | 122151897 | rs2278865 | -0.964 | -0.49 | -1.535 | 0.149 | 0.015 | -0.03 | -0.03 | 2.124 | 0.35 | 0.75 | 0.51 | 0.43 | 0.34 | 0.21 |
| 4 | 122156142 | rs7671286 | -1.07 | -0.831 | -1.523 | -0.472 | 0.011 | 0.005 | 0.005 | 0 | 0.53 | 0.72 | 0.61 | 0.44 | 0.48 | 0.38 |
| 4 | 122161207 | rs11938619 | -0.365 | -0.946 | -0.644 | -0.704 | 0.001 | -0.109 | -0.109 | 0.5 | 0.24 | 0.31 | 0.13 | 0.04 | 0.2 | 0.22 |
| 4 | 122163552 | rs12650479 | 0.552 | -0.944 | -0.745 | -0.453 | 0.004 | 0.029 | 0.029 | 0 | 0.06 | 0.06 | 0.06 | 0.14 | 0.11 | 0.05 |
| 4 | 122170912 | rs7663303 | -1.461 | -1.115 | -0.171 | -0.624 | 0.042 | 0.075 | 0.075 | 0 | 0.47 | 0.69 | 0.54 | 0.27 | 0.37 | 0.33 |
| 4 | 122176770 | rs9993738 | -1.611 | -1.021 | -0.336 | -0.714 | 0.044 | 0.086 | 0.086 | 0 | 0.5 | 0.69 | 0.56 | 0.31 | 0.4 | 0.38 |
| 4 | 122181277 | rs9997417 | -0.243 | -1.319 | -0.06 | -0.596 | 0.02 | -0.012 | -0.012 | 0 | 0.2 | 0.14 | 0.4 | 0.25 | 0.14 | 0.11 |
| 4 | 122184678 | rs17051324 | 0.352 | -1.126 | -0.217 | -0.166 | -0.028 | -0.03 | -0.03 | 0 | 0.06 | 0 | 0.06 | 0.13 | 0.12 | 0.06 |
| 4 | 122184750 | rs7693881 | -1.651 | -1.613 | 0.373 | -0.461 | -0.009 | -0.005 | -0.005 | 0 | 0.32 | 0.38 | 0.28 | 0.2 | 0.31 | 0.25 |
| 4 | 122187618 | rs7654307 | 0.25 | -1.61 | 0.59 | -0.467 | -0.022 | -0.027 | -0.027 | 0.016 | 0.15 | 0.16 | 0.15 | 0.04 | 0.08 | 0.09 |
| 4 | 122187933 | rs13128635 | -0.429 | -1.562 | 0.455 | -0.458 | 0.002 | -0.009 | -0.009 | 0 | 0.07 | 0.1 | 0.06 | 0.02 | 0.11 | 0.09 |
| 4 | 122193376 | rs4833711 | 0.743 | -1.999 | 0.018 | -0.614 | 0.018 | -0.039 | -0.039 | 0 | 0.11 | 0.03 | 0.12 | 0.22 | 0.08 | 0.05 |
| 4 | 122197904 | rs1986022 | 0.743 | -1.999 | 0.018 | -0.579 | 0.018 | -0.04 | -0.04 | 0 | 0.11 | 0.03 | 0.12 | 0.22 | 0.08 | 0.05 |
| 4 | 122201508 | rs7664875 | -0.022 | -2.181 | 0.335 | -0.699 | 0.019 | -0.022 | -0.022 | 0.97 | 0.07 | 0.15 | 0.05 | 0.03 | 0.08 | 0.07 |
| 4 | 122203163 | rs17051336 | 0.66 | -2.128 | -0.066 | -0.633 | 0.02 | -0.046 | -0.046 | 0 | 0.12 | 0.02 | 0.12 | 0.22 | 0.08 | 0.06 |
| 4 | 122205406 | rs12647412 | -1.668 | -1.918 | 0.165 | -0.595 | 0.047 | -0.006 | -0.006 | 0.014 | 0.48 | 0.63 | 0.42 | 0.42 | 0.41 | 0.33 |
| 4 | 122219233 | rs12642727 | -0.942 | -1.923 | 0.089 | -0.611 | 0.009 | 0.008 | 0.008 | 0 | 0.42 | 0.57 | 0.36 | 0.29 | 0.29 | 0.27 |
| 4 | 122222233 | rs4642269 | -0.779 | -1.952 | 0.176 | -0.613 | -0.02 | -0.01 | -0.01 | 0 | 0.39 | 0.55 | 0.32 | 0.27 | 0.23 | 0.23 |
| 4 | 122226998 | rs13119531 | -0.179 | -1.628 | -0.571 | -0.383 | -0.034 | -0.037 | -0.037 | 0 | 0.09 | 0.03 | 0.1 | 0.16 | 0.18 | 0.1 |
| 4 | 122231301 | rs12711093 | -1.431 | -1.106 | -0.566 | -0.622 | -0.023 | -0.071 | -0.071 | 0 | 0.58 | 0.85 | 0.49 | 0.44 | 0.48 | 0.41 |
| 4 | 122235791 | rs10222821 | -1.19 | -0.57 | 0.122 | -0.814 | 0.01 | -0.063 | -0.063 | 0.859 | 0.24 | 0.35 | 0.15 | 0.14 | 0.27 | 0.22 |
| 4 | 122241609 | rs4642270 | 0.852 | -0.471 | -1.232 | -1.374 | -0.017 | 0.098 | 0.098 | 0 | 0.36 | 0.19 | 0.29 | 0.14 | 0.36 | 0.2 |
| 4 | 122244597 | rs12639850 | -0.096 | 0.632 | -0.996 | -1.222 | -0.012 | 0.023 | 0.023 | 0 | 0.36 | 0.27 | 0.55 | 0.75 | 0.43 | 0.59 |
| 4 | 122249743 | rs11721375 | -0.09 | 0.641 | -0.991 | -1.217 | -0.013 | 0.023 | 0.023 | 0 | 0.36 | 0.27 | 0.55 | 0.75 | 0.43 | 0.59 |

| | | | | | | | | | | | | | | | | |
|---|---|---|---|---|---|---|---|---|---|---|---|---|---|---|---|---|
| 4 | 122250734 | rs2302310 | 1.313 | 0.786 | -1.019 | -0.584 | -0.001 | 0.025 | 0.025 | 0.602 | 0.17 | 0.16 | 0.41 | 0.54 | 0.17 | 0.41 |
| 4 | 122257538 | rs4273489 | 1.923 | 0.716 | -1.071 | -0.697 | -0.018 | 0.083 | 0.083 | 0 | 0.36 | 0.18 | 0.29 | 0.16 | 0.35 | 0.2 |
| 4 | 122258063 | rs4490481 | -0.405 | 0.651 | -1.139 | -0.766 | -0.036 | -0.019 | -0.019 | 0 | 0.29 | 0.18 | 0.56 | 0.73 | 0.47 | 0.58 |
| 4 | 122258950 | rs35623795 | 1.336 | 0.755 | -1.08 | -0.725 | -0.004 | 0.003 | 0.003 | 0 | 0.1 | 0.08 | 0.11 | 0.06 | 0.22 | 0.08 |
| 4 | 122268413 | rs13110738 | -0.062 | 0.494 | -0.968 | -0.814 | -0.055 | -0.045 | -0.045 | 0.419 | 0.69 | 0.44 | 0.88 | 0.9 | 0.83 | 0.82 |
| 4 | 122275299 | rs11733055 | 1.668 | 0.935 | -1.107 | -0.574 | -0.022 | 0.086 | 0.086 | 0 | 0.36 | 0.18 | 0.29 | 0.16 | 0.35 | 0.2 |
| 4 | 122281107 | rs13114721 | 1.236 | 1.511 | -0.448 | 0.443 | 0.002 | 0.084 | 0.084 | 0.214 | 0.08 | 0.08 | 0.34 | 0.52 | 0.14 | 0.41 |
| 4 | 122291131 | rs2013534 | 0.999 | 1.445 | -0.115 | -0.282 | 0.009 | -0.03 | -0.03 | 0 | 0.64 | 0.65 | 0.45 | 0.17 | 0.52 | 0.48 |
| 4 | 122296807 | rs691712 | -0.924 | 1.457 | -0.106 | -0.155 | 0 | 0 | 0 | 0 | 0.28 | 0.26 | 0.2 | 0.18 | 0.33 | 0.12 |
| 4 | 122300413 | rs12642360 | 1.161 | 1.314 | 0.116 | -0.308 | -0.006 | 0.015 | 0.015 | 0.68 | 0.71 | 0.42 | 0.57 | 0.35 | 0.75 | 0.55 |
| 4 | 122301622 | rs17438900 | 0.83 | 1.306 | 0.042 | -0.416 | -0.047 | -0.018 | -0.018 | 0 | 0.09 | 0.02 | 0.15 | 0.07 | 0.23 | 0.08 |
| 4 | 122308022 | rs692496 | -1.162 | 0.812 | 0.239 | -0.414 | -0.066 | -0.033 | -0.033 | 0 | 0.25 | 0.45 | 0.08 | 0.11 | 0.13 | 0.09 |
| 4 | 122309865 | rs535465 | -0.62 | 0.875 | 0.251 | -0.56 | -0.056 | -0.149 | -0.149 | 0 | 0.2 | 0.44 | 0.07 | 0 | 0.11 | 0.06 |
| 4 | 122310963 | rs691684 | 0.619 | 0.987 | 0.048 | -0.479 | -0.016 | -0.06 | -0.06 | 0.009 | 0.13 | 0.15 | 0.04 | 0 | 0.09 | 0.01 |
| 4 | 122315742 | rs6857701 | 0.305 | 1.279 | -0.287 | -0.661 | -0.001 | 0.014 | 0.014 | 3.022 | 0.19 | 0.09 | 0.08 | 0 | 0.14 | 0.02 |
| 4 | 122315993 | rs17439459 | 1.153 | 1.558 | -0.289 | -0.554 | -0.055 | -0.03 | -0.03 | 0.006 | 0.08 | 0.01 | 0.15 | 0.07 | 0.22 | 0.07 |
| 4 | 122324112 | rs4425401 | 0.598 | 1.454 | -0.265 | -0.567 | -0.007 | 0.028 | 0.028 | 0 | 0.64 | 0.76 | 0.54 | 0.22 | 0.65 | 0.54 |
| 4 | 122343345 | rs9884280 | -0.6 | 1.284 | -0.146 | -0.408 | 0.022 | 0.034 | 0.034 | 0.648 | 0.22 | 0.39 | 0.19 | 0.19 | 0.27 | 0.14 |
| 4 | 122358044 | rs691345 | -1.177 | 0.88 | -0.309 | -0.415 | 0.008 | 0.072 | 0.072 | 0 | 0.48 | 0.36 | 0.6 | 0.84 | 0.53 | 0.56 |
| 4 | 122361299 | rs10010505 | 0.564 | 0.795 | -0.311 | 0.043 | -0.006 | -0.026 | -0.026 | 0 | 0.06 | 0.08 | 0.03 | 0 | 0.06 | 0.03 |
| 4 | 122361381 | rs10010429 | -0.799 | 0.707 | -0.614 | -0.087 | 0.006 | 0.018 | 0.018 | 0 | 0.74 | 0.78 | 0.84 | 0.93 | 0.78 | 0.68 |
| 4 | 122361659 | rs17051353 | -0.136 | 1.21 | -0.275 | 1.136 | -0.002 | 0.069 | 0.069 | 0.165 | 0.09 | 0.1 | 0.26 | 0.56 | 0.12 | 0.32 |
| 4 | 122369141 | rs12647248 | -0.266 | 0.798 | 0.129 | 0.885 | -0.028 | 0.095 | 0.095 | 0 | 0.14 | 0.25 | 0.04 | 0.22 | 0.07 | 0.09 |
| 4 | 122369974 | rs1022251 | 0.15 | 0.793 | -0.078 | 0.883 | 0.005 | -0.079 | -0.079 | 0 | 0.12 | 0.01 | 0.04 | 0.22 | 0.06 | 0.08 |
| 4 | 122370550 | rs577737 | -0.181 | 1.796 | -0.512 | 0.619 | -0.077 | 0.007 | 0.007 | 0 | 0.13 | 0.04 | 0.18 | 0.05 | 0.31 | 0.08 |
| 4 | 122370729 | rs692380 | -1.542 | 1.63 | -0.067 | 0.416 | 0.032 | 0.022 | 0.022 | 0 | 0.32 | 0.48 | 0.26 | 0.27 | 0.43 | 0.19 |
| 4 | 122377715 | rs12510951 | -1.305 | 1.158 | -0.478 | -0.15 | 0.044 | 0.07 | 0.07 | 0 | 0.39 | 0.77 | 0.46 | 0.73 | 0.29 | 0.45 |
| 4 | 122392108 | rs1513702 | -0.632 | 1.709 | 0.013 | -0.663 | 0.035 | -0.039 | -0.039 | 0 | 0.17 | 0.23 | 0.18 | 0.04 | 0.38 | 0.08 |
| 4 | 122394649 | rs10031780 | 0.026 | 1.466 | 0.475 | -0.498 | -0.032 | 0.119 | 0.119 | 0.509 | 0.19 | 0.35 | 0.29 | 0.57 | 0.15 | 0.32 |
| 4 | 122395119 | rs13137076 | -2.251 | 1.431 | -0.161 | -0.563 | 0.03 | 0.222 | 0.222 | 0 | 0.5 | 0.84 | 0.52 | 0.75 | 0.59 | 0.47 |
| 4 | 122401640 | rs6842940 | -0.832 | 1.97 | -0.088 | -0.321 | -0.027 | 0.017 | 0.017 | 0 | 0.17 | 0.16 | 0.18 | 0.04 | 0.38 | 0.08 |
| 4 | 122421393 | rs1513707 | -1.366 | 1.872 | -0.113 | -0.299 | -0.074 | 0.026 | 0.026 | 0 | 0.29 | 0.18 | 0.21 | 0.18 | 0.44 | 0.15 |
| 4 | 122423129 | rs13124630 | -0.39 | 2.113 | -0.019 | -0.302 | -0.043 | 0.03 | 0.03 | 0 | 0.14 | 0.09 | 0.14 | 0 | 0.34 | 0.08 |
| 4 | 122426180 | rs10015843 | 0.972 | 2.021 | 0.591 | -0.338 | 0.061 | -0.021 | -0.021 | 0 | 0.74 | 0.51 | 0.8 | 0.83 | 0.65 | 0.84 |
| 4 | 122426239 | rs17382098 | -0.717 | 2.194 | 0.662 | -0.394 | 0.036 | -0.047 | -0.047 | 0 | 0.16 | 0.19 | 0.18 | 0.02 | 0.34 | 0.09 |
| 4 | 122426864 | rs9993990 | 0.44 | 1.285 | 0.312 | -0.701 | 0.031 | 0.08 | 0.08 | 0 | 0.56 | 0.37 | 0.56 | 0.27 | 0.67 | 0.59 |
| 4 | 122429378 | rs4833736 | -0.236 | 1.23 | 0.397 | -0.66 | -0.004 | 0.011 | 0.011 | 0 | 0.15 | 0.2 | 0.09 | 0.19 | 0.1 | 0.11 |
| 4 | 122431338 | rs7698292 | -0.489 | 0.974 | -0.112 | -1.093 | 0.002 | -0.004 | -0.004 | 0 | 0.27 | 0.22 | 0.29 | 0.26 | 0.24 | 0.28 |
| 4 | 122431579 | rs883029 | 1.173 | 0.992 | 0.04 | -1.139 | -0.012 | 0.032 | 0.032 | 0 | 0.28 | 0.29 | 0.34 | 0.54 | 0.23 | 0.3 |
| 4 | 122436261 | rs6828751 | -0.771 | 0.997 | 0.024 | -1.144 | -0.003 | 0.015 | 0.015 | 0 | 0.55 | 0.53 | 0.63 | 0.81 | 0.46 | 0.58 |
| 4 | 122437789 | rs17051364 | 0.231 | 0.994 | -0.015 | -1.144 | 0.011 | -0.04 | -0.04 | 1.073 | 0.22 | 0.08 | 0.24 | 0.25 | 0.19 | 0.26 |
| 4 | 122438484 | rs1016802 | -0.335 | 0.999 | -0.02 | -1.144 | 0.008 | 0.005 | 0.005 | 0 | 0.5 | 0.41 | 0.58 | 0.79 | 0.42 | 0.56 |
| 4 | 122440829 | rs12648087 | 0.084 | 0.805 | -0.619 | -0.847 | 0.011 | -0.042 | -0.042 | 0 | 0.22 | 0.08 | 0.23 | 0.24 | 0.19 | 0.26 |
| 4 | 122446250 | rs4833741 | -0.027 | 0.96 | -0.32 | -0.861 | -0.008 | -0.024 | -0.024 | 0 | 0.6 | 0.61 | 0.58 | 0.28 | 0.69 | 0.62 |
| 4 | 122448971 | rs12501209 | 1.198 | 0.979 | -0.987 | -1.093 | -0.027 | 0.023 | 0.023 | 0 | 0.1 | 0.01 | 0.16 | 0.04 | 0.12 | 0.05 |
| 4 | 122449393 | rs4833218 | 1.436 | 0.752 | -0.796 | -0.821 | -0.003 | -0.052 | -0.052 | 0 | 0.09 | 0.01 | 0.03 | 0.15 | 0.06 | 0.07 |
| 4 | 122449490 | rs4833219 | -0.393 | 0.984 | -0.709 | -0.92 | -0.005 | 0.005 | 0.005 | 0 | 0.35 | 0.37 | 0.33 | 0.19 | 0.44 | 0.23 |
| 4 | 122450475 | rs6840582 | 0.888 | 0.975 | -0.871 | -0.925 | 0.005 | 0.012 | 0.012 | 0 | 0.35 | 0.31 | 0.29 | 0.25 | 0.3 | 0.46 |
| 4 | 122450852 | rs7683438 | -0.032 | 1.355 | -0.377 | -0.75 | 0.035 | -0.091 | -0.091 | 0 | 0.27 | 0.41 | 0.3 | 0.04 | 0.39 | 0.16 |
| 4 | 122451829 | rs9994817 | 0.128 | 0.954 | -0.463 | 0.208 | -0.001 | 0.001 | 0.001 | 0.528 | 0.7 | 0.73 | 0.62 | 0.39 | 0.74 | 0.68 |
| 4 | 122463069 | rs4370152 | -0.569 | 1.152 | -0.312 | 0.429 | -0.004 | -0.006 | -0.006 | 0 | 0.44 | 0.49 | 0.44 | 0.63 | 0.41 | 0.5 |
| 4 | 122473563 | rs12504183 | -0.36 | 1.354 | -0.14 | 0.015 | 0.004 | -0.053 | -0.053 | 0 | 0.12 | 0.15 | 0.09 | 0.01 | 0.12 | 0.14 |
| 4 | 122475832 | rs982887 | 0.794 | 1.39 | -0.162 | 0.034 | -0.007 | 0.011 | 0.011 | 0 | 0.44 | 0.32 | 0.47 | 0.36 | 0.47 | 0.37 |
| 4 | 122480241 | rs4565102 | 0.652 | 1.226 | 0.501 | 0.002 | 0.016 | -0.091 | -0.091 | 1.406 | 0.86 | 0.66 | 0.9 | 0.99 | 0.88 | 0.87 |
| 4 | 122482578 | rs1480296 | 1.045 | 1.17 | 0.548 | -0.074 | -0.027 | 0.009 | 0.009 | 0 | 0.42 | 0.23 | 0.48 | 0.36 | 0.49 | 0.36 |
| 4 | 122487647 | rs6534295 | 0.743 | 1.097 | 0.314 | 0.261 | 0.005 | -0.099 | -0.099 | 0 | 0.26 | 0.04 | 0.15 | 0.41 | 0.19 | 0.29 |
| 4 | 122494838 | rs1159702 | 0.585 | 1.098 | 0.489 | 0.285 | 0 | 0.011 | 0.011 | 0 | 0.08 | 0.08 | 0.18 | 0.22 | 0.11 | 0.18 |
| 4 | 122499665 | rs11731548 | 1.162 | 1.044 | 0.481 | 0.23 | 0.028 | 0.037 | 0.037 | 0 | 0.5 | 0.31 | 0.67 | 0.57 | 0.6 | 0.54 |
| 4 | 122502810 | rs7690087 | 0.77 | 0.983 | 0.63 | 0.233 | 0.016 | 0.007 | 0.007 | 0 | 0.55 | 0.38 | 0.72 | 0.57 | 0.69 | 0.57 |
| 4 | 122508654 | rs6534300 | 0.99 | 1.127 | 0.652 | 0.149 | 0.038 | -0.106 | -0.106 | 0 | 0.86 | 0.52 | 0.89 | 1 | 0.88 | 0.91 |
| 4 | 122510701 | rs1480276 | 0.592 | 1.437 | 0.342 | 0.232 | -0.037 | -0.042 | -0.042 | 0 | 0.08 | 0.02 | 0.19 | 0.21 | 0.13 | 0.18 |
| 4 | 122511492 | rs17051373 | -1.07 | 1.427 | 0.771 | 0.152 | 0.036 | -0.104 | -0.104 | 0 | 0.14 | 0.39 | 0.11 | 0 | 0.13 | 0.1 |
| 4 | 122511622 | rs10022795 | 0.793 | 1.381 | 0.718 | 0.187 | 0.024 | -0.069 | -0.069 | 0.082 | 0.27 | 0.1 | 0.15 | 0.42 | 0.19 | 0.29 |
| 4 | 122512853 | rs10518393 | -0.719 | 1.389 | 0.775 | 0.226 | 0.038 | -0.073 | -0.073 | 0 | 0.13 | 0.31 | 0.1 | 0.02 | 0.13 | 0.1 |
| 4 | 122515534 | rs13114712 | 0.675 | 1.432 | 0.741 | 0.201 | 0.006 | 0.019 | 0.019 | 0 | 0.54 | 0.59 | 0.57 | 0.33 | 0.56 | 0.46 |
| 4 | 122520868 | rs6534302 | -0.437 | 1.28 | 0.55 | 0.333 | -0.021 | -0.044 | -0.044 | 0 | 0.33 | 0.42 | 0.19 | 0.26 | 0.24 | 0.29 |
| 4 | 122525783 | rs9685971 | -1.367 | 1.345 | 0.687 | 0.373 | 0.018 | -0.004 | -0.004 | 0 | 0.23 | 0.45 | 0.17 | 0.16 | 0.21 | 0.2 |
| 4 | 122526728 | rs7684836 | 1.364 | 1.33 | 0.655 | 0.386 | 0.025 | -0.029 | -0.029 | 0 | 0.19 | 0.03 | 0.09 | 0.26 | 0.11 | 0.19 |
| 4 | 122529718 | rs4490484 | -0.46 | 1.372 | 0.725 | 0.375 | 0.037 | 0.116 | 0.116 | 0 | 0.91 | 0.72 | 0.8 | 0.77 | 0.88 | 0.83 |

| | | | | | | | | | | | | | | | | |
|---|---|---|---|---|---|---|---|---|---|---|---|---|---|---|---|---|
| 4 | 122531359 | rs3811745 | 2.045 | 1.29 | 0.604 | 0.373 | -0.018 | 0.011 | 0.011 | 0 | 0.4 | 0.2 | 0.46 | 0.33 | 0.43 | 0.36 |
| 4 | 122532114 | rs7654348 | -1.364 | 1.25 | 0.792 | 0.442 | -0.003 | 0 | 0 | 0 | 0.43 | 0.48 | 0.39 | 0.51 | 0.4 | 0.47 |
| 4 | 122532992 | rs6840518 | 1.172 | 1.264 | 0.571 | 0.432 | 0.025 | -0.035 | -0.035 | 0 | 0.19 | 0.03 | 0.09 | 0.28 | 0.12 | 0.22 |
| 4 | 122533936 | rs17442542 | -1.11 | 1.223 | 0.822 | 0.533 | 0.04 | 0.025 | 0.025 | 0 | 0.18 | 0.26 | 0.28 | 0.23 | 0.26 | 0.24 |
| 4 | 122535735 | rs4460015 | -0.875 | 1.249 | 0.805 | 0.59 | 0.066 | 0.064 | 0.064 | 0 | 0.15 | 0.35 | 0.25 | 0.23 | 0.22 | 0.17 |
| 4 | 122544756 | rs7675417 | -1.695 | 1.537 | 0.924 | 0.606 | 0.046 | 0.06 | 0.06 | 0 | 0.28 | 0.49 | 0.37 | 0.39 | 0.37 | 0.32 |
| 4 | 122550874 | rs4833753 | 1.197 | 2.018 | 0.782 | 0.531 | -0.002 | -0.005 | -0.005 | 0.543 | 0.49 | 0.48 | 0.49 | 0.35 | 0.44 | 0.38 |
| 4 | 122551505 | rs4522886 | -1.285 | 2.024 | 0.734 | 0.682 | 0.025 | 0.056 | 0.056 | 0 | 0.23 | 0.3 | 0.32 | 0.38 | 0.28 | 0.31 |
| 4 | 122552373 | rs11735972 | -0.503 | 1.868 | 0.594 | 0.5 | 0.002 | -0.002 | -0.002 | 0.704 | 0.16 | 0.21 | 0.13 | 0.13 | 0.14 | 0.15 |
| 4 | 122555682 | rs4460016 | 1.359 | 1.697 | 0.676 | 0.386 | -0.003 | -0.015 | -0.015 | 0 | 0.94 | 0.67 | 0.96 | 1 | 0.97 | 0.95 |
| 4 | 122557052 | rs11722834 | 0.841 | 1.869 | 0.57 | 0.488 | 0.015 | 0.005 | 0.005 | 0 | 0.6 | 0.5 | 0.59 | 0.52 | 0.51 | 0.52 |
| 4 | 122557607 | rs12499696 | 0.841 | 1.869 | 0.57 | 0.488 | 0.015 | 0.005 | 0.005 | 0 | 0.6 | 0.5 | 0.59 | 0.52 | 0.51 | 0.52 |
| 4 | 122559883 | rs5020375 | -0.942 | 2.01 | -0.017 | 0.556 | -0.018 | -0.009 | -0.009 | 0 | 0.38 | 0.28 | 0.39 | 0.48 | 0.49 | 0.45 |
| 4 | 122565377 | rs2390235 | 0.007 | 1.811 | -0.438 | 0.753 | 0.007 | 0.019 | 0.019 | 0 | 0.08 | 0.1 | 0.19 | 0.21 | 0.13 | 0.18 |
| 4 | 122565698 | rs17051378 | 0.022 | 1.414 | -0.617 | 0.403 | -0.021 | -0.003 | -0.003 | 0 | 0.25 | 0.1 | 0.17 | 0.28 | 0.28 | 0.27 |
| 4 | 122567739 | rs11937669 | -0.402 | 0.943 | -0.202 | -0.292 | -0.002 | -0.012 | -0.012 | 1.442 | 0.42 | 0.35 | 0.21 | 0.24 | 0.33 | 0.32 |
| 4 | 122568322 | rs725882 | 1.609 | 1.538 | 0.639 | -0.2 | -0.001 | 0.004 | 0.004 | 0 | 0.74 | 0.23 | 0.76 | 0.82 | 0.77 | 0.74 |
| 4 | 122571226 | rs10022259 | -0.047 | 1.515 | 1.457 | -0.254 | 0.01 | -0.023 | -0.023 | 0 | 0.05 | 0.17 | 0.02 | 0 | 0.02 | 0.01 |
| 4 | 122572851 | rs10003585 | 0.325 | 1.532 | 1.979 | -0.215 | 0.047 | 0.086 | 0.086 | 0 | 0.91 | 0.74 | 0.87 | 0.82 | 0.9 | 0.78 |
| 4 | 122573054 | rs6823721 | 1.407 | 1.8 | 1.609 | -0.358 | 0 | -0.001 | -0.001 | 0.315 | 0.6 | 0.44 | 0.57 | 0.63 | 0.51 | 0.63 |
| 4 | 122574626 | rs11937561 | -0.657 | 1.785 | 1.237 | -0.35 | -0.003 | 0.001 | 0.001 | 0 | 0.26 | 0.25 | 0.28 | 0.19 | 0.37 | 0.14 |
| 4 | 122580875 | rs17384008 | -0.627 | 1.82 | 1.297 | -0.601 | 0.013 | 0.008 | 0.008 | 0 | 0.13 | 0.17 | 0.16 | 0.16 | 0.17 | 0.17 |
| 4 | 122583666 | rs11098633 | 2.182 | 1.957 | 1.223 | -0.583 | -0.005 | 0.063 | 0.063 | 0 | 0.52 | 0.07 | 0.51 | 0.7 | 0.44 | 0.59 |
| 4 | 122589283 | rs17718 | 0.28 | 1.968 | 1.848 | -0.602 | 0.039 | -0.001 | -0.001 | 0 | 0.05 | 0.24 | 0.02 | 0 | 0.02 | 0.01 |
| 4 | 122591475 | rs2306420 | 0.046 | 1.998 | 1.687 | -0.6 | -0.06 | 0.019 | 0.019 | 0 | 0.19 | 0.04 | 0.24 | 0.1 | 0.31 | 0.13 |
| 4 | 122600471 | rs11098637 | 2.208 | 1.954 | 1.629 | -0.594 | -0.001 | 0.063 | 0.063 | 0 | 0.52 | 0.07 | 0.5 | 0.7 | 0.44 | 0.59 |
| 4 | 122607728 | rs17449954 | 0.201 | 2.035 | 1.654 | -0.647 | 0.009 | 0.008 | 0.008 | 0 | 0.09 | 0.04 | 0.05 | 0 | 0.06 | 0.12 |
| 4 | 122607848 | rs10518391 | -0.064 | 2.002 | 1.811 | -0.578 | -0.001 | -0.001 | -0.001 | 0.34 | 0.11 | 0.11 | 0.13 | 0.17 | 0.11 | 0.16 |
| 4 | 122609929 | rs6534309 | -0.308 | 2.017 | 1.96 | -0.598 | 0.068 | -0.026 | -0.026 | 0 | 0.07 | 0.24 | 0.05 | 0 | 0.07 | 0.02 |
| 4 | 122611285 | rs6857766 | -1.296 | 2.013 | 1.936 | -0.595 | 0.032 | 0.002 | 0.002 | 0 | 0.18 | 0.35 | 0.18 | 0.17 | 0.18 | 0.18 |
| 4 | 122614755 | rs6830321 | 2.154 | 2.02 | 1.121 | -0.591 | -0.018 | 0.07 | 0.07 | 0 | 0.55 | 0.16 | 0.52 | 0.73 | 0.48 | 0.6 |
| 4 | 122616166 | rs13145977 | -0.246 | 1.912 | 0.865 | -0.326 | -0.042 | 0.023 | 0.023 | 0 | 0.2 | 0.06 | 0.25 | 0.1 | 0.31 | 0.13 |
| 4 | 122617688 | rs2306416 | -0.711 | 1.992 | 0.931 | -0.267 | 0.018 | 0.047 | 0.047 | 0 | 0.13 | 0.27 | 0.15 | 0.19 | 0.14 | 0.17 |
| 4 | 122624065 | rs7680787 | -0.804 | 1.779 | 0.453 | -0.493 | 0.011 | -0.026 | -0.026 | 0 | 0.28 | 0.37 | 0.32 | 0.11 | 0.38 | 0.27 |
| 4 | 122628680 | rs9997310 | -0.408 | 1.72 | 0.697 | -0.393 | 0.054 | 0.008 | 0.008 | 0 | 0.06 | 0.21 | 0.06 | 0.04 | 0.1 | 0.01 |
| 4 | 122633535 | rs4833757 | 0.547 | 1.246 | 0.243 | -0.449 | 0.005 | -0.027 | -0.027 | 0 | 0.25 | 0.19 | 0.48 | 0.55 | 0.22 | 0.23 |
| 4 | 122639133 | rs1383228 | -0.11 | 1.174 | 0.737 | -0.723 | -0.001 | 0 | 0 | 0 | 0.88 | 0.81 | 0.93 | 0.89 | 0.92 | 0.86 |
| 4 | 122644567 | rs2892966 | 0.342 | 1.199 | 0.896 | -1.002 | 0 | 0.007 | 0.007 | 0 | 0.57 | 0.55 | 0.69 | 0.82 | 0.56 | 0.59 |
| 4 | 122647904 | rs1031923 | -1.42 | 1.176 | 0.965 | -0.978 | 0.123 | 0.083 | 0.083 | 4.267 | 0.09 | 0.41 | 0.08 | 0.09 | 0.1 | 0.08 |
| 4 | 122653098 | rs17051396 | -1.541 | 1.159 | 0.724 | -1.016 | 0.112 | 0.062 | 0.062 | 4.55 | 0.09 | 0.47 | 0.08 | 0.09 | 0.1 | 0.09 |
| 4 | 122653300 | rs11732306 | 0.735 | 1.214 | 0.363 | -0.969 | 0.021 | 0.113 | 0.113 | 0 | 0.5 | 0.27 | 0.62 | 0.74 | 0.46 | 0.5 |
| 4 | 122656144 | rs6838346 | 0.535 | 0.885 | 0.108 | -0.823 | 0 | -0.004 | -0.004 | 0 | 0.53 | 0.45 | 0.66 | 0.69 | 0.48 | 0.5 |
| 4 | 122656714 | rs2390227 | 0.325 | 0.903 | -0.308 | -0.924 | -0.01 | 0.048 | 0.048 | 0 | 0.61 | 0.63 | 0.71 | 0.79 | 0.57 | 0.66 |
| 4 | 122657875 | rs4833765 | 0.567 | 0.572 | 0.021 | -1.043 | -0.006 | -0.088 | -0.088 | 0 | 0.64 | 0.5 | 0.79 | 0.87 | 0.69 | 0.59 |
| 4 | 122659257 | rs17051398 | -0.717 | 0.157 | 0.516 | -0.581 | -0.023 | -0.016 | -0.016 | 0 | 0.11 | 0.28 | 0.07 | 0.05 | 0.06 | 0.15 |
| 4 | 122659350 | rs2706783 | -0.332 | 0.296 | 0.225 | -0.927 | -0.013 | 0.036 | 0.036 | 0 | 0.16 | 0.06 | 0.12 | 0.08 | 0.19 | 0.11 |
| 4 | 122659704 | rs2706784 | 1.154 | -0.058 | 0.887 | -0.454 | -0.058 | 0.038 | 0.038 | 1.571 | 0.82 | 0.34 | 0.87 | 0.77 | 0.93 | 0.77 |
| 4 | 122664323 | rs28532673 | -0.858 | 0.294 | 0.214 | -0.272 | -0.006 | 0.016 | 0.016 | 0 | 0.43 | 0.71 | 0.31 | 0.39 | 0.43 | 0.52 |
| 4 | 122669747 | rs4629472 | -0.441 | 0.305 | -0.013 | -0.25 | -0.004 | -0.018 | -0.018 | 0 | 0.84 | 0.87 | 0.87 | 0.7 | 0.83 | 0.85 |
| 4 | 122677514 | rs11934307 | 1.258 | 0.272 | -0.138 | -0.275 | 0.012 | -0.034 | -0.034 | 0 | 0.42 | 0.19 | 0.57 | 0.5 | 0.39 | 0.37 |
| 4 | 122680075 | rs11937861 | -0.144 | 0.337 | 0.128 | -0.381 | -0.018 | -0.049 | -0.049 | 0 | 0.09 | 0.23 | 0.07 | 0 | 0.06 | 0.12 |
| 4 | 122683007 | rs4370153 | -0.118 | 0.569 | -0.174 | -0.272 | -0.01 | 0.012 | 0.012 | 0 | 0.85 | 0.93 | 0.89 | 0.88 | 0.83 | 0.89 |
| 4 | 122688744 | rs4833233 | 0.736 | 0.387 | 0.11 | -0.232 | 0.002 | -0.009 | -0.009 | 0 | 0.56 | 0.59 | 0.58 | 0.42 | 0.46 | 0.49 |
| 4 | 122692023 | rs6534321 | -0.278 | 0.365 | 0.018 | -0.05 | -0.005 | -0.03 | -0.03 | 0.002 | 0.84 | 0.87 | 0.9 | 0.64 | 0.83 | 0.86 |
| 4 | 122702459 | rs4519820 | -0.445 | 0.408 | -0.184 | -0.185 | -0.043 | -0.026 | -0.026 | 0.173 | 0.86 | 0.96 | 0.9 | 0.86 | 0.83 | 0.9 |
| 4 | 122720999 | rs6838153 | 0.137 | 0.345 | -0.204 | -0.251 | -0.086 | 0.03 | 0.03 | 0 | 0.26 | 0.1 | 0.3 | 0.2 | 0.34 | 0.36 |
| 4 | 122722368 | rs3828485 | 0.067 | 0.32 | -0.217 | -0.255 | -0.085 | 0.03 | 0.03 | 0 | 0.27 | 0.1 | 0.3 | 0.2 | 0.34 | 0.37 |
| 4 | 122726073 | rs6836544 | -1.109 | 0.068 | 0.617 | -0.379 | 0.03 | 0.026 | 0.026 | 2.166 | 0.07 | 0.43 | 0.06 | 0.02 | 0.01 | 0.01 |
| 4 | 122735142 | rs1803183 | -0.367 | 0.062 | 0.813 | -0.381 | 0.033 | 0.035 | 0.035 | 0.059 | 0.05 | 0.26 | 0.05 | 0.02 | 0.01 | 0.01 |
| 4 | 122740162 | rs2071486 | -0.743 | 0.119 | 0.978 | -0.443 | 0 | -0.005 | -0.005 | 0 | 0.34 | 0.62 | 0.37 | 0.22 | 0.35 | 0.37 |
| 4 | 122741807 | rs769243 | -1.243 | 0.12 | 1.024 | -0.441 | 0.028 | 0.024 | 0.024 | 2.166 | 0.07 | 0.42 | 0.06 | 0.02 | 0.01 | 0.01 |
| 4 | 122748996 | rs1048433 | 1.972 | -0.362 | 0.614 | -0.729 | -0.081 | -0.177 | -0.177 | 0.046 | 0.34 | 0.02 | 0.45 | 0.45 | 0.36 | 0.28 |
| 4 | 122749541 | rs1507994 | -0.168 | -0.348 | 0.623 | -0.727 | 0.02 | -0.049 | -0.049 | 1.028 | 0.33 | 0.53 | 0.36 | 0.22 | 0.35 | 0.37 |
| 4 | 122750079 | rs13145213 | 1.862 | -0.311 | -0.014 | -0.709 | -0.007 | -0.065 | -0.065 | 0 | 0.36 | 0.12 | 0.44 | 0.43 | 0.36 | 0.27 |
| 4 | 122774917 | rs12499222 | 0.461 | -0.231 | -0.091 | -0.484 | -0.021 | -0.05 | -0.05 | 0 | 0.72 | 0.91 | 0.81 | 0.65 | 0.71 | 0.7 |
| 4 | 122781263 | rs6838198 | 0.449 | -0.205 | -0.078 | -0.652 | -0.008 | 0.073 | 0.073 | 0 | 0.72 | 0.88 | 0.82 | 0.86 | 0.71 | 0.74 |
| 4 | 122806228 | rs11726196 | 0.914 | -0.291 | 0.271 | -0.886 | -0.011 | -0.039 | -0.039 | 0 | 0.6 | 0.47 | 0.72 | 0.82 | 0.67 | 0.51 |
| 4 | 122813459 | rs12502635 | 0.955 | -0.094 | -0.027 | -0.93 | -0.08 | -0.017 | -0.017 | 0 | 0.18 | 0.05 | 0.27 | 0.18 | 0.34 | 0.2 |
| 4 | 122816181 | rs2135976 | 0.992 | -0.146 | 0.062 | -0.956 | 0.003 | 0.01 | 0.01 | 0 | 0.09 | 0.05 | 0.07 | 0.03 | 0.05 | 0.17 |

| | | | | | | | | | | | | | | | | |
|---|---|---|---|---|---|---|---|---|---|---|---|---|---|---|---|---|
| 4 | 122824052 | rs11732666 | 2.153 | -0.264 | 0.054 | -0.874 | -0.025 | -0.092 | -0.092 | 0.282 | 0.34 | 0.1 | 0.4 | 0.41 | 0.31 | 0.27 |
| 4 | 122824591 | rs17517624 | -0.206 | -0.352 | 0.342 | -0.897 | 0 | 0.055 | 0.055 | 0 | 0.28 | 0.13 | 0.18 | 0.13 | 0.28 | 0.28 |
| 4 | 122825364 | rs3762839 | -0.713 | -0.353 | 0.054 | -0.901 | -0.025 | 0.023 | 0.023 | 0 | 0.16 | 0.33 | 0.11 | 0.24 | 0.14 | 0.09 |
| 4 | 122830406 | rs884701 | 1.765 | -0.215 | 0.152 | -0.736 | 0 | -0.062 | -0.062 | 0 | 0.35 | 0.16 | 0.4 | 0.41 | 0.31 | 0.26 |
| 4 | 122830800 | rs13127488 | 2.042 | -0.232 | -0.125 | -0.744 | -0.079 | -0.165 | -0.165 | 0.225 | 0.33 | 0.02 | 0.39 | 0.41 | 0.31 | 0.26 |
| 4 | 122833314 | rs906496 | 1.507 | -0.204 | -0.005 | -0.638 | 0.007 | -0.013 | -0.013 | 2.836 | 0.37 | 0.32 | 0.33 | 0.39 | 0.31 | 0.26 |
| 4 | 122842267 | rs4292355 | 0.782 | -0.297 | 0.303 | -0.606 | 0.002 | 0.003 | 0.003 | 0 | 0.62 | 0.64 | 0.65 | 0.81 | 0.66 | 0.52 |
| 4 | 122857568 | rs6841843 | 0.295 | -0.134 | 0.296 | -0.536 | -0.001 | -0.012 | -0.012 | 0 | 0.72 | 0.69 | 0.72 | 0.85 | 0.7 | 0.68 |
| 4 | 122861426 | rs4833779 | 0.753 | -0.029 | 0.022 | -0.669 | 0 | 0.002 | 0.002 | 0 | 0.52 | 0.57 | 0.6 | 0.59 | 0.55 | 0.44 |
| 4 | 122864241 | rs950574 | 1.019 | 0.107 | -0.46 | -0.547 | -0.002 | 0.039 | 0.039 | 0 | 0.46 | 0.24 | 0.55 | 0.59 | 0.54 | 0.44 |
| 4 | 122871632 | rs970349 | 0.355 | 0.679 | -0.3 | -0.334 | 0.001 | -0.012 | -0.012 | 0 | 0.76 | 0.64 | 0.7 | 0.82 | 0.74 | 0.7 |
| 4 | 122886240 | rs4001038 | 0.034 | 1.065 | -0.676 | -0.622 | -0.005 | -0.002 | -0.002 | 0 | 0.79 | 0.81 | 0.85 | 0.98 | 0.7 | 0.73 |
| 4 | 122896734 | rs1358229 | 0.424 | 0.964 | -1.351 | -0.19 | 0.018 | 0.032 | 0.032 | 0 | 0.45 | 0.21 | 0.39 | 0.37 | 0.46 | 0.33 |
| 4 | 122899165 | rs6838639 | 0.069 | 1.054 | -0.862 | 0.054 | -0.007 | -0.017 | -0.017 | 0 | 0.26 | 0.17 | 0.43 | 0.34 | 0.24 | 0.34 |
| 4 | 122900529 | rs13109219 | -0.719 | 0.763 | -0.41 | -0.204 | 0.029 | -0.093 | -0.093 | 0.114 | 0.14 | 0.35 | 0.13 | 0.01 | 0.19 | 0.07 |
| 4 | 122902084 | rs9999992 | -0.384 | 0.149 | -1.215 | 0.246 | 0.016 | -0.001 | -0.001 | 0 | 0.17 | 0.1 | 0.11 | 0.16 | 0.13 | 0.21 |
| 4 | 122902923 | rs6817255 | 0.049 | -0.575 | -2.271 | 0.274 | 0.041 | -0.195 | -0.195 | 0 | 0.3 | 0.01 | 0.37 | 0.38 | 0.2 | 0.29 |
| 4 | 122903090 | rs11098653 | -0.035 | 0.054 | -2.103 | -0.22 | -0.02 | 0.019 | 0.019 | 0.129 | 0.35 | 0.69 | 0.45 | 0.28 | 0.5 | 0.39 |
| 4 | 122903206 | rs13130390 | 0.408 | 1.091 | -1.65 | -0.257 | -0.023 | 0.011 | 0.011 | 1.864 | 0.17 | 0.13 | 0.32 | 0.18 | 0.37 | 0.26 |
| 4 | 122909415 | rs11098654 | -0.506 | 0.789 | -1.584 | 0.08 | 0.053 | 0.111 | 0.111 | 0 | 0.95 | 0.84 | 0.94 | 0.79 | 0.92 | 0.95 |
| 4 | 122924236 | rs17596104 | 1.162 | 0.55 | -2.037 | -0.187 | -0.028 | 0.03 | 0.03 | 0 | 0.21 | 0.01 | 0.09 | 0.12 | 0.18 | 0.21 |
| 4 | 122933691 | rs4833241 | 0.776 | 1.174 | -1.458 | -0.026 | -0.011 | -0.005 | -0.005 | 0 | 0.6 | 0.26 | 0.5 | 0.5 | 0.42 | 0.48 |
| 4 | 122947939 | rs1965582 | 0.521 | 1.365 | -0.38 | 0.379 | 0.043 | 0.101 | 0.101 | 0.338 | 0.95 | 0.79 | 0.95 | 0.82 | 0.95 | 0.9 |
| 4 | 122949052 | rs10004294 | -0.816 | 1.593 | 0.011 | -0.017 | 0.042 | -0.023 | -0.023 | 0.329 | 0.28 | 0.47 | 0.34 | 0.24 | 0.4 | 0.28 |
| 4 | 122949327 | rs10016562 | -0.172 | 1.688 | 0.563 | 0.019 | 0.006 | -0.001 | -0.001 | 0 | 0.2 | 0.26 | 0.31 | 0.24 | 0.38 | 0.27 |
| 4 | 122959398 | rs4833798 | 1.262 | 1.81 | 0.423 | -0.157 | 0.116 | 0.083 | 0.083 | 0.93 | 0.86 | 0.49 | 0.84 | 0.77 | 0.82 | 0.85 |
| 4 | 122962378 | rs6850993 | 1.282 | 1.807 | 0.427 | -0.158 | 0.119 | 0.085 | 0.085 | 0.966 | 0.86 | 0.5 | 0.84 | 0.76 | 0.82 | 0.85 |
| 4 | 122971505 | rs13114230 | -0.687 | 1.503 | 0.597 | -0.403 | 0.058 | -0.066 | -0.066 | 0.937 | 0.25 | 0.39 | 0.33 | 0.12 | 0.39 | 0.26 |
| 4 | 122974607 | rs10516615 | 0.49 | 2.088 | 0.268 | -0.506 | -0.016 | 0.047 | 0.047 | 0.525 | 0.11 | 0.03 | 0.06 | 0 | 0.13 | 0.12 |
| 4 | 122974644 | rs10516616 | -0.097 | 2.382 | 0.325 | -0.489 | 0.021 | -0.007 | -0.007 | 0 | 0.07 | 0.11 | 0.09 | 0.03 | 0.1 | 0.04 |
| 4 | 122978844 | rs6840372 | 1.744 | 2.25 | 0.563 | -0.515 | -0.008 | 0.017 | 0.017 | 0 | 0.61 | 0.36 | 0.64 | 0.67 | 0.45 | 0.63 |
| 4 | 122982259 | rs7661311 | -0.147 | 2.069 | 0.195 | -0.394 | 0.017 | -0.006 | -0.006 | 0 | 0.06 | 0.11 | 0.08 | 0.03 | 0.1 | 0.04 |
| 4 | 122987188 | rs1880864 | 2.129 | 2.34 | -0.533 | -0.734 | 0.085 | 0.06 | 0.06 | 0.228 | 0.52 | 0.05 | 0.55 | 0.6 | 0.34 | 0.6 |
| 4 | 122989473 | rs13116766 | -0.7 | 2.714 | 0.06 | -0.806 | 0.056 | -0.111 | -0.111 | 0 | 0.2 | 0.49 | 0.18 | 0.05 | 0.27 | 0.11 |
| 4 | 122992499 | rs4833808 | 0.931 | 2.407 | -0.965 | -0.733 | -0.024 | -0.114 | -0.114 | 1.193 | 0.16 | 0.03 | 0.43 | 0.47 | 0.13 | 0.2 |
| 4 | 122993170 | rs10009575 | 2.04 | 2.891 | -0.371 | -0.533 | -0.024 | 0.077 | 0.077 | 0 | 0.56 | 0.11 | 0.62 | 0.63 | 0.43 | 0.68 |
| 4 | 122994218 | rs1526926 | 2.12 | 2.877 | -0.388 | -0.537 | -0.022 | 0.082 | 0.082 | 0 | 0.55 | 0.09 | 0.61 | 0.63 | 0.41 | 0.64 |
| 4 | 123001769 | rs941130 | 0.593 | 2.851 | 0.006 | -0.288 | 0.046 | 0.034 | 0.034 | 0.044 | 0.84 | 0.71 | 0.82 | 0.68 | 0.72 | 0.76 |
| 4 | 123009174 | rs4279237 | 0.028 | 2.372 | -0.203 | -0.267 | 0 | -0.001 | -0.001 | 0.18 | 0.11 | 0.09 | 0.13 | 0.12 | 0.24 | 0.13 |
| 4 | 123010587 | rs13132933 | -0.186 | 2.144 | 0.037 | -0.33 | 0.045 | -0.06 | -0.06 | 0.255 | 0.14 | 0.46 | 0.1 | 0.05 | 0.16 | 0.1 |
| 4 | 123016755 | rs11936230 | 0.572 | 2.43 | -0.683 | -0.02 | -0.017 | -0.013 | -0.013 | 0 | 0.83 | 0.91 | 0.76 | 0.68 | 0.61 | 0.76 |
| 4 | 123018390 | rs4833810 | 1.65 | 2.119 | -1.048 | -0.17 | 0.012 | 0.032 | 0.032 | 0.01 | 0.54 | 0.05 | 0.59 | 0.59 | 0.36 | 0.61 |
| 4 | 123021560 | rs3108396 | 0.937 | 2.485 | -0.217 | 0.056 | 0.089 | 0.033 | 0.033 | 0 | 0.82 | 0.73 | 0.75 | 0.7 | 0.6 | 0.77 |
| 4 | 123026869 | rs6534338 | 0.451 | 2.753 | -0.352 | 0.049 | 0.028 | 0.005 | 0.005 | 0 | 0.89 | 0.89 | 0.81 | 0.68 | 0.67 | 0.82 |
| 4 | 123027723 | rs4833811 | 1.829 | 1.901 | -0.803 | 0.345 | 0.169 | 0.082 | 0.082 | 0.023 | 0.49 | 0.13 | 0.42 | 0.36 | 0.28 | 0.52 |
| 4 | 123073009 | rs11938795 | -0.529 | 2.296 | 0.291 | 0.335 | 0.004 | -0.082 | -0.082 | 0 | 0.21 | 0.35 | 0.15 | 0.06 | 0.24 | 0.12 |
| 4 | 123115502 | rs13151961 | 0.851 | 2.296 | 0.353 | 0.299 | -0.025 | 0.042 | 0.042 | 0.004 | 0.11 | 0.02 | 0.06 | 0 | 0.16 | 0.11 |
| 4 | 123146225 | rs7664452 | -0.96 | 2.57 | 0.273 | 0.436 | -0.064 | -0.067 | -0.067 | 2.509 | 0.15 | 0.08 | 0.44 | 0.43 | 0.35 | 0.3 |
| 4 | 123150286 | rs6848868 | 0.807 | 2.493 | 0.328 | 0.373 | 0.002 | 0.013 | 0.013 | 0 | 0.07 | 0.03 | 0.07 | 0 | 0.08 | 0.01 |
| 4 | 123228113 | rs11734090 | -1.086 | 2.467 | 0.421 | 0.367 | -0.002 | -0.083 | -0.083 | 0.004 | 0.22 | 0.34 | 0.16 | 0.06 | 0.24 | 0.12 |
| 4 | 123250384 | rs10027161 | 0.382 | 2.402 | 0.374 | 0.265 | 0.023 | -0.041 | -0.041 | 0 | 0.08 | 0.25 | 0.06 | 0 | 0.08 | 0.04 |
| 4 | 123280860 | rs1127348 | 0.865 | 2.827 | 0.152 | 0.309 | -0.044 | -0.036 | -0.036 | 0 | 0.07 | 0.01 | 0.13 | 0.12 | 0.24 | 0.15 |
| 4 | 123281293 | rs17005693 | 1.854 | 1.314 | -0.662 | -0.269 | 0.113 | 0.032 | 0.032 | 0 | 0.24 | 0.02 | 0.08 | 0.15 | 0.12 | 0.27 |
| 4 | 123306223 | rs10032704 | 1.802 | 2.894 | 0.527 | 0.197 | -0.001 | 0 | 0 | 1.483 | 0.64 | 0.66 | 0.39 | 0.38 | 0.38 | 0.57 |
| 4 | 123329362 | rs17388568 | 0.264 | 3.012 | 0.518 | 0.212 | -0.047 | -0.027 | -0.027 | 6.006 | 0.09 | 0.02 | 0.16 | 0.12 | 0.29 | 0.15 |
| 4 | 123333177 | rs12499753 | -0.826 | 3.001 | 0.526 | 0.225 | -0.051 | -0.05 | -0.05 | 0.24 | 0.14 | 0.08 | 0.44 | 0.46 | 0.34 | 0.29 |
| 4 | 123341159 | rs7684187 | 0.744 | 3.001 | 0.534 | 0.225 | 0.001 | -0.009 | -0.009 | 2.929 | 0.78 | 0.77 | 0.83 | 0.84 | 0.73 | 0.86 |
| 4 | 123342616 | rs1383043 | 1.86 | 2.998 | 0.53 | 0.224 | -0.001 | 0 | 0 | 1.328 | 0.64 | 0.65 | 0.39 | 0.38 | 0.38 | 0.57 |
| 4 | 123348345 | rs11732095 | 0.687 | 2.983 | 0.515 | 0.218 | 0.003 | 0.015 | 0.015 | 0 | 0.07 | 0.03 | 0.07 | 0 | 0.09 | 0.01 |
| 4 | 123368516 | rs10027390 | 0.642 | 2.977 | 0.506 | 0.234 | -0.001 | -0.001 | -0.001 | 0 | 0.79 | 0.8 | 0.83 | 0.84 | 0.73 | 0.86 |
| 4 | 123371049 | rs11575812 | -0.874 | 2.977 | 0.497 | 0.235 | -0.009 | 0.026 | 0.026 | 0 | 0.21 | 0.12 | 0.16 | 0.16 | 0.27 | 0.14 |
| 4 | 123377482 | rs2069763 | -0.895 | 3.162 | 0.537 | 0.358 | -0.051 | -0.05 | -0.05 | 0 | 0.14 | 0.08 | 0.44 | 0.46 | 0.34 | 0.29 |
| 4 | 123395041 | rs6848139 | -0.586 | 2.587 | 0.651 | 0.117 | 0.048 | -0.058 | -0.058 | 0 | 0.11 | 0.47 | 0.08 | 0 | 0.09 | 0.07 |
| 4 | 123506056 | rs1512973 | -1.51 | 2.825 | -0.55 | 0.313 | 0.014 | 0.072 | 0.072 | 0.004 | 0.26 | 0.38 | 0.36 | 0.48 | 0.32 | 0.29 |
| 4 | 123507914 | rs11098666 | -0.258 | 2.849 | -0.638 | 0.299 | -0.02 | -0.003 | -0.003 | 0.99 | 0.87 | 0.91 | 0.81 | 0.89 | 0.66 | 0.83 |
| 4 | 123508501 | rs12642902 | -1.387 | 2.767 | -0.614 | 0.32 | 0.007 | 0.037 | 0.037 | 0 | 0.25 | 0.32 | 0.35 | 0.48 | 0.32 | 0.29 |
| 4 | 123509421 | rs6822844 | 1.097 | 2.804 | -0.719 | 0.309 | -0.035 | 0.04 | 0.04 | 0 | 0.09 | 0.01 | 0.06 | 0 | 0.15 | 0.07 |
| 4 | 123521180 | rs975405 | -1.802 | 2.707 | -0.716 | 0.333 | 0.012 | 0.034 | 0.034 | 0 | 0.32 | 0.44 | 0.41 | 0.49 | 0.4 | 0.35 |
| 4 | 123533820 | rs17886348 | 0.799 | 2.884 | -1.379 | 0.396 | -0.024 | 0.033 | 0.033 | 0 | 0.06 | 0 | 0.04 | 0 | 0.08 | 0.04 |

| | | | | | | | | | | | | | | | | |
|---|---|---|---|---|---|---|---|---|---|---|---|---|---|---|---|---|
| 4 | 123536963 | rs4833837 | -0.334 | 3.02 | -1.39 | 0.4 | -0.042 | -0.014 | -0.014 | 0.116 | 0.88 | 0.95 | 0.82 | 0.89 | 0.66 | 0.84 |
| 4 | 123549699 | rs17005934 | 0.207 | 2.655 | -1.228 | 0.65 | 0 | 0.197 | 0.197 | 0 | 0.05 | 0.11 | 0.18 | 0.33 | 0.05 | 0.15 |
| 4 | 123554707 | rs6840978 | -0.294 | 2.378 | -1.076 | -0.037 | -0.003 | 0.001 | 0.001 | 0 | 0.13 | 0.13 | 0.1 | 0.16 | 0.18 | 0.13 |
| 4 | 123558254 | rs2137497 | -0.955 | 1.853 | -1.175 | -0.091 | -0.015 | 0.035 | 0.035 | 0 | 0.38 | 0.25 | 0.33 | 0.29 | 0.59 | 0.3 |
| 4 | 123558330 | rs2390352 | 0.902 | 1.483 | -0.258 | 0.072 | 0.058 | -0.025 | -0.025 | 1.614 | 0.92 | 0.39 | 0.93 | 1 | 0.95 | 0.94 |
| 4 | 123563078 | rs1022234 | 0.316 | 1.991 | -0.774 | 0.327 | 0.015 | -0.003 | -0.003 | 0 | 0.82 | 0.81 | 0.83 | 0.79 | 0.69 | 0.86 |
| 4 | 123564458 | rs1533236 | -0.004 | 1.047 | -0.963 | 0.234 | -0.007 | 0.012 | 0.012 | 0 | 0.46 | 0.36 | 0.51 | 0.49 | 0.53 | 0.47 |
| 4 | 123580354 | rs10518405 | 0.06 | -0.996 | -1.121 | -0.932 | 0.023 | 0.063 | 0.063 | 0 | 0.14 | 0.05 | 0.05 | 0 | 0.09 | 0.11 |
| 4 | 123581235 | rs6844999 | 0.591 | -2.171 | -1.384 | -1.093 | -0.013 | 0.008 | 0.008 | 0 | 0.27 | 0.24 | 0.2 | 0.33 | 0.12 | 0.36 |
| 4 | 123582426 | rs10518400 | 1.203 | -1.935 | -1.509 | -0.974 | -0.001 | 0.012 | 0.012 | 1.041 | 0.1 | 0.05 | 0.14 | 0.08 | 0.08 | 0.09 |
| 4 | 123601697 | rs6836610 | -0.638 | -2.107 | -1.143 | -1.086 | 0.081 | -0.037 | -0.037 | 1.273 | 0.55 | 0.85 | 0.46 | 0.42 | 0.28 | 0.56 |
| 4 | 123614762 | rs309398 | -0.499 | -1.88 | -0.026 | -1.647 | 0.025 | -0.013 | -0.013 | 0 | 0.73 | 0.64 | 0.57 | 0.84 | 0.52 | 0.78 |
| 4 | 123617561 | rs13101575 | 1.231 | -2.019 | 0.011 | -1.888 | -0.011 | -0.005 | -0.005 | 0 | 0.17 | 0.1 | 0.13 | 0 | 0.24 | 0.11 |
| 4 | 123620912 | rs309414 | -1.834 | -2.038 | 0.198 | -1.899 | 0.036 | -0.011 | -0.011 | 0 | 0.83 | 0.75 | 0.71 | 0.95 | 0.56 | 0.87 |
| 4 | 123623893 | rs1026157 | 0.87 | -2.376 | 0.288 | -1.82 | 0.048 | 0.012 | 0.012 | 0 | 0.39 | 0.24 | 0.23 | 0.29 | 0.21 | 0.4 |
| 4 | 123629427 | rs416556 | -1.128 | -2.674 | 1.139 | -1.453 | 0.012 | 0.002 | 0.002 | 0 | 0.43 | 0.39 | 0.46 | 0.66 | 0.33 | 0.47 |
| 4 | 123631308 | rs1512974 | -0.17 | -2.674 | 0.969 | -1.403 | 0.078 | -0.082 | -0.082 | 0.159 | 0.25 | 0.15 | 0.32 | 0.66 | 0.08 | 0.35 |
| 4 | 123652580 | rs13138426 | 0.964 | -2.477 | 1.105 | -1.426 | -0.05 | 0.08 | 0.08 | 0 | 0.17 | 0.05 | 0.11 | 0 | 0.21 | 0.11 |
| 4 | 123664204 | rs309370 | -1.43 | -2.597 | 1.361 | -1.436 | 0.003 | 0 | 0 | 0 | 0.45 | 0.58 | 0.47 | 0.66 | 0.32 | 0.46 |
| 4 | 123664446 | rs13135778 | 0.811 | -2.623 | 1.482 | -1.436 | -0.043 | 0.075 | 0.075 | 0 | 0.17 | 0.05 | 0.11 | 0 | 0.21 | 0.11 |
| 4 | 123664919 | rs13102440 | 0.999 | -2.623 | 1.471 | -1.436 | -0.05 | 0.08 | 0.08 | 0 | 0.17 | 0.05 | 0.11 | 0 | 0.21 | 0.11 |
| 4 | 123665428 | rs4833843 | -0.456 | -2.635 | 1.512 | -1.435 | -0.042 | 0.025 | 0.025 | 0 | 0.27 | 0.38 | 0.35 | 0.66 | 0.09 | 0.35 |
| 4 | 123668544 | rs4833844 | -0.192 | -2.625 | 1.468 | -1.419 | 0.04 | -0.039 | -0.039 | 2.16 | 0.25 | 0.22 | 0.33 | 0.66 | 0.08 | 0.35 |
| 4 | 123671551 | rs6811183 | -1.387 | -2.601 | 1.384 | -1.409 | -0.006 | -0.002 | -0.002 | 0 | 0.45 | 0.55 | 0.48 | 0.66 | 0.32 | 0.46 |
| 4 | 123679002 | rs13142203 | 0.201 | -1.958 | 1.799 | -2.195 | 0.018 | -0.082 | -0.082 | 0 | 0.19 | 0.25 | 0.14 | 0 | 0.24 | 0.11 |
| 4 | 123705387 | rs921301 | -0.65 | -2.073 | 1.579 | -2.128 | -0.061 | -0.008 | -0.008 | 1.275 | 0.15 | 0.33 | 0.15 | 0.17 | 0.03 | 0.21 |
| 4 | 123715055 | rs10005301 | -1.438 | -2.125 | 1.513 | -2.2 | 0.012 | 0.035 | 0.035 | 0 | 0.36 | 0.75 | 0.32 | 0.17 | 0.28 | 0.35 |
| 4 | 123715828 | rs13113055 | 0.576 | -1.817 | 1.482 | -1.892 | -0.073 | -0.013 | -0.013 | 0 | 0.84 | 0.64 | 0.84 | 0.83 | 0.97 | 0.77 |
| 4 | 123738546 | rs308447 | -1.252 | -1.682 | 0.565 | -2.252 | 0.068 | 0.241 | 0.241 | 0 | 0.5 | 0.83 | 0.37 | 0.13 | 0.36 | 0.42 |
| 4 | 123746883 | rs308394 | -1.804 | -1.4 | 1.578 | -1.177 | -0.028 | -0.029 | -0.029 | 0 | 0.05 | 0.12 | 0.01 | 0.01 | 0 | 0.16 |
| 4 | 123748086 | rs1449683 | -0.862 | -1.409 | 1.784 | -1.202 | 0.012 | -0.006 | -0.006 | 0 | 0.11 | 0.3 | 0.19 | 0.15 | 0.1 | 0.12 |
| 4 | 123752017 | rs308412 | -2.113 | -1.496 | 1.207 | -1.257 | -0.01 | -0.028 | -0.028 | 0 | 0.31 | 0.65 | 0.25 | 0.16 | 0.16 | 0.36 |
| 4 | 123767943 | rs308420 | -1.374 | -1.021 | 0.694 | -0.977 | -0.004 | -0.007 | -0.007 | 0 | 0.14 | 0.18 | 0.17 | 0.16 | 0.09 | 0.28 |
| 4 | 123771828 | rs308434 | -1.387 | -0.909 | 1.191 | -1.206 | -0.033 | -0.134 | -0.134 | 0.791 | 0.23 | 0.6 | 0.11 | 0.01 | 0.14 | 0.29 |
| 4 | 123772018 | rs308435 | -1.808 | -0.975 | 1.278 | -1.315 | -0.016 | -0.086 | -0.086 | 1.045 | 0.27 | 0.64 | 0.17 | 0.02 | 0.19 | 0.3 |
| 4 | 123772887 | rs308438 | -1.341 | -0.768 | 1.441 | -1.072 | -0.011 | -0.033 | -0.033 | 0.067 | 0.07 | 0.18 | 0.02 | 0.01 | 0.02 | 0.11 |
| 4 | 123773579 | rs308439 | -1.563 | -0.745 | 1.759 | -1.093 | 0.008 | -0.035 | -0.035 | 0.336 | 0.07 | 0.25 | 0.05 | 0.01 | 0.05 | 0.12 |
| 4 | 123774065 | rs308441 | -1.875 | -0.872 | 1.419 | -1.25 | 0.006 | 0.008 | 0.008 | 0 | 0.3 | 0.71 | 0.29 | 0.09 | 0.23 | 0.26 |
| 4 | 123774832 | rs17472986 | -0.381 | -0.895 | 1.009 | -1.14 | 0.011 | 0.023 | 0.023 | 0.074 | 0.15 | 0.07 | 0.18 | 0.08 | 0.09 | 0.1 |
| 4 | 123775767 | rs308443 | -1.574 | -0.856 | 1.426 | -0.919 | -0.006 | -0.033 | -0.033 | 0.922 | 0.08 | 0.3 | 0.03 | 0.01 | 0.03 | 0.11 |
| 4 | 123779341 | rs17407577 | -0.553 | -0.4 | 0.635 | -0.906 | -0.005 | 0.002 | 0.002 | 0 | 0.05 | 0.03 | 0.04 | 0 | 0.06 | 0.02 |
| 4 | 123782568 | rs1960669 | -1.11 | -0.468 | 0.822 | -1.07 | 0.019 | -0.054 | -0.054 | 0 | 0.12 | 0.19 | 0.09 | 0 | 0.16 | 0.07 |
| 4 | 123787079 | rs308388 | -0.765 | -1.281 | 0.4 | -0.816 | 0.005 | 0.005 | 0.005 | 0.298 | 0.48 | 0.48 | 0.33 | 0.46 | 0.36 | 0.62 |
| 4 | 123788495 | rs7694627 | 0.056 | -1.547 | 0.134 | -1.098 | -0.007 | 0.008 | 0.008 | 0 | 0.17 | 0.12 | 0.11 | 0 | 0.17 | 0.08 |
| 4 | 123799987 | rs10028204 | -1.191 | -2.214 | 0.26 | -1.53 | -0.02 | -0.025 | -0.025 | 0.012 | 0.82 | 0.8 | 0.84 | 0.95 | 0.94 | 0.92 |
| 4 | 123800617 | rs17408557 | 0.689 | -2.167 | 0.179 | -1.518 | -0.003 | 0.014 | 0.014 | 0 | 0.15 | 0.1 | 0.09 | 0 | 0.13 | 0.07 |
| 4 | 123810328 | rs10452197 | 0.683 | -2.187 | 0.167 | -1.528 | -0.001 | -0.001 | -0.001 | 0 | 0.15 | 0.12 | 0.09 | 0 | 0.14 | 0.07 |
| 4 | 123814900 | rs3804158 | 1.604 | -2.226 | 0.013 | -1.508 | -0.185 | -0.176 | -0.176 | 1.361 | 0.29 | 0.03 | 0.47 | 0.43 | 0.54 | 0.24 |
| 4 | 123816707 | rs6854081 | 0.688 | -2.253 | 0.044 | -1.56 | 0 | 0 | 0 | 0.101 | 0.15 | 0.12 | 0.09 | 0 | 0.14 | 0.07 |
| 4 | 123818074 | rs3747676 | 1.603 | -2.234 | 0.059 | -1.546 | -0.174 | -0.16 | -0.16 | 0.018 | 0.3 | 0.05 | 0.48 | 0.49 | 0.56 | 0.3 |
| 4 | 123826780 | rs11737764 | 0.107 | -2.603 | 0.764 | -1.731 | -0.067 | -0.083 | -0.083 | 0 | 0.7 | 0.48 | 0.77 | 0.94 | 0.91 | 0.75 |
| 4 | 123835879 | rs1458758 | 0.855 | -2.623 | 0.74 | -1.747 | 0.062 | 0.076 | 0.076 | 2.728 | 0.54 | 0.35 | 0.68 | 0.94 | 0.77 | 0.67 |
| 4 | 123838758 | rs12648093 | 1.187 | -2.661 | 0.879 | -1.77 | 0.117 | 0.238 | 0.238 | 2.186 | 0.51 | 0.2 | 0.64 | 0.94 | 0.74 | 0.65 |
| 4 | 123840064 | rs898091 | -0.979 | -2.64 | 1.097 | -1.741 | -0.055 | -0.072 | -0.072 | 0.007 | 0.8 | 0.64 | 0.83 | 0.95 | 0.93 | 0.92 |
| 4 | 123843716 | rs13134412 | -0.192 | -2.651 | 1.054 | -1.743 | -0.022 | -0.047 | -0.047 | 0.048 | 0.75 | 0.34 | 0.77 | 0.95 | 0.89 | 0.9 |
| 4 | 123852184 | rs10222773 | 0.108 | -2.747 | 0.948 | -1.767 | -0.008 | -0.038 | -0.038 | 0 | 0.37 | 0.17 | 0.24 | 0.45 | 0.31 | 0.43 |
| 4 | 123873534 | rs13106595 | 1.762 | -2.72 | 1 | -1.755 | -0.113 | -0.104 | -0.104 | 0.922 | 0.29 | 0.07 | 0.49 | 0.49 | 0.56 | 0.29 |
| 4 | 123881864 | rs906833 | 0.047 | -2.714 | 1.009 | -1.732 | -0.014 | 0.003 | 0.003 | 0 | 0.57 | 0.5 | 0.64 | 0.5 | 0.73 | 0.55 |
| 4 | 123885815 | rs7673580 | -0.728 | -2.597 | 1.093 | -1.639 | -0.048 | -0.076 | -0.076 | 0 | 0.79 | 0.6 | 0.8 | 0.95 | 0.91 | 0.9 |
| 4 | 123890773 | rs1467994 | -0.469 | -2.599 | 1.044 | -1.641 | -0.064 | -0.102 | -0.102 | 0.967 | 0.78 | 0.52 | 0.79 | 0.95 | 0.91 | 0.9 |
| 4 | 123949556 | rs1472949 | -1.298 | -2.661 | 0.979 | -1.675 | -0.037 | -0.039 | -0.039 | 0 | 0.82 | 0.76 | 0.84 | 0.95 | 0.95 | 0.92 |
| 4 | 123976639 | rs11731720 | 1.771 | -2.574 | 0.678 | -1.226 | -0.173 | -0.106 | -0.106 | 0 | 0.28 | 0.04 | 0.48 | 0.37 | 0.55 | 0.29 |
| 4 | 123984242 | rs12650805 | 1.679 | -2.633 | 0.717 | -1.67 | -0.171 | -0.136 | -0.136 | 0.074 | 0.29 | 0.05 | 0.5 | 0.51 | 0.57 | 0.3 |
| 4 | 123989201 | rs12650569 | 0.034 | -2.762 | 1.175 | -1.64 | -0.022 | -0.069 | -0.069 | 0 | 0.75 | 0.34 | 0.77 | 0.97 | 0.89 | 0.91 |
| 4 | 124005579 | rs11937761 | 0.679 | -2.668 | 0.873 | -1.652 | -0.003 | 0.03 | 0.03 | 0 | 0.15 | 0.09 | 0.08 | 0 | 0.13 | 0.07 |
| 4 | 124008458 | rs433220 | 0.286 | -2.791 | 1.042 | -1.719 | 0.008 | -0.03 | -0.03 | 0 | 0.65 | 0.7 | 0.74 | 0.97 | 0.78 | 0.84 |
| 4 | 124015188 | rs412925 | 0.625 | -2.803 | 1.07 | -1.727 | -0.008 | -0.099 | -0.099 | 0 | 0.62 | 0.58 | 0.7 | 0.97 | 0.76 | 0.82 |
| 4 | 124018404 | rs10518410 | 0.687 | -2.653 | 0.872 | -1.622 | -0.003 | 0.031 | 0.031 | 0 | 0.16 | 0.09 | 0.08 | 0 | 0.13 | 0.07 |
| 4 | 124066948 | rs303084 | 0.813 | -2.68 | 0.65 | -1.635 | 0.071 | 0.091 | 0.091 | 0.795 | 0.57 | 0.3 | 0.7 | 0.97 | 0.79 | 0.74 |

| | | | | | | | | | | | | | | | | |
|---|---|---|---|---|---|---|---|---|---|---|---|---|---|---|---|---|
| 4 | 124080712 | rs304653 | 0.977 | -2.636 | 0.466 | -1.652 | 0.094 | 0.34 | 0.34 | 2.291 | 0.53 | 0.12 | 0.66 | 0.96 | 0.75 | 0.71 |
| 4 | 124083918 | rs304650 | 0.849 | -2.388 | 0.637 | -1.196 | 0.012 | -0.023 | -0.023 | 0 | 0.39 | 0.21 | 0.57 | 0.52 | 0.72 | 0.39 |
| 4 | 124107641 | rs303130 | 0.697 | -2.425 | 0.968 | -1.197 | -0.006 | 0.002 | 0.002 | 0.13 | 0.4 | 0.43 | 0.57 | 0.52 | 0.71 | 0.37 |
| 4 | 124108270 | rs304658 | -0.187 | -2.562 | 0.753 | -1.204 | 0.082 | -0.017 | -0.017 | 0 | 0.61 | 0.78 | 0.7 | 0.53 | 0.87 | 0.46 |
| 4 | 124119834 | rs11944380 | -0.649 | -2.575 | 0.739 | -1.191 | 0.063 | -0.028 | -0.028 | 1.167 | 0.66 | 0.8 | 0.75 | 0.55 | 0.88 | 0.58 |
| 4 | 124132528 | rs303100 | 0.498 | -2.409 | 0.81 | -1.161 | 0.034 | 0.008 | 0.008 | 0.496 | 0.44 | 0.24 | 0.62 | 0.54 | 0.72 | 0.5 |
| 4 | 124140719 | rs2561151 | -2.033 | -2.254 | 0.913 | -1.201 | 0.065 | -0.011 | -0.011 | 0 | 0.08 | 0.53 | 0.07 | 0.01 | 0.05 | 0.02 |
| 4 | 124154217 | rs4479722 | 0.557 | -2.274 | 0.845 | -1.083 | 0.001 | -0.001 | -0.001 | 0 | 0.43 | 0.56 | 0.72 | 0.53 | 0.71 | 0.39 |
| 4 | 124171702 | rs11940736 | -0.421 | -2.426 | 0.77 | -1.013 | 0.022 | -0.014 | -0.014 | 2.699 | 0.64 | 0.66 | 0.83 | 0.54 | 0.87 | 0.58 |
| 4 | 124173635 | rs991183 | -0.482 | -2.429 | 1.172 | -1.178 | -0.012 | -0.098 | -0.098 | 0 | 0.24 | 0.34 | 0.14 | 0.03 | 0.17 | 0.2 |
| 4 | 124185679 | rs11735364 | -0.076 | -2.516 | 1.376 | -1.138 | -0.009 | 0.001 | 0.001 | 0.01 | 0.59 | 0.42 | 0.79 | 0.55 | 0.83 | 0.6 |
| 4 | 124192894 | rs1031135 | -0.517 | -2.574 | 1.342 | -1.163 | 0.029 | -0.015 | -0.015 | 0.11 | 0.64 | 0.68 | 0.84 | 0.56 | 0.88 | 0.62 |
| 4 | 124206187 | rs17006770 | -0.217 | -2.612 | 1.439 | -1.214 | -0.018 | -0.1 | -0.1 | 0.224 | 0.22 | 0.32 | 0.12 | 0.01 | 0.14 | 0.18 |
| 4 | 124211262 | rs13109534 | -0.155 | -2.539 | 1.277 | -1.144 | -0.011 | 0.002 | 0.002 | 0 | 0.61 | 0.54 | 0.8 | 0.54 | 0.83 | 0.61 |
| 4 | 124239525 | rs6832894 | 0.392 | -2.521 | 1.232 | -1.163 | 0.026 | -0.002 | -0.002 | 0 | 0.55 | 0.37 | 0.78 | 0.54 | 0.82 | 0.49 |
| 4 | 124246965 | rs2291934 | -0.516 | -2.501 | 0.664 | -1.133 | 0.031 | -0.016 | -0.016 | 1.072 | 0.64 | 0.71 | 0.84 | 0.56 | 0.88 | 0.61 |
| 4 | 124256835 | rs300559 | -1.371 | -2.738 | 0.473 | -1.15 | 0.039 | -0.071 | -0.071 | 1.85 | 0.8 | 0.9 | 0.89 | 0.59 | 0.92 | 0.68 |
| 4 | 124257091 | rs7677059 | 0.4 | -2.46 | 0.647 | -1.058 | -0.008 | -0.001 | -0.001 | 0.028 | 0.46 | 0.45 | 0.74 | 0.56 | 0.74 | 0.55 |
| 4 | 124267444 | rs871648 | 0.138 | -2.417 | 0.223 | -0.818 | 0 | -0.002 | -0.002 | 0 | 0.57 | 0.51 | 0.75 | 0.51 | 0.81 | 0.51 |
| 4 | 124275630 | rs7655411 | -0.458 | -2.26 | -0.55 | -0.636 | 0.067 | -0.024 | -0.024 | 0 | 0.63 | 0.79 | 0.79 | 0.61 | 0.82 | 0.63 |
| 4 | 124282603 | rs879905 | -0.397 | -1.436 | -0.965 | -0.455 | 0.03 | -0.002 | -0.002 | 1.023 | 0.57 | 0.75 | 0.71 | 0.61 | 0.67 | 0.57 |
| 4 | 124283138 | rs1027374 | 0.366 | -0.08 | -0.979 | 0.232 | -0.104 | -0.116 | -0.116 | 1.642 | 0.28 | 0.09 | 0.56 | 0.51 | 0.5 | 0.33 |
| 4 | 124285735 | rs6845023 | -0.795 | -0.398 | -0.088 | 0.174 | -0.01 | -0.019 | -0.019 | 2.596 | 0.08 | 0.14 | 0.07 | 0.05 | 0.06 | 0.11 |
| 4 | 124288412 | rs7659671 | -1.66 | -0.475 | 0.303 | 0.804 | -0.016 | 0.095 | 0.095 | 0.463 | 0.17 | 0.6 | 0.35 | 0.38 | 0.15 | 0.25 |
| 4 | 124291946 | rs300550 | 1.204 | -0.429 | 0.625 | 0.645 | -0.024 | 0.146 | 0.146 | 0.063 | 0.85 | 0.6 | 0.66 | 0.63 | 0.89 | 0.72 |
| 4 | 124297264 | rs11098708 | 0.034 | -0.204 | -0.565 | 0.254 | -0.033 | -0.059 | -0.059 | 0.911 | 0.38 | 0.15 | 0.45 | 0.48 | 0.57 | 0.38 |
| 4 | 124305017 | rs300556 | 0.442 | -0.169 | 0.296 | 0.664 | -0.007 | 0.237 | 0.237 | 0.858 | 0.86 | 0.63 | 0.63 | 0.58 | 0.9 | 0.75 |
| 4 | 124315710 | rs6836221 | 0.814 | -0.996 | 0.234 | 0.011 | -0.027 | -0.036 | -0.036 | 0.822 | 0.35 | 0.55 | 0.17 | 0.16 | 0.24 | 0.33 |
| 4 | 124319144 | rs300569 | 1.125 | -0.959 | 0.016 | -0.14 | -0.007 | 0.042 | 0.042 | 0.071 | 0.15 | 0.05 | 0.07 | 0.01 | 0.11 | 0.03 |
| 4 | 124323738 | rs300574 | -0.082 | -1.021 | 0.163 | -0.061 | 0.102 | 0.04 | 0.04 | 0.217 | 0.51 | 0.22 | 0.48 | 0.41 | 0.67 | 0.4 |
| 4 | 124329732 | rs7665238 | 0.686 | -1.9 | 0.357 | -0.059 | -0.018 | 0.028 | 0.028 | 0 | 0.67 | 0.5 | 0.54 | 0.56 | 0.76 | 0.69 |
| 4 | 124334279 | rs923982 | 0.469 | -1.434 | -0.281 | -0.309 | -0.032 | 0 | 0 | 0.005 | 0.35 | 0.24 | 0.37 | 0.36 | 0.45 | 0.38 |
| 4 | 124337522 | rs12505913 | 1.075 | -1.302 | -0.572 | -0.383 | -0.075 | -0.039 | -0.039 | 0 | 0.25 | 0.09 | 0.34 | 0.35 | 0.38 | 0.33 |
| 4 | 124340706 | rs13124930 | 0.859 | -1.282 | -0.569 | -0.366 | -0.024 | -0.009 | -0.009 | 0.044 | 0.27 | 0.2 | 0.35 | 0.35 | 0.39 | 0.33 |
| 4 | 124342410 | rs2390522 | -0.435 | -1.436 | -0.561 | -0.465 | -0.012 | -0.025 | -0.025 | 0 | 0.19 | 0.13 | 0.41 | 0.44 | 0.25 | 0.23 |
| 4 | 124343118 | rs10857095 | 0.965 | -1.365 | -0.771 | -0.424 | -0.026 | -0.009 | -0.009 | 0 | 0.27 | 0.2 | 0.35 | 0.35 | 0.39 | 0.33 |
| 4 | 124343951 | rs6534397 | 1.428 | -2.128 | -1.081 | -0.83 | 0.02 | -0.036 | -0.036 | 0 | 0.19 | 0.07 | 0.07 | 0.2 | 0.14 | 0.33 |
| 4 | 124344717 | rs12233855 | 0.408 | -2.124 | -1.075 | -0.836 | -0.003 | 0.003 | 0.003 | 0.247 | 0.45 | 0.27 | 0.42 | 0.55 | 0.53 | 0.66 |
| 4 | 124345346 | rs10021762 | 0.063 | -1.846 | -0.86 | -0.809 | 0.022 | 0.033 | 0.033 | 0.7 | 0.42 | 0.29 | 0.75 | 0.73 | 0.62 | 0.54 |
| 4 | 124353489 | rs11098710 | 0.002 | -1.686 | -0.716 | -0.349 | -0.001 | 0.003 | 0.003 | 0 | 0.48 | 0.47 | 0.5 | 0.35 | 0.53 | 0.31 |
| 4 | 124359022 | rs17006926 | 0.93 | -1.743 | -0.773 | -0.322 | 0.058 | -0.043 | -0.043 | 0 | 0.23 | 0.08 | 0.06 | 0.27 | 0.1 | 0.41 |
| 4 | 124359046 | rs6534401 | 0.134 | -0.929 | -0.156 | -0.318 | 0.028 | 0.034 | 0.034 | 0 | 0.69 | 0.56 | 0.56 | 0.62 | 0.63 | 0.73 |
| 4 | 124362035 | rs17006928 | -0.182 | -1.43 | -0.263 | -0.884 | -0.009 | -0.037 | -0.037 | 0 | 0.19 | 0.16 | 0.06 | 0 | 0.1 | 0.06 |
| 4 | 124368370 | rs3860074 | 0.494 | -0.301 | -0.551 | -0.312 | -0.001 | -0.001 | -0.001 | 0 | 0.75 | 0.73 | 0.85 | 0.84 | 0.68 | 0.78 |
| 4 | 124371328 | rs11734331 | 0.821 | -0.349 | -1.318 | -0.374 | 0.004 | 0.027 | 0.027 | 0.45 | 0.58 | 0.25 | 0.63 | 0.7 | 0.57 | 0.62 |
| 4 | 124372645 | rs13139048 | 0.928 | -0.087 | -1.945 | -0.233 | 0.023 | 0.085 | 0.085 | 0.116 | 0.51 | 0.15 | 0.59 | 0.7 | 0.55 | 0.61 |
| 4 | 124375248 | rs7677413 | 0.235 | 0.908 | -1.252 | -0.187 | -0.011 | 0.005 | 0.005 | 0.15 | 0.84 | 0.93 | 0.89 | 1 | 0.8 | 0.95 |
| 4 | 124387408 | rs11732507 | 0.738 | 0.531 | -1.425 | 0.166 | 0.006 | 0.005 | 0.005 | 0.111 | 0.8 | 0.89 | 0.89 | 0.84 | 0.85 | 0.6 |
| 4 | 124387459 | rs17807333 | 0.175 | 0.455 | -1.32 | -1.27 | -0.002 | -0.101 | -0.101 | 0 | 0.58 | 0.51 | 0.61 | 0.92 | 0.57 | 0.8 |
| 4 | 124387479 | rs10776501 | -0.005 | 0.076 | -0.653 | -1.682 | 0.015 | -0.068 | -0.068 | 0 | 0.84 | 0.68 | 0.87 | 0.98 | 0.84 | 0.97 |
| 4 | 124387560 | rs7697652 | -0.312 | 0.111 | -0.857 | -1.798 | -0.001 | 0.012 | 0.012 | 0 | 0.36 | 0.61 | 0.31 | 0.21 | 0.39 | 0.32 |
| 4 | 124389966 | rs10029753 | 0.324 | 0.297 | -1.429 | -2.278 | -0.019 | 0.099 | 0.099 | 0 | 0.67 | 0.81 | 0.7 | 0.9 | 0.64 | 0.7 |
| 4 | 124393598 | rs17745658 | 1.022 | 0.403 | -1.136 | -1.833 | -0.002 | 0 | 0 | 0 | 0.1 | 0.07 | 0.1 | 0.13 | 0.1 | 0.06 |
| 4 | 124395024 | rs6534407 | 0.748 | 0.129 | -1.318 | -0.972 | 0.023 | -0.027 | -0.027 | 0 | 0.2 | 0.1 | 0.12 | 0.26 | 0.1 | 0.23 |
| 4 | 124397309 | rs1452731 | -0.796 | 0.596 | -1.194 | -0.66 | 0.006 | 0.001 | 0.001 | 0 | 0.93 | 0.95 | 0.89 | 0.91 | 0.84 | 0.82 |
| 4 | 124397732 | rs12233795 | 1.243 | 0.53 | -1.282 | -0.776 | 0.004 | 0.011 | 0.011 | 0 | 0.12 | 0.1 | 0.07 | 0.03 | 0.09 | 0.15 |
| 4 | 124399060 | rs1349683 | -1.412 | 0.531 | -1.285 | -0.78 | 0.002 | 0.014 | 0.014 | 0 | 0.87 | 0.91 | 0.94 | 0.97 | 0.9 | 0.84 |
| 4 | 124399672 | rs737010 | -1.037 | 0.492 | -1.295 | -0.766 | -0.002 | -0.004 | -0.004 | 0 | 0.93 | 0.96 | 0.89 | 0.91 | 0.84 | 0.82 |
| 4 | 124399996 | rs737009 | 0.451 | 0.223 | -0.16 | -0.719 | 0.004 | 0.002 | 0.002 | 0 | 0.76 | 0.47 | 0.79 | 0.79 | 0.76 | 0.58 |
| 4 | 124402023 | rs1901694 | 0.442 | 0.212 | -0.197 | -0.715 | 0.021 | 0.003 | 0.003 | 0 | 0.76 | 0.51 | 0.78 | 0.79 | 0.75 | 0.58 |
| 4 | 124403102 | rs6831251 | 0.251 | 0.73 | -0.504 | 0.364 | -0.013 | 0.014 | 0.014 | 0.057 | 0.09 | 0.07 | 0.17 | 0.06 | 0.12 | 0.08 |
| 4 | 124410268 | rs10471022 | 0.181 | 0.603 | -0.312 | 0.412 | -0.005 | 0.018 | 0.018 | 0 | 0.67 | 0.37 | 0.72 | 0.58 | 0.73 | 0.51 |
| 4 | 124417151 | rs896678 | -0.195 | 0.519 | 0.01 | 0.442 | -0.007 | 0.027 | 0.027 | 0 | 0.7 | 0.55 | 0.75 | 0.67 | 0.75 | 0.6 |
| 4 | 124417350 | rs896676 | 0.488 | 0.648 | -0.409 | 0.336 | 0.001 | -0.005 | -0.005 | 0 | 0.61 | 0.34 | 0.66 | 0.57 | 0.68 | 0.36 |
| 4 | 124423491 | rs1947372 | 0.012 | 0.89 | -0.275 | 0.049 | 0.001 | 0 | 0 | 0 | 0.81 | 0.8 | 0.75 | 0.8 | 0.77 | 0.73 |
| 4 | 124426163 | rs7688738 | 1.368 | 0.834 | -0.277 | -0.422 | 0.006 | -0.025 | -0.025 | 0 | 0.58 | 0.46 | 0.62 | 0.79 | 0.56 | 0.6 |
| 4 | 124430104 | rs4833925 | 0.161 | 0.912 | -0.21 | -0.384 | 0.005 | -0.028 | -0.028 | 0 | 0.71 | 0.57 | 0.8 | 0.85 | 0.73 | 0.72 |
| 4 | 124437413 | rs17007028 | 1.376 | 0.549 | -0.584 | -0.143 | 0.023 | 0.017 | 0.017 | 0.368 | 0.15 | 0.06 | 0.1 | 0.12 | 0.08 | 0.07 |
| 4 | 124437842 | rs1993191 | 0.442 | 1.269 | -0.005 | -0.034 | 0.021 | -0.006 | -0.006 | 0 | 0.74 | 0.66 | 0.71 | 0.78 | 0.63 | 0.57 |

| | | | | | | | | | | | | | | | | |
|---|---|---|---|---|---|---|---|---|---|---|---|---|---|---|---|---|
| 4 | 124439866 | rs6534410 | -0.158 | 1.31 | 0.011 | -0.041 | 0.026 | -0.005 | -0.005 | 0 | 0.79 | 0.7 | 0.77 | 0.84 | 0.71 | 0.65 |
| 4 | 124443473 | rs17007051 | 1.515 | 1.252 | -0.228 | 0.011 | -0.005 | 0.001 | 0.001 | 0 | 0.05 | 0.03 | 0.06 | 0.06 | 0.08 | 0.09 |
| 4 | 124450148 | rs10461205 | 1.098 | 1.036 | -0.052 | -0.066 | 0.01 | -0.009 | -0.009 | 0 | 0.66 | 0.39 | 0.58 | 0.74 | 0.58 | 0.49 |
| 4 | 124469200 | rs7670704 | 0.781 | 0.33 | -0.081 | 0.313 | 0.001 | 0.006 | 0.006 | 0 | 0.67 | 0.58 | 0.63 | 0.64 | 0.67 | 0.61 |
| 4 | 124473935 | rs10011590 | -1.075 | 0.353 | -0.465 | 0.363 | -0.044 | -0.027 | -0.027 | 0 | 0.85 | 0.95 | 0.87 | 0.84 | 0.8 | 0.8 |
| 4 | 124496348 | rs4833930 | 0.963 | 0.445 | -0.269 | 0.31 | 0 | -0.001 | -0.001 | 0 | 0.63 | 0.38 | 0.61 | 0.64 | 0.65 | 0.59 |
| 4 | 124509674 | rs1806533 | 1.2 | 0.216 | -0.682 | 0.57 | -0.091 | -0.061 | -0.061 | 0 | 0.2 | 0.08 | 0.34 | 0.33 | 0.41 | 0.27 |
| 4 | 124512827 | rs6534418 | 0.082 | 0.165 | -0.574 | 0.528 | 0.027 | -0.008 | -0.008 | 0.037 | 0.75 | 0.55 | 0.78 | 0.78 | 0.73 | 0.7 |
| 4 | 124519432 | rs2553364 | 0.672 | 0.095 | -0.163 | 0.666 | -0.034 | -0.006 | -0.006 | 0 | 0.78 | 0.44 | 0.74 | 0.79 | 0.84 | 0.79 |
| 4 | 124527958 | rs752499 | 0.786 | 0.349 | -0.589 | 0.77 | -0.094 | -0.061 | -0.061 | 2.774 | 0.2 | 0.08 | 0.34 | 0.33 | 0.4 | 0.26 |
| 4 | 124528377 | rs752501 | 0.167 | -0.644 | 0.128 | 0.563 | 0.005 | -0.005 | -0.005 | 0 | 0.12 | 0.1 | 0.16 | 0.14 | 0.07 | 0.11 |
| 4 | 124530321 | rs10017009 | 0.857 | -0.844 | -0.002 | 0.537 | -0.038 | -0.017 | -0.017 | 0 | 0.21 | 0.28 | 0.13 | 0.15 | 0.08 | 0.2 |
| 4 | 124531422 | rs2589850 | 0.033 | -0.19 | -0.604 | 0.704 | 0.024 | 0.011 | 0.011 | 0 | 0.59 | 0.33 | 0.63 | 0.65 | 0.77 | 0.58 |
| 4 | 124545508 | rs13115814 | -1.091 | 1.435 | -0.892 | 1.306 | -0.021 | -0.028 | -0.028 | 0.373 | 0.14 | 0.07 | 0.15 | 0.29 | 0.23 | 0.3 |
| 4 | 124550693 | rs2589864 | 0.492 | 1.204 | -1.698 | 0.888 | 0.012 | 0.056 | 0.056 | 0 | 0.42 | 0.18 | 0.35 | 0.37 | 0.37 | 0.26 |
| 4 | 124564632 | rs17753395 | 0.374 | 1.233 | -1.822 | 0.827 | -0.029 | 0.001 | 0.001 | 0 | 0.08 | 0 | 0.08 | 0.06 | 0.14 | 0.12 |
| 4 | 124565964 | rs1425419 | 0.937 | 0.866 | -2.167 | 0.701 | 0.009 | -0.118 | -0.118 | 0 | 0.24 | 0.02 | 0.33 | 0.31 | 0.16 | 0.39 |
| 4 | 124570408 | rs7695956 | 0.977 | 0.914 | -2.09 | 0.726 | 0.009 | -0.107 | -0.107 | 0 | 0.24 | 0.02 | 0.35 | 0.31 | 0.2 | 0.4 |
| 4 | 124570507 | rs2589858 | 0.287 | 0.987 | -1.101 | 0.584 | 0.003 | 0.003 | 0.003 | 0 | 0.5 | 0.46 | 0.39 | 0.35 | 0.38 | 0.3 |
| 4 | 124574008 | rs6534420 | 0.733 | 1.393 | -0.292 | 0.759 | 0.127 | 0.062 | 0.062 | 0.427 | 0.77 | 0.55 | 0.75 | 0.66 | 0.61 | 0.7 |
| 4 | 124590869 | rs2553355 | 1.148 | 1.141 | -0.749 | 0.257 | -0.039 | 0.105 | 0.105 | 0.061 | 0.26 | 0.01 | 0.11 | 0.12 | 0.2 | 0.1 |
| 4 | 124592367 | rs12509315 | -0.275 | 1.665 | -0.251 | 0.597 | -0.005 | 0.001 | 0.001 | 0.123 | 0.09 | 0.09 | 0.13 | 0.24 | 0.16 | 0.18 |
| 4 | 124592723 | rs2589838 | -0.647 | 1.49 | 0.041 | 0.478 | 0.062 | -0.045 | -0.045 | 0.302 | 0.9 | 0.85 | 0.91 | 0.98 | 0.79 | 0.96 |
| 4 | 124608621 | rs2589845 | -0.645 | 1.487 | 0.156 | 0.477 | 0.061 | -0.044 | -0.044 | 0 | 0.9 | 0.79 | 0.9 | 0.98 | 0.79 | 0.96 |
| 4 | 124621650 | rs890180 | -0.377 | 1.192 | 0.133 | 0.556 | -0.002 | -0.001 | -0.001 | 0.004 | 0.6 | 0.64 | 0.53 | 0.61 | 0.57 | 0.42 |
| 4 | 124623291 | rs12500929 | 0.019 | 1.439 | 0.366 | 0.547 | 0.095 | -0.03 | -0.03 | 0 | 0.87 | 0.72 | 0.89 | 0.92 | 0.77 | 0.84 |
| 4 | 124627790 | rs11723044 | -0.042 | 1.549 | -0.027 | 0.778 | 0.009 | 0.031 | 0.031 | 0.051 | 0.05 | 0.09 | 0.1 | 0.21 | 0.13 | 0.14 |
| 4 | 124628029 | rs13117812 | -0.29 | 1.544 | 0.124 | 0.594 | 0.045 | -0.01 | -0.01 | 0 | 0.14 | 0.17 | 0.12 | 0.11 | 0.25 | 0.14 |
| 4 | 124628278 | rs1425416 | -0.055 | 1.04 | -0.072 | 0.451 | 0.002 | 0.006 | 0.006 | 0 | 0.5 | 0.53 | 0.4 | 0.37 | 0.42 | 0.24 |
| 4 | 124650005 | rs1834378 | 0.103 | 0.279 | 0.243 | 0.595 | 0.038 | 0.011 | 0.011 | 0 | 0.48 | 0.62 | 0.51 | 0.41 | 0.36 | 0.5 |
| 4 | 124652559 | rs11098717 | 0.103 | 0.279 | 0.243 | 0.595 | 0.038 | 0.011 | 0.011 | 0 | 0.48 | 0.62 | 0.5 | 0.41 | 0.36 | 0.5 |
| 4 | 124654530 | rs4833940 | 0.899 | 0.146 | 0.649 | 0.529 | -0.013 | -0.001 | -0.001 | 0 | 0.36 | 0.32 | 0.39 | 0.36 | 0.22 | 0.45 |
| 4 | 124654693 | rs4833288 | 1.039 | 0.104 | 0.61 | 0.508 | 0.066 | -0.023 | -0.023 | 0.286 | 0.34 | 0.12 | 0.37 | 0.36 | 0.22 | 0.45 |
| 4 | 124659199 | rs17007240 | -0.444 | 0.696 | 0.594 | 0.511 | -0.003 | 0.003 | 0.003 | 0 | 0.44 | 0.43 | 0.31 | 0.41 | 0.52 | 0.22 |
| 4 | 124672450 | rs4833947 | -0.071 | 0.783 | 0.612 | 0.545 | 0 | -0.009 | -0.009 | 0.693 | 0.12 | 0.21 | 0.1 | 0.05 | 0.14 | 0.05 |
| 4 | 124673938 | rs2194865 | 0.05 | 0.918 | 0.848 | 0.665 | 0.037 | 0.054 | 0.054 | 0.048 | 0.92 | 0.77 | 0.89 | 0.85 | 0.89 | 0.84 |
| 4 | 124679431 | rs6828406 | -0.055 | 0.812 | 0.805 | 0.841 | 0.001 | -0.012 | -0.012 | 0.983 | 0.12 | 0.22 | 0.1 | 0.05 | 0.14 | 0.05 |
| 4 | 124683891 | rs1425406 | -0.521 | 0.781 | 0.909 | 0.827 | 0.001 | -0.025 | -0.025 | 0 | 0.13 | 0.34 | 0.12 | 0.05 | 0.14 | 0.05 |
| 4 | 124704038 | rs3097925 | -0.756 | 0.685 | 0.76 | 0.792 | -0.015 | 0.002 | 0.002 | 0.764 | 0.67 | 0.74 | 0.79 | 0.65 | 0.61 | 0.82 |
| 4 | 124706056 | rs1433219 | 0.348 | 0.692 | 0.557 | 0.785 | -0.001 | 0.006 | 0.006 | 0 | 0.11 | 0.13 | 0.08 | 0.05 | 0.13 | 0.04 |
| 4 | 124707000 | rs1433218 | -0.892 | 0.67 | 0.58 | 0.775 | -0.039 | -0.004 | -0.004 | 0 | 0.68 | 0.83 | 0.8 | 0.65 | 0.61 | 0.82 |
| 4 | 124710584 | rs7692299 | -0.086 | 0.573 | 0.46 | 0.749 | -0.003 | -0.013 | -0.013 | 1.134 | 0.12 | 0.23 | 0.1 | 0.05 | 0.13 | 0.05 |
| 4 | 124711445 | rs4833953 | 1.283 | 0.38 | 0.139 | 0.737 | 0.054 | 0.02 | 0.02 | 0 | 0.33 | 0.15 | 0.34 | 0.31 | 0.23 | 0.44 |
| 4 | 124713216 | rs6821590 | 0.309 | 0.548 | -0.106 | 0.801 | 0 | 0.008 | 0.008 | 0 | 0.11 | 0.12 | 0.09 | 0.05 | 0.13 | 0.04 |
| 4 | 124718116 | rs2390586 | 1.397 | 0.318 | -0.33 | 0.835 | 0.001 | -0.048 | -0.048 | 0 | 0.29 | 0.03 | 0.33 | 0.31 | 0.21 | 0.42 |
| 4 | 124718998 | rs3097935 | -0.136 | 0.517 | 0.089 | 0.93 | 0.014 | 0.025 | 0.025 | 0 | 0.08 | 0.11 | 0.12 | 0.14 | 0.11 | 0.16 |
| 4 | 124725097 | rs4833956 | 1.323 | 0.341 | -0.15 | 1.143 | 0.026 | -0.122 | -0.122 | 0 | 0.3 | 0.07 | 0.37 | 0.55 | 0.22 | 0.45 |
| 4 | 124729590 | rs3097914 | 0.467 | 0.258 | 0.236 | 0.944 | 0.015 | -0.005 | -0.005 | 0 | 0.39 | 0.28 | 0.46 | 0.41 | 0.33 | 0.57 |
| 4 | 124731732 | rs10028801 | 0.321 | 0.239 | 0.561 | 0.735 | -0.002 | -0.028 | -0.028 | 0 | 0.13 | 0.14 | 0.12 | 0.03 | 0.11 | 0.12 |
| 4 | 124733305 | rs1433216 | 0.306 | 0.209 | 0.881 | 0.699 | 0.008 | -0.002 | -0.002 | 0 | 0.46 | 0.34 | 0.54 | 0.46 | 0.45 | 0.6 |
| 4 | 124741086 | rs3113387 | -0.033 | 0.214 | 0.959 | 0.527 | 0.055 | 0.047 | 0.047 | 0 | 0.1 | 0.28 | 0.14 | 0.09 | 0.12 | 0.16 |
| 4 | 124742689 | rs9991633 | 0.059 | 0.194 | 0.961 | 0.501 | -0.007 | -0.052 | -0.052 | 0 | 0.15 | 0.23 | 0.13 | 0.03 | 0.12 | 0.12 |
| 4 | 124743259 | rs1347188 | -0.9 | 0.452 | 1.049 | 0.56 | 0.023 | 0.014 | 0.014 | 0.104 | 0.18 | 0.33 | 0.22 | 0.14 | 0.24 | 0.19 |
| 4 | 124745047 | rs1991839 | -0.559 | 0.512 | 0.691 | 0.633 | -0.002 | 0 | 0 | 0.699 | 0.62 | 0.59 | 0.7 | 0.61 | 0.59 | 0.78 |
| 4 | 124749377 | rs978534 | -0.695 | 0.538 | 0.633 | 0.616 | -0.015 | 0.002 | 0.002 | 0.659 | 0.64 | 0.68 | 0.71 | 0.6 | 0.59 | 0.79 |
| 4 | 124752974 | rs1433212 | -0.602 | 0.69 | 0.427 | 0.558 | -0.012 | 0.005 | 0.005 | 0.432 | 0.15 | 0.12 | 0.18 | 0.09 | 0.2 | 0.16 |
| 4 | 124754548 | rs17491036 | 0.113 | 0.634 | 0.229 | 0.465 | -0.036 | 0.045 | 0.045 | 0.41 | 0.11 | 0 | 0.12 | 0.02 | 0.11 | 0.06 |
| 4 | 124757420 | rs1347184 | 1.78 | 0.733 | 0.035 | 0.507 | 0.012 | -0.044 | -0.044 | 0 | 0.33 | 0.03 | 0.36 | 0.31 | 0.27 | 0.42 |
| 4 | 124763465 | rs1433211 | -1.722 | 0.752 | 0.643 | 0.556 | 0.002 | 0.002 | 0.002 | 2.459 | 0.33 | 0.6 | 0.37 | 0.33 | 0.37 | 0.37 |
| 4 | 124766115 | rs3113390 | -1.422 | 0.759 | 0.864 | 0.495 | 0.018 | 0.011 | 0.011 | 1.353 | 0.31 | 0.5 | 0.36 | 0.28 | 0.37 | 0.37 |
| 4 | 124767640 | rs1036253 | -0.952 | 0.657 | 0.941 | 0.476 | -0.039 | -0.016 | -0.016 | 0 | 0.68 | 0.84 | 0.75 | 0.6 | 0.64 | 0.78 |
| 4 | 124770843 | rs4449426 | -0.454 | 0.346 | 0.592 | 0.283 | -0.012 | 0.002 | 0.002 | 0 | 0.23 | 0.36 | 0.25 | 0.2 | 0.25 | 0.21 |
| 4 | 124770865 | rs4340803 | 1.454 | 0.343 | 0.681 | 0.297 | 0.005 | 0 | 0 | 0 | 0.07 | 0.1 | 0.07 | 0.05 | 0.12 | 0.02 |
| 4 | 124775267 | rs11731103 | 0.91 | 0.318 | 0.552 | 0.298 | 0.012 | -0.01 | -0.01 | 0 | 0.15 | 0.05 | 0.15 | 0.15 | 0.13 | 0.19 |
| 4 | 124775472 | rs3113392 | -1.707 | 0.319 | 0.551 | 0.3 | -0.002 | 0.006 | 0.006 | 2.177 | 0.41 | 0.61 | 0.43 | 0.3 | 0.45 | 0.38 |
| 4 | 124784586 | rs3113396 | -1.09 | 0.359 | 0.625 | 0.292 | -0.002 | -0.001 | -0.001 | 0 | 0.66 | 0.64 | 0.72 | 0.6 | 0.63 | 0.73 |
| 4 | 124791509 | rs1560738 | 0.483 | 0.905 | 0.808 | 0.363 | 0.008 | -0.011 | -0.011 | 1.437 | 0.15 | 0.04 | 0.15 | 0.14 | 0.13 | 0.18 |
| 4 | 124794203 | rs986345 | -0.401 | 0.99 | 0.926 | 0.384 | 0.006 | -0.007 | -0.007 | 0 | 0.18 | 0.22 | 0.18 | 0.11 | 0.2 | 0.17 |
| 4 | 124797438 | rs17491837 | 1.074 | 0.99 | 0.769 | 0.477 | -0.022 | -0.014 | -0.014 | 0 | 0.06 | 0.01 | 0.06 | 0.05 | 0.12 | 0.02 |

| | | | | | | | | | | | | | | | | |
|---|---|---|---|---|---|---|---|---|---|---|---|---|---|---|---|---|
| 4 | 124802131 | rs1433225 | -1.064 | 1.25 | 1.291 | 0.413 | -0.013 | -0.017 | -0.017 | 1.314 | 0.26 | 0.46 | 0.26 | 0.19 | 0.28 | 0.24 |
| 4 | 124802903 | rs1347189 | -2.323 | 1.364 | 1.174 | 0.331 | -0.019 | 0.034 | 0.034 | 0.306 | 0.43 | 0.75 | 0.44 | 0.3 | 0.48 | 0.4 |
| 4 | 124807935 | rs17007553 | 0.901 | 1.333 | 1.404 | 0.343 | 0.002 | -0.003 | -0.003 | 1.085 | 0.07 | 0.1 | 0.07 | 0.05 | 0.11 | 0.02 |
| 4 | 124809136 | rs9991120 | -0.229 | 1.343 | 1.236 | 0.334 | -0.004 | -0.004 | -0.004 | 0.088 | 0.19 | 0.24 | 0.18 | 0.15 | 0.17 | 0.22 |
| 4 | 124810078 | rs1433223 | -0.037 | 1.344 | 1.012 | 0.335 | 0.006 | 0.005 | 0.005 | 0.159 | 0.19 | 0.15 | 0.18 | 0.15 | 0.17 | 0.19 |
| 4 | 124812897 | rs17492375 | -0.14 | 1.408 | 1.029 | 0.344 | 0 | 0.001 | 0.001 | 0 | 0.15 | 0.14 | 0.16 | 0.1 | 0.19 | 0.17 |
| 4 | 124819109 | rs7691701 | -1.613 | 1.338 | 0.981 | 0.352 | 0.001 | -0.009 | -0.009 | 0.573 | 0.33 | 0.39 | 0.29 | 0.19 | 0.35 | 0.34 |
| 4 | 124819486 | rs1857 | 1.65 | 1.342 | 0.462 | 0.434 | -0.016 | -0.019 | -0.019 | 0.852 | 0.1 | 0.07 | 0.12 | 0.11 | 0.13 | 0.07 |
| 4 | 124820655 | rs1469717 | 1.773 | 1.347 | 0.373 | 0.437 | -0.038 | -0.04 | -0.04 | 0 | 0.1 | 0.01 | 0.11 | 0.11 | 0.13 | 0.07 |
| 4 | 124833001 | rs1433220 | -1.871 | 1.435 | 0.625 | 0.433 | -0.005 | -0.001 | -0.001 | 0.235 | 0.34 | 0.74 | 0.37 | 0.3 | 0.38 | 0.38 |
| 4 | 124834717 | rs3113378 | -1.849 | 1.493 | 0.597 | 0.435 | -0.009 | -0.002 | -0.002 | 1.52 | 0.34 | 0.74 | 0.37 | 0.31 | 0.38 | 0.4 |
| 4 | 124844059 | rs1519242 | 0.223 | 1.526 | 0.819 | 0.485 | 0.008 | -0.005 | -0.005 | 0.229 | 0.13 | 0.16 | 0.16 | 0.08 | 0.17 | 0.15 |
| 4 | 124846679 | rs17007605 | 1.547 | 1.83 | 0.868 | 0.044 | 0.021 | -0.011 | -0.011 | 0 | 0.61 | 0.5 | 0.62 | 0.7 | 0.54 | 0.53 |
| 4 | 124847477 | rs10034152 | -0.827 | 1.834 | 0.842 | 0.046 | 0.008 | -0.005 | -0.005 | 0.043 | 0.87 | 0.84 | 0.84 | 0.92 | 0.83 | 0.85 |
| 4 | 124850901 | rs13151047 | -0.12 | 1.707 | 0.247 | 0.11 | -0.003 | -0.006 | -0.006 | 0.465 | 0.3 | 0.28 | 0.29 | 0.36 | 0.32 | 0.15 |
| 4 | 124851510 | rs6838706 | 1.565 | 1.961 | 0.273 | 0.084 | -0.002 | -0.013 | -0.013 | 0 | 0.56 | 0.52 | 0.58 | 0.7 | 0.46 | 0.52 |
| 4 | 124851726 | rs1048174 | 0.332 | 2.052 | 0.184 | 0.195 | -0.001 | 0 | 0 | 0 | 0.24 | 0.27 | 0.24 | 0.36 | 0.24 | 0.14 |
| 4 | 124855529 | rs11946288 | -0.691 | 2.145 | 0.307 | 0.294 | -0.001 | -0.001 | -0.001 | 0 | 0.23 | 0.17 | 0.2 | 0.21 | 0.28 | 0.23 |
| 4 | 124859063 | rs1022373 | -1.439 | 2.175 | 0.05 | 0.244 | 0.013 | -0.004 | -0.004 | 4.147 | 0.89 | 0.84 | 0.85 | 0.91 | 0.84 | 0.87 |
| 4 | 124861428 | rs10027482 | 1.697 | 2.259 | -0.045 | 0.268 | 0.046 | 0.042 | 0.042 | 2.213 | 0.43 | 0.15 | 0.37 | 0.33 | 0.33 | 0.48 |
| 4 | 124865191 | rs2390610 | 1.727 | 2.255 | -0.045 | 0.286 | 0.043 | 0.035 | 0.035 | 2.06 | 0.43 | 0.15 | 0.37 | 0.33 | 0.33 | 0.47 |
| 4 | 124871513 | rs1996580 | -0.946 | 2.408 | -0.047 | 0.302 | 0.006 | -0.007 | -0.007 | 8.087 | 0.85 | 0.82 | 0.84 | 0.9 | 0.82 | 0.78 |
| 4 | 124874115 | rs4833296 | 0.993 | 2.132 | -0.427 | 0.488 | -0.02 | 0 | 0 | 0 | 0.18 | 0.01 | 0.11 | 0.14 | 0.17 | 0.04 |
| 4 | 124875936 | rs17007675 | 0.638 | 2.203 | -0.136 | 0.537 | -0.013 | -0.02 | -0.02 | 0 | 0.09 | 0.03 | 0.12 | 0.18 | 0.13 | 0.16 |
| 4 | 124876173 | rs1519238 | 0.638 | 2.203 | -0.136 | 0.537 | -0.013 | -0.02 | -0.02 | 0 | 0.09 | 0.03 | 0.12 | 0.18 | 0.13 | 0.16 |
| 4 | 124888965 | rs9998439 | 2.398 | 2.147 | 0.037 | 0.542 | 0.032 | -0.009 | -0.009 | 0 | 0.31 | 0.21 | 0.34 | 0.33 | 0.23 | 0.38 |
| 4 | 124890766 | rs1474147 | 1.251 | 2.172 | -0.006 | 0.534 | 0 | 0.012 | 0.012 | 0.002 | 0.08 | 0.02 | 0.06 | 0.02 | 0.07 | 0.11 |
| 4 | 124891317 | rs4833961 | 1.628 | 2.252 | 0.16 | 0.634 | 0.022 | -0.026 | -0.026 | 0 | 0.4 | 0.28 | 0.47 | 0.51 | 0.35 | 0.54 |
| 4 | 124905690 | rs9307535 | 1.262 | 2.257 | 0.499 | 0.852 | 0.034 | 0.008 | 0.008 | 0 | 0.39 | 0.27 | 0.41 | 0.35 | 0.3 | 0.49 |
| 4 | 124910861 | rs1519225 | -0.289 | 2.165 | 0.486 | 0.845 | 0.006 | 0.012 | 0.012 | 0.21 | 0.66 | 0.56 | 0.67 | 0.6 | 0.63 | 0.85 |
| 4 | 124926477 | rs2893068 | 0.264 | 2.272 | -0.13 | 0.936 | -0.007 | -0.011 | -0.011 | 0 | 0.3 | 0.26 | 0.31 | 0.38 | 0.34 | 0.15 |
| 4 | 124929001 | rs12641524 | 0.134 | 2.301 | -0.043 | 0.812 | 0.005 | -0.008 | -0.008 | 1.543 | 0.17 | 0.2 | 0.16 | 0.11 | 0.2 | 0.23 |
| 4 | 124955670 | rs17756952 | 0.078 | 2.323 | 0.144 | 1.278 | -0.006 | -0.011 | -0.011 | 0 | 0.3 | 0.26 | 0.31 | 0.38 | 0.34 | 0.15 |
| 4 | 124965262 | rs1850756 | 0.087 | 2.421 | 0.149 | 1.282 | -0.002 | -0.011 | -0.011 | 0 | 0.3 | 0.26 | 0.3 | 0.38 | 0.32 | 0.14 |
| 4 | 124966525 | rs10003267 | 0.542 | 2.284 | 0.444 | 1.692 | 0.002 | 0 | 0 | 0 | 0.49 | 0.4 | 0.53 | 0.52 | 0.49 | 0.63 |
| 4 | 124974957 | rs17813612 | -0.605 | 2.418 | -0.031 | 1.451 | 0.012 | 0.041 | 0.041 | 0.01 | 0.44 | 0.34 | 0.3 | 0.28 | 0.38 | 0.28 |
| 4 | 124992645 | rs10007602 | -1.15 | 1.842 | 0.756 | 1.302 | 0.072 | 0.02 | 0.02 | 0.213 | 0.16 | 0.41 | 0.32 | 0.14 | 0.2 | 0.14 |
| 4 | 125013856 | rs4371629 | 0.48 | 1.863 | 0.784 | 1.251 | 0.004 | -0.003 | -0.003 | 0.018 | 0.05 | 0.07 | 0.23 | 0.02 | 0.09 | 0.03 |
| 4 | 125017034 | rs12647636 | -0.105 | 1.884 | 0.735 | 1.617 | -0.001 | 0.003 | 0.003 | 3.662 | 0.93 | 0.94 | 0.89 | 0.91 | 0.94 | 0.92 |
| 4 | 125020390 | rs10029877 | -0.105 | 1.884 | 0.735 | 1.617 | -0.001 | 0.003 | 0.003 | 3.662 | 0.93 | 0.94 | 0.89 | 0.91 | 0.94 | 0.92 |
| 4 | 125037794 | rs4421000 | -0.914 | 1.518 | 0.532 | 1.377 | -0.016 | -0.018 | -0.018 | 0 | 0.59 | 0.73 | 0.57 | 0.48 | 0.52 | 0.66 |
| 4 | 125046178 | rs11945388 | 0.823 | 1.493 | 0.525 | 1.287 | 0 | -0.019 | -0.019 | 0.133 | 0.07 | 0.04 | 0.04 | 0 | 0.07 | 0.07 |
| 4 | 125078861 | rs893558 | -0.15 | 1.519 | 0.553 | 1.297 | -0.016 | -0.002 | -0.002 | 0.265 | 0.1 | 0.04 | 0.23 | 0.06 | 0.14 | 0.11 |
| 4 | 125095213 | rs11933201 | -0.936 | 1.496 | 0.58 | 1.294 | -0.019 | 0.014 | 0.014 | 0 | 0.16 | 0.07 | 0.27 | 0.06 | 0.21 | 0.18 |
| 4 | 125104155 | rs7664648 | -1.498 | 1.432 | 1.011 | 1.362 | 0.055 | 0.019 | 0.019 | 0 | 0.2 | 0.41 | 0.32 | 0.19 | 0.25 | 0.22 |
| 4 | 125109084 | rs6834028 | -0.589 | 1.157 | 0.695 | 1.232 | -0.001 | 0 | 0 | 0 | 0.53 | 0.44 | 0.53 | 0.44 | 0.49 | 0.56 |
| 4 | 125120257 | rs960925 | -0.604 | 1.143 | 0.686 | 1.225 | -0.001 | -0.001 | -0.001 | 0 | 0.54 | 0.43 | 0.53 | 0.44 | 0.49 | 0.57 |
| 4 | 125127477 | rs17007911 | -1.311 | 1.205 | 0.72 | 1.295 | 0.035 | 0.013 | 0.013 | 0 | 0.2 | 0.33 | 0.31 | 0.21 | 0.25 | 0.22 |
| 4 | 125133465 | rs11937163 | 1.116 | 1.151 | 0.592 | 0.784 | -0.002 | 0 | 0 | 0 | 0.06 | 0.03 | 0.05 | 0 | 0.07 | 0.07 |
| 4 | 125141551 | rs1356837 | 0.775 | 1.191 | 0.618 | 0.828 | 0.002 | 0.005 | 0.005 | 0 | 0.05 | 0.06 | 0.11 | 0.06 | 0.05 | 0.08 |
| 4 | 125157264 | rs1123772 | 0.277 | 1.178 | 0.573 | 0.786 | 0.045 | 0.032 | 0.032 | 0.692 | 0.42 | 0.24 | 0.3 | 0.38 | 0.35 | 0.45 |
| 4 | 125188048 | rs725001 | -1.135 | 1.178 | 0.877 | 0.767 | -0.018 | -0.02 | -0.02 | 1.076 | 0.61 | 0.75 | 0.58 | 0.46 | 0.52 | 0.63 |
| 4 | 125209067 | rs13145092 | -0.204 | 1.196 | 0.829 | 0.785 | 0.013 | -0.006 | -0.006 | 0 | 0.07 | 0.33 | 0.05 | 0.02 | 0.03 | 0.06 |
| 4 | 125214511 | rs17759829 | -0.053 | 1.22 | 0.796 | 0.775 | 0.006 | -0.001 | -0.001 | 2 | 0.11 | 0.1 | 0.23 | 0.06 | 0.15 | 0.11 |
| 4 | 125220846 | rs11735374 | -0.924 | 1.283 | 0.741 | 0.852 | -0.002 | -0.002 | -0.002 | 0 | 0.59 | 0.63 | 0.57 | 0.46 | 0.52 | 0.63 |
| 4 | 125225084 | rs10518427 | -0.16 | 1.223 | 0.5 | 0.879 | 0.011 | 0.03 | 0.03 | 0 | 0.09 | 0.14 | 0.08 | 0.17 | 0.11 | 0.11 |
| 4 | 125225107 | rs17816583 | 0.938 | 1.222 | 0.291 | 0.712 | 0 | -0.013 | -0.013 | 0.287 | 0.06 | 0.06 | 0.05 | 0 | 0.07 | 0.07 |
| 4 | 125241747 | rs13137088 | -0.431 | 1.232 | 0.316 | 0.84 | 0.004 | 0.003 | 0.003 | 0.237 | 0.56 | 0.49 | 0.55 | 0.47 | 0.49 | 0.56 |
| 4 | 125249002 | rs2168692 | 0.872 | 1.191 | 0.09 | 0.792 | -0.031 | 0.029 | 0.029 | 0 | 0.06 | 0 | 0.13 | 0 | 0.09 | 0.04 |
| 4 | 125265313 | rs2125000 | 0.27 | 0.945 | -0.5 | 0.667 | -0.003 | -0.044 | -0.044 | 0 | 0.3 | 0.03 | 0.24 | 0.29 | 0.3 | 0.27 |
| 4 | 125266797 | rs672858 | 1.286 | 0.871 | -0.17 | 0.742 | -0.014 | -0.013 | -0.013 | 0 | 0.46 | 0.23 | 0.43 | 0.49 | 0.49 | 0.52 |
| 4 | 125268531 | rs593371 | 1.413 | 1.169 | 0.856 | 0.615 | 0.015 | 0.011 | 0.011 | 0.29 | 0.77 | 0.36 | 0.68 | 0.77 | 0.78 | 0.78 |
| 4 | 125280852 | rs640246 | 0.88 | 1.133 | 0.632 | 0.501 | 0.031 | 0.009 | 0.009 | 0.55 | 0.83 | 0.55 | 0.8 | 0.82 | 0.83 | 0.87 |
| 4 | 125295432 | rs4833978 | 0.603 | 1.126 | -0.047 | 0.257 | -0.003 | 0.01 | 0.01 | 0.703 | 0.25 | 0.15 | 0.27 | 0.22 | 0.28 | 0.08 |
| 4 | 125296033 | rs9999775 | 0.133 | 0.954 | 0.319 | 0.316 | 0.014 | 0.066 | 0.066 | 0 | 0.94 | 0.8 | 0.94 | 0.84 | 0.95 | 0.96 |
| 4 | 125298963 | rs1159819 | 1.057 | 0.997 | 0.305 | 0.374 | 0.017 | 0.062 | 0.062 | 0.055 | 0.89 | 0.71 | 0.84 | 0.78 | 0.9 | 0.87 |
| 4 | 125303951 | rs2390684 | 1.341 | 0.974 | -0.268 | 0.249 | -0.03 | -0.111 | -0.111 | 0 | 0.39 | 0.11 | 0.33 | 0.44 | 0.41 | 0.62 |
| 4 | 125318885 | rs1523504 | 1.064 | 1.107 | 0.457 | 0.262 | -0.07 | 0.005 | 0.005 | 0.967 | 0.56 | 0.12 | 0.53 | 0.54 | 0.57 | 0.7 |
| 4 | 125320611 | rs17008117 | -0.789 | 1.223 | 0.766 | 0.419 | 0.016 | 0.056 | 0.056 | 0 | 0.07 | 0.15 | 0.14 | 0.22 | 0.08 | 0.12 |

| | | | | | | | | | | | | | | | | |
|---|---|---|---|---|---|---|---|---|---|---|---|---|---|---|---|---|
| 4 | 125326845 | rs1851689 | -0.418 | 1.329 | 0.677 | 0.429 | -0.01 | 0.018 | 0.018 | 0.188 | 0.32 | 0.2 | 0.38 | 0.27 | 0.35 | 0.2 |
| 4 | 125332916 | rs7691579 | 0.038 | 1.312 | 0.596 | 0.448 | -0.004 | 0.021 | 0.021 | 0.329 | 0.28 | 0.16 | 0.28 | 0.22 | 0.3 | 0.1 |
| 4 | 125355040 | rs7661156 | 0.237 | 1.361 | 0.411 | 0.553 | 0.017 | 0.011 | 0.011 | 0 | 0.91 | 0.73 | 0.87 | 0.94 | 0.93 | 0.9 |
| 4 | 125361767 | rs7655016 | 0.456 | 1.295 | -0.283 | 0.516 | -0.002 | 0.014 | 0.014 | 1.812 | 0.6 | 0.28 | 0.54 | 0.56 | 0.61 | 0.77 |
| 4 | 125371187 | rs12506469 | -0.517 | 1.238 | -0.321 | 0.374 | 0.008 | 0.031 | 0.031 | 1.273 | 0.33 | 0.21 | 0.3 | 0.28 | 0.33 | 0.19 |
| 4 | 125379709 | rs6839627 | -0.537 | 1.149 | -0.471 | 0.23 | -0.001 | -0.001 | -0.001 | 0 | 0.23 | 0.21 | 0.24 | 0.28 | 0.25 | 0.18 |
| 4 | 125380922 | rs11936470 | 1.248 | 1.661 | 0.217 | 0.432 | 0.003 | 0.002 | 0.002 | 3.085 | 0.54 | 0.53 | 0.54 | 0.43 | 0.51 | 0.75 |
| 4 | 125391283 | rs17008220 | -0.039 | 1.67 | 0.509 | 0.334 | 0.005 | -0.038 | -0.038 | 0.112 | 0.17 | 0.27 | 0.15 | 0.09 | 0.22 | 0.16 |
| 4 | 125399775 | rs13119813 | 0.88 | 1.764 | 0.887 | 0.464 | 0.001 | -0.005 | -0.005 | 1.555 | 0.61 | 0.42 | 0.63 | 0.63 | 0.57 | 0.77 |
| 4 | 125404239 | rs962939 | -0.121 | 1.552 | 0.634 | 0.382 | 0.003 | 0.011 | 0.011 | 0.23 | 0.27 | 0.25 | 0.2 | 0.09 | 0.29 | 0.17 |
| 4 | 125423945 | rs10857100 | 0.836 | 1.701 | 0.985 | 0.509 | 0 | 0 | 0 | 0 | 0.6 | 0.44 | 0.66 | 0.62 | 0.55 | 0.72 |
| 4 | 125426113 | rs6848373 | 0.096 | 1.678 | 1.244 | 0.475 | -0.002 | -0.023 | -0.023 | 0 | 0.26 | 0.24 | 0.19 | 0.1 | 0.29 | 0.16 |
| 4 | 125427661 | rs12641839 | 0.216 | 1.692 | 1.248 | 0.491 | 0.042 | 0.094 | 0.094 | 0 | 0.87 | 0.68 | 0.86 | 0.73 | 0.84 | 0.88 |
| 4 | 125435781 | rs101927 | 0.819 | 1.671 | 1.461 | 0.476 | 0.003 | -0.008 | -0.008 | 1.4 | 0.61 | 0.48 | 0.66 | 0.63 | 0.56 | 0.73 |
| 4 | 125437668 | rs312508 | -0.006 | 1.746 | 1.398 | 0.542 | 0.01 | 0.005 | 0.005 | 0 | 0.88 | 0.71 | 0.87 | 0.89 | 0.87 | 0.92 |
| 4 | 125440570 | rs17008280 | 0.424 | 1.756 | 1.333 | 0.547 | 0.01 | 0.04 | 0.04 | 0 | 0.09 | 0.12 | 0.11 | 0.27 | 0.13 | 0.07 |
| 4 | 125445182 | rs17008303 | 1.448 | 1.66 | 0.453 | 0.439 | 0.008 | 0.017 | 0.017 | 0 | 0.26 | 0.17 | 0.23 | 0.2 | 0.19 | 0.33 |
| 4 | 125448257 | rs312492 | -0.45 | 1.611 | 0.535 | 0.382 | -0.022 | -0.094 | -0.094 | 0 | 0.72 | 0.49 | 0.78 | 0.9 | 0.71 | 0.84 |
| 4 | 125448288 | rs312493 | 0.345 | 1.469 | 0.471 | 0.261 | -0.001 | 0.001 | 0.001 | 0.841 | 0.6 | 0.35 | 0.65 | 0.63 | 0.55 | 0.72 |
| 4 | 125451364 | rs10029005 | -0.272 | 1.404 | 0.419 | 0.36 | -0.001 | -0.001 | -0.001 | 0.239 | 0.36 | 0.41 | 0.31 | 0.37 | 0.41 | 0.23 |
| 4 | 125455038 | rs1509524 | 1.118 | 1.267 | 0.002 | 0.23 | 0.008 | 0.016 | 0.016 | 0 | 0.26 | 0.17 | 0.23 | 0.2 | 0.2 | 0.33 |
| 4 | 125457073 | rs11946186 | -0.828 | 1.473 | 0.522 | 0.52 | -0.007 | -0.021 | -0.021 | 3.779 | 0.3 | 0.61 | 0.25 | 0.26 | 0.31 | 0.2 |
| 4 | 125462793 | rs975107 | -0.302 | 1.507 | 0.28 | 0.602 | -0.023 | -0.038 | -0.038 | 0 | 0.94 | 0.99 | 0.9 | 0.8 | 0.94 | 0.98 |
| 4 | 125463533 | rs7666278 | -1.012 | 1.284 | 0.605 | -0.012 | -0.006 | -0.015 | -0.015 | 0 | 0.3 | 0.63 | 0.25 | 0.26 | 0.31 | 0.2 |
| 4 | 125465163 | rs1567396 | -0.977 | 1.331 | 0.306 | 0.011 | -0.007 | -0.02 | -0.02 | 0 | 0.3 | 0.57 | 0.25 | 0.28 | 0.31 | 0.2 |
| 4 | 125479073 | rs558208 | -0.398 | 1.461 | -0.257 | -0.376 | -0.007 | 0.012 | 0.012 | 0 | 0.19 | 0.09 | 0.13 | 0.01 | 0.22 | 0.15 |
| 4 | 125480782 | rs1092825 | -0.724 | 1.435 | 0.072 | -0.427 | 0.005 | -0.165 | -0.165 | 0 | 0.25 | 0.49 | 0.19 | 0.01 | 0.24 | 0.15 |
| 4 | 125484598 | rs491645 | -0.8 | 1.495 | 0.033 | -0.484 | 0.002 | -0.077 | -0.077 | 0 | 0.28 | 0.31 | 0.19 | 0.01 | 0.3 | 0.19 |
| 4 | 125494504 | rs508410 | 0.309 | 1.925 | -1.105 | 0.909 | -0.067 | -0.081 | -0.081 | 0.857 | 0.82 | 0.95 | 0.79 | 0.81 | 0.69 | 0.82 |
| 4 | 125500592 | rs7677431 | 0.222 | 1.553 | -1.203 | 1.269 | 0.004 | 0.001 | 0.001 | 0.18 | 0.51 | 0.42 | 0.47 | 0.47 | 0.44 | 0.61 |
| 4 | 125511115 | rs17008385 | 0.647 | 1.593 | -1.251 | 0.924 | 0.003 | -0.01 | -0.01 | 0.633 | 0.09 | 0.07 | 0.07 | 0.02 | 0.12 | 0.06 |
| 4 | 125512711 | rs2048265 | -0.536 | 1.577 | -1.767 | 0.486 | 0.023 | 0.035 | 0.035 | 0 | 0.66 | 0.46 | 0.56 | 0.47 | 0.57 | 0.64 |
| 4 | 125513521 | rs4833990 | 0.236 | 1.377 | -1.786 | 0.354 | 0.001 | 0.023 | 0.023 | 0 | 0.09 | 0.08 | 0.16 | 0.21 | 0.13 | 0.22 |
| 4 | 125518999 | rs7689192 | -0.791 | 1.363 | -1.692 | 0.825 | 0.027 | 0.013 | 0.013 | 0 | 0.66 | 0.48 | 0.62 | 0.53 | 0.57 | 0.65 |
| 4 | 125519380 | rs4833993 | 0.701 | 0.997 | -1.932 | 0.521 | 0.031 | -0.007 | -0.007 | 0.462 | 0.37 | 0.08 | 0.35 | 0.28 | 0.28 | 0.53 |
| 4 | 125523042 | rs1390079 | 1.312 | 1.048 | -1.351 | 0.713 | 0.009 | 0.015 | 0.015 | 0.157 | 0.15 | 0.07 | 0.15 | 0.06 | 0.13 | 0.03 |
| 4 | 125531787 | rs6846249 | 1.26 | 1.055 | -1.356 | 0.561 | 0.008 | 0.041 | 0.041 | 0.179 | 0.15 | 0.06 | 0.09 | 0 | 0.12 | 0.03 |
| 4 | 125544726 | rs9996039 | -0.984 | 1.215 | -0.9 | 0.762 | 0.034 | 0.052 | 0.052 | 0 | 0.2 | 0.37 | 0.26 | 0.24 | 0.25 | 0.28 |
| 4 | 125554113 | rs870741 | 0.696 | 1.063 | -0.989 | 0.938 | -0.003 | -0.018 | -0.018 | 0 | 0.08 | 0.13 | 0.04 | 0.02 | 0.06 | 0.06 |
| 4 | 125582871 | rs10006893 | -1.294 | 1.076 | 0.014 | 1.104 | 0.007 | 0.08 | 0.08 | 0 | 0.18 | 0.39 | 0.21 | 0.26 | 0.2 | 0.28 |
| 4 | 125584268 | rs7675483 | 0.967 | 0.942 | -0.627 | 0.935 | -0.046 | 0.047 | 0.047 | 0 | 0.25 | 0.04 | 0.21 | 0.15 | 0.29 | 0.27 |
| 4 | 125587561 | rs3172524 | -0.592 | 0.968 | -0.421 | 1.101 | 0.014 | 0.086 | 0.086 | 0.003 | 0.17 | 0.35 | 0.21 | 0.26 | 0.2 | 0.28 |
| 4 | 125590478 | rs1874748 | 0.099 | 0.969 | -0.42 | 1.102 | 0.039 | 0.131 | 0.131 | 0 | 0.09 | 0.23 | 0.18 | 0.24 | 0.16 | 0.24 |
| 4 | 125591687 | rs3733471 | -1.06 | 0.97 | -0.476 | 1.1 | 0.014 | 0.077 | 0.077 | 0 | 0.18 | 0.39 | 0.22 | 0.26 | 0.21 | 0.29 |
| 4 | 125592440 | rs3733472 | 1.258 | 0.964 | -0.429 | 1.089 | 0 | 0.001 | 0.001 | 1.325 | 0.07 | 0.05 | 0.04 | 0.02 | 0.05 | 0.05 |
| 4 | 125602470 | rs7656458 | 1.21 | 0.712 | -0.031 | 1.018 | -0.008 | -0.004 | -0.004 | 0 | 0.34 | 0.24 | 0.3 | 0.39 | 0.3 | 0.32 |
| 4 | 125610859 | rs13110120 | -0.584 | 0.652 | 0.397 | 1.342 | 0.048 | 0.154 | 0.154 | 0 | 0.11 | 0.35 | 0.19 | 0.24 | 0.16 | 0.24 |
| 4 | 125619501 | rs13435256 | 1.176 | 0.65 | 0.377 | 1.33 | 0 | 0 | 0 | 0 | 0.07 | 0.05 | 0.03 | 0.02 | 0.04 | 0.04 |
| 4 | 125620031 | rs2036204 | -1.103 | 0.659 | 0.347 | 1.334 | 0.012 | 0.082 | 0.082 | 0 | 0.18 | 0.39 | 0.21 | 0.26 | 0.2 | 0.28 |
| 4 | 125636426 | rs10025996 | -0.923 | 0.135 | 0.074 | 1.14 | -0.015 | 0.01 | 0.01 | 0.068 | 0.2 | 0.3 | 0.27 | 0.24 | 0.13 | 0.15 |
| 4 | 125637940 | rs17008503 | -0.159 | 0.199 | -0.062 | 1.233 | 0.001 | -0.002 | -0.002 | 0 | 0.12 | 0.17 | 0.24 | 0.22 | 0.09 | 0.11 |
| 4 | 125642349 | rs11728675 | 1.59 | -0.074 | -0.197 | 1.284 | 0.009 | 0.07 | 0.07 | 0.193 | 0.13 | 0.02 | 0.08 | 0 | 0.09 | 0.02 |
| 4 | 125649024 | rs11098769 | 1.27 | -0.084 | -0.789 | 0.923 | -0.035 | 0.068 | 0.068 | 2.444 | 0.28 | 0.07 | 0.24 | 0.14 | 0.26 | 0.3 |
| 4 | 125649621 | rs9993784 | 0.526 | -0.182 | -0.703 | 0.895 | -0.004 | 0.078 | 0.078 | 1.567 | 0.36 | 0.16 | 0.28 | 0.16 | 0.3 | 0.35 |
| 4 | 125652879 | rs10518438 | -0.11 | -0.102 | -0.243 | 1.142 | 0.006 | 0.049 | 0.049 | 0.155 | 0.53 | 0.38 | 0.46 | 0.24 | 0.39 | 0.42 |
| 4 | 125653594 | rs10518436 | 1.06 | 0.471 | -0.824 | 1.437 | -0.02 | -0.024 | -0.024 | 0 | 0.28 | 0.07 | 0.23 | 0.24 | 0.3 | 0.28 |
| 4 | 125659471 | rs17772791 | 0.464 | 1.102 | -0.205 | 0.968 | 0.066 | 0.018 | 0.018 | 0 | 0.1 | 0.2 | 0.18 | 0.13 | 0.25 | 0.15 |
| 4 | 125659601 | rs10008933 | 0.457 | 0.461 | -0.806 | 0.79 | -0.009 | 0.008 | 0.008 | 1.522 | 0.28 | 0.17 | 0.33 | 0.22 | 0.26 | 0.35 |
| 4 | 125666014 | rs13102814 | 0.438 | -0.328 | -0.487 | 1.019 | -0.003 | 0.061 | 0.061 | 0 | 0.05 | 0.09 | 0.15 | 0.21 | 0.07 | 0.09 |
| 4 | 125666719 | rs6841569 | 0.645 | -0.47 | -0.293 | 0.912 | -0.01 | 0.045 | 0.045 | 0 | 0.05 | 0.13 | 0.08 | 0.13 | 0.07 | 0.04 |
| 4 | 125668245 | rs10020316 | 0.577 | -0.911 | -0.163 | 0.613 | 0 | -0.005 | -0.005 | 0 | 0.31 | 0.34 | 0.32 | 0.2 | 0.25 | 0.32 |
| 4 | 125670010 | rs11722399 | -1.055 | -1.099 | -0.849 | -0.104 | -0.086 | 0.127 | 0.127 | 0.121 | 0.76 | 0.96 | 0.76 | 1 | 0.66 | 0.89 |
| 4 | 125675055 | rs2036203 | -0.456 | -1.112 | -0.315 | -0.18 | 0.012 | -0.037 | -0.037 | 0.26 | 0.69 | 0.59 | 0.66 | 0.87 | 0.57 | 0.86 |
| 4 | 125683296 | rs1506551 | 1.144 | -1.085 | -0.697 | 0.095 | 0.002 | -0.086 | -0.086 | 0 | 0.25 | 0.1 | 0.22 | 0.56 | 0.24 | 0.34 |
| 4 | 125686468 | rs17008575 | 0.049 | -0.937 | -0.285 | 0.047 | 0 | 0 | 0 | 0 | 0.13 | 0.1 | 0.11 | 0.1 | 0.07 | 0.19 |
| 4 | 125687748 | rs17008580 | 0.008 | -0.937 | -0.273 | 0.047 | -0.005 | -0.004 | -0.004 | 0 | 0.13 | 0.12 | 0.11 | 0.1 | 0.08 | 0.19 |
| 4 | 125707202 | rs10518450 | -0.748 | -0.957 | -0.063 | -0.122 | -0.016 | 0.049 | 0.049 | 0 | 0.08 | 0.14 | 0.04 | 0.14 | 0.03 | 0.12 |
| 4 | 125707389 | rs17008594 | -0.778 | -0.935 | -0.045 | -0.116 | -0.013 | 0.055 | 0.055 | 0 | 0.08 | 0.16 | 0.04 | 0.14 | 0.03 | 0.12 |
| 4 | 125712060 | rs993704 | 0.219 | -1.221 | 0.233 | -0.319 | -0.011 | 0.008 | 0.008 | 0 | 0.37 | 0.43 | 0.34 | 0.63 | 0.31 | 0.46 |

| | | | | | | | | | | | | | | | | |
|---|---|---|---|---|---|---|---|---|---|---|---|---|---|---|---|---|
| 4 | 125720431 | rs11931944 | 0.142 | -0.779 | 0.149 | -0.494 | -0.013 | 0 | 0 | 1.361 | 0.15 | 0.13 | 0.25 | 0.13 | 0.32 | 0.1 |
| 4 | 125724482 | rs6829814 | 0.16 | -0.884 | 0.662 | -0.571 | 0.001 | -0.047 | -0.047 | 0 | 0.13 | 0.24 | 0.14 | 0.07 | 0.1 | 0.17 |
| 4 | 125726204 | rs10518444 | -0.064 | -1.456 | 0.644 | -0.324 | -0.014 | 0.007 | 0.007 | 0 | 0.11 | 0.18 | 0.09 | 0.15 | 0.08 | 0.16 |
| 4 | 125728477 | rs10518443 | -0.134 | -1.556 | 0.871 | -0.24 | -0.024 | 0.185 | 0.185 | 0 | 0.09 | 0.28 | 0.11 | 0.25 | 0.07 | 0.06 |
| 4 | 125732734 | rs10029225 | -1.524 | -1.659 | 0.72 | -0.32 | 0.036 | -0.032 | -0.032 | 0 | 0.33 | 0.71 | 0.32 | 0.48 | 0.21 | 0.36 |
| 4 | 125746220 | rs1543936 | -1.756 | -1.523 | 0.873 | -0.431 | 0 | -0.001 | -0.001 | 0 | 0.4 | 0.57 | 0.47 | 0.36 | 0.46 | 0.39 |
| 4 | 125755019 | rs1848056 | -1.393 | -2.071 | 0.735 | -0.59 | -0.012 | 0.054 | 0.054 | 0.794 | 0.53 | 0.68 | 0.53 | 0.36 | 0.55 | 0.41 |
| 4 | 125763629 | rs9997902 | -1.338 | -2.084 | 0.724 | -0.579 | -0.007 | 0.038 | 0.038 | 0.639 | 0.53 | 0.64 | 0.52 | 0.35 | 0.55 | 0.42 |
| 4 | 125767985 | rs1395330 | -0.537 | -2.109 | 0.565 | -0.598 | -0.042 | -0.026 | -0.026 | 0 | 0.22 | 0.36 | 0.13 | 0.14 | 0.16 | 0.22 |
| 4 | 125769617 | rs13124340 | 0.474 | -2.091 | 0.372 | -0.58 | -0.012 | 0.014 | 0.014 | 0 | 0.1 | 0.12 | 0.08 | 0.14 | 0.1 | 0.13 |
| 4 | 125771540 | rs1395331 | -1.534 | -2.077 | -0.031 | -0.542 | -0.05 | -0.041 | -0.041 | 0 | 0.55 | 0.8 | 0.56 | 0.48 | 0.55 | 0.43 |
| 4 | 125773999 | rs17008684 | 0.702 | -1.949 | -0.177 | -0.441 | 0.011 | -0.028 | -0.028 | 0 | 0.1 | 0.04 | 0.1 | 0.16 | 0.07 | 0.06 |
| 4 | 125781529 | rs17786141 | 0.55 | -2.179 | -0.005 | -0.699 | 0.013 | -0.021 | -0.021 | 0 | 0.26 | 0.38 | 0.25 | 0.12 | 0.33 | 0.11 |
| 4 | 125784021 | rs2390740 | -0.695 | -2.434 | -0.304 | -0.865 | -0.012 | 0.008 | 0.008 | 0 | 0.84 | 0.86 | 0.82 | 0.8 | 0.86 | 0.79 |
| 4 | 125786036 | rs7695725 | 1.087 | -2.389 | -0.417 | -0.865 | 0.005 | 0.021 | 0.021 | 0.229 | 0.09 | 0.03 | 0.05 | 0.03 | 0.07 | 0.07 |
| 4 | 125788673 | rs4833310 | 2.505 | -2.395 | -0.588 | -0.839 | -0.059 | -0.048 | -0.048 | 0 | 0.16 | 0.01 | 0.26 | 0.22 | 0.16 | 0.21 |
| 4 | 125790657 | rs13148187 | 0.886 | -2.589 | 0.148 | -0.872 | 0.006 | -0.004 | -0.004 | 1.457 | 0.09 | 0.01 | 0.03 | 0.09 | 0.09 | 0.12 |
| 4 | 125791252 | rs2086765 | 2.237 | -2.565 | 0.177 | -0.946 | -0.092 | -0.077 | -0.077 | 0 | 0.35 | 0.09 | 0.43 | 0.63 | 0.39 | 0.52 |
| 4 | 125796096 | rs10034686 | 0.884 | -2.188 | 0.556 | -0.726 | -0.027 | -0.016 | -0.016 | 0 | 0.72 | 0.65 | 0.75 | 0.79 | 0.8 | 0.7 |
| 4 | 125804479 | rs1395326 | 0.114 | -2.159 | 0.322 | -0.644 | -0.032 | 0.066 | 0.066 | 0 | 0.81 | 0.69 | 0.76 | 0.68 | 0.85 | 0.76 |
| 4 | 125807868 | rs9993168 | 1.395 | -1.983 | 0.433 | -0.686 | -0.006 | 0.02 | 0.02 | 0 | 0.48 | 0.33 | 0.49 | 0.64 | 0.48 | 0.57 |
| 4 | 125812575 | rs1995626 | 1.012 | -2.001 | 0.279 | -0.709 | -0.024 | 0.013 | 0.013 | 0 | 0.66 | 0.63 | 0.72 | 0.6 | 0.78 | 0.62 |
| 4 | 125813232 | rs10033902 | 1.133 | -1.865 | 0.321 | -0.874 | -0.057 | 0.071 | 0.071 | 0.595 | 0.18 | 0.09 | 0.22 | 0 | 0.31 | 0.06 |
| 4 | 125813774 | rs16997819 | 2.25 | -1.884 | 0.298 | -0.874 | -0.058 | -0.056 | -0.056 | 0.041 | 0.38 | 0.15 | 0.44 | 0.61 | 0.41 | 0.5 |
| 4 | 125815138 | rs9990572 | 0.762 | -1.908 | 0.017 | -0.776 | -0.046 | 0.039 | 0.039 | 0.026 | 0.73 | 0.64 | 0.76 | 0.66 | 0.84 | 0.71 |
| 4 | 125831214 | rs6838702 | 1.113 | -1.509 | -0.633 | -0.664 | -0.048 | 0 | 0 | 0.349 | 0.21 | 0.09 | 0.24 | 0.14 | 0.31 | 0.09 |
| 4 | 125835783 | rs7698052 | -1.055 | -1.56 | -0.631 | -0.683 | 0.015 | 0.024 | 0.024 | 0 | 0.89 | 0.95 | 0.91 | 0.94 | 0.94 | 0.88 |
| 4 | 125840550 | rs17008804 | 0.493 | -1.593 | -0.485 | -0.68 | -0.008 | 0.075 | 0.075 | 0 | 0.12 | 0.33 | 0.12 | 0.13 | 0.1 | 0.14 |
| 4 | 125841892 | rs7356273 | 0.509 | -1.587 | -0.64 | -0.723 | -0.032 | 0.062 | 0.062 | 0.238 | 0.6 | 0.83 | 0.63 | 0.77 | 0.56 | 0.73 |
| 4 | 125842481 | rs1909100 | 0.641 | -1.599 | -0.582 | -0.729 | -0.004 | 0.054 | 0.054 | 0 | 0.58 | 0.71 | 0.62 | 0.76 | 0.56 | 0.73 |
| 4 | 125842539 | rs10470922 | 0.79 | -1.501 | -0.867 | -0.695 | 0 | -0.008 | -0.008 | 0 | 0.08 | 0.11 | 0.06 | 0.01 | 0.05 | 0.08 |
| 4 | 125842552 | rs10518455 | 0.717 | -1.493 | -0.87 | -0.689 | 0.004 | 0.012 | 0.012 | 0 | 0.07 | 0.03 | 0.03 | 0.03 | 0.07 | 0.06 |
| 4 | 125842641 | rs1483594 | 0.864 | -1.173 | -0.716 | -0.528 | -0.048 | 0.004 | 0.004 | 0.182 | 0.22 | 0.09 | 0.24 | 0.14 | 0.31 | 0.09 |
| 4 | 125844550 | rs931651 | 0.935 | -1.37 | 0.133 | -0.336 | 0.018 | 0.026 | 0.026 | 0 | 0.8 | 0.84 | 0.85 | 0.92 | 0.89 | 0.8 |
| 4 | 125846011 | rs7662742 | -0.008 | -1.235 | -0.143 | -0.249 | 0.018 | 0.027 | 0.027 | 0 | 0.88 | 0.95 | 0.91 | 0.94 | 0.95 | 0.89 |
| 4 | 125849986 | rs6847509 | -0.706 | -1.023 | 0.068 | -0.12 | -0.047 | -0.004 | -0.004 | 0.498 | 0.2 | 0.45 | 0.17 | 0.14 | 0.15 | 0.22 |
| 4 | 125853748 | rs1483599 | 0.194 | -0.69 | -1.005 | -0.057 | 0.011 | 0.03 | 0.03 | 0 | 0.84 | 0.93 | 0.88 | 0.91 | 0.88 | 0.83 |
| 4 | 125856465 | rs6852823 | -0.779 | -0.78 | -0.484 | 0.021 | -0.035 | -0.003 | -0.003 | 0.004 | 0.19 | 0.56 | 0.16 | 0.14 | 0.15 | 0.23 |
| 4 | 125863974 | rs10022194 | -0.684 | -0.468 | -0.79 | 0.019 | -0.002 | -0.009 | -0.009 | 0 | 0.13 | 0.14 | 0.12 | 0.06 | 0.07 | 0.12 |
| 4 | 125864653 | rs2170743 | 0.639 | -0.314 | -1.259 | 0.158 | 0.019 | -0.003 | -0.003 | 0 | 0.53 | 0.2 | 0.48 | 0.54 | 0.56 | 0.59 |
| 4 | 125865096 | rs1021629 | -0.033 | -0.496 | -1.062 | -0.248 | 0.018 | 0.03 | 0.03 | 0 | 0.41 | 0.36 | 0.37 | 0.78 | 0.36 | 0.49 |
| 4 | 125865319 | rs1021631 | -0.146 | -1.012 | -0.724 | -0.024 | 0.022 | 0.001 | 0.001 | 0 | 0.46 | 0.32 | 0.39 | 0.56 | 0.42 | 0.54 |
| 4 | 125874154 | rs7684522 | 1.209 | -1.146 | -1.252 | -0.274 | -0.072 | -0.114 | -0.114 | 0.089 | 0.27 | 0.01 | 0.25 | 0.38 | 0.34 | 0.36 |
| 4 | 125875734 | rs1498049 | 0.238 | -1.172 | -0.326 | -0.173 | 0.024 | 0.017 | 0.017 | 0 | 0.47 | 0.31 | 0.35 | 0.59 | 0.46 | 0.46 |
| 4 | 125876427 | rs13133741 | 0.735 | -0.751 | 0.021 | -0.28 | -0.008 | 0.015 | 0.015 | 0.241 | 0.31 | 0.22 | 0.38 | 0.29 | 0.29 | 0.33 |
| 4 | 125878081 | rs17008912 | 0.261 | -0.838 | 0.236 | -0.244 | 0.016 | 0.014 | 0.014 | 0 | 0.51 | 0.4 | 0.42 | 0.61 | 0.47 | 0.55 |
| 4 | 125880959 | rs11723954 | 0.325 | -0.839 | 0.351 | -0.258 | 0.003 | 0.003 | 0.003 | 0 | 0.47 | 0.42 | 0.4 | 0.62 | 0.39 | 0.53 |
| 4 | 125886011 | rs6852293 | 0.873 | -0.804 | 0.277 | -0.253 | 0.025 | 0.003 | 0.003 | 0.007 | 0.42 | 0.3 | 0.31 | 0.59 | 0.37 | 0.44 |
| 4 | 125892629 | rs12640447 | -0.821 | -0.424 | 0.557 | -0.163 | 0.054 | -0.055 | -0.055 | 0.049 | 0.85 | 0.77 | 0.84 | 0.97 | 0.76 | 0.89 |
| 4 | 125894102 | rs12331172 | -0.162 | -0.434 | 0.498 | -0.147 | 0.101 | -0.018 | -0.018 | 0 | 0.82 | 0.55 | 0.78 | 0.91 | 0.74 | 0.89 |
| 4 | 125907632 | rs4834012 | 0.242 | -0.376 | 0.011 | -0.479 | 0.03 | -0.027 | -0.027 | 0 | 0.73 | 0.66 | 0.74 | 0.83 | 0.62 | 0.83 |
| 4 | 125914639 | rs11939336 | 0.776 | -0.626 | -0.098 | -0.5 | -0.001 | -0.001 | -0.001 | 0 | 0.47 | 0.48 | 0.39 | 0.65 | 0.4 | 0.57 |
| 4 | 125931345 | rs6534454 | 0.371 | -0.365 | -0.046 | -0.249 | -0.007 | -0.009 | -0.009 | 0 | 0.09 | 0.09 | 0.1 | 0.15 | 0.14 | 0.05 |
| 4 | 125933427 | rs1498047 | -0.896 | -0.631 | -0.097 | -0.003 | 0.008 | -0.004 | -0.004 | 0 | 0.42 | 0.6 | 0.48 | 0.43 | 0.26 | 0.39 |
| 4 | 125933472 | rs1498046 | -0.896 | -0.631 | -0.097 | -0.003 | 0.008 | -0.004 | -0.004 | 0 | 0.42 | 0.6 | 0.48 | 0.43 | 0.26 | 0.4 |
| 4 | 125937256 | rs1498040 | -1.482 | -0.313 | 0.13 | 0.19 | 0.01 | 0.017 | 0.017 | 0.901 | 0.48 | 0.62 | 0.58 | 0.58 | 0.43 | 0.46 |
| 4 | 125941623 | rs1532492 | -0.973 | -0.223 | 0.147 | 0.224 | -0.004 | 0.025 | 0.025 | 0.068 | 0.07 | 0.14 | 0.13 | 0.17 | 0.06 | 0.12 |
| 4 | 125946668 | rs1874753 | -1.998 | -0.071 | -0.001 | 0.122 | -0.06 | 0.052 | 0.052 | 0 | 0.91 | 0.99 | 0.88 | 0.99 | 0.81 | 0.95 |
| 4 | 125956403 | rs6851045 | -0.502 | -0.961 | 0.576 | 0.567 | -0.034 | 0.108 | 0.108 | 0.711 | 0.33 | 0.12 | 0.35 | 0.16 | 0.34 | 0.32 |
| 4 | 125958645 | rs6534456 | -1.365 | -0.906 | 0.789 | 0.632 | -0.002 | -0.006 | -0.006 | 0 | 0.42 | 0.53 | 0.54 | 0.37 | 0.4 | 0.43 |
| 4 | 125960784 | rs10009302 | 0.289 | -0.36 | 0.456 | 1.213 | -0.013 | 0.083 | 0.083 | 0 | 0.05 | 0.1 | 0.03 | 0.22 | 0.02 | 0.03 |
| 4 | 125973337 | rs2647130 | -1.199 | -0.479 | 0.534 | 1.215 | 0.001 | -0.001 | -0.001 | 0 | 0.33 | 0.32 | 0.48 | 0.61 | 0.26 | 0.43 |
| 4 | 125973565 | rs1391397 | -0.719 | -0.525 | 0.454 | 1.028 | 0.004 | -0.013 | -0.013 | 0.464 | 0.28 | 0.27 | 0.32 | 0.38 | 0.21 | 0.34 |
| 4 | 125976652 | rs6534458 | -0.285 | -0.419 | 0.284 | 1.166 | -0.012 | 0.15 | 0.15 | 0 | 0.94 | 0.83 | 0.96 | 0.78 | 0.97 | 0.96 |
| 4 | 125978009 | rs9784420 | 1.029 | -0.362 | 0.31 | 0.887 | 0.001 | -0.002 | -0.002 | 0 | 0.61 | 0.61 | 0.57 | 0.47 | 0.65 | 0.6 |
| 4 | 125996360 | rs12512644 | 0.932 | -0.031 | -0.003 | 1.195 | -0.001 | 0.001 | 0.001 | 0.163 | 0.64 | 0.63 | 0.7 | 0.47 | 0.66 | 0.65 |
| 4 | 126002667 | rs9685753 | -0.074 | 0.288 | -0.187 | 1.126 | -0.006 | 0.003 | 0.003 | 0 | 0.07 | 0.05 | 0.06 | 0 | 0.11 | 0.03 |
| 4 | 126003832 | rs17791437 | -0.485 | 0.247 | -0.392 | 1.511 | -0.012 | -0.044 | -0.044 | 0 | 0.14 | 0.05 | 0.15 | 0.26 | 0.16 | 0.21 |
| 4 | 126007059 | rs2390797 | -0.702 | 0.155 | -0.217 | 1.691 | -0.001 | -0.025 | -0.025 | 0 | 0.15 | 0.13 | 0.21 | 0.38 | 0.16 | 0.2 |

| | | | | | | | | | | | | | | | | |
|---|---|---|---|---|---|---|---|---|---|---|---|---|---|---|---|---|
| 4 | 126007196 | rs2390798 | -1.792 | -0.071 | 0.272 | 2.006 | -0.004 | 0.023 | 0.023 | 0.113 | 0.22 | 0.27 | 0.3 | 0.56 | 0.22 | 0.33 |
| 4 | 126007788 | rs12645899 | -0.767 | -0.155 | 0.55 | 2.108 | -0.012 | 0.038 | 0.038 | 1.477 | 0.07 | 0.14 | 0.05 | 0.13 | 0.05 | 0.05 |
| 4 | 126010408 | rs13124640 | -0.645 | -0.144 | 0.568 | 2.113 | -0.012 | 0.062 | 0.062 | 0.785 | 0.07 | 0.18 | 0.05 | 0.12 | 0.05 | 0.05 |
| 4 | 126011991 | rs4395523 | -0.681 | -0.144 | 0.543 | 2.114 | -0.011 | 0.038 | 0.038 | 1.227 | 0.07 | 0.14 | 0.04 | 0.12 | 0.06 | 0.05 |
| 4 | 126013202 | rs2390771 | 2.497 | -0.271 | 1.216 | 1.917 | -0.024 | 0.039 | 0.039 | 0 | 0.7 | 0.44 | 0.66 | 0.45 | 0.75 | 0.66 |
| 4 | 126028567 | rs13102452 | -0.429 | -0.157 | 1.091 | 1.798 | -0.009 | 0.051 | 0.051 | 0 | 0.12 | 0.15 | 0.14 | 0.26 | 0.11 | 0.16 |
| 4 | 126036776 | rs7675988 | -0.892 | -0.19 | 1.124 | 1.893 | -0.01 | 0.102 | 0.102 | 0 | 0.13 | 0.21 | 0.19 | 0.41 | 0.12 | 0.27 |
| 4 | 126039335 | rs3956574 | 1.541 | -1.001 | 1.161 | 1.555 | -0.033 | 0.017 | 0.017 | 0 | 0.65 | 0.48 | 0.57 | 0.46 | 0.75 | 0.62 |
| 4 | 126053047 | rs4240275 | -0.396 | -0.974 | 1.05 | 1.229 | 0.002 | 0 | 0 | 0 | 0.94 | 0.94 | 0.96 | 0.89 | 0.95 | 0.95 |
| 4 | 126056711 | rs6826117 | -0.214 | -0.964 | 1.085 | 1.354 | -0.005 | 0.07 | 0.07 | 0 | 0.12 | 0.21 | 0.19 | 0.28 | 0.11 | 0.18 |
| 4 | 126059221 | rs12506923 | -0.434 | -0.925 | 0.998 | 1.134 | -0.006 | -0.001 | -0.001 | 0.044 | 0.2 | 0.18 | 0.23 | 0.13 | 0.13 | 0.11 |
| 4 | 126064719 | rs13119112 | -1.574 | -0.798 | 1.026 | 1.199 | -0.034 | 0.116 | 0.116 | 0 | 0.11 | 0.26 | 0.1 | 0.26 | 0.06 | 0.14 |
| 4 | 126066704 | rs10213602 | -0.099 | -0.728 | 0.958 | 0.948 | 0.001 | -0.049 | -0.049 | 0.093 | 0.09 | 0.15 | 0.06 | 0 | 0.08 | 0.03 |
| 4 | 126081446 | rs4129246 | 0.794 | -0.601 | 0.854 | 1.199 | -0.052 | 0.026 | 0.026 | 0.351 | 0.75 | 0.61 | 0.69 | 0.71 | 0.84 | 0.74 |
| 4 | 126084755 | rs13136491 | -0.606 | -0.616 | 0.903 | 1.242 | -0.009 | 0.049 | 0.049 | 1.019 | 0.07 | 0.15 | 0.06 | 0.14 | 0.08 | 0.06 |
| 4 | 126087650 | rs17791550 | 1.282 | -0.198 | 0.143 | 1.034 | -0.049 | 0.003 | 0.003 | 0 | 0.12 | 0.01 | 0.1 | 0.09 | 0.14 | 0.19 |
| 4 | 126093508 | rs7690379 | 0.528 | -0.187 | 0.627 | 1.104 | 0.011 | 0 | 0 | 0 | 0.15 | 0.12 | 0.23 | 0.14 | 0.07 | 0.19 |
| 4 | 126103475 | rs13145439 | 0.886 | -0.224 | 0.718 | 1.104 | -0.065 | 0.033 | 0.033 | 0.568 | 0.75 | 0.59 | 0.69 | 0.71 | 0.84 | 0.75 |
| 4 | 126103526 | rs11098796 | -0.02 | 0.069 | 0.411 | 1.169 | 0.003 | -0.007 | -0.007 | 0.231 | 0.06 | 0.04 | 0.05 | 0.13 | 0.07 | 0.06 |
| 4 | 126119697 | rs12512915 | -0.249 | 0.604 | 0.379 | 1.12 | -0.003 | -0.002 | -0.002 | 0 | 0.07 | 0.06 | 0.08 | 0.11 | 0.13 | 0.07 |
| 4 | 126122232 | rs6534467 | 1.038 | 0.431 | 0.409 | 1.169 | -0.001 | 0 | 0 | 0.874 | 0.8 | 0.8 | 0.73 | 0.71 | 0.74 | 0.77 |
| 4 | 126122584 | rs7675694 | 0.603 | 0.591 | 0.22 | 1.086 | 0.01 | 0 | 0 | 0 | 0.15 | 0.09 | 0.26 | 0.14 | 0.12 | 0.21 |
| 4 | 126126219 | rs12499358 | 0.063 | 0.431 | 0.222 | 1.118 | 0 | -0.002 | -0.002 | 0 | 0.12 | 0.1 | 0.19 | 0.18 | 0.12 | 0.16 |
| 4 | 126138152 | rs7685639 | -1.415 | 0.422 | 0.372 | 1.087 | -0.018 | 0.018 | 0.018 | 0.145 | 0.35 | 0.55 | 0.53 | 0.45 | 0.34 | 0.42 |
| 4 | 126144198 | rs17009332 | -0.489 | 0.96 | -0.214 | 1.192 | 0.002 | -0.045 | -0.045 | 0 | 0.23 | 0.11 | 0.41 | 0.35 | 0.21 | 0.29 |
| 4 | 126159605 | rs11935229 | -1.412 | 1.178 | 0.609 | 1.045 | 0.065 | 0.144 | 0.144 | 0.762 | 0.05 | 0.22 | 0.1 | 0.12 | 0.06 | 0.04 |
| 4 | 126163379 | rs10518459 | -1.94 | 1.398 | 0.721 | 0.918 | 0.051 | 0.025 | 0.025 | 6.117 | 0.2 | 0.5 | 0.25 | 0.22 | 0.3 | 0.22 |
| 4 | 126165609 | rs7667073 | -0.831 | 1.724 | 0.538 | 0.716 | 0.041 | -0.013 | -0.013 | 0 | 0.22 | 0.42 | 0.37 | 0.2 | 0.28 | 0.19 |
| 4 | 126169900 | rs7676378 | -0.546 | 1.983 | 0.102 | 0.951 | 0.007 | 0.07 | 0.07 | 0 | 0.81 | 0.74 | 0.81 | 0.54 | 0.79 | 0.64 |
| 4 | 126170152 | rs9998852 | 1.001 | 1.751 | -0.213 | 0.526 | 0.027 | 0.031 | 0.031 | 0 | 0.47 | 0.36 | 0.45 | 0.35 | 0.37 | 0.36 |
| 4 | 126172651 | rs1039811 | -0.736 | 1.645 | -0.014 | 0.301 | 0.012 | -0.023 | -0.023 | 0 | 0.35 | 0.43 | 0.37 | 0.21 | 0.42 | 0.29 |
| 4 | 126176069 | rs1396688 | 0.966 | 1.666 | 0.015 | 0.223 | 0 | -0.034 | -0.034 | 0.447 | 0.14 | 0.15 | 0.19 | 0 | 0.12 | 0.06 |
| 4 | 126182392 | rs7696578 | 1.553 | 1.139 | -0.414 | -0.063 | -0.009 | -0.022 | -0.022 | 0 | 0.18 | 0.28 | 0.1 | 0.05 | 0.12 | 0.09 |
| 4 | 126186133 | rs1396696 | -0.805 | 1.785 | -0.062 | 0.319 | 0.013 | -0.029 | -0.029 | 0 | 0.35 | 0.45 | 0.38 | 0.21 | 0.43 | 0.29 |
| 4 | 126196021 | rs1509306 | 0.542 | 1.283 | 0.024 | 0.382 | 0.004 | -0.014 | -0.014 | 0 | 0.15 | 0.33 | 0.2 | 0.1 | 0.13 | 0.17 |
| 4 | 126198201 | rs2663289 | -1.05 | 1.562 | -0.216 | 0.726 | -0.005 | -0.018 | -0.018 | 0.19 | 0.81 | 0.89 | 0.77 | 0.55 | 0.77 | 0.65 |
| 4 | 126199915 | rs6534472 | 0.334 | 1.822 | -0.325 | 0.513 | 0.009 | -0.006 | -0.006 | 0 | 0.19 | 0.21 | 0.19 | 0.12 | 0.3 | 0.15 |
| 4 | 126205746 | rs2663265 | 0.814 | 1.84 | -0.341 | 0.484 | 0.008 | -0.011 | -0.011 | 0 | 0.15 | 0.27 | 0.21 | 0.09 | 0.14 | 0.18 |
| 4 | 126205766 | rs2663266 | -1.258 | 1.58 | -0.899 | 0.63 | -0.016 | -0.042 | -0.042 | 0 | 0.82 | 0.92 | 0.78 | 0.53 | 0.77 | 0.69 |
| 4 | 126208377 | rs2940781 | -0.8 | 1.707 | -0.717 | 0.372 | 0.02 | -0.023 | -0.023 | 0 | 0.35 | 0.54 | 0.39 | 0.21 | 0.44 | 0.32 |
| 4 | 126215371 | rs11941590 | 0.19 | 1.038 | -1.611 | 0.464 | -0.003 | 0.049 | 0.049 | 0 | 0.37 | 0.17 | 0.45 | 0.25 | 0.33 | 0.39 |
| 4 | 126216229 | rs1355167 | -0.791 | 0.879 | -1.603 | 0.463 | 0.006 | 0.002 | 0.002 | 0.393 | 0.57 | 0.37 | 0.6 | 0.52 | 0.61 | 0.7 |
| 4 | 126216963 | rs4834025 | 0.583 | 1.048 | -1.638 | 0.293 | -0.009 | 0.005 | 0.005 | 0 | 0.16 | 0.13 | 0.13 | 0.11 | 0.23 | 0.2 |
| 4 | 126217989 | rs11735586 | -0.053 | 0.936 | -1.44 | 0.347 | -0.014 | 0.004 | 0.004 | 0 | 0.38 | 0.22 | 0.47 | 0.3 | 0.38 | 0.42 |
| 4 | 126219619 | rs1912045 | -0.123 | 0.994 | -1.411 | 0.281 | -0.012 | 0.003 | 0.003 | 0 | 0.38 | 0.22 | 0.47 | 0.32 | 0.38 | 0.42 |
| 4 | 126234727 | rs2710572 | 0.278 | 0.618 | -1.358 | -0.234 | 0.014 | 0.022 | 0.022 | 1.975 | 0.23 | 0.18 | 0.08 | 0.02 | 0.15 | 0.11 |
| 4 | 126239986 | rs1039808 | -0.089 | 0.817 | -1.241 | -0.14 | 0 | -0.002 | -0.002 | 0 | 0.36 | 0.61 | 0.41 | 0.23 | 0.42 | 0.39 |
| 4 | 126240510 | rs2940779 | -1.227 | 0.863 | -1.65 | 0.26 | -0.04 | -0.094 | -0.094 | 0 | 0.78 | 0.93 | 0.73 | 0.46 | 0.71 | 0.72 |
| 4 | 126241871 | rs2710555 | -1.193 | 0.847 | -1.493 | 0.275 | -0.025 | -0.069 | -0.069 | 1.347 | 0.78 | 0.9 | 0.73 | 0.45 | 0.71 | 0.72 |
| 4 | 126252650 | rs2710534 | -0.543 | 0.675 | -1.252 | 0.187 | -0.024 | -0.045 | -0.045 | 0 | 0.72 | 0.82 | 0.7 | 0.38 | 0.66 | 0.69 |
| 4 | 126257065 | rs7692092 | 1.162 | 0.685 | -1.364 | -0.033 | -0.022 | 0.011 | 0.011 | 0.146 | 0.15 | 0.07 | 0.14 | 0.1 | 0.22 | 0.15 |
| 4 | 126261961 | rs17800919 | 0.493 | 0.293 | -1.39 | -0.325 | -0.013 | 0.087 | 0.087 | 0 | 0.22 | 0.07 | 0.25 | 0 | 0.24 | 0.2 |
| 4 | 126262415 | rs1355164 | 1.303 | 0.171 | -0.459 | 0.061 | -0.001 | 0.132 | 0.132 | 0 | 0.1 | 0.33 | 0.2 | 0.19 | 0.06 | 0.15 |
| 4 | 126267588 | rs11737230 | 0.124 | 0.36 | -0.599 | 0.032 | 0.002 | -0.007 | -0.007 | 0 | 0.57 | 0.51 | 0.57 | 0.28 | 0.5 | 0.45 |
| 4 | 126267997 | rs17009497 | 1.4 | 0.473 | -0.671 | 0.282 | -0.021 | -0.021 | -0.021 | 0 | 0.08 | 0.02 | 0.06 | 0.09 | 0.13 | 0.11 |
| 4 | 126270773 | rs7676058 | 1.059 | 0.306 | -0.583 | 0.033 | -0.023 | -0.019 | -0.019 | 0 | 0.1 | 0.23 | 0.17 | 0.05 | 0.06 | 0.09 |
| 4 | 126271899 | rs2710540 | -1.308 | 0.374 | -0.645 | 0.499 | -0.046 | -0.084 | -0.084 | 0 | 0.76 | 0.93 | 0.76 | 0.5 | 0.7 | 0.71 |
| 4 | 126272004 | rs9307560 | 1.05 | 0.29 | -0.619 | 0.372 | -0.023 | -0.019 | -0.019 | 1.358 | 0.1 | 0.23 | 0.17 | 0.05 | 0.06 | 0.09 |
| 4 | 126273052 | rs11098809 | 0.086 | 0.445 | -0.679 | 0.465 | 0.002 | -0.009 | -0.009 | 0 | 0.56 | 0.5 | 0.57 | 0.28 | 0.5 | 0.45 |
| 4 | 126275373 | rs2710562 | 1.709 | 0.379 | -0.649 | 0.765 | 0.004 | -0.009 | -0.009 | 0 | 0.08 | 0.08 | 0.11 | 0.2 | 0.07 | 0.16 |
| 4 | 126277208 | rs17009521 | 1.499 | 0.448 | -0.647 | 0.595 | -0.002 | 0.002 | 0.002 | 0 | 0.08 | 0.06 | 0.04 | 0 | 0.08 | 0.08 |
| 4 | 126280006 | rs2710560 | -0.274 | 0.486 | -0.364 | 0.454 | 0.004 | -0.018 | -0.018 | 0.05 | 0.43 | 0.5 | 0.65 | 0.59 | 0.5 | 0.54 |
| 4 | 126281130 | rs17193280 | 1.112 | 0.441 | -0.656 | 0.566 | -0.11 | -0.034 | -0.034 | 5.226 | 0.26 | 0.02 | 0.35 | 0.19 | 0.37 | 0.21 |
| 4 | 126281438 | rs992122 | -0.229 | 0.414 | 0.05 | 0.858 | 0.053 | 0.188 | 0.188 | 0.176 | 0.07 | 0.29 | 0.18 | 0.13 | 0.06 | 0.08 |
| 4 | 126282962 | rs1396686 | 1.965 | 0.36 | -0.141 | 0.787 | 0.007 | -0.014 | -0.014 | 0 | 0.09 | 0.07 | 0.11 | 0.26 | 0.07 | 0.24 |
| 4 | 126306884 | rs12650847 | 0.31 | 0.401 | 0.636 | 0.62 | 0.018 | 0.014 | 0.014 | 0 | 0.13 | 0.32 | 0.08 | 0.07 | 0.12 | 0.13 |
| 4 | 126308835 | rs1509289 | 0.585 | 0.375 | 0.359 | 0.405 | -0.015 | -0.021 | -0.021 | 0 | 0.26 | 0.12 | 0.22 | 0.41 | 0.27 | 0.3 |
| 4 | 126313149 | rs1912044 | -0.899 | 0.399 | 0.426 | 0.38 | 0.003 | 0.059 | 0.059 | 0.244 | 0.5 | 0.61 | 0.56 | 0.81 | 0.47 | 0.7 |
| 4 | 126318368 | rs7666354 | 0.245 | 0.414 | 0.496 | 0.476 | 0.065 | 0.176 | 0.176 | 0.007 | 0.05 | 0.26 | 0.04 | 0.09 | 0.05 | 0.04 |

| | | | | | | | | | | | | | | | | |
|---|---|---|---|---|---|---|---|---|---|---|---|---|---|---|---|---|
| 4 | 126319772 | rs11736643 | 2.085 | 0.359 | -0.047 | 0.156 | 0.007 | 0.016 | 0.016 | 4.218 | 0.2 | 0.09 | 0.05 | 0 | 0.12 | 0.07 |
| 4 | 126351995 | rs17009641 | 1.185 | 0.627 | -0.288 | 0.225 | -0.027 | -0.083 | -0.083 | 0 | 0.18 | 0.03 | 0.18 | 0.41 | 0.18 | 0.25 |
| 4 | 126352482 | rs13141820 | 0.265 | 0.7 | -0.096 | 0.189 | 0.013 | -0.002 | -0.002 | 0.006 | 0.35 | 0.53 | 0.42 | 0.26 | 0.45 | 0.26 |
| 4 | 126354307 | rs1395233 | 0.342 | 0.706 | -0.039 | 0.192 | 0.013 | -0.001 | -0.001 | 0 | 0.34 | 0.42 | 0.41 | 0.26 | 0.45 | 0.26 |
| 4 | 126364565 | rs9968420 | -0.816 | 0.636 | -0.57 | 0.344 | 0.007 | -0.004 | -0.004 | 1.481 | 0.81 | 0.81 | 0.72 | 0.77 | 0.86 | 0.71 |
| 4 | 126366934 | rs17009673 | 0.27 | 0.403 | -1.208 | 0.064 | 0.021 | 0.027 | 0.027 | 0 | 0.29 | 0.16 | 0.31 | 0.23 | 0.2 | 0.29 |
| 4 | 126372742 | rs1567047 | 0.703 | 0.861 | -1.052 | 0.411 | -0.05 | -0.123 | -0.123 | 0 | 0.21 | 0.02 | 0.25 | 0.39 | 0.3 | 0.26 |
| 4 | 126376877 | rs13128061 | 0.835 | 0.75 | -0.594 | 0.379 | 0.01 | 0.087 | 0.087 | 0.436 | 0.51 | 0.28 | 0.43 | 0.26 | 0.5 | 0.44 |
| 4 | 126379019 | rs7696523 | 0.594 | 0.738 | -0.518 | 0.479 | 0.009 | 0.017 | 0.017 | 0.361 | 0.55 | 0.35 | 0.46 | 0.34 | 0.55 | 0.48 |
| 4 | 126380367 | rs10518466 | 0.603 | 0.739 | -0.518 | 0.474 | 0.009 | 0.021 | 0.021 | 0.377 | 0.55 | 0.35 | 0.46 | 0.32 | 0.55 | 0.48 |
| 4 | 126384595 | rs4146009 | 0.646 | 0.986 | -0.583 | 0.352 | -0.001 | -0.047 | -0.047 | 0 | 0.45 | 0.45 | 0.35 | 0.06 | 0.4 | 0.26 |
| 4 | 126389047 | rs6813101 | 0.381 | 1.709 | -0.307 | 0.956 | -0.001 | -0.031 | -0.031 | 0 | 0.25 | 0.28 | 0.27 | 0.06 | 0.24 | 0.17 |
| 4 | 126391828 | rs6828096 | 1.021 | 1.658 | -0.076 | 0.956 | -0.003 | 0.026 | 0.026 | 0 | 0.43 | 0.35 | 0.27 | 0.28 | 0.39 | 0.38 |
| 4 | 126398824 | rs1506360 | -0.637 | 1.691 | -0.338 | 1.152 | -0.003 | -0.016 | -0.016 | 0 | 0.75 | 0.67 | 0.72 | 0.85 | 0.76 | 0.74 |
| 4 | 126402618 | rs10026069 | 0.331 | 1.698 | -0.338 | 1.148 | -0.002 | -0.031 | -0.031 | 0 | 0.25 | 0.28 | 0.27 | 0.06 | 0.24 | 0.17 |
| 4 | 126402890 | rs6824160 | 1.299 | 1.78 | -0.541 | 1.16 | -0.042 | -0.04 | -0.04 | 0 | 0.07 | 0.01 | 0.11 | 0.26 | 0.13 | 0.2 |
| 4 | 126409835 | rs17009848 | 0.419 | 1.143 | -0.247 | 1.659 | -0.046 | -0.104 | -0.104 | 0.226 | 0.21 | 0.02 | 0.21 | 0.35 | 0.29 | 0.27 |
| 4 | 126412575 | rs1014866 | -0.447 | 1.16 | -0.104 | 1.141 | -0.004 | -0.018 | -0.018 | 0 | 0.75 | 0.65 | 0.72 | 0.85 | 0.76 | 0.74 |
| 4 | 126414605 | rs10031677 | 0.838 | 0.842 | -0.164 | 1.125 | -0.006 | -0.07 | -0.07 | 0 | 0.2 | 0.23 | 0.23 | 0 | 0.18 | 0.13 |
| 4 | 126415914 | rs4314296 | 0.932 | 1.011 | -0.414 | 1.946 | 0.016 | -0.034 | -0.034 | 1.137 | 0.12 | 0.01 | 0.09 | 0.17 | 0.09 | 0.17 |
| 4 | 126419726 | rs10518469 | 0.079 | 1.052 | -0.255 | 1.365 | -0.028 | -0.104 | -0.104 | 0 | 0.23 | 0.07 | 0.39 | 0.59 | 0.3 | 0.32 |
| 4 | 126423687 | rs1395227 | -0.256 | 0.533 | 0.547 | 1.006 | 0.015 | 0.024 | 0.024 | 0 | 0.9 | 0.65 | 0.92 | 0.9 | 0.92 | 0.87 |
| 4 | 126426695 | rs12639834 | 0.304 | 0.918 | 0.143 | 1.336 | -0.037 | -0.045 | -0.045 | 0 | 0.25 | 0.1 | 0.26 | 0.38 | 0.4 | 0.42 |
| 4 | 126431919 | rs950063 | -0.578 | 0.909 | 0.057 | 1.298 | -0.062 | -0.032 | -0.032 | 0 | 0.75 | 0.91 | 0.74 | 0.73 | 0.6 | 0.66 |
| 4 | 126433214 | rs1156232 | 0.339 | 0.795 | 0.472 | 1.494 | -0.012 | 0.049 | 0.049 | 0 | 0.1 | 0.27 | 0.24 | 0.19 | 0.06 | 0.11 |
| 4 | 126434087 | rs1532883 | 0.831 | 0.517 | 0.664 | 1.51 | -0.005 | 0.03 | 0.03 | 0.277 | 0.59 | 0.3 | 0.45 | 0.32 | 0.53 | 0.43 |
| 4 | 126435279 | rs1532881 | 1.053 | 0.51 | 0.657 | 1.462 | -0.02 | 0.032 | 0.032 | 0 | 0.56 | 0.23 | 0.45 | 0.41 | 0.49 | 0.51 |
| 4 | 126447482 | rs1567049 | -1.942 | 0.141 | 1.711 | 1.561 | 0.019 | 0.265 | 0.265 | 0.166 | 0.09 | 0.44 | 0.06 | 0.22 | 0.05 | 0.14 |
| 4 | 126447899 | rs10518470 | 0.1 | 0.038 | 1.351 | 1.34 | -0.04 | 0.053 | 0.053 | 0.15 | 0.19 | 0.01 | 0.25 | 0.07 | 0.21 | 0.24 |
| 4 | 126456859 | rs4834056 | -1.693 | -0.005 | 1.57 | 1.728 | -0.039 | 0.1 | 0.1 | 0.734 | 0.17 | 0.68 | 0.29 | 0.41 | 0.07 | 0.23 |
| 4 | 126462071 | rs11098817 | -1.321 | 0.149 | 1.209 | 1.364 | -0.04 | 0.174 | 0.174 | 0.573 | 0.15 | 0.56 | 0.27 | 0.32 | 0.07 | 0.15 |
| 4 | 126476695 | rs12500867 | -0.526 | 0.396 | 0.285 | 1.195 | -0.011 | 0.04 | 0.04 | 0 | 0.11 | 0.28 | 0.25 | 0.19 | 0.08 | 0.11 |
| 4 | 126480268 | rs6834632 | -0.432 | 0.581 | -0.071 | 0.791 | 0.002 | -0.009 | -0.009 | 0 | 0.22 | 0.18 | 0.27 | 0.23 | 0.21 | 0.29 |
| 4 | 126482267 | rs13138789 | 0.884 | 0.818 | 0.77 | 0.526 | 0.053 | -0.056 | -0.056 | 0.174 | 0.86 | 0.57 | 0.86 | 0.88 | 0.83 | 0.94 |
| 4 | 126490303 | rs6818629 | 1.316 | 0.385 | 0.299 | 0.63 | 0.01 | -0.007 | -0.007 | 0 | 0.66 | 0.31 | 0.56 | 0.6 | 0.67 | 0.6 |
| 4 | 126494752 | rs10016783 | 1.221 | 0.473 | 0.428 | 0.659 | -0.001 | 0 | 0 | 0 | 0.67 | 0.35 | 0.57 | 0.6 | 0.67 | 0.61 |
| 4 | 126500482 | rs1429901 | 1.432 | 0.583 | 0.809 | 0.397 | -0.004 | -0.04 | -0.04 | 0.032 | 0.76 | 0.45 | 0.64 | 0.88 | 0.78 | 0.68 |
| 4 | 126504601 | rs6534494 | -0.498 | 0.648 | 1.142 | 0.36 | 0.012 | -0.015 | -0.015 | 0.746 | 0.08 | 0.18 | 0.24 | 0.03 | 0.08 | 0.09 |
| 4 | 126505272 | rs13126049 | 0.828 | 0.563 | 1.07 | 0.444 | 0.004 | -0.043 | -0.043 | 0 | 0.89 | 0.86 | 0.9 | 1 | 0.87 | 0.94 |
| 4 | 126524825 | rs1864363 | 0.574 | 0.768 | 0.413 | 0.507 | 0.005 | -0.001 | -0.001 | 0 | 0.77 | 0.69 | 0.82 | 0.76 | 0.78 | 0.84 |
| 4 | 126535503 | rs10001030 | -1.188 | 0.858 | 0.125 | 0.318 | 0.001 | -0.009 | -0.009 | 0 | 0.19 | 0.22 | 0.09 | 0.11 | 0.17 | 0.23 |
| 4 | 126535676 | rs1346715 | -1.254 | 0.759 | 0.257 | 0.684 | 0 | 0.01 | 0.01 | 0 | 0.2 | 0.27 | 0.09 | 0.22 | 0.17 | 0.38 |
| 4 | 126538037 | rs12331173 | -0.909 | 0.762 | 0.259 | 0.623 | 0.025 | -0.01 | -0.01 | 0.949 | 0.14 | 0.24 | 0.06 | 0.11 | 0.14 | 0.21 |
| 4 | 126540305 | rs6534495 | -0.225 | 0.782 | -0.715 | 0.746 | -0.053 | -0.081 | -0.081 | 0.019 | 0.67 | 0.91 | 0.57 | 0.58 | 0.64 | 0.74 |
| 4 | 126553888 | rs9307573 | -0.219 | 0.879 | -0.243 | 0.53 | -0.032 | -0.078 | -0.078 | 2.7 | 0.68 | 0.86 | 0.55 | 0.53 | 0.63 | 0.72 |
| 4 | 126556868 | rs6836731 | 1.207 | 0.578 | 0.056 | 0.468 | 0 | 0.001 | 0.001 | 2.202 | 0.47 | 0.5 | 0.45 | 0.37 | 0.46 | 0.37 |
| 4 | 126561331 | rs12643523 | -0.143 | 0.65 | -0.107 | 0.431 | -0.037 | -0.069 | -0.069 | 0 | 0.71 | 0.87 | 0.56 | 0.51 | 0.63 | 0.69 |
| 4 | 126561925 | rs13140903 | 0.85 | 0.544 | 0.218 | 0.525 | 0 | -0.004 | -0.004 | 0 | 0.56 | 0.52 | 0.49 | 0.35 | 0.55 | 0.52 |
| 4 | 126572145 | rs4834060 | -0.259 | -0.121 | -0.654 | -0.298 | -0.001 | -0.001 | -0.001 | 0 | 0.37 | 0.35 | 0.4 | 0.49 | 0.43 | 0.49 |
| 4 | 126576950 | rs34095747 | 0.834 | -0.106 | -0.685 | -0.288 | -0.078 | -0.082 | -0.082 | 3.118 | 0.25 | 0.07 | 0.35 | 0.34 | 0.36 | 0.33 |
| 4 | 126594533 | rs17010235 | -0.015 | 0.015 | -0.738 | -0.347 | -0.009 | -0.026 | -0.026 | 0 | 0.45 | 0.55 | 0.2 | 0.17 | 0.29 | 0.37 |
| 4 | 126598111 | rs10434043 | 0.866 | 0.074 | -0.974 | -0.143 | -0.077 | -0.085 | -0.085 | 0.252 | 0.22 | 0.06 | 0.33 | 0.34 | 0.33 | 0.31 |
| 4 | 126599698 | rs17010239 | 0.427 | 0.102 | -0.715 | -0.056 | 0.029 | 0.041 | 0.041 | 0 | 0.25 | 0.34 | 0.35 | 0.34 | 0.32 | 0.31 |
| 4 | 126603140 | rs6844035 | 0.843 | 0.044 | -0.543 | -0.067 | 0.002 | -0.003 | -0.003 | 0 | 0.1 | 0.09 | 0.05 | 0.16 | 0.07 | 0.17 |
| 4 | 126613855 | rs1367066 | 0.908 | -0.025 | -0.381 | 0.09 | 0.017 | 0.024 | 0.024 | 0.013 | 0.31 | 0.3 | 0.15 | 0.14 | 0.22 | 0.21 |
| 4 | 126629426 | rs6831614 | -0.214 | 0.042 | -0.412 | 0.201 | -0.01 | -0.003 | -0.003 | 0.286 | 0.45 | 0.32 | 0.5 | 0.51 | 0.46 | 0.57 |
| 4 | 126654773 | rs9307576 | -0.338 | 0.14 | -1.227 | 0.281 | -0.001 | -0.001 | -0.001 | 0 | 0.47 | 0.56 | 0.5 | 0.49 | 0.46 | 0.57 |
| 4 | 126683716 | rs10018249 | -0.481 | 0.179 | -1.349 | 0.275 | -0.003 | -0.002 | -0.002 | 0 | 0.48 | 0.63 | 0.51 | 0.49 | 0.46 | 0.56 |
| 4 | 126688743 | rs4446326 | -0.85 | 0.074 | -1.483 | 0.219 | -0.044 | -0.017 | -0.017 | 0.078 | 0.74 | 0.81 | 0.6 | 0.7 | 0.56 | 0.66 |
| 4 | 126694369 | rs10013424 | 0.407 | -0.205 | -1.592 | 0.128 | -0.005 | 0.011 | 0.011 | 0.371 | 0.35 | 0.52 | 0.44 | 0.39 | 0.3 | 0.46 |
| 4 | 126696308 | rs4394015 | 0.208 | -0.183 | -2.011 | 0.149 | 0.038 | 0.031 | 0.031 | 0.903 | 0.38 | 0.29 | 0.16 | 0.31 | 0.27 | 0.19 |
| 4 | 126697893 | rs4591618 | 0.84 | -0.153 | -1.968 | 0.105 | 0.004 | 0.002 | 0.002 | 0.117 | 0.28 | 0.29 | 0.14 | 0.16 | 0.21 | 0.12 |
| 4 | 126709492 | rs7678929 | 0.454 | 0.323 | -1.853 | 0.096 | 0.038 | 0.062 | 0.062 | 0 | 0.41 | 0.31 | 0.2 | 0.17 | 0.3 | 0.19 |
| 4 | 126709616 | rs10034133 | -0.175 | 1.303 | -1.946 | 0.633 | -0.127 | -0.067 | -0.067 | 0.017 | 0.74 | 0.92 | 0.63 | 0.69 | 0.57 | 0.66 |
| 4 | 126714120 | rs9307578 | -0.153 | 1.354 | -1.954 | 0.656 | -0.115 | -0.06 | -0.06 | 0 | 0.74 | 0.93 | 0.63 | 0.69 | 0.57 | 0.66 |
| 4 | 126736372 | rs6841080 | -0.549 | 1.231 | -1.883 | 0.8 | -0.023 | -0.02 | -0.02 | 0 | 0.48 | 0.75 | 0.66 | 0.49 | 0.55 | 0.28 |
| 4 | 126741907 | rs10222842 | -0.113 | 1.119 | -1.184 | 0.699 | -0.026 | 0.07 | 0.07 | 0 | 0.24 | 0.35 | 0.25 | 0.42 | 0.2 | 0.15 |
| 4 | 126742571 | rs9998979 | -1.045 | 1.121 | -1.257 | 0.712 | -0.019 | -0.016 | -0.016 | 4.005 | 0.48 | 0.77 | 0.67 | 0.49 | 0.55 | 0.29 |
| 4 | 126747317 | rs6854255 | -0.472 | 1.306 | -1.122 | 0.842 | 0.012 | 0 | 0 | 0 | 0.35 | 0.44 | 0.59 | 0.36 | 0.47 | 0.22 |

| | | | | | | | | | | | | | | | | |
|---|---|---|---|---|---|---|---|---|---|---|---|---|---|---|---|---|
| 4 | 126749816 | rs4541524 | -0.833 | 1.519 | -1.894 | 0.616 | -0.058 | -0.025 | -0.025 | 0 | 0.8 | 0.95 | 0.84 | 0.77 | 0.78 | 0.68 |
| 4 | 126749905 | rs4077738 | 0.533 | 1.944 | -1.88 | 0.35 | -0.052 | -0.004 | -0.004 | 0 | 0.12 | 0 | 0.29 | 0.07 | 0.18 | 0.08 |
| 4 | 126752842 | rs12506206 | 0.152 | 2.074 | -1.387 | 0.692 | 0.013 | 0.051 | 0.051 | 0.012 | 0.09 | 0.14 | 0.34 | 0.33 | 0.12 | 0.31 |
| 4 | 126754145 | rs4590050 | -0.383 | 1.732 | -1.176 | 0.015 | 0.005 | 0.018 | 0.018 | 0 | 0.52 | 0.63 | 0.44 | 0.62 | 0.57 | 0.5 |
| 4 | 126761599 | rs7660904 | -0.715 | 1.83 | -0.851 | -0.108 | -0.018 | 0.079 | 0.079 | 0 | 0.55 | 0.64 | 0.69 | 0.87 | 0.52 | 0.69 |
| 4 | 126763987 | rs12513118 | 0.194 | 1.54 | -0.955 | 0.548 | -0.002 | -0.011 | -0.011 | 0 | 0.19 | 0.16 | 0.4 | 0.6 | 0.18 | 0.36 |
| 4 | 126774098 | rs1911014 | 1.501 | 1.552 | -0.904 | 0.446 | -0.028 | -0.003 | -0.003 | 0 | 0.15 | 0.06 | 0.07 | 0.11 | 0.17 | 0.09 |
| 4 | 126780071 | rs10857108 | 1.455 | 1.592 | -0.672 | 0.418 | -0.011 | 0.015 | 0.015 | 0.503 | 0.18 | 0.08 | 0.07 | 0.11 | 0.17 | 0.09 |
| 4 | 126799654 | rs12645313 | 1.265 | 1.531 | -0.706 | 0.595 | -0.005 | -0.006 | -0.006 | 0 | 0.07 | 0.05 | 0.29 | 0.23 | 0.08 | 0.26 |
| 4 | 126808685 | rs3956582 | 0.979 | 1.476 | -0.257 | 0.052 | -0.022 | 0.137 | 0.137 | 0 | 0.42 | 0.23 | 0.27 | 0.03 | 0.44 | 0.27 |
| 4 | 126811649 | rs10518477 | -0.441 | 1.396 | -0.038 | 0.75 | -0.008 | 0.037 | 0.037 | 0 | 0.82 | 0.78 | 0.63 | 0.51 | 0.87 | 0.69 |
| 4 | 126812573 | rs1506169 | -0.064 | 1.503 | -0.327 | 0.728 | 0.007 | -0.061 | -0.061 | 0 | 0.2 | 0.1 | 0.37 | 0.49 | 0.17 | 0.35 |
| 4 | 126838888 | rs6534520 | -0.013 | 1.35 | 0.315 | 0.511 | 0.048 | 0.081 | 0.081 | 0.152 | 0.93 | 0.73 | 0.92 | 0.88 | 0.92 | 0.92 |
| 4 | 126840854 | rs11721858 | 1.139 | 1.415 | 0.088 | 0.43 | -0.013 | 0.018 | 0.018 | 0.588 | 0.18 | 0.06 | 0.07 | 0.1 | 0.17 | 0.1 |
| 4 | 126864721 | rs4348108 | 0.613 | 1.08 | 0.087 | 0.443 | 0 | -0.002 | -0.002 | 0 | 0.23 | 0.25 | 0.13 | 0.22 | 0.22 | 0.17 |
| 4 | 126875119 | rs17224022 | 0.693 | 1.499 | -0.255 | 0.383 | 0.038 | -0.02 | -0.02 | 0 | 0.08 | 0.12 | 0.08 | 0.01 | 0.19 | 0.14 |
| 4 | 126880304 | rs7691677 | 0.189 | 0.484 | -0.386 | 0.581 | -0.008 | 0.004 | 0.004 | 0.512 | 0.59 | 0.5 | 0.73 | 0.52 | 0.67 | 0.77 |
| 4 | 126882844 | rs12505759 | 1.454 | 0.45 | -0.494 | 0.631 | 0.022 | -0.065 | -0.065 | 0.792 | 0.1 | 0 | 0.07 | 0.26 | 0.06 | 0.05 |
| 4 | 126884861 | rs10019346 | 0.086 | 0.584 | -0.315 | 0.509 | -0.008 | 0.004 | 0.004 | 0.512 | 0.59 | 0.5 | 0.73 | 0.52 | 0.67 | 0.77 |
| 4 | 126891842 | rs13128559 | 0.974 | 0.505 | -0.534 | 0.205 | -0.006 | 0.027 | 0.027 | 0 | 0.23 | 0.16 | 0.13 | 0.09 | 0.21 | 0.12 |
| 4 | 126892733 | rs7665514 | -0.962 | 0.065 | -1.245 | -0.427 | -0.011 | 0.12 | 0.12 | 0.059 | 0.8 | 0.97 | 0.87 | 0.98 | 0.8 | 0.9 |
| 4 | 126896243 | rs11098847 | 0.778 | -0.028 | -0.613 | 0.069 | -0.002 | 0.151 | 0.151 | 0 | 0.14 | 0.29 | 0.19 | 0.36 | 0.11 | 0.23 |
| 4 | 126896321 | rs1506149 | 0.55 | -0.096 | -0.474 | 0.003 | 0.005 | -0.009 | -0.009 | 0 | 0.33 | 0.38 | 0.47 | 0.29 | 0.42 | 0.49 |
| 4 | 126899057 | rs12512463 | 1.39 | 0.116 | -0.62 | 0.033 | -0.005 | -0.002 | -0.002 | 0 | 0.07 | 0.04 | 0.28 | 0.28 | 0.09 | 0.25 |
| 4 | 126902635 | rs12641642 | 0.64 | -0.08 | -0.805 | -0.088 | 0.019 | 0.005 | 0.005 | 0 | 0.3 | 0.04 | 0.2 | 0.25 | 0.26 | 0.16 |
| 4 | 126910178 | rs12651652 | 1.329 | -0.111 | -0.671 | -0.256 | 0.001 | -0.001 | -0.001 | 0 | 0.21 | 0.13 | 0.37 | 0.28 | 0.22 | 0.32 |
| 4 | 126915614 | rs1021940 | -0.053 | -0.154 | -0.23 | -0.338 | 0.048 | 0.108 | 0.108 | 0 | 0.25 | 0.52 | 0.29 | 0.38 | 0.31 | 0.4 |
| 4 | 126925575 | rs17010573 | 2.474 | -0.125 | -0.333 | -0.424 | 0.01 | 0.011 | 0.011 | 0 | 0.14 | 0.08 | 0.09 | 0.01 | 0.13 | 0.07 |
| 4 | 126931413 | rs2390928 | 1.375 | -0.126 | -0.337 | -0.281 | 0.001 | -0.005 | -0.005 | 0.519 | 0.21 | 0.13 | 0.41 | 0.3 | 0.22 | 0.32 |
| 4 | 126931623 | rs7434822 | 1.254 | -0.01 | -0.199 | -0.314 | 0.031 | -0.035 | -0.035 | 0.682 | 0.1 | 0.17 | 0.09 | 0 | 0.19 | 0.17 |
| 4 | 126932102 | rs10026567 | 0.725 | -0.53 | -0.2 | -0.206 | -0.002 | 0.01 | 0.01 | 0 | 0.44 | 0.36 | 0.54 | 0.4 | 0.43 | 0.45 |
| 4 | 126937174 | rs6534526 | 1.657 | -0.555 | -0.275 | -0.167 | -0.003 | 0.011 | 0.011 | 0 | 0.07 | 0.05 | 0.32 | 0.29 | 0.09 | 0.26 |
| 4 | 126939153 | rs4495049 | -1.374 | -0.66 | -0.391 | -0.599 | 0.006 | 0.103 | 0.103 | 0 | 0.79 | 0.96 | 0.87 | 0.98 | 0.8 | 0.9 |
| 4 | 126941560 | rs2893160 | -1.564 | -0.714 | -0.486 | -0.662 | -0.011 | 0.037 | 0.037 | 0 | 0.92 | 1 | 0.92 | 0.98 | 0.9 | 0.95 |
| 4 | 126942485 | rs34044861 | 2.405 | -0.77 | -0.473 | -0.734 | -0.027 | 0.045 | 0.045 | 0 | 0.21 | 0.07 | 0.12 | 0.1 | 0.21 | 0.13 |
| 4 | 126943520 | rs2390939 | -0.854 | -0.8 | -0.434 | -0.457 | 0 | -0.001 | -0.001 | 0 | 0.69 | 0.73 | 0.5 | 0.61 | 0.68 | 0.63 |
| 4 | 126947359 | rs12644234 | 2.476 | -0.806 | -0.493 | -0.406 | 0.012 | -0.057 | -0.057 | 0 | 0.09 | 0 | 0.03 | 0.21 | 0.06 | 0.05 |
| 4 | 126949007 | rs4834082 | 0.182 | -0.705 | -0.318 | -0.401 | 0.024 | 0.012 | 0.012 | 0.019 | 0.67 | 0.5 | 0.63 | 0.68 | 0.6 | 0.56 |
| 4 | 126950447 | rs2390944 | -1.149 | -0.493 | -0.138 | -0.053 | 0.003 | -0.013 | -0.013 | 0.104 | 0.92 | 0.94 | 0.89 | 0.86 | 0.9 | 0.95 |
| 4 | 126950568 | rs2390945 | 0.996 | -0.364 | 0.099 | 0.046 | 0.005 | -0.006 | -0.006 | 0 | 0.14 | 0.11 | 0.06 | 0 | 0.1 | 0.05 |
| 4 | 126959315 | rs9985873 | 0.393 | -0.108 | 0.157 | 0.393 | -0.002 | 0.006 | 0.006 | 0 | 0.22 | 0.2 | 0.41 | 0.37 | 0.22 | 0.32 |
| 4 | 126975201 | rs7656535 | -0.62 | -0.257 | -0.24 | 0.122 | -0.023 | 0.125 | 0.125 | 0.088 | 0.6 | 0.88 | 0.8 | 0.82 | 0.64 | 0.78 |
| 4 | 126976466 | rs4279227 | 0.206 | -0.046 | 0.003 | 0.409 | 0.004 | -0.023 | -0.023 | 0.18 | 0.23 | 0.15 | 0.42 | 0.39 | 0.23 | 0.34 |
| 4 | 126976898 | rs12499838 | 0.288 | -0.041 | -0.027 | 0.411 | 0.003 | -0.029 | -0.029 | 0 | 0.22 | 0.14 | 0.42 | 0.39 | 0.23 | 0.34 |
| 4 | 126986023 | rs7693642 | -0.861 | -0.074 | 0.01 | 0.302 | 0.002 | 0 | 0 | 0 | 0.81 | 0.79 | 0.85 | 0.9 | 0.82 | 0.9 |
| 4 | 126989652 | rs2271873 | 0.857 | -0.266 | 0.294 | 0.425 | -0.02 | 0.004 | 0.004 | 0.559 | 0.3 | 0.23 | 0.39 | 0.25 | 0.39 | 0.42 |
| 4 | 127000579 | rs4586960 | -0.323 | -0.431 | 0.659 | 0.528 | -0.018 | 0.053 | 0.053 | 0 | 0.27 | 0.41 | 0.39 | 0.64 | 0.23 | 0.36 |
| 4 | 127006583 | rs17010666 | -0.113 | -0.358 | 0.559 | 0.568 | -0.003 | 0.035 | 0.035 | 0 | 0.24 | 0.31 | 0.36 | 0.49 | 0.23 | 0.35 |
| 4 | 127008222 | rs9994809 | 0.453 | -0.521 | 0.749 | 0.573 | -0.018 | 0.037 | 0.037 | 0 | 0.71 | 0.54 | 0.57 | 0.33 | 0.72 | 0.6 |
| 4 | 127009851 | rs10518483 | 0.614 | -0.421 | 0.327 | 0.646 | 0.002 | 0.081 | 0.081 | 0 | 0.07 | 0.05 | 0.23 | 0.4 | 0.09 | 0.29 |
| 4 | 127018677 | rs4530647 | 0.614 | -0.467 | 0.505 | 0.538 | 0.006 | -0.001 | -0.001 | 0 | 0.18 | 0.19 | 0.14 | 0.18 | 0.15 | 0.08 |
| 4 | 127027927 | rs9686033 | -0.254 | -0.454 | 0.728 | 0.563 | -0.017 | 0.043 | 0.043 | 0 | 0.26 | 0.41 | 0.34 | 0.65 | 0.23 | 0.27 |
| 4 | 127030561 | rs10857109 | -0.2 | -0.461 | 0.689 | 0.577 | -0.004 | 0.02 | 0.02 | 0 | 0.24 | 0.27 | 0.33 | 0.58 | 0.23 | 0.27 |
| 4 | 127037075 | rs11721735 | 0.717 | -0.489 | 0.599 | 0.528 | 0.013 | 0.001 | 0.001 | 0 | 0.19 | 0.12 | 0.14 | 0.18 | 0.19 | 0.12 |
| 4 | 127041554 | rs7685518 | -0.39 | -0.479 | 0.673 | 0.456 | -0.01 | 0.027 | 0.027 | 0 | 0.28 | 0.43 | 0.36 | 0.66 | 0.28 | 0.32 |
| 4 | 127043282 | rs11098856 | 0.93 | -0.504 | 0.052 | 0.084 | -0.099 | -0.015 | -0.015 | 0 | 0.29 | 0.07 | 0.37 | 0.23 | 0.37 | 0.49 |
| 4 | 127045418 | rs2062209 | 0.095 | -0.444 | 0.116 | 0.199 | -0.002 | 0.015 | 0.015 | 0 | 0.27 | 0.29 | 0.34 | 0.58 | 0.28 | 0.32 |
| 4 | 127048641 | rs13146295 | 1.608 | -0.61 | -0.658 | -0.28 | -0.074 | 0.02 | 0.02 | 0 | 0.2 | 0.01 | 0.15 | 0.1 | 0.22 | 0.12 |
| 4 | 127072596 | rs561662 | -0.562 | -0.522 | -0.109 | -0.352 | 0.009 | 0.05 | 0.05 | 0.08 | 0.61 | 0.72 | 0.77 | 0.89 | 0.68 | 0.84 |
| 4 | 127077168 | rs1351732 | 0.447 | -0.424 | 0.273 | -0.094 | -0.019 | -0.057 | -0.057 | 0 | 0.21 | 0.37 | 0.14 | 0.09 | 0.19 | 0.11 |
| 4 | 127093759 | rs1017863 | 0.765 | -0.001 | 0.016 | 0.737 | 0.001 | 0.098 | 0.098 | 0 | 0.05 | 0.03 | 0.19 | 0.4 | 0.08 | 0.19 |
| 4 | 127096872 | rs17010784 | 0.765 | -0.001 | 0.016 | 0.737 | 0.001 | 0.098 | 0.098 | 0 | 0.05 | 0.03 | 0.19 | 0.4 | 0.08 | 0.19 |
| 4 | 127113427 | rs7696522 | -0.124 | -0.332 | 0.074 | 0.172 | -0.016 | -0.007 | -0.007 | 0 | 0.31 | 0.24 | 0.33 | 0.29 | 0.36 | 0.48 |
| 4 | 127116244 | rs6838566 | -0.043 | -0.344 | 0.032 | -0.094 | -0.023 | 0.042 | 0.042 | 0 | 0.29 | 0.19 | 0.31 | 0.11 | 0.36 | 0.47 |
| 4 | 127123816 | rs4144814 | 0.601 | -0.598 | 0.071 | -0.469 | -0.014 | 0.185 | 0.185 | 0.338 | 0.54 | 0.2 | 0.61 | 0.86 | 0.57 | 0.49 |
| 4 | 127134711 | rs6829892 | 0.241 | -0.556 | -0.032 | -0.452 | -0.032 | 0.053 | 0.053 | 0 | 0.27 | 0.16 | 0.29 | 0.08 | 0.34 | 0.49 |
| 4 | 127136369 | rs4560423 | 0.157 | -0.535 | 0.132 | -0.231 | -0.02 | 0 | 0 | 1.942 | 0.8 | 0.31 | 0.87 | 0.79 | 0.92 | 0.94 |
| 4 | 127157577 | rs2390994 | -0.907 | -0.461 | -0.075 | -0.259 | -0.018 | -0.055 | -0.055 | 0 | 0.86 | 0.82 | 0.95 | 1 | 0.94 | 0.96 |
| 4 | 127171396 | rs17835005 | 0.679 | -0.375 | -0.752 | -0.214 | -0.016 | 0.121 | 0.121 | 0.219 | 0.51 | 0.12 | 0.57 | 0.71 | 0.57 | 0.43 |

| | | | | | | | | | | | | | | | | |
|---|---|---|---|---|---|---|---|---|---|---|---|---|---|---|---|---|
| 4 | 127182592 | rs989576 | 0.157 | 0.164 | -0.09 | -0.323 | 0.001 | -0.02 | -0.02 | 0 | 0.26 | 0.29 | 0.25 | 0.08 | 0.28 | 0.45 |
| 4 | 127196618 | rs6855524 | -0.253 | 0.168 | -0.277 | -0.428 | 0 | -0.001 | -0.001 | 0 | 0.61 | 0.64 | 0.76 | 0.85 | 0.67 | 0.86 |
| 4 | 127201582 | rs7683349 | 1.256 | 0.125 | -0.195 | -0.522 | -0.002 | 0.011 | 0.011 | 0 | 0.21 | 0.14 | 0.24 | 0.08 | 0.22 | 0.43 |
| 4 | 127201933 | rs7666758 | -0.323 | 0.309 | -0.764 | -0.424 | 0.005 | 0.017 | 0.017 | 0 | 0.61 | 0.67 | 0.76 | 0.85 | 0.67 | 0.86 |
| 4 | 127204248 | rs1548111 | 0.394 | 0.273 | -0.92 | -0.462 | -0.007 | -0.067 | -0.067 | 1.952 | 0.29 | 0.5 | 0.3 | 0.11 | 0.27 | 0.46 |
| 4 | 127207959 | rs13130903 | 0.061 | 0.123 | -1.639 | -0.48 | -0.004 | -0.009 | -0.009 | 0 | 0.18 | 0.18 | 0.08 | 0.16 | 0.13 | 0.09 |
| 4 | 127208717 | rs2128782 | 0.054 | 0.12 | -1.665 | -0.49 | 0.022 | 0.038 | 0.038 | 0 | 0.6 | 0.76 | 0.78 | 0.71 | 0.69 | 0.84 |
| 4 | 127208746 | rs13132177 | 1.166 | 0.167 | -1.76 | -0.489 | 0.001 | -0.006 | -0.006 | 0.521 | 0.11 | 0.09 | 0.05 | 0 | 0.1 | 0.04 |
| 4 | 127213903 | rs10011584 | -1.456 | 0.189 | -1.899 | -0.469 | -0.008 | -0.005 | -0.005 | 0.977 | 0.78 | 0.94 | 0.85 | 0.88 | 0.82 | 0.93 |
| 4 | 127220695 | rs11098860 | -0.917 | 0.759 | -1.394 | 0.435 | -0.074 | 0.016 | 0.016 | 0 | 0.73 | 0.94 | 0.73 | 0.83 | 0.64 | 0.79 |
| 4 | 127224641 | rs10213170 | 0.815 | 0.684 | -1.184 | -0.368 | -0.003 | -0.016 | -0.016 | 0 | 0.45 | 0.49 | 0.38 | 0.2 | 0.4 | 0.5 |
| 4 | 127232483 | rs11725332 | 0.248 | 1.138 | -1.872 | -0.471 | -0.002 | 0.029 | 0.029 | 0 | 0.11 | 0.04 | 0.05 | 0 | 0.1 | 0.04 |
| 4 | 127235715 | rs1320244 | 0.154 | 1.043 | -1.574 | 0.075 | -0.02 | -0.006 | -0.006 | 0 | 0.65 | 0.72 | 0.48 | 0.36 | 0.55 | 0.64 |
| 4 | 127242389 | rs2893181 | -0.104 | 1.325 | -1.887 | -0.182 | -0.131 | -0.063 | -0.063 | 0 | 0.25 | 0.09 | 0.57 | 0.65 | 0.47 | 0.43 |
| 4 | 127242422 | rs17240023 | 0.339 | 0.166 | -2.806 | -1.065 | -0.006 | 0.102 | 0.102 | 0 | 0.19 | 0.01 | 0.08 | 0.02 | 0.14 | 0.23 |
| 4 | 127243397 | rs6852950 | -0.15 | -0.741 | -3.111 | -1.663 | 0.042 | 0.094 | 0.094 | 0.035 | 0.35 | 0.13 | 0.2 | 0.13 | 0.27 | 0.36 |
| 4 | 127247924 | rs17836083 | 0.938 | -0.326 | -2.985 | -1.728 | -0.078 | -0.044 | -0.044 | 0 | 0.24 | 0.11 | 0.53 | 0.65 | 0.4 | 0.42 |
| 4 | 127249062 | rs6534545 | -0.774 | -0.734 | -3.035 | -1.977 | 0.029 | 0.071 | 0.071 | 0.293 | 0.65 | 0.26 | 0.78 | 0.79 | 0.73 | 0.83 |
| 4 | 127249276 | rs6534548 | -0.814 | -0.805 | -2.22 | -2.011 | -0.001 | -0.008 | -0.008 | 0 | 0.66 | 0.41 | 0.8 | 0.79 | 0.73 | 0.83 |
| 4 | 127251381 | rs2391031 | -0.263 | -0.811 | -2.252 | -2.01 | 0.054 | 0.149 | 0.149 | 0 | 0.42 | 0.17 | 0.26 | 0.14 | 0.33 | 0.41 |
| 4 | 127255447 | rs11938961 | 0.496 | -1.263 | -1.743 | -2.332 | -0.049 | -0.026 | -0.026 | 0.33 | 0.24 | 0.64 | 0.1 | 0.19 | 0.09 | 0.09 |
| 4 | 127256413 | rs6846321 | 1.169 | -1.681 | -1.405 | -2.555 | -0.038 | -0.085 | -0.085 | 0.134 | 0.13 | 0.37 | 0.05 | 0 | 0.04 | 0.03 |
| 4 | 127256427 | rs6534549 | -0.033 | -1.613 | -1.551 | -2.502 | -0.059 | -0.024 | -0.024 | 0.009 | 0.23 | 0.64 | 0.09 | 0.19 | 0.07 | 0.08 |
| 4 | 127257660 | rs6827629 | 0.013 | -1.403 | -1.661 | -2.547 | -0.014 | 0.101 | 0.101 | 0 | 0.46 | 0.73 | 0.62 | 0.84 | 0.48 | 0.51 |
| 4 | 127259366 | rs9991404 | -0.031 | -1.332 | -1.35 | -2.29 | -0.076 | -0.037 | -0.037 | 0 | 0.22 | 0.47 | 0.08 | 0.19 | 0.09 | 0.09 |
| 4 | 127260284 | rs6534550 | -1.342 | -0.31 | -1.267 | -1.769 | 0.016 | -0.036 | -0.036 | 0 | 0.89 | 0.89 | 0.88 | 0.98 | 0.82 | 0.92 |
| 4 | 127264759 | rs12500408 | 0.643 | -0.543 | -1.673 | -1.58 | -0.004 | -0.003 | -0.003 | 0 | 0.16 | 0.06 | 0.12 | 0.12 | 0.13 | 0.14 |
| 4 | 127266906 | rs4538495 | -0.496 | -0.187 | -0.983 | -1.463 | 0.03 | -0.14 | -0.14 | 1.068 | 0.76 | 0.56 | 0.78 | 0.96 | 0.7 | 0.82 |
| 4 | 127267782 | rs12509279 | -0.084 | 0.033 | -1.033 | -0.617 | -0.027 | 0.005 | 0.005 | 0 | 0.34 | 0.24 | 0.54 | 0.64 | 0.41 | 0.45 |
| 4 | 127273449 | rs1318464 | -0.333 | 0.218 | -1.23 | -0.887 | -0.006 | 0.059 | 0.059 | 0 | 0.39 | 0.3 | 0.55 | 0.83 | 0.42 | 0.5 |
| 4 | 127281188 | rs13135378 | -0.766 | -0.053 | -1.033 | -0.378 | 0.011 | -0.001 | -0.001 | 0.157 | 0.13 | 0.11 | 0.05 | 0.14 | 0.09 | 0.12 |
| 4 | 127283300 | rs6534554 | -1.053 | 0.295 | -1.644 | 0.057 | -0.044 | -0.051 | -0.051 | 0 | 0.58 | 0.85 | 0.65 | 0.53 | 0.54 | 0.79 |
| 4 | 127287894 | rs12508982 | 0.403 | 0.194 | -1.989 | -0.196 | -0.015 | 0.007 | 0.007 | 0 | 0.2 | 0.05 | 0.33 | 0.16 | 0.18 | 0.3 |
| 4 | 127289047 | rs13103146 | 0.348 | 0.162 | -1.211 | -0.325 | 0.002 | -0.017 | -0.017 | 0 | 0.11 | 0.17 | 0.17 | 0.04 | 0.13 | 0.27 |
| 4 | 127290078 | rs13110774 | -0.556 | -0.069 | -0.294 | -0.288 | -0.006 | -0.025 | -0.025 | 1.248 | 0.2 | 0.32 | 0.12 | 0.14 | 0.21 | 0.15 |
| 4 | 127296342 | rs7694254 | -0.083 | -0.233 | -0.334 | -0.209 | 0.008 | 0.016 | 0.016 | 0 | 0.76 | 0.88 | 0.66 | 0.85 | 0.83 | 0.77 |
| 4 | 127297701 | rs13124314 | 0.364 | -0.204 | -0.361 | -0.265 | 0.001 | 0.031 | 0.031 | 0.459 | 0.09 | 0.02 | 0.07 | 0.02 | 0.09 | 0.22 |
| 4 | 127303936 | rs7699436 | 0.746 | -0.593 | -0.161 | -0.446 | 0.021 | 0.021 | 0.021 | 0 | 0.58 | 0.75 | 0.54 | 0.69 | 0.72 | 0.55 |
| 4 | 127309605 | rs17011033 | 1.99 | -0.311 | 0.015 | -0.31 | -0.071 | -0.058 | -0.058 | 0 | 0.39 | 0.2 | 0.34 | 0.44 | 0.5 | 0.21 |
| 4 | 127309862 | rs553145 | -0.525 | -0.002 | 0.954 | -0.229 | 0.117 | 0.201 | 0.201 | 0.005 | 0.07 | 0.35 | 0.1 | 0.24 | 0.12 | 0.08 |
| 4 | 127313065 | rs592813 | -1.694 | -0.053 | 1.053 | -0.25 | -0.005 | 0.01 | 0.01 | 0 | 0.32 | 0.51 | 0.22 | 0.39 | 0.31 | 0.52 |
| 4 | 127313506 | rs545314 | -1.683 | -0.053 | 1.051 | -0.25 | -0.004 | 0.008 | 0.008 | 0 | 0.32 | 0.47 | 0.22 | 0.39 | 0.31 | 0.52 |
| 4 | 127314275 | rs13120906 | 0.198 | -0.055 | 1.079 | -0.272 | 0.002 | 0.004 | 0.004 | 0.348 | 0.09 | 0.1 | 0.07 | 0.02 | 0.1 | 0.22 |
| 4 | 127315346 | rs473695 | -1.501 | -0.033 | 0.986 | -0.248 | -0.002 | -0.061 | -0.061 | 2.477 | 0.31 | 0.47 | 0.21 | 0.14 | 0.3 | 0.48 |
| 4 | 127315595 | rs624738 | -1.219 | -0.013 | 0.5 | -0.23 | 0.021 | 0.031 | 0.031 | 2.978 | 0.19 | 0.57 | 0.19 | 0.26 | 0.23 | 0.33 |
| 4 | 127316370 | rs6812090 | -0.57 | 0.018 | -0.018 | -0.205 | 0.004 | 0.003 | 0.003 | 0 | 0.17 | 0.12 | 0.06 | 0.12 | 0.1 | 0.22 |
| 4 | 127321888 | rs12331943 | -0.691 | 0.2 | -0.088 | -0.452 | 0 | -0.008 | -0.008 | 0 | 0.27 | 0.26 | 0.19 | 0.11 | 0.25 | 0.28 |
| 4 | 127327343 | rs666205 | -0.208 | -0.187 | 0.103 | -0.585 | -0.004 | -0.01 | -0.01 | 0 | 0.21 | 0.19 | 0.08 | 0.11 | 0.13 | 0.19 |
| 4 | 127329737 | rs549638 | -0.264 | -0.148 | 0.254 | -0.6 | -0.02 | -0.046 | -0.046 | 0 | 0.22 | 0.3 | 0.09 | 0.11 | 0.13 | 0.19 |
| 4 | 127335358 | rs634701 | 0.142 | -0.117 | -0.046 | -0.887 | -0.003 | -0.01 | -0.01 | 0.024 | 0.76 | 0.79 | 0.84 | 0.89 | 0.82 | 0.71 |
| 4 | 127336047 | rs12647099 | 1.983 | -0.131 | 0.117 | -0.389 | -0.042 | -0.019 | -0.019 | 0 | 0.43 | 0.21 | 0.64 | 0.49 | 0.52 | 0.4 |
| 4 | 127337839 | rs636729 | -0.083 | -0.142 | -0.042 | -0.366 | 0.009 | 0.013 | 0.013 | 0 | 0.78 | 0.84 | 0.86 | 0.9 | 0.83 | 0.76 |
| 4 | 127343002 | rs2391054 | 0.664 | 0.142 | -0.358 | -0.222 | 0.03 | 0.005 | 0.005 | 0 | 0.53 | 0.41 | 0.74 | 0.51 | 0.68 | 0.55 |
| 4 | 127351139 | rs6534562 | -0.441 | 0.761 | -0.449 | -0.153 | 0.01 | -0.021 | -0.021 | 0.255 | 0.1 | 0.1 | 0.11 | 0.01 | 0.17 | 0.14 |
| 4 | 127353144 | rs1396531 | -0.617 | 0.111 | -0.821 | 0.177 | 0.001 | -0.013 | -0.013 | 1.804 | 0.2 | 0.02 | 0.12 | 0.13 | 0.14 | 0.25 |
| 4 | 127354073 | rs1396529 | 0.594 | 0.043 | -0.214 | 0.04 | 0.01 | -0.035 | -0.035 | 0.215 | 0.78 | 0.62 | 0.87 | 0.84 | 0.82 | 0.7 |
| 4 | 127360862 | rs519699 | 0.396 | 0.213 | -0.265 | 0.016 | 0 | -0.006 | -0.006 | 0.599 | 0.9 | 0.88 | 0.85 | 0.96 | 0.9 | 0.83 |
| 4 | 127363097 | rs12498806 | -0.091 | -0.088 | 0.155 | 0.47 | -0.017 | 0.093 | 0.093 | 0 | 0.06 | 0.17 | 0.02 | 0.24 | 0.02 | 0.1 |
| 4 | 127365329 | rs4833346 | 1.891 | -0.761 | -0.249 | 0.544 | 0.076 | 0.044 | 0.044 | 1.359 | 0.5 | 0.13 | 0.52 | 0.4 | 0.64 | 0.35 |
| 4 | 127365899 | rs484051 | -1.14 | -0.745 | 0.725 | 0.446 | -0.033 | 0.02 | 0.02 | 0.696 | 0.21 | 0.6 | 0.11 | 0.3 | 0.13 | 0.24 |
| 4 | 127373021 | rs1508983 | -0.915 | -0.469 | 0.652 | 0.425 | 0 | 0.023 | 0.023 | 0 | 0.13 | 0.26 | 0.13 | 0.25 | 0.12 | 0.2 |
| 4 | 127374038 | rs1508986 | -0.968 | -0.067 | 0.281 | -0.184 | 0.007 | -0.022 | -0.022 | 0.153 | 0.35 | 0.42 | 0.25 | 0.27 | 0.38 | 0.29 |
| 4 | 127378114 | rs10016992 | 0.667 | -0.007 | -0.031 | -0.171 | -0.01 | 0.002 | 0.002 | 0.435 | 0.22 | 0.16 | 0.12 | 0.02 | 0.27 | 0.08 |
| 4 | 127378765 | rs12499737 | -0.264 | 0.144 | 0.018 | 0.299 | 0.007 | 0.031 | 0.031 | 0 | 0.12 | 0.24 | 0.12 | 0.25 | 0.12 | 0.2 |
| 4 | 127382084 | rs12647338 | -0.627 | 0.047 | -0.24 | 0.329 | 0.001 | -0.002 | -0.002 | 0 | 0.17 | 0.21 | 0.13 | 0.24 | 0.14 | 0.24 |
| 4 | 127399765 | rs6822951 | -0.764 | 0.27 | -0.434 | 0.433 | -0.001 | -0.001 | -0.001 | 0.784 | 0.88 | 0.86 | 0.94 | 0.85 | 0.9 | 0.87 |
| 4 | 127404615 | rs7699051 | -0.686 | 0.141 | -0.554 | 0.587 | -0.016 | -0.026 | -0.026 | 0 | 0.57 | 0.75 | 0.57 | 0.47 | 0.53 | 0.61 |
| 4 | 127419470 | rs7673418 | -0.61 | -1.136 | -0.264 | 0.425 | 0.058 | -0.009 | -0.009 | 0 | 0.36 | 0.2 | 0.38 | 0.6 | 0.21 | 0.38 |
| 4 | 127429942 | rs6820022 | -0.608 | -1.685 | -0.281 | 0.41 | -0.018 | -0.012 | -0.012 | 0 | 0.28 | 0.38 | 0.35 | 0.19 | 0.15 | 0.3 |

| | | | | | | | | | | | | | | | | |
|---|---|---|---|---|---|---|---|---|---|---|---|---|---|---|---|---|
| 4 | 127431975 | rs7698081 | 1.502 | -1.348 | -0.699 | 0.121 | 0.022 | 0.028 | 0.028 | 1.308 | 0.44 | 0.13 | 0.46 | 0.38 | 0.64 | 0.43 |
| 4 | 127438435 | rs17011171 | -0.474 | -1.498 | -0.77 | 0.158 | -0.033 | 0.118 | 0.118 | 1.333 | 0.14 | 0.28 | 0.05 | 0.3 | 0.1 | 0.11 |
| 4 | 127440009 | rs1397676 | -0.556 | -1.644 | -0.675 | 0.087 | -0.037 | 0.063 | 0.063 | 0.093 | 0.25 | 0.36 | 0.13 | 0.39 | 0.13 | 0.32 |
| 4 | 127440570 | rs17011180 | 0.757 | -1.844 | -0.873 | 0.14 | 0.024 | -0.064 | -0.064 | 0.559 | 0.19 | 0.01 | 0.28 | 0.22 | 0.12 | 0.15 |
| 4 | 127440659 | rs990364 | 0.086 | -1.425 | -0.388 | 0.295 | 0.01 | 0.022 | 0.022 | 0 | 0.19 | 0.12 | 0.09 | 0.1 | 0.11 | 0.26 |
| 4 | 127442885 | rs9994335 | -0.151 | -1.245 | -0.266 | 0.475 | -0.009 | 0.305 | 0.305 | 4.288 | 0.06 | 0.25 | 0.04 | 0.29 | 0.02 | 0.06 |
| 4 | 127448140 | rs1877169 | -1.17 | -1.325 | -0.454 | 0.259 | -0.029 | 0.021 | 0.021 | 0.415 | 0.34 | 0.4 | 0.14 | 0.4 | 0.21 | 0.38 |
| 4 | 127450408 | rs17197167 | 0.68 | -1.353 | -0.652 | -0.104 | 0.009 | 0.014 | 0.014 | 0 | 0.14 | 0.05 | 0.06 | 0.1 | 0.07 | 0.17 |
| 4 | 127458699 | rs12650680 | 1.655 | -1.262 | -0.736 | -0.176 | 0.004 | 0.002 | 0.002 | 0.243 | 0.33 | 0.33 | 0.37 | 0.41 | 0.49 | 0.37 |
| 4 | 127458938 | rs1543000 | -0.262 | -1.484 | -0.614 | -0.366 | 0.002 | -0.006 | -0.006 | 0.474 | 0.66 | 0.69 | 0.55 | 0.52 | 0.77 | 0.71 |
| 4 | 127467854 | rs992181 | 0.439 | -1.459 | -0.621 | -0.385 | -0.013 | 0.002 | 0.002 | 0 | 0.56 | 0.5 | 0.46 | 0.52 | 0.65 | 0.64 |
| 4 | 127473539 | rs17257398 | 0.801 | -0.566 | -0.966 | -0.514 | -0.015 | 0.043 | 0.043 | 0.512 | 0.34 | 0.07 | 0.41 | 0.2 | 0.33 | 0.23 |
| 4 | 127485146 | rs313096 | 1.709 | -0.457 | -0.804 | -0.501 | 0.014 | 0.026 | 0.026 | 0.039 | 0.27 | 0.34 | 0.3 | 0.41 | 0.36 | 0.38 |
| 4 | 127485170 | rs313097 | 1.722 | -0.456 | -0.808 | -0.501 | 0.011 | 0.019 | 0.019 | 0.054 | 0.27 | 0.31 | 0.3 | 0.41 | 0.35 | 0.38 |
| 4 | 127485630 | rs313098 | 1.722 | -0.456 | -0.808 | -0.501 | 0.011 | 0.019 | 0.019 | 0.054 | 0.27 | 0.31 | 0.3 | 0.41 | 0.36 | 0.38 |
| 4 | 127493571 | rs885600 | 0.138 | -0.596 | -0.834 | -0.57 | 0.003 | -0.026 | -0.026 | 0 | 0.24 | 0.31 | 0.16 | 0.11 | 0.21 | 0.3 |
| 4 | 127516859 | rs1511138 | 1.01 | -0.538 | -0.969 | -0.456 | 0.009 | -0.014 | -0.014 | 0.153 | 0.1 | 0.07 | 0.11 | 0.29 | 0.08 | 0.08 |
| 4 | 127519940 | rs12508410 | 1.126 | -0.649 | -1.28 | -0.585 | -0.033 | 0.02 | 0.02 | 1.998 | 0.32 | 0.03 | 0.4 | 0.21 | 0.31 | 0.22 |
| 4 | 127525092 | rs17258452 | -0.078 | -0.523 | -0.92 | -0.556 | -0.001 | -0.001 | -0.001 | 0.12 | 0.26 | 0.26 | 0.17 | 0.11 | 0.24 | 0.24 |
| 4 | 127525235 | rs7695327 | 0.386 | -0.482 | -1.046 | -0.398 | 0.007 | -0.011 | -0.011 | 0 | 0.12 | 0.08 | 0.11 | 0.29 | 0.1 | 0.1 |
| 4 | 127529033 | rs2054981 | 1.072 | -0.503 | -0.507 | -0.574 | -0.021 | -0.047 | -0.047 | 0 | 0.14 | 0.25 | 0.07 | 0.07 | 0.07 | 0.17 |
| 4 | 127529360 | rs13434700 | 0.447 | -0.642 | -0.997 | -0.678 | -0.002 | 0.068 | 0.068 | 3.687 | 0.41 | 0.12 | 0.43 | 0.25 | 0.4 | 0.28 |
| 4 | 127535445 | rs1511144 | 1.351 | -0.238 | -0.739 | -0.772 | 0.012 | 0.042 | 0.042 | 0 | 0.28 | 0.42 | 0.29 | 0.39 | 0.33 | 0.36 |
| 4 | 127535865 | rs7681514 | -0.052 | -0.309 | -0.841 | -0.897 | 0 | 0.006 | 0.006 | 2.443 | 0.53 | 0.3 | 0.61 | 0.6 | 0.48 | 0.5 |
| 4 | 127536521 | rs17258848 | 1.247 | -0.333 | -1.08 | -0.884 | -0.003 | 0.023 | 0.023 | 1.843 | 0.3 | 0.07 | 0.49 | 0.23 | 0.32 | 0.28 |
| 4 | 127548421 | rs1511140 | 0.52 | -0.298 | -0.861 | -0.897 | -0.008 | -0.003 | -0.003 | 0 | 0.65 | 0.43 | 0.81 | 0.63 | 0.74 | 0.65 |
| 4 | 127551274 | rs447507 | -0.313 | -0.288 | -0.922 | -0.961 | -0.002 | -0.102 | -0.102 | 0 | 0.3 | 0.4 | 0.16 | 0.07 | 0.24 | 0.34 |
| 4 | 127551516 | rs17011345 | 1.355 | -0.048 | -0.928 | -0.696 | -0.029 | -0.041 | -0.041 | 0.315 | 0.27 | 0.03 | 0.46 | 0.26 | 0.29 | 0.26 |
| 4 | 127557937 | rs10518503 | 2.047 | 0.119 | -0.703 | -0.633 | -0.021 | 0.03 | 0.03 | 1.153 | 0.08 | 0.03 | 0.05 | 0 | 0.09 | 0.01 |
| 4 | 127562447 | rs313045 | 0.257 | -0.082 | -0.601 | -0.632 | -0.001 | 0 | 0 | 0.19 | 0.14 | 0.17 | 0.08 | 0.27 | 0.17 | 0.13 |
| 4 | 127565540 | rs313040 | -0.804 | -0.18 | -0.924 | -0.854 | -0.013 | 0.02 | 0.02 | 0 | 0.82 | 0.88 | 0.85 | 1 | 0.78 | 0.94 |
| 4 | 127565650 | rs6534573 | 1.278 | -0.132 | -0.837 | -0.715 | -0.022 | 0.019 | 0.019 | 0 | 0.06 | 0.03 | 0.05 | 0 | 0.12 | 0.05 |
| 4 | 127574191 | rs170197 | 0.018 | -0.185 | -0.542 | -0.638 | 0.031 | 0.051 | 0.051 | 0 | 0.16 | 0.37 | 0.11 | 0.27 | 0.17 | 0.12 |
| 4 | 127580432 | rs386871 | -1.275 | -0.428 | -0.561 | -0.625 | -0.017 | 0.021 | 0.021 | 0 | 0.86 | 0.91 | 0.88 | 1 | 0.83 | 0.97 |
| 4 | 127597069 | rs313055 | 0.448 | -0.641 | -0.782 | -0.725 | 0.01 | 0.005 | 0.005 | 0 | 0.36 | 0.44 | 0.33 | 0.39 | 0.46 | 0.42 |
| 4 | 127605685 | rs398578 | -0.495 | -0.937 | -0.8 | -0.892 | 0.014 | 0.041 | 0.041 | 0.983 | 0.59 | 0.7 | 0.46 | 0.74 | 0.59 | 0.71 |
| 4 | 127605711 | rs12513097 | 1.456 | -1.013 | -0.823 | -0.881 | 0.027 | -0.014 | -0.014 | 0 | 0.25 | 0.1 | 0.42 | 0.26 | 0.24 | 0.27 |
| 4 | 127607021 | rs17011412 | 0.574 | -0.948 | -0.694 | -0.83 | -0.022 | -0.047 | -0.047 | 0.826 | 0.23 | 0.24 | 0.12 | 0.08 | 0.13 | 0.27 |
| 4 | 127609379 | rs1552665 | 0.629 | -0.816 | -0.871 | -0.805 | -0.037 | 0.039 | 0.039 | 0 | 0.23 | 0.37 | 0.13 | 0.35 | 0.13 | 0.28 |
| 4 | 127610903 | rs313071 | 0.665 | -0.721 | -1.118 | -0.801 | 0.05 | 0.074 | 0.074 | 0 | 0.55 | 0.18 | 0.71 | 0.65 | 0.67 | 0.66 |
| 4 | 127622803 | rs17011442 | 1.665 | -0.718 | -1.294 | -0.775 | 0.014 | -0.05 | -0.05 | 0 | 0.23 | 0.03 | 0.4 | 0.26 | 0.22 | 0.25 |
| 4 | 127624935 | rs313048 | 0.474 | -0.36 | -0.529 | -0.694 | -0.006 | -0.01 | -0.01 | 0 | 0.6 | 0.49 | 0.73 | 0.66 | 0.68 | 0.68 |
| 4 | 127638432 | rs7697724 | -1.025 | 0.165 | -0.119 | -0.496 | 0.001 | 0 | 0 | 2.39 | 0.23 | 0.74 | 0.18 | 0.24 | 0.18 | 0.04 |
| 4 | 127644271 | rs2654715 | -0.702 | 0.389 | 0.261 | -0.517 | 0.055 | 0.034 | 0.034 | 0.065 | 0.2 | 0.43 | 0.18 | 0.23 | 0.22 | 0.09 |
| 4 | 127646765 | rs2969612 | -1.217 | 0.395 | 0.248 | -0.601 | -0.004 | -0.011 | -0.011 | 0.126 | 0.87 | 0.92 | 0.88 | 1 | 0.84 | 0.98 |
| 4 | 127649306 | rs6846559 | 1.391 | 0.036 | -0.126 | -0.812 | 0.058 | -0.004 | -0.004 | 0.768 | 0.3 | 0.05 | 0.42 | 0.26 | 0.23 | 0.31 |
| 4 | 127655701 | rs10857117 | 0.871 | 0.541 | -0.217 | -1.164 | -0.004 | -0.026 | -0.026 | 0 | 0.56 | 0.34 | 0.67 | 0.99 | 0.54 | 0.69 |
| 4 | 127656069 | rs1167190 | 0.536 | 0.515 | -0.045 | -1.202 | -0.011 | 0.026 | 0.026 | 0 | 0.23 | 0.15 | 0.14 | 0.01 | 0.25 | 0.23 |
| 4 | 127657442 | rs1388240 | 1.034 | 0.554 | -0.334 | -1.079 | 0.013 | 0.029 | 0.029 | 0.071 | 0.26 | 0.27 | 0.28 | 0.48 | 0.34 | 0.42 |
| 4 | 127660479 | rs17011528 | -0.009 | 0.504 | 0.069 | -1.013 | 0.004 | -0.099 | -0.099 | 0.982 | 0.27 | 0.42 | 0.17 | 0.01 | 0.25 | 0.23 |
| 4 | 127662110 | rs11098892 | 0.568 | 0.49 | -0.287 | -0.991 | -0.002 | -0.05 | -0.05 | 0 | 0.6 | 0.35 | 0.71 | 0.99 | 0.59 | 0.75 |
| 4 | 127706053 | rs1503626 | -0.217 | 0.087 | -0.176 | -0.859 | -0.032 | -0.139 | -0.139 | 0.077 | 0.73 | 0.55 | 0.85 | 0.99 | 0.81 | 0.8 |
| 4 | 127712289 | rs4834142 | 0.746 | 0.111 | -0.365 | -0.9 | 0.001 | 0.23 | 0.23 | 0.662 | 0.59 | 0.14 | 0.71 | 0.99 | 0.65 | 0.77 |
| 4 | 127714708 | rs12507245 | 0.939 | 0.044 | -0.253 | -0.792 | 0.03 | -0.021 | -0.021 | 0 | 0.4 | 0.29 | 0.51 | 0.51 | 0.3 | 0.36 |
| 4 | 127720918 | rs12508795 | 1.687 | 0.14 | -0.387 | -0.784 | -0.03 | 0.036 | 0.036 | 1.149 | 0.11 | 0.05 | 0.11 | 0 | 0.16 | 0.02 |
| 4 | 127727154 | rs12642632 | 0.889 | 0.038 | -0.189 | -0.749 | 0.016 | -0.011 | -0.011 | 0 | 0.4 | 0.27 | 0.5 | 0.52 | 0.3 | 0.36 |
| 4 | 127745441 | rs13130797 | 0.885 | 0.058 | -0.181 | -0.727 | -0.005 | -0.02 | -0.02 | 0 | 0.56 | 0.47 | 0.64 | 0.74 | 0.6 | 0.76 |
| 4 | 127754030 | rs6829454 | -0.226 | 0.064 | -0.257 | -0.792 | -0.02 | -0.095 | -0.095 | 6.067 | 0.72 | 0.41 | 0.84 | 0.99 | 0.8 | 0.8 |
| 4 | 127768849 | rs1503609 | 0.553 | 0.096 | -0.339 | -0.715 | -0.031 | -0.047 | -0.047 | 0.268 | 0.26 | 0.22 | 0.26 | 0.48 | 0.37 | 0.41 |
| 4 | 127770015 | rs2134642 | 0.88 | 0.002 | -0.363 | -0.701 | 0.053 | -0.062 | -0.062 | 0.651 | 0.39 | 0.12 | 0.5 | 0.51 | 0.31 | 0.39 |
| 4 | 127773649 | rs4834144 | -0.036 | -0.027 | 0.034 | -0.531 | -0.017 | -0.072 | -0.072 | 0.501 | 0.72 | 0.4 | 0.82 | 0.96 | 0.79 | 0.8 |
| 4 | 127774985 | rs17272221 | 1.065 | -0.056 | -0.205 | -0.502 | -0.035 | 0.113 | 0.113 | 0.187 | 0.44 | 0.06 | 0.45 | 0.23 | 0.41 | 0.4 |
| 4 | 127779997 | rs13114412 | -0.162 | 0.108 | 0.091 | -0.714 | -0.03 | -0.127 | -0.127 | 0.134 | 0.73 | 0.53 | 0.81 | 0.96 | 0.77 | 0.8 |
| 4 | 127788899 | rs4270599 | 0.836 | 0.337 | -0.063 | -0.211 | -0.017 | -0.026 | -0.026 | 1.244 | 0.21 | 0.21 | 0.38 | 0.57 | 0.33 | 0.51 |
| 4 | 127799124 | rs2391097 | 0.858 | 0.314 | -0.241 | -0.286 | -0.066 | -0.104 | -0.104 | 0.888 | 0.21 | 0.12 | 0.38 | 0.56 | 0.33 | 0.51 |
| 4 | 127822133 | rs4833358 | 0.66 | 0.015 | -0.397 | -0.371 | -0.104 | -0.148 | -0.148 | 0 | 0.27 | 0.1 | 0.39 | 0.54 | 0.39 | 0.54 |
| 4 | 127825185 | rs6534591 | -0.894 | -0.224 | 0.168 | -0.538 | 0.017 | 0.182 | 0.182 | 1.545 | 0.55 | 0.24 | 0.64 | 0.89 | 0.65 | 0.75 |
| 4 | 127831937 | rs1510677 | -0.975 | -0.19 | 0.164 | -0.619 | -0.003 | -0.036 | -0.036 | 2.145 | 0.56 | 0.56 | 0.67 | 0.89 | 0.65 | 0.75 |
| 4 | 127862511 | rs1510694 | -1.378 | -0.138 | 0.208 | -0.583 | -0.005 | 0.1 | 0.1 | 0.466 | 0.61 | 0.85 | 0.69 | 0.88 | 0.65 | 0.75 |

| | | | | | | | | | | | | | | | | |
|---|---|---|---|---|---|---|---|---|---|---|---|---|---|---|---|---|
| 4 | 127874789 | rs10016117 | -0.172 | -0.006 | 0.186 | -0.553 | -0.004 | 0.008 | 0.008 | 1.157 | 0.35 | 0.38 | 0.51 | 0.85 | 0.41 | 0.58 |
| 4 | 127925448 | rs4834162 | 0.253 | -0.098 | 0.272 | -0.435 | 0 | -0.001 | -0.001 | 0.111 | 0.69 | 0.69 | 0.73 | 0.65 | 0.72 | 0.77 |
| 4 | 127973615 | rs340441 | 0.118 | 0.356 | -0.015 | -0.373 | -0.072 | -0.113 | -0.113 | 0 | 0.21 | 0.55 | 0.14 | 0.03 | 0.13 | 0.12 |
| 4 | 127977716 | rs340446 | -0.143 | 0.378 | -0.166 | -0.361 | -0.065 | -0.111 | -0.111 | 0 | 0.23 | 0.57 | 0.15 | 0.03 | 0.15 | 0.16 |
| 4 | 127978786 | rs6834065 | 0.47 | 0.946 | -0.269 | -0.094 | -0.002 | 0.001 | 0.001 | 0 | 0.24 | 0.25 | 0.38 | 0.54 | 0.35 | 0.51 |
| 4 | 127981170 | rs1383522 | -0.09 | 0.839 | -0.635 | -0.434 | -0.001 | -0.006 | -0.006 | 0 | 0.92 | 0.9 | 0.95 | 1 | 0.91 | 0.96 |
| 4 | 127991638 | rs17278402 | 1.029 | 0.642 | -1.11 | -0.71 | 0.02 | 0.168 | 0.168 | 0 | 0.34 | 0.03 | 0.25 | 0.08 | 0.27 | 0.27 |
| 4 | 127992106 | rs12646249 | 0.577 | 0.881 | -0.705 | -0.49 | -0.008 | -0.01 | -0.01 | 0 | 0.25 | 0.23 | 0.38 | 0.56 | 0.37 | 0.51 |
| 4 | 127993106 | rs7670859 | 0.325 | 0.697 | -0.36 | -0.599 | 0.016 | 0.021 | 0.021 | 0 | 0.21 | 0.19 | 0.13 | 0.03 | 0.18 | 0.21 |
| 4 | 128005362 | rs6852817 | -0.108 | 0.769 | 0.32 | -0.58 | 0.045 | -0.038 | -0.038 | 0.076 | 0.91 | 0.73 | 0.94 | 1 | 0.92 | 0.96 |
| 4 | 128023339 | rs1871738 | 0.723 | 0.777 | 0.456 | -0.486 | 0.003 | -0.008 | -0.008 | 0 | 0.13 | 0.19 | 0.24 | 0.04 | 0.14 | 0.01 |
| 4 | 128046856 | rs13107241 | -0.465 | 0.875 | 0.305 | -0.532 | -0.004 | -0.002 | -0.002 | 0.069 | 0.44 | 0.42 | 0.35 | 0.57 | 0.52 | 0.69 |
| 4 | 128055169 | rs17012111 | 0.583 | 0.862 | 0.286 | -0.568 | 0.017 | 0.016 | 0.016 | 0 | 0.2 | 0.18 | 0.12 | 0.03 | 0.17 | 0.18 |
| 4 | 128063104 | rs4385081 | 1.129 | 0.952 | 0.213 | -0.451 | -0.01 | 0.011 | 0.011 | 0.227 | 0.12 | 0.09 | 0.23 | 0.04 | 0.13 | 0.01 |
| 4 | 128064562 | rs1155135 | 0.164 | 0.961 | 0.209 | -0.403 | 0.046 | 0.191 | 0.191 | 0.943 | 0.9 | 0.64 | 0.85 | 0.69 | 0.91 | 0.95 |
| 4 | 128072643 | rs12503951 | 1.497 | 0.902 | 0.078 | -0.411 | -0.052 | 0 | 0 | 0 | 0.11 | 0.01 | 0.21 | 0.04 | 0.13 | 0.01 |
| 4 | 128072800 | rs1026824 | 0.208 | 1.088 | 0.177 | -0.407 | 0.057 | 0.088 | 0.088 | 1.144 | 0.78 | 0.51 | 0.62 | 0.65 | 0.77 | 0.93 |
| 4 | 128079327 | rs6837753 | -0.476 | 1.077 | 0.072 | -0.461 | -0.002 | -0.002 | -0.002 | 0 | 0.44 | 0.44 | 0.35 | 0.56 | 0.5 | 0.69 |
| 4 | 128096615 | rs1471556 | -0.174 | 1.196 | 0.149 | -0.42 | 0.027 | -0.03 | -0.03 | 0.01 | 0.81 | 0.72 | 0.69 | 0.96 | 0.76 | 0.95 |
| 4 | 128097963 | rs11726994 | 0.866 | 1.161 | -0.177 | -0.233 | -0.095 | -0.157 | -0.157 | 0 | 0.21 | 0.06 | 0.18 | 0.53 | 0.31 | 0.46 |
| 4 | 128113424 | rs6831679 | 1.046 | 1.003 | 0.454 | -0.207 | -0.017 | 0.002 | 0.002 | 0 | 0.36 | 0.3 | 0.35 | 0.39 | 0.26 | 0.27 |
| 4 | 128125715 | rs1383518 | -0.18 | 1.183 | 0.432 | -0.243 | 0.027 | -0.034 | -0.034 | 0 | 0.92 | 0.78 | 0.91 | 1 | 0.89 | 0.95 |
| 4 | 128132034 | rs12498324 | -0.588 | 1.084 | 0.358 | -0.268 | 0.014 | 0.016 | 0.016 | 0 | 0.56 | 0.37 | 0.55 | 0.6 | 0.63 | 0.69 |
| 4 | 128140708 | rs12501497 | 0.17 | 1.111 | 0.297 | -0.442 | 0.003 | 0.026 | 0.026 | 0 | 0.31 | 0.26 | 0.35 | 0.07 | 0.3 | 0.18 |
| 4 | 128149355 | rs9307596 | -0.47 | 1.098 | 0.602 | -0.142 | 0.01 | -0.014 | -0.014 | 0.009 | 0.93 | 0.86 | 0.92 | 1 | 0.89 | 0.95 |
| 4 | 128158126 | rs4997057 | -0.714 | 0.849 | 0.766 | -0.079 | 0.002 | 0.106 | 0.106 | 0 | 0.6 | 0.78 | 0.66 | 0.91 | 0.64 | 0.71 |
| 4 | 128160430 | rs967532 | -0.681 | 0.909 | 0.987 | -0.102 | 0.003 | -0.006 | -0.006 | 0 | 0.59 | 0.62 | 0.65 | 0.91 | 0.67 | 0.72 |
| 4 | 128167093 | rs1899548 | -0.342 | 0.901 | 0.836 | 0.064 | 0.032 | 0.02 | 0.02 | 0 | 0.54 | 0.34 | 0.55 | 0.59 | 0.65 | 0.69 |
| 4 | 128228213 | rs13122922 | -0.847 | 0.793 | 0.783 | -0.295 | 0 | 0.036 | 0.036 | 0 | 0.63 | 0.73 | 0.65 | 0.91 | 0.64 | 0.74 |
| 4 | 128248863 | rs1372965 | -0.088 | 0.702 | 0.872 | 0.042 | 0.039 | 0.092 | 0.092 | 0 | 0.51 | 0.28 | 0.53 | 0.7 | 0.6 | 0.71 |
| 4 | 128258252 | rs1838081 | -0.796 | 0.856 | 0.912 | 0.024 | 0.009 | 0.086 | 0.086 | 0 | 0.61 | 0.76 | 0.63 | 0.9 | 0.65 | 0.73 |
| 4 | 128274696 | rs1443178 | 1.1 | 1.342 | 0.657 | 0.563 | -0.055 | -0.049 | -0.049 | 0 | 0.09 | 0.01 | 0.09 | 0.17 | 0.17 | 0.15 |
| 4 | 128289824 | rs2120087 | -0.311 | 0.585 | 0.897 | 0.145 | 0.037 | 0.065 | 0.065 | 0 | 0.57 | 0.28 | 0.55 | 0.68 | 0.63 | 0.73 |
| 4 | 128290263 | rs10428477 | -0.83 | 0.628 | 0.955 | 0.094 | 0.012 | 0.077 | 0.077 | 0 | 0.63 | 0.76 | 0.66 | 0.89 | 0.64 | 0.74 |
| 4 | 128313686 | rs13141037 | -0.318 | 0.279 | 0.989 | 0.119 | 0.051 | 0.066 | 0.066 | 0 | 0.57 | 0.27 | 0.55 | 0.69 | 0.66 | 0.73 |
| 4 | 128337411 | rs2034077 | 0.55 | 0.548 | 0.317 | 0.433 | -0.049 | 0.158 | 0.158 | 0 | 0.44 | 0.08 | 0.52 | 0.17 | 0.45 | 0.27 |
| 4 | 128342120 | rs724950 | -0.024 | 0.775 | 0.25 | 0.509 | -0.031 | 0.031 | 0.031 | 0 | 0.56 | 0.72 | 0.67 | 0.37 | 0.49 | 0.33 |
| 4 | 128367991 | rs17012435 | -0.372 | 0.855 | 0.487 | 0.646 | 0.037 | -0.029 | -0.029 | 0 | 0.08 | 0.25 | 0.06 | 0 | 0.08 | 0.06 |
| 4 | 128377396 | rs7660782 | -1.067 | 0.857 | 0.335 | 0.688 | 0.017 | 0.048 | 0.048 | 0 | 0.14 | 0.32 | 0.15 | 0.2 | 0.11 | 0.12 |
| 4 | 128391591 | rs11724136 | 0.025 | 0.539 | -0.616 | 0.663 | 0.006 | -0.046 | -0.046 | 0.124 | 0.59 | 0.8 | 0.53 | 0.53 | 0.4 | 0.59 |
| 4 | 128392771 | rs7679411 | 0.646 | 0.818 | -0.271 | 0.52 | 0 | 0 | 0 | 0 | 0.69 | 0.66 | 0.76 | 0.71 | 0.72 | 0.76 |
| 4 | 128400330 | rs1443171 | 0.191 | 0.753 | -0.331 | 0.467 | -0.008 | 0.091 | 0.091 | 0 | 0.77 | 0.89 | 0.88 | 0.95 | 0.8 | 0.84 |
| 4 | 128410916 | rs2002543 | 0.273 | 0.754 | -0.211 | 0.497 | 0.001 | -0.008 | -0.008 | 0 | 0.76 | 0.81 | 0.8 | 0.71 | 0.79 | 0.82 |
| 4 | 128411344 | rs6534622 | 0.383 | 0.729 | -0.04 | 0.484 | 0 | -0.002 | -0.002 | 0 | 0.75 | 0.74 | 0.79 | 0.71 | 0.79 | 0.82 |
| 4 | 128419398 | rs2391184 | 0.334 | 0.605 | -0.008 | 0.528 | -0.088 | -0.097 | -0.097 | 5.13 | 0.24 | 0.08 | 0.33 | 0.4 | 0.41 | 0.29 |
| 4 | 128419456 | rs7696963 | 0.524 | 0.562 | 0.333 | 0.469 | -0.001 | -0.031 | -0.031 | 0.077 | 0.07 | 0.14 | 0.03 | 0 | 0.07 | 0.06 |
| 4 | 128425000 | rs925656 | 0.653 | 0.559 | 0.417 | 0.467 | -0.001 | -0.021 | -0.021 | 0 | 0.07 | 0.09 | 0.03 | 0 | 0.07 | 0.06 |
| 4 | 128427778 | rs1355924 | -0.986 | 0.513 | 0.349 | 0.48 | -0.028 | 0.05 | 0.05 | 0 | 0.84 | 0.93 | 0.89 | 0.95 | 0.81 | 0.88 |
| 4 | 128432062 | rs7653949 | -0.579 | 0.493 | 0.423 | 0.526 | 0.014 | 0.035 | 0.035 | 0 | 0.82 | 0.68 | 0.79 | 0.71 | 0.8 | 0.86 |
| 4 | 128441567 | rs17012484 | -0.811 | 0.398 | 0.594 | 0.504 | -0.03 | 0.138 | 0.138 | 0.271 | 0.1 | 0.37 | 0.14 | 0.24 | 0.03 | 0.06 |
| 4 | 128446769 | rs1399215 | 0.515 | 0.473 | 0.547 | 0.49 | -0.005 | 0.005 | 0.005 | 0 | 0.74 | 0.56 | 0.75 | 0.71 | 0.78 | 0.82 |
| 4 | 128454610 | rs12509921 | 1.684 | 0.323 | 0.872 | 0.414 | 0.031 | 0.032 | 0.032 | 0 | 0.42 | 0.32 | 0.39 | 0.31 | 0.3 | 0.46 |
| 4 | 128463930 | rs11730901 | -0.108 | 0.486 | 0.823 | 0.447 | -0.063 | -0.058 | -0.058 | 0.289 | 0.3 | 0.15 | 0.35 | 0.4 | 0.47 | 0.36 |
| 4 | 128467881 | rs17012511 | 1.428 | -0.06 | 1.178 | 0.439 | 0.007 | 0.007 | 0.007 | 0 | 0.42 | 0.36 | 0.4 | 0.31 | 0.3 | 0.46 |
| 4 | 128467975 | rs12505574 | -0.234 | 0.098 | 0.696 | 0.577 | 0.008 | -0.004 | -0.004 | 0 | 0.08 | 0.08 | 0.12 | 0.24 | 0.03 | 0.06 |
| 4 | 128474999 | rs10007344 | 0.115 | 0.032 | 0.547 | 0.572 | -0.022 | -0.081 | -0.081 | 0 | 0.61 | 0.88 | 0.63 | 0.56 | 0.41 | 0.59 |
| 4 | 128476186 | rs11933223 | 0.232 | 0.026 | 0.869 | 0.52 | -0.003 | -0.03 | -0.03 | 0 | 0.06 | 0.17 | 0.05 | 0 | 0.05 | 0.02 |
| 4 | 128478408 | rs10005550 | 0.244 | 0.016 | 0.868 | 0.578 | -0.003 | 0.104 | 0.104 | 0 | 0.06 | 0.17 | 0.14 | 0.24 | 0.06 | 0.03 |
| 4 | 128487450 | rs1008995 | 1.4 | -0.093 | 0.98 | 0.472 | 0.029 | 0.031 | 0.031 | 0 | 0.43 | 0.31 | 0.44 | 0.32 | 0.31 | 0.52 |
| 4 | 128488553 | rs4834203 | 0.099 | -0.055 | 0.887 | 0.392 | -0.001 | -0.051 | -0.051 | 0 | 0.6 | 0.82 | 0.63 | 0.56 | 0.41 | 0.59 |
| 4 | 128495654 | rs1355925 | 0.437 | -0.029 | 0.927 | 0.422 | 0.049 | -0.024 | -0.024 | 1.396 | 0.54 | 0.8 | 0.61 | 0.56 | 0.4 | 0.55 |
| 4 | 128506962 | rs6534623 | 1.09 | -0.058 | 1.117 | 0.344 | -0.002 | -0.025 | -0.025 | 0 | 0.44 | 0.47 | 0.34 | 0.17 | 0.3 | 0.51 |
| 4 | 128528617 | rs10434049 | 0.414 | 0.319 | 0.36 | 0.608 | -0.018 | -0.038 | -0.038 | 0 | 0.61 | 0.8 | 0.64 | 0.58 | 0.45 | 0.59 |
| 4 | 128534796 | rs10857124 | 0.794 | 0.051 | 0.752 | 0.28 | 0.047 | 0.119 | 0.119 | 0 | 0.54 | 0.28 | 0.37 | 0.16 | 0.36 | 0.57 |
| 4 | 128554404 | rs2255457 | 1.247 | 0.495 | 1.235 | 0.494 | 0 | -0.009 | -0.009 | 1.68 | 0.6 | 0.43 | 0.45 | 0.27 | 0.51 | 0.67 |
| 4 | 128565000 | rs3749507 | 0.569 | 0.485 | 1.592 | 0.791 | 0.001 | 0 | 0 | 0.082 | 0.67 | 0.67 | 0.58 | 0.55 | 0.54 | 0.72 |
| 4 | 128569904 | rs6847397 | 1.233 | 0.556 | 1.217 | 0.706 | -0.047 | 0.106 | 0.106 | 0 | 0.59 | 0.24 | 0.44 | 0.28 | 0.5 | 0.67 |
| 4 | 128607832 | rs2125573 | 0.959 | 0.856 | 1.507 | 0.843 | 0.035 | 0.03 | 0.03 | 0 | 0.66 | 0.59 | 0.58 | 0.55 | 0.55 | 0.71 |
| 4 | 128668806 | rs13147065 | 1.333 | 2.103 | 0.601 | 1.632 | -0.037 | -0.013 | -0.013 | 0.755 | 0.06 | 0 | 0.06 | 0.25 | 0.1 | 0.12 |

| | | | | | | | | | | | | | | | | |
|---|---|---|---|---|---|---|---|---|---|---|---|---|---|---|---|---|
| 4 | 128674030 | rs9995738 | 1.105 | 2.632 | 1.248 | 1.519 | 0.009 | 0.003 | 0.003 | 0 | 0.81 | 0.77 | 0.69 | 0.41 | 0.66 | 0.77 |
| 4 | 128697519 | rs11098937 | 0.254 | 2.752 | 1.093 | 1.505 | -0.062 | -0.055 | -0.055 | 0.01 | 0.09 | 0.01 | 0.15 | 0.29 | 0.2 | 0.19 |
| 4 | 128723042 | rs1380154 | 0.794 | 2.663 | 1.838 | 1.445 | 0.014 | 0.02 | 0.02 | 0 | 0.82 | 0.8 | 0.77 | 0.68 | 0.69 | 0.78 |
| 4 | 128732926 | rs2305957 | 1.192 | 2.625 | 1.906 | 1.419 | 0.098 | 0.126 | 0.126 | 0 | 0.8 | 0.62 | 0.77 | 0.67 | 0.68 | 0.78 |
| 4 | 128761940 | rs17314566 | 0.817 | 2.411 | 1.339 | 1.006 | -0.005 | 0.039 | 0.039 | 0 | 0.09 | 0.03 | 0.04 | 0 | 0.09 | 0.04 |
| 4 | 128766074 | rs1824331 | 0.875 | 2.448 | 1.113 | 0.936 | -0.05 | -0.029 | -0.029 | 0 | 0.12 | 0 | 0.21 | 0.16 | 0.15 | 0.14 |
| 4 | 128803159 | rs3811741 | 0.679 | 2.459 | 1.351 | 1.063 | 0.031 | 0.056 | 0.056 | 0 | 0.85 | 0.81 | 0.78 | 0.68 | 0.73 | 0.78 |
| 4 | 128814964 | rs17012739 | 0.817 | 2.357 | 1.339 | 1.06 | 0.015 | 0.022 | 0.022 | 0 | 0.82 | 0.8 | 0.77 | 0.67 | 0.68 | 0.77 |
| 4 | 128832078 | rs12503257 | 1.408 | 2.22 | 1.392 | 0.984 | 0.092 | 0.12 | 0.12 | 0 | 0.78 | 0.59 | 0.76 | 0.67 | 0.68 | 0.75 |
| 4 | 128843480 | rs1602084 | 1.882 | 0.717 | 0.407 | 0.152 | 0.06 | 0.084 | 0.084 | 0.325 | 0.15 | 0 | 0.04 | 0.03 | 0.06 | 0.17 |
| 4 | 128901402 | rs748630 | 0.34 | 1.144 | 0.705 | 0.321 | 0.024 | 0.032 | 0.032 | 0 | 0.81 | 0.78 | 0.76 | 0.67 | 0.68 | 0.77 |
| 4 | 128940313 | rs28687198 | -0.705 | 2.07 | 0.404 | 0.688 | 0.024 | 0.086 | 0.086 | 0 | 0.15 | 0.21 | 0.22 | 0.38 | 0.23 | 0.25 |
| 4 | 128954647 | rs17315466 | 0.744 | 0.661 | -0.238 | 0.227 | -0.052 | 0.152 | 0.152 | 0.507 | 0.44 | 0.13 | 0.39 | 0.28 | 0.44 | 0.48 |
| 4 | 128960569 | rs11098945 | 0.716 | 0.627 | -0.238 | 0.227 | -0.054 | 0.156 | 0.156 | 0.479 | 0.44 | 0.13 | 0.39 | 0.27 | 0.44 | 0.46 |
| 4 | 128981336 | rs4834224 | 0.268 | 0.926 | -0.103 | 0.403 | 0.09 | 0.119 | 0.119 | 0 | 0.78 | 0.58 | 0.75 | 0.68 | 0.67 | 0.74 |
| 4 | 128986874 | rs4834226 | 0.292 | 0.983 | -0.058 | 0.049 | 0.108 | 0.151 | 0.151 | 0 | 0.8 | 0.57 | 0.72 | 0.53 | 0.68 | 0.76 |
| 4 | 129012181 | rs2306054 | 0.292 | 0.983 | -0.058 | 0.049 | 0.108 | 0.151 | 0.151 | 0 | 0.8 | 0.57 | 0.72 | 0.53 | 0.68 | 0.76 |
| 4 | 129024273 | rs4834232 | 1.033 | 0.866 | -0.298 | -0.06 | 0.046 | 0.133 | 0.133 | 0 | 0.31 | 0.03 | 0.23 | 0.14 | 0.23 | 0.26 |
| 4 | 129045317 | rs10857129 | 0.259 | 1.721 | 0.606 | 0.646 | 0.032 | 0.047 | 0.047 | 0.104 | 0.84 | 0.79 | 0.75 | 0.53 | 0.68 | 0.76 |
| 4 | 129082491 | rs10000748 | -1.519 | 1.432 | -0.496 | 0.298 | -0.056 | 0.001 | 0.001 | 0 | 0.85 | 0.99 | 0.88 | 0.88 | 0.84 | 0.73 |
| 4 | 129104537 | rs2172813 | 0.751 | 1.646 | 0.198 | 0.693 | 0.1 | 0.132 | 0.132 | 0 | 0.79 | 0.55 | 0.71 | 0.52 | 0.68 | 0.76 |
| 4 | 129113495 | rs6856312 | 0.917 | 1.623 | 0.022 | 0.657 | 0.044 | 0.059 | 0.059 | 0.173 | 0.78 | 0.36 | 0.68 | 0.52 | 0.68 | 0.76 |
| 4 | 129160316 | rs4975300 | 0.207 | 1.604 | 0.039 | 0.766 | 0.113 | 0.252 | 0.252 | 0 | 0.84 | 0.56 | 0.75 | 0.47 | 0.76 | 0.76 |
| 4 | 129180759 | rs13133067 | 1.542 | 0.377 | -1.414 | -0.108 | 0.003 | 0.166 | 0.166 | 0.102 | 0.37 | 0.06 | 0.26 | 0.14 | 0.29 | 0.28 |
| 4 | 129189040 | rs11726595 | 1.34 | -0.155 | -1.21 | 0.063 | -0.066 | 0.004 | 0.004 | 0.236 | 0.14 | 0.01 | 0.1 | 0.13 | 0.17 | 0.22 |
| 4 | 129213745 | rs4975221 | 1.7 | -0.167 | -0.818 | 0.246 | -0.017 | -0.012 | -0.012 | 0.147 | 0.07 | 0.01 | 0.2 | 0.1 | 0.1 | 0.05 |
| 4 | 129218126 | rs1391980 | -0.084 | -0.339 | -0.286 | 0.102 | 0.046 | 0.053 | 0.053 | 0 | 0.68 | 0.53 | 0.65 | 0.55 | 0.57 | 0.68 |
| 4 | 129221901 | rs6534677 | 1.36 | -0.533 | -0.435 | 0.011 | 0.029 | 0.056 | 0.056 | 1.113 | 0.44 | 0.27 | 0.34 | 0.37 | 0.36 | 0.49 |
| 4 | 129223349 | rs10001857 | -0.223 | -0.605 | 0.004 | 0.164 | -0.013 | -0.008 | -0.008 | 1.251 | 0.23 | 0.27 | 0.31 | 0.18 | 0.21 | 0.2 |
| 4 | 129239583 | rs10019366 | 1.501 | -0.687 | -0.26 | -0.015 | 0.031 | 0.045 | 0.045 | 0 | 0.35 | 0.28 | 0.26 | 0.16 | 0.21 | 0.25 |
| 4 | 129244451 | rs11098956 | 0.554 | -0.736 | -0.328 | -0.07 | 0.059 | 0.105 | 0.105 | 0 | 0.5 | 0.36 | 0.48 | 0.29 | 0.37 | 0.4 |
| 4 | 129247395 | rs4975181 | -0.035 | -0.636 | -0.292 | 0.364 | 0.007 | -0.022 | -0.022 | 0 | 0.59 | 0.74 | 0.56 | 0.49 | 0.4 | 0.44 |
| 4 | 129254598 | rs17337491 | 0.702 | -0.643 | -0.651 | 0.33 | -0.027 | 0.025 | 0.025 | 0.432 | 0.17 | 0.01 | 0.22 | 0.13 | 0.18 | 0.15 |
| 4 | 129261079 | rs290003 | 0.489 | -0.211 | -0.548 | 0.523 | -0.182 | -0.166 | -0.166 | 0.277 | 0.26 | 0.01 | 0.32 | 0.41 | 0.45 | 0.31 |
| 4 | 129281445 | rs11723210 | 1.17 | -0.846 | 0.344 | 0.574 | 0.036 | 0.046 | 0.046 | 0 | 0.34 | 0.18 | 0.29 | 0.24 | 0.21 | 0.29 |
| 4 | 129282956 | rs1982904 | 0.523 | -0.854 | 0.501 | 0.692 | 0.062 | 0.009 | 0.009 | 0 | 0.41 | 0.76 | 0.34 | 0.36 | 0.22 | 0.3 |
| 4 | 129303169 | rs7695173 | 0.712 | -0.569 | 0.171 | 0.862 | -0.013 | 0.004 | 0.004 | 0 | 0.38 | 0.41 | 0.31 | 0.55 | 0.22 | 0.33 |
| 4 | 129309601 | rs3853107 | 1.331 | -1.015 | 0.412 | 0.779 | 0.039 | 0.039 | 0.039 | 0.052 | 0.3 | 0.2 | 0.24 | 0.18 | 0.16 | 0.32 |
| 4 | 129311823 | rs3870129 | -0.348 | -1.049 | -0.217 | 0.652 | 0.099 | 0.036 | 0.036 | 0 | 0.54 | 0.85 | 0.49 | 0.65 | 0.32 | 0.45 |
| 4 | 129314562 | rs17013261 | 1.078 | -0.884 | -0.133 | 0.588 | 0.018 | -0.026 | -0.026 | 1.551 | 0.05 | 0.09 | 0.05 | 0.01 | 0.08 | 0.06 |
| 4 | 129314791 | rs10022598 | 1.453 | -0.994 | 0.213 | 0.756 | -0.046 | 0.004 | 0.004 | 0 | 0.31 | 0.37 | 0.27 | 0.29 | 0.16 | 0.32 |
| 4 | 129314887 | rs10020014 | -0.155 | -0.814 | 0.076 | 0.749 | 0.057 | 0.008 | 0.008 | 0.063 | 0.56 | 0.78 | 0.5 | 0.64 | 0.34 | 0.46 |
| 4 | 129336928 | rs17013269 | 0.774 | -0.567 | -0.134 | 0.659 | 0.003 | 0.008 | 0.008 | 0 | 0.18 | 0.17 | 0.2 | 0.11 | 0.15 | 0.11 |
| 4 | 129345343 | rs12508336 | 0.07 | -0.494 | -0.104 | 0.735 | 0.04 | 0.018 | 0.018 | 0 | 0.54 | 0.69 | 0.49 | 0.64 | 0.33 | 0.47 |
| 4 | 129346970 | rs11098966 | 0.487 | -0.443 | 0.107 | 0.842 | 0.037 | 0.048 | 0.048 | 0.234 | 0.51 | 0.42 | 0.45 | 0.29 | 0.32 | 0.42 |
| 4 | 129349449 | rs10021346 | 0.208 | -0.408 | -0.012 | 0.799 | 0.032 | 0.014 | 0.014 | 0.072 | 0.55 | 0.69 | 0.49 | 0.65 | 0.33 | 0.47 |
| 4 | 129369697 | rs4305531 | 0.746 | -0.167 | -0.479 | 0.414 | -0.041 | 0.048 | 0.048 | 0.075 | 0.19 | 0.01 | 0.21 | 0.1 | 0.19 | 0.16 |
| 4 | 129370768 | rs11728419 | 0.969 | 0.481 | -0.193 | 0.592 | -0.028 | 0.011 | 0.011 | 0 | 0.11 | 0.03 | 0.13 | 0.11 | 0.18 | 0.2 |
| 4 | 129373425 | rs17013315 | 0.768 | -0.081 | -0.465 | 0.293 | 0.039 | 0.042 | 0.042 | 0.054 | 0.33 | 0.25 | 0.25 | 0.19 | 0.17 | 0.31 |
| 4 | 129375881 | rs9307607 | 0.636 | -0.057 | -0.5 | 0.339 | 0.017 | 0.016 | 0.016 | 0.001 | 0.34 | 0.33 | 0.25 | 0.19 | 0.16 | 0.31 |
| 4 | 129382079 | rs6534682 | -0.128 | 1.267 | -0.837 | 0.856 | -0.041 | -0.015 | -0.015 | 5.613 | 0.16 | 0.1 | 0.18 | 0.15 | 0.24 | 0.2 |
| 4 | 129382308 | rs6534683 | -0.841 | 1.679 | -0.237 | 1.322 | 0.034 | 0.062 | 0.062 | 0.106 | 0.22 | 0.39 | 0.21 | 0.41 | 0.32 | 0.22 |
| 4 | 129382634 | rs17013332 | 1.379 | 1.261 | -0.809 | 1.211 | 0.025 | -0.008 | -0.008 | 0 | 0.27 | 0.14 | 0.24 | 0.26 | 0.19 | 0.25 |
| 4 | 129389738 | rs6840580 | -1.345 | 1.489 | -1.555 | 0.845 | -0.082 | 0.013 | 0.013 | 0.576 | 0.72 | 0.99 | 0.67 | 0.77 | 0.71 | 0.68 |
| 4 | 129392060 | rs10000490 | 0.813 | 1.598 | -1.168 | 1.013 | -0.029 | -0.023 | -0.023 | 0 | 0.13 | 0.07 | 0.14 | 0.16 | 0.21 | 0.11 |
| 4 | 129401741 | rs10012794 | 0.034 | 1.383 | -1.116 | 0.901 | 0.014 | 0.002 | 0.002 | 0 | 0.44 | 0.32 | 0.41 | 0.59 | 0.43 | 0.44 |
| 4 | 129409513 | rs7690289 | 0.351 | 1.654 | -1.283 | 0.894 | -0.073 | -0.042 | -0.042 | 0 | 0.17 | 0.06 | 0.17 | 0.18 | 0.32 | 0.15 |
| 4 | 129412788 | rs13134269 | 0.291 | 1.315 | -0.697 | 0.733 | 0.013 | -0.028 | -0.028 | 0 | 0.2 | 0.29 | 0.21 | 0.1 | 0.24 | 0.19 |
| 4 | 129414681 | rs17013357 | -0.754 | 1.147 | -0.278 | 0.833 | 0.063 | 0.273 | 0.273 | 4.66 | 0.05 | 0.33 | 0.03 | 0.08 | 0.01 | 0.02 |
| 4 | 129431298 | rs4549394 | 0.428 | 1.185 | -0.747 | 0.741 | -0.004 | -0.031 | -0.031 | 0 | 0.19 | 0.31 | 0.18 | 0.1 | 0.16 | 0.18 |
| 4 | 129436625 | rs7441245 | -1.386 | 0.984 | -1.126 | 0.784 | -0.094 | 0.044 | 0.044 | 0.381 | 0.73 | 0.99 | 0.67 | 0.79 | 0.71 | 0.69 |
| 4 | 129439313 | rs6821705 | 0.705 | 1.086 | -1.046 | 0.458 | -0.047 | -0.041 | -0.041 | 1.544 | 0.16 | 0.06 | 0.14 | 0.18 | 0.24 | 0.13 |
| 4 | 129441798 | rs6534689 | -0.698 | 1.065 | -0.783 | 0.45 | 0.019 | -0.008 | -0.008 | 0 | 0.37 | 0.47 | 0.36 | 0.27 | 0.48 | 0.32 |
| 4 | 129442692 | rs6843137 | -1.438 | 0.86 | -0.842 | 0.413 | -0.028 | -0.024 | -0.024 | 0 | 0.47 | 0.92 | 0.42 | 0.53 | 0.53 | 0.43 |
| 4 | 129445452 | rs4975234 | 0.052 | 0.794 | -0.824 | 0.344 | -0.001 | -0.008 | -0.008 | 0 | 0.49 | 0.5 | 0.46 | 0.27 | 0.42 | 0.45 |
| 4 | 129445759 | rs7695069 | 0.422 | 0.797 | -0.925 | 0.362 | 0.017 | -0.043 | -0.043 | 0 | 0.21 | 0.36 | 0.21 | 0.1 | 0.24 | 0.19 |
| 4 | 129447957 | rs4975235 | 1.413 | 0.686 | -1.376 | 0.401 | 0.063 | 0.064 | 0.064 | 0 | 0.27 | 0.07 | 0.25 | 0.18 | 0.18 | 0.26 |
| 4 | 129450677 | rs1603886 | -0.807 | 1.263 | -0.948 | 0.73 | -0.088 | -0.083 | -0.083 | 2.246 | 0.74 | 0.98 | 0.66 | 0.72 | 0.71 | 0.69 |
| 4 | 129458258 | rs10518533 | -0.404 | 1.236 | -0.498 | 0.814 | -0.001 | -0.002 | -0.002 | 0 | 0.65 | 0.66 | 0.62 | 0.72 | 0.68 | 0.67 |

| | | | | | | | | | | | | | | | | |
|---|---|---|---|---|---|---|---|---|---|---|---|---|---|---|---|---|
| 4 | 129462245 | rs6835518 | -1.072 | 1.499 | 0.246 | 1.324 | 0.051 | 0.015 | 0.015 | 0.071 | 0.23 | 0.49 | 0.18 | 0.19 | 0.27 | 0.2 |
| 4 | 129465489 | rs1032673 | -1.081 | 1.423 | 0.182 | 1.214 | 0.057 | 0.125 | 0.125 | 0 | 0.25 | 0.49 | 0.36 | 0.44 | 0.33 | 0.27 |
| 4 | 129471494 | rs7678706 | -0.317 | 0.418 | 0.085 | 0.807 | -0.008 | -0.027 | -0.027 | 0 | 0.23 | 0.36 | 0.19 | 0.05 | 0.17 | 0.11 |
| 4 | 129472039 | rs2173502 | -0.926 | 0.498 | 0.27 | 1.229 | 0.006 | -0.002 | -0.002 | 2.123 | 0.3 | 0.72 | 0.38 | 0.35 | 0.24 | 0.2 |
| 4 | 129477404 | rs9991179 | 0.979 | 0.433 | -0.294 | 0.811 | 0.001 | -0.011 | -0.011 | 0 | 0.33 | 0.27 | 0.21 | 0.12 | 0.26 | 0.33 |
| 4 | 129477529 | rs12510203 | 1.001 | 0.404 | -0.4 | 1.153 | -0.133 | -0.025 | -0.025 | 0 | 0.31 | 0.02 | 0.27 | 0.2 | 0.4 | 0.38 |
| 4 | 129483230 | rs6858471 | 0.687 | -0.017 | 0.096 | -0.53 | 0.001 | -0.001 | -0.001 | 0 | 0.33 | 0.26 | 0.23 | 0.12 | 0.29 | 0.35 |
| 4 | 129483448 | rs2036237 | -1.078 | -0.346 | 0.714 | 0.276 | -0.016 | -0.035 | -0.035 | 3.787 | 0.24 | 0.38 | 0.2 | 0.05 | 0.17 | 0.1 |
| 4 | 129494050 | rs17013481 | -1.35 | -0.221 | 0.887 | 0.832 | -0.018 | -0.005 | -0.005 | 0 | 0.09 | 0.18 | 0.1 | 0.05 | 0.06 | 0.11 |
| 4 | 129495183 | rs17013484 | -0.71 | -0.232 | 0.803 | 0.76 | 0.031 | -0.011 | -0.011 | 2.528 | 0.05 | 0.2 | 0.03 | 0 | 0.03 | 0.06 |
| 4 | 129495243 | rs17013486 | -1.133 | -0.268 | 0.834 | 0.741 | 0.023 | -0.018 | -0.018 | 0.032 | 0.06 | 0.25 | 0.04 | 0 | 0.04 | 0.05 |
| 4 | 129495459 | rs1216370 | 2.024 | -0.383 | 0.971 | 0.774 | -0.035 | -0.027 | -0.027 | 0.166 | 0.79 | 0.36 | 0.82 | 0.88 | 0.88 | 0.81 |
| 4 | 129499923 | rs1216378 | 1.874 | -0.219 | 1.127 | 0.905 | -0.031 | -0.015 | -0.015 | 0.183 | 0.82 | 0.35 | 0.83 | 0.88 | 0.91 | 0.85 |
| 4 | 129503678 | rs1216383 | 1.106 | -0.103 | 0.946 | 0.924 | -0.014 | 0 | 0 | 3.2 | 0.89 | 0.63 | 0.87 | 0.95 | 0.94 | 0.91 |
| 4 | 129509265 | rs1216394 | 1.005 | -0.118 | 0.158 | 1.175 | -0.028 | 0.089 | 0.089 | 2.23 | 0.84 | 0.43 | 0.84 | 0.76 | 0.91 | 0.84 |
| 4 | 129522518 | rs904477 | 0.186 | 0.013 | -1.374 | 0.845 | -0.02 | 0.003 | 0.003 | 0 | 0.31 | 0.06 | 0.26 | 0.29 | 0.29 | 0.35 |
| 4 | 129526565 | rs10518534 | -0.577 | -0.459 | -1.254 | 1.106 | 0.035 | -0.014 | -0.014 | 0 | 0.43 | 0.19 | 0.35 | 0.43 | 0.36 | 0.49 |
| 4 | 129527656 | rs1216405 | 1.166 | -0.225 | -1.922 | 1.006 | -0.167 | -0.111 | -0.111 | 0 | 0.36 | 0.03 | 0.49 | 0.38 | 0.47 | 0.44 |
| 4 | 129529446 | rs1216411 | -0.402 | -0.46 | -1.127 | 0.868 | 0.002 | 0.003 | 0.003 | 0 | 0.59 | 0.61 | 0.68 | 0.62 | 0.62 | 0.58 |
| 4 | 129529604 | rs1216412 | -0.861 | -0.504 | -1.327 | 0.671 | 0.011 | 0.028 | 0.028 | 0 | 0.65 | 0.7 | 0.73 | 0.8 | 0.71 | 0.64 |
| 4 | 129531333 | rs1216416 | 0.869 | 0.216 | -1.831 | 0.844 | -0.147 | -0.104 | -0.104 | 0 | 0.35 | 0.04 | 0.49 | 0.4 | 0.47 | 0.44 |
| 4 | 129543559 | rs1216440 | 0.981 | -0.032 | -1.284 | 0.95 | -0.011 | -0.068 | -0.068 | 0.023 | 0.76 | 0.11 | 0.71 | 0.6 | 0.8 | 0.82 |
| 4 | 129557924 | rs1216348 | 0.865 | -0.173 | -1.603 | 0.248 | -0.066 | 0.131 | 0.131 | 0 | 0.38 | 0.14 | 0.31 | 0.13 | 0.5 | 0.54 |
| 4 | 129558393 | rs7670660 | -0.773 | -0.411 | -0.513 | 0.396 | 0.039 | 0.018 | 0.018 | 0.148 | 0.2 | 0.49 | 0.15 | 0.18 | 0.2 | 0.1 |
| 4 | 129568096 | rs1615523 | -0.138 | -0.775 | -0.517 | 0.849 | 0.077 | 0.083 | 0.083 | 0 | 0.58 | 0.81 | 0.7 | 0.63 | 0.68 | 0.62 |
| 4 | 129574927 | rs17013522 | -0.47 | -0.084 | 0.836 | 1.547 | 0.048 | 0.077 | 0.077 | 0 | 0.09 | 0.23 | 0.08 | 0.18 | 0.13 | 0.06 |
| 4 | 129581554 | rs2391291 | 0.466 | -0.88 | 0.344 | 1.644 | -0.02 | -0.057 | -0.057 | 0.261 | 0.19 | 0.01 | 0.17 | 0.32 | 0.19 | 0.15 |
| 4 | 129585163 | rs11098979 | -2.366 | -0.703 | 0.9 | 1.821 | -0.014 | 0.311 | 0.311 | 0 | 0.13 | 0.44 | 0.34 | 0.44 | 0.08 | 0.16 |
| 4 | 129587598 | rs1553143 | -1.535 | -0.696 | 0.984 | 1.843 | -0.012 | 0.313 | 0.313 | 0 | 0.08 | 0.23 | 0.29 | 0.44 | 0.04 | 0.13 |
| 4 | 129593111 | rs17013593 | -2.214 | -0.559 | 0.696 | 1.477 | 0.01 | 0.242 | 0.242 | 0 | 0.13 | 0.41 | 0.39 | 0.55 | 0.09 | 0.19 |
| 4 | 129598078 | rs7700079 | -2.214 | -0.559 | 0.685 | 1.477 | 0.01 | 0.242 | 0.242 | 0 | 0.13 | 0.41 | 0.39 | 0.55 | 0.09 | 0.19 |
| 4 | 129611736 | rs6856260 | 0.704 | -0.527 | -0.044 | 0.524 | -0.001 | -0.024 | -0.024 | 0 | 0.66 | 0.63 | 0.51 | 0.11 | 0.67 | 0.67 |
| 4 | 129611773 | rs12512245 | -0.846 | -0.471 | -0.033 | 0.767 | -0.004 | 0.021 | 0.021 | 0 | 0.07 | 0.07 | 0.29 | 0.65 | 0.07 | 0.1 |
| 4 | 129612729 | rs6837389 | 0.292 | -0.554 | 0.117 | 0.748 | -0.006 | -0.006 | -0.006 | 0.592 | 0.72 | 0.69 | 0.8 | 0.76 | 0.75 | 0.78 |
| 4 | 129646870 | rs13147397 | 1.16 | -0.514 | -0.351 | 0.814 | 0.095 | 0.163 | 0.163 | 0 | 0.53 | 0.12 | 0.4 | 0.2 | 0.67 | 0.39 |
| 4 | 129651071 | rs6534699 | 1.394 | 0.019 | 0.96 | 1.979 | -0.015 | 0.053 | 0.053 | 0 | 0.74 | 0.46 | 0.83 | 0.58 | 0.77 | 0.61 |
| 4 | 129654251 | rs1425619 | -1.739 | 0.363 | 0.786 | 1.999 | 0 | 0.009 | 0.009 | 0.095 | 0.23 | 0.27 | 0.44 | 0.56 | 0.24 | 0.34 |
| 4 | 129675024 | rs6820206 | 1.243 | 0.63 | 0.572 | 1.139 | 0.038 | -0.031 | -0.031 | 0.811 | 0.83 | 0.71 | 0.87 | 0.88 | 0.77 | 0.73 |
| 4 | 129676284 | rs2033052 | 1.273 | 0.787 | 0.756 | 1.384 | 0.076 | -0.004 | -0.004 | 0 | 0.87 | 0.65 | 0.89 | 0.88 | 0.81 | 0.77 |
| 4 | 129679066 | rs17013657 | -1.482 | 0.146 | -0.101 | 1.571 | 0.014 | -0.021 | -0.021 | 0 | 0.18 | 0.11 | 0.14 | 0.26 | 0.16 | 0.14 |
| 4 | 129682775 | rs4435757 | 1.417 | 0.562 | 0.451 | 1.218 | 0.04 | -0.024 | -0.024 | 0 | 0.9 | 0.74 | 0.93 | 0.94 | 0.87 | 0.79 |
| 4 | 129685158 | rs11723731 | 1.453 | -0.005 | -0.709 | 1.403 | 0.024 | -0.01 | -0.01 | 2.81 | 0.54 | 0.06 | 0.47 | 0.39 | 0.64 | 0.46 |
| 4 | 129692038 | rs11941279 | -0.184 | -0.156 | -0.432 | 1.426 | -0.009 | -0.008 | -0.008 | 0 | 0.07 | 0.09 | 0.03 | 0.03 | 0.04 | 0.1 |
| 4 | 129694008 | rs17013678 | -0.68 | -0.288 | -0.142 | 1.879 | -0.004 | 0.017 | 0.017 | 0 | 0.11 | 0.17 | 0.19 | 0.21 | 0.08 | 0.16 |
| 4 | 129694554 | rs4975253 | 1.633 | -0.241 | 0.295 | 2.019 | 0.004 | -0.012 | -0.012 | 1.287 | 0.63 | 0.33 | 0.6 | 0.54 | 0.68 | 0.56 |
| 4 | 129696911 | rs10017881 | 1.149 | -0.268 | 0.181 | 2.013 | -0.028 | 0.067 | 0.067 | 0 | 0.83 | 0.66 | 0.71 | 0.64 | 0.89 | 0.74 |
| 4 | 129698469 | rs1863337 | -0.713 | -0.148 | -0.12 | 1.883 | -0.004 | 0.017 | 0.017 | 0 | 0.11 | 0.17 | 0.19 | 0.21 | 0.08 | 0.16 |
| 4 | 129700043 | rs11098986 | 1.522 | 0.031 | -0.383 | 1.275 | 0.047 | 0.02 | 0.02 | 0.783 | 0.55 | 0.13 | 0.49 | 0.39 | 0.64 | 0.46 |
| 4 | 129708024 | rs4975260 | 0.893 | 0 | 0.578 | 1.004 | -0.046 | -0.112 | -0.112 | 0 | 0.82 | 0.48 | 0.9 | 0.99 | 0.9 | 0.89 |
| 4 | 129709109 | rs13101730 | 1.404 | 0.468 | 0.297 | 1.343 | 0.007 | 0.032 | 0.032 | 2.532 | 0.62 | 0.2 | 0.6 | 0.37 | 0.65 | 0.59 |
| 4 | 129709244 | rs6813608 | -1.825 | 0.399 | 1.067 | 2.274 | 0.023 | 0.402 | 0.402 | 0.637 | 0.08 | 0.35 | 0.16 | 0.39 | 0.05 | 0.05 |
| 4 | 129713511 | rs9991981 | 1.071 | 0.562 | 0.591 | 1.901 | 0.009 | 0.033 | 0.033 | 0.312 | 0.74 | 0.61 | 0.77 | 0.63 | 0.72 | 0.72 |
| 4 | 129721687 | rs1031691 | 1.433 | 0.366 | -0.062 | 2.06 | -0.004 | -0.027 | -0.027 | 0.007 | 0.63 | 0.17 | 0.59 | 0.46 | 0.65 | 0.69 |
| 4 | 129729412 | rs13143771 | 1.469 | 0.488 | -0.383 | 1.678 | 0.004 | 0.041 | 0.041 | 0.447 | 0.69 | 0.15 | 0.69 | 0.7 | 0.75 | 0.83 |
| 4 | 129737089 | rs10017436 | -1.781 | -0.907 | -1.665 | 0.768 | -0.011 | 0.033 | 0.033 | 0.044 | 0.56 | 0.93 | 0.48 | 0.38 | 0.4 | 0.32 |
| 4 | 129746735 | rs17013734 | -0.111 | -0.978 | -1.336 | 0.646 | 0.051 | 0.051 | 0.051 | 0 | 0.27 | 0.13 | 0.19 | 0.09 | 0.18 | 0.24 |
| 4 | 129757215 | rs4975270 | 1.819 | -0.971 | -1.365 | 0.658 | -0.022 | 0.025 | 0.025 | 0.151 | 0.43 | 0.07 | 0.51 | 0.62 | 0.59 | 0.67 |
| 4 | 129774375 | rs6847753 | 0.367 | -1.141 | -0.15 | 0.586 | 0.051 | 0.049 | 0.049 | 0.165 | 0.85 | 0.48 | 0.78 | 0.72 | 0.86 | 0.93 |
| 4 | 129779076 | rs2162126 | 0.546 | -1.225 | 0.759 | 0.539 | -0.003 | -0.008 | -0.008 | 1.44 | 0.78 | 0.44 | 0.78 | 0.73 | 0.83 | 0.93 |
| 4 | 129793559 | rs11933240 | -0.492 | -1.592 | 0.479 | 0.35 | 0.034 | 0.044 | 0.044 | 0 | 0.26 | 0.12 | 0.2 | 0.09 | 0.21 | 0.26 |
| 4 | 129803970 | rs11940298 | -0.468 | -1.597 | 0.411 | 0.345 | 0.025 | 0.041 | 0.041 | 0 | 0.26 | 0.12 | 0.2 | 0.09 | 0.22 | 0.25 |
| 4 | 129825629 | rs186444 | -1.405 | -0.634 | -0.974 | -0.062 | -0.01 | -0.006 | -0.006 | 0 | 0.77 | 0.78 | 0.77 | 0.71 | 0.68 | 0.6 |
| 4 | 129836255 | rs318543 | 0.002 | -0.723 | -0.939 | -0.064 | -0.001 | -0.006 | -0.006 | 0.415 | 0.22 | 0.52 | 0.22 | 0.27 | 0.18 | 0.07 |
| 4 | 129838879 | rs318549 | -1.51 | -0.674 | -1.096 | -0.049 | -0.033 | -0.018 | -0.018 | 1.541 | 0.78 | 0.85 | 0.79 | 0.72 | 0.68 | 0.6 |
| 4 | 129840234 | rs11722253 | 0.262 | -0.775 | -1.15 | -0.081 | 0.03 | 0.045 | 0.045 | 0 | 0.25 | 0.09 | 0.19 | 0.1 | 0.21 | 0.25 |
| 4 | 129860963 | rs6840353 | 1.675 | -0.7 | -1.202 | 0.002 | -0.005 | -0.032 | -0.032 | 0 | 0.28 | 0.13 | 0.35 | 0.33 | 0.29 | 0.28 |
| 4 | 129862073 | rs2034497 | 0.017 | -0.656 | -1.286 | -0.054 | -0.024 | 0.006 | 0.006 | 0 | 0.55 | 0.8 | 0.64 | 0.63 | 0.51 | 0.36 |
| 4 | 129866452 | rs7682145 | -0.794 | -0.418 | -1.374 | -0.039 | -0.024 | 0.033 | 0.033 | 0 | 0.76 | 0.9 | 0.85 | 0.86 | 0.71 | 0.61 |
| 4 | 129867280 | rs10028124 | -1.015 | -0.337 | -1.455 | -0.113 | -0.083 | -0.055 | -0.055 | 1.52 | 0.77 | 0.96 | 0.76 | 0.7 | 0.67 | 0.64 |

| | | | | | | | | | | | | | | | | |
|---|---|---|---|---|---|---|---|---|---|---|---|---|---|---|---|---|
| 4 | 129873147 | rs4975288 | -0.958 | -0.361 | -1.459 | 0.015 | -0.082 | -0.054 | -0.054 | 0.262 | 0.77 | 0.96 | 0.76 | 0.71 | 0.67 | 0.62 |
| 4 | 129879744 | rs921377 | 1.977 | -0.401 | -1.773 | -0.118 | -0.004 | -0.031 | -0.031 | 0 | 0.32 | 0.16 | 0.36 | 0.32 | 0.32 | 0.33 |
| 4 | 129913345 | rs13151322 | 1.303 | -0.532 | -1.957 | -0.171 | -0.001 | -0.029 | -0.029 | 0.031 | 0.17 | 0.02 | 0.13 | 0.1 | 0.15 | 0.2 |
| 4 | 129914268 | rs280601 | 0.897 | -0.563 | -1.935 | -0.189 | 0.041 | 0.045 | 0.045 | 0.188 | 0.55 | 0.15 | 0.58 | 0.61 | 0.64 | 0.71 |
| 4 | 129924977 | rs3113487 | 1.002 | -0.557 | -1.851 | -0.217 | 0.034 | 0.042 | 0.042 | 0.105 | 0.56 | 0.22 | 0.61 | 0.64 | 0.64 | 0.71 |
| 4 | 129927804 | rs2044523 | 2.07 | -0.553 | -1.902 | -0.218 | -0.001 | -0.031 | -0.031 | 0.744 | 0.32 | 0.14 | 0.36 | 0.33 | 0.32 | 0.33 |
| 4 | 129999071 | rs789982 | 0.315 | -0.561 | -1.786 | -0.221 | -0.002 | -0.01 | -0.01 | 0 | 0.72 | 0.32 | 0.72 | 0.7 | 0.76 | 0.85 |
| 4 | 130003237 | rs1699388 | -0.325 | -0.577 | -1.628 | -0.234 | -0.008 | -0.03 | -0.03 | 0.07 | 0.75 | 0.39 | 0.75 | 0.75 | 0.81 | 0.92 |
| 4 | 130013447 | rs11728895 | 1.586 | -0.552 | -1.668 | -0.211 | 0.015 | -0.003 | -0.003 | 0.2 | 0.15 | 0.01 | 0.11 | 0.06 | 0.12 | 0.14 |
| 4 | 130019889 | rs13136220 | 0.685 | -0.577 | -1.655 | -0.21 | -0.003 | -0.001 | -0.001 | 0 | 0.2 | 0.17 | 0.16 | 0.12 | 0.17 | 0.21 |
| 4 | 130025037 | rs1030831 | -0.32 | -0.589 | -1.694 | -0.226 | -0.008 | -0.023 | -0.023 | 2.203 | 0.75 | 0.36 | 0.73 | 0.72 | 0.81 | 0.92 |
| 4 | 130025117 | rs1030830 | 2.044 | -0.568 | -1.749 | -0.203 | -0.005 | -0.04 | -0.04 | 0 | 0.32 | 0.12 | 0.36 | 0.32 | 0.32 | 0.33 |
| 4 | 130030644 | rs2271570 | 0.788 | -0.614 | -1.685 | -0.266 | 0.01 | 0.006 | 0.006 | 0 | 0.19 | 0.12 | 0.15 | 0.11 | 0.19 | 0.21 |
| 4 | 130030652 | rs337277 | -0.35 | -0.646 | -1.66 | -0.28 | -0.007 | -0.029 | -0.029 | 0 | 0.75 | 0.39 | 0.73 | 0.74 | 0.81 | 0.92 |
| 4 | 130030944 | rs337276 | 1.032 | -0.403 | -1.695 | -0.294 | 0.02 | 0.032 | 0.032 | 0 | 0.55 | 0.27 | 0.57 | 0.63 | 0.59 | 0.69 |
| 4 | 130032119 | rs1867951 | 0.838 | -0.388 | -1.737 | -0.315 | 0.01 | 0.008 | 0.008 | 0 | 0.19 | 0.11 | 0.16 | 0.11 | 0.21 | 0.22 |
| 4 | 130042232 | rs337270 | -0.074 | -0.431 | -1.685 | -0.369 | -0.005 | -0.011 | -0.011 | 0.012 | 0.74 | 0.32 | 0.73 | 0.72 | 0.81 | 0.92 |
| 4 | 130051748 | rs12500557 | 1.952 | -0.395 | -1.804 | -0.357 | 0.011 | -0.016 | -0.016 | 0.093 | 0.32 | 0.19 | 0.34 | 0.31 | 0.27 | 0.31 |
| 4 | 130056276 | rs234264 | 0.726 | -0.389 | -1.877 | -0.35 | -0.131 | -0.101 | -0.101 | 0 | 0.24 | 0.05 | 0.25 | 0.35 | 0.37 | 0.45 |
| 4 | 130057766 | rs17014243 | 1.347 | -0.497 | -2.047 | -0.476 | -0.019 | -0.048 | -0.048 | 0 | 0.13 | 0.01 | 0.15 | 0.25 | 0.09 | 0.05 |
| 4 | 130058367 | rs1849762 | 1.059 | -0.426 | -1.793 | -0.375 | 0.018 | 0.001 | 0.001 | 0 | 0.5 | 0.35 | 0.5 | 0.41 | 0.43 | 0.47 |
| 4 | 130058670 | rs1757923 | -0.167 | -0.43 | -1.708 | -0.399 | -0.005 | -0.032 | -0.032 | 0.504 | 0.75 | 0.41 | 0.75 | 0.75 | 0.81 | 0.92 |
| 4 | 130060230 | rs11946237 | 0.954 | -0.351 | -1.346 | -0.33 | -0.016 | -0.014 | -0.014 | 0.063 | 0.18 | 0.18 | 0.16 | 0.1 | 0.16 | 0.16 |
| 4 | 130072652 | rs4616770 | 1.218 | -0.205 | -1.22 | -0.179 | 0.006 | -0.018 | -0.018 | 0.061 | 0.35 | 0.24 | 0.37 | 0.34 | 0.31 | 0.38 |
| 4 | 130074947 | rs13123966 | 0.258 | 0.68 | -1.263 | -0.301 | -0.001 | -0.001 | -0.001 | 2.011 | 0.56 | 0.43 | 0.6 | 0.47 | 0.6 | 0.55 |
| 4 | 130077663 | rs11099001 | 1.802 | 0.801 | -1.289 | -0.242 | 0.007 | -0.006 | -0.006 | 0.115 | 0.34 | 0.28 | 0.37 | 0.36 | 0.29 | 0.32 |
| 4 | 130083679 | rs13132155 | 0.331 | 0.786 | -1.266 | -0.224 | 0 | -0.003 | -0.003 | 0 | 0.54 | 0.42 | 0.55 | 0.38 | 0.56 | 0.48 |
| 4 | 130089052 | rs11726808 | 0.197 | 0.774 | -1.159 | -0.249 | -0.001 | 0 | 0 | 0 | 0.55 | 0.48 | 0.58 | 0.43 | 0.57 | 0.49 |
| 4 | 130089540 | rs11735242 | 1.589 | 0.843 | -1.2 | -0.187 | -0.021 | -0.02 | -0.02 | 0 | 0.14 | 0.01 | 0.13 | 0.07 | 0.17 | 0.14 |
| 4 | 130093303 | rs1113473 | 0.896 | 0.959 | -1.036 | -0.18 | 0.027 | -0.022 | -0.022 | 0 | 0.05 | 0.11 | 0.05 | 0 | 0.09 | 0.02 |
| 4 | 130095095 | rs2037073 | 0.697 | 0.733 | -1.329 | -0.47 | -0.012 | 0.04 | 0.04 | 0 | 0.29 | 0.22 | 0.25 | 0.12 | 0.33 | 0.19 |
| 4 | 130096106 | rs17790511 | 1.603 | 0.93 | -1.098 | -0.251 | 0.001 | 0.006 | 0.006 | 0 | 0.17 | 0.15 | 0.16 | 0.06 | 0.19 | 0.14 |
| 4 | 130112548 | rs12511477 | 0.266 | 1.039 | -1.058 | -0.216 | 0.007 | -0.004 | -0.004 | 0.007 | 0.24 | 0.27 | 0.25 | 0.17 | 0.35 | 0.24 |
| 4 | 130114848 | rs10518545 | 0.882 | 1.06 | -1.112 | -0.21 | -0.006 | 0.005 | 0.005 | 0 | 0.21 | 0.22 | 0.22 | 0.12 | 0.29 | 0.17 |
| 4 | 130121328 | rs10857138 | 0.639 | 1.017 | -0.88 | -0.096 | 0.03 | -0.001 | -0.001 | 0 | 0.5 | 0.25 | 0.55 | 0.39 | 0.46 | 0.47 |
| 4 | 130122117 | rs6857889 | -0.511 | 1.161 | -0.94 | -0.041 | -0.001 | 0.002 | 0.002 | 0 | 0.6 | 0.52 | 0.65 | 0.5 | 0.62 | 0.57 |
| 4 | 130123251 | rs7659741 | 0.273 | 0.827 | -1.138 | -0.103 | 0.009 | -0.002 | -0.002 | 0 | 0.52 | 0.41 | 0.56 | 0.39 | 0.46 | 0.46 |
| 4 | 130130312 | rs488685 | 0.384 | 0.758 | -0.714 | -0.252 | -0.001 | 0 | 0 | 0.164 | 0.47 | 0.51 | 0.62 | 0.65 | 0.49 | 0.64 |
| 4 | 130130625 | rs512552 | 0.001 | 0.412 | -0.687 | -0.562 | -0.031 | -0.032 | -0.032 | 0 | 0.29 | 0.43 | 0.14 | 0.26 | 0.16 | 0.15 |
| 4 | 130130914 | rs17014455 | 1.155 | 0.369 | -0.806 | -0.58 | -0.001 | -0.025 | -0.025 | 0 | 0.13 | 0.14 | 0.04 | 0 | 0.09 | 0.08 |
| 4 | 130133372 | rs477926 | 1.349 | 0.684 | -0.752 | -0.578 | -0.001 | -0.002 | -0.002 | 0 | 0.41 | 0.42 | 0.56 | 0.59 | 0.39 | 0.61 |
| 4 | 130136452 | rs7675783 | 1.253 | 0.869 | -0.715 | -0.148 | -0.018 | 0.006 | 0.006 | 0.467 | 0.15 | 0.25 | 0.09 | 0.26 | 0.07 | 0.07 |
| 4 | 130136648 | rs908103 | -0.937 | 0.952 | -0.786 | -0.279 | -0.012 | 0.024 | 0.024 | 0.04 | 0.74 | 0.85 | 0.76 | 0.91 | 0.68 | 0.8 |
| 4 | 130137058 | rs908102 | 0.222 | 0.9 | -0.873 | -0.217 | 0.001 | 0.002 | 0.002 | 0.054 | 0.59 | 0.6 | 0.65 | 0.59 | 0.61 | 0.72 |
| 4 | 130140333 | rs1699376 | -1.061 | 0.68 | -1.095 | -0.16 | -0.012 | 0.002 | 0.002 | 0 | 0.78 | 0.98 | 0.81 | 0.92 | 0.77 | 0.82 |
| 4 | 130146176 | rs1757939 | 0.044 | 0.914 | -0.456 | 0.219 | -0.002 | 0.001 | 0.001 | 0 | 0.58 | 0.59 | 0.48 | 0.62 | 0.57 | 0.66 |
| 4 | 130155315 | rs971446 | 0.953 | 1.143 | -0.51 | 0.361 | 0.025 | 0.051 | 0.051 | 0 | 0.07 | 0.21 | 0.18 | 0.14 | 0.08 | 0.11 |
| 4 | 130155588 | rs6842443 | 0.292 | 1.127 | -0.328 | 0.325 | 0.001 | -0.001 | -0.001 | 0.535 | 0.55 | 0.47 | 0.32 | 0.49 | 0.49 | 0.54 |
| 4 | 130158317 | rs7682166 | -1.47 | 1.118 | -0.468 | 0.315 | 0.017 | 0.039 | 0.039 | 0.786 | 0.77 | 0.91 | 0.8 | 0.92 | 0.8 | 0.81 |
| 4 | 130160706 | rs1514572 | -1.462 | 1.187 | -0.686 | -0.021 | 0.022 | 0.016 | 0.016 | 0 | 0.38 | 0.47 | 0.57 | 0.43 | 0.47 | 0.43 |
| 4 | 130160851 | rs1355962 | -0.171 | 1.315 | -0.651 | -0.095 | -0.011 | 0.012 | 0.012 | 0.053 | 0.24 | 0.15 | 0.14 | 0 | 0.28 | 0.21 |
| 4 | 130161004 | rs1514570 | -1.102 | 1.302 | -0.637 | 0.026 | 0.019 | -0.029 | -0.029 | 0.57 | 0.33 | 0.44 | 0.35 | 0.21 | 0.38 | 0.35 |
| 4 | 130161349 | rs7677291 | 1.058 | 1.195 | -0.643 | 0.123 | -0.007 | 0.001 | 0.001 | 0 | 0.4 | 0.46 | 0.23 | 0.5 | 0.34 | 0.4 |
| 4 | 130163564 | rs17014505 | 1.529 | 1.33 | -0.917 | 0.121 | -0.005 | 0.011 | 0.011 | 0.028 | 0.06 | 0.05 | 0.24 | 0.22 | 0.12 | 0.09 |
| 4 | 130163823 | rs17432813 | 0.201 | 1.252 | -0.958 | -0.118 | 0.003 | 0.001 | 0.001 | 0.014 | 0.17 | 0.12 | 0.1 | 0 | 0.17 | 0.17 |
| 4 | 130167235 | rs17367253 | 1.841 | 1.35 | -1.162 | -0.062 | -0.016 | 0.013 | 0.013 | 0 | 0.07 | 0.03 | 0.06 | 0 | 0.14 | 0.11 |
| 4 | 130167642 | rs11099004 | -1.028 | 1.303 | -1.172 | 0.035 | -0.009 | 0.066 | 0.066 | 0 | 0.7 | 0.86 | 0.74 | 0.92 | 0.68 | 0.79 |
| 4 | 130171981 | rs6824516 | 1.183 | 0.91 | -1.048 | 0.017 | -0.002 | 0.001 | 0.001 | 0 | 0.37 | 0.37 | 0.19 | 0.49 | 0.3 | 0.37 |
| 4 | 130178102 | rs6829641 | -0.957 | 0.946 | -0.504 | -0.178 | 0.002 | -0.04 | -0.04 | 0.073 | 0.26 | 0.4 | 0.28 | 0.15 | 0.26 | 0.31 |
| 4 | 130184243 | rs6534714 | -1.338 | 0.783 | -1.339 | -0.056 | 0.015 | -0.019 | -0.019 | 0.932 | 0.57 | 0.84 | 0.69 | 0.6 | 0.59 | 0.64 |
| 4 | 130185974 | rs9994767 | -0.818 | 0.749 | -1.336 | -0.111 | 0.007 | 0.025 | 0.025 | 0.442 | 0.39 | 0.78 | 0.37 | 0.4 | 0.34 | 0.43 |
| 4 | 130187875 | rs4132289 | -0.118 | 0.795 | -0.37 | -0.014 | 0.053 | 0.087 | 0.087 | 0 | 0.11 | 0.32 | 0.22 | 0.18 | 0.12 | 0.16 |
| 4 | 130192103 | rs9917851 | -0.217 | -0.203 | -1.015 | -0.155 | 0.004 | -0.05 | -0.05 | 0 | 0.66 | 0.79 | 0.74 | 0.57 | 0.68 | 0.65 |
| 4 | 130194362 | rs13119203 | 1.884 | -0.13 | -1.351 | -0.187 | -0.09 | -0.022 | -0.022 | 1.569 | 0.27 | 0.03 | 0.38 | 0.19 | 0.33 | 0.24 |
| 4 | 130196647 | rs4604070 | 1.101 | -0.475 | -0.299 | 0.145 | -0.025 | 0.01 | 0.01 | 0 | 0.43 | 0.23 | 0.61 | 0.36 | 0.57 | 0.45 |
| 4 | 130200129 | rs4484315 | -0.804 | -0.497 | 0.25 | 0.138 | -0.011 | -0.089 | -0.089 | 0.671 | 0.22 | 0.48 | 0.14 | 0.05 | 0.16 | 0.11 |
| 4 | 130200748 | rs4975208 | -1.04 | -0.449 | -0.208 | 0.393 | 0.009 | -0.1 | -0.1 | 0.086 | 0.8 | 0.96 | 0.83 | 0.6 | 0.82 | 0.73 |
| 4 | 130205744 | rs6825050 | 1.27 | -0.35 | 0.003 | 0.286 | 0.019 | -0.005 | -0.005 | 0.621 | 0.07 | 0.11 | 0.08 | 0.03 | 0.12 | 0.03 |
| 4 | 130216263 | rs4426799 | -0.761 | -0.64 | 0.093 | 0.384 | 0.016 | -0.036 | -0.036 | 0 | 0.78 | 0.85 | 0.83 | 0.6 | 0.84 | 0.77 |

| | | | | | | | | | | | | | | | | |
|---|---|---|---|---|---|---|---|---|---|---|---|---|---|---|---|---|
| 4 | 130221251 | rs6849054 | -0.418 | -0.803 | 0.323 | 0.214 | 0 | -0.008 | -0.008 | 1.781 | 0.26 | 0.26 | 0.15 | 0.23 | 0.22 | 0.24 |
| 4 | 130222259 | rs10857139 | 0.908 | -0.889 | 0.353 | 0.268 | 0.007 | 0.023 | 0.023 | 0 | 0.45 | 0.28 | 0.7 | 0.36 | 0.61 | 0.48 |
| 4 | 130230383 | rs4355401 | 1.272 | -0.818 | 0.228 | 0.357 | -0.051 | 0.043 | 0.043 | 0 | 0.4 | 0.22 | 0.6 | 0.29 | 0.53 | 0.46 |
| 4 | 130236239 | rs6856123 | 0.028 | -0.836 | 0.527 | 0.472 | -0.006 | 0.004 | 0.004 | 0 | 0.76 | 0.71 | 0.83 | 0.71 | 0.77 | 0.78 |
| 4 | 130245233 | rs4334769 | 0.997 | -0.98 | 0.691 | 0.303 | 0 | 0 | 0 | 0 | 0.46 | 0.44 | 0.6 | 0.31 | 0.5 | 0.52 |
| 4 | 130245888 | rs6813691 | 0.447 | -0.871 | 0.705 | 0.078 | 0.012 | -0.005 | -0.005 | 0.052 | 0.14 | 0.26 | 0.12 | 0.18 | 0.15 | 0.08 |
| 4 | 130247519 | rs9994470 | -0.113 | -0.814 | 0.531 | 0.138 | 0 | -0.002 | -0.002 | 4.041 | 0.78 | 0.82 | 0.86 | 0.71 | 0.78 | 0.76 |
| 4 | 130260879 | rs11099007 | 1.07 | -0.923 | 0.551 | 0.04 | -0.005 | 0.011 | 0.011 | 0 | 0.4 | 0.35 | 0.49 | 0.12 | 0.47 | 0.33 |
| 4 | 130264384 | rs10031400 | -0.487 | -0.895 | 0.442 | 0.118 | 0.001 | 0 | 0 | 0 | 0.38 | 0.35 | 0.32 | 0.61 | 0.25 | 0.43 |
| 4 | 130267046 | rs4975217 | 0.008 | -0.897 | 0.176 | 0.093 | 0.001 | -0.003 | -0.003 | 0.76 | 0.71 | 0.67 | 0.79 | 0.73 | 0.67 | 0.75 |
| 4 | 130267085 | rs4975218 | 1.775 | -0.836 | -0.36 | 0.113 | -0.019 | 0.039 | 0.039 | 0 | 0.33 | 0.25 | 0.46 | 0.12 | 0.43 | 0.32 |
| 4 | 130267509 | rs17014612 | 0.537 | -0.41 | -0.981 | 0.511 | -0.022 | -0.134 | -0.134 | 1.117 | 0.23 | 0.03 | 0.32 | 0.57 | 0.18 | 0.35 |
| 4 | 130277165 | rs6820523 | 0.385 | -0.036 | -0.506 | 0.464 | -0.024 | 0.012 | 0.012 | 0 | 0.71 | 0.81 | 0.79 | 0.73 | 0.66 | 0.72 |
| 4 | 130308015 | rs10014575 | 0.307 | -0.084 | 0.286 | 0.657 | 0.015 | -0.002 | -0.002 | 0 | 0.14 | 0.29 | 0.12 | 0.12 | 0.15 | 0.09 |
| 4 | 130310751 | rs11933154 | -1.381 | -0.497 | 0.002 | 0.484 | -0.005 | 0.152 | 0.152 | 0 | 0.53 | 0.81 | 0.5 | 0.77 | 0.43 | 0.61 |
| 4 | 130314764 | rs4376156 | -1.431 | -0.509 | 0.015 | 0.478 | -0.017 | 0.156 | 0.156 | 0.007 | 0.54 | 0.87 | 0.51 | 0.77 | 0.43 | 0.61 |
| 4 | 130315957 | rs11942882 | -1.118 | -0.822 | 0.426 | 1.187 | -0.065 | 0.108 | 0.108 | 0.021 | 0.16 | 0.43 | 0.26 | 0.25 | 0.08 | 0.16 |
| 4 | 130316519 | rs1433378 | 0.278 | -0.806 | 0.353 | 1.181 | -0.019 | -0.086 | -0.086 | 0 | 0.22 | 0.05 | 0.12 | 0.39 | 0.21 | 0.37 |
| 4 | 130317282 | rs6845051 | -0.304 | -0.854 | 0.553 | 1.239 | -0.015 | 0.042 | 0.042 | 0 | 0.09 | 0.22 | 0.21 | 0.17 | 0.05 | 0.05 |
| 4 | 130318033 | rs13139447 | 0.35 | -0.83 | 0.287 | 1.251 | 0.003 | -0.046 | -0.046 | 0 | 0.08 | 0.01 | 0.19 | 0.17 | 0.05 | 0.05 |
| 4 | 130318128 | rs1030696 | 0.196 | -0.773 | 0.336 | 1.186 | -0.004 | -0.051 | -0.051 | 0.229 | 0.23 | 0.08 | 0.13 | 0.39 | 0.21 | 0.37 |
| 4 | 130325201 | rs10001834 | -1.601 | -0.725 | 0.936 | 1.438 | -0.068 | 0.138 | 0.138 | 0 | 0.16 | 0.48 | 0.26 | 0.28 | 0.08 | 0.16 |
| 4 | 130328109 | rs17014740 | -1.418 | -0.638 | 0.047 | 1.906 | -0.043 | 0.207 | 0.207 | 0.222 | 0.14 | 0.27 | 0.26 | 0.53 | 0.08 | 0.28 |
| 4 | 130332644 | rs6837783 | -1.789 | -0.294 | -0.907 | 1.417 | -0.018 | -0.01 | -0.01 | 0 | 0.49 | 0.92 | 0.34 | 0.63 | 0.43 | 0.43 |
| 4 | 130333958 | rs2043372 | 0.179 | -0.318 | -0.902 | 1.588 | 0.008 | -0.024 | -0.024 | 0.567 | 0.08 | 0.04 | 0.04 | 0.3 | 0.05 | 0.16 |
| 4 | 130338059 | rs1433375 | -1.545 | -0.233 | -0.78 | 1.087 | -0.001 | 0.04 | 0.04 | 0.546 | 0.48 | 0.88 | 0.34 | 0.66 | 0.43 | 0.44 |
| 4 | 130338172 | rs11730148 | 0.273 | -0.204 | -0.914 | 0.968 | -0.045 | 0.067 | 0.067 | 0 | 0.12 | 0.01 | 0.09 | 0 | 0.13 | 0.04 |
| 4 | 130340880 | rs17014760 | -0.387 | -0.274 | -0.517 | 0.998 | -0.004 | 0.044 | 0.044 | 0.071 | 0.29 | 0.66 | 0.21 | 0.53 | 0.27 | 0.28 |
| 4 | 130347005 | rs1816429 | -0.938 | -0.141 | -1.002 | 1.195 | -0.003 | -0.033 | -0.033 | 0 | 0.44 | 0.83 | 0.32 | 0.55 | 0.42 | 0.35 |
| 4 | 130364131 | rs995638 | -0.919 | 0.013 | -1.177 | 1.082 | 0.002 | 0.001 | 0.001 | 0.375 | 0.45 | 0.8 | 0.31 | 0.6 | 0.41 | 0.38 |
| 4 | 130364383 | rs995637 | -0.982 | 0.143 | -1.481 | 0.946 | 0.001 | 0.037 | 0.037 | 0.428 | 0.47 | 0.89 | 0.34 | 0.65 | 0.42 | 0.43 |
| 4 | 130367473 | rs13125482 | -0.244 | 0.335 | -1.158 | 0.512 | 0.015 | 0.04 | 0.04 | 0 | 0.48 | 0.4 | 0.35 | 0.25 | 0.43 | 0.4 |
| 4 | 130373727 | rs1865178 | -0.038 | -0.115 | -0.997 | 0.707 | 0.004 | -0.001 | -0.001 | 0 | 0.36 | 0.38 | 0.25 | 0.42 | 0.28 | 0.31 |
| 4 | 130375150 | rs1433384 | -0.863 | 0.409 | -0.569 | 0.883 | 0 | 0 | 0 | 0 | 0.7 | 0.65 | 0.7 | 0.74 | 0.64 | 0.8 |
| 4 | 130375622 | rs1865176 | 0.466 | 0.433 | -0.831 | 1.131 | 0.003 | 0.002 | 0.002 | 0 | 0.36 | 0.38 | 0.24 | 0.34 | 0.27 | 0.29 |
| 4 | 130378087 | rs17370136 | 1.491 | 0.723 | -1.307 | 1.398 | -0.036 | -0.048 | -0.048 | 0 | 0.09 | 0.01 | 0.18 | 0.15 | 0.13 | 0.04 |
| 4 | 130378616 | rs621740 | 0.958 | 0.597 | -1.322 | 1.226 | -0.057 | -0.003 | -0.003 | 0 | 0.17 | 0.01 | 0.11 | 0.12 | 0.23 | 0.16 |
| 4 | 130384715 | rs623865 | -0.136 | 0.49 | -0.784 | 1.254 | -0.004 | -0.011 | -0.011 | 0.113 | 0.31 | 0.24 | 0.33 | 0.34 | 0.37 | 0.23 |
| 4 | 130390365 | rs7666727 | 0.703 | 0.437 | -0.984 | 1.17 | -0.051 | 0.062 | 0.062 | 0 | 0.32 | 0.11 | 0.34 | 0.14 | 0.36 | 0.38 |
| 4 | 130396639 | rs592988 | -0.866 | 0.429 | -0.74 | 1.196 | -0.024 | -0.004 | -0.004 | 0 | 0.67 | 0.81 | 0.58 | 0.69 | 0.64 | 0.53 |
| 4 | 130403749 | rs3097948 | 0.679 | 0.287 | -1.183 | 0.879 | -0.095 | -0.024 | -0.024 | 0.265 | 0.22 | 0.06 | 0.18 | 0.2 | 0.35 | 0.37 |
| 4 | 130412286 | rs17050480 | 0.62 | 0.076 | -1.032 | 0.895 | -0.004 | -0.023 | -0.023 | 1.079 | 0.18 | 0.06 | 0.25 | 0.22 | 0.17 | 0.24 |
| 4 | 130412508 | rs17050482 | 0.686 | 0.073 | -1.034 | 0.874 | -0.005 | -0.002 | -0.002 | 0 | 0.17 | 0.05 | 0.2 | 0.17 | 0.16 | 0.21 |
| 4 | 130417557 | rs650483 | -1.501 | 0.07 | -0.722 | 0.797 | 0.008 | 0.001 | 0.001 | 0 | 0.86 | 0.83 | 0.8 | 0.83 | 0.79 | 0.95 |
| 4 | 130421169 | rs600121 | -0.247 | 0.027 | -0.688 | 0.783 | -0.002 | -0.018 | -0.018 | 0 | 0.69 | 0.75 | 0.47 | 0.53 | 0.63 | 0.66 |
| 4 | 130439039 | rs650052 | 2.39 | 0.515 | -1.119 | 0.351 | -0.043 | 0.046 | 0.046 | 0 | 0.08 | 0.01 | 0.06 | 0 | 0.12 | 0.07 |
| 4 | 130444832 | rs724156 | 0.844 | -0.034 | -0.761 | 0.223 | 0.059 | 0.036 | 0.036 | 0 | 0.4 | 0.22 | 0.24 | 0.34 | 0.28 | 0.32 |
| 4 | 130450256 | rs685845 | 0.069 | 0.159 | 0.215 | 0.717 | 0 | 0.002 | 0.002 | 0 | 0.67 | 0.53 | 0.53 | 0.62 | 0.63 | 0.77 |
| 4 | 130453157 | rs666306 | 0.501 | 0.177 | -0.132 | 0.602 | 0.008 | 0.034 | 0.034 | 0 | 0.5 | 0.43 | 0.41 | 0.28 | 0.49 | 0.54 |
| 4 | 130458147 | rs659322 | -0.342 | 0.308 | -0.243 | 0.622 | -0.003 | 0 | 0 | 0 | 0.67 | 0.78 | 0.55 | 0.63 | 0.61 | 0.77 |
| 4 | 130464486 | rs1074762 | 1.328 | 0.281 | -0.572 | 0.567 | -0.001 | 0.009 | 0.009 | 0.122 | 0.15 | 0.05 | 0.07 | 0.11 | 0.16 | 0.1 |
| 4 | 130469043 | rs2117060 | 0.769 | 0.101 | -0.353 | 0.68 | -0.007 | 0.026 | 0.026 | 0 | 0.15 | 0.15 | 0.11 | 0.33 | 0.12 | 0.23 |
| 4 | 130469784 | rs17372420 | 1.464 | 0.285 | -0.473 | 0.609 | -0.057 | -0.028 | -0.028 | 0 | 0.13 | 0.01 | 0.18 | 0.15 | 0.2 | 0.05 |
| 4 | 130474564 | rs10857140 | 0.141 | 0.183 | -0.405 | 0.63 | 0.006 | -0.011 | -0.011 | 0 | 0.31 | 0.2 | 0.18 | 0.46 | 0.28 | 0.33 |
| 4 | 130482494 | rs1356342 | 0.387 | 0.181 | -0.137 | 0.572 | 0 | 0 | 0 | 0 | 0.54 | 0.52 | 0.61 | 0.4 | 0.52 | 0.63 |
| 4 | 130508349 | rs11725348 | 1.28 | 0.656 | -0.621 | 0.483 | -0.062 | 0.012 | 0.012 | 0 | 0.1 | 0.02 | 0.1 | 0.03 | 0.2 | 0.07 |
| 4 | 130523279 | rs628527 | -0.391 | -0.66 | -0.149 | 0.758 | -0.018 | 0.02 | 0.02 | 0.043 | 0.26 | 0.65 | 0.25 | 0.42 | 0.15 | 0.36 |
| 4 | 130530633 | rs12649182 | 1.802 | -0.787 | -0.374 | 0.654 | -0.011 | -0.001 | -0.001 | 0.001 | 0.15 | 0.02 | 0.07 | 0.13 | 0.16 | 0.1 |
| 4 | 130535115 | rs3106876 | -0.506 | -0.546 | 0.343 | 0.891 | -0.018 | 0.058 | 0.058 | 0 | 0.29 | 0.35 | 0.34 | 0.5 | 0.26 | 0.28 |
| 4 | 130536283 | rs10518552 | 2.043 | -0.48 | 0.682 | 1.008 | 0.003 | 0.046 | 0.046 | 0.188 | 0.07 | 0.22 | 0.13 | 0.1 | 0.05 | 0.14 |
| 4 | 130539982 | rs6849510 | 0.492 | -0.308 | 0.742 | 1.159 | -0.014 | 0.029 | 0.029 | 0 | 0.15 | 0.15 | 0.11 | 0.32 | 0.1 | 0.23 |
| 4 | 130543768 | rs17439675 | -0.805 | -0.398 | 0.455 | 1.107 | -0.105 | 0.064 | 0.064 | 0 | 0.62 | 0.87 | 0.73 | 0.79 | 0.53 | 0.59 |
| 4 | 130547353 | rs10001198 | -0.707 | -0.441 | 1.179 | 1.467 | -0.06 | 0.104 | 0.104 | 0 | 0.18 | 0.39 | 0.12 | 0.33 | 0.1 | 0.23 |
| 4 | 130552827 | rs10017568 | -0.455 | -0.4 | 1.028 | 1.488 | -0.047 | 0.09 | 0.09 | 0 | 0.17 | 0.3 | 0.12 | 0.33 | 0.1 | 0.23 |
| 4 | 130562576 | rs7674283 | -0.209 | 0.128 | 0.895 | 1.103 | 0.019 | 0.062 | 0.062 | 0.018 | 0.11 | 0.39 | 0.13 | 0.11 | 0.09 | 0.19 |
| 4 | 130568677 | rs17050583 | 0.09 | 0.237 | 0.442 | 0.879 | 0.002 | 0.022 | 0.022 | 0 | 0.57 | 0.39 | 0.55 | 0.28 | 0.57 | 0.33 |
| 4 | 130569024 | rs9307614 | -0.277 | 0.393 | 0.22 | 1.145 | 0.003 | 0.005 | 0.005 | 0 | 0.63 | 0.62 | 0.7 | 0.45 | 0.58 | 0.47 |
| 4 | 130580317 | rs1907454 | -0.334 | 0.389 | 0.184 | 1.132 | -0.014 | -0.02 | -0.02 | 0 | 0.63 | 0.75 | 0.71 | 0.44 | 0.58 | 0.47 |
| 4 | 130591851 | rs11931950 | -0.45 | 0.818 | 0.205 | 0.731 | 0.029 | -0.039 | -0.039 | 0 | 0.27 | 0.48 | 0.23 | 0.16 | 0.32 | 0.12 |

| | | | | | | | | | | | | | | | | |
|---|---|---|---|---|---|---|---|---|---|---|---|---|---|---|---|---|
| 4 | 130592857 | rs17050588 | 0.365 | 0.616 | -0.207 | 1.156 | -0.004 | -0.021 | -0.021 | 5.434 | 0.33 | 0.22 | 0.44 | 0.45 | 0.35 | 0.45 |
| 4 | 130593382 | rs2728765 | -1.021 | 1.317 | -0.023 | 1.651 | 0.011 | 0.011 | 0.011 | 2.368 | 0.33 | 0.61 | 0.37 | 0.57 | 0.41 | 0.43 |
| 4 | 130601796 | rs2728740 | 0.639 | 0.803 | -0.291 | 1.588 | 0.014 | 0.027 | 0.027 | 0 | 0.53 | 0.65 | 0.5 | 0.4 | 0.46 | 0.57 |
| 4 | 130606797 | rs4864109 | 0.384 | 1.059 | -0.452 | 1.594 | 0.005 | 0.007 | 0.007 | 0.306 | 0.18 | 0.25 | 0.23 | 0.3 | 0.27 | 0.17 |
| 4 | 130619114 | rs2728760 | -0.185 | 0.907 | -0.71 | 1.441 | -0.03 | 0.001 | 0.001 | 0.063 | 0.72 | 0.94 | 0.79 | 0.79 | 0.73 | 0.77 |
| 4 | 130626785 | rs2637613 | -0.278 | 0.889 | -0.762 | 1.469 | -0.03 | -0.057 | -0.057 | 0 | 0.73 | 0.94 | 0.72 | 0.7 | 0.72 | 0.74 |
| 4 | 130645957 | rs13115271 | 1.585 | 0.888 | -0.231 | 1.504 | 0.009 | 0.029 | 0.029 | 0 | 0.42 | 0.3 | 0.45 | 0.26 | 0.4 | 0.4 |
| 4 | 130646527 | rs2728752 | -0.229 | 0.781 | 0.248 | 1.52 | -0.015 | 0.031 | 0.031 | 0.762 | 0.06 | 0.21 | 0.01 | 0.07 | 0 | 0.07 |
| 4 | 130650949 | rs2728748 | -0.083 | 0.782 | 0.264 | 1.521 | -0.015 | 0.03 | 0.03 | 0.388 | 0.06 | 0.2 | 0.01 | 0.07 | 0 | 0.07 |
| 4 | 130654405 | rs10022354 | 0.107 | 0.851 | 0.404 | 1.474 | 0.013 | -0.026 | -0.026 | 0 | 0.06 | 0.12 | 0.05 | 0 | 0.06 | 0.06 |
| 4 | 130655673 | rs10020857 | -0.02 | 0.818 | 0.359 | 1.617 | 0.011 | -0.012 | -0.012 | 0 | 0.67 | 0.77 | 0.72 | 0.62 | 0.72 | 0.66 |
| 4 | 130655964 | rs10011452 | -0.37 | 0.808 | 0.458 | 1.69 | 0.011 | -0.001 | -0.001 | 0 | 0.23 | 0.27 | 0.19 | 0.21 | 0.32 | 0.18 |
| 4 | 130656378 | rs2637592 | 0.083 | 0.785 | 0.448 | 1.696 | -0.022 | 0.042 | 0.042 | 0.149 | 0.06 | 0.15 | 0.04 | 0.09 | 0 | 0.08 |
| 4 | 130658868 | rs2728743 | -1.092 | 0.82 | 0.386 | 1.693 | -0.027 | -0.003 | -0.003 | 0 | 0.76 | 0.98 | 0.86 | 0.85 | 0.77 | 0.84 |
| 4 | 130661821 | rs11726911 | 0.584 | 0.854 | 0.423 | 1.676 | -0.012 | -0.006 | -0.006 | 0 | 0.17 | 0.15 | 0.15 | 0.2 | 0.26 | 0.12 |
| 4 | 130664077 | rs2035793 | 0.023 | 0.778 | 0.423 | 1.678 | 0.007 | -0.008 | -0.008 | 0 | 0.67 | 0.75 | 0.72 | 0.62 | 0.72 | 0.66 |
| 4 | 130667787 | rs6827640 | 1.743 | 0.828 | 0.491 | 1.689 | 0.006 | 0.032 | 0.032 | 0 | 0.42 | 0.3 | 0.45 | 0.25 | 0.4 | 0.4 |
| 4 | 130672736 | rs13126098 | 1.643 | 0.818 | 0.511 | 1.677 | 0.007 | 0.009 | 0.009 | 0 | 0.43 | 0.33 | 0.51 | 0.41 | 0.4 | 0.43 |
| 4 | 130678673 | rs961410 | -0.734 | 0.914 | 0.567 | 1.721 | 0.04 | 0.008 | 0.008 | 0 | 0.25 | 0.44 | 0.21 | 0.27 | 0.32 | 0.24 |
| 4 | 130680257 | rs2391481 | 1.523 | 0.863 | 0.563 | 1.737 | 0.006 | 0.05 | 0.05 | 0.836 | 0.86 | 0.6 | 0.8 | 0.84 | 0.89 | 0.73 |
| 4 | 130680689 | rs4608819 | 1.432 | 0.859 | 0.528 | 1.745 | -0.006 | 0.07 | 0.07 | 1.98 | 0.84 | 0.54 | 0.79 | 0.77 | 0.89 | 0.72 |
| 4 | 130680706 | rs10010057 | 0.353 | 0.837 | 0.475 | 1.747 | -0.002 | 0.097 | 0.097 | 0.09 | 0.91 | 0.72 | 0.85 | 0.84 | 0.95 | 0.79 |
| 4 | 130681587 | rs17050638 | 0.87 | 0.854 | 0.236 | 1.756 | -0.04 | -0.025 | -0.025 | 0 | 0.16 | 0.07 | 0.14 | 0.2 | 0.25 | 0.12 |
| 4 | 130684221 | rs4417988 | 0.07 | 0.481 | 0.199 | 1.822 | -0.008 | 0.063 | 0.063 | 0 | 0.92 | 0.84 | 0.86 | 0.84 | 0.95 | 0.84 |
| 4 | 130691794 | rs10518556 | 0.963 | 0.482 | 0.094 | 1.837 | 0.002 | -0.001 | -0.001 | 0 | 0.58 | 0.4 | 0.59 | 0.44 | 0.64 | 0.52 |
| 4 | 130696787 | rs4317218 | 0.844 | 0.468 | 0.071 | 1.773 | -0.016 | 0.126 | 0.126 | 0.018 | 0.9 | 0.82 | 0.8 | 0.59 | 0.94 | 0.79 |
| 4 | 130701197 | rs2391560 | 0.948 | 0.49 | 0.026 | 1.778 | 0.001 | -0.001 | -0.001 | 0 | 0.58 | 0.4 | 0.59 | 0.44 | 0.64 | 0.52 |
| 4 | 130707281 | rs17690833 | 0.717 | 0.49 | 0 | 1.759 | -0.029 | 0 | 0 | 0 | 0.24 | 0.02 | 0.14 | 0.15 | 0.23 | 0.16 |
| 4 | 130707396 | rs7692863 | 0.167 | 0.499 | 0.199 | 1.657 | -0.005 | 0.068 | 0.068 | 0 | 0.92 | 0.85 | 0.86 | 0.82 | 0.94 | 0.84 |
| 4 | 130710218 | rs11099019 | 0.39 | 0.556 | -0.179 | 1.238 | 0.001 | -0.001 | -0.001 | 0 | 0.58 | 0.4 | 0.59 | 0.44 | 0.64 | 0.52 |
| 4 | 130714768 | rs6534743 | 0.912 | 0.14 | -0.979 | 0.776 | 0.001 | 0.064 | 0.064 | 0.026 | 0.37 | 0.05 | 0.39 | 0.23 | 0.3 | 0.38 |
| 4 | 130724902 | rs11099020 | -0.311 | 0.326 | -0.78 | 0.793 | 0.06 | 0.062 | 0.062 | 1.676 | 0.44 | 0.15 | 0.42 | 0.24 | 0.63 | 0.44 |
| 4 | 130731072 | rs2391504 | -1.23 | -0.12 | -0.37 | 0.934 | 0.047 | 0.075 | 0.075 | 0 | 0.59 | 0.77 | 0.77 | 0.79 | 0.68 | 0.63 |
| 4 | 130756174 | rs2391535 | -1.163 | -0.212 | -0.294 | 0.904 | 0.038 | 0.047 | 0.047 | 0 | 0.61 | 0.76 | 0.79 | 0.82 | 0.73 | 0.64 |
| 4 | 130756415 | rs12233773 | -1.185 | -0.195 | 0.427 | 1.337 | -0.026 | 0.05 | 0.05 | 0 | 0.12 | 0.28 | 0.18 | 0.21 | 0.06 | 0.28 |
| 4 | 130758713 | rs2952862 | -0.006 | -0.226 | 0.399 | 1.446 | 0.053 | 0.097 | 0.097 | 2.244 | 0.44 | 0.09 | 0.42 | 0.23 | 0.65 | 0.3 |
| 4 | 130764605 | rs2860199 | -0.941 | -0.225 | 0.87 | 1.458 | 0.017 | 0.012 | 0.012 | 0.511 | 0.55 | 0.62 | 0.64 | 0.67 | 0.71 | 0.48 |
| 4 | 130765713 | rs10022903 | -0.149 | -0.072 | 0.996 | 1.522 | 0.025 | 0.019 | 0.019 | 0 | 0.07 | 0.22 | 0.04 | 0.05 | 0.05 | 0.16 |
| 4 | 130767230 | rs10028795 | -1.481 | -0.075 | 0.828 | 1.528 | 0.008 | 0.352 | 0.352 | 2.49 | 0.11 | 0.53 | 0.22 | 0.44 | 0.07 | 0.18 |
| 4 | 130767559 | rs4863907 | 1.145 | 1.054 | 0.506 | 1.266 | -0.06 | 0.037 | 0.037 | 0 | 0.11 | 0.03 | 0.2 | 0.02 | 0.3 | 0.15 |
| 4 | 130767899 | rs17050689 | -1.342 | 0.75 | 0.574 | 1.316 | 0.008 | 0.352 | 0.352 | 2.49 | 0.11 | 0.53 | 0.22 | 0.44 | 0.07 | 0.18 |
| 4 | 130784545 | rs2968999 | -1.295 | 0.602 | 0.254 | 1.019 | 0.033 | 0.015 | 0.015 | 0.755 | 0.58 | 0.69 | 0.65 | 0.66 | 0.73 | 0.53 |
| 4 | 130824678 | rs3111741 | -1.211 | 0.695 | 0.413 | 1.06 | 0.001 | 0.001 | 0.001 | 0 | 0.57 | 0.63 | 0.76 | 0.77 | 0.69 | 0.51 |
| 4 | 130827136 | rs2968959 | -0.45 | 0.662 | 0.469 | 1.19 | 0.11 | 0.048 | 0.048 | 1.602 | 0.51 | 0.21 | 0.58 | 0.41 | 0.68 | 0.49 |
| 4 | 130827478 | rs4447877 | 1.199 | 1.084 | 0.37 | 1.127 | -0.059 | 0.019 | 0.019 | 0 | 0.11 | 0.03 | 0.2 | 0.06 | 0.28 | 0.16 |
| 4 | 130839795 | rs2952871 | -1.653 | 0.891 | 0.689 | 1.212 | 0.037 | 0.041 | 0.041 | 0 | 0.64 | 0.8 | 0.79 | 0.82 | 0.74 | 0.67 |
| 4 | 130850180 | rs12644225 | -1.538 | 0.867 | 0.901 | 1.337 | -0.021 | 0.389 | 0.389 | 3.558 | 0.08 | 0.32 | 0.32 | 0.5 | 0.03 | 0.14 |
| 4 | 130857829 | rs2968955 | -0.097 | 1.045 | 0.776 | 1.246 | 0.048 | 0.121 | 0.121 | 1.243 | 0.46 | 0.13 | 0.43 | 0.24 | 0.65 | 0.32 |
| 4 | 130866817 | rs10025344 | -1.524 | 0.908 | 0.9 | 1.333 | -0.021 | 0.384 | 0.384 | 3.97 | 0.08 | 0.32 | 0.32 | 0.49 | 0.03 | 0.14 |
| 4 | 130876818 | rs6825946 | -0.129 | 1.191 | 0.137 | 1.148 | 0.032 | 0.128 | 0.128 | 1.487 | 0.53 | 0.87 | 0.58 | 0.74 | 0.38 | 0.69 |
| 4 | 130887651 | rs1995720 | -0.378 | 1.163 | -0.199 | 1.307 | 0.056 | 0.1 | 0.1 | 1.232 | 0.48 | 0.18 | 0.56 | 0.36 | 0.63 | 0.46 |
| 4 | 130892616 | rs17388944 | 1.388 | 1.533 | -0.515 | 0.968 | -0.037 | 0.054 | 0.054 | 0 | 0.12 | 0.02 | 0.12 | 0 | 0.18 | 0.03 |
| 4 | 130903511 | rs17455138 | 1.67 | 0.148 | -0.674 | 1.045 | -0.02 | 0.009 | 0.009 | 0 | 0.28 | 0.01 | 0.13 | 0.15 | 0.22 | 0.3 |
| 4 | 130908815 | rs2968933 | -0.214 | 0.529 | -0.243 | 1.245 | 0.043 | 0.143 | 0.143 | 0.978 | 0.47 | 0.22 | 0.43 | 0.24 | 0.62 | 0.35 |
| 4 | 130912560 | rs2391577 | 0.606 | 0.487 | -0.642 | 1.108 | -0.074 | 0.069 | 0.069 | 0.073 | 0.29 | 0.11 | 0.21 | 0.14 | 0.37 | 0.12 |
| 4 | 130913879 | rs2940273 | 0.884 | 0.56 | -0.665 | 1.012 | -0.031 | 0.046 | 0.046 | 0 | 0.17 | 0.06 | 0.2 | 0.09 | 0.25 | 0.2 |
| 4 | 130918048 | rs2940288 | -0.288 | -0.098 | -0.126 | 1.361 | 0.013 | 0.01 | 0.01 | 0.482 | 0.53 | 0.43 | 0.59 | 0.45 | 0.64 | 0.48 |
| 4 | 130928325 | rs1567192 | -1.012 | -0.026 | -0.921 | 1.001 | 0.034 | 0.051 | 0.051 | 0 | 0.62 | 0.82 | 0.81 | 0.82 | 0.73 | 0.66 |
| 4 | 130948872 | rs17050809 | 0.268 | 0.503 | -1.36 | 0.635 | -0.013 | -0.005 | -0.005 | 0 | 0.05 | 0.02 | 0.02 | 0.05 | 0.08 | 0.16 |
| 4 | 130949366 | rs7661528 | -0.294 | 0.368 | -1.324 | 0.583 | 0.02 | 0.086 | 0.086 | 0.742 | 0.46 | 0.2 | 0.42 | 0.32 | 0.6 | 0.34 |
| 4 | 130956277 | rs2135927 | 0.338 | 0.635 | -1.026 | 0.421 | 0.008 | 0.083 | 0.083 | 0.321 | 0.37 | 0.16 | 0.2 | 0.16 | 0.28 | 0.15 |
| 4 | 130962263 | rs1567193 | -0.202 | 0.451 | -0.427 | 1.063 | 0.045 | 0.029 | 0.029 | 0 | 0.52 | 0.74 | 0.49 | 0.59 | 0.32 | 0.3 |
| 4 | 130962301 | rs7665451 | 0.305 | 0.408 | -0.355 | 1.195 | 0.036 | -0.026 | -0.026 | 0 | 0.44 | 0.21 | 0.4 | 0.51 | 0.3 | 0.25 |
| 4 | 130966165 | rs1022009 | 0.004 | 0.39 | -0.233 | 1.339 | 0.015 | 0.004 | 0.004 | 0 | 0.52 | 0.65 | 0.46 | 0.52 | 0.35 | 0.24 |
| 4 | 130978989 | rs6534762 | -0.552 | 0.783 | -0.374 | 1.42 | -0.071 | 0.039 | 0.039 | 0 | 0.6 | 0.9 | 0.55 | 0.67 | 0.44 | 0.41 |
| 4 | 130980955 | rs4864311 | -1.313 | 0.743 | -0.18 | 1.441 | 0.056 | 0.094 | 0.094 | 5.667 | 0.12 | 0.63 | 0.15 | 0.18 | 0.11 | 0.17 |
| 4 | 130983055 | rs6844491 | -1.03 | 0.753 | 0.204 | 1.434 | 0.119 | 0.15 | 0.15 | 2.256 | 0.1 | 0.53 | 0.13 | 0.15 | 0.11 | 0.16 |
| 4 | 130983605 | rs12499245 | -0.422 | 0.993 | 0.034 | 1.476 | -0.059 | 0.031 | 0.031 | 0 | 0.61 | 0.9 | 0.57 | 0.67 | 0.47 | 0.42 |
| 4 | 130984088 | rs4863950 | -1.377 | 0.904 | -0.148 | 1.389 | 0.036 | -0.014 | -0.014 | 0.851 | 0.78 | 0.97 | 0.9 | 0.79 | 0.83 | 0.61 |

| | | | | | | | | | | | | | | | | |
|---|---|---|---|---|---|---|---|---|---|---|---|---|---|---|---|---|
| 4 | 130997758 | rs13134698 | 0.245 | 1.044 | 0.076 | 1.404 | -0.001 | 0.027 | 0.027 | 0.242 | 0.52 | 0.64 | 0.28 | 0.27 | 0.39 | 0.28 |
| 4 | 131003605 | rs11934226 | -0.717 | 1.178 | 0.153 | 1.419 | 0.059 | 0.033 | 0.033 | 0.011 | 0.05 | 0.14 | 0.05 | 0.08 | 0.09 | 0.11 |
| 4 | 131013140 | rs11933661 | -0.081 | 1.076 | 0.197 | 1.489 | -0.004 | 0.051 | 0.051 | 0 | 0.56 | 0.83 | 0.56 | 0.64 | 0.45 | 0.39 |
| 4 | 131016112 | rs1507787 | 0.142 | 1.11 | 0.286 | 1.518 | 0.009 | 0.039 | 0.039 | 0.285 | 0.53 | 0.76 | 0.51 | 0.6 | 0.44 | 0.33 |
| 4 | 131017136 | rs6827509 | 0.162 | 0.948 | 0.554 | 1.452 | -0.02 | 0.034 | 0.034 | 2.043 | 0.07 | 0.29 | 0.05 | 0.07 | 0.01 | 0.04 |
| 4 | 131018271 | rs2391598 | 0.301 | 1.174 | 0.586 | 1.553 | -0.001 | -0.006 | -0.006 | 0.098 | 0.49 | 0.58 | 0.32 | 0.27 | 0.36 | 0.27 |
| 4 | 131021262 | rs11099035 | 0.028 | 1.17 | 0.46 | 1.545 | -0.007 | 0.026 | 0.026 | 0.003 | 0.53 | 0.71 | 0.37 | 0.31 | 0.39 | 0.33 |
| 4 | 131061059 | rs2391603 | -0.667 | 1.211 | 0.175 | 1.578 | -0.007 | 0.015 | 0.015 | 0 | 0.67 | 0.7 | 0.68 | 0.44 | 0.72 | 0.51 |
| 4 | 131062487 | rs13109471 | 0.089 | 1.309 | 0.181 | 1.609 | 0 | 0.001 | 0.001 | 0 | 0.61 | 0.42 | 0.55 | 0.32 | 0.63 | 0.34 |
| 4 | 131063868 | rs6853869 | -0.375 | 1.15 | -0.068 | 1.272 | 0 | 0.006 | 0.006 | 0 | 0.54 | 0.66 | 0.38 | 0.32 | 0.4 | 0.35 |
| 4 | 131071699 | rs1845224 | -1.06 | 0.693 | 0.159 | 1.361 | 0.001 | 0.071 | 0.071 | 0 | 0.08 | 0.25 | 0.1 | 0.13 | 0.05 | 0.16 |
| 4 | 131083338 | rs17050863 | -0.653 | 0.463 | 0.098 | 1.298 | -0.049 | 0.036 | 0.036 | 0.877 | 0.11 | 0.33 | 0.04 | 0.14 | 0.01 | 0.05 |
| 4 | 131091376 | rs7656732 | -0.691 | 0.538 | -0.142 | 1.316 | -0.009 | 0.024 | 0.024 | 1.241 | 0.14 | 0.55 | 0.09 | 0.13 | 0.06 | 0.07 |
| 4 | 131093532 | rs6826948 | -0.273 | 0.721 | -0.718 | 1.446 | -0.11 | 0.117 | 0.117 | 0.028 | 0.56 | 0.94 | 0.54 | 0.69 | 0.39 | 0.34 |
| 4 | 131093554 | rs6827503 | -1.011 | 0.835 | -0.244 | 1.985 | 0.01 | 0.366 | 0.366 | 0 | 0.06 | 0.24 | 0.26 | 0.44 | 0.05 | 0.18 |
| 4 | 131097850 | rs10003688 | -0.144 | 1.116 | -0.914 | 0.739 | 0.001 | 0.007 | 0.007 | 0 | 0.49 | 0.43 | 0.46 | 0.58 | 0.4 | 0.25 |
| 4 | 131100838 | rs17407415 | -0.871 | 1.638 | -1.528 | 0.326 | -0.065 | 0.06 | 0.06 | 0.691 | 0.19 | 0.03 | 0.32 | 0.07 | 0.29 | 0.25 |
| 4 | 131101141 | rs13107934 | -1.16 | 0.894 | -1.232 | 0.545 | -0.007 | 0.001 | 0.001 | 0 | 0.1 | 0.13 | 0.05 | 0.07 | 0.07 | 0.11 |
| 4 | 131103736 | rs17050881 | -0.539 | 0.658 | -1.817 | 0.726 | 0.007 | -0.025 | -0.025 | 0 | 0.23 | 0.12 | 0.28 | 0.29 | 0.2 | 0.39 |
| 4 | 131104359 | rs13106127 | 0.428 | 0.017 | -1.183 | 1.483 | -0.025 | 0.139 | 0.139 | 0.873 | 0.89 | 0.76 | 0.85 | 0.7 | 0.94 | 0.83 |
| 4 | 131107527 | rs1979689 | 0.381 | 0.064 | -1.873 | 0.906 | -0.003 | 0.017 | 0.017 | 0 | 0.44 | 0.64 | 0.38 | 0.39 | 0.43 | 0.23 |
| 4 | 131107545 | rs1979690 | 0.512 | -0.239 | -0.984 | 0.757 | -0.001 | 0.001 | 0.001 | 0 | 0.41 | 0.43 | 0.37 | 0.39 | 0.43 | 0.23 |
| 4 | 131108390 | rs2893315 | 0.802 | 0.049 | -1.294 | 0.292 | -0.001 | -0.002 | -0.002 | 0 | 0.25 | 0.3 | 0.27 | 0.23 | 0.25 | 0.14 |
| 4 | 131109747 | rs2220754 | -1.09 | -0.152 | -1.205 | 0.166 | 0.007 | 0.051 | 0.051 | 0 | 0.66 | 0.79 | 0.77 | 0.86 | 0.68 | 0.74 |
| 4 | 131110436 | rs6837104 | 0.546 | -0.256 | -1.166 | 0.253 | -0.01 | -0.01 | -0.01 | 0 | 0.27 | 0.52 | 0.28 | 0.24 | 0.26 | 0.14 |
| 4 | 131116188 | rs1873867 | 1.31 | -0.233 | -1.32 | 0.448 | -0.054 | -0.064 | -0.064 | 0 | 0.19 | 0.01 | 0.26 | 0.25 | 0.2 | 0.36 |
| 4 | 131116943 | rs1120962 | -0.037 | -0.429 | -1.11 | -0.015 | -0.003 | -0.123 | -0.123 | 0.014 | 0.29 | 0.53 | 0.22 | 0.06 | 0.26 | 0.13 |
| 4 | 131117354 | rs7690486 | -1.56 | -0.34 | -1.108 | 0.081 | -0.035 | 0.072 | 0.072 | 0 | 0.71 | 0.88 | 0.79 | 0.89 | 0.66 | 0.77 |
| 4 | 131118067 | rs1602816 | 0.287 | -0.542 | -0.577 | 0.627 | -0.002 | 0.028 | 0.028 | 0 | 0.15 | 0.19 | 0.17 | 0.27 | 0.15 | 0.09 |
| 4 | 131119614 | rs10001806 | -0.418 | -0.508 | -0.572 | 0.773 | -0.004 | 0.015 | 0.015 | 0 | 0.39 | 0.46 | 0.46 | 0.57 | 0.39 | 0.56 |
| 4 | 131121249 | rs12504035 | 0.022 | -0.096 | -0.609 | 1.017 | 0 | 0.086 | 0.086 | 0 | 0.14 | 0.18 | 0.27 | 0.46 | 0.15 | 0.14 |
| 4 | 131122852 | rs13149325 | 0.304 | -0.23 | -0.798 | 0.559 | 0.02 | 0.002 | 0.002 | 0 | 0.34 | 0.19 | 0.5 | 0.65 | 0.28 | 0.48 |
| 4 | 131125809 | rs6819699 | 1.233 | -0.218 | -0.987 | 0.174 | -0.085 | 0.042 | 0.042 | 0.19 | 0.25 | 0.1 | 0.31 | 0.16 | 0.42 | 0.18 |
| 4 | 131127703 | rs17408481 | 0.913 | -0.163 | -0.613 | 0.433 | -0.003 | -0.027 | -0.027 | 0 | 0.08 | 0.11 | 0.11 | 0 | 0.07 | 0.05 |
| 4 | 131131416 | rs11099040 | -0.503 | -0.544 | -0.001 | 0.385 | -0.032 | 0.023 | 0.023 | 0.766 | 0.25 | 0.46 | 0.21 | 0.32 | 0.23 | 0.22 |
| 4 | 131132900 | rs13113376 | 1.411 | -0.57 | -0.244 | 0.598 | -0.003 | -0.098 | -0.098 | 1.077 | 0.24 | 0.07 | 0.29 | 0.43 | 0.18 | 0.35 |
| 4 | 131137761 | rs1024020 | -0.516 | -0.65 | -0.087 | 0.342 | -0.015 | 0.013 | 0.013 | 0 | 0.66 | 0.63 | 0.67 | 0.5 | 0.8 | 0.61 |
| 4 | 131167056 | rs4449432 | -1.399 | -0.565 | -0.232 | 0.409 | 0.068 | -0.068 | -0.068 | 0.088 | 0.36 | 0.91 | 0.42 | 0.46 | 0.26 | 0.25 |
| 4 | 131169265 | rs7437554 | -0.829 | -0.615 | 0.385 | 0.429 | -0.026 | 0.009 | 0.009 | 3.71 | 0.31 | 0.57 | 0.31 | 0.36 | 0.25 | 0.22 |
| 4 | 131174088 | rs12506063 | -0.464 | -0.613 | 0.127 | 0.437 | -0.012 | 0.005 | 0.005 | 0.124 | 0.26 | 0.36 | 0.28 | 0.31 | 0.25 | 0.2 |
| 4 | 131174498 | rs11099042 | -0.413 | -0.51 | 0.182 | 0.488 | -0.011 | 0.009 | 0.009 | 0.125 | 0.25 | 0.36 | 0.28 | 0.31 | 0.24 | 0.2 |
| 4 | 131175071 | rs6833836 | -0.306 | -0.508 | 0.183 | 0.489 | -0.011 | 0.01 | 0.01 | 0.207 | 0.25 | 0.36 | 0.28 | 0.31 | 0.24 | 0.2 |
| 4 | 131194571 | rs4383635 | -1.267 | -0.446 | -0.823 | 0.358 | -0.103 | 0.256 | 0.256 | 1.003 | 0.59 | 0.95 | 0.72 | 0.92 | 0.49 | 0.67 |
| 4 | 131198147 | rs11944858 | -1.59 | -0.071 | -0.195 | 0.827 | 0.055 | -0.068 | -0.068 | 0 | 0.34 | 0.9 | 0.43 | 0.46 | 0.26 | 0.26 |
| 4 | 131202954 | rs7697995 | -1.553 | -0.502 | -0.179 | 0.842 | 0.053 | -0.062 | -0.062 | 0 | 0.33 | 0.87 | 0.42 | 0.46 | 0.27 | 0.26 |
| 4 | 131206901 | rs17050940 | -0.165 | -0.635 | 0.142 | 0.555 | -0.07 | -0.086 | -0.086 | 0.153 | 0.13 | 0.46 | 0.05 | 0 | 0.02 | 0.04 |
| 4 | 131207582 | rs10024383 | -0.523 | -0.65 | -0.017 | 0.547 | -0.083 | -0.098 | -0.098 | 0.307 | 0.14 | 0.54 | 0.06 | 0 | 0.02 | 0.04 |
| 4 | 131215475 | rs1033018 | -1.61 | -0.508 | -0.73 | 0.858 | 0.04 | -0.076 | -0.076 | 0.113 | 0.32 | 0.86 | 0.43 | 0.43 | 0.27 | 0.21 |
| 4 | 131238094 | rs1022079 | -0.124 | 0.172 | -0.553 | 0.891 | -0.012 | -0.015 | -0.015 | 0 | 0.15 | 0.1 | 0.26 | 0.23 | 0.24 | 0.13 |
| 4 | 131242185 | rs9307624 | 0.04 | -0.629 | -0.516 | 0.733 | -0.018 | -0.001 | -0.001 | 0 | 0.42 | 0.53 | 0.48 | 0.59 | 0.27 | 0.49 |
| 4 | 131242827 | rs9307625 | -0.211 | -0.435 | -0.242 | 0.776 | 0.004 | 0.008 | 0.008 | 0 | 0.34 | 0.5 | 0.58 | 0.32 | 0.4 | 0.26 |
| 4 | 131250760 | rs7662168 | 1.196 | -0.508 | -0.6 | 0.818 | 0.001 | -0.058 | -0.058 | 0 | 0.28 | 0.05 | 0.19 | 0.43 | 0.24 | 0.38 |
| 4 | 131250944 | rs4146036 | -0.39 | -0.549 | -0.517 | 0.789 | 0.003 | -0.015 | -0.015 | 0 | 0.38 | 0.75 | 0.61 | 0.37 | 0.39 | 0.26 |
| 4 | 131252392 | rs10032131 | -1.444 | -0.69 | -0.612 | 0.704 | -0.113 | 0.112 | 0.112 | 0 | 0.71 | 0.98 | 0.84 | 0.9 | 0.64 | 0.73 |
| 4 | 131253096 | rs6815723 | -0.95 | -0.74 | -0.613 | 0.683 | -0.021 | 0.045 | 0.045 | 0 | 0.67 | 0.81 | 0.82 | 0.81 | 0.64 | 0.65 |
| 4 | 131253173 | rs6838581 | -0.95 | -0.74 | -0.613 | 0.686 | -0.021 | 0.044 | 0.044 | 0 | 0.67 | 0.81 | 0.82 | 0.8 | 0.64 | 0.65 |
| 4 | 131253435 | rs6838871 | -1.004 | -0.738 | -0.612 | 0.685 | -0.021 | 0.043 | 0.043 | 0 | 0.67 | 0.81 | 0.82 | 0.81 | 0.64 | 0.65 |
| 4 | 131254621 | rs7688496 | -0.527 | -0.677 | -0.547 | 0.669 | 0.005 | -0.015 | -0.015 | 0.524 | 0.38 | 0.75 | 0.62 | 0.38 | 0.4 | 0.26 |
| 4 | 131263678 | rs1396766 | 0.511 | -0.59 | -0.247 | 0.748 | -0.016 | 0.013 | 0.013 | 0.077 | 0.36 | 0.51 | 0.24 | 0.53 | 0.25 | 0.48 |
| 4 | 131264117 | rs13143102 | 0.09 | -0.519 | -0.558 | 0.762 | -0.041 | 0.018 | 0.018 | 0 | 0.3 | 0.15 | 0.36 | 0.22 | 0.38 | 0.23 |
| 4 | 131265231 | rs9307626 | -0.691 | -0.316 | -0.239 | 0.779 | -0.021 | 0.021 | 0.021 | 0 | 0.67 | 0.85 | 0.64 | 0.76 | 0.63 | 0.72 |
| 4 | 131269444 | rs6827936 | -1.027 | -0.294 | -0.809 | 0.731 | -0.111 | 0.024 | 0.024 | 0.011 | 0.69 | 0.98 | 0.65 | 0.82 | 0.63 | 0.73 |
| 4 | 131272587 | rs2310878 | -1.005 | -0.196 | -0.757 | 0.324 | -0.122 | 0.117 | 0.117 | 0 | 0.71 | 0.98 | 0.84 | 0.9 | 0.64 | 0.74 |
| 4 | 131280307 | rs17050983 | -1.083 | -0.248 | -0.09 | 0.647 | 0 | 0.178 | 0.178 | 3.668 | 0.08 | 0.46 | 0.07 | 0.15 | 0.01 | 0.1 |
| 4 | 131282678 | rs1912101 | -0.959 | 0.11 | -1.087 | 0.198 | -0.132 | 0.106 | 0.106 | 0.131 | 0.73 | 0.98 | 0.84 | 0.9 | 0.63 | 0.7 |
| 4 | 131282886 | rs1509446 | -0.695 | -0.502 | -0.581 | 0.344 | -0.009 | -0.009 | -0.009 | 2.696 | 0.39 | 0.57 | 0.4 | 0.33 | 0.4 | 0.22 |
| 4 | 131286529 | rs16998043 | -0.219 | -0.31 | -0.7 | 0.72 | -0.038 | -0.072 | -0.072 | 0.309 | 0.17 | 0.06 | 0.33 | 0.43 | 0.31 | 0.22 |
| 4 | 131290993 | rs11737037 | -0.44 | -0.712 | 0.021 | 0.472 | -0.003 | -0.009 | -0.009 | 0.004 | 0.05 | 0.16 | 0.05 | 0.01 | 0.04 | 0.05 |
| 4 | 131296975 | rs1581037 | -0.681 | -0.754 | 0.042 | 0.451 | -0.012 | -0.018 | -0.018 | 0 | 0.06 | 0.15 | 0.05 | 0.01 | 0.04 | 0.05 |

| | | | | | | | | | | | | | | | | |
|---|---|---|---|---|---|---|---|---|---|---|---|---|---|---|---|---|
| 4 | 131297208 | rs12509540 | -1.1 | -0.8 | 0.474 | 0.425 | 0.013 | 0.08 | 0.08 | 0 | 0.25 | 0.33 | 0.39 | 0.44 | 0.36 | 0.27 |
| 4 | 131297764 | rs4458467 | -1.64 | -1.166 | 0.163 | -0.371 | -0.034 | 0.031 | 0.031 | 0.024 | 0.73 | 0.82 | 0.83 | 0.98 | 0.64 | 0.87 |
| 4 | 131300750 | rs10033555 | 0.427 | -1.166 | 0.152 | -0.338 | -0.039 | -0.071 | -0.071 | 0 | 0.18 | 0.06 | 0.34 | 0.43 | 0.32 | 0.22 |
| 4 | 131301036 | rs1396757 | 0.06 | -1.544 | 0.246 | -0.223 | -0.014 | 0.062 | 0.062 | 0.129 | 0.11 | 0.23 | 0.09 | 0.27 | 0.05 | 0.2 |
| 4 | 131304008 | rs6534789 | -0.364 | -1.502 | 0.248 | -0.196 | 0.017 | 0.005 | 0.005 | 0 | 0.5 | 0.6 | 0.26 | 0.48 | 0.28 | 0.59 |
| 4 | 131309152 | rs11931985 | 1.604 | -1.895 | 0.14 | -0.483 | 0.027 | 0.015 | 0.015 | 0 | 0.23 | 0.1 | 0.09 | 0.2 | 0.15 | 0.33 |
| 4 | 131311267 | rs11099051 | 0.005 | -1.747 | 0.209 | -0.402 | 0.003 | 0.022 | 0.022 | 0 | 0.23 | 0.2 | 0.39 | 0.44 | 0.36 | 0.27 |
| 4 | 131314946 | rs12331990 | 0.221 | -1.779 | 0.086 | -0.546 | 0.018 | 0.021 | 0.021 | 0 | 0.39 | 0.37 | 0.17 | 0.22 | 0.23 | 0.39 |
| 4 | 131340652 | rs7675553 | -1.936 | -1.594 | -0.174 | -0.569 | -0.057 | 0.075 | 0.075 | 0.042 | 0.73 | 0.87 | 0.84 | 0.98 | 0.64 | 0.87 |
| 4 | 131342779 | rs1355187 | -0.912 | -1.654 | 0.01 | -0.505 | -0.004 | 0.004 | 0.004 | 0 | 0.62 | 0.66 | 0.75 | 0.71 | 0.59 | 0.66 |
| 4 | 131353634 | rs7657931 | 0.804 | -1.62 | -0.213 | -0.484 | -0.002 | 0.023 | 0.023 | 0 | 0.08 | 0.09 | 0.08 | 0.27 | 0.05 | 0.2 |
| 4 | 131354219 | rs4863816 | -1.915 | -1.54 | -0.347 | -0.511 | -0.087 | 0.131 | 0.131 | 0.048 | 0.73 | 0.91 | 0.83 | 0.98 | 0.65 | 0.87 |
| 4 | 131381117 | rs7665833 | 1.659 | -1.567 | -0.342 | -0.297 | 0.018 | -0.035 | -0.035 | 0 | 0.21 | 0.01 | 0.07 | 0.21 | 0.14 | 0.33 |
| 4 | 131389611 | rs10026321 | -0.003 | -1.325 | 0.371 | -0.298 | 0.036 | 0.134 | 0.134 | 1.053 | 0.21 | 0.39 | 0.55 | 0.5 | 0.32 | 0.23 |
| 4 | 131390997 | rs10020348 | -0.554 | -1.375 | 0.458 | -0.348 | 0.014 | 0.086 | 0.086 | 0.319 | 0.25 | 0.52 | 0.58 | 0.5 | 0.35 | 0.27 |
| 4 | 131392589 | rs6813929 | -0.561 | -1.376 | 0.431 | -0.376 | 0.013 | 0.081 | 0.081 | 0.294 | 0.25 | 0.53 | 0.59 | 0.51 | 0.35 | 0.27 |
| 4 | 131392651 | rs10001417 | 0.165 | -1.186 | 0.313 | -0.331 | 0.002 | 0.02 | 0.02 | 0.462 | 0.19 | 0.16 | 0.34 | 0.43 | 0.32 | 0.22 |
| 4 | 131415182 | rs6853908 | -0.165 | -1.538 | 0.369 | -0.273 | -0.009 | 0.002 | 0.002 | 0.003 | 0.4 | 0.55 | 0.43 | 0.5 | 0.24 | 0.56 |
| 4 | 131417196 | rs2926419 | -0.162 | -1.632 | -0.049 | -0.317 | 0.095 | -0.037 | -0.037 | 0.042 | 0.42 | 0.78 | 0.43 | 0.56 | 0.23 | 0.55 |
| 4 | 131438739 | rs6817368 | 0.38 | -1.526 | -0.917 | -0.242 | 0.047 | -0.028 | -0.028 | 0.187 | 0.36 | 0.21 | 0.38 | 0.48 | 0.23 | 0.54 |
| 4 | 131438987 | rs2926412 | 0.077 | -1.178 | -1.079 | -0.427 | 0.017 | 0.06 | 0.06 | 0 | 0.14 | 0.04 | 0.07 | 0 | 0.09 | 0.07 |
| 4 | 131451250 | rs7664079 | -0.663 | -1.253 | -1.311 | -0.68 | -0.079 | 0.206 | 0.206 | 0 | 0.61 | 0.86 | 0.74 | 0.98 | 0.52 | 0.73 |
| 4 | 131451365 | rs12502992 | 0.591 | -1.498 | -1.507 | -0.147 | -0.074 | -0.116 | -0.116 | 0.021 | 0.16 | 0.03 | 0.33 | 0.39 | 0.28 | 0.21 |
| 4 | 131453828 | rs7677917 | 0.041 | -1.62 | -1.138 | -0.28 | 0.081 | -0.035 | -0.035 | 0.03 | 0.42 | 0.71 | 0.41 | 0.54 | 0.23 | 0.54 |
| 4 | 131456285 | rs2969152 | 0.151 | -1.387 | -1.322 | -0.37 | -0.02 | -0.079 | -0.079 | 0.054 | 0.69 | 0.85 | 0.61 | 0.58 | 0.63 | 0.71 |
| 4 | 131457148 | rs2969153 | 0.17 | -1.392 | -1.325 | -0.373 | -0.019 | -0.079 | -0.079 | 0 | 0.69 | 0.85 | 0.61 | 0.58 | 0.63 | 0.71 |
| 4 | 131459940 | rs1459308 | 0.11 | -1.67 | -1.121 | -0.141 | -0.005 | 0.017 | 0.017 | 0 | 0.1 | 0.1 | 0.27 | 0.31 | 0.05 | 0.14 |
| 4 | 131468686 | rs13105686 | 0.996 | -1.781 | -0.555 | -0.162 | -0.005 | -0.008 | -0.008 | 0 | 0.29 | 0.26 | 0.12 | 0.17 | 0.18 | 0.38 |
| 4 | 131473531 | rs4864001 | -0.632 | -1.56 | -0.856 | -0.029 | 0.065 | -0.028 | -0.028 | 0.168 | 0.57 | 0.81 | 0.3 | 0.49 | 0.33 | 0.61 |
| 4 | 131484721 | rs11099064 | 1.633 | -1.389 | -0.816 | -0.103 | -0.074 | 0.079 | 0.079 | 0 | 0.27 | 0.12 | 0.37 | 0.1 | 0.39 | 0.19 |
| 4 | 131494414 | rs11945492 | -1.015 | -1.471 | -0.892 | -0.011 | 0.004 | -0.087 | -0.087 | 0.137 | 0.6 | 0.91 | 0.5 | 0.56 | 0.36 | 0.61 |
| 4 | 131496373 | rs10518572 | 1.103 | -1.528 | -0.903 | -0.092 | -0.001 | 0.073 | 0.073 | 0 | 0.12 | 0 | 0.05 | 0 | 0.09 | 0.07 |
| 4 | 131498335 | rs12642857 | 0.494 | -1.68 | -0.624 | -0.13 | -0.025 | 0.002 | 0.002 | 0 | 0.37 | 0.38 | 0.22 | 0.44 | 0.25 | 0.5 |
| 4 | 131499709 | rs2169718 | -0.048 | -1.436 | -1.495 | -0.105 | 0.093 | -0.012 | -0.012 | 0 | 0.45 | 0.79 | 0.27 | 0.5 | 0.27 | 0.54 |
| 4 | 131501118 | rs17051049 | 0.754 | -0.965 | -1.227 | 0.405 | -0.008 | 0.023 | 0.023 | 0 | 0.12 | 0 | 0.23 | 0.07 | 0.09 | 0.07 |
| 4 | 131503899 | rs4273485 | -0.403 | -0.637 | -0.846 | 0.695 | -0.005 | -0.001 | -0.001 | 0 | 0.85 | 0.84 | 0.69 | 0.59 | 0.77 | 0.8 |
| 4 | 131508305 | rs13125488 | 0.564 | -0.913 | -0.872 | 0.156 | 0.029 | 0.023 | 0.023 | 0 | 0.23 | 0.12 | 0.09 | 0.17 | 0.16 | 0.34 |
| 4 | 131518894 | rs4479720 | -0.28 | -0.766 | -0.746 | 0.568 | -0.001 | 0 | 0 | 0 | 0.85 | 0.84 | 0.77 | 0.86 | 0.83 | 0.82 |
| 4 | 131524184 | rs899131 | -0.209 | -0.848 | -0.586 | 0.78 | -0.006 | 0.028 | 0.028 | 0 | 0.12 | 0.11 | 0.09 | 0.28 | 0.08 | 0.09 |
| 4 | 131528197 | rs1459288 | 0.674 | -0.816 | -0.622 | 0.688 | -0.006 | 0.024 | 0.024 | 0 | 0.74 | 0.73 | 0.67 | 0.58 | 0.74 | 0.73 |
| 4 | 131528824 | rs10518574 | 1.345 | -0.736 | -1.147 | -0.099 | -0.083 | 0.136 | 0.136 | 0.016 | 0.3 | 0.06 | 0.21 | 0.05 | 0.39 | 0.22 |
| 4 | 131532350 | rs2125639 | 0.725 | -1.037 | -0.414 | 0.125 | 0.007 | -0.026 | -0.026 | 1.319 | 0.2 | 0.24 | 0.09 | 0.13 | 0.22 | 0.26 |
| 4 | 131536250 | rs2085678 | 0.943 | -1.054 | -0.398 | 0.117 | -0.023 | 0.04 | 0.04 | 0 | 0.19 | 0.1 | 0.08 | 0.13 | 0.21 | 0.26 |
| 4 | 131539495 | rs2871751 | 0.255 | -1.217 | -0.163 | 0.26 | 0 | -0.024 | -0.024 | 0 | 0.23 | 0.29 | 0.29 | 0.19 | 0.23 | 0.28 |
| 4 | 131540018 | rs7668962 | 1.747 | -1.433 | -0.544 | -0.045 | -0.132 | 0.193 | 0.193 | 0 | 0.34 | 0.05 | 0.22 | 0.06 | 0.44 | 0.24 |
| 4 | 131541099 | rs7675752 | 1.218 | -1.398 | -0.452 | 0.058 | -0.02 | 0.039 | 0.039 | 0 | 0.18 | 0.1 | 0.09 | 0.13 | 0.21 | 0.26 |
| 4 | 131545432 | rs17051056 | -0.237 | -1.71 | -0.071 | 0.385 | -0.081 | 0.074 | 0.074 | 1.883 | 0.26 | 0.48 | 0.2 | 0.58 | 0.12 | 0.28 |
| 4 | 131551052 | rs7659897 | 0.362 | -1.702 | 0.25 | 0.737 | -0.014 | 0.01 | 0.01 | 0.166 | 0.18 | 0.23 | 0.13 | 0.29 | 0.08 | 0.21 |
| 4 | 131558064 | rs1948210 | 0.423 | -1.348 | 0.12 | 0.565 | 0.005 | -0.008 | -0.008 | 1.934 | 0.6 | 0.53 | 0.43 | 0.62 | 0.56 | 0.53 |
| 4 | 131561136 | rs2054931 | 0.39 | -1.24 | 0.207 | 0.739 | -0.001 | -0.001 | -0.001 | 0.148 | 0.23 | 0.23 | 0.3 | 0.29 | 0.23 | 0.28 |
| 4 | 131568230 | rs6534808 | 0.428 | -1.463 | 0.216 | 0.626 | -0.017 | 0.032 | 0.032 | 0 | 0.77 | 0.59 | 0.53 | 0.76 | 0.77 | 0.76 |
| 4 | 131569148 | rs1304183 | 1.459 | -1.472 | -0.502 | 0.516 | 0.049 | 0.138 | 0.138 | 0.528 | 0.46 | 0.19 | 0.33 | 0.3 | 0.6 | 0.48 |
| 4 | 131578668 | rs7687787 | 1.614 | -1.378 | -0.633 | 0.411 | -0.123 | 0.144 | 0.144 | 0.062 | 0.36 | 0.06 | 0.24 | 0.13 | 0.45 | 0.29 |
| 4 | 131589008 | rs1387452 | -0.283 | -1.346 | 0.162 | 0.771 | -0.034 | 0.019 | 0.019 | 0 | 0.26 | 0.42 | 0.17 | 0.26 | 0.15 | 0.24 |
| 4 | 131589420 | rs1032272 | -0.634 | -1.23 | 0.152 | 0.9 | 0.002 | 0.012 | 0.012 | 0 | 0.83 | 0.81 | 0.73 | 0.71 | 0.81 | 0.77 |
| 4 | 131594623 | rs1994541 | 1.762 | -1.265 | 0.052 | 0.879 | -0.003 | 0.003 | 0.003 | 0 | 0.15 | 0.15 | 0.09 | 0.16 | 0.18 | 0.21 |
| 4 | 131596752 | rs10006622 | -0.16 | -1.301 | 0.196 | 1.366 | 0.009 | 0.027 | 0.027 | 0 | 0.82 | 0.75 | 0.71 | 0.71 | 0.8 | 0.76 |
| 4 | 131599180 | rs4864014 | -0.618 | -1.045 | -0.255 | 1.265 | 0.015 | -0.016 | -0.016 | 0.526 | 0.24 | 0.24 | 0.15 | 0.27 | 0.14 | 0.23 |
| 4 | 131606179 | rs1844327 | 1 | -0.713 | -0.147 | 0.015 | -0.002 | -0.025 | -0.025 | 1.161 | 0.73 | 0.67 | 0.7 | 0.86 | 0.71 | 0.71 |
| 4 | 131611683 | rs7674705 | 0.587 | -0.602 | -0.656 | 0.048 | -0.024 | 0.041 | 0.041 | 0 | 0.76 | 0.86 | 0.7 | 0.88 | 0.7 | 0.78 |
| 4 | 131629960 | rs2311780 | -0.847 | -0.944 | -0.398 | 0.019 | -0.005 | -0.005 | -0.005 | 0 | 0.37 | 0.54 | 0.49 | 0.58 | 0.35 | 0.35 |
| 4 | 131634859 | rs931418 | 0.291 | -1.236 | -0.45 | -0.049 | -0.022 | 0.004 | 0.004 | 2.582 | 0.2 | 0.25 | 0.08 | 0.22 | 0.09 | 0.21 |
| 4 | 131638465 | rs6811023 | -0.48 | -0.852 | 0.123 | -0.119 | -0.016 | -0.039 | -0.039 | 0 | 0.15 | 0.35 | 0.13 | 0.08 | 0.12 | 0.11 |
| 4 | 131641319 | rs10009557 | 0.058 | -0.875 | -0.183 | -0.262 | 0.005 | 0.013 | 0.013 | 0 | 0.11 | 0.08 | 0.06 | 0 | 0.13 | 0.11 |
| 4 | 131641387 | rs306344 | 0.932 | -0.907 | -0.217 | -0.471 | 0.019 | 0.08 | 0.08 | 0 | 0.5 | 0.34 | 0.45 | 0.34 | 0.57 | 0.52 |
| 4 | 131647396 | rs385027 | -0.685 | -0.893 | -0.047 | -0.406 | -0.003 | -0.014 | -0.014 | 0.003 | 0.93 | 0.89 | 0.92 | 0.95 | 0.96 | 0.97 |
| 4 | 131649821 | rs306369 | 0.991 | -0.89 | 0.02 | -0.421 | 0.014 | 0.052 | 0.052 | 0 | 0.5 | 0.38 | 0.45 | 0.34 | 0.57 | 0.52 |
| 4 | 131650867 | rs17051076 | 1.123 | -0.883 | -0.149 | -0.291 | -0.021 | -0.018 | -0.018 | 0 | 0.05 | 0 | 0.03 | 0.09 | 0.05 | 0.07 |
| 4 | 131652041 | rs306364 | 1.19 | -0.851 | 0.184 | -0.262 | 0.002 | 0.005 | 0.005 | 0 | 0.5 | 0.47 | 0.46 | 0.34 | 0.57 | 0.52 |

| | | | | | | | | | | | | | | | | |
|---|---|---|---|---|---|---|---|---|---|---|---|---|---|---|---|---|
| 4 | 131652629 | rs17051078 | 0.375 | -0.896 | -0.238 | -0.22 | 0 | -0.001 | -0.001 | 0.095 | 0.12 | 0.12 | 0.09 | 0.04 | 0.14 | 0.11 |
| 4 | 131662183 | rs306353 | 0.7 | -0.686 | -0.059 | -0.012 | -0.008 | -0.01 | -0.01 | 0 | 0.62 | 0.54 | 0.54 | 0.38 | 0.71 | 0.63 |
| 4 | 131662676 | rs990046 | 0.376 | -0.487 | -0.715 | 0.348 | -0.04 | -0.113 | -0.113 | 0 | 0.18 | 0.02 | 0.31 | 0.47 | 0.18 | 0.22 |
| 4 | 131671137 | rs17051084 | -0.02 | -0.99 | -0.634 | -0.551 | 0.027 | 0.016 | 0.016 | 4.192 | 0.25 | 0.18 | 0.09 | 0 | 0.17 | 0.12 |
| 4 | 131672725 | rs11938827 | -0.031 | -0.967 | -0.422 | -0.505 | 0.019 | 0.025 | 0.025 | 3.213 | 0.25 | 0.19 | 0.13 | 0.03 | 0.18 | 0.14 |
| 4 | 131680834 | rs4146375 | 0.511 | -0.721 | -0.124 | 0.352 | 0.002 | 0.067 | 0.067 | 0 | 0.17 | 0.2 | 0.35 | 0.43 | 0.15 | 0.23 |
| 4 | 131695431 | rs3113500 | 0.12 | -0.555 | -0.15 | -0.046 | -0.004 | -0.001 | -0.001 | 0 | 0.7 | 0.65 | 0.73 | 0.63 | 0.75 | 0.67 |
| 4 | 131695570 | rs3103398 | -0.045 | -0.558 | -0.223 | -0.128 | -0.011 | -0.003 | -0.003 | 0.002 | 0.76 | 0.69 | 0.76 | 0.72 | 0.81 | 0.74 |
| 4 | 131696764 | rs424918 | -0.688 | -0.578 | -0.223 | -0.129 | -0.02 | -0.031 | -0.031 | 0.155 | 0.85 | 0.72 | 0.94 | 0.93 | 0.93 | 0.91 |
| 4 | 131700214 | rs306377 | 1.714 | -0.367 | -0.336 | -0.075 | 0.027 | -0.005 | -0.005 | 0 | 0.26 | 0.28 | 0.24 | 0.21 | 0.37 | 0.25 |
| 4 | 131701952 | rs12505414 | -0.004 | -0.427 | -0.763 | 0.082 | 0 | -0.036 | -0.036 | 0.294 | 0.27 | 0.16 | 0.39 | 0.51 | 0.21 | 0.36 |
| 4 | 131714357 | rs11099081 | 0.791 | -0.113 | -0.841 | -0.058 | -0.029 | -0.03 | -0.03 | 0 | 0.07 | 0 | 0.15 | 0.16 | 0.13 | 0.17 |
| 4 | 131723626 | rs306393 | 0.687 | -0.376 | -0.582 | -0.327 | 0.019 | 0.077 | 0.077 | 0 | 0.48 | 0.3 | 0.62 | 0.67 | 0.56 | 0.56 |
| 4 | 131724116 | rs11099082 | -0.263 | -0.542 | -0.577 | -0.421 | -0.011 | -0.025 | -0.025 | 2.093 | 0.14 | 0.06 | 0.19 | 0.28 | 0.16 | 0.2 |
| 4 | 131727377 | rs2312337 | -1.063 | -0.781 | -0.147 | -0.598 | -0.014 | -0.035 | -0.035 | 0 | 0.86 | 0.79 | 0.95 | 0.96 | 0.93 | 0.91 |
| 4 | 131727827 | rs1824748 | 1.592 | -0.759 | -0.71 | 0.007 | -0.048 | -0.061 | -0.061 | 0 | 0.15 | 0.01 | 0.12 | 0.16 | 0.14 | 0.12 |
| 4 | 131727847 | rs1478075 | 0.269 | -0.858 | -0.305 | -0.292 | 0.034 | 0.05 | 0.05 | 0.419 | 0.6 | 0.27 | 0.77 | 0.92 | 0.71 | 0.76 |
| 4 | 131747478 | rs12651109 | 0.787 | -0.609 | -0.236 | 0.191 | -0.001 | 0.006 | 0.006 | 0 | 0.06 | 0.07 | 0.15 | 0.2 | 0.09 | 0.14 |
| 4 | 131747961 | rs12645823 | 0.787 | -0.609 | -0.236 | 0.191 | -0.001 | 0.006 | 0.006 | 0 | 0.06 | 0.07 | 0.15 | 0.2 | 0.09 | 0.14 |
| 4 | 131750641 | rs6854661 | -0.154 | -0.657 | 0.157 | 0.166 | 0.005 | 0.054 | 0.054 | 0.272 | 0.08 | 0.21 | 0.16 | 0.2 | 0.09 | 0.14 |
| 4 | 131750954 | rs6811623 | 0.133 | -0.638 | 0.045 | 0.176 | -0.001 | 0.024 | 0.024 | 0.196 | 0.08 | 0.15 | 0.16 | 0.2 | 0.09 | 0.14 |
| 4 | 131772771 | rs6836232 | 0.742 | -0.607 | 0.023 | 0.186 | 0.001 | 0.016 | 0.016 | 0 | 0.06 | 0.12 | 0.15 | 0.18 | 0.09 | 0.13 |
| 4 | 131778067 | rs1382171 | 0.128 | -0.825 | 0.479 | -0.031 | -0.021 | -0.011 | -0.011 | 0 | 0.8 | 0.56 | 0.79 | 0.78 | 0.85 | 0.79 |
| 4 | 131779157 | rs7440566 | 1.061 | -0.97 | 0.241 | -0.367 | 0.026 | 0.111 | 0.111 | 0 | 0.48 | 0.24 | 0.62 | 0.7 | 0.56 | 0.58 |
| 4 | 131782781 | rs13133329 | 0.985 | -0.997 | 0.245 | -0.362 | 0.036 | 0.124 | 0.124 | 0 | 0.49 | 0.23 | 0.62 | 0.7 | 0.56 | 0.58 |
| 4 | 131787782 | rs2035213 | -0.916 | -1.062 | 0.657 | -0.387 | -0.014 | -0.04 | -0.04 | 3.149 | 0.86 | 0.77 | 0.95 | 0.96 | 0.93 | 0.91 |
| 4 | 131788216 | rs6824018 | 0.333 | -1.107 | 0.685 | -0.46 | -0.008 | -0.055 | -0.055 | 0 | 0.62 | 0.37 | 0.77 | 0.93 | 0.71 | 0.76 |
| 4 | 131790501 | rs1478091 | -0.542 | -1.083 | 0.721 | -0.377 | -0.037 | -0.058 | -0.058 | 0 | 0.84 | 0.54 | 0.9 | 0.93 | 0.92 | 0.91 |
| 4 | 131792029 | rs6850860 | -0.145 | -1.003 | 0.315 | 0.263 | -0.003 | 0.005 | 0.005 | 0 | 0.79 | 0.76 | 0.85 | 0.71 | 0.84 | 0.82 |
| 4 | 131804232 | rs1478095 | 0.48 | -1.089 | 0.273 | 0.15 | 0.01 | 0.013 | 0.013 | 0.125 | 0.58 | 0.61 | 0.73 | 0.73 | 0.63 | 0.75 |
| 4 | 131804615 | rs2126823 | -0.428 | -1.08 | 0.16 | 0.204 | 0.007 | 0.028 | 0.028 | 0.01 | 0.35 | 0.26 | 0.28 | 0.25 | 0.35 | 0.32 |
| 4 | 131809519 | rs12711163 | 2.183 | -1.013 | -0.027 | 0.266 | -0.039 | -0.035 | -0.035 | 0 | 0.25 | 0.12 | 0.4 | 0.34 | 0.34 | 0.36 |
| 4 | 131811462 | rs13116382 | 1.096 | -1.011 | -0.088 | 0.328 | -0.007 | -0.002 | -0.002 | 0 | 0.06 | 0.01 | 0.11 | 0.25 | 0.09 | 0.08 |
| 4 | 131814499 | rs6847750 | 1.258 | -1.021 | 0.082 | 0.319 | -0.015 | -0.013 | -0.013 | 0 | 0.42 | 0.27 | 0.5 | 0.42 | 0.49 | 0.5 |
| 4 | 131824376 | rs1382177 | 0.961 | -1.016 | -0.274 | 0.16 | 0.016 | 0.077 | 0.077 | 0.251 | 0.21 | 0.1 | 0.12 | 0 | 0.21 | 0.14 |
| 4 | 131834405 | rs4864021 | 0.925 | -0.896 | -0.017 | 0.723 | 0.003 | -0.005 | -0.005 | 0.247 | 0.64 | 0.37 | 0.63 | 0.46 | 0.7 | 0.59 |
| 4 | 131838432 | rs11941740 | 0.426 | -0.897 | -0.017 | 0.74 | -0.015 | 0.028 | 0.028 | 0.12 | 0.7 | 0.43 | 0.78 | 0.57 | 0.77 | 0.76 |
| 4 | 131838796 | rs10010210 | 1.123 | -0.628 | -0.255 | 0.672 | -0.002 | -0.003 | -0.003 | 0.355 | 0.06 | 0.03 | 0.12 | 0.13 | 0.07 | 0.17 |
| 4 | 131856379 | rs11724421 | 0.772 | -0.593 | -0.21 | 0.597 | -0.004 | 0.002 | 0.002 | 0 | 0.08 | 0.05 | 0.15 | 0.11 | 0.12 | 0.22 |
| 4 | 131857035 | rs7658306 | 0.375 | -1.084 | -0.059 | 0.645 | -0.003 | 0.032 | 0.032 | 0 | 0.48 | 0.64 | 0.58 | 0.33 | 0.54 | 0.39 |
| 4 | 131866484 | rs1846790 | -1.241 | -1.072 | -0.429 | 0.809 | -0.005 | 0.019 | 0.019 | 0.149 | 0.89 | 0.89 | 0.89 | 0.78 | 0.93 | 0.94 |
| 4 | 131866685 | rs12498616 | 0.916 | -0.99 | -0.599 | 0.426 | 0.015 | -0.046 | -0.046 | 0 | 0.19 | 0.07 | 0.31 | 0.27 | 0.17 | 0.18 |
| 4 | 131866907 | rs9994110 | 1.54 | -0.928 | -0.116 | 0.247 | 0.024 | -0.085 | -0.085 | 0.638 | 0.24 | 0.29 | 0.21 | 0.03 | 0.37 | 0.21 |
| 4 | 131868112 | rs2055392 | 1.164 | -0.908 | -0.206 | 0.249 | 0.029 | -0.12 | -0.12 | 0.345 | 0.27 | 0.37 | 0.22 | 0.03 | 0.38 | 0.28 |
| 4 | 131880992 | rs12508862 | 0.847 | -1.05 | -0.601 | 0.675 | 0.019 | -0.045 | -0.045 | 0.607 | 0.16 | 0.07 | 0.29 | 0.28 | 0.13 | 0.15 |
| 4 | 131884313 | rs6830509 | -0.682 | -0.89 | -0.666 | 0.575 | -0.01 | -0.023 | -0.023 | 0 | 0.64 | 0.78 | 0.6 | 0.56 | 0.63 | 0.58 |
| 4 | 131886381 | rs2312546 | 0.53 | -0.97 | -0.349 | 0.482 | 0.017 | -0.063 | -0.063 | 0 | 0.36 | 0.45 | 0.25 | 0.17 | 0.41 | 0.31 |
| 4 | 131887689 | rs2872072 | 1.777 | -0.803 | -0.504 | 0.32 | 0.021 | -0.093 | -0.093 | 1.493 | 0.23 | 0.26 | 0.19 | 0.01 | 0.33 | 0.21 |
| 4 | 131897140 | rs1948720 | -0.286 | -0.906 | -0.484 | 0.679 | 0 | -0.002 | -0.002 | 0 | 0.54 | 0.59 | 0.6 | 0.47 | 0.44 | 0.54 |
| 4 | 131905081 | rs17445005 | 0.973 | -0.837 | -0.832 | 0.79 | -0.006 | -0.015 | -0.015 | 0 | 0.09 | 0.02 | 0.04 | 0.11 | 0.08 | 0.12 |
| 4 | 131916429 | rs1500515 | -0.397 | -0.87 | -0.753 | 0.736 | -0.001 | 0.001 | 0.001 | 1.783 | 0.53 | 0.56 | 0.61 | 0.48 | 0.45 | 0.55 |
| 4 | 131916948 | rs11941696 | 1.321 | -1.071 | -1.109 | 0.56 | 0.01 | -0.021 | -0.021 | 0 | 0.11 | 0.07 | 0.28 | 0.29 | 0.09 | 0.13 |
| 4 | 131920334 | rs12647096 | 0.667 | -0.95 | -1.175 | 0.066 | -0.027 | 0.091 | 0.091 | 0.612 | 0.32 | 0.16 | 0.27 | 0.08 | 0.31 | 0.37 |
| 4 | 131938086 | rs9996563 | 1.38 | -0.808 | -1.198 | 0.129 | 0.016 | -0.031 | -0.031 | 0 | 0.32 | 0.34 | 0.21 | 0.23 | 0.39 | 0.3 |
| 4 | 131939529 | rs12642294 | 0.807 | -0.828 | -1.264 | 0.097 | -0.027 | 0.09 | 0.09 | 0.661 | 0.32 | 0.16 | 0.27 | 0.08 | 0.31 | 0.37 |
| 4 | 131941333 | rs10006329 | 0.62 | -0.84 | -1.234 | 0.183 | -0.001 | -0.001 | -0.001 | 0 | 0.42 | 0.35 | 0.27 | 0.35 | 0.49 | 0.42 |
| 4 | 131946375 | rs6822842 | -0.153 | -0.349 | -1.52 | 0.272 | -0.043 | 0.103 | 0.103 | 0 | 0.42 | 0.19 | 0.32 | 0.21 | 0.41 | 0.5 |
| 4 | 131963445 | rs17051142 | 1.151 | -0.031 | -1.301 | 0.903 | -0.027 | -0.029 | -0.029 | 0.075 | 0.06 | 0 | 0.06 | 0.23 | 0.05 | 0.05 |
| 4 | 131965131 | rs3104157 | -0.342 | -0.451 | -1.687 | 0.755 | -0.021 | 0.038 | 0.038 | 1.651 | 0.62 | 0.8 | 0.63 | 0.71 | 0.61 | 0.59 |
| 4 | 131969350 | rs3104159 | -1.508 | -0.778 | -1.864 | 0.588 | 0.008 | -0.01 | -0.01 | 0.011 | 0.9 | 0.93 | 0.9 | 0.8 | 0.92 | 0.93 |
| 4 | 131972377 | rs13145260 | 0.608 | -1.151 | -1.296 | 0.47 | 0.004 | -0.001 | -0.001 | 0 | 0.42 | 0.4 | 0.29 | 0.43 | 0.48 | 0.43 |
| 4 | 131983567 | rs2220898 | 1.157 | -1.06 | -1.128 | 0.595 | 0.023 | -0.028 | -0.028 | 0 | 0.15 | 0.08 | 0.31 | 0.24 | 0.13 | 0.16 |
| 4 | 131985143 | rs440783 | 0.725 | -0.79 | -1.019 | -0.091 | -0.003 | 0.001 | 0.001 | 0 | 0.36 | 0.6 | 0.27 | 0.33 | 0.38 | 0.3 |
| 4 | 131998959 | rs17446401 | 1.398 | -0.786 | -1.041 | -0.122 | -0.032 | -0.041 | -0.041 | 0 | 0.09 | 0.01 | 0.05 | 0.13 | 0.11 | 0.12 |
| 4 | 132008409 | rs7692456 | -0.37 | -0.69 | -0.963 | -0.084 | -0.008 | -0.004 | -0.004 | 0 | 0.54 | 0.39 | 0.68 | 0.56 | 0.51 | 0.58 |
| 4 | 132024484 | rs4864029 | -0.381 | -0.662 | -0.894 | 0.074 | 0.006 | 0.048 | 0.048 | 3.59 | 0.88 | 0.71 | 0.92 | 0.86 | 0.89 | 0.89 |
| 4 | 132051076 | rs10031771 | -0.923 | -0.586 | -1.132 | -0.286 | 0.005 | -0.013 | -0.013 | 0 | 0.59 | 0.48 | 0.73 | 0.71 | 0.55 | 0.64 |
| 4 | 132057086 | rs12651384 | -1.439 | -0.609 | -0.998 | -1.163 | 0.003 | -0.114 | -0.114 | 2.18 | 0.69 | 0.49 | 0.82 | 0.96 | 0.67 | 0.73 |
| 4 | 132080596 | rs7690099 | -0.543 | -0.631 | -1.075 | -1.096 | 0.004 | -0.015 | -0.015 | 0.24 | 0.61 | 0.47 | 0.76 | 0.8 | 0.56 | 0.61 |

| | | | | | | | | | | | | | | | | |
|---|---|---|---|---|---|---|---|---|---|---|---|---|---|---|---|---|
| 4 | 132082388 | rs12509451 | 1.406 | -0.602 | -1.129 | -0.925 | -0.034 | -0.044 | -0.044 | 0 | 0.07 | 0 | 0.05 | 0.13 | 0.1 | 0.12 |
| 4 | 132118585 | rs17447684 | 1.839 | -0.676 | -1.079 | -0.968 | -0.001 | -0.012 | -0.012 | 0 | 0.06 | 0.05 | 0.03 | 0 | 0.04 | 0.11 |
| 4 | 132125839 | rs500131 | -0.279 | -0.904 | -0.712 | -1.294 | -0.008 | 0.036 | 0.036 | 0 | 0.56 | 0.69 | 0.32 | 0.2 | 0.55 | 0.55 |
| 4 | 132126271 | rs10018513 | 0.391 | -0.937 | -0.706 | -1.348 | -0.001 | -0.001 | -0.001 | 0.173 | 0.16 | 0.18 | 0.09 | 0.13 | 0.16 | 0.16 |
| 4 | 132129741 | rs490321 | 1.614 | -0.955 | -0.845 | -1.379 | 0.012 | -0.101 | -0.101 | 0.066 | 0.3 | 0.42 | 0.18 | 0.04 | 0.34 | 0.26 |
| 4 | 132134734 | rs9999452 | -0.846 | -0.938 | -1.122 | -1.315 | -0.048 | 0.017 | 0.017 | 0 | 0.64 | 0.84 | 0.45 | 0.25 | 0.62 | 0.79 |
| 4 | 132143726 | rs6811871 | 1.113 | -0.808 | -1.006 | -1.29 | 0.001 | -0.067 | -0.067 | 0 | 0.36 | 0.4 | 0.31 | 0.09 | 0.4 | 0.5 |
| 4 | 132157647 | rs10021892 | 0.103 | -0.687 | -1.197 | -0.717 | 0.006 | -0.005 | -0.005 | 0 | 0.16 | 0.08 | 0.09 | 0.13 | 0.16 | 0.16 |
| 4 | 132169789 | rs6855567 | -0.725 | -0.638 | -1.39 | -0.772 | -0.011 | -0.044 | -0.044 | 1.047 | 0.76 | 0.9 | 0.56 | 0.27 | 0.77 | 0.74 |
| 4 | 132177261 | rs12507587 | 0.249 | -0.564 | -1.331 | -0.743 | 0.001 | -0.007 | -0.007 | 0 | 0.15 | 0.06 | 0.08 | 0.13 | 0.16 | 0.16 |
| 4 | 132177937 | rs17051169 | 0.725 | -0.68 | -1.166 | -1.072 | 0.009 | 0.075 | 0.075 | 0 | 0.53 | 0.63 | 0.44 | 0.15 | 0.57 | 0.56 |
| 4 | 132179341 | rs11938816 | -0.048 | -0.369 | -1.016 | -0.919 | -0.025 | -0.074 | -0.074 | 0 | 0.14 | 0.24 | 0.07 | 0 | 0.1 | 0.06 |
| 4 | 132179527 | rs6534832 | 0.782 | -0.764 | -0.756 | -1.133 | -0.001 | 0 | 0 | 0 | 0.42 | 0.43 | 0.43 | 0.14 | 0.54 | 0.44 |
| 4 | 132183047 | rs11943774 | -0.986 | -0.788 | -0.692 | -0.83 | 0 | -0.001 | -0.001 | 0 | 0.27 | 0.27 | 0.1 | 0.1 | 0.19 | 0.22 |
| 4 | 132189098 | rs6534833 | -0.685 | -0.491 | -0.822 | -0.89 | -0.001 | 0.004 | 0.004 | 3.236 | 0.77 | 0.8 | 0.75 | 0.84 | 0.77 | 0.79 |
| 4 | 132192738 | rs9992696 | 0.327 | -0.486 | -0.882 | -0.976 | 0.016 | 0.062 | 0.062 | 10.016 | 0.64 | 0.78 | 0.66 | 0.84 | 0.67 | 0.74 |
| 4 | 132196533 | rs11728768 | 1.141 | 0.109 | -0.912 | -0.814 | -0.056 | -0.028 | -0.028 | 0.631 | 0.11 | 0 | 0.09 | 0.08 | 0.17 | 0.14 |
| 4 | 132240434 | rs10000209 | -0.094 | -0.094 | -0.3 | -0.158 | -0.004 | 0.042 | 0.042 | 0 | 0.31 | 0.46 | 0.38 | 0.44 | 0.32 | 0.37 |
| 4 | 132242896 | rs4133013 | -1.436 | 0.279 | -0.624 | -0.255 | 0.006 | -0.02 | -0.02 | 0 | 0.61 | 0.73 | 0.57 | 0.52 | 0.68 | 0.57 |
| 4 | 132252355 | rs7676822 | 0.047 | 0.291 | -0.703 | -0.391 | 0.006 | 0.042 | 0.042 | 0.919 | 0.33 | 0.21 | 0.35 | 0.17 | 0.31 | 0.3 |
| 4 | 132257490 | rs1039782 | -1.229 | 0.269 | -0.436 | -0.142 | -0.004 | -0.002 | -0.002 | 0 | 0.55 | 0.54 | 0.55 | 0.53 | 0.65 | 0.55 |
| 4 | 132288496 | rs1318943 | 0.983 | 0.044 | -0.587 | -0.002 | -0.056 | -0.028 | -0.028 | 0.072 | 0.13 | 0.01 | 0.09 | 0.09 | 0.21 | 0.12 |
| 4 | 132298239 | rs1911877 | -0.461 | -0.093 | -0.379 | -0.205 | -0.004 | -0.024 | -0.024 | 0 | 0.37 | 0.37 | 0.38 | 0.16 | 0.34 | 0.32 |
| 4 | 132305574 | rs4864039 | -1.853 | -0.319 | -0.417 | -0.249 | -0.014 | -0.014 | -0.014 | 1 | 0.58 | 0.7 | 0.52 | 0.54 | 0.58 | 0.62 |
| 4 | 132310610 | rs6534838 | -0.054 | -0.372 | -0.352 | -0.33 | 0.004 | 0.056 | 0.056 | 0 | 0.34 | 0.21 | 0.34 | 0.17 | 0.32 | 0.37 |
| 4 | 132328214 | rs6811541 | -1.293 | -0.507 | -0.495 | -0.35 | -0.005 | 0.059 | 0.059 | 0 | 0.51 | 0.66 | 0.39 | 0.34 | 0.52 | 0.45 |
| 4 | 132347889 | rs17051193 | 1.048 | -0.453 | -0.612 | -0.333 | 0.01 | -0.041 | -0.041 | 0.338 | 0.07 | 0 | 0.13 | 0.2 | 0.06 | 0.17 |
| 4 | 132351485 | rs17457419 | 1.038 | -0.221 | -0.503 | -0.538 | -0.053 | 0.018 | 0.018 | 0.176 | 0.14 | 0.02 | 0.09 | 0.04 | 0.21 | 0.12 |
| 4 | 132351655 | rs7690600 | -0.444 | -0.429 | -0.015 | -0.384 | -0.003 | -0.122 | -0.122 | 0 | 0.31 | 0.32 | 0.25 | 0.02 | 0.27 | 0.22 |
| 4 | 132360407 | rs7441235 | -1.133 | -0.418 | -0.133 | -0.268 | 0.013 | 0.015 | 0.015 | 0.02 | 0.37 | 0.66 | 0.31 | 0.31 | 0.3 | 0.33 |
| 4 | 132363754 | rs6823370 | -0.725 | -0.558 | 0.139 | -0.257 | -0.008 | 0.201 | 0.201 | 1.494 | 0.08 | 0.3 | 0.08 | 0.29 | 0.04 | 0.11 |
| 4 | 132371344 | rs1280680 | -0.956 | -0.344 | -0.134 | -0.26 | -0.007 | -0.006 | -0.006 | 1.446 | 0.36 | 0.54 | 0.28 | 0.31 | 0.28 | 0.32 |
| 4 | 132381454 | rs2662065 | 1.682 | 0.088 | -0.481 | -0.056 | -0.058 | 0.042 | 0.042 | 0.246 | 0.1 | 0.01 | 0.15 | 0.02 | 0.18 | 0.05 |
| 4 | 132392727 | rs971005 | 0.077 | -0.346 | -0.275 | -0.272 | -0.001 | -0.048 | -0.048 | 0.84 | 0.23 | 0.18 | 0.19 | 0.03 | 0.22 | 0.19 |
| 4 | 132399049 | rs1280687 | -1.138 | -0.371 | -0.842 | -0.213 | 0.017 | 0.009 | 0.009 | 6.891 | 0.39 | 0.65 | 0.31 | 0.34 | 0.31 | 0.34 |
| 4 | 132408562 | rs1280699 | -0.72 | -0.412 | -0.754 | -0.202 | 0.002 | -0.001 | -0.001 | 0 | 0.34 | 0.58 | 0.26 | 0.34 | 0.26 | 0.33 |
| 4 | 132409029 | rs351098 | -0.275 | -0.429 | -0.691 | -0.46 | -0.022 | -0.102 | -0.102 | 0 | 0.3 | 0.46 | 0.21 | 0.05 | 0.23 | 0.27 |
| 4 | 132409579 | rs351096 | -0.912 | -0.576 | -1.176 | -0.035 | 0.029 | -0.002 | -0.002 | 0.087 | 0.45 | 0.66 | 0.43 | 0.56 | 0.37 | 0.48 |
| 4 | 132413729 | rs17458198 | 0.722 | 0.087 | -1.282 | 0.066 | -0.052 | 0.02 | 0.02 | 0 | 0.11 | 0.02 | 0.09 | 0.04 | 0.2 | 0.11 |
| 4 | 132424332 | rs17458385 | -0.016 | -0.285 | -1.671 | 0.043 | -0.048 | 0.017 | 0.017 | 0.049 | 0.37 | 0.11 | 0.46 | 0.36 | 0.39 | 0.46 |
| 4 | 132426530 | rs12506711 | -0.019 | -0.298 | -1.677 | 0.114 | -0.048 | -0.003 | -0.003 | 0 | 0.37 | 0.11 | 0.46 | 0.39 | 0.39 | 0.46 |
| 4 | 132435333 | rs1911889 | -0.214 | -0.398 | -0.842 | 0.154 | -0.001 | 0.004 | 0.004 | 0.203 | 0.39 | 0.36 | 0.47 | 0.39 | 0.39 | 0.47 |
| 4 | 132443909 | rs7653972 | -0.978 | -0.49 | -0.921 | -0.131 | -0.003 | 0.013 | 0.013 | 1.218 | 0.77 | 0.85 | 0.84 | 0.98 | 0.76 | 0.8 |
| 4 | 132455666 | rs2036891 | 0.603 | -0.326 | -0.462 | 0.569 | 0.006 | 0.145 | 0.145 | 0 | 0.08 | 0.15 | 0.07 | 0.28 | 0.06 | 0.07 |
| 4 | 132455995 | rs1548174 | -1.059 | -0.687 | -0.501 | 0.088 | -0.002 | 0.019 | 0.019 | 0 | 0.61 | 0.72 | 0.77 | 0.92 | 0.62 | 0.51 |
| 4 | 132456338 | rs1705709 | 1.029 | -0.718 | -0.747 | 0.019 | -0.025 | 0.047 | 0.047 | 0 | 0.16 | 0.08 | 0.1 | 0.03 | 0.22 | 0.11 |
| 4 | 132457822 | rs1548157 | 0.944 | -1.11 | -0.923 | 0.107 | -0.079 | -0.132 | -0.132 | 3.57 | 0.22 | 0.03 | 0.44 | 0.4 | 0.27 | 0.2 |
| 4 | 132460604 | rs11099104 | 1.128 | -0.75 | -0.326 | 0.052 | -0.024 | 0.01 | 0.01 | 0 | 0.13 | 0.18 | 0.15 | 0.21 | 0.06 | 0.13 |
| 4 | 132465027 | rs4864048 | 1.734 | -0.496 | -0.704 | 0.129 | -0.087 | -0.024 | -0.024 | 0 | 0.15 | 0.01 | 0.14 | 0.11 | 0.25 | 0.13 |
| 4 | 132474137 | rs2062586 | 1.036 | -0.381 | -0.309 | 0.394 | 0.005 | 0.139 | 0.139 | 0 | 0.07 | 0.12 | 0.07 | 0.3 | 0.06 | 0.07 |
| 4 | 132479903 | rs11737708 | 0.706 | -0.248 | -0.537 | 0.118 | -0.025 | 0.052 | 0.052 | 0 | 0.17 | 0.08 | 0.14 | 0.03 | 0.2 | 0.1 |
| 4 | 132485551 | rs350959 | 0.964 | -0.439 | -0.401 | 0.151 | 0.014 | -0.008 | -0.008 | 0 | 0.6 | 0.34 | 0.67 | 0.45 | 0.66 | 0.66 |
| 4 | 132489231 | rs350995 | 1.204 | -0.369 | -0.459 | 0.119 | 0.015 | 0.064 | 0.064 | 0 | 0.46 | 0.27 | 0.53 | 0.34 | 0.42 | 0.56 |
| 4 | 132496097 | rs350993 | 0.761 | -0.321 | -0.24 | 0.369 | -0.003 | 0.003 | 0.003 | 0 | 0.66 | 0.37 | 0.71 | 0.45 | 0.69 | 0.69 |
| 4 | 132496237 | rs9997012 | 0.829 | 0.295 | -0.244 | 0.362 | 0.023 | -0.039 | -0.039 | 0 | 0.12 | 0.24 | 0.12 | 0.03 | 0.17 | 0.1 |
| 4 | 132500347 | rs13118882 | 0.691 | 0.227 | -0.498 | 0.71 | 0.009 | 0.134 | 0.134 | 0 | 0.07 | 0.13 | 0.08 | 0.28 | 0.07 | 0.07 |
| 4 | 132525794 | rs979640 | -0.444 | 0.221 | -1.191 | -0.097 | -0.03 | 0.087 | 0.087 | 0 | 0.7 | 0.82 | 0.81 | 0.91 | 0.66 | 0.74 |
| 4 | 132526933 | rs1511502 | 0.472 | 0.225 | -1.275 | -0.183 | -0.003 | -0.001 | -0.001 | 0.331 | 0.15 | 0.16 | 0.12 | 0.03 | 0.17 | 0.11 |
| 4 | 132535129 | rs350951 | 0.913 | -0.441 | -1.045 | -0.163 | 0.003 | -0.005 | -0.005 | 0.168 | 0.45 | 0.47 | 0.63 | 0.36 | 0.5 | 0.42 |
| 4 | 132535239 | rs17051227 | 1.592 | -0.595 | -1.191 | 0.103 | 0.019 | -0.054 | -0.054 | 0 | 0.09 | 0 | 0.07 | 0.23 | 0.07 | 0.16 |
| 4 | 132540869 | rs13121110 | -0.139 | -0.534 | -1.087 | -0.129 | 0.025 | -0.045 | -0.045 | 0.006 | 0.27 | 0.17 | 0.2 | 0.54 | 0.19 | 0.31 |
| 4 | 132548856 | rs13136106 | 1.706 | -0.451 | -1.04 | -0.153 | -0.019 | 0.022 | 0.022 | 0 | 0.11 | 0.06 | 0.12 | 0.03 | 0.17 | 0.12 |
| 4 | 132551302 | rs10003774 | 1.628 | -0.457 | -0.989 | -0.272 | -0.016 | 0 | 0 | 0 | 0.28 | 0.36 | 0.45 | 0.25 | 0.23 | 0.3 |
| 4 | 132556879 | rs2088797 | -1.037 | -0.43 | -1.025 | -0.31 | -0.001 | -0.013 | -0.013 | 0 | 0.7 | 0.65 | 0.81 | 0.9 | 0.66 | 0.73 |
| 4 | 132565470 | rs1500780 | -0.814 | -0.427 | -1.039 | -0.308 | -0.006 | 0.008 | 0.008 | 0 | 0.7 | 0.69 | 0.81 | 0.9 | 0.66 | 0.73 |
| 4 | 132574038 | rs1500777 | -2.118 | -0.602 | -1.33 | -0.203 | -0.009 | 0.055 | 0.055 | 0 | 0.83 | 0.92 | 0.93 | 0.94 | 0.83 | 0.86 |
| 4 | 132575313 | rs11931589 | -0.161 | -0.597 | -1.426 | -0.105 | 0.016 | -0.027 | -0.027 | 0 | 0.28 | 0.21 | 0.21 | 0.54 | 0.19 | 0.32 |
| 4 | 132587682 | rs2173805 | -2.311 | -0.184 | -1.451 | -0.192 | -0.038 | 0.084 | 0.084 | 0 | 0.85 | 0.99 | 0.94 | 0.94 | 0.84 | 0.86 |
| 4 | 132595111 | rs1909854 | -2.233 | -0.132 | -1.468 | -0.205 | -0.041 | 0.084 | 0.084 | 0 | 0.85 | 0.98 | 0.93 | 0.95 | 0.82 | 0.85 |

| | | | | | | | | | | | | | | | | |
|---|---|---|---|---|---|---|---|---|---|---|---|---|---|---|---|---|
| 4 | 132610422 | rs350694 | -0.056 | -0.253 | -1.435 | -0.099 | 0.005 | -0.007 | -0.007 | 0 | 0.66 | 0.78 | 0.78 | 0.6 | 0.68 | 0.69 |
| 4 | 132612328 | rs1909852 | -0.304 | 0.015 | -1.36 | 0.173 | 0.013 | 0.019 | 0.019 | 0 | 0.43 | 0.27 | 0.32 | 0.64 | 0.38 | 0.44 |
| 4 | 132689843 | rs13122986 | -0.606 | -0.246 | -1.524 | 0.315 | -0.009 | 0.008 | 0.008 | 0.474 | 0.73 | 0.66 | 0.57 | 0.74 | 0.77 | 0.7 |
| 4 | 132773067 | rs2863673 | -0.652 | -0.012 | -1.94 | 0.343 | 0.006 | -0.04 | -0.04 | 0 | 0.13 | 0.06 | 0.12 | 0.33 | 0.11 | 0.15 |
| 4 | 132902055 | rs349293 | -0.418 | 0.29 | -1.751 | -0.392 | 0.001 | 0.033 | 0.033 | 0.214 | 0.71 | 0.45 | 0.62 | 0.36 | 0.77 | 0.57 |
| 4 | 132914907 | rs1367832 | 0.346 | 0.229 | -1.993 | -0.874 | -0.02 | 0.02 | 0.02 | 1.024 | 0.57 | 0.21 | 0.45 | 0.31 | 0.61 | 0.48 |
| 4 | 132919103 | rs11734516 | 0.189 | 0.799 | -1.945 | -0.509 | 0.003 | -0.032 | -0.032 | 0.052 | 0.12 | 0.18 | 0.16 | 0.03 | 0.15 | 0.08 |
| 4 | 132921851 | rs17468174 | 0.917 | 0.311 | -2.261 | -0.411 | 0.002 | 0.011 | 0.011 | 0.981 | 0.22 | 0.03 | 0.15 | 0.12 | 0.22 | 0.14 |
| 4 | 132932804 | rs17547161 | -0.008 | 0.202 | -1.515 | 0.2 | 0.001 | -0.005 | -0.005 | 0 | 0.12 | 0.12 | 0.06 | 0.15 | 0.12 | 0.14 |
| 4 | 132932856 | rs4863859 | 0.192 | 0.206 | -1.558 | -0.317 | 0.02 | 0.152 | 0.152 | 0 | 0.51 | 0.21 | 0.53 | 0.19 | 0.6 | 0.38 |
| 4 | 132961872 | rs6848554 | -0.95 | -0.158 | -0.886 | -0.081 | 0.006 | 0.006 | 0.006 | 0.017 | 0.15 | 0.18 | 0.11 | 0.08 | 0.07 | 0.12 |
| 4 | 132962490 | rs6849829 | -0.95 | -0.158 | -0.886 | -0.081 | 0.006 | 0.006 | 0.006 | 0.017 | 0.15 | 0.18 | 0.11 | 0.08 | 0.07 | 0.12 |
| 4 | 132964641 | rs1346962 | -0.106 | 0.084 | -0.657 | -0.486 | -0.009 | 0.008 | 0.008 | 0 | 0.29 | 0.27 | 0.38 | 0.17 | 0.38 | 0.22 |
| 4 | 132980813 | rs1431686 | -1.23 | -0.074 | 0.132 | -0.604 | 0 | -0.014 | -0.014 | 0 | 0.45 | 0.51 | 0.5 | 0.22 | 0.46 | 0.31 |
| 4 | 132988240 | rs11726701 | 0.206 | -0.176 | 0.209 | -0.705 | -0.022 | 0.005 | 0.005 | 0 | 0.26 | 0.19 | 0.25 | 0.02 | 0.37 | 0.2 |
| 4 | 132999062 | rs1505413 | -0.182 | -0.356 | 0.828 | -0.676 | -0.005 | -0.027 | -0.027 | 0 | 0.89 | 0.74 | 0.92 | 0.96 | 0.92 | 0.86 |
| 4 | 133001527 | rs10018031 | 0.287 | -0.244 | 0.562 | -0.727 | -0.003 | -0.043 | -0.043 | 0 | 0.27 | 0.28 | 0.26 | 0.01 | 0.34 | 0.18 |
| 4 | 133003374 | rs1431688 | -1.031 | -0.374 | 0.709 | -0.607 | 0.004 | -0.022 | -0.022 | 0.021 | 0.43 | 0.47 | 0.48 | 0.23 | 0.45 | 0.31 |
| 4 | 133005854 | rs4864099 | -0.792 | -0.382 | 0.704 | -0.611 | 0.007 | -0.021 | -0.021 | 0 | 0.41 | 0.43 | 0.48 | 0.23 | 0.45 | 0.31 |
| 4 | 133016451 | rs13108543 | 0.1 | -0.444 | 0.438 | 0.13 | -0.008 | 0.008 | 0.008 | 0 | 0.3 | 0.23 | 0.26 | 0.22 | 0.34 | 0.28 |
| 4 | 133028269 | rs1910873 | 0.1 | -0.48 | 0.411 | 0.123 | -0.015 | 0.014 | 0.014 | 0 | 0.3 | 0.21 | 0.25 | 0.22 | 0.34 | 0.28 |
| 4 | 133047300 | rs9996530 | -0.85 | -0.293 | 0.327 | -0.055 | 0.001 | 0 | 0 | 1.949 | 0.43 | 0.45 | 0.36 | 0.39 | 0.34 | 0.35 |
| 4 | 133061307 | rs11943359 | 0.281 | -0.036 | 0.363 | -0.213 | -0.015 | 0.01 | 0.01 | 2.284 | 0.27 | 0.21 | 0.24 | 0.01 | 0.31 | 0.2 |
| 4 | 133062685 | rs10461213 | -0.438 | -0.241 | 0.18 | -0.06 | 0.009 | -0.029 | -0.029 | 0.322 | 0.26 | 0.2 | 0.34 | 0.48 | 0.22 | 0.29 |
| 4 | 133071142 | rs10028352 | -1.046 | -0.065 | 0.227 | -0.011 | -0.001 | -0.001 | -0.001 | 0 | 0.54 | 0.57 | 0.52 | 0.5 | 0.49 | 0.48 |
| 4 | 133084358 | rs6828393 | 0.206 | 0.19 | 0.342 | -0.252 | -0.003 | -0.027 | -0.027 | 2.562 | 0.27 | 0.23 | 0.19 | 0.02 | 0.28 | 0.2 |
| 4 | 133084604 | rs4479723 | 0.041 | 0.473 | 0.109 | 0.178 | -0.002 | 0.001 | 0.001 | 1.17 | 0.1 | 0.09 | 0.11 | 0.25 | 0.15 | 0.18 |
| 4 | 133101094 | rs13137045 | 0.377 | 0.519 | -0.502 | -0.145 | -0.019 | 0.032 | 0.032 | 0 | 0.25 | 0.14 | 0.18 | 0.01 | 0.28 | 0.21 |
| 4 | 133118600 | rs10021585 | -1.207 | 0.216 | -0.291 | 0.241 | 0.002 | 0.005 | 0.005 | 0.889 | 0.64 | 0.54 | 0.62 | 0.59 | 0.62 | 0.64 |
| 4 | 133134553 | rs348071 | -0.675 | 0.157 | -0.342 | 0.324 | -0.002 | -0.001 | -0.001 | 0 | 0.54 | 0.43 | 0.52 | 0.45 | 0.5 | 0.5 |
| 4 | 133145345 | rs13150182 | 1.054 | 0.321 | -0.509 | 0.044 | -0.014 | 0.042 | 0.042 | 0 | 0.23 | 0.1 | 0.14 | 0 | 0.26 | 0.17 |
| 4 | 133151211 | rs17557587 | 0.792 | 0.251 | -0.479 | 0.09 | -0.01 | -0.017 | -0.017 | 0 | 0.08 | 0.03 | 0.09 | 0.14 | 0.12 | 0.14 |
| 4 | 133156160 | rs1603924 | -0.48 | -0.036 | -0.346 | 0.282 | -0.018 | 0.021 | 0.021 | 0.025 | 0.9 | 0.85 | 0.83 | 0.82 | 0.94 | 0.95 |
| 4 | 133157721 | rs6811696 | 0.398 | 0.013 | -0.421 | 0.289 | -0.01 | -0.055 | -0.055 | 3.081 | 0.16 | 0.04 | 0.15 | 0.25 | 0.15 | 0.2 |
| 4 | 133165898 | rs1499328 | 0.959 | 0.034 | -0.32 | 0.277 | -0.006 | -0.003 | -0.003 | 0 | 0.46 | 0.62 | 0.48 | 0.55 | 0.5 | 0.5 |
| 4 | 133173784 | rs13434717 | 0.918 | 0.035 | -0.32 | 0.277 | -0.005 | -0.002 | -0.002 | 0.279 | 0.47 | 0.64 | 0.48 | 0.55 | 0.5 | 0.5 |
| 4 | 133189144 | rs17048863 | 0.333 | 0.139 | -0.463 | 0.384 | -0.024 | -0.064 | -0.064 | 0 | 0.16 | 0.04 | 0.17 | 0.27 | 0.16 | 0.22 |
| 4 | 133196593 | rs904516 | -0.164 | 0.021 | -0.36 | 0.264 | 0 | 0.025 | 0.025 | 0 | 0.66 | 0.73 | 0.65 | 0.81 | 0.66 | 0.75 |
| 4 | 133199152 | rs967914 | -0.304 | 0.005 | -0.382 | 0.249 | -0.013 | 0.006 | 0.006 | 0 | 0.88 | 0.84 | 0.81 | 0.82 | 0.93 | 0.93 |
| 4 | 133199870 | rs1499319 | 1.255 | -0.082 | -0.355 | 0.279 | 0.001 | -0.001 | -0.001 | 0 | 0.46 | 0.49 | 0.48 | 0.56 | 0.51 | 0.54 |
| 4 | 133205896 | rs1392015 | 0.18 | 0.067 | -0.492 | 0.365 | -0.015 | -0.048 | -0.048 | 0 | 0.17 | 0.07 | 0.17 | 0.26 | 0.16 | 0.22 |
| 4 | 133208543 | rs17049065 | 0.329 | 0.086 | -0.51 | 0.374 | -0.024 | -0.059 | -0.059 | 0.186 | 0.16 | 0.04 | 0.17 | 0.26 | 0.16 | 0.22 |
| 4 | 133223152 | rs13103069 | 0.874 | 0.085 | -0.433 | 0.173 | -0.023 | 0.034 | 0.034 | 0 | 0.22 | 0.12 | 0.15 | 0.01 | 0.26 | 0.18 |
| 4 | 133227636 | rs6857942 | 0.407 | -0.219 | -0.118 | 0.507 | -0.032 | 0.019 | 0.019 | 0.205 | 0.88 | 0.77 | 0.81 | 0.82 | 0.93 | 0.94 |
| 4 | 133236522 | rs13121719 | 1.151 | -0.042 | -0.018 | 0.564 | -0.003 | 0.019 | 0.019 | 0 | 0.65 | 0.59 | 0.47 | 0.51 | 0.68 | 0.62 |
| 4 | 133236937 | rs2062467 | 0.151 | -0.074 | -0.086 | 0.591 | -0.025 | 0.015 | 0.015 | 0.009 | 0.16 | 0.1 | 0.23 | 0.11 | 0.22 | 0.2 |
| 4 | 133251547 | rs17049500 | 0.263 | -0.005 | 0.291 | 0.344 | 0.009 | 0.009 | 0.009 | 0 | 0.12 | 0.23 | 0.07 | 0.11 | 0.14 | 0.11 |
| 4 | 133254836 | rs17559165 | 0.652 | 0.199 | 0.034 | 0.238 | -0.064 | -0.001 | -0.001 | 0 | 0.14 | 0.01 | 0.19 | 0.09 | 0.19 | 0.17 |
| 4 | 133255316 | rs17559291 | 1.47 | 0.301 | -0.485 | -0.107 | -0.02 | 0.082 | 0.082 | 0 | 0.3 | 0.12 | 0.18 | 0.04 | 0.32 | 0.3 |
| 4 | 133261123 | rs2062464 | 0.361 | 0.553 | 0.001 | 0.626 | -0.031 | 0.015 | 0.015 | 0 | 0.15 | 0.05 | 0.19 | 0.09 | 0.19 | 0.18 |
| 4 | 133278971 | rs1499330 | -0.606 | 0.535 | -0.015 | 0.807 | -0.045 | -0.06 | -0.06 | 0 | 0.21 | 0.09 | 0.34 | 0.33 | 0.25 | 0.33 |
| 4 | 133284503 | rs10006396 | -0.662 | 0.53 | 0.113 | 0.804 | -0.017 | -0.023 | -0.023 | 0 | 0.21 | 0.15 | 0.35 | 0.32 | 0.25 | 0.32 |
| 4 | 133287048 | rs2088629 | -0.131 | 0.421 | 0.283 | 0.835 | -0.029 | 0.038 | 0.038 | 1.119 | 0.91 | 0.79 | 0.83 | 0.83 | 0.94 | 0.96 |
| 4 | 133287497 | rs6831400 | -0.638 | 0.534 | 0.091 | 0.768 | -0.017 | -0.023 | -0.023 | 0.888 | 0.21 | 0.15 | 0.35 | 0.32 | 0.25 | 0.32 |
| 4 | 133289007 | rs17559561 | 0.401 | 0.518 | -0.022 | 0.686 | -0.029 | 0.015 | 0.015 | 0 | 0.15 | 0.05 | 0.19 | 0.09 | 0.2 | 0.17 |
| 4 | 133353530 | rs1499348 | 1.207 | 0.4 | 0.238 | 0.78 | 0.004 | 0.039 | 0.039 | 0 | 0.69 | 0.61 | 0.47 | 0.54 | 0.68 | 0.62 |
| 4 | 133354687 | rs1499349 | -0.098 | 0.348 | 0.167 | 0.761 | -0.023 | 0.038 | 0.038 | 0.813 | 0.91 | 0.78 | 0.82 | 0.86 | 0.94 | 0.96 |
| 4 | 133397984 | rs1909456 | 1.442 | 0.316 | 0.215 | 0.929 | 0.004 | 0.018 | 0.018 | 0 | 0.65 | 0.52 | 0.44 | 0.41 | 0.65 | 0.56 |
| 4 | 133407553 | rs6812055 | -0.974 | 0.378 | 0.247 | 0.983 | 0.008 | 0.019 | 0.019 | 0 | 0.25 | 0.29 | 0.38 | 0.44 | 0.29 | 0.37 |
| 4 | 133431421 | rs2013016 | 0.223 | -0.028 | 0.367 | 0.791 | -0.005 | 0.003 | 0.003 | 0 | 0.88 | 0.83 | 0.82 | 0.86 | 0.88 | 0.86 |
| 4 | 133441627 | rs13101445 | 0.317 | 0.185 | 0.13 | 0.613 | -0.046 | 0.035 | 0.035 | 0.248 | 0.21 | 0.1 | 0.27 | 0.09 | 0.33 | 0.26 |
| 4 | 133441732 | rs9307675 | 1.083 | 0.088 | 0.111 | 0.608 | 0.012 | -0.007 | -0.007 | 0.18 | 0.62 | 0.56 | 0.5 | 0.67 | 0.52 | 0.51 |
| 4 | 133447771 | rs1580810 | 0.226 | 0.021 | -0.15 | 0.343 | -0.018 | 0.036 | 0.036 | 0 | 0.86 | 0.98 | 0.92 | 0.94 | 0.85 | 0.84 |
| 4 | 133458239 | rs13147558 | 1.295 | -0.161 | 0.437 | 0.584 | -0.008 | 0.015 | 0.015 | 0.706 | 0.55 | 0.33 | 0.35 | 0.38 | 0.48 | 0.43 |
| 4 | 133463042 | rs11935217 | -0.636 | -0.178 | 0.603 | 0.654 | 0.025 | 0.047 | 0.047 | 0 | 0.33 | 0.51 | 0.6 | 0.56 | 0.38 | 0.39 |
| 4 | 133464597 | rs4498140 | 0.123 | -0.141 | 0.669 | 0.587 | -0.002 | 0.007 | 0.007 | 0 | 0.85 | 0.87 | 0.9 | 0.93 | 0.84 | 0.79 |
| 4 | 133466826 | rs7695551 | 0.08 | -0.135 | 0.729 | 0.587 | -0.003 | 0.006 | 0.006 | 0 | 0.86 | 0.87 | 0.9 | 0.94 | 0.84 | 0.85 |
| 4 | 133466974 | rs13149899 | 0.08 | -0.135 | 0.729 | 0.587 | -0.003 | 0.006 | 0.006 | 0 | 0.86 | 0.87 | 0.91 | 0.94 | 0.84 | 0.85 |
| 4 | 133469080 | rs6849769 | 0.372 | -0.152 | 0.777 | 0.578 | 0.008 | -0.042 | -0.042 | 0 | 0.83 | 0.66 | 0.89 | 0.94 | 0.83 | 0.85 |

| | | | | | | | | | | | | | | | | |
|---|---|---|---|---|---|---|---|---|---|---|---|---|---|---|---|---|
| 4 | 133476393 | rs13150301 | 2.218 | -0.857 | -0.028 | -0.026 | -0.082 | 0.199 | 0.199 | 0 | 0.3 | 0.03 | 0.18 | 0.01 | 0.33 | 0.21 |
| 4 | 133478128 | rs1499346 | 0.081 | -0.43 | 0.868 | 0.99 | -0.024 | -0.022 | -0.022 | 0 | 0.27 | 0.2 | 0.42 | 0.36 | 0.37 | 0.28 |
| 4 | 133479165 | rs6856692 | 0.522 | -0.476 | 0.982 | 1.009 | -0.01 | -0.049 | -0.049 | 0 | 0.82 | 0.62 | 0.88 | 0.93 | 0.84 | 0.79 |
| 4 | 133480032 | rs6814930 | 1.301 | -0.426 | 0.757 | 0.947 | 0.004 | 0.017 | 0.017 | 0.452 | 0.54 | 0.33 | 0.31 | 0.37 | 0.45 | 0.46 |
| 4 | 133492317 | rs1816512 | 0.5 | -0.849 | 0.97 | 0.997 | -0.003 | 0.006 | 0.006 | 0 | 0.83 | 0.75 | 0.76 | 0.74 | 0.82 | 0.74 |
| 4 | 133508013 | rs726332 | 0.964 | -0.836 | 0.688 | 1.004 | -0.015 | 0.027 | 0.027 | 0.102 | 0.8 | 0.55 | 0.74 | 0.74 | 0.82 | 0.74 |
| 4 | 133513991 | rs10518615 | 1.063 | -0.694 | 0.423 | 1.128 | 0 | 0.05 | 0.05 | 0.544 | 0.52 | 0.29 | 0.3 | 0.29 | 0.46 | 0.42 |
| 4 | 133519291 | rs11732021 | -0.169 | -0.248 | 0.574 | 1.244 | -0.008 | -0.001 | -0.001 | 0 | 0.27 | 0.27 | 0.47 | 0.33 | 0.36 | 0.27 |
| 4 | 133519301 | rs11723884 | -0.495 | -0.566 | 0.45 | 1.442 | 0 | 0.011 | 0.011 | 0 | 0.11 | 0.21 | 0.07 | 0.13 | 0.11 | 0.26 |
| 4 | 133529212 | rs4864123 | 0.117 | -0.76 | 0.287 | 1.163 | -0.049 | 0.021 | 0.021 | 0 | 0.31 | 0.18 | 0.52 | 0.31 | 0.48 | 0.32 |
| 4 | 133530587 | rs13107972 | 1.052 | -1.265 | 0.731 | 1.344 | 0.025 | -0.014 | -0.014 | 0 | 0.48 | 0.23 | 0.37 | 0.49 | 0.43 | 0.38 |
| 4 | 133530621 | rs6844375 | -0.065 | -1.3 | 1.139 | 1.357 | -0.002 | 0.006 | 0.006 | 0 | 0.85 | 0.81 | 0.92 | 0.82 | 0.88 | 0.71 |
| 4 | 133534583 | rs1946687 | 0.381 | -1.332 | 1.603 | 1.361 | -0.023 | 0.04 | 0.04 | 0 | 0.82 | 0.64 | 0.9 | 0.8 | 0.88 | 0.68 |
| 4 | 133535054 | rs12510292 | 1.226 | -1.359 | 1.651 | 1.353 | 0.021 | -0.013 | -0.013 | 0.935 | 0.47 | 0.23 | 0.37 | 0.47 | 0.43 | 0.35 |
| 4 | 133537182 | rs17563797 | 0.125 | -1.336 | 1.723 | 1.321 | 0.002 | -0.002 | -0.002 | 2.57 | 0.33 | 0.35 | 0.54 | 0.33 | 0.45 | 0.31 |
| 4 | 133560170 | rs11944559 | -0.89 | -1.262 | 1.545 | 1.168 | 0.006 | -0.002 | -0.002 | 4.669 | 0.05 | 0.28 | 0.02 | 0 | 0 | 0.02 |
| 4 | 133561788 | rs13142383 | 1.283 | -1.313 | 1.067 | 1.582 | 0.01 | 0.139 | 0.139 | 0.021 | 0.44 | 0.15 | 0.19 | 0.17 | 0.41 | 0.31 |
| 4 | 133562149 | rs1435313 | -0.053 | -1.087 | 0.716 | 1.694 | -0.008 | -0.006 | -0.006 | 0 | 0.34 | 0.32 | 0.7 | 0.63 | 0.46 | 0.35 |
| 4 | 133589514 | rs2419533 | 1.272 | -1.311 | 0.701 | 1.692 | 0.005 | 0.003 | 0.003 | 0.773 | 0.69 | 0.34 | 0.54 | 0.31 | 0.82 | 0.54 |
| 4 | 133597653 | rs11099203 | 1 | -1.459 | 0.245 | 1.377 | -0.093 | 0.013 | 0.013 | 0 | 0.23 | 0.07 | 0.33 | 0.22 | 0.41 | 0.22 |
| 4 | 133601521 | rs11737152 | 0.87 | -1.473 | 0.271 | 1.371 | -0.065 | 0.015 | 0.015 | 0 | 0.24 | 0.12 | 0.33 | 0.22 | 0.41 | 0.22 |
| 4 | 133603296 | rs7689868 | 0.305 | -1.527 | 0.425 | 1.509 | -0.04 | -0.002 | -0.002 | 3.041 | 0.31 | 0.2 | 0.37 | 0.34 | 0.45 | 0.29 |
| 4 | 133609486 | rs1435305 | 1.906 | -1.857 | -0.065 | 0.995 | -0.071 | 0.203 | 0.203 | 0 | 0.3 | 0.02 | 0.15 | 0 | 0.33 | 0.24 |
| 4 | 133622871 | rs13143170 | 0.593 | -0.824 | 1.109 | 1.693 | -0.025 | 0.044 | 0.044 | 0.047 | 0.81 | 0.63 | 0.72 | 0.75 | 0.87 | 0.68 |
| 4 | 133628547 | rs12504144 | -1.039 | -0.78 | 0.782 | 1.793 | -0.005 | 0.087 | 0.087 | 0 | 0.07 | 0.07 | 0.14 | 0.32 | 0.04 | 0.18 |
| 4 | 133637568 | rs13117639 | 0.738 | -0.876 | 0.914 | 1.651 | -0.006 | -0.023 | -0.023 | 0 | 0.55 | 0.49 | 0.26 | 0.27 | 0.42 | 0.41 |
| 4 | 133642296 | rs13102108 | 0.687 | -0.884 | 0.927 | 1.663 | -0.006 | -0.025 | -0.025 | 0 | 0.56 | 0.49 | 0.27 | 0.27 | 0.43 | 0.42 |
| 4 | 133652752 | rs7691446 | 0.583 | -0.883 | 0.986 | 1.668 | -0.002 | 0.002 | 0.002 | 0 | 0.57 | 0.53 | 0.4 | 0.5 | 0.44 | 0.46 |
| 4 | 133671899 | rs12645565 | -0.374 | -0.869 | 1.013 | 1.674 | -0.013 | -0.011 | -0.011 | 0.302 | 0.37 | 0.26 | 0.58 | 0.42 | 0.55 | 0.41 |
| 4 | 133680303 | rs9999397 | 0.032 | -0.828 | 0.94 | 1.657 | -0.009 | 0.071 | 0.071 | 0 | 0.11 | 0.17 | 0.19 | 0.35 | 0.08 | 0.08 |
| 4 | 133685637 | rs13119882 | 1.213 | -0.896 | 1.05 | 1.632 | 0.066 | 0.031 | 0.031 | 0 | 0.46 | 0.26 | 0.33 | 0.38 | 0.37 | 0.41 |
| 4 | 133689782 | rs11943009 | -1.006 | -0.896 | 1.106 | 1.625 | -0.029 | 0.061 | 0.061 | 0 | 0.16 | 0.44 | 0.09 | 0.2 | 0.09 | 0.18 |
| 4 | 133690420 | rs2419540 | 0.686 | -0.845 | 0.989 | 1.629 | -0.001 | 0 | 0 | 0 | 0.09 | 0.08 | 0.05 | 0.12 | 0.07 | 0.05 |
| 4 | 133694117 | rs11930973 | 0.531 | -0.859 | 1.052 | 1.485 | 0.003 | 0.019 | 0.019 | 0.211 | 0.57 | 0.35 | 0.26 | 0.35 | 0.44 | 0.56 |
| 4 | 133694135 | rs13132952 | 1.01 | -0.821 | 0.863 | 1.491 | 0.06 | 0.186 | 0.186 | 0.06 | 0.45 | 0.19 | 0.2 | 0.15 | 0.35 | 0.38 |
| 4 | 133703268 | rs7665789 | -0.551 | 0.273 | 0.232 | 1.651 | 0.002 | 0.008 | 0.008 | 0 | 0.13 | 0.16 | 0.26 | 0.19 | 0.13 | 0.18 |
| 4 | 133707298 | rs12641171 | 0.237 | 0.184 | -0.052 | 1.721 | -0.084 | -0.003 | -0.003 | 0 | 0.23 | 0.08 | 0.31 | 0.24 | 0.41 | 0.22 |
| 4 | 133714445 | rs9996230 | 0.79 | 0.182 | 0.13 | 1.784 | 0.005 | 0.001 | 0.001 | 0 | 0.6 | 0.55 | 0.43 | 0.58 | 0.48 | 0.59 |
| 4 | 133729234 | rs11099215 | -0.075 | 0.542 | -0.894 | 1.683 | -0.09 | 0.021 | 0.021 | 0 | 0.27 | 0.09 | 0.31 | 0.23 | 0.42 | 0.23 |
| 4 | 133741753 | rs1374944 | -0.731 | 0.019 | -0.37 | 1.602 | 0 | -0.003 | -0.003 | 0.261 | 0.1 | 0.09 | 0.25 | 0.2 | 0.08 | 0.17 |
| 4 | 133750858 | rs1020440 | -0.581 | -0.64 | 0.328 | 1.696 | -0.009 | 0.034 | 0.034 | 0 | 0.19 | 0.23 | 0.25 | 0.6 | 0.11 | 0.29 |
| 4 | 133752631 | rs1037177 | -0.641 | -0.65 | 0.015 | 0.686 | -0.001 | -0.004 | -0.004 | 0 | 0.56 | 0.53 | 0.52 | 0.72 | 0.45 | 0.6 |
| 4 | 133764742 | rs10518621 | 0.059 | -0.714 | 0.162 | 0.669 | -0.032 | 0.121 | 0.121 | 0.034 | 0.2 | 0.37 | 0.26 | 0.6 | 0.11 | 0.3 |
| 4 | 133768010 | rs11942306 | 0.589 | -0.845 | 0.131 | 0.627 | -0.008 | 0.053 | 0.053 | 0 | 0.41 | 0.31 | 0.35 | 0.19 | 0.55 | 0.37 |
| 4 | 133774665 | rs17509240 | 0.527 | -0.745 | -0.086 | 0.293 | -0.033 | 0.205 | 0.205 | 0.11 | 0.33 | 0.06 | 0.25 | 0.06 | 0.32 | 0.3 |
| 4 | 133786573 | rs12501689 | 0.242 | -0.562 | 0.042 | 0.505 | -0.003 | 0.046 | 0.046 | 0 | 0.43 | 0.34 | 0.35 | 0.19 | 0.56 | 0.37 |
| 4 | 133814532 | rs10518627 | 0.258 | -0.429 | 0.503 | 1.06 | -0.005 | -0.009 | -0.009 | 0 | 0.62 | 0.53 | 0.64 | 0.64 | 0.65 | 0.62 |
| 4 | 133821810 | rs1349050 | 0.258 | -0.429 | 0.503 | 1.06 | -0.005 | -0.009 | -0.009 | 0 | 0.62 | 0.53 | 0.65 | 0.64 | 0.65 | 0.61 |
| 4 | 133826719 | rs1447996 | -0.822 | -0.468 | 0.633 | 1.236 | -0.002 | 0.209 | 0.209 | 0.299 | 0.11 | 0.26 | 0.29 | 0.37 | 0.07 | 0.13 |
| 4 | 133827600 | rs12510100 | 0.309 | -0.416 | -0.05 | 0.683 | -0.034 | 0.115 | 0.115 | 0 | 0.28 | 0.07 | 0.26 | 0.04 | 0.31 | 0.22 |
| 4 | 133839342 | rs1448019 | -0.375 | -0.476 | 0.263 | 0.739 | -0.01 | 0.084 | 0.084 | 0 | 0.37 | 0.21 | 0.31 | 0.07 | 0.35 | 0.33 |
| 4 | 133840998 | rs1972970 | -1.335 | -0.758 | 0.839 | 1.16 | 0.015 | 0.036 | 0.036 | 0 | 0.47 | 0.65 | 0.64 | 0.7 | 0.4 | 0.51 |
| 4 | 133842910 | rs7683176 | -1.246 | -0.749 | 0.849 | 1.495 | 0.007 | 0.015 | 0.015 | 0 | 0.46 | 0.64 | 0.64 | 0.63 | 0.4 | 0.51 |
| 4 | 133858930 | rs11099230 | -1.287 | -0.673 | 0.912 | 1.382 | 0.001 | -0.009 | -0.009 | 0 | 0.44 | 0.47 | 0.6 | 0.71 | 0.43 | 0.51 |
| 4 | 133862954 | rs12641896 | -0.983 | -0.67 | 0.855 | 1.384 | -0.011 | 0.016 | 0.016 | 0 | 0.41 | 0.33 | 0.58 | 0.68 | 0.42 | 0.5 |
| 4 | 133876492 | rs1448009 | -0.552 | -0.664 | 0.833 | 1.33 | -0.028 | 0.088 | 0.088 | 0 | 0.36 | 0.21 | 0.3 | 0.16 | 0.38 | 0.33 |
| 4 | 133892106 | rs7691971 | -1.039 | -0.568 | 0.889 | 1.528 | 0.002 | 0.382 | 0.382 | 0.114 | 0.08 | 0.26 | 0.3 | 0.54 | 0.05 | 0.18 |
| 4 | 133906801 | rs10518624 | -0.116 | -0.427 | 0.299 | 0.702 | -0.02 | 0.053 | 0.053 | 0 | 0.3 | 0.22 | 0.28 | 0.12 | 0.34 | 0.22 |
| 4 | 133918746 | rs10002224 | -1.017 | -0.485 | 0.139 | 0.821 | 0.005 | -0.003 | -0.003 | 0 | 0.61 | 0.57 | 0.6 | 0.66 | 0.55 | 0.64 |
| 4 | 133955962 | rs7675150 | -1.308 | -0.465 | 1.051 | 0.867 | -0.029 | 0.102 | 0.102 | 3.022 | 0.12 | 0.5 | 0.14 | 0.21 | 0.05 | 0.02 |
| 4 | 133961722 | rs10026639 | -1.002 | -0.47 | 1.255 | 0.875 | -0.003 | 0.144 | 0.144 | 3.152 | 0.1 | 0.42 | 0.13 | 0.17 | 0.04 | 0.01 |
| 4 | 133981517 | rs7699528 | -0.934 | -0.645 | 0.616 | 0.268 | 0 | -0.022 | -0.022 | 0.419 | 0.46 | 0.47 | 0.47 | 0.22 | 0.45 | 0.33 |
| 4 | 133985941 | rs6832145 | -1.091 | -0.676 | 0.639 | 0.599 | -0.001 | -0.004 | -0.004 | 0 | 0.48 | 0.49 | 0.54 | 0.34 | 0.46 | 0.42 |
| 4 | 133987761 | rs6840712 | -1.091 | -0.676 | 0.639 | 0.599 | -0.001 | -0.004 | -0.004 | 1.959 | 0.48 | 0.49 | 0.54 | 0.34 | 0.46 | 0.42 |
| 4 | 134030542 | rs17585807 | -0.26 | -0.613 | 0.525 | 0.439 | -0.087 | 0.233 | 0.233 | 0 | 0.39 | 0.11 | 0.34 | 0.04 | 0.43 | 0.32 |
| 4 | 134038614 | rs12650564 | -0.571 | -0.637 | 0.684 | 0.456 | -0.01 | 0.135 | 0.135 | 3.433 | 0.1 | 0.45 | 0.13 | 0.18 | 0.04 | 0.01 |
| 4 | 134046930 | rs4404561 | -0.44 | -0.585 | 0.258 | 0.469 | -0.066 | 0.112 | 0.112 | 0 | 0.4 | 0.13 | 0.42 | 0.17 | 0.45 | 0.41 |
| 4 | 134062016 | rs4864175 | 0.72 | -0.544 | 0.457 | 0.491 | -0.018 | 0.083 | 0.083 | 0 | 0.14 | 0.01 | 0.09 | 0 | 0.18 | 0.09 |
| 4 | 134069362 | rs28565197 | -0.615 | -0.64 | 0.664 | 0.508 | -0.042 | -0.05 | -0.05 | 0.052 | 0.11 | 0.55 | 0.04 | 0.03 | 0.02 | 0.01 |

| | | | | | | | | | | | | | | | | |
|---|---|---|---|---|---|---|---|---|---|---|---|---|---|---|---|---|
| 4 | 134074058 | rs3749589 | -1.626 | -0.253 | 0.207 | 0.557 | 0.029 | 0.038 | 0.038 | 0.042 | 0.5 | 0.67 | 0.53 | 0.38 | 0.44 | 0.39 |
| 4 | 134083486 | rs9307688 | -0.634 | -0.345 | 0.483 | -0.063 | 0.001 | -0.031 | -0.031 | 0 | 0.24 | 0.27 | 0.14 | 0.01 | 0.22 | 0.13 |
| 4 | 134083809 | rs10084852 | 0.393 | 0.386 | 0.396 | 0.825 | 0.002 | 0.005 | 0.005 | 0 | 0.93 | 0.95 | 0.95 | 0.86 | 0.91 | 0.92 |
| 4 | 134120109 | rs6846610 | 1.127 | -0.198 | -0.97 | 0.37 | -0.002 | -0.001 | -0.001 | 0.534 | 0.56 | 0.53 | 0.69 | 0.58 | 0.58 | 0.51 |
| 4 | 134120138 | rs2420155 | -0.024 | -0.185 | -1.915 | -0.16 | 0.019 | -0.002 | -0.002 | 0 | 0.88 | 0.93 | 0.97 | 0.88 | 0.93 | 0.85 |
| 4 | 134123827 | rs7663268 | -0.18 | 0.375 | -1.663 | -0.112 | -0.002 | 0.01 | 0.01 | 0 | 0.14 | 0.09 | 0.13 | 0.02 | 0.16 | 0.17 |
| 4 | 134124586 | rs971869 | -0.633 | 0.493 | -2.133 | 0.044 | -0.016 | -0.022 | -0.022 | 0 | 0.3 | 0.11 | 0.29 | 0.29 | 0.28 | 0.23 |
| 4 | 134127276 | rs13122589 | 0.221 | 0.506 | -2.115 | 0.074 | -0.009 | -0.066 | -0.066 | 0 | 0.21 | 0.06 | 0.21 | 0.29 | 0.18 | 0.16 |
| 4 | 134134331 | rs6534937 | 0.703 | 0.441 | -2.085 | 0.022 | 0.011 | -0.01 | -0.01 | 0.312 | 0.63 | 0.73 | 0.65 | 0.61 | 0.67 | 0.68 |
| 4 | 134139650 | rs7676691 | -0.04 | 0.498 | -1.827 | 0.195 | 0.003 | 0.009 | 0.009 | 0 | 0.84 | 0.84 | 0.86 | 0.89 | 0.84 | 0.84 |
| 4 | 134153787 | rs976259 | -0.349 | 0.495 | -1.691 | 0.519 | -0.004 | 0.044 | 0.044 | 0 | 0.51 | 0.35 | 0.48 | 0.38 | 0.5 | 0.49 |
| 4 | 134169341 | rs7675918 | -0.467 | 0.507 | -2.189 | 0.586 | -0.048 | 0.044 | 0.044 | 0.658 | 0.51 | 0.96 | 0.46 | 0.62 | 0.42 | 0.53 |
| 4 | 134201194 | rs10213461 | -0.378 | -0.216 | -0.369 | 0.896 | 0.001 | 0.022 | 0.022 | 0.058 | 0.39 | 0.51 | 0.36 | 0.45 | 0.37 | 0.4 |
| 4 | 134203735 | rs7677931 | -0.07 | -0.218 | -0.767 | 0.839 | -0.015 | -0.121 | -0.121 | 0 | 0.17 | 0.02 | 0.12 | 0.35 | 0.15 | 0.15 |
| 4 | 134221580 | rs13115954 | 0.431 | -0.325 | -0.707 | 0.755 | -0.038 | 0.046 | 0.046 | 0 | 0.15 | 0.05 | 0.18 | 0 | 0.19 | 0.21 |
| 4 | 134223023 | rs7684286 | 0.208 | -0.33 | -0.541 | 0.759 | -0.011 | 0.012 | 0.012 | 0 | 0.16 | 0.15 | 0.19 | 0.09 | 0.19 | 0.24 |
| 4 | 134223969 | rs1346065 | -1.361 | -0.412 | -0.547 | 0.732 | 0 | 0.001 | 0.001 | 0.24 | 0.39 | 0.42 | 0.36 | 0.45 | 0.37 | 0.42 |
| 4 | 134238544 | rs17022723 | -0.014 | -0.374 | -0.627 | 1.041 | -0.003 | 0.007 | 0.007 | 0.205 | 0.1 | 0.07 | 0.1 | 0.28 | 0.11 | 0.07 |
| 4 | 134266978 | rs1895778 | -1.425 | -0.705 | 0.343 | 0.929 | -0.009 | 0.074 | 0.074 | 0.243 | 0.18 | 0.29 | 0.13 | 0.31 | 0.16 | 0.14 |
| 4 | 134274037 | rs13101921 | 0.426 | -0.744 | -0.085 | 0.708 | -0.062 | 0.083 | 0.083 | 3.946 | 0.14 | 0.01 | 0.16 | 0 | 0.16 | 0.16 |
| 4 | 134291887 | rs6818278 | -1.674 | -0.763 | 0.134 | 0.734 | 0 | 0 | 0 | 0.2 | 0.36 | 0.37 | 0.3 | 0.32 | 0.33 | 0.32 |
| 4 | 134293912 | rs10518633 | -0.558 | -0.634 | -0.157 | 0.872 | -0.012 | -0.007 | -0.007 | 0 | 0.07 | 0.12 | 0.02 | 0.04 | 0.05 | 0.08 |
| 4 | 134298899 | rs10031161 | -2.169 | -0.618 | -0.47 | 0.854 | -0.098 | 0.063 | 0.063 | 0.954 | 0.53 | 0.97 | 0.52 | 0.66 | 0.43 | 0.53 |
| 4 | 134303084 | rs6857817 | -0.008 | -0.456 | -0.286 | 0.605 | 0 | 0.036 | 0.036 | 0.334 | 0.11 | 0.11 | 0.1 | 0.28 | 0.11 | 0.07 |
| 4 | 134312838 | rs1425565 | -1.955 | -0.599 | 0.011 | 0.558 | 0.02 | -0.009 | -0.009 | 0 | 0.51 | 0.85 | 0.37 | 0.48 | 0.41 | 0.47 |
| 4 | 134319928 | rs2009274 | -2.246 | -0.617 | 0.042 | 0.515 | 0.041 | 0.012 | 0.012 | 0.268 | 0.5 | 0.8 | 0.36 | 0.38 | 0.4 | 0.43 |
| 4 | 134322289 | rs1425562 | 0.214 | -0.605 | -0.034 | 0.557 | 0 | 0.055 | 0.055 | 0.99 | 0.08 | 0.07 | 0.09 | 0.28 | 0.09 | 0.06 |
| 4 | 134326078 | rs1834407 | -0.112 | -0.661 | -0.03 | 0.283 | 0.002 | 0.002 | 0.002 | 0 | 0.08 | 0.07 | 0.02 | 0.03 | 0.05 | 0.08 |
| 4 | 134330928 | rs10518635 | -0.599 | -0.341 | 0.057 | 0.243 | -0.001 | -0.132 | -0.132 | 0 | 0.28 | 0.52 | 0.23 | 0.07 | 0.25 | 0.27 |
| 4 | 134335871 | rs1389331 | -1.192 | 0.526 | -1.003 | 0.29 | -0.044 | -0.046 | -0.046 | 0 | 0.56 | 0.85 | 0.29 | 0.43 | 0.47 | 0.49 |
| 4 | 134340488 | rs1493499 | -1.035 | 0.464 | -0.955 | 0.249 | -0.005 | 0.008 | 0.008 | 0.998 | 0.5 | 0.76 | 0.28 | 0.4 | 0.43 | 0.42 |
| 4 | 134343880 | rs7679087 | -1.024 | 1.057 | -0.742 | -0.085 | -0.01 | 0.009 | 0.009 | 0 | 0.24 | 0.21 | 0.15 | 0.14 | 0.3 | 0.15 |
| 4 | 134346117 | rs13127497 | -0.359 | 0.749 | -0.252 | -0.693 | 0.001 | -0.061 | -0.061 | 0 | 0.21 | 0.23 | 0.24 | 0 | 0.22 | 0.15 |
| 4 | 134352803 | rs17024567 | -0.146 | 0.306 | 0.267 | -0.88 | -0.031 | -0.033 | -0.033 | 2.187 | 0.06 | 0.18 | 0.03 | 0 | 0.01 | 0.01 |
| 4 | 134352903 | rs1032439 | -1.493 | 0.449 | 0.011 | -0.242 | -0.02 | 0.062 | 0.062 | 0 | 0.26 | 0.4 | 0.15 | 0.5 | 0.23 | 0.19 |
| 4 | 134357855 | rs11099240 | -1.095 | 0.561 | 0.099 | -0.151 | -0.021 | -0.034 | -0.034 | 3.538 | 0.16 | 0.35 | 0.13 | 0.08 | 0.14 | 0.08 |
| 4 | 134358917 | rs13107301 | 0.161 | 0.561 | 0.078 | -0.185 | 0.001 | -0.048 | -0.048 | 0 | 0.22 | 0.21 | 0.24 | 0.01 | 0.23 | 0.15 |
| 4 | 134359051 | rs17024656 | -1.224 | 0.549 | 0.162 | -0.171 | -0.013 | -0.029 | -0.029 | 0 | 0.16 | 0.31 | 0.12 | 0.08 | 0.15 | 0.08 |
| 4 | 134376495 | rs2420407 | -0.318 | 0.535 | 0.16 | -0.165 | -0.003 | -0.075 | -0.075 | 0 | 0.25 | 0.31 | 0.24 | 0.01 | 0.22 | 0.16 |
| 4 | 134379548 | rs13149700 | -2.4 | 0.359 | 0.161 | -0.025 | 0.017 | 0 | 0 | 0.001 | 0.48 | 0.64 | 0.4 | 0.49 | 0.45 | 0.36 |
| 4 | 134384370 | rs7375925 | -2.496 | 0.253 | 0.089 | 0.202 | 0.027 | 0.005 | 0.005 | 0 | 0.44 | 0.68 | 0.36 | 0.44 | 0.38 | 0.36 |
| 4 | 134384791 | rs17024794 | -0.209 | 0.234 | 0.077 | 0.231 | -0.014 | -0.037 | -0.037 | 0 | 0.13 | 0.04 | 0.1 | 0.37 | 0.18 | 0.12 |
| 4 | 134389485 | rs7662704 | -2.308 | 0.282 | 0.001 | 0.27 | 0.027 | 0.014 | 0.014 | 0 | 0.44 | 0.68 | 0.36 | 0.42 | 0.38 | 0.35 |
| 4 | 134391195 | rs1493508 | -1.705 | 0.406 | 0.067 | 0.296 | -0.006 | 0.009 | 0.009 | 0 | 0.35 | 0.53 | 0.34 | 0.42 | 0.36 | 0.31 |
| 4 | 134392080 | rs1908125 | -1.065 | 0.296 | -0.215 | 0.035 | 0 | -0.005 | -0.005 | 0 | 0.23 | 0.23 | 0.22 | 0.07 | 0.16 | 0.21 |
| 4 | 134394882 | rs11935955 | -1.125 | 0.343 | -0.257 | 0.058 | 0.018 | 0.027 | 0.027 | 0 | 0.62 | 0.71 | 0.6 | 0.76 | 0.72 | 0.61 |
| 4 | 134396454 | rs11732664 | -1.331 | 0.446 | -0.247 | -0.016 | 0.024 | 0.045 | 0.045 | 0 | 0.65 | 0.77 | 0.69 | 0.81 | 0.73 | 0.61 |
| 4 | 134397039 | rs12505494 | -0.936 | 0.494 | -0.307 | -0.026 | 0.004 | 0.006 | 0.006 | 1.361 | 0.12 | 0.09 | 0.15 | 0.08 | 0.12 | 0.09 |
| 4 | 134398990 | rs1493493 | 0.768 | 0.522 | -0.247 | 0.015 | 0.008 | 0.009 | 0.009 | 0 | 0.39 | 0.35 | 0.42 | 0.26 | 0.29 | 0.42 |
| 4 | 134415283 | rs7696123 | 0.823 | 0.514 | -0.332 | 0.54 | 0.023 | -0.012 | -0.012 | 0.017 | 0.39 | 0.24 | 0.42 | 0.56 | 0.33 | 0.45 |
| 4 | 134425824 | rs6839762 | 1.339 | 0.559 | -0.358 | 0.457 | 0.035 | 0.042 | 0.042 | 0 | 0.34 | 0.21 | 0.3 | 0.19 | 0.23 | 0.4 |
| 4 | 134439150 | rs4593189 | -0.692 | 0.898 | -0.248 | 0.162 | -0.001 | -0.005 | -0.005 | 0 | 0.22 | 0.25 | 0.22 | 0.08 | 0.27 | 0.12 |
| 4 | 134448546 | rs12500334 | -0.109 | 0.997 | -0.047 | 0.245 | 0.013 | 0.007 | 0.007 | 0 | 0.6 | 0.53 | 0.54 | 0.56 | 0.55 | 0.54 |
| 4 | 134458405 | rs6836197 | -0.055 | 0.431 | -0.035 | 0.253 | 0.016 | 0.007 | 0.007 | 0 | 0.4 | 0.48 | 0.45 | 0.45 | 0.48 | 0.46 |
| 4 | 134463193 | rs1032438 | -0.132 | 0.244 | 0.284 | -0.362 | 0.008 | -0.033 | -0.033 | 0 | 0.42 | 0.47 | 0.46 | 0.71 | 0.52 | 0.45 |
| 4 | 134471862 | rs1493517 | -1.055 | 0.136 | 0.289 | -0.453 | 0.003 | 0.004 | 0.004 | 0 | 0.56 | 0.59 | 0.55 | 0.71 | 0.65 | 0.49 |
| 4 | 134478128 | rs1565733 | -1.335 | 0.147 | 0.137 | -0.448 | 0.016 | 0.067 | 0.067 | 0 | 0.59 | 0.75 | 0.56 | 0.71 | 0.65 | 0.49 |
| 4 | 134482149 | rs903092 | -0.097 | 0.153 | 0.137 | -0.454 | 0.004 | -0.024 | -0.024 | 0 | 0.43 | 0.5 | 0.47 | 0.71 | 0.52 | 0.44 |
| 4 | 134482324 | rs903094 | -0.107 | 0.153 | 0.137 | -0.455 | 0.004 | -0.024 | -0.024 | 0.268 | 0.43 | 0.5 | 0.47 | 0.71 | 0.52 | 0.44 |
| 4 | 134489656 | rs13106321 | -0.678 | 0.469 | 0.198 | -0.404 | -0.012 | -0.071 | -0.071 | 0.339 | 0.16 | 0.26 | 0.1 | 0 | 0.13 | 0.07 |
| 4 | 134514218 | rs12508742 | 1.191 | 0.314 | -0.087 | -0.369 | -0.004 | 0.078 | 0.078 | 0.525 | 0.4 | 0.14 | 0.39 | 0.28 | 0.37 | 0.49 |
| 4 | 134522725 | rs10518641 | 0.948 | 0.445 | 0.068 | -0.411 | 0.021 | 0.079 | 0.079 | 0.096 | 0.44 | 0.26 | 0.41 | 0.27 | 0.37 | 0.5 |
| 4 | 134530386 | rs925084 | 0.928 | 0.454 | 0.039 | -0.41 | 0.018 | 0.05 | 0.05 | 0 | 0.46 | 0.36 | 0.42 | 0.27 | 0.37 | 0.49 |
| 4 | 134535101 | rs906372 | 1.397 | 0.427 | -0.7 | -0.374 | -0.006 | 0.095 | 0.095 | 0.788 | 0.4 | 0.1 | 0.33 | 0.28 | 0.36 | 0.49 |
| 4 | 134538656 | rs13101958 | 1.188 | 0.84 | -0.782 | -0.291 | -0.032 | 0.061 | 0.061 | 0 | 0.1 | 0.01 | 0.06 | 0 | 0.13 | 0.03 |
| 4 | 134540393 | rs2713289 | -0.193 | 0.767 | -0.298 | -0.285 | -0.006 | 0.005 | 0.005 | 0 | 0.64 | 0.66 | 0.47 | 0.28 | 0.52 | 0.58 |
| 4 | 134553039 | rs7676258 | 0.9 | 1.195 | -0.327 | -0.071 | -0.031 | 0.015 | 0.015 | 0 | 0.17 | 0.08 | 0.13 | 0.11 | 0.3 | 0.12 |
| 4 | 134570882 | rs7700155 | 0.404 | 0.637 | -0.08 | 0.07 | 0.028 | 0.049 | 0.049 | 0 | 0.51 | 0.62 | 0.37 | 0.28 | 0.37 | 0.55 |
| 4 | 134572146 | rs1948964 | 0.341 | 0.96 | -0.263 | 0.03 | 0.012 | 0 | 0 | 1.03 | 0.49 | 0.44 | 0.47 | 0.5 | 0.38 | 0.61 |

| | | | | | | | | | | | | | | | | |
|---|---|---|---|---|---|---|---|---|---|---|---|---|---|---|---|---|
| 4 | 134576263 | rs4864206 | 0.482 | 1.001 | -0.446 | -0.016 | -0.052 | 0.078 | 0.078 | 0 | 0.31 | 0.16 | 0.22 | 0.1 | 0.45 | 0.17 |
| 4 | 134578828 | rs1567176 | -0.374 | 0.369 | -0.436 | 0.118 | 0.022 | -0.001 | -0.001 | 0 | 0.48 | 0.38 | 0.47 | 0.5 | 0.41 | 0.6 |
| 4 | 134603887 | rs1395763 | -0.282 | 0.049 | -0.704 | 0.172 | 0.002 | 0 | 0 | 0 | 0.5 | 0.49 | 0.48 | 0.5 | 0.41 | 0.61 |
| 4 | 134626218 | rs2724267 | -0.156 | -0.112 | -2.125 | 0.356 | 0.074 | -0.059 | -0.059 | 0.303 | 0.38 | 0.08 | 0.35 | 0.35 | 0.29 | 0.55 |
| 4 | 134627285 | rs2660733 | -0.248 | -0.037 | -2.018 | 0.295 | 0.065 | -0.043 | -0.043 | 0 | 0.39 | 0.07 | 0.36 | 0.35 | 0.31 | 0.57 |
| 4 | 134639697 | rs2724272 | 0.165 | -0.295 | -2.169 | -0.693 | 0.045 | 0.145 | 0.145 | 0 | 0.25 | 0.03 | 0.07 | 0 | 0.14 | 0.18 |
| 4 | 134640333 | rs10518642 | 0.391 | -0.148 | -2.77 | -1.508 | -0.019 | 0.111 | 0.111 | 0.259 | 0.25 | 0.05 | 0.32 | 0.02 | 0.25 | 0.23 |
| 4 | 134670981 | rs12508593 | -0.513 | -0.483 | -1.304 | -1.583 | -0.067 | -0.161 | -0.161 | 0 | 0.71 | 0.52 | 0.88 | 1 | 0.86 | 0.82 |
| 4 | 134702716 | rs12646011 | 0.83 | 0.328 | -0.887 | -0.841 | -0.03 | -0.047 | -0.047 | 0 | 0.3 | 0.21 | 0.22 | 0.42 | 0.41 | 0.28 |
| 4 | 134706510 | rs13135185 | 0.928 | 0.11 | -0.658 | -0.921 | 0.027 | 0.021 | 0.021 | 0 | 0.25 | 0.12 | 0.09 | 0 | 0.14 | 0.18 |
| 4 | 134710417 | rs7668810 | 0.256 | 0.627 | -0.188 | -0.636 | 0.007 | -0.006 | -0.006 | 0 | 0.13 | 0.17 | 0.21 | 0.04 | 0.18 | 0.21 |
| 4 | 134714897 | rs4613620 | -0.278 | 0.061 | 0.284 | -0.572 | -0.005 | 0.016 | 0.016 | 0.444 | 0.71 | 0.7 | 0.56 | 0.45 | 0.74 | 0.68 |
| 4 | 134733250 | rs4505885 | -0.801 | -0.204 | 0.492 | -0.838 | -0.042 | -0.158 | -0.158 | 0 | 0.7 | 0.59 | 0.88 | 1 | 0.83 | 0.8 |
| 4 | 134748873 | rs10006443 | -0.329 | 0.08 | 0.863 | -0.874 | -0.008 | 0.022 | 0.022 | 0 | 0.37 | 0.44 | 0.61 | 0.59 | 0.36 | 0.51 |
| 4 | 134754555 | rs10034320 | 0.239 | 0.059 | 0.773 | -0.866 | 0 | -0.002 | -0.002 | 0 | 0.08 | 0.14 | 0.16 | 0.04 | 0.09 | 0.18 |
| 4 | 134763851 | rs4541580 | -1.481 | -0.018 | 0.769 | -0.926 | 0.028 | 0.122 | 0.122 | 0 | 0.74 | 0.91 | 0.9 | 1 | 0.83 | 0.8 |
| 4 | 134766022 | rs10084961 | 0.299 | -0.03 | 0.936 | -0.898 | 0.013 | 0.008 | 0.008 | 0 | 0.38 | 0.49 | 0.29 | 0.41 | 0.47 | 0.3 |
| 4 | 134768896 | rs4293839 | -0.737 | -0.166 | 0.836 | -0.917 | 0.028 | 0.104 | 0.104 | 0.078 | 0.66 | 0.83 | 0.74 | 0.96 | 0.74 | 0.61 |
| 4 | 134775136 | rs12499235 | 0.829 | -0.082 | 1.071 | -0.93 | -0.002 | 0.079 | 0.079 | 0.603 | 0.24 | 0.07 | 0.3 | 0.01 | 0.25 | 0.25 |
| 4 | 134777686 | rs4629515 | -0.379 | -0.196 | 1.343 | -0.944 | 0.004 | -0.004 | -0.004 | 0.092 | 0.62 | 0.65 | 0.6 | 0.43 | 0.73 | 0.55 |
| 4 | 134780470 | rs7699345 | 0.494 | -0.211 | 1.371 | -0.942 | -0.045 | -0.027 | -0.027 | 0.084 | 0.36 | 0.25 | 0.28 | 0.41 | 0.47 | 0.3 |
| 4 | 134782403 | rs4582208 | -0.632 | -0.136 | 1.303 | -0.959 | 0.007 | -0.008 | -0.008 | 0 | 0.64 | 0.68 | 0.58 | 0.43 | 0.72 | 0.54 |
| 4 | 134784379 | rs6811342 | -0.416 | -0.103 | 1.098 | -0.942 | -0.016 | 0.011 | 0.011 | 0 | 0.62 | 0.57 | 0.57 | 0.43 | 0.72 | 0.54 |
| 4 | 134788400 | rs17611459 | 0.4 | -0.066 | 0.791 | -0.914 | -0.044 | -0.025 | -0.025 | 0 | 0.36 | 0.26 | 0.26 | 0.39 | 0.47 | 0.29 |
| 4 | 134815036 | rs4415013 | -0.32 | 0.022 | 0.766 | -1.184 | -0.023 | -0.091 | -0.091 | 0 | 0.61 | 0.51 | 0.66 | 0.95 | 0.72 | 0.61 |
| 4 | 134847697 | rs17540695 | 0.275 | 0.206 | 0.829 | -0.74 | 0.001 | 0.037 | 0.037 | 0.367 | 0.25 | 0.13 | 0.27 | 0.02 | 0.25 | 0.23 |
| 4 | 134848916 | rs11099261 | -0.62 | 0.117 | 1.285 | -0.81 | -0.007 | -0.1 | -0.1 | 3.558 | 0.34 | 0.5 | 0.52 | 0.07 | 0.34 | 0.39 |
| 4 | 134860496 | rs10001384 | -0.359 | -0.02 | 1.412 | -0.35 | -0.019 | 0.055 | 0.055 | 0 | 0.31 | 0.48 | 0.42 | 0.52 | 0.26 | 0.31 |
| 4 | 134865463 | rs7340759 | -1.199 | -0.029 | 1.374 | -0.362 | 0.014 | -0.023 | -0.023 | 0 | 0.39 | 0.74 | 0.65 | 0.58 | 0.35 | 0.46 |
| 4 | 134869724 | rs6840096 | -0.865 | -0.044 | 1.381 | -0.322 | 0.004 | 0.362 | 0.362 | 0.355 | 0.08 | 0.34 | 0.15 | 0.5 | 0.03 | 0.09 |
| 4 | 134888185 | rs11731250 | 0.338 | 0.147 | 0.789 | -0.758 | 0.003 | 0.048 | 0.048 | 0.966 | 0.23 | 0.13 | 0.27 | 0.02 | 0.22 | 0.22 |
| 4 | 134889067 | rs12508354 | -0.487 | 0.209 | 0.628 | -0.734 | -0.005 | -0.076 | -0.076 | 0 | 0.31 | 0.39 | 0.51 | 0.08 | 0.32 | 0.37 |
| 4 | 134924076 | rs17613678 | 0.041 | 0.5 | -0.023 | -0.273 | -0.024 | -0.014 | -0.014 | 0 | 0.05 | 0.01 | 0.22 | 0.05 | 0.09 | 0.15 |
| 4 | 134930037 | rs17031068 | -0.172 | 0.445 | 0.142 | -0.304 | 0 | -0.001 | -0.001 | 0 | 0.06 | 0.07 | 0.12 | 0.05 | 0.09 | 0.15 |
| 4 | 134940358 | rs17613932 | -0.444 | 0.155 | 0.052 | -0.836 | 0.002 | 0.041 | 0.041 | 0 | 0.31 | 0.22 | 0.39 | 0.07 | 0.32 | 0.38 |
| 4 | 134947801 | rs1494978 | 0.047 | 0.438 | -0.024 | -1.551 | -0.018 | -0.053 | -0.053 | 0.376 | 0.36 | 0.59 | 0.3 | 0.92 | 0.27 | 0.39 |
| 4 | 134964519 | rs11099266 | 0.834 | 1.071 | 0.229 | -1.183 | 0.027 | -0.038 | -0.038 | 0 | 0.2 | 0.11 | 0.11 | 0.42 | 0.13 | 0.22 |
| 4 | 134964598 | rs6839208 | -1.02 | 1.03 | 0.111 | -1.282 | 0.02 | 0.057 | 0.057 | 0 | 0.61 | 0.75 | 0.75 | 0.94 | 0.68 | 0.65 |
| 4 | 134972554 | rs12642075 | -0.571 | 0.657 | 0.176 | -1.081 | -0.018 | -0.055 | -0.055 | 0 | 0.36 | 0.59 | 0.3 | 0.92 | 0.27 | 0.38 |
| 4 | 134984561 | rs1126293 | 0.379 | 1.011 | 0.397 | -0.76 | 0.008 | 0.17 | 0.17 | 0 | 0.12 | 0.24 | 0.17 | 0.5 | 0.14 | 0.17 |
| 4 | 135019322 | rs1390022 | 1.177 | 1.073 | -0.239 | -1.229 | -0.053 | 0.106 | 0.106 | 1.484 | 0.21 | 0.04 | 0.27 | 0.01 | 0.26 | 0.24 |
| 4 | 135019690 | rs1494989 | -1.823 | 0.92 | -0.384 | -1.57 | 0.016 | 0.131 | 0.131 | 0.009 | 0.71 | 0.93 | 0.89 | 1 | 0.77 | 0.8 |
| 4 | 135020910 | rs1390021 | -1.684 | 0.232 | -0.227 | -1.54 | -0.069 | 0.105 | 0.105 | 0 | 0.68 | 0.89 | 0.83 | 1 | 0.59 | 0.79 |
| 4 | 135046236 | rs1494984 | -1.006 | 0.154 | -0.043 | -1.448 | 0.086 | 0.149 | 0.149 | 0 | 0.46 | 0.79 | 0.55 | 0.99 | 0.33 | 0.55 |
| 4 | 135062735 | rs3980965 | -1.095 | 0.153 | -0.058 | -1.362 | 0.106 | 0.218 | 0.218 | 0 | 0.47 | 0.83 | 0.55 | 0.96 | 0.33 | 0.55 |
| 4 | 135090280 | rs2088124 | -1.718 | 0.199 | -0.072 | -1.42 | -0.063 | 0.09 | 0.09 | 0 | 0.68 | 0.87 | 0.83 | 1 | 0.59 | 0.79 |
| 4 | 135121066 | rs11099273 | -1.034 | 0.102 | 0.168 | -1.364 | 0.054 | 0.065 | 0.065 | 0 | 0.45 | 0.69 | 0.54 | 0.98 | 0.33 | 0.56 |
| 4 | 135121368 | rs2044116 | -0.975 | 0.103 | 0.16 | -1.363 | 0.063 | 0.087 | 0.087 | 0 | 0.45 | 0.72 | 0.54 | 0.98 | 0.33 | 0.55 |
| 4 | 135121601 | rs10009368 | -0.965 | 0.094 | 0.156 | -1.367 | 0.062 | 0.086 | 0.086 | 0 | 0.45 | 0.72 | 0.54 | 0.98 | 0.33 | 0.55 |
| 4 | 135126327 | rs17033938 | 1.11 | -0.179 | -0.177 | -1.356 | 0.043 | -0.112 | -0.112 | 1.003 | 0.17 | 0.03 | 0.12 | 0.42 | 0.12 | 0.23 |
| 4 | 135137960 | rs4337777 | -0.36 | -0.125 | 0.358 | -1.643 | -0.038 | -0.135 | -0.135 | 0.014 | 0.33 | 0.46 | 0.41 | 0.97 | 0.24 | 0.4 |
| 4 | 135142988 | rs1017507 | -0.836 | -0.07 | 0.151 | -1.654 | -0.058 | 0.172 | 0.172 | 0.045 | 0.6 | 0.87 | 0.74 | 0.98 | 0.5 | 0.64 |
| 4 | 135144904 | rs17034156 | 0.593 | -0.113 | 0.114 | -1.689 | 0.001 | -0.008 | -0.008 | 0.491 | 0.09 | 0.07 | 0.1 | 0.01 | 0.09 | 0.15 |
| 4 | 135149801 | rs2901383 | -0.498 | -0.143 | 0.142 | -1.946 | 0.004 | -0.047 | -0.047 | 0 | 0.4 | 0.61 | 0.47 | 0.97 | 0.32 | 0.42 |
| 4 | 135176705 | rs6822094 | -0.808 | -0.011 | 0.274 | -1.963 | 0 | -0.08 | -0.08 | 0.174 | 0.55 | 0.53 | 0.69 | 0.98 | 0.51 | 0.64 |
| 4 | 135183337 | rs10015286 | -1.281 | -0.024 | 0.345 | -1.983 | 0.085 | 0.289 | 0.289 | 0 | 0.51 | 0.84 | 0.46 | 0.97 | 0.4 | 0.55 |
| 4 | 135202678 | rs10019779 | 0.321 | -0.045 | 0.659 | -1.658 | 0.007 | 0.3 | 0.3 | 0.136 | 0.15 | 0.41 | 0.29 | 0.57 | 0.11 | 0.17 |
| 4 | 135206722 | rs11733152 | -2.058 | 0.004 | 0.6 | -1.718 | -0.108 | 0.169 | 0.169 | 0 | 0.69 | 0.94 | 0.84 | 1 | 0.6 | 0.79 |
| 4 | 135211409 | rs11935714 | -1.734 | 0.018 | 0.635 | -1.711 | -0.017 | -0.01 | -0.01 | 0 | 0.67 | 0.71 | 0.81 | 1 | 0.6 | 0.79 |
| 4 | 135250225 | rs17553221 | 0.46 | -0.045 | 0.555 | -1.649 | 0.003 | 0.01 | 0.01 | 0 | 0.16 | 0.11 | 0.12 | 0.01 | 0.14 | 0.16 |
| 4 | 135252930 | rs12649164 | -0.991 | -0.346 | 0.512 | -1.862 | 0.038 | 0.093 | 0.093 | 0 | 0.5 | 0.67 | 0.57 | 0.98 | 0.42 | 0.59 |
| 4 | 135278000 | rs1486983 | -0.039 | -0.295 | 0.869 | -1.754 | -0.005 | 0.028 | 0.028 | 0 | 0.25 | 0.29 | 0.39 | 0.57 | 0.21 | 0.34 |
| 4 | 135288420 | rs1386363 | -0.395 | -0.271 | 0.639 | -1.743 | -0.028 | 0.107 | 0.107 | 0 | 0.27 | 0.49 | 0.41 | 0.57 | 0.21 | 0.34 |
| 4 | 135294098 | rs1601648 | -1.029 | -0.352 | 0.607 | -1.798 | 0.049 | 0.167 | 0.167 | 0 | 0.51 | 0.72 | 0.59 | 0.99 | 0.42 | 0.58 |
| 4 | 135297475 | rs1486994 | 0.984 | -0.305 | 0.325 | -1.724 | 0.004 | -0.099 | -0.099 | 0 | 0.21 | 0.06 | 0.16 | 0.42 | 0.21 | 0.24 |
| 4 | 135310068 | rs17036736 | 0.979 | -0.297 | 0.322 | -1.721 | 0.007 | -0.077 | -0.077 | 1.719 | 0.21 | 0.07 | 0.16 | 0.42 | 0.21 | 0.24 |
| 4 | 135312908 | rs12644853 | -1.008 | -0.184 | 0.671 | -2.234 | 0.001 | -0.083 | -0.083 | 3.117 | 0.49 | 0.49 | 0.55 | 0.99 | 0.42 | 0.58 |
| 4 | 135317096 | rs2421334 | -0.918 | -0.168 | 0.565 | -2.226 | 0.025 | 0.039 | 0.039 | 0 | 0.51 | 0.64 | 0.57 | 0.99 | 0.42 | 0.58 |
| 4 | 135332361 | rs12510007 | 0.877 | -0.135 | 0.368 | -2.26 | 0 | -0.05 | -0.05 | 0 | 0.18 | 0.19 | 0.28 | 0.01 | 0.22 | 0.21 |

| | | | | | | | | | | | | | | | | |
|---|---|---|---|---|---|---|---|---|---|---|---|---|---|---|---|---|
| 4 | 135339629 | rs1486978 | -2.081 | -0.126 | 0.367 | -2.328 | -0.034 | 0.028 | 0.028 | 0 | 0.7 | 0.82 | 0.8 | 1 | 0.61 | 0.79 |
| 4 | 135345377 | rs17037260 | 0.831 | -0.19 | 0.161 | -1.934 | 0.017 | -0.092 | -0.092 | 0 | 0.23 | 0.07 | 0.21 | 0.57 | 0.18 | 0.25 |
| 4 | 135358154 | rs11729282 | 0.507 | 0.314 | 0.675 | -2.627 | 0 | -0.045 | -0.045 | 0 | 0.18 | 0.19 | 0.28 | 0.01 | 0.22 | 0.25 |
| 4 | 135385287 | rs11730209 | 0.424 | 0.312 | 0.679 | -2.628 | 0 | -0.086 | -0.086 | 0.748 | 0.19 | 0.25 | 0.28 | 0.01 | 0.22 | 0.25 |
| 4 | 135410583 | rs983035 | -1.306 | 0.465 | 0.155 | -2.576 | -0.023 | -0.006 | -0.006 | 0.792 | 0.69 | 0.75 | 0.68 | 1 | 0.59 | 0.78 |
| 4 | 135420199 | rs13129759 | 0.995 | 0.451 | -0.204 | -2.539 | -0.057 | 0.078 | 0.078 | 0 | 0.3 | 0.13 | 0.2 | 0 | 0.39 | 0.21 |
| 4 | 135421707 | rs10006423 | -0.038 | 0.321 | 0.515 | -1.882 | -0.008 | -0.146 | -0.146 | 0 | 0.79 | 0.52 | 0.71 | 0.98 | 0.78 | 0.76 |
| 4 | 135428455 | rs4388143 | 0.662 | 0.788 | -0.146 | -2.3 | 0.003 | -0.121 | -0.121 | 0 | 0.57 | 0.46 | 0.55 | 0.98 | 0.55 | 0.58 |
| 4 | 135430486 | rs13140370 | 0.167 | 0.784 | -0.302 | -2.467 | -0.018 | 0.018 | 0.018 | 0 | 0.21 | 0.13 | 0.17 | 0 | 0.27 | 0.12 |
| 4 | 135438724 | rs11099284 | 0.145 | 0.292 | -0.146 | -1.942 | -0.008 | -0.08 | -0.08 | 0 | 0.78 | 0.64 | 0.72 | 0.98 | 0.82 | 0.7 |
| 4 | 135454396 | rs6844188 | 0.015 | 0.862 | -0.212 | -1.646 | -0.024 | 0.027 | 0.027 | 0 | 0.13 | 0.08 | 0.11 | 0 | 0.2 | 0.08 |
| 4 | 135457628 | rs6535025 | 0.015 | 0.862 | -0.15 | -1.646 | -0.017 | 0.012 | 0.012 | 0 | 0.13 | 0.12 | 0.11 | 0 | 0.2 | 0.08 |
| 4 | 135474690 | rs11934048 | 0.886 | 0.468 | 0.26 | -2.066 | 0.004 | -0.105 | -0.105 | 0.207 | 0.69 | 0.53 | 0.79 | 0.99 | 0.68 | 0.8 |
| 4 | 135514757 | rs4479759 | -0.61 | 0.467 | 0.868 | -2.064 | 0.04 | -0.075 | -0.075 | 0 | 0.18 | 0.4 | 0.19 | 0.03 | 0.24 | 0.14 |
| 4 | 135523007 | rs4452492 | 1.663 | 0.053 | 0.657 | -1.951 | 0.002 | 0.004 | 0.004 | 0 | 0.6 | 0.31 | 0.66 | 0.84 | 0.65 | 0.62 |
| 4 | 135537719 | rs6535028 | -0.627 | 0.187 | 0.578 | -1.865 | 0.001 | -0.016 | -0.016 | 0.926 | 0.86 | 0.86 | 0.96 | 1 | 0.91 | 0.96 |
| 4 | 135547294 | rs10010236 | -0.684 | -0.012 | 0.984 | -1.931 | 0.05 | -0.124 | -0.124 | 0.349 | 0.2 | 0.48 | 0.26 | 0.03 | 0.24 | 0.15 |
| 4 | 135580589 | rs4475200 | 0.77 | -0.297 | 0.805 | -2.008 | 0.009 | -0.006 | -0.006 | 0 | 0.63 | 0.33 | 0.67 | 0.96 | 0.74 | 0.77 |
| 4 | 135598459 | rs10020543 | -0.824 | -0.355 | 0.819 | -2.061 | -0.018 | -0.112 | -0.112 | 0 | 0.36 | 0.53 | 0.32 | 0.04 | 0.26 | 0.25 |
| 4 | 135610277 | rs2404308 | 0.144 | -0.402 | 0.984 | -2.099 | 0.009 | -0.062 | -0.062 | 0.118 | 0.19 | 0.26 | 0.22 | 0.04 | 0.19 | 0.15 |
| 4 | 135623679 | rs16998192 | 0.409 | -0.35 | 0.731 | -2.076 | 0.033 | 0.044 | 0.044 | 3.556 | 0.16 | 0.11 | 0.1 | 0.01 | 0.08 | 0.09 |
| 4 | 135623942 | rs7672011 | 0.03 | -0.277 | 0.721 | -2.042 | 0.019 | -0.107 | -0.107 | 0 | 0.2 | 0.39 | 0.23 | 0.04 | 0.19 | 0.16 |
| 4 | 135643583 | rs6535038 | -0.753 | -0.2 | 0.345 | -1.934 | 0.015 | -0.001 | -0.001 | 0 | 0.4 | 0.65 | 0.34 | 0.05 | 0.26 | 0.3 |
| 4 | 135659888 | rs586090 | -1.491 | 0.002 | -0.644 | -1.635 | -0.02 | -0.013 | -0.013 | 0 | 0.83 | 0.97 | 0.86 | 0.79 | 0.8 | 0.85 |
| 4 | 135678228 | rs11938483 | 0.57 | -0.281 | -0.261 | -1.902 | 0.001 | -0.017 | -0.017 | 0 | 0.19 | 0.16 | 0.12 | 0.01 | 0.1 | 0.17 |
| 4 | 135685643 | rs1506621 | 0.728 | -0.289 | -0.276 | -1.906 | 0.025 | 0.03 | 0.03 | 7.584 | 0.18 | 0.09 | 0.12 | 0.01 | 0.1 | 0.17 |
| 4 | 135686477 | rs548172 | 0.775 | -0.081 | -0.592 | -1.821 | -0.015 | -0.004 | -0.004 | 0 | 0.16 | 0.1 | 0.08 | 0.19 | 0.18 | 0.26 |
| 4 | 135687493 | rs644708 | -0.742 | -0.17 | -0.233 | -1.856 | -0.024 | -0.053 | -0.053 | 0.522 | 0.39 | 0.56 | 0.22 | 0.19 | 0.28 | 0.43 |
| 4 | 135697955 | rs7693807 | 0.025 | -0.193 | 0.028 | -1.581 | -0.049 | -0.108 | -0.108 | 0 | 0.24 | 0.46 | 0.16 | 0.01 | 0.13 | 0.17 |
| 4 | 135698868 | rs4863918 | -0.693 | -0.076 | -0.122 | -1.169 | 0.006 | -0.001 | -0.001 | 0 | 0.45 | 0.66 | 0.52 | 0.42 | 0.4 | 0.37 |
| 4 | 135709542 | rs514587 | -1.799 | -0.572 | -0.203 | -1.198 | -0.041 | -0.095 | -0.095 | 0 | 0.77 | 0.97 | 0.78 | 0.65 | 0.71 | 0.77 |
| 4 | 135726024 | rs1354750 | -0.255 | -1.226 | 0.601 | -0.855 | -0.017 | -0.054 | -0.054 | 0 | 0.23 | 0.33 | 0.15 | 0.01 | 0.13 | 0.18 |
| 4 | 135735591 | rs13144694 | -1.259 | -1.032 | 0.421 | -0.824 | 0.012 | -0.001 | -0.001 | 0 | 0.45 | 0.65 | 0.49 | 0.42 | 0.33 | 0.39 |
| 4 | 135740001 | rs11934675 | 0.052 | -0.517 | -0.749 | -0.719 | 0.018 | 0.017 | 0.017 | 3.238 | 0.21 | 0.16 | 0.13 | 0.01 | 0.13 | 0.18 |
| 4 | 135741863 | rs7667372 | -0.128 | 0.342 | -0.604 | 0.078 | -0.002 | 0.007 | 0.007 | 0 | 0.18 | 0.15 | 0.33 | 0.41 | 0.19 | 0.19 |
| 4 | 135766067 | rs2055671 | -0.493 | 0.192 | -0.427 | -0.058 | -0.002 | -0.004 | -0.004 | 0 | 0.23 | 0.19 | 0.45 | 0.34 | 0.27 | 0.27 |
| 4 | 135766089 | rs2055670 | -1.221 | -0.399 | -0.247 | -0.562 | -0.002 | -0.004 | -0.004 | 1.064 | 0.44 | 0.38 | 0.56 | 0.35 | 0.4 | 0.38 |
| 4 | 135768224 | rs17573425 | 0.412 | -0.4 | -0.484 | -0.588 | 0.012 | 0.095 | 0.095 | 0 | 0.18 | 0.03 | 0.1 | 0.01 | 0.12 | 0.11 |
| 4 | 135772152 | rs13119012 | -1.201 | -0.462 | 0.062 | -0.589 | 0.001 | 0.002 | 0.002 | 0 | 0.45 | 0.43 | 0.57 | 0.35 | 0.4 | 0.38 |
| 4 | 135775238 | rs10013441 | -0.981 | -0.514 | 0.004 | -1.608 | -0.003 | -0.053 | -0.053 | 0 | 0.4 | 0.4 | 0.42 | 0.09 | 0.31 | 0.39 |
| 4 | 135778261 | rs12505192 | -0.759 | -0.519 | 0.047 | -1.607 | 0.008 | -0.003 | -0.003 | 0.157 | 0.38 | 0.3 | 0.41 | 0.09 | 0.3 | 0.39 |
| 4 | 135779831 | rs7684927 | -1.312 | -0.389 | 0.237 | -1.698 | -0.013 | 0.021 | 0.021 | 0 | 0.57 | 0.65 | 0.51 | 0.11 | 0.45 | 0.53 |
| 4 | 135787960 | rs13130652 | 1.013 | -0.38 | 0.043 | -1.666 | -0.001 | -0.035 | -0.035 | 0 | 0.43 | 0.4 | 0.49 | 0.89 | 0.55 | 0.47 |
| 4 | 135788408 | rs4412039 | -0.097 | -0.317 | -0.015 | -0.844 | 0.001 | -0.032 | -0.032 | 0 | 0.83 | 0.83 | 0.7 | 0.91 | 0.83 | 0.75 |
| 4 | 135792739 | rs4864233 | -0.586 | -0.493 | -0.159 | -0.878 | 0.02 | 0.077 | 0.077 | 0.003 | 0.47 | 0.37 | 0.34 | 0.18 | 0.36 | 0.5 |
| 4 | 135793327 | rs6853230 | 0.844 | -0.45 | 0.002 | -0.431 | -0.001 | 0.006 | 0.006 | 0 | 0.46 | 0.55 | 0.56 | 0.65 | 0.55 | 0.49 |
| 4 | 135795762 | rs2896920 | 0.67 | -0.418 | -0.201 | -0.419 | -0.003 | 0.008 | 0.008 | 0.058 | 0.05 | 0.16 | 0.17 | 0.06 | 0.05 | 0.05 |
| 4 | 135796837 | rs10034996 | 0.378 | -0.532 | -0.896 | -0.667 | 0.005 | 0.015 | 0.015 | 1.375 | 0.2 | 0.16 | 0.09 | 0.01 | 0.15 | 0.1 |
| 4 | 135798890 | rs6856596 | 0.24 | -0.352 | -0.64 | 0.176 | 0.003 | 0.069 | 0.069 | 1.257 | 0.92 | 0.86 | 0.89 | 0.8 | 0.9 | 0.96 |
| 4 | 135803220 | rs6813668 | -0.354 | -0.375 | -0.827 | -0.495 | -0.005 | 0.075 | 0.075 | 0 | 0.25 | 0.08 | 0.23 | 0.15 | 0.21 | 0.36 |
| 4 | 135818060 | rs10010200 | -1.414 | -0.156 | 0.085 | -0.122 | -0.003 | -0.023 | -0.023 | 0 | 0.15 | 0.26 | 0.08 | 0.08 | 0.14 | 0.1 |
| 4 | 135829981 | rs10008735 | -1.404 | -0.723 | 0.169 | -0.477 | -0.01 | -0.013 | -0.013 | 0.079 | 0.16 | 0.23 | 0.05 | 0.08 | 0.12 | 0.13 |
| 4 | 135831516 | rs11737779 | 1.203 | -0.919 | -0.019 | -0.187 | 0.064 | 0.005 | 0.005 | 0 | 0.56 | 0.2 | 0.5 | 0.39 | 0.65 | 0.59 |
| 4 | 135832268 | rs11722290 | 1.203 | -0.919 | -0.014 | -0.187 | 0.063 | 0.005 | 0.005 | 0 | 0.56 | 0.21 | 0.5 | 0.39 | 0.65 | 0.59 |
| 4 | 135838960 | rs9998198 | 0.18 | -0.922 | 0.11 | -0.184 | -0.009 | 0.032 | 0.032 | 0 | 0.71 | 0.4 | 0.54 | 0.47 | 0.75 | 0.72 |
| 4 | 135860798 | rs4290941 | -0.881 | -1.099 | 0.822 | -0.285 | 0.019 | -0.026 | -0.026 | 0 | 0.36 | 0.27 | 0.44 | 0.53 | 0.29 | 0.36 |
| 4 | 135886041 | rs7692242 | 1.408 | -0.926 | 0.089 | -0.216 | 0.073 | 0.027 | 0.027 | 0 | 0.51 | 0.27 | 0.47 | 0.38 | 0.63 | 0.59 |
| 4 | 135911869 | rs7668080 | 0.308 | -0.799 | 0.102 | -0.149 | -0.012 | -0.025 | -0.025 | 0 | 0.1 | 0.11 | 0.02 | 0 | 0.04 | 0.09 |
| 4 | 135912595 | rs13140134 | 1.212 | -0.008 | 0.435 | 0.33 | 0.018 | -0.017 | -0.017 | 0.036 | 0.61 | 0.28 | 0.5 | 0.39 | 0.68 | 0.67 |
| 4 | 135916528 | rs4343778 | -1.361 | 0.029 | 0.386 | -0.089 | -0.004 | -0.036 | -0.036 | 0 | 0.11 | 0.16 | 0.06 | 0 | 0.08 | 0.05 |
| 4 | 135918927 | rs4361440 | -0.817 | 0.445 | 0.098 | 0.098 | -0.019 | 0.032 | 0.032 | 0 | 0.18 | 0.06 | 0.11 | 0.13 | 0.19 | 0.14 |
| 4 | 135928708 | rs12509818 | -0.163 | 0.896 | 0.496 | 0.436 | 0.013 | 0.04 | 0.04 | 0 | 0.27 | 0.4 | 0.57 | 0.43 | 0.29 | 0.25 |
| 4 | 135939520 | rs4527530 | -0.733 | 0.631 | 0.262 | 0.42 | -0.001 | -0.001 | -0.001 | 0 | 0.44 | 0.48 | 0.68 | 0.57 | 0.47 | 0.37 |
| 4 | 135949225 | rs4507423 | -0.768 | 0.888 | -0.527 | 0.561 | 0 | 0 | 0 | 0 | 0.46 | 0.4 | 0.69 | 0.57 | 0.51 | 0.39 |
| 4 | 135953351 | rs6535065 | -0.357 | 1.117 | -0.643 | 0.715 | -0.01 | -0.014 | -0.014 | 4.465 | 0.35 | 0.3 | 0.64 | 0.54 | 0.4 | 0.29 |
| 4 | 135954321 | rs4565135 | -0.688 | 1.12 | -0.759 | 0.67 | 0 | -0.001 | -0.001 | 0.602 | 0.45 | 0.57 | 0.62 | 0.36 | 0.44 | 0.34 |
| 4 | 135954367 | rs4563542 | -0.422 | 1.141 | -0.417 | 0.632 | -0.005 | -0.01 | -0.01 | 0 | 0.28 | 0.41 | 0.44 | 0.25 | 0.25 | 0.23 |
| 4 | 135954865 | rs4292380 | 0.431 | 1.551 | -1.166 | 0.249 | -0.072 | 0.013 | 0.013 | 0.023 | 0.18 | 0.06 | 0.14 | 0.14 | 0.31 | 0.16 |
| 4 | 135966148 | rs4530692 | -0.121 | 0.33 | -1.592 | -0.037 | -0.001 | -0.001 | -0.001 | 0 | 0.61 | 0.61 | 0.71 | 0.67 | 0.57 | 0.68 |

| | | | | | | | | | | | | | | | | |
|---|---|---|---|---|---|---|---|---|---|---|---|---|---|---|---|---|
| 4 | 135966595 | rs28568985 | 0.794 | 0.135 | -1.952 | 0.161 | 0.017 | 0.066 | 0.066 | 0 | 0.27 | 0.5 | 0.47 | 0.37 | 0.27 | 0.22 |
| 4 | 135966954 | rs28703919 | -1.369 | -0.035 | -1.706 | 0.03 | 0.001 | 0.034 | 0.034 | 0 | 0.93 | 0.81 | 0.97 | 0.83 | 0.95 | 0.95 |
| 4 | 135974322 | rs12509986 | -0.431 | -0.243 | -1.579 | -0.09 | 0.028 | -0.013 | -0.013 | 0 | 0.73 | 0.61 | 0.82 | 0.74 | 0.66 | 0.81 |
| 4 | 135976246 | rs11730178 | 0.597 | -0.349 | -1.726 | -0.061 | 0.074 | 0.002 | 0.002 | 0 | 0.4 | 0.13 | 0.32 | 0.37 | 0.29 | 0.44 |
| 4 | 135987864 | rs6856823 | -1.084 | -0.299 | -1.787 | -0.166 | -0.005 | 0.017 | 0.017 | 0 | 0.92 | 0.8 | 0.97 | 0.85 | 0.94 | 0.95 |
| 4 | 135989375 | rs7664664 | -0.07 | -0.414 | -1.833 | -0.065 | 0.001 | 0.002 | 0.002 | 0 | 0.41 | 0.57 | 0.54 | 0.53 | 0.43 | 0.43 |
| 4 | 136008847 | rs7687865 | -0.242 | -0.715 | -1.947 | -0.233 | -0.006 | -0.006 | -0.006 | 2.401 | 0.59 | 0.65 | 0.69 | 0.56 | 0.57 | 0.56 |
| 4 | 136012580 | rs12647065 | 1.372 | -0.87 | -1.858 | -0.154 | 0.014 | -0.102 | -0.102 | 0.37 | 0.2 | 0.02 | 0.15 | 0.34 | 0.15 | 0.3 |
| 4 | 136020662 | rs11099317 | 1.398 | -0.868 | -1.858 | -0.189 | 0.009 | -0.103 | -0.103 | 0 | 0.2 | 0.02 | 0.15 | 0.33 | 0.15 | 0.31 |
| 4 | 136022178 | rs4133331 | 0.235 | -0.747 | -1.767 | -0.15 | -0.001 | 0.004 | 0.004 | 0 | 0.47 | 0.63 | 0.64 | 0.4 | 0.46 | 0.37 |
| 4 | 136024521 | rs4128296 | 0.331 | -0.311 | -1.754 | -0.459 | -0.116 | -0.006 | -0.006 | 0 | 0.27 | 0.05 | 0.18 | 0.26 | 0.39 | 0.31 |
| 4 | 136034687 | rs9997836 | 0.562 | -0.067 | -0.869 | -0.554 | 0.002 | -0.119 | -0.119 | 0 | 0.24 | 0.45 | 0.2 | 0.05 | 0.21 | 0.18 |
| 4 | 136037410 | rs4330400 | 0.575 | -0.178 | -1.404 | -0.793 | 0.006 | 0.012 | 0.012 | 0 | 0.42 | 0.4 | 0.59 | 0.7 | 0.39 | 0.5 |
| 4 | 136042022 | rs4452495 | 0.042 | 0.464 | -1.729 | -0.098 | -0.067 | 0.046 | 0.046 | 0 | 0.71 | 0.88 | 0.82 | 0.75 | 0.61 | 0.7 |
| 4 | 136046765 | rs6815438 | 0.039 | -0.674 | -1.062 | 0.098 | 0.003 | 0.004 | 0.004 | 0 | 0.29 | 0.25 | 0.2 | 0.2 | 0.18 | 0.2 |
| 4 | 136057950 | rs4128811 | 0.248 | -0.001 | -1.092 | 0.345 | -0.006 | -0.005 | -0.005 | 0 | 0.56 | 0.6 | 0.49 | 0.46 | 0.49 | 0.62 |
| 4 | 136064120 | rs4501276 | 0.979 | -0.579 | -1.673 | 0.091 | 0.018 | 0.009 | 0.009 | 0 | 0.18 | 0.1 | 0.21 | 0.16 | 0.13 | 0.3 |
| 4 | 136068696 | rs11099320 | 0.299 | -1.107 | -1.919 | -0.275 | 0.077 | 0.093 | 0.093 | 0 | 0.38 | 0.17 | 0.35 | 0.21 | 0.26 | 0.42 |
| 4 | 136074901 | rs2404639 | -1.07 | -1.494 | -2.335 | -0.577 | -0.052 | -0.022 | -0.022 | 0.376 | 0.71 | 0.93 | 0.8 | 0.78 | 0.68 | 0.74 |
| 4 | 136078993 | rs11099322 | -0.17 | -1.207 | -2.263 | -0.355 | 0.122 | 0.171 | 0.171 | 0.039 | 0.49 | 0.21 | 0.37 | 0.22 | 0.33 | 0.52 |
| 4 | 136079340 | rs12505219 | 1.463 | -0.76 | -2.075 | -0.143 | 0.018 | 0.025 | 0.025 | 0 | 0.18 | 0.1 | 0.2 | 0.13 | 0.13 | 0.29 |
| 4 | 136084138 | rs11734733 | 1.206 | -0.225 | -1.457 | 0.178 | 0.07 | 0.087 | 0.087 | 0 | 0.09 | 0.1 | 0.35 | 0.44 | 0.31 | 0.16 |
| 4 | 136087584 | rs10031835 | 1.28 | -0.213 | -1.459 | 0.178 | 0.061 | 0.079 | 0.079 | 0 | 0.09 | 0.09 | 0.36 | 0.44 | 0.31 | 0.15 |
| 4 | 136094675 | rs10009456 | -0.624 | -0.845 | -1.174 | -0.384 | 0.006 | 0.014 | 0.014 | 0 | 0.69 | 0.76 | 0.83 | 0.83 | 0.72 | 0.75 |
| 4 | 136111450 | rs7658327 | 0.065 | -1.15 | -0.779 | -0.718 | -0.003 | -0.005 | -0.005 | 0 | 0.63 | 0.55 | 0.76 | 0.77 | 0.67 | 0.66 |
| 4 | 136111898 | rs10519350 | 1.563 | -1.539 | -1.047 | -0.958 | 0.015 | 0.054 | 0.054 | 0.541 | 0.21 | 0.09 | 0.21 | 0.11 | 0.15 | 0.28 |
| 4 | 136120558 | rs10014016 | -1.791 | -1.695 | -1.175 | -1.084 | -0.011 | 0.049 | 0.049 | 0 | 0.8 | 0.98 | 0.88 | 0.94 | 0.8 | 0.81 |
| 4 | 136123827 | rs13101517 | 0.449 | -1.679 | -0.598 | -1.056 | -0.017 | 0.007 | 0.007 | 0 | 0.39 | 0.29 | 0.54 | 0.66 | 0.49 | 0.37 |
| 4 | 136125132 | rs1841409 | 0.51 | -1.628 | -0.47 | -1.03 | -0.003 | 0.001 | 0.001 | 0 | 0.38 | 0.34 | 0.54 | 0.66 | 0.49 | 0.37 |
| 4 | 136126570 | rs17637542 | 0.754 | -1.651 | -0.39 | -1.042 | -0.024 | 0.004 | 0.004 | 0 | 0.37 | 0.26 | 0.53 | 0.66 | 0.49 | 0.37 |
| 4 | 136131907 | rs10519351 | 1.262 | -1.457 | -0.293 | -0.902 | 0 | -0.02 | -0.02 | 0 | 0.08 | 0.05 | 0.06 | 0.17 | 0.1 | 0.08 |
| 4 | 136147584 | rs11099332 | -0.756 | -1.517 | -0.117 | -0.98 | -0.009 | -0.006 | -0.006 | 0 | 0.69 | 0.61 | 0.8 | 0.77 | 0.72 | 0.73 |
| 4 | 136148697 | rs1824483 | 0.527 | -1.592 | -0.232 | -1.033 | 0.051 | 0.044 | 0.044 | 0 | 0.19 | 0.23 | 0.39 | 0.61 | 0.37 | 0.24 |
| 4 | 136151551 | rs2219587 | -0.061 | -1.656 | -0.298 | -1.057 | 0.097 | 0.068 | 0.068 | 0 | 0.25 | 0.35 | 0.41 | 0.61 | 0.39 | 0.25 |
| 4 | 136164162 | rs17638685 | 1.108 | -1.777 | -0.493 | -1.129 | 0.068 | 0.159 | 0.159 | 0.145 | 0.38 | 0.11 | 0.36 | 0.16 | 0.29 | 0.4 |
| 4 | 136176700 | rs4356958 | -0.487 | -1.599 | -0.192 | -1 | 0.072 | 0.034 | 0.034 | 0 | 0.3 | 0.44 | 0.42 | 0.63 | 0.43 | 0.33 |
| 4 | 136179916 | rs2054294 | 0.476 | -1.479 | -0.324 | -0.934 | 0.07 | 0.058 | 0.058 | 0 | 0.21 | 0.27 | 0.39 | 0.62 | 0.37 | 0.25 |
| 4 | 136192112 | rs17639159 | 1.165 | -1.76 | -0.566 | -1.11 | 0.058 | 0.156 | 0.156 | 0 | 0.37 | 0.1 | 0.36 | 0.15 | 0.29 | 0.38 |
| 4 | 136199802 | rs1460689 | 0.683 | -1.686 | -0.299 | -1.104 | 0.029 | 0.023 | 0.023 | 0 | 0.2 | 0.23 | 0.39 | 0.62 | 0.37 | 0.24 |
| 4 | 136211250 | rs1460686 | -0.455 | -1.779 | -0.214 | -1.136 | 0.084 | 0.047 | 0.047 | 0 | 0.29 | 0.45 | 0.42 | 0.62 | 0.4 | 0.32 |
| 4 | 136217865 | rs1460687 | -0.798 | -1.814 | -0.221 | -1.153 | 0.003 | 0.004 | 0.004 | 0 | 0.67 | 0.67 | 0.79 | 0.77 | 0.69 | 0.7 |
| 4 | 136223378 | rs1460682 | 0.547 | -1.592 | -0.644 | -1 | 0.052 | 0.043 | 0.043 | 0 | 0.2 | 0.29 | 0.43 | 0.62 | 0.37 | 0.26 |
| 4 | 136232574 | rs9307731 | 0.141 | -1.619 | -0.46 | -0.966 | -0.005 | -0.05 | -0.05 | 0 | 0.09 | 0.23 | 0.03 | 0 | 0.03 | 0.08 |
| 4 | 136242130 | rs6844518 | 0.355 | -1.573 | -1.463 | -0.899 | -0.013 | 0.035 | 0.035 | 0.07 | 0.39 | 0.21 | 0.61 | 0.74 | 0.54 | 0.52 |
| 4 | 136257232 | rs1460691 | -1.776 | -1.675 | -1.914 | -1.127 | 0.016 | 0.104 | 0.104 | 0.084 | 0.78 | 0.96 | 0.83 | 0.95 | 0.8 | 0.86 |
| 4 | 136258488 | rs13147783 | 0.029 | -1.169 | -1.52 | -0.584 | -0.018 | 0.015 | 0.015 | 0.059 | 0.39 | 0.24 | 0.57 | 0.71 | 0.52 | 0.51 |
| 4 | 136288768 | rs2035469 | 0.1 | -1.123 | -1.618 | -0.589 | -0.006 | 0.038 | 0.038 | 1.641 | 0.41 | 0.18 | 0.57 | 0.71 | 0.55 | 0.5 |
| 4 | 136294099 | rs6848293 | -1.415 | -1.659 | -2.031 | -0.947 | 0.016 | 0.157 | 0.157 | 0.063 | 0.71 | 0.96 | 0.79 | 0.96 | 0.77 | 0.83 |
| 4 | 136302270 | rs4864269 | -1.448 | -1.668 | -1.69 | -0.961 | 0.038 | 0.105 | 0.105 | 0.009 | 0.7 | 0.86 | 0.79 | 0.96 | 0.77 | 0.83 |
| 4 | 136302474 | rs12510384 | 0.635 | 0.175 | -1.571 | -0.883 | -0.017 | -0.015 | -0.015 | 0 | 0.17 | 0.11 | 0.19 | 0.23 | 0.26 | 0.26 |
| 4 | 136311803 | rs11099340 | 0.216 | 0.087 | -1.414 | -0.766 | -0.035 | 0.046 | 0.046 | 1.655 | 0.41 | 0.2 | 0.54 | 0.73 | 0.47 | 0.5 |
| 4 | 136316187 | rs1905461 | 1.352 | 0.526 | -1.297 | -0.232 | -0.001 | -0.001 | -0.001 | 0 | 0.09 | 0.07 | 0.04 | 0.09 | 0.07 | 0.09 |
| 4 | 136319647 | rs10519357 | 0.345 | 0.919 | -1.686 | -0.428 | -0.043 | 0.009 | 0.009 | 0.158 | 0.21 | 0.06 | 0.14 | 0.18 | 0.24 | 0.16 |
| 4 | 136339396 | rs13135756 | 0.7 | 1.069 | -0.997 | 0.491 | 0.059 | 0.082 | 0.082 | 0 | 0.09 | 0.18 | 0.27 | 0.26 | 0.21 | 0.09 |
| 4 | 136354391 | rs316693 | 1.955 | 0.36 | -1.481 | -0.137 | 0.037 | 0.092 | 0.092 | 0 | 0.23 | 0.08 | 0.17 | 0.05 | 0.17 | 0.14 |
| 4 | 136374928 | rs13112126 | 0.121 | 0.263 | -1.353 | -0.163 | -0.021 | 0.006 | 0.006 | 0.159 | 0.3 | 0.21 | 0.26 | 0.23 | 0.38 | 0.3 |
| 4 | 136381862 | rs17579742 | 1.782 | 0.09 | -1.049 | -0.238 | 0.041 | 0.058 | 0.058 | 0 | 0.26 | 0.16 | 0.18 | 0.05 | 0.18 | 0.16 |
| 4 | 136406835 | rs984160 | 0.083 | 0.199 | -0.636 | 0.029 | -0.001 | 0.005 | 0.005 | 2.159 | 0.58 | 0.43 | 0.44 | 0.28 | 0.56 | 0.46 |
| 4 | 136415590 | rs17641646 | 1.557 | 0.153 | -0.831 | 0.229 | -0.021 | -0.026 | -0.026 | 2.919 | 0.15 | 0.09 | 0.31 | 0.27 | 0.26 | 0.08 |
| 4 | 136417496 | rs11099346 | 0.091 | 0.039 | -0.488 | -0.004 | 0.007 | 0.045 | 0.045 | 0 | 0.51 | 0.38 | 0.42 | 0.26 | 0.52 | 0.47 |
| 4 | 136422205 | rs13110301 | -0.484 | 0.139 | -0.144 | 0.215 | 0.011 | -0.009 | -0.009 | 0 | 0.71 | 0.77 | 0.77 | 0.59 | 0.79 | 0.59 |
| 4 | 136426582 | rs1382998 | 1.027 | 0.97 | -0.005 | 0.129 | 0.025 | 0.006 | 0.006 | 0.333 | 0.07 | 0.09 | 0.12 | 0.08 | 0.14 | 0.12 |
| 4 | 136445508 | rs2404866 | 0.009 | 0.519 | -0.048 | -0.233 | -0.005 | -0.004 | -0.004 | 0 | 0.38 | 0.33 | 0.4 | 0.42 | 0.38 | 0.59 |
| 4 | 136453566 | rs1479776 | -0.36 | -0.032 | 0.26 | -0.282 | -0.001 | -0.004 | -0.004 | 0 | 0.56 | 0.58 | 0.53 | 0.68 | 0.55 | 0.66 |
| 4 | 136455685 | rs1871490 | 0.551 | 0.211 | 0.168 | 0.005 | -0.001 | -0.001 | -0.001 | 0 | 0.26 | 0.25 | 0.37 | 0.28 | 0.3 | 0.17 |
| 4 | 136464719 | rs17047924 | 1.132 | 0.288 | -0.019 | 0.155 | 0.005 | -0.002 | -0.002 | 0.408 | 0.11 | 0.09 | 0.18 | 0.37 | 0.09 | 0.29 |
| 4 | 136466850 | rs12503254 | -0.684 | 0.389 | 0.27 | -0.128 | -0.005 | -0.009 | -0.009 | 0 | 0.55 | 0.59 | 0.55 | 0.7 | 0.59 | 0.67 |
| 4 | 136467679 | rs17047931 | -0.673 | 0.409 | 0.354 | -0.118 | 0.004 | 0.008 | 0.008 | 0 | 0.55 | 0.49 | 0.54 | 0.7 | 0.59 | 0.67 |
| 4 | 136468089 | rs17581218 | 1.114 | 0.748 | -0.028 | -0.169 | -0.048 | 0.034 | 0.034 | 0.545 | 0.16 | 0.08 | 0.21 | 0.05 | 0.24 | 0.18 |

| | | | | | | | | | | | | | | | | |
|---|---|---|---|---|---|---|---|---|---|---|---|---|---|---|---|---|
| 4 | 136482197 | rs7340923 | 1.302 | 0.643 | 0.16 | -0.015 | -0.01 | 0.078 | 0.078 | 0 | 0.11 | 0.17 | 0.19 | 0.38 | 0.09 | 0.28 |
| 4 | 136485458 | rs6857087 | 0.751 | 0.443 | 0.165 | -0.274 | 0.001 | -0.001 | -0.001 | 0.869 | 0.35 | 0.29 | 0.33 | 0.41 | 0.27 | 0.46 |
| 4 | 136486203 | rs17047940 | 1.399 | 0.408 | -0.03 | -0.382 | 0 | -0.003 | -0.003 | 0 | 0.1 | 0.09 | 0.04 | 0.03 | 0.1 | 0.03 |
| 4 | 136490362 | rs17648258 | 0.241 | 0.577 | -0.068 | -0.407 | 0.001 | 0.002 | 0.002 | 0.132 | 0.28 | 0.25 | 0.4 | 0.43 | 0.34 | 0.49 |
| 4 | 136491544 | rs931121 | 1.365 | 0.596 | -0.224 | -0.367 | -0.034 | 0.03 | 0.03 | 0.956 | 0.17 | 0.11 | 0.21 | 0.05 | 0.25 | 0.2 |
| 4 | 136494967 | rs2101018 | 1.51 | 0.557 | -0.181 | -0.377 | -0.001 | 0 | 0 | 0 | 0.17 | 0.2 | 0.11 | 0.25 | 0.16 | 0.13 |
| 4 | 136496620 | rs11935796 | 1.347 | 0.688 | -0.584 | -0.418 | -0.03 | -0.067 | -0.067 | 0 | 0.19 | 0.06 | 0.3 | 0.27 | 0.23 | 0.18 |
| 4 | 136499081 | rs1382988 | 1.228 | 0.615 | -0.556 | -0.483 | -0.049 | 0.04 | 0.04 | 0.913 | 0.17 | 0.08 | 0.21 | 0.05 | 0.24 | 0.18 |
| 4 | 136503631 | rs4146972 | -1.742 | 0.44 | -0.408 | -0.509 | -0.022 | -0.083 | -0.083 | 0 | 0.75 | 0.68 | 0.85 | 0.97 | 0.82 | 0.83 |
| 4 | 136510557 | rs2086327 | -0.015 | 0.597 | -1.407 | -0.401 | -0.068 | -0.02 | -0.02 | 0.456 | 0.37 | 0.18 | 0.49 | 0.34 | 0.46 | 0.37 |
| 4 | 136511399 | rs13117043 | -1.998 | 0.547 | -1.318 | -0.432 | 0.01 | 0.014 | 0.014 | 0 | 0.77 | 0.85 | 0.87 | 0.97 | 0.82 | 0.83 |
| 4 | 136512755 | rs1038151 | -0.44 | 0.856 | -1.499 | 0.072 | -0.068 | -0.021 | -0.021 | 0.345 | 0.37 | 0.18 | 0.49 | 0.34 | 0.46 | 0.37 |
| 4 | 136519891 | rs2404867 | 0.649 | 0.782 | -1.615 | 0.088 | -0.017 | -0.065 | -0.065 | 0 | 0.2 | 0.07 | 0.27 | 0.31 | 0.22 | 0.19 |
| 4 | 136521106 | rs11737486 | 0.737 | 0.808 | -1.758 | 0.101 | -0.03 | -0.082 | -0.082 | 0 | 0.19 | 0.05 | 0.27 | 0.31 | 0.22 | 0.19 |
| 4 | 136525023 | rs10029804 | 0.676 | 0.082 | -1.559 | 0.029 | -0.033 | 0.04 | 0.04 | 0.969 | 0.17 | 0.1 | 0.17 | 0.04 | 0.23 | 0.12 |
| 4 | 136543136 | rs7685332 | 0.76 | 0.164 | -1.568 | 0.26 | 0.037 | -0.064 | -0.064 | 0.105 | 0.17 | 0.07 | 0.28 | 0.33 | 0.13 | 0.17 |
| 4 | 136543541 | rs905772 | 0.746 | 0.131 | -1.462 | 0.259 | 0.009 | -0.012 | -0.012 | 0.742 | 0.17 | 0.15 | 0.3 | 0.33 | 0.13 | 0.17 |
| 4 | 136549624 | rs7666345 | 0.516 | 0.555 | -1.797 | -0.087 | -0.025 | 0.005 | 0.005 | 0 | 0.38 | 0.16 | 0.34 | 0.36 | 0.41 | 0.49 |
| 4 | 136561308 | rs17649956 | -0.13 | 0.97 | -1.699 | -0.524 | -0.04 | 0.062 | 0.062 | 0.038 | 0.24 | 0.14 | 0.19 | 0.07 | 0.29 | 0.16 |
| 4 | 136566098 | rs1610224 | -0.423 | 0.214 | -1.33 | -0.46 | 0.018 | 0.046 | 0.046 | 0 | 0.36 | 0.26 | 0.54 | 0.86 | 0.32 | 0.6 |
| 4 | 136566650 | rs1610225 | 0.407 | -0.029 | -1.283 | 0.056 | 0.004 | -0.041 | -0.041 | 0 | 0.67 | 0.22 | 0.69 | 0.43 | 0.74 | 0.57 |
| 4 | 136570472 | rs17048026 | 0.105 | -0.595 | -1.314 | 0.011 | 0.025 | -0.03 | -0.03 | 0 | 0.16 | 0.1 | 0.35 | 0.27 | 0.11 | 0.16 |
| 4 | 136572585 | rs958632 | 0.133 | -0.664 | -0.93 | -0.031 | 0.018 | -0.017 | -0.017 | 0.021 | 0.16 | 0.15 | 0.36 | 0.27 | 0.11 | 0.16 |
| 4 | 136586509 | rs1354539 | -0.266 | -0.505 | -0.724 | -0.539 | 0 | 0.002 | 0.002 | 0 | 0.47 | 0.48 | 0.58 | 0.37 | 0.48 | 0.38 |
| 4 | 136591498 | rs1504865 | -0.135 | -0.525 | -0.769 | -0.547 | -0.001 | 0.013 | 0.013 | 0.022 | 0.45 | 0.39 | 0.57 | 0.37 | 0.48 | 0.38 |
| 4 | 136599787 | rs12505927 | 0.903 | -0.484 | -0.79 | -0.528 | 0.01 | -0.026 | -0.026 | 0 | 0.29 | 0.36 | 0.23 | 0.1 | 0.36 | 0.23 |
| 4 | 136606270 | rs1566874 | 0.222 | -0.515 | -0.365 | -0.585 | 0.004 | 0.053 | 0.053 | 0 | 0.5 | 0.26 | 0.38 | 0.63 | 0.49 | 0.6 |
| 4 | 136613685 | rs11734150 | 0.952 | -0.513 | -0.338 | -0.516 | 0.008 | -0.02 | -0.02 | 0 | 0.29 | 0.35 | 0.22 | 0.11 | 0.36 | 0.23 |
| 4 | 136614519 | rs1394542 | -0.311 | -0.101 | -0.703 | -0.153 | 0.004 | 0.045 | 0.045 | 0 | 0.5 | 0.26 | 0.38 | 0.62 | 0.49 | 0.61 |
| 4 | 136618495 | rs13127387 | 0.165 | 0.012 | -1.124 | 0.694 | -0.091 | -0.111 | -0.111 | 0.135 | 0.26 | 0.04 | 0.24 | 0.36 | 0.33 | 0.46 |
| 4 | 136633424 | rs2174391 | -0.719 | -0.522 | -0.437 | 0.825 | -0.023 | 0.111 | 0.111 | 0 | 0.1 | 0.13 | 0.06 | 0.47 | 0.05 | 0.39 |
| 4 | 136649383 | rs963645 | -0.302 | -0.741 | -0.609 | 0.492 | 0.007 | -0.018 | -0.018 | 0 | 0.16 | 0.09 | 0.34 | 0.33 | 0.16 | 0.17 |
| 4 | 136650042 | rs2134985 | 0.74 | -1.03 | -0.24 | 0.296 | -0.021 | -0.033 | -0.033 | 0.444 | 0.06 | 0.13 | 0.03 | 0 | 0.02 | 0 |
| 4 | 136651244 | rs6811227 | 0.175 | -1.207 | 0.075 | 0.294 | -0.028 | 0.142 | 0.142 | 0 | 0.9 | 0.84 | 0.95 | 0.51 | 0.95 | 0.61 |
| 4 | 136653594 | rs13111781 | 1.016 | -0.866 | -0.459 | -0.403 | -0.072 | 0.104 | 0.104 | 0.178 | 0.26 | 0.11 | 0.26 | 0.07 | 0.34 | 0.24 |
| 4 | 136660767 | rs7655183 | 1.156 | -0.88 | -0.444 | -0.339 | -0.1 | 0.137 | 0.137 | 0.558 | 0.37 | 0.07 | 0.27 | 0.11 | 0.42 | 0.2 |
| 4 | 136667668 | rs2134987 | 0.763 | -0.897 | 0.294 | -0.143 | -0.013 | -0.032 | -0.032 | 0.087 | 0.07 | 0.16 | 0.03 | 0 | 0.03 | 0 |
| 4 | 136681841 | rs1354538 | 0.541 | -0.972 | 0.412 | 0.044 | -0.039 | 0.135 | 0.135 | 0.001 | 0.82 | 0.69 | 0.9 | 0.53 | 0.9 | 0.66 |
| 4 | 136688436 | rs1604945 | -0.14 | -1.076 | 0.507 | 0.066 | 0 | 0 | 0 | 0 | 0.18 | 0.16 | 0.32 | 0.36 | 0.17 | 0.22 |
| 4 | 136694463 | rs2018601 | -0.628 | -1.005 | 0.371 | -0.201 | 0.001 | 0.025 | 0.025 | 0 | 0.49 | 0.61 | 0.63 | 0.44 | 0.49 | 0.44 |
| 4 | 136698656 | rs17048197 | -0.387 | -1.303 | 1.128 | 0.569 | 0.012 | -0.03 | -0.03 | 0 | 0.18 | 0.09 | 0.31 | 0.36 | 0.17 | 0.22 |
| 4 | 136711280 | rs6535115 | 1.661 | -1.23 | 1.418 | 0.549 | 0.008 | 0.008 | 0.008 | 0.152 | 0.63 | 0.34 | 0.57 | 0.17 | 0.74 | 0.46 |
| 4 | 136716868 | rs10028711 | -1.484 | -1.008 | 1.741 | 0.807 | -0.044 | 0.022 | 0.022 | 0 | 0.25 | 0.51 | 0.35 | 0.33 | 0.18 | 0.2 |
| 4 | 136724563 | rs12500395 | -1.902 | -1.192 | 1.959 | 0.779 | -0.057 | 0.188 | 0.188 | 0.348 | 0.19 | 0.43 | 0.12 | 0.5 | 0.11 | 0.33 |
| 4 | 136727025 | rs7657021 | 1.895 | -1.248 | 2.022 | 0.751 | -0.054 | 0.09 | 0.09 | 3.567 | 0.77 | 0.4 | 0.87 | 0.5 | 0.89 | 0.67 |
| 4 | 136727303 | rs12513116 | -0.373 | -1.17 | 1.913 | 0.79 | 0.006 | 0.023 | 0.023 | 0 | 0.1 | 0.06 | 0.07 | 0.5 | 0.08 | 0.33 |
| 4 | 136728487 | rs4311361 | 0.133 | -1.161 | 1.955 | 0.683 | -0.017 | -0.037 | -0.037 | 0.041 | 0.08 | 0.27 | 0.04 | 0 | 0.02 | 0 |
| 4 | 136730024 | rs11099363 | -1.745 | -0.871 | 1.498 | 0.653 | -0.001 | -0.006 | -0.006 | 0 | 0.29 | 0.3 | 0.39 | 0.83 | 0.28 | 0.52 |
| 4 | 136737730 | rs6535121 | 1.508 | -0.9 | 1.033 | 0.861 | -0.005 | 0.001 | 0.001 | 0.408 | 0.71 | 0.36 | 0.62 | 0.64 | 0.77 | 0.79 |
| 4 | 136741799 | rs2068179 | -0.441 | -0.792 | 0.738 | 0.951 | -0.006 | -0.016 | -0.016 | 0.021 | 0.17 | 0.12 | 0.33 | 0.31 | 0.2 | 0.21 |
| 4 | 136750399 | rs13139177 | 1.622 | -0.833 | 0.427 | 0.596 | 0.033 | 0.068 | 0.068 | 1.025 | 0.59 | 0.29 | 0.5 | 0.17 | 0.68 | 0.46 |
| 4 | 136754367 | rs6852986 | 1.525 | -0.908 | 0.337 | 1.018 | 0.021 | -0.022 | -0.022 | 0 | 0.73 | 0.24 | 0.77 | 0.45 | 0.84 | 0.69 |
| 4 | 136754442 | rs6853180 | 1.867 | -0.903 | 0.295 | 1.001 | 0.029 | -0.026 | -0.026 | 0.112 | 0.71 | 0.24 | 0.76 | 0.45 | 0.8 | 0.68 |
| 4 | 136754993 | rs10005321 | -1.658 | -0.94 | 0.502 | 1.226 | -0.014 | 0.016 | 0.016 | 2.484 | 0.25 | 0.67 | 0.2 | 0.43 | 0.13 | 0.29 |
| 4 | 136755136 | rs4443343 | -2.073 | -0.931 | 0.474 | 1.199 | 0.014 | -0.014 | -0.014 | 0 | 0.28 | 0.69 | 0.23 | 0.55 | 0.2 | 0.32 |
| 4 | 136755552 | rs4302513 | 2.023 | -0.931 | 0.443 | 0.926 | 0.052 | 0.074 | 0.074 | 0.818 | 0.58 | 0.13 | 0.48 | 0.29 | 0.68 | 0.48 |
| 4 | 136760894 | rs6535122 | -1.87 | -0.9 | 1.359 | 0.755 | 0.023 | 0.085 | 0.085 | 1.553 | 0.1 | 0.42 | 0.06 | 0.12 | 0.07 | 0.03 |
| 4 | 136763493 | rs7672503 | 0.968 | -0.993 | 1.337 | 0.696 | -0.017 | -0.124 | -0.124 | 0 | 0.77 | 0.53 | 0.85 | 0.99 | 0.84 | 0.8 |
| 4 | 136764416 | rs12645397 | 1.792 | -0.964 | 0.765 | 0.735 | 0.034 | 0.113 | 0.113 | 1.496 | 0.67 | 0.1 | 0.79 | 0.87 | 0.76 | 0.77 |
| 4 | 136778939 | rs4610390 | 1.497 | -0.831 | 1.487 | 1.037 | -0.03 | 0.014 | 0.014 | 1.609 | 0.68 | 0.43 | 0.55 | 0.36 | 0.76 | 0.55 |
| 4 | 136828780 | rs11722042 | -1.445 | -0.319 | 1.145 | 0.45 | -0.001 | 0.029 | 0.029 | 0 | 0.53 | 0.6 | 0.68 | 0.8 | 0.57 | 0.58 |
| 4 | 136837128 | rs4863945 | 1.542 | -0.326 | 0.999 | 0.447 | -0.06 | 0.191 | 0.191 | 0 | 0.45 | 0.1 | 0.3 | 0.2 | 0.42 | 0.42 |
| 4 | 136844538 | rs11734469 | -1.575 | -0.318 | 1.035 | 0.451 | -0.025 | 0.163 | 0.163 | 0 | 0.54 | 0.77 | 0.68 | 0.8 | 0.58 | 0.58 |
| 4 | 136847562 | rs4339269 | -1.221 | -0.376 | 1.406 | 0.405 | -0.009 | 0.089 | 0.089 | 0 | 0.47 | 0.31 | 0.63 | 0.8 | 0.56 | 0.56 |
| 4 | 136866722 | rs12509309 | 0.221 | -0.376 | 1.245 | 0.557 | -0.001 | 0.029 | 0.029 | 0 | 0.61 | 0.66 | 0.67 | 0.81 | 0.55 | 0.69 |
| 4 | 136871982 | rs12507949 | 1.541 | -0.27 | 0.454 | 0.617 | -0.048 | 0.189 | 0.189 | 0 | 0.45 | 0.12 | 0.32 | 0.19 | 0.43 | 0.42 |
| 4 | 136882276 | rs7679231 | 0.309 | -0.233 | 0.508 | 0.898 | -0.014 | -0.009 | -0.009 | 0 | 0.09 | 0.02 | 0.3 | 0.61 | 0.11 | 0.26 |
| 4 | 136910271 | rs4318703 | -1.474 | -0.429 | 0.787 | 0.723 | -0.045 | 0.022 | 0.022 | 2.959 | 0.31 | 0.56 | 0.46 | 0.64 | 0.24 | 0.38 |
| 4 | 136910907 | rs4455480 | -0.633 | 0.294 | 0.721 | 0.963 | 0.004 | 0 | 0 | 0 | 0.77 | 0.76 | 0.77 | 0.79 | 0.67 | 0.81 |

| | | | | | | | | | | | | | | | | |
|---|---|---|---|---|---|---|---|---|---|---|---|---|---|---|---|---|
| 4 | 136919049 | rs11733579 | -0.654 | 0.126 | 0.622 | 0.778 | -0.003 | 0.002 | 0.002 | 0 | 0.4 | 0.38 | 0.58 | 0.66 | 0.46 | 0.5 |
| 4 | 136940740 | rs4580703 | -0.975 | 0.227 | 0.899 | 0.777 | -0.031 | 0.007 | 0.007 | 0 | 0.12 | 0.25 | 0.16 | 0.09 | 0.11 | 0.15 |
| 4 | 136958364 | rs6535135 | -0.699 | 1.128 | 0.444 | 0.806 | -0.01 | -0.001 | -0.001 | 0 | 0.34 | 0.28 | 0.51 | 0.69 | 0.44 | 0.47 |
| 4 | 137003101 | rs4145876 | -0.551 | 1.326 | 1.043 | 1.599 | 0.002 | 0.012 | 0.012 | 0.349 | 0.25 | 0.21 | 0.43 | 0.5 | 0.29 | 0.44 |
| 4 | 137016357 | rs17048466 | -0.598 | 1.173 | 0.993 | 1.412 | -0.001 | 0 | 0 | 0 | 0.07 | 0.07 | 0.11 | 0.03 | 0.07 | 0.13 |
| 4 | 137018385 | rs1344033 | 1.318 | 1.351 | 0.827 | 1.747 | -0.05 | -0.06 | -0.06 | 0.281 | 0.1 | 0 | 0.32 | 0.25 | 0.15 | 0.14 |
| 4 | 137021516 | rs12511521 | -1.916 | 0.929 | 1.288 | 1.362 | -0.021 | 0.126 | 0.126 | 1.257 | 0.16 | 0.6 | 0.19 | 0.32 | 0.13 | 0.18 |
| 4 | 137043545 | rs7675695 | 1.163 | 1.76 | 0.825 | 1.102 | -0.04 | 0.028 | 0.028 | 0.469 | 0.06 | 0.01 | 0.07 | 0 | 0.13 | 0.03 |
| 4 | 137071335 | rs1451109 | -0.228 | 1.531 | 1.199 | 1.386 | 0.023 | 0.046 | 0.046 | 0.986 | 0.24 | 0.31 | 0.37 | 0.47 | 0.32 | 0.42 |
| 4 | 137074119 | rs1724563 | 1.666 | 0.674 | 1.669 | 1.274 | -0.027 | 0.19 | 0.19 | 0.714 | 0.87 | 0.54 | 0.83 | 0.73 | 0.89 | 0.85 |
| 4 | 137088592 | rs1283049 | 1.811 | 0.616 | 1.29 | 1.264 | -0.044 | 0.158 | 0.158 | 1.554 | 0.86 | 0.46 | 0.83 | 0.73 | 0.9 | 0.85 |
| 4 | 137105336 | rs2897153 | -0.732 | 0.627 | 0.86 | 1.267 | -0.022 | 0.034 | 0.034 | 0 | 0.1 | 0.16 | 0.12 | 0.26 | 0.07 | 0.14 |
| 4 | 137116996 | rs2034535 | 1.819 | 1.011 | 1.442 | 1.244 | -0.011 | 0.111 | 0.111 | 0.015 | 0.82 | 0.42 | 0.78 | 0.67 | 0.83 | 0.81 |
| 4 | 137138361 | rs12506698 | -0.272 | 0.797 | 0.9 | 0.58 | -0.048 | -0.031 | -0.031 | 0 | 0.35 | 0.19 | 0.45 | 0.56 | 0.5 | 0.56 |
| 4 | 137154796 | rs2053896 | 0.25 | 0.725 | 0.959 | 1.047 | -0.026 | -0.065 | -0.065 | 0 | 0.13 | 0.01 | 0.07 | 0.23 | 0.13 | 0.29 |
| 4 | 137181847 | rs968639 | 1.627 | 0.397 | 1.609 | 0.624 | 0.029 | 0.287 | 0.287 | 2.698 | 0.91 | 0.46 | 0.89 | 0.7 | 0.94 | 0.92 |
| 4 | 137196244 | rs1451111 | 1.807 | 0.304 | 1.072 | 0.546 | -0.012 | 0.007 | 0.007 | 0.959 | 0.81 | 0.26 | 0.76 | 0.69 | 0.87 | 0.83 |
| 4 | 137213250 | rs11723419 | 0.9 | -0.411 | 0.318 | -0.216 | -0.004 | 0.166 | 0.166 | 1.52 | 0.32 | 0.05 | 0.21 | 0.12 | 0.24 | 0.2 |
| 4 | 137213987 | rs6857747 | -1.311 | -0.43 | 1.478 | 0.189 | -0.052 | 0.082 | 0.082 | 2.245 | 0.19 | 0.5 | 0.23 | 0.29 | 0.11 | 0.15 |
| 4 | 137223663 | rs11099379 | -1.5 | -0.395 | 1.19 | 0.194 | -0.036 | 0.058 | 0.058 | 0.034 | 0.21 | 0.47 | 0.26 | 0.29 | 0.16 | 0.19 |
| 4 | 137233717 | rs2110186 | 0.45 | -0.271 | 0.873 | -0.503 | 0.032 | -0.029 | -0.029 | 0 | 0.91 | 0.59 | 0.93 | 1 | 0.95 | 0.96 |
| 4 | 137241357 | rs2192201 | 1.432 | -0.293 | 0.848 | -0.536 | -0.024 | -0.078 | -0.078 | 3.297 | 0.86 | 0.35 | 0.89 | 0.97 | 0.93 | 0.91 |
| 4 | 137246038 | rs1558514 | 0.735 | -0.249 | 0.723 | -0.506 | 0.02 | -0.049 | -0.049 | 0 | 0.89 | 0.54 | 0.9 | 0.97 | 0.93 | 0.91 |
| 4 | 137247266 | rs10026312 | 1.094 | -0.234 | -0.387 | -0.484 | 0.018 | 0.028 | 0.028 | 1.831 | 0.72 | 0.23 | 0.71 | 0.91 | 0.82 | 0.8 |
| 4 | 137252090 | rs10007624 | 0.419 | 0.014 | -0.745 | -0.284 | -0.001 | -0.001 | -0.001 | 0 | 0.33 | 0.26 | 0.41 | 0.36 | 0.37 | 0.36 |
| 4 | 137253127 | rs11947164 | 0.367 | -0.017 | -0.995 | -0.533 | -0.03 | 0.066 | 0.066 | 2.204 | 0.11 | 0.02 | 0.13 | 0 | 0.12 | 0.1 |
| 4 | 137258380 | rs6858670 | -0.467 | -0.127 | -0.557 | -0.585 | -0.001 | -0.001 | -0.001 | 0.021 | 0.47 | 0.41 | 0.56 | 0.36 | 0.49 | 0.46 |
| 4 | 137259372 | rs13102201 | -0.312 | -0.118 | -0.54 | -0.589 | -0.002 | -0.043 | -0.043 | 0.254 | 0.14 | 0.15 | 0.14 | 0 | 0.12 | 0.1 |
| 4 | 137259828 | rs10013817 | -0.76 | -0.373 | -0.899 | -0.364 | 0.01 | -0.009 | -0.009 | 0 | 0.6 | 0.43 | 0.64 | 0.69 | 0.59 | 0.76 |
| 4 | 137299152 | rs17048626 | 0.359 | -0.527 | -1.045 | -0.301 | 0.016 | -0.092 | -0.092 | 0.513 | 0.14 | 0.01 | 0.07 | 0.34 | 0.1 | 0.32 |
| 4 | 137318397 | rs13110422 | 0.158 | -0.684 | -1.25 | -0.967 | -0.007 | 0.03 | 0.03 | 3.617 | 0.39 | 0.26 | 0.33 | 0.08 | 0.4 | 0.31 |
| 4 | 137323344 | rs4864340 | -0.654 | -0.747 | -0.972 | -0.806 | -0.015 | 0.02 | 0.02 | 0.003 | 0.84 | 0.75 | 0.89 | 0.65 | 0.9 | 0.68 |
| 4 | 137324597 | rs7682599 | 1.108 | -0.448 | -1.18 | -0.863 | -0.023 | 0.031 | 0.031 | 0 | 0.12 | 0.06 | 0.11 | 0 | 0.19 | 0.06 |
| 4 | 137328242 | rs17048646 | 0.111 | -0.339 | -1.201 | -0.685 | -0.02 | -0.046 | -0.046 | 0.653 | 0.21 | 0.14 | 0.19 | 0.3 | 0.28 | 0.11 |
| 4 | 137332146 | rs10489022 | 1.666 | -0.391 | -1.271 | -0.928 | -0.006 | -0.01 | -0.01 | 0 | 0.27 | 0.16 | 0.37 | 0.34 | 0.31 | 0.29 |
| 4 | 137333550 | rs10489023 | 1.339 | -0.449 | -1.317 | -0.888 | 0.005 | -0.068 | -0.068 | 0 | 0.1 | 0 | 0.03 | 0.26 | 0.08 | 0.25 |
| 4 | 137335125 | rs17719661 | 1.877 | -0.398 | -1.331 | -1.077 | -0.064 | 0.074 | 0.074 | 3.899 | 0.18 | 0.03 | 0.15 | 0.07 | 0.2 | 0.17 |
| 4 | 137335180 | rs13132705 | -0.137 | -0.427 | -1.164 | -1.053 | 0.01 | 0.084 | 0.084 | 0.475 | 0.77 | 0.44 | 0.78 | 0.72 | 0.77 | 0.91 |
| 4 | 137336902 | rs13146292 | -0.626 | -0.393 | -1.127 | -1.027 | 0.005 | 0.054 | 0.054 | 0.505 | 0.5 | 0.28 | 0.42 | 0.38 | 0.47 | 0.62 |
| 4 | 137349221 | rs11732083 | 1.4 | -0.697 | -1.124 | -0.917 | -0.004 | -0.005 | -0.005 | 0.361 | 0.19 | 0.12 | 0.15 | 0.07 | 0.2 | 0.17 |
| 4 | 137355816 | rs982315 | 0.179 | -0.264 | -1.176 | -0.629 | 0.025 | 0.063 | 0.063 | 0.378 | 0.77 | 0.54 | 0.78 | 0.73 | 0.74 | 0.91 |
| 4 | 137358857 | rs2016184 | 1.032 | -0.253 | -1.408 | -0.648 | -0.005 | -0.009 | -0.009 | 0 | 0.31 | 0.23 | 0.26 | 0.38 | 0.29 | 0.45 |
| 4 | 137359917 | rs13106779 | 0.049 | -0.344 | -1.441 | -0.366 | 0.03 | 0.088 | 0.088 | 0 | 0.18 | 0.05 | 0.14 | 0.01 | 0.14 | 0.16 |
| 4 | 137364012 | rs6848260 | 1.014 | -0.355 | -1.364 | -0.38 | -0.003 | -0.001 | -0.001 | 0 | 0.33 | 0.32 | 0.28 | 0.38 | 0.31 | 0.46 |
| 4 | 137364820 | rs1476909 | -0.083 | -0.364 | -1.372 | -0.389 | 0.001 | 0.039 | 0.039 | 0.051 | 0.51 | 0.37 | 0.43 | 0.38 | 0.45 | 0.61 |
| 4 | 137376628 | rs2215746 | 0.727 | -0.151 | -1.4 | -0.356 | -0.033 | -0.048 | -0.048 | 0 | 0.13 | 0.05 | 0.15 | 0.27 | 0.21 | 0.06 |
| 4 | 137382422 | rs17721005 | -0.094 | -0.115 | -1.355 | -0.309 | 0.032 | 0.103 | 0.103 | 0 | 0.19 | 0.05 | 0.14 | 0.01 | 0.15 | 0.14 |
| 4 | 137396283 | rs6848692 | 0.547 | 0.35 | -0.805 | 0.39 | -0.003 | 0.007 | 0.007 | 0 | 0.43 | 0.32 | 0.34 | 0.4 | 0.4 | 0.52 |
| 4 | 137397046 | rs6854954 | -0.421 | 0.395 | 0.059 | 0.289 | 0.003 | -0.002 | -0.002 | 0 | 0.29 | 0.33 | 0.39 | 0.34 | 0.32 | 0.33 |
| 4 | 137408872 | rs6535149 | 0.177 | 0.126 | -0.521 | 0.639 | 0.01 | 0 | 0 | 0 | 0.57 | 0.41 | 0.76 | 0.46 | 0.71 | 0.62 |
| 4 | 137409145 | rs7662012 | -0.212 | -0.315 | 0.024 | 0.086 | 0 | -0.049 | -0.049 | 0 | 0.25 | 0.21 | 0.1 | 0 | 0.2 | 0.17 |
| 4 | 137413277 | rs10028445 | -0.064 | -0.427 | -0.008 | 0.012 | 0 | -0.053 | -0.053 | 0.563 | 0.21 | 0.21 | 0.1 | 0 | 0.19 | 0.16 |
| 4 | 137414375 | rs6853939 | 1.183 | -1.421 | -0.901 | -0.944 | -0.106 | 0.151 | 0.151 | 0 | 0.32 | 0.1 | 0.52 | 0.11 | 0.58 | 0.3 |
| 4 | 137417231 | rs12505589 | 0.209 | -2.181 | -0.678 | -0.741 | 0.015 | 0.008 | 0.008 | 0 | 0.23 | 0.14 | 0.08 | 0.17 | 0.12 | 0.24 |
| 4 | 137423151 | rs6855815 | 0.537 | -2.124 | -0.62 | -0.788 | -0.001 | -0.033 | -0.033 | 2.802 | 0.24 | 0.2 | 0.1 | 0 | 0.22 | 0.16 |
| 4 | 137423956 | rs7678967 | 0.207 | -2.251 | -0.381 | 0.279 | -0.032 | 0.026 | 0.026 | 0.053 | 0.72 | 0.64 | 0.88 | 0.53 | 0.89 | 0.63 |
| 4 | 137429284 | rs10000852 | 0.433 | -2.326 | -0.409 | 0.052 | -0.009 | -0.072 | -0.072 | 0.761 | 0.17 | 0.21 | 0.06 | 0 | 0.09 | 0.07 |
| 4 | 137429980 | rs13104804 | 0.148 | -2.49 | -0.509 | -0.043 | 0.011 | 0.034 | 0.034 | 0.113 | 0.3 | 0.15 | 0.13 | 0.17 | 0.24 | 0.32 |
| 4 | 137445190 | rs12503505 | 0.263 | -2.489 | -0.356 | -0.045 | 0.005 | 0.015 | 0.015 | 0.033 | 0.3 | 0.22 | 0.14 | 0.18 | 0.24 | 0.33 |
| 4 | 137445988 | rs6535158 | -0.805 | -2.486 | -0.396 | -0.113 | -0.012 | -0.088 | -0.088 | 0.269 | 0.83 | 0.71 | 0.94 | 1 | 0.91 | 0.93 |
| 4 | 137454023 | rs4315856 | -1.426 | -2.614 | 0.146 | -0.06 | 0.014 | -0.028 | -0.028 | 0 | 0.56 | 0.75 | 0.26 | 0.56 | 0.35 | 0.52 |
| 4 | 137474142 | rs9685146 | 0.157 | -2.463 | 0.366 | -0.15 | -0.017 | -0.092 | -0.092 | 0.018 | 0.18 | 0.3 | 0.07 | 0 | 0.09 | 0.06 |
| 4 | 137479869 | rs9683947 | -0.573 | -2.842 | 0.662 | -0.188 | 0.005 | 0.002 | 0.002 | 0 | 0.42 | 0.39 | 0.18 | 0.13 | 0.29 | 0.39 |
| 4 | 137480493 | rs11733040 | -1.331 | -3.027 | 0.679 | -0.061 | 0.019 | -0.01 | -0.01 | 0 | 0.55 | 0.7 | 0.26 | 0.5 | 0.35 | 0.52 |
| 4 | 137483215 | rs4607283 | -1.018 | -3.008 | 0.773 | -0.052 | -0.052 | 0.052 | 0.052 | 0 | 0.24 | 0.51 | 0.11 | 0.37 | 0.12 | 0.18 |
| 4 | 137485929 | rs10519392 | -0.329 | -2.957 | 0.804 | -0.015 | -0.025 | 0.193 | 0.193 | 0 | 0.14 | 0.45 | 0.08 | 0.37 | 0.07 | 0.13 |
| 4 | 137493086 | rs1343837 | -0.346 | -3.066 | 0.748 | -0.051 | 0.021 | 0.061 | 0.061 | 0 | 0.41 | 0.25 | 0.18 | 0.13 | 0.29 | 0.39 |
| 4 | 137493839 | rs1317842 | 0.402 | -3.008 | 0.812 | -0.019 | -0.016 | 0.134 | 0.134 | 0 | 0.12 | 0.25 | 0.07 | 0.36 | 0.07 | 0.14 |
| 4 | 137504762 | rs7695493 | -1.437 | -3.047 | 0.608 | -0.002 | 0.003 | -0.047 | -0.047 | 0.575 | 0.57 | 0.87 | 0.28 | 0.49 | 0.36 | 0.53 |

| | | | | | | | | | | | | | | | | |
|---|---|---|---|---|---|---|---|---|---|---|---|---|---|---|---|---|
| 4 | 137518143 | rs7682676 | -1.438 | -3.072 | 0.596 | -0.001 | 0.003 | -0.047 | -0.047 | 0 | 0.57 | 0.87 | 0.28 | 0.49 | 0.36 | 0.53 |
| 4 | 137524479 | rs4304024 | -0.606 | -3.126 | 0.727 | -0.033 | 0.001 | -0.009 | -0.009 | 0 | 0.43 | 0.4 | 0.18 | 0.13 | 0.29 | 0.39 |
| 4 | 137524981 | rs13115947 | 0.544 | -3.061 | 0.553 | 0.01 | 0.01 | 0.05 | 0.05 | 0.573 | 0.3 | 0.16 | 0.14 | 0.13 | 0.23 | 0.35 |
| 4 | 137526751 | rs10012428 | -0.878 | -2.704 | 0.836 | 0.248 | -0.045 | 0.145 | 0.145 | 0.544 | 0.16 | 0.6 | 0.1 | 0.37 | 0.07 | 0.15 |
| 4 | 137529812 | rs17048799 | -0.299 | -2.135 | 0.549 | -0.13 | -0.027 | -0.084 | -0.084 | 0 | 0.17 | 0.28 | 0.06 | 0.05 | 0.09 | 0.11 |
| 4 | 137533307 | rs11099401 | 0.836 | -0.712 | -0.264 | -0.077 | -0.012 | -0.009 | -0.009 | 0 | 0.26 | 0.23 | 0.38 | 0.3 | 0.38 | 0.22 |
| 4 | 137533710 | rs12509153 | 0.578 | 0.229 | -0.33 | -0.755 | 0.03 | -0.055 | -0.055 | 0 | 0.12 | 0.22 | 0.18 | 0.01 | 0.18 | 0.09 |
| 4 | 137534902 | rs12509396 | -0.419 | 1.005 | -0.603 | -0.632 | 0.005 | -0.015 | -0.015 | 0 | 0.83 | 0.82 | 0.88 | 0.9 | 0.82 | 0.76 |
| 4 | 137535800 | rs2406020 | 0.756 | 0.775 | -1.102 | -0.785 | 0.02 | 0.091 | 0.091 | 0 | 0.3 | 0.08 | 0.31 | 0.12 | 0.23 | 0.34 |
| 4 | 137535918 | rs4864355 | -0.234 | 0.793 | -0.559 | -0.645 | 0.005 | 0.025 | 0.025 | 0 | 0.51 | 0.56 | 0.55 | 0.78 | 0.58 | 0.4 |
| 4 | 137539440 | rs6815580 | 0.006 | 0.955 | 0.406 | -0.031 | -0.001 | 0 | 0 | 0.014 | 0.52 | 0.51 | 0.59 | 0.47 | 0.48 | 0.55 |
| 4 | 137542931 | rs2405826 | -1.84 | 0.899 | 0.577 | -0.347 | 0.018 | -0.021 | -0.021 | 0 | 0.24 | 0.5 | 0.2 | 0.28 | 0.23 | 0.29 |
| 4 | 137551442 | rs13125562 | -0.658 | 1.07 | 0.053 | -0.038 | 0.04 | 0.076 | 0.076 | 0.115 | 0.07 | 0.21 | 0.09 | 0.22 | 0.09 | 0.1 |
| 4 | 137567044 | rs2406055 | 0.747 | 0.967 | -0.514 | -0.277 | -0.02 | 0.051 | 0.051 | 0 | 0.41 | 0.21 | 0.61 | 0.68 | 0.43 | 0.41 |
| 4 | 137573355 | rs17615017 | -0.219 | 1.53 | -0.398 | -0.156 | 0.016 | -0.012 | -0.012 | 0 | 0.09 | 0.16 | 0.09 | 0.06 | 0.13 | 0.11 |
| 4 | 137573579 | rs7685386 | -0.602 | 1.45 | -0.905 | -0.355 | -0.05 | 0.141 | 0.141 | 1.497 | 0.72 | 0.96 | 0.85 | 0.95 | 0.7 | 0.7 |
| 4 | 137581282 | rs6812959 | 1.022 | 1.377 | -0.734 | -0.143 | 0.014 | 0.079 | 0.079 | 0 | 0.43 | 0.17 | 0.52 | 0.74 | 0.38 | 0.45 |
| 4 | 137589137 | rs11728206 | -0.702 | 1.429 | -0.521 | -0.228 | -0.007 | 0.099 | 0.099 | 0.059 | 0.71 | 0.86 | 0.85 | 0.95 | 0.7 | 0.7 |
| 4 | 137590858 | rs2406041 | -0.845 | 1.709 | -0.745 | 0.104 | -0.021 | -0.003 | -0.003 | 0.967 | 0.13 | 0.07 | 0.19 | 0.14 | 0.23 | 0.21 |
| 4 | 137590890 | rs2406040 | -1.192 | 1.66 | -0.47 | 0.3 | 0.038 | 0.043 | 0.043 | 0 | 0.17 | 0.28 | 0.21 | 0.32 | 0.23 | 0.21 |
| 4 | 137591468 | rs2406037 | -0.488 | 1.649 | -0.404 | 0.29 | -0.005 | -0.045 | -0.045 | 0 | 0.59 | 0.47 | 0.72 | 0.88 | 0.61 | 0.65 |
| 4 | 137593405 | rs12512934 | 0.237 | 1.663 | -0.478 | 0.271 | 0 | 0.063 | 0.063 | 0.016 | 0.5 | 0.23 | 0.6 | 0.62 | 0.51 | 0.55 |
| 4 | 137593654 | rs12513015 | 0.136 | 1.684 | -0.507 | 0.28 | -0.009 | 0.007 | 0.007 | 0 | 0.08 | 0.05 | 0.09 | 0.06 | 0.13 | 0.11 |
| 4 | 137599574 | rs7658823 | 0.405 | 1.233 | -0.565 | 0.55 | 0.013 | -0.008 | -0.008 | 0 | 0.42 | 0.18 | 0.51 | 0.57 | 0.38 | 0.44 |
| 4 | 137614143 | rs17048874 | -1.439 | 1.472 | 0.05 | 1.094 | 0.061 | 0.123 | 0.123 | 0 | 0.09 | 0.31 | 0.12 | 0.22 | 0.11 | 0.11 |
| 4 | 137626639 | rs2125883 | 0.031 | 1.205 | -0.6 | 0.675 | -0.004 | 0.006 | 0.006 | 0 | 0.68 | 0.71 | 0.73 | 0.78 | 0.67 | 0.62 |
| 4 | 137632211 | rs2612181 | 0.017 | 1.227 | -0.578 | 0.711 | -0.009 | 0.049 | 0.049 | 0 | 0.72 | 0.81 | 0.85 | 0.88 | 0.72 | 0.68 |
| 4 | 137634247 | rs7658630 | -0.535 | 0.328 | -0.322 | 0.065 | -0.042 | 0.016 | 0.016 | 0 | 0.25 | 0.39 | 0.24 | 0.24 | 0.19 | 0.28 |
| 4 | 137634874 | rs1460160 | 0.307 | 1.26 | -0.903 | 0.944 | -0.047 | -0.023 | -0.023 | 0 | 0.08 | 0.01 | 0.23 | 0.43 | 0.18 | 0.1 |
| 4 | 137638537 | rs6832309 | -0.516 | 0.773 | -0.693 | 0.442 | 0.011 | 0.019 | 0.019 | 0 | 0.47 | 0.38 | 0.26 | 0.07 | 0.41 | 0.47 |
| 4 | 137643932 | rs13109827 | -0.371 | 0.363 | -0.462 | 0.861 | 0.013 | 0.015 | 0.015 | 0 | 0.19 | 0.1 | 0.2 | 0.12 | 0.17 | 0.25 |
| 4 | 137647418 | rs6831651 | -0.908 | 0.377 | 0.209 | 0.995 | -0.023 | 0.107 | 0.107 | 0.315 | 0.07 | 0.3 | 0.04 | 0.11 | 0.02 | 0.03 |
| 4 | 137650213 | rs9996792 | -0.737 | 0.426 | 0.576 | 1.189 | 0.005 | 0.114 | 0.114 | 0 | 0.12 | 0.31 | 0.11 | 0.25 | 0.12 | 0.12 |
| 4 | 137652477 | rs10003357 | -0.715 | 0.427 | 0.576 | 1.189 | 0.006 | 0.116 | 0.116 | 0 | 0.12 | 0.31 | 0.11 | 0.25 | 0.12 | 0.12 |
| 4 | 137654086 | rs13120314 | 0.113 | -0.211 | 0.237 | 0.017 | 0.015 | -0.011 | -0.011 | 0 | 0.32 | 0.28 | 0.14 | 0.02 | 0.21 | 0.16 |
| 4 | 137654857 | rs10009978 | 1.238 | -0.948 | 0.059 | -0.2 | -0.001 | 0 | 0 | 0.051 | 0.16 | 0.13 | 0.15 | 0.12 | 0.1 | 0.28 |
| 4 | 137657271 | rs17617045 | 0.948 | -0.078 | -0.398 | -0.13 | -0.022 | 0.03 | 0.03 | 0 | 0.19 | 0.04 | 0.13 | 0.12 | 0.18 | 0.26 |
| 4 | 137662425 | rs1460155 | -0.808 | -0.298 | 0.327 | 0.524 | 0.006 | 0.186 | 0.186 | 0 | 0.88 | 0.67 | 0.89 | 0.67 | 0.88 | 0.89 |
| 4 | 137662604 | rs7692053 | -0.258 | 0.807 | 0.086 | 0.734 | -0.014 | -0.021 | -0.021 | 0.023 | 0.44 | 0.26 | 0.5 | 0.55 | 0.47 | 0.66 |
| 4 | 137665438 | rs6833955 | 1.131 | 0.809 | 0.019 | 0.732 | -0.022 | 0.03 | 0.03 | 0 | 0.19 | 0.04 | 0.13 | 0.12 | 0.18 | 0.3 |
| 4 | 137669227 | rs2406056 | 0.89 | 1.385 | 0.077 | 0.732 | 0.008 | 0.006 | 0.006 | 0 | 0.07 | 0.12 | 0.06 | 0.08 | 0.09 | 0.08 |
| 4 | 137671615 | rs736339 | -0.689 | 1.368 | 0.044 | 0.739 | 0.014 | 0.02 | 0.02 | 0 | 0.49 | 0.39 | 0.7 | 0.65 | 0.55 | 0.71 |
| 4 | 137673414 | rs17673757 | 0.467 | 1.448 | -0.082 | 0.688 | 0.017 | 0.027 | 0.027 | 0 | 0.35 | 0.23 | 0.28 | 0.23 | 0.28 | 0.54 |
| 4 | 137676856 | rs1460138 | -0.052 | 0.853 | -0.51 | 0.306 | -0.002 | -0.004 | -0.004 | 0 | 0.34 | 0.32 | 0.34 | 0.36 | 0.36 | 0.2 |
| 4 | 137677024 | rs17048925 | 1.716 | 0.975 | -0.628 | 0.275 | -0.034 | 0.005 | 0.005 | 0 | 0.18 | 0.02 | 0.12 | 0.12 | 0.17 | 0.3 |
| 4 | 137683836 | rs17048948 | 0.115 | 1.522 | -0.579 | 0.343 | 0.02 | 0.04 | 0.04 | 0 | 0.1 | 0.2 | 0.21 | 0.2 | 0.1 | 0.08 |
| 4 | 137684462 | rs1460157 | 1.02 | 1.413 | -0.618 | 0.338 | 0.021 | 0.037 | 0.037 | 0 | 0.34 | 0.15 | 0.27 | 0.24 | 0.28 | 0.54 |
| 4 | 137689373 | rs2634904 | -0.795 | 1.2 | -0.388 | 0.446 | 0.009 | 0.024 | 0.024 | 0 | 0.7 | 0.49 | 0.63 | 0.59 | 0.66 | 0.75 |
| 4 | 137691499 | rs6842883 | 1.683 | 0.703 | -0.46 | 0.291 | 0.003 | 0.003 | 0.003 | 0 | 0.18 | 0.13 | 0.19 | 0.12 | 0.16 | 0.25 |
| 4 | 137692712 | rs2011114 | -0.887 | 0.575 | -0.454 | 0.136 | 0.002 | -0.007 | -0.007 | 0 | 0.63 | 0.37 | 0.71 | 0.88 | 0.76 | 0.68 |
| 4 | 137693116 | rs17680173 | 0.461 | 0.959 | -0.606 | 0.149 | -0.131 | -0.037 | -0.037 | 0 | 0.32 | 0.06 | 0.54 | 0.68 | 0.46 | 0.5 |
| 4 | 137694429 | rs11733819 | 0.639 | 0.926 | -0.574 | 0.273 | -0.026 | -0.005 | -0.005 | 0 | 0.09 | 0.03 | 0.23 | 0.54 | 0.16 | 0.17 |
| 4 | 137697208 | rs10016963 | 0.779 | 0.908 | -0.543 | 0.302 | -0.026 | 0.065 | 0.065 | 0 | 0.4 | 0.15 | 0.48 | 0.25 | 0.44 | 0.57 |
| 4 | 137704034 | rs1841293 | 0.198 | 0.66 | -0.508 | 0.372 | 0 | -0.002 | -0.002 | 0 | 0.31 | 0.3 | 0.17 | 0.21 | 0.3 | 0.18 |
| 4 | 137706263 | rs17680592 | 1.547 | 0.486 | -0.553 | 0.272 | 0.004 | 0.042 | 0.042 | 0 | 0.16 | 0.06 | 0.06 | 0.02 | 0.14 | 0.1 |
| 4 | 137707853 | rs1460148 | 0.054 | 0.442 | -0.433 | 0.534 | -0.004 | -0.015 | -0.015 | 0.772 | 0.35 | 0.3 | 0.34 | 0.51 | 0.37 | 0.27 |
| 4 | 137712561 | rs10857199 | 0.083 | 0.396 | -0.202 | 0.545 | -0.001 | -0.009 | -0.009 | 0 | 0.35 | 0.25 | 0.32 | 0.43 | 0.36 | 0.23 |
| 4 | 137713519 | rs17619001 | 2.033 | 0.736 | -0.567 | 0.315 | -0.075 | -0.014 | -0.014 | 0 | 0.21 | 0.02 | 0.28 | 0.16 | 0.2 | 0.3 |
| 4 | 137713685 | rs9990891 | 1.374 | 0.592 | -0.164 | 0.306 | 0.02 | -0.009 | -0.009 | 0 | 0.26 | 0.34 | 0.36 | 0.23 | 0.29 | 0.35 |
| 4 | 137714170 | rs2634888 | -2.237 | 0.478 | -0.384 | 0.182 | 0.024 | 0.037 | 0.037 | 0 | 0.84 | 0.95 | 0.91 | 0.92 | 0.86 | 0.87 |
| 4 | 137720486 | rs17829074 | 1.998 | 0.506 | -0.307 | 0.172 | 0.002 | 0.012 | 0.012 | 0.073 | 0.2 | 0.21 | 0.21 | 0.12 | 0.21 | 0.26 |
| 4 | 137727040 | rs2612144 | -3.652 | 0.438 | -0.351 | 0.139 | 0.016 | 0.015 | 0.015 | 0 | 0.9 | 0.94 | 0.96 | 1 | 0.94 | 0.95 |
| 4 | 137727533 | rs2169147 | 0.38 | 0.493 | -0.337 | 0.494 | 0 | 0 | 0 | 0 | 0.32 | 0.3 | 0.52 | 0.29 | 0.37 | 0.39 |
| 4 | 137727894 | rs13132581 | 0.56 | 0.609 | -0.393 | 0.43 | -0.014 | 0.025 | 0.025 | 0 | 0.31 | 0.15 | 0.17 | 0.28 | 0.34 | 0.21 |
| 4 | 137727959 | rs7662945 | 0.125 | 0.403 | -0.216 | 0.347 | -0.011 | -0.009 | -0.009 | 0.472 | 0.28 | 0.45 | 0.43 | 0.24 | 0.3 | 0.34 |
| 4 | 137731357 | rs17049031 | 1.46 | 0.628 | -0.746 | 0.471 | -0.009 | -0.04 | -0.04 | 0.236 | 0.07 | 0 | 0.19 | 0.13 | 0.08 | 0.08 |
| 4 | 137733154 | rs4864365 | 0.463 | 0.659 | -0.962 | 0.253 | 0 | -0.001 | -0.001 | 1.698 | 0.36 | 0.31 | 0.22 | 0.43 | 0.35 | 0.32 |
| 4 | 137733596 | rs2634900 | 0.663 | 0.4 | -1.271 | 0.052 | -0.052 | -0.046 | -0.046 | 1.027 | 0.42 | 0.16 | 0.52 | 0.44 | 0.5 | 0.54 |
| 4 | 137733835 | rs2612134 | -0.429 | 0.393 | -1.049 | -0.019 | 0.044 | 0.041 | 0.041 | 0 | 0.49 | 0.25 | 0.71 | 0.57 | 0.58 | 0.63 |

| | | | | | | | | | | | | | | | | |
|---|---|---|---|---|---|---|---|---|---|---|---|---|---|---|---|---|
| 4 | 137734912 | rs1551706 | 1.934 | 0.387 | -1.616 | -0.211 | 0.01 | 0.072 | 0.072 | 0 | 0.16 | 0.03 | 0.06 | 0.01 | 0.12 | 0.11 |
| 4 | 137735357 | rs1551705 | 1.566 | 0.635 | -1.499 | 0.082 | -0.062 | 0.027 | 0.027 | 0 | 0.17 | 0.01 | 0.24 | 0.13 | 0.17 | 0.23 |
| 4 | 137736095 | rs13101322 | 1.144 | 0.633 | -1.495 | 0.085 | -0.084 | 0.045 | 0.045 | 0 | 0.2 | 0.02 | 0.27 | 0.13 | 0.25 | 0.23 |
| 4 | 137740596 | rs898427 | -0.672 | 0.527 | -1.087 | 0.12 | 0.002 | 0.002 | 0.002 | 0 | 0.52 | 0.58 | 0.75 | 0.57 | 0.58 | 0.59 |
| 4 | 137743560 | rs2612121 | -2.97 | 0.667 | -1.71 | -0.009 | 0.02 | 0.055 | 0.055 | 0 | 0.9 | 0.98 | 0.97 | 1 | 0.93 | 0.95 |
| 4 | 137745250 | rs7658921 | 0.595 | 1.145 | -1.493 | 0.299 | -0.087 | 0.009 | 0.009 | 0 | 0.22 | 0.07 | 0.31 | 0.19 | 0.29 | 0.26 |
| 4 | 137746765 | rs9884859 | 1.015 | 0.951 | -0.912 | 0.775 | 0.001 | -0.03 | -0.03 | 0 | 0.2 | 0.09 | 0.21 | 0.25 | 0.2 | 0.31 |
| 4 | 137755834 | rs1380627 | 1.062 | 0.917 | -0.827 | 0.665 | 0.001 | -0.024 | -0.024 | 0.529 | 0.19 | 0.07 | 0.2 | 0.24 | 0.2 | 0.32 |
| 4 | 137757203 | rs2634862 | 1.037 | 0.916 | -0.801 | 0.664 | 0.004 | -0.01 | -0.01 | 0 | 0.19 | 0.1 | 0.2 | 0.24 | 0.19 | 0.32 |
| 4 | 137757692 | rs2612131 | 0.732 | 1.135 | -0.763 | 0.674 | -0.01 | -0.01 | -0.01 | 0 | 0.22 | 0.12 | 0.26 | 0.29 | 0.28 | 0.35 |
| 4 | 137760899 | rs2009775 | 0.99 | 0.939 | -1.083 | 0.303 | -0.005 | 0.101 | 0.101 | 0 | 0.3 | 0.15 | 0.29 | 0.12 | 0.25 | 0.28 |
| 4 | 137761486 | rs2612165 | -0.415 | 1.278 | -0.255 | 1.174 | -0.014 | 0.05 | 0.05 | 0 | 0.9 | 0.87 | 0.78 | 0.83 | 0.9 | 0.96 |
| 4 | 137775757 | rs6842060 | -0.277 | 1.149 | -0.344 | 0.895 | -0.007 | -0.006 | -0.006 | 0 | 0.37 | 0.34 | 0.26 | 0.47 | 0.46 | 0.37 |
| 4 | 137778900 | rs10519401 | 0.23 | 0.846 | -0.351 | 0.816 | -0.005 | 0.014 | 0.014 | 0 | 0.08 | 0.1 | 0.09 | 0.12 | 0.09 | 0.04 |
| 4 | 137804904 | rs2590762 | -0.506 | 1.076 | -0.491 | 0.748 | -0.001 | 0 | 0 | 0 | 0.92 | 0.9 | 0.92 | 0.83 | 0.91 | 0.88 |
| 4 | 137808928 | rs7698161 | -0.512 | 1.219 | -0.313 | 0.526 | 0.001 | 0.007 | 0.007 | 0 | 0.36 | 0.37 | 0.41 | 0.55 | 0.39 | 0.38 |
| 4 | 137814303 | rs7685312 | -1.151 | 1.107 | -0.202 | 0.363 | -0.003 | 0.013 | 0.013 | 0.156 | 0.46 | 0.51 | 0.47 | 0.74 | 0.49 | 0.52 |
| 4 | 137816124 | rs11099419 | 0.072 | 1.059 | -0.17 | 0.707 | 0.004 | -0.032 | -0.032 | 0.405 | 0.24 | 0.15 | 0.14 | 0.48 | 0.24 | 0.32 |
| 4 | 137822979 | rs1992694 | 1.124 | 1.038 | -1.02 | 0.189 | -0.021 | 0.017 | 0.017 | 2.883 | 0.14 | 0.03 | 0.1 | 0.12 | 0.15 | 0.12 |
| 4 | 137824601 | rs10007934 | 0.965 | 1.019 | -0.573 | 0.422 | 0 | -0.009 | -0.009 | 0 | 0.08 | 0.11 | 0.05 | 0 | 0.11 | 0.01 |
| 4 | 137828866 | rs1445523 | -0.745 | 1.047 | -0.635 | 0.804 | 0 | 0.003 | 0.003 | 0.241 | 0.33 | 0.31 | 0.22 | 0.64 | 0.33 | 0.44 |
| 4 | 137832558 | rs12331904 | -0.384 | 0.861 | -1.056 | 0.789 | 0.006 | -0.02 | -0.02 | 0.828 | 0.12 | 0.04 | 0.06 | 0.17 | 0.1 | 0.22 |
| 4 | 137834818 | rs994911 | -0.189 | 1.321 | -1.138 | 0.206 | -0.02 | 0.032 | 0.032 | 0.307 | 0.23 | 0.15 | 0.16 | 0.08 | 0.28 | 0.13 |
| 4 | 137834834 | rs13124866 | 0.802 | 1.131 | -1.243 | 0.492 | 0.01 | 0.011 | 0.011 | 5.561 | 0.53 | 0.26 | 0.36 | 0.64 | 0.44 | 0.72 |
| 4 | 137834967 | rs994912 | 0.568 | 1.261 | -1.182 | 0.398 | -0.008 | 0.018 | 0.018 | 0.262 | 0.2 | 0.13 | 0.13 | 0.08 | 0.22 | 0.11 |
| 4 | 137838411 | rs7692485 | 0.411 | 1.188 | -0.15 | 0.623 | 0.063 | 0.135 | 0.135 | 0 | 0.11 | 0.28 | 0.3 | 0.24 | 0.16 | 0.14 |
| 4 | 137842889 | rs7677705 | -0.411 | 1.189 | -0.081 | 0.746 | 0.023 | 0.016 | 0.016 | 1.047 | 0.31 | 0.37 | 0.48 | 0.36 | 0.39 | 0.21 |
| 4 | 137845624 | rs17049140 | 0.681 | 1.23 | -0.193 | 0.886 | -0.021 | 0.024 | 0.024 | 0.261 | 0.2 | 0.1 | 0.17 | 0.12 | 0.23 | 0.11 |
| 4 | 137851689 | rs12500811 | 0.681 | 1.216 | -0.251 | 0.865 | -0.021 | 0.039 | 0.039 | 0.261 | 0.2 | 0.1 | 0.17 | 0.08 | 0.23 | 0.11 |
| 4 | 137851858 | rs1373895 | -0.266 | 0.6 | -1.11 | 0.144 | 0.043 | -0.014 | -0.014 | 0 | 0.43 | 0.27 | 0.3 | 0.56 | 0.38 | 0.53 |
| 4 | 137852484 | rs693365 | 0.315 | 1.016 | -0.506 | 0.48 | -0.033 | -0.018 | -0.018 | 0 | 0.67 | 0.27 | 0.49 | 0.66 | 0.61 | 0.82 |
| 4 | 137874783 | rs1400918 | -1.184 | 0.838 | -0.266 | 0.091 | -0.005 | 0.011 | 0.011 | 0.182 | 0.43 | 0.54 | 0.55 | 0.32 | 0.48 | 0.29 |
| 4 | 137875449 | rs10519404 | -0.459 | 0.946 | -0.281 | 0.101 | -0.001 | -0.038 | -0.038 | 0 | 0.34 | 0.36 | 0.26 | 0.19 | 0.34 | 0.17 |
| 4 | 137878063 | rs17049170 | 0.091 | 1.39 | -0.576 | 0.424 | -0.019 | 0.039 | 0.039 | 0 | 0.23 | 0.1 | 0.18 | 0.11 | 0.25 | 0.15 |
| 4 | 137886834 | rs7695609 | -0.292 | 1.395 | -0.372 | 0.85 | 0.018 | 0.006 | 0.006 | 0 | 0.1 | 0.13 | 0.28 | 0.14 | 0.14 | 0.16 |
| 4 | 137900265 | rs17049203 | -0.197 | 1.013 | -0.354 | 1.216 | -0.007 | 0.171 | 0.171 | 0 | 0.07 | 0.1 | 0.06 | 0.55 | 0.06 | 0.15 |
| 4 | 137912755 | rs1517955 | 0.442 | 1.095 | -0.942 | 0.444 | 0.006 | 0.086 | 0.086 | 0.393 | 0.49 | 0.31 | 0.55 | 0.3 | 0.47 | 0.55 |
| 4 | 137912903 | rs1517956 | 1.271 | 1.12 | -1.207 | 0.263 | 0.027 | 0.164 | 0.164 | 0.832 | 0.37 | 0.11 | 0.26 | 0.07 | 0.32 | 0.34 |
| 4 | 137914565 | rs1996452 | 1.266 | 1.146 | -1.183 | 0.244 | 0.029 | 0.153 | 0.153 | 0.659 | 0.37 | 0.14 | 0.26 | 0.07 | 0.32 | 0.34 |
| 4 | 137923777 | rs679959 | 0.297 | 1.175 | -1.573 | 0.344 | -0.105 | 0.016 | 0.016 | 1.535 | 0.28 | 0.03 | 0.21 | 0.16 | 0.32 | 0.14 |
| 4 | 137937395 | rs1517944 | 0.315 | 1.011 | -0.903 | 0.678 | -0.03 | 0.203 | 0.203 | 0 | 0.15 | 0.4 | 0.17 | 0.46 | 0.12 | 0.23 |
| 4 | 137938336 | rs1517945 | 0.778 | 0.98 | -1.206 | -0.03 | 0.027 | 0.017 | 0.017 | 1.387 | 0.49 | 0.44 | 0.4 | 0.61 | 0.4 | 0.56 |
| 4 | 137944935 | rs529654 | -0.314 | 1.309 | -1.303 | -0.267 | 0.006 | -0.067 | -0.067 | 0.808 | 0.78 | 0.71 | 0.84 | 0.96 | 0.73 | 0.88 |
| 4 | 137947710 | rs572260 | 1.054 | 1.716 | -1.509 | -0.325 | -0.089 | 0.006 | 0.006 | 1.743 | 0.17 | 0.04 | 0.16 | 0.11 | 0.22 | 0.13 |
| 4 | 137953807 | rs515800 | 0.958 | 1.486 | -1.009 | -0.434 | 0.012 | -0.016 | -0.016 | 0 | 0.64 | 0.48 | 0.55 | 0.72 | 0.58 | 0.66 |
| 4 | 137954283 | rs534612 | 0.412 | 0.957 | -0.732 | -0.134 | 0.002 | 0.009 | 0.009 | 0.429 | 0.31 | 0.43 | 0.31 | 0.64 | 0.3 | 0.32 |
| 4 | 137964269 | rs545844 | 0.942 | 0.916 | -0.459 | 0.354 | 0.005 | -0.058 | -0.058 | 0.779 | 0.77 | 0.71 | 0.84 | 0.96 | 0.73 | 0.87 |
| 4 | 137966680 | rs481565 | 0.442 | 0.948 | -0.419 | 0.501 | -0.002 | -0.005 | -0.005 | 0 | 0.85 | 0.81 | 0.89 | 0.89 | 0.86 | 0.74 |
| 4 | 137968302 | rs17049291 | -0.678 | 0.96 | -0.521 | 0.644 | 0 | -0.005 | -0.005 | 0 | 0.14 | 0.12 | 0.09 | 0.04 | 0.17 | 0.04 |
| 4 | 137984231 | rs11735377 | -0.991 | 0.827 | -0.181 | 1.332 | 0.005 | 0.053 | 0.053 | 0.074 | 0.17 | 0.27 | 0.18 | 0.33 | 0.14 | 0.3 |
| 4 | 137985807 | rs17049311 | -0.203 | 0.828 | -0.131 | 1.394 | 0.008 | 0.101 | 0.101 | 0 | 0.07 | 0.12 | 0.12 | 0.26 | 0.07 | 0.14 |
| 4 | 137986252 | rs7692042 | -1.463 | 0.652 | -0.235 | 0.968 | 0.004 | -0.012 | -0.012 | 0 | 0.42 | 0.75 | 0.34 | 0.42 | 0.37 | 0.44 |
| 4 | 137986727 | rs10049824 | -1.035 | 0.894 | -0.371 | 1.286 | 0.004 | 0.068 | 0.068 | 0.105 | 0.16 | 0.27 | 0.18 | 0.37 | 0.14 | 0.3 |
| 4 | 137989392 | rs598364 | 0.519 | 0.405 | -0.352 | 0.973 | 0.007 | 0.007 | 0.007 | 0.874 | 0.6 | 0.38 | 0.69 | 0.65 | 0.66 | 0.61 |
| 4 | 137990451 | rs12511969 | -0.763 | 0.311 | 0.27 | 1.099 | -0.009 | 0.026 | 0.026 | 0.069 | 0.29 | 0.53 | 0.31 | 0.37 | 0.21 | 0.42 |
| 4 | 137997347 | rs720733 | -1.077 | 0.337 | -0.434 | 0.782 | -0.004 | -0.009 | -0.009 | 0.654 | 0.65 | 0.71 | 0.54 | 0.58 | 0.62 | 0.78 |
| 4 | 137997845 | rs12510407 | 0.176 | 0.327 | 0.329 | 0.768 | -0.01 | 0.024 | 0.024 | 0 | 0.25 | 0.3 | 0.28 | 0.36 | 0.18 | 0.36 |
| 4 | 137999526 | rs12647735 | 0.054 | 0.337 | 0.354 | 0.918 | -0.002 | 0.007 | 0.007 | 0 | 0.4 | 0.4 | 0.26 | 0.28 | 0.44 | 0.42 |
| 4 | 138002681 | rs1456266 | -1.43 | 0.032 | 0.114 | 0.722 | -0.001 | -0.001 | -0.001 | 0.033 | 0.81 | 0.85 | 0.61 | 0.77 | 0.72 | 0.87 |
| 4 | 138004950 | rs10018409 | 0.522 | -0.011 | 0.01 | 0.714 | -0.011 | 0.012 | 0.012 | 0 | 0.25 | 0.32 | 0.28 | 0.31 | 0.18 | 0.36 |
| 4 | 138010057 | rs10519416 | 0.919 | -0.017 | -0.086 | 0.643 | -0.01 | 0.033 | 0.033 | 0 | 0.33 | 0.2 | 0.23 | 0.2 | 0.39 | 0.31 |
| 4 | 138016196 | rs1542389 | 1.391 | 0.229 | 0.01 | 0.858 | 0.017 | -0.019 | -0.019 | 0.044 | 0.16 | 0.06 | 0.06 | 0.19 | 0.1 | 0.09 |
| 4 | 138038058 | rs1118368 | -0.942 | 0.173 | 0.175 | 0.78 | 0.046 | 0.002 | 0.002 | 0.023 | 0.48 | 0.79 | 0.39 | 0.57 | 0.33 | 0.57 |
| 4 | 138042250 | rs7692372 | 1.403 | 0.175 | 0.028 | 0.822 | 0.013 | -0.03 | -0.03 | 0 | 0.15 | 0.06 | 0.06 | 0.22 | 0.1 | 0.09 |
| 4 | 138044886 | rs10519415 | 0.277 | 0.19 | 0.356 | 0.83 | -0.006 | 0.006 | 0.006 | 0 | 0.26 | 0.32 | 0.29 | 0.31 | 0.18 | 0.36 |
| 4 | 138048759 | rs17642711 | 1.413 | 0.048 | -0.085 | 0.655 | -0.105 | -0.051 | -0.051 | 0 | 0.2 | 0.01 | 0.17 | 0.2 | 0.34 | 0.13 |
| 4 | 138049464 | rs12505431 | 0.089 | -0.336 | 0.655 | 0.697 | 0.004 | -0.008 | -0.008 | 0.06 | 0.24 | 0.22 | 0.28 | 0.31 | 0.18 | 0.41 |
| 4 | 138052684 | rs17049362 | 0.322 | -0.184 | 0.451 | 0.699 | 0.003 | -0.006 | -0.006 | 0.006 | 0.2 | 0.21 | 0.09 | 0.24 | 0.15 | 0.15 |
| 4 | 138080761 | rs989519 | -0.745 | -0.147 | -0.322 | 0.631 | 0.049 | -0.02 | -0.02 | 2.351 | 0.47 | 0.76 | 0.37 | 0.52 | 0.32 | 0.55 |

| | | | | | | | | | | | | | | | | |
|---|---|---|---|---|---|---|---|---|---|---|---|---|---|---|---|---|
| 4 | 138087831 | rs10010440 | 0.012 | -0.384 | 0.224 | 0.881 | -0.015 | 0.058 | 0.058 | 0 | 0.07 | 0.17 | 0.09 | 0.16 | 0.03 | 0.17 |
| 4 | 138103313 | rs1913606 | -0.328 | -0.841 | 0.451 | 0.909 | -0.037 | 0.035 | 0.035 | 0.226 | 0.2 | 0.65 | 0.19 | 0.38 | 0.1 | 0.26 |
| 4 | 138111659 | rs7660328 | -0.297 | -0.857 | 0.804 | 0.902 | -0.052 | 0.056 | 0.056 | 0.178 | 0.2 | 0.59 | 0.18 | 0.38 | 0.1 | 0.26 |
| 4 | 138115250 | rs9307765 | -0.769 | -1.798 | 0.528 | 0.295 | 0.071 | 0.015 | 0.015 | 0 | 0.37 | 0.72 | 0.22 | 0.37 | 0.17 | 0.41 |
| 4 | 138118905 | rs13148112 | -1.171 | -1.263 | 0.087 | 0.193 | 0.007 | -0.018 | -0.018 | 4.672 | 0.79 | 0.75 | 0.63 | 0.87 | 0.75 | 0.8 |
| 4 | 138119828 | rs4577637 | 1.844 | -1.307 | -0.357 | 0.246 | 0.009 | -0.011 | -0.011 | 0 | 0.11 | 0.07 | 0.05 | 0.2 | 0.07 | 0.08 |
| 4 | 138121754 | rs1124481 | -0.377 | -1.322 | -0.098 | 0.222 | -0.003 | -0.003 | -0.003 | 0 | 0.66 | 0.41 | 0.55 | 0.66 | 0.68 | 0.72 |
| 4 | 138126978 | rs10008805 | 0.056 | -1.403 | -0.585 | 0.24 | 0.07 | 0.049 | 0.049 | 0 | 0.32 | 0.14 | 0.12 | 0.2 | 0.16 | 0.25 |
| 4 | 138129929 | rs10519412 | 0.056 | -1.398 | -0.627 | 0.259 | 0.07 | 0.049 | 0.049 | 0 | 0.32 | 0.14 | 0.12 | 0.2 | 0.16 | 0.25 |
| 4 | 138145125 | rs7696692 | 0.252 | -1.157 | -1.102 | -0.002 | 0.003 | -0.003 | -0.003 | 0 | 0.3 | 0.68 | 0.44 | 0.3 | 0.34 | 0.34 |
| 4 | 138157047 | rs2081521 | -1.409 | -0.38 | -1.678 | 0.098 | -0.018 | -0.024 | -0.024 | 0 | 0.89 | 0.97 | 0.92 | 0.89 | 0.88 | 0.87 |
| 4 | 138160754 | rs2406716 | -0.804 | -0.768 | -1.641 | -0.001 | -0.142 | -0.118 | -0.118 | 2.317 | 0.66 | 0.93 | 0.6 | 0.51 | 0.52 | 0.67 |
| 4 | 138161364 | rs4561979 | -0.296 | -0.764 | -1.546 | -0.048 | -0.042 | 0.026 | 0.026 | 1.092 | 0.6 | 0.8 | 0.52 | 0.36 | 0.5 | 0.53 |
| 4 | 138161923 | rs17701651 | 0.968 | -0.923 | -1.816 | -0.076 | 0.063 | 0.081 | 0.081 | 0 | 0.17 | 0.04 | 0.04 | 0 | 0.06 | 0.15 |
| 4 | 138164950 | rs1425979 | -0.226 | -0.348 | -1.68 | 0.191 | -0.063 | -0.049 | -0.049 | 1.328 | 0.58 | 0.9 | 0.61 | 0.51 | 0.47 | 0.61 |
| 4 | 138167524 | rs1365138 | 0.067 | -0.593 | -1.707 | 0.411 | 0.007 | -0.038 | -0.038 | 0 | 0.51 | 0.85 | 0.61 | 0.56 | 0.44 | 0.64 |
| 4 | 138168552 | rs1425982 | 0.127 | -0.304 | -1.048 | 0.988 | -0.009 | 0.022 | 0.022 | 1.261 | 0.27 | 0.37 | 0.22 | 0.42 | 0.22 | 0.16 |
| 4 | 138168682 | rs1425983 | -0.763 | -0.146 | -0.96 | 0.936 | -0.005 | 0.008 | 0.008 | 0 | 0.77 | 0.84 | 0.84 | 0.84 | 0.71 | 0.74 |
| 4 | 138171595 | rs6816675 | 0.645 | -0.165 | -0.942 | 0.973 | 0 | 0.004 | 0.004 | 0.486 | 0.5 | 0.46 | 0.6 | 0.41 | 0.49 | 0.58 |
| 4 | 138185006 | rs7681206 | -0.328 | 0.008 | -0.966 | 0.734 | -0.002 | -0.002 | -0.002 | 0.041 | 0.38 | 0.43 | 0.42 | 0.68 | 0.32 | 0.37 |
| 4 | 138187352 | rs736733 | 0.938 | 0.398 | -1.088 | 0.886 | -0.01 | -0.008 | -0.008 | 0 | 0.21 | 0.06 | 0.18 | 0.18 | 0.21 | 0.1 |
| 4 | 138190394 | rs6535206 | -1.558 | 0.399 | -1.297 | 0.854 | 0 | 0 | 0 | 0.222 | 0.87 | 0.87 | 0.91 | 0.87 | 0.85 | 0.87 |
| 4 | 138206940 | rs12505076 | 0.85 | 0.291 | -1.045 | 0.849 | 0.01 | -0.008 | -0.008 | 0 | 0.2 | 0.1 | 0.17 | 0.18 | 0.18 | 0.1 |
| 4 | 138213225 | rs7670687 | 1.234 | 0.263 | -0.783 | 0.355 | -0.002 | 0.035 | 0.035 | 0.404 | 0.35 | 0.26 | 0.39 | 0.15 | 0.34 | 0.35 |
| 4 | 138218981 | rs6832097 | -0.578 | -0.278 | 0.138 | 0.902 | -0.016 | -0.002 | -0.002 | 0 | 0.34 | 0.5 | 0.38 | 0.65 | 0.28 | 0.37 |
| 4 | 138221687 | rs6847311 | -1.747 | -0.406 | -0.014 | 0.737 | 0.044 | 0.082 | 0.082 | 0.135 | 0.48 | 0.78 | 0.45 | 0.64 | 0.39 | 0.41 |
| 4 | 138224531 | rs1425969 | 0.716 | -0.406 | 0.263 | 1.078 | -0.004 | 0.019 | 0.019 | 0.061 | 0.18 | 0.2 | 0.19 | 0.23 | 0.16 | 0.12 |
| 4 | 138226204 | rs1365132 | 0.351 | -0.399 | 0.204 | 1.006 | 0.005 | -0.06 | -0.06 | 4.528 | 0.13 | 0.2 | 0.07 | 0 | 0.13 | 0.04 |
| 4 | 138227609 | rs987360 | -1.902 | -0.345 | -0.308 | 1.091 | 0.039 | 0.072 | 0.072 | 0 | 0.48 | 0.71 | 0.44 | 0.64 | 0.39 | 0.41 |
| 4 | 138227898 | rs768728 | -0.316 | -0.37 | -0.291 | 1.307 | -0.011 | 0.032 | 0.032 | 0 | 0.3 | 0.34 | 0.36 | 0.45 | 0.26 | 0.37 |
| 4 | 138229547 | rs13129710 | -0.507 | 0.152 | -0.752 | 1.1 | -0.008 | 0.019 | 0.019 | 0 | 0.35 | 0.54 | 0.27 | 0.43 | 0.29 | 0.16 |
| 4 | 138229727 | rs17049516 | 0.992 | 0.486 | -0.967 | 0.956 | -0.011 | -0.035 | -0.035 | 2.274 | 0.07 | 0.01 | 0.05 | 0.16 | 0.07 | 0.09 |
| 4 | 138238306 | rs1365133 | -0.023 | 0.573 | -0.911 | 0.675 | -0.001 | -0.003 | -0.003 | 0 | 0.36 | 0.34 | 0.39 | 0.5 | 0.38 | 0.2 |
| 4 | 138247146 | rs10857203 | -0.155 | 0.532 | -0.964 | 0.429 | 0.011 | -0.053 | -0.053 | 0.952 | 0.39 | 0.13 | 0.39 | 0.5 | 0.36 | 0.3 |
| 4 | 138247502 | rs1425993 | -0.53 | -0.183 | -1.18 | -0.189 | 0.033 | 0.035 | 0.035 | 0 | 0.64 | 0.5 | 0.6 | 0.55 | 0.54 | 0.55 |
| 4 | 138261366 | rs7672454 | -1.919 | 0.28 | -1.4 | -0.118 | -0.034 | -0.023 | -0.023 | 0 | 0.93 | 1 | 0.88 | 0.89 | 0.87 | 0.84 |
| 4 | 138279041 | rs1991260 | -0.322 | 0.122 | -0.962 | -0.218 | 0.014 | 0.034 | 0.034 | 0 | 0.69 | 0.77 | 0.77 | 0.88 | 0.74 | 0.6 |
| 4 | 138279845 | rs10519423 | 0.743 | 0.005 | -1.17 | -0.184 | 0.021 | 0.036 | 0.036 | 0.421 | 0.17 | 0.03 | 0.14 | 0.07 | 0.14 | 0.17 |
| 4 | 138283405 | rs6811274 | -0.015 | 0.04 | -0.779 | -0.109 | 0.025 | 0.003 | 0.003 | 0.027 | 0.49 | 0.67 | 0.56 | 0.55 | 0.44 | 0.32 |
| 4 | 138287439 | rs7692730 | -0.05 | 0.054 | -0.805 | -0.119 | 0.031 | 0.017 | 0.017 | 0.049 | 0.5 | 0.72 | 0.48 | 0.5 | 0.44 | 0.31 |
| 4 | 138293584 | rs11730485 | 0.134 | -0.125 | -0.952 | -0.197 | -0.026 | -0.042 | -0.042 | 1.073 | 0.39 | 0.19 | 0.27 | 0.34 | 0.37 | 0.45 |
| 4 | 138299994 | rs6831492 | 0.689 | -0.11 | -0.225 | -0.103 | -0.003 | -0.027 | -0.027 | 0 | 0.08 | 0.35 | 0.05 | 0 | 0.03 | 0.01 |
| 4 | 138301340 | rs11929773 | 0.501 | 0.028 | -0.531 | -0.044 | 0 | -0.007 | -0.007 | 3.584 | 0.41 | 0.35 | 0.42 | 0.5 | 0.41 | 0.28 |
| 4 | 138313173 | rs10519420 | 1.044 | 0.122 | -0.497 | -0.138 | -0.068 | -0.09 | -0.09 | 0.323 | 0.2 | 0.09 | 0.21 | 0.33 | 0.29 | 0.3 |
| 4 | 138328077 | rs13118887 | -0.013 | -0.29 | -0.452 | -0.272 | -0.026 | -0.028 | -0.028 | 0 | 0.62 | 0.71 | 0.53 | 0.54 | 0.5 | 0.42 |
| 4 | 138333658 | rs5020420 | 0.599 | -0.381 | -0.392 | -0.366 | 0.001 | -0.026 | -0.026 | 0 | 0.19 | 0.17 | 0.09 | 0 | 0.09 | 0.14 |
| 4 | 138346339 | rs7699794 | 0.144 | -0.216 | -0.276 | -0.177 | 0.006 | 0.006 | 0.006 | 0 | 0.61 | 0.58 | 0.52 | 0.54 | 0.5 | 0.43 |
| 4 | 138348864 | rs17049579 | 0.311 | -0.097 | -0.039 | -0.491 | 0.01 | -0.05 | -0.05 | 0.526 | 0.09 | 0.22 | 0.05 | 0 | 0.1 | 0.08 |
| 4 | 138349799 | rs6831421 | 1.106 | 0.1 | -0.241 | -0.451 | 0.003 | 0.011 | 0.011 | 0 | 0.4 | 0.23 | 0.38 | 0.42 | 0.41 | 0.31 |
| 4 | 138352416 | rs11936879 | 0.368 | -0.3 | -0.35 | -0.737 | 0 | -0.001 | -0.001 | 0 | 0.58 | 0.43 | 0.46 | 0.43 | 0.5 | 0.41 |
| 4 | 138355102 | rs6811783 | 0.825 | -0.444 | -0.345 | -0.725 | -0.014 | -0.051 | -0.051 | 0 | 0.16 | 0.2 | 0.08 | 0 | 0.08 | 0.1 |
| 4 | 138356364 | rs17207630 | 1.483 | -0.019 | -0.448 | -0.428 | 0.023 | 0.012 | 0.012 | 0 | 0.05 | 0.08 | 0.12 | 0.11 | 0.13 | 0.11 |
| 4 | 138367052 | rs7698992 | -0.967 | -0.402 | -0.596 | -0.462 | -0.004 | 0.005 | 0.005 | 0 | 0.17 | 0.55 | 0.13 | 0.24 | 0.16 | 0.13 |
| 4 | 138367353 | rs2406747 | -1.051 | -0.637 | -0.863 | -0.561 | 0.041 | 0.033 | 0.033 | 0 | 0.36 | 0.81 | 0.21 | 0.38 | 0.26 | 0.24 |
| 4 | 138367672 | rs17049611 | 1.679 | -0.588 | -0.843 | -0.537 | -0.024 | -0.03 | -0.03 | 1.025 | 0.05 | 0.02 | 0.07 | 0.17 | 0.1 | 0.06 |
| 4 | 138367713 | rs4398579 | -1.139 | -0.939 | -1.399 | -1.012 | 0.034 | 0.072 | 0.072 | 0.206 | 0.54 | 0.84 | 0.33 | 0.43 | 0.41 | 0.36 |
| 4 | 138369786 | rs10519425 | -0.061 | -0.689 | -1.01 | -0.778 | -0.003 | 0.006 | 0.006 | 0 | 0.17 | 0.55 | 0.13 | 0.24 | 0.16 | 0.13 |
| 4 | 138369854 | rs17049637 | -0.41 | -0.535 | -1.902 | -0.683 | 0.051 | 0.042 | 0.042 | 0.006 | 0.37 | 0.8 | 0.22 | 0.38 | 0.27 | 0.24 |
| 4 | 138370960 | rs6535218 | 1.067 | 0.325 | -1.833 | -0.478 | -0.035 | 0.044 | 0.044 | 0 | 0.22 | 0.04 | 0.24 | 0.12 | 0.27 | 0.16 |
| 4 | 138372112 | rs7660985 | -0.944 | 1.08 | -1.54 | 0.59 | -0.019 | -0.007 | -0.007 | 0.11 | 0.93 | 0.97 | 0.89 | 0.72 | 0.83 | 0.83 |
| 4 | 138374116 | rs4864420 | 0.536 | 0.517 | -1.5 | 0.105 | 0.004 | 0.015 | 0.015 | 0.045 | 0.47 | 0.29 | 0.65 | 0.53 | 0.53 | 0.5 |
| 4 | 138380321 | rs7679076 | -0.491 | 0.31 | -0.998 | -0.145 | -0.001 | 0 | 0 | 0 | 0.64 | 0.66 | 0.73 | 0.59 | 0.65 | 0.56 |
| 4 | 138381268 | rs12498882 | 0.532 | 0.051 | -0.927 | 0.026 | -0.01 | 0.021 | 0.021 | 0 | 0.46 | 0.32 | 0.62 | 0.54 | 0.51 | 0.45 |
| 4 | 138382556 | rs12500066 | -0.042 | 0.235 | 0.027 | 0.548 | 0.002 | 0.016 | 0.016 | 0 | 0.09 | 0.08 | 0.1 | 0.27 | 0.16 | 0.23 |
| 4 | 138390066 | rs2406756 | 1.517 | 0.068 | 0.08 | 0.495 | 0.006 | -0.02 | -0.02 | 0 | 0.47 | 0.22 | 0.54 | 0.49 | 0.55 | 0.48 |
| 4 | 138392703 | rs1320341 | -0.569 | -0.061 | 0.395 | 0.453 | -0.01 | -0.017 | -0.017 | 0.1 | 0.75 | 0.68 | 0.82 | 0.89 | 0.84 | 0.84 |
| 4 | 138393839 | rs17049672 | 0.575 | -0.151 | 0.494 | 0.417 | -0.005 | -0.017 | -0.017 | 0.12 | 0.08 | 0.16 | 0.08 | 0.05 | 0.08 | 0.11 |
| 4 | 138395132 | rs2406613 | -1.154 | -0.119 | 0.441 | 0.461 | -0.003 | 0.005 | 0.005 | 0 | 0.27 | 0.35 | 0.27 | 0.39 | 0.29 | 0.36 |
| 4 | 138395782 | rs12643295 | 1.167 | 0.063 | -0.029 | 0.553 | 0.008 | -0.005 | -0.005 | 0 | 0.48 | 0.36 | 0.54 | 0.49 | 0.55 | 0.48 |

| | | | | | | | | | | | | | | | | |
|---|---|---|---|---|---|---|---|---|---|---|---|---|---|---|---|---|
| 4 | 138397770 | rs10032588 | 0.247 | -0.012 | -0.334 | 0.769 | -0.023 | -0.048 | -0.048 | 0 | 0.13 | 0.03 | 0.18 | 0.32 | 0.2 | 0.23 |
| 4 | 138413244 | rs13112145 | -1.477 | -0.308 | -0.21 | 0.185 | 0.03 | -0.004 | -0.004 | 0 | 0.48 | 0.69 | 0.45 | 0.5 | 0.42 | 0.45 |
| 4 | 138415363 | rs9685021 | 0.192 | -0.126 | 0.362 | 0.352 | -0.04 | -0.016 | -0.016 | 0.625 | 0.2 | 0.51 | 0.17 | 0.12 | 0.1 | 0.07 |
| 4 | 138433801 | rs34680700 | 0.024 | 0.17 | -0.173 | 0.32 | -0.001 | -0.034 | -0.034 | 0 | 0.07 | 0.15 | 0.06 | 0 | 0.08 | 0.08 |
| 4 | 138434955 | rs9330352 | -1.345 | 0.229 | -0.238 | 0.437 | -0.001 | -0.014 | -0.014 | 0.556 | 0.92 | 0.96 | 0.82 | 0.78 | 0.93 | 0.88 |
| 4 | 138449244 | rs35678237 | 0.504 | 0.568 | -0.229 | 0.355 | 0 | -0.01 | -0.01 | 0 | 0.07 | 0.1 | 0.05 | 0 | 0.07 | 0.08 |
| 4 | 138449411 | rs13132611 | 0.504 | 0.568 | -0.229 | 0.355 | 0 | -0.01 | -0.01 | 0 | 0.07 | 0.1 | 0.05 | 0 | 0.07 | 0.08 |
| 4 | 138449683 | rs10018837 | -1.53 | 0.571 | -0.462 | 0.368 | 0.021 | -0.067 | -0.067 | 0 | 0.46 | 0.9 | 0.45 | 0.51 | 0.42 | 0.44 |
| 4 | 138455637 | rs28433055 | -0.007 | 1.194 | -0.142 | 0.912 | 0.013 | 0.056 | 0.056 | 0 | 0.13 | 0.19 | 0.2 | 0.35 | 0.21 | 0.19 |
| 4 | 138459997 | rs6841487 | 0.53 | 1.454 | -0.603 | 0.787 | -0.062 | 0.027 | 0.027 | 0 | 0.11 | 0.02 | 0.09 | 0.05 | 0.2 | 0.12 |
| 4 | 138468166 | rs9307912 | -1.082 | 0.798 | 0.165 | 1.192 | 0.016 | 0.144 | 0.144 | 0 | 0.18 | 0.38 | 0.21 | 0.38 | 0.22 | 0.26 |
| 4 | 138469564 | rs6605362 | -1.591 | 0.728 | 0.16 | 1.148 | -0.003 | 0.01 | 0.01 | 0 | 0.35 | 0.54 | 0.35 | 0.51 | 0.33 | 0.35 |
| 4 | 138470185 | rs10028163 | -0.496 | 1.146 | 0.049 | 1.316 | 0.024 | 0.095 | 0.095 | 0 | 0.14 | 0.26 | 0.2 | 0.36 | 0.21 | 0.19 |
| 4 | 138477450 | rs6824771 | -0.346 | 1.046 | -0.844 | 0.74 | 0.002 | 0 | 0 | 0.95 | 0.17 | 0.18 | 0.14 | 0.13 | 0.11 | 0.09 |
| 4 | 138482973 | rs9761983 | -0.937 | 1.618 | -0.841 | 1.114 | 0.016 | 0.05 | 0.05 | 0 | 0.2 | 0.26 | 0.23 | 0.49 | 0.32 | 0.28 |
| 4 | 138493361 | rs4282237 | 0.37 | 1.021 | -0.699 | 0.381 | -0.008 | -0.052 | -0.052 | 0 | 0.85 | 0.87 | 0.91 | 1 | 0.86 | 0.96 |
| 4 | 138496793 | rs10023753 | 0.387 | 1.013 | -0.703 | 0.377 | -0.008 | -0.052 | -0.052 | 0 | 0.85 | 0.87 | 0.91 | 1 | 0.86 | 0.96 |
| 4 | 138498854 | rs4478233 | 1.348 | 1.531 | -0.842 | 0.491 | 0.001 | 0 | 0 | 0 | 0.57 | 0.54 | 0.61 | 0.39 | 0.44 | 0.58 |
| 4 | 138506420 | rs6842329 | -0.748 | 1.791 | -0.814 | 0.511 | 0.024 | -0.033 | -0.033 | 0 | 0.17 | 0.22 | 0.14 | 0.1 | 0.24 | 0.18 |
| 4 | 138511495 | rs4554127 | -1.527 | 1.417 | -1.425 | 0.193 | -0.007 | -0.002 | -0.002 | 0 | 0.62 | 0.39 | 0.46 | 0.71 | 0.72 | 0.63 |
| 4 | 138512291 | rs4327536 | 0.453 | 1.486 | -1.41 | 0.198 | 0.01 | -0.047 | -0.047 | 1.932 | 0.08 | 0.2 | 0.05 | 0 | 0.07 | 0.07 |
| 4 | 138514969 | rs3922840 | -0.005 | 1.636 | -1.701 | 0.508 | -0.023 | -0.07 | -0.07 | 0.615 | 0.22 | 0.11 | 0.22 | 0.5 | 0.31 | 0.25 |
| 4 | 138520936 | rs4391086 | 1.231 | 1.634 | -1.71 | 0.502 | 0.036 | 0.052 | 0.052 | 0.749 | 0.46 | 0.75 | 0.6 | 0.27 | 0.34 | 0.44 |
| 4 | 138528410 | rs4346697 | -0.096 | 1.77 | -1.742 | 0.602 | -0.036 | -0.094 | -0.094 | 0 | 0.22 | 0.08 | 0.22 | 0.51 | 0.31 | 0.25 |
| 4 | 138534164 | rs12645301 | -0.116 | 1.18 | -1.546 | 0.443 | -0.075 | -0.071 | -0.071 | 0.219 | 0.31 | 0.12 | 0.31 | 0.63 | 0.46 | 0.37 |
| 4 | 138535434 | rs10021969 | 0.18 | 0.677 | -0.82 | 0.362 | 0 | -0.002 | -0.002 | 0.034 | 0.78 | 0.84 | 0.91 | 0.9 | 0.83 | 0.81 |
| 4 | 138541971 | rs6841316 | -0.943 | 1.027 | -0.273 | 1.018 | 0.009 | 0.047 | 0.047 | 0.625 | 0.22 | 0.37 | 0.3 | 0.38 | 0.21 | 0.24 |
| 4 | 138542282 | rs9995514 | -0.757 | 1.258 | 0.152 | 1.087 | 0.018 | 0.012 | 0.012 | 10.895 | 0.21 | 0.36 | 0.27 | 0.23 | 0.24 | 0.24 |
| 4 | 138545488 | rs2114927 | -1.041 | 1.529 | -0.126 | 1.414 | -0.044 | -0.007 | -0.007 | 0 | 0.11 | 0.03 | 0.11 | 0.1 | 0.17 | 0.16 |
| 4 | 138563488 | rs1110119 | -1.194 | 1.192 | 0.112 | 1.297 | -0.003 | 0.005 | 0.005 | 0 | 0.13 | 0.1 | 0.11 | 0.09 | 0.15 | 0.14 |
| 4 | 138571663 | rs7666878 | -1.489 | 1.11 | 0.013 | 1.493 | -0.002 | 0.014 | 0.014 | 0 | 0.34 | 0.41 | 0.33 | 0.56 | 0.34 | 0.47 |
| 4 | 138576003 | rs17049735 | 0.481 | 1.022 | -0.086 | 1.5 | -0.052 | -0.081 | -0.081 | 0 | 0.12 | 0.01 | 0.18 | 0.44 | 0.18 | 0.3 |
| 4 | 138577151 | rs1976929 | 1.243 | 1.013 | -0.813 | 0.784 | -0.016 | -0.044 | -0.044 | 0 | 0.39 | 0.41 | 0.18 | 0.15 | 0.28 | 0.27 |
| 4 | 138578594 | rs6839772 | -1.512 | 1.06 | -0.627 | 0.792 | 0.01 | 0.011 | 0.011 | 0 | 0.38 | 0.53 | 0.36 | 0.6 | 0.42 | 0.55 |
| 4 | 138582711 | rs17049741 | 0.945 | 1.008 | -0.787 | 0.928 | -0.052 | -0.081 | -0.081 | 0 | 0.12 | 0.01 | 0.18 | 0.44 | 0.18 | 0.29 |
| 4 | 138593451 | rs2434137 | 0.502 | 0.738 | -0.481 | 0.323 | -0.015 | -0.056 | -0.056 | 0 | 0.8 | 0.59 | 0.84 | 0.91 | 0.84 | 0.85 |
| 4 | 138597155 | rs2462870 | 0.698 | 1.032 | -1.138 | 0.358 | -0.079 | 0.019 | 0.019 | 0 | 0.22 | 0.05 | 0.34 | 0.14 | 0.3 | 0.17 |
| 4 | 138598424 | rs4618363 | -0.05 | 0.186 | -1.011 | 0.345 | 0.008 | -0.02 | -0.02 | 0.361 | 0.2 | 0.13 | 0.19 | 0.6 | 0.15 | 0.31 |
| 4 | 138602702 | rs2166356 | 0.447 | 0.622 | -1.358 | -0.441 | -0.074 | 0.061 | 0.061 | 0 | 0.26 | 0.13 | 0.34 | 0.11 | 0.35 | 0.26 |
| 4 | 138603311 | rs1949905 | 0.392 | 0.052 | -0.628 | -0.569 | -0.044 | 0.046 | 0.046 | 4.624 | 0.11 | 0.31 | 0.14 | 0.14 | 0.02 | 0.12 |
| 4 | 138604190 | rs1596795 | -0.612 | -0.221 | -0.587 | -0.739 | -0.001 | 0.002 | 0.002 | 1.392 | 0.41 | 0.6 | 0.5 | 0.24 | 0.38 | 0.3 |
| 4 | 138609373 | rs2559897 | -0.601 | -0.259 | -0.761 | -0.872 | -0.014 | 0.008 | 0.008 | 0 | 0.9 | 0.94 | 0.89 | 0.94 | 0.85 | 0.89 |
| 4 | 138609462 | rs3843891 | 0.993 | -0.237 | -0.74 | -0.872 | 0.011 | 0.043 | 0.043 | 0.442 | 0.49 | 0.32 | 0.39 | 0.71 | 0.48 | 0.58 |
| 4 | 138611802 | rs2693550 | 0.741 | -0.037 | -0.588 | -0.674 | -0.068 | 0.055 | 0.055 | 0 | 0.25 | 0.13 | 0.32 | 0.1 | 0.34 | 0.18 |
| 4 | 138611810 | rs11737372 | 2.244 | 0.118 | -0.65 | -0.746 | -0.057 | 0.061 | 0.061 | 0 | 0.13 | 0.01 | 0.18 | 0.01 | 0.17 | 0.08 |
| 4 | 138616339 | rs11946875 | 0.168 | -0.315 | -0.172 | -0.574 | -0.059 | 0.062 | 0.062 | 5.36 | 0.12 | 0.43 | 0.15 | 0.12 | 0.01 | 0.04 |
| 4 | 138633459 | rs17049806 | 1.166 | -0.128 | -0.847 | -0.616 | 0.033 | -0.023 | -0.023 | 0 | 0.08 | 0.02 | 0.13 | 0.09 | 0.01 | 0.04 |
| 4 | 138633828 | rs1526237 | 0.724 | 0.09 | -0.988 | -0.872 | -0.067 | 0.053 | 0.053 | 0 | 0.25 | 0.14 | 0.33 | 0.1 | 0.33 | 0.17 |
| 4 | 138636149 | rs13132522 | 0.724 | 0.09 | -0.988 | -0.872 | -0.067 | 0.053 | 0.053 | 0 | 0.25 | 0.14 | 0.33 | 0.1 | 0.33 | 0.17 |
| 4 | 138644879 | rs1526242 | 0.764 | 0.075 | -0.967 | -0.879 | -0.085 | 0.056 | 0.056 | 0 | 0.25 | 0.12 | 0.32 | 0.1 | 0.33 | 0.17 |
| 4 | 138663042 | rs17049850 | 1.33 | -0.098 | -1.264 | -0.992 | -0.008 | -0.024 | -0.024 | 0 | 0.15 | 0.09 | 0.22 | 0.61 | 0.21 | 0.28 |
| 4 | 138663262 | rs13142558 | 1.167 | -0.061 | -1.27 | -0.995 | -0.094 | 0.053 | 0.053 | 0 | 0.24 | 0.1 | 0.32 | 0.1 | 0.33 | 0.17 |
| 4 | 138664579 | rs1880774 | 0.026 | -0.196 | -0.752 | -0.999 | -0.008 | 0.019 | 0.019 | 0 | 0.4 | 0.36 | 0.55 | 0.71 | 0.54 | 0.46 |
| 4 | 138678024 | rs12374402 | -0.965 | -0.572 | -1.016 | -1.342 | -0.011 | 0.036 | 0.036 | 0 | 0.84 | 0.95 | 0.86 | 0.94 | 0.81 | 0.85 |
| 4 | 138680839 | rs10049648 | 1.782 | -0.512 | -0.971 | -1.272 | 0.033 | -0.023 | -0.023 | 0 | 0.08 | 0.02 | 0.13 | 0.09 | 0.01 | 0.04 |
| 4 | 138688076 | rs9308023 | -0.326 | -0.426 | -0.836 | -1.328 | 0.018 | 0.057 | 0.057 | 0.523 | 0.51 | 0.43 | 0.69 | 0.79 | 0.56 | 0.5 |
| 4 | 138694227 | rs7681896 | 0.459 | -0.431 | -1.125 | -1.266 | 0.069 | -0.001 | -0.001 | 1.32 | 0.17 | 0.04 | 0.22 | 0.11 | 0.07 | 0.13 |
| 4 | 138694514 | rs7672176 | -1.034 | -0.59 | -1.118 | -1.486 | -0.008 | 0.005 | 0.005 | 0 | 0.67 | 0.62 | 0.61 | 0.86 | 0.53 | 0.74 |
| 4 | 138701890 | rs2089615 | -0.917 | -0.664 | -0.951 | -1.383 | 0.042 | -0.002 | -0.002 | 0 | 0.34 | 0.14 | 0.44 | 0.74 | 0.29 | 0.48 |
| 4 | 138702796 | rs2175072 | 1.351 | -0.573 | -0.927 | -1.333 | -0.029 | -0.078 | -0.078 | 0 | 0.16 | 0.05 | 0.22 | 0.62 | 0.22 | 0.35 |
| 4 | 138712304 | rs1509268 | 1.494 | -0.827 | -0.54 | -2.061 | -0.013 | -0.041 | -0.041 | 0 | 0.27 | 0.31 | 0.14 | 0.13 | 0.21 | 0.23 |
| 4 | 138714027 | rs1027876 | 1.444 | -0.688 | -0.58 | -1.987 | -0.026 | -0.061 | -0.061 | 0.17 | 0.16 | 0.06 | 0.34 | 0.72 | 0.22 | 0.35 |
| 4 | 138717331 | rs7670760 | -1.043 | -0.719 | -0.496 | -2.166 | 0 | -0.008 | -0.008 | 1.78 | 0.62 | 0.56 | 0.59 | 0.86 | 0.49 | 0.71 |
| 4 | 138726604 | rs4863722 | 1.531 | -0.603 | -0.795 | -2.013 | -0.028 | -0.056 | -0.056 | 2.099 | 0.15 | 0.05 | 0.34 | 0.73 | 0.21 | 0.28 |
| 4 | 138733886 | rs11940232 | 1.168 | -0.549 | -0.689 | -2.071 | -0.013 | -0.055 | -0.055 | 0 | 0.29 | 0.31 | 0.14 | 0.1 | 0.21 | 0.24 |
| 4 | 138739406 | rs13132156 | 1.03 | -0.082 | -1.044 | -1.87 | -0.112 | 0.096 | 0.096 | 0 | 0.32 | 0.09 | 0.35 | 0.1 | 0.38 | 0.23 |
| 4 | 138763411 | rs6815259 | 0.754 | 0.518 | -0.603 | -2.146 | -0.019 | 0.007 | 0.007 | 0 | 0.16 | 0.11 | 0.19 | 0.09 | 0.19 | 0.12 |
| 4 | 138764286 | rs6823023 | -1.346 | 0.25 | -0.46 | -2.544 | 0 | -0.019 | -0.019 | 0 | 0.49 | 0.57 | 0.61 | 0.87 | 0.46 | 0.62 |
| 4 | 138783682 | rs11930640 | -0.55 | -0.045 | -0.319 | -2.578 | -0.011 | -0.049 | -0.049 | 0.846 | 0.36 | 0.47 | 0.26 | 0.09 | 0.28 | 0.27 |

| | | | | | | | | | | | | | | | | |
|---|---|---|---|---|---|---|---|---|---|---|---|---|---|---|---|---|
| 4 | 138794021 | rs7679412 | 1.546 | -0.048 | -0.351 | -2.594 | 0.008 | 0.083 | 0.083 | 0.509 | 0.45 | 0.29 | 0.34 | 0.06 | 0.41 | 0.26 |
| 4 | 138801711 | rs12511738 | 2.043 | -0.146 | -0.175 | -2.718 | 0 | -0.04 | -0.04 | 0 | 0.22 | 0.22 | 0.23 | 0.01 | 0.2 | 0.09 |
| 4 | 138802565 | rs6855385 | -0.131 | 0.077 | 0.297 | -2.219 | -0.069 | -0.091 | -0.091 | 0.01 | 0.2 | 0.34 | 0.07 | 0.03 | 0.06 | 0.19 |
| 4 | 138803834 | rs12503655 | 0.603 | 0.619 | -0.344 | -1.889 | -0.027 | 0.013 | 0.013 | 0 | 0.17 | 0.12 | 0.2 | 0.07 | 0.21 | 0.14 |
| 4 | 138825270 | rs13117595 | 0.785 | 0.757 | -0.36 | -1.831 | -0.039 | 0.028 | 0.028 | 0 | 0.19 | 0.09 | 0.19 | 0.07 | 0.21 | 0.14 |
| 4 | 138829787 | rs1509255 | -0.191 | 0.72 | -0.405 | -1.85 | -0.046 | 0.058 | 0.058 | 0 | 0.34 | 0.21 | 0.54 | 0.84 | 0.43 | 0.55 |
| 4 | 138831486 | rs1960161 | -0.284 | 0.346 | -0.208 | -1.574 | 0.001 | 0.001 | 0.001 | 1.055 | 0.84 | 0.84 | 0.9 | 0.91 | 0.85 | 0.85 |
| 4 | 138840995 | rs13128867 | 0.838 | 0.537 | -0.341 | -1.54 | -0.018 | -0.032 | -0.032 | 0 | 0.14 | 0.06 | 0.33 | 0.73 | 0.19 | 0.38 |
| 4 | 138852972 | rs1519324 | -0.217 | 0.304 | -0.106 | -1.654 | 0.019 | -0.036 | -0.036 | 0.346 | 0.88 | 0.74 | 0.9 | 0.95 | 0.84 | 0.87 |
| 4 | 138853076 | rs4640713 | 1.302 | 0.387 | -0.064 | -1.943 | 0.001 | -0.035 | -0.035 | 0 | 0.48 | 0.42 | 0.38 | 0.07 | 0.45 | 0.28 |
| 4 | 138862113 | rs1589210 | 0.035 | 0.271 | 0.345 | -1.668 | 0.016 | -0.039 | -0.039 | 0.064 | 0.88 | 0.71 | 0.9 | 0.95 | 0.84 | 0.87 |
| 4 | 138873552 | rs6851757 | 0.995 | 0.28 | 0.223 | -1.683 | -0.046 | 0.008 | 0.008 | 0 | 0.18 | 0.03 | 0.18 | 0.11 | 0.19 | 0.14 |
| 4 | 138877026 | rs7667668 | 0.753 | 0.258 | 0.307 | -1.666 | -0.005 | -0.034 | -0.034 | 0 | 0.62 | 0.42 | 0.69 | 0.8 | 0.64 | 0.68 |
| 4 | 138881933 | rs17231411 | 0.293 | 0.231 | 0.105 | -1.792 | 0 | 0.004 | 0.004 | 1.289 | 0.51 | 0.43 | 0.64 | 0.77 | 0.51 | 0.6 |
| 4 | 138895657 | rs2139320 | -0.79 | 0.252 | 0.028 | -1.718 | 0.003 | 0.013 | 0.013 | 0.051 | 0.54 | 0.46 | 0.33 | 0.18 | 0.43 | 0.47 |
| 4 | 138898310 | rs938025 | -0.181 | 0.354 | 0.15 | -1.608 | 0.011 | 0.004 | 0.004 | 0 | 0.88 | 0.66 | 0.9 | 0.9 | 0.88 | 0.93 |
| 4 | 138900717 | rs13131913 | 0.187 | 0.638 | -0.061 | -1.589 | -0.038 | 0.118 | 0.118 | 0.526 | 0.37 | 0.12 | 0.26 | 0.14 | 0.37 | 0.32 |
| 4 | 138913554 | rs4863533 | -0.008 | 0.703 | -0.208 | -1.626 | -0.003 | 0.069 | 0.069 | 0.363 | 0.39 | 0.24 | 0.27 | 0.15 | 0.37 | 0.34 |
| 4 | 138939662 | rs2056275 | 0.177 | 0.7 | -0.025 | -1.455 | 0.006 | -0.036 | -0.036 | 0.255 | 0.66 | 0.57 | 0.65 | 0.79 | 0.65 | 0.7 |
| 4 | 138954084 | rs6834471 | 0.947 | 0.759 | 0.298 | -1.194 | 0.01 | 0.008 | 0.008 | 0 | 0.09 | 0.05 | 0.03 | 0 | 0.06 | 0.1 |
| 4 | 138956770 | rs1807894 | 0.119 | 0.695 | 0.259 | -1.305 | 0.001 | 0.019 | 0.019 | 0 | 0.5 | 0.6 | 0.58 | 0.77 | 0.48 | 0.53 |
| 4 | 138959728 | rs1607529 | -0.25 | 0.596 | 0.211 | -1.262 | 0.004 | -0.03 | -0.03 | 0.294 | 0.88 | 0.82 | 0.92 | 0.95 | 0.86 | 0.87 |
| 4 | 138976805 | rs9284677 | 0.601 | 0.677 | 0.185 | -1.293 | -0.028 | 0.088 | 0.088 | 0 | 0.37 | 0.13 | 0.33 | 0.16 | 0.39 | 0.32 |
| 4 | 138977173 | rs1439502 | -0.169 | 0.688 | 0.207 | -1.288 | -0.003 | 0.119 | 0.119 | 0 | 0.46 | 0.2 | 0.36 | 0.16 | 0.45 | 0.42 |
| 4 | 138983809 | rs6842285 | 0.297 | 0.607 | 0.156 | -1.274 | 0.009 | 0.052 | 0.052 | 0 | 0.5 | 0.63 | 0.58 | 0.77 | 0.47 | 0.53 |
| 4 | 139005538 | rs11734678 | -0.091 | 0.835 | 0.471 | -0.755 | 0.002 | -0.005 | -0.005 | 0.461 | 0.59 | 0.59 | 0.61 | 0.68 | 0.54 | 0.63 |
| 4 | 139014474 | rs11100776 | 0.35 | 0.731 | 0.465 | -0.461 | 0.002 | 0.008 | 0.008 | 0 | 0.5 | 0.57 | 0.59 | 0.61 | 0.52 | 0.59 |
| 4 | 139019356 | rs1982203 | 0.527 | 0.907 | 0.111 | -0.459 | -0.06 | 0.003 | 0.003 | 0 | 0.36 | 0.06 | 0.32 | 0.25 | 0.36 | 0.33 |
| 4 | 139064115 | rs1344976 | 0.261 | 0.978 | 0.373 | -0.459 | 0.004 | 0.022 | 0.022 | 0 | 0.5 | 0.62 | 0.59 | 0.61 | 0.52 | 0.53 |
| 4 | 139079966 | rs4863765 | 0.08 | 0.908 | 0.432 | -0.427 | 0.003 | 0.023 | 0.023 | 0 | 0.5 | 0.6 | 0.59 | 0.63 | 0.52 | 0.53 |
| 4 | 139084238 | rs11942121 | -0.492 | 0.873 | 0.413 | -0.431 | 0.001 | 0.02 | 0.02 | 0 | 0.49 | 0.39 | 0.4 | 0.37 | 0.47 | 0.45 |
| 4 | 139084657 | rs7688870 | -0.447 | 0.955 | 0.801 | -0.372 | 0 | -0.002 | -0.002 | 0 | 0.52 | 0.51 | 0.43 | 0.37 | 0.49 | 0.46 |
| 4 | 139089463 | rs7674870 | -0.076 | 1.406 | 0.794 | -0.041 | -0.018 | 0.039 | 0.039 | 0 | 0.42 | 0.21 | 0.38 | 0.29 | 0.45 | 0.41 |
| 4 | 139090804 | rs13144383 | -0.123 | 1.818 | 1.238 | 1.098 | -0.043 | -0.065 | -0.065 | 0 | 0.1 | 0.01 | 0.29 | 0.57 | 0.16 | 0.31 |
| 4 | 139110577 | rs6537201 | 1.373 | 1.58 | 0.925 | 0.624 | 0.11 | 0.049 | 0.049 | 1.276 | 0.92 | 0.56 | 0.9 | 0.94 | 0.88 | 0.95 |
| 4 | 139123748 | rs4131888 | -0.409 | 1.531 | -0.102 | 0.752 | -0.013 | -0.03 | -0.03 | 0 | 0.17 | 0.07 | 0.25 | 0.2 | 0.18 | 0.12 |
| 4 | 139125041 | rs4602539 | -1.144 | 1.803 | -0.008 | 1.78 | -0.021 | -0.045 | -0.045 | 0.042 | 0.23 | 0.17 | 0.35 | 0.54 | 0.27 | 0.29 |
| 4 | 139150283 | rs7666936 | 1.8 | 1.599 | 1.086 | 1.704 | 0.034 | 0.254 | 0.254 | 0.018 | 0.84 | 0.43 | 0.81 | 0.53 | 0.84 | 0.73 |
| 4 | 139155804 | rs7685529 | -1.66 | 2.374 | 0.954 | 1.264 | -0.024 | 0.022 | 0.022 | 0.41 | 0.31 | 0.23 | 0.58 | 0.88 | 0.46 | 0.47 |
| 4 | 139175695 | rs6537290 | -1.594 | 1.502 | 1.159 | 0.892 | 0.012 | 0.051 | 0.051 | 0.147 | 0.49 | 0.61 | 0.57 | 0.72 | 0.5 | 0.45 |
| 4 | 139178894 | rs11737641 | 1.275 | 1.647 | 0.914 | 0.762 | -0.009 | 0.072 | 0.072 | 0.319 | 0.53 | 0.23 | 0.41 | 0.32 | 0.52 | 0.52 |
| 4 | 139179074 | rs6816398 | -1.071 | 1.691 | 0.914 | 0.949 | 0.009 | -0.015 | -0.015 | 0.227 | 0.39 | 0.46 | 0.31 | 0.33 | 0.39 | 0.31 |
| 4 | 139183123 | rs17300990 | 1.729 | 1.663 | 0.252 | 0.789 | -0.021 | -0.006 | -0.006 | 0 | 0.46 | 0.07 | 0.38 | 0.35 | 0.47 | 0.53 |
| 4 | 139184664 | rs6850624 | -1.104 | 1.614 | 0.593 | 0.667 | 0.014 | 0.107 | 0.107 | 0 | 0.13 | 0.22 | 0.28 | 0.39 | 0.12 | 0.15 |
| 4 | 139184781 | rs2203291 | 0.171 | 1.767 | 0.415 | 0.416 | 0 | -0.038 | -0.038 | 0 | 0.17 | 0.22 | 0.18 | 0.03 | 0.17 | 0.21 |
| 4 | 139185003 | rs7687419 | 0.176 | 1.755 | 0.046 | 0.48 | -0.009 | 0.005 | 0.005 | 0.08 | 0.07 | 0.04 | 0.05 | 0.02 | 0.1 | 0.02 |
| 4 | 139188359 | rs11726463 | 0.4 | 1.629 | 0.008 | 0.544 | -0.003 | -0.018 | -0.018 | 0.089 | 0.15 | 0.15 | 0.15 | 0.03 | 0.18 | 0.2 |
| 4 | 139192424 | rs7657480 | 0.477 | 1.101 | 0.258 | 1 | 0.003 | 0.004 | 0.004 | 1.963 | 0.11 | 0.14 | 0.1 | 0.12 | 0.14 | 0.09 |
| 4 | 139193151 | rs1513765 | 0.88 | 0.971 | 0.347 | 1.13 | -0.025 | 0.076 | 0.076 | 1.106 | 0.77 | 0.4 | 0.64 | 0.56 | 0.83 | 0.84 |
| 4 | 139194027 | rs9308184 | -1.134 | 0.867 | -0.037 | 1.263 | -0.01 | 0.062 | 0.062 | 0 | 0.15 | 0.17 | 0.3 | 0.37 | 0.09 | 0.15 |
| 4 | 139195152 | rs4241941 | 0.603 | 0.928 | -0.597 | 1.118 | 0.005 | -0.021 | -0.021 | 0 | 0.62 | 0.27 | 0.52 | 0.44 | 0.66 | 0.75 |
| 4 | 139203950 | rs10021045 | -0.043 | -0.124 | -0.716 | 1.183 | 0.014 | -0.007 | -0.007 | 0 | 0.7 | 0.8 | 0.74 | 0.65 | 0.83 | 0.57 |
| 4 | 139209701 | rs6817763 | 0.557 | 0.566 | -0.983 | 0.941 | 0 | 0 | 0 | 1.284 | 0.5 | 0.58 | 0.56 | 0.54 | 0.51 | 0.44 |
| 4 | 139211038 | rs13119725 | -0.013 | 0.669 | -1.03 | 1.415 | -0.008 | -0.006 | -0.006 | 0 | 0.22 | 0.24 | 0.33 | 0.34 | 0.33 | 0.16 |
| 4 | 139212420 | rs4441808 | -0.84 | 0.267 | -1.193 | 1.053 | -0.003 | -0.005 | -0.005 | 0 | 0.89 | 0.94 | 0.8 | 0.82 | 0.88 | 0.81 |
| 4 | 139212919 | rs10519453 | 0.548 | 0.511 | -1.058 | 1.001 | -0.074 | 0.015 | 0.015 | 1.235 | 0.17 | 0.06 | 0.16 | 0.11 | 0.32 | 0.13 |
| 4 | 139223367 | rs6537323 | -0.115 | -0.422 | -1.019 | 0.93 | 0.038 | -0.026 | -0.026 | 10.783 | 0.72 | 0.85 | 0.75 | 0.65 | 0.84 | 0.57 |
| 4 | 139229237 | rs7657780 | -0.057 | 0.444 | -0.758 | 0.444 | -0.001 | -0.001 | -0.001 | 0 | 0.34 | 0.37 | 0.28 | 0.3 | 0.3 | 0.32 |
| 4 | 139234254 | rs13120633 | -0.317 | 1.248 | -0.755 | 0.515 | 0.005 | 0 | 0 | 0 | 0.1 | 0.12 | 0.16 | 0.15 | 0.17 | 0.13 |
| 4 | 139238565 | rs6810458 | -0.191 | -0.427 | -0.567 | -0.116 | 0.038 | 0.044 | 0.044 | 0.178 | 0.54 | 0.64 | 0.67 | 0.81 | 0.73 | 0.56 |
| 4 | 139243656 | rs7670671 | 0.273 | -0.19 | -0.139 | 0.128 | -0.011 | 0.003 | 0.003 | 0 | 0.66 | 0.43 | 0.69 | 0.62 | 0.69 | 0.49 |
| 4 | 139249761 | rs6849956 | 1.005 | -0.304 | -0.35 | 0.051 | -0.017 | 0.031 | 0.031 | 0 | 0.13 | 0.02 | 0.05 | 0.05 | 0.14 | 0.15 |
| 4 | 139253392 | rs7669522 | -0.068 | 0.02 | 0.229 | -0.131 | -0.021 | 0.07 | 0.07 | 1.686 | 0.79 | 0.5 | 0.76 | 0.67 | 0.84 | 0.65 |
| 4 | 139253454 | rs7669668 | -0.358 | 0.015 | 0.538 | -0.131 | -0.006 | -0.006 | -0.006 | 0.106 | 0.82 | 0.73 | 0.79 | 0.85 | 0.88 | 0.65 |
| 4 | 139258276 | rs7682572 | 0.488 | -0.139 | -0.016 | -0.147 | 0.021 | 0.137 | 0.137 | 3.146 | 0.45 | 0.19 | 0.38 | 0.22 | 0.58 | 0.29 |
| 4 | 139260411 | rs3096659 | -0.393 | 0.195 | 0.42 | 0.383 | 0.005 | 0.003 | 0.003 | 3.932 | 0.84 | 0.8 | 0.82 | 0.87 | 0.89 | 0.76 |
| 4 | 139266486 | rs298469 | 0.17 | 1.102 | -0.016 | 0.469 | 0.005 | 0.005 | 0.005 | 1.357 | 0.22 | 0.23 | 0.22 | 0.32 | 0.29 | 0.12 |
| 4 | 139267650 | rs298472 | -0.489 | 1.069 | 0.102 | -0.132 | 0.024 | -0.044 | -0.044 | 0 | 0.76 | 0.6 | 0.74 | 0.87 | 0.72 | 0.86 |
| 4 | 139268507 | rs7687008 | -0.139 | 0.943 | 0.359 | 0.119 | 0.034 | 0.053 | 0.053 | 0 | 0.28 | 0.4 | 0.49 | 0.56 | 0.31 | 0.35 |

| | | | | | | | | | | | | | | | | |
|---|---|---|---|---|---|---|---|---|---|---|---|---|---|---|---|---|
| 4 | 139271665 | rs7657271 | 0.732 | 0.969 | 0.316 | 0.068 | 0.007 | -0.001 | -0.001 | 0 | 0.22 | 0.24 | 0.18 | 0.23 | 0.26 | 0.11 |
| 4 | 139274999 | rs448469 | 0.073 | 0.961 | 0.428 | 0.068 | 0.015 | -0.005 | -0.005 | 1.79 | 0.67 | 0.45 | 0.5 | 0.34 | 0.67 | 0.67 |
| 4 | 139282314 | rs417342 | 0.701 | 0.909 | 0.37 | 0.059 | 0.074 | 0.154 | 0.154 | 0 | 0.48 | 0.2 | 0.29 | 0.25 | 0.41 | 0.54 |
| 4 | 139285096 | rs17531534 | 0.26 | 0.93 | 0.367 | 0.088 | 0.007 | 0.213 | 0.213 | 0 | 0.06 | 0.11 | 0.27 | 0.43 | 0.05 | 0.17 |
| 4 | 139285676 | rs17050321 | 0.26 | 0.93 | 0.366 | 0.099 | 0.006 | 0.195 | 0.195 | 0 | 0.06 | 0.11 | 0.27 | 0.4 | 0.05 | 0.17 |
| 4 | 139285764 | rs426029 | 0.81 | 0.925 | 0.306 | -0.1 | 0.066 | 0.182 | 0.182 | 0 | 0.47 | 0.18 | 0.28 | 0.21 | 0.41 | 0.54 |
| 4 | 139286917 | rs11944210 | -0.034 | 1.015 | 0.497 | 0.206 | 0.012 | 0.02 | 0.02 | 0.173 | 0.69 | 0.54 | 0.53 | 0.4 | 0.69 | 0.78 |
| 4 | 139291850 | rs17242939 | 1.184 | 0.489 | -0.439 | -0.067 | 0.002 | 0.051 | 0.051 | 0 | 0.16 | 0.04 | 0.09 | 0.05 | 0.17 | 0.17 |
| 4 | 139300684 | rs1398925 | -0.731 | 0.956 | 0.19 | 0.527 | 0.012 | 0.035 | 0.035 | 0 | 0.92 | 0.75 | 0.86 | 0.86 | 0.94 | 0.88 |
| 4 | 139300738 | rs6837219 | 0.294 | 0.935 | -0.304 | 0.407 | 0.001 | -0.011 | -0.011 | 0 | 0.27 | 0.28 | 0.2 | 0.19 | 0.27 | 0.17 |
| 4 | 139301744 | rs12505439 | 1.463 | 0.9 | -0.997 | 0.37 | 0.031 | 0.079 | 0.079 | 0 | 0.37 | 0.18 | 0.2 | 0.2 | 0.31 | 0.46 |
| 4 | 139302720 | rs298462 | 0.879 | 0.927 | -1.024 | 0.352 | -0.04 | -0.056 | -0.056 | 2.144 | 0.1 | 0 | 0.29 | 0.38 | 0.14 | 0.1 |
| 4 | 139302835 | rs298461 | -0.302 | 0.966 | -0.94 | 0.359 | -0.007 | -0.015 | -0.015 | 0 | 0.3 | 0.23 | 0.45 | 0.5 | 0.36 | 0.25 |
| 4 | 139302875 | rs7678098 | 0.991 | 1.016 | -1.272 | -0.172 | -0.001 | -0.001 | -0.001 | 0 | 0.17 | 0.17 | 0.15 | 0.11 | 0.21 | 0.15 |
| 4 | 139303058 | rs6537378 | -0.778 | 1.013 | -1.267 | -0.172 | 0.013 | 0.165 | 0.165 | 0.396 | 0.8 | 0.61 | 0.55 | 0.47 | 0.8 | 0.78 |
| 4 | 139304119 | rs6842871 | 0.939 | 1.134 | -1.573 | -0.254 | -0.002 | 0 | 0 | 0.019 | 0.17 | 0.17 | 0.15 | 0.12 | 0.21 | 0.15 |
| 4 | 139304278 | rs11935389 | 0.863 | 1.13 | -1.571 | -0.308 | -0.011 | 0.016 | 0.016 | 0 | 0.22 | 0.17 | 0.19 | 0.13 | 0.27 | 0.14 |
| 4 | 139305143 | rs13108768 | 0.872 | 1.236 | -1.504 | -0.201 | -0.002 | 0 | 0 | 0.044 | 0.17 | 0.17 | 0.15 | 0.13 | 0.21 | 0.15 |
| 4 | 139305156 | rs12506913 | 0.984 | 1.216 | -1.598 | -0.215 | 0.033 | 0.092 | 0.092 | 0 | 0.37 | 0.18 | 0.21 | 0.17 | 0.31 | 0.46 |
| 4 | 139305290 | rs2052750 | 0.082 | 1.085 | -1.177 | 0.356 | -0.01 | 0.021 | 0.021 | 0 | 0.25 | 0.19 | 0.19 | 0.13 | 0.27 | 0.14 |
| 4 | 139308745 | rs6857347 | 0.027 | 1.074 | -1.44 | 0.161 | -0.005 | 0.039 | 0.039 | 0 | 0.54 | 0.36 | 0.36 | 0.29 | 0.52 | 0.61 |
| 4 | 139312114 | rs7356153 | 0.036 | 1.177 | -2.137 | -0.008 | -0.071 | 0.054 | 0.054 | 0.002 | 0.67 | 0.97 | 0.75 | 0.85 | 0.62 | 0.77 |
| 4 | 139312812 | rs6848489 | 0.601 | 1.362 | -1.094 | 0.418 | 0 | -0.013 | -0.013 | 0 | 0.52 | 0.53 | 0.46 | 0.33 | 0.47 | 0.52 |
| 4 | 139324474 | rs6825410 | -0.409 | 1.364 | -0.777 | 0.335 | -0.017 | 0.071 | 0.071 | 0 | 0.7 | 0.91 | 0.77 | 0.9 | 0.69 | 0.83 |
| 4 | 139325031 | rs17050362 | 0.029 | 1.413 | -0.384 | 0.667 | 0.078 | 0.413 | 0.413 | 0.041 | 0.06 | 0.23 | 0.24 | 0.46 | 0.1 | 0.2 |
| 4 | 139326271 | rs1544891 | -0.511 | 1.425 | -0.446 | 0.666 | -0.06 | 0.089 | 0.089 | 0 | 0.71 | 0.98 | 0.78 | 0.93 | 0.7 | 0.83 |
| 4 | 139327019 | rs4863797 | 0.661 | 1.412 | -0.08 | 0.663 | 0 | -0.001 | -0.001 | 1.617 | 0.54 | 0.48 | 0.47 | 0.38 | 0.51 | 0.56 |
| 4 | 139331299 | rs2314413 | -0.613 | 1.418 | -0.082 | 0.702 | -0.032 | -0.05 | -0.05 | 0 | 0.12 | 0.04 | 0.26 | 0.52 | 0.18 | 0.26 |
| 4 | 139335602 | rs6816442 | 0.114 | 1.421 | -0.023 | -0.234 | 0.029 | -0.083 | -0.083 | 0 | 0.23 | 0.41 | 0.31 | 0.05 | 0.28 | 0.3 |
| 4 | 139336708 | rs2976499 | -0.519 | 1.25 | 0.133 | -0.425 | -0.003 | -0.001 | -0.001 | 0 | 0.63 | 0.69 | 0.51 | 0.45 | 0.56 | 0.51 |
| 4 | 139336851 | rs11732157 | 1.37 | 1.119 | 0.01 | -0.492 | 0.034 | 0.031 | 0.031 | 1.798 | 0.31 | 0.14 | 0.16 | 0.3 | 0.21 | 0.15 |
| 4 | 139342080 | rs17050391 | 0.498 | 1.408 | -0.036 | -0.385 | 0.008 | 0.02 | 0.02 | 0 | 0.06 | 0.12 | 0.06 | 0.07 | 0.07 | 0.14 |
| 4 | 139345618 | rs2912467 | -0.731 | 1.562 | -0.04 | -0.416 | -0.009 | 0.076 | 0.076 | 0 | 0.67 | 0.82 | 0.73 | 0.92 | 0.64 | 0.71 |
| 4 | 139350368 | rs17050398 | 0.402 | 1.675 | 0.065 | -0.237 | 0.012 | 0.02 | 0.02 | 0 | 0.06 | 0.12 | 0.06 | 0.07 | 0.07 | 0.13 |
| 4 | 139361172 | rs2912470 | -0.924 | 1.899 | -0.216 | -0.24 | -0.012 | 0.137 | 0.137 | 0 | 0.67 | 0.91 | 0.77 | 0.89 | 0.69 | 0.8 |
| 4 | 139366615 | rs10519466 | 0.033 | 2.182 | -0.187 | -0.032 | -0.004 | 0.088 | 0.088 | 0 | 0.07 | 0.05 | 0.33 | 0.49 | 0.16 | 0.22 |
| 4 | 139367276 | rs2976490 | 0.842 | 2.213 | 0.013 | -0.139 | 0.051 | -0.032 | -0.032 | 0.128 | 0.12 | 0.31 | 0.17 | 0.06 | 0.16 | 0.23 |
| 4 | 139367951 | rs17245609 | 1.049 | 2.214 | -0.374 | -0.165 | -0.012 | -0.019 | -0.019 | 0 | 0.07 | 0.03 | 0.04 | 0.09 | 0.08 | 0.06 |
| 4 | 139376233 | rs971950 | 0.15 | 2.244 | -0.332 | -0.229 | 0.006 | 0.032 | 0.032 | 0.14 | 0.53 | 0.7 | 0.4 | 0.38 | 0.43 | 0.55 |
| 4 | 139376425 | rs1426520 | -0.977 | 2.04 | -0.746 | -0.773 | -0.016 | 0.137 | 0.137 | 0 | 0.67 | 0.89 | 0.78 | 0.93 | 0.67 | 0.82 |
| 4 | 139377761 | rs1834624 | -0.499 | 2.023 | -0.571 | -0.625 | -0.021 | 0.14 | 0.14 | 0.028 | 0.6 | 0.84 | 0.74 | 0.85 | 0.58 | 0.75 |
| 4 | 139379432 | rs2912475 | 0.317 | 1.982 | 0.182 | -0.295 | 0.036 | -0.03 | -0.03 | 0.126 | 0.11 | 0.24 | 0.16 | 0.06 | 0.16 | 0.22 |
| 4 | 139387054 | rs755954 | 0.187 | 2.175 | -0.086 | -0.398 | 0.011 | -0.034 | -0.034 | 0 | 0.08 | 0.11 | 0.13 | 0 | 0.15 | 0.04 |
| 4 | 139387890 | rs1346214 | -0.075 | 2.951 | 0.152 | 0.138 | 0.062 | 0.222 | 0.222 | 0 | 0.05 | 0.14 | 0.28 | 0.51 | 0.13 | 0.21 |
| 4 | 139388475 | rs6818743 | -0.857 | 2.31 | -0.509 | -0.897 | 0.023 | 0.139 | 0.139 | 0.006 | 0.64 | 0.89 | 0.81 | 0.87 | 0.7 | 0.74 |
| 4 | 139391659 | rs2912473 | 0.058 | 2.339 | -0.462 | -0.977 | 0 | 0.123 | 0.123 | 0 | 0.52 | 0.71 | 0.62 | 0.81 | 0.48 | 0.56 |
| 4 | 139396007 | rs9308208 | 0.11 | 2.341 | -0.706 | -1.133 | 0.016 | 0.124 | 0.124 | 1.685 | 0.46 | 0.74 | 0.55 | 0.78 | 0.4 | 0.48 |
| 4 | 139396693 | rs13148881 | 0.473 | 2.162 | -0.265 | -1.116 | 0 | 0.006 | 0.006 | 0.045 | 0.37 | 0.31 | 0.43 | 0.66 | 0.31 | 0.3 |
| 4 | 139398829 | rs2976477 | -0.511 | 1.964 | -0.477 | -1.03 | 0.023 | 0.037 | 0.037 | 0 | 0.54 | 0.32 | 0.68 | 0.71 | 0.62 | 0.55 |
| 4 | 139399972 | rs13108353 | -0.242 | 1.692 | -0.283 | -0.805 | -0.007 | 0.024 | 0.024 | 0.056 | 0.45 | 0.32 | 0.55 | 0.66 | 0.46 | 0.33 |
| 4 | 139403484 | rs17050442 | 0.163 | 1.06 | -0.269 | -1.002 | 0.034 | 0.036 | 0.036 | 0 | 0.05 | 0.24 | 0.11 | 0.07 | 0.06 | 0.16 |
| 4 | 139406421 | rs12644806 | 0.017 | 1.166 | -0.61 | -1.082 | -0.009 | 0.028 | 0.028 | 1.156 | 0.16 | 0.1 | 0.11 | 0.06 | 0.18 | 0.05 |
| 4 | 139407719 | rs10519463 | 0.761 | 1.285 | -0.669 | -1.16 | -0.003 | 0 | 0 | 2.235 | 0.05 | 0.05 | 0.11 | 0.06 | 0.09 | 0.17 |
| 4 | 139409081 | rs10018963 | -0.838 | 0.871 | -0.446 | -1.647 | -0.006 | -0.016 | -0.016 | 0 | 0.71 | 0.65 | 0.68 | 0.82 | 0.72 | 0.6 |
| 4 | 139409291 | rs17306529 | 0.848 | 0.859 | -0.54 | -1.614 | -0.01 | 0.022 | 0.022 | 0.112 | 0.14 | 0.1 | 0.11 | 0.06 | 0.18 | 0.04 |
| 4 | 139431355 | rs299113 | -0.46 | 0.856 | -0.272 | -1.634 | -0.001 | -0.008 | -0.008 | 0 | 0.43 | 0.44 | 0.43 | 0.24 | 0.45 | 0.44 |
| 4 | 139434076 | rs299120 | -0.069 | 0.947 | -0.061 | -1.731 | 0 | -0.007 | -0.007 | 0 | 0.38 | 0.34 | 0.32 | 0.12 | 0.37 | 0.26 |
| 4 | 139443162 | rs1450432 | 0.064 | 0.368 | 0.387 | -1.69 | 0.042 | 0.049 | 0.049 | 1.882 | 0.65 | 0.16 | 0.63 | 0.76 | 0.71 | 0.58 |
| 4 | 139460404 | rs299160 | 1.103 | 0.209 | 0.786 | -1.409 | -0.003 | -0.008 | -0.008 | 0 | 0.24 | 0.24 | 0.21 | 0.12 | 0.2 | 0.24 |
| 4 | 139462440 | rs17050484 | -1.007 | 0.219 | 0.764 | -1.511 | -0.025 | -0.02 | -0.02 | 0.136 | 0.74 | 0.48 | 0.77 | 0.8 | 0.8 | 0.76 |
| 4 | 139464920 | rs13135708 | -0.893 | 0.211 | 0.506 | -1.494 | -0.034 | -0.023 | -0.023 | 0 | 0.73 | 0.47 | 0.77 | 0.8 | 0.81 | 0.75 |
| 4 | 139482703 | rs11730882 | -0.076 | 0.418 | -0.072 | -1.62 | 0.003 | 0.002 | 0.002 | 0 | 0.53 | 0.6 | 0.44 | 0.19 | 0.42 | 0.51 |
| 4 | 139482958 | rs7694744 | 0.331 | 0.691 | -0.609 | -1.382 | 0.016 | 0.074 | 0.074 | 0 | 0.4 | 0.25 | 0.24 | 0.1 | 0.33 | 0.32 |
| 4 | 139485229 | rs1376082 | -0.506 | 0.983 | -0.346 | -0.826 | 0.005 | 0.003 | 0.003 | 0.009 | 0.82 | 0.87 | 0.65 | 0.26 | 0.75 | 0.64 |
| 4 | 139486064 | rs1530798 | 0.245 | -0.082 | -0.488 | -1.025 | 0.029 | 0.039 | 0.039 | 0 | 0.15 | 0.07 | 0.19 | 0.08 | 0.1 | 0.2 |
| 4 | 139486920 | rs6831698 | 1.065 | -0.318 | -0.651 | -1.082 | 0.003 | 0.011 | 0.011 | 0 | 0.24 | 0.1 | 0.16 | 0.15 | 0.23 | 0.23 |
| 4 | 139488136 | rs1817031 | -0.817 | -0.301 | -0.218 | -0.675 | 0.006 | -0.033 | -0.033 | 0 | 0.61 | 0.55 | 0.7 | 0.82 | 0.57 | 0.75 |
| 4 | 139493398 | rs1450439 | -0.07 | -0.338 | -0.063 | -0.597 | -0.023 | -0.094 | -0.094 | 0.005 | 0.29 | 0.43 | 0.29 | 0.1 | 0.22 | 0.25 |
| 4 | 139494811 | rs1376086 | 0.906 | -0.127 | -0.275 | -0.298 | -0.004 | 0.038 | 0.038 | 0 | 0.09 | 0.04 | 0.25 | 0.57 | 0.12 | 0.26 |

| | | | | | | | | | | | | | | | | |
|---|---|---|---|---|---|---|---|---|---|---|---|---|---|---|---|---|
| 4 | 139495121 | rs1376088 | 0.475 | -0.144 | -0.293 | -0.395 | -0.013 | 0.039 | 0.039 | 0.1 | 0.33 | 0.12 | 0.41 | 0.72 | 0.34 | 0.51 |
| 4 | 139495763 | rs13142307 | -0.47 | -0.468 | 0.314 | -0.369 | -0.001 | -0.004 | -0.004 | 0 | 0.59 | 0.48 | 0.71 | 0.69 | 0.62 | 0.7 |
| 4 | 139496147 | rs1376091 | -0.023 | -0.19 | 0.688 | -0.116 | -0.035 | -0.004 | -0.004 | 0 | 0.81 | 0.62 | 0.76 | 0.78 | 0.86 | 0.77 |
| 4 | 139497265 | rs7435327 | 0.286 | -0.037 | 0.51 | -0.054 | -0.007 | 0 | 0 | 0 | 0.17 | 0.2 | 0.23 | 0.23 | 0.14 | 0.15 |
| 4 | 139500215 | rs13131639 | 1.229 | 0.105 | 0.513 | 0.268 | -0.013 | -0.023 | -0.023 | 0.022 | 0.23 | 0.15 | 0.15 | 0.4 | 0.27 | 0.33 |
| 4 | 139503481 | rs12643388 | 1.378 | 0.047 | 0.795 | 0.322 | 0.002 | 0.046 | 0.046 | 0 | 0.07 | 0.17 | 0.03 | 0.13 | 0.05 | 0.15 |
| 4 | 139506065 | rs6841254 | -0.339 | 0.002 | 0.762 | 0.208 | -0.002 | -0.029 | -0.029 | 0 | 0.27 | 0.39 | 0.4 | 0.17 | 0.31 | 0.19 |
| 4 | 139507392 | rs12504508 | 0.096 | -0.004 | 0.821 | 0.195 | 0.009 | -0.038 | -0.038 | 0 | 0.17 | 0.25 | 0.18 | 0.08 | 0.23 | 0.15 |
| 4 | 139507777 | rs12644755 | 1.666 | 0.001 | 0.728 | 0.204 | -0.003 | -0.003 | -0.003 | 1.449 | 0.06 | 0.1 | 0.02 | 0.08 | 0.05 | 0.11 |
| 4 | 139510249 | rs9307846 | -0.814 | 0.005 | 0.602 | 0.219 | -0.006 | -0.051 | -0.051 | 0.818 | 0.34 | 0.5 | 0.42 | 0.25 | 0.36 | 0.29 |
| 4 | 139513580 | rs9685123 | 1.413 | 0.021 | 0.521 | 0.235 | -0.038 | -0.063 | -0.063 | 0 | 0.23 | 0.08 | 0.15 | 0.39 | 0.27 | 0.33 |
| 4 | 139514099 | rs4863565 | -1.728 | -0.054 | 0.533 | 0.122 | 0.009 | 0.006 | 0.006 | 0 | 0.71 | 0.82 | 0.64 | 0.8 | 0.76 | 0.72 |
| 4 | 139519532 | rs1450453 | -1.434 | -0.052 | 0.475 | 0.095 | 0.01 | 0.062 | 0.062 | 0 | 0.49 | 0.74 | 0.5 | 0.36 | 0.49 | 0.39 |
| 4 | 139528388 | rs17050529 | 1.284 | -0.057 | 0.419 | 0.124 | 0 | 0.001 | 0.001 | 0 | 0.14 | 0.15 | 0.2 | 0.13 | 0.1 | 0.13 |
| 4 | 139530255 | rs7679163 | -0.547 | -0.017 | 0.456 | 0.151 | -0.001 | -0.002 | -0.002 | 0 | 0.55 | 0.57 | 0.51 | 0.62 | 0.54 | 0.52 |
| 4 | 139530310 | rs7680585 | 1.535 | 0.042 | 0.124 | 0.072 | 0 | -0.031 | -0.031 | 0 | 0.13 | 0.17 | 0.07 | 0 | 0.12 | 0.05 |
| 4 | 139532860 | rs12639806 | -1.493 | 0.219 | -0.424 | 0.343 | 0.006 | 0.003 | 0.003 | 0 | 0.72 | 0.85 | 0.65 | 0.8 | 0.76 | 0.72 |
| 4 | 139537855 | rs2198030 | 1.384 | 0.248 | -1.157 | -0.014 | -0.003 | -0.013 | -0.013 | 2.947 | 0.09 | 0.09 | 0.1 | 0 | 0.17 | 0.04 |
| 4 | 139539662 | rs975477 | -1.215 | 0.042 | -1.458 | 0.367 | 0.006 | 0.032 | 0.032 | 0 | 0.72 | 0.92 | 0.64 | 0.83 | 0.75 | 0.72 |
| 4 | 139540219 | rs6851978 | 0.057 | -0.034 | -1.429 | 0.337 | -0.014 | 0.026 | 0.026 | 0.087 | 0.18 | 0.23 | 0.29 | 0.36 | 0.16 | 0.2 |
| 4 | 139541397 | rs12233744 | 0.589 | 0.092 | -1.328 | -0.179 | 0.009 | -0.154 | -0.154 | 0 | 0.31 | 0.56 | 0.2 | 0.02 | 0.33 | 0.15 |
| 4 | 139550075 | rs11941340 | -0.651 | -0.35 | -1.379 | 0.734 | -0.016 | 0 | 0 | 0 | 0.87 | 0.84 | 0.82 | 0.81 | 0.93 | 0.91 |
| 4 | 139552686 | rs719922 | -0.319 | 0.393 | -1.703 | 0.796 | 0.004 | -0.036 | -0.036 | 0 | 0.32 | 0.14 | 0.41 | 0.38 | 0.32 | 0.39 |
| 4 | 139555962 | rs1450454 | 0.098 | 0.093 | -1.96 | 0.382 | 0.108 | 0.088 | 0.088 | 0 | 0.51 | 0.17 | 0.52 | 0.65 | 0.59 | 0.57 |
| 4 | 139557368 | rs4863572 | 0.093 | 0.193 | -1.827 | 0.011 | 0.049 | 0.039 | 0.039 | 1.91 | 0.58 | 0.29 | 0.66 | 0.73 | 0.69 | 0.61 |
| 4 | 139567521 | rs11721545 | 0.262 | 0.499 | -1.13 | -0.074 | 0.031 | 0.04 | 0.04 | 1.797 | 0.55 | 0.32 | 0.65 | 0.74 | 0.63 | 0.66 |
| 4 | 139568754 | rs12502933 | 0.186 | 0.623 | -1.082 | -0.074 | 0.025 | 0.046 | 0.046 | 2.366 | 0.54 | 0.32 | 0.64 | 0.73 | 0.61 | 0.6 |
| 4 | 139570038 | rs4863578 | 0.123 | 0.588 | -0.423 | -0.092 | 0.006 | 0.011 | 0.011 | 1.903 | 0.55 | 0.42 | 0.65 | 0.73 | 0.61 | 0.6 |
| 4 | 139571883 | rs6535613 | 0.216 | 0.565 | -0.435 | -0.093 | 0.026 | 0.048 | 0.048 | 2.248 | 0.54 | 0.32 | 0.64 | 0.73 | 0.62 | 0.6 |
| 4 | 139574585 | rs1450449 | 0.265 | 0.557 | -0.372 | -0.077 | 0.039 | 0.074 | 0.074 | 2.644 | 0.53 | 0.23 | 0.64 | 0.72 | 0.61 | 0.6 |
| 4 | 139575076 | rs12504582 | 0.165 | 0.548 | -0.387 | -0.081 | 0.025 | 0.045 | 0.045 | 2.405 | 0.54 | 0.31 | 0.65 | 0.72 | 0.61 | 0.6 |
| 4 | 139575494 | rs12501198 | 0.275 | 0.571 | -0.553 | -0.07 | 0.039 | 0.081 | 0.081 | 2.685 | 0.53 | 0.16 | 0.63 | 0.72 | 0.61 | 0.6 |
| 4 | 139580179 | rs10017576 | 0.218 | 0.611 | -0.093 | -0.167 | 0.019 | 0.043 | 0.043 | 3.438 | 0.56 | 0.23 | 0.66 | 0.73 | 0.62 | 0.6 |
| 4 | 139580945 | rs2122509 | 0.247 | 0.746 | 0.361 | -0.129 | 0.001 | 0.019 | 0.019 | 1.509 | 0.56 | 0.37 | 0.62 | 0.7 | 0.59 | 0.63 |
| 4 | 139581543 | rs9685238 | -0.233 | 0.961 | 0.671 | -0.085 | -0.001 | 0.017 | 0.017 | 1.074 | 0.42 | 0.32 | 0.38 | 0.3 | 0.41 | 0.33 |
| 4 | 139582664 | rs13135384 | -0.759 | 0.942 | 0.137 | -0.236 | 0.008 | -0.012 | -0.012 | 0 | 0.66 | 0.73 | 0.77 | 0.4 | 0.75 | 0.6 |
| 4 | 139584101 | rs6835813 | -0.72 | 0.921 | 0.144 | -0.601 | 0.005 | 0 | 0 | 0 | 0.66 | 0.7 | 0.77 | 0.31 | 0.75 | 0.62 |
| 4 | 139590199 | rs7692637 | -0.192 | 1.241 | 0.179 | -0.61 | 0.036 | -0.057 | -0.057 | 1.121 | 0.31 | 0.44 | 0.44 | 0.11 | 0.47 | 0.32 |
| 4 | 139605165 | rs4863585 | -1.132 | 0.728 | 0.415 | -0.328 | 0.076 | 0.17 | 0.17 | 5.71 | 0.06 | 0.47 | 0.15 | 0.07 | 0.01 | 0.07 |
| 4 | 139605950 | rs10004565 | -0.059 | 0.603 | -0.586 | -0.575 | 0.04 | 0.043 | 0.043 | 4.11 | 0.4 | 0.76 | 0.45 | 0.26 | 0.25 | 0.27 |
| 4 | 139609402 | rs7685323 | -0.199 | 1.515 | -0.435 | -0.688 | 0 | -0.018 | -0.018 | 1.762 | 0.33 | 0.4 | 0.46 | 0.09 | 0.47 | 0.32 |
| 4 | 139612743 | rs7693492 | -1.059 | 1.39 | 0.131 | 0.152 | 0.004 | -0.011 | -0.011 | 0 | 0.27 | 0.38 | 0.41 | 0.75 | 0.29 | 0.31 |
| 4 | 139624155 | rs7667792 | -1.089 | 1.064 | 0.526 | 0.413 | 0.001 | 0.022 | 0.022 | 0 | 0.24 | 0.41 | 0.4 | 0.71 | 0.26 | 0.28 |
| 4 | 139626820 | rs9790693 | -0.614 | 0.768 | 0.528 | 1.187 | -0.004 | 0.092 | 0.092 | 2.364 | 0.15 | 0.21 | 0.29 | 0.51 | 0.16 | 0.17 |
| 4 | 139629908 | rs7655082 | -1.666 | 0.435 | -0.325 | -0.05 | -0.001 | 0.164 | 0.164 | 0.368 | 0.54 | 0.85 | 0.67 | 0.84 | 0.48 | 0.6 |
| 4 | 139631377 | rs905998 | -1.085 | 0.306 | 0.497 | -0.006 | 0.001 | 0.001 | 0.001 | 0 | 0.31 | 0.32 | 0.5 | 0.78 | 0.28 | 0.51 |
| 4 | 139636653 | rs4863590 | -1.007 | 0.277 | 0.6 | 0.083 | -0.003 | 0.004 | 0.004 | 0 | 0.29 | 0.34 | 0.5 | 0.69 | 0.28 | 0.35 |
| 4 | 139637374 | rs6851223 | 0.231 | 0.137 | 1.111 | -0.002 | -0.008 | -0.04 | -0.04 | 0.255 | 0.18 | 0.23 | 0.09 | 0.03 | 0.14 | 0.11 |
| 4 | 139646534 | rs2135287 | -0.659 | -0.002 | 1.252 | -0.013 | -0.045 | -0.039 | -0.039 | 0.831 | 0.21 | 0.44 | 0.12 | 0.14 | 0.13 | 0.23 |
| 4 | 139648420 | rs6824816 | -2.228 | 0.463 | 0.996 | -0.059 | -0.078 | 0.179 | 0.179 | 0 | 0.54 | 0.93 | 0.66 | 0.83 | 0.49 | 0.64 |
| 4 | 139651607 | rs6853043 | 0.31 | 0.354 | 1.403 | -0.074 | -0.015 | -0.056 | -0.056 | 0 | 0.16 | 0.29 | 0.09 | 0.03 | 0.12 | 0.07 |
| 4 | 139653067 | rs6812423 | -1.479 | 0.305 | 1.374 | -0.051 | 0.004 | 0.003 | 0.003 | 0 | 0.31 | 0.7 | 0.19 | 0.16 | 0.2 | 0.31 |
| 4 | 139655744 | rs1505866 | 0.248 | 0.344 | 1.169 | -0.024 | 0 | -0.004 | -0.004 | 0 | 0.08 | 0.08 | 0.05 | 0.02 | 0.06 | 0.07 |
| 4 | 139655759 | rs1505867 | 0.248 | 0.344 | 1.151 | -0.024 | -0.013 | -0.026 | -0.026 | 0.253 | 0.08 | 0.13 | 0.06 | 0.02 | 0.06 | 0.07 |
| 4 | 139658216 | rs6821816 | -2.226 | 0.398 | 0.943 | 0.141 | -0.039 | 0.152 | 0.152 | 0 | 0.52 | 0.89 | 0.64 | 0.8 | 0.47 | 0.62 |
| 4 | 139687856 | rs10020229 | -0.97 | 0.201 | 1.26 | 0.061 | -0.036 | -0.091 | -0.091 | 0.208 | 0.27 | 0.46 | 0.17 | 0.08 | 0.19 | 0.17 |
| 4 | 139690422 | rs7686170 | -0.712 | 0.207 | 1.269 | 0.069 | -0.026 | -0.062 | -0.062 | 0 | 0.13 | 0.3 | 0.09 | 0.04 | 0.09 | 0.11 |
| 4 | 139692069 | rs11099746 | -0.367 | 0.47 | 0.458 | 0.071 | -0.044 | -0.054 | -0.054 | 0.117 | 0.24 | 0.13 | 0.45 | 0.64 | 0.32 | 0.34 |
| 4 | 139692813 | rs1540744 | -2.188 | 0.459 | 0.298 | 0.069 | -0.081 | 0.053 | 0.053 | 0 | 0.55 | 0.92 | 0.66 | 0.72 | 0.51 | 0.51 |
| 4 | 139695631 | rs6535696 | -2.185 | 0.461 | 0.3 | -0.027 | -0.081 | 0.124 | 0.124 | 0 | 0.55 | 0.93 | 0.66 | 0.78 | 0.51 | 0.5 |
| 4 | 139696731 | rs11736543 | 0.421 | 0.195 | 0.887 | 0.043 | -0.01 | -0.045 | -0.045 | 0.029 | 0.14 | 0.21 | 0.09 | 0.04 | 0.1 | 0.05 |
| 4 | 139698522 | rs6855592 | -1.12 | 0.28 | 0.18 | 0.097 | -0.025 | -0.029 | -0.029 | 0 | 0.29 | 0.55 | 0.18 | 0.16 | 0.2 | 0.32 |
| 4 | 139708493 | rs9996437 | -1.074 | 0.274 | -0.284 | 0.121 | 0.015 | 0.015 | 0.015 | 0 | 0.31 | 0.68 | 0.18 | 0.15 | 0.19 | 0.3 |
| 4 | 139710480 | rs1948898 | -1.237 | 0.586 | -0.367 | 0.512 | -0.007 | -0.004 | -0.004 | 0 | 0.37 | 0.34 | 0.55 | 0.67 | 0.39 | 0.43 |
| 4 | 139711242 | rs9307869 | 0.785 | 0.502 | -0.046 | 0.648 | -0.011 | 0.007 | 0.007 | 0 | 0.73 | 0.66 | 0.54 | 0.32 | 0.76 | 0.58 |
| 4 | 139711726 | rs13136052 | 1.81 | 0.769 | -0.788 | -0.112 | -0.102 | 0.268 | 0.268 | 0 | 0.41 | 0.08 | 0.29 | 0.05 | 0.48 | 0.27 |
| 4 | 139716412 | rs1505869 | -0.529 | 0.598 | -0.621 | 0.683 | -0.02 | -0.066 | -0.066 | 0 | 0.87 | 0.99 | 0.72 | 0.7 | 0.85 | 0.86 |
| 4 | 139717020 | rs1995598 | 0.201 | 0.704 | -0.227 | -0.03 | -0.006 | -0.01 | -0.01 | 0 | 0.7 | 0.54 | 0.47 | 0.3 | 0.72 | 0.56 |
| 4 | 139727944 | rs6844434 | 0.93 | 0.846 | -0.622 | -0.236 | -0.031 | 0.116 | 0.116 | 0 | 0.44 | 0.24 | 0.32 | 0.21 | 0.5 | 0.46 |

| | | | | | | | | | | | | | | | | |
|---|---|---|---|---|---|---|---|---|---|---|---|---|---|---|---|---|
| 4 | 139729522 | rs7657235 | 0.995 | 1.321 | -0.724 | -0.242 | -0.013 | 0.084 | 0.084 | 0 | 0.42 | 0.14 | 0.32 | 0.27 | 0.41 | 0.39 |
| 4 | 139729690 | rs6535707 | 0.344 | 0.674 | -0.209 | -0.243 | 0.003 | -0.078 | -0.078 | 0.06 | 0.21 | 0.33 | 0.12 | 0.07 | 0.18 | 0.1 |
| 4 | 139731065 | rs1505864 | -0.013 | 0.221 | -0.354 | 0.107 | -0.003 | 0.003 | 0.003 | 0.068 | 0.74 | 0.68 | 0.55 | 0.34 | 0.72 | 0.59 |
| 4 | 139734961 | rs13118554 | 0.115 | -0.066 | -1.137 | -0.605 | 0.063 | 0.091 | 0.091 | 0.783 | 0.53 | 0.3 | 0.48 | 0.19 | 0.66 | 0.33 |
| 4 | 139739456 | rs11728868 | 0.836 | 0.63 | -1.763 | -0.508 | -0.079 | 0.05 | 0.05 | 0.068 | 0.26 | 0.1 | 0.26 | 0.14 | 0.38 | 0.19 |
| 4 | 139745929 | rs2102309 | 0.005 | 0.007 | -1.516 | -0.426 | 0.048 | 0.072 | 0.072 | 0.004 | 0.55 | 0.67 | 0.46 | 0.23 | 0.64 | 0.33 |
| 4 | 139748640 | rs4481278 | -0.149 | -0.275 | -1.987 | 0.115 | 0.045 | -0.022 | -0.022 | 0 | 0.61 | 0.87 | 0.49 | 0.35 | 0.67 | 0.48 |
| 4 | 139753601 | rs10006363 | -0.369 | -0.495 | -2.146 | 0.13 | 0.025 | -0.065 | -0.065 | 0 | 0.68 | 0.93 | 0.49 | 0.37 | 0.71 | 0.56 |
| 4 | 139755093 | rs10022416 | -0.752 | -0.37 | -2.112 | 0.888 | -0.047 | -0.091 | -0.091 | 0.249 | 0.86 | 0.97 | 0.57 | 0.58 | 0.79 | 0.78 |
| 4 | 139764049 | rs6854489 | -0.155 | -0.388 | -2.593 | 0.376 | -0.118 | 0.129 | 0.129 | 2.143 | 0.6 | 0.91 | 0.31 | 0.2 | 0.43 | 0.5 |
| 4 | 139768109 | rs17050640 | 0.522 | 0.01 | -2.366 | 0.803 | -0.041 | -0.089 | -0.089 | 6.217 | 0.15 | 0.03 | 0.36 | 0.4 | 0.19 | 0.2 |
| 4 | 139771078 | rs10001269 | 0.581 | -0.521 | -1.528 | 0.108 | -0.021 | -0.035 | -0.035 | 0 | 0.19 | 0.28 | 0.09 | 0.1 | 0.12 | 0.19 |
| 4 | 139771441 | rs2135289 | -0.325 | 0.043 | -1.852 | 0.373 | -0.024 | -0.021 | -0.021 | 0 | 0.59 | 0.77 | 0.34 | 0.42 | 0.44 | 0.53 |
| 4 | 139773223 | rs7669416 | -0.35 | 0.067 | -2.037 | 0.374 | -0.026 | -0.02 | -0.02 | 0 | 0.6 | 0.76 | 0.33 | 0.43 | 0.44 | 0.53 |
| 4 | 139779568 | rs11728621 | 0.424 | 1.003 | -2.017 | -0.026 | -0.036 | 0.021 | 0.021 | 4.361 | 0.15 | 0.07 | 0.11 | 0.08 | 0.24 | 0.1 |
| 4 | 139789454 | rs6844950 | -0.543 | 0.867 | -2.071 | -0.204 | -0.058 | 0.078 | 0.078 | 0.835 | 0.74 | 0.89 | 0.73 | 0.92 | 0.61 | 0.75 |
| 4 | 139790626 | rs7689448 | 0.276 | 0.299 | -1.978 | -0.054 | 0.064 | 0.094 | 0.094 | 4.739 | 0.51 | 0.71 | 0.39 | 0.77 | 0.32 | 0.48 |
| 4 | 139794345 | rs4074855 | 0.059 | -0.032 | -2.033 | -0.172 | 0.107 | 0.031 | 0.031 | 8.873 | 0.49 | 0.81 | 0.3 | 0.42 | 0.3 | 0.45 |
| 4 | 139794516 | rs4074856 | -0.124 | 0.383 | -1.668 | 0.495 | -0.106 | -0.128 | -0.128 | 0.461 | 0.26 | 0.05 | 0.44 | 0.49 | 0.37 | 0.29 |
| 4 | 139799012 | rs6535748 | -0.299 | 0.628 | -1.312 | 0.082 | -0.016 | 0.103 | 0.103 | 0 | 0.5 | 0.78 | 0.62 | 0.66 | 0.49 | 0.5 |
| 4 | 139814341 | rs7684118 | 0.016 | 0.503 | -1.291 | 0.109 | 0.001 | 0.09 | 0.09 | 0 | 0.5 | 0.23 | 0.39 | 0.35 | 0.52 | 0.53 |
| 4 | 139826720 | rs4639023 | 0.154 | 0.86 | -1.122 | 0.24 | 0.005 | 0.054 | 0.054 | 0 | 0.52 | 0.3 | 0.38 | 0.41 | 0.53 | 0.62 |
| 4 | 139833891 | rs13103414 | 0.93 | 0.711 | 0.166 | 0.134 | 0.044 | 0.051 | 0.051 | 0 | 0.94 | 0.7 | 0.96 | 0.96 | 0.98 | 0.95 |
| 4 | 139837873 | rs10004074 | 1.228 | 1.417 | -0.123 | -0.093 | 0.046 | -0.022 | -0.022 | 0.192 | 0.86 | 0.74 | 0.91 | 0.92 | 0.85 | 0.81 |
| 4 | 139839354 | rs13146053 | -0.282 | 1.974 | -0.794 | -0.989 | -0.053 | 0.089 | 0.089 | 8.756 | 0.22 | 0.07 | 0.27 | 0.01 | 0.32 | 0.16 |
| 4 | 139842608 | rs4241923 | 1.449 | 1.584 | -0.531 | -1.101 | 0.009 | -0.048 | -0.048 | 0 | 0.61 | 0.6 | 0.65 | 0.9 | 0.53 | 0.66 |
| 4 | 139847316 | rs10028549 | -1.217 | 2.162 | 0.546 | -1.303 | -0.001 | -0.008 | -0.008 | 0 | 0.29 | 0.24 | 0.28 | 0.09 | 0.33 | 0.28 |
| 4 | 139848044 | rs6858505 | 0.599 | 2.136 | 0.344 | -1.229 | -0.033 | 0.033 | 0.033 | 0.157 | 0.06 | 0 | 0.06 | 0 | 0.13 | 0.08 |
| 4 | 139856728 | rs6839690 | 0.061 | 2.114 | 0.548 | -1.341 | -0.029 | -0.075 | -0.075 | 0 | 0.83 | 0.66 | 0.87 | 0.98 | 0.86 | 0.93 |
| 4 | 139860920 | rs6848189 | 0.128 | 2.169 | 0.546 | -1.343 | -0.007 | -0.076 | -0.076 | 0 | 0.82 | 0.65 | 0.85 | 0.98 | 0.83 | 0.92 |
| 4 | 139861084 | rs9790390 | 2.16 | 1.325 | 0.755 | -1.758 | 0.024 | 0.16 | 0.16 | 1.393 | 0.49 | 0.23 | 0.58 | 0.84 | 0.48 | 0.56 |
| 4 | 139861368 | rs7697018 | -1.177 | 1.516 | 0.754 | -1.703 | 0.035 | -0.038 | -0.038 | 0 | 0.25 | 0.3 | 0.26 | 0.08 | 0.37 | 0.3 |
| 4 | 139861418 | rs11731798 | 0.361 | 1.446 | 0.421 | -1.548 | -0.003 | -0.017 | -0.017 | 0 | 0.14 | 0.16 | 0.11 | 0.02 | 0.14 | 0.07 |
| 4 | 139876605 | rs4863620 | 1.135 | 1.425 | 0.348 | -1.567 | -0.012 | 0.017 | 0.017 | 0.01 | 0.64 | 0.31 | 0.67 | 0.91 | 0.63 | 0.71 |
| 4 | 139884321 | rs11723586 | 1.025 | 1.127 | -0.42 | -0.816 | 0.008 | 0.082 | 0.082 | 0 | 0.51 | 0.29 | 0.51 | 0.36 | 0.53 | 0.38 |
| 4 | 139899443 | rs6858120 | -0.851 | 0.515 | -0.135 | -1.038 | 0.001 | 0.001 | 0.001 | 0 | 0.45 | 0.42 | 0.49 | 0.73 | 0.4 | 0.55 |
| 4 | 139906122 | rs7659060 | -0.108 | 0.393 | 0.084 | -0.796 | -0.006 | 0.021 | 0.021 | 0.181 | 0.24 | 0.29 | 0.36 | 0.56 | 0.19 | 0.22 |
| 4 | 139907427 | rs13101244 | -0.205 | 0.353 | -0.055 | -0.88 | -0.001 | -0.013 | -0.013 | 0.143 | 0.19 | 0.18 | 0.11 | 0.06 | 0.19 | 0.12 |
| 4 | 139908503 | rs4309784 | 1.285 | 0.425 | -0.22 | -0.911 | -0.01 | 0.168 | 0.168 | 0.015 | 0.51 | 0.11 | 0.45 | 0.27 | 0.56 | 0.62 |
| 4 | 139915883 | rs6817458 | 1.152 | 0.345 | -0.103 | -0.842 | 0.004 | 0.151 | 0.151 | 0 | 0.5 | 0.15 | 0.45 | 0.29 | 0.55 | 0.61 |
| 4 | 139925351 | rs6848587 | -0.043 | 0.292 | 0.156 | -0.62 | -0.005 | 0.019 | 0.019 | 0.22 | 0.24 | 0.28 | 0.37 | 0.57 | 0.19 | 0.24 |
| 4 | 139930032 | rs13116075 | 0.407 | 0.522 | 0.215 | -0.652 | -0.035 | 0.052 | 0.052 | 0.027 | 0.11 | 0.02 | 0.09 | 0 | 0.18 | 0.08 |
| 4 | 139956970 | rs1112828 | -0.635 | 0.804 | -0.162 | -0.579 | -0.028 | -0.032 | -0.032 | 0 | 0.13 | 0.06 | 0.48 | 0.63 | 0.2 | 0.18 |
| 4 | 139981918 | rs3796637 | 1.446 | 1.042 | -0.366 | -0.387 | -0.072 | 0.091 | 0.091 | 1.478 | 0.25 | 0.02 | 0.14 | 0.07 | 0.25 | 0.18 |
| 4 | 140006680 | rs2292837 | -0.249 | 1.208 | 0.348 | -0.352 | 0.002 | 0.073 | 0.073 | 0.213 | 0.51 | 0.35 | 0.32 | 0.12 | 0.52 | 0.42 |
| 4 | 140046907 | rs11725721 | 1.263 | 1.183 | 0.434 | -0.339 | -0.013 | 0.094 | 0.094 | 0.05 | 0.29 | 0.11 | 0.16 | 0.08 | 0.26 | 0.29 |
| 4 | 140065768 | rs13134328 | -0.148 | 1.605 | 1.083 | -0.107 | 0.001 | -0.025 | -0.025 | 0 | 0.23 | 0.24 | 0.17 | 0.06 | 0.26 | 0.13 |
| 4 | 140075052 | rs2046661 | 1.426 | 1.52 | 1.148 | -0.05 | -0.011 | -0.012 | -0.012 | 0 | 0.3 | 0.37 | 0.18 | 0.21 | 0.27 | 0.3 |
| 4 | 140093565 | rs4597768 | 1.68 | 1.551 | 0.73 | -0.061 | 0.056 | 0.001 | 0.001 | 0 | 0.28 | 0.07 | 0.13 | 0.21 | 0.21 | 0.29 |
| 4 | 140099768 | rs1471928 | -0.122 | 1.605 | 0.749 | -0.087 | 0.001 | -0.039 | -0.039 | 0 | 0.23 | 0.24 | 0.17 | 0.05 | 0.26 | 0.13 |
| 4 | 140100756 | rs4586876 | 1.796 | 1.609 | 0.769 | -0.02 | 0.041 | -0.031 | -0.031 | 0 | 0.26 | 0.01 | 0.12 | 0.21 | 0.2 | 0.29 |
| 4 | 140115744 | rs11941589 | -0.477 | 1.68 | 0.836 | 0.001 | 0.001 | 0.001 | 0.001 | 0 | 0.6 | 0.35 | 0.62 | 0.59 | 0.61 | 0.6 |
| 4 | 140117588 | rs6825915 | -0.947 | 1.777 | 0.811 | 0.043 | -0.024 | 0.012 | 0.012 | 0.151 | 0.94 | 0.98 | 0.82 | 0.95 | 0.88 | 0.92 |
| 4 | 140128447 | rs12648234 | 1.018 | 1.584 | 0.715 | 0.031 | -0.01 | 0.002 | 0.002 | 0 | 0.33 | 0.62 | 0.21 | 0.36 | 0.28 | 0.31 |
| 4 | 140134659 | rs6820313 | 1.107 | 1.427 | 0.697 | 0.057 | 0.016 | -0.022 | -0.022 | 0 | 0.66 | 0.41 | 0.61 | 0.75 | 0.55 | 0.75 |
| 4 | 140151736 | rs7660661 | -0.696 | 1.12 | 0.8 | 0.005 | 0.035 | -0.053 | -0.053 | 0.011 | 0.27 | 0.45 | 0.21 | 0.19 | 0.33 | 0.16 |
| 4 | 140159339 | rs13143516 | -0.643 | 0.986 | 0.214 | 0.017 | 0.002 | 0.032 | 0.032 | 0 | 0.55 | 0.81 | 0.36 | 0.38 | 0.5 | 0.51 |
| 4 | 140165770 | rs4863639 | 1.095 | 0.834 | 0.148 | 0.17 | -0.054 | -0.022 | -0.022 | 0.947 | 0.28 | 0.57 | 0.16 | 0.22 | 0.19 | 0.33 |
| 4 | 140166713 | rs6835286 | -0.317 | 1.017 | 0.138 | 0.179 | 0.058 | 0.071 | 0.071 | 0 | 0.51 | 0.79 | 0.32 | 0.37 | 0.45 | 0.47 |
| 4 | 140167394 | rs6841480 | 0.025 | 1.14 | -0.014 | 0.24 | 0.001 | -0.005 | -0.005 | 1.12 | 0.22 | 0.22 | 0.16 | 0.16 | 0.26 | 0.15 |
| 4 | 140170092 | rs2029288 | 0.04 | 1.329 | -0.65 | 0.615 | -0.046 | -0.034 | -0.034 | 0.021 | 0.89 | 0.95 | 0.69 | 0.5 | 0.75 | 0.82 |
| 4 | 140177399 | rs17050728 | 1.536 | 0.154 | -0.472 | 0.177 | 0.053 | -0.096 | -0.096 | 0 | 0.23 | 0.01 | 0.1 | 0.24 | 0.16 | 0.31 |
| 4 | 140177562 | rs6536020 | -0.545 | 0.364 | -0.339 | 0.161 | 0.019 | -0.024 | -0.024 | 0.053 | 0.53 | 0.85 | 0.32 | 0.42 | 0.45 | 0.48 |
| 4 | 140186178 | rs13103115 | -0.937 | 0.583 | 0.449 | 0.077 | 0.002 | -0.053 | -0.053 | 0 | 0.27 | 0.54 | 0.19 | 0.16 | 0.26 | 0.17 |
| 4 | 140189613 | rs13120574 | -0.735 | 0.593 | 1.108 | 0.018 | 0.034 | -0.047 | -0.047 | 0 | 0.24 | 0.53 | 0.18 | 0.19 | 0.27 | 0.17 |
| 4 | 140199813 | rs13127307 | -0.088 | 0.61 | 1.113 | 0.007 | -0.028 | 0.066 | 0.066 | 0 | 0.17 | 0.04 | 0.1 | 0.06 | 0.2 | 0.15 |
| 4 | 140216412 | rs6816703 | -0.987 | 0.615 | 1.623 | 0.003 | 0.009 | -0.023 | -0.023 | 0 | 0.22 | 0.36 | 0.17 | 0.16 | 0.25 | 0.18 |
| 4 | 140225539 | rs11729130 | 1.081 | 0.486 | 1.282 | -0.056 | -0.117 | -0.108 | -0.108 | 2.874 | 0.39 | 0.06 | 0.48 | 0.51 | 0.45 | 0.31 |
| 4 | 140228808 | rs13147688 | 0.205 | 0.521 | 1.277 | -0.033 | -0.034 | 0.068 | 0.068 | 0 | 0.17 | 0.04 | 0.1 | 0.06 | 0.2 | 0.15 |

| | | | | | | | | | | | | | | | | |
|---|---|---|---|---|---|---|---|---|---|---|---|---|---|---|---|---|
| 4 | 140235620 | rs1901173 | 0.998 | 0.455 | 0.834 | -0.076 | -0.039 | 0.004 | 0.004 | 0 | 0.3 | 0.34 | 0.14 | 0.26 | 0.2 | 0.41 |
| 4 | 140243755 | rs12645529 | -0.097 | 0.817 | 0.866 | -0.156 | -0.032 | 0.016 | 0.016 | 0 | 0.18 | 0.07 | 0.15 | 0.17 | 0.25 | 0.18 |
| 4 | 140254518 | rs747004 | -0.993 | 0.804 | 0.943 | -0.162 | 0.031 | -0.048 | -0.048 | 0 | 0.23 | 0.49 | 0.18 | 0.19 | 0.26 | 0.19 |
| 4 | 140259458 | rs6820950 | -0.678 | 0.833 | 0.748 | -0.143 | -0.008 | 0.006 | 0.006 | 0 | 0.94 | 0.96 | 0.8 | 0.94 | 0.9 | 0.92 |
| 4 | 140289285 | rs13117654 | -0.91 | 0.788 | 0.814 | -0.15 | 0.014 | -0.024 | -0.024 | 0 | 0.23 | 0.39 | 0.18 | 0.19 | 0.26 | 0.18 |
| 4 | 140330090 | rs7687436 | -0.853 | 0.789 | 0.727 | -0.15 | 0.018 | -0.029 | -0.029 | 0 | 0.23 | 0.4 | 0.17 | 0.19 | 0.26 | 0.18 |
| 4 | 140370149 | rs13128827 | -0.846 | 0.727 | 0.661 | -0.153 | 0.019 | -0.029 | -0.029 | 0.692 | 0.22 | 0.4 | 0.17 | 0.19 | 0.26 | 0.19 |
| 4 | 140373081 | rs1142627 | -0.43 | 0.725 | 0.635 | -0.154 | -0.034 | 0.018 | 0.018 | 0 | 0.19 | 0.1 | 0.15 | 0.19 | 0.26 | 0.18 |
| 4 | 140395583 | rs4074940 | -0.177 | 0.758 | 0.344 | -0.139 | -0.029 | 0.016 | 0.016 | 0 | 0.18 | 0.08 | 0.15 | 0.17 | 0.25 | 0.18 |
| 4 | 140396931 | rs3088313 | -0.662 | 0.759 | 0.383 | -0.138 | 0.006 | -0.016 | -0.016 | 0.198 | 0.22 | 0.33 | 0.16 | 0.17 | 0.25 | 0.18 |
| 4 | 140406637 | rs3967770 | 0.477 | 0.612 | -0.01 | -0.019 | -0.012 | -0.056 | -0.056 | 0 | 0.45 | 0.12 | 0.69 | 0.51 | 0.54 | 0.41 |
| 4 | 140414333 | rs7666900 | 1.556 | 1.27 | 1.359 | -0.044 | 0.028 | -0.067 | -0.067 | 0 | 0.88 | 0.6 | 0.77 | 0.99 | 0.89 | 0.87 |
| 4 | 140420250 | rs2592966 | 1.561 | 1.438 | 1.163 | -0.11 | 0 | -0.022 | -0.022 | 0 | 0.74 | 0.31 | 0.73 | 0.88 | 0.73 | 0.77 |
| 4 | 140423134 | rs17050782 | -1.468 | 2.007 | 0.973 | 0.133 | 0.004 | 0.024 | 0.024 | 0 | 0.12 | 0.12 | 0.13 | 0.39 | 0.2 | 0.11 |
| 4 | 140425004 | rs720257 | 1.923 | 2.018 | 0.982 | 0.035 | 0.001 | 0.007 | 0.007 | 1.013 | 0.76 | 0.32 | 0.72 | 0.65 | 0.75 | 0.77 |
| 4 | 140429555 | rs1026048 | -1.65 | 1.63 | 1.406 | -0.356 | 0.004 | -0.126 | -0.126 | 0.454 | 0.19 | 0.42 | 0.14 | 0.04 | 0.17 | 0.19 |
| 4 | 140430625 | rs895600 | 1.435 | 1.607 | 1.442 | -0.345 | -0.002 | -0.13 | -0.13 | 0.673 | 0.8 | 0.58 | 0.87 | 0.96 | 0.83 | 0.81 |
| 4 | 140431931 | rs17050790 | -1.759 | 1.61 | 1.506 | -0.326 | 0.02 | -0.005 | -0.005 | 0 | 0.05 | 0.21 | 0.04 | 0.02 | 0.03 | 0.01 |
| 4 | 140436646 | rs2725773 | 1.461 | 1.606 | 1.58 | -0.34 | 0.01 | -0.143 | -0.143 | 0 | 0.81 | 0.51 | 0.86 | 0.96 | 0.83 | 0.8 |
| 4 | 140442145 | rs2592980 | 1.221 | 1.767 | 1.555 | -0.28 | 0.024 | -0.105 | -0.105 | 0 | 0.82 | 0.66 | 0.87 | 0.95 | 0.8 | 0.83 |
| 4 | 140447105 | rs7680948 | -1.901 | 2.475 | 1.533 | 0.137 | 0.018 | 0.099 | 0.099 | 0.158 | 0.22 | 0.37 | 0.27 | 0.49 | 0.25 | 0.29 |
| 4 | 140449420 | rs2725779 | 2.734 | 2.262 | 1.196 | -0.057 | -0.052 | -0.03 | -0.03 | 0.056 | 0.57 | 0.16 | 0.58 | 0.47 | 0.54 | 0.49 |
| 4 | 140449824 | rs4863658 | -1.994 | 2.258 | 1.247 | -0.146 | 0.047 | -0.098 | -0.098 | 1.054 | 0.25 | 0.44 | 0.27 | 0.12 | 0.29 | 0.36 |
| 4 | 140458672 | rs10025309 | -0.957 | 2.214 | 1.231 | -0.026 | 0.031 | -0.001 | -0.001 | 0 | 0.1 | 0.28 | 0.16 | 0.1 | 0.08 | 0.16 |
| 4 | 140460303 | rs2725790 | 1.212 | 2.706 | 1.225 | 0.074 | 0.004 | 0.026 | 0.026 | 0 | 0.85 | 0.8 | 0.83 | 0.56 | 0.78 | 0.86 |
| 4 | 140466311 | rs1995103 | -2.831 | 2.716 | 1.241 | -0.091 | -0.016 | -0.01 | -0.01 | 0 | 0.43 | 0.75 | 0.43 | 0.56 | 0.49 | 0.56 |
| 4 | 140466604 | rs6828434 | -1.512 | 2.419 | 1.181 | 0.406 | 0.018 | 0.125 | 0.125 | 0 | 0.17 | 0.3 | 0.26 | 0.49 | 0.22 | 0.27 |
| 4 | 140470212 | rs2592984 | -2.101 | 2.534 | 1.153 | 0.359 | 0.049 | 0.119 | 0.119 | 0.022 | 0.24 | 0.44 | 0.33 | 0.54 | 0.3 | 0.34 |
| 4 | 140476214 | rs706308 | -2.238 | 3.164 | 1.73 | -0.326 | 0.161 | -0.042 | -0.042 | 0.162 | 0.16 | 0.46 | 0.16 | 0.14 | 0.24 | 0.32 |
| 4 | 140478478 | rs706309 | -2.659 | 2.935 | 1.554 | -0.551 | 0.023 | -0.055 | -0.055 | 0 | 0.26 | 0.61 | 0.23 | 0.13 | 0.3 | 0.44 |
| 4 | 140481510 | rs706310 | -1.967 | 3.046 | 1.486 | -0.434 | 0.081 | -0.014 | -0.014 | 0.464 | 0.1 | 0.22 | 0.11 | 0.08 | 0.14 | 0.18 |
| 4 | 140495051 | rs795617 | -2.232 | 2.865 | 0.694 | -0.787 | -0.01 | 0.031 | 0.031 | 2.542 | 0.44 | 0.64 | 0.32 | 0.23 | 0.49 | 0.57 |
| 4 | 140497312 | rs11726931 | -2.096 | 2.833 | 0.25 | -0.442 | 0.05 | -0.021 | -0.021 | 0 | 0.26 | 0.41 | 0.28 | 0.19 | 0.38 | 0.33 |
| 4 | 140505632 | rs2166596 | -1.126 | 3.304 | 0.248 | -0.297 | 0.169 | -0.014 | -0.014 | 0.831 | 0.12 | 0.35 | 0.19 | 0.06 | 0.25 | 0.2 |
| 4 | 140511212 | rs706328 | 0.01 | 1.944 | -0.652 | -0.207 | 0.022 | 0.018 | 0.018 | 0 | 0.12 | 0.05 | 0.03 | 0.07 | 0.05 | 0.13 |
| 4 | 140513877 | rs17282576 | -0.818 | 2.407 | -0.588 | -0.087 | -0.044 | -0.028 | -0.028 | 0 | 0.09 | 0.02 | 0.12 | 0.14 | 0.18 | 0.09 |
| 4 | 140517700 | rs706333 | -1.338 | 1.861 | -0.673 | -0.494 | -0.005 | 0.081 | 0.081 | 2.375 | 0.47 | 0.71 | 0.44 | 0.22 | 0.57 | 0.56 |
| 4 | 140518308 | rs7689415 | 0.078 | 1.927 | -0.247 | -0.756 | -0.001 | -0.04 | -0.04 | 0 | 0.18 | 0.17 | 0.09 | 0 | 0.16 | 0.15 |
| 4 | 140519467 | rs795377 | -1.091 | 1.961 | 0.275 | -0.493 | 0.025 | -0.009 | -0.009 | 0.032 | 0.27 | 0.4 | 0.31 | 0.22 | 0.39 | 0.4 |
| 4 | 140522504 | rs706336 | -0.915 | 1.935 | 0.399 | -0.369 | 0.074 | 0.063 | 0.063 | 0.115 | 0.17 | 0.45 | 0.27 | 0.2 | 0.28 | 0.21 |
| 4 | 140523889 | rs706337 | -0.603 | 1.996 | 0.006 | -0.352 | 0.057 | 0.047 | 0.047 | 0 | 0.16 | 0.34 | 0.27 | 0.2 | 0.28 | 0.21 |
| 4 | 140528564 | rs2602248 | -0.875 | 0.876 | -1.139 | -1.228 | 0.049 | 0.145 | 0.145 | 1.374 | 0.58 | 0.75 | 0.8 | 0.9 | 0.68 | 0.62 |
| 4 | 140528612 | rs13112005 | -0.042 | 0.961 | -1.428 | -0.313 | -0.003 | -0.003 | -0.003 | 0.198 | 0.3 | 0.28 | 0.45 | 0.25 | 0.32 | 0.22 |
| 4 | 140531714 | rs6536296 | -0.933 | 0.543 | -1.648 | -0.525 | 0.018 | 0.123 | 0.123 | 0.973 | 0.72 | 0.93 | 0.86 | 0.9 | 0.79 | 0.68 |
| 4 | 140534118 | rs13114510 | -0.931 | 0.543 | -1.646 | -0.53 | 0.018 | 0.127 | 0.127 | 0.929 | 0.72 | 0.93 | 0.86 | 0.91 | 0.79 | 0.68 |
| 4 | 140537127 | rs2602234 | 0.336 | 1.358 | -0.778 | 0.563 | 0.019 | 0.027 | 0.027 | 0.191 | 0.8 | 0.82 | 0.71 | 0.49 | 0.63 | 0.75 |
| 4 | 140537671 | rs11936429 | -0.263 | 0.928 | -0.758 | 0.673 | -0.015 | 0.057 | 0.057 | 0 | 0.07 | 0.12 | 0.14 | 0.22 | 0.04 | 0.03 |
| 4 | 140542332 | rs13111220 | 0.001 | 0.742 | -1.362 | 0.222 | -0.018 | 0.073 | 0.073 | 0.71 | 0.14 | 0.02 | 0.06 | 0.04 | 0.15 | 0.2 |
| 4 | 140542427 | rs10027416 | -0.279 | 0.817 | -0.935 | 0.242 | 0.001 | -0.023 | -0.023 | 0 | 0.16 | 0.19 | 0.11 | 0.09 | 0.18 | 0.22 |
| 4 | 140544895 | rs17355469 | 1.469 | 0.568 | -1.29 | 0.015 | 0.047 | 0.254 | 0.254 | 0.678 | 0.5 | 0.12 | 0.43 | 0.18 | 0.4 | 0.5 |
| 4 | 140545348 | rs706341 | 0.432 | 1.312 | -0.66 | 0.725 | -0.018 | -0.018 | -0.018 | 0 | 0.8 | 0.83 | 0.7 | 0.44 | 0.62 | 0.73 |
| 4 | 140549569 | rs17050838 | 0.743 | 0.458 | -0.64 | 0.432 | -0.002 | 0.023 | 0.023 | 0.67 | 0.52 | 0.36 | 0.56 | 0.35 | 0.47 | 0.48 |
| 4 | 140549622 | rs706345 | -0.94 | 0.78 | -0.189 | 0.91 | 0 | -0.001 | -0.001 | 0 | 0.26 | 0.21 | 0.27 | 0.33 | 0.28 | 0.23 |
| 4 | 140552120 | rs4241931 | -0.294 | 0.957 | 0.006 | 1.166 | 0.025 | 0.015 | 0.015 | 0 | 0.1 | 0.17 | 0.17 | 0.11 | 0.15 | 0.1 |
| 4 | 140552399 | rs1027472 | 0.562 | 0.642 | 0.264 | 1.229 | 0.001 | 0.017 | 0.017 | 2.806 | 0.6 | 0.57 | 0.71 | 0.45 | 0.52 | 0.57 |
| 4 | 140560388 | rs885156 | 0.339 | 0.91 | 0.069 | 1.275 | 0.005 | 0.002 | 0.002 | 0 | 0.08 | 0.09 | 0.13 | 0.09 | 0.12 | 0.06 |
| 4 | 140563222 | rs11100193 | 1.61 | 0.902 | 0.142 | 1.252 | 0.008 | 0.026 | 0.026 | 0.767 | 0.39 | 0.24 | 0.5 | 0.3 | 0.33 | 0.36 |
| 4 | 140563620 | rs10023253 | 0.029 | 0.478 | 0.358 | 1.38 | 0.005 | 0.002 | 0.002 | 0 | 0.08 | 0.09 | 0.13 | 0.09 | 0.12 | 0.06 |
| 4 | 140573599 | rs1316503 | -1.356 | 0.341 | 0.457 | 1.566 | 0.014 | -0.008 | -0.008 | 0.091 | 0.44 | 0.63 | 0.28 | 0.54 | 0.37 | 0.54 |
| 4 | 140575134 | rs2602219 | -0.981 | 0.212 | 0.77 | 1.644 | -0.007 | 0.011 | 0.011 | 1.052 | 0.36 | 0.43 | 0.19 | 0.46 | 0.34 | 0.49 |
| 4 | 140575501 | rs2602218 | -0.062 | 0.213 | 0.939 | 1.547 | 0.056 | -0.032 | -0.032 | 0 | 0.06 | 0.2 | 0.06 | 0 | 0.14 | 0.05 |
| 4 | 140576200 | rs1078764 | 0.045 | 0.007 | 0.825 | 1.562 | 0.006 | 0.022 | 0.022 | 0.489 | 0.13 | 0.16 | 0.14 | 0.24 | 0.13 | 0.2 |
| 4 | 140577920 | rs3755996 | 0.238 | -0.025 | 0.841 | 1.566 | 0.026 | -0.026 | -0.026 | 0 | 0.17 | 0.09 | 0.06 | 0.28 | 0.1 | 0.28 |
| 4 | 140578500 | rs17050847 | -0.948 | -0.002 | 1.027 | 1.583 | 0.013 | 0.026 | 0.026 | 0.979 | 0.16 | 0.33 | 0.16 | 0.15 | 0.15 | 0.11 |
| 4 | 140579042 | rs3755992 | 0.35 | -0.065 | 0.553 | 1.593 | -0.001 | 0.009 | 0.009 | 0.351 | 0.12 | 0.15 | 0.06 | 0.18 | 0.11 | 0.16 |
| 4 | 140579455 | rs924199 | 0.284 | -0.085 | 0.543 | 1.604 | 0.026 | -0.028 | -0.028 | 0.047 | 0.17 | 0.09 | 0.06 | 0.29 | 0.09 | 0.28 |
| 4 | 140580748 | rs12640152 | -0.044 | 0.033 | 0.282 | 1.576 | 0.006 | 0.019 | 0.019 | 0.475 | 0.14 | 0.17 | 0.14 | 0.24 | 0.13 | 0.2 |
| 4 | 140581932 | rs795602 | -2.03 | 0.077 | 0.261 | 1.531 | -0.005 | 0.046 | 0.046 | 1.579 | 0.52 | 0.76 | 0.35 | 0.61 | 0.48 | 0.6 |
| 4 | 140585031 | rs8191997 | -1.313 | 0.076 | 0.312 | 1.466 | -0.006 | 0.014 | 0.014 | 0 | 0.19 | 0.28 | 0.15 | 0.25 | 0.13 | 0.21 |

| | | | | | | | | | | | | | | | | |
|---|---|---|---|---|---|---|---|---|---|---|---|---|---|---|---|---|
| 4 | 140585089 | rs8191998 | -0.37 | 0.351 | 0.136 | 1.379 | -0.002 | 0.006 | 0.006 | 0.283 | 0.13 | 0.16 | 0.06 | 0.17 | 0.11 | 0.17 |
| 4 | 140586701 | rs8192004 | -0.25 | 0.5 | -0.157 | 1.798 | -0.013 | -0.011 | -0.011 | 0 | 0.08 | 0.05 | 0.22 | 0.29 | 0.09 | 0.11 |
| 4 | 140587414 | rs3822045 | -1.491 | 0.865 | 0.17 | 1.82 | 0.02 | 0.034 | 0.034 | 0.001 | 0.17 | 0.22 | 0.29 | 0.42 | 0.28 | 0.2 |
| 4 | 140587985 | rs1000222 | -1.448 | -0.264 | 0.515 | 1.376 | -0.002 | -0.003 | -0.003 | 2.354 | 0.47 | 0.48 | 0.41 | 0.7 | 0.4 | 0.43 |
| 4 | 140589998 | rs795596 | -2.142 | -0.644 | 0.185 | 1.287 | -0.052 | 0.143 | 0.143 | 0.139 | 0.59 | 0.84 | 0.48 | 0.87 | 0.47 | 0.55 |
| 4 | 140590595 | rs795594 | -2.142 | -0.644 | 0.185 | 1.287 | -0.049 | 0.143 | 0.143 | 0.139 | 0.59 | 0.84 | 0.48 | 0.87 | 0.47 | 0.55 |
| 4 | 140590947 | rs8192018 | -2.145 | -0.654 | 0.18 | 1.269 | -0.05 | 0.145 | 0.145 | 0.169 | 0.59 | 0.84 | 0.48 | 0.87 | 0.47 | 0.55 |
| 4 | 140592641 | rs8192024 | -1.934 | -0.762 | 0.427 | 1.252 | 0.004 | 0.021 | 0.021 | 0 | 0.29 | 0.65 | 0.36 | 0.5 | 0.32 | 0.28 |
| 4 | 140593075 | rs8192031 | -0.258 | -0.765 | 0.382 | 1.252 | -0.037 | 0.021 | 0.021 | 0 | 0.08 | 0.23 | 0.05 | 0.08 | 0.02 | 0.04 |
| 4 | 140593308 | rs2602238 | -2.109 | -0.549 | -0.388 | 1.298 | -0.108 | 0.149 | 0.149 | 0.231 | 0.6 | 0.92 | 0.49 | 0.86 | 0.48 | 0.56 |
| 4 | 140594551 | rs2602239 | -0.535 | 0.305 | 0.058 | 1.651 | 0.059 | 0.07 | 0.07 | 0 | 0.06 | 0.28 | 0.04 | 0.09 | 0.05 | 0.08 |
| 4 | 140596699 | rs8192047 | -0.613 | 0.281 | -0.759 | 1.671 | -0.042 | -0.045 | -0.045 | 0 | 0.14 | 0.04 | 0.28 | 0.34 | 0.27 | 0.15 |
| 4 | 140597052 | rs8192049 | -0.024 | -0.259 | -0.759 | 1.354 | -0.006 | -0.039 | -0.039 | 0 | 0.07 | 0 | 0.03 | 0.25 | 0.05 | 0.1 |
| 4 | 140599053 | rs8192060 | 0.977 | -0.26 | -0.98 | 0.246 | 0.017 | 0.064 | 0.064 | 0.364 | 0.11 | 0.01 | 0.05 | 0 | 0.06 | 0.05 |
| 4 | 140600125 | rs795589 | -0.768 | -0.152 | -0.494 | -0.017 | 0.001 | 0.001 | 0.001 | 1.355 | 0.48 | 0.49 | 0.42 | 0.77 | 0.43 | 0.51 |
| 4 | 140600307 | rs706349 | -0.774 | -0.267 | -0.427 | -0.051 | -0.001 | -0.006 | -0.006 | 0 | 0.47 | 0.47 | 0.39 | 0.74 | 0.36 | 0.49 |
| 4 | 140600397 | rs1027473 | -0.868 | 0.353 | -0.524 | -0.277 | 0.012 | 0.04 | 0.04 | 2.082 | 0.51 | 0.43 | 0.65 | 0.82 | 0.46 | 0.62 |
| 4 | 140600437 | rs706350 | -0.891 | 0.302 | -0.364 | 0.074 | 0.013 | 0.02 | 0.02 | 1.238 | 0.51 | 0.6 | 0.42 | 0.73 | 0.42 | 0.59 |
| 4 | 140601036 | rs4241932 | 0.119 | -0.255 | -0.382 | 0.687 | -0.009 | 0.029 | 0.029 | 0 | 0.26 | 0.3 | 0.27 | 0.46 | 0.2 | 0.28 |
| 4 | 140601791 | rs1587265 | -1.842 | 0.561 | -1.094 | 0.181 | 0.027 | 0.192 | 0.192 | 2.218 | 0.55 | 0.85 | 0.76 | 0.85 | 0.6 | 0.54 |
| 4 | 140601934 | rs2262377 | -1.918 | 0.845 | -0.924 | 0.23 | 0.052 | -0.009 | -0.009 | 0.013 | 0.28 | 0.53 | 0.45 | 0.26 | 0.36 | 0.22 |
| 4 | 140602972 | rs2132847 | -1.431 | 0.977 | -0.49 | 0.158 | 0.028 | -0.062 | -0.062 | 0 | 0.18 | 0.41 | 0.34 | 0.06 | 0.19 | 0.1 |
| 4 | 140605684 | rs8192075 | -0.132 | 0.738 | -0.399 | 0.165 | -0.001 | -0.027 | -0.027 | 0 | 0.05 | 0.12 | 0.03 | 0 | 0.04 | 0.03 |
| 4 | 140606181 | rs8192081 | -0.132 | 0.738 | -0.399 | 0.165 | -0.001 | -0.027 | -0.027 | 0 | 0.05 | 0.12 | 0.03 | 0 | 0.04 | 0.03 |
| 4 | 140606425 | rs7664138 | -0.117 | 0.54 | -0.293 | 0.397 | -0.034 | 0.065 | 0.065 | 0 | 0.29 | 0.45 | 0.32 | 0.49 | 0.19 | 0.29 |
| 4 | 140608002 | rs795593 | -2.847 | 0.355 | -0.843 | -0.143 | 0.011 | 0.094 | 0.094 | 2.088 | 0.85 | 0.99 | 0.93 | 1 | 0.9 | 0.95 |
| 4 | 140608645 | rs795592 | -0.26 | 1.311 | 0.998 | 1.628 | -0.017 | -0.033 | -0.033 | 0 | 0.06 | 0.13 | 0.04 | 0 | 0.03 | 0.02 |
| 4 | 140612059 | rs2602243 | -2.645 | 1.26 | 1.333 | 2.459 | -0.012 | 0.028 | 0.028 | 0 | 0.27 | 0.6 | 0.29 | 0.47 | 0.2 | 0.26 |
| 4 | 140614711 | rs4147590 | -2.248 | 1.408 | 1.203 | 2.348 | -0.011 | 0.024 | 0.024 | 0.067 | 0.23 | 0.68 | 0.29 | 0.35 | 0.18 | 0.17 |
| 4 | 140616983 | rs8192100 | -0.952 | 1.489 | 0.938 | 2.263 | -0.006 | 0.008 | 0.008 | 0 | 0.12 | 0.08 | 0.12 | 0.09 | 0.15 | 0.12 |
| 4 | 140626105 | rs345978 | 0.8 | 1.513 | 0.879 | 2.281 | -0.004 | 0.003 | 0.003 | 0.076 | 0.88 | 0.9 | 0.87 | 0.91 | 0.85 | 0.88 |
| 4 | 140627667 | rs345976 | 0.812 | 1.527 | 0.928 | 2.283 | -0.005 | 0.004 | 0.004 | 0.17 | 0.88 | 0.9 | 0.87 | 0.91 | 0.84 | 0.88 |
| 4 | 140637703 | rs11944843 | -1.628 | 1.593 | 0.969 | 2.295 | -0.002 | 0.007 | 0.007 | 0 | 0.16 | 0.14 | 0.13 | 0.16 | 0.16 | 0.19 |
| 4 | 140637815 | rs13112141 | 2.233 | 1.581 | 1.131 | 2.596 | 0.042 | 0.064 | 0.064 | 0.824 | 0.49 | 0.2 | 0.26 | 0.41 | 0.43 | 0.5 |
| 4 | 140638963 | rs345983 | 0.725 | 1.993 | 1.692 | 3.078 | 0.024 | 0.021 | 0.021 | 0 | 0.71 | 0.49 | 0.5 | 0.7 | 0.69 | 0.72 |
| 4 | 140640283 | rs10519504 | -1.31 | 2.002 | 1.439 | 2.954 | 0.002 | -0.001 | -0.001 | 0 | 0.13 | 0.15 | 0.1 | 0.16 | 0.14 | 0.18 |
| 4 | 140643322 | rs4863507 | -1.259 | 2.058 | 1.468 | 3.067 | -0.003 | 0.02 | 0.02 | 0.31 | 0.33 | 0.57 | 0.42 | 0.44 | 0.32 | 0.29 |
| 4 | 140644144 | rs6835548 | -1.152 | 1.938 | 1.183 | 2.976 | -0.034 | -0.036 | -0.036 | 0.249 | 0.15 | 0.07 | 0.19 | 0.26 | 0.23 | 0.2 |
| 4 | 140650730 | rs2246759 | 2.125 | 1.843 | 1.096 | 2.944 | 0.027 | 0.185 | 0.185 | 1.258 | 0.51 | 0.18 | 0.28 | 0.25 | 0.45 | 0.46 |
| 4 | 140650787 | rs17050893 | -0.293 | 1.517 | 1.328 | 2.98 | -0.004 | 0.04 | 0.04 | 0.299 | 0.22 | 0.31 | 0.25 | 0.29 | 0.22 | 0.21 |
| 4 | 140663621 | rs7674828 | 1.948 | 1.321 | 1.443 | 2.656 | -0.007 | 0.103 | 0.103 | 0 | 0.73 | 0.53 | 0.68 | 0.46 | 0.77 | 0.67 |
| 4 | 140668374 | rs7678266 | -2.178 | 1.445 | 1.411 | 2.383 | -0.006 | 0.008 | 0.008 | 1.034 | 0.41 | 0.57 | 0.65 | 0.74 | 0.47 | 0.43 |
| 4 | 140670043 | rs12641953 | -0.906 | 1.394 | 1.366 | 2.34 | -0.002 | 0.013 | 0.013 | 0 | 0.23 | 0.3 | 0.2 | 0.22 | 0.25 | 0.24 |
| 4 | 140676385 | rs4863683 | 0.971 | 1.416 | 1.358 | 2.128 | 0.019 | 0.099 | 0.099 | 0.134 | 0.77 | 0.41 | 0.57 | 0.43 | 0.8 | 0.72 |
| 4 | 140677733 | rs17050895 | -0.528 | 0.868 | 0.848 | 1.304 | -0.011 | -0.007 | -0.007 | 0 | 0.19 | 0.23 | 0.12 | 0.12 | 0.19 | 0.15 |
| 4 | 140678281 | rs4863687 | -0.415 | 1.014 | 0.576 | 1.266 | 0.004 | 0.011 | 0.011 | 0.011 | 0.2 | 0.19 | 0.32 | 0.23 | 0.27 | 0.19 |
| 4 | 140679983 | rs10857325 | -1.284 | 1.131 | 0.165 | 0.421 | 0.007 | 0.03 | 0.03 | 0.413 | 0.43 | 0.63 | 0.4 | 0.83 | 0.41 | 0.42 |
| 4 | 140684195 | rs4863510 | 0.501 | 1.096 | -0.036 | 0.653 | 0.014 | 0.047 | 0.047 | 3.414 | 0.44 | 0.28 | 0.41 | 0.33 | 0.44 | 0.52 |
| 4 | 140684567 | rs10021952 | 1.279 | 1.106 | 0.897 | 1.654 | 0.013 | 0.114 | 0.114 | 0 | 0.87 | 0.76 | 0.72 | 0.52 | 0.82 | 0.88 |
| 4 | 140686645 | rs907320 | -1.736 | 1.01 | 0.926 | 1.385 | 0.012 | 0.091 | 0.091 | 0 | 0.16 | 0.29 | 0.32 | 0.46 | 0.2 | 0.1 |
| 4 | 140687323 | rs4073315 | 1.268 | 0.632 | 0.995 | 0.222 | -0.013 | 0.096 | 0.096 | 0.356 | 0.92 | 0.63 | 0.97 | 0.92 | 0.99 | 0.96 |
| 4 | 140693020 | rs6835277 | 0.601 | 1.181 | 0.34 | -0.002 | 0.018 | -0.012 | -0.012 | 0.002 | 0.69 | 0.57 | 0.59 | 0.26 | 0.65 | 0.69 |
| 4 | 140693868 | rs6815550 | 0.358 | 0.605 | 0.143 | -0.787 | -0.018 | -0.062 | -0.062 | 0 | 0.68 | 0.49 | 0.82 | 0.9 | 0.73 | 0.77 |
| 4 | 140693870 | rs6816973 | 0.358 | 0.605 | 0.143 | -0.787 | -0.018 | -0.062 | -0.062 | 0 | 0.68 | 0.5 | 0.82 | 0.9 | 0.73 | 0.77 |
| 4 | 140694305 | rs11721831 | -0.329 | 0.37 | 0.27 | -0.88 | 0.013 | 0.016 | 0.016 | 0 | 0.23 | 0.16 | 0.13 | 0.09 | 0.2 | 0.18 |
| 4 | 140694724 | rs7690424 | -1.333 | 0.71 | -0.182 | -1.427 | -0.011 | 0.041 | 0.041 | 0 | 0.59 | 0.63 | 0.67 | 0.87 | 0.55 | 0.6 |
| 4 | 140694991 | rs6848976 | 0.131 | 1 | -0.227 | -0.784 | -0.004 | -0.028 | -0.028 | 3.65 | 0.07 | 0.2 | 0.03 | 0.01 | 0.02 | 0.04 |
| 4 | 140696130 | rs932012 | -0.218 | 1.307 | -0.688 | -0.6 | -0.025 | -0.01 | -0.01 | 0 | 0.23 | 0.17 | 0.41 | 0.71 | 0.33 | 0.35 |
| 4 | 140696703 | rs7679592 | -0.711 | 0.541 | -0.743 | -1.022 | -0.015 | -0.059 | -0.059 | 0.15 | 0.14 | 0.14 | 0.05 | 0.01 | 0.08 | 0.1 |
| 4 | 140697617 | rs11100269 | 0.612 | 1.224 | -2.139 | -1.249 | -0.183 | 0.034 | 0.034 | 0.284 | 0.28 | 0.01 | 0.38 | 0.12 | 0.44 | 0.28 |
| 4 | 140700785 | rs7695171 | -0.648 | 0.668 | -1.72 | -0.873 | -0.004 | 0.023 | 0.023 | 0 | 0.37 | 0.34 | 0.42 | 0.28 | 0.41 | 0.29 |
| 4 | 140702406 | rs6839816 | -0.139 | 0.625 | -1.917 | -0.92 | 0.004 | -0.001 | -0.001 | 0.001 | 0.17 | 0.23 | 0.16 | 0.18 | 0.24 | 0.16 |
| 4 | 140705510 | rs6812244 | -0.927 | 0.639 | -1.903 | -0.899 | 0.005 | 0.017 | 0.017 | 0 | 0.34 | 0.32 | 0.39 | 0.27 | 0.35 | 0.27 |
| 4 | 140709444 | rs2175458 | 0.627 | 1.241 | -2.2 | -1.131 | -0.081 | 0.116 | 0.116 | 2.994 | 0.35 | 0.15 | 0.47 | 0.11 | 0.48 | 0.33 |
| 4 | 140709905 | rs6823459 | 0.538 | 0.721 | -1.379 | -0.921 | -0.018 | -0.042 | -0.042 | 0 | 0.88 | 0.83 | 0.96 | 1 | 0.94 | 0.86 |
| 4 | 140713471 | rs795998 | -0.322 | 1.495 | -1.353 | -0.3 | 0 | -0.001 | -0.001 | 0 | 0.14 | 0.13 | 0.17 | 0.2 | 0.24 | 0.08 |
| 4 | 140721565 | rs6849438 | 0.347 | 1.239 | -0.772 | -0.555 | -0.001 | 0.006 | 0.006 | 0 | 0.32 | 0.32 | 0.29 | 0.57 | 0.26 | 0.32 |
| 4 | 140724600 | rs2604931 | 0.349 | 0.956 | -0.462 | -0.519 | -0.002 | 0.001 | 0.001 | 0.05 | 0.85 | 0.81 | 0.84 | 0.83 | 0.82 | 0.9 |
| 4 | 140728731 | rs7670072 | 0.08 | 1.142 | -0.194 | -0.64 | 0.047 | -0.029 | -0.029 | 0 | 0.05 | 0.1 | 0.04 | 0 | 0.1 | 0.07 |

| | | | | | | | | | | | | | | | | |
|---|---|---|---|---|---|---|---|---|---|---|---|---|---|---|---|---|
| 4 | 140729346 | rs795989 | 0.265 | 0.562 | -0.119 | -0.932 | -0.005 | -0.007 | -0.007 | 1.096 | 0.36 | 0.43 | 0.34 | 0.59 | 0.33 | 0.48 |
| 4 | 140736222 | rs795992 | 0.434 | 0.758 | -0.129 | -0.811 | 0.006 | -0.004 | -0.004 | 0 | 0.8 | 0.71 | 0.8 | 0.83 | 0.72 | 0.83 |
| 4 | 140744599 | rs1350038 | 0.409 | 0.497 | -0.353 | -0.898 | 0.015 | 0.036 | 0.036 | 0 | 0.49 | 0.43 | 0.48 | 0.24 | 0.39 | 0.37 |
| 4 | 140747153 | rs795988 | -0.096 | 0.416 | -0.396 | -0.861 | 0.003 | 0.004 | 0.004 | 0 | 0.39 | 0.68 | 0.35 | 0.58 | 0.34 | 0.49 |
| 4 | 140749745 | rs13112683 | 0.353 | -0.086 | -0.168 | -1.442 | 0.016 | 0.032 | 0.032 | 0.197 | 0.48 | 0.37 | 0.46 | 0.19 | 0.37 | 0.35 |
| 4 | 140762420 | rs6840907 | -0.355 | -0.626 | 0.003 | -1.108 | 0.01 | 0.018 | 0.018 | 0.022 | 0.39 | 0.28 | 0.36 | 0.64 | 0.33 | 0.5 |
| 4 | 140769923 | rs11100308 | 0.474 | -0.739 | -0.105 | -1.165 | -0.01 | 0.009 | 0.009 | 0.575 | 0.3 | 0.14 | 0.33 | 0.64 | 0.31 | 0.47 |
| 4 | 140776128 | rs795971 | 0.965 | -0.929 | -0.162 | -1.469 | -0.001 | -0.082 | -0.082 | 0 | 0.42 | 0.51 | 0.34 | 0.08 | 0.41 | 0.34 |
| 4 | 140778960 | rs2667370 | -0.128 | -0.568 | -1.016 | -1.234 | 0.004 | -0.001 | -0.001 | 0 | 0.81 | 0.78 | 0.7 | 0.73 | 0.74 | 0.84 |
| 4 | 140782082 | rs2667364 | 0.365 | -0.498 | -1.86 | -1.247 | -0.04 | 0.274 | 0.274 | 1.66 | 0.55 | 0.91 | 0.39 | 0.08 | 0.43 | 0.38 |
| 4 | 140782542 | rs17367750 | 0.623 | -0.499 | -1.868 | -1.225 | -0.051 | -0.033 | -0.033 | 0 | 0.28 | 0.04 | 0.33 | 0.63 | 0.31 | 0.46 |
| 4 | 140783603 | rs11100314 | 0.231 | -0.539 | -1.76 | -1.316 | -0.04 | 0.274 | 0.274 | 1.646 | 0.55 | 0.91 | 0.39 | 0.08 | 0.43 | 0.38 |
| 4 | 140793181 | rs2320535 | 0.065 | -0.56 | -1.609 | -0.861 | -0.05 | -0.022 | -0.022 | 0 | 0.79 | 0.94 | 0.64 | 0.72 | 0.74 | 0.73 |
| 4 | 140795327 | rs1531070 | 1.082 | -0.79 | -1.723 | -1.026 | -0.103 | 0.043 | 0.043 | 4.251 | 0.3 | 0.06 | 0.24 | 0.13 | 0.41 | 0.28 |
| 4 | 140801949 | rs9999030 | -0.384 | -0.88 | -1.429 | -0.918 | 0.025 | 0.043 | 0.043 | 0 | 0.1 | 0.02 | 0.03 | 0 | 0.04 | 0.07 |
| 4 | 140802689 | rs795981 | -1.675 | -0.974 | -0.89 | -1.155 | -0.008 | -0.049 | -0.049 | 0 | 0.2 | 0.65 | 0.15 | 0.01 | 0.09 | 0.11 |
| 4 | 140806021 | rs17050970 | -1.132 | -0.837 | -0.343 | -0.932 | -0.002 | -0.025 | -0.025 | 0 | 0.08 | 0.13 | 0.09 | 0.01 | 0.05 | 0.08 |
| 4 | 140808139 | rs4863700 | 0.204 | -1.206 | -0.092 | -0.691 | -0.001 | 0.014 | 0.014 | 0.482 | 0.32 | 0.32 | 0.34 | 0.56 | 0.28 | 0.45 |
| 4 | 140809656 | rs6839194 | 0.569 | -1.036 | 0.101 | -0.669 | 0.011 | -0.025 | -0.025 | 0 | 0.36 | 0.34 | 0.31 | 0.12 | 0.44 | 0.27 |
| 4 | 140812788 | rs11726067 | 0.617 | -1.307 | 0.635 | -0.32 | -0.075 | 0.061 | 0.061 | 0 | 0.77 | 0.46 | 0.71 | 0.67 | 0.88 | 0.8 |
| 4 | 140814922 | rs12509463 | 1.365 | -1.316 | 0.456 | -0.382 | 0.054 | -0.015 | -0.015 | 0 | 0.68 | 0.21 | 0.61 | 0.66 | 0.81 | 0.69 |
| 4 | 140815184 | rs12505461 | -0.133 | -1.165 | 0.103 | -0.307 | 0.024 | 0.037 | 0.037 | 0 | 0.15 | 0.06 | 0.26 | 0.02 | 0.07 | 0.17 |
| 4 | 140815925 | rs17050982 | -0.786 | -1.148 | 0.141 | -0.299 | -0.051 | -0.122 | -0.122 | 1.25 | 0.21 | 0.35 | 0.28 | 0.03 | 0.1 | 0.24 |
| 4 | 140816174 | rs10026813 | 1.231 | -1.09 | 0.162 | -0.26 | -0.036 | 0.019 | 0.019 | 0 | 0.71 | 0.4 | 0.62 | 0.66 | 0.81 | 0.69 |
| 4 | 140816848 | rs10029731 | 1.008 | -1.138 | -0.126 | -0.139 | -0.052 | 0.027 | 0.027 | 0 | 0.73 | 0.41 | 0.7 | 0.66 | 0.84 | 0.73 |
| 4 | 140821391 | rs9997981 | 0.519 | -0.23 | -0.974 | 0.379 | -0.005 | 0.075 | 0.075 | 0 | 0.89 | 0.75 | 0.89 | 0.69 | 0.89 | 0.92 |
| 4 | 140823857 | rs11724190 | 0.485 | -0.206 | -1.004 | 0.39 | -0.004 | 0.059 | 0.059 | 0 | 0.89 | 0.76 | 0.89 | 0.69 | 0.89 | 0.92 |
| 4 | 140824428 | rs11100354 | 0.155 | -0.233 | -0.965 | 0.476 | -0.008 | 0.122 | 0.122 | 0 | 0.91 | 0.72 | 0.9 | 0.69 | 0.91 | 0.94 |
| 4 | 140825430 | rs17051001 | -0.357 | 0.441 | -1.805 | 0.262 | -0.119 | -0.161 | -0.161 | 0 | 0.38 | 0.04 | 0.42 | 0.54 | 0.38 | 0.54 |
| 4 | 140834736 | rs17005223 | -0.512 | -0.027 | -1.008 | 0.346 | -0.003 | 0.002 | 0.002 | 0 | 0.35 | 0.39 | 0.34 | 0.36 | 0.31 | 0.43 |
| 4 | 140835297 | rs17370238 | -0.335 | 0.009 | -0.92 | -0.133 | 0.027 | 0.02 | 0.02 | 0 | 0.42 | 0.26 | 0.53 | 0.62 | 0.33 | 0.53 |
| 4 | 140835853 | rs1378473 | -0.103 | 0.102 | -1.061 | -0.086 | 0.003 | -0.014 | -0.014 | 0.476 | 0.35 | 0.18 | 0.53 | 0.6 | 0.33 | 0.5 |
| 4 | 140839169 | rs769654 | -0.691 | 0.373 | -0.992 | -0.539 | -0.009 | 0.067 | 0.067 | 0.134 | 0.62 | 0.8 | 0.7 | 0.89 | 0.6 | 0.79 |
| 4 | 140849333 | rs13127289 | -0.07 | 0.321 | -0.573 | -0.219 | 0.021 | 0.018 | 0.018 | 0.068 | 0.45 | 0.26 | 0.56 | 0.6 | 0.33 | 0.51 |
| 4 | 140873893 | rs4863708 | 0.034 | -0.376 | -0.346 | -0.641 | 0.045 | 0.012 | 0.012 | 0.857 | 0.43 | 0.22 | 0.53 | 0.59 | 0.3 | 0.47 |
| 4 | 140882750 | rs769673 | 0.224 | -0.325 | 0.171 | -0.929 | 0.042 | 0.016 | 0.016 | 0 | 0.24 | 0.57 | 0.18 | 0.31 | 0.31 | 0.32 |
| 4 | 140887838 | rs17313183 | 1.187 | -0.453 | -0.302 | -1.174 | -0.092 | 0.021 | 0.021 | 0.306 | 0.23 | 0.01 | 0.21 | 0.11 | 0.31 | 0.17 |
| 4 | 140888081 | rs13119079 | -0.044 | -0.495 | 0.704 | -1.001 | -0.005 | -0.002 | -0.002 | 5.639 | 0.49 | 0.53 | 0.57 | 0.57 | 0.31 | 0.55 |
| 4 | 140891385 | rs4863711 | 0.408 | -0.493 | 0.563 | -1.011 | 0.042 | -0.001 | -0.001 | 0.837 | 0.44 | 0.26 | 0.56 | 0.57 | 0.31 | 0.47 |
| 4 | 140892233 | rs17313646 | 0.278 | -0.491 | 0.609 | -1.01 | 0.026 | 0.006 | 0.006 | 0.472 | 0.46 | 0.37 | 0.57 | 0.57 | 0.31 | 0.47 |
| 4 | 140901754 | rs10519515 | -0.232 | -0.953 | 0.631 | -1.088 | 0 | -0.009 | -0.009 | 0.036 | 0.62 | 0.37 | 0.72 | 0.89 | 0.6 | 0.67 |
| 4 | 140907254 | rs4863713 | -0.976 | -0.973 | 0.704 | -1.012 | -0.001 | -0.004 | -0.004 | 0 | 0.69 | 0.66 | 0.75 | 0.9 | 0.63 | 0.75 |
| 4 | 140909793 | rs12504594 | 0.897 | -0.674 | 0.523 | -0.96 | -0.124 | -0.094 | -0.094 | 0 | 0.2 | 0.03 | 0.21 | 0.33 | 0.36 | 0.22 |
| 4 | 140919083 | rs7656823 | -0.366 | -1.402 | 0.478 | -1.053 | 0.005 | 0.004 | 0.004 | 0 | 0.49 | 0.51 | 0.64 | 0.6 | 0.46 | 0.5 |
| 4 | 140929111 | rs2048382 | -0.393 | -1.377 | 0.132 | -1.049 | 0.017 | 0.017 | 0.017 | 0.09 | 0.5 | 0.55 | 0.65 | 0.61 | 0.46 | 0.5 |
| 4 | 140939653 | rs1521491 | -0.514 | -1.329 | -0.36 | -1.086 | 0.029 | 0.047 | 0.047 | 1.202 | 0.65 | 0.27 | 0.84 | 0.93 | 0.82 | 0.63 |
| 4 | 140942859 | rs6851797 | -0.03 | -0.933 | -0.463 | -0.827 | -0.025 | 0.027 | 0.027 | 0.001 | 0.66 | 0.75 | 0.41 | 0.18 | 0.57 | 0.6 |
| 4 | 140946828 | rs17314804 | 1.139 | -0.909 | -0.566 | -0.695 | -0.103 | -0.081 | -0.081 | 0.406 | 0.19 | 0.03 | 0.19 | 0.33 | 0.35 | 0.22 |
| 4 | 140948284 | rs6838273 | -0.231 | -1.033 | -0.497 | -0.684 | -0.024 | 0.092 | 0.092 | 0 | 0.85 | 0.79 | 0.61 | 0.51 | 0.92 | 0.83 |
| 4 | 140949552 | rs6850591 | -0.162 | -0.841 | -0.404 | -0.722 | -0.013 | 0.011 | 0.011 | 0 | 0.67 | 0.74 | 0.41 | 0.18 | 0.56 | 0.6 |
| 4 | 140952110 | rs11100449 | -0.336 | -1.029 | -1.098 | -1.044 | -0.012 | 0.153 | 0.153 | 0 | 0.47 | 0.28 | 0.65 | 0.9 | 0.48 | 0.61 |
| 4 | 140952449 | rs17374508 | 1.213 | -1.578 | -1.411 | -0.955 | -0.009 | 0.043 | 0.043 | 0 | 0.15 | 0.06 | 0.08 | 0 | 0.13 | 0.09 |
| 4 | 140954867 | rs13116165 | 0.024 | -1.515 | -1.403 | -0.593 | 0.026 | -0.102 | -0.102 | 1.758 | 0.15 | 0.02 | 0.43 | 0.51 | 0.1 | 0.16 |
| 4 | 140959594 | rs6851301 | -0.416 | -1.126 | -0.986 | -1.434 | -0.001 | -0.05 | -0.05 | 0.966 | 0.43 | 0.56 | 0.35 | 0.06 | 0.39 | 0.28 |
| 4 | 140960502 | rs17374883 | -0.052 | -1.017 | -1.788 | -1.378 | 0.013 | 0.139 | 0.139 | 0 | 0.33 | 0.08 | 0.24 | 0.11 | 0.3 | 0.3 |
| 4 | 140961792 | rs17315705 | 0.879 | -0.204 | -0.858 | -0.864 | 0.005 | -0.04 | -0.04 | 0.937 | 0.12 | 0.12 | 0.08 | 0 | 0.12 | 0.07 |
| 4 | 140968980 | rs6828897 | -0.029 | -0.287 | -1.399 | -0.341 | -0.005 | -0.001 | -0.001 | 1.379 | 0.27 | 0.3 | 0.3 | 0.25 | 0.21 | 0.19 |
| 4 | 140975013 | rs1474246 | -0.181 | -0.058 | -1.697 | -0.24 | -0.021 | 0.099 | 0.099 | 0 | 0.2 | 0.06 | 0.24 | 0.05 | 0.24 | 0.17 |
| 4 | 140981966 | rs10519522 | -0.447 | -0.13 | -0.848 | 0.19 | -0.024 | -0.03 | -0.03 | 0 | 0.06 | 0.16 | 0.02 | 0 | 0.02 | 0.04 |
| 4 | 140983576 | rs10440457 | 0.162 | -0.166 | -2.072 | 0.522 | -0.012 | -0.04 | -0.04 | 0.387 | 0.25 | 0.06 | 0.28 | 0.33 | 0.28 | 0.29 |
| 4 | 140985905 | rs1402669 | 0.745 | -0.5 | -1.973 | 0.944 | 0.058 | 0.018 | 0.018 | 1.805 | 0.55 | 0.23 | 0.56 | 0.43 | 0.62 | 0.53 |
| 4 | 140996427 | rs11100475 | -0.338 | -0.189 | -1.994 | 1.123 | 0.01 | -0.032 | -0.032 | 0 | 0.32 | 0.16 | 0.33 | 0.38 | 0.31 | 0.39 |
| 4 | 141000337 | rs10017961 | 0.219 | -0.195 | -2.04 | 1.135 | 0.005 | -0.023 | -0.023 | 0.007 | 0.24 | 0.13 | 0.27 | 0.33 | 0.26 | 0.27 |
| 4 | 141009952 | rs12648900 | -0.014 | 0.326 | -2.239 | 1.117 | -0.097 | -0.077 | -0.077 | 0.674 | 0.4 | 0.05 | 0.33 | 0.36 | 0.44 | 0.42 |
| 4 | 141012918 | rs7691693 | -0.426 | 1.133 | -1.513 | 1.064 | -0.02 | -0.028 | -0.028 | 2.001 | 0.26 | 0.2 | 0.43 | 0.5 | 0.36 | 0.37 |
| 4 | 141019884 | rs6817684 | -1.221 | 0.411 | -0.687 | 0.51 | -0.016 | -0.018 | -0.018 | 0 | 0.09 | 0.25 | 0.05 | 0.03 | 0.07 | 0.04 |
| 4 | 141029512 | rs1546716 | 0.715 | 0.41 | -1.038 | 0.217 | 0.003 | 0.008 | 0.008 | 3.049 | 0.76 | 0.28 | 0.77 | 0.91 | 0.82 | 0.75 |
| 4 | 141030652 | rs6820015 | -0.197 | 0.662 | -1.249 | 0.487 | 0.003 | -0.022 | -0.022 | 0.003 | 0.57 | 0.73 | 0.66 | 0.44 | 0.58 | 0.5 |
| 4 | 141035847 | rs11100496 | 0.039 | 1.058 | -1.679 | 0.876 | -0.094 | -0.041 | -0.041 | 0 | 0.24 | 0.02 | 0.22 | 0.3 | 0.28 | 0.3 |

| | | | | | | | | | | | | | | | | |
|---|---|---|---|---|---|---|---|---|---|---|---|---|---|---|---|---|
| 4 | 141035921 | rs2139926 | 0.341 | 0.711 | -0.919 | 0.987 | 0.003 | 0 | 0 | 1.483 | 0.6 | 0.49 | 0.63 | 0.4 | 0.57 | 0.57 |
| 4 | 141048606 | rs1464435 | -0.996 | 0.17 | -0.899 | 1.253 | 0.001 | -0.02 | -0.02 | 0.039 | 0.21 | 0.17 | 0.23 | 0.33 | 0.2 | 0.25 |
| 4 | 141049040 | rs1464434 | -1.062 | 0.187 | -0.588 | 1.17 | -0.004 | 0.026 | 0.026 | 0 | 0.22 | 0.33 | 0.23 | 0.3 | 0.2 | 0.25 |
| 4 | 141053044 | rs2280500 | -0.468 | -0.214 | -0.628 | 0.994 | -0.01 | 0.057 | 0.057 | 0 | 0.16 | 0.28 | 0.28 | 0.27 | 0.13 | 0.21 |
| 4 | 141053848 | rs6858531 | -0.585 | -0.284 | -0.339 | 0.959 | -0.017 | 0.095 | 0.095 | 0 | 0.17 | 0.43 | 0.27 | 0.27 | 0.13 | 0.2 |
| 4 | 141055896 | rs6822207 | 1.045 | -0.107 | -1.303 | 0.581 | -0.159 | -0.035 | -0.035 | 0 | 0.25 | 0.04 | 0.31 | 0.23 | 0.43 | 0.23 |
| 4 | 141060111 | rs7692035 | 0.939 | -1.171 | -0.787 | 0.802 | 0.035 | 0.063 | 0.063 | 0.013 | 0.43 | 0.18 | 0.4 | 0.31 | 0.61 | 0.4 |
| 4 | 141062977 | rs13112302 | -0.512 | -1.59 | -0.627 | 1.032 | -0.018 | 0.001 | 0.001 | 0 | 0.4 | 0.49 | 0.33 | 0.43 | 0.26 | 0.37 |
| 4 | 141069147 | rs1521496 | -0.329 | -0.916 | -0.531 | 1.807 | 0.024 | -0.036 | -0.036 | 1.072 | 0.19 | 0.12 | 0.22 | 0.39 | 0.15 | 0.2 |
| 4 | 141069646 | rs6536733 | -1.008 | -0.081 | 0 | 1.546 | -0.03 | 0.063 | 0.063 | 0 | 0.13 | 0.23 | 0.21 | 0.26 | 0.06 | 0.24 |
| 4 | 141109029 | rs17005617 | -1.382 | 0.461 | 0.174 | 1.06 | 0.004 | 0.002 | 0.002 | 0 | 0.23 | 0.3 | 0.26 | 0.27 | 0.2 | 0.42 |
| 4 | 141112167 | rs13102559 | 1.759 | -0.105 | 1.261 | 0.766 | -0.01 | -0.011 | -0.011 | 0 | 0.7 | 0.42 | 0.68 | 0.72 | 0.77 | 0.57 |
| 4 | 141155942 | rs11733305 | -1.076 | 0.239 | 1.231 | 0.617 | -0.016 | 0.04 | 0.04 | 0.049 | 0.11 | 0.02 | 0.07 | 0.01 | 0.15 | 0.15 |
| 4 | 141172413 | rs7659292 | -2.307 | -0.045 | 2.386 | 0.987 | -0.011 | 0.041 | 0.041 | 0 | 0.19 | 0.62 | 0.22 | 0.3 | 0.11 | 0.27 |
| 4 | 141183514 | rs12502106 | -1.924 | -0.167 | 2.789 | 0.937 | -0.024 | -0.07 | -0.07 | 0.028 | 0.16 | 0.41 | 0.09 | 0.04 | 0.07 | 0.13 |
| 4 | 141194324 | rs6831812 | -2.969 | -0.106 | 2.847 | 1.138 | -0.001 | 0.081 | 0.081 | 0.165 | 0.14 | 0.64 | 0.27 | 0.32 | 0.08 | 0.24 |
| 4 | 141205306 | rs17005684 | -2.352 | -0.167 | 3.042 | 0.888 | -0.003 | -0.056 | -0.056 | 0.877 | 0.11 | 0.55 | 0.07 | 0.02 | 0.03 | 0.1 |
| 4 | 141219784 | rs868653 | -2.161 | -0.144 | 2.718 | 0.684 | 0 | -0.034 | -0.034 | 0.864 | 0.16 | 0.21 | 0.07 | 0.03 | 0.11 | 0.21 |
| 4 | 141227577 | rs1877410 | -3.318 | 0.019 | 3.142 | 1.007 | -0.001 | -0.002 | -0.002 | 0 | 0.32 | 0.75 | 0.36 | 0.41 | 0.29 | 0.46 |
| 4 | 141231258 | rs17005729 | -1.674 | 0.062 | 3.297 | 1.007 | 0.007 | -0.061 | -0.061 | 0 | 0.13 | 0.25 | 0.08 | 0.01 | 0.13 | 0.13 |
| 4 | 141234930 | rs13128278 | -1.85 | 0.097 | 3.385 | 1.05 | 0.014 | 0.142 | 0.142 | 0.006 | 0.1 | 0.3 | 0.23 | 0.35 | 0.08 | 0.17 |
| 4 | 141236020 | rs4956333 | -2.458 | 0.027 | 3.308 | 1.675 | 0.003 | 0.038 | 0.038 | 0 | 0.22 | 0.32 | 0.36 | 0.52 | 0.21 | 0.31 |
| 4 | 141237134 | rs7678137 | -1.569 | 0.033 | 3.302 | 1.349 | 0.067 | -0.007 | -0.007 | 0 | 0.06 | 0.26 | 0.06 | 0 | 0.08 | 0.05 |
| 4 | 141246800 | rs2137267 | -2.135 | -0.715 | 2.247 | 1.496 | 0.035 | -0.032 | -0.032 | 0.591 | 0.42 | 0.77 | 0.4 | 0.56 | 0.28 | 0.46 |
| 4 | 141249813 | rs11100615 | -1.572 | -0.683 | 2.451 | 1.466 | -0.006 | 0.027 | 0.027 | 0 | 0.18 | 0.28 | 0.23 | 0.22 | 0.13 | 0.2 |
| 4 | 141253396 | rs2874445 | -1.705 | -0.647 | 2.583 | 1.438 | -0.011 | 0.024 | 0.024 | 0 | 0.21 | 0.38 | 0.15 | 0.22 | 0.14 | 0.23 |
| 4 | 141262661 | rs1039987 | 1.124 | -0.883 | 2.398 | 1.146 | -0.015 | -0.013 | -0.013 | 2.076 | 0.76 | 0.64 | 0.73 | 0.75 | 0.83 | 0.71 |
| 4 | 141262836 | rs1039988 | 1.015 | -0.885 | 2.425 | 1.144 | 0.007 | 0.006 | 0.006 | 0.038 | 0.77 | 0.82 | 0.73 | 0.75 | 0.83 | 0.71 |
| 4 | 141265134 | rs996116 | 0.861 | -0.885 | 2.41 | 1.144 | 0.005 | 0.006 | 0.006 | 0.35 | 0.78 | 0.84 | 0.83 | 0.76 | 0.83 | 0.72 |
| 4 | 141267864 | rs11945466 | -1.777 | -1.189 | 2.449 | 1.028 | -0.006 | -0.001 | -0.001 | 0.008 | 0.25 | 0.29 | 0.16 | 0.22 | 0.18 | 0.33 |
| 4 | 141276725 | rs10009892 | 0.951 | -1.153 | 2.499 | 1.031 | -0.019 | -0.006 | -0.006 | 0.284 | 0.78 | 0.56 | 0.74 | 0.76 | 0.87 | 0.79 |
| 4 | 141284111 | rs4956366 | -0.149 | -1.156 | 2.3 | 0.999 | 0.002 | 0.082 | 0.082 | 3.604 | 0.87 | 0.98 | 0.96 | 1 | 0.95 | 0.91 |
| 4 | 141284198 | rs7663838 | -1.263 | -1.154 | 2.481 | 1.01 | 0.041 | 0.133 | 0.133 | 0 | 0.07 | 0.24 | 0.21 | 0.24 | 0.07 | 0.11 |
| 4 | 141286226 | rs10024214 | 1.006 | -1.154 | 2.492 | 1.001 | -0.012 | 0.001 | 0.001 | 0.204 | 0.79 | 0.66 | 0.74 | 0.75 | 0.87 | 0.79 |
| 4 | 141298142 | rs358308 | -0.145 | -1.233 | 2.266 | 0.924 | 0.028 | 0.074 | 0.074 | 3.171 | 0.14 | 0.04 | 0.04 | 0 | 0.04 | 0.1 |
| 4 | 141298709 | rs358311 | -0.145 | -1.233 | 2.266 | 0.924 | 0.028 | 0.074 | 0.074 | 3.171 | 0.14 | 0.04 | 0.04 | 0 | 0.04 | 0.1 |
| 4 | 141304666 | rs4362897 | 2.784 | -1.191 | 2.478 | 0.936 | 0.009 | 0.001 | 0.001 | 0 | 0.54 | 0.39 | 0.59 | 0.53 | 0.69 | 0.47 |
| 4 | 141310226 | rs3805317 | -1.102 | -1.162 | 2.409 | 0.962 | 0.014 | 0.079 | 0.079 | 0 | 0.07 | 0.16 | 0.2 | 0.25 | 0.08 | 0.11 |
| 4 | 141319793 | rs2175562 | 2.044 | -1.181 | 2.466 | 0.954 | -0.003 | 0.012 | 0.012 | 0 | 0.69 | 0.64 | 0.64 | 0.53 | 0.73 | 0.59 |
| 4 | 141320021 | rs2175563 | -0.218 | -1.21 | 2.401 | 0.922 | 0.024 | 0.026 | 0.026 | 2.521 | 0.14 | 0.11 | 0.05 | 0 | 0.04 | 0.1 |
| 4 | 141322236 | rs7681627 | -1.462 | -1.225 | 2.517 | 0.917 | 0.034 | 0.132 | 0.132 | 0 | 0.07 | 0.2 | 0.21 | 0.25 | 0.08 | 0.11 |
| 4 | 141323162 | rs2567241 | -0.354 | -1.262 | 2.162 | 0.884 | 0.026 | 0.029 | 0.029 | 3.743 | 0.15 | 0.11 | 0.05 | 0 | 0.04 | 0.1 |
| 4 | 141325738 | rs17005845 | -1.631 | -1.307 | 2.104 | 0.977 | -0.016 | -0.034 | -0.034 | 0 | 0.15 | 0.04 | 0.29 | 0.3 | 0.17 | 0.25 |
| 4 | 141341312 | rs10519554 | -0.315 | -1.281 | 2.159 | 0.947 | 0.008 | -0.007 | -0.007 | 0 | 0.15 | 0.15 | 0.05 | 0 | 0.04 | 0.1 |
| 4 | 141343133 | rs10012138 | 3.098 | -1.332 | 2.252 | 0.93 | 0.062 | 0.015 | 0.015 | 0 | 0.52 | 0.25 | 0.58 | 0.53 | 0.68 | 0.47 |
| 4 | 141350602 | rs1367705 | 0.241 | -1.181 | 2.139 | 0.851 | -0.023 | -0.058 | -0.058 | 0 | 0.85 | 0.76 | 0.95 | 1 | 0.96 | 0.89 |
| 4 | 141358232 | rs2060074 | 0.156 | -1.174 | 1.809 | 0.912 | 0.004 | -0.007 | -0.007 | 0 | 0.86 | 0.86 | 0.96 | 0.98 | 0.96 | 0.88 |
| 4 | 141364288 | rs17005918 | -1.919 | -1.183 | 1.593 | 0.993 | 0.001 | -0.005 | -0.005 | 0 | 0.24 | 0.16 | 0.16 | 0.22 | 0.2 | 0.29 |
| 4 | 141371764 | rs6830503 | 2.264 | -1.094 | 1.566 | 1.066 | 0.02 | 0.007 | 0.007 | 0 | 0.55 | 0.38 | 0.59 | 0.54 | 0.7 | 0.47 |
| 4 | 141372158 | rs1897280 | 2.011 | -1.167 | 1.553 | 0.96 | -0.011 | -0.016 | -0.016 | 0 | 0.6 | 0.43 | 0.78 | 0.78 | 0.76 | 0.58 |
| 4 | 141379314 | rs6836265 | -1.85 | -0.909 | 1.228 | 1.057 | 0.002 | 0.001 | 0.001 | 0 | 0.23 | 0.2 | 0.24 | 0.24 | 0.15 | 0.26 |
| 4 | 141381635 | rs12509826 | -2.62 | -0.967 | 1.238 | 1.056 | 0.006 | 0.005 | 0.005 | 0 | 0.3 | 0.22 | 0.32 | 0.29 | 0.22 | 0.37 |
| 4 | 141388317 | rs1431346 | -1.017 | -0.785 | 1.243 | 1.183 | 0.011 | 0.042 | 0.042 | 0 | 0.06 | 0.1 | 0.2 | 0.21 | 0.09 | 0.13 |
| 4 | 141391868 | rs6818155 | -0.839 | -0.701 | 1.335 | 1.205 | 0 | 0 | 0 | 0 | 0.09 | 0.11 | 0.1 | 0.06 | 0.08 | 0.11 |
| 4 | 141403999 | rs2321267 | -1.349 | -0.71 | 1.42 | 1.262 | 0.014 | 0.055 | 0.055 | 0 | 0.07 | 0.15 | 0.2 | 0.2 | 0.09 | 0.15 |
| 4 | 141404389 | rs2874478 | -2.481 | -0.728 | 1.474 | 1.672 | 0.01 | 0.124 | 0.124 | 0 | 0.16 | 0.31 | 0.31 | 0.5 | 0.17 | 0.29 |
| 4 | 141409324 | rs10519560 | -1.932 | -0.719 | 1.455 | 1.684 | 0.013 | 0.181 | 0.181 | 0 | 0.08 | 0.2 | 0.21 | 0.43 | 0.09 | 0.17 |
| 4 | 141410251 | rs7670320 | -2.119 | -0.747 | 1.795 | 1.759 | 0.04 | 0.293 | 0.293 | 0 | 0.08 | 0.28 | 0.22 | 0.44 | 0.09 | 0.17 |
| 4 | 141414355 | rs17389716 | -0.829 | -0.76 | 1.821 | 1.752 | 0 | 0 | 0 | 0.874 | 0.09 | 0.11 | 0.09 | 0.06 | 0.08 | 0.11 |
| 4 | 141416100 | rs12508462 | -2.751 | -0.759 | 1.814 | 1.74 | 0.018 | 0.203 | 0.203 | 0 | 0.17 | 0.39 | 0.31 | 0.5 | 0.17 | 0.28 |
| 4 | 141428878 | rs9790415 | -2.062 | -0.802 | 1.644 | 1.521 | 0.015 | 0.02 | 0.02 | 0 | 0.13 | 0.23 | 0.27 | 0.2 | 0.18 | 0.21 |
| 4 | 141436854 | rs1241046 | 2.913 | -0.966 | 1.996 | 1.448 | -0.006 | 0.058 | 0.058 | 0 | 0.69 | 0.47 | 0.61 | 0.48 | 0.71 | 0.6 |
| 4 | 141438865 | rs1123003 | -1.995 | -0.963 | 1.988 | 1.449 | 0.015 | 0.02 | 0.02 | 0 | 0.13 | 0.23 | 0.27 | 0.2 | 0.18 | 0.21 |
| 4 | 141445816 | rs11945078 | -2.907 | -0.964 | 2 | 1.441 | 0.014 | 0.131 | 0.131 | 0 | 0.23 | 0.44 | 0.37 | 0.5 | 0.26 | 0.35 |
| 4 | 141449275 | rs3104107 | 1.178 | -1.156 | 2.343 | 1.265 | -0.037 | -0.088 | -0.088 | 1.379 | 0.83 | 0.64 | 0.93 | 0.97 | 0.93 | 0.87 |
| 4 | 141459925 | rs17006095 | -2.714 | -0.937 | 2.314 | 1.279 | 0.008 | -0.025 | -0.025 | 0 | 0.23 | 0.51 | 0.33 | 0.21 | 0.24 | 0.3 |
| 4 | 141465244 | rs2321272 | 3.029 | -1.008 | 2.219 | 1.363 | 0.016 | -0.03 | -0.03 | 0 | 0.6 | 0.25 | 0.57 | 0.48 | 0.67 | 0.52 |
| 4 | 141468569 | rs6818701 | -0.618 | -0.983 | 2.18 | 1.062 | 0.006 | 0.032 | 0.032 | 0 | 0.06 | 0.08 | 0.2 | 0.2 | 0.09 | 0.14 |
| 4 | 141481581 | rs6536991 | -2.635 | -0.991 | 2.094 | 1.065 | -0.014 | -0.029 | -0.029 | 1.594 | 0.31 | 0.61 | 0.33 | 0.23 | 0.25 | 0.34 |

| | | | | | | | | | | | | | | | | |
|---|---|---|---|---|---|---|---|---|---|---|---|---|---|---|---|---|
| 4 | 141487748 | rs6822807 | -2.377 | -0.748 | 1.511 | 1.255 | 0.01 | 0.015 | 0.015 | 0 | 0.22 | 0.34 | 0.36 | 0.27 | 0.26 | 0.33 |
| 4 | 141489996 | rs10011540 | -0.698 | -1.032 | 0.37 | 0.645 | 0.003 | 0.002 | 0.002 | 0 | 0.09 | 0.18 | 0.09 | 0.07 | 0.08 | 0.18 |
| 4 | 141494104 | rs2043125 | -2.015 | -0.962 | 0.363 | 1.132 | 0.003 | -0.018 | -0.018 | 0 | 0.3 | 0.73 | 0.41 | 0.47 | 0.24 | 0.39 |
| 4 | 141496369 | rs12645959 | -2.281 | -1.031 | 0.24 | 1.014 | 0.004 | -0.039 | -0.039 | 0 | 0.36 | 0.75 | 0.46 | 0.47 | 0.33 | 0.44 |
| 4 | 141499558 | rs17006132 | -2.018 | -1.045 | 0.404 | 1.25 | -0.01 | -0.014 | -0.014 | 0 | 0.2 | 0.06 | 0.2 | 0.19 | 0.22 | 0.32 |
| 4 | 141500395 | rs3851422 | -0.603 | -1.095 | 0.978 | 1.255 | 0.012 | 0.025 | 0.025 | 0 | 0.13 | 0.35 | 0.24 | 0.15 | 0.11 | 0.09 |
| 4 | 141509633 | rs12509905 | -0.76 | -1.151 | 0.314 | 1.044 | -0.02 | 0.008 | 0.008 | 0 | 0.12 | 0.01 | 0.13 | 0.06 | 0.1 | 0.25 |
| 4 | 141513385 | rs2195645 | -2.676 | -0.082 | 0.289 | 0.998 | -0.015 | -0.006 | -0.006 | 0 | 0.4 | 0.81 | 0.4 | 0.37 | 0.38 | 0.5 |
| 4 | 141518965 | rs12640614 | -1.35 | 0.099 | 0.396 | 1.018 | 0.026 | -0.015 | -0.015 | 0.656 | 0.22 | 0.59 | 0.21 | 0.2 | 0.26 | 0.14 |
| 4 | 141520308 | rs4956454 | -1.191 | 0.106 | 0.3 | 0.916 | 0.042 | -0.115 | -0.115 | 0.071 | 0.21 | 0.5 | 0.21 | 0.06 | 0.26 | 0.13 |
| 4 | 141525704 | rs4956323 | -1.428 | 0.101 | -0.559 | 1.22 | 0 | -0.034 | -0.034 | 0.914 | 0.16 | 0.1 | 0.17 | 0.3 | 0.12 | 0.31 |
| 4 | 141533137 | rs4956324 | -0.999 | 0.182 | -1.496 | 1.075 | -0.013 | -0.14 | -0.14 | 0.116 | 0.79 | 0.98 | 0.83 | 0.64 | 0.81 | 0.88 |
| 4 | 141533832 | rs12503379 | 1.148 | 0.724 | -1.944 | 0.809 | -0.062 | -0.083 | -0.083 | 0.624 | 0.26 | 0.1 | 0.42 | 0.44 | 0.29 | 0.29 |
| 4 | 141537771 | rs4956325 | -0.527 | 0.091 | -1.589 | 0.582 | 0.023 | -0.023 | -0.023 | 0 | 0.63 | 0.8 | 0.71 | 0.63 | 0.67 | 0.71 |
| 4 | 141548569 | rs3816933 | 0.18 | -0.013 | -1.157 | 0.316 | -0.03 | -0.078 | -0.078 | 0.779 | 0.22 | 0.53 | 0.15 | 0.14 | 0.15 | 0.18 |
| 4 | 141549393 | rs2291711 | -0.666 | -0.072 | -1.159 | 0.329 | 0.02 | -0.018 | -0.018 | 0 | 0.62 | 0.76 | 0.7 | 0.63 | 0.67 | 0.71 |
| 4 | 141551799 | rs1430581 | 0.576 | -0.062 | -1.194 | 0.331 | -0.039 | -0.041 | -0.041 | 0 | 0.39 | 0.2 | 0.54 | 0.49 | 0.52 | 0.53 |
| 4 | 141554145 | rs4956327 | 0.227 | -0.054 | -1.136 | 0.33 | -0.028 | -0.073 | -0.073 | 0.196 | 0.22 | 0.55 | 0.16 | 0.14 | 0.15 | 0.18 |
| 4 | 141554949 | rs10002459 | 0.49 | 0.069 | -1.518 | 0.408 | -0.048 | -0.051 | -0.051 | 0 | 0.39 | 0.17 | 0.54 | 0.49 | 0.51 | 0.53 |
| 4 | 141561353 | rs6813507 | -0.225 | 0.075 | -0.921 | 0.209 | -0.033 | -0.072 | -0.072 | 0 | 0.22 | 0.56 | 0.15 | 0.13 | 0.15 | 0.19 |
| 4 | 141562798 | rs35060427 | 0.184 | 0.73 | -1.91 | 0.479 | -0.058 | 0.045 | 0.045 | 0.139 | 0.41 | 0.15 | 0.51 | 0.33 | 0.49 | 0.39 |
| 4 | 141563992 | rs6537003 | -0.614 | 1.215 | -1.851 | 0.833 | -0.027 | 0 | 0 | 0 | 0.76 | 0.88 | 0.79 | 0.82 | 0.71 | 0.8 |
| 4 | 141569847 | rs17392415 | -0.278 | 1.456 | -1.952 | 1.034 | -0.024 | -0.027 | -0.027 | 0 | 0.1 | 0.03 | 0.12 | 0.19 | 0.19 | 0.24 |
| 4 | 141570672 | rs1430580 | -0.702 | 1.406 | -2.052 | 1.011 | -0.025 | 0.019 | 0.019 | 0 | 0.75 | 0.87 | 0.78 | 0.83 | 0.7 | 0.79 |
| 4 | 141573742 | rs13109751 | 0.859 | 1.438 | -1.148 | 1.25 | 0 | -0.003 | -0.003 | 0 | 0.54 | 0.52 | 0.48 | 0.43 | 0.42 | 0.38 |
| 4 | 141574613 | rs4956471 | -1.455 | 1.134 | -0.962 | 1.403 | 0.029 | 0.068 | 0.068 | 0 | 0.19 | 0.29 | 0.28 | 0.38 | 0.28 | 0.31 |
| 4 | 141576673 | rs2163322 | 1.065 | 0.111 | -1.72 | 0.636 | -0.106 | -0.057 | -0.057 | 0.204 | 0.45 | 0.04 | 0.28 | 0.44 | 0.42 | 0.36 |
| 4 | 141578155 | rs7683748 | -0.209 | -0.044 | -0.934 | 0.222 | 0.003 | 0.003 | 0.003 | 0 | 0.66 | 0.58 | 0.57 | 0.65 | 0.54 | 0.59 |
| 4 | 141580561 | rs3886094 | -0.774 | -0.331 | -0.871 | -0.289 | -0.005 | -0.02 | -0.02 | 0.025 | 0.14 | 0.16 | 0.17 | 0.04 | 0.1 | 0.12 |
| 4 | 141582368 | rs12640364 | 0.86 | -0.322 | -1.447 | -0.163 | -0.099 | -0.013 | -0.013 | 0 | 0.46 | 0.04 | 0.35 | 0.43 | 0.43 | 0.43 |
| 4 | 141589301 | rs6845527 | -0.024 | -0.686 | -0.837 | -0.339 | -0.012 | -0.006 | -0.006 | 0.008 | 0.63 | 0.7 | 0.55 | 0.47 | 0.51 | 0.6 |
| 4 | 141593715 | rs12643256 | 1.098 | -0.709 | -1.047 | -0.361 | -0.108 | -0.014 | -0.014 | 0.035 | 0.47 | 0.04 | 0.37 | 0.44 | 0.46 | 0.44 |
| 4 | 141596785 | rs7688253 | 0.288 | -0.574 | -0.383 | -0.377 | -0.014 | -0.041 | -0.041 | 0 | 0.07 | 0.18 | 0.06 | 0 | 0.05 | 0.01 |
| 4 | 141597913 | rs1469539 | -0.537 | -0.365 | -0.167 | -0.221 | -0.001 | 0 | 0 | 0 | 0.7 | 0.69 | 0.61 | 0.56 | 0.67 | 0.62 |
| 4 | 141598290 | rs2303912 | -0.362 | -0.37 | 0.024 | -0.225 | 0.007 | 0.001 | 0.001 | 0 | 0.09 | 0.1 | 0.07 | 0.09 | 0.16 | 0.08 |
| 4 | 141600320 | rs4956329 | -0.449 | -0.246 | 0.063 | -0.172 | -0.001 | -0.001 | -0.001 | 0 | 0.7 | 0.69 | 0.61 | 0.57 | 0.67 | 0.62 |
| 4 | 141601420 | rs6813209 | -0.429 | -0.243 | -0.116 | -0.172 | -0.001 | 0.012 | 0.012 | 0 | 0.7 | 0.66 | 0.61 | 0.56 | 0.67 | 0.62 |
| 4 | 141616979 | rs11736541 | -0.029 | -0.191 | -0.279 | -0.414 | -0.003 | 0.06 | 0.06 | 0 | 0.32 | 0.47 | 0.42 | 0.45 | 0.33 | 0.38 |
| 4 | 141620931 | rs13149166 | 1.038 | -0.045 | -1.345 | -0.882 | -0.099 | 0.031 | 0.031 | 0.224 | 0.46 | 0.05 | 0.37 | 0.36 | 0.47 | 0.42 |
| 4 | 141625217 | rs11731292 | -0.084 | 0.049 | -1.209 | -0.875 | 0.017 | -0.026 | -0.026 | 0 | 0.36 | 0.78 | 0.44 | 0.57 | 0.31 | 0.44 |
| 4 | 141629176 | rs923307 | -0.351 | 0.179 | -1.19 | -0.912 | -0.029 | 0.009 | 0.009 | 0 | 0.91 | 0.96 | 0.92 | 0.95 | 0.83 | 0.92 |
| 4 | 141629826 | rs10009007 | 0.852 | -0.448 | -1.105 | -0.89 | 0.004 | 0.008 | 0.008 | 0 | 0.06 | 0.03 | 0.04 | 0 | 0.06 | 0.01 |
| 4 | 141633234 | rs10004266 | 0.08 | -0.461 | -1.038 | -0.897 | -0.009 | -0.03 | -0.03 | 0.546 | 0.07 | 0.13 | 0.05 | 0.01 | 0.06 | 0.05 |
| 4 | 141637275 | rs2306815 | -0.1 | -0.489 | -1.151 | -0.882 | 0.008 | -0.014 | -0.014 | 0 | 0.34 | 0.69 | 0.42 | 0.56 | 0.3 | 0.4 |
| 4 | 141639072 | rs901687 | -0.69 | -0.132 | -1.362 | -1.057 | -0.042 | 0.037 | 0.037 | 0 | 0.85 | 0.92 | 0.87 | 0.96 | 0.73 | 0.91 |
| 4 | 141642216 | rs7688945 | 0.506 | -0.209 | -1.453 | -1.071 | -0.031 | 0.048 | 0.048 | 0 | 0.75 | 0.83 | 0.82 | 0.95 | 0.67 | 0.86 |
| 4 | 141652922 | rs17006401 | 0.494 | -0.265 | -1.534 | -1.069 | 0.012 | -0.029 | -0.029 | 0 | 0.31 | 0.74 | 0.39 | 0.56 | 0.25 | 0.33 |
| 4 | 141656467 | rs1532013 | 0.257 | -0.163 | -1.613 | -1.018 | -0.01 | 0.045 | 0.045 | 0 | 0.53 | 0.18 | 0.47 | 0.39 | 0.48 | 0.59 |
| 4 | 141658697 | rs4555724 | 0.586 | -0.175 | -1.467 | -0.79 | -0.034 | -0.014 | -0.014 | 0 | 0.44 | 0.07 | 0.42 | 0.39 | 0.4 | 0.55 |
| 4 | 141682299 | rs6843792 | -0.113 | -0.182 | -1.14 | -0.729 | 0.003 | -0.007 | -0.007 | 0.025 | 0.34 | 0.63 | 0.5 | 0.6 | 0.32 | 0.41 |
| 4 | 141688766 | rs12331218 | -0.369 | -0.296 | -1.251 | -0.852 | 0.016 | 0 | 0 | 0 | 0.4 | 0.63 | 0.5 | 0.58 | 0.35 | 0.44 |
| 4 | 141710146 | rs11930453 | 0.273 | -0.116 | -1.445 | -0.83 | -0.012 | 0.036 | 0.036 | 0 | 0.63 | 0.67 | 0.69 | 0.91 | 0.57 | 0.76 |
| 4 | 141716419 | rs6810701 | 0.572 | -0.059 | -1.792 | -0.247 | -0.021 | -0.044 | -0.044 | 0 | 0.38 | 0.06 | 0.31 | 0.37 | 0.33 | 0.46 |
| 4 | 141720070 | rs11727409 | 0.648 | 0.055 | -1.67 | -0.148 | -0.016 | -0.034 | -0.034 | 0.021 | 0.37 | 0.09 | 0.32 | 0.36 | 0.33 | 0.45 |
| 4 | 141723577 | rs2171412 | -0.575 | -0.226 | -0.362 | 0.596 | -0.004 | 0.004 | 0.004 | 0 | 0.33 | 0.35 | 0.38 | 0.57 | 0.28 | 0.41 |
| 4 | 141765064 | rs9993702 | -0.55 | -0.137 | -0.734 | 0.945 | 0.012 | -0.086 | -0.086 | 0 | 0.22 | 0.11 | 0.27 | 0.54 | 0.2 | 0.24 |
| 4 | 141770572 | rs6836292 | -1.092 | -0.615 | 0.111 | 0.603 | -0.019 | 0.021 | 0.021 | 0.919 | 0.33 | 0.6 | 0.34 | 0.54 | 0.23 | 0.26 |
| 4 | 141776759 | rs7696023 | -1.179 | -0.7 | 1.121 | 0.341 | -0.007 | -0.019 | -0.019 | 0 | 0.14 | 0.38 | 0.15 | 0.12 | 0.11 | 0.18 |
| 4 | 141777368 | rs1155206 | 1.409 | -0.981 | 1.123 | 0.522 | -0.011 | 0.01 | 0.01 | 0.497 | 0.76 | 0.33 | 0.76 | 0.68 | 0.84 | 0.73 |
| 4 | 141779593 | rs1487565 | 0.947 | -1.113 | 1.321 | 0.367 | -0.013 | 0.038 | 0.038 | 0 | 0.77 | 0.63 | 0.77 | 0.71 | 0.82 | 0.81 |
| 4 | 141783289 | rs17421358 | 0.206 | -1.458 | 0.052 | -0.076 | 0.02 | 0.042 | 0.042 | 0 | 0.17 | 0.02 | 0.17 | 0.08 | 0.15 | 0.24 |
| 4 | 141783627 | rs3733478 | 0.176 | -1.634 | -0.038 | 0.512 | -0.005 | 0.159 | 0.159 | 0 | 0.05 | 0.09 | 0.08 | 0.39 | 0.05 | 0.1 |
| 4 | 141783990 | rs3733480 | 0.149 | -1.633 | -0.038 | 0.512 | -0.005 | 0.159 | 0.159 | 0 | 0.05 | 0.09 | 0.08 | 0.39 | 0.05 | 0.1 |
| 4 | 141784596 | rs7655676 | 1.454 | -1.683 | -0.147 | 0.098 | 0.056 | 0.225 | 0.225 | 2.106 | 0.47 | 0.12 | 0.46 | 0.19 | 0.6 | 0.38 |
| 4 | 141786545 | rs998013 | 0.646 | -1.195 | 0.411 | 0.133 | -0.001 | -0.007 | -0.007 | 0 | 0.65 | 0.34 | 0.71 | 0.84 | 0.72 | 0.71 |
| 4 | 141787735 | rs3796699 | 0.919 | -0.366 | -0.504 | -0.049 | -0.017 | 0.009 | 0.009 | 0 | 0.27 | 0.18 | 0.29 | 0.23 | 0.35 | 0.4 |
| 4 | 141795525 | rs11730663 | 0.953 | -0.09 | -0.407 | -0.556 | -0.078 | 0.111 | 0.111 | 0.061 | 0.31 | 0.15 | 0.27 | 0.04 | 0.46 | 0.45 |
| 4 | 141804520 | rs6829449 | 0.228 | -0.256 | -0.624 | 0.245 | -0.004 | 0.033 | 0.033 | 0.271 | 0.15 | 0.16 | 0.21 | 0.38 | 0.12 | 0.12 |
| 4 | 141812185 | rs1863305 | 0.301 | -0.607 | -0.624 | 0.016 | 0.013 | 0.024 | 0.024 | 0 | 0.36 | 0.29 | 0.17 | 0.16 | 0.28 | 0.27 |

| | | | | | | | | | | | | | | | | |
|---|---|---|---|---|---|---|---|---|---|---|---|---|---|---|---|---|
| 4 | 141812645 | rs1863307 | -0.322 | -0.245 | -0.71 | 0.31 | 0.002 | -0.014 | -0.014 | 0 | 0.56 | 0.69 | 0.42 | 0.54 | 0.42 | 0.41 |
| 4 | 141816657 | rs767761 | 0.125 | 0.271 | -1.115 | 0.402 | 0.004 | 0.034 | 0.034 | 1.418 | 0.51 | 0.19 | 0.4 | 0.47 | 0.44 | 0.45 |
| 4 | 141820214 | rs3844179 | -0.463 | 0.422 | -0.649 | -0.403 | 0 | -0.017 | -0.017 | 1.705 | 0.64 | 0.38 | 0.72 | 0.83 | 0.59 | 0.62 |
| 4 | 141826780 | rs1030356 | -0.113 | 0.106 | -1.224 | -0.289 | -0.025 | 0.077 | 0.077 | 3.483 | 0.27 | 0.03 | 0.17 | 0.14 | 0.21 | 0.11 |
| 4 | 141831660 | rs1863301 | -0.097 | 0.107 | -0.683 | -0.227 | 0.037 | 0.051 | 0.051 | 0 | 0.55 | 0.4 | 0.43 | 0.27 | 0.62 | 0.37 |
| 4 | 141833025 | rs874776 | -0.475 | 0.472 | -0.226 | 0.006 | 0.027 | 0.039 | 0.039 | 0 | 0.49 | 0.44 | 0.59 | 0.26 | 0.6 | 0.35 |
| 4 | 141833103 | rs874775 | 0.228 | 0.611 | 0.137 | -0.064 | 0.017 | 0.034 | 0.034 | 0 | 0.5 | 0.43 | 0.42 | 0.73 | 0.43 | 0.64 |
| 4 | 141833535 | rs1834382 | -0.569 | 0.486 | 0.129 | 0.062 | -0.009 | 0.014 | 0.014 | 2.75 | 0.81 | 0.88 | 0.72 | 0.85 | 0.82 | 0.89 |
| 4 | 141834589 | rs17338560 | 1.054 | 0.249 | 0.348 | 0.105 | -0.028 | 0.023 | 0.023 | 0 | 0.26 | 0.25 | 0.31 | 0.1 | 0.43 | 0.26 |
| 4 | 141836576 | rs6537018 | -1.048 | 0.305 | 0.092 | 0.133 | 0.087 | -0.096 | -0.096 | 0 | 0.72 | 0.5 | 0.66 | 0.9 | 0.56 | 0.74 |
| 4 | 141845421 | rs1425446 | 0.634 | 0.124 | 0.163 | 0.127 | 0.016 | 0.016 | 0.016 | 0 | 0.43 | 0.33 | 0.32 | 0.69 | 0.31 | 0.54 |
| 4 | 141851355 | rs13118519 | -0.599 | 0.592 | 0.374 | 0.934 | 0.073 | 0.049 | 0.049 | 0 | 0.69 | 0.48 | 0.58 | 0.59 | 0.53 | 0.66 |
| 4 | 141853440 | rs7658326 | -0.476 | -0.419 | 0.519 | 1.404 | -0.02 | 0.074 | 0.074 | 0 | 0.15 | 0.26 | 0.14 | 0.31 | 0.1 | 0.17 |
| 4 | 141865303 | rs6537023 | 0.141 | -0.544 | 0.49 | 1.103 | 0.005 | 0.005 | 0.005 | 0 | 0.52 | 0.55 | 0.65 | 0.36 | 0.62 | 0.43 |
| 4 | 141868515 | rs1001487 | 0.102 | -0.505 | 0.474 | 1.038 | 0.037 | 0.043 | 0.043 | 0 | 0.49 | 0.38 | 0.61 | 0.33 | 0.6 | 0.44 |
| 4 | 141870648 | rs4956494 | 0.539 | -0.424 | 0.402 | 0.922 | 0 | -0.013 | -0.013 | 0 | 0.43 | 0.52 | 0.48 | 0.19 | 0.51 | 0.34 |
| 4 | 141872192 | rs11724213 | 0.651 | -0.428 | -0.191 | 0.944 | -0.003 | 0.019 | 0.019 | 0 | 0.42 | 0.37 | 0.48 | 0.19 | 0.51 | 0.35 |
| 4 | 141875683 | rs2321689 | 0.72 | -0.398 | -0.749 | 0.95 | -0.022 | 0.077 | 0.077 | 0 | 0.39 | 0.27 | 0.46 | 0.19 | 0.5 | 0.34 |
| 4 | 141877688 | rs17424664 | 0.296 | -0.508 | -0.562 | 1.062 | 0.064 | 0.055 | 0.055 | 0 | 0.47 | 0.28 | 0.59 | 0.36 | 0.6 | 0.44 |
| 4 | 141879419 | rs7689080 | 0.12 | -0.261 | -0.829 | 0.988 | 0.025 | -0.005 | -0.005 | 0 | 0.34 | 0.24 | 0.25 | 0.36 | 0.29 | 0.4 |
| 4 | 141880666 | rs6814533 | -0.978 | -0.161 | -0.543 | 0.982 | -0.003 | 0.002 | 0.002 | 0 | 0.2 | 0.24 | 0.3 | 0.24 | 0.18 | 0.2 |
| 4 | 141880770 | rs13127505 | -0.675 | 0.099 | -0.792 | 0.959 | -0.031 | -0.03 | -0.03 | 0 | 0.09 | 0.02 | 0.14 | 0.14 | 0.11 | 0.1 |
| 4 | 141881059 | rs10003541 | -0.31 | -0.023 | -1.098 | 0.593 | -0.03 | 0.073 | 0.073 | 0 | 0.6 | 0.76 | 0.52 | 0.75 | 0.48 | 0.52 |
| 4 | 141881246 | rs1425443 | -0.682 | -0.51 | -0.101 | 1.324 | 0.002 | 0.08 | 0.08 | 0 | 0.17 | 0.29 | 0.34 | 0.32 | 0.2 | 0.15 |
| 4 | 141888451 | rs11724521 | 0.691 | -0.769 | -0.231 | 0.458 | -0.111 | 0.013 | 0.013 | 0 | 0.29 | 0.12 | 0.46 | 0.21 | 0.44 | 0.29 |
| 4 | 141890683 | rs1364880 | -0.342 | -1.249 | 0.26 | 0.221 | -0.023 | 0.044 | 0.044 | 0 | 0.59 | 0.27 | 0.47 | 0.75 | 0.51 | 0.66 |
| 4 | 141891566 | rs1425431 | -0.52 | -1.244 | 0.406 | 0.22 | 0.007 | -0.012 | -0.012 | 0.218 | 0.62 | 0.45 | 0.48 | 0.79 | 0.52 | 0.68 |
| 4 | 141896970 | rs1364881 | 0.681 | -1.855 | -0.176 | -0.283 | -0.003 | 0.077 | 0.077 | 0 | 0.19 | 0.09 | 0.2 | 0.03 | 0.18 | 0.09 |
| 4 | 141903402 | rs6820265 | -0.152 | -1.859 | -0.126 | -0.177 | 0.029 | 0.025 | 0.025 | 0 | 0.38 | 0.2 | 0.27 | 0.72 | 0.34 | 0.57 |
| 4 | 141907543 | rs6537024 | 0.438 | -1.453 | 0.055 | -0.158 | 0.001 | -0.013 | -0.013 | 0 | 0.21 | 0.2 | 0.21 | 0.02 | 0.18 | 0.09 |
| 4 | 141920037 | rs11726692 | 0.353 | -2.203 | 0.072 | 0.382 | -0.022 | 0.021 | 0.021 | 0 | 0.19 | 0.24 | 0.15 | 0.39 | 0.11 | 0.4 |
| 4 | 141929562 | rs12643749 | 0.535 | -2.299 | -0.125 | 0.415 | -0.015 | -0.054 | -0.054 | 0 | 0.14 | 0.05 | 0.17 | 0.35 | 0.17 | 0.15 |
| 4 | 141933097 | rs7687054 | -0.481 | -2.629 | 0.31 | -0.265 | -0.035 | -0.093 | -0.093 | 0.54 | 0.84 | 0.72 | 0.93 | 1 | 0.92 | 0.93 |
| 4 | 141937337 | rs3851424 | 0.8 | -2.613 | 0.245 | -0.263 | 0.001 | -0.005 | -0.005 | 0 | 0.22 | 0.21 | 0.22 | 0.07 | 0.19 | 0.11 |
| 4 | 141942359 | rs2059499 | 0.788 | -2.814 | -0.213 | -0.262 | 0.03 | 0.031 | 0.031 | 0.44 | 0.43 | 0.3 | 0.64 | 0.61 | 0.62 | 0.43 |
| 4 | 141943052 | rs17425634 | -0.3 | -2.473 | -0.574 | -0.568 | 0.035 | 0.146 | 0.146 | 0 | 0.34 | 0.15 | 0.18 | 0 | 0.27 | 0.16 |
| 4 | 141945926 | rs13151459 | 0.431 | -2.164 | -0.139 | 0.137 | -0.007 | 0.019 | 0.019 | 3.657 | 0.12 | 0.06 | 0.16 | 0.07 | 0.16 | 0.11 |
| 4 | 141953830 | rs1907159 | -0.411 | -2.856 | -0.172 | 0.413 | 0.054 | -0.043 | -0.043 | 0.125 | 0.7 | 0.82 | 0.81 | 0.52 | 0.87 | 0.5 |
| 4 | 141957064 | rs11100699 | 0.498 | -2.469 | -0.22 | 0.206 | -0.001 | -0.003 | -0.003 | 0 | 0.12 | 0.1 | 0.16 | 0.07 | 0.14 | 0.1 |
| 4 | 141957740 | rs9997698 | 0.976 | -2.416 | -0.387 | 0.118 | 0.012 | -0.001 | -0.001 | 0 | 0.14 | 0.11 | 0.07 | 0 | 0.09 | 0.03 |
| 4 | 141959896 | rs3851427 | 1.273 | -2.475 | -0.227 | 0.316 | -0.006 | 0.002 | 0.002 | 0 | 0.34 | 0.23 | 0.53 | 0.26 | 0.57 | 0.29 |
| 4 | 141960841 | rs9993321 | 0.555 | -2.532 | 0.134 | 0.406 | 0.033 | 0.001 | 0.001 | 0 | 0.51 | 0.38 | 0.59 | 0.45 | 0.67 | 0.36 |
| 4 | 141963286 | rs7687232 | -0.376 | -2.342 | 0.191 | 0.556 | 0.04 | -0.029 | -0.029 | 0 | 0.67 | 0.78 | 0.73 | 0.53 | 0.8 | 0.47 |
| 4 | 141965158 | rs9991795 | 0.403 | -1.953 | 0.145 | 0.178 | 0.015 | 0.018 | 0.018 | 1.543 | 0.27 | 0.19 | 0.22 | 0.07 | 0.21 | 0.19 |
| 4 | 141968129 | rs12505705 | 0.094 | -2.044 | 0.046 | 0.2 | 0.035 | -0.004 | -0.004 | 0 | 0.59 | 0.33 | 0.68 | 0.34 | 0.76 | 0.46 |
| 4 | 141970192 | rs12650097 | 0.372 | -1.809 | -0.203 | -0.537 | 0.015 | 0.021 | 0.021 | 1.667 | 0.27 | 0.19 | 0.22 | 0.07 | 0.2 | 0.19 |
| 4 | 141972002 | rs13116646 | 0.016 | -1.444 | -0.041 | 0.287 | -0.032 | -0.022 | -0.022 | 0.35 | 0.69 | 0.64 | 0.72 | 0.74 | 0.81 | 0.56 |
| 4 | 141975349 | rs2042630 | 0.004 | -1.441 | -0.25 | 0.288 | 0.005 | 0.004 | 0.004 | 0 | 0.69 | 0.75 | 0.74 | 0.74 | 0.82 | 0.56 |
| 4 | 141978179 | rs3863129 | 0.084 | -1.64 | -0.029 | 0.251 | -0.038 | -0.026 | -0.026 | 0 | 0.69 | 0.61 | 0.73 | 0.75 | 0.82 | 0.56 |
| 4 | 141980240 | rs4956344 | 0.447 | -1.128 | -0.944 | -0.088 | -0.078 | -0.053 | -0.053 | 0.02 | 0.33 | 0.13 | 0.43 | 0.3 | 0.52 | 0.31 |
| 4 | 141982395 | rs4361465 | 0.012 | -1.478 | -0.339 | -0.343 | 0.012 | 0.012 | 0.012 | 0 | 0.42 | 0.38 | 0.43 | 0.31 | 0.59 | 0.38 |
| 4 | 141995467 | rs2321859 | -0.663 | -1.844 | -0.117 | -0.072 | -0.033 | 0.1 | 0.1 | 0 | 0.77 | 0.6 | 0.79 | 0.51 | 0.86 | 0.61 |
| 4 | 141996416 | rs3851432 | 0.235 | -1.929 | -0.133 | -0.793 | -0.005 | -0.001 | -0.001 | 0.832 | 0.27 | 0.25 | 0.3 | 0.23 | 0.24 | 0.17 |
| 4 | 141999286 | rs3851433 | 0.546 | -1.592 | 0.397 | -1.133 | -0.052 | -0.111 | -0.111 | 0 | 0.84 | 0.6 | 0.93 | 1 | 0.91 | 0.9 |
| 4 | 142007952 | rs6537029 | 0.067 | -1.466 | -0.252 | -1.047 | -0.03 | -0.017 | -0.017 | 0 | 0.35 | 0.43 | 0.29 | 0.76 | 0.21 | 0.49 |
| 4 | 142009392 | rs11100704 | 0.862 | -0.695 | -1.143 | -1.338 | -0.111 | 0.024 | 0.024 | 0.103 | 0.27 | 0.08 | 0.49 | 0.21 | 0.46 | 0.28 |
| 4 | 142011361 | rs6836603 | -0.715 | -0.786 | -1.005 | -1.394 | -0.003 | -0.08 | -0.08 | 0 | 0.69 | 0.37 | 0.82 | 1 | 0.76 | 0.85 |
| 4 | 142011941 | rs11734545 | -0.344 | -0.518 | -1.195 | -0.599 | -0.054 | -0.006 | -0.006 | 0 | 0.66 | 0.5 | 0.76 | 0.37 | 0.8 | 0.51 |
| 4 | 142016173 | rs10454797 | -0.272 | 1.022 | -1.686 | -0.414 | -0.092 | 0.018 | 0.018 | 0 | 0.29 | 0.1 | 0.43 | 0.23 | 0.46 | 0.32 |
| 4 | 142021835 | rs1507179 | 0.044 | 0.768 | -1.996 | -0.848 | -0.16 | 0.074 | 0.074 | 0 | 0.33 | 0.06 | 0.52 | 0.15 | 0.48 | 0.31 |
| 4 | 142024322 | rs4629541 | -0.366 | 0.037 | -0.998 | -1.157 | -0.016 | -0.05 | -0.05 | 4.686 | 0.81 | 0.56 | 0.86 | 0.89 | 0.81 | 0.84 |
| 4 | 142043379 | rs4956517 | 0.068 | -0.208 | -0.546 | -1.463 | -0.019 | -0.088 | -0.088 | 0 | 0.75 | 0.52 | 0.83 | 0.88 | 0.76 | 0.81 |
| 4 | 142061946 | rs6853785 | -0.167 | -0.205 | -0.397 | -1.498 | -0.001 | -0.013 | -0.013 | 0.299 | 0.23 | 0.26 | 0.14 | 0.12 | 0.24 | 0.18 |
| 4 | 142066755 | rs4956520 | 0.36 | -0.578 | -0.037 | -1.859 | -0.002 | -0.007 | -0.007 | 0 | 0.42 | 0.39 | 0.3 | 0.8 | 0.28 | 0.43 |
| 4 | 142068023 | rs241867 | 0.062 | -0.708 | 0.169 | -1.806 | -0.045 | -0.17 | -0.17 | 0.501 | 0.74 | 0.48 | 0.87 | 0.99 | 0.81 | 0.82 |
| 4 | 142068473 | rs4956349 | 0.239 | -0.394 | 0.402 | -1.889 | 0.007 | 0.092 | 0.092 | 0 | 0.42 | 0.27 | 0.35 | 0.82 | 0.37 | 0.5 |
| 4 | 142070069 | rs354936 | 0.664 | -0.152 | 0.47 | -1.691 | -0.133 | 0.069 | 0.069 | 0 | 0.33 | 0.04 | 0.5 | 0.17 | 0.45 | 0.36 |
| 4 | 142074259 | rs10014312 | -0.487 | -0.207 | 0.859 | -1.718 | 0.012 | -0.038 | -0.038 | 0 | 0.62 | 0.55 | 0.47 | 0.81 | 0.55 | 0.63 |
| 4 | 142081109 | rs4956353 | -0.624 | -0.196 | 1.012 | -1.802 | -0.005 | 0.002 | 0.002 | 0 | 0.59 | 0.39 | 0.45 | 0.82 | 0.5 | 0.64 |

| | | | | | | | | | | | | | | | | |
|---|---|---|---|---|---|---|---|---|---|---|---|---|---|---|---|---|
| 4 | 142092124 | rs7663005 | 0.746 | -0.192 | 0.676 | -1.802 | -0.128 | 0.13 | 0.13 | 0 | 0.36 | 0.06 | 0.51 | 0.16 | 0.51 | 0.37 |
| 4 | 142095763 | rs7666238 | 0.196 | -0.452 | 0.366 | -1.363 | 0.011 | 0.082 | 0.082 | 0 | 0.31 | 0.09 | 0.22 | 0.19 | 0.24 | 0.35 |
| 4 | 142112913 | rs3851435 | 0.122 | -0.402 | 0.569 | -1.247 | 0.006 | -0.01 | -0.01 | 0 | 0.27 | 0.17 | 0.23 | 0.64 | 0.25 | 0.32 |
| 4 | 142117367 | rs4526055 | 0.066 | -0.261 | 0.772 | -1.302 | 0.001 | 0.036 | 0.036 | 0 | 0.32 | 0.21 | 0.25 | 0.2 | 0.26 | 0.31 |
| 4 | 142117424 | rs1550057 | 0.207 | -0.25 | 0.707 | -1.343 | 0.008 | -0.023 | -0.023 | 0 | 0.28 | 0.18 | 0.23 | 0.62 | 0.25 | 0.32 |
| 4 | 142119839 | rs2117576 | 0.02 | -0.364 | 1.037 | -1.322 | -0.003 | -0.038 | -0.038 | 0 | 0.32 | 0.38 | 0.26 | 0.2 | 0.25 | 0.31 |
| 4 | 142125380 | rs6822047 | -0.967 | -0.337 | 0.994 | -1.29 | 0.014 | -0.045 | -0.045 | 0 | 0.61 | 0.53 | 0.47 | 0.81 | 0.49 | 0.61 |
| 4 | 142129233 | rs13105932 | -0.867 | -0.271 | 1.043 | -1.3 | -0.012 | 0.044 | 0.044 | 0 | 0.59 | 0.34 | 0.46 | 0.81 | 0.5 | 0.61 |
| 4 | 142129314 | rs13129840 | 0.179 | -0.256 | 0.667 | -1.259 | 0.005 | 0.064 | 0.064 | 0 | 0.31 | 0.13 | 0.24 | 0.2 | 0.25 | 0.31 |
| 4 | 142137165 | rs6837873 | -0.176 | -0.267 | 0.709 | -1.249 | -0.002 | -0.01 | -0.01 | 0 | 0.34 | 0.36 | 0.26 | 0.2 | 0.25 | 0.31 |
| 4 | 142155214 | rs3822128 | 0.511 | -0.29 | -0.099 | -1.27 | -0.056 | 0.028 | 0.028 | 0 | 0.23 | 0.01 | 0.17 | 0.14 | 0.21 | 0.16 |
| 4 | 142155518 | rs3733484 | -0.093 | -0.077 | 0.26 | -1.014 | -0.02 | 0.018 | 0.018 | 0 | 0.31 | 0.52 | 0.26 | 0.62 | 0.24 | 0.3 |
| 4 | 142159511 | rs1435127 | -0.032 | -0.102 | 0.18 | -1.002 | -0.023 | 0.025 | 0.025 | 0 | 0.3 | 0.49 | 0.26 | 0.62 | 0.24 | 0.3 |
| 4 | 142165741 | rs7672861 | -0.711 | 0.134 | -0.095 | -1.075 | -0.032 | 0.049 | 0.049 | 0 | 0.64 | 0.78 | 0.5 | 0.82 | 0.49 | 0.61 |
| 4 | 142166712 | rs6537038 | -0.721 | 0.24 | 0.028 | -0.997 | 0.004 | -0.014 | -0.014 | 0.271 | 0.7 | 0.7 | 0.6 | 0.82 | 0.62 | 0.72 |
| 4 | 142168903 | rs1012864 | 0.285 | -0.083 | -0.528 | -1.237 | -0.019 | -0.054 | -0.054 | 0.012 | 0.13 | 0.23 | 0.06 | 0.01 | 0.04 | 0.07 |
| 4 | 142169704 | rs10002917 | -0.139 | -0.07 | -0.86 | -0.913 | 0.009 | 0.133 | 0.133 | 0.086 | 0.51 | 0.18 | 0.48 | 0.75 | 0.55 | 0.68 |
| 4 | 142177058 | rs978157 | -0.208 | 0.097 | -0.81 | -0.702 | -0.001 | 0.051 | 0.051 | 0 | 0.43 | 0.25 | 0.4 | 0.73 | 0.42 | 0.66 |
| 4 | 142187853 | rs1304390 | -0.522 | 0.282 | -0.947 | -0.632 | 0.005 | 0.003 | 0.003 | 0 | 0.44 | 0.56 | 0.42 | 0.57 | 0.36 | 0.35 |
| 4 | 142188804 | rs354843 | 1.296 | 0.339 | -1.039 | -0.707 | -0.046 | 0.043 | 0.043 | 0 | 0.31 | 0.14 | 0.41 | 0.22 | 0.4 | 0.26 |
| 4 | 142197917 | rs6840028 | -0.664 | 0.296 | -0.921 | -0.616 | 0.007 | -0.011 | -0.011 | 0 | 0.75 | 0.71 | 0.83 | 0.79 | 0.75 | 0.6 |
| 4 | 142198497 | rs13102572 | 0.582 | 0.188 | -0.984 | -0.735 | -0.038 | 0.064 | 0.064 | 0 | 0.35 | 0.2 | 0.46 | 0.22 | 0.45 | 0.28 |
| 4 | 142203334 | rs3114024 | -0.143 | 0.512 | -1.308 | -0.173 | -0.014 | 0.063 | 0.063 | 0 | 0.53 | 0.76 | 0.39 | 0.27 | 0.48 | 0.29 |
| 4 | 142204581 | rs3098027 | -0.162 | 0.5 | -0.666 | 0.174 | -0.016 | 0.113 | 0.113 | 0 | 0.79 | 0.69 | 0.7 | 0.45 | 0.83 | 0.59 |
| 4 | 142212781 | rs354850 | -0.54 | 0.303 | -0.417 | 0.048 | -0.002 | 0 | 0 | 0 | 0.33 | 0.27 | 0.29 | 0.26 | 0.33 | 0.25 |
| 4 | 142218748 | rs6857575 | 0.138 | 0.418 | 0.084 | 0.013 | -0.004 | -0.003 | -0.003 | 0 | 0.36 | 0.39 | 0.22 | 0.26 | 0.36 | 0.36 |
| 4 | 142220324 | rs189564 | 0.711 | 0.603 | 0.178 | -0.154 | -0.03 | 0.151 | 0.151 | 0 | 0.43 | 0.09 | 0.39 | 0.18 | 0.43 | 0.5 |
| 4 | 142230739 | rs6818727 | 0.045 | 0.495 | 0.414 | -0.015 | -0.011 | -0.003 | -0.003 | 0 | 0.38 | 0.51 | 0.24 | 0.32 | 0.38 | 0.36 |
| 4 | 142246430 | rs1369970 | 0.255 | 0.718 | 0.573 | 0.015 | 0.034 | 0.021 | 0.021 | 0 | 0.44 | 0.36 | 0.23 | 0.33 | 0.38 | 0.37 |
| 4 | 142247487 | rs1946731 | -0.287 | 0.848 | 0.981 | 0.021 | 0.029 | -0.04 | -0.04 | 0.231 | 0.11 | 0.45 | 0.14 | 0.08 | 0.06 | 0.04 |
| 4 | 142250415 | rs10002181 | 0.137 | 0.844 | 1.174 | -0.007 | 0.01 | -0.041 | -0.041 | 0 | 0.09 | 0.27 | 0.12 | 0.08 | 0.06 | 0.04 |
| 4 | 142255485 | rs7663070 | -0.349 | 0.843 | 1.175 | -0.008 | 0.029 | -0.045 | -0.045 | 0.004 | 0.11 | 0.45 | 0.14 | 0.08 | 0.06 | 0.04 |
| 4 | 142266475 | rs10003789 | -0.268 | 0.845 | 1.167 | -0.007 | 0.021 | -0.046 | -0.046 | 0 | 0.1 | 0.39 | 0.14 | 0.08 | 0.06 | 0.04 |
| 4 | 142268897 | rs4956372 | 0.129 | 0.871 | 1.002 | 0.006 | 0.011 | -0.041 | -0.041 | 0 | 0.09 | 0.27 | 0.12 | 0.08 | 0.06 | 0.04 |
| 4 | 142275621 | rs746484 | 0.09 | 0.851 | 0.973 | 0.004 | 0.004 | -0.043 | -0.043 | 0 | 0.09 | 0.27 | 0.12 | 0.08 | 0.06 | 0.04 |
| 4 | 142280519 | rs893037 | -1.171 | 1.19 | 0.865 | 0.201 | -0.018 | 0.045 | 0.045 | 0 | 0.94 | 0.97 | 0.78 | 0.62 | 0.87 | 0.9 |
| 4 | 142291367 | rs6838972 | 0.012 | -0.112 | 1.099 | -0.54 | 0.006 | -0.042 | -0.042 | 0 | 0.09 | 0.26 | 0.12 | 0.07 | 0.05 | 0.03 |
| 4 | 142291626 | rs9884442 | -0.157 | -0.112 | 0.841 | -0.54 | 0.029 | -0.035 | -0.035 | 0 | 0.1 | 0.39 | 0.13 | 0.07 | 0.05 | 0.03 |
| 4 | 142296923 | rs6825690 | -0.125 | -0.111 | 0.788 | -0.539 | 0.024 | -0.037 | -0.037 | 1.38 | 0.09 | 0.36 | 0.13 | 0.07 | 0.05 | 0.03 |
| 4 | 142322777 | rs1561152 | -1.16 | 0.043 | -0.013 | -0.56 | -0.054 | -0.01 | -0.01 | 0 | 0.65 | 0.84 | 0.43 | 0.39 | 0.54 | 0.49 |
| 4 | 142326350 | rs17007248 | 0.462 | -0.067 | -0.14 | -0.309 | -0.004 | -0.039 | -0.039 | 0.002 | 0.13 | 0.14 | 0.06 | 0 | 0.09 | 0.06 |
| 4 | 142328977 | rs17007252 | 0.257 | 0.504 | -0.367 | 0.226 | -0.019 | -0.061 | -0.061 | 0 | 0.1 | 0.01 | 0.09 | 0.27 | 0.14 | 0.17 |
| 4 | 142331044 | rs10519598 | 0.276 | 0.439 | -0.246 | 0.048 | -0.002 | -0.07 | -0.07 | 0 | 0.48 | 0.54 | 0.24 | 0.08 | 0.35 | 0.31 |
| 4 | 142331924 | rs6828736 | 0.194 | 0.441 | -0.3 | 0.05 | 0.01 | -0.036 | -0.036 | 3.784 | 0.49 | 0.58 | 0.24 | 0.08 | 0.35 | 0.31 |
| 4 | 142333286 | rs1079115 | -0.344 | 0.503 | -0.637 | 0.529 | 0 | 0.01 | 0.01 | 0 | 0.11 | 0.12 | 0.1 | 0.27 | 0.14 | 0.17 |
| 4 | 142336683 | rs6831004 | -0.123 | 0.282 | -0.536 | -0.548 | -0.005 | -0.075 | -0.075 | 0 | 0.48 | 0.54 | 0.24 | 0.08 | 0.34 | 0.31 |
| 4 | 142352369 | rs1057260 | -0.72 | -0.041 | -0.741 | 0.273 | 0.016 | -0.042 | -0.042 | 0 | 0.34 | 0.23 | 0.47 | 0.49 | 0.32 | 0.34 |
| 4 | 142353425 | rs10519597 | -0.415 | -0.038 | -0.824 | 0.46 | 0.003 | 0.139 | 0.139 | 0 | 0.31 | 0.09 | 0.24 | 0.12 | 0.31 | 0.25 |
| 4 | 142354175 | rs6828927 | -0.525 | -0.053 | -0.793 | 0.457 | 0.018 | 0.111 | 0.111 | 0 | 0.31 | 0.14 | 0.24 | 0.12 | 0.3 | 0.25 |
| 4 | 142376136 | rs12510810 | -0.966 | -0.467 | -0.25 | 0.526 | -0.034 | -0.02 | -0.02 | 0.012 | 0.28 | 0.45 | 0.32 | 0.22 | 0.19 | 0.15 |
| 4 | 142419423 | rs13140249 | -0.676 | -0.499 | -0.23 | 1.144 | 0.001 | 0 | 0 | 0 | 0.25 | 0.26 | 0.29 | 0.22 | 0.19 | 0.16 |
| 4 | 142423973 | rs7693458 | -0.609 | -0.605 | 0.002 | 1.125 | -0.009 | 0.043 | 0.043 | 0 | 0.07 | 0.13 | 0.13 | 0.17 | 0.04 | 0.03 |
| 4 | 142424409 | rs7698747 | -1.323 | -0.657 | -0.135 | 1.11 | -0.04 | -0.032 | -0.032 | 0 | 0.28 | 0.48 | 0.32 | 0.22 | 0.19 | 0.16 |
| 4 | 142432407 | rs12644130 | 1.4 | 0.449 | -0.968 | 0.648 | -0.124 | -0.076 | -0.076 | 0 | 0.2 | 0.02 | 0.26 | 0.22 | 0.37 | 0.38 |
| 4 | 142466804 | rs13132036 | -0.668 | 0.136 | -0.753 | 0.536 | -0.039 | 0.096 | 0.096 | 0 | 0.25 | 0.48 | 0.35 | 0.37 | 0.17 | 0.19 |
| 4 | 142478548 | rs6857587 | -0.795 | -0.366 | -1.126 | -0.492 | -0.009 | 0.117 | 0.117 | 0 | 0.67 | 0.86 | 0.72 | 0.85 | 0.69 | 0.76 |
| 4 | 142484038 | rs1879685 | -0.806 | -0.366 | -1.104 | -0.485 | -0.026 | 0.137 | 0.137 | 0 | 0.67 | 0.91 | 0.72 | 0.85 | 0.69 | 0.76 |
| 4 | 142487363 | rs2322183 | -0.554 | -0.45 | -1.159 | -0.532 | -0.011 | 0.14 | 0.14 | 0 | 0.63 | 0.85 | 0.71 | 0.85 | 0.66 | 0.76 |
| 4 | 142501053 | rs4371699 | -1.738 | -0.732 | -1.365 | -0.802 | 0.009 | 0.064 | 0.064 | 0 | 0.82 | 0.94 | 0.87 | 0.99 | 0.79 | 0.89 |
| 4 | 142533402 | rs6835391 | -0.456 | -0.721 | -1.222 | -0.89 | 0.007 | 0.042 | 0.042 | 0 | 0.7 | 0.8 | 0.8 | 0.99 | 0.72 | 0.81 |
| 4 | 142545451 | rs2139383 | 0.716 | -0.354 | -1.021 | -1.015 | 0 | -0.015 | -0.015 | 0 | 0.43 | 0.46 | 0.53 | 0.87 | 0.47 | 0.65 |
| 4 | 142570472 | rs1519551 | 0.247 | -0.339 | -0.864 | -0.919 | 0.002 | 0.004 | 0.004 | 0.363 | 0.53 | 0.46 | 0.5 | 0.85 | 0.55 | 0.75 |
| 4 | 142575266 | rs2322262 | -0.657 | -0.338 | -0.869 | -0.951 | 0.001 | -0.006 | -0.006 | 0 | 0.7 | 0.74 | 0.8 | 0.99 | 0.71 | 0.81 |
| 4 | 142579816 | rs957077 | 0.902 | 0.06 | -1.044 | -0.782 | -0.002 | 0.014 | 0.014 | 0 | 0.12 | 0.01 | 0.26 | 0.08 | 0.1 | 0.07 |
| 4 | 142590103 | rs12504148 | -0.029 | 0.076 | -0.964 | -1.08 | -0.009 | 0.001 | 0.001 | 0 | 0.12 | 0.07 | 0.07 | 0.06 | 0.13 | 0.08 |
| 4 | 142601496 | rs12510514 | 0.768 | 0.078 | -1.003 | -1.08 | -0.045 | 0.002 | 0.002 | 0 | 0.09 | 0 | 0.05 | 0.04 | 0.11 | 0.07 |
| 4 | 142635648 | rs1607595 | -0.391 | 0.024 | -0.879 | -1.108 | 0.014 | -0.008 | -0.008 | 0 | 0.14 | 0.23 | 0.09 | 0.06 | 0.13 | 0.08 |
| 4 | 142636313 | rs2322301 | 0.949 | -0.182 | -0.997 | -0.975 | 0.004 | 0.004 | 0.004 | 0 | 0.41 | 0.47 | 0.41 | 0.53 | 0.46 | 0.62 |
| 4 | 142637101 | rs10519610 | -0.472 | -0.179 | -0.994 | -0.983 | 0.02 | -0.013 | -0.013 | 0 | 0.15 | 0.32 | 0.09 | 0.06 | 0.14 | 0.08 |

| | | | | | | | | | | | | | | | | |
|---|---|---|---|---|---|---|---|---|---|---|---|---|---|---|---|---|
| 4 | 142637305 | rs6850492 | -0.295 | -0.184 | -0.767 | -0.696 | -0.007 | 0.09 | 0.09 | 0 | 0.56 | 0.7 | 0.74 | 0.93 | 0.57 | 0.74 |
| 4 | 142649655 | rs3775597 | 0.533 | -0.147 | -0.84 | -0.647 | -0.001 | 0.036 | 0.036 | 0 | 0.13 | 0.12 | 0.33 | 0.4 | 0.11 | 0.12 |
| 4 | 142650413 | rs13122930 | 0.481 | -0.108 | -0.852 | -0.643 | 0 | 0 | 0 | 0 | 0.42 | 0.57 | 0.41 | 0.53 | 0.46 | 0.62 |
| 4 | 142653701 | rs10519612 | 0.799 | -0.08 | -1.114 | -0.495 | -0.004 | -0.074 | -0.074 | 0 | 0.12 | 0.02 | 0.33 | 0.4 | 0.11 | 0.11 |
| 4 | 142654547 | rs10833 | -1.18 | -0.171 | -1.241 | -0.704 | -0.03 | 0.109 | 0.109 | 0.623 | 0.69 | 0.91 | 0.79 | 0.93 | 0.65 | 0.82 |
| 4 | 142657057 | rs1493025 | 0.465 | -0.018 | -1.018 | -0.647 | 0.002 | 0.002 | 0.002 | 0 | 0.42 | 0.55 | 0.41 | 0.53 | 0.46 | 0.62 |
| 4 | 142675412 | rs9990563 | -0.516 | 0.076 | -1.205 | -0.384 | -0.004 | -0.084 | -0.084 | 0 | 0.23 | 0.08 | 0.36 | 0.4 | 0.18 | 0.2 |
| 4 | 142677084 | rs10004459 | 0.602 | 0.542 | -0.909 | -0.484 | -0.005 | 0.025 | 0.025 | 0 | 0.11 | 0.06 | 0.04 | 0 | 0.08 | 0.09 |
| 4 | 142682770 | rs10519629 | 0.845 | 0.483 | -0.872 | -0.425 | 0.006 | 0.004 | 0.004 | 0.023 | 0.18 | 0.08 | 0.11 | 0.21 | 0.16 | 0.23 |
| 4 | 142686873 | rs9884645 | 0.535 | 0.57 | -0.822 | -0.55 | 0 | -0.004 | -0.004 | 0.501 | 0.13 | 0.1 | 0.04 | 0 | 0.09 | 0.12 |
| 4 | 142687795 | rs4956410 | 0.081 | 0.543 | -0.678 | -0.386 | -0.001 | 0.155 | 0.155 | 0 | 0.12 | 0.23 | 0.37 | 0.46 | 0.09 | 0.14 |
| 4 | 142688390 | rs4254850 | 0.322 | 0.636 | -0.587 | -0.519 | 0.014 | 0.011 | 0.011 | 0 | 0.37 | 0.48 | 0.35 | 0.44 | 0.45 | 0.57 |
| 4 | 142688981 | rs7663689 | 0.852 | 0.444 | -0.882 | -0.638 | 0 | 0.008 | 0.008 | 0 | 0.21 | 0.08 | 0.18 | 0.28 | 0.19 | 0.26 |
| 4 | 142691963 | rs1493015 | -1.217 | 0.366 | -0.843 | -0.663 | 0.007 | 0.082 | 0.082 | 0.6 | 0.64 | 0.81 | 0.77 | 0.91 | 0.64 | 0.82 |
| 4 | 142696583 | rs7680696 | -1.322 | 0.31 | -1.02 | -0.587 | -0.061 | 0.141 | 0.141 | 0 | 0.64 | 0.89 | 0.79 | 0.91 | 0.62 | 0.77 |
| 4 | 142699341 | rs9996010 | -0.444 | 0.169 | -0.724 | -0.487 | -0.036 | 0.089 | 0.089 | 0 | 0.23 | 0.38 | 0.36 | 0.4 | 0.14 | 0.22 |
| 4 | 142705418 | rs1389098 | 0.164 | 0.208 | -0.703 | -0.479 | 0.004 | 0.003 | 0.003 | 0 | 0.41 | 0.45 | 0.42 | 0.52 | 0.48 | 0.55 |
| 4 | 142710442 | rs1493023 | 1.565 | 0.381 | -0.659 | -0.376 | 0.044 | 0.052 | 0.052 | 0 | 0.18 | 0.36 | 0.22 | 0.24 | 0.26 | 0.29 |
| 4 | 142711243 | rs17701271 | 0.474 | 0.429 | -1.078 | -0.541 | 0.009 | 0.026 | 0.026 | 0 | 0.26 | 0.13 | 0.21 | 0.29 | 0.24 | 0.27 |
| 4 | 142715747 | rs6537072 | -1.368 | 0.334 | -1.103 | -0.575 | -0.05 | 0.118 | 0.118 | 0.468 | 0.67 | 0.86 | 0.79 | 0.92 | 0.63 | 0.78 |
| 4 | 142715821 | rs10013727 | 0.234 | 0.492 | -0.604 | -0.198 | -0.007 | -0.048 | -0.048 | 2.073 | 0.12 | 0.16 | 0.04 | 0 | 0.07 | 0.09 |
| 4 | 142723797 | rs4956412 | 0.875 | 0.754 | -0.72 | 0.029 | 0.05 | 0.026 | 0.026 | 0 | 0.23 | 0.43 | 0.23 | 0.24 | 0.3 | 0.28 |
| 4 | 142725468 | rs12511035 | 1.014 | 0.903 | -0.964 | 0.209 | -0.011 | 0.01 | 0.01 | 0 | 0.1 | 0.06 | 0.4 | 0.46 | 0.12 | 0.15 |
| 4 | 142728396 | rs7691866 | -0.67 | 0.648 | -0.947 | -0.296 | 0.01 | 0.055 | 0.055 | 0.045 | 0.46 | 0.64 | 0.71 | 0.75 | 0.51 | 0.55 |
| 4 | 142728951 | rs10018731 | 0.068 | 0.416 | -1.151 | -0.625 | 0.011 | 0.037 | 0.037 | 0.213 | 0.28 | 0.21 | 0.13 | 0.12 | 0.24 | 0.29 |
| 4 | 142729438 | rs17015015 | -1.094 | 0.394 | -1 | -0.588 | -0.003 | -0.002 | -0.002 | 0 | 0.45 | 0.54 | 0.45 | 0.64 | 0.38 | 0.49 |
| 4 | 142732715 | rs10033645 | -0.839 | 0.605 | -1.106 | -0.483 | 0.012 | -0.032 | -0.032 | 0.005 | 0.6 | 0.76 | 0.56 | 0.49 | 0.62 | 0.71 |
| 4 | 142733937 | rs4956305 | -1.71 | 0.588 | -1.091 | -0.509 | 0.032 | 0.052 | 0.052 | 1.573 | 0.72 | 0.83 | 0.87 | 0.99 | 0.78 | 0.89 |
| 4 | 142746336 | rs12506931 | 0.249 | 0.924 | -0.782 | 0.603 | -0.002 | 0.001 | 0.001 | 0 | 0.07 | 0.08 | 0.06 | 0.11 | 0.09 | 0.11 |
| 4 | 142762276 | rs10009998 | 0.6 | 0.908 | -0.824 | 0.581 | -0.019 | 0.041 | 0.041 | 0 | 0.08 | 0.02 | 0.03 | 0 | 0.06 | 0.09 |
| 4 | 142763570 | rs2099884 | 0.408 | 0.941 | -0.768 | 0.689 | -0.018 | -0.019 | -0.019 | 0.226 | 0.12 | 0.07 | 0.31 | 0.5 | 0.15 | 0.18 |
| 4 | 142775313 | rs6823281 | -0.012 | 0.965 | -0.774 | 0.675 | 0.011 | 0.117 | 0.117 | 0 | 0.15 | 0.25 | 0.32 | 0.52 | 0.15 | 0.2 |
| 4 | 142785171 | rs12502051 | 0.364 | 0.842 | -0.952 | 0.83 | -0.008 | -0.008 | -0.008 | 0 | 0.06 | 0.05 | 0.06 | 0.11 | 0.09 | 0.11 |
| 4 | 142792211 | rs17015119 | -0.531 | 0.48 | -0.744 | 0.696 | 0.003 | 0.042 | 0.042 | 0.005 | 0.26 | 0.38 | 0.36 | 0.52 | 0.22 | 0.3 |
| 4 | 142797440 | rs11944032 | -0.534 | 0.483 | -0.742 | 0.706 | 0.002 | 0.042 | 0.042 | 0.015 | 0.27 | 0.37 | 0.36 | 0.52 | 0.22 | 0.3 |
| 4 | 142808503 | rs12503750 | -0.645 | 0.622 | -0.66 | 0.564 | -0.001 | 0 | 0 | 0 | 0.42 | 0.46 | 0.45 | 0.63 | 0.38 | 0.49 |
| 4 | 142811058 | rs17015143 | -1.145 | 0.177 | 0.131 | 0.68 | -0.037 | 0.024 | 0.024 | 0.019 | 0.17 | 0.51 | 0.12 | 0.24 | 0.11 | 0.11 |
| 4 | 142812073 | rs1500840 | 1.143 | 0.371 | -1.003 | 0.588 | -0.064 | -0.093 | -0.093 | 0 | 0.42 | 0.11 | 0.45 | 0.5 | 0.51 | 0.48 |
| 4 | 142820445 | rs713281 | -1.304 | 0.458 | -1.046 | 0.395 | 0.01 | -0.018 | -0.018 | 0 | 0.34 | 0.68 | 0.36 | 0.41 | 0.27 | 0.33 |
| 4 | 142833862 | rs1392773 | -1.432 | 0.58 | -1.009 | 0.413 | 0.014 | -0.048 | -0.048 | 0 | 0.34 | 0.8 | 0.37 | 0.41 | 0.27 | 0.32 |
| 4 | 142834152 | rs9993798 | -0.214 | 0.552 | -0.878 | 0.561 | 0.005 | 0.007 | 0.007 | 0 | 0.15 | 0.16 | 0.23 | 0.14 | 0.16 | 0.21 |
| 4 | 142836161 | rs2133926 | 1.705 | 0.527 | -0.464 | 0.466 | -0.007 | -0.04 | -0.04 | 0.537 | 0.43 | 0.23 | 0.47 | 0.51 | 0.5 | 0.49 |
| 4 | 142849228 | rs11731062 | 1.516 | 0.807 | -0.368 | 0.307 | -0.001 | 0.013 | 0.013 | 0 | 0.49 | 0.42 | 0.52 | 0.56 | 0.55 | 0.53 |
| 4 | 142853017 | rs1392779 | -0.508 | 0.995 | -0.376 | 0.175 | -0.007 | 0.056 | 0.056 | 0 | 0.8 | 0.95 | 0.86 | 0.92 | 0.83 | 0.85 |
| 4 | 142874553 | rs6537091 | 0.828 | 0.908 | 0.044 | 0.098 | -0.016 | 0.037 | 0.037 | 0 | 0.65 | 0.77 | 0.64 | 0.82 | 0.67 | 0.62 |
| 4 | 142880833 | rs4455499 | 0.839 | 0.896 | 0.081 | 0.093 | -0.007 | 0.021 | 0.021 | 0 | 0.64 | 0.7 | 0.64 | 0.82 | 0.67 | 0.62 |
| 4 | 142898552 | rs1604236 | -0.433 | 0.288 | 0.709 | -0.033 | -0.001 | 0.012 | 0.012 | 0 | 0.23 | 0.15 | 0.19 | 0.01 | 0.22 | 0.15 |
| 4 | 142916958 | rs1995960 | 2.336 | 0.079 | 0.549 | 0.208 | 0.055 | 0.033 | 0.033 | 0.003 | 0.5 | 0.08 | 0.51 | 0.64 | 0.63 | 0.52 |
| 4 | 142917074 | rs1995959 | 2.365 | -0.239 | 1.197 | 0.668 | 0.044 | -0.039 | -0.039 | 2.788 | 0.72 | 0.17 | 0.68 | 0.65 | 0.84 | 0.71 |
| 4 | 142944080 | rs11736202 | 2.262 | -0.221 | 1.356 | 0.409 | 0.025 | -0.014 | -0.014 | 0 | 0.73 | 0.25 | 0.79 | 0.67 | 0.84 | 0.7 |
| 4 | 142945451 | rs336361 | 0.233 | -0.162 | 1.45 | 0.456 | 0 | 0.002 | 0.002 | 0 | 0.92 | 0.91 | 0.9 | 0.9 | 0.92 | 0.83 |
| 4 | 142952946 | rs336355 | -0.806 | -1.266 | 1.816 | 0.414 | -0.006 | -0.027 | -0.027 | 0 | 0.05 | 0.17 | 0.04 | 0 | 0.03 | 0.02 |
| 4 | 142954285 | rs4245935 | 1.818 | -1.29 | 2.071 | 0.408 | -0.054 | 0.035 | 0.035 | 0.993 | 0.83 | 0.41 | 0.9 | 0.76 | 0.93 | 0.87 |
| 4 | 142955593 | rs336350 | 0.85 | -1.124 | 1.627 | 0.526 | -0.025 | 0.073 | 0.073 | 0.579 | 0.9 | 0.74 | 0.96 | 0.76 | 0.96 | 0.89 |
| 4 | 142964158 | rs2645810 | 1.097 | -1.311 | 1.283 | 0.078 | -0.07 | -0.073 | -0.073 | 0.505 | 0.77 | 0.49 | 0.88 | 0.91 | 0.91 | 0.9 |
| 4 | 142965814 | rs2645813 | 0.746 | -1.376 | 1.609 | 0.046 | -0.062 | -0.058 | -0.058 | 0.283 | 0.81 | 0.55 | 0.9 | 0.91 | 0.92 | 0.9 |
| 4 | 142968081 | rs3775605 | 1.815 | -2.189 | 1.101 | 0.168 | 0.149 | 0.012 | 0.012 | 4.521 | 0.6 | 0.13 | 0.75 | 0.66 | 0.84 | 0.79 |
| 4 | 142969773 | rs2667096 | 2.4 | -1.998 | 0.837 | 0.16 | 0.138 | -0.006 | -0.006 | 0.349 | 0.49 | 0.12 | 0.63 | 0.56 | 0.7 | 0.55 |
| 4 | 142974842 | rs1511252 | 0.583 | -1.782 | 1.515 | 0.25 | -0.073 | -0.07 | -0.07 | 0 | 0.8 | 0.5 | 0.89 | 0.91 | 0.92 | 0.9 |
| 4 | 142976073 | rs2667100 | 0.563 | -1.765 | 1.524 | 0.259 | -0.069 | -0.065 | -0.065 | 0 | 0.81 | 0.5 | 0.89 | 0.91 | 0.92 | 0.9 |
| 4 | 142984889 | rs4956433 | 0.711 | -1.669 | 1.244 | 0.264 | 0.012 | 0.018 | 0.018 | 0 | 0.11 | 0.06 | 0.07 | 0.02 | 0.07 | 0.05 |
| 4 | 142994344 | rs12501475 | 0.383 | -1.546 | 1.398 | 0.363 | -0.003 | 0.004 | 0.004 | 4.787 | 0.14 | 0.14 | 0.08 | 0.25 | 0.06 | 0.08 |
| 4 | 143013388 | rs336292 | 1.594 | -1.661 | 1.52 | 0.294 | 0.025 | -0.011 | -0.011 | 0 | 0.66 | 0.32 | 0.76 | 0.63 | 0.78 | 0.67 |
| 4 | 143014875 | rs336297 | 2.026 | -1.759 | 1.389 | 0.215 | 0.099 | -0.02 | -0.02 | 0 | 0.55 | 0.17 | 0.68 | 0.57 | 0.69 | 0.62 |
| 4 | 143019415 | rs336304 | 0.736 | -1.677 | 1.684 | 0.223 | 0.071 | 0.117 | 0.117 | 0.26 | 0.94 | 0.59 | 0.97 | 0.93 | 0.99 | 0.98 |
| 4 | 143020388 | rs17468315 | -0.303 | -1.617 | 1.119 | 0.206 | -0.037 | 0.003 | 0.003 | 0 | 0.1 | 0 | 0.11 | 0.08 | 0.14 | 0.22 |
| 4 | 143045599 | rs978752 | -0.938 | -1.624 | 1.233 | 0.216 | 0.001 | 0.007 | 0.007 | 0 | 0.21 | 0.21 | 0.18 | 0.13 | 0.21 | 0.25 |
| 4 | 143049620 | rs3775648 | 0.439 | -1.701 | 1.02 | 0.335 | 0.025 | -0.037 | -0.037 | 2.71 | 0.13 | 0.06 | 0.08 | 0.23 | 0.06 | 0.08 |
| 4 | 143049980 | rs3775652 | -0.72 | -1.73 | 1.1 | 0.316 | -0.012 | 0.009 | 0.009 | 1.467 | 0.18 | 0.21 | 0.12 | 0.24 | 0.11 | 0.09 |

| | | | | | | | | | | | | | | | | |
|---|---|---|---|---|---|---|---|---|---|---|---|---|---|---|---|---|
| 4 | 143050454 | rs3775654 | -0.299 | -1.704 | 0.846 | 0.322 | 0.014 | -0.015 | -0.015 | 0.597 | 0.17 | 0.11 | 0.1 | 0.24 | 0.09 | 0.09 |
| 4 | 143052584 | rs3775658 | -0.281 | -1.703 | 0.758 | 0.329 | 0.025 | -0.026 | -0.026 | 0.004 | 0.17 | 0.1 | 0.1 | 0.23 | 0.09 | 0.09 |
| 4 | 143054921 | rs336407 | 1.842 | -2.026 | 0.308 | 0.343 | 0.136 | -0.092 | -0.092 | 0 | 0.57 | 0.07 | 0.7 | 0.54 | 0.73 | 0.77 |
| 4 | 143055538 | rs336408 | 1.004 | -2.004 | 1.402 | -0.483 | -0.056 | -0.051 | -0.051 | 0.086 | 0.67 | 0.45 | 0.82 | 0.8 | 0.82 | 0.84 |
| 4 | 143064439 | rs13148456 | -0.756 | -2.051 | 1.109 | -0.573 | -0.01 | 0.016 | 0.016 | 1.434 | 0.12 | 0.01 | 0.11 | 0.06 | 0.12 | 0.22 |
| 4 | 143066199 | rs10519631 | 0.749 | -2.113 | 1.504 | -0.578 | -0.03 | -0.022 | -0.022 | 0 | 0.13 | 0.16 | 0.06 | 0.06 | 0.05 | 0.07 |
| 4 | 143076406 | rs336391 | 2.181 | -2.058 | 1.456 | -0.605 | 0.136 | 0.149 | 0.149 | 0.802 | 0.55 | 0.1 | 0.69 | 0.73 | 0.71 | 0.61 |
| 4 | 143077401 | rs336394 | 1.102 | -2.039 | 1.772 | -0.614 | 0.05 | 0.058 | 0.058 | 1.386 | 0.67 | 0.21 | 0.81 | 0.9 | 0.82 | 0.86 |
| 4 | 143080615 | rs11724936 | 0.576 | -1.995 | 1.793 | -0.591 | 0.007 | 0.031 | 0.031 | 0.007 | 0.14 | 0.09 | 0.09 | 0.07 | 0.12 | 0.07 |
| 4 | 143089849 | rs336384 | 1.245 | -2.248 | 2.108 | -0.635 | -0.015 | -0.003 | -0.003 | 0 | 0.66 | 0.35 | 0.82 | 0.76 | 0.83 | 0.81 |
| 4 | 143094698 | rs1390993 | 0.421 | -2.249 | 2.182 | -0.638 | -0.049 | -0.059 | -0.059 | 0 | 0.15 | 0.24 | 0.06 | 0.03 | 0.05 | 0.07 |
| 4 | 143096594 | rs1219273 | -0.14 | -2.255 | 2.256 | -0.638 | -0.024 | -0.001 | -0.001 | 0 | 0.83 | 0.75 | 0.9 | 0.79 | 0.89 | 0.88 |
| 4 | 143098083 | rs2636683 | 2.206 | -2.25 | 2.2 | -0.574 | -0.025 | -0.016 | -0.016 | 0 | 0.57 | 0.49 | 0.72 | 0.69 | 0.71 | 0.57 |
| 4 | 143105706 | rs2055212 | -1.679 | -2.219 | 2.12 | -0.556 | -0.012 | 0.016 | 0.016 | 0.034 | 0.15 | 0.18 | 0.14 | 0.21 | 0.13 | 0.3 |
| 4 | 143114548 | rs992257 | 1.521 | -2.261 | 2.116 | -0.717 | 0.156 | 0.1 | 0.1 | 5.462 | 0.62 | 0.1 | 0.8 | 0.76 | 0.83 | 0.81 |
| 4 | 143138844 | rs2627804 | 2.924 | -1.951 | 1.192 | -1.355 | -0.122 | 0.017 | 0.017 | 0.276 | 0.38 | 0.03 | 0.55 | 0.7 | 0.5 | 0.46 |
| 4 | 143153264 | rs13109869 | -1.56 | -2.279 | 1.848 | -0.869 | -0.011 | 0.014 | 0.014 | 0.014 | 0.15 | 0.18 | 0.13 | 0.24 | 0.12 | 0.3 |
| 4 | 143156127 | rs2627801 | 0.687 | -2.275 | 1.802 | -0.89 | -0.041 | -0.009 | -0.009 | 0.123 | 0.79 | 0.38 | 0.89 | 0.78 | 0.93 | 0.85 |
| 4 | 143208349 | rs2627813 | 2.433 | -2.189 | 1.35 | -0.86 | 0.167 | 0.157 | 0.157 | 1.255 | 0.51 | 0.08 | 0.69 | 0.72 | 0.7 | 0.54 |
| 4 | 143211964 | rs2200997 | -0.715 | -2.014 | 1.57 | -0.592 | -0.005 | 0.037 | 0.037 | 0.132 | 0.06 | 0.09 | 0.03 | 0.19 | 0.03 | 0.06 |
| 4 | 143214649 | rs2636632 | 0.861 | -2.02 | 1.412 | -0.668 | 0.006 | -0.004 | -0.004 | 0.35 | 0.72 | 0.7 | 0.86 | 0.95 | 0.84 | 0.85 |
| 4 | 143217391 | rs2172023 | 0.75 | -1.946 | 0.923 | -0.628 | -0.07 | -0.102 | -0.102 | 3.465 | 0.7 | 0.49 | 0.85 | 0.95 | 0.84 | 0.85 |
| 4 | 143220613 | rs10002445 | -0.417 | -1.056 | 0.703 | -0.021 | -0.002 | 0.001 | 0.001 | 0 | 0.07 | 0.09 | 0.11 | 0.07 | 0.08 | 0.19 |
| 4 | 143224296 | rs3822152 | -1.992 | -1.165 | 0.855 | 0.025 | -0.01 | 0.083 | 0.083 | 0.628 | 0.14 | 0.37 | 0.17 | 0.27 | 0.12 | 0.26 |
| 4 | 143224971 | rs3775692 | -1.746 | -1.185 | -0.394 | -0.008 | 0.082 | 0.013 | 0.013 | 5.465 | 0.3 | 0.82 | 0.25 | 0.3 | 0.17 | 0.34 |
| 4 | 143226496 | rs17015882 | -0.599 | -0.115 | 0.102 | 0.226 | -0.001 | 0 | 0 | 0 | 0.06 | 0.06 | 0.11 | 0.08 | 0.08 | 0.2 |
| 4 | 143227742 | rs1907108 | 1.739 | -0.101 | 0.618 | 0.034 | 0.016 | -0.027 | -0.027 | 0.973 | 0.85 | 0.64 | 0.76 | 0.92 | 0.82 | 0.79 |
| 4 | 143228914 | rs3775696 | -0.788 | 0.113 | 0.368 | -0.265 | 0 | 0.018 | 0.018 | 0 | 0.29 | 0.21 | 0.2 | 0.2 | 0.27 | 0.29 |
| 4 | 143229634 | rs1907107 | 1.43 | 0.274 | -0.334 | -0.495 | -0.059 | 0.002 | 0.002 | 0 | 0.37 | 0.08 | 0.47 | 0.63 | 0.45 | 0.52 |
| 4 | 143230028 | rs2636638 | 0.615 | 0.191 | 0.315 | -0.611 | -0.001 | -0.005 | -0.005 | 0 | 0.51 | 0.49 | 0.61 | 0.73 | 0.63 | 0.62 |
| 4 | 143232702 | rs1872294 | -0.535 | 0.127 | 0.656 | -0.597 | -0.014 | -0.062 | -0.062 | 0 | 0.28 | 0.53 | 0.29 | 0.19 | 0.23 | 0.15 |
| 4 | 143234593 | rs2636670 | 0.426 | 0.017 | 1.009 | -0.218 | 0.013 | -0.004 | -0.004 | 0.003 | 0.63 | 0.55 | 0.61 | 0.41 | 0.6 | 0.57 |
| 4 | 143234663 | rs10024947 | 0.688 | -0.004 | 1.158 | -0.213 | -0.038 | -0.045 | -0.045 | 0.743 | 0.18 | 0.31 | 0.18 | 0.07 | 0.09 | 0.05 |
| 4 | 143234955 | rs2636671 | 1.446 | -0.054 | 0.95 | -0.252 | -0.13 | 0.024 | 0.024 | 0 | 0.41 | 0.05 | 0.4 | 0.34 | 0.5 | 0.51 |
| 4 | 143236158 | rs960678 | 0.53 | -0.025 | 1.176 | -0.172 | 0.059 | 0.014 | 0.014 | 0 | 0.1 | 0.27 | 0.1 | 0.13 | 0.15 | 0.09 |
| 4 | 143240651 | rs17015920 | 0.66 | -0.049 | 0.912 | -0.22 | 0.005 | 0.003 | 0.003 | 0.063 | 0.18 | 0.18 | 0.17 | 0.06 | 0.09 | 0.07 |
| 4 | 143243556 | rs2635429 | 1.601 | 0.446 | 1.018 | -0.01 | -0.093 | 0.023 | 0.023 | 0 | 0.32 | 0.02 | 0.34 | 0.24 | 0.39 | 0.45 |
| 4 | 143247240 | rs6844681 | 0.443 | 0.357 | 1.197 | -0.001 | 0 | 0 | 0 | 0.012 | 0.47 | 0.46 | 0.46 | 0.44 | 0.51 | 0.52 |
| 4 | 143257940 | rs2646091 | 1.688 | 0.504 | 0.63 | -0.04 | -0.097 | 0.016 | 0.016 | 0 | 0.29 | 0.02 | 0.23 | 0.23 | 0.35 | 0.41 |
| 4 | 143259305 | rs7666932 | -0.665 | 0.422 | 0.885 | 0.038 | 0 | 0.001 | 0.001 | 0 | 0.37 | 0.38 | 0.38 | 0.61 | 0.37 | 0.39 |
| 4 | 143268702 | rs1908968 | 0.267 | 0.46 | 0.55 | 0.08 | -0.065 | -0.142 | -0.142 | 1.153 | 0.25 | 0.03 | 0.27 | 0.45 | 0.25 | 0.33 |
| 4 | 143272497 | rs716762 | 1.333 | 0.413 | 0.571 | 0.038 | -0.061 | 0.069 | 0.069 | 0 | 0.34 | 0.09 | 0.26 | 0.23 | 0.39 | 0.48 |
| 4 | 143286268 | rs10519644 | 0.854 | 0.425 | 0.633 | 0.041 | 0 | -0.001 | -0.001 | 0.041 | 0.1 | 0.1 | 0.09 | 0.16 | 0.12 | 0.06 |
| 4 | 143290199 | rs11938760 | -0.413 | 0.435 | 0.627 | 0.004 | -0.008 | -0.038 | -0.038 | 0 | 0.7 | 0.57 | 0.65 | 0.84 | 0.75 | 0.81 |
| 4 | 143297732 | rs17016027 | 0.762 | 0.363 | 0.522 | -0.066 | 0 | -0.002 | -0.002 | 0 | 0.24 | 0.26 | 0.31 | 0.14 | 0.19 | 0.11 |
| 4 | 143304592 | rs1497389 | 1.628 | 0.302 | 0.459 | 0.185 | -0.08 | -0.028 | -0.028 | 0 | 0.29 | 0.02 | 0.23 | 0.29 | 0.35 | 0.42 |
| 4 | 143305002 | rs11930321 | 0.278 | 0.286 | 0.647 | 0.208 | -0.001 | 0.014 | 0.014 | 0 | 0.14 | 0.38 | 0.12 | 0.15 | 0.14 | 0.06 |
| 4 | 143309895 | rs1391097 | -0.972 | 0.281 | 0.523 | 0.219 | -0.004 | -0.038 | -0.038 | 0 | 0.73 | 0.61 | 0.65 | 0.86 | 0.77 | 0.87 |
| 4 | 143318673 | rs10006795 | 0.115 | 0.302 | 0.418 | 0.247 | 0.021 | 0.085 | 0.085 | 0.034 | 0.59 | 0.17 | 0.53 | 0.7 | 0.63 | 0.82 |
| 4 | 143324036 | rs1982965 | 0.055 | 0.318 | 0.185 | 0.225 | 0.02 | 0.093 | 0.093 | 0.033 | 0.59 | 0.16 | 0.53 | 0.7 | 0.63 | 0.82 |
| 4 | 143324094 | rs1982966 | 1.349 | 0.388 | 0.426 | 0.116 | -0.085 | -0.028 | -0.028 | 0 | 0.3 | 0.03 | 0.24 | 0.29 | 0.35 | 0.42 |
| 4 | 143331154 | rs1489577 | 0.956 | 0.579 | 0.557 | 0.238 | -0.014 | 0.024 | 0.024 | 0 | 0.34 | 0.18 | 0.25 | 0.3 | 0.38 | 0.49 |
| 4 | 143339542 | rs1497388 | -0.535 | 0.648 | 0.597 | 0.409 | -0.004 | -0.04 | -0.04 | 0 | 0.73 | 0.59 | 0.65 | 0.87 | 0.76 | 0.88 |
| 4 | 143343605 | rs6820463 | 1.156 | 0.716 | 0.552 | 0.437 | 0.001 | -0.004 | -0.004 | 0 | 0.28 | 0.24 | 0.16 | 0.04 | 0.29 | 0.27 |
| 4 | 143343632 | rs12499068 | -0.179 | 0.787 | -0.037 | 0.488 | -0.004 | -0.004 | -0.004 | 0 | 0.37 | 0.3 | 0.41 | 0.57 | 0.4 | 0.38 |
| 4 | 143346040 | rs1497391 | -0.501 | 0.884 | -0.293 | 1.357 | 0 | -0.004 | -0.004 | 0 | 0.34 | 0.29 | 0.44 | 0.4 | 0.32 | 0.34 |
| 4 | 143360851 | rs979511 | -1.116 | 0.878 | 0.296 | 2.077 | -0.004 | 0.068 | 0.068 | 0 | 0.13 | 0.19 | 0.38 | 0.39 | 0.11 | 0.23 |
| 4 | 143369780 | rs1489581 | -1.897 | 0.457 | 1.079 | 1.843 | -0.056 | 0.105 | 0.105 | 0 | 0.25 | 0.47 | 0.45 | 0.42 | 0.15 | 0.29 |
| 4 | 143373910 | rs1391099 | -2.12 | 0.367 | 1.202 | 1.798 | -0.014 | 0.014 | 0.014 | 0 | 0.27 | 0.68 | 0.46 | 0.42 | 0.16 | 0.28 |
| 4 | 143375877 | rs1013397 | -0.951 | 0.335 | 1.231 | 1.932 | -0.003 | 0.031 | 0.031 | 0 | 0.14 | 0.16 | 0.28 | 0.35 | 0.1 | 0.19 |
| 4 | 143393572 | rs11930849 | -0.266 | 0.329 | 1.364 | 1.907 | -0.001 | -0.006 | -0.006 | 0 | 0.07 | 0.05 | 0.05 | 0.01 | 0.04 | 0.06 |
| 4 | 143394613 | rs3913163 | -2.136 | 0.315 | 1.437 | 2.214 | -0.017 | 0.015 | 0.015 | 0 | 0.23 | 0.3 | 0.44 | 0.54 | 0.16 | 0.3 |
| 4 | 143397125 | rs7695918 | -1.316 | 0.32 | 1.263 | 2.054 | -0.019 | 0.074 | 0.074 | 0 | 0.16 | 0.26 | 0.39 | 0.41 | 0.11 | 0.23 |
| 4 | 143414537 | rs1593360 | -2.701 | 1.397 | 0.83 | 1.91 | -0.024 | 0.002 | 0.002 | 0 | 0.36 | 0.74 | 0.62 | 0.53 | 0.41 | 0.4 |
| 4 | 143418354 | rs17016248 | -0.735 | 0.193 | 1.137 | 2.021 | 0.003 | -0.005 | -0.005 | 0 | 0.13 | 0.1 | 0.28 | 0.36 | 0.14 | 0.2 |
| 4 | 143421175 | rs1425533 | -1.856 | 0.225 | 1.103 | 1.946 | 0 | 0.007 | 0.007 | 0 | 0.21 | 0.24 | 0.34 | 0.42 | 0.18 | 0.26 |
| 4 | 143426031 | rs1425531 | -2.216 | 0.197 | 0.849 | 1.453 | -0.012 | 0.015 | 0.015 | 0 | 0.27 | 0.69 | 0.47 | 0.36 | 0.19 | 0.29 |
| 4 | 143437219 | rs1364920 | -1.223 | 0.763 | 0.087 | 1.446 | -0.002 | 0.019 | 0.019 | 0 | 0.14 | 0.17 | 0.29 | 0.29 | 0.14 | 0.2 |
| 4 | 143452535 | rs10857395 | 1.444 | 0.645 | 1.418 | 0.528 | -0.003 | 0.024 | 0.024 | 0.649 | 0.86 | 0.47 | 0.85 | 0.87 | 0.89 | 0.8 |

| | | | | | | | | | | | | | | | | |
|---|---|---|---|---|---|---|---|---|---|---|---|---|---|---|---|---|
| 4 | 143474601 | rs1425520 | 0.01 | 0.361 | 0.113 | -0.922 | 0.031 | 0.232 | 0.232 | 0 | 0.52 | 0.14 | 0.43 | 0.18 | 0.64 | 0.6 |
| 4 | 143476181 | rs6814775 | 0.28 | -0.036 | 0.592 | -1.062 | 0 | 0.002 | 0.002 | 0 | 0.31 | 0.32 | 0.44 | 0.78 | 0.24 | 0.28 |
| 4 | 143488095 | rs1473222 | 0.988 | -0.507 | -0.47 | -1.663 | -0.022 | 0.114 | 0.114 | 0 | 0.58 | 0.07 | 0.68 | 0.77 | 0.66 | 0.61 |
| 4 | 143493559 | rs1425523 | -0.491 | -0.562 | 0.136 | -1.719 | 0.004 | 0.005 | 0.005 | 0 | 0.8 | 0.89 | 0.88 | 0.84 | 0.83 | 0.8 |
| 4 | 143534216 | rs6856817 | 0.163 | -0.682 | 1.043 | -1.783 | 0 | -0.03 | -0.03 | 0 | 0.1 | 0.15 | 0.11 | 0.02 | 0.11 | 0.16 |
| 4 | 143562086 | rs7685002 | 0.118 | -0.159 | 1.414 | -1.779 | -0.019 | -0.039 | -0.039 | 0 | 0.76 | 0.56 | 0.85 | 0.83 | 0.81 | 0.81 |
| 4 | 143578523 | rs13119090 | 0.894 | -0.156 | 1.536 | -1.778 | -0.003 | -0.013 | -0.013 | 0 | 0.66 | 0.39 | 0.72 | 0.79 | 0.71 | 0.66 |
| 4 | 143607404 | rs6537123 | 0.893 | -0.194 | 1.542 | -1.802 | -0.003 | -0.013 | -0.013 | 0.213 | 0.67 | 0.39 | 0.72 | 0.79 | 0.71 | 0.66 |
| 4 | 143608324 | rs10020322 | 0.177 | -0.192 | 1.543 | -1.805 | 0.003 | -0.033 | -0.033 | 0 | 0.1 | 0.15 | 0.11 | 0.02 | 0.1 | 0.16 |
| 4 | 143610293 | rs13133181 | 1.315 | -0.571 | 1.189 | -1.97 | 0.004 | 0.039 | 0.039 | 0 | 0.59 | 0.33 | 0.7 | 0.77 | 0.65 | 0.58 |
| 4 | 143613959 | rs13150837 | -0.087 | -0.573 | 1.258 | -1.949 | -0.016 | -0.05 | -0.05 | 0 | 0.77 | 0.56 | 0.85 | 0.84 | 0.82 | 0.81 |
| 4 | 143617099 | rs12650777 | -0.072 | -0.573 | 1.291 | -1.949 | -0.017 | -0.048 | -0.048 | 0 | 0.77 | 0.57 | 0.85 | 0.84 | 0.82 | 0.84 |
| 4 | 143617304 | rs11943397 | 1.374 | -0.518 | 1.021 | -2.03 | 0.002 | 0.045 | 0.045 | 0 | 0.59 | 0.33 | 0.68 | 0.78 | 0.62 | 0.55 |
| 4 | 143626567 | rs11942593 | -0.385 | -0.488 | 0.774 | -2.017 | 0 | -0.001 | -0.001 | 0 | 0.78 | 0.83 | 0.86 | 0.83 | 0.81 | 0.84 |
| 4 | 143629934 | rs10009093 | -0.758 | -0.493 | 0.821 | -1.988 | 0.005 | -0.02 | -0.02 | 0.82 | 0.11 | 0.21 | 0.11 | 0.04 | 0.12 | 0.24 |
| 4 | 143634746 | rs12644329 | 1.803 | -0.646 | 0.73 | -2.072 | 0.018 | 0.142 | 0.142 | 0 | 0.54 | 0.16 | 0.68 | 0.75 | 0.62 | 0.52 |
| 4 | 143639211 | rs4690698 | 0.402 | -0.494 | 0.58 | -1.988 | -0.001 | -0.013 | -0.013 | 0 | 0.7 | 0.69 | 0.76 | 0.79 | 0.68 | 0.59 |
| 4 | 143655174 | rs986081 | -0.585 | -0.54 | 0.321 | -1.889 | 0.012 | -0.001 | -0.001 | 0 | 0.06 | 0.1 | 0.05 | 0.03 | 0.06 | 0.06 |
| 4 | 143677136 | rs17016749 | -0.907 | -0.457 | 0.354 | -1.911 | 0 | -0.012 | -0.012 | 0.866 | 0.09 | 0.12 | 0.02 | 0.01 | 0.07 | 0.07 |
| 4 | 143683303 | rs17016755 | -1.055 | -0.47 | 0.353 | -1.917 | 0 | -0.026 | -0.026 | 0.049 | 0.1 | 0.16 | 0.03 | 0.01 | 0.07 | 0.07 |
| 4 | 143691110 | rs17732348 | 1.174 | -0.506 | 0.219 | -1.961 | 0.027 | 0.036 | 0.036 | 0 | 0.14 | 0.03 | 0.04 | 0 | 0.09 | 0.03 |
| 4 | 143708921 | rs12504681 | -1.072 | -0.263 | 0.441 | -1.742 | 0.003 | -0.03 | -0.03 | 0 | 0.09 | 0.16 | 0.03 | 0.01 | 0.07 | 0.07 |
| 4 | 143723381 | rs12509337 | -0.837 | -0.426 | 0.27 | -1.733 | -0.02 | -0.026 | -0.026 | 0 | 0.09 | 0.13 | 0.01 | 0.01 | 0.02 | 0.06 |
| 4 | 143735741 | rs1014464 | -1.52 | 0.306 | -0.587 | -2.333 | -0.134 | 0.159 | 0.159 | 0 | 0.73 | 0.98 | 0.69 | 0.95 | 0.61 | 0.71 |
| 4 | 143737159 | rs1992418 | -0.431 | -0.054 | 0.451 | -2.271 | 0.006 | -0.013 | -0.013 | 0 | 0.28 | 0.3 | 0.24 | 0.2 | 0.34 | 0.37 |
| 4 | 143747354 | rs1444949 | -0.39 | -0.216 | 0.578 | -2.286 | -0.012 | 0.012 | 0.012 | 0 | 0.25 | 0.22 | 0.2 | 0.2 | 0.32 | 0.37 |
| 4 | 143772767 | rs2218240 | -1.699 | -0.5 | 1.825 | -2.47 | 0 | -0.029 | -0.029 | 0 | 0.39 | 0.49 | 0.28 | 0.26 | 0.38 | 0.5 |
| 4 | 143779613 | rs6537129 | 0.042 | -0.368 | 1.591 | -2.44 | -0.006 | 0.009 | 0.009 | 0 | 0.23 | 0.17 | 0.18 | 0.21 | 0.28 | 0.36 |
| 4 | 143795347 | rs6852412 | -1.422 | -0.644 | 1.748 | -2.537 | 0.011 | 0.028 | 0.028 | 0 | 0.38 | 0.72 | 0.25 | 0.23 | 0.29 | 0.3 |
| 4 | 143796487 | rs2874907 | -0.965 | -0.664 | 2.031 | -2.546 | -0.005 | -0.031 | -0.031 | 0.013 | 0.34 | 0.5 | 0.24 | 0.23 | 0.28 | 0.3 |
| 4 | 143804147 | rs1372241 | -1.459 | -0.686 | 1.873 | -2.549 | 0.012 | 0.03 | 0.03 | 0 | 0.38 | 0.73 | 0.26 | 0.23 | 0.29 | 0.3 |
| 4 | 143823839 | rs1441413 | -1.87 | -0.836 | 1.817 | -2.72 | 0.038 | 0.129 | 0.129 | 0 | 0.43 | 0.91 | 0.29 | 0.2 | 0.34 | 0.36 |
| 4 | 143835306 | rs716878 | -0.575 | -0.797 | 1.995 | -2.652 | 0 | 0.037 | 0.037 | 0 | 0.29 | 0.18 | 0.19 | 0.15 | 0.26 | 0.25 |
| 4 | 143842458 | rs732601 | -0.742 | -0.794 | 2.18 | -2.651 | -0.002 | -0.012 | -0.012 | 0 | 0.32 | 0.34 | 0.21 | 0.16 | 0.26 | 0.25 |
| 4 | 143861293 | rs6834860 | -0.575 | -0.968 | 1.43 | -2.647 | 0.008 | 0.034 | 0.034 | 0 | 0.3 | 0.21 | 0.18 | 0.17 | 0.25 | 0.26 |
| 4 | 143861348 | rs6834625 | -1.341 | -1.039 | 1.577 | -2.703 | 0.039 | 0.074 | 0.074 | 0 | 0.37 | 0.8 | 0.22 | 0.15 | 0.25 | 0.26 |
| 4 | 143866615 | rs13115045 | -0.623 | -0.948 | 1.426 | -2.698 | 0.01 | 0.033 | 0.033 | 0 | 0.3 | 0.22 | 0.19 | 0.15 | 0.26 | 0.26 |
| 4 | 143882545 | rs1510136 | -1.49 | -0.949 | 1.015 | -2.668 | 0.053 | 0.11 | 0.11 | 0 | 0.38 | 0.83 | 0.23 | 0.15 | 0.27 | 0.26 |
| 4 | 143899483 | rs17678540 | -0.692 | -0.797 | 0.858 | -2.637 | 0.008 | 0.046 | 0.046 | 0 | 0.3 | 0.19 | 0.2 | 0.13 | 0.26 | 0.27 |
| 4 | 143907480 | rs1588408 | -0.637 | -0.761 | 0.434 | -2.631 | -0.008 | 0.071 | 0.071 | 0 | 0.29 | 0.11 | 0.19 | 0.13 | 0.26 | 0.26 |
| 4 | 143927609 | rs6817062 | 0.321 | -1.209 | 0.494 | -2.736 | 0.003 | 0.002 | 0.002 | 0 | 0.56 | 0.41 | 0.5 | 0.27 | 0.68 | 0.56 |
| 4 | 143936903 | rs1104696 | 0.339 | -1.213 | 0.573 | -2.738 | 0.015 | 0.018 | 0.018 | 0 | 0.56 | 0.39 | 0.5 | 0.27 | 0.68 | 0.56 |
| 4 | 143960787 | rs13136216 | -0.919 | -1.217 | 0.829 | -2.751 | 0.006 | -0.025 | -0.025 | 0 | 0.09 | 0.17 | 0.07 | 0.01 | 0.08 | 0.06 |
| 4 | 143971806 | rs10001212 | -1.38 | -1.235 | 0.819 | -2.759 | 0.007 | -0.029 | -0.029 | 0 | 0.09 | 0.18 | 0.07 | 0.01 | 0.08 | 0.06 |
| 4 | 143972745 | rs4690760 | 0.527 | -1.285 | 0.896 | -2.778 | -0.009 | -0.003 | -0.003 | 0.569 | 0.34 | 0.37 | 0.43 | 0.72 | 0.24 | 0.38 |
| 4 | 143998282 | rs13126430 | 0.501 | -1.325 | 0.859 | -2.725 | 0.1 | 0.129 | 0.129 | 0 | 0.53 | 0.15 | 0.5 | 0.26 | 0.68 | 0.56 |
| 4 | 144003159 | rs7686660 | 0.299 | -1.298 | 0.901 | -2.71 | -0.038 | -0.015 | -0.015 | 0 | 0.36 | 0.45 | 0.42 | 0.73 | 0.24 | 0.39 |
| 4 | 144027729 | rs10032938 | -0.135 | -0.907 | 1.076 | -2.378 | -0.017 | 0.014 | 0.014 | 0 | 0.06 | 0.02 | 0.05 | 0.01 | 0.08 | 0.06 |
| 4 | 144033656 | rs10471103 | 0.21 | -0.184 | 0.731 | -2.44 | 0.003 | -0.001 | -0.001 | 0 | 0.58 | 0.59 | 0.65 | 0.82 | 0.64 | 0.52 |
| 4 | 144043466 | rs11100769 | 0.492 | -0.15 | 0.883 | -2.498 | 0.038 | 0.073 | 0.073 | 0 | 0.57 | 0.31 | 0.63 | 0.82 | 0.67 | 0.58 |
| 4 | 144051108 | rs4690772 | 0.904 | -0.092 | 1.148 | -2.494 | 0.061 | -0.002 | -0.002 | 1.845 | 0.92 | 0.69 | 0.93 | 0.99 | 0.92 | 0.94 |
| 4 | 144055140 | rs11933189 | -0.783 | -0.091 | 1.37 | -2.494 | 0.021 | -0.02 | -0.02 | 0 | 0.08 | 0.17 | 0.06 | 0.01 | 0.08 | 0.06 |
| 4 | 144060464 | rs7666785 | 0.662 | -0.149 | 1.409 | -2.545 | -0.003 | -0.022 | -0.022 | 2.173 | 0.55 | 0.53 | 0.6 | 0.81 | 0.6 | 0.49 |
| 4 | 144073912 | rs11100775 | -0.122 | -0.148 | 1.379 | -2.52 | 0.018 | 0.108 | 0.108 | 0 | 0.36 | 0.16 | 0.33 | 0.18 | 0.32 | 0.45 |
| 4 | 144086419 | rs13109025 | -1.782 | -0.222 | 1.235 | -2.631 | -0.013 | 0.056 | 0.056 | 0 | 0.86 | 0.97 | 0.82 | 0.96 | 0.87 | 0.74 |
| 4 | 144104906 | rs11100780 | 0.652 | -0.109 | 1.552 | -2.513 | -0.001 | -0.01 | -0.01 | 0 | 0.55 | 0.56 | 0.6 | 0.81 | 0.6 | 0.49 |
| 4 | 144114413 | rs13140326 | -0.756 | -0.123 | 1.526 | -2.442 | 0.014 | -0.018 | -0.018 | 0.386 | 0.08 | 0.14 | 0.06 | 0.01 | 0.08 | 0.06 |
| 4 | 144130750 | rs7670426 | -0.858 | -0.123 | 1.687 | -2.465 | 0.018 | -0.022 | -0.022 | 0.168 | 0.08 | 0.17 | 0.06 | 0.01 | 0.08 | 0.06 |
| 4 | 144135718 | rs4285045 | -0.204 | -0.075 | 1.459 | -2.461 | 0.012 | 0.108 | 0.108 | 0 | 0.64 | 0.84 | 0.67 | 0.82 | 0.68 | 0.56 |
| 4 | 144136193 | rs10000911 | -0.194 | -0.05 | 1.46 | -2.461 | 0.009 | 0.108 | 0.108 | 0 | 0.64 | 0.84 | 0.67 | 0.82 | 0.68 | 0.56 |
| 4 | 144174773 | rs300942 | -2.064 | -0.045 | 1.65 | -2.383 | 0.033 | -0.047 | -0.047 | 1.151 | 0.12 | 0.41 | 0.09 | 0.01 | 0.09 | 0.08 |
| 4 | 144182073 | rs10519678 | 1.005 | 0.025 | 1.426 | -2.471 | 0.025 | 0.101 | 0.101 | 0 | 0.49 | 0.34 | 0.57 | 0.81 | 0.58 | 0.43 |
| 4 | 144187380 | rs893929 | 1.518 | 0.107 | 1.441 | -2.488 | -0.02 | 0.07 | 0.07 | 0 | 0.44 | 0.24 | 0.51 | 0.78 | 0.48 | 0.33 |
| 4 | 144193117 | rs763918 | 0.297 | 0.103 | 1.652 | -2.528 | 0.003 | -0.005 | -0.005 | 1.851 | 0.57 | 0.53 | 0.65 | 0.82 | 0.65 | 0.49 |
| 4 | 144193695 | rs13114280 | 0.297 | 0.103 | 1.652 | -2.528 | 0.003 | -0.005 | -0.005 | 0 | 0.57 | 0.53 | 0.65 | 0.82 | 0.65 | 0.49 |
| 4 | 144198204 | rs300948 | -0.741 | 0.005 | 1.466 | -2.633 | -0.001 | -0.024 | -0.024 | 0 | 0.14 | 0.14 | 0.09 | 0.01 | 0.12 | 0.12 |
| 4 | 144201383 | rs300951 | -0.324 | 0.147 | 1.597 | -2.53 | 0.016 | 0.059 | 0.059 | 0 | 0.59 | 0.69 | 0.64 | 0.84 | 0.63 | 0.49 |
| 4 | 144206948 | rs12643013 | 1.39 | 0.212 | 1.566 | -2.52 | -0.013 | 0.044 | 0.044 | 0 | 0.43 | 0.31 | 0.54 | 0.83 | 0.51 | 0.37 |
| 4 | 144207686 | rs17686361 | 1.883 | 0.167 | 1.539 | -2.518 | -0.017 | 0.045 | 0.045 | 0 | 0.37 | 0.23 | 0.47 | 0.8 | 0.41 | 0.27 |

| | | | | | | | | | | | | | | | | |
|---|---|---|---|---|---|---|---|---|---|---|---|---|---|---|---|---|
| 4 | 144220578 | rs522027 | -0.821 | -0.506 | 1.474 | -2.441 | -0.003 | -0.018 | -0.018 | 0 | 0.14 | 0.16 | 0.09 | 0.01 | 0.08 | 0.11 |
| 4 | 144230902 | rs6537153 | 1.661 | -0.29 | 1.248 | -2.354 | -0.012 | 0.035 | 0.035 | 0 | 0.4 | 0.26 | 0.53 | 0.84 | 0.51 | 0.35 |
| 4 | 144240473 | rs300905 | 1.247 | -0.264 | 1.14 | -2.042 | 0.004 | 0.061 | 0.061 | 0 | 0.44 | 0.25 | 0.53 | 0.73 | 0.55 | 0.39 |
| 4 | 144261486 | rs6825973 | -0.455 | -0.255 | 1.415 | -1.968 | -0.026 | -0.053 | -0.053 | 0 | 0.1 | 0.24 | 0.05 | 0 | 0.04 | 0.06 |
| 4 | 144273358 | rs17017629 | -0.485 | -0.252 | 1.413 | -1.967 | -0.018 | -0.049 | -0.049 | 0 | 0.1 | 0.32 | 0.05 | 0 | 0.04 | 0.06 |
| 4 | 144276643 | rs300913 | -0.386 | -0.363 | 1.243 | -2.133 | 0.005 | 0.033 | 0.033 | 0 | 0.63 | 0.78 | 0.65 | 0.73 | 0.63 | 0.52 |
| 4 | 144276717 | rs300912 | 1.342 | -0.368 | 1.167 | -2.161 | 0.008 | 0.082 | 0.082 | 0 | 0.46 | 0.21 | 0.57 | 0.73 | 0.55 | 0.41 |
| 4 | 144282005 | rs17738628 | 1.522 | -0.327 | 1.098 | -2.056 | -0.019 | 0.048 | 0.048 | 0 | 0.17 | 0.01 | 0.11 | 0.06 | 0.15 | 0.2 |
| 4 | 144289305 | rs1472873 | 0.059 | -0.153 | 1.407 | -1.714 | 0.018 | 0.027 | 0.027 | 0 | 0.36 | 0.25 | 0.34 | 0.27 | 0.32 | 0.51 |
| 4 | 144289502 | rs3805253 | -1.471 | -0.199 | 1.268 | -1.72 | 0.02 | 0.065 | 0.065 | 0 | 0.52 | 0.72 | 0.44 | 0.28 | 0.41 | 0.64 |
| 4 | 144293053 | rs6537155 | -1.471 | -0.199 | 1.268 | -1.72 | 0.02 | 0.065 | 0.065 | 0 | 0.52 | 0.72 | 0.44 | 0.28 | 0.41 | 0.64 |
| 4 | 144312520 | rs7680959 | -1.265 | -0.177 | 0.973 | -1.702 | 0.015 | 0.023 | 0.023 | 0 | 0.48 | 0.57 | 0.42 | 0.28 | 0.38 | 0.6 |
| 4 | 144319745 | rs1397528 | 0.001 | 0.007 | 0.277 | -1.578 | -0.001 | 0.033 | 0.033 | 0.055 | 0.36 | 0.2 | 0.36 | 0.29 | 0.34 | 0.5 |
| 4 | 144323529 | rs17739332 | -1.477 | -0.034 | 0.547 | -1.324 | 0.005 | 0.035 | 0.035 | 0.061 | 0.11 | 0.08 | 0.06 | 0.01 | 0.09 | 0.15 |
| 4 | 144335735 | rs2666350 | -0.239 | 0.189 | 0.174 | -1.223 | 0.002 | -0.032 | -0.032 | 0 | 0.06 | 0.19 | 0.03 | 0 | 0.05 | 0.05 |
| 4 | 144340560 | rs1995931 | 0.082 | 0.381 | -0.32 | -1.535 | -0.018 | 0.058 | 0.058 | 0 | 0.28 | 0.16 | 0.24 | 0.12 | 0.29 | 0.31 |
| 4 | 144357737 | rs3805236 | -1.283 | 0.541 | -1.084 | -1.701 | -0.067 | -0.034 | -0.034 | 0 | 0.66 | 0.99 | 0.6 | 0.28 | 0.66 | 0.72 |
| 4 | 144381401 | rs3795246 | -0.864 | 0.201 | 0.217 | -1.706 | 0.013 | 0.048 | 0.048 | 0.009 | 0.41 | 0.72 | 0.31 | 0.11 | 0.33 | 0.49 |
| 4 | 144388923 | rs7696619 | -0.829 | 0.173 | 0.237 | -1.691 | 0.012 | 0.044 | 0.044 | 0.077 | 0.41 | 0.73 | 0.31 | 0.11 | 0.33 | 0.48 |
| 4 | 144398793 | rs11947756 | -0.013 | 0.255 | 0.116 | -1.632 | -0.003 | -0.049 | -0.049 | 2.044 | 0.3 | 0.27 | 0.19 | 0.08 | 0.22 | 0.23 |
| 4 | 144401524 | rs11735967 | -1.024 | 0.182 | 0.241 | -1.741 | 0.024 | 0.073 | 0.073 | 0 | 0.42 | 0.76 | 0.29 | 0.1 | 0.32 | 0.52 |
| 4 | 144406061 | rs17017810 | -0.991 | 0.415 | 0.644 | -0.902 | -0.002 | -0.053 | -0.053 | 1.353 | 0.12 | 0.44 | 0.1 | 0.02 | 0.1 | 0.29 |
| 4 | 144412846 | rs6818844 | -0.357 | 0.74 | 0.176 | -0.709 | 0.006 | -0.022 | -0.022 | 1.3 | 0.37 | 0.41 | 0.42 | 0.77 | 0.42 | 0.44 |
| 4 | 144426798 | rs11730153 | 0.057 | 0.828 | 0.289 | -0.271 | 0 | 0.039 | 0.039 | 0 | 0.19 | 0.25 | 0.26 | 0.6 | 0.21 | 0.17 |
| 4 | 144451912 | rs11944532 | -0.285 | 0.579 | 0.423 | -0.507 | 0.003 | -0.007 | -0.007 | 0 | 0.19 | 0.18 | 0.16 | 0.15 | 0.21 | 0.3 |
| 4 | 144458198 | rs3924845 | 0.898 | 0.506 | 0.737 | -0.53 | 0.005 | -0.02 | -0.02 | 0 | 0.62 | 0.49 | 0.57 | 0.25 | 0.58 | 0.53 |
| 4 | 144466050 | rs7692376 | 0.898 | 0.506 | 0.737 | -0.53 | 0.005 | -0.02 | -0.02 | 3.084 | 0.62 | 0.49 | 0.57 | 0.25 | 0.58 | 0.53 |
| 4 | 144477601 | rs11732122 | 1.212 | 0.69 | 0.802 | -0.794 | -0.004 | 0.059 | 0.059 | 0 | 0.53 | 0.39 | 0.47 | 0.19 | 0.44 | 0.39 |
| 4 | 144479917 | rs924232 | 0.761 | 0.597 | 0.954 | -0.843 | -0.001 | 0.001 | 0.001 | 0 | 0.61 | 0.4 | 0.57 | 0.25 | 0.58 | 0.54 |
| 4 | 144480013 | rs924233 | 0.78 | 0.595 | 0.971 | -0.844 | -0.001 | 0.001 | 0.001 | 0 | 0.61 | 0.4 | 0.57 | 0.25 | 0.58 | 0.54 |
| 4 | 144480231 | rs17017907 | 0.567 | 0.733 | 0.453 | -0.77 | 0 | 0.026 | 0.026 | 0 | 0.18 | 0.22 | 0.26 | 0.61 | 0.21 | 0.19 |
| 4 | 144493185 | rs2323187 | -0.462 | 0.601 | 0.361 | -0.928 | 0.006 | -0.011 | -0.011 | 0 | 0.36 | 0.39 | 0.42 | 0.76 | 0.42 | 0.46 |
| 4 | 144496477 | rs17690723 | -0.273 | 0.19 | 0.011 | -0.613 | -0.01 | -0.006 | -0.006 | 0.178 | 0.08 | 0.04 | 0.1 | 0.06 | 0.14 | 0.13 |
| 4 | 144522627 | rs10519696 | -0.091 | -0.127 | -0.01 | -1.149 | 0.002 | -0.003 | -0.003 | 0 | 0.49 | 0.44 | 0.43 | 0.14 | 0.44 | 0.45 |
| 4 | 144524582 | rs1497614 | 0.291 | -0.371 | 0.092 | -1.256 | -0.048 | 0.232 | 0.232 | 0.251 | 0.43 | 0.12 | 0.39 | 0.09 | 0.43 | 0.45 |
| 4 | 144526199 | rs13149438 | -0.012 | -0.378 | 0.241 | -1.26 | 0.005 | 0.066 | 0.066 | 0.004 | 0.47 | 0.33 | 0.41 | 0.14 | 0.44 | 0.46 |
| 4 | 144539138 | rs1018092 | 0.237 | -0.472 | -0.01 | -1.342 | -0.055 | 0.28 | 0.28 | 0.274 | 0.46 | 0.11 | 0.39 | 0.09 | 0.44 | 0.46 |
| 4 | 144543099 | rs1391187 | -0.246 | -0.53 | 0.751 | -1.379 | 0.001 | 0.018 | 0.018 | 0 | 0.53 | 0.57 | 0.45 | 0.15 | 0.45 | 0.46 |
| 4 | 144546275 | rs13143261 | 0.294 | -0.599 | 0.474 | -1.124 | -0.006 | -0.013 | -0.013 | 0 | 0.14 | 0.06 | 0.12 | 0.2 | 0.13 | 0.24 |
| 4 | 144550211 | rs1497608 | 0.642 | -0.606 | 0.481 | -1.29 | -0.051 | 0.264 | 0.264 | 0.303 | 0.46 | 0.12 | 0.41 | 0.08 | 0.47 | 0.47 |
| 4 | 144559530 | rs1450237 | -0.49 | -0.599 | 0.666 | -1.263 | 0 | 0.053 | 0.053 | 0 | 0.48 | 0.4 | 0.54 | 0.85 | 0.53 | 0.53 |
| 4 | 144574139 | rs1826927 | -1.144 | -0.592 | 0.165 | -1.415 | -0.007 | 0.169 | 0.169 | 0 | 0.64 | 0.87 | 0.64 | 0.95 | 0.67 | 0.66 |
| 4 | 144575873 | rs1349325 | -0.138 | -0.769 | 0.532 | -1.113 | 0.018 | 0.093 | 0.093 | 0 | 0.51 | 0.28 | 0.49 | 0.74 | 0.56 | 0.53 |
| 4 | 144576742 | rs1450242 | 0.32 | -0.622 | 0.5 | -0.923 | -0.008 | -0.003 | -0.003 | 0 | 0.15 | 0.05 | 0.12 | 0.18 | 0.14 | 0.24 |
| 4 | 144578932 | rs1375988 | -1.399 | -0.618 | 0.574 | -0.978 | -0.007 | 0.093 | 0.093 | 0 | 0.72 | 0.87 | 0.68 | 0.95 | 0.72 | 0.76 |
| 4 | 144586417 | rs11100801 | -0.339 | -0.687 | 0.625 | -0.911 | 0.002 | 0.021 | 0.021 | 0 | 0.61 | 0.32 | 0.53 | 0.74 | 0.62 | 0.6 |
| 4 | 144616366 | rs7676614 | 1.179 | -0.507 | 0.308 | -0.828 | -0.062 | 0.24 | 0.24 | 0.053 | 0.36 | 0.03 | 0.33 | 0.04 | 0.35 | 0.32 |
| 4 | 144617842 | rs1545437 | 0.857 | -0.531 | 0.704 | -0.84 | 0.01 | 0.056 | 0.056 | 0 | 0.39 | 0.28 | 0.35 | 0.04 | 0.35 | 0.32 |
| 4 | 144635129 | rs2019159 | 0.674 | -0.234 | 0.728 | -0.328 | -0.016 | -0.024 | -0.024 | 0 | 0.22 | 0.16 | 0.35 | 0.6 | 0.35 | 0.26 |
| 4 | 144669996 | rs9685306 | 1.251 | -0.458 | 0.202 | -0.474 | -0.039 | 0.253 | 0.253 | 0.259 | 0.38 | 0.06 | 0.33 | 0.04 | 0.34 | 0.31 |
| 4 | 144676269 | rs35243309 | 1.006 | -0.507 | 0.466 | -0.427 | 0.018 | 0.053 | 0.053 | 0.013 | 0.41 | 0.27 | 0.41 | 0.22 | 0.35 | 0.32 |
| 4 | 144680954 | rs7687915 | 0.996 | -0.507 | 0.466 | -0.427 | 0.018 | 0.054 | 0.054 | 0.008 | 0.41 | 0.27 | 0.41 | 0.22 | 0.35 | 0.32 |
| 4 | 144687660 | rs1580002 | 1.267 | -0.481 | 0.221 | -0.465 | -0.039 | 0.253 | 0.253 | 0.259 | 0.38 | 0.05 | 0.33 | 0.04 | 0.34 | 0.31 |
| 4 | 144727743 | rs13101541 | 1.445 | -0.462 | 0.219 | -0.45 | -0.061 | 0.241 | 0.241 | 0.274 | 0.36 | 0.03 | 0.33 | 0.04 | 0.34 | 0.31 |
| 4 | 144740199 | rs13105102 | 1.253 | -0.448 | 0.333 | -0.443 | -0.007 | 0.203 | 0.203 | 0.275 | 0.38 | 0.1 | 0.33 | 0.04 | 0.34 | 0.31 |
| 4 | 144746002 | rs17018500 | 1.046 | -0.472 | 0.549 | -0.405 | 0.01 | 0.026 | 0.026 | 0.058 | 0.41 | 0.32 | 0.41 | 0.22 | 0.35 | 0.32 |
| 4 | 144782841 | rs7698581 | -0.151 | -0.435 | 0.419 | -0.417 | -0.012 | 0.001 | 0.001 | 1.369 | 0.31 | 0.34 | 0.23 | 0.35 | 0.19 | 0.29 |
| 4 | 144784844 | rs1822842 | 0.044 | -0.42 | 0.617 | -0.456 | 0.022 | 0.009 | 0.009 | 0.208 | 0.31 | 0.4 | 0.43 | 0.61 | 0.47 | 0.39 |
| 4 | 144792505 | rs4240334 | -1.831 | -0.422 | 0.568 | -0.466 | -0.059 | 0.242 | 0.242 | 0.211 | 0.64 | 0.97 | 0.67 | 0.96 | 0.66 | 0.69 |
| 4 | 144792959 | rs1808991 | -0.132 | -0.463 | 0.62 | -0.44 | -0.012 | 0.002 | 0.002 | 1.244 | 0.3 | 0.34 | 0.22 | 0.35 | 0.19 | 0.29 |
| 4 | 144825754 | rs9997931 | -0.956 | -0.37 | 0.682 | -0.465 | 0 | -0.004 | -0.004 | 0.776 | 0.57 | 0.45 | 0.57 | 0.78 | 0.65 | 0.68 |
| 4 | 144829906 | rs10857415 | -1.199 | -0.288 | 0.502 | -0.593 | 0.008 | 0.003 | 0.003 | 0 | 0.6 | 0.67 | 0.64 | 0.96 | 0.65 | 0.69 |
| 4 | 144869008 | rs2323413 | 1.341 | -0.252 | 0.097 | -0.583 | -0.051 | 0.241 | 0.241 | 0.183 | 0.36 | 0.04 | 0.33 | 0.04 | 0.34 | 0.31 |
| 4 | 144878737 | rs1912745 | -1.353 | -0.094 | 0.506 | -0.506 | 0.007 | -0.013 | -0.013 | 0 | 0.59 | 0.66 | 0.64 | 0.96 | 0.66 | 0.69 |
| 4 | 144882744 | rs4613514 | -1.346 | -0.102 | 0.542 | -0.509 | 0 | -0.053 | -0.053 | 2.698 | 0.59 | 0.59 | 0.64 | 0.96 | 0.66 | 0.69 |
| 4 | 144894260 | rs9799404 | 0.028 | -0.045 | 0.185 | -0.132 | 0.009 | 0.007 | 0.007 | 0.002 | 0.28 | 0.28 | 0.42 | 0.57 | 0.47 | 0.37 |
| 4 | 144949348 | rs34617896 | -0.016 | -0.206 | -0.421 | -0.04 | 0.004 | -0.007 | -0.007 | 0 | 0.28 | 0.07 | 0.14 | 0.23 | 0.19 | 0.31 |
| 4 | 144994903 | rs7663170 | -0.902 | 0.275 | 0.099 | 0.812 | 0.014 | 0.187 | 0.187 | 6.467 | 0.07 | 0.55 | 0.1 | 0.16 | 0.01 | 0.01 |
| 4 | 145009802 | rs7657795 | -0.28 | 1.002 | -0.846 | 0.91 | -0.027 | -0.046 | -0.046 | 0 | 0.78 | 0.85 | 0.65 | 0.46 | 0.66 | 0.72 |

| | | | | | | | | | | | | | | | | |
|---|---|---|---|---|---|---|---|---|---|---|---|---|---|---|---|---|
| 4 | 145024799 | rs6844670 | 0.339 | 0.29 | -0.587 | 0.814 | 0 | -0.001 | -0.001 | 0 | 0.5 | 0.45 | 0.61 | 0.59 | 0.38 | 0.46 |
| 4 | 145029870 | rs3936169 | -1.128 | -1.161 | 0.451 | 0.162 | -0.068 | -0.069 | -0.069 | 1.975 | 0.23 | 0.44 | 0.13 | 0.11 | 0.07 | 0.06 |
| 4 | 145029887 | rs3936168 | 1.005 | -1.137 | 0.216 | 0.174 | -0.051 | -0.054 | -0.054 | 0 | 0.75 | 0.32 | 0.86 | 0.89 | 0.92 | 0.94 |
| 4 | 145030701 | rs11728240 | 1.024 | -1.136 | 0.217 | 0.174 | -0.051 | -0.054 | -0.054 | 0 | 0.75 | 0.32 | 0.86 | 0.89 | 0.92 | 0.94 |
| 4 | 145033532 | rs4425323 | 1.323 | -0.527 | -0.705 | 0.352 | -0.117 | -0.027 | -0.027 | 0 | 0.43 | 0.04 | 0.5 | 0.46 | 0.47 | 0.54 |
| 4 | 145033810 | rs4374581 | -0.406 | -0.575 | -0.291 | 0.409 | -0.047 | -0.028 | -0.028 | 0 | 0.88 | 0.68 | 0.95 | 0.91 | 0.95 | 0.97 |
| 4 | 145035788 | rs6857303 | 0.112 | -0.441 | -0.434 | 0.512 | -0.007 | -0.007 | -0.007 | 0 | 0.32 | 0.25 | 0.35 | 0.43 | 0.45 | 0.4 |
| 4 | 145056487 | rs13103731 | 0.073 | -0.47 | -0.55 | 0.573 | -0.026 | -0.028 | -0.028 | 0 | 0.28 | 0.21 | 0.28 | 0.42 | 0.45 | 0.39 |
| 4 | 145076356 | rs7688971 | 0.011 | -0.906 | -0.001 | 0.238 | -0.012 | -0.02 | -0.02 | 0 | 0.07 | 0.11 | 0.07 | 0.02 | 0.02 | 0.03 |
| 4 | 145088692 | rs4402988 | 1.203 | -1.094 | -0.4 | 0 | 0.005 | 0.004 | 0.004 | 1.087 | 0.72 | 0.26 | 0.73 | 0.89 | 0.85 | 0.9 |
| 4 | 145092857 | rs4539990 | -1.677 | -0.917 | -1.035 | -0.391 | 0.036 | 0.146 | 0.146 | 0.822 | 0.67 | 0.86 | 0.81 | 0.94 | 0.79 | 0.59 |
| 4 | 145102264 | rs6812128 | 1.217 | -0.463 | -0.789 | 0.748 | -0.026 | 0.261 | 0.261 | 1.112 | 0.47 | 0.07 | 0.38 | 0.12 | 0.42 | 0.47 |
| 4 | 145111329 | rs3923603 | 0.229 | -0.184 | -0.413 | 1.901 | -0.014 | -0.014 | -0.014 | 0 | 0.2 | 0.14 | 0.31 | 0.43 | 0.36 | 0.12 |
| 4 | 145123453 | rs4599356 | -0.201 | -0.213 | -0.429 | 1.832 | -0.01 | -0.01 | -0.01 | 0 | 0.78 | 0.82 | 0.58 | 0.36 | 0.62 | 0.87 |
| 4 | 145135525 | rs4303930 | -1.145 | -0.192 | 0.027 | 1.719 | 0.007 | -0.005 | -0.005 | 0.861 | 0.48 | 0.63 | 0.49 | 0.51 | 0.57 | 0.44 |
| 4 | 145141405 | rs4423828 | 1.415 | 0.093 | 0.306 | 1.92 | -0.018 | 0.213 | 0.213 | 1.635 | 0.9 | 0.68 | 0.85 | 0.71 | 0.96 | 0.95 |
| 4 | 145142144 | rs4425324 | -0.829 | 0.19 | -0.201 | 1.662 | -0.013 | -0.01 | -0.01 | 0 | 0.39 | 0.67 | 0.44 | 0.46 | 0.5 | 0.39 |
| 4 | 145151018 | rs7658605 | 1.109 | 0.357 | -1.15 | 1.166 | -0.063 | 0.058 | 0.058 | 0.822 | 0.43 | 0.02 | 0.47 | 0.25 | 0.4 | 0.51 |
| 4 | 145177682 | rs11100843 | -0.513 | 0.787 | -0.286 | 1.062 | -0.029 | -0.014 | -0.014 | 0.009 | 0.35 | 0.32 | 0.34 | 0.44 | 0.44 | 0.17 |
| 4 | 145195649 | rs7689179 | -0.889 | 0.795 | -0.468 | 1.049 | -0.003 | -0.003 | -0.003 | 0 | 0.44 | 0.68 | 0.39 | 0.49 | 0.46 | 0.19 |
| 4 | 145203640 | rs7681569 | 0.711 | 0.519 | -0.855 | 0.978 | -0.002 | 0.085 | 0.085 | 0.174 | 0.49 | 0.18 | 0.39 | 0.42 | 0.5 | 0.49 |
| 4 | 145210834 | rs13146588 | -0.008 | 0.344 | -0.414 | 1.13 | -0.011 | 0.031 | 0.031 | 0.659 | 0.32 | 0.21 | 0.18 | 0.16 | 0.37 | 0.19 |
| 4 | 145213954 | rs12711416 | -0.356 | 0.498 | -0.224 | 1.139 | 0 | 0.009 | 0.009 | 0.026 | 0.41 | 0.46 | 0.48 | 0.45 | 0.45 | 0.48 |
| 4 | 145214922 | rs7674504 | -0.384 | 0.391 | -0.029 | 1.234 | 0.014 | 0.11 | 0.11 | 0 | 0.07 | 0.26 | 0.11 | 0.13 | 0.05 | 0.03 |
| 4 | 145225998 | rs6821055 | -0.465 | 0.514 | -0.347 | 1.152 | -0.007 | 0.013 | 0.013 | 0.004 | 0.41 | 0.5 | 0.46 | 0.45 | 0.42 | 0.47 |
| 4 | 145226422 | rs11935505 | -1.268 | 0.497 | 0.228 | 1.11 | 0.014 | 0.108 | 0.108 | 0.081 | 0.09 | 0.37 | 0.12 | 0.13 | 0.06 | 0.03 |
| 4 | 145227600 | rs13118083 | 0.222 | 0.545 | -1.288 | 0.778 | -0.052 | -0.021 | -0.021 | 0.653 | 0.49 | 0.1 | 0.5 | 0.49 | 0.51 | 0.6 |
| 4 | 145240011 | rs4256191 | 0.354 | 0.228 | -0.837 | 0.486 | 0.012 | 0.007 | 0.007 | 0 | 0.62 | 0.39 | 0.73 | 0.69 | 0.71 | 0.83 |
| 4 | 145254685 | rs7377575 | 0.247 | 0.048 | 0.163 | 0.398 | -0.006 | -0.003 | -0.003 | 0 | 0.64 | 0.48 | 0.75 | 0.69 | 0.71 | 0.83 |
| 4 | 145258807 | rs10030023 | -0.235 | -0.048 | 0.072 | 0.353 | -0.037 | -0.057 | -0.057 | 0.45 | 0.73 | 0.92 | 0.79 | 0.69 | 0.72 | 0.85 |
| 4 | 145261999 | rs6852276 | -0.792 | 0.352 | -0.453 | 0.482 | -0.004 | -0.006 | -0.006 | 0 | 0.33 | 0.36 | 0.45 | 0.43 | 0.4 | 0.38 |
| 4 | 145263286 | rs2719341 | -0.201 | 0.408 | -0.552 | 0.44 | -0.012 | -0.01 | -0.01 | 0.191 | 0.12 | 0.11 | 0.22 | 0.23 | 0.2 | 0.13 |
| 4 | 145264014 | rs6840871 | 0.064 | 0.554 | -0.214 | 0.336 | 0.014 | 0.014 | 0.014 | 0 | 0.72 | 0.62 | 0.75 | 0.69 | 0.72 | 0.87 |
| 4 | 145275039 | rs6537279 | -1.592 | 0.705 | -0.289 | 0.202 | -0.062 | 0.087 | 0.087 | 0.948 | 0.86 | 1 | 0.87 | 1 | 0.8 | 0.96 |
| 4 | 145278837 | rs17766287 | 2.165 | 0.276 | -0.153 | 0.389 | -0.003 | 0.052 | 0.052 | 0 | 0.35 | 0.06 | 0.28 | 0.23 | 0.32 | 0.49 |
| 4 | 145282779 | rs2657799 | 0.366 | 0.24 | 0.13 | 0.375 | -0.003 | 0.012 | 0.012 | 0 | 0.2 | 0.16 | 0.22 | 0.22 | 0.2 | 0.25 |
| 4 | 145293039 | rs970022 | -1.135 | 0.244 | 0.115 | 0.407 | 0.009 | 0.022 | 0.022 | 0 | 0.36 | 0.53 | 0.47 | 0.45 | 0.4 | 0.38 |
| 4 | 145302155 | rs1394998 | 2.115 | 0.223 | 0.219 | 0.392 | 0.026 | 0.075 | 0.075 | 0 | 0.36 | 0.11 | 0.28 | 0.23 | 0.32 | 0.49 |
| 4 | 145334169 | rs17019341 | 0.515 | 0.341 | 0.265 | 0.396 | -0.069 | -0.064 | -0.064 | 0 | 0.22 | 0.05 | 0.22 | 0.3 | 0.28 | 0.11 |
| 4 | 145347980 | rs12512146 | 0.81 | 0.383 | 0.576 | 0.475 | -0.007 | -0.011 | -0.011 | 2.824 | 0.66 | 0.64 | 0.56 | 0.57 | 0.6 | 0.62 |
| 4 | 145363247 | rs1602238 | -0.73 | 0.006 | -0.022 | 0.095 | -0.04 | 0.025 | 0.025 | 0 | 0.82 | 0.98 | 0.8 | 0.81 | 0.8 | 0.75 |
| 4 | 145419510 | rs7684769 | 0.848 | 0.137 | 0.176 | 0.228 | -0.001 | -0.001 | -0.001 | 0 | 0.66 | 0.6 | 0.58 | 0.65 | 0.68 | 0.62 |
| 4 | 145424843 | rs12641251 | 1.091 | 0.098 | 0.024 | 0.209 | 0.008 | 0.008 | 0.008 | 0.011 | 0.63 | 0.29 | 0.55 | 0.65 | 0.68 | 0.62 |
| 4 | 145429056 | rs11935246 | 1.362 | 0.064 | -0.138 | 0.209 | 0.034 | 0.033 | 0.033 | 0.008 | 0.6 | 0.24 | 0.55 | 0.65 | 0.68 | 0.6 |
| 4 | 145430297 | rs6827794 | -0.641 | 0.101 | -0.319 | 0.13 | -0.043 | 0.048 | 0.048 | 0 | 0.8 | 0.98 | 0.8 | 0.84 | 0.78 | 0.73 |
| 4 | 145431497 | rs1512282 | 1.428 | 0.241 | -0.089 | 0.442 | -0.03 | 0.009 | 0.009 | 0 | 0.37 | 0.12 | 0.32 | 0.32 | 0.39 | 0.49 |
| 4 | 145434688 | rs1032296 | -0.547 | 0.018 | -0.12 | 0.406 | -0.045 | -0.051 | -0.051 | 0.708 | 0.65 | 0.88 | 0.68 | 0.64 | 0.59 | 0.87 |
| 4 | 145434744 | rs1032297 | 1.587 | -0.166 | -0.192 | 0.397 | -0.097 | -0.029 | -0.029 | 0 | 0.34 | 0.08 | 0.32 | 0.32 | 0.43 | 0.46 |
| 4 | 145437319 | rs7677035 | -0.3 | -0.222 | 0.062 | 0.402 | -0.019 | -0.012 | -0.012 | 0 | 0.6 | 0.78 | 0.66 | 0.64 | 0.56 | 0.85 |
| 4 | 145441759 | rs6854832 | -0.364 | -0.191 | 0.559 | 0.365 | -0.021 | 0.014 | 0.014 | 0 | 0.2 | 0.36 | 0.31 | 0.32 | 0.13 | 0.39 |
| 4 | 145452783 | rs6845536 | -0.112 | -0.154 | 0.48 | 0.39 | -0.003 | 0 | 0 | 0 | 0.57 | 0.74 | 0.65 | 0.64 | 0.56 | 0.83 |
| 4 | 145474473 | rs1489759 | 1.081 | -0.16 | 0.636 | 0.373 | 0.01 | -0.009 | -0.009 | 0 | 0.37 | 0.49 | 0.33 | 0.31 | 0.43 | 0.44 |
| 4 | 145485738 | rs1980057 | 1.537 | -0.16 | 0.311 | 0.373 | -0.123 | -0.043 | -0.043 | 0 | 0.33 | 0.04 | 0.3 | 0.31 | 0.42 | 0.44 |
| 4 | 145485915 | rs7655625 | 1.325 | -0.123 | 0.487 | 0.391 | 0.002 | -0.001 | -0.001 | 0 | 0.36 | 0.32 | 0.32 | 0.31 | 0.42 | 0.44 |
| 4 | 145499204 | rs17019499 | 0.261 | 0.211 | -0.504 | 0.726 | 0.009 | -0.006 | -0.006 | 4.425 | 0.14 | 0.12 | 0.16 | 0.23 | 0.1 | 0.32 |
| 4 | 145511875 | rs7670758 | -0.362 | 0.321 | -0.839 | 0.548 | -0.044 | -0.039 | -0.039 | 0.008 | 0.63 | 0.91 | 0.67 | 0.65 | 0.59 | 0.86 |
| 4 | 145513053 | rs7677662 | -0.732 | 0.233 | -0.482 | 0.445 | -0.008 | -0.004 | -0.004 | 0 | 0.2 | 0.28 | 0.18 | 0.21 | 0.11 | 0.34 |
| 4 | 145520608 | rs4835180 | 0.683 | 0.676 | -1.042 | 0.62 | -0.026 | -0.026 | -0.026 | 4.679 | 0.57 | 0.23 | 0.59 | 0.47 | 0.48 | 0.77 |
| 4 | 145544836 | rs17776795 | 1.201 | 0.99 | -0.979 | 0.382 | 0.021 | 0.101 | 0.101 | 0 | 0.45 | 0.18 | 0.43 | 0.21 | 0.41 | 0.5 |
| 4 | 145546540 | rs2639582 | -0.435 | 0.834 | -0.301 | 0.638 | -0.029 | 0.042 | 0.042 | 0 | 0.36 | 0.2 | 0.3 | 0.16 | 0.48 | 0.19 |
| 4 | 145548933 | rs2575570 | -0.429 | 0.784 | 0.357 | 0.711 | -0.019 | 0.194 | 0.194 | 0 | 0.89 | 0.7 | 0.85 | 0.61 | 0.92 | 0.92 |
| 4 | 145552729 | rs2575572 | -0.389 | 0.787 | 0.397 | 0.671 | -0.04 | 0.057 | 0.057 | 0 | 0.36 | 0.17 | 0.3 | 0.16 | 0.48 | 0.19 |
| 4 | 145562457 | rs1398244 | -0.257 | 1.244 | 0.246 | 0.554 | -0.021 | 0.018 | 0.018 | 0 | 0.28 | 0.16 | 0.22 | 0.16 | 0.37 | 0.16 |
| 4 | 145565826 | rs6845999 | 2.155 | 1.195 | -0.014 | 0.526 | -0.013 | 0.102 | 0.102 | 0 | 0.44 | 0.09 | 0.43 | 0.2 | 0.41 | 0.5 |
| 4 | 145568352 | rs7689420 | 0.044 | 1.165 | 0.11 | 0.553 | -0.008 | 0.059 | 0.059 | 0 | 0.81 | 0.7 | 0.77 | 0.61 | 0.82 | 0.88 |
| 4 | 145582536 | rs2306924 | 0.555 | 1.047 | 0.177 | 0.292 | -0.016 | -0.009 | -0.009 | 0.471 | 0.58 | 0.68 | 0.6 | 0.45 | 0.49 | 0.76 |
| 4 | 145586900 | rs982902 | 1.503 | 1.062 | 0.156 | 0.26 | -0.015 | 0.108 | 0.108 | 0 | 0.45 | 0.12 | 0.43 | 0.21 | 0.41 | 0.5 |
| 4 | 145605033 | rs6819412 | 1.6 | 1.113 | 0.151 | 0.183 | -0.08 | 0.139 | 0.139 | 0 | 0.45 | 0.1 | 0.43 | 0.17 | 0.47 | 0.49 |
| 4 | 145605075 | rs6537309 | 0.526 | 1.135 | 0.534 | 0.204 | -0.004 | -0.006 | -0.006 | 0 | 0.59 | 0.64 | 0.63 | 0.45 | 0.55 | 0.8 |

| | | | | | | | | | | | | | | | | |
|---|---|---|---|---|---|---|---|---|---|---|---|---|---|---|---|---|
| 4 | 145619413 | rs1492819 | 1.21 | 1.1 | 0.614 | 0.205 | -0.012 | 0.003 | 0.003 | 0 | 0.51 | 0.25 | 0.58 | 0.45 | 0.5 | 0.76 |
| 4 | 145623107 | rs6838704 | 0.49 | 1.131 | 0.704 | 0.217 | -0.001 | 0 | 0 | 0.732 | 0.59 | 0.59 | 0.63 | 0.45 | 0.55 | 0.79 |
| 4 | 145625730 | rs7654947 | 0.532 | 1.068 | 0.488 | 0.23 | 0.001 | 0.007 | 0.007 | 0 | 0.59 | 0.5 | 0.63 | 0.45 | 0.57 | 0.77 |
| 4 | 145643079 | rs6854783 | 0.658 | 1.221 | 0.379 | 0.356 | 0 | 0 | 0 | 0 | 0.58 | 0.47 | 0.68 | 0.5 | 0.57 | 0.77 |
| 4 | 145646554 | rs1492818 | 0.677 | 1.22 | 0.406 | 0.355 | -0.001 | -0.002 | -0.002 | 0 | 0.58 | 0.4 | 0.68 | 0.5 | 0.57 | 0.77 |
| 4 | 145650021 | rs1492820 | 1.06 | 1.25 | 0.376 | 0.366 | 0.004 | 0.01 | 0.01 | 0 | 0.52 | 0.33 | 0.64 | 0.5 | 0.53 | 0.74 |
| 4 | 145655532 | rs1472174 | 0.642 | 1.267 | 0.251 | 0.403 | -0.001 | -0.002 | -0.002 | 0 | 0.58 | 0.41 | 0.68 | 0.51 | 0.57 | 0.77 |
| 4 | 145659064 | rs11727676 | 1.483 | 0.979 | -0.048 | 0.235 | -0.012 | 0.037 | 0.037 | 0 | 0.07 | 0 | 0.05 | 0 | 0.08 | 0.01 |
| 4 | 145670871 | rs11944404 | 1.133 | 0.821 | 0.184 | 0.386 | -0.014 | 0.159 | 0.159 | 0 | 0.48 | 0.15 | 0.44 | 0.17 | 0.47 | 0.53 |
| 4 | 145687147 | rs4835186 | -0.563 | 0.932 | 0.736 | 0.896 | 0.046 | 0.246 | 0.246 | 0 | 0.08 | 0.31 | 0.22 | 0.31 | 0.06 | 0.23 |
| 4 | 145697040 | rs2353934 | -1.243 | 0.919 | 0.773 | 0.858 | 0.025 | 0.17 | 0.17 | 0 | 0.12 | 0.37 | 0.26 | 0.31 | 0.1 | 0.26 |
| 4 | 145707662 | rs7442087 | 1.446 | 0.881 | 0.824 | 0.838 | 0.012 | 0.153 | 0.153 | 0 | 0.86 | 0.61 | 0.74 | 0.68 | 0.9 | 0.74 |
| 4 | 145770167 | rs12651029 | 0.943 | 1.195 | 0.367 | 0.015 | -0.07 | -0.015 | -0.015 | 0.344 | 0.21 | 0.05 | 0.19 | 0.25 | 0.27 | 0.18 |
| 4 | 145771221 | rs13102609 | 0.563 | 0.922 | 0.432 | 0.19 | -0.001 | 0.056 | 0.056 | 0 | 0.07 | 0.05 | 0.25 | 0.45 | 0.08 | 0.26 |
| 4 | 145782963 | rs4555661 | -1.741 | 0.689 | 0.887 | 0.072 | -0.037 | 0.108 | 0.108 | 1.599 | 0.2 | 0.57 | 0.32 | 0.45 | 0.12 | 0.29 |
| 4 | 145822014 | rs11100870 | -1.904 | 0.451 | 0.757 | -0.2 | -0.029 | 0.022 | 0.022 | 0 | 0.57 | 0.85 | 0.63 | 0.75 | 0.53 | 0.55 |
| 4 | 145858473 | rs813139 | -0.019 | 0.478 | 0.78 | -0.478 | -0.057 | 0.04 | 0.04 | 1.137 | 0.34 | 0.16 | 0.3 | 0.3 | 0.41 | 0.26 |
| 4 | 145889835 | rs789344 | -1.359 | 0.615 | 0.914 | -0.328 | -0.029 | 0.192 | 0.192 | 0.564 | 0.19 | 0.44 | 0.31 | 0.47 | 0.13 | 0.26 |
| 4 | 145914748 | rs813138 | -1.359 | 0.665 | 0.914 | -0.328 | -0.014 | 0.192 | 0.192 | 0.564 | 0.19 | 0.45 | 0.31 | 0.47 | 0.13 | 0.26 |
| 4 | 145920099 | rs1086103 | 0.755 | 0.587 | 0.887 | -0.336 | 0.004 | 0.034 | 0.034 | 0 | 0.06 | 0 | 0.03 | 0 | 0.04 | 0.03 |
| 4 | 145961040 | rs2007345 | -1.353 | 0.587 | 0.922 | -0.333 | -0.029 | 0.156 | 0.156 | 0.718 | 0.2 | 0.52 | 0.32 | 0.47 | 0.13 | 0.26 |
| 4 | 145974688 | rs11734845 | -0.247 | 0.11 | 0.528 | -0.635 | -0.008 | 0.019 | 0.019 | 1.023 | 0.45 | 0.33 | 0.36 | 0.4 | 0.48 | 0.48 |
| 4 | 145978910 | rs10032067 | 0.78 | 0.112 | 0.519 | -0.634 | -0.042 | 0.108 | 0.108 | 0.944 | 0.79 | 0.44 | 0.68 | 0.53 | 0.87 | 0.74 |
| 4 | 145984839 | rs17781702 | 1.94 | 0.12 | 0.493 | -0.662 | -0.099 | 0.033 | 0.033 | 0 | 0.33 | 0.02 | 0.31 | 0.13 | 0.39 | 0.26 |
| 4 | 145989190 | rs12331496 | 1.43 | 0.187 | 0.554 | -0.649 | 0.002 | 0.011 | 0.011 | 0 | 0.1 | 0.14 | 0.06 | 0.1 | 0.08 | 0.19 |
| 4 | 145996754 | rs13115967 | -0.622 | -0.016 | 0.488 | -0.598 | -0.016 | 0.144 | 0.144 | 0.058 | 0.17 | 0.3 | 0.3 | 0.47 | 0.13 | 0.26 |
| 4 | 146013265 | rs13136718 | 0.736 | 0.013 | 0.479 | -0.579 | -0.002 | 0.061 | 0.061 | 0 | 0.07 | 0.04 | 0.25 | 0.47 | 0.08 | 0.23 |
| 4 | 146036388 | rs34149094 | 0.888 | 0.135 | 0.312 | -0.575 | 0 | 0.069 | 0.069 | 0 | 0.12 | 0.01 | 0.06 | 0 | 0.09 | 0.04 |
| 4 | 146037320 | rs34330278 | 1.782 | -0.006 | 0.178 | -0.709 | -0.006 | 0.026 | 0.026 | 0 | 0.05 | 0 | 0.05 | 0 | 0.05 | 0.01 |
| 4 | 146055117 | rs11545157 | 0.689 | 0.168 | 0.224 | -0.475 | -0.017 | 0.041 | 0.041 | 0 | 0.12 | 0.01 | 0.09 | 0.04 | 0.12 | 0.08 |
| 4 | 146061560 | rs11731583 | 0.627 | 0.456 | 0.812 | -0.166 | -0.001 | 0.01 | 0.01 | 0.543 | 0.47 | 0.42 | 0.37 | 0.39 | 0.48 | 0.49 |
| 4 | 146068652 | rs12502109 | -0.171 | 0.454 | 0.814 | -0.16 | -0.023 | 0.157 | 0.157 | 0.261 | 0.18 | 0.37 | 0.31 | 0.47 | 0.13 | 0.26 |
| 4 | 146081096 | rs11734405 | 0.676 | 0.549 | 0.161 | -0.117 | -0.002 | 0.016 | 0.016 | 0.676 | 0.46 | 0.39 | 0.37 | 0.39 | 0.48 | 0.49 |
| 4 | 146125343 | rs2355233 | -0.097 | -0.055 | -0.586 | 0.073 | -0.019 | 0.198 | 0.198 | 0.072 | 0.17 | 0.39 | 0.31 | 0.47 | 0.13 | 0.25 |
| 4 | 146139896 | rs11736535 | 0.797 | 0.086 | -1.307 | 0.133 | -0.062 | 0.005 | 0.005 | 0 | 0.3 | 0.02 | 0.3 | 0.14 | 0.35 | 0.24 |
| 4 | 146167657 | rs6844904 | -0.083 | 0.201 | -1.195 | 0.187 | -0.028 | 0.107 | 0.107 | 0.522 | 0.77 | 0.47 | 0.65 | 0.53 | 0.83 | 0.69 |
| 4 | 146173197 | rs10013728 | -0.168 | 0.846 | -1.271 | -0.018 | -0.002 | 0.026 | 0.026 | 0.788 | 0.45 | 0.32 | 0.34 | 0.39 | 0.48 | 0.47 |
| 4 | 146184565 | rs11100872 | 0.024 | 0.503 | -1.124 | -0.641 | 0.022 | 0.067 | 0.067 | 0 | 0.49 | 0.32 | 0.48 | 0.84 | 0.4 | 0.54 |
| 4 | 146188270 | rs4835001 | -0.584 | 0.816 | -1.718 | -1.062 | -0.157 | 0.128 | 0.128 | 0.306 | 0.77 | 0.98 | 0.67 | 0.97 | 0.6 | 0.84 |
| 4 | 146188881 | rs13141761 | 0.238 | 0.404 | -1.357 | -1.239 | -0.065 | 0.1 | 0.1 | 0.703 | 0.65 | 0.83 | 0.56 | 0.84 | 0.47 | 0.61 |
| 4 | 146193967 | rs6833910 | -1.133 | 0.178 | -1.365 | -0.929 | 0.007 | 0.01 | 0.01 | 0 | 0.39 | 0.71 | 0.34 | 0.28 | 0.32 | 0.44 |
| 4 | 146194938 | rs6822403 | 0.427 | 0.207 | -1.492 | -0.797 | -0.001 | -0.001 | -0.001 | 0 | 0.1 | 0.14 | 0.09 | 0.15 | 0.13 | 0.25 |
| 4 | 146197201 | rs17019992 | 0.38 | 0.187 | -1.478 | -0.747 | -0.001 | 0 | 0 | 0 | 0.1 | 0.14 | 0.09 | 0.16 | 0.13 | 0.25 |
| 4 | 146201203 | rs2355620 | 0.818 | 0.07 | -1.568 | -0.67 | -0.025 | -0.045 | -0.045 | 0 | 0.09 | 0.02 | 0.13 | 0.17 | 0.1 | 0.13 |
| 4 | 146205494 | rs10006257 | -1.343 | 0.13 | -1.125 | -0.816 | 0.003 | 0.005 | 0.005 | 0 | 0.43 | 0.71 | 0.35 | 0.34 | 0.38 | 0.46 |
| 4 | 146210435 | rs2355617 | -0.996 | 0.438 | -1.69 | -1.02 | -0.077 | 0.083 | 0.083 | 0.275 | 0.77 | 0.93 | 0.68 | 0.95 | 0.66 | 0.81 |
| 4 | 146211177 | rs13109698 | 0.318 | -0.178 | -0.797 | -0.446 | -0.009 | -0.04 | -0.04 | 3.514 | 0.13 | 0.16 | 0.07 | 0 | 0.07 | 0.04 |
| 4 | 146213427 | rs6825120 | -0.707 | 0.317 | -0.794 | 0.167 | -0.01 | -0.007 | -0.007 | 0 | 0.14 | 0.12 | 0.12 | 0.17 | 0.2 | 0.26 |
| 4 | 146220802 | rs4373182 | -0.283 | -0.464 | -0.117 | 0.492 | -0.05 | 0.026 | 0.026 | 0.074 | 0.17 | 0.38 | 0.2 | 0.19 | 0.1 | 0.16 |
| 4 | 146220888 | rs11930564 | -0.242 | -0.423 | -0.438 | 1.244 | 0 | 0.072 | 0.072 | 0 | 0.05 | 0.1 | 0.02 | 0.29 | 0.04 | 0.05 |
| 4 | 146226787 | rs2705575 | -1.379 | -1.033 | -0.822 | 1.076 | 0.105 | -0.023 | -0.023 | 0.76 | 0.41 | 0.85 | 0.35 | 0.63 | 0.28 | 0.33 |
| 4 | 146232373 | rs11728665 | -0.803 | -0.159 | -0.424 | 1.352 | -0.011 | 0.026 | 0.026 | 0 | 0.21 | 0.26 | 0.13 | 0.45 | 0.18 | 0.17 |
| 4 | 146243587 | rs28606996 | -0.989 | 0.573 | -0.496 | 1.686 | 0.013 | 0.072 | 0.072 | 0 | 0.15 | 0.26 | 0.24 | 0.61 | 0.19 | 0.3 |
| 4 | 146249655 | rs6537340 | -0.886 | 1.198 | -1.329 | 0.259 | -0.024 | 0.005 | 0.005 | 0 | 0.74 | 0.81 | 0.84 | 0.99 | 0.7 | 0.82 |
| 4 | 146254791 | rs11931253 | 0.226 | 0.357 | -1.762 | 0.193 | 0.03 | 0.018 | 0.018 | 0.051 | 0.2 | 0.08 | 0.11 | 0.13 | 0.15 | 0.16 |
| 4 | 146256922 | rs1489152 | 0.454 | -0.128 | -2.181 | 0.309 | -0.026 | -0.073 | -0.073 | 0.146 | 0.29 | 0.14 | 0.42 | 0.47 | 0.29 | 0.35 |
| 4 | 146257067 | rs4240328 | 0.172 | 0.163 | -2.09 | 0.191 | 0.014 | 0.011 | 0.011 | 0 | 0.19 | 0.08 | 0.13 | 0.15 | 0.17 | 0.17 |
| 4 | 146258152 | rs1596480 | -0.822 | -0.367 | -1.77 | -0.241 | -0.003 | -0.028 | -0.028 | 0.112 | 0.2 | 0.14 | 0.35 | 0.4 | 0.18 | 0.43 |
| 4 | 146260382 | rs7668650 | -0.492 | -0.717 | -2.059 | -1.513 | -0.008 | 0.076 | 0.076 | 0.245 | 0.55 | 0.75 | 0.69 | 0.75 | 0.51 | 0.7 |
| 4 | 146262747 | rs568752 | 0.091 | -0.57 | -2.013 | -1.689 | -0.047 | 0.077 | 0.077 | 0 | 0.64 | 0.75 | 0.52 | 0.81 | 0.56 | 0.61 |
| 4 | 146270748 | rs11731479 | 0.999 | -0.667 | -1.797 | -1.706 | -0.023 | -0.091 | -0.091 | 0 | 0.33 | 0.08 | 0.33 | 0.58 | 0.34 | 0.33 |
| 4 | 146270827 | rs514422 | -1.295 | -0.391 | -1.586 | -1.719 | -0.077 | 0.096 | 0.096 | 0 | 0.79 | 0.93 | 0.84 | 0.98 | 0.72 | 0.92 |
| 4 | 146273558 | rs7673272 | 0.251 | -0.524 | -1.271 | -1.522 | 0.024 | -0.008 | -0.008 | 0.003 | 0.34 | 0.23 | 0.27 | 0.33 | 0.26 | 0.47 |
| 4 | 146276195 | rs920840 | 0.385 | -0.489 | -1.309 | -1.517 | 0.032 | -0.02 | -0.02 | 1.005 | 0.33 | 0.17 | 0.26 | 0.33 | 0.27 | 0.47 |
| 4 | 146281857 | rs3849041 | 0.096 | -0.59 | -0.659 | -2.159 | 0.003 | -0.116 | -0.116 | 0 | 0.61 | 0.45 | 0.73 | 0.92 | 0.66 | 0.81 |
| 4 | 146285738 | rs12513059 | -0.796 | -0.502 | -0.603 | -2.115 | 0.025 | -0.069 | -0.069 | 0 | 0.75 | 0.65 | 0.8 | 0.92 | 0.7 | 0.87 |
| 4 | 146297328 | rs2044276 | 0.194 | -0.174 | -0.12 | -1.483 | 0.011 | 0.001 | 0.001 | 0 | 0.33 | 0.2 | 0.3 | 0.3 | 0.31 | 0.43 |
| 4 | 146307650 | rs6842280 | -0.134 | 0.219 | -0.643 | -1.52 | -0.054 | -0.044 | -0.044 | 0 | 0.21 | 0.29 | 0.19 | 0.12 | 0.11 | 0.22 |
| 4 | 146309232 | rs17728991 | -0.845 | 0.508 | -0.943 | -1.478 | 0.012 | 0.066 | 0.066 | 0 | 0.55 | 0.7 | 0.63 | 0.31 | 0.44 | 0.61 |

| | | | | | | | | | | | | | | | | |
|---|---|---|---|---|---|---|---|---|---|---|---|---|---|---|---|---|
| 4 | 146314357 | rs10016954 | -1.553 | 0.537 | -1.025 | -1.32 | -0.015 | -0.022 | -0.022 | 0 | 0.71 | 0.87 | 0.77 | 0.38 | 0.67 | 0.75 |
| 4 | 146314863 | rs1444086 | -1.081 | 1.002 | -0.782 | -0.828 | 0.005 | 0.015 | 0.015 | 0 | 0.56 | 0.61 | 0.69 | 0.38 | 0.59 | 0.68 |
| 4 | 146315063 | rs7674130 | -0.322 | 0.825 | -1.11 | -0.441 | 0.003 | 0.013 | 0.013 | 0 | 0.27 | 0.21 | 0.25 | 0.19 | 0.25 | 0.26 |
| 4 | 146334754 | rs1992598 | -0.951 | 1.088 | -0.941 | -0.495 | -0.004 | 0.007 | 0.007 | 0 | 0.2 | 0.15 | 0.19 | 0.07 | 0.21 | 0.17 |
| 4 | 146347867 | rs13113847 | -0.722 | 1.15 | -0.973 | -0.815 | -0.031 | 0.044 | 0.044 | 0 | 0.36 | 0.25 | 0.32 | 0.19 | 0.49 | 0.38 |
| 4 | 146350184 | rs9684285 | -0.116 | 0.581 | 0.092 | -0.925 | 0.019 | -0.002 | -0.002 | 2.422 | 0.07 | 0.31 | 0.07 | 0.02 | 0.05 | 0.19 |
| 4 | 146358876 | rs499629 | -1.137 | 0.245 | 0.598 | -0.995 | -0.019 | -0.027 | -0.027 | 0 | 0.11 | 0.56 | 0.11 | 0.09 | 0.06 | 0.19 |
| 4 | 146369756 | rs13109195 | -0.128 | 0.28 | 0.345 | -1.178 | 0.01 | 0.008 | 0.008 | 0 | 0.13 | 0.1 | 0.08 | 0.09 | 0.07 | 0.13 |
| 4 | 146402592 | rs6537355 | -1.456 | 0.125 | 0.769 | -1.242 | -0.019 | -0.046 | -0.046 | 2.787 | 0.16 | 0.48 | 0.13 | 0.11 | 0.11 | 0.26 |
| 4 | 146407524 | rs6816306 | -1.302 | 0.13 | 1.443 | -1.154 | 0.06 | 0.124 | 0.124 | 3.699 | 0.06 | 0.44 | 0.05 | 0.09 | 0.01 | 0.03 |
| 4 | 146411000 | rs6823513 | -1.178 | 0.123 | 1.5 | -1.153 | 0.062 | 0.087 | 0.087 | 3.131 | 0.05 | 0.41 | 0.05 | 0.06 | 0.01 | 0.03 |
| 4 | 146414267 | rs11944685 | -1.964 | -0.207 | 0.82 | -1.559 | 0 | 0.157 | 0.157 | 0.953 | 0.49 | 0.92 | 0.47 | 0.31 | 0.57 | 0.62 |
| 4 | 146430293 | rs7694604 | -0.94 | -0.116 | 1.056 | -1.515 | -0.066 | -0.027 | -0.027 | 0.936 | 0.15 | 0.64 | 0.12 | 0.16 | 0.04 | 0.17 |
| 4 | 146450970 | rs11100883 | -0.119 | -0.007 | 1.021 | -1.613 | -0.137 | 0.054 | 0.054 | 1.126 | 0.23 | 0.05 | 0.28 | 0.1 | 0.45 | 0.34 |
| 4 | 146452773 | rs6852200 | -0.855 | -0.142 | 1.691 | -1.581 | -0.078 | -0.019 | -0.019 | 0 | 0.15 | 0.5 | 0.12 | 0.16 | 0.04 | 0.17 |
| 4 | 146453921 | rs7670486 | 0.228 | -0.108 | 1.712 | -1.565 | -0.036 | -0.008 | -0.008 | 0 | 0.1 | 0.22 | 0.07 | 0.11 | 0.04 | 0.13 |
| 4 | 146461534 | rs3756021 | -0.555 | -0.131 | 1.447 | -1.578 | -0.063 | -0.01 | -0.01 | 0.704 | 0.13 | 0.53 | 0.13 | 0.16 | 0.04 | 0.19 |
| 4 | 146464773 | rs7662543 | -0.845 | -0.031 | 1.321 | -1.612 | -0.066 | 0.064 | 0.064 | 0.177 | 0.29 | 0.16 | 0.34 | 0.13 | 0.51 | 0.4 |
| 4 | 146478467 | rs6829932 | -0.728 | -0.088 | 1.355 | -1.541 | -0.075 | -0.021 | -0.021 | 0 | 0.15 | 0.54 | 0.13 | 0.16 | 0.04 | 0.19 |
| 4 | 146478779 | rs1016792 | -1.017 | -0.907 | -0.151 | -0.895 | -0.024 | -0.139 | -0.139 | 0 | 0.76 | 0.98 | 0.62 | 0.63 | 0.79 | 0.74 |
| 4 | 146482020 | rs13120931 | -0.292 | -0.446 | 0.336 | -0.986 | -0.104 | 0.069 | 0.069 | 1.257 | 0.26 | 0.09 | 0.33 | 0.13 | 0.51 | 0.36 |
| 4 | 146484997 | rs6848760 | -0.543 | -0.54 | 0.626 | -0.899 | -0.088 | -0.021 | -0.021 | 0.024 | 0.16 | 0.52 | 0.07 | 0.16 | 0.05 | 0.22 |
| 4 | 146488217 | rs11944100 | -0.186 | -0.365 | 0.544 | -0.942 | -0.064 | 0.038 | 0.038 | 0.617 | 0.24 | 0.12 | 0.3 | 0.12 | 0.45 | 0.33 |
| 4 | 146488327 | rs11939979 | -1.482 | -0.404 | 0.477 | -0.944 | 0.007 | -0.019 | -0.019 | 0.648 | 0.4 | 0.61 | 0.37 | 0.29 | 0.5 | 0.55 |
| 4 | 146495355 | rs7662930 | 0.322 | -0.095 | 0.021 | -0.831 | -0.151 | 0.016 | 0.016 | 0.585 | 0.19 | 0.02 | 0.25 | 0.1 | 0.4 | 0.27 |
| 4 | 146501892 | rs6842568 | -0.829 | -0.793 | 0.699 | -0.384 | -0.078 | 0.01 | 0.01 | 0.124 | 0.16 | 0.45 | 0.06 | 0.19 | 0.05 | 0.22 |
| 4 | 146509970 | rs1874572 | 0.265 | -0.179 | 0.223 | -0.447 | -0.011 | -0.013 | -0.013 | 0 | 0.38 | 0.41 | 0.27 | 0.28 | 0.32 | 0.25 |
| 4 | 146539630 | rs1497125 | -0.489 | 0.762 | 0.027 | -0.207 | 0.017 | -0.007 | -0.007 | 0.057 | 0.47 | 0.6 | 0.38 | 0.49 | 0.37 | 0.34 |
| 4 | 146543677 | rs17020432 | -1.215 | 0.468 | -0.132 | -0.39 | 0.008 | 0.033 | 0.033 | 0 | 0.27 | 0.24 | 0.18 | 0.11 | 0.23 | 0.27 |
| 4 | 146545470 | rs17008702 | 0.624 | 0.43 | -0.331 | -0.465 | 0.019 | 0.024 | 0.024 | 0 | 0.09 | 0.05 | 0.03 | 0 | 0.04 | 0.06 |
| 4 | 146546701 | rs1390981 | -1.966 | 0.325 | 0.3 | -0.505 | -0.024 | -0.074 | -0.074 | 0 | 0.3 | 0.56 | 0.21 | 0.11 | 0.24 | 0.27 |
| 4 | 146551432 | rs13130076 | 1.064 | 0.637 | 0.202 | -0.63 | 0 | 0 | 0 | 0 | 0.09 | 0.07 | 0.21 | 0.07 | 0.1 | 0.08 |
| 4 | 146559160 | rs17020467 | 1.174 | 0.211 | 0.11 | -0.533 | -0.007 | 0.001 | 0.001 | 0 | 0.06 | 0.06 | 0.06 | 0.01 | 0.1 | 0.04 |
| 4 | 146560236 | rs4835011 | -0.437 | 0.119 | 0.062 | -0.513 | 0.021 | 0.054 | 0.054 | 0 | 0.14 | 0.05 | 0.05 | 0 | 0.09 | 0.06 |
| 4 | 146566650 | rs4835014 | 0.667 | 0.154 | 0.102 | -0.479 | 0.019 | 0.022 | 0.022 | 0 | 0.09 | 0.05 | 0.03 | 0 | 0.04 | 0.06 |
| 4 | 146567584 | rs17014946 | 0.602 | 0.061 | 0.226 | -0.413 | -0.004 | 0.023 | 0.023 | 0.077 | 0.29 | 0.23 | 0.27 | 0.48 | 0.24 | 0.27 |
| 4 | 146579452 | rs17020500 | 0.649 | 0.066 | -0.018 | -0.412 | 0.005 | -0.05 | -0.05 | 0.002 | 0.28 | 0.15 | 0.27 | 0.48 | 0.24 | 0.26 |
| 4 | 146586214 | rs9308191 | -2.033 | 0.179 | 0.098 | -0.396 | -0.036 | 0.013 | 0.013 | 0 | 0.63 | 0.78 | 0.66 | 0.66 | 0.53 | 0.61 |
| 4 | 146589780 | rs6831899 | 1.275 | 0.12 | 0.112 | -0.286 | -0.002 | 0.019 | 0.019 | 0.013 | 0.23 | 0.2 | 0.25 | 0.47 | 0.19 | 0.25 |
| 4 | 146592982 | rs7688920 | -0.313 | -0.006 | 0.075 | -0.434 | 0.014 | 0.038 | 0.038 | 0 | 0.21 | 0.17 | 0.12 | 0.01 | 0.18 | 0.09 |
| 4 | 146595347 | rs17744367 | -1.209 | -0.014 | 0.327 | -0.428 | 0.022 | 0.026 | 0.026 | 0 | 0.52 | 0.68 | 0.44 | 0.59 | 0.42 | 0.5 |
| 4 | 146601349 | rs17020528 | -0.177 | -0.114 | 0.539 | -0.595 | 0.02 | 0.073 | 0.073 | 0 | 0.08 | 0.32 | 0.08 | 0.1 | 0.05 | 0.16 |
| 4 | 146601701 | rs7682360 | -1.322 | -0.147 | 0.481 | -0.71 | 0.006 | 0.019 | 0.019 | 0 | 0.33 | 0.34 | 0.35 | 0.08 | 0.29 | 0.19 |
| 4 | 146602241 | rs17020536 | 0.922 | -0.087 | 0.343 | -0.657 | 0.004 | -0.009 | -0.009 | 0 | 0.11 | 0.15 | 0.23 | 0.07 | 0.12 | 0.1 |
| 4 | 146603460 | rs7678910 | -1.727 | -0.025 | 0.226 | -0.612 | 0.004 | 0.007 | 0.007 | 0.195 | 0.41 | 0.65 | 0.42 | 0.18 | 0.34 | 0.35 |
| 4 | 146614073 | rs2132778 | -0.675 | 0.139 | 0.089 | -0.025 | 0.011 | 0.013 | 0.013 | 0 | 0.5 | 0.54 | 0.56 | 0.38 | 0.4 | 0.4 |
| 4 | 146628662 | rs10084798 | -1.724 | 0.165 | -0.313 | -0.009 | -0.073 | -0.086 | -0.086 | 2.659 | 0.64 | 0.88 | 0.69 | 0.5 | 0.55 | 0.62 |
| 4 | 146635806 | rs6537368 | 0.085 | -0.088 | 0.429 | 0.048 | -0.037 | -0.079 | -0.079 | 0.068 | 0.14 | 0.22 | 0.05 | 0 | 0.07 | 0.06 |
| 4 | 146638036 | rs9999510 | -0.991 | 0.049 | 0.159 | 0.247 | 0.04 | 0.003 | 0.003 | 0 | 0.51 | 0.73 | 0.4 | 0.46 | 0.4 | 0.52 |
| 4 | 146639774 | rs10003327 | 0.076 | -0.074 | 0.379 | -0.132 | 0.01 | -0.008 | -0.008 | 0 | 0.17 | 0.2 | 0.08 | 0 | 0.1 | 0.07 |
| 4 | 146642017 | rs9997749 | -0.496 | 0.1 | 0.176 | 0.151 | -0.005 | -0.012 | -0.012 | 0.63 | 0.43 | 0.58 | 0.36 | 0.26 | 0.33 | 0.35 |
| 4 | 146651446 | rs3935131 | -0.534 | 0.273 | 0.407 | 0.841 | -0.002 | 0.053 | 0.053 | 0.025 | 0.18 | 0.14 | 0.25 | 0.38 | 0.14 | 0.25 |
| 4 | 146656795 | rs6841736 | -0.213 | 0.206 | 0.693 | 0.933 | -0.003 | 0.196 | 0.196 | 0 | 0.14 | 0.24 | 0.24 | 0.38 | 0.1 | 0.23 |
| 4 | 146662794 | rs7690184 | -0.276 | 0.122 | 0.658 | 1.116 | -0.003 | 0.155 | 0.155 | 0 | 0.14 | 0.25 | 0.18 | 0.35 | 0.1 | 0.26 |
| 4 | 146663390 | rs11934305 | 1.419 | -0.054 | 0.063 | -0.013 | -0.045 | 0.125 | 0.125 | 1.765 | 0.42 | 0.2 | 0.32 | 0.18 | 0.53 | 0.29 |
| 4 | 146669242 | rs4591626 | 0.164 | 0.108 | 0.368 | -0.051 | 0.002 | 0 | 0 | 0.522 | 0.61 | 0.37 | 0.58 | 0.83 | 0.71 | 0.73 |
| 4 | 146685889 | rs11939704 | 1.372 | 0.058 | 0.139 | -0.108 | -0.038 | 0.162 | 0.162 | 2.895 | 0.43 | 0.18 | 0.32 | 0.14 | 0.53 | 0.31 |
| 4 | 146697205 | rs4835257 | 1.274 | 0.042 | 0.142 | -0.109 | -0.024 | 0.176 | 0.176 | 3.629 | 0.44 | 0.18 | 0.33 | 0.14 | 0.55 | 0.3 |
| 4 | 146709412 | rs4521364 | -0.102 | 0.225 | 0.479 | 0.311 | 0.006 | 0.092 | 0.092 | 0 | 0.17 | 0.23 | 0.25 | 0.66 | 0.16 | 0.43 |
| 4 | 146717169 | rs10010535 | -0.293 | -0.062 | 0.355 | -0.005 | 0.001 | -0.004 | -0.004 | 0 | 0.33 | 0.39 | 0.33 | 0.13 | 0.24 | 0.21 |
| 4 | 146722152 | rs6854834 | -0.043 | 0.293 | 0.164 | 0.231 | -0.002 | -0.031 | -0.031 | 0.541 | 0.16 | 0.09 | 0.25 | 0.64 | 0.16 | 0.43 |
| 4 | 146730687 | rs1396055 | -0.776 | -0.108 | 0.273 | -0.267 | 0.006 | 0.033 | 0.033 | 0.146 | 0.35 | 0.32 | 0.2 | 0.08 | 0.29 | 0.22 |
| 4 | 146732654 | rs17806780 | -0.571 | -0.135 | 0.165 | -0.24 | -0.002 | 0.045 | 0.045 | 0.667 | 0.21 | 0.13 | 0.13 | 0.09 | 0.2 | 0.18 |
| 4 | 146737421 | rs7656212 | -0.679 | -0.079 | 0.716 | 0.385 | -0.022 | 0.067 | 0.067 | 0 | 0.19 | 0.31 | 0.24 | 0.61 | 0.13 | 0.36 |
| 4 | 146745835 | rs9790518 | -1.404 | 0.018 | 0.397 | -0.122 | 0.039 | 0.118 | 0.118 | 4.278 | 0.5 | 0.75 | 0.55 | 0.75 | 0.38 | 0.58 |
| 4 | 146746960 | rs11100901 | -0.43 | 0.186 | 0.634 | -0.186 | -0.001 | -0.026 | -0.026 | 0 | 0.27 | 0.29 | 0.16 | 0.05 | 0.22 | 0.21 |
| 4 | 146747838 | rs10011020 | -0.718 | 0.883 | 0.939 | 0.47 | 0.017 | 0.041 | 0.041 | 0 | 0.22 | 0.27 | 0.43 | 0.68 | 0.24 | 0.38 |
| 4 | 146748876 | rs17745264 | 0.049 | -0.391 | 0.823 | 0.018 | -0.017 | 0.068 | 0.068 | 0 | 0.28 | 0.19 | 0.18 | 0.08 | 0.29 | 0.2 |
| 4 | 146750966 | rs13149290 | -1.119 | -0.36 | 0.804 | -0.026 | 0.01 | 0.009 | 0.009 | 0 | 0.28 | 0.22 | 0.47 | 0.8 | 0.23 | 0.5 |

| | | | | | | | | | | | | | | | | |
|---|---|---|---|---|---|---|---|---|---|---|---|---|---|---|---|---|
| 4 | 146751207 | rs17020662 | 0.207 | -0.308 | 0.922 | -0.015 | -0.009 | 0.033 | 0.033 | 0 | 0.27 | 0.22 | 0.17 | 0.08 | 0.3 | 0.2 |
| 4 | 146764719 | rs17020691 | 0.225 | -0.313 | 1.179 | 0.047 | 0.025 | -0.008 | -0.008 | 0 | 0.05 | 0.16 | 0.04 | 0 | 0.05 | 0.02 |
| 4 | 146764883 | rs988163 | -1.524 | -0.493 | 1.233 | 0.226 | -0.001 | 0.021 | 0.021 | 0 | 0.52 | 0.54 | 0.65 | 0.71 | 0.52 | 0.54 |
| 4 | 146768394 | rs4544728 | -0.771 | -0.508 | 0.961 | 0.234 | -0.007 | -0.023 | -0.023 | 0 | 0.37 | 0.48 | 0.24 | 0.25 | 0.36 | 0.39 |
| 4 | 146775428 | rs17303543 | 0.238 | -0.463 | 0.548 | 0.179 | -0.015 | 0.055 | 0.055 | 0 | 0.27 | 0.2 | 0.18 | 0.08 | 0.3 | 0.2 |
| 4 | 146776836 | rs13128681 | 0.122 | -0.406 | 0.626 | 0.284 | -0.01 | 0.052 | 0.052 | 0 | 0.31 | 0.27 | 0.22 | 0.08 | 0.34 | 0.22 |
| 4 | 146781747 | rs4835261 | -0.652 | -0.358 | 0.056 | 0.488 | 0.014 | -0.039 | -0.039 | 0.126 | 0.2 | 0.1 | 0.42 | 0.63 | 0.17 | 0.31 |
| 4 | 146788825 | rs925136 | -0.104 | -0.276 | -0.514 | 0.546 | -0.001 | 0.036 | 0.036 | 0 | 0.16 | 0.11 | 0.41 | 0.42 | 0.16 | 0.26 |
| 4 | 146794621 | rs4547811 | -0.106 | -0.233 | -0.697 | 0.568 | -0.001 | 0.084 | 0.084 | 0 | 0.16 | 0.17 | 0.42 | 0.42 | 0.16 | 0.27 |
| 4 | 146795226 | rs11100902 | -1.675 | -0.341 | -1.153 | 0.397 | -0.094 | 0.163 | 0.163 | 0 | 0.6 | 0.87 | 0.68 | 0.86 | 0.52 | 0.71 |
| 4 | 146800922 | rs17020769 | -1.209 | 0.397 | -0.977 | 0.932 | -0.001 | 0 | 0 | 0 | 0.42 | 0.33 | 0.54 | 0.42 | 0.44 | 0.43 |
| 4 | 146818215 | rs9790707 | -0.408 | 0.839 | -1.681 | 1.223 | 0.006 | -0.033 | -0.033 | 0 | 0.31 | 0.2 | 0.19 | 0.4 | 0.3 | 0.37 |
| 4 | 146824778 | rs4835267 | -0.749 | 1.113 | -2.077 | 1.77 | -0.058 | -0.072 | -0.072 | 0 | 0.11 | 0.01 | 0.33 | 0.36 | 0.16 | 0.18 |
| 4 | 146825650 | rs2135961 | -0.882 | -0.064 | -2.309 | 1.522 | 0.014 | -0.017 | -0.017 | 0 | 0.35 | 0.32 | 0.33 | 0.47 | 0.28 | 0.45 |
| 4 | 146826299 | rs4835268 | -0.327 | -0.127 | -2.838 | 0.9 | -0.001 | 0.033 | 0.033 | 0 | 0.21 | 0.09 | 0.09 | 0.12 | 0.18 | 0.17 |
| 4 | 146826444 | rs4835024 | -0.991 | 0.393 | -3.369 | 0.785 | -0.183 | 0.039 | 0.039 | 0 | 0.74 | 0.97 | 0.7 | 0.86 | 0.54 | 0.8 |
| 4 | 146829530 | rs12332040 | -0.264 | -0.201 | -3.105 | 0.607 | -0.008 | -0.027 | -0.027 | 0 | 0.1 | 0.09 | 0.04 | 0 | 0.06 | 0.06 |
| 4 | 146840077 | rs1567224 | -0.479 | -0.549 | -3.452 | 0.782 | -0.066 | -0.024 | -0.024 | 0.009 | 0.59 | 0.96 | 0.47 | 0.67 | 0.41 | 0.76 |
| 4 | 146842502 | rs12641158 | 0.298 | -0.764 | -2.969 | 0.861 | 0.032 | 0.034 | 0.034 | 0 | 0.44 | 0.66 | 0.22 | 0.26 | 0.29 | 0.57 |
| 4 | 146849295 | rs12331851 | -0.048 | -0.44 | -2.877 | 0.888 | -0.02 | -0.026 | -0.026 | 0.122 | 0.07 | 0.17 | 0.02 | 0.04 | 0.02 | 0.04 |
| 4 | 146849715 | rs1472866 | -1.208 | -0.059 | -3.148 | 1.018 | -0.191 | 0.063 | 0.063 | 0.018 | 0.7 | 0.98 | 0.67 | 0.86 | 0.52 | 0.77 |
| 4 | 146862215 | rs9996164 | 0.416 | -0.546 | -2.96 | 1.075 | 0.007 | 0.03 | 0.03 | 0 | 0.23 | 0.11 | 0.09 | 0.12 | 0.21 | 0.2 |
| 4 | 146866150 | rs7659884 | 0.666 | -0.517 | -2.924 | 0.824 | -0.01 | -0.004 | -0.004 | 0 | 0.29 | 0.67 | 0.23 | 0.14 | 0.13 | 0.37 |
| 4 | 146867376 | rs6537382 | -0.188 | -0.128 | -2.967 | 0.673 | -0.012 | 0.015 | 0.015 | 0 | 0.57 | 0.93 | 0.56 | 0.67 | 0.39 | 0.68 |
| 4 | 146870683 | rs6537383 | -0.304 | -0.062 | -1.965 | 0.875 | 0.033 | -0.017 | -0.017 | 1.454 | 0.42 | 0.68 | 0.38 | 0.43 | 0.29 | 0.45 |
| 4 | 146875551 | rs723794 | -0.035 | -0.276 | -1.405 | 0.41 | 0.021 | -0.006 | -0.006 | 0.657 | 0.54 | 0.42 | 0.71 | 0.43 | 0.7 | 0.53 |
| 4 | 146883385 | rs7698461 | 0.394 | -0.987 | -1.093 | 0.426 | -0.004 | 0 | 0 | 0.122 | 0.2 | 0.16 | 0.13 | 0.38 | 0.09 | 0.3 |
| 4 | 146889338 | rs10025229 | 0.342 | -1.205 | -0.843 | 0.288 | -0.04 | 0.008 | 0.008 | 0 | 0.14 | 0.27 | 0.06 | 0.14 | 0.06 | 0.1 |
| 4 | 146890261 | rs9968429 | 0.136 | -0.63 | -1.51 | -0.305 | 0.005 | -0.002 | -0.002 | 0.285 | 0.46 | 0.45 | 0.38 | 0.46 | 0.4 | 0.3 |
| 4 | 146893493 | rs7657882 | 1.65 | -0.771 | -1.925 | -0.754 | -0.007 | 0.078 | 0.078 | 0.032 | 0.18 | 0.05 | 0.07 | 0 | 0.16 | 0.03 |
| 4 | 146893952 | rs4543136 | -1.956 | -0.721 | -1.925 | -0.748 | -0.047 | 0.021 | 0.021 | 0 | 0.9 | 0.99 | 0.92 | 0.98 | 0.85 | 0.96 |
| 4 | 146894530 | rs1579388 | 1.16 | -0.727 | -1.899 | -0.738 | -0.046 | 0.008 | 0.008 | 0 | 0.14 | 0.28 | 0.06 | 0.14 | 0.06 | 0.1 |
| 4 | 146897773 | rs6820440 | 0.904 | -0.255 | -1.517 | -0.631 | 0.019 | -0.025 | -0.025 | 0 | 0.24 | 0.28 | 0.39 | 0.11 | 0.36 | 0.36 |
| 4 | 146901801 | rs7660035 | -0.388 | -0.267 | -1.518 | -0.627 | 0.007 | -0.022 | -0.022 | 0 | 0.65 | 0.61 | 0.52 | 0.84 | 0.49 | 0.6 |
| 4 | 146903211 | rs719091 | 1.066 | -0.266 | -1.545 | -0.595 | -0.032 | -0.085 | -0.085 | 1.207 | 0.14 | 0.05 | 0.25 | 0.33 | 0.18 | 0.16 |
| 4 | 146904304 | rs11100904 | 0.821 | -0.142 | -1.44 | -0.579 | 0.024 | -0.033 | -0.033 | 0.656 | 0.25 | 0.3 | 0.39 | 0.11 | 0.36 | 0.36 |
| 4 | 146931256 | rs17020816 | 1.709 | -0.384 | -1.548 | -0.647 | 0.005 | -0.006 | -0.006 | 0 | 0.16 | 0.11 | 0.13 | 0.37 | 0.08 | 0.28 |
| 4 | 146932089 | rs10434119 | 1.422 | -0.271 | -1.623 | -0.28 | -0.034 | -0.033 | -0.033 | 0 | 0.06 | 0.01 | 0.22 | 0.19 | 0.12 | 0.09 |
| 4 | 146933473 | rs6537391 | 0.909 | -0.294 | -1.439 | -0.342 | 0.02 | -0.036 | -0.036 | 0 | 0.25 | 0.29 | 0.27 | 0.08 | 0.37 | 0.37 |
| 4 | 146945603 | rs4835275 | -0.745 | -0.191 | -1.408 | -0.381 | 0.004 | -0.008 | -0.008 | 0.682 | 0.71 | 0.79 | 0.75 | 0.6 | 0.74 | 0.66 |
| 4 | 146946713 | rs17809246 | 1.646 | 0.478 | -1.41 | -0.714 | -0.008 | 0.003 | 0.003 | 0.036 | 0.09 | 0.05 | 0.06 | 0.01 | 0.12 | 0.18 |
| 4 | 146948669 | rs17020895 | 1.524 | 0.162 | -1.647 | -0.727 | 0.014 | 0.04 | 0.04 | 0 | 0.23 | 0.13 | 0.09 | 0 | 0.18 | 0.03 |
| 4 | 146958602 | rs17020919 | 1.34 | 0.64 | -1.144 | -0.364 | 0.035 | 0.052 | 0.052 | 0.041 | 0.32 | 0.19 | 0.12 | 0.17 | 0.21 | 0.14 |
| 4 | 146961679 | rs13125215 | 0.29 | 0.772 | -1.104 | -0.437 | -0.001 | -0.008 | -0.008 | 0 | 0.56 | 0.47 | 0.45 | 0.25 | 0.54 | 0.49 |
| 4 | 146968904 | rs9991712 | 0.811 | 1.619 | -0.373 | -0.184 | -0.027 | 0.093 | 0.093 | 0 | 0.1 | 0.26 | 0.15 | 0.17 | 0.06 | 0.07 |
| 4 | 146974543 | rs13142208 | -0.521 | 1.76 | -0.513 | -0.225 | 0.012 | -0.008 | -0.008 | 1.13 | 0.34 | 0.54 | 0.48 | 0.27 | 0.39 | 0.42 |
| 4 | 146983240 | rs1426888 | 0.997 | 2.16 | -0.465 | 0.251 | 0.058 | -0.006 | -0.006 | 0 | 0.75 | 0.58 | 0.66 | 0.82 | 0.62 | 0.79 |
| 4 | 146985627 | rs12642634 | -0.747 | 2.249 | -0.472 | 0.241 | 0.068 | 0.041 | 0.041 | 3.013 | 0.16 | 0.4 | 0.26 | 0.17 | 0.23 | 0.16 |
| 4 | 146987110 | rs10007440 | -0.324 | 1.826 | -0.373 | 0.234 | -0.026 | -0.016 | -0.016 | 0 | 0.86 | 0.91 | 0.72 | 0.83 | 0.71 | 0.91 |
| 4 | 146987500 | rs11932033 | 0.809 | 1.827 | -0.542 | 0.235 | 0.079 | -0.007 | -0.007 | 3.63 | 0.75 | 0.53 | 0.65 | 0.82 | 0.62 | 0.79 |
| 4 | 146994618 | rs6845844 | -0.755 | 1.747 | -1.093 | 0.162 | 0.018 | 0.009 | 0.009 | 0.021 | 0.37 | 0.76 | 0.44 | 0.28 | 0.31 | 0.36 |
| 4 | 146999111 | rs2241521 | 0.09 | 1.286 | -1.067 | 0.179 | -0.001 | -0.001 | -0.001 | 0 | 0.26 | 0.27 | 0.36 | 0.22 | 0.22 | 0.23 |
| 4 | 147002718 | rs6821225 | 0.011 | 1.007 | -1.653 | 0.139 | 0.035 | -0.002 | -0.002 | 0 | 0.49 | 0.18 | 0.28 | 0.56 | 0.4 | 0.56 |
| 4 | 147012038 | rs11100914 | 0.538 | 1.19 | -1.254 | -0.085 | -0.011 | 0.045 | 0.045 | 0.018 | 0.83 | 0.94 | 0.82 | 0.92 | 0.79 | 0.85 |
| 4 | 147012650 | rs7666409 | 0.782 | 1.233 | -1.312 | -0.113 | -0.014 | 0.049 | 0.049 | 0.949 | 0.81 | 0.93 | 0.81 | 0.91 | 0.76 | 0.85 |
| 4 | 147015936 | rs10012073 | 1.179 | 1.279 | -0.245 | -0.223 | 0.016 | -0.075 | -0.075 | 0 | 0.76 | 0.53 | 0.75 | 0.88 | 0.71 | 0.78 |
| 4 | 147019583 | rs10857425 | 1.132 | 1.274 | -0.238 | -0.229 | 0.016 | -0.077 | -0.077 | 0 | 0.76 | 0.53 | 0.75 | 0.89 | 0.72 | 0.78 |
| 4 | 147028405 | rs4835030 | -0.803 | 1.462 | -0.351 | -0.882 | 0.006 | -0.046 | -0.046 | 0.847 | 0.32 | 0.39 | 0.27 | 0.08 | 0.37 | 0.17 |
| 4 | 147029511 | rs10017509 | -0.519 | 1.504 | -0.348 | -0.9 | -0.044 | 0.08 | 0.08 | 0.881 | 0.26 | 0.09 | 0.21 | 0.08 | 0.31 | 0.15 |
| 4 | 147037222 | rs11734320 | -0.554 | 1.446 | -0.386 | -0.928 | -0.045 | 0.087 | 0.087 | 0.759 | 0.27 | 0.08 | 0.21 | 0.08 | 0.31 | 0.15 |
| 4 | 147037235 | rs11722991 | -0.711 | 1.44 | -0.313 | -0.93 | -0.017 | 0.037 | 0.037 | 0.234 | 0.28 | 0.17 | 0.22 | 0.08 | 0.31 | 0.15 |
| 4 | 147042373 | rs11721815 | -0.777 | 1.371 | -0.212 | -1.065 | -0.005 | -0.003 | -0.003 | 0.079 | 0.31 | 0.31 | 0.24 | 0.09 | 0.33 | 0.15 |
| 4 | 147051347 | rs13141665 | 0.373 | 2.01 | -0.949 | -1.171 | -0.069 | -0.008 | -0.008 | 0 | 0.13 | 0.02 | 0.22 | 0.06 | 0.2 | 0.15 |
| 4 | 147054644 | rs2007366 | -0.247 | 1.978 | -0.883 | -1.205 | -0.042 | 0.091 | 0.091 | 2.755 | 0.29 | 0.16 | 0.22 | 0.09 | 0.33 | 0.14 |
| 4 | 147059435 | rs13101861 | 0.18 | 1.938 | -0.182 | -1.065 | 0.003 | -0.001 | -0.001 | 0 | 0.13 | 0.11 | 0.23 | 0.06 | 0.2 | 0.16 |
| 4 | 147071801 | rs6841228 | -1.32 | 1.28 | -0.038 | -1.162 | -0.001 | -0.014 | -0.014 | 0 | 0.07 | 0.34 | 0.03 | 0 | 0.02 | 0.01 |
| 4 | 147090211 | rs7356451 | -1.046 | 1.24 | 0.108 | -1.171 | 0.005 | -0.006 | -0.006 | 0 | 0.06 | 0.3 | 0.03 | 0 | 0.02 | 0.01 |
| 4 | 147097293 | rs6815527 | -0.58 | 1.294 | -0.097 | -1.195 | -0.011 | 0.035 | 0.035 | 1.279 | 0.31 | 0.23 | 0.23 | 0.11 | 0.35 | 0.16 |
| 4 | 147104263 | rs919483 | -0.104 | 1.292 | -0.185 | -1.194 | -0.055 | 0.088 | 0.088 | 2.341 | 0.27 | 0.11 | 0.21 | 0.11 | 0.34 | 0.15 |

| | | | | | | | | | | | | | | | | |
|---|---|---|---|---|---|---|---|---|---|---|---|---|---|---|---|---|
| 4 | 147112495 | rs11100918 | -0.017 | 1.314 | -0.45 | -1.414 | -0.028 | 0.031 | 0.031 | 0 | 0.94 | 1 | 0.92 | 1 | 0.9 | 0.94 |
| 4 | 147118845 | rs6844381 | 0.125 | 1.084 | -0.391 | -1.374 | -0.07 | 0.088 | 0.088 | 2.281 | 0.26 | 0.1 | 0.21 | 0.11 | 0.34 | 0.15 |
| 4 | 147120075 | rs1865532 | -0.797 | 0.499 | -0.463 | -1.013 | 0 | -0.006 | -0.006 | 0 | 0.38 | 0.43 | 0.29 | 0.26 | 0.4 | 0.23 |
| 4 | 147127987 | rs6821444 | -0.324 | 0.516 | -0.452 | -1.067 | -0.003 | -0.006 | -0.006 | 0.583 | 0.32 | 0.37 | 0.24 | 0.11 | 0.35 | 0.16 |
| 4 | 147135668 | rs12374409 | -0.272 | -0.292 | -1.071 | -0.602 | 0.023 | 0.043 | 0.043 | 5.261 | 0.38 | 0.16 | 0.21 | 0.27 | 0.32 | 0.28 |
| 4 | 147138193 | rs16998675 | -0.259 | -0.292 | -1.138 | -0.602 | 0.016 | 0.04 | 0.04 | 5.387 | 0.38 | 0.13 | 0.21 | 0.27 | 0.32 | 0.28 |
| 4 | 147143572 | rs1864372 | -0.617 | -0.428 | -0.267 | -0.345 | -0.05 | -0.012 | -0.012 | 0.423 | 0.19 | 0.22 | 0.05 | 0.15 | 0.08 | 0.2 |
| 4 | 147151858 | rs2278471 | -0.399 | -0.538 | -0.196 | -0.123 | -0.007 | -0.009 | -0.009 | 0 | 0.43 | 0.41 | 0.27 | 0.3 | 0.37 | 0.22 |
| 4 | 147159504 | rs7676643 | -0.152 | 0.009 | -0.298 | 0.619 | -0.003 | -0.006 | -0.006 | 0 | 0.64 | 0.71 | 0.74 | 0.47 | 0.57 | 0.72 |
| 4 | 147163132 | rs874851 | 0.508 | -0.175 | -0.193 | 0.597 | -0.001 | 0.038 | 0.038 | 2.733 | 0.51 | 0.39 | 0.65 | 0.35 | 0.47 | 0.61 |
| 4 | 147165756 | rs6857756 | 0.68 | -0.221 | -0.258 | 0.569 | -0.002 | 0.005 | 0.005 | 0 | 0.11 | 0.15 | 0.07 | 0.12 | 0.1 | 0.12 |
| 4 | 147168885 | rs11729554 | -0.787 | -0.167 | -0.37 | 0.854 | -0.015 | 0.002 | 0.002 | 4.302 | 0.68 | 0.87 | 0.8 | 0.66 | 0.64 | 0.83 |
| 4 | 147169761 | rs7656598 | 0.001 | 0.107 | -0.436 | 0.751 | 0.019 | 0.009 | 0.009 | 1.019 | 0.5 | 0.72 | 0.61 | 0.49 | 0.51 | 0.58 |
| 4 | 147170097 | rs10032233 | 0.377 | 0.302 | -0.425 | 0.682 | 0 | -0.001 | -0.001 | 0 | 0.39 | 0.57 | 0.54 | 0.38 | 0.43 | 0.46 |
| 4 | 147171388 | rs13122077 | 0.761 | 0.151 | -0.832 | 0.417 | -0.003 | 0.021 | 0.021 | 0.315 | 0.34 | 0.29 | 0.46 | 0.16 | 0.32 | 0.34 |
| 4 | 147174997 | rs7676356 | 0.747 | -0.464 | -0.956 | 0.508 | -0.011 | -0.002 | -0.002 | 0.308 | 0.19 | 0.23 | 0.23 | 0.15 | 0.11 | 0.13 |
| 4 | 147176603 | rs1036016 | -1.352 | 0.237 | -1.631 | 0.555 | -0.057 | -0.039 | -0.039 | 0.471 | 0.91 | 0.99 | 0.89 | 0.88 | 0.79 | 0.94 |
| 4 | 147177341 | rs1057560 | 1.225 | -0.165 | -2.135 | 0.393 | -0.144 | 0.007 | 0.007 | 0.733 | 0.37 | 0.04 | 0.42 | 0.34 | 0.44 | 0.38 |
| 4 | 147177468 | rs11930848 | -0.137 | -0.803 | -1.38 | 0.467 | -0.024 | -0.003 | -0.003 | 0 | 0.35 | 0.58 | 0.33 | 0.27 | 0.22 | 0.25 |
| 4 | 147177822 | rs28406249 | -0.266 | -0.018 | -1.159 | 0.451 | 0.071 | 0.123 | 0.123 | 0.158 | 0.49 | 0.8 | 0.47 | 0.26 | 0.39 | 0.43 |
| 4 | 147180527 | rs2356940 | -0.111 | -0.506 | -0.632 | 0.105 | -0.003 | 0.009 | 0.009 | 0.319 | 0.59 | 0.62 | 0.71 | 0.83 | 0.51 | 0.76 |
| 4 | 147198667 | rs1960018 | 0.551 | -0.371 | -0.514 | 0.245 | -0.004 | -0.003 | -0.003 | 0 | 0.45 | 0.43 | 0.61 | 0.57 | 0.45 | 0.47 |
| 4 | 147215026 | rs10032088 | 0.704 | -0.183 | -0.523 | 0.296 | 0.008 | 0.002 | 0.002 | 0 | 0.13 | 0.12 | 0.09 | 0.08 | 0.1 | 0.12 |
| 4 | 147216089 | rs9784437 | -0.517 | -0.167 | -0.294 | 0.205 | -0.001 | -0.032 | -0.032 | 0 | 0.19 | 0.33 | 0.14 | 0.09 | 0.19 | 0.16 |
| 4 | 147216213 | rs9784489 | -1.867 | -0.248 | -0.595 | 0.06 | -0.06 | 0.042 | 0.042 | 1.228 | 0.79 | 0.97 | 0.85 | 0.9 | 0.71 | 0.91 |
| 4 | 147217867 | rs17021355 | 1.211 | -0.518 | -0.776 | 0.147 | 0.04 | -0.038 | -0.038 | 0 | 0.13 | 0.02 | 0.09 | 0.15 | 0.07 | 0.18 |
| 4 | 147220412 | rs17021362 | 0.859 | -0.541 | -0.58 | 0.271 | -0.015 | 0.01 | 0.01 | 0 | 0.14 | 0.14 | 0.11 | 0.23 | 0.07 | 0.21 |
| 4 | 147221103 | rs2101345 | -0.519 | -0.493 | -0.451 | 0.283 | 0.008 | -0.006 | -0.006 | 0.304 | 0.64 | 0.78 | 0.75 | 0.65 | 0.64 | 0.68 |
| 4 | 147230765 | rs10212747 | 0.866 | -0.533 | -0.474 | 0.253 | 0.004 | 0.004 | 0.004 | 0 | 0.44 | 0.5 | 0.61 | 0.56 | 0.45 | 0.51 |
| 4 | 147246721 | rs6844812 | 0.936 | -0.506 | -0.524 | -0.054 | -0.002 | 0 | 0 | 0 | 0.44 | 0.45 | 0.62 | 0.63 | 0.45 | 0.55 |
| 4 | 147247694 | rs4835294 | 0.618 | -0.515 | -0.447 | -0.07 | 0.001 | 0.003 | 0.003 | 0 | 0.14 | 0.14 | 0.09 | 0.09 | 0.13 | 0.15 |
| 4 | 147252599 | rs9991483 | 0.901 | -0.494 | -0.468 | -0.025 | -0.001 | 0 | 0 | 0 | 0.44 | 0.46 | 0.62 | 0.63 | 0.45 | 0.55 |
| 4 | 147260476 | rs4835297 | 0.035 | -0.46 | -0.421 | -0.03 | -0.001 | -0.032 | -0.032 | 0 | 0.19 | 0.3 | 0.13 | 0.09 | 0.19 | 0.15 |
| 4 | 147269334 | rs6828612 | -0.007 | -0.495 | -0.534 | -0.047 | 0.005 | -0.026 | -0.026 | 0 | 0.58 | 0.53 | 0.71 | 0.78 | 0.52 | 0.75 |
| 4 | 147280685 | rs10009867 | -0.918 | -0.545 | -0.486 | 0.03 | -0.026 | -0.018 | -0.018 | 0 | 0.35 | 0.54 | 0.24 | 0.24 | 0.26 | 0.35 |
| 4 | 147295128 | rs2679144 | -0.883 | -0.694 | -0.434 | -0.113 | -0.025 | -0.02 | -0.02 | 0.162 | 0.38 | 0.44 | 0.23 | 0.24 | 0.27 | 0.34 |
| 4 | 147314034 | rs3849034 | 0.351 | -0.307 | -0.317 | -0.326 | 0 | 0.002 | 0.002 | 0.101 | 0.54 | 0.55 | 0.72 | 0.8 | 0.51 | 0.73 |
| 4 | 147320219 | rs2356820 | 0.459 | -0.311 | -0.214 | -0.328 | 0.004 | 0.03 | 0.03 | 0.039 | 0.53 | 0.43 | 0.71 | 0.8 | 0.51 | 0.73 |
| 4 | 147323331 | rs6826676 | -0.462 | -0.382 | -0.049 | -0.27 | -0.018 | -0.056 | -0.056 | 0 | 0.24 | 0.35 | 0.13 | 0.08 | 0.2 | 0.16 |
| 4 | 147325925 | rs2679141 | -0.531 | -0.358 | -0.067 | -0.272 | -0.028 | -0.087 | -0.087 | 1.274 | 0.25 | 0.42 | 0.14 | 0.08 | 0.2 | 0.16 |
| 4 | 147326974 | rs6858794 | -0.309 | -0.329 | -0.269 | -0.258 | -0.005 | -0.024 | -0.024 | 0 | 0.23 | 0.27 | 0.12 | 0.08 | 0.2 | 0.16 |
| 4 | 147336656 | rs2679157 | -0.65 | -0.32 | -0.525 | -0.261 | -0.024 | -0.083 | -0.083 | 0 | 0.25 | 0.43 | 0.14 | 0.1 | 0.21 | 0.2 |
| 4 | 147339907 | rs7699206 | 0.403 | -0.194 | -0.499 | -0.331 | -0.001 | -0.015 | -0.015 | 0 | 0.54 | 0.5 | 0.69 | 0.79 | 0.52 | 0.7 |
| 4 | 147351628 | rs10017463 | -1.845 | 0.031 | -0.86 | -0.208 | -0.049 | 0.031 | 0.031 | 1.816 | 0.79 | 0.96 | 0.84 | 0.89 | 0.73 | 0.91 |
| 4 | 147365170 | rs13105153 | -1.759 | 0.048 | -0.831 | -0.174 | -0.031 | 0.029 | 0.029 | 1.583 | 0.78 | 0.92 | 0.83 | 0.89 | 0.73 | 0.91 |
| 4 | 147370156 | rs6537420 | 0.793 | -0.331 | -0.947 | -0.204 | -0.006 | 0.006 | 0.006 | 0.395 | 0.42 | 0.59 | 0.67 | 0.65 | 0.46 | 0.56 |
| 4 | 147374979 | rs11944523 | -0.644 | -0.329 | -0.892 | -0.39 | 0.035 | 0.046 | 0.046 | 0 | 0.36 | 0.26 | 0.15 | 0.23 | 0.27 | 0.33 |
| 4 | 147376576 | rs7663212 | -0.56 | -0.327 | -0.919 | -0.389 | 0.044 | 0.061 | 0.061 | 0 | 0.35 | 0.21 | 0.15 | 0.23 | 0.27 | 0.33 |
| 4 | 147391468 | rs11734649 | -1.49 | -0.101 | -1.124 | -0.361 | -0.009 | 0.009 | 0.009 | 1.656 | 0.77 | 0.82 | 0.83 | 0.95 | 0.72 | 0.9 |
| 4 | 147412986 | rs12711436 | -0.426 | -0.354 | -1.3 | -0.471 | 0.006 | 0.079 | 0.079 | 1.55 | 0.63 | 0.82 | 0.8 | 0.83 | 0.65 | 0.71 |
| 4 | 147416669 | rs13149914 | 1.205 | 0.81 | -1.432 | 0.472 | -0.03 | 0.017 | 0.017 | 0 | 0.07 | 0.03 | 0.36 | 0.46 | 0.16 | 0.13 |
| 4 | 147424678 | rs4835040 | -1.775 | 0.523 | -1.462 | 0.1 | -0.018 | 0.033 | 0.033 | 1.7 | 0.77 | 0.87 | 0.83 | 0.96 | 0.72 | 0.91 |
| 4 | 147426105 | rs6849958 | -1.688 | 0.531 | -1.462 | 0.105 | -0.008 | 0.005 | 0.005 | 1.513 | 0.77 | 0.83 | 0.83 | 0.95 | 0.72 | 0.88 |
| 4 | 147429316 | rs13152024 | 0.028 | 0.708 | -1.459 | 0.348 | 0.045 | 0.109 | 0.109 | 0 | 0.17 | 0.23 | 0.45 | 0.54 | 0.27 | 0.2 |
| 4 | 147438019 | rs3914628 | 0.441 | 0.248 | -1.82 | 0.173 | 0.009 | 0.023 | 0.023 | 0 | 0.16 | 0.1 | 0.07 | 0.1 | 0.14 | 0.17 |
| 4 | 147439250 | rs4550954 | 1.455 | 0.069 | -2.025 | -0.205 | -0.042 | -0.047 | -0.047 | 0 | 0.26 | 0.02 | 0.25 | 0.2 | 0.24 | 0.28 |
| 4 | 147447028 | rs2884092 | 0.802 | 0.472 | -1.899 | -0.132 | -0.048 | -0.037 | -0.037 | 0.086 | 0.41 | 0.03 | 0.3 | 0.31 | 0.36 | 0.48 |
| 4 | 147454992 | rs7679939 | -0.059 | 0.507 | -0.786 | -0.14 | -0.004 | -0.032 | -0.032 | 0 | 0.14 | 0.2 | 0.06 | 0.03 | 0.11 | 0.1 |
| 4 | 147464245 | rs4835042 | -0.182 | 0.83 | -0.354 | 0.545 | 0.029 | 0.116 | 0.116 | 0 | 0.16 | 0.25 | 0.41 | 0.41 | 0.21 | 0.23 |
| 4 | 147469133 | rs1504355 | -1.564 | 0.831 | -0.199 | 0.03 | 0.026 | -0.031 | -0.031 | 1.124 | 0.86 | 0.76 | 0.9 | 0.94 | 0.85 | 0.94 |
| 4 | 147473314 | rs9996620 | -0.118 | 0.363 | 0.167 | 0.023 | 0.023 | -0.045 | -0.045 | 0 | 0.11 | 0.26 | 0.09 | 0.03 | 0.13 | 0.1 |
| 4 | 147481601 | rs17770092 | 1.033 | -0.142 | -0.123 | -0.216 | 0.01 | 0.009 | 0.009 | 0 | 0.18 | 0.12 | 0.07 | 0.1 | 0.12 | 0.18 |
| 4 | 147495449 | rs1504359 | 0.89 | -0.044 | -0.211 | 0.01 | -0.008 | -0.003 | -0.003 | 1.419 | 0.12 | 0.12 | 0.32 | 0.39 | 0.19 | 0.19 |
| 4 | 147499642 | rs17832354 | 1.126 | -0.026 | -0.055 | 0.117 | 0.009 | 0.009 | 0.009 | 0 | 0.17 | 0.12 | 0.07 | 0.1 | 0.12 | 0.18 |
| 4 | 147505431 | rs10005397 | -1.406 | 0.003 | 0.235 | 0.238 | -0.028 | 0.009 | 0.009 | 0.131 | 0.24 | 0.58 | 0.21 | 0.29 | 0.18 | 0.23 |
| 4 | 147507651 | rs7680105 | -0.186 | 0.172 | -0.391 | -0.006 | -0.002 | 0.006 | 0.006 | 0 | 0.16 | 0.14 | 0.11 | 0.03 | 0.17 | 0.12 |
| 4 | 147514051 | rs10014502 | 0.295 | 0.213 | -0.192 | 0.234 | 0.001 | -0.009 | -0.009 | 0 | 0.11 | 0.14 | 0.07 | 0.04 | 0.13 | 0.1 |
| 4 | 147522465 | rs17021624 | -0.553 | 0.137 | 0.082 | 0.183 | 0.04 | -0.029 | -0.029 | 0.858 | 0.08 | 0.36 | 0.07 | 0 | 0.06 | 0.03 |
| 4 | 147529209 | rs6835151 | -0.79 | 0.195 | -0.214 | 0.213 | -0.004 | -0.039 | -0.039 | 0.675 | 0.16 | 0.3 | 0.1 | 0.05 | 0.14 | 0.11 |

| | | | | | | | | | | | | | | | | |
|---|---|---|---|---|---|---|---|---|---|---|---|---|---|---|---|---|
| 4 | 147535168 | rs2134852 | 0.329 | -0.295 | -0.683 | 0.143 | 0.019 | -0.044 | -0.044 | 0 | 0.34 | 0.21 | 0.39 | 0.49 | 0.32 | 0.38 |
| 4 | 147538064 | rs1875844 | 1.732 | 0.11 | -0.902 | 0.466 | -0.092 | 0.002 | 0.002 | 1.78 | 0.2 | 0.01 | 0.18 | 0.1 | 0.25 | 0.11 |
| 4 | 147545471 | rs13138020 | -0.215 | -0.379 | 0.221 | 1.053 | -0.001 | -0.002 | -0.002 | 0 | 0.09 | 0.13 | 0.04 | 0.04 | 0.08 | 0.08 |
| 4 | 147551074 | rs970674 | 1.056 | -0.402 | 0.228 | 1.101 | -0.001 | 0.025 | 0.025 | 0 | 0.06 | 0.08 | 0.1 | 0.24 | 0.05 | 0.06 |
| 4 | 147551241 | rs9994157 | -1.618 | -0.11 | 0.27 | 0.512 | -0.027 | -0.066 | -0.066 | 0 | 0.16 | 0.39 | 0.09 | 0.04 | 0.1 | 0.08 |
| 4 | 147560411 | rs13152799 | -1.094 | -0.029 | -0.254 | 0.575 | -0.011 | -0.023 | -0.023 | 0 | 0.13 | 0.19 | 0.07 | 0.04 | 0.1 | 0.08 |
| 4 | 147566988 | rs6537429 | -1.407 | 0.131 | -0.236 | 0.629 | -0.012 | -0.046 | -0.046 | 0 | 0.13 | 0.31 | 0.08 | 0.04 | 0.1 | 0.07 |
| 4 | 147573688 | rs7689436 | -0.137 | 0.424 | -1.051 | 0.808 | -0.002 | 0.008 | 0.008 | 0 | 0.07 | 0.07 | 0.04 | 0.03 | 0.08 | 0.06 |
| 4 | 147580638 | rs17021755 | -0.387 | 0.471 | -1.336 | 1.2 | -0.022 | -0.056 | -0.056 | 0 | 0.21 | 0.05 | 0.33 | 0.32 | 0.22 | 0.25 |
| 4 | 147591367 | rs1857721 | 1.519 | 0.344 | -1.777 | 0.867 | -0.138 | 0.112 | 0.112 | 0 | 0.33 | 0.02 | 0.26 | 0.11 | 0.38 | 0.3 |
| 4 | 147619089 | rs17772451 | 0.37 | -0.458 | -1.226 | 0.77 | 0.005 | 0.002 | 0.002 | 0 | 0.17 | 0.14 | 0.07 | 0.04 | 0.13 | 0.13 |
| 4 | 147633134 | rs1394272 | 0.02 | -0.601 | -0.532 | 1.479 | -0.029 | 0.081 | 0.081 | 0 | 0.8 | 0.36 | 0.78 | 0.68 | 0.85 | 0.83 |
| 4 | 147636699 | rs1504354 | 0.322 | -0.408 | -0.876 | 1.399 | -0.061 | -0.103 | -0.103 | 0.077 | 0.22 | 0.03 | 0.37 | 0.37 | 0.25 | 0.28 |
| 4 | 147641087 | rs2713878 | -0.012 | -0.587 | -0.554 | 1.484 | -0.028 | 0.031 | 0.031 | 0 | 0.81 | 0.54 | 0.8 | 0.81 | 0.86 | 0.83 |
| 4 | 147642738 | rs4835336 | 0.612 | -0.597 | -0.398 | 1.466 | 0 | 0.017 | 0.017 | 0.196 | 0.1 | 0.07 | 0.07 | 0.03 | 0.1 | 0.05 |
| 4 | 147648358 | rs4476620 | 0.156 | -0.581 | -0.444 | 1.477 | -0.018 | 0.045 | 0.045 | 0.011 | 0.82 | 0.54 | 0.8 | 0.81 | 0.86 | 0.9 |
| 4 | 147651538 | rs4484319 | 1.644 | -0.19 | -0.991 | 1.486 | -0.062 | -0.054 | -0.054 | 0 | 0.36 | 0.08 | 0.29 | 0.37 | 0.4 | 0.47 |
| 4 | 147658082 | rs10027612 | 0.473 | -0.195 | -0.029 | 1.764 | -0.005 | 0.034 | 0.034 | 0 | 0.67 | 0.47 | 0.49 | 0.46 | 0.68 | 0.69 |
| 4 | 147664762 | rs1509332 | 0.45 | -0.313 | 0.176 | 1.682 | 0.012 | -0.024 | -0.024 | 0 | 0.68 | 0.74 | 0.49 | 0.57 | 0.7 | 0.65 |
| 4 | 147666797 | rs7671336 | -0.981 | -0.329 | 0.417 | 1.66 | -0.01 | 0.011 | 0.011 | 0.385 | 0.34 | 0.55 | 0.54 | 0.43 | 0.31 | 0.36 |
| 4 | 147688226 | rs10519819 | 1.719 | -0.588 | -0.189 | 1.604 | 0.009 | -0.018 | -0.018 | 0 | 0.07 | 0 | 0.16 | 0.04 | 0.04 | 0.04 |
| 4 | 147701619 | rs952639 | -0.554 | -0.601 | -0.146 | 1.644 | 0.007 | -0.035 | -0.035 | 0.394 | 0.25 | 0.11 | 0.33 | 0.33 | 0.24 | 0.32 |
| 4 | 147717009 | rs10446756 | -0.426 | -0.606 | -0.113 | 1.635 | 0.013 | -0.035 | -0.035 | 0 | 0.24 | 0.11 | 0.33 | 0.33 | 0.22 | 0.3 |
| 4 | 147720217 | rs7674628 | 1.371 | -0.695 | -0.219 | 1.561 | -0.032 | 0.038 | 0.038 | 0 | 0.09 | 0.03 | 0.08 | 0 | 0.16 | 0.03 |
| 4 | 147734508 | rs12504096 | -1.313 | -0.852 | 0.512 | 1.62 | -0.02 | 0.004 | 0.004 | 0 | 0.34 | 0.56 | 0.53 | 0.38 | 0.27 | 0.35 |
| 4 | 147751943 | rs6857108 | 1.531 | -0.848 | 0.628 | 1.627 | 0.009 | -0.001 | -0.001 | 0.115 | 0.08 | 0.06 | 0.17 | 0.05 | 0.05 | 0.05 |
| 4 | 147755975 | rs2036728 | -0.502 | -0.769 | 0.736 | 2.001 | -0.025 | 0.022 | 0.022 | 0 | 0.25 | 0.5 | 0.37 | 0.33 | 0.23 | 0.3 |
| 4 | 147757175 | rs1509339 | -2.138 | -0.891 | 0.228 | 1.717 | -0.087 | 0.063 | 0.063 | 0 | 0.62 | 0.92 | 0.75 | 0.78 | 0.6 | 0.67 |
| 4 | 147761369 | rs17022009 | 0.873 | -0.997 | 0.927 | 1.769 | -0.014 | 0.027 | 0.027 | 0 | 0.09 | 0.3 | 0.18 | 0.05 | 0.05 | 0.05 |
| 4 | 147768045 | rs17022031 | 1.432 | -0.965 | 0.505 | 1.786 | 0.007 | -0.001 | -0.001 | 0 | 0.08 | 0.05 | 0.16 | 0.05 | 0.05 | 0.04 |
| 4 | 147768206 | rs6855967 | 0.381 | -0.96 | 0.676 | 1.926 | -0.009 | 0.04 | 0.04 | 0 | 0.11 | 0.2 | 0.19 | 0.18 | 0.08 | 0.14 |
| 4 | 147783128 | rs17610019 | 0.499 | -0.963 | 0.639 | 1.921 | -0.005 | 0.019 | 0.019 | 0 | 0.11 | 0.14 | 0.19 | 0.18 | 0.08 | 0.14 |
| 4 | 147783160 | rs10018635 | -0.397 | -1.488 | 0.689 | 1.86 | -0.022 | 0 | 0 | 0 | 0.34 | 0.52 | 0.41 | 0.41 | 0.3 | 0.47 |
| 4 | 147785578 | rs12507364 | 0.644 | -1.481 | 0.766 | 1.857 | -0.008 | 0.064 | 0.064 | 0 | 0.11 | 0.22 | 0.2 | 0.18 | 0.08 | 0.14 |
| 4 | 147787745 | rs1396716 | -2.889 | -1.635 | 0.473 | 1.737 | -0.17 | -0.019 | -0.019 | 3.833 | 0.66 | 0.98 | 0.71 | 0.77 | 0.53 | 0.68 |
| 4 | 147788709 | rs10013280 | -0.245 | -1.485 | 0.611 | 1.795 | -0.013 | 0.001 | 0.001 | 0 | 0.34 | 0.45 | 0.4 | 0.41 | 0.3 | 0.47 |
| 4 | 147797214 | rs10001365 | -1.306 | -1.451 | 0.341 | 1.712 | 0.025 | 0 | 0 | 0 | 0.45 | 0.67 | 0.6 | 0.58 | 0.38 | 0.61 |
| 4 | 147809311 | rs7656258 | -1.99 | -1.732 | 0.266 | 1.659 | -0.071 | 0.046 | 0.046 | 0 | 0.6 | 0.92 | 0.7 | 0.76 | 0.5 | 0.66 |
| 4 | 147820477 | rs984696 | -0.023 | -1.754 | 0.484 | 1.895 | 0.006 | -0.021 | -0.021 | 0.915 | 0.31 | 0.13 | 0.38 | 0.38 | 0.3 | 0.44 |
| 4 | 147824864 | rs17610219 | -1.169 | -1.773 | 0.669 | 1.911 | 0.026 | -0.011 | -0.011 | 0 | 0.44 | 0.66 | 0.59 | 0.55 | 0.36 | 0.56 |
| 4 | 147834758 | rs13122121 | 0.629 | -1.856 | 0.585 | 1.908 | 0.007 | 0.001 | 0.001 | 0 | 0.24 | 0.13 | 0.33 | 0.26 | 0.23 | 0.27 |
| 4 | 147839631 | rs17022118 | 0.664 | -1.837 | 0.707 | 1.922 | -0.023 | 0.145 | 0.145 | 0.021 | 0.11 | 0.41 | 0.21 | 0.17 | 0.06 | 0.11 |
| 4 | 147850007 | rs1509333 | -1.166 | -1.777 | 0.522 | 1.847 | 0.029 | -0.004 | -0.004 | 0 | 0.44 | 0.66 | 0.57 | 0.5 | 0.36 | 0.56 |
| 4 | 147867307 | rs3827595 | 0.251 | -1.542 | 0.825 | 1.486 | -0.018 | 0.002 | 0.002 | 0 | 0.27 | 0.48 | 0.37 | 0.33 | 0.25 | 0.25 |
| 4 | 147897673 | rs7663340 | -1.44 | -1.642 | 0.761 | 1.465 | 0.045 | 0.064 | 0.064 | 0 | 0.51 | 0.71 | 0.68 | 0.71 | 0.45 | 0.56 |
| 4 | 147905897 | rs1347655 | -0.492 | -1.733 | 0.81 | 1.493 | -0.001 | -0.001 | -0.001 | 0 | 0.24 | 0.24 | 0.29 | 0.31 | 0.19 | 0.27 |
| 4 | 147908124 | rs9918010 | 0.137 | -1.724 | 0.829 | 1.512 | -0.002 | 0.004 | 0.004 | 0 | 0.16 | 0.16 | 0.12 | 0.28 | 0.14 | 0.2 |
| 4 | 147909409 | rs1595309 | 0.623 | -1.715 | 0.832 | 1.519 | -0.001 | -0.051 | -0.051 | 0.038 | 0.25 | 0.13 | 0.36 | 0.4 | 0.25 | 0.29 |
| 4 | 147917821 | rs4467588 | -1.089 | -1.766 | 0.843 | 1.413 | -0.038 | 0.016 | 0.016 | 0.403 | 0.24 | 0.41 | 0.15 | 0.28 | 0.17 | 0.24 |
| 4 | 147919958 | rs17022256 | 0.362 | -1.703 | 0.833 | 1.343 | -0.01 | -0.033 | -0.033 | 0 | 0.06 | 0.1 | 0.01 | 0 | 0.02 | 0.04 |
| 4 | 147925900 | rs974278 | -0.653 | -1.76 | 0.872 | 1.387 | -0.002 | -0.001 | -0.001 | 0 | 0.41 | 0.53 | 0.56 | 0.43 | 0.34 | 0.39 |
| 4 | 147927227 | rs12498817 | 0.376 | -1.736 | 0.791 | 1.248 | -0.011 | -0.018 | -0.018 | 0 | 0.13 | 0.14 | 0.18 | 0.03 | 0.07 | 0.1 |
| 4 | 147935557 | rs1370299 | -0.032 | -1.805 | 0.617 | 1.37 | -0.014 | 0.014 | 0.014 | 0 | 0.16 | 0.21 | 0.12 | 0.27 | 0.12 | 0.23 |
| 4 | 147940034 | rs6537444 | -1.66 | -1.869 | 0.381 | 1.38 | 0.062 | -0.028 | -0.028 | 1.152 | 0.52 | 0.84 | 0.6 | 0.52 | 0.4 | 0.56 |
| 4 | 147941013 | rs6813768 | -0.109 | -1.899 | 0.947 | 1.373 | -0.055 | 0.036 | 0.036 | 0.016 | 0.13 | 0.48 | 0.08 | 0.11 | 0.06 | 0.2 |
| 4 | 147941253 | rs6537445 | -0.441 | -1.95 | 0.969 | 1.365 | -0.005 | 0.003 | 0.003 | 0 | 0.37 | 0.6 | 0.42 | 0.49 | 0.33 | 0.49 |
| 4 | 147941480 | rs1370300 | 0.695 | -2.016 | 0.648 | 1.351 | -0.005 | -0.059 | -0.059 | 0 | 0.12 | 0.01 | 0.09 | 0.23 | 0.11 | 0.17 |
| 4 | 147945451 | rs7665036 | 1.131 | -2.016 | 0.644 | 1.36 | -0.01 | -0.04 | -0.04 | 0 | 0.22 | 0.12 | 0.33 | 0.38 | 0.25 | 0.26 |
| 4 | 147946604 | rs10019569 | -1.496 | -1.957 | 0.439 | 1.375 | 0.1 | -0.015 | -0.015 | 0 | 0.49 | 0.83 | 0.59 | 0.52 | 0.37 | 0.54 |
| 4 | 147947622 | rs1370301 | -2.491 | -1.871 | 0.415 | 1.339 | -0.111 | -0.009 | -0.009 | 0 | 0.64 | 0.94 | 0.69 | 0.76 | 0.51 | 0.73 |
| 4 | 147964863 | rs1561223 | -1.822 | -1.721 | 0.506 | 0.892 | 0.002 | -0.021 | -0.021 | 0 | 0.55 | 0.83 | 0.51 | 0.63 | 0.47 | 0.51 |
| 4 | 147964933 | rs11734148 | 0.849 | -1.673 | 0.615 | 0.639 | -0.029 | -0.041 | -0.041 | 0 | 0.09 | 0.21 | 0.03 | 0.02 | 0.05 | 0.02 |
| 4 | 147970022 | rs10008795 | -1.275 | -1.713 | 0.53 | 0.704 | -0.004 | -0.002 | -0.002 | 0 | 0.31 | 0.31 | 0.28 | 0.28 | 0.21 | 0.26 |
| 4 | 147970692 | rs4835363 | -1.77 | -1.382 | 0.553 | 0.952 | -0.032 | -0.059 | -0.059 | 0 | 0.61 | 0.84 | 0.65 | 0.57 | 0.49 | 0.48 |
| 4 | 147972073 | rs10446782 | -0.418 | -1.587 | 0.862 | 0.716 | -0.016 | -0.02 | -0.02 | 0 | 0.41 | 0.27 | 0.48 | 0.54 | 0.47 | 0.42 |
| 4 | 147979721 | rs7674049 | -0.443 | -1.667 | 0.906 | 0.723 | 0 | 0.007 | 0.007 | 0 | 0.42 | 0.64 | 0.6 | 0.4 | 0.45 | 0.36 |
| 4 | 147980595 | rs4835063 | -0.055 | -1.637 | 0.666 | 0.722 | 0.009 | -0.039 | -0.039 | 0 | 0.2 | 0.01 | 0.11 | 0.26 | 0.16 | 0.21 |
| 4 | 147983095 | rs4835367 | -1.452 | -1.602 | 0.961 | 0.737 | 0.016 | 0.013 | 0.013 | 0 | 0.62 | 0.55 | 0.69 | 0.66 | 0.6 | 0.57 |
| 4 | 147991697 | rs6855501 | -0.061 | -1.701 | 0.665 | 0.526 | 0.017 | -0.012 | -0.012 | 0.299 | 0.81 | 0.91 | 0.89 | 0.72 | 0.85 | 0.8 |

| | | | | | | | | | | | | | | | | |
|---|---|---|---|---|---|---|---|---|---|---|---|---|---|---|---|---|
| 4 | 147993702 | rs1507500 | 0.654 | -1.849 | 0.773 | 0.541 | -0.01 | -0.003 | -0.003 | 0.676 | 0.16 | 0.2 | 0.21 | 0.14 | 0.11 | 0.12 |
| 4 | 147994381 | rs10034640 | 1.003 | -1.758 | 0.866 | 0.537 | -0.004 | -0.002 | -0.002 | 0.78 | 0.74 | 0.76 | 0.87 | 0.71 | 0.84 | 0.76 |
| 4 | 147998571 | rs7667695 | 1.117 | -1.771 | 0.901 | 0.481 | -0.03 | -0.018 | -0.018 | 0 | 0.73 | 0.64 | 0.86 | 0.72 | 0.84 | 0.76 |
| 4 | 147998958 | rs4571308 | -0.083 | -1.535 | 0.308 | 0.65 | -0.006 | -0.047 | -0.047 | 0 | 0.81 | 0.99 | 0.9 | 0.72 | 0.86 | 0.8 |
| 4 | 148001517 | rs17022539 | 0.521 | -2.066 | 0.538 | 0.517 | -0.021 | -0.008 | -0.008 | 0 | 0.17 | 0.25 | 0.22 | 0.14 | 0.12 | 0.15 |
| 4 | 148001572 | rs10212894 | 0.668 | -2.037 | 0.619 | 0.475 | -0.015 | -0.004 | -0.004 | 0 | 0.62 | 0.54 | 0.66 | 0.57 | 0.75 | 0.67 |
| 4 | 148005660 | rs4383597 | 0.284 | -1.453 | 0.734 | 0.371 | 0.001 | -0.002 | -0.002 | 0 | 0.41 | 0.52 | 0.33 | 0.29 | 0.41 | 0.49 |
| 4 | 148006860 | rs4379045 | 0.071 | -1.391 | 0.214 | 0.363 | 0.02 | 0.016 | 0.016 | 0 | 0.82 | 0.98 | 0.9 | 0.81 | 0.88 | 0.83 |
| 4 | 148011063 | rs4414935 | 0.078 | -1.388 | 0.219 | 0.343 | 0.023 | 0.023 | 0.023 | 0 | 0.82 | 0.97 | 0.9 | 0.81 | 0.89 | 0.83 |
| 4 | 148012476 | rs1979956 | -0.19 | -1.253 | 1.27 | 0.661 | 0.002 | -0.004 | -0.004 | 0 | 0.41 | 0.53 | 0.33 | 0.29 | 0.42 | 0.5 |
| 4 | 148021144 | rs1911588 | 0.213 | -0.45 | 0.128 | 0.955 | -0.066 | -0.046 | -0.046 | 0.034 | 0.28 | 0.08 | 0.27 | 0.29 | 0.36 | 0.42 |
| 4 | 148025559 | rs4543091 | 0.957 | -1.424 | 1.029 | 1.32 | -0.017 | 0.076 | 0.076 | 0.329 | 0.71 | 0.5 | 0.75 | 0.54 | 0.78 | 0.5 |
| 4 | 148040980 | rs4835378 | 0.958 | -1.429 | 0.528 | 1.129 | -0.029 | 0.006 | 0.006 | 0 | 0.69 | 0.43 | 0.78 | 0.63 | 0.8 | 0.62 |
| 4 | 148051452 | rs4485803 | 0.702 | -1.192 | -0.651 | 1.078 | 0.04 | -0.021 | -0.021 | 0 | 0.87 | 0.95 | 0.92 | 0.81 | 0.95 | 0.89 |
| 4 | 148066627 | rs12499925 | 0.818 | -1.106 | -0.693 | 1.258 | -0.022 | -0.069 | -0.069 | 0 | 0.11 | 0.01 | 0.17 | 0.25 | 0.12 | 0.22 |
| 4 | 148069540 | rs2174801 | -1.329 | -1.864 | -1.025 | 1.444 | 0.086 | -0.056 | -0.056 | 0 | 0.24 | 0.07 | 0.13 | 0.29 | 0.12 | 0.22 |
| 4 | 148075340 | rs1354884 | -1.334 | -1.887 | -0.971 | 1.427 | 0.059 | -0.018 | -0.018 | 0 | 0.25 | 0.17 | 0.12 | 0.29 | 0.12 | 0.22 |
| 4 | 148077418 | rs2357604 | -1.369 | -1.902 | -0.818 | 1.426 | 0.032 | -0.007 | -0.007 | 0 | 0.25 | 0.22 | 0.12 | 0.29 | 0.12 | 0.22 |
| 4 | 148081634 | rs952688 | -0.341 | -1.973 | -0.871 | 1.317 | 0.03 | -0.04 | -0.04 | 0.373 | 0.11 | 0.04 | 0.08 | 0.22 | 0.06 | 0.14 |
| 4 | 148084305 | rs4399964 | -0.015 | -1.975 | -0.354 | 1.422 | 0.054 | 0.161 | 0.161 | 0.074 | 0.86 | 0.62 | 0.82 | 0.72 | 0.86 | 0.69 |
| 4 | 148093804 | rs17022766 | -0.096 | -1.994 | -0.756 | 1.309 | 0.014 | 0.015 | 0.015 | 0 | 0.12 | 0.1 | 0.03 | 0.04 | 0.06 | 0.08 |
| 4 | 148099831 | rs17022775 | 0.001 | -1.997 | -0.76 | 1.308 | 0.035 | 0.033 | 0.033 | 0 | 0.12 | 0.04 | 0.03 | 0.04 | 0.06 | 0.08 |
| 4 | 148131784 | rs4835382 | 2.134 | -2.143 | -1.096 | 1.081 | -0.089 | 0.053 | 0.053 | 0.515 | 0.41 | 0.03 | 0.46 | 0.22 | 0.57 | 0.24 |
| 4 | 148162962 | rs7656015 | -0.011 | -2.323 | -0.197 | 1.043 | 0.03 | -0.03 | -0.03 | 0 | 0.9 | 0.94 | 0.92 | 0.79 | 0.95 | 0.9 |
| 4 | 148171265 | rs7658280 | 0.508 | -2.2 | -0.137 | 1.066 | 0.09 | -0.03 | -0.03 | 0 | 0.78 | 0.92 | 0.9 | 0.75 | 0.91 | 0.8 |
| 4 | 148174084 | rs4835074 | 1.819 | -2.275 | 0.157 | 1.063 | 0.017 | -0.01 | -0.01 | 0 | 0.6 | 0.36 | 0.7 | 0.46 | 0.77 | 0.5 |
| 4 | 148200205 | rs12644017 | -0.676 | -2.208 | 0.045 | 1.083 | 0.022 | 0.105 | 0.105 | 0 | 0.16 | 0.32 | 0.19 | 0.29 | 0.13 | 0.31 |
| 4 | 148209824 | rs6841482 | 0.745 | -1.999 | -0.757 | 1.05 | 0.069 | -0.017 | -0.017 | 0.308 | 0.78 | 0.87 | 0.9 | 0.76 | 0.91 | 0.81 |
| 4 | 148229316 | rs2714886 | 1.058 | -2.105 | -0.546 | 1.217 | -0.003 | -0.011 | -0.011 | 0.791 | 0.75 | 0.71 | 0.89 | 0.55 | 0.91 | 0.78 |
| 4 | 148268027 | rs2163012 | 1.687 | -2.378 | -0.861 | 0.867 | 0.125 | -0.076 | -0.076 | 3.871 | 0.57 | 0.11 | 0.65 | 0.46 | 0.79 | 0.51 |
| 4 | 148277011 | rs4835078 | 0.365 | -2.542 | -0.675 | 0.886 | 0.006 | -0.005 | -0.005 | 0 | 0.85 | 0.85 | 0.91 | 0.78 | 0.96 | 0.92 |
| 4 | 148280874 | rs17023122 | -0.596 | -2.442 | 0.409 | 0.778 | 0.108 | 0.007 | 0.007 | 0.299 | 0.06 | 0.31 | 0.06 | 0 | 0.06 | 0.02 |
| 4 | 148289389 | rs1429142 | -1.632 | -2.52 | 0.113 | 0.788 | 0.017 | -0.016 | -0.016 | 0 | 0.26 | 0.79 | 0.26 | 0.33 | 0.16 | 0.39 |
| 4 | 148291697 | rs1560228 | 0.181 | -2.386 | 0.089 | 0.417 | 0.023 | 0.014 | 0.014 | 0 | 0.9 | 0.95 | 0.94 | 0.91 | 0.97 | 0.94 |
| 4 | 148301706 | rs1429102 | -1.7 | -2.467 | 0.353 | 0.096 | -0.008 | -0.001 | -0.001 | 3.565 | 0.36 | 0.55 | 0.34 | 0.36 | 0.31 | 0.4 |
| 4 | 148306058 | rs11936867 | 1.958 | -2.691 | 0.196 | -0.03 | 0.012 | 0.006 | 0.006 | 0 | 0.54 | 0.44 | 0.6 | 0.57 | 0.66 | 0.46 |
| 4 | 148308126 | rs7669258 | -1.419 | -2.761 | 0.929 | -0.122 | -0.004 | 0.017 | 0.017 | 0 | 0.25 | 0.28 | 0.23 | 0.35 | 0.19 | 0.35 |
| 4 | 148309369 | rs13140736 | 0.609 | -2.71 | 0.812 | -0.097 | -0.006 | 0.037 | 0.037 | 0 | 0.08 | 0.04 | 0.09 | 0 | 0.12 | 0.05 |
| 4 | 148311822 | rs17674732 | -0.965 | -3.067 | 0.747 | -0.262 | 0.035 | -0.058 | -0.058 | 0 | 0.21 | 0.09 | 0.21 | 0.35 | 0.15 | 0.38 |
| 4 | 148318048 | rs7671190 | 1.954 | -3.008 | 0.866 | -0.207 | 0.01 | 0.002 | 0.002 | 0 | 0.58 | 0.41 | 0.66 | 0.57 | 0.74 | 0.46 |
| 4 | 148322387 | rs1429107 | 0.725 | -3.266 | 1.141 | -0.397 | -0.067 | -0.06 | -0.06 | 0.072 | 0.82 | 0.72 | 0.9 | 0.92 | 0.96 | 0.87 |
| 4 | 148328556 | rs2195469 | 0.147 | -3.2 | 0.954 | -0.364 | 0.016 | 0.016 | 0.016 | 0 | 0.86 | 0.92 | 0.92 | 0.92 | 0.97 | 0.87 |
| 4 | 148329646 | rs17023204 | -1.357 | -3.049 | 0.952 | -0.278 | -0.014 | 0.019 | 0.019 | 0.18 | 0.27 | 0.4 | 0.25 | 0.33 | 0.22 | 0.38 |
| 4 | 148337747 | rs1429115 | 1.083 | -2.84 | 0.765 | -0.525 | -0.003 | -0.005 | -0.005 | 0 | 0.6 | 0.6 | 0.66 | 0.69 | 0.65 | 0.54 |
| 4 | 148349756 | rs1594085 | 2.573 | -3.053 | 0.545 | -0.55 | -0.079 | -0.059 | -0.059 | 2.454 | 0.41 | 0.06 | 0.53 | 0.58 | 0.59 | 0.42 |
| 4 | 148353423 | rs4835398 | 0.949 | -3.06 | 0.631 | -0.554 | -0.002 | -0.004 | -0.004 | 0 | 0.61 | 0.62 | 0.66 | 0.69 | 0.65 | 0.54 |
| 4 | 148353940 | rs1429123 | 2.216 | -3 | -0.092 | -0.527 | -0.083 | -0.063 | -0.063 | 0 | 0.4 | 0.06 | 0.53 | 0.58 | 0.6 | 0.42 |
| 4 | 148362457 | rs17023262 | 1.129 | -2.889 | 0.424 | -0.499 | 0.013 | 0.02 | 0.02 | 0.019 | 0.59 | 0.32 | 0.64 | 0.69 | 0.65 | 0.54 |
| 4 | 148362953 | rs1560231 | 1.092 | -2.928 | 0.65 | -0.52 | 0.007 | 0.001 | 0.001 | 2.32 | 0.67 | 0.27 | 0.71 | 0.69 | 0.75 | 0.58 |
| 4 | 148370419 | rs2063936 | 1.705 | -2.857 | 0.516 | -0.705 | 0.113 | 0.074 | 0.074 | 0.824 | 0.5 | 0.15 | 0.72 | 0.64 | 0.72 | 0.57 |
| 4 | 148380805 | rs1400558 | 1.579 | -2.9 | 1.514 | -0.725 | -0.005 | -0.002 | -0.002 | 0 | 0.52 | 0.38 | 0.74 | 0.64 | 0.72 | 0.57 |
| 4 | 148396927 | rs4333164 | -0.694 | -3.133 | 1.537 | -0.604 | -0.018 | 0 | 0 | 1.508 | 0.13 | 0.21 | 0.04 | 0.11 | 0.05 | 0.17 |
| 4 | 148400819 | rs6842241 | -1.016 | -3.277 | 1.668 | -0.667 | -0.073 | -0.034 | -0.034 | 0.71 | 0.26 | 0.45 | 0.17 | 0.23 | 0.15 | 0.25 |
| 4 | 148402369 | rs1801708 | -2.524 | -3.566 | 1.088 | -1.092 | 0.108 | 0.049 | 0.049 | 0.15 | 0.54 | 0.94 | 0.33 | 0.34 | 0.32 | 0.46 |
| 4 | 148402737 | rs6855875 | -1.782 | -3.494 | 1.328 | -1.039 | -0.051 | -0.009 | -0.009 | 0 | 0.37 | 0.51 | 0.21 | 0.34 | 0.19 | 0.38 |
| 4 | 148414651 | rs17612742 | -0.45 | -3.271 | 1.085 | -1.035 | 0.014 | 0.005 | 0.005 | 0 | 0.22 | 0.19 | 0.15 | 0.23 | 0.14 | 0.21 |
| 4 | 148422625 | rs6848108 | -1.634 | -3.104 | 0.455 | -1.091 | 0.038 | 0.039 | 0.039 | 0 | 0.51 | 0.58 | 0.32 | 0.32 | 0.33 | 0.44 |
| 4 | 148429102 | rs4591581 | -1.397 | -3.042 | 0.851 | -1.23 | -0.05 | -0.048 | -0.048 | 1.181 | 0.21 | 0.59 | 0.09 | 0.1 | 0.11 | 0.12 |
| 4 | 148434688 | rs10305895 | -2.043 | -3.293 | 0.573 | -1.375 | -0.031 | -0.016 | -0.016 | 0 | 0.38 | 0.43 | 0.32 | 0.3 | 0.24 | 0.39 |
| 4 | 148435446 | rs6827096 | -1.62 | -3.243 | -0.022 | -1.367 | 0.019 | 0.027 | 0.027 | 0.846 | 0.24 | 0.8 | 0.22 | 0.1 | 0.14 | 0.14 |
| 4 | 148440625 | rs1878404 | -2.202 | -3.238 | -0.902 | -1.289 | 0.138 | 0.042 | 0.042 | 0.181 | 0.42 | 0.91 | 0.22 | 0.3 | 0.21 | 0.39 |
| 4 | 148441139 | rs4835412 | -0.767 | -2.782 | -0.696 | -1.112 | 0.041 | -0.037 | -0.037 | 0 | 0.18 | 0.03 | 0.12 | 0.2 | 0.1 | 0.27 |
| 4 | 148445429 | rs10008744 | -1.84 | -2.901 | -0.738 | -1.223 | 0.101 | 0.041 | 0.041 | 0 | 0.39 | 0.89 | 0.33 | 0.29 | 0.2 | 0.41 |
| 4 | 148448870 | rs4639051 | -0.271 | -2.764 | -0.198 | -1.4 | -0.004 | -0.064 | -0.064 | 0 | 0.21 | 0.38 | 0.21 | 0.07 | 0.15 | 0.11 |
| 4 | 148461037 | rs5333 | -1.328 | -2.662 | -0.463 | -1.283 | 0.019 | 0.023 | 0.023 | 0 | 0.39 | 0.65 | 0.34 | 0.24 | 0.22 | 0.37 |
| 4 | 148461073 | rs5334 | -1.328 | -2.662 | -0.463 | -1.283 | 0.019 | 0.023 | 0.023 | 0 | 0.39 | 0.65 | 0.34 | 0.24 | 0.22 | 0.37 |
| 4 | 148464992 | rs5343 | 0.778 | -1.029 | -1.063 | -1.115 | 0.015 | 0.077 | 0.077 | 0 | 0.37 | 0.11 | 0.37 | 0.23 | 0.3 | 0.35 |
| 4 | 148486299 | rs17613180 | -0.555 | -1.591 | -0.659 | -1.143 | -0.002 | 0 | 0 | 0.033 | 0.6 | 0.62 | 0.44 | 0.44 | 0.36 | 0.67 |
| 4 | 148488545 | rs2090422 | 0.508 | -1.344 | -1.308 | -1.034 | 0.015 | 0.001 | 0.001 | 0 | 0.17 | 0.11 | 0.04 | 0.17 | 0.05 | 0.23 |

| | | | | | | | | | | | | | | | | |
|---|---|---|---|---|---|---|---|---|---|---|---|---|---|---|---|---|
| 4 | 148499607 | rs4110 | 0.584 | -0.629 | -1.235 | -1.097 | 0.012 | 0.079 | 0.079 | 0 | 0.37 | 0.1 | 0.37 | 0.23 | 0.31 | 0.38 |
| 4 | 148509519 | rs4835088 | -1.835 | 0.508 | -0.507 | -0.746 | -0.039 | 0.159 | 0.159 | 0.009 | 0.5 | 0.92 | 0.45 | 0.32 | 0.47 | 0.46 |
| 4 | 148510474 | rs10028794 | -0.393 | 0.901 | 0.087 | -0.888 | 0.005 | -0.04 | -0.04 | 0.338 | 0.25 | 0.47 | 0.28 | 0.21 | 0.25 | 0.21 |
| 4 | 148510725 | rs10519904 | -0.88 | 0.862 | -0.063 | -0.418 | -0.007 | 0.007 | 0.007 | 0 | 0.18 | 0.13 | 0.1 | 0.03 | 0.18 | 0.09 |
| 4 | 148522294 | rs4835438 | 0.957 | 0.495 | 0.385 | -0.518 | 0.001 | -0.136 | -0.136 | 0 | 0.73 | 0.64 | 0.86 | 0.99 | 0.79 | 0.85 |
| 4 | 148529844 | rs6848237 | -0.841 | -0.052 | 1.087 | 0.452 | 0 | 0.011 | 0.011 | 0 | 0.05 | 0.03 | 0.01 | 0.23 | 0.01 | 0.06 |
| 4 | 148550590 | rs10519911 | -1.28 | -0.099 | 1.292 | 0.433 | -0.004 | 0.006 | 0.006 | 0 | 0.06 | 0.06 | 0.02 | 0.24 | 0.01 | 0.06 |
| 4 | 148553461 | rs13119467 | 2.206 | -0.442 | 1.509 | 0.717 | 0.009 | -0.009 | -0.009 | 0.056 | 0.77 | 0.21 | 0.85 | 0.67 | 0.9 | 0.77 |
| 4 | 148554421 | rs2654937 | 2.219 | -0.413 | 1.807 | 0.623 | -0.048 | 0.137 | 0.137 | 1.229 | 0.89 | 0.52 | 0.95 | 0.76 | 0.99 | 0.94 |
| 4 | 148566463 | rs17475359 | 0.562 | -0.043 | 1.197 | 0.486 | -0.027 | 0.027 | 0.027 | 0 | 0.05 | 0.01 | 0.03 | 0 | 0.08 | 0.04 |
| 4 | 148612488 | rs11731298 | 0.463 | -0.607 | 1.214 | 0.825 | -0.006 | -0.018 | -0.018 | 0 | 0.07 | 0 | 0.04 | 0.11 | 0.08 | 0.08 |
| 4 | 148631527 | rs4835446 | 1.737 | -0.534 | 1.757 | 0.777 | -0.06 | 0.036 | 0.036 | 1.387 | 0.8 | 0.54 | 0.92 | 0.72 | 0.9 | 0.81 |
| 4 | 148648861 | rs12502667 | 1.994 | -0.564 | 0.929 | 0.761 | 0.044 | -0.011 | -0.011 | 0 | 0.75 | 0.15 | 0.83 | 0.7 | 0.89 | 0.74 |
| 4 | 148654651 | rs6836509 | 1.937 | -0.591 | 0.953 | 0.625 | 0.028 | 0.007 | 0.007 | 6.025 | 0.76 | 0.17 | 0.84 | 0.78 | 0.89 | 0.75 |
| 4 | 148658108 | rs7675343 | 0.993 | -0.715 | 0.388 | 0.847 | -0.072 | -0.103 | -0.103 | 0 | 0.39 | 0.05 | 0.32 | 0.45 | 0.36 | 0.39 |
| 4 | 148659137 | rs17023732 | -1.599 | -1.039 | 1.933 | 1.081 | -0.015 | 0.063 | 0.063 | 0 | 0.07 | 0.17 | 0.03 | 0.15 | 0.01 | 0.05 |
| 4 | 148687610 | rs6845560 | 2.013 | -1.27 | 1.324 | 0.719 | 0.01 | 0.012 | 0.012 | 6.674 | 0.78 | 0.18 | 0.86 | 0.88 | 0.93 | 0.77 |
| 4 | 148694658 | rs6535561 | 1.787 | -1.15 | 1.412 | 0.809 | -0.002 | 0.004 | 0.004 | 7.446 | 0.79 | 0.18 | 0.86 | 0.85 | 0.94 | 0.78 |
| 4 | 148695956 | rs964170 | 1.787 | -1.15 | 1.412 | 0.809 | -0.002 | 0.004 | 0.004 | 7.446 | 0.79 | 0.18 | 0.86 | 0.85 | 0.94 | 0.79 |
| 4 | 148698365 | rs13136202 | 1.585 | -1.19 | 1.794 | 0.788 | -0.067 | -0.051 | -0.051 | 0.209 | 0.82 | 0.4 | 0.88 | 0.89 | 0.94 | 0.79 |
| 4 | 148722083 | rs6837391 | 1.914 | -1.17 | 1.621 | 0.849 | -0.018 | -0.004 | -0.004 | 0.343 | 0.79 | 0.21 | 0.86 | 0.85 | 0.93 | 0.78 |
| 4 | 148727355 | rs17023845 | -1.718 | -1.08 | 2.369 | 0.874 | 0.043 | 0.097 | 0.097 | 0 | 0.06 | 0.38 | 0.04 | 0.07 | 0.01 | 0.04 |
| 4 | 148730825 | rs747097 | 2.447 | -1.051 | 1.93 | 0.939 | 0.042 | 0.004 | 0.004 | 6.495 | 0.75 | 0.14 | 0.81 | 0.82 | 0.86 | 0.74 |
| 4 | 148733764 | rs4835457 | 2.189 | -1.043 | 2.065 | 0.825 | -0.002 | -0.003 | -0.003 | 0 | 0.78 | 0.26 | 0.82 | 0.88 | 0.86 | 0.76 |
| 4 | 148734627 | rs7683103 | -2.634 | -0.927 | 2.877 | 0.963 | 0.015 | 0.084 | 0.084 | 0 | 0.12 | 0.49 | 0.15 | 0.14 | 0.09 | 0.12 |
| 4 | 148735644 | rs10519917 | -1.572 | -0.936 | 2.736 | 0.937 | 0.021 | 0.011 | 0.011 | 0 | 0.08 | 0.29 | 0.06 | 0.07 | 0.08 | 0.05 |
| 4 | 148740219 | rs2043644 | 2.799 | -1.119 | 2.374 | 1.251 | 0.005 | -0.004 | -0.004 | 0 | 0.83 | 0.19 | 0.81 | 0.79 | 0.91 | 0.78 |
| 4 | 148753310 | rs9996108 | -1.489 | -1.108 | 2.946 | 1.267 | 0.002 | 0.068 | 0.068 | 0 | 0.09 | 0.21 | 0.07 | 0.16 | 0.08 | 0.06 |
| 4 | 148760097 | rs10461235 | 3.026 | -0.835 | 2.825 | 1.252 | 0.004 | -0.002 | -0.002 | 0 | 0.81 | 0.19 | 0.79 | 0.8 | 0.87 | 0.8 |
| 4 | 148762097 | rs6846899 | -3.202 | -0.905 | 2.724 | 1.194 | -0.013 | 0.055 | 0.055 | 0.213 | 0.18 | 0.53 | 0.11 | 0.21 | 0.12 | 0.14 |
| 4 | 148763687 | rs11737346 | 3.346 | -0.988 | 2.232 | 1.398 | 0.039 | -0.05 | -0.05 | 0.419 | 0.77 | 0.1 | 0.78 | 0.7 | 0.86 | 0.81 |
| 4 | 148783402 | rs1013842 | 2.673 | -0.882 | 2.629 | 1.489 | -0.017 | 0.012 | 0.012 | 0.973 | 0.82 | 0.29 | 0.87 | 0.79 | 0.88 | 0.86 |
| 4 | 148787669 | rs4240341 | 2.58 | -1.318 | 2.664 | 1.393 | -0.042 | 0.039 | 0.039 | 0.712 | 0.83 | 0.3 | 0.82 | 0.7 | 0.92 | 0.87 |
| 4 | 148789141 | rs12331562 | -1.985 | -1.306 | 2.803 | 1.377 | -0.027 | 0.13 | 0.13 | 0 | 0.12 | 0.28 | 0.15 | 0.26 | 0.08 | 0.06 |
| 4 | 148798090 | rs11734619 | 1.317 | -1.568 | 1.866 | 1.209 | 0.096 | 0.003 | 0.003 | 0.658 | 0.5 | 0.06 | 0.57 | 0.44 | 0.62 | 0.53 |
| 4 | 148798473 | rs10026985 | -1.615 | -1.45 | 2.235 | 1.339 | -0.021 | 0.162 | 0.162 | 0 | 0.11 | 0.3 | 0.14 | 0.26 | 0.08 | 0.06 |
| 4 | 148799613 | rs6840771 | -2.143 | -1.454 | 2.263 | 1.3 | -0.033 | 0.176 | 0.176 | 0.718 | 0.13 | 0.5 | 0.2 | 0.27 | 0.08 | 0.08 |
| 4 | 148815028 | rs17676080 | -0.907 | -1.437 | 2.259 | 1.284 | -0.01 | 0.093 | 0.093 | 0 | 0.09 | 0.14 | 0.18 | 0.27 | 0.08 | 0.06 |
| 4 | 148820632 | rs893033 | 2.346 | -1.449 | 2.185 | 1.261 | -0.047 | 0.133 | 0.133 | 0.028 | 0.85 | 0.4 | 0.78 | 0.71 | 0.92 | 0.86 |
| 4 | 148822783 | rs3857095 | 1.866 | -1.494 | 1.536 | 1.094 | -0.027 | -0.016 | -0.016 | 0 | 0.93 | 0.67 | 0.97 | 0.98 | 0.99 | 0.92 |
| 4 | 148827747 | rs12509618 | 1.861 | -1.445 | 0.629 | 1.187 | 0.054 | -0.058 | -0.058 | 0.017 | 0.79 | 0.12 | 0.73 | 0.65 | 0.91 | 0.84 |
| 4 | 148828922 | rs12643550 | -1.531 | -1.276 | 0.674 | 1.26 | -0.047 | 0.18 | 0.18 | 0.124 | 0.16 | 0.59 | 0.24 | 0.34 | 0.09 | 0.15 |
| 4 | 148830472 | rs1529935 | 0.657 | -1.01 | -0.655 | 1.048 | 0.094 | 0.051 | 0.051 | 0.659 | 0.48 | 0.06 | 0.58 | 0.37 | 0.61 | 0.52 |
| 4 | 148838054 | rs7655265 | -1.772 | -0.783 | 0.8 | 1.323 | -0.028 | 0.277 | 0.277 | 0.038 | 0.1 | 0.42 | 0.19 | 0.32 | 0.01 | 0.08 |
| 4 | 148839717 | rs4027115 | -1.271 | -0.784 | 0.957 | 1.499 | 0.024 | 0.228 | 0.228 | 0.042 | 0.06 | 0.38 | 0.16 | 0.21 | 0.01 | 0.05 |
| 4 | 148848283 | rs4835111 | -0.338 | -0.825 | 0.373 | 0.548 | 0 | 0.023 | 0.023 | 0 | 0.23 | 0.14 | 0.14 | 0 | 0.17 | 0.15 |
| 4 | 148856057 | rs187005 | 1.298 | -0.988 | 0.944 | 0.897 | 0.001 | -0.006 | -0.006 | 0 | 0.93 | 0.76 | 0.96 | 0.98 | 0.96 | 0.96 |
| 4 | 148866526 | rs2635266 | 1.727 | -0.989 | 0.823 | 0.909 | -0.038 | 0.07 | 0.07 | 0.971 | 0.84 | 0.28 | 0.78 | 0.66 | 0.94 | 0.87 |
| 4 | 148873538 | rs2262881 | 1.813 | -1.221 | 1.411 | 0.519 | -0.043 | -0.025 | -0.025 | 0.432 | 0.82 | 0.32 | 0.86 | 0.9 | 0.93 | 0.8 |
| 4 | 148874351 | rs2635259 | 1.785 | -1.223 | 1.627 | 0.504 | -0.057 | -0.035 | -0.035 | 0 | 0.83 | 0.37 | 0.87 | 0.9 | 0.93 | 0.8 |
| 4 | 148891027 | rs17024235 | -1.979 | -1.314 | 1.872 | 0.364 | -0.035 | -0.049 | -0.049 | 0 | 0.14 | 0.32 | 0.06 | 0.08 | 0.08 | 0.13 |
| 4 | 148897030 | rs9684012 | -2.766 | -1.408 | 1.788 | 0.332 | -0.053 | -0.045 | -0.045 | 0.208 | 0.21 | 0.63 | 0.14 | 0.1 | 0.1 | 0.24 |
| 4 | 148899082 | rs11099673 | 0.44 | -1.273 | 1.182 | 0.117 | -0.003 | -0.007 | -0.007 | 0 | 0.64 | 0.62 | 0.62 | 0.72 | 0.71 | 0.78 |
| 4 | 148903388 | rs13129809 | 1.282 | -1.447 | 0.541 | 0.058 | 0.042 | 0.035 | 0.035 | 0.01 | 0.49 | 0.09 | 0.49 | 0.63 | 0.65 | 0.58 |
| 4 | 148916437 | rs10519933 | -1.628 | -1.179 | 1.047 | -0.047 | -0.048 | -0.039 | -0.039 | 0.249 | 0.14 | 0.34 | 0.09 | 0.05 | 0.07 | 0.16 |
| 4 | 148928002 | rs10025443 | -0.157 | -0.621 | 0.326 | -0.434 | -0.002 | 0.004 | 0.004 | 0 | 0.65 | 0.68 | 0.69 | 0.87 | 0.68 | 0.82 |
| 4 | 148928213 | rs10003349 | 0.084 | -0.155 | 0.114 | -0.305 | -0.001 | 0.021 | 0.021 | 0.03 | 0.51 | 0.58 | 0.5 | 0.71 | 0.59 | 0.46 |
| 4 | 148928372 | rs12643170 | 0.412 | -0.321 | -0.066 | -0.113 | 0.001 | 0 | 0 | 0.851 | 0.44 | 0.2 | 0.51 | 0.63 | 0.61 | 0.44 |
| 4 | 148936414 | rs10213376 | -0.675 | -0.35 | 0.87 | -0.009 | 0.011 | 0.013 | 0.013 | 0 | 0.07 | 0.31 | 0.05 | 0.06 | 0.06 | 0.02 |
| 4 | 148938598 | rs6815554 | -0.675 | -0.35 | 0.87 | -0.009 | 0.011 | 0.013 | 0.013 | 0 | 0.07 | 0.31 | 0.05 | 0.06 | 0.06 | 0.02 |
| 4 | 148940093 | rs9991394 | -1.901 | -0.763 | 0.732 | 0.023 | -0.069 | -0.002 | -0.002 | 3.214 | 0.22 | 0.55 | 0.1 | 0.2 | 0.1 | 0.27 |
| 4 | 148950412 | rs13111299 | -0.532 | -0.724 | 0.685 | 0.043 | 0.02 | -0.012 | -0.012 | 0 | 0.13 | 0.05 | 0.03 | 0.14 | 0.04 | 0.23 |
| 4 | 148950873 | rs7668569 | -1.768 | -0.724 | 0.643 | 0.043 | -0.077 | 0.011 | 0.011 | 0 | 0.21 | 0.46 | 0.09 | 0.2 | 0.1 | 0.33 |
| 4 | 148952982 | rs6839459 | -0.821 | -0.476 | 0.594 | 0.04 | 0.022 | 0.028 | 0.028 | 0.016 | 0.08 | 0.37 | 0.05 | 0.06 | 0.06 | 0.02 |
| 4 | 148974602 | rs6845865 | -1.242 | -0.018 | 0.348 | 0.06 | -0.034 | 0.003 | 0.003 | 0 | 0.22 | 0.51 | 0.23 | 0.19 | 0.16 | 0.28 |
| 4 | 148979174 | rs1021956 | -0.896 | -0.051 | -0.762 | -0.487 | -0.013 | 0.074 | 0.074 | 0 | 0.75 | 0.93 | 0.87 | 0.9 | 0.8 | 0.78 |
| 4 | 148979700 | rs10857228 | -0.236 | -0.056 | -0.169 | -0.648 | -0.012 | -0.031 | -0.031 | 0 | 0.64 | 0.41 | 0.77 | 0.85 | 0.72 | 0.76 |
| 4 | 148983139 | rs13109093 | 1.397 | 0.37 | -1.244 | -1.183 | -0.074 | -0.087 | -0.087 | 0 | 0.25 | 0.05 | 0.44 | 0.65 | 0.36 | 0.38 |
| 4 | 148983376 | rs10519931 | 1.118 | 0.204 | -1.085 | -1.47 | 0.003 | 0.058 | 0.058 | 0.012 | 0.18 | 0.07 | 0.08 | 0 | 0.15 | 0.08 |

| | | | | | | | | | | | | | | | | |
|---|---|---|---|---|---|---|---|---|---|---|---|---|---|---|---|---|
| 4 | 148984321 | rs2276932 | 1.485 | 0.196 | -1.205 | -1.422 | -0.015 | -0.024 | -0.024 | 0.832 | 0.06 | 0.01 | 0.05 | 0.06 | 0.07 | 0.03 |
| 4 | 148987187 | rs12650033 | -1.121 | 0.114 | -0.566 | -1.477 | -0.022 | -0.055 | -0.055 | 0 | 0.65 | 0.48 | 0.78 | 0.85 | 0.73 | 0.75 |
| 4 | 148987972 | rs7660368 | 0.001 | 0.099 | -0.97 | -1.467 | -0.007 | 0.122 | 0.122 | 0.318 | 0.44 | 0.17 | 0.66 | 0.83 | 0.58 | 0.66 |
| 4 | 148990995 | rs6824449 | -0.442 | 0.106 | -0.961 | -1.392 | 0.001 | 0.079 | 0.079 | 0.602 | 0.61 | 0.2 | 0.75 | 0.83 | 0.62 | 0.72 |
| 4 | 148997402 | rs11933380 | -0.453 | 0.098 | -0.965 | -1.314 | 0.022 | 0.083 | 0.083 | 0.577 | 0.61 | 0.17 | 0.78 | 0.83 | 0.67 | 0.73 |
| 4 | 148998566 | rs7696997 | -1.158 | 0.061 | -0.388 | -1.333 | -0.007 | -0.047 | -0.047 | 0 | 0.68 | 0.56 | 0.81 | 0.85 | 0.69 | 0.77 |
| 4 | 149000024 | rs2871 | -1.05 | -0.011 | -0.399 | -1.344 | -0.005 | -0.046 | -0.046 | 0 | 0.67 | 0.56 | 0.8 | 0.83 | 0.68 | 0.79 |
| 4 | 149001085 | rs5534 | 0.236 | -0.05 | -1.097 | -1.441 | -0.005 | 0.146 | 0.146 | 0.263 | 0.46 | 0.17 | 0.72 | 0.83 | 0.54 | 0.68 |
| 4 | 149016541 | rs9993133 | -0.064 | -0.33 | -0.652 | -1.267 | -0.023 | -0.038 | -0.038 | 0.054 | 0.26 | 0.6 | 0.12 | 0.11 | 0.19 | 0.16 |
| 4 | 149018311 | rs6840127 | -0.544 | -0.356 | -0.856 | -1.28 | 0.026 | 0.033 | 0.033 | 2.934 | 0.29 | 0.75 | 0.14 | 0.11 | 0.19 | 0.17 |
| 4 | 149022906 | rs13137823 | 0.29 | 0.464 | -1.331 | -1.276 | -0.041 | 0.007 | 0.007 | 0 | 0.18 | 0.02 | 0.32 | 0.13 | 0.21 | 0.24 |
| 4 | 149024260 | rs17024387 | 0.285 | 0.088 | -1.515 | -1.469 | 0.016 | 0.049 | 0.049 | 0 | 0.7 | 0.82 | 0.62 | 0.78 | 0.71 | 0.75 |
| 4 | 149024413 | rs11737660 | 1.597 | 0.956 | -1.485 | -1.105 | -0.077 | -0.106 | -0.106 | 0 | 0.14 | 0.01 | 0.3 | 0.57 | 0.21 | 0.26 |
| 4 | 149025406 | rs1879827 | -0.104 | 0.616 | -0.825 | -1.466 | 0.033 | -0.05 | -0.05 | 0 | 0.19 | 0.48 | 0.14 | 0.1 | 0.22 | 0.21 |
| 4 | 149030027 | rs13142954 | 0.143 | 0.88 | -1.255 | -1.299 | -0.067 | -0.009 | -0.009 | 0.314 | 0.19 | 0.01 | 0.29 | 0.13 | 0.22 | 0.25 |
| 4 | 149035449 | rs1879829 | -0.393 | 1.026 | -0.354 | -1.105 | 0.043 | -0.003 | -0.003 | 0 | 0.32 | 0.47 | 0.46 | 0.68 | 0.44 | 0.5 |
| 4 | 149040160 | rs17483391 | 0.428 | 1.036 | 0.018 | -1.039 | 0.035 | -0.052 | -0.052 | 0 | 0.09 | 0.15 | 0.06 | 0 | 0.14 | 0.04 |
| 4 | 149042293 | rs1403142 | 0.76 | 0.851 | -0.212 | -1.14 | 0.025 | 0.104 | 0.104 | 0 | 0.53 | 0.2 | 0.5 | 0.72 | 0.54 | 0.46 |
| 4 | 149042334 | rs1403143 | 0.545 | 0.853 | -0.198 | -1.139 | 0.007 | 0.025 | 0.025 | 0 | 0.54 | 0.34 | 0.51 | 0.72 | 0.54 | 0.47 |
| 4 | 149055437 | rs6857011 | 1.903 | 0.754 | -0.517 | -1.205 | 0.002 | 0.119 | 0.119 | 0.183 | 0.32 | 0.06 | 0.15 | 0.1 | 0.27 | 0.21 |
| 4 | 149060307 | rs17024404 | -0.107 | 0.674 | 0.18 | -1.224 | -0.003 | 0.035 | 0.035 | 0.385 | 0.12 | 0.43 | 0.09 | 0.08 | 0.11 | 0.2 |
| 4 | 149063275 | rs11736161 | -0.368 | 0.715 | -0.226 | -1.25 | -0.051 | 0.071 | 0.071 | 0 | 0.68 | 0.94 | 0.61 | 0.8 | 0.65 | 0.67 |
| 4 | 149064158 | rs12651282 | 0.559 | 0.926 | -0.154 | -1.205 | -0.03 | -0.043 | -0.043 | 1.499 | 0.2 | 0.14 | 0.34 | 0.61 | 0.27 | 0.26 |
| 4 | 149073555 | rs2272089 | -0.084 | 0.555 | -0.004 | -1.227 | 0.018 | 0.03 | 0.03 | 0 | 0.71 | 0.79 | 0.67 | 0.76 | 0.73 | 0.67 |
| 4 | 149081621 | rs6844155 | -0.08 | 0.559 | -0.002 | -1.078 | 0.018 | 0.007 | 0.007 | 0 | 0.71 | 0.79 | 0.67 | 0.72 | 0.73 | 0.66 |
| 4 | 149081808 | rs17024437 | 0.226 | 0.823 | 0.163 | -0.903 | 0.002 | 0.017 | 0.017 | 0 | 0.09 | 0.2 | 0.06 | 0.08 | 0.09 | 0.17 |
| 4 | 149084112 | rs6810951 | 0.339 | 0.933 | -0.057 | -0.899 | -0.056 | -0.073 | -0.073 | 0 | 0.26 | 0.13 | 0.43 | 0.54 | 0.37 | 0.25 |
| 4 | 149087127 | rs6856803 | -0.262 | 0.896 | 0.103 | -0.932 | 0.024 | 0.036 | 0.036 | 0 | 0.32 | 0.57 | 0.48 | 0.55 | 0.38 | 0.3 |
| 4 | 149090088 | rs6535584 | 0.296 | 0.913 | -0.366 | -1.225 | -0.006 | -0.001 | -0.001 | 0.1 | 0.63 | 0.67 | 0.59 | 0.69 | 0.57 | 0.47 |
| 4 | 149090326 | rs4835490 | 1.732 | 0.734 | -0.787 | -1.244 | 0.029 | 0.148 | 0.148 | 0.402 | 0.31 | 0.07 | 0.13 | 0.07 | 0.22 | 0.16 |
| 4 | 149098559 | rs10519942 | 0.869 | 1.246 | -0.414 | -0.789 | 0.007 | 0.021 | 0.021 | 0 | 0.07 | 0.14 | 0.04 | 0.08 | 0.09 | 0.22 |
| 4 | 149100252 | rs17024456 | 0.598 | 1.741 | -0.509 | -0.726 | 0.084 | -0.062 | -0.062 | 0 | 0.1 | 0.29 | 0.09 | 0 | 0.16 | 0.04 |
| 4 | 149103454 | rs11099681 | -0.394 | 1.408 | -1.153 | -0.677 | -0.074 | -0.077 | -0.077 | 0 | 0.76 | 0.96 | 0.7 | 0.75 | 0.7 | 0.69 |
| 4 | 149116534 | rs3910054 | -0.083 | 1.317 | -1.093 | -0.704 | -0.093 | -0.048 | -0.048 | 0 | 0.72 | 0.97 | 0.66 | 0.77 | 0.65 | 0.7 |
| 4 | 149117955 | rs3846302 | 0.874 | 1.405 | -0.737 | -0.698 | 0.098 | -0.045 | -0.045 | 0.089 | 0.09 | 0.3 | 0.08 | 0 | 0.14 | 0.03 |
| 4 | 149131264 | rs3846306 | 0.772 | 1.178 | -0.842 | -0.584 | -0.014 | -0.044 | -0.044 | 0.048 | 0.19 | 0.13 | 0.48 | 0.6 | 0.22 | 0.26 |
| 4 | 149132756 | rs6535594 | 0.734 | 1.784 | -0.36 | -0.401 | 0.022 | -0.027 | -0.027 | 0 | 0.61 | 0.53 | 0.57 | 0.76 | 0.49 | 0.65 |
| 4 | 149134594 | rs7699349 | 0.437 | 1.276 | -0.54 | -0.512 | -0.004 | -0.046 | -0.046 | 0.171 | 0.75 | 0.66 | 0.85 | 0.9 | 0.75 | 0.87 |
| 4 | 149135547 | rs10050229 | -0.472 | 1.466 | -0.974 | -0.378 | 0.004 | -0.021 | -0.021 | 0 | 0.3 | 0.57 | 0.2 | 0.12 | 0.36 | 0.39 |
| 4 | 149136414 | rs17483979 | -0.016 | 1.745 | -1.443 | 0.213 | -0.074 | -0.045 | -0.045 | 4.761 | 0.14 | 0.01 | 0.26 | 0.13 | 0.21 | 0.22 |
| 4 | 149136440 | rs3857080 | -0.275 | 0.9 | -1.143 | 0.099 | 0.01 | 0.019 | 0.019 | 0 | 0.85 | 0.91 | 0.93 | 0.93 | 0.89 | 0.94 |
| 4 | 149139067 | rs13116347 | 0.399 | 1.142 | -0.146 | 0.519 | 0.037 | 0.082 | 0.082 | 0 | 0.89 | 0.65 | 0.92 | 0.84 | 0.86 | 0.75 |
| 4 | 149139764 | rs12641471 | -0.657 | 1.031 | -0.352 | 0.548 | -0.001 | 0.002 | 0.002 | 1.138 | 0.47 | 0.45 | 0.55 | 0.52 | 0.55 | 0.61 |
| 4 | 149147823 | rs3910044 | 0.016 | 1.241 | -0.217 | 0.233 | 0.001 | 0.018 | 0.018 | 0 | 0.63 | 0.48 | 0.51 | 0.48 | 0.59 | 0.46 |
| 4 | 149148578 | rs2883917 | 0.129 | 1.124 | -0.411 | 0.066 | 0.003 | -0.049 | -0.049 | 0 | 0.12 | 0.2 | 0.07 | 0 | 0.12 | 0.07 |
| 4 | 149157085 | rs982076 | 0.586 | 1.096 | -1.551 | 0.52 | -0.024 | -0.152 | -0.152 | 0 | 0.28 | 0.04 | 0.23 | 0.49 | 0.24 | 0.26 |
| 4 | 149159119 | rs3846313 | 0.708 | 1.087 | -1.659 | -0.649 | -0.006 | 0.06 | 0.06 | 0 | 0.2 | 0.04 | 0.1 | 0 | 0.18 | 0.06 |
| 4 | 149164905 | rs17484063 | 1.504 | 1.855 | -1.685 | -0.243 | -0.041 | -0.037 | -0.037 | 0 | 0.07 | 0 | 0.21 | 0.18 | 0.2 | 0.17 |
| 4 | 149165351 | rs3846317 | -0.278 | 1.775 | -1.158 | -0.299 | 0.022 | 0.02 | 0.02 | 0 | 0.27 | 0.37 | 0.5 | 0.4 | 0.39 | 0.39 |
| 4 | 149165388 | rs3846318 | -0.47 | 1.424 | -1.306 | -0.454 | 0.025 | 0.121 | 0.121 | 0 | 0.52 | 0.73 | 0.72 | 0.85 | 0.6 | 0.67 |
| 4 | 149166943 | rs3846321 | 1.466 | 1.314 | -1.542 | -0.395 | -0.048 | -0.142 | -0.142 | 0 | 0.21 | 0.03 | 0.18 | 0.46 | 0.2 | 0.27 |
| 4 | 149173629 | rs3846322 | 0.376 | 1.235 | -1.129 | -0.455 | 0 | 0 | 0 | 0 | 0.22 | 0.15 | 0.29 | 0.22 | 0.24 | 0.23 |
| 4 | 149183374 | rs6855341 | 0.723 | 1.213 | -1.022 | -0.428 | -0.001 | 0.002 | 0.002 | 0 | 0.19 | 0.2 | 0.28 | 0.22 | 0.2 | 0.22 |
| 4 | 149188327 | rs6836191 | -0.359 | 1.262 | -0.804 | -0.617 | 0.009 | 0.02 | 0.02 | 0 | 0.49 | 0.43 | 0.7 | 0.85 | 0.62 | 0.67 |
| 4 | 149196991 | rs6812904 | -0.713 | 1.168 | -0.302 | -0.643 | -0.022 | 0.003 | 0.003 | 0.676 | 0.38 | 0.75 | 0.56 | 0.4 | 0.43 | 0.42 |
| 4 | 149198415 | rs17484357 | 1.244 | 0.672 | -0.224 | -0.827 | 0.022 | 0.1 | 0.1 | 0.003 | 0.27 | 0.05 | 0.1 | 0 | 0.19 | 0.06 |
| 4 | 149198522 | rs10012553 | 0.116 | 0.74 | -0.054 | -0.7 | -0.004 | -0.044 | -0.044 | 0 | 0.09 | 0.34 | 0.05 | 0 | 0.04 | 0.04 |
| 4 | 149203945 | rs3846323 | 0.415 | 0.77 | -0.547 | -0.611 | 0.024 | 0.074 | 0.074 | 0 | 0.37 | 0.16 | 0.24 | 0.15 | 0.3 | 0.31 |
| 4 | 149209008 | rs17484454 | 0.388 | 0.713 | -0.208 | -0.572 | 0.024 | 0.044 | 0.044 | 0 | 0.37 | 0.21 | 0.22 | 0.18 | 0.29 | 0.34 |
| 4 | 149210677 | rs3910047 | 0.387 | 0.844 | -0.002 | -0.496 | 0.001 | 0.005 | 0.005 | 0 | 0.2 | 0.24 | 0.3 | 0.25 | 0.2 | 0.18 |
| 4 | 149215821 | rs13147775 | 1.533 | 0.957 | -0.322 | -0.5 | -0.067 | -0.155 | -0.155 | 0 | 0.25 | 0.03 | 0.24 | 0.42 | 0.25 | 0.25 |
| 4 | 149221110 | rs2883929 | -1.5 | 1.224 | 0.142 | -0.419 | 0.031 | 0.004 | 0.004 | 0 | 0.89 | 0.79 | 0.79 | 0.84 | 0.76 | 0.8 |
| 4 | 149229338 | rs3846329 | 0.422 | 0.701 | 0.131 | -0.518 | -0.001 | -0.001 | -0.001 | 0 | 0.19 | 0.16 | 0.28 | 0.26 | 0.21 | 0.18 |
| 4 | 149245293 | rs10434128 | 0.82 | 0.338 | 0.242 | -0.777 | 0 | 0 | 0 | 0 | 0.12 | 0.11 | 0.12 | 0.15 | 0.15 | 0.11 |
| 4 | 149251809 | rs6851669 | 0.85 | 0.184 | 0.317 | -0.799 | -0.009 | 0.06 | 0.06 | 0 | 0.12 | 0.27 | 0.13 | 0.17 | 0.09 | 0.27 |
| 4 | 149251850 | rs6826213 | 0.894 | 0.187 | 0.215 | -0.798 | -0.008 | 0.032 | 0.032 | 0 | 0.12 | 0.2 | 0.12 | 0.17 | 0.1 | 0.27 |
| 4 | 149264255 | rs1546531 | 1.75 | 0.201 | -0.28 | -0.859 | -0.086 | -0.168 | -0.168 | 0.32 | 0.29 | 0.03 | 0.46 | 0.56 | 0.32 | 0.4 |
| 4 | 149264399 | rs1546529 | 0.618 | 0.571 | 0.412 | -0.52 | -0.002 | 0.014 | 0.014 | 0.169 | 0.73 | 0.39 | 0.72 | 0.67 | 0.76 | 0.62 |
| 4 | 149264663 | rs17024631 | 0.299 | 0.571 | 0.316 | -0.577 | -0.008 | -0.018 | -0.018 | 2.577 | 0.08 | 0.15 | 0.02 | 0 | 0.04 | 0.04 |

| | | | | | | | | | | | | | | | | |
|---|---|---|---|---|---|---|---|---|---|---|---|---|---|---|---|---|
| 4 | 149265166 | rs2137335 | -0.507 | 0.643 | 0.348 | -0.538 | 0.004 | 0.056 | 0.056 | 0.11 | 0.24 | 0.5 | 0.26 | 0.32 | 0.23 | 0.37 |
| 4 | 149265483 | rs10034789 | -0.157 | 0.843 | 0.576 | -1.172 | -0.03 | 0.083 | 0.083 | 0 | 0.32 | 0.16 | 0.21 | 0.11 | 0.39 | 0.17 |
| 4 | 149268136 | rs17484671 | 0.515 | 0.911 | 0.559 | -1.195 | -0.004 | 0.002 | 0.002 | 0 | 0.13 | 0.13 | 0.14 | 0.11 | 0.24 | 0.12 |
| 4 | 149268146 | rs17484678 | 0.929 | 0.943 | 0.417 | -1.178 | 0 | 0.082 | 0.082 | 0 | 0.2 | 0.04 | 0.07 | 0 | 0.15 | 0.05 |
| 4 | 149269636 | rs992556 | 1.559 | 0.944 | 0.453 | -1.171 | -0.026 | 0.12 | 0.12 | 0 | 0.43 | 0.1 | 0.59 | 0.74 | 0.43 | 0.67 |
| 4 | 149284080 | rs2063555 | 0.583 | 0.91 | 0.735 | -1.103 | 0.012 | 0.004 | 0.004 | 0 | 0.11 | 0.16 | 0.12 | 0.15 | 0.14 | 0.11 |
| 4 | 149284425 | rs2137330 | 0.844 | 0.913 | 0.718 | -1.101 | 0.008 | 0.003 | 0.003 | 0 | 0.1 | 0.11 | 0.12 | 0.15 | 0.14 | 0.11 |
| 4 | 149287807 | rs10434100 | 0.675 | 0.906 | 0.715 | -1.101 | 0.006 | 0.002 | 0.002 | 0 | 0.11 | 0.12 | 0.12 | 0.15 | 0.14 | 0.09 |
| 4 | 149291532 | rs13109808 | 1.49 | 0.89 | 0.819 | -1.249 | 0.003 | 0.027 | 0.027 | 0 | 0.44 | 0.36 | 0.61 | 0.73 | 0.42 | 0.68 |
| 4 | 149297675 | rs10519963 | 0.704 | 0.967 | 0.327 | -1.101 | 0.005 | 0 | 0 | 0 | 0.11 | 0.15 | 0.12 | 0.13 | 0.14 | 0.1 |
| 4 | 149299071 | rs17581884 | 0.885 | 0.869 | 0.315 | -1.147 | 0.002 | 0.068 | 0.068 | 0 | 0.2 | 0.05 | 0.08 | 0 | 0.16 | 0.05 |
| 4 | 149302984 | rs907620 | -1.09 | 0.876 | 0.437 | -1.147 | 0 | -0.039 | -0.039 | 0 | 0.25 | 0.63 | 0.21 | 0.13 | 0.27 | 0.17 |
| 4 | 149311728 | rs11725509 | 0.834 | 0.828 | 0.372 | -1.13 | -0.019 | -0.032 | -0.032 | 1.622 | 0.07 | 0.15 | 0.03 | 0 | 0.02 | 0.04 |
| 4 | 149313878 | rs6831212 | 1.672 | 0.861 | 0.29 | -1.153 | -0.032 | 0.129 | 0.129 | 0 | 0.44 | 0.11 | 0.59 | 0.74 | 0.44 | 0.65 |
| 4 | 149321346 | rs1490453 | -0.743 | 0.913 | 0.404 | -1.117 | 0.007 | -0.049 | -0.049 | 0.031 | 0.2 | 0.45 | 0.12 | 0.07 | 0.16 | 0.09 |
| 4 | 149322379 | rs1512327 | 1.722 | 0.907 | 0.135 | -1.126 | -0.028 | 0.129 | 0.129 | 0 | 0.44 | 0.11 | 0.59 | 0.74 | 0.44 | 0.64 |
| 4 | 149326597 | rs6834935 | 0.329 | 1.011 | 0.882 | -0.129 | -0.003 | 0 | 0 | 0 | 0.09 | 0.09 | 0.07 | 0.06 | 0.13 | 0.05 |
| 4 | 149330922 | rs1994624 | 1.422 | 1.097 | 0.455 | -0.115 | 0.01 | -0.025 | -0.025 | 0.191 | 0.65 | 0.44 | 0.7 | 0.7 | 0.61 | 0.71 |
| 4 | 149332071 | rs7698307 | -0.405 | 2.045 | 0.403 | 0.477 | 0.009 | 0 | 0 | 0.227 | 0.13 | 0.24 | 0.08 | 0.11 | 0.13 | 0.05 |
| 4 | 149340536 | rs17024708 | -0.857 | 1.759 | -0.082 | 0.534 | 0.044 | 0.005 | 0.005 | 0 | 0.13 | 0.25 | 0.16 | 0.14 | 0.23 | 0.13 |
| 4 | 149347854 | rs10026568 | -0.711 | 1.596 | -0.856 | 0.65 | -0.016 | -0.023 | -0.023 | 0.091 | 0.37 | 0.22 | 0.31 | 0.41 | 0.45 | 0.33 |
| 4 | 149350527 | rs4635799 | -0.439 | 1.577 | -1.438 | -0.207 | -0.025 | 0.054 | 0.054 | 0 | 0.39 | 0.11 | 0.54 | 0.75 | 0.49 | 0.68 |
| 4 | 149352962 | rs2172002 | -0.231 | 1.633 | -1.241 | -0.266 | -0.001 | 0.001 | 0.001 | 0 | 0.91 | 0.93 | 0.83 | 0.84 | 0.91 | 0.9 |
| 4 | 149357475 | rs5522 | -0.167 | 1.534 | -0.306 | -0.319 | -0.001 | 0 | 0 | 0 | 0.91 | 0.93 | 0.83 | 0.85 | 0.92 | 0.9 |
| 4 | 149379537 | rs6535611 | 0.406 | 1.791 | -0.138 | -0.649 | -0.022 | 0.062 | 0.062 | 0 | 0.58 | 0.76 | 0.32 | 0.1 | 0.47 | 0.26 |
| 4 | 149384030 | rs6535612 | -0.181 | 1.84 | -0.061 | -0.511 | 0.008 | 0.008 | 0.008 | 0 | 0.91 | 0.9 | 0.81 | 0.86 | 0.87 | 0.9 |
| 4 | 149386704 | rs1490463 | -0.64 | 1.766 | 0.158 | -0.458 | 0.046 | 0.022 | 0.022 | 0 | 0.06 | 0.21 | 0.06 | 0.03 | 0.08 | 0.05 |
| 4 | 149398778 | rs1472061 | 0.024 | 1.659 | -0.686 | -0.447 | -0.025 | 0.024 | 0.024 | 0 | 0.32 | 0.11 | 0.49 | 0.76 | 0.4 | 0.63 |
| 4 | 149403565 | rs4324532 | -0.843 | 1.292 | -0.435 | -0.993 | -0.004 | -0.079 | -0.079 | 0 | 0.84 | 0.77 | 0.9 | 1 | 0.85 | 0.96 |
| 4 | 149425641 | rs17485372 | 0.719 | 0.909 | -1.392 | -1.379 | 0.005 | 0.065 | 0.065 | 0.01 | 0.16 | 0.05 | 0.07 | 0.03 | 0.12 | 0.06 |
| 4 | 149447366 | rs875758 | -0.292 | 1.655 | -1.781 | -1.311 | -0.039 | 0.042 | 0.042 | 0 | 0.35 | 0.22 | 0.65 | 0.78 | 0.49 | 0.64 |
| 4 | 149469901 | rs2130644 | -0.124 | 1.455 | -1.735 | -0.608 | -0.008 | 0.021 | 0.021 | 0 | 0.9 | 0.83 | 0.94 | 0.82 | 0.92 | 0.91 |
| 4 | 149522696 | rs4475130 | 0.814 | 1.543 | -2.088 | -0.532 | 0.005 | -0.008 | -0.008 | 0.278 | 0.06 | 0.04 | 0.02 | 0.16 | 0.05 | 0.06 |
| 4 | 149537415 | rs12644406 | 0.635 | 2.184 | -2.242 | 0.267 | -0.058 | -0.043 | -0.043 | 0.007 | 0.1 | 0.01 | 0.26 | 0.49 | 0.24 | 0.37 |
| 4 | 149555789 | rs4835529 | -0.566 | 1.061 | -2.142 | -0.999 | -0.057 | 0.161 | 0.161 | 0 | 0.67 | 0.98 | 0.85 | 0.93 | 0.71 | 0.8 |
| 4 | 149562277 | rs17582780 | 0.009 | 1.102 | -2.007 | -0.806 | -0.012 | 0.065 | 0.065 | 0.304 | 0.35 | 0.08 | 0.16 | 0.27 | 0.31 | 0.28 |
| 4 | 149562638 | rs17582787 | 0.538 | 0.752 | -2.121 | -0.914 | 0.007 | -0.003 | -0.003 | 0.116 | 0.16 | 0.03 | 0.07 | 0.16 | 0.13 | 0.17 |
| 4 | 149570069 | rs10452224 | -0.627 | 1.288 | -1.814 | -1.134 | -0.026 | 0.087 | 0.087 | 0 | 0.41 | 0.24 | 0.23 | 0.29 | 0.42 | 0.29 |
| 4 | 149574788 | rs7437928 | 0.045 | 1.329 | -1.349 | -1.566 | 0.003 | -0.021 | -0.021 | 0 | 0.57 | 0.47 | 0.35 | 0.24 | 0.53 | 0.3 |
| 4 | 149590581 | rs2358469 | -0.279 | 1.42 | -1.643 | -1.33 | -0.057 | 0.081 | 0.081 | 0.101 | 0.43 | 0.13 | 0.3 | 0.4 | 0.54 | 0.49 |
| 4 | 149598573 | rs6840553 | 0.048 | 0.922 | -0.667 | -1.706 | -0.022 | -0.084 | -0.084 | 0 | 0.23 | 0.53 | 0.16 | 0.15 | 0.21 | 0.17 |
| 4 | 149598635 | rs6840754 | -0.315 | 0.785 | -0.965 | -2.084 | -0.033 | 0.017 | 0.017 | 0 | 0.71 | 0.89 | 0.43 | 0.32 | 0.64 | 0.39 |
| 4 | 149600181 | rs17024901 | 0.679 | 0.656 | -0.603 | -2.121 | 0.018 | 0.055 | 0.055 | 0 | 0.37 | 0.25 | 0.21 | 0.06 | 0.31 | 0.12 |
| 4 | 149601351 | rs1316785 | 0.916 | 0.739 | -0.597 | -1.985 | -0.057 | 0.055 | 0.055 | 0 | 0.23 | 0.09 | 0.17 | 0.12 | 0.29 | 0.12 |
| 4 | 149605569 | rs6826228 | 1.347 | 0.633 | -0.695 | -1.994 | 0.008 | 0.092 | 0.092 | 0 | 0.26 | 0.07 | 0.13 | 0.06 | 0.22 | 0.11 |
| 4 | 149614024 | rs7666326 | -0.1 | 0.756 | -0.767 | -1.939 | -0.032 | 0.188 | 0.188 | 0 | 0.54 | 0.87 | 0.35 | 0.16 | 0.48 | 0.23 |
| 4 | 149616993 | rs4835530 | -0.013 | 0.761 | -0.709 | -1.937 | -0.018 | 0.175 | 0.175 | 0 | 0.54 | 0.84 | 0.35 | 0.16 | 0.48 | 0.23 |
| 4 | 149619238 | rs7674926 | 0.56 | 0.614 | -0.496 | -1.949 | -0.005 | -0.047 | -0.047 | 0 | 0.3 | 0.68 | 0.19 | 0.06 | 0.23 | 0.1 |
| 4 | 149619375 | rs13137679 | 0.345 | 0.432 | -0.502 | -1.791 | -0.036 | 0.109 | 0.109 | 0.161 | 0.3 | 0.11 | 0.19 | 0.1 | 0.29 | 0.1 |
| 4 | 149624705 | rs17008423 | 0.9 | 0.324 | 0.449 | -1.839 | 0.001 | -0.046 | -0.046 | 0.445 | 0.21 | 0.34 | 0.15 | 0.08 | 0.19 | 0.15 |
| 4 | 149625757 | rs10008892 | 0.43 | 0.307 | 0.583 | -1.854 | -0.004 | -0.076 | -0.076 | 0.918 | 0.24 | 0.64 | 0.18 | 0.08 | 0.2 | 0.17 |
| 4 | 149635025 | rs10519980 | 0.929 | 0.336 | 0.512 | -1.855 | 0.001 | -0.048 | -0.048 | 0 | 0.21 | 0.33 | 0.13 | 0.08 | 0.19 | 0.15 |
| 4 | 149637571 | rs7682301 | -0.509 | 0.338 | 0.085 | -1.908 | -0.037 | 0.103 | 0.103 | 0 | 0.63 | 0.84 | 0.35 | 0.17 | 0.52 | 0.3 |
| 4 | 149641824 | rs2358649 | 1.143 | 0.357 | 0.026 | -1.812 | 0.005 | 0.037 | 0.037 | 0 | 0.27 | 0.18 | 0.14 | 0.09 | 0.23 | 0.12 |
| 4 | 149643143 | rs7681551 | 0.461 | 0.326 | 0.123 | -1.896 | -0.006 | -0.093 | -0.093 | 0 | 0.25 | 0.6 | 0.16 | 0.08 | 0.2 | 0.16 |
| 4 | 149648035 | rs17583283 | -0.165 | 0.33 | 0.069 | -1.883 | -0.025 | 0.239 | 0.239 | 0 | 0.53 | 0.9 | 0.31 | 0.17 | 0.43 | 0.29 |
| 4 | 149650033 | rs17025017 | 0.3 | 0.658 | 0.647 | -1.338 | 0.002 | -0.085 | -0.085 | 0.286 | 0.23 | 0.46 | 0.15 | 0.08 | 0.2 | 0.15 |
| 4 | 149652478 | rs11935941 | -0.041 | 0.638 | 0.715 | -1.347 | -0.006 | -0.059 | -0.059 | 4.568 | 0.26 | 0.7 | 0.17 | 0.08 | 0.2 | 0.17 |
| 4 | 149652582 | rs11945082 | 0.093 | 0.65 | 0.694 | -1.342 | -0.002 | -0.111 | -0.111 | 0 | 0.24 | 0.55 | 0.16 | 0.08 | 0.2 | 0.17 |
| 4 | 149692497 | rs549653 | 0.564 | 0.785 | 0.177 | -0.626 | -0.001 | 0.012 | 0.012 | 0 | 0.18 | 0.18 | 0.21 | 0.33 | 0.16 | 0.15 |
| 4 | 149697086 | rs6848368 | 0.995 | 0.827 | 0.004 | -0.689 | -0.041 | 0.089 | 0.089 | 0 | 0.19 | 0.05 | 0.11 | 0.07 | 0.19 | 0.15 |
| 4 | 149700674 | rs558244 | -0.1 | 0.755 | 0.458 | -0.586 | -0.035 | 0.093 | 0.093 | 0 | 0.22 | 0.52 | 0.25 | 0.33 | 0.16 | 0.15 |
| 4 | 149710794 | rs495515 | 0.546 | 1.114 | -0.514 | -0.403 | 0.002 | -0.043 | -0.043 | 0 | 0.17 | 0.1 | 0.2 | 0.34 | 0.17 | 0.15 |
| 4 | 149712041 | rs10519990 | 0.809 | 1.165 | -0.806 | -0.507 | -0.052 | 0.102 | 0.102 | 0 | 0.2 | 0.03 | 0.11 | 0.07 | 0.2 | 0.16 |
| 4 | 149712613 | rs517151 | -0.681 | 1.145 | -0.699 | -0.513 | -0.001 | -0.001 | -0.001 | 0 | 0.39 | 0.35 | 0.34 | 0.41 | 0.37 | 0.3 |
| 4 | 149713872 | rs2204206 | 0.315 | 1.623 | -1.929 | -0.405 | -0.004 | 0.001 | 0.001 | 0 | 0.19 | 0.17 | 0.45 | 0.5 | 0.31 | 0.52 |
| 4 | 149724882 | rs13134311 | 0.106 | 1.271 | -1.911 | -0.203 | -0.051 | 0.103 | 0.103 | 0 | 0.2 | 0.03 | 0.12 | 0.07 | 0.2 | 0.16 |
| 4 | 149729151 | rs525368 | -0.354 | 1.047 | -1.386 | 0.562 | -0.018 | -0.065 | -0.065 | 0 | 0.17 | 0.03 | 0.2 | 0.28 | 0.19 | 0.14 |
| 4 | 149733739 | rs486116 | 0.203 | 0.996 | -1.062 | 0.108 | -0.001 | -0.007 | -0.007 | 0.013 | 0.76 | 0.8 | 0.87 | 0.87 | 0.77 | 0.88 |

| | | | | | | | | | | | | | | | | |
|---|---|---|---|---|---|---|---|---|---|---|---|---|---|---|---|---|
| 4 | 149734385 | rs559072 | 0.119 | 0.584 | -0.414 | -0.13 | 0 | -0.025 | -0.025 | 0 | 0.09 | 0.25 | 0.05 | 0.03 | 0.06 | 0.08 |
| 4 | 149736293 | rs17487044 | 0.302 | 0.53 | -0.739 | -0.172 | 0.015 | 0 | 0 | 0 | 0.16 | 0.01 | 0.04 | 0.1 | 0.11 | 0.07 |
| 4 | 149742079 | rs4282147 | 0.62 | 0.406 | -0.998 | -0.339 | -0.002 | 0.01 | 0.01 | 0 | 0.43 | 0.4 | 0.6 | 0.65 | 0.56 | 0.7 |
| 4 | 149742690 | rs11099710 | 0.269 | 0.114 | -1.279 | -0.551 | 0.024 | -0.096 | -0.096 | 0.135 | 0.23 | 0.03 | 0.19 | 0.31 | 0.15 | 0.12 |
| 4 | 149743531 | rs552274 | 0.045 | 0.521 | -1.437 | 0.005 | -0.003 | 0.038 | 0.038 | 0 | 0.19 | 0.26 | 0.28 | 0.31 | 0.17 | 0.2 |
| 4 | 149744201 | rs17623254 | 1.409 | 0.524 | -1.458 | 0.033 | -0.038 | -0.061 | -0.061 | 5.295 | 0.1 | 0 | 0.23 | 0.29 | 0.11 | 0.14 |
| 4 | 149744518 | rs471967 | -0.439 | 0.461 | -1.072 | -0.056 | -0.002 | -0.01 | -0.01 | 2.643 | 0.43 | 0.47 | 0.43 | 0.32 | 0.39 | 0.3 |
| 4 | 149750527 | rs6854659 | 1.043 | 0.493 | -1.258 | -0.095 | -0.001 | -0.039 | -0.039 | 0 | 0.24 | 0.18 | 0.15 | 0 | 0.23 | 0.1 |
| 4 | 149762698 | rs9307850 | 0.12 | 0.513 | -0.939 | 0.045 | -0.002 | 0.046 | 0.046 | 0 | 0.19 | 0.33 | 0.28 | 0.32 | 0.17 | 0.2 |
| 4 | 149765628 | rs853721 | -1.418 | 0.794 | -1.569 | -0.385 | -0.121 | -0.101 | -0.101 | 0 | 0.8 | 0.98 | 0.8 | 0.65 | 0.67 | 0.46 |
| 4 | 149771033 | rs862657 | -0.522 | 0.964 | -1.278 | -0.375 | -0.017 | 0.018 | 0.018 | 0.245 | 0.71 | 0.73 | 0.75 | 0.82 | 0.6 | 0.56 |
| 4 | 149771314 | rs12498869 | 1.073 | 0.927 | -1.574 | -0.191 | -0.034 | -0.067 | -0.067 | 0 | 0.12 | 0 | 0.12 | 0.2 | 0.14 | 0.16 |
| 4 | 149772371 | rs17778070 | 1.296 | 0.966 | -1.247 | -0.448 | 0 | -0.043 | -0.043 | 1.909 | 0.24 | 0.18 | 0.15 | 0 | 0.22 | 0.09 |
| 4 | 149774113 | rs1357440 | 0.183 | 0.975 | -1.208 | -0.42 | 0.005 | 0.032 | 0.032 | 0 | 0.26 | 0.62 | 0.37 | 0.34 | 0.26 | 0.26 |
| 4 | 149775941 | rs1634070 | 0.269 | 0.971 | -1.204 | -0.422 | 0.012 | 0.049 | 0.049 | 0 | 0.26 | 0.57 | 0.36 | 0.34 | 0.25 | 0.27 |
| 4 | 149794284 | rs10003214 | 1.267 | 0.934 | -1.247 | -0.467 | -0.012 | -0.096 | -0.096 | 1.277 | 0.25 | 0.24 | 0.16 | 0 | 0.21 | 0.09 |
| 4 | 149802894 | rs886148 | -1.576 | 1.043 | -1.502 | -0.28 | -0.089 | -0.05 | -0.05 | 1.294 | 0.84 | 0.98 | 0.83 | 0.76 | 0.76 | 0.61 |
| 4 | 149817052 | rs2520496 | -0.122 | 0.992 | -1.564 | -0.312 | -0.074 | 0.043 | 0.043 | 0 | 0.51 | 0.93 | 0.54 | 0.34 | 0.47 | 0.36 |
| 4 | 149832511 | rs1010178 | -0.319 | 0.935 | -1.228 | -0.402 | 0.001 | -0.033 | -0.033 | 0 | 0.56 | 0.58 | 0.66 | 0.75 | 0.54 | 0.53 |
| 4 | 149841708 | rs12374234 | 0.36 | 0.9 | -1.485 | -0.373 | -0.012 | -0.015 | -0.015 | 0.248 | 0.55 | 0.19 | 0.43 | 0.42 | 0.5 | 0.35 |
| 4 | 149846648 | rs12233660 | 0.772 | 0.916 | -1.585 | -0.205 | -0.03 | -0.158 | -0.158 | 0 | 0.31 | 0.02 | 0.28 | 0.42 | 0.29 | 0.25 |
| 4 | 149871390 | rs6535646 | -0.454 | 1.118 | -1.185 | -0.16 | -0.025 | 0.065 | 0.065 | 0 | 0.62 | 0.79 | 0.69 | 0.73 | 0.56 | 0.55 |
| 4 | 149872845 | rs17025520 | 0.982 | 1.248 | -1.222 | -0.051 | -0.076 | -0.051 | -0.051 | 0 | 0.15 | 0.03 | 0.15 | 0.24 | 0.22 | 0.36 |
| 4 | 149878147 | rs12644174 | -0.222 | 1.146 | -0.696 | -0.036 | 0.011 | 0.029 | 0.029 | 0.001 | 0.31 | 0.47 | 0.43 | 0.59 | 0.35 | 0.57 |
| 4 | 149894321 | rs13129108 | 0.588 | 0.893 | -0.74 | -0.309 | -0.002 | -0.002 | -0.002 | 0 | 0.57 | 0.36 | 0.48 | 0.42 | 0.55 | 0.34 |
| 4 | 149901085 | rs724043 | 0.381 | 0.872 | -0.685 | -0.317 | 0.006 | 0.006 | 0.006 | 0.522 | 0.59 | 0.42 | 0.49 | 0.42 | 0.55 | 0.34 |
| 4 | 149902599 | rs10489052 | 0.995 | 0.834 | -1.106 | -0.596 | -0.015 | -0.099 | -0.099 | 0 | 0.35 | 0.07 | 0.29 | 0.42 | 0.31 | 0.26 |
| 4 | 149903798 | rs756862 | -1.445 | 0.534 | -0.988 | -0.795 | 0 | 0.003 | 0.003 | 0 | 0.88 | 0.89 | 0.92 | 1 | 0.89 | 0.92 |
| 4 | 149903982 | rs17025566 | 1.122 | 0.564 | -0.926 | -0.763 | 0.003 | -0.104 | -0.104 | 0 | 0.22 | 0.28 | 0.18 | 0 | 0.23 | 0.07 |
| 4 | 149920515 | rs7691878 | 0.959 | 0.552 | -0.763 | -0.496 | -0.004 | 0.077 | 0.077 | 0 | 0.14 | 0.31 | 0.28 | 0.28 | 0.12 | 0.19 |
| 4 | 149929686 | rs2520494 | 1.193 | 0.719 | -1.783 | -0.27 | -0.103 | -0.104 | -0.104 | 0 | 0.18 | 0.02 | 0.32 | 0.41 | 0.28 | 0.39 |
| 4 | 149933235 | rs17025612 | 1.728 | 0.627 | -1.773 | -0.26 | -0.027 | -0.073 | -0.073 | 0 | 0.11 | 0.01 | 0.25 | 0.28 | 0.12 | 0.19 |
| 4 | 149936868 | rs10001495 | -1.576 | 0.626 | -1.762 | -0.316 | -0.008 | -0.089 | -0.089 | 0 | 0.7 | 0.98 | 0.87 | 0.72 | 0.75 | 0.78 |
| 4 | 149943840 | rs6839286 | 0.686 | 1.038 | -1.127 | -0.589 | -0.003 | -0.033 | -0.033 | 0.257 | 0.11 | 0.16 | 0.06 | 0 | 0.09 | 0.06 |
| 4 | 149948850 | rs6824983 | 0.578 | 1.244 | -1.284 | -0.325 | -0.134 | -0.105 | -0.105 | 0 | 0.2 | 0.03 | 0.36 | 0.41 | 0.36 | 0.41 |
| 4 | 149954364 | rs11946353 | 0.233 | 0.757 | -0.254 | -0.69 | -0.003 | -0.018 | -0.018 | 0 | 0.24 | 0.18 | 0.19 | 0 | 0.26 | 0.1 |
| 4 | 150009235 | rs10434101 | -0.591 | 0.313 | -0.411 | 0.082 | 0.013 | 0.048 | 0.048 | 0 | 0.49 | 0.38 | 0.58 | 0.67 | 0.44 | 0.48 |
| 4 | 150060844 | rs1438768 | -0.955 | -0.02 | -0.453 | -0.005 | -0.087 | 0.039 | 0.039 | 0 | 0.56 | 0.95 | 0.65 | 0.65 | 0.45 | 0.52 |
| 4 | 150070596 | rs4522846 | -0.942 | -0.05 | -0.338 | -0.003 | -0.049 | 0.058 | 0.058 | 0 | 0.56 | 0.9 | 0.64 | 0.65 | 0.45 | 0.52 |
| 4 | 150074544 | rs6842668 | 0.262 | -0.375 | 0.157 | 0.24 | -0.034 | -0.025 | -0.025 | 0 | 0.29 | 0.49 | 0.22 | 0.2 | 0.22 | 0.16 |
| 4 | 150089373 | rs979227 | -1.406 | -0.338 | -0.102 | 0.263 | -0.088 | 0.034 | 0.034 | 0 | 0.55 | 0.97 | 0.64 | 0.63 | 0.44 | 0.51 |
| 4 | 150092178 | rs1438781 | -0.133 | -0.497 | 0.11 | 0.172 | 0.031 | 0.019 | 0.019 | 0 | 0.33 | 0.82 | 0.25 | 0.23 | 0.23 | 0.19 |
| 4 | 150108607 | rs11099718 | -1.371 | -0.283 | -0.245 | 0.206 | -0.117 | 0.04 | 0.04 | 0 | 0.56 | 0.96 | 0.64 | 0.62 | 0.45 | 0.5 |
| 4 | 150122796 | rs7679738 | -1.535 | -0.294 | -0.414 | 0.084 | -0.131 | 0.021 | 0.021 | 0 | 0.61 | 0.97 | 0.66 | 0.65 | 0.5 | 0.7 |
| 4 | 150122817 | rs10520008 | 0.128 | -0.356 | -0.308 | 0.119 | -0.011 | -0.033 | -0.033 | 0 | 0.26 | 0.13 | 0.42 | 0.38 | 0.26 | 0.34 |
| 4 | 150129324 | rs562329 | 1.423 | -0.295 | -0.528 | 0.14 | -0.131 | 0.021 | 0.021 | 0 | 0.39 | 0.03 | 0.34 | 0.35 | 0.5 | 0.29 |
| 4 | 150129861 | rs1438765 | 0.918 | -0.428 | -0.076 | -0.062 | 0.007 | -0.012 | -0.012 | 0 | 0.22 | 0.19 | 0.15 | 0.23 | 0.18 | 0.33 |
| 4 | 150135657 | rs17025936 | 1.32 | -0.396 | 0.073 | 0.003 | 0.004 | 0.004 | 0.004 | 0 | 0.08 | 0.1 | 0.04 | 0.04 | 0.06 | 0.02 |
| 4 | 150156579 | rs11721563 | 0.438 | -0.271 | 0.233 | 0.064 | 0.003 | 0.002 | 0.002 | 0 | 0.34 | 0.4 | 0.22 | 0.27 | 0.25 | 0.36 |
| 4 | 150167413 | rs6828493 | 0.793 | -0.182 | 0.052 | 0.137 | 0.004 | 0.001 | 0.001 | 0 | 0.31 | 0.29 | 0.27 | 0.26 | 0.22 | 0.33 |
| 4 | 150178430 | rs580399 | 0.931 | -0.083 | -0.284 | 0.027 | -0.092 | 0.055 | 0.055 | 0 | 0.39 | 0.11 | 0.37 | 0.32 | 0.5 | 0.29 |
| 4 | 150181884 | rs3106916 | -0.237 | -0.114 | 0.444 | 0.217 | 0.006 | 0.083 | 0.083 | 0 | 0.77 | 0.65 | 0.77 | 0.6 | 0.75 | 0.62 |
| 4 | 150184393 | rs3106921 | 1.15 | -0.268 | 0.02 | 0.101 | -0.153 | 0.05 | 0.05 | 0.274 | 0.41 | 0.03 | 0.43 | 0.31 | 0.49 | 0.26 |
| 4 | 150200706 | rs546978 | 0.546 | -0.316 | 0.428 | 0.093 | -0.007 | 0.03 | 0.03 | 0.042 | 0.45 | 0.36 | 0.46 | 0.33 | 0.5 | 0.29 |
| 4 | 150202672 | rs4835570 | 1.564 | -0.385 | 0.279 | 0.052 | 0.005 | -0.006 | -0.006 | 0 | 0.26 | 0.13 | 0.26 | 0.23 | 0.2 | 0.19 |
| 4 | 150205408 | rs9307855 | 1.041 | -0.223 | -0.247 | 0.113 | -0.004 | -0.002 | -0.002 | 0 | 0.32 | 0.33 | 0.29 | 0.27 | 0.23 | 0.33 |
| 4 | 150208355 | rs538844 | 0.722 | -0.129 | -0.588 | -0.135 | -0.096 | 0.066 | 0.066 | 0.375 | 0.41 | 0.1 | 0.43 | 0.32 | 0.5 | 0.26 |
| 4 | 150233544 | rs13124924 | 0.067 | 0.311 | -0.343 | -0.321 | 0 | -0.003 | -0.003 | 0 | 0.3 | 0.33 | 0.27 | 0.2 | 0.31 | 0.13 |
| 4 | 150242751 | rs7668712 | 0.021 | 0.458 | -0.408 | -0.401 | 0.01 | -0.038 | -0.038 | 0.047 | 0.08 | 0.26 | 0.04 | 0.01 | 0.06 | 0.04 |
| 4 | 150253761 | rs2359333 | 1.379 | 0.479 | -0.43 | -0.137 | 0.009 | -0.006 | -0.006 | 4.768 | 0.45 | 0.33 | 0.43 | 0.47 | 0.42 | 0.47 |
| 4 | 150267639 | rs6535675 | 1.19 | 0.127 | -0.44 | -0.005 | 0 | 0 | 0 | 0.491 | 0.5 | 0.5 | 0.52 | 0.48 | 0.47 | 0.52 |
| 4 | 150280975 | rs17624486 | 1.392 | 0.088 | -0.674 | -0.074 | -0.049 | 0.008 | 0.008 | 2.606 | 0.22 | 0.03 | 0.22 | 0.19 | 0.25 | 0.09 |
| 4 | 150283932 | rs8180206 | -0.081 | 0.04 | -0.573 | 0.116 | -0.003 | -0.008 | -0.008 | 0 | 0.19 | 0.11 | 0.17 | 0.3 | 0.2 | 0.34 |
| 4 | 150288111 | rs7653957 | -0.173 | 0.181 | -0.345 | 0.202 | -0.001 | 0.016 | 0.016 | 1.113 | 0.3 | 0.29 | 0.27 | 0.2 | 0.3 | 0.13 |
| 4 | 150419759 | rs17624848 | -0.084 | -0.346 | -0.162 | 0.813 | 0.002 | 0.011 | 0.011 | 0 | 0.15 | 0.11 | 0.25 | 0.31 | 0.14 | 0.37 |
| 4 | 150433523 | rs13144096 | 0.692 | -0.276 | -0.26 | 0.681 | 0.003 | 0 | 0 | 1.886 | 0.59 | 0.49 | 0.49 | 0.5 | 0.55 | 0.54 |
| 4 | 150457508 | rs1394028 | 0.811 | -0.247 | -0.363 | 0.689 | -0.055 | 0.009 | 0.009 | 4.712 | 0.22 | 0.03 | 0.23 | 0.19 | 0.3 | 0.08 |
| 4 | 150459436 | rs10019047 | -0.236 | -0.36 | -0.404 | 0.846 | -0.003 | -0.001 | -0.001 | 0.014 | 0.73 | 0.83 | 0.76 | 0.75 | 0.67 | 0.73 |
| 4 | 150473832 | rs2654804 | 0.847 | -0.355 | -0.409 | 0.821 | -0.055 | 0.009 | 0.009 | 4.712 | 0.22 | 0.03 | 0.23 | 0.19 | 0.3 | 0.08 |

| | | | | | | | | | | | | | | | | |
|---|---|---|---|---|---|---|---|---|---|---|---|---|---|---|---|---|
| 4 | 150479466 | rs11099736 | -0.921 | -0.355 | -0.398 | 0.821 | -0.012 | 0.026 | 0.026 | 4.685 | 0.77 | 0.86 | 0.76 | 0.81 | 0.69 | 0.91 |
| 4 | 150480129 | rs17625032 | 0.148 | -0.377 | -0.486 | 0.83 | -0.004 | -0.01 | -0.01 | 0.009 | 0.15 | 0.08 | 0.25 | 0.31 | 0.14 | 0.37 |
| 4 | 150486553 | rs1503697 | 0.916 | -0.342 | -0.522 | 0.837 | -0.001 | -0.002 | -0.002 | 0 | 0.57 | 0.58 | 0.49 | 0.45 | 0.53 | 0.37 |
| 4 | 150554591 | rs6814124 | -1.534 | -0.351 | -0.7 | 0.834 | 0 | 0 | 0 | 0 | 0.47 | 0.57 | 0.53 | 0.57 | 0.48 | 0.64 |
| 4 | 150585770 | rs6835892 | 0.864 | 0.023 | -1.472 | 1.038 | -0.068 | -0.007 | -0.007 | 0 | 0.2 | 0.03 | 0.22 | 0.2 | 0.3 | 0.06 |
| 4 | 150613311 | rs7659523 | 0.717 | 0.034 | -1.466 | 1.083 | -0.057 | -0.044 | -0.044 | 0.107 | 0.21 | 0.03 | 0.22 | 0.23 | 0.3 | 0.06 |
| 4 | 150616733 | rs17687220 | -0.304 | 0.289 | -1.287 | 0.527 | 0.001 | 0.007 | 0.007 | 0 | 0.16 | 0.11 | 0.26 | 0.3 | 0.15 | 0.38 |
| 4 | 150632186 | rs1394048 | -0.114 | 0.368 | -1.379 | 0.486 | -0.025 | 0.008 | 0.008 | 2.714 | 0.26 | 0.08 | 0.24 | 0.22 | 0.32 | 0.26 |
| 4 | 150632409 | rs1354363 | -0.103 | 0.327 | -1.377 | 0.489 | -0.022 | 0.008 | 0.008 | 2.732 | 0.26 | 0.08 | 0.24 | 0.22 | 0.32 | 0.26 |
| 4 | 150634191 | rs905576 | -0.147 | 0.287 | -0.624 | 0.556 | 0.005 | 0.026 | 0.026 | 0 | 0.55 | 0.64 | 0.55 | 0.38 | 0.6 | 0.5 |
| 4 | 150646588 | rs7684095 | -1.146 | -0.211 | 0.027 | 1.121 | -0.024 | 0.031 | 0.031 | 0.346 | 0.29 | 0.42 | 0.43 | 0.51 | 0.23 | 0.37 |
| 4 | 150648381 | rs6832401 | -1.517 | -0.376 | 0.038 | 1.026 | 0.001 | -0.003 | -0.003 | 0.212 | 0.37 | 0.69 | 0.52 | 0.55 | 0.29 | 0.51 |
| 4 | 150655652 | rs17502597 | 1.415 | 0.442 | -0.689 | 1.041 | -0.016 | 0.006 | 0.006 | 2.024 | 0.54 | 0.18 | 0.28 | 0.42 | 0.53 | 0.42 |
| 4 | 150659552 | rs1394050 | 0.389 | 0.296 | -0.255 | 0.574 | -0.028 | 0.029 | 0.029 | 0 | 0.86 | 0.94 | 0.81 | 0.99 | 0.82 | 0.92 |
| 4 | 150672514 | rs17026425 | -0.727 | -0.319 | 0.294 | 1.267 | 0.01 | -0.006 | -0.006 | 0 | 0.08 | 0.06 | 0.06 | 0.07 | 0.06 | 0.15 |
| 4 | 150678416 | rs10021126 | -1.426 | -0.338 | 0.389 | 1.386 | -0.043 | 0.065 | 0.065 | 2.061 | 0.15 | 0.57 | 0.23 | 0.21 | 0.11 | 0.2 |
| 4 | 150684793 | rs1394843 | -1.274 | -0.26 | 0.36 | 1.804 | -0.024 | 0.07 | 0.07 | 0.72 | 0.24 | 0.58 | 0.44 | 0.42 | 0.21 | 0.28 |
| 4 | 150688687 | rs17631073 | -0.752 | -0.39 | 0.229 | 2.032 | -0.005 | 0.027 | 0.027 | 0 | 0.13 | 0.17 | 0.11 | 0.2 | 0.11 | 0.2 |
| 4 | 150690457 | rs1503748 | -0.319 | -0.525 | 0.38 | 2.623 | 0.006 | -0.02 | -0.02 | 0 | 0.14 | 0.05 | 0.2 | 0.16 | 0.16 | 0.11 |
| 4 | 150692380 | rs17687541 | -0.721 | -0.547 | 0.392 | 2.968 | -0.005 | 0.049 | 0.049 | 0 | 0.13 | 0.17 | 0.11 | 0.34 | 0.13 | 0.2 |
| 4 | 150711342 | rs1876069 | -0.449 | -0.543 | 0.844 | 2.926 | -0.002 | 0 | 0 | 0.022 | 0.15 | 0.17 | 0.19 | 0.16 | 0.15 | 0.11 |
| 4 | 150714094 | rs17026442 | -0.605 | -0.559 | 1.496 | 2.876 | 0.02 | 0.248 | 0.248 | 1.863 | 0.06 | 0.34 | 0.22 | 0.25 | 0.01 | 0.13 |
| 4 | 150737037 | rs4835597 | -0.78 | -0.596 | 1.478 | 2.745 | 0.017 | 0.019 | 0.019 | 1.397 | 0.07 | 0.33 | 0.05 | 0.04 | 0.01 | 0.05 |
| 4 | 150741301 | rs11936885 | -1.767 | -0.743 | 1.763 | 2.76 | -0.007 | 0.04 | 0.04 | 1.05 | 0.11 | 0.61 | 0.1 | 0.19 | 0.07 | 0.1 |
| 4 | 150756808 | rs6849686 | -2.788 | -0.872 | 1.423 | 3.306 | 0.01 | -0.07 | -0.07 | 0.128 | 0.39 | 0.93 | 0.36 | 0.63 | 0.34 | 0.43 |
| 4 | 150766017 | rs17631265 | -0.523 | -0.801 | 1.554 | 3.219 | 0.005 | -0.026 | -0.026 | 0.525 | 0.15 | 0.1 | 0.12 | 0.24 | 0.16 | 0.12 |
| 4 | 150798489 | rs7655916 | -2.007 | -0.821 | 1.389 | 3.198 | 0.019 | -0.031 | -0.031 | 2.836 | 0.26 | 0.82 | 0.22 | 0.43 | 0.23 | 0.22 |
| 4 | 150818662 | rs6852150 | -2.103 | -0.847 | 1.713 | 3.223 | -0.021 | 0.053 | 0.053 | 2.079 | 0.12 | 0.7 | 0.11 | 0.22 | 0.07 | 0.11 |
| 4 | 150827830 | rs12642812 | -0.33 | -0.866 | 1.397 | 3.284 | -0.003 | -0.05 | -0.05 | 0 | 0.14 | 0.03 | 0.11 | 0.22 | 0.15 | 0.11 |
| 4 | 150844720 | rs4835604 | 2.198 | -0.657 | 0.364 | 1.996 | -0.141 | 0.144 | 0.144 | 1.489 | 0.46 | 0.03 | 0.37 | 0.19 | 0.47 | 0.44 |
| 4 | 150845428 | rs6830133 | -0.07 | -0.399 | 0.521 | 1.92 | -0.061 | -0.022 | -0.022 | 0 | 0.82 | 0.98 | 0.68 | 0.83 | 0.8 | 0.77 |
| 4 | 150852782 | rs905900 | 1.468 | -0.325 | 0.757 | 1.77 | -0.001 | -0.025 | -0.025 | 0.858 | 0.69 | 0.33 | 0.56 | 0.81 | 0.72 | 0.67 |
| 4 | 150853668 | rs17631619 | -0.113 | -0.389 | 0.376 | 1.874 | 0.033 | -0.038 | -0.038 | 0 | 0.15 | 0.02 | 0.15 | 0.18 | 0.07 | 0.18 |
| 4 | 150854267 | rs17587757 | -2.138 | -0.548 | 0.071 | 1.305 | 0.06 | 0.173 | 0.173 | 3.143 | 0.47 | 0.85 | 0.42 | 0.79 | 0.38 | 0.51 |
| 4 | 150854889 | rs1505473 | 1.159 | -0.373 | 0.844 | 1.24 | 0.094 | -0.038 | -0.038 | 0 | 0.91 | 0.64 | 0.8 | 0.98 | 0.87 | 0.93 |
| 4 | 150856693 | rs11726953 | -2.161 | -1.007 | 0.339 | 1.106 | 0.082 | 0.129 | 0.129 | 2.404 | 0.43 | 0.9 | 0.45 | 0.77 | 0.34 | 0.53 |
| 4 | 150858865 | rs1505474 | -1.604 | -0.816 | 0.639 | 1.242 | -0.005 | -0.007 | -0.007 | 0 | 0.36 | 0.31 | 0.39 | 0.74 | 0.33 | 0.46 |
| 4 | 150862026 | rs17631722 | -0.989 | -0.805 | 0.665 | 1.434 | 0.013 | -0.028 | -0.028 | 0 | 0.23 | 0.13 | 0.29 | 0.43 | 0.17 | 0.33 |
| 4 | 150884878 | rs12642120 | -0.389 | -0.498 | 0.628 | 1.445 | 0 | 0.027 | 0.027 | 0 | 0.12 | 0.2 | 0.09 | 0.16 | 0.15 | 0.13 |
| 4 | 150901440 | rs9759379 | -0.124 | -0.701 | 0.502 | 1.595 | -0.027 | 0.01 | 0.01 | 0 | 0.14 | 0.27 | 0.07 | 0.13 | 0.08 | 0.23 |
| 4 | 150901757 | rs11930273 | -0.979 | -0.755 | -0.09 | 1.767 | 0.002 | 0.004 | 0.004 | 0.32 | 0.09 | 0.1 | 0.14 | 0.28 | 0.07 | 0.08 |
| 4 | 150902638 | rs10034329 | 0.073 | -0.809 | -0.21 | 1.716 | 0.005 | 0.012 | 0.012 | 0 | 0.06 | 0.04 | 0.12 | 0.28 | 0.06 | 0.08 |
| 4 | 150904037 | rs11099749 | 0.313 | -0.86 | -0.212 | 1.656 | 0.019 | -0.018 | -0.018 | 0.649 | 0.13 | 0.06 | 0.06 | 0.15 | 0.08 | 0.23 |
| 4 | 150915414 | rs10002187 | -0.144 | -0.785 | -0.126 | 1.332 | 0.001 | 0.001 | 0.001 | 0 | 0.17 | 0.29 | 0.13 | 0.16 | 0.18 | 0.15 |
| 4 | 150918798 | rs3923657 | -1.021 | -0.713 | -0.171 | 1.357 | -0.001 | 0.034 | 0.034 | 0 | 0.24 | 0.34 | 0.33 | 0.45 | 0.26 | 0.2 |
| 4 | 150919730 | rs6535702 | -1.457 | -0.441 | -0.968 | 0.967 | 0.057 | 0.16 | 0.16 | 3.705 | 0.46 | 0.89 | 0.45 | 0.77 | 0.39 | 0.5 |
| 4 | 150924437 | rs4501203 | -0.881 | -0.09 | -0.306 | 1.013 | -0.004 | 0.129 | 0.129 | 0 | 0.08 | 0.19 | 0.2 | 0.3 | 0.07 | 0.08 |
| 4 | 150930752 | rs4260523 | 0.085 | 0.567 | -1.439 | -0.477 | -0.1 | 0.173 | 0.173 | 0 | 0.27 | 0.03 | 0.22 | 0 | 0.35 | 0.11 |
| 4 | 150946383 | rs4317185 | 0.244 | -0.373 | -0.332 | 0.265 | -0.055 | 0.009 | 0.009 | 0 | 0.77 | 0.45 | 0.87 | 0.77 | 0.86 | 0.76 |
| 4 | 150951692 | rs6833562 | -0.716 | -0.501 | -0.621 | -0.155 | -0.001 | -0.002 | -0.002 | 0 | 0.26 | 0.28 | 0.11 | 0.07 | 0.13 | 0.26 |
| 4 | 150954229 | rs4634199 | -0.047 | -0.072 | -1.569 | -0.986 | -0.054 | 0.152 | 0.152 | 0 | 0.27 | 0.04 | 0.18 | 0 | 0.3 | 0.07 |
| 4 | 150955500 | rs7655571 | 0.687 | -0.78 | -0.918 | -1.483 | -0.008 | -0.046 | -0.046 | 0.052 | 0.59 | 0.53 | 0.76 | 0.92 | 0.69 | 0.6 |
| 4 | 150960142 | rs10857241 | 0.498 | -0.824 | -1.236 | -1.495 | 0.028 | 0.157 | 0.157 | 0.427 | 0.62 | 0.87 | 0.78 | 0.93 | 0.69 | 0.66 |
| 4 | 150960542 | rs10017432 | -0.498 | -0.75 | -1.069 | -1.289 | 0.021 | 0.023 | 0.023 | 0.365 | 0.83 | 0.92 | 0.92 | 0.93 | 0.87 | 0.71 |
| 4 | 150968276 | rs13130722 | 0.355 | -0.449 | -1.614 | -0.785 | 0.011 | 0.025 | 0.025 | 0 | 0.5 | 0.16 | 0.44 | 0.44 | 0.58 | 0.33 |
| 4 | 150974325 | rs4835616 | -0.9 | -0.545 | -1.603 | -0.025 | 0.037 | -0.061 | -0.061 | 0 | 0.83 | 0.94 | 0.72 | 0.68 | 0.9 | 0.6 |
| 4 | 150976000 | rs4835617 | -0.441 | -0.651 | -1.624 | 0.114 | 0.038 | -0.058 | -0.058 | 0 | 0.8 | 0.88 | 0.61 | 0.6 | 0.88 | 0.51 |
| 4 | 150984857 | rs4423856 | 0.89 | -0.742 | -1.885 | -0.153 | -0.067 | 0.053 | 0.053 | 0 | 0.17 | 0.05 | 0.1 | 0.07 | 0.21 | 0.11 |
| 4 | 150995468 | rs4270569 | 0.464 | -0.942 | -1.438 | -0.37 | -0.008 | -0.005 | -0.005 | 0.39 | 0.4 | 0.34 | 0.38 | 0.48 | 0.51 | 0.32 |
| 4 | 150997068 | rs6818273 | -0.096 | -0.991 | -1.128 | -0.423 | 0.006 | -0.06 | -0.06 | 0.01 | 0.43 | 0.52 | 0.35 | 0.16 | 0.41 | 0.29 |
| 4 | 151005404 | rs11099754 | 1.725 | -1.011 | -1.175 | -0.183 | -0.005 | -0.008 | -0.008 | 0.021 | 0.27 | 0.23 | 0.24 | 0.38 | 0.37 | 0.23 |
| 4 | 151005467 | rs9307865 | -0.011 | -0.906 | -0.613 | -0.017 | -0.013 | 0.032 | 0.032 | 0 | 0.4 | 0.37 | 0.34 | 0.16 | 0.41 | 0.27 |
| 4 | 151030964 | rs6858003 | -1.088 | -1.055 | -0.675 | 0.347 | 0.048 | -0.069 | -0.069 | 0.323 | 0.84 | 0.95 | 0.73 | 0.63 | 0.92 | 0.58 |
| 4 | 151047967 | rs4274825 | 0.137 | -0.67 | -0.202 | -0.023 | -0.02 | -0.074 | -0.074 | 1.092 | 0.24 | 0.38 | 0.24 | 0.09 | 0.19 | 0.15 |
| 4 | 151049574 | rs12506111 | 1.153 | -0.625 | -0.337 | -0.135 | -0.021 | 0.084 | 0.084 | 0 | 0.14 | 0.02 | 0.08 | 0 | 0.13 | 0.04 |
| 4 | 151050375 | rs7667342 | -0.175 | -0.585 | -0.289 | -0.099 | -0.019 | -0.072 | -0.072 | 0 | 0.24 | 0.38 | 0.24 | 0.09 | 0.19 | 0.15 |
| 4 | 151063849 | rs10857243 | 0.458 | -0.009 | -1.01 | 0.29 | 0.024 | 0.018 | 0.018 | 0.008 | 0.48 | 0.14 | 0.55 | 0.57 | 0.49 | 0.71 |
| 4 | 151063917 | rs11099756 | 1.235 | -0.351 | -1.143 | 0.734 | 0.041 | -0.055 | -0.055 | 0 | 0.15 | 0.04 | 0.24 | 0.28 | 0.09 | 0.41 |
| 4 | 151070611 | rs4240358 | -1.09 | -0.069 | -0.56 | 0.726 | 0.005 | -0.005 | -0.005 | 0 | 0.8 | 0.73 | 0.84 | 0.8 | 0.81 | 0.94 |

| | | | | | | | | | | | | | | | | |
|---|---|---|---|---|---|---|---|---|---|---|---|---|---|---|---|---|
| 4 | 151080475 | rs7340876 | 0.255 | -0.015 | -0.538 | 0.944 | 0.003 | -0.007 | -0.007 | 0 | 0.09 | 0.09 | 0.14 | 0.15 | 0.08 | 0.13 |
| 4 | 151082561 | rs4240359 | 0.116 | -0.176 | -0.337 | 1.04 | 0.037 | 0.054 | 0.054 | 0 | 0.34 | 0.26 | 0.37 | 0.21 | 0.24 | 0.5 |
| 4 | 151091007 | rs9683765 | 1.36 | 0.475 | -0.516 | 1.117 | -0.045 | -0.01 | -0.01 | 0 | 0.3 | 0.07 | 0.28 | 0.26 | 0.39 | 0.26 |
| 4 | 151101413 | rs6535715 | 0.347 | 0.878 | 0.917 | 1.814 | 0.006 | 0.022 | 0.022 | 0.001 | 0.69 | 0.53 | 0.75 | 0.66 | 0.71 | 0.78 |
| 4 | 151103731 | rs10520059 | 0.424 | 0.983 | 0.501 | 1.752 | -0.018 | -0.007 | -0.007 | 0 | 0.19 | 0.16 | 0.12 | 0.19 | 0.21 | 0.1 |
| 4 | 151117651 | rs7658837 | 0.68 | 0.841 | 0.842 | 1.992 | 0.014 | 0.19 | 0.19 | 0 | 0.93 | 0.82 | 0.84 | 0.65 | 0.94 | 0.87 |
| 4 | 151126725 | rs12374344 | 0.197 | 1.289 | 0.88 | 1.996 | -0.016 | 0.001 | 0.001 | 0 | 0.21 | 0.19 | 0.16 | 0.16 | 0.29 | 0.1 |
| 4 | 151133778 | rs17631799 | 0.799 | 0.605 | 0.885 | 2.063 | 0.026 | -0.017 | -0.017 | 0.351 | 0.13 | 0.05 | 0.27 | 0.22 | 0.08 | 0.51 |
| 4 | 151157195 | rs1459744 | 1.556 | 0.832 | 1.411 | 1.991 | 0.06 | 0.357 | 0.357 | 3.807 | 0.91 | 0.49 | 0.81 | 0.65 | 0.93 | 0.93 |
| 4 | 151158031 | rs1459748 | 0.38 | 0.926 | 0.842 | 2.033 | 0.024 | 0.207 | 0.207 | 0.036 | 0.94 | 0.81 | 0.84 | 0.65 | 0.94 | 0.93 |
| 4 | 151164724 | rs1993656 | 0.765 | 1.047 | 0.778 | 1.791 | -0.005 | 0.006 | 0.006 | 0 | 0.61 | 0.36 | 0.39 | 0.25 | 0.59 | 0.57 |
| 4 | 151166237 | rs1531335 | 0.113 | 1.033 | 1.097 | 1.973 | 0.065 | 0.079 | 0.079 | 0 | 0.74 | 0.6 | 0.68 | 0.47 | 0.66 | 0.8 |
| 4 | 151168929 | rs2359940 | 0.607 | 0.754 | 0.691 | 1.811 | 0.032 | -0.02 | -0.02 | 2.03 | 0.14 | 0.02 | 0.28 | 0.23 | 0.09 | 0.34 |
| 4 | 151170387 | rs1459750 | 1.083 | 0.929 | 0.82 | 1.625 | -0.01 | 0.024 | 0.024 | 0 | 0.57 | 0.34 | 0.38 | 0.25 | 0.54 | 0.54 |
| 4 | 151182255 | rs6844989 | -0.077 | 0.941 | 1.113 | 1.997 | -0.028 | 0.201 | 0.201 | 0 | 0.82 | 0.55 | 0.59 | 0.51 | 0.88 | 0.7 |
| 4 | 151185651 | rs11059 | 0.468 | 1.233 | 1.3 | 2.131 | 0.028 | 0.034 | 0.034 | 0.021 | 0.74 | 0.37 | 0.7 | 0.48 | 0.65 | 0.82 |
| 4 | 151190804 | rs13134607 | 0.39 | 1.233 | 1.46 | 2.124 | 0.061 | 0.076 | 0.076 | 0 | 0.74 | 0.45 | 0.7 | 0.48 | 0.65 | 0.82 |
| 4 | 151199080 | rs2290846 | 0.667 | 1.287 | 1.287 | 2.1 | -0.109 | -0.076 | -0.076 | 0.418 | 0.16 | 0.01 | 0.13 | 0.24 | 0.27 | 0.1 |
| 4 | 151202952 | rs11099759 | 0.519 | 1.132 | 1.456 | 2.13 | 0.055 | 0.298 | 0.298 | 0.71 | 0.95 | 0.77 | 0.88 | 0.72 | 0.96 | 0.94 |
| 4 | 151204693 | rs11944163 | 1.02 | 1.078 | 1.355 | 2.078 | -0.02 | 0.019 | 0.019 | 0 | 0.62 | 0.33 | 0.42 | 0.26 | 0.57 | 0.57 |
| 4 | 151207127 | rs3749574 | 0.376 | 1.116 | 1.174 | 2.028 | -0.115 | -0.065 | -0.065 | 0 | 0.18 | 0.04 | 0.16 | 0.24 | 0.3 | 0.12 |
| 4 | 151210761 | rs7657457 | 0.104 | 1.117 | 1.287 | 2.028 | 0.044 | 0.011 | 0.011 | 0 | 0.2 | 0.26 | 0.17 | 0.24 | 0.3 | 0.12 |
| 4 | 151214038 | rs10007367 | 1.045 | 0.597 | 0.995 | 2.034 | 0.02 | -0.032 | -0.032 | 0 | 0.11 | 0.05 | 0.27 | 0.21 | 0.08 | 0.25 |
| 4 | 151216870 | rs4696254 | -0.615 | 0.522 | 1.021 | 2.005 | 0.021 | -0.035 | -0.035 | 0 | 0.84 | 0.9 | 0.64 | 0.66 | 0.9 | 0.54 |
| 4 | 151222266 | rs1960918 | 1.211 | 0.81 | 1.203 | 1.906 | 0 | -0.017 | -0.017 | 2.296 | 0.61 | 0.41 | 0.45 | 0.26 | 0.62 | 0.41 |
| 4 | 151229489 | rs11734677 | 0.282 | 0.778 | 1.105 | 1.933 | -0.021 | -0.011 | -0.011 | 0 | 0.18 | 0.15 | 0.15 | 0.22 | 0.26 | 0.1 |
| 4 | 151231371 | rs1813134 | 0.766 | 0.764 | 1.196 | 1.96 | 0 | 0.001 | 0.001 | 0 | 0.64 | 0.58 | 0.48 | 0.44 | 0.65 | 0.43 |
| 4 | 151233934 | rs13151875 | 1.492 | 0.438 | 0.642 | 1.575 | -0.058 | 0.197 | 0.197 | 0 | 0.4 | 0.1 | 0.24 | 0.1 | 0.42 | 0.32 |
| 4 | 151238436 | rs962626 | 0.63 | 0.588 | 0.755 | 1.691 | -0.001 | 0.002 | 0.002 | 0 | 0.63 | 0.53 | 0.48 | 0.44 | 0.65 | 0.43 |
| 4 | 151252297 | rs17588629 | 0.458 | 0.811 | 0.36 | 1.605 | -0.034 | -0.019 | -0.019 | 0 | 0.18 | 0.12 | 0.14 | 0.22 | 0.25 | 0.1 |
| 4 | 151271758 | rs17588741 | 0.586 | 0.323 | 0.526 | 1.435 | 0.001 | 0.001 | 0.001 | 0.173 | 0.22 | 0.2 | 0.19 | 0.16 | 0.2 | 0.08 |
| 4 | 151272102 | rs11945847 | 1.323 | 0.241 | 0.575 | 1.258 | 0.003 | 0.051 | 0.051 | 0.192 | 0.39 | 0.28 | 0.32 | 0.12 | 0.37 | 0.31 |
| 4 | 151275634 | rs13136928 | -0.704 | 0.175 | 0.999 | 1.57 | -0.011 | 0.017 | 0.017 | 0.009 | 0.83 | 0.8 | 0.71 | 0.68 | 0.9 | 0.53 |
| 4 | 151275797 | rs4458434 | -0.378 | 0.168 | 1.036 | 1.566 | -0.041 | 0.058 | 0.058 | 0.024 | 0.81 | 0.66 | 0.7 | 0.68 | 0.9 | 0.52 |
| 4 | 151284982 | rs7657948 | -0.378 | 0.19 | 0.545 | 1.578 | -0.041 | 0.058 | 0.058 | 0.024 | 0.81 | 0.66 | 0.7 | 0.68 | 0.9 | 0.52 |
| 4 | 151301638 | rs7699001 | 0.394 | 0.897 | 0.603 | 1.533 | 0.006 | 0.009 | 0.009 | 0.13 | 0.2 | 0.13 | 0.11 | 0.11 | 0.16 | 0.08 |
| 4 | 151312761 | rs6848343 | 0.067 | 0.73 | 0.843 | 1.561 | -0.022 | 0.028 | 0.028 | 0.738 | 0.89 | 0.65 | 0.86 | 0.88 | 0.95 | 0.72 |
| 4 | 151331191 | rs11736546 | 0.404 | 1.005 | 0.579 | 1.714 | -0.08 | -0.026 | -0.026 | 0 | 0.19 | 0.07 | 0.15 | 0.2 | 0.33 | 0.11 |
| 4 | 151337243 | rs7686589 | 1.482 | 0.945 | 0.702 | 1.828 | 0.02 | 0.056 | 0.056 | 0 | 0.46 | 0.23 | 0.49 | 0.34 | 0.43 | 0.46 |
| 4 | 151343482 | rs1979920 | 0.68 | 0.761 | 0.744 | 1.635 | 0.013 | 0.025 | 0.025 | 0.24 | 0.19 | 0.06 | 0.11 | 0.11 | 0.16 | 0.07 |
| 4 | 151349664 | rs7689850 | 0.036 | 0.74 | 0.994 | 1.763 | -0.017 | 0.044 | 0.044 | 0.913 | 0.89 | 0.61 | 0.84 | 0.86 | 0.95 | 0.72 |
| 4 | 151356081 | rs4696352 | 0.656 | 0.676 | 0.867 | 1.654 | 0.053 | -0.037 | -0.037 | 0.061 | 0.65 | 0.17 | 0.69 | 0.6 | 0.77 | 0.61 |
| 4 | 151368816 | rs7698239 | 1.127 | 0.724 | 0.907 | 1.634 | 0.009 | 0.006 | 0.006 | 0 | 0.49 | 0.37 | 0.6 | 0.57 | 0.47 | 0.52 |
| 4 | 151369107 | rs11733926 | -0.603 | 0.743 | 0.834 | 1.634 | -0.049 | 0.012 | 0.012 | 0.001 | 0.38 | 0.18 | 0.26 | 0.31 | 0.48 | 0.19 |
| 4 | 151376718 | rs11727758 | 0.447 | 0.828 | 0.754 | 1.699 | -0.088 | -0.017 | -0.017 | 0 | 0.18 | 0.06 | 0.16 | 0.16 | 0.33 | 0.12 |
| 4 | 151385976 | rs1875899 | -0.882 | 0.78 | 0.897 | 1.679 | -0.002 | -0.004 | -0.004 | 0.329 | 0.41 | 0.43 | 0.3 | 0.46 | 0.51 | 0.23 |
| 4 | 151388101 | rs1504781 | 0.261 | 0.707 | 0.937 | 1.641 | -0.034 | 0.027 | 0.027 | 1.079 | 0.87 | 0.57 | 0.84 | 0.88 | 0.95 | 0.71 |
| 4 | 151397759 | rs2036552 | 0.822 | 0.755 | 0.916 | 1.656 | 0.005 | 0.008 | 0.008 | 0 | 0.19 | 0.13 | 0.1 | 0.1 | 0.16 | 0.07 |
| 4 | 151399246 | rs4696435 | 0.711 | 0.753 | 0.908 | 1.682 | -0.039 | 0.1 | 0.1 | 6.595 | 0.84 | 0.36 | 0.81 | 0.69 | 0.92 | 0.67 |
| 4 | 151403684 | rs7688410 | 1.557 | 0.768 | 0.87 | 1.678 | 0.016 | 0.034 | 0.034 | 0.163 | 0.46 | 0.14 | 0.54 | 0.43 | 0.44 | 0.48 |
| 4 | 151406414 | rs1504786 | 0.669 | 0.753 | 0.882 | 1.678 | 0.044 | -0.035 | -0.035 | 0 | 0.65 | 0.21 | 0.7 | 0.59 | 0.77 | 0.6 |
| 4 | 151413637 | rs13102532 | 0.689 | 0.753 | 0.868 | 1.677 | 0.051 | -0.045 | -0.045 | 3.842 | 0.65 | 0.18 | 0.69 | 0.58 | 0.77 | 0.6 |
| 4 | 151420304 | rs7688842 | -0.568 | 0.806 | 0.848 | 1.629 | -0.036 | 0.019 | 0.019 | 0 | 0.38 | 0.22 | 0.26 | 0.26 | 0.48 | 0.19 |
| 4 | 151444619 | rs11931161 | -0.159 | 0.813 | 0.838 | 1.596 | -0.015 | -0.013 | -0.013 | 2.718 | 0.23 | 0.25 | 0.19 | 0.35 | 0.35 | 0.16 |
| 4 | 151444833 | rs11931175 | -0.068 | 0.773 | 0.93 | 1.575 | -0.031 | 0.014 | 0.014 | 0.303 | 0.88 | 0.66 | 0.86 | 0.88 | 0.95 | 0.71 |
| 4 | 151446347 | rs13150736 | 1.669 | 0.751 | 0.73 | 1.543 | -0.041 | -0.02 | -0.02 | 1.201 | 0.45 | 0.04 | 0.54 | 0.43 | 0.44 | 0.48 |
| 4 | 151454015 | rs952225 | 0.572 | 0.607 | 0.87 | 1.503 | 0.04 | -0.009 | -0.009 | 0 | 0.68 | 0.52 | 0.69 | 0.72 | 0.62 | 0.59 |
| 4 | 151479467 | rs12649589 | 1.316 | 0.883 | 0.698 | 1.49 | 0 | 0.028 | 0.028 | 0.144 | 0.48 | 0.26 | 0.59 | 0.43 | 0.49 | 0.48 |
| 4 | 151496759 | rs17589398 | 0.474 | 0.876 | 0.558 | 1.283 | -0.095 | -0.033 | -0.033 | 0 | 0.18 | 0.02 | 0.13 | 0.16 | 0.29 | 0.11 |
| 4 | 151499440 | rs10022844 | 1.309 | 0.759 | 0.633 | 1.314 | -0.004 | 0.014 | 0.014 | 0.037 | 0.47 | 0.26 | 0.59 | 0.45 | 0.49 | 0.46 |
| 4 | 151502267 | rs9993229 | 1.183 | 0.748 | 0.372 | 1.414 | -0.017 | 0.014 | 0.014 | 0 | 0.48 | 0.23 | 0.6 | 0.61 | 0.51 | 0.5 |
| 4 | 151519232 | rs4435730 | 0.856 | 0.845 | 0.47 | 1.239 | 0.005 | -0.006 | -0.006 | 0.245 | 0.83 | 0.18 | 0.87 | 0.74 | 0.92 | 0.67 |
| 4 | 151534545 | rs9999828 | 1.524 | 0.903 | 0.373 | 1.232 | -0.042 | -0.046 | -0.046 | 0 | 0.45 | 0.04 | 0.55 | 0.46 | 0.45 | 0.43 |
| 4 | 151547243 | rs6847725 | 0.761 | 0.305 | 0.786 | 1.286 | -0.048 | -0.026 | -0.026 | 0 | 0.85 | 0.34 | 0.91 | 0.92 | 0.95 | 0.68 |
| 4 | 151632108 | rs10857249 | 0.759 | 0.269 | 0.898 | 1.4 | -0.013 | 0.147 | 0.147 | 1.027 | 0.92 | 0.79 | 0.85 | 0.68 | 0.94 | 0.66 |
| 4 | 151637154 | rs17027030 | -0.047 | 0.213 | 0.998 | 1.335 | -0.018 | -0.051 | -0.051 | 0 | 0.1 | 0.56 | 0.1 | 0 | 0.05 | 0.01 |
| 4 | 151637268 | rs6535746 | -0.346 | 0.463 | 0.982 | 1.265 | -0.036 | 0.01 | 0.01 | 0 | 0.36 | 0.22 | 0.32 | 0.27 | 0.45 | 0.22 |
| 4 | 151659605 | rs7690485 | 0.631 | 0.41 | 0.992 | 1.217 | -0.012 | 0.151 | 0.151 | 1.183 | 0.92 | 0.79 | 0.85 | 0.69 | 0.94 | 0.65 |
| 4 | 151665097 | rs1451634 | 0.624 | 0.234 | 1.264 | 1.232 | -0.039 | 0.013 | 0.013 | 0.333 | 0.86 | 0.71 | 0.83 | 0.87 | 0.92 | 0.68 |

| | | | | | | | | | | | | | | | | |
|---|---|---|---|---|---|---|---|---|---|---|---|---|---|---|---|---|
| 4 | 151730224 | rs716473 | 0.743 | 0.615 | 0.82 | 1.248 | -0.084 | -0.047 | -0.047 | 0.429 | 0.14 | 0.01 | 0.11 | 0.15 | 0.26 | 0.1 |
| 4 | 151730255 | rs716474 | 0.074 | 0.464 | 0.966 | 1.25 | 0.004 | -0.033 | -0.033 | 0 | 0.09 | 0.44 | 0.08 | 0 | 0.05 | 0.01 |
| 4 | 151761599 | rs6846361 | 0.356 | 0.534 | 0.763 | 1.259 | -0.096 | -0.029 | -0.029 | 0 | 0.17 | 0.02 | 0.13 | 0.15 | 0.3 | 0.13 |
| 4 | 151765243 | rs186080 | 0.687 | 0.469 | 0.972 | 1.072 | -0.031 | 0.041 | 0.041 | 1.737 | 0.86 | 0.56 | 0.81 | 0.87 | 0.92 | 0.69 |
| 4 | 151772410 | rs10012720 | 1.404 | 0.467 | 0.351 | 0.897 | -0.053 | 0.036 | 0.036 | 0.825 | 0.46 | 0.07 | 0.44 | 0.4 | 0.44 | 0.44 |
| 4 | 151790540 | rs2407549 | 1.551 | -0.212 | 1.039 | 0.931 | 0.03 | -0.019 | -0.019 | 0 | 0.79 | 0.15 | 0.75 | 0.72 | 0.89 | 0.64 |
| 4 | 151794207 | rs17027158 | -0.036 | -0.148 | 1.406 | 0.895 | 0.002 | -0.031 | -0.031 | 0 | 0.09 | 0.36 | 0.07 | 0 | 0.05 | 0.01 |
| 4 | 151815783 | rs1201214 | -0.635 | -0.286 | 1.368 | 0.997 | -0.01 | 0.051 | 0.051 | 0.198 | 0.09 | 0.12 | 0.14 | 0.28 | 0.06 | 0.34 |
| 4 | 151829824 | rs1201208 | -0.903 | -0.311 | 1.798 | 0.921 | -0.042 | -0.013 | -0.013 | 0.033 | 0.14 | 0.25 | 0.14 | 0.1 | 0.07 | 0.29 |
| 4 | 151852107 | rs1148642 | 0.351 | -0.317 | 2.149 | 0.89 | -0.011 | -0.039 | -0.039 | 0 | 0.9 | 0.5 | 0.93 | 1 | 0.96 | 0.99 |
| 4 | 151859674 | rs1201198 | -0.643 | -0.311 | 2.06 | 0.978 | -0.011 | 0.057 | 0.057 | 0.138 | 0.09 | 0.13 | 0.12 | 0.28 | 0.06 | 0.33 |
| 4 | 151887280 | rs6839352 | 0.85 | -0.334 | 2.204 | 0.803 | -0.045 | -0.022 | -0.022 | 0 | 0.82 | 0.28 | 0.82 | 0.9 | 0.93 | 0.7 |
| 4 | 151948111 | rs4696648 | 0.291 | -0.462 | 2.016 | 0.951 | -0.041 | -0.014 | -0.014 | 0.044 | 0.86 | 0.74 | 0.86 | 0.9 | 0.93 | 0.67 |
| 4 | 151978131 | rs6819599 | 0.185 | -0.333 | 2.035 | 0.913 | -0.036 | -0.035 | -0.035 | 0 | 0.86 | 0.74 | 0.86 | 0.93 | 0.92 | 0.71 |
| 4 | 151998622 | rs7697992 | 0.212 | -0.338 | 2.121 | 0.922 | -0.055 | -0.055 | -0.055 | 0 | 0.84 | 0.56 | 0.85 | 0.93 | 0.92 | 0.71 |
| 4 | 152000891 | rs12647989 | 1.149 | -0.393 | 2.093 | 1.07 | -0.006 | 0 | 0 | 0.674 | 0.8 | 0.22 | 0.8 | 0.73 | 0.89 | 0.66 |
| 4 | 152014653 | rs13145829 | 1.11 | -0.393 | 2.093 | 1.07 | -0.007 | 0 | 0 | 3.621 | 0.8 | 0.22 | 0.8 | 0.73 | 0.89 | 0.66 |
| 4 | 152015704 | rs11733964 | 1.151 | -0.393 | 2.093 | 1.07 | -0.006 | 0 | 0 | 0 | 0.8 | 0.22 | 0.8 | 0.73 | 0.89 | 0.66 |
| 4 | 152015776 | rs13101417 | 1.11 | -0.392 | 2.094 | 1.07 | -0.007 | 0 | 0 | 3.621 | 0.8 | 0.22 | 0.8 | 0.73 | 0.89 | 0.66 |
| 4 | 152016931 | rs2897659 | 0.35 | -0.384 | 2.224 | 1.075 | -0.045 | 0.103 | 0.103 | 0.112 | 0.84 | 0.53 | 0.82 | 0.73 | 0.9 | 0.66 |
| 4 | 152020229 | rs4696226 | 0.197 | -0.382 | 2.207 | 1.076 | -0.042 | -0.041 | -0.041 | 1.579 | 0.86 | 0.67 | 0.85 | 0.93 | 0.92 | 0.71 |
| 4 | 152053302 | rs7659146 | 0.179 | -0.392 | 2.113 | 1.059 | -0.043 | -0.036 | -0.036 | 1.846 | 0.85 | 0.66 | 0.84 | 0.92 | 0.92 | 0.71 |
| 4 | 152055073 | rs11938266 | 0.246 | -0.392 | 2.135 | 1.059 | -0.064 | -0.055 | -0.055 | 0.579 | 0.84 | 0.46 | 0.83 | 0.92 | 0.92 | 0.71 |
| 4 | 152065684 | rs6818900 | 0.216 | -0.19 | 1.177 | 1.173 | -0.122 | -0.07 | -0.07 | 1.923 | 0.35 | 0.02 | 0.25 | 0.29 | 0.42 | 0.14 |
| 4 | 152066906 | rs7669180 | 0.744 | -0.151 | 1.462 | 1.098 | -0.009 | 0.001 | 0.001 | 3.643 | 0.8 | 0.21 | 0.8 | 0.73 | 0.89 | 0.66 |
| 4 | 152095256 | rs11734586 | -0.093 | -0.18 | 1.548 | 1.078 | -0.059 | -0.042 | -0.042 | 0.048 | 0.84 | 0.6 | 0.85 | 0.92 | 0.92 | 0.71 |
| 4 | 152097962 | rs11934792 | 0.936 | -0.128 | 0.991 | 1.064 | 0.019 | -0.012 | -0.012 | 0.047 | 0.78 | 0.18 | 0.78 | 0.72 | 0.87 | 0.63 |
| 4 | 152105400 | rs12647306 | 0.887 | -0.113 | 1.001 | 1.077 | 0.016 | -0.011 | -0.011 | 0 | 0.78 | 0.18 | 0.78 | 0.72 | 0.87 | 0.63 |
| 4 | 152109552 | rs13120989 | 1.191 | -0.053 | 0.715 | 1.177 | -0.096 | 0.032 | 0.032 | 0 | 0.38 | 0.07 | 0.43 | 0.37 | 0.44 | 0.44 |
| 4 | 152144869 | rs2407406 | 0.417 | -0.072 | 1.347 | 1.392 | -0.072 | -0.058 | -0.058 | 0 | 0.84 | 0.45 | 0.83 | 0.92 | 0.92 | 0.7 |
| 4 | 152147262 | rs10520101 | 0.805 | -0.041 | 0.958 | 1.426 | -0.033 | 0.023 | 0.023 | 1.205 | 0.79 | 0.32 | 0.79 | 0.72 | 0.87 | 0.63 |
| 4 | 152147441 | rs10520102 | 0.805 | -0.041 | 0.929 | 1.426 | -0.032 | 0.021 | 0.021 | 1.205 | 0.79 | 0.32 | 0.79 | 0.72 | 0.87 | 0.63 |
| 4 | 152164786 | rs12509302 | -1.204 | 1.212 | 0.865 | 1.937 | 0.115 | 0.17 | 0.17 | 0 | 0.14 | 0.37 | 0.22 | 0.27 | 0.23 | 0.12 |
| 4 | 152172980 | rs2407167 | 0.056 | 0.49 | 1.493 | 2.154 | 0.007 | 0.219 | 0.219 | 0.275 | 0.92 | 0.71 | 0.87 | 0.74 | 0.94 | 0.69 |
| 4 | 152175168 | rs7663659 | 0 | 0.107 | 0.149 | 1.668 | -0.095 | 0.087 | 0.087 | 0.807 | 0.37 | 0.04 | 0.25 | 0.18 | 0.42 | 0.19 |
| 4 | 152179311 | rs10520104 | 0.079 | -0.114 | 0.259 | 1.924 | -0.008 | 0.012 | 0.012 | 0 | 0.12 | 0.12 | 0.14 | 0.2 | 0.05 | 0.04 |
| 4 | 152192808 | rs10016704 | -0.023 | -0.28 | 0.383 | 1.829 | -0.043 | 0.127 | 0.127 | 0.014 | 0.41 | 0.15 | 0.27 | 0.18 | 0.44 | 0.23 |
| 4 | 152196218 | rs6832149 | -0.908 | -0.323 | 0.203 | 1.814 | -0.01 | 0 | 0 | 0 | 0.53 | 0.27 | 0.41 | 0.38 | 0.49 | 0.26 |
| 4 | 152197483 | rs10031902 | 0.105 | -0.32 | 0.173 | 1.798 | -0.07 | 0.107 | 0.107 | 0.018 | 0.38 | 0.08 | 0.25 | 0.18 | 0.43 | 0.19 |
| 4 | 152205278 | rs13112985 | 1.421 | -0.361 | 0.347 | 1.71 | -0.073 | 0.036 | 0.036 | 0 | 0.37 | 0.13 | 0.44 | 0.36 | 0.45 | 0.43 |
| 4 | 152212486 | rs13126069 | 1.42 | -0.361 | 0.347 | 1.71 | -0.073 | 0.036 | 0.036 | 0 | 0.37 | 0.13 | 0.44 | 0.36 | 0.45 | 0.43 |
| 4 | 152213113 | rs11734518 | -0.045 | -0.226 | -0.025 | 1.524 | -0.042 | 0.046 | 0.046 | 0.004 | 0.41 | 0.15 | 0.34 | 0.27 | 0.44 | 0.23 |
| 4 | 152221451 | rs6852223 | -0.335 | -0.19 | 0.264 | 1.545 | -0.008 | 0.001 | 0.001 | 0 | 0.12 | 0.12 | 0.07 | 0.11 | 0.05 | 0.04 |
| 4 | 152227582 | rs6535787 | -1.064 | -0.208 | 0.069 | 1.375 | 0.006 | 0.023 | 0.023 | 0 | 0.53 | 0.78 | 0.51 | 0.56 | 0.47 | 0.56 |
| 4 | 152229767 | rs11935905 | -0.315 | -0.529 | 0.225 | 1.281 | -0.002 | -0.063 | -0.063 | 0 | 0.33 | 0.21 | 0.3 | 0.42 | 0.34 | 0.24 |
| 4 | 152234938 | rs12646254 | -1.706 | -0.625 | -0.475 | 1.033 | -0.017 | 0.017 | 0.017 | 0 | 0.65 | 0.78 | 0.58 | 0.79 | 0.56 | 0.76 |
| 4 | 152256782 | rs2407171 | -1.16 | -0.378 | -0.115 | 1.157 | -0.001 | 0.003 | 0.003 | 0 | 0.55 | 0.62 | 0.55 | 0.54 | 0.55 | 0.63 |
| 4 | 152280646 | rs10033020 | 0.12 | -0.422 | 0.426 | 1.205 | -0.027 | 0.033 | 0.033 | 0.077 | 0.09 | 0.28 | 0.11 | 0.09 | 0.04 | 0.04 |
| 4 | 152294729 | rs6535791 | 0.348 | -0.255 | 0.532 | 1.252 | 0.022 | 0.015 | 0.015 | 0.66 | 0.09 | 0.19 | 0.07 | 0.13 | 0.1 | 0.03 |
| 4 | 152297368 | rs9968292 | -0.954 | -0.374 | 0.822 | 1.261 | 0.005 | -0.002 | -0.002 | 0.054 | 0.44 | 0.45 | 0.42 | 0.37 | 0.52 | 0.35 |
| 4 | 152306809 | rs6535797 | 0.526 | -0.347 | 0.762 | 1.264 | 0.024 | 0.016 | 0.016 | 0.632 | 0.09 | 0.2 | 0.07 | 0.13 | 0.1 | 0.03 |
| 4 | 152321881 | rs2724553 | -1.947 | -0.497 | 0.483 | 1.232 | -0.103 | -0.09 | -0.09 | 0 | 0.6 | 0.97 | 0.56 | 0.52 | 0.57 | 0.64 |
| 4 | 152355268 | rs2709828 | -0.904 | -0.451 | 0.816 | 1.241 | -0.001 | 0.003 | 0.003 | 0 | 0.45 | 0.56 | 0.43 | 0.37 | 0.53 | 0.36 |
| 4 | 152375803 | rs6845067 | 0.814 | -0.446 | 0.844 | 1.266 | 0.017 | 0.015 | 0.015 | 0.572 | 0.08 | 0.15 | 0.06 | 0.16 | 0.1 | 0.03 |
| 4 | 152379820 | rs11941136 | 0.448 | -0.382 | 0.203 | 1.241 | -0.073 | -0.048 | -0.048 | 0 | 0.11 | 0.01 | 0.18 | 0.13 | 0.24 | 0.24 |
| 4 | 152401909 | rs11732407 | -0.291 | -0.515 | 0.474 | 1.243 | -0.032 | 0.009 | 0.009 | 0.29 | 0.35 | 0.21 | 0.36 | 0.23 | 0.43 | 0.33 |
| 4 | 152417635 | rs17027640 | 0.02 | -0.556 | 0.657 | 1.244 | -0.033 | 0.006 | 0.006 | 0.337 | 0.14 | 0.24 | 0.11 | 0.15 | 0.04 | 0.29 |
| 4 | 152428258 | rs11947054 | -0.438 | -0.352 | 0.472 | 1.186 | -0.048 | 0.006 | 0.006 | 0.381 | 0.35 | 0.17 | 0.36 | 0.24 | 0.43 | 0.32 |
| 4 | 152450126 | rs1443088 | -1.074 | -0.358 | 0.678 | 1.211 | -0.001 | 0 | 0 | 0 | 0.44 | 0.46 | 0.43 | 0.37 | 0.53 | 0.35 |
| 4 | 152455339 | rs1946999 | -1.185 | -0.716 | 0.756 | 1.184 | -0.008 | 0.01 | 0.01 | 0 | 0.44 | 0.63 | 0.4 | 0.34 | 0.45 | 0.33 |
| 4 | 152486018 | rs11946622 | -0.007 | -0.624 | 0.559 | 1.355 | -0.054 | -0.054 | -0.054 | 0 | 0.3 | 0.07 | 0.33 | 0.31 | 0.34 | 0.3 |
| 4 | 152488075 | rs7695412 | -0.381 | -0.634 | 0.771 | 1.082 | 0 | -0.001 | -0.001 | 0 | 0.33 | 0.29 | 0.32 | 0.19 | 0.35 | 0.29 |
| 4 | 152498625 | rs17275536 | -0.388 | -0.784 | 0.773 | 1.076 | 0 | -0.001 | -0.001 | 0 | 0.33 | 0.29 | 0.32 | 0.19 | 0.34 | 0.29 |
| 4 | 152511721 | rs1473305 | -0.129 | -0.785 | 0.802 | 1.076 | -0.009 | 0.006 | 0.006 | 0.207 | 0.31 | 0.18 | 0.31 | 0.19 | 0.34 | 0.29 |
| 4 | 152523327 | rs17275584 | 0.84 | -0.793 | 0.916 | 1.124 | 0.011 | 0.01 | 0.01 | 0.468 | 0.08 | 0.14 | 0.07 | 0.11 | 0.1 | 0.03 |
| 4 | 152543783 | rs6825241 | -1.032 | -0.802 | 0.868 | 1.087 | 0.002 | -0.003 | -0.003 | 0 | 0.42 | 0.55 | 0.39 | 0.3 | 0.45 | 0.32 |
| 4 | 152557786 | rs12651635 | -0.016 | -0.759 | 0.529 | 1.098 | -0.055 | -0.011 | -0.011 | 0 | 0.3 | 0.07 | 0.31 | 0.19 | 0.34 | 0.29 |
| 4 | 152567239 | rs9631757 | 0.072 | -1.003 | 0.525 | 1.186 | -0.039 | 0.022 | 0.022 | 0.001 | 0.13 | 0.23 | 0.04 | 0.2 | 0.04 | 0.31 |
| 4 | 152568478 | rs17275598 | -0.005 | -0.905 | -0.208 | 1.265 | -0.096 | -0.031 | -0.031 | 0 | 0.29 | 0.02 | 0.3 | 0.18 | 0.34 | 0.27 |

| | | | | | | | | | | | | | | | | |
|---|---|---|---|---|---|---|---|---|---|---|---|---|---|---|---|---|
| 4 | 152569864 | rs12509425 | 1.144 | -0.759 | 0.099 | 1.311 | 0.01 | -0.04 | -0.04 | 0 | 0.07 | 0 | 0.02 | 0.13 | 0.04 | 0.05 |
| 4 | 152575079 | rs7699787 | -0.077 | -0.581 | 0.451 | 1.087 | 0.034 | 0.022 | 0.022 | 1.223 | 0.1 | 0.31 | 0.08 | 0.11 | 0.1 | 0.03 |
| 4 | 152588541 | rs1366910 | 0.554 | -0.564 | 0.385 | 1.097 | 0.015 | 0.009 | 0.009 | 0.42 | 0.08 | 0.17 | 0.06 | 0.11 | 0.1 | 0.03 |
| 4 | 152590337 | rs17027780 | -0.048 | -0.575 | 0.4 | 1.09 | 0.027 | 0.017 | 0.017 | 1.154 | 0.1 | 0.29 | 0.07 | 0.11 | 0.11 | 0.03 |
| 4 | 152592191 | rs4571 | -1.56 | -0.875 | -0.218 | 1.06 | -0.086 | -0.054 | -0.054 | 0.121 | 0.56 | 0.93 | 0.44 | 0.48 | 0.49 | 0.6 |
| 4 | 152596508 | rs3749563 | -1.527 | -0.868 | -0.228 | 0.936 | -0.088 | -0.059 | -0.059 | 0 | 0.55 | 0.93 | 0.44 | 0.48 | 0.49 | 0.6 |
| 4 | 152609304 | rs7669761 | -0.957 | -0.9 | 0.104 | 0.924 | 0.001 | 0 | 0 | 0 | 0.44 | 0.54 | 0.35 | 0.35 | 0.38 | 0.58 |
| 4 | 152610318 | rs6833277 | 0.563 | -0.87 | 0.206 | 0.888 | 0.02 | 0.013 | 0.013 | 0.085 | 0.09 | 0.22 | 0.08 | 0.12 | 0.11 | 0.03 |
| 4 | 152610894 | rs9996600 | -0.142 | -0.887 | 0.266 | 0.879 | 0.061 | 0.046 | 0.046 | 0.794 | 0.11 | 0.46 | 0.1 | 0.12 | 0.11 | 0.03 |
| 4 | 152621150 | rs4696288 | -0.017 | -1.093 | -0.172 | 0.905 | -0.046 | 0.007 | 0.007 | 0.381 | 0.29 | 0.07 | 0.16 | 0.16 | 0.33 | 0.27 |
| 4 | 152625446 | rs11732441 | 0.543 | -0.854 | 0.088 | 0.867 | 0.042 | 0.031 | 0.031 | 0.217 | 0.09 | 0.32 | 0.09 | 0.12 | 0.11 | 0.03 |
| 4 | 152631472 | rs4696290 | -1.115 | -1.007 | 0.074 | 0.892 | 0.007 | 0.001 | 0.001 | 0 | 0.45 | 0.52 | 0.31 | 0.34 | 0.37 | 0.59 |
| 4 | 152640305 | rs6832872 | -2.307 | -1.354 | -0.557 | 0.894 | -0.169 | 0.063 | 0.063 | 0 | 0.69 | 0.99 | 0.56 | 0.88 | 0.6 | 0.82 |
| 4 | 152650882 | rs6535818 | -1.945 | -0.733 | 0.277 | 1.291 | -0.082 | -0.077 | -0.077 | 0.018 | 0.56 | 0.92 | 0.4 | 0.46 | 0.48 | 0.62 |
| 4 | 152672882 | rs7684350 | -1.897 | -0.715 | 0.305 | 1.3 | -0.034 | -0.031 | -0.031 | 0 | 0.55 | 0.8 | 0.39 | 0.46 | 0.48 | 0.62 |
| 4 | 152677580 | rs17275724 | 0.279 | -0.838 | 0.109 | 1.146 | -0.095 | 0.022 | 0.022 | 0 | 0.21 | 0.01 | 0.11 | 0.09 | 0.28 | 0.26 |
| 4 | 152681952 | rs1429559 | -1.56 | -0.807 | 0.439 | 1.071 | 0.017 | 0.007 | 0.007 | 0 | 0.46 | 0.55 | 0.31 | 0.33 | 0.38 | 0.59 |
| 4 | 152682805 | rs3762906 | 0.415 | -0.805 | 0.446 | 1.129 | 0.035 | 0.041 | 0.041 | 0 | 0.09 | 0.25 | 0.08 | 0.13 | 0.1 | 0.03 |
| 4 | 152701085 | rs2048433 | -0.505 | -0.949 | 0.709 | 1.009 | -0.066 | 0.008 | 0.008 | 0.321 | 0.18 | 0.39 | 0.16 | 0.17 | 0.08 | 0.33 |
| 4 | 152701656 | rs2175409 | -1.553 | -0.74 | 0.446 | 0.541 | 0.031 | 0.015 | 0.015 | 0 | 0.47 | 0.61 | 0.31 | 0.33 | 0.38 | 0.6 |
| 4 | 152710943 | rs13137669 | 1.692 | -0.675 | 0.308 | 0.517 | -0.015 | -0.015 | -0.015 | 0.582 | 0.44 | 0.28 | 0.62 | 0.54 | 0.52 | 0.38 |
| 4 | 152727299 | rs11099829 | 1.024 | -0.633 | 0.27 | 0.387 | 0.004 | 0.002 | 0.002 | 0 | 0.54 | 0.44 | 0.69 | 0.67 | 0.63 | 0.42 |
| 4 | 152730201 | rs6846809 | 1.646 | -0.532 | 0.288 | -0.409 | 0.002 | 0.015 | 0.015 | 0 | 0.46 | 0.36 | 0.54 | 0.37 | 0.54 | 0.31 |
| 4 | 152740233 | rs2136959 | -0.499 | 0.069 | 0.429 | -0.184 | -0.028 | -0.059 | -0.059 | 0.174 | 0.83 | 0.91 | 0.64 | 0.44 | 0.75 | 0.69 |
| 4 | 152744583 | rs17027878 | 0.589 | -0.25 | 0.358 | -0.229 | 0.005 | -0.006 | -0.006 | 0.369 | 0.14 | 0.12 | 0.06 | 0 | 0.09 | 0.05 |
| 4 | 152746079 | rs2136961 | 0.481 | -0.19 | 0.548 | -0.175 | 0.013 | 0.029 | 0.029 | 0 | 0.73 | 0.62 | 0.59 | 0.44 | 0.68 | 0.69 |
| 4 | 152752413 | rs1397684 | 1.056 | -0.049 | -0.19 | 0.055 | 0.004 | 0.085 | 0.085 | 0 | 0.48 | 0.18 | 0.49 | 0.36 | 0.54 | 0.35 |
| 4 | 152761633 | rs361191 | 0.322 | 0.267 | 0.103 | -0.438 | 0.009 | 0.044 | 0.044 | 0 | 0.6 | 0.32 | 0.41 | 0.17 | 0.61 | 0.58 |
| 4 | 152764822 | rs6843959 | 1.171 | 0.534 | -0.455 | -0.382 | -0.054 | 0.216 | 0.216 | 0 | 0.45 | 0.11 | 0.35 | 0.11 | 0.5 | 0.3 |
| 4 | 152779838 | rs7657092 | -1.474 | 0.374 | 0.872 | 0.304 | -0.016 | -0.009 | -0.009 | 1.134 | 0.18 | 0.5 | 0.09 | 0.19 | 0.14 | 0.31 |
| 4 | 152782639 | rs17027958 | -1.677 | 0.259 | 0.995 | 0.003 | -0.028 | -0.097 | -0.097 | 0.049 | 0.17 | 0.39 | 0.15 | 0.07 | 0.1 | 0.28 |
| 4 | 152784064 | rs1682083 | 1.35 | 0.505 | 1.531 | 0.044 | 0.011 | -0.013 | -0.013 | 0 | 0.89 | 0.72 | 0.92 | 0.93 | 0.87 | 0.88 |
| 4 | 152784669 | rs1872513 | -0.719 | 0.948 | 1.252 | -0.105 | -0.004 | 0 | 0 | 2.356 | 0.22 | 0.2 | 0.4 | 0.76 | 0.27 | 0.38 |
| 4 | 152801922 | rs715947 | -0.487 | 1.192 | 1.486 | 0.198 | -0.004 | -0.003 | -0.003 | 0.504 | 0.15 | 0.13 | 0.35 | 0.7 | 0.21 | 0.25 |
| 4 | 152804140 | rs901207 | -0.236 | 1.196 | 1.46 | 0.101 | -0.002 | 0.002 | 0.002 | 0.063 | 0.09 | 0.06 | 0.12 | 0.06 | 0.12 | 0.13 |
| 4 | 152807559 | rs11723144 | -0.17 | 1.198 | 1.35 | 0.103 | -0.024 | -0.002 | -0.002 | 0.837 | 0.09 | 0.02 | 0.11 | 0.06 | 0.12 | 0.14 |
| 4 | 152832903 | rs360919 | 1.434 | 0.688 | 1.923 | -0.3 | -0.037 | -0.048 | -0.048 | 0 | 0.81 | 0.51 | 0.85 | 0.88 | 0.86 | 0.75 |
| 4 | 152833119 | rs10517561 | -1.234 | 0.771 | 2.079 | -0.367 | -0.01 | 0.019 | 0.019 | 0.617 | 0.27 | 0.18 | 0.49 | 0.83 | 0.31 | 0.51 |
| 4 | 152841392 | rs361110 | 1.18 | 0.767 | 2.164 | -0.226 | -0.002 | -0.013 | -0.013 | 0 | 0.87 | 0.69 | 0.92 | 0.92 | 0.87 | 0.88 |
| 4 | 152851437 | rs1631928 | 1.366 | 0.769 | 2.197 | -0.227 | -0.018 | -0.016 | -0.016 | 0 | 0.82 | 0.66 | 0.86 | 0.86 | 0.87 | 0.75 |
| 4 | 152854159 | rs189672 | 2.281 | 1.124 | 2.125 | 0.021 | 0.014 | -0.006 | -0.006 | 1.889 | 0.72 | 0.49 | 0.62 | 0.76 | 0.73 | 0.69 |
| 4 | 152854578 | rs360939 | 2.469 | 1.117 | 1.953 | 0.018 | 0.001 | -0.005 | -0.005 | 3.696 | 0.71 | 0.32 | 0.61 | 0.76 | 0.72 | 0.69 |
| 4 | 152855624 | rs19412 | 2.176 | 1.047 | 1.668 | 0.179 | 0.001 | 0.003 | 0.003 | 0.263 | 0.66 | 0.34 | 0.6 | 0.66 | 0.68 | 0.62 |
| 4 | 152861685 | rs165579 | 1.902 | 0.611 | 1.698 | 0.186 | -0.009 | -0.052 | -0.052 | 0 | 0.6 | 0.45 | 0.54 | 0.13 | 0.6 | 0.37 |
| 4 | 152872349 | rs12645594 | -0.152 | 0.71 | 1.555 | 0.328 | -0.004 | -0.023 | -0.023 | 1.493 | 0.18 | 0.11 | 0.28 | 0.66 | 0.22 | 0.33 |
| 4 | 152887879 | rs360915 | 1.083 | 0.765 | 1.323 | 0.478 | 0.004 | -0.012 | -0.012 | 0.11 | 0.82 | 0.8 | 0.86 | 0.72 | 0.84 | 0.7 |
| 4 | 152895947 | rs168629 | 0.703 | 0.923 | 1.144 | 0.507 | 0 | -0.013 | -0.013 | 2.21 | 0.85 | 0.9 | 0.86 | 0.72 | 0.84 | 0.7 |
| 4 | 152899554 | rs975820 | -0.877 | 0.941 | 1.307 | 0.621 | -0.01 | 0.111 | 0.111 | 1.267 | 0.22 | 0.57 | 0.31 | 0.6 | 0.22 | 0.33 |
| 4 | 152900692 | rs6535830 | -1.175 | 0.88 | 1.303 | 0.617 | -0.001 | 0.065 | 0.065 | 0.049 | 0.27 | 0.58 | 0.34 | 0.6 | 0.29 | 0.34 |
| 4 | 152933610 | rs10015937 | -0.999 | 0.765 | 1.02 | 0.209 | -0.004 | 0.003 | 0.003 | 0 | 0.3 | 0.48 | 0.32 | 0.6 | 0.32 | 0.42 |
| 4 | 152936693 | rs17028387 | -1.165 | 0.576 | 0.916 | -0.03 | 0.002 | -0.004 | -0.004 | 0 | 0.15 | 0.18 | 0.22 | 0.15 | 0.13 | 0.22 |
| 4 | 152943626 | rs17028403 | -1.371 | 0.819 | 1.237 | -0.072 | 0.083 | 0.167 | 0.167 | 0 | 0.06 | 0.35 | 0.14 | 0.22 | 0.09 | 0.07 |
| 4 | 152944433 | rs17028407 | -0.248 | 0.752 | 1.037 | -0.083 | -0.007 | -0.009 | -0.009 | 0.442 | 0.08 | 0.05 | 0.18 | 0.14 | 0.06 | 0.2 |
| 4 | 152945315 | rs7667919 | 0.762 | 0.854 | 0.535 | -0.549 | -0.008 | 0.014 | 0.014 | 0 | 0.11 | 0.04 | 0.07 | 0 | 0.13 | 0.04 |
| 4 | 152948812 | rs12505581 | -1.276 | 0.498 | 0.918 | -0.258 | 0.023 | 0.096 | 0.096 | 0.109 | 0.1 | 0.29 | 0.14 | 0.24 | 0.12 | 0.08 |
| 4 | 152957069 | rs7664595 | 0.119 | 0.495 | 0.997 | -0.66 | -0.004 | 0.023 | 0.023 | 0 | 0.77 | 0.77 | 0.73 | 0.43 | 0.76 | 0.66 |
| 4 | 152962166 | rs1559695 | 0.526 | 0.43 | 0.793 | -0.781 | -0.003 | 0.012 | 0.012 | 0 | 0.7 | 0.81 | 0.55 | 0.3 | 0.7 | 0.43 |
| 4 | 152967911 | rs7655280 | 1.424 | 0.341 | 0.671 | -1.357 | 0 | -0.061 | -0.061 | 0 | 0.59 | 0.52 | 0.42 | 0.09 | 0.58 | 0.34 |
| 4 | 152983232 | rs16998854 | 0.332 | 0.487 | 0.659 | -0.941 | -0.003 | -0.027 | -0.027 | 0 | 0.06 | 0.09 | 0.02 | 0 | 0.05 | 0.02 |
| 4 | 152988228 | rs11737270 | -0.692 | 0.61 | 0.658 | -0.772 | 0 | 0 | 0 | 0 | 0.19 | 0.19 | 0.36 | 0.77 | 0.26 | 0.39 |
| 4 | 153001662 | rs1318845 | -0.016 | 0.697 | -0.226 | -0.603 | -0.041 | -0.077 | -0.077 | 0 | 0.15 | 0.03 | 0.28 | 0.65 | 0.2 | 0.31 |
| 4 | 153005716 | rs4696314 | 0.376 | 0.648 | 0.016 | -1.793 | 0.002 | -0.028 | -0.028 | 0 | 0.53 | 0.6 | 0.36 | 0.04 | 0.55 | 0.33 |
| 4 | 153010765 | rs6535835 | 0.182 | 0.271 | 0.119 | -1.675 | -0.001 | -0.007 | -0.007 | 0 | 0.7 | 0.7 | 0.48 | 0.16 | 0.7 | 0.39 |
| 4 | 153012499 | rs9992175 | 0.396 | -0.054 | 0.341 | -2.063 | 0.011 | 0.019 | 0.019 | 4.208 | 0.26 | 0.19 | 0.19 | 0.13 | 0.17 | 0.11 |
| 4 | 153016797 | rs17361160 | 0.761 | -0.057 | 0.7 | -2.133 | -0.016 | 0.129 | 0.129 | 0.458 | 0.44 | 0.25 | 0.28 | 0.03 | 0.53 | 0.26 |
| 4 | 153017452 | rs747110 | -0.572 | -0.054 | 0.441 | -2.044 | 0.002 | 0.002 | 0.002 | 0.231 | 0.77 | 0.76 | 0.61 | 0.23 | 0.8 | 0.44 |
| 4 | 153022375 | rs13140099 | 0.228 | -0.071 | 0.596 | -2.329 | 0.004 | 0.091 | 0.091 | 1.367 | 0.3 | 0.11 | 0.29 | 0.13 | 0.25 | 0.17 |
| 4 | 153023402 | rs6535838 | -0.134 | 0.298 | 0.469 | -2.458 | -0.004 | 0.028 | 0.028 | 0 | 0.7 | 0.75 | 0.58 | 0.15 | 0.7 | 0.41 |
| 4 | 153029463 | rs17361216 | 0.421 | 0.525 | 0.131 | -2.526 | 0.002 | 0.081 | 0.081 | 1.48 | 0.3 | 0.17 | 0.31 | 0.09 | 0.26 | 0.13 |

| | | | | | | | | | | | | | | | | |
|---|---|---|---|---|---|---|---|---|---|---|---|---|---|---|---|---|
| 4 | 153035743 | rs10517562 | 0.929 | 0.586 | 0.289 | -2.449 | -0.002 | -0.055 | -0.055 | 0 | 0.38 | 0.34 | 0.24 | 0.02 | 0.42 | 0.26 |
| 4 | 153055930 | rs13126953 | 1.009 | 0.608 | -0.369 | -2.367 | -0.02 | 0.078 | 0.078 | 0.728 | 0.37 | 0.18 | 0.22 | 0.02 | 0.42 | 0.26 |
| 4 | 153075234 | rs11727862 | 0.296 | 0.473 | -0.615 | -2.199 | -0.006 | 0.119 | 0.119 | 1.841 | 0.3 | 0.08 | 0.33 | 0.1 | 0.27 | 0.15 |
| 4 | 153091323 | rs2897464 | 0.992 | 0.915 | -0.291 | -2.103 | -0.012 | 0.016 | 0.016 | 1.07 | 0.33 | 0.18 | 0.25 | 0.03 | 0.38 | 0.24 |
| 4 | 153092536 | rs2001705 | 0.272 | 0.848 | -0.28 | -2.163 | -0.001 | -0.07 | -0.07 | 0 | 0.42 | 0.41 | 0.31 | 0.04 | 0.49 | 0.27 |
| 4 | 153093492 | rs4466023 | -0.081 | 1.133 | -1.14 | -1.948 | -0.001 | -0.011 | -0.011 | 0 | 0.07 | 0.11 | 0.03 | 0 | 0.1 | 0.02 |
| 4 | 153097096 | rs17361412 | 0.83 | 0.416 | -1.244 | -1.819 | -0.062 | 0.094 | 0.094 | 2.712 | 0.34 | 0.18 | 0.26 | 0.05 | 0.45 | 0.25 |
| 4 | 153100692 | rs11732677 | -0.334 | 0.028 | -1.323 | -1.038 | 0.028 | 0.065 | 0.065 | 4.258 | 0.35 | 0.13 | 0.36 | 0.17 | 0.3 | 0.14 |
| 4 | 153102015 | rs2897462 | 0.402 | -0.071 | -0.967 | -1.045 | 0.035 | 0.05 | 0.05 | 4.548 | 0.55 | 0.65 | 0.74 | 0.81 | 0.69 | 0.88 |
| 4 | 153109365 | rs13119810 | 0.832 | 0.243 | -1.44 | -1.499 | -0.051 | 0.135 | 0.135 | 0.068 | 0.33 | 0.13 | 0.19 | 0.04 | 0.39 | 0.22 |
| 4 | 153110256 | rs13127295 | 0.805 | 0.234 | -1.384 | -1.504 | -0.044 | 0.125 | 0.125 | 0 | 0.33 | 0.14 | 0.2 | 0.04 | 0.4 | 0.22 |
| 4 | 153111134 | rs13109486 | -0.354 | -0.162 | -0.784 | -1.143 | 0.006 | 0.007 | 0.007 | 0 | 0.69 | 0.63 | 0.49 | 0.34 | 0.65 | 0.42 |
| 4 | 153121818 | rs6856727 | -0.585 | -0.589 | -0.241 | -0.594 | -0.029 | 0.009 | 0.009 | 0 | 0.28 | 0.44 | 0.25 | 0.27 | 0.18 | 0.15 |
| 4 | 153134959 | rs11727661 | -0.477 | -0.356 | 0.223 | -0.224 | 0.005 | 0.032 | 0.032 | 0 | 0.13 | 0.24 | 0.07 | 0.12 | 0.1 | 0.07 |
| 4 | 153135952 | rs11728805 | -0.013 | -0.356 | 0.212 | -0.234 | -0.002 | 0.045 | 0.045 | 0 | 0.1 | 0.17 | 0.05 | 0.09 | 0.08 | 0.07 |
| 4 | 153136602 | rs11737308 | -0.698 | -0.355 | 0.174 | -0.272 | 0.002 | 0.011 | 0.011 | 0 | 0.14 | 0.18 | 0.09 | 0.12 | 0.12 | 0.08 |
| 4 | 153143877 | rs3749541 | -0.913 | -0.38 | 0.214 | -0.541 | 0 | -0.003 | -0.003 | 0 | 0.29 | 0.34 | 0.24 | 0.22 | 0.25 | 0.1 |
| 4 | 153149284 | rs6847949 | -0.484 | -0.366 | 0.235 | -0.494 | 0.006 | 0.012 | 0.012 | 0 | 0.25 | 0.22 | 0.22 | 0.19 | 0.22 | 0.1 |
| 4 | 153151602 | rs6856075 | -0.949 | -0.242 | 0.377 | -0.46 | -0.021 | -0.051 | -0.051 | 0.206 | 0.29 | 0.44 | 0.24 | 0.23 | 0.23 | 0.1 |
| 4 | 153156044 | rs6810751 | -0.298 | -0.259 | 0.609 | 0.327 | -0.011 | 0.074 | 0.074 | 1.173 | 0.12 | 0.24 | 0.4 | 0.27 | 0.1 | 0.19 |
| 4 | 153156200 | rs6836275 | -1.104 | -0.194 | 0.154 | 0.552 | -0.014 | -0.037 | -0.037 | 0 | 0.32 | 0.5 | 0.16 | 0.26 | 0.27 | 0.1 |
| 4 | 153162845 | rs2406624 | -0.296 | -0.263 | -0.008 | 1.092 | -0.003 | 0.037 | 0.037 | 0 | 0.22 | 0.34 | 0.2 | 0.3 | 0.21 | 0.1 |
| 4 | 153167171 | rs4554046 | -0.689 | -0.245 | -0.048 | 1.137 | 0.008 | 0.037 | 0.037 | 0.05 | 0.24 | 0.36 | 0.22 | 0.33 | 0.24 | 0.1 |
| 4 | 153184316 | rs2406623 | -1.539 | 0.387 | -0.519 | 0.99 | -0.005 | -0.002 | -0.002 | 0 | 0.14 | 0.14 | 0.2 | 0.04 | 0.16 | 0.06 |
| 4 | 153188678 | rs10517563 | -0.918 | 0.294 | -0.603 | 1.036 | -0.026 | 0.027 | 0.027 | 0 | 0.08 | 0.01 | 0.06 | 0.04 | 0.1 | 0.03 |
| 4 | 153206597 | rs3923993 | -1.011 | 0.088 | -0.494 | 1.381 | -0.003 | 0.025 | 0.025 | 0 | 0.09 | 0.11 | 0.23 | 0.27 | 0.08 | 0.16 |
| 4 | 153211026 | rs6838430 | 0.795 | 0.254 | -0.911 | 0.766 | -0.034 | 0.062 | 0.062 | 1.93 | 0.42 | 0.17 | 0.37 | 0.23 | 0.51 | 0.36 |
| 4 | 153219231 | rs4416456 | -0.419 | 0.558 | -0.076 | 0.943 | -0.009 | -0.009 | -0.009 | 0 | 0.05 | 0.04 | 0.13 | 0.15 | 0.07 | 0.04 |
| 4 | 153220331 | rs17276848 | -1 | 0.547 | -0.07 | 0.89 | -0.013 | -0.014 | -0.014 | 0 | 0.07 | 0.06 | 0.15 | 0.18 | 0.09 | 0.05 |
| 4 | 153229868 | rs7678272 | 1.133 | 0.25 | 0.53 | 0.91 | 0.041 | 0.082 | 0.082 | 1.606 | 0.93 | 0.66 | 0.86 | 0.85 | 0.95 | 0.97 |
| 4 | 153252061 | rs10033601 | -1.085 | -0.682 | 0.14 | 0.78 | 0.035 | -0.012 | -0.012 | 0 | 0.41 | 0.71 | 0.45 | 0.45 | 0.3 | 0.41 |
| 4 | 153294875 | rs2714805 | 0.964 | -1.071 | 0.604 | 0.681 | -0.001 | -0.001 | -0.001 | 0 | 0.65 | 0.34 | 0.56 | 0.59 | 0.72 | 0.66 |
| 4 | 153309538 | rs2255137 | 1.753 | -1.157 | 0.006 | 0.534 | -0.108 | -0.119 | -0.119 | 0 | 0.44 | 0.03 | 0.45 | 0.43 | 0.55 | 0.38 |
| 4 | 153332380 | rs12644477 | -0.968 | -0.763 | 1.223 | 0.823 | -0.02 | 0.102 | 0.102 | 1.578 | 0.1 | 0.23 | 0.21 | 0.28 | 0.06 | 0.15 |
| 4 | 153363612 | rs17028899 | -0.702 | -0.824 | 1.583 | 0.753 | -0.045 | -0.098 | -0.098 | 0 | 0.18 | 0.3 | 0.09 | 0 | 0.09 | 0.03 |
| 4 | 153373680 | rs2628570 | -1.667 | -0.563 | 1.052 | 0.974 | 0.054 | -0.072 | -0.072 | 0.538 | 0.44 | 0.86 | 0.49 | 0.51 | 0.33 | 0.44 |
| 4 | 153380801 | rs1516820 | -1.671 | -0.527 | 1.034 | 0.974 | 0.046 | -0.084 | -0.084 | 0.028 | 0.44 | 0.88 | 0.49 | 0.51 | 0.33 | 0.44 |
| 4 | 153415654 | rs17028932 | -1.375 | -0.603 | 1.049 | 0.972 | 0.018 | -0.011 | -0.011 | 0.304 | 0.42 | 0.66 | 0.47 | 0.46 | 0.33 | 0.41 |
| 4 | 153455628 | rs1484879 | -1.652 | -0.593 | 0.75 | 0.976 | 0.054 | -0.069 | -0.069 | 0 | 0.44 | 0.86 | 0.49 | 0.51 | 0.33 | 0.44 |
| 4 | 153456102 | rs1351903 | -1.664 | -0.599 | 0.646 | 1.029 | 0.05 | -0.073 | -0.073 | 0.049 | 0.44 | 0.87 | 0.49 | 0.52 | 0.33 | 0.44 |
| 4 | 153482738 | rs1072789 | 0.301 | -0.281 | 0.54 | 0.812 | 0.005 | -0.006 | -0.006 | 0 | 0.05 | 0.01 | 0.18 | 0.25 | 0.04 | 0.12 |
| 4 | 153485919 | rs2054757 | -0.734 | -0.1 | 0.698 | 0.566 | -0.011 | 0 | 0 | 0 | 0.41 | 0.53 | 0.44 | 0.42 | 0.3 | 0.41 |
| 4 | 153495515 | rs10011174 | -0.807 | 0.079 | 0.559 | 0.574 | -0.009 | 0.004 | 0.004 | 0.049 | 0.42 | 0.48 | 0.45 | 0.47 | 0.34 | 0.44 |
| 4 | 153519781 | rs10020796 | -0.803 | -0.174 | 0.563 | 0.644 | -0.014 | -0.001 | -0.001 | 0 | 0.44 | 0.51 | 0.44 | 0.48 | 0.3 | 0.42 |
| 4 | 153532240 | rs7695877 | -1.698 | 0.455 | 1.183 | 0.739 | -0.01 | 0.063 | 0.063 | 0 | 0.19 | 0.47 | 0.31 | 0.3 | 0.18 | 0.21 |
| 4 | 153537671 | rs10032212 | -1.706 | 0.113 | 1.018 | 0.696 | -0.025 | 0.012 | 0.012 | 0.078 | 0.21 | 0.7 | 0.3 | 0.29 | 0.15 | 0.19 |
| 4 | 153548037 | rs3922934 | 2.149 | 0.732 | 1.415 | 0.312 | 0.049 | 0.03 | 0.03 | 0.375 | 0.92 | 0.64 | 0.95 | 0.92 | 0.94 | 0.94 |
| 4 | 153554397 | rs4696329 | 2.552 | 1.168 | 1.266 | 0.21 | 0.038 | -0.034 | -0.034 | 0 | 0.85 | 0.62 | 0.76 | 0.9 | 0.85 | 0.93 |
| 4 | 153557795 | rs12642676 | 2.043 | 1.307 | 1.23 | 0.035 | 0.071 | -0.045 | -0.045 | 0 | 0.88 | 0.61 | 0.76 | 0.97 | 0.87 | 0.95 |
| 4 | 153558041 | rs6813566 | -1.723 | 0.975 | 0.882 | 0.24 | 0.005 | -0.019 | -0.019 | 0 | 0.11 | 0.28 | 0.08 | 0.09 | 0.08 | 0.03 |
| 4 | 153559468 | rs4696332 | -1.606 | 1.002 | 0.569 | 0.225 | -0.001 | -0.024 | -0.024 | 0.274 | 0.11 | 0.23 | 0.07 | 0.07 | 0.08 | 0.03 |
| 4 | 153565838 | rs3935569 | -0.31 | 1.125 | -0.241 | -0.086 | -0.007 | 0.027 | 0.027 | 0 | 0.43 | 0.36 | 0.32 | 0.24 | 0.51 | 0.47 |
| 4 | 153565969 | rs4443258 | 0.204 | 1.083 | -0.671 | -0.418 | 0 | 0.063 | 0.063 | 0.015 | 0.56 | 0.37 | 0.41 | 0.24 | 0.61 | 0.57 |
| 4 | 153569094 | rs10034000 | -0.385 | 0.956 | -0.721 | -0.147 | 0.01 | 0.022 | 0.022 | 0 | 0.52 | 0.62 | 0.68 | 0.39 | 0.54 | 0.41 |
| 4 | 153571180 | rs4508863 | -0.377 | 0.931 | -0.65 | -0.142 | 0.013 | 0.02 | 0.02 | 0 | 0.52 | 0.6 | 0.68 | 0.4 | 0.56 | 0.41 |
| 4 | 153573491 | rs6849505 | -0.09 | 0.644 | -0.164 | -0.324 | -0.026 | -0.048 | -0.048 | 0.305 | 0.08 | 0.19 | 0.05 | 0 | 0.05 | 0.01 |
| 4 | 153575339 | rs6814401 | -0.296 | 0.77 | -0.777 | -0.208 | -0.008 | 0.018 | 0.018 | 0 | 0.45 | 0.58 | 0.64 | 0.34 | 0.49 | 0.35 |
| 4 | 153583949 | rs11721759 | -0.302 | 0.961 | -1.012 | -0.226 | -0.02 | -0.015 | -0.015 | 0 | 0.06 | 0 | 0.04 | 0.06 | 0.06 | 0.06 |
| 4 | 153586001 | rs4437231 | -0.571 | 0.885 | -1.124 | -0.114 | 0.03 | 0.019 | 0.019 | 0 | 0.49 | 0.76 | 0.65 | 0.44 | 0.57 | 0.43 |
| 4 | 153588500 | rs4447840 | 0.868 | 0.869 | -0.536 | -0.119 | 0.005 | -0.037 | -0.037 | 0.798 | 0.9 | 0.79 | 0.83 | 0.99 | 0.91 | 0.95 |
| 4 | 153590063 | rs10010565 | -0.883 | 0.767 | 0.508 | -0.204 | 0.061 | 0.012 | 0.012 | 0.162 | 0.08 | 0.33 | 0.07 | 0.03 | 0.08 | 0.03 |
| 4 | 153601716 | rs12650067 | 1.614 | 0.744 | 0.42 | -0.224 | -0.005 | -0.055 | -0.055 | 0 | 0.87 | 0.4 | 0.89 | 0.95 | 0.91 | 0.93 |
| 4 | 153610164 | rs6857876 | 0.306 | 1.092 | -1.105 | -0.296 | -0.07 | -0.097 | -0.097 | 0 | 0.45 | 0.07 | 0.44 | 0.52 | 0.49 | 0.47 |
| 4 | 153626114 | rs4513549 | 0.606 | 0.915 | -0.158 | -0.96 | 0.002 | -0.108 | -0.108 | 0.015 | 0.68 | 0.45 | 0.78 | 0.91 | 0.7 | 0.8 |
| 4 | 153629791 | rs13124365 | 0.765 | 0.488 | 0.504 | -0.822 | -0.041 | -0.085 | -0.085 | 0 | 0.84 | 0.51 | 0.89 | 0.98 | 0.92 | 0.94 |
| 4 | 153635369 | rs11930234 | -0.255 | 0.893 | 0.237 | -0.981 | -0.006 | 0.007 | 0.007 | 1.039 | 0.19 | 0.16 | 0.21 | 0.08 | 0.22 | 0.13 |
| 4 | 153641782 | rs4128306 | 0.014 | 0.282 | -0.219 | -0.47 | -0.001 | 0 | 0 | 0.446 | 0.56 | 0.4 | 0.72 | 0.67 | 0.65 | 0.69 |
| 4 | 153644873 | rs4696345 | -0.411 | 0.194 | 0.188 | -0.288 | -0.001 | -0.001 | -0.001 | 0 | 0.63 | 0.37 | 0.73 | 0.66 | 0.67 | 0.73 |
| 4 | 153656788 | rs4077297 | -0.032 | -0.009 | 0.23 | -0.324 | 0.028 | 0.027 | 0.027 | 0 | 0.56 | 0.35 | 0.73 | 0.7 | 0.67 | 0.69 |

| | | | | | | | | | | | | | | | | |
|---|---|---|---|---|---|---|---|---|---|---|---|---|---|---|---|---|
| 4 | 153658324 | rs7340869 | 0.339 | -0.051 | 0.581 | -0.322 | -0.011 | -0.004 | -0.004 | 0 | 0.34 | 0.37 | 0.21 | 0.3 | 0.28 | 0.25 |
| 4 | 153658895 | rs10003999 | -0.716 | -0.386 | 0.587 | -0.514 | 0.002 | 0.001 | 0.001 | 1.519 | 0.22 | 0.2 | 0.12 | 0.1 | 0.14 | 0.07 |
| 4 | 153660407 | rs10009955 | -1.122 | 0.135 | 0.602 | -0.595 | -0.018 | 0.028 | 0.028 | 0 | 0.63 | 0.73 | 0.76 | 0.73 | 0.61 | 0.76 |
| 4 | 153669008 | rs12500918 | 0.129 | -0.015 | 0.621 | -0.264 | 0 | 0.001 | 0.001 | 0.922 | 0.39 | 0.52 | 0.52 | 0.53 | 0.45 | 0.63 |
| 4 | 153669404 | rs4540026 | -1.518 | -0.017 | 0.626 | -0.254 | -0.017 | 0.025 | 0.025 | 0 | 0.64 | 0.73 | 0.76 | 0.73 | 0.62 | 0.76 |
| 4 | 153681139 | rs11732223 | 0.805 | -0.03 | 1.047 | -0.243 | -0.006 | -0.035 | -0.035 | 0 | 0.32 | 0.2 | 0.47 | 0.51 | 0.4 | 0.58 |
| 4 | 153685314 | rs11099852 | -1.36 | -0.028 | 1.106 | -0.266 | -0.017 | 0.029 | 0.029 | 0 | 0.62 | 0.69 | 0.75 | 0.73 | 0.61 | 0.75 |
| 4 | 153688776 | rs7674876 | 0.6 | -0.034 | 1.329 | -0.239 | -0.001 | 0.004 | 0.004 | 0.244 | 0.34 | 0.34 | 0.48 | 0.48 | 0.4 | 0.58 |
| 4 | 153689963 | rs6839340 | 0.507 | -0.071 | 1.308 | -0.257 | -0.001 | 0.002 | 0.002 | 0.402 | 0.35 | 0.34 | 0.48 | 0.48 | 0.4 | 0.58 |
| 4 | 153695501 | rs4359884 | -0.696 | -0.036 | 1.356 | -0.239 | 0.002 | 0 | 0 | 0.202 | 0.27 | 0.26 | 0.27 | 0.22 | 0.21 | 0.17 |
| 4 | 153695779 | rs4276265 | 0.332 | 0.087 | 1.308 | -0.265 | -0.003 | 0.019 | 0.019 | 0.378 | 0.35 | 0.43 | 0.49 | 0.48 | 0.4 | 0.58 |
| 4 | 153696635 | rs6850467 | 0.48 | 0.104 | 1.144 | -0.256 | -0.001 | 0.005 | 0.005 | 0.244 | 0.34 | 0.34 | 0.48 | 0.48 | 0.4 | 0.58 |
| 4 | 153701130 | rs4619875 | 0.797 | -0.053 | 1.049 | -0.321 | -0.027 | -0.053 | -0.053 | 0 | 0.3 | 0.18 | 0.55 | 0.48 | 0.41 | 0.48 |
| 4 | 153701640 | rs10028182 | -0.273 | 0.061 | 1.35 | -0.222 | -0.012 | -0.026 | -0.026 | 0 | 0.18 | 0.25 | 0.09 | 0.09 | 0.13 | 0.05 |
| 4 | 153703306 | rs11099853 | 0.812 | 0.037 | 1.296 | -0.214 | -0.03 | -0.055 | -0.055 | 0 | 0.34 | 0.19 | 0.57 | 0.48 | 0.43 | 0.4 |
| 4 | 153720692 | rs7699456 | 1.22 | -0.017 | 1.478 | -0.223 | -0.014 | -0.013 | -0.013 | 0 | 0.73 | 0.45 | 0.8 | 0.81 | 0.81 | 0.68 |
| 4 | 153729627 | rs28769919 | -0.157 | -0.014 | 1.387 | -0.242 | -0.002 | -0.006 | -0.006 | 0.079 | 0.18 | 0.19 | 0.09 | 0.09 | 0.13 | 0.08 |
| 4 | 153732359 | rs4504228 | 0.93 | -0.014 | 1.389 | -0.222 | -0.006 | 0.006 | 0.006 | 0 | 0.39 | 0.32 | 0.24 | 0.34 | 0.39 | 0.29 |
| 4 | 153765098 | rs10007643 | 0.1 | -0.034 | 1.326 | -0.254 | -0.005 | -0.006 | -0.006 | 0 | 0.15 | 0.18 | 0.08 | 0.09 | 0.1 | 0.08 |
| 4 | 153779819 | rs13102607 | -0.464 | 0.042 | 1.376 | -0.184 | -0.006 | -0.003 | -0.003 | 0 | 0.21 | 0.24 | 0.15 | 0.17 | 0.15 | 0.15 |
| 4 | 153793179 | rs10013266 | 0.013 | 0.044 | 1.258 | -0.185 | 0.004 | 0.007 | 0.007 | 0 | 0.17 | 0.15 | 0.08 | 0.09 | 0.13 | 0.08 |
| 4 | 153800029 | rs4254753 | -1.042 | 0.034 | 1.074 | -0.179 | -0.024 | -0.025 | -0.025 | 0.184 | 0.27 | 0.48 | 0.19 | 0.19 | 0.18 | 0.32 |
| 4 | 153829996 | rs7664303 | 0.761 | 0.156 | 0.122 | -0.112 | -0.034 | -0.061 | -0.061 | 0 | 0.34 | 0.18 | 0.57 | 0.48 | 0.43 | 0.4 |
| 4 | 153860134 | rs2004794 | -0.772 | 0.103 | 0.059 | 0.058 | 0 | -0.001 | -0.001 | 0 | 0.61 | 0.56 | 0.75 | 0.66 | 0.61 | 0.71 |
| 4 | 153863214 | rs1474028 | -0.477 | 0.668 | -0.344 | 0.372 | -0.003 | 0.003 | 0.003 | 0 | 0.16 | 0.15 | 0.1 | 0.09 | 0.17 | 0.08 |
| 4 | 153872443 | rs6836054 | -0.18 | 0.379 | -0.773 | 0.524 | 0.003 | 0.002 | 0.002 | 0 | 0.54 | 0.36 | 0.65 | 0.55 | 0.45 | 0.58 |
| 4 | 153878480 | rs867322 | -0.435 | 0.775 | -1.106 | 0.702 | -0.002 | 0.003 | 0.003 | 0 | 0.46 | 0.29 | 0.55 | 0.54 | 0.42 | 0.58 |
| 4 | 153879878 | rs11737767 | -0.467 | 0.744 | -1.093 | 0.773 | 0.004 | 0.001 | 0.001 | 0 | 0.47 | 0.48 | 0.56 | 0.54 | 0.38 | 0.66 |
| 4 | 153881835 | rs3811834 | -0.167 | 0.937 | -1.56 | 1.056 | -0.037 | -0.098 | -0.098 | 0 | 0.3 | 0.09 | 0.41 | 0.5 | 0.34 | 0.45 |
| 4 | 153888640 | rs1018316 | -0.304 | 0.796 | -1.234 | 0.725 | -0.013 | 0.011 | 0.011 | 0 | 0.66 | 0.75 | 0.76 | 0.69 | 0.63 | 0.69 |
| 4 | 153890646 | rs10008050 | 0.658 | 0.247 | -0.671 | 1.228 | -0.017 | 0.039 | 0.039 | 1.354 | 0.76 | 0.73 | 0.7 | 0.58 | 0.83 | 0.63 |
| 4 | 153893354 | rs11099864 | 1.252 | 0.052 | -0.582 | 1.103 | 0.045 | -0.062 | -0.062 | 0 | 0.64 | 0.14 | 0.6 | 0.53 | 0.77 | 0.51 |
| 4 | 153894243 | rs908526 | 0.612 | 0.032 | 0.442 | 1.07 | -0.027 | 0.044 | 0.044 | 1.406 | 0.76 | 0.6 | 0.68 | 0.62 | 0.83 | 0.61 |
| 4 | 153898178 | rs4696370 | -0.206 | -0.081 | 1.413 | 0.903 | 0.022 | -0.042 | -0.042 | 0.359 | 0.92 | 0.83 | 0.95 | 1 | 0.93 | 0.99 |
| 4 | 153899573 | rs1059354 | 0.027 | -0.084 | 1.548 | 0.894 | 0.022 | -0.05 | -0.05 | 0 | 0.91 | 0.8 | 0.94 | 1 | 0.93 | 0.97 |
| 4 | 153899658 | rs1059355 | -0.226 | -0.075 | 1.45 | 0.899 | 0.022 | -0.046 | -0.046 | 0.515 | 0.92 | 0.83 | 0.95 | 1 | 0.93 | 0.99 |
| 4 | 153901275 | rs7681249 | -0.876 | -0.183 | 1.524 | 0.987 | -0.008 | 0.015 | 0.015 | 0 | 0.18 | 0.14 | 0.28 | 0.36 | 0.11 | 0.35 |
| 4 | 153902011 | rs12512990 | 0.58 | -0.377 | 1.187 | 0.551 | 0.002 | -0.001 | -0.001 | 0.505 | 0.09 | 0.07 | 0.22 | 0.02 | 0.05 | 0.03 |
| 4 | 153902939 | rs6851633 | 1.786 | 0.316 | 1.323 | 0.627 | -0.001 | -0.009 | -0.009 | 0 | 0.66 | 0.29 | 0.61 | 0.55 | 0.69 | 0.52 |
| 4 | 153903766 | rs6535886 | -0.98 | 0.352 | 1.224 | 0.834 | 0 | 0.014 | 0.014 | 0 | 0.2 | 0.17 | 0.28 | 0.37 | 0.2 | 0.37 |
| 4 | 153915003 | rs13131114 | -1.294 | 0.36 | 0.037 | 0.914 | -0.002 | -0.003 | -0.003 | 0 | 0.3 | 0.21 | 0.28 | 0.33 | 0.25 | 0.44 |
| 4 | 153916577 | rs11931660 | 0.687 | -0.188 | -0.655 | 1.036 | 0.012 | -0.013 | -0.013 | 1.646 | 0.19 | 0.16 | 0.18 | 0.26 | 0.14 | 0.18 |
| 4 | 153917822 | rs13120414 | 1.488 | 0.262 | -1.247 | 0.957 | -0.093 | 0.01 | 0.01 | 3.925 | 0.37 | 0.06 | 0.32 | 0.35 | 0.45 | 0.31 |
| 4 | 153919578 | rs1878284 | 0.758 | 0.258 | -1.183 | 1.011 | -0.008 | -0.016 | -0.016 | 0 | 0.61 | 0.27 | 0.52 | 0.62 | 0.62 | 0.53 |
| 4 | 153920579 | rs12504106 | 0.688 | 0.217 | -0.845 | 1.077 | 0.008 | -0.009 | -0.009 | 0 | 0.19 | 0.19 | 0.19 | 0.27 | 0.14 | 0.17 |
| 4 | 153928731 | rs6811075 | 0.636 | 0.813 | -0.478 | 0.976 | 0.029 | 0.01 | 0.01 | 0 | 0.69 | 0.5 | 0.58 | 0.67 | 0.69 | 0.51 |
| 4 | 153930963 | rs6837953 | -0.51 | 0.749 | 0.261 | 0.96 | 0.084 | 0.039 | 0.039 | 0.401 | 0.09 | 0.43 | 0.08 | 0.08 | 0.08 | 0.12 |
| 4 | 153934639 | rs11731715 | 1.553 | 0.72 | -0.502 | 1.034 | -0.034 | -0.025 | -0.025 | 0.017 | 0.37 | 0.15 | 0.29 | 0.42 | 0.39 | 0.29 |
| 4 | 153936488 | rs4696379 | -0.912 | 0.138 | -0.335 | 1.034 | -0.02 | 0.01 | 0.01 | 0 | 0.07 | 0.23 | 0.03 | 0.07 | 0.02 | 0.09 |
| 4 | 153938095 | rs6535888 | -0.291 | 0.373 | -0.498 | 0.974 | 0.001 | -0.002 | -0.002 | 1.997 | 0.24 | 0.18 | 0.37 | 0.22 | 0.26 | 0.3 |
| 4 | 153940203 | rs11722167 | -0.13 | 0.875 | 0.246 | 1.221 | 0.066 | 0.046 | 0.046 | 0 | 0.07 | 0.28 | 0.06 | 0.08 | 0.08 | 0.12 |
| 4 | 153941622 | rs11723393 | 0.688 | 0.71 | -0.267 | 1.267 | -0.003 | -0.001 | -0.001 | 0.119 | 0.62 | 0.42 | 0.63 | 0.64 | 0.68 | 0.59 |
| 4 | 153942865 | rs11722890 | -0.081 | 0.828 | -0.628 | 1.369 | 0.002 | -0.041 | -0.041 | 0 | 0.22 | 0.08 | 0.23 | 0.3 | 0.22 | 0.15 |
| 4 | 153943149 | rs1533458 | 0.262 | 0.579 | -0.065 | 1.368 | 0.027 | -0.024 | -0.024 | 0 | 0.14 | 0.05 | 0.17 | 0.19 | 0.1 | 0.1 |
| 4 | 153943571 | rs6818398 | -0.23 | 1.224 | 0.522 | 1.493 | -0.001 | 0.076 | 0.076 | 0 | 0.12 | 0.22 | 0.2 | 0.25 | 0.11 | 0.24 |
| 4 | 153945320 | rs17029499 | 0.095 | 1.792 | 0.652 | 1.032 | 0.006 | -0.007 | -0.007 | 0 | 0.48 | 0.31 | 0.41 | 0.45 | 0.45 | 0.52 |
| 4 | 153947145 | rs6814889 | 0.789 | 2.057 | 1.369 | 0.949 | 0.126 | -0.001 | -0.001 | 0 | 0.88 | 0.57 | 0.87 | 0.95 | 0.81 | 0.82 |
| 4 | 153949047 | rs1111986 | 0.938 | 2.026 | 1.045 | 0.955 | 0.033 | -0.019 | -0.019 | 0 | 0.33 | 0.07 | 0.41 | 0.4 | 0.3 | 0.23 |
| 4 | 153951159 | rs2406136 | 1.463 | 1.995 | 1.125 | 0.964 | 0.039 | -0.003 | -0.003 | 0 | 0.81 | 0.35 | 0.82 | 0.84 | 0.75 | 0.75 |
| 4 | 153957777 | rs1607074 | 0.672 | 1.906 | 1.331 | 0.95 | 0.011 | 0.006 | 0.006 | 8.759 | 0.36 | 0.27 | 0.45 | 0.39 | 0.33 | 0.23 |
| 4 | 153958198 | rs17029520 | 0.808 | 1.907 | 1.235 | 0.958 | 0.035 | 0.004 | 0.004 | 8.47 | 0.34 | 0.17 | 0.42 | 0.39 | 0.3 | 0.23 |
| 4 | 153958505 | rs1516691 | 1.126 | 1.879 | 1.058 | 0.903 | 0.097 | 0.023 | 0.023 | 2.661 | 0.83 | 0.55 | 0.86 | 0.85 | 0.77 | 0.74 |
| 4 | 153960368 | rs1024148 | 0.591 | 1.852 | 0.696 | 0.919 | 0.01 | -0.001 | -0.001 | 0.739 | 0.45 | 0.66 | 0.42 | 0.42 | 0.4 | 0.49 |
| 4 | 153963155 | rs1516692 | 0.463 | 1.843 | 0.658 | 0.937 | 0.004 | 0.006 | 0.006 | 1.1 | 0.47 | 0.72 | 0.45 | 0.4 | 0.43 | 0.49 |
| 4 | 153968142 | rs908535 | 0.869 | 1.894 | 0.159 | 0.9 | 0.029 | -0.022 | -0.022 | 0.16 | 0.34 | 0.07 | 0.43 | 0.39 | 0.31 | 0.24 |
| 4 | 153969414 | rs10030391 | 1.334 | 2.155 | 0.833 | 0.777 | 0.006 | -0.001 | -0.001 | 0 | 0.86 | 0.88 | 0.9 | 0.91 | 0.78 | 0.8 |
| 4 | 153970416 | rs4696161 | 1.479 | 2.114 | 0.451 | 0.773 | 0.039 | 0.006 | 0.006 | 1.814 | 0.79 | 0.47 | 0.85 | 0.83 | 0.75 | 0.74 |
| 4 | 153970964 | rs4696164 | 1.486 | 2.115 | 0.437 | 0.773 | 0.039 | 0.006 | 0.006 | 1.831 | 0.79 | 0.48 | 0.85 | 0.83 | 0.75 | 0.74 |
| 4 | 153977593 | rs2056168 | 0.321 | 2.072 | -0.44 | 0.597 | -0.01 | 0.016 | 0.016 | 0.409 | 0.88 | 0.92 | 0.92 | 0.97 | 0.85 | 0.84 |

| | | | | | | | | | | | | | | | |
|---|---|---|---|---|---|---|---|---|---|---|---|---|---|---|---|
| 4 | 153979941 | rs12651351 | 0.874 | 1.825 | -0.803 | 0.762 | 0.021 | -0.1 | -0.1 | 0 | 0.39 | 0.07 | 0.41 | 0.42 | 0.29 | 0.43 |
| 4 | 153980805 | rs4696392 | 1.642 | 2.224 | 0.592 | 0.735 | -0.026 | -0.008 | -0.008 | 0 | 0.67 | 0.25 | 0.55 | 0.73 | 0.63 | 0.7 |
| 4 | 153981236 | rs6535893 | 1.067 | 2.562 | 0.525 | 0.972 | 0.009 | 0.005 | 0.005 | 0 | 0.82 | 0.74 | 0.8 | 0.81 | 0.78 | 0.79 |
| 4 | 153984283 | rs7683508 | -1.279 | 1.992 | -0.228 | 0.924 | -0.003 | -0.001 | -0.001 | 0.031 | 0.19 | 0.21 | 0.29 | 0.17 | 0.22 | 0.11 |
| 4 | 153986325 | rs2090380 | 1.378 | 1.558 | 0.412 | 1.321 | 0.037 | 0.196 | 0.196 | 0.032 | 0.91 | 0.66 | 0.81 | 0.78 | 0.9 | 0.84 |
| 4 | 153991639 | rs6858521 | -0.014 | 1.548 | -0.076 | 0.74 | -0.01 | 0.007 | 0.007 | 0.018 | 0.61 | 0.66 | 0.5 | 0.34 | 0.5 | 0.39 |
| 4 | 153999303 | rs12641081 | -0.068 | 1.203 | -0.527 | 0.472 | -0.01 | -0.037 | -0.037 | 0 | 0.09 | 0 | 0.05 | 0.17 | 0.08 | 0.06 |
| 4 | 154000240 | rs4696394 | 0.117 | 0.692 | -0.217 | -0.1 | -0.008 | -0.008 | -0.008 | 0.916 | 0.14 | 0.4 | 0.25 | 0.07 | 0.1 | 0.08 |
| 4 | 154000267 | rs4696395 | -1.24 | 0.247 | -0.506 | -0.185 | -0.029 | -0.002 | -0.002 | 0 | 0.32 | 0.55 | 0.34 | 0.31 | 0.23 | 0.24 |
| 4 | 154000968 | rs4696396 | -0.104 | 0.313 | -0.743 | -0.574 | -0.003 | -0.036 | -0.036 | 0 | 0.39 | 0.57 | 0.39 | 0.08 | 0.32 | 0.17 |
| 4 | 154003295 | rs7691951 | -0.998 | 0.475 | -0.528 | -0.609 | -0.002 | 0.015 | 0.015 | 0.423 | 0.56 | 0.64 | 0.48 | 0.33 | 0.45 | 0.34 |
| 4 | 154005032 | rs1516684 | 0.065 | 0.468 | -0.186 | -0.617 | -0.006 | -0.051 | -0.051 | 2.415 | 0.38 | 0.41 | 0.38 | 0.08 | 0.32 | 0.17 |
| 4 | 154006090 | rs12648638 | -1.167 | 0.438 | 0.091 | -0.385 | 0.006 | 0.004 | 0.004 | 0.088 | 0.25 | 0.16 | 0.12 | 0.26 | 0.2 | 0.24 |
| 4 | 154008453 | rs7683638 | 1.453 | 0.947 | -0.652 | -0.043 | -0.063 | -0.107 | -0.107 | 0 | 0.13 | 0.01 | 0.17 | 0.48 | 0.19 | 0.22 |
| 4 | 154008962 | rs12651091 | -0.34 | 0.498 | -0.224 | -0.014 | -0.009 | 0.005 | 0.005 | 0 | 0.18 | 0.26 | 0.1 | 0.24 | 0.13 | 0.16 |
| 4 | 154011656 | rs13112096 | 0.756 | 0.43 | 0.083 | -0.098 | -0.012 | -0.001 | -0.001 | 0 | 0.12 | 0.22 | 0.23 | 0.08 | 0.1 | 0.07 |
| 4 | 154012751 | rs1878285 | -1.436 | 0.541 | -0.081 | -0.116 | 0.007 | 0 | 0 | 0 | 0.4 | 0.65 | 0.39 | 0.42 | 0.3 | 0.32 |
| 4 | 154012784 | rs2406241 | 0.234 | 0.219 | -0.101 | -0.164 | -0.002 | -0.042 | -0.042 | 0 | 0.09 | 0.2 | 0.05 | 0.02 | 0.07 | 0.08 |
| 4 | 154014175 | rs4696402 | -1.024 | 0.214 | -0.419 | 0.013 | 0.007 | -0.002 | -0.002 | 0 | 0.39 | 0.63 | 0.38 | 0.42 | 0.24 | 0.3 |
| 4 | 154015564 | rs13130102 | 0.082 | 0.465 | -0.616 | -0.251 | 0 | 0.001 | 0.001 | 2.275 | 0.29 | 0.29 | 0.36 | 0.15 | 0.29 | 0.31 |
| 4 | 154016744 | rs4696404 | 0.423 | 0.242 | -0.802 | 0.325 | -0.02 | 0.004 | 0.004 | 0 | 0.28 | 0.28 | 0.19 | 0.32 | 0.19 | 0.29 |
| 4 | 154016973 | rs17029682 | 0.491 | 0.175 | -0.793 | 0.442 | -0.017 | -0.022 | -0.022 | 0.027 | 0.27 | 0.27 | 0.14 | 0.2 | 0.18 | 0.28 |
| 4 | 154017805 | rs2176304 | 0.909 | 0.491 | -0.815 | 0.501 | -0.005 | -0.031 | -0.031 | 0.493 | 0.25 | 0.16 | 0.28 | 0.47 | 0.27 | 0.3 |
| 4 | 154017964 | rs7662116 | -0.191 | 0.532 | -0.62 | 0.389 | -0.013 | 0.001 | 0.001 | 1.421 | 0.63 | 0.38 | 0.55 | 0.67 | 0.69 | 0.68 |
| 4 | 154020068 | rs1516689 | -0.357 | 0.624 | -0.618 | -0.175 | -0.014 | -0.027 | -0.027 | 0 | 0.64 | 0.4 | 0.59 | 0.79 | 0.69 | 0.69 |
| 4 | 154022065 | rs1588982 | -0.67 | 1.107 | -1.041 | -0.029 | 0 | 0.001 | 0.001 | 0.889 | 0.73 | 0.74 | 0.64 | 0.83 | 0.72 | 0.7 |
| 4 | 154035960 | rs11725217 | 0.179 | 0.712 | -1.394 | -0.291 | -0.013 | 0.103 | 0.103 | 1.481 | 0.3 | 0.12 | 0.16 | 0.09 | 0.3 | 0.33 |
| 4 | 154040215 | rs6849116 | -0.686 | 0.338 | -0.518 | 0.265 | 0.01 | -0.014 | -0.014 | 0 | 0.39 | 0.45 | 0.36 | 0.31 | 0.42 | 0.36 |
| 4 | 154044043 | rs10032068 | 0.751 | -0.079 | -0.29 | 0.347 | -0.031 | 0.005 | 0.005 | 0 | 0.15 | 0.18 | 0.07 | 0.16 | 0.11 | 0.12 |
| 4 | 154053258 | rs4323085 | -1.043 | -0.109 | -0.856 | 0.436 | -0.006 | -0.001 | -0.001 | 0 | 0.46 | 0.8 | 0.34 | 0.45 | 0.47 | 0.41 |
| 4 | 154055176 | rs10001638 | 0.095 | 0.113 | -0.631 | 0.411 | 0.001 | 0.034 | 0.034 | 0.282 | 0.17 | 0.25 | 0.14 | 0.27 | 0.16 | 0.21 |
| 4 | 154058319 | rs880972 | -0.586 | 0.33 | -0.381 | 0.354 | 0 | -0.037 | -0.037 | 0 | 0.29 | 0.54 | 0.2 | 0.18 | 0.31 | 0.2 |
| 4 | 154066142 | rs4075881 | -1.524 | 0.577 | -1.473 | 0.457 | -0.032 | 0.05 | 0.05 | 1.72 | 0.68 | 0.84 | 0.72 | 0.76 | 0.65 | 0.55 |
| 4 | 154066319 | rs4075879 | 0.213 | 0.631 | -1.492 | 0.196 | -0.033 | 0.036 | 0.036 | 0 | 0.13 | 0.09 | 0.1 | 0.02 | 0.21 | 0.09 |
| 4 | 154075881 | rs34205168 | 0.585 | 0.393 | -1.473 | 0.486 | -0.013 | 0.129 | 0.129 | 0.567 | 0.43 | 0.23 | 0.37 | 0.23 | 0.55 | 0.43 |
| 4 | 154080813 | rs6817112 | 0.158 | 0.23 | -1.282 | 0.702 | 0.002 | -0.017 | -0.017 | 0.164 | 0.25 | 0.21 | 0.28 | 0.34 | 0.21 | 0.25 |
| 4 | 154087520 | rs4696428 | 0.725 | 0.354 | -1.609 | 0.528 | -0.006 | 0.086 | 0.086 | 0.444 | 0.44 | 0.25 | 0.37 | 0.29 | 0.56 | 0.55 |
| 4 | 154087943 | rs4696433 | -1.214 | 0.142 | -1.159 | 0.865 | 0.04 | -0.038 | -0.038 | 0 | 0.87 | 0.97 | 0.93 | 0.84 | 0.9 | 0.89 |
| 4 | 154094001 | rs10023596 | -0.857 | 1.009 | -0.449 | 0.778 | -0.003 | -0.002 | -0.002 | 0 | 0.37 | 0.56 | 0.49 | 0.43 | 0.47 | 0.34 |
| 4 | 154110795 | rs6535904 | -0.674 | 0.672 | 0.067 | 0.627 | 0.008 | 0.003 | 0.003 | 0.023 | 0.2 | 0.43 | 0.29 | 0.21 | 0.18 | 0.31 |
| 4 | 154118168 | rs6817587 | 0.461 | 0.798 | -0.369 | 0.241 | 0.006 | 0.03 | 0.03 | 0.069 | 0.44 | 0.36 | 0.27 | 0.27 | 0.39 | 0.4 |
| 4 | 154125998 | rs10019436 | -0.453 | 0.405 | 0.429 | 0.305 | -0.031 | -0.027 | -0.027 | 0 | 0.16 | 0.42 | 0.2 | 0.08 | 0.07 | 0.09 |
| 4 | 154135457 | rs9997214 | -0.287 | 0.335 | 0.209 | 0.421 | -0.001 | 0.001 | 0.001 | 0 | 0.17 | 0.15 | 0.06 | 0.17 | 0.15 | 0.2 |
| 4 | 154135588 | rs7686229 | 0.859 | 0.875 | 0.391 | -0.015 | 0.031 | -0.008 | -0.008 | 0 | 0.65 | 0.48 | 0.7 | 0.72 | 0.5 | 0.65 |
| 4 | 154138086 | rs6535908 | -0.514 | 1.093 | 0.114 | -0.024 | -0.054 | 0.008 | 0.008 | 0 | 0.86 | 0.95 | 0.81 | 0.89 | 0.68 | 0.86 |
| 4 | 154138758 | rs11723584 | 1.67 | 0.784 | 0.103 | -0.084 | 0.025 | -0.03 | -0.03 | 0.087 | 0.46 | 0.18 | 0.5 | 0.6 | 0.38 | 0.52 |
| 4 | 154139583 | rs7687844 | 0.154 | 1.109 | 0.596 | -0.263 | -0.029 | 0.015 | 0.015 | 0 | 0.82 | 0.87 | 0.77 | 0.89 | 0.59 | 0.84 |
| 4 | 154151988 | rs4696443 | -0.554 | 0.212 | 1.047 | -0.307 | -0.016 | -0.001 | -0.001 | 1.256 | 0.17 | 0.53 | 0.24 | 0.1 | 0.13 | 0.09 |
| 4 | 154155013 | rs4596222 | -1.745 | -0.078 | -0.291 | -0.565 | -0.02 | 0.008 | 0.008 | 0.511 | 0.83 | 1 | 0.9 | 0.9 | 0.85 | 0.82 |
| 4 | 154159547 | rs6813648 | -1.421 | -0.155 | 0.602 | -0.45 | -0.015 | -0.012 | -0.012 | 0 | 0.23 | 0.7 | 0.28 | 0.13 | 0.17 | 0.13 |
| 4 | 154159965 | rs4696175 | 0.704 | -0.174 | 0.64 | -0.383 | -0.003 | -0.066 | -0.066 | 0 | 0.11 | 0.32 | 0.19 | 0.01 | 0.06 | 0.07 |
| 4 | 154163394 | rs10009522 | -0.784 | -0.08 | 0.059 | -0.349 | -0.03 | -0.015 | -0.015 | 0 | 0.21 | 0.6 | 0.25 | 0.13 | 0.13 | 0.12 |
| 4 | 154176379 | rs10033643 | -1.068 | 0.189 | -1 | -0.511 | -0.014 | 0.053 | 0.053 | 0 | 0.57 | 0.82 | 0.49 | 0.27 | 0.58 | 0.55 |
| 4 | 154191710 | rs1470104 | 0.558 | 0.496 | -1.234 | -0.813 | -0.033 | 0.043 | 0.043 | 0 | 0.26 | 0.03 | 0.15 | 0.13 | 0.27 | 0.38 |
| 4 | 154196082 | rs6833828 | 0.004 | 0.365 | -1.023 | -0.863 | -0.006 | -0.044 | -0.044 | 0 | 0.2 | 0.7 | 0.25 | 0.06 | 0.17 | 0.12 |
| 4 | 154200320 | rs1455840 | -1.336 | 0.424 | -1.415 | -0.811 | -0.035 | 0.094 | 0.094 | 0 | 0.56 | 0.86 | 0.48 | 0.23 | 0.56 | 0.54 |
| 4 | 154200347 | rs1455841 | 0.789 | 0.481 | -1.408 | -0.78 | -0.04 | -0.008 | -0.008 | 0 | 0.09 | 0 | 0.07 | 0.05 | 0.14 | 0.05 |
| 4 | 154216710 | rs893805 | -0.666 | 0.38 | -1.271 | -0.776 | 0.001 | 0.073 | 0.073 | 0.046 | 0.46 | 0.65 | 0.4 | 0.18 | 0.46 | 0.51 |
| 4 | 154232522 | rs2033880 | -1.14 | 0.484 | -1.805 | -0.488 | -0.021 | 0.031 | 0.031 | 0 | 0.57 | 0.84 | 0.49 | 0.29 | 0.57 | 0.56 |
| 4 | 154240476 | rs961358 | -1.096 | 0.002 | -1.64 | 0.025 | 0.008 | -0.017 | -0.017 | 1.132 | 0.93 | 1 | 0.97 | 0.91 | 0.97 | 0.96 |
| 4 | 154244315 | rs17029874 | -0.72 | 0.388 | -1.061 | -0.018 | 0 | -0.001 | -0.001 | 0 | 0.3 | 0.27 | 0.18 | 0.13 | 0.3 | 0.39 |
| 4 | 154244953 | rs10517569 | -0.253 | -0.209 | -2.06 | 0.231 | -0.014 | 0.03 | 0.03 | 0 | 0.23 | 0.01 | 0.09 | 0.1 | 0.17 | 0.29 |
| 4 | 154262551 | rs6535914 | -0.755 | -0.081 | -1.764 | 0.363 | 0.026 | 0.048 | 0.048 | 3.092 | 0.53 | 0.87 | 0.41 | 0.31 | 0.36 | 0.54 |
| 4 | 154265347 | rs10025256 | -0.159 | -0.731 | -0.821 | 0.277 | 0 | -0.004 | -0.004 | 0.427 | 0.23 | 0.4 | 0.26 | 0.23 | 0.16 | 0.25 |
| 4 | 154274664 | rs6841435 | -1.573 | -0.827 | -1.018 | 0.098 | -0.04 | -0.04 | -0.04 | 0 | 0.56 | 0.86 | 0.45 | 0.39 | 0.42 | 0.57 |
| 4 | 154283051 | rs13132311 | 0.18 | -0.863 | -1.031 | 0.023 | -0.021 | 0.034 | 0.034 | 0 | 0.07 | 0.01 | 0.06 | 0 | 0.12 | 0.05 |
| 4 | 154284935 | rs17029903 | -0.444 | -1.106 | -0.528 | -0.001 | -0.021 | -0.035 | -0.035 | 0.696 | 0.3 | 0.35 | 0.15 | 0.11 | 0.24 | 0.25 |
| 4 | 154288126 | rs17029907 | 0.193 | -1.39 | 0.128 | -0.163 | 0.022 | 0.025 | 0.025 | 0 | 0.16 | 0.28 | 0.21 | 0.23 | 0.13 | 0.11 |
| 4 | 154292316 | rs12499126 | -0.117 | -1.347 | 0.133 | -0.103 | 0.012 | 0.042 | 0.042 | 0 | 0.18 | 0.28 | 0.23 | 0.29 | 0.15 | 0.16 |

| | | | | | | | | | | | | | | | | |
|---|---|---|---|---|---|---|---|---|---|---|---|---|---|---|---|---|
| 4 | 154296368 | rs12503471 | -0.117 | -1.344 | 0.286 | -0.102 | 0.012 | 0.042 | 0.042 | 0 | 0.18 | 0.28 | 0.23 | 0.29 | 0.15 | 0.16 |
| 4 | 154299829 | rs7699617 | 0.387 | -1.415 | 0.373 | -0.151 | -0.028 | -0.029 | -0.029 | 0 | 0.07 | 0.17 | 0.04 | 0.01 | 0.02 | 0.16 |
| 4 | 154311774 | rs6535916 | -1.096 | -1.477 | 0.266 | -0.202 | 0.002 | -0.008 | -0.008 | 0.833 | 0.37 | 0.58 | 0.2 | 0.1 | 0.26 | 0.38 |
| 4 | 154314931 | rs10517571 | -2.017 | -1.44 | 0.235 | -0.166 | -0.032 | -0.032 | -0.032 | 0 | 0.56 | 0.84 | 0.44 | 0.39 | 0.41 | 0.54 |
| 4 | 154329842 | rs10016021 | -0.146 | -1.377 | 0.448 | 0.08 | -0.032 | -0.034 | -0.034 | 0.025 | 0.08 | 0.2 | 0.05 | 0.01 | 0.02 | 0.16 |
| 4 | 154334786 | rs12510542 | -1.308 | -1.389 | 0.425 | 0.075 | -0.006 | 0.047 | 0.047 | 0.601 | 0.25 | 0.48 | 0.28 | 0.31 | 0.17 | 0.32 |
| 4 | 154339482 | rs10030294 | -0.229 | -1.454 | 0.137 | 0.184 | -0.03 | -0.031 | -0.031 | 0 | 0.07 | 0.18 | 0.05 | 0.01 | 0.02 | 0.16 |
| 4 | 154354566 | rs17029966 | -0.88 | -1.355 | 0.022 | 0.259 | -0.016 | 0.006 | 0.006 | 0 | 0.09 | 0.13 | 0.06 | 0.09 | 0.04 | 0.2 |
| 4 | 154355313 | rs17029977 | -0.306 | -1.365 | -0.027 | -0.083 | 0.033 | 0.041 | 0.041 | 2.201 | 0.26 | 0.14 | 0.11 | 0.09 | 0.13 | 0.33 |
| 4 | 154357175 | rs17029990 | -0.924 | -1.09 | 0.301 | 0.129 | -0.001 | 0.006 | 0.006 | 0 | 0.07 | 0.14 | 0.04 | 0.13 | 0.05 | 0.04 |
| 4 | 154363772 | rs13113769 | -0.582 | -0.936 | 0.382 | -0.047 | 0 | -0.015 | -0.015 | 0 | 0.08 | 0.08 | 0.07 | 0 | 0.12 | 0.04 |
| 4 | 154369706 | rs13105829 | -0.992 | -1.416 | 0.817 | 0.203 | -0.019 | -0.011 | -0.011 | 0 | 0.38 | 0.47 | 0.21 | 0.31 | 0.32 | 0.43 |
| 4 | 154371229 | rs11941933 | -0.314 | -1.474 | 1.34 | 0.181 | -0.024 | 0.007 | 0.007 | 0 | 0.3 | 0.37 | 0.13 | 0.31 | 0.2 | 0.39 |
| 4 | 154373196 | rs11728644 | 0.063 | -1.485 | 1.009 | 0.193 | 0.018 | -0.044 | -0.044 | 0 | 0.27 | 0.07 | 0.11 | 0.31 | 0.2 | 0.39 |
| 4 | 154374984 | rs6817170 | -0.922 | -1.514 | 1.089 | 0.152 | -0.016 | -0.006 | -0.006 | 0 | 0.37 | 0.57 | 0.19 | 0.32 | 0.27 | 0.47 |
| 4 | 154376222 | rs11732163 | -1.686 | -1.525 | 0.875 | 0.181 | 0.021 | 0.018 | 0.018 | 0 | 0.45 | 0.63 | 0.25 | 0.33 | 0.36 | 0.5 |
| 4 | 154378572 | rs1560940 | 0.009 | -1.38 | 0.365 | 0.348 | 0.007 | -0.036 | -0.036 | 0 | 0.27 | 0.09 | 0.13 | 0.31 | 0.22 | 0.41 |
| 4 | 154389653 | rs1836401 | -0.314 | -1.45 | 0.738 | 0.486 | 0.022 | -0.008 | -0.008 | 0 | 0.33 | 0.19 | 0.16 | 0.33 | 0.27 | 0.43 |
| 4 | 154391492 | rs11936132 | -1.902 | -1.361 | 1.35 | 0.462 | 0.063 | 0.053 | 0.053 | 4.931 | 0.05 | 0.38 | 0.03 | 0 | 0 | 0.01 |
| 4 | 154401015 | rs2405712 | 0.044 | -1.9 | 0.141 | 0.695 | 0.038 | -0.048 | -0.048 | 0 | 0.13 | 0 | 0.23 | 0.23 | 0.07 | 0.09 |
| 4 | 154408454 | rs713248 | 0.181 | -2.143 | 0.218 | 0.738 | 0.02 | -0.02 | -0.02 | 0 | 0.32 | 0.17 | 0.15 | 0.34 | 0.27 | 0.44 |
| 4 | 154415184 | rs13119846 | -0.957 | -1.366 | 0.392 | 0.679 | 0.002 | -0.003 | -0.003 | 0 | 0.55 | 0.39 | 0.46 | 0.58 | 0.45 | 0.6 |
| 4 | 154418561 | rs3816086 | -0.579 | -1.191 | 0.3 | 0.667 | 0.013 | 0.04 | 0.04 | 1.695 | 0.46 | 0.31 | 0.23 | 0.31 | 0.4 | 0.44 |
| 4 | 154427319 | rs10030193 | -1.47 | -1.168 | 0.358 | 0.965 | -0.036 | 0.099 | 0.099 | 1.989 | 0.13 | 0.45 | 0.26 | 0.24 | 0.05 | 0.08 |
| 4 | 154437863 | rs13125518 | -0.458 | -1.263 | -0.667 | 0.966 | -0.074 | -0.035 | -0.035 | 0 | 0.43 | 0.03 | 0.25 | 0.36 | 0.42 | 0.43 |
| 4 | 154448872 | rs7660240 | -2.158 | -0.55 | 0.796 | 1.137 | 0.004 | -0.007 | -0.007 | 1.548 | 0.22 | 0.33 | 0.35 | 0.22 | 0.24 | 0.27 |
| 4 | 154452248 | rs6850598 | -1.281 | -0.454 | 0.645 | 1.165 | 0.003 | -0.003 | -0.003 | 0 | 0.14 | 0.18 | 0.13 | 0.07 | 0.19 | 0.14 |
| 4 | 154460415 | rs4466024 | -0.769 | -0.05 | 0.491 | 1.226 | -0.016 | 0.003 | 0.003 | 0 | 0.11 | 0.09 | 0.11 | 0.07 | 0.19 | 0.15 |
| 4 | 154461884 | rs6810447 | -2.305 | -0.38 | 0.77 | 1.483 | 0.001 | -0.011 | -0.011 | 0 | 0.29 | 0.74 | 0.27 | 0.38 | 0.25 | 0.35 |
| 4 | 154465994 | rs6535924 | -2.233 | -0.249 | 0.515 | 1.578 | -0.003 | 0.004 | 0.004 | 1.014 | 0.25 | 0.72 | 0.31 | 0.2 | 0.24 | 0.15 |
| 4 | 154466939 | rs3864176 | -2.418 | -0.247 | 0.294 | 1.607 | -0.002 | -0.091 | -0.091 | 0 | 0.36 | 0.85 | 0.43 | 0.54 | 0.29 | 0.42 |
| 4 | 154468420 | rs11724474 | -1.564 | -0.271 | 0.81 | 1.726 | -0.046 | 0.061 | 0.061 | 1.23 | 0.2 | 0.45 | 0.3 | 0.34 | 0.1 | 0.24 |
| 4 | 154468648 | rs11736782 | -1.519 | -0.257 | 0.794 | 1.442 | -0.049 | 0.108 | 0.108 | 1.739 | 0.22 | 0.48 | 0.3 | 0.43 | 0.11 | 0.3 |
| 4 | 154469214 | rs6535925 | -1.123 | -0.043 | 0.789 | 1.386 | 0.085 | 0.012 | 0.012 | 0.032 | 0.14 | 0.39 | 0.13 | 0.11 | 0.18 | 0.12 |
| 4 | 154474250 | rs10011687 | -0.894 | -0.152 | 0.262 | 1.394 | 0.028 | 0.002 | 0.002 | 0 | 0.13 | 0.28 | 0.12 | 0.1 | 0.17 | 0.12 |
| 4 | 154474931 | rs10034657 | -1.617 | -0.251 | -0.553 | 1.075 | -0.002 | -0.12 | -0.12 | 0 | 0.35 | 0.89 | 0.3 | 0.52 | 0.27 | 0.48 |
| 4 | 154478222 | rs10001845 | -1.418 | -0.652 | 0.142 | 1.063 | -0.022 | 0.011 | 0.011 | 0.042 | 0.35 | 0.51 | 0.44 | 0.52 | 0.2 | 0.38 |
| 4 | 154479430 | rs6848033 | -1.474 | -0.705 | -0.339 | 0.647 | 0.084 | 0.071 | 0.071 | 0.01 | 0.5 | 0.83 | 0.62 | 0.58 | 0.34 | 0.51 |
| 4 | 154479728 | rs6848629 | -0.562 | -0.013 | 0.173 | 0.541 | -0.06 | 0.016 | 0.016 | 0.171 | 0.15 | 0.02 | 0.14 | 0.06 | 0.18 | 0.17 |
| 4 | 154486552 | rs10010874 | -0.935 | -0.133 | 1.496 | 0.651 | 0.025 | 0 | 0 | 0 | 0.07 | 0.31 | 0.04 | 0 | 0.02 | 0.01 |
| 4 | 154492573 | rs10517573 | -1.855 | -0.178 | 1.591 | 0.629 | -0.016 | -0.043 | -0.043 | 0 | 0.1 | 0.46 | 0.05 | 0 | 0.01 | 0.01 |
| 4 | 154502068 | rs7673291 | -1.962 | -0.168 | 1.146 | 0.628 | -0.053 | -0.079 | -0.079 | 1.254 | 0.12 | 0.45 | 0.05 | 0 | 0.02 | 0.02 |
| 4 | 154510843 | rs10517574 | -0.678 | 0.526 | 0.271 | 1.148 | -0.049 | -0.049 | -0.049 | 0 | 0.22 | 0.06 | 0.21 | 0.22 | 0.31 | 0.23 |
| 4 | 154513627 | rs7669418 | -0.743 | 0.515 | 0.28 | 1.137 | -0.011 | -0.011 | -0.011 | 0 | 0.23 | 0.15 | 0.21 | 0.24 | 0.31 | 0.24 |
| 4 | 154513722 | rs7669741 | -0.743 | 0.515 | 0.28 | 1.137 | -0.011 | -0.011 | -0.011 | 0 | 0.23 | 0.15 | 0.21 | 0.24 | 0.31 | 0.24 |
| 4 | 154514965 | rs17370297 | -0.743 | 0.515 | 0.28 | 1.137 | -0.011 | -0.011 | -0.011 | 1.143 | 0.23 | 0.15 | 0.22 | 0.24 | 0.31 | 0.24 |
| 4 | 154520071 | rs17030216 | -1.48 | 0.226 | 0.573 | 1.098 | 0.012 | 0.008 | 0.008 | 1.006 | 0.06 | 0.34 | 0.02 | 0 | 0.01 | 0.01 |
| 4 | 154525458 | rs1063151 | -0.964 | 0.859 | -0.544 | 1.912 | -0.03 | 0.022 | 0.022 | 0 | 0.05 | 0.01 | 0.22 | 0.33 | 0.12 | 0.21 |
| 4 | 154530639 | rs6821970 | -0.416 | 0.67 | -0.745 | 1.177 | 0.016 | 0.009 | 0.009 | 0.884 | 0.49 | 0.58 | 0.54 | 0.49 | 0.56 | 0.52 |
| 4 | 154532047 | rs13135353 | -0.384 | 0.578 | -1.202 | 1.114 | 0.021 | -0.001 | -0.001 | 0 | 0.52 | 0.83 | 0.56 | 0.49 | 0.58 | 0.58 |
| 4 | 154540056 | rs6535930 | -0.603 | 0.265 | -1.81 | 0.81 | -0.072 | -0.154 | -0.154 | 0.122 | 0.63 | 0.96 | 0.59 | 0.44 | 0.61 | 0.62 |
| 4 | 154541296 | rs11099891 | -0.153 | 0.215 | -1.655 | 0.609 | -0.001 | -0.019 | -0.019 | 2.459 | 0.47 | 0.15 | 0.38 | 0.44 | 0.53 | 0.48 |
| 4 | 154544267 | rs2289319 | -0.431 | 0.765 | -0.973 | 0.51 | -0.007 | -0.009 | -0.009 | 0.449 | 0.53 | 0.68 | 0.55 | 0.41 | 0.47 | 0.43 |
| 4 | 154549663 | rs6535932 | -0.693 | 0.852 | -0.494 | 0.059 | 0.006 | 0.015 | 0.015 | 0.194 | 0.5 | 0.52 | 0.34 | 0.22 | 0.44 | 0.3 |
| 4 | 154550603 | rs2083288 | -0.624 | 0.595 | -0.081 | 0.258 | -0.015 | -0.012 | -0.012 | 3.913 | 0.27 | 0.35 | 0.19 | 0.21 | 0.2 | 0.21 |
| 4 | 154558964 | rs7687239 | 0.49 | 0.853 | -0.104 | 0.638 | -0.004 | 0.043 | 0.043 | 0.085 | 0.68 | 0.44 | 0.72 | 0.46 | 0.72 | 0.65 |
| 4 | 154559151 | rs2033831 | 0.922 | 1.246 | -0.022 | 0.531 | -0.003 | -0.001 | -0.001 | 0 | 0.84 | 0.85 | 0.8 | 0.79 | 0.81 | 0.83 |
| 4 | 154562769 | rs6835748 | -1.241 | 0.795 | 0.679 | 0.384 | 0.005 | -0.059 | -0.059 | 1.306 | 0.1 | 0.27 | 0.06 | 0 | 0.07 | 0.06 |
| 4 | 154563897 | rs1025748 | 1.009 | 1.437 | 0.841 | 0.179 | 0.046 | -0.009 | -0.009 | 0 | 0.79 | 0.54 | 0.86 | 0.78 | 0.74 | 0.84 |
| 4 | 154571036 | rs2165084 | -2.027 | 1.094 | 1.155 | 0.055 | -0.029 | -0.031 | -0.031 | 0 | 0.07 | 0.66 | 0.05 | 0 | 0.01 | 0.03 |
| 4 | 154571570 | rs7656224 | -1.782 | 0.663 | 0.237 | 0.034 | -0.018 | -0.01 | -0.01 | 0.464 | 0.4 | 0.79 | 0.29 | 0.39 | 0.42 | 0.31 |
| 4 | 154573663 | rs10030403 | 1.235 | 0.493 | 0.993 | -0.106 | -0.013 | -0.099 | -0.099 | 0 | 0.74 | 0.51 | 0.87 | 0.9 | 0.77 | 0.86 |
| 4 | 154581208 | rs6853866 | 0.239 | 0.277 | 0.728 | -0.099 | -0.007 | 0.015 | 0.015 | 0 | 0.82 | 0.86 | 0.9 | 0.88 | 0.78 | 0.86 |
| 4 | 154584591 | rs1595625 | 2.126 | -0.308 | -0.006 | -0.301 | -0.045 | -0.04 | -0.04 | 0 | 0.47 | 0.13 | 0.7 | 0.56 | 0.55 | 0.67 |
| 4 | 154585922 | rs11727442 | -0.472 | -0.388 | 0.724 | -0.307 | 0.027 | 0.195 | 0.195 | 0 | 0.16 | 0.4 | 0.13 | 0.31 | 0.14 | 0.14 |
| 4 | 154589724 | rs17370673 | 0.228 | -0.4 | 0.424 | -0.297 | -0.044 | -0.081 | -0.081 | 0.076 | 0.13 | 0.01 | 0.08 | 0.27 | 0.14 | 0.13 |
| 4 | 154590349 | rs6835636 | 0.224 | -0.113 | 0.033 | -0.346 | -0.038 | 0.064 | 0.064 | 1.238 | 0.66 | 0.81 | 0.76 | 0.76 | 0.57 | 0.78 |
| 4 | 154599622 | rs13150331 | 1.946 | -0.144 | 0.002 | -0.357 | -0.039 | -0.071 | -0.071 | 0.03 | 0.38 | 0.12 | 0.6 | 0.45 | 0.41 | 0.64 |
| 4 | 154605745 | rs7696323 | 0.158 | -0.178 | 0.503 | -0.314 | -0.044 | 0.007 | 0.007 | 1.625 | 0.25 | 0.1 | 0.2 | 0.27 | 0.37 | 0.15 |

| | | | | | | | | | | | | | | | | |
|---|---|---|---|---|---|---|---|---|---|---|---|---|---|---|---|---|
| 4 | 154609523 | rs1816702 | -0.083 | -0.327 | 0.485 | -0.563 | -0.029 | -0.135 | -0.135 | 1.192 | 0.82 | 0.57 | 0.84 | 1 | 0.88 | 0.91 |
| 4 | 154621946 | rs11938228 | 2.176 | -0.062 | 0.094 | -0.34 | -0.007 | -0.048 | -0.048 | 0 | 0.34 | 0.13 | 0.51 | 0.42 | 0.34 | 0.59 |
| 4 | 154624656 | rs3804099 | -1.606 | 0.034 | 0.453 | -0.22 | 0.001 | 0.026 | 0.026 | 0 | 0.48 | 0.64 | 0.33 | 0.28 | 0.44 | 0.29 |
| 4 | 154625409 | rs3804100 | 0.821 | 0.043 | 0.434 | -0.097 | 0.005 | -0.014 | -0.014 | 0 | 0.11 | 0.05 | 0.06 | 0.24 | 0.06 | 0.14 |
| 4 | 154627655 | rs7656411 | -1.242 | 0.127 | 0.303 | 0.227 | -0.001 | -0.001 | -0.001 | 3.312 | 0.39 | 0.62 | 0.24 | 0.48 | 0.27 | 0.25 |
| 4 | 154632838 | rs17030340 | 0.059 | 0.006 | 0.212 | 0.135 | -0.002 | -0.054 | -0.054 | 0 | 0.22 | 0.22 | 0.1 | 0 | 0.19 | 0.02 |
| 4 | 154639667 | rs7654018 | 2.14 | -0.031 | -0.28 | 0.249 | -0.035 | 0.036 | 0.036 | 0.238 | 0.3 | 0.11 | 0.46 | 0.23 | 0.33 | 0.51 |
| 4 | 154641489 | rs1439166 | -0.738 | -0.279 | -0.12 | 0.283 | 0.015 | 0.02 | 0.02 | 0 | 0.51 | 0.4 | 0.39 | 0.31 | 0.64 | 0.27 |
| 4 | 154643503 | rs1869947 | -0.242 | -0.133 | 0.041 | 0.511 | -0.043 | 0.311 | 0.311 | 0.204 | 0.88 | 0.65 | 0.88 | 0.53 | 0.96 | 0.79 |
| 4 | 154644537 | rs10517577 | 0.475 | 0.429 | -0.818 | -0.021 | -0.096 | 0.101 | 0.101 | 0 | 0.16 | 0.01 | 0.16 | 0 | 0.22 | 0.12 |
| 4 | 154645471 | rs10440446 | 1.649 | 0.167 | -0.465 | 0.286 | 0.001 | 0.025 | 0.025 | 0.031 | 0.31 | 0.21 | 0.46 | 0.26 | 0.31 | 0.52 |
| 4 | 154664697 | rs11099896 | -0.298 | 0.143 | 0.012 | 0.64 | -0.007 | -0.019 | -0.019 | 0 | 0.27 | 0.32 | 0.24 | 0.81 | 0.26 | 0.33 |
| 4 | 154667382 | rs1598632 | -1.047 | 0.258 | -0.421 | 0.337 | -0.033 | 0.116 | 0.116 | 1.983 | 0.42 | 0.18 | 0.39 | 0.81 | 0.5 | 0.43 |
| 4 | 154670813 | rs17030417 | 0.294 | 0.194 | -0.453 | 0.891 | -0.026 | 0.173 | 0.173 | 0 | 0.09 | 0.18 | 0.09 | 0.44 | 0.04 | 0.2 |
| 4 | 154673127 | rs17299041 | 0.523 | 0.261 | -0.535 | 0.561 | -0.102 | 0.101 | 0.101 | 0 | 0.16 | 0.01 | 0.16 | 0 | 0.24 | 0.12 |
| 4 | 154675860 | rs6838237 | 1.172 | 0.097 | 0.032 | 0.548 | -0.002 | 0.005 | 0.005 | 1.503 | 0.56 | 0.61 | 0.59 | 0.19 | 0.49 | 0.55 |
| 4 | 154685712 | rs1020702 | 0.523 | -0.177 | -0.142 | 0.583 | -0.005 | 0.004 | 0.004 | 0 | 0.69 | 0.7 | 0.78 | 0.78 | 0.71 | 0.74 |
| 4 | 154686183 | rs11099905 | 1.054 | -0.15 | -0.093 | 0.71 | -0.015 | -0.052 | -0.052 | 0 | 0.17 | 0.07 | 0.15 | 0.39 | 0.22 | 0.15 |
| 4 | 154689140 | rs6857738 | 0.595 | -0.243 | -0.113 | 0.655 | -0.003 | 0.003 | 0.003 | 0 | 0.67 | 0.69 | 0.78 | 0.78 | 0.7 | 0.74 |
| 4 | 154700374 | rs12639885 | 0.999 | -0.231 | -0.188 | 0.698 | -0.015 | -0.052 | -0.052 | 2.489 | 0.17 | 0.07 | 0.15 | 0.39 | 0.22 | 0.15 |
| 4 | 154706016 | rs6853435 | 1.207 | -0.244 | -0.147 | 0.608 | 0.011 | 0.019 | 0.019 | 0.021 | 0.49 | 0.6 | 0.63 | 0.4 | 0.49 | 0.61 |
| 4 | 154709480 | rs3810765 | 1.216 | -0.244 | -0.147 | 0.623 | 0.011 | 0.018 | 0.018 | 0.018 | 0.49 | 0.61 | 0.63 | 0.42 | 0.49 | 0.61 |
| 4 | 154709854 | rs4643790 | 0.908 | -0.644 | -0.548 | 0.214 | -0.009 | -0.085 | -0.085 | 0 | 0.19 | 0.28 | 0.11 | 0.01 | 0.19 | 0.11 |
| 4 | 154711186 | rs11937424 | 2.416 | -0.657 | -0.595 | 0.296 | -0.01 | 0.038 | 0.038 | 0.01 | 0.28 | 0.12 | 0.46 | 0.21 | 0.3 | 0.43 |
| 4 | 154725119 | rs732681 | -0.949 | -0.464 | -0.354 | 0.732 | -0.044 | 0.027 | 0.027 | 0.002 | 0.41 | 0.23 | 0.34 | 0.39 | 0.47 | 0.27 |
| 4 | 154726634 | rs1105491 | 0.552 | -0.465 | -0.311 | 0.724 | 0.02 | -0.012 | -0.012 | 0.104 | 0.72 | 0.54 | 0.74 | 0.79 | 0.74 | 0.81 |
| 4 | 154727070 | rs10026531 | 1.135 | -0.357 | -0.736 | 0.751 | -0.024 | -0.066 | -0.066 | 0 | 0.17 | 0.05 | 0.14 | 0.39 | 0.22 | 0.15 |
| 4 | 154733070 | rs11723258 | -1.618 | -0.418 | -0.696 | 0.634 | 0.02 | 0.042 | 0.042 | 0.039 | 0.5 | 0.28 | 0.36 | 0.6 | 0.51 | 0.39 |
| 4 | 154743191 | rs17030494 | 0.635 | -0.395 | -0.634 | 0.603 | -0.022 | -0.033 | -0.033 | 0.045 | 0.08 | 0.17 | 0.04 | 0 | 0.02 | 0 |
| 4 | 154746806 | rs10031057 | 1.666 | -0.445 | -0.558 | 0.583 | 0.003 | 0.046 | 0.046 | 0.009 | 0.45 | 0.34 | 0.56 | 0.21 | 0.48 | 0.56 |
| 4 | 154760600 | rs6843172 | -0.168 | -0.278 | -0.957 | 0.695 | 0.002 | -0.071 | -0.071 | 0 | 0.26 | 0.1 | 0.16 | 0.58 | 0.26 | 0.27 |
| 4 | 154768011 | rs994598 | 0.44 | -0.263 | -0.887 | 0.596 | -0.002 | 0 | 0 | 2.856 | 0.61 | 0.6 | 0.63 | 0.38 | 0.52 | 0.66 |
| 4 | 154769052 | rs1125228 | -0.132 | -0.107 | -1.657 | 0.436 | 0.005 | -0.081 | -0.081 | 2.62 | 0.29 | 0.07 | 0.17 | 0.61 | 0.27 | 0.27 |
| 4 | 154769597 | rs7673508 | 0.853 | -0.058 | -1.363 | -0.137 | -0.01 | 0.107 | 0.107 | 0 | 0.4 | 0.19 | 0.52 | 0.16 | 0.43 | 0.52 |
| 4 | 154771776 | rs954368 | -0.607 | -0.099 | -1.522 | 0.218 | -0.014 | -0.015 | -0.015 | 0.306 | 0.12 | 0.05 | 0.08 | 0.14 | 0.1 | 0.29 |
| 4 | 154771820 | rs9968342 | 0.068 | -0.37 | -1.651 | -0.552 | 0.006 | -0.007 | -0.007 | 0.012 | 0.61 | 0.4 | 0.59 | 0.29 | 0.52 | 0.54 |
| 4 | 154772185 | rs11099916 | 0.219 | -1.592 | -2.409 | -0.566 | 0.078 | -0.017 | -0.017 | 0 | 0.28 | 0.1 | 0.32 | 0.26 | 0.13 | 0.26 |
| 4 | 154787090 | rs1378933 | 0.82 | -1.22 | -1.832 | -0.565 | 0.006 | -0.006 | -0.006 | 0 | 0.4 | 0.56 | 0.43 | 0.63 | 0.45 | 0.44 |
| 4 | 154791497 | rs5009912 | 0.44 | -1.327 | -1.792 | -0.579 | -0.015 | 0.01 | 0.01 | 0 | 0.44 | 0.64 | 0.45 | 0.61 | 0.47 | 0.44 |
| 4 | 154805474 | rs7669875 | 0.501 | -1.558 | -1.911 | -0.833 | -0.055 | 0.146 | 0.146 | 0 | 0.32 | 0.07 | 0.19 | 0.11 | 0.35 | 0.26 |
| 4 | 154818522 | rs17030719 | 0.77 | -1.926 | -1.793 | -0.371 | -0.009 | -0.026 | -0.026 | 0 | 0.09 | 0.48 | 0.04 | 0 | 0.01 | 0.06 |
| 4 | 154824112 | rs1993423 | 1.05 | -1.937 | -2.03 | -0.4 | 0 | -0.001 | -0.001 | 0 | 0.51 | 0.54 | 0.4 | 0.52 | 0.55 | 0.44 |
| 4 | 154824331 | rs17299833 | 0.737 | -1.485 | -2.562 | -0.461 | -0.035 | 0.05 | 0.05 | 0 | 0.18 | 0.06 | 0.14 | 0.1 | 0.22 | 0.2 |
| 4 | 154827630 | rs6831196 | 0.782 | -1.487 | -2.542 | -0.438 | -0.033 | 0.04 | 0.04 | 0 | 0.18 | 0.07 | 0.18 | 0.11 | 0.22 | 0.2 |
| 4 | 154827974 | rs4696521 | -0.986 | -1.889 | -2.668 | -0.556 | -0.034 | -0.019 | -0.019 | 4.234 | 0.61 | 0.3 | 0.62 | 0.45 | 0.56 | 0.64 |
| 4 | 154828366 | rs9995821 | 0.066 | -2.053 | -2.782 | -0.683 | 0.026 | 0.023 | 0.023 | 0 | 0.28 | 0.2 | 0.35 | 0.22 | 0.22 | 0.32 |
| 4 | 154829156 | rs9307914 | 1.335 | -2 | -2.559 | -0.312 | 0.001 | 0.001 | 0.001 | 0 | 0.36 | 0.58 | 0.37 | 0.47 | 0.43 | 0.34 |
| 4 | 154831899 | rs2045323 | 1.348 | -2.096 | -2.525 | -0.211 | 0.03 | -0.029 | -0.029 | 0.198 | 0.16 | 0.03 | 0.27 | 0.2 | 0.09 | 0.23 |
| 4 | 154834707 | rs10031184 | 0.783 | -1.83 | -2.194 | -0.119 | -0.016 | -0.014 | -0.014 | 0 | 0.41 | 0.65 | 0.39 | 0.47 | 0.47 | 0.38 |
| 4 | 154838816 | rs11933412 | 1.837 | -1.128 | -2.088 | 0.108 | -0.014 | 0.086 | 0.086 | 0 | 0.14 | 0 | 0.05 | 0 | 0.12 | 0.06 |
| 4 | 154850532 | rs11724021 | 1.383 | -1.164 | -2.091 | 0.243 | -0.174 | -0.142 | -0.142 | 0.012 | 0.31 | 0.04 | 0.32 | 0.46 | 0.44 | 0.33 |
| 4 | 154850835 | rs11099924 | 1.188 | -0.985 | -0.898 | 0.77 | 0.016 | 0.021 | 0.021 | 0 | 0.52 | 0.57 | 0.49 | 0.65 | 0.61 | 0.49 |
| 4 | 154855886 | rs4235233 | 1.204 | -0.989 | -0.924 | 0.832 | 0.015 | 0.015 | 0.015 | 0 | 0.52 | 0.57 | 0.48 | 0.62 | 0.61 | 0.49 |
| 4 | 154863536 | rs6835798 | 0.46 | -0.882 | -0.487 | 0.96 | -0.001 | -0.003 | -0.003 | 0 | 0.08 | 0.09 | 0.12 | 0.12 | 0.07 | 0.16 |
| 4 | 154863623 | rs6835664 | 0.526 | -0.88 | -0.501 | 0.96 | -0.002 | -0.005 | -0.005 | 0.378 | 0.08 | 0.07 | 0.12 | 0.12 | 0.07 | 0.16 |
| 4 | 154864750 | rs13135284 | 0.839 | -0.841 | -0.46 | 0.912 | -0.005 | -0.002 | -0.002 | 0 | 0.67 | 0.6 | 0.54 | 0.65 | 0.73 | 0.54 |
| 4 | 154867538 | rs10024897 | -0.077 | -0.063 | -0.58 | 0.783 | 0.002 | -0.024 | -0.024 | 0 | 0.05 | 0.09 | 0.06 | 0 | 0.09 | 0.07 |
| 4 | 154873677 | rs4128249 | -0.691 | -0.198 | -0.628 | 0.972 | 0.041 | 0.007 | 0.007 | 0 | 0.43 | 0.29 | 0.27 | 0.41 | 0.33 | 0.47 |
| 4 | 154883493 | rs6818192 | 0.329 | 0.43 | -0.599 | 1.125 | 0.002 | -0.024 | -0.024 | 0 | 0.22 | 0.15 | 0.14 | 0.34 | 0.24 | 0.24 |
| 4 | 154883600 | rs7661078 | 0.431 | 0.255 | -0.462 | 1.34 | 0.017 | -0.013 | -0.013 | 3.351 | 0.63 | 0.71 | 0.6 | 0.43 | 0.68 | 0.57 |
| 4 | 154884393 | rs4696523 | -0.237 | 0.204 | -0.377 | 1.317 | 0.003 | -0.004 | -0.004 | 0.015 | 0.85 | 0.88 | 0.74 | 0.76 | 0.92 | 0.81 |
| 4 | 154885560 | rs10026016 | 0.854 | 0.572 | -0.461 | 0.645 | 0.012 | 0.052 | 0.052 | 0 | 0.53 | 0.69 | 0.5 | 0.31 | 0.52 | 0.44 |
| 4 | 154887604 | rs1878449 | -1.353 | 0.843 | 0.11 | 0.39 | 0.005 | 0.048 | 0.048 | 0.111 | 0.49 | 0.63 | 0.54 | 0.7 | 0.47 | 0.56 |
| 4 | 154888777 | rs12505957 | 0.342 | 0.856 | 0.222 | 0.507 | -0.003 | 0.001 | 0.001 | 0 | 0.15 | 0.18 | 0.27 | 0.17 | 0.13 | 0.18 |
| 4 | 154890126 | rs4640650 | -0.706 | 0.855 | 0.363 | 0.554 | 0.003 | 0.004 | 0.004 | 0.01 | 0.41 | 0.35 | 0.44 | 0.63 | 0.39 | 0.44 |
| 4 | 154890983 | rs9790432 | -0.606 | 0.853 | 0.364 | 0.548 | 0.018 | 0.061 | 0.061 | 0 | 0.21 | 0.34 | 0.39 | 0.63 | 0.25 | 0.39 |
| 4 | 154891251 | rs1878452 | 0.09 | 0.876 | 0.502 | 0.514 | 0.005 | 0.01 | 0.01 | 0.277 | 0.07 | 0.08 | 0.06 | 0.12 | 0.09 | 0.07 |
| 4 | 154891356 | rs9790702 | -0.131 | 0.907 | 0.385 | 0.618 | 0.037 | 0.227 | 0.227 | 0 | 0.07 | 0.15 | 0.13 | 0.46 | 0.13 | 0.21 |
| 4 | 154891741 | rs11947700 | 1.099 | 0.521 | 0.099 | 0.248 | 0.016 | 0.116 | 0.116 | 0 | 0.18 | 0 | 0.05 | 0 | 0.13 | 0.05 |

| | | | | | | | | | | | | | | | | |
|---|---|---|---|---|---|---|---|---|---|---|---|---|---|---|---|---|
| 4 | 154896241 | rs12508706 | 0.244 | 0.62 | 0.259 | 0.402 | -0.004 | 0.003 | 0.003 | 0 | 0.14 | 0.17 | 0.26 | 0.17 | 0.12 | 0.18 |
| 4 | 154908045 | rs12508232 | 0.082 | 0.624 | 0.252 | 0.369 | -0.01 | 0.005 | 0.005 | 0 | 0.15 | 0.21 | 0.26 | 0.17 | 0.12 | 0.18 |
| 4 | 154908572 | rs7663313 | 0.096 | 0.629 | 0.264 | 0.371 | -0.01 | 0.006 | 0.006 | 0 | 0.15 | 0.23 | 0.26 | 0.17 | 0.12 | 0.18 |
| 4 | 154909285 | rs12505740 | -0.134 | 0.912 | 0.465 | 0.592 | -0.011 | -0.024 | -0.024 | 0 | 0.41 | 0.47 | 0.41 | 0.23 | 0.33 | 0.35 |
| 4 | 154930494 | rs6813540 | -0.686 | 0.878 | 0.207 | 0.638 | -0.018 | -0.017 | -0.017 | 0 | 0.6 | 0.54 | 0.76 | 0.69 | 0.66 | 0.66 |
| 4 | 154931179 | rs13105395 | -0.053 | 1.865 | 0.18 | 1.193 | 0 | 0.028 | 0.028 | 0 | 0.13 | 0.11 | 0.25 | 0.41 | 0.18 | 0.26 |
| 4 | 154932704 | rs17030932 | -0.138 | 1.696 | 0.267 | 0.776 | 0.013 | 0.101 | 0.101 | 0 | 0.13 | 0.19 | 0.25 | 0.5 | 0.15 | 0.25 |
| 4 | 154937685 | rs2897074 | -0.279 | 1.149 | 0.047 | 0.079 | -0.008 | 0.04 | 0.04 | 0 | 0.45 | 0.38 | 0.35 | 0.25 | 0.51 | 0.44 |
| 4 | 154938328 | rs12641646 | -0.451 | 1.323 | -0.742 | 0.164 | -0.001 | 0.033 | 0.033 | 0 | 0.48 | 0.64 | 0.37 | 0.25 | 0.51 | 0.44 |
| 4 | 154938646 | rs7663305 | -0.571 | 1.694 | -0.864 | 0.6 | -0.025 | -0.001 | -0.001 | 0 | 0.08 | 0.07 | 0.1 | 0.06 | 0.15 | 0.15 |
| 4 | 154950464 | rs12512005 | -0.699 | 1.099 | -0.7 | 0.486 | -0.001 | -0.002 | -0.002 | 0.022 | 0.16 | 0.15 | 0.3 | 0.12 | 0.15 | 0.15 |
| 4 | 154951725 | rs17031027 | 0.11 | 0.784 | -0.93 | 0.426 | 0.005 | 0.008 | 0.008 | 0 | 0.35 | 0.27 | 0.26 | 0.27 | 0.31 | 0.43 |
| 4 | 154951914 | rs17031035 | 0.153 | 0.762 | -0.829 | 0.394 | 0.013 | 0.011 | 0.011 | 0 | 0.34 | 0.24 | 0.25 | 0.26 | 0.29 | 0.42 |
| 4 | 154960908 | rs4696526 | 0.275 | 0.483 | 0.018 | 0.445 | -0.015 | 0.008 | 0.008 | 0.116 | 0.24 | 0.3 | 0.21 | 0.27 | 0.18 | 0.25 |
| 4 | 154962558 | rs1320100 | 0.484 | 0.226 | -0.431 | 0.581 | 0.052 | 0.026 | 0.026 | 0.025 | 0.41 | 0.23 | 0.25 | 0.36 | 0.3 | 0.5 |
| 4 | 154968120 | rs17372382 | 0.031 | 0.551 | -0.111 | 0.453 | 0.006 | 0.004 | 0.004 | 0 | 0.53 | 0.37 | 0.33 | 0.48 | 0.43 | 0.63 |
| 4 | 154968875 | rs10517583 | 1.175 | -0.014 | -0.825 | -0.078 | 0.034 | 0.047 | 0.047 | 0.039 | 0.22 | 0.12 | 0.05 | 0.01 | 0.13 | 0.13 |
| 4 | 154972357 | rs6835823 | -0.427 | 0.347 | -1.164 | -0.35 | -0.003 | 0.024 | 0.024 | 0 | 0.47 | 0.35 | 0.41 | 0.62 | 0.47 | 0.63 |
| 4 | 154974131 | rs11099935 | 0.931 | 0.364 | -1.044 | -0.238 | -0.004 | 0.009 | 0.009 | 0.124 | 0.15 | 0.13 | 0.22 | 0.44 | 0.2 | 0.4 |
| 4 | 154974203 | rs11099936 | 0.418 | 0.211 | -0.922 | 0 | 0.006 | -0.011 | -0.011 | 0 | 0.38 | 0.25 | 0.28 | 0.44 | 0.32 | 0.53 |
| 4 | 154975869 | rs9999890 | 0.075 | 0.304 | -0.335 | 0.11 | 0.035 | 0.07 | 0.07 | 0 | 0.09 | 0.33 | 0.07 | 0.08 | 0.08 | 0.1 |
| 4 | 154988567 | rs2404921 | -0.088 | 0.245 | -1.208 | -0.035 | -0.009 | 0.004 | 0.004 | 0 | 0.28 | 0.2 | 0.45 | 0.24 | 0.27 | 0.22 |
| 4 | 154995223 | rs2404917 | -0.733 | -0.489 | -1.339 | -0.105 | 0.022 | 0.049 | 0.049 | 0.15 | 0.85 | 0.94 | 0.96 | 1 | 0.92 | 0.96 |
| 4 | 154995463 | rs2404916 | -0.477 | -0.849 | -1.171 | 0.127 | -0.011 | 0.055 | 0.055 | 1.051 | 0.32 | 0.49 | 0.31 | 0.49 | 0.31 | 0.57 |
| 4 | 154997713 | rs6827515 | 1.214 | -1.176 | -1.495 | 0.043 | 0.05 | 0.008 | 0.008 | 0.82 | 0.2 | 0.07 | 0.12 | 0.2 | 0.11 | 0.09 |
| 4 | 155000585 | rs9307919 | -1.34 | -1.104 | -1.16 | 0.137 | 0.019 | -0.017 | -0.017 | 0.467 | 0.65 | 0.46 | 0.45 | 0.71 | 0.49 | 0.66 |
| 4 | 155007625 | rs10005886 | 0.81 | -1.475 | -1.052 | 0.191 | 0.044 | 0.018 | 0.018 | 1.3 | 0.24 | 0.14 | 0.32 | 0.21 | 0.1 | 0.15 |
| 4 | 155008496 | rs17031131 | 0.063 | -1.08 | -0.996 | 0.387 | 0 | -0.007 | -0.007 | 1.613 | 0.22 | 0.17 | 0.21 | 0.5 | 0.24 | 0.43 |
| 4 | 155009671 | rs7667589 | 1.171 | -1.079 | -1.128 | 0.082 | 0.004 | -0.009 | -0.009 | 0 | 0.37 | 0.44 | 0.54 | 0.28 | 0.54 | 0.37 |
| 4 | 155013194 | rs17031138 | 0.7 | -0.972 | -1.146 | 0.487 | -0.027 | -0.078 | -0.078 | 0 | 0.13 | 0.02 | 0.15 | 0.3 | 0.11 | 0.36 |
| 4 | 155018810 | rs4266272 | 0.139 | -0.75 | -0.884 | 0.5 | -0.008 | 0.046 | 0.046 | 0 | 0.24 | 0.33 | 0.21 | 0.39 | 0.23 | 0.42 |
| 4 | 155027927 | rs1878448 | -0.2 | -0.318 | -0.907 | 0.703 | -0.006 | 0 | 0 | 0 | 0.62 | 0.64 | 0.74 | 0.6 | 0.7 | 0.54 |
| 4 | 155042470 | rs17031151 | -0.111 | -0.388 | -1.13 | 1.009 | 0.011 | -0.015 | -0.015 | 0 | 0.24 | 0.17 | 0.19 | 0.33 | 0.16 | 0.15 |
| 4 | 155047033 | rs4484280 | -0.479 | 0.441 | -0.898 | 0.908 | -0.001 | -0.016 | -0.016 | 0 | 0.27 | 0.23 | 0.41 | 0.5 | 0.3 | 0.34 |
| 4 | 155047531 | rs2014131 | 0.792 | 0.824 | -1.12 | -0.198 | 0.005 | 0.037 | 0.037 | 0 | 0.43 | 0.33 | 0.27 | 0.06 | 0.38 | 0.24 |
| 4 | 155049484 | rs938716 | -0.209 | 0.982 | -1.157 | 0.752 | -0.077 | -0.094 | -0.094 | 0 | 0.18 | 0.01 | 0.31 | 0.29 | 0.26 | 0.25 |
| 4 | 155055498 | rs17372674 | 0.786 | 0.759 | -0.671 | 0.27 | -0.031 | 0.05 | 0.05 | 0 | 0.57 | 0.25 | 0.44 | 0.69 | 0.56 | 0.72 |
| 4 | 155057837 | rs1574232 | -1.703 | 0.141 | 0.649 | 0.416 | 0.011 | 0.002 | 0.002 | 0 | 0.2 | 0.31 | 0.16 | 0.21 | 0.19 | 0.11 |
| 4 | 155062069 | rs10028852 | 1.963 | -0.254 | 1.454 | 0.137 | -0.021 | -0.052 | -0.052 | 0 | 0.72 | 0.58 | 0.88 | 0.85 | 0.83 | 0.87 |
| 4 | 155071496 | rs2404903 | 0.498 | -0.281 | 1.844 | 0.125 | -0.027 | -0.059 | -0.059 | 0 | 0.89 | 0.63 | 0.97 | 1 | 0.98 | 1 |
| 4 | 155071789 | rs4349583 | -1.818 | -0.269 | 1.778 | 0.129 | -0.002 | 0.005 | 0.005 | 0.843 | 0.22 | 0.21 | 0.12 | 0.15 | 0.17 | 0.12 |
| 4 | 155080330 | rs4432731 | -1.55 | -0.903 | 2.118 | 0.293 | -0.014 | 0.011 | 0.011 | 1.524 | 0.1 | 0.26 | 0.07 | 0.1 | 0.06 | 0.07 |
| 4 | 155083506 | rs6813896 | 1.969 | -0.913 | 1.999 | 0.377 | 0.005 | 0 | 0 | 4.696 | 0.78 | 0.2 | 0.85 | 0.8 | 0.9 | 0.93 |
| 4 | 155084367 | rs17031193 | -0.706 | -0.734 | 2.142 | 0.373 | -0.012 | 0.024 | 0.024 | 0.06 | 0.08 | 0.25 | 0.05 | 0.09 | 0.04 | 0 |
| 4 | 155103453 | rs4235235 | 1.442 | -0.943 | 1.799 | -0.02 | -0.086 | -0.157 | -0.157 | 0.821 | 0.77 | 0.5 | 0.93 | 1 | 0.93 | 0.99 |
| 4 | 155123244 | rs4393979 | 1.434 | -0.86 | 2.725 | 0.426 | -0.049 | -0.004 | -0.004 | 0 | 0.84 | 0.52 | 0.7 | 0.89 | 0.92 | 0.98 |
| 4 | 155132707 | rs11099942 | 1.377 | -0.875 | 2.989 | 0.433 | -0.051 | -0.085 | -0.085 | 0 | 0.85 | 0.54 | 0.91 | 0.96 | 0.92 | 0.99 |
| 4 | 155135688 | rs4385038 | 2.199 | -1.252 | 2.727 | 0.754 | 0.018 | -0.009 | -0.009 | 0.111 | 0.78 | 0.2 | 0.65 | 0.74 | 0.91 | 0.88 |
| 4 | 155156207 | rs7655799 | 1.791 | -1.256 | 2.744 | 0.695 | -0.023 | 0.007 | 0.007 | 4.583 | 0.82 | 0.26 | 0.65 | 0.77 | 0.91 | 0.89 |
| 4 | 155166327 | rs6821649 | 1.932 | -1.25 | 2.817 | 0.502 | -0.052 | -0.055 | -0.055 | 0.403 | 0.8 | 0.38 | 0.69 | 0.92 | 0.92 | 0.96 |
| 4 | 155182784 | rs3828478 | -0.426 | -1.028 | 1.618 | 0.671 | 0.01 | 0 | 0 | 0.714 | 0.46 | 0.62 | 0.51 | 0.54 | 0.37 | 0.43 |
| 4 | 155193398 | rs4312742 | 0.739 | -0.936 | 1.264 | 0.742 | 0.011 | -0.015 | -0.015 | 0.404 | 0.44 | 0.23 | 0.45 | 0.46 | 0.57 | 0.52 |
| 4 | 155197891 | rs6823540 | -0.465 | -1.094 | 1.318 | 0.883 | -0.003 | -0.001 | -0.001 | 0 | 0.31 | 0.67 | 0.19 | 0.26 | 0.24 | 0.31 |
| 4 | 155205880 | rs4447842 | -0.442 | -0.752 | 1.03 | 1.018 | -0.019 | -0.073 | -0.073 | 0 | 0.7 | 0.89 | 0.7 | 0.54 | 0.68 | 0.58 |
| 4 | 155209141 | rs12508287 | -0.436 | -0.841 | 2.131 | 0.969 | -0.001 | 0.041 | 0.041 | 0 | 0.35 | 0.43 | 0.35 | 0.47 | 0.33 | 0.42 |
| 4 | 155219361 | rs17031387 | -0.174 | -1.118 | 1.999 | 0.818 | 0.021 | 0.037 | 0.037 | 0.077 | 0.23 | 0.17 | 0.11 | 0.15 | 0.17 | 0.24 |
| 4 | 155229324 | rs4696546 | 0.374 | -0.975 | 1.86 | 0.91 | 0 | -0.027 | -0.027 | 0 | 0.61 | 0.37 | 0.61 | 0.53 | 0.59 | 0.57 |
| 4 | 155241572 | rs11935573 | 0.44 | -1.026 | 1.587 | 1.156 | -0.038 | -0.048 | -0.048 | 0 | 0.28 | 0.06 | 0.3 | 0.36 | 0.32 | 0.36 |
| 4 | 155244475 | rs12500437 | 1.984 | -1.064 | 2.012 | 1.15 | -0.011 | 0.059 | 0.059 | 1.631 | 0.89 | 0.45 | 0.91 | 0.88 | 0.91 | 0.93 |
| 4 | 155248664 | rs1388082 | 1.44 | -1.048 | 2.014 | 1.159 | 0.025 | 0.122 | 0.122 | 0.016 | 0.92 | 0.62 | 0.94 | 0.9 | 0.93 | 0.94 |
| 4 | 155251015 | rs10517595 | 0.692 | -1.071 | 1.527 | 1.053 | -0.014 | -0.008 | -0.008 | 0 | 0.66 | 0.33 | 0.6 | 0.74 | 0.74 | 0.68 |
| 4 | 155251785 | rs2130710 | 0.38 | -0.948 | 1.114 | 0.999 | -0.029 | -0.04 | -0.04 | 0 | 0.29 | 0.1 | 0.3 | 0.37 | 0.32 | 0.37 |
| 4 | 155252934 | rs4696548 | 1.654 | -0.832 | 0.819 | 1.056 | -0.148 | 0.009 | 0.009 | 0 | 0.33 | 0.04 | 0.28 | 0.27 | 0.42 | 0.25 |
| 4 | 155257709 | rs1021381 | 0.01 | -0.828 | 1.271 | 1.044 | 0 | 0 | 0 | 0 | 0.63 | 0.41 | 0.63 | 0.61 | 0.62 | 0.62 |
| 4 | 155258749 | rs10517592 | -0.523 | -0.761 | 0.749 | 1.025 | -0.007 | 0.053 | 0.053 | 0 | 0.07 | 0.23 | 0.06 | 0.12 | 0.02 | 0.24 |
| 4 | 155269311 | rs11726534 | -0.043 | -0.587 | -0.069 | 0.926 | 0.002 | 0.003 | 0.003 | 0 | 0.34 | 0.3 | 0.35 | 0.38 | 0.37 | 0.32 |
| 4 | 155269388 | rs4696207 | 0.989 | -0.005 | -1.124 | 0.652 | -0.14 | 0.049 | 0.049 | 0 | 0.35 | 0.03 | 0.28 | 0.25 | 0.45 | 0.26 |
| 4 | 155270460 | rs7681342 | 0.974 | -0.009 | -1.127 | 0.649 | -0.146 | 0.05 | 0.05 | 0 | 0.35 | 0.03 | 0.28 | 0.25 | 0.45 | 0.26 |
| 4 | 155273743 | rs17031489 | -0.049 | -0.012 | -0.791 | 0.955 | -0.024 | -0.049 | -0.049 | 0 | 0.27 | 0.17 | 0.45 | 0.43 | 0.29 | 0.33 |

| | | | | | | | | | | | | | | | | |
|---|---|---|---|---|---|---|---|---|---|---|---|---|---|---|---|---|
| 4 | 155281772 | rs7687041 | -0.176 | -0.753 | -0.288 | 0.628 | -0.02 | -0.016 | -0.016 | 0 | 0.79 | 0.79 | 0.9 | 0.89 | 0.87 | 0.94 |
| 4 | 155285052 | rs951728 | 1.275 | -0.796 | -0.464 | 0.431 | -0.078 | 0.003 | 0.003 | 0 | 0.38 | 0.1 | 0.36 | 0.37 | 0.46 | 0.46 |
| 4 | 155287535 | rs10017772 | 0.836 | -0.971 | 0.295 | 0.398 | -0.011 | 0.005 | 0.005 | 0 | 0.45 | 0.39 | 0.41 | 0.45 | 0.49 | 0.58 |
| 4 | 155294563 | rs11099951 | 1.157 | -1.032 | -0.061 | 0.375 | -0.077 | 0.015 | 0.015 | 0 | 0.43 | 0.13 | 0.36 | 0.39 | 0.48 | 0.44 |
| 4 | 155295076 | rs11721758 | 1.161 | -1.046 | -0.048 | 0.368 | -0.075 | 0.015 | 0.015 | 0 | 0.43 | 0.14 | 0.36 | 0.39 | 0.49 | 0.44 |
| 4 | 155296207 | rs13111929 | 0.577 | -1.051 | 0.629 | 0.118 | -0.002 | 0.048 | 0.048 | 0 | 0.54 | 0.38 | 0.34 | 0.24 | 0.56 | 0.59 |
| 4 | 155297245 | rs6536003 | -0.375 | -1.136 | 0.647 | 0.02 | -0.006 | -0.018 | -0.018 | 0 | 0.83 | 0.79 | 0.93 | 0.9 | 0.86 | 0.91 |
| 4 | 155299483 | rs13134088 | 1.682 | -1.144 | 0.9 | 0.063 | -0.063 | 0.174 | 0.174 | 0 | 0.47 | 0.11 | 0.28 | 0.2 | 0.49 | 0.43 |
| 4 | 155300448 | rs4696556 | -0.379 | -1.084 | 0.927 | 0.085 | -0.005 | -0.017 | -0.017 | 0 | 0.83 | 0.8 | 0.93 | 0.9 | 0.86 | 0.91 |
| 4 | 155300938 | rs1388070 | -0.157 | -0.979 | 0.713 | 0.104 | 0.011 | 0.019 | 0.019 | 0 | 0.27 | 0.37 | 0.59 | 0.65 | 0.3 | 0.32 |
| 4 | 155319043 | rs7686473 | 0.865 | -1.022 | 0.422 | -0.045 | -0.02 | 0.011 | 0.011 | 0.099 | 0.75 | 0.4 | 0.85 | 0.7 | 0.8 | 0.76 |
| 4 | 155329567 | rs11099953 | -1.013 | -1.075 | 0.713 | -0.145 | -0.024 | 0.027 | 0.027 | 0 | 0.24 | 0.4 | 0.13 | 0.3 | 0.2 | 0.22 |
| 4 | 155331774 | rs4550900 | -1.615 | -1.119 | 0.126 | -0.226 | 0.03 | 0.005 | 0.005 | 1.662 | 0.36 | 0.68 | 0.18 | 0.34 | 0.25 | 0.38 |
| 4 | 155333714 | rs6851087 | -1.708 | -1.153 | -0.393 | -0.734 | -0.029 | 0.188 | 0.188 | 0 | 0.56 | 0.78 | 0.73 | 0.98 | 0.55 | 0.7 |
| 4 | 155339195 | rs902455 | 1.259 | -1.204 | 0.072 | -0.555 | -0.063 | -0.021 | -0.021 | 0.047 | 0.85 | 0.42 | 0.93 | 0.85 | 0.95 | 0.82 |
| 4 | 155343176 | rs2896868 | 0.968 | -1.021 | 0.272 | -0.447 | 0.02 | 0.091 | 0.091 | 1.106 | 0.91 | 0.51 | 0.93 | 0.85 | 0.96 | 0.82 |
| 4 | 155354916 | rs4696572 | 0.115 | -0.635 | -1.011 | -0.406 | -0.066 | -0.063 | -0.063 | 0 | 0.22 | 0.07 | 0.36 | 0.69 | 0.31 | 0.33 |
| 4 | 155364122 | rs13146084 | 1.402 | -0.147 | -1.547 | -0.859 | -0.007 | 0.067 | 0.067 | 0 | 0.42 | 0.09 | 0.47 | 0.25 | 0.37 | 0.45 |
| 4 | 155364486 | rs13122913 | 0.961 | 0.056 | -0.869 | -0.884 | -0.005 | 0.013 | 0.013 | 0 | 0.58 | 0.33 | 0.59 | 0.35 | 0.55 | 0.68 |
| 4 | 155371246 | rs4696576 | 0.135 | 0.482 | -0.871 | -0.724 | -0.071 | -0.084 | -0.084 | 0 | 0.18 | 0.02 | 0.28 | 0.64 | 0.27 | 0.25 |
| 4 | 155379883 | rs11731813 | 0.388 | 0.53 | -1.017 | -0.755 | -0.019 | -0.047 | -0.047 | 0.169 | 0.16 | 0.07 | 0.3 | 0.64 | 0.23 | 0.24 |
| 4 | 155397517 | rs4696584 | -0.901 | 0.42 | -0.851 | -1.428 | -0.017 | -0.028 | -0.028 | 0 | 0.73 | 0.62 | 0.67 | 0.85 | 0.79 | 0.69 |
| 4 | 155400984 | rs10517593 | 0.436 | 0.664 | -1.039 | -1.788 | 0 | -0.006 | -0.006 | 0 | 0.52 | 0.48 | 0.5 | 0.83 | 0.52 | 0.56 |
| 4 | 155403593 | rs9284660 | 1.485 | 0.579 | -0.852 | -1.793 | -0.001 | 0.038 | 0.038 | 5.291 | 0.34 | 0.2 | 0.18 | 0.19 | 0.3 | 0.26 |
| 4 | 155408038 | rs2404478 | -0.155 | 0.681 | -0.931 | -1.992 | 0.015 | -0.143 | -0.143 | 0.127 | 0.76 | 0.52 | 0.81 | 0.99 | 0.75 | 0.79 |
| 4 | 155411065 | rs7656522 | -0.399 | 0.67 | -0.921 | -1.997 | 0.013 | -0.118 | -0.118 | 0 | 0.77 | 0.6 | 0.81 | 0.99 | 0.75 | 0.79 |
| 4 | 155413362 | rs1873367 | 1.51 | 0.619 | -1.132 | -1.917 | -0.013 | 0.018 | 0.018 | 0 | 0.23 | 0.04 | 0.31 | 0.15 | 0.22 | 0.24 |
| 4 | 155413510 | rs13132637 | -0.398 | 0.644 | -0.947 | -1.96 | -0.004 | 0.066 | 0.066 | 0 | 0.45 | 0.36 | 0.49 | 0.17 | 0.46 | 0.44 |
| 4 | 155414179 | rs13112066 | 0.358 | 0.917 | -0.692 | -1.832 | -0.007 | -0.013 | -0.013 | 0.002 | 0.21 | 0.2 | 0.17 | 0.01 | 0.24 | 0.21 |
| 4 | 155414923 | rs17031739 | 0.099 | 0.968 | -0.695 | -1.786 | 0.013 | -0.098 | -0.098 | 0.614 | 0.2 | 0.32 | 0.14 | 0.01 | 0.21 | 0.2 |
| 4 | 155420487 | rs10014500 | 1.055 | 0.775 | -0.606 | -1.842 | -0.001 | -0.037 | -0.037 | 0.113 | 0.28 | 0.32 | 0.39 | 0.15 | 0.27 | 0.28 |
| 4 | 155421710 | rs4345160 | 0.082 | 0.572 | -0.439 | -1.435 | 0.001 | -0.043 | -0.043 | 0.329 | 0.46 | 0.48 | 0.53 | 0.16 | 0.48 | 0.47 |
| 4 | 155424231 | rs4323084 | -0.66 | 0.436 | -1.513 | -1.404 | 0.006 | -0.004 | -0.004 | 0.397 | 0.8 | 0.85 | 0.71 | 0.27 | 0.77 | 0.77 |
| 4 | 155427505 | rs10019863 | 0.166 | 0.471 | -1.479 | -1.446 | -0.007 | 0.098 | 0.098 | 0 | 0.48 | 0.67 | 0.55 | 0.16 | 0.47 | 0.47 |
| 4 | 155430666 | rs6536017 | 0.622 | 0.811 | -1.657 | -1.234 | -0.027 | 0.107 | 0.107 | 0.793 | 0.31 | 0.13 | 0.16 | 0.11 | 0.31 | 0.33 |
| 4 | 155438339 | rs4235247 | -0.557 | 0.182 | -1.441 | -0.666 | -0.001 | -0.068 | -0.068 | 0 | 0.84 | 0.94 | 0.84 | 0.49 | 0.83 | 0.85 |
| 4 | 155442246 | rs11737226 | 0.671 | -0.08 | -0.919 | -1.124 | -0.008 | 0.002 | 0.002 | 0 | 0.65 | 0.6 | 0.67 | 0.43 | 0.68 | 0.64 |
| 4 | 155446305 | rs4482740 | 0.681 | 0.129 | -1.1 | -1.45 | -0.014 | 0.011 | 0.011 | 0 | 0.31 | 0.2 | 0.47 | 0.3 | 0.35 | 0.31 |
| 4 | 155448553 | rs10008078 | 0.253 | -0.006 | -0.581 | -1.569 | -0.007 | 0.046 | 0.046 | 0 | 0.21 | 0.24 | 0.2 | 0.54 | 0.2 | 0.2 |
| 4 | 155449629 | rs7689945 | -0.236 | 0.012 | -0.513 | -1.828 | 0.008 | -0.022 | -0.022 | 1.844 | 0.6 | 0.49 | 0.38 | 0.68 | 0.53 | 0.57 |
| 4 | 155449832 | rs13435101 | -0.236 | 0.012 | -0.513 | -1.824 | 0.008 | -0.023 | -0.023 | 0.339 | 0.6 | 0.49 | 0.38 | 0.68 | 0.53 | 0.57 |
| 4 | 155451187 | rs6819508 | -0.57 | 0.061 | -0.479 | -1.818 | 0.046 | -0.066 | -0.066 | 0 | 0.87 | 0.61 | 0.84 | 0.95 | 0.85 | 0.87 |
| 4 | 155451527 | rs11099958 | 0.962 | 0.002 | -0.494 | -1.866 | -0.003 | 0.139 | 0.139 | 0.976 | 0.38 | 0.15 | 0.18 | 0.14 | 0.33 | 0.37 |
| 4 | 155452233 | rs4622984 | 0.211 | -0.154 | -0.316 | -1.75 | 0.006 | 0.015 | 0.015 | 0 | 0.68 | 0.5 | 0.65 | 0.42 | 0.65 | 0.67 |
| 4 | 155456608 | rs7654425 | 0.925 | -0.159 | -0.719 | -1.7 | 0.017 | 0.139 | 0.139 | 1.717 | 0.38 | 0.15 | 0.19 | 0.14 | 0.32 | 0.36 |
| 4 | 155460459 | rs12642770 | 0.364 | -0.07 | -0.636 | -1.422 | -0.011 | -0.049 | -0.049 | 0.1 | 0.19 | 0.1 | 0.2 | 0.54 | 0.21 | 0.22 |
| 4 | 155474222 | rs12642469 | 0.672 | -0.152 | -0.728 | -1.404 | -0.009 | -0.065 | -0.065 | 0 | 0.17 | 0.07 | 0.18 | 0.54 | 0.19 | 0.21 |
| 4 | 155481289 | rs4508864 | 1.021 | -0.131 | -0.69 | -1.589 | -0.001 | 0 | 0 | 0 | 0.22 | 0.17 | 0.16 | 0.22 | 0.22 | 0.14 |
| 4 | 155484015 | rs1800787 | 1.164 | -0.138 | -0.712 | -1.592 | -0.012 | 0 | 0 | 0 | 0.21 | 0.09 | 0.15 | 0.22 | 0.22 | 0.14 |
| 4 | 155486381 | rs2227401 | 1.181 | -0.132 | -0.709 | -1.568 | -0.012 | 0 | 0 | 2.393 | 0.21 | 0.09 | 0.16 | 0.22 | 0.22 | 0.14 |
| 4 | 155488821 | rs6056 | 1.206 | -0.168 | -0.823 | -1.573 | -0.007 | -0.001 | -0.001 | 0 | 0.2 | 0.08 | 0.15 | 0.22 | 0.2 | 0.14 |
| 4 | 155490832 | rs4681 | 1.28 | -0.243 | -0.768 | -1.597 | -0.052 | -0.039 | -0.039 | 0.006 | 0.19 | 0.02 | 0.15 | 0.22 | 0.2 | 0.14 |
| 4 | 155491759 | rs4220 | 1.22 | -0.238 | -0.757 | -1.595 | -0.007 | -0.002 | -0.002 | 0.001 | 0.2 | 0.08 | 0.15 | 0.22 | 0.2 | 0.14 |
| 4 | 155492224 | rs2227421 | 1.016 | -0.506 | -0.931 | -1.73 | 0.004 | 0.142 | 0.142 | 1.641 | 0.37 | 0.11 | 0.18 | 0.14 | 0.33 | 0.33 |
| 4 | 155493352 | rs1044291 | 1.023 | -0.496 | -0.926 | -1.725 | 0.007 | 0.142 | 0.142 | 1.706 | 0.37 | 0.11 | 0.18 | 0.14 | 0.33 | 0.33 |
| 4 | 155495533 | rs4463047 | 0.445 | -0.289 | -0.624 | -0.979 | 0.007 | -0.004 | -0.004 | 0 | 0.17 | 0.15 | 0.17 | 0.43 | 0.12 | 0.2 |
| 4 | 155504038 | rs28673871 | 0.896 | -0.274 | -0.702 | -0.981 | -0.027 | 0.013 | 0.013 | 0 | 0.14 | 0.02 | 0.34 | 0.11 | 0.18 | 0.24 |
| 4 | 155504948 | rs2070022 | 0.885 | -0.257 | -0.581 | -0.973 | -0.002 | 0.012 | 0.012 | 0 | 0.15 | 0.11 | 0.33 | 0.11 | 0.18 | 0.24 |
| 4 | 155508627 | rs2070018 | -1.043 | -0.179 | -0.454 | -0.983 | 0.004 | -0.001 | -0.001 | 0.088 | 0.92 | 0.92 | 0.85 | 0.93 | 0.87 | 0.85 |
| 4 | 155510314 | rs2070016 | 1.476 | -0.417 | -0.703 | -1.175 | -0.014 | -0.018 | -0.018 | 0.92 | 0.19 | 0.03 | 0.11 | 0.22 | 0.16 | 0.13 |
| 4 | 155511897 | rs2070011 | 0.414 | -0.448 | 0.086 | -0.809 | -0.01 | -0.034 | -0.034 | 0 | 0.68 | 0.79 | 0.64 | 0.5 | 0.64 | 0.61 |
| 4 | 155514879 | rs13109457 | -0.106 | -0.649 | 0.48 | -0.614 | -0.007 | 0.036 | 0.036 | 0 | 0.23 | 0.27 | 0.21 | 0.43 | 0.23 | 0.25 |
| 4 | 155520930 | rs7659024 | 0.067 | -0.064 | 0.395 | -0.216 | -0.009 | 0.085 | 0.085 | 0.383 | 0.21 | 0.34 | 0.21 | 0.44 | 0.22 | 0.24 |
| 4 | 155525276 | rs2066865 | 0.067 | -0.064 | 0.397 | -0.237 | -0.009 | 0.086 | 0.086 | 0.383 | 0.21 | 0.35 | 0.21 | 0.44 | 0.22 | 0.24 |
| 4 | 155525970 | rs1049636 | -0.342 | -0.026 | 0.34 | -0.303 | -0.007 | 0.005 | 0.005 | 0.355 | 0.77 | 0.81 | 0.52 | 0.81 | 0.68 | 0.6 |
| 4 | 155534408 | rs1800792 | 1.385 | -0.287 | -1.2 | -0.32 | -0.098 | 0.081 | 0.081 | 0.139 | 0.45 | 0.09 | 0.28 | 0.37 | 0.45 | 0.36 |
| 4 | 155537321 | rs12644950 | -0.22 | 1.21 | -0.769 | 0.368 | -0.006 | 0.067 | 0.067 | 0.336 | 0.2 | 0.3 | 0.21 | 0.44 | 0.22 | 0.24 |
| 4 | 155541289 | rs12648395 | 0.036 | 1.305 | -0.996 | 0.326 | -0.032 | 0.019 | 0.019 | 0.309 | 0.77 | 0.87 | 0.52 | 0.81 | 0.68 | 0.6 |
| 4 | 155545072 | rs7654093 | -0.732 | 0.78 | -0.345 | 1.037 | -0.008 | 0.091 | 0.091 | 0.382 | 0.2 | 0.35 | 0.21 | 0.45 | 0.22 | 0.24 |

| | | | | | | | | | | | | | | | | |
|---|---|---|---|---|---|---|---|---|---|---|---|---|---|---|---|---|
| 4 | 155548453 | rs13137121 | -0.294 | 1.057 | -0.646 | 0.898 | 0.005 | 0.006 | 0.006 | 0 | 0.48 | 0.55 | 0.6 | 0.42 | 0.42 | 0.6 |
| 4 | 155552487 | rs9790537 | -1.053 | 2.285 | -0.911 | 1.357 | -0.134 | -0.155 | -0.155 | 0.324 | 0.23 | 0.04 | 0.3 | 0.45 | 0.36 | 0.29 |
| 4 | 155559499 | rs4386585 | -1.75 | 1.913 | -0.93 | 1.106 | -0.081 | 0.024 | 0.024 | 0 | 0.38 | 0.15 | 0.34 | 0.66 | 0.47 | 0.43 |
| 4 | 155580418 | rs6828439 | -0.907 | 1.68 | -0.269 | 0.993 | 0.013 | 0.048 | 0.048 | 0 | 0.07 | 0.23 | 0.34 | 0.14 | 0.07 | 0.24 |
| 4 | 155609285 | rs17031902 | -0.024 | 1.791 | 0.427 | 0.969 | -0.028 | 0.002 | 0.002 | 0 | 0.11 | 0.01 | 0.05 | 0.09 | 0.1 | 0.12 |
| 4 | 155627814 | rs10517596 | -1.139 | 1.697 | 0.753 | 0.921 | 0.043 | 0.095 | 0.095 | 0.194 | 0.09 | 0.53 | 0.36 | 0.14 | 0.07 | 0.24 |
| 4 | 155632606 | rs1546370 | 1.411 | 1.881 | 0.139 | 0.941 | -0.073 | 0.023 | 0.023 | 0 | 0.61 | 0.83 | 0.65 | 0.35 | 0.53 | 0.58 |
| 4 | 155651896 | rs17031957 | -1.027 | 1.68 | 0.604 | 0.9 | 0.001 | -0.045 | -0.045 | 2.486 | 0.09 | 0.16 | 0.37 | 0.03 | 0.09 | 0.25 |
| 4 | 155652184 | rs17031960 | -0.643 | 1.975 | 0.413 | 1.121 | -0.046 | -0.101 | -0.101 | 0 | 0.15 | 0.02 | 0.22 | 0.43 | 0.24 | 0.2 |
| 4 | 155668710 | rs201824 | -1.554 | 1.966 | 0.307 | 1.104 | -0.025 | 0.005 | 0.005 | 0.231 | 0.31 | 0.2 | 0.31 | 0.66 | 0.41 | 0.36 |
| 4 | 155671626 | rs12507608 | -0.821 | 1.414 | 0.776 | 0.801 | -0.003 | 0.012 | 0.012 | 0 | 0.08 | 0.05 | 0.36 | 0.03 | 0.08 | 0.25 |
| 4 | 155672958 | rs17032000 | -1.192 | 1.374 | 0.835 | 0.799 | 0.003 | -0.048 | -0.048 | 0 | 0.09 | 0.19 | 0.37 | 0.03 | 0.08 | 0.24 |
| 4 | 155677315 | rs156500 | -1.213 | 1.485 | 0.052 | 0.911 | 0.014 | 0.041 | 0.041 | 0 | 0.14 | 0.19 | 0.08 | 0.23 | 0.15 | 0.16 |
| 4 | 155678345 | rs156501 | -1.457 | 1.502 | 0.084 | 0.956 | 0.008 | 0.011 | 0.011 | 0 | 0.21 | 0.27 | 0.43 | 0.25 | 0.23 | 0.39 |
| 4 | 155681012 | rs156553 | -1.412 | 1.515 | -0.289 | 0.955 | -0.016 | -0.013 | -0.013 | 0 | 0.21 | 0.17 | 0.43 | 0.26 | 0.24 | 0.4 |
| 4 | 155686380 | rs7662464 | -0.646 | 0.792 | -0.798 | 0.419 | -0.026 | 0.043 | 0.043 | 0.66 | 0.27 | 0.11 | 0.21 | 0.13 | 0.27 | 0.28 |
| 4 | 155701520 | rs11940892 | -0.668 | 0.792 | -0.797 | 0.419 | -0.025 | 0.045 | 0.045 | 0.68 | 0.27 | 0.11 | 0.21 | 0.13 | 0.26 | 0.28 |
| 4 | 155705144 | rs156550 | -1.739 | 0.275 | -0.772 | 0.427 | -0.006 | 0.21 | 0.21 | 0 | 0.52 | 0.92 | 0.68 | 0.75 | 0.42 | 0.58 |
| 4 | 155720274 | rs156502 | -1.739 | 0.275 | -0.772 | 0.427 | -0.006 | 0.21 | 0.21 | 0 | 0.52 | 0.92 | 0.68 | 0.75 | 0.42 | 0.58 |
| 4 | 155720349 | rs12651031 | -0.459 | 0.175 | 0.222 | 0.951 | 0.003 | 0.146 | 0.146 | 0 | 0.06 | 0.3 | 0.05 | 0.17 | 0.03 | 0.05 |
| 4 | 155739051 | rs7691720 | -0.173 | 0.122 | 0.348 | 0.947 | -0.01 | 0.075 | 0.075 | 0 | 0.15 | 0.17 | 0.38 | 0.44 | 0.12 | 0.24 |
| 4 | 155742578 | rs166682 | -1.823 | 0.209 | -0.196 | 0.905 | 0.019 | 0.21 | 0.21 | 0 | 0.52 | 0.89 | 0.68 | 0.75 | 0.42 | 0.58 |
| 4 | 155742865 | rs156569 | -0.615 | 0.622 | -0.127 | 0.952 | 0 | 0.019 | 0.019 | 0.707 | 0.28 | 0.21 | 0.22 | 0.13 | 0.27 | 0.28 |
| 4 | 155761481 | rs12054563 | -1.018 | -0.008 | -0.274 | 1.163 | -0.013 | 0.019 | 0.019 | 0 | 0.1 | 0.17 | 0.07 | 0.22 | 0.06 | 0.11 |
| 4 | 155764635 | rs17302908 | -0.326 | 0.872 | -0.725 | 1.09 | -0.148 | -0.037 | -0.037 | 0 | 0.27 | 0.05 | 0.31 | 0.24 | 0.44 | 0.35 |
| 4 | 155766258 | rs4449396 | -1.341 | 0.608 | -0.274 | 0.736 | 0.003 | 0.005 | 0.005 | 0.033 | 0.49 | 0.55 | 0.73 | 0.69 | 0.56 | 0.6 |
| 4 | 155772689 | rs12498643 | -1.025 | 0.314 | -0.042 | 1.689 | -0.027 | 0.128 | 0.128 | 0 | 0.14 | 0.23 | 0.3 | 0.48 | 0.09 | 0.13 |
| 4 | 155773993 | rs4696608 | 0.158 | 0.587 | -0.426 | 1.082 | 0.008 | 0.001 | 0.001 | 1.711 | 0.5 | 0.47 | 0.39 | 0.55 | 0.44 | 0.5 |
| 4 | 155775417 | rs987196 | -0.4 | 0.679 | -1.028 | 1.076 | -0.047 | -0.013 | -0.013 | 0.507 | 0.7 | 0.93 | 0.61 | 0.73 | 0.62 | 0.81 |
| 4 | 155777542 | rs6536031 | -0.027 | -0.174 | 0.446 | 1.092 | -0.004 | -0.009 | -0.009 | 0.366 | 0.64 | 0.54 | 0.55 | 0.71 | 0.61 | 0.8 |
| 4 | 155778844 | rs1559689 | -0.421 | -0.222 | 0.755 | 1.11 | -0.001 | 0.006 | 0.006 | 1.068 | 0.07 | 0.39 | 0.04 | 0.01 | 0.01 | 0.01 |
| 4 | 155780165 | rs1392554 | -0.573 | -0.224 | 0.785 | 1.11 | -0.011 | -0.02 | -0.02 | 0 | 0.69 | 0.85 | 0.57 | 0.6 | 0.62 | 0.76 |
| 4 | 155786387 | rs1500387 | 0.236 | -0.203 | 0.714 | 1.132 | -0.007 | -0.001 | -0.001 | 0 | 0.61 | 0.45 | 0.53 | 0.61 | 0.6 | 0.74 |
| 4 | 155790646 | rs17032149 | 0.914 | -0.785 | 0.328 | 1.4 | -0.006 | 0.008 | 0.008 | 0 | 0.11 | 0.14 | 0.11 | 0.18 | 0.05 | 0.08 |
| 4 | 155794220 | rs17375199 | 0.238 | 0.179 | 0.426 | 0.505 | -0.026 | -0.004 | -0.004 | 0 | 0.15 | 0.07 | 0.18 | 0.12 | 0.22 | 0.31 |
| 4 | 155794261 | rs17032154 | 0.779 | 0.042 | 0.428 | 0.447 | -0.009 | -0.032 | -0.032 | 0.273 | 0.43 | 0.21 | 0.35 | 0.49 | 0.38 | 0.43 |
| 4 | 155794591 | rs6813070 | -0.662 | 0.144 | 0.737 | 0.456 | 0.02 | -0.023 | -0.023 | 0 | 0.22 | 0.41 | 0.21 | 0.13 | 0.23 | 0.32 |
| 4 | 155796464 | rs1500383 | 0.1 | -0.143 | 0.719 | 0.512 | 0.005 | 0 | 0 | 0.098 | 0.48 | 0.45 | 0.38 | 0.6 | 0.39 | 0.49 |
| 4 | 155803664 | rs17032180 | 0.284 | -0.503 | 0.68 | 0.354 | -0.032 | 0 | 0 | 0 | 0.08 | 0.14 | 0.08 | 0.09 | 0.03 | 0.07 |
| 4 | 155811101 | rs1343937 | -0.164 | -0.577 | 0.666 | 0.17 | 0.06 | 0.01 | 0.01 | 0 | 0.48 | 0.62 | 0.35 | 0.58 | 0.28 | 0.47 |
| 4 | 155811145 | rs2404198 | 0.702 | -0.654 | 0.757 | 0.138 | 0.043 | -0.036 | -0.036 | 6.253 | 0.37 | 0.25 | 0.25 | 0.49 | 0.25 | 0.4 |
| 4 | 155820246 | rs17032207 | 0.012 | -0.546 | 0.627 | 0.171 | -0.04 | 0.005 | 0.005 | 0.001 | 0.1 | 0.25 | 0.09 | 0.1 | 0.03 | 0.07 |
| 4 | 155821855 | rs13101382 | -0.11 | 0.057 | -0.008 | -0.02 | -0.049 | 0.005 | 0.005 | 1.806 | 0.23 | 0.15 | 0.28 | 0.15 | 0.4 | 0.34 |
| 4 | 155826061 | rs10033629 | -1.465 | -0.284 | -0.081 | -0.054 | -0.064 | -0.01 | -0.01 | 0.001 | 0.73 | 0.93 | 0.63 | 0.73 | 0.64 | 0.8 |
| 4 | 155830634 | rs10002121 | 0.41 | -0.222 | 0.102 | -0.004 | 0.006 | 0 | 0 | 0 | 0.46 | 0.54 | 0.35 | 0.57 | 0.28 | 0.43 |
| 4 | 155833011 | rs2896784 | -0.309 | 0.275 | -0.145 | 0.199 | -0.05 | 0.002 | 0.002 | 1.924 | 0.22 | 0.12 | 0.26 | 0.15 | 0.34 | 0.35 |
| 4 | 155835510 | rs6536038 | -0.998 | 0.26 | -0.168 | 0.205 | -0.013 | 0.003 | 0.003 | 2.95 | 0.3 | 0.23 | 0.34 | 0.24 | 0.37 | 0.43 |
| 4 | 155849300 | rs4691385 | 0.509 | 0.926 | -0.67 | 0.301 | -0.022 | -0.013 | -0.013 | 0.083 | 0.05 | 0.01 | 0.12 | 0.05 | 0.16 | 0.04 |
| 4 | 155854976 | rs12498703 | -0.594 | 0.353 | -0.82 | 0.302 | -0.083 | -0.009 | -0.009 | 0 | 0.26 | 0.1 | 0.36 | 0.17 | 0.37 | 0.38 |
| 4 | 155861540 | rs4691329 | -0.967 | -0.191 | -1.184 | 0.007 | -0.011 | 0.059 | 0.059 | 0.053 | 0.44 | 0.28 | 0.45 | 0.62 | 0.48 | 0.66 |
| 4 | 155861812 | rs17032275 | 0.546 | -0.442 | -1.315 | -0.121 | -0.07 | -0.013 | -0.013 | 0.745 | 0.13 | 0.51 | 0.12 | 0.11 | 0.04 | 0.1 |
| 4 | 155867729 | rs2124656 | -1.215 | -0.244 | -1.861 | -0.102 | 0 | 0.002 | 0.002 | 0.116 | 0.54 | 0.57 | 0.55 | 0.74 | 0.52 | 0.75 |
| 4 | 155886241 | rs17032316 | 0.731 | -0.004 | -3.167 | 0.194 | 0.007 | -0.011 | -0.011 | 0 | 0.18 | 0.15 | 0.08 | 0.44 | 0.11 | 0.28 |
| 4 | 155891502 | rs4691555 | -1.005 | 0.407 | -3.747 | -0.05 | -0.097 | 0.073 | 0.073 | 0 | 0.61 | 0.96 | 0.57 | 0.72 | 0.51 | 0.75 |
| 4 | 155900720 | rs1456457 | -0.334 | 0.632 | -3.232 | 0.359 | -0.078 | -0.039 | -0.039 | 0 | 0.19 | 0.01 | 0.22 | 0.13 | 0.23 | 0.33 |
| 4 | 155903741 | rs1378965 | -0.294 | 0.377 | -2.605 | 0.178 | -0.022 | 0.037 | 0.037 | 0 | 0.35 | 0.08 | 0.38 | 0.27 | 0.36 | 0.24 |
| 4 | 155914904 | rs11729306 | -0.107 | 0.53 | -2.447 | 0.008 | -0.062 | 0.089 | 0.089 | 0 | 0.3 | 0.05 | 0.22 | 0.19 | 0.33 | 0.22 |
| 4 | 155922530 | rs6845282 | -0.234 | -0.724 | -2.361 | 0.56 | 0.097 | -0.07 | -0.07 | 0 | 0.42 | 0.89 | 0.28 | 0.48 | 0.24 | 0.42 |
| 4 | 155927800 | rs1456459 | 0.274 | -1.336 | -0.886 | 0.043 | -0.009 | -0.013 | -0.013 | 0 | 0.77 | 0.76 | 0.9 | 0.85 | 0.84 | 0.75 |
| 4 | 155932934 | rs6829756 | 0.995 | -2.081 | -0.522 | 0.259 | -0.029 | 0.017 | 0.017 | 0 | 0.62 | 0.52 | 0.63 | 0.51 | 0.72 | 0.66 |
| 4 | 155933753 | rs11099970 | 1.1 | -2.215 | -0.914 | 0.046 | -0.008 | -0.007 | -0.007 | 1.248 | 0.33 | 0.32 | 0.48 | 0.41 | 0.52 | 0.4 |
| 4 | 155938012 | rs10009064 | 0.864 | -2.095 | -0.805 | 0.07 | -0.016 | -0.004 | -0.004 | 0.023 | 0.13 | 0.19 | 0.07 | 0.18 | 0.1 | 0.07 |
| 4 | 155947269 | rs9998561 | 0.705 | -2.062 | -0.881 | -0.197 | 0 | -0.01 | -0.01 | 0 | 0.11 | 0.11 | 0.03 | 0.01 | 0.07 | 0.09 |
| 4 | 155947430 | rs7436102 | -1.489 | -1.918 | -1.105 | -0.093 | -0.013 | 0.012 | 0.012 | 0 | 0.9 | 0.99 | 0.94 | 0.92 | 0.9 | 0.84 |
| 4 | 155951509 | rs7656217 | 0.695 | -2.134 | -0.659 | -0.063 | -0.008 | -0.015 | -0.015 | 0 | 0.3 | 0.3 | 0.15 | 0.1 | 0.21 | 0.26 |
| 4 | 155957348 | rs10020550 | -0.564 | -1.245 | -0.592 | 0.306 | -0.003 | -0.008 | -0.008 | 0 | 0.65 | 0.73 | 0.5 | 0.58 | 0.43 | 0.57 |
| 4 | 155960819 | rs1378964 | 0.011 | -1.137 | -0.428 | 0.438 | 0.011 | 0.003 | 0.003 | 0 | 0.51 | 0.59 | 0.44 | 0.51 | 0.33 | 0.5 |
| 4 | 155967231 | rs10085000 | 0.01 | -1.194 | -0.403 | 0.421 | 0.037 | 0.008 | 0.008 | 0 | 0.51 | 0.65 | 0.44 | 0.51 | 0.32 | 0.5 |
| 4 | 155967748 | rs10013268 | -0.36 | -1.288 | -0.324 | 0.464 | -0.001 | 0 | 0 | 0 | 0.63 | 0.63 | 0.3 | 0.39 | 0.43 | 0.55 |

| | | | | | | | | | | | | | | | | |
|---|---|---|---|---|---|---|---|---|---|---|---|---|---|---|---|---|
| 4 | 155968185 | rs876450 | -0.479 | -2.356 | -0.596 | 0.095 | -0.076 | -0.041 | -0.041 | 0 | 0.34 | 0.52 | 0.14 | 0.29 | 0.17 | 0.26 |
| 4 | 155968437 | rs17032387 | -0.092 | -1.044 | -0.896 | 0.719 | 0.023 | -0.098 | -0.098 | 0 | 0.26 | 0.11 | 0.13 | 0.49 | 0.23 | 0.26 |
| 4 | 155969377 | rs1869965 | -0.53 | -0.518 | -0.305 | 0.94 | 0.146 | 0.099 | 0.099 | 0.369 | 0.79 | 0.62 | 0.42 | 0.7 | 0.63 | 0.66 |
| 4 | 155969710 | rs17032390 | 0.405 | -0.804 | -0.838 | 0.84 | 0.016 | -0.012 | -0.012 | 0 | 0.1 | 0.09 | 0.04 | 0.2 | 0.04 | 0.14 |
| 4 | 155979144 | rs1869966 | -0.262 | -0.24 | -1.105 | 0.555 | -0.018 | 0.014 | 0.014 | 0 | 0.26 | 0.22 | 0.15 | 0.15 | 0.32 | 0.26 |
| 4 | 155985107 | rs7668866 | 0.702 | -0.237 | -1.293 | 0.498 | -0.074 | 0.04 | 0.04 | 0 | 0.14 | 0.03 | 0.11 | 0.05 | 0.24 | 0.22 |
| 4 | 155989391 | rs6832923 | -0.918 | -0.237 | -0.932 | 0.743 | 0.007 | 0.003 | 0.003 | 0 | 0.65 | 0.55 | 0.45 | 0.57 | 0.65 | 0.55 |
| 4 | 155995373 | rs11735554 | 1.198 | -0.604 | -1.415 | 0.68 | -0.04 | -0.113 | -0.113 | 2.103 | 0.2 | 0.03 | 0.19 | 0.32 | 0.22 | 0.14 |
| 4 | 155995695 | rs11727277 | 0.43 | -0.586 | -1.372 | 0.722 | -0.022 | -0.087 | -0.087 | 0 | 0.28 | 0.06 | 0.22 | 0.32 | 0.29 | 0.15 |
| 4 | 156002908 | rs4690860 | 0.882 | -0.431 | -1.121 | 0.588 | -0.033 | -0.139 | -0.139 | 0 | 0.22 | 0.03 | 0.2 | 0.42 | 0.22 | 0.14 |
| 4 | 156007879 | rs17032433 | -0.754 | -0.46 | -0.66 | 0.683 | -0.001 | 0 | 0 | 0 | 0.64 | 0.41 | 0.44 | 0.63 | 0.65 | 0.55 |
| 4 | 156008037 | rs10008264 | 0.185 | -0.417 | -0.695 | 0.656 | -0.01 | 0.066 | 0.066 | 0 | 0.4 | 0.26 | 0.23 | 0.21 | 0.43 | 0.4 |
| 4 | 156016684 | rs6813291 | -0.416 | -0.266 | -0.431 | 0.881 | 0 | -0.003 | -0.003 | 0 | 0.65 | 0.63 | 0.46 | 0.58 | 0.64 | 0.52 |
| 4 | 156018884 | rs6826870 | -0.044 | 0.121 | -0.621 | 0.891 | -0.001 | -0.002 | -0.002 | 0 | 0.61 | 0.49 | 0.39 | 0.52 | 0.58 | 0.55 |
| 4 | 156024958 | rs6831797 | 0.152 | 0.136 | -0.652 | 1.107 | -0.03 | 0.064 | 0.064 | 0.539 | 0.85 | 0.6 | 0.86 | 0.79 | 0.87 | 0.86 |
| 4 | 156029149 | rs1378957 | 0.009 | 0.876 | -1.611 | 1.286 | -0.014 | -0.011 | -0.011 | 0 | 0.23 | 0.22 | 0.48 | 0.32 | 0.29 | 0.27 |
| 4 | 156038542 | rs12650515 | -0.132 | 0.251 | -1.821 | 1.034 | 0.002 | -0.072 | -0.072 | 0.295 | 0.2 | 0.04 | 0.15 | 0.31 | 0.16 | 0.1 |
| 4 | 156045227 | rs6848512 | -0.07 | 0.384 | -1.815 | 0.595 | 0.004 | 0.041 | 0.041 | 0 | 0.51 | 0.78 | 0.54 | 0.51 | 0.51 | 0.46 |
| 4 | 156051299 | rs17304378 | 0.331 | -0.188 | -1.312 | 0.126 | 0.006 | 0.021 | 0.021 | 4.732 | 0.09 | 0.03 | 0.04 | 0 | 0.08 | 0.02 |
| 4 | 156055565 | rs17032501 | 0.49 | -0.212 | -0.729 | 0.133 | 0.011 | 0.018 | 0.018 | 0 | 0.09 | 0.08 | 0.17 | 0.14 | 0.09 | 0.13 |
| 4 | 156055961 | rs10027655 | -0.861 | -0.453 | -0.721 | -0.773 | -0.017 | 0.13 | 0.13 | 0 | 0.55 | 0.8 | 0.68 | 0.98 | 0.52 | 0.69 |
| 4 | 156057352 | rs4323056 | -1.634 | -0.532 | -0.767 | -0.807 | -0.148 | 0.233 | 0.233 | 0.032 | 0.66 | 0.96 | 0.73 | 0.98 | 0.59 | 0.71 |
| 4 | 156062435 | rs17032524 | 0.185 | -1.069 | -0.453 | -0.361 | 0 | 0.004 | 0.004 | 0 | 0.13 | 0.16 | 0.12 | 0.27 | 0.11 | 0.24 |
| 4 | 156069879 | rs2880418 | -1.185 | -1.273 | -0.067 | -0.505 | -0.002 | -0.052 | -0.052 | 1.419 | 0.55 | 0.71 | 0.68 | 0.98 | 0.52 | 0.7 |
| 4 | 156074074 | rs13138293 | 0.842 | -1.423 | -0.317 | -0.534 | -0.066 | -0.128 | -0.128 | 0.337 | 0.26 | 0.07 | 0.36 | 0.56 | 0.32 | 0.32 |
| 4 | 156081090 | rs1316749 | 1.381 | -1.373 | -0.053 | -0.603 | -0.001 | 0 | 0 | 0 | 0.09 | 0.06 | 0.16 | 0.14 | 0.09 | 0.13 |
| 4 | 156081675 | rs9790645 | -0.717 | -1.409 | -0.117 | -0.64 | 0.015 | 0.089 | 0.089 | 0 | 0.42 | 0.35 | 0.48 | 0.83 | 0.41 | 0.56 |
| 4 | 156083037 | rs17376560 | 0.781 | -1.279 | -0.515 | -0.542 | -0.062 | -0.121 | -0.121 | 0 | 0.25 | 0.07 | 0.34 | 0.57 | 0.29 | 0.33 |
| 4 | 156087185 | rs10030872 | -1.225 | -1.281 | -0.23 | -0.35 | 0.005 | 0.004 | 0.004 | 0.66 | 0.28 | 0.73 | 0.26 | 0.14 | 0.19 | 0.16 |
| 4 | 156091935 | rs1317400 | -0.973 | -1.295 | -0.057 | -0.34 | -0.058 | 0.022 | 0.022 | 2.41 | 0.24 | 0.45 | 0.18 | 0.26 | 0.2 | 0.25 |
| 4 | 156093651 | rs2880414 | -1.454 | -1.202 | -0.446 | -0.346 | 0.031 | -0.012 | -0.012 | 0.272 | 0.36 | 0.76 | 0.37 | 0.41 | 0.3 | 0.39 |
| 4 | 156103755 | rs6854846 | -0.245 | -0.832 | -0.455 | -0.404 | -0.019 | -0.057 | -0.057 | 2.094 | 0.17 | 0.24 | 0.07 | 0 | 0.11 | 0.02 |
| 4 | 156111912 | rs13111894 | 1.226 | -0.373 | -0.977 | -0.049 | -0.028 | -0.032 | -0.032 | 0 | 0.06 | 0 | 0.04 | 0.14 | 0.06 | 0.03 |
| 4 | 156112853 | rs10517606 | 0.045 | -0.523 | -0.727 | -0.343 | 0.005 | -0.008 | -0.008 | 0 | 0.36 | 0.38 | 0.51 | 0.69 | 0.38 | 0.46 |
| 4 | 156115288 | rs13152353 | -0.531 | -0.491 | -0.701 | -0.78 | 0.002 | -0.088 | -0.088 | 1.058 | 0.45 | 0.43 | 0.63 | 0.93 | 0.52 | 0.78 |
| 4 | 156123035 | rs13120088 | -0.149 | -0.018 | -0.963 | -0.159 | 0.057 | 0.006 | 0.006 | 0.013 | 0.51 | 0.7 | 0.45 | 0.52 | 0.39 | 0.38 |
| 4 | 156124223 | rs6849115 | -0.033 | 0.033 | -0.978 | -0.183 | 0.009 | 0.069 | 0.069 | 0.8 | 0.15 | 0.24 | 0.29 | 0.41 | 0.23 | 0.42 |
| 4 | 156125333 | rs12649641 | -0.466 | -0.292 | -0.72 | 0.049 | 0.058 | 0.006 | 0.006 | 0 | 0.51 | 0.7 | 0.45 | 0.52 | 0.39 | 0.38 |
| 4 | 156130948 | rs17376826 | -0.21 | -0.157 | -0.48 | -0.155 | -0.011 | -0.032 | -0.032 | 0 | 0.1 | 0.14 | 0.07 | 0 | 0.07 | 0.02 |
| 4 | 156133712 | rs1574175 | -0.328 | 0.186 | -0.607 | -0.146 | 0 | -0.008 | -0.008 | 3.276 | 0.12 | 0.15 | 0.07 | 0.06 | 0.15 | 0.18 |
| 4 | 156135676 | rs1047214 | -1.174 | 0.388 | -1.673 | -0.582 | -0.057 | 0.172 | 0.172 | 0.158 | 0.56 | 0.94 | 0.5 | 0.81 | 0.54 | 0.56 |
| 4 | 156138372 | rs17304901 | 1.196 | 0.323 | -2.18 | -1.003 | -0.047 | 0.119 | 0.119 | 0 | 0.25 | 0.02 | 0.21 | 0.05 | 0.27 | 0.19 |
| 4 | 156139467 | rs6846703 | 0.517 | 0.45 | -1.044 | 0.368 | -0.003 | 0.109 | 0.109 | 0 | 0.11 | 0.16 | 0.21 | 0.52 | 0.09 | 0.18 |
| 4 | 156143159 | rs6832844 | 0.294 | 0.421 | -0.088 | 0.078 | 0.08 | 0.036 | 0.036 | 0 | 0.19 | 0.33 | 0.14 | 0.19 | 0.22 | 0.18 |
| 4 | 156149811 | rs2342654 | 0.496 | 0.351 | 0.038 | 0.028 | 0.011 | -0.034 | -0.034 | 0 | 0.12 | 0.23 | 0.08 | 0.06 | 0.15 | 0.18 |
| 4 | 156154314 | rs2342649 | 0.372 | 0.399 | 0.077 | 0.059 | 0.042 | 0.017 | 0.017 | 0 | 0.18 | 0.26 | 0.13 | 0.19 | 0.22 | 0.18 |
| 4 | 156163904 | rs1822806 | -0.388 | 0.133 | -0.083 | -0.006 | -0.004 | -0.036 | -0.036 | 0 | 0.23 | 0.24 | 0.15 | 0.06 | 0.23 | 0.2 |
| 4 | 156174594 | rs11099996 | 0.375 | 0.135 | 0.016 | 0.372 | -0.003 | 0.066 | 0.066 | 0 | 0.11 | 0.16 | 0.26 | 0.64 | 0.09 | 0.19 |
| 4 | 156185574 | rs17032732 | 0.034 | -0.094 | 0.32 | 0.303 | -0.033 | -0.013 | -0.013 | 0 | 0.15 | 0.26 | 0.09 | 0.09 | 0.09 | 0.07 |
| 4 | 156189220 | rs17032744 | 0.326 | -0.116 | 0.037 | 0.57 | -0.055 | -0.059 | -0.059 | 0 | 0.17 | 0.02 | 0.13 | 0.21 | 0.22 | 0.29 |
| 4 | 156189953 | rs991097 | -1.394 | -0.152 | 0.101 | 0.429 | -0.031 | 0.035 | 0.035 | 0 | 0.38 | 0.84 | 0.42 | 0.72 | 0.34 | 0.31 |
| 4 | 156200509 | rs35125259 | 0.257 | 0.158 | 0.482 | 0.576 | 0.079 | 0.025 | 0.025 | 0 | 0.17 | 0.45 | 0.13 | 0.15 | 0.2 | 0.16 |
| 4 | 156216726 | rs1078275 | -0.543 | 0.535 | 0.426 | 0.898 | 0.015 | 0.019 | 0.019 | 0 | 0.22 | 0.28 | 0.18 | 0.34 | 0.33 | 0.37 |
| 4 | 156217344 | rs17377092 | 0.869 | 0.417 | 0.376 | 0.774 | 0.029 | 0.048 | 0.048 | 0.879 | 0.13 | 0.04 | 0.07 | 0 | 0.09 | 0.02 |
| 4 | 156227224 | rs17032793 | -0.315 | 0.469 | 0.386 | 0.793 | 0.04 | 0.001 | 0.001 | 0 | 0.19 | 0.54 | 0.13 | 0.15 | 0.2 | 0.16 |
| 4 | 156237313 | rs11726054 | -1.067 | 0.424 | 0.411 | 0.829 | 0.017 | 0.005 | 0.005 | 0 | 0.35 | 0.52 | 0.26 | 0.36 | 0.44 | 0.44 |
| 4 | 156238370 | rs1822804 | -2.251 | 0.413 | 0.328 | 0.804 | -0.086 | 0.308 | 0.308 | 0 | 0.56 | 0.96 | 0.55 | 0.93 | 0.56 | 0.57 |
| 4 | 156239090 | rs17032814 | 0.825 | 0.437 | 0.926 | 0.858 | 0 | 0.009 | 0.009 | 0 | 0.13 | 0.16 | 0.08 | 0.15 | 0.12 | 0.13 |
| 4 | 156245641 | rs750068 | -0.533 | 0.107 | 0.975 | 0.888 | 0.024 | 0.08 | 0.08 | 0 | 0.22 | 0.61 | 0.32 | 0.59 | 0.22 | 0.2 |
| 4 | 156264133 | rs10034144 | 0.714 | 0.147 | 0.576 | 0.932 | 0.004 | 0 | 0 | 0 | 0.15 | 0.14 | 0.11 | 0.15 | 0.2 | 0.16 |
| 4 | 156265409 | rs6817266 | 0.714 | 0.147 | 0.576 | 0.932 | 0.017 | 0.005 | 0.005 | 0 | 0.15 | 0.19 | 0.11 | 0.15 | 0.2 | 0.16 |
| 4 | 156273768 | rs2305050 | -1.776 | 0.094 | 0.27 | 0.871 | 0.016 | 0.124 | 0.124 | 0 | 0.52 | 0.78 | 0.53 | 0.79 | 0.56 | 0.52 |
| 4 | 156285170 | rs6842102 | -1.776 | 0.154 | 0.292 | 0.942 | 0.016 | 0.124 | 0.124 | 0 | 0.52 | 0.78 | 0.53 | 0.79 | 0.56 | 0.52 |
| 4 | 156289900 | rs3733390 | -0.675 | 0.204 | -0.307 | 0.982 | -0.013 | -0.077 | -0.077 | 0.078 | 0.31 | 0.16 | 0.4 | 0.64 | 0.35 | 0.36 |
| 4 | 156289917 | rs3733391 | 0.184 | 0.758 | 0.188 | 1.315 | -0.066 | -0.058 | -0.058 | 0 | 0.16 | 0.02 | 0.13 | 0.21 | 0.24 | 0.29 |
| 4 | 156297060 | rs17377679 | 0.094 | 0.477 | 0.762 | 1.079 | 0.009 | 0.002 | 0.002 | 0 | 0.15 | 0.15 | 0.11 | 0.15 | 0.2 | 0.16 |
| 4 | 156300040 | rs3775255 | -1.137 | 0.503 | 0.3 | 0.953 | -0.007 | -0.022 | -0.022 | 0 | 0.24 | 0.75 | 0.15 | 0.28 | 0.2 | 0.21 |
| 4 | 156311510 | rs7663620 | -0.561 | -0.191 | -0.098 | -0.049 | -0.072 | 0.099 | 0.099 | 0 | 0.36 | 0.11 | 0.43 | 0.82 | 0.44 | 0.58 |
| 4 | 156321133 | rs6847136 | -1.119 | -0.367 | 1.161 | -0.136 | 0.039 | 0.034 | 0.034 | 5.487 | 0.06 | 0.47 | 0.04 | 0 | 0 | 0 |

| | | | | | | | | | | | | | | | | |
|---|---|---|---|---|---|---|---|---|---|---|---|---|---|---|---|---|
| 4 | 156322989 | rs11100000 | -0.784 | -0.267 | 1.151 | -0.142 | -0.004 | 0.019 | 0.019 | 0 | 0.38 | 0.33 | 0.44 | 0.86 | 0.42 | 0.56 |
| 4 | 156325539 | rs1460128 | -0.967 | -0.268 | 1.029 | -0.275 | 0 | -0.101 | -0.101 | 1.275 | 0.42 | 0.49 | 0.45 | 0.91 | 0.43 | 0.6 |
| 4 | 156346580 | rs9991561 | -2.047 | -0.385 | 1.604 | 0.224 | -0.044 | 0.114 | 0.114 | 2.196 | 0.1 | 0.54 | 0.05 | 0.11 | 0.01 | 0.05 |
| 4 | 156353245 | rs4690966 | -1.429 | -0.33 | 1.773 | 0.84 | -0.012 | -0.028 | -0.028 | 0 | 0.39 | 0.78 | 0.26 | 0.41 | 0.39 | 0.48 |
| 4 | 156355515 | rs4691631 | -0.138 | -0.183 | 1.35 | 0.929 | -0.095 | -0.044 | -0.044 | 0 | 0.26 | 0.05 | 0.19 | 0.31 | 0.37 | 0.43 |
| 4 | 156370726 | rs9996409 | -1.605 | -0.181 | 1.673 | 0.936 | -0.017 | 0.134 | 0.134 | 1.204 | 0.08 | 0.34 | 0.04 | 0.11 | 0.01 | 0.05 |
| 4 | 156378568 | rs4691643 | -0.511 | -0.117 | 1.121 | 0.86 | -0.033 | -0.038 | -0.038 | 0.772 | 0.28 | 0.21 | 0.21 | 0.38 | 0.37 | 0.48 |
| 4 | 156383288 | rs12643194 | -1.263 | -0.166 | 0.464 | 1.124 | -0.004 | 0.112 | 0.112 | 0 | 0.08 | 0.08 | 0.25 | 0.5 | 0.08 | 0.16 |
| 4 | 156393197 | rs1517293 | -0.579 | -0.039 | -0.064 | 0.272 | -0.006 | -0.018 | -0.018 | 0 | 0.31 | 0.26 | 0.52 | 0.62 | 0.34 | 0.4 |
| 4 | 156401693 | rs17033050 | 0.957 | 0.242 | -0.538 | 0.224 | -0.042 | -0.052 | -0.052 | 0 | 0.09 | 0.01 | 0.11 | 0.24 | 0.17 | 0.23 |
| 4 | 156410972 | rs10013797 | 0.253 | 0.178 | -0.436 | 0.08 | -0.025 | -0.058 | -0.058 | 0 | 0.7 | 0.87 | 0.46 | 0.39 | 0.66 | 0.6 |
| 4 | 156414715 | rs1155097 | 0.111 | -0.347 | -0.154 | -0.229 | -0.001 | 0.033 | 0.033 | 0 | 0.16 | 0.23 | 0.12 | 0.24 | 0.18 | 0.26 |
| 4 | 156415404 | rs717695 | 0.914 | -0.249 | -0.261 | -0.308 | -0.016 | 0.077 | 0.077 | 0 | 0.56 | 0.7 | 0.33 | 0.16 | 0.48 | 0.34 |
| 4 | 156417909 | rs6837085 | 1.375 | -0.474 | 0.428 | -0.455 | 0.001 | 0.037 | 0.037 | 0.081 | 0.5 | 0.43 | 0.31 | 0.13 | 0.48 | 0.34 |
| 4 | 156434164 | rs7440928 | -0.706 | -0.421 | -0.08 | -0.493 | 0.016 | -0.077 | -0.077 | 0 | 0.84 | 0.95 | 0.76 | 0.49 | 0.89 | 0.79 |
| 4 | 156438083 | rs1517294 | 2.126 | -0.579 | 0.08 | -0.812 | -0.02 | 0.161 | 0.161 | 0.549 | 0.42 | 0.18 | 0.26 | 0.14 | 0.42 | 0.28 |
| 4 | 156442666 | rs13124618 | 0.609 | -0.586 | 0.492 | -0.709 | -0.002 | 0 | 0 | 0 | 0.63 | 0.62 | 0.58 | 0.39 | 0.67 | 0.49 |
| 4 | 156443877 | rs1400623 | 0.577 | -0.538 | 0.403 | -0.699 | -0.018 | 0.022 | 0.022 | 0 | 0.18 | 0.07 | 0.13 | 0.1 | 0.22 | 0.31 |
| 4 | 156446231 | rs17033109 | 0.563 | -0.426 | 0.568 | -0.539 | -0.047 | -0.037 | -0.037 | 0.151 | 0.14 | 0.05 | 0.29 | 0.22 | 0.23 | 0.2 |
| 4 | 156449830 | rs17312460 | 1.89 | -0.918 | 0.027 | -1.282 | -0.026 | 0.112 | 0.112 | 0.026 | 0.34 | 0.15 | 0.22 | 0.14 | 0.33 | 0.23 |
| 4 | 156450200 | rs7436887 | -1.333 | -0.728 | 0.051 | -1.31 | -0.007 | 0.087 | 0.087 | 0 | 0.5 | 0.8 | 0.51 | 0.33 | 0.54 | 0.56 |
| 4 | 156455542 | rs2138741 | -0.373 | -0.609 | 0.239 | -1.23 | 0.021 | 0.058 | 0.058 | 0.304 | 0.22 | 0.48 | 0.35 | 0.28 | 0.23 | 0.22 |
| 4 | 156460954 | rs7671138 | 0.373 | -0.488 | 0.133 | -1.532 | -0.002 | -0.013 | -0.013 | 0.047 | 0.27 | 0.27 | 0.18 | 0.06 | 0.32 | 0.37 |
| 4 | 156470555 | rs10016993 | 0.926 | -0.91 | 0.405 | -1.714 | -0.001 | -0.09 | -0.09 | 0 | 0.35 | 0.41 | 0.22 | 0.05 | 0.31 | 0.22 |
| 4 | 156472766 | rs6850722 | 0.141 | -0.687 | 0.053 | -1.616 | -0.022 | 0.109 | 0.109 | 0 | 0.46 | 0.74 | 0.5 | 0.25 | 0.46 | 0.32 |
| 4 | 156482330 | rs7659237 | 0.802 | -0.573 | 0.031 | -1.477 | 0.002 | 0.003 | 0.003 | 0 | 0.11 | 0.15 | 0.26 | 0.21 | 0.15 | 0.13 |
| 4 | 156484198 | rs2879909 | 0.62 | -0.576 | 0.075 | -1.482 | 0.012 | 0.019 | 0.019 | 0.043 | 0.12 | 0.22 | 0.26 | 0.21 | 0.15 | 0.13 |
| 4 | 156492983 | rs6854265 | 0.758 | -0.824 | 0.409 | -1.393 | -0.013 | 0.023 | 0.023 | 0 | 0.4 | 0.38 | 0.45 | 0.23 | 0.44 | 0.32 |
| 4 | 156499985 | rs1032051 | 0.415 | -0.824 | 0.283 | -1.379 | -0.009 | 0.033 | 0.033 | 0 | 0.45 | 0.62 | 0.48 | 0.24 | 0.45 | 0.32 |
| 4 | 156503440 | rs1483041 | -0.483 | -0.493 | 0.127 | -1.247 | 0 | 0.041 | 0.041 | 0 | 0.5 | 0.4 | 0.51 | 0.73 | 0.53 | 0.61 |
| 4 | 156511302 | rs1483037 | -0.379 | -0.595 | -0.322 | -0.994 | -0.006 | 0.011 | 0.011 | 0 | 0.18 | 0.18 | 0.26 | 0.63 | 0.1 | 0.22 |
| 4 | 156515995 | rs2625247 | -0.5 | -0.218 | -0.725 | -0.956 | -0.014 | 0.05 | 0.05 | 0.156 | 0.84 | 0.8 | 0.72 | 0.36 | 0.89 | 0.8 |
| 4 | 156517289 | rs1466516 | -0.11 | 0.261 | -1.267 | -1.348 | -0.034 | 0.093 | 0.093 | 0 | 0.33 | 0.13 | 0.24 | 0.11 | 0.42 | 0.41 |
| 4 | 156518902 | rs2625248 | 0.488 | 0.407 | -1.134 | -1.612 | -0.002 | 0.063 | 0.063 | 2.052 | 0.31 | 0.25 | 0.21 | 0.02 | 0.29 | 0.15 |
| 4 | 156519123 | rs2625249 | 0.194 | 0.252 | -0.837 | -1.31 | 0.024 | 0.145 | 0.145 | 0 | 0.51 | 0.71 | 0.54 | 0.23 | 0.55 | 0.38 |
| 4 | 156519199 | rs9998255 | -0.442 | 0.213 | -0.2 | -0.847 | -0.001 | 0.001 | 0.001 | 0 | 0.51 | 0.52 | 0.44 | 0.2 | 0.46 | 0.48 |
| 4 | 156523316 | rs13110106 | -1.495 | -0.136 | 0.257 | 0.306 | 0.009 | -0.019 | -0.019 | 0 | 0.38 | 0.67 | 0.29 | 0.58 | 0.35 | 0.56 |
| 4 | 156524118 | rs6813207 | -0.329 | -0.004 | 0.445 | 0.095 | 0.001 | -0.013 | -0.013 | 0 | 0.08 | 0.17 | 0.11 | 0.05 | 0.07 | 0.09 |
| 4 | 156524415 | rs4691787 | -1.294 | -0.054 | 0.64 | 0.039 | -0.006 | -0.034 | -0.034 | 0 | 0.25 | 0.25 | 0.19 | 0.07 | 0.19 | 0.31 |
| 4 | 156526246 | rs4691022 | -1.544 | -0.047 | 0.609 | 0.046 | -0.017 | -0.111 | -0.111 | 0 | 0.28 | 0.45 | 0.22 | 0.07 | 0.24 | 0.35 |
| 4 | 156540258 | rs7680331 | -0.769 | 0.228 | 0.246 | 0.595 | -0.005 | 0.121 | 0.121 | 0 | 0.13 | 0.23 | 0.13 | 0.53 | 0.12 | 0.2 |
| 4 | 156547554 | rs2062177 | -0.448 | 0.101 | -0.294 | 0.293 | 0.013 | 0.032 | 0.032 | 0 | 0.21 | 0.09 | 0.11 | 0.02 | 0.14 | 0.3 |
| 4 | 156550942 | rs727773 | -0.996 | 0.247 | -0.136 | 0.514 | -0.001 | 0.002 | 0.002 | 0 | 0.35 | 0.32 | 0.26 | 0.56 | 0.32 | 0.51 |
| 4 | 156555556 | rs7692799 | -1.001 | 0.036 | -0.232 | 0.419 | -0.005 | 0.003 | 0.003 | 0 | 0.4 | 0.53 | 0.23 | 0.54 | 0.29 | 0.52 |
| 4 | 156555835 | rs7693386 | -0.914 | 0.273 | -0.548 | 0.569 | 0.005 | -0.015 | -0.015 | 0.236 | 0.37 | 0.15 | 0.21 | 0.63 | 0.31 | 0.59 |
| 4 | 156557255 | rs12642995 | -0.589 | 0.564 | -0.55 | -0.346 | 0 | -0.024 | -0.024 | 0.252 | 0.16 | 0.15 | 0.18 | 0.01 | 0.17 | 0.12 |
| 4 | 156558695 | rs2705434 | 0.476 | 0.177 | -1.049 | -0.716 | 0.016 | 0.039 | 0.039 | 0 | 0.31 | 0.18 | 0.24 | 0.14 | 0.31 | 0.21 |
| 4 | 156561632 | rs4691036 | -0.295 | 0.7 | -0.901 | -0.536 | 0 | -0.009 | -0.009 | 0 | 0.21 | 0.21 | 0.4 | 0.06 | 0.2 | 0.29 |
| 4 | 156563407 | rs2128650 | -0.424 | 0.533 | -0.816 | -0.135 | 0.026 | 0.102 | 0.102 | 0 | 0.8 | 0.6 | 0.79 | 0.51 | 0.76 | 0.73 |
| 4 | 156569802 | rs2705462 | -0.182 | 0.636 | -1.598 | -0.757 | -0.088 | 0.094 | 0.094 | 0.13 | 0.24 | 0.06 | 0.39 | 0.06 | 0.29 | 0.33 |
| 4 | 156575786 | rs6841458 | 0.634 | 0.44 | -1.1 | -0.766 | 0.011 | 0.013 | 0.013 | 0 | 0.55 | 0.64 | 0.42 | 0.33 | 0.47 | 0.41 |
| 4 | 156580768 | rs2705441 | -0.26 | 0.166 | -1.214 | -1.03 | -0.022 | 0.07 | 0.07 | 0.758 | 0.55 | 0.8 | 0.67 | 0.22 | 0.5 | 0.51 |
| 4 | 156584397 | rs11722434 | 1.262 | 0.172 | -1.328 | -0.988 | -0.011 | 0.009 | 0.009 | 0 | 0.19 | 0.06 | 0.11 | 0.19 | 0.19 | 0.21 |
| 4 | 156586804 | rs3806776 | 0.756 | 0.352 | -1.312 | -1.081 | -0.008 | -0.033 | -0.033 | 0 | 0.16 | 0.23 | 0.25 | 0.04 | 0.07 | 0.24 |
| 4 | 156586949 | rs3806777 | 1.467 | 0.501 | -1.757 | -1.049 | 0.023 | -0.009 | -0.009 | 0.618 | 0.24 | 0.11 | 0.22 | 0.16 | 0.21 | 0.17 |
| 4 | 156597290 | rs4691841 | 1.036 | 0.721 | -1.973 | -0.948 | 0.002 | 0.024 | 0.024 | 0 | 0.14 | 0.08 | 0.23 | 0.04 | 0.07 | 0.24 |
| 4 | 156598008 | rs4691047 | -0.035 | 0.938 | -1.695 | -0.584 | 0.005 | 0.06 | 0.06 | 1.513 | 0.57 | 0.28 | 0.56 | 0.82 | 0.56 | 0.67 |
| 4 | 156598020 | rs4691048 | -0.035 | 0.959 | -1.719 | -0.584 | 0.003 | 0.061 | 0.061 | 1.513 | 0.57 | 0.28 | 0.56 | 0.82 | 0.56 | 0.67 |
| 4 | 156598081 | rs4691842 | -0.035 | 0.959 | -1.719 | -0.584 | 0.005 | 0.061 | 0.061 | 1.513 | 0.57 | 0.28 | 0.56 | 0.82 | 0.56 | 0.67 |
| 4 | 156598385 | rs7685675 | 0.23 | 1.838 | -1.505 | 0.105 | -0.07 | -0.136 | -0.136 | 0 | 0.22 | 0.03 | 0.2 | 0.59 | 0.3 | 0.23 |
| 4 | 156600197 | rs2625276 | -0.598 | 1.384 | -0.304 | -0.599 | 0.049 | -0.099 | -0.099 | 0 | 0.17 | 0.34 | 0.2 | 0.01 | 0.23 | 0.16 |
| 4 | 156607143 | rs7658967 | -0.381 | 1.027 | -1.117 | -0.117 | -0.056 | -0.015 | -0.015 | 0.014 | 0.33 | 0.16 | 0.28 | 0.37 | 0.39 | 0.27 |
| 4 | 156614954 | rs2101180 | -0.314 | 0.486 | -0.097 | -0.559 | -0.001 | 0.002 | 0.002 | 0 | 0.26 | 0.23 | 0.12 | 0.2 | 0.24 | 0.21 |
| 4 | 156617994 | rs2306557 | 0.103 | 0.099 | -0.587 | -0.941 | -0.007 | 0.027 | 0.027 | 0 | 0.27 | 0.12 | 0.25 | 0.23 | 0.29 | 0.26 |
| 4 | 156619204 | rs13143871 | 0.012 | 0.033 | -0.536 | -0.973 | -0.001 | 0.03 | 0.03 | 0 | 0.28 | 0.16 | 0.26 | 0.23 | 0.29 | 0.26 |
| 4 | 156627416 | rs11945064 | 0.594 | -0.273 | -0.003 | -1.421 | 0.003 | -0.001 | -0.001 | 0 | 0.18 | 0.12 | 0.25 | 0.06 | 0.12 | 0.22 |
| 4 | 156630713 | rs7689264 | 0.411 | -0.41 | 0.059 | -1.454 | 0.036 | 0.156 | 0.156 | 0 | 0.5 | 0.19 | 0.59 | 0.74 | 0.62 | 0.57 |
| 4 | 156633186 | rs10019835 | -0.411 | -0.289 | 0.119 | -1.572 | -0.002 | -0.046 | -0.046 | 0 | 0.75 | 0.73 | 0.72 | 0.94 | 0.8 | 0.74 |
| 4 | 156633270 | rs17033385 | 0.468 | -0.288 | 0.127 | -1.571 | 0.001 | 0.01 | 0.01 | 0 | 0.22 | 0.15 | 0.27 | 0.06 | 0.19 | 0.26 |

| | | | | | | | | | | | | | | | | |
|---|---|---|---|---|---|---|---|---|---|---|---|---|---|---|---|---|
| 4 | 156635885 | rs3796588 | -0.139 | -0.38 | -0.043 | -1.683 | 0 | -0.02 | -0.02 | 1.032 | 0.7 | 0.73 | 0.69 | 0.94 | 0.74 | 0.67 |
| 4 | 156638573 | rs2306556 | 1.458 | -0.343 | 0.392 | -1.716 | -0.015 | -0.005 | -0.005 | 0 | 0.18 | 0.06 | 0.21 | 0.21 | 0.2 | 0.18 |
| 4 | 156639174 | rs3796585 | 0.1 | -0.348 | 0.404 | -1.713 | -0.005 | -0.005 | -0.005 | 0 | 0.43 | 0.48 | 0.34 | 0.41 | 0.38 | 0.35 |
| 4 | 156641244 | rs6854751 | -0.174 | -0.222 | 0.452 | -1.655 | 0.002 | -0.007 | -0.007 | 0 | 0.69 | 0.74 | 0.69 | 0.94 | 0.74 | 0.67 |
| 4 | 156643068 | rs3816763 | 0.947 | -0.27 | 0.555 | -1.599 | 0.018 | 0.029 | 0.029 | 0 | 0.07 | 0.16 | 0.03 | 0.12 | 0.07 | 0.03 |
| 4 | 156644655 | rs9993460 | 1.352 | -0.381 | 0.418 | -1.657 | 0.009 | 0.017 | 0.017 | 0.321 | 0.23 | 0.28 | 0.36 | 0.53 | 0.36 | 0.31 |
| 4 | 156648354 | rs3796579 | -0.496 | -0.646 | 0.312 | -1.654 | 0.028 | 0.116 | 0.116 | 0 | 0.53 | 0.77 | 0.4 | 0.26 | 0.36 | 0.43 |
| 4 | 156654735 | rs7435347 | 0.499 | -0.449 | 0.09 | -1.437 | -0.005 | -0.003 | -0.003 | 0.007 | 0.27 | 0.28 | 0.11 | 0.2 | 0.17 | 0.17 |
| 4 | 156663407 | rs11722059 | 0.666 | -0.295 | -0.396 | -1.45 | -0.018 | 0.006 | 0.006 | 0 | 0.2 | 0.07 | 0.22 | 0.21 | 0.22 | 0.26 |
| 4 | 156666845 | rs6814688 | 0.614 | -0.376 | -0.287 | -1.562 | 0.009 | 0.012 | 0.012 | 0 | 0.26 | 0.19 | 0.12 | 0.19 | 0.18 | 0.18 |
| 4 | 156671888 | rs4691073 | 0.46 | -0.485 | -0.747 | -1.823 | -0.014 | 0.114 | 0.114 | 0 | 0.44 | 0.25 | 0.59 | 0.76 | 0.59 | 0.49 |
| 4 | 156672159 | rs6838513 | -0.878 | -0.479 | -0.421 | -1.766 | -0.023 | -0.027 | -0.027 | 0.346 | 0.72 | 0.62 | 0.85 | 0.81 | 0.82 | 0.82 |
| 4 | 156675369 | rs6838946 | -0.043 | -0.322 | 0.105 | -1.656 | -0.013 | -0.108 | -0.108 | 0.239 | 0.22 | 0.38 | 0.23 | 0.05 | 0.17 | 0.26 |
| 4 | 156677592 | rs10517621 | -0.201 | -0.173 | -0.3 | -1.361 | -0.001 | 0.031 | 0.031 | 0 | 0.25 | 0.16 | 0.27 | 0.21 | 0.26 | 0.25 |
| 4 | 156683281 | rs2219538 | -0.45 | 0.018 | -0.498 | -1.378 | -0.003 | 0.082 | 0.082 | 0.004 | 0.35 | 0.17 | 0.22 | 0.17 | 0.34 | 0.31 |
| 4 | 156687170 | rs10012242 | -0.88 | -0.488 | -0.16 | -1.493 | -0.036 | -0.039 | -0.039 | 0 | 0.19 | 0.36 | 0.21 | 0.13 | 0.12 | 0.12 |
| 4 | 156688928 | rs3796564 | -1.53 | 0.499 | -1.022 | -1.367 | -0.084 | -0.083 | -0.083 | 0 | 0.81 | 0.95 | 0.61 | 0.35 | 0.61 | 0.71 |
| 4 | 156692818 | rs1563806 | -0.637 | 0.375 | -0.322 | -1.578 | 0.093 | 0.056 | 0.056 | 0.235 | 0.73 | 0.58 | 0.53 | 0.35 | 0.51 | 0.69 |
| 4 | 156699636 | rs3796570 | 0.691 | -1.695 | -0.967 | -2.145 | 0.003 | 0.004 | 0.004 | 0 | 0.29 | 0.25 | 0.16 | 0.12 | 0.11 | 0.17 |
| 4 | 156703084 | rs3775087 | -0.079 | -1.084 | -1.435 | -1.965 | -0.03 | 0.099 | 0.099 | 0 | 0.24 | 0.06 | 0.14 | 0.09 | 0.23 | 0.21 |
| 4 | 156713031 | rs7661128 | -1.512 | 0.02 | -1.292 | -1.486 | -0.047 | -0.042 | -0.042 | 0.032 | 0.79 | 0.87 | 0.61 | 0.33 | 0.6 | 0.73 |
| 4 | 156721410 | rs3796574 | -0.172 | -0.619 | -1.414 | -1.98 | -0.006 | 0.031 | 0.031 | 0 | 0.6 | 0.64 | 0.49 | 0.18 | 0.44 | 0.52 |
| 4 | 156722774 | rs4560354 | -1.728 | -0.275 | -1.569 | -1.839 | -0.027 | -0.021 | -0.021 | 0.001 | 0.79 | 0.86 | 0.61 | 0.3 | 0.62 | 0.74 |
| 4 | 156727377 | rs17033572 | 0.202 | -0.419 | -2.113 | -2.025 | 0.038 | 0.08 | 0.08 | 0 | 0.33 | 0.19 | 0.19 | 0.07 | 0.25 | 0.36 |
| 4 | 156731162 | rs5024165 | 0.049 | -0.522 | -2.097 | -2.013 | 0.003 | 0.078 | 0.078 | 0 | 0.35 | 0.24 | 0.24 | 0.07 | 0.34 | 0.38 |
| 4 | 156731952 | rs922134 | 0.718 | -1.107 | -2.279 | -1.91 | 0.025 | 0.013 | 0.013 | 0.201 | 0.18 | 0.01 | 0.21 | 0.1 | 0.1 | 0.14 |
| 4 | 156739530 | rs17313835 | -0.553 | -0.853 | -1.482 | -1.76 | 0.002 | -0.005 | -0.005 | 0 | 0.17 | 0.21 | 0.14 | 0.12 | 0.19 | 0.16 |
| 4 | 156741858 | rs10032771 | 0.495 | -1.442 | -1.518 | -1.699 | 0.017 | 0.023 | 0.023 | 0 | 0.66 | 0.78 | 0.83 | 0.86 | 0.76 | 0.65 |
| 4 | 156745450 | rs12499875 | -0.199 | -1.045 | -2.182 | -1.853 | 0.026 | 0.031 | 0.031 | 0 | 0.19 | 0.05 | 0.18 | 0.09 | 0.12 | 0.17 |
| 4 | 156745915 | rs6536096 | -0.806 | -0.629 | -2.059 | -1.647 | 0.046 | 0.249 | 0.249 | 0.154 | 0.46 | 0.86 | 0.32 | 0.12 | 0.3 | 0.3 |
| 4 | 156758741 | rs9992550 | -0.881 | -0.756 | -0.69 | -1.188 | -0.023 | -0.008 | -0.008 | 0 | 0.11 | 0.28 | 0.07 | 0.07 | 0.05 | 0.17 |
| 4 | 156762493 | rs13148223 | 1.046 | -1.003 | -1.575 | -1.944 | -0.085 | 0.112 | 0.112 | 0.421 | 0.39 | 0.08 | 0.55 | 0.79 | 0.56 | 0.48 |
| 4 | 156772468 | rs1563804 | 1.193 | -1.169 | -0.696 | -2.05 | 0.016 | 0.023 | 0.023 | 0 | 0.65 | 0.29 | 0.65 | 0.82 | 0.76 | 0.64 |
| 4 | 156780170 | rs6817090 | 0.72 | -1.25 | -0.081 | -2.092 | 0.014 | -0.016 | -0.016 | 2.316 | 0.81 | 0.54 | 0.79 | 0.91 | 0.84 | 0.87 |
| 4 | 156784671 | rs17033623 | 0.815 | -1.258 | -0.038 | -2.072 | -0.028 | -0.016 | -0.016 | 0 | 0.06 | 0.13 | 0.02 | 0.03 | 0.01 | 0.13 |
| 4 | 156787340 | rs6848883 | 0.859 | -1.091 | 0.07 | -2.017 | 0.029 | 0.03 | 0.03 | 0.013 | 0.9 | 0.57 | 0.92 | 0.96 | 0.95 | 0.95 |
| 4 | 156787854 | rs17033637 | 0.194 | -1.189 | -0.498 | -2.075 | 0 | -0.006 | -0.006 | 0 | 0.17 | 0.13 | 0.07 | 0.02 | 0.17 | 0.09 |
| 4 | 156810755 | rs17314234 | -0.102 | -1.021 | -0.313 | -1.571 | -0.013 | 0.006 | 0.006 | 0 | 0.1 | 0.04 | 0.13 | 0.05 | 0.1 | 0.08 |
| 4 | 156823379 | rs3775085 | -1.249 | -1.294 | 0.472 | -1.53 | -0.018 | -0.025 | -0.025 | 0.013 | 0.16 | 0.37 | 0.18 | 0.1 | 0.11 | 0.16 |
| 4 | 156831444 | rs10517626 | -1.43 | -1.653 | 0.694 | -1.651 | -0.064 | -0.06 | -0.06 | 0 | 0.2 | 0.38 | 0.16 | 0.12 | 0.12 | 0.21 |
| 4 | 156832841 | rs12502494 | 1.706 | -1.687 | 0.796 | -1.708 | -0.075 | -0.081 | -0.081 | 0 | 0.75 | 0.4 | 0.82 | 0.88 | 0.87 | 0.79 |
| 4 | 156834028 | rs10020753 | -0.818 | -1.416 | 0.465 | -1.595 | -0.005 | -0.01 | -0.01 | 0.048 | 0.13 | 0.14 | 0.14 | 0.07 | 0.11 | 0.14 |
| 4 | 156839084 | rs13128945 | 0.807 | -1.13 | 0.686 | -1.581 | 0.006 | -0.03 | -0.03 | 0.458 | 0.91 | 0.78 | 0.95 | 0.98 | 0.92 | 0.97 |
| 4 | 156839953 | rs6856558 | 0.717 | -1.581 | 0.334 | -2.285 | 0.042 | 0.07 | 0.07 | 0 | 0.46 | 0.34 | 0.7 | 0.82 | 0.64 | 0.52 |
| 4 | 156844396 | rs12509679 | 0.435 | -1.594 | 0.52 | -2.28 | 0.027 | 0.014 | 0.014 | 1.22 | 0.94 | 0.67 | 0.97 | 1 | 0.98 | 1 |
| 4 | 156846087 | rs9044 | 1.304 | -1.518 | 0.097 | -2.276 | -0.021 | 0.083 | 0.083 | 0 | 0.38 | 0.21 | 0.62 | 0.78 | 0.54 | 0.46 |
| 4 | 156847230 | rs1543108 | 1.321 | -1.746 | 0.317 | -2.247 | -0.023 | -0.029 | -0.029 | 0 | 0.81 | 0.33 | 0.86 | 0.91 | 0.89 | 0.86 |
| 4 | 156855433 | rs12510557 | 0.864 | -0.754 | -0.363 | -2.415 | -0.032 | 0.108 | 0.108 | 0 | 0.41 | 0.2 | 0.65 | 0.81 | 0.48 | 0.52 |
| 4 | 156859190 | rs1355875 | 1.316 | -0.838 | -0.463 | -2.34 | -0.072 | 0.045 | 0.045 | 0 | 0.34 | 0.15 | 0.58 | 0.74 | 0.42 | 0.44 |
| 4 | 156860391 | rs7684248 | 1.697 | -1.035 | -0.442 | -2.324 | -0.037 | 0.002 | 0.002 | 0 | 0.3 | 0.15 | 0.52 | 0.71 | 0.36 | 0.36 |
| 4 | 156862104 | rs11100024 | 1.634 | -1.033 | -0.443 | -2.325 | -0.044 | 0.014 | 0.014 | 0 | 0.31 | 0.15 | 0.53 | 0.73 | 0.37 | 0.37 |
| 4 | 156863252 | rs13128318 | 0.461 | -1.018 | -0.384 | -2.489 | 0.054 | 0.128 | 0.128 | 0 | 0.36 | 0.15 | 0.13 | 0.01 | 0.25 | 0.25 |
| 4 | 156865486 | rs10019161 | 0.596 | -1.001 | -0.133 | -2.479 | 0.031 | 0.005 | 0.005 | 0 | 0.11 | 0.17 | 0.14 | 0.15 | 0.23 | 0.17 |
| 4 | 156865988 | rs7693476 | 0.649 | -1.041 | -0.264 | -2.489 | 0.087 | 0.221 | 0.221 | 0 | 0.47 | 0.2 | 0.76 | 0.91 | 0.68 | 0.59 |
| 4 | 156871631 | rs13108152 | 1.57 | -1.063 | 0.054 | -2.229 | -0.029 | 0.16 | 0.16 | 0 | 0.51 | 0.16 | 0.57 | 0.73 | 0.5 | 0.5 |
| 4 | 156872831 | rs10002427 | -0.369 | -0.804 | 0.634 | -2.049 | -0.034 | 0.005 | 0.005 | 0 | 0.13 | 0.28 | 0.15 | 0.15 | 0.08 | 0.23 |
| 4 | 156874028 | rs10019975 | 1.495 | -0.789 | 0.278 | -2.053 | -0.002 | 0 | 0 | 0.166 | 0.71 | 0.32 | 0.66 | 0.73 | 0.63 | 0.68 |
| 4 | 156885169 | rs13108041 | -0.253 | -1.121 | 0.544 | -2.005 | -0.021 | -0.045 | -0.045 | 1.037 | 0.32 | 0.38 | 0.13 | 0.08 | 0.18 | 0.28 |
| 4 | 156886528 | rs4691182 | 1.243 | -0.854 | 0.319 | -1.918 | -0.039 | -0.018 | -0.018 | 0 | 0.08 | 0.02 | 0.13 | 0.09 | 0.23 | 0.08 |
| 4 | 156902320 | rs17033879 | 0.544 | -1.252 | 0.434 | -2.025 | 0.072 | 0.098 | 0.098 | 0 | 0.24 | 0.06 | 0.07 | 0 | 0.14 | 0.16 |
| 4 | 156906461 | rs1355874 | 1.767 | -1.277 | 0.411 | -2.059 | -0.076 | 0.084 | 0.084 | 0 | 0.43 | 0.13 | 0.56 | 0.72 | 0.48 | 0.39 |
| 4 | 156906856 | rs13106367 | 0.081 | -1.211 | 0.391 | -1.947 | -0.001 | -0.001 | -0.001 | 0.042 | 0.07 | 0.11 | 0.04 | 0.08 | 0.03 | 0.24 |
| 4 | 156916490 | rs10029718 | -0.664 | -1.208 | 0.433 | -1.955 | -0.001 | 0 | 0 | 6.662 | 0.1 | 0.21 | 0.09 | 0.11 | 0.06 | 0.29 |
| 4 | 156918475 | rs4358343 | 0.692 | -1.118 | 0.614 | -1.907 | -0.022 | -0.049 | -0.049 | 0.012 | 0.57 | 0.44 | 0.8 | 0.89 | 0.78 | 0.56 |
| 4 | 156930971 | rs11939710 | -0.853 | -0.927 | 0.26 | -1.879 | -0.037 | -0.043 | -0.043 | 1.771 | 0.92 | 0.81 | 0.97 | 1 | 0.98 | 1 |
| 4 | 156931181 | rs11939830 | 0.551 | -0.845 | 0.249 | -1.921 | 0.044 | -0.03 | -0.03 | 0.348 | 0.76 | 0.49 | 0.72 | 0.83 | 0.67 | 0.82 |
| 4 | 156937210 | rs11100027 | -0.673 | -0.534 | -0.624 | -1.564 | 0 | -0.025 | -0.025 | 0.222 | 0.46 | 0.53 | 0.36 | 0.16 | 0.47 | 0.37 |
| 4 | 156947188 | rs7683170 | 0.571 | -0.052 | -1.432 | -0.151 | -0.058 | -0.087 | -0.087 | 1.568 | 0.15 | 0.04 | 0.33 | 0.4 | 0.24 | 0.17 |
| 4 | 156950378 | rs11941518 | -0.59 | -1.015 | -1.63 | -0.353 | 0.009 | 0.009 | 0.009 | 0.393 | 0.3 | 0.67 | 0.24 | 0.17 | 0.16 | 0.23 |

| | | | | | | | | | | | | | | | | |
|---|---|---|---|---|---|---|---|---|---|---|---|---|---|---|---|---|
| 4 | 156954890 | rs4691233 | 0.031 | -0.878 | -1.715 | -0.263 | -0.064 | -0.095 | -0.095 | 0.012 | 0.23 | 0.08 | 0.39 | 0.56 | 0.32 | 0.32 |
| 4 | 156963692 | rs4691238 | 1.198 | -1.156 | -2.309 | -0.442 | -0.118 | -0.079 | -0.079 | 0 | 0.25 | 0.06 | 0.35 | 0.36 | 0.42 | 0.15 |
| 4 | 156964349 | rs6841250 | 1.271 | -0.956 | -1.336 | -0.276 | 0.017 | 0.06 | 0.06 | 0.051 | 0.15 | 0.14 | 0.34 | 0.4 | 0.24 | 0.17 |
| 4 | 156967385 | rs4691240 | 1.271 | -0.956 | -1.331 | -0.276 | 0.018 | 0.063 | 0.063 | 0.051 | 0.15 | 0.14 | 0.34 | 0.4 | 0.24 | 0.17 |
| 4 | 156974697 | rs6826026 | -0.478 | -1.212 | -1.664 | -0.415 | -0.001 | -0.024 | -0.024 | 0 | 0.38 | 0.72 | 0.53 | 0.41 | 0.35 | 0.29 |
| 4 | 156986004 | rs4605601 | 0.577 | -1.539 | -1.677 | -0.492 | 0.079 | 0.081 | 0.081 | 0.167 | 0.64 | 0.18 | 0.76 | 0.9 | 0.83 | 0.64 |
| 4 | 156988133 | rs6845616 | 0.47 | -1.52 | -1.116 | -0.351 | 0 | -0.004 | -0.004 | 0 | 0.42 | 0.39 | 0.44 | 0.52 | 0.41 | 0.49 |
| 4 | 156991316 | rs13149635 | 0.241 | -1.065 | -0.544 | -0.136 | -0.045 | -0.104 | -0.104 | 0 | 0.81 | 0.46 | 0.8 | 0.99 | 0.9 | 0.89 |
| 4 | 156994724 | rs11722256 | 0.846 | -0.56 | -0.904 | 0.098 | -0.037 | -0.037 | -0.037 | 0.087 | 0.12 | 0.06 | 0.23 | 0.35 | 0.35 | 0.12 |
| 4 | 156998599 | rs4690843 | 0.876 | -1.044 | -1.247 | 0.191 | 0.141 | 0.098 | 0.098 | 0.289 | 0.52 | 0.1 | 0.62 | 0.75 | 0.77 | 0.51 |
| 4 | 157004706 | rs4428239 | -0.772 | -1.132 | -0.828 | 0.175 | -0.01 | -0.007 | -0.007 | 0.198 | 0.79 | 0.77 | 0.87 | 0.89 | 0.88 | 0.81 |
| 4 | 157006346 | rs4417922 | 1.223 | -0.751 | -0.97 | 0.308 | -0.062 | -0.06 | -0.06 | 0 | 0.11 | 0.04 | 0.23 | 0.36 | 0.35 | 0.13 |
| 4 | 157006954 | rs11100040 | 0.449 | -0.879 | -0.838 | 0.263 | 0.061 | 0.038 | 0.038 | 0.089 | 0.59 | 0.3 | 0.63 | 0.74 | 0.79 | 0.62 |
| 4 | 157008429 | rs13117709 | 1.608 | -0.883 | -0.922 | 0.217 | -0.068 | -0.032 | -0.032 | 0.159 | 0.41 | 0.04 | 0.37 | 0.39 | 0.42 | 0.39 |
| 4 | 157011923 | rs10030044 | 0.702 | -0.281 | -0.25 | 0.632 | 0.088 | 0.082 | 0.082 | 0 | 0.76 | 0.53 | 0.63 | 0.58 | 0.6 | 0.73 |
| 4 | 157012721 | rs6836081 | 0.154 | -0.184 | -0.975 | 0.686 | 0.012 | 0.011 | 0.011 | 0 | 0.8 | 0.74 | 0.66 | 0.58 | 0.62 | 0.73 |
| 4 | 157014586 | rs4555581 | 0.106 | -0.173 | -1.013 | 0.574 | -0.006 | -0.003 | -0.003 | 0.072 | 0.8 | 0.77 | 0.67 | 0.54 | 0.62 | 0.72 |
| 4 | 157021404 | rs4437204 | -0.322 | -0.385 | -0.843 | 0.483 | -0.013 | -0.008 | -0.008 | 4.031 | 0.93 | 0.8 | 0.91 | 0.96 | 0.98 | 0.93 |
| 4 | 157022492 | rs13143174 | 1.249 | -0.498 | -1.23 | 0.315 | -0.029 | 0.094 | 0.094 | 1.857 | 0.46 | 0.08 | 0.38 | 0.37 | 0.44 | 0.37 |
| 4 | 157030680 | rs4558821 | 0.792 | -0.369 | -0.953 | 0.356 | 0.015 | 0.066 | 0.066 | 0 | 0.52 | 0.68 | 0.44 | 0.37 | 0.46 | 0.36 |
| 4 | 157034642 | rs4690853 | -0.054 | 0.02 | -1.048 | 0.649 | -0.011 | -0.007 | -0.007 | 0.455 | 0.16 | 0.12 | 0.23 | 0.43 | 0.27 | 0.24 |
| 4 | 157050382 | rs4259017 | 0.382 | -0.724 | -1.189 | -0.055 | 0.071 | 0.071 | 0.071 | 1.829 | 0.73 | 0.16 | 0.81 | 0.91 | 0.87 | 0.86 |
| 4 | 157065076 | rs7661151 | -0.176 | 0.462 | -1.831 | 0.009 | 0.004 | 0.085 | 0.085 | 0.444 | 0.38 | 0.17 | 0.4 | 0.17 | 0.37 | 0.32 |
| 4 | 157068585 | rs10016298 | 0.1 | 0.156 | -1.647 | 0.526 | -0.03 | 0.056 | 0.056 | 0 | 0.23 | 0.58 | 0.25 | 0.4 | 0.15 | 0.22 |
| 4 | 157068808 | rs10004557 | 0.1 | 0.156 | -1.625 | 0.545 | -0.03 | 0.055 | 0.055 | 0 | 0.23 | 0.58 | 0.26 | 0.39 | 0.15 | 0.22 |
| 4 | 157074296 | rs12331916 | -0.7 | 0.925 | -2.044 | 0.547 | -0.016 | -0.014 | -0.014 | 0 | 0.85 | 0.95 | 0.82 | 0.82 | 0.82 | 0.75 |
| 4 | 157077635 | rs11100049 | 0.802 | 0.709 | -2.413 | 0.429 | -0.001 | -0.004 | -0.004 | 0.055 | 0.71 | 0.74 | 0.52 | 0.66 | 0.71 | 0.57 |
| 4 | 157086546 | rs17314974 | 1.891 | 1.378 | -2.597 | 0.383 | -0.032 | -0.024 | -0.024 | 0.051 | 0.05 | 0 | 0.06 | 0.09 | 0.11 | 0.07 |
| 4 | 157089145 | rs17380673 | 0.638 | 0.955 | -2.443 | 0.194 | -0.017 | -0.016 | -0.016 | 0 | 0.15 | 0.05 | 0.18 | 0.18 | 0.18 | 0.25 |
| 4 | 157089605 | rs5027023 | -0.552 | 0.935 | -2.436 | 0.186 | -0.01 | -0.016 | -0.016 | 0.019 | 0.22 | 0.13 | 0.23 | 0.26 | 0.26 | 0.31 |
| 4 | 157090452 | rs12499949 | 1.622 | 0.931 | -2.451 | 0.198 | -0.006 | -0.03 | -0.03 | 0.072 | 0.05 | 0.01 | 0.24 | 0.09 | 0.03 | 0.12 |
| 4 | 157094509 | rs11721807 | 1.187 | 0.873 | -2.774 | 0.011 | -0.059 | 0.112 | 0.112 | 0.019 | 0.28 | 0.02 | 0.12 | 0.08 | 0.29 | 0.19 |
| 4 | 157096990 | rs4129759 | -1.189 | 0.715 | -2.433 | 0.306 | -0.005 | -0.036 | -0.036 | 0.062 | 0.28 | 0.19 | 0.48 | 0.35 | 0.29 | 0.43 |
| 4 | 157098758 | rs7676050 | -0.263 | 0.72 | -2.451 | 0.311 | -0.008 | -0.035 | -0.035 | 0 | 0.21 | 0.1 | 0.43 | 0.27 | 0.21 | 0.37 |
| 4 | 157100233 | rs17380799 | -0.227 | 0.928 | -2.443 | 0.325 | -0.024 | -0.039 | -0.039 | 0.361 | 0.11 | 0.05 | 0.3 | 0.17 | 0.14 | 0.18 |
| 4 | 157100986 | rs17034170 | 0.173 | 0.804 | -2.431 | 0.347 | -0.012 | -0.043 | -0.043 | 0.106 | 0.1 | 0.03 | 0.27 | 0.17 | 0.08 | 0.18 |
| 4 | 157104688 | rs12511271 | -1.032 | 0.817 | -2.359 | 0.331 | -0.001 | -0.001 | -0.001 | 0 | 0.28 | 0.27 | 0.48 | 0.37 | 0.29 | 0.34 |
| 4 | 157109135 | rs12186255 | -0.97 | 0.856 | -2.363 | 0.335 | -0.003 | -0.016 | -0.016 | 0.061 | 0.28 | 0.23 | 0.48 | 0.35 | 0.29 | 0.33 |
| 4 | 157109267 | rs4296721 | -1.367 | 0.727 | -1.862 | 0.081 | 0 | -0.01 | -0.01 | 1.071 | 0.92 | 0.93 | 0.96 | 1 | 0.94 | 0.96 |
| 4 | 157116213 | rs17034196 | -0.561 | 0.82 | -2.192 | -0.005 | 0.027 | 0.059 | 0.059 | 0 | 0.62 | 0.78 | 0.85 | 0.83 | 0.7 | 0.64 |
| 4 | 157118100 | rs4691308 | 0.084 | 0.972 | -2.087 | 0.346 | -0.016 | -0.017 | -0.017 | 0.048 | 0.15 | 0.06 | 0.18 | 0.2 | 0.18 | 0.16 |
| 4 | 157119346 | rs10014673 | 0.103 | 1.014 | -2.102 | 0.355 | 0.009 | -0.014 | -0.014 | 0 | 0.4 | 0.74 | 0.49 | 0.58 | 0.37 | 0.34 |
| 4 | 157124728 | rs13103675 | -0.034 | 0.863 | -1.522 | 0.492 | 0.013 | 0.004 | 0.004 | 0.121 | 0.48 | 0.4 | 0.64 | 0.54 | 0.65 | 0.34 |
| 4 | 157128226 | rs11726343 | -0.84 | 0.441 | -1.2 | 0.42 | 0.001 | -0.005 | -0.005 | 0 | 0.67 | 0.67 | 0.72 | 0.68 | 0.71 | 0.51 |
| 4 | 157129218 | rs2340252 | 1.23 | 1.086 | -1.043 | 0.881 | -0.039 | 0.027 | 0.027 | 0 | 0.35 | 0.21 | 0.31 | 0.21 | 0.46 | 0.25 |
| 4 | 157129639 | rs10029573 | -0.707 | 0.99 | -0.902 | 0.856 | -0.018 | 0.028 | 0.028 | 0 | 0.31 | 0.37 | 0.4 | 0.48 | 0.24 | 0.27 |
| 4 | 157130289 | rs17315339 | 1.569 | 1.282 | -0.991 | 0.787 | -0.015 | 0.015 | 0.015 | 0 | 0.32 | 0.2 | 0.27 | 0.21 | 0.37 | 0.24 |
| 4 | 157131028 | rs13102097 | -1.315 | 1.282 | -1.031 | 0.737 | -0.003 | -0.023 | -0.023 | 0 | 0.69 | 0.67 | 0.8 | 0.83 | 0.75 | 0.57 |
| 4 | 157144661 | rs4600965 | -0.235 | 0.965 | -0.994 | 0.745 | 0.025 | 0.01 | 0.01 | 0 | 0.55 | 0.67 | 0.75 | 0.62 | 0.72 | 0.42 |
| 4 | 157145352 | rs10517634 | 1.475 | 0.888 | -1.104 | 0.78 | -0.019 | -0.001 | -0.001 | 0 | 0.33 | 0.27 | 0.34 | 0.29 | 0.44 | 0.32 |
| 4 | 157146395 | rs4691312 | 0.613 | 0.787 | -0.986 | 0.763 | -0.008 | 0.044 | 0.044 | 0.588 | 0.14 | 0.22 | 0.32 | 0.33 | 0.15 | 0.09 |
| 4 | 157150175 | rs13112531 | 0.49 | 0.794 | -1.087 | 0.761 | 0.015 | -0.004 | -0.004 | 0 | 0.43 | 0.39 | 0.49 | 0.44 | 0.61 | 0.39 |
| 4 | 157152021 | rs6833995 | -0.028 | 0.565 | -1.15 | 0.296 | -0.004 | -0.001 | -0.001 | 0 | 0.54 | 0.47 | 0.8 | 0.77 | 0.76 | 0.48 |
| 4 | 157158799 | rs2879724 | 0.632 | 0.712 | -1.159 | 0.097 | 0 | 0 | 0 | 0 | 0.45 | 0.45 | 0.65 | 0.62 | 0.53 | 0.41 |
| 4 | 157160921 | rs6536158 | -1.132 | 0.682 | -0.999 | 0.235 | 0.038 | 0.101 | 0.101 | 0 | 0.74 | 0.92 | 0.89 | 0.91 | 0.84 | 0.67 |
| 4 | 157162514 | rs10006636 | 0.555 | 0.762 | -0.782 | 0.351 | 0.001 | 0.004 | 0.004 | 0 | 0.5 | 0.47 | 0.67 | 0.62 | 0.59 | 0.42 |
| 4 | 157163691 | rs6830845 | -0.498 | 0.328 | -1.021 | -0.005 | 0.045 | 0.102 | 0.102 | 0 | 0.72 | 0.91 | 0.89 | 0.9 | 0.84 | 0.65 |
| 4 | 157166746 | rs11728161 | -0.718 | 1.02 | -0.647 | 0.445 | -0.034 | -0.041 | -0.041 | 0.007 | 0.93 | 1 | 0.97 | 0.82 | 0.93 | 0.93 |
| 4 | 157167891 | rs7700147 | 1.628 | 0.604 | -0.018 | 0.461 | -0.001 | 0 | 0 | 0 | 0.68 | 0.34 | 0.83 | 0.56 | 0.77 | 0.64 |
| 4 | 157168163 | rs10517633 | -0.143 | 0.555 | 0.22 | 0.462 | 0.022 | -0.015 | -0.015 | 0 | 0.16 | 0.09 | 0.08 | 0.24 | 0.13 | 0.27 |
| 4 | 157169331 | rs13434949 | -1.394 | 0.543 | 0.73 | 0.438 | 0.028 | 0.006 | 0.006 | 0.123 | 0.06 | 0.22 | 0.04 | 0.04 | 0.07 | 0.09 |
| 4 | 157171078 | rs4440278 | -1.083 | 0.537 | 0.753 | 0.421 | -0.024 | 0.003 | 0.003 | 0 | 0.2 | 0.28 | 0.11 | 0.24 | 0.16 | 0.27 |
| 4 | 157171201 | rs2340254 | 0.219 | 0.557 | 0.841 | 0.289 | 0.006 | 0.031 | 0.031 | 0 | 0.9 | 0.81 | 0.96 | 0.8 | 0.92 | 0.92 |
| 4 | 157174024 | rs11734958 | 1.091 | 0.917 | 0.707 | -0.01 | 0.032 | -0.04 | -0.04 | 0 | 0.74 | 0.53 | 0.81 | 0.8 | 0.77 | 0.84 |
| 4 | 157176193 | rs10517632 | -0.161 | 0.619 | 0.174 | 0.208 | 0.025 | -0.032 | -0.032 | 0 | 0.16 | 0.08 | 0.14 | 0.29 | 0.13 | 0.28 |
| 4 | 157181209 | rs2340277 | 2.281 | 0.941 | 0.234 | -0.321 | 0.002 | -0.055 | -0.055 | 1.007 | 0.43 | 0.38 | 0.22 | 0.03 | 0.38 | 0.28 |
| 4 | 157185914 | rs17034315 | 0.056 | 0.831 | 0.332 | 0.103 | 0.025 | -0.038 | -0.038 | 0.203 | 0.15 | 0.05 | 0.13 | 0.27 | 0.12 | 0.27 |
| 4 | 157190788 | rs7668315 | 1.454 | 1.044 | 0.466 | 0.097 | -0.005 | 0.02 | 0.02 | 0.054 | 0.69 | 0.38 | 0.71 | 0.52 | 0.73 | 0.65 |
| 4 | 157197295 | rs6818678 | 0.598 | 1.292 | -0.107 | 0 | -0.01 | 0.015 | 0.015 | 0 | 0.15 | 0.1 | 0.12 | 0.01 | 0.14 | 0.09 |

| | | | | | | | | | | | | | | | | |
|---|---|---|---|---|---|---|---|---|---|---|---|---|---|---|---|---|
| 4 | 157197353 | rs4690859 | -0.179 | 1.123 | -0.073 | 0.194 | -0.044 | 0.033 | 0.033 | 0.128 | 0.15 | 0.24 | 0.14 | 0.23 | 0.08 | 0.18 |
| 4 | 157213766 | rs2220585 | -0.382 | 1.811 | -0.479 | 0.361 | -0.055 | 0.01 | 0.01 | 0.84 | 0.91 | 0.99 | 0.89 | 0.93 | 0.86 | 0.92 |
| 4 | 157221159 | rs6825273 | -0.814 | 1.413 | -0.508 | 0.366 | 0.003 | -0.041 | -0.041 | 0 | 0.25 | 0.11 | 0.22 | 0.32 | 0.22 | 0.4 |
| 4 | 157232731 | rs10517639 | -0.467 | 1.493 | -0.104 | 0.282 | 0.001 | -0.018 | -0.018 | 0 | 0.19 | 0.26 | 0.13 | 0.06 | 0.17 | 0.2 |
| 4 | 157235050 | rs10030326 | 1.419 | 1.638 | 0.344 | 0.274 | 0.036 | -0.044 | -0.044 | 0 | 0.68 | 0.47 | 0.72 | 0.75 | 0.67 | 0.73 |
| 4 | 157244229 | rs1352703 | -0.083 | 1.653 | 0.454 | 0.296 | 0.025 | 0.007 | 0.007 | 0.084 | 0.89 | 0.86 | 0.87 | 0.81 | 0.83 | 0.92 |
| 4 | 157249803 | rs13103662 | -0.454 | 1.588 | 0.451 | 0.252 | 0.002 | -0.045 | -0.045 | 0.151 | 0.2 | 0.34 | 0.14 | 0.06 | 0.17 | 0.2 |
| 4 | 157252132 | rs1873358 | 0.536 | 1.639 | 0.462 | 0.512 | 0.043 | 0.009 | 0.009 | 0.03 | 0.87 | 0.77 | 0.86 | 0.81 | 0.84 | 0.92 |
| 4 | 157264625 | rs2046876 | 0.761 | 1.626 | 0.535 | 0.482 | 0.056 | 0.008 | 0.008 | 0 | 0.86 | 0.68 | 0.85 | 0.82 | 0.84 | 0.92 |
| 4 | 157267440 | rs10021338 | 0.208 | 1.466 | 0.245 | 0.644 | -0.014 | -0.012 | -0.012 | 0 | 0.12 | 0.05 | 0.37 | 0.45 | 0.19 | 0.26 |
| 4 | 157279663 | rs1532154 | -0.175 | 1.532 | 0.08 | 0.631 | -0.007 | -0.008 | -0.008 | 0 | 0.14 | 0.08 | 0.38 | 0.47 | 0.2 | 0.27 |
| 4 | 157295801 | rs2340289 | -0.886 | 1.489 | 0.161 | 0.06 | -0.011 | -0.015 | -0.015 | 0 | 0.26 | 0.37 | 0.51 | 0.74 | 0.27 | 0.45 |
| 4 | 157304598 | rs10004475 | -0.045 | 1.672 | 0.08 | -0.127 | 0 | -0.014 | -0.014 | 0.101 | 0.17 | 0.22 | 0.13 | 0.02 | 0.14 | 0.11 |
| 4 | 157306101 | rs6536171 | -1.479 | 1.641 | -0.103 | -0.038 | 0.001 | 0.009 | 0.009 | 0.32 | 0.44 | 0.6 | 0.64 | 0.75 | 0.42 | 0.56 |
| 4 | 157306780 | rs1352705 | -0.146 | 2.064 | -0.305 | 0.343 | -0.053 | -0.038 | -0.038 | 0 | 0.91 | 0.99 | 0.88 | 0.82 | 0.86 | 0.92 |
| 4 | 157309545 | rs17034504 | 0.475 | 1.857 | -0.088 | 0.098 | 0.009 | -0.026 | -0.026 | 1.176 | 0.06 | 0.07 | 0.06 | 0 | 0.08 | 0.01 |
| 4 | 157317266 | rs17034515 | -1.077 | 1.905 | 0.203 | 0.739 | 0.031 | 0.216 | 0.216 | 0.227 | 0.1 | 0.21 | 0.22 | 0.39 | 0.1 | 0.28 |
| 4 | 157328376 | rs7670864 | 0.578 | 1.757 | -0.209 | 0.456 | -0.002 | 0.004 | 0.004 | 0.105 | 0.64 | 0.43 | 0.51 | 0.47 | 0.69 | 0.66 |
| 4 | 157332033 | rs6813766 | -0.163 | 1.873 | -0.75 | 0.008 | -0.051 | 0.065 | 0.065 | 0 | 0.22 | 0.11 | 0.15 | 0.06 | 0.29 | 0.12 |
| 4 | 157342073 | rs10517637 | -0.257 | 1.48 | -0.098 | 0.744 | 0.007 | 0.109 | 0.109 | 0.465 | 0.07 | 0.13 | 0.09 | 0.3 | 0.07 | 0.28 |
| 4 | 157345022 | rs6818354 | 1 | 1.486 | -0.043 | 0.589 | -0.001 | 0 | 0 | 4.042 | 0.59 | 0.43 | 0.36 | 0.38 | 0.63 | 0.64 |
| 4 | 157345834 | rs7667504 | 0.55 | 1.42 | -0.51 | 0.755 | -0.002 | -0.025 | -0.025 | 0.002 | 0.67 | 0.81 | 0.61 | 0.59 | 0.69 | 0.72 |
| 4 | 157349944 | rs17034575 | 0.35 | 1.617 | -0.24 | 1.082 | -0.024 | -0.032 | -0.032 | 0.036 | 0.06 | 0 | 0.18 | 0.23 | 0.09 | 0.08 |
| 4 | 157351242 | rs10517636 | 1.385 | 1.68 | 0.019 | 0.603 | 0.005 | -0.006 | -0.006 | 0 | 0.47 | 0.59 | 0.37 | 0.14 | 0.43 | 0.36 |
| 4 | 157352057 | rs6536176 | -0.427 | 1.827 | -0.16 | 0.83 | 0 | 0.007 | 0.007 | 0.544 | 0.25 | 0.25 | 0.37 | 0.47 | 0.23 | 0.46 |
| 4 | 157354625 | rs7669612 | 0.323 | 1.668 | -0.129 | 0.909 | -0.013 | 0.011 | 0.011 | 0 | 0.1 | 0.06 | 0.07 | 0.04 | 0.16 | 0.05 |
| 4 | 157363345 | rs13141045 | 0.792 | 1.487 | -0.077 | 0.944 | -0.001 | 0.001 | 0.001 | 0 | 0.07 | 0.08 | 0.03 | 0.04 | 0.08 | 0.04 |
| 4 | 157363709 | rs7669823 | 0.08 | 1.433 | 0.281 | 1.001 | 0.004 | 0.022 | 0.022 | 0.156 | 0.77 | 0.77 | 0.76 | 0.65 | 0.74 | 0.86 |
| 4 | 157365096 | rs6834515 | -0.511 | 1.388 | 0.16 | 1.116 | 0.003 | 0.035 | 0.035 | 0.023 | 0.23 | 0.28 | 0.36 | 0.48 | 0.2 | 0.41 |
| 4 | 157371613 | rs11938052 | 1.404 | 1.294 | -0.19 | 1.123 | -0.01 | 0.101 | 0.101 | 0.022 | 0.51 | 0.12 | 0.44 | 0.33 | 0.51 | 0.46 |
| 4 | 157372570 | rs9997210 | -0.402 | 1.473 | -0.044 | 1.067 | -0.004 | -0.006 | -0.006 | 0.201 | 0.81 | 0.86 | 0.82 | 0.74 | 0.84 | 0.64 |
| 4 | 157380782 | rs1352701 | -1.67 | 0.962 | -0.015 | 0.62 | -0.007 | 0.034 | 0.034 | 0 | 0.49 | 0.76 | 0.57 | 0.61 | 0.51 | 0.48 |
| 4 | 157381223 | rs1490571 | -0.214 | 0.976 | 0.133 | 0.651 | -0.001 | -0.001 | -0.001 | 0.386 | 0.11 | 0.13 | 0.06 | 0.11 | 0.13 | 0.17 |
| 4 | 157381961 | rs4691343 | -1.211 | 0.976 | 0.73 | 0.555 | -0.005 | 0.011 | 0.011 | 0 | 0.35 | 0.51 | 0.42 | 0.61 | 0.31 | 0.43 |
| 4 | 157383426 | rs6833736 | 2.211 | 0.941 | 0.654 | 0.386 | -0.031 | 0.133 | 0.133 | 0 | 0.36 | 0.12 | 0.3 | 0.14 | 0.39 | 0.33 |
| 4 | 157383920 | rs1586690 | -1.008 | 1.01 | 0.68 | 0.563 | 0 | 0.003 | 0.003 | 2.194 | 0.88 | 0.89 | 0.89 | 0.79 | 0.9 | 0.87 |
| 4 | 157385298 | rs10021375 | -1.217 | 1.139 | 0.91 | 0.758 | -0.002 | 0.026 | 0.026 | 0 | 0.17 | 0.31 | 0.2 | 0.3 | 0.17 | 0.19 |
| 4 | 157398759 | rs937068 | -1.596 | 1.354 | 0.456 | 0.648 | -0.022 | -0.012 | -0.012 | 0 | 0.36 | 0.72 | 0.52 | 0.43 | 0.47 | 0.42 |
| 4 | 157403920 | rs6824856 | -1.552 | 1.429 | 0.461 | 0.581 | -0.012 | 0.038 | 0.038 | 0 | 0.45 | 0.71 | 0.55 | 0.65 | 0.48 | 0.51 |
| 4 | 157405224 | rs13435328 | 0.985 | 1.519 | 0.753 | 0.616 | 0.02 | -0.065 | -0.065 | 0 | 0.13 | 0.17 | 0.15 | 0 | 0.2 | 0.04 |
| 4 | 157409735 | rs1123714 | -1.135 | 1.186 | 0.864 | 1.087 | -0.011 | 0.019 | 0.019 | 0 | 0.32 | 0.49 | 0.4 | 0.65 | 0.28 | 0.46 |
| 4 | 157416571 | rs7671552 | -1.081 | 1.212 | 0.77 | 1.101 | -0.009 | 0.02 | 0.02 | 0 | 0.31 | 0.48 | 0.4 | 0.65 | 0.28 | 0.46 |
| 4 | 157420901 | rs13106670 | 0.229 | 1.409 | 0.326 | 1.318 | -0.002 | -0.064 | -0.064 | 0.973 | 0.18 | 0.08 | 0.21 | 0.33 | 0.13 | 0.23 |
| 4 | 157433363 | rs4328961 | -0.261 | 1.219 | 0.379 | 0.986 | -0.001 | 0.002 | 0.002 | 1.822 | 0.87 | 0.88 | 0.88 | 0.79 | 0.89 | 0.88 |
| 4 | 157437970 | rs7694359 | -0.156 | 1.133 | 0.583 | 0.971 | 0.028 | 0.051 | 0.051 | 0 | 0.07 | 0.24 | 0.05 | 0.11 | 0.09 | 0.14 |
| 4 | 157444701 | rs11100060 | -0.557 | 1.176 | -0.241 | 1.024 | 0 | 0.005 | 0.005 | 0.057 | 0.24 | 0.29 | 0.36 | 0.41 | 0.22 | 0.36 |
| 4 | 157448161 | rs34455702 | -0.182 | 1.011 | -0.594 | 0.447 | -0.01 | 0.012 | 0.012 | 0 | 0.31 | 0.2 | 0.35 | 0.25 | 0.33 | 0.2 |
| 4 | 157449015 | rs1402787 | 1.276 | 0.978 | -0.317 | 0.425 | 0.004 | 0.095 | 0.095 | 0 | 0.42 | 0.24 | 0.36 | 0.3 | 0.41 | 0.37 |
| 4 | 157449136 | rs1402788 | -0.499 | 0.961 | -0.421 | 0.289 | -0.001 | 0.001 | 0.001 | 0 | 0.09 | 0.11 | 0.18 | 0.25 | 0.12 | 0.12 |
| 4 | 157450822 | rs10517643 | 0.19 | 0.944 | -0.229 | 0.312 | -0.004 | 0.069 | 0.069 | 0 | 0.72 | 0.48 | 0.72 | 0.56 | 0.75 | 0.58 |
| 4 | 157458306 | rs952244 | 0.301 | 0.954 | 0.343 | 0.312 | -0.019 | 0.091 | 0.091 | 0 | 0.22 | 0.33 | 0.23 | 0.41 | 0.13 | 0.31 |
| 4 | 157458846 | rs17034885 | 0.46 | 1.02 | -0.321 | 0.339 | -0.005 | -0.012 | -0.012 | 0 | 0.67 | 0.32 | 0.58 | 0.49 | 0.66 | 0.47 |
| 4 | 157459397 | rs6832535 | 1.947 | 0.973 | -0.376 | 0.286 | 0.014 | 0.156 | 0.156 | 0 | 0.33 | 0.08 | 0.18 | 0.06 | 0.29 | 0.25 |
| 4 | 157466506 | rs1521710 | 0.59 | 1.09 | -0.236 | 0.38 | -0.004 | -0.01 | -0.01 | 0.554 | 0.66 | 0.33 | 0.58 | 0.49 | 0.66 | 0.47 |
| 4 | 157469352 | rs1521713 | -0.507 | 1.342 | -0.243 | 0.538 | -0.015 | 0.005 | 0.005 | 0 | 0.34 | 0.25 | 0.38 | 0.28 | 0.46 | 0.32 |
| 4 | 157484641 | rs6536193 | 0.374 | 1.148 | -0.324 | 0.78 | -0.003 | -0.013 | -0.013 | 0 | 0.66 | 0.7 | 0.65 | 0.58 | 0.66 | 0.65 |
| 4 | 157492586 | rs1608012 | 0.912 | 0.82 | -0.642 | 0.51 | -0.023 | -0.021 | -0.021 | 0 | 0.38 | 0.23 | 0.42 | 0.46 | 0.47 | 0.31 |
| 4 | 157512342 | rs28648624 | 0.157 | 0.634 | -0.705 | 0.685 | -0.016 | -0.026 | -0.026 | 0.478 | 0.12 | 0.08 | 0.26 | 0.39 | 0.13 | 0.21 |
| 4 | 157517071 | rs4569797 | 1.105 | 0.629 | -0.215 | 0.618 | -0.001 | -0.007 | -0.007 | 0.043 | 0.37 | 0.33 | 0.39 | 0.46 | 0.41 | 0.31 |
| 4 | 157518336 | rs10517648 | 0.536 | 0.557 | -0.453 | 0.457 | -0.006 | 0.03 | 0.03 | 0 | 0.06 | 0 | 0.01 | 0 | 0.03 | 0.03 |
| 4 | 157530667 | rs10028138 | 1.316 | 0.435 | 0.661 | 0.432 | 0.003 | 0.012 | 0.012 | 0 | 0.36 | 0.42 | 0.39 | 0.48 | 0.42 | 0.29 |
| 4 | 157533294 | rs1521717 | -0.727 | 0.189 | 0.587 | 0.299 | 0.001 | 0 | 0 | 0 | 0.37 | 0.38 | 0.38 | 0.32 | 0.29 | 0.32 |
| 4 | 157535527 | rs6811244 | 0.537 | 0.157 | 0.53 | 0.17 | -0.008 | -0.008 | -0.008 | 0 | 0.83 | 0.8 | 0.88 | 0.92 | 0.9 | 0.85 |
| 4 | 157545843 | rs1521724 | -1.126 | 0.117 | 0.742 | 0.21 | -0.001 | -0.001 | -0.001 | 0 | 0.48 | 0.53 | 0.5 | 0.47 | 0.48 | 0.54 |
| 4 | 157548520 | rs8180255 | 0.112 | 0.12 | 0.829 | 0.185 | 0.01 | -0.017 | -0.017 | 0 | 0.19 | 0.23 | 0.13 | 0.15 | 0.22 | 0.27 |
| 4 | 157548539 | rs2342141 | 0.185 | 0.092 | 0.834 | 0.205 | 0 | -0.003 | -0.003 | 0.03 | 0.28 | 0.29 | 0.36 | 0.32 | 0.25 | 0.26 |
| 4 | 157551268 | rs6817409 | 0.626 | 0.134 | 0.96 | 0.186 | -0.015 | -0.024 | -0.024 | 0 | 0.81 | 0.74 | 0.86 | 0.92 | 0.89 | 0.83 |
| 4 | 157554917 | rs12499279 | -0.159 | 0.174 | 0.834 | 0.232 | -0.009 | -0.004 | -0.004 | 0 | 0.12 | 0.16 | 0.11 | 0.08 | 0.08 | 0.14 |
| 4 | 157555252 | rs12507707 | 0.333 | 0.2 | 0.638 | 0.246 | 0.003 | -0.002 | -0.002 | 0 | 0.1 | 0.07 | 0.11 | 0.08 | 0.08 | 0.14 |

| | | | | | | | | | | | | | | | | |
|---|---|---|---|---|---|---|---|---|---|---|---|---|---|---|---|---|
| 4 | 157557522 | rs4690871 | -1.485 | 0.21 | 0.607 | 0.232 | -0.003 | -0.003 | -0.003 | 0.088 | 0.56 | 0.51 | 0.6 | 0.54 | 0.55 | 0.67 |
| 4 | 157557726 | rs11943855 | 1.195 | 0.283 | 0.169 | 0.218 | 0.04 | 0.004 | 0.004 | 0 | 0.51 | 0.28 | 0.49 | 0.6 | 0.64 | 0.57 |
| 4 | 157560957 | rs12498803 | -0.939 | 0.047 | 0.068 | 0.148 | 0.018 | 0.016 | 0.016 | 0.034 | 0.21 | 0.13 | 0.14 | 0.1 | 0.14 | 0.21 |
| 4 | 157563908 | rs17035016 | -0.082 | 0.041 | -1.086 | 0.215 | -0.096 | -0.084 | -0.084 | 0 | 0.38 | 0.12 | 0.46 | 0.45 | 0.44 | 0.46 |
| 4 | 157568858 | rs9998752 | -1.321 | -0.015 | -0.956 | 0.17 | 0.013 | -0.001 | -0.001 | 0 | 0.56 | 0.34 | 0.6 | 0.53 | 0.55 | 0.62 |
| 4 | 157577439 | rs7698424 | -0.445 | 0.213 | -1.345 | 0.252 | 0 | 0.003 | 0.003 | 0 | 0.16 | 0.11 | 0.16 | 0.11 | 0.13 | 0.18 |
| 4 | 157583132 | rs17317061 | 1.725 | 0.298 | -1.694 | 0.278 | -0.026 | -0.04 | -0.04 | 0 | 0.34 | 0.15 | 0.34 | 0.45 | 0.4 | 0.28 |
| 4 | 157586380 | rs1521716 | -0.609 | 0.897 | -1.705 | 0.284 | -0.071 | -0.042 | -0.042 | 0 | 0.38 | 0.11 | 0.44 | 0.4 | 0.43 | 0.45 |
| 4 | 157589751 | rs4691370 | 0.367 | 0.668 | -0.984 | 0.354 | -0.001 | -0.017 | -0.017 | 1.62 | 0.92 | 0.92 | 0.97 | 0.98 | 0.94 | 0.95 |
| 4 | 157593703 | rs4691372 | 0.107 | 0.817 | -1.372 | 0.604 | -0.032 | -0.048 | -0.048 | 0 | 0.78 | 0.91 | 0.67 | 0.73 | 0.77 | 0.78 |
| 4 | 157594988 | rs7684999 | -0.971 | 0.418 | -0.892 | 0.187 | -0.006 | -0.033 | -0.033 | 0 | 0.08 | 0.12 | 0.04 | 0 | 0.06 | 0.08 |
| 4 | 157601193 | rs10030015 | 0.111 | 0.063 | -1.567 | -0.265 | 0.022 | 0.085 | 0.085 | 0 | 0.42 | 0.23 | 0.36 | 0.2 | 0.37 | 0.39 |
| 4 | 157615583 | rs1996770 | -0.947 | -0.336 | -1.521 | 0.165 | -0.005 | -0.039 | -0.039 | 0.054 | 0.6 | 0.76 | 0.39 | 0.46 | 0.41 | 0.66 |
| 4 | 157622193 | rs6847708 | -0.347 | -0.705 | -1.362 | 0.005 | -0.006 | -0.014 | -0.014 | 0.017 | 0.05 | 0.08 | 0.05 | 0 | 0.03 | 0.05 |
| 4 | 157626640 | rs9307949 | -0.083 | -1.295 | -0.791 | -0.081 | -0.017 | 0.052 | 0.052 | 0 | 0.15 | 0.18 | 0.08 | 0.29 | 0.09 | 0.26 |
| 4 | 157629331 | rs7665232 | 0.412 | -1.698 | -0.3 | -0.723 | -0.008 | -0.027 | -0.027 | 0 | 0.24 | 0.34 | 0.2 | 0.15 | 0.16 | 0.3 |
| 4 | 157633839 | rs9307950 | -1.5 | -1.754 | -0.331 | -0.625 | 0.06 | 0.06 | 0.06 | 0 | 0.52 | 0.74 | 0.32 | 0.36 | 0.35 | 0.57 |
| 4 | 157642163 | rs1469245 | -1.999 | -1.708 | -0.378 | -0.6 | 0.015 | 0.018 | 0.018 | 0 | 0.57 | 0.83 | 0.37 | 0.36 | 0.38 | 0.62 |
| 4 | 157643986 | rs2194868 | -0.759 | -1.629 | -0.27 | -0.538 | -0.038 | -0.019 | -0.019 | 0 | 0.28 | 0.35 | 0.12 | 0.21 | 0.19 | 0.27 |
| 4 | 157655811 | rs28403897 | -0.437 | -1.741 | -0.366 | -0.501 | -0.02 | 0.006 | 0.006 | 0 | 0.19 | 0.21 | 0.12 | 0.21 | 0.1 | 0.3 |
| 4 | 157662111 | rs10012388 | -1.283 | -1.882 | -0.088 | -0.572 | 0.015 | 0.007 | 0.007 | 0.067 | 0.35 | 0.63 | 0.2 | 0.29 | 0.21 | 0.37 |
| 4 | 157670537 | rs6855363 | -1.948 | -1.887 | -0.227 | -0.609 | 0.024 | 0.034 | 0.034 | 0 | 0.58 | 0.83 | 0.37 | 0.33 | 0.36 | 0.63 |
| 4 | 157683685 | rs1425486 | -1.961 | -1.905 | -0.212 | -0.599 | 0.02 | 0.044 | 0.044 | 0 | 0.58 | 0.84 | 0.37 | 0.32 | 0.36 | 0.63 |
| 4 | 157694651 | rs4691377 | 0.472 | -1.839 | -0.06 | -0.594 | 0.005 | 0.022 | 0.022 | 0 | 0.22 | 0.17 | 0.17 | 0.06 | 0.16 | 0.26 |
| 4 | 157695392 | rs983473 | -1.443 | -1.873 | 0.484 | -0.586 | -0.018 | -0.025 | -0.025 | 9.831 | 0.43 | 0.51 | 0.3 | 0.23 | 0.25 | 0.56 |
| 4 | 157711526 | rs6834932 | -1.632 | -1.881 | 0.351 | -0.581 | 0.087 | 0.117 | 0.117 | 0 | 0.46 | 0.78 | 0.32 | 0.23 | 0.26 | 0.56 |
| 4 | 157715062 | rs17035257 | 0.147 | -1.853 | 0.315 | -0.557 | 0.005 | 0.01 | 0.01 | 0 | 0.23 | 0.21 | 0.16 | 0.17 | 0.17 | 0.42 |
| 4 | 157722648 | rs10023148 | -0.723 | -1.862 | 0.625 | -0.544 | -0.042 | -0.042 | -0.042 | 0 | 0.2 | 0.29 | 0.14 | 0.07 | 0.09 | 0.14 |
| 4 | 157730274 | rs4691381 | -1.693 | -1.92 | 0.552 | -0.687 | -0.011 | -0.018 | -0.018 | 0.153 | 0.44 | 0.53 | 0.3 | 0.23 | 0.26 | 0.57 |
| 4 | 157733990 | rs10517653 | 0.536 | -1.922 | 0.504 | -0.713 | 0.003 | 0.044 | 0.044 | 0 | 0.21 | 0.13 | 0.16 | 0.06 | 0.16 | 0.26 |
| 4 | 157734312 | rs10033375 | -0.713 | -1.848 | 0.687 | -0.593 | -0.047 | -0.061 | -0.061 | 0.194 | 0.2 | 0.32 | 0.14 | 0.06 | 0.09 | 0.15 |
| 4 | 157765129 | rs10008978 | -0.247 | -1.881 | 0.868 | -0.473 | -0.011 | -0.008 | -0.008 | 0 | 0.14 | 0.14 | 0.11 | 0.08 | 0.07 | 0.13 |
| 4 | 157771062 | rs2974254 | -1.526 | -2.145 | 0.698 | -0.536 | 0.013 | 0.003 | 0.003 | 0.035 | 0.32 | 0.66 | 0.2 | 0.29 | 0.18 | 0.41 |
| 4 | 157776293 | rs7682857 | 0.767 | -1.849 | 1.054 | -0.402 | -0.021 | -0.005 | -0.005 | 0 | 0.12 | 0.21 | 0.06 | 0.11 | 0.1 | 0.07 |
| 4 | 157783398 | rs2911941 | 0.11 | -1.908 | 1.097 | -0.428 | -0.011 | -0.005 | -0.005 | 0 | 0.12 | 0.15 | 0.11 | 0.07 | 0.07 | 0.11 |
| 4 | 157785454 | rs6811964 | 0.737 | -1.913 | 1.105 | -0.391 | -0.021 | -0.004 | -0.004 | 4.701 | 0.13 | 0.22 | 0.06 | 0.11 | 0.1 | 0.06 |
| 4 | 157786028 | rs10517657 | 0.178 | -2.226 | 0.975 | -0.583 | 0.01 | -0.003 | -0.003 | 0 | 0.14 | 0.08 | 0.13 | 0.16 | 0.09 | 0.44 |
| 4 | 157786225 | rs17035367 | 0.457 | -2.128 | 0.9 | -0.562 | -0.018 | -0.01 | -0.01 | 4.111 | 0.14 | 0.23 | 0.06 | 0.11 | 0.11 | 0.08 |
| 4 | 157789987 | rs342317 | -1.15 | -2.165 | 0.867 | -0.583 | -0.02 | -0.009 | -0.009 | 0 | 0.3 | 0.38 | 0.26 | 0.24 | 0.16 | 0.55 |
| 4 | 157795071 | rs1469243 | 1.101 | -2.208 | 0.612 | -0.686 | 0.009 | 0.035 | 0.035 | 4.672 | 0.1 | 0.05 | 0.04 | 0 | 0.06 | 0.01 |
| 4 | 157797342 | rs342308 | -2.203 | -2.116 | 0.769 | -0.603 | 0.071 | 0.047 | 0.047 | 0.019 | 0.53 | 0.79 | 0.37 | 0.35 | 0.32 | 0.63 |
| 4 | 157815887 | rs6812101 | 0.718 | -2.099 | 1.019 | -0.541 | -0.017 | 0.015 | 0.015 | 0 | 0.12 | 0.33 | 0.07 | 0.11 | 0.09 | 0.06 |
| 4 | 157818132 | rs17231377 | 1.442 | -1.246 | 0.475 | -0.437 | 0.003 | 0.023 | 0.023 | 0 | 0.49 | 0.57 | 0.65 | 0.61 | 0.59 | 0.34 |
| 4 | 157825397 | rs11728198 | 1.247 | -1.278 | 0.262 | -0.464 | 0.001 | 0.025 | 0.025 | 0 | 0.52 | 0.57 | 0.65 | 0.7 | 0.59 | 0.37 |
| 4 | 157836610 | rs4691383 | 2.437 | -0.736 | -0.951 | -0.244 | -0.093 | -0.188 | -0.188 | 0 | 0.28 | 0.02 | 0.29 | 0.49 | 0.31 | 0.21 |
| 4 | 157837205 | rs17231573 | -0.365 | -1.542 | -0.213 | 0.112 | -0.065 | 0.132 | 0.132 | 0 | 0.3 | 0.12 | 0.23 | 0.05 | 0.33 | 0.35 |
| 4 | 157842396 | rs10517660 | 2.094 | -1.461 | 0.018 | 0.233 | -0.011 | 0.033 | 0.033 | 0 | 0.45 | 0.22 | 0.61 | 0.62 | 0.59 | 0.34 |
| 4 | 157843781 | rs10010472 | -1.606 | -1.751 | 0.128 | 0.387 | -0.014 | -0.024 | -0.024 | 0 | 0.4 | 0.55 | 0.25 | 0.3 | 0.29 | 0.54 |
| 4 | 157845568 | rs6811767 | -1.234 | -1.525 | 0.789 | 0.724 | 0.006 | 0.121 | 0.121 | 0 | 0.07 | 0.36 | 0.02 | 0.11 | 0.01 | 0.03 |
| 4 | 157849689 | rs17035447 | -0.958 | -1.503 | 0.8 | 0.735 | 0.027 | 0.148 | 0.148 | 0 | 0.06 | 0.37 | 0.02 | 0.11 | 0.01 | 0.01 |
| 4 | 157856084 | rs4382082 | -0.873 | -1.535 | 0.48 | 0.722 | -0.033 | 0.054 | 0.054 | 0 | 0.29 | 0.13 | 0.42 | 0.16 | 0.31 | 0.35 |
| 4 | 157884105 | rs6845322 | -1.515 | -1.251 | 0.813 | 0.999 | -0.007 | 0.007 | 0.007 | 0 | 0.39 | 0.59 | 0.47 | 0.46 | 0.32 | 0.55 |
| 4 | 157886145 | rs6811213 | -1.615 | -1.483 | 1.306 | 0.918 | 0.034 | 0.157 | 0.157 | 0 | 0.06 | 0.35 | 0.02 | 0.12 | 0.01 | 0.01 |
| 4 | 157897554 | rs1443237 | 1.53 | -1.999 | 0.785 | 0.807 | -0.001 | -0.001 | -0.001 | 0 | 0.61 | 0.37 | 0.79 | 0.6 | 0.77 | 0.62 |
| 4 | 157906976 | rs6833575 | -0.897 | -2.111 | 1.002 | 0.553 | -0.013 | -0.035 | -0.035 | 0.187 | 0.13 | 0.16 | 0.07 | 0.03 | 0.09 | 0.1 |
| 4 | 157914261 | rs1015540 | 2.117 | -2.246 | 1.348 | 0.67 | -0.012 | 0.002 | 0.002 | 0.031 | 0.61 | 0.4 | 0.8 | 0.61 | 0.76 | 0.61 |
| 4 | 157916511 | rs11723576 | -1.003 | -1.894 | 1.019 | 0.819 | -0.001 | -0.008 | -0.008 | 0 | 0.17 | 0.15 | 0.1 | 0.27 | 0.13 | 0.24 |
| 4 | 157921480 | rs1443230 | 2.239 | -2.254 | 0.999 | 0.195 | 0.102 | 0.057 | 0.057 | 2.828 | 0.53 | 0.19 | 0.73 | 0.65 | 0.72 | 0.6 |
| 4 | 157931688 | rs716680 | 1.768 | -2.409 | 1.385 | 0.121 | 0.026 | -0.002 | -0.002 | 0 | 0.66 | 0.28 | 0.79 | 0.66 | 0.81 | 0.65 |
| 4 | 157936643 | rs11100086 | -0.811 | -2.413 | 1.801 | 0.127 | -0.003 | 0.02 | 0.02 | 0 | 0.17 | 0.25 | 0.14 | 0.22 | 0.13 | 0.19 |
| 4 | 157940550 | rs765986 | -1.799 | -2.46 | 1.828 | -0.102 | -0.003 | -0.028 | -0.028 | 0 | 0.18 | 0.2 | 0.09 | 0.03 | 0.14 | 0.16 |
| 4 | 157959931 | rs4691389 | 1.283 | -2.37 | 1.968 | -0.008 | -0.068 | 0.006 | 0.006 | 0.074 | 0.89 | 0.67 | 0.96 | 0.91 | 0.99 | 0.94 |
| 4 | 157959959 | rs7664666 | -0.413 | -2.287 | 1.684 | 0.001 | -0.003 | -0.013 | -0.013 | 0.369 | 0.07 | 0.08 | 0.03 | 0 | 0.03 | 0.04 |
| 4 | 157964285 | rs4691391 | 0.826 | -2.136 | 1.367 | 0.268 | -0.03 | -0.005 | -0.005 | 1.641 | 0.07 | 0.01 | 0.36 | 0.55 | 0.1 | 0.27 |
| 4 | 157965809 | rs2343742 | 2.463 | -2.226 | 2.064 | -0.412 | 0.032 | 0.045 | 0.045 | 2.342 | 0.59 | 0.33 | 0.74 | 0.76 | 0.73 | 0.61 |
| 4 | 157965879 | rs11100087 | -0.874 | -2.166 | 1.931 | -0.402 | -0.017 | -0.023 | -0.023 | 0.062 | 0.18 | 0.04 | 0.15 | 0.21 | 0.17 | 0.2 |
| 4 | 157966554 | rs2343744 | -2.451 | -2.169 | 2.093 | -0.393 | -0.007 | -0.015 | -0.015 | 0 | 0.34 | 0.4 | 0.23 | 0.24 | 0.26 | 0.34 |
| 4 | 157980872 | rs2343753 | 1.99 | -2.154 | 2.276 | -0.426 | -0.04 | -0.029 | -0.029 | 0 | 0.82 | 0.58 | 0.92 | 0.88 | 0.9 | 0.84 |
| 4 | 157991592 | rs6812324 | 2.124 | -2.229 | 2.316 | -0.386 | -0.024 | -0.006 | -0.006 | 0 | 0.63 | 0.4 | 0.78 | 0.69 | 0.8 | 0.68 |

| | | | | | | | | | | | | | | | | |
|---|---|---|---|---|---|---|---|---|---|---|---|---|---|---|---|---|
| 4 | 157993607 | rs2880691 | -1.546 | -2.071 | 2.264 | -0.346 | 0.001 | 0.001 | 0.001 | 0.332 | 0.24 | 0.23 | 0.14 | 0.21 | 0.16 | 0.26 |
| 4 | 157994894 | rs4690875 | 1.763 | -1.967 | 2.372 | -0.358 | -0.035 | 0.018 | 0.018 | 4.335 | 0.67 | 0.5 | 0.8 | 0.55 | 0.79 | 0.59 |
| 4 | 158002896 | rs6852066 | -2.118 | -1.996 | 2.043 | -0.347 | 0.001 | -0.001 | -0.001 | 0 | 0.36 | 0.35 | 0.18 | 0.45 | 0.21 | 0.39 |
| 4 | 158018180 | rs17035648 | -0.834 | -1.859 | 1.846 | -0.31 | 0.021 | -0.005 | -0.005 | 1.546 | 0.21 | 0.11 | 0.14 | 0.3 | 0.14 | 0.27 |
| 4 | 158035818 | rs11100094 | -1.31 | -1.772 | 2.219 | -0.365 | -0.016 | -0.007 | -0.007 | 0 | 0.25 | 0.32 | 0.17 | 0.3 | 0.17 | 0.29 |
| 4 | 158036110 | rs11100096 | -0.789 | -1.771 | 2.203 | -0.364 | 0.015 | -0.005 | -0.005 | 0.97 | 0.21 | 0.11 | 0.14 | 0.3 | 0.15 | 0.28 |
| 4 | 158051385 | rs7655209 | -2.713 | -1.632 | 2.169 | -0.425 | -0.011 | -0.013 | -0.013 | 0 | 0.26 | 0.63 | 0.15 | 0.17 | 0.17 | 0.19 |
| 4 | 158064484 | rs3775721 | -3.077 | -1.568 | 2.017 | -0.531 | 0.054 | 0.02 | 0.02 | 0.305 | 0.48 | 0.76 | 0.3 | 0.48 | 0.32 | 0.47 |
| 4 | 158073913 | rs1801154 | -0.821 | -0.625 | 2.504 | -0.914 | 0.002 | 0.006 | 0.006 | 0 | 0.09 | 0.08 | 0.02 | 0.03 | 0.04 | 0.11 |
| 4 | 158074503 | rs17035810 | -1.551 | -0.62 | 2.552 | -0.917 | 0 | 0.001 | 0.001 | 0 | 0.12 | 0.14 | 0.07 | 0.05 | 0.08 | 0.14 |
| 4 | 158074816 | rs11945868 | -2.586 | -0.59 | 2.294 | -0.93 | 0.063 | 0.011 | 0.011 | 3.345 | 0.07 | 0.44 | 0.05 | 0 | 0.04 | 0.01 |
| 4 | 158088464 | rs17035816 | -1.288 | -0.567 | 1.229 | -0.782 | 0.02 | -0.017 | -0.017 | 0.181 | 0.07 | 0.32 | 0.04 | 0.01 | 0.04 | 0.02 |
| 4 | 158093061 | rs10517662 | -1.038 | -0.58 | 1.081 | -0.789 | -0.009 | -0.017 | -0.017 | 0.135 | 0.11 | 0.15 | 0.03 | 0.04 | 0.04 | 0.13 |
| 4 | 158093737 | rs17035827 | -1.546 | -0.211 | 0.702 | -0.721 | 0.017 | -0.036 | -0.036 | 0 | 0.1 | 0.31 | 0.08 | 0.02 | 0.08 | 0.05 |
| 4 | 158105424 | rs10517663 | -0.924 | -0.208 | -0.309 | -0.462 | -0.024 | -0.024 | -0.024 | 0 | 0.05 | 0 | 0.03 | 0.14 | 0.07 | 0.04 |
| 4 | 158115848 | rs6831272 | -1.211 | -0.822 | -0.114 | -1.2 | -0.01 | -0.036 | -0.036 | 0 | 0.4 | 0.56 | 0.24 | 0.19 | 0.3 | 0.26 |
| 4 | 158131579 | rs10025251 | 0.723 | -1.007 | 0.25 | -1.316 | -0.002 | -0.003 | -0.003 | 0.219 | 0.8 | 0.75 | 0.86 | 0.8 | 0.83 | 0.82 |
| 4 | 158143886 | rs6536221 | -0.897 | -1.297 | 0.676 | -1.602 | -0.001 | -0.04 | -0.04 | 0.499 | 0.21 | 0.24 | 0.08 | 0.01 | 0.17 | 0.08 |
| 4 | 158155690 | rs17035889 | -0.781 | -1.328 | 0.79 | -1.609 | 0.053 | 0.049 | 0.049 | 0.92 | 0.08 | 0.23 | 0.12 | 0.2 | 0.14 | 0.06 |
| 4 | 158166689 | rs17035903 | -1.621 | -1.412 | 0.822 | -1.655 | -0.011 | -0.023 | -0.023 | 0 | 0.21 | 0.36 | 0.14 | 0.2 | 0.17 | 0.18 |
| 4 | 158168547 | rs10461267 | -0.822 | -1.38 | 0.769 | -1.683 | 0.016 | 0.051 | 0.051 | 0.477 | 0.2 | 0.12 | 0.07 | 0.01 | 0.16 | 0.08 |
| 4 | 158186342 | rs9993365 | -2.492 | -1.388 | 0.869 | -1.795 | -0.008 | -0.04 | -0.04 | 1.513 | 0.42 | 0.51 | 0.22 | 0.21 | 0.34 | 0.26 |
| 4 | 158199532 | rs7682101 | -0.79 | -1.371 | 0.911 | -1.779 | 0.061 | 0.054 | 0.054 | 1.005 | 0.08 | 0.25 | 0.12 | 0.2 | 0.14 | 0.06 |
| 4 | 158205538 | rs6844775 | -0.415 | -1.474 | 0.981 | -1.828 | 0.002 | -0.015 | -0.015 | 0 | 0.13 | 0.11 | 0.02 | 0 | 0.04 | 0.12 |
| 4 | 158221234 | rs10008950 | -2.417 | -1.457 | 0.839 | -1.82 | -0.01 | -0.046 | -0.046 | 0.325 | 0.42 | 0.51 | 0.22 | 0.21 | 0.34 | 0.26 |
| 4 | 158222905 | rs6536229 | -1.616 | -1.437 | 0.884 | -1.786 | -0.012 | -0.035 | -0.035 | 0 | 0.22 | 0.44 | 0.15 | 0.2 | 0.18 | 0.18 |
| 4 | 158242540 | rs4475186 | 2.036 | -1.155 | 0.742 | -1.86 | -0.003 | -0.027 | -0.027 | 0 | 0.56 | 0.48 | 0.75 | 0.79 | 0.63 | 0.73 |
| 4 | 158252243 | rs17244653 | -0.647 | -0.954 | 0.611 | -1.746 | -0.014 | 0.053 | 0.053 | 0 | 0.11 | 0.03 | 0.07 | 0 | 0.11 | 0.03 |
| 4 | 158276014 | rs6536234 | -2.074 | -0.632 | 0.642 | -1.601 | -0.007 | 0.001 | 0.001 | 0.504 | 0.33 | 0.77 | 0.31 | 0.36 | 0.34 | 0.16 |
| 4 | 158279954 | rs10012124 | -0.751 | -0.541 | 0.841 | -1.564 | 0.081 | 0.207 | 0.207 | 1.001 | 0.08 | 0.47 | 0.13 | 0.17 | 0.09 | 0.09 |
| 4 | 158292824 | rs4550983 | 2.437 | -0.664 | 0.379 | -1.668 | -0.045 | 0.091 | 0.091 | 0.612 | 0.49 | 0.08 | 0.63 | 0.62 | 0.54 | 0.67 |
| 4 | 158322542 | rs4691398 | 1.923 | -0.738 | 0.474 | -1.795 | 0.01 | 0.04 | 0.04 | 0 | 0.54 | 0.56 | 0.75 | 0.78 | 0.6 | 0.69 |
| 4 | 158324379 | rs11725238 | 2.202 | -0.813 | 0.483 | -1.855 | -0.004 | -0.007 | -0.007 | 0 | 0.74 | 0.25 | 0.71 | 0.64 | 0.72 | 0.75 |
| 4 | 158327705 | rs6823909 | 1.1 | -0.813 | 0.489 | -1.855 | 0.001 | -0.003 | -0.003 | 0 | 0.83 | 0.75 | 0.84 | 0.79 | 0.81 | 0.85 |
| 4 | 158333646 | rs4234911 | -0.426 | -0.798 | 0.578 | -1.825 | 0.102 | 0.238 | 0.238 | 0.306 | 0.08 | 0.45 | 0.13 | 0.17 | 0.09 | 0.1 |
| 4 | 158336110 | rs4691400 | -0.868 | -0.798 | 0.572 | -1.825 | 0.056 | 0.169 | 0.169 | 1.405 | 0.09 | 0.5 | 0.13 | 0.17 | 0.09 | 0.1 |
| 4 | 158346878 | rs9992749 | -0.611 | -0.796 | 0.669 | -1.824 | 0.095 | 0.23 | 0.23 | 0.508 | 0.08 | 0.46 | 0.13 | 0.17 | 0.09 | 0.1 |
| 4 | 158349609 | rs12502016 | 2.999 | -0.619 | 0.474 | -1.872 | -0.122 | 0.042 | 0.042 | 0 | 0.45 | 0.04 | 0.61 | 0.63 | 0.49 | 0.6 |
| 4 | 158359767 | rs1865353 | 3.006 | -0.621 | 0.515 | -1.872 | -0.127 | 0.033 | 0.033 | 0 | 0.45 | 0.04 | 0.61 | 0.62 | 0.49 | 0.6 |
| 4 | 158361729 | rs10029781 | -2.836 | -0.607 | 0.561 | -1.791 | -0.041 | 0.064 | 0.064 | 0 | 0.53 | 0.85 | 0.39 | 0.39 | 0.51 | 0.4 |
| 4 | 158363688 | rs6536237 | -2.881 | -0.588 | 0.494 | -1.834 | -0.037 | 0.059 | 0.059 | 0 | 0.53 | 0.86 | 0.39 | 0.39 | 0.51 | 0.41 |
| 4 | 158366623 | rs1369028 | -2.286 | -0.635 | 0.617 | -1.81 | -0.004 | -0.009 | -0.009 | 2.24 | 0.26 | 0.75 | 0.29 | 0.37 | 0.28 | 0.24 |
| 4 | 158367001 | rs10213312 | -1.871 | -0.64 | 0.663 | -1.814 | 0.003 | 0.008 | 0.008 | 0 | 0.36 | 0.62 | 0.23 | 0.2 | 0.32 | 0.26 |
| 4 | 158375017 | rs6842542 | -1.448 | -0.745 | 0.572 | -1.866 | 0.002 | -0.005 | -0.005 | 0 | 0.17 | 0.24 | 0.16 | 0.2 | 0.19 | 0.14 |
| 4 | 158375048 | rs11731394 | -0.957 | -0.734 | 0.451 | -1.858 | 0.003 | 0.099 | 0.099 | 0 | 0.26 | 0.1 | 0.1 | 0.01 | 0.23 | 0.16 |
| 4 | 158382194 | rs17186879 | -2.303 | -0.68 | 0.539 | -1.785 | -0.001 | 0.13 | 0.13 | 0 | 0.44 | 0.28 | 0.25 | 0.21 | 0.41 | 0.3 |
| 4 | 158383828 | rs17246641 | -2.248 | -0.671 | 0.543 | -1.783 | -0.002 | 0.13 | 0.13 | 0 | 0.44 | 0.28 | 0.25 | 0.21 | 0.41 | 0.3 |
| 4 | 158391794 | rs12647438 | 1.339 | -0.677 | 0.997 | -1.49 | -0.012 | -0.011 | -0.011 | 0 | 0.73 | 0.34 | 0.81 | 0.81 | 0.79 | 0.82 |
| 4 | 158416500 | rs17036216 | -1.002 | -0.634 | 1.133 | -1.596 | 0.026 | -0.058 | -0.058 | 0 | 0.11 | 0.22 | 0.09 | 0.01 | 0.14 | 0.07 |
| 4 | 158419053 | rs6820047 | -0.852 | -0.627 | 1.169 | -1.543 | 0.137 | 0.426 | 0.426 | 0 | 0.05 | 0.34 | 0.13 | 0.35 | 0.08 | 0.1 |
| 4 | 158419205 | rs4691413 | 1.453 | -0.612 | 0.491 | -1.538 | -0.007 | 0.001 | 0.001 | 0 | 0.71 | 0.34 | 0.77 | 0.63 | 0.77 | 0.8 |
| 4 | 158426633 | rs1156225 | 1.861 | -0.48 | -0.022 | -1.481 | -0.017 | -0.001 | -0.001 | 0 | 0.61 | 0.12 | 0.67 | 0.61 | 0.63 | 0.73 |
| 4 | 158430547 | rs12507865 | 1.602 | -0.287 | 0.1 | -1.403 | -0.019 | -0.006 | -0.006 | 0 | 0.65 | 0.18 | 0.68 | 0.62 | 0.63 | 0.77 |
| 4 | 158443622 | rs725037 | 1.219 | -0.492 | 0.856 | -1.716 | -0.025 | -0.034 | -0.034 | 0 | 0.81 | 0.46 | 0.88 | 0.85 | 0.87 | 0.87 |
| 4 | 158444071 | rs1550017 | -1.178 | -0.492 | 1.136 | -1.717 | 0.075 | 0.08 | 0.08 | 0 | 0.06 | 0.27 | 0.08 | 0.14 | 0.1 | 0.11 |
| 4 | 158444947 | rs12054636 | -0.938 | -0.483 | 1.068 | -1.712 | 0.062 | 0.066 | 0.066 | 0 | 0.06 | 0.22 | 0.08 | 0.14 | 0.1 | 0.11 |
| 4 | 158447249 | rs6536246 | 1.721 | -0.585 | 0.422 | -1.857 | -0.001 | -0.012 | -0.012 | 0.282 | 0.66 | 0.32 | 0.82 | 0.85 | 0.76 | 0.72 |
| 4 | 158452144 | rs4475188 | 0.909 | -0.355 | 0.353 | -1.38 | -0.035 | -0.028 | -0.028 | 0 | 0.82 | 0.51 | 0.89 | 0.85 | 0.89 | 0.87 |
| 4 | 158455363 | rs1025554 | 0.865 | -0.057 | -1.139 | -1.456 | -0.124 | 0.038 | 0.038 | 0 | 0.37 | 0.07 | 0.63 | 0.75 | 0.48 | 0.57 |
| 4 | 158462362 | rs1594869 | -1.273 | -0.022 | 0.113 | -1.625 | -0.018 | -0.093 | -0.093 | 0.017 | 0.18 | 0.42 | 0.09 | 0.03 | 0.14 | 0.17 |
| 4 | 158467400 | rs6827325 | -0.049 | 0.165 | -0.019 | -1.514 | 0 | -0.024 | -0.024 | 0 | 0.07 | 0.12 | 0.04 | 0 | 0.06 | 0.03 |
| 4 | 158468178 | rs4234914 | 0.484 | -0.048 | -0.082 | -1.751 | -0.007 | -0.002 | -0.002 | 0 | 0.77 | 0.76 | 0.81 | 0.79 | 0.67 | 0.68 |
| 4 | 158475705 | rs17036351 | -0.689 | 0.088 | 0.297 | -2.057 | 0.037 | -0.059 | -0.059 | 0 | 0.1 | 0.33 | 0.06 | 0 | 0.08 | 0.06 |
| 4 | 158479475 | rs12331558 | 0.557 | -0.352 | 0.031 | -2.362 | 0.009 | 0.002 | 0.002 | 0 | 0.5 | 0.46 | 0.69 | 0.86 | 0.56 | 0.6 |
| 4 | 158479507 | rs1434384 | 0.583 | -0.362 | -0.027 | -2.366 | 0.002 | -0.019 | -0.019 | 0.009 | 0.52 | 0.47 | 0.7 | 0.86 | 0.56 | 0.6 |
| 4 | 158492974 | rs1036342 | -1.222 | -0.348 | 0.328 | -2.064 | -0.011 | -0.024 | -0.024 | 2.42 | 0.28 | 0.64 | 0.2 | 0.18 | 0.26 | 0.31 |
| 4 | 158503661 | rs10857301 | -0.191 | -0.477 | 0.619 | -2.448 | 0.015 | -0.074 | -0.074 | 0 | 0.13 | 0.25 | 0.11 | 0 | 0.13 | 0.05 |
| 4 | 158503818 | rs10857302 | 0.178 | -0.435 | 0.673 | -2.398 | 0.05 | -0.043 | -0.043 | 0.134 | 0.08 | 0.25 | 0.08 | 0 | 0.09 | 0.01 |
| 4 | 158524662 | rs728492 | -1.123 | -0.561 | 0.787 | -2.245 | -0.012 | -0.03 | -0.03 | 7.462 | 0.28 | 0.62 | 0.2 | 0.18 | 0.26 | 0.31 |

| | | | | | | | | | | | | | | | | |
|---|---|---|---|---|---|---|---|---|---|---|---|---|---|---|---|---|
| 4 | 158530437 | rs1982845 | -0.86 | -0.464 | 0.345 | -2.198 | -0.006 | -0.041 | -0.041 | 0.481 | 0.24 | 0.32 | 0.17 | 0.13 | 0.25 | 0.29 |
| 4 | 158538338 | rs11724621 | 1.468 | -0.251 | -0.302 | -2.479 | -0.124 | 0.247 | 0.247 | 0.191 | 0.43 | 0.03 | 0.66 | 0.89 | 0.54 | 0.51 |
| 4 | 158540416 | rs1907089 | 0.587 | -0.518 | 0.537 | -2.669 | 0.007 | 0.052 | 0.052 | 0.269 | 0.59 | 0.69 | 0.76 | 0.93 | 0.63 | 0.57 |
| 4 | 158541687 | rs7697239 | -0.847 | -0.556 | 1.274 | -2.688 | -0.006 | -0.058 | -0.058 | 0.008 | 0.1 | 0.24 | 0.07 | 0 | 0.07 | 0.04 |
| 4 | 158541777 | rs1490530 | -0.235 | -0.552 | 1.207 | -2.784 | -0.014 | 0.15 | 0.15 | 0 | 0.72 | 0.91 | 0.86 | 1 | 0.75 | 0.81 |
| 4 | 158542093 | rs6856493 | 1.726 | -0.522 | 1.518 | -2.748 | -0.017 | 0.176 | 0.176 | 0.123 | 0.45 | 0.16 | 0.67 | 0.89 | 0.55 | 0.51 |
| 4 | 158550241 | rs17036576 | -0.609 | -0.606 | 1.899 | -2.445 | -0.004 | -0.001 | -0.001 | 0 | 0.08 | 0.19 | 0.06 | 0.07 | 0.05 | 0.04 |
| 4 | 158562269 | rs1352695 | 1.73 | -0.603 | 1.44 | -2.47 | -0.009 | 0.183 | 0.183 | 0.011 | 0.46 | 0.12 | 0.67 | 0.82 | 0.57 | 0.52 |
| 4 | 158562472 | rs1352696 | 1.778 | -0.599 | 1.349 | -2.469 | -0.037 | 0.2 | 0.2 | 0.036 | 0.45 | 0.07 | 0.67 | 0.81 | 0.57 | 0.52 |
| 4 | 158574310 | rs6821900 | -1.507 | -0.345 | 1.917 | -2.084 | -0.028 | -0.075 | -0.075 | 0.151 | 0.13 | 0.38 | 0.07 | 0.01 | 0.06 | 0.05 |
| 4 | 158576241 | rs6835594 | -0.079 | -0.48 | 2.085 | -2.189 | -0.003 | -0.061 | -0.061 | 0 | 0.77 | 0.63 | 0.86 | 0.92 | 0.79 | 0.87 |
| 4 | 158577288 | rs10017629 | -0.041 | -0.484 | 2.084 | -2.207 | -0.004 | -0.063 | -0.063 | 0 | 0.77 | 0.63 | 0.86 | 0.93 | 0.79 | 0.87 |
| 4 | 158577315 | rs9995432 | -0.079 | -0.48 | 2.086 | -2.205 | -0.003 | -0.063 | -0.063 | 0.253 | 0.77 | 0.63 | 0.86 | 0.93 | 0.79 | 0.87 |
| 4 | 158580884 | rs10005357 | -1.353 | -0.567 | 2.094 | -2.27 | -0.016 | -0.043 | -0.043 | 0 | 0.2 | 0.3 | 0.15 | 0.09 | 0.14 | 0.25 |
| 4 | 158584331 | rs1490541 | -0.117 | -0.543 | 1.97 | -2.252 | -0.003 | -0.063 | -0.063 | 0.323 | 0.77 | 0.63 | 0.86 | 0.93 | 0.79 | 0.87 |
| 4 | 158584939 | rs10001392 | -0.117 | -0.543 | 1.97 | -2.252 | -0.003 | -0.063 | -0.063 | 0.112 | 0.77 | 0.63 | 0.85 | 0.93 | 0.79 | 0.87 |
| 4 | 158586638 | rs6839727 | 2.027 | -0.551 | 1.478 | -2.261 | 0.001 | 0.237 | 0.237 | 0 | 0.51 | 0.04 | 0.68 | 0.82 | 0.62 | 0.59 |
| 4 | 158591563 | rs1352693 | 0.648 | -0.48 | 1.995 | -2.145 | -0.001 | -0.018 | -0.018 | 0 | 0.86 | 0.65 | 0.88 | 0.91 | 0.89 | 0.77 |
| 4 | 158593927 | rs10517673 | -0.274 | -0.467 | 1.696 | -2.132 | -0.008 | -0.028 | -0.028 | 0.229 | 0.08 | 0.1 | 0.04 | 0 | 0.03 | 0.02 |
| 4 | 158600677 | rs2220582 | 0.583 | -0.491 | 1.873 | -2.075 | 0.015 | 0.057 | 0.057 | 1.204 | 0.85 | 0.55 | 0.84 | 0.83 | 0.87 | 0.75 |
| 4 | 158602153 | rs10024159 | -0.549 | -0.523 | 2.009 | -2.096 | -0.017 | -0.044 | -0.044 | 0.01 | 0.09 | 0.17 | 0.04 | 0 | 0.03 | 0.02 |
| 4 | 158605184 | rs17036640 | -0.7 | -0.518 | 2.147 | -2.092 | 0.008 | 0.042 | 0.042 | 0.466 | 0.14 | 0.29 | 0.14 | 0.17 | 0.13 | 0.25 |
| 4 | 158608616 | rs9998812 | -0.817 | -0.631 | 1.957 | -2.234 | 0.007 | -0.029 | -0.029 | 1.618 | 0.3 | 0.3 | 0.15 | 0 | 0.23 | 0.13 |
| 4 | 158609602 | rs4484348 | 1.66 | -0.638 | 1.94 | -2.213 | -0.02 | -0.018 | -0.018 | 0 | 0.76 | 0.4 | 0.8 | 0.83 | 0.83 | 0.73 |
| 4 | 158620303 | rs11100128 | 2.222 | -0.692 | 1.385 | -2.224 | 0.043 | 0.042 | 0.042 | 1.547 | 0.7 | 0.13 | 0.78 | 0.82 | 0.81 | 0.7 |
| 4 | 158622875 | rs4691434 | 1.91 | -0.448 | 1.533 | -2.234 | 0.021 | 0.277 | 0.277 | 0 | 0.52 | 0.13 | 0.72 | 0.89 | 0.62 | 0.6 |
| 4 | 158634769 | rs10030915 | -1.408 | -0.438 | 1.349 | -2.157 | 0 | -0.013 | -0.013 | 0 | 0.36 | 0.65 | 0.22 | 0.1 | 0.26 | 0.17 |
| 4 | 158663069 | rs1904949 | 1.282 | -0.318 | 1.358 | -1.981 | -0.009 | -0.054 | -0.054 | 0 | 0.83 | 0.39 | 0.87 | 0.95 | 0.88 | 0.74 |
| 4 | 158669702 | rs1383707 | -1.039 | -0.316 | 0.968 | -2.162 | -0.009 | 0.013 | 0.013 | 0 | 0.34 | 0.29 | 0.21 | 0.12 | 0.35 | 0.32 |
| 4 | 158672216 | rs11100132 | 1.535 | -0.351 | 0.567 | -2.142 | -0.049 | 0.18 | 0.18 | 0 | 0.55 | 0.05 | 0.72 | 0.83 | 0.64 | 0.63 |
| 4 | 158689938 | rs756334 | 2.082 | -0.079 | 0.245 | -2.392 | -0.142 | 0.187 | 0.187 | 0 | 0.44 | 0.03 | 0.64 | 0.84 | 0.53 | 0.55 |
| 4 | 158697324 | rs12648731 | 1.754 | 0.166 | 0.577 | -2.266 | 0.001 | -0.007 | -0.007 | 0 | 0.76 | 0.27 | 0.81 | 0.95 | 0.79 | 0.67 |
| 4 | 158704610 | rs1383704 | 0.976 | 0.372 | 0.524 | -1.471 | 0.027 | 0.041 | 0.041 | 0.807 | 0.86 | 0.65 | 0.89 | 0.88 | 0.88 | 0.74 |
| 4 | 158706314 | rs985731 | 0.557 | 0.401 | 0.364 | -1.294 | 0.02 | 0.175 | 0.175 | 1.239 | 0.94 | 0.7 | 0.96 | 0.88 | 0.97 | 0.94 |
| 4 | 158711676 | rs7671734 | -0.784 | 0.675 | -0.785 | -1.583 | -0.041 | 0.107 | 0.107 | 0 | 0.29 | 0.13 | 0.18 | 0.07 | 0.31 | 0.19 |
| 4 | 158715149 | rs7667778 | -0.497 | 0.689 | -0.698 | -1.551 | -0.025 | 0.066 | 0.066 | 0 | 0.27 | 0.15 | 0.13 | 0.08 | 0.27 | 0.22 |
| 4 | 158715453 | rs1531713 | 1.467 | 0.344 | -0.57 | -1.596 | 0.031 | 0.032 | 0.032 | 0.072 | 0.72 | 0.22 | 0.79 | 0.84 | 0.81 | 0.8 |
| 4 | 158724848 | rs727755 | -0.626 | 1.073 | -0.338 | -1.573 | -0.048 | 0.058 | 0.058 | 0 | 0.26 | 0.11 | 0.18 | 0 | 0.34 | 0.15 |
| 4 | 158730857 | rs13150049 | -0.728 | 1.016 | 0.202 | -1.473 | 0.011 | 0.012 | 0.012 | 0.657 | 0.09 | 0.23 | 0.1 | 0.07 | 0.07 | 0.1 |
| 4 | 158731019 | rs7434819 | -0.867 | 1.049 | 0.504 | -1.526 | 0.03 | -0.138 | -0.138 | 0 | 0.29 | 0.36 | 0.19 | 0 | 0.36 | 0.16 |
| 4 | 158737934 | rs6812217 | -1.651 | 0.808 | 0.858 | -0.775 | 0.004 | 0.052 | 0.052 | 0.39 | 0.14 | 0.4 | 0.13 | 0.16 | 0.1 | 0.1 |
| 4 | 158742582 | rs7677854 | -1.536 | 2.045 | 0.751 | -1.553 | 0.073 | -0.105 | -0.105 | 0.167 | 0.19 | 0.33 | 0.17 | 0.01 | 0.29 | 0.22 |
| 4 | 158760439 | rs7654999 | -0.446 | 0.959 | 0.604 | -1.861 | -0.005 | -0.048 | -0.048 | 2.008 | 0.38 | 0.48 | 0.45 | 0.83 | 0.33 | 0.61 |
| 4 | 158765677 | rs6536284 | -0.973 | 0.879 | 0.228 | -2.435 | -0.002 | 0.004 | 0.004 | 0 | 0.82 | 0.86 | 0.71 | 0.9 | 0.81 | 0.83 |
| 4 | 158771076 | rs6536285 | -1.084 | 0.875 | 0.239 | -2.437 | -0.012 | 0.016 | 0.016 | 0 | 0.83 | 0.93 | 0.72 | 0.9 | 0.81 | 0.83 |
| 4 | 158778839 | rs7693369 | 0.093 | 0.787 | 0.28 | -2.445 | -0.005 | -0.054 | -0.054 | 0.006 | 0.39 | 0.48 | 0.45 | 0.84 | 0.34 | 0.61 |
| 4 | 158782030 | rs9991982 | 0.121 | 1.057 | 0.433 | -2.551 | 0.006 | -0.05 | -0.05 | 0 | 0.41 | 0.48 | 0.49 | 0.84 | 0.43 | 0.62 |
| 4 | 158790324 | rs7698656 | 0.268 | -0.114 | 1.422 | -2.384 | -0.016 | -0.043 | -0.043 | 0.001 | 0.74 | 0.41 | 0.67 | 0.86 | 0.79 | 0.75 |
| 4 | 158793663 | rs1481223 | 1.184 | -0.408 | 1.186 | -2.605 | 0.012 | 0.034 | 0.034 | 1.254 | 0.62 | 0.29 | 0.61 | 0.84 | 0.67 | 0.69 |
| 4 | 158813228 | rs1481231 | 0.137 | -0.383 | 1.215 | -2.573 | -0.01 | -0.026 | -0.026 | 1.878 | 0.73 | 0.37 | 0.66 | 0.86 | 0.79 | 0.75 |
| 4 | 158813324 | rs6857469 | 0.114 | -0.255 | 0.909 | -2.531 | 0.002 | 0.025 | 0.025 | 0 | 0.11 | 0.09 | 0.06 | 0.02 | 0.12 | 0.06 |
| 4 | 158815510 | rs1448451 | 0.456 | -0.207 | 1.068 | -2.489 | -0.008 | -0.071 | -0.071 | 0 | 0.66 | 0.47 | 0.63 | 0.87 | 0.67 | 0.76 |
| 4 | 158840495 | rs11939514 | -1.319 | -0.411 | 1.764 | -2.147 | -0.013 | 0.013 | 0.013 | 0.025 | 0.07 | 0.39 | 0.04 | 0.03 | 0 | 0.06 |
| 4 | 158842148 | rs1375138 | -1.131 | -0.405 | 1.118 | -2.336 | -0.02 | 0.036 | 0.036 | 0 | 0.78 | 0.9 | 0.68 | 0.92 | 0.72 | 0.84 |
| 4 | 158857135 | rs1448450 | -0.192 | -0.467 | 1.194 | -2.472 | -0.006 | 0.023 | 0.023 | 0 | 0.68 | 0.76 | 0.64 | 0.88 | 0.65 | 0.79 |
| 4 | 158863930 | rs2725476 | -1.203 | -0.456 | 0.974 | -2.326 | -0.017 | 0.024 | 0.024 | 0 | 0.78 | 0.86 | 0.68 | 0.9 | 0.72 | 0.85 |
| 4 | 158866939 | rs721604 | 0.971 | -1.087 | 0.988 | -2.596 | 0.007 | 0.123 | 0.123 | 0 | 0.48 | 0.22 | 0.52 | 0.78 | 0.52 | 0.62 |
| 4 | 158867973 | rs10002472 | -0.379 | -1.01 | 0.996 | -2.524 | -0.026 | 0.055 | 0.055 | 0 | 0.67 | 0.82 | 0.61 | 0.85 | 0.6 | 0.81 |
| 4 | 158898321 | rs10212653 | 1.483 | -0.924 | 0.588 | -2.432 | -0.01 | -0.043 | -0.043 | 0 | 0.22 | 0.06 | 0.35 | 0.68 | 0.2 | 0.51 |
| 4 | 158898891 | rs4447905 | -1.099 | -1.458 | 1.168 | -2.445 | -0.028 | 0.016 | 0.016 | 0 | 0.92 | 0.99 | 0.93 | 0.97 | 0.88 | 0.96 |
| 4 | 158909445 | rs10517684 | 0.795 | -1.471 | 1.253 | -2.396 | -0.025 | 0.019 | 0.019 | 0 | 0.23 | 0.11 | 0.31 | 0.18 | 0.28 | 0.13 |
| 4 | 158917997 | rs10857308 | 0.885 | -1.523 | 1.25 | -2.396 | -0.028 | 0.019 | 0.019 | 0 | 0.22 | 0.08 | 0.31 | 0.18 | 0.28 | 0.13 |
| 4 | 158924505 | rs10461245 | 0.689 | -1.631 | 1.853 | -2.453 | -0.044 | -0.067 | -0.067 | 0.018 | 0.76 | 0.6 | 0.87 | 0.94 | 0.84 | 0.82 |
| 4 | 158932876 | rs6847800 | 0.944 | -1.62 | 1.968 | -2.438 | 0.004 | 0.01 | 0.01 | 0 | 0.52 | 0.41 | 0.54 | 0.77 | 0.56 | 0.69 |
| 4 | 158983480 | rs4569796 | -0.056 | -1.598 | 1.908 | -2.42 | -0.072 | -0.056 | -0.056 | 3.906 | 0.16 | 0.33 | 0.06 | 0.03 | 0.04 | 0.15 |
| 4 | 158988325 | rs13144391 | -0.625 | -1.599 | 1.9 | -2.413 | -0.005 | -0.012 | -0.012 | 0.106 | 0.39 | 0.43 | 0.37 | 0.21 | 0.32 | 0.28 |
| 4 | 158991908 | rs13116234 | 0.066 | -1.518 | 1.9 | -2.361 | -0.055 | -0.041 | -0.041 | 0 | 0.14 | 0.27 | 0.05 | 0.03 | 0.04 | 0.15 |
| 4 | 159014453 | rs4388135 | 0.768 | -1.486 | 1.86 | -2.344 | -0.003 | -0.01 | -0.01 | 0 | 0.54 | 0.43 | 0.55 | 0.76 | 0.58 | 0.69 |
| 4 | 159033327 | rs10517685 | 0.471 | -1.458 | 1.807 | -2.303 | 0.003 | -0.015 | -0.015 | 0.482 | 0.08 | 0.12 | 0.08 | 0.03 | 0.11 | 0.03 |

| | | | | | | | | | | | | | | | | |
|---|---|---|---|---|---|---|---|---|---|---|---|---|---|---|---|---|
| 4 | 159043610 | rs1320370 | -0.383 | -1.517 | 1.914 | -2.328 | -0.055 | -0.044 | -0.044 | 0 | 0.86 | 0.72 | 0.95 | 0.98 | 0.96 | 0.86 |
| 4 | 159045843 | rs7384 | 1 | -1.539 | 1.887 | -2.361 | -0.036 | 0.009 | 0.009 | 0 | 0.22 | 0.06 | 0.3 | 0.18 | 0.28 | 0.14 |
| 4 | 159045966 | rs8370 | 0.571 | -1.508 | 1.72 | -2.333 | -0.041 | -0.064 | -0.064 | 0.048 | 0.77 | 0.61 | 0.87 | 0.95 | 0.85 | 0.83 |
| 4 | 159048217 | rs4691455 | 0.411 | -1.504 | 1.67 | -2.331 | -0.026 | -0.043 | -0.043 | 0 | 0.78 | 0.67 | 0.88 | 0.95 | 0.85 | 0.83 |
| 4 | 159049771 | rs4501266 | 0.562 | -1.502 | 1.617 | -2.33 | -0.041 | -0.064 | -0.064 | 0 | 0.77 | 0.61 | 0.87 | 0.95 | 0.85 | 0.83 |
| 4 | 159055292 | rs7692254 | 0.903 | -1.504 | 1.616 | -2.334 | 0.033 | 0.1 | 0.1 | 0 | 0.52 | 0.27 | 0.55 | 0.77 | 0.57 | 0.67 |
| 4 | 159055765 | rs4691458 | -0.264 | -1.414 | 1.617 | -2.222 | -0.078 | -0.065 | -0.065 | 0.183 | 0.85 | 0.63 | 0.94 | 0.98 | 0.96 | 0.86 |
| 4 | 159064672 | rs6833295 | 0.886 | -1.491 | 1.083 | -2.288 | 0.05 | 0.139 | 0.139 | 0 | 0.48 | 0.23 | 0.54 | 0.8 | 0.57 | 0.67 |
| 4 | 159065799 | rs2346784 | 0.74 | -1.486 | 1.4 | -2.263 | -0.072 | -0.101 | -0.101 | 0 | 0.76 | 0.53 | 0.87 | 0.95 | 0.85 | 0.81 |
| 4 | 159076826 | rs6830702 | 0.757 | -1.477 | 1.221 | -2.259 | -0.07 | -0.099 | -0.099 | 0.219 | 0.76 | 0.51 | 0.86 | 0.95 | 0.85 | 0.81 |
| 4 | 159083696 | rs4690893 | -0.688 | -1.483 | 1.252 | -2.262 | -0.035 | -0.026 | -0.026 | 0 | 0.87 | 0.8 | 0.96 | 0.98 | 0.96 | 0.86 |
| 4 | 159086166 | rs17037201 | 0.421 | -1.446 | 0.954 | -2.244 | 0.005 | -0.026 | -0.026 | 0 | 0.09 | 0.17 | 0.08 | 0.03 | 0.11 | 0.05 |
| 4 | 159089861 | rs6849902 | -0.394 | -1.41 | 0.851 | -2.212 | -0.066 | -0.05 | -0.05 | 0.037 | 0.86 | 0.69 | 0.94 | 0.98 | 0.96 | 0.86 |
| 4 | 159092633 | rs2881373 | 0.003 | -1.152 | 0.012 | -2.078 | -0.071 | -0.099 | -0.099 | 0.307 | 0.75 | 0.5 | 0.86 | 0.95 | 0.85 | 0.82 |
| 4 | 159098784 | rs13126694 | 0.912 | -0.953 | -0.61 | -1.932 | -0.02 | 0.065 | 0.065 | 0 | 0.4 | 0.2 | 0.52 | 0.72 | 0.52 | 0.59 |
| 4 | 159101703 | rs10517688 | -0.221 | -0.11 | -0.215 | -1.311 | 0 | 0.013 | 0.013 | 0 | 0.28 | 0.23 | 0.32 | 0.17 | 0.29 | 0.14 |
| 4 | 159127462 | rs7662398 | 0.091 | -0.504 | -0.111 | -1.622 | 0.02 | 0.014 | 0.014 | 0.088 | 0.66 | 0.23 | 0.73 | 0.94 | 0.74 | 0.86 |
| 4 | 159128631 | rs13126054 | -0.111 | -0.642 | 0.705 | -1.689 | -0.012 | -0.07 | -0.07 | 0 | 0.3 | 0.37 | 0.24 | 0.06 | 0.26 | 0.14 |
| 4 | 159129407 | rs11100154 | -0.015 | -0.567 | 0.023 | -1.673 | 0.008 | -0.013 | -0.013 | 0.354 | 0.67 | 0.25 | 0.73 | 0.94 | 0.74 | 0.86 |
| 4 | 159132582 | rs10019333 | 0.201 | -0.856 | -0.267 | -1.739 | 0.086 | 0.13 | 0.13 | 0.053 | 0.61 | 0.17 | 0.72 | 0.94 | 0.74 | 0.83 |
| 4 | 159158275 | rs2272050 | -0.145 | 0.85 | -0.37 | -0.798 | -0.062 | -0.023 | -0.023 | 0.029 | 0.12 | 0.03 | 0.1 | 0.1 | 0.24 | 0.13 |
| 4 | 159158677 | rs12504074 | 0.152 | 0.965 | -0.662 | -0.955 | 0 | 0.041 | 0.041 | 0.823 | 0.15 | 0.05 | 0.15 | 0.04 | 0.14 | 0.05 |
| 4 | 159160422 | rs12510198 | -0.153 | 0.925 | -0.545 | -0.974 | -0.003 | -0.019 | -0.019 | 0.617 | 0.17 | 0.14 | 0.15 | 0.04 | 0.14 | 0.07 |
| 4 | 159164640 | rs3805144 | 0.528 | 0.821 | -0.505 | -0.855 | 0.013 | -0.037 | -0.037 | 0.092 | 0.67 | 0.53 | 0.7 | 0.85 | 0.62 | 0.79 |
| 4 | 159173102 | rs1879656 | 0.528 | 0.821 | -0.59 | -0.869 | 0.011 | -0.033 | -0.033 | 0 | 0.67 | 0.54 | 0.7 | 0.85 | 0.63 | 0.79 |
| 4 | 159176109 | rs1047725 | -0.151 | 0.744 | -1.341 | -0.749 | -0.02 | 0.043 | 0.043 | 1.633 | 0.4 | 0.19 | 0.37 | 0.26 | 0.41 | 0.36 |
| 4 | 159179862 | rs1410789 | 0.57 | 0.338 | -1.091 | -1.373 | 0.015 | -0.019 | -0.019 | 0 | 0.69 | 0.44 | 0.66 | 0.81 | 0.71 | 0.74 |
| 4 | 159185633 | rs1879658 | -0.172 | 0.224 | -0.944 | -1.409 | -0.046 | -0.092 | -0.092 | 0.013 | 0.87 | 0.46 | 0.92 | 1 | 0.95 | 0.98 |
| 4 | 159185770 | rs10857310 | -0.072 | 0.219 | -1.433 | -1.674 | 0.066 | 0.104 | 0.104 | 1.859 | 0.54 | 0.26 | 0.66 | 0.88 | 0.67 | 0.8 |
| 4 | 159190290 | rs1328925 | 0.082 | 0.275 | -1.486 | -1.407 | -0.009 | 0.077 | 0.077 | 0.954 | 0.47 | 0.24 | 0.6 | 0.77 | 0.49 | 0.73 |
| 4 | 159194559 | rs17037324 | 0.646 | 0.675 | -1.701 | -1.385 | -0.044 | -0.078 | -0.078 | 0.066 | 0.19 | 0.06 | 0.35 | 0.6 | 0.33 | 0.33 |
| 4 | 159206103 | rs11932623 | -0.829 | -0.138 | -0.624 | -1.594 | -0.074 | -0.031 | -0.031 | 0 | 0.18 | 0.54 | 0.25 | 0.12 | 0.06 | 0.2 |
| 4 | 159224413 | rs1574386 | -0.686 | -0.118 | -0.11 | -1.707 | -0.035 | -0.009 | -0.009 | 4.526 | 0.14 | 0.32 | 0.23 | 0.11 | 0.06 | 0.25 |
| 4 | 159224455 | rs1887891 | -0.583 | -0.128 | -0.106 | -1.71 | -0.036 | -0.005 | -0.005 | 0 | 0.13 | 0.31 | 0.23 | 0.13 | 0.04 | 0.24 |
| 4 | 159227601 | rs4691475 | -0.609 | -0.145 | 0.112 | -1.722 | -0.039 | 0.002 | 0.002 | 0 | 0.14 | 0.37 | 0.23 | 0.11 | 0.04 | 0.24 |
| 4 | 159235840 | rs11100160 | -0.564 | -0.144 | 0.121 | -1.721 | -0.038 | 0.001 | 0.001 | 0 | 0.14 | 0.36 | 0.23 | 0.11 | 0.04 | 0.24 |
| 4 | 159241440 | rs4577629 | -0.061 | -0.192 | -0.261 | -1.876 | 0 | -0.015 | -0.015 | 0 | 0.17 | 0.17 | 0.11 | 0.03 | 0.19 | 0.16 |
| 4 | 159255213 | rs4364309 | -1.651 | -0.343 | -0.563 | -1.892 | -0.038 | -0.087 | -0.087 | 0 | 0.85 | 0.97 | 0.79 | 0.4 | 0.78 | 0.72 |
| 4 | 159255270 | rs4379102 | 0.886 | -0.792 | -0.196 | -2.31 | 0.004 | -0.02 | -0.02 | 0 | 0.07 | 0.01 | 0.19 | 0.11 | 0.04 | 0.23 |
| 4 | 159266567 | rs6536306 | -0.355 | -0.761 | 0.003 | -2.519 | -0.003 | -0.052 | -0.052 | 0 | 0.18 | 0.24 | 0.12 | 0.04 | 0.17 | 0.16 |
| 4 | 159275588 | rs7435789 | -0.949 | -0.736 | -0.108 | -2.501 | -0.013 | -0.061 | -0.061 | 0 | 0.26 | 0.38 | 0.26 | 0.1 | 0.21 | 0.39 |
| 4 | 159280560 | rs12508350 | -0.858 | -0.892 | -0.199 | -2.768 | -0.029 | -0.084 | -0.084 | 0 | 0.36 | 0.57 | 0.28 | 0.06 | 0.21 | 0.39 |
| 4 | 159295359 | rs7441062 | -0.868 | -0.892 | -0.201 | -2.768 | -0.027 | -0.081 | -0.081 | 0.911 | 0.36 | 0.58 | 0.29 | 0.06 | 0.22 | 0.39 |
| 4 | 159314091 | rs6536309 | -1.502 | -0.719 | -0.46 | -2.988 | -0.055 | 0.155 | 0.155 | 0 | 0.69 | 0.93 | 0.77 | 0.89 | 0.7 | 0.79 |
| 4 | 159317442 | rs10857312 | 0.301 | -0.994 | 0.7 | -2.122 | -0.002 | -0.033 | -0.033 | 0 | 0.74 | 0.72 | 0.87 | 0.98 | 0.85 | 0.84 |
| 4 | 159319980 | rs2219191 | -0.557 | -0.528 | 0.421 | -1.626 | -0.013 | -0.033 | -0.033 | 0 | 0.1 | 0.25 | 0.16 | 0.04 | 0.05 | 0.23 |
| 4 | 159331657 | rs4691493 | -0.75 | -0.343 | 0.188 | -1.716 | -0.006 | -0.099 | -0.099 | 0 | 0.19 | 0.42 | 0.21 | 0.02 | 0.17 | 0.22 |
| 4 | 159367847 | rs6847255 | 0.948 | -0.614 | -0.72 | -1.684 | 0.067 | 0.146 | 0.146 | 0 | 0.59 | 0.21 | 0.67 | 0.95 | 0.7 | 0.59 |
| 4 | 159399146 | rs10517691 | -0.114 | -0.638 | -0.426 | -1.508 | 0.015 | 0.01 | 0.01 | 0 | 0.2 | 0.17 | 0.1 | 0.03 | 0.13 | 0.19 |
| 4 | 159412248 | rs7375480 | 1.204 | -0.843 | -0.473 | -1.83 | 0.081 | 0.191 | 0.191 | 0 | 0.59 | 0.16 | 0.67 | 0.95 | 0.71 | 0.59 |
| 4 | 159430701 | rs7377646 | 0.244 | -0.913 | 1.075 | -1.853 | -0.072 | -0.127 | -0.127 | 0.588 | 0.75 | 0.53 | 0.88 | 0.97 | 0.87 | 0.8 |
| 4 | 159433657 | rs13117144 | 0.37 | -1.355 | 1.028 | -2.143 | 0.019 | 0.005 | 0.005 | 0 | 0.71 | 0.76 | 0.88 | 0.98 | 0.85 | 0.69 |
| 4 | 159433681 | rs13143911 | 1.091 | -1.136 | 0.678 | -2.667 | 0.006 | -0.026 | -0.026 | 0.221 | 0.17 | 0.19 | 0.18 | 0.09 | 0.17 | 0.08 |
| 4 | 159434669 | rs10517692 | 1.298 | -1.135 | 0.647 | -2.666 | -0.021 | 0.036 | 0.036 | 0 | 0.16 | 0.06 | 0.17 | 0.09 | 0.16 | 0.07 |
| 4 | 159440817 | rs6536350 | -0.238 | -1.074 | 0.652 | -2.688 | 0.02 | 0.008 | 0.008 | 0 | 0.71 | 0.75 | 0.88 | 0.98 | 0.81 | 0.69 |
| 4 | 159441020 | rs1455454 | 0.533 | -1.109 | 0.706 | -2.655 | -0.003 | -0.039 | -0.039 | 0.572 | 0.55 | 0.57 | 0.7 | 0.89 | 0.65 | 0.62 |
| 4 | 159441539 | rs7678831 | -0.208 | -0.939 | 0.692 | -2.65 | 0.017 | 0.008 | 0.008 | 0 | 0.71 | 0.75 | 0.87 | 0.98 | 0.79 | 0.69 |
| 4 | 159442046 | rs1455456 | -0.154 | -0.935 | 0.597 | -2.648 | -0.029 | -0.098 | -0.098 | 0 | 0.71 | 0.64 | 0.86 | 0.98 | 0.79 | 0.7 |
| 4 | 159442057 | rs7679484 | -0.041 | -0.714 | 0.964 | -2.382 | -0.002 | -0.085 | -0.085 | 0.01 | 0.15 | 0.39 | 0.2 | 0.02 | 0.13 | 0.2 |
| 4 | 159445028 | rs13139892 | 0.905 | -0.541 | 0.54 | -2.073 | -0.008 | 0.009 | 0.009 | 0 | 0.15 | 0.07 | 0.17 | 0.14 | 0.15 | 0.08 |
| 4 | 159460722 | rs6845779 | -1.035 | -0.821 | 1.483 | -2.308 | 0.009 | -0.098 | -0.098 | 0 | 0.15 | 0.48 | 0.23 | 0.02 | 0.13 | 0.17 |
| 4 | 159481490 | rs6832186 | -0.588 | -0.862 | 1.802 | -2.329 | 0.008 | -0.062 | -0.062 | 3.146 | 0.13 | 0.31 | 0.21 | 0.02 | 0.13 | 0.17 |
| 4 | 159500966 | rs11940491 | -1.423 | -0.958 | 1.631 | -2.376 | -0.015 | -0.088 | -0.088 | 0 | 0.17 | 0.65 | 0.24 | 0.02 | 0.13 | 0.17 |
| 4 | 159503774 | rs16999138 | -1.091 | -0.95 | 1.621 | -2.372 | 0.004 | -0.099 | -0.099 | 0 | 0.15 | 0.49 | 0.22 | 0.02 | 0.13 | 0.17 |
| 4 | 159588731 | rs3210749 | 1.028 | -0.464 | 0.105 | -2.613 | -0.087 | 0.234 | 0.234 | 0 | 0.46 | 0.06 | 0.56 | 0.85 | 0.51 | 0.56 |
| 4 | 159619426 | rs12640862 | 1.103 | -0.342 | 1.531 | -1.973 | -0.008 | -0.026 | -0.026 | 0.889 | 0.91 | 0.74 | 0.95 | 0.96 | 0.95 | 0.97 |
| 4 | 159620035 | rs7679753 | -0.916 | -0.905 | 1.211 | -2.525 | 0.006 | -0.032 | -0.032 | 0 | 0.36 | 0.38 | 0.17 | 0.16 | 0.36 | 0.34 |
| 4 | 159621250 | rs11737481 | 1.309 | -1.271 | 1.065 | -2.643 | 0.013 | 0.151 | 0.151 | 0.341 | 0.6 | 0.13 | 0.81 | 0.83 | 0.7 | 0.62 |
| 4 | 159629814 | rs17843966 | 0.1 | -1.28 | 1.266 | -2.695 | -0.019 | 0.021 | 0.021 | 0.004 | 0.22 | 0.14 | 0.1 | 0.13 | 0.25 | 0.28 |

| | | | | | | | | | | | | | | | | |
|---|---|---|---|---|---|---|---|---|---|---|---|---|---|---|---|---|
| 4 | 159630817 | rs8396 | -0.405 | -1.243 | 1.195 | -2.672 | 0.013 | -0.036 | -0.036 | 0 | 0.27 | 0.35 | 0.14 | 0.15 | 0.29 | 0.32 |
| 4 | 159635244 | rs12505475 | 1.023 | -1.135 | 0.473 | -2.629 | 0.013 | 0.164 | 0.164 | 0.2 | 0.61 | 0.15 | 0.82 | 0.87 | 0.7 | 0.62 |
| 4 | 159672059 | rs4690912 | 0.192 | -0.272 | 0.153 | -2.137 | 0.013 | 0.09 | 0.09 | 0.135 | 0.53 | 0.35 | 0.67 | 0.89 | 0.57 | 0.68 |
| 4 | 159677712 | rs7666007 | 0.082 | -0.425 | 0.534 | -2.216 | 0 | -0.037 | -0.037 | 0 | 0.38 | 0.39 | 0.27 | 0.05 | 0.38 | 0.23 |
| 4 | 159682676 | rs11100196 | 0.178 | -0.416 | 0.468 | -2.21 | -0.022 | -0.027 | -0.027 | 0 | 0.08 | 0.16 | 0.05 | 0.05 | 0.05 | 0.09 |
| 4 | 159688477 | rs4234923 | 0.201 | -0.316 | 0.166 | -2.146 | 0.013 | 0.085 | 0.085 | 0.192 | 0.53 | 0.36 | 0.67 | 0.89 | 0.56 | 0.68 |
| 4 | 159729794 | rs10857319 | 1.482 | -0.241 | -0.491 | -2.096 | -0.093 | 0.031 | 0.031 | 0 | 0.3 | 0.03 | 0.47 | 0.81 | 0.36 | 0.39 |
| 4 | 159732671 | rs11725623 | -0.283 | -0.172 | -0.01 | -2.21 | 0.003 | 0.007 | 0.007 | 0.224 | 0.47 | 0.57 | 0.32 | 0.14 | 0.44 | 0.33 |
| 4 | 159779241 | rs6828666 | 0.628 | -0.073 | -0.189 | -2.213 | -0.024 | 0.067 | 0.067 | 0.769 | 0.2 | 0.1 | 0.18 | 0.04 | 0.2 | 0.26 |
| 4 | 159791519 | rs11946928 | 0.53 | -0.188 | 0.324 | -2.023 | -0.028 | -0.025 | -0.025 | 0.116 | 0.05 | 0.16 | 0.01 | 0.03 | 0.01 | 0.02 |
| 4 | 159804265 | rs4600878 | 0.071 | -0.171 | 0.036 | -2.037 | -0.031 | 0.062 | 0.062 | 0 | 0.37 | 0.21 | 0.29 | 0.07 | 0.44 | 0.24 |
| 4 | 159827313 | rs10589 | 0.518 | -0.457 | 0.095 | -1.906 | -0.033 | 0.061 | 0.061 | 0 | 0.3 | 0.14 | 0.25 | 0.05 | 0.37 | 0.22 |
| 4 | 159834341 | rs4690915 | 0.664 | -0.519 | 0.205 | -1.928 | 0.012 | 0.1 | 0.1 | 0.297 | 0.52 | 0.36 | 0.67 | 0.87 | 0.57 | 0.68 |
| 4 | 159835412 | rs9998332 | -0.774 | -0.454 | 0.088 | -1.917 | -0.026 | 0.057 | 0.057 | 0 | 0.68 | 0.83 | 0.75 | 0.95 | 0.63 | 0.77 |
| 4 | 159839313 | rs10010963 | -0.713 | -0.404 | 0.092 | -1.917 | -0.028 | 0.064 | 0.064 | 0 | 0.68 | 0.82 | 0.75 | 0.95 | 0.63 | 0.78 |
| 4 | 159860293 | rs4146954 | 0.22 | -0.171 | 0.512 | -1.869 | 0.008 | -0.038 | -0.038 | 0 | 0.72 | 0.71 | 0.75 | 0.91 | 0.67 | 0.76 |
| 4 | 159863158 | rs11724252 | 0.532 | -0.218 | 0.719 | -1.895 | 0.003 | -0.103 | -0.103 | 0 | 0.66 | 0.54 | 0.73 | 0.87 | 0.66 | 0.74 |
| 4 | 159873196 | rs1820510 | -1.116 | -0.377 | 1.085 | -1.708 | -0.052 | -0.062 | -0.062 | 0.107 | 0.13 | 0.42 | 0.06 | 0.06 | 0.05 | 0.11 |
| 4 | 159876379 | rs6536380 | -0.992 | -0.408 | 0.681 | -1 | -0.001 | -0.01 | -0.01 | 0 | 0.6 | 0.46 | 0.54 | 0.67 | 0.63 | 0.78 |
| 4 | 159877388 | rs10003821 | -0.203 | -0.292 | 0.412 | -0.654 | 0 | -0.002 | -0.002 | 0.05 | 0.34 | 0.34 | 0.35 | 0.38 | 0.35 | 0.51 |
| 4 | 159881479 | rs9784569 | 1.146 | -0.225 | 0.244 | -0.631 | -0.019 | -0.003 | -0.003 | 0 | 0.2 | 0.12 | 0.2 | 0.24 | 0.27 | 0.2 |
| 4 | 159881943 | rs11722868 | -1.495 | -0.395 | 0.719 | -0.685 | -0.004 | -0.007 | -0.007 | 0 | 0.64 | 0.63 | 0.58 | 0.68 | 0.68 | 0.82 |
| 4 | 159882696 | rs7698077 | 0.63 | -0.505 | 0.67 | -0.822 | -0.012 | -0.053 | -0.053 | 0.34 | 0.11 | 0.33 | 0.05 | 0.04 | 0.08 | 0.18 |
| 4 | 159885770 | rs714462 | -0.973 | -0.263 | 0.367 | -0.519 | 0.026 | 0.061 | 0.061 | 0 | 0.53 | 0.29 | 0.53 | 0.64 | 0.61 | 0.63 |
| 4 | 159888222 | rs1971087 | 0.402 | -0.366 | 0.188 | -0.313 | 0.048 | -0.072 | -0.072 | 0 | 0.19 | 0.1 | 0.2 | 0.36 | 0.12 | 0.18 |
| 4 | 159890417 | rs13136816 | -0.635 | -0.182 | 0.228 | -0.349 | -0.004 | -0.09 | -0.09 | 0 | 0.33 | 0.14 | 0.33 | 0.4 | 0.33 | 0.43 |
| 4 | 159894391 | rs17037864 | 0.639 | -0.067 | 0.269 | -0.22 | -0.036 | 0.018 | 0.018 | 0 | 0.06 | 0 | 0.06 | 0.33 | 0.13 | 0.27 |
| 4 | 159912840 | rs109119 | 0.397 | -0.181 | 0.835 | -0.305 | -0.009 | 0.011 | 0.011 | 0 | 0.64 | 0.53 | 0.64 | 0.6 | 0.67 | 0.57 |
| 4 | 159920048 | rs12510910 | -0.507 | -0.078 | 0.313 | -0.279 | 0.013 | -0.045 | -0.045 | 0 | 0.34 | 0.23 | 0.34 | 0.4 | 0.33 | 0.43 |
| 4 | 159927765 | rs999411 | -0.545 | -0.065 | 0.363 | -0.272 | 0.012 | -0.03 | -0.03 | 0 | 0.34 | 0.27 | 0.34 | 0.4 | 0.33 | 0.43 |
| 4 | 159931261 | rs1549850 | 1.046 | 0.022 | 0.009 | -0.402 | -0.033 | 0.004 | 0.004 | 0 | 0.38 | 0.21 | 0.42 | 0.39 | 0.39 | 0.33 |
| 4 | 159944851 | rs3929142 | 0.453 | 1.002 | 0.705 | -0.558 | 0.014 | -0.039 | -0.039 | 0 | 0.17 | 0.19 | 0.17 | 0.02 | 0.23 | 0.16 |
| 4 | 159948180 | rs7660477 | -0.855 | 0.956 | 0.651 | -0.202 | 0.025 | -0.017 | -0.017 | 0 | 0.36 | 0.5 | 0.48 | 0.25 | 0.44 | 0.42 |
| 4 | 159950990 | rs6849580 | -0.746 | 1.057 | 0.539 | -0.099 | 0.047 | 0.074 | 0.074 | 0 | 0.11 | 0.32 | 0.12 | 0.19 | 0.16 | 0.17 |
| 4 | 159952353 | rs1559887 | 0.05 | 1.035 | -0.108 | -0.184 | -0.024 | 0.056 | 0.056 | 0 | 0.25 | 0.11 | 0.26 | 0.07 | 0.29 | 0.25 |
| 4 | 159954244 | rs601196 | -0.952 | 1.048 | -0.444 | 0.185 | -0.041 | -0.085 | -0.085 | 0 | 0.62 | 0.83 | 0.61 | 0.49 | 0.61 | 0.53 |
| 4 | 159958150 | rs2053346 | 0.052 | 1.323 | 0.179 | -0.479 | -0.011 | 0.046 | 0.046 | 0 | 0.46 | 0.67 | 0.49 | 0.74 | 0.48 | 0.61 |
| 4 | 159961231 | rs6834079 | 0.684 | 1.385 | 0.618 | -0.413 | 0.005 | -0.001 | -0.001 | 0 | 0.38 | 0.34 | 0.37 | 0.6 | 0.33 | 0.46 |
| 4 | 159963042 | rs12331691 | -0.399 | 1.527 | 0.49 | -0.251 | 0.114 | 0.07 | 0.07 | 0.114 | 0.08 | 0.3 | 0.12 | 0.14 | 0.15 | 0.15 |
| 4 | 159976179 | rs588811 | 0.736 | 1.256 | -0.036 | -0.344 | 0.005 | -0.02 | -0.02 | 0.041 | 0.36 | 0.19 | 0.36 | 0.6 | 0.33 | 0.45 |
| 4 | 159981640 | rs581849 | 0.822 | 0.982 | 0.534 | -0.565 | -0.014 | -0.008 | -0.008 | 0 | 0.38 | 0.33 | 0.37 | 0.56 | 0.47 | 0.44 |
| 4 | 160019492 | rs2014078 | -0.258 | 0.739 | 0.69 | -0.909 | 0.004 | -0.029 | -0.029 | 0.033 | 0.34 | 0.47 | 0.41 | 0.24 | 0.36 | 0.39 |
| 4 | 160026546 | rs2306651 | 0.209 | 0.699 | 0.407 | -0.925 | 0.055 | 0.04 | 0.04 | 0 | 0.28 | 0.13 | 0.22 | 0.2 | 0.17 | 0.18 |
| 4 | 160053187 | rs10003940 | -1.692 | 0.748 | 0.436 | -0.889 | -0.052 | -0.033 | -0.033 | 1.152 | 0.64 | 0.82 | 0.65 | 0.44 | 0.53 | 0.56 |
| 4 | 160079961 | rs13148447 | 0.538 | 0.65 | 0.362 | -0.943 | -0.024 | 0.051 | 0.051 | 0 | 0.25 | 0.15 | 0.26 | 0.08 | 0.29 | 0.23 |
| 4 | 160084403 | rs7681750 | 0.163 | 0.521 | 0.427 | -1.174 | 0.038 | 0.029 | 0.029 | 0 | 0.28 | 0.16 | 0.22 | 0.2 | 0.17 | 0.18 |
| 4 | 160088137 | rs7656697 | -1.224 | 0.635 | 0.705 | -1.119 | 0.006 | 0.003 | 0.003 | 0 | 0.61 | 0.6 | 0.63 | 0.44 | 0.53 | 0.56 |
| 4 | 160102354 | rs4410485 | -1.76 | 0.046 | 0.297 | -1.751 | -0.013 | 0.002 | 0.002 | 0 | 0.66 | 0.9 | 0.66 | 0.29 | 0.68 | 0.59 |
| 4 | 160118458 | rs9997385 | -0.713 | 0.389 | 1.229 | -1.722 | -0.002 | 0.005 | 0.005 | 0.032 | 0.35 | 0.64 | 0.4 | 0.23 | 0.39 | 0.39 |
| 4 | 160124407 | rs10050099 | -0.168 | 0.384 | 1.302 | -1.779 | -0.001 | 0 | 0 | 0 | 0.29 | 0.29 | 0.34 | 0.11 | 0.35 | 0.25 |
| 4 | 160132990 | rs7685028 | -0.673 | 0.155 | 0.674 | -1.718 | -0.008 | 0.031 | 0.031 | 0.111 | 0.36 | 0.73 | 0.4 | 0.23 | 0.39 | 0.39 |
| 4 | 160143615 | rs12502799 | -0.073 | -0.189 | 1.15 | -2.037 | 0.041 | -0.053 | -0.053 | 0 | 0.27 | 0.36 | 0.29 | 0.08 | 0.37 | 0.3 |
| 4 | 160147159 | rs11730876 | -0.042 | -0.199 | 1.187 | -2.071 | 0.055 | -0.066 | -0.066 | 0 | 0.27 | 0.37 | 0.3 | 0.08 | 0.37 | 0.29 |
| 4 | 160151176 | rs13139002 | -1.324 | -0.553 | 0.691 | -2.616 | 0.024 | 0.163 | 0.163 | 0 | 0.57 | 0.76 | 0.54 | 0.12 | 0.58 | 0.42 |
| 4 | 160173041 | rs6536404 | -1.327 | -0.525 | 0.712 | -2.662 | 0.025 | 0.16 | 0.16 | 0 | 0.57 | 0.76 | 0.55 | 0.12 | 0.6 | 0.42 |
| 4 | 160173942 | rs4690933 | -0.452 | -0.607 | 0.7 | -2.437 | 0.017 | 0.049 | 0.049 | 0 | 0.25 | 0.15 | 0.2 | 0.08 | 0.19 | 0.11 |
| 4 | 160184658 | rs11100213 | -1.045 | -0.555 | 1.232 | -2.335 | -0.031 | -0.082 | -0.082 | 0 | 0.29 | 0.52 | 0.28 | 0.09 | 0.2 | 0.1 |
| 4 | 160185669 | rs13145422 | 0.44 | -0.448 | 1.114 | -2.551 | -0.074 | 0.116 | 0.116 | 0 | 0.28 | 0.17 | 0.28 | 0.04 | 0.35 | 0.32 |
| 4 | 160186284 | rs10428456 | -0.868 | -0.417 | 1.303 | -2.54 | -0.032 | -0.106 | -0.106 | 0 | 0.29 | 0.47 | 0.28 | 0.09 | 0.22 | 0.1 |
| 4 | 160208630 | rs4396948 | -1.593 | -0.451 | 1.401 | -2.536 | -0.003 | 0.164 | 0.164 | 0 | 0.59 | 0.83 | 0.59 | 0.15 | 0.57 | 0.43 |
| 4 | 160209816 | rs6840293 | -0.879 | -0.458 | 1.416 | -2.544 | -0.027 | -0.083 | -0.083 | 0.037 | 0.29 | 0.52 | 0.28 | 0.07 | 0.2 | 0.1 |
| 4 | 160210862 | rs4295222 | 0.241 | -0.466 | 1.283 | -2.517 | -0.039 | 0.065 | 0.065 | 0 | 0.3 | 0.24 | 0.29 | 0.07 | 0.36 | 0.33 |
| 4 | 160223891 | rs9799685 | -0.321 | -0.486 | 0.954 | -2.503 | 0.007 | 0.048 | 0.048 | 0 | 0.24 | 0.14 | 0.26 | 0.07 | 0.21 | 0.1 |
| 4 | 160240899 | rs7678811 | -1.425 | -0.66 | 0.892 | -2.434 | 0.002 | 0.149 | 0.149 | 0 | 0.59 | 0.8 | 0.59 | 0.15 | 0.57 | 0.43 |
| 4 | 160249722 | rs10022399 | -0.064 | -0.738 | 1.527 | -2.081 | -0.011 | -0.004 | -0.004 | 0 | 0.34 | 0.33 | 0.33 | 0.07 | 0.43 | 0.35 |
| 4 | 160261053 | rs13134868 | -1.139 | -0.855 | 1.096 | -2.133 | 0.047 | 0.221 | 0.221 | 0 | 0.53 | 0.85 | 0.6 | 0.13 | 0.56 | 0.44 |
| 4 | 160267334 | rs11723634 | 0.33 | -0.781 | 0.937 | -2.271 | -0.128 | 0.152 | 0.152 | 0 | 0.32 | 0.14 | 0.31 | 0.04 | 0.42 | 0.33 |
| 4 | 160278606 | rs10025560 | -0.144 | -0.674 | 1.091 | -2.212 | 0.001 | 0.014 | 0.014 | 0 | 0.16 | 0.14 | 0.24 | 0.05 | 0.13 | 0.11 |
| 4 | 160283091 | rs1969578 | 0.453 | -0.853 | 0.468 | -2.255 | -0.129 | 0.155 | 0.155 | 0 | 0.32 | 0.13 | 0.32 | 0.05 | 0.43 | 0.33 |

| | | | | | | | | | | | | | | | |
|---|---|---|---|---|---|---|---|---|---|---|---|---|---|---|---|
| 4 | 160284623 | rs2016061 | -0.5 | -0.827 | 0.477 | -2.094 | 0.049 | 0.214 | 0.214 | 0 | 0.46 | 0.23 | 0.55 | 0.1 | 0.55 | 0.42 |
| 4 | 160299086 | rs12505843 | -1 | -0.712 | 0.23 | -1.978 | 0.045 | 0.181 | 0.181 | 0 | 0.54 | 0.85 | 0.6 | 0.11 | 0.58 | 0.44 |
| 4 | 160327856 | rs4440173 | -0.509 | -0.634 | 0.409 | -2.044 | 0.025 | 0.22 | 0.22 | 0 | 0.46 | 0.21 | 0.51 | 0.1 | 0.53 | 0.41 |
| 4 | 160331540 | rs1125409 | 0.11 | -0.558 | 0.3 | -2.142 | 0 | 0.013 | 0.013 | 0 | 0.14 | 0.1 | 0.2 | 0.06 | 0.13 | 0.09 |
| 4 | 160338309 | rs11730206 | 1.461 | -0.803 | 0.291 | -2.102 | -0.023 | 0.062 | 0.062 | 0 | 0.35 | 0.09 | 0.36 | 0.8 | 0.39 | 0.51 |
| 4 | 160341351 | rs2348164 | 0.669 | -0.762 | 0.463 | -2.138 | -0.043 | -0.018 | -0.018 | 0 | 0.82 | 0.32 | 0.88 | 0.9 | 0.92 | 0.91 |
| 4 | 160344788 | rs6812880 | 0.504 | -0.759 | 0.727 | -2.136 | -0.085 | -0.03 | -0.03 | 0.045 | 0.83 | 0.49 | 0.89 | 0.9 | 0.92 | 0.91 |
| 4 | 160347129 | rs2699023 | -0.396 | -0.629 | 0.472 | -2.159 | -0.014 | -0.009 | -0.009 | 0 | 0.88 | 0.88 | 0.93 | 0.92 | 0.92 | 0.93 |
| 4 | 160349432 | rs2699027 | 0.261 | -0.662 | 0.16 | -2.248 | -0.041 | 0.074 | 0.074 | 0 | 0.35 | 0.25 | 0.32 | 0.07 | 0.41 | 0.33 |
| 4 | 160350425 | rs2699032 | -0.021 | -0.404 | 0.158 | -1.889 | -0.041 | -0.027 | -0.027 | 0.96 | 0.86 | 0.7 | 0.91 | 0.92 | 0.92 | 0.93 |
| 4 | 160355746 | rs4690942 | 0.65 | 0.279 | -2.202 | -1.509 | -0.019 | 0.056 | 0.056 | 0 | 0.34 | 0.1 | 0.38 | 0.79 | 0.37 | 0.5 |
| 4 | 160357877 | rs4591547 | -0.293 | 0.863 | -1.888 | -1.681 | -0.105 | 0.139 | 0.139 | 0 | 0.32 | 0.1 | 0.31 | 0.09 | 0.41 | 0.46 |
| 4 | 160361747 | rs11100229 | 0.554 | 0.884 | -2.195 | -1.892 | 0.006 | 0.202 | 0.202 | 0.275 | 0.44 | 0.1 | 0.54 | 0.78 | 0.38 | 0.42 |
| 4 | 160364016 | rs4691567 | -0.015 | 0.167 | -0.857 | -2.029 | -0.051 | -0.038 | -0.038 | 0 | 0.87 | 0.69 | 0.96 | 0.93 | 0.96 | 0.94 |
| 4 | 160369165 | rs2707687 | 0.746 | 0.777 | -0.097 | -2.283 | 0.003 | 0.028 | 0.028 | 0 | 0.59 | 0.4 | 0.78 | 0.83 | 0.6 | 0.54 |
| 4 | 160374255 | rs11947461 | -0.303 | 0.85 | 0.204 | -2.229 | 0.015 | -0.094 | -0.094 | 3.91 | 0.31 | 0.52 | 0.21 | 0.1 | 0.36 | 0.4 |
| 4 | 160377484 | rs7675279 | 0.684 | 0.87 | 0.466 | -2.08 | -0.007 | -0.03 | -0.03 | 0 | 0.66 | 0.58 | 0.83 | 0.83 | 0.72 | 0.6 |
| 4 | 160381253 | rs2699006 | -0.229 | 0.786 | 0.501 | -2.11 | -0.002 | 0 | 0 | 0 | 0.25 | 0.26 | 0.17 | 0.1 | 0.25 | 0.34 |
| 4 | 160382933 | rs5008449 | 0.785 | 0.742 | 0.56 | -1.97 | -0.004 | -0.013 | -0.013 | 0 | 0.65 | 0.6 | 0.79 | 0.76 | 0.71 | 0.61 |
| 4 | 160384694 | rs6833365 | 0.785 | 0.742 | 0.56 | -1.97 | -0.004 | -0.013 | -0.013 | 0 | 0.65 | 0.6 | 0.79 | 0.76 | 0.71 | 0.61 |
| 4 | 160388504 | rs13111198 | 0.847 | 0.671 | 0.555 | -1.992 | -0.009 | -0.022 | -0.022 | 0 | 0.64 | 0.57 | 0.79 | 0.77 | 0.72 | 0.61 |
| 4 | 160406190 | rs2851820 | 0.945 | 0.782 | 0.535 | -2.054 | 0.003 | 0.015 | 0.015 | 0 | 0.63 | 0.66 | 0.78 | 0.77 | 0.68 | 0.63 |
| 4 | 160412034 | rs13144517 | 1.097 | 0.758 | 0.499 | -2.065 | -0.003 | -0.015 | -0.015 | 0 | 0.62 | 0.58 | 0.78 | 0.77 | 0.68 | 0.65 |
| 4 | 160412665 | rs2851828 | -0.25 | 0.766 | 0.474 | -2.052 | 0.025 | -0.028 | -0.028 | 0 | 0.92 | 0.99 | 0.98 | 0.87 | 0.96 | 0.94 |
| 4 | 160419271 | rs1922645 | 1.053 | 0.753 | 0.49 | -2.068 | -0.003 | -0.015 | -0.015 | 0 | 0.63 | 0.58 | 0.78 | 0.77 | 0.68 | 0.64 |
| 4 | 160421579 | rs2851836 | 0.914 | 0.694 | 0.39 | -2.043 | 0.002 | 0.045 | 0.045 | 0 | 0.65 | 0.81 | 0.8 | 0.76 | 0.68 | 0.64 |
| 4 | 160423997 | rs1922642 | 0.084 | 0.79 | 0.347 | -1.993 | -0.03 | 0.058 | 0.058 | 0 | 0.75 | 0.9 | 0.83 | 0.89 | 0.72 | 0.7 |
| 4 | 160425090 | rs2851769 | 0.105 | 0.84 | 1.076 | -1.981 | -0.039 | 0.06 | 0.06 | 0 | 0.75 | 0.91 | 0.83 | 0.88 | 0.72 | 0.73 |
| 4 | 160426520 | rs4621382 | 0.373 | 0.507 | 1.069 | -1.988 | -0.002 | 0.028 | 0.028 | 0 | 0.7 | 0.78 | 0.82 | 0.86 | 0.71 | 0.7 |
| 4 | 160426970 | rs1317333 | 0.222 | 0.539 | 0.928 | -1.974 | -0.026 | 0.063 | 0.063 | 0 | 0.73 | 0.9 | 0.83 | 0.86 | 0.71 | 0.7 |
| 4 | 160442645 | rs11724165 | 0.168 | 0.556 | 0.821 | -2.082 | -0.007 | 0.031 | 0.031 | 0 | 0.2 | 0.08 | 0.11 | 0.13 | 0.19 | 0.28 |
| 4 | 160446640 | rs2851798 | 0.921 | 0.174 | 1.537 | -1.818 | -0.028 | -0.015 | -0.015 | 0 | 0.84 | 0.72 | 0.9 | 0.86 | 0.91 | 0.92 |
| 4 | 160456060 | rs6824835 | 1.536 | 0.156 | 1.428 | -1.833 | -0.038 | -0.047 | -0.047 | 1.924 | 0.79 | 0.44 | 0.86 | 0.86 | 0.87 | 0.91 |
| 4 | 160474408 | rs1893953 | 0.258 | 0.571 | 0.628 | -2.216 | -0.003 | 0.002 | 0.002 | 0 | 0.59 | 0.65 | 0.72 | 0.71 | 0.51 | 0.59 |
| 4 | 160477057 | rs11100238 | 0.184 | 0.516 | 0.906 | -2.241 | -0.041 | 0.069 | 0.069 | 0 | 0.61 | 0.76 | 0.73 | 0.72 | 0.51 | 0.59 |
| 4 | 160483950 | rs6823356 | 1.076 | -0.038 | 1.103 | -2.6 | -0.004 | -0.023 | -0.023 | 0 | 0.4 | 0.46 | 0.57 | 0.67 | 0.41 | 0.36 |
| 4 | 160485488 | rs11945017 | 0.99 | -0.007 | 1.162 | -2.66 | 0.01 | -0.027 | -0.027 | 1.03 | 0.05 | 0.08 | 0.04 | 0 | 0.05 | 0.03 |
| 4 | 160486937 | rs4691577 | 0.392 | -0.055 | 1.178 | -2.678 | 0.012 | 0.054 | 0.054 | 0 | 0.46 | 0.62 | 0.68 | 0.69 | 0.46 | 0.38 |
| 4 | 160488212 | rs1393576 | 0.567 | 0.004 | 0.895 | -2.771 | -0.056 | 0.096 | 0.096 | 0 | 0.34 | 0.14 | 0.23 | 0.14 | 0.44 | 0.4 |
| 4 | 160494830 | rs998888 | 1.065 | 0.029 | 0.934 | -2.727 | 0 | -0.002 | -0.002 | 0 | 0.11 | 0.13 | 0.07 | 0.04 | 0.05 | 0.17 |
| 4 | 160497236 | rs2348414 | 1.106 | 0.077 | 0.964 | -2.711 | -0.004 | -0.024 | -0.024 | 0 | 0.41 | 0.48 | 0.63 | 0.67 | 0.41 | 0.36 |
| 4 | 160512123 | rs6858329 | 1.919 | 0.01 | 0.965 | -2.738 | -0.001 | 0.017 | 0.017 | 0 | 0.32 | 0.2 | 0.57 | 0.67 | 0.36 | 0.33 |
| 4 | 160512801 | rs924678 | -0.528 | 0.013 | 0.969 | -2.749 | -0.004 | 0.003 | 0.003 | 0 | 0.92 | 0.87 | 0.96 | 0.86 | 0.95 | 0.94 |
| 4 | 160514021 | rs6848958 | 1.982 | -0.036 | 0.932 | -2.768 | -0.014 | 0.015 | 0.015 | 0 | 0.32 | 0.13 | 0.57 | 0.67 | 0.36 | 0.34 |
| 4 | 160515218 | rs9994725 | -0.594 | 0.051 | 1.175 | -2.767 | -0.047 | -0.134 | -0.134 | 0 | 0.19 | 0.42 | 0.09 | 0.01 | 0.1 | 0.12 |
| 4 | 160515337 | rs961787 | -1.515 | 0.066 | 0.819 | -2.769 | 0.005 | 0.09 | 0.09 | 0 | 0.6 | 0.74 | 0.39 | 0.19 | 0.59 | 0.55 |
| 4 | 160516090 | rs17239575 | 0.397 | 0.165 | 0.563 | -2.698 | -0.034 | 0.062 | 0.062 | 0 | 0.35 | 0.2 | 0.26 | 0.17 | 0.45 | 0.4 |
| 4 | 160521480 | rs2000886 | -0.828 | 0.181 | 0.387 | -2.66 | 0.013 | 0.143 | 0.143 | 0 | 0.49 | 0.69 | 0.35 | 0.18 | 0.54 | 0.45 |
| 4 | 160523363 | rs1502739 | -0.377 | 0.163 | 0.507 | -2.069 | -0.004 | -0.019 | -0.019 | 0 | 0.06 | 0.08 | 0.02 | 0 | 0.05 | 0.01 |
| 4 | 160526077 | rs13125739 | -1.341 | -0.009 | 0.393 | -2.199 | -0.018 | -0.054 | -0.054 | 0.122 | 0.28 | 0.58 | 0.15 | 0.1 | 0.16 | 0.2 |
| 4 | 160526796 | rs17318405 | 0.615 | 0.803 | 0.149 | -1.56 | -0.017 | -0.014 | -0.014 | 0.082 | 0.06 | 0.02 | 0.11 | 0.11 | 0.14 | 0.1 |
| 4 | 160558972 | rs4383572 | 0.723 | 0.285 | 0.177 | -1.784 | -0.008 | 0.063 | 0.063 | 0 | 0.44 | 0.3 | 0.67 | 0.75 | 0.47 | 0.47 |
| 4 | 160560872 | rs11933100 | 0.478 | 0.12 | 0.301 | -1.943 | -0.013 | 0.134 | 0.134 | 0 | 0.37 | 0.15 | 0.22 | 0.09 | 0.39 | 0.39 |
| 4 | 160563128 | rs12331504 | -0.374 | 0.263 | 0.305 | -1.717 | 0.003 | 0.034 | 0.034 | 0 | 0.5 | 0.53 | 0.3 | 0.15 | 0.48 | 0.5 |
| 4 | 160574450 | rs1502752 | 0.573 | 0.286 | 0.272 | -1.702 | 0.017 | 0.034 | 0.034 | 0.782 | 0.94 | 0.78 | 0.96 | 0.9 | 0.95 | 0.96 |
| 4 | 160578021 | rs10517704 | -0.741 | 0.285 | 0.097 | -1.758 | 0.001 | 0.008 | 0.008 | 0 | 0.5 | 0.47 | 0.29 | 0.15 | 0.48 | 0.5 |
| 4 | 160591060 | rs9985638 | -1.373 | 0.179 | 0.603 | -1.463 | -0.001 | -0.038 | -0.038 | 0 | 0.16 | 0.36 | 0.08 | 0.07 | 0.1 | 0.09 |
| 4 | 160594757 | rs6536452 | 0.789 | -0.085 | 0.183 | -0.719 | -0.017 | -0.012 | -0.012 | 0.184 | 0.36 | 0.22 | 0.59 | 0.6 | 0.41 | 0.52 |
| 4 | 160596412 | rs10517706 | -0.43 | -0.142 | 0.291 | -0.729 | 0.002 | 0.058 | 0.058 | 0 | 0.52 | 0.63 | 0.67 | 0.67 | 0.53 | 0.61 |
| 4 | 160604277 | rs1118040 | 0.657 | -0.397 | 0.196 | -0.79 | -0.038 | 0.086 | 0.086 | 0 | 0.4 | 0.18 | 0.29 | 0.25 | 0.45 | 0.36 |
| 4 | 160607399 | rs1502736 | -0.022 | -0.383 | 0.75 | -0.63 | 0.003 | 0.004 | 0.004 | 0 | 0.45 | 0.55 | 0.62 | 0.6 | 0.43 | 0.54 |
| 4 | 160609723 | rs1580882 | 0.228 | -0.386 | 0.715 | -0.631 | -0.004 | -0.005 | -0.005 | 0 | 0.42 | 0.44 | 0.61 | 0.6 | 0.41 | 0.51 |
| 4 | 160630642 | rs9762089 | 1.664 | -0.275 | 0.14 | -0.458 | -0.033 | -0.074 | -0.074 | 0 | 0.2 | 0.06 | 0.51 | 0.55 | 0.29 | 0.38 |
| 4 | 160640315 | rs2348616 | -0.103 | -0.326 | 0.27 | -0.531 | 0.009 | 0.043 | 0.043 | 0 | 0.53 | 0.38 | 0.36 | 0.32 | 0.56 | 0.45 |
| 4 | 160645543 | rs2348620 | -1.159 | -0.266 | 0.665 | -0.439 | -0.01 | -0.051 | -0.051 | 0.602 | 0.16 | 0.36 | 0.09 | 0.06 | 0.12 | 0.07 |
| 4 | 160647109 | rs17319860 | 0.013 | -0.266 | 0.641 | -0.44 | 0.004 | 0.011 | 0.011 | 0.163 | 0.11 | 0.04 | 0.05 | 0.06 | 0.09 | 0.07 |
| 4 | 160654172 | rs1224511 | -1.667 | -0.241 | 0.809 | -0.258 | -0.001 | 0.001 | 0.001 | 0.145 | 0.18 | 0.4 | 0.11 | 0.14 | 0.14 | 0.09 |
| 4 | 160661776 | rs6536467 | 1.421 | -0.291 | 1.181 | -0.273 | 0.006 | -0.019 | -0.019 | 0 | 0.85 | 0.49 | 0.91 | 0.9 | 0.89 | 0.94 |
| 4 | 160665557 | rs13114723 | 0.039 | -0.252 | 0.778 | -0.124 | -0.006 | 0.067 | 0.067 | 0 | 0.5 | 0.24 | 0.32 | 0.36 | 0.51 | 0.44 |

| | | | | | | | | | | | | | | | | |
|---|---|---|---|---|---|---|---|---|---|---|---|---|---|---|---|---|
| 4 | 160700862 | rs6814758 | 1.561 | -0.002 | 0.967 | 0.146 | 0.062 | 0.042 | 0.042 | 1.196 | 0.94 | 0.63 | 0.96 | 0.96 | 0.97 | 0.99 |
| 4 | 160708061 | rs10020212 | 0.973 | -0.081 | 0.658 | 0.189 | -0.011 | 0.006 | 0.006 | 0 | 0.84 | 0.75 | 0.92 | 0.81 | 0.89 | 0.91 |
| 4 | 160708104 | rs13118321 | 0.579 | 0.302 | 0.214 | 0.287 | -0.05 | -0.062 | -0.062 | 0.255 | 0.1 | 0 | 0.12 | 0.23 | 0.15 | 0.27 |
| 4 | 160715001 | rs1457806 | 0.802 | -0.03 | 0.68 | 0.026 | 0.008 | 0.079 | 0.079 | 1.011 | 0.85 | 0.58 | 0.91 | 0.79 | 0.89 | 0.92 |
| 4 | 160715660 | rs17038866 | 0.796 | -0.198 | -0.302 | -0.161 | 0.015 | 0.006 | 0.006 | 0 | 0.2 | 0.12 | 0.14 | 0.16 | 0.15 | 0.17 |
| 4 | 160715825 | rs6831972 | -0.16 | 0 | 0.184 | 0.223 | -0.003 | -0.014 | -0.014 | 0 | 0.33 | 0.25 | 0.52 | 0.59 | 0.36 | 0.32 |
| 4 | 160719567 | rs9307996 | 1.32 | -0.231 | 0.767 | 0.096 | -0.008 | 0.028 | 0.028 | 0.19 | 0.84 | 0.55 | 0.9 | 0.86 | 0.9 | 0.93 |
| 4 | 160742791 | rs6811354 | 1.257 | -0.232 | 0.629 | 0.116 | -0.023 | 0.004 | 0.004 | 0.744 | 0.82 | 0.57 | 0.91 | 0.86 | 0.89 | 0.91 |
| 4 | 160751151 | rs13150069 | 1.414 | -0.244 | 0.588 | 0.103 | -0.031 | -0.009 | -0.009 | 0.023 | 0.81 | 0.57 | 0.91 | 0.86 | 0.89 | 0.9 |
| 4 | 160753524 | rs13115100 | 0.132 | -0.069 | 0.095 | 0.225 | -0.042 | -0.06 | -0.06 | 0 | 0.1 | 0 | 0.09 | 0.16 | 0.15 | 0.22 |
| 4 | 160766276 | rs4690958 | 1.129 | -0.018 | 0.275 | 0.23 | -0.007 | 0.038 | 0.038 | 0 | 0.92 | 0.72 | 0.95 | 0.91 | 0.97 | 0.97 |
| 4 | 160815681 | rs7672637 | 1.551 | 0.239 | -0.361 | 0.17 | -0.004 | -0.009 | -0.009 | 0 | 0.56 | 0.54 | 0.72 | 0.69 | 0.56 | 0.55 |
| 4 | 160816351 | rs11945058 | -0.326 | 0.293 | -0.038 | 0.832 | 0.002 | 0.009 | 0.009 | 0 | 0.13 | 0.15 | 0.12 | 0.17 | 0.17 | 0.3 |
| 4 | 160817334 | rs4690963 | 0.145 | 0.448 | -0.325 | 0.748 | 0.006 | 0.004 | 0.004 | 0.049 | 0.89 | 0.93 | 0.96 | 0.93 | 0.92 | 0.93 |
| 4 | 160822830 | rs4119125 | 0.745 | 0.47 | -0.216 | 0.824 | -0.016 | 0.024 | 0.024 | 0 | 0.68 | 0.52 | 0.8 | 0.69 | 0.71 | 0.57 |
| 4 | 160823974 | rs6849301 | 0.751 | 0.562 | -0.004 | 1.071 | -0.005 | 0.053 | 0.053 | 0 | 0.75 | 0.59 | 0.55 | 0.37 | 0.76 | 0.67 |
| 4 | 160828500 | rs13133802 | 0.091 | -0.342 | 0.445 | 0.561 | -0.032 | -0.047 | -0.047 | 0 | 0.83 | 0.68 | 0.83 | 0.89 | 0.88 | 0.9 |
| 4 | 160834831 | rs7659695 | 0.867 | -0.347 | -0.817 | 0.135 | -0.057 | 0.022 | 0.022 | 0 | 0.21 | 0.01 | 0.11 | 0.12 | 0.2 | 0.23 |
| 4 | 160869705 | rs4271965 | -0.403 | -1.271 | -0.379 | -0.008 | 0.007 | 0 | 0 | 0.49 | 0.64 | 0.57 | 0.65 | 0.65 | 0.61 | 0.44 |
| 4 | 160884054 | rs6814814 | -0.393 | -1.488 | -0.824 | -0.462 | -0.007 | -0.032 | -0.032 | 0 | 0.6 | 0.55 | 0.72 | 0.77 | 0.65 | 0.69 |
| 4 | 160885494 | rs2349006 | -0.367 | -1.502 | -0.874 | -0.469 | 0.001 | 0.006 | 0.006 | 2.372 | 0.6 | 0.65 | 0.72 | 0.77 | 0.65 | 0.68 |
| 4 | 160889694 | rs6536489 | -0.234 | -1.585 | -1.02 | -0.55 | 0.009 | 0.066 | 0.066 | 1.671 | 0.53 | 0.69 | 0.69 | 0.78 | 0.58 | 0.69 |
| 4 | 160904953 | rs13142965 | -0.172 | -1.647 | -0.992 | -0.652 | -0.052 | -0.09 | -0.09 | 0 | 0.24 | 0.48 | 0.21 | 0.1 | 0.18 | 0.1 |
| 4 | 160905205 | rs12650598 | 0.476 | -1.841 | -1.526 | -0.627 | -0.135 | 0.022 | 0.022 | 0.194 | 0.44 | 0.04 | 0.37 | 0.28 | 0.46 | 0.55 |
| 4 | 160907341 | rs4690967 | -0.707 | -1.782 | -1.007 | -1.169 | -0.002 | 0.02 | 0.02 | 0.023 | 0.68 | 0.83 | 0.85 | 0.74 | 0.71 | 0.72 |
| 4 | 160930806 | rs1035895 | -0.09 | -1.315 | -1.078 | -0.883 | 0.002 | 0.002 | 0.002 | 0.01 | 0.2 | 0.19 | 0.18 | 0.12 | 0.18 | 0.15 |
| 4 | 160937416 | rs17324206 | 0.663 | -1.235 | -1.316 | -0.874 | -0.007 | 0.016 | 0.016 | 0.324 | 0.15 | 0.01 | 0.15 | 0.09 | 0.13 | 0.1 |
| 4 | 160965723 | rs1428412 | -0.19 | -1.141 | -1.384 | -0.897 | 0.017 | 0.082 | 0.082 | 0 | 0.25 | 0.07 | 0.21 | 0.13 | 0.22 | 0.16 |
| 4 | 160974552 | rs2711273 | 0.996 | -1.269 | -1.276 | -0.985 | 0.001 | -0.002 | -0.002 | 0.301 | 0.22 | 0.25 | 0.12 | 0.18 | 0.21 | 0.17 |
| 4 | 160979988 | rs2708233 | -1.014 | -1.518 | -1.514 | -1.23 | -0.011 | 0.014 | 0.014 | 0.022 | 0.58 | 0.65 | 0.4 | 0.31 | 0.51 | 0.39 |
| 4 | 160981899 | rs7689243 | 0.129 | -1.511 | -1.462 | -1.249 | 0.027 | 0.07 | 0.07 | 0 | 0.26 | 0.15 | 0.22 | 0.13 | 0.22 | 0.17 |
| 4 | 160983529 | rs2708232 | -0.562 | -1.533 | -1.475 | -1.211 | 0.019 | 0.063 | 0.063 | 0.786 | 0.51 | 0.64 | 0.36 | 0.27 | 0.43 | 0.34 |
| 4 | 160991913 | rs1366345 | 0.411 | -1.53 | -1.969 | -1.173 | -0.01 | 0.014 | 0.014 | 0.332 | 0.32 | 0.13 | 0.15 | 0.23 | 0.29 | 0.21 |
| 4 | 160992265 | rs7674336 | -0.075 | -1.375 | -1.798 | -1.195 | 0.028 | 0.071 | 0.071 | 0 | 0.26 | 0.14 | 0.22 | 0.13 | 0.22 | 0.17 |
| 4 | 160993666 | rs17326052 | 0.491 | -1.341 | -1.844 | -1.177 | -0.004 | 0.053 | 0.053 | 0 | 0.21 | 0.01 | 0.18 | 0.11 | 0.17 | 0.14 |
| 4 | 160999691 | rs17326199 | -0.478 | -1.328 | -1.898 | -1.323 | 0.015 | 0.12 | 0.12 | 0 | 0.31 | 0.1 | 0.26 | 0.13 | 0.28 | 0.2 |
| 4 | 161000801 | rs10031516 | -0.621 | -1.305 | -1.841 | -1.312 | 0.028 | 0.092 | 0.092 | 0 | 0.33 | 0.18 | 0.26 | 0.13 | 0.28 | 0.2 |
| 4 | 161001529 | rs6839487 | -0.514 | -1.495 | -1.965 | -1.243 | 0.046 | 0.101 | 0.101 | 0 | 0.3 | 0.14 | 0.23 | 0.13 | 0.23 | 0.15 |
| 4 | 161004631 | rs7655275 | -0.352 | -1.407 | -1.981 | -1.182 | 0.042 | 0.1 | 0.1 | 0 | 0.28 | 0.13 | 0.22 | 0.12 | 0.23 | 0.15 |
| 4 | 161010407 | rs1593935 | 0.397 | -1.132 | -2.268 | -1.111 | -0.017 | 0.02 | 0.02 | 0 | 0.31 | 0.1 | 0.14 | 0.21 | 0.28 | 0.23 |
| 4 | 161014288 | rs1651215 | -1.145 | -0.637 | -2.173 | -0.794 | 0.035 | 0.012 | 0.012 | 0 | 0.72 | 0.65 | 0.61 | 0.37 | 0.61 | 0.47 |
| 4 | 161019327 | rs1651214 | -0.115 | -0.656 | -2.193 | -0.866 | 0.057 | 0.067 | 0.067 | 0 | 0.33 | 0.05 | 0.34 | 0.24 | 0.27 | 0.25 |
| 4 | 161024158 | rs1612279 | -0.961 | -0.301 | -2.048 | -0.673 | 0.036 | 0.022 | 0.022 | 0 | 0.71 | 0.63 | 0.6 | 0.42 | 0.61 | 0.45 |
| 4 | 161041088 | rs2464855 | -0.281 | 0.079 | -2.319 | -0.193 | 0.05 | 0.155 | 0.155 | 0 | 0.41 | 0.04 | 0.37 | 0.22 | 0.34 | 0.3 |
| 4 | 161062144 | rs1013340 | -0.658 | 0.922 | -1.401 | 0.33 | -0.022 | 0.028 | 0.028 | 0 | 0.05 | 0 | 0.05 | 0 | 0.09 | 0.07 |
| 4 | 161066459 | rs11723182 | 0.906 | 1.124 | -2.223 | 0.583 | -0.028 | -0.03 | -0.03 | 0 | 0.71 | 0.05 | 0.64 | 0.81 | 0.75 | 0.78 |
| 4 | 161069027 | rs10857326 | 0.283 | 0.932 | -0.733 | 0.652 | 0.031 | 0.143 | 0.143 | 0 | 0.93 | 0.66 | 0.87 | 0.91 | 0.98 | 0.9 |
| 4 | 161083178 | rs6852530 | -1.061 | 1.614 | 0.663 | 0.617 | 0.061 | -0.036 | -0.036 | 1.429 | 0.19 | 0.5 | 0.13 | 0.09 | 0.23 | 0.12 |
| 4 | 161083206 | rs17262799 | -0.627 | 1.532 | 1.216 | 0.577 | 0.004 | -0.001 | -0.001 | 0.476 | 0.15 | 0.17 | 0.1 | 0.09 | 0.23 | 0.12 |
| 4 | 161086241 | rs12331818 | -1.497 | 1.528 | 1.39 | 0.596 | 0.041 | -0.02 | -0.02 | 0.353 | 0.19 | 0.42 | 0.13 | 0.1 | 0.23 | 0.12 |
| 4 | 161094977 | rs11930198 | -1.853 | 2.227 | 0.979 | 1.518 | 0.046 | 0.053 | 0.053 | 0.052 | 0.3 | 0.47 | 0.37 | 0.52 | 0.43 | 0.34 |
| 4 | 161095976 | rs10017226 | -0.974 | 2.309 | 0.781 | 1.523 | 0.008 | 0.04 | 0.04 | 0 | 0.15 | 0.22 | 0.36 | 0.43 | 0.22 | 0.25 |
| 4 | 161097392 | rs9998533 | -0.611 | 2.304 | 0.796 | 1.521 | 0.013 | 0.055 | 0.055 | 0 | 0.13 | 0.2 | 0.26 | 0.43 | 0.2 | 0.22 |
| 4 | 161102726 | rs1922875 | -1.01 | 2.335 | 0.789 | 1.543 | 0.007 | 0.041 | 0.041 | 0 | 0.15 | 0.21 | 0.35 | 0.43 | 0.21 | 0.25 |
| 4 | 161104701 | rs10004177 | -2.101 | 2.316 | 0.833 | 1.533 | 0.003 | 0.004 | 0.004 | 0 | 0.32 | 0.66 | 0.55 | 0.52 | 0.45 | 0.36 |
| 4 | 161105967 | rs17329555 | -0.456 | 2.397 | 0.406 | 1.503 | 0.004 | -0.002 | -0.002 | 6.376 | 0.16 | 0.18 | 0.1 | 0.09 | 0.23 | 0.12 |
| 4 | 161113226 | rs12506633 | -0.579 | 2.417 | 0.322 | 1.668 | 0.012 | 0.056 | 0.056 | 0 | 0.12 | 0.18 | 0.44 | 0.42 | 0.21 | 0.25 |
| 4 | 161114136 | rs11731668 | -0.118 | 2.471 | 0.022 | 1.714 | -0.053 | -0.066 | -0.066 | 0 | 0.11 | 0.02 | 0.42 | 0.42 | 0.21 | 0.25 |
| 4 | 161114166 | rs1922869 | -0.713 | 2.569 | 0.367 | 1.254 | 0.039 | -0.005 | -0.005 | 0 | 0.13 | 0.21 | 0.1 | 0.09 | 0.27 | 0.16 |
| 4 | 161115744 | rs12648235 | -0.922 | 2.124 | 0.647 | 2.316 | 0.013 | 0.205 | 0.205 | 0 | 0.14 | 0.31 | 0.42 | 0.46 | 0.16 | 0.23 |
| 4 | 161116677 | rs1279186 | 0.479 | 1.918 | 0.305 | 2.101 | -0.001 | -0.003 | -0.003 | 0 | 0.88 | 0.89 | 0.83 | 0.79 | 0.87 | 0.88 |
| 4 | 161117083 | rs2349537 | 0.29 | 1.343 | -0.068 | 1.28 | 0.016 | 0.032 | 0.032 | 0 | 0.26 | 0.15 | 0.11 | 0.13 | 0.21 | 0.11 |
| 4 | 161129045 | rs960665 | 0.478 | 1.315 | -0.296 | 1.332 | -0.022 | 0.066 | 0.066 | 0 | 0.44 | 0.21 | 0.5 | 0.35 | 0.49 | 0.4 |
| 4 | 161129696 | rs17039338 | 0.356 | 1.286 | 0.352 | 1.519 | -0.012 | 0.052 | 0.052 | 0 | 0.16 | 0.31 | 0.1 | 0.25 | 0.14 | 0.16 |
| 4 | 161130442 | rs17039340 | 0.559 | 1.518 | 0.143 | 1.316 | 0.001 | -0.003 | -0.003 | 0.213 | 0.15 | 0.15 | 0.09 | 0.09 | 0.19 | 0.1 |
| 4 | 161130933 | rs1922876 | 0.123 | 1.473 | 0.402 | 1.539 | 0.015 | 0.034 | 0.034 | 1.094 | 0.17 | 0.29 | 0.11 | 0.22 | 0.19 | 0.11 |
| 4 | 161131305 | rs10461271 | 1.211 | 1.254 | -0.135 | 1.721 | -0.038 | -0.085 | -0.085 | 0 | 0.27 | 0.03 | 0.41 | 0.38 | 0.3 | 0.25 |
| 4 | 161134770 | rs17039360 | -0.116 | 1.209 | 0.566 | 1.513 | 0.014 | -0.002 | -0.002 | 1.233 | 0.44 | 0.61 | 0.32 | 0.37 | 0.36 | 0.22 |
| 4 | 161135028 | rs13138661 | 1.332 | 0.661 | 0.56 | 1.282 | -0.015 | -0.013 | -0.013 | 0 | 0.2 | 0.22 | 0.09 | 0.11 | 0.13 | 0.1 |

| | | | | | | | | | | | | | | | | |
|---|---|---|---|---|---|---|---|---|---|---|---|---|---|---|---|---|
| 4 | 161137084 | rs13152810 | -0.484 | 0.696 | 0.518 | 1.297 | 0.013 | -0.001 | -0.001 | 0 | 0.44 | 0.6 | 0.31 | 0.37 | 0.36 | 0.22 |
| 4 | 161141895 | rs7686798 | -0.042 | 0.745 | 0.587 | 1.224 | 0.004 | 0.018 | 0.018 | 0 | 0.23 | 0.35 | 0.14 | 0.23 | 0.23 | 0.12 |
| 4 | 161149134 | rs17039372 | 2.452 | 0.765 | 0.155 | 1.253 | -0.008 | -0.031 | -0.031 | 0 | 0.05 | 0 | 0.02 | 0.12 | 0.05 | 0.01 |
| 4 | 161149981 | rs17039374 | -0.33 | 0.761 | 0.281 | 1.24 | 0.007 | 0 | 0 | 0 | 0.43 | 0.56 | 0.23 | 0.36 | 0.36 | 0.22 |
| 4 | 161156298 | rs11725459 | 1.46 | 0.774 | 0.002 | 1.086 | -0.028 | 0.048 | 0.048 | 2.92 | 0.09 | 0 | 0.06 | 0 | 0.11 | 0.06 |
| 4 | 161159633 | rs17039386 | 0.528 | 0.806 | -0.009 | 1.14 | 0.031 | -0.016 | -0.016 | 0 | 0.46 | 0.23 | 0.69 | 0.51 | 0.54 | 0.53 |
| 4 | 161160240 | rs6835451 | -1.046 | 0.671 | 0.257 | 1.108 | -0.003 | -0.01 | -0.01 | 0 | 0.84 | 0.77 | 0.9 | 0.86 | 0.84 | 0.71 |
| 4 | 161179694 | rs295240 | -1.532 | 0.952 | -0.219 | 1.124 | -0.033 | 0.013 | 0.013 | 2.94 | 0.67 | 0.85 | 0.56 | 0.66 | 0.63 | 0.73 |
| 4 | 161206712 | rs13145112 | 2.171 | 0.491 | -0.212 | 0.791 | 0.037 | 0.047 | 0.047 | 0 | 0.17 | 0.06 | 0.07 | 0 | 0.11 | 0.09 |
| 4 | 161211232 | rs295251 | -0.745 | 0.7 | 0.05 | 1.062 | -0.021 | 0.015 | 0.015 | 3.095 | 0.61 | 0.49 | 0.69 | 0.53 | 0.69 | 0.69 |
| 4 | 161213118 | rs1895182 | -0.353 | 0.773 | 0.201 | 1.062 | 0.002 | 0.004 | 0.004 | 0.336 | 0.48 | 0.43 | 0.62 | 0.4 | 0.55 | 0.45 |
| 4 | 161213904 | rs17271770 | 2.409 | 1.151 | 0.126 | 1.182 | -0.03 | -0.023 | -0.023 | 0 | 0.07 | 0.04 | 0.12 | 0.09 | 0.12 | 0.09 |
| 4 | 161214912 | rs953014 | -0.46 | 1.191 | 0.209 | 1.098 | 0.005 | 0.004 | 0.004 | 2.218 | 0.35 | 0.33 | 0.5 | 0.43 | 0.46 | 0.49 |
| 4 | 161215280 | rs953012 | -1.43 | 1.123 | 0.257 | 1.258 | -0.012 | -0.004 | -0.004 | 0 | 0.68 | 0.6 | 0.73 | 0.65 | 0.74 | 0.71 |
| 4 | 161221634 | rs10017605 | 0.954 | 1.179 | -0.12 | 1.233 | 0.004 | 0.013 | 0.013 | 0 | 0.15 | 0.21 | 0.16 | 0.22 | 0.16 | 0.11 |
| 4 | 161224554 | rs12374311 | 1.355 | 1.289 | -0.626 | 1.17 | -0.049 | 0.015 | 0.015 | 0 | 0.26 | 0.07 | 0.39 | 0.21 | 0.3 | 0.25 |
| 4 | 161225387 | rs7694810 | -0.234 | 1.012 | -0.4 | 1.2 | 0.031 | 0.021 | 0.021 | 0.005 | 0.51 | 0.29 | 0.62 | 0.52 | 0.59 | 0.47 |
| 4 | 161225712 | rs1874347 | 1.528 | 1.116 | -1.019 | 1.012 | -0.013 | 0.057 | 0.057 | 1.408 | 0.28 | 0.04 | 0.22 | 0.21 | 0.26 | 0.24 |
| 4 | 161227697 | rs12500560 | 0.38 | 1.112 | -1.019 | 1.255 | -0.088 | -0.02 | -0.02 | 0 | 0.33 | 0.14 | 0.52 | 0.33 | 0.42 | 0.45 |
| 4 | 161229808 | rs11939980 | 0.46 | 1.145 | -1.157 | 1.077 | -0.094 | 0.005 | 0.005 | 0.984 | 0.42 | 0.08 | 0.59 | 0.42 | 0.5 | 0.46 |
| 4 | 161234438 | rs2112267 | -0.197 | 1.063 | -0.197 | 1.287 | -0.002 | -0.004 | -0.004 | 0.043 | 0.56 | 0.5 | 0.68 | 0.68 | 0.57 | 0.54 |
| 4 | 161239313 | rs1495603 | -0.428 | 1.246 | -0.229 | 1.299 | 0.055 | 0.061 | 0.061 | 1.288 | 0.8 | 0.64 | 0.59 | 0.69 | 0.7 | 0.73 |
| 4 | 161240066 | rs2349960 | 1.34 | 0.974 | -0.589 | 0.975 | 0.003 | 0 | 0 | 0 | 0.11 | 0.09 | 0.04 | 0.12 | 0.07 | 0.12 |
| 4 | 161240911 | rs1532391 | 0.993 | 1.107 | -0.478 | 0.784 | 0.008 | 0.083 | 0.083 | 0.998 | 0.34 | 0.18 | 0.28 | 0.2 | 0.32 | 0.3 |
| 4 | 161249029 | rs13123632 | 1.062 | 0.667 | -0.083 | 0.761 | 0.001 | 0.028 | 0.028 | 3.227 | 0.09 | 0.15 | 0.08 | 0.15 | 0.1 | 0.06 |
| 4 | 161255092 | rs6846953 | 0.562 | 0.649 | -0.051 | 0.616 | 0.001 | -0.015 | -0.015 | 0 | 0.3 | 0.34 | 0.25 | 0.14 | 0.31 | 0.37 |
| 4 | 161257623 | rs6818044 | -0.275 | 0.438 | 0.013 | 0.907 | -0.015 | 0.091 | 0.091 | 0.097 | 0.82 | 0.61 | 0.87 | 0.71 | 0.85 | 0.87 |
| 4 | 161267468 | rs13115184 | 0.658 | 0.447 | -0.156 | 0.508 | -0.004 | -0.023 | -0.023 | 0 | 0.32 | 0.38 | 0.24 | 0.13 | 0.28 | 0.35 |
| 4 | 161270471 | rs12503435 | 0.707 | 0.847 | -0.679 | 0.344 | -0.002 | 0.001 | 0.001 | 0 | 0.14 | 0.12 | 0.21 | 0.17 | 0.16 | 0.08 |
| 4 | 161274885 | rs3910586 | 1.119 | 0.57 | -0.476 | 0.271 | 0.038 | 0.014 | 0.014 | 0 | 0.11 | 0.23 | 0.17 | 0.13 | 0.14 | 0.23 |
| 4 | 161280429 | rs294514 | -0.651 | 0.41 | -0.65 | 0.241 | -0.019 | -0.043 | -0.043 | 0 | 0.64 | 0.52 | 0.79 | 0.8 | 0.68 | 0.78 |
| 4 | 161287144 | rs17272220 | 0.438 | 0.759 | -0.699 | 0.503 | 0.025 | -0.019 | -0.019 | 0 | 0.2 | 0.31 | 0.26 | 0.17 | 0.24 | 0.29 |
| 4 | 161287754 | rs17272241 | 0.922 | 0.421 | -1.838 | 0.332 | -0.024 | 0.02 | 0.02 | 0.022 | 0.18 | 0.01 | 0.08 | 0.07 | 0.15 | 0.16 |
| 4 | 161291816 | rs2350244 | 0.823 | 1.186 | -1.718 | 0.758 | -0.086 | -0.063 | -0.063 | 0 | 0.14 | 0.01 | 0.3 | 0.23 | 0.22 | 0.23 |
| 4 | 161292814 | rs3846243 | 0.274 | 0.876 | -1.581 | 0.731 | -0.047 | -0.038 | -0.038 | 0 | 0.22 | 0.11 | 0.33 | 0.32 | 0.3 | 0.3 |
| 4 | 161294866 | rs294526 | -0.221 | -0.344 | -1.136 | 0.469 | 0.079 | 0.085 | 0.085 | 0 | 0.53 | 0.72 | 0.81 | 0.8 | 0.67 | 0.66 |
| 4 | 161300559 | rs9884424 | -0.824 | -0.557 | -0.52 | 0.475 | -0.051 | -0.02 | -0.02 | 0 | 0.76 | 0.66 | 0.88 | 0.83 | 0.86 | 0.86 |
| 4 | 161300792 | rs9308003 | -0.376 | 0.076 | -0.686 | 0.82 | 0.001 | -0.007 | -0.007 | 0 | 0.46 | 0.49 | 0.44 | 0.31 | 0.49 | 0.37 |
| 4 | 161300938 | rs17332185 | 0.112 | 0.59 | -1.054 | 0.975 | -0.067 | 0 | 0 | 0 | 0.19 | 0.08 | 0.25 | 0.19 | 0.29 | 0.24 |
| 4 | 161301370 | rs17039504 | 0.306 | 0.41 | -0.932 | 1.369 | -0.014 | -0.024 | -0.024 | 0.123 | 0.09 | 0.03 | 0.21 | 0.38 | 0.1 | 0.25 |
| 4 | 161306164 | rs6835941 | -0.813 | 0.149 | -0.34 | 1.114 | -0.007 | 0.021 | 0.021 | 0 | 0.26 | 0.33 | 0.33 | 0.48 | 0.22 | 0.35 |
| 4 | 161307463 | rs1946867 | -1.961 | 0.285 | -0.203 | 0.665 | 0.01 | 0.134 | 0.134 | 0.645 | 0.5 | 0.67 | 0.64 | 0.86 | 0.54 | 0.59 |
| 4 | 161308423 | rs6536518 | -1.521 | 0.304 | -0.233 | 0.643 | -0.011 | 0.078 | 0.078 | 0.105 | 0.44 | 0.66 | 0.59 | 0.86 | 0.45 | 0.53 |
| 4 | 161320477 | rs1583732 | -1.036 | 0.085 | 0.099 | 0.772 | 0.003 | 0.022 | 0.022 | 0 | 0.36 | 0.45 | 0.33 | 0.44 | 0.34 | 0.25 |
| 4 | 161323274 | rs17039540 | -0.809 | 0.142 | 0.299 | 0.837 | 0.009 | 0.013 | 0.013 | 0.37 | 0.24 | 0.28 | 0.3 | 0.39 | 0.32 | 0.22 |
| 4 | 161336215 | rs7678343 | -0.159 | 0.049 | 0.255 | 1.043 | 0.002 | 0.131 | 0.131 | 0 | 0.12 | 0.27 | 0.28 | 0.42 | 0.14 | 0.3 |
| 4 | 161337411 | rs13124867 | 0.555 | -0.212 | 0.262 | 1.022 | -0.034 | 0.037 | 0.037 | 0.41 | 0.76 | 0.4 | 0.8 | 0.65 | 0.86 | 0.85 |
| 4 | 161343388 | rs7376552 | -1.058 | 0.165 | -0.063 | 1.098 | -0.052 | -0.049 | -0.049 | 0 | 0.32 | 0.17 | 0.49 | 0.58 | 0.45 | 0.45 |
| 4 | 161344370 | rs4691655 | 0.087 | 0.197 | 0.111 | 1.117 | 0 | 0 | 0 | 0.443 | 0.07 | 0.07 | 0.09 | 0.07 | 0.1 | 0.06 |
| 4 | 161366230 | rs1595663 | 0.134 | -0.023 | 0.758 | 1.175 | -0.05 | 0.008 | 0.008 | 0 | 0.17 | 0.4 | 0.13 | 0.12 | 0.09 | 0.07 |
| 4 | 161366623 | rs12640159 | -0.268 | 0.372 | 0.027 | 1.305 | -0.061 | -0.103 | -0.103 | 0 | 0.21 | 0.09 | 0.39 | 0.5 | 0.29 | 0.38 |
| 4 | 161379828 | rs1347903 | -0.883 | 0.061 | -0.01 | 1.283 | -0.067 | -0.037 | -0.037 | 0.043 | 0.26 | 0.15 | 0.5 | 0.69 | 0.35 | 0.44 |
| 4 | 161381482 | rs1371251 | 1.679 | -0.105 | 0.479 | 1.129 | -0.008 | 0.059 | 0.059 | 1.153 | 0.56 | 0.3 | 0.34 | 0.19 | 0.54 | 0.48 |
| 4 | 161386383 | rs1946869 | 0.366 | 0.706 | 0.133 | 1.685 | -0.036 | -0.075 | -0.075 | 0 | 0.13 | 0.04 | 0.37 | 0.31 | 0.17 | 0.27 |
| 4 | 161402299 | rs1898871 | -1 | 0.72 | 0.184 | 1.622 | -0.079 | -0.078 | -0.078 | 0.052 | 0.24 | 0.13 | 0.47 | 0.56 | 0.38 | 0.45 |
| 4 | 161406478 | rs17039615 | 0.183 | 0.688 | 0.189 | 1.643 | -0.031 | -0.074 | -0.074 | 0 | 0.15 | 0.06 | 0.38 | 0.3 | 0.19 | 0.29 |
| 4 | 161411394 | rs13151634 | 1.222 | 0.596 | 0.647 | 1.561 | -0.033 | 0.019 | 0.019 | 0 | 0.62 | 0.7 | 0.43 | 0.38 | 0.54 | 0.49 |
| 4 | 161413994 | rs12498920 | 0.195 | 0.666 | 0.184 | 1.539 | -0.031 | -0.077 | -0.077 | 0 | 0.15 | 0.06 | 0.37 | 0.31 | 0.19 | 0.29 |
| 4 | 161421752 | rs4525928 | 1.174 | 0.579 | 0.274 | 1.178 | 0.006 | -0.017 | -0.017 | 0 | 0.59 | 0.5 | 0.37 | 0.31 | 0.52 | 0.48 |
| 4 | 161425366 | rs10857329 | 0.476 | 0.413 | -0.376 | 1.326 | -0.007 | -0.057 | -0.057 | 0 | 0.17 | 0.1 | 0.41 | 0.3 | 0.18 | 0.35 |
| 4 | 161427827 | rs9995630 | 0.112 | 0.367 | -0.677 | 0.567 | -0.014 | -0.013 | -0.013 | 0 | 0.35 | 0.47 | 0.26 | 0.29 | 0.3 | 0.29 |
| 4 | 161428827 | rs4690982 | 0.016 | 0.763 | -0.923 | 0.771 | -0.032 | -0.035 | -0.035 | 0 | 0.28 | 0.18 | 0.53 | 0.56 | 0.38 | 0.51 |
| 4 | 161429238 | rs4690983 | 0.049 | 0.773 | -1.109 | 0.775 | -0.084 | -0.09 | -0.09 | 0 | 0.28 | 0.13 | 0.52 | 0.56 | 0.38 | 0.51 |
| 4 | 161436797 | rs1898872 | 1.227 | 0.286 | -0.979 | 0.637 | 0.016 | 0.009 | 0.009 | 0 | 0.14 | 0.06 | 0.07 | 0.05 | 0.11 | 0.07 |
| 4 | 161441599 | rs1595667 | -1.266 | 0.238 | -1.12 | 0.792 | 0.002 | -0.075 | -0.075 | 0 | 0.8 | 0.95 | 0.6 | 0.67 | 0.82 | 0.63 |
| 4 | 161442245 | rs7681545 | 1.399 | 0.192 | -1.126 | 0.782 | 0.019 | 0.007 | 0.007 | 0 | 0.14 | 0.06 | 0.07 | 0.06 | 0.11 | 0.07 |
| 4 | 161443164 | rs1561492 | 1.334 | 0.036 | -1.022 | 0.326 | 0.011 | 0.018 | 0.018 | 0 | 0.33 | 0.28 | 0.19 | 0.15 | 0.29 | 0.21 |
| 4 | 161444504 | rs17039652 | 1.795 | 0.309 | -1.133 | 0.514 | -0.066 | -0.052 | -0.052 | 0 | 0.11 | 0.02 | 0.11 | 0.26 | 0.2 | 0.12 |
| 4 | 161447090 | rs17039654 | 0.198 | 0.128 | -0.711 | 0.489 | 0.017 | -0.023 | -0.023 | 0 | 0.35 | 0.28 | 0.49 | 0.38 | 0.29 | 0.41 |

| | | | | | | | | | | | | | | | | |
|---|---|---|---|---|---|---|---|---|---|---|---|---|---|---|---|---|
| 4 | 161451545 | rs11930824 | 0.223 | 0.105 | -0.623 | 0.249 | 0.003 | 0.006 | 0.006 | 0 | 0.36 | 0.43 | 0.58 | 0.54 | 0.37 | 0.55 |
| 4 | 161453186 | rs10032921 | -1.368 | 0.074 | -0.688 | 0.181 | 0.014 | 0.031 | 0.031 | 0 | 0.63 | 0.7 | 0.78 | 0.85 | 0.67 | 0.74 |
| 4 | 161464002 | rs1439395 | 0.222 | 0 | -0.724 | 0.247 | 0.004 | 0.02 | 0.02 | 0 | 0.5 | 0.57 | 0.35 | 0.41 | 0.53 | 0.35 |
| 4 | 161468822 | rs11941490 | 0.063 | -0.132 | -0.475 | 0.229 | -0.005 | -0.026 | -0.026 | 0.089 | 0.34 | 0.49 | 0.28 | 0.23 | 0.34 | 0.32 |
| 4 | 161472909 | rs17355669 | 1.77 | -0.279 | -0.514 | 0.248 | -0.006 | -0.006 | -0.006 | 0 | 0.13 | 0 | 0.08 | 0.05 | 0.12 | 0.07 |
| 4 | 161473319 | rs10018218 | 0.748 | -0.39 | -0.477 | 0.208 | 0.03 | -0.011 | -0.011 | 0 | 0.19 | 0.5 | 0.2 | 0.17 | 0.2 | 0.12 |
| 4 | 161477815 | rs17039750 | 0.266 | -0.266 | -0.565 | 0.174 | -0.003 | -0.001 | -0.001 | 0 | 0.27 | 0.25 | 0.23 | 0.32 | 0.33 | 0.2 |
| 4 | 161479466 | rs17277682 | 1.451 | -0.133 | -0.475 | 0.275 | 0.04 | -0.025 | -0.025 | 0 | 0.16 | 0.36 | 0.17 | 0.09 | 0.18 | 0.08 |
| 4 | 161482994 | rs4691681 | 1.609 | -0.081 | -0.93 | 0.328 | -0.075 | -0.06 | -0.06 | 0 | 0.11 | 0.01 | 0.14 | 0.27 | 0.2 | 0.13 |
| 4 | 161489245 | rs13140082 | -0.508 | -1.495 | -1.086 | 0.079 | 0.074 | 0.05 | 0.05 | 0 | 0.64 | 0.84 | 0.83 | 0.8 | 0.77 | 0.68 |
| 4 | 161495175 | rs10031651 | -1.424 | -1.516 | -1.061 | 0.083 | 0.015 | 0.012 | 0.012 | 0 | 0.77 | 0.84 | 0.94 | 0.86 | 0.9 | 0.77 |
| 4 | 161501719 | rs12511322 | 1.542 | -1.509 | -1.111 | 0.186 | 0.017 | -0.07 | -0.07 | 0 | 0.2 | 0.05 | 0.39 | 0.35 | 0.19 | 0.38 |
| 4 | 161508641 | rs11938367 | -0.076 | -1.34 | -0.97 | 0.11 | 0.001 | -0.001 | -0.001 | 0.259 | 0.4 | 0.61 | 0.57 | 0.52 | 0.36 | 0.5 |
| 4 | 161517400 | rs2350469 | 0.244 | -1.224 | -1.106 | -0.081 | -0.01 | -0.025 | -0.025 | 0 | 0.32 | 0.56 | 0.29 | 0.22 | 0.3 | 0.19 |
| 4 | 161522527 | rs7667168 | -1.536 | -1.286 | -1.395 | 0.223 | 0.056 | -0.013 | -0.013 | 0 | 0.77 | 0.92 | 0.93 | 0.77 | 0.87 | 0.75 |
| 4 | 161524114 | rs4443235 | 0.224 | -1.156 | -1.282 | 0.192 | -0.009 | -0.021 | -0.021 | 0 | 0.32 | 0.57 | 0.29 | 0.22 | 0.3 | 0.19 |
| 4 | 161534644 | rs6847636 | 1.237 | -1.123 | -1.027 | 0.222 | 0.031 | 0.066 | 0.066 | 0 | 0.18 | 0.48 | 0.18 | 0.22 | 0.18 | 0.12 |
| 4 | 161548582 | rs1992108 | 1.676 | -1.022 | -1.146 | 0.257 | -0.037 | 0.014 | 0.014 | 0 | 0.24 | 0.21 | 0.24 | 0.28 | 0.36 | 0.2 |
| 4 | 161552712 | rs11945623 | -0.195 | -1.181 | -1.18 | 0.162 | 0.066 | 0.001 | 0.001 | 0 | 0.56 | 0.82 | 0.53 | 0.5 | 0.68 | 0.39 |
| 4 | 161567725 | rs11100301 | 1.387 | -1.213 | -1.085 | 0.382 | 0.016 | -0.052 | -0.052 | 0.113 | 0.21 | 0.05 | 0.39 | 0.27 | 0.19 | 0.36 |
| 4 | 161571071 | rs6536530 | 0.786 | -1.443 | -1.172 | 0.188 | 0.008 | 0.004 | 0.004 | 0 | 0.46 | 0.6 | 0.42 | 0.49 | 0.57 | 0.35 |
| 4 | 161575183 | rs4691690 | -1.607 | -1.498 | -1.181 | 0.19 | 0.011 | -0.004 | -0.004 | 0 | 0.8 | 0.86 | 0.93 | 0.77 | 0.88 | 0.76 |
| 4 | 161575686 | rs1436944 | 1.342 | -1.25 | -1.007 | 0.365 | -0.004 | 0.01 | 0.01 | 0 | 0.21 | 0.2 | 0.41 | 0.28 | 0.19 | 0.35 |
| 4 | 161578427 | rs17359034 | 0.595 | -1.159 | -1.057 | 0.335 | 0.002 | 0.001 | 0.001 | 0 | 0.46 | 0.59 | 0.42 | 0.49 | 0.57 | 0.35 |
| 4 | 161578634 | rs17039966 | 1.491 | -0.76 | -1.43 | 0.415 | -0.101 | 0.052 | 0.052 | 0 | 0.27 | 0.13 | 0.27 | 0.22 | 0.39 | 0.23 |
| 4 | 161579050 | rs17039968 | 1.484 | -0.748 | -1.432 | 0.413 | -0.099 | 0.053 | 0.053 | 0 | 0.27 | 0.13 | 0.27 | 0.22 | 0.38 | 0.23 |
| 4 | 161590091 | rs11100304 | 0.778 | -1.05 | -1.158 | 0.646 | 0.009 | -0.021 | -0.021 | 0 | 0.21 | 0.13 | 0.4 | 0.28 | 0.19 | 0.35 |
| 4 | 161595023 | rs1436942 | -1.646 | -1.057 | -1.22 | 0.487 | -0.007 | -0.083 | -0.083 | 1.234 | 0.86 | 0.98 | 0.62 | 0.59 | 0.86 | 0.76 |
| 4 | 161595152 | rs17039992 | 0.591 | -1.225 | -0.688 | 0.152 | 0.034 | -0.014 | -0.014 | 0 | 0.19 | 0.39 | 0.14 | 0.09 | 0.22 | 0.11 |
| 4 | 161597298 | rs4690990 | 0.1 | -1.614 | -1.285 | -0.127 | 0.048 | 0.048 | 0.048 | 0.016 | 0.49 | 0.64 | 0.46 | 0.29 | 0.63 | 0.37 |
| 4 | 161598173 | rs2164631 | 1.692 | -1.129 | -1.608 | -0.373 | -0.11 | 0.039 | 0.039 | 0.031 | 0.25 | 0.08 | 0.3 | 0.2 | 0.38 | 0.22 |
| 4 | 161598367 | rs1036471 | -1.52 | -0.958 | -1.212 | 0.042 | 0 | -0.074 | -0.074 | 0.326 | 0.85 | 0.98 | 0.62 | 0.59 | 0.86 | 0.76 |
| 4 | 161617009 | rs6843805 | 0.366 | -1.167 | -1.06 | -0.24 | 0.012 | 0.008 | 0.008 | 0 | 0.2 | 0.13 | 0.07 | 0.16 | 0.13 | 0.17 |
| 4 | 161620939 | rs2350477 | -0.508 | -1.238 | -0.443 | -0.78 | 0 | 0.001 | 0.001 | 0 | 0.75 | 0.64 | 0.85 | 0.76 | 0.75 | 0.74 |
| 4 | 161623467 | rs9308012 | 0.688 | -1.69 | -0.378 | -0.987 | 0.001 | 0.018 | 0.018 | 0 | 0.45 | 0.4 | 0.38 | 0.32 | 0.51 | 0.4 |
| 4 | 161632032 | rs1370171 | -0.503 | -1.469 | -0.546 | -0.882 | 0 | -0.001 | -0.001 | 0.107 | 0.75 | 0.74 | 0.86 | 0.75 | 0.75 | 0.76 |
| 4 | 161636751 | rs6536531 | -0.413 | -1.452 | -0.588 | -0.875 | 0.001 | 0 | 0 | 0 | 0.74 | 0.75 | 0.86 | 0.74 | 0.74 | 0.74 |
| 4 | 161646978 | rs4691696 | 0.242 | -1.462 | -0.725 | -0.654 | 0.018 | -0.054 | -0.054 | 0 | 0.24 | 0.14 | 0.45 | 0.44 | 0.21 | 0.32 |
| 4 | 161656017 | rs6843165 | -0.494 | -1.434 | -0.448 | -0.793 | -0.001 | 0.014 | 0.014 | 0 | 0.48 | 0.53 | 0.63 | 0.61 | 0.5 | 0.58 |
| 4 | 161657993 | rs6857015 | 0.363 | -1.278 | -0.395 | -0.886 | 0.014 | 0.064 | 0.064 | 0 | 0.47 | 0.35 | 0.31 | 0.31 | 0.42 | 0.37 |
| 4 | 161659251 | rs4081678 | 0.425 | -1.252 | -0.353 | -0.869 | 0.006 | 0.032 | 0.032 | 0 | 0.46 | 0.42 | 0.32 | 0.3 | 0.42 | 0.37 |
| 4 | 161662308 | rs3846247 | 0.737 | -1.268 | -0.415 | -0.8 | -0.001 | -0.028 | -0.028 | 0.374 | 0.23 | 0.13 | 0.46 | 0.33 | 0.24 | 0.3 |
| 4 | 161666367 | rs10011041 | -0.672 | -1.707 | -0.629 | -1.334 | -0.019 | -0.022 | -0.022 | 0.004 | 0.43 | 0.47 | 0.18 | 0.18 | 0.27 | 0.25 |
| 4 | 161676056 | rs13130301 | 0.653 | -1.826 | -0.711 | -1.4 | 0.012 | 0.009 | 0.009 | 0 | 0.22 | 0.14 | 0.08 | 0.14 | 0.17 | 0.16 |
| 4 | 161683458 | rs3846250 | 0.531 | -1.618 | -0.415 | -1.335 | -0.03 | -0.016 | -0.016 | 0 | 0.34 | 0.22 | 0.52 | 0.6 | 0.41 | 0.52 |
| 4 | 161685302 | rs10517723 | 1.387 | -1.626 | -0.364 | -1.378 | -0.004 | -0.021 | -0.021 | 0 | 0.09 | 0.1 | 0.06 | 0 | 0.07 | 0.03 |
| 4 | 161698345 | rs12499554 | 2.177 | -1.38 | -0.591 | -1.214 | -0.056 | -0.087 | -0.087 | 0 | 0.12 | 0.01 | 0.39 | 0.34 | 0.18 | 0.26 |
| 4 | 161703150 | rs6853708 | -2.571 | -1.799 | 0.359 | -2.026 | -0.032 | 0.008 | 0.008 | 0 | 0.84 | 0.92 | 0.78 | 0.86 | 0.77 | 0.82 |
| 4 | 161707186 | rs1420877 | 0.966 | -1.856 | 0.357 | -2.035 | -0.054 | 0.015 | 0.015 | 0 | 0.37 | 0.19 | 0.57 | 0.68 | 0.5 | 0.57 |
| 4 | 161713693 | rs10031243 | -0.972 | -1.882 | 0.374 | -2.042 | -0.027 | -0.031 | -0.031 | 0 | 0.44 | 0.5 | 0.18 | 0.18 | 0.28 | 0.25 |
| 4 | 161718544 | rs3909235 | -1.661 | -1.124 | 0.686 | -1.595 | 0.024 | 0.003 | 0.003 | 0 | 0.82 | 0.75 | 0.75 | 0.79 | 0.77 | 0.82 |
| 4 | 161733077 | rs4488900 | -0.258 | -1.383 | 0.602 | -1.694 | 0.006 | 0.005 | 0.005 | 0 | 0.22 | 0.16 | 0.09 | 0.13 | 0.16 | 0.17 |
| 4 | 161736289 | rs4691702 | -0.671 | -0.947 | 0.895 | -1.75 | -0.032 | 0.028 | 0.028 | 0 | 0.76 | 0.85 | 0.7 | 0.86 | 0.66 | 0.75 |
| 4 | 161736792 | rs4691703 | -0.681 | -0.942 | 0.898 | -1.747 | -0.032 | 0.028 | 0.028 | 0 | 0.76 | 0.85 | 0.7 | 0.86 | 0.66 | 0.75 |
| 4 | 161741448 | rs3909239 | -2.337 | -1.623 | 2.055 | -1.199 | -0.035 | -0.043 | -0.043 | 0 | 0.31 | 0.48 | 0.17 | 0.2 | 0.23 | 0.24 |
| 4 | 161742881 | rs2349301 | -1.711 | -1.614 | 1.81 | -1.194 | 0 | 0.002 | 0.002 | 0 | 0.14 | 0.12 | 0.11 | 0.13 | 0.16 | 0.16 |
| 4 | 161745970 | rs13112276 | -1.317 | -1.626 | 1.635 | -1.205 | -0.009 | -0.003 | -0.003 | 0.004 | 0.18 | 0.18 | 0.09 | 0.17 | 0.14 | 0.15 |
| 4 | 161756548 | rs7667388 | -2.474 | -1.479 | 1.145 | -1.129 | 0.092 | 0.115 | 0.115 | 0 | 0.42 | 0.79 | 0.23 | 0.2 | 0.29 | 0.27 |
| 4 | 161761676 | rs17040246 | -1.186 | -1.168 | 1.101 | -0.985 | -0.007 | -0.053 | -0.053 | 0 | 0.12 | 0.22 | 0.09 | 0.02 | 0.1 | 0.06 |
| 4 | 161763818 | rs17040262 | -1.345 | -0.932 | 1.095 | -0.812 | 0.019 | 0.014 | 0.014 | 0 | 0.16 | 0.08 | 0.06 | 0.13 | 0.12 | 0.14 |
| 4 | 161764629 | rs1420884 | -1.993 | -1.009 | 1.585 | -0.856 | -0.073 | -0.064 | -0.064 | 0.136 | 0.25 | 0.42 | 0.1 | 0.14 | 0.17 | 0.18 |
| 4 | 161771838 | rs4594673 | -2.306 | -0.978 | 1.495 | -0.833 | -0.036 | -0.048 | -0.048 | 0 | 0.32 | 0.43 | 0.14 | 0.19 | 0.24 | 0.25 |
| 4 | 161777781 | rs1362705 | -2.158 | -0.687 | 1.552 | -0.644 | -0.01 | -0.015 | -0.015 | 0 | 0.26 | 0.3 | 0.14 | 0.18 | 0.22 | 0.23 |
| 4 | 161778204 | rs4295220 | -1.523 | -0.337 | 1.632 | -0.373 | 0.027 | -0.066 | -0.066 | 0 | 0.14 | 0.38 | 0.08 | 0.04 | 0.13 | 0.09 |
| 4 | 161783422 | rs6840597 | -2.625 | -0.315 | 1.482 | -0.448 | -0.025 | -0.049 | -0.049 | 1.013 | 0.27 | 0.52 | 0.14 | 0.18 | 0.2 | 0.23 |
| 4 | 161785031 | rs13130758 | -0.327 | -0.317 | 1.381 | -0.447 | 0.017 | -0.035 | -0.035 | 0.014 | 0.06 | 0.01 | 0.02 | 0.12 | 0.03 | 0.12 |
| 4 | 161785599 | rs9991811 | -1.317 | -0.318 | 1.43 | -0.45 | 0.016 | -0.059 | -0.059 | 0 | 0.14 | 0.33 | 0.07 | 0.04 | 0.13 | 0.09 |
| 4 | 161785781 | rs6854952 | -1.621 | -0.452 | 0.789 | -0.433 | -0.003 | -0.001 | -0.001 | 0 | 0.13 | 0.17 | 0.06 | 0.14 | 0.07 | 0.14 |
| 4 | 161790472 | rs4273418 | -2.597 | -0.448 | 0.944 | -0.454 | 0.014 | 0.019 | 0.019 | 0 | 0.29 | 0.74 | 0.18 | 0.18 | 0.2 | 0.25 |

| | | | | | | | | | | | | | | | | |
|---|---|---|---|---|---|---|---|---|---|---|---|---|---|---|---|---|
| 4 | 161790594 | rs7694824 | -1.134 | 0.19 | 0.39 | -0.435 | 0.009 | -0.018 | -0.018 | 0.015 | 0.1 | 0.16 | 0.08 | 0.05 | 0.14 | 0.07 |
| 4 | 161795832 | rs988569 | -1.134 | -0.151 | 0.157 | -0.129 | -0.039 | -0.049 | -0.049 | 0 | 0.1 | 0.22 | 0.03 | 0 | 0.03 | 0.04 |
| 4 | 161799400 | rs17040351 | -0.415 | 0.42 | -0.317 | 0.102 | -0.002 | 0 | 0 | 0 | 0.09 | 0.09 | 0.05 | 0.08 | 0.12 | 0.03 |
| 4 | 161808171 | rs9997826 | -0.061 | 0.563 | -0.938 | -0.976 | -0.005 | 0.041 | 0.041 | 0 | 0.22 | 0.11 | 0.13 | 0.01 | 0.21 | 0.2 |
| 4 | 161808304 | rs4463025 | -0.868 | 0.721 | -0.924 | -1.113 | 0.002 | -0.002 | -0.002 | 0 | 0.8 | 0.82 | 0.67 | 0.85 | 0.76 | 0.87 |
| 4 | 161809149 | rs11737365 | 1.17 | 0.479 | -1.665 | -0.409 | -0.027 | -0.053 | -0.053 | 0 | 0.26 | 0.07 | 0.29 | 0.32 | 0.27 | 0.33 |
| 4 | 161818208 | rs4399939 | 0.597 | 0.242 | -1.589 | -0.384 | -0.019 | -0.084 | -0.084 | 0.48 | 0.18 | 0.06 | 0.15 | 0.42 | 0.17 | 0.34 |
| 4 | 161828952 | rs4613522 | 0.29 | -0.12 | -0.624 | -0.249 | 0.001 | 0.009 | 0.009 | 0 | 0.52 | 0.5 | 0.46 | 0.39 | 0.46 | 0.53 |
| 4 | 161837715 | rs4691713 | -0.002 | 0.057 | -0.228 | 0.062 | 0.071 | -0.002 | -0.002 | 0 | 0.79 | 0.56 | 0.78 | 0.76 | 0.69 | 0.81 |
| 4 | 161840651 | rs9992795 | 0.384 | -0.585 | -0.699 | -0.276 | -0.019 | -0.015 | -0.015 | 0 | 0.31 | 0.43 | 0.36 | 0.23 | 0.29 | 0.21 |
| 4 | 161846866 | rs4444780 | 1.677 | -0.858 | -1.16 | -0.424 | 0.039 | -0.08 | -0.08 | 0 | 0.26 | 0.07 | 0.29 | 0.4 | 0.17 | 0.49 |
| 4 | 161848243 | rs10050122 | 0.421 | -0.854 | -0.782 | -0.522 | -0.011 | -0.017 | -0.017 | 0.045 | 0.34 | 0.53 | 0.39 | 0.23 | 0.3 | 0.25 |
| 4 | 161851303 | rs7695380 | 1.727 | -0.87 | -0.728 | -0.516 | 0.034 | -0.054 | -0.054 | 0.207 | 0.26 | 0.14 | 0.29 | 0.4 | 0.17 | 0.49 |
| 4 | 161856084 | rs9993721 | -0.136 | -0.929 | -0.723 | -0.621 | 0.001 | 0.003 | 0.003 | 0 | 0.47 | 0.55 | 0.48 | 0.36 | 0.48 | 0.29 |
| 4 | 161856343 | rs9993952 | 1.233 | -0.958 | -0.747 | -0.663 | -0.007 | -0.011 | -0.011 | 1.305 | 0.23 | 0.31 | 0.32 | 0.19 | 0.21 | 0.14 |
| 4 | 161861640 | rs7436001 | -0.622 | -0.854 | -0.868 | -0.584 | 0.001 | -0.001 | -0.001 | 2.659 | 0.74 | 0.73 | 0.78 | 0.75 | 0.65 | 0.78 |
| 4 | 161863893 | rs13122745 | -2.483 | -0.465 | -0.894 | -0.706 | -0.014 | 0.011 | 0.011 | 0 | 0.89 | 0.92 | 0.92 | 0.99 | 0.83 | 0.95 |
| 4 | 161864128 | rs6858095 | 0.308 | -0.509 | -0.81 | -0.624 | -0.008 | -0.011 | -0.011 | 0.223 | 0.34 | 0.56 | 0.4 | 0.24 | 0.3 | 0.25 |
| 4 | 161875145 | rs4529002 | 0.949 | -0.421 | -0.927 | -0.578 | 0.003 | 0.002 | 0.002 | 0 | 0.15 | 0.16 | 0.14 | 0.24 | 0.19 | 0.17 |
| 4 | 161878326 | rs11735987 | 0.499 | -0.54 | -1.015 | -0.574 | -0.005 | -0.008 | -0.008 | 0 | 0.33 | 0.39 | 0.38 | 0.23 | 0.3 | 0.25 |
| 4 | 161889983 | rs12503470 | 0.674 | -0.482 | -1.547 | -0.454 | 0.035 | -0.063 | -0.063 | 0 | 0.4 | 0.16 | 0.38 | 0.52 | 0.35 | 0.53 |
| 4 | 161907084 | rs13132985 | 0.331 | -0.409 | -1.204 | -0.434 | -0.055 | -0.086 | -0.086 | 0 | 0.28 | 0.08 | 0.21 | 0.35 | 0.29 | 0.27 |
| 4 | 161912988 | rs4345122 | -1.148 | 0.288 | -1.078 | -0.467 | -0.031 | -0.029 | -0.029 | 0 | 0.62 | 0.89 | 0.66 | 0.59 | 0.62 | 0.5 |
| 4 | 161924832 | rs11100327 | -0.382 | 0.284 | -0.609 | -0.482 | -0.001 | 0 | 0 | 0 | 0.47 | 0.55 | 0.56 | 0.54 | 0.47 | 0.41 |
| 4 | 161935342 | rs1982202 | 1.439 | 0.288 | -0.586 | -0.324 | -0.004 | -0.026 | -0.026 | 0 | 0.15 | 0.12 | 0.24 | 0.35 | 0.13 | 0.23 |
| 4 | 161936611 | rs4379019 | -1.598 | 0.239 | -0.57 | -0.374 | -0.001 | -0.015 | -0.015 | 0 | 0.87 | 0.88 | 0.9 | 0.79 | 0.88 | 0.77 |
| 4 | 161940430 | rs4467511 | -0.139 | 0.291 | -0.596 | -0.483 | -0.002 | 0.001 | 0.001 | 0 | 0.48 | 0.45 | 0.46 | 0.46 | 0.48 | 0.54 |
| 4 | 161940921 | rs9991854 | 0.568 | 0.287 | -0.708 | -0.474 | 0.001 | -0.012 | -0.012 | 2.437 | 0.37 | 0.24 | 0.37 | 0.41 | 0.33 | 0.45 |
| 4 | 161945302 | rs1439260 | -0.268 | 0.275 | -0.479 | -0.474 | 0 | 0.044 | 0.044 | 0 | 0.7 | 0.53 | 0.57 | 0.6 | 0.7 | 0.62 |
| 4 | 161946368 | rs4234935 | -1.497 | 0.286 | -0.482 | -0.449 | -0.001 | 0.006 | 0.006 | 0.071 | 0.86 | 0.83 | 0.9 | 0.79 | 0.88 | 0.76 |
| 4 | 161952228 | rs6536560 | -0.883 | 0.244 | -0.572 | -0.523 | 0.002 | 0.003 | 0.003 | 0 | 0.65 | 0.75 | 0.77 | 0.72 | 0.68 | 0.54 |
| 4 | 161953729 | rs17301015 | 2.183 | 0.359 | -0.419 | -0.514 | 0.009 | -0.019 | -0.019 | 0 | 0.11 | 0.17 | 0.08 | 0.05 | 0.15 | 0.08 |
| 4 | 161968079 | rs2028307 | -0.835 | 0.332 | -0.704 | -0.512 | -0.016 | -0.019 | -0.019 | 0 | 0.64 | 0.86 | 0.79 | 0.65 | 0.66 | 0.67 |
| 4 | 161982414 | rs17040579 | 0.25 | -0.067 | -0.458 | -0.496 | 0.004 | -0.016 | -0.016 | 0.053 | 0.29 | 0.44 | 0.42 | 0.24 | 0.33 | 0.22 |
| 4 | 161987342 | rs2218107 | 2.039 | 0.008 | -0.722 | -0.412 | 0.001 | 0.013 | 0.013 | 0 | 0.21 | 0.14 | 0.13 | 0.07 | 0.2 | 0.22 |
| 4 | 161996191 | rs17040617 | 1.471 | 0.157 | -0.33 | 0.139 | 0 | 0.005 | 0.005 | 0 | 0.16 | 0.2 | 0.33 | 0.19 | 0.18 | 0.14 |
| 4 | 162002577 | rs1523558 | -0.959 | 0.127 | -0.719 | 0.21 | 0.015 | -0.007 | -0.007 | 0 | 0.65 | 0.78 | 0.81 | 0.65 | 0.7 | 0.71 |
| 4 | 162002921 | rs17375159 | 1.895 | 0.084 | -0.807 | 0.218 | -0.004 | 0.045 | 0.045 | 0.096 | 0.24 | 0.1 | 0.12 | 0.14 | 0.22 | 0.09 |
| 4 | 162003624 | rs4591550 | -1.588 | 0.224 | -0.8 | 0.385 | 0.005 | -0.011 | -0.011 | 0 | 0.89 | 0.89 | 0.94 | 0.79 | 0.92 | 0.8 |
| 4 | 162009525 | rs12644817 | 1.24 | 0.441 | -0.889 | 0.262 | -0.044 | -0.077 | -0.077 | 7.069 | 0.15 | 0.03 | 0.25 | 0.34 | 0.17 | 0.26 |
| 4 | 162024164 | rs12645681 | 0.406 | 0.341 | -0.691 | 0.167 | -0.007 | 0.003 | 0.003 | 0.652 | 0.46 | 0.22 | 0.39 | 0.49 | 0.46 | 0.38 |
| 4 | 162027911 | rs10011336 | 0.723 | 0.388 | -0.53 | 0.234 | 0 | -0.005 | -0.005 | 0 | 0.24 | 0.22 | 0.28 | 0.35 | 0.24 | 0.29 |
| 4 | 162028591 | rs10014253 | 0.723 | 0.388 | -0.53 | 0.234 | 0 | -0.005 | -0.005 | 0.073 | 0.24 | 0.22 | 0.28 | 0.35 | 0.24 | 0.29 |
| 4 | 162029797 | rs2313008 | -0.035 | 0.429 | -0.198 | 0.141 | 0.007 | -0.002 | -0.002 | 0 | 0.38 | 0.52 | 0.38 | 0.41 | 0.42 | 0.39 |
| 4 | 162059629 | rs6813133 | -1.033 | 0.355 | -0.325 | -0.013 | 0.011 | -0.027 | -0.027 | 2.186 | 0.89 | 0.91 | 0.71 | 0.62 | 0.92 | 0.79 |
| 4 | 162065899 | rs13132807 | -0.114 | 0.442 | 0.056 | -0.126 | 0 | 0.062 | 0.062 | 0.056 | 0.82 | 0.72 | 0.64 | 0.62 | 0.84 | 0.73 |
| 4 | 162076088 | rs10517736 | 1.433 | 0.544 | 0.245 | -0.224 | 0.024 | -0.026 | -0.026 | 0.093 | 0.12 | 0.21 | 0.1 | 0.06 | 0.18 | 0.1 |
| 4 | 162080425 | rs10212879 | 1.168 | 0.133 | 0.682 | 0.115 | -0.001 | 0.005 | 0.005 | 0 | 0.19 | 0.13 | 0.15 | 0.07 | 0.18 | 0.25 |
| 4 | 162082401 | rs17040713 | 0.831 | 0.15 | 0.732 | 0.084 | 0.016 | -0.034 | -0.034 | 0 | 0.09 | 0.14 | 0.06 | 0.01 | 0.1 | 0.09 |
| 4 | 162088272 | rs17378274 | -0.137 | 0.06 | 0.511 | 0.216 | 0 | 0.036 | 0.036 | 0 | 0.44 | 0.29 | 0.65 | 0.73 | 0.43 | 0.55 |
| 4 | 162105557 | rs6536577 | 0.819 | -0.237 | 0.303 | 0.235 | 0 | -0.029 | -0.029 | 0 | 0.39 | 0.54 | 0.47 | 0.23 | 0.41 | 0.19 |
| 4 | 162109687 | rs17040749 | 0.545 | -0.258 | 0.2 | 0.253 | 0.009 | -0.039 | -0.039 | 0 | 0.13 | 0.2 | 0.09 | 0.03 | 0.12 | 0.14 |
| 4 | 162115247 | rs13148588 | 1.055 | -0.127 | -0.533 | 0.372 | 0 | 0.008 | 0.008 | 0 | 0.2 | 0.15 | 0.15 | 0.07 | 0.18 | 0.25 |
| 4 | 162129844 | rs13110266 | 0.327 | -0.127 | -0.243 | 0.321 | -0.02 | -0.047 | -0.047 | 0.195 | 0.39 | 0.18 | 0.41 | 0.52 | 0.41 | 0.49 |
| 4 | 162147112 | rs719880 | -0.095 | 0.749 | -1.037 | -0.318 | -0.014 | 0.067 | 0.067 | 0 | 0.35 | 0.23 | 0.22 | 0.06 | 0.4 | 0.18 |
| 4 | 162153936 | rs992955 | -0.134 | 0.737 | -0.859 | 0.704 | -0.003 | -0.023 | -0.023 | 0 | 0.72 | 0.74 | 0.56 | 0.42 | 0.7 | 0.55 |
| 4 | 162155361 | rs6831434 | 0.669 | 0.691 | -0.114 | 0.468 | -0.001 | -0.008 | -0.008 | 0 | 0.54 | 0.51 | 0.7 | 0.71 | 0.57 | 0.6 |
| 4 | 162158370 | rs10517739 | -1.134 | 0.541 | -0.007 | 0.513 | -0.01 | 0.004 | 0.004 | 0 | 0.3 | 0.43 | 0.29 | 0.31 | 0.23 | 0.17 |
| 4 | 162158922 | rs17314203 | -0.124 | 0.634 | 0.041 | 0.554 | -0.018 | 0.027 | 0.027 | 0.234 | 0.31 | 0.11 | 0.33 | 0.28 | 0.35 | 0.28 |
| 4 | 162163105 | rs1444285 | -0.873 | 0.765 | -0.01 | 0.664 | 0.018 | 0.062 | 0.062 | 0.668 | 0.17 | 0.36 | 0.22 | 0.29 | 0.15 | 0.12 |
| 4 | 162168945 | rs10026817 | -0.258 | 0.808 | -0.164 | 0.457 | 0.004 | -0.029 | -0.029 | 0 | 0.13 | 0.13 | 0.06 | 0.02 | 0.15 | 0.11 |
| 4 | 162192797 | rs958247 | 0.195 | 0.606 | -0.208 | 0.511 | -0.004 | 0.02 | 0.02 | 0.278 | 0.28 | 0.07 | 0.3 | 0.26 | 0.27 | 0.21 |
| 4 | 162193568 | rs1348530 | -1.583 | 0.548 | 0.03 | 0.538 | -0.004 | 0.004 | 0.004 | 0.131 | 0.55 | 0.69 | 0.61 | 0.58 | 0.53 | 0.47 |
| 4 | 162194539 | rs7665845 | -1.422 | 0.556 | -0.122 | 0.541 | 0.003 | 0.007 | 0.007 | 0 | 0.49 | 0.44 | 0.57 | 0.57 | 0.51 | 0.4 |
| 4 | 162202272 | rs10517742 | -0.814 | 0.61 | -0.196 | 0.507 | 0.019 | 0.03 | 0.03 | 0 | 0.22 | 0.35 | 0.27 | 0.31 | 0.24 | 0.19 |
| 4 | 162205711 | rs10029508 | -1.434 | 0.718 | -0.801 | 0.43 | -0.007 | 0.002 | 0.002 | 0.228 | 0.55 | 0.77 | 0.61 | 0.58 | 0.53 | 0.47 |
| 4 | 162207646 | rs9308024 | -0.325 | 0.196 | -1.214 | 0.025 | -0.02 | -0.041 | -0.041 | 0 | 0.08 | 0.12 | 0.04 | 0.01 | 0.03 | 0.03 |
| 4 | 162207662 | rs13102687 | -0.531 | 0.667 | -1.614 | 0.397 | -0.009 | 0.015 | 0.015 | 0.997 | 0.27 | 0.04 | 0.3 | 0.26 | 0.27 | 0.21 |
| 4 | 162219569 | rs7667597 | -0.557 | 0.843 | -0.813 | 0.502 | -0.003 | -0.014 | -0.014 | 0 | 0.1 | 0.21 | 0.08 | 0.06 | 0.08 | 0.02 |

| | | | | | | | | | | | | | | | | |
|---|---|---|---|---|---|---|---|---|---|---|---|---|---|---|---|---|
| 4 | 162221987 | rs13106146 | -1.218 | 1.061 | -0.88 | 0.57 | 0 | 0.025 | 0.025 | 0 | 0.48 | 0.73 | 0.55 | 0.6 | 0.46 | 0.43 |
| 4 | 162222374 | rs13107435 | -0.996 | 1.102 | -0.391 | 0.491 | 0.009 | -0.057 | -0.057 | 1.281 | 0.22 | 0.5 | 0.19 | 0.13 | 0.21 | 0.19 |
| 4 | 162227134 | rs7679790 | -1.259 | 0.848 | -0.543 | 0.784 | -0.003 | 0.01 | 0.01 | 0.516 | 0.26 | 0.73 | 0.21 | 0.35 | 0.22 | 0.19 |
| 4 | 162227234 | rs4691748 | -1.838 | 0.974 | -0.682 | 0.676 | -0.001 | 0.014 | 0.014 | 0 | 0.45 | 0.74 | 0.39 | 0.36 | 0.43 | 0.32 |
| 4 | 162228270 | rs7687260 | 0.289 | 1.051 | -0.941 | 0.606 | -0.06 | 0.112 | 0.112 | 0 | 0.17 | 0.01 | 0.15 | 0.01 | 0.2 | 0.12 |
| 4 | 162234554 | rs11730185 | -0.124 | 1.119 | -0.673 | 0.993 | -0.003 | -0.001 | -0.001 | 0 | 0.08 | 0.05 | 0.1 | 0.1 | 0.11 | 0.15 |
| 4 | 162235785 | rs5010472 | -1.519 | 1.548 | -0.793 | 0.837 | 0 | 0.016 | 0.016 | 0 | 0.52 | 0.6 | 0.6 | 0.73 | 0.51 | 0.47 |
| 4 | 162235993 | rs5010468 | -1.363 | 1.342 | -0.413 | 1.346 | -0.022 | 0.115 | 0.115 | 0.022 | 0.24 | 0.52 | 0.25 | 0.46 | 0.2 | 0.2 |
| 4 | 162236018 | rs34018667 | -0.103 | 1.446 | -0.782 | 0.908 | -0.001 | -0.001 | -0.001 | 0 | 0.07 | 0.05 | 0.09 | 0.1 | 0.1 | 0.14 |
| 4 | 162236867 | rs1530425 | -0.799 | 1.344 | -0.855 | 0.439 | 0.006 | -0.021 | -0.021 | 0.746 | 0.43 | 0.51 | 0.54 | 0.79 | 0.48 | 0.4 |
| 4 | 162241086 | rs1017543 | -0.193 | 0.933 | -1.014 | 0.585 | 0.006 | 0.005 | 0.005 | 0 | 0.45 | 0.38 | 0.43 | 0.37 | 0.45 | 0.4 |
| 4 | 162243468 | rs2313748 | -0.212 | 0.788 | -0.75 | 0.775 | 0.008 | 0 | 0 | 0 | 0.5 | 0.44 | 0.7 | 0.43 | 0.59 | 0.59 |
| 4 | 162248546 | rs12331118 | -0.407 | 1.171 | -0.404 | 0.997 | 0.01 | 0.015 | 0.015 | 0 | 0.26 | 0.31 | 0.21 | 0.32 | 0.31 | 0.22 |
| 4 | 162249966 | rs2313565 | 1.545 | 1.174 | -0.429 | 0.98 | -0.008 | -0.003 | -0.003 | 0.033 | 0.12 | 0.1 | 0.15 | 0.09 | 0.19 | 0.06 |
| 4 | 162250538 | rs10003505 | -1.51 | 1.255 | -0.055 | 1.06 | 0 | 0 | 0 | 0 | 0.63 | 0.62 | 0.68 | 0.61 | 0.66 | 0.63 |
| 4 | 162252857 | rs11100365 | -0.079 | 1.037 | 0.501 | 1.215 | -0.005 | 0.004 | 0.004 | 0 | 0.22 | 0.2 | 0.19 | 0.13 | 0.27 | 0.2 |
| 4 | 162255770 | rs12512282 | 1.011 | 0.818 | 0.715 | 1.303 | 0.033 | 0.211 | 0.211 | 0 | 0.91 | 0.67 | 0.87 | 0.72 | 0.92 | 0.83 |
| 4 | 162257301 | rs4691007 | 1.552 | 0.347 | 0.513 | 1.285 | -0.021 | 0.009 | 0.009 | 0 | 0.07 | 0.07 | 0.14 | 0.11 | 0.03 | 0.13 |
| 4 | 162260229 | rs6857927 | -0.164 | 0.19 | 0.139 | 0.921 | -0.031 | -0.017 | -0.017 | 0.303 | 0.7 | 0.78 | 0.6 | 0.68 | 0.52 | 0.62 |
| 4 | 162260938 | rs4446272 | -0.06 | -0.104 | 0.177 | 0.636 | -0.018 | -0.008 | -0.008 | 0 | 0.7 | 0.66 | 0.73 | 0.8 | 0.89 | 0.72 |
| 4 | 162261170 | rs4327441 | 2.44 | -0.234 | 0.126 | 0.599 | -0.031 | -0.002 | -0.002 | 0 | 0.3 | 0.06 | 0.21 | 0.29 | 0.28 | 0.2 |
| 4 | 162262333 | rs7438721 | -0.328 | -0.159 | 0.397 | 0.589 | 0.011 | 0.012 | 0.012 | 0 | 0.37 | 0.39 | 0.43 | 0.46 | 0.56 | 0.4 |
| 4 | 162265796 | rs2314137 | -1.458 | -0.074 | 0.208 | 0.735 | 0.032 | -0.035 | -0.035 | 0 | 0.61 | 0.84 | 0.65 | 0.54 | 0.69 | 0.63 |
| 4 | 162268088 | rs17040932 | -0.18 | -0.197 | 0.276 | 0.681 | -0.026 | 0.006 | 0.006 | 0 | 0.11 | 0.27 | 0.17 | 0.11 | 0.09 | 0.14 |
| 4 | 162272259 | rs10007083 | 1.465 | -0.062 | 0.161 | 0.797 | -0.008 | -0.008 | -0.008 | 0 | 0.36 | 0.27 | 0.27 | 0.44 | 0.36 | 0.23 |
| 4 | 162273772 | rs10010638 | -0.069 | -0.084 | 0.211 | 0.859 | 0.051 | 0.148 | 0.148 | 0 | 0.05 | 0.24 | 0.04 | 0.15 | 0.05 | 0.03 |
| 4 | 162274024 | rs6832320 | 2.027 | -0.075 | -0.218 | 0.81 | -0.056 | 0.003 | 0.003 | 0 | 0.31 | 0.07 | 0.23 | 0.29 | 0.31 | 0.2 |
| 4 | 162276769 | rs17336367 | -0.229 | -0.068 | -0.424 | 0.088 | 0.023 | 0.017 | 0.017 | 0 | 0.26 | 0.16 | 0.23 | 0.22 | 0.22 | 0.39 |
| 4 | 162281464 | rs16999316 | 1.743 | -0.07 | -0.424 | 0.303 | 0.033 | -0.031 | -0.031 | 0.365 | 0.06 | 0.01 | 0.13 | 0.22 | 0.04 | 0.17 |
| 4 | 162281985 | rs2314141 | -1.272 | 0.043 | -0.207 | 0.04 | -0.001 | 0.042 | 0.042 | 0 | 0.49 | 0.75 | 0.5 | 0.4 | 0.51 | 0.59 |
| 4 | 162284626 | rs12643813 | -0.081 | 0.073 | -0.957 | 0.012 | -0.007 | 0.002 | 0.002 | 0 | 0.35 | 0.23 | 0.36 | 0.31 | 0.37 | 0.41 |
| 4 | 162285736 | rs17336722 | 1.751 | -0.4 | -0.846 | 0.069 | -0.041 | 0.048 | 0.048 | 0 | 0.09 | 0.01 | 0.08 | 0 | 0.13 | 0.03 |
| 4 | 162288572 | rs13142107 | 0.141 | -0.435 | -0.479 | 0.163 | 0.008 | -0.036 | -0.036 | 0 | 0.09 | 0.13 | 0.03 | 0.03 | 0.11 | 0.08 |
| 4 | 162301366 | rs17040974 | 1.107 | -0.618 | -0.822 | 0.368 | 0.04 | -0.04 | -0.04 | 0 | 0.07 | 0.01 | 0.13 | 0.17 | 0.03 | 0.13 |
| 4 | 162303020 | rs2313567 | 0.25 | -0.658 | -0.348 | 0.23 | -0.004 | 0.003 | 0.003 | 0 | 0.78 | 0.64 | 0.67 | 0.78 | 0.79 | 0.73 |
| 4 | 162303613 | rs6823286 | 1.858 | -0.626 | -1.336 | 0.099 | 0.049 | -0.028 | -0.028 | 0 | 0.34 | 0.15 | 0.17 | 0.38 | 0.22 | 0.18 |
| 4 | 162307000 | rs17040982 | 1.894 | -0.528 | -1.294 | 0.14 | 0.028 | -0.043 | -0.043 | 0 | 0.07 | 0 | 0.16 | 0.18 | 0.03 | 0.15 |
| 4 | 162307060 | rs17040984 | 1.903 | -0.546 | -1.368 | 0.07 | 0.037 | -0.053 | -0.053 | 0.281 | 0.33 | 0.04 | 0.17 | 0.32 | 0.23 | 0.2 |
| 4 | 162311913 | rs12374210 | 1.376 | 0.089 | -0.837 | 0.253 | -0.002 | 0.001 | 0.001 | 0 | 0.1 | 0.08 | 0.09 | 0.05 | 0.14 | 0.07 |
| 4 | 162320903 | rs11725954 | 1.459 | -0.597 | -1.365 | -0.165 | -0.011 | 0.126 | 0.126 | 0 | 0.25 | 0.07 | 0.14 | 0.05 | 0.19 | 0.22 |
| 4 | 162324395 | rs6820464 | 0.54 | -0.223 | -1.154 | -0.112 | 0.005 | -0.01 | -0.01 | 0.26 | 0.09 | 0.08 | 0.13 | 0.07 | 0.13 | 0.1 |
| 4 | 162324909 | rs12500921 | 1.846 | -0.089 | -1.335 | 0.009 | -0.031 | -0.036 | -0.036 | 0 | 0.06 | 0.01 | 0.12 | 0.13 | 0.12 | 0.22 |
| 4 | 162327265 | rs4547755 | -0.616 | 0.064 | -0.759 | -0.273 | 0.012 | -0.03 | -0.03 | 0.167 | 0.6 | 0.55 | 0.69 | 0.78 | 0.56 | 0.72 |
| 4 | 162328955 | rs12639796 | 1.949 | -0.041 | -1.015 | -0.074 | 0.031 | -0.047 | -0.047 | 0 | 0.08 | 0 | 0.17 | 0.19 | 0.04 | 0.15 |
| 4 | 162329244 | rs17338054 | 1.753 | -0.058 | -1.001 | -0.106 | -0.002 | 0.001 | 0.001 | 0 | 0.11 | 0.09 | 0.09 | 0.05 | 0.13 | 0.07 |
| 4 | 162334309 | rs2872530 | 1.188 | -0.02 | -1.228 | -0.106 | 0.024 | -0.039 | -0.039 | 0.037 | 0.31 | 0.23 | 0.29 | 0.47 | 0.26 | 0.41 |
| 4 | 162350385 | rs4518193 | 0.589 | 0.064 | -1.533 | -0.256 | -0.002 | -0.041 | -0.041 | 0 | 0.19 | 0.04 | 0.26 | 0.24 | 0.18 | 0.23 |
| 4 | 162355688 | rs7694864 | 0.54 | 0.37 | -1.3 | -0.31 | -0.011 | 0.062 | 0.062 | 0 | 0.3 | 0.16 | 0.25 | 0.12 | 0.27 | 0.29 |
| 4 | 162361217 | rs7669737 | 0.618 | 0.348 | -1.419 | -0.14 | -0.015 | 0.019 | 0.019 | 0 | 0.4 | 0.23 | 0.41 | 0.6 | 0.47 | 0.41 |
| 4 | 162362473 | rs4691761 | 0.805 | 0.174 | -1.592 | -0.536 | -0.028 | 0.064 | 0.064 | 0 | 0.55 | 0.12 | 0.55 | 0.67 | 0.43 | 0.66 |
| 4 | 162367270 | rs11945487 | 0.634 | 0.508 | -1.427 | -0.388 | -0.065 | -0.043 | -0.043 | 0 | 0.22 | 0.12 | 0.22 | 0.31 | 0.37 | 0.24 |
| 4 | 162370089 | rs7681811 | -0.245 | 0.595 | -0.691 | -0.469 | 0.055 | -0.001 | -0.001 | 0.246 | 0.91 | 0.69 | 0.89 | 0.95 | 0.91 | 0.89 |
| 4 | 162376761 | rs12650551 | 1.033 | 0.289 | -1.547 | -0.598 | -0.019 | 0.067 | 0.067 | 0 | 0.23 | 0.05 | 0.12 | 0.15 | 0.18 | 0.3 |
| 4 | 162379746 | rs16999322 | 1.315 | 0.145 | -1.547 | -0.476 | 0.009 | -0.022 | -0.022 | 0 | 0.11 | 0.06 | 0.12 | 0.14 | 0.11 | 0.2 |
| 4 | 162380016 | rs17041092 | 1.035 | -0.033 | -1.814 | -0.517 | 0.045 | -0.103 | -0.103 | 0.46 | 0.33 | 0.09 | 0.42 | 0.49 | 0.25 | 0.35 |
| 4 | 162382240 | rs6536595 | -1.659 | -0.041 | -1.66 | -0.555 | -0.008 | 0.023 | 0.023 | 0 | 0.77 | 0.83 | 0.84 | 0.95 | 0.78 | 0.8 |
| 4 | 162383466 | rs17041096 | 0.806 | 0.006 | -1.387 | -0.256 | 0.05 | -0.099 | -0.099 | 0 | 0.33 | 0.09 | 0.42 | 0.49 | 0.25 | 0.35 |
| 4 | 162384818 | rs4358350 | 1.534 | 0.061 | -1.351 | -0.265 | -0.012 | -0.043 | -0.043 | 0 | 0.11 | 0 | 0.12 | 0.14 | 0.11 | 0.19 |
| 4 | 162389854 | rs11937911 | 0.7 | 0.036 | -0.714 | -0.33 | 0.045 | 0.03 | 0.03 | 0 | 0.27 | 0.43 | 0.31 | 0.34 | 0.38 | 0.24 |
| 4 | 162390574 | rs6839543 | -0.377 | 0.764 | -0.469 | -0.486 | -0.024 | 0.109 | 0.109 | 0 | 0.63 | 0.82 | 0.76 | 0.8 | 0.64 | 0.64 |
| 4 | 162396765 | rs6832611 | -0.044 | 0.297 | -0.502 | -0.22 | -0.004 | 0.048 | 0.048 | 0 | 0.55 | 0.63 | 0.65 | 0.76 | 0.55 | 0.48 |
| 4 | 162406039 | rs6858725 | 0.064 | 0.335 | -0.833 | -0.167 | 0 | -0.018 | -0.018 | 0.486 | 0.54 | 0.51 | 0.64 | 0.75 | 0.55 | 0.48 |
| 4 | 162415916 | rs6842030 | 0.242 | 0.282 | -1.364 | 0.177 | -0.013 | -0.016 | -0.016 | 0.036 | 0.51 | 0.22 | 0.55 | 0.46 | 0.47 | 0.63 |
| 4 | 162419531 | rs11100371 | 0.683 | 0.033 | -0.624 | 0.208 | -0.033 | 0.025 | 0.025 | 0 | 0.17 | 0.34 | 0.18 | 0.17 | 0.12 | 0.19 |
| 4 | 162424868 | rs17339916 | 0.604 | 0.028 | -0.76 | 0.124 | -0.052 | 0.026 | 0.026 | 0 | 0.24 | 0.13 | 0.24 | 0.21 | 0.38 | 0.08 |
| 4 | 162426149 | rs17399436 | 2.005 | 0.192 | -0.677 | 0.041 | -0.077 | -0.048 | -0.048 | 0.009 | 0.12 | 0.01 | 0.14 | 0.21 | 0.26 | 0.05 |
| 4 | 162426629 | rs4518198 | 0.997 | -0.525 | -1.072 | -0.174 | -0.094 | 0.047 | 0.047 | 0 | 0.29 | 0.06 | 0.25 | 0.25 | 0.33 | 0.45 |
| 4 | 162428196 | rs17340154 | 0.774 | -0.019 | -0.863 | 0.004 | -0.043 | 0.026 | 0.026 | 0 | 0.24 | 0.13 | 0.23 | 0.21 | 0.37 | 0.08 |
| 4 | 162439632 | rs4434196 | 1.042 | -0.237 | -0.838 | 0.006 | 0.012 | -0.015 | -0.015 | 0.113 | 0.26 | 0.18 | 0.31 | 0.38 | 0.17 | 0.28 |

| | | | | | | | | | | | | | | | | |
|---|---|---|---|---|---|---|---|---|---|---|---|---|---|---|---|---|
| 4 | 162447206 | rs13104132 | 1.07 | 0.14 | -1.056 | -0.196 | -0.105 | 0.094 | 0.094 | 0 | 0.21 | 0.02 | 0.22 | 0.04 | 0.34 | 0.07 |
| 4 | 162458688 | rs4355345 | 0.422 | -0.045 | -0.011 | -0.318 | 0.032 | -0.106 | -0.106 | 2.292 | 0.27 | 0.39 | 0.24 | 0.04 | 0.34 | 0.09 |
| 4 | 162460563 | rs16999330 | 0.702 | -0.442 | -0.619 | -0.058 | 0.033 | -0.06 | -0.06 | 0 | 0.27 | 0.12 | 0.33 | 0.56 | 0.21 | 0.34 |
| 4 | 162461996 | rs12512822 | 1.05 | -0.731 | -0.797 | -0.217 | 0.026 | -0.042 | -0.042 | 0 | 0.15 | 0.08 | 0.15 | 0.31 | 0.08 | 0.22 |
| 4 | 162473868 | rs13141240 | -1.383 | -0.743 | -0.523 | -0.224 | -0.007 | 0.014 | 0.014 | 0 | 0.85 | 0.86 | 0.85 | 0.69 | 0.92 | 0.77 |
| 4 | 162477082 | rs4501178 | 1.476 | -0.781 | -0.783 | -0.418 | 0.018 | -0.041 | -0.041 | 0 | 0.22 | 0.12 | 0.32 | 0.4 | 0.18 | 0.26 |
| 4 | 162482595 | rs9996901 | 0.117 | -0.587 | -0.616 | -0.298 | 0.03 | -0.076 | -0.076 | 1.646 | 0.18 | 0.07 | 0.16 | 0.48 | 0.12 | 0.32 |
| 4 | 162489958 | rs4330312 | 0.024 | -0.23 | -0.136 | -0.173 | -0.001 | -0.002 | -0.002 | 0 | 0.49 | 0.37 | 0.55 | 0.44 | 0.53 | 0.35 |
| 4 | 162501251 | rs11100375 | 0.132 | -0.595 | -0.375 | -0.286 | 0.011 | -0.002 | -0.002 | 0 | 0.51 | 0.39 | 0.57 | 0.45 | 0.62 | 0.38 |
| 4 | 162505383 | rs10003025 | -0.122 | -0.657 | -0.221 | -0.468 | 0.025 | -0.02 | -0.02 | 0 | 0.67 | 0.41 | 0.57 | 0.78 | 0.47 | 0.73 |
| 4 | 162511370 | rs6858225 | 0.493 | -1.094 | -0.498 | -0.631 | -0.001 | 0.003 | 0.003 | 0.275 | 0.57 | 0.37 | 0.57 | 0.73 | 0.47 | 0.54 |
| 4 | 162514902 | rs10003547 | -0.668 | -1.383 | -0.854 | -0.556 | 0.048 | 0.027 | 0.027 | 0 | 0.5 | 0.61 | 0.48 | 0.58 | 0.31 | 0.57 |
| 4 | 162516553 | rs4355346 | -1.29 | -1.045 | -0.917 | -0.567 | -0.005 | 0.008 | 0.008 | 0.127 | 0.61 | 0.65 | 0.72 | 0.76 | 0.55 | 0.76 |
| 4 | 162521710 | rs10517746 | 1.904 | -1.013 | -0.948 | -0.544 | -0.054 | -0.031 | -0.031 | 0 | 0.11 | 0.04 | 0.25 | 0.18 | 0.24 | 0.19 |
| 4 | 162522247 | rs13149906 | 1.902 | -0.951 | -0.915 | -0.532 | 0.011 | 0.019 | 0.019 | 1.28 | 0.23 | 0.15 | 0.15 | 0.18 | 0.2 | 0.17 |
| 4 | 162525511 | rs10857338 | -0.751 | -0.475 | -1.471 | -0.088 | -0.021 | 0 | 0 | 0.76 | 0.71 | 0.79 | 0.76 | 0.75 | 0.63 | 0.72 |
| 4 | 162530770 | rs11100377 | 0.01 | -0.687 | -1.457 | -0.082 | 0.015 | -0.052 | -0.052 | 0 | 0.34 | 0.14 | 0.42 | 0.55 | 0.29 | 0.37 |
| 4 | 162548816 | rs17343587 | 0.62 | -0.284 | -1.098 | -0.123 | 0.007 | 0 | 0 | 0.06 | 0.15 | 0.18 | 0.27 | 0.21 | 0.26 | 0.24 |
| 4 | 162552883 | rs17403181 | 0.043 | -0.557 | -1.252 | -0.315 | 0.003 | 0.019 | 0.019 | 0 | 0.5 | 0.56 | 0.44 | 0.37 | 0.47 | 0.55 |
| 4 | 162556751 | rs17343847 | 1.443 | -0.051 | -1.737 | -0.044 | -0.045 | -0.023 | -0.023 | 0.13 | 0.11 | 0.05 | 0.25 | 0.18 | 0.26 | 0.18 |
| 4 | 162577430 | rs17638339 | 0.435 | -0.774 | -1.668 | 0.16 | 0.008 | -0.059 | -0.059 | 2.906 | 0.21 | 0.08 | 0.3 | 0.33 | 0.16 | 0.15 |
| 4 | 162595918 | rs9308035 | -1.554 | -0.343 | -1.401 | 0.144 | 0.01 | -0.012 | -0.012 | 0 | 0.85 | 0.81 | 0.87 | 0.95 | 0.76 | 0.93 |
| 4 | 162600018 | rs4532194 | 0.151 | -1.992 | -1.584 | 0.324 | 0.012 | -0.016 | -0.016 | 0 | 0.41 | 0.33 | 0.45 | 0.5 | 0.35 | 0.29 |
| 4 | 162602209 | rs17638464 | 0.388 | -1.773 | -1.96 | 0.113 | 0.024 | -0.027 | -0.027 | 0 | 0.12 | 0.07 | 0.07 | 0.18 | 0.05 | 0.18 |
| 4 | 162612565 | rs11100382 | 0.366 | -1.649 | -2.201 | 0.047 | 0.035 | -0.009 | -0.009 | 0 | 0.43 | 0.09 | 0.64 | 0.67 | 0.67 | 0.35 |
| 4 | 162633257 | rs7442468 | -0.123 | -1.55 | -1.294 | -0.623 | 0 | -0.019 | -0.019 | 0 | 0.37 | 0.58 | 0.25 | 0.13 | 0.27 | 0.28 |
| 4 | 162637121 | rs12500089 | 0.473 | -2.507 | -1.864 | -0.363 | 0.128 | 0.048 | 0.048 | 0.332 | 0.45 | 0.1 | 0.66 | 0.67 | 0.69 | 0.47 |
| 4 | 162644069 | rs10013537 | 0.54 | -2.881 | -1.468 | -0.314 | 0.117 | 0.041 | 0.041 | 0 | 0.47 | 0.23 | 0.66 | 0.64 | 0.69 | 0.55 |
| 4 | 162646894 | rs1982121 | 0.39 | -3.036 | -1.523 | -0.39 | 0.164 | 0.06 | 0.06 | 0 | 0.49 | 0.18 | 0.66 | 0.64 | 0.7 | 0.58 |
| 4 | 162647126 | rs6820280 | 0.695 | -3.398 | -0.998 | -0.398 | -0.006 | 0.01 | 0.01 | 0 | 0.13 | 0.13 | 0.12 | 0.23 | 0.13 | 0.13 |
| 4 | 162650185 | rs13119737 | 0.759 | -3.343 | -0.986 | -0.503 | -0.007 | 0.005 | 0.005 | 0 | 0.13 | 0.14 | 0.12 | 0.18 | 0.13 | 0.13 |
| 4 | 162650988 | rs10517750 | -0.398 | -3.483 | -0.623 | -0.479 | -0.031 | -0.012 | -0.012 | 0 | 0.06 | 0.1 | 0.07 | 0.03 | 0.01 | 0.19 |
| 4 | 162651017 | rs17596009 | 0.732 | -3.052 | -0.601 | -0.433 | 0 | 0.004 | 0.004 | 0 | 0.13 | 0.14 | 0.13 | 0.19 | 0.15 | 0.13 |
| 4 | 162651104 | rs1347353 | -0.096 | -3.291 | -0.659 | -0.577 | -0.065 | -0.046 | -0.046 | 0 | 0.68 | 0.44 | 0.81 | 0.88 | 0.87 | 0.8 |
| 4 | 162657759 | rs986704 | 1.076 | -3.207 | -0.985 | -0.535 | 0.157 | 0.072 | 0.072 | 0 | 0.49 | 0.16 | 0.61 | 0.66 | 0.71 | 0.58 |
| 4 | 162663106 | rs10517752 | 0.248 | -3.432 | -0.648 | -0.712 | -0.028 | -0.019 | -0.019 | 0 | 0.21 | 0.3 | 0.14 | 0.09 | 0.1 | 0.12 |
| 4 | 162666856 | rs1434614 | 1.096 | -3.201 | -1.384 | -0.603 | 0.162 | 0.059 | 0.059 | 0 | 0.48 | 0.1 | 0.6 | 0.65 | 0.7 | 0.58 |
| 4 | 162669309 | rs4691025 | -1.015 | -3.272 | -1.116 | -0.68 | 0.126 | 0.13 | 0.13 | 0 | 0.45 | 0.73 | 0.28 | 0.14 | 0.23 | 0.31 |
| 4 | 162678311 | rs11100385 | -0.701 | -3.425 | -1.032 | -0.752 | -0.022 | -0.011 | -0.011 | 0.318 | 0.28 | 0.66 | 0.13 | 0.15 | 0.1 | 0.17 |
| 4 | 162687669 | rs11942761 | -0.449 | -3.396 | -1.076 | -0.715 | -0.081 | -0.034 | -0.034 | 0.135 | 0.26 | 0.58 | 0.15 | 0.15 | 0.07 | 0.16 |
| 4 | 162709000 | rs7690085 | 0.831 | -3.63 | -0.899 | -0.681 | 0.082 | 0.032 | 0.032 | 0.149 | 0.58 | 0.25 | 0.6 | 0.65 | 0.72 | 0.67 |
| 4 | 162710868 | rs7668203 | 1.585 | -2.805 | -1.563 | -1.051 | -0.105 | -0.13 | -0.13 | 0 | 0.23 | 0.05 | 0.29 | 0.56 | 0.35 | 0.36 |
| 4 | 162714672 | rs1560926 | 0.384 | -3.018 | -0.918 | -0.652 | -0.084 | -0.007 | -0.007 | 0 | 0.16 | 0.4 | 0.12 | 0.13 | 0.04 | 0.08 |
| 4 | 162718372 | rs13130537 | 0.029 | -2.057 | -0.97 | -0.557 | -0.016 | -0.006 | -0.006 | 1.928 | 0.74 | 0.68 | 0.72 | 0.76 | 0.76 | 0.76 |
| 4 | 162719690 | rs17041550 | 0.185 | -1.13 | -0.577 | -0.02 | -0.032 | -0.024 | -0.024 | 0.021 | 0.1 | 0.18 | 0.04 | 0.04 | 0.03 | 0.08 |
| 4 | 162733762 | rs2082669 | 0.275 | -1.041 | -0.703 | 0.108 | -0.027 | 0.006 | 0.006 | 0 | 0.77 | 0.67 | 0.76 | 0.74 | 0.83 | 0.76 |
| 4 | 162733928 | rs2014158 | 0.167 | -0.521 | -1.047 | 0.573 | 0.004 | 0.001 | 0.001 | 0 | 0.39 | 0.43 | 0.35 | 0.44 | 0.45 | 0.46 |
| 4 | 162734547 | rs12504254 | -0.86 | -0.482 | -1.491 | 0.794 | 0.039 | -0.03 | -0.03 | 0 | 0.18 | 0.02 | 0.25 | 0.19 | 0.11 | 0.17 |
| 4 | 162734835 | rs359147 | -1.123 | 0.808 | -1.347 | 1.268 | -0.019 | -0.021 | -0.021 | 0 | 0.18 | 0.14 | 0.42 | 0.37 | 0.29 | 0.25 |
| 4 | 162735000 | rs359144 | -0.822 | 1.068 | -1.109 | 1.671 | 0.015 | 0.016 | 0.016 | 0 | 0.12 | 0.15 | 0.16 | 0.27 | 0.24 | 0.27 |
| 4 | 162737097 | rs359142 | -0.989 | 1.26 | -0.866 | 1.897 | 0.022 | 0.053 | 0.053 | 0.095 | 0.11 | 0.19 | 0.15 | 0.29 | 0.2 | 0.26 |
| 4 | 162738654 | rs1946656 | 1.496 | 0.904 | -0.59 | 1.185 | -0.001 | -0.003 | -0.003 | 0 | 0.48 | 0.41 | 0.51 | 0.53 | 0.47 | 0.42 |
| 4 | 162741975 | rs17596748 | -0.167 | 0.616 | -0.719 | 0.909 | 0.019 | 0.042 | 0.042 | 0.674 | 0.2 | 0.05 | 0.11 | 0.14 | 0.14 | 0.17 |
| 4 | 162763646 | rs9308039 | 0.901 | 0.396 | -0.361 | 0.727 | -0.005 | -0.02 | -0.02 | 0 | 0.7 | 0.62 | 0.83 | 0.84 | 0.77 | 0.72 |
| 4 | 162764014 | rs1440588 | -0.603 | 0.415 | -0.25 | 0.782 | -0.002 | 0.001 | 0.001 | 0 | 0.24 | 0.19 | 0.29 | 0.17 | 0.29 | 0.24 |
| 4 | 162770250 | rs11938253 | 2.25 | 0.447 | -0.194 | 0.844 | -0.003 | -0.005 | -0.005 | 0.466 | 0.45 | 0.39 | 0.5 | 0.53 | 0.45 | 0.37 |
| 4 | 162782993 | rs16999366 | -1.064 | 0.347 | -0.212 | 0.816 | -0.008 | -0.017 | -0.017 | 0 | 0.25 | 0.3 | 0.14 | 0.15 | 0.2 | 0.22 |
| 4 | 162783324 | rs732224 | -2.364 | 0.25 | -0.225 | 0.852 | 0 | -0.004 | -0.004 | 0 | 0.52 | 0.45 | 0.44 | 0.31 | 0.51 | 0.48 |
| 4 | 162785412 | rs6831077 | -1.922 | 0.71 | 0.072 | 1.176 | -0.005 | -0.011 | -0.011 | 0.023 | 0.38 | 0.27 | 0.44 | 0.46 | 0.45 | 0.55 |
| 4 | 162787495 | rs11732750 | 2.162 | 0.657 | 0.137 | 1.144 | -0.008 | -0.006 | -0.006 | 0.159 | 0.55 | 0.36 | 0.51 | 0.53 | 0.47 | 0.4 |
| 4 | 162801373 | rs4691032 | -1.214 | 0.664 | 0.201 | 1.148 | -0.023 | 0.04 | 0.04 | 0 | 0.29 | 0.09 | 0.3 | 0.17 | 0.31 | 0.32 |
| 4 | 162814045 | rs2053430 | 1.305 | 0.712 | 0.337 | 1.272 | 0.072 | 0.193 | 0.193 | 1.176 | 0.84 | 0.5 | 0.68 | 0.67 | 0.79 | 0.71 |
| 4 | 162816953 | rs17536725 | -0.015 | 0.748 | 0.047 | 1.1 | -0.01 | 0.093 | 0.093 | 0 | 0.17 | 0.02 | 0.09 | 0 | 0.18 | 0.14 |
| 4 | 162817423 | rs1440604 | 2.07 | 0.632 | 0.153 | 1.176 | 0.002 | 0 | 0 | 0 | 0.61 | 0.46 | 0.59 | 0.67 | 0.57 | 0.51 |
| 4 | 162827306 | rs7666181 | 2.398 | 0.935 | 0.552 | 1.483 | 0.017 | 0.018 | 0.018 | 0.239 | 0.67 | 0.42 | 0.55 | 0.52 | 0.53 | 0.47 |
| 4 | 162837458 | rs1348065 | 1.93 | 0.953 | 0.58 | 1.67 | 0.031 | 0.146 | 0.146 | 0.319 | 0.88 | 0.59 | 0.87 | 0.81 | 0.89 | 0.81 |
| 4 | 162840766 | rs1549828 | 1.245 | 0.907 | 0.621 | 1.692 | 0.017 | 0.21 | 0.21 | 0.261 | 0.93 | 0.71 | 0.9 | 0.81 | 0.96 | 0.84 |
| 4 | 162841653 | rs359502 | -0.903 | 0.801 | 0.733 | 1.644 | 0.008 | 0.025 | 0.025 | 0.225 | 0.06 | 0.25 | 0.04 | 0.03 | 0.01 | 0.11 |
| 4 | 162844090 | rs1440610 | 2.265 | 0.881 | 0.588 | 1.646 | 0.034 | 0.055 | 0.055 | 0 | 0.73 | 0.57 | 0.65 | 0.65 | 0.63 | 0.55 |

| | | | | | | | | | | | | | | | | |
|---|---|---|---|---|---|---|---|---|---|---|---|---|---|---|---|---|
| 4 | 162871521 | rs2067907 | -0.165 | 0.645 | -0.061 | 1.463 | -0.117 | 0.043 | 0.043 | 0.396 | 0.28 | 0.08 | 0.34 | 0.16 | 0.44 | 0.3 |
| 4 | 162876772 | rs994654 | -0.803 | 0.58 | -0.009 | 1.904 | -0.092 | -0.053 | -0.053 | 0 | 0.35 | 0.18 | 0.41 | 0.37 | 0.51 | 0.38 |
| 4 | 162881087 | rs359512 | -1.935 | 0.542 | 0.207 | 1.874 | -0.003 | -0.004 | -0.004 | 0 | 0.45 | 0.54 | 0.51 | 0.38 | 0.62 | 0.5 |
| 4 | 162885218 | rs1440587 | -0.56 | 0.55 | 0.225 | 1.944 | -0.075 | -0.024 | -0.024 | 0 | 0.32 | 0.16 | 0.42 | 0.38 | 0.55 | 0.35 |
| 4 | 162891609 | rs1842985 | 1.474 | 0.546 | 0.333 | 1.933 | 0.003 | 0.004 | 0.004 | 0 | 0.55 | 0.43 | 0.49 | 0.6 | 0.37 | 0.5 |
| 4 | 162894103 | rs45426 | 0.147 | 0.544 | 0.307 | 1.905 | -0.059 | -0.057 | -0.057 | 7.618 | 0.7 | 0.83 | 0.63 | 0.63 | 0.56 | 0.65 |
| 4 | 162900278 | rs466478 | -1.294 | 0.557 | 0.319 | 1.787 | -0.009 | -0.035 | -0.035 | 0.261 | 0.12 | 0.3 | 0.09 | 0.03 | 0.08 | 0.15 |
| 4 | 162904449 | rs10028812 | -0.079 | 0.557 | 0.217 | 1.762 | -0.117 | 0.045 | 0.045 | 0.54 | 0.28 | 0.06 | 0.34 | 0.16 | 0.44 | 0.3 |
| 4 | 162904939 | rs1483549 | 2.027 | 0.553 | 0.293 | 1.795 | 0.03 | 0.048 | 0.048 | 0.392 | 0.83 | 0.54 | 0.83 | 0.77 | 0.82 | 0.83 |
| 4 | 162907833 | rs359486 | 0.379 | 0.682 | 0.219 | 1.852 | -0.024 | -0.002 | -0.002 | 0 | 0.67 | 0.72 | 0.57 | 0.63 | 0.45 | 0.65 |
| 4 | 162912081 | rs17041860 | 0.795 | 0.528 | 0.259 | 1.819 | 0.027 | 0.021 | 0.021 | 0 | 0.62 | 0.5 | 0.56 | 0.61 | 0.49 | 0.53 |
| 4 | 162923324 | rs10022486 | 0.252 | 0.604 | -0.102 | 1.592 | -0.027 | 0.029 | 0.029 | 0 | 0.22 | 0.14 | 0.16 | 0.06 | 0.35 | 0.16 |
| 4 | 162925724 | rs1483550 | -0.809 | 0.742 | 0.044 | 2.509 | 0.004 | 0.039 | 0.039 | 0 | 0.12 | 0.16 | 0.3 | 0.44 | 0.21 | 0.25 |
| 4 | 162934834 | rs7658962 | -0.506 | 0.402 | -0.344 | 1.98 | 0.002 | 0.025 | 0.025 | 0.136 | 0.15 | 0.05 | 0.09 | 0.09 | 0.13 | 0.1 |
| 4 | 162944335 | rs1905363 | 0.571 | 0.179 | -0.425 | 1.636 | -0.025 | 0.029 | 0.029 | 0.179 | 0.22 | 0.14 | 0.17 | 0.1 | 0.32 | 0.17 |
| 4 | 162953565 | rs1471693 | 1.322 | 0.016 | -0.198 | 1.663 | 0.017 | 0.04 | 0.04 | 0 | 0.48 | 0.38 | 0.42 | 0.24 | 0.38 | 0.41 |
| 4 | 162967006 | rs11942587 | 1.121 | 0.18 | -0.311 | 1.883 | 0.019 | 0.052 | 0.052 | 0 | 0.52 | 0.58 | 0.44 | 0.19 | 0.4 | 0.4 |
| 4 | 162967558 | rs1483557 | 0.408 | 0.27 | -0.346 | 2.231 | -0.06 | 0.003 | 0.003 | 0.124 | 0.24 | 0.12 | 0.25 | 0.3 | 0.39 | 0.24 |
| 4 | 162985367 | rs1492457 | 1.174 | 0.056 | -0.32 | 2.327 | -0.009 | 0.016 | 0.016 | 0 | 0.06 | 0.05 | 0.13 | 0.12 | 0.03 | 0.12 |
| 4 | 162985402 | rs17459954 | 0.608 | 0.06 | -0.352 | 2.327 | -0.052 | 0.075 | 0.075 | 0.116 | 0.22 | 0.09 | 0.17 | 0.11 | 0.3 | 0.2 |
| 4 | 162991888 | rs2101216 | -1.46 | 0.041 | -0.353 | 2.329 | -0.023 | 0.077 | 0.077 | 0 | 0.44 | 0.25 | 0.4 | 0.32 | 0.46 | 0.43 |
| 4 | 162992246 | rs2086754 | 0.599 | 0.404 | 0.078 | 2.627 | -0.007 | 0.007 | 0.007 | 0 | 0.07 | 0.12 | 0.14 | 0.12 | 0.03 | 0.12 |
| 4 | 162995995 | rs10030663 | 0.318 | 0.431 | 0.055 | 2.54 | -0.039 | 0.061 | 0.061 | 0.029 | 0.23 | 0.11 | 0.17 | 0.11 | 0.3 | 0.21 |
| 4 | 163005050 | rs6814301 | -0.343 | 0.244 | 0.093 | 2.493 | -0.031 | -0.023 | -0.023 | 0 | 0.75 | 0.79 | 0.75 | 0.66 | 0.64 | 0.73 |
| 4 | 163007304 | rs971300 | 0.341 | 0.242 | 0.103 | 2.363 | 0.01 | 0.018 | 0.018 | 0.143 | 0.68 | 0.59 | 0.6 | 0.47 | 0.61 | 0.55 |
| 4 | 163008238 | rs973516 | -0.042 | 0.425 | 0.156 | 2.221 | 0.024 | 0.032 | 0.032 | 0.124 | 0.73 | 0.61 | 0.73 | 0.58 | 0.63 | 0.66 |
| 4 | 163011663 | rs10005652 | -0.335 | 0.37 | -0.188 | 2.046 | -0.041 | -0.052 | -0.052 | 1.13 | 0.28 | 0.19 | 0.29 | 0.52 | 0.39 | 0.39 |
| 4 | 163012287 | rs9991417 | -0.642 | -0.705 | 0.403 | 1.677 | -0.029 | -0.01 | -0.01 | 0.12 | 0.23 | 0.35 | 0.14 | 0.21 | 0.18 | 0.2 |
| 4 | 163019507 | rs1388834 | 0.627 | -1.055 | 0.248 | 1.688 | -0.012 | 0.002 | 0.002 | 0 | 0.68 | 0.63 | 0.81 | 0.58 | 0.85 | 0.67 |
| 4 | 163030012 | rs1492468 | -0.026 | -1.092 | 0.196 | 1.66 | -0.008 | -0.008 | -0.008 | 0 | 0.78 | 0.73 | 0.88 | 0.87 | 0.89 | 0.81 |
| 4 | 163038241 | rs17538472 | 0.152 | -0.721 | -0.055 | 1.672 | -0.055 | -0.017 | -0.017 | 0.136 | 0.09 | 0 | 0.09 | 0.08 | 0.23 | 0.12 |
| 4 | 163038864 | rs11931941 | -0.99 | -0.796 | -0.013 | 1.779 | -0.021 | -0.071 | -0.071 | 0 | 0.19 | 0.08 | 0.15 | 0.38 | 0.27 | 0.26 |
| 4 | 163039228 | rs17599262 | 0.653 | -0.806 | -0.055 | 1.775 | 0.029 | -0.055 | -0.055 | 0 | 0.09 | 0.01 | 0.06 | 0.29 | 0.04 | 0.13 |
| 4 | 163041211 | rs13134589 | -1.04 | -0.751 | 0.057 | 1.441 | 0.002 | 0.02 | 0.02 | 0 | 0.32 | 0.25 | 0.21 | 0.22 | 0.34 | 0.31 |
| 4 | 163043184 | rs13110363 | -0.486 | -0.879 | 0.91 | 1.757 | -0.054 | 0.01 | 0.01 | 0 | 0.23 | 0.4 | 0.14 | 0.24 | 0.11 | 0.2 |
| 4 | 163043362 | rs13138416 | 0.025 | -0.909 | 0.855 | 1.724 | 0.021 | 0.018 | 0.018 | 0 | 0.2 | 0.13 | 0.09 | 0.13 | 0.09 | 0.17 |
| 4 | 163044149 | rs17041987 | -0.857 | -0.372 | 1.338 | 2.117 | 0.047 | 0.039 | 0.039 | 0 | 0.12 | 0.2 | 0.13 | 0.18 | 0.25 | 0.16 |
| 4 | 163046421 | rs6833160 | -1.77 | -0.371 | 1.765 | 2.332 | 0.099 | 0.29 | 0.29 | 0.073 | 0.07 | 0.36 | 0.09 | 0.23 | 0.07 | 0.13 |
| 4 | 163048238 | rs1032394 | 0.56 | -0.725 | 1.613 | 2.171 | -0.01 | 0 | 0 | 0 | 0.68 | 0.64 | 0.81 | 0.59 | 0.83 | 0.7 |
| 4 | 163048530 | rs4441714 | -0.851 | -0.728 | 1.6 | 2.17 | 0.003 | -0.004 | -0.004 | 0 | 0.32 | 0.34 | 0.18 | 0.41 | 0.17 | 0.3 |
| 4 | 163055350 | rs2872800 | 2.148 | -0.541 | 1.48 | 2.144 | 0.003 | 0.142 | 0.142 | 0.715 | 0.49 | 0.16 | 0.58 | 0.26 | 0.49 | 0.42 |
| 4 | 163057553 | rs17042032 | -1.046 | -0.506 | 1.513 | 2.127 | 0.029 | 0.025 | 0.025 | 0 | 0.14 | 0.17 | 0.17 | 0.22 | 0.28 | 0.19 |
| 4 | 163060499 | rs13109373 | 0.177 | -0.552 | 1.517 | 2.118 | -0.002 | 0 | 0 | 0 | 0.21 | 0.21 | 0.12 | 0.13 | 0.13 | 0.16 |
| 4 | 163061051 | rs17460693 | 0.199 | -0.463 | 1.439 | 2.082 | -0.055 | -0.015 | -0.015 | 0 | 0.09 | 0 | 0.09 | 0.08 | 0.23 | 0.12 |
| 4 | 163062608 | rs7375984 | 0.199 | -0.463 | 1.439 | 2.082 | -0.055 | -0.015 | -0.015 | 0 | 0.09 | 0 | 0.09 | 0.08 | 0.23 | 0.12 |
| 4 | 163073169 | rs1388840 | -1.182 | -0.467 | 1.515 | 2.11 | 0.029 | 0.025 | 0.025 | 0 | 0.14 | 0.17 | 0.17 | 0.22 | 0.28 | 0.19 |
| 4 | 163074444 | rs6818975 | 1.701 | -0.402 | 1.584 | 2.118 | 0.012 | 0.013 | 0.013 | 0.086 | 0.54 | 0.38 | 0.64 | 0.39 | 0.54 | 0.51 |
| 4 | 163079683 | rs1492463 | 1.165 | -0.644 | 1.552 | 2.128 | 0.011 | -0.012 | -0.012 | 0 | 0.61 | 0.67 | 0.73 | 0.53 | 0.6 | 0.58 |
| 4 | 163092108 | rs11731180 | 1.443 | -0.223 | 1.842 | 2.361 | -0.004 | 0.002 | 0.002 | 0 | 0.61 | 0.47 | 0.69 | 0.43 | 0.57 | 0.55 |
| 4 | 163092479 | rs13112114 | 0.219 | -0.355 | 1.851 | 2.415 | 0 | 0 | 0 | 0 | 0.18 | 0.16 | 0.11 | 0.13 | 0.13 | 0.16 |
| 4 | 163094997 | rs12646105 | -1.101 | -0.382 | 1.842 | 2.352 | -0.045 | 0.051 | 0.051 | 0 | 0.31 | 0.39 | 0.22 | 0.51 | 0.2 | 0.34 |
| 4 | 163099369 | rs13135573 | -0.883 | -0.329 | 1.868 | 2.38 | 0.019 | -0.015 | -0.015 | 0 | 0.29 | 0.22 | 0.19 | 0.41 | 0.18 | 0.31 |
| 4 | 163100289 | rs13142621 | -1.003 | -0.302 | 1.506 | 1.96 | -0.045 | 0.054 | 0.054 | 0 | 0.31 | 0.39 | 0.22 | 0.51 | 0.2 | 0.34 |
| 4 | 163110932 | rs6839078 | -0.946 | -0.249 | 1.481 | 1.909 | -0.022 | 0.021 | 0.021 | 0 | 0.3 | 0.33 | 0.27 | 0.56 | 0.2 | 0.34 |
| 4 | 163135702 | rs4574362 | -1.855 | -0.184 | 1.157 | 1.878 | -0.001 | -0.003 | -0.003 | 0.95 | 0.43 | 0.54 | 0.41 | 0.64 | 0.49 | 0.46 |
| 4 | 163149785 | rs13101933 | 0.02 | -0.365 | 1.062 | 1.796 | -0.04 | 0.043 | 0.043 | 0 | 0.2 | 0.34 | 0.15 | 0.23 | 0.13 | 0.18 |
| 4 | 163152054 | rs4292288 | -1.006 | -0.242 | 0.657 | 1.659 | -0.022 | 0.013 | 0.013 | 0 | 0.36 | 0.55 | 0.33 | 0.53 | 0.25 | 0.35 |
| 4 | 163159563 | rs4234947 | -1.441 | -0.091 | -0.189 | 1.171 | 0.034 | 0.132 | 0.132 | 0 | 0.46 | 0.85 | 0.45 | 0.77 | 0.48 | 0.54 |
| 4 | 163178092 | rs13108616 | -0.37 | 0.256 | 0.662 | 1.382 | -0.008 | 0.037 | 0.037 | 0 | 0.17 | 0.25 | 0.13 | 0.24 | 0.14 | 0.19 |
| 4 | 163179938 | rs10009342 | -1.517 | 0.595 | 0.992 | 1.034 | 0.035 | 0.051 | 0.051 | 7.969 | 0.08 | 0.68 | 0.11 | 0.14 | 0.11 | 0.04 |
| 4 | 163186335 | rs7672069 | -0.086 | 0.744 | 0.009 | 0.041 | -0.004 | -0.002 | -0.002 | 0 | 0.77 | 0.83 | 0.86 | 1 | 0.74 | 0.9 |
| 4 | 163191154 | rs4306919 | 0.345 | 1.45 | 0.044 | -0.22 | -0.008 | -0.018 | -0.018 | 1.153 | 0.72 | 0.76 | 0.82 | 0.99 | 0.65 | 0.79 |
| 4 | 163195242 | rs4429685 | 0.355 | 1.359 | -0.297 | 0.028 | 0 | -0.018 | -0.018 | 0 | 0.27 | 0.21 | 0.19 | 0.42 | 0.27 | 0.25 |
| 4 | 163200863 | rs6536647 | 0.57 | 1.152 | 0.229 | 0.748 | 0.012 | 0.012 | 0.012 | 0 | 0.4 | 0.37 | 0.43 | 0.33 | 0.31 | 0.46 |
| 4 | 163211495 | rs4691041 | -0.271 | 1.046 | 0.076 | 0.41 | 0 | -0.053 | -0.053 | 0.813 | 0.67 | 0.69 | 0.82 | 0.96 | 0.62 | 0.79 |
| 4 | 163216127 | rs4343679 | 0.205 | 0.908 | -0.203 | 0.542 | 0.001 | -0.007 | -0.007 | 0.221 | 0.59 | 0.48 | 0.59 | 0.71 | 0.53 | 0.72 |
| 4 | 163221194 | rs4355349 | 0.642 | 0.846 | -0.317 | 0.642 | -0.005 | 0.001 | 0.001 | 0 | 0.34 | 0.42 | 0.37 | 0.42 | 0.24 | 0.52 |
| 4 | 163230830 | rs13141291 | 1.001 | 0.604 | -0.891 | 0.458 | -0.066 | -0.091 | -0.091 | 1.472 | 0.43 | 0.08 | 0.46 | 0.47 | 0.4 | 0.41 |
| 4 | 163237694 | rs4398485 | -1.258 | 0.914 | -1.001 | 0.34 | -0.043 | 0.116 | 0.116 | 0.686 | 0.8 | 0.98 | 0.89 | 1 | 0.76 | 0.91 |

| | | | | | | | | | | | | | | | | |
|---|---|---|---|---|---|---|---|---|---|---|---|---|---|---|---|---|
| 4 | 163237790 | rs6852854 | 0.804 | 0.816 | -0.841 | 0.484 | 0.024 | -0.035 | -0.035 | 0 | 0.13 | 0.34 | 0.12 | 0.05 | 0.13 | 0.15 |
| 4 | 163237809 | rs4413364 | 0.471 | 0.934 | -1.521 | 0.834 | -0.01 | -0.008 | -0.008 | 0 | 0.64 | 0.75 | 0.63 | 0.58 | 0.57 | 0.59 |
| 4 | 163239474 | rs4405972 | -0.063 | 0.953 | -1.618 | 0.241 | -0.019 | 0.058 | 0.058 | 0 | 0.74 | 0.9 | 0.82 | 0.89 | 0.68 | 0.88 |
| 4 | 163243053 | rs4440177 | -0.443 | 0.441 | -1.776 | 0.559 | 0.049 | -0.012 | -0.012 | 0 | 0.42 | 0.73 | 0.42 | 0.44 | 0.35 | 0.43 |
| 4 | 163243244 | rs4533716 | -0.198 | 0.998 | -1.637 | 0.68 | -0.018 | 0.001 | 0.001 | 0 | 0.41 | 0.3 | 0.53 | 0.42 | 0.52 | 0.38 |
| 4 | 163245174 | rs7684439 | -0.805 | 1.309 | -0.984 | 0.981 | 0.017 | 0.017 | 0.017 | 0 | 0.13 | 0.19 | 0.14 | 0.17 | 0.17 | 0.17 |
| 4 | 163248122 | rs11941679 | -0.63 | 1.317 | -0.973 | 0.937 | 0.046 | 0.009 | 0.009 | 0.184 | 0.08 | 0.17 | 0.11 | 0.09 | 0.13 | 0.13 |
| 4 | 163256188 | rs7697133 | 0.41 | 1.29 | -1.272 | 0.848 | -0.014 | 0.006 | 0.006 | 0 | 0.4 | 0.3 | 0.48 | 0.39 | 0.43 | 0.35 |
| 4 | 163263038 | rs13138079 | 0.095 | 0.992 | -0.816 | 0.342 | 0 | -0.005 | -0.005 | 0 | 0.78 | 0.77 | 0.86 | 0.91 | 0.8 | 0.8 |
| 4 | 163264951 | rs6857753 | 0.244 | 0.882 | -0.576 | 0.298 | -0.017 | -0.053 | -0.053 | 0.156 | 0.76 | 0.64 | 0.83 | 0.92 | 0.81 | 0.8 |
| 4 | 163265313 | rs1993024 | 0.388 | 1.047 | -0.538 | 0.556 | -0.012 | 0.031 | 0.031 | 0.002 | 0.75 | 0.61 | 0.6 | 0.67 | 0.74 | 0.78 |
| 4 | 163267782 | rs12508239 | 0.738 | 0.887 | -0.295 | 0.996 | -0.026 | 0.074 | 0.074 | 0.667 | 0.72 | 0.51 | 0.59 | 0.55 | 0.74 | 0.7 |
| 4 | 163269056 | rs10517764 | 1.295 | 1.079 | -0.565 | 0.725 | -0.042 | 0.068 | 0.068 | 0 | 0.36 | 0.2 | 0.23 | 0.17 | 0.38 | 0.32 |
| 4 | 163281547 | rs921344 | 0.088 | 0.77 | -0.404 | 0.949 | -0.093 | 0.05 | 0.05 | 0.019 | 0.39 | 0.13 | 0.37 | 0.32 | 0.51 | 0.45 |
| 4 | 163282170 | rs1118864 | -0.357 | 0.65 | -0.02 | 0.737 | -0.011 | -0.088 | -0.088 | 0 | 0.82 | 0.68 | 0.87 | 0.98 | 0.84 | 0.84 |
| 4 | 163283456 | rs11100437 | 0.327 | 0.66 | 0.107 | 0.824 | -0.028 | -0.061 | -0.061 | 0 | 0.76 | 0.62 | 0.83 | 0.91 | 0.81 | 0.8 |
| 4 | 163284691 | rs7656923 | 1.854 | 0.638 | 0.194 | 0.886 | -0.002 | -0.06 | -0.06 | 0 | 0.3 | 0.34 | 0.14 | 0.1 | 0.23 | 0.19 |
| 4 | 163288053 | rs11936687 | 1.529 | 0.709 | 0.435 | 0.827 | 0.013 | -0.002 | -0.002 | 0 | 0.35 | 0.41 | 0.4 | 0.34 | 0.34 | 0.27 |
| 4 | 163302616 | rs11731765 | 0.824 | 0.694 | 0.422 | 0.882 | -0.022 | 0.262 | 0.262 | 2.933 | 0.87 | 0.69 | 0.65 | 0.57 | 0.91 | 0.9 |
| 4 | 163314790 | rs1450384 | 0.017 | 0.912 | -0.251 | 0.546 | -0.004 | -0.006 | -0.006 | 0 | 0.15 | 0.15 | 0.12 | 0.02 | 0.18 | 0.15 |
| 4 | 163318230 | rs1597469 | -0.466 | 0.875 | -0.118 | 0.665 | 0 | -0.001 | -0.001 | 0 | 0.42 | 0.45 | 0.45 | 0.58 | 0.46 | 0.5 |
| 4 | 163318348 | rs13140705 | 0.513 | 0.938 | -0.917 | 0.346 | -0.064 | 0.078 | 0.078 | 0 | 0.2 | 0.01 | 0.18 | 0.1 | 0.23 | 0.22 |
| 4 | 163324016 | rs2597444 | -1.165 | 0.59 | -0.323 | 0.691 | -0.001 | -0.001 | -0.001 | 0 | 0.58 | 0.49 | 0.73 | 0.6 | 0.66 | 0.6 |
| 4 | 163324468 | rs6845152 | 0.262 | 1.155 | 0.001 | 0.748 | -0.004 | 0.011 | 0.011 | 0 | 0.1 | 0.05 | 0.06 | 0 | 0.12 | 0.09 |
| 4 | 163326947 | rs1901120 | 1.831 | 0.994 | -0.113 | 0.586 | -0.031 | 0.163 | 0.163 | 0 | 0.36 | 0.08 | 0.16 | 0.11 | 0.3 | 0.3 |
| 4 | 163329336 | rs17600575 | 0.381 | 0.952 | 0.107 | 0.946 | -0.006 | -0.017 | -0.017 | 0 | 0.07 | 0.03 | 0.25 | 0.25 | 0.08 | 0.07 |
| 4 | 163332671 | rs2060670 | -0.086 | 0.797 | 0.168 | 0.778 | -0.013 | 0.034 | 0.034 | 0 | 0.65 | 0.71 | 0.66 | 0.91 | 0.56 | 0.71 |
| 4 | 163334309 | rs2712675 | 1.623 | 0.751 | 0.434 | 0.97 | 0.004 | -0.018 | -0.018 | 0 | 0.44 | 0.23 | 0.3 | 0.57 | 0.36 | 0.49 |
| 4 | 163337191 | rs11100443 | -1.066 | 0.769 | 0.801 | 0.923 | -0.003 | 0.093 | 0.093 | 1.313 | 0.12 | 0.26 | 0.28 | 0.25 | 0.11 | 0.12 |
| 4 | 163337355 | rs12645341 | 0.349 | 0.817 | 0.405 | 0.958 | 0.001 | 0.012 | 0.012 | 0 | 0.07 | 0.05 | 0.25 | 0.23 | 0.08 | 0.06 |
| 4 | 163339290 | rs2712683 | 0.037 | 0.805 | 0.439 | 0.357 | -0.042 | 0.125 | 0.125 | 0 | 0.65 | 0.83 | 0.66 | 0.91 | 0.56 | 0.71 |
| 4 | 163353217 | rs13129429 | -0.86 | 0.771 | 0.494 | 0.753 | 0.003 | -0.018 | -0.018 | 0 | 0.25 | 0.21 | 0.43 | 0.53 | 0.29 | 0.23 |
| 4 | 163356033 | rs1597472 | 0.898 | 0.96 | 0.554 | 0.457 | -0.006 | 0.024 | 0.024 | 0 | 0.59 | 0.33 | 0.63 | 0.85 | 0.55 | 0.63 |
| 4 | 163366516 | rs2712674 | -0.325 | 1.023 | 0.713 | 0.491 | -0.035 | 0.065 | 0.065 | 0 | 0.73 | 0.85 | 0.73 | 0.91 | 0.65 | 0.74 |
| 4 | 163367829 | rs2597484 | 1.967 | 0.755 | 0.656 | 0.695 | -0.019 | -0.009 | -0.009 | 0.337 | 0.39 | 0.1 | 0.22 | 0.36 | 0.33 | 0.41 |
| 4 | 163370862 | rs11100444 | 0.115 | 1.328 | 0.657 | -0.132 | -0.064 | 0.104 | 0.104 | 0 | 0.7 | 0.88 | 0.72 | 0.91 | 0.58 | 0.73 |
| 4 | 163376243 | rs10029930 | -1.362 | 1.228 | 1.324 | -0.032 | 0.044 | 0.068 | 0.068 | 0 | 0.08 | 0.51 | 0.07 | 0.05 | 0.04 | 0.09 |
| 4 | 163385544 | rs1450395 | -0.048 | 1.451 | 0.319 | 0.25 | -0.027 | -0.106 | -0.106 | 0.172 | 0.22 | 0.06 | 0.44 | 0.49 | 0.3 | 0.3 |
| 4 | 163385682 | rs1450394 | -1.049 | 1.386 | 0.651 | 0.168 | 0.043 | 0.066 | 0.066 | 0.013 | 0.07 | 0.39 | 0.06 | 0.05 | 0.04 | 0.09 |
| 4 | 163390849 | rs17462361 | -0.214 | 1.586 | 0.484 | 0.083 | 0.015 | -0.018 | -0.018 | 0.022 | 0.12 | 0.2 | 0.1 | 0.06 | 0.19 | 0.13 |
| 4 | 163390933 | rs4139 | -0.995 | 1.712 | 0.368 | -0.068 | -0.03 | 0.079 | 0.079 | 0 | 0.79 | 0.95 | 0.8 | 0.94 | 0.76 | 0.8 |
| 4 | 163392037 | rs10517768 | -0.308 | 1.636 | 1.24 | 0.798 | 0.013 | -0.017 | -0.017 | 2.123 | 0.1 | 0.13 | 0.09 | 0.03 | 0.18 | 0.1 |
| 4 | 163403791 | rs10021001 | -1.336 | 1.395 | 1.085 | 0.817 | 0.009 | -0.005 | -0.005 | 0 | 0.38 | 0.58 | 0.56 | 0.62 | 0.46 | 0.51 |
| 4 | 163406113 | rs4521302 | -1.015 | 1.517 | 0.281 | 0.87 | -0.005 | -0.004 | -0.004 | 0 | 0.34 | 0.33 | 0.52 | 0.59 | 0.43 | 0.44 |
| 4 | 163417093 | rs10003033 | -1.652 | 1.23 | 0.321 | 0.875 | 0.007 | 0.009 | 0.009 | 0 | 0.17 | 0.23 | 0.16 | 0.25 | 0.22 | 0.16 |
| 4 | 163419055 | rs6839249 | -1.216 | 1.146 | 0.844 | 0.813 | 0.052 | 0.128 | 0.128 | 0 | 0.09 | 0.3 | 0.12 | 0.24 | 0.15 | 0.12 |
| 4 | 163432830 | rs2045631 | 1.973 | 1.057 | 0.492 | 0.888 | 0.002 | 0.018 | 0.018 | 0 | 0.79 | 0.69 | 0.83 | 0.71 | 0.78 | 0.78 |
| 4 | 163443708 | rs1471260 | -0.351 | 1.134 | 0.358 | 0.927 | 0.012 | 0.038 | 0.038 | 0 | 0.07 | 0.13 | 0.11 | 0.23 | 0.15 | 0.12 |
| 4 | 163444684 | rs1380722 | -1.164 | 0.748 | 0.316 | 0.477 | -0.002 | -0.026 | -0.026 | 0.355 | 0.11 | 0.1 | 0.05 | 0.01 | 0.07 | 0.04 |
| 4 | 163475257 | rs10428343 | -0.009 | 0.744 | -0.031 | 0.18 | 0.001 | 0.001 | 0.001 | 0.001 | 0.3 | 0.25 | 0.21 | 0.11 | 0.27 | 0.26 |
| 4 | 163484868 | rs6856230 | -1.44 | 0.918 | -0.227 | 0.294 | -0.004 | -0.037 | -0.037 | 0 | 0.76 | 0.81 | 0.8 | 0.55 | 0.76 | 0.75 |
| 4 | 163487232 | rs11100450 | -0.81 | 0.696 | 0.252 | 0.139 | -0.004 | 0.022 | 0.022 | 0 | 0.68 | 0.6 | 0.76 | 0.52 | 0.73 | 0.69 |
| 4 | 163494122 | rs1870602 | 1.536 | 0.636 | 0.38 | 0.11 | -0.016 | -0.068 | -0.068 | 0 | 0.23 | 0.11 | 0.18 | 0.44 | 0.24 | 0.26 |
| 4 | 163494546 | rs6536667 | 0.251 | 0.622 | 0.441 | 0.067 | -0.001 | -0.001 | -0.001 | 0 | 0.38 | 0.37 | 0.55 | 0.42 | 0.45 | 0.44 |
| 4 | 163495571 | rs13147036 | 1.318 | 0.706 | 0.45 | 0.247 | -0.008 | -0.05 | -0.05 | 0.23 | 0.23 | 0.16 | 0.19 | 0.47 | 0.24 | 0.26 |
| 4 | 163497956 | rs4691857 | 0.152 | 0.665 | 0.206 | -0.025 | -0.015 | -0.047 | -0.047 | 0 | 0.92 | 0.77 | 0.95 | 0.99 | 0.97 | 0.95 |
| 4 | 163523865 | rs195898 | -0.021 | 0.669 | 0.143 | -0.023 | -0.013 | -0.043 | -0.043 | 0 | 0.08 | 0.15 | 0.04 | 0.01 | 0.04 | 0.05 |
| 4 | 163525235 | rs2872973 | 1.335 | 0.72 | -0.442 | 0.173 | -0.04 | -0.139 | -0.139 | 2.166 | 0.23 | 0.04 | 0.23 | 0.52 | 0.23 | 0.25 |
| 4 | 163525786 | rs7688628 | 1.349 | 0.75 | -0.442 | 0.159 | -0.043 | -0.147 | -0.147 | 2.635 | 0.23 | 0.04 | 0.23 | 0.49 | 0.23 | 0.25 |
| 4 | 163527435 | rs195894 | -0.761 | 0.974 | -0.345 | -0.021 | -0.02 | -0.015 | -0.015 | 0.119 | 0.71 | 0.83 | 0.57 | 0.65 | 0.63 | 0.7 |
| 4 | 163532717 | rs195891 | -0.988 | 0.869 | 0.013 | 0.331 | -0.001 | 0.038 | 0.038 | 0 | 0.15 | 0.48 | 0.14 | 0.14 | 0.11 | 0.14 |
| 4 | 163533054 | rs6810504 | 0.984 | 1.04 | -0.132 | 0.485 | -0.04 | -0.147 | -0.147 | 0.01 | 0.23 | 0.04 | 0.22 | 0.48 | 0.23 | 0.25 |
| 4 | 163534998 | rs182016 | -1.09 | 0.917 | 0.796 | -0.001 | 0.039 | 0.133 | 0.133 | 0.984 | 0.1 | 0.48 | 0.11 | 0.13 | 0.08 | 0.1 |
| 4 | 163540150 | rs1903291 | -0.301 | 0.992 | 0.463 | -0.817 | -0.001 | -0.099 | -0.099 | 0 | 0.28 | 0.38 | 0.17 | 0.01 | 0.29 | 0.26 |
| 4 | 163543945 | rs6849368 | 1.502 | 1.414 | 0.2 | -0.765 | -0.005 | -0.007 | -0.007 | 0 | 0.3 | 0.11 | 0.53 | 0.71 | 0.28 | 0.35 |
| 4 | 163564805 | rs195982 | -0.333 | 1.395 | 0.566 | -1.088 | -0.002 | 0 | 0 | 0 | 0.64 | 0.68 | 0.77 | 0.84 | 0.59 | 0.71 |
| 4 | 163565725 | rs10489050 | 0.987 | 1.466 | 0.474 | -1.154 | -0.022 | 0.03 | 0.03 | 0 | 0.06 | 0.01 | 0.05 | 0.01 | 0.07 | 0.09 |
| 4 | 163573991 | rs195913 | -0.828 | 1.371 | 0.723 | -1.109 | -0.008 | 0.064 | 0.064 | 0.016 | 0.72 | 0.89 | 0.84 | 0.9 | 0.69 | 0.81 |
| 4 | 163580271 | rs195937 | 1.192 | 1.347 | 0.777 | -0.952 | 0.002 | 0.009 | 0.009 | 0 | 0.33 | 0.21 | 0.58 | 0.71 | 0.3 | 0.39 |

| | | | | | | | | | | | | | | | |
|---|---|---|---|---|---|---|---|---|---|---|---|---|---|---|---|
| 4 | 163605363 | rs195921 | 0.059 | 1.453 | 0.796 | -0.746 | -0.002 | 0.003 | 0.003 | 0 | 0.63 | 0.67 | 0.77 | 0.81 | 0.59 | 0.71 |
| 4 | 163650534 | rs195559 | -0.166 | 1.433 | 0.767 | -0.773 | -0.002 | 0.017 | 0.017 | 0.126 | 0.65 | 0.73 | 0.78 | 0.89 | 0.62 | 0.76 |
| 4 | 163658871 | rs195557 | -0.145 | 1.435 | 0.768 | -0.772 | -0.002 | 0.015 | 0.015 | 0.128 | 0.65 | 0.74 | 0.78 | 0.89 | 0.62 | 0.76 |
| 4 | 163671744 | rs17042829 | -1.369 | 1.349 | 0.92 | -0.482 | 0.03 | 0.114 | 0.114 | 0.033 | 0.06 | 0.21 | 0.04 | 0.18 | 0.05 | 0.15 |
| 4 | 163703018 | rs2320181 | 0.331 | 1.276 | 0.946 | -0.91 | 0.005 | -0.063 | -0.063 | 0.06 | 0.7 | 0.48 | 0.83 | 0.85 | 0.71 | 0.66 |
| 4 | 163707598 | rs6839846 | 0.268 | 1.257 | 0.422 | -0.953 | 0.004 | 0.03 | 0.03 | 0 | 0.3 | 0.2 | 0.17 | 0.19 | 0.31 | 0.3 |
| 4 | 163713049 | rs10517773 | 0.051 | 1.388 | 0.283 | -0.924 | 0.014 | -0.134 | -0.134 | 9.662 | 0.25 | 0.41 | 0.15 | 0.01 | 0.27 | 0.21 |
| 4 | 163718686 | rs13130233 | 0.314 | 1.297 | -0.122 | -0.923 | -0.005 | 0.044 | 0.044 | 0 | 0.28 | 0.13 | 0.17 | 0.18 | 0.3 | 0.28 |
| 4 | 163725002 | rs995302 | -0.483 | 1.201 | -0.251 | -0.514 | 0.006 | 0.031 | 0.031 | 0 | 0.56 | 0.67 | 0.35 | 0.26 | 0.6 | 0.55 |
| 4 | 163731187 | rs10857349 | 0.319 | 1.272 | -0.384 | -0.401 | 0.002 | 0.02 | 0.02 | 0 | 0.65 | 0.29 | 0.72 | 0.71 | 0.69 | 0.7 |
| 4 | 163736954 | rs17042954 | 0.002 | 1.442 | -0.3 | -0.655 | 0.001 | -0.007 | -0.007 | 0 | 0.14 | 0.16 | 0.06 | 0.12 | 0.14 | 0.11 |
| 4 | 163739342 | rs11729764 | 0.285 | 1.28 | -0.486 | -0.83 | 0.006 | 0.013 | 0.013 | 0.675 | 0.63 | 0.35 | 0.7 | 0.81 | 0.69 | 0.72 |
| 4 | 163749607 | rs11723207 | 0.265 | 1.37 | -0.2 | -0.898 | -0.003 | -0.011 | -0.011 | 8.957 | 0.63 | 0.42 | 0.71 | 0.81 | 0.69 | 0.72 |
| 4 | 163758401 | rs4234951 | 1.654 | 1.582 | -0.446 | -0.575 | -0.024 | -0.079 | -0.079 | 0.006 | 0.41 | 0.06 | 0.49 | 0.48 | 0.41 | 0.42 |
| 4 | 163758431 | rs4144993 | -0.779 | 1.106 | 0.452 | -0.953 | 0.017 | -0.059 | -0.059 | 0 | 0.86 | 0.76 | 0.88 | 0.99 | 0.81 | 0.95 |
| 4 | 163758757 | rs745351 | 0.553 | 1.1 | 0.723 | -0.985 | 0.006 | -0.086 | -0.086 | 0.528 | 0.71 | 0.51 | 0.81 | 0.9 | 0.71 | 0.84 |
| 4 | 163766726 | rs10033012 | -0.654 | 1.099 | 0.772 | -0.947 | -0.001 | 0 | 0 | 0 | 0.2 | 0.28 | 0.19 | 0.23 | 0.18 | 0.23 |
| 4 | 163770664 | rs17470149 | 0.385 | 1.119 | 0.353 | -0.925 | -0.001 | -0.012 | -0.012 | 0 | 0.06 | 0.04 | 0.11 | 0.1 | 0.05 | 0.12 |
| 4 | 163771109 | rs17470184 | 0.875 | 1.133 | 0.327 | -0.922 | -0.004 | 0.014 | 0.014 | 0.067 | 0.05 | 0.02 | 0.1 | 0.02 | 0.05 | 0.05 |
| 4 | 163781864 | rs2054210 | 1.539 | 0.97 | 0.314 | -0.907 | -0.007 | 0.084 | 0.084 | 0.169 | 0.49 | 0.27 | 0.62 | 0.66 | 0.53 | 0.59 |
| 4 | 163782459 | rs1458140 | 1.552 | 0.967 | 0.278 | -1.264 | -0.012 | 0.152 | 0.152 | 0.209 | 0.49 | 0.23 | 0.62 | 0.73 | 0.53 | 0.59 |
| 4 | 163793257 | rs4691874 | 0.336 | 0.987 | 0.031 | -1.246 | 0 | -0.002 | -0.002 | 0.768 | 0.14 | 0.14 | 0.07 | 0.14 | 0.13 | 0.11 |
| 4 | 163795102 | rs2012358 | 0.251 | 0.776 | -0.157 | -1.043 | -0.009 | -0.012 | -0.012 | 0 | 0.16 | 0.17 | 0.07 | 0.09 | 0.1 | 0.14 |
| 4 | 163795426 | rs10007144 | 0.322 | 1.005 | -0.627 | -0.689 | 0.003 | 0.11 | 0.11 | 0.218 | 0.5 | 0.35 | 0.63 | 0.74 | 0.54 | 0.63 |
| 4 | 163799376 | rs4691060 | -0.03 | 0.663 | -0.767 | -0.864 | -0.011 | 0.094 | 0.094 | 0.035 | 0.31 | 0.12 | 0.24 | 0.08 | 0.32 | 0.34 |
| 4 | 163802929 | rs7694367 | 0.19 | 0.375 | 0.558 | -0.711 | -0.007 | -0.043 | -0.043 | 0 | 0.63 | 0.47 | 0.72 | 0.85 | 0.67 | 0.61 |
| 4 | 163804334 | rs11100465 | 0.507 | 0.051 | 0.953 | -0.787 | 0.021 | 0.052 | 0.052 | 0.251 | 0.59 | 0.27 | 0.7 | 0.8 | 0.67 | 0.52 |
| 4 | 163808442 | rs2315664 | 0.317 | 0.044 | 1.058 | -0.79 | 0 | -0.003 | -0.003 | 0 | 0.61 | 0.35 | 0.71 | 0.85 | 0.67 | 0.61 |
| 4 | 163817354 | rs4361343 | 0.478 | 0.062 | 1.603 | -0.796 | 0.026 | 0.071 | 0.071 | 0.26 | 0.59 | 0.25 | 0.7 | 0.8 | 0.67 | 0.53 |
| 4 | 163830516 | rs12501673 | 0.192 | 0.361 | 0.593 | -0.903 | -0.019 | 0.088 | 0.088 | 0 | 0.3 | 0.12 | 0.28 | 0.09 | 0.33 | 0.35 |
| 4 | 163836677 | rs7673703 | -0.469 | 0.367 | 0.913 | -0.766 | -0.007 | 0.011 | 0.011 | 0.442 | 0.34 | 0.2 | 0.51 | 0.71 | 0.41 | 0.4 |
| 4 | 163843547 | rs6841751 | 0.062 | 0.33 | 0.881 | -0.798 | -0.005 | 0.01 | 0.01 | 0.864 | 0.66 | 0.81 | 0.49 | 0.29 | 0.59 | 0.6 |
| 4 | 163843642 | rs6842513 | 0.046 | 0.326 | 0.88 | -0.8 | -0.006 | 0.009 | 0.009 | 0.363 | 0.67 | 0.81 | 0.49 | 0.29 | 0.59 | 0.61 |
| 4 | 163854178 | rs6852825 | 0.411 | 0.183 | 1.682 | -0.617 | 0.013 | 0.024 | 0.024 | 0.21 | 0.59 | 0.78 | 0.5 | 0.3 | 0.59 | 0.57 |
| 4 | 163868041 | rs11100469 | 0.836 | 0.213 | 2.311 | -0.657 | 0.003 | 0.095 | 0.095 | 0.463 | 0.53 | 0.27 | 0.41 | 0.27 | 0.52 | 0.53 |
| 4 | 163879822 | rs11938430 | 0.95 | 0.303 | 2.54 | -0.593 | -0.002 | 0.059 | 0.059 | 0 | 0.56 | 0.31 | 0.42 | 0.28 | 0.53 | 0.53 |
| 4 | 163882204 | rs11100472 | 0.659 | 0.302 | 2.572 | -0.592 | 0 | -0.001 | -0.001 | 0 | 0.58 | 0.42 | 0.44 | 0.26 | 0.54 | 0.53 |
| 4 | 163896310 | rs4263350 | 0.509 | 0.329 | 2.48 | -0.575 | 0.002 | -0.03 | -0.03 | 0 | 0.59 | 0.49 | 0.46 | 0.27 | 0.55 | 0.53 |
| 4 | 163906288 | rs17470878 | 0.928 | 0.331 | 1.919 | -0.58 | -0.007 | 0.071 | 0.071 | 0 | 0.56 | 0.27 | 0.41 | 0.26 | 0.53 | 0.52 |
| 4 | 163913951 | rs4546199 | 0.247 | 0.495 | 1.535 | -0.563 | -0.005 | -0.005 | -0.005 | 1.212 | 0.64 | 0.81 | 0.49 | 0.35 | 0.63 | 0.61 |
| 4 | 163934608 | rs6536694 | 0.487 | 0.66 | 1.198 | -0.519 | -0.007 | -0.019 | -0.019 | 0.734 | 0.62 | 0.45 | 0.46 | 0.27 | 0.65 | 0.6 |
| 4 | 163938611 | rs7662233 | 0.662 | 0.718 | 0.726 | -0.466 | -0.003 | 0.008 | 0.008 | 0 | 0.57 | 0.39 | 0.43 | 0.31 | 0.54 | 0.6 |
| 4 | 163939622 | rs6849055 | 0.402 | 0.664 | 0.247 | -0.365 | -0.005 | -0.001 | -0.001 | 0 | 0.66 | 0.5 | 0.5 | 0.3 | 0.68 | 0.66 |
| 4 | 163966691 | rs4301068 | 0.38 | 0.575 | -1.026 | -0.462 | 0.042 | 0.119 | 0.119 | 0.513 | 0.48 | 0.24 | 0.38 | 0.21 | 0.58 | 0.45 |
| 4 | 163969466 | rs4691888 | 0.354 | 0.589 | -1.484 | -0.215 | -0.005 | 0.064 | 0.064 | 0.11 | 0.44 | 0.21 | 0.34 | 0.26 | 0.41 | 0.44 |
| 4 | 163974516 | rs6846180 | -0.359 | 0.738 | -1.594 | -0.055 | -0.017 | -0.015 | -0.015 | 0.1 | 0.68 | 0.8 | 0.45 | 0.33 | 0.61 | 0.56 |
| 4 | 163980423 | rs4691889 | -0.267 | 0.665 | -1.727 | -0.024 | -0.03 | -0.026 | -0.026 | 0 | 0.66 | 0.82 | 0.44 | 0.34 | 0.6 | 0.56 |
| 4 | 163990008 | rs2320331 | -0.688 | 0.21 | -0.903 | 0.281 | -0.004 | -0.001 | -0.001 | 0 | 0.26 | 0.73 | 0.16 | 0.18 | 0.21 | 0.21 |
| 4 | 164000953 | rs2086241 | -0.321 | 0.254 | -0.994 | 0.606 | 0.006 | -0.009 | -0.009 | 0 | 0.39 | 0.33 | 0.48 | 0.55 | 0.36 | 0.55 |
| 4 | 164007992 | rs7676823 | 0.341 | 0.247 | -1.362 | 0.012 | 0.012 | 0.035 | 0.035 | 0 | 0.61 | 0.28 | 0.64 | 0.72 | 0.64 | 0.68 |
| 4 | 164010468 | rs11100478 | -0.542 | 0.201 | -0.234 | -0.176 | 0.015 | -0.053 | -0.053 | 0 | 0.09 | 0.27 | 0.08 | 0 | 0.1 | 0.07 |
| 4 | 164012901 | rs11100479 | -0.142 | 0.194 | 0.531 | -0.14 | 0.003 | 0.007 | 0.007 | 0 | 0.74 | 0.81 | 0.78 | 0.79 | 0.76 | 0.81 |
| 4 | 164013880 | rs11941112 | -0.311 | 0.222 | 0.577 | -0.167 | 0.004 | 0.01 | 0.01 | 0 | 0.77 | 0.85 | 0.8 | 0.83 | 0.79 | 0.85 |
| 4 | 164025528 | rs17574812 | 0.292 | 0.218 | 0.688 | -0.203 | -0.002 | -0.005 | -0.005 | 0 | 0.05 | 0.04 | 0.06 | 0 | 0.09 | 0.03 |
| 4 | 164045822 | rs1055263 | -0.188 | 0.127 | 1.006 | 0.004 | 0 | 0 | 0 | 0 | 0.76 | 0.81 | 0.79 | 0.79 | 0.8 | 0.85 |
| 4 | 164049916 | rs1061075 | 0.407 | 0.711 | 0.46 | -0.64 | 0.017 | -0.081 | -0.081 | 0 | 0.23 | 0.42 | 0.21 | 0.04 | 0.25 | 0.23 |
| 4 | 164066710 | rs7693533 | -0.516 | 0.641 | 0.402 | -0.62 | 0 | 0 | 0 | 0 | 0.74 | 0.79 | 0.78 | 0.79 | 0.76 | 0.84 |
| 4 | 164083616 | rs6536704 | 0.942 | 0.53 | 0.526 | -0.534 | 0.007 | 0.026 | 0.026 | 0.17 | 0.43 | 0.12 | 0.46 | 0.64 | 0.39 | 0.54 |
| 4 | 164101614 | rs10001148 | 0.557 | 0.577 | 0.571 | -0.663 | -0.003 | -0.014 | -0.014 | 0 | 0.45 | 0.43 | 0.5 | 0.75 | 0.41 | 0.54 |
| 4 | 164114844 | rs12644622 | -0.773 | 0.704 | 0.798 | -0.726 | 0.054 | 0.031 | 0.031 | 0.368 | 0.09 | 0.24 | 0.12 | 0.1 | 0.12 | 0.13 |
| 4 | 164123026 | rs6536707 | 0.427 | 0.778 | 0.741 | -0.868 | 0.004 | -0.059 | -0.059 | 0 | 0.25 | 0.38 | 0.2 | 0.04 | 0.26 | 0.3 |
| 4 | 164126538 | rs2127214 | -0.813 | 0.651 | 0.493 | -0.852 | 0.005 | 0.017 | 0.017 | 0 | 0.77 | 0.84 | 0.78 | 0.84 | 0.81 | 0.88 |
| 4 | 164134572 | rs4293752 | 0.768 | 0.66 | 0.431 | -0.52 | 0.002 | 0.043 | 0.043 | 0 | 0.41 | 0.1 | 0.45 | 0.7 | 0.39 | 0.44 |
| 4 | 164158859 | rs11732470 | -0.95 | 0.728 | 0.955 | -0.202 | 0.076 | 0 | 0 | 0 | 0.08 | 0.31 | 0.09 | 0.03 | 0.09 | 0.1 |
| 4 | 164191370 | rs11932242 | -0.281 | 0.503 | 0.002 | -0.444 | -0.011 | 0.025 | 0.025 | 0 | 0.16 | 0.08 | 0.1 | 0.04 | 0.23 | 0.11 |
| 4 | 164200171 | rs4642195 | 0.615 | -0.393 | 0.143 | -0.05 | 0.02 | -0.017 | -0.017 | 0 | 0.7 | 0.27 | 0.75 | 0.38 | 0.83 | 0.69 |
| 4 | 164203507 | rs4128839 | 1.902 | 0.115 | -0.679 | -0.517 | -0.062 | 0.185 | 0.185 | 0 | 0.32 | 0.04 | 0.26 | 0.02 | 0.33 | 0.12 |
| 4 | 164211883 | rs3967761 | -0.325 | -0.407 | 0.62 | 0.053 | -0.036 | -0.091 | -0.091 | 0 | 0.81 | 0.57 | 0.86 | 0.97 | 0.89 | 0.84 |
| 4 | 164221286 | rs7688754 | -0.463 | -0.111 | 0.708 | 0.249 | -0.011 | 0.055 | 0.055 | 0 | 0.26 | 0.34 | 0.21 | 0.62 | 0.28 | 0.24 |

| | | | | | | | | | | | | | | | | |
|---|---|---|---|---|---|---|---|---|---|---|---|---|---|---|---|---|
| 4 | 164225132 | rs4057791 | 1.795 | -0.182 | 1.014 | 0.2 | 0.023 | 0.007 | 0.007 | 0 | 0.58 | 0.27 | 0.66 | 0.35 | 0.62 | 0.64 |
| 4 | 164230540 | rs6828820 | -0.594 | -0.098 | 1.181 | 0.211 | -0.002 | 0.047 | 0.047 | 0 | 0.27 | 0.35 | 0.22 | 0.59 | 0.31 | 0.25 |
| 4 | 164242092 | rs1574637 | -0.381 | -0.122 | 1.386 | 0.179 | -0.032 | -0.107 | -0.107 | 0.106 | 0.84 | 0.43 | 0.89 | 0.99 | 0.91 | 0.91 |
| 4 | 164245405 | rs9764 | -0.518 | -0.007 | 1.198 | 0.256 | 0 | 0.044 | 0.044 | 0 | 0.27 | 0.32 | 0.22 | 0.59 | 0.31 | 0.24 |
| 4 | 164245854 | rs10033119 | 0.382 | -0.027 | 1.192 | 0.96 | -0.01 | 0.141 | 0.141 | 0 | 0.06 | 0.12 | 0.02 | 0.35 | 0.07 | 0.13 |
| 4 | 164249485 | rs4691075 | -0.36 | 0.024 | 1.17 | 0.578 | -0.006 | -0.049 | -0.049 | 0 | 0.86 | 0.63 | 0.87 | 0.96 | 0.9 | 0.85 |
| 4 | 164250436 | rs12510421 | -0.407 | 0.024 | 1.176 | 0.578 | -0.004 | -0.049 | -0.049 | 0 | 0.86 | 0.63 | 0.87 | 0.96 | 0.9 | 0.85 |
| 4 | 164250797 | rs7687423 | 1.522 | -0.019 | 0.947 | 0.5 | 0.005 | 0.004 | 0.004 | 0 | 0.59 | 0.34 | 0.65 | 0.38 | 0.59 | 0.61 |
| 4 | 164252839 | rs11100490 | -0.272 | -0.254 | 1.052 | 0.334 | -0.01 | -0.072 | -0.072 | 0 | 0.87 | 0.66 | 0.92 | 0.99 | 0.92 | 0.91 |
| 4 | 164259120 | rs6852829 | 0.202 | -0.188 | 0.882 | 0.326 | 0.003 | -0.077 | -0.077 | 0 | 0.86 | 0.66 | 0.91 | 0.99 | 0.91 | 0.92 |
| 4 | 164259613 | rs10022662 | 0.155 | -0.187 | 0.819 | 0.326 | 0 | -0.068 | -0.068 | 0 | 0.87 | 0.7 | 0.91 | 0.99 | 0.91 | 0.92 |
| 4 | 164260276 | rs4234955 | 1.426 | -0.602 | 0.274 | 0.833 | 0.033 | 0.007 | 0.007 | 0.047 | 0.63 | 0.3 | 0.65 | 0.63 | 0.77 | 0.58 |
| 4 | 164264776 | rs4314240 | 1.039 | -0.533 | 0.307 | 1.335 | -0.001 | 0.012 | 0.012 | 0 | 0.87 | 0.89 | 0.8 | 0.7 | 0.91 | 0.8 |
| 4 | 164270253 | rs11100494 | 0.652 | -0.334 | 0.341 | 1.466 | 0.001 | 0.01 | 0.01 | 0 | 0.06 | 0.07 | 0.12 | 0.15 | 0.06 | 0.19 |
| 4 | 164276942 | rs4443234 | -1.434 | -1.017 | 0.828 | 1.684 | -0.025 | 0.023 | 0.023 | 3.111 | 0.33 | 0.42 | 0.27 | 0.45 | 0.22 | 0.42 |
| 4 | 164278765 | rs7662762 | -1.562 | -1.059 | 1.046 | 1.666 | -0.036 | 0.026 | 0.026 | 3.207 | 0.34 | 0.45 | 0.28 | 0.45 | 0.22 | 0.42 |
| 4 | 164282763 | rs13151717 | -1.061 | -1.07 | 1.291 | 1.661 | -0.049 | 0.001 | 0.001 | 2.763 | 0.28 | 0.44 | 0.16 | 0.29 | 0.16 | 0.21 |
| 4 | 164283085 | rs6851444 | -0.777 | -1.12 | 1.481 | 1.833 | 0.012 | 0.012 | 0.012 | 3.917 | 0.26 | 0.2 | 0.1 | 0.13 | 0.13 | 0.17 |
| 4 | 164283347 | rs6831008 | 1.264 | -1.175 | 1.27 | 1.734 | 0.033 | 0.036 | 0.036 | 0.812 | 0.11 | 0.03 | 0.01 | 0.05 | 0.02 | 0.07 |
| 4 | 164284929 | rs12503838 | 1.123 | -1.101 | 1.293 | 1.787 | 0.003 | 0.02 | 0.02 | 0 | 0.06 | 0.08 | 0.12 | 0.15 | 0.06 | 0.19 |
| 4 | 164287042 | rs4234956 | 1.473 | -1.136 | 1.175 | 1.714 | -0.031 | 0.039 | 0.039 | 0 | 0.07 | 0.01 | 0.06 | 0 | 0.08 | 0.03 |
| 4 | 164288523 | rs10031046 | -0.229 | -1.12 | 1.444 | 2.301 | -0.004 | 0.048 | 0.048 | 0 | 0.11 | 0.15 | 0.07 | 0.29 | 0.1 | 0.13 |
| 4 | 164293345 | rs7658224 | 2.221 | -1.271 | 1.456 | 2.237 | 0.028 | 0.018 | 0.018 | 0.581 | 0.61 | 0.36 | 0.71 | 0.3 | 0.76 | 0.49 |
| 4 | 164310785 | rs11938071 | 0.068 | -1.172 | 1.156 | 1.859 | -0.015 | 0.004 | 0.004 | 0 | 0.84 | 0.92 | 0.81 | 0.85 | 0.84 | 0.74 |
| 4 | 164313181 | rs10517780 | -0.473 | -1.198 | 1.259 | 2.025 | 0 | 0.046 | 0.046 | 0 | 0.1 | 0.12 | 0.11 | 0.41 | 0.1 | 0.12 |
| 4 | 164325064 | rs1441716 | 0.748 | -1.232 | 1.189 | 2.016 | 0 | 0.029 | 0.029 | 0 | 0.78 | 0.87 | 0.83 | 0.87 | 0.83 | 0.74 |
| 4 | 164328368 | rs4691082 | 2.873 | -0.501 | 1.335 | 1.903 | 0.012 | 0.037 | 0.037 | 1.544 | 0.52 | 0.38 | 0.67 | 0.34 | 0.63 | 0.44 |
| 4 | 164333380 | rs6536728 | -1.458 | -0.72 | 1.349 | 1.924 | -0.058 | 0.111 | 0.111 | 0.03 | 0.24 | 0.49 | 0.15 | 0.53 | 0.14 | 0.29 |
| 4 | 164340073 | rs11937784 | -1.178 | -0.528 | 1.421 | 1.985 | 0.017 | 0.341 | 0.341 | 0 | 0.12 | 0.4 | 0.12 | 0.41 | 0.1 | 0.16 |
| 4 | 164350705 | rs9992637 | -0.231 | -0.497 | 1.074 | 1.953 | -0.002 | 0.108 | 0.108 | 0 | 0.09 | 0.11 | 0.26 | 0.43 | 0.1 | 0.12 |
| 4 | 164357974 | rs1441720 | 0.855 | -0.4 | 0.874 | 1.681 | -0.019 | 0.018 | 0.018 | 0 | 0.77 | 0.89 | 0.82 | 0.82 | 0.8 | 0.65 |
| 4 | 164362850 | rs10014739 | -0.523 | -0.705 | 0.925 | 1.456 | -0.001 | 0.008 | 0.008 | 0 | 0.22 | 0.18 | 0.29 | 0.49 | 0.15 | 0.21 |
| 4 | 164366036 | rs6536732 | -0.289 | -0.601 | 0.893 | 1.347 | -0.004 | -0.003 | -0.003 | 0 | 0.9 | 0.93 | 0.9 | 0.86 | 0.92 | 0.76 |
| 4 | 164377038 | rs993264 | -2.97 | -0.429 | 2.116 | 1.597 | 0.054 | 0.126 | 0.126 | 0.374 | 0.07 | 0.38 | 0.02 | 0.07 | 0.02 | 0.1 |
| 4 | 164377063 | rs2028488 | 1.647 | -0.288 | 1.235 | 0.988 | -0.098 | 0.072 | 0.072 | 0.113 | 0.17 | 0.03 | 0.32 | 0.03 | 0.35 | 0.22 |
| 4 | 164378054 | rs2320767 | 0.403 | -0.569 | 1.042 | 0.838 | -0.025 | 0 | 0 | 0 | 0.33 | 0.34 | 0.44 | 0.32 | 0.54 | 0.37 |
| 4 | 164379003 | rs1545226 | -1.978 | -0.644 | 0.745 | 0.794 | -0.008 | 0.144 | 0.144 | 0 | 0.64 | 0.88 | 0.74 | 0.86 | 0.69 | 0.66 |
| 4 | 164380632 | rs13121637 | 1.062 | -0.426 | 1.183 | 0.968 | 0.022 | 0.036 | 0.036 | 0 | 0.13 | 0.15 | 0.08 | 0.29 | 0.19 | 0.13 |
| 4 | 164383199 | rs12642830 | 1.36 | -0.509 | 1.054 | 0.967 | -0.048 | -0.054 | -0.054 | 0 | 0.12 | 0.03 | 0.08 | 0.28 | 0.19 | 0.13 |
| 4 | 164384209 | rs10005483 | -0.591 | -0.575 | 1.378 | 1.087 | 0.051 | 0.071 | 0.071 | 0 | 0.23 | 0.44 | 0.29 | 0.61 | 0.28 | 0.32 |
| 4 | 164387952 | rs10017973 | -0.832 | -0.585 | 1.397 | 1.082 | 0.024 | 0.036 | 0.036 | 0 | 0.24 | 0.52 | 0.29 | 0.61 | 0.28 | 0.32 |
| 4 | 164389550 | rs4234960 | 1.089 | -0.468 | 1.283 | 0.927 | 0.005 | -0.021 | -0.021 | 0 | 0.22 | 0.28 | 0.38 | 0.05 | 0.38 | 0.25 |
| 4 | 164392225 | rs9308060 | 0.265 | -0.634 | 1.481 | 1.147 | 0.021 | 0.038 | 0.038 | 0 | 0.17 | 0.21 | 0.09 | 0.29 | 0.2 | 0.13 |
| 4 | 164398884 | rs2119677 | 0.326 | -0.617 | 1.204 | 1.166 | 0.004 | 0.183 | 0.183 | 0 | 0.08 | 0.21 | 0.19 | 0.29 | 0.08 | 0.13 |
| 4 | 164407600 | rs2320618 | -1.754 | -0.562 | 0.882 | 0.683 | 0.008 | 0.109 | 0.109 | 0 | 0.61 | 0.79 | 0.73 | 0.89 | 0.68 | 0.66 |
| 4 | 164408033 | rs2119678 | 1.078 | -0.292 | 0.822 | 0.801 | 0.009 | -0.026 | -0.026 | 0 | 0.2 | 0.26 | 0.37 | 0.05 | 0.35 | 0.22 |
| 4 | 164409556 | rs1073547 | -0.444 | -0.792 | 1.005 | 1.362 | -0.03 | 0.067 | 0.067 | 0 | 0.21 | 0.27 | 0.26 | 0.48 | 0.11 | 0.22 |
| 4 | 164411073 | rs10034495 | -0.935 | -0.72 | 0.995 | 1.067 | -0.002 | -0.001 | -0.001 | 0 | 0.56 | 0.68 | 0.75 | 0.55 | 0.58 | 0.49 |
| 4 | 164415792 | rs2276978 | 0.23 | -0.575 | 1.185 | 1.227 | 0.005 | 0.002 | 0.002 | 0 | 0.15 | 0.13 | 0.12 | 0.14 | 0.08 | 0.09 |
| 4 | 164415930 | rs2276977 | 0.165 | -0.577 | 1.385 | 1.226 | -0.007 | -0.003 | -0.003 | 0 | 0.15 | 0.23 | 0.12 | 0.14 | 0.08 | 0.09 |
| 4 | 164417245 | rs11730351 | 1.363 | -0.598 | 1 | 1.288 | -0.044 | -0.075 | -0.075 | 0 | 0.11 | 0 | 0.06 | 0.35 | 0.15 | 0.21 |
| 4 | 164418356 | rs7657886 | 0.535 | -0.734 | 1.482 | 1.208 | -0.01 | -0.003 | -0.003 | 0.367 | 0.33 | 0.27 | 0.61 | 0.34 | 0.48 | 0.35 |
| 4 | 164436103 | rs12500221 | -1.223 | -0.813 | 1.643 | 1.133 | -0.013 | -0.016 | -0.016 | 0.2 | 0.58 | 0.79 | 0.75 | 0.54 | 0.58 | 0.48 |
| 4 | 164447976 | rs1044722 | -0.994 | -0.697 | 1.663 | 1.261 | 0.023 | 0.018 | 0.018 | 0 | 0.54 | 0.66 | 0.75 | 0.57 | 0.61 | 0.47 |
| 4 | 164454771 | rs9993575 | -0.815 | -0.559 | 1.163 | 1.28 | -0.001 | -0.001 | -0.001 | 0 | 0.67 | 0.7 | 0.39 | 0.6 | 0.5 | 0.61 |
| 4 | 164457232 | rs2044052 | -0.613 | -0.616 | 1.007 | 1.128 | 0.014 | -0.004 | -0.004 | 0 | 0.51 | 0.34 | 0.72 | 0.49 | 0.56 | 0.42 |
| 4 | 164460645 | rs17473073 | 1.782 | -0.612 | 0.879 | 1.085 | -0.011 | -0.017 | -0.017 | 0.24 | 0.08 | 0.01 | 0.03 | 0.09 | 0.05 | 0.06 |
| 4 | 164463778 | rs17473108 | 0.008 | -0.615 | 0.903 | 1.081 | -0.003 | -0.004 | -0.004 | 0.98 | 0.15 | 0.15 | 0.13 | 0.17 | 0.11 | 0.11 |
| 4 | 164465746 | rs12508980 | -0.929 | -0.613 | 0.998 | 1.118 | -0.003 | -0.001 | -0.001 | 0 | 0.54 | 0.51 | 0.75 | 0.53 | 0.61 | 0.47 |
| 4 | 164466824 | rs13130399 | 0.503 | -0.41 | 0.736 | 0.987 | -0.009 | -0.004 | -0.004 | 0 | 0.32 | 0.28 | 0.6 | 0.35 | 0.46 | 0.34 |
| 4 | 164470301 | rs11100507 | -0.395 | -0.27 | 0.55 | 0.99 | 0.014 | 0.046 | 0.046 | 0 | 0.51 | 0.36 | 0.51 | 0.41 | 0.55 | 0.47 |
| 4 | 164471823 | rs6815345 | -1.299 | -0.245 | 0.34 | 0.518 | 0.007 | 0.003 | 0.003 | 0 | 0.66 | 0.72 | 0.81 | 0.91 | 0.72 | 0.7 |
| 4 | 164474468 | rs2320771 | 0.002 | 0.039 | -0.008 | 0.765 | -0.015 | -0.003 | -0.003 | 0 | 0.39 | 0.35 | 0.67 | 0.42 | 0.53 | 0.39 |
| 4 | 164474591 | rs17473171 | 0.405 | 0.058 | -0.09 | 0.616 | 0.001 | -0.001 | -0.001 | 0 | 0.11 | 0.1 | 0.04 | 0.08 | 0.07 | 0.13 |
| 4 | 164475425 | rs11100508 | -0.38 | 0.403 | -0.156 | 0.693 | 0.027 | 0.055 | 0.055 | 0 | 0.25 | 0.47 | 0.24 | 0.32 | 0.23 | 0.33 |
| 4 | 164476240 | rs6843683 | -0.115 | 0.187 | -0.57 | 0.239 | 0.017 | 0.063 | 0.063 | 0 | 0.67 | 0.8 | 0.82 | 0.93 | 0.73 | 0.7 |
| 4 | 164478874 | rs11100509 | 1.033 | 0.471 | -0.052 | 1.541 | 0.002 | 0.057 | 0.057 | 0 | 0.84 | 0.8 | 0.67 | 0.59 | 0.84 | 0.79 |
| 4 | 164479967 | rs10029699 | -1.911 | 0.201 | 0.97 | 1.527 | 0.004 | 0.143 | 0.143 | 0 | 0.12 | 0.42 | 0.14 | 0.27 | 0.07 | 0.17 |
| 4 | 164487447 | rs17473283 | 0.392 | 0.4 | 0.332 | 1.172 | -0.058 | 0.07 | 0.07 | 0 | 0.32 | 0.08 | 0.29 | 0.15 | 0.38 | 0.3 |

| | | | | | | | | | | | | | | | | |
|---|---|---|---|---|---|---|---|---|---|---|---|---|---|---|---|---|
| 4 | 164489038 | rs13115712 | 0.689 | 0.194 | 1.183 | 1.227 | 0.019 | -0.004 | -0.004 | 0 | 0.49 | 0.32 | 0.52 | 0.52 | 0.47 | 0.42 |
| 4 | 164490018 | rs1877314 | -0.475 | 0.262 | 1.125 | 1.21 | -0.004 | -0.004 | -0.004 | 0.333 | 0.95 | 0.96 | 0.95 | 0.94 | 0.93 | 0.92 |
| 4 | 164492395 | rs10001689 | 0.881 | 0.069 | 1.562 | 1.226 | 0.009 | -0.009 | -0.009 | 0.038 | 0.46 | 0.33 | 0.51 | 0.54 | 0.46 | 0.37 |
| 4 | 164499999 | rs7693363 | -0.163 | 0.089 | 1.811 | 1.271 | 0.013 | 0.126 | 0.126 | 0 | 0.06 | 0.22 | 0.1 | 0.21 | 0.04 | 0.02 |
| 4 | 164500909 | rs6821802 | -1.476 | 0.08 | 1.847 | 1.224 | 0.004 | 0.081 | 0.081 | 0 | 0.09 | 0.25 | 0.13 | 0.27 | 0.08 | 0.15 |
| 4 | 164514168 | rs7693970 | 0.52 | 0.039 | 1.65 | 1.169 | -0.037 | 0.052 | 0.052 | 0 | 0.33 | 0.1 | 0.3 | 0.16 | 0.38 | 0.32 |
| 4 | 164521066 | rs10213166 | 0.604 | -0.043 | 1.372 | 1.465 | 0.012 | 0.012 | 0.012 | 0 | 0.51 | 0.44 | 0.52 | 0.44 | 0.47 | 0.36 |
| 4 | 164522666 | rs7692553 | -0.175 | 0.134 | 0.99 | 1.443 | -0.005 | -0.006 | -0.006 | 0.19 | 0.4 | 0.3 | 0.44 | 0.5 | 0.48 | 0.51 |
| 4 | 164528452 | rs1511779 | -0.435 | 0.358 | 0.642 | 1.46 | -0.033 | -0.011 | -0.011 | 0 | 0.94 | 0.99 | 0.95 | 0.95 | 0.92 | 0.9 |
| 4 | 164529581 | rs17473521 | 0.326 | 0.376 | 0.71 | 1.455 | -0.002 | 0 | 0 | 0 | 0.36 | 0.28 | 0.34 | 0.15 | 0.42 | 0.37 |
| 4 | 164529829 | rs950565 | 1.319 | -0.076 | 0.383 | 1.635 | -0.022 | 0.03 | 0.03 | 0.014 | 0.8 | 0.5 | 0.74 | 0.74 | 0.9 | 0.83 |
| 4 | 164532325 | rs907435 | -0.461 | 0.307 | -0.871 | 1.79 | -0.024 | -0.004 | -0.004 | 0.078 | 0.94 | 0.99 | 0.94 | 0.95 | 0.91 | 0.9 |
| 4 | 164532973 | rs3792614 | 0.147 | 0.055 | -0.704 | 1.854 | -0.016 | 0.028 | 0.028 | 0 | 0.07 | 0.15 | 0.18 | 0.2 | 0.01 | 0.05 |
| 4 | 164536663 | rs4413365 | 0.312 | 0.049 | -0.819 | 1.88 | -0.027 | -0.013 | -0.013 | 0.026 | 0.44 | 0.19 | 0.39 | 0.58 | 0.48 | 0.46 |
| 4 | 164540801 | rs10030322 | -0.083 | 0.009 | -0.877 | 1.795 | -0.001 | 0.098 | 0.098 | 0.03 | 0.51 | 0.35 | 0.57 | 0.79 | 0.5 | 0.46 |
| 4 | 164542625 | rs7686154 | -1.091 | -0.011 | -1.188 | 1.903 | 0.003 | 0.134 | 0.134 | 0 | 0.05 | 0.13 | 0.11 | 0.35 | 0.06 | 0.16 |
| 4 | 164547230 | rs2036905 | 0.385 | 0.075 | -1.52 | 1.795 | -0.057 | -0.027 | -0.027 | 0 | 0.4 | 0.08 | 0.29 | 0.33 | 0.43 | 0.3 |
| 4 | 164547492 | rs2175490 | 0.488 | 0.022 | -1.53 | 1.569 | -0.073 | 0.018 | 0.018 | 0.48 | 0.39 | 0.05 | 0.27 | 0.24 | 0.42 | 0.3 |
| 4 | 164561013 | rs11725683 | -0.127 | 0.188 | -1.376 | 1.704 | 0.024 | -0.003 | -0.003 | 0 | 0.47 | 0.18 | 0.45 | 0.59 | 0.44 | 0.46 |
| 4 | 164562638 | rs13149834 | 0.241 | 0.536 | -1.503 | 1.237 | -0.031 | 0.046 | 0.046 | 0.137 | 0.39 | 0.11 | 0.28 | 0.23 | 0.42 | 0.3 |
| 4 | 164570672 | rs13105536 | -0.514 | 0.628 | -1.582 | 0.042 | -0.05 | 0.11 | 0.11 | 0 | 0.21 | 0.02 | 0.15 | 0.03 | 0.24 | 0.15 |
| 4 | 164571149 | rs17044007 | -0.106 | 0.472 | -1.201 | -0.352 | -0.004 | -0.049 | -0.049 | 0 | 0.13 | 0.14 | 0.05 | 0 | 0.1 | 0.03 |
| 4 | 164571285 | rs17473710 | -0.237 | 0.599 | -1.826 | -0.704 | -0.029 | 0.052 | 0.052 | 0.085 | 0.21 | 0.07 | 0.21 | 0.07 | 0.25 | 0.17 |
| 4 | 164572108 | rs7655593 | 0.326 | 0.3 | -2.016 | -0.428 | 0.016 | -0.021 | -0.021 | 0 | 0.31 | 0.29 | 0.39 | 0.46 | 0.24 | 0.24 |
| 4 | 164575991 | rs1533151 | 0.608 | -0.305 | -1.921 | -0.451 | -0.011 | 0.012 | 0.012 | 0 | 0.2 | 0.22 | 0.31 | 0.44 | 0.11 | 0.13 |
| 4 | 164582457 | rs1511776 | 0.74 | 0.329 | -2.296 | -0.673 | -0.122 | 0.112 | 0.112 | 0.844 | 0.23 | 0.01 | 0.22 | 0.05 | 0.31 | 0.18 |
| 4 | 164595552 | rs6536747 | 0.215 | -0.542 | -1.049 | -0.548 | -0.086 | 0.091 | 0.091 | 0 | 0.24 | 0.61 | 0.34 | 0.39 | 0.13 | 0.18 |
| 4 | 164595666 | rs17473787 | 0.243 | -0.282 | -1.391 | -0.58 | -0.171 | 0.132 | 0.132 | 0 | 0.32 | 0.02 | 0.29 | 0.08 | 0.41 | 0.3 |
| 4 | 164602209 | rs6826691 | -0.945 | -0.55 | -0.395 | -0.294 | 0.001 | -0.017 | -0.017 | 3.535 | 0.63 | 0.73 | 0.67 | 0.52 | 0.62 | 0.51 |
| 4 | 164611944 | rs998117 | -0.299 | -0.612 | 0.304 | -0.419 | 0.001 | -0.021 | -0.021 | 1.604 | 0.41 | 0.47 | 0.4 | 0.16 | 0.52 | 0.33 |
| 4 | 164621883 | rs9308064 | -0.253 | -0.624 | 0.404 | -0.398 | -0.006 | 0.021 | 0.021 | 0 | 0.41 | 0.41 | 0.39 | 0.16 | 0.52 | 0.34 |
| 4 | 164623738 | rs17577307 | 0.155 | -0.653 | 0.374 | -0.453 | -0.044 | 0.083 | 0.083 | 0 | 0.35 | 0.22 | 0.33 | 0.07 | 0.49 | 0.32 |
| 4 | 164632127 | rs1511783 | 0.627 | -0.553 | 0.208 | -0.479 | -0.168 | 0.154 | 0.154 | 0 | 0.3 | 0.04 | 0.3 | 0.06 | 0.43 | 0.27 |
| 4 | 164634372 | rs1565840 | -0.24 | -0.674 | 0.683 | 0.28 | 0.011 | 0.005 | 0.005 | 0 | 0.4 | 0.5 | 0.56 | 0.36 | 0.49 | 0.39 |
| 4 | 164655112 | rs6814421 | 0.803 | -1.098 | 1.481 | 0.372 | 0.002 | 0.009 | 0.009 | 0 | 0.41 | 0.3 | 0.25 | 0.45 | 0.41 | 0.46 |
| 4 | 164661559 | rs7667981 | 0.224 | -1.101 | 1.289 | 0.236 | -0.024 | 0.036 | 0.036 | 0 | 0.37 | 0.33 | 0.36 | 0.11 | 0.47 | 0.32 |
| 4 | 164668080 | rs9308066 | -1.377 | -1.037 | 1.451 | 0.508 | 0.002 | -0.003 | -0.003 | 0.034 | 0.84 | 0.85 | 0.83 | 0.73 | 0.91 | 0.86 |
| 4 | 164669761 | rs6857627 | -0.836 | -1.043 | 1.202 | 0.195 | 0.016 | 0.065 | 0.065 | 0 | 0.07 | 0.28 | 0.06 | 0.1 | 0.04 | 0.04 |
| 4 | 164675031 | rs2131861 | 1.263 | -1.38 | 0.931 | 0.22 | -0.03 | 0.023 | 0.023 | 0 | 0.32 | 0.13 | 0.18 | 0.34 | 0.35 | 0.37 |
| 4 | 164679594 | rs10517786 | 0.023 | -0.485 | 1.267 | -0.054 | -0.004 | -0.007 | -0.007 | 1.173 | 0.36 | 0.4 | 0.34 | 0.11 | 0.43 | 0.31 |
| 4 | 164690560 | rs4256198 | 0.724 | -0.678 | 1.339 | 0.041 | -0.023 | -0.046 | -0.046 | 0 | 0.84 | 0.65 | 0.89 | 0.95 | 0.9 | 0.91 |
| 4 | 164694796 | rs1389706 | 0.683 | -0.734 | 1.426 | 0.015 | -0.036 | -0.032 | -0.032 | 0.673 | 0.87 | 0.7 | 0.92 | 0.96 | 0.95 | 0.93 |
| 4 | 164701659 | rs1038975 | 0.694 | -0.741 | 1.109 | -0.094 | 0.008 | 0.025 | 0.025 | 0 | 0.49 | 0.44 | 0.54 | 0.83 | 0.43 | 0.58 |
| 4 | 164715878 | rs8180235 | 1.014 | -0.741 | 1.145 | 0.298 | -0.008 | -0.005 | -0.005 | 0.895 | 0.48 | 0.18 | 0.39 | 0.58 | 0.45 | 0.52 |
| 4 | 164720383 | rs12500778 | 0.936 | -0.68 | 0.964 | 0.067 | -0.028 | -0.022 | -0.022 | 0 | 0.88 | 0.59 | 0.93 | 0.96 | 0.95 | 0.98 |
| 4 | 164721357 | rs12645753 | 0.7 | -0.673 | 0.893 | 0.072 | 0.001 | 0.011 | 0.011 | 0 | 0.52 | 0.49 | 0.57 | 0.84 | 0.47 | 0.6 |
| 4 | 164732179 | rs1908290 | 1.457 | -0.614 | 0.862 | 0.127 | -0.031 | -0.011 | -0.011 | 1.63 | 0.8 | 0.54 | 0.83 | 0.86 | 0.86 | 0.88 |
| 4 | 164748523 | rs2131862 | 0.323 | -0.496 | 0.503 | 0.046 | -0.102 | 0.177 | 0.177 | 0 | 0.32 | 0.11 | 0.29 | 0.02 | 0.41 | 0.26 |
| 4 | 164752195 | rs17044211 | 0.239 | -0.447 | 0.641 | 0.056 | -0.001 | -0.011 | -0.011 | 0.891 | 0.08 | 0.1 | 0.06 | 0.01 | 0.05 | 0.06 |
| 4 | 164753387 | rs1874094 | 1.193 | -0.573 | 0.728 | 0.198 | 0.011 | -0.001 | -0.001 | 0.228 | 0.44 | 0.23 | 0.33 | 0.57 | 0.41 | 0.52 |
| 4 | 164758471 | rs11939053 | -0.369 | -0.588 | 0.748 | 0.347 | -0.012 | -0.011 | -0.011 | 0 | 0.65 | 0.69 | 0.54 | 0.71 | 0.58 | 0.66 |
| 4 | 164758636 | rs7681115 | -0.363 | -0.572 | 0.76 | 0.352 | -0.007 | -0.007 | -0.007 | 0 | 0.64 | 0.65 | 0.54 | 0.7 | 0.58 | 0.66 |
| 4 | 164759917 | rs2047160 | 0.968 | -0.599 | 0.993 | 0.307 | -0.024 | 0.012 | 0.012 | 0 | 0.85 | 0.62 | 0.87 | 0.88 | 0.9 | 0.96 |
| 4 | 164761718 | rs1565842 | 1.279 | -0.373 | 0.932 | 0.458 | 0.008 | 0.003 | 0.003 | 0 | 0.88 | 0.77 | 0.87 | 0.9 | 0.89 | 0.89 |
| 4 | 164765518 | rs7670349 | 0.363 | -0.461 | 0.587 | 0.479 | 0.01 | 0.007 | 0.007 | 0 | 0.5 | 0.37 | 0.39 | 0.6 | 0.48 | 0.52 |
| 4 | 164766062 | rs7693115 | 0.7 | -0.415 | 0.696 | 0.862 | -0.006 | 0.339 | 0.339 | 0.74 | 0.91 | 0.54 | 0.74 | 0.76 | 0.96 | 0.89 |
| 4 | 164772977 | rs11100520 | -0.05 | 0.032 | -0.424 | -0.214 | -0.027 | 0.103 | 0.103 | 0 | 0.4 | 0.22 | 0.34 | 0.12 | 0.49 | 0.29 |
| 4 | 164777197 | rs923913 | 0.007 | 0.304 | -0.56 | -0.079 | -0.002 | 0.028 | 0.028 | 0 | 0.56 | 0.64 | 0.63 | 0.86 | 0.5 | 0.7 |
| 4 | 164779320 | rs10517791 | -0.536 | 0.055 | -0.749 | 0.375 | 0.016 | 0.008 | 0.008 | 0 | 0.33 | 0.34 | 0.33 | 0.32 | 0.28 | 0.33 |
| 4 | 164780120 | rs978402 | -0.631 | 0.849 | -0.684 | 0.705 | 0.004 | 0.014 | 0.014 | 0 | 0.49 | 0.7 | 0.42 | 0.41 | 0.49 | 0.44 |
| 4 | 164782515 | rs7655259 | 0.427 | 1.337 | 0.41 | 1.262 | 0.054 | 0.142 | 0.142 | 0.27 | 0.83 | 0.47 | 0.81 | 0.63 | 0.8 | 0.84 |
| 4 | 164790801 | rs17044246 | 0.2 | 1.487 | 0.307 | 1.035 | 0.012 | -0.02 | -0.02 | 0.114 | 0.09 | 0.15 | 0.08 | 0.04 | 0.1 | 0.05 |
| 4 | 164795529 | rs12504215 | 1.048 | 1.341 | 0.658 | 1.233 | 0.003 | 0.009 | 0.009 | 2.296 | 0.74 | 0.29 | 0.76 | 0.56 | 0.69 | 0.78 |
| 4 | 164796501 | rs12645283 | 1.093 | 1.315 | 0.644 | 1.229 | 0.001 | 0.006 | 0.006 | 2.042 | 0.73 | 0.28 | 0.73 | 0.55 | 0.69 | 0.78 |
| 4 | 164797386 | rs1494284 | 1.248 | 1.27 | 0.015 | 0.653 | 0.022 | -0.026 | -0.026 | 0 | 0.52 | 0.19 | 0.54 | 0.51 | 0.44 | 0.43 |
| 4 | 164797407 | rs17474536 | 0.428 | 1.283 | -0.047 | 0.522 | -0.007 | 0.021 | 0.021 | 0 | 0.21 | 0.11 | 0.2 | 0.05 | 0.25 | 0.34 |
| 4 | 164799338 | rs17474585 | 0.542 | 1.342 | -0.048 | 0.545 | -0.009 | 0.031 | 0.031 | 0 | 0.2 | 0.09 | 0.18 | 0.05 | 0.23 | 0.35 |
| 4 | 164801854 | rs6821574 | 0.914 | 0.856 | 0.695 | 0.873 | 0.005 | 0.019 | 0.019 | 1.862 | 0.55 | 0.36 | 0.58 | 0.7 | 0.45 | 0.44 |
| 4 | 164820243 | rs6536757 | -0.931 | 0.83 | 0.831 | 0.679 | -0.001 | -0.006 | -0.006 | 0.592 | 0.42 | 0.44 | 0.39 | 0.35 | 0.54 | 0.48 |

| | | | | | | | | | | | | | | | | |
|---|---|---|---|---|---|---|---|---|---|---|---|---|---|---|---|---|
| 4 | 164829751 | rs3890761 | 0.376 | 1.084 | 1.425 | 1.027 | -0.01 | -0.003 | -0.003 | 0 | 0.21 | 0.12 | 0.2 | 0.26 | 0.23 | 0.35 |
| 4 | 164830114 | rs6844489 | 0.701 | 1.26 | 1.659 | 1.277 | -0.023 | -0.025 | -0.025 | 0 | 0.64 | 0.32 | 0.67 | 0.51 | 0.62 | 0.51 |
| 4 | 164832145 | rs12501935 | 0.623 | 1.294 | 1.614 | 1.256 | -0.059 | 0.086 | 0.086 | 0 | 0.19 | 0.03 | 0.17 | 0.04 | 0.22 | 0.34 |
| 4 | 164841877 | rs10008162 | -0.493 | 1.264 | 1.727 | 1.333 | 0.082 | 0.321 | 0.321 | 1.001 | 0.14 | 0.43 | 0.14 | 0.41 | 0.15 | 0.13 |
| 4 | 164845121 | rs11100528 | -0.381 | 1.249 | 1.743 | 1.47 | 0.003 | -0.013 | -0.013 | 1.42 | 0.77 | 0.73 | 0.81 | 0.88 | 0.78 | 0.63 |
| 4 | 164855278 | rs4691940 | 0.366 | 1.248 | 1.942 | 1.443 | 0.004 | -0.012 | -0.012 | 0.448 | 0.68 | 0.64 | 0.74 | 0.86 | 0.66 | 0.58 |
| 4 | 164858736 | rs11725690 | 0.418 | 1.305 | 1.85 | 1.407 | -0.005 | 0.003 | 0.003 | 0 | 0.1 | 0.09 | 0.08 | 0.06 | 0.11 | 0.11 |
| 4 | 164865947 | rs10857365 | 0.336 | 1.31 | 1.804 | 1.41 | 0.023 | -0.02 | -0.02 | 0 | 0.86 | 0.71 | 0.89 | 0.93 | 0.86 | 0.89 |
| 4 | 164870129 | rs4449375 | 0.574 | -0.05 | 0.94 | 1.305 | 0.07 | 0.015 | 0.015 | 0 | 0.19 | 0.05 | 0.21 | 0.16 | 0.15 | 0.13 |
| 4 | 164872277 | rs12505619 | 0.707 | 0.473 | 1.209 | 1.52 | -0.069 | 0.074 | 0.074 | 0 | 0.22 | 0.04 | 0.17 | 0.08 | 0.24 | 0.36 |
| 4 | 164873243 | rs4270550 | 0.069 | 0.458 | 1.262 | 1.536 | 0.021 | -0.042 | -0.042 | 0.677 | 0.67 | 0.55 | 0.73 | 0.85 | 0.66 | 0.58 |
| 4 | 164876016 | rs6536764 | 0.404 | 0.474 | 0.93 | 1.533 | -0.032 | -0.029 | -0.029 | 1.109 | 0.63 | 0.24 | 0.68 | 0.47 | 0.63 | 0.51 |
| 4 | 164884894 | rs3967463 | 0.364 | 0.411 | 0.904 | 1.575 | -0.016 | -0.019 | -0.019 | 0 | 0.63 | 0.32 | 0.68 | 0.47 | 0.63 | 0.52 |
| 4 | 164889372 | rs17578295 | 0.736 | 0.474 | -0.447 | 1.095 | -0.076 | 0.082 | 0.082 | 0 | 0.22 | 0.02 | 0.17 | 0.08 | 0.24 | 0.36 |
| 4 | 164889887 | rs9308069 | 0.144 | 0.182 | -0.302 | 1.352 | -0.03 | -0.046 | -0.046 | 0 | 0.62 | 0.24 | 0.68 | 0.54 | 0.63 | 0.5 |
| 4 | 164892274 | rs17578337 | 1.121 | 0.072 | -1.117 | 0.557 | -0.066 | -0.011 | -0.011 | 0 | 0.25 | 0.03 | 0.21 | 0.22 | 0.28 | 0.25 |
| 4 | 164905719 | rs3792673 | -0.064 | 0.283 | -0.389 | 0.411 | 0.016 | -0.065 | -0.065 | 0.115 | 0.64 | 0.59 | 0.73 | 0.89 | 0.64 | 0.58 |
| 4 | 164922532 | rs12511868 | 0.197 | -0.239 | -0.632 | 1.462 | 0.007 | 0.023 | 0.023 | 0 | 0.06 | 0.03 | 0.17 | 0.31 | 0.04 | 0.07 |
| 4 | 164934874 | rs10007601 | 0.244 | -1.135 | -0.163 | 0.986 | -0.05 | -0.034 | -0.034 | 0.06 | 0.71 | 0.57 | 0.83 | 0.87 | 0.76 | 0.84 |
| 4 | 164941060 | rs7687757 | 0.699 | -0.97 | -0.804 | 0.49 | 0.001 | -0.017 | -0.017 | 0 | 0.12 | 0.04 | 0.05 | 0.15 | 0.08 | 0.23 |
| 4 | 164959804 | rs4533720 | -1.353 | -1.429 | -0.774 | 0.028 | -0.006 | 0.013 | 0.013 | 0 | 0.62 | 0.74 | 0.47 | 0.6 | 0.46 | 0.52 |
| 4 | 164962875 | rs4560357 | -0.682 | -1.44 | -0.584 | -0.006 | 0.023 | 0.001 | 0.001 | 0 | 0.48 | 0.6 | 0.42 | 0.46 | 0.38 | 0.29 |
| 4 | 164968164 | rs2001227 | -1.34 | -1.442 | -0.591 | 0.057 | -0.007 | 0.012 | 0.012 | 0 | 0.62 | 0.74 | 0.47 | 0.6 | 0.46 | 0.52 |
| 4 | 164968546 | rs10026455 | -0.05 | -1.916 | -0.441 | 0.074 | -0.11 | 0.061 | 0.061 | 0 | 0.31 | 0.5 | 0.29 | 0.45 | 0.15 | 0.22 |
| 4 | 164972162 | rs6536779 | -0.493 | -1.33 | -0.312 | 0.107 | 0.006 | -0.002 | -0.002 | 0.792 | 0.84 | 0.74 | 0.9 | 0.86 | 0.86 | 0.76 |
| 4 | 164972256 | rs6536780 | -0.743 | -1.619 | -0.587 | 0.016 | -0.011 | 0 | 0 | 0 | 0.43 | 0.46 | 0.36 | 0.46 | 0.32 | 0.26 |
| 4 | 164972450 | rs13123163 | 0.433 | -0.949 | -0.972 | -0.295 | -0.01 | 0.032 | 0.032 | 0 | 0.14 | 0.04 | 0.08 | 0.01 | 0.16 | 0.06 |
| 4 | 164976680 | rs17475275 | 0.912 | -1.417 | -1.749 | -0.452 | -0.096 | 0.096 | 0.096 | 1.223 | 0.35 | 0.03 | 0.55 | 0.21 | 0.58 | 0.5 |
| 4 | 164982057 | rs891577 | -0.035 | -2.293 | -0.456 | -0.114 | -0.012 | -0.001 | -0.001 | 0 | 0.62 | 0.62 | 0.69 | 0.49 | 0.78 | 0.71 |
| 4 | 164991575 | rs1430987 | 0.232 | -2.36 | 0.175 | -0.125 | -0.008 | 0 | 0 | 0.427 | 0.59 | 0.43 | 0.68 | 0.44 | 0.77 | 0.7 |
| 4 | 164992016 | rs971945 | -0.226 | -2.565 | 0.27 | -0.227 | 0.019 | -0.022 | -0.022 | 0 | 0.75 | 0.79 | 0.75 | 0.49 | 0.86 | 0.73 |
| 4 | 164996728 | rs2321210 | -0.191 | -2.593 | 0.319 | -0.188 | 0.017 | -0.02 | -0.02 | 0 | 0.75 | 0.79 | 0.75 | 0.49 | 0.86 | 0.72 |
| 4 | 165013565 | rs919574 | -0.021 | -2.623 | 0.447 | -0.003 | -0.043 | 0.038 | 0.038 | 0.012 | 0.76 | 0.67 | 0.77 | 0.49 | 0.89 | 0.74 |
| 4 | 165037553 | rs4691956 | 0.517 | -2.84 | 0.567 | -0.108 | -0.049 | -0.002 | -0.002 | 0.124 | 0.61 | 0.41 | 0.71 | 0.47 | 0.81 | 0.72 |
| 4 | 165049116 | rs7691235 | 0.523 | -2.84 | 0.568 | -0.09 | -0.05 | -0.002 | -0.002 | 0.314 | 0.61 | 0.41 | 0.71 | 0.48 | 0.81 | 0.72 |
| 4 | 165052223 | rs4691961 | 1.131 | -2.419 | 0.283 | -0.253 | -0.001 | -0.001 | -0.001 | 0 | 0.42 | 0.39 | 0.56 | 0.44 | 0.54 | 0.58 |
| 4 | 165052304 | rs17579243 | 0.775 | -2.319 | -0.137 | -0.405 | -0.075 | 0.099 | 0.099 | 0.2 | 0.19 | 0.03 | 0.16 | 0.03 | 0.27 | 0.14 |
| 4 | 165057027 | rs2195697 | 1.571 | -2.305 | 0.063 | -0.403 | -0.077 | 0.093 | 0.093 | 0 | 0.35 | 0.18 | 0.49 | 0.24 | 0.47 | 0.45 |
| 4 | 165060898 | rs982257 | 0.571 | -2.494 | 0.799 | -0.32 | -0.007 | -0.002 | -0.002 | 0 | 0.67 | 0.35 | 0.52 | 0.34 | 0.82 | 0.62 |
| 4 | 165076871 | rs1430986 | 0.84 | -2.551 | 0.649 | -0.317 | 0.13 | 0.131 | 0.131 | 0.344 | 0.53 | 0.25 | 0.47 | 0.26 | 0.73 | 0.59 |
| 4 | 165087375 | rs1430997 | 1.773 | -2.457 | 0.556 | -0.316 | -0.087 | 0.086 | 0.086 | 0 | 0.33 | 0.15 | 0.31 | 0.22 | 0.46 | 0.44 |
| 4 | 165093156 | rs17044711 | -0.013 | -2.45 | 0.957 | -0.067 | 0.012 | 0.079 | 0.079 | 0.096 | 0.07 | 0.18 | 0.23 | 0.17 | 0.08 | 0.14 |
| 4 | 165108381 | rs2163395 | -0.399 | -2.678 | 1.11 | -0.069 | 0.027 | -0.029 | -0.029 | 0 | 0.77 | 0.82 | 0.78 | 0.5 | 0.9 | 0.74 |
| 4 | 165108975 | rs1430976 | 0.086 | -2.79 | 1.251 | -0.131 | -0.003 | -0.003 | -0.003 | 0 | 0.64 | 0.65 | 0.72 | 0.5 | 0.82 | 0.72 |
| 4 | 165109476 | rs28710221 | 1.718 | -2.37 | 1.18 | 0.03 | 0.044 | 0.044 | 0.044 | 0.183 | 0.12 | 0.06 | 0.04 | 0 | 0.08 | 0.02 |
| 4 | 165111673 | rs1835507 | -0.906 | -2.337 | 1.09 | 0.14 | 0.007 | 0.1 | 0.1 | 1.089 | 0.09 | 0.3 | 0.24 | 0.17 | 0.08 | 0.2 |
| 4 | 165117009 | rs1430974 | -0.962 | -2.342 | 1.087 | 0.137 | 0.004 | 0.096 | 0.096 | 1.092 | 0.1 | 0.3 | 0.23 | 0.17 | 0.08 | 0.2 |
| 4 | 165118652 | rs2288675 | 1.912 | -2.405 | 0.411 | -0.046 | -0.127 | 0.107 | 0.107 | 0.255 | 0.33 | 0.11 | 0.31 | 0.22 | 0.47 | 0.42 |
| 4 | 165129362 | rs4502640 | -0.154 | -2.434 | 0.58 | 0.34 | -0.025 | 0.002 | 0.002 | 0.39 | 0.23 | 0.3 | 0.32 | 0.23 | 0.16 | 0.23 |
| 4 | 165131350 | rs6814420 | -0.069 | -2.37 | 0.04 | 0.961 | -0.036 | -0.05 | -0.05 | 0 | 0.67 | 0.92 | 0.77 | 0.65 | 0.68 | 0.71 |
| 4 | 165139274 | rs9968537 | 1.12 | -2.083 | 0.342 | 0.909 | -0.001 | 0 | 0 | 0.802 | 0.41 | 0.45 | 0.42 | 0.29 | 0.47 | 0.47 |
| 4 | 165145428 | rs7668234 | -0.318 | -2.412 | 0.355 | 1.237 | 0.016 | -0.02 | -0.02 | 0.368 | 0.87 | 0.95 | 0.93 | 0.83 | 0.94 | 0.9 |
| 4 | 165153650 | rs2321373 | -0.38 | -2.411 | 0.344 | 0.95 | 0.016 | 0 | 0 | 0 | 0.87 | 0.96 | 0.93 | 0.88 | 0.94 | 0.9 |
| 4 | 165161269 | rs17676292 | -0.455 | -2.514 | 0.554 | 0.938 | -0.006 | 0.01 | 0.01 | 0.068 | 0.3 | 0.34 | 0.45 | 0.39 | 0.22 | 0.33 |
| 4 | 165168921 | rs4561891 | 0.608 | -2.678 | 0.487 | 0.978 | -0.049 | -0.045 | -0.045 | 0 | 0.73 | 0.66 | 0.86 | 0.88 | 0.85 | 0.89 |
| 4 | 165170408 | rs4496540 | -0.211 | -2.49 | 0.191 | 1.081 | 0.02 | -0.002 | -0.002 | 0.144 | 0.87 | 0.96 | 0.94 | 0.88 | 0.94 | 0.9 |
| 4 | 165173550 | rs10222965 | -0.332 | -2.471 | 0.2 | 1.09 | 0.018 | -0.003 | -0.003 | 0 | 0.87 | 0.97 | 0.94 | 0.88 | 0.94 | 0.9 |
| 4 | 165202419 | rs1985065 | 0.525 | -2.323 | 0.404 | 1.064 | -0.024 | 0.01 | 0.01 | 0 | 0.61 | 0.48 | 0.48 | 0.48 | 0.76 | 0.59 |
| 4 | 165202724 | rs2321377 | 1.258 | -2.412 | 0.41 | 0.804 | 0.096 | 0.073 | 0.073 | 0.05 | 0.46 | 0.16 | 0.37 | 0.35 | 0.67 | 0.54 |
| 4 | 165202782 | rs2321376 | 0.6 | -2.355 | 0.38 | 0.837 | -0.022 | -0.012 | -0.012 | 0 | 0.6 | 0.48 | 0.45 | 0.35 | 0.74 | 0.55 |
| 4 | 165209309 | rs17614438 | 0.911 | -2.123 | -0.423 | 0.638 | 0.108 | 0.027 | 0.027 | 0.223 | 0.51 | 0.22 | 0.4 | 0.44 | 0.68 | 0.56 |
| 4 | 165214649 | rs11941402 | 0.455 | -1.556 | -0.315 | 0.891 | -0.016 | 0.008 | 0.008 | 0.291 | 0.62 | 0.6 | 0.48 | 0.44 | 0.75 | 0.57 |
| 4 | 165221971 | rs6829588 | -1.103 | -1.812 | -0.43 | 0.948 | 0.064 | -0.01 | -0.01 | 0 | 0.29 | 0.76 | 0.25 | 0.24 | 0.13 | 0.12 |
| 4 | 165222183 | rs6829484 | 1.063 | -1.022 | 0.263 | 1.103 | -0.022 | 0.021 | 0.021 | 0 | 0.73 | 0.44 | 0.74 | 0.72 | 0.8 | 0.9 |
| 4 | 165232273 | rs17044895 | -0.274 | -0.719 | 0.781 | 0.707 | 0.001 | -0.027 | -0.027 | 0 | 0.1 | 0.08 | 0.1 | 0.01 | 0.12 | 0.07 |
| 4 | 165234545 | rs7675835 | -1.058 | -1.175 | 1.132 | 1.371 | -0.028 | 0.147 | 0.147 | 0 | 0.12 | 0.22 | 0.33 | 0.4 | 0.07 | 0.31 |
| 4 | 165234678 | rs7675571 | -0.798 | -1.239 | 0.705 | 0.693 | 0.011 | -0.005 | -0.005 | 0 | 0.59 | 0.65 | 0.8 | 0.58 | 0.67 | 0.65 |
| 4 | 165236998 | rs4434204 | 0.98 | -1.275 | 0.814 | 0.647 | -0.087 | 0.122 | 0.122 | 0 | 0.36 | 0.14 | 0.35 | 0.16 | 0.48 | 0.26 |
| 4 | 165237207 | rs4622958 | -0.771 | -1.262 | 1.168 | 0.665 | 0.018 | -0.016 | -0.016 | 0 | 0.6 | 0.71 | 0.8 | 0.58 | 0.67 | 0.65 |

| | | | | | | | | | | | | | | | | |
|---|---|---|---|---|---|---|---|---|---|---|---|---|---|---|---|---|
| 4 | 165239116 | rs7680916 | -0.367 | -1.08 | 0.828 | 0.786 | -0.028 | 0.149 | 0.149 | 0 | 0.12 | 0.22 | 0.33 | 0.4 | 0.07 | 0.32 |
| 4 | 165254752 | rs7669453 | 0.425 | -1.161 | 0.818 | 0.961 | -0.02 | 0.008 | 0.008 | 0 | 0.39 | 0.31 | 0.37 | 0.34 | 0.51 | 0.35 |
| 4 | 165262655 | rs7662252 | -0.385 | -1.159 | 0.739 | 0.983 | 0.003 | 0.009 | 0.009 | 0 | 0.49 | 0.43 | 0.69 | 0.6 | 0.56 | 0.56 |
| 4 | 165284574 | rs4431170 | -0.18 | -1.088 | 0.204 | 0.935 | 0.013 | -0.014 | -0.014 | 0.608 | 0.09 | 0.06 | 0.31 | 0.26 | 0.05 | 0.22 |
| 4 | 165305889 | rs7658861 | -0.56 | -1.293 | 0.181 | 0.04 | 0.02 | 0.06 | 0.06 | 0 | 0.48 | 0.65 | 0.62 | 0.78 | 0.39 | 0.65 |
| 4 | 165307934 | rs1027755 | -0.73 | -1.22 | -0.306 | -0.075 | 0.001 | 0.053 | 0.053 | 0 | 0.86 | 0.97 | 0.96 | 0.99 | 0.88 | 0.92 |
| 4 | 165314229 | rs17483750 | 0.759 | -1.272 | -0.341 | 0.189 | -0.123 | -0.06 | -0.06 | 0 | 0.38 | 0.12 | 0.33 | 0.37 | 0.5 | 0.35 |
| 4 | 165315996 | rs4691974 | 0.895 | -1.265 | -0.346 | 0.267 | -0.123 | 0.04 | 0.04 | 0 | 0.36 | 0.12 | 0.33 | 0.25 | 0.5 | 0.27 |
| 4 | 165317260 | rs12510747 | 0.914 | -1.665 | -0.234 | 0.143 | 0.02 | -0.07 | -0.07 | 0.109 | 0.25 | 0.13 | 0.39 | 0.47 | 0.16 | 0.46 |
| 4 | 165317732 | rs6835711 | 0.747 | -1.713 | 0.602 | 0.134 | 0.006 | 0.005 | 0.005 | 0.045 | 0.42 | 0.47 | 0.39 | 0.49 | 0.51 | 0.37 |
| 4 | 165322933 | rs9994016 | 0.99 | -1.701 | 0.384 | 0.118 | -0.026 | 0.016 | 0.016 | 0 | 0.4 | 0.34 | 0.38 | 0.38 | 0.5 | 0.27 |
| 4 | 165324215 | rs11100552 | 0.827 | -1.485 | 0.172 | 0.123 | 0.024 | 0.039 | 0.039 | 0.64 | 0.61 | 0.33 | 0.73 | 0.72 | 0.66 | 0.72 |
| 4 | 165329158 | rs1605057 | -0.438 | -1.777 | 0.229 | -0.141 | 0.028 | 0.014 | 0.014 | 0 | 0.43 | 0.64 | 0.52 | 0.59 | 0.32 | 0.54 |
| 4 | 165329655 | rs10004121 | -0.269 | -1.761 | 0.461 | -0.133 | -0.002 | -0.001 | -0.001 | 0 | 0.41 | 0.51 | 0.52 | 0.59 | 0.32 | 0.54 |
| 4 | 165351210 | rs1566934 | 0.235 | -2.164 | -0.253 | -0.539 | -0.009 | 0.002 | 0.002 | 3.354 | 0.37 | 0.43 | 0.41 | 0.39 | 0.51 | 0.4 |
| 4 | 165351616 | rs4691118 | 1.22 | -2.351 | -0.288 | -0.58 | 0.016 | -0.052 | -0.052 | 0 | 0.26 | 0.15 | 0.4 | 0.51 | 0.18 | 0.4 |
| 4 | 165353749 | rs1505383 | -0.375 | -2.35 | -0.282 | -0.585 | 0.076 | 0.092 | 0.092 | 0.453 | 0.5 | 0.86 | 0.53 | 0.4 | 0.64 | 0.43 |
| 4 | 165355188 | rs6536816 | 1.502 | -2.305 | -0.15 | -0.577 | 0.006 | -0.04 | -0.04 | 0.39 | 0.1 | 0.09 | 0.1 | 0.01 | 0.12 | 0.03 |
| 4 | 165361556 | rs11931372 | 0.633 | -2.314 | -0.317 | -0.568 | 0.053 | 0.022 | 0.022 | 0.395 | 0.13 | 0.39 | 0.12 | 0.09 | 0.13 | 0.09 |
| 4 | 165364934 | rs2874568 | -0.242 | -2.337 | -0.362 | -0.584 | -0.02 | -0.07 | -0.07 | 0.46 | 0.64 | 0.58 | 0.81 | 0.9 | 0.69 | 0.8 |
| 4 | 165370126 | rs1857798 | -0.48 | -2.181 | -0.872 | -0.511 | 0.068 | -0.016 | -0.016 | 0 | 0.51 | 0.89 | 0.53 | 0.48 | 0.64 | 0.5 |
| 4 | 165386097 | rs2089223 | 1.552 | -0.466 | -0.988 | -0.753 | -0.024 | 0.033 | 0.033 | 0 | 0.11 | 0.04 | 0.11 | 0.02 | 0.17 | 0.07 |
| 4 | 165392008 | rs4691121 | -0.312 | -0.745 | -0.78 | -0.95 | -0.004 | -0.063 | -0.063 | 0 | 0.63 | 0.62 | 0.82 | 0.95 | 0.67 | 0.81 |
| 4 | 165395936 | rs1960980 | 0.557 | -0.843 | -0.936 | -0.867 | 0.005 | 0.035 | 0.035 | 0 | 0.15 | 0.29 | 0.15 | 0.2 | 0.13 | 0.12 |
| 4 | 165400016 | rs1394807 | 1.356 | -0.595 | -1.989 | -0.733 | 0.005 | -0.135 | -0.135 | 1.304 | 0.27 | 0.09 | 0.41 | 0.51 | 0.2 | 0.43 |
| 4 | 165403992 | rs7690372 | 0.438 | -0.608 | -1.993 | -0.748 | -0.03 | -0.064 | -0.064 | 0 | 0.39 | 0.14 | 0.52 | 0.52 | 0.36 | 0.52 |
| 4 | 165406393 | rs12650464 | 0.98 | -0.576 | -1.944 | -0.446 | 0 | -0.001 | -0.001 | 0 | 0.21 | 0.23 | 0.25 | 0.26 | 0.34 | 0.28 |
| 4 | 165409295 | rs6834683 | 0.988 | -0.552 | -2.409 | -0.292 | 0.01 | -0.14 | -0.14 | 0.069 | 0.27 | 0.09 | 0.41 | 0.5 | 0.2 | 0.45 |
| 4 | 165417351 | rs1028212 | 1.09 | -0.733 | -2.382 | -0.337 | 0.058 | -0.129 | -0.129 | 0 | 0.26 | 0.08 | 0.4 | 0.5 | 0.16 | 0.42 |
| 4 | 165422101 | rs1505392 | -0.702 | -0.059 | -1.748 | 0.176 | 0.027 | 0.004 | 0.004 | 0.029 | 0.75 | 0.82 | 0.88 | 0.76 | 0.84 | 0.8 |
| 4 | 165425204 | rs1505390 | 0.124 | 1.368 | -1.421 | 0.429 | -0.034 | -0.008 | -0.008 | 0 | 0.19 | 0.11 | 0.21 | 0.22 | 0.32 | 0.21 |
| 4 | 165434864 | rs13128973 | -0.006 | 1.768 | -1.478 | 0.663 | -0.056 | -0.002 | -0.002 | 0.174 | 0.25 | 0.13 | 0.3 | 0.26 | 0.42 | 0.28 |
| 4 | 165440926 | rs7654643 | -0.392 | 1.246 | -0.596 | 1.544 | 0.061 | 0.215 | 0.215 | 0 | 0.12 | 0.27 | 0.37 | 0.36 | 0.18 | 0.26 |
| 4 | 165446252 | rs12648459 | 0.234 | -0.348 | -0.587 | 0.793 | -0.015 | -0.013 | -0.013 | 0 | 0.2 | 0.21 | 0.08 | 0.16 | 0.11 | 0.14 |
| 4 | 165448658 | rs6839763 | 0.062 | -1.506 | -0.912 | -0.152 | 0.016 | 0.021 | 0.021 | 0 | 0.48 | 0.67 | 0.57 | 0.57 | 0.41 | 0.47 |
| 4 | 165452112 | rs4597781 | -0.485 | -2.135 | -0.833 | -0.224 | -0.033 | 0.053 | 0.053 | 0 | 0.63 | 0.83 | 0.63 | 0.74 | 0.5 | 0.59 |
| 4 | 165452601 | rs12650712 | 0.864 | -3.933 | -1.21 | -0.647 | 0.017 | 0.021 | 0.021 | 1.243 | 0.21 | 0.14 | 0.15 | 0.17 | 0.16 | 0.17 |
| 4 | 165452969 | rs10001210 | -1.163 | -3.393 | -1.116 | -0.487 | -0.006 | 0 | 0 | 0.909 | 0.74 | 0.77 | 0.67 | 0.74 | 0.58 | 0.71 |
| 4 | 165456150 | rs12508710 | 0.102 | -3.24 | -1.25 | -0.469 | 0.007 | -0.004 | -0.004 | 0.305 | 0.36 | 0.3 | 0.43 | 0.54 | 0.31 | 0.42 |
| 4 | 165458952 | rs4079693 | 0.12 | -2.106 | 0.478 | 0.231 | -0.008 | 0.238 | 0.238 | 0 | 0.08 | 0.17 | 0.28 | 0.37 | 0.05 | 0.19 |
| 4 | 165471242 | rs6536824 | -0.859 | -1.798 | 0.072 | -0.536 | -0.042 | 0.005 | 0.005 | 0 | 0.77 | 0.89 | 0.7 | 0.81 | 0.6 | 0.72 |
| 4 | 165483847 | rs7692840 | 0.261 | -2.766 | 0.997 | -0.19 | -0.047 | 0.004 | 0.004 | 0 | 0.32 | 0.45 | 0.39 | 0.56 | 0.18 | 0.41 |
| 4 | 165484115 | rs7693270 | 0.518 | -2.768 | 1.128 | -0.24 | 0 | -0.011 | -0.011 | 0 | 0.7 | 0.72 | 0.82 | 0.84 | 0.72 | 0.88 |
| 4 | 165491839 | rs1110968 | 0.409 | -2.323 | 0.704 | -0.034 | 0.003 | 0.003 | 0.003 | 1.256 | 0.45 | 0.43 | 0.48 | 0.31 | 0.64 | 0.5 |
| 4 | 165492082 | rs1528698 | 0.448 | -2.304 | 1.251 | 0.534 | 0.01 | 0.01 | 0.01 | 0.129 | 0.53 | 0.44 | 0.53 | 0.31 | 0.74 | 0.51 |
| 4 | 165498202 | rs17045152 | -0.628 | -1.565 | 2.021 | 1.252 | -0.004 | 0.002 | 0.002 | 1.057 | 0.19 | 0.2 | 0.09 | 0.14 | 0.09 | 0.09 |
| 4 | 165499278 | rs10517809 | 0.179 | -1.548 | 1.586 | 1.257 | 0.044 | 0.213 | 0.213 | 0.071 | 0.93 | 0.48 | 0.93 | 0.75 | 0.97 | 0.94 |
| 4 | 165513998 | rs1534577 | 0.677 | -1.425 | 0.687 | 1.179 | 0.059 | -0.015 | -0.015 | 0 | 0.72 | 0.17 | 0.83 | 0.6 | 0.88 | 0.86 |
| 4 | 165514302 | rs1534578 | 1.115 | -1.387 | 0.613 | 1.228 | 0.077 | -0.075 | -0.075 | 0.2 | 0.64 | 0.08 | 0.77 | 0.49 | 0.77 | 0.73 |
| 4 | 165515604 | rs13114673 | 0.813 | -1.635 | 1.005 | 0.913 | 0.059 | -0.01 | -0.01 | 0 | 0.72 | 0.17 | 0.83 | 0.62 | 0.88 | 0.87 |
| 4 | 165560063 | rs9790550 | -1.084 | -1.285 | 0.243 | 1.042 | -0.04 | 0.006 | 0.006 | 0.374 | 0.23 | 0.38 | 0.17 | 0.28 | 0.12 | 0.17 |
| 4 | 165561062 | rs17045285 | -0.7 | -1.467 | 0.304 | 1.3 | -0.013 | -0.001 | -0.001 | 0 | 0.16 | 0.22 | 0.06 | 0.15 | 0.03 | 0.11 |
| 4 | 165561662 | rs9991808 | -1.266 | -2.545 | 0.078 | 1.257 | -0.044 | 0.015 | 0.015 | 0.044 | 0.22 | 0.57 | 0.19 | 0.29 | 0.13 | 0.11 |
| 4 | 165562187 | rs17678153 | -0.63 | -2.44 | -0.332 | 1.906 | -0.001 | 0.101 | 0.101 | 0 | 0.07 | 0.06 | 0.09 | 0.43 | 0.12 | 0.19 |
| 4 | 165564515 | rs6824404 | -1.226 | -2.283 | -0.343 | 0.584 | 0 | -0.025 | -0.025 | 0 | 0.45 | 0.46 | 0.45 | 0.86 | 0.49 | 0.27 |
| 4 | 165564711 | rs17678175 | -0.647 | -2.249 | -0.484 | 0.794 | -0.023 | -0.069 | -0.069 | 0 | 0.14 | 0.03 | 0.23 | 0.69 | 0.22 | 0.17 |
| 4 | 165564737 | rs11936514 | -1.773 | -2.353 | -0.591 | 0.547 | 0.003 | 0.037 | 0.037 | 0.021 | 0.52 | 0.6 | 0.47 | 0.89 | 0.52 | 0.43 |
| 4 | 165576740 | rs12499842 | 1.167 | -2.469 | -0.543 | 0.894 | 0.006 | -0.017 | -0.017 | 1.367 | 0.74 | 0.68 | 0.76 | 0.41 | 0.74 | 0.69 |
| 4 | 165578100 | rs12651422 | -0.338 | -2.475 | -0.579 | 0.93 | -0.017 | -0.078 | -0.078 | 0 | 0.13 | 0.03 | 0.18 | 0.55 | 0.2 | 0.15 |
| 4 | 165579891 | rs1406365 | -0.148 | -2.476 | -0.456 | 0.87 | -0.001 | -0.051 | -0.051 | 0 | 0.27 | 0.41 | 0.21 | 0.17 | 0.26 | 0.1 |
| 4 | 165582170 | rs11932404 | -0.987 | -2.431 | -0.449 | 0.89 | -0.017 | -0.042 | -0.042 | 0 | 0.39 | 0.49 | 0.23 | 0.19 | 0.29 | 0.2 |
| 4 | 165585096 | rs1528690 | 0.643 | -2.367 | -0.438 | 0.939 | 0 | 0.069 | 0.069 | 0 | 0.83 | 0.73 | 0.77 | 0.43 | 0.77 | 0.8 |
| 4 | 165585563 | rs12507345 | 1.267 | -2.336 | -0.093 | 0.766 | 0.002 | 0.075 | 0.075 | 0.043 | 0.48 | 0.29 | 0.55 | 0.24 | 0.48 | 0.6 |
| 4 | 165605068 | rs10001932 | 0.186 | -3.182 | 0.019 | 1.038 | 0.003 | -0.01 | -0.01 | 0 | 0.65 | 0.74 | 0.7 | 0.47 | 0.66 | 0.72 |
| 4 | 165607255 | rs7680144 | 0.924 | -3.594 | 0.435 | 0.906 | 0.006 | 0.005 | 0.005 | 0 | 0.52 | 0.6 | 0.63 | 0.35 | 0.58 | 0.55 |
| 4 | 165607558 | rs11724238 | 1.527 | -3.831 | 0.611 | 0.944 | 0.006 | 0.026 | 0.026 | 0 | 0.43 | 0.44 | 0.58 | 0.25 | 0.52 | 0.54 |
| 4 | 165609979 | rs4691129 | 0.724 | -3.845 | 0.63 | 0.984 | -0.012 | -0.007 | -0.007 | 0 | 0.57 | 0.61 | 0.65 | 0.38 | 0.6 | 0.7 |
| 4 | 165613783 | rs6536843 | 1.399 | -3.485 | 0.766 | 0.961 | -0.001 | -0.007 | -0.007 | 0 | 0.43 | 0.53 | 0.57 | 0.25 | 0.51 | 0.52 |
| 4 | 165614362 | rs11935111 | -0.101 | -3.686 | 0.859 | 1.095 | 0 | -0.001 | -0.001 | 0 | 0.3 | 0.25 | 0.21 | 0.49 | 0.3 | 0.24 |

| | | | | | | | | | | | | | | | | |
|---|---|---|---|---|---|---|---|---|---|---|---|---|---|---|---|---|
| 4 | 165614650 | rs6536847 | 0.948 | -3.677 | 0.859 | 0.979 | 0.016 | 0.01 | 0.01 | 0 | 0.51 | 0.63 | 0.62 | 0.37 | 0.57 | 0.54 |
| 4 | 165617850 | rs11939098 | 0.021 | -3.637 | 0.706 | 0.996 | -0.018 | 0.03 | 0.03 | 0 | 0.09 | 0.1 | 0.05 | 0.12 | 0.06 | 0.02 |
| 4 | 165618268 | rs7671059 | 0.021 | -3.637 | 0.706 | 0.996 | -0.018 | 0.03 | 0.03 | 0 | 0.09 | 0.1 | 0.05 | 0.12 | 0.06 | 0.02 |
| 4 | 165618371 | rs17582753 | 2.352 | -3.886 | 0.691 | 0.821 | -0.084 | 0.042 | 0.042 | 0 | 0.35 | 0.09 | 0.53 | 0.24 | 0.5 | 0.52 |
| 4 | 165621955 | rs10000041 | 0.598 | -3.875 | 0.913 | 0.68 | -0.053 | -0.051 | -0.051 | 1.452 | 0.73 | 0.62 | 0.79 | 0.84 | 0.86 | 0.76 |
| 4 | 165626856 | rs6830070 | 0.796 | -3.582 | 0.84 | 0.823 | -0.045 | 0.042 | 0.042 | 0 | 0.87 | 0.71 | 0.93 | 0.85 | 0.93 | 0.98 |
| 4 | 165627020 | rs6536849 | 0.796 | -3.582 | 0.84 | 0.823 | -0.045 | 0.042 | 0.042 | 0 | 0.87 | 0.71 | 0.93 | 0.85 | 0.93 | 0.98 |
| 4 | 165628168 | rs17045377 | -0.459 | -3.917 | 0.262 | -0.287 | -0.01 | -0.046 | -0.046 | 0 | 0.4 | 0.5 | 0.29 | 0.12 | 0.31 | 0.36 |
| 4 | 165628814 | rs1980391 | 1.415 | -4.076 | 0.172 | -0.085 | 0.05 | 0.037 | 0.037 | 0 | 0.44 | 0.2 | 0.64 | 0.68 | 0.63 | 0.62 |
| 4 | 165629466 | rs9308087 | 0.269 | -4.208 | 0.326 | -0.225 | -0.025 | -0.121 | -0.121 | 0 | 0.58 | 0.44 | 0.73 | 0.95 | 0.72 | 0.76 |
| 4 | 165630021 | rs12640091 | 2.708 | -4.64 | -0.555 | -0.126 | -0.178 | -0.07 | -0.07 | 0 | 0.24 | 0.02 | 0.53 | 0.24 | 0.48 | 0.41 |
| 4 | 165634055 | rs13120623 | 1.055 | -4.51 | -0.342 | 0.118 | -0.017 | -0.102 | -0.102 | 6.581 | 0.15 | 0.01 | 0.07 | 0.38 | 0.13 | 0.2 |
| 4 | 165638228 | rs10001077 | 0.131 | -4.272 | 0.283 | 0.041 | -0.019 | 0.138 | 0.138 | 0 | 0.07 | 0.16 | 0.04 | 0.14 | 0.05 | 0.02 |
| 4 | 165642091 | rs7677517 | 0.028 | -4.483 | 0.18 | -0.194 | -0.016 | -0.013 | -0.013 | 0.436 | 0.27 | 0.42 | 0.16 | 0.14 | 0.23 | 0.07 |
| 4 | 165644827 | rs7696505 | 0.998 | -4.448 | 0.02 | -0.25 | -0.001 | -0.034 | -0.034 | 0 | 0.2 | 0.19 | 0.12 | 0 | 0.18 | 0.05 |
| 4 | 165646228 | rs17622247 | -1.003 | -4.269 | 0.057 | 0.1 | 0.024 | -0.038 | -0.038 | 0 | 0.12 | 0.03 | 0.09 | 0.27 | 0.1 | 0.14 |
| 4 | 165646778 | rs1113272 | -0.983 | -4.265 | 0.057 | 0.101 | 0.022 | -0.037 | -0.037 | 1.517 | 0.12 | 0.03 | 0.09 | 0.27 | 0.1 | 0.14 |
| 4 | 165647164 | rs1881523 | 0.346 | -4.507 | 0.221 | -0.245 | -0.007 | -0.089 | -0.089 | 0.232 | 0.34 | 0.43 | 0.23 | 0.01 | 0.26 | 0.23 |
| 4 | 165647923 | rs4692007 | 1.917 | -4.048 | 0.321 | 0.452 | -0.025 | 0.002 | 0.002 | 2.575 | 0.26 | 0.15 | 0.52 | 0.22 | 0.47 | 0.4 |
| 4 | 165649008 | rs17045434 | 0.863 | -4.084 | 0.339 | 0.447 | 0.011 | -0.001 | -0.001 | 0.395 | 0.06 | 0.02 | 0.03 | 0.14 | 0.05 | 0.02 |
| 4 | 165649271 | rs2056759 | 1.192 | -4.093 | 0.344 | 0.469 | -0.023 | -0.014 | -0.014 | 0 | 0.32 | 0.18 | 0.6 | 0.37 | 0.53 | 0.49 |
| 4 | 165649300 | rs2056760 | 1.924 | -3.937 | 0.323 | 0.448 | -0.024 | -0.002 | -0.002 | 2.302 | 0.25 | 0.15 | 0.55 | 0.25 | 0.46 | 0.37 |
| 4 | 165656200 | rs4691131 | 0.751 | -3.983 | 0.612 | 0.43 | 0.013 | 0.005 | 0.005 | 2.542 | 0.34 | 0.44 | 0.62 | 0.37 | 0.53 | 0.49 |
| 4 | 165658642 | rs12502646 | 1.233 | -3.839 | 0.029 | 0.493 | -0.028 | -0.03 | -0.03 | 0 | 0.31 | 0.17 | 0.57 | 0.39 | 0.51 | 0.38 |
| 4 | 165669160 | rs4254724 | -0.498 | -3.525 | 0.127 | 0.504 | 0.002 | 0.012 | 0.012 | 0 | 0.08 | 0.18 | 0.09 | 0.12 | 0.08 | 0.13 |
| 4 | 165682010 | rs10006449 | 0.369 | -3.459 | 0.156 | 0.451 | -0.017 | -0.031 | -0.031 | 1.459 | 0.39 | 0.27 | 0.6 | 0.43 | 0.53 | 0.42 |
| 4 | 165683452 | rs4593086 | -0.275 | -3.47 | 0.195 | 0.446 | -0.002 | -0.001 | -0.001 | 0 | 0.47 | 0.54 | 0.69 | 0.56 | 0.61 | 0.55 |
| 4 | 165683917 | rs17045455 | -0.562 | -3.232 | -0.038 | 0.484 | 0.03 | 0.048 | 0.048 | 0 | 0.08 | 0.26 | 0.09 | 0.12 | 0.09 | 0.13 |
| 4 | 165686723 | rs11732101 | 1.612 | -3.071 | -0.751 | 0.35 | -0.115 | -0.03 | -0.03 | 0.432 | 0.24 | 0.06 | 0.53 | 0.25 | 0.46 | 0.37 |
| 4 | 165692104 | rs11100576 | 0.266 | -2.66 | -0.416 | 0.284 | 0.001 | -0.001 | -0.001 | 0 | 0.45 | 0.48 | 0.67 | 0.47 | 0.58 | 0.52 |
| 4 | 165697040 | rs10155303 | 0.595 | -0.293 | -0.438 | 0.282 | -0.001 | -0.001 | -0.001 | 0 | 0.57 | 0.53 | 0.39 | 0.58 | 0.48 | 0.62 |
| 4 | 165698208 | rs13113465 | 0.634 | -0.422 | -0.298 | 0.237 | -0.001 | 0 | 0 | 0 | 0.57 | 0.52 | 0.4 | 0.59 | 0.47 | 0.62 |
| 4 | 165700679 | rs6841926 | -1.014 | -1.32 | -0.999 | -0.239 | 0.103 | 0.086 | 0.086 | 1.663 | 0.31 | 0.89 | 0.17 | 0.08 | 0.12 | 0.18 |
| 4 | 165707728 | rs4539996 | 0.46 | -0.048 | -1.228 | -0.178 | -0.032 | -0.029 | -0.029 | 2.154 | 0.56 | 0.83 | 0.48 | 0.5 | 0.53 | 0.49 |
| 4 | 165710503 | rs7668032 | -0.002 | 0.18 | -0.564 | 0.015 | 0 | 0.052 | 0.052 | 0.593 | 0.76 | 0.68 | 0.6 | 0.56 | 0.73 | 0.73 |
| 4 | 165721118 | rs13152512 | 0.239 | 0.231 | -0.445 | -0.013 | -0.003 | 0.104 | 0.104 | 2.047 | 0.74 | 0.5 | 0.58 | 0.56 | 0.73 | 0.75 |
| 4 | 165726037 | rs17045582 | -0.789 | 0.343 | -0.218 | -0.872 | 0.003 | -0.051 | -0.051 | 0 | 0.14 | 0.22 | 0.13 | 0.03 | 0.14 | 0.08 |
| 4 | 165726126 | rs13126014 | 0.125 | 0.462 | -0.47 | -0.987 | -0.003 | 0.061 | 0.061 | 0 | 0.26 | 0.11 | 0.14 | 0.08 | 0.21 | 0.26 |
| 4 | 165726289 | rs17486421 | -0.401 | 0.642 | -0.481 | -0.995 | -0.02 | 0.023 | 0.023 | 0 | 0.05 | 0.04 | 0.04 | 0 | 0.07 | 0.03 |
| 4 | 165726891 | rs4572830 | 1.743 | 0.838 | -0.43 | -0.852 | -0.024 | -0.004 | -0.004 | 0 | 0.43 | 0.15 | 0.6 | 0.61 | 0.47 | 0.48 |
| 4 | 165728860 | rs4518194 | 1.124 | 0.98 | -0.28 | -0.794 | 0.01 | 0.032 | 0.032 | 0.099 | 0.48 | 0.27 | 0.65 | 0.6 | 0.57 | 0.51 |
| 4 | 165730192 | rs10776539 | -0.836 | 0.984 | 0.069 | -0.77 | 0.002 | 0.075 | 0.075 | 1.713 | 0.83 | 0.72 | 0.87 | 0.71 | 0.83 | 0.83 |
| 4 | 165737589 | rs7683083 | -1.09 | 1.063 | 0.042 | -0.816 | 0 | -0.028 | -0.028 | 0 | 0.38 | 0.49 | 0.27 | 0.11 | 0.36 | 0.34 |
| 4 | 165739878 | rs6536867 | -0.755 | 0.808 | 0.029 | -0.546 | -0.001 | -0.005 | -0.005 | 0.166 | 0.33 | 0.4 | 0.21 | 0.09 | 0.29 | 0.31 |
| 4 | 165740749 | rs6536868 | -0.345 | 0.812 | -0.018 | -0.497 | -0.002 | -0.016 | -0.016 | 0 | 0.27 | 0.36 | 0.16 | 0.09 | 0.23 | 0.28 |
| 4 | 165742640 | rs13139212 | 0.108 | 0.685 | -0.377 | -0.453 | 0.028 | 0.045 | 0.045 | 0 | 0.23 | 0.12 | 0.11 | 0.08 | 0.15 | 0.24 |
| 4 | 165742801 | rs11723570 | 0.433 | 0.941 | -0.335 | -0.204 | 0.002 | 0.031 | 0.031 | 0 | 0.5 | 0.28 | 0.66 | 0.62 | 0.54 | 0.51 |
| 4 | 165744136 | rs4339166 | 0.607 | 1.118 | -0.966 | -0.285 | -0.007 | 0.037 | 0.037 | 0 | 0.48 | 0.3 | 0.66 | 0.66 | 0.54 | 0.52 |
| 4 | 165752701 | rs4234980 | -0.143 | 1.183 | -0.969 | -0.516 | -0.025 | 0.043 | 0.043 | 0 | 0.89 | 0.98 | 0.93 | 1 | 0.85 | 0.99 |
| 4 | 165756691 | rs6827249 | -0.629 | 0.943 | -0.711 | -0.359 | 0.002 | -0.011 | -0.011 | 0 | 0.44 | 0.64 | 0.38 | 0.26 | 0.53 | 0.34 |
| 4 | 165767472 | rs4593087 | -0.786 | 0.958 | -0.976 | -0.637 | -0.008 | 0.058 | 0.058 | 0 | 0.46 | 0.65 | 0.38 | 0.17 | 0.54 | 0.33 |
| 4 | 165771791 | rs4346606 | -0.753 | 1.041 | -1.413 | -0.121 | -0.032 | -0.005 | -0.005 | 0 | 0.23 | 0.08 | 0.21 | 0.21 | 0.24 | 0.24 |
| 4 | 165788675 | rs17679574 | -0.41 | 0.784 | -1.059 | 0.011 | 0.001 | -0.001 | -0.001 | 0 | 0.07 | 0.07 | 0.02 | 0.1 | 0.04 | 0.08 |
| 4 | 165789042 | rs17045668 | -0.203 | 0.75 | -1.232 | -0.106 | 0 | 0.008 | 0.008 | 2.976 | 0.15 | 0.13 | 0.06 | 0.08 | 0.13 | 0.03 |
| 4 | 165793001 | rs4459954 | -0.818 | 0.589 | -0.784 | -0.17 | -0.035 | -0.047 | -0.047 | 0.001 | 0.28 | 0.46 | 0.11 | 0.16 | 0.18 | 0.14 |
| 4 | 165798986 | rs12500546 | -0.044 | 0.54 | -1.094 | -0.003 | -0.021 | -0.005 | -0.005 | 0 | 0.39 | 0.27 | 0.41 | 0.38 | 0.44 | 0.5 |
| 4 | 165800154 | rs4541465 | -0.772 | 0.799 | -1.063 | 0.415 | 0.002 | -0.005 | -0.005 | 0.131 | 0.59 | 0.69 | 0.48 | 0.54 | 0.55 | 0.64 |
| 4 | 165808470 | rs11728450 | -0.497 | 1.024 | -0.986 | 0.533 | 0.002 | -0.002 | -0.002 | 0 | 0.31 | 0.3 | 0.37 | 0.19 | 0.33 | 0.24 |
| 4 | 165814359 | rs7691785 | -0.616 | 0.855 | -1.188 | 0.815 | -0.004 | -0.021 | -0.021 | 0 | 0.29 | 0.21 | 0.22 | 0.46 | 0.29 | 0.33 |
| 4 | 165814775 | rs13152225 | -0.386 | 0.244 | -0.701 | 0.673 | -0.021 | -0.033 | -0.033 | 0 | 0.27 | 0.56 | 0.16 | 0.16 | 0.19 | 0.16 |
| 4 | 165822659 | rs6846439 | -0.749 | 0.261 | -1.03 | 0.812 | 0.024 | -0.007 | -0.007 | 0 | 0.42 | 0.66 | 0.27 | 0.52 | 0.28 | 0.3 |
| 4 | 165827749 | rs6854993 | -1.027 | 0.241 | -0.337 | -0.108 | -0.001 | -0.081 | -0.081 | 0 | 0.25 | 0.49 | 0.17 | 0.1 | 0.21 | 0.17 |
| 4 | 165836060 | rs4429695 | -1.465 | 0.901 | -0.989 | -0.522 | -0.057 | 0.117 | 0.117 | 0.973 | 0.42 | 0.82 | 0.5 | 0.18 | 0.42 | 0.36 |
| 4 | 165837436 | rs17487218 | 0.232 | 0.895 | -0.949 | -0.491 | -0.084 | -0.004 | -0.004 | 0 | 0.14 | 0.02 | 0.31 | 0.08 | 0.2 | 0.18 |
| 4 | 165837631 | rs17584342 | 0.232 | 0.918 | -0.949 | -0.491 | -0.085 | -0.004 | -0.004 | 0 | 0.14 | 0.02 | 0.31 | 0.08 | 0.2 | 0.18 |
| 4 | 165839378 | rs6852326 | -0.92 | 0.843 | -0.686 | -0.754 | -0.011 | 0.048 | 0.048 | 0 | 0.25 | 0.15 | 0.35 | 0.08 | 0.25 | 0.24 |
| 4 | 165845582 | rs6856935 | -1.261 | 0.525 | -0.386 | -0.96 | 0 | -0.006 | -0.006 | 0 | 0.31 | 0.3 | 0.38 | 0.09 | 0.27 | 0.29 |
| 4 | 165849934 | rs10006119 | -0.708 | 0.423 | -0.668 | -1.242 | 0.017 | -0.036 | -0.036 | 0.473 | 0.64 | 0.48 | 0.63 | 0.74 | 0.54 | 0.67 |
| 4 | 165851057 | rs10011691 | 0.999 | 0.138 | -0.734 | -1.203 | 0 | 0.008 | 0.008 | 0 | 0.05 | 0.05 | 0.04 | 0 | 0.05 | 0.08 |

| | | | | | | | | | | | | | | | | |
|---|---|---|---|---|---|---|---|---|---|---|---|---|---|---|---|---|
| 4 | 165852918 | rs3889248 | 0.9 | 0.33 | -0.808 | -1.15 | -0.058 | -0.036 | -0.036 | 0 | 0.39 | 0.06 | 0.3 | 0.67 | 0.36 | 0.42 |
| 4 | 165853130 | rs3889246 | -0.874 | 0.316 | -0.65 | -1.229 | 0.004 | -0.02 | -0.02 | 0 | 0.67 | 0.43 | 0.65 | 0.74 | 0.6 | 0.68 |
| 4 | 165855463 | rs4690795 | -0.609 | 0.251 | -0.562 | -1.266 | -0.001 | 0.005 | 0.005 | 0 | 0.63 | 0.33 | 0.63 | 0.74 | 0.56 | 0.66 |
| 4 | 165859951 | rs4359860 | -0.899 | 0.249 | -0.337 | -1.227 | 0.003 | -0.066 | -0.066 | 0.051 | 0.28 | 0.41 | 0.35 | 0.09 | 0.26 | 0.27 |
| 4 | 165869823 | rs10020204 | -0.967 | 0.194 | -0.339 | -1.235 | -0.004 | -0.067 | -0.067 | 0 | 0.29 | 0.42 | 0.35 | 0.1 | 0.27 | 0.27 |
| 4 | 165869919 | rs10020142 | -0.416 | 0.213 | -0.451 | -1.212 | -0.001 | -0.017 | -0.017 | 0 | 0.25 | 0.22 | 0.32 | 0.09 | 0.22 | 0.25 |
| 4 | 165875359 | rs28544252 | -0.416 | 0.213 | -0.451 | -1.212 | -0.001 | -0.018 | -0.018 | 0 | 0.25 | 0.22 | 0.32 | 0.1 | 0.22 | 0.25 |
| 4 | 165875608 | rs6834073 | 0.768 | 0.138 | -0.452 | -1.147 | -0.012 | -0.006 | -0.006 | 0 | 0.37 | 0.17 | 0.31 | 0.66 | 0.35 | 0.44 |
| 4 | 165877963 | rs6848637 | -1.21 | -1.09 | 0.319 | -1.218 | -0.002 | -0.067 | -0.067 | 6.408 | 0.33 | 0.5 | 0.39 | 0.09 | 0.32 | 0.28 |
| 4 | 165877990 | rs9762579 | -1.538 | -1.337 | 0.334 | -1.292 | -0.015 | 0.044 | 0.044 | 1.513 | 0.6 | 0.73 | 0.57 | 0.75 | 0.47 | 0.66 |
| 4 | 165878335 | rs3733418 | 1.187 | -1.288 | 0.179 | -1.256 | 0.058 | -0.103 | -0.103 | 2.932 | 0.26 | 0.06 | 0.17 | 0.65 | 0.16 | 0.39 |
| 4 | 165879007 | rs10034083 | -0.599 | -1.304 | 0.348 | -1.275 | -0.002 | 0.01 | 0.01 | 0 | 0.29 | 0.17 | 0.34 | 0.09 | 0.29 | 0.28 |
| 4 | 165879381 | rs10001548 | -0.88 | -1.327 | 0.146 | -1.249 | 0 | -0.075 | -0.075 | 0 | 0.33 | 0.42 | 0.38 | 0.1 | 0.32 | 0.28 |
| 4 | 165880958 | rs8180162 | 1.082 | -1.193 | -0.316 | -1.118 | 0.057 | -0.105 | -0.105 | 0.587 | 0.26 | 0.06 | 0.17 | 0.65 | 0.16 | 0.39 |
| 4 | 165889612 | rs13134940 | 1.075 | -1.186 | -0.313 | -1.137 | 0.058 | -0.103 | -0.103 | 0.978 | 0.26 | 0.06 | 0.17 | 0.65 | 0.16 | 0.39 |
| 4 | 165905118 | rs13120990 | 1.015 | -1.169 | -0.264 | -0.993 | 0.065 | -0.103 | -0.103 | 0 | 0.27 | 0.08 | 0.17 | 0.63 | 0.16 | 0.39 |
| 4 | 165908721 | rs4691139 | -1.184 | -0.942 | -0.112 | -1.093 | -0.014 | 0.024 | 0.024 | 2.453 | 0.61 | 0.68 | 0.57 | 0.75 | 0.48 | 0.68 |
| 4 | 165926286 | rs17584656 | 0.286 | -1.299 | -0.248 | -0.732 | 0.002 | 0.004 | 0.004 | 0 | 0.15 | 0.07 | 0.24 | 0.08 | 0.1 | 0.15 |
| 4 | 165930168 | rs11942693 | 0.405 | -1.24 | 0.172 | -0.613 | -0.002 | -0.002 | -0.002 | 0 | 0.32 | 0.33 | 0.23 | 0.67 | 0.18 | 0.4 |
| 4 | 165930299 | rs11942750 | 0.415 | -1.241 | 0.171 | -0.613 | -0.002 | -0.001 | -0.001 | 0 | 0.32 | 0.33 | 0.23 | 0.67 | 0.18 | 0.4 |
| 4 | 165944299 | rs7684974 | -1.181 | -1.264 | -0.12 | -0.783 | 0.013 | 0.097 | 0.097 | 5.199 | 0.57 | 0.76 | 0.55 | 0.77 | 0.38 | 0.62 |
| 4 | 165948306 | rs6847301 | -1.079 | -1.038 | -0.109 | -0.754 | -0.001 | -0.005 | -0.005 | 0.616 | 0.22 | 0.26 | 0.32 | 0.1 | 0.19 | 0.2 |
| 4 | 165950675 | rs6536885 | -0.696 | -0.945 | -0.258 | -0.724 | 0.002 | -0.016 | -0.016 | 0 | 0.2 | 0.2 | 0.3 | 0.08 | 0.19 | 0.21 |
| 4 | 165951703 | rs12505213 | -0.696 | -0.945 | -0.258 | -0.724 | 0.002 | -0.016 | -0.016 | 0.097 | 0.2 | 0.19 | 0.3 | 0.08 | 0.19 | 0.21 |
| 4 | 165954865 | rs10000216 | 0.768 | -1.37 | -0.45 | -0.667 | -0.008 | 0.042 | 0.042 | 0 | 0.08 | 0.02 | 0.06 | 0 | 0.09 | 0.14 |
| 4 | 165956628 | rs13110766 | 0.206 | -1.696 | -0.23 | -0.595 | -0.067 | -0.002 | -0.002 | 0.033 | 0.37 | 0.47 | 0.25 | 0.68 | 0.19 | 0.4 |
| 4 | 165964290 | rs7654142 | 0.308 | -1.493 | -0.315 | -0.724 | -0.002 | -0.001 | -0.001 | 3.543 | 0.11 | 0.07 | 0.22 | 0.08 | 0.07 | 0.09 |
| 4 | 165965120 | rs12503351 | -0.081 | -1.946 | -0.292 | -0.51 | -0.083 | -0.013 | -0.013 | 0 | 0.25 | 0.53 | 0.11 | 0.16 | 0.09 | 0.08 |
| 4 | 165968037 | rs6536887 | -1.51 | -1.921 | -1.127 | -0.585 | 0.137 | 0.06 | 0.06 | 0.644 | 0.45 | 0.86 | 0.39 | 0.25 | 0.24 | 0.26 |
| 4 | 165971999 | rs6828376 | -0.058 | -1.341 | -0.744 | -0.622 | 0.004 | 0.006 | 0.006 | 0 | 0.37 | 0.32 | 0.3 | 0.17 | 0.34 | 0.26 |
| 4 | 165975122 | rs6848914 | 0.084 | -0.813 | -0.643 | -0.216 | -0.002 | -0.001 | -0.001 | 0 | 0.11 | 0.07 | 0.23 | 0.08 | 0.07 | 0.09 |
| 4 | 165978659 | rs6826711 | -0.489 | -0.88 | -0.484 | -0.122 | -0.002 | 0 | 0 | 0 | 0.1 | 0.05 | 0.09 | 0.08 | 0.07 | 0.12 |
| 4 | 165981142 | rs6819917 | -0.969 | -0.928 | -0.284 | -0.149 | -0.031 | -0.005 | -0.005 | 0 | 0.14 | 0.29 | 0.11 | 0.08 | 0.07 | 0.11 |
| 4 | 165982775 | rs4691148 | -0.265 | -0.624 | -0.927 | -0.267 | -0.069 | 0.091 | 0.091 | 0 | 0.38 | 0.1 | 0.49 | 0.21 | 0.43 | 0.33 |
| 4 | 165991632 | rs13146344 | 0.2 | -0.815 | 0.499 | 0.158 | -0.072 | -0.047 | -0.047 | 0 | 0.83 | 0.46 | 0.92 | 0.96 | 0.95 | 0.91 |
| 4 | 165995106 | rs10027850 | 1.294 | -0.305 | 1.169 | 0.419 | -0.035 | -0.017 | -0.017 | 0.893 | 0.86 | 0.57 | 0.88 | 0.94 | 0.93 | 0.9 |
| 4 | 166000195 | rs9999037 | 1.414 | -0.269 | 1.284 | 0.442 | -0.036 | -0.022 | -0.022 | 0 | 0.85 | 0.58 | 0.89 | 0.94 | 0.92 | 0.9 |
| 4 | 166010815 | rs12498760 | -0.7 | -0.03 | 0.706 | 0.7 | -0.036 | -0.035 | -0.035 | 0 | 0.22 | 0.09 | 0.43 | 0.24 | 0.3 | 0.27 |
| 4 | 166014765 | rs4285049 | 1.254 | -0.134 | 1.393 | 0.626 | -0.044 | -0.052 | -0.052 | 0 | 0.86 | 0.67 | 0.91 | 0.95 | 0.91 | 0.94 |
| 4 | 166014766 | rs4643773 | 1.043 | -0.121 | 1.356 | 0.639 | -0.032 | -0.028 | -0.028 | 0 | 0.85 | 0.59 | 0.89 | 0.94 | 0.9 | 0.9 |
| 4 | 166023258 | rs12642069 | -0.729 | 0.213 | 0.954 | 1.32 | -0.012 | -0.029 | -0.029 | 0 | 0.1 | 0.06 | 0.11 | 0.3 | 0.12 | 0.09 |
| 4 | 166026033 | rs4690801 | 1.899 | 0.044 | 1.214 | 1.203 | -0.004 | 0.006 | 0.006 | 0.731 | 0.73 | 0.37 | 0.76 | 0.63 | 0.81 | 0.8 |
| 4 | 166030858 | rs17045969 | -0.903 | -0.024 | 1.347 | 1.018 | -0.035 | -0.02 | -0.02 | 0.922 | 0.1 | 0.3 | 0.1 | 0.04 | 0.05 | 0.05 |
| 4 | 166035174 | rs2056255 | 1.275 | 0.26 | 0.841 | 1.078 | -0.007 | 0.006 | 0.006 | 0.087 | 0.73 | 0.42 | 0.66 | 0.71 | 0.68 | 0.61 |
| 4 | 166040374 | rs7661552 | -1.453 | 0.139 | 0.626 | 1.338 | 0.009 | 0.084 | 0.084 | 0 | 0.1 | 0.24 | 0.11 | 0.22 | 0.11 | 0.08 |
| 4 | 166052268 | rs1877536 | -0.995 | -0.039 | 0.096 | 0.702 | 0 | 0.032 | 0.032 | 0.022 | 0.11 | 0.02 | 0.15 | 0.05 | 0.13 | 0.33 |
| 4 | 166052741 | rs4691153 | -0.926 | 0.024 | 0.146 | 0.688 | -0.004 | 0.023 | 0.023 | 0 | 0.1 | 0.06 | 0.15 | 0.05 | 0.14 | 0.31 |
| 4 | 166055516 | rs17623971 | -1.489 | 0.085 | 0.917 | 1.276 | 0.004 | 0.256 | 0.256 | 0 | 0.11 | 0.29 | 0.13 | 0.38 | 0.09 | 0.1 |
| 4 | 166070974 | rs10023696 | -1.971 | -0.554 | 1.094 | -0.034 | -0.04 | -0.051 | -0.051 | 0 | 0.16 | 0.69 | 0.08 | 0.04 | 0.05 | 0.05 |
| 4 | 166072693 | rs4691168 | -0.352 | -0.22 | 0.811 | -0.052 | -0.033 | 0.01 | 0.01 | 0 | 0.09 | 0.01 | 0.14 | 0.05 | 0.13 | 0.3 |
| 4 | 166080237 | rs9308098 | -1.074 | -0.296 | 1.385 | -0.036 | 0.008 | -0.018 | -0.018 | 0.017 | 0.11 | 0.24 | 0.16 | 0.06 | 0.13 | 0.37 |
| 4 | 166089111 | rs10014286 | -2.245 | -0.543 | 1.331 | -0.16 | 0.005 | 0.011 | 0.011 | 0 | 0.18 | 0.17 | 0.18 | 0.1 | 0.18 | 0.4 |
| 4 | 166090564 | rs4234891 | -1.426 | -0.448 | 1.758 | -0.107 | 0.028 | -0.021 | -0.021 | 1.791 | 0.13 | 0.41 | 0.17 | 0.06 | 0.14 | 0.36 |
| 4 | 166099190 | rs11731244 | -1.508 | -0.191 | 1.207 | -0.427 | -0.004 | -0.031 | -0.031 | 0 | 0.19 | 0.26 | 0.12 | 0.11 | 0.15 | 0.13 |
| 4 | 166112950 | rs4605609 | -1.747 | -0.253 | 1.42 | -0.466 | -0.016 | -0.073 | -0.073 | 0.105 | 0.2 | 0.49 | 0.14 | 0.08 | 0.13 | 0.14 |
| 4 | 166114231 | rs17585333 | -0.843 | -0.165 | 0.889 | -0.423 | -0.001 | -0.002 | -0.002 | 0.131 | 0.15 | 0.16 | 0.11 | 0.08 | 0.13 | 0.08 |
| 4 | 166123245 | rs10010241 | -1.91 | -0.367 | 0.602 | -0.419 | 0.027 | 0.126 | 0.126 | 0.913 | 0.36 | 0.85 | 0.32 | 0.22 | 0.28 | 0.44 |
| 4 | 166144008 | rs17585396 | 0.463 | -0.094 | 0.303 | -0.862 | -0.047 | 0.044 | 0.044 | 2.026 | 0.18 | 0.06 | 0.13 | 0.02 | 0.3 | 0.07 |
| 4 | 166170390 | rs10517833 | 0.31 | -0.102 | 0.646 | -0.705 | -0.023 | 0.015 | 0.015 | 1.173 | 0.19 | 0.11 | 0.14 | 0.02 | 0.31 | 0.1 |
| 4 | 166199390 | rs10016586 | -0.846 | -0.195 | 0.711 | -0.58 | -0.023 | -0.008 | -0.008 | 3.433 | 0.16 | 0.23 | 0.09 | 0.19 | 0.09 | 0.11 |
| 4 | 166211057 | rs11726531 | -0.745 | -0.607 | 0.619 | -0.831 | 0.009 | 0.075 | 0.075 | 1.13 | 0.5 | 0.65 | 0.28 | 0.26 | 0.46 | 0.21 |
| 4 | 166223992 | rs11729242 | -0.57 | -0.383 | 0.849 | -0.681 | -0.007 | 0.007 | 0.007 | 0 | 0.05 | 0.19 | 0.03 | 0.05 | 0.02 | 0.01 |
| 4 | 166237389 | rs6536903 | -0.636 | -0.379 | 0.448 | -0.439 | 0 | 0 | 0 | 1.918 | 0.46 | 0.56 | 0.28 | 0.29 | 0.46 | 0.19 |
| 4 | 166248202 | rs10009994 | -0.45 | -0.288 | -0.101 | -0.399 | 0 | 0.044 | 0.044 | 3.456 | 0.43 | 0.32 | 0.24 | 0.17 | 0.42 | 0.14 |
| 4 | 166252950 | rs17686351 | -0.221 | -0.734 | -0.77 | 0.825 | 0.037 | -0.118 | -0.118 | 2.124 | 0.22 | 0.04 | 0.16 | 0.51 | 0.16 | 0.16 |
| 4 | 166253765 | rs1370689 | -1.412 | -0.757 | 0.42 | 0.861 | 0.021 | 0.14 | 0.14 | 3.57 | 0.07 | 0.59 | 0.13 | 0.14 | 0 | 0.09 |
| 4 | 166261800 | rs1550270 | 0.336 | -0.884 | -0.087 | 0.502 | 0.006 | 0.002 | 0.002 | 0 | 0.35 | 0.26 | 0.5 | 0.35 | 0.32 | 0.48 |
| 4 | 166267119 | rs7667236 | 0.081 | -0.43 | 0.11 | 0.022 | 0.014 | 0 | 0 | 0.125 | 0.39 | 0.26 | 0.29 | 0.6 | 0.33 | 0.49 |
| 4 | 166267383 | rs17686529 | 0.977 | -0.115 | -0.077 | -0.031 | 0.032 | -0.009 | -0.009 | 0.114 | 0.33 | 0.14 | 0.47 | 0.35 | 0.26 | 0.39 |

| | | | | | | | | | | | | | | | | |
|---|---|---|---|---|---|---|---|---|---|---|---|---|---|---|---|---|
| 4 | 166270327 | rs12641145 | 0.486 | -0.11 | -0.163 | 0.32 | -0.005 | 0.004 | 0.004 | 0 | 0.37 | 0.45 | 0.48 | 0.42 | 0.33 | 0.46 |
| 4 | 166279952 | rs6828152 | -0.056 | -0.199 | -0.206 | 0.367 | -0.023 | -0.005 | -0.005 | 0 | 0.42 | 0.41 | 0.3 | 0.6 | 0.34 | 0.45 |
| 4 | 166281891 | rs17046348 | -0.046 | -0.233 | 0.004 | 0.784 | -0.004 | 0.053 | 0.053 | 0.008 | 0.14 | 0.35 | 0.11 | 0.18 | 0.11 | 0.19 |
| 4 | 166282004 | rs12513198 | 1.131 | -0.371 | -0.144 | 0.686 | 0.017 | -0.053 | -0.053 | 3.455 | 0.31 | 0.13 | 0.47 | 0.4 | 0.25 | 0.38 |
| 4 | 166283914 | rs17488814 | 1.424 | -0.391 | -0.189 | 0.575 | -0.092 | 0.087 | 0.087 | 0 | 0.19 | 0.07 | 0.15 | 0.02 | 0.33 | 0.06 |
| 4 | 166286171 | rs4621390 | -0.486 | -0.448 | 0.204 | 0.674 | 0.02 | 0.002 | 0.002 | 0 | 0.48 | 0.57 | 0.35 | 0.56 | 0.42 | 0.55 |
| 4 | 166287429 | rs6837095 | -0.096 | -0.53 | 0.36 | 0.701 | -0.01 | 0.055 | 0.055 | 0.341 | 0.14 | 0.43 | 0.12 | 0.18 | 0.11 | 0.18 |
| 4 | 166289408 | rs12501589 | 1.074 | -0.509 | -0.154 | 0.731 | 0.018 | -0.05 | -0.05 | 0 | 0.3 | 0.13 | 0.47 | 0.39 | 0.23 | 0.38 |
| 4 | 166296379 | rs1370682 | 1.079 | -0.488 | -0.199 | 0.717 | 0.049 | -0.041 | -0.041 | 0.64 | 0.31 | 0.08 | 0.19 | 0.36 | 0.24 | 0.29 |
| 4 | 166297180 | rs1438113 | 0.065 | -0.291 | 0.039 | 0.513 | 0.004 | -0.01 | -0.01 | 0 | 0.65 | 0.37 | 0.69 | 0.75 | 0.49 | 0.67 |
| 4 | 166298451 | rs1583645 | 0.056 | -0.31 | 0.039 | 0.513 | 0.005 | -0.01 | -0.01 | 0 | 0.65 | 0.37 | 0.69 | 0.76 | 0.49 | 0.67 |
| 4 | 166301778 | rs10022413 | -0.624 | 0.105 | 0.123 | 0.858 | 0.001 | 0.016 | 0.016 | 0.807 | 0.51 | 0.61 | 0.63 | 0.61 | 0.48 | 0.62 |
| 4 | 166304172 | rs6850000 | -0.781 | 0.087 | 0.175 | 0.881 | 0.035 | 0.036 | 0.036 | 0.266 | 0.18 | 0.48 | 0.17 | 0.25 | 0.21 | 0.27 |
| 4 | 166307302 | rs12509669 | 0.855 | 0.033 | -0.17 | 0.846 | 0.015 | -0.045 | -0.045 | 0 | 0.33 | 0.13 | 0.46 | 0.37 | 0.27 | 0.35 |
| 4 | 166309213 | rs9997245 | 0.915 | 0.065 | 0.23 | 1.057 | 0.019 | -0.056 | -0.056 | 0.039 | 0.32 | 0.14 | 0.24 | 0.42 | 0.28 | 0.32 |
| 4 | 166310679 | rs2322291 | -0.698 | 0.16 | 0.354 | 1.091 | 0.011 | 0.044 | 0.044 | 0.157 | 0.5 | 0.75 | 0.45 | 0.59 | 0.49 | 0.54 |
| 4 | 166311553 | rs934813 | -0.393 | 0.184 | 0.578 | 1.051 | 0.01 | 0.021 | 0.021 | 0 | 0.13 | 0.23 | 0.16 | 0.15 | 0.15 | 0.13 |
| 4 | 166315725 | rs6536912 | -0.393 | 0.185 | 0.579 | 1.052 | 0.01 | 0.02 | 0.02 | 0 | 0.13 | 0.23 | 0.16 | 0.15 | 0.15 | 0.13 |
| 4 | 166319702 | rs1543915 | -0.916 | 0.029 | 0.271 | 0.99 | -0.002 | 0.044 | 0.044 | 1.745 | 0.54 | 0.81 | 0.47 | 0.65 | 0.49 | 0.47 |
| 4 | 166323057 | rs6811043 | 0.1 | 0.329 | 0.473 | 1.318 | 0.019 | 0.101 | 0.101 | 0 | 0.08 | 0.24 | 0.11 | 0.15 | 0.1 | 0.05 |
| 4 | 166327010 | rs934819 | -0.488 | 0.101 | 0.324 | 1.442 | 0.01 | -0.005 | -0.005 | 0 | 0.54 | 0.62 | 0.37 | 0.46 | 0.44 | 0.39 |
| 4 | 166334446 | rs9992011 | 0.612 | 0.037 | 0.035 | 1.087 | 0.019 | 0.018 | 0.018 | 0 | 0.38 | 0.24 | 0.23 | 0.31 | 0.31 | 0.3 |
| 4 | 166334840 | rs1866224 | 0.978 | -0.085 | -0.053 | 1.021 | 0.002 | -0.005 | -0.005 | 0 | 0.34 | 0.07 | 0.21 | 0.3 | 0.3 | 0.29 |
| 4 | 166342375 | rs13125242 | 0.737 | 0.007 | 0.304 | 1.269 | 0.007 | -0.005 | -0.005 | 0 | 0.37 | 0.32 | 0.23 | 0.36 | 0.3 | 0.3 |
| 4 | 166358437 | rs7692208 | -0.75 | 0.216 | 0.403 | 1.329 | -0.025 | 0.049 | 0.049 | 1.228 | 0.6 | 0.75 | 0.46 | 0.76 | 0.54 | 0.51 |
| 4 | 166358532 | rs10517849 | 0.32 | 0.221 | 0.626 | 1.329 | 0.002 | 0.001 | 0.001 | 0.007 | 0.44 | 0.44 | 0.28 | 0.35 | 0.35 | 0.32 |
| 4 | 166359421 | rs4481204 | 0.768 | 0.25 | 0.478 | 1.391 | 0.004 | 0.001 | 0.001 | 0 | 0.41 | 0.23 | 0.26 | 0.35 | 0.35 | 0.32 |
| 4 | 166365138 | rs2083159 | -0.646 | 0.24 | 0.412 | 1.575 | 0.004 | 0.122 | 0.122 | 0.021 | 0.16 | 0.3 | 0.19 | 0.4 | 0.19 | 0.19 |
| 4 | 166369281 | rs6836074 | -0.489 | 0.235 | 0.44 | 1.597 | -0.013 | 0.037 | 0.037 | 0 | 0.15 | 0.28 | 0.16 | 0.23 | 0.13 | 0.12 |
| 4 | 166369661 | rs6841638 | -0.489 | 0.235 | 0.44 | 1.597 | -0.013 | 0.037 | 0.037 | 0 | 0.15 | 0.28 | 0.16 | 0.23 | 0.13 | 0.12 |
| 4 | 166370111 | rs6842167 | -0.663 | 0.232 | 0.444 | 1.485 | 0.003 | 0.12 | 0.12 | 0.017 | 0.16 | 0.3 | 0.19 | 0.39 | 0.19 | 0.19 |
| 4 | 166370156 | rs1438114 | -0.393 | 0.137 | 0.41 | 1.258 | -0.015 | 0.076 | 0.076 | 0.236 | 0.55 | 0.73 | 0.43 | 0.76 | 0.51 | 0.5 |
| 4 | 166371512 | rs10021007 | -1.016 | 0.121 | 0.515 | 1.587 | -0.002 | 0.072 | 0.072 | 0 | 0.17 | 0.29 | 0.16 | 0.35 | 0.18 | 0.18 |
| 4 | 166386944 | rs3775311 | 0.877 | 0.603 | 0.043 | 1.247 | -0.027 | 0.004 | 0.004 | 0 | 0.12 | 0.01 | 0.21 | 0.09 | 0.15 | 0.06 |
| 4 | 166389558 | rs1530042 | -1.429 | 0.442 | 0.74 | 1.415 | 0.009 | 0.052 | 0.052 | 1.007 | 0.13 | 0.47 | 0.15 | 0.24 | 0.1 | 0.13 |
| 4 | 166390220 | rs1370686 | 0.864 | 0.601 | 0.246 | 1.409 | -0.007 | 0.008 | 0.008 | 0 | 0.41 | 0.24 | 0.28 | 0.33 | 0.4 | 0.38 |
| 4 | 166398831 | rs11100614 | -0.778 | 0.528 | 0.576 | 1.434 | 0.001 | 0.024 | 0.024 | 0 | 0.22 | 0.43 | 0.21 | 0.37 | 0.26 | 0.13 |
| 4 | 166399738 | rs11730342 | 0.115 | 0.54 | 0.423 | 1.542 | 0.016 | 0.04 | 0.04 | 0 | 0.54 | 0.27 | 0.4 | 0.59 | 0.62 | 0.5 |
| 4 | 166408782 | rs3822062 | 0.198 | 0.633 | 0.789 | 1.576 | -0.002 | -0.002 | -0.002 | 0.42 | 0.43 | 0.6 | 0.35 | 0.42 | 0.43 | 0.47 |
| 4 | 166416643 | rs7692951 | -1.991 | 0.951 | 1.298 | 1.793 | 0.011 | 0.142 | 0.142 | 2.233 | 0.13 | 0.63 | 0.18 | 0.37 | 0.12 | 0.12 |
| 4 | 166419478 | rs6822718 | -1.484 | 0.888 | 1.191 | 1.364 | 0.019 | 0.044 | 0.044 | 0.845 | 0.1 | 0.39 | 0.14 | 0.16 | 0.05 | 0.06 |
| 4 | 166431563 | rs17046580 | -0.514 | 0.709 | 0.778 | 1.195 | -0.015 | -0.018 | -0.018 | 0 | 0.07 | 0.12 | 0.1 | 0.06 | 0.04 | 0.03 |
| 4 | 166433829 | rs6536916 | -0.578 | 0.464 | 1.048 | 1.355 | -0.004 | 0.063 | 0.063 | 0 | 0.05 | 0.14 | 0.01 | 0.08 | 0.01 | 0.05 |
| 4 | 166436027 | rs7656395 | 1.208 | 0.45 | 0.488 | 1.372 | 0.001 | -0.011 | -0.011 | 0.595 | 0.42 | 0.13 | 0.55 | 0.36 | 0.39 | 0.45 |
| 4 | 166438457 | rs7699278 | 0.356 | 0.485 | 0.551 | 1.335 | 0.002 | 0.005 | 0.005 | 0 | 0.55 | 0.66 | 0.7 | 0.42 | 0.46 | 0.48 |
| 4 | 166444375 | rs2033765 | 0.262 | 0.554 | 0.646 | 1.51 | -0.007 | 0.006 | 0.006 | 0 | 0.56 | 0.61 | 0.72 | 0.62 | 0.51 | 0.55 |
| 4 | 166444924 | rs2033766 | -0.779 | 0.928 | 0.304 | 1.399 | -0.12 | -0.052 | -0.052 | 0.858 | 0.7 | 0.95 | 0.78 | 0.7 | 0.62 | 0.65 |
| 4 | 166448333 | rs1837172 | 0.641 | 0.523 | 0.642 | 1.02 | 0.024 | -0.006 | -0.006 | 1.23 | 0.51 | 0.59 | 0.6 | 0.43 | 0.41 | 0.5 |
| 4 | 166448674 | rs1438118 | 0.949 | 0.19 | 0.02 | 0.843 | 0.045 | 0.029 | 0.029 | 0 | 0.4 | 0.21 | 0.47 | 0.25 | 0.31 | 0.33 |
| 4 | 166455839 | rs2322122 | -2.094 | 0.004 | 0.337 | 1.048 | 0.001 | 0.05 | 0.05 | 0.294 | 0.48 | 0.87 | 0.39 | 0.69 | 0.43 | 0.48 |
| 4 | 166456376 | rs17503083 | 0.394 | 0.112 | 0.191 | 0.919 | -0.045 | -0.025 | -0.025 | 0 | 0.17 | 0.03 | 0.12 | 0.17 | 0.17 | 0.23 |
| 4 | 166457130 | rs6835679 | -0.916 | 0.088 | 0.738 | 0.719 | 0.016 | -0.091 | -0.091 | 0.307 | 0.13 | 0.42 | 0.06 | 0 | 0.1 | 0.05 |
| 4 | 166468810 | rs17046700 | -0.66 | 0.091 | 0.615 | 0.769 | -0.009 | -0.049 | -0.049 | 0.01 | 0.14 | 0.26 | 0.07 | 0.06 | 0.11 | 0.05 |
| 4 | 166469496 | rs17587610 | 0.504 | 0.13 | 0.313 | 0.809 | -0.06 | -0.021 | -0.021 | 0 | 0.17 | 0.01 | 0.12 | 0.14 | 0.17 | 0.22 |
| 4 | 166469526 | rs9990545 | -0.4 | -0.003 | 0.737 | 0.864 | -0.005 | 0.02 | 0.02 | 0 | 0.1 | 0.27 | 0.11 | 0.11 | 0.07 | 0.08 |
| 4 | 166471139 | rs7654319 | -2.232 | -0.059 | 0.782 | 1.117 | 0.014 | -0.021 | -0.021 | 0 | 0.32 | 0.72 | 0.25 | 0.51 | 0.26 | 0.3 |
| 4 | 166472187 | rs17687965 | 0.164 | -0.009 | 0.433 | 1.186 | -0.007 | 0.001 | 0.001 | 0 | 0.06 | 0.03 | 0.06 | 0.32 | 0.08 | 0.15 |
| 4 | 166472744 | rs10028219 | -0.394 | 0.065 | 0.489 | 1.076 | -0.006 | 0.038 | 0.038 | 0 | 0.1 | 0.27 | 0.11 | 0.13 | 0.07 | 0.1 |
| 4 | 166473571 | rs7697374 | -1.732 | 0.088 | 0.013 | 1.081 | 0.013 | -0.017 | -0.017 | 0.083 | 0.32 | 0.71 | 0.25 | 0.49 | 0.26 | 0.28 |
| 4 | 166476045 | rs17587680 | 0.642 | 0.1 | -0.088 | 1.029 | -0.059 | -0.023 | -0.023 | 0 | 0.16 | 0.01 | 0.11 | 0.14 | 0.16 | 0.22 |
| 4 | 166481918 | rs1914839 | -2.202 | -0.218 | 0.75 | 1.254 | 0.021 | -0.031 | -0.031 | 0 | 0.32 | 0.74 | 0.25 | 0.51 | 0.26 | 0.3 |
| 4 | 166488615 | rs10017297 | -0.142 | -0.169 | 0.676 | 1.305 | 0.016 | 0.09 | 0.09 | 0 | 0.07 | 0.31 | 0.03 | 0.1 | 0.02 | 0.05 |
| 4 | 166489134 | rs1320227 | -1.399 | 0.281 | 0.406 | 1.197 | -0.01 | 0.049 | 0.049 | 0 | 0.21 | 0.34 | 0.13 | 0.38 | 0.19 | 0.2 |
| 4 | 166494614 | rs6843136 | -1.327 | 0.389 | 0.439 | 0.531 | 0 | -0.074 | -0.074 | 0 | 0.16 | 0.32 | 0.14 | 0.03 | 0.16 | 0.12 |
| 4 | 166495142 | rs7662154 | -1.413 | 0.526 | 0.432 | 0.827 | -0.006 | 0.004 | 0.004 | 0 | 0.29 | 0.62 | 0.13 | 0.29 | 0.23 | 0.31 |
| 4 | 166495319 | rs4691205 | 0.239 | -0.08 | 0.176 | 0.428 | -0.024 | -0.029 | -0.029 | 0 | 0.66 | 0.53 | 0.71 | 0.82 | 0.79 | 0.65 |
| 4 | 166497317 | rs10006198 | -0.406 | 0.295 | 0.663 | 0.423 | 0.054 | -0.021 | -0.021 | 0.119 | 0.05 | 0.2 | 0.1 | 0.03 | 0.1 | 0.03 |
| 4 | 166497693 | rs12331357 | -0.787 | 0.251 | 0.897 | 0.757 | 0.08 | 0.09 | 0.09 | 0 | 0.06 | 0.28 | 0.1 | 0.12 | 0.1 | 0.03 |
| 4 | 166508518 | rs1320225 | -0.294 | 0.442 | 0.506 | 0.653 | -0.012 | -0.014 | -0.014 | 1.616 | 0.35 | 0.29 | 0.61 | 0.48 | 0.41 | 0.52 |

| | | | | | | | | | | | | | | | |
|---|---|---|---|---|---|---|---|---|---|---|---|---|---|---|---|
| 4 | 166519866 | rs17046824 | -0.377 | 0.539 | 0.846 | 0.61 | 0.051 | -0.019 | -0.019 | 0 | 0.06 | 0.24 | 0.09 | 0.03 | 0.08 | 0.02 |
| 4 | 166532454 | rs9308109 | -1.545 | 0.409 | 1.13 | 0.85 | 0.064 | 0.249 | 0.249 | 0.049 | 0.09 | 0.49 | 0.11 | 0.27 | 0.09 | 0.09 |
| 4 | 166547323 | rs11721568 | -0.55 | 0.3 | 0.301 | 0.463 | 0 | 0.001 | 0.001 | 0 | 0.38 | 0.42 | 0.6 | 0.53 | 0.41 | 0.49 |
| 4 | 166547603 | rs1459506 | -1.334 | 0.009 | -0.606 | -0.022 | 0.044 | -0.059 | -0.059 | 0 | 0.56 | 0.86 | 0.77 | 0.53 | 0.61 | 0.55 |
| 4 | 166562938 | rs11938725 | 1.222 | -0.165 | -0.505 | 0.239 | 0.003 | 0.021 | 0.021 | 0 | 0.08 | 0.07 | 0.02 | 0.31 | 0.04 | 0.11 |
| 4 | 166568732 | rs2714834 | -0.013 | -0.188 | -0.53 | 0.094 | 0 | 0.008 | 0.008 | 0 | 0.84 | 0.85 | 0.86 | 0.68 | 0.84 | 0.85 |
| 4 | 166574651 | rs2611212 | -0.776 | -0.222 | -0.552 | 0.115 | -0.004 | 0.013 | 0.013 | 0 | 0.88 | 0.89 | 0.94 | 0.69 | 0.94 | 0.87 |
| 4 | 166575465 | rs2611210 | 0.482 | -0.228 | -0.805 | -0.112 | 0 | -0.015 | -0.015 | 0 | 0.14 | 0.16 | 0.09 | 0 | 0.13 | 0.05 |
| 4 | 166575815 | rs1380348 | 0.6 | -0.227 | -0.81 | -0.111 | 0 | -0.012 | -0.012 | 0 | 0.14 | 0.15 | 0.08 | 0 | 0.13 | 0.05 |
| 4 | 166578236 | rs2611209 | -1.186 | -0.103 | -0.799 | -0.44 | -0.004 | -0.008 | -0.008 | 0.86 | 0.43 | 0.45 | 0.3 | 0.34 | 0.39 | 0.3 |
| 4 | 166585445 | rs4691212 | 0.918 | -0.268 | -1.172 | -0.511 | 0.077 | 0.055 | 0.055 | 0 | 0.55 | 0.29 | 0.73 | 0.68 | 0.68 | 0.69 |
| 4 | 166588048 | rs2322264 | 2.343 | -0.093 | -1.548 | -0.568 | -0.077 | 0.056 | 0.056 | 0 | 0.29 | 0.02 | 0.16 | 0.14 | 0.32 | 0.3 |
| 4 | 166588085 | rs2322265 | -0.009 | -0.059 | -1.333 | -0.592 | -0.009 | 0.112 | 0.112 | 0.022 | 0.27 | 0.1 | 0.18 | 0 | 0.26 | 0.18 |
| 4 | 166588960 | rs11932029 | 0.792 | -0.113 | -0.605 | 0.049 | -0.033 | 0.114 | 0.114 | 0 | 0.11 | 0.17 | 0.02 | 0.32 | 0.04 | 0.12 |
| 4 | 166591321 | rs2714807 | 0.107 | -0.183 | -0.944 | 0.232 | -0.043 | -0.009 | -0.009 | 0 | 0.37 | 0.26 | 0.57 | 0.38 | 0.41 | 0.33 |
| 4 | 166592304 | rs17688688 | 0.426 | -0.366 | -1.074 | 0.327 | -0.041 | 0.154 | 0.154 | 0 | 0.12 | 0.26 | 0.03 | 0.32 | 0.04 | 0.12 |
| 4 | 166592849 | rs17504069 | 0.429 | 0.12 | -2.121 | -0.448 | -0.017 | 0.063 | 0.063 | 0 | 0.41 | 0.16 | 0.27 | 0.28 | 0.43 | 0.51 |
| 4 | 166594188 | rs17588573 | 0.115 | 0.404 | -2.33 | -0.623 | -0.096 | 0.056 | 0.056 | 0 | 0.28 | 0.03 | 0.21 | 0.14 | 0.35 | 0.36 |
| 4 | 166598495 | rs2611246 | -0.4 | 0.071 | -2.318 | -0.32 | 0.032 | -0.002 | -0.002 | 0 | 0.46 | 0.78 | 0.34 | 0.4 | 0.4 | 0.33 |
| 4 | 166602111 | rs1459509 | 1.147 | 0.468 | -2.105 | -0.021 | -0.024 | -0.028 | -0.028 | 0.764 | 0.06 | 0 | 0.11 | 0.11 | 0.06 | 0.11 |
| 4 | 166602585 | rs1380355 | -0.376 | 0.207 | -2.298 | -0.328 | 0.016 | 0.17 | 0.17 | 0 | 0.5 | 0.84 | 0.66 | 0.75 | 0.4 | 0.66 |
| 4 | 166604001 | rs2611248 | -0.615 | 0.115 | -1.883 | -0.408 | 0.009 | 0.08 | 0.08 | 0 | 0.63 | 0.78 | 0.77 | 0.9 | 0.65 | 0.71 |
| 4 | 166605363 | rs1870478 | 0.705 | -0.053 | -1.095 | -0.285 | -0.006 | 0.041 | 0.041 | 0 | 0.34 | 0.54 | 0.47 | 0.51 | 0.3 | 0.41 |
| 4 | 166606298 | rs2611249 | 1.013 | 0.357 | -1.537 | 0.399 | -0.068 | -0.015 | -0.015 | 0 | 0.16 | 0.02 | 0.11 | 0.16 | 0.25 | 0.05 |
| 4 | 166606890 | rs4691215 | -0.345 | -0.041 | -0.772 | 0.524 | -0.007 | 0.016 | 0.016 | 0 | 0.56 | 0.64 | 0.6 | 0.64 | 0.46 | 0.61 |
| 4 | 166607504 | rs1013356 | -1.391 | -0.065 | 0.515 | 0.863 | -0.032 | 0.098 | 0.098 | 0.101 | 0.12 | 0.45 | 0.08 | 0.13 | 0.05 | 0.15 |
| 4 | 166616007 | rs17046936 | -1.248 | 0.151 | 0.112 | 0.935 | -0.017 | 0.023 | 0.023 | 0 | 0.09 | 0.16 | 0.07 | 0.12 | 0.05 | 0.17 |
| 4 | 166619952 | rs2611259 | 0.443 | 0.08 | 0.626 | 1.454 | 0.002 | 0.088 | 0.088 | 0 | 0.81 | 0.51 | 0.82 | 0.74 | 0.82 | 0.81 |
| 4 | 166627996 | rs17688831 | 0.231 | 0.528 | 0.052 | 1.375 | -0.043 | -0.017 | -0.017 | 0 | 0.09 | 0.01 | 0.09 | 0.11 | 0.1 | 0.13 |
| 4 | 166629269 | rs2611251 | 1.446 | 0.723 | 0.32 | 1.423 | -0.006 | 0.03 | 0.03 | 0 | 0.71 | 0.34 | 0.73 | 0.61 | 0.73 | 0.63 |
| 4 | 166631681 | rs11936747 | 0.688 | 0.622 | 0.272 | 1.648 | -0.002 | -0.035 | -0.035 | 0 | 0.07 | 0.01 | 0.03 | 0.16 | 0.04 | 0.06 |
| 4 | 166642590 | rs17632444 | 0.378 | 0.718 | 0.546 | 1.698 | -0.005 | 0.036 | 0.036 | 1.745 | 0.08 | 0.15 | 0.06 | 0.17 | 0.04 | 0.06 |
| 4 | 166646557 | rs17046950 | -0.276 | 0.81 | 0.472 | 1.562 | 0.019 | 0.023 | 0.023 | 0 | 0.07 | 0.13 | 0.1 | 0.11 | 0.13 | 0.12 |
| 4 | 166655128 | rs6816303 | -0.347 | 0.464 | 0.509 | 1.457 | -0.025 | 0.011 | 0.011 | 0 | 0.06 | 0.08 | 0.01 | 0.1 | 0.01 | 0.17 |
| 4 | 166659698 | rs716399 | 0.314 | 0.93 | -0.27 | 1.542 | -0.016 | -0.016 | -0.016 | 0 | 0.15 | 0.1 | 0.41 | 0.3 | 0.19 | 0.3 |
| 4 | 166661136 | rs10517866 | -0.455 | 0.858 | -0.03 | 1.565 | -0.005 | 0.018 | 0.018 | 0 | 0.09 | 0.15 | 0.07 | 0.13 | 0.11 | 0.1 |
| 4 | 166662954 | rs17589098 | -0.293 | 1.288 | -0.279 | 1.773 | -0.035 | -0.024 | -0.024 | 0 | 0.08 | 0.03 | 0.1 | 0.11 | 0.13 | 0.11 |
| 4 | 166663108 | rs9308115 | -1.91 | 1.19 | -0.276 | 2.016 | -0.001 | 0 | 0 | 0 | 0.23 | 0.24 | 0.37 | 0.41 | 0.28 | 0.28 |
| 4 | 166663278 | rs17504644 | 1.072 | 0.393 | -0.879 | 1.392 | 0.04 | 0.075 | 0.075 | 0.037 | 0.37 | 0.18 | 0.33 | 0.26 | 0.29 | 0.41 |
| 4 | 166664618 | rs13111857 | -0.322 | 0.562 | -0.945 | 1.583 | 0.01 | 0.041 | 0.041 | 0 | 0.15 | 0.22 | 0.14 | 0.29 | 0.16 | 0.21 |
| 4 | 166665840 | rs2714819 | 0.895 | 0.505 | -1.479 | 1.433 | -0.143 | -0.079 | -0.079 | 0 | 0.3 | 0.05 | 0.38 | 0.3 | 0.44 | 0.19 |
| 4 | 166666480 | rs881821 | 1.135 | -0.087 | -0.97 | 1.256 | 0.001 | 0.001 | 0.001 | 0 | 0.68 | 0.38 | 0.73 | 0.37 | 0.74 | 0.6 |
| 4 | 166667602 | rs922096 | 1.517 | -0.68 | -1.797 | 1.082 | 0.025 | -0.063 | -0.063 | 0 | 0.22 | 0.02 | 0.1 | 0.23 | 0.16 | 0.27 |
| 4 | 166667833 | rs10517867 | 0.591 | -0.367 | -1.94 | 1.059 | -0.078 | -0.034 | -0.034 | 0.454 | 0.12 | 0.01 | 0.15 | 0.12 | 0.21 | 0.17 |
| 4 | 166671434 | rs7700172 | 1.059 | -0.623 | -2.405 | 0.961 | -0.11 | -0.006 | -0.006 | 0 | 0.33 | 0.09 | 0.38 | 0.28 | 0.42 | 0.2 |
| 4 | 166671834 | rs355196 | -1.86 | -0.676 | -1.867 | 0.91 | 0.011 | -0.023 | -0.023 | 0 | 0.94 | 0.81 | 0.95 | 1 | 0.96 | 1 |
| 4 | 166674512 | rs191644 | 1.015 | -0.622 | -1.947 | 0.968 | -0.102 | 0 | 0 | 0 | 0.33 | 0.1 | 0.38 | 0.28 | 0.42 | 0.2 |
| 4 | 166678420 | rs1460053 | -0.599 | -0.61 | -1.465 | 0.785 | 0 | 0.005 | 0.005 | 0 | 0.38 | 0.53 | 0.42 | 0.44 | 0.37 | 0.52 |
| 4 | 166689405 | rs355171 | 0.76 | -0.3 | -2.149 | 0.802 | -0.063 | -0.009 | -0.009 | 0 | 0.33 | 0.16 | 0.42 | 0.33 | 0.43 | 0.21 |
| 4 | 166691492 | rs355168 | 0.197 | -0.266 | -2.455 | 0.879 | -0.013 | -0.112 | -0.112 | 0 | 0.41 | 0.05 | 0.5 | 0.45 | 0.61 | 0.38 |
| 4 | 166693657 | rs2611830 | 0.997 | -0.299 | -2.323 | 0.769 | -0.145 | -0.048 | -0.048 | 0 | 0.29 | 0.02 | 0.36 | 0.25 | 0.4 | 0.21 |
| 4 | 166698701 | rs12331967 | 0.789 | -0.867 | -1.681 | 0.925 | 0.004 | -0.014 | -0.014 | 0 | 0.58 | 0.33 | 0.55 | 0.43 | 0.61 | 0.52 |
| 4 | 166699421 | rs1563831 | 0.344 | -1.581 | -1.875 | 0.903 | 0.037 | -0.021 | -0.021 | 3.156 | 0.19 | 0.11 | 0.22 | 0.35 | 0.12 | 0.28 |
| 4 | 166705990 | rs7694605 | -1.626 | -1.219 | -2.173 | 0.542 | -0.145 | -0.001 | -0.001 | 0 | 0.74 | 0.95 | 0.6 | 0.82 | 0.57 | 0.88 |
| 4 | 166708955 | rs17504826 | 0.145 | -1.144 | -1.874 | 0.302 | -0.011 | 0.007 | 0.007 | 0 | 0.12 | 0.12 | 0.12 | 0 | 0.18 | 0.14 |
| 4 | 166709866 | rs17632649 | 1.012 | -1.616 | -1.946 | 0.545 | 0.017 | -0.026 | -0.026 | 0 | 0.2 | 0.06 | 0.07 | 0.26 | 0.14 | 0.19 |
| 4 | 166717673 | rs7691783 | 0.952 | -1.515 | -1.468 | 0.522 | -0.013 | 0.004 | 0.004 | 0 | 0.08 | 0.14 | 0.04 | 0.1 | 0.03 | 0.08 |
| 4 | 166721110 | rs10517871 | 0.173 | -1.467 | -1.386 | 0.398 | 0 | -0.023 | -0.023 | 0 | 0.13 | 0.16 | 0.13 | 0 | 0.18 | 0.14 |
| 4 | 166726468 | rs12648621 | 0.566 | -2.197 | -1.09 | 0.994 | 0.01 | -0.024 | -0.024 | 0.001 | 0.19 | 0.03 | 0.23 | 0.21 | 0.14 | 0.29 |
| 4 | 166734132 | rs1380582 | -0.102 | -2.199 | -0.937 | 1.024 | 0.023 | -0.002 | -0.002 | 0.477 | 0.27 | 0.19 | 0.28 | 0.31 | 0.18 | 0.37 |
| 4 | 166734530 | rs7687480 | 0.644 | -2.191 | -0.185 | 1.074 | -0.04 | 0.019 | 0.019 | 0.84 | 0.22 | 0.3 | 0.09 | 0.28 | 0.11 | 0.18 |
| 4 | 166747763 | rs11100620 | -1.025 | -2.094 | -0.492 | 1.012 | 0 | 0.064 | 0.064 | 0 | 0.56 | 0.75 | 0.62 | 0.73 | 0.43 | 0.62 |
| 4 | 166752233 | rs2874792 | 0.415 | -2.993 | 0.115 | 1.136 | -0.003 | -0.002 | -0.002 | 0 | 0.23 | 0.25 | 0.38 | 0.22 | 0.18 | 0.27 |
| 4 | 166763264 | rs17632760 | 0.824 | -2.848 | 0.245 | 1.46 | -0.007 | -0.023 | -0.023 | 0.614 | 0.11 | 0.05 | 0.12 | 0.15 | 0.12 | 0.12 |
| 4 | 166764043 | rs6816331 | -0.717 | -2.946 | 0.3 | 1.314 | -0.005 | 0.005 | 0.005 | 3.716 | 0.4 | 0.41 | 0.55 | 0.51 | 0.33 | 0.47 |
| 4 | 166764589 | rs6844820 | -1.17 | -2.499 | 0.535 | 0.966 | -0.005 | 0.053 | 0.053 | 0.545 | 0.61 | 0.75 | 0.6 | 0.81 | 0.39 | 0.7 |
| 4 | 166772855 | rs12642989 | -1.441 | -2.29 | 0.459 | 1.056 | -0.05 | 0.061 | 0.061 | 0.681 | 0.65 | 0.81 | 0.59 | 0.82 | 0.42 | 0.73 |
| 4 | 166785331 | rs6536934 | -1.173 | -2.291 | 0.713 | 1.356 | -0.014 | 0.012 | 0.012 | 0.043 | 0.62 | 0.69 | 0.61 | 0.75 | 0.42 | 0.66 |
| 4 | 166785679 | rs6811456 | 0.2 | -2.009 | 1.067 | 1.46 | -0.01 | 0.008 | 0.008 | 0 | 0.08 | 0.22 | 0.03 | 0.09 | 0.03 | 0.08 |

| | | | | | | | | | | | | | | | |
|---|---|---|---|---|---|---|---|---|---|---|---|---|---|---|---|
| 4 | 166793201 | rs719753 | -0.472 | -2.104 | 0.98 | 1.704 | -0.006 | 0.12 | 0.12 | 0 | 0.14 | 0.28 | 0.15 | 0.37 | 0.11 | 0.14 |
| 4 | 166794829 | rs1460072 | -1.488 | -2.218 | 0.937 | 1.611 | -0.032 | 0.097 | 0.097 | 0.192 | 0.24 | 0.59 | 0.18 | 0.46 | 0.13 | 0.25 |
| 4 | 166796028 | rs1460073 | 0.384 | -1.987 | 0.532 | 1.516 | -0.004 | -0.024 | -0.024 | 0.115 | 0.11 | 0.06 | 0.11 | 0.17 | 0.1 | 0.11 |
| 4 | 166797660 | rs2322443 | -1.388 | -2.168 | 0.35 | 1.168 | -0.055 | 0.073 | 0.073 | 0.359 | 0.64 | 0.79 | 0.37 | 0.86 | 0.45 | 0.75 |
| 4 | 166803528 | rs9637646 | -0.196 | -2.342 | 1.019 | 1.502 | 0.001 | 0 | 0 | 0 | 0.45 | 0.38 | 0.46 | 0.5 | 0.33 | 0.55 |
| 4 | 166806601 | rs2125862 | -1.129 | -2.408 | 1.013 | 1.306 | -0.004 | 0.004 | 0.004 | 0 | 0.61 | 0.62 | 0.61 | 0.75 | 0.44 | 0.67 |
| 4 | 166808248 | rs17633011 | -0.631 | -1.876 | 1.658 | 1.947 | -0.015 | 0.105 | 0.105 | 0 | 0.15 | 0.31 | 0.13 | 0.29 | 0.1 | 0.15 |
| 4 | 166815429 | rs1460060 | -0.143 | -2.117 | 1.535 | 1.858 | 0.002 | -0.006 | -0.006 | 0.161 | 0.31 | 0.29 | 0.47 | 0.53 | 0.25 | 0.51 |
| 4 | 166827370 | rs10025830 | -0.486 | -2.115 | 1.333 | 1.852 | -0.014 | 0.086 | 0.086 | 0 | 0.15 | 0.27 | 0.12 | 0.29 | 0.1 | 0.15 |
| 4 | 166827464 | rs11100626 | -0.369 | -2.116 | 1.425 | 1.849 | -0.016 | 0.034 | 0.034 | 0.104 | 0.33 | 0.4 | 0.48 | 0.57 | 0.25 | 0.51 |
| 4 | 166830580 | rs9990589 | -1.337 | -1.949 | 1.018 | 1.576 | 0.004 | -0.001 | -0.001 | 0 | 0.46 | 0.46 | 0.59 | 0.86 | 0.35 | 0.66 |
| 4 | 166833808 | rs13136952 | 0.689 | -1.911 | 0.343 | 1.544 | -0.023 | -0.102 | -0.102 | 1.681 | 0.21 | 0.02 | 0.37 | 0.41 | 0.18 | 0.43 |
| 4 | 166837654 | rs7658883 | -1.488 | -1.741 | 0.776 | 1.16 | 0.018 | 0.062 | 0.062 | 0.03 | 0.54 | 0.69 | 0.64 | 0.98 | 0.37 | 0.78 |
| 4 | 166838505 | rs1566750 | -1.292 | -1.68 | 0.362 | 1.216 | 0.004 | -0.02 | -0.02 | 0 | 0.52 | 0.52 | 0.62 | 0.93 | 0.37 | 0.78 |
| 4 | 166848001 | rs1018139 | -1.856 | -1.028 | 1.258 | 2.417 | -0.039 | 0.188 | 0.188 | 0 | 0.19 | 0.41 | 0.13 | 0.42 | 0.09 | 0.29 |
| 4 | 166850443 | rs10517873 | -0.595 | -0.56 | 0.475 | 2.278 | -0.004 | 0.068 | 0.068 | 0 | 0.1 | 0.12 | 0.08 | 0.26 | 0.07 | 0.18 |
| 4 | 166854139 | rs11942650 | -0.533 | -0.379 | 0.368 | 1.971 | -0.01 | -0.004 | -0.004 | 0.016 | 0.05 | 0.04 | 0.09 | 0.15 | 0.05 | 0.07 |
| 4 | 166854660 | rs7654070 | -1.969 | -0.583 | 1.327 | 1.983 | 0.044 | 0.229 | 0.229 | 0.328 | 0.1 | 0.42 | 0.12 | 0.2 | 0.06 | 0.08 |
| 4 | 166867875 | rs1503292 | -1.814 | -1.057 | 1.132 | 1.402 | -0.063 | -0.029 | -0.029 | 0 | 0.12 | 0.57 | 0.06 | 0.05 | 0.02 | 0.03 |
| 4 | 166882179 | rs7696087 | 0.135 | -0.708 | -0.358 | 1.865 | -0.078 | -0.114 | -0.114 | 0.185 | 0.75 | 0.98 | 0.83 | 0.54 | 0.73 | 0.57 |
| 4 | 166905263 | rs1503294 | 0.236 | -0.734 | 0.577 | 1.847 | -0.007 | -0.016 | -0.016 | 0.051 | 0.75 | 0.86 | 0.81 | 0.58 | 0.72 | 0.59 |
| 4 | 166921482 | rs13123379 | 0.876 | -1.32 | -0.127 | 1.324 | 0.033 | 0.007 | 0.007 | 0.921 | 0.55 | 0.27 | 0.69 | 0.33 | 0.65 | 0.45 |
| 4 | 166928627 | rs17505658 | -0.132 | -0.519 | -0.659 | 1.05 | -0.065 | -0.007 | -0.007 | 0 | 0.15 | 0.02 | 0.1 | 0.16 | 0.21 | 0.1 |
| 4 | 166933854 | rs1393856 | -0.581 | -0.591 | -0.688 | 0.366 | 0.039 | 0.021 | 0.021 | 0 | 0.51 | 0.69 | 0.38 | 0.55 | 0.43 | 0.47 |
| 4 | 166939154 | rs17689952 | 0.278 | -0.651 | -0.613 | 0.358 | 0.016 | -0.065 | -0.065 | 0 | 0.23 | 0.08 | 0.14 | 0.37 | 0.19 | 0.29 |
| 4 | 166944581 | rs10049682 | -0.638 | -0.581 | 0.154 | 0.281 | 0.034 | 0.078 | 0.078 | 2.032 | 0.09 | 0.47 | 0.09 | 0.13 | 0.04 | 0.04 |
| 4 | 166945293 | rs10003271 | -0.722 | -0.576 | 0.148 | 0.314 | 0.028 | 0.099 | 0.099 | 1.819 | 0.1 | 0.48 | 0.09 | 0.15 | 0.04 | 0.04 |
| 4 | 166945866 | rs7675062 | -0.798 | -0.673 | -0.327 | 0.125 | 0.015 | -0.012 | -0.012 | 0.274 | 0.39 | 0.66 | 0.28 | 0.47 | 0.26 | 0.32 |
| 4 | 166952047 | rs17689976 | -0.386 | 0.127 | -0.119 | -0.18 | 0.001 | 0.017 | 0.017 | 0.1 | 0.1 | 0.1 | 0.1 | 0.26 | 0.13 | 0.1 |
| 4 | 166956652 | rs17047207 | -0.123 | 0.074 | 0.506 | -0.497 | 0.016 | -0.05 | -0.05 | 0 | 0.14 | 0.3 | 0.1 | 0.07 | 0.12 | 0.15 |
| 4 | 166960363 | rs17590344 | -1.444 | -0.351 | 0.208 | -1.039 | -0.001 | 0.052 | 0.052 | 0 | 0.55 | 0.74 | 0.39 | 0.38 | 0.47 | 0.49 |
| 4 | 166965201 | rs6842100 | 0.064 | -0.249 | 0.332 | -0.894 | 0.008 | -0.055 | -0.055 | 0.103 | 0.14 | 0.27 | 0.1 | 0.06 | 0.12 | 0.15 |
| 4 | 166967504 | rs17505781 | -0.485 | -0.386 | -0.139 | -1.052 | 0.02 | 0.017 | 0.017 | 0 | 0.34 | 0.19 | 0.22 | 0.3 | 0.3 | 0.32 |
| 4 | 166971075 | rs9991292 | -0.516 | -0.03 | 0.287 | -1.183 | -0.019 | -0.094 | -0.094 | 0 | 0.41 | 0.5 | 0.23 | 0.12 | 0.32 | 0.38 |
| 4 | 166980214 | rs11730398 | -0.949 | 0.629 | 0.298 | -0.522 | 0.005 | -0.004 | -0.004 | 0 | 0.4 | 0.68 | 0.33 | 0.49 | 0.34 | 0.19 |
| 4 | 166981852 | rs7693478 | -0.972 | 0.581 | 0.262 | -0.545 | 0.011 | -0.004 | -0.004 | 0 | 0.41 | 0.72 | 0.34 | 0.49 | 0.34 | 0.19 |
| 4 | 166996347 | rs2292082 | -1.299 | 0.542 | 0.206 | -0.432 | 0.015 | -0.017 | -0.017 | 0 | 0.43 | 0.83 | 0.38 | 0.48 | 0.36 | 0.26 |
| 4 | 166996414 | rs2292083 | -0.781 | 0.543 | 0.575 | -0.422 | -0.003 | -0.001 | -0.001 | 0 | 0.37 | 0.43 | 0.3 | 0.42 | 0.33 | 0.2 |
| 4 | 167008090 | rs17505880 | 0.251 | 1.023 | 0.054 | -0.169 | -0.026 | 0.062 | 0.062 | 0.024 | 0.13 | 0.02 | 0.1 | 0.02 | 0.12 | 0.09 |
| 4 | 167010685 | rs1503293 | 0.92 | 0.97 | 0.454 | -0.077 | 0.01 | 0.001 | 0.001 | 0.257 | 0.67 | 0.29 | 0.68 | 0.62 | 0.76 | 0.75 |
| 4 | 167014137 | rs1995126 | 0.151 | 1.488 | 0.322 | 0.52 | 0.013 | -0.052 | -0.052 | 0 | 0.78 | 0.73 | 0.83 | 0.91 | 0.74 | 0.74 |
| 4 | 167016664 | rs4690833 | 0.505 | 1.219 | -0.062 | 0.588 | 0.023 | 0.042 | 0.042 | 0.123 | 0.72 | 0.55 | 0.71 | 0.56 | 0.69 | 0.68 |
| 4 | 167019072 | rs1826956 | -0.71 | 0.66 | -0.215 | 0.523 | -0.015 | 0.004 | 0.004 | 0 | 0.32 | 0.49 | 0.23 | 0.63 | 0.3 | 0.27 |
| 4 | 167020324 | rs10018177 | -0.569 | 0.878 | -0.428 | -0.4 | -0.012 | 0.041 | 0.041 | 0 | 0.34 | 0.25 | 0.39 | 0.07 | 0.32 | 0.37 |
| 4 | 167020644 | rs2291822 | 0.003 | 0.541 | 0.066 | 0.52 | -0.015 | 0.275 | 0.275 | 0 | 0.07 | 0.19 | 0.06 | 0.43 | 0.04 | 0.06 |
| 4 | 167022359 | rs2279723 | 0.969 | 0.191 | 0.703 | 0.287 | 0.006 | 0 | 0 | 0.28 | 0.53 | 0.33 | 0.4 | 0.5 | 0.57 | 0.54 |
| 4 | 167023760 | rs1909083 | -0.001 | 0.179 | 0.706 | 0.276 | -0.018 | 0.261 | 0.261 | 0 | 0.07 | 0.19 | 0.06 | 0.43 | 0.04 | 0.06 |
| 4 | 167024715 | rs1497949 | 0.866 | 0.139 | 0.68 | 0.257 | 0.003 | -0.005 | -0.005 | 0 | 0.54 | 0.34 | 0.41 | 0.5 | 0.58 | 0.55 |
| 4 | 167027707 | rs1391334 | 0.217 | 0.207 | 0.58 | 0.132 | 0.003 | -0.096 | -0.096 | 0 | 0.63 | 0.54 | 0.59 | 0.92 | 0.67 | 0.62 |
| 4 | 167040095 | rs12649030 | -0.057 | 0.202 | 0.743 | 0.458 | 0.006 | 0.001 | 0.001 | 0.93 | 0.7 | 0.66 | 0.65 | 0.79 | 0.66 | 0.71 |
| 4 | 167044948 | rs904246 | -0.686 | 0.227 | 0.865 | 0.609 | -0.012 | 0.173 | 0.173 | 0 | 0.94 | 0.84 | 0.97 | 0.83 | 0.98 | 0.95 |
| 4 | 167046795 | rs17690647 | 0.471 | 0.224 | 0.788 | 0.603 | -0.016 | 0.07 | 0.07 | 0 | 0.06 | 0.1 | 0.03 | 0.17 | 0.03 | 0.05 |
| 4 | 167049629 | rs1391332 | 0.441 | -0.079 | 0.35 | -0.26 | 0.019 | 0.037 | 0.037 | 11.686 | 0.3 | 0.21 | 0.21 | 0.03 | 0.26 | 0.21 |
| 4 | 167054170 | rs6825989 | 0.747 | -0.078 | 0.571 | -0.173 | -0.012 | 0.278 | 0.278 | 0.122 | 0.89 | 0.67 | 0.8 | 0.58 | 0.92 | 0.93 |
| 4 | 167057609 | rs6826343 | -0.561 | -0.217 | 0.182 | -1.131 | -0.004 | 0.036 | 0.036 | 1.221 | 0.57 | 0.67 | 0.53 | 0.08 | 0.55 | 0.43 |
| 4 | 167069396 | rs12650012 | 0.51 | 0.163 | 0.588 | -0.639 | 0.006 | -0.056 | -0.056 | 0.053 | 0.58 | 0.51 | 0.49 | 0.16 | 0.52 | 0.43 |
| 4 | 167074691 | rs2874823 | 0.625 | 0.232 | 0.355 | -0.602 | -0.024 | 0.028 | 0.028 | 0.495 | 0.21 | 0.16 | 0.31 | 0.04 | 0.3 | 0.23 |
| 4 | 167079178 | rs1039178 | 0.408 | 0.398 | 0.321 | -0.563 | -0.009 | 0.043 | 0.043 | 6.007 | 0.55 | 0.38 | 0.47 | 0.16 | 0.52 | 0.44 |
| 4 | 167079944 | rs10517878 | 0.548 | 0.405 | 0.176 | -0.572 | -0.002 | 0.039 | 0.039 | 6.049 | 0.54 | 0.37 | 0.36 | 0.16 | 0.48 | 0.42 |
| 4 | 167081486 | rs13112070 | 0.344 | 0.446 | -0.364 | -0.471 | -0.008 | -0.004 | -0.004 | 6.96 | 0.27 | 0.27 | 0.34 | 0.21 | 0.32 | 0.29 |
| 4 | 167082048 | rs1497961 | -0.169 | 0.554 | -0.995 | -1.188 | -0.016 | 0.12 | 0.12 | 6.838 | 0.56 | 0.28 | 0.51 | 0.16 | 0.53 | 0.46 |
| 4 | 167082557 | rs6536948 | 0.316 | 0.336 | -1.187 | -0.992 | -0.016 | 0.179 | 0.179 | 2.139 | 0.4 | 0.16 | 0.27 | 0.12 | 0.42 | 0.36 |
| 4 | 167095645 | rs2221058 | 0.099 | 0.531 | -0.736 | -0.65 | -0.005 | 0.001 | 0.001 | 3.755 | 0.56 | 0.41 | 0.48 | 0.15 | 0.52 | 0.42 |
| 4 | 167099687 | rs6838584 | -0.204 | 0.331 | -0.566 | 0.052 | -0.003 | 0.014 | 0.014 | 0 | 0.12 | 0.17 | 0.19 | 0.21 | 0.13 | 0.22 |
| 4 | 167099916 | rs17047336 | -0.362 | -0.07 | -0.456 | -0.362 | -0.013 | -0.011 | -0.011 | 0 | 0.35 | 0.52 | 0.3 | 0.38 | 0.32 | 0.6 |
| 4 | 167102631 | rs28415748 | -0.868 | -0.743 | -0.873 | -0.807 | -0.001 | -0.002 | -0.002 | 0 | 0.49 | 0.55 | 0.35 | 0.4 | 0.37 | 0.65 |
| 4 | 167104872 | rs10028492 | -0.877 | -0.759 | -0.878 | -0.784 | -0.001 | -0.001 | -0.001 | 0 | 0.5 | 0.55 | 0.35 | 0.41 | 0.38 | 0.65 |
| 4 | 167106550 | rs10012204 | -0.877 | -0.759 | -0.878 | -0.784 | -0.001 | -0.001 | -0.001 | 1.429 | 0.5 | 0.55 | 0.35 | 0.41 | 0.38 | 0.65 |
| 4 | 167109588 | rs1497956 | 1.039 | -0.903 | -0.861 | -0.645 | -0.053 | 0.053 | 0.053 | 0 | 0.33 | 0.15 | 0.49 | 0.14 | 0.52 | 0.28 |

| | | | | | | | | | | | | | | | | |
|---|---|---|---|---|---|---|---|---|---|---|---|---|---|---|---|---|
| 4 | 167110433 | rs1497957 | -0.35 | -0.847 | -0.909 | -0.66 | -0.001 | -0.001 | -0.001 | 0.202 | 0.5 | 0.55 | 0.35 | 0.41 | 0.38 | 0.65 |
| 4 | 167118729 | rs833609 | -0.14 | -0.963 | -0.736 | -0.663 | 0.023 | 0.023 | 0.023 | 0 | 0.5 | 0.62 | 0.35 | 0.41 | 0.38 | 0.65 |
| 4 | 167124766 | rs833605 | 0.584 | -0.864 | -0.756 | -0.608 | 0.006 | 0.071 | 0.071 | 0 | 0.42 | 0.21 | 0.64 | 0.26 | 0.64 | 0.49 |
| 4 | 167125430 | rs353867 | 0.858 | -1.021 | -0.255 | -0.17 | -0.045 | 0.02 | 0.02 | 0 | 0.66 | 0.58 | 0.77 | 0.45 | 0.82 | 0.87 |
| 4 | 167143385 | rs17047381 | -0.29 | -0.99 | 0.404 | -0.178 | -0.066 | -0.08 | -0.08 | 0 | 0.18 | 0.27 | 0.07 | 0.02 | 0.05 | 0.05 |
| 4 | 167146175 | rs11940089 | -0.06 | -0.988 | 0.399 | -0.177 | -0.047 | -0.058 | -0.058 | 1.19 | 0.17 | 0.24 | 0.07 | 0.02 | 0.05 | 0.05 |
| 4 | 167163302 | rs435881 | 0.863 | -0.887 | 0.507 | 0.005 | -0.054 | 0.055 | 0.055 | 0 | 0.69 | 0.61 | 0.76 | 0.51 | 0.84 | 0.89 |
| 4 | 167174293 | rs1459835 | 0.207 | -0.742 | 0.388 | 0.021 | 0.001 | -0.015 | -0.015 | 0.034 | 0.12 | 0.09 | 0.07 | 0 | 0.09 | 0.04 |
| 4 | 167179581 | rs382628 | 0.621 | -0.815 | 0.603 | 0.124 | -0.006 | 0.066 | 0.066 | 0.96 | 0.81 | 0.62 | 0.75 | 0.27 | 0.86 | 0.77 |
| 4 | 167204445 | rs1380497 | 1.142 | -0.929 | 0.506 | -0.075 | -0.046 | -0.025 | -0.025 | 0 | 0.66 | 0.57 | 0.73 | 0.26 | 0.83 | 0.73 |
| 4 | 167222523 | rs895035 | 1.005 | -0.999 | 0.457 | -0.074 | -0.022 | -0.013 | -0.013 | 0 | 0.67 | 0.63 | 0.73 | 0.27 | 0.78 | 0.71 |
| 4 | 167229729 | rs11726963 | 1.204 | -0.998 | 0.477 | -0.084 | -0.045 | -0.027 | -0.027 | 0.438 | 0.64 | 0.54 | 0.72 | 0.27 | 0.78 | 0.71 |
| 4 | 167232997 | rs425222 | -1.006 | -1.028 | 0.569 | 0.094 | -0.019 | 0.018 | 0.018 | 0.259 | 0.31 | 0.28 | 0.22 | 0.48 | 0.17 | 0.11 |
| 4 | 167237947 | rs368067 | 1.08 | -0.937 | 0.602 | 0.136 | 0.016 | 0.146 | 0.146 | 0 | 0.95 | 0.82 | 0.95 | 0.75 | 0.95 | 0.83 |
| 4 | 167249784 | rs384968 | -1.184 | -0.927 | 0.561 | 0.13 | -0.02 | 0.022 | 0.022 | 0.073 | 0.3 | 0.28 | 0.22 | 0.48 | 0.17 | 0.11 |
| 4 | 167258089 | rs7436496 | 1.164 | -0.975 | 0.605 | 0.121 | -0.024 | -0.011 | -0.011 | 0 | 0.66 | 0.63 | 0.73 | 0.28 | 0.78 | 0.71 |
| 4 | 167265509 | rs12506355 | 1.261 | -0.926 | 0.638 | 0.198 | -0.044 | -0.016 | -0.016 | 0 | 0.66 | 0.58 | 0.73 | 0.3 | 0.78 | 0.83 |
| 4 | 167284116 | rs7653953 | -0.385 | -0.732 | 0.569 | 0.455 | -0.026 | -0.006 | -0.006 | 0 | 0.18 | 0.24 | 0.07 | 0.13 | 0.1 | 0.07 |
| 4 | 167287685 | rs1982582 | 1.065 | 0.008 | 0.1 | 0.035 | -0.066 | 0.071 | 0.071 | 0.179 | 0.13 | 0.01 | 0.28 | 0 | 0.16 | 0.15 |
| 4 | 167300345 | rs1080788 | 1.369 | -0.523 | 0.726 | 0.435 | -0.043 | 0.04 | 0.04 | 0.169 | 0.7 | 0.57 | 0.76 | 0.44 | 0.8 | 0.88 |
| 4 | 167309607 | rs2639302 | 1.008 | -0.335 | 0.751 | 0.611 | 0.045 | 0.255 | 0.255 | 2.177 | 0.87 | 0.66 | 0.8 | 0.48 | 0.85 | 0.92 |
| 4 | 167310371 | rs1492533 | -0.534 | -0.309 | 0.894 | 0.533 | 0 | -0.041 | -0.041 | 0 | 0.89 | 0.89 | 0.94 | 1 | 0.89 | 0.95 |
| 4 | 167321777 | rs1565650 | 0.656 | -0.318 | 1.089 | 0.603 | 0.023 | 0.226 | 0.226 | 1.758 | 0.84 | 0.61 | 0.81 | 0.46 | 0.85 | 0.91 |
| 4 | 167329523 | rs10440459 | -1.8 | -0.435 | 1.077 | 0.606 | 0.059 | 0.128 | 0.128 | 1.192 | 0.06 | 0.28 | 0.04 | 0.11 | 0.05 | 0.03 |
| 4 | 167351495 | rs1388879 | 1.04 | -0.686 | 1.106 | 0.429 | -0.076 | 0.033 | 0.033 | 0.206 | 0.7 | 0.58 | 0.76 | 0.36 | 0.85 | 0.73 |
| 4 | 167352346 | rs1492523 | 1.634 | -0.545 | 0.79 | -0.174 | -0.003 | 0.042 | 0.042 | 0.035 | 0.34 | 0.3 | 0.22 | 0.17 | 0.35 | 0.39 |
| 4 | 167356733 | rs1602681 | -0.382 | -0.563 | 0.859 | -0.11 | -0.012 | 0.153 | 0.153 | 0.14 | 0.87 | 0.78 | 0.83 | 0.57 | 0.91 | 0.93 |
| 4 | 167366409 | rs6845532 | 0.193 | -0.565 | 0.676 | -0.202 | -0.033 | -0.001 | -0.001 | 0 | 0.35 | 0.24 | 0.62 | 0.63 | 0.5 | 0.35 |
| 4 | 167366750 | rs11100648 | 1.576 | -0.581 | 0.689 | -0.211 | 0 | 0.044 | 0.044 | 0.247 | 0.35 | 0.32 | 0.22 | 0.17 | 0.35 | 0.39 |
| 4 | 167368812 | rs4596200 | 0.392 | -0.5 | 0.575 | -0.159 | -0.089 | 0.084 | 0.084 | 1.043 | 0.33 | 0.12 | 0.51 | 0.15 | 0.45 | 0.31 |
| 4 | 167372087 | rs13141694 | 0.179 | -0.821 | 0.546 | -0.396 | -0.033 | -0.002 | -0.002 | 0 | 0.35 | 0.24 | 0.62 | 0.63 | 0.5 | 0.36 |
| 4 | 167383599 | rs1492539 | -1.01 | -0.712 | 0.347 | -0.311 | 0 | 0.025 | 0.025 | 0 | 0.89 | 0.97 | 0.94 | 1 | 0.91 | 0.94 |
| 4 | 167400016 | rs10857379 | 1.274 | -0.503 | -0.003 | -0.056 | -0.002 | 0.053 | 0.053 | 0.059 | 0.34 | 0.29 | 0.22 | 0.17 | 0.35 | 0.39 |
| 4 | 167407097 | rs993097 | -0.498 | -0.243 | 0.562 | 0.823 | -0.006 | 0.003 | 0.003 | 0 | 0.38 | 0.34 | 0.66 | 0.64 | 0.51 | 0.39 |
| 4 | 167408477 | rs11100652 | -0.509 | -0.614 | 0.576 | 1.143 | -0.031 | 0.11 | 0.11 | 2.946 | 0.43 | 0.25 | 0.56 | 0.2 | 0.46 | 0.35 |
| 4 | 167421242 | rs992021 | -0.263 | -0.688 | 0.487 | 1.18 | -0.044 | 0.104 | 0.104 | 2.432 | 0.4 | 0.22 | 0.54 | 0.19 | 0.44 | 0.33 |
| 4 | 167422116 | rs978230 | -0.903 | -0.936 | 0.899 | 1.479 | -0.001 | -0.004 | -0.004 | 0 | 0.52 | 0.52 | 0.7 | 0.67 | 0.47 | 0.38 |
| 4 | 167423956 | rs12504728 | -0.247 | -0.72 | 0.193 | 1.57 | -0.091 | 0.118 | 0.118 | 2.743 | 0.39 | 0.13 | 0.54 | 0.19 | 0.44 | 0.33 |
| 4 | 167437817 | rs10034660 | -1.491 | -0.789 | 0.441 | 1.329 | -0.04 | 0.067 | 0.067 | 0.068 | 0.6 | 0.82 | 0.75 | 0.75 | 0.5 | 0.59 |
| 4 | 167448941 | rs875441 | -1.284 | -1.034 | 0.92 | 1.569 | -0.061 | 0.16 | 0.16 | 0.954 | 0.19 | 0.59 | 0.21 | 0.5 | 0.07 | 0.2 |
| 4 | 167450233 | rs6826162 | -0.146 | -1.082 | 1.212 | 2.216 | -0.026 | 0.179 | 0.179 | 0.398 | 0.08 | 0.24 | 0.14 | 0.28 | 0.02 | 0.04 |
| 4 | 167455493 | rs11941399 | -1.65 | -1.092 | 0.864 | 2.174 | -0.041 | 0.07 | 0.07 | 1.736 | 0.2 | 0.68 | 0.21 | 0.5 | 0.07 | 0.2 |
| 4 | 167460285 | rs2131252 | -1.593 | -1.033 | 0.686 | 1.769 | -0.076 | 0.088 | 0.088 | 0.039 | 0.61 | 0.88 | 0.76 | 0.76 | 0.51 | 0.59 |
| 4 | 167488592 | rs7439050 | -1.57 | -1.134 | 1.237 | 1.966 | -0.06 | 0.078 | 0.078 | 0.152 | 0.18 | 0.53 | 0.11 | 0.27 | 0.07 | 0.19 |
| 4 | 167514725 | rs4691299 | -0.984 | -0.887 | 1.26 | 1.649 | -0.001 | 0.012 | 0.012 | 0.155 | 0.51 | 0.53 | 0.69 | 0.78 | 0.46 | 0.57 |
| 4 | 167528623 | rs6536987 | -1.323 | -0.939 | 1.311 | 1.623 | -0.005 | 0.025 | 0.025 | 0 | 0.55 | 0.61 | 0.71 | 0.78 | 0.46 | 0.57 |
| 4 | 167529635 | rs28840815 | -1.474 | -1.014 | 1.172 | 1.587 | -0.036 | 0.09 | 0.09 | 0.021 | 0.57 | 0.82 | 0.74 | 0.78 | 0.46 | 0.57 |
| 4 | 167540460 | rs7673384 | 0.181 | -0.914 | 1.143 | 1.888 | -0.05 | 0.065 | 0.065 | 0.02 | 0.61 | 0.89 | 0.47 | 0.75 | 0.61 | 0.62 |
| 4 | 167547429 | rs7673076 | 0.307 | -0.966 | 1.508 | 2.131 | -0.021 | -0.045 | -0.045 | 0 | 0.61 | 0.78 | 0.36 | 0.52 | 0.61 | 0.68 |
| 4 | 167550669 | rs10001950 | 0.943 | -1.056 | 2.069 | 2.127 | 0.009 | 0.007 | 0.007 | 1.471 | 0.53 | 0.3 | 0.33 | 0.46 | 0.6 | 0.6 |
| 4 | 167550780 | rs10034869 | 0.943 | -1.056 | 2.069 | 2.127 | 0.009 | 0.007 | 0.007 | 1.471 | 0.53 | 0.3 | 0.33 | 0.46 | 0.6 | 0.6 |
| 4 | 167552158 | rs6822910 | -1.676 | -1.041 | 1.955 | 1.443 | 0.003 | 0.068 | 0.068 | 6.87 | 0.08 | 0.48 | 0.04 | 0.06 | 0.01 | 0.09 |
| 4 | 167571068 | rs28713613 | -0.317 | -1.086 | 0.989 | 1.412 | -0.049 | 0.137 | 0.137 | 0 | 0.33 | 0.15 | 0.49 | 0.07 | 0.34 | 0.27 |
| 4 | 167594972 | rs10032330 | -1.311 | -1.108 | 1.613 | 2.415 | 0.034 | -0.043 | -0.043 | 0 | 0.35 | 0.79 | 0.32 | 0.46 | 0.25 | 0.33 |
| 4 | 167602105 | rs3924566 | -0.658 | -1.08 | 1.997 | 2.148 | -0.019 | -0.038 | -0.038 | 0 | 0.28 | 0.41 | 0.25 | 0.18 | 0.22 | 0.24 |
| 4 | 167607492 | rs7692681 | -0.669 | -1.049 | 1.845 | 1.731 | -0.019 | -0.038 | -0.038 | 0 | 0.28 | 0.41 | 0.25 | 0.18 | 0.22 | 0.24 |
| 4 | 167607726 | rs7661107 | -0.723 | -1.05 | 1.887 | 1.73 | -0.024 | -0.048 | -0.048 | 0 | 0.29 | 0.46 | 0.25 | 0.18 | 0.22 | 0.24 |
| 4 | 167622160 | rs6553485 | -0.66 | -0.991 | 0.791 | 1.772 | -0.017 | -0.032 | -0.032 | 0 | 0.28 | 0.39 | 0.25 | 0.18 | 0.22 | 0.24 |
| 4 | 167625748 | rs7674577 | -1.904 | -1.13 | -0.162 | 1.87 | 0.072 | 0.023 | 0.023 | 2.611 | 0.45 | 0.95 | 0.42 | 0.74 | 0.3 | 0.55 |
| 4 | 167634706 | rs12503564 | -0.585 | -0.479 | 0.228 | 1.404 | 0.004 | 0.009 | 0.009 | 0 | 0.26 | 0.19 | 0.22 | 0.18 | 0.22 | 0.24 |
| 4 | 167635674 | rs2198738 | -0.601 | -0.468 | 0.138 | 1.41 | -0.001 | -0.001 | -0.001 | 0 | 0.27 | 0.24 | 0.23 | 0.18 | 0.22 | 0.24 |
| 4 | 167644235 | rs17052538 | -1.349 | -0.344 | 0.604 | 2.077 | -0.047 | 0.313 | 0.313 | 0.858 | 0.13 | 0.49 | 0.12 | 0.42 | 0.07 | 0.18 |
| 4 | 167645029 | rs13109075 | -0.774 | 0.22 | 0.029 | 1.657 | 0.008 | 0.012 | 0.012 | 0 | 0.34 | 0.39 | 0.43 | 0.41 | 0.36 | 0.43 |
| 4 | 167647323 | rs4860006 | -0.488 | 0.458 | -0.436 | 1.242 | -0.003 | 0.011 | 0.011 | 0 | 0.54 | 0.64 | 0.64 | 0.59 | 0.54 | 0.5 |
| 4 | 167648055 | rs4859971 | -0.341 | 0.61 | -0.484 | 0.951 | 0.002 | -0.01 | -0.01 | 1.704 | 0.44 | 0.69 | 0.57 | 0.51 | 0.41 | 0.43 |
| 4 | 167649165 | rs2198737 | 0.238 | 0.639 | 0.049 | 1.281 | 0 | -0.007 | -0.007 | 0.761 | 0.34 | 0.26 | 0.51 | 0.34 | 0.34 | 0.35 |
| 4 | 167652786 | rs11943562 | -0.937 | 0.616 | 0.112 | 1.176 | 0.022 | -0.127 | -0.127 | 0 | 0.2 | 0.45 | 0.13 | 0.05 | 0.18 | 0.12 |
| 4 | 167655482 | rs3762244 | 0.297 | 0.596 | 0.089 | 1.207 | 0 | -0.007 | -0.007 | 0.001 | 0.34 | 0.27 | 0.51 | 0.34 | 0.34 | 0.35 |
| 4 | 167655803 | rs7695294 | -0.014 | 0.638 | -0.023 | 1.063 | 0.001 | -0.001 | -0.001 | 0 | 0.39 | 0.37 | 0.55 | 0.36 | 0.4 | 0.36 |

| | | | | | | | | | | | | | | | | |
|---|---|---|---|---|---|---|---|---|---|---|---|---|---|---|---|---|
| 4 | 167659338 | rs7681289 | -1.086 | 0.4 | -0.595 | 0.502 | 0.001 | -0.019 | -0.019 | 0.208 | 0.8 | 0.86 | 0.78 | 0.54 | 0.81 | 0.83 |
| 4 | 167661584 | rs7678726 | -0.255 | 0.409 | -0.585 | 0.42 | 0 | -0.017 | -0.017 | 1.215 | 0.45 | 0.52 | 0.27 | 0.21 | 0.47 | 0.48 |
| 4 | 167663664 | rs7691894 | 1.05 | 0.39 | -0.591 | 0.516 | 0 | -0.009 | -0.009 | 0 | 0.34 | 0.28 | 0.51 | 0.35 | 0.35 | 0.35 |
| 4 | 167664591 | rs2318485 | -0.841 | 0.442 | -0.517 | 0.549 | 0.001 | 0.001 | 0.001 | 0 | 0.79 | 0.81 | 0.78 | 0.55 | 0.81 | 0.83 |
| 4 | 167685213 | rs7689440 | 1.197 | 0.625 | -0.656 | 0.536 | -0.039 | 0.086 | 0.086 | 0 | 0.25 | 0.06 | 0.14 | 0.07 | 0.28 | 0.34 |
| 4 | 167695661 | rs897511 | -0.002 | 0.739 | -0.484 | 0.61 | 0.002 | -0.042 | -0.042 | 0 | 0.39 | 0.55 | 0.21 | 0.12 | 0.4 | 0.45 |
| 4 | 167698248 | rs6817936 | -0.079 | 0.51 | -0.725 | 0.945 | -0.006 | -0.025 | -0.025 | 1.639 | 0.44 | 0.26 | 0.55 | 0.46 | 0.46 | 0.44 |
| 4 | 167698563 | rs17052594 | 0.247 | 0.39 | 0.017 | 0.833 | 0.011 | -0.034 | -0.034 | 1.251 | 0.09 | 0.16 | 0.05 | 0.05 | 0.08 | 0.08 |
| 4 | 167699942 | rs7698457 | -0.846 | 0.179 | -0.342 | 1.427 | -0.004 | 0.014 | 0.014 | 0 | 0.28 | 0.33 | 0.27 | 0.52 | 0.24 | 0.26 |
| 4 | 167700120 | rs7698841 | -0.843 | 0.111 | -0.317 | 1.447 | -0.007 | 0.022 | 0.022 | 0 | 0.28 | 0.34 | 0.28 | 0.51 | 0.23 | 0.26 |
| 4 | 167711854 | rs10517903 | -0.785 | 0.367 | -1.003 | 1.255 | -0.024 | -0.068 | -0.068 | 0.551 | 0.28 | 0.1 | 0.18 | 0.35 | 0.27 | 0.26 |
| 4 | 167727136 | rs1551491 | -1.202 | -0.164 | -0.205 | 1.238 | -0.008 | -0.01 | -0.01 | 0.558 | 0.33 | 0.6 | 0.33 | 0.72 | 0.26 | 0.36 |
| 4 | 167738944 | rs1023875 | 0.545 | -0.341 | 0.076 | 1.085 | -0.015 | -0.034 | -0.034 | 0 | 0.13 | 0.18 | 0.03 | 0 | 0.02 | 0.09 |
| 4 | 167745216 | rs921856 | 0.157 | -0.324 | -0.308 | 0.577 | 0.01 | 0.013 | 0.013 | 0.158 | 0.67 | 0.7 | 0.64 | 0.79 | 0.76 | 0.79 |
| 4 | 167749783 | rs17701990 | 0.794 | -0.174 | 0.094 | 0.952 | 0.004 | 0.059 | 0.059 | 0.026 | 0.08 | 0.07 | 0.08 | 0.28 | 0.1 | 0.19 |
| 4 | 167760175 | rs7698485 | 1.923 | -0.344 | -0.272 | 0.638 | -0.145 | 0.266 | 0.266 | 0 | 0.44 | 0.04 | 0.31 | 0.09 | 0.49 | 0.51 |
| 4 | 167776873 | rs897514 | 0.463 | -0.342 | -0.18 | 0.712 | 0.01 | -0.019 | -0.019 | 0 | 0.62 | 0.7 | 0.59 | 0.54 | 0.67 | 0.61 |
| 4 | 167785513 | rs4859997 | 0.741 | -0.388 | 0.185 | 0.774 | -0.018 | -0.001 | -0.001 | 1.23 | 0.62 | 0.45 | 0.46 | 0.36 | 0.69 | 0.73 |
| 4 | 167789207 | rs7440269 | 0.209 | -0.382 | 0.331 | 0.976 | -0.001 | 0.115 | 0.115 | 0 | 0.06 | 0.09 | 0.19 | 0.37 | 0.07 | 0.06 |
| 4 | 167790958 | rs7672200 | 1.379 | -0.383 | 0.356 | 0.878 | 0.004 | 0.005 | 0.005 | 0.158 | 0.55 | 0.41 | 0.41 | 0.35 | 0.59 | 0.7 |
| 4 | 167817905 | rs10517908 | 0.883 | -0.407 | 0.209 | 0.687 | -0.011 | -0.031 | -0.031 | 0 | 0.61 | 0.45 | 0.58 | 0.72 | 0.66 | 0.75 |
| 4 | 167822014 | rs12649447 | 0.563 | -0.167 | -0.25 | 0.702 | 0.003 | -0.006 | -0.006 | 0 | 0.09 | 0.13 | 0.06 | 0.08 | 0.09 | 0.05 |
| 4 | 167837678 | rs17645839 | 0.438 | 0.086 | -0.728 | 0.716 | -0.043 | -0.067 | -0.067 | 1.388 | 0.11 | 0 | 0.1 | 0.27 | 0.15 | 0.22 |
| 4 | 167837707 | rs6846723 | 1.232 | -0.047 | -0.497 | -0.339 | -0.014 | 0.094 | 0.094 | 0 | 0.43 | 0.25 | 0.35 | 0.11 | 0.46 | 0.48 |
| 4 | 167843608 | rs10020692 | -0.592 | 0.256 | -1.035 | -0.814 | -0.029 | 0.033 | 0.033 | 0 | 0.61 | 0.8 | 0.47 | 0.71 | 0.54 | 0.64 |
| 4 | 167844096 | rs8180142 | -0.389 | -0.068 | -0.475 | -0.608 | -0.002 | -0.002 | -0.002 | 0 | 0.4 | 0.59 | 0.33 | 0.68 | 0.3 | 0.34 |
| 4 | 167853560 | rs6828808 | -0.291 | 0.527 | -0.211 | -0.941 | 0.041 | -0.073 | -0.073 | 0.126 | 0.79 | 0.67 | 0.83 | 0.94 | 0.75 | 0.84 |
| 4 | 167857369 | rs6851188 | 0.78 | 0.415 | -0.795 | -1.018 | -0.04 | 0.089 | 0.089 | 0.65 | 0.64 | 0.19 | 0.75 | 0.81 | 0.65 | 0.71 |
| 4 | 167864930 | rs12644678 | 1.038 | 0.075 | -1.355 | -1.388 | -0.061 | 0.005 | 0.005 | 0.04 | 0.37 | 0.12 | 0.37 | 0.63 | 0.45 | 0.38 |
| 4 | 167866712 | rs6823924 | -0.468 | -0.02 | 0.205 | -0.94 | 0.007 | -0.043 | -0.043 | 0.006 | 0.14 | 0.37 | 0.08 | 0.05 | 0.1 | 0.1 |
| 4 | 167867095 | rs7682844 | 0.332 | -0.032 | 0.523 | -0.908 | 0 | -0.004 | -0.004 | 0 | 0.08 | 0.16 | 0.04 | 0.05 | 0.06 | 0.08 |
| 4 | 167899470 | rs1896479 | -0.529 | -0.126 | 0.329 | -1.254 | 0.02 | -0.03 | -0.03 | 0 | 0.23 | 0.36 | 0.17 | 0.16 | 0.23 | 0.25 |
| 4 | 167902573 | rs1593770 | 1.071 | -0.164 | 0.22 | -1.276 | -0.007 | 0.119 | 0.119 | 0 | 0.2 | 0.03 | 0.1 | 0.01 | 0.15 | 0.22 |
| 4 | 167907991 | rs2703841 | 0.673 | -0.102 | 0.566 | -1.23 | 0.005 | -0.021 | -0.021 | 2.087 | 0.75 | 0.36 | 0.8 | 0.83 | 0.75 | 0.74 |
| 4 | 167914437 | rs1863782 | 0.872 | -0.125 | 0.696 | -1.242 | -0.002 | 0.001 | 0.001 | 0.924 | 0.74 | 0.29 | 0.8 | 0.83 | 0.75 | 0.73 |
| 4 | 167925272 | rs17646122 | 1.813 | -0.261 | 0.251 | -1.166 | 0.007 | -0.011 | -0.011 | 0.01 | 0.07 | 0.04 | 0.26 | 0.18 | 0.06 | 0.07 |
| 4 | 167933010 | rs6553490 | -0.942 | -0.06 | 0.487 | -1.357 | 0.001 | -0.012 | -0.012 | 0 | 0.26 | 0.66 | 0.2 | 0.16 | 0.24 | 0.25 |
| 4 | 167938604 | rs2703842 | 1.066 | -0.112 | 0.379 | -1.416 | 0.003 | 0.045 | 0.045 | 0 | 0.54 | 0.37 | 0.71 | 0.83 | 0.62 | 0.52 |
| 4 | 167943152 | rs7685624 | -0.942 | -0.05 | 0.454 | -1.343 | 0.025 | -0.05 | -0.05 | 0.011 | 0.25 | 0.55 | 0.19 | 0.17 | 0.25 | 0.26 |
| 4 | 167953146 | rs1427624 | 1.136 | -0.042 | 0.382 | -1.309 | -0.028 | 0.203 | 0.203 | 0.012 | 0.55 | 0.16 | 0.69 | 0.83 | 0.61 | 0.53 |
| 4 | 167956606 | rs6818745 | 0.517 | 0.433 | 0.67 | -0.728 | 0.072 | -0.007 | -0.007 | 2.028 | 0.13 | 0.38 | 0.12 | 0.11 | 0.14 | 0.14 |
| 4 | 167959369 | rs7660050 | -0.527 | 0.342 | 0.249 | -0.719 | 0.02 | -0.029 | -0.029 | 0 | 0.2 | 0.55 | 0.16 | 0.16 | 0.19 | 0.22 |
| 4 | 167982046 | rs13137632 | 0.779 | -0.118 | -0.265 | -1.096 | 0.039 | 0.072 | 0.072 | 0 | 0.52 | 0.36 | 0.54 | 0.77 | 0.64 | 0.58 |
| 4 | 167995689 | rs6829378 | -1.243 | -0.429 | -0.348 | -1.122 | -0.006 | -0.004 | -0.004 | 0.708 | 0.3 | 0.62 | 0.36 | 0.23 | 0.22 | 0.2 |
| 4 | 167999350 | rs1427635 | 0.005 | -0.479 | -0.172 | -1.136 | -0.031 | 0.001 | 0.001 | 0 | 0.18 | 0.3 | 0.3 | 0.19 | 0.13 | 0.12 |
| 4 | 168003155 | rs17599740 | 1.721 | -0.465 | -0.282 | -1.171 | -0.006 | 0.103 | 0.103 | 0 | 0.16 | 0.02 | 0.07 | 0.01 | 0.11 | 0.21 |
| 4 | 168008751 | rs7688165 | -0.402 | -0.323 | 0.079 | -0.912 | -0.049 | 0.002 | 0.002 | 0 | 0.19 | 0.42 | 0.31 | 0.19 | 0.13 | 0.12 |
| 4 | 168022694 | rs11939479 | 0.803 | 0.078 | -0.388 | -0.977 | -0.043 | -0.035 | -0.035 | 0 | 0.13 | 0.03 | 0.16 | 0.15 | 0.17 | 0.1 |
| 4 | 168029135 | rs1579404 | -0.895 | -0.001 | -0.321 | -1.029 | 0.001 | -0.001 | -0.001 | 1.061 | 0.24 | 0.22 | 0.21 | 0.19 | 0.26 | 0.18 |
| 4 | 168034257 | rs815213 | 0.094 | 0.027 | -0.43 | -1.021 | -0.061 | -0.023 | -0.023 | 0 | 0.17 | 0.03 | 0.18 | 0.16 | 0.22 | 0.13 |
| 4 | 168036224 | rs815212 | -1.746 | -0.069 | -0.255 | -1.126 | 0.006 | 0.014 | 0.014 | 0 | 0.43 | 0.61 | 0.52 | 0.32 | 0.39 | 0.31 |
| 4 | 168038730 | rs815216 | -0.225 | -0.007 | -0.267 | -1.06 | -0.002 | -0.006 | -0.006 | 0 | 0.2 | 0.09 | 0.18 | 0.19 | 0.2 | 0.17 |
| 4 | 168039224 | rs10517917 | -0.028 | -0.038 | -0.211 | -0.988 | -0.024 | -0.038 | -0.038 | 0 | 0.17 | 0.32 | 0.31 | 0.12 | 0.14 | 0.1 |
| 4 | 168044726 | rs7435823 | 0.984 | -0.052 | -1.074 | -0.937 | 0.004 | 0.003 | 0.003 | 0 | 0.12 | 0.01 | 0.27 | 0.12 | 0.11 | 0.09 |
| 4 | 168052793 | rs13125626 | 0.88 | 0.395 | -0.725 | -0.758 | 0.096 | 0.024 | 0.024 | 0.488 | 0.13 | 0.41 | 0.18 | 0.14 | 0.19 | 0.2 |
| 4 | 168055022 | rs12498755 | 1.978 | 0.039 | -1.159 | -0.678 | -0.075 | -0.151 | -0.151 | 0.833 | 0.24 | 0.02 | 0.23 | 0.51 | 0.29 | 0.31 |
| 4 | 168056478 | rs12640303 | 1.035 | -0.135 | -0.64 | -0.579 | -0.022 | -0.015 | -0.015 | 0 | 0.62 | 0.51 | 0.5 | 0.66 | 0.69 | 0.73 |
| 4 | 168080117 | rs17703374 | -0.825 | -0.436 | -0.448 | -0.3 | -0.007 | -0.002 | -0.002 | 0.43 | 0.22 | 0.22 | 0.31 | 0.22 | 0.15 | 0.15 |
| 4 | 168084157 | rs10017958 | -1.489 | -0.418 | -0.378 | -0.394 | -0.047 | -0.053 | -0.053 | 0.155 | 0.29 | 0.52 | 0.37 | 0.22 | 0.21 | 0.18 |
| 4 | 168098935 | rs10029237 | -1.471 | -0.404 | -0.348 | -0.389 | -0.046 | -0.052 | -0.052 | 0 | 0.28 | 0.5 | 0.37 | 0.22 | 0.21 | 0.18 |
| 4 | 168100489 | rs7667846 | -1.455 | -0.407 | -0.348 | -0.382 | -0.05 | -0.069 | -0.069 | 0 | 0.28 | 0.49 | 0.36 | 0.18 | 0.2 | 0.16 |
| 4 | 168131137 | rs13143783 | -0.638 | -0.101 | -0.879 | -0.202 | 0.001 | -0.028 | -0.028 | 0 | 0.08 | 0.15 | 0.1 | 0 | 0.08 | 0.06 |
| 4 | 168131448 | rs10022799 | -1.436 | -0.224 | -0.85 | -0.248 | -0.029 | -0.022 | -0.022 | 1.918 | 0.23 | 0.41 | 0.35 | 0.22 | 0.18 | 0.1 |
| 4 | 168136671 | rs9996196 | -0.712 | -0.188 | -0.963 | -0.202 | -0.016 | -0.003 | -0.003 | 0 | 0.18 | 0.23 | 0.26 | 0.22 | 0.14 | 0.05 |
| 4 | 168139380 | rs9312517 | 0.044 | -0.207 | -0.923 | -0.155 | -0.031 | 0.012 | 0.012 | 0 | 0.14 | 0.22 | 0.24 | 0.22 | 0.1 | 0.04 |
| 4 | 168147524 | rs10517927 | 0.948 | 0.155 | -1.623 | 0.112 | -0.079 | -0.177 | -0.177 | 0.119 | 0.32 | 0.01 | 0.24 | 0.4 | 0.36 | 0.35 |
| 4 | 168150023 | rs6850485 | -1.013 | -0.073 | -0.682 | 0.066 | -0.029 | -0.022 | -0.022 | 0 | 0.23 | 0.4 | 0.35 | 0.22 | 0.18 | 0.1 |
| 4 | 168154181 | rs7657608 | -1.168 | 0.379 | -0.927 | -0.177 | 0.008 | -0.026 | -0.026 | 0 | 0.25 | 0.34 | 0.21 | 0.11 | 0.27 | 0.31 |
| 4 | 168162185 | rs17600543 | -0.082 | 0.483 | -1.059 | -0.2 | -0.026 | 0.018 | 0.018 | 0 | 0.15 | 0.06 | 0.1 | 0.1 | 0.17 | 0.23 |

| | | | | | | | | | | | | | | | | |
|---|---|---|---|---|---|---|---|---|---|---|---|---|---|---|---|---|
| 4 | 168166789 | rs6833251 | -1.506 | 0.459 | -1.049 | -0.246 | 0.002 | -0.014 | -0.014 | 0.071 | 0.27 | 0.33 | 0.22 | 0.12 | 0.3 | 0.31 |
| 4 | 168169259 | rs17053127 | 1.065 | 0.484 | -1.006 | -0.22 | 0.013 | -0.029 | -0.029 | 0.043 | 0.07 | 0.12 | 0.04 | 0 | 0.1 | 0.02 |
| 4 | 168175697 | rs7694278 | -1.85 | 0.297 | -1.623 | -0.077 | -0.094 | -0.143 | -0.143 | 0 | 0.71 | 0.99 | 0.76 | 0.58 | 0.62 | 0.68 |
| 4 | 168180681 | rs10517930 | -1.579 | 0.466 | -0.75 | 0.175 | 0.001 | -0.015 | -0.015 | 0 | 0.26 | 0.33 | 0.21 | 0.13 | 0.27 | 0.31 |
| 4 | 168187191 | rs4266319 | -1.66 | 0.447 | -0.749 | 0.164 | 0.002 | -0.018 | -0.018 | 0 | 0.25 | 0.33 | 0.21 | 0.13 | 0.27 | 0.31 |
| 4 | 168191092 | rs13117708 | -0.463 | 0.251 | -0.936 | 0.179 | -0.034 | 0.016 | 0.016 | 0 | 0.15 | 0.05 | 0.1 | 0.11 | 0.17 | 0.23 |
| 4 | 168204564 | rs35520897 | -0.436 | 0.178 | -0.907 | 0.177 | -0.023 | 0.018 | 0.018 | 0 | 0.14 | 0.05 | 0.1 | 0.1 | 0.16 | 0.23 |
| 4 | 168211052 | rs10004837 | -1.564 | 0.168 | -0.242 | 0.322 | 0.006 | 0.002 | 0.002 | 0 | 0.34 | 0.53 | 0.48 | 0.4 | 0.33 | 0.33 |
| 4 | 168214408 | rs2319265 | 0.291 | -0.081 | -0.408 | 0.162 | 0.001 | -0.03 | -0.03 | 4.262 | 0.06 | 0.11 | 0.03 | 0 | 0.06 | 0.02 |
| 4 | 168236901 | rs10008779 | 0.697 | -0.505 | -0.401 | 0.251 | -0.003 | 0.039 | 0.039 | 0.062 | 0.47 | 0.32 | 0.35 | 0.3 | 0.53 | 0.5 |
| 4 | 168240912 | rs7665794 | 0.035 | -0.854 | -0.874 | 0.641 | -0.099 | -0.074 | -0.074 | 0 | 0.25 | 0.05 | 0.25 | 0.26 | 0.33 | 0.33 |
| 4 | 168252561 | rs2319140 | 1.369 | -1.021 | -0.608 | 0.527 | -0.016 | 0.051 | 0.051 | 0.003 | 0.45 | 0.26 | 0.33 | 0.3 | 0.52 | 0.46 |
| 4 | 168256202 | rs11942633 | -0.711 | -1.196 | 0 | 0.905 | -0.008 | -0.015 | -0.015 | 0 | 0.34 | 0.27 | 0.5 | 0.4 | 0.37 | 0.42 |
| 4 | 168256989 | rs11132983 | -0.624 | -1.286 | 0.501 | 1.025 | -0.027 | 0.068 | 0.068 | 1.056 | 0.09 | 0.22 | 0.27 | 0.17 | 0.04 | 0.1 |
| 4 | 168259395 | rs17703652 | -0.094 | -1.25 | 0.777 | 0.891 | -0.015 | 0.029 | 0.029 | 2.789 | 0.14 | 0.2 | 0.09 | 0.25 | 0.08 | 0.08 |
| 4 | 168260187 | rs4385095 | 0.079 | -1.014 | 0.658 | 0.751 | -0.022 | 0.006 | 0.006 | 0 | 0.92 | 0.78 | 0.93 | 0.96 | 0.97 | 0.96 |
| 4 | 168272305 | rs17053176 | -1.006 | -1.093 | 1.056 | 0.64 | 0.013 | -0.008 | -0.008 | 0 | 0.08 | 0.39 | 0.04 | 0 | 0.02 | 0.03 |
| 4 | 168273744 | rs17053183 | -0.595 | -1.089 | 1.025 | 0.642 | -0.001 | -0.019 | -0.019 | 0.835 | 0.07 | 0.31 | 0.05 | 0 | 0.02 | 0.03 |
| 4 | 168284744 | rs2319270 | -0.086 | -1.243 | 0.696 | 0.882 | 0.008 | -0.034 | -0.034 | 0.138 | 0.86 | 0.95 | 0.87 | 0.8 | 0.9 | 0.88 |
| 4 | 168288328 | rs2319250 | 1.555 | -1.489 | 1.061 | 0.942 | -0.011 | 0.009 | 0.009 | 0.565 | 0.69 | 0.36 | 0.58 | 0.49 | 0.84 | 0.75 |
| 4 | 168288641 | rs17053204 | -1.449 | -1.451 | 1.21 | 0.942 | -0.063 | 0.145 | 0.145 | 0.749 | 0.16 | 0.45 | 0.28 | 0.32 | 0.06 | 0.14 |
| 4 | 168292265 | rs2319248 | -1.521 | -1.461 | 0.254 | 0.773 | -0.002 | -0.054 | -0.054 | 0.57 | 0.44 | 0.78 | 0.55 | 0.53 | 0.4 | 0.45 |
| 4 | 168292438 | rs2319247 | 0.451 | -1.369 | 0.122 | 0.732 | -0.092 | -0.017 | -0.017 | 0 | 0.23 | 0.05 | 0.23 | 0.21 | 0.32 | 0.28 |
| 4 | 168294218 | rs17601255 | 1.783 | -1.338 | 0.101 | 0.688 | -0.115 | 0.049 | 0.049 | 0.43 | 0.42 | 0.07 | 0.31 | 0.28 | 0.5 | 0.44 |
| 4 | 168301771 | rs2252135 | -0.129 | -1.39 | 0.722 | 0.864 | -0.016 | 0.116 | 0.116 | 1.61 | 0.07 | 0.21 | 0.24 | 0.2 | 0.02 | 0.12 |
| 4 | 168307924 | rs2952429 | 0.979 | -1.479 | 0.823 | 0.804 | 0.012 | -0.013 | -0.013 | 0.877 | 0.73 | 0.77 | 0.69 | 0.6 | 0.83 | 0.74 |
| 4 | 168308058 | rs17601548 | 1.666 | -1.457 | 0.681 | 0.778 | -0.021 | 0.049 | 0.049 | 0 | 0.44 | 0.27 | 0.42 | 0.28 | 0.51 | 0.43 |
| 4 | 168316748 | rs2706826 | 0.823 | -1.408 | 0.6 | 0.814 | 0.032 | -0.044 | -0.044 | 0 | 0.74 | 0.86 | 0.69 | 0.6 | 0.83 | 0.74 |
| 4 | 168316848 | rs2706825 | -0.032 | -1.104 | 0.809 | 0.896 | 0.023 | 0.025 | 0.025 | 0 | 0.26 | 0.39 | 0.25 | 0.31 | 0.31 | 0.3 |
| 4 | 168327233 | rs3860620 | -0.661 | -1.666 | 0.357 | 0.289 | -0.024 | 0 | 0 | 0 | 0.42 | 0.51 | 0.32 | 0.43 | 0.26 | 0.36 |
| 4 | 168334028 | rs2706836 | -1.416 | -1.208 | 0.416 | -0.304 | 0.018 | -0.001 | -0.001 | 0 | 0.92 | 0.86 | 0.91 | 0.96 | 0.87 | 0.94 |
| 4 | 168338506 | rs2952482 | -0.121 | -1.436 | 0.452 | -0.406 | -0.032 | -0.019 | -0.019 | 0 | 0.33 | 0.52 | 0.19 | 0.23 | 0.22 | 0.24 |
| 4 | 168348929 | rs4419497 | 1.336 | -1.112 | 0.108 | -0.417 | -0.031 | 0.074 | 0.074 | 0.376 | 0.42 | 0.23 | 0.43 | 0.26 | 0.46 | 0.43 |
| 4 | 168351130 | rs6553778 | 1.213 | -1.058 | 0.275 | -0.451 | 0.036 | 0.12 | 0.12 | 0 | 0.15 | 0.27 | 0.27 | 0.51 | 0.22 | 0.35 |
| 4 | 168353430 | rs13136714 | -0.476 | -1.138 | 0.47 | -0.36 | 0.011 | 0.016 | 0.016 | 0.551 | 0.25 | 0.16 | 0.19 | 0.08 | 0.17 | 0.13 |
| 4 | 168362952 | rs2706822 | -0.781 | -1.125 | 0.445 | -0.236 | -0.001 | -0.001 | -0.001 | 0 | 0.81 | 0.82 | 0.82 | 0.95 | 0.77 | 0.92 |
| 4 | 168364638 | rs2319242 | 1.675 | -1.131 | 0.426 | -0.229 | -0.031 | 0.007 | 0.007 | 0 | 0.26 | 0.16 | 0.32 | 0.22 | 0.32 | 0.31 |
| 4 | 168366518 | rs11133037 | 0.189 | -1.357 | 0.819 | 0.304 | -0.033 | -0.069 | -0.069 | 0 | 0.29 | 0.18 | 0.36 | 0.55 | 0.36 | 0.45 |
| 4 | 168375863 | rs6854438 | 0.477 | -1.41 | 0.832 | 0.306 | -0.016 | -0.011 | -0.011 | 0 | 0.12 | 0.13 | 0.08 | 0.08 | 0.06 | 0.09 |
| 4 | 168390470 | rs17704183 | 0.186 | -1.286 | 0.475 | 0.41 | -0.019 | -0.047 | -0.047 | 0.14 | 0.3 | 0.23 | 0.36 | 0.55 | 0.36 | 0.45 |
| 4 | 168391112 | rs17053293 | 0.302 | -1.322 | 0.543 | 0.404 | -0.023 | -0.018 | -0.018 | 0.096 | 0.13 | 0.18 | 0.08 | 0.08 | 0.07 | 0.1 |
| 4 | 168391333 | rs6817078 | -0.591 | -0.827 | 0.177 | 0.555 | -0.06 | 0.036 | 0.036 | 2.352 | 0.9 | 0.78 | 0.98 | 0.9 | 0.99 | 0.97 |
| 4 | 168401003 | rs2169736 | 0.026 | -0.051 | -0.572 | 0.33 | 0.011 | 0.019 | 0.019 | 0 | 0.49 | 0.47 | 0.55 | 0.29 | 0.58 | 0.43 |
| 4 | 168414411 | rs1600106 | -0.418 | -0.915 | -0.576 | 0.12 | 0.005 | 0.004 | 0.004 | 0 | 0.35 | 0.33 | 0.33 | 0.33 | 0.29 | 0.37 |
| 4 | 168431604 | rs10517942 | -1.007 | -1.476 | -1.075 | 0.38 | 0.065 | 0 | 0 | 0.097 | 0.47 | 0.26 | 0.4 | 0.49 | 0.38 | 0.42 |
| 4 | 168434661 | rs2219758 | 0.62 | -1.504 | -1.103 | 0.274 | 0.013 | -0.048 | -0.048 | 0 | 0.09 | 0.02 | 0.07 | 0.16 | 0.07 | 0.04 |
| 4 | 168442929 | rs17053332 | 0.88 | -1.892 | -0.85 | -0.128 | -0.032 | -0.063 | -0.063 | 0 | 0.18 | 0.44 | 0.16 | 0.08 | 0.13 | 0.06 |
| 4 | 168448425 | rs1600103 | -0.375 | -2.053 | -0.982 | -0.312 | -0.018 | 0.007 | 0.007 | 0.474 | 0.84 | 0.81 | 0.91 | 0.84 | 0.89 | 0.96 |
| 4 | 168450615 | rs1518203 | 0.301 | -1.991 | -0.778 | -0.111 | 0.029 | 0.022 | 0.022 | 0 | 0.48 | 0.64 | 0.59 | 0.62 | 0.58 | 0.58 |
| 4 | 168455386 | rs1817389 | 0.256 | -2.059 | -0.617 | -0.155 | 0.031 | 0.025 | 0.025 | 0 | 0.49 | 0.62 | 0.59 | 0.63 | 0.58 | 0.58 |
| 4 | 168470029 | rs1478241 | -0.473 | -2.378 | -0.549 | -0.639 | -0.009 | -0.021 | -0.021 | 0.062 | 0.76 | 0.71 | 0.78 | 0.84 | 0.81 | 0.8 |
| 4 | 168472046 | rs7668194 | 0.686 | -2.569 | -0.49 | -0.737 | -0.032 | -0.059 | -0.059 | 0 | 0.18 | 0.44 | 0.16 | 0.08 | 0.13 | 0.07 |
| 4 | 168472510 | rs1878685 | -0.029 | -2.562 | -0.724 | -0.731 | -0.01 | -0.022 | -0.022 | 0 | 0.74 | 0.68 | 0.78 | 0.85 | 0.81 | 0.8 |
| 4 | 168472966 | rs6816291 | 1.16 | -3.109 | -1.347 | -0.873 | 0.031 | -0.005 | -0.005 | 0.087 | 0.4 | 0.09 | 0.52 | 0.58 | 0.63 | 0.55 |
| 4 | 168473019 | rs6845819 | 0.017 | -3.668 | -0.842 | -1.822 | -0.029 | -0.017 | -0.017 | 1.605 | 0.79 | 0.72 | 0.82 | 0.84 | 0.92 | 0.84 |
| 4 | 168478645 | rs7672872 | -0.084 | -3.763 | -0.372 | -2.069 | -0.015 | -0.036 | -0.036 | 0 | 0.89 | 0.89 | 0.96 | 0.98 | 0.92 | 0.9 |
| 4 | 168485751 | rs992771 | 1.017 | -4.312 | 0.247 | -2.105 | -0.054 | -0.01 | -0.01 | 0.042 | 0.68 | 0.39 | 0.81 | 0.78 | 0.83 | 0.79 |
| 4 | 168490955 | rs1914621 | 1.085 | -4.34 | 0.233 | -2.084 | -0.052 | -0.004 | -0.004 | 0 | 0.68 | 0.37 | 0.8 | 0.75 | 0.83 | 0.79 |
| 4 | 168508173 | rs17704359 | 1.417 | -3.983 | 0.133 | -2.149 | 0.048 | 0.009 | 0.009 | 0 | 0.44 | 0.26 | 0.56 | 0.61 | 0.61 | 0.6 |
| 4 | 168515495 | rs1401014 | 1.515 | -3.6 | 0.247 | -1.976 | 0.048 | 0.01 | 0.01 | 0 | 0.46 | 0.26 | 0.56 | 0.59 | 0.61 | 0.59 |
| 4 | 168520452 | rs17053380 | -0.971 | -3.546 | 0.667 | -2.109 | -0.076 | 0.016 | 0.016 | 0 | 0.18 | 0.41 | 0.1 | 0.17 | 0.09 | 0.07 |
| 4 | 168523786 | rs7657818 | 0.865 | -3.495 | 0.728 | -2.419 | -0.013 | -0.041 | -0.041 | 0 | 0.67 | 0.36 | 0.8 | 0.94 | 0.79 | 0.77 |
| 4 | 168536364 | rs7675475 | -0.313 | -3.074 | 0.562 | -2.499 | -0.017 | -0.074 | -0.074 | 2.248 | 0.82 | 0.76 | 0.92 | 1 | 0.87 | 0.92 |
| 4 | 168536686 | rs1401022 | -0.328 | -3.073 | 0.452 | -2.498 | -0.007 | -0.05 | -0.05 | 0.182 | 0.82 | 0.82 | 0.92 | 1 | 0.87 | 0.92 |
| 4 | 168539717 | rs13140381 | -0.522 | -3.23 | 0.511 | -2.495 | -0.011 | 0.002 | 0.002 | 0 | 0.14 | 0.15 | 0.12 | 0.17 | 0.1 | 0.08 |
| 4 | 168545366 | rs2169735 | 1.016 | -3.227 | 0.514 | -2.494 | -0.033 | -0.023 | -0.023 | 0 | 0.67 | 0.63 | 0.8 | 0.83 | 0.77 | 0.85 |
| 4 | 168545610 | rs1356511 | -0.18 | -2.627 | 0.378 | -2.759 | -0.009 | -0.055 | -0.055 | 0 | 0.82 | 0.82 | 0.92 | 1 | 0.87 | 0.92 |
| 4 | 168564107 | rs1533547 | -1.492 | -1.746 | 0.261 | -3.001 | 0.014 | 0.115 | 0.115 | 0 | 0.51 | 0.34 | 0.39 | 0.12 | 0.49 | 0.26 |
| 4 | 168576098 | rs12645719 | -0.66 | -1.797 | 0.152 | -3.055 | -0.058 | 0.118 | 0.118 | 0 | 0.32 | 0.08 | 0.3 | 0.11 | 0.37 | 0.18 |

| | | | | | | | | | | | | | | | | |
|---|---|---|---|---|---|---|---|---|---|---|---|---|---|---|---|---|
| 4 | 168576599 | rs6850404 | 0.818 | -1.906 | 0.405 | -3.142 | -0.061 | -0.156 | -0.156 | 1.288 | 0.71 | 0.5 | 0.86 | 0.98 | 0.81 | 0.83 |
| 4 | 168583227 | rs17524463 | 2.058 | -1.883 | 0.422 | -3.128 | 0.007 | -0.036 | -0.036 | 0 | 0.39 | 0.41 | 0.56 | 0.87 | 0.44 | 0.66 |
| 4 | 168590008 | rs7683919 | 0.473 | -1.862 | 0.354 | -3.107 | -0.025 | -0.09 | -0.09 | 2.202 | 0.75 | 0.64 | 0.87 | 0.98 | 0.81 | 0.83 |
| 4 | 168593397 | rs1533551 | -1.697 | -1.878 | 0.28 | -3.139 | 0.005 | 0.041 | 0.041 | 0 | 0.52 | 0.39 | 0.39 | 0.11 | 0.5 | 0.26 |
| 4 | 168593787 | rs7664708 | -0.672 | -1.867 | 0.263 | -2.976 | -0.031 | 0.095 | 0.095 | 0 | 0.29 | 0.07 | 0.29 | 0.1 | 0.33 | 0.16 |
| 4 | 168593961 | rs2203769 | -1.567 | -1.795 | 0.348 | -2.958 | 0.029 | 0.132 | 0.132 | 0 | 0.47 | 0.29 | 0.36 | 0.1 | 0.45 | 0.23 |
| 4 | 168594710 | rs1518186 | 1.324 | -0.997 | 0.924 | -2.552 | -0.001 | 0.013 | 0.013 | 0 | 0.71 | 0.76 | 0.72 | 0.87 | 0.7 | 0.72 |
| 4 | 168598025 | rs17053475 | -0.77 | -0.694 | 0.812 | -2.413 | -0.009 | -0.039 | -0.039 | 0 | 0.1 | 0.19 | 0.04 | 0.02 | 0.07 | 0.13 |
| 4 | 168602424 | rs6552196 | -0.312 | -0.737 | 0.815 | -2.449 | 0.003 | -0.018 | -0.018 | 0 | 0.85 | 0.86 | 0.92 | 1 | 0.89 | 0.93 |
| 4 | 168610661 | rs6816694 | 2.111 | -0.648 | 0.909 | -2.502 | -0.007 | -0.132 | -0.132 | 2.071 | 0.6 | 0.47 | 0.78 | 0.97 | 0.65 | 0.72 |
| 4 | 168611058 | rs10517954 | -0.736 | -0.585 | 0.787 | -2.467 | -0.009 | -0.038 | -0.038 | 0 | 0.1 | 0.19 | 0.04 | 0.02 | 0.07 | 0.13 |
| 4 | 168613796 | rs17053510 | -0.114 | -0.574 | 0.815 | -2.463 | 0.005 | -0.016 | -0.016 | 0 | 0.16 | 0.15 | 0.08 | 0 | 0.11 | 0.07 |
| 4 | 168615218 | rs12647151 | 1.264 | -0.414 | 0.521 | -2.386 | 0.005 | -0.077 | -0.077 | 0 | 0.77 | 0.71 | 0.86 | 0.98 | 0.76 | 0.78 |
| 4 | 168616720 | rs7677082 | 0.092 | -0.141 | -0.031 | -1.921 | -0.014 | 0.004 | 0.004 | 0 | 0.88 | 0.95 | 0.91 | 0.99 | 0.83 | 0.91 |
| 4 | 168617513 | rs9968440 | 0.627 | -0.351 | -0.09 | -1.632 | -0.018 | -0.137 | -0.137 | 0 | 0.65 | 0.48 | 0.81 | 0.97 | 0.69 | 0.72 |
| 4 | 168621364 | rs7690929 | -1.172 | -0.743 | 0.104 | -1.295 | -0.057 | -0.119 | -0.119 | 0 | 0.24 | 0.62 | 0.11 | 0.05 | 0.09 | 0.13 |
| 4 | 168626439 | rs11730375 | -1.218 | -0.791 | 0.174 | -1.329 | -0.067 | -0.13 | -0.13 | 0 | 0.24 | 0.59 | 0.1 | 0.05 | 0.09 | 0.13 |
| 4 | 168628323 | rs10471169 | -1.039 | -0.781 | 0.696 | -1.318 | -0.052 | -0.11 | -0.11 | 0 | 0.2 | 0.37 | 0.09 | 0.01 | 0.07 | 0.1 |
| 4 | 168631913 | rs10016865 | -1.06 | -0.794 | 0.69 | -1.324 | -0.052 | -0.11 | -0.11 | 0.02 | 0.2 | 0.37 | 0.09 | 0.01 | 0.07 | 0.1 |
| 4 | 168637332 | rs959169 | -1.034 | -0.987 | 1.356 | -1.442 | -0.052 | -0.095 | -0.095 | 2.874 | 0.14 | 0.33 | 0.04 | 0.01 | 0.04 | 0.08 |
| 4 | 168656374 | rs4331802 | -1.155 | -0.731 | 1.538 | -1.539 | -0.01 | -0.014 | -0.014 | 0 | 0.34 | 0.62 | 0.41 | 0.67 | 0.22 | 0.43 |
| 4 | 168669808 | rs13128223 | 0.743 | 0.123 | 0.696 | -2.081 | -0.059 | -0.055 | -0.055 | 1.26 | 0.13 | 0.03 | 0.18 | 0.22 | 0.29 | 0.21 |
| 4 | 168671673 | rs6848110 | -1.606 | -0.234 | 0.686 | -2.355 | 0.001 | 0.147 | 0.147 | 0 | 0.54 | 0.76 | 0.61 | 0.89 | 0.57 | 0.69 |
| 4 | 168679351 | rs6552243 | -1.596 | -0.234 | 0.686 | -2.355 | 0.001 | 0.148 | 0.148 | 0 | 0.54 | 0.76 | 0.61 | 0.89 | 0.57 | 0.69 |
| 4 | 168679456 | rs6834993 | -0.942 | -0.311 | 1.203 | -2.394 | -0.01 | 0.015 | 0.015 | 0 | 0.44 | 0.36 | 0.55 | 0.88 | 0.51 | 0.64 |
| 4 | 168705038 | rs10002325 | 1.456 | -0.564 | 1.269 | -2.42 | -0.009 | -0.035 | -0.035 | 0 | 0.92 | 0.74 | 0.97 | 1 | 0.97 | 0.96 |
| 4 | 168721503 | rs13124475 | -1.344 | -0.38 | 1.322 | -2.372 | 0.002 | -0.026 | -0.026 | 0.755 | 0.49 | 0.57 | 0.6 | 0.86 | 0.57 | 0.66 |
| 4 | 168726851 | rs3885749 | 1.04 | -0.407 | 1.298 | -2.41 | 0.009 | 0.008 | 0.008 | 0 | 0.49 | 0.4 | 0.39 | 0.14 | 0.41 | 0.31 |
| 4 | 168737612 | rs1971169 | 0.493 | -0.415 | 1.62 | -2.37 | -0.011 | 0.073 | 0.073 | 0 | 0.58 | 0.68 | 0.44 | 0.15 | 0.46 | 0.36 |
| 4 | 168764988 | rs2007133 | 1.234 | -0.933 | 0.562 | -3.329 | 0.016 | 0.045 | 0.045 | 0 | 0.29 | 0.18 | 0.27 | 0.1 | 0.22 | 0.24 |
| 4 | 168768607 | rs17525095 | 1.67 | -1.292 | 0.152 | -3.634 | -0.008 | 0.115 | 0.115 | 0 | 0.22 | 0.01 | 0.21 | 0.06 | 0.15 | 0.21 |
| 4 | 168793863 | rs304327 | 0.297 | -1.433 | 0.391 | -3.666 | 0.034 | 0.048 | 0.048 | 0.206 | 0.61 | 0.31 | 0.81 | 0.94 | 0.74 | 0.84 |
| 4 | 168798179 | rs407213 | -0.797 | -1.268 | 0.617 | -3.618 | 0 | -0.002 | -0.002 | 0 | 0.49 | 0.46 | 0.35 | 0.28 | 0.54 | 0.39 |
| 4 | 168807727 | rs1847688 | 0.288 | -1.248 | 0.484 | -3.563 | -0.001 | -0.011 | -0.011 | 0 | 0.53 | 0.55 | 0.67 | 0.76 | 0.49 | 0.65 |
| 4 | 168808600 | rs12500312 | 1.659 | -1.274 | 0.427 | -3.586 | -0.045 | -0.041 | -0.041 | 1.039 | 0.19 | 0.11 | 0.38 | 0.66 | 0.24 | 0.38 |
| 4 | 168810105 | rs1562143 | -0.059 | -1.146 | 0.648 | -3.294 | 0.002 | -0.017 | -0.017 | 0 | 0.25 | 0.32 | 0.21 | 0.19 | 0.3 | 0.25 |
| 4 | 168815347 | rs9994795 | 0.535 | -1.101 | 0.398 | -3.268 | 0 | -0.008 | -0.008 | 0 | 0.54 | 0.55 | 0.67 | 0.75 | 0.49 | 0.67 |
| 4 | 168818443 | rs304351 | 0.221 | -1.311 | 0.651 | -3.143 | -0.026 | -0.101 | -0.101 | 0.664 | 0.68 | 0.58 | 0.84 | 0.95 | 0.78 | 0.9 |
| 4 | 168818710 | rs17525374 | 0.794 | -0.852 | 0.016 | -2.897 | -0.009 | 0.042 | 0.042 | 0 | 0.18 | 0.06 | 0.11 | 0.05 | 0.19 | 0.07 |
| 4 | 168824533 | rs10022932 | -0.362 | -0.922 | -0.111 | -2.978 | 0.018 | 0.101 | 0.101 | 0.975 | 0.3 | 0.13 | 0.12 | 0.05 | 0.24 | 0.11 |
| 4 | 168829106 | rs7682960 | -0.469 | -1.024 | 0.033 | -3.013 | -0.001 | -0.003 | -0.003 | 0 | 0.61 | 0.61 | 0.57 | 0.72 | 0.53 | 0.44 |
| 4 | 168832046 | rs6817742 | -0.219 | -0.641 | -0.642 | -2.913 | -0.001 | -0.062 | -0.062 | 0 | 0.29 | 0.39 | 0.17 | 0.1 | 0.26 | 0.12 |
| 4 | 168841128 | rs2166186 | -1.513 | -0.612 | 0.203 | -2.333 | 0.009 | -0.011 | -0.011 | 0.009 | 0.09 | 0.41 | 0.05 | 0.04 | 0.03 | 0.04 |
| 4 | 168850375 | rs13107574 | -0.248 | 0.008 | -0.193 | -2.39 | 0.024 | -0.074 | -0.074 | 2.511 | 0.14 | 0.25 | 0.11 | 0 | 0.19 | 0.06 |
| 4 | 168853211 | rs17525479 | -0.554 | 0.238 | -0.267 | -2.445 | -0.001 | -0.046 | -0.046 | 0.492 | 0.16 | 0.19 | 0.1 | 0 | 0.16 | 0.07 |
| 4 | 168857462 | rs13140875 | -0.616 | -0.136 | -0.354 | -1.337 | 0.014 | -0.005 | -0.005 | 0.748 | 0.18 | 0.16 | 0.07 | 0 | 0.12 | 0.06 |
| 4 | 168871037 | rs17610208 | 0.151 | -0.133 | -0.382 | -1.279 | 0.027 | 0.058 | 0.058 | 0 | 0.14 | 0.04 | 0.11 | 0 | 0.1 | 0.04 |
| 4 | 168893805 | rs304341 | -1.38 | -0.061 | -0.288 | -1.182 | 0.01 | 0.04 | 0.04 | 0 | 0.44 | 0.66 | 0.54 | 0.63 | 0.44 | 0.37 |
| 4 | 168935601 | rs11944180 | -0.906 | -0.363 | 0.043 | -1.028 | -0.05 | -0.107 | -0.107 | 1.041 | 0.17 | 0.32 | 0.14 | 0 | 0.1 | 0.03 |
| 4 | 168945264 | rs13140793 | -0.239 | -0.202 | -0.953 | -0.95 | 0.027 | 0.07 | 0.07 | 0.177 | 0.14 | 0.04 | 0.11 | 0 | 0.1 | 0.04 |
| 4 | 168965733 | rs11934907 | -0.03 | 0.165 | -0.463 | -0.587 | -0.004 | -0.005 | -0.005 | 0 | 0.54 | 0.47 | 0.55 | 0.62 | 0.4 | 0.5 |
| 4 | 168967476 | rs10006894 | -0.928 | -0.403 | 0.378 | -0.959 | -0.018 | -0.038 | -0.038 | 0.032 | 0.09 | 0.41 | 0.05 | 0 | 0.05 | 0.04 |
| 4 | 168968218 | rs11728448 | 0.381 | 0.312 | 0.16 | -1.322 | -0.012 | -0.002 | -0.002 | 1.18 | 0.31 | 0.29 | 0.31 | 0.34 | 0.37 | 0.43 |
| 4 | 168972967 | rs11132000 | 1.442 | -0.294 | -0.367 | -1.646 | 0.004 | -0.039 | -0.039 | 0.383 | 0.35 | 0.04 | 0.4 | 0.64 | 0.24 | 0.44 |
| 4 | 168977085 | rs9999798 | -0.663 | -0.404 | 0.369 | -1.507 | -0.024 | -0.101 | -0.101 | 0 | 0.17 | 0.34 | 0.16 | 0 | 0.15 | 0.07 |
| 4 | 168980017 | rs7678466 | -1.267 | -0.074 | 0.026 | -1.466 | -0.005 | -0.005 | -0.005 | 0 | 0.87 | 0.91 | 0.87 | 0.98 | 0.74 | 0.95 |
| 4 | 168987754 | rs11132008 | 0.169 | -0.148 | -0.049 | -1.459 | 0.002 | 0.002 | 0.002 | 0 | 0.54 | 0.56 | 0.51 | 0.63 | 0.4 | 0.54 |
| 4 | 168994309 | rs4443240 | 1.277 | -0.09 | -0.752 | -1.362 | -0.022 | -0.029 | -0.029 | 0 | 0.35 | 0.06 | 0.4 | 0.63 | 0.3 | 0.46 |
| 4 | 169008885 | rs7662898 | -1.531 | -0.036 | -0.048 | -0.748 | 0.012 | -0.043 | -0.043 | 0.122 | 0.62 | 0.83 | 0.58 | 0.42 | 0.66 | 0.51 |
| 4 | 169017774 | rs12646377 | 1.258 | -0.069 | 0.778 | -0.947 | 0 | -0.01 | -0.01 | 0 | 0.12 | 0.09 | 0.11 | 0.01 | 0.22 | 0.05 |
| 4 | 169017909 | rs9799627 | -1.039 | -0.07 | 1.008 | -0.957 | 0.025 | -0.005 | -0.005 | 0 | 0.55 | 0.32 | 0.53 | 0.43 | 0.64 | 0.51 |
| 4 | 169019867 | rs10517976 | -0.93 | -0.096 | 0.965 | -1.009 | 0.041 | 0.013 | 0.013 | 0 | 0.5 | 0.3 | 0.48 | 0.42 | 0.58 | 0.45 |
| 4 | 169020566 | rs2319707 | 0.973 | 0.091 | 0.57 | -0.986 | -0.076 | -0.09 | -0.09 | 0.215 | 0.15 | 0.02 | 0.16 | 0.29 | 0.18 | 0.2 |
| 4 | 169032268 | rs4692853 | -1.098 | -0.015 | 0.889 | -0.915 | 0.023 | 0.002 | 0.002 | 0 | 0.53 | 0.35 | 0.49 | 0.42 | 0.6 | 0.45 |
| 4 | 169035968 | rs7653928 | -1.667 | -0.025 | 0.745 | -0.86 | 0.012 | -0.023 | -0.023 | 2.674 | 0.6 | 0.8 | 0.56 | 0.42 | 0.64 | 0.49 |
| 4 | 169041179 | rs9991416 | -1.419 | -0.029 | 0.743 | -0.862 | 0.018 | -0.013 | -0.013 | 0 | 0.57 | 0.79 | 0.52 | 0.42 | 0.6 | 0.45 |
| 4 | 169042167 | rs4405980 | -1.917 | 0.006 | 0.425 | -0.839 | -0.025 | -0.1 | -0.1 | 0 | 0.63 | 0.94 | 0.59 | 0.42 | 0.66 | 0.51 |
| 4 | 169046172 | rs7662687 | -2.068 | 0.058 | 0.501 | -0.855 | -0.005 | -0.043 | -0.043 | 2.876 | 0.65 | 0.87 | 0.59 | 0.39 | 0.66 | 0.51 |
| 4 | 169049597 | rs7699520 | -1.174 | 0.163 | 0.498 | -1.087 | 0.016 | 0.023 | 0.023 | 0 | 0.54 | 0.38 | 0.48 | 0.3 | 0.59 | 0.44 |

| | | | | | | | | | | | | | | | | |
|---|---|---|---|---|---|---|---|---|---|---|---|---|---|---|---|---|
| 4 | 169056777 | rs6832786 | -2.097 | 0.167 | 0.412 | -1.084 | -0.029 | -0.029 | -0.029 | 0 | 0.65 | 0.94 | 0.57 | 0.31 | 0.65 | 0.5 |
| 4 | 169059063 | rs6841013 | 0.707 | 0.3 | 0.554 | -1.06 | 0.005 | -0.032 | -0.032 | 1.345 | 0.17 | 0.17 | 0.14 | 0.04 | 0.25 | 0.07 |
| 4 | 169062297 | rs4518204 | -0.688 | 0.418 | 0.079 | -0.946 | -0.024 | 0.033 | 0.033 | 0 | 0.38 | 0.26 | 0.34 | 0.27 | 0.34 | 0.37 |
| 4 | 169072888 | rs1317046 | -1.836 | 0.528 | 0.146 | -0.738 | -0.037 | -0.025 | -0.025 | 0 | 0.59 | 0.89 | 0.64 | 0.37 | 0.59 | 0.52 |
| 4 | 169076192 | rs11936562 | -1.567 | 0.52 | 0.572 | -0.744 | 0.01 | 0.013 | 0.013 | 0.108 | 0.57 | 0.74 | 0.62 | 0.37 | 0.6 | 0.54 |
| 4 | 169083694 | rs6836994 | -1.552 | 0.514 | 0.581 | -0.7 | 0.011 | 0.066 | 0.066 | 0.114 | 0.56 | 0.74 | 0.47 | 0.24 | 0.59 | 0.53 |
| 4 | 169086441 | rs4405979 | -1.508 | 0.499 | 0.577 | -0.705 | 0.011 | 0.069 | 0.069 | 0.461 | 0.56 | 0.74 | 0.47 | 0.24 | 0.59 | 0.53 |
| 4 | 169090279 | rs978726 | -1.648 | 0.511 | 0.549 | -0.693 | 0.003 | 0.009 | 0.009 | 0.286 | 0.57 | 0.8 | 0.63 | 0.37 | 0.6 | 0.54 |
| 4 | 169098914 | rs477897 | 0.633 | 0.616 | 0.9 | -0.739 | -0.001 | 0.004 | 0.004 | 0 | 0.15 | 0.11 | 0.13 | 0.05 | 0.23 | 0.07 |
| 4 | 169098984 | rs10024092 | -0.855 | 0.617 | 1.45 | -0.754 | 0.07 | 0.192 | 0.192 | 0 | 0.06 | 0.37 | 0.24 | 0.13 | 0.05 | 0.03 |
| 4 | 169107212 | rs642729 | -0.228 | 0.613 | 1.609 | -0.77 | 0.002 | -0.028 | -0.028 | 0 | 0.2 | 0.23 | 0.18 | 0.05 | 0.27 | 0.09 |
| 4 | 169108921 | rs17053750 | 0.308 | 0.615 | 1.771 | -0.769 | 0 | -0.022 | -0.022 | 0 | 0.17 | 0.21 | 0.14 | 0.05 | 0.23 | 0.06 |
| 4 | 169112742 | rs670905 | -0.17 | 0.619 | 1.667 | -0.765 | 0.003 | -0.043 | -0.043 | 0 | 0.2 | 0.31 | 0.19 | 0.05 | 0.27 | 0.1 |
| 4 | 169123534 | rs644911 | -0.665 | 0.601 | 1.536 | -0.77 | -0.007 | 0.001 | 0.001 | 0 | 0.23 | 0.57 | 0.37 | 0.17 | 0.28 | 0.1 |
| 4 | 169126043 | rs602731 | -0.384 | 0.661 | 1.416 | -0.793 | 0 | -0.055 | -0.055 | 0 | 0.21 | 0.36 | 0.2 | 0.05 | 0.28 | 0.1 |
| 4 | 169134541 | rs9685229 | -1.635 | 0.852 | 0.811 | -0.654 | -0.015 | 0.041 | 0.041 | 0 | 0.43 | 0.75 | 0.47 | 0.31 | 0.38 | 0.45 |
| 4 | 169138291 | rs17611096 | 0.222 | 1.212 | 0.384 | -0.594 | -0.072 | -0.017 | -0.017 | 0 | 0.13 | 0.02 | 0.11 | 0.12 | 0.17 | 0.17 |
| 4 | 169140553 | rs17611110 | -0.757 | 0.938 | 0.775 | -0.605 | -0.018 | 0.025 | 0.025 | 0.008 | 0.32 | 0.23 | 0.23 | 0.17 | 0.32 | 0.44 |
| 4 | 169158471 | rs6552559 | -1.343 | 0.711 | 1.49 | -0.44 | 0.049 | 0.151 | 0.151 | 0 | 0.08 | 0.44 | 0.24 | 0.13 | 0.07 | 0.04 |
| 4 | 169161977 | rs12510170 | -1.372 | 0.709 | 1.489 | -0.441 | 0.047 | 0.148 | 0.148 | 0 | 0.08 | 0.43 | 0.24 | 0.13 | 0.07 | 0.04 |
| 4 | 169174299 | rs6857534 | -1.433 | 0.704 | 1.49 | -0.444 | 0.045 | 0.146 | 0.146 | 0 | 0.08 | 0.43 | 0.24 | 0.13 | 0.07 | 0.04 |
| 4 | 169175069 | rs17053871 | -0.928 | 0.717 | 1.559 | -0.449 | 0.008 | -0.034 | -0.034 | 0 | 0.07 | 0.25 | 0.07 | 0 | 0.06 | 0.03 |
| 4 | 169181574 | rs1671317 | -1.287 | 0.814 | 1.2 | -0.589 | -0.001 | 0.04 | 0.04 | 0.029 | 0.39 | 0.67 | 0.27 | 0.22 | 0.32 | 0.44 |
| 4 | 169185207 | rs958769 | -0.875 | 0.913 | 1.521 | -0.616 | 0.005 | -0.032 | -0.032 | 0.018 | 0.35 | 0.45 | 0.23 | 0.22 | 0.3 | 0.42 |
| 4 | 169191536 | rs593342 | -1.389 | 0.947 | 1.09 | -0.63 | -0.013 | 0.036 | 0.036 | 0.32 | 0.4 | 0.67 | 0.4 | 0.26 | 0.4 | 0.42 |
| 4 | 169202696 | rs665375 | -1.868 | 1.028 | -0.305 | -0.567 | -0.081 | -0.067 | -0.067 | 0 | 0.57 | 0.98 | 0.53 | 0.38 | 0.6 | 0.56 |
| 4 | 169236274 | rs2684370 | -1.231 | 1.301 | 0.761 | -0.476 | -0.001 | -0.005 | -0.005 | 0 | 0.29 | 0.6 | 0.31 | 0.23 | 0.25 | 0.38 |
| 4 | 169237664 | rs497179 | -1.244 | 1.308 | 0.797 | -0.499 | 0.003 | -0.011 | -0.011 | 0 | 0.29 | 0.58 | 0.35 | 0.23 | 0.3 | 0.41 |
| 4 | 169243273 | rs6843318 | -0.869 | 1.087 | 0.882 | -0.567 | 0.041 | -0.036 | -0.036 | 0.584 | 0.09 | 0.33 | 0.07 | 0 | 0.06 | 0.01 |
| 4 | 169244726 | rs522085 | 1.189 | 1.099 | 0.023 | -0.532 | 0.003 | 0.021 | 0.021 | 0 | 0.67 | 0.34 | 0.78 | 0.84 | 0.73 | 0.67 |
| 4 | 169255850 | rs559203 | 0.264 | 1.258 | -0.306 | -0.395 | -0.015 | 0.022 | 0.022 | 0.209 | 0.78 | 0.88 | 0.86 | 0.84 | 0.8 | 0.69 |
| 4 | 169279364 | rs11736382 | -0.227 | 1.919 | -0.556 | -0.494 | -0.021 | 0.035 | 0.035 | 0 | 0.07 | 0 | 0.03 | 0 | 0.07 | 0.01 |
| 4 | 169293182 | rs13145897 | 1.211 | 1.723 | 0.155 | -0.786 | 0.005 | -0.056 | -0.056 | 0.289 | 0.68 | 0.48 | 0.79 | 0.84 | 0.73 | 0.69 |
| 4 | 169297132 | rs12509086 | 1.9 | 1.679 | -0.674 | -0.857 | -0.09 | -0.059 | -0.059 | 0 | 0.42 | 0.03 | 0.49 | 0.6 | 0.43 | 0.47 |
| 4 | 169297397 | rs17540046 | 0.316 | 1.785 | -0.758 | -0.935 | -0.055 | 0.008 | 0.008 | 0.725 | 0.15 | 0 | 0.08 | 0.09 | 0.14 | 0.2 |
| 4 | 169298000 | rs2046770 | -0.715 | 1.612 | -0.357 | -0.845 | 0.005 | -0.014 | -0.014 | 0 | 0.3 | 0.49 | 0.32 | 0.26 | 0.33 | 0.23 |
| 4 | 169306106 | rs902003 | 1.768 | 1.808 | 0.004 | -0.752 | -0.014 | 0.037 | 0.037 | 0 | 0.58 | 0.36 | 0.6 | 0.72 | 0.56 | 0.62 |
| 4 | 169307654 | rs3762853 | 0.716 | 1.868 | 0.327 | -0.762 | 0.047 | 0.027 | 0.027 | 0.039 | 0.06 | 0.12 | 0.17 | 0.13 | 0.14 | 0.12 |
| 4 | 169311333 | rs1994530 | 0.038 | 1.891 | 0.215 | -0.841 | -0.069 | -0.015 | -0.015 | 0 | 0.16 | 0.01 | 0.12 | 0.15 | 0.19 | 0.17 |
| 4 | 169315867 | rs17612144 | 0.641 | 1.803 | 0.667 | -0.687 | 0.003 | 0.004 | 0.004 | 0 | 0.07 | 0.04 | 0.03 | 0 | 0.05 | 0.02 |
| 4 | 169316026 | rs3828494 | 2.049 | 1.832 | 0.587 | -0.663 | 0.009 | -0.013 | -0.013 | 0 | 0.58 | 0.47 | 0.61 | 0.62 | 0.56 | 0.56 |
| 4 | 169323978 | rs9284933 | -1.115 | 1.856 | 0.716 | -0.534 | 0.007 | 0.015 | 0.015 | 0 | 0.19 | 0.24 | 0.14 | 0.28 | 0.19 | 0.29 |
| 4 | 169324016 | rs6853665 | 2.09 | 1.758 | 0.609 | -0.431 | -0.008 | -0.002 | -0.002 | 0 | 0.62 | 0.35 | 0.65 | 0.6 | 0.59 | 0.56 |
| 4 | 169328289 | rs17540220 | 1.672 | 2.264 | -0.257 | -0.765 | -0.046 | 0.06 | 0.06 | 0 | 0.1 | 0.01 | 0.09 | 0 | 0.15 | 0.07 |
| 4 | 169329321 | rs1387330 | -1.588 | 1.891 | 0.534 | -0.294 | 0.014 | 0.019 | 0.019 | 0 | 0.27 | 0.34 | 0.3 | 0.4 | 0.35 | 0.43 |
| 4 | 169329930 | rs17612290 | 0.011 | 1.881 | -0.109 | -0.509 | -0.021 | 0.043 | 0.043 | 0 | 0.08 | 0 | 0.04 | 0 | 0.06 | 0.02 |
| 4 | 169331408 | rs11942034 | 1.107 | 1.761 | 0.299 | -0.333 | 0.05 | 0.002 | 0.002 | 0 | 0.86 | 0.66 | 0.8 | 0.88 | 0.79 | 0.85 |
| 4 | 169337313 | rs3762848 | -1.016 | 1.497 | -0.276 | -0.181 | -0.008 | 0.041 | 0.041 | 0.336 | 0.36 | 0.68 | 0.31 | 0.2 | 0.36 | 0.23 |
| 4 | 169337400 | rs1565266 | 1.242 | 1.416 | -0.135 | -0.2 | -0.002 | 0.038 | 0.038 | 0 | 0.59 | 0.31 | 0.69 | 0.72 | 0.6 | 0.69 |
| 4 | 169337545 | rs1565265 | 0.879 | 1.538 | -0.429 | -0.032 | -0.034 | -0.033 | -0.033 | 0 | 0.06 | 0.01 | 0.16 | 0.14 | 0.16 | 0.14 |
| 4 | 169338127 | rs6848102 | -0.076 | 1.568 | 0.056 | -0.209 | 0.017 | -0.084 | -0.084 | 0 | 0.15 | 0.35 | 0.14 | 0.01 | 0.16 | 0.1 |
| 4 | 169338131 | rs6552758 | -0.654 | 1.474 | -0.082 | -0.116 | 0.006 | -0.052 | -0.052 | 0 | 0.31 | 0.52 | 0.26 | 0.14 | 0.35 | 0.22 |
| 4 | 169338520 | rs1963569 | 0.895 | 1.222 | -0.327 | -0.121 | -0.028 | 0.049 | 0.049 | 0.07 | 0.61 | 0.28 | 0.57 | 0.75 | 0.49 | 0.65 |
| 4 | 169340639 | rs1021318 | 1.167 | 1.275 | -0.534 | -0.244 | 0.002 | 0.027 | 0.027 | 0 | 0.51 | 0.22 | 0.51 | 0.58 | 0.43 | 0.55 |
| 4 | 169343522 | rs1000775 | -1.431 | 1.249 | 0.142 | -0.446 | 0.003 | -0.056 | -0.056 | 0.319 | 0.1 | 0.28 | 0.05 | 0 | 0.07 | 0.01 |
| 4 | 169346189 | rs1021317 | -0.628 | 1.184 | -0.288 | -0.954 | -0.001 | -0.016 | -0.016 | 0 | 0.22 | 0.25 | 0.12 | 0.01 | 0.25 | 0.19 |
| 4 | 169346949 | rs2246498 | -0.812 | 1.168 | -0.311 | -0.981 | 0.001 | -0.034 | -0.034 | 0 | 0.07 | 0.23 | 0.04 | 0 | 0.05 | 0.02 |
| 4 | 169347712 | rs2712119 | -1.796 | 1.135 | -0.384 | -0.965 | -0.001 | -0.122 | -0.122 | 0 | 0.33 | 0.54 | 0.22 | 0.02 | 0.32 | 0.24 |
| 4 | 169348435 | rs2712120 | -1.986 | 1.215 | -0.417 | -0.885 | -0.01 | -0.108 | -0.108 | 0.198 | 0.17 | 0.54 | 0.14 | 0.01 | 0.13 | 0.09 |
| 4 | 169349568 | rs17540470 | 0.233 | 1.132 | -0.712 | -0.791 | -0.051 | 0.087 | 0.087 | 0 | 0.16 | 0.01 | 0.07 | 0.01 | 0.2 | 0.17 |
| 4 | 169358017 | rs2319851 | -0.218 | 1.267 | -0.723 | -0.43 | 0.037 | 0.022 | 0.022 | 0 | 0.24 | 0.43 | 0.21 | 0.3 | 0.28 | 0.4 |
| 4 | 169361897 | rs11728342 | 0.919 | 1.262 | -0.853 | -0.421 | -0.048 | -0.051 | -0.051 | 1.821 | 0.16 | 0.11 | 0.17 | 0.28 | 0.24 | 0.26 |
| 4 | 169362557 | rs13151700 | 2.013 | 1.243 | -1.134 | -0.473 | -0.029 | -0.047 | -0.047 | 0.211 | 0.39 | 0.05 | 0.52 | 0.61 | 0.34 | 0.3 |
| 4 | 169369920 | rs12507582 | 2.02 | 1.238 | -1.144 | -0.52 | -0.041 | -0.055 | -0.055 | 0 | 0.39 | 0.02 | 0.52 | 0.62 | 0.34 | 0.3 |
| 4 | 169372709 | rs2634951 | 0.624 | 1.164 | -0.995 | -0.604 | -0.001 | 0.01 | 0.01 | 0 | 0.81 | 0.84 | 0.83 | 0.91 | 0.8 | 0.83 |
| 4 | 169382991 | rs17612630 | 0.818 | 1.227 | -1.175 | -0.574 | -0.057 | 0.087 | 0.087 | 0 | 0.16 | 0.01 | 0.06 | 0.01 | 0.19 | 0.14 |
| 4 | 169387941 | rs1387324 | 1.088 | 1.195 | -1.034 | -0.584 | -0.016 | -0.062 | -0.062 | 0.493 | 0.76 | 0.54 | 0.8 | 0.89 | 0.8 | 0.83 |
| 4 | 169390717 | rs10004447 | -0.544 | 1.257 | -1.131 | -0.508 | 0.028 | -0.024 | -0.024 | 0 | 0.37 | 0.44 | 0.27 | 0.28 | 0.46 | 0.54 |
| 4 | 169394695 | rs7678260 | 0.992 | 1.312 | -1.814 | -0.228 | -0.121 | -0.022 | -0.022 | 0 | 0.55 | 0.03 | 0.59 | 0.62 | 0.54 | 0.46 |

| | | | | | | | | | | | | | | | | |
|---|---|---|---|---|---|---|---|---|---|---|---|---|---|---|---|---|
| 4 | 169396512 | rs2029385 | 1.631 | 1.31 | -1.831 | -0.289 | -0.034 | -0.078 | -0.078 | 0 | 0.39 | 0.02 | 0.49 | 0.66 | 0.33 | 0.32 |
| 4 | 169396757 | rs17054232 | 0.311 | 1.448 | -1.481 | -0.279 | 0.037 | 0.017 | 0.017 | 0 | 0.25 | 0.45 | 0.24 | 0.22 | 0.32 | 0.43 |
| 4 | 169397830 | rs10012902 | -0.031 | 1.1 | -1.905 | -0.379 | -0.005 | 0.068 | 0.068 | 0 | 0.2 | 0.13 | 0.13 | 0.04 | 0.2 | 0.19 |
| 4 | 169398762 | rs2634959 | 0.636 | 1.353 | -1.731 | -0.07 | -0.055 | -0.03 | -0.03 | 0 | 0.61 | 0.21 | 0.62 | 0.63 | 0.54 | 0.5 |
| 4 | 169401199 | rs13353710 | 0.051 | 1.535 | -1.541 | 0.045 | 0.028 | 0.019 | 0.019 | 0 | 0.27 | 0.53 | 0.27 | 0.3 | 0.34 | 0.45 |
| 4 | 169407947 | rs2029386 | -0.339 | 1.285 | -1.721 | 0.344 | -0.019 | 0.023 | 0.023 | 0 | 0.43 | 0.67 | 0.34 | 0.3 | 0.5 | 0.57 |
| 4 | 169408390 | rs2243724 | 0.638 | 1.415 | -0.973 | 0.762 | 0.008 | -0.004 | -0.004 | 0 | 0.89 | 0.84 | 0.88 | 0.92 | 0.85 | 0.96 |
| 4 | 169413936 | rs17650332 | -1.28 | 1.088 | -1.314 | 1.262 | -0.004 | -0.055 | -0.055 | 0 | 0.09 | 0.02 | 0.05 | 0.28 | 0.07 | 0.41 |
| 4 | 169414429 | rs6819953 | -1.168 | 1.323 | -1.225 | 0.817 | 0 | -0.006 | -0.006 | 0 | 0.08 | 0.11 | 0.03 | 0.03 | 0.11 | 0.05 |
| 4 | 169417232 | rs2712134 | 0.203 | 1.458 | -1.981 | 0.272 | -0.042 | 0.044 | 0.044 | 0 | 0.84 | 0.98 | 0.88 | 0.9 | 0.81 | 0.83 |
| 4 | 169419849 | rs2712135 | 0.815 | 1.109 | -1.425 | -0.242 | 0.001 | 0 | 0 | 0 | 0.4 | 0.37 | 0.53 | 0.6 | 0.29 | 0.28 |
| 4 | 169422705 | rs2712149 | 0.057 | 1.288 | -1.115 | -0.331 | 0 | 0.002 | 0.002 | 0 | 0.69 | 0.68 | 0.62 | 0.76 | 0.69 | 0.59 |
| 4 | 169425016 | rs2002727 | 1.14 | 0.682 | -0.959 | -0.23 | 0.013 | 0.008 | 0.008 | 0 | 0.43 | 0.32 | 0.37 | 0.6 | 0.37 | 0.31 |
| 4 | 169425339 | rs4692943 | 0.586 | 0.52 | -1.244 | -0.55 | 0.005 | 0.066 | 0.066 | 0 | 0.52 | 0.28 | 0.47 | 0.67 | 0.55 | 0.48 |
| 4 | 169425778 | rs10517996 | 1.401 | 0.314 | -0.913 | -0.699 | 0.018 | 0.012 | 0.012 | 0 | 0.39 | 0.2 | 0.36 | 0.62 | 0.33 | 0.31 |
| 4 | 169426262 | rs17054263 | -0.31 | -0.186 | -0.557 | -0.926 | -0.036 | -0.014 | -0.014 | 0.069 | 0.11 | 0.25 | 0.06 | 0.07 | 0.04 | 0.19 |
| 4 | 169428546 | rs1962363 | 1.414 | 0.163 | -0.708 | -0.642 | -0.015 | 0.005 | 0.005 | 0.072 | 0.41 | 0.2 | 0.42 | 0.6 | 0.47 | 0.39 |
| 4 | 169429657 | rs10518001 | 1.691 | 0.777 | -1.021 | 0.072 | -0.038 | -0.038 | -0.038 | 0 | 0.11 | 0.02 | 0.05 | 0.17 | 0.12 | 0.07 |
| 4 | 169429928 | rs6857497 | -0.928 | 0.651 | -0.235 | -0.168 | -0.01 | -0.004 | -0.004 | 0.218 | 0.27 | 0.44 | 0.19 | 0.25 | 0.23 | 0.15 |
| 4 | 169430365 | rs10440507 | -0.581 | 0.616 | -0.351 | -0.201 | -0.006 | 0.003 | 0.003 | 0 | 0.26 | 0.23 | 0.18 | 0.19 | 0.33 | 0.24 |
| 4 | 169433333 | rs7673220 | -0.566 | 0.619 | -0.355 | -0.197 | -0.005 | 0.002 | 0.002 | 0 | 0.26 | 0.24 | 0.18 | 0.19 | 0.33 | 0.24 |
| 4 | 169434602 | rs6841109 | 0.942 | 0.479 | -0.061 | -0.045 | -0.001 | 0.004 | 0.004 | 0.028 | 0.06 | 0.19 | 0.04 | 0.05 | 0.03 | 0.01 |
| 4 | 169439108 | rs11735275 | 1.521 | 0.258 | -0.486 | -0.164 | 0.006 | 0.006 | 0.006 | 0.021 | 0.14 | 0.11 | 0.07 | 0.07 | 0.09 | 0.24 |
| 4 | 169439280 | rs11939452 | -0.565 | 0.658 | -0.732 | -0.358 | -0.016 | 0.041 | 0.041 | 0 | 0.75 | 0.82 | 0.64 | 0.86 | 0.71 | 0.7 |
| 4 | 169439902 | rs11132283 | -1.772 | 0.578 | -0.914 | -0.331 | -0.058 | 0.023 | 0.023 | 0 | 0.89 | 0.96 | 0.71 | 0.93 | 0.8 | 0.93 |
| 4 | 169448628 | rs9312333 | -0.758 | 0.31 | -0.315 | -0.217 | 0.001 | -0.013 | -0.013 | 0 | 0.74 | 0.73 | 0.64 | 0.86 | 0.71 | 0.7 |
| 4 | 169454317 | rs17054309 | 0.518 | 0.279 | -0.322 | -0.252 | 0.004 | 0.009 | 0.009 | 0 | 0.16 | 0.21 | 0.14 | 0.19 | 0.17 | 0.12 |
| 4 | 169457246 | rs13145788 | 2.043 | 0.152 | -0.766 | -0.175 | -0.013 | -0.071 | -0.071 | 0.963 | 0.41 | 0.08 | 0.38 | 0.57 | 0.36 | 0.3 |
| 4 | 169457744 | rs1986369 | -0.748 | 0.341 | -0.328 | 0.037 | -0.023 | 0.048 | 0.048 | 2.053 | 0.2 | 0.55 | 0.16 | 0.24 | 0.15 | 0.07 |
| 4 | 169459736 | rs10023820 | -1.738 | -0.071 | -0.316 | 0.151 | -0.015 | 0.009 | 0.009 | 0 | 0.3 | 0.57 | 0.23 | 0.28 | 0.28 | 0.39 |
| 4 | 169474972 | rs6857016 | 0.33 | 0.277 | -0.661 | -0.155 | 0.01 | -0.027 | -0.027 | 0 | 0.61 | 0.57 | 0.54 | 0.82 | 0.48 | 0.53 |
| 4 | 169475264 | rs17707180 | 1.704 | 0.318 | -1.024 | 0.183 | -0.039 | -0.061 | -0.061 | 1.934 | 0.15 | 0.04 | 0.18 | 0.28 | 0.15 | 0.07 |
| 4 | 169476590 | rs11132322 | -1.732 | 0.172 | -0.851 | -0.118 | -0.004 | 0.007 | 0.007 | 1.586 | 0.28 | 0.2 | 0.22 | 0.19 | 0.32 | 0.43 |
| 4 | 169482396 | rs10518011 | 1.059 | 0.054 | -0.721 | -0.086 | 0.001 | -0.004 | -0.004 | 0 | 0.11 | 0.14 | 0.07 | 0.1 | 0.08 | 0.07 |
| 4 | 169484303 | rs10022111 | 1.478 | -0.058 | -0.768 | -0.148 | 0.016 | 0.011 | 0.011 | 0 | 0.44 | 0.37 | 0.43 | 0.66 | 0.37 | 0.3 |
| 4 | 169485237 | rs10015043 | 0.755 | -0.216 | -0.816 | -0.18 | 0 | -0.009 | -0.009 | 0.433 | 0.55 | 0.52 | 0.5 | 0.76 | 0.44 | 0.37 |
| 4 | 169491370 | rs10004025 | 0.461 | -0.083 | -0.693 | -0.7 | -0.002 | 0.006 | 0.006 | 0 | 0.57 | 0.49 | 0.52 | 0.79 | 0.59 | 0.53 |
| 4 | 169492567 | rs1962022 | 1.126 | -0.03 | -0.743 | -0.703 | 0.014 | 0.074 | 0.074 | 0 | 0.49 | 0.4 | 0.45 | 0.78 | 0.44 | 0.36 |
| 4 | 169494427 | rs7693542 | -0.576 | 0.076 | -0.692 | -0.797 | 0.032 | -0.077 | -0.077 | 0 | 0.71 | 0.63 | 0.64 | 0.9 | 0.64 | 0.72 |
| 4 | 169499240 | rs7668720 | 0.85 | -0.086 | -0.87 | -0.714 | -0.009 | 0.019 | 0.019 | 1.056 | 0.09 | 0.04 | 0.07 | 0.01 | 0.15 | 0.17 |
| 4 | 169499863 | rs4458418 | 1.464 | -0.063 | -0.936 | -0.577 | -0.076 | -0.081 | -0.081 | 0.456 | 0.15 | 0.02 | 0.18 | 0.27 | 0.18 | 0.08 |
| 4 | 169500402 | rs17650886 | 0.239 | -0.075 | -0.705 | -0.758 | 0.006 | 0.002 | 0.002 | 0.149 | 0.13 | 0.07 | 0.12 | 0.1 | 0.08 | 0.17 |
| 4 | 169503531 | rs6815330 | 0.086 | 0.151 | -0.533 | -0.935 | 0.037 | -0.092 | -0.092 | 0 | 0.65 | 0.54 | 0.58 | 0.9 | 0.53 | 0.63 |
| 4 | 169504186 | rs17650892 | 0.459 | 0.306 | -0.614 | -0.603 | -0.034 | 0.051 | 0.051 | 0 | 0.1 | 0.01 | 0.18 | 0.02 | 0.12 | 0.04 |
| 4 | 169506877 | rs2874112 | -0.315 | 0.06 | -0.027 | -0.247 | -0.004 | -0.004 | -0.004 | 1.493 | 0.27 | 0.21 | 0.34 | 0.38 | 0.25 | 0.23 |
| 4 | 169509103 | rs17707379 | 0.892 | -0.238 | -0.087 | -0.437 | 0.026 | -0.024 | -0.024 | 0 | 0.34 | 0.31 | 0.25 | 0.53 | 0.31 | 0.41 |
| 4 | 169509436 | rs2319909 | 0.947 | 0.071 | -0.136 | -0.299 | -0.056 | -0.075 | -0.075 | 0.006 | 0.17 | 0.05 | 0.19 | 0.33 | 0.18 | 0.11 |
| 4 | 169517130 | rs4144994 | 0.267 | 0.04 | -0.179 | -0.319 | 0.012 | 0 | 0 | 0 | 0.44 | 0.41 | 0.4 | 0.58 | 0.38 | 0.53 |
| 4 | 169523335 | rs4132492 | -0.515 | 0.275 | -0.826 | -0.456 | -0.041 | 0.084 | 0.084 | 0 | 0.64 | 0.84 | 0.57 | 0.9 | 0.57 | 0.64 |
| 4 | 169526903 | rs4371580 | -0.546 | 0.315 | -0.817 | -0.37 | -0.041 | 0.072 | 0.072 | 0 | 0.65 | 0.84 | 0.57 | 0.88 | 0.57 | 0.64 |
| 4 | 169530339 | rs4389538 | -0.196 | 0.405 | -1.556 | -0.121 | -0.013 | 0.176 | 0.176 | 0 | 0.49 | 0.21 | 0.43 | 0.8 | 0.51 | 0.39 |
| 4 | 169532292 | rs12647503 | 0.523 | 0.142 | -1.512 | 0.17 | 0.033 | -0.102 | -0.102 | 0 | 0.31 | 0.12 | 0.24 | 0.49 | 0.33 | 0.28 |
| 4 | 169532319 | rs12650216 | 0.986 | -0.083 | -1.403 | 0.34 | 0.06 | -0.103 | -0.103 | 0 | 0.24 | 0.1 | 0.18 | 0.47 | 0.19 | 0.18 |
| 4 | 169537189 | rs17707568 | 0.602 | 0.545 | -0.969 | 0.061 | 0.046 | -0.032 | -0.032 | 0 | 0.39 | 0.24 | 0.34 | 0.43 | 0.33 | 0.5 |
| 4 | 169538426 | rs2723687 | 0.425 | 1.411 | -0.366 | -0.086 | 0.01 | 0.011 | 0.011 | 0 | 0.61 | 0.54 | 0.53 | 0.46 | 0.58 | 0.55 |
| 4 | 169538443 | rs2710851 | 0.632 | 1.467 | -0.419 | -0.112 | -0.011 | -0.022 | -0.022 | 0 | 0.58 | 0.72 | 0.52 | 0.46 | 0.49 | 0.52 |
| 4 | 169539216 | rs2723688 | -0.201 | 2.303 | 0.102 | -0.051 | 0.018 | 0.009 | 0.009 | 0 | 0.79 | 0.68 | 0.88 | 0.76 | 0.77 | 0.67 |
| 4 | 169549646 | rs3109799 | -1.303 | 2.319 | -0.032 | -0.141 | 0.009 | -0.046 | -0.046 | 0.044 | 0.21 | 0.65 | 0.15 | 0.06 | 0.23 | 0.14 |
| 4 | 169551542 | rs2723696 | -1.873 | 2.482 | -0.112 | 0.349 | 0.071 | 0.003 | 0.003 | 0 | 0.21 | 0.46 | 0.31 | 0.24 | 0.28 | 0.09 |
| 4 | 169558119 | rs2723698 | -0.903 | 2.435 | -0.135 | 0.247 | 0.081 | -0.061 | -0.061 | 8.082 | 0.1 | 0.36 | 0.13 | 0 | 0.12 | 0.02 |
| 4 | 169567817 | rs12643131 | -1.027 | 2.207 | -0.965 | 0.79 | -0.031 | -0.031 | -0.031 | 0 | 0.09 | 0.05 | 0.18 | 0.24 | 0.16 | 0.06 |
| 4 | 169569729 | rs2723704 | 0.454 | 1.531 | -0.776 | -0.065 | -0.001 | -0.047 | -0.047 | 0 | 0.72 | 0.63 | 0.84 | 0.88 | 0.77 | 0.66 |
| 4 | 169571481 | rs17614077 | -0.127 | 0.634 | -1.996 | -0.603 | -0.001 | 0.085 | 0.085 | 0 | 0.26 | 0.04 | 0.22 | 0.11 | 0.21 | 0.25 |
| 4 | 169577535 | rs10010321 | -0.659 | 0.342 | -2.352 | -0.758 | -0.006 | 0.089 | 0.089 | 0 | 0.31 | 0.07 | 0.29 | 0.17 | 0.3 | 0.21 |
| 4 | 169579967 | rs12642267 | -0.457 | -0.259 | -1.66 | -1.093 | -0.001 | -0.016 | -0.016 | 9.016 | 0.19 | 0.2 | 0.11 | 0.02 | 0.13 | 0.13 |
| 4 | 169580501 | rs17054449 | -1.213 | -0.186 | -1.341 | -1.121 | -0.008 | -0.031 | -0.031 | 1.839 | 0.25 | 0.28 | 0.25 | 0.08 | 0.2 | 0.21 |
| 4 | 169584060 | rs4314247 | -1.356 | -0.408 | -1.749 | -0.974 | -0.075 | 0.004 | 0.004 | 0 | 0.62 | 0.83 | 0.63 | 0.38 | 0.52 | 0.55 |
| 4 | 169585678 | rs6839585 | -0.021 | -0.168 | -1.243 | -0.751 | -0.002 | -0.019 | -0.019 | 0.895 | 0.19 | 0.2 | 0.11 | 0.02 | 0.13 | 0.16 |
| 4 | 169586731 | rs6846708 | -0.627 | 0.176 | 0.177 | -0.198 | 0.044 | -0.007 | -0.007 | 0 | 0.09 | 0.26 | 0.06 | 0.06 | 0.07 | 0.15 |

| | | | | | | | | | | | | | | | | |
|---|---|---|---|---|---|---|---|---|---|---|---|---|---|---|---|---|
| 4 | 169588252 | rs12649186 | -0.261 | 0.191 | 0.141 | -0.134 | 0.015 | -0.005 | -0.005 | 0 | 0.07 | 0.19 | 0.15 | 0.06 | 0.07 | 0.08 |
| 4 | 169589214 | rs2723705 | -0.989 | -0.074 | 0.465 | -0.071 | 0.001 | 0.011 | 0.011 | 0 | 0.41 | 0.61 | 0.55 | 0.29 | 0.45 | 0.38 |
| 4 | 169592669 | rs2723708 | -1.117 | -0.298 | 0.556 | 0.07 | -0.01 | 0.041 | 0.041 | 0 | 0.4 | 0.77 | 0.52 | 0.32 | 0.46 | 0.39 |
| 4 | 169595421 | rs11933816 | -0.66 | -0.447 | 0.822 | 0.038 | -0.025 | -0.056 | -0.056 | 8.059 | 0.33 | 0.5 | 0.25 | 0.17 | 0.3 | 0.29 |
| 4 | 169596133 | rs2710824 | -1.05 | -0.298 | 0.344 | 0.204 | -0.015 | 0.025 | 0.025 | 0.217 | 0.36 | 0.78 | 0.45 | 0.31 | 0.41 | 0.38 |
| 4 | 169596649 | rs13137200 | 0.993 | -0.159 | 0.371 | 0.521 | 0.005 | -0.043 | -0.043 | 0.45 | 0.57 | 0.17 | 0.52 | 0.48 | 0.58 | 0.58 |
| 4 | 169599633 | rs6832582 | -1.85 | -0.39 | 1.637 | 0.888 | -0.012 | 0.039 | 0.039 | 0.113 | 0.21 | 0.67 | 0.27 | 0.44 | 0.15 | 0.29 |
| 4 | 169605174 | rs17054465 | -1.111 | -0.232 | 1.877 | 0.543 | 0.037 | 0.012 | 0.012 | 0.034 | 0.14 | 0.48 | 0.25 | 0.14 | 0.13 | 0.1 |
| 4 | 169605918 | rs4692947 | -0.855 | -0.008 | 1.791 | 0.694 | 0.103 | 0.093 | 0.093 | 0.036 | 0.05 | 0.31 | 0.22 | 0.08 | 0.08 | 0.08 |
| 4 | 169612528 | rs4260495 | -1.5 | -0.188 | 1.351 | 0.699 | -0.019 | 0.14 | 0.14 | 0 | 0.2 | 0.44 | 0.26 | 0.4 | 0.18 | 0.31 |
| 4 | 169618128 | rs4425335 | -1.333 | -0.358 | 1.146 | 0.422 | -0.009 | 0.015 | 0.015 | 1.337 | 0.31 | 0.64 | 0.3 | 0.45 | 0.26 | 0.17 |
| 4 | 169618430 | rs6842732 | 0.095 | -0.38 | 1.451 | 0.49 | -0.007 | 0.041 | 0.041 | 0.795 | 0.15 | 0.22 | 0.05 | 0.33 | 0.1 | 0.04 |
| 4 | 169618568 | rs4280700 | -0.629 | -0.364 | 1.333 | 0.477 | 0.016 | -0.03 | -0.03 | 0 | 0.14 | 0.35 | 0.24 | 0.09 | 0.15 | 0.13 |
| 4 | 169622650 | rs12649675 | -0.752 | -0.431 | 1.458 | 0.764 | -0.023 | 0.081 | 0.081 | 0.004 | 0.18 | 0.35 | 0.07 | 0.36 | 0.12 | 0.21 |
| 4 | 169634434 | rs4635780 | -0.254 | 0.034 | 0.719 | 0.567 | 0.008 | 0.055 | 0.055 | 0 | 0.47 | 0.67 | 0.62 | 0.66 | 0.48 | 0.57 |
| 4 | 169639326 | rs7670597 | 0.449 | 0.15 | 0.4 | 0.614 | -0.019 | 0.015 | 0.015 | 0 | 0.37 | 0.22 | 0.41 | 0.3 | 0.41 | 0.38 |
| 4 | 169641425 | rs9884230 | -0.157 | 0.299 | 0.555 | 0.548 | -0.008 | 0.077 | 0.077 | 0 | 0.55 | 0.72 | 0.51 | 0.7 | 0.56 | 0.72 |
| 4 | 169641509 | rs4599370 | -1.07 | 0.487 | 0.236 | 0.498 | -0.052 | 0.025 | 0.025 | 0 | 0.87 | 0.99 | 0.7 | 0.94 | 0.81 | 0.89 |
| 4 | 169644844 | rs11132434 | -0.45 | 0.332 | 1.142 | 0.925 | 0.015 | 0.163 | 0.163 | 0 | 0.15 | 0.25 | 0.12 | 0.47 | 0.15 | 0.24 |
| 4 | 169644989 | rs17542430 | 0.527 | 0.209 | 1.219 | 0.868 | -0.009 | 0.024 | 0.024 | 0 | 0.35 | 0.23 | 0.35 | 0.23 | 0.39 | 0.31 |
| 4 | 169657096 | rs1500800 | -0.402 | 0.303 | 0.575 | 0.955 | -0.013 | 0.036 | 0.036 | 0 | 0.46 | 0.3 | 0.43 | 0.29 | 0.5 | 0.62 |
| 4 | 169663615 | rs6811238 | -0.764 | 0.334 | 0.569 | 0.835 | 0 | 0.024 | 0.024 | 0 | 0.49 | 0.55 | 0.47 | 0.62 | 0.5 | 0.63 |
| 4 | 169664679 | rs7656460 | 0.522 | 0.341 | -0.206 | 0.818 | 0 | -0.003 | -0.003 | 0 | 0.1 | 0.09 | 0.09 | 0.04 | 0.09 | 0.14 |
| 4 | 169667766 | rs6857155 | -0.58 | 0.53 | -0.734 | 0.691 | -0.006 | 0.064 | 0.064 | 0 | 0.52 | 0.66 | 0.49 | 0.62 | 0.51 | 0.63 |
| 4 | 169670530 | rs6817535 | -0.193 | 1.058 | -1.296 | 0.422 | -0.021 | 0.007 | 0.007 | 0 | 0.33 | 0.17 | 0.34 | 0.26 | 0.39 | 0.33 |
| 4 | 169676032 | rs6852874 | -0.363 | 1.02 | -1.478 | 0.404 | -0.029 | 0.022 | 0.022 | 0 | 0.38 | 0.21 | 0.37 | 0.26 | 0.43 | 0.35 |
| 4 | 169685036 | rs17542654 | -0.627 | 1.029 | -2.119 | -0.449 | -0.069 | 0.127 | 0.127 | 1.22 | 0.2 | 0.02 | 0.11 | 0.01 | 0.18 | 0.11 |
| 4 | 169685812 | rs12499664 | 0.081 | 0.688 | -1.914 | -0.237 | -0.001 | 0 | 0 | 0 | 0.37 | 0.34 | 0.52 | 0.56 | 0.31 | 0.46 |
| 4 | 169689516 | rs17542688 | 0.986 | 0.55 | -1.718 | -0.295 | -0.017 | 0.019 | 0.019 | 0 | 0.28 | 0.21 | 0.28 | 0.13 | 0.41 | 0.29 |
| 4 | 169692342 | rs17708307 | 0.481 | 0.287 | -1.266 | -0.432 | -0.008 | 0.007 | 0.007 | 0 | 0.38 | 0.55 | 0.54 | 0.55 | 0.29 | 0.45 |
| 4 | 169692626 | rs4692948 | 0.066 | 0.209 | -1.1 | -0.453 | -0.007 | 0.008 | 0.008 | 0 | 0.67 | 0.75 | 0.81 | 0.66 | 0.7 | 0.74 |
| 4 | 169692809 | rs12510663 | 0.524 | 0.208 | -1.099 | -0.453 | -0.008 | 0.007 | 0.007 | 0 | 0.38 | 0.55 | 0.54 | 0.55 | 0.29 | 0.45 |
| 4 | 169693353 | rs7679564 | 1.033 | 0.238 | -1.09 | -0.439 | -0.013 | 0.021 | 0.021 | 0 | 0.29 | 0.22 | 0.28 | 0.11 | 0.42 | 0.29 |
| 4 | 169694079 | rs2247733 | -1.468 | 0.189 | -1.128 | -0.251 | 0 | -0.003 | -0.003 | 0.008 | 0.89 | 0.96 | 0.94 | 0.88 | 0.9 | 0.87 |
| 4 | 169698873 | rs999958 | -0.51 | 0.187 | -1.105 | -0.261 | -0.031 | 0.047 | 0.047 | 0 | 0.61 | 0.78 | 0.67 | 0.77 | 0.49 | 0.57 |
| 4 | 169703162 | rs7688994 | -0.114 | -0.014 | -1.004 | -0.394 | 0.033 | 0.053 | 0.053 | 0 | 0.51 | 0.39 | 0.39 | 0.3 | 0.61 | 0.41 |
| 4 | 169703442 | rs17614733 | 0.583 | 0.122 | -1.137 | -0.377 | -0.075 | 0.105 | 0.105 | 0.34 | 0.21 | 0.02 | 0.11 | 0.04 | 0.19 | 0.12 |
| 4 | 169717641 | rs1542869 | -0.442 | 0.009 | -0.285 | -0.31 | -0.011 | -0.015 | -0.015 | 0 | 0.88 | 0.79 | 0.93 | 0.9 | 0.93 | 0.88 |
| 4 | 169718700 | rs9312367 | 0.238 | -0.262 | -0.379 | -0.426 | 0.01 | 0.151 | 0.151 | 0 | 0.45 | 0.2 | 0.37 | 0.13 | 0.58 | 0.44 |
| 4 | 169718848 | rs11733873 | 1.332 | -0.367 | -0.741 | 0.068 | -0.047 | -0.084 | -0.084 | 1.724 | 0.12 | 0.01 | 0.23 | 0.42 | 0.19 | 0.23 |
| 4 | 169722992 | rs17614822 | 0.738 | -0.177 | -0.583 | -0.125 | -0.044 | 0.061 | 0.061 | 0 | 0.18 | 0.07 | 0.2 | 0.05 | 0.19 | 0.12 |
| 4 | 169725009 | rs1875297 | -1.544 | -0.242 | -0.547 | -0.151 | -0.001 | 0 | 0 | 0 | 0.92 | 0.94 | 0.94 | 0.91 | 0.92 | 0.88 |
| 4 | 169725161 | rs1875296 | 0.655 | -0.205 | -0.479 | 0.009 | -0.026 | -0.018 | -0.018 | 0 | 0.18 | 0.09 | 0.21 | 0.23 | 0.19 | 0.12 |
| 4 | 169725605 | rs7681510 | -0.842 | 0.228 | -0.343 | -0.209 | -0.007 | 0.029 | 0.029 | 0 | 0.64 | 0.7 | 0.7 | 0.83 | 0.56 | 0.66 |
| 4 | 169726027 | rs13129779 | -0.672 | -0.016 | -0.333 | -0.16 | -0.013 | 0.026 | 0.026 | 0 | 0.61 | 0.67 | 0.66 | 0.83 | 0.51 | 0.65 |
| 4 | 169728142 | rs6822195 | 1.325 | -0.058 | -0.538 | -0.273 | -0.031 | 0.039 | 0.039 | 0 | 0.26 | 0.16 | 0.27 | 0.09 | 0.39 | 0.19 |
| 4 | 169728236 | rs6854137 | 0.044 | -0.144 | -0.321 | -0.267 | -0.001 | -0.031 | -0.031 | 0 | 0.43 | 0.46 | 0.48 | 0.14 | 0.55 | 0.3 |
| 4 | 169732171 | rs1566499 | 0.542 | -0.261 | -0.238 | -0.332 | -0.007 | 0.004 | 0.004 | 0 | 0.37 | 0.62 | 0.35 | 0.09 | 0.45 | 0.32 |
| 4 | 169732320 | rs1566498 | -0.99 | -1.114 | -1.007 | 0.272 | -0.036 | -0.131 | -0.131 | 0 | 0.72 | 0.99 | 0.73 | 0.58 | 0.72 | 0.72 |
| 4 | 169733334 | rs4692949 | 0.377 | -0.37 | -0.442 | 0.097 | 0.031 | 0.029 | 0.029 | 0 | 0.45 | 0.33 | 0.52 | 0.33 | 0.61 | 0.45 |
| 4 | 169740931 | rs10005789 | 0.379 | -0.355 | -0.933 | -0.074 | 0.04 | 0.132 | 0.132 | 0 | 0.44 | 0.24 | 0.5 | 0.14 | 0.6 | 0.45 |
| 4 | 169745078 | rs1392750 | -0.265 | 0.106 | -1.194 | 0.043 | 0.013 | -0.002 | -0.002 | 0.395 | 0.59 | 0.68 | 0.61 | 0.43 | 0.67 | 0.64 |
| 4 | 169754830 | rs2133911 | 0.134 | 0.703 | -1.928 | -0.449 | -0.042 | 0.128 | 0.128 | 0.022 | 0.38 | 0.1 | 0.38 | 0.15 | 0.43 | 0.4 |
| 4 | 169762454 | rs867901 | -0.058 | 0.952 | -1.664 | -1.312 | -0.014 | -0.024 | -0.024 | 0 | 0.3 | 0.3 | 0.24 | 0 | 0.39 | 0.35 |
| 4 | 169763252 | rs973990 | -0.573 | 0.175 | -1.525 | -1.439 | 0.001 | -0.033 | -0.033 | 0.005 | 0.49 | 0.47 | 0.33 | 0.09 | 0.52 | 0.55 |
| 4 | 169764312 | rs12643033 | 1.444 | 0.145 | -2.3 | -1.092 | 0.022 | -0.008 | -0.008 | 0 | 0.14 | 0.03 | 0.13 | 0.15 | 0.11 | 0.09 |
| 4 | 169764489 | rs1318822 | -1.057 | 0.025 | -1.972 | -1.17 | -0.007 | 0.024 | 0.024 | 0.653 | 0.68 | 0.8 | 0.49 | 0.24 | 0.65 | 0.64 |
| 4 | 169765728 | rs3811758 | 0.974 | -0.383 | -1.594 | -0.612 | 0.021 | -0.014 | -0.014 | 8.475 | 0.14 | 0.03 | 0.13 | 0.17 | 0.11 | 0.09 |
| 4 | 169766595 | rs7688533 | -1.047 | -0.795 | -2.186 | -0.626 | -0.04 | -0.109 | -0.109 | 0 | 0.61 | 0.98 | 0.5 | 0.65 | 0.6 | 0.43 |
| 4 | 169766963 | rs1392751 | 0.108 | -1.902 | -1.104 | -1.476 | 0.009 | 0.029 | 0.029 | 0 | 0.2 | 0.12 | 0.1 | 0.07 | 0.15 | 0.21 |
| 4 | 169769793 | rs4692553 | 0.846 | -1.905 | -0.712 | -1.731 | 0.038 | 0.141 | 0.141 | 1.549 | 0.54 | 0.27 | 0.69 | 0.91 | 0.64 | 0.63 |
| 4 | 169771612 | rs13114906 | -1.562 | -2.123 | 0.609 | -1.27 | -0.054 | -0.091 | -0.091 | 0 | 0.18 | 0.53 | 0.12 | 0.03 | 0.09 | 0.19 |
| 4 | 169779025 | rs7682426 | -1.599 | -2.305 | 0.41 | -1.273 | 0.083 | 0.172 | 0.172 | 1.452 | 0.37 | 0.85 | 0.19 | 0.1 | 0.23 | 0.28 |
| 4 | 169781253 | rs867632 | -0.32 | -2.298 | 1.083 | -1.265 | 0.005 | -0.089 | -0.089 | 0 | 0.8 | 0.64 | 0.82 | 0.94 | 0.81 | 0.88 |
| 4 | 169788148 | rs7690353 | 1.526 | -2.366 | 1.421 | -1.334 | -0.011 | -0.073 | -0.073 | 0 | 0.61 | 0.45 | 0.66 | 0.91 | 0.66 | 0.73 |
| 4 | 169801952 | rs12643097 | 1.982 | -2.391 | 1.308 | -0.88 | 0.037 | -0.09 | -0.09 | 0.441 | 0.56 | 0.1 | 0.57 | 0.54 | 0.64 | 0.66 |
| 4 | 169812782 | rs1875299 | 2.036 | -2.394 | 1.491 | -0.908 | -0.023 | 0.058 | 0.058 | 0 | 0.74 | 0.38 | 0.63 | 0.6 | 0.78 | 0.77 |
| 4 | 169815472 | rs2047633 | 0.151 | -2.344 | 1.511 | -2.449 | 0.019 | -0.048 | -0.048 | 0 | 0.91 | 0.75 | 0.84 | 1 | 0.89 | 0.85 |
| 4 | 169819067 | rs1566496 | 1.614 | -2.341 | 1.589 | -2.448 | -0.005 | -0.077 | -0.077 | 0.548 | 0.78 | 0.66 | 0.72 | 0.97 | 0.78 | 0.8 |

| | | | | | | | | | | | | | | | | |
|---|---|---|---|---|---|---|---|---|---|---|---|---|---|---|---|---|
| 4 | 169827618 | rs6819031 | 0.431 | -2.366 | 1.475 | -2.441 | -0.021 | -0.069 | -0.069 | 0.392 | 0.18 | 0.31 | 0.07 | 0.07 | 0.14 | 0.1 |
| 4 | 169828862 | rs1500795 | -1.314 | -2.367 | 1.478 | -2.442 | -0.011 | -0.035 | -0.035 | 0 | 0.14 | 0.21 | 0.13 | 0.03 | 0.1 | 0.04 |
| 4 | 169845720 | rs4396955 | 0.558 | -2.352 | 1.085 | -2.432 | -0.018 | -0.063 | -0.063 | 0 | 0.17 | 0.26 | 0.07 | 0.06 | 0.13 | 0.1 |
| 4 | 169863970 | rs1115717 | 1.92 | -2.459 | 0.989 | -2.486 | 0.069 | 0.268 | 0.268 | 0.847 | 0.54 | 0.15 | 0.64 | 0.9 | 0.65 | 0.7 |
| 4 | 169874986 | rs17543378 | 0.194 | -2.415 | 1.013 | -2.492 | 0.003 | 0.018 | 0.018 | 0 | 0.07 | 0.02 | 0.06 | 0.03 | 0.07 | 0.04 |
| 4 | 169889068 | rs4446276 | 2.058 | -2.416 | 0.97 | -2.507 | 0.024 | 0.309 | 0.309 | 0.706 | 0.54 | 0.08 | 0.6 | 0.9 | 0.63 | 0.65 |
| 4 | 169899023 | rs10518037 | 2.023 | -2.29 | 1.161 | -2.361 | -0.016 | -0.08 | -0.08 | 0 | 0.74 | 0.36 | 0.7 | 0.97 | 0.78 | 0.83 |
| 4 | 169909593 | rs17054604 | 0.663 | -2.275 | 1.186 | -2.357 | -0.011 | -0.042 | -0.042 | 0 | 0.17 | 0.18 | 0.05 | 0.06 | 0.13 | 0.08 |
| 4 | 169917245 | rs898002 | 1.939 | -2.159 | 1.104 | -2.459 | -0.018 | -0.104 | -0.104 | 0 | 0.75 | 0.42 | 0.7 | 0.97 | 0.79 | 0.84 |
| 4 | 169928842 | rs2877380 | 1.939 | -2.151 | 1.108 | -2.455 | -0.019 | -0.1 | -0.1 | 0.806 | 0.75 | 0.42 | 0.7 | 0.97 | 0.79 | 0.81 |
| 4 | 169931752 | rs1551612 | -0.151 | -2.195 | 0.562 | -1.985 | -0.007 | -0.021 | -0.021 | 0 | 0.95 | 0.93 | 0.95 | 1 | 0.97 | 1 |
| 4 | 169934087 | rs17615362 | -0.299 | -2.211 | 0.542 | -2.004 | 0 | 0.007 | 0.007 | 0 | 0.08 | 0.04 | 0.07 | 0.03 | 0.08 | 0.04 |
| 4 | 169934459 | rs17054653 | 0.74 | -2.278 | 0.287 | -2.079 | 0.001 | 0.073 | 0.073 | 0.025 | 0.14 | 0.03 | 0.05 | 0.02 | 0.12 | 0.07 |
| 4 | 169935881 | rs1458292 | 0.952 | -2.14 | 0.603 | -1.978 | 0.014 | -0.002 | -0.002 | 0 | 0.64 | 0.37 | 0.78 | 0.95 | 0.75 | 0.86 |
| 4 | 169936671 | rs2331453 | 0.919 | -1.172 | 0.695 | -0.967 | -0.008 | -0.028 | -0.028 | 0 | 0.93 | 0.89 | 0.94 | 1 | 0.97 | 1 |
| 4 | 169947762 | rs17054666 | -1.117 | -1.093 | 0.838 | -0.868 | -0.009 | -0.04 | -0.04 | 1.279 | 0.09 | 0.12 | 0.06 | 0 | 0.08 | 0.05 |
| 4 | 169948102 | rs13434426 | -1.308 | -1.363 | 1.552 | -1.02 | -0.012 | -0.024 | -0.024 | 0 | 0.1 | 0.41 | 0.05 | 0.03 | 0.03 | 0.17 |
| 4 | 169951395 | rs2331452 | 2.585 | -1.569 | 1.435 | -1.159 | 0.05 | 0.081 | 0.081 | 1.592 | 0.73 | 0.19 | 0.78 | 0.96 | 0.84 | 0.66 |
| 4 | 169960188 | rs13147728 | -1.045 | -1.234 | 1.764 | -0.975 | 0 | -0.013 | -0.013 | 0 | 0.09 | 0.34 | 0.05 | 0.02 | 0.03 | 0.17 |
| 4 | 169962604 | rs2331548 | 1.13 | -0.966 | 1.321 | -0.789 | 0.026 | -0.026 | -0.026 | 1.693 | 0.92 | 0.7 | 0.89 | 1 | 0.95 | 0.88 |
| 4 | 169967051 | rs4692691 | 0.997 | -1.068 | 0.656 | -0.841 | -0.02 | -0.091 | -0.091 | 0 | 0.87 | 0.65 | 0.9 | 0.99 | 0.91 | 0.95 |
| 4 | 169969076 | rs7664949 | 1.208 | -1.318 | -1.046 | -0.704 | 0.101 | 0.096 | 0.096 | 0.279 | 0.6 | 0.11 | 0.61 | 0.76 | 0.74 | 0.52 |
| 4 | 169971917 | rs752625 | 1.217 | -1.306 | -1.097 | -0.68 | 0.097 | 0.082 | 0.082 | 0.75 | 0.6 | 0.1 | 0.61 | 0.75 | 0.73 | 0.49 |
| 4 | 169972311 | rs10009678 | 0.868 | -0.851 | 0.048 | -0.473 | -0.01 | -0.002 | -0.002 | 0.167 | 0.78 | 0.73 | 0.77 | 0.78 | 0.8 | 0.68 |
| 4 | 169972982 | rs7673669 | 0.802 | -0.749 | -0.481 | -0.479 | 0.008 | -0.006 | -0.006 | 0.022 | 0.79 | 0.91 | 0.7 | 0.79 | 0.8 | 0.65 |
| 4 | 169974756 | rs7686225 | 1.335 | -0.345 | 0.572 | -0.557 | 0.022 | 0.001 | 0.001 | 0.368 | 0.88 | 0.51 | 0.94 | 0.95 | 0.92 | 0.95 |
| 4 | 169977902 | rs12498384 | 0.756 | 0.196 | 0.337 | -1.078 | -0.005 | -0.085 | -0.085 | 0.981 | 0.68 | 0.49 | 0.73 | 0.9 | 0.74 | 0.65 |
| 4 | 169979856 | rs17054702 | -0.346 | 0.308 | -0.029 | -1.118 | -0.002 | -0.018 | -0.018 | 0 | 0.14 | 0.21 | 0.09 | 0.06 | 0.11 | 0.22 |
| 4 | 169981893 | rs17626424 | -0.507 | 0.354 | -0.464 | -0.998 | -0.041 | 0.074 | 0.074 | 0 | 0.16 | 0 | 0.1 | 0.04 | 0.13 | 0.14 |
| 4 | 169984410 | rs17543834 | -1.376 | 1.049 | -0.32 | -0.474 | 0 | 0 | 0 | 0.639 | 0.18 | 0.19 | 0.25 | 0.19 | 0.23 | 0.12 |
| 4 | 169985016 | rs4692693 | -0.914 | 1.032 | -0.179 | -0.356 | 0.003 | 0.006 | 0.006 | 0 | 0.13 | 0.19 | 0.22 | 0.18 | 0.15 | 0.11 |
| 4 | 169990206 | rs4692561 | -1.507 | 1.261 | -0.738 | -0.163 | 0.059 | 0.039 | 0.039 | 0 | 0.49 | 0.75 | 0.54 | 0.43 | 0.57 | 0.64 |
| 4 | 169991158 | rs4692694 | -1.13 | 1.848 | -0.705 | -0.235 | -0.002 | 0.001 | 0.001 | 0.892 | 0.31 | 0.66 | 0.4 | 0.22 | 0.39 | 0.41 |
| 4 | 169992206 | rs4692695 | 1.364 | 2.25 | -0.155 | 0.134 | -0.019 | -0.004 | -0.004 | 0 | 0.7 | 0.24 | 0.58 | 0.62 | 0.64 | 0.69 |
| 4 | 169992966 | rs12505635 | 1.869 | 2.46 | 0.588 | -0.223 | 0.01 | -0.028 | -0.028 | 0 | 0.72 | 0.41 | 0.73 | 0.8 | 0.75 | 0.57 |
| 4 | 169993004 | rs10012798 | 1.849 | 2.56 | 0.581 | -0.182 | 0.02 | 0.001 | 0.001 | 0 | 0.78 | 0.73 | 0.68 | 0.79 | 0.63 | 0.74 |
| 4 | 169994701 | rs11736883 | -1.079 | 2.291 | 0.294 | -0.147 | -0.005 | -0.002 | -0.002 | 0 | 0.08 | 0.05 | 0.06 | 0.07 | 0.1 | 0.2 |
| 4 | 169998230 | rs17658335 | 1.971 | 1.859 | -0.052 | -0.26 | -0.076 | 0.059 | 0.059 | 0.01 | 0.58 | 0.06 | 0.5 | 0.68 | 0.53 | 0.49 |
| 4 | 170008947 | rs6844814 | -1.898 | 1.14 | 1.425 | 0.007 | 0.004 | 0.11 | 0.11 | 0 | 0.09 | 0.18 | 0.28 | 0.24 | 0.1 | 0.12 |
| 4 | 170013910 | rs17626617 | -1.563 | 1.722 | 1.068 | -0.129 | -0.017 | 0.013 | 0.013 | 0 | 0.09 | 0.06 | 0.08 | 0.04 | 0.15 | 0.22 |
| 4 | 170014786 | rs2608852 | 2.241 | 1.286 | 0.767 | -0.514 | -0.035 | 0.032 | 0.032 | 0.009 | 0.59 | 0.13 | 0.49 | 0.63 | 0.59 | 0.53 |
| 4 | 170015977 | rs13139803 | -1.334 | 1.377 | 0.739 | -0.482 | -0.033 | 0.019 | 0.019 | 0 | 0.09 | 0.03 | 0.08 | 0.04 | 0.15 | 0.21 |
| 4 | 170019039 | rs6553418 | -1.001 | 0.979 | 1.393 | -0.422 | -0.01 | 0.006 | 0.006 | 0 | 0.22 | 0.28 | 0.21 | 0.26 | 0.14 | 0.17 |
| 4 | 170037323 | rs10017472 | -2.588 | 0.905 | 1.531 | -0.601 | 0 | 0.027 | 0.027 | 1.65 | 0.39 | 0.74 | 0.32 | 0.3 | 0.36 | 0.48 |
| 4 | 170037572 | rs3811813 | -0.708 | 0.964 | 1.263 | -0.74 | -0.033 | -0.011 | -0.011 | 1.032 | 0.06 | 0.01 | 0.06 | 0.08 | 0.14 | 0.19 |
| 4 | 170043589 | rs3811815 | -2.701 | 0.932 | 1.929 | -0.751 | 0.032 | -0.003 | -0.003 | 0 | 0.13 | 0.24 | 0.12 | 0.13 | 0.19 | 0.23 |
| 4 | 170059119 | rs7655930 | -0.593 | 0.939 | 1.865 | -0.748 | -0.035 | -0.015 | -0.015 | 0 | 0.06 | 0 | 0.06 | 0.08 | 0.14 | 0.19 |
| 4 | 170059252 | rs6553420 | -2.739 | 0.92 | 1.958 | -0.757 | 0.031 | -0.008 | -0.008 | 0 | 0.14 | 0.27 | 0.12 | 0.13 | 0.19 | 0.23 |
| 4 | 170064574 | rs9684601 | -1.183 | 0.922 | 1.994 | -0.752 | -0.009 | -0.017 | -0.017 | 0 | 0.24 | 0.26 | 0.18 | 0.15 | 0.16 | 0.23 |
| 4 | 170068750 | rs10866333 | -2.42 | 0.868 | 1.901 | -0.734 | 0 | -0.01 | -0.01 | 0.766 | 0.37 | 0.51 | 0.3 | 0.28 | 0.35 | 0.47 |
| 4 | 170072418 | rs7660681 | -1.859 | 0.86 | 2.028 | -0.759 | 0.006 | -0.075 | -0.075 | 0.658 | 0.31 | 0.4 | 0.2 | 0.09 | 0.31 | 0.29 |
| 4 | 170072881 | rs7666351 | -1.173 | 0.861 | 2.035 | -0.729 | 0 | 0.096 | 0.096 | 0 | 0.06 | 0.14 | 0.1 | 0.18 | 0.03 | 0.17 |
| 4 | 170077789 | rs10518039 | -1.945 | 0.979 | 1.751 | -1.005 | 0.007 | -0.076 | -0.076 | 0 | 0.3 | 0.4 | 0.2 | 0.09 | 0.31 | 0.29 |
| 4 | 170081113 | rs7689533 | -1.968 | 0.971 | 1.747 | -1.009 | 0.006 | -0.076 | -0.076 | 0.299 | 0.31 | 0.4 | 0.2 | 0.09 | 0.31 | 0.29 |
| 4 | 170092033 | rs13118066 | -2.036 | 0.913 | 1.811 | -0.962 | -0.03 | 0.007 | 0.007 | 3.13 | 0.07 | 0.22 | 0.03 | 0.04 | 0.04 | 0.02 |
| 4 | 170093506 | rs9991235 | -1.703 | 1.061 | 1.625 | -0.999 | 0.015 | -0.103 | -0.103 | 0 | 0.28 | 0.39 | 0.2 | 0.05 | 0.28 | 0.26 |
| 4 | 170109896 | rs17593695 | -0.372 | 0.63 | 1.24 | -1 | 0.009 | 0.037 | 0.037 | 0 | 0.18 | 0.09 | 0.09 | 0 | 0.13 | 0.13 |
| 4 | 170115171 | rs2706740 | 1.83 | 0.415 | 0.97 | -1.473 | 0.033 | 0.129 | 0.129 | 0 | 0.49 | 0.23 | 0.58 | 0.78 | 0.52 | 0.48 |
| 4 | 170133911 | rs2660416 | -0.654 | 0.442 | 0.957 | -1.478 | 0 | -0.003 | -0.003 | 0 | 0.06 | 0.08 | 0.08 | 0.06 | 0.05 | 0.1 |
| 4 | 170145050 | rs7698807 | 0.264 | 0.251 | 1.102 | -1.555 | -0.016 | 0.044 | 0.044 | 0 | 0.05 | 0.09 | 0.15 | 0.11 | 0.02 | 0.05 |
| 4 | 170146632 | rs1876398 | -0.194 | 0.209 | 1.069 | -1.586 | -0.016 | 0.035 | 0.035 | 0 | 0.21 | 0.15 | 0.14 | 0.11 | 0.32 | 0.33 |
| 4 | 170149701 | rs2660404 | 1.965 | 0.198 | 1.164 | -1.58 | 0.032 | 0.099 | 0.099 | 0 | 0.5 | 0.28 | 0.58 | 0.78 | 0.52 | 0.5 |
| 4 | 170153297 | rs2331608 | 0.123 | 0.535 | 0.704 | -1.712 | -0.003 | 0.021 | 0.021 | 0 | 0.17 | 0.13 | 0.11 | 0.08 | 0.26 | 0.1 |
| 4 | 170158097 | rs6814821 | -0.702 | 0.477 | 0.89 | -1.284 | 0.005 | -0.017 | -0.017 | 0 | 0.2 | 0.18 | 0.1 | 0 | 0.13 | 0.13 |
| 4 | 170182897 | rs13130049 | -0.497 | 0.88 | -0.457 | -0.664 | -0.015 | 0.017 | 0.017 | 0.498 | 0.06 | 0.01 | 0.08 | 0.03 | 0.07 | 0.23 |
| 4 | 170198039 | rs17710102 | -0.737 | 0.91 | -0.432 | -0.512 | -0.015 | -0.019 | -0.019 | 1.887 | 0.07 | 0.03 | 0.07 | 0.11 | 0.12 | 0.07 |
| 4 | 170203367 | rs7683893 | -1.519 | 0.989 | -0.036 | -1.098 | 0.003 | -0.025 | -0.025 | 0 | 0.43 | 0.47 | 0.32 | 0.22 | 0.49 | 0.62 |
| 4 | 170203848 | rs13112591 | -1.485 | 0.902 | 0.124 | -1.126 | 0.002 | -0.013 | -0.013 | 2.893 | 0.42 | 0.43 | 0.3 | 0.22 | 0.49 | 0.61 |
| 4 | 170208954 | rs992939 | 2.014 | 0.196 | 0.012 | -1.401 | -0.054 | 0.123 | 0.123 | 0.231 | 0.46 | 0.11 | 0.65 | 0.81 | 0.47 | 0.37 |

| | | | | | | | | | | | | | | | | |
|---|---|---|---|---|---|---|---|---|---|---|---|---|---|---|---|---|
| 4 | 170212576 | rs6841955 | 0.689 | 0.322 | 0.265 | -1.548 | -0.002 | -0.06 | -0.06 | 0.343 | 0.67 | 0.65 | 0.81 | 0.97 | 0.68 | 0.53 |
| 4 | 170212887 | rs10009607 | -1.426 | 0.398 | 0.708 | -1.481 | 0.053 | 0.05 | 0.05 | 0 | 0.16 | 0.37 | 0.13 | 0.15 | 0.21 | 0.16 |
| 4 | 170228549 | rs4692709 | 1.531 | 0.312 | 0.66 | -1.473 | -0.028 | 0.037 | 0.037 | 0.032 | 0.6 | 0.27 | 0.69 | 0.81 | 0.54 | 0.41 |
| 4 | 170230174 | rs4295223 | -0.246 | 0.184 | 1.009 | -1.612 | 0.008 | 0.04 | 0.04 | 0 | 0.93 | 0.86 | 0.93 | 0.9 | 0.9 | 0.94 |
| 4 | 170230419 | rs12511773 | 1.579 | 0.203 | 1.006 | -1.685 | -0.004 | -0.012 | -0.012 | 0.857 | 0.72 | 0.64 | 0.85 | 0.85 | 0.73 | 0.79 |
| 4 | 170235438 | rs11736716 | 0.283 | 0.166 | 0.751 | -1.711 | 0.021 | 0.005 | 0.005 | 0 | 0.81 | 0.97 | 0.92 | 0.89 | 0.83 | 0.88 |
| 4 | 170237660 | rs10017932 | 1.846 | 0.218 | 0.934 | -1.727 | -0.004 | 0.04 | 0.04 | 0 | 0.52 | 0.6 | 0.71 | 0.84 | 0.49 | 0.38 |
| 4 | 170245260 | rs10002073 | -0.826 | 0.379 | 0.998 | -1.55 | 0.04 | 0.001 | 0.001 | 0 | 0.09 | 0.21 | 0.06 | 0.04 | 0.11 | 0.1 |
| 4 | 170252904 | rs13435077 | -1.178 | 0.35 | 0.856 | -1.564 | 0.034 | -0.012 | -0.012 | 0 | 0.11 | 0.3 | 0.07 | 0.04 | 0.11 | 0.1 |
| 4 | 170262104 | rs7671546 | 0.368 | 0.347 | 0.078 | -1.468 | 0.023 | 0.012 | 0.012 | 0 | 0.81 | 0.92 | 0.92 | 0.89 | 0.83 | 0.88 |
| 4 | 170263443 | rs4535301 | 2.03 | 0.445 | -0.004 | -1.581 | -0.058 | 0.165 | 0.165 | 0.29 | 0.47 | 0.1 | 0.62 | 0.82 | 0.47 | 0.42 |
| 4 | 170266486 | rs13106653 | -1.162 | 0.706 | 0.353 | -1.448 | 0.005 | -0.042 | -0.042 | 0 | 0.31 | 0.66 | 0.26 | 0.07 | 0.35 | 0.47 |
| 4 | 170286096 | rs6845753 | 1 | 0.449 | 0.512 | -1.28 | -0.007 | -0.035 | -0.035 | 0 | 0.63 | 0.5 | 0.72 | 0.78 | 0.61 | 0.56 |
| 4 | 170291213 | rs4331740 | -0.885 | 0.079 | 0.544 | -1.612 | 0.003 | 0.018 | 0.018 | 0 | 0.27 | 0.21 | 0.18 | 0.03 | 0.26 | 0.4 |
| 4 | 170297709 | rs11132803 | 2.149 | 0.064 | 1.168 | -1.719 | 0.001 | 0.028 | 0.028 | 0.151 | 0.49 | 0.37 | 0.66 | 0.75 | 0.51 | 0.4 |
| 4 | 170302498 | rs10866338 | 2.412 | -0.024 | 1.116 | -1.69 | -0.103 | 0.111 | 0.111 | 0.653 | 0.47 | 0.03 | 0.62 | 0.72 | 0.51 | 0.4 |
| 4 | 170305035 | rs4349564 | 2.229 | 0.029 | 1.619 | -1.687 | -0.007 | 0.11 | 0.11 | 0.267 | 0.48 | 0.22 | 0.65 | 0.73 | 0.51 | 0.4 |
| 4 | 170310446 | rs4263355 | -0.4 | 0.26 | 1.343 | -1.587 | 0.002 | -0.033 | -0.033 | 0 | 0.89 | 0.96 | 0.92 | 0.8 | 0.89 | 0.92 |
| 4 | 170312589 | rs4544675 | -1.812 | 0.292 | 1.504 | -1.673 | 0.004 | -0.12 | -0.12 | 0 | 0.36 | 0.56 | 0.24 | 0.07 | 0.34 | 0.5 |
| 4 | 170314951 | rs13212 | 0.232 | 0.282 | 1.481 | -1.678 | -0.06 | 0.107 | 0.107 | 0 | 0.19 | 0.03 | 0.15 | 0.04 | 0.23 | 0.35 |
| 4 | 170315166 | rs1129694 | -1.804 | 0.287 | 1.507 | -1.668 | 0.004 | -0.12 | -0.12 | 1.728 | 0.36 | 0.56 | 0.24 | 0.07 | 0.35 | 0.51 |
| 4 | 170319471 | rs7687276 | -1.684 | 0.277 | 1.507 | -1.673 | 0.004 | -0.091 | -0.091 | 0 | 0.36 | 0.49 | 0.23 | 0.07 | 0.34 | 0.51 |
| 4 | 170338453 | rs12644838 | 1.284 | 0.244 | 1.529 | -1.591 | -0.002 | 0.005 | 0.005 | 0 | 0.79 | 0.37 | 0.82 | 0.79 | 0.83 | 0.81 |
| 4 | 170345893 | rs7655924 | -0.461 | 0.251 | 1.577 | -1.588 | 0.043 | 0.001 | 0.001 | 0.093 | 0.07 | 0.19 | 0.05 | 0.03 | 0.07 | 0.11 |
| 4 | 170362151 | rs4334719 | -0.619 | 0.239 | 1.665 | -1.589 | 0 | 0.075 | 0.075 | 0 | 0.14 | 0.43 | 0.13 | 0.18 | 0.1 | 0.08 |
| 4 | 170367235 | rs4353861 | -0.65 | 0.244 | 1.596 | -1.625 | -0.069 | 0.108 | 0.108 | 0 | 0.82 | 0.98 | 0.85 | 0.96 | 0.77 | 0.65 |
| 4 | 170448464 | rs6553441 | 2.272 | 0.272 | 1.585 | -1.659 | -0.009 | 0.167 | 0.167 | 1.005 | 0.51 | 0.12 | 0.64 | 0.74 | 0.58 | 0.43 |
| 4 | 170451435 | rs6856378 | -1.475 | 0.241 | 1.595 | -1.651 | 0.01 | 0.023 | 0.023 | 0 | 0.19 | 0.37 | 0.15 | 0.21 | 0.17 | 0.19 |
| 4 | 170452770 | rs11941560 | 0.091 | 0.246 | 1.241 | -1.648 | 0.003 | -0.015 | -0.015 | 4.62 | 0.11 | 0.08 | 0.09 | 0.18 | 0.1 | 0.08 |
| 4 | 170462979 | rs11722114 | -1.862 | 0.243 | 1.14 | -1.644 | -0.005 | -0.001 | -0.001 | 0 | 0.24 | 0.63 | 0.18 | 0.22 | 0.17 | 0.21 |
| 4 | 170471183 | rs10001274 | -2.119 | 0.243 | 1.14 | -1.644 | 0.001 | 0.001 | 0.001 | 0 | 0.27 | 0.71 | 0.2 | 0.22 | 0.2 | 0.24 |
| 4 | 170486710 | rs10032225 | -1.867 | 0.255 | 1.12 | -1.63 | -0.005 | -0.001 | -0.001 | 0 | 0.24 | 0.63 | 0.18 | 0.22 | 0.17 | 0.21 |
| 4 | 170511551 | rs4235024 | 2.067 | 0.281 | 0.865 | -1.579 | -0.029 | 0.172 | 0.172 | 1.119 | 0.51 | 0.1 | 0.64 | 0.74 | 0.57 | 0.43 |
| 4 | 170540000 | rs11935607 | -1.605 | 0.573 | 1.103 | -1.537 | -0.003 | -0.003 | -0.003 | 0 | 0.23 | 0.61 | 0.2 | 0.21 | 0.2 | 0.21 |
| 4 | 170541041 | rs3797041 | 1.077 | 0.523 | 1.27 | -1.509 | 0.022 | -0.008 | -0.008 | 0 | 0.92 | 0.73 | 0.95 | 0.97 | 0.93 | 0.89 |
| 4 | 170549443 | rs7682861 | -0.439 | 0.578 | 1.168 | -1.482 | 0.014 | -0.004 | -0.004 | 0 | 0.06 | 0.13 | 0.04 | 0.03 | 0.07 | 0.12 |
| 4 | 170549935 | rs11937417 | -1.134 | 0.904 | 1.319 | -1.629 | 0.005 | -0.017 | -0.017 | 0 | 0.34 | 0.34 | 0.24 | 0.22 | 0.37 | 0.44 |
| 4 | 170558032 | rs10023330 | 0.095 | 0.937 | 1.372 | -1.653 | 0.019 | -0.081 | -0.081 | 0 | 0.19 | 0.24 | 0.14 | 0.03 | 0.25 | 0.34 |
| 4 | 170559440 | rs17659581 | 1.133 | 1.175 | 0.695 | -1.205 | -0.045 | -0.048 | -0.048 | 0 | 0.1 | 0.01 | 0.2 | 0.49 | 0.19 | 0.17 |
| 4 | 170568789 | rs10866341 | 0.636 | 0.607 | 1.186 | -1.969 | 0.022 | -0.112 | -0.112 | 0 | 0.72 | 0.51 | 0.81 | 0.93 | 0.68 | 0.53 |
| 4 | 170579002 | rs17627811 | 2.015 | 0.485 | 1.165 | -2.023 | -0.007 | 0.112 | 0.112 | 0.727 | 0.51 | 0.23 | 0.68 | 0.75 | 0.53 | 0.4 |
| 4 | 170579585 | rs17627829 | -0.096 | 0.521 | 1.01 | -2.004 | -0.001 | -0.011 | -0.011 | 0.817 | 0.15 | 0.11 | 0.1 | 0.18 | 0.13 | 0.09 |
| 4 | 170586793 | rs35899341 | 0.535 | 0.526 | 0.969 | -1.961 | -0.077 | 0.089 | 0.089 | 0 | 0.18 | 0.02 | 0.11 | 0.04 | 0.25 | 0.34 |
| 4 | 170590389 | rs12511877 | 2.073 | 0.525 | 1.043 | -1.94 | -0.023 | 0.148 | 0.148 | 1.452 | 0.49 | 0.14 | 0.67 | 0.74 | 0.52 | 0.4 |
| 4 | 170590989 | rs13435853 | -1.052 | 0.465 | 1.138 | -1.857 | 0.076 | 0.014 | 0.014 | 0.166 | 0.1 | 0.41 | 0.07 | 0.03 | 0.09 | 0.12 |
| 4 | 170598148 | rs10222938 | -1.082 | 0.464 | 1.138 | -1.85 | 0.075 | 0.024 | 0.024 | 0.16 | 0.1 | 0.41 | 0.07 | 0.04 | 0.09 | 0.12 |
| 4 | 170626552 | rs10520163 | 2.063 | 0.399 | 0.686 | -1.994 | -0.016 | 0.118 | 0.118 | 0.515 | 0.48 | 0.17 | 0.66 | 0.75 | 0.5 | 0.39 |
| 4 | 170634408 | rs1010713 | -1.357 | 0.109 | 1.322 | -1.496 | 0.07 | 0.014 | 0.014 | 0.245 | 0.1 | 0.41 | 0.08 | 0.03 | 0.09 | 0.13 |
| 4 | 170642781 | rs13550 | -1.002 | -0.119 | 0.957 | -1.334 | 0.002 | 0 | 0 | 2.068 | 0.17 | 0.2 | 0.11 | 0.18 | 0.13 | 0.12 |
| 4 | 170651811 | rs4692742 | 1.887 | -0.033 | 0.614 | -1.866 | 0.035 | 0.192 | 0.192 | 0.067 | 0.53 | 0.15 | 0.68 | 0.82 | 0.61 | 0.46 |
| 4 | 170658028 | rs6850430 | -1.747 | -0.02 | 1.862 | -1.451 | 0.052 | 0.052 | 0.052 | 0.005 | 0.09 | 0.38 | 0.07 | 0.06 | 0.06 | 0.04 |
| 4 | 170658731 | rs17055114 | -1.604 | -0.048 | 1.486 | -1.499 | -0.002 | -0.017 | -0.017 | 0 | 0.24 | 0.33 | 0.19 | 0.13 | 0.21 | 0.15 |
| 4 | 170662275 | rs7684296 | -0.782 | -0.226 | 1.498 | -1.668 | 0 | -0.015 | -0.015 | 0 | 0.17 | 0.21 | 0.14 | 0.07 | 0.16 | 0.13 |
| 4 | 170663235 | rs1047642 | 0.67 | -0.2 | 1.367 | -1.649 | -0.004 | -0.031 | -0.031 | 0.106 | 0.79 | 0.67 | 0.85 | 0.91 | 0.82 | 0.75 |
| 4 | 170668031 | rs2331721 | 0.442 | -0.282 | 0.852 | -1.62 | -0.004 | -0.027 | -0.027 | 0 | 0.79 | 0.73 | 0.87 | 0.91 | 0.83 | 0.8 |
| 4 | 170670592 | rs10020331 | 1.293 | -0.184 | -0.147 | -1.57 | 0.035 | 0.062 | 0.062 | 1.014 | 0.67 | 0.2 | 0.79 | 0.9 | 0.81 | 0.72 |
| 4 | 170675646 | rs17055122 | -0.526 | -0.055 | -0.157 | -1.454 | -0.009 | -0.027 | -0.027 | 0 | 0.05 | 0.14 | 0.02 | 0 | 0.02 | 0.03 |
| 4 | 170676328 | rs13122912 | 0.998 | -0.016 | -0.624 | -1.452 | 0.055 | 0.108 | 0.108 | 0.446 | 0.64 | 0.17 | 0.78 | 0.9 | 0.8 | 0.71 |
| 4 | 170688979 | rs9312454 | 1.164 | 0.33 | -0.505 | -1.454 | 0.007 | 0.134 | 0.134 | 1.476 | 0.6 | 0.2 | 0.71 | 0.97 | 0.67 | 0.77 |
| 4 | 170690690 | rs7673018 | 0.659 | 0.318 | 0.577 | -1.636 | 0.063 | -0.088 | -0.088 | 0 | 0.86 | 0.71 | 0.86 | 1 | 0.81 | 0.84 |
| 4 | 170693902 | rs4692749 | 1.784 | 0.284 | 0.512 | -1.61 | 0.008 | 0.135 | 0.135 | 1.633 | 0.6 | 0.2 | 0.71 | 0.96 | 0.67 | 0.77 |
| 4 | 170695748 | rs4692750 | 1.657 | 0.375 | 0.503 | -1.643 | 0.007 | 0.114 | 0.114 | 0.408 | 0.62 | 0.2 | 0.73 | 0.97 | 0.69 | 0.84 |
| 4 | 170706770 | rs4692751 | 0.928 | 0.337 | 1.147 | -1.656 | 0.08 | -0.058 | -0.058 | 0.222 | 0.83 | 0.62 | 0.84 | 0.97 | 0.76 | 0.82 |
| 4 | 170713166 | rs6553459 | -1.62 | 0.168 | 1.637 | -1.618 | 0.023 | 0.042 | 0.042 | 0.273 | 0.07 | 0.31 | 0.06 | 0.03 | 0.03 | 0.02 |
| 4 | 170717008 | rs11132816 | 1.792 | 0.223 | 1.209 | -1.531 | -0.004 | 0.132 | 0.132 | 1.15 | 0.59 | 0.2 | 0.7 | 0.94 | 0.63 | 0.76 |
| 4 | 170719215 | rs6835612 | -1.918 | 0.254 | 1.443 | -1.583 | -0.003 | 0.009 | 0.009 | 1.925 | 0.39 | 0.66 | 0.28 | 0.03 | 0.37 | 0.23 |
| 4 | 170737119 | rs1995305 | 1.619 | -0.626 | 1.998 | -1.082 | 0.06 | 0.072 | 0.072 | 0.168 | 0.94 | 0.69 | 0.96 | 0.97 | 0.97 | 0.98 |
| 4 | 170740266 | rs4072504 | -1.671 | -0.742 | 1.43 | -1.16 | -0.06 | -0.093 | -0.093 | 0 | 0.23 | 0.54 | 0.13 | 0.03 | 0.11 | 0.06 |
| 4 | 170754412 | rs10866343 | 1.463 | -0.721 | 1.197 | -1.15 | -0.05 | -0.076 | -0.076 | 0.633 | 0.75 | 0.39 | 0.87 | 0.97 | 0.89 | 0.94 |

| | | | | | | | | | | | | | | | | |
|---|---|---|---|---|---|---|---|---|---|---|---|---|---|---|---|---|
| 4 | 170770486 | rs13110366 | 1.46 | -0.871 | 0.863 | -1.022 | -0.062 | -0.088 | -0.088 | 0.483 | 0.74 | 0.4 | 0.88 | 0.97 | 0.9 | 0.94 |
| 4 | 170773712 | rs4692758 | -1.354 | -0.735 | 0.122 | -0.886 | -0.009 | -0.024 | -0.024 | 0.656 | 0.28 | 0.29 | 0.25 | 0.07 | 0.18 | 0.19 |
| 4 | 170778068 | rs7659128 | -0.811 | -0.613 | 0.051 | -0.301 | -0.011 | 0.013 | 0.013 | 0 | 0.08 | 0.13 | 0.06 | 0.14 | 0.06 | 0.04 |
| 4 | 170778702 | rs6810718 | 0.881 | -1.332 | -0.104 | -0.448 | -0.007 | -0.001 | -0.001 | 0 | 0.61 | 0.38 | 0.66 | 0.63 | 0.72 | 0.64 |
| 4 | 170782760 | rs6811785 | -0.593 | -1.181 | -0.484 | -0.346 | 0.014 | 0.022 | 0.022 | 0 | 0.55 | 0.34 | 0.4 | 0.63 | 0.45 | 0.48 |
| 4 | 170787254 | rs6840086 | -0.88 | -1.321 | -0.485 | -0.38 | 0.01 | 0.01 | 0.01 | 0.12 | 0.19 | 0.2 | 0.11 | 0.13 | 0.13 | 0.09 |
| 4 | 170801265 | rs10004128 | 0.947 | -1.3 | 0.082 | -0.667 | -0.014 | 0.033 | 0.033 | 0 | 0.42 | 0.31 | 0.53 | 0.17 | 0.44 | 0.39 |
| 4 | 170801990 | rs11132826 | 0.669 | -1.339 | 0.758 | -0.556 | -0.008 | 0.007 | 0.007 | 0.684 | 0.79 | 0.6 | 0.85 | 0.7 | 0.78 | 0.79 |
| 4 | 170807597 | rs4077273 | 0.572 | -1.458 | 0.693 | -0.67 | -0.001 | -0.083 | -0.083 | 0 | 0.3 | 0.05 | 0.28 | 0.53 | 0.28 | 0.35 |
| 4 | 170814998 | rs11132829 | 0.384 | -1.51 | 0.846 | -0.721 | -0.001 | -0.034 | -0.034 | 0 | 0.81 | 0.69 | 0.88 | 0.88 | 0.78 | 0.88 |
| 4 | 170818615 | rs4076940 | 0.723 | -1.526 | 1.199 | -0.733 | 0.001 | 0.001 | 0.001 | 0 | 0.48 | 0.45 | 0.55 | 0.16 | 0.5 | 0.44 |
| 4 | 170826315 | rs10002607 | 0.064 | -1.632 | 1.244 | -0.744 | -0.006 | 0.016 | 0.016 | 0 | 0.92 | 0.73 | 0.92 | 0.89 | 0.95 | 0.93 |
| 4 | 170832177 | rs10004720 | 1.065 | -1.617 | 1.182 | -0.841 | 0 | -0.015 | -0.015 | 0 | 0.44 | 0.45 | 0.53 | 0.16 | 0.44 | 0.4 |
| 4 | 170834427 | rs4634183 | 0.215 | -1.568 | 1.242 | -0.775 | -0.003 | -0.004 | -0.004 | 0.575 | 0.57 | 0.57 | 0.61 | 0.37 | 0.62 | 0.55 |
| 4 | 170842930 | rs2002221 | 0.197 | -1.609 | 0.817 | -0.986 | -0.063 | -0.022 | -0.022 | 0.207 | 0.33 | 0.03 | 0.34 | 0.68 | 0.37 | 0.33 |
| 4 | 170848562 | rs2955269 | -0.447 | -1.419 | 1.172 | -0.822 | -0.005 | 0.029 | 0.029 | 0 | 0.07 | 0.27 | 0.05 | 0.11 | 0.03 | 0.06 |
| 4 | 170857504 | rs10010614 | 1.53 | -1.624 | 1.113 | -1.01 | -0.011 | 0.017 | 0.017 | 0 | 0.43 | 0.31 | 0.46 | 0.04 | 0.43 | 0.42 |
| 4 | 170859150 | rs7684116 | 0.14 | -1.694 | 1.156 | -0.989 | -0.001 | -0.025 | -0.025 | 0.006 | 0.63 | 0.6 | 0.57 | 0.23 | 0.6 | 0.63 |
| 4 | 170867341 | rs4692771 | -1.466 | -1.572 | 0.781 | -0.775 | 0.002 | -0.004 | -0.004 | 0.639 | 0.14 | 0.19 | 0.09 | 0.2 | 0.11 | 0.32 |
| 4 | 170872226 | rs10008105 | 0.901 | -1.828 | 1.278 | -0.956 | 0.001 | -0.001 | -0.001 | 0 | 0.51 | 0.46 | 0.54 | 0.25 | 0.53 | 0.61 |
| 4 | 170875915 | rs2955267 | -1.056 | -1.789 | 1.48 | -0.879 | -0.016 | -0.013 | -0.013 | 0 | 0.13 | 0.3 | 0.09 | 0.11 | 0.08 | 0.06 |
| 4 | 170877933 | rs6553477 | 0.393 | -2.009 | 1.904 | -1.032 | 0 | 0.012 | 0.012 | 0.065 | 0.62 | 0.67 | 0.57 | 0.15 | 0.58 | 0.38 |
| 4 | 170880049 | rs6821170 | -0.693 | -2.016 | 1.914 | -0.872 | 0.001 | 0.009 | 0.009 | 1.695 | 0.74 | 0.76 | 0.66 | 0.39 | 0.68 | 0.71 |
| 4 | 170880883 | rs11723530 | 1.864 | -2.07 | 1.913 | -0.92 | -0.015 | 0.225 | 0.225 | 0 | 0.46 | 0.16 | 0.47 | 0.04 | 0.5 | 0.32 |
| 4 | 170883505 | rs13137856 | 0.608 | -2.009 | 1.943 | -0.802 | -0.001 | -0.022 | -0.022 | 0 | 0.61 | 0.64 | 0.57 | 0.16 | 0.59 | 0.39 |
| 4 | 170887367 | rs4692775 | -0.303 | -1.892 | 1.86 | -0.983 | -0.018 | -0.059 | -0.059 | 0 | 0.1 | 0.01 | 0.06 | 0.21 | 0.09 | 0.32 |
| 4 | 170892618 | rs3797042 | 0.299 | -2.001 | 1.988 | -0.956 | 0.001 | -0.003 | -0.003 | 0 | 0.65 | 0.63 | 0.56 | 0.33 | 0.6 | 0.63 |
| 4 | 170894451 | rs13118628 | 0.682 | -2.006 | 1.931 | -1.129 | -0.004 | -0.027 | -0.027 | 0.01 | 0.62 | 0.52 | 0.55 | 0.28 | 0.6 | 0.61 |
| 4 | 170895510 | rs4588426 | 0.78 | -2.008 | 1.939 | -1.13 | -0.002 | -0.006 | -0.006 | 0 | 0.61 | 0.43 | 0.54 | 0.28 | 0.6 | 0.61 |
| 4 | 170896626 | rs6553482 | 0.478 | -1.923 | 1.761 | -1.064 | -0.008 | -0.097 | -0.097 | 0.515 | 0.65 | 0.5 | 0.61 | 0.07 | 0.63 | 0.47 |
| 4 | 170897935 | rs13114528 | -0.687 | -1.787 | 1.679 | -1.01 | -0.009 | -0.01 | -0.01 | 0.016 | 0.76 | 0.85 | 0.67 | 0.3 | 0.69 | 0.72 |
| 4 | 170900925 | rs17055338 | -1.387 | -2.463 | 1.651 | -1.023 | -0.001 | -0.003 | -0.003 | 0 | 0.43 | 0.56 | 0.45 | 0.72 | 0.36 | 0.41 |
| 4 | 170901019 | rs10033615 | 0.266 | -2.155 | 1.073 | -0.583 | -0.005 | 0.008 | 0.008 | 1.265 | 0.23 | 0.21 | 0.3 | 0.64 | 0.22 | 0.24 |
| 4 | 170904165 | rs9684599 | 0.298 | -2.152 | 0.779 | -0.558 | 0.005 | -0.011 | -0.011 | 0 | 0.23 | 0.16 | 0.3 | 0.64 | 0.22 | 0.25 |
| 4 | 170906940 | rs17055345 | -0.812 | -2.16 | 0.952 | -0.847 | -0.043 | -0.085 | -0.085 | 0 | 0.16 | 0.34 | 0.06 | 0 | 0.1 | 0.04 |
| 4 | 170908445 | rs6553483 | -0.429 | -2.252 | 0.862 | -0.932 | -0.011 | -0.036 | -0.036 | 0 | 0.07 | 0.19 | 0.05 | 0.01 | 0.04 | 0.03 |
| 4 | 170910322 | rs4320103 | -0.429 | -2.431 | 0.617 | -0.817 | -0.011 | -0.036 | -0.036 | 0 | 0.07 | 0.19 | 0.05 | 0.01 | 0.04 | 0.03 |
| 4 | 170912278 | rs7661337 | -2.105 | -2.547 | 0.133 | -0.964 | 0.067 | 0.058 | 0.058 | 4.243 | 0.38 | 0.77 | 0.21 | 0.25 | 0.24 | 0.37 |
| 4 | 170919666 | rs6553484 | -1.922 | -2.59 | -0.249 | -1.199 | -0.01 | 0.18 | 0.18 | 0.359 | 0.61 | 0.86 | 0.51 | 0.94 | 0.47 | 0.65 |
| 4 | 170924737 | rs7694808 | -1.612 | -2.721 | -0.667 | -1.369 | 0.085 | 0.294 | 0.294 | 0.047 | 0.52 | 0.91 | 0.5 | 0.92 | 0.4 | 0.63 |
| 4 | 170927723 | rs6841912 | -0.96 | -2.812 | 0.157 | -1.216 | -0.017 | 0.097 | 0.097 | 1.921 | 0.13 | 0.47 | 0.1 | 0.2 | 0.08 | 0.32 |
| 4 | 170931125 | rs17055385 | -0.485 | -2.718 | 0.315 | -1.207 | -0.026 | -0.066 | -0.066 | 0 | 0.12 | 0.22 | 0.04 | 0 | 0.07 | 0.03 |
| 4 | 170933203 | rs6842442 | -1.717 | -2.675 | 0.212 | -1.197 | 0.001 | -0.001 | -0.001 | 0 | 0.47 | 0.49 | 0.49 | 0.74 | 0.39 | 0.42 |
| 4 | 170949341 | rs3806826 | -0.606 | -2.294 | -0.319 | -1.065 | -0.004 | -0.018 | -0.018 | 0 | 0.1 | 0.13 | 0.08 | 0.02 | 0.07 | 0.06 |
| 4 | 170959988 | rs6821231 | -0.36 | -2.526 | -0.574 | -0.669 | 0.015 | -0.032 | -0.032 | 0.643 | 0.26 | 0.09 | 0.27 | 0.66 | 0.26 | 0.27 |
| 4 | 170960062 | rs2727525 | -2.646 | -3.071 | -0.852 | -1.112 | -0.014 | 0.263 | 0.263 | 0 | 0.58 | 0.9 | 0.53 | 0.95 | 0.44 | 0.69 |
| 4 | 170975204 | rs12650424 | -0.48 | -2.975 | -0.781 | -0.98 | 0.029 | 0.002 | 0.002 | 0 | 0.28 | 0.09 | 0.27 | 0.7 | 0.26 | 0.29 |
| 4 | 170981783 | rs34202278 | 0.23 | -2.764 | -0.648 | -0.977 | 0 | -0.002 | -0.002 | 9.165 | 0.05 | 0.03 | 0.05 | 0 | 0.06 | 0.03 |
| 4 | 170993771 | rs2466990 | -2.003 | -2.645 | -0.112 | -1.23 | 0.052 | 0.19 | 0.19 | 0 | 0.53 | 0.78 | 0.52 | 0.98 | 0.43 | 0.66 |
| 4 | 170995066 | rs963660 | 0.594 | -2.839 | -0.303 | -0.003 | 0.012 | -0.071 | -0.071 | 0 | 0.13 | 0.01 | 0.14 | 0.46 | 0.1 | 0.17 |
| 4 | 170999860 | rs2622068 | -2.152 | -2.579 | -0.187 | -0.842 | 0.057 | 0.201 | 0.201 | 0 | 0.52 | 0.81 | 0.53 | 0.98 | 0.42 | 0.65 |
| 4 | 171009360 | rs10213032 | -1.241 | -2.755 | -0.261 | -0.586 | -0.003 | 0.001 | 0.001 | 0 | 0.15 | 0.19 | 0.1 | 0.27 | 0.1 | 0.3 |
| 4 | 171013833 | rs2955256 | -0.556 | -2.589 | -0.026 | -1.415 | -0.006 | -0.027 | -0.027 | 0 | 0.33 | 0.41 | 0.39 | 0.71 | 0.32 | 0.36 |
| 4 | 171017638 | rs2727526 | -2.45 | -2.403 | -0.201 | -1.459 | -0.05 | 0.251 | 0.251 | 0 | 0.57 | 0.87 | 0.57 | 0.99 | 0.49 | 0.75 |
| 4 | 171031170 | rs11736939 | -0.062 | -2.275 | 0.167 | -1.283 | -0.009 | -0.051 | -0.051 | 0 | 0.1 | 0.17 | 0.08 | 0 | 0.07 | 0.04 |
| 4 | 171034742 | rs10000291 | -0.843 | -2.215 | 0.446 | -1.244 | -0.001 | 0.002 | 0.002 | 0 | 0.33 | 0.29 | 0.44 | 0.69 | 0.35 | 0.38 |
| 4 | 171044796 | rs12510576 | -0.366 | -2.203 | 0.384 | -1.277 | -0.006 | -0.014 | -0.014 | 0 | 0.11 | 0.16 | 0.06 | 0.07 | 0.08 | 0.14 |
| 4 | 171049915 | rs2101065 | 0.532 | -2.222 | 0.066 | -1.26 | -0.054 | -0.043 | -0.043 | 0 | 0.21 | 0.04 | 0.33 | 0.69 | 0.27 | 0.34 |
| 4 | 171055510 | rs519160 | -2.193 | -2.163 | 0.074 | -1.588 | -0.041 | 0.227 | 0.227 | 0 | 0.55 | 0.84 | 0.57 | 0.99 | 0.49 | 0.75 |
| 4 | 171062113 | rs540256 | -1.528 | -2.074 | 0.48 | -1.216 | -0.009 | 0.02 | 0.02 | 0.072 | 0.22 | 0.32 | 0.23 | 0.31 | 0.18 | 0.24 |
| 4 | 171064513 | rs548846 | -1.9 | -1.516 | -0.801 | -2.028 | -0.064 | 0.209 | 0.209 | 0 | 0.64 | 0.91 | 0.64 | 0.98 | 0.61 | 0.77 |
| 4 | 171076562 | rs3846272 | -0.832 | -1.593 | -0.499 | -2.005 | 0.016 | 0.096 | 0.096 | 0.33 | 0.48 | 0.75 | 0.47 | 0.94 | 0.44 | 0.65 |
| 4 | 171082172 | rs1947859 | 1.647 | -1.411 | -0.588 | -1.794 | -0.016 | 0.035 | 0.035 | 0 | 0.07 | 0.01 | 0.07 | 0 | 0.08 | 0.01 |
| 4 | 171089568 | rs532160 | -1.251 | -1.346 | -0.496 | -1.779 | -0.002 | -0.01 | -0.01 | 0.484 | 0.92 | 0.93 | 0.92 | 0.96 | 0.93 | 0.93 |
| 4 | 171092557 | rs13150580 | 0.93 | -1.4 | -0.257 | -1.781 | -0.006 | 0.019 | 0.019 | 0.14 | 0.09 | 0.04 | 0.08 | 0.04 | 0.09 | 0.07 |
| 4 | 171095168 | rs11132838 | -1.296 | -1.394 | -0.513 | -2.026 | -0.045 | 0.218 | 0.218 | 0 | 0.55 | 0.87 | 0.55 | 0.97 | 0.51 | 0.65 |
| 4 | 171098146 | rs512592 | 1.073 | -1.373 | -0.515 | -2.015 | -0.072 | 0.295 | 0.295 | 0 | 0.46 | 0.1 | 0.45 | 0.03 | 0.49 | 0.35 |
| 4 | 171101240 | rs13125979 | 0.526 | -1.371 | -0.236 | -2.005 | 0.004 | -0.032 | -0.032 | 0 | 0.1 | 0.16 | 0.1 | 0.04 | 0.09 | 0.07 |
| 4 | 171110103 | rs1522605 | 1.719 | -1.378 | -0.3 | -2.026 | -0.068 | 0.226 | 0.226 | 0 | 0.37 | 0.08 | 0.36 | 0.03 | 0.4 | 0.3 |

| | | | | | | | | | | | | | | | | |
|---|---|---|---|---|---|---|---|---|---|---|---|---|---|---|---|---|
| 4 | 171123515 | rs1949621 | -1.134 | -1.371 | -0.189 | -1.944 | -0.012 | 0.075 | 0.075 | 0 | 0.53 | 0.76 | 0.53 | 0.97 | 0.51 | 0.65 |
| 4 | 171124116 | rs1581421 | -1.292 | -1.37 | -0.199 | -1.944 | -0.025 | 0.148 | 0.148 | 0 | 0.54 | 0.81 | 0.54 | 0.97 | 0.51 | 0.65 |
| 4 | 171127756 | rs1718800 | 1.09 | -1.37 | -0.2 | -1.944 | -0.041 | 0.218 | 0.218 | 0 | 0.46 | 0.15 | 0.45 | 0.03 | 0.5 | 0.35 |
| 4 | 171148608 | rs7692357 | -1.111 | -1.232 | -0.126 | -1.938 | -0.013 | 0.056 | 0.056 | 0 | 0.55 | 0.75 | 0.53 | 0.97 | 0.51 | 0.68 |
| 4 | 171149711 | rs1522600 | 0.648 | -1.372 | 0.246 | -1.92 | -0.008 | 0.02 | 0.02 | 0.902 | 0.05 | 0.21 | 0.04 | 0.06 | 0.03 | 0.06 |
| 4 | 171152392 | rs1915509 | -0.618 | -1.259 | 0.124 | -2.008 | 0.01 | 0.033 | 0.033 | 0 | 0.46 | 0.7 | 0.45 | 0.93 | 0.42 | 0.61 |
| 4 | 171163418 | rs17629627 | 1.637 | -1.249 | -0.239 | -2.092 | -0.014 | 0.027 | 0.027 | 0 | 0.07 | 0 | 0.03 | 0.02 | 0.06 | 0.04 |
| 4 | 171163569 | rs1718781 | 0.649 | -1.325 | -0.111 | -1.694 | -0.018 | 0.073 | 0.073 | 0 | 0.47 | 0.22 | 0.53 | 0.31 | 0.5 | 0.47 |
| 4 | 171167671 | rs10030257 | 1.507 | -1.189 | -0.149 | -1.691 | -0.001 | -0.032 | -0.032 | 3.847 | 0.13 | 0.21 | 0.06 | 0.04 | 0.14 | 0.09 |
| 4 | 171167789 | rs10030422 | 1.287 | -1.364 | 0.096 | -1.749 | 0 | 0.027 | 0.027 | 0 | 0.25 | 0.31 | 0.31 | 0.6 | 0.26 | 0.36 |
| 4 | 171168272 | rs12647798 | 0.101 | -1.244 | 0.087 | -1.961 | 0 | -0.005 | -0.005 | 0.668 | 0.35 | 0.34 | 0.4 | 0.7 | 0.35 | 0.4 |
| 4 | 171168353 | rs10023223 | 1.113 | -1.353 | 0.192 | -2.025 | -0.012 | 0.027 | 0.027 | 0 | 0.16 | 0.27 | 0.12 | 0.28 | 0.12 | 0.21 |
| 4 | 171172831 | rs12648095 | -1.298 | -1.427 | 0.198 | -2.172 | -0.007 | 0.043 | 0.043 | 0 | 0.55 | 0.66 | 0.59 | 0.95 | 0.48 | 0.67 |
| 4 | 171175255 | rs9991425 | -0.653 | -1.439 | 0.205 | -2.173 | 0.012 | 0.018 | 0.018 | 0 | 0.45 | 0.36 | 0.5 | 0.92 | 0.39 | 0.65 |
| 4 | 171175320 | rs17661170 | 1.386 | -1.45 | 0.225 | -2.178 | -0.002 | -0.013 | -0.013 | 0 | 0.26 | 0.15 | 0.32 | 0.67 | 0.26 | 0.37 |
| 4 | 171175522 | rs10028534 | -0.644 | -1.441 | 0.218 | -2.141 | 0.011 | 0.039 | 0.039 | 0 | 0.44 | 0.35 | 0.44 | 0.85 | 0.39 | 0.65 |
| 4 | 171179337 | rs10520181 | 0.215 | -1.357 | 0.047 | -2.133 | 0.007 | 0.028 | 0.028 | 0 | 0.15 | 0.08 | 0.1 | 0 | 0.13 | 0.08 |
| 4 | 171180273 | rs4692812 | -0.95 | -1.388 | 0.333 | -1.844 | 0.004 | 0.03 | 0.03 | 0 | 0.49 | 0.59 | 0.47 | 0.71 | 0.47 | 0.48 |
| 4 | 171193885 | rs7699981 | 1.684 | -1.018 | 0.307 | -2.431 | 0 | -0.008 | -0.008 | 0.569 | 0.13 | 0.13 | 0.05 | 0.03 | 0.12 | 0.08 |
| 4 | 171203364 | rs17055545 | 0.834 | -1.023 | 0.314 | -2.398 | -0.014 | -0.037 | -0.037 | 0.081 | 0.08 | 0.12 | 0.03 | 0 | 0.03 | 0.03 |
| 4 | 171204964 | rs12647693 | 0.694 | -0.576 | -0.025 | -2.085 | -0.001 | 0.025 | 0.025 | 0 | 0.27 | 0.15 | 0.32 | 0.79 | 0.29 | 0.45 |
| 4 | 171206696 | rs12649329 | 0.249 | -0.578 | -0.079 | -1.876 | 0 | -0.007 | -0.007 | 0 | 0.36 | 0.37 | 0.41 | 0.82 | 0.37 | 0.47 |
| 4 | 171207036 | rs1522603 | -0.365 | -0.447 | -0.231 | -1.715 | 0 | -0.018 | -0.018 | 0 | 0.49 | 0.53 | 0.46 | 0.85 | 0.49 | 0.56 |
| 4 | 171215850 | rs1717072 | -0.624 | 0.078 | 0.038 | -1.425 | 0.006 | 0.072 | 0.072 | 0 | 0.51 | 0.39 | 0.46 | 0.19 | 0.53 | 0.33 |
| 4 | 171215956 | rs11132841 | 0.502 | 0.053 | 0.287 | -1.437 | -0.001 | -0.013 | -0.013 | 0 | 0.48 | 0.49 | 0.52 | 0.81 | 0.47 | 0.67 |
| 4 | 171224046 | rs17629990 | 0.648 | 0.075 | -0.222 | -1.473 | 0.017 | -0.003 | -0.003 | 0 | 0.22 | 0.14 | 0.09 | 0.19 | 0.16 | 0.13 |
| 4 | 171234457 | rs17548053 | -0.615 | 0.279 | -0.785 | -1.492 | -0.014 | 0.074 | 0.074 | 0 | 0.43 | 0.25 | 0.36 | 0.19 | 0.44 | 0.28 |
| 4 | 171234790 | rs4692821 | 0.072 | 1.04 | -0.81 | -1.427 | -0.014 | 0.034 | 0.034 | 0 | 0.06 | 0.03 | 0.08 | 0 | 0.09 | 0.05 |
| 4 | 171244684 | rs6814090 | -0.146 | 0.912 | -0.414 | -1.247 | 0.003 | -0.044 | -0.044 | 0 | 0.22 | 0.19 | 0.29 | 0 | 0.28 | 0.15 |
| 4 | 171269504 | rs1406514 | -0.107 | -0.322 | -0.319 | -1.606 | -0.008 | -0.068 | -0.068 | 0.116 | 0.63 | 0.6 | 0.71 | 0.09 | 0.69 | 0.43 |
| 4 | 171271368 | rs12506448 | 0.022 | -0.481 | -0.193 | -1.689 | -0.014 | -0.091 | -0.091 | 0 | 0.6 | 0.52 | 0.7 | 0.08 | 0.69 | 0.4 |
| 4 | 171288473 | rs4692824 | 0.071 | -0.541 | -0.245 | -1.452 | -0.004 | -0.06 | -0.06 | 0.416 | 0.6 | 0.59 | 0.69 | 0.09 | 0.68 | 0.4 |
| 4 | 171292771 | rs1879500 | -0.185 | -0.55 | -0.189 | -1.416 | 0.011 | -0.013 | -0.013 | 0 | 0.95 | 0.79 | 0.96 | 1 | 0.97 | 0.97 |
| 4 | 171297661 | rs1919502 | 0.573 | -0.603 | -0.435 | -1.427 | -0.026 | 0.045 | 0.045 | 0 | 0.3 | 0.05 | 0.18 | 0.75 | 0.29 | 0.44 |
| 4 | 171307586 | rs10020695 | 0.057 | -0.635 | -0.303 | -1.543 | -0.016 | -0.11 | -0.11 | 0.014 | 0.62 | 0.52 | 0.66 | 0.08 | 0.7 | 0.41 |
| 4 | 171312561 | rs2141473 | -0.438 | -2.04 | -0.677 | -1.385 | 0.033 | 0.01 | 0.01 | 0 | 0.3 | 0.26 | 0.19 | 0.28 | 0.16 | 0.42 |
| 4 | 171318582 | rs1919506 | 0.035 | -1.952 | -0.586 | -1.326 | 0.025 | -0.005 | -0.005 | 0 | 0.25 | 0.17 | 0.15 | 0.27 | 0.12 | 0.36 |
| 4 | 171327181 | rs7670352 | -0.993 | -1.517 | -0.211 | -1.377 | -0.008 | 0.007 | 0.007 | 0 | 0.62 | 0.68 | 0.65 | 0.32 | 0.48 | 0.54 |
| 4 | 171339965 | rs2612406 | 0.864 | -2.22 | 0.633 | -1.353 | -0.045 | -0.055 | -0.055 | 0.889 | 0.77 | 0.43 | 0.68 | 0.85 | 0.8 | 0.81 |
| 4 | 171340897 | rs1609380 | -0.329 | -2.609 | 0.713 | -1.844 | 0.013 | 0.02 | 0.02 | 0 | 0.68 | 0.76 | 0.68 | 0.85 | 0.81 | 0.75 |
| 4 | 171351306 | rs1919510 | 0.927 | -2.4 | 1.187 | -1.553 | -0.069 | -0.058 | -0.058 | 0 | 0.82 | 0.4 | 0.75 | 0.85 | 0.92 | 0.84 |
| 4 | 171351736 | rs2877520 | -0.221 | -2.408 | 1.375 | -1.516 | 0 | 0.062 | 0.062 | 0 | 0.34 | 0.16 | 0.37 | 0.18 | 0.31 | 0.27 |
| 4 | 171371794 | rs1528945 | 0.271 | -2.347 | 1.798 | -1.664 | -0.017 | -0.06 | -0.06 | 0.68 | 0.89 | 0.58 | 0.83 | 0.99 | 0.94 | 0.96 |
| 4 | 171386268 | rs1534659 | 0.008 | -2.398 | 1.441 | -1.735 | 0.005 | 0.042 | 0.042 | 0 | 0.63 | 0.73 | 0.72 | 0.86 | 0.66 | 0.72 |
| 4 | 171402225 | rs1581676 | 0.847 | -2.442 | 1.728 | -1.76 | 0.017 | 0.037 | 0.037 | 0 | 0.52 | 0.37 | 0.44 | 0.85 | 0.6 | 0.7 |
| 4 | 171403760 | rs13134352 | -0.243 | -2.525 | 1.55 | -1.749 | 0.022 | 0.152 | 0.152 | 0.148 | 0.41 | 0.18 | 0.28 | 0.14 | 0.34 | 0.29 |
| 4 | 171418411 | rs2020059 | 0.442 | -2.322 | 1.437 | -1.599 | 0.001 | -0.017 | -0.017 | 0 | 0.5 | 0.48 | 0.44 | 0.83 | 0.6 | 0.69 |
| 4 | 171420400 | rs2661081 | 1.227 | -2.153 | 0.807 | -1.482 | -0.017 | 0.091 | 0.091 | 0 | 0.4 | 0.21 | 0.37 | 0.8 | 0.54 | 0.62 |
| 4 | 171423826 | rs2661079 | 1.114 | -2.171 | 0.659 | -1.479 | -0.023 | 0.116 | 0.116 | 0 | 0.4 | 0.15 | 0.57 | 0.8 | 0.55 | 0.62 |
| 4 | 171426045 | rs7662606 | -0.906 | -1.697 | 2.03 | -0.86 | 0.001 | 0.012 | 0.012 | 0 | 0.36 | 0.36 | 0.27 | 0.26 | 0.3 | 0.26 |
| 4 | 171448717 | rs2661071 | 0.581 | -2.11 | 2.146 | -0.844 | -0.01 | -0.022 | -0.022 | 0 | 0.66 | 0.47 | 0.72 | 0.72 | 0.73 | 0.67 |
| 4 | 171471531 | rs371100 | 1.509 | -2.179 | 2.076 | -1.143 | 0.044 | 0.17 | 0.17 | 0.072 | 0.57 | 0.19 | 0.44 | 0.82 | 0.67 | 0.66 |
| 4 | 171481511 | rs403730 | 0.501 | -2.196 | 2.25 | -1.153 | -0.009 | -0.042 | -0.042 | 0 | 0.67 | 0.48 | 0.71 | 0.83 | 0.73 | 0.7 |
| 4 | 171484438 | rs424540 | 1.555 | -2.129 | 1.661 | -1.118 | 0.027 | 0.203 | 0.203 | 1.192 | 0.55 | 0.08 | 0.43 | 0.82 | 0.67 | 0.66 |
| 4 | 171499216 | rs446973 | 1.574 | -2.092 | 1.728 | -1.074 | 0.045 | 0.212 | 0.212 | 0.493 | 0.55 | 0.1 | 0.47 | 0.84 | 0.68 | 0.66 |
| 4 | 171505217 | rs2055717 | -0.608 | -2.166 | 2.332 | -1.121 | 0.001 | -0.001 | -0.001 | 0 | 0.3 | 0.27 | 0.23 | 0.14 | 0.24 | 0.19 |
| 4 | 171509449 | rs413812 | 2.763 | -2.158 | 2.37 | -1.115 | -0.036 | -0.086 | -0.086 | 0.004 | 0.86 | 0.4 | 0.71 | 0.98 | 0.92 | 0.85 |
| 4 | 171510501 | rs406542 | 0.979 | -2.165 | 2.659 | -1.119 | 0.039 | 0.005 | 0.005 | 0 | 0.94 | 0.65 | 0.76 | 0.99 | 0.97 | 0.98 |
| 4 | 171513737 | rs368771 | 1.705 | -2.173 | 2.607 | -1.125 | 0.053 | 0.201 | 0.201 | 0.19 | 0.56 | 0.13 | 0.48 | 0.84 | 0.68 | 0.66 |
| 4 | 171525130 | rs13129109 | -0.607 | -2.063 | 2.424 | -1.068 | 0.004 | 0.012 | 0.012 | 0 | 0.3 | 0.24 | 0.23 | 0.14 | 0.24 | 0.19 |
| 4 | 171537901 | rs4692612 | -1.381 | -1.68 | 2.406 | -0.96 | 0.043 | 0.008 | 0.008 | 0 | 0.06 | 0.34 | 0.24 | 0.01 | 0.03 | 0.02 |
| 4 | 171544366 | rs10021531 | -2.096 | -1.677 | 1.86 | -0.959 | -0.012 | -0.069 | -0.069 | 0 | 0.12 | 0.54 | 0.28 | 0.02 | 0.08 | 0.1 |
| 4 | 171576294 | rs1596248 | -1.57 | -1.64 | 1.337 | -0.996 | 0.059 | 0.136 | 0.136 | 0.086 | 0.44 | 0.78 | 0.48 | 0.09 | 0.28 | 0.24 |
| 4 | 171579204 | rs10028123 | -0.824 | -1.699 | 1.804 | -1.006 | -0.008 | -0.056 | -0.056 | 0 | 0.34 | 0.37 | 0.22 | 0.07 | 0.25 | 0.19 |
| 4 | 171590438 | rs10030592 | -1.149 | -1.718 | 1.802 | -0.808 | -0.027 | -0.024 | -0.024 | 0 | 0.14 | 0.26 | 0.32 | 0.08 | 0.11 | 0.05 |
| 4 | 171592467 | rs7676680 | -1.117 | -1.718 | 1.816 | -0.808 | -0.026 | -0.023 | -0.023 | 0 | 0.14 | 0.26 | 0.32 | 0.08 | 0.1 | 0.05 |
| 4 | 171592684 | rs7665640 | -0.637 | -1.884 | 2.045 | 0.107 | -0.024 | 0.096 | 0.096 | 0 | 0.28 | 0.48 | 0.18 | 0.47 | 0.18 | 0.24 |
| 4 | 171596102 | rs7681130 | -2.085 | -1.905 | 1.844 | 0.092 | 0.072 | -0.04 | -0.04 | 0.001 | 0.45 | 0.93 | 0.53 | 0.55 | 0.29 | 0.31 |
| 4 | 171603479 | rs10012640 | -0.575 | -1.776 | 2.233 | 0.165 | -0.01 | 0.044 | 0.044 | 0 | 0.27 | 0.34 | 0.17 | 0.47 | 0.17 | 0.24 |

| | | | | | | | | | | | | | | | | |
|---|---|---|---|---|---|---|---|---|---|---|---|---|---|---|---|---|
| 4 | 171619313 | rs11935380 | -0.894 | -1.785 | 2.336 | 0.117 | -0.018 | 0.035 | 0.035 | 0 | 0.29 | 0.58 | 0.15 | 0.4 | 0.18 | 0.24 |
| 4 | 171620514 | rs11132859 | -1.425 | -1.892 | 2.338 | 0.008 | -0.012 | 0.023 | 0.023 | 0 | 0.37 | 0.52 | 0.26 | 0.47 | 0.29 | 0.3 |
| 4 | 171634702 | rs9312487 | -1.378 | -1.795 | 1.818 | -0.137 | 0.009 | -0.034 | -0.034 | 0.046 | 0.36 | 0.69 | 0.36 | 0.03 | 0.23 | 0.17 |
| 4 | 171635106 | rs7658788 | -1.381 | -1.801 | 1.816 | -0.14 | 0.009 | -0.034 | -0.034 | 0 | 0.36 | 0.69 | 0.36 | 0.03 | 0.23 | 0.17 |
| 4 | 171637914 | rs9312489 | -1.67 | -1.913 | 1.543 | 0.042 | 0.039 | -0.029 | -0.029 | 0 | 0.39 | 0.71 | 0.25 | 0.48 | 0.27 | 0.3 |
| 4 | 171644884 | rs1533837 | -2.142 | -1.913 | 1.295 | 0.041 | 0.091 | -0.109 | -0.109 | 0 | 0.46 | 0.95 | 0.47 | 0.49 | 0.32 | 0.3 |
| 4 | 171645207 | rs1533835 | -1.977 | -1.4 | 2.169 | 0.724 | 0.032 | 0.553 | 0.553 | 1.516 | 0.07 | 0.41 | 0.06 | 0.41 | 0.03 | 0.12 |
| 4 | 171651895 | rs9312490 | -2.24 | -1.42 | 1.837 | 0.746 | -0.037 | 0.269 | 0.269 | 1.784 | 0.13 | 0.54 | 0.27 | 0.41 | 0.07 | 0.13 |
| 4 | 171654737 | rs2177183 | -0.39 | -1.433 | 1.695 | 0.716 | -0.002 | -0.002 | -0.002 | 0 | 0.11 | 0.13 | 0.11 | 0.06 | 0.1 | 0.06 |
| 4 | 171663745 | rs7697640 | -2.196 | -1.207 | 1.771 | 0.882 | -0.023 | 0.342 | 0.342 | 1.07 | 0.12 | 0.47 | 0.26 | 0.41 | 0.07 | 0.12 |
| 4 | 171666694 | rs17730512 | -1.174 | -1.053 | 1.068 | 1.015 | -0.006 | 0.302 | 0.302 | 0 | 0.07 | 0.23 | 0.23 | 0.41 | 0.05 | 0.12 |
| 4 | 171672038 | rs2332027 | 0.178 | -0.585 | 0.463 | 0.061 | 0.007 | -0.012 | -0.012 | 0 | 0.6 | 0.43 | 0.64 | 0.65 | 0.5 | 0.77 |
| 4 | 171673765 | rs11931207 | -1.172 | -1.68 | 1.627 | 0.311 | -0.046 | 0.064 | 0.064 | 0.035 | 0.13 | 0.43 | 0.09 | 0.18 | 0.06 | 0.11 |
| 4 | 171680223 | rs10084822 | -2.118 | -1.802 | 1.761 | 0.278 | -0.046 | 0.002 | 0.002 | 0.564 | 0.18 | 0.59 | 0.3 | 0.22 | 0.11 | 0.15 |
| 4 | 171692801 | rs10007705 | -2.582 | -1.977 | 1.372 | 0.252 | 0.03 | -0.008 | -0.008 | 0 | 0.23 | 0.8 | 0.34 | 0.3 | 0.12 | 0.18 |
| 4 | 171694360 | rs1589560 | 0.353 | -1.904 | 1.61 | 0.161 | -0.029 | -0.059 | -0.059 | 0 | 0.89 | 0.78 | 0.95 | 0.99 | 0.95 | 0.99 |
| 4 | 171713039 | rs4615138 | -1.576 | -1.775 | 1.062 | 0.233 | -0.001 | -0.01 | -0.01 | 0 | 0.06 | 0.16 | 0.21 | 0.06 | 0.04 | 0.03 |
| 4 | 171722438 | rs10012933 | -1.653 | -1.786 | 1.036 | 0.228 | 0.005 | -0.005 | -0.005 | 0 | 0.06 | 0.2 | 0.21 | 0.06 | 0.04 | 0.03 |
| 4 | 171743216 | rs2877558 | -2.185 | -1.954 | 0.678 | 0.198 | 0.072 | -0.014 | -0.014 | 0.052 | 0.24 | 0.85 | 0.34 | 0.31 | 0.11 | 0.2 |
| 4 | 171751059 | rs10001385 | -2 | -2 | 0.629 | 0.236 | 0.03 | -0.006 | -0.006 | 0.907 | 0.24 | 0.81 | 0.35 | 0.31 | 0.11 | 0.21 |
| 4 | 171760509 | rs1432095 | -1.936 | -2.091 | -0.151 | 0.005 | 0.128 | -0.005 | -0.005 | 0 | 0.46 | 0.94 | 0.46 | 0.4 | 0.27 | 0.34 |
| 4 | 171762217 | rs1432094 | -0.755 | -1.432 | 0.551 | 0.273 | 0 | -0.011 | -0.011 | 1.625 | 0.06 | 0.19 | 0.2 | 0.05 | 0.03 | 0.03 |
| 4 | 171769237 | rs10011172 | -1.626 | -1.805 | 0.125 | 0.253 | 0.049 | -0.01 | -0.01 | 0 | 0.35 | 0.75 | 0.42 | 0.38 | 0.23 | 0.24 |
| 4 | 171769714 | rs10520196 | -1.523 | -2.06 | 0.79 | 0.246 | -0.026 | 0.023 | 0.023 | 0.003 | 0.28 | 0.42 | 0.35 | 0.38 | 0.18 | 0.25 |
| 4 | 171773560 | rs10520197 | -0.848 | -1.998 | 0.593 | 0.141 | 0 | 0.007 | 0.007 | 0 | 0.18 | 0.17 | 0.08 | 0.09 | 0.17 | 0.11 |
| 4 | 171775253 | rs17056275 | -1.752 | -2.11 | 1.331 | 0.406 | -0.012 | 0.015 | 0.015 | 0 | 0.26 | 0.35 | 0.17 | 0.33 | 0.2 | 0.26 |
| 4 | 171788150 | rs1560566 | -1.578 | -2.1 | 1.17 | 0.429 | 0.01 | -0.028 | -0.028 | 0 | 0.24 | 0.2 | 0.15 | 0.33 | 0.2 | 0.27 |
| 4 | 171790788 | rs1432089 | -2.715 | -2.307 | 1.674 | 0.86 | 0.023 | -0.016 | -0.016 | 0 | 0.39 | 0.72 | 0.25 | 0.49 | 0.26 | 0.3 |
| 4 | 171807535 | rs7673440 | -0.242 | -3.089 | 0.814 | 0.187 | 0.036 | 0.046 | 0.046 | 1.984 | 0.25 | 0.14 | 0.13 | 0.09 | 0.17 | 0.1 |
| 4 | 171808256 | rs7695819 | -1.752 | -3.183 | 0.924 | 0.19 | 0.023 | 0.006 | 0.006 | 6.271 | 0.41 | 0.71 | 0.27 | 0.35 | 0.22 | 0.28 |
| 4 | 171813412 | rs1432083 | -2.203 | -3.277 | 0.812 | 0.098 | 0.064 | 0 | 0 | 0 | 0.5 | 0.72 | 0.32 | 0.49 | 0.31 | 0.32 |
| 4 | 171821091 | rs6821882 | -1.197 | -3.3 | 0.924 | 0.207 | -0.031 | 0.049 | 0.049 | 0 | 0.1 | 0.23 | 0.26 | 0.25 | 0.05 | 0.16 |
| 4 | 171821267 | rs2332074 | 0.255 | -3.321 | 0.248 | -0.353 | 0.011 | 0.134 | 0.134 | 1.713 | 0.32 | 0.1 | 0.17 | 0.06 | 0.24 | 0.18 |
| 4 | 171824549 | rs2082317 | -2.659 | -3.481 | -0.494 | -0.825 | -0.172 | -0.192 | -0.192 | 3.773 | 0.69 | 0.98 | 0.6 | 0.49 | 0.47 | 0.52 |
| 4 | 171828587 | rs6822509 | -1.246 | -3.538 | 0.119 | -0.757 | 0.026 | 0.022 | 0.022 | 6.576 | 0.48 | 0.51 | 0.45 | 0.32 | 0.3 | 0.37 |
| 4 | 171842726 | rs6850897 | -1.058 | -3.554 | 0.04 | -0.769 | 0.033 | 0.026 | 0.026 | 3.61 | 0.47 | 0.48 | 0.42 | 0.32 | 0.3 | 0.37 |
| 4 | 171852518 | rs4392476 | -1.828 | -2.896 | 0.217 | -0.415 | 0 | 0 | 0 | 0.287 | 0.59 | 0.68 | 0.52 | 0.45 | 0.41 | 0.47 |
| 4 | 171853059 | rs28666968 | -0.874 | -2.81 | 0.106 | -0.474 | -0.023 | 0.039 | 0.039 | 1.005 | 0.1 | 0.21 | 0.25 | 0.25 | 0.04 | 0.19 |
| 4 | 171860973 | rs11725347 | -1.824 | -2.774 | 0.165 | -0.485 | 0.003 | 0.006 | 0.006 | 3.857 | 0.59 | 0.63 | 0.51 | 0.44 | 0.41 | 0.44 |
| 4 | 171872498 | rs13121066 | -0.634 | -2.754 | -0.004 | -0.631 | 0.018 | 0.065 | 0.065 | 0.829 | 0.45 | 0.35 | 0.24 | 0.2 | 0.36 | 0.24 |
| 4 | 171884710 | rs17245674 | 0.38 | -2.883 | -0.126 | -0.714 | -0.025 | 0.141 | 0.141 | 0.156 | 0.33 | 0.13 | 0.18 | 0.08 | 0.28 | 0.22 |
| 4 | 171888656 | rs7674455 | -0.565 | -2.844 | -0.057 | -0.647 | 0.015 | 0.005 | 0.005 | 0 | 0.14 | 0.1 | 0.06 | 0.13 | 0.09 | 0.04 |
| 4 | 171889916 | rs7656121 | -0.565 | -2.844 | -0.057 | -0.647 | 0.015 | 0.007 | 0.007 | 0 | 0.14 | 0.1 | 0.06 | 0.13 | 0.09 | 0.04 |
| 4 | 171890298 | rs7658356 | -0.73 | -2.886 | -0.078 | -0.67 | 0.018 | 0.007 | 0.007 | 0 | 0.15 | 0.1 | 0.06 | 0.13 | 0.09 | 0.04 |
| 4 | 171914470 | rs7682721 | -0.358 | -2.93 | -0.027 | -0.737 | 0.005 | 0.031 | 0.031 | 0.435 | 0.64 | 0.7 | 0.79 | 0.92 | 0.7 | 0.75 |
| 4 | 171919389 | rs13126363 | 0.157 | -2.873 | -0.117 | -0.71 | -0.018 | 0.154 | 0.154 | 0.755 | 0.35 | 0.13 | 0.19 | 0.08 | 0.29 | 0.2 |
| 4 | 171936344 | rs1508508 | 0.686 | -2.824 | -0.128 | -0.673 | 0.014 | 0.044 | 0.044 | 1.287 | 0.52 | 0.59 | 0.74 | 0.79 | 0.62 | 0.72 |
| 4 | 171957000 | rs1949007 | -1.153 | -2.915 | -0.116 | -0.714 | 0 | 0 | 0 | 0 | 0.49 | 0.56 | 0.46 | 0.35 | 0.33 | 0.39 |
| 4 | 171962965 | rs17056407 | -0.409 | -2.914 | -0.144 | -0.701 | 0.016 | -0.011 | -0.011 | 0 | 0.09 | 0.1 | 0.24 | 0.27 | 0.03 | 0.2 |
| 4 | 171963233 | rs10006864 | -1.931 | -2.849 | -0.085 | -0.674 | 0 | 0.002 | 0.002 | 3.456 | 0.6 | 0.69 | 0.51 | 0.44 | 0.41 | 0.48 |
| 4 | 172002025 | rs4450884 | -1.558 | -3.12 | -0.353 | -0.567 | 0.004 | 0.001 | 0.001 | 1.518 | 0.52 | 0.52 | 0.46 | 0.38 | 0.33 | 0.41 |
| 4 | 172006998 | rs11731647 | 0.533 | -2.995 | -0.356 | -0.474 | 0 | 0.014 | 0.014 | 0 | 0.1 | 0.04 | 0.04 | 0.06 | 0.07 | 0.02 |
| 4 | 172027644 | rs13136443 | -1.398 | -3.042 | -0.423 | -0.45 | 0.003 | 0 | 0 | 2.867 | 0.51 | 0.52 | 0.45 | 0.41 | 0.3 | 0.32 |
| 4 | 172031172 | rs17247383 | -0.192 | -2.152 | -0.959 | -0.715 | -0.002 | 0.124 | 0.124 | 0.784 | 0.29 | 0.1 | 0.15 | 0.06 | 0.21 | 0.18 |
| 4 | 172033270 | rs13135657 | -0.813 | -1.92 | -0.523 | -0.536 | 0.01 | 0.02 | 0.02 | 0 | 0.39 | 0.28 | 0.47 | 0.35 | 0.32 | 0.33 |
| 4 | 172033347 | rs13137432 | -0.821 | -1.922 | -0.525 | -0.537 | 0.01 | 0.02 | 0.02 | 0 | 0.39 | 0.28 | 0.47 | 0.35 | 0.32 | 0.36 |
| 4 | 172034838 | rs7671049 | -0.618 | -2.087 | 0.193 | -0.754 | -0.047 | -0.033 | -0.033 | 0 | 0.13 | 0.28 | 0.05 | 0.08 | 0.05 | 0.1 |
| 4 | 172036039 | rs17056485 | 0.027 | -2.176 | 0.172 | -0.842 | -0.009 | -0.034 | -0.034 | 0.716 | 0.06 | 0.16 | 0.04 | 0 | 0.04 | 0.05 |
| 4 | 172041735 | rs1508511 | -2.288 | -2.447 | 0.289 | -0.997 | -0.086 | -0.107 | -0.107 | 0.426 | 0.27 | 0.61 | 0.13 | 0.08 | 0.11 | 0.16 |
| 4 | 172041974 | rs17056509 | -1.499 | -2.321 | 0.328 | -0.952 | -0.034 | -0.071 | -0.071 | 0 | 0.13 | 0.3 | 0.08 | 0 | 0.06 | 0.06 |
| 4 | 172050379 | rs2063213 | -0.976 | -2.23 | 0.271 | -0.887 | 0.024 | 0.013 | 0.013 | 0 | 0.44 | 0.37 | 0.47 | 0.41 | 0.3 | 0.4 |
| 4 | 172052739 | rs10004389 | -1.092 | -2.125 | 0.702 | -1.673 | -0.03 | -0.029 | -0.029 | 0 | 0.14 | 0.24 | 0.06 | 0.09 | 0.08 | 0.11 |
| 4 | 172055275 | rs9997273 | -0.809 | -1.754 | 0.788 | -1.565 | 0.037 | 0.088 | 0.088 | 0 | 0.43 | 0.27 | 0.23 | 0.14 | 0.28 | 0.31 |
| 4 | 172056196 | rs11725022 | -1.272 | -1.68 | 0.833 | -1.499 | 0.005 | -0.02 | -0.02 | 0 | 0.52 | 0.59 | 0.28 | 0.14 | 0.34 | 0.32 |
| 4 | 172078453 | rs2055733 | -1.429 | -1.68 | 1.113 | -1.562 | -0.002 | -0.077 | -0.077 | 0 | 0.48 | 0.54 | 0.26 | 0.09 | 0.33 | 0.28 |
| 4 | 172083089 | rs7673719 | -1.443 | -1.764 | 1.209 | -1.547 | 0.039 | 0.118 | 0.118 | 1.469 | 0.52 | 0.72 | 0.28 | 0.09 | 0.33 | 0.35 |
| 4 | 172089287 | rs1508294 | -1.587 | -1.791 | 1.247 | -1.496 | 0.008 | -0.008 | -0.008 | 0 | 0.05 | 0.25 | 0.03 | 0 | 0.02 | 0.04 |
| 4 | 172104856 | rs10520201 | 0.109 | -1.649 | 1.114 | -1.618 | 0.018 | 0.092 | 0.092 | 0 | 0.31 | 0.16 | 0.18 | 0.04 | 0.25 | 0.2 |
| 4 | 172108672 | rs715990 | -1.618 | -1.472 | 1.243 | -1.403 | -0.006 | -0.023 | -0.023 | 0 | 0.07 | 0.32 | 0.04 | 0 | 0.02 | 0.04 |

| | | | | | | | | | | | | | | | | |
|---|---|---|---|---|---|---|---|---|---|---|---|---|---|---|---|---|
| 4 | 172112864 | rs10033316 | -2.003 | -1.487 | 1.289 | -1.433 | -0.021 | -0.11 | -0.11 | 0 | 0.2 | 0.54 | 0.12 | 0.05 | 0.13 | 0.15 |
| 4 | 172124768 | rs6841957 | -1.04 | -1.329 | 0.746 | -1.361 | 0.003 | -0.047 | -0.047 | 0.647 | 0.14 | 0.23 | 0.08 | 0.05 | 0.12 | 0.07 |
| 4 | 172132826 | rs7681110 | -0.042 | -1.286 | 0.416 | -1.29 | -0.012 | 0.02 | 0.02 | 0 | 0.89 | 0.96 | 0.94 | 0.93 | 0.89 | 0.94 |
| 4 | 172133199 | rs11132876 | 0.88 | -1.265 | 0.408 | -1.28 | -0.002 | -0.047 | -0.047 | 0 | 0.84 | 0.73 | 0.9 | 0.93 | 0.87 | 0.93 |
| 4 | 172147796 | rs6830239 | 0.426 | -1.148 | 0.159 | -1.2 | 0 | -0.012 | -0.012 | 1.463 | 0.87 | 0.85 | 0.93 | 0.93 | 0.89 | 0.93 |
| 4 | 172155235 | rs7685133 | 0.426 | -1.148 | 0.159 | -1.192 | 0 | -0.008 | -0.008 | 2.068 | 0.87 | 0.85 | 0.93 | 0.89 | 0.89 | 0.93 |
| 4 | 172167031 | rs6841319 | -0.506 | -1.351 | 0.222 | -1.429 | -0.027 | -0.148 | -0.148 | 0.762 | 0.38 | 0.49 | 0.23 | 0.04 | 0.26 | 0.28 |
| 4 | 172167890 | rs13145065 | 0.764 | -1.113 | 0.027 | -1.197 | 0.02 | 0.056 | 0.056 | 0.699 | 0.49 | 0.35 | 0.7 | 0.88 | 0.62 | 0.65 |
| 4 | 172186360 | rs1491332 | -1.033 | -1.608 | -0.25 | -0.306 | 0.058 | 0.032 | 0.032 | 0.023 | 0.43 | 0.76 | 0.51 | 0.34 | 0.28 | 0.42 |
| 4 | 172199176 | rs13135493 | -0.215 | -2.173 | -0.193 | -0.394 | 0.073 | 0.128 | 0.128 | 0.035 | 0.28 | 0.12 | 0.1 | 0.06 | 0.15 | 0.23 |
| 4 | 172200213 | rs7681428 | -1.183 | -2.02 | 0.109 | -0.37 | 0.037 | 0.019 | 0.019 | 0 | 0.41 | 0.73 | 0.49 | 0.35 | 0.27 | 0.42 |
| 4 | 172203864 | rs11132893 | -1.192 | -2.019 | 0.11 | -0.37 | 0.037 | 0.02 | 0.02 | 0 | 0.41 | 0.73 | 0.49 | 0.35 | 0.28 | 0.42 |
| 4 | 172203928 | rs12503744 | -0.937 | -1.517 | 1.076 | 0.294 | 0.052 | 0.391 | 0.391 | 0 | 0.05 | 0.35 | 0.31 | 0.29 | 0.03 | 0.13 |
| 4 | 172227195 | rs11132894 | -0.607 | -2.088 | 0.251 | 0.337 | 0.041 | -0.011 | -0.011 | 0 | 0.45 | 0.34 | 0.47 | 0.52 | 0.28 | 0.4 |
| 4 | 172236706 | rs13135574 | -0.169 | -2.372 | 0.647 | 0.32 | 0.077 | 0.098 | 0.098 | 0.004 | 0.38 | 0.17 | 0.18 | 0.16 | 0.21 | 0.31 |
| 4 | 172239480 | rs1491355 | -0.169 | -2.372 | 0.647 | 0.32 | 0.077 | 0.098 | 0.098 | 0.004 | 0.38 | 0.17 | 0.18 | 0.16 | 0.21 | 0.31 |
| 4 | 172243508 | rs333766 | 0.576 | -2.292 | 0.635 | 0.431 | 0.003 | 0 | 0 | 0 | 0.52 | 0.41 | 0.51 | 0.48 | 0.72 | 0.6 |
| 4 | 172263476 | rs333736 | 0.698 | -2.739 | 0.533 | 0.776 | 0.124 | 0.01 | 0.01 | 1.327 | 0.54 | 0.2 | 0.55 | 0.56 | 0.78 | 0.65 |
| 4 | 172267887 | rs952008 | 0.752 | -2.848 | 0.755 | 0.871 | 0.054 | -0.004 | -0.004 | 0 | 0.56 | 0.29 | 0.53 | 0.49 | 0.76 | 0.63 |
| 4 | 172269952 | rs7664958 | -0.585 | -2.891 | 0.963 | 0.906 | -0.005 | 0 | 0 | 0 | 0.4 | 0.62 | 0.19 | 0.18 | 0.17 | 0.27 |
| 4 | 172274229 | rs2877640 | -1.036 | -2.886 | 0.913 | 0.907 | 0.088 | -0.005 | -0.005 | 6.816 | 0.45 | 0.78 | 0.47 | 0.51 | 0.24 | 0.37 |
| 4 | 172275212 | rs9994171 | -1.164 | -2.911 | 0.906 | 0.894 | 0.136 | -0.014 | -0.014 | 0 | 0.46 | 0.84 | 0.48 | 0.51 | 0.24 | 0.38 |
| 4 | 172278868 | rs6830090 | -1.029 | -2.905 | 0.892 | 0.897 | 0.088 | -0.005 | -0.005 | 0 | 0.45 | 0.78 | 0.47 | 0.51 | 0.24 | 0.37 |
| 4 | 172311051 | rs17056840 | -1.779 | -2.784 | 1.286 | 0.897 | 0.004 | -0.023 | -0.023 | 0 | 0.09 | 0.38 | 0.06 | 0 | 0.03 | 0.09 |
| 4 | 172328758 | rs11132900 | 0.879 | -2.575 | 0.477 | 0.69 | 0.104 | 0.138 | 0.138 | 1.566 | 0.49 | 0.14 | 0.72 | 0.74 | 0.71 | 0.68 |
| 4 | 172331641 | rs6832175 | 0.01 | -2.761 | 0.923 | 0.544 | -0.003 | -0.008 | -0.008 | 0.606 | 0.6 | 0.6 | 0.76 | 0.79 | 0.71 | 0.71 |
| 4 | 172338295 | rs3913189 | 0.064 | -2.772 | 0.923 | 0.535 | 0.014 | 0.023 | 0.023 | 6.811 | 0.6 | 0.67 | 0.77 | 0.79 | 0.72 | 0.71 |
| 4 | 172338748 | rs17056855 | -0.965 | -2.89 | 0.824 | 0.471 | 0.034 | 0.022 | 0.022 | 0.836 | 0.09 | 0.3 | 0.07 | 0.1 | 0.06 | 0.08 |
| 4 | 172347176 | rs3913195 | 0.525 | -2.893 | 1.028 | 0.478 | 0.049 | 0.074 | 0.074 | 4.469 | 0.54 | 0.31 | 0.74 | 0.8 | 0.71 | 0.68 |
| 4 | 172350228 | rs7694458 | 0.027 | -2.894 | 0.836 | 0.477 | 0.011 | 0.017 | 0.017 | 6.274 | 0.6 | 0.64 | 0.77 | 0.8 | 0.72 | 0.71 |
| 4 | 172352348 | rs13120580 | 0.698 | -2.822 | -0.353 | 0.516 | 0.101 | 0.163 | 0.163 | 5.395 | 0.52 | 0.16 | 0.72 | 0.8 | 0.71 | 0.68 |
| 4 | 172354182 | rs2332216 | 1.719 | -2.5 | -0.852 | 0.604 | -0.152 | -0.048 | -0.048 | 0 | 0.31 | 0.02 | 0.42 | 0.3 | 0.59 | 0.46 |
| 4 | 172367126 | rs7662060 | 0.503 | -2.435 | -0.472 | 0.065 | 0.057 | 0.08 | 0.08 | 4.732 | 0.56 | 0.35 | 0.79 | 0.87 | 0.75 | 0.7 |
| 4 | 172367540 | rs4692883 | -1.248 | -2.509 | -0.548 | 0.023 | -0.001 | 0.06 | 0.06 | 3.837 | 0.73 | 0.99 | 0.87 | 0.91 | 0.81 | 0.83 |
| 4 | 172373388 | rs4692885 | 0.4 | -2.308 | -0.413 | 0.118 | 0.058 | 0.083 | 0.083 | 4.732 | 0.56 | 0.35 | 0.79 | 0.87 | 0.75 | 0.7 |
| 4 | 172388728 | rs7664076 | -2.038 | -1.518 | 0.033 | 1.163 | 0.069 | -0.093 | -0.093 | 0.592 | 0.36 | 0.96 | 0.42 | 0.58 | 0.19 | 0.36 |
| 4 | 172394198 | rs6831239 | -1.104 | -1.596 | 0.738 | 1.131 | -0.012 | 0.02 | 0.02 | 0 | 0.23 | 0.25 | 0.33 | 0.56 | 0.14 | 0.25 |
| 4 | 172405095 | rs4144862 | -2.829 | -0.826 | 0.812 | 1.334 | 0 | -0.121 | -0.121 | 0 | 0.33 | 0.96 | 0.36 | 0.55 | 0.22 | 0.37 |
| 4 | 172406840 | rs10014387 | -1.839 | -0.713 | 1.859 | 1.439 | -0.017 | 0.08 | 0.08 | 1.869 | 0.22 | 0.62 | 0.31 | 0.53 | 0.15 | 0.28 |
| 4 | 172425609 | rs1590121 | 1.288 | -0.595 | 2.331 | 1.34 | -0.009 | -0.026 | -0.026 | 0 | 0.8 | 0.49 | 0.88 | 0.84 | 0.85 | 0.85 |
| 4 | 172432296 | rs1358596 | 0.777 | -0.343 | 2.513 | 1.013 | 0.006 | 0.066 | 0.066 | 0.195 | 0.84 | 0.76 | 0.71 | 0.41 | 0.82 | 0.74 |
| 4 | 172451166 | rs6857184 | -1.183 | -0.22 | 2.195 | 0.611 | 0.004 | -0.031 | -0.031 | 0 | 0.11 | 0.17 | 0.05 | 0.01 | 0.09 | 0.13 |
| 4 | 172455594 | rs6553575 | -2.157 | -0.254 | 2.126 | 0.528 | 0 | -0.103 | -0.103 | 0 | 0.18 | 0.54 | 0.1 | 0.02 | 0.13 | 0.15 |
| 4 | 172473735 | rs1569214 | 0.697 | -0.181 | 1.294 | 1.062 | 0 | 0.034 | 0.034 | 0 | 0.84 | 0.77 | 0.73 | 0.46 | 0.83 | 0.74 |
| 4 | 172481191 | rs10025346 | 2.362 | 0.41 | 1.177 | 0.734 | -0.048 | -0.038 | -0.038 | 0 | 0.62 | 0.17 | 0.62 | 0.47 | 0.62 | 0.58 |
| 4 | 172487231 | rs11939052 | -1.997 | 0.309 | 2.008 | 0.508 | 0.025 | -0.095 | -0.095 | 0.041 | 0.21 | 0.37 | 0.12 | 0.02 | 0.22 | 0.16 |
| 4 | 172491592 | rs13109214 | 2.56 | 0.19 | 1.52 | 0.591 | -0.028 | -0.007 | -0.007 | 0 | 0.61 | 0.16 | 0.64 | 0.43 | 0.64 | 0.62 |
| 4 | 172492100 | rs12643448 | 2.529 | 0.212 | 1.532 | 0.585 | -0.021 | 0 | 0 | 0.009 | 0.61 | 0.2 | 0.64 | 0.42 | 0.64 | 0.62 |
| 4 | 172496760 | rs12500769 | 2.463 | 0.228 | 1.55 | 0.593 | -0.016 | 0 | 0 | 0 | 0.62 | 0.23 | 0.64 | 0.42 | 0.64 | 0.62 |
| 4 | 172496926 | rs1406510 | 0.38 | 0.302 | 1.548 | 0.625 | -0.003 | -0.015 | -0.015 | 0 | 0.87 | 0.9 | 0.79 | 0.44 | 0.86 | 0.77 |
| 4 | 172498522 | rs10009454 | -2.136 | 0.304 | 1.727 | 0.556 | 0.029 | -0.115 | -0.115 | 0.586 | 0.22 | 0.48 | 0.14 | 0.02 | 0.22 | 0.16 |
| 4 | 172504661 | rs1919495 | 0.63 | 0.299 | 1.145 | 0.751 | -0.002 | -0.015 | -0.015 | 0 | 0.87 | 0.9 | 0.79 | 0.44 | 0.86 | 0.77 |
| 4 | 172509162 | rs7654240 | -0.645 | 0.33 | 1.146 | 0.602 | -0.001 | -0.013 | -0.013 | 0.004 | 0.07 | 0.09 | 0.02 | 0.01 | 0.05 | 0.03 |
| 4 | 172509887 | rs10020087 | -1.915 | 0.342 | 1.682 | 0.55 | 0.016 | -0.062 | -0.062 | 4.122 | 0.13 | 0.43 | 0.07 | 0.01 | 0.08 | 0.05 |
| 4 | 172516685 | rs7683440 | -1.535 | 0.336 | 1.684 | 0.553 | 0.008 | -0.033 | -0.033 | 1.61 | 0.1 | 0.43 | 0.06 | 0.01 | 0.05 | 0.03 |
| 4 | 172516878 | rs10005145 | 2.801 | 0.319 | 1.853 | 0.58 | -0.016 | -0.006 | -0.006 | 0.013 | 0.62 | 0.24 | 0.6 | 0.42 | 0.64 | 0.62 |
| 4 | 172525210 | rs17057154 | -1.631 | 0.382 | 1.833 | 0.443 | 0.029 | -0.062 | -0.062 | 3.393 | 0.12 | 0.21 | 0.08 | 0.01 | 0.14 | 0.11 |
| 4 | 172537391 | rs6833580 | -1.353 | 0.286 | 1.561 | 0.158 | 0.004 | -0.079 | -0.079 | 0 | 0.18 | 0.23 | 0.1 | 0.01 | 0.18 | 0.14 |
| 4 | 172540259 | rs6847702 | 0.183 | 0.312 | 1.565 | 0.374 | -0.008 | -0.024 | -0.024 | 0.532 | 0.88 | 0.93 | 0.8 | 0.46 | 0.87 | 0.76 |
| 4 | 172542936 | rs1406506 | -0.73 | 0.32 | 1.576 | 0.355 | 0.048 | 0.026 | 0.026 | 0.489 | 0.05 | 0.21 | 0.02 | 0.01 | 0.02 | 0.09 |
| 4 | 172545173 | rs9991140 | -1.774 | 0.529 | 1.295 | 0.327 | 0.016 | -0.055 | -0.055 | 0 | 0.12 | 0.33 | 0.05 | 0.01 | 0.08 | 0.04 |
| 4 | 172551496 | rs4692898 | 2.275 | 0.489 | 1.176 | 0.352 | 0.003 | -0.003 | -0.003 | 0 | 0.6 | 0.5 | 0.65 | 0.43 | 0.61 | 0.58 |
| 4 | 172557879 | rs9312509 | 1.827 | 0.714 | 0.681 | 0.638 | 0.009 | 0.032 | 0.032 | 0 | 0.71 | 0.6 | 0.68 | 0.46 | 0.69 | 0.65 |
| 4 | 172558394 | rs9312510 | 1.083 | 1.162 | 0.184 | 0.078 | -0.003 | 0 | 0 | 0.067 | 0.77 | 0.74 | 0.54 | 0.21 | 0.72 | 0.62 |
| 4 | 172559687 | rs2172009 | 1.427 | 0.768 | -0.244 | -0.393 | -0.019 | 0.13 | 0.13 | 1.071 | 0.57 | 0.14 | 0.67 | 0.78 | 0.58 | 0.74 |
| 4 | 172560014 | rs17057224 | -1.181 | 1.421 | 0.747 | -0.136 | 0.053 | -0.034 | -0.034 | 0 | 0.09 | 0.26 | 0.11 | 0.02 | 0.15 | 0.08 |
| 4 | 172566190 | rs12644946 | 1.651 | 1.259 | 0.744 | -0.16 | 0.043 | -0.054 | -0.054 | 0 | 0.77 | 0.58 | 0.77 | 0.9 | 0.69 | 0.79 |
| 4 | 172568268 | rs13136852 | 1.49 | 1.18 | 0.598 | -0.133 | 0.032 | -0.04 | -0.04 | 4.697 | 0.78 | 0.69 | 0.78 | 0.9 | 0.69 | 0.8 |
| 4 | 172590031 | rs6844440 | 0.173 | 0.16 | 0.014 | 0.071 | 0 | -0.007 | -0.007 | 0 | 0.9 | 0.99 | 0.92 | 0.92 | 0.92 | 0.89 |

| | | | | | | | | | | | | | | | | |
|---|---|---|---|---|---|---|---|---|---|---|---|---|---|---|---|---|
| 4 | 172597919 | rs6553586 | -0.687 | 0.411 | 0.234 | -0.337 | -0.003 | -0.027 | -0.027 | 0 | 0.83 | 0.85 | 0.85 | 1 | 0.77 | 0.94 |
| 4 | 172609720 | rs1907084 | -1.975 | -0.622 | 1.55 | 0.14 | -0.034 | -0.076 | -0.076 | 1.154 | 0.16 | 0.51 | 0.13 | 0.05 | 0.13 | 0.15 |
| 4 | 172633978 | rs9993484 | -1.798 | -0.6 | 1.816 | 0.534 | -0.02 | 0.034 | 0.034 | 0.037 | 0.15 | 0.47 | 0.11 | 0.13 | 0.12 | 0.08 |
| 4 | 172651944 | rs768709 | -0.2 | -0.346 | 1.156 | 0.73 | -0.056 | -0.117 | -0.117 | 0 | 0.19 | 0.02 | 0.15 | 0.39 | 0.24 | 0.27 |
| 4 | 172661677 | rs836028 | 2.006 | -0.665 | 2.271 | 0.215 | -0.028 | -0.005 | -0.005 | 0 | 0.84 | 0.28 | 0.87 | 0.85 | 0.9 | 0.9 |
| 4 | 172675143 | rs836022 | 0.859 | -0.562 | 2.182 | 0.245 | 0.032 | 0.033 | 0.033 | 0 | 0.92 | 0.63 | 0.91 | 0.96 | 0.95 | 0.98 |
| 4 | 172699232 | rs17057429 | -0.5 | -0.615 | 1.514 | 0.276 | 0.011 | -0.017 | -0.017 | 0 | 0.05 | 0.02 | 0.02 | 0.12 | 0.04 | 0.07 |
| 4 | 172705236 | rs10003818 | 1.26 | -0.633 | 1.716 | 0.271 | -0.054 | 0.026 | 0.026 | 0.281 | 0.84 | 0.55 | 0.88 | 0.84 | 0.9 | 0.92 |
| 4 | 172713976 | rs1495288 | 1.064 | -0.454 | 0.885 | 0.286 | -0.034 | 0.003 | 0.003 | 0 | 0.87 | 0.63 | 0.92 | 0.88 | 0.91 | 0.92 |
| 4 | 172733788 | rs3733432 | 1.067 | -0.535 | 0.19 | 0.259 | -0.053 | -0.012 | -0.012 | 0 | 0.81 | 0.37 | 0.87 | 0.84 | 0.9 | 0.88 |
| 4 | 172740734 | rs901236 | 0.607 | -0.182 | -0.747 | -0.552 | 0.025 | 0.146 | 0.146 | 0.01 | 0.55 | 0.29 | 0.72 | 0.87 | 0.6 | 0.7 |
| 4 | 172742577 | rs494770 | 0.698 | -0.775 | -0.233 | -0.346 | 0.005 | 0.005 | 0.005 | 0.124 | 0.68 | 0.37 | 0.84 | 0.82 | 0.82 | 0.61 |
| 4 | 172747852 | rs10027752 | -0.881 | -0.701 | 0.079 | -0.401 | -0.035 | -0.034 | -0.034 | 0.673 | 0.32 | 0.53 | 0.16 | 0.18 | 0.21 | 0.4 |
| 4 | 172748274 | rs570686 | -0.071 | -0.717 | -0.598 | -0.612 | -0.006 | 0.01 | 0.01 | 0 | 0.28 | 0.26 | 0.28 | 0.19 | 0.35 | 0.11 |
| 4 | 172750974 | rs580091 | -1.705 | -0.711 | -0.747 | -0.623 | -0.081 | -0.03 | -0.03 | 0.504 | 0.6 | 0.9 | 0.44 | 0.37 | 0.56 | 0.51 |
| 4 | 172751111 | rs558129 | -0.208 | -1.468 | -0.465 | -0.506 | -0.013 | 0.018 | 0.018 | 0 | 0.28 | 0.24 | 0.28 | 0.19 | 0.35 | 0.11 |
| 4 | 172758442 | rs494219 | -0.012 | -1.836 | -0.401 | -0.459 | 0 | -0.005 | -0.005 | 0 | 0.26 | 0.3 | 0.24 | 0.2 | 0.28 | 0.1 |
| 4 | 172762249 | rs2046482 | 1.178 | -2.128 | -0.153 | 0.312 | 0.101 | -0.027 | -0.027 | 0 | 0.48 | 0.13 | 0.59 | 0.46 | 0.63 | 0.54 |
| 4 | 172763249 | rs2877674 | 0.439 | -2.277 | 0.267 | 0.304 | -0.107 | 0.014 | 0.014 | 0 | 0.73 | 0.5 | 0.86 | 0.69 | 0.9 | 0.65 |
| 4 | 172764187 | rs11132917 | 1.119 | -2.203 | -0.106 | 0.349 | 0.09 | 0.024 | 0.024 | 0 | 0.5 | 0.25 | 0.63 | 0.57 | 0.63 | 0.54 |
| 4 | 172776138 | rs7676508 | -0.072 | -2.081 | 0.031 | 0.384 | 0.002 | 0.006 | 0.006 | 0 | 0.74 | 0.76 | 0.73 | 0.77 | 0.73 | 0.89 |
| 4 | 172786927 | rs6818860 | 0.976 | -1.811 | -0.318 | 0.516 | 0.121 | -0.029 | -0.029 | 0 | 0.51 | 0.15 | 0.6 | 0.49 | 0.64 | 0.53 |
| 4 | 172789426 | rs12503485 | 0.782 | -1.786 | -0.008 | 0.435 | 0.044 | 0.014 | 0.014 | 0 | 0.53 | 0.36 | 0.67 | 0.6 | 0.65 | 0.54 |
| 4 | 172790557 | rs7666914 | -0.923 | -2.174 | 0.353 | 0.573 | -0.071 | -0.002 | -0.002 | 0.073 | 0.24 | 0.53 | 0.12 | 0.23 | 0.1 | 0.36 |
| 4 | 172797779 | rs10031590 | -1.045 | -1.908 | -0.009 | 0.883 | 0.009 | 0.007 | 0.007 | 0 | 0.55 | 0.63 | 0.35 | 0.57 | 0.44 | 0.46 |
| 4 | 172801023 | rs13101636 | -1.095 | -1.931 | 0.008 | 0.938 | 0.003 | 0.001 | 0.001 | 0 | 0.56 | 0.78 | 0.39 | 0.58 | 0.45 | 0.46 |
| 4 | 172808394 | rs6553597 | -1.178 | -2.013 | -0.114 | 0.842 | -0.006 | -0.007 | -0.007 | 0 | 0.59 | 0.68 | 0.37 | 0.57 | 0.47 | 0.46 |
| 4 | 172812489 | rs4124044 | -1.18 | -2 | -0.208 | 0.922 | -0.023 | -0.032 | -0.032 | 0 | 0.6 | 0.79 | 0.41 | 0.56 | 0.47 | 0.46 |
| 4 | 172816200 | rs718913 | 0.076 | -2.077 | 0.294 | 1.298 | 0.006 | 0.019 | 0.019 | 0 | 0.33 | 0.23 | 0.25 | 0.2 | 0.3 | 0.15 |
| 4 | 172832692 | rs6845054 | -0.947 | -2.158 | 0.74 | 1.244 | -0.016 | 0.002 | 0.002 | 0 | 0.31 | 0.64 | 0.17 | 0.36 | 0.2 | 0.34 |
| 4 | 172842419 | rs4692643 | -1.634 | -1.751 | -0.087 | 1.456 | -0.087 | -0.003 | -0.003 | 0 | 0.59 | 0.92 | 0.41 | 0.65 | 0.47 | 0.45 |
| 4 | 172849152 | rs10005702 | -1.637 | -1.795 | -0.044 | 1.457 | -0.07 | 0.011 | 0.011 | 0.011 | 0.6 | 0.91 | 0.41 | 0.65 | 0.47 | 0.45 |
| 4 | 172849554 | rs11931247 | -0.445 | -1.799 | 0.494 | 1.618 | -0.023 | 0.015 | 0.015 | 0 | 0.23 | 0.34 | 0.12 | 0.26 | 0.19 | 0.29 |
| 4 | 172864640 | rs332995 | 0.493 | -1.64 | 0.377 | 1.534 | -0.025 | 0.02 | 0.02 | 0 | 0.74 | 0.43 | 0.83 | 0.66 | 0.82 | 0.69 |
| 4 | 172870192 | rs17057610 | 0.096 | -1.821 | 0.319 | 1.497 | 0.029 | 0.02 | 0.02 | 0.521 | 0.33 | 0.18 | 0.24 | 0.25 | 0.28 | 0.15 |
| 4 | 172870885 | rs6553603 | -0.849 | -1.979 | 0.46 | 1.587 | -0.025 | 0.03 | 0.03 | 0 | 0.27 | 0.6 | 0.2 | 0.4 | 0.19 | 0.29 |
| 4 | 172871125 | rs333003 | 0.77 | -1.22 | 0.641 | 1.056 | -0.009 | 0.069 | 0.069 | 0 | 0.41 | 0.13 | 0.55 | 0.31 | 0.54 | 0.56 |
| 4 | 172872173 | rs405831 | -0.562 | -1.237 | 0.661 | 0.959 | 0.005 | 0.012 | 0.012 | 0 | 0.55 | 0.59 | 0.44 | 0.67 | 0.46 | 0.44 |
| 4 | 172880449 | rs332969 | -0.519 | -1.006 | 0.342 | 0.927 | 0.02 | 0.018 | 0.018 | 0.158 | 0.53 | 0.57 | 0.37 | 0.59 | 0.44 | 0.44 |
| 4 | 172886425 | rs4692907 | 0.581 | -0.706 | -0.361 | 0.935 | -0.037 | 0.053 | 0.053 | 2.395 | 0.44 | 0.07 | 0.56 | 0.34 | 0.57 | 0.59 |
| 4 | 172893271 | rs409190 | -0.253 | -0.606 | -0.213 | 1.007 | 0.016 | 0.004 | 0.004 | 0.521 | 0.46 | 0.32 | 0.35 | 0.54 | 0.43 | 0.39 |
| 4 | 172896473 | rs230 | 0.214 | -0.46 | -0.065 | 1.159 | 0.062 | 0.113 | 0.113 | 1.907 | 0.49 | 0.17 | 0.57 | 0.33 | 0.59 | 0.59 |
| 4 | 172902011 | rs184564 | 0.878 | 0.413 | 0.284 | 1.09 | 0.045 | 0.11 | 0.11 | 0.46 | 0.95 | 0.62 | 0.94 | 0.93 | 1 | 0.98 |
| 4 | 172920376 | rs2135196 | -0.777 | 1.459 | -1.029 | 1.281 | -0.106 | -0.092 | -0.092 | 0 | 0.18 | 0.01 | 0.16 | 0.24 | 0.25 | 0.27 |
| 4 | 172929844 | rs1979931 | 0.041 | 0.284 | 0.067 | 0.944 | -0.056 | -0.001 | -0.001 | 0.098 | 0.73 | 0.58 | 0.85 | 0.76 | 0.8 | 0.71 |
| 4 | 172945557 | rs12645248 | -0.248 | 0.186 | 1.348 | 0.85 | 0.001 | -0.007 | -0.007 | 0 | 0.4 | 0.41 | 0.57 | 0.5 | 0.42 | 0.55 |
| 4 | 172946287 | rs983560 | -0.604 | 0.603 | 1.337 | 0.418 | 0.01 | 0.056 | 0.056 | 0.206 | 0.57 | 0.76 | 0.75 | 0.76 | 0.59 | 0.68 |
| 4 | 172950660 | rs6843490 | -0.133 | 0.566 | 1.531 | 0.526 | 0.008 | 0.016 | 0.016 | 0.402 | 0.45 | 0.7 | 0.67 | 0.66 | 0.44 | 0.61 |
| 4 | 172953080 | rs12646854 | 0.336 | 0.594 | 1.198 | 0.537 | 0.001 | -0.017 | -0.017 | 0 | 0.39 | 0.36 | 0.57 | 0.49 | 0.42 | 0.56 |
| 4 | 172955918 | rs10780085 | 0.449 | 0.683 | 0.22 | 0.479 | -0.014 | 0.087 | 0.087 | 0.095 | 0.43 | 0.12 | 0.23 | 0.24 | 0.41 | 0.32 |
| 4 | 172957428 | rs4146317 | 0.311 | 0.596 | 0.138 | 0.547 | -0.024 | -0.068 | -0.068 | 0 | 0.34 | 0.15 | 0.56 | 0.49 | 0.42 | 0.55 |
| 4 | 172965481 | rs13134541 | 0.248 | 0.766 | -0.319 | 0.492 | -0.029 | -0.036 | -0.036 | 0 | 0.36 | 0.12 | 0.41 | 0.66 | 0.36 | 0.55 |
| 4 | 172968055 | rs7654286 | 0.285 | 0.869 | 0.444 | 0.034 | 0.024 | -0.063 | -0.063 | 1.347 | 0.78 | 0.59 | 0.82 | 0.88 | 0.79 | 0.85 |
| 4 | 172979002 | rs1394860 | 0.153 | 0.568 | 0.433 | 0.502 | -0.003 | 0.029 | 0.029 | 0 | 0.8 | 0.66 | 0.8 | 0.75 | 0.84 | 0.88 |
| 4 | 172985760 | rs1394859 | -0.242 | 0.376 | 0.135 | 0.179 | 0 | -0.01 | -0.01 | 0 | 0.51 | 0.48 | 0.51 | 0.21 | 0.51 | 0.35 |
| 4 | 172997356 | rs13139421 | -0.721 | 0.064 | 0.31 | 0.527 | -0.013 | 0.086 | 0.086 | 0 | 0.89 | 0.76 | 0.89 | 0.76 | 0.93 | 0.94 |
| 4 | 172999916 | rs2877689 | 1.162 | 0.14 | 0.061 | 0.411 | -0.076 | -0.12 | -0.12 | 0 | 0.32 | 0.09 | 0.35 | 0.54 | 0.39 | 0.55 |
| 4 | 173001831 | rs13136240 | -0.113 | 0.015 | 0.519 | -0.197 | 0 | -0.015 | -0.015 | 0 | 0.51 | 0.49 | 0.51 | 0.21 | 0.51 | 0.34 |
| 4 | 173005057 | rs17058017 | 0.738 | -0.057 | 0.273 | -0.312 | -0.026 | 0.031 | 0.031 | 0.078 | 0.12 | 0.02 | 0.23 | 0.08 | 0.11 | 0.12 |
| 4 | 173005461 | rs9990764 | -0.524 | -0.025 | 0.48 | -0.381 | -0.008 | 0.051 | 0.051 | 0 | 0.56 | 0.62 | 0.53 | 0.21 | 0.53 | 0.35 |
| 4 | 173013636 | rs1354631 | 0.43 | -0.116 | 0.72 | -0.293 | 0.001 | 0.027 | 0.027 | 0 | 0.4 | 0.35 | 0.3 | 0.17 | 0.42 | 0.26 |
| 4 | 173015421 | rs2131002 | 0.464 | 0.113 | 1.016 | -0.054 | 0.026 | -0.014 | -0.014 | 0 | 0.92 | 0.68 | 0.96 | 1 | 0.96 | 0.99 |
| 4 | 173024088 | rs6846244 | 0.127 | -0.064 | 0.541 | -0.36 | -0.011 | -0.053 | -0.053 | 0 | 0.42 | 0.53 | 0.28 | 0.13 | 0.38 | 0.27 |
| 4 | 173025475 | rs17219104 | 0.254 | 0.011 | 0.161 | -0.295 | -0.002 | 0.015 | 0.015 | 0 | 0.1 | 0.04 | 0.23 | 0.07 | 0.09 | 0.14 |
| 4 | 173035260 | rs2201113 | 1.622 | 0.12 | 0.03 | 0.187 | -0.088 | -0.099 | -0.099 | 0 | 0.31 | 0.07 | 0.32 | 0.51 | 0.41 | 0.53 |
| 4 | 173040504 | rs17219493 | 0.434 | 0.343 | 0.171 | 0.116 | -0.002 | 0.004 | 0.004 | 0.018 | 0.07 | 0.03 | 0.09 | 0.04 | 0.08 | 0.15 |
| 4 | 173047549 | rs10016776 | 0.216 | 0.074 | 0.42 | -0.06 | 0.003 | 0.024 | 0.024 | 0 | 0.41 | 0.39 | 0.41 | 0.14 | 0.4 | 0.27 |
| 4 | 173052098 | rs6553608 | -0.137 | 0.042 | 0.589 | -0.071 | 0.004 | 0.007 | 0.007 | 0.103 | 0.45 | 0.61 | 0.44 | 0.14 | 0.4 | 0.27 |
| 4 | 173068658 | rs6553610 | -0.145 | 0.042 | 0.593 | -0.067 | 0.003 | -0.001 | -0.001 | 0 | 0.45 | 0.6 | 0.43 | 0.14 | 0.4 | 0.27 |

| | | | | | | | | | | | | | | | | |
|---|---|---|---|---|---|---|---|---|---|---|---|---|---|---|---|---|
| 4 | 173084108 | rs13110849 | -0.524 | 0.005 | 0.635 | -0.038 | 0.019 | 0.066 | 0.066 | 0.177 | 0.42 | 0.7 | 0.43 | 0.14 | 0.38 | 0.21 |
| 4 | 173089205 | rs13124593 | 0.21 | 0.044 | 0.623 | 0.263 | -0.002 | 0.058 | 0.058 | 0 | 0.77 | 0.68 | 0.72 | 0.63 | 0.8 | 0.79 |
| 4 | 173109636 | rs2202281 | -0.314 | 0.057 | 0.768 | -0.334 | 0.002 | 0.006 | 0.006 | 0 | 0.37 | 0.38 | 0.37 | 0.12 | 0.36 | 0.22 |
| 4 | 173125742 | rs6825169 | 0.2 | 0.097 | 0.542 | -0.345 | -0.013 | 0.036 | 0.036 | 0 | 0.15 | 0.08 | 0.08 | 0.03 | 0.13 | 0.08 |
| 4 | 173129218 | rs836306 | 0.501 | 0.06 | 0.891 | -0.202 | 0.013 | 0.101 | 0.101 | 0 | 0.54 | 0.26 | 0.56 | 0.85 | 0.59 | 0.76 |
| 4 | 173142594 | rs9997484 | -0.456 | 0.115 | 0.927 | -0.176 | -0.001 | -0.028 | -0.028 | 0 | 0.42 | 0.47 | 0.41 | 0.14 | 0.42 | 0.23 |
| 4 | 173152095 | rs836325 | 0.835 | 0.108 | 0.794 | -0.153 | -0.029 | 0.192 | 0.192 | 0.423 | 0.49 | 0.12 | 0.47 | 0.79 | 0.52 | 0.68 |
| 4 | 173152405 | rs10012505 | -0.477 | 0.106 | 0.957 | -0.176 | -0.001 | -0.012 | -0.012 | 0 | 0.42 | 0.43 | 0.4 | 0.14 | 0.42 | 0.24 |
| 4 | 173164661 | rs836321 | 0.431 | 0.12 | 0.845 | -0.169 | 0.009 | 0.057 | 0.057 | 0 | 0.55 | 0.32 | 0.57 | 0.85 | 0.58 | 0.76 |
| 4 | 173167398 | rs836322 | 0.563 | 0.117 | 0.636 | -0.171 | 0.019 | 0.134 | 0.134 | 0 | 0.53 | 0.23 | 0.55 | 0.85 | 0.58 | 0.77 |
| 4 | 173172113 | rs10049623 | -0.502 | -0.046 | 0.994 | -0.051 | 0.003 | 0.046 | 0.046 | 0 | 0.38 | 0.34 | 0.36 | 0.15 | 0.38 | 0.26 |
| 4 | 173175173 | rs1354356 | -1.483 | -0.125 | 0.903 | 0.389 | -0.037 | -0.044 | -0.044 | 0 | 0.58 | 0.82 | 0.56 | 0.45 | 0.48 | 0.36 |
| 4 | 173175931 | rs6843010 | -1.041 | -0.141 | 1.046 | 0.361 | -0.001 | -0.001 | -0.001 | 0 | 0.53 | 0.45 | 0.52 | 0.44 | 0.48 | 0.35 |
| 4 | 173177403 | rs905571 | -1.266 | -0.122 | 1.056 | 0.37 | -0.002 | -0.002 | -0.002 | 0 | 0.56 | 0.61 | 0.54 | 0.44 | 0.48 | 0.36 |
| 4 | 173178085 | rs10002142 | 0.673 | -0.085 | 0.538 | 0.401 | -0.003 | 0.019 | 0.019 | 0.998 | 0.15 | 0.13 | 0.16 | 0.29 | 0.1 | 0.1 |
| 4 | 173179915 | rs12505892 | 0.694 | -0.082 | 0.526 | 0.402 | -0.002 | 0.017 | 0.017 | 1.07 | 0.15 | 0.13 | 0.17 | 0.29 | 0.1 | 0.1 |
| 4 | 173185572 | rs10021776 | -1.378 | 0.01 | -0.043 | 0.429 | -0.036 | -0.043 | -0.043 | 2.564 | 0.58 | 0.82 | 0.56 | 0.44 | 0.48 | 0.36 |
| 4 | 173206315 | rs9991827 | -0.837 | 0.376 | 0.04 | 0.028 | -0.001 | -0.01 | -0.01 | 1.595 | 0.43 | 0.47 | 0.45 | 0.21 | 0.44 | 0.34 |
| 4 | 173208191 | rs9312523 | -0.803 | 0.451 | -0.661 | 0.11 | 0.028 | 0.112 | 0.112 | 0.098 | 0.41 | 0.8 | 0.44 | 0.21 | 0.32 | 0.26 |
| 4 | 173213263 | rs1955282 | 0.795 | 0.974 | 0.207 | 0.697 | 0.004 | 0.007 | 0.007 | 0 | 0.69 | 0.65 | 0.75 | 0.64 | 0.65 | 0.84 |
| 4 | 173215669 | rs6839113 | -0.17 | 1.101 | 0.272 | 0.621 | 0.004 | -0.035 | -0.035 | 0.601 | 0.15 | 0.16 | 0.09 | 0.01 | 0.19 | 0.06 |
| 4 | 173227094 | rs2654774 | -0.1 | 0.867 | 1.003 | 0.496 | -0.016 | -0.004 | -0.004 | 1.324 | 0.81 | 0.65 | 0.85 | 0.8 | 0.84 | 0.84 |
| 4 | 173227165 | rs17289279 | 2.071 | 0.911 | 0.698 | 0.49 | -0.058 | -0.002 | -0.002 | 0.588 | 0.47 | 0.09 | 0.56 | 0.43 | 0.49 | 0.67 |
| 4 | 173238445 | rs2054684 | 0.259 | 0.486 | 0.986 | 0.672 | 0.014 | -0.002 | -0.002 | 0 | 0.15 | 0.07 | 0.12 | 0.16 | 0.11 | 0.13 |
| 4 | 173245389 | rs333384 | -0.236 | 0.48 | 0.898 | 0.808 | 0 | 0.016 | 0.016 | 0 | 0.17 | 0.19 | 0.16 | 0.35 | 0.16 | 0.1 |
| 4 | 173271397 | rs4692917 | 1.325 | 0.486 | 0.932 | 0.562 | 0.019 | -0.011 | -0.011 | 0.248 | 0.6 | 0.21 | 0.66 | 0.46 | 0.68 | 0.73 |
| 4 | 173287422 | rs1484019 | 1.471 | 0.4 | 1.241 | 0.526 | 0.028 | -0.022 | -0.022 | 0.077 | 0.58 | 0.14 | 0.66 | 0.46 | 0.68 | 0.73 |
| 4 | 173302198 | rs1508257 | -0.042 | 0.526 | 1.511 | 0.754 | 0.023 | 0.107 | 0.107 | 1.322 | 0.84 | 0.59 | 0.86 | 0.81 | 0.88 | 0.87 |
| 4 | 173323928 | rs335998 | -0.581 | 0.703 | 1.443 | 0.841 | 0.008 | 0.034 | 0.034 | 0.019 | 0.88 | 0.83 | 0.89 | 0.82 | 0.88 | 0.89 |
| 4 | 173324955 | rs1508260 | -0.425 | 0.631 | 1.366 | 0.817 | -0.004 | 0.006 | 0.006 | 0 | 0.23 | 0.28 | 0.19 | 0.35 | 0.21 | 0.15 |
| 4 | 173331787 | rs6852501 | -0.579 | 0.625 | 1.363 | 0.814 | -0.019 | 0.015 | 0.015 | 0.021 | 0.25 | 0.36 | 0.2 | 0.35 | 0.21 | 0.15 |
| 4 | 173333056 | rs335983 | -0.037 | 0.849 | 1.489 | 0.923 | 0.017 | 0.07 | 0.07 | 0.005 | 0.87 | 0.72 | 0.87 | 0.82 | 0.88 | 0.89 |
| 4 | 173340994 | rs663643 | -0.374 | 0.885 | 1.138 | 0.928 | 0.007 | 0.05 | 0.05 | 0 | 0.88 | 0.84 | 0.88 | 0.75 | 0.88 | 0.89 |
| 4 | 173353213 | rs335972 | 0.308 | 0.863 | 0.912 | 0.9 | 0.021 | 0.142 | 0.142 | 0.074 | 0.85 | 0.71 | 0.88 | 0.74 | 0.88 | 0.87 |
| 4 | 173354354 | rs10024431 | 1.156 | 1.233 | 0.847 | 0.665 | 0.015 | 0.011 | 0.011 | 0 | 0.7 | 0.43 | 0.77 | 0.73 | 0.69 | 0.82 |
| 4 | 173356104 | rs163185 | -0.627 | 1.148 | 0.922 | 0.744 | 0.011 | 0.066 | 0.066 | 0 | 0.89 | 0.87 | 0.89 | 0.74 | 0.88 | 0.9 |
| 4 | 173358095 | rs336003 | -0.145 | 1.162 | 0.893 | 0.732 | 0.022 | 0.111 | 0.111 | 0 | 0.87 | 0.78 | 0.88 | 0.74 | 0.88 | 0.87 |
| 4 | 173360981 | rs12644293 | -0.187 | 0.992 | 0.54 | 0.552 | -0.028 | 0.035 | 0.035 | 0 | 0.23 | 0.37 | 0.2 | 0.36 | 0.16 | 0.13 |
| 4 | 173399986 | rs338021 | -0.875 | 1.258 | 0.638 | 0.287 | 0.004 | 0.001 | 0.001 | 0 | 0.25 | 0.27 | 0.2 | 0.22 | 0.33 | 0.15 |
| 4 | 173404269 | rs13142236 | 0.476 | 1.285 | 0.528 | 0.275 | -0.022 | 0.01 | 0.01 | 0 | 0.12 | 0.08 | 0.09 | 0 | 0.2 | 0.06 |
| 4 | 173405278 | rs10019934 | 1.194 | 1.2 | 0.546 | 0.281 | 0.023 | 0.005 | 0.005 | 0 | 0.7 | 0.42 | 0.76 | 0.77 | 0.67 | 0.82 |
| 4 | 173418304 | rs6553634 | -0.039 | 1.113 | 0.465 | 0.37 | -0.022 | 0.01 | 0.01 | 0 | 0.21 | 0.35 | 0.18 | 0.26 | 0.14 | 0.09 |
| 4 | 173423366 | rs1551446 | -0.694 | 1.557 | 0.374 | 0.067 | 0.045 | -0.084 | -0.084 | 2.074 | 0.15 | 0.27 | 0.1 | 0 | 0.2 | 0.06 |
| 4 | 173424803 | rs2124204 | 0.664 | 1.538 | 0.383 | 0.062 | 0.049 | -0.099 | -0.099 | 0.014 | 0.83 | 0.65 | 0.89 | 0.99 | 0.8 | 0.92 |
| 4 | 173440948 | rs17233636 | -0.682 | 1.655 | 0.099 | 0.113 | 0.046 | -0.086 | -0.086 | 0 | 0.15 | 0.27 | 0.1 | 0 | 0.2 | 0.05 |
| 4 | 173466872 | rs13130915 | -0.424 | 1.752 | -0.426 | 0.101 | -0.08 | 0.076 | 0.076 | 0.034 | 0.15 | 0.04 | 0.11 | 0 | 0.28 | 0.08 |
| 4 | 173476675 | rs6819214 | 0.097 | 0.689 | 0.021 | 0.758 | -0.026 | -0.039 | -0.039 | 0.037 | 0.89 | 0.94 | 0.88 | 0.78 | 0.86 | 0.9 |
| 4 | 173478710 | rs2332519 | 0.543 | 0.671 | 0.084 | 0.731 | 0.005 | 0.055 | 0.055 | 0.41 | 0.84 | 0.75 | 0.85 | 0.72 | 0.86 | 0.82 |
| 4 | 173491064 | rs13102637 | 0.591 | 0.559 | -0.448 | 0.136 | -0.04 | 0.012 | 0.012 | 0 | 0.42 | 0.19 | 0.56 | 0.66 | 0.55 | 0.51 |
| 4 | 173495158 | rs17234582 | 0.382 | 0.389 | -0.102 | 0.074 | -0.013 | 0.021 | 0.021 | 0 | 0.44 | 0.27 | 0.56 | 0.66 | 0.55 | 0.61 |
| 4 | 173513908 | rs10434280 | 0.139 | 0.064 | 0.466 | -0.177 | -0.001 | 0.02 | 0.02 | 0 | 0.53 | 0.42 | 0.63 | 0.75 | 0.55 | 0.74 |
| 4 | 173521239 | rs6858309 | 0.939 | 0.076 | 0.316 | -0.097 | -0.067 | -0.003 | -0.003 | 0 | 0.42 | 0.16 | 0.56 | 0.66 | 0.54 | 0.53 |
| 4 | 173535503 | rs1455129 | -0.182 | 0.21 | 0.681 | -0.317 | 0.002 | -0.064 | -0.064 | 0 | 0.44 | 0.46 | 0.25 | 0.01 | 0.36 | 0.22 |
| 4 | 173541374 | rs1551447 | -0.263 | 0.226 | 0.701 | -0.301 | -0.007 | -0.121 | -0.121 | 0 | 0.45 | 0.52 | 0.25 | 0.01 | 0.36 | 0.22 |
| 4 | 173543874 | rs2332544 | 0.586 | 0.295 | 0.756 | -0.235 | 0.014 | 0.079 | 0.079 | 0 | 0.44 | 0.29 | 0.69 | 0.93 | 0.63 | 0.63 |
| 4 | 173562564 | rs4695771 | 1.043 | 0.309 | 0.491 | -0.131 | -0.1 | -0.027 | -0.027 | 0 | 0.39 | 0.12 | 0.42 | 0.67 | 0.52 | 0.56 |
| 4 | 173597016 | rs1437842 | -1.206 | 0.372 | 0.677 | -0.138 | -0.017 | 0.011 | 0.011 | 0 | 0.59 | 0.76 | 0.56 | 0.33 | 0.48 | 0.42 |
| 4 | 173598033 | rs10520242 | -0.897 | 0.105 | 0.948 | 0.187 | 0.043 | 0.171 | 0.171 | 0 | 0.06 | 0.14 | 0.27 | 0.26 | 0.1 | 0.06 |
| 4 | 173614720 | rs6848466 | 0.118 | -0.046 | 0.972 | 0.076 | -0.031 | -0.012 | -0.012 | 0 | 0.09 | 0.15 | 0.05 | 0.05 | 0.01 | 0.12 |
| 4 | 173632325 | rs1946797 | 1.311 | 0.009 | 1.326 | 0.146 | 0.042 | 0.191 | 0.191 | 0 | 0.91 | 0.76 | 0.71 | 0.74 | 0.88 | 0.91 |
| 4 | 173637709 | rs9312527 | 1.643 | -0.022 | 1.432 | 0.131 | 0.054 | 0.235 | 0.235 | 0 | 0.9 | 0.66 | 0.7 | 0.73 | 0.88 | 0.91 |
| 4 | 173643640 | rs17058802 | -1.346 | -0.023 | 1.483 | 0.13 | 0.018 | 0.116 | 0.116 | 0 | 0.09 | 0.17 | 0.27 | 0.27 | 0.12 | 0.09 |
| 4 | 173646842 | rs11132959 | 1.57 | -0.138 | 1.453 | 0.07 | -0.03 | 0.137 | 0.137 | 0 | 0.8 | 0.52 | 0.65 | 0.68 | 0.87 | 0.79 |
| 4 | 173653680 | rs1437845 | -0.15 | -0.126 | 1.202 | -0.581 | -0.001 | -0.079 | -0.079 | 0.585 | 0.39 | 0.42 | 0.23 | 0.02 | 0.32 | 0.22 |
| 4 | 173663511 | rs2083136 | -0.777 | -0.089 | 1.15 | -0.555 | 0.003 | -0.042 | -0.042 | 0 | 0.48 | 0.56 | 0.28 | 0.07 | 0.33 | 0.34 |
| 4 | 173664444 | rs716647 | -0.777 | -0.089 | 1.15 | -0.555 | 0.003 | -0.042 | -0.042 | 1.098 | 0.48 | 0.56 | 0.28 | 0.07 | 0.33 | 0.34 |
| 4 | 173684873 | rs12642442 | 1.213 | 0.018 | 0.673 | -0.366 | -0.071 | 0.05 | 0.05 | 0.073 | 0.41 | 0.13 | 0.42 | 0.71 | 0.54 | 0.57 |
| 4 | 173693980 | rs9997085 | 0.454 | 0.064 | 0.841 | -0.343 | 0 | 0.044 | 0.044 | 0 | 0.52 | 0.38 | 0.48 | 0.77 | 0.55 | 0.72 |
| 4 | 173720434 | rs10019400 | -0.211 | -0.54 | 0.825 | -0.532 | 0.012 | 0.04 | 0.04 | 0 | 0.77 | 0.83 | 0.88 | 0.96 | 0.83 | 0.87 |

| | | | | | | | | | | | | | | | | |
|---|---|---|---|---|---|---|---|---|---|---|---|---|---|---|---|---|
| 4 | 173731762 | rs750376 | -1.353 | -0.469 | 1.384 | -0.212 | 0.067 | 0.198 | 0.198 | 0 | 0.1 | 0.37 | 0.28 | 0.26 | 0.12 | 0.12 |
| 4 | 173739244 | rs1459157 | 1.487 | -0.741 | 0.965 | -0.447 | 0.042 | 0.073 | 0.073 | 0.491 | 0.53 | 0.05 | 0.51 | 0.69 | 0.68 | 0.6 |
| 4 | 173739355 | rs9884518 | 0.264 | -0.728 | 1.186 | -0.459 | -0.027 | -0.015 | -0.015 | 0.057 | 0.11 | 0.13 | 0.05 | 0.04 | 0.01 | 0.12 |
| 4 | 173741357 | rs10007423 | -0.901 | -0.731 | 1.276 | -0.466 | -0.036 | -0.084 | -0.084 | 0.353 | 0.36 | 0.46 | 0.2 | 0.09 | 0.2 | 0.25 |
| 4 | 173742700 | rs1551664 | -1.482 | -0.279 | 1.641 | 0.119 | 0.035 | 0.104 | 0.104 | 0 | 0.09 | 0.24 | 0.27 | 0.22 | 0.12 | 0.12 |
| 4 | 173753757 | rs10033472 | -1.052 | -0.325 | 1.385 | 0.012 | -0.089 | -0.073 | -0.073 | 0 | 0.16 | 0.41 | 0.22 | 0.07 | 0.04 | 0.19 |
| 4 | 173757115 | rs17058946 | -1.033 | -0.325 | 1.368 | 0.013 | -0.088 | -0.068 | -0.068 | 0.067 | 0.16 | 0.4 | 0.22 | 0.07 | 0.03 | 0.19 |
| 4 | 173758268 | rs10001933 | -1.453 | -0.242 | 0.393 | 0.133 | 0.013 | -0.005 | -0.005 | 0 | 0.22 | 0.76 | 0.34 | 0.26 | 0.14 | 0.21 |
| 4 | 173764577 | rs1824347 | 0.405 | 0.504 | -0.855 | -0.87 | -0.056 | 0.192 | 0.192 | 0.217 | 0.41 | 0.11 | 0.62 | 0.89 | 0.54 | 0.66 |
| 4 | 173770121 | rs17058964 | 0.347 | 0.048 | -0.174 | -1.485 | -0.018 | -0.051 | -0.051 | 0.929 | 0.35 | 0.41 | 0.21 | 0.1 | 0.23 | 0.19 |
| 4 | 173770940 | rs6823952 | 0.285 | 0.018 | 0.139 | -1.589 | -0.024 | -0.062 | -0.062 | 0 | 0.14 | 0.15 | 0.07 | 0 | 0.06 | 0.07 |
| 4 | 173777083 | rs6833917 | 0.395 | 0.03 | 0.119 | -1.556 | 0.13 | 0.243 | 0.243 | 0.047 | 0.49 | 0.18 | 0.69 | 0.89 | 0.72 | 0.74 |
| 4 | 173786855 | rs6818634 | 0.818 | 0.318 | -0.151 | -0.73 | 0.004 | 0.021 | 0.021 | 0.05 | 0.45 | 0.14 | 0.51 | 0.6 | 0.6 | 0.61 |
| 4 | 173797032 | rs7660673 | 0.472 | 0.03 | -0.107 | -1.146 | -0.016 | -0.044 | -0.044 | 0.222 | 0.33 | 0.39 | 0.19 | 0.1 | 0.21 | 0.17 |
| 4 | 173797669 | rs12512152 | 0.513 | 0.467 | -0.529 | -0.932 | -0.009 | 0.198 | 0.198 | 0.608 | 0.44 | 0.12 | 0.65 | 0.88 | 0.58 | 0.68 |
| 4 | 173797856 | rs1903126 | 0.154 | 0.119 | -0.284 | -0.829 | 0.068 | 0.094 | 0.094 | 0.181 | 0.52 | 0.32 | 0.72 | 0.9 | 0.74 | 0.76 |
| 4 | 173803914 | rs6810619 | -0.823 | 1.575 | -0.284 | -0.511 | -0.004 | 0.001 | 0.001 | 0 | 0.3 | 0.26 | 0.34 | 0.14 | 0.41 | 0.29 |
| 4 | 173820365 | rs1993616 | 0.072 | 1.641 | -0.107 | 0.042 | -0.005 | -0.004 | -0.004 | 0 | 0.14 | 0.13 | 0.21 | 0.21 | 0.21 | 0.17 |
| 4 | 173822571 | rs2653824 | -1.608 | 1.548 | 0.731 | -0.083 | 0.037 | -0.05 | -0.05 | 0.154 | 0.16 | 0.58 | 0.12 | 0.06 | 0.17 | 0.19 |
| 4 | 173849497 | rs2653831 | -1.444 | 1.361 | 1.113 | -0.104 | 0.058 | -0.07 | -0.07 | 0.767 | 0.13 | 0.43 | 0.09 | 0.02 | 0.13 | 0.14 |
| 4 | 173868291 | rs17254079 | -0.631 | 1.395 | 0.672 | 0.009 | -0.013 | 0.008 | 0.008 | 0.623 | 0.08 | 0.04 | 0.05 | 0.05 | 0.11 | 0.11 |
| 4 | 173872313 | rs13150573 | 0.763 | 0.986 | 0.159 | 0.166 | 0 | 0 | 0 | 0 | 0.53 | 0.47 | 0.53 | 0.53 | 0.51 | 0.58 |
| 4 | 173872401 | rs2653841 | 0.33 | 1.027 | -0.45 | 0.216 | -0.052 | -0.046 | -0.046 | 0 | 0.58 | 0.81 | 0.57 | 0.55 | 0.53 | 0.62 |
| 4 | 173874976 | rs17254336 | 0.6 | 1.053 | -0.588 | 0.432 | -0.063 | -0.01 | -0.01 | 0 | 0.25 | 0.01 | 0.26 | 0.17 | 0.26 | 0.14 |
| 4 | 173885229 | rs4696036 | -0.502 | 1.21 | -0.156 | 0.335 | -0.019 | -0.021 | -0.021 | 0 | 0.42 | 0.29 | 0.44 | 0.46 | 0.47 | 0.38 |
| 4 | 173889191 | rs10520251 | -0.431 | 1.566 | 0.277 | 0.292 | 0.006 | 0 | 0 | 0 | 0.08 | 0.09 | 0.07 | 0.06 | 0.14 | 0.1 |
| 4 | 173914254 | rs6811225 | 0.289 | 1.431 | 0.005 | 0.21 | -0.036 | -0.053 | -0.053 | 0 | 0.13 | 0 | 0.29 | 0.19 | 0.12 | 0.11 |
| 4 | 173916231 | rs4574383 | -1.436 | 1.33 | 0.431 | -0.002 | 0.024 | -0.024 | -0.024 | 0 | 0.14 | 0.26 | 0.11 | 0.06 | 0.2 | 0.13 |
| 4 | 173930848 | rs17059082 | -0.598 | 1.051 | 0.83 | -0.106 | -0.01 | -0.003 | -0.003 | 0.032 | 0.18 | 0.27 | 0.18 | 0.2 | 0.15 | 0.13 |
| 4 | 173935056 | rs6832095 | -1.363 | 1.243 | 1.012 | -0.293 | 0.025 | -0.032 | -0.032 | 0 | 0.34 | 0.54 | 0.43 | 0.29 | 0.39 | 0.18 |
| 4 | 173944939 | rs7659038 | -1.103 | 1.233 | 0.915 | -0.758 | 0.02 | -0.012 | -0.012 | 0 | 0.05 | 0.21 | 0.03 | 0 | 0.03 | 0.01 |
| 4 | 173945730 | rs10020676 | -1.474 | 1.27 | 1.059 | -0.822 | 0.023 | -0.092 | -0.092 | 0 | 0.34 | 0.53 | 0.39 | 0.09 | 0.37 | 0.16 |
| 4 | 173949124 | rs4331761 | -0.725 | 1.346 | 0.474 | -0.81 | -0.018 | 0.025 | 0.025 | 0 | 0.27 | 0.23 | 0.35 | 0.07 | 0.34 | 0.15 |
| 4 | 173949815 | rs10005134 | -1.51 | 0.956 | 0.308 | -0.479 | 0 | -0.028 | -0.028 | 0 | 0.05 | 0.14 | 0.03 | 0 | 0.03 | 0.01 |
| 4 | 173962425 | rs2019683 | 0.537 | 0.505 | -0.103 | -1.042 | 0.022 | 0.031 | 0.031 | 0 | 0.61 | 0.36 | 0.73 | 0.95 | 0.7 | 0.84 |
| 4 | 173963006 | rs11132977 | 0.784 | 0.324 | -0.887 | -0.937 | -0.137 | -0.048 | -0.048 | 0 | 0.36 | 0.06 | 0.55 | 0.35 | 0.48 | 0.37 |
| 4 | 173966463 | rs6817670 | -0.009 | 0.29 | -0.835 | -0.95 | 0.022 | 0.031 | 0.031 | 0 | 0.61 | 0.36 | 0.73 | 0.95 | 0.7 | 0.84 |
| 4 | 173991635 | rs4309818 | 0.671 | 0.354 | -0.778 | -0.9 | -0.098 | -0.025 | -0.025 | 0.021 | 0.37 | 0.1 | 0.56 | 0.34 | 0.48 | 0.37 |
| 4 | 173995332 | rs7667977 | 1.064 | 0.126 | -0.753 | -0.992 | 0.001 | -0.003 | -0.003 | 0.304 | 0.22 | 0.19 | 0.17 | 0.6 | 0.22 | 0.46 |
| 4 | 173995383 | rs4293766 | -0.293 | 0.059 | -0.392 | -1.099 | 0.021 | 0.013 | 0.013 | 0 | 0.66 | 0.76 | 0.79 | 0.96 | 0.73 | 0.87 |
| 4 | 173996293 | rs11725123 | 1.09 | -0.115 | -0.515 | -1.237 | -0.001 | 0.022 | 0.022 | 0 | 0.16 | 0.09 | 0.07 | 0 | 0.15 | 0.08 |
| 4 | 174007048 | rs10018524 | 0.799 | 0.048 | -0.658 | -1.061 | -0.103 | -0.017 | -0.017 | 0 | 0.37 | 0.09 | 0.55 | 0.32 | 0.48 | 0.38 |
| 4 | 174016641 | rs4048413 | -1.247 | -0.073 | -0.546 | -1.144 | 0.001 | -0.009 | -0.009 | 0 | 0.83 | 0.86 | 0.85 | 0.96 | 0.87 | 0.97 |
| 4 | 174020403 | rs11935600 | 0.839 | 0.843 | -0.874 | -0.984 | -0.079 | -0.016 | -0.016 | 0 | 0.35 | 0.08 | 0.46 | 0.31 | 0.33 | 0.37 |
| 4 | 174035061 | rs6856402 | 0.713 | 0.067 | -0.89 | -0.998 | -0.107 | 0.026 | 0.026 | 0 | 0.35 | 0.08 | 0.55 | 0.23 | 0.49 | 0.37 |
| 4 | 174046343 | rs3966013 | 0.259 | -0.088 | -0.448 | -1.176 | -0.001 | -0.043 | -0.043 | 0.61 | 0.23 | 0.24 | 0.1 | 0 | 0.17 | 0.08 |
| 4 | 174048218 | rs13138296 | 0.234 | -0.129 | -0.453 | -1.182 | -0.001 | -0.042 | -0.042 | 0 | 0.23 | 0.24 | 0.1 | 0 | 0.17 | 0.08 |
| 4 | 174050460 | rs10520253 | 1.058 | -0.18 | -0.465 | -0.755 | 0.003 | -0.008 | -0.008 | 0.206 | 0.21 | 0.17 | 0.16 | 0.66 | 0.2 | 0.46 |
| 4 | 174051114 | rs6818771 | -0.716 | -0.006 | -0.298 | -0.967 | 0.016 | 0.018 | 0.018 | 0.038 | 0.83 | 0.9 | 0.85 | 0.94 | 0.87 | 0.96 |
| 4 | 174053361 | rs4356895 | 0.309 | 0.669 | 0.141 | -1.237 | -0.009 | 0.012 | 0.012 | 0 | 0.32 | 0.28 | 0.45 | 0.14 | 0.39 | 0.29 |
| 4 | 174057435 | rs3966014 | 0.381 | 0.76 | -0.085 | -1.501 | 0.013 | 0.102 | 0.102 | 0 | 0.47 | 0.18 | 0.32 | 0.74 | 0.44 | 0.6 |
| 4 | 174061455 | rs6829566 | 0.079 | 0.817 | 0.554 | -1.581 | 0.002 | -0.097 | -0.097 | 0 | 0.18 | 0.3 | 0.13 | 0 | 0.18 | 0.06 |
| 4 | 174062064 | rs716357 | -0.592 | 0.865 | 0.663 | -1.555 | -0.001 | -0.001 | -0.001 | 0 | 0.66 | 0.67 | 0.48 | 0.81 | 0.62 | 0.67 |
| 4 | 174062281 | rs4639046 | -0.528 | 0.746 | 0.569 | -1.679 | -0.004 | -0.017 | -0.017 | 0 | 0.79 | 0.81 | 0.8 | 0.93 | 0.81 | 0.85 |
| 4 | 174063974 | rs6822683 | -0.46 | 0.97 | 0.796 | -1.225 | 0.019 | 0.009 | 0.009 | 0 | 0.7 | 0.58 | 0.48 | 0.72 | 0.64 | 0.67 |
| 4 | 174072292 | rs6553679 | -0.579 | 0.975 | 0.786 | -1.232 | -0.005 | 0.001 | 0.001 | 0 | 0.72 | 0.74 | 0.49 | 0.77 | 0.64 | 0.69 |
| 4 | 174072755 | rs7689781 | -0.247 | 0.92 | 1.089 | -1.393 | 0.008 | -0.069 | -0.069 | 0 | 0.2 | 0.39 | 0.25 | 0.02 | 0.22 | 0.1 |
| 4 | 174075363 | rs11734882 | 0.758 | 0.867 | 1.038 | -1.369 | 0.036 | 0.052 | 0.052 | 0 | 0.49 | 0.19 | 0.23 | 0.69 | 0.41 | 0.56 |
| 4 | 174081834 | rs7669267 | -0.357 | 0.585 | 1.12 | -1.356 | 0.003 | -0.049 | -0.049 | 0 | 0.2 | 0.35 | 0.24 | 0.02 | 0.21 | 0.1 |
| 4 | 174083622 | rs4419455 | -0.69 | 0.766 | 1.101 | -1.256 | -0.001 | 0 | 0 | 1.605 | 0.74 | 0.75 | 0.53 | 0.77 | 0.7 | 0.69 |
| 4 | 174089080 | rs1054496 | -0.403 | 0.734 | 1.195 | -1.233 | 0.006 | 0.017 | 0.017 | 0 | 0.72 | 0.58 | 0.52 | 0.72 | 0.7 | 0.66 |
| 4 | 174092791 | rs1006003 | 0.382 | 0.821 | 1.096 | -1.346 | 0.01 | -0.012 | -0.012 | 0 | 0.58 | 0.51 | 0.39 | 0.72 | 0.5 | 0.58 |
| 4 | 174098017 | rs722827 | 0.197 | 0.551 | 1.199 | -1.369 | -0.012 | 0.024 | 0.024 | 0 | 0.16 | 0.11 | 0.14 | 0 | 0.18 | 0.13 |
| 4 | 174101050 | rs11942280 | -0.793 | 0.528 | 1.228 | -1.325 | 0.004 | -0.067 | -0.067 | 0 | 0.2 | 0.36 | 0.23 | 0.05 | 0.2 | 0.16 |
| 4 | 174107537 | rs13151627 | -0.92 | 0.523 | 1.26 | -1.319 | 0.002 | -0.039 | -0.039 | 0 | 0.21 | 0.46 | 0.23 | 0.11 | 0.2 | 0.19 |
| 4 | 174119049 | rs11732929 | 0.8 | 0.314 | 0.814 | -1.531 | -0.001 | 0.039 | 0.039 | 0 | 0.14 | 0.07 | 0.06 | 0 | 0.13 | 0.06 |
| 4 | 174127031 | rs4535319 | 0.789 | 0.106 | 1.187 | -1.281 | -0.009 | 0.037 | 0.037 | 0 | 0.53 | 0.25 | 0.3 | 0.68 | 0.51 | 0.5 |
| 4 | 174129740 | rs10213525 | 0.754 | 0.103 | 1.186 | -1.283 | -0.009 | 0.034 | 0.034 | 0 | 0.53 | 0.26 | 0.3 | 0.68 | 0.51 | 0.5 |
| 4 | 174130503 | rs11737748 | 0.904 | 0.071 | 1.163 | -1.3 | -0.049 | 0.016 | 0.016 | 0.273 | 0.23 | 0.07 | 0.38 | 0.15 | 0.3 | 0.28 |

| | | | | | | | | | | | | | | | | |
|---|---|---|---|---|---|---|---|---|---|---|---|---|---|---|---|---|
| 4 | 174132146 | rs10001613 | 0.675 | 0.036 | 1.396 | -1.304 | 0 | -0.005 | -0.005 | 0 | 0.56 | 0.43 | 0.37 | 0.79 | 0.52 | 0.56 |
| 4 | 174136952 | rs9995556 | 0.846 | 0.059 | 1.271 | -1.295 | -0.01 | -0.057 | -0.057 | 0.022 | 0.77 | 0.42 | 0.69 | 0.89 | 0.81 | 0.8 |
| 4 | 174146299 | rs4608791 | 0.631 | 0.049 | 1.104 | -1.303 | -0.012 | -0.094 | -0.094 | 0.004 | 0.78 | 0.51 | 0.76 | 0.94 | 0.81 | 0.83 |
| 4 | 174156383 | rs4696054 | 1.155 | 0.037 | 0.586 | -1.262 | -0.006 | -0.019 | -0.019 | 0 | 0.74 | 0.28 | 0.68 | 0.88 | 0.8 | 0.76 |
| 4 | 174156538 | rs12639929 | 0.537 | -0.209 | 0.739 | -0.798 | -0.049 | -0.029 | -0.029 | 0 | 0.23 | 0.07 | 0.38 | 0.21 | 0.3 | 0.27 |
| 4 | 174165126 | rs7672108 | -0.016 | -0.179 | 0.855 | -0.976 | -0.003 | -0.011 | -0.011 | 0.799 | 0.69 | 0.38 | 0.47 | 0.77 | 0.69 | 0.64 |
| 4 | 174186322 | rs10471199 | -1.975 | -0.286 | 1.624 | -0.713 | -0.024 | 0.074 | 0.074 | 1.511 | 0.12 | 0.56 | 0.1 | 0.11 | 0.06 | 0.14 |
| 4 | 174190222 | rs13128039 | -0.036 | -0.607 | 0.903 | -1.024 | -0.048 | 0.201 | 0.201 | 0 | 0.38 | 0.07 | 0.3 | 0.12 | 0.4 | 0.39 |
| 4 | 174199490 | rs6816923 | -0.549 | -0.293 | 1.328 | -0.673 | -0.008 | -0.009 | -0.009 | 0 | 0.05 | 0.24 | 0.04 | 0.01 | 0.03 | 0.08 |
| 4 | 174205734 | rs6553691 | -1.405 | -0.327 | 1.204 | -0.675 | -0.019 | 0.017 | 0.017 | 0.357 | 0.08 | 0.38 | 0.06 | 0.04 | 0.03 | 0.06 |
| 4 | 174210358 | rs6825703 | 0.922 | -0.38 | 0.935 | -0.725 | 0.007 | 0.1 | 0.1 | 0.152 | 0.49 | 0.05 | 0.53 | 0.71 | 0.57 | 0.43 |
| 4 | 174222981 | rs3775591 | -2.113 | -0.274 | 1.365 | -0.621 | -0.049 | 0.03 | 0.03 | 1.576 | 0.13 | 0.64 | 0.09 | 0.12 | 0.06 | 0.12 |
| 4 | 174232666 | rs6842693 | -1.889 | -0.261 | 1.371 | -0.619 | -0.039 | 0.045 | 0.045 | 1.493 | 0.13 | 0.53 | 0.07 | 0.11 | 0.06 | 0.12 |
| 4 | 174235624 | rs7658148 | 0.933 | -0.215 | 0.812 | -0.583 | 0.054 | 0.119 | 0.119 | 0 | 0.5 | 0.12 | 0.56 | 0.69 | 0.57 | 0.44 |
| 4 | 174243631 | rs3775590 | -0.589 | 0.069 | 0.636 | -0.258 | -0.007 | 0.061 | 0.061 | 0 | 0.05 | 0.16 | 0.02 | 0.1 | 0.02 | 0.07 |
| 4 | 174262102 | rs12498911 | -0.181 | 0.215 | 0.159 | -0.56 | -0.001 | 0.036 | 0.036 | 0 | 0.67 | 0.79 | 0.64 | 0.83 | 0.66 | 0.58 |
| 4 | 174271800 | rs4696060 | 0.609 | 0.646 | 0.413 | -0.195 | 0.072 | 0.031 | 0.031 | 0 | 0.95 | 0.75 | 0.93 | 0.97 | 0.93 | 0.94 |
| 4 | 174301717 | rs12498508 | 0.422 | 0.41 | -0.954 | 0.22 | -0.05 | -0.04 | -0.04 | 0.01 | 0.43 | 0.1 | 0.42 | 0.6 | 0.48 | 0.38 |
| 4 | 174315913 | rs2101092 | 0.587 | 0.463 | -0.916 | 0.383 | -0.076 | -0.075 | -0.075 | 0 | 0.43 | 0.1 | 0.34 | 0.57 | 0.49 | 0.3 |
| 4 | 174323228 | rs13148735 | 0.743 | 0.369 | -0.204 | 0.32 | -0.109 | -0.092 | -0.092 | 0 | 0.41 | 0.08 | 0.34 | 0.56 | 0.5 | 0.29 |
| 4 | 174327618 | rs7655786 | 0.143 | -0.034 | 0.611 | 0.099 | -0.003 | 0.004 | 0.004 | 0 | 0.54 | 0.35 | 0.43 | 0.57 | 0.55 | 0.4 |
| 4 | 174344177 | rs13147839 | 1.137 | -0.001 | 0.171 | 0.2 | -0.113 | -0.066 | -0.066 | 0 | 0.42 | 0.08 | 0.3 | 0.62 | 0.53 | 0.31 |
| 4 | 174347475 | rs2086477 | 1.099 | 0 | 0.226 | 0.201 | -0.082 | -0.041 | -0.041 | 0 | 0.42 | 0.12 | 0.3 | 0.62 | 0.53 | 0.3 |
| 4 | 174348397 | rs1481121 | -0.477 | 0.215 | 0.585 | 0.552 | -0.011 | 0.013 | 0.013 | 0 | 0.44 | 0.26 | 0.58 | 0.35 | 0.43 | 0.55 |
| 4 | 174353610 | rs35033382 | -0.723 | 0.206 | 0.645 | 0.596 | -0.017 | 0.005 | 0.005 | 0 | 0.45 | 0.26 | 0.62 | 0.35 | 0.44 | 0.55 |
| 4 | 174366240 | rs1946968 | 1.266 | 0.095 | 0.994 | 0.749 | 0.023 | -0.016 | -0.016 | 0.181 | 0.56 | 0.29 | 0.59 | 0.61 | 0.65 | 0.39 |
| 4 | 174369546 | rs17059455 | -0.081 | 0.127 | 0.629 | 0.794 | -0.008 | -0.003 | -0.003 | 0 | 0.05 | 0.03 | 0.06 | 0.06 | 0.08 | 0.07 |
| 4 | 174369908 | rs1561792 | 0.057 | -0.066 | 0.615 | 0.638 | -0.012 | 0.066 | 0.066 | 1.204 | 0.23 | 0.09 | 0.15 | 0 | 0.2 | 0.19 |
| 4 | 174371557 | rs17059458 | -0.344 | 0.243 | 0.65 | 0.875 | 0.02 | 0.038 | 0.038 | 0 | 0.06 | 0.14 | 0.07 | 0.1 | 0.09 | 0.08 |
| 4 | 174376198 | rs7659018 | -0.691 | -0.075 | 0.788 | 0.782 | -0.002 | 0.001 | 0.001 | 0 | 0.36 | 0.37 | 0.32 | 0.32 | 0.26 | 0.52 |
| 4 | 174392507 | rs6419984 | -0.886 | 0.018 | 0.781 | 0.616 | 0 | 0 | 0 | 0.454 | 0.38 | 0.37 | 0.32 | 0.29 | 0.3 | 0.52 |
| 4 | 174398324 | rs10520259 | 0.814 | 0.076 | 0.402 | 0.645 | 0.034 | 0.016 | 0.016 | 0 | 0.59 | 0.25 | 0.64 | 0.7 | 0.7 | 0.47 |
| 4 | 174401970 | rs4496557 | 0.712 | 0.323 | 0.079 | 0.766 | -0.016 | -0.006 | -0.006 | 0 | 0.61 | 0.48 | 0.66 | 0.71 | 0.7 | 0.5 |
| 4 | 174425236 | rs13141534 | 0.416 | 0.727 | -0.596 | 0.878 | -0.033 | 0.006 | 0.006 | 0 | 0.66 | 0.49 | 0.73 | 0.71 | 0.77 | 0.53 |
| 4 | 174426144 | rs1020231 | 0.431 | 0.613 | -0.611 | 0.85 | -0.009 | 0.018 | 0.018 | 0 | 0.64 | 0.48 | 0.66 | 0.66 | 0.7 | 0.5 |
| 4 | 174437070 | rs7661281 | -0.002 | 0.419 | -1.617 | 0.713 | -0.056 | -0.028 | -0.028 | 0.012 | 0.25 | 0.01 | 0.18 | 0.28 | 0.23 | 0.21 |
| 4 | 174438227 | rs6820150 | -0.824 | 0.292 | -0.965 | 0.736 | -0.006 | -0.007 | -0.007 | 0.409 | 0.19 | 0.39 | 0.17 | 0.17 | 0.17 | 0.15 |
| 4 | 174438897 | rs7655560 | 0.022 | 0.257 | -0.983 | 0.626 | 0 | -0.015 | -0.015 | 0 | 0.08 | 0.07 | 0.04 | 0 | 0.07 | 0.06 |
| 4 | 174442919 | rs12644553 | 0.455 | -0.064 | -1.463 | 0.48 | -0.02 | 0.031 | 0.031 | 0 | 0.29 | 0.03 | 0.17 | 0.26 | 0.21 | 0.23 |
| 4 | 174443496 | rs17059527 | -0.097 | 0.105 | -1.133 | 0.645 | 0.006 | 0.003 | 0.003 | 0 | 0.07 | 0.09 | 0.05 | 0.1 | 0.1 | 0.06 |
| 4 | 174447349 | rs10520260 | 0.297 | 0.755 | -1.207 | 1.029 | -0.019 | -0.016 | -0.016 | 0 | 0.16 | 0.13 | 0.41 | 0.22 | 0.28 | 0.24 |
| 4 | 174447362 | rs13141356 | 1.042 | 0.319 | -1.648 | 1.115 | -0.009 | -0.084 | -0.084 | 0 | 0.19 | 0.04 | 0.1 | 0.2 | 0.18 | 0.25 |
| 4 | 174454083 | rs6813810 | -0.477 | 0.199 | -0.686 | 1.198 | -0.015 | 0.043 | 0.043 | 0 | 0.74 | 0.61 | 0.84 | 0.57 | 0.79 | 0.66 |
| 4 | 174455297 | rs2332691 | -1.382 | 0.412 | -0.501 | 1.448 | -0.001 | -0.005 | -0.005 | 0.015 | 0.85 | 0.84 | 0.89 | 0.79 | 0.84 | 0.79 |
| 4 | 174455685 | rs2877766 | -0.786 | 0.656 | 0.501 | 1.395 | 0.027 | 0.01 | 0.01 | 0 | 0.29 | 0.41 | 0.52 | 0.28 | 0.41 | 0.35 |
| 4 | 174456068 | rs13114604 | -0.794 | 0.288 | 0.584 | 1.46 | 0.018 | 0.017 | 0.017 | 0.553 | 0.28 | 0.43 | 0.5 | 0.28 | 0.36 | 0.32 |
| 4 | 174456843 | rs4695845 | -0.883 | 0.29 | 0.599 | 1.431 | 0.018 | 0.05 | 0.05 | 0.423 | 0.28 | 0.43 | 0.51 | 0.37 | 0.36 | 0.39 |
| 4 | 174458901 | rs13131421 | -1.714 | 0.271 | 0.52 | 1.376 | -0.009 | 0 | 0 | 0 | 0.47 | 0.64 | 0.6 | 0.48 | 0.52 | 0.53 |
| 4 | 174459823 | rs744724 | -1.499 | -0.834 | 0.883 | 0.904 | -0.003 | -0.002 | -0.002 | 0 | 0.34 | 0.27 | 0.19 | 0.3 | 0.31 | 0.41 |
| 4 | 174461343 | rs6820651 | -0.78 | -0.541 | 0.464 | 1.003 | -0.009 | 0.018 | 0.018 | 0.209 | 0.22 | 0.34 | 0.11 | 0.26 | 0.2 | 0.31 |
| 4 | 174471671 | rs4695849 | 0.827 | -0.268 | 0.421 | 1.212 | -0.028 | 0.009 | 0.009 | 4.317 | 0.83 | 0.63 | 0.88 | 0.87 | 0.9 | 0.82 |
| 4 | 174476098 | rs11132992 | -1.319 | 0.048 | -0.364 | 1.505 | -0.004 | 0.02 | 0.02 | 0.369 | 0.21 | 0.26 | 0.19 | 0.46 | 0.16 | 0.3 |
| 4 | 174484121 | rs4695853 | -1.224 | 0.263 | -0.222 | 1.031 | 0.009 | 0.078 | 0.078 | 0 | 0.07 | 0.23 | 0.14 | 0.17 | 0.05 | 0.1 |
| 4 | 174497836 | rs4490433 | -0.072 | 0.223 | -1.32 | 0.184 | 0.046 | 0.137 | 0.137 | 1.439 | 0.47 | 0.15 | 0.6 | 0.76 | 0.62 | 0.62 |
| 4 | 174502457 | rs4695857 | 0.913 | 0.415 | -1.44 | 0.41 | -0.005 | -0.04 | -0.04 | 0.018 | 0.74 | 0.22 | 0.49 | 0.63 | 0.74 | 0.73 |
| 4 | 174507822 | rs7681766 | 0.747 | 0.257 | -1.035 | -0.194 | 0.001 | 0.007 | 0.007 | 0 | 0.49 | 0.43 | 0.41 | 0.62 | 0.56 | 0.46 |
| 4 | 174509660 | rs17328110 | 0.801 | 0.441 | -1.534 | -0.197 | -0.05 | -0.043 | -0.043 | 0 | 0.11 | 0.01 | 0.33 | 0.15 | 0.16 | 0.13 |
| 4 | 174509911 | rs4160 | -0.261 | 0.289 | -1.425 | -0.288 | 0.016 | 0.081 | 0.081 | 0 | 0.35 | 0.1 | 0.25 | 0.22 | 0.27 | 0.37 |
| 4 | 174513213 | rs13136539 | -0.194 | 0.28 | -1.378 | -0.415 | 0.006 | 0.077 | 0.077 | 0.1 | 0.34 | 0.11 | 0.25 | 0.2 | 0.27 | 0.37 |
| 4 | 174515682 | rs12646107 | 0.837 | 0.445 | -1.162 | -0.299 | -0.017 | -0.019 | -0.019 | 0 | 0.1 | 0.04 | 0.1 | 0.13 | 0.14 | 0.12 |
| 4 | 174516590 | rs11132998 | 0.875 | 0.453 | -1.227 | -0.292 | -0.037 | -0.038 | -0.038 | 0 | 0.1 | 0.01 | 0.09 | 0.13 | 0.14 | 0.12 |
| 4 | 174519735 | rs11724478 | -0.417 | 0.109 | -0.894 | -0.547 | 0.01 | -0.009 | -0.009 | 0.331 | 0.34 | 0.11 | 0.27 | 0.32 | 0.27 | 0.34 |
| 4 | 174521315 | rs4639045 | 0.442 | 0.144 | -0.784 | -0.601 | -0.006 | 0.008 | 0.008 | 0 | 0.13 | 0.07 | 0.08 | 0.01 | 0.14 | 0.14 |
| 4 | 174530500 | rs13110724 | -0.421 | 0.097 | -0.844 | -0.595 | 0.013 | -0.012 | -0.012 | 0 | 0.34 | 0.11 | 0.26 | 0.32 | 0.27 | 0.35 |
| 4 | 174530652 | rs11945239 | -0.421 | 0.097 | -0.844 | -0.595 | 0.013 | -0.012 | -0.012 | 0 | 0.34 | 0.11 | 0.26 | 0.32 | 0.27 | 0.35 |
| 4 | 174530927 | rs11731252 | 1.17 | 0.025 | -0.154 | -0.637 | 0.009 | 0.03 | 0.03 | 0 | 0.49 | 0.42 | 0.63 | 0.67 | 0.59 | 0.52 |
| 4 | 174540510 | rs13143445 | -0.356 | 0.058 | -0.323 | -0.603 | 0.014 | -0.014 | -0.014 | 0 | 0.34 | 0.11 | 0.26 | 0.32 | 0.27 | 0.35 |
| 4 | 174555529 | rs13111144 | 0.2 | 0.393 | -0.205 | -0.483 | -0.002 | -0.012 | -0.012 | 0.47 | 0.13 | 0.1 | 0.08 | 0.01 | 0.14 | 0.13 |
| 4 | 174562083 | rs4359880 | 1.003 | 0.146 | -0.446 | -0.56 | -0.001 | 0.016 | 0.016 | 1.514 | 0.48 | 0.44 | 0.39 | 0.66 | 0.53 | 0.45 |

| | | | | | | | | | | | | | | | | |
|---|---|---|---|---|---|---|---|---|---|---|---|---|---|---|---|---|
| 4 | 174565021 | rs4282145 | 0.67 | 0.294 | -0.89 | -0.455 | 0.015 | 0.021 | 0.021 | 0 | 0.62 | 0.32 | 0.43 | 0.67 | 0.68 | 0.62 |
| 4 | 174571498 | rs11725485 | 0.234 | 0.672 | -1.129 | -0.399 | -0.004 | -0.004 | -0.004 | 0.008 | 0.1 | 0.11 | 0.08 | 0.07 | 0.08 | 0.09 |
| 4 | 174573033 | rs17263178 | 0.334 | 0.946 | -1.481 | -0.66 | -0.035 | 0.044 | 0.044 | 0 | 0.1 | 0.01 | 0.08 | 0.01 | 0.12 | 0.09 |
| 4 | 174578566 | rs11723222 | -0.203 | 0.752 | -1.194 | -0.43 | 0.002 | 0.003 | 0.003 | 0.017 | 0.13 | 0.11 | 0.11 | 0.07 | 0.13 | 0.1 |
| 4 | 174578616 | rs11728325 | -0.165 | 0.765 | -1.429 | -0.424 | 0.008 | 0.022 | 0.022 | 0.013 | 0.13 | 0.06 | 0.11 | 0.07 | 0.13 | 0.1 |
| 4 | 174579326 | rs17263395 | 0.196 | 1.118 | -1.138 | -0.2 | -0.011 | 0.002 | 0.002 | 0 | 0.14 | 0.07 | 0.38 | 0.18 | 0.18 | 0.17 |
| 4 | 174580179 | rs12647892 | 0.03 | 1.321 | -1.049 | 0.127 | -0.034 | -0.019 | -0.019 | 0.115 | 0.09 | 0.01 | 0.07 | 0.13 | 0.13 | 0.1 |
| 4 | 174591579 | rs6841699 | 0.016 | 1.109 | -0.962 | 0.005 | -0.045 | -0.025 | -0.025 | 3.424 | 0.32 | 0.13 | 0.5 | 0.37 | 0.39 | 0.41 |
| 4 | 174596609 | rs10018807 | 0.975 | 0.776 | -0.368 | -0.413 | 0.003 | -0.011 | -0.011 | 0.372 | 0.41 | 0.33 | 0.25 | 0.08 | 0.33 | 0.29 |
| 4 | 174599459 | rs4615152 | 0.012 | 0.625 | -0.045 | -0.129 | -0.002 | -0.007 | -0.007 | 0 | 0.8 | 0.8 | 0.83 | 0.49 | 0.77 | 0.75 |
| 4 | 174602048 | rs11727767 | -0.378 | 0.6 | 0.052 | -0.621 | 0.003 | 0.024 | 0.024 | 0 | 0.49 | 0.59 | 0.64 | 0.37 | 0.51 | 0.56 |
| 4 | 174603650 | rs17263985 | 0.991 | 0.542 | -0.117 | -0.538 | -0.058 | -0.005 | -0.005 | 0 | 0.11 | 0.01 | 0.1 | 0.09 | 0.19 | 0.12 |
| 4 | 174604104 | rs7698692 | -0.193 | 0.534 | 0.013 | -0.505 | -0.028 | 0.285 | 0.285 | 0 | 0.91 | 0.71 | 0.87 | 0.57 | 0.96 | 0.88 |
| 4 | 174605859 | rs17264076 | 1.822 | 0.486 | -0.071 | -0.543 | -0.002 | 0.095 | 0.095 | 0.775 | 0.3 | 0.07 | 0.18 | 0.12 | 0.26 | 0.2 |
| 4 | 174605978 | rs13142579 | 0.329 | 0.314 | 0.437 | -0.387 | 0.01 | -0.063 | -0.063 | 1.134 | 0.13 | 0.24 | 0.09 | 0.01 | 0.14 | 0.15 |
| 4 | 174606591 | rs12649191 | 1.017 | 0.153 | 0.197 | -0.447 | -0.079 | -0.07 | -0.07 | 0.009 | 0.28 | 0.08 | 0.48 | 0.36 | 0.36 | 0.41 |
| 4 | 174609530 | rs6813832 | 1.063 | 0.118 | 0.676 | -0.188 | 0.013 | -0.045 | -0.045 | 0.097 | 0.09 | 0.16 | 0.06 | 0 | 0.09 | 0.09 |
| 4 | 174609693 | rs13132889 | 0.398 | -0.003 | 0.413 | 0.127 | -0.026 | 0.328 | 0.328 | 0.052 | 0.9 | 0.66 | 0.86 | 0.57 | 0.96 | 0.88 |
| 4 | 174620399 | rs9312547 | 0.2 | 0.284 | -0.13 | -0.847 | -0.004 | 0.076 | 0.076 | 0 | 0.44 | 0.25 | 0.31 | 0.21 | 0.45 | 0.33 |
| 4 | 174621493 | rs12499670 | -0.493 | 0.454 | -0.25 | -0.805 | 0 | -0.02 | -0.02 | 0 | 0.55 | 0.56 | 0.72 | 0.86 | 0.61 | 0.7 |
| 4 | 174623476 | rs4695873 | -0.025 | 0.61 | -0.291 | -0.627 | -0.003 | 0.02 | 0.02 | 0 | 0.45 | 0.4 | 0.63 | 0.78 | 0.46 | 0.58 |
| 4 | 174630567 | rs11133012 | 0.809 | 0.669 | -0.724 | -0.557 | 0.003 | 0.119 | 0.119 | 0.018 | 0.5 | 0.19 | 0.65 | 0.69 | 0.54 | 0.5 |
| 4 | 174631643 | rs4401434 | 0.653 | 0.496 | -0.115 | -0.812 | -0.013 | 0.001 | 0.001 | 0.347 | 0.14 | 0.25 | 0.11 | 0.14 | 0.08 | 0.07 |
| 4 | 174631819 | rs4434222 | -0.123 | 0.563 | 0.458 | -1.006 | 0.004 | -0.055 | -0.055 | 0 | 0.75 | 0.6 | 0.84 | 0.92 | 0.71 | 0.74 |
| 4 | 174632969 | rs4631014 | -0.163 | 0.563 | 0.38 | -1.002 | 0.003 | -0.002 | -0.002 | 2.015 | 0.19 | 0.33 | 0.17 | 0.22 | 0.15 | 0.2 |
| 4 | 174635674 | rs4364235 | 1.716 | 0.303 | -0.236 | -0.942 | -0.052 | -0.002 | -0.002 | 0.51 | 0.4 | 0.12 | 0.55 | 0.62 | 0.44 | 0.43 |
| 4 | 174642677 | rs6816095 | 0.674 | 0.355 | 0.192 | -1.158 | 0.006 | -0.063 | -0.063 | 0 | 0.71 | 0.63 | 0.82 | 0.92 | 0.7 | 0.68 |
| 4 | 174643499 | rs6553719 | -0.607 | 0.366 | 0.226 | -1.11 | 0.001 | -0.057 | -0.057 | 1.443 | 0.81 | 0.76 | 0.9 | 1 | 0.8 | 0.86 |
| 4 | 174655914 | rs4521323 | 1.232 | 0.24 | -0.017 | -1.008 | 0 | 0.034 | 0.034 | 0 | 0.57 | 0.32 | 0.67 | 0.77 | 0.54 | 0.55 |
| 4 | 174657554 | rs7692617 | -1.077 | 0.122 | -0.071 | -1.014 | -0.012 | -0.058 | -0.058 | 0.149 | 0.32 | 0.42 | 0.22 | 0.12 | 0.25 | 0.33 |
| 4 | 174657851 | rs7694565 | 0.772 | -0.112 | -0.446 | -1.019 | -0.001 | 0 | 0 | 0 | 0.33 | 0.31 | 0.48 | 0.66 | 0.32 | 0.36 |
| 4 | 174659094 | rs17264373 | 1.086 | -0.077 | -0.074 | -0.924 | -0.016 | 0.026 | 0.026 | 0.005 | 0.1 | 0.04 | 0.03 | 0 | 0.1 | 0.04 |
| 4 | 174659937 | rs10520263 | -0.341 | -0.08 | -0.064 | -0.928 | 0.004 | -0.078 | -0.078 | 0 | 0.18 | 0.37 | 0.12 | 0.06 | 0.13 | 0.21 |
| 4 | 174667332 | rs6830691 | -1.592 | 0.047 | -0.21 | -0.895 | -0.008 | 0.024 | 0.024 | 0 | 0.7 | 0.76 | 0.75 | 0.92 | 0.66 | 0.82 |
| 4 | 174670065 | rs6813034 | -0.191 | 0.077 | -0.65 | -0.834 | -0.002 | 0.054 | 0.054 | 0 | 0.53 | 0.38 | 0.63 | 0.86 | 0.53 | 0.61 |
| 4 | 174679198 | rs13108803 | -0.302 | -0.06 | -0.402 | -0.272 | 0.002 | -0.013 | -0.013 | 0 | 0.6 | 0.49 | 0.65 | 0.76 | 0.54 | 0.65 |
| 4 | 174684932 | rs11929802 | -0.471 | -0.088 | -0.787 | -0.046 | -0.006 | -0.005 | -0.005 | 0 | 0.25 | 0.33 | 0.18 | 0.23 | 0.2 | 0.37 |
| 4 | 174686346 | rs4695885 | 0.725 | -0.183 | -1.53 | 0.132 | -0.036 | 0.017 | 0.017 | 0 | 0.3 | 0.1 | 0.43 | 0.33 | 0.32 | 0.21 |
| 4 | 174686458 | rs6553727 | -0.015 | -0.441 | -0.802 | 0.325 | 0.004 | -0.007 | -0.007 | 0.191 | 0.17 | 0.14 | 0.12 | 0.18 | 0.14 | 0.19 |
| 4 | 174701186 | rs12506204 | -0.063 | -0.449 | -1.066 | -0.03 | -0.009 | 0.029 | 0.029 | 0 | 0.32 | 0.09 | 0.24 | 0.18 | 0.3 | 0.36 |
| 4 | 174713351 | rs3922692 | -0.762 | -0.468 | -0.609 | 0.035 | 0.042 | 0.056 | 0.056 | 0.083 | 0.51 | 0.67 | 0.4 | 0.33 | 0.38 | 0.57 |
| 4 | 174723835 | rs4695886 | -1.106 | -0.803 | -0.403 | -0.225 | 0.042 | 0.082 | 0.082 | 0 | 0.47 | 0.72 | 0.33 | 0.17 | 0.37 | 0.42 |
| 4 | 174729270 | rs7689307 | 0.013 | -0.612 | -0.99 | -0.129 | -0.011 | 0.03 | 0.03 | 0.224 | 0.31 | 0.06 | 0.24 | 0.17 | 0.31 | 0.36 |
| 4 | 174731952 | rs7683033 | -1.04 | -0.397 | 0.152 | 0.08 | -0.013 | -0.053 | -0.053 | 0 | 0.1 | 0.25 | 0.05 | 0 | 0.06 | 0.05 |
| 4 | 174741938 | rs4615153 | 0.977 | -0.612 | 0.295 | 0.365 | -0.051 | 0.007 | 0.007 | 0 | 0.79 | 0.52 | 0.84 | 0.82 | 0.84 | 0.79 |
| 4 | 174751009 | rs6833126 | 0.71 | -0.561 | -0.352 | 0.193 | 0.041 | 0.092 | 0.092 | 0 | 0.49 | 0.24 | 0.57 | 0.72 | 0.55 | 0.42 |
| 4 | 174760546 | rs4544692 | 0.512 | -0.489 | -0.7 | 0.016 | -0.042 | 0.025 | 0.025 | 0.018 | 0.21 | 0.04 | 0.18 | 0.16 | 0.28 | 0.19 |
| 4 | 174772568 | rs6813541 | 1.166 | -0.778 | -0.159 | 0.848 | 0.017 | 0.038 | 0.038 | 0 | 0.49 | 0.33 | 0.63 | 0.44 | 0.54 | 0.38 |
| 4 | 174777061 | rs7700171 | -1.212 | -0.762 | 0.2 | 1.373 | -0.005 | 0.285 | 0.285 | 1.53 | 0.09 | 0.31 | 0.05 | 0.33 | 0.03 | 0.1 |
| 4 | 174777578 | rs13138013 | 0.18 | -0.562 | -0.563 | 1.035 | 0.001 | -0.02 | -0.02 | 0 | 0.17 | 0.04 | 0.14 | 0.14 | 0.15 | 0.31 |
| 4 | 174794014 | rs6553733 | -0.256 | -0.486 | -0.886 | 0.555 | 0.064 | 0.108 | 0.108 | 0 | 0.45 | 0.76 | 0.32 | 0.26 | 0.38 | 0.42 |
| 4 | 174800374 | rs6827215 | -1.025 | 0.247 | -1.193 | 0.778 | -0.038 | -0.035 | -0.035 | 0 | 0.66 | 0.81 | 0.5 | 0.42 | 0.64 | 0.62 |
| 4 | 174802918 | rs4600896 | 0.204 | 0.205 | -1.259 | 0.528 | -0.044 | -0.024 | -0.024 | 0.11 | 0.19 | 0.05 | 0.2 | 0.16 | 0.25 | 0.22 |
| 4 | 174803040 | rs9997597 | 0.216 | 0.003 | -0.95 | 0.02 | 0.024 | 0.02 | 0.02 | 0 | 0.41 | 0.66 | 0.27 | 0.1 | 0.33 | 0.36 |
| 4 | 174803471 | rs10471202 | -0.695 | 0.122 | -0.788 | 0.766 | -0.014 | -0.015 | -0.015 | 0 | 0.28 | 0.2 | 0.25 | 0.58 | 0.33 | 0.38 |
| 4 | 174817063 | rs4358372 | -0.857 | -0.124 | 0.383 | 0.553 | -0.018 | 0.05 | 0.05 | 0 | 0.1 | 0.35 | 0.07 | 0.16 | 0.06 | 0.16 |
| 4 | 174817710 | rs4695894 | -0.203 | 0 | 0.043 | 0.339 | -0.001 | 0 | 0 | 0 | 0.47 | 0.57 | 0.58 | 0.48 | 0.46 | 0.55 |
| 4 | 174819012 | rs4333160 | 0.016 | 0.153 | -0.353 | 0.226 | 0.004 | -0.025 | -0.025 | 0 | 0.86 | 0.79 | 0.9 | 0.94 | 0.88 | 0.89 |
| 4 | 174819200 | rs4398502 | 0.639 | 0.171 | -0.808 | 0.255 | -0.054 | 0.018 | 0.018 | 0 | 0.37 | 0.19 | 0.51 | 0.32 | 0.41 | 0.47 |
| 4 | 174819487 | rs4615154 | 0.18 | 0.209 | -0.51 | 0.519 | 0.037 | 0.039 | 0.039 | 0 | 0.5 | 0.28 | 0.6 | 0.4 | 0.53 | 0.47 |
| 4 | 174820018 | rs6824200 | -0.04 | 0.639 | -0.613 | 0.586 | 0 | -0.001 | -0.001 | 0.018 | 0.56 | 0.55 | 0.68 | 0.55 | 0.55 | 0.65 |
| 4 | 174822905 | rs7657936 | 0.323 | 0.842 | -1.119 | 0.345 | 0.004 | -0.011 | -0.011 | 0 | 0.16 | 0.12 | 0.35 | 0.2 | 0.13 | 0.17 |
| 4 | 174824576 | rs10033133 | -0.427 | 0.91 | -1.163 | 0.213 | -0.005 | 0.003 | 0.003 | 0.145 | 0.44 | 0.5 | 0.47 | 0.49 | 0.42 | 0.32 |
| 4 | 174826120 | rs4330337 | -0.665 | 1.293 | -1.34 | 0.591 | -0.038 | -0.071 | -0.071 | 0 | 0.79 | 0.86 | 0.69 | 0.59 | 0.7 | 0.77 |
| 4 | 174826157 | rs3887907 | 0.133 | 0.937 | -1.112 | 0.677 | -0.002 | -0.002 | -0.002 | 0 | 0.27 | 0.25 | 0.13 | 0.27 | 0.25 | 0.12 |
| 4 | 174831241 | rs7700127 | -0.102 | 0.67 | -1.02 | 0.44 | -0.004 | -0.007 | -0.007 | 0.337 | 0.45 | 0.49 | 0.49 | 0.32 | 0.36 | 0.59 |
| 4 | 174837887 | rs4695895 | -0.229 | 0.575 | -1.489 | 0.456 | -0.029 | 0.057 | 0.057 | 0.067 | 0.55 | 0.8 | 0.48 | 0.64 | 0.48 | 0.51 |
| 4 | 174850559 | rs15884 | 0.321 | 0.542 | -1.345 | 1.046 | -0.012 | -0.009 | -0.009 | 2.558 | 0.24 | 0.16 | 0.28 | 0.37 | 0.32 | 0.19 |
| 4 | 174852035 | rs13107942 | 0.568 | -0.131 | -2.099 | 0.88 | -0.114 | 0.04 | 0.04 | 0 | 0.38 | 0.06 | 0.28 | 0.27 | 0.49 | 0.31 |

| | | | | | | | | | | | | | | | | |
|---|---|---|---|---|---|---|---|---|---|---|---|---|---|---|---|---|
| 4 | 174853695 | rs9999393 | 0.566 | -0.793 | -0.242 | 1.286 | -0.014 | 0.106 | 0.106 | 1.408 | 0.93 | 0.64 | 0.92 | 0.89 | 1 | 0.9 |
| 4 | 174854581 | rs7670348 | -0.418 | -0.78 | -0.366 | 1.314 | 0.002 | 0.003 | 0.003 | 0 | 0.2 | 0.09 | 0.27 | 0.17 | 0.17 | 0.36 |
| 4 | 174858993 | rs12500903 | -1.379 | -0.935 | 0.566 | 1.326 | -0.058 | 0.145 | 0.145 | 1.249 | 0.16 | 0.41 | 0.24 | 0.34 | 0.08 | 0.22 |
| 4 | 174868363 | rs6858266 | -0.11 | -0.665 | 0.693 | 1.168 | 0.02 | -0.008 | -0.008 | 0 | 0.16 | 0.28 | 0.25 | 0.12 | 0.17 | 0.23 |
| 4 | 174873969 | rs7657182 | -0.849 | -0.902 | 0.373 | 0.925 | 0.011 | -0.027 | -0.027 | 0 | 0.23 | 0.2 | 0.26 | 0.36 | 0.18 | 0.29 |
| 4 | 174894345 | rs6845317 | 0.303 | -0.821 | 0.715 | 0.859 | 0.002 | -0.046 | -0.046 | 0 | 0.79 | 0.5 | 0.74 | 0.9 | 0.81 | 0.77 |
| 4 | 174900688 | rs11934296 | -0.194 | -0.839 | 1.035 | 0.849 | 0.018 | -0.023 | -0.023 | 0 | 0.17 | 0.34 | 0.29 | 0.1 | 0.17 | 0.23 |
| 4 | 174906286 | rs7659538 | 0.317 | -0.838 | 1.029 | 0.85 | 0.008 | -0.049 | -0.049 | 0 | 0.8 | 0.52 | 0.69 | 0.9 | 0.81 | 0.77 |
| 4 | 174908521 | rs13119610 | 0.793 | -1.072 | 0.071 | 0.781 | -0.111 | 0.119 | 0.119 | 4.86 | 0.45 | 0.06 | 0.3 | 0.19 | 0.52 | 0.29 |
| 4 | 174920657 | rs4495020 | 0.559 | -1.152 | 0.387 | 0.852 | -0.013 | 0.108 | 0.108 | 3.385 | 0.48 | 0.22 | 0.32 | 0.22 | 0.52 | 0.33 |
| 4 | 174928071 | rs4235203 | -0.211 | -1.201 | 0.807 | 1.169 | -0.003 | -0.005 | -0.005 | 0 | 0.42 | 0.4 | 0.56 | 0.62 | 0.44 | 0.61 |
| 4 | 174937446 | rs6823093 | 1.597 | -1.105 | 0.572 | 1.197 | -0.012 | -0.032 | -0.032 | 0 | 0.2 | 0.12 | 0.18 | 0.36 | 0.22 | 0.23 |
| 4 | 174945087 | rs13144752 | 1.699 | -1.237 | 0.272 | 0.907 | -0.12 | 0.008 | 0.008 | 7.964 | 0.28 | 0.01 | 0.24 | 0.13 | 0.36 | 0.21 |
| 4 | 174947684 | rs6812370 | -0.653 | -1.299 | 1.213 | 1.472 | -0.003 | 0.136 | 0.136 | 0.645 | 0.06 | 0.21 | 0.04 | 0.15 | 0.02 | 0.02 |
| 4 | 174956412 | rs10520267 | 0.133 | -1.316 | 0.989 | 1.401 | -0.024 | 0.007 | 0.007 | 0.974 | 0.65 | 0.47 | 0.66 | 0.31 | 0.72 | 0.59 |
| 4 | 174961164 | rs11133026 | 1.038 | -1.242 | 0.183 | 1.394 | -0.145 | 0.1 | 0.1 | 1.938 | 0.44 | 0.04 | 0.35 | 0.2 | 0.51 | 0.32 |
| 4 | 174963224 | rs4695759 | -0.131 | -1.251 | 0.58 | 1.406 | -0.018 | 0.022 | 0.022 | 0 | 0.35 | 0.54 | 0.33 | 0.52 | 0.3 | 0.4 |
| 4 | 174964203 | rs17060079 | -1.12 | -1.336 | 0.275 | 1.355 | -0.055 | 0.067 | 0.067 | 1.067 | 0.13 | 0.56 | 0.14 | 0.17 | 0.04 | 0.16 |
| 4 | 174965215 | rs13128730 | 1.049 | -1.324 | -0.899 | 1.323 | -0.112 | 0.111 | 0.111 | 4.41 | 0.44 | 0.07 | 0.35 | 0.21 | 0.51 | 0.29 |
| 4 | 174966483 | rs1540448 | -1.298 | -1.375 | -0.909 | 1.416 | -0.046 | -0.048 | -0.048 | 0 | 0.77 | 0.97 | 0.81 | 0.65 | 0.75 | 0.74 |
| 4 | 174969224 | rs10020606 | 0.486 | -1.348 | -0.791 | 1.474 | -0.01 | -0.006 | -0.006 | 1.008 | 0.14 | 0.1 | 0.26 | 0.12 | 0.16 | 0.27 |
| 4 | 174984230 | rs9996330 | -1.096 | -1.316 | -1.034 | 1.41 | 0.048 | -0.102 | -0.102 | 3.729 | 0.31 | 0.86 | 0.44 | 0.44 | 0.21 | 0.43 |
| 4 | 174987671 | rs7670262 | -1.031 | -1.328 | -0.906 | 1.175 | -0.016 | 0.109 | 0.109 | 2.691 | 0.53 | 0.79 | 0.54 | 0.76 | 0.47 | 0.7 |
| 4 | 174991468 | rs6553765 | 0.66 | -0.792 | -0.714 | 1.333 | -0.032 | -0.022 | -0.022 | 2.44 | 0.24 | 0.11 | 0.21 | 0.37 | 0.26 | 0.28 |
| 4 | 174996402 | rs2196368 | -0.155 | -0.974 | -0.706 | 1.461 | 0.002 | -0.043 | -0.043 | 0.94 | 0.83 | 0.91 | 0.89 | 0.73 | 0.84 | 0.61 |
| 4 | 175000272 | rs13118468 | -0.216 | -0.895 | -0.358 | 1.205 | -0.025 | -0.041 | -0.041 | 0 | 0.58 | 0.79 | 0.68 | 0.39 | 0.58 | 0.34 |
| 4 | 175002057 | rs2332756 | 0.303 | -0.798 | -0.369 | 1.244 | -0.027 | -0.017 | -0.017 | 0.266 | 0.37 | 0.18 | 0.31 | 0.48 | 0.38 | 0.51 |
| 4 | 175006016 | rs877019 | -0.592 | -0.789 | -0.574 | 0.857 | -0.103 | 0.019 | 0.019 | 0 | 0.61 | 0.88 | 0.46 | 0.74 | 0.51 | 0.69 |
| 4 | 175008790 | rs4695907 | -0.254 | -0.555 | 0.022 | 0.917 | -0.001 | -0.001 | -0.001 | 0.147 | 0.48 | 0.55 | 0.6 | 0.38 | 0.55 | 0.52 |
| 4 | 175017835 | rs11133030 | 0.792 | -0.661 | 0.349 | 1.214 | -0.057 | 0.041 | 0.041 | 0.109 | 0.81 | 0.39 | 0.89 | 0.72 | 0.9 | 0.83 |
| 4 | 175022693 | rs6834670 | -0.584 | -0.026 | 0.013 | 1.139 | 0.001 | 0.004 | 0.004 | 0 | 0.43 | 0.46 | 0.38 | 0.39 | 0.46 | 0.39 |
| 4 | 175025442 | rs11946672 | 0.532 | 0.194 | -0.099 | 1.104 | -0.001 | 0 | 0 | 0 | 0.44 | 0.4 | 0.38 | 0.5 | 0.43 | 0.42 |
| 4 | 175031632 | rs2196387 | 0.769 | 0.687 | 0.334 | 1.56 | -0.006 | 0.049 | 0.049 | 0 | 0.77 | 0.58 | 0.69 | 0.69 | 0.83 | 0.66 |
| 4 | 175033463 | rs6841986 | 1.54 | 0.88 | 0.183 | 1.855 | 0.002 | 0.051 | 0.051 | 0 | 0.68 | 0.47 | 0.59 | 0.52 | 0.69 | 0.43 |
| 4 | 175033659 | rs1992021 | 0.252 | 0.9 | 0.191 | 2.042 | -0.02 | -0.03 | -0.03 | 0 | 0.09 | 0.04 | 0.1 | 0.18 | 0.11 | 0.23 |
| 4 | 175035169 | rs10015990 | -1.193 | 0.881 | 0.5 | 1.977 | 0.009 | 0.018 | 0.018 | 0 | 0.17 | 0.27 | 0.3 | 0.2 | 0.2 | 0.21 |
| 4 | 175035811 | rs7657736 | 1.037 | 0.859 | 0.506 | 1.988 | -0.056 | 0.041 | 0.041 | 0.025 | 0.83 | 0.53 | 0.78 | 0.79 | 0.94 | 0.76 |
| 4 | 175039764 | rs2332758 | 0.945 | 0.929 | 0.377 | 1.874 | 0.009 | 0.028 | 0.028 | 0 | 0.9 | 0.81 | 0.89 | 0.83 | 0.85 | 0.78 |
| 4 | 175041257 | rs6838064 | 1.136 | 0.955 | 0.359 | 1.805 | 0.007 | 0.015 | 0.015 | 0 | 0.82 | 0.77 | 0.79 | 0.73 | 0.75 | 0.67 |
| 4 | 175045002 | rs6838591 | -0.985 | 0.861 | 0.255 | 1.807 | -0.023 | 0.086 | 0.086 | 0.905 | 0.11 | 0.4 | 0.05 | 0.19 | 0.04 | 0.2 |
| 4 | 175046985 | rs10520270 | 0.273 | 0.915 | 0.081 | 1.622 | -0.027 | -0.019 | -0.019 | 0.207 | 0.06 | 0.01 | 0.1 | 0.07 | 0.14 | 0.1 |
| 4 | 175053051 | rs10002205 | 0.887 | 0.614 | 0.263 | 1.586 | 0.001 | 0.056 | 0.056 | 0 | 0.68 | 0.52 | 0.75 | 0.54 | 0.68 | 0.45 |
| 4 | 175057033 | rs17060221 | 0.318 | 0.487 | -0.042 | 1.72 | 0 | 0.006 | 0.006 | 0.009 | 0.11 | 0.1 | 0.17 | 0.35 | 0.09 | 0.07 |
| 4 | 175057725 | rs2196381 | -0.59 | 0.714 | 0.078 | 1.055 | 0 | -0.003 | -0.003 | 0 | 0.83 | 0.79 | 0.68 | 0.91 | 0.77 | 0.87 |
| 4 | 175059383 | rs4695912 | -1.291 | 0.736 | 0.316 | 0.941 | 0.003 | -0.003 | -0.003 | 0.448 | 0.57 | 0.54 | 0.75 | 0.84 | 0.67 | 0.73 |
| 4 | 175060620 | rs6827803 | -0.848 | 0.739 | 0.58 | 1.197 | 0.019 | 0.002 | 0.002 | 0 | 0.35 | 0.43 | 0.42 | 0.67 | 0.41 | 0.42 |
| 4 | 175061541 | rs17334566 | 2.454 | 0.53 | 0.274 | 0.959 | -0.033 | 0.165 | 0.165 | 0 | 0.32 | 0.02 | 0.19 | 0.06 | 0.27 | 0.18 |
| 4 | 175062207 | rs1992017 | 0.812 | 0.449 | 0.373 | 1.077 | -0.044 | -0.001 | -0.001 | 0.013 | 0.2 | 0.01 | 0.33 | 0.1 | 0.26 | 0.13 |
| 4 | 175063224 | rs7662489 | -0.398 | 0.601 | 0.751 | 1.244 | -0.003 | -0.023 | -0.023 | 0.028 | 0.33 | 0.22 | 0.54 | 0.57 | 0.37 | 0.37 |
| 4 | 175063668 | rs1992016 | 0.838 | 0.478 | 1.195 | 1.39 | -0.004 | 0.028 | 0.028 | 1.355 | 0.89 | 0.51 | 0.94 | 0.91 | 0.94 | 0.91 |
| 4 | 175065347 | rs17060286 | -0.797 | 0.691 | 1.141 | 1.578 | 0.025 | 0.02 | 0.02 | 0 | 0.25 | 0.39 | 0.21 | 0.29 | 0.29 | 0.36 |
| 4 | 175065821 | rs2196380 | -1.594 | 0.592 | 1.255 | 1.721 | 0.002 | 0.006 | 0.006 | 0 | 0.52 | 0.56 | 0.57 | 0.61 | 0.59 | 0.73 |
| 4 | 175066056 | rs4342175 | -0.262 | 0.654 | 0.595 | 1.764 | -0.019 | -0.018 | -0.018 | 0.443 | 0.23 | 0.18 | 0.19 | 0.29 | 0.29 | 0.36 |
| 4 | 175066104 | rs10520272 | -1.038 | 0.616 | 0.583 | 1.741 | -0.011 | -0.014 | -0.014 | 0.024 | 0.31 | 0.3 | 0.29 | 0.62 | 0.36 | 0.4 |
| 4 | 175067059 | rs6851647 | -1.525 | 0.614 | 0.59 | 1.753 | -0.013 | -0.001 | -0.001 | 0 | 0.36 | 0.36 | 0.45 | 0.7 | 0.4 | 0.45 |
| 4 | 175067665 | rs2217855 | -0.629 | 0.669 | 0.844 | 1.771 | 0.043 | 0.036 | 0.036 | 0 | 0.25 | 0.51 | 0.22 | 0.32 | 0.3 | 0.37 |
| 4 | 175067806 | rs10520273 | 0.975 | 0.645 | -0.315 | 1.765 | 0.018 | 0.003 | 0.003 | 1.624 | 0.6 | 0.25 | 0.5 | 0.36 | 0.62 | 0.57 |
| 4 | 175068695 | rs749493 | 0.178 | 0.66 | -0.317 | 1.723 | 0 | 0.03 | 0.03 | 0 | 0.1 | 0.13 | 0.1 | 0.33 | 0.07 | 0.05 |
| 4 | 175068856 | rs1865656 | 2.08 | 0.769 | -1.012 | 1.605 | -0.019 | 0.198 | 0.198 | 0 | 0.37 | 0.05 | 0.31 | 0.06 | 0.33 | 0.22 |
| 4 | 175069861 | rs873372 | 0.44 | 1.033 | -0.694 | 2.072 | -0.085 | -0.059 | -0.059 | 0.436 | 0.15 | 0.02 | 0.13 | 0.21 | 0.23 | 0.32 |
| 4 | 175070243 | rs6836227 | 0.97 | 0.712 | -0.188 | 2.008 | 0.017 | -0.037 | -0.037 | 0 | 0.64 | 0.27 | 0.66 | 0.43 | 0.66 | 0.62 |
| 4 | 175072660 | rs6855522 | -0.189 | 1.006 | -0.714 | 1.464 | 0.006 | -0.022 | -0.022 | 0 | 0.22 | 0.1 | 0.19 | 0.2 | 0.26 | 0.3 |
| 4 | 175074766 | rs4695772 | 0.085 | 1.109 | -0.796 | 1.393 | 0.002 | 0.007 | 0.007 | 0.705 | 0.75 | 0.92 | 0.84 | 0.83 | 0.72 | 0.68 |
| 4 | 175076940 | rs11733331 | 1.657 | 1.247 | -0.838 | 1.474 | 0.019 | 0.11 | 0.11 | 0 | 0.45 | 0.21 | 0.37 | 0.21 | 0.4 | 0.29 |
| 4 | 175077652 | rs11734349 | 0.207 | 1.276 | -1.021 | 1.552 | -0.043 | -0.007 | -0.007 | 0 | 0.23 | 0.02 | 0.15 | 0.11 | 0.27 | 0.14 |
| 4 | 175080533 | rs17276390 | -0.201 | 1.369 | -0.685 | 1.904 | -0.024 | -0.018 | -0.018 | 0 | 0.2 | 0.19 | 0.31 | 0.3 | 0.26 | 0.33 |
| 4 | 175084416 | rs2196379 | -0.058 | 1.122 | -0.252 | 1.69 | 0.079 | 0.337 | 0.337 | 0.365 | 0.92 | 0.58 | 0.85 | 0.68 | 0.93 | 0.95 |
| 4 | 175086079 | rs10003923 | 1.395 | 1.155 | -0.648 | 1.653 | 0.015 | 0.093 | 0.093 | 0 | 0.46 | 0.27 | 0.38 | 0.21 | 0.41 | 0.28 |
| 4 | 175086260 | rs4695913 | -1.193 | 1.351 | -1.408 | 1.439 | -0.014 | -0.018 | -0.018 | 0.022 | 0.46 | 0.31 | 0.46 | 0.47 | 0.53 | 0.67 |

| | | | | | | | | | | | | | | | | |
|---|---|---|---|---|---|---|---|---|---|---|---|---|---|---|---|---|
| 4 | 175089750 | rs2164380 | -0.355 | 1.022 | -1.758 | 0.967 | -0.035 | 0.001 | 0.001 | 0 | 0.25 | 0.01 | 0.23 | 0.13 | 0.29 | 0.16 |
| 4 | 175094250 | rs7657652 | 0.763 | 1.904 | -0.832 | 1.593 | 0.006 | 0.014 | 0.014 | 0 | 0.77 | 0.77 | 0.69 | 0.54 | 0.64 | 0.71 |
| 4 | 175097637 | rs7661496 | 0.631 | 1.848 | -0.73 | 1.208 | -0.001 | 0 | 0 | 0 | 0.92 | 0.92 | 0.95 | 0.93 | 0.92 | 0.93 |
| 4 | 175103590 | rs6828239 | -0.913 | 1.617 | -0.577 | 0.967 | 0.004 | -0.029 | -0.029 | 0 | 0.37 | 0.47 | 0.54 | 0.24 | 0.44 | 0.32 |
| 4 | 175105239 | rs6553774 | -0.895 | 1.78 | -0.737 | 1.483 | 0.004 | 0.039 | 0.039 | 0 | 0.3 | 0.45 | 0.23 | 0.46 | 0.36 | 0.22 |
| 4 | 175112520 | rs11942884 | -0.928 | 1.984 | -1.143 | 1.275 | -0.03 | 0.014 | 0.014 | 0 | 0.07 | 0.02 | 0.11 | 0.02 | 0.09 | 0.2 |
| 4 | 175113110 | rs17060350 | -0.485 | 1.858 | -1.313 | 1.498 | -0.024 | -0.033 | -0.033 | 0 | 0.08 | 0.02 | 0.1 | 0.12 | 0.12 | 0.06 |
| 4 | 175114675 | rs1467003 | -0.941 | 1.326 | -0.844 | 1.295 | -0.006 | 0.005 | 0.005 | 0 | 0.41 | 0.42 | 0.51 | 0.51 | 0.44 | 0.3 |
| 4 | 175115254 | rs1467005 | 0.174 | 0.913 | -0.605 | 1.437 | -0.031 | 0.046 | 0.046 | 0.405 | 0.76 | 0.59 | 0.61 | 0.63 | 0.77 | 0.83 |
| 4 | 175120513 | rs2174904 | 0.14 | 1.056 | -1.118 | 1.188 | 0.002 | 0.048 | 0.048 | 0 | 0.1 | 0.15 | 0.08 | 0.22 | 0.09 | 0.07 |
| 4 | 175121923 | rs12651020 | -0.474 | 1.183 | -1.139 | 1.009 | -0.006 | 0.011 | 0.011 | 0.072 | 0.34 | 0.62 | 0.49 | 0.51 | 0.36 | 0.36 |
| 4 | 175122454 | rs11947112 | 0.324 | 1.801 | -1.444 | 0.544 | -0.058 | 0.041 | 0.041 | 0 | 0.8 | 0.92 | 0.71 | 0.89 | 0.72 | 0.72 |
| 4 | 175125617 | rs4235204 | 0.322 | 1.218 | -1.103 | 0.555 | 0.003 | -0.005 | -0.005 | 0 | 0.59 | 0.48 | 0.43 | 0.64 | 0.59 | 0.55 |
| 4 | 175126378 | rs12651414 | -0.592 | 0.181 | -1.26 | 0.44 | 0.01 | -0.019 | -0.019 | 0 | 0.35 | 0.32 | 0.32 | 0.47 | 0.23 | 0.5 |
| 4 | 175133901 | rs1032988 | -1 | -0.053 | -1.841 | -0.013 | 0.047 | -0.055 | -0.055 | 0 | 0.49 | 0.9 | 0.43 | 0.49 | 0.35 | 0.62 |
| 4 | 175135244 | rs4695914 | -0.049 | -0.574 | -1.478 | -0.545 | -0.023 | 0.001 | 0.001 | 0.17 | 0.67 | 0.65 | 0.72 | 0.3 | 0.77 | 0.72 |
| 4 | 175135624 | rs970096 | 1.082 | -0.442 | -1.605 | -0.997 | -0.139 | 0.185 | 0.185 | 0.515 | 0.32 | 0.09 | 0.38 | 0.02 | 0.51 | 0.31 |
| 4 | 175139380 | rs954926 | -0.202 | -1.09 | -0.408 | -1.003 | -0.021 | -0.014 | -0.014 | 0.087 | 0.34 | 0.4 | 0.31 | 0.74 | 0.27 | 0.36 |
| 4 | 175143493 | rs13150997 | 1.742 | -0.179 | -0.697 | -1.226 | -0.055 | 0.088 | 0.088 | 0.295 | 0.26 | 0.11 | 0.2 | 0.04 | 0.37 | 0.23 |
| 4 | 175150481 | rs11736418 | 0.392 | -0.179 | -0.56 | -1.17 | 0.055 | 0.124 | 0.124 | 1.839 | 0.06 | 0.21 | 0.12 | 0.15 | 0.06 | 0.12 |
| 4 | 175157743 | rs17357120 | 0.777 | -0.548 | -0.37 | -1.496 | -0.004 | -0.03 | -0.03 | 0 | 0.38 | 0.37 | 0.28 | 0.02 | 0.48 | 0.28 |
| 4 | 175158412 | rs14278 | 1.295 | -0.577 | -0.412 | -1.502 | -0.022 | 0.033 | 0.033 | 0 | 0.06 | 0.01 | 0.05 | 0.02 | 0.09 | 0.06 |
| 4 | 175162767 | rs7664501 | 0.245 | -0.587 | -0.283 | -1.486 | -0.001 | -0.005 | -0.005 | 0.254 | 0.09 | 0.1 | 0.06 | 0.04 | 0.11 | 0.08 |
| 4 | 175167253 | rs17060428 | 1.422 | -0.706 | -0.351 | -1.39 | 0.01 | -0.034 | -0.034 | 0 | 0.22 | 0.12 | 0.24 | 0.59 | 0.18 | 0.13 |
| 4 | 175168083 | rs6824643 | 2.191 | -0.661 | -0.305 | -1.367 | -0.075 | 0.126 | 0.126 | 0.622 | 0.26 | 0.07 | 0.19 | 0.02 | 0.37 | 0.21 |
| 4 | 175168153 | rs6851312 | 1.869 | -0.604 | -0.273 | -1.34 | -0.014 | 0.018 | 0.018 | 0.842 | 0.28 | 0.24 | 0.21 | 0.02 | 0.37 | 0.21 |
| 4 | 175170526 | rs13109634 | 0.012 | -0.54 | -0.185 | -1.272 | -0.003 | -0.015 | -0.015 | 0 | 0.65 | 0.61 | 0.54 | 0.76 | 0.72 | 0.64 |
| 4 | 175183493 | rs10520276 | 1.015 | -0.508 | -0.58 | -1.295 | -0.067 | 0.02 | 0.02 | 0 | 0.21 | 0.01 | 0.29 | 0.09 | 0.17 | 0.19 |
| 4 | 175184671 | rs17060434 | 0.311 | -0.477 | -0.478 | -1.244 | -0.001 | 0 | 0 | 0 | 0.05 | 0.09 | 0.02 | 0.14 | 0.05 | 0.23 |
| 4 | 175202610 | rs11730346 | 1.012 | -0.493 | -0.564 | -1.305 | -0.067 | 0.02 | 0.02 | 0 | 0.21 | 0.01 | 0.29 | 0.09 | 0.17 | 0.19 |
| 4 | 175203977 | rs7685970 | 0.228 | -0.565 | -0.377 | -1.212 | -0.001 | 0 | 0 | 0 | 0.06 | 0.09 | 0.02 | 0.13 | 0.05 | 0.23 |
| 4 | 175204676 | rs11555602 | -0.269 | -0.687 | -0.452 | -1.347 | 0.001 | 0 | 0 | 0 | 0.15 | 0.1 | 0.1 | 0.01 | 0.14 | 0.11 |
| 4 | 175219813 | rs13101580 | 0.834 | -0.725 | -0.127 | -1.196 | -0.013 | 0.054 | 0.054 | 1.176 | 0.4 | 0.33 | 0.28 | 0.03 | 0.48 | 0.29 |
| 4 | 175225452 | rs4695918 | 0.07 | -0.701 | -0.281 | -1.142 | 0 | -0.026 | -0.026 | 2.435 | 0.48 | 0.58 | 0.32 | 0.16 | 0.55 | 0.52 |
| 4 | 175230896 | rs17060455 | -0.381 | -0.666 | -0.235 | -1.119 | -0.004 | 0.007 | 0.007 | 0 | 0.07 | 0.12 | 0.03 | 0.13 | 0.07 | 0.23 |
| 4 | 175234130 | rs4695920 | 0.02 | -0.657 | -0.441 | -1.082 | -0.013 | 0.151 | 0.151 | 0.017 | 0.52 | 0.78 | 0.34 | 0.16 | 0.56 | 0.52 |
| 4 | 175237408 | rs1553669 | -0.321 | -0.038 | -0.389 | -0.639 | 0 | -0.006 | -0.006 | 0 | 0.68 | 0.7 | 0.56 | 0.74 | 0.74 | 0.64 |
| 4 | 175246203 | rs6553781 | 0.176 | 0.221 | -0.7 | -0.359 | 0.01 | -0.004 | -0.004 | 1.876 | 0.4 | 0.49 | 0.36 | 0.59 | 0.48 | 0.33 |
| 4 | 175249433 | rs17060468 | 0.329 | -0.08 | -1.706 | -0.61 | 0.045 | 0.055 | 0.055 | 0 | 0.29 | 0.16 | 0.13 | 0.16 | 0.21 | 0.27 |
| 4 | 175270804 | rs11934405 | 0.697 | 0.676 | -1.228 | -0.205 | -0.117 | 0.139 | 0.139 | 1.883 | 0.26 | 0.01 | 0.21 | 0.02 | 0.38 | 0.21 |
| 4 | 175273562 | rs4695924 | -0.31 | 0.791 | -1.045 | 0.252 | -0.039 | -0.056 | -0.056 | 0.087 | 0.37 | 0.17 | 0.34 | 0.46 | 0.46 | 0.31 |
| 4 | 175282806 | rs4467538 | 0.319 | 0.713 | -0.927 | -0.109 | -0.011 | 0.047 | 0.047 | 1.85 | 0.31 | 0.19 | 0.23 | 0.02 | 0.39 | 0.22 |
| 4 | 175300338 | rs6553785 | -0.237 | 1.169 | -0.718 | 0.739 | 0.007 | 0.097 | 0.097 | 0.114 | 0.09 | 0.09 | 0.16 | 0.52 | 0.13 | 0.11 |
| 4 | 175340298 | rs4518223 | -0.524 | 0.675 | -0.609 | 0.298 | -0.007 | 0.001 | 0.001 | 0 | 0.47 | 0.66 | 0.46 | 0.54 | 0.53 | 0.33 |
| 4 | 175345619 | rs4499680 | 0.361 | 0.991 | -0.285 | -0.059 | -0.002 | -0.014 | -0.014 | 0 | 0.21 | 0.22 | 0.15 | 0.01 | 0.3 | 0.17 |
| 4 | 175347444 | rs4484269 | 0.193 | 0.811 | 0.011 | -0.043 | 0.008 | 0.047 | 0.047 | 0.199 | 0.79 | 0.69 | 0.77 | 0.34 | 0.81 | 0.68 |
| 4 | 175351787 | rs7664475 | 1.03 | 0.727 | 0.515 | -0.071 | -0.004 | 0.011 | 0.011 | 0.021 | 0.57 | 0.38 | 0.6 | 0.34 | 0.51 | 0.5 |
| 4 | 175351902 | rs7669690 | -0.954 | 0.716 | 0.577 | 0.252 | 0.006 | 0.072 | 0.072 | 0 | 0.92 | 0.82 | 0.95 | 0.86 | 0.95 | 0.81 |
| 4 | 175352929 | rs6833543 | 0.197 | 1.062 | 0.876 | 0.149 | -0.004 | 0.01 | 0.01 | 0 | 0.66 | 0.33 | 0.75 | 0.85 | 0.61 | 0.63 |
| 4 | 175354852 | rs13135838 | 0.525 | 1.1 | 0.683 | 0.239 | -0.013 | -0.014 | -0.014 | 0 | 0.12 | 0.04 | 0.18 | 0.53 | 0.14 | 0.13 |
| 4 | 175358647 | rs12498292 | 0.99 | 0.98 | 0.795 | 0.038 | -0.005 | 0.047 | 0.047 | 0.238 | 0.54 | 0.26 | 0.56 | 0.32 | 0.47 | 0.5 |
| 4 | 175372799 | rs1035804 | 0.441 | 0.584 | 0.543 | -0.307 | -0.007 | -0.02 | -0.02 | 0 | 0.57 | 0.6 | 0.51 | 0.3 | 0.66 | 0.51 |
| 4 | 175375824 | rs12649767 | 1.689 | 0.54 | 0.396 | -0.389 | -0.022 | 0.019 | 0.019 | 0 | 0.31 | 0.12 | 0.29 | 0.29 | 0.33 | 0.34 |
| 4 | 175376440 | rs2555622 | 0.453 | 0.613 | 0.393 | -0.363 | -0.003 | -0.01 | -0.01 | 0 | 0.57 | 0.65 | 0.5 | 0.3 | 0.65 | 0.51 |
| 4 | 175384927 | rs1426936 | 0.262 | 0.527 | 0.181 | -0.157 | -0.006 | -0.014 | -0.014 | 0 | 0.31 | 0.23 | 0.38 | 0.5 | 0.42 | 0.29 |
| 4 | 175388125 | rs3846297 | 0.865 | 0.371 | 0.111 | -0.028 | 0.027 | 0.071 | 0.071 | 0 | 0.23 | 0.06 | 0.22 | 0.07 | 0.16 | 0.17 |
| 4 | 175389785 | rs6827776 | 0.928 | 0.529 | 0.258 | 0.171 | -0.01 | -0.001 | -0.001 | 0 | 0.1 | 0.04 | 0.21 | 0.49 | 0.12 | 0.13 |
| 4 | 175390534 | rs1426946 | -0.291 | 0.591 | 0.564 | 0.07 | -0.014 | -0.017 | -0.017 | 0.501 | 0.64 | 0.5 | 0.68 | 0.79 | 0.75 | 0.62 |
| 4 | 175411953 | rs9312555 | 0.263 | 0.407 | 0.264 | 0.154 | -0.004 | -0.001 | -0.001 | 1.42 | 0.14 | 0.11 | 0.32 | 0.53 | 0.16 | 0.15 |
| 4 | 175412477 | rs8752 | 0.548 | 0.52 | 0.384 | 0.018 | 0.005 | 0.089 | 0.089 | 0 | 0.49 | 0.31 | 0.36 | 0.28 | 0.58 | 0.45 |
| 4 | 175415991 | rs13106936 | 0.329 | 0.46 | 0.58 | 0.029 | 0 | 0.024 | 0.024 | 0 | 0.14 | 0.13 | 0.31 | 0.53 | 0.16 | 0.15 |
| 4 | 175416331 | rs3857075 | -0.742 | 0.394 | 0.466 | 0.02 | -0.026 | -0.026 | -0.026 | 0 | 0.67 | 0.52 | 0.71 | 0.81 | 0.77 | 0.61 |
| 4 | 175421464 | rs11133041 | 0.496 | 0.501 | -0.928 | 0.07 | -0.014 | 0.106 | 0.106 | 0 | 0.48 | 0.2 | 0.36 | 0.29 | 0.58 | 0.45 |
| 4 | 175422289 | rs2612656 | -0.972 | 0.568 | -1.4 | 0.118 | 0.017 | 0.016 | 0.016 | 0 | 0.69 | 0.82 | 0.74 | 0.81 | 0.77 | 0.61 |
| 4 | 175424172 | rs7349744 | 0.9 | 1.101 | -1.832 | -0.152 | -0.031 | -0.019 | -0.019 | 0.102 | 0.3 | 0.07 | 0.21 | 0.27 | 0.3 | 0.31 |
| 4 | 175428732 | rs7673685 | 0.381 | 1.296 | -1.038 | -0.047 | -0.028 | 0.04 | 0.04 | 0 | 0.16 | 0.09 | 0.13 | 0.01 | 0.27 | 0.15 |
| 4 | 175429808 | rs3775977 | -0.229 | 0.603 | -1.223 | 0.413 | 0.012 | -0.024 | -0.024 | 0 | 0.32 | 0.23 | 0.32 | 0.68 | 0.26 | 0.35 |
| 4 | 175433338 | rs2612659 | -0.128 | 1.055 | -0.755 | -0.66 | 0.006 | -0.094 | -0.094 | 0 | 0.29 | 0.47 | 0.19 | 0.02 | 0.3 | 0.11 |
| 4 | 175434074 | rs11722919 | 0.675 | 1.101 | -0.872 | -0.705 | -0.045 | 0.106 | 0.106 | 0 | 0.22 | 0.07 | 0.11 | 0.01 | 0.25 | 0.1 |

| | | | | | | | | | | | | | | | | |
|---|---|---|---|---|---|---|---|---|---|---|---|---|---|---|---|---|
| 4 | 175434162 | rs2555632 | -0.997 | 1.034 | -0.68 | -1.294 | 0.003 | -0.061 | -0.061 | 0 | 0.73 | 0.67 | 0.68 | 0.94 | 0.72 | 0.61 |
| 4 | 175434520 | rs3797012 | 2.019 | 0.743 | -0.335 | -0.696 | -0.006 | 0.024 | 0.024 | 0.032 | 0.12 | 0.17 | 0.13 | 0.23 | 0.1 | 0.18 |
| 4 | 175437117 | rs2081654 | 0.36 | 0.689 | -0.704 | -0.47 | 0.002 | 0.04 | 0.04 | 0 | 0.24 | 0.27 | 0.3 | 0.65 | 0.23 | 0.35 |
| 4 | 175437621 | rs2555645 | 0.395 | 0.5 | -0.658 | -0.201 | -0.019 | 0.014 | 0.014 | 0.133 | 0.25 | 0.33 | 0.23 | 0.27 | 0.2 | 0.14 |
| 4 | 175443156 | rs1050145 | -0.11 | 1.065 | -1.225 | -0.236 | -0.049 | -0.036 | -0.036 | 0 | 0.35 | 0.26 | 0.32 | 0.46 | 0.41 | 0.27 |
| 4 | 175444382 | rs1426945 | -0.147 | 1.175 | -1.28 | -0.572 | -0.013 | 0.043 | 0.043 | 2.51 | 0.57 | 0.23 | 0.48 | 0.66 | 0.54 | 0.41 |
| 4 | 175454164 | rs17361009 | 0.523 | 1.568 | -1.04 | -0.285 | -0.008 | 0.014 | 0.014 | 0 | 0.06 | 0.02 | 0.04 | 0 | 0.06 | 0.06 |
| 4 | 175457629 | rs12645863 | 0.466 | 1.982 | -1.138 | 0.245 | -0.039 | -0.017 | -0.017 | 0 | 0.09 | 0.02 | 0.17 | 0.48 | 0.15 | 0.14 |
| 4 | 175458289 | rs2251316 | -1.009 | 2.395 | -0.59 | 0.028 | 0.083 | 0.138 | 0.138 | 0 | 0.17 | 0.38 | 0.28 | 0.48 | 0.31 | 0.16 |
| 4 | 175460403 | rs7657391 | 0.275 | 2.495 | -0.752 | -0.106 | -0.032 | -0.022 | -0.022 | 0 | 0.13 | 0.08 | 0.15 | 0.16 | 0.19 | 0.1 |
| 4 | 175461517 | rs17295603 | 0.602 | 2.525 | -0.659 | -0.098 | -0.007 | 0 | 0 | 0 | 0.07 | 0.04 | 0.07 | 0.04 | 0.08 | 0.08 |
| 4 | 175461530 | rs2555639 | -0.256 | 2.204 | -0.323 | -0.381 | 0.039 | 0.023 | 0.023 | 0 | 0.27 | 0.46 | 0.32 | 0.29 | 0.35 | 0.29 |
| 4 | 175462630 | rs9990951 | 0.384 | 1.926 | -0.254 | -0.33 | 0.027 | 0.03 | 0.03 | 0 | 0.22 | 0.43 | 0.25 | 0.25 | 0.27 | 0.22 |
| 4 | 175466101 | rs2555674 | 0.121 | 1.999 | -0.64 | -0.317 | 0.014 | 0.016 | 0.016 | 0.011 | 0.23 | 0.61 | 0.27 | 0.25 | 0.28 | 0.21 |
| 4 | 175470669 | rs1365625 | -0.448 | 1.84 | -0.944 | -0.62 | 0.002 | 0.011 | 0.011 | 0 | 0.61 | 0.69 | 0.58 | 0.79 | 0.61 | 0.65 |
| 4 | 175475048 | rs17060632 | 0.831 | 1.894 | -1.086 | -0.472 | -0.015 | -0.012 | -0.012 | 0.559 | 0.09 | 0.04 | 0.09 | 0.11 | 0.12 | 0.1 |
| 4 | 175475745 | rs10520286 | 0.664 | 2.012 | -1.179 | -0.546 | -0.045 | 0.005 | 0.005 | 0 | 0.13 | 0.05 | 0.15 | 0.12 | 0.2 | 0.1 |
| 4 | 175476462 | rs6837393 | -0.436 | 1.118 | -0.776 | -0.553 | -0.005 | -0.004 | -0.004 | 0 | 0.37 | 0.36 | 0.38 | 0.41 | 0.44 | 0.43 |
| 4 | 175479449 | rs2555665 | -1.367 | 0.704 | -0.949 | -0.494 | -0.001 | -0.008 | -0.008 | 1.068 | 0.74 | 0.78 | 0.69 | 0.9 | 0.79 | 0.75 |
| 4 | 175479552 | rs7671048 | 1.38 | 1.146 | -1.739 | -0.292 | -0.015 | 0.011 | 0.011 | 0 | 0.16 | 0.22 | 0.13 | 0.18 | 0.12 | 0.23 |
| 4 | 175485146 | rs7657168 | -0.931 | 1.865 | -2.199 | 0.057 | 0.092 | 0.076 | 0.076 | 0.001 | 0.54 | 0.21 | 0.56 | 0.72 | 0.67 | 0.52 |
| 4 | 175485318 | rs7687916 | -0.117 | 1.397 | -2.002 | 0.29 | -0.104 | -0.045 | -0.045 | 0 | 0.19 | 0.03 | 0.27 | 0.23 | 0.33 | 0.2 |
| 4 | 175486220 | rs1365621 | -0.334 | 0.962 | -1.433 | -0.12 | 0.04 | 0.104 | 0.104 | 0.062 | 0.5 | 0.2 | 0.45 | 0.72 | 0.53 | 0.49 |
| 4 | 175492358 | rs2332895 | 0.239 | 0.825 | -0.931 | -0.338 | -0.001 | -0.011 | -0.011 | 0.119 | 0.68 | 0.39 | 0.69 | 0.88 | 0.72 | 0.64 |
| 4 | 175496482 | rs2555655 | -0.247 | 0.75 | -0.495 | -0.559 | 0.009 | 0.021 | 0.021 | 0 | 0.23 | 0.19 | 0.2 | 0.07 | 0.19 | 0.26 |
| 4 | 175497431 | rs1820525 | 0.451 | 1.033 | -0.89 | 0.178 | -0.02 | -0.053 | -0.053 | 0 | 0.16 | 0.07 | 0.18 | 0.34 | 0.2 | 0.22 |
| 4 | 175499645 | rs2612680 | -0.033 | 0.727 | -1.052 | -0.423 | -0.003 | 0.142 | 0.142 | 0 | 0.56 | 0.75 | 0.4 | 0.12 | 0.43 | 0.42 |
| 4 | 175500748 | rs1426938 | -0.871 | 0.725 | -0.999 | -0.354 | -0.023 | -0.037 | -0.037 | 0 | 0.71 | 0.79 | 0.58 | 0.45 | 0.63 | 0.65 |
| 4 | 175503770 | rs7662981 | -0.122 | 0.545 | -0.63 | -0.386 | 0.081 | 0.039 | 0.039 | 0 | 0.1 | 0.4 | 0.11 | 0.05 | 0.08 | 0.06 |
| 4 | 175523040 | rs2555675 | 0.372 | 0.517 | -0.91 | -0.575 | 0.025 | 0.051 | 0.051 | 0 | 0.46 | 0.33 | 0.3 | 0.08 | 0.37 | 0.36 |
| 4 | 175523073 | rs4695935 | 0.365 | 0.627 | -1.101 | -0.434 | -0.063 | -0.102 | -0.102 | 0 | 0.21 | 0.04 | 0.25 | 0.37 | 0.27 | 0.28 |
| 4 | 175526690 | rs2332838 | 0.9 | 0.433 | -1.144 | -0.565 | -0.054 | -0.103 | -0.103 | 0 | 0.26 | 0.1 | 0.39 | 0.55 | 0.33 | 0.39 |
| 4 | 175526746 | rs12642720 | 0.237 | 0.677 | -1.03 | -0.415 | -0.076 | -0.108 | -0.108 | 0.8 | 0.22 | 0.04 | 0.26 | 0.38 | 0.29 | 0.28 |
| 4 | 175528204 | rs2555678 | 0.316 | 0.675 | -0.572 | -0.056 | 0.011 | 0.017 | 0.017 | 0 | 0.54 | 0.35 | 0.52 | 0.38 | 0.45 | 0.5 |
| 4 | 175535392 | rs4315752 | 0.05 | 0.821 | -0.887 | 0.147 | -0.04 | -0.057 | -0.057 | 0 | 0.24 | 0.1 | 0.26 | 0.37 | 0.31 | 0.23 |
| 4 | 175535408 | rs12505186 | 0.166 | 0.807 | -0.69 | 0.093 | -0.004 | 0 | 0 | 0 | 0.57 | 0.73 | 0.59 | 0.41 | 0.45 | 0.56 |
| 4 | 175543263 | rs6846164 | 0.098 | 0.59 | -1.142 | 0.155 | -0.062 | -0.124 | -0.124 | 0 | 0.27 | 0.04 | 0.31 | 0.46 | 0.32 | 0.24 |
| 4 | 175543957 | rs9685165 | 1.206 | 0.687 | -0.962 | 0.179 | -0.02 | -0.009 | -0.009 | 0 | 0.19 | 0.1 | 0.13 | 0.21 | 0.24 | 0.28 |
| 4 | 175552622 | rs4695937 | 0.192 | 0.561 | -0.592 | 0.105 | 0.001 | 0.001 | 0.001 | 0 | 0.48 | 0.45 | 0.51 | 0.32 | 0.41 | 0.48 |
| 4 | 175556222 | rs9999152 | -0.094 | -0.079 | -1.18 | -0.302 | 0 | -0.001 | -0.001 | 0 | 0.59 | 0.76 | 0.65 | 0.54 | 0.61 | 0.71 |
| 4 | 175562084 | rs6849847 | -0.493 | -0.196 | -1.7 | -0.287 | 0.064 | 0.017 | 0.017 | 0 | 0.49 | 0.77 | 0.65 | 0.42 | 0.41 | 0.53 |
| 4 | 175565245 | rs4235212 | 0.382 | -0.316 | -1.669 | -0.366 | 0.034 | -0.047 | -0.047 | 0 | 0.36 | 0.12 | 0.52 | 0.39 | 0.33 | 0.34 |
| 4 | 175570846 | rs9998793 | 0.306 | -0.3 | -1.526 | -0.452 | 0.118 | 0.032 | 0.032 | 0.013 | 0.07 | 0.43 | 0.09 | 0.03 | 0.06 | 0.18 |
| 4 | 175571589 | rs4695945 | 0.63 | 0.221 | -3.142 | -0.34 | -0.066 | -0.052 | -0.052 | 0 | 0.25 | 0.06 | 0.22 | 0.36 | 0.29 | 0.18 |
| 4 | 175579442 | rs12511536 | -0.347 | -0.181 | -2.294 | -0.215 | 0.031 | -0.048 | -0.048 | 0 | 0.39 | 0.08 | 0.52 | 0.37 | 0.34 | 0.35 |
| 4 | 175582984 | rs4425356 | -0.347 | -0.203 | -2.294 | -0.215 | 0.035 | -0.048 | -0.048 | 4.587 | 0.39 | 0.08 | 0.52 | 0.37 | 0.34 | 0.35 |
| 4 | 175585803 | rs7686453 | -1.052 | 0.204 | -1.274 | 0.057 | 0.004 | -0.014 | -0.014 | 0 | 0.18 | 0.72 | 0.13 | 0.03 | 0.1 | 0.24 |
| 4 | 175601668 | rs12647358 | 0.216 | 0.553 | -1.572 | 0.914 | -0.035 | -0.084 | -0.084 | 1.615 | 0.12 | 0.01 | 0.22 | 0.35 | 0.17 | 0.14 |
| 4 | 175602273 | rs13142152 | 0.188 | 1.019 | -1.691 | 1.056 | -0.006 | -0.006 | -0.006 | 0 | 0.13 | 0.1 | 0.21 | 0.38 | 0.2 | 0.15 |
| 4 | 175606405 | rs1352058 | 1.024 | 0.842 | -1.837 | -0.241 | 0.011 | 0.175 | 0.175 | 0 | 0.47 | 0.23 | 0.39 | 0.04 | 0.44 | 0.36 |
| 4 | 175610550 | rs1471817 | 0.56 | 0.821 | -1.6 | -0.399 | 0.002 | -0.003 | -0.003 | 0 | 0.1 | 0.08 | 0.05 | 0 | 0.05 | 0.14 |
| 4 | 175615500 | rs6553806 | 0.049 | 0.753 | -0.529 | -0.313 | 0.001 | 0.002 | 0.002 | 0 | 0.69 | 0.63 | 0.55 | 0.25 | 0.62 | 0.68 |
| 4 | 175621502 | rs4695952 | 0.758 | 0.516 | -0.919 | -0.81 | 0.003 | 0.158 | 0.158 | 0 | 0.5 | 0.27 | 0.41 | 0.04 | 0.46 | 0.5 |
| 4 | 175631986 | rs4695954 | 0.671 | 0.397 | -1.17 | -1.299 | 0.012 | 0.077 | 0.077 | 0 | 0.29 | 0.13 | 0.18 | 0 | 0.24 | 0.34 |
| 4 | 175632405 | rs1485945 | 1.096 | 0.317 | -0.699 | -0.99 | 0.012 | 0.04 | 0.04 | 0.035 | 0.13 | 0.2 | 0.12 | 0.18 | 0.13 | 0.14 |
| 4 | 175637324 | rs10024014 | 1.72 | 0.072 | -0.86 | -1.146 | 0.003 | 0.067 | 0.067 | 0 | 0.21 | 0.07 | 0.13 | 0 | 0.2 | 0.22 |
| 4 | 175641906 | rs4594702 | 1.694 | 0.163 | -0.931 | -1.012 | -0.012 | -0.019 | -0.019 | 0 | 0.13 | 0.09 | 0.21 | 0.35 | 0.2 | 0.15 |
| 4 | 175644564 | rs10000984 | 0.102 | 0.266 | -0.837 | -0.963 | -0.003 | 0.02 | 0.02 | 0 | 0.42 | 0.34 | 0.4 | 0.36 | 0.46 | 0.49 |
| 4 | 175644689 | rs12650824 | 0.978 | 0.091 | -1.198 | -1.198 | 0.016 | -0.09 | -0.09 | 0 | 0.24 | 0.08 | 0.23 | 0.45 | 0.21 | 0.16 |
| 4 | 175649941 | rs10000456 | 1.757 | 0.208 | -1.156 | -1.499 | 0.006 | 0.035 | 0.035 | 0 | 0.2 | 0.08 | 0.12 | 0 | 0.18 | 0.21 |
| 4 | 175650116 | rs9312559 | -0.555 | 0.151 | -1.126 | -1.506 | 0.001 | 0.004 | 0.004 | 0.445 | 0.6 | 0.38 | 0.62 | 0.37 | 0.66 | 0.7 |
| 4 | 175652512 | rs11133053 | 1.11 | 0.277 | -1.501 | -1.528 | -0.033 | 0.053 | 0.053 | 0.405 | 0.18 | 0.04 | 0.22 | 0.07 | 0.21 | 0.22 |
| 4 | 175654447 | rs9999658 | 0.742 | 0.374 | -0.983 | -1.19 | 0 | 0.097 | 0.097 | 0 | 0.22 | 0.05 | 0.12 | 0 | 0.2 | 0.23 |
| 4 | 175658175 | rs4695959 | -0.858 | 1.082 | -0.076 | -0.988 | 0.054 | -0.066 | -0.066 | 0.277 | 0.19 | 0.36 | 0.25 | 0.07 | 0.27 | 0.21 |
| 4 | 175662891 | rs6827308 | -1.235 | 1.281 | 0.396 | -1.178 | 0.033 | -0.042 | -0.042 | 0 | 0.1 | 0.37 | 0.1 | 0.02 | 0.1 | 0.08 |
| 4 | 175667791 | rs7685430 | -0.206 | 1.133 | 0.898 | -1.265 | 0.086 | -0.034 | -0.034 | 0 | 0.87 | 0.73 | 0.8 | 0.94 | 0.74 | 0.87 |
| 4 | 175674144 | rs13113891 | 2.242 | 0.844 | 0.552 | -0.548 | -0.023 | -0.026 | -0.026 | 0.163 | 0.62 | 0.25 | 0.51 | 0.51 | 0.58 | 0.56 |
| 4 | 175675480 | rs2029570 | -1.255 | 0.98 | 0.582 | -0.481 | -0.001 | 0.001 | 0.001 | 0 | 0.26 | 0.28 | 0.35 | 0.47 | 0.37 | 0.33 |
| 4 | 175685891 | rs1352804 | 1.072 | 0.961 | 0.47 | -0.389 | 0.007 | 0.028 | 0.028 | 0 | 0.08 | 0.03 | 0.13 | 0.39 | 0.05 | 0.15 |

| | | | | | | | | | | | | | | | | |
|---|---|---|---|---|---|---|---|---|---|---|---|---|---|---|---|---|
| 4 | 175686589 | rs2029569 | -0.305 | 1.01 | 0.468 | -0.429 | 0.018 | -0.006 | -0.006 | 0 | 0.16 | 0.21 | 0.2 | 0.08 | 0.31 | 0.17 |
| 4 | 175688141 | rs12651268 | 1.05 | 1.088 | -0.447 | -1.079 | 0.019 | -0.075 | -0.075 | 1.257 | 0.44 | 0.11 | 0.32 | 0.52 | 0.37 | 0.43 |
| 4 | 175690792 | rs6858127 | -1.259 | 1.002 | -0.034 | -1.051 | 0.001 | -0.105 | -0.105 | 0.431 | 0.75 | 0.65 | 0.73 | 1 | 0.74 | 0.83 |
| 4 | 175696544 | rs1385838 | -1.243 | 0.91 | -0.034 | -1.035 | 0.005 | -0.101 | -0.101 | 0 | 0.74 | 0.65 | 0.73 | 1 | 0.72 | 0.79 |
| 4 | 175699276 | rs2200457 | 0.945 | 0.661 | -0.355 | -0.909 | 0.069 | -0.052 | -0.052 | 0.129 | 0.43 | 0.16 | 0.29 | 0.51 | 0.33 | 0.42 |
| 4 | 175707292 | rs9992190 | -0.403 | 1.231 | 0.093 | -0.748 | 0.016 | -0.018 | -0.018 | 0 | 0.17 | 0.24 | 0.18 | 0.1 | 0.27 | 0.14 |
| 4 | 175709806 | rs10049967 | -0.322 | 1.164 | 0.107 | -0.709 | 0.013 | -0.014 | -0.014 | 0 | 0.15 | 0.23 | 0.17 | 0.1 | 0.24 | 0.12 |
| 4 | 175711800 | rs10021711 | 0.509 | 0.792 | 0.468 | -0.547 | -0.006 | 0.273 | 0.273 | 0.132 | 0.1 | 0.21 | 0.17 | 0.42 | 0.07 | 0.14 |
| 4 | 175713278 | rs4695963 | -1 | 0.376 | 0.036 | -1.012 | 0.031 | -0.033 | -0.033 | 0 | 0.61 | 0.78 | 0.55 | 0.51 | 0.68 | 0.69 |
| 4 | 175716041 | rs11934209 | 0.896 | 0.825 | 0.388 | -1.301 | -0.005 | 0.04 | 0.04 | 2.06 | 0.28 | 0.23 | 0.18 | 0.02 | 0.31 | 0.24 |
| 4 | 175716626 | rs1994313 | -0.403 | 0.858 | 0.385 | -1.363 | -0.007 | 0.031 | 0.031 | 1.036 | 0.43 | 0.62 | 0.33 | 0.05 | 0.45 | 0.42 |
| 4 | 175723995 | rs6825046 | -0.839 | 0.646 | 0.396 | -1.06 | 0.023 | -0.044 | -0.044 | 0 | 0.16 | 0.28 | 0.14 | 0.05 | 0.15 | 0.16 |
| 4 | 175724056 | rs1485938 | 0.309 | 0.547 | -0.076 | -1.3 | -0.006 | 0.04 | 0.04 | 0 | 0.16 | 0.07 | 0.11 | 0.04 | 0.15 | 0.1 |
| 4 | 175734620 | rs1352062 | -0.799 | 0.449 | 0.277 | -1.138 | 0.029 | -0.046 | -0.046 | 0 | 0.16 | 0.32 | 0.14 | 0.05 | 0.15 | 0.16 |
| 4 | 175740569 | rs6824735 | -0.255 | -0.072 | 0.168 | -0.928 | -0.022 | 0.009 | 0.009 | 0.377 | 0.61 | 0.45 | 0.54 | 0.46 | 0.69 | 0.69 |
| 4 | 175743525 | rs6840133 | -0.493 | 0.11 | -0.529 | -0.747 | -0.004 | 0.015 | 0.015 | 0 | 0.37 | 0.36 | 0.3 | 0.18 | 0.35 | 0.29 |
| 4 | 175753571 | rs13116721 | 0.077 | 0.124 | -0.581 | -0.889 | -0.012 | 0.084 | 0.084 | 0 | 0.37 | 0.22 | 0.23 | 0.03 | 0.43 | 0.31 |
| 4 | 175753871 | rs7437645 | 0.888 | 0.263 | -0.732 | -0.813 | 0.005 | 0.023 | 0.023 | 4.057 | 0.11 | 0.07 | 0.05 | 0.02 | 0.1 | 0.07 |
| 4 | 175761435 | rs17300435 | 0.984 | 0.477 | -0.446 | -0.329 | -0.041 | 0.081 | 0.081 | 0.213 | 0.23 | 0.08 | 0.17 | 0.01 | 0.33 | 0.24 |
| 4 | 175764593 | rs1074475 | -0.505 | 0.296 | -0.122 | 0.193 | -0.024 | 0.063 | 0.063 | 0.013 | 0.75 | 0.62 | 0.6 | 0.48 | 0.79 | 0.76 |
| 4 | 175777871 | rs1491411 | -0.465 | 0.314 | -0.245 | 0.124 | -0.026 | 0.061 | 0.061 | 0.002 | 0.75 | 0.59 | 0.61 | 0.49 | 0.79 | 0.75 |
| 4 | 175792412 | rs992363 | -0.361 | 0.603 | -0.597 | -0.107 | 0.001 | 0.008 | 0.008 | 0.183 | 0.48 | 0.36 | 0.41 | 0.48 | 0.46 | 0.52 |
| 4 | 175796630 | rs1121336 | 0.422 | 0.807 | -0.462 | -0.133 | -0.001 | 0 | 0 | 0 | 0.26 | 0.28 | 0.19 | 0.16 | 0.27 | 0.18 |
| 4 | 175798347 | rs7667865 | 1.233 | 0.555 | -0.59 | -0.479 | -0.019 | 0.055 | 0.055 | 0.283 | 0.27 | 0.15 | 0.19 | 0.01 | 0.34 | 0.24 |
| 4 | 175801325 | rs2029566 | 0.536 | 0.593 | -0.721 | -0.51 | -0.024 | 0.006 | 0.006 | 0 | 0.08 | 0.02 | 0.11 | 0.03 | 0.08 | 0.09 |
| 4 | 175816129 | rs6858430 | -1.193 | 0.334 | -0.162 | -0.471 | -0.002 | 0.004 | 0.004 | 0 | 0.74 | 0.7 | 0.6 | 0.49 | 0.78 | 0.75 |
| 4 | 175816341 | rs11933901 | -0.263 | 0.495 | -0.376 | -0.54 | -0.013 | -0.036 | -0.036 | 2.15 | 0.2 | 0.08 | 0.21 | 0.31 | 0.18 | 0.33 |
| 4 | 175817225 | rs10003605 | 0.929 | 0.613 | -0.382 | -0.634 | -0.016 | 0.046 | 0.046 | 0.318 | 0.26 | 0.16 | 0.18 | 0.01 | 0.33 | 0.24 |
| 4 | 175828036 | rs7697216 | 0.029 | 0.472 | 0.304 | -0.226 | 0.018 | 0.126 | 0.126 | 0 | 0.86 | 0.53 | 0.81 | 0.77 | 0.89 | 0.79 |
| 4 | 175831859 | rs4388065 | 0.353 | 0.495 | -0.033 | -0.652 | -0.026 | 0.093 | 0.093 | 0.058 | 0.51 | 0.19 | 0.49 | 0.72 | 0.47 | 0.39 |
| 4 | 175836048 | rs6553847 | -0.863 | 0.402 | 0.083 | -0.695 | -0.014 | 0.017 | 0.017 | 0.327 | 0.67 | 0.75 | 0.68 | 0.95 | 0.57 | 0.61 |
| 4 | 175838816 | rs6553849 | 0.393 | 0.4 | 0.054 | -0.617 | -0.018 | 0.081 | 0.081 | 0.047 | 0.51 | 0.19 | 0.49 | 0.71 | 0.46 | 0.4 |
| 4 | 175839432 | rs1104945 | -0.411 | 0.401 | 0.09 | -0.612 | -0.013 | 0.084 | 0.084 | 0 | 0.16 | 0.51 | 0.19 | 0.24 | 0.11 | 0.21 |
| 4 | 175842230 | rs12512467 | 0.028 | 0.442 | -0.407 | -0.536 | -0.014 | 0.056 | 0.056 | 0 | 0.54 | 0.3 | 0.51 | 0.71 | 0.47 | 0.4 |
| 4 | 175854023 | rs12647292 | 0.733 | 0.441 | -0.785 | -0.32 | 0.003 | -0.127 | -0.127 | 0 | 0.24 | 0.09 | 0.35 | 0.55 | 0.22 | 0.21 |
| 4 | 175858796 | rs4695984 | 0.632 | 0.538 | -0.128 | -0.319 | 0.014 | 0.112 | 0.112 | 0 | 0.11 | 0.27 | 0.18 | 0.25 | 0.11 | 0.22 |
| 4 | 175859612 | rs13147656 | 0.602 | 0.445 | -0.525 | -0.613 | -0.006 | 0.051 | 0.051 | 0 | 0.28 | 0.17 | 0.16 | 0.15 | 0.24 | 0.19 |
| 4 | 175860563 | rs11934547 | 0.969 | 0.572 | -0.502 | -0.741 | -0.04 | 0.108 | 0.108 | 0.637 | 0.28 | 0.1 | 0.19 | 0.02 | 0.35 | 0.29 |
| 4 | 175862049 | rs13136874 | 0.67 | 0.589 | 0.329 | -0.055 | 0.014 | 0.104 | 0.104 | 0 | 0.8 | 0.44 | 0.7 | 0.71 | 0.82 | 0.68 |
| 4 | 175876473 | rs13130150 | -0.673 | 0.643 | 0.243 | -0.125 | 0 | 0.005 | 0.005 | 0 | 0.42 | 0.55 | 0.35 | 0.44 | 0.35 | 0.45 |
| 4 | 175878038 | rs7661012 | -0.289 | 0.562 | 0.183 | -0.241 | -0.009 | -0.053 | -0.053 | 0 | 0.34 | 0.54 | 0.2 | 0.18 | 0.25 | 0.25 |
| 4 | 175890127 | rs4128922 | -0.99 | 0.614 | 0.434 | -0.229 | 0.002 | 0.005 | 0.005 | 0 | 0.45 | 0.64 | 0.41 | 0.45 | 0.41 | 0.47 |
| 4 | 175891945 | rs4129839 | -0.162 | 0.516 | 0.002 | -0.611 | -0.046 | -0.067 | -0.067 | 2.643 | 0.2 | 0.03 | 0.23 | 0.31 | 0.22 | 0.31 |
| 4 | 175894380 | rs6845431 | -1.018 | 0.43 | 0.259 | -0.661 | -0.01 | 0.009 | 0.009 | 0 | 0.27 | 0.48 | 0.28 | 0.31 | 0.22 | 0.31 |
| 4 | 175908720 | rs28833210 | 0.123 | 0.455 | 0.365 | -0.711 | -0.027 | -0.077 | -0.077 | 0 | 0.12 | 0.32 | 0.05 | 0 | 0.05 | 0.03 |
| 4 | 175913798 | rs10026079 | 0.595 | 0.41 | -0.023 | -0.695 | -0.011 | 0.036 | 0.036 | 0 | 0.17 | 0.09 | 0.14 | 0.08 | 0.18 | 0.13 |
| 4 | 175923101 | rs12502984 | 0.84 | 0.344 | -0.113 | -0.611 | -0.001 | -0.002 | -0.002 | 0.165 | 0.55 | 0.44 | 0.59 | 0.62 | 0.59 | 0.56 |
| 4 | 175926862 | rs11734965 | 0.826 | 0.348 | -0.11 | -0.613 | -0.001 | -0.003 | -0.003 | 0.075 | 0.56 | 0.44 | 0.59 | 0.62 | 0.59 | 0.56 |
| 4 | 175928058 | rs10461333 | 0.829 | 0.374 | -0.488 | -0.6 | 0.002 | 0.017 | 0.017 | 0 | 0.55 | 0.36 | 0.6 | 0.62 | 0.59 | 0.56 |
| 4 | 175929930 | rs2127292 | 0.642 | 0.005 | -0.914 | -0.704 | 0.01 | 0.004 | 0.004 | 0.021 | 0.15 | 0.11 | 0.19 | 0 | 0.09 | 0.06 |
| 4 | 175936490 | rs6553853 | 0.151 | 0.425 | -0.77 | -0.08 | -0.005 | -0.012 | -0.012 | 1.447 | 0.87 | 0.88 | 0.84 | 0.74 | 0.85 | 0.75 |
| 4 | 175939626 | rs12513233 | -0.006 | 0.432 | -1.076 | -0.076 | -0.048 | -0.066 | -0.066 | 0.49 | 0.12 | 0.02 | 0.15 | 0.26 | 0.14 | 0.25 |
| 4 | 175947638 | rs4382019 | 0.247 | 0.607 | -1.501 | 0.173 | -0.059 | -0.043 | -0.043 | 0.526 | 0.36 | 0.11 | 0.35 | 0.42 | 0.38 | 0.28 |
| 4 | 175948807 | rs12509972 | 0.274 | 1.003 | -1.625 | 0.357 | -0.048 | -0.066 | -0.066 | 0 | 0.11 | 0.02 | 0.15 | 0.26 | 0.14 | 0.25 |
| 4 | 175950601 | rs11932717 | -0.049 | 0.495 | -0.837 | -0.102 | 0.012 | 0.033 | 0.033 | 0 | 0.49 | 0.37 | 0.47 | 0.32 | 0.51 | 0.48 |
| 4 | 175954666 | rs11941955 | 0.178 | 0.571 | -0.745 | 0.071 | 0.006 | 0.014 | 0.014 | 0 | 0.44 | 0.33 | 0.43 | 0.32 | 0.45 | 0.47 |
| 4 | 175964181 | rs1842178 | -1.055 | 0.301 | -0.084 | -0.445 | -0.018 | -0.104 | -0.104 | 0 | 0.24 | 0.42 | 0.17 | 0.02 | 0.17 | 0.18 |
| 4 | 175967160 | rs1471480 | 0.374 | 0.578 | -0.077 | -0.581 | -0.007 | 0.012 | 0.012 | 0.528 | 0.31 | 0.23 | 0.28 | 0.22 | 0.39 | 0.4 |
| 4 | 175968691 | rs1479380 | 0.051 | 0.442 | 0.124 | -0.523 | -0.001 | -0.02 | -0.02 | 0 | 0.08 | 0.09 | 0.1 | 0.01 | 0.07 | 0.07 |
| 4 | 175970464 | rs7356528 | -0.53 | 0.414 | 0.374 | -0.04 | 0.04 | 0.33 | 0.33 | 0.003 | 0.93 | 0.78 | 0.81 | 0.41 | 0.94 | 0.79 |
| 4 | 175987359 | rs1491413 | 0.233 | 0.396 | 0.071 | -0.328 | -0.001 | -0.02 | -0.02 | 0 | 0.08 | 0.09 | 0.1 | 0 | 0.07 | 0.06 |
| 4 | 176009525 | rs11733561 | 0.647 | 0.491 | 0.177 | -0.368 | -0.039 | 0.035 | 0.035 | 1.737 | 0.29 | 0.14 | 0.29 | 0.2 | 0.4 | 0.4 |
| 4 | 176023652 | rs330502 | -0.916 | 0.486 | -0.151 | -0.053 | -0.015 | 0.084 | 0.084 | 0 | 0.56 | 0.76 | 0.49 | 0.29 | 0.58 | 0.53 |
| 4 | 176027111 | rs1352806 | 0.847 | 0.241 | 0.243 | 0.167 | -0.015 | 0.059 | 0.059 | 0 | 0.76 | 0.57 | 0.72 | 0.38 | 0.82 | 0.47 |
| 4 | 176031901 | rs1602491 | 0.836 | 0.243 | 0.244 | 0.168 | -0.015 | 0.06 | 0.06 | 0 | 0.76 | 0.56 | 0.72 | 0.38 | 0.82 | 0.47 |
| 4 | 176036504 | rs2219824 | -0.707 | 0.236 | 0.177 | -0.39 | -0.006 | 0.13 | 0.13 | 1.512 | 0.46 | 0.26 | 0.42 | 0.18 | 0.5 | 0.39 |
| 4 | 176055295 | rs6849435 | 0.738 | 0.088 | 0.191 | -0.018 | -0.007 | 0.018 | 0.018 | 3.581 | 0.33 | 0.21 | 0.39 | 0.24 | 0.39 | 0.23 |
| 4 | 176056502 | rs6811622 | 1.428 | 0.207 | 0.374 | -0.019 | -0.008 | 0.044 | 0.044 | 0.064 | 0.4 | 0.32 | 0.37 | 0.2 | 0.41 | 0.23 |
| 4 | 176066263 | rs4695809 | 0.519 | 0.389 | 0.239 | 0.056 | 0.002 | 0.021 | 0.021 | 3.818 | 0.34 | 0.21 | 0.39 | 0.24 | 0.37 | 0.24 |

| | | | | | | | | | | | | | | | | |
|---|---|---|---|---|---|---|---|---|---|---|---|---|---|---|---|---|
| 4 | 176090779 | rs162449 | -0.369 | 0.3 | 0.571 | 0.211 | 0.012 | 0.165 | 0.165 | 0 | 0.89 | 0.73 | 0.86 | 0.38 | 0.91 | 0.61 |
| 4 | 176104892 | rs162471 | -0.369 | 0.249 | 0.85 | 0.179 | 0.012 | 0.162 | 0.162 | 0 | 0.89 | 0.72 | 0.86 | 0.38 | 0.91 | 0.62 |
| 4 | 176109168 | rs1491416 | -0.335 | 0.215 | 0.684 | -0.219 | -0.012 | 0.135 | 0.135 | 0.006 | 0.42 | 0.19 | 0.41 | 0.17 | 0.43 | 0.37 |
| 4 | 176113346 | rs11133085 | 1.998 | 0.209 | 0.565 | -0.238 | 0.004 | -0.004 | -0.004 | 0 | 0.7 | 0.31 | 0.74 | 0.37 | 0.77 | 0.45 |
| 4 | 176116299 | rs13142619 | 1.214 | 0.415 | 0.487 | -0.352 | -0.007 | 0.042 | 0.042 | 0 | 0.44 | 0.36 | 0.43 | 0.2 | 0.46 | 0.23 |
| 4 | 176122398 | rs13134673 | 0.372 | 0.545 | 0.075 | 0.755 | -0.019 | -0.14 | -0.14 | 0.041 | 0.34 | 0.05 | 0.37 | 0.47 | 0.33 | 0.34 |
| 4 | 176122493 | rs13134743 | -0.186 | 0.499 | 0.169 | 0.816 | -0.013 | -0.073 | -0.073 | 0 | 0.4 | 0.15 | 0.42 | 0.47 | 0.4 | 0.37 |
| 4 | 176124452 | rs2637795 | -0.243 | 0.635 | 0.69 | 1.012 | 0.05 | 0.284 | 0.284 | 0 | 0.92 | 0.71 | 0.91 | 0.68 | 0.92 | 0.74 |
| 4 | 176126370 | rs647003 | 1.023 | 0.602 | 0.046 | -0.054 | 0.023 | -0.066 | -0.066 | 0 | 0.23 | 0.3 | 0.3 | 0.08 | 0.29 | 0.1 |
| 4 | 176126818 | rs6812564 | -0.181 | -0.354 | -0.416 | 0.2 | 0.062 | 0.005 | 0.005 | 0.164 | 0.43 | 0.22 | 0.4 | 0.44 | 0.37 | 0.38 |
| 4 | 176161041 | rs10014039 | 1.147 | -0.249 | -1.577 | -0.29 | -0.005 | 0.068 | 0.068 | 0.177 | 0.36 | 0.2 | 0.23 | 0.08 | 0.36 | 0.22 |
| 4 | 176164722 | rs4146 | 0.141 | 0.156 | -2.268 | 0.273 | -0.065 | -0.089 | -0.089 | 0 | 0.35 | 0.07 | 0.34 | 0.53 | 0.36 | 0.53 |
| 4 | 176170783 | rs13132343 | -0.141 | 0.403 | -1.462 | 0.439 | -0.015 | -0.028 | -0.028 | 0.062 | 0.1 | 0.04 | 0.3 | 0.18 | 0.15 | 0.12 |
| 4 | 176176194 | rs493293 | -0.424 | 0.464 | -1.588 | 0.305 | -0.012 | 0.031 | 0.031 | 0 | 0.76 | 0.44 | 0.86 | 0.63 | 0.8 | 0.78 |
| 4 | 176181659 | rs13146658 | -0.494 | 0.68 | -1.749 | 1.266 | -0.025 | -0.044 | -0.044 | 0 | 0.11 | 0.07 | 0.32 | 0.17 | 0.17 | 0.14 |
| 4 | 176184572 | rs4696017 | -0.371 | 0.747 | -1.373 | 1.221 | -0.056 | -0.117 | -0.117 | 0.776 | 0.23 | 0.07 | 0.35 | 0.46 | 0.29 | 0.49 |
| 4 | 176195262 | rs4696019 | 0.273 | 0.189 | -0.489 | 0.863 | -0.015 | 0.008 | 0.008 | 3.937 | 0.93 | 0.9 | 0.95 | 0.92 | 0.99 | 1 |
| 4 | 176200230 | rs7673394 | 0.968 | 0.209 | -0.684 | 0.676 | 0.005 | -0.003 | -0.003 | 0 | 0.78 | 0.84 | 0.64 | 0.8 | 0.8 | 0.8 |
| 4 | 176214028 | rs10520296 | 0.154 | 0.433 | -1.093 | 0.641 | -0.095 | -0.063 | -0.063 | 0 | 0.18 | 0.02 | 0.24 | 0.26 | 0.27 | 0.19 |
| 4 | 176226337 | rs1711988 | -1.899 | -0.614 | 0.455 | 0.405 | -0.038 | -0.058 | -0.058 | 0 | 0.15 | 0.32 | 0.07 | 0.05 | 0.06 | 0.07 |
| 4 | 176227974 | rs4417946 | -1.255 | -0.555 | 0.481 | 0.442 | -0.02 | -0.027 | -0.027 | 0 | 0.11 | 0.2 | 0.04 | 0.05 | 0.05 | 0.01 |
| 4 | 176252966 | rs13133504 | -0.943 | -0.525 | 0.416 | 0.446 | 0 | -0.002 | -0.002 | 0 | 0.09 | 0.1 | 0.03 | 0.04 | 0.05 | 0.01 |
| 4 | 176257561 | rs13135411 | -1.115 | -0.532 | 0.533 | 0.439 | 0.005 | -0.007 | -0.007 | 0 | 0.1 | 0.09 | 0.04 | 0 | 0.05 | 0 |
| 4 | 176285962 | rs2333166 | -0.426 | -1.047 | 1.708 | 0.406 | -0.031 | -0.026 | -0.026 | 0 | 0.16 | 0 | 0.22 | 0.13 | 0.18 | 0.21 |
| 4 | 176289800 | rs1879629 | 2.271 | -0.836 | 2.4 | 0.435 | -0.043 | -0.02 | -0.02 | 0.216 | 0.81 | 0.34 | 0.89 | 0.87 | 0.93 | 0.88 |
| 4 | 176344294 | rs9994051 | -1.998 | -0.675 | 2.564 | 0.441 | -0.044 | 0.019 | 0.019 | 1.298 | 0.14 | 0.42 | 0.07 | 0.11 | 0.06 | 0.03 |
| 4 | 176349829 | rs9684098 | 1.892 | -0.69 | 2.574 | 0.433 | -0.058 | -0.011 | -0.011 | 0 | 0.84 | 0.43 | 0.93 | 0.88 | 0.94 | 0.97 |
| 4 | 176366005 | rs9990657 | 1.444 | -0.688 | 2.283 | 0.44 | 0.044 | 0.126 | 0.126 | 0.925 | 0.95 | 0.7 | 0.98 | 0.93 | 1 | 0.99 |
| 4 | 176366314 | rs13434491 | -1.181 | -0.7 | 1.451 | 0.418 | -0.011 | -0.033 | -0.033 | 2.41 | 0.1 | 0.14 | 0.04 | 0 | 0.05 | 0.01 |
| 4 | 176370331 | rs13149814 | 1.503 | -0.736 | 1.421 | 0.402 | -0.006 | 0 | 0 | 0.078 | 0.81 | 0.27 | 0.85 | 0.88 | 0.93 | 0.92 |
| 4 | 176393194 | rs7699269 | 1.263 | -0.586 | 0.396 | 0.339 | 0 | 0.001 | 0.001 | 0.522 | 0.83 | 0.27 | 0.86 | 0.9 | 0.91 | 0.88 |
| 4 | 176399587 | rs13107001 | 1.203 | -0.581 | 0.363 | 0.341 | 0.008 | 0.008 | 0.008 | 0.59 | 0.83 | 0.24 | 0.86 | 0.9 | 0.91 | 0.89 |
| 4 | 176410587 | rs6829020 | 1.306 | -1.154 | -0.545 | 0.526 | 0.09 | -0.003 | -0.003 | 0 | 0.6 | 0.13 | 0.55 | 0.66 | 0.73 | 0.61 |
| 4 | 176432982 | rs10015016 | -1.196 | -1.082 | 1.556 | 0.667 | 0.05 | 0.031 | 0.031 | 0 | 0.05 | 0.3 | 0.05 | 0 | 0.03 | 0.02 |
| 4 | 176451803 | rs4622948 | 0.968 | -1.063 | 1.566 | 0.729 | -0.041 | 0.068 | 0.068 | 0.763 | 0.9 | 0.55 | 0.96 | 0.89 | 0.99 | 0.94 |
| 4 | 176480544 | rs9992999 | -0.783 | -1 | 0.945 | 0.808 | -0.038 | 0.028 | 0.028 | 0.475 | 0.08 | 0.18 | 0.01 | 0.13 | 0.01 | 0.03 |
| 4 | 176487623 | rs4311268 | -0.653 | -0.193 | 0.399 | 1.244 | -0.002 | -0.001 | -0.001 | 4.515 | 0.22 | 0.21 | 0.39 | 0.45 | 0.22 | 0.31 |
| 4 | 176491850 | rs11735559 | 1.292 | -0.546 | -1.028 | 0.693 | -0.131 | -0.025 | -0.025 | 0 | 0.39 | 0.04 | 0.42 | 0.3 | 0.49 | 0.5 |
| 4 | 176502740 | rs12502137 | 0.197 | -0.268 | -1.056 | 0.675 | -0.004 | -0.032 | -0.032 | 2.596 | 0.2 | 0.12 | 0.37 | 0.44 | 0.17 | 0.26 |
| 4 | 176513040 | rs13111738 | -0.456 | -0.455 | -0.613 | 0.621 | -0.016 | -0.013 | -0.013 | 0 | 0.42 | 0.29 | 0.52 | 0.5 | 0.41 | 0.36 |
| 4 | 176514624 | rs4690379 | 0.711 | -0.647 | -0.061 | 0.52 | -0.048 | 0.138 | 0.138 | 0 | 0.88 | 0.51 | 0.9 | 0.81 | 0.95 | 0.9 |
| 4 | 176538654 | rs2333233 | -0.637 | -0.691 | -0.082 | 0.525 | -0.005 | -0.003 | -0.003 | 0 | 0.43 | 0.38 | 0.49 | 0.49 | 0.42 | 0.36 |
| 4 | 176541897 | rs10027928 | -0.333 | -0.715 | -0.17 | 0.513 | -0.039 | -0.019 | -0.019 | 0 | 0.41 | 0.18 | 0.47 | 0.47 | 0.42 | 0.36 |
| 4 | 176545324 | rs4690535 | 1.952 | -0.444 | -0.413 | 0.424 | -0.115 | 0.055 | 0.055 | 0 | 0.4 | 0.05 | 0.34 | 0.26 | 0.48 | 0.48 |
| 4 | 176546320 | rs10006242 | -0.913 | -0.36 | -0.171 | 0.419 | -0.004 | -0.005 | -0.005 | 0 | 0.44 | 0.33 | 0.46 | 0.49 | 0.39 | 0.34 |
| 4 | 176547576 | rs11728358 | 0.108 | -0.491 | -0.243 | 0.894 | -0.058 | 0.006 | 0.006 | 0.006 | 0.31 | 0.1 | 0.35 | 0.29 | 0.33 | 0.23 |
| 4 | 176548782 | rs1014381 | 1.15 | 0.076 | -0.252 | 0.527 | 0.016 | 0.066 | 0.066 | 0 | 0.49 | 0.35 | 0.38 | 0.25 | 0.47 | 0.45 |
| 4 | 176549722 | rs2333243 | 0.067 | 0.034 | 0.017 | 0.573 | -0.001 | 0.031 | 0.031 | 0.512 | 0.91 | 0.84 | 0.93 | 0.79 | 0.91 | 0.84 |
| 4 | 176555585 | rs3733398 | 0.206 | 0.114 | -0.337 | 0.614 | -0.017 | 0.087 | 0.087 | 0.922 | 0.86 | 0.57 | 0.88 | 0.79 | 0.88 | 0.83 |
| 4 | 176560091 | rs10520301 | 0.06 | 0.184 | -1.399 | 0.499 | -0.02 | -0.031 | -0.031 | 0 | 0.33 | 0.18 | 0.41 | 0.46 | 0.33 | 0.34 |
| 4 | 176560547 | rs17313669 | 1.337 | 0.532 | -1.499 | 0.55 | -0.137 | 0.08 | 0.08 | 0.046 | 0.42 | 0.04 | 0.37 | 0.25 | 0.51 | 0.47 |
| 4 | 176566551 | rs2333251 | 1.11 | 0.492 | -0.388 | 1.005 | 0.003 | 0.06 | 0.06 | 2.529 | 0.81 | 0.36 | 0.79 | 0.74 | 0.79 | 0.8 |
| 4 | 176566962 | rs7678146 | 0.965 | 0.163 | -0.356 | 1.018 | -0.021 | 0.094 | 0.094 | 1.899 | 0.82 | 0.36 | 0.8 | 0.7 | 0.84 | 0.8 |
| 4 | 176571011 | rs4429679 | 0.221 | 0.663 | -1.428 | 0.026 | -0.025 | 0.093 | 0.093 | 0.247 | 0.41 | 0.19 | 0.31 | 0.14 | 0.42 | 0.31 |
| 4 | 176582766 | rs2333255 | 1.032 | 0.409 | -1.83 | 0.127 | -0.143 | 0.167 | 0.167 | 1.176 | 0.44 | 0.04 | 0.38 | 0.22 | 0.49 | 0.32 |
| 4 | 176587929 | rs11940383 | -0.184 | 0.3 | -1.829 | -0.407 | -0.056 | 0.095 | 0.095 | 2.359 | 0.22 | 0.05 | 0.14 | 0 | 0.25 | 0.05 |
| 4 | 176591669 | rs35424424 | 0.661 | 0.352 | -1.848 | -0.411 | -0.071 | 0.075 | 0.075 | 2.115 | 0.12 | 0.01 | 0.11 | 0 | 0.21 | 0.03 |
| 4 | 176601113 | rs10213204 | 0.357 | -0.243 | -0.219 | 0.156 | -0.029 | 0.078 | 0.078 | 0.268 | 0.83 | 0.47 | 0.83 | 0.71 | 0.89 | 0.81 |
| 4 | 176608039 | rs10520302 | 0.068 | -0.28 | -0.71 | 0.723 | -0.004 | -0.019 | -0.019 | 0.867 | 0.07 | 0 | 0.04 | 0.34 | 0.04 | 0.13 |
| 4 | 176609256 | rs2333257 | 0.255 | -0.207 | -0.669 | 0.499 | -0.06 | 0.076 | 0.076 | 1.258 | 0.15 | 0.03 | 0.13 | 0.01 | 0.24 | 0.17 |
| 4 | 176610969 | rs17061779 | 0.719 | -0.292 | -0.652 | 0.519 | 0.013 | 0.028 | 0.028 | 0.025 | 0.09 | 0.02 | 0.03 | 0 | 0.06 | 0.02 |
| 4 | 176612351 | rs17061785 | 0.747 | -0.28 | -0.646 | 0.525 | 0.012 | 0.028 | 0.028 | 0.08 | 0.09 | 0.02 | 0.03 | 0 | 0.06 | 0.02 |
| 4 | 176626272 | rs12640626 | 1.503 | -0.271 | -0.579 | 0.507 | -0.02 | 0.12 | 0.12 | 0.808 | 0.55 | 0.16 | 0.66 | 0.7 | 0.59 | 0.62 |
| 4 | 176626693 | rs12512924 | 2.3 | -0.267 | -0.724 | 0.597 | -0.132 | 0.285 | 0.285 | 1.475 | 0.44 | 0.05 | 0.49 | 0.08 | 0.51 | 0.3 |
| 4 | 176632858 | rs10022458 | 0.104 | -0.243 | -0.345 | 0.684 | -0.012 | 0.129 | 0.129 | 0.042 | 0.07 | 0.17 | 0.09 | 0.28 | 0.06 | 0.08 |
| 4 | 176641467 | rs2581761 | 1.536 | -0.307 | -0.044 | 0.581 | 0.008 | 0.031 | 0.031 | 0.924 | 0.53 | 0.31 | 0.6 | 0.35 | 0.57 | 0.39 |
| 4 | 176641650 | rs2644005 | 1.158 | -0.273 | -0.019 | 0.598 | 0 | 0 | 0 | 0.517 | 0.62 | 0.33 | 0.63 | 0.35 | 0.63 | 0.41 |
| 4 | 176644662 | rs2644007 | -0.563 | 0.063 | 0.074 | 0.59 | 0.006 | -0.028 | -0.028 | 0.118 | 0.95 | 0.88 | 0.96 | 0.99 | 0.95 | 0.84 |
| 4 | 176644821 | rs1472328 | 0.776 | -0.004 | 0.094 | 0.705 | -0.004 | 0.072 | 0.072 | 0 | 0.86 | 0.7 | 0.83 | 0.71 | 0.88 | 0.72 |

| | | | | | | | | | | | | | | | |
|---|---|---|---|---|---|---|---|---|---|---|---|---|---|---|---|
| 4 | 176648332 | rs9284954 | 1.589 | -0.069 | 0.122 | 0.766 | -0.001 | -0.001 | -0.001 | 1.422 | 0.54 | 0.46 | 0.55 | 0.44 | 0.55 | 0.46 |
| 4 | 176648393 | rs9312583 | 1.099 | -0.015 | 0.132 | 0.801 | 0.002 | 0.01 | 0.01 | 0.018 | 0.63 | 0.48 | 0.58 | 0.44 | 0.61 | 0.48 |
| 4 | 176651926 | rs7685659 | -1.249 | -0.04 | 0.371 | 1.368 | -0.008 | 0.071 | 0.071 | 0 | 0.1 | 0.18 | 0.14 | 0.64 | 0.07 | 0.21 |
| 4 | 176652429 | rs2644002 | -0.342 | -0.029 | 0.391 | 1.316 | -0.01 | 0.17 | 0.17 | 0 | 0.06 | 0.18 | 0.1 | 0.28 | 0.04 | 0.06 |
| 4 | 176660420 | rs13152426 | -0.039 | 0.229 | -0.153 | 1.34 | -0.004 | -0.004 | -0.004 | 0 | 0.18 | 0.14 | 0.2 | 0.27 | 0.26 | 0.1 |
| 4 | 176662797 | rs1845732 | -1.267 | 0.168 | -0.148 | 1.296 | 0 | -0.004 | -0.004 | 0.177 | 0.35 | 0.29 | 0.37 | 0.55 | 0.37 | 0.32 |
| 4 | 176666186 | rs13144140 | -0.471 | 0.777 | -0.013 | 1.776 | -0.004 | -0.004 | -0.004 | 0 | 0.18 | 0.14 | 0.2 | 0.27 | 0.26 | 0.1 |
| 4 | 176667918 | rs2643988 | 1.099 | 0.373 | 0.312 | 1.685 | 0.011 | 0.301 | 0.301 | 0.081 | 0.9 | 0.61 | 0.92 | 0.63 | 0.95 | 0.79 |
| 4 | 176670796 | rs1472324 | -0.856 | 0.785 | -0.786 | 1.028 | -0.003 | -0.002 | -0.002 | 0.737 | 0.34 | 0.25 | 0.47 | 0.63 | 0.36 | 0.29 |
| 4 | 176671191 | rs2643990 | -0.059 | 1.136 | -0.617 | 0.256 | 0.003 | -0.02 | -0.02 | 0 | 0.73 | 0.72 | 0.68 | 0.9 | 0.71 | 0.85 |
| 4 | 176672144 | rs10011322 | -0.236 | 0.42 | -0.654 | 0.734 | -0.003 | 0.009 | 0.009 | 1.542 | 0.75 | 0.73 | 0.79 | 0.49 | 0.81 | 0.57 |
| 4 | 176673554 | rs2047246 | 0.837 | 0.838 | -1.071 | 0.459 | -0.019 | -0.015 | -0.015 | 0 | 0.22 | 0.16 | 0.32 | 0.24 | 0.28 | 0.29 |
| 4 | 176674283 | rs2643991 | -1.224 | 0.942 | -1.138 | 1.228 | 0.013 | -0.028 | -0.028 | 0 | 0.81 | 0.88 | 0.84 | 0.62 | 0.87 | 0.67 |
| 4 | 176676303 | rs17658370 | -0.227 | 1.3 | -0.913 | 0.899 | 0 | 0.01 | 0.01 | 0.076 | 0.07 | 0.11 | 0.09 | 0.12 | 0.1 | 0.1 |
| 4 | 176677770 | rs722825 | 0.295 | 1.054 | -0.69 | 0.472 | -0.042 | 0.035 | 0.035 | 0 | 0.36 | 0.22 | 0.37 | 0.26 | 0.43 | 0.25 |
| 4 | 176690799 | rs17061991 | -0.662 | 1.257 | 0.056 | 0.246 | 0.006 | -0.026 | -0.026 | 0.825 | 0.05 | 0.1 | 0.02 | 0 | 0.04 | 0.03 |
| 4 | 176698383 | rs2643998 | -0.288 | 1.472 | -0.287 | 0.028 | -0.003 | 0.008 | 0.008 | 0.106 | 0.82 | 0.83 | 0.73 | 0.9 | 0.75 | 0.87 |
| 4 | 176706135 | rs1390343 | 1.028 | 1.828 | -0.305 | -0.329 | -0.047 | 0.081 | 0.081 | 0.112 | 0.63 | 0.2 | 0.53 | 0.88 | 0.46 | 0.75 |
| 4 | 176707862 | rs17062019 | -0.379 | 1.976 | -0.014 | -0.298 | 0.034 | -0.036 | -0.036 | 0 | 0.07 | 0.15 | 0.09 | 0 | 0.14 | 0.03 |
| 4 | 176711191 | rs13140287 | 0.994 | 1.438 | -0.351 | -0.272 | -0.014 | 0.074 | 0.074 | 0.566 | 0.63 | 0.2 | 0.58 | 0.88 | 0.58 | 0.77 |
| 4 | 176716522 | rs13113026 | 1.051 | 1.108 | -0.082 | -0.153 | -0.019 | -0.064 | -0.064 | 0 | 0.83 | 0.38 | 0.89 | 0.96 | 0.9 | 0.92 |
| 4 | 176728576 | rs1495716 | -0.37 | 0.701 | -0.723 | 0.218 | -0.046 | -0.055 | -0.055 | 0.018 | 0.38 | 0.16 | 0.36 | 0.42 | 0.4 | 0.4 |
| 4 | 176728614 | rs13142920 | 1.203 | 0.695 | -0.831 | 0.219 | 0.009 | -0.12 | -0.12 | 0.025 | 0.2 | 0 | 0.08 | 0.36 | 0.14 | 0.23 |
| 4 | 176730094 | rs2291761 | 0.448 | 0.713 | -0.192 | 0.38 | -0.003 | -0.006 | -0.006 | 0 | 0.54 | 0.4 | 0.51 | 0.48 | 0.52 | 0.5 |
| 4 | 176745747 | rs2132339 | 1.151 | 0.808 | 0.171 | 0.232 | 0.119 | 0.134 | 0.134 | 0.182 | 0.94 | 0.54 | 0.88 | 0.89 | 0.93 | 0.89 |
| 4 | 176747173 | rs6838421 | 1.162 | 0.806 | 0.17 | 0.231 | 0.117 | 0.131 | 0.131 | 0.106 | 0.94 | 0.54 | 0.88 | 0.89 | 0.93 | 0.89 |
| 4 | 176754093 | rs11133116 | -0.73 | 0.767 | -0.434 | 0.404 | -0.012 | 0.001 | 0.001 | 0.406 | 0.55 | 0.36 | 0.52 | 0.58 | 0.55 | 0.65 |
| 4 | 176754675 | rs17062130 | 0.799 | 0.726 | -0.29 | 0.424 | 0.004 | 0.014 | 0.014 | 0 | 0.23 | 0.25 | 0.35 | 0.39 | 0.26 | 0.16 |
| 4 | 176766796 | rs10033784 | 0.194 | 0.393 | -0.125 | 0.443 | -0.001 | 0.006 | 0.006 | 0 | 0.33 | 0.36 | 0.41 | 0.36 | 0.34 | 0.33 |
| 4 | 176768164 | rs10520312 | 0.204 | 0.396 | -0.124 | 0.433 | 0 | 0.006 | 0.006 | 0 | 0.33 | 0.36 | 0.41 | 0.37 | 0.34 | 0.33 |
| 4 | 176782054 | rs11940577 | 1.058 | 0.244 | -0.605 | 0.452 | 0.026 | -0.016 | -0.016 | 0 | 0.23 | 0.17 | 0.09 | 0.28 | 0.19 | 0.31 |
| 4 | 176792194 | rs17062188 | 0.348 | 0.951 | -0.446 | 0.6 | -0.007 | -0.022 | -0.022 | 0 | 0.3 | 0.19 | 0.39 | 0.37 | 0.32 | 0.17 |
| 4 | 176795898 | rs4306906 | -0.435 | 0.855 | -0.57 | 0.405 | 0.029 | 0.104 | 0.104 | 0 | 0.49 | 0.31 | 0.66 | 0.69 | 0.56 | 0.43 |
| 4 | 176804052 | rs1826763 | -0.315 | 0.436 | 0.035 | 0.516 | -0.005 | -0.008 | -0.008 | 1.315 | 0.47 | 0.53 | 0.65 | 0.71 | 0.59 | 0.48 |
| 4 | 176820049 | rs11133119 | -0.381 | 0.441 | 0.098 | 0.767 | 0.01 | 0.005 | 0.005 | 0 | 0.33 | 0.34 | 0.57 | 0.58 | 0.54 | 0.4 |
| 4 | 176820095 | rs10049819 | 0.988 | -0.41 | -0.279 | -0.12 | 0.036 | 0.034 | 0.034 | 0 | 0.23 | 0.13 | 0.07 | 0 | 0.13 | 0.04 |
| 4 | 176823651 | rs7436322 | -0.268 | 0.012 | 0.55 | 0.651 | 0.059 | 0.028 | 0.028 | 0 | 0.69 | 0.58 | 0.44 | 0.67 | 0.53 | 0.61 |
| 4 | 176825221 | rs10033742 | -0.008 | 0.002 | 1.091 | 1.116 | -0.001 | 0.025 | 0.025 | 0 | 0.71 | 0.67 | 0.72 | 0.45 | 0.73 | 0.67 |
| 4 | 176827467 | rs903650 | 1.825 | -0.112 | 1.487 | 1.314 | -0.006 | 0.099 | 0.099 | 0 | 0.94 | 0.81 | 0.94 | 0.72 | 0.95 | 0.77 |
| 4 | 176831865 | rs10520316 | -1.936 | -0.2 | 1.698 | 0.912 | 0.005 | 0.003 | 0.003 | 0.566 | 0.05 | 0.3 | 0.03 | 0 | 0 | 0.04 |
| 4 | 176832899 | rs17062252 | -0.519 | -0.389 | 1.666 | 0.912 | -0.008 | 0.001 | 0.001 | 0 | 0.21 | 0.24 | 0.12 | 0.34 | 0.18 | 0.49 |
| 4 | 176837789 | rs1495714 | 1.843 | -0.509 | 1.378 | 0.993 | 0.054 | -0.038 | -0.038 | 0.267 | 0.6 | 0.12 | 0.7 | 0.56 | 0.69 | 0.28 |
| 4 | 176838690 | rs7438439 | 0.828 | -0.494 | 2.107 | 0.933 | -0.036 | -0.01 | -0.01 | 0 | 0.75 | 0.37 | 0.84 | 0.71 | 0.83 | 0.49 |
| 4 | 176839335 | rs9686024 | -1.582 | -0.287 | 2.081 | 0.95 | 0.009 | 0.004 | 0.004 | 0 | 0.14 | 0.22 | 0.13 | 0.12 | 0.16 | 0.2 |
| 4 | 176840816 | rs6818081 | -0.996 | -0.209 | 2.117 | 0.994 | 0.014 | 0.014 | 0.014 | 0 | 0.1 | 0.16 | 0.11 | 0.12 | 0.14 | 0.17 |
| 4 | 176850196 | rs17062274 | -0.087 | -0.288 | 2.24 | 0.924 | -0.015 | -0.007 | -0.007 | 0 | 0.17 | 0.29 | 0.08 | 0.17 | 0.12 | 0.27 |
| 4 | 176861301 | rs1106568 | 1.249 | -0.097 | 1.993 | 0.914 | 0.004 | 0.018 | 0.018 | 0 | 0.79 | 0.66 | 0.79 | 0.73 | 0.75 | 0.58 |
| 4 | 176869131 | rs34044173 | 0.452 | -0.142 | 1.896 | 0.92 | 0.004 | 0.005 | 0.005 | 0.283 | 0.15 | 0.08 | 0.07 | 0.16 | 0.13 | 0.25 |
| 4 | 176872920 | rs4690647 | 2.349 | -0.206 | 1.806 | 0.818 | 0.022 | 0.011 | 0.011 | 0.094 | 0.56 | 0.19 | 0.67 | 0.56 | 0.6 | 0.28 |
| 4 | 176880351 | rs10520320 | 0.427 | -0.254 | 1.824 | 0.466 | 0.001 | 0.008 | 0.008 | 0.278 | 0.14 | 0.06 | 0.07 | 0.15 | 0.13 | 0.26 |
| 4 | 176882871 | rs7658712 | -0.872 | -0.253 | 1.917 | 0.466 | -0.037 | -0.054 | -0.054 | 0.512 | 0.22 | 0.43 | 0.11 | 0.15 | 0.14 | 0.31 |
| 4 | 176902231 | rs17062345 | -0.064 | 0.02 | 1.043 | 0.708 | 0.001 | -0.002 | -0.002 | 0.663 | 0.17 | 0.13 | 0.07 | 0.23 | 0.14 | 0.23 |
| 4 | 176918741 | rs7675676 | -1.502 | 0.356 | 1.057 | 0.579 | 0.014 | -0.045 | -0.045 | 0 | 0.33 | 0.48 | 0.26 | 0.3 | 0.35 | 0.45 |
| 4 | 176922278 | rs6553899 | 1.464 | 0.436 | 1.02 | 0.798 | 0.018 | 0.003 | 0.003 | 0.157 | 0.75 | 0.69 | 0.74 | 0.7 | 0.69 | 0.53 |
| 4 | 176924716 | rs1471797 | 1.392 | 0.395 | 1.006 | 1.152 | 0.022 | 0.062 | 0.062 | 0.574 | 0.81 | 0.68 | 0.73 | 0.65 | 0.75 | 0.59 |
| 4 | 176926946 | rs1471798 | 1.056 | 0.361 | 0.989 | 1.168 | -0.007 | 0.184 | 0.184 | 0 | 0.92 | 0.71 | 0.9 | 0.76 | 0.95 | 0.71 |
| 4 | 176936038 | rs997779 | -0.711 | 0.483 | 0.549 | 1.086 | -0.066 | -0.018 | -0.018 | 0 | 0.11 | 0.01 | 0.17 | 0.11 | 0.2 | 0.12 |
| 4 | 176949352 | rs11936134 | -1.588 | 0.149 | 0.556 | 0.719 | -0.003 | 0.002 | 0.002 | 0 | 0.43 | 0.52 | 0.34 | 0.52 | 0.4 | 0.65 |
| 4 | 176950029 | rs11133123 | 1.454 | 0.151 | -0.113 | 0.706 | -0.002 | -0.031 | -0.031 | 6.166 | 0.53 | 0.11 | 0.61 | 0.48 | 0.58 | 0.35 |
| 4 | 176957224 | rs12499459 | 1.657 | 0.084 | -0.24 | 0.912 | -0.011 | -0.002 | -0.002 | 0 | 0.49 | 0.08 | 0.6 | 0.4 | 0.58 | 0.34 |
| 4 | 176957395 | rs6823563 | -1.761 | 0.089 | 0.146 | 0.971 | 0.003 | 0.012 | 0.012 | 0 | 0.24 | 0.29 | 0.27 | 0.36 | 0.27 | 0.43 |
| 4 | 176970155 | rs17669562 | -0.514 | 0.634 | -0.079 | 0.842 | -0.072 | -0.043 | -0.043 | 0 | 0.12 | 0.02 | 0.2 | 0.15 | 0.23 | 0.13 |
| 4 | 176971792 | rs973732 | 0.617 | 0.459 | 0.449 | 0.723 | -0.001 | -0.001 | -0.001 | 0 | 0.65 | 0.66 | 0.71 | 0.57 | 0.64 | 0.65 |
| 4 | 176973932 | rs6553905 | 0.789 | 0.926 | 0.128 | 0.754 | -0.002 | -0.001 | -0.001 | 0.412 | 0.6 | 0.44 | 0.64 | 0.57 | 0.56 | 0.65 |
| 4 | 176975408 | rs13146736 | 0.071 | 0.845 | -0.1 | 0.553 | -0.009 | -0.024 | -0.024 | 0 | 0.61 | 0.51 | 0.46 | 0.73 | 0.65 | 0.74 |
| 4 | 176976075 | rs7692309 | 0.64 | 1.458 | -0.037 | 0.658 | 0.005 | 0.021 | 0.021 | 0 | 0.51 | 0.34 | 0.36 | 0.6 | 0.46 | 0.68 |
| 4 | 176978086 | rs7657735 | -0.469 | 1.479 | -0.078 | 0.667 | 0.027 | 0.013 | 0.013 | 0 | 0.1 | 0.17 | 0.09 | 0.13 | 0.19 | 0.06 |
| 4 | 176980809 | rs2333391 | 0.66 | 1.403 | -0.028 | 0.639 | 0.007 | 0.028 | 0.028 | 0 | 0.51 | 0.3 | 0.36 | 0.6 | 0.46 | 0.68 |
| 4 | 176993039 | rs1385614 | 0.308 | 0.952 | -0.137 | 0.536 | 0.036 | 0.019 | 0.019 | 0 | 0.54 | 0.29 | 0.4 | 0.6 | 0.61 | 0.67 |

| | | | | | | | | | | | | | | | | |
|---|---|---|---|---|---|---|---|---|---|---|---|---|---|---|---|---|
| 4 | 177006043 | rs12641442 | -0.553 | 0.636 | -0.127 | 0.252 | -0.001 | 0.001 | 0.001 | 0 | 0.31 | 0.32 | 0.3 | 0.44 | 0.39 | 0.42 |
| 4 | 177016992 | rs17624517 | -0.552 | 0.593 | 0.151 | 0.231 | 0 | 0.002 | 0.002 | 0 | 0.31 | 0.32 | 0.3 | 0.44 | 0.39 | 0.42 |
| 4 | 177045953 | rs10009701 | -0.918 | 0.637 | -0.066 | -0.124 | 0 | 0.002 | 0.002 | 0 | 0.65 | 0.68 | 0.48 | 0.72 | 0.67 | 0.75 |
| 4 | 177061748 | rs7678897 | 0.805 | 0.534 | 0.006 | -0.209 | -0.014 | -0.051 | -0.051 | 0 | 0.33 | 0.33 | 0.14 | 0.16 | 0.25 | 0.27 |
| 4 | 177079374 | rs2128422 | -0.643 | 0.402 | -0.051 | -0.242 | 0.023 | 0.034 | 0.034 | 0 | 0.62 | 0.76 | 0.5 | 0.73 | 0.68 | 0.76 |
| 4 | 177085016 | rs2170576 | 0.565 | 0.332 | -0.054 | -0.348 | -0.006 | 0.002 | 0.002 | 0 | 0.61 | 0.4 | 0.63 | 0.39 | 0.55 | 0.47 |
| 4 | 177091829 | rs4475089 | 0.081 | 0.192 | -0.123 | -0.43 | 0.003 | 0.001 | 0.001 | 0.051 | 0.69 | 0.41 | 0.64 | 0.39 | 0.56 | 0.52 |
| 4 | 177095908 | rs4690665 | 0.428 | 0.264 | -0.023 | -0.703 | 0.006 | 0.005 | 0.005 | 0 | 0.3 | 0.28 | 0.42 | 0.26 | 0.26 | 0.16 |
| 4 | 177098285 | rs17625943 | 0.711 | 0.181 | -0.037 | -0.736 | 0.002 | 0 | 0 | 0 | 0.27 | 0.25 | 0.42 | 0.26 | 0.26 | 0.16 |
| 4 | 177100644 | rs11736872 | 1.447 | 0.237 | -0.002 | -0.735 | -0.022 | 0.098 | 0.098 | 0 | 0.3 | 0.14 | 0.21 | 0.13 | 0.28 | 0.31 |
| 4 | 177101060 | rs7699964 | 0.741 | 0.308 | -0.029 | -0.684 | 0.001 | 0 | 0 | 0 | 0.27 | 0.28 | 0.42 | 0.26 | 0.26 | 0.16 |
| 4 | 177105388 | rs11133135 | -0.778 | 0.286 | 0.197 | -0.695 | -0.009 | -0.005 | -0.005 | 0.99 | 0.72 | 0.54 | 0.56 | 0.74 | 0.74 | 0.84 |
| 4 | 177107335 | rs2333402 | 0.698 | 0.375 | 0.162 | -0.489 | 0.003 | 0.044 | 0.044 | 0 | 0.34 | 0.29 | 0.22 | 0.13 | 0.28 | 0.31 |
| 4 | 177113779 | rs12502935 | -0.162 | 0.826 | 0.089 | -0.341 | 0.003 | 0.008 | 0.008 | 0 | 0.26 | 0.27 | 0.24 | 0.61 | 0.31 | 0.38 |
| 4 | 177113836 | rs6832177 | 1.115 | 0.896 | -0.055 | -0.727 | -0.013 | 0.025 | 0.025 | 0 | 0.34 | 0.22 | 0.36 | 0.26 | 0.37 | 0.24 |
| 4 | 177113956 | rs2291244 | 0.895 | 0.908 | -0.125 | 0.011 | 0 | 0.015 | 0.015 | 0 | 0.16 | 0.16 | 0.16 | 0.38 | 0.18 | 0.26 |
| 4 | 177114889 | rs6820936 | -0.947 | 0.843 | -0.029 | -0.181 | -0.003 | -0.051 | -0.051 | 0 | 0.66 | 0.62 | 0.63 | 0.86 | 0.69 | 0.63 |
| 4 | 177117349 | rs1905184 | 0.174 | 1.102 | -0.416 | -0.012 | -0.002 | -0.003 | -0.003 | 0 | 0.39 | 0.38 | 0.36 | 0.46 | 0.46 | 0.37 |
| 4 | 177119860 | rs10004500 | -0.745 | 1.154 | -0.527 | -0.168 | 0 | -0.001 | -0.001 | 0 | 0.53 | 0.54 | 0.55 | 0.46 | 0.63 | 0.42 |
| 4 | 177121983 | rs728846 | 0.73 | 1.25 | -0.402 | -0.519 | -0.002 | 0.003 | 0.003 | 0.245 | 0.34 | 0.32 | 0.34 | 0.24 | 0.37 | 0.25 |
| 4 | 177124254 | rs4626145 | -0.379 | 1.362 | -0.634 | 0.026 | 0.003 | 0.018 | 0.018 | 0 | 0.27 | 0.34 | 0.28 | 0.49 | 0.3 | 0.33 |
| 4 | 177126245 | rs2333406 | 0.348 | 1.459 | -1.109 | 0.322 | -0.029 | -0.07 | -0.07 | 0 | 0.2 | 0.09 | 0.21 | 0.49 | 0.25 | 0.29 |
| 4 | 177127021 | rs11133136 | -0.474 | 0.894 | -0.835 | -0.039 | -0.007 | -0.008 | -0.008 | 0 | 0.68 | 0.64 | 0.79 | 0.73 | 0.77 | 0.57 |
| 4 | 177131268 | rs17062621 | -0.688 | 1.491 | -0.499 | 0.589 | 0.059 | 0.05 | 0.05 | 3.406 | 0.21 | 0.57 | 0.25 | 0.3 | 0.29 | 0.27 |
| 4 | 177134283 | rs13120122 | 1.304 | 1.33 | -1.44 | -0.24 | -0.07 | 0.132 | 0.132 | 0 | 0.2 | 0.01 | 0.19 | 0 | 0.22 | 0.07 |
| 4 | 177135620 | rs10003543 | -1.686 | 0.708 | -1.812 | -0.312 | 0.012 | -0.003 | -0.003 | 0 | 0.86 | 0.92 | 0.89 | 0.83 | 0.92 | 0.73 |
| 4 | 177136149 | rs13109244 | 1.665 | 0.805 | -1.787 | -0.287 | -0.069 | 0.133 | 0.133 | 0 | 0.2 | 0.01 | 0.19 | 0 | 0.22 | 0.07 |
| 4 | 177136512 | rs922849 | -1.557 | 0.456 | -1.722 | -0.395 | 0.012 | 0.003 | 0.003 | 0 | 0.83 | 0.88 | 0.89 | 0.83 | 0.92 | 0.69 |
| 4 | 177137373 | rs17062628 | 1.592 | 0.037 | -1.928 | -0.253 | 0.006 | -0.007 | -0.007 | 0 | 0.08 | 0.03 | 0.06 | 0.11 | 0.07 | 0.06 |
| 4 | 177137988 | rs6827525 | 0.112 | 0.3 | -1.473 | -0.105 | 0.003 | 0.003 | 0.003 | 0 | 0.5 | 0.52 | 0.55 | 0.57 | 0.48 | 0.47 |
| 4 | 177142776 | rs10032098 | -1.928 | 0.289 | -1.473 | -0.105 | 0.012 | 0.003 | 0.003 | 0 | 0.83 | 0.88 | 0.89 | 0.83 | 0.91 | 0.69 |
| 4 | 177143727 | rs6830358 | 1.246 | 0.266 | -1.384 | -0.129 | 0.096 | 0.123 | 0.123 | 0.402 | 0.12 | 0.28 | 0.15 | 0.26 | 0.21 | 0.15 |
| 4 | 177144364 | rs6814870 | -1.967 | 0.213 | -1.376 | -0.056 | 0.001 | -0.001 | -0.001 | 0 | 0.83 | 0.82 | 0.92 | 0.82 | 0.94 | 0.71 |
| 4 | 177146627 | rs10013975 | 0.596 | 0.397 | -1.415 | -0.045 | 0 | 0 | 0 | 0 | 0.32 | 0.29 | 0.34 | 0.26 | 0.44 | 0.22 |
| 4 | 177149518 | rs17673171 | 0.677 | 0.172 | -1.55 | -0.435 | -0.009 | 0.009 | 0.009 | 0 | 0.14 | 0.1 | 0.3 | 0.12 | 0.23 | 0.18 |
| 4 | 177150491 | rs1351632 | -0.429 | -0.42 | -1.619 | -0.788 | -0.002 | -0.023 | -0.023 | 0 | 0.52 | 0.48 | 0.59 | 0.23 | 0.57 | 0.37 |
| 4 | 177150925 | rs2046262 | 0.011 | -0.465 | -1.817 | -1.074 | -0.006 | 0.036 | 0.036 | 0 | 0.45 | 0.62 | 0.42 | 0.7 | 0.47 | 0.45 |
| 4 | 177154323 | rs13117768 | 0.644 | -0.389 | -1.835 | -1.063 | -0.072 | 0.142 | 0.142 | 0 | 0.34 | 0.12 | 0.48 | 0.12 | 0.46 | 0.25 |
| 4 | 177157830 | rs7684092 | 0.897 | -0.33 | -1.808 | -1.067 | -0.024 | 0.071 | 0.071 | 0 | 0.33 | 0.2 | 0.44 | 0.09 | 0.38 | 0.19 |
| 4 | 177160262 | rs7688636 | 1.61 | -0.221 | -2.125 | -0.994 | -0.017 | 0.014 | 0.014 | 0 | 0.08 | 0.02 | 0.25 | 0.09 | 0.16 | 0.12 |
| 4 | 177162117 | rs9991242 | -1.386 | -0.383 | -2.042 | -0.929 | 0.005 | 0.001 | 0.001 | 0.018 | 0.78 | 0.8 | 0.87 | 0.79 | 0.86 | 0.64 |
| 4 | 177163414 | rs9312595 | -1.385 | -0.353 | -2.049 | -0.752 | 0.006 | -0.005 | -0.005 | 0.153 | 0.78 | 0.8 | 0.87 | 0.72 | 0.85 | 0.63 |
| 4 | 177164064 | rs9790840 | -0.075 | -0.374 | -2.096 | -0.825 | -0.015 | 0.045 | 0.045 | 0 | 0.45 | 0.66 | 0.43 | 0.63 | 0.47 | 0.45 |
| 4 | 177165556 | rs4690671 | -1.966 | -0.331 | -1.918 | -0.322 | 0.029 | -0.024 | -0.024 | 0 | 0.84 | 0.91 | 0.92 | 0.75 | 0.93 | 0.69 |
| 4 | 177167740 | rs17679314 | 1.967 | 0.17 | -1.656 | -0.348 | 0.005 | -0.013 | -0.013 | 0 | 0.06 | 0.09 | 0.05 | 0.03 | 0.08 | 0.06 |
| 4 | 177180575 | rs10520340 | 1.845 | 0.347 | -1.925 | -0.316 | -0.036 | 0.002 | 0.002 | 0 | 0.06 | 0 | 0.15 | 0.09 | 0.13 | 0.12 |
| 4 | 177184276 | rs10005873 | -0.394 | 0.198 | -1.927 | -0.419 | 0.001 | 0.012 | 0.012 | 0 | 0.64 | 0.55 | 0.71 | 0.47 | 0.67 | 0.49 |
| 4 | 177186882 | rs1454167 | -0.828 | -0.075 | -2.046 | -0.583 | 0.003 | 0.012 | 0.012 | 0 | 0.67 | 0.61 | 0.72 | 0.47 | 0.67 | 0.5 |
| 4 | 177194007 | rs12649379 | -0.438 | -0.076 | -2.09 | -0.583 | 0 | 0.009 | 0.009 | 0 | 0.63 | 0.56 | 0.72 | 0.47 | 0.67 | 0.49 |
| 4 | 177195145 | rs6820078 | 1.155 | 0.188 | -2.212 | -0.386 | 0.093 | 0.102 | 0.102 | 0.191 | 0.12 | 0.27 | 0.18 | 0.28 | 0.24 | 0.18 |
| 4 | 177202275 | rs13129947 | 1.654 | -0.306 | -3.002 | -1.127 | -0.061 | 0.155 | 0.155 | 0 | 0.24 | 0.02 | 0.28 | 0 | 0.26 | 0.08 |
| 4 | 177202400 | rs4690681 | -1.122 | -0.275 | -2.296 | -0.393 | 0.04 | -0.011 | -0.011 | 0.093 | 0.79 | 0.89 | 0.9 | 0.75 | 0.91 | 0.69 |
| 4 | 177202549 | rs4690682 | -0.146 | 0.522 | -1.984 | -0.332 | -0.067 | 0.064 | 0.064 | 1.043 | 0.17 | 0.01 | 0.21 | 0.06 | 0.24 | 0.18 |
| 4 | 177212135 | rs11133142 | -0.121 | -0.096 | -1.393 | 0.022 | 0.018 | 0.015 | 0.015 | 0 | 0.58 | 0.69 | 0.76 | 0.59 | 0.7 | 0.48 |
| 4 | 177224449 | rs13115859 | -0.579 | 0.046 | -1.473 | 0.09 | -0.011 | -0.024 | -0.024 | 0 | 0.66 | 0.76 | 0.5 | 0.58 | 0.57 | 0.6 |
| 4 | 177227998 | rs12507341 | 0.844 | 0.096 | -1.091 | 0.133 | 0 | -0.002 | -0.002 | 0 | 0.27 | 0.27 | 0.13 | 0.25 | 0.24 | 0.44 |
| 4 | 177232708 | rs6553943 | -0.753 | 0.113 | -1.332 | -0.342 | -0.006 | -0.012 | -0.012 | 0 | 0.58 | 0.74 | 0.48 | 0.58 | 0.54 | 0.59 |
| 4 | 177233904 | rs7670905 | -0.178 | 0.101 | -1.347 | -0.474 | 0.015 | 0.002 | 0.002 | 0.935 | 0.47 | 0.64 | 0.31 | 0.58 | 0.36 | 0.55 |
| 4 | 177234129 | rs13115857 | -2.409 | 0.162 | -1.39 | -0.259 | -0.011 | -0.005 | -0.005 | 0 | 0.84 | 0.9 | 0.62 | 0.84 | 0.7 | 0.84 |
| 4 | 177234906 | rs723820 | 0.379 | -0.248 | -0.935 | 0.062 | 0 | 0.017 | 0.017 | 0 | 0.3 | 0.35 | 0.23 | 0.58 | 0.27 | 0.5 |
| 4 | 177235286 | rs9992247 | 0.183 | -0.022 | -1.44 | 0.043 | -0.009 | 0.024 | 0.024 | 0 | 0.36 | 0.59 | 0.24 | 0.52 | 0.27 | 0.49 |
| 4 | 177235586 | rs13130930 | -0.177 | 0.536 | -1.516 | 0.206 | -0.044 | 0.052 | 0.052 | 0 | 0.1 | 0.01 | 0.17 | 0 | 0.18 | 0.04 |
| 4 | 177237359 | rs11735748 | -0.434 | 0.104 | -1.729 | 0.453 | -0.045 | -0.061 | -0.061 | 0 | 0.31 | 0.08 | 0.21 | 0.33 | 0.3 | 0.34 |
| 4 | 177237631 | rs17062749 | -0.392 | 0.08 | -0.788 | 0.606 | 0.004 | 0.077 | 0.077 | 0 | 0.11 | 0.21 | 0.13 | 0.24 | 0.09 | 0.14 |
| 4 | 177239269 | rs1454162 | -0.506 | -0.022 | -0.805 | 0.201 | 0.001 | -0.031 | -0.031 | 0 | 0.13 | 0.14 | 0.12 | 0 | 0.15 | 0.05 |
| 4 | 177239338 | rs1454160 | -0.492 | -0.013 | -0.782 | 0.205 | 0.001 | -0.034 | -0.034 | 0 | 0.12 | 0.15 | 0.12 | 0 | 0.15 | 0.05 |
| 4 | 177242959 | rs4690686 | -1.294 | -0.257 | -0.888 | 0.208 | 0.031 | 0.063 | 0.063 | 0 | 0.48 | 0.3 | 0.25 | 0.32 | 0.39 | 0.32 |
| 4 | 177244326 | rs17681035 | 0.348 | -0.097 | -0.975 | 0.088 | -0.004 | -0.005 | -0.005 | 0 | 0.08 | 0.1 | 0.04 | 0 | 0.07 | 0.06 |
| 4 | 177250491 | rs17629999 | 0.747 | -0.078 | -1.015 | 0.1 | -0.034 | 0.031 | 0.031 | 0 | 0.06 | 0.02 | 0.08 | 0 | 0.12 | 0.02 |

| | | | | | | | | | | | | | | | | |
|---|---|---|---|---|---|---|---|---|---|---|---|---|---|---|---|---|
| 4 | 177255407 | rs13117427 | 0.451 | -0.074 | -0.99 | 0.107 | -0.01 | 0.01 | 0.01 | 0 | 0.08 | 0.06 | 0.04 | 0 | 0.08 | 0.07 |
| 4 | 177256699 | rs10027831 | -1.458 | -0.307 | -0.462 | 0.215 | 0.004 | 0.007 | 0.007 | 0.599 | 0.55 | 0.66 | 0.35 | 0.53 | 0.43 | 0.42 |
| 4 | 177259635 | rs4690687 | -0.453 | -0.323 | -0.945 | 0.306 | 0.021 | 0.007 | 0.007 | 0 | 0.37 | 0.13 | 0.16 | 0.3 | 0.29 | 0.3 |
| 4 | 177263983 | rs4690429 | -0.601 | -0.246 | -0.882 | -0.042 | 0.051 | -0.034 | -0.034 | 0 | 0.41 | 0.21 | 0.23 | 0.53 | 0.3 | 0.4 |
| 4 | 177285894 | rs7694145 | -1.556 | 0.454 | 0.461 | -0.232 | 0.009 | -0.147 | -0.147 | 0 | 0.22 | 0.57 | 0.18 | 0 | 0.19 | 0.08 |
| 4 | 177290945 | rs1454158 | -1.163 | 0.849 | 0.138 | 0.297 | -0.002 | 0.006 | 0.006 | 1.461 | 0.24 | 0.36 | 0.21 | 0.31 | 0.26 | 0.22 |
| 4 | 177292032 | rs13144122 | -1.156 | 1.322 | -0.291 | -0.063 | -0.005 | -0.024 | -0.024 | 0.457 | 0.23 | 0.25 | 0.19 | 0.03 | 0.32 | 0.16 |
| 4 | 177296975 | rs11944672 | 0.88 | 0.796 | -0.753 | -0.256 | 0.005 | 0.3 | 0.3 | 0 | 0.48 | 0.15 | 0.43 | 0.09 | 0.55 | 0.35 |
| 4 | 177298249 | rs7682500 | 0 | 0.632 | 0.244 | -0.147 | -0.032 | -0.07 | -0.07 | 0 | 0.88 | 0.77 | 0.94 | 1 | 0.92 | 0.95 |
| 4 | 177307459 | rs11932699 | -0.31 | 0.641 | 0.471 | -0.15 | -0.03 | -0.078 | -0.078 | 0.053 | 0.12 | 0.26 | 0.06 | 0 | 0.08 | 0.05 |
| 4 | 177308667 | rs6553951 | 1.319 | 0.625 | 0.469 | -0.177 | -0.021 | -0.123 | -0.123 | 0 | 0.68 | 0.56 | 0.83 | 0.9 | 0.74 | 0.75 |
| 4 | 177308956 | rs13139292 | -0.18 | 1.36 | -0.062 | 0.343 | -0.012 | 0.02 | 0.02 | 0 | 0.05 | 0.01 | 0.02 | 0.01 | 0.05 | 0.06 |
| 4 | 177310481 | rs17631173 | -1.274 | 1.441 | 0.094 | 0.53 | -0.04 | 0.041 | 0.041 | 0 | 0.11 | 0.03 | 0.12 | 0 | 0.2 | 0.08 |
| 4 | 177324650 | rs10018184 | 1.5 | 0.44 | 0.726 | 1.595 | -0.035 | 0.087 | 0.087 | 0 | 0.8 | 0.59 | 0.64 | 0.45 | 0.88 | 0.67 |
| 4 | 177328296 | rs2123861 | 1.831 | 0.468 | 0.873 | 1.541 | -0.034 | 0.08 | 0.08 | 0 | 0.76 | 0.52 | 0.6 | 0.45 | 0.82 | 0.66 |
| 4 | 177334892 | rs6832586 | 0.74 | 0.58 | 0.894 | 2.17 | -0.011 | 0.134 | 0.134 | 0 | 0.89 | 0.69 | 0.78 | 0.67 | 0.92 | 0.85 |
| 4 | 177336067 | rs6553957 | 0.885 | 0.593 | 0.73 | 2.178 | -0.01 | 0.166 | 0.166 | 0 | 0.88 | 0.63 | 0.78 | 0.67 | 0.91 | 0.85 |
| 4 | 177337800 | rs12331128 | -1.625 | 0.842 | 0.625 | 1.96 | 0.01 | 0.239 | 0.239 | 0 | 0.09 | 0.25 | 0.17 | 0.33 | 0.07 | 0.27 |
| 4 | 177339963 | rs11133152 | -1.094 | 1.529 | -0.012 | 1.476 | 0.003 | 0 | 0 | 0.001 | 0.09 | 0.08 | 0.25 | 0.07 | 0.14 | 0.11 |
| 4 | 177340086 | rs4690437 | -0.251 | 1.191 | -0.364 | 1.55 | 0 | -0.009 | -0.009 | 0 | 0.4 | 0.39 | 0.48 | 0.49 | 0.36 | 0.39 |
| 4 | 177340623 | rs4146092 | -0.476 | 1.584 | -1.13 | 1.616 | 0.001 | -0.033 | -0.033 | 0 | 0.53 | 0.84 | 0.61 | 0.51 | 0.51 | 0.43 |
| 4 | 177343885 | rs6553961 | -0.371 | 2.133 | -0.1 | 1.713 | 0.093 | 0.049 | 0.049 | 0 | 0.12 | 0.48 | 0.09 | 0.15 | 0.13 | 0.07 |
| 4 | 177350693 | rs2167094 | -0.283 | 1.887 | -0.678 | 1.764 | -0.001 | -0.019 | -0.019 | 0 | 0.18 | 0.13 | 0.17 | 0.28 | 0.21 | 0.39 |
| 4 | 177355622 | rs10009750 | 0.869 | 2.109 | -0.204 | 1.542 | -0.018 | -0.096 | -0.096 | 2.089 | 0.37 | 0.45 | 0.32 | 0.13 | 0.29 | 0.25 |
| 4 | 177358612 | rs4690442 | 0.529 | 2.161 | -0.227 | 1.77 | -0.023 | -0.027 | -0.027 | 0.133 | 0.15 | 0.04 | 0.16 | 0.19 | 0.2 | 0.31 |
| 4 | 177359397 | rs28562735 | 1.25 | 2.164 | -0.284 | 1.87 | 0.034 | 0.022 | 0.022 | 0 | 0.69 | 0.52 | 0.57 | 0.41 | 0.61 | 0.52 |
| 4 | 177360669 | rs921691 | -0.503 | 2.143 | -0.278 | 1.87 | 0 | 0 | 0 | 0 | 0.83 | 0.82 | 0.83 | 0.82 | 0.79 | 0.69 |
| 4 | 177366729 | rs1377841 | 0.515 | 2.163 | -0.265 | 2.044 | 0.005 | -0.001 | -0.001 | 0.001 | 0.78 | 0.74 | 0.77 | 0.77 | 0.7 | 0.64 |
| 4 | 177371366 | rs17062990 | 0.759 | 2.164 | -0.263 | 2.056 | 0.026 | 0.196 | 0.196 | 0 | 0.09 | 0.21 | 0.2 | 0.36 | 0.09 | 0.12 |
| 4 | 177372753 | rs2167096 | 1.64 | 2.14 | -0.336 | 2.023 | -0.043 | 0.022 | 0.022 | 0 | 0.29 | 0.06 | 0.23 | 0.18 | 0.31 | 0.18 |
| 4 | 177373302 | rs2167095 | -0.321 | 2.11 | -0.333 | 1.918 | 0.012 | -0.004 | -0.004 | 0 | 0.06 | 0.08 | 0.06 | 0.05 | 0.09 | 0.06 |
| 4 | 177374643 | rs309753 | -0.894 | 2.083 | -0.309 | 2.073 | -0.011 | -0.023 | -0.023 | 0.233 | 0.85 | 0.91 | 0.72 | 0.8 | 0.79 | 0.7 |
| 4 | 177378346 | rs6837598 | 0.79 | 1.999 | -0.213 | 1.927 | -0.015 | -0.043 | -0.043 | 0.307 | 0.4 | 0.51 | 0.34 | 0.23 | 0.3 | 0.33 |
| 4 | 177391167 | rs365051 | 0.236 | 2.145 | -0.227 | 1.953 | 0.001 | 0.02 | 0.02 | 0 | 0.46 | 0.59 | 0.41 | 0.27 | 0.42 | 0.4 |
| 4 | 177394679 | rs309732 | -0.672 | 2.249 | -0.281 | 2.147 | -0.022 | -0.017 | -0.017 | 0.175 | 0.58 | 0.85 | 0.51 | 0.62 | 0.51 | 0.55 |
| 4 | 177396008 | rs309729 | -0.765 | 2.229 | -0.437 | 2.044 | -0.018 | 0.028 | 0.028 | 0.06 | 0.58 | 0.86 | 0.63 | 0.67 | 0.53 | 0.57 |
| 4 | 177397059 | rs13132587 | 1.674 | 2.239 | -0.526 | 2.045 | -0.063 | -0.004 | -0.004 | 0 | 0.28 | 0.02 | 0.22 | 0.18 | 0.31 | 0.18 |
| 4 | 177400257 | rs4146433 | 0.258 | 2.247 | -0.315 | 2.059 | 0.028 | 0.19 | 0.19 | 0 | 0.1 | 0.2 | 0.2 | 0.46 | 0.12 | 0.18 |
| 4 | 177401671 | rs309719 | -0.505 | 2.303 | -0.171 | 1.898 | 0.046 | 0.005 | 0.005 | 0 | 0.06 | 0.12 | 0.06 | 0.07 | 0.1 | 0.07 |
| 4 | 177406393 | rs4690445 | -1.71 | 2.229 | -0.339 | 1.869 | -0.026 | 0.026 | 0.026 | 0 | 0.71 | 0.92 | 0.78 | 0.82 | 0.69 | 0.82 |
| 4 | 177406476 | rs10866385 | -0.819 | 2.247 | -0.275 | 2.055 | -0.004 | 0.008 | 0.008 | 0 | 0.61 | 0.69 | 0.72 | 0.67 | 0.59 | 0.74 |
| 4 | 177407175 | rs1449727 | 0.329 | 2.163 | -0.292 | 2.027 | 0.019 | 0.029 | 0.029 | 0 | 0.1 | 0.23 | 0.06 | 0.15 | 0.09 | 0.08 |
| 4 | 177410613 | rs11133155 | 0.75 | 2.174 | -0.324 | 2.047 | -0.008 | -0.01 | -0.01 | 0 | 0.12 | 0.05 | 0.15 | 0.15 | 0.17 | 0.25 |
| 4 | 177419908 | rs4690351 | 0.414 | 2.159 | -0.267 | 1.916 | -0.001 | -0.001 | -0.001 | 0 | 0.13 | 0.1 | 0.16 | 0.15 | 0.17 | 0.25 |
| 4 | 177420945 | rs1375738 | 0.547 | 2.181 | -0.289 | 1.882 | 0.03 | 0.042 | 0.042 | 0 | 0.09 | 0.23 | 0.06 | 0.15 | 0.09 | 0.08 |
| 4 | 177422510 | rs403742 | -0.781 | 2.107 | -0.301 | 1.786 | -0.012 | -0.06 | -0.06 | 2.185 | 0.57 | 0.44 | 0.64 | 0.85 | 0.69 | 0.68 |
| 4 | 177429154 | rs309703 | -1.147 | 2.111 | -0.482 | 1.957 | 0.011 | 0.035 | 0.035 | 0 | 0.2 | 0.19 | 0.36 | 0.52 | 0.28 | 0.43 |
| 4 | 177430724 | rs448336 | -1.162 | 2.266 | -0.691 | 2.053 | 0 | 0.049 | 0.049 | 0 | 0.1 | 0.08 | 0.2 | 0.4 | 0.14 | 0.19 |
| 4 | 177436430 | rs377413 | -0.337 | 1.829 | -0.619 | 1.718 | -0.016 | 0.019 | 0.019 | 0 | 0.58 | 0.67 | 0.56 | 0.41 | 0.55 | 0.64 |
| 4 | 177440967 | rs416203 | 0.583 | 1.682 | -0.538 | 1.779 | 0.025 | 0.134 | 0.134 | 0 | 0.85 | 0.73 | 0.8 | 0.58 | 0.86 | 0.8 |
| 4 | 177442955 | rs12500053 | 0.552 | 1.382 | -0.541 | 1.2 | 0 | 0.002 | 0.002 | 0 | 0.38 | 0.36 | 0.37 | 0.24 | 0.35 | 0.36 |
| 4 | 177444335 | rs425571 | -0.308 | 1.349 | -1.112 | 1.293 | -0.014 | 0.062 | 0.062 | 0 | 0.55 | 0.7 | 0.5 | 0.37 | 0.5 | 0.63 |
| 4 | 177460096 | rs13115445 | 1.581 | 1.406 | -1.805 | 1.187 | -0.05 | 0.072 | 0.072 | 0 | 0.28 | 0.05 | 0.26 | 0.09 | 0.33 | 0.09 |
| 4 | 177461365 | rs1375749 | 1.612 | 1.406 | -1.808 | 1.187 | -0.063 | 0.067 | 0.067 | 0 | 0.28 | 0.05 | 0.26 | 0.1 | 0.33 | 0.09 |
| 4 | 177462810 | rs309780 | 0.502 | 1.424 | -1.642 | 1.048 | 0.01 | 0.042 | 0.042 | 0 | 0.43 | 0.63 | 0.4 | 0.36 | 0.35 | 0.37 |
| 4 | 177464940 | rs309789 | 0.304 | 1.472 | -1.614 | 1.27 | 0.011 | 0.077 | 0.077 | 0 | 0.85 | 0.72 | 0.76 | 0.58 | 0.83 | 0.76 |
| 4 | 177466128 | rs309794 | 0.076 | 1.52 | -1.719 | 1.455 | -0.003 | 0.021 | 0.021 | 0 | 0.92 | 0.89 | 0.83 | 0.73 | 0.94 | 0.85 |
| 4 | 177466797 | rs309798 | -0.975 | 1.32 | -1.758 | 0.499 | -0.042 | 0.013 | 0.013 | 0 | 0.65 | 0.81 | 0.68 | 0.74 | 0.56 | 0.83 |
| 4 | 177466989 | rs11133161 | 0.809 | 1.364 | -1.703 | 0.534 | 0.022 | 0.037 | 0.037 | 0 | 0.08 | 0.13 | 0.07 | 0.16 | 0.11 | 0.09 |
| 4 | 177467220 | rs13133225 | 1.455 | 1.134 | -1.812 | 0.452 | -0.066 | 0.061 | 0.061 | 0 | 0.27 | 0.04 | 0.26 | 0.1 | 0.33 | 0.08 |
| 4 | 177467880 | rs309800 | 0.549 | 0.642 | -1.334 | -0.338 | 0 | 0 | 0 | 0 | 0.77 | 0.76 | 0.82 | 0.78 | 0.77 | 0.62 |
| 4 | 177469582 | rs13148526 | 0.391 | 0.545 | -1.571 | -1.081 | -0.065 | 0.289 | 0.289 | 0.129 | 0.45 | 0.09 | 0.27 | 0.08 | 0.5 | 0.39 |
| 4 | 177478482 | rs2034861 | 0.343 | 0.054 | -1.032 | -1.186 | -0.003 | -0.007 | -0.007 | 0 | 0.59 | 0.42 | 0.45 | 0.17 | 0.51 | 0.42 |
| 4 | 177482558 | rs4690449 | 0.368 | -0.215 | -0.949 | -1.231 | -0.026 | -0.032 | -0.032 | 0 | 0.14 | 0.31 | 0.07 | 0.1 | 0.09 | 0.17 |
| 4 | 177490659 | rs17688977 | 0.439 | -0.196 | -1.265 | -1.117 | -0.095 | 0.033 | 0.033 | 0.331 | 0.29 | 0.04 | 0.22 | 0.19 | 0.37 | 0.35 |
| 4 | 177497169 | rs10023049 | -1.331 | -0.185 | -0.744 | -1.021 | 0.003 | 0.028 | 0.028 | 0 | 0.55 | 0.72 | 0.49 | 0.47 | 0.58 | 0.65 |
| 4 | 177497711 | rs6811268 | -1.176 | -0.373 | -0.335 | -1.099 | -0.03 | -0.015 | -0.015 | 0.358 | 0.12 | 0.42 | 0.16 | 0.09 | 0.06 | 0.05 |
| 4 | 177507768 | rs13119438 | 0.757 | 0.021 | -0.36 | -1.134 | 0.013 | 0.029 | 0.029 | 0 | 0.39 | 0.3 | 0.22 | 0.16 | 0.32 | 0.2 |
| 4 | 177508930 | rs1449749 | 0.757 | 0.021 | -0.324 | -1.134 | 0.008 | 0.005 | 0.005 | 0 | 0.39 | 0.33 | 0.22 | 0.16 | 0.32 | 0.2 |

| | | | | | | | | | | | | | | | | |
|---|---|---|---|---|---|---|---|---|---|---|---|---|---|---|---|---|
| 4 | 177514623 | rs11725315 | 0.803 | -0.442 | -0.559 | -1.235 | -0.127 | 0.133 | 0.133 | 0 | 0.25 | 0.04 | 0.22 | 0.04 | 0.4 | 0.32 |
| 4 | 177522973 | rs12644889 | 0.32 | -0.782 | -0.237 | -1.036 | -0.005 | 0.001 | 0.001 | 0 | 0.08 | 0.2 | 0.05 | 0.14 | 0.05 | 0.08 |
| 4 | 177529489 | rs1017912 | -0.264 | -0.874 | -0.044 | -0.955 | -0.008 | 0.019 | 0.019 | 0 | 0.43 | 0.37 | 0.42 | 0.32 | 0.52 | 0.56 |
| 4 | 177529663 | rs13131892 | 1.464 | -0.87 | -0.204 | -0.967 | 0.01 | -0.029 | -0.029 | 0.073 | 0.32 | 0.22 | 0.15 | 0.02 | 0.24 | 0.11 |
| 4 | 177530812 | rs7660760 | -0.851 | -0.676 | -0.376 | -0.724 | 0.019 | 0.039 | 0.039 | 0 | 0.52 | 0.65 | 0.46 | 0.45 | 0.56 | 0.63 |
| 4 | 177531472 | rs6553973 | -1.829 | -0.51 | -1.035 | -0.626 | -0.046 | -0.089 | -0.089 | 0.763 | 0.84 | 0.95 | 0.63 | 0.49 | 0.81 | 0.74 |
| 4 | 177531891 | rs6824464 | 0.031 | -0.861 | -0.707 | -0.932 | 0.107 | 0.238 | 0.238 | 0 | 0.44 | 0.84 | 0.24 | 0.17 | 0.3 | 0.2 |
| 4 | 177533410 | rs17063414 | 0.189 | -0.121 | -0.022 | -0.084 | 0 | -0.002 | -0.002 | 0 | 0.06 | 0.11 | 0.04 | 0.08 | 0.05 | 0.07 |
| 4 | 177539619 | rs7663611 | -0.484 | -0.437 | 0.089 | -0.102 | -0.021 | -0.005 | -0.005 | 0 | 0.08 | 0.16 | 0.04 | 0.09 | 0.04 | 0.07 |
| 4 | 177542973 | rs13111989 | 1.091 | -0.8 | -0.437 | -0.521 | 0.07 | 0.196 | 0.196 | 0 | 0.38 | 0.07 | 0.18 | 0.12 | 0.29 | 0.31 |
| 4 | 177544609 | rs2062416 | 0.006 | -0.558 | -0.387 | -0.012 | -0.001 | -0.002 | -0.002 | 2.239 | 0.67 | 0.31 | 0.59 | 0.54 | 0.6 | 0.58 |
| 4 | 177548598 | rs1002976 | -0.592 | -0.582 | -0.421 | -0.131 | -0.014 | -0.013 | -0.013 | 0.599 | 0.35 | 0.28 | 0.41 | 0.49 | 0.44 | 0.51 |
| 4 | 177555700 | rs17063436 | 0.44 | -1.753 | -0.918 | 0.147 | 0.069 | -0.089 | -0.089 | 0 | 0.42 | 0.12 | 0.66 | 0.39 | 0.65 | 0.54 |
| 4 | 177561651 | rs13144350 | 0.677 | -1.513 | -0.849 | 0.145 | 0.06 | 0.182 | 0.182 | 0 | 0.38 | 0.07 | 0.18 | 0.12 | 0.29 | 0.28 |
| 4 | 177563237 | rs13135363 | -0.325 | -1.528 | -0.604 | 0.15 | 0.021 | 0.015 | 0.015 | 0 | 0.49 | 0.47 | 0.24 | 0.17 | 0.34 | 0.36 |
| 4 | 177566170 | rs6852384 | -0.298 | -1.207 | -0.134 | 0.57 | -0.023 | -0.027 | -0.027 | 0 | 0.11 | 0.39 | 0.05 | 0.05 | 0.05 | 0.08 |
| 4 | 177571116 | rs7671044 | -0.304 | -1.112 | -0.327 | 0.6 | -0.025 | -0.029 | -0.029 | 0 | 0.11 | 0.39 | 0.06 | 0.05 | 0.05 | 0.08 |
| 4 | 177572182 | rs17063470 | -0.456 | -0.359 | -1.185 | 1.048 | -0.016 | 0.086 | 0.086 | 0 | 0.06 | 0.09 | 0.09 | 0.41 | 0.01 | 0.09 |
| 4 | 177573314 | rs6842237 | -0.105 | -0.201 | -1.514 | 0.615 | 0.083 | -0.019 | -0.019 | 0 | 0.55 | 0.73 | 0.33 | 0.58 | 0.34 | 0.46 |
| 4 | 177575052 | rs1471810 | -0.387 | 0.864 | -1.645 | 1.016 | -0.044 | -0.059 | -0.059 | 0 | 0.19 | 0.07 | 0.2 | 0.39 | 0.24 | 0.22 |
| 4 | 177575129 | rs10011748 | 0.949 | 0.755 | -1.871 | 0.442 | -0.03 | -0.005 | -0.005 | 0 | 0.58 | 0.74 | 0.52 | 0.55 | 0.56 | 0.67 |
| 4 | 177575300 | rs12505396 | 0.523 | 0.857 | -1.736 | 0.001 | 0.007 | -0.022 | -0.022 | 0 | 0.08 | 0.1 | 0.08 | 0 | 0.1 | 0.02 |
| 4 | 177580725 | rs11133168 | 0.058 | 0.753 | -2.082 | 0.253 | -0.058 | -0.103 | -0.103 | 0 | 0.25 | 0.06 | 0.23 | 0.72 | 0.28 | 0.34 |
| 4 | 177583933 | rs4690358 | -0.479 | 0.266 | -0.658 | 0.369 | -0.01 | 0.094 | 0.094 | 0 | 0.2 | 0.32 | 0.35 | 0.26 | 0.16 | 0.22 |
| 4 | 177587894 | rs10520358 | 0.569 | 0.239 | -0.363 | 0.312 | 0.01 | -0.023 | -0.023 | 0 | 0.08 | 0.1 | 0.08 | 0 | 0.1 | 0.02 |
| 4 | 177591869 | rs6826173 | -1.793 | 0.189 | -0.363 | 0.289 | 0.002 | -0.006 | -0.006 | 0 | 0.35 | 0.68 | 0.45 | 0.26 | 0.27 | 0.24 |
| 4 | 177605696 | rs2877961 | 0.842 | 0.268 | -0.569 | 0.363 | -0.038 | 0.061 | 0.061 | 0 | 0.74 | 0.5 | 0.63 | 0.74 | 0.83 | 0.77 |
| 4 | 177607731 | rs1485762 | 1.256 | 0.33 | -0.655 | 0.389 | -0.003 | 0.007 | 0.007 | 0 | 0.66 | 0.45 | 0.56 | 0.74 | 0.73 | 0.75 |
| 4 | 177608707 | rs7664413 | -0.761 | 0.341 | -0.668 | 0.876 | -0.008 | 0.083 | 0.083 | 0 | 0.2 | 0.32 | 0.35 | 0.26 | 0.16 | 0.22 |
| 4 | 177610884 | rs1485766 | -0.382 | 0.693 | -1.618 | 0.532 | -0.114 | -0.101 | -0.101 | 0 | 0.4 | 0.08 | 0.37 | 0.5 | 0.48 | 0.45 |
| 4 | 177611211 | rs3775202 | -0.753 | 0.499 | -0.641 | 0.39 | 0 | -0.001 | -0.001 | 0 | 0.47 | 0.55 | 0.44 | 0.6 | 0.46 | 0.47 |
| 4 | 177612566 | rs3775198 | -1.511 | 0.133 | 0.411 | 0.085 | -0.016 | -0.051 | -0.051 | 0.306 | 0.15 | 0.57 | 0.22 | 0.09 | 0.1 | 0.1 |
| 4 | 177621711 | rs10007002 | -0.529 | 0.112 | 0.496 | 0.139 | -0.009 | -0.004 | -0.004 | 0 | 0.07 | 0.13 | 0.14 | 0.09 | 0.02 | 0.08 |
| 4 | 177632236 | rs1531956 | 1.883 | -0.248 | 0.728 | -0.091 | -0.037 | -0.036 | -0.036 | 0 | 0.88 | 0.75 | 0.84 | 0.94 | 0.98 | 0.87 |
| 4 | 177634522 | rs1564922 | 2.575 | -0.449 | 1.074 | -0.254 | 0.044 | 0.064 | 0.064 | 4.651 | 0.78 | 0.15 | 0.75 | 0.94 | 0.89 | 0.86 |
| 4 | 177645549 | rs6820170 | 2.448 | -0.449 | 1.493 | -0.254 | -0.006 | -0.022 | -0.022 | 4.371 | 0.79 | 0.24 | 0.75 | 0.93 | 0.89 | 0.86 |
| 4 | 177647977 | rs3775189 | -2.249 | -0.464 | 1.768 | -0.257 | -0.013 | -0.007 | -0.007 | 0 | 0.07 | 0.23 | 0.16 | 0.07 | 0.02 | 0.13 |
| 4 | 177651791 | rs1485767 | -2.213 | -0.476 | 2.282 | -0.262 | 0.022 | 0.028 | 0.028 | 0.08 | 0.07 | 0.35 | 0.16 | 0.07 | 0.02 | 0.13 |
| 4 | 177653598 | rs6838834 | -0.717 | -0.47 | 2.26 | -0.276 | -0.021 | -0.029 | -0.029 | 0.011 | 0.06 | 0.11 | 0.02 | 0 | 0.01 | 0 |
| 4 | 177656790 | rs475106 | 0.77 | -0.476 | 2.296 | -0.266 | 0.026 | -0.034 | -0.034 | 0.078 | 0.92 | 0.71 | 0.93 | 1 | 0.92 | 0.99 |
| 4 | 177662092 | rs13122901 | 2.626 | -0.483 | 2.313 | -0.253 | -0.041 | -0.073 | -0.073 | 0 | 0.8 | 0.42 | 0.81 | 0.92 | 0.89 | 0.86 |
| 4 | 177667073 | rs964812 | 2.67 | -0.497 | 2.298 | -0.258 | 0.025 | 0.032 | 0.032 | 4.77 | 0.79 | 0.2 | 0.79 | 0.92 | 0.89 | 0.86 |
| 4 | 177689133 | rs4557213 | 1.94 | -0.466 | 2.246 | -0.271 | -0.041 | -0.116 | -0.116 | 1.333 | 0.84 | 0.55 | 0.93 | 1 | 0.91 | 0.99 |
| 4 | 177720359 | rs11131764 | 0.068 | -0.384 | 2.277 | -0.248 | 0.061 | -0.007 | -0.007 | 0.504 | 0.94 | 0.73 | 0.94 | 1 | 0.92 | 0.99 |
| 4 | 177726720 | rs20533 | -2.935 | -0.436 | 2.332 | -0.251 | -0.068 | -0.09 | -0.09 | 0 | 0.15 | 0.57 | 0.16 | 0.01 | 0.03 | 0.13 |
| 4 | 177737136 | rs9992272 | 2.014 | -0.204 | 1.931 | -0.496 | -0.027 | -0.09 | -0.09 | 0 | 0.76 | 0.32 | 0.78 | 0.99 | 0.83 | 0.79 |
| 4 | 177738885 | rs11734535 | 1.924 | -0.569 | 1.101 | -0.517 | 0.016 | 0.123 | 0.123 | 0 | 0.56 | 0.11 | 0.51 | 0.76 | 0.61 | 0.55 |
| 4 | 177739471 | rs17063777 | -0.291 | -0.846 | 1.919 | -0.479 | -0.015 | -0.046 | -0.046 | 0.077 | 0.1 | 0.34 | 0.14 | 0.01 | 0.07 | 0.2 |
| 4 | 177740692 | rs4690363 | 1.056 | -0.796 | 1.843 | -0.487 | 0.092 | -0.018 | -0.018 | 0.206 | 0.91 | 0.59 | 0.92 | 1 | 0.89 | 0.99 |
| 4 | 177741229 | rs7654391 | 0.068 | -0.811 | 1.295 | -0.381 | 0.003 | -0.003 | -0.003 | 0 | 0.08 | 0.1 | 0.12 | 0.01 | 0.07 | 0.2 |
| 4 | 177743630 | rs17063785 | -0.061 | -0.732 | 1.251 | -0.315 | -0.009 | -0.013 | -0.013 | 0 | 0.06 | 0.05 | 0.01 | 0 | 0.01 | 0 |
| 4 | 177761552 | rs2333571 | 0.934 | -0.422 | 1.034 | -0.051 | -0.006 | -0.021 | -0.021 | 0.01 | 0.74 | 0.81 | 0.76 | 0.69 | 0.74 | 0.55 |
| 4 | 177776703 | rs10006747 | 1.318 | -0.474 | 1.534 | -0.077 | -0.003 | 0.005 | 0.005 | 0 | 0.71 | 0.58 | 0.74 | 0.69 | 0.74 | 0.54 |
| 4 | 177785568 | rs6552198 | 1.91 | -0.508 | 1.803 | -0.153 | 0.001 | 0.01 | 0.01 | 1.11 | 0.63 | 0.26 | 0.67 | 0.69 | 0.64 | 0.53 |
| 4 | 177789086 | rs6850178 | -2.119 | -0.943 | 1.835 | -0.174 | -0.012 | -0.064 | -0.064 | 0 | 0.12 | 0.7 | 0.11 | 0 | 0.11 | 0.02 |
| 4 | 177802943 | rs10034876 | 1.313 | -0.882 | 1.645 | -0.116 | -0.012 | 0.002 | 0.002 | 0.193 | 0.67 | 0.49 | 0.69 | 0.68 | 0.68 | 0.56 |
| 4 | 177804746 | rs7668750 | 1.443 | -0.874 | 1.497 | -0.112 | 0 | -0.001 | -0.001 | 0 | 0.66 | 0.3 | 0.68 | 0.68 | 0.68 | 0.56 |
| 4 | 177814863 | rs7700025 | 1.151 | -0.798 | 1.253 | -0.039 | -0.003 | 0.008 | 0.008 | 2.744 | 0.69 | 0.37 | 0.63 | 0.63 | 0.68 | 0.59 |
| 4 | 177815859 | rs13105194 | 1.419 | -0.793 | 0.751 | -0.036 | 0.017 | -0.012 | -0.012 | 0 | 0.65 | 0.24 | 0.61 | 0.63 | 0.68 | 0.55 |
| 4 | 177841949 | rs1836829 | 1.544 | -1.908 | 0.88 | -0.585 | -0.07 | -0.09 | -0.09 | 0.313 | 0.68 | 0.42 | 0.77 | 0.96 | 0.84 | 0.69 |
| 4 | 177842009 | rs1836828 | 1.544 | -1.888 | 0.928 | -0.587 | -0.077 | -0.107 | -0.107 | 0 | 0.68 | 0.45 | 0.77 | 0.96 | 0.83 | 0.66 |
| 4 | 177842267 | rs13152811 | 1.361 | -1.942 | 1.007 | -0.615 | -0.072 | -0.103 | -0.103 | 0.65 | 0.74 | 0.61 | 0.83 | 0.99 | 0.9 | 0.7 |
| 4 | 177844568 | rs2574904 | 1.702 | -1.836 | 0.526 | -0.555 | 0.082 | 0.092 | 0.092 | 0.138 | 0.7 | 0.18 | 0.75 | 0.96 | 0.83 | 0.67 |
| 4 | 177854130 | rs1545089 | 0.976 | -1.517 | 0.642 | -0.304 | -0.035 | -0.028 | -0.028 | 0 | 0.87 | 0.79 | 0.87 | 0.96 | 0.95 | 0.76 |
| 4 | 177862546 | rs1436197 | 1.785 | -1.339 | 0.224 | -0.206 | -0.047 | -0.031 | -0.031 | 0.535 | 0.9 | 0.32 | 0.94 | 0.98 | 0.98 | 0.82 |
| 4 | 177870509 | rs2724754 | 0.973 | -1.1 | -0.86 | -0.25 | -0.117 | -0.123 | -0.123 | 0 | 0.42 | 0.04 | 0.45 | 0.61 | 0.54 | 0.48 |
| 4 | 177870554 | rs873632 | -0.184 | -1.198 | -0.111 | -0.347 | -0.002 | 0.021 | 0.021 | 0 | 0.64 | 0.51 | 0.65 | 0.62 | 0.65 | 0.62 |
| 4 | 177871569 | rs873350 | -1.047 | -0.955 | 0.177 | -0.455 | -0.043 | -0.094 | -0.094 | 0.042 | 0.16 | 0.36 | 0.19 | 0.01 | 0.11 | 0.13 |
| 4 | 177880120 | rs17063948 | -1.171 | -0.951 | 0.052 | -0.442 | -0.043 | -0.079 | -0.079 | 0 | 0.17 | 0.32 | 0.22 | 0.03 | 0.11 | 0.14 |

| | | | | | | | | | | | | | | | |
|---|---|---|---|---|---|---|---|---|---|---|---|---|---|---|---|
| 4 | 177885655 | rs12507320 | -0.29 | -0.376 | -0.193 | -0.209 | 0.011 | -0.031 | -0.031 | 0 | 0.06 | 0.13 | 0.06 | 0 | 0.07 | 0 |
| 4 | 177890893 | rs6812865 | -0.232 | -0.452 | -0.368 | -0.054 | -0.01 | 0.011 | 0.011 | 0 | 0.68 | 0.78 | 0.65 | 0.73 | 0.67 | 0.72 |
| 4 | 177891850 | rs7662445 | 0.56 | -1.022 | -0.426 | -0.022 | 0.018 | 0.021 | 0.021 | 0 | 0.62 | 0.26 | 0.56 | 0.69 | 0.64 | 0.62 |
| 4 | 177895429 | rs13104005 | 0.268 | -0.927 | -1.059 | 0.045 | -0.005 | 0.007 | 0.007 | 0 | 0.37 | 0.33 | 0.45 | 0.27 | 0.38 | 0.4 |
| 4 | 177897523 | rs6837144 | -0.669 | -0.859 | -0.587 | 0.002 | 0.038 | 0.145 | 0.145 | 0.233 | 0.06 | 0.42 | 0.09 | 0.08 | 0.01 | 0.06 |
| 4 | 177900078 | rs12648879 | 0.514 | -0.845 | -0.746 | 0.024 | -0.052 | -0.094 | -0.094 | 0 | 0.33 | 0.03 | 0.23 | 0.42 | 0.3 | 0.31 |
| 4 | 177907949 | rs13434489 | -0.648 | -0.733 | -0.437 | -0.383 | 0.047 | 0.153 | 0.153 | 0.114 | 0.06 | 0.38 | 0.09 | 0.08 | 0.01 | 0.06 |
| 4 | 177916874 | rs13110230 | 0.751 | -0.414 | -1.265 | -0.408 | -0.039 | -0.037 | -0.037 | 0 | 0.07 | 0.01 | 0.06 | 0.13 | 0.11 | 0.11 |
| 4 | 177928041 | rs7655591 | -0.445 | -0.83 | -1.039 | -0.25 | 0.015 | 0.037 | 0.037 | 0 | 0.66 | 0.82 | 0.78 | 0.66 | 0.71 | 0.64 |
| 4 | 177952118 | rs7663143 | -0.552 | -0.803 | -1.351 | -0.308 | 0.015 | 0.006 | 0.006 | 0 | 0.72 | 0.89 | 0.81 | 0.66 | 0.76 | 0.73 |
| 4 | 177964148 | rs10025425 | -0.126 | -0.395 | -1.056 | -0.362 | -0.001 | 0.01 | 0.01 | 0 | 0.47 | 0.43 | 0.49 | 0.23 | 0.44 | 0.36 |
| 4 | 177966548 | rs17720291 | 0.507 | 0.265 | -1.824 | 0.021 | -0.074 | 0.036 | 0.036 | 0 | 0.31 | 0.09 | 0.43 | 0.23 | 0.38 | 0.23 |
| 4 | 177968578 | rs4282116 | 0.109 | 0.344 | -1.193 | 0.351 | 0.008 | -0.028 | -0.028 | 0.032 | 0.77 | 0.9 | 0.77 | 0.59 | 0.78 | 0.68 |
| 4 | 177974062 | rs13112548 | -0.44 | 0.399 | -1.689 | -0.566 | 0.004 | 0.115 | 0.115 | 0.504 | 0.26 | 0.1 | 0.18 | 0 | 0.25 | 0.18 |
| 4 | 177974857 | rs13119914 | -0.765 | 0.856 | -1.592 | 0.347 | -0.009 | -0.024 | -0.024 | 0 | 0.17 | 0.1 | 0.28 | 0.28 | 0.24 | 0.19 |
| 4 | 177974921 | rs17064100 | -0.139 | 1.032 | -1.664 | 0.564 | -0.005 | -0.032 | -0.032 | 0 | 0.23 | 0.15 | 0.29 | 0.41 | 0.24 | 0.29 |
| 4 | 177975350 | rs17064106 | 0.11 | 0.437 | -1.236 | -0.235 | -0.003 | 0.001 | 0.001 | 0 | 0.25 | 0.26 | 0.24 | 0.21 | 0.33 | 0.35 |
| 4 | 177976662 | rs10004643 | 0.271 | -0.265 | -0.245 | -0.75 | -0.006 | -0.016 | -0.016 | 0 | 0.59 | 0.53 | 0.48 | 0.27 | 0.68 | 0.59 |
| 4 | 177982668 | rs10780058 | -0.012 | -0.915 | 0.492 | -0.344 | -0.032 | 0.146 | 0.146 | 0 | 0.85 | 0.65 | 0.76 | 0.65 | 0.89 | 0.86 |
| 4 | 177993233 | rs2583194 | 0.214 | -1.304 | 0.61 | -0.757 | 0.01 | -0.001 | -0.001 | 0 | 0.57 | 0.43 | 0.55 | 0.56 | 0.68 | 0.7 |
| 4 | 178001276 | rs3114031 | -0.76 | -1.278 | 0.75 | -0.774 | -0.024 | -0.07 | -0.07 | 0 | 0.71 | 0.67 | 0.78 | 0.91 | 0.79 | 0.81 |
| 4 | 178008182 | rs1866505 | 0.515 | -1.03 | 0.839 | -0.905 | 0.041 | 0.107 | 0.107 | 0 | 0.51 | 0.36 | 0.42 | 0.15 | 0.61 | 0.54 |
| 4 | 178013213 | rs12639884 | -0.624 | -1.054 | 0.97 | -0.788 | 0.005 | 0.153 | 0.153 | 0 | 0.13 | 0.22 | 0.21 | 0.49 | 0.1 | 0.2 |
| 4 | 178016011 | rs4398480 | 0.14 | -1.034 | 1 | -0.774 | 0.02 | 0.209 | 0.209 | 0 | 0.07 | 0.11 | 0.13 | 0.41 | 0.07 | 0.16 |
| 4 | 178019148 | rs6846031 | 0.055 | -0.909 | 0.885 | -1.057 | -0.006 | -0.006 | -0.006 | 0 | 0.45 | 0.42 | 0.39 | 0.44 | 0.46 | 0.39 |
| 4 | 178028186 | rs1898854 | -0.098 | -0.916 | 1.004 | -1.033 | 0.003 | 0.01 | 0.01 | 0 | 0.51 | 0.49 | 0.47 | 0.55 | 0.52 | 0.43 |
| 4 | 178029963 | rs2580062 | 0.775 | -0.942 | 0.993 | -1.321 | -0.016 | 0.061 | 0.061 | 0 | 0.35 | 0.24 | 0.32 | 0.13 | 0.38 | 0.43 |
| 4 | 178041034 | rs17064315 | 0.468 | -0.898 | 0.835 | -1.117 | 0.006 | 0.131 | 0.131 | 0 | 0.07 | 0.06 | 0.14 | 0.43 | 0.08 | 0.17 |
| 4 | 178043001 | rs13130688 | 0.394 | -0.926 | 0.848 | -1.248 | 0.01 | -0.014 | -0.014 | 0 | 0.14 | 0.1 | 0.21 | 0.31 | 0.1 | 0.11 |
| 4 | 178045600 | rs12501567 | 0.681 | -1.132 | 0.7 | -1.53 | 0.026 | 0.069 | 0.069 | 0.043 | 0.32 | 0.17 | 0.27 | 0.03 | 0.28 | 0.38 |
| 4 | 178049288 | rs2580069 | 1.116 | -0.864 | 0.855 | -1.444 | -0.008 | 0.054 | 0.054 | 0.078 | 0.45 | 0.36 | 0.3 | 0.09 | 0.46 | 0.3 |
| 4 | 178052785 | rs2635835 | 1.62 | -0.895 | 0.747 | -1.451 | -0.012 | 0.023 | 0.023 | 0 | 0.25 | 0.17 | 0.17 | 0.09 | 0.28 | 0.21 |
| 4 | 178056849 | rs9996167 | 0.483 | -0.881 | 0.736 | -1.408 | -0.011 | -0.012 | -0.012 | 0 | 0.24 | 0.23 | 0.27 | 0.5 | 0.33 | 0.26 |
| 4 | 178061968 | rs1439343 | 0.529 | -1.033 | 0.538 | -1.392 | 0.002 | -0.002 | -0.002 | 0 | 0.15 | 0.16 | 0.22 | 0.3 | 0.11 | 0.14 |
| 4 | 178069059 | rs7656045 | 1.527 | -1.044 | 0.574 | -1.377 | -0.013 | 0.015 | 0.015 | 0.045 | 0.17 | 0.15 | 0.13 | 0.07 | 0.24 | 0.14 |
| 4 | 178069165 | rs13125748 | 0.513 | -1.15 | 0.13 | -1.38 | 0.045 | 0.016 | 0.016 | 0 | 0.34 | 0.18 | 0.33 | 0.31 | 0.28 | 0.22 |
| 4 | 178072504 | rs6838906 | 1.619 | -1.202 | 0.152 | -1.371 | -0.001 | -0.006 | -0.006 | 0.256 | 0.05 | 0 | 0.05 | 0.07 | 0.04 | 0.03 |
| 4 | 178084093 | rs4299533 | 1.406 | -1.151 | 0.255 | -1.367 | 0.014 | -0.027 | -0.027 | 2.934 | 0.17 | 0.24 | 0.14 | 0.06 | 0.24 | 0.12 |
| 4 | 178089529 | rs2583221 | -1.986 | -0.931 | -0.288 | -1.171 | -0.001 | -0.04 | -0.04 | 0 | 0.81 | 0.85 | 0.89 | 1 | 0.85 | 0.92 |
| 4 | 178092059 | rs2583165 | 0.785 | -0.338 | 0.263 | -0.234 | -0.02 | 0.032 | 0.032 | 0 | 0.24 | 0.16 | 0.18 | 0.08 | 0.3 | 0.21 |
| 4 | 178093317 | rs1489958 | -0.571 | -0.187 | 0.18 | 0.072 | 0.001 | 0.015 | 0.015 | 0 | 0.44 | 0.37 | 0.46 | 0.47 | 0.45 | 0.38 |
| 4 | 178100729 | rs2246360 | 0.785 | -0.421 | 0.209 | 1.494 | -0.013 | 0.041 | 0.041 | 0 | 0.88 | 0.84 | 0.78 | 0.71 | 0.9 | 0.89 |
| 4 | 178106459 | rs966023 | -0.534 | -0.259 | -0.327 | 1.301 | 0.001 | -0.002 | -0.002 | 0 | 0.09 | 0.06 | 0.13 | 0.03 | 0.08 | 0.26 |
| 4 | 178116819 | rs13105496 | -1.917 | 0.083 | -0.269 | 1.786 | 0.007 | 0.022 | 0.022 | 0 | 0.21 | 0.11 | 0.36 | 0.77 | 0.19 | 0.27 |
| 4 | 178123945 | rs4690374 | -0.927 | 0.43 | -0.082 | 2.154 | -0.007 | -0.013 | -0.013 | 0 | 0.11 | 0.05 | 0.15 | 0.44 | 0.09 | 0.14 |
| 4 | 178145355 | rs1992264 | -0.837 | 0.478 | -0.055 | 2.021 | -0.005 | 0 | 0 | 0.367 | 0.11 | 0.07 | 0.08 | 0.38 | 0.09 | 0.14 |
| 4 | 178151435 | rs1009523 | 1.006 | 0.282 | -0.119 | 0.607 | -0.023 | -0.017 | -0.017 | 0 | 0.74 | 0.43 | 0.61 | 0.28 | 0.83 | 0.71 |
| 4 | 178152242 | rs4690376 | -0.329 | -0.34 | -1.231 | 0.669 | 0.024 | -0.014 | -0.014 | 0 | 0.3 | 0.11 | 0.2 | 0.26 | 0.2 | 0.28 |
| 4 | 178152876 | rs6821871 | -0.504 | -0.222 | -1.202 | 0.647 | 0.012 | 0.009 | 0.009 | 0 | 0.46 | 0.32 | 0.36 | 0.44 | 0.41 | 0.48 |
| 4 | 178153009 | rs6821735 | 0.337 | -0.43 | -1.79 | 0.566 | 0.059 | -0.055 | -0.055 | 0 | 0.6 | 0.08 | 0.66 | 0.52 | 0.72 | 0.53 |
| 4 | 178160453 | rs11942838 | 0.498 | 0.191 | -1.814 | 0.499 | 0.002 | -0.061 | -0.061 | 0 | 0.59 | 0.09 | 0.63 | 0.53 | 0.65 | 0.52 |
| 4 | 178160858 | rs11734226 | 0.007 | 0.41 | -1.725 | 0.501 | -0.039 | -0.008 | -0.008 | 29.433 | 0.06 | 0.02 | 0.06 | 0.09 | 0.14 | 0.13 |
| 4 | 178167074 | rs2118922 | 0.862 | 0.085 | -1.087 | -0.014 | 0.003 | 0.006 | 0.006 | 0 | 0.58 | 0.46 | 0.46 | 0.48 | 0.53 | 0.46 |
| 4 | 178168551 | rs17064500 | 0.216 | 0.075 | -0.431 | -0.001 | -0.004 | 0.029 | 0.029 | 0 | 0.21 | 0.28 | 0.23 | 0.38 | 0.19 | 0.3 |
| 4 | 178169500 | rs3828517 | 1.29 | -0.2 | 0.347 | -0.466 | -0.002 | -0.02 | -0.02 | 0 | 0.59 | 0.35 | 0.6 | 0.81 | 0.6 | 0.54 |
| 4 | 178170776 | rs10005919 | -0.552 | -0.193 | 0.452 | -0.159 | 0.03 | -0.016 | -0.016 | 0 | 0.07 | 0.3 | 0.05 | 0 | 0.04 | 0 |
| 4 | 178173007 | rs9761093 | -0.223 | -0.181 | -0.304 | -0.167 | -0.009 | 0.095 | 0.095 | 0.017 | 0.21 | 0.06 | 0.1 | 0.07 | 0.2 | 0.33 |
| 4 | 178178160 | rs1542722 | 1.179 | -0.091 | -0.016 | -0.174 | 0.007 | 0.014 | 0.014 | 0.002 | 0.75 | 0.23 | 0.6 | 0.84 | 0.77 | 0.85 |
| 4 | 178181681 | rs9784497 | 2.438 | 0.284 | -0.022 | -0.201 | -0.029 | 0.122 | 0.122 | 0.035 | 0.39 | 0.16 | 0.24 | 0.1 | 0.37 | 0.26 |
| 4 | 178184819 | rs13125007 | 2.034 | 0.356 | 0.002 | 0.487 | -0.022 | -0.014 | -0.014 | 0 | 0.47 | 0.21 | 0.33 | 0.45 | 0.46 | 0.38 |
| 4 | 178186997 | rs17725302 | 1.019 | 0.484 | -0.208 | 0.313 | -0.047 | -0.013 | -0.013 | 0 | 0.1 | 0.01 | 0.28 | 0.14 | 0.16 | 0.1 |
| 4 | 178194441 | rs9684452 | -0.115 | -0.02 | 0.567 | 0.641 | 0.008 | 0.081 | 0.081 | 0 | 0.21 | 0.06 | 0.1 | 0.06 | 0.19 | 0.37 |
| 4 | 178197237 | rs1567111 | 0.4 | 0.317 | 1.25 | 1.03 | 0.003 | -0.049 | -0.049 | 0.004 | 0.91 | 0.65 | 0.94 | 1 | 0.93 | 1 |
| 4 | 178199981 | rs17064538 | 0.504 | 0.343 | 0.765 | 1.29 | -0.017 | -0.047 | -0.047 | 0.251 | 0.1 | 0.02 | 0.11 | 0.36 | 0.14 | 0.14 |
| 4 | 178201257 | rs4690504 | 1.718 | 0.418 | 1.113 | 1.314 | -0.006 | 0.152 | 0.152 | 1.608 | 0.82 | 0.45 | 0.74 | 0.67 | 0.86 | 0.87 |
| 4 | 178203463 | rs10520376 | -0.256 | 0.74 | 0.619 | 1.51 | -0.01 | 0.035 | 0.035 | 0 | 0.08 | 0.04 | 0.2 | 0.33 | 0.12 | 0.15 |
| 4 | 178206711 | rs13144731 | 0.219 | 0.054 | 0.995 | 1.225 | 0.075 | 0.022 | 0.022 | 0.349 | 0.94 | 0.65 | 0.95 | 1 | 0.95 | 1 |
| 4 | 178209089 | rs1844483 | 0.26 | 0.082 | 0.745 | 1.699 | 0.02 | 0.274 | 0.274 | 0 | 0.93 | 0.73 | 0.83 | 0.71 | 0.94 | 0.93 |
| 4 | 178213319 | rs11131791 | -0.163 | 0.12 | -0.011 | 1.38 | -0.006 | -0.022 | -0.022 | 0 | 0.13 | 0.08 | 0.13 | 0.39 | 0.16 | 0.19 |

| | | | | | | | | | | | | | | | | |
|---|---|---|---|---|---|---|---|---|---|---|---|---|---|---|---|---|
| 4 | 178214726 | rs17064564 | -1.015 | 0.035 | 0.054 | 1.132 | 0.01 | -0.017 | -0.017 | 0.723 | 0.37 | 0.3 | 0.34 | 0.58 | 0.34 | 0.55 |
| 4 | 178221198 | rs17064578 | -1.29 | 0.429 | 0.601 | 1.157 | 0.012 | 0.105 | 0.105 | 0.013 | 0.11 | 0.31 | 0.25 | 0.3 | 0.09 | 0.11 |
| 4 | 178224638 | rs2068197 | -1.507 | 0.187 | 1.142 | 0.711 | 0.049 | 0.04 | 0.04 | 1.511 | 0.05 | 0.41 | 0.04 | 0 | 0.01 | 0.01 |
| 4 | 178225675 | rs2135617 | 0.384 | -0.598 | 0.558 | 0.272 | 0.007 | 0.007 | 0.007 | 0.207 | 0.53 | 0.41 | 0.41 | 0.22 | 0.6 | 0.52 |
| 4 | 178229612 | rs17726200 | -0.073 | 0.308 | -0.134 | -0.343 | -0.001 | -0.005 | -0.005 | 0 | 0.32 | 0.42 | 0.22 | 0.21 | 0.28 | 0.24 |
| 4 | 178229925 | rs10009807 | 0.982 | 0.802 | 0.222 | -0.477 | 0.009 | -0.016 | -0.016 | 0 | 0.16 | 0.16 | 0.21 | 0.03 | 0.26 | 0.23 |
| 4 | 178231152 | rs10013040 | -0.218 | 0.21 | -0.324 | 0.044 | 0.01 | 0.049 | 0.049 | 0 | 0.27 | 0.14 | 0.17 | 0.14 | 0.22 | 0.19 |
| 4 | 178235334 | rs6817959 | -1.437 | 0.117 | -0.391 | -0.055 | 0 | -0.011 | -0.011 | 0 | 0.54 | 0.78 | 0.52 | 0.47 | 0.51 | 0.51 |
| 4 | 178236402 | rs6850861 | -1.372 | 0.054 | -0.278 | -0.013 | 0.018 | 0.29 | 0.29 | 0.936 | 0.1 | 0.47 | 0.17 | 0.31 | 0.04 | 0.09 |
| 4 | 178236896 | rs6856628 | -1.488 | 0.053 | -0.279 | -0.013 | 0.013 | 0.281 | 0.281 | 1.302 | 0.1 | 0.48 | 0.17 | 0.32 | 0.04 | 0.1 |
| 4 | 178242398 | rs6830266 | -0.592 | 0.083 | 0.036 | -0.034 | -0.06 | 0 | 0 | 0.834 | 0.31 | 0.52 | 0.26 | 0.32 | 0.16 | 0.17 |
| 4 | 178242537 | rs6834946 | -0.799 | 0.094 | 0.095 | -0.022 | 0.035 | 0.273 | 0.273 | 0.266 | 0.08 | 0.4 | 0.14 | 0.29 | 0.04 | 0.07 |
| 4 | 178247400 | rs12642885 | -0.482 | 0.149 | -0.487 | -0.049 | 0.004 | -0.003 | -0.003 | 0 | 0.29 | 0.27 | 0.27 | 0.34 | 0.15 | 0.18 |
| 4 | 178255747 | rs2048076 | -1.28 | 0.219 | -0.897 | -0.176 | 0 | -0.011 | -0.011 | 0 | 0.53 | 0.77 | 0.52 | 0.48 | 0.5 | 0.51 |
| 4 | 178259934 | rs17064661 | -0.224 | -0.017 | -0.999 | 0.091 | 0.025 | 0.024 | 0.024 | 0.081 | 0.22 | 0.12 | 0.12 | 0.03 | 0.12 | 0.08 |
| 4 | 178260872 | rs12645561 | -0.293 | 0.685 | -0.795 | 0.927 | -0.028 | -0.009 | -0.009 | 0.484 | 0.07 | 0.02 | 0.15 | 0.33 | 0.11 | 0.17 |
| 4 | 178261895 | rs11940019 | 0.077 | 0.276 | -0.799 | 0.06 | -0.001 | 0.013 | 0.013 | 0 | 0.18 | 0.14 | 0.13 | 0.11 | 0.2 | 0.14 |
| 4 | 178268663 | rs1983130 | 0.367 | 0.079 | -0.328 | -0.284 | -0.005 | 0.008 | 0.008 | 0 | 0.59 | 0.4 | 0.61 | 0.31 | 0.6 | 0.46 |
| 4 | 178268897 | rs2877985 | -0.437 | -0.841 | 0.096 | -0.038 | -0.013 | 0.024 | 0.024 | 0 | 0.14 | 0.23 | 0.28 | 0.22 | 0.14 | 0.19 |
| 4 | 178269253 | rs12501016 | -0.551 | -0.904 | 0.641 | -0.103 | -0.029 | 0.063 | 0.063 | 0 | 0.14 | 0.29 | 0.27 | 0.23 | 0.11 | 0.19 |
| 4 | 178269670 | rs9637678 | -0.318 | -0.458 | 0.781 | 0.12 | -0.005 | 0.067 | 0.067 | 0 | 0.09 | 0.17 | 0.22 | 0.23 | 0.09 | 0.17 |
| 4 | 178270916 | rs2048073 | -0.234 | -0.554 | 1.089 | 0.158 | 0.005 | -0.011 | -0.011 | 0.002 | 0.78 | 0.8 | 0.73 | 0.45 | 0.83 | 0.65 |
| 4 | 178272916 | rs3805169 | -0.122 | -0.498 | 1.15 | 0.236 | -0.004 | 0.04 | 0.04 | 0 | 0.09 | 0.17 | 0.22 | 0.16 | 0.09 | 0.13 |
| 4 | 178273030 | rs3828488 | -0.973 | -0.742 | 1.141 | 0.085 | -0.033 | 0.055 | 0.055 | 0.244 | 0.17 | 0.36 | 0.31 | 0.24 | 0.12 | 0.21 |
| 4 | 178274694 | rs10007075 | 0.952 | -0.622 | 0.904 | 0.108 | -0.01 | 0.001 | 0.001 | 0.002 | 0.65 | 0.44 | 0.61 | 0.38 | 0.76 | 0.57 |
| 4 | 178274750 | rs13112358 | 0.506 | -0.318 | 0.185 | 0.274 | -0.01 | 0.007 | 0.007 | 0 | 0.7 | 0.67 | 0.63 | 0.38 | 0.77 | 0.58 |
| 4 | 178274835 | rs13112390 | 0.054 | 0.023 | 0.208 | 0.408 | -0.008 | 0.04 | 0.04 | 0 | 0.77 | 0.71 | 0.71 | 0.45 | 0.8 | 0.64 |
| 4 | 178275220 | rs13140743 | 0.035 | 0.038 | -0.045 | 0.415 | -0.002 | 0.013 | 0.013 | 0 | 0.77 | 0.75 | 0.71 | 0.45 | 0.8 | 0.64 |
| 4 | 178280473 | rs12507296 | -0.295 | -0.37 | -0.022 | -0.384 | 0.002 | -0.014 | -0.014 | 0 | 0.19 | 0.14 | 0.12 | 0.02 | 0.08 | 0.15 |
| 4 | 178283967 | rs1055678 | -0.413 | -0.526 | 0.489 | -0.42 | -0.041 | -0.069 | -0.069 | 0.095 | 0.21 | 0.27 | 0.14 | 0.02 | 0.08 | 0.15 |
| 4 | 178287162 | rs7667508 | 1.034 | -0.227 | -0.726 | -1.327 | -0.073 | 0.296 | 0.296 | 0 | 0.46 | 0.1 | 0.53 | 0.08 | 0.52 | 0.43 |
| 4 | 178288939 | rs17090614 | 0.412 | -0.26 | -0.606 | -1.277 | 0.016 | 0.007 | 0.007 | 0 | 0.19 | 0.13 | 0.12 | 0.14 | 0.08 | 0.21 |
| 4 | 178290368 | rs6846587 | 0.41 | 0.223 | -0.301 | -1.218 | -0.003 | 0.011 | 0.011 | 0 | 0.18 | 0.23 | 0.23 | 0.68 | 0.18 | 0.27 |
| 4 | 178291257 | rs6830385 | -0.122 | 0.102 | -1.028 | -1.251 | 0.003 | -0.007 | -0.007 | 0.026 | 0.67 | 0.36 | 0.67 | 0.22 | 0.6 | 0.63 |
| 4 | 178294146 | rs1506811 | 0.348 | 0.559 | -1.813 | -1.172 | -0.057 | 0.25 | 0.25 | 0.026 | 0.39 | 0.05 | 0.37 | 0.04 | 0.44 | 0.23 |
| 4 | 178295429 | rs17739370 | -0.421 | 0.939 | -1.306 | -0.717 | -0.005 | 0.003 | 0.003 | 0.004 | 0.35 | 0.35 | 0.38 | 0.22 | 0.42 | 0.36 |
| 4 | 178295725 | rs17064756 | -0.47 | 0.32 | -1.532 | -0.012 | 0.026 | -0.018 | -0.018 | 0 | 0.39 | 0.32 | 0.31 | 0.51 | 0.31 | 0.25 |
| 4 | 178295766 | rs4690508 | -0.274 | 0.627 | -1.302 | 0.462 | 0.006 | -0.003 | -0.003 | 0.109 | 0.41 | 0.36 | 0.46 | 0.5 | 0.42 | 0.38 |
| 4 | 178296863 | rs9312207 | -0.243 | -0.315 | -0.539 | 0.595 | -0.048 | 0.004 | 0.004 | 0 | 0.25 | 0.61 | 0.3 | 0.3 | 0.11 | 0.23 |
| 4 | 178297365 | rs11131792 | 0.831 | 0.366 | 0.212 | 1.151 | 0.001 | 0.14 | 0.14 | 0 | 0.81 | 0.55 | 0.75 | 0.6 | 0.86 | 0.79 |
| 4 | 178297393 | rs1395480 | 0.828 | 0.735 | 0.741 | 1.206 | 0.056 | 0.166 | 0.166 | 0.303 | 0.84 | 0.45 | 0.77 | 0.67 | 0.86 | 0.81 |
| 4 | 178298024 | rs17677649 | 0.284 | 0.449 | -0.02 | 0.355 | 0.023 | 0.05 | 0.05 | 0 | 0.26 | 0.11 | 0.16 | 0.17 | 0.17 | 0.34 |
| 4 | 178299038 | rs10866214 | 0.354 | 0.669 | -0.039 | -0.146 | -0.005 | -0.031 | -0.031 | 0.005 | 0.54 | 0.53 | 0.6 | 0.82 | 0.63 | 0.56 |
| 4 | 178299498 | rs7671847 | 1.033 | 0.95 | 0.256 | 0.286 | -0.001 | -0.021 | -0.021 | 0 | 0.72 | 0.53 | 0.6 | 0.82 | 0.72 | 0.83 |
| 4 | 178301110 | rs17677777 | -0.146 | 1.01 | -0.459 | 0.126 | -0.025 | 0.065 | 0.065 | 0.714 | 0.13 | 0.02 | 0.21 | 0 | 0.17 | 0.08 |
| 4 | 178309091 | rs6832432 | 0.89 | 0.629 | 0.043 | 0.401 | -0.002 | -0.003 | -0.003 | 0 | 0.85 | 0.83 | 0.7 | 0.93 | 0.85 | 0.92 |
| 4 | 178310307 | rs5004774 | 1.524 | 0.453 | 0.698 | 0.314 | -0.021 | -0.031 | -0.031 | 0 | 0.81 | 0.58 | 0.67 | 0.93 | 0.85 | 0.92 |
| 4 | 178310766 | rs13108563 | 0.571 | 0.543 | 0.45 | 0.374 | 0 | 0.012 | 0.012 | 0 | 0.9 | 0.81 | 0.84 | 0.93 | 0.91 | 0.96 |
| 4 | 178313274 | rs17064782 | -0.962 | 0.492 | 0.598 | 0.454 | -0.01 | -0.007 | -0.007 | 0.034 | 0.13 | 0.24 | 0.23 | 0.08 | 0.11 | 0.06 |
| 4 | 178313450 | rs17064783 | -1.101 | 0.6 | -0.15 | 0.128 | 0.001 | 0.012 | 0.012 | 0 | 0.49 | 0.5 | 0.53 | 0.25 | 0.42 | 0.33 |
| 4 | 178313514 | rs17739994 | 0.029 | 0.4 | -0.86 | 0.1 | 0.002 | 0.003 | 0.003 | 0 | 0.09 | 0.05 | 0.2 | 0.05 | 0.09 | 0.05 |
| 4 | 178314542 | rs2135615 | 1.07 | 0.541 | -1.341 | 0.413 | -0.042 | -0.127 | -0.127 | 0 | 0.37 | 0.05 | 0.24 | 0.49 | 0.38 | 0.48 |
| 4 | 178317692 | rs1506807 | 0.595 | 0.726 | -0.938 | 0.811 | -0.087 | -0.063 | -0.063 | 0 | 0.23 | 0.01 | 0.16 | 0.22 | 0.28 | 0.23 |
| 4 | 178321406 | rs17064813 | -0.177 | 0.736 | -0.088 | 1.012 | -0.007 | 0.052 | 0.052 | 0.264 | 0.11 | 0.19 | 0.19 | 0.19 | 0.08 | 0.1 |
| 4 | 178321418 | rs1395483 | 0.737 | 0.742 | -0.233 | 0.844 | -0.01 | 0.006 | 0.006 | 0 | 0.51 | 0.2 | 0.46 | 0.52 | 0.54 | 0.61 |
| 4 | 178327053 | rs4690513 | -1.222 | 0.546 | 1.043 | 1.411 | 0.034 | 0.091 | 0.091 | 0.581 | 0.12 | 0.3 | 0.26 | 0.22 | 0.14 | 0.12 |
| 4 | 178327855 | rs6552231 | 1.196 | 0.391 | 1.527 | 1.295 | -0.017 | -0.015 | -0.015 | 0 | 0.9 | 0.67 | 0.79 | 0.96 | 0.96 | 0.94 |
| 4 | 178328596 | rs17064824 | 0.926 | 0.19 | 1.402 | 1.533 | -0.015 | 0.09 | 0.09 | 0 | 0.82 | 0.69 | 0.89 | 0.71 | 0.85 | 0.78 |
| 4 | 178328886 | rs17064825 | -0.893 | 0.498 | 0.908 | 1.695 | 0.039 | 0.192 | 0.192 | 0 | 0.06 | 0.18 | 0.25 | 0.23 | 0.08 | 0.1 |
| 4 | 178335508 | rs1911239 | 0.655 | -0.093 | 0.103 | 0.939 | 0.042 | 0.006 | 0.006 | 0.073 | 0.58 | 0.32 | 0.57 | 0.63 | 0.7 | 0.61 |
| 4 | 178337145 | rs1588051 | 0.755 | 0.027 | 0.445 | 0.855 | -0.015 | -0.005 | -0.005 | 0.564 | 0.56 | 0.54 | 0.57 | 0.63 | 0.67 | 0.62 |
| 4 | 178340126 | rs4690516 | 1.071 | 0.098 | 0.242 | 0.883 | 0.05 | -0.019 | -0.019 | 1.785 | 0.5 | 0.19 | 0.46 | 0.52 | 0.6 | 0.54 |
| 4 | 178351295 | rs12641389 | 0.082 | 0.026 | 0.823 | 0.783 | 0.05 | 0.056 | 0.056 | 0.978 | 0.93 | 0.71 | 0.79 | 0.92 | 0.93 | 0.93 |
| 4 | 178351803 | rs3749480 | 0.28 | -0.205 | 0.352 | 0.918 | -0.041 | -0.029 | -0.029 | 0 | 0.68 | 0.54 | 0.66 | 0.78 | 0.79 | 0.79 |
| 4 | 178353473 | rs3828487 | -0.182 | 0.159 | -0.468 | 1.127 | -0.009 | -0.033 | -0.033 | 0 | 0.15 | 0.12 | 0.12 | 0.27 | 0.16 | 0.21 |
| 4 | 178354886 | rs2279932 | 1.168 | 0.158 | -0.64 | 0.976 | 0.051 | -0.009 | -0.009 | 0.441 | 0.51 | 0.18 | 0.47 | 0.5 | 0.63 | 0.54 |
| 4 | 178356930 | rs12499096 | 0.188 | 0.036 | -1.356 | 0.644 | 0.03 | -0.059 | -0.059 | 0 | 0.35 | 0.09 | 0.29 | 0.57 | 0.28 | 0.58 |
| 4 | 178359679 | rs2271100 | 1.027 | 0.209 | -1.62 | 0.156 | 0 | 0.086 | 0.086 | 0 | 0.27 | 0.1 | 0.16 | 0.14 | 0.26 | 0.12 |
| 4 | 178361051 | rs4690522 | 0.331 | 0.788 | -1.244 | 0.068 | -0.023 | -0.009 | -0.009 | 0 | 0.22 | 0.16 | 0.32 | 0.25 | 0.34 | 0.2 |

| | | | | | | | | | | | | | | | | |
|---|---|---|---|---|---|---|---|---|---|---|---|---|---|---|---|---|
| 4 | 178361162 | rs4690523 | -0.653 | 0.58 | -0.41 | 0.035 | 0.042 | -0.009 | -0.009 | 0 | 0.34 | 0.41 | 0.47 | 0.28 | 0.47 | 0.28 |
| 4 | 178369766 | rs7680586 | -1.159 | 0.176 | -0.176 | -0.066 | -0.01 | -0.039 | -0.039 | 0 | 0.69 | 0.6 | 0.82 | 0.85 | 0.74 | 0.87 |
| 4 | 178374869 | rs6847279 | 0.488 | 0.553 | -0.151 | 0.418 | 0.042 | -0.02 | -0.02 | 0 | 0.38 | 0.23 | 0.28 | 0.4 | 0.28 | 0.58 |
| 4 | 178377527 | rs17064851 | 1.795 | 0.203 | -0.598 | 0.175 | -0.004 | 0.028 | 0.028 | 0 | 0.05 | 0 | 0.02 | 0 | 0.03 | 0.05 |
| 4 | 178381669 | rs17064860 | 0.31 | 0.379 | -0.449 | 0.319 | 0.043 | -0.018 | -0.018 | 1.311 | 0.38 | 0.24 | 0.29 | 0.4 | 0.28 | 0.58 |
| 4 | 178384657 | rs12504521 | 0.32 | 0.379 | -0.437 | 0.319 | 0.041 | -0.018 | -0.018 | 0 | 0.38 | 0.24 | 0.28 | 0.4 | 0.28 | 0.58 |
| 4 | 178387902 | rs7693160 | 1.59 | 0.33 | -0.442 | 0.249 | -0.009 | 0.076 | 0.076 | 0 | 0.27 | 0.08 | 0.15 | 0.14 | 0.26 | 0.13 |
| 4 | 178388432 | rs6852945 | 0.377 | 0.355 | -0.443 | 0.264 | 0.043 | -0.018 | -0.018 | 0 | 0.38 | 0.23 | 0.28 | 0.4 | 0.28 | 0.58 |
| 4 | 178393335 | rs35955682 | 0.711 | 0.454 | -0.537 | 0.241 | -0.14 | -0.129 | -0.129 | 0 | 0.22 | 0.04 | 0.35 | 0.41 | 0.37 | 0.23 |
| 4 | 178394279 | rs6823035 | 0.149 | 0.377 | -0.15 | 0.275 | 0.002 | -0.002 | -0.002 | 0.252 | 0.39 | 0.38 | 0.34 | 0.41 | 0.28 | 0.6 |
| 4 | 178405429 | rs6814553 | 0.212 | 0.394 | -0.298 | 0.283 | 0.023 | -0.009 | -0.009 | 0.978 | 0.39 | 0.31 | 0.33 | 0.41 | 0.28 | 0.6 |
| 4 | 178419101 | rs17742216 | 0.97 | 0.638 | -0.187 | 0.224 | 0.003 | -0.018 | -0.018 | 0 | 0.07 | 0.07 | 0.07 | 0.01 | 0.09 | 0.03 |
| 4 | 178426799 | rs11944393 | 0.158 | 0.533 | -0.326 | 0.27 | -0.002 | -0.011 | -0.011 | 0 | 0.46 | 0.47 | 0.41 | 0.36 | 0.49 | 0.28 |
| 4 | 178438637 | rs17742420 | 1.104 | 0.352 | -0.199 | 0.457 | -0.031 | 0.077 | 0.077 | 0.013 | 0.4 | 0.09 | 0.29 | 0.24 | 0.39 | 0.23 |
| 4 | 178449352 | rs1882160 | 0.667 | 0.408 | 0.058 | 0.921 | 0.001 | 0.012 | 0.012 | 0.085 | 0.51 | 0.52 | 0.44 | 0.35 | 0.5 | 0.27 |
| 4 | 178454290 | rs12510932 | -0.081 | -0.558 | 0.572 | 0.974 | 0.059 | -0.031 | -0.031 | 0 | 0.25 | 0.08 | 0.17 | 0.29 | 0.15 | 0.46 |
| 4 | 178471300 | rs4047774 | 0.198 | 0.306 | -0.094 | 0.501 | -0.033 | -0.044 | -0.044 | 0 | 0.66 | 0.51 | 0.69 | 0.76 | 0.77 | 0.52 |
| 4 | 178477452 | rs1920975 | 0.438 | 1.006 | 0.364 | 0.91 | -0.01 | 0 | 0 | 0.197 | 0.53 | 0.33 | 0.52 | 0.55 | 0.51 | 0.4 |
| 4 | 178479822 | rs1316294 | -0.35 | 0.867 | 0.352 | 0.844 | -0.113 | -0.082 | -0.082 | 2.773 | 0.42 | 0.09 | 0.42 | 0.45 | 0.49 | 0.6 |
| 4 | 178491443 | rs4690536 | 0.131 | 0.567 | 0.372 | 0.355 | 0.007 | 0.069 | 0.069 | 0 | 0.7 | 0.53 | 0.65 | 0.56 | 0.67 | 0.69 |
| 4 | 178498527 | rs10015045 | -0.126 | 0.537 | 0.25 | 0.44 | 0.175 | 0.257 | 0.257 | 0.702 | 0.07 | 0.45 | 0.13 | 0.18 | 0.08 | 0.1 |
| 4 | 178500973 | rs1920969 | 0.377 | 0.768 | -0.872 | 0.301 | -0.072 | -0.088 | -0.088 | 0 | 0.39 | 0.11 | 0.48 | 0.43 | 0.41 | 0.57 |
| 4 | 178501916 | rs12509926 | -1.224 | 0.987 | -0.302 | 0.677 | -0.021 | -0.057 | -0.057 | 0.865 | 0.35 | 0.79 | 0.41 | 0.46 | 0.4 | 0.33 |
| 4 | 178503993 | rs13116233 | -0.221 | 0.99 | -0.357 | 0.647 | -0.009 | -0.017 | -0.017 | 0 | 0.26 | 0.14 | 0.25 | 0.25 | 0.31 | 0.22 |
| 4 | 178506363 | rs17065006 | 0.623 | 0.913 | -0.766 | 0.466 | -0.023 | -0.058 | -0.058 | 0 | 0.65 | 0.19 | 0.59 | 0.57 | 0.6 | 0.68 |
| 4 | 178509336 | rs17743886 | 0.786 | 0.693 | -0.832 | 0.125 | 0.011 | 0.017 | 0.017 | 0 | 0.11 | 0.01 | 0.08 | 0.04 | 0.09 | 0.14 |
| 4 | 178510236 | rs11732864 | 0.534 | 1.227 | -0.68 | 0.054 | 0.017 | -0.004 | -0.004 | 0 | 0.07 | 0.1 | 0.14 | 0.03 | 0.21 | 0.08 |
| 4 | 178511146 | rs6816548 | -0.077 | 0.975 | -0.713 | -0.086 | 0 | 0 | 0 | 0 | 0.65 | 0.65 | 0.51 | 0.48 | 0.65 | 0.48 |
| 4 | 178516328 | rs13114753 | -0.084 | 0.966 | -0.749 | -0.09 | -0.005 | -0.019 | -0.019 | 0 | 0.65 | 0.71 | 0.52 | 0.48 | 0.65 | 0.48 |
| 4 | 178517058 | rs6834239 | 1.558 | 0.749 | -1.116 | -0.137 | -0.035 | 0.099 | 0.099 | 0.097 | 0.37 | 0.05 | 0.23 | 0.23 | 0.33 | 0.23 |
| 4 | 178520166 | rs17749802 | 1.527 | 0.845 | -1.298 | -0.339 | -0.041 | 0.05 | 0.05 | 0 | 0.13 | 0.05 | 0.12 | 0.01 | 0.2 | 0.12 |
| 4 | 178524155 | rs10866216 | -0.766 | 0.701 | -1.254 | -0.262 | 0.014 | -0.057 | -0.057 | 0 | 0.8 | 0.9 | 0.65 | 0.55 | 0.85 | 0.66 |
| 4 | 178536315 | rs7661921 | 0.148 | 0.731 | -1.211 | -0.222 | -0.028 | -0.074 | -0.074 | 0 | 0.65 | 0.82 | 0.52 | 0.54 | 0.63 | 0.54 |
| 4 | 178538777 | rs10520382 | 1.645 | 0.787 | -1.29 | -0.196 | -0.035 | 0.103 | 0.103 | 0.107 | 0.37 | 0.05 | 0.22 | 0.23 | 0.33 | 0.23 |
| 4 | 178556723 | rs13102468 | -0.308 | 0.611 | -0.462 | 0.004 | 0.006 | -0.002 | -0.002 | 0.015 | 0.28 | 0.43 | 0.28 | 0.24 | 0.33 | 0.27 |
| 4 | 178564250 | rs10019475 | -0.228 | 0.585 | -0.465 | 0.328 | 0.002 | -0.02 | -0.02 | 0 | 0.85 | 0.92 | 0.69 | 0.54 | 0.88 | 0.71 |
| 4 | 178579037 | rs13147867 | -0.086 | 0.539 | -0.739 | -0.037 | -0.007 | -0.02 | -0.02 | 4.79 | 0.39 | 0.31 | 0.55 | 0.49 | 0.41 | 0.45 |
| 4 | 178593268 | rs13149928 | -1.222 | 0.793 | -0.421 | -0.068 | -0.001 | -0.007 | -0.007 | 0 | 0.75 | 0.75 | 0.79 | 0.78 | 0.77 | 0.82 |
| 4 | 178594906 | rs4443228 | 0.215 | 0.685 | -0.085 | 0.256 | 0.065 | 0.123 | 0.123 | 4.213 | 0.94 | 0.57 | 0.97 | 0.92 | 1 | 0.99 |
| 4 | 178596794 | rs4690549 | 0.093 | 0.71 | -0.232 | 0.354 | 0.067 | 0.16 | 0.16 | 3.794 | 0.95 | 0.57 | 0.97 | 0.87 | 1 | 0.99 |
| 4 | 178597709 | rs9684887 | 0.38 | 0.819 | -0.27 | 0.089 | 0.018 | -0.033 | -0.033 | 0 | 0.14 | 0.27 | 0.19 | 0.04 | 0.22 | 0.16 |
| 4 | 178598308 | rs6845027 | 0.174 | 0.688 | -0.253 | 0.406 | -0.005 | -0.012 | -0.012 | 0 | 0.37 | 0.35 | 0.55 | 0.43 | 0.41 | 0.45 |
| 4 | 178599771 | rs4690554 | 0.707 | 0.751 | 0.314 | 0.447 | 0.076 | 0.072 | 0.072 | 3.001 | 0.95 | 0.58 | 0.97 | 1 | 1 | 1 |
| 4 | 178607009 | rs12502947 | 0.707 | 0.751 | 0.314 | 0.447 | 0.076 | 0.072 | 0.072 | 3.001 | 0.95 | 0.58 | 0.97 | 1 | 1 | 1 |
| 4 | 178617740 | rs4432701 | -0.671 | 1.297 | -0.851 | 0.645 | -0.019 | -0.003 | -0.003 | 0.877 | 0.77 | 0.89 | 0.8 | 0.75 | 0.7 | 0.79 |
| 4 | 178620683 | rs1993279 | -0.248 | 1.158 | -1.155 | 0.582 | -0.065 | 0.016 | 0.016 | 0.058 | 0.66 | 0.88 | 0.77 | 0.73 | 0.6 | 0.75 |
| 4 | 178621004 | rs1993280 | 0.588 | 0.884 | -0.662 | 0.94 | 0.015 | 0.01 | 0.01 | 0 | 0.44 | 0.37 | 0.31 | 0.32 | 0.34 | 0.35 |
| 4 | 178621097 | rs1993281 | -0.543 | 0.956 | -0.772 | 1.068 | -0.058 | -0.018 | -0.018 | 0 | 0.66 | 0.88 | 0.76 | 0.61 | 0.61 | 0.74 |
| 4 | 178644817 | rs1454212 | 0.363 | 0.043 | 0.665 | 1.196 | -0.027 | 0.015 | 0.015 | 1.145 | 0.81 | 0.3 | 0.86 | 0.74 | 0.91 | 0.85 |
| 4 | 178646542 | rs1454213 | 0.845 | 0.102 | 0.913 | 1.212 | 0.098 | 0.195 | 0.195 | 1.58 | 0.95 | 0.61 | 0.95 | 0.89 | 0.98 | 0.99 |
| 4 | 178649091 | rs3943470 | 0.097 | -0.159 | 0.118 | 0.977 | 0.064 | -0.019 | -0.019 | 0.335 | 0.6 | 0.21 | 0.68 | 0.57 | 0.69 | 0.67 |
| 4 | 178651027 | rs11730575 | 1.378 | 0.124 | 0.022 | 0.7 | -0.002 | 0.02 | 0.02 | 0 | 0.19 | 0.09 | 0.17 | 0.16 | 0.21 | 0.17 |
| 4 | 178662910 | rs4690560 | 0.11 | 0.4 | 0.183 | 0.953 | 0.056 | 0.005 | 0.005 | 1.103 | 0.76 | 0.5 | 0.51 | 0.76 | 0.69 | 0.57 |
| 4 | 178669888 | rs9312212 | 0.138 | 0.387 | 0.61 | 0.969 | -0.003 | -0.007 | -0.007 | 0 | 0.55 | 0.53 | 0.32 | 0.62 | 0.42 | 0.4 |
| 4 | 178674077 | rs4690563 | -0.399 | 0.018 | 0.23 | 0.941 | 0.055 | 0.003 | 0.003 | 1.214 | 0.53 | 0.22 | 0.7 | 0.44 | 0.69 | 0.64 |
| 4 | 178678485 | rs6829056 | 1.813 | -0.174 | -0.023 | 1.232 | -0.064 | -0.033 | -0.033 | 0 | 0.27 | 0.02 | 0.16 | 0.24 | 0.23 | 0.14 |
| 4 | 178681212 | rs10222887 | -0.02 | 0.394 | 0.459 | 1.365 | 0.004 | 0.009 | 0.009 | 0 | 0.59 | 0.4 | 0.7 | 0.65 | 0.59 | 0.63 |
| 4 | 178682034 | rs6851156 | 0.135 | 0.404 | 0.445 | 1.338 | -0.009 | -0.009 | -0.009 | 0 | 0.55 | 0.39 | 0.32 | 0.48 | 0.46 | 0.41 |
| 4 | 178682882 | rs7698744 | 1.024 | 0.361 | 0.698 | 1.32 | 0 | 0.001 | 0.001 | 0 | 0.61 | 0.36 | 0.45 | 0.57 | 0.56 | 0.44 |
| 4 | 178683194 | rs7677777 | 1.762 | 0.354 | 0.675 | 1.304 | 0 | -0.005 | -0.005 | 0 | 0.21 | 0.15 | 0.2 | 0.15 | 0.25 | 0.16 |
| 4 | 178683325 | rs11131820 | 1.151 | 0.424 | 0.29 | 1.327 | 0.03 | -0.019 | -0.019 | 0.528 | 0.4 | 0.18 | 0.25 | 0.42 | 0.3 | 0.28 |
| 4 | 178686626 | rs17700785 | 0.533 | 0.528 | 0.046 | 1.34 | -0.022 | 0 | 0 | 0.752 | 0.14 | 0.03 | 0.06 | 0.06 | 0.15 | 0.14 |
| 4 | 178689242 | rs2085067 | 0.144 | 0.61 | 0.241 | 1.352 | -0.006 | -0.007 | -0.007 | 0 | 0.2 | 0.21 | 0.45 | 0.37 | 0.29 | 0.42 |
| 4 | 178692400 | rs7691749 | 0.454 | 0.666 | -0.111 | 1.275 | -0.059 | -0.031 | -0.031 | 0 | 0.18 | 0.06 | 0.44 | 0.21 | 0.28 | 0.35 |
| 4 | 178701283 | rs13140117 | -1.029 | 0.535 | 0.465 | 1.302 | -0.006 | -0.003 | -0.003 | 0 | 0.74 | 0.57 | 0.77 | 0.71 | 0.75 | 0.76 |
| 4 | 178709040 | rs12506671 | -0.802 | 0.619 | 0.681 | 1.318 | 0.017 | 0.001 | 0.001 | 0 | 0.33 | 0.43 | 0.53 | 0.27 | 0.43 | 0.46 |
| 4 | 178710704 | rs1902019 | 0.866 | 0.199 | 0.833 | 1.647 | 0.006 | 0.129 | 0.129 | 0.396 | 0.82 | 0.52 | 0.88 | 0.75 | 0.84 | 0.71 |
| 4 | 178710940 | rs12508860 | -0.274 | 0.332 | 0.379 | 1.4 | -0.032 | -0.047 | -0.047 | 0 | 0.11 | 0.03 | 0.41 | 0.24 | 0.16 | 0.43 |
| 4 | 178711450 | rs10011916 | 0.617 | 0.1 | 0.888 | 1.143 | 0.011 | 0.054 | 0.054 | 0 | 0.84 | 0.66 | 0.6 | 0.79 | 0.85 | 0.67 |

| | | | | | | | | | | | | | | | | |
|---|---|---|---|---|---|---|---|---|---|---|---|---|---|---|---|---|
| 4 | 178713104 | rs1454218 | -0.013 | 0.057 | 0.197 | 1.165 | 0 | 0.004 | 0.004 | 0 | 0.88 | 0.89 | 0.64 | 0.79 | 0.88 | 0.65 |
| 4 | 178715833 | rs17751904 | 0.389 | 0.023 | -0.211 | 1.054 | -0.056 | 0.032 | 0.032 | 0 | 0.48 | 0.12 | 0.27 | 0.39 | 0.45 | 0.32 |
| 4 | 178718914 | rs17752004 | 1.125 | 0.128 | -0.21 | 0.891 | -0.072 | 0.014 | 0.014 | 0.005 | 0.31 | 0.08 | 0.28 | 0.21 | 0.38 | 0.23 |
| 4 | 178721539 | rs10017412 | -0.186 | 0.163 | -0.093 | 0.908 | -0.029 | -0.022 | -0.022 | 0 | 0.56 | 0.77 | 0.34 | 0.58 | 0.49 | 0.41 |
| 4 | 178731036 | rs975438 | 0.233 | -0.234 | -0.035 | 0.629 | -0.02 | 0.046 | 0.046 | 0 | 0.42 | 0.33 | 0.33 | 0.21 | 0.47 | 0.37 |
| 4 | 178734746 | rs11723791 | -0.262 | 0 | 0.039 | 0.965 | 0.009 | 0.061 | 0.061 | 0 | 0.89 | 0.72 | 0.67 | 0.81 | 0.88 | 0.65 |
| 4 | 178740648 | rs4997950 | 0.401 | -0.065 | 0.132 | 0.932 | -0.01 | 0.057 | 0.057 | 0.071 | 0.85 | 0.41 | 0.64 | 0.81 | 0.88 | 0.65 |
| 4 | 178752567 | rs4574348 | 1.105 | -0.053 | -1.111 | 0.622 | -0.141 | -0.011 | -0.011 | 0 | 0.32 | 0.04 | 0.3 | 0.21 | 0.42 | 0.28 |
| 4 | 178754587 | rs10049757 | -0.145 | 0.289 | -0.922 | 1.023 | -0.002 | 0.089 | 0.089 | 0 | 0.88 | 0.8 | 0.65 | 0.62 | 0.88 | 0.64 |
| 4 | 178755843 | rs6827227 | -0.223 | 0.25 | -1.381 | 0.673 | 0.002 | 0.028 | 0.028 | 0 | 0.46 | 0.17 | 0.28 | 0.41 | 0.43 | 0.36 |
| 4 | 178759663 | rs6843725 | 0.094 | 0.118 | -1.64 | 0.627 | -0.025 | -0.015 | -0.015 | 0.779 | 0.36 | 0.12 | 0.28 | 0.39 | 0.39 | 0.25 |
| 4 | 178759888 | rs17065284 | 0.042 | -0.328 | -1.219 | 0.366 | 0.006 | 0.002 | 0.002 | 0 | 0.16 | 0.12 | 0.06 | 0.03 | 0.08 | 0.1 |
| 4 | 178760454 | rs1869511 | -0.438 | -0.207 | -1.126 | 0.892 | -0.001 | -0.002 | -0.002 | 0.455 | 0.18 | 0.2 | 0.21 | 0.34 | 0.16 | 0.37 |
| 4 | 178760909 | rs11131829 | -0.555 | -0.384 | -0.643 | 0.792 | -0.001 | -0.005 | -0.005 | 0 | 0.39 | 0.29 | 0.54 | 0.58 | 0.42 | 0.56 |
| 4 | 178762043 | rs2378775 | 1.558 | -0.678 | -0.778 | 0.424 | -0.083 | 0.049 | 0.049 | 0 | 0.36 | 0.15 | 0.35 | 0.21 | 0.48 | 0.27 |
| 4 | 178770448 | rs2168041 | -0.101 | -0.748 | -0.392 | 0.431 | -0.061 | -0.023 | -0.023 | 0 | 0.24 | 0.44 | 0.09 | 0.2 | 0.11 | 0.16 |
| 4 | 178772728 | rs12643723 | 0.568 | -0.775 | -0.535 | 0.265 | -0.034 | -0.049 | -0.049 | 0 | 0.13 | 0.02 | 0.12 | 0.19 | 0.15 | 0.34 |
| 4 | 178773349 | rs11131831 | 0.309 | -0.955 | -0.082 | 0.321 | -0.006 | 0.043 | 0.043 | 1.567 | 0.27 | 0.29 | 0.42 | 0.57 | 0.28 | 0.25 |
| 4 | 178775388 | rs17065363 | 0.175 | -0.954 | -0.237 | 0.35 | 0.001 | -0.025 | -0.025 | 0 | 0.21 | 0.23 | 0.08 | 0.02 | 0.11 | 0.12 |
| 4 | 178776502 | rs6849372 | -0.102 | -0.089 | 0.003 | 0.984 | -0.009 | 0.051 | 0.051 | 1.056 | 0.31 | 0.48 | 0.44 | 0.57 | 0.3 | 0.27 |
| 4 | 178793689 | rs4690581 | -0.458 | -0.078 | -0.523 | 1.253 | -0.02 | -0.009 | -0.009 | 0 | 0.63 | 0.77 | 0.48 | 0.54 | 0.54 | 0.69 |
| 4 | 178794478 | rs2168040 | 0.244 | -0.102 | -0.525 | 1.231 | -0.071 | 0.053 | 0.053 | 0 | 0.35 | 0.16 | 0.49 | 0.23 | 0.43 | 0.24 |
| 4 | 178797990 | rs4690582 | -0.248 | -0.402 | -0.603 | 1.15 | 0.046 | 0.022 | 0.022 | 0.738 | 0.52 | 0.21 | 0.63 | 0.58 | 0.6 | 0.57 |
| 4 | 178799980 | rs11722051 | 0.637 | -0.352 | -0.699 | 1.078 | -0.089 | -0.06 | -0.06 | 0 | 0.34 | 0.12 | 0.47 | 0.39 | 0.44 | 0.23 |
| 4 | 178801018 | rs6824106 | 0.6 | -0.708 | -0.443 | 0.796 | 0.004 | -0.003 | -0.003 | 0 | 0.58 | 0.39 | 0.64 | 0.44 | 0.64 | 0.4 |
| 4 | 178803423 | rs11725712 | 1.465 | -0.675 | -0.415 | 0.758 | -0.006 | -0.043 | -0.043 | 0 | 0.24 | 0.27 | 0.17 | 0.05 | 0.2 | 0.17 |
| 4 | 178805083 | rs6851593 | 2.031 | -0.752 | -0.543 | 0.721 | -0.031 | 0.082 | 0.082 | 0 | 0.2 | 0.02 | 0.14 | 0.05 | 0.19 | 0.17 |
| 4 | 178806432 | rs4277728 | -1.004 | -0.921 | -0.082 | 0.747 | -0.016 | -0.017 | -0.017 | 0 | 0.81 | 0.64 | 0.81 | 0.88 | 0.84 | 0.83 |
| 4 | 178806449 | rs4306905 | -0.228 | -0.828 | 0.076 | 0.884 | -0.023 | 0.075 | 0.075 | 0 | 0.76 | 0.57 | 0.8 | 0.56 | 0.8 | 0.59 |
| 4 | 178812009 | rs2219692 | -0.02 | -0.628 | 0.136 | 1.134 | -0.004 | -0.003 | -0.003 | 0.369 | 0.43 | 0.42 | 0.34 | 0.34 | 0.36 | 0.38 |
| 4 | 178813276 | rs13110833 | 0.335 | -0.646 | 0.129 | 1.142 | 0.019 | 0.007 | 0.007 | 0 | 0.4 | 0.23 | 0.3 | 0.35 | 0.35 | 0.38 |
| 4 | 178817591 | rs13118487 | -0.26 | -0.598 | -0.157 | 0.631 | -0.004 | 0.005 | 0.005 | 0 | 0.72 | 0.67 | 0.8 | 0.58 | 0.77 | 0.68 |
| 4 | 178820545 | rs2378799 | -0.198 | -0.569 | -0.168 | 0.266 | -0.005 | -0.016 | -0.016 | 0 | 0.43 | 0.42 | 0.33 | 0.18 | 0.36 | 0.45 |
| 4 | 178821372 | rs11939187 | 0.239 | -0.407 | -0.535 | 0.235 | 0 | 0.045 | 0.045 | 0 | 0.33 | 0.18 | 0.29 | 0.13 | 0.35 | 0.43 |
| 4 | 178827897 | rs17065447 | 0.514 | -0.793 | -0.413 | 0.642 | 0 | -0.001 | -0.001 | 0 | 0.22 | 0.25 | 0.21 | 0.26 | 0.19 | 0.25 |
| 4 | 178836429 | rs17703021 | 0.904 | -0.699 | -0.732 | 0.621 | 0 | 0.026 | 0.026 | 0 | 0.25 | 0.03 | 0.1 | 0.16 | 0.19 | 0.28 |
| 4 | 178841353 | rs12645056 | 0.664 | -0.683 | -0.682 | 0.582 | 0.001 | 0 | 0 | 0 | 0.23 | 0.25 | 0.22 | 0.24 | 0.19 | 0.24 |
| 4 | 178873922 | rs2126506 | 0.461 | -0.142 | 0.253 | 0.919 | 0.009 | -0.009 | -0.009 | 0 | 0.16 | 0.08 | 0.11 | 0.2 | 0.11 | 0.23 |
| 4 | 178883188 | rs7673686 | 0.616 | -0.039 | -0.114 | 0.91 | 0.036 | 0.052 | 0.052 | 0 | 0.47 | 0.26 | 0.32 | 0.39 | 0.38 | 0.56 |
| 4 | 178888659 | rs8180172 | 0.811 | 0.212 | -0.468 | 0.989 | -0.086 | -0.073 | -0.073 | 0 | 0.25 | 0.1 | 0.46 | 0.38 | 0.42 | 0.18 |
| 4 | 178891411 | rs10049915 | -0.203 | 0.379 | -0.243 | 0.662 | 0 | -0.009 | -0.009 | 0 | 0.55 | 0.54 | 0.4 | 0.39 | 0.47 | 0.6 |
| 4 | 178910307 | rs10006596 | -0.551 | 0.917 | -0.006 | 0.796 | 0.05 | 0.087 | 0.087 | 0 | 0.29 | 0.48 | 0.28 | 0.47 | 0.37 | 0.38 |
| 4 | 178910702 | rs1599605 | 1.034 | 0.108 | -0.615 | 0.214 | 0.055 | 0.101 | 0.101 | 2.59 | 0.22 | 0.02 | 0.04 | 0.05 | 0.09 | 0.18 |
| 4 | 178911822 | rs17065494 | 0.307 | 0.13 | -0.148 | 0.481 | 0.017 | 0.124 | 0.124 | 0 | 0.06 | 0.17 | 0.09 | 0.2 | 0.05 | 0.09 |
| 4 | 178911873 | rs17065497 | 1.072 | -0.136 | -0.267 | 0.437 | 0.019 | 0.007 | 0.007 | 0 | 0.21 | 0.16 | 0.14 | 0.17 | 0.16 | 0.17 |
| 4 | 178915050 | rs1462513 | -1.15 | -0.159 | 0.369 | 0.347 | -0.001 | -0.001 | -0.001 | 0 | 0.72 | 0.75 | 0.45 | 0.69 | 0.62 | 0.72 |
| 4 | 178918074 | rs12641431 | 1.477 | -0.08 | 0.06 | 0.287 | -0.058 | -0.064 | -0.064 | 0.182 | 0.22 | 0.04 | 0.48 | 0.31 | 0.3 | 0.25 |
| 4 | 178922921 | rs1462514 | 0.161 | -0.269 | 0.155 | 0.485 | 0.047 | 0.021 | 0.021 | 0 | 0.25 | 0.45 | 0.21 | 0.27 | 0.32 | 0.34 |
| 4 | 178933641 | rs1711390 | -0.179 | -0.252 | -0.3 | 0.489 | 0 | 0.019 | 0.019 | 0.696 | 0.45 | 0.36 | 0.25 | 0.32 | 0.4 | 0.5 |
| 4 | 178943989 | rs2054380 | 0.705 | -0.141 | -0.994 | 0.566 | -0.101 | -0.071 | -0.071 | 0 | 0.19 | 0.05 | 0.16 | 0.24 | 0.31 | 0.31 |
| 4 | 178944245 | rs2054381 | 0.681 | -0.135 | -0.986 | 0.569 | -0.08 | -0.054 | -0.054 | 0.384 | 0.2 | 0.08 | 0.16 | 0.24 | 0.31 | 0.31 |
| 4 | 178951714 | rs1676169 | -0.487 | -0.44 | -1.025 | 0.619 | -0.034 | 0.016 | 0.016 | 0 | 0.44 | 0.27 | 0.39 | 0.61 | 0.51 | 0.53 |
| 4 | 178955488 | rs1711376 | 0.407 | -0.193 | -1.061 | 0.642 | 0 | -0.007 | -0.007 | 0 | 0.26 | 0.26 | 0.26 | 0.38 | 0.27 | 0.22 |
| 4 | 178962336 | rs2045757 | 0.54 | 0.102 | -1.188 | 0.603 | -0.001 | -0.001 | -0.001 | 0 | 0.19 | 0.19 | 0.16 | 0.23 | 0.25 | 0.27 |
| 4 | 178978140 | rs1676187 | -0.353 | 0.194 | -1.18 | 0.598 | 0.015 | -0.002 | -0.002 | 0 | 0.46 | 0.33 | 0.3 | 0.48 | 0.39 | 0.56 |
| 4 | 178979494 | rs1381698 | 0.243 | 0.07 | -1.582 | 0.494 | 0.043 | 0.013 | 0.013 | 0 | 0.28 | 0.14 | 0.13 | 0.25 | 0.14 | 0.29 |
| 4 | 178981852 | rs1676196 | 0.243 | 0.08 | -1.573 | 0.503 | 0.042 | 0.013 | 0.013 | 0 | 0.28 | 0.14 | 0.14 | 0.26 | 0.15 | 0.29 |
| 4 | 178988148 | rs1711386 | -0.013 | 1.271 | -1.317 | 0.694 | -0.04 | -0.005 | -0.005 | 0 | 0.7 | 0.85 | 0.5 | 0.72 | 0.59 | 0.72 |
| 4 | 179002061 | rs1462505 | 0.414 | 0.864 | -0.896 | 0.439 | 0.001 | 0.001 | 0.001 | 0.995 | 0.85 | 0.86 | 0.91 | 0.93 | 0.89 | 0.8 |
| 4 | 179004868 | rs1711391 | -0.725 | 1.518 | -0.941 | 1.074 | -0.014 | -0.017 | -0.017 | 0 | 0.12 | 0.11 | 0.34 | 0.22 | 0.18 | 0.05 |
| 4 | 179005829 | rs10520402 | -0.772 | 1.099 | 0.065 | 1.032 | 0.045 | 0.095 | 0.095 | 0.189 | 0.05 | 0.31 | 0.05 | 0.1 | 0.03 | 0.06 |
| 4 | 179010647 | rs1632770 | -0.774 | 1.248 | -0.054 | 1.131 | 0.004 | 0.007 | 0.007 | 0 | 0.12 | 0.16 | 0.34 | 0.18 | 0.17 | 0.05 |
| 4 | 179011288 | rs1676157 | -1.525 | 1.39 | -0.111 | 0.885 | 0.009 | 0.01 | 0.01 | 0 | 0.19 | 0.27 | 0.39 | 0.29 | 0.28 | 0.08 |
| 4 | 179012585 | rs17756250 | -0.935 | 1.396 | -0.502 | 0.781 | -0.016 | -0.027 | -0.027 | 2.331 | 0.09 | 0.02 | 0.11 | 0.13 | 0.09 | 0.12 |
| 4 | 179023206 | rs533025 | -1.007 | 1.552 | 0.046 | 1.28 | 0.011 | 0.043 | 0.043 | 0 | 0.24 | 0.38 | 0.16 | 0.36 | 0.23 | 0.22 |
| 4 | 179032994 | rs672041 | -0.195 | 1.523 | -0.048 | 1.515 | -0.039 | -0.044 | -0.044 | 1.777 | 0.13 | 0.08 | 0.12 | 0.23 | 0.21 | 0.15 |
| 4 | 179046619 | rs553372 | -1.418 | 1.484 | 0.13 | 1.482 | 0.004 | 0.019 | 0.019 | 0 | 0.22 | 0.26 | 0.15 | 0.36 | 0.23 | 0.22 |
| 4 | 179046640 | rs11131851 | 0.53 | 1.05 | -0.828 | 0.917 | -0.003 | -0.011 | -0.011 | 0 | 0.46 | 0.15 | 0.55 | 0.44 | 0.43 | 0.44 |
| 4 | 179055616 | rs9993819 | 0.768 | 0.634 | -0.164 | 0.363 | -0.009 | 0.018 | 0.018 | 1.794 | 0.28 | 0.21 | 0.27 | 0.2 | 0.34 | 0.34 |

| | | | | | | | | | | | | | | | | |
|---|---|---|---|---|---|---|---|---|---|---|---|---|---|---|---|---|
| 4 | 179067086 | rs604363 | -0.218 | 0.672 | 0.065 | 0.394 | -0.003 | -0.006 | -0.006 | 0.092 | 0.62 | 0.45 | 0.69 | 0.67 | 0.64 | 0.59 |
| 4 | 179069109 | rs493668 | 0.594 | 0.276 | -0.584 | 0.372 | -0.055 | -0.123 | -0.123 | 0.324 | 0.23 | 0.08 | 0.18 | 0.52 | 0.23 | 0.29 |
| 4 | 179071004 | rs664886 | -0.365 | 0.056 | -0.189 | 0.189 | -0.001 | -0.002 | -0.002 | 0 | 0.62 | 0.43 | 0.68 | 0.69 | 0.63 | 0.58 |
| 4 | 179107663 | rs1458719 | 0.601 | -0.196 | -0.342 | 0.003 | -0.057 | 0 | 0 | 0.202 | 0.29 | 0.07 | 0.25 | 0.27 | 0.36 | 0.38 |
| 4 | 179114110 | rs1458717 | 0.212 | -0.414 | -0.219 | -0.239 | -0.073 | 0.006 | 0.006 | 0.072 | 0.41 | 0.16 | 0.54 | 0.59 | 0.47 | 0.47 |
| 4 | 179119001 | rs1545672 | 0.938 | -0.601 | -0.141 | -0.365 | 0.005 | 0.017 | 0.017 | 0 | 0.15 | 0.08 | 0.11 | 0.09 | 0.14 | 0.1 |
| 4 | 179125887 | rs13132202 | -0.48 | -0.564 | 0.232 | -0.455 | -0.004 | -0.025 | -0.025 | 0 | 0.62 | 0.55 | 0.7 | 0.73 | 0.63 | 0.58 |
| 4 | 179135822 | rs17758367 | 0.571 | -0.516 | -0.245 | -0.916 | -0.016 | 0.101 | 0.101 | 0 | 0.34 | 0.19 | 0.2 | 0.11 | 0.36 | 0.4 |
| 4 | 179176878 | rs2063396 | 0.299 | -0.43 | -0.386 | -0.785 | -0.058 | 0.116 | 0.116 | 0 | 0.42 | 0.14 | 0.62 | 0.75 | 0.49 | 0.48 |
| 4 | 179184331 | rs12650801 | 0.353 | -0.421 | -0.462 | -0.866 | -0.075 | 0.136 | 0.136 | 0.019 | 0.42 | 0.1 | 0.62 | 0.79 | 0.49 | 0.48 |
| 4 | 179185906 | rs6818908 | -1.48 | -0.517 | 0.05 | -1.156 | -0.001 | -0.085 | -0.085 | 0 | 0.75 | 0.7 | 0.83 | 0.98 | 0.73 | 0.7 |
| 4 | 179194375 | rs6552287 | 0.317 | 0.133 | -0.278 | -0.996 | -0.014 | 0.11 | 0.11 | 0 | 0.43 | 0.21 | 0.63 | 0.78 | 0.44 | 0.49 |
| 4 | 179198970 | rs4690620 | -0.092 | 0.075 | -0.058 | -1.174 | 0 | 0.071 | 0.071 | 1.572 | 0.59 | 0.31 | 0.73 | 0.86 | 0.57 | 0.56 |
| 4 | 179199152 | rs17708346 | 0.823 | -0.188 | 0.011 | -1.219 | -0.022 | 0.118 | 0.118 | 0.151 | 0.32 | 0.1 | 0.17 | 0.1 | 0.35 | 0.41 |
| 4 | 179200338 | rs12648022 | -0.077 | -0.125 | 0.016 | -1.212 | 0.022 | 0.05 | 0.05 | 1.314 | 0.6 | 0.32 | 0.77 | 0.86 | 0.65 | 0.57 |
| 4 | 179208760 | rs6552290 | 0.043 | -0.043 | -0.07 | -1.126 | 0.022 | -0.04 | -0.04 | 0 | 0.18 | 0.27 | 0.17 | 0.13 | 0.22 | 0.11 |
| 4 | 179227598 | rs1567475 | 0.368 | -0.043 | -0.078 | -0.874 | -0.028 | 0.109 | 0.109 | 0.102 | 0.43 | 0.14 | 0.62 | 0.72 | 0.43 | 0.45 |
| 4 | 179227880 | rs1567474 | -0.454 | -0.038 | 0.179 | -0.872 | 0.002 | 0.032 | 0.032 | 0 | 0.54 | 0.6 | 0.36 | 0.28 | 0.56 | 0.55 |
| 4 | 179247331 | rs925373 | 0.347 | 0.423 | 0.157 | -1.036 | -0.006 | -0.12 | -0.12 | 0 | 0.28 | 0.58 | 0.14 | 0.02 | 0.22 | 0.15 |
| 4 | 179251931 | rs6837438 | 0.751 | 0.557 | -0.163 | -0.558 | 0.029 | -0.022 | -0.022 | 0 | 0.41 | 0.23 | 0.59 | 0.45 | 0.34 | 0.44 |
| 4 | 179266260 | rs2666044 | -0.756 | 0.608 | -0.233 | -0.45 | -0.045 | 0.034 | 0.034 | 0 | 0.69 | 0.81 | 0.73 | 0.78 | 0.57 | 0.65 |
| 4 | 179275567 | rs1473016 | 0.598 | 0.624 | -0.27 | -0.398 | -0.014 | 0.02 | 0.02 | 4.675 | 0.28 | 0.45 | 0.13 | 0.33 | 0.22 | 0.21 |
| 4 | 179277574 | rs2063393 | 1.124 | 0.598 | -0.627 | -0.456 | 0.013 | -0.079 | -0.079 | 0 | 0.34 | 0.09 | 0.55 | 0.45 | 0.31 | 0.41 |
| 4 | 179283569 | rs3109075 | -0.566 | 1.027 | -0.442 | -0.314 | 0.048 | 0.014 | 0.014 | 0 | 0.17 | 0.47 | 0.17 | 0.16 | 0.19 | 0.1 |
| 4 | 179286562 | rs12643657 | 0.733 | 1.269 | -0.267 | -0.192 | -0.012 | -0.008 | -0.008 | 0 | 0.11 | 0.04 | 0.14 | 0.16 | 0.18 | 0.1 |
| 4 | 179290393 | rs2705988 | 0.078 | 0.828 | -0.337 | -0.021 | 0.011 | -0.039 | -0.039 | 4.179 | 0.38 | 0.48 | 0.26 | 0.21 | 0.39 | 0.25 |
| 4 | 179311419 | rs1510631 | 0.099 | 0.907 | -0.625 | 0.025 | 0.013 | -0.041 | -0.041 | 0 | 0.38 | 0.46 | 0.26 | 0.22 | 0.39 | 0.25 |
| 4 | 179323762 | rs1473017 | -1.094 | 0.826 | -0.766 | 0.026 | -0.007 | -0.023 | -0.023 | 0 | 0.78 | 0.78 | 0.82 | 0.65 | 0.73 | 0.65 |
| 4 | 179326362 | rs17065950 | 0.105 | 0.684 | -0.7 | 0.334 | 0.017 | -0.01 | -0.01 | 0 | 0.43 | 0.35 | 0.6 | 0.47 | 0.4 | 0.49 |
| 4 | 179334721 | rs2263116 | -0.319 | 1.321 | -0.41 | 0.119 | 0.026 | -0.029 | -0.029 | 1.782 | 0.31 | 0.44 | 0.5 | 0.21 | 0.33 | 0.23 |
| 4 | 179335261 | rs9991342 | -0.343 | 0.363 | -0.116 | -0.121 | 0.008 | -0.02 | -0.02 | 0 | 0.42 | 0.48 | 0.53 | 0.23 | 0.48 | 0.34 |
| 4 | 179337999 | rs2715399 | 0.175 | 0.156 | -0.129 | 0.281 | -0.005 | 0.015 | 0.015 | 0.169 | 0.36 | 0.51 | 0.27 | 0.44 | 0.32 | 0.39 |
| 4 | 179339480 | rs1567477 | -0.573 | -0.261 | -0.522 | -0.12 | 0.002 | 0.065 | 0.065 | 0 | 0.68 | 0.82 | 0.79 | 0.91 | 0.71 | 0.71 |
| 4 | 179340650 | rs2666052 | 1.518 | -0.307 | -0.552 | 0.118 | 0.008 | -0.01 | -0.01 | 0 | 0.14 | 0.11 | 0.07 | 0.2 | 0.1 | 0.2 |
| 4 | 179341460 | rs2715402 | 0.11 | -0.345 | -0.58 | 0.142 | 0.013 | 0.062 | 0.062 | 0 | 0.52 | 0.67 | 0.38 | 0.75 | 0.44 | 0.6 |
| 4 | 179346683 | rs1606021 | 0.931 | -0.368 | -0.198 | 0.357 | -0.001 | 0.001 | 0.001 | 0 | 0.36 | 0.38 | 0.29 | 0.38 | 0.34 | 0.39 |
| 4 | 179350247 | rs2666062 | -0.582 | -0.237 | -0.416 | 0.13 | -0.003 | 0.053 | 0.053 | 0 | 0.69 | 0.84 | 0.8 | 0.83 | 0.71 | 0.7 |
| 4 | 179359572 | rs2705977 | 1.186 | 0.015 | -0.434 | 0.322 | -0.002 | 0.008 | 0.008 | 0 | 0.1 | 0.11 | 0.07 | 0.15 | 0.1 | 0.12 |
| 4 | 179362277 | rs6819722 | 1.063 | 0.052 | -0.543 | -0.018 | -0.083 | -0.051 | -0.051 | 0.062 | 0.16 | 0.02 | 0.38 | 0.16 | 0.27 | 0.11 |
| 4 | 179363159 | rs2706018 | 1.269 | 0.048 | -0.51 | -0.035 | -0.003 | -0.009 | -0.009 | 0 | 0.1 | 0.13 | 0.08 | 0.06 | 0.1 | 0.11 |
| 4 | 179365152 | rs2702386 | -0.088 | 0.018 | -0.493 | -0.043 | -0.017 | -0.01 | -0.01 | 0 | 0.26 | 0.16 | 0.47 | 0.31 | 0.36 | 0.23 |
| 4 | 179374122 | rs736591 | 1.093 | 0.223 | -0.516 | -0.051 | -0.079 | -0.038 | -0.038 | 0 | 0.15 | 0.01 | 0.34 | 0.14 | 0.26 | 0.11 |
| 4 | 179378497 | rs2702383 | -1.442 | 0.1 | -0.014 | 0.091 | 0.011 | 0.217 | 0.217 | 0 | 0.9 | 0.7 | 0.87 | 0.62 | 0.91 | 0.8 |
| 4 | 179380391 | rs11944902 | 0.988 | 0.043 | 0.272 | 0.026 | -0.026 | 0.022 | 0.022 | 0 | 0.15 | 0.3 | 0.11 | 0.18 | 0.09 | 0.24 |
| 4 | 179380432 | rs11940603 | 0.988 | 0.043 | 0.272 | 0.026 | -0.026 | 0.022 | 0.022 | 0 | 0.15 | 0.3 | 0.11 | 0.18 | 0.09 | 0.24 |
| 4 | 179380467 | rs1903524 | 0.24 | -0.145 | -0.129 | -0.168 | -0.001 | -0.014 | -0.014 | 0.002 | 0.52 | 0.52 | 0.42 | 0.17 | 0.55 | 0.45 |
| 4 | 179383805 | rs2702385 | 1.3 | -0.113 | -0.382 | -0.013 | -0.007 | 0.033 | 0.033 | 0 | 0.17 | 0.2 | 0.13 | 0.38 | 0.1 | 0.2 |
| 4 | 179385177 | rs2256732 | -1.359 | -0.128 | -0.373 | -0.188 | -0.01 | -0.056 | -0.056 | 0.009 | 0.84 | 0.72 | 0.88 | 0.95 | 0.87 | 0.89 |
| 4 | 179385208 | rs13120364 | 1.764 | 0.098 | -0.565 | -0.277 | -0.03 | 0.048 | 0.048 | 0.298 | 0.23 | 0.05 | 0.14 | 0.1 | 0.24 | 0.2 |
| 4 | 179385635 | rs10471173 | 0.191 | 0.19 | -0.618 | -0.21 | 0 | 0 | 0 | 3.003 | 0.35 | 0.39 | 0.48 | 0.33 | 0.4 | 0.36 |
| 4 | 179387496 | rs2249887 | -0.207 | 0.112 | -1.06 | 0.002 | -0.026 | -0.026 | -0.026 | 0 | 0.7 | 0.57 | 0.78 | 0.76 | 0.76 | 0.65 |
| 4 | 179387659 | rs2126272 | -0.079 | 0.173 | -1.816 | 0.087 | 0.012 | 0.023 | 0.023 | 0 | 0.58 | 0.31 | 0.65 | 0.62 | 0.63 | 0.53 |
| 4 | 179388852 | rs2614609 | -0.69 | 0.294 | -1.281 | 1.019 | -0.013 | 0.002 | 0.002 | 0 | 0.32 | 0.34 | 0.23 | 0.69 | 0.2 | 0.46 |
| 4 | 179389332 | rs12501890 | -0.203 | 0.912 | -1.409 | 0.955 | -0.004 | 0.01 | 0.01 | 0 | 0.15 | 0.14 | 0.09 | 0.31 | 0.1 | 0.25 |
| 4 | 179389470 | rs12501924 | -0.203 | 0.853 | -1.409 | 0.918 | -0.005 | 0.009 | 0.009 | 0 | 0.15 | 0.14 | 0.09 | 0.3 | 0.1 | 0.25 |
| 4 | 179390635 | rs2715373 | -0.638 | 1.623 | -1.576 | 0.407 | -0.055 | 0.001 | 0.001 | 0 | 0.2 | 0.07 | 0.41 | 0.2 | 0.34 | 0.14 |
| 4 | 179391633 | rs2250361 | -0.122 | 1.054 | -1.031 | 0.119 | -0.003 | -0.014 | -0.014 | 0.398 | 0.62 | 0.47 | 0.45 | 0.28 | 0.66 | 0.62 |
| 4 | 179392878 | rs6840687 | -0.328 | 1.651 | -1.366 | -0.34 | -0.031 | 0.037 | 0.037 | 0 | 0.21 | 0.09 | 0.17 | 0.06 | 0.32 | 0.19 |
| 4 | 179395117 | rs7696617 | -0.121 | 1.387 | -1.097 | -0.896 | 0 | -0.025 | -0.025 | 0 | 0.28 | 0.3 | 0.16 | 0.06 | 0.32 | 0.09 |
| 4 | 179395250 | rs17713627 | -0.114 | 1.865 | -1.442 | -1.119 | -0.033 | 0.037 | 0.037 | 0 | 0.1 | 0.02 | 0.09 | 0.02 | 0.16 | 0.12 |
| 4 | 179398668 | rs1870793 | -0.173 | 1.456 | -0.901 | -0.502 | 0 | -0.017 | -0.017 | 0 | 0.33 | 0.47 | 0.21 | 0.22 | 0.34 | 0.15 |
| 4 | 179399523 | rs2702414 | 0.5 | 1.193 | -1.003 | 0.283 | 0.001 | 0.026 | 0.026 | 0 | 0.08 | 0.06 | 0.36 | 0.39 | 0.08 | 0.15 |
| 4 | 179401433 | rs2702449 | 0.779 | 0.394 | -1.208 | -0.325 | 0.008 | -0.022 | -0.022 | 0 | 0.38 | 0.25 | 0.31 | 0.48 | 0.34 | 0.45 |
| 4 | 179401563 | rs2702450 | 0.869 | 0.262 | -0.907 | 0.32 | 0.021 | 0.087 | 0.087 | 0.016 | 0.76 | 0.51 | 0.51 | 0.54 | 0.71 | 0.62 |
| 4 | 179402364 | rs2614589 | -0.267 | 0.022 | -1.45 | 0.43 | 0.013 | -0.079 | -0.079 | 0 | 0.21 | 0.04 | 0.22 | 0.32 | 0.18 | 0.34 |
| 4 | 179404023 | rs2465532 | 0.006 | 0.141 | -1.332 | 0.084 | -0.028 | -0.086 | -0.086 | 0 | 0.22 | 0.06 | 0.44 | 0.41 | 0.25 | 0.3 |
| 4 | 179407255 | rs1903532 | 0.598 | 0.178 | -1.271 | 0.175 | 0.008 | -0.044 | -0.044 | 0 | 0.12 | 0.04 | 0.19 | 0.47 | 0.12 | 0.26 |
| 4 | 179408291 | rs2715388 | 0.361 | 0.333 | -0.438 | -0.283 | -0.005 | -0.067 | -0.067 | 1.664 | 0.74 | 0.61 | 0.82 | 0.9 | 0.76 | 0.72 |
| 4 | 179410247 | rs1903535 | 0.049 | 0.006 | -0.103 | 0.694 | -0.032 | 0.094 | 0.094 | 0 | 0.82 | 0.66 | 0.68 | 0.66 | 0.88 | 0.8 |

| | | | | | | | | | | | | | | | | |
|---|---|---|---|---|---|---|---|---|---|---|---|---|---|---|---|---|
| 4 | 179410273 | rs1903536 | 0.225 | 0.25 | -0.913 | -0.037 | -0.004 | -0.002 | -0.002 | 0.026 | 0.64 | 0.64 | 0.58 | 0.65 | 0.71 | 0.71 |
| 4 | 179420957 | rs7669485 | 0.176 | 0.976 | -0.688 | -0.682 | 0.007 | -0.018 | -0.018 | 0 | 0.16 | 0.27 | 0.11 | 0.06 | 0.24 | 0.17 |
| 4 | 179423869 | rs2702432 | -0.421 | 0.625 | -1.374 | -0.529 | 0.02 | 0.048 | 0.048 | 0 | 0.5 | 0.63 | 0.46 | 0.32 | 0.62 | 0.52 |
| 4 | 179424481 | rs2702431 | 1.404 | 0.672 | -1.784 | -0.819 | -0.057 | 0.122 | 0.122 | 0 | 0.3 | 0.1 | 0.26 | 0.07 | 0.37 | 0.22 |
| 4 | 179426861 | rs2702430 | 0.123 | 0.506 | -1.455 | -0.746 | -0.004 | 0.147 | 0.147 | 0 | 0.44 | 0.2 | 0.35 | 0.14 | 0.6 | 0.39 |
| 4 | 179442000 | rs13140529 | 1.479 | 0.843 | -1.344 | -0.722 | -0.048 | 0.109 | 0.109 | 0 | 0.18 | 0.02 | 0.11 | 0 | 0.17 | 0.11 |
| 4 | 179447164 | rs998257 | -0.848 | 0.873 | -0.341 | -0.429 | 0.041 | -0.026 | -0.026 | 0.09 | 0.18 | 0.61 | 0.21 | 0.12 | 0.24 | 0.12 |
| 4 | 179448086 | rs2715500 | -0.999 | 0.851 | -0.438 | -0.145 | 0.019 | 0.014 | 0.014 | 1.969 | 0.19 | 0.68 | 0.17 | 0.17 | 0.24 | 0.17 |
| 4 | 179451516 | rs13146234 | -0.407 | 0.697 | -0.472 | -0.598 | 0.011 | 0.039 | 0.039 | 0 | 0.49 | 0.6 | 0.34 | 0.1 | 0.61 | 0.38 |
| 4 | 179454416 | rs2614633 | 0.769 | 0.728 | -0.721 | 0.099 | -0.013 | -0.052 | -0.052 | 3.104 | 0.27 | 0.15 | 0.19 | 0.35 | 0.27 | 0.33 |
| 4 | 179466683 | rs2246528 | 0.14 | 0.342 | -0.368 | -0.015 | -0.004 | 0.001 | 0.001 | 0 | 0.62 | 0.56 | 0.45 | 0.36 | 0.68 | 0.54 |
| 4 | 179467290 | rs2614651 | -0.43 | 0.268 | -0.485 | -0.028 | -0.003 | -0.021 | -0.021 | 0.838 | 0.69 | 0.64 | 0.79 | 0.89 | 0.71 | 0.68 |
| 4 | 179471262 | rs17066329 | 1.169 | 0.108 | -0.497 | -0.042 | 0.037 | 0.031 | 0.031 | 0.416 | 0.21 | 0.13 | 0.11 | 0.08 | 0.12 | 0.23 |
| 4 | 179475641 | rs1903558 | 0.487 | 0.14 | -0.453 | -0.151 | 0.049 | -0.052 | -0.052 | 0 | 0.27 | 0.19 | 0.45 | 0.62 | 0.14 | 0.37 |
| 4 | 179477407 | rs1903557 | -0.728 | 0.725 | -0.223 | -0.137 | 0.059 | -0.043 | -0.043 | 0 | 0.9 | 0.81 | 0.89 | 0.97 | 0.8 | 0.91 |
| 4 | 179479829 | rs13108016 | 0.444 | 0.19 | -0.01 | 0.02 | -0.004 | 0.001 | 0.001 | 0 | 0.3 | 0.32 | 0.19 | 0.27 | 0.26 | 0.39 |
| 4 | 179481211 | rs1903552 | 0.119 | 0.347 | 0.176 | -0.014 | 0.05 | 0.01 | 0.01 | 0 | 0.92 | 0.77 | 0.93 | 0.91 | 0.89 | 0.93 |
| 4 | 179481298 | rs1903551 | 0.399 | 0.2 | -0.144 | -0.165 | -0.038 | 0.036 | 0.036 | 0 | 0.3 | 0.37 | 0.48 | 0.58 | 0.19 | 0.35 |
| 4 | 179481522 | rs2702464 | -0.376 | 0.544 | -0.032 | -0.229 | 0.065 | 0.018 | 0.018 | 0 | 0.93 | 0.77 | 0.93 | 0.91 | 0.89 | 0.93 |
| 4 | 179482210 | rs3108797 | 0.155 | 0.502 | -0.29 | -0.308 | -0.013 | -0.025 | -0.025 | 0 | 0.38 | 0.32 | 0.29 | 0.01 | 0.47 | 0.21 |
| 4 | 179486578 | rs4861888 | 0.658 | 0.481 | -0.476 | -0.255 | -0.084 | 0.132 | 0.132 | 0 | 0.32 | 0.14 | 0.26 | 0.06 | 0.44 | 0.19 |
| 4 | 179488838 | rs6811144 | -0.127 | 0.41 | -0.287 | 0.053 | -0.009 | -0.007 | -0.007 | 0 | 0.42 | 0.39 | 0.63 | 0.55 | 0.51 | 0.31 |
| 4 | 179492948 | rs880240 | 1.066 | 0.252 | 0.373 | 0.432 | 0 | -0.04 | -0.04 | 0.057 | 0.07 | 0.16 | 0.04 | 0 | 0.06 | 0.04 |
| 4 | 179493721 | rs2715544 | 1.001 | 0.363 | -0.018 | 0.354 | -0.02 | 0.171 | 0.171 | 0 | 0.52 | 0.11 | 0.32 | 0.2 | 0.57 | 0.47 |
| 4 | 179498010 | rs2614555 | 0.817 | 0.886 | 0.521 | 0.448 | 0.036 | -0.008 | -0.008 | 0.403 | 0.82 | 0.65 | 0.85 | 0.81 | 0.77 | 0.81 |
| 4 | 179503140 | rs2126270 | 0.404 | 1.133 | 0.657 | 0.331 | 0.063 | -0.017 | -0.017 | 0 | 0.87 | 0.68 | 0.89 | 0.9 | 0.81 | 0.88 |
| 4 | 179511788 | rs2715518 | -1.09 | 1.437 | 0.2 | 0.74 | 0.133 | 0.108 | 0.108 | 0.018 | 0.09 | 0.27 | 0.13 | 0.15 | 0.17 | 0.15 |
| 4 | 179514388 | rs2715520 | -0.532 | 0.564 | 0.025 | 0.54 | -0.015 | -0.014 | -0.014 | 0 | 0.8 | 0.74 | 0.88 | 0.92 | 0.88 | 0.79 |
| 4 | 179521436 | rs4607266 | 0.82 | 0.555 | -0.118 | 0.596 | 0.006 | 0.001 | 0.001 | 0.008 | 0.65 | 0.36 | 0.72 | 0.68 | 0.69 | 0.57 |
| 4 | 179525874 | rs4541553 | -0.57 | 0.678 | -0.074 | 0.572 | 0.115 | 0.11 | 0.11 | 0 | 0.94 | 0.74 | 0.91 | 0.88 | 0.9 | 0.9 |
| 4 | 179530781 | rs1434035 | 0.358 | 0.651 | -0.13 | 0.555 | -0.011 | -0.012 | -0.012 | 0 | 0.73 | 0.46 | 0.79 | 0.8 | 0.78 | 0.69 |
| 4 | 179535767 | rs1036313 | 0.059 | 0.198 | -0.558 | 0.732 | -0.019 | -0.018 | -0.018 | 0.02 | 0.2 | 0.27 | 0.12 | 0.09 | 0.12 | 0.22 |
| 4 | 179535956 | rs17066480 | -0.344 | 0.656 | -0.968 | 1.381 | 0.003 | 0.013 | 0.013 | 0 | 0.11 | 0.09 | 0.19 | 0.5 | 0.09 | 0.18 |
| 4 | 179539753 | rs5012200 | -0.416 | 0.647 | -1.042 | 0.804 | 0.004 | 0.031 | 0.031 | 0 | 0.52 | 0.42 | 0.4 | 0.14 | 0.48 | 0.41 |
| 4 | 179541008 | rs2715490 | -0.356 | 0.662 | -1.043 | 0.331 | 0.007 | -0.003 | -0.003 | 0 | 0.49 | 0.45 | 0.31 | 0.05 | 0.43 | 0.34 |
| 4 | 179541036 | rs2715491 | 0.451 | 0.651 | -1.05 | 0.203 | -0.002 | -0.039 | -0.039 | 1.84 | 0.27 | 0.25 | 0.15 | 0.02 | 0.22 | 0.23 |
| 4 | 179541387 | rs2715493 | 0.047 | 1.4 | -0.856 | 0.993 | 0.007 | -0.018 | -0.018 | 0 | 0.57 | 0.52 | 0.51 | 0.7 | 0.54 | 0.58 |
| 4 | 179545940 | rs12644125 | -0.868 | 1.756 | -1.067 | 2.125 | -0.016 | -0.026 | -0.026 | 0 | 0.16 | 0.09 | 0.36 | 0.34 | 0.21 | 0.26 |
| 4 | 179547537 | rs2890028 | -0.987 | 1.772 | -1.022 | 2.076 | -0.012 | -0.019 | -0.019 | 0 | 0.17 | 0.13 | 0.38 | 0.41 | 0.22 | 0.34 |
| 4 | 179549418 | rs2570725 | -0.598 | 1.393 | -0.466 | 1.468 | 0.007 | 0.013 | 0.013 | 0.425 | 0.28 | 0.45 | 0.35 | 0.32 | 0.3 | 0.19 |
| 4 | 179549653 | rs10434287 | -1.139 | 1.501 | 0.098 | 2.311 | 0.048 | 0.331 | 0.331 | 0 | 0.09 | 0.25 | 0.27 | 0.51 | 0.11 | 0.19 |
| 4 | 179550103 | rs4484343 | -0.043 | 0.923 | -0.223 | 1.275 | -0.011 | -0.019 | -0.019 | 0 | 0.62 | 0.6 | 0.57 | 0.26 | 0.69 | 0.65 |
| 4 | 179550271 | rs7661561 | -0.17 | 1.297 | 0.024 | 1.009 | 0.007 | 0.002 | 0.002 | 0.456 | 0.21 | 0.28 | 0.26 | 0.22 | 0.23 | 0.16 |
| 4 | 179557134 | rs12498819 | 0.969 | 0.94 | -0.38 | 0.432 | -0.093 | 0.201 | 0.201 | 3.84 | 0.37 | 0.11 | 0.29 | 0.01 | 0.49 | 0.42 |
| 4 | 179563805 | rs1583358 | -0.489 | 0.594 | 0.302 | 1.336 | -0.013 | 0.015 | 0.015 | 1.545 | 0.63 | 0.58 | 0.55 | 0.51 | 0.71 | 0.66 |
| 4 | 179564248 | rs1897760 | -1.088 | 0.561 | 0.574 | 2.151 | -0.004 | 0.005 | 0.005 | 0.398 | 0.28 | 0.61 | 0.39 | 0.71 | 0.23 | 0.36 |
| 4 | 179573368 | rs1434046 | 0.794 | 0.621 | 0.376 | 2.118 | -0.05 | -0.027 | -0.027 | 0 | 0.13 | 0.02 | 0.15 | 0.14 | 0.17 | 0.06 |
| 4 | 179576074 | rs10007324 | -0.36 | 0.605 | 0.525 | 2.201 | 0.012 | 0.087 | 0.087 | 0.017 | 0.19 | 0.31 | 0.25 | 0.45 | 0.19 | 0.16 |
| 4 | 179584676 | rs2570731 | -0.003 | 0.616 | 0.528 | 2.161 | -0.006 | 0.001 | 0.001 | 1.03 | 0.57 | 0.56 | 0.63 | 0.46 | 0.68 | 0.58 |
| 4 | 179594357 | rs10014516 | -0.019 | 0.532 | 0.592 | 2.102 | 0.023 | 0.024 | 0.024 | 0.004 | 0.17 | 0.31 | 0.25 | 0.21 | 0.19 | 0.15 |
| 4 | 179604212 | rs346101 | 1.409 | 0.405 | 0.245 | 1.656 | -0.032 | 0.022 | 0.022 | 3.161 | 0.33 | 0.2 | 0.29 | 0.01 | 0.46 | 0.29 |
| 4 | 179609480 | rs118806 | -0.553 | 0.205 | 0.113 | 1.968 | 0.034 | 0.056 | 0.056 | 1.4 | 0.62 | 0.77 | 0.79 | 0.77 | 0.69 | 0.71 |
| 4 | 179612525 | rs427796 | 0.051 | 0.202 | 0.191 | 2.123 | 0.014 | 0.011 | 0.011 | 0.854 | 0.54 | 0.63 | 0.65 | 0.67 | 0.68 | 0.57 |
| 4 | 179615916 | rs17066635 | 0.104 | 0.182 | 0.241 | 2.294 | -0.015 | 0.023 | 0.023 | 0.103 | 0.08 | 0.17 | 0.16 | 0.11 | 0.02 | 0.14 |
| 4 | 179619524 | rs379519 | 0.551 | 0.078 | 0.292 | 2.2 | -0.001 | 0.001 | 0.001 | 1.092 | 0.46 | 0.55 | 0.48 | 0.34 | 0.49 | 0.55 |
| 4 | 179620557 | rs390455 | 0.385 | 0.088 | 0.299 | 2.097 | -0.001 | 0.002 | 0.002 | 1.212 | 0.47 | 0.56 | 0.56 | 0.58 | 0.5 | 0.57 |
| 4 | 179620725 | rs17066644 | 0.491 | 0.216 | 0.373 | 2.107 | 0.016 | 0.013 | 0.013 | 0 | 0.15 | 0.21 | 0.23 | 0.21 | 0.2 | 0.15 |
| 4 | 179627762 | rs10520412 | 1.703 | 0.504 | -0.405 | 1.473 | -0.109 | 0.23 | 0.23 | 1.708 | 0.32 | 0.02 | 0.23 | 0.01 | 0.37 | 0.3 |
| 4 | 179629091 | rs17731793 | 1.511 | 0.114 | -0.256 | 1.545 | -0.071 | 0.142 | 0.142 | 1.661 | 0.34 | 0.08 | 0.28 | 0.01 | 0.45 | 0.31 |
| 4 | 179633509 | rs7666980 | 0.08 | -0.673 | -0.487 | 1.713 | -0.002 | -0.068 | -0.068 | 0 | 0.2 | 0.01 | 0.11 | 0.22 | 0.12 | 0.25 |
| 4 | 179640780 | rs17066688 | 0.118 | -0.412 | -0.059 | 1.685 | 0.003 | -0.02 | -0.02 | 0.013 | 0.11 | 0.08 | 0.06 | 0.2 | 0.11 | 0.16 |
| 4 | 179661007 | rs4861516 | 0.899 | -0.559 | 0.299 | 1.925 | -0.031 | 0.131 | 0.131 | 0.126 | 0.81 | 0.44 | 0.84 | 0.57 | 0.86 | 0.73 |
| 4 | 179665370 | rs418063 | 0.224 | -0.535 | 0.598 | 1.964 | -0.009 | 0.088 | 0.088 | 0 | 0.86 | 0.79 | 0.93 | 0.63 | 0.87 | 0.84 |
| 4 | 179677801 | rs17732937 | 0.935 | -0.518 | -0.618 | 1.464 | -0.004 | 0.144 | 0.144 | 0.155 | 0.49 | 0.12 | 0.35 | 0.23 | 0.59 | 0.49 |
| 4 | 179681882 | rs4862126 | -0.381 | -0.318 | -0.569 | 1.531 | -0.085 | -0.092 | -0.092 | 0 | 0.44 | 0.04 | 0.39 | 0.43 | 0.39 | 0.36 |
| 4 | 179683403 | rs346119 | 0.82 | -0.267 | -0.294 | 1.417 | 0.002 | 0.056 | 0.056 | 0 | 0.48 | 0.34 | 0.4 | 0.3 | 0.58 | 0.57 |
| 4 | 179692090 | rs1448958 | 1.757 | -0.311 | -1.371 | 0.635 | -0.141 | 0.168 | 0.168 | 0.337 | 0.34 | 0.03 | 0.4 | 0.08 | 0.46 | 0.41 |
| 4 | 179698359 | rs4862147 | -0.566 | -0.099 | -1.097 | 0.894 | -0.006 | 0.076 | 0.076 | 0 | 0.43 | 0.22 | 0.42 | 0.7 | 0.38 | 0.54 |
| 4 | 179699093 | rs7679972 | 0.351 | 0.013 | -1.894 | 0.98 | -0.009 | 0.031 | 0.031 | 0 | 0.3 | 0.01 | 0.14 | 0.18 | 0.19 | 0.27 |

| | | | | | | | | | | | | | | | | |
|---|---|---|---|---|---|---|---|---|---|---|---|---|---|---|---|---|
| 4 | 179702044 | rs12508820 | 1.688 | 0.169 | -2.262 | 0.764 | -0.009 | -0.022 | -0.022 | 0.151 | 0.05 | 0 | 0.05 | 0.22 | 0.05 | 0.02 |
| 4 | 179704645 | rs11937114 | 0.049 | 0.175 | -2.271 | 0.846 | -0.053 | -0.062 | -0.062 | 2.025 | 0.41 | 0.06 | 0.36 | 0.59 | 0.36 | 0.36 |
| 4 | 179704824 | rs12504394 | 0.688 | 0.273 | -1.988 | 0.124 | 0.004 | 0.011 | 0.011 | 0 | 0.09 | 0.08 | 0.04 | 0.01 | 0.1 | 0.05 |
| 4 | 179705308 | rs1947202 | -0.413 | 0.437 | -1.793 | 0.542 | -0.071 | 0.167 | 0.167 | 0 | 0.46 | 0.07 | 0.41 | 0.81 | 0.42 | 0.38 |
| 4 | 179729425 | rs4862165 | -1.908 | -0.035 | 0.251 | 2.006 | -0.043 | 0.205 | 0.205 | 0.577 | 0.15 | 0.45 | 0.29 | 0.37 | 0.09 | 0.19 |
| 4 | 179759509 | rs7684851 | -1.068 | -0.07 | 0.474 | 1.57 | -0.008 | -0.028 | -0.028 | 0 | 0.05 | 0.18 | 0.05 | 0 | 0.04 | 0 |
| 4 | 179802170 | rs1348365 | -2.456 | -0.031 | 0.612 | 1.721 | -0.01 | -0.025 | -0.025 | 0 | 0.1 | 0.48 | 0.09 | 0.01 | 0.04 | 0.02 |
| 4 | 179833516 | rs12502810 | 1.02 | 0.366 | 0.142 | 1.471 | 0.006 | -0.015 | -0.015 | 0 | 0.59 | 0.41 | 0.72 | 0.73 | 0.5 | 0.75 |
| 4 | 179837718 | rs6847330 | -0.632 | 0.628 | -0.195 | 0.944 | -0.007 | -0.026 | -0.026 | 0 | 0.05 | 0.12 | 0.05 | 0 | 0.04 | 0 |
| 4 | 179865606 | rs11733087 | 1.838 | 0.178 | -1.224 | 0.793 | -0.068 | 0.23 | 0.23 | 0 | 0.45 | 0.02 | 0.36 | 0.14 | 0.4 | 0.47 |
| 4 | 179872123 | rs11929794 | -0.051 | 0.334 | -1.148 | 0.89 | -0.182 | 0.014 | 0.014 | 0.313 | 0.29 | 0.04 | 0.18 | 0.19 | 0.43 | 0.2 |
| 4 | 179894901 | rs10014264 | -1.397 | 0.114 | -0.56 | 1.004 | 0 | -0.004 | -0.004 | 0.053 | 0.44 | 0.51 | 0.37 | 0.35 | 0.43 | 0.33 |
| 4 | 179896707 | rs9998128 | -0.598 | 0.105 | -0.516 | 0.755 | 0.001 | -0.016 | -0.016 | 0.294 | 0.34 | 0.41 | 0.19 | 0.16 | 0.34 | 0.21 |
| 4 | 179899458 | rs10033034 | -1.074 | 0.112 | -0.539 | 0.762 | 0.001 | -0.015 | -0.015 | 0.03 | 0.39 | 0.45 | 0.23 | 0.2 | 0.39 | 0.22 |
| 4 | 179899843 | rs1348368 | -1.407 | 0.136 | -0.53 | 0.796 | 0 | -0.006 | -0.006 | 1.508 | 0.44 | 0.51 | 0.27 | 0.31 | 0.43 | 0.32 |
| 4 | 179917392 | rs930972 | 1.294 | 0.237 | -0.594 | 0.616 | 0.012 | 0.138 | 0.138 | 0.031 | 0.52 | 0.23 | 0.46 | 0.19 | 0.56 | 0.52 |
| 4 | 179921499 | rs2044868 | 1.225 | 0.361 | -0.321 | 0.652 | 0.008 | 0.075 | 0.075 | 0.006 | 0.53 | 0.32 | 0.47 | 0.19 | 0.55 | 0.52 |
| 4 | 179925680 | rs7689898 | 0.088 | 0.279 | -0.317 | 0.65 | -0.06 | 0.22 | 0.22 | 0 | 0.83 | 0.56 | 0.64 | 0.42 | 0.9 | 0.74 |
| 4 | 179926087 | rs9995115 | -0.374 | 0.454 | -1.272 | 0.688 | -0.018 | 0.018 | 0.018 | 0 | 0.3 | 0.23 | 0.17 | 0.23 | 0.34 | 0.22 |
| 4 | 179929126 | rs17736809 | -0.038 | 0.219 | -1.478 | 0.446 | -0.1 | 0.02 | 0.02 | 0 | 0.24 | 0.03 | 0.13 | 0.17 | 0.29 | 0.2 |
| 4 | 179929454 | rs6833369 | 0.466 | 0.523 | -1.077 | 0.076 | 0.014 | 0.077 | 0.077 | 0.494 | 0.57 | 0.28 | 0.48 | 0.19 | 0.58 | 0.55 |
| 4 | 179932194 | rs13144575 | 0.466 | 0.633 | -1.638 | 0.159 | 0.027 | 0.154 | 0.154 | 1.55 | 0.54 | 0.14 | 0.46 | 0.19 | 0.58 | 0.55 |
| 4 | 179937829 | rs7687736 | -0.675 | 0.737 | -1.253 | 0.26 | -0.029 | 0.074 | 0.074 | 1.806 | 0.29 | 0.13 | 0.31 | 0.08 | 0.3 | 0.3 |
| 4 | 179940301 | rs13137879 | -0.731 | 0.304 | 0.468 | 1.179 | -0.005 | -0.025 | -0.025 | 0 | 0.31 | 0.23 | 0.42 | 0.43 | 0.31 | 0.4 |
| 4 | 179942389 | rs2167373 | 1.035 | 0.286 | 0.857 | 0.924 | -0.002 | -0.001 | -0.001 | 0.774 | 0.48 | 0.42 | 0.35 | 0.48 | 0.51 | 0.48 |
| 4 | 179944405 | rs17799333 | -0.455 | 0.376 | 0.302 | 0.715 | -0.052 | 0.143 | 0.143 | 3.876 | 0.34 | 0.11 | 0.34 | 0.11 | 0.4 | 0.26 |
| 4 | 179944540 | rs7698185 | -0.109 | 0.363 | 0.477 | 0.95 | 0.002 | -0.036 | -0.036 | 0.476 | 0.08 | 0.13 | 0.05 | 0 | 0.07 | 0.03 |
| 4 | 179947803 | rs17066974 | -0.976 | 0.097 | 0.754 | 1.208 | -0.008 | 0.105 | 0.105 | 0 | 0.07 | 0.11 | 0.25 | 0.41 | 0.03 | 0.21 |
| 4 | 179962359 | rs9312239 | 0.854 | 0.151 | 1.526 | 0.962 | -0.004 | 0.002 | 0.002 | 2.489 | 0.55 | 0.49 | 0.4 | 0.48 | 0.58 | 0.5 |
| 4 | 179973430 | rs1823603 | -2.353 | 0.395 | 2.448 | 2.128 | 0.012 | 0.247 | 0.247 | 1.246 | 0.08 | 0.35 | 0.11 | 0.27 | 0.04 | 0.19 |
| 4 | 179974064 | rs11736034 | -0.757 | 0.391 | 2.124 | 1.726 | 0.014 | -0.001 | -0.001 | 0 | 0.11 | 0.06 | 0.11 | 0.11 | 0.08 | 0.08 |
| 4 | 179983413 | rs7671216 | -2.36 | 0.685 | 2.785 | 1.825 | 0.031 | -0.018 | -0.018 | 0 | 0.27 | 0.45 | 0.26 | 0.24 | 0.32 | 0.26 |
| 4 | 179992133 | rs10520417 | -1.085 | 0.745 | 2.503 | 1.835 | 0.028 | -0.001 | -0.001 | 0 | 0.13 | 0.17 | 0.13 | 0.13 | 0.25 | 0.16 |
| 4 | 180000268 | rs1993095 | 2.587 | 0.552 | 2.431 | 2.254 | -0.008 | -0.037 | -0.037 | 0.714 | 0.65 | 0.23 | 0.58 | 0.47 | 0.63 | 0.62 |
| 4 | 180002879 | rs11724611 | -2.405 | 0.674 | 2.278 | 2.3 | -0.004 | 0.002 | 0.002 | 1.614 | 0.29 | 0.3 | 0.29 | 0.25 | 0.36 | 0.26 |
| 4 | 180005125 | rs17801260 | 0.37 | 0.843 | 1.849 | 2.133 | -0.027 | 0.026 | 0.026 | 0 | 0.05 | 0.01 | 0.06 | 0 | 0.14 | 0.03 |
| 4 | 180014056 | rs11131908 | 2.132 | 0.706 | 2.108 | 2.222 | 0.021 | 0.059 | 0.059 | 2.771 | 0.69 | 0.54 | 0.61 | 0.47 | 0.63 | 0.59 |
| 4 | 180014143 | rs1451407 | 2.275 | 0.708 | 1.935 | 2.223 | 0.007 | 0.027 | 0.027 | 4.444 | 0.68 | 0.43 | 0.6 | 0.47 | 0.63 | 0.59 |
| 4 | 180014954 | rs17067096 | -1.064 | -0.038 | 1.1 | 1.975 | 0.017 | 0.003 | 0.003 | 0 | 0.13 | 0.1 | 0.12 | 0.12 | 0.08 | 0.1 |
| 4 | 180024042 | rs1975651 | 1.666 | 0.014 | 1.283 | 2.308 | -0.009 | 0.051 | 0.051 | 0 | 0.7 | 0.53 | 0.57 | 0.56 | 0.74 | 0.58 |
| 4 | 180032926 | rs7699104 | -0.815 | -0.141 | 1.233 | 2.847 | 0.006 | -0.002 | -0.002 | 2.56 | 0.08 | 0.08 | 0.02 | 0.09 | 0.04 | 0.09 |
| 4 | 180036877 | rs13126556 | -1.7 | -0.051 | 1.658 | 3.099 | 0 | -0.001 | -0.001 | 0.568 | 0.16 | 0.13 | 0.19 | 0.1 | 0.16 | 0.21 |
| 4 | 180039967 | rs2028492 | -3.202 | -0.168 | 1.854 | 3.624 | -0.007 | -0.006 | -0.006 | 0 | 0.32 | 0.33 | 0.52 | 0.7 | 0.28 | 0.58 |
| 4 | 180040368 | rs17067100 | -0.464 | -0.151 | 1.708 | 3.878 | -0.001 | 0.065 | 0.065 | 0 | 0.06 | 0.04 | 0.25 | 0.36 | 0.08 | 0.16 |
| 4 | 180045846 | rs17801853 | -0.995 | -0.159 | 1.688 | 3.845 | -0.043 | -0.022 | -0.022 | 0.239 | 0.12 | 0.01 | 0.17 | 0.09 | 0.13 | 0.19 |
| 4 | 180047257 | rs10004699 | -1.298 | -0.162 | 1.815 | 3.803 | -0.017 | -0.003 | -0.003 | 0 | 0.07 | 0.14 | 0.03 | 0.05 | 0.03 | 0.09 |
| 4 | 180048362 | rs17067123 | -2.167 | -0.161 | 1.792 | 3.824 | -0.002 | 0.026 | 0.026 | 0 | 0.13 | 0.15 | 0.27 | 0.41 | 0.11 | 0.25 |
| 4 | 180053797 | rs13145669 | -2.552 | -0.19 | 1.762 | 3.808 | 0 | -0.003 | -0.003 | 0 | 0.2 | 0.17 | 0.26 | 0.28 | 0.18 | 0.3 |
| 4 | 180055500 | rs17067153 | -3.804 | -0.322 | 2.514 | 3.954 | -0.01 | -0.014 | -0.014 | 2.325 | 0.34 | 0.55 | 0.54 | 0.73 | 0.28 | 0.52 |
| 4 | 180056687 | rs1451420 | -2.18 | -0.287 | 2.397 | 3.902 | -0.001 | 0.002 | 0.002 | 0 | 0.17 | 0.16 | 0.19 | 0.08 | 0.16 | 0.17 |
| 4 | 180063431 | rs10011636 | -1.958 | -0.303 | 2.297 | 3.994 | -0.009 | 0.03 | 0.03 | 0 | 0.1 | 0.11 | 0.09 | 0.24 | 0.05 | 0.21 |
| 4 | 180068417 | rs1036767 | -2.005 | -0.304 | 2.337 | 4.023 | -0.003 | 0.327 | 0.327 | 0.012 | 0.13 | 0.44 | 0.29 | 0.45 | 0.1 | 0.22 |
| 4 | 180072512 | rs1016186 | -2.136 | -0.057 | 2.019 | 3.471 | -0.02 | 0.016 | 0.016 | 0 | 0.18 | 0.06 | 0.19 | 0.09 | 0.18 | 0.2 |
| 4 | 180072951 | rs1016184 | -2.457 | -0.085 | 2.04 | 3.418 | -0.003 | -0.029 | -0.029 | 0 | 0.21 | 0.14 | 0.26 | 0.27 | 0.19 | 0.3 |
| 4 | 180085436 | rs11939017 | -2.95 | -0.203 | 2.189 | 2.73 | -0.013 | 0.007 | 0.007 | 0 | 0.34 | 0.53 | 0.25 | 0.34 | 0.29 | 0.42 |
| 4 | 180085873 | rs17067173 | -1.702 | 0.33 | 2.276 | 2.466 | 0.005 | -0.006 | -0.006 | 0 | 0.12 | 0.2 | 0.13 | 0.09 | 0.12 | 0.1 |
| 4 | 180095990 | rs2383273 | 2.649 | 0.341 | 2.153 | 2.562 | 0 | 0.003 | 0.003 | 0.019 | 0.71 | 0.31 | 0.73 | 0.75 | 0.77 | 0.67 |
| 4 | 180106072 | rs7671333 | -2.247 | 0.701 | 1.616 | 2.256 | 0.024 | -0.001 | -0.001 | 0 | 0.23 | 0.41 | 0.21 | 0.21 | 0.23 | 0.23 |
| 4 | 180116455 | rs17746595 | -1.941 | 0.897 | 0.741 | 1.968 | 0.018 | -0.018 | -0.018 | 1.364 | 0.21 | 0.29 | 0.19 | 0.14 | 0.23 | 0.19 |
| 4 | 180121844 | rs11934681 | -1.715 | 0.673 | 0.861 | 2.468 | 0.001 | 0.187 | 0.187 | 0.007 | 0.17 | 0.35 | 0.36 | 0.49 | 0.13 | 0.23 |
| 4 | 180126785 | rs1350327 | 1.993 | 0.93 | 1.748 | 2.722 | 0.075 | 0.364 | 0.364 | 1.022 | 0.89 | 0.6 | 0.76 | 0.57 | 0.88 | 0.77 |
| 4 | 180131863 | rs1379410 | 1.803 | 1.244 | 1.641 | 2.434 | 0.001 | 0.002 | 0.002 | 0 | 0.69 | 0.37 | 0.55 | 0.44 | 0.63 | 0.52 |
| 4 | 180133742 | rs1457402 | -0.431 | 1.144 | 1.633 | 2.23 | 0.005 | 0.012 | 0.012 | 0.083 | 0.06 | 0.07 | 0.13 | 0.13 | 0.09 | 0.16 |
| 4 | 180137681 | rs9991397 | -0.614 | 0.498 | 1.626 | 1.4 | -0.003 | 0.003 | 0.003 | 0 | 0.47 | 0.57 | 0.55 | 0.43 | 0.5 | 0.38 |
| 4 | 180137800 | rs4621480 | -0.569 | 0.516 | 1.588 | 1.394 | 0 | -0.002 | -0.002 | 0.043 | 0.46 | 0.5 | 0.53 | 0.38 | 0.49 | 0.37 |
| 4 | 180147019 | rs17803697 | 1.179 | -0.003 | 1.224 | 1.329 | -0.06 | 0.056 | 0.056 | 0 | 0.38 | 0.11 | 0.27 | 0.23 | 0.4 | 0.35 |
| 4 | 180148587 | rs2383278 | 0.428 | -0.117 | 1.34 | 1.076 | -0.011 | 0.024 | 0.024 | 2.014 | 0.35 | 0.25 | 0.25 | 0.01 | 0.4 | 0.16 |
| 4 | 180157413 | rs12641267 | 0.042 | 0.183 | 0.127 | 1.284 | -0.002 | 0.01 | 0.01 | 0.042 | 0.56 | 0.38 | 0.5 | 0.61 | 0.53 | 0.69 |
| 4 | 180165656 | rs17747864 | 0.783 | 0.339 | -0.633 | 1.16 | -0.041 | 0.059 | 0.059 | 0 | 0.4 | 0.14 | 0.27 | 0.24 | 0.41 | 0.39 |

| | | | | | | | | | | | | | | | | |
|---|---|---|---|---|---|---|---|---|---|---|---|---|---|---|---|---|
| 4 | 180168625 | rs10000174 | -0.475 | 0.321 | 0.009 | 0.723 | -0.001 | 0.014 | 0.014 | 0 | 0.52 | 0.43 | 0.48 | 0.39 | 0.51 | 0.36 |
| 4 | 180173268 | rs7682192 | -0.193 | 0.406 | 0.057 | 0.65 | 0 | 0 | 0 | 0.117 | 0.45 | 0.4 | 0.37 | 0.34 | 0.41 | 0.19 |
| 4 | 180173785 | rs13108774 | -1.046 | 0.425 | 0.11 | 0.68 | 0 | -0.001 | -0.001 | 0.023 | 0.58 | 0.54 | 0.54 | 0.45 | 0.58 | 0.47 |
| 4 | 180182483 | rs10005160 | -1.059 | 0.414 | 0.199 | 0.637 | -0.001 | -0.002 | -0.002 | 0 | 0.58 | 0.59 | 0.57 | 0.47 | 0.58 | 0.47 |
| 4 | 180186551 | rs17067290 | 1.221 | 0.264 | -0.174 | 0.392 | -0.032 | 0.057 | 0.057 | 1.286 | 0.23 | 0.04 | 0.18 | 0.15 | 0.22 | 0.08 |
| 4 | 180187024 | rs6850582 | 0.139 | 0.072 | 0.23 | 0.42 | -0.004 | -0.003 | -0.003 | 1.236 | 0.21 | 0.24 | 0.17 | 0.19 | 0.19 | 0.11 |
| 4 | 180188121 | rs1457413 | -0.733 | 0.189 | 0.291 | 0.586 | -0.003 | -0.004 | -0.004 | 0.664 | 0.59 | 0.64 | 0.57 | 0.47 | 0.58 | 0.47 |
| 4 | 180191273 | rs12500931 | -0.438 | 0.095 | 0.376 | 0.652 | -0.004 | 0.004 | 0.004 | 0 | 0.54 | 0.58 | 0.52 | 0.45 | 0.51 | 0.37 |
| 4 | 180193914 | rs6828433 | -0.965 | -0.072 | 0.451 | 0.25 | -0.031 | -0.062 | -0.062 | 0.149 | 0.11 | 0.3 | 0.08 | 0 | 0.03 | 0.06 |
| 4 | 180203808 | rs6812195 | -0.929 | -0.053 | 0.427 | 0.245 | -0.031 | -0.06 | -0.06 | 0 | 0.11 | 0.26 | 0.07 | 0 | 0.03 | 0.04 |
| 4 | 180206478 | rs7670049 | -1.808 | -0.178 | 0.885 | 0.577 | 0 | -0.001 | -0.001 | 0 | 0.29 | 0.64 | 0.27 | 0.31 | 0.22 | 0.27 |
| 4 | 180207412 | rs10010601 | -0.664 | -0.058 | 1.23 | 0.481 | -0.022 | -0.06 | -0.06 | 0 | 0.11 | 0.32 | 0.09 | 0 | 0.03 | 0.04 |
| 4 | 180240950 | rs13136851 | 0.645 | 0.016 | -0.323 | 0.24 | -0.109 | 0.067 | 0.067 | 0 | 0.4 | 0.04 | 0.25 | 0.18 | 0.42 | 0.41 |
| 4 | 180248838 | rs6552356 | 1.602 | -0.151 | 0.966 | 0.91 | -0.022 | 0.187 | 0.187 | 0 | 0.88 | 0.69 | 0.77 | 0.49 | 0.94 | 0.69 |
| 4 | 180250564 | rs17817727 | -0.758 | -0.227 | 0.728 | 0.681 | -0.015 | 0.013 | 0.013 | 0 | 0.11 | 0.05 | 0.1 | 0.04 | 0.17 | 0.19 |
| 4 | 180250778 | rs13146351 | 0.92 | -0.477 | 0.553 | 0.054 | 0 | -0.001 | -0.001 | 0 | 0.54 | 0.49 | 0.53 | 0.43 | 0.56 | 0.42 |
| 4 | 180251982 | rs921839 | 0.069 | -0.413 | 0.686 | 0.095 | -0.003 | 0.009 | 0.009 | 0 | 0.66 | 0.63 | 0.65 | 0.45 | 0.73 | 0.62 |
| 4 | 180253471 | rs7665006 | 0.404 | -0.263 | 0.505 | -0.062 | 0.001 | -0.004 | -0.004 | 0 | 0.61 | 0.62 | 0.45 | 0.4 | 0.66 | 0.53 |
| 4 | 180257370 | rs6826259 | 1.373 | -0.389 | 1.051 | -0.176 | -0.013 | 0.036 | 0.036 | 0 | 0.48 | 0.34 | 0.33 | 0.36 | 0.49 | 0.35 |
| 4 | 180263649 | rs7654367 | -0.491 | -0.39 | 1.076 | -0.143 | 0.002 | -0.026 | -0.026 | 0.186 | 0.37 | 0.25 | 0.54 | 0.6 | 0.34 | 0.47 |
| 4 | 180267949 | rs1017671 | 0.8 | -0.381 | 1.386 | -0.117 | 0.011 | -0.043 | -0.043 | 0 | 0.84 | 0.48 | 0.86 | 0.96 | 0.84 | 0.81 |
| 4 | 180269114 | rs11933503 | 0.752 | -0.371 | 1.286 | -0.094 | 0.024 | -0.021 | -0.021 | 0 | 0.84 | 0.55 | 0.87 | 0.93 | 0.84 | 0.79 |
| 4 | 180271128 | rs12511057 | 0.302 | -0.254 | 1.236 | -0.142 | 0.005 | -0.024 | -0.024 | 0.531 | 0.88 | 0.82 | 0.88 | 0.98 | 0.84 | 0.91 |
| 4 | 180275389 | rs10019140 | 1.01 | -0.367 | 1.283 | -0.079 | -0.01 | 0.282 | 0.282 | 0.347 | 0.89 | 0.67 | 0.62 | 0.42 | 0.9 | 0.8 |
| 4 | 180279328 | rs17768364 | 1.568 | -0.5 | 0.679 | -0.528 | -0.034 | 0.021 | 0.021 | 0.176 | 0.41 | 0.25 | 0.3 | 0.32 | 0.46 | 0.29 |
| 4 | 180282887 | rs980165 | -1.32 | -0.338 | 0.785 | -0.431 | 0 | -0.004 | -0.004 | 0 | 0.53 | 0.47 | 0.66 | 0.68 | 0.51 | 0.65 |
| 4 | 180284473 | rs6828941 | -0.095 | -0.366 | 0.39 | -0.77 | 0.008 | 0.134 | 0.134 | 0 | 0.28 | 0.12 | 0.16 | 0.02 | 0.19 | 0.24 |
| 4 | 180293616 | rs4861773 | -0.141 | -0.036 | 0.596 | -0.464 | -0.001 | -0.07 | -0.07 | 1.885 | 0.75 | 0.64 | 0.78 | 0.96 | 0.79 | 0.74 |
| 4 | 180298293 | rs17090630 | 0.375 | -0.142 | 0.458 | 0.051 | 0.004 | -0.016 | -0.016 | 0 | 0.66 | 0.34 | 0.56 | 0.46 | 0.72 | 0.52 |
| 4 | 180302451 | rs1378554 | 0.094 | -0.025 | 0.585 | -0.713 | -0.003 | -0.046 | -0.046 | 0 | 0.68 | 0.43 | 0.58 | 0.9 | 0.73 | 0.64 |
| 4 | 180305067 | rs10002760 | -0.254 | 0.224 | -0.443 | -0.865 | -0.003 | -0.038 | -0.038 | 0 | 0.25 | 0.32 | 0.16 | 0.07 | 0.24 | 0.2 |
| 4 | 180306130 | rs1455698 | -0.642 | 0.37 | -1.118 | -0.365 | -0.004 | 0.119 | 0.119 | 0 | 0.53 | 0.7 | 0.55 | 0.16 | 0.52 | 0.55 |
| 4 | 180313511 | rs6841508 | -0.227 | 0.519 | -1.102 | -0.057 | 0.001 | 0.009 | 0.009 | 0 | 0.22 | 0.14 | 0.13 | 0.12 | 0.17 | 0.26 |
| 4 | 180316651 | rs4861775 | -0.327 | 0.751 | -1.711 | 0.606 | -0.048 | -0.095 | -0.095 | 0.788 | 0.17 | 0.02 | 0.22 | 0.29 | 0.21 | 0.1 |
| 4 | 180319376 | rs6841214 | 0.341 | 0.542 | -1.268 | 0.609 | -0.007 | 0.006 | 0.006 | 0 | 0.88 | 0.8 | 0.93 | 0.88 | 0.91 | 0.76 |
| 4 | 180319752 | rs7688643 | 0.811 | 0.4 | -0.191 | 0.478 | -0.003 | 0.033 | 0.033 | 0 | 0.85 | 0.57 | 0.91 | 0.88 | 0.91 | 0.75 |
| 4 | 180320695 | rs12508123 | -0.222 | 0.438 | -0.995 | 0.505 | -0.05 | -0.056 | -0.056 | 0 | 0.2 | 0.03 | 0.21 | 0.26 | 0.24 | 0.1 |
| 4 | 180324614 | rs17090633 | 0.351 | 0.459 | -1.006 | 0.479 | -0.029 | -0.057 | -0.057 | 0 | 0.14 | 0.04 | 0.18 | 0.27 | 0.18 | 0.07 |
| 4 | 180330521 | rs6837023 | 0.561 | -0.421 | -0.657 | 0.002 | -0.04 | -0.03 | -0.03 | 0 | 0.15 | 0.32 | 0.05 | 0.07 | 0.06 | 0.14 |
| 4 | 180332717 | rs9312251 | -0.557 | -0.795 | -1.031 | -0.469 | 0.033 | 0.045 | 0.045 | 0 | 0.67 | 0.78 | 0.79 | 0.83 | 0.74 | 0.63 |
| 4 | 180334106 | rs13122924 | 2.803 | -0.639 | -1.128 | -0.74 | -0.054 | 0.057 | 0.057 | 0 | 0.2 | 0.08 | 0.26 | 0.02 | 0.33 | 0.18 |
| 4 | 180334183 | rs1401916 | -1.143 | -0.953 | -1.178 | -0.773 | 0.004 | 0.028 | 0.028 | 0 | 0.72 | 0.94 | 0.81 | 0.83 | 0.74 | 0.64 |
| 4 | 180336832 | rs10009088 | 0.19 | -0.847 | -0.68 | -0.622 | 0.006 | 0.11 | 0.11 | 0 | 0.5 | 0.78 | 0.59 | 0.74 | 0.54 | 0.5 |
| 4 | 180337114 | rs1455659 | -0.254 | -0.947 | -0.678 | -0.604 | 0.031 | 0.063 | 0.063 | 0 | 0.56 | 0.78 | 0.63 | 0.73 | 0.61 | 0.58 |
| 4 | 180337189 | rs10031494 | 0.015 | -0.841 | -0.271 | -0.241 | 0.042 | -0.022 | -0.022 | 0.401 | 0.06 | 0.2 | 0.06 | 0 | 0.06 | 0.04 |
| 4 | 180339842 | rs11939068 | 0.057 | -0.959 | -0.317 | -0.185 | -0.036 | -0.064 | -0.064 | 4.069 | 0.21 | 0.33 | 0.11 | 0.07 | 0.12 | 0.17 |
| 4 | 180344099 | rs1455661 | 0.229 | -0.988 | -0.483 | -0.051 | -0.071 | 0.275 | 0.275 | 0 | 0.17 | 0.48 | 0.26 | 0.56 | 0.09 | 0.24 |
| 4 | 180347106 | rs4861401 | 0.463 | -0.979 | -0.839 | -0.218 | -0.061 | -0.076 | -0.076 | 0 | 0.2 | 0.03 | 0.2 | 0.26 | 0.23 | 0.15 |
| 4 | 180347365 | rs7658508 | -0.582 | -0.994 | -0.467 | -0.005 | -0.055 | 0.131 | 0.131 | 0 | 0.24 | 0.52 | 0.31 | 0.55 | 0.15 | 0.25 |
| 4 | 180348448 | rs17821431 | 1.718 | -0.5 | -0.86 | -0.275 | -0.002 | 0.011 | 0.011 | 0 | 0.07 | 0.03 | 0.05 | 0 | 0.06 | 0.03 |
| 4 | 180351563 | rs6552362 | -0.88 | -0.536 | -0.566 | -0.494 | -0.004 | 0.033 | 0.033 | 0 | 0.47 | 0.57 | 0.55 | 0.82 | 0.49 | 0.46 |
| 4 | 180352288 | rs4861781 | 2.193 | -0.778 | -0.937 | -0.947 | -0.064 | 0.075 | 0.075 | 0 | 0.22 | 0.09 | 0.23 | 0.01 | 0.32 | 0.2 |
| 4 | 180356846 | rs1378552 | -0.151 | -0.863 | -0.782 | -0.81 | -0.006 | -0.012 | -0.012 | 0 | 0.38 | 0.54 | 0.42 | 0.82 | 0.33 | 0.32 |
| 4 | 180362145 | rs2605050 | -1.672 | -1.015 | -0.837 | -0.968 | -0.006 | -0.028 | -0.028 | 0 | 0.93 | 0.92 | 0.95 | 1 | 0.94 | 0.99 |
| 4 | 180365721 | rs12643739 | 1.245 | -1.21 | -1.332 | -0.669 | -0.02 | -0.069 | -0.069 | 0.724 | 0.2 | 0.02 | 0.11 | 0.23 | 0.17 | 0.14 |
| 4 | 180365895 | rs17770817 | 1.49 | -1.312 | -1.406 | -0.758 | -0.032 | 0.037 | 0.037 | 0 | 0.14 | 0 | 0.14 | 0.08 | 0.16 | 0.21 |
| 4 | 180383312 | rs2716816 | -0.083 | -1.085 | -0.425 | -0.598 | 0.022 | -0.121 | -0.121 | 0.286 | 0.84 | 0.6 | 0.89 | 1 | 0.85 | 0.9 |
| 4 | 180387187 | rs767002 | 0.901 | -1.274 | -0.303 | -0.299 | -0.013 | -0.018 | -0.018 | 0 | 0.38 | 0.4 | 0.48 | 0.56 | 0.51 | 0.42 |
| 4 | 180389661 | rs13107405 | 1.721 | -1.439 | -0.209 | -0.106 | -0.024 | -0.027 | -0.027 | 0 | 0.31 | 0.32 | 0.46 | 0.54 | 0.45 | 0.34 |
| 4 | 180393970 | rs6823606 | -0.655 | -1.435 | -0.229 | -0.252 | 0.021 | 0.026 | 0.026 | 0 | 0.72 | 0.81 | 0.75 | 0.84 | 0.81 | 0.61 |
| 4 | 180412136 | rs6826419 | 1.262 | -1.098 | -0.143 | -0.209 | -0.018 | -0.018 | -0.018 | 0 | 0.37 | 0.37 | 0.48 | 0.55 | 0.51 | 0.38 |
| 4 | 180417262 | rs1455693 | 0.67 | -1.156 | -0.165 | -0.514 | 0.019 | 0.04 | 0.04 | 0 | 0.27 | 0.17 | 0.25 | 0.16 | 0.21 | 0.39 |
| 4 | 180418475 | rs2254490 | -0.141 | -1.173 | -0.13 | -0.16 | -0.015 | 0.016 | 0.016 | 0 | 0.64 | 0.54 | 0.73 | 0.63 | 0.72 | 0.76 |
| 4 | 180419383 | rs13129106 | 1.771 | -1.052 | -0.2 | -0.11 | -0.012 | -0.007 | -0.007 | 0 | 0.32 | 0.33 | 0.45 | 0.47 | 0.43 | 0.33 |
| 4 | 180420134 | rs2605070 | -1.333 | -1.071 | -0.429 | -0.052 | 0.006 | -0.059 | -0.059 | 0 | 0.79 | 0.91 | 0.89 | 0.69 | 0.83 | 0.82 |
| 4 | 180433430 | rs2681358 | 0.938 | -0.981 | -0.532 | -0.16 | 0.016 | 0.023 | 0.023 | 0 | 0.23 | 0.14 | 0.25 | 0.16 | 0.16 | 0.34 |
| 4 | 180436364 | rs2681357 | 0.638 | -0.935 | -0.264 | -0.136 | -0.001 | -0.001 | -0.001 | 0 | 0.55 | 0.54 | 0.71 | 0.65 | 0.59 | 0.67 |
| 4 | 180442301 | rs1464292 | -0.003 | -0.901 | -0.372 | -0.241 | 0.007 | 0.035 | 0.035 | 0 | 0.32 | 0.24 | 0.27 | 0.13 | 0.28 | 0.43 |
| 4 | 180446118 | rs4861784 | 0.972 | -0.938 | -0.401 | -0.197 | 0.007 | 0.014 | 0.014 | 0 | 0.22 | 0.17 | 0.24 | 0.13 | 0.16 | 0.34 |

| | | | | | | | | | | | | | | | | |
|---|---|---|---|---|---|---|---|---|---|---|---|---|---|---|---|---|
| 4 | 180466309 | rs10017041 | 0.077 | -1.147 | -0.394 | -0.284 | 0.018 | 0.033 | 0.033 | 0 | 0.33 | 0.23 | 0.25 | 0.13 | 0.23 | 0.41 |
| 4 | 180466423 | rs10005349 | 0.494 | -1.121 | -0.392 | -0.252 | 0.014 | 0.025 | 0.025 | 0 | 0.27 | 0.19 | 0.24 | 0.13 | 0.2 | 0.38 |
| 4 | 180466511 | rs6852751 | 2.005 | -0.954 | -0.136 | -0.226 | 0.002 | -0.032 | -0.032 | 0.051 | 0.06 | 0.16 | 0.06 | 0 | 0.06 | 0.01 |
| 4 | 180467353 | rs2584396 | 1.807 | -1.041 | -0.338 | -0.086 | 0.004 | -0.01 | -0.01 | 1.408 | 0.08 | 0.04 | 0.12 | 0.1 | 0.09 | 0.04 |
| 4 | 180467437 | rs3911604 | 0.804 | -0.858 | -0.11 | 0.244 | -0.001 | -0.086 | -0.086 | 0.264 | 0.21 | 0.07 | 0.11 | 0.3 | 0.16 | 0.21 |
| 4 | 180472146 | rs3907136 | 1.136 | -0.934 | -0.015 | -0.002 | 0.023 | 0.049 | 0.049 | 0 | 0.3 | 0.18 | 0.28 | 0.13 | 0.25 | 0.38 |
| 4 | 180472174 | rs3911610 | -0.342 | -1.032 | 0.344 | 0.047 | -0.037 | 0.259 | 0.259 | 0.895 | 0.83 | 0.44 | 0.68 | 0.46 | 0.89 | 0.83 |
| 4 | 180472339 | rs2716771 | 0.761 | -0.887 | -0.011 | 0.077 | -0.009 | -0.018 | -0.018 | 0 | 0.11 | 0.07 | 0.24 | 0.44 | 0.16 | 0.13 |
| 4 | 180472353 | rs2716772 | -1.883 | -0.725 | 0.137 | -0.005 | -0.006 | -0.011 | -0.011 | 0 | 0.72 | 0.74 | 0.77 | 0.97 | 0.61 | 0.74 |
| 4 | 180476903 | rs17067795 | 0.516 | -0.872 | 0.024 | 0.389 | 0.001 | -0.042 | -0.042 | 1.346 | 0.22 | 0.08 | 0.11 | 0.28 | 0.17 | 0.2 |
| 4 | 180493646 | rs3911612 | 0.388 | -0.964 | 0.063 | 0.386 | 0.033 | 0.007 | 0.007 | 0 | 0.66 | 0.25 | 0.64 | 0.28 | 0.77 | 0.68 |
| 4 | 180497015 | rs2333688 | 0.562 | -0.853 | -0.103 | 0.386 | 0.009 | 0.068 | 0.068 | 0 | 0.29 | 0.16 | 0.29 | 0.14 | 0.26 | 0.37 |
| 4 | 180498452 | rs2716785 | 0.659 | -0.88 | 0.329 | 0.564 | -0.009 | 0.199 | 0.199 | 0.129 | 0.79 | 0.5 | 0.66 | 0.46 | 0.81 | 0.79 |
| 4 | 180499234 | rs2333689 | 0.47 | -0.844 | 0.263 | 0.559 | -0.031 | -0.037 | -0.037 | 3.411 | 0.11 | 0.01 | 0.12 | 0.1 | 0.12 | 0.09 |
| 4 | 180500797 | rs2584414 | 0.453 | -1.86 | 0.551 | 0.758 | -0.027 | 0.252 | 0.252 | 0.867 | 0.82 | 0.53 | 0.67 | 0.46 | 0.85 | 0.83 |
| 4 | 180501487 | rs2611020 | -0.148 | -1.81 | 0.566 | 0.776 | -0.008 | 0.272 | 0.272 | 4.54 | 0.89 | 0.6 | 0.79 | 0.57 | 0.92 | 0.87 |
| 4 | 180504324 | rs6552374 | 0.885 | -1.791 | 0.259 | 0.73 | 0.003 | -0.014 | -0.014 | 0.094 | 0.08 | 0.04 | 0.12 | 0.1 | 0.08 | 0.04 |
| 4 | 180505632 | rs2920105 | 0.099 | -1.902 | 0.297 | 0.585 | -0.008 | 0.008 | 0.008 | 0.037 | 0.6 | 0.44 | 0.5 | 0.42 | 0.65 | 0.48 |
| 4 | 180509267 | rs2716798 | -0.939 | -1.982 | 0.534 | 0.632 | -0.009 | 0.191 | 0.191 | 1.312 | 0.91 | 0.76 | 0.8 | 0.57 | 0.92 | 0.87 |
| 4 | 180510815 | rs2716800 | 0.611 | -2.062 | -0.203 | 0.463 | -0.003 | -0.079 | -0.079 | 0.988 | 0.21 | 0.03 | 0.12 | 0.28 | 0.17 | 0.18 |
| 4 | 180512607 | rs2584430 | 0.203 | -1.996 | 0.038 | 0.961 | -0.012 | -0.005 | -0.005 | 0 | 0.6 | 0.47 | 0.49 | 0.33 | 0.62 | 0.47 |
| 4 | 180512644 | rs10009213 | 0.488 | -1.79 | -0.506 | 1.093 | -0.017 | 0.002 | 0.002 | 0 | 0.28 | 0.1 | 0.27 | 0.26 | 0.26 | 0.4 |
| 4 | 180514510 | rs3902658 | 0.081 | -1.751 | -0.615 | 0.983 | -0.028 | 0.052 | 0.052 | 0 | 0.33 | 0.07 | 0.35 | 0.21 | 0.31 | 0.43 |
| 4 | 180518801 | rs2584434 | 0.9 | -1.67 | -0.533 | 0.613 | -0.018 | 0.053 | 0.053 | 0 | 0.25 | 0.07 | 0.26 | 0.16 | 0.25 | 0.36 |
| 4 | 180519077 | rs2716805 | -0.03 | -1.679 | 0.033 | 0.703 | -0.021 | 0.012 | 0.012 | 0 | 0.7 | 0.53 | 0.66 | 0.31 | 0.76 | 0.66 |
| 4 | 180521957 | rs4345226 | 1.273 | -1.648 | 0.363 | 1.017 | -0.009 | -0.009 | -0.009 | 0 | 0.34 | 0.26 | 0.45 | 0.5 | 0.43 | 0.35 |
| 4 | 180522174 | rs2584437 | 0.664 | -1.609 | 0.021 | 1.011 | 0.009 | -0.003 | -0.003 | 0.034 | 0.29 | 0.06 | 0.28 | 0.27 | 0.26 | 0.38 |
| 4 | 180524520 | rs6827730 | -0.047 | -1.427 | 0.267 | 0.367 | 0 | 0.002 | 0.002 | 0 | 0.1 | 0.14 | 0.12 | 0.08 | 0.11 | 0.07 |
| 4 | 180528017 | rs17831521 | 0.5 | -1.445 | 0.065 | 0.321 | -0.014 | -0.026 | -0.026 | 0 | 0.08 | 0.01 | 0.05 | 0.1 | 0.07 | 0.04 |
| 4 | 180528263 | rs2611043 | 0.815 | -1.478 | 0.36 | 0.125 | -0.002 | -0.042 | -0.042 | 2.736 | 0.41 | 0.42 | 0.28 | 0.05 | 0.49 | 0.31 |
| 4 | 180531692 | rs6816822 | -0.046 | -1.34 | 0.374 | 0.533 | 0.002 | -0.012 | -0.012 | 0 | 0.9 | 0.74 | 0.95 | 0.94 | 0.93 | 0.96 |
| 4 | 180536545 | rs10520430 | 0.724 | -1.252 | -0.343 | 0.446 | -0.01 | 0.106 | 0.106 | 0 | 0.29 | 0.07 | 0.21 | 0.13 | 0.28 | 0.37 |
| 4 | 180537212 | rs17067866 | 0.172 | -1.132 | -0.306 | 0.729 | 0.009 | -0.029 | -0.029 | 0 | 0.16 | 0.12 | 0.09 | 0.29 | 0.11 | 0.13 |
| 4 | 180537663 | rs13146175 | 0.577 | -1.058 | -0.16 | 0.689 | 0 | 0.001 | 0.001 | 0 | 0.71 | 0.67 | 0.79 | 0.59 | 0.76 | 0.78 |
| 4 | 180542136 | rs3849602 | 1.452 | -0.984 | -0.883 | 0.094 | -0.086 | 0.106 | 0.106 | 0.544 | 0.26 | 0.11 | 0.25 | 0.04 | 0.41 | 0.22 |
| 4 | 180542333 | rs3849603 | 0.116 | -0.696 | -0.148 | 0.667 | 0.006 | 0.001 | 0.001 | 0 | 0.56 | 0.45 | 0.41 | 0.39 | 0.63 | 0.46 |
| 4 | 180548076 | rs3849606 | 0.263 | -0.522 | 0.122 | 0.338 | -0.02 | -0.066 | -0.066 | 0 | 0.8 | 0.66 | 0.89 | 0.95 | 0.85 | 0.8 |
| 4 | 180551100 | rs4526010 | 0.707 | -0.447 | -0.477 | 0.244 | 0.007 | -0.001 | -0.001 | 0 | 0.86 | 0.81 | 0.93 | 0.99 | 0.93 | 0.84 |
| 4 | 180564205 | rs6854193 | 0.194 | -0.391 | -0.203 | 0.79 | 0.004 | 0.04 | 0.04 | 0 | 0.2 | 0.36 | 0.16 | 0.23 | 0.15 | 0.27 |
| 4 | 180564815 | rs10520432 | -0.025 | -0.214 | -0.872 | 0.736 | 0.007 | 0.004 | 0.004 | 0 | 0.55 | 0.57 | 0.47 | 0.34 | 0.65 | 0.51 |
| 4 | 180570932 | rs7659991 | 0.785 | 0.292 | -1.433 | 0.899 | -0.109 | 0.115 | 0.115 | 0 | 0.32 | 0.09 | 0.29 | 0.1 | 0.46 | 0.24 |
| 4 | 180582498 | rs12509486 | -0.461 | 0.208 | -1.103 | 1.129 | 0.011 | -0.017 | -0.017 | 0 | 0.33 | 0.18 | 0.35 | 0.51 | 0.3 | 0.31 |
| 4 | 180586598 | rs6840953 | -0.679 | 0.256 | -1.091 | 1.044 | 0.011 | -0.025 | -0.025 | 0 | 0.38 | 0.22 | 0.25 | 0.52 | 0.35 | 0.39 |
| 4 | 180588881 | rs10032248 | 0.261 | -0.337 | -1.053 | 0.785 | 0.03 | 0.017 | 0.017 | 0 | 0.78 | 0.87 | 0.93 | 0.86 | 0.89 | 0.81 |
| 4 | 180589341 | rs17067909 | -0.767 | 0.15 | -1.186 | 1.288 | -0.011 | -0.019 | -0.019 | 0 | 0.12 | 0.11 | 0.3 | 0.3 | 0.15 | 0.17 |
| 4 | 180591910 | rs2584393 | 0.828 | -0.975 | -0.782 | 0.298 | -0.068 | 0.009 | 0.009 | 0 | 0.71 | 0.47 | 0.78 | 0.68 | 0.84 | 0.58 |
| 4 | 180596542 | rs3849612 | 0.913 | -1.213 | 0.11 | 0.169 | 0.002 | -0.013 | -0.013 | 0.124 | 0.67 | 0.69 | 0.7 | 0.91 | 0.76 | 0.73 |
| 4 | 180608477 | rs6552386 | 1.805 | -1.434 | 0.439 | 0.076 | 0.041 | 0.08 | 0.08 | 0 | 0.46 | 0.22 | 0.52 | 0.66 | 0.63 | 0.47 |
| 4 | 180608655 | rs982088 | -1.83 | -1.619 | 0.7 | -0.012 | -0.015 | -0.017 | -0.017 | 0.78 | 0.45 | 0.47 | 0.44 | 0.33 | 0.34 | 0.39 |
| 4 | 180613345 | rs12500438 | 1.508 | -1.848 | 0.429 | -0.309 | -0.001 | 0.001 | 0.001 | 0 | 0.55 | 0.48 | 0.57 | 0.67 | 0.7 | 0.63 |
| 4 | 180615657 | rs17498828 | 0.13 | -1.712 | 0.388 | -0.15 | 0.035 | 0.025 | 0.025 | 0 | 0.12 | 0.05 | 0.04 | 0.07 | 0.04 | 0.16 |
| 4 | 180619408 | rs1020237 | 0.518 | -1.789 | 0.457 | -0.201 | 0.021 | 0.022 | 0.022 | 0 | 0.81 | 0.86 | 0.92 | 0.93 | 0.9 | 0.79 |
| 4 | 180624051 | rs4861798 | -0.97 | -1.132 | -0.117 | -0.854 | 0.006 | 0.011 | 0.011 | 0 | 0.47 | 0.63 | 0.54 | 0.9 | 0.43 | 0.62 |
| 4 | 180627328 | rs4373197 | 1.054 | -1.192 | -0.107 | -0.731 | -0.013 | -0.017 | -0.017 | 0 | 0.22 | 0.21 | 0.34 | 0.57 | 0.3 | 0.24 |
| 4 | 180635834 | rs10084772 | -0.509 | -1.246 | -0.08 | -0.772 | -0.002 | -0.009 | -0.009 | 0 | 0.4 | 0.58 | 0.42 | 0.68 | 0.35 | 0.54 |
| 4 | 180652005 | rs6852457 | -0.094 | -1.071 | 0.032 | -0.719 | -0.025 | -0.049 | -0.049 | 0 | 0.1 | 0.29 | 0.06 | 0.02 | 0.07 | 0.17 |
| 4 | 180652782 | rs13122669 | 0.579 | -0.995 | -0.169 | -0.535 | 0.014 | 0.02 | 0.02 | 0 | 0.24 | 0.32 | 0.36 | 0.62 | 0.3 | 0.29 |
| 4 | 180659401 | rs12502245 | -0.453 | -0.795 | -0.359 | -0.49 | 0.002 | 0.007 | 0.007 | 0.557 | 0.21 | 0.23 | 0.25 | 0.09 | 0.17 | 0.1 |
| 4 | 180659403 | rs17068069 | -0.055 | -0.915 | 0.065 | -0.708 | 0.051 | -0.013 | -0.013 | 0 | 0.08 | 0.35 | 0.05 | 0 | 0.04 | 0.05 |
| 4 | 180661580 | rs1380000 | 1.003 | -0.583 | -1.145 | -0.934 | -0.008 | -0.037 | -0.037 | 0.541 | 0.37 | 0.17 | 0.33 | 0.6 | 0.38 | 0.47 |
| 4 | 180663940 | rs10050025 | -1.287 | -0.568 | -1.196 | -0.979 | -0.008 | 0.014 | 0.014 | 0 | 0.66 | 0.7 | 0.63 | 0.84 | 0.61 | 0.68 |
| 4 | 180664123 | rs13113541 | 0.71 | -0.455 | -1.377 | -0.134 | -0.031 | -0.124 | -0.124 | 0 | 0.37 | 0.13 | 0.31 | 0.47 | 0.38 | 0.47 |
| 4 | 180667647 | rs17090640 | 0.359 | -0.082 | -0.267 | -0.161 | 0.055 | -0.011 | -0.011 | 0 | 0.07 | 0.33 | 0.04 | 0 | 0.05 | 0.05 |
| 4 | 180668187 | rs11724099 | 0.746 | 0.542 | -1.375 | 0.512 | -0.011 | -0.009 | -0.009 | 0 | 0.11 | 0.11 | 0.11 | 0.11 | 0.15 | 0.1 |
| 4 | 180669418 | rs10520433 | 0.263 | 0.198 | -1.38 | 0.327 | 0.014 | 0.01 | 0.01 | 0 | 0.15 | 0.09 | 0.21 | 0.1 | 0.1 | 0.14 |
| 4 | 180671987 | rs17746764 | 1.174 | 0.552 | -1.537 | 0.092 | -0.021 | 0.04 | 0.04 | 0 | 0.09 | 0.04 | 0.06 | 0 | 0.11 | 0.01 |
| 4 | 180673271 | rs2100686 | -1.055 | -0.541 | -0.572 | -0.012 | -0.001 | -0.001 | -0.001 | 0 | 0.34 | 0.66 | 0.38 | 0.29 | 0.27 | 0.19 |
| 4 | 180673823 | rs11736914 | -1.102 | -1.214 | -1.098 | 0.256 | 0.084 | 0.034 | 0.034 | 0.149 | 0.51 | 0.76 | 0.53 | 0.56 | 0.35 | 0.39 |
| 4 | 180678164 | rs1903032 | -1.778 | -0.218 | -1.638 | -0.146 | -0.027 | 0.044 | 0.044 | 0.142 | 0.78 | 0.86 | 0.73 | 0.89 | 0.68 | 0.68 |

| | | | | | | | | | | | | | | | | |
|---|---|---|---|---|---|---|---|---|---|---|---|---|---|---|---|---|
| 4 | 180683092 | rs17068136 | 1.515 | -0.38 | -1.366 | 0.172 | 0.008 | 0.021 | 0.021 | 0 | 0.17 | 0.08 | 0.14 | 0.08 | 0.15 | 0.11 |
| 4 | 180683541 | rs17068140 | 2.619 | -0.529 | -1.407 | 0.154 | 0.002 | -0.087 | -0.087 | 0 | 0.13 | 0 | 0.14 | 0.26 | 0.08 | 0.2 |
| 4 | 180684491 | rs17803891 | 1.263 | -0.109 | -1.356 | 0.142 | -0.019 | 0.028 | 0.028 | 0.044 | 0.18 | 0.08 | 0.16 | 0.08 | 0.22 | 0.12 |
| 4 | 180687498 | rs11131944 | 0.862 | -0.553 | -1.178 | 0.293 | 0.003 | -0.004 | -0.004 | 0 | 0.21 | 0.22 | 0.29 | 0.3 | 0.16 | 0.16 |
| 4 | 180688000 | rs6844851 | -0.57 | -0.273 | -1.441 | 0.373 | 0.078 | 0.035 | 0.035 | 0.048 | 0.5 | 0.75 | 0.52 | 0.56 | 0.36 | 0.39 |
| 4 | 180693794 | rs17747662 | 1.242 | 0.256 | -1.353 | 0.554 | -0.033 | -0.051 | -0.051 | 0 | 0.23 | 0.09 | 0.32 | 0.3 | 0.28 | 0.32 |
| 4 | 180696728 | rs7678609 | -0.117 | -0.603 | -0.72 | 0.624 | 0.001 | -0.004 | -0.004 | 0 | 0.39 | 0.56 | 0.41 | 0.52 | 0.34 | 0.29 |
| 4 | 180700264 | rs1017710 | -1.073 | -0.534 | -0.787 | 0.779 | 0 | 0.008 | 0.008 | 0 | 0.94 | 0.89 | 0.97 | 0.85 | 0.92 | 0.73 |
| 4 | 180702862 | rs12510224 | 1.542 | -0.644 | -0.935 | 0.625 | -0.011 | 0.016 | 0.016 | 0 | 0.33 | 0.21 | 0.44 | 0.25 | 0.37 | 0.3 |
| 4 | 180704767 | rs7674308 | 0.133 | -0.824 | -0.359 | 0.656 | 0.006 | -0.013 | -0.013 | 0 | 0.38 | 0.42 | 0.4 | 0.34 | 0.36 | 0.26 |
| 4 | 180704918 | rs11495908 | 0.376 | -0.676 | -0.926 | 0.159 | -0.003 | 0.049 | 0.049 | 0 | 0.2 | 0.08 | 0.11 | 0.07 | 0.2 | 0.14 |
| 4 | 180711597 | rs17068192 | 1.313 | -0.766 | -0.919 | 0.213 | 0.013 | 0.011 | 0.011 | 0 | 0.13 | 0.08 | 0.05 | 0.07 | 0.12 | 0.11 |
| 4 | 180714806 | rs10035040 | 0.978 | -0.742 | -0.427 | 0.169 | 0.004 | -0.081 | -0.081 | 0 | 0.27 | 0.28 | 0.19 | 0.05 | 0.26 | 0.14 |
| 4 | 180715010 | rs1903027 | 0.022 | -0.517 | -1.163 | 0.455 | -0.004 | 0.035 | 0.035 | 0 | 0.54 | 0.31 | 0.55 | 0.32 | 0.57 | 0.44 |
| 4 | 180715058 | rs17068194 | 0.911 | -0.048 | -1.144 | 0.482 | -0.019 | -0.021 | -0.021 | 0 | 0.05 | 0.02 | 0.03 | 0.16 | 0.08 | 0.27 |
| 4 | 180716739 | rs13115756 | 0.712 | -0.179 | -1.024 | 0.039 | -0.022 | 0.063 | 0.063 | 0 | 0.33 | 0.16 | 0.35 | 0.16 | 0.37 | 0.28 |
| 4 | 180719585 | rs11729049 | 1.12 | 0.144 | -0.732 | 0.17 | -0.035 | 0.064 | 0.064 | 0 | 0.3 | 0.11 | 0.33 | 0.15 | 0.36 | 0.27 |
| 4 | 180745286 | rs13101669 | 0.877 | -0.122 | -0.638 | 0.145 | -0.029 | 0.064 | 0.064 | 0 | 0.29 | 0.11 | 0.32 | 0.14 | 0.32 | 0.27 |
| 4 | 180745339 | rs17068207 | 0.982 | -0.192 | -0.362 | -0.305 | -0.002 | -0.034 | -0.034 | 9.796 | 0.21 | 0.21 | 0.15 | 0.02 | 0.24 | 0.15 |
| 4 | 180747463 | rs1458669 | 0.421 | -0.336 | 0.038 | 0.245 | -0.011 | 0.046 | 0.046 | 0 | 0.78 | 0.62 | 0.66 | 0.38 | 0.8 | 0.56 |
| 4 | 180757043 | rs1379983 | 0.649 | -0.309 | 0.041 | 0.073 | 0.004 | 0.025 | 0.025 | 0 | 0.28 | 0.17 | 0.17 | 0.06 | 0.26 | 0.18 |
| 4 | 180758188 | rs7685478 | 0.077 | -0.303 | 0.313 | 0.75 | 0.004 | 0.118 | 0.118 | 0.019 | 0.07 | 0.18 | 0.12 | 0.26 | 0.08 | 0.24 |
| 4 | 180764869 | rs6552396 | 0.559 | -0.417 | 0.476 | 0.445 | 0.005 | 0.008 | 0.008 | 0 | 0.53 | 0.44 | 0.64 | 0.32 | 0.57 | 0.45 |
| 4 | 180764928 | rs10520438 | 0.003 | -0.363 | 0.362 | 0.185 | 0.01 | 0.032 | 0.032 | 0 | 0.36 | 0.22 | 0.21 | 0.06 | 0.33 | 0.21 |
| 4 | 180770819 | rs17068232 | -0.039 | -0.452 | 0.361 | 0.154 | 0.001 | -0.077 | -0.077 | 0.091 | 0.33 | 0.48 | 0.25 | 0.16 | 0.32 | 0.23 |
| 4 | 180772704 | rs986247 | -0.929 | -0.376 | 0.089 | 0.124 | 0.008 | 0.02 | 0.02 | 0 | 0.73 | 0.81 | 0.84 | 0.93 | 0.76 | 0.86 |
| 4 | 180774423 | rs6832008 | 0.822 | -0.291 | 0.431 | 0.402 | 0.028 | -0.029 | -0.029 | 0 | 0.23 | 0.42 | 0.21 | 0.18 | 0.27 | 0.34 |
| 4 | 180790098 | rs12651279 | 1.653 | -0.543 | 0.22 | 0.693 | -0.045 | 0.043 | 0.043 | 0 | 0.27 | 0.11 | 0.43 | 0.16 | 0.33 | 0.3 |
| 4 | 180794840 | rs17068292 | -0.058 | -0.566 | 0.359 | 0.831 | 0.005 | 0.033 | 0.033 | 0 | 0.11 | 0.09 | 0.11 | 0.48 | 0.06 | 0.18 |
| 4 | 180798886 | rs7654672 | -1.542 | -0.558 | 0.386 | 0.505 | 0.003 | -0.007 | -0.007 | 0 | 0.91 | 0.9 | 0.95 | 1 | 0.91 | 0.97 |
| 4 | 180798964 | rs7655189 | -0.239 | -0.449 | 0.362 | 0.569 | -0.012 | -0.039 | -0.039 | 0.93 | 0.66 | 0.47 | 0.66 | 0.77 | 0.68 | 0.62 |
| 4 | 180799005 | rs10520440 | 1.35 | -0.43 | -0.127 | 0.745 | -0.079 | 0.054 | 0.054 | 0 | 0.29 | 0.06 | 0.43 | 0.16 | 0.33 | 0.3 |
| 4 | 180801466 | rs6826314 | 0.088 | -0.453 | 0.351 | 0.647 | 0 | -0.004 | -0.004 | 0 | 0.48 | 0.45 | 0.45 | 0.29 | 0.52 | 0.47 |
| 4 | 180802030 | rs17068316 | -0.276 | -0.691 | 0.201 | 1.349 | -0.005 | 0.019 | 0.019 | 0 | 0.1 | 0.11 | 0.02 | 0.23 | 0.05 | 0.17 |
| 4 | 180803490 | rs1072776 | 1.345 | -0.505 | -0.124 | 1.075 | -0.075 | 0.05 | 0.05 | 0 | 0.31 | 0.1 | 0.31 | 0.19 | 0.36 | 0.36 |
| 4 | 180818705 | rs1458664 | 0.875 | -1.112 | 0.578 | 0.709 | -0.042 | 0.079 | 0.079 | 0 | 0.76 | 0.42 | 0.86 | 0.59 | 0.83 | 0.66 |
| 4 | 180819627 | rs6552399 | 0.431 | -1.095 | 0.537 | 0.519 | -0.001 | 0.03 | 0.03 | 0 | 0.39 | 0.28 | 0.32 | 0.11 | 0.4 | 0.22 |
| 4 | 180819809 | rs17068350 | 0.575 | -1.114 | 0.676 | 0.587 | -0.018 | 0 | 0 | 0 | 0.09 | 0.17 | 0.12 | 0.1 | 0.06 | 0.19 |
| 4 | 180822313 | rs1947622 | 0.368 | -1.03 | 0.477 | 0.552 | -0.001 | 0.045 | 0.045 | 0 | 0.4 | 0.27 | 0.32 | 0.11 | 0.41 | 0.22 |
| 4 | 180823827 | rs11934076 | 1.118 | -0.977 | 0.254 | 0.461 | -0.009 | 0.041 | 0.041 | 0 | 0.24 | 0.12 | 0.19 | 0.07 | 0.26 | 0.08 |
| 4 | 180824638 | rs1870347 | 1.086 | -0.398 | 0.477 | 0.834 | -0.039 | -0.033 | -0.033 | 0 | 0.33 | 0.23 | 0.45 | 0.4 | 0.39 | 0.39 |
| 4 | 180834630 | rs1545671 | -0.004 | -0.745 | 0.068 | 0.724 | 0.009 | 0.011 | 0.011 | 0 | 0.54 | 0.63 | 0.62 | 0.67 | 0.6 | 0.65 |
| 4 | 180844991 | rs12512992 | -0.628 | -0.631 | -0.207 | 0.767 | 0.001 | -0.003 | -0.003 | 1.247 | 0.89 | 0.91 | 0.94 | 0.83 | 0.89 | 0.83 |
| 4 | 180846512 | rs13112173 | 1.284 | -0.656 | -0.492 | 0.724 | -0.177 | -0.135 | -0.135 | 0 | 0.3 | 0.03 | 0.42 | 0.36 | 0.4 | 0.37 |
| 4 | 180856616 | rs10780062 | 0.798 | -0.737 | -0.267 | 0.673 | -0.079 | -0.036 | -0.036 | 0 | 0.36 | 0.18 | 0.44 | 0.36 | 0.42 | 0.4 |
| 4 | 180857874 | rs9991082 | 0.413 | -0.888 | -0.05 | 0.604 | -0.003 | -0.015 | -0.015 | 0 | 0.35 | 0.34 | 0.33 | 0.16 | 0.3 | 0.19 |
| 4 | 180859149 | rs10000486 | 0.71 | -0.841 | 0.122 | 0.687 | 0.003 | 0.006 | 0.006 | 0 | 0.32 | 0.29 | 0.33 | 0.19 | 0.3 | 0.26 |
| 4 | 180862232 | rs17068418 | 0.618 | -0.687 | 0.075 | 0.803 | 0.01 | 0.014 | 0.014 | 0 | 0.35 | 0.26 | 0.32 | 0.16 | 0.29 | 0.17 |
| 4 | 180867238 | rs2125431 | 0.409 | -0.685 | 0.028 | 0.812 | -0.043 | -0.008 | -0.008 | 0 | 0.42 | 0.18 | 0.49 | 0.36 | 0.51 | 0.46 |
| 4 | 180875233 | rs6836502 | 1.525 | -0.553 | 0.143 | 0.803 | -0.008 | 0.035 | 0.035 | 0 | 0.07 | 0 | 0.04 | 0 | 0.09 | 0.01 |
| 4 | 180877617 | rs7675742 | -0.307 | -0.508 | 0.293 | 0.943 | -0.01 | 0.117 | 0.117 | 0 | 0.77 | 0.45 | 0.81 | 0.52 | 0.8 | 0.64 |
| 4 | 180880956 | rs4861822 | 0.917 | -0.443 | -0.095 | 0.847 | -0.006 | 0.02 | 0.02 | 0 | 0.17 | 0.2 | 0.15 | 0.28 | 0.15 | 0.12 |
| 4 | 180886822 | rs7675700 | 0.025 | -0.457 | -0.132 | 0.69 | 0.005 | 0.005 | 0.005 | 0 | 0.52 | 0.46 | 0.49 | 0.61 | 0.58 | 0.57 |
| 4 | 180893766 | rs6810891 | -0.292 | -0.441 | -0.503 | 0.576 | 0 | 0 | 0 | 0 | 0.74 | 0.75 | 0.71 | 0.76 | 0.75 | 0.84 |
| 4 | 180894794 | rs12510169 | -1.343 | -0.432 | -0.505 | 0.575 | -0.001 | 0.003 | 0.003 | 0 | 0.82 | 0.8 | 0.78 | 0.76 | 0.85 | 0.87 |
| 4 | 180896929 | rs7674405 | 0.41 | -0.463 | -0.679 | 0.474 | -0.008 | -0.025 | -0.025 | 0 | 0.61 | 0.54 | 0.63 | 0.76 | 0.65 | 0.81 |
| 4 | 180899782 | rs1994813 | 1.473 | -0.136 | -1.074 | -0.027 | -0.123 | -0.1 | -0.1 | 2.478 | 0.24 | 0.06 | 0.29 | 0.33 | 0.35 | 0.33 |
| 4 | 180901257 | rs17068502 | 1.334 | -0.086 | -0.987 | -0.037 | -0.002 | -0.004 | -0.004 | 0.405 | 0.08 | 0.06 | 0.08 | 0 | 0.1 | 0.03 |
| 4 | 180902914 | rs6836406 | 1.104 | -0.743 | -0.906 | 0.077 | -0.1 | -0.04 | -0.04 | 0 | 0.31 | 0.14 | 0.33 | 0.33 | 0.45 | 0.35 |
| 4 | 180903852 | rs4861824 | 0.593 | -0.505 | -0.009 | 0.456 | -0.014 | 0.017 | 0.017 | 0.008 | 0.67 | 0.53 | 0.7 | 0.61 | 0.71 | 0.69 |
| 4 | 180909740 | rs13111060 | -0.133 | -0.309 | -0.236 | 0.648 | -0.001 | 0.01 | 0.01 | 0 | 0.79 | 0.76 | 0.78 | 0.62 | 0.83 | 0.75 |
| 4 | 180911019 | rs17068534 | -0.528 | -0.394 | -0.007 | 0.28 | -0.004 | -0.01 | -0.01 | 0 | 0.11 | 0.17 | 0.17 | 0.07 | 0.12 | 0.18 |
| 4 | 180911596 | rs6552405 | 0.084 | -0.587 | 0.826 | 0.188 | 0.003 | -0.06 | -0.06 | 0 | 0.9 | 0.77 | 0.93 | 1 | 0.91 | 0.99 |
| 4 | 180915780 | rs9991368 | -0.515 | -0.67 | 0.078 | -0.207 | -0.005 | -0.011 | -0.011 | 0 | 0.62 | 0.67 | 0.61 | 0.48 | 0.59 | 0.53 |
| 4 | 180916219 | rs9991928 | -0.512 | -0.626 | 0.114 | -0.21 | -0.005 | -0.008 | -0.008 | 0 | 0.62 | 0.65 | 0.61 | 0.48 | 0.59 | 0.53 |
| 4 | 180917828 | rs17821516 | 1.227 | -0.697 | 0.045 | -0.211 | 0.003 | 0.018 | 0.018 | 0 | 0.08 | 0.04 | 0.05 | 0 | 0.09 | 0.01 |
| 4 | 180920056 | rs6858037 | -0.515 | -0.738 | 0.182 | -0.165 | -0.004 | -0.01 | -0.01 | 0.565 | 0.61 | 0.66 | 0.6 | 0.49 | 0.6 | 0.53 |
| 4 | 180920998 | rs4861826 | 1.317 | -0.852 | -0.573 | -0.591 | -0.035 | 0.124 | 0.124 | 0 | 0.25 | 0.06 | 0.16 | 0.04 | 0.28 | 0.09 |
| 4 | 180922702 | rs10019132 | 0.088 | -1.473 | -0.361 | -0.226 | -0.078 | 0.181 | 0.181 | 1.372 | 0.83 | 0.48 | 0.88 | 0.57 | 0.94 | 0.76 |

| | | | | | | | | | | | | | | | | |
|---|---|---|---|---|---|---|---|---|---|---|---|---|---|---|---|---|
| 4 | 180926735 | rs7698405 | 1.071 | -1.42 | -0.519 | -0.3 | 0.059 | 0.017 | 0.017 | 1.493 | 0.52 | 0.24 | 0.49 | 0.4 | 0.64 | 0.46 |
| 4 | 180929339 | rs6835516 | 0.779 | -1.476 | -0.565 | -0.321 | -0.042 | -0.019 | -0.019 | 0 | 0.36 | 0.21 | 0.38 | 0.36 | 0.46 | 0.37 |
| 4 | 180929612 | rs6552407 | 0.805 | -1.342 | -0.378 | -0.154 | 0.042 | -0.017 | -0.017 | 0 | 0.6 | 0.24 | 0.53 | 0.4 | 0.72 | 0.46 |
| 4 | 180931381 | rs4241716 | 0.086 | -1.218 | -0.286 | -0.037 | -0.001 | 0.015 | 0.015 | 0.094 | 0.85 | 0.81 | 0.78 | 0.75 | 0.87 | 0.68 |
| 4 | 180933174 | rs2309139 | 0.764 | -1.165 | -0.409 | -0.066 | -0.015 | -0.012 | -0.012 | 0 | 0.16 | 0.24 | 0.16 | 0.11 | 0.08 | 0.12 |
| 4 | 180933347 | rs6837073 | 1.473 | -1.285 | -0.569 | -0.241 | -0.02 | 0.152 | 0.152 | 0.093 | 0.3 | 0.06 | 0.18 | 0.03 | 0.32 | 0.17 |
| 4 | 180934921 | rs4521383 | -0.765 | -1.427 | -0.295 | -0.15 | -0.029 | 0.02 | 0.02 | 0.2 | 0.33 | 0.66 | 0.4 | 0.47 | 0.16 | 0.33 |
| 4 | 180935476 | rs13114766 | 0.335 | -1.408 | -0.473 | -0.127 | -0.009 | -0.004 | -0.004 | 0 | 0.2 | 0.25 | 0.31 | 0.18 | 0.12 | 0.22 |
| 4 | 180942234 | rs2871129 | 1.342 | -1.366 | -0.978 | -0.121 | -0.162 | -0.109 | -0.109 | 0 | 0.28 | 0.06 | 0.3 | 0.37 | 0.42 | 0.37 |
| 4 | 180942572 | rs1316917 | 0.759 | -1.699 | -0.738 | 0.002 | -0.045 | 0.038 | 0.038 | 0 | 0.11 | 0.3 | 0.08 | 0.16 | 0.04 | 0.1 |
| 4 | 180942678 | rs4254817 | -2.134 | -1.688 | -0.736 | -0.017 | -0.006 | 0 | 0 | 1.057 | 0.95 | 0.93 | 0.83 | 0.93 | 0.97 | 0.9 |
| 4 | 180951070 | rs10013059 | -0.519 | -1.622 | -0.741 | -0.011 | -0.04 | 0.019 | 0.019 | 0.131 | 0.29 | 0.65 | 0.26 | 0.38 | 0.12 | 0.27 |
| 4 | 180953886 | rs2309129 | 0.542 | -1.654 | -0.713 | 0 | -0.045 | 0.11 | 0.11 | 0 | 0.12 | 0.32 | 0.09 | 0.28 | 0.04 | 0.11 |
| 4 | 180954083 | rs13327938 | -0.126 | -1.385 | -1.068 | -0.006 | -0.035 | 0.019 | 0.019 | 0.131 | 0.29 | 0.65 | 0.26 | 0.39 | 0.13 | 0.27 |
| 4 | 180955602 | rs17771724 | 1.313 | -1.315 | -1.201 | -0.189 | -0.106 | 0.175 | 0.175 | 0 | 0.29 | 0.04 | 0.21 | 0.03 | 0.39 | 0.16 |
| 4 | 180958428 | rs2309124 | -0.206 | -1.12 | -0.876 | 0.073 | -0.043 | -0.001 | -0.001 | 0.076 | 0.28 | 0.63 | 0.26 | 0.27 | 0.12 | 0.26 |
| 4 | 180959787 | rs1112857 | 0.636 | -0.165 | -1.311 | 0.435 | -0.123 | -0.12 | -0.12 | 0 | 0.24 | 0.04 | 0.26 | 0.37 | 0.35 | 0.34 |
| 4 | 180962074 | rs6829907 | -0.293 | -0.739 | -0.416 | 0.281 | 0.004 | 0 | 0 | 0.053 | 0.11 | 0.1 | 0.11 | 0.13 | 0.09 | 0.14 |
| 4 | 180968431 | rs1994816 | -0.91 | -0.782 | -0.07 | 0.362 | -0.062 | 0.014 | 0.014 | 0.103 | 0.26 | 0.51 | 0.31 | 0.32 | 0.17 | 0.37 |
| 4 | 180982097 | rs1665841 | 0.84 | -0.433 | 0.52 | 0.234 | 0.018 | 0 | 0 | 0 | 0.93 | 0.68 | 0.83 | 0.99 | 0.97 | 0.94 |
| 4 | 180988794 | rs7690743 | -0.272 | -0.107 | -0.138 | 0.057 | 0.002 | 0.004 | 0.004 | 0 | 0.43 | 0.42 | 0.45 | 0.44 | 0.45 | 0.45 |
| 4 | 180998616 | rs1665831 | 0.47 | 0.371 | -0.164 | 0.246 | 0.003 | 0.041 | 0.041 | 0 | 0.71 | 0.52 | 0.61 | 0.64 | 0.71 | 0.65 |
| 4 | 181008167 | rs1689004 | -0.047 | 0.533 | -0.317 | 0.561 | 0 | 0.022 | 0.022 | 0 | 0.8 | 0.75 | 0.79 | 0.65 | 0.81 | 0.75 |
| 4 | 181011016 | rs6830355 | 0.66 | 0.248 | -0.575 | 0.395 | -0.001 | 0 | 0 | 0 | 0.66 | 0.63 | 0.71 | 0.5 | 0.64 | 0.62 |
| 4 | 181022656 | rs12499924 | -0.651 | 0.081 | -0.707 | 0.229 | -0.018 | 0.018 | 0.018 | 0 | 0.35 | 0.31 | 0.37 | 0.78 | 0.46 | 0.45 |
| 4 | 181027678 | rs7656301 | 0.769 | -0.344 | -0.51 | 0.645 | -0.005 | 0.028 | 0.028 | 0 | 0.52 | 0.35 | 0.43 | 0.31 | 0.5 | 0.36 |
| 4 | 181028368 | rs7679909 | 0.34 | -0.363 | -0.354 | 0.951 | 0.003 | 0.006 | 0.006 | 0.164 | 0.61 | 0.47 | 0.54 | 0.49 | 0.59 | 0.5 |
| 4 | 181031805 | rs720778 | -1.004 | 0.29 | -0.401 | 0.936 | 0.007 | 0.072 | 0.072 | 0.286 | 0.07 | 0.16 | 0.11 | 0.21 | 0.07 | 0.09 |
| 4 | 181032789 | rs13148326 | 0.541 | 0.344 | -1.231 | 0.377 | -0.076 | 0.131 | 0.131 | 0 | 0.26 | 0.03 | 0.21 | 0.06 | 0.3 | 0.19 |
| 4 | 181038957 | rs1689020 | 0.956 | 0.168 | -0.038 | 1.104 | 0.013 | 0.248 | 0.248 | 1.112 | 0.9 | 0.54 | 0.89 | 0.76 | 0.96 | 0.88 |
| 4 | 181039757 | rs1665844 | 0.984 | 0.185 | -0.617 | 1.078 | -0.004 | 0.014 | 0.014 | 0 | 0.61 | 0.44 | 0.54 | 0.46 | 0.62 | 0.51 |
| 4 | 181045298 | rs13146117 | 1.216 | 0.117 | -1.211 | 0.48 | -0.127 | 0.116 | 0.116 | 0.033 | 0.29 | 0.02 | 0.24 | 0.09 | 0.34 | 0.19 |
| 4 | 181046018 | rs9990459 | -0.659 | -0.275 | -0.919 | 0.458 | 0.05 | 0.097 | 0.097 | 0.394 | 0.65 | 0.24 | 0.77 | 0.88 | 0.75 | 0.78 |
| 4 | 181046865 | rs1689014 | 0.676 | -0.319 | -0.69 | 0.466 | -0.02 | -0.049 | -0.049 | 0 | 0.33 | 0.56 | 0.32 | 0.11 | 0.25 | 0.23 |
| 4 | 181048589 | rs13135628 | -0.179 | -0.314 | -0.819 | 0.468 | 0.08 | 0.166 | 0.166 | 0.529 | 0.6 | 0.14 | 0.64 | 0.86 | 0.72 | 0.76 |
| 4 | 181048849 | rs1665851 | -0.001 | -0.244 | -0.586 | 0.545 | 0.028 | 0.025 | 0.025 | 0.357 | 0.95 | 0.74 | 0.97 | 0.97 | 0.97 | 0.98 |
| 4 | 181051943 | rs10520442 | 0.977 | -0.169 | -0.909 | 0.705 | -0.064 | -0.134 | -0.134 | 0.012 | 0.35 | 0.07 | 0.39 | 0.48 | 0.39 | 0.3 |
| 4 | 181056626 | rs1689021 | 1.665 | -0.789 | -0.239 | 1.085 | -0.016 | 0.123 | 0.123 | 0.135 | 0.86 | 0.5 | 0.76 | 0.78 | 0.91 | 0.82 |
| 4 | 181066033 | rs1447317 | 0.888 | -0.781 | -0.187 | 0.976 | -0.031 | -0.03 | -0.03 | 0 | 0.89 | 0.64 | 0.83 | 0.95 | 0.94 | 0.96 |
| 4 | 181075687 | rs1447320 | 0.535 | -0.774 | -0.247 | 0.98 | -0.029 | -0.028 | -0.028 | 0 | 0.9 | 0.69 | 0.83 | 0.95 | 0.94 | 0.96 |
| 4 | 181076187 | rs13129209 | -0.018 | -0.863 | -0.71 | 0.927 | 0.026 | 0.02 | 0.02 | 0.148 | 0.6 | 0.1 | 0.57 | 0.7 | 0.68 | 0.61 |
| 4 | 181084084 | rs13121605 | 0.649 | -0.744 | -0.565 | 0.972 | -0.024 | -0.023 | -0.023 | 0 | 0.89 | 0.64 | 0.84 | 0.95 | 0.94 | 0.96 |
| 4 | 181098462 | rs7677721 | 0.279 | -0.641 | -1.296 | 1.151 | 0.07 | -0.015 | -0.015 | 1.076 | 0.52 | 0.09 | 0.52 | 0.56 | 0.63 | 0.56 |
| 4 | 181101509 | rs17068825 | 0.388 | -0.389 | -1.725 | 1.241 | -0.015 | -0.058 | -0.058 | 0 | 0.14 | 0.02 | 0.11 | 0.21 | 0.12 | 0.22 |
| 4 | 181108510 | rs6836062 | -0.491 | -0.567 | -1.525 | 0.765 | 0.022 | 0.02 | 0.02 | 1.288 | 0.68 | 0.21 | 0.69 | 0.75 | 0.73 | 0.71 |
| 4 | 181118760 | rs10027059 | 0.144 | -0.407 | -1.576 | 0.799 | 0.015 | -0.008 | -0.008 | 2.142 | 0.11 | 0.1 | 0.17 | 0.16 | 0.08 | 0.07 |
| 4 | 181119137 | rs17068866 | 0.16 | -0.068 | -1.616 | 0.638 | 0.084 | -0.016 | -0.016 | 0 | 0.53 | 0.13 | 0.52 | 0.54 | 0.63 | 0.62 |
| 4 | 181124250 | rs1868056 | -0.549 | 0.747 | -0.722 | 0.417 | 0.003 | -0.017 | -0.017 | 0.852 | 0.37 | 0.43 | 0.55 | 0.74 | 0.4 | 0.57 |
| 4 | 181124269 | rs6855724 | 0.509 | 0.619 | -0.739 | 0.333 | -0.009 | 0.028 | 0.028 | 0.936 | 0.56 | 0.4 | 0.39 | 0.25 | 0.5 | 0.37 |
| 4 | 181125964 | rs11734697 | 0.43 | -0.126 | -0.137 | 0.913 | -0.027 | 0.071 | 0.071 | 0.382 | 0.77 | 0.59 | 0.8 | 0.71 | 0.84 | 0.81 |
| 4 | 181126498 | rs10007350 | -0.005 | -1.376 | -0.531 | 0.504 | 0.012 | 0.013 | 0.013 | 0 | 0.24 | 0.21 | 0.11 | 0.13 | 0.11 | 0.17 |
| 4 | 181126541 | rs9312259 | -0.509 | -1.178 | -0.602 | 0.58 | 0.007 | -0.042 | -0.042 | 0.196 | 0.83 | 0.9 | 0.86 | 0.71 | 0.86 | 0.83 |
| 4 | 181136629 | rs1447325 | -0.491 | -1.197 | -0.563 | 0.57 | 0.006 | -0.034 | -0.034 | 0 | 0.82 | 0.88 | 0.86 | 0.7 | 0.87 | 0.8 |
| 4 | 181137134 | rs7655191 | -0.326 | -1.444 | -0.32 | 0.445 | -0.096 | -0.079 | -0.079 | 0 | 0.3 | 0.48 | 0.17 | 0.13 | 0.15 | 0.19 |
| 4 | 181137418 | rs17068948 | 0.647 | -1.262 | -0.426 | 0.543 | 0.027 | 0.036 | 0.036 | 0.611 | 0.08 | 0.18 | 0.11 | 0.1 | 0.1 | 0.09 |
| 4 | 181138753 | rs2053745 | -0.72 | -1.385 | -0.499 | 0.566 | 0.007 | -0.042 | -0.042 | 0 | 0.82 | 0.9 | 0.86 | 0.7 | 0.87 | 0.8 |
| 4 | 181139421 | rs17068958 | 0.564 | -1.096 | -0.517 | 0.453 | 0.063 | 0.002 | 0.002 | 0 | 0.52 | 0.37 | 0.68 | 0.57 | 0.72 | 0.62 |
| 4 | 181148678 | rs7663206 | 0.182 | -0.634 | -0.054 | 0.793 | -0.002 | 0.073 | 0.073 | 0.399 | 0.79 | 0.6 | 0.81 | 0.72 | 0.84 | 0.83 |
| 4 | 181152554 | rs1447323 | -0.163 | -0.509 | -0.559 | 0.434 | 0.011 | -0.001 | -0.001 | 0 | 0.39 | 0.28 | 0.51 | 0.64 | 0.36 | 0.5 |
| 4 | 181152961 | rs17068995 | 0.361 | -0.749 | -1.299 | 0.192 | 0.018 | -0.036 | -0.036 | 0.008 | 0.11 | 0.02 | 0.06 | 0.18 | 0.09 | 0.1 |
| 4 | 181153885 | rs10020349 | -0.098 | -0.592 | -1.168 | -0.047 | 0 | -0.009 | -0.009 | 0 | 0.76 | 0.84 | 0.79 | 0.72 | 0.76 | 0.8 |
| 4 | 181154351 | rs7697719 | 0.672 | -0.263 | -0.992 | -0.071 | 0.01 | 0.027 | 0.027 | 0 | 0.07 | 0.13 | 0.04 | 0.08 | 0.07 | 0.15 |
| 4 | 181157746 | rs4861844 | 1.658 | -0.216 | -1.285 | 0.131 | -0.051 | -0.099 | -0.099 | 0 | 0.14 | 0.02 | 0.23 | 0.39 | 0.18 | 0.22 |
| 4 | 181161490 | rs17069022 | -0.278 | 0.02 | -0.333 | -0.105 | 0.013 | 0.038 | 0.038 | 0.063 | 0.09 | 0.25 | 0.05 | 0.08 | 0.07 | 0.15 |
| 4 | 181169328 | rs17069038 | 0.329 | 0.355 | -0.888 | -0.235 | 0.017 | -0.001 | -0.001 | 0 | 0.1 | 0.14 | 0.06 | 0.06 | 0.13 | 0.16 |
| 4 | 181181237 | rs34136869 | 0.665 | 0.407 | -1.821 | -0.603 | -0.118 | 0.139 | 0.139 | 0 | 0.21 | 0.01 | 0.19 | 0 | 0.31 | 0.14 |
| 4 | 181186334 | rs11131981 | -0.608 | 0.911 | -1.379 | -0.07 | -0.049 | 0.012 | 0.012 | 0 | 0.35 | 0.17 | 0.34 | 0.37 | 0.41 | 0.27 |
| 4 | 181193110 | rs7667016 | 1.128 | -0.271 | -0.632 | 0.289 | -0.006 | 0 | 0 | 0.348 | 0.61 | 0.42 | 0.65 | 0.57 | 0.66 | 0.56 |
| 4 | 181196155 | rs17069051 | -0.727 | -0.06 | -1.349 | 0.834 | 0.007 | -0.012 | -0.012 | 0.063 | 0.25 | 0.19 | 0.26 | 0.41 | 0.21 | 0.28 |

| | | | | | | | | | | | | | | | | |
|---|---|---|---|---|---|---|---|---|---|---|---|---|---|---|---|---|
| 4 | 181205991 | rs10018538 | -0.437 | -0.288 | -1.636 | 0.703 | 0.03 | -0.005 | -0.005 | 0 | 0.21 | 0.13 | 0.27 | 0.26 | 0.16 | 0.22 |
| 4 | 181207294 | rs17777325 | 0.132 | 0.59 | -2.322 | 0.42 | -0.093 | -0.003 | -0.003 | 0.183 | 0.27 | 0.05 | 0.24 | 0.23 | 0.33 | 0.31 |
| 4 | 181210711 | rs17069068 | 0.461 | 0.343 | -1.829 | 0.905 | -0.002 | -0.005 | -0.005 | 0 | 0.07 | 0.03 | 0.04 | 0.23 | 0.06 | 0.13 |
| 4 | 181211523 | rs7696117 | 0.101 | 0.132 | -1.436 | 0.226 | -0.007 | -0.012 | -0.012 | 6.246 | 0.1 | 0.12 | 0.09 | 0.05 | 0.09 | 0.18 |
| 4 | 181214528 | rs17069075 | 0.065 | -0.023 | -1.336 | 0.225 | -0.012 | -0.007 | -0.007 | 0 | 0.13 | 0.14 | 0.16 | 0.09 | 0.08 | 0.12 |
| 4 | 181215239 | rs7671393 | 0.215 | 0.659 | -1.294 | 0.674 | 0.012 | 0.019 | 0.019 | 0.323 | 0.13 | 0.17 | 0.26 | 0.28 | 0.18 | 0.16 |
| 4 | 181215806 | rs6834475 | 0.422 | -0.404 | -1.115 | -0.005 | -0.012 | -0.052 | -0.052 | 0 | 0.59 | 0.54 | 0.44 | 0.19 | 0.62 | 0.4 |
| 4 | 181221012 | rs12501188 | 1.378 | 0.192 | -1.798 | -0.77 | -0.085 | 0.103 | 0.103 | 0 | 0.17 | 0.01 | 0.23 | 0 | 0.21 | 0.13 |
| 4 | 181222881 | rs13109268 | -0.438 | -0.859 | -0.857 | -0.909 | 0 | -0.001 | -0.001 | 0 | 0.65 | 0.37 | 0.61 | 0.36 | 0.68 | 0.48 |
| 4 | 181224923 | rs17069165 | 1.586 | -0.918 | -1.018 | -0.644 | -0.019 | -0.082 | -0.082 | 0.458 | 0.12 | 0.02 | 0.07 | 0.47 | 0.11 | 0.3 |
| 4 | 181225524 | rs13130442 | 0.859 | -0.828 | -1.562 | -1.968 | -0.125 | 0.131 | 0.131 | 0 | 0.27 | 0.05 | 0.41 | 0.07 | 0.37 | 0.18 |
| 4 | 181226698 | rs17069175 | 0.768 | -0.888 | -1.364 | -2.328 | -0.035 | -0.03 | -0.03 | 0 | 0.37 | 0.47 | 0.27 | 0.2 | 0.25 | 0.24 |
| 4 | 181227321 | rs4991519 | -1.502 | -0.752 | -1.245 | -2.244 | 0.015 | 0.019 | 0.019 | 0 | 0.77 | 0.71 | 0.81 | 0.36 | 0.72 | 0.57 |
| 4 | 181231080 | rs10866234 | 0.314 | -0.767 | -1.357 | -2.262 | 0.014 | 0.019 | 0.019 | 0 | 0.47 | 0.45 | 0.45 | 0.28 | 0.4 | 0.29 |
| 4 | 181235156 | rs13106873 | -0.709 | -0.663 | -1.236 | -2.13 | 0.013 | -0.025 | -0.025 | 0 | 0.64 | 0.51 | 0.68 | 0.28 | 0.61 | 0.42 |
| 4 | 181239345 | rs2871184 | 1.303 | -0.705 | -1.038 | -2.069 | 0.002 | -0.03 | -0.03 | 0 | 0.16 | 0.36 | 0.28 | 0.09 | 0.14 | 0.22 |
| 4 | 181239467 | rs13150189 | 1.322 | -1.269 | -1.519 | -2.445 | 0.004 | 0.042 | 0.042 | 0 | 0.2 | 0.11 | 0.13 | 0.11 | 0.2 | 0.08 |
| 4 | 181239651 | rs4290930 | 0.739 | -1.475 | -1.475 | -2.563 | 0.002 | -0.025 | -0.025 | 3.544 | 0.32 | 0.33 | 0.49 | 0.09 | 0.35 | 0.35 |
| 4 | 181243214 | rs6839892 | 1.534 | -1.444 | -1.422 | -2.551 | -0.005 | -0.034 | -0.034 | 0 | 0.15 | 0.31 | 0.27 | 0.08 | 0.13 | 0.21 |
| 4 | 181247001 | rs17069214 | 2.33 | -1.633 | -1.558 | -2.605 | 0.002 | -0.042 | -0.042 | 0 | 0.14 | 0.01 | 0.13 | 0.16 | 0.15 | 0.12 |
| 4 | 181247685 | rs6847807 | 0.609 | -1.336 | -1.411 | -2.493 | 0.002 | -0.026 | -0.026 | 6.276 | 0.32 | 0.33 | 0.49 | 0.09 | 0.35 | 0.34 |
| 4 | 181251312 | rs2309330 | -0.71 | -1.525 | -1.227 | -2.581 | -0.054 | -0.036 | -0.036 | 0.043 | 0.63 | 0.49 | 0.77 | 0.79 | 0.77 | 0.8 |
| 4 | 181253016 | rs6552415 | 0.786 | -1.1 | -1.045 | -2.278 | -0.03 | -0.062 | -0.062 | 0.481 | 0.24 | 0.37 | 0.31 | 0.09 | 0.16 | 0.24 |
| 4 | 181253362 | rs7688867 | -0.352 | -0.491 | -0.895 | -1.847 | 0.006 | -0.003 | -0.003 | 0 | 0.67 | 0.68 | 0.71 | 0.29 | 0.56 | 0.52 |
| 4 | 181259377 | rs4861849 | 0.596 | -0.724 | -0.874 | -2.39 | -0.036 | -0.07 | -0.07 | 0.944 | 0.17 | 0.24 | 0.07 | 0 | 0.08 | 0.07 |
| 4 | 181263217 | rs6830593 | -1.333 | 0.039 | -0.94 | -2.442 | 0.019 | -0.026 | -0.026 | 0 | 0.84 | 0.79 | 0.88 | 0.94 | 0.76 | 0.88 |
| 4 | 181266001 | rs13113937 | -0.053 | -0.141 | -1.141 | -2.357 | 0 | -0.019 | -0.019 | 0.465 | 0.32 | 0.4 | 0.43 | 0.71 | 0.31 | 0.44 |
| 4 | 181271289 | rs17779404 | 0.646 | -0.134 | -1.217 | -2.373 | 0.004 | 0.028 | 0.028 | 0 | 0.22 | 0.14 | 0.22 | 0 | 0.2 | 0.17 |
| 4 | 181271864 | rs2309337 | 0.631 | -0.135 | -1.218 | -2.374 | 0.005 | 0.028 | 0.028 | 0 | 0.23 | 0.14 | 0.22 | 0 | 0.2 | 0.16 |
| 4 | 181273127 | rs4241719 | -0.811 | -0.003 | -1.324 | -2.181 | 0 | -0.01 | -0.01 | 0.829 | 0.56 | 0.55 | 0.65 | 0.73 | 0.52 | 0.63 |
| 4 | 181275147 | rs13141204 | -0.185 | 0.503 | -1.092 | -1.342 | 0.001 | -0.005 | -0.005 | 0 | 0.27 | 0.33 | 0.41 | 0.72 | 0.27 | 0.41 |
| 4 | 181289806 | rs13136634 | 0.507 | 0.178 | -1.516 | -1.164 | -0.018 | -0.052 | -0.052 | 0 | 0.17 | 0.03 | 0.16 | 0.21 | 0.16 | 0.18 |
| 4 | 181290296 | rs7667245 | -0.079 | 0.493 | -1.714 | -1.345 | -0.097 | -0.052 | -0.052 | 0.236 | 0.25 | 0.05 | 0.26 | 0.26 | 0.35 | 0.23 |
| 4 | 181292279 | rs4560455 | 0.054 | 0.146 | -1.421 | -1.551 | 0.021 | -0.004 | -0.004 | 0 | 0.36 | 0.67 | 0.45 | 0.62 | 0.28 | 0.51 |
| 4 | 181294474 | rs28627320 | -0.086 | 0.21 | -1.285 | -1.448 | 0.047 | 0.16 | 0.16 | 1.895 | 0.51 | 0.71 | 0.6 | 0.8 | 0.43 | 0.7 |
| 4 | 181295223 | rs7699693 | -0.69 | 0.465 | -0.969 | -1.099 | -0.019 | 0.042 | 0.042 | 0.135 | 0.75 | 0.84 | 0.82 | 0.91 | 0.69 | 0.89 |
| 4 | 181295874 | rs11934651 | 0.254 | 0.344 | -0.55 | -1.203 | -0.019 | 0.053 | 0.053 | 0 | 0.58 | 0.64 | 0.66 | 0.89 | 0.48 | 0.76 |
| 4 | 181297475 | rs12649854 | 0.496 | 0.479 | -0.559 | -1.162 | 0.036 | 0.058 | 0.058 | 0 | 0.47 | 0.58 | 0.51 | 0.82 | 0.32 | 0.68 |
| 4 | 181301107 | rs2309241 | -0.093 | 0.417 | -0.255 | -1.079 | -0.001 | 0.049 | 0.049 | 0.965 | 0.25 | 0.43 | 0.33 | 0.62 | 0.26 | 0.34 |
| 4 | 181303512 | rs10032868 | 0.704 | 0.743 | -0.53 | -1.554 | 0.026 | 0.038 | 0.038 | 0 | 0.46 | 0.57 | 0.5 | 0.82 | 0.32 | 0.67 |
| 4 | 181306647 | rs10020466 | -0.271 | 0.594 | -0.791 | -1.429 | -0.002 | 0.055 | 0.055 | 1.798 | 0.7 | 0.77 | 0.76 | 0.89 | 0.73 | 0.83 |
| 4 | 181310122 | rs10013756 | -0.385 | 0.293 | -0.258 | -0.75 | -0.005 | -0.071 | -0.071 | 0 | 0.67 | 0.64 | 0.77 | 0.96 | 0.67 | 0.81 |
| 4 | 181310204 | rs10780063 | 0.57 | 0.091 | -0.471 | -0.649 | 0.024 | 0.042 | 0.042 | 0 | 0.52 | 0.41 | 0.59 | 0.87 | 0.6 | 0.56 |
| 4 | 181310570 | rs4437296 | -0.003 | 1.132 | -0.73 | -0.337 | 0 | -0.026 | -0.026 | 0.002 | 0.68 | 0.7 | 0.77 | 0.96 | 0.67 | 0.81 |
| 4 | 181312837 | rs6811892 | -0.655 | 1.834 | -0.664 | 1.211 | 0.015 | 0.073 | 0.073 | 0 | 0.1 | 0.14 | 0.2 | 0.41 | 0.23 | 0.2 |
| 4 | 181315369 | rs4473690 | -0.145 | 1.396 | -1.104 | 0.506 | 0.025 | 0.07 | 0.07 | 0.007 | 0.53 | 0.28 | 0.46 | 0.67 | 0.58 | 0.5 |
| 4 | 181316366 | rs6858814 | -0.129 | 1.013 | -0.355 | 0.409 | -0.023 | -0.015 | -0.015 | 2.88 | 0.84 | 0.74 | 0.88 | 0.84 | 0.89 | 0.89 |
| 4 | 181316450 | rs6833224 | 0.435 | 1.007 | 0.331 | 0.322 | -0.028 | -0.029 | -0.029 | 0.02 | 0.8 | 0.47 | 0.72 | 0.8 | 0.86 | 0.86 |
| 4 | 181318838 | rs7659432 | 0.297 | 0.886 | 0.578 | 0.495 | -0.003 | 0.003 | 0.003 | 0.024 | 0.32 | 0.42 | 0.43 | 0.36 | 0.32 | 0.5 |
| 4 | 181320766 | rs2871153 | 0.825 | 1.317 | -0.318 | -0.252 | 0.007 | -0.043 | -0.043 | 0.172 | 0.34 | 0.17 | 0.25 | 0.48 | 0.34 | 0.28 |
| 4 | 181321511 | rs17069334 | 0.742 | 0.903 | -0.709 | -0.21 | -0.013 | 0.07 | 0.07 | 0 | 0.14 | 0.15 | 0.14 | 0.38 | 0.09 | 0.2 |
| 4 | 181321999 | rs13127344 | 0.918 | 0.913 | -0.732 | -0.241 | -0.009 | 0.033 | 0.033 | 0.613 | 0.49 | 0.32 | 0.41 | 0.4 | 0.54 | 0.54 |
| 4 | 181327110 | rs17069342 | 1.377 | 0.865 | -0.828 | -0.274 | 0.003 | 0.026 | 0.026 | 0 | 0.07 | 0.04 | 0.06 | 0.02 | 0.07 | 0.03 |
| 4 | 181327429 | rs2309244 | -1.032 | 0.855 | -0.528 | -0.236 | 0.027 | 0.096 | 0.096 | 1.184 | 0.92 | 0.66 | 0.93 | 0.85 | 0.96 | 0.93 |
| 4 | 181327662 | rs2309246 | 1.047 | 0.933 | -0.398 | -0.21 | -0.004 | -0.001 | -0.001 | 0 | 0.34 | 0.49 | 0.46 | 0.34 | 0.34 | 0.5 |
| 4 | 181328600 | rs2871159 | -0.461 | 0.906 | -0.547 | -0.245 | 0.041 | 0.009 | 0.009 | 0.068 | 0.48 | 0.21 | 0.33 | 0.51 | 0.43 | 0.33 |
| 4 | 181329037 | rs6552422 | -0.777 | 1.078 | -0.574 | -0.051 | 0.034 | -0.021 | -0.021 | 0.017 | 0.82 | 0.76 | 0.8 | 0.85 | 0.77 | 0.83 |
| 4 | 181342591 | rs12509071 | 0.966 | 0.751 | -0.26 | 0.129 | 0.003 | 0.262 | 0.262 | 0 | 0.06 | 0.18 | 0.12 | 0.51 | 0.05 | 0.19 |
| 4 | 181343571 | rs2677008 | 0.737 | 0.958 | -0.242 | -0.299 | -0.007 | 0.02 | 0.02 | 0 | 0.45 | 0.42 | 0.44 | 0.32 | 0.48 | 0.57 |
| 4 | 181346824 | rs9990567 | 0.038 | 1.076 | -0.418 | -0.254 | 0.009 | 0.15 | 0.15 | 0 | 0.1 | 0.21 | 0.18 | 0.51 | 0.11 | 0.22 |
| 4 | 181347530 | rs17793705 | 1.317 | 1.066 | -0.964 | -0.429 | -0.063 | 0.046 | 0.046 | 0 | 0.21 | 0.02 | 0.11 | 0.11 | 0.25 | 0.08 |
| 4 | 181350463 | rs1320084 | 0.078 | 1.057 | -0.669 | -0.386 | 0.001 | 0.038 | 0.038 | 0 | 0.36 | 0.25 | 0.34 | 0.17 | 0.38 | 0.16 |
| 4 | 181350614 | rs1320083 | 0.522 | 0.785 | -0.655 | -0.088 | -0.013 | 0.204 | 0.204 | 0.196 | 0.8 | 0.56 | 0.76 | 0.49 | 0.86 | 0.74 |
| 4 | 181351303 | rs2677005 | 0.23 | 1.019 | -1.19 | -0.419 | 0.005 | 0.023 | 0.023 | 0 | 0.37 | 0.26 | 0.34 | 0.2 | 0.33 | 0.41 |
| 4 | 181351635 | rs7678786 | -0.214 | 1.272 | -1.689 | -0.83 | 0.002 | 0.029 | 0.029 | 0 | 0.39 | 0.35 | 0.33 | 0.14 | 0.38 | 0.25 |
| 4 | 181353732 | rs2309341 | 0.357 | 0.858 | -2.745 | -0.475 | -0.067 | -0.012 | -0.012 | 0 | 0.5 | 0.07 | 0.41 | 0.64 | 0.51 | 0.45 |
| 4 | 181353789 | rs4597870 | 0.152 | 0.837 | -2.184 | -0.493 | -0.017 | 0.006 | 0.006 | 0 | 0.6 | 0.34 | 0.48 | 0.68 | 0.57 | 0.52 |
| 4 | 181356882 | rs4241721 | -0.921 | 1.233 | -1.98 | -1.214 | -0.009 | 0.001 | 0.001 | 0 | 0.35 | 0.3 | 0.25 | 0.03 | 0.42 | 0.26 |
| 4 | 181358136 | rs17069404 | -0.619 | 0.31 | -1.764 | -1.003 | -0.002 | -0.037 | -0.037 | 0 | 0.28 | 0.29 | 0.17 | 0.03 | 0.25 | 0.15 |

| | | | | | | | | | | | | | | | | |
|---|---|---|---|---|---|---|---|---|---|---|---|---|---|---|---|---|
| 4 | 181358718 | rs17794036 | 0.33 | 0.492 | -1.605 | -0.351 | 0.005 | 0.005 | 0.005 | 0.095 | 0.09 | 0.06 | 0.09 | 0.04 | 0.05 | 0.06 |
| 4 | 181361148 | rs2727427 | 0.133 | 0.517 | -1.837 | -0.235 | 0.011 | 0.032 | 0.032 | 0 | 0.59 | 0.78 | 0.37 | 0.76 | 0.47 | 0.43 |
| 4 | 181361180 | rs13127074 | 0.011 | 0.738 | -1.195 | -0.451 | 0.04 | -0.031 | -0.031 | 0 | 0.7 | 0.6 | 0.57 | 0.83 | 0.62 | 0.73 |
| 4 | 181366289 | rs4552520 | 0.471 | -0.008 | -1.008 | 0.277 | 0.006 | -0.002 | -0.002 | 0.111 | 0.59 | 0.31 | 0.63 | 0.62 | 0.62 | 0.54 |
| 4 | 181368684 | rs2546589 | -0.37 | 0.31 | -0.958 | 0.142 | 0.029 | -0.012 | -0.012 | 0 | 0.23 | 0.26 | 0.28 | 0.19 | 0.33 | 0.22 |
| 4 | 181370064 | rs2546588 | 1.843 | -0.224 | -1.412 | 0.372 | -0.06 | 0.142 | 0.142 | 0 | 0.37 | 0.08 | 0.26 | 0.13 | 0.41 | 0.22 |
| 4 | 181375630 | rs7674341 | 0.935 | 0.367 | -0.926 | 0.446 | -0.02 | 0.011 | 0.011 | 0 | 0.13 | 0.09 | 0.1 | 0 | 0.18 | 0.12 |
| 4 | 181380227 | rs2727438 | 0.837 | 0.427 | -0.816 | 0.589 | 0.033 | 0.025 | 0.025 | 0 | 0.09 | 0.15 | 0.18 | 0.19 | 0.14 | 0.1 |
| 4 | 181382631 | rs2546581 | 1.184 | 0.101 | -0.858 | 0.537 | 0.006 | 0.021 | 0.021 | 0 | 0.53 | 0.24 | 0.46 | 0.43 | 0.48 | 0.42 |
| 4 | 181386404 | rs2727439 | 1.607 | 0.066 | -1.103 | 0.532 | -0.021 | -0.015 | -0.015 | 0 | 0.44 | 0.09 | 0.37 | 0.43 | 0.45 | 0.3 |
| 4 | 181399345 | rs17069436 | 0.134 | 0.05 | -0.958 | 1.024 | 0.004 | -0.015 | -0.015 | 0 | 0.08 | 0.02 | 0.15 | 0.32 | 0.06 | 0.26 |
| 4 | 181401306 | rs2727444 | 0.696 | 0.183 | -0.844 | 0.918 | 0.029 | 0.025 | 0.025 | 0 | 0.09 | 0.14 | 0.17 | 0.19 | 0.15 | 0.09 |
| 4 | 181405666 | rs2546575 | -1.332 | -0.232 | -0.27 | 0.836 | 0.018 | 0.004 | 0.004 | 0 | 0.4 | 0.48 | 0.57 | 0.42 | 0.44 | 0.5 |
| 4 | 181411378 | rs7687661 | -0.658 | -0.437 | 0.032 | 0.951 | -0.047 | -0.011 | -0.011 | 0 | 0.81 | 0.64 | 0.89 | 0.85 | 0.87 | 0.76 |
| 4 | 181421221 | rs2546571 | 1.464 | -0.089 | -0.456 | 1.023 | -0.033 | -0.04 | -0.04 | 0 | 0.39 | 0.12 | 0.32 | 0.44 | 0.41 | 0.29 |
| 4 | 181426359 | rs2546614 | -0.233 | 0.288 | -0.109 | 1.157 | 0.011 | 0.026 | 0.026 | 0 | 0.13 | 0.17 | 0.25 | 0.24 | 0.18 | 0.22 |
| 4 | 181437203 | rs2546606 | -1.566 | 0.037 | -0.014 | 1.052 | 0.002 | 0.006 | 0.006 | 0 | 0.56 | 0.7 | 0.64 | 0.56 | 0.58 | 0.7 |
| 4 | 181438804 | rs6829918 | -1.965 | -0.142 | -0.102 | 1.178 | -0.013 | 0.052 | 0.052 | 0.023 | 0.63 | 0.77 | 0.69 | 0.79 | 0.61 | 0.78 |
| 4 | 181440620 | rs2546595 | -1.223 | -0.145 | 0.518 | 1.197 | 0 | -0.003 | -0.003 | 0 | 0.48 | 0.43 | 0.6 | 0.51 | 0.52 | 0.62 |
| 4 | 181440776 | rs6843673 | -0.662 | -0.128 | 0.53 | 1.276 | 0.005 | 0.028 | 0.028 | 0.689 | 0.4 | 0.32 | 0.41 | 0.25 | 0.37 | 0.51 |
| 4 | 181441363 | rs6845332 | -0.899 | -0.335 | 0.801 | 1.262 | -0.039 | 0.01 | 0.01 | 0 | 0.29 | 0.39 | 0.29 | 0.3 | 0.19 | 0.43 |
| 4 | 181441802 | rs2059117 | -1.125 | -0.312 | 0.821 | 1.272 | -0.024 | 0.056 | 0.056 | 0 | 0.32 | 0.52 | 0.45 | 0.5 | 0.29 | 0.46 |
| 4 | 181441904 | rs2059116 | -0.253 | -0.373 | 0.437 | 1.157 | -0.027 | 0.002 | 0.002 | 0 | 0.22 | 0.27 | 0.25 | 0.25 | 0.13 | 0.35 |
| 4 | 181448254 | rs13134793 | 0.657 | -0.103 | 0.328 | 1.212 | -0.006 | 0.009 | 0.009 | 0 | 0.07 | 0.09 | 0.09 | 0.11 | 0.05 | 0.15 |
| 4 | 181449355 | rs11943526 | 0.972 | -0.123 | 0.192 | 1.221 | 0.027 | -0.002 | -0.002 | 0 | 0.14 | 0.1 | 0.06 | 0.12 | 0.1 | 0.21 |
| 4 | 181450318 | rs3113753 | -0.795 | 0.304 | 0.315 | 1.427 | 0.006 | 0.028 | 0.028 | 0.055 | 0.92 | 0.89 | 0.71 | 0.73 | 0.84 | 0.89 |
| 4 | 181450417 | rs1422738 | 1.21 | 0.125 | 0.31 | 1.347 | 0.007 | 0.003 | 0.003 | 0 | 0.58 | 0.48 | 0.39 | 0.43 | 0.51 | 0.57 |
| 4 | 181451934 | rs17069618 | 0.694 | 0.098 | -0.14 | 1.363 | 0.004 | 0.004 | 0.004 | 0 | 0.07 | 0.05 | 0.04 | 0.23 | 0.03 | 0.08 |
| 4 | 181454233 | rs17180827 | 0.703 | 0.214 | -0.225 | 1.237 | -0.016 | -0.026 | -0.026 | 0 | 0.05 | 0 | 0.04 | 0.06 | 0.06 | 0.08 |
| 4 | 181456588 | rs17795466 | 0.612 | 0.321 | -0.456 | 0.898 | -0.097 | 0.122 | 0.122 | 0.709 | 0.19 | 0.01 | 0.23 | 0 | 0.25 | 0.17 |
| 4 | 181460127 | rs6853596 | -1.637 | -0.257 | 0.128 | 0.935 | 0.015 | 0.015 | 0.015 | 0 | 0.5 | 0.67 | 0.4 | 0.42 | 0.44 | 0.54 |
| 4 | 181461281 | rs9684754 | -0.098 | -0.272 | 0.595 | 1.081 | 0.008 | 0.074 | 0.074 | 0 | 0.07 | 0.22 | 0.04 | 0.11 | 0.05 | 0.07 |
| 4 | 181462101 | rs7662339 | -0.02 | -0.294 | 0.635 | 1.063 | -0.031 | 0.031 | 0.031 | 0 | 0.25 | 0.14 | 0.27 | 0.11 | 0.31 | 0.25 |
| 4 | 181462408 | rs12502732 | -1.45 | -0.374 | 0.773 | 1.016 | -0.003 | -0.002 | -0.002 | 0 | 0.46 | 0.51 | 0.38 | 0.41 | 0.43 | 0.51 |
| 4 | 181463189 | rs7669248 | -2.204 | -0.423 | 0.797 | 1.078 | -0.024 | -0.019 | -0.019 | 0.793 | 0.57 | 0.75 | 0.45 | 0.52 | 0.5 | 0.67 |
| 4 | 181463308 | rs7669469 | 0.68 | -0.246 | 0.759 | 1.003 | -0.02 | -0.002 | -0.002 | 0.284 | 0.15 | 0.21 | 0.07 | 0.12 | 0.1 | 0.22 |
| 4 | 181468667 | rs17069681 | -0.112 | -0.079 | 0.61 | 1.054 | -0.022 | 0.022 | 0.022 | 0 | 0.25 | 0.17 | 0.28 | 0.13 | 0.31 | 0.25 |
| 4 | 181468882 | rs3095972 | -0.65 | 0.074 | 0.627 | 1.296 | 0.014 | 0.043 | 0.043 | 0.171 | 0.91 | 0.87 | 0.7 | 0.72 | 0.84 | 0.89 |
| 4 | 181468920 | rs3095971 | 2.269 | -0.101 | 0.683 | 1.303 | 0.003 | 0.041 | 0.041 | 0 | 0.41 | 0.2 | 0.29 | 0.31 | 0.4 | 0.37 |
| 4 | 181469160 | rs998760 | -0.063 | -0.102 | 0.686 | 1.313 | -0.047 | 0.022 | 0.022 | 0 | 0.21 | 0.34 | 0.12 | 0.27 | 0.12 | 0.27 |
| 4 | 181470273 | rs3113741 | 0.803 | 0.24 | 0.535 | 1.083 | -0.006 | -0.002 | -0.002 | 0 | 0.64 | 0.65 | 0.42 | 0.55 | 0.54 | 0.64 |
| 4 | 181471913 | rs6851088 | -0.906 | 0.377 | 0.552 | 0.81 | 0.005 | -0.024 | -0.024 | 2.119 | 0.29 | 0.38 | 0.28 | 0.14 | 0.31 | 0.25 |
| 4 | 181472405 | rs12331673 | -0.313 | 0.451 | 0.371 | 0.848 | -0.008 | 0.009 | 0.009 | 0 | 0.25 | 0.21 | 0.27 | 0.13 | 0.29 | 0.24 |
| 4 | 181474231 | rs11943826 | 0.436 | 0.447 | -0.128 | 0.812 | -0.018 | -0.001 | -0.001 | 0 | 0.15 | 0.24 | 0.07 | 0.11 | 0.11 | 0.19 |
| 4 | 181474291 | rs17796537 | 0.75 | 0.541 | -0.285 | 0.57 | -0.076 | 0.11 | 0.11 | 1.163 | 0.17 | 0.01 | 0.21 | 0 | 0.22 | 0.09 |
| 4 | 181476804 | rs1422506 | -1.17 | 0.506 | -0.076 | 0.623 | 0.001 | 0.011 | 0.011 | 0.04 | 0.42 | 0.38 | 0.34 | 0.29 | 0.41 | 0.44 |
| 4 | 181479147 | rs1559103 | -0.827 | 0.699 | -0.044 | 0.637 | 0 | 0.001 | 0.001 | 0.024 | 0.32 | 0.36 | 0.32 | 0.28 | 0.32 | 0.42 |
| 4 | 181479706 | rs17069770 | 0.164 | 0.769 | -0.305 | 0.79 | -0.032 | -0.032 | -0.032 | 0 | 0.07 | 0.01 | 0.23 | 0.14 | 0.15 | 0.16 |
| 4 | 181479865 | rs1030188 | -0.657 | 0.898 | -0.184 | 0.216 | -0.005 | -0.002 | -0.002 | 0.966 | 0.3 | 0.31 | 0.33 | 0.25 | 0.37 | 0.35 |
| 4 | 181480238 | rs13134598 | -0.814 | 0.846 | -0.114 | 0.159 | 0.027 | 0.002 | 0.002 | 0 | 0.32 | 0.45 | 0.37 | 0.25 | 0.4 | 0.39 |
| 4 | 181485416 | rs17796970 | 0.338 | 0.882 | -0.533 | -0.072 | -0.086 | 0.135 | 0.135 | 0 | 0.2 | 0.01 | 0.19 | 0 | 0.28 | 0.12 |
| 4 | 181489069 | rs7656346 | -0.971 | 0.747 | -0.229 | 0.03 | 0.021 | -0.011 | -0.011 | 0 | 0.37 | 0.52 | 0.42 | 0.27 | 0.45 | 0.42 |
| 4 | 181492183 | rs3095968 | 0.506 | 0.39 | -0.447 | -0.216 | 0.003 | 0.028 | 0.028 | 0 | 0.54 | 0.37 | 0.47 | 0.14 | 0.55 | 0.35 |
| 4 | 181492677 | rs3113749 | 0.273 | -0.02 | -0.582 | -0.1 | -0.013 | -0.004 | -0.004 | 0 | 0.55 | 0.43 | 0.61 | 0.35 | 0.67 | 0.56 |
| 4 | 181493355 | rs6552431 | 0.361 | -0.091 | -0.699 | -0.223 | -0.027 | 0.005 | 0.005 | 0 | 0.6 | 0.48 | 0.71 | 0.35 | 0.75 | 0.57 |
| 4 | 181493608 | rs6846354 | 0.958 | -0.091 | -0.647 | -0.163 | 0.001 | -0.001 | -0.001 | 1.153 | 0.13 | 0.11 | 0.07 | 0.12 | 0.13 | 0.2 |
| 4 | 181494406 | rs749463 | 0.789 | -0.069 | -0.536 | -0.185 | 0.023 | 0.028 | 0.028 | 0 | 0.48 | 0.34 | 0.64 | 0.23 | 0.62 | 0.38 |
| 4 | 181497819 | rs13114558 | 0.383 | 0.36 | -0.605 | -0.248 | 0.031 | 0.03 | 0.03 | 0 | 0.07 | 0.1 | 0.18 | 0.1 | 0.13 | 0.1 |
| 4 | 181498188 | rs7669125 | -0.701 | -0.015 | -0.73 | -0.471 | -0.009 | -0.016 | -0.016 | 0 | 0.88 | 0.83 | 0.92 | 0.91 | 0.9 | 0.93 |
| 4 | 181499684 | rs6851035 | 0.654 | -0.151 | -0.75 | -0.158 | -0.037 | 0.012 | 0.012 | 0 | 0.17 | 0.28 | 0.07 | 0.23 | 0.05 | 0.13 |
| 4 | 181501862 | rs3095966 | 0.347 | 0.809 | -0.966 | -0.223 | -0.003 | 0.002 | 0.002 | 0 | 0.59 | 0.61 | 0.56 | 0.7 | 0.47 | 0.56 |
| 4 | 181504039 | rs1422505 | 0.619 | 0.428 | -0.871 | 0.403 | -0.003 | -0.01 | -0.01 | 0 | 0.86 | 0.84 | 0.93 | 0.91 | 0.89 | 0.93 |
| 4 | 181518101 | rs1035423 | -0.887 | 0.551 | -1.233 | 0.58 | -0.003 | -0.001 | -0.001 | 0.03 | 0.34 | 0.23 | 0.24 | 0.28 | 0.39 | 0.33 |
| 4 | 181519492 | rs3113752 | -1.056 | 0.74 | -0.589 | 0.878 | 0.016 | 0.007 | 0.007 | 0 | 0.27 | 0.4 | 0.4 | 0.36 | 0.32 | 0.21 |
| 4 | 181520702 | rs3113757 | 0.451 | 0.208 | -0.923 | 0.772 | -0.004 | 0 | 0 | 0 | 0.75 | 0.75 | 0.78 | 0.71 | 0.84 | 0.71 |
| 4 | 181529714 | rs17798302 | -0.61 | 0.483 | -1.146 | 0.168 | 0.002 | 0.035 | 0.035 | 0 | 0.31 | 0.19 | 0.26 | 0.15 | 0.28 | 0.22 |
| 4 | 181535808 | rs3113765 | -1.057 | 0.771 | -0.91 | 0.139 | 0.006 | 0.024 | 0.024 | 0 | 0.4 | 0.33 | 0.29 | 0.16 | 0.34 | 0.26 |
| 4 | 181536607 | rs2059096 | -0.822 | 0.793 | -1.098 | -0.001 | -0.02 | 0.057 | 0.057 | 0 | 0.17 | 0.05 | 0.12 | 0.04 | 0.18 | 0.16 |
| 4 | 181539774 | rs2309360 | -1.061 | 0.565 | -0.67 | -0.035 | 0.001 | 0.006 | 0.006 | 0.007 | 0.37 | 0.32 | 0.29 | 0.16 | 0.32 | 0.21 |

| | | | | | | | | | | | | | | | | |
|---|---|---|---|---|---|---|---|---|---|---|---|---|---|---|---|---|
| 4 | 181540757 | rs3113773 | -0.221 | 0.226 | -0.908 | -0.016 | -0.007 | 0.095 | 0.095 | 0 | 0.19 | 0.03 | 0.06 | 0.04 | 0.16 | 0.08 |
| 4 | 181546303 | rs2446802 | -0.452 | 0.02 | -0.657 | 0.61 | -0.011 | 0.013 | 0.013 | 0.115 | 0.77 | 0.73 | 0.72 | 0.61 | 0.82 | 0.82 |
| 4 | 181555711 | rs10000265 | 0.846 | 0.546 | -0.892 | 0.279 | 0.013 | 0.037 | 0.037 | 0 | 0.44 | 0.79 | 0.48 | 0.28 | 0.41 | 0.56 |
| 4 | 181559236 | rs2546286 | 0.13 | 0.489 | 0.103 | 0.551 | -0.006 | 0.016 | 0.016 | 0 | 0.69 | 0.59 | 0.55 | 0.32 | 0.7 | 0.66 |
| 4 | 181563321 | rs2613041 | 0.05 | 0.565 | 0.149 | 0.558 | 0.002 | 0.013 | 0.013 | 0 | 0.72 | 0.72 | 0.59 | 0.32 | 0.7 | 0.66 |
| 4 | 181571062 | rs4861441 | 1.327 | 0.301 | -0.382 | 0.41 | -0.091 | 0.163 | 0.163 | 0.05 | 0.41 | 0.13 | 0.43 | 0.14 | 0.46 | 0.47 |
| 4 | 181571097 | rs2613047 | -0.356 | 0.186 | 0.099 | 0.53 | -0.001 | 0.003 | 0.003 | 0 | 0.3 | 0.32 | 0.27 | 0.34 | 0.3 | 0.28 |
| 4 | 181571542 | rs1422513 | -0.259 | 0.169 | -0.013 | 0.522 | 0.001 | -0.011 | -0.011 | 0 | 0.3 | 0.26 | 0.27 | 0.34 | 0.3 | 0.29 |
| 4 | 181572734 | rs10032205 | 1.357 | 0.133 | 0.032 | 0.62 | -0.103 | 0.15 | 0.15 | 0 | 0.4 | 0.13 | 0.43 | 0.12 | 0.46 | 0.46 |
| 4 | 181572822 | rs2546267 | -0.472 | 0.228 | 0.245 | 0.733 | -0.001 | -0.001 | -0.001 | 0.143 | 0.31 | 0.34 | 0.29 | 0.29 | 0.3 | 0.28 |
| 4 | 181574201 | rs1422511 | -0.709 | 0.23 | 0.158 | 0.709 | -0.002 | -0.013 | -0.013 | 0 | 0.34 | 0.48 | 0.34 | 0.28 | 0.34 | 0.3 |
| 4 | 181576122 | rs2613055 | -0.447 | 0.217 | 0.115 | 0.62 | 0.007 | -0.046 | -0.046 | 3.358 | 0.31 | 0.48 | 0.33 | 0.21 | 0.32 | 0.19 |
| 4 | 181576712 | rs10049679 | 0.943 | 0.215 | -0.1 | 0.755 | -0.011 | -0.008 | -0.008 | 0 | 0.46 | 0.29 | 0.61 | 0.4 | 0.52 | 0.66 |
| 4 | 181577045 | rs2194101 | 0.116 | -0.124 | 0.16 | 0.862 | -0.052 | 0.053 | 0.053 | 0.118 | 0.7 | 0.49 | 0.76 | 0.5 | 0.84 | 0.72 |
| 4 | 181577800 | rs2112580 | 1.186 | 0.371 | -0.015 | 0.734 | -0.018 | 0.003 | 0.003 | 0 | 0.34 | 0.51 | 0.4 | 0.3 | 0.35 | 0.56 |
| 4 | 181580617 | rs13114018 | 0.621 | 0.671 | -0.287 | 0.41 | -0.028 | 0.061 | 0.061 | 0.05 | 0.26 | 0.13 | 0.16 | 0.04 | 0.31 | 0.16 |
| 4 | 181582856 | rs17069961 | 1.449 | 0.446 | -0.489 | 0.345 | 0.006 | -0.029 | -0.029 | 0 | 0.05 | 0.09 | 0.02 | 0 | 0.04 | 0.04 |
| 4 | 181583727 | rs11132009 | 2.236 | 0.336 | -1.041 | 0.336 | -0.034 | 0.063 | 0.063 | 0 | 0.29 | 0.11 | 0.36 | 0.12 | 0.33 | 0.37 |
| 4 | 181584314 | rs2546260 | -0.289 | 0.738 | -0.743 | 0.122 | 0 | 0.017 | 0.017 | 0 | 0.33 | 0.36 | 0.33 | 0.53 | 0.31 | 0.27 |
| 4 | 181585352 | rs1422509 | -0.089 | 0.882 | -1.196 | -0.458 | -0.079 | 0.136 | 0.136 | 0.194 | 0.25 | 0.03 | 0.15 | 0.04 | 0.3 | 0.15 |
| 4 | 181591556 | rs17806047 | 0.426 | 0.803 | -0.736 | -0.14 | 0 | -0.003 | -0.003 | 0 | 0.15 | 0.16 | 0.09 | 0.06 | 0.17 | 0.12 |
| 4 | 181598141 | rs2613072 | -0.439 | 0.952 | -1.077 | -0.086 | -0.022 | 0.032 | 0.032 | 0 | 0.69 | 0.84 | 0.75 | 0.79 | 0.69 | 0.82 |
| 4 | 181600151 | rs11132010 | 1.109 | 0.71 | -0.245 | -0.243 | 0.007 | 0.012 | 0.012 | 0 | 0.61 | 0.33 | 0.74 | 0.68 | 0.65 | 0.69 |
| 4 | 181608225 | rs2161326 | 1.166 | -0.131 | -0.112 | -0.42 | 0.029 | 0.017 | 0.017 | 0 | 0.62 | 0.33 | 0.78 | 0.7 | 0.75 | 0.66 |
| 4 | 181608833 | rs7667149 | -0.082 | -0.017 | 0.533 | -0.384 | -0.007 | 0.007 | 0.007 | 0 | 0.22 | 0.25 | 0.13 | 0.29 | 0.2 | 0.22 |
| 4 | 181612806 | rs2216624 | -0.342 | 0.311 | 1.062 | -0.076 | -0.034 | -0.079 | -0.079 | 0 | 0.13 | 0.3 | 0.06 | 0.01 | 0.06 | 0.09 |
| 4 | 181636440 | rs1363207 | 1.712 | 0.65 | 0.568 | -0.175 | -0.003 | 0.024 | 0.024 | 0 | 0.54 | 0.36 | 0.71 | 0.65 | 0.56 | 0.59 |
| 4 | 181637915 | rs2545308 | 1.331 | 0.607 | 0.848 | -0.074 | 0.003 | -0.038 | -0.038 | 0 | 0.58 | 0.49 | 0.75 | 0.83 | 0.59 | 0.68 |
| 4 | 181638467 | rs6552438 | -0.935 | 0.602 | 0.892 | -0.117 | 0.014 | 0.022 | 0.022 | 0 | 0.26 | 0.32 | 0.19 | 0.34 | 0.3 | 0.27 |
| 4 | 181639976 | rs17807368 | -0.329 | 0.6 | 0.653 | -0.129 | -0.081 | 0.013 | 0.013 | 0 | 0.21 | 0.04 | 0.14 | 0.16 | 0.28 | 0.19 |
| 4 | 181647206 | rs1421998 | 1.5 | 1.44 | 1.176 | 0.102 | 0.029 | -0.072 | -0.072 | 0 | 0.63 | 0.5 | 0.74 | 0.85 | 0.57 | 0.71 |
| 4 | 181648742 | rs2545288 | 1.499 | 1.44 | 1.177 | -0.048 | 0.029 | -0.089 | -0.089 | 0.433 | 0.63 | 0.51 | 0.74 | 0.89 | 0.57 | 0.71 |
| 4 | 181663616 | rs12643977 | 0.033 | 1.972 | 1.013 | 0.305 | -0.05 | 0.006 | 0.006 | 0 | 0.14 | 0.04 | 0.13 | 0.11 | 0.25 | 0.15 |
| 4 | 181680688 | rs11734848 | -0.964 | 1.984 | 1.053 | 0.567 | 0.009 | 0.047 | 0.047 | 1.312 | 0.12 | 0.15 | 0.17 | 0.27 | 0.16 | 0.22 |
| 4 | 181682943 | rs2613007 | 0.882 | 1.941 | 1.147 | 0.539 | 0.032 | 0.191 | 0.191 | 0.137 | 0.87 | 0.59 | 0.81 | 0.74 | 0.85 | 0.85 |
| 4 | 181691738 | rs918402 | -0.73 | 1.954 | 0.842 | 0.509 | 0.012 | 0.079 | 0.079 | 2.318 | 0.12 | 0.18 | 0.1 | 0.28 | 0.14 | 0.21 |
| 4 | 181699644 | rs3101265 | 1.331 | 1.853 | 0.868 | 0.396 | 0.029 | 0.016 | 0.016 | 1.365 | 0.65 | 0.45 | 0.63 | 0.61 | 0.51 | 0.58 |
| 4 | 181707386 | rs2024530 | -1.634 | 1.936 | 0.59 | 0.753 | 0.061 | 0.058 | 0.058 | 0.749 | 0.29 | 0.48 | 0.33 | 0.39 | 0.41 | 0.36 |
| 4 | 181720229 | rs2545306 | -1.208 | 1.104 | -0.369 | 0.495 | 0.004 | -0.002 | -0.002 | 0.029 | 0.46 | 0.57 | 0.59 | 0.46 | 0.6 | 0.54 |
| 4 | 181720516 | rs2250756 | -0.238 | 1.271 | -0.411 | 0.506 | -0.069 | 0.037 | 0.037 | 0 | 0.33 | 0.12 | 0.38 | 0.23 | 0.43 | 0.34 |
| 4 | 181727899 | rs2613029 | -1.421 | 1.362 | -0.004 | 0.772 | -0.005 | -0.007 | -0.007 | 0 | 0.43 | 0.61 | 0.59 | 0.45 | 0.58 | 0.53 |
| 4 | 181729352 | rs2613032 | -0.266 | 1.542 | -1.109 | 0.358 | -0.074 | -0.028 | -0.028 | 0 | 0.28 | 0.13 | 0.47 | 0.26 | 0.44 | 0.36 |
| 4 | 181730895 | rs7693824 | -0.913 | 0.88 | -0.402 | 0.179 | -0.001 | -0.001 | -0.001 | 0 | 0.09 | 0.08 | 0.04 | 0 | 0.08 | 0.06 |
| 4 | 181734640 | rs4861444 | -0.798 | 0.334 | -0.085 | 0.376 | -0.062 | -0.013 | -0.013 | 0 | 0.2 | 0.37 | 0.1 | 0.21 | 0.11 | 0.14 |
| 4 | 181748551 | rs17070118 | -0.188 | 1.501 | -0.695 | 0.749 | -0.03 | -0.048 | -0.048 | 4.052 | 0.1 | 0.04 | 0.34 | 0.21 | 0.18 | 0.2 |
| 4 | 181752254 | rs17203326 | -0.134 | 1.175 | -0.742 | 0.309 | 0.005 | -0.011 | -0.011 | 0 | 0.14 | 0.07 | 0.04 | 0 | 0.07 | 0.03 |
| 4 | 181756792 | rs10866239 | -0.665 | 1.963 | -0.671 | 0.912 | -0.047 | -0.05 | -0.05 | 2.766 | 0.17 | 0.06 | 0.38 | 0.24 | 0.32 | 0.23 |
| 4 | 181757422 | rs7657018 | -1.093 | 1.584 | -0.098 | 0.607 | 0.002 | -0.004 | -0.004 | 0 | 0.08 | 0.08 | 0.06 | 0.02 | 0.15 | 0.03 |
| 4 | 181782574 | rs12648100 | -1.201 | 0.368 | 0.399 | 1.652 | -0.028 | 0.264 | 0.264 | 0 | 0.17 | 0.43 | 0.27 | 0.46 | 0.1 | 0.18 |
| 4 | 181782616 | rs12644875 | -0.925 | 1.227 | 0.098 | 2.291 | -0.008 | -0.003 | -0.003 | 0.119 | 0.17 | 0.16 | 0.25 | 0.44 | 0.26 | 0.4 |
| 4 | 181782692 | rs6832395 | -0.067 | 0.611 | 0.199 | 1.106 | -0.023 | -0.077 | -0.077 | 0 | 0.27 | 0.45 | 0.31 | 0.13 | 0.21 | 0.22 |
| 4 | 181783984 | rs17070257 | -0.941 | 0.952 | 0.253 | 1.901 | 0.019 | 0.141 | 0.141 | 0 | 0.08 | 0.12 | 0.09 | 0.37 | 0.12 | 0.19 |
| 4 | 181785074 | rs11722768 | -0.211 | 0.947 | -0.072 | 1.225 | -0.019 | 0.036 | 0.036 | 0 | 0.26 | 0.16 | 0.26 | 0.12 | 0.31 | 0.19 |
| 4 | 181785938 | rs6854060 | 0.468 | 0.273 | 0.284 | 0.876 | -0.003 | -0.024 | -0.024 | 0.168 | 0.55 | 0.54 | 0.63 | 0.78 | 0.57 | 0.6 |
| 4 | 181786172 | rs6854130 | 1.087 | 0.439 | -0.259 | 0.541 | -0.021 | 0.008 | 0.008 | 0 | 0.2 | 0.1 | 0.16 | 0.2 | 0.26 | 0.17 |
| 4 | 181797596 | rs7659727 | -0.18 | -0.142 | 0.13 | 0.439 | 0.002 | 0.018 | 0.018 | 0 | 0.57 | 0.42 | 0.5 | 0.67 | 0.62 | 0.53 |
| 4 | 181801052 | rs7692001 | 0.833 | -0.058 | -0.222 | 0.207 | -0.019 | 0.041 | 0.041 | 0.083 | 0.26 | 0.14 | 0.23 | 0.13 | 0.28 | 0.19 |
| 4 | 181808235 | rs6850503 | -0.08 | -0.089 | -0.204 | 0.286 | 0.004 | 0.008 | 0.008 | 0 | 0.5 | 0.52 | 0.64 | 0.38 | 0.53 | 0.46 |
| 4 | 181810029 | rs10866240 | -0.15 | -0.292 | -0.261 | -0.048 | 0.002 | 0.002 | 0.002 | 0 | 0.56 | 0.56 | 0.73 | 0.72 | 0.62 | 0.63 |
| 4 | 181810622 | rs13122971 | -1.086 | -0.018 | -0.279 | 0.509 | -0.013 | -0.013 | -0.013 | 0.086 | 0.85 | 0.84 | 0.94 | 0.91 | 0.92 | 0.82 |
| 4 | 181813489 | rs17070275 | -0.38 | 0.616 | -0.33 | 0.983 | -0.012 | -0.016 | -0.016 | 0 | 0.29 | 0.21 | 0.46 | 0.58 | 0.34 | 0.44 |
| 4 | 181814242 | rs12504749 | 0.236 | 0.945 | -0.544 | 1.088 | -0.045 | -0.03 | -0.03 | 0 | 0.22 | 0.06 | 0.37 | 0.27 | 0.26 | 0.28 |
| 4 | 181814803 | rs2309396 | 0.867 | 1.038 | -0.124 | 0.732 | -0.017 | -0.003 | -0.003 | 0 | 0.65 | 0.34 | 0.55 | 0.39 | 0.57 | 0.51 |
| 4 | 181814917 | rs6813148 | -0.76 | 1.181 | 0.668 | 0.885 | 0.052 | 0.08 | 0.08 | 0 | 0.27 | 0.45 | 0.38 | 0.49 | 0.34 | 0.33 |
| 4 | 181821983 | rs1589580 | -0.5 | 1.274 | 0.439 | 1.151 | 0 | 0.022 | 0.022 | 0 | 0.17 | 0.13 | 0.28 | 0.46 | 0.2 | 0.33 |
| 4 | 181825165 | rs1533873 | -0.608 | 0.967 | 0.062 | 0.253 | -0.004 | 0.058 | 0.058 | 0 | 0.48 | 0.72 | 0.5 | 0.42 | 0.49 | 0.33 |
| 4 | 181826016 | rs12507955 | -0.852 | 0.805 | 0.523 | 0.101 | -0.009 | -0.03 | -0.03 | 0 | 0.68 | 0.46 | 0.74 | 0.77 | 0.68 | 0.82 |
| 4 | 181850257 | rs17070354 | 0.153 | 0.71 | 0.655 | 0.379 | -0.005 | -0.006 | -0.006 | 0 | 0.14 | 0.13 | 0.32 | 0.25 | 0.17 | 0.1 |
| 4 | 181851323 | rs10015408 | 0.39 | 0.63 | 1.066 | 0.243 | 0.015 | -0.029 | -0.029 | 0 | 0.92 | 0.68 | 0.95 | 0.99 | 0.94 | 0.97 |

| | | | | | | | | | | | | | | | | |
|---|---|---|---|---|---|---|---|---|---|---|---|---|---|---|---|---|
| 4 | 181867512 | rs10866243 | 0.362 | 0.893 | 0.653 | 0.092 | 0.045 | 0.024 | 0.024 | 0 | 0.49 | 0.16 | 0.38 | 0.5 | 0.43 | 0.68 |
| 4 | 181868940 | rs6820051 | 0.314 | 0.876 | 0.997 | 0.118 | 0.032 | -0.018 | -0.018 | 0 | 0.06 | 0.21 | 0.05 | 0 | 0.06 | 0.02 |
| 4 | 181885905 | rs13115400 | 0.447 | 1.003 | 0.72 | 0.185 | 0.015 | -0.02 | -0.02 | 0 | 0.3 | 0.34 | 0.39 | 0.24 | 0.36 | 0.14 |
| 4 | 181888903 | rs10018261 | 0.497 | 0.962 | 0.697 | 0.185 | 0.019 | -0.038 | -0.038 | 0.207 | 0.31 | 0.37 | 0.4 | 0.2 | 0.36 | 0.16 |
| 4 | 181892753 | rs11727972 | 0.018 | 0.899 | 0.271 | 0.407 | 0.006 | -0.015 | -0.015 | 0.113 | 0.63 | 0.46 | 0.44 | 0.66 | 0.62 | 0.64 |
| 4 | 181895905 | rs11931741 | -0.841 | 1.267 | -0.018 | 0.609 | 0.005 | 0.068 | 0.068 | 1.765 | 0.28 | 0.44 | 0.28 | 0.44 | 0.32 | 0.43 |
| 4 | 181898870 | rs13435468 | -0.283 | 1.701 | -0.314 | 0.293 | 0 | 0 | 0 | 0 | 0.79 | 0.83 | 0.73 | 0.75 | 0.79 | 0.79 |
| 4 | 181904385 | rs7688750 | 0.892 | 1.73 | -0.253 | -0.087 | 0.012 | -0.008 | -0.008 | 5.742 | 0.47 | 0.43 | 0.26 | 0.07 | 0.36 | 0.22 |
| 4 | 181905572 | rs2056490 | 1.009 | 1.72 | -0.259 | -0.093 | 0.007 | -0.018 | -0.018 | 1.047 | 0.46 | 0.43 | 0.26 | 0.07 | 0.36 | 0.2 |
| 4 | 181907769 | rs10016729 | 0.307 | 1.969 | 0.194 | 0.571 | 0.041 | 0.094 | 0.094 | 0 | 0.09 | 0.21 | 0.08 | 0.26 | 0.12 | 0.17 |
| 4 | 181910926 | rs1916109 | 0.209 | 1.942 | 0.343 | 0.54 | 0.023 | 0.108 | 0.108 | 0 | 0.79 | 0.68 | 0.61 | 0.48 | 0.73 | 0.69 |
| 4 | 181912806 | rs1969399 | 0.696 | 1.901 | 0.23 | 0.491 | -0.001 | 0 | 0 | 0 | 0.71 | 0.73 | 0.55 | 0.46 | 0.64 | 0.53 |
| 4 | 181919047 | rs11132035 | 1.011 | 1.76 | 0.154 | -0.023 | -0.005 | -0.002 | -0.002 | 0 | 0.58 | 0.59 | 0.4 | 0.13 | 0.5 | 0.37 |
| 4 | 181920297 | rs11942320 | 1.397 | 1.27 | -0.984 | -0.855 | -0.018 | 0.115 | 0.115 | 0 | 0.28 | 0.03 | 0.22 | 0.07 | 0.23 | 0.2 |
| 4 | 181920483 | rs11132037 | 1.272 | 0.946 | -0.473 | -0.891 | 0.001 | -0.04 | -0.04 | 0 | 0.23 | 0.21 | 0.12 | 0.01 | 0.2 | 0.16 |
| 4 | 181923958 | rs13353673 | 0.587 | 1.016 | -0.325 | -0.687 | 0.01 | -0.017 | -0.017 | 0 | 0.37 | 0.29 | 0.26 | 0.4 | 0.27 | 0.3 |
| 4 | 181925065 | rs6843038 | 0.332 | 0.939 | -0.301 | -0.726 | -0.007 | 0.023 | 0.023 | 0.094 | 0.36 | 0.46 | 0.29 | 0.37 | 0.32 | 0.33 |
| 4 | 181926518 | rs11946898 | -0.593 | 1.502 | -0.708 | -0.411 | -0.017 | -0.056 | -0.056 | 0.289 | 0.78 | 0.87 | 0.63 | 0.52 | 0.71 | 0.7 |
| 4 | 181927704 | rs11132039 | 1.255 | 1.54 | -0.743 | -0.42 | -0.059 | 0.056 | 0.056 | 1.831 | 0.14 | 0.03 | 0.22 | 0.03 | 0.21 | 0.21 |
| 4 | 181928515 | rs10022243 | 0.625 | 1.287 | -0.002 | 0.074 | 0.006 | 0.218 | 0.218 | 0.263 | 0.16 | 0.43 | 0.2 | 0.37 | 0.16 | 0.17 |
| 4 | 181929211 | rs17070458 | 0.817 | 1.543 | -0.816 | 0.374 | -0.044 | -0.061 | -0.061 | 0.058 | 0.1 | 0.01 | 0.22 | 0.26 | 0.11 | 0.19 |
| 4 | 181933056 | rs12642493 | 0.125 | 1.367 | -0.757 | 0.117 | -0.014 | -0.048 | -0.048 | 3.442 | 0.76 | 0.85 | 0.62 | 0.52 | 0.7 | 0.69 |
| 4 | 181934106 | rs7663613 | 0.501 | 1.065 | -0.517 | -0.288 | -0.001 | -0.07 | -0.07 | 0.048 | 0.27 | 0.3 | 0.17 | 0 | 0.25 | 0.16 |
| 4 | 181935405 | rs1357353 | -0.273 | 0.872 | -0.776 | 0.491 | -0.001 | -0.004 | -0.004 | 1.535 | 0.45 | 0.49 | 0.5 | 0.31 | 0.39 | 0.4 |
| 4 | 181941847 | rs17070467 | 0.044 | 1.023 | -1.21 | 1.166 | 0 | -0.004 | -0.004 | 0 | 0.1 | 0.05 | 0.2 | 0.44 | 0.09 | 0.22 |
| 4 | 181942406 | rs17070468 | -0.139 | 1.128 | -1.205 | 0.554 | -0.003 | -0.007 | -0.007 | 0.013 | 0.27 | 0.2 | 0.37 | 0.24 | 0.25 | 0.34 |
| 4 | 181944160 | rs9997006 | -0.057 | 1.083 | -0.721 | 0.517 | 0.006 | -0.004 | -0.004 | 0 | 0.37 | 0.45 | 0.64 | 0.29 | 0.39 | 0.48 |
| 4 | 181945777 | rs7659915 | 0.145 | 1.615 | -0.807 | 0.518 | 0.004 | 0.024 | 0.024 | 0 | 0.89 | 0.87 | 0.93 | 0.73 | 0.83 | 0.87 |
| 4 | 181957456 | rs13121678 | 0.359 | 1.38 | -1.026 | 0.219 | -0.032 | 0.036 | 0.036 | 0.039 | 0.62 | 0.83 | 0.78 | 0.7 | 0.55 | 0.62 |
| 4 | 181982520 | rs10019828 | -0.5 | 1.429 | -0.466 | -0.083 | -0.021 | 0.027 | 0.027 | 0 | 0.17 | 0.08 | 0.2 | 0.03 | 0.2 | 0.16 |
| 4 | 181987292 | rs6856731 | 0.262 | 0.948 | 0.185 | 0.068 | -0.01 | -0.016 | -0.016 | 0.303 | 0.76 | 0.68 | 0.83 | 0.84 | 0.76 | 0.69 |
| 4 | 182007048 | rs12645608 | 0.868 | 0.691 | 0.615 | 0.05 | 0.009 | 0.008 | 0.008 | 0.946 | 0.7 | 0.29 | 0.78 | 0.74 | 0.75 | 0.59 |
| 4 | 182009132 | rs7659755 | 0.048 | 0.755 | 1.145 | -0.129 | 0.001 | -0.068 | -0.068 | 0 | 0.78 | 0.7 | 0.77 | 0.97 | 0.81 | 0.85 |
| 4 | 182024045 | rs9312270 | 1.327 | 0.743 | 0.952 | -0.114 | 0.014 | 0.162 | 0.162 | 0 | 0.52 | 0.17 | 0.57 | 0.71 | 0.56 | 0.43 |
| 4 | 182029163 | rs11132044 | -0.53 | 0.66 | 1.451 | -0.051 | -0.027 | -0.026 | -0.026 | 0 | 0.26 | 0.56 | 0.19 | 0.24 | 0.25 | 0.33 |
| 4 | 182032424 | rs13117129 | 0.293 | 0.646 | 1.732 | -0.09 | 0.003 | 0.006 | 0.006 | 0 | 0.18 | 0.13 | 0.11 | 0.18 | 0.18 | 0.26 |
| 4 | 182033941 | rs4861905 | -1.188 | 0.738 | 1.477 | -0.094 | 0.015 | 0.047 | 0.047 | 0 | 0.41 | 0.66 | 0.36 | 0.23 | 0.33 | 0.44 |
| 4 | 182046294 | rs7435210 | -0.131 | 0.806 | 1.618 | -0.162 | -0.006 | -0.066 | -0.066 | 1.837 | 0.79 | 0.73 | 0.77 | 0.97 | 0.84 | 0.81 |
| 4 | 182053380 | rs4488971 | -0.206 | 0.746 | 1.732 | -0.106 | -0.026 | -0.042 | -0.042 | 0.36 | 0.22 | 0.39 | 0.13 | 0.16 | 0.19 | 0.33 |
| 4 | 182054829 | rs998399 | 1.383 | 0.837 | 1.045 | -0.122 | 0.036 | 0.156 | 0.156 | 0 | 0.52 | 0.22 | 0.58 | 0.77 | 0.56 | 0.46 |
| 4 | 182061996 | rs6852438 | 0.601 | 0.873 | 1.004 | -0.1 | -0.019 | -0.086 | -0.086 | 0.341 | 0.73 | 0.58 | 0.7 | 0.89 | 0.76 | 0.73 |
| 4 | 182064722 | rs4861466 | 1.307 | 0.94 | 0.015 | -0.093 | 0.033 | 0.206 | 0.206 | 0 | 0.5 | 0.13 | 0.56 | 0.74 | 0.56 | 0.42 |
| 4 | 182065258 | rs6824538 | 1.006 | 1.181 | 0.526 | 0.935 | -0.003 | 0.016 | 0.016 | 0 | 0.7 | 0.46 | 0.51 | 0.7 | 0.73 | 0.71 |
| 4 | 182065314 | rs6825217 | 1.079 | 1.184 | 0.504 | 0.938 | -0.002 | 0.012 | 0.012 | 0 | 0.69 | 0.46 | 0.51 | 0.7 | 0.73 | 0.71 |
| 4 | 182065498 | rs4349637 | -0.657 | 1.41 | -0.652 | 0.211 | -0.004 | 0.038 | 0.038 | 0 | 0.21 | 0.09 | 0.23 | 0.03 | 0.19 | 0.16 |
| 4 | 182066869 | rs6552486 | -1.219 | 2.124 | -0.379 | 1.063 | 0.043 | 0.028 | 0.028 | 0 | 0.05 | 0.06 | 0.19 | 0.1 | 0.17 | 0.09 |
| 4 | 182067359 | rs12498222 | -1.797 | 2.168 | 0.076 | 1.303 | 0.083 | 0.075 | 0.075 | 0 | 0.09 | 0.24 | 0.27 | 0.16 | 0.2 | 0.14 |
| 4 | 182068097 | rs6552487 | 0.196 | 1.755 | -0.933 | 1.082 | -0.076 | -0.001 | -0.001 | 0 | 0.37 | 0.12 | 0.37 | 0.36 | 0.41 | 0.44 |
| 4 | 182068819 | rs17070655 | -0.858 | 1.108 | 0.341 | 1.461 | 0.052 | 0.168 | 0.168 | 0.056 | 0.05 | 0.24 | 0.04 | 0.13 | 0.04 | 0.15 |
| 4 | 182072142 | rs10520467 | 0.062 | 0.339 | -0.398 | 1.227 | 0.002 | -0.014 | -0.014 | 0 | 0.3 | 0.23 | 0.27 | 0.24 | 0.33 | 0.21 |
| 4 | 182078896 | rs12507155 | 0.399 | 0.096 | -0.207 | 0.984 | 0.035 | -0.001 | -0.001 | 0.118 | 0.48 | 0.31 | 0.62 | 0.54 | 0.59 | 0.67 |
| 4 | 182089409 | rs7688965 | 0.783 | 0.018 | -0.219 | 0.972 | -0.013 | 0.011 | 0.011 | 3.872 | 0.43 | 0.2 | 0.41 | 0.42 | 0.5 | 0.47 |
| 4 | 182092437 | rs6844876 | 0.65 | -0.511 | -0.267 | 0.929 | 0.025 | -0.005 | -0.005 | 0 | 0.22 | 0.13 | 0.15 | 0.27 | 0.11 | 0.15 |
| 4 | 182094811 | rs1993327 | 1.222 | -0.457 | -0.376 | 0.941 | -0.043 | -0.005 | -0.005 | 0.003 | 0.42 | 0.13 | 0.4 | 0.4 | 0.49 | 0.46 |
| 4 | 182095358 | rs17828403 | 1.23 | -0.899 | -0.524 | 0.553 | -0.008 | -0.006 | -0.006 | 2.411 | 0.08 | 0 | 0.05 | 0.04 | 0.07 | 0.09 |
| 4 | 182098564 | rs4861919 | 0.296 | -0.739 | 0.173 | 0.657 | 0.015 | 0.067 | 0.067 | 0 | 0.92 | 0.68 | 0.79 | 0.89 | 0.9 | 0.83 |
| 4 | 182101689 | rs10018288 | -0.063 | -0.789 | -0.373 | 0.521 | -0.05 | 0.018 | 0.018 | 0 | 0.68 | 0.46 | 0.64 | 0.62 | 0.78 | 0.64 |
| 4 | 182101765 | rs17828571 | -0.266 | -0.393 | -0.837 | 0.26 | 0.026 | 0.046 | 0.046 | 2.711 | 0.23 | 0.12 | 0.13 | 0.05 | 0.18 | 0.11 |
| 4 | 182104055 | rs9991738 | 0.431 | 0.076 | -1.133 | 0.545 | -0.03 | 0.008 | 0.008 | 0 | 0.41 | 0.24 | 0.44 | 0.41 | 0.44 | 0.43 |
| 4 | 182111358 | rs1454711 | 0.112 | -0.087 | -1.462 | 0.219 | 0.031 | 0.093 | 0.093 | 1.207 | 0.23 | 0.1 | 0.14 | 0.07 | 0.16 | 0.08 |
| 4 | 182117057 | rs1902112 | -0.531 | -0.304 | -0.885 | 0.854 | -0.011 | 0.003 | 0.003 | 1.091 | 0.78 | 0.75 | 0.88 | 0.67 | 0.88 | 0.8 |
| 4 | 182118509 | rs6845037 | -0.426 | 0.376 | -0.778 | 0.652 | -0.002 | 0.002 | 0.002 | 0.052 | 0.43 | 0.58 | 0.61 | 0.28 | 0.51 | 0.41 |
| 4 | 182120631 | rs9999847 | -0.779 | 0.581 | 0.016 | 0.969 | 0.008 | 0.085 | 0.085 | 0 | 0.17 | 0.28 | 0.36 | 0.26 | 0.16 | 0.3 |
| 4 | 182126122 | rs5019445 | -0.01 | 0.715 | 0.002 | 0.395 | 0 | -0.036 | -0.036 | 0.098 | 0.71 | 0.68 | 0.75 | 0.96 | 0.72 | 0.89 |
| 4 | 182131530 | rs9995325 | 0.988 | 0.707 | 0.046 | 0.505 | 0 | 0 | 0 | 0 | 0.56 | 0.58 | 0.38 | 0.65 | 0.55 | 0.6 |
| 4 | 182133522 | rs10520469 | 0.709 | 0.457 | -0.873 | 0.609 | 0.019 | -0.043 | -0.043 | 0 | 0.13 | 0.04 | 0.05 | 0.32 | 0.1 | 0.13 |
| 4 | 182135395 | rs1869606 | -0.727 | 0.938 | -0.462 | 0.561 | -0.062 | 0.018 | 0.018 | 0.166 | 0.9 | 0.99 | 0.84 | 0.96 | 0.84 | 0.93 |
| 4 | 182137188 | rs6836317 | -0.454 | 0.921 | -0.09 | 0.565 | -0.001 | -0.064 | -0.064 | 0.684 | 0.35 | 0.51 | 0.31 | 0.04 | 0.31 | 0.12 |
| 4 | 182140400 | rs9994253 | 0.568 | 0.876 | -0.678 | 0.756 | -0.006 | 0.003 | 0.003 | 0 | 0.57 | 0.4 | 0.39 | 0.67 | 0.55 | 0.61 |

| | | | | | | | | | | | | | | | | |
|---|---|---|---|---|---|---|---|---|---|---|---|---|---|---|---|---|
| 4 | 182153124 | rs4861928 | -0.238 | 1.011 | -0.875 | 1.298 | -0.015 | -0.023 | -0.023 | 0 | 0.12 | 0.07 | 0.26 | 0.42 | 0.17 | 0.29 |
| 4 | 182154578 | rs17070824 | -0.562 | 0.577 | -0.633 | 0.437 | -0.004 | 0 | 0 | 0.018 | 0.18 | 0.25 | 0.23 | 0.23 | 0.16 | 0.23 |
| 4 | 182155026 | rs10012888 | -0.094 | 0.673 | -0.681 | -0.31 | -0.003 | -0.085 | -0.085 | 0 | 0.72 | 0.69 | 0.77 | 1 | 0.76 | 0.94 |
| 4 | 182159150 | rs17070835 | -0.485 | 0.584 | -0.837 | 0.752 | 0.001 | -0.007 | -0.007 | 0 | 0.18 | 0.14 | 0.24 | 0.3 | 0.17 | 0.3 |
| 4 | 182163907 | rs10024557 | 0.255 | 0.264 | -0.695 | 0.292 | -0.007 | -0.016 | -0.016 | 0 | 0.87 | 0.83 | 0.73 | 0.94 | 0.91 | 0.94 |
| 4 | 182164515 | rs4861929 | 0.188 | 0.309 | -0.791 | 0.384 | 0.001 | -0.006 | -0.006 | 0.549 | 0.07 | 0.06 | 0.07 | 0.14 | 0.05 | 0.15 |
| 4 | 182167822 | rs9992143 | 0.978 | 0.745 | -1.295 | 0.776 | -0.031 | -0.053 | -0.053 | 0 | 0.15 | 0.04 | 0.13 | 0.3 | 0.22 | 0.12 |
| 4 | 182170424 | rs17070839 | 0.559 | 0.586 | -1.535 | 0.708 | -0.037 | -0.006 | -0.006 | 0.017 | 0.19 | 0.05 | 0.14 | 0.2 | 0.22 | 0.17 |
| 4 | 182178058 | rs7669856 | 0.801 | 0.299 | -2.033 | -0.507 | 0.017 | 0.05 | 0.05 | 0 | 0.34 | 0.26 | 0.28 | 0.17 | 0.3 | 0.31 |
| 4 | 182179570 | rs11727860 | 0.197 | 0.128 | -1.538 | -0.544 | -0.015 | -0.015 | -0.015 | 0.674 | 0.3 | 0.24 | 0.52 | 0.52 | 0.4 | 0.42 |
| 4 | 182179788 | rs17070889 | 1.007 | -0.054 | -1.358 | -0.669 | -0.039 | -0.004 | -0.004 | 0 | 0.3 | 0.11 | 0.19 | 0.34 | 0.33 | 0.3 |
| 4 | 182181119 | rs7438836 | 1.33 | 0.088 | -1.205 | -0.492 | 0.012 | 0.052 | 0.052 | 0 | 0.16 | 0.19 | 0.2 | 0.45 | 0.22 | 0.29 |
| 4 | 182182652 | rs6849417 | -0.101 | -0.057 | -0.913 | -0.52 | 0.028 | 0.029 | 0.029 | 0 | 0.33 | 0.52 | 0.57 | 0.52 | 0.41 | 0.42 |
| 4 | 182183915 | rs10520473 | 1.387 | -0.061 | -1.224 | -0.385 | -0.021 | -0.048 | -0.048 | 0 | 0.1 | 0.01 | 0.05 | 0.21 | 0.1 | 0.13 |
| 4 | 182185950 | rs6844811 | 1.034 | -0.194 | -0.76 | -0.413 | 0 | 0.024 | 0.024 | 3.109 | 0.46 | 0.33 | 0.34 | 0.37 | 0.44 | 0.4 |
| 4 | 182187373 | rs1031500 | 0.826 | 0.107 | -0.867 | -0.489 | -0.011 | 0.034 | 0.034 | 0 | 0.15 | 0.06 | 0.33 | 0.06 | 0.2 | 0.13 |
| 4 | 182189284 | rs17240768 | 1.634 | -0.11 | -0.755 | -0.492 | 0.011 | 0.013 | 0.013 | 0 | 0.25 | 0.11 | 0.19 | 0.2 | 0.19 | 0.23 |
| 4 | 182189790 | rs7441242 | -0.611 | -0.102 | -0.354 | -0.404 | 0.012 | 0.007 | 0.007 | 0 | 0.4 | 0.57 | 0.62 | 0.42 | 0.5 | 0.49 |
| 4 | 182192995 | rs7666029 | -0.561 | -0.178 | -0.46 | -0.439 | 0.001 | 0.052 | 0.052 | 0 | 0.56 | 0.71 | 0.45 | 0.74 | 0.53 | 0.59 |
| 4 | 182195518 | rs11930417 | 1.023 | 0.27 | 0.084 | -0.163 | 0.041 | 0.135 | 0.135 | 0 | 0.17 | 0.37 | 0.23 | 0.37 | 0.2 | 0.31 |
| 4 | 182197947 | rs1454694 | 1.045 | 0.053 | -0.223 | -0.15 | -0.007 | 0.01 | 0.01 | 1.778 | 0.24 | 0.1 | 0.13 | 0.22 | 0.25 | 0.15 |
| 4 | 182199391 | rs1563213 | 1.101 | 0.016 | -0.253 | -0.223 | 0.031 | -0.006 | -0.006 | 0 | 0.35 | 0.14 | 0.22 | 0.35 | 0.26 | 0.36 |
| 4 | 182199523 | rs1031501 | 1.733 | 0.293 | 0.153 | 0.369 | -0.002 | 0.06 | 0.06 | 0 | 0.06 | 0.09 | 0.04 | 0.23 | 0.03 | 0.12 |
| 4 | 182204531 | rs4861931 | 0.149 | 0.289 | -0.183 | -0.223 | -0.001 | 0.012 | 0.012 | 1.927 | 0.25 | 0.17 | 0.2 | 0.2 | 0.25 | 0.15 |
| 4 | 182210944 | rs4861938 | 0.636 | 0.651 | -0.115 | -0.046 | -0.005 | 0.003 | 0.003 | 2.159 | 0.16 | 0.16 | 0.16 | 0.2 | 0.15 | 0.06 |
| 4 | 182211159 | rs1454696 | -1.23 | 0.683 | -0.15 | -0.139 | 0 | -0.023 | -0.023 | 0 | 0.92 | 0.82 | 0.93 | 1 | 0.9 | 0.94 |
| 4 | 182213030 | rs7681232 | 0.545 | 0.523 | -0.109 | 0.003 | -0.01 | 0.012 | 0.012 | 2.324 | 0.17 | 0.22 | 0.16 | 0.2 | 0.15 | 0.06 |
| 4 | 182214915 | rs9996848 | 0.909 | 0.481 | -0.497 | 0.168 | -0.014 | -0.016 | -0.016 | 0 | 0.06 | 0.03 | 0.09 | 0.16 | 0.11 | 0.1 |
| 4 | 182216904 | rs6817589 | 0.565 | 0.315 | -0.202 | -0.04 | -0.004 | -0.013 | -0.013 | 0.035 | 0.75 | 0.35 | 0.73 | 0.78 | 0.73 | 0.81 |
| 4 | 182221687 | rs17071008 | 0.779 | 0.059 | -1.302 | 0.096 | 0.043 | 0.008 | 0.008 | 0 | 0.31 | 0.09 | 0.25 | 0.28 | 0.21 | 0.34 |
| 4 | 182222033 | rs2309470 | 0.024 | 0.303 | -0.953 | 0.174 | 0.032 | 0.006 | 0.006 | 0 | 0.49 | 0.38 | 0.39 | 0.56 | 0.36 | 0.56 |
| 4 | 182222287 | rs2309467 | 0.416 | 0.995 | -1.067 | -0.212 | -0.011 | 0.007 | 0.007 | 0 | 0.8 | 0.82 | 0.66 | 0.92 | 0.7 | 0.89 |
| 4 | 182222727 | rs17183114 | -0.259 | 0.803 | -0.794 | 0.14 | 0.004 | -0.007 | -0.007 | 0 | 0.09 | 0.09 | 0.1 | 0.03 | 0.16 | 0.02 |
| 4 | 182226071 | rs11930570 | -0.887 | 0.379 | -1.155 | 0.233 | 0.006 | -0.002 | -0.002 | 0 | 0.25 | 0.16 | 0.16 | 0.23 | 0.24 | 0.18 |
| 4 | 182226435 | rs10027292 | 0.281 | 0.427 | -0.827 | 0.145 | 0.039 | -0.01 | -0.01 | 0 | 0.3 | 0.43 | 0.41 | 0.25 | 0.39 | 0.35 |
| 4 | 182231482 | rs10520475 | 0.266 | 0.212 | -1.162 | 0.028 | 0.032 | -0.054 | -0.054 | 0 | 0.13 | 0.05 | 0.05 | 0.21 | 0.07 | 0.12 |
| 4 | 182231885 | rs9312276 | 0.193 | -0.025 | -0.968 | -0.622 | 0.01 | 0.042 | 0.042 | 0 | 0.49 | 0.56 | 0.51 | 0.23 | 0.53 | 0.45 |
| 4 | 182234393 | rs1115988 | 0.694 | 0.061 | -0.957 | -0.733 | 0.004 | -0.008 | -0.008 | 0 | 0.26 | 0.19 | 0.16 | 0.01 | 0.24 | 0.14 |
| 4 | 182234443 | rs17071059 | 1.272 | -0.015 | -0.905 | -0.67 | -0.001 | 0.004 | 0.004 | 0 | 0.16 | 0.1 | 0.15 | 0.15 | 0.17 | 0.17 |
| 4 | 182235447 | rs9994198 | 1.73 | 0.258 | -0.763 | -0.715 | 0.016 | -0.001 | -0.001 | 0 | 0.22 | 0.29 | 0.33 | 0.22 | 0.26 | 0.31 |
| 4 | 182237515 | rs17071066 | 1.738 | 0.281 | -0.818 | -0.609 | 0.005 | -0.013 | -0.013 | 0 | 0.16 | 0.13 | 0.08 | 0 | 0.11 | 0.12 |
| 4 | 182239040 | rs7654697 | 0.597 | 0.411 | -0.802 | -0.604 | 0.015 | 0.015 | 0.015 | 0 | 0.25 | 0.16 | 0.14 | 0 | 0.2 | 0.13 |
| 4 | 182247565 | rs1584937 | -0.194 | 0.582 | -0.734 | -0.488 | 0.001 | 0.055 | 0.055 | 0 | 0.35 | 0.26 | 0.21 | 0.13 | 0.35 | 0.24 |
| 4 | 182249501 | rs6552489 | 0.37 | 0.512 | -0.73 | -0.463 | 0.023 | 0.051 | 0.051 | 0 | 0.29 | 0.17 | 0.15 | 0.12 | 0.23 | 0.22 |
| 4 | 182256121 | rs1563219 | -1.358 | 0.404 | -0.476 | -0.424 | 0.016 | -0.015 | -0.015 | 0 | 0.69 | 0.6 | 0.61 | 0.78 | 0.63 | 0.68 |
| 4 | 182259455 | rs17071132 | 0.306 | 0.511 | -0.521 | -0.343 | 0.009 | 0.03 | 0.03 | 0 | 0.31 | 0.25 | 0.16 | 0.13 | 0.26 | 0.23 |
| 4 | 182262320 | rs11723033 | 1.177 | 0.588 | -0.529 | -0.162 | 0.001 | -0.029 | -0.029 | 0 | 0.08 | 0.15 | 0.06 | 0 | 0.09 | 0.01 |
| 4 | 182266682 | rs7675654 | -0.952 | 0.629 | -0.5 | -0.103 | 0 | 0.002 | 0.002 | 0 | 0.53 | 0.54 | 0.54 | 0.66 | 0.57 | 0.48 |
| 4 | 182271880 | rs7674604 | 0.866 | 0.321 | -0.436 | 0.584 | -0.001 | 0 | 0 | 0 | 0.08 | 0.09 | 0.17 | 0.15 | 0.11 | 0.07 |
| 4 | 182276030 | rs11729166 | -0.844 | -0.099 | -0.262 | 0.709 | 0.008 | -0.002 | -0.002 | 0.019 | 0.48 | 0.43 | 0.44 | 0.56 | 0.44 | 0.44 |
| 4 | 182278111 | rs2309460 | 0.15 | -0.246 | 0.083 | 0.733 | 0.014 | 0.008 | 0.008 | 0 | 0.6 | 0.37 | 0.65 | 0.64 | 0.68 | 0.66 |
| 4 | 182278636 | rs1987162 | 0.924 | -0.325 | 0.216 | 0.704 | -0.004 | 0 | 0 | 0 | 0.24 | 0.24 | 0.25 | 0.28 | 0.19 | 0.28 |
| 4 | 182279787 | rs17071170 | 1.174 | -0.287 | -0.01 | 0.466 | -0.012 | -0.013 | -0.013 | 0.05 | 0.41 | 0.35 | 0.45 | 0.45 | 0.45 | 0.54 |
| 4 | 182280205 | rs12647115 | 0.346 | -0.29 | -0.046 | 0.557 | -0.016 | -0.059 | -0.059 | 0 | 0.16 | 0.06 | 0.18 | 0.42 | 0.18 | 0.29 |
| 4 | 182280270 | rs4861945 | -0.184 | -0.299 | 0.036 | 0.522 | -0.017 | -0.02 | -0.02 | 0.069 | 0.84 | 0.66 | 0.91 | 0.9 | 0.87 | 0.94 |
| 4 | 182280855 | rs17185924 | 2.035 | -0.244 | -0.159 | 0.546 | -0.03 | -0.018 | -0.018 | 0 | 0.1 | 0.01 | 0.2 | 0.13 | 0.13 | 0.07 |
| 4 | 182280863 | rs4621476 | -0.018 | -0.033 | 0.193 | 0.644 | 0.029 | 0.048 | 0.048 | 0 | 0.5 | 0.37 | 0.57 | 0.64 | 0.57 | 0.64 |
| 4 | 182282929 | rs17071183 | 0.464 | -0.034 | 0.143 | 0.643 | 0.004 | 0.103 | 0.103 | 1.476 | 0.08 | 0.16 | 0.06 | 0.22 | 0.06 | 0.18 |
| 4 | 182283087 | rs4283709 | 0.908 | -0.091 | 0.265 | 0.622 | -0.012 | 0.001 | 0.001 | 0 | 0.73 | 0.59 | 0.78 | 0.72 | 0.75 | 0.84 |
| 4 | 182284224 | rs4861947 | 0.725 | -0.051 | 0.255 | 0.434 | 0.011 | 0.027 | 0.027 | 0.09 | 0.32 | 0.18 | 0.32 | 0.26 | 0.29 | 0.3 |
| 4 | 182284314 | rs6829606 | 0.2 | 0.013 | 0.113 | 0.485 | -0.001 | 0.007 | 0.007 | 1.33 | 0.17 | 0.15 | 0.18 | 0.4 | 0.18 | 0.28 |
| 4 | 182285215 | rs7657136 | 0.447 | -0.02 | 0.144 | 0.472 | -0.008 | 0.049 | 0.049 | 0.512 | 0.76 | 0.69 | 0.78 | 0.51 | 0.78 | 0.68 |
| 4 | 182285422 | rs6836888 | -0.575 | 0.025 | 0.045 | 0.487 | 0.006 | 0.032 | 0.032 | 0 | 0.5 | 0.39 | 0.51 | 0.67 | 0.48 | 0.59 |
| 4 | 182286222 | rs17071211 | 0.239 | 0.089 | -0.182 | 0.497 | -0.003 | 0.002 | 0.002 | 1.511 | 0.17 | 0.16 | 0.2 | 0.41 | 0.2 | 0.28 |
| 4 | 182287004 | rs7692027 | 0.203 | 0.097 | -0.203 | 0.1 | -0.012 | 0.034 | 0.034 | 0 | 0.41 | 0.34 | 0.43 | 0.24 | 0.46 | 0.35 |
| 4 | 182288767 | rs6552495 | 0.528 | -0.057 | 0.134 | 0.495 | -0.022 | 0.033 | 0.033 | 0 | 0.92 | 0.78 | 0.96 | 0.92 | 0.96 | 0.96 |
| 4 | 182290596 | rs12641677 | 0.091 | 0.072 | 0.052 | 0.327 | -0.005 | -0.016 | -0.016 | 0 | 0.91 | 0.79 | 0.91 | 0.98 | 0.92 | 0.98 |
| 4 | 182292227 | rs7682400 | -0.46 | -0.78 | -0.08 | -0.046 | -0.045 | -0.049 | -0.049 | 0 | 0.39 | 0.46 | 0.27 | 0.23 | 0.23 | 0.2 |
| 4 | 182292713 | rs10520479 | 1.127 | -0.15 | -1.179 | 0.27 | -0.125 | -0.129 | -0.129 | 0 | 0.25 | 0.04 | 0.32 | 0.32 | 0.33 | 0.34 |

| | | | | | | | | | | | | | | | | |
|---|---|---|---|---|---|---|---|---|---|---|---|---|---|---|---|---|
| 4 | 182294897 | rs17071237 | 1.011 | -0.892 | -1.164 | -0.004 | 0.043 | 0.014 | 0.014 | 0.083 | 0.19 | 0.05 | 0.17 | 0.13 | 0.11 | 0.09 |
| 4 | 182295274 | rs6834709 | 0.911 | -0.436 | -0.821 | 0.211 | -0.058 | -0.048 | -0.048 | 0.113 | 0.27 | 0.12 | 0.35 | 0.32 | 0.37 | 0.35 |
| 4 | 182295676 | rs10520478 | 0.372 | -0.548 | -0.423 | 0.136 | 0.043 | 0.04 | 0.04 | 0.522 | 0.36 | 0.23 | 0.23 | 0.2 | 0.19 | 0.18 |
| 4 | 182297666 | rs13145131 | 0.89 | -0.382 | -0.672 | 0.216 | -0.122 | -0.105 | -0.105 | 2.711 | 0.27 | 0.08 | 0.35 | 0.31 | 0.37 | 0.35 |
| 4 | 182300733 | rs2309472 | -0.508 | -0.099 | 0.022 | 0.347 | 0.004 | 0.015 | 0.015 | 0 | 0.66 | 0.47 | 0.64 | 0.52 | 0.62 | 0.53 |
| 4 | 182307648 | rs6838270 | -0.609 | -0.37 | 0.18 | 0.238 | 0.013 | 0.006 | 0.006 | 0 | 0.07 | 0.25 | 0.08 | 0.03 | 0.06 | 0.06 |
| 4 | 182311928 | rs4479746 | 0.353 | -1.015 | -0.455 | -0.028 | 0.097 | 0.079 | 0.079 | 0 | 0.56 | 0.28 | 0.66 | 0.71 | 0.75 | 0.73 |
| 4 | 182317147 | rs13129340 | 0.447 | -0.61 | -0.556 | 0.51 | 0.086 | 0 | 0 | 0.017 | 0.51 | 0.17 | 0.47 | 0.45 | 0.66 | 0.65 |
| 4 | 182318445 | rs1030790 | -0.529 | -0.862 | 0.037 | 0.488 | -0.036 | -0.002 | -0.002 | 0.455 | 0.78 | 0.7 | 0.79 | 0.71 | 0.88 | 0.8 |
| 4 | 182322805 | rs11132055 | -0.752 | -0.845 | -0.214 | 0.246 | -0.002 | -0.015 | -0.015 | 0 | 0.8 | 0.82 | 0.92 | 0.95 | 0.89 | 0.88 |
| 4 | 182327465 | rs17071292 | 0.303 | 0.082 | -0.926 | 0.428 | -0.066 | -0.074 | -0.074 | 0 | 0.24 | 0.06 | 0.32 | 0.27 | 0.29 | 0.28 |
| 4 | 182327838 | rs724658 | 0.878 | 0.084 | -0.942 | 0.337 | -0.047 | -0.081 | -0.081 | 0 | 0.18 | 0.06 | 0.26 | 0.27 | 0.21 | 0.26 |
| 4 | 182327868 | rs724657 | -0.523 | 0.017 | -1.467 | 0.13 | -0.047 | -0.083 | -0.083 | 0 | 0.34 | 0.1 | 0.36 | 0.34 | 0.36 | 0.52 |
| 4 | 182327943 | rs724659 | 0.605 | -0.252 | -0.706 | 0.214 | -0.02 | 0.025 | 0.025 | 0 | 0.37 | 0.33 | 0.38 | 0.23 | 0.46 | 0.28 |
| 4 | 182329791 | rs6857563 | -0.56 | -0.335 | -0.385 | 0.218 | -0.001 | -0.001 | -0.001 | 0 | 0.6 | 0.38 | 0.51 | 0.55 | 0.58 | 0.66 |
| 4 | 182329856 | rs6857599 | 0.36 | -0.245 | -0.246 | 0.351 | 0.001 | 0.003 | 0.003 | 0 | 0.1 | 0.04 | 0.07 | 0.06 | 0.11 | 0.2 |
| 4 | 182338549 | rs7698202 | 0.825 | -0.135 | 0.043 | 0.411 | -0.02 | 0.06 | 0.06 | 0 | 0.9 | 0.74 | 0.94 | 0.81 | 0.93 | 0.79 |
| 4 | 182342069 | rs6552500 | 1.407 | 0.365 | 0.504 | 0.446 | 0.046 | 0.097 | 0.097 | 0 | 0.81 | 0.52 | 0.75 | 0.66 | 0.76 | 0.74 |
| 4 | 182342394 | rs1369479 | 0.085 | 0.314 | 0.412 | 0.472 | -0.019 | 0.028 | 0.028 | 0 | 0.9 | 0.83 | 0.95 | 0.81 | 0.93 | 0.8 |
| 4 | 182345272 | rs7692661 | 0.715 | 0.062 | 0.135 | 0.372 | 0.021 | 0.005 | 0.005 | 0 | 0.28 | 0.18 | 0.25 | 0.3 | 0.19 | 0.17 |
| 4 | 182347670 | rs4861475 | 0.025 | 0.241 | -0.032 | 0.252 | -0.005 | 0.047 | 0.047 | 0 | 0.46 | 0.69 | 0.45 | 0.35 | 0.53 | 0.34 |
| 4 | 182347745 | rs17195559 | 1.541 | 0.199 | -0.057 | 0.256 | -0.005 | -0.011 | -0.011 | 0 | 0.07 | 0.02 | 0.05 | 0.14 | 0.08 | 0.04 |
| 4 | 182348386 | rs13112319 | 0.017 | 0.382 | -0.086 | 0.234 | 0.032 | 0.048 | 0.048 | 0 | 0.52 | 0.24 | 0.59 | 0.59 | 0.57 | 0.51 |
| 4 | 182350247 | rs892862 | -0.489 | 0.375 | -0.078 | 0.156 | 0 | 0.002 | 0.002 | 0 | 0.64 | 0.7 | 0.68 | 0.63 | 0.66 | 0.6 |
| 4 | 182350796 | rs4549433 | -0.388 | 0.239 | 0.4 | -0.007 | -0.001 | -0.031 | -0.031 | 2.485 | 0.58 | 0.48 | 0.72 | 0.87 | 0.6 | 0.52 |
| 4 | 182353549 | rs17195902 | 0.899 | 0.298 | -0.05 | 0.068 | -0.031 | 0.033 | 0.033 | 0 | 0.12 | 0.01 | 0.17 | 0.06 | 0.15 | 0.13 |
| 4 | 182354145 | rs6827352 | 1.125 | 0.271 | 0.428 | 0.096 | -0.001 | -0.028 | -0.028 | 0.001 | 0.3 | 0.29 | 0.18 | 0.07 | 0.28 | 0.24 |
| 4 | 182362615 | rs6828912 | 1.174 | 0.209 | 0.446 | 0.096 | -0.001 | -0.015 | -0.015 | 0 | 0.31 | 0.29 | 0.21 | 0.07 | 0.32 | 0.24 |
| 4 | 182363381 | rs11723828 | 0.597 | 0.173 | 0.344 | 0.366 | -0.033 | -0.05 | -0.05 | 0 | 0.24 | 0.1 | 0.32 | 0.27 | 0.29 | 0.32 |
| 4 | 182366956 | rs17071491 | 1.257 | 0.095 | 0.543 | 0.415 | 0.04 | -0.039 | -0.039 | 4.145 | 0.25 | 0.11 | 0.2 | 0.45 | 0.12 | 0.18 |
| 4 | 182367808 | rs1991989 | 0.012 | 0.235 | 0.689 | 0.396 | -0.008 | -0.009 | -0.009 | 0 | 0.66 | 0.66 | 0.53 | 0.53 | 0.58 | 0.52 |
| 4 | 182370185 | rs10002388 | 0.743 | 0.245 | 0.464 | 0.388 | -0.007 | -0.002 | -0.002 | 0 | 0.1 | 0.08 | 0.1 | 0.01 | 0.13 | 0.05 |
| 4 | 182371433 | rs17071527 | 1.668 | -0.129 | 0.446 | 0.469 | 0.035 | -0.017 | -0.017 | 0 | 0.22 | 0.1 | 0.2 | 0.31 | 0.12 | 0.17 |
| 4 | 182372333 | rs4861960 | -0.042 | -0.055 | 0.77 | 0.487 | 0.077 | 0.053 | 0.053 | 0 | 0.08 | 0.24 | 0.2 | 0.14 | 0.17 | 0.06 |
| 4 | 182374325 | rs4861962 | -0.441 | -0.159 | 0.52 | 0.497 | -0.037 | -0.084 | -0.084 | 0 | 0.73 | 0.87 | 0.68 | 0.67 | 0.69 | 0.56 |
| 4 | 182376357 | rs1435499 | 0.842 | -0.241 | 0.716 | 0.441 | -0.022 | 0.03 | 0.03 | 0.869 | 0.36 | 0.3 | 0.37 | 0.21 | 0.49 | 0.34 |
| 4 | 182377628 | rs10027214 | -0.555 | -0.169 | 0.256 | 0.5 | -0.014 | -0.084 | -0.084 | 0 | 0.73 | 0.87 | 0.71 | 0.67 | 0.74 | 0.57 |
| 4 | 182383645 | rs1435501 | -0.259 | -0.178 | 0.295 | 0.414 | 0.06 | 0.09 | 0.09 | 0 | 0.52 | 0.76 | 0.51 | 0.36 | 0.62 | 0.4 |
| 4 | 182384993 | rs7676405 | -0.305 | -0.065 | 0.228 | 0.573 | -0.012 | -0.066 | -0.066 | 0.245 | 0.72 | 0.87 | 0.58 | 0.43 | 0.73 | 0.49 |
| 4 | 182387603 | rs7695194 | -0.019 | -0.26 | 0.471 | 0.635 | 0.03 | -0.009 | -0.009 | 0 | 0.61 | 0.68 | 0.68 | 0.48 | 0.74 | 0.64 |
| 4 | 182391631 | rs1991988 | 0.025 | -0.267 | 0.598 | 0.632 | -0.012 | 0.001 | 0.001 | 0 | 0.6 | 0.56 | 0.67 | 0.47 | 0.74 | 0.62 |
| 4 | 182391672 | rs1991987 | 0.672 | -0.296 | 0.615 | 0.632 | -0.042 | 0.092 | 0.092 | 0 | 0.89 | 0.72 | 0.84 | 0.67 | 0.97 | 0.74 |
| 4 | 182391939 | rs28437883 | 0.768 | -0.189 | 0.363 | 0.563 | 0.02 | 0.036 | 0.036 | 0 | 0.29 | 0.16 | 0.17 | 0.19 | 0.23 | 0.13 |
| 4 | 182396586 | rs7687690 | 1.129 | -0.136 | 0.427 | 0.491 | 0 | 0.029 | 0.029 | 0.101 | 0.32 | 0.23 | 0.18 | 0.08 | 0.3 | 0.26 |
| 4 | 182398593 | rs4861967 | 0.426 | -0.326 | 0.597 | 0.59 | -0.002 | -0.003 | -0.003 | 0.118 | 0.41 | 0.57 | 0.35 | 0.33 | 0.32 | 0.39 |
| 4 | 182399556 | rs981847 | 0.505 | -0.243 | 0.115 | 0.625 | -0.133 | -0.142 | -0.142 | 1.11 | 0.25 | 0.05 | 0.47 | 0.4 | 0.44 | 0.35 |
| 4 | 182403395 | rs17251860 | 1.575 | -0.313 | 0.1 | 0.571 | -0.024 | 0.015 | 0.015 | 0 | 0.06 | 0 | 0.04 | 0.04 | 0.08 | 0.03 |
| 4 | 182403824 | rs1561000 | 0.02 | -0.071 | 0.39 | 0.757 | 0.036 | 0.036 | 0.036 | 0.325 | 0.52 | 0.24 | 0.6 | 0.56 | 0.62 | 0.44 |
| 4 | 182405735 | rs6822070 | 0.722 | 0.243 | -0.102 | 0.687 | -0.009 | 0.016 | 0.016 | 0.578 | 0.35 | 0.25 | 0.34 | 0.33 | 0.36 | 0.37 |
| 4 | 182407529 | rs17252128 | 1.491 | 0.057 | -0.384 | 0.456 | 0.044 | 0.059 | 0.059 | 0 | 0.2 | 0.11 | 0.08 | 0.07 | 0.11 | 0.06 |
| 4 | 182407681 | rs4861970 | 0.283 | 0.582 | -0.008 | 0.674 | 0 | -0.004 | -0.004 | 0 | 0.39 | 0.39 | 0.35 | 0.25 | 0.44 | 0.35 |
| 4 | 182410954 | rs12649512 | 0.532 | 0.277 | -0.288 | 0.671 | -0.003 | 0.029 | 0.029 | 0.015 | 0.08 | 0.13 | 0.05 | 0.15 | 0.09 | 0.23 |
| 4 | 182411580 | rs4861477 | 1.112 | 0.419 | -0.65 | 0.705 | -0.013 | 0.021 | 0.021 | 0 | 0.29 | 0.21 | 0.32 | 0.1 | 0.33 | 0.26 |
| 4 | 182412992 | rs10520489 | 0.101 | 0.047 | -0.249 | 1.224 | 0 | -0.01 | -0.01 | 1.527 | 0.62 | 0.65 | 0.51 | 0.47 | 0.59 | 0.71 |
| 4 | 182415937 | rs4861973 | 0.11 | 0.044 | -0.25 | 1.223 | 0 | -0.01 | -0.01 | 0 | 0.62 | 0.65 | 0.51 | 0.47 | 0.59 | 0.72 |
| 4 | 182416097 | rs4861975 | 0.537 | 0.447 | -0.365 | 0.854 | 0.007 | 0.031 | 0.031 | 0 | 0.47 | 0.31 | 0.43 | 0.32 | 0.5 | 0.44 |
| 4 | 182420303 | rs10006195 | 0.603 | 1.062 | -0.079 | 1.164 | 0.016 | 0.009 | 0.009 | 0 | 0.9 | 0.79 | 0.88 | 0.92 | 0.88 | 0.96 |
| 4 | 182421815 | rs7654623 | 0.798 | 1.054 | -0.222 | 1.447 | 0.012 | 0.111 | 0.111 | 0 | 0.73 | 0.48 | 0.55 | 0.49 | 0.7 | 0.61 |
| 4 | 182422900 | rs7661191 | -0.342 | 1.498 | -0.851 | 1.7 | -0.018 | -0.043 | -0.043 | 0 | 0.17 | 0.07 | 0.15 | 0.41 | 0.19 | 0.26 |
| 4 | 182426033 | rs930308 | 0.049 | 1.611 | -1.126 | 1.224 | 0.004 | 0.003 | 0.003 | 0.014 | 0.68 | 0.63 | 0.49 | 0.53 | 0.66 | 0.64 |
| 4 | 182429303 | rs11731430 | -0.026 | 1.698 | -1.163 | 1.222 | 0.015 | 0.019 | 0.019 | 0 | 0.78 | 0.69 | 0.6 | 0.62 | 0.76 | 0.83 |
| 4 | 182431113 | rs920083 | -0.304 | 1.742 | -0.813 | 1.213 | 0.005 | 0.014 | 0.014 | 0 | 0.81 | 0.74 | 0.62 | 0.64 | 0.81 | 0.83 |
| 4 | 182432295 | rs13107304 | 0.429 | 1.934 | -0.445 | 0.415 | 0.029 | -0.087 | -0.087 | 0 | 0.14 | 0.33 | 0.12 | 0.01 | 0.17 | 0.04 |
| 4 | 182433238 | rs4861977 | 0.788 | 0.905 | -0.872 | 0.128 | -0.036 | 0.024 | 0.024 | 0 | 0.22 | 0.27 | 0.08 | 0.27 | 0.12 | 0.33 |
| 4 | 182433814 | rs10520484 | 1.855 | 0.74 | -1.401 | 0.192 | -0.072 | 0.022 | 0.022 | 0 | 0.23 | 0.02 | 0.15 | 0.1 | 0.28 | 0.19 |
| 4 | 182434469 | rs1435497 | -0.2 | 1.011 | -1.058 | 0.316 | 0.006 | -0.007 | -0.007 | 0.005 | 0.34 | 0.43 | 0.36 | 0.27 | 0.36 | 0.3 |
| 4 | 182438604 | rs17071683 | 0.492 | 0.742 | -0.922 | 0.42 | 0.016 | 0.019 | 0.019 | 0.029 | 0.51 | 0.27 | 0.33 | 0.35 | 0.49 | 0.67 |
| 4 | 182444131 | rs11939305 | 1.31 | 0.876 | -0.982 | 0.554 | -0.01 | -0.022 | -0.022 | 0.31 | 0.15 | 0.06 | 0.11 | 0.2 | 0.14 | 0.13 |
| 4 | 182447323 | rs974578 | -1.011 | 0.836 | -1.119 | 0.472 | 0.004 | 0.003 | 0.003 | 0 | 0.78 | 0.76 | 0.58 | 0.65 | 0.73 | 0.86 |

| | | | | | | | | | | | | | | | | |
|---|---|---|---|---|---|---|---|---|---|---|---|---|---|---|---|---|
| 4 | 182450706 | rs1439276 | 1.393 | 0.791 | -1.242 | 0.562 | -0.023 | -0.043 | -0.043 | 0 | 0.14 | 0.04 | 0.1 | 0.2 | 0.14 | 0.13 |
| 4 | 182453413 | rs4861981 | 0.897 | 0.602 | -1.35 | 0.27 | -0.017 | -0.016 | -0.016 | 0 | 0.37 | 0.1 | 0.21 | 0.26 | 0.4 | 0.48 |
| 4 | 182458937 | rs17071724 | 0.082 | 0.185 | -0.751 | 0.292 | 0.017 | -0.005 | -0.005 | 0 | 0.3 | 0.14 | 0.21 | 0.29 | 0.19 | 0.36 |
| 4 | 182459990 | rs12650613 | 0.641 | 0.106 | -0.163 | 0.513 | -0.037 | 0.065 | 0.065 | 0.002 | 0.14 | 0.34 | 0.37 | 0.29 | 0.08 | 0.06 |
| 4 | 182461424 | rs7680020 | -0.579 | 0.416 | 0.03 | 0.242 | 0.054 | 0.004 | 0.004 | 0 | 0.52 | 0.72 | 0.64 | 0.59 | 0.33 | 0.45 |
| 4 | 182461858 | rs7663547 | -1.284 | 0.505 | -0.108 | 0.243 | -0.024 | -0.025 | -0.025 | 0 | 0.83 | 0.89 | 0.84 | 0.68 | 0.7 | 0.68 |
| 4 | 182463503 | rs7697865 | 1.546 | 0.482 | 0.175 | 0.262 | -0.019 | -0.02 | -0.02 | 0 | 0.06 | 0.13 | 0.03 | 0.01 | 0.01 | 0.02 |
| 4 | 182463971 | rs9998972 | 0.961 | 0.444 | 0.397 | 0.242 | -0.003 | -0.007 | -0.007 | 0 | 0.08 | 0.18 | 0.16 | 0.09 | 0.06 | 0.17 |
| 4 | 182465917 | rs10005266 | -0.561 | 0.503 | 0.136 | 0.11 | -0.052 | -0.045 | -0.045 | 0 | 0.18 | 0.38 | 0.15 | 0.11 | 0.09 | 0.21 |
| 4 | 182467304 | rs11132072 | 0.376 | -0.074 | -0.22 | 0.128 | -0.016 | -0.007 | -0.007 | 0 | 0.11 | 0.17 | 0.11 | 0.1 | 0.07 | 0.19 |
| 4 | 182467989 | rs17071811 | -0.362 | 0.301 | -0.842 | 0.206 | 0.018 | -0.007 | -0.007 | 0 | 0.54 | 0.66 | 0.66 | 0.59 | 0.38 | 0.46 |
| 4 | 182470191 | rs11132079 | 0.121 | 0.304 | -0.647 | 0.284 | -0.014 | 0.005 | 0.005 | 0 | 0.65 | 0.48 | 0.69 | 0.64 | 0.67 | 0.55 |
| 4 | 182470210 | rs17071826 | 0.197 | 0.226 | -0.666 | -0.019 | 0.003 | -0.003 | -0.003 | 0 | 0.1 | 0.12 | 0.17 | 0.14 | 0.09 | 0.14 |
| 4 | 182470792 | rs11944460 | -0.563 | -0.609 | -0.654 | -0.31 | 0.009 | 0.008 | 0.008 | 0.028 | 0.42 | 0.58 | 0.29 | 0.29 | 0.27 | 0.35 |
| 4 | 182471822 | rs13140378 | -0.042 | -0.493 | -0.249 | -0.335 | -0.023 | -0.03 | -0.03 | 0 | 0.22 | 0.35 | 0.11 | 0.17 | 0.14 | 0.24 |
| 4 | 182472171 | rs4861482 | -1.263 | -0.688 | -0.497 | -0.739 | -0.025 | -0.064 | -0.064 | 0 | 0.4 | 0.44 | 0.22 | 0.17 | 0.29 | 0.29 |
| 4 | 182472791 | rs4362864 | -0.135 | -0.679 | -0.572 | -0.723 | 0.022 | 0.017 | 0.017 | 0 | 0.2 | 0.09 | 0.1 | 0 | 0.15 | 0.12 |
| 4 | 182473726 | rs13103500 | -0.641 | -0.482 | -0.426 | -0.605 | -0.005 | -0.02 | -0.02 | 0.219 | 0.37 | 0.38 | 0.2 | 0.17 | 0.28 | 0.21 |
| 4 | 182473823 | rs13103727 | -0.908 | -0.468 | -0.468 | -0.597 | -0.026 | -0.065 | -0.065 | 0 | 0.39 | 0.44 | 0.22 | 0.17 | 0.28 | 0.26 |
| 4 | 182477095 | rs11934219 | 0.326 | -0.602 | -0.282 | -0.558 | -0.013 | -0.058 | -0.058 | 1.608 | 0.15 | 0.16 | 0.09 | 0 | 0.13 | 0.05 |
| 4 | 182477386 | rs2043893 | 0.051 | 0.257 | -0.438 | -0.037 | -0.002 | -0.003 | -0.003 | 0 | 0.66 | 0.61 | 0.56 | 0.39 | 0.62 | 0.43 |
| 4 | 182480385 | rs1025758 | 0.466 | -0.407 | -0.306 | -0.345 | 0.023 | 0.015 | 0.015 | 0 | 0.2 | 0.14 | 0.09 | 0.17 | 0.1 | 0.16 |
| 4 | 182482693 | rs2309504 | -0.236 | -0.823 | -0.463 | -0.598 | 0.003 | 0.006 | 0.006 | 0 | 0.5 | 0.49 | 0.37 | 0.36 | 0.41 | 0.43 |
| 4 | 182482973 | rs7440035 | -0.526 | -1.034 | -0.527 | -0.723 | -0.043 | -0.064 | -0.064 | 0.261 | 0.4 | 0.47 | 0.33 | 0.19 | 0.25 | 0.27 |
| 4 | 182484769 | rs1439283 | 0.018 | -0.708 | -0.778 | -0.524 | 0.024 | 0.083 | 0.083 | 0 | 0.46 | 0.66 | 0.49 | 0.2 | 0.44 | 0.26 |
| 4 | 182487135 | rs6845687 | 0.014 | -1.059 | -0.264 | -0.284 | -0.001 | -0.002 | -0.002 | 0 | 0.22 | 0.23 | 0.11 | 0.15 | 0.14 | 0.21 |
| 4 | 182490447 | rs6845256 | -0.022 | -1.121 | 0.102 | -0.159 | -0.044 | -0.024 | -0.024 | 0.564 | 0.22 | 0.36 | 0.13 | 0.21 | 0.13 | 0.16 |
| 4 | 182490475 | rs6845276 | 0.28 | -1.029 | -0.438 | -0.114 | 0.004 | 0.003 | 0.003 | 0 | 0.21 | 0.21 | 0.11 | 0.21 | 0.12 | 0.16 |
| 4 | 182493718 | rs4861485 | -0.339 | -0.506 | -0.286 | 0.201 | -0.031 | -0.011 | -0.011 | 0 | 0.26 | 0.52 | 0.24 | 0.22 | 0.23 | 0.38 |
| 4 | 182507266 | rs12511842 | 0.956 | -0.448 | -0.799 | 0.169 | 0.003 | 0.003 | 0.003 | 0 | 0.15 | 0.1 | 0.13 | 0.07 | 0.11 | 0.07 |
| 4 | 182514392 | rs10520496 | 0.638 | -0.052 | -1.358 | 0.444 | -0.1 | -0.073 | -0.073 | 0 | 0.23 | 0.1 | 0.4 | 0.41 | 0.4 | 0.21 |
| 4 | 182520502 | rs6849949 | -0.452 | -0.932 | -1.44 | 0.12 | 0.027 | 0.01 | 0.01 | 0.382 | 0.66 | 0.28 | 0.68 | 0.75 | 0.72 | 0.64 |
| 4 | 182522272 | rs11132083 | -0.359 | -0.576 | -0.813 | 0.231 | -0.001 | 0.005 | 0.005 | 0 | 0.42 | 0.45 | 0.55 | 0.51 | 0.52 | 0.47 |
| 4 | 182523766 | rs17210704 | 0.059 | -0.154 | -1.119 | 0.342 | -0.001 | -0.011 | -0.011 | 1.973 | 0.06 | 0.11 | 0.08 | 0.03 | 0.05 | 0.04 |
| 4 | 182528430 | rs1439279 | 0.746 | 0.029 | -1.338 | 0.547 | -0.092 | -0.065 | -0.065 | 0 | 0.22 | 0.1 | 0.4 | 0.42 | 0.4 | 0.23 |
| 4 | 182537718 | rs9799857 | -0.241 | -0.868 | -0.889 | 0.209 | 0.036 | 0.034 | 0.034 | 0 | 0.58 | 0.63 | 0.66 | 0.72 | 0.65 | 0.53 |
| 4 | 182538969 | rs17071911 | 0.439 | -1.089 | -0.38 | 0.606 | 0.007 | -0.011 | -0.011 | 0 | 0.42 | 0.36 | 0.57 | 0.55 | 0.55 | 0.36 |
| 4 | 182539137 | rs17071912 | 0.769 | -1.124 | -0.193 | 0.598 | -0.002 | 0.013 | 0.013 | 0 | 0.14 | 0.13 | 0.07 | 0.18 | 0.1 | 0.21 |
| 4 | 182543945 | rs10002408 | -1.435 | -1.904 | -0.351 | 0.162 | -0.021 | -0.05 | -0.05 | 0 | 0.83 | 0.77 | 0.92 | 0.98 | 0.89 | 0.87 |
| 4 | 182544354 | rs11721527 | 1.154 | -1.885 | -0.379 | 0.199 | -0.007 | -0.04 | -0.04 | 0 | 0.13 | 0.05 | 0.06 | 0.17 | 0.1 | 0.2 |
| 4 | 182544544 | rs745704 | 1.176 | -1.948 | -0.443 | 0.204 | 0.033 | 0.003 | 0.003 | 0 | 0.23 | 0.13 | 0.13 | 0.21 | 0.19 | 0.17 |
| 4 | 182544594 | rs745705 | 0.809 | -1.423 | -0.29 | 0.48 | 0.001 | -0.012 | -0.012 | 0 | 0.4 | 0.35 | 0.55 | 0.53 | 0.53 | 0.34 |
| 4 | 182548076 | rs715745 | -0.132 | -1.132 | -0.3 | 0.521 | -0.01 | 0.028 | 0.028 | 0 | 0.17 | 0.24 | 0.16 | 0.21 | 0.12 | 0.21 |
| 4 | 182553043 | rs6815916 | 1.069 | -1.045 | -0.636 | 0.558 | -0.001 | 0.014 | 0.014 | 0 | 0.09 | 0.1 | 0.05 | 0.13 | 0.07 | 0.12 |
| 4 | 182564825 | rs17211911 | 1.693 | -1.03 | -1.006 | 0.555 | -0.011 | -0.048 | -0.048 | 0 | 0.15 | 0.01 | 0.1 | 0.17 | 0.12 | 0.07 |
| 4 | 182565693 | rs7662677 | 0.079 | -1.307 | -0.981 | 0.533 | -0.003 | 0 | 0 | 0 | 0.41 | 0.41 | 0.56 | 0.44 | 0.52 | 0.44 |
| 4 | 182566129 | rs1971822 | -0.822 | -1.349 | -0.955 | 0.49 | 0.007 | 0.003 | 0.003 | 0 | 0.59 | 0.59 | 0.71 | 0.64 | 0.68 | 0.61 |
| 4 | 182569100 | rs17212220 | -0.276 | 0.232 | -1.411 | 0.241 | 0.008 | -0.003 | -0.003 | 0.352 | 0.25 | 0.24 | 0.2 | 0.12 | 0.36 | 0.27 |
| 4 | 182571325 | rs17212276 | 1.609 | 0.104 | -1.632 | 0.362 | -0.024 | -0.011 | -0.011 | 0.003 | 0.11 | 0.01 | 0.05 | 0.08 | 0.1 | 0.15 |
| 4 | 182572102 | rs10520501 | -0.528 | -0.03 | -1.67 | 0.419 | -0.002 | -0.028 | -0.028 | 0 | 0.36 | 0.2 | 0.36 | 0.38 | 0.32 | 0.47 |
| 4 | 182572180 | rs11727308 | 1.061 | 0.272 | -1.486 | 0.315 | -0.023 | -0.009 | -0.009 | 0 | 0.11 | 0.01 | 0.05 | 0.08 | 0.1 | 0.16 |
| 4 | 182576586 | rs6552517 | -0.557 | 0.029 | -1.245 | 0.319 | 0.055 | -0.004 | -0.004 | 0 | 0.47 | 0.32 | 0.57 | 0.49 | 0.59 | 0.65 |
| 4 | 182578680 | rs4590086 | -0.182 | 0.264 | -1.309 | -0.019 | -0.041 | 0.038 | 0.038 | 0 | 0.23 | 0.15 | 0.15 | 0.11 | 0.31 | 0.17 |
| 4 | 182581686 | rs6856771 | 0.432 | -0.4 | -1.29 | 0.122 | -0.006 | 0.003 | 0.003 | 0.244 | 0.43 | 0.57 | 0.58 | 0.44 | 0.42 | 0.42 |
| 4 | 182584177 | rs4861488 | -0.096 | -1.182 | -1.4 | -0.101 | -0.011 | 0.01 | 0.01 | 3.781 | 0.63 | 0.53 | 0.54 | 0.46 | 0.65 | 0.58 |
| 4 | 182584331 | rs1439272 | 0.83 | -1.175 | -1.177 | -0.056 | -0.011 | 0.04 | 0.04 | 0 | 0.11 | 0.29 | 0.2 | 0.12 | 0.1 | 0.1 |
| 4 | 182585855 | rs6835945 | 1.201 | -0.838 | -1.233 | -0.238 | 0.044 | -0.026 | -0.026 | 0 | 0.24 | 0.37 | 0.3 | 0.12 | 0.35 | 0.3 |
| 4 | 182588504 | rs1439273 | 1.497 | -0.991 | -1.284 | -0.291 | -0.013 | -0.016 | -0.016 | 0 | 0.26 | 0.35 | 0.23 | 0.2 | 0.23 | 0.14 |
| 4 | 182589046 | rs10471178 | -0.793 | -1.11 | -1.386 | -0.326 | 0.006 | -0.069 | -0.069 | 0.505 | 0.68 | 0.84 | 0.59 | 0.46 | 0.67 | 0.58 |
| 4 | 182591864 | rs13144599 | -0.402 | -1.007 | -1.372 | -0.292 | 0.002 | -0.014 | -0.014 | 0 | 0.44 | 0.5 | 0.36 | 0.26 | 0.46 | 0.44 |
| 4 | 182592079 | rs1439281 | 1.113 | -0.695 | -1.466 | -0.233 | -0.001 | -0.001 | -0.001 | 0 | 0.13 | 0.15 | 0.06 | 0.13 | 0.09 | 0.14 |
| 4 | 182592541 | rs17279486 | 1.312 | -0.372 | -1.459 | -0.131 | -0.002 | 0.001 | 0.001 | 0 | 0.23 | 0.22 | 0.28 | 0.13 | 0.35 | 0.28 |
| 4 | 182594449 | rs13133342 | -0.185 | -0.776 | -1.614 | 0.267 | 0.026 | -0.023 | -0.023 | 0.044 | 0.58 | 0.77 | 0.8 | 0.47 | 0.67 | 0.54 |
| 4 | 182594688 | rs4861998 | 0.428 | -0.742 | -1.532 | 0.177 | 0.027 | 0.048 | 0.048 | 0.021 | 0.53 | 0.77 | 0.61 | 0.34 | 0.59 | 0.44 |
| 4 | 182597939 | rs17072040 | 1.127 | -0.635 | -1.194 | 0.553 | -0.012 | -0.029 | -0.029 | 0 | 0.09 | 0 | 0.04 | 0.12 | 0.08 | 0.04 |
| 4 | 182598039 | rs6847130 | -0.433 | -0.622 | -1.131 | 0.81 | 0 | 0.025 | 0.025 | 0 | 0.78 | 0.72 | 0.79 | 0.66 | 0.82 | 0.69 |
| 4 | 182602940 | rs6844975 | -0.029 | -0.96 | -1.183 | 0.606 | -0.031 | -0.012 | -0.012 | 0.107 | 0.34 | 0.5 | 0.29 | 0.34 | 0.26 | 0.18 |
| 4 | 182606762 | rs7666018 | -0.863 | -0.7 | -1.333 | 0.495 | 0.009 | 0.038 | 0.038 | 0 | 0.81 | 0.72 | 0.88 | 0.71 | 0.83 | 0.75 |
| 4 | 182611537 | rs4527522 | 1.48 | -0.829 | -1.431 | 0.476 | -0.124 | 0.114 | 0.114 | 0.635 | 0.39 | 0.08 | 0.28 | 0.19 | 0.5 | 0.46 |

| | | | | | | | | | | | | | | | | |
|---|---|---|---|---|---|---|---|---|---|---|---|---|---|---|---|---|
| 4 | 182612164 | rs12503151 | -0.553 | -0.141 | -0.814 | 0.903 | 0.01 | 0.041 | 0.041 | 0 | 0.81 | 0.72 | 0.88 | 0.71 | 0.83 | 0.74 |
| 4 | 182614559 | rs2309573 | 0.781 | -0.059 | -1.293 | 0.854 | -0.009 | 0.089 | 0.089 | 0.062 | 0.52 | 0.11 | 0.36 | 0.34 | 0.57 | 0.55 |
| 4 | 182616401 | rs17072125 | 0.056 | 0.22 | -0.903 | 0.891 | 0 | 0 | 0 | 0.02 | 0.28 | 0.3 | 0.53 | 0.4 | 0.26 | 0.2 |
| 4 | 182621264 | rs7689930 | 0.056 | 0.572 | -0.608 | 0.452 | 0.025 | 0.052 | 0.052 | 0 | 0.79 | 0.69 | 0.86 | 0.68 | 0.78 | 0.73 |
| 4 | 182621558 | rs17072140 | 1.07 | 1.216 | -1.113 | 0.315 | -0.064 | -0.099 | -0.099 | 0.1 | 0.16 | 0.01 | 0.26 | 0.36 | 0.19 | 0.12 |
| 4 | 182622759 | rs17280931 | 1.15 | 1.06 | -0.819 | -0.774 | -0.035 | 0.006 | 0.006 | 0 | 0.41 | 0.19 | 0.55 | 0.62 | 0.49 | 0.43 |
| 4 | 182626599 | rs10520505 | -1.796 | 1.296 | -0.489 | -0.826 | 0.015 | -0.003 | -0.003 | 0 | 0.93 | 0.84 | 0.94 | 0.94 | 0.92 | 0.93 |
| 4 | 182626706 | rs10520506 | -0.489 | 0.809 | 0.065 | -1.162 | 0 | 0.001 | 0.001 | 0 | 0.36 | 0.34 | 0.31 | 0.33 | 0.35 | 0.43 |
| 4 | 182631262 | rs2309534 | -1.884 | 0.949 | -0.014 | -1.095 | 0.015 | -0.004 | -0.004 | 0.256 | 0.93 | 0.84 | 0.94 | 0.94 | 0.92 | 0.93 |
| 4 | 182632276 | rs17215232 | 2.583 | 1.274 | -0.196 | -1.2 | -0.031 | 0.029 | 0.029 | 0 | 0.06 | 0.01 | 0.07 | 0 | 0.17 | 0.04 |
| 4 | 182635289 | rs17281245 | 0.809 | 0.024 | 0.421 | -0.948 | -0.001 | -0.023 | -0.023 | 0.154 | 0.13 | 0.07 | 0.03 | 0 | 0.07 | 0.08 |
| 4 | 182637590 | rs13143181 | -1.191 | -0.129 | 0.545 | -1.054 | -0.037 | -0.062 | -0.062 | 0 | 0.27 | 0.4 | 0.15 | 0.12 | 0.18 | 0.18 |
| 4 | 182639404 | rs17215414 | 1.639 | 0.025 | 0.121 | -1.071 | -0.03 | 0.038 | 0.038 | 0 | 0.07 | 0 | 0.02 | 0 | 0.08 | 0.03 |
| 4 | 182641214 | rs12502114 | -0.05 | 0.042 | 0.288 | -0.945 | -0.022 | 0.001 | 0.001 | 0.173 | 0.7 | 0.74 | 0.71 | 0.73 | 0.66 | 0.61 |
| 4 | 182643207 | rs2309541 | 1.196 | -0.006 | 0.443 | -0.982 | 0.002 | -0.034 | -0.034 | 0 | 0.72 | 0.4 | 0.81 | 0.81 | 0.77 | 0.76 |
| 4 | 182646450 | rs13113072 | -0.081 | -0.031 | 0.361 | -0.988 | 0.012 | 0.027 | 0.027 | 0 | 0.3 | 0.19 | 0.26 | 0.19 | 0.27 | 0.31 |
| 4 | 182648949 | rs2309544 | 1.401 | 0.075 | 0.436 | -0.904 | 0.021 | 0.047 | 0.047 | 0 | 0.64 | 0.31 | 0.78 | 0.82 | 0.74 | 0.75 |
| 4 | 182649354 | rs2309545 | -0.243 | 0.394 | 0.61 | -0.722 | 0.072 | 0.027 | 0.027 | 1.656 | 0.87 | 0.65 | 0.88 | 0.81 | 0.83 | 0.82 |
| 4 | 182655999 | rs17072184 | 0.581 | 0.44 | -0.08 | -0.59 | -0.029 | -0.009 | -0.009 | 0 | 0.23 | 0.14 | 0.24 | 0.2 | 0.26 | 0.31 |
| 4 | 182662286 | rs11934518 | 0.466 | 0.325 | -0.018 | -0.631 | -0.012 | -0.018 | -0.018 | 0.23 | 0.18 | 0.26 | 0.11 | 0.12 | 0.13 | 0.11 |
| 4 | 182669719 | rs6855392 | 1.567 | 0.496 | -0.357 | -0.55 | -0.026 | -0.021 | -0.021 | 4.855 | 0.38 | 0.2 | 0.51 | 0.6 | 0.44 | 0.42 |
| 4 | 182669892 | rs6552521 | -0.828 | 1.292 | -0.371 | -0.019 | 0.02 | -0.003 | -0.003 | 0 | 0.92 | 0.9 | 0.94 | 0.93 | 0.91 | 0.94 |
| 4 | 182673224 | rs2309550 | -1.001 | 0.975 | -0.771 | -0.01 | 0.001 | 0.001 | 0.001 | 0 | 0.42 | 0.49 | 0.34 | 0.33 | 0.35 | 0.41 |
| 4 | 182677604 | rs7656889 | -0.915 | 0.849 | -1.243 | 0.194 | -0.002 | 0.008 | 0.008 | 0 | 0.4 | 0.41 | 0.35 | 0.33 | 0.39 | 0.42 |
| 4 | 182686437 | rs10520510 | -0.962 | 0.394 | -1.694 | 0.383 | -0.067 | -0.019 | -0.019 | 2.227 | 0.28 | 0.11 | 0.28 | 0.24 | 0.31 | 0.36 |
| 4 | 182686491 | rs10520509 | -0.059 | 0.125 | -1.448 | 0.553 | -0.023 | -0.039 | -0.039 | 2.76 | 0.13 | 0.06 | 0.21 | 0.17 | 0.15 | 0.23 |
| 4 | 182689025 | rs17072241 | 0.661 | -0.399 | -1.132 | 0.39 | -0.003 | 0.006 | 0.006 | 3.532 | 0.09 | 0.11 | 0.14 | 0.13 | 0.08 | 0.1 |
| 4 | 182691071 | rs17072249 | -0.154 | -0.955 | -1.475 | 0.283 | 0.034 | 0 | 0 | 0 | 0.14 | 0.06 | 0.05 | 0.11 | 0.07 | 0.11 |
| 4 | 182692368 | rs12508055 | 0.436 | -0.753 | -1.461 | 0.307 | -0.05 | -0.025 | -0.025 | 0 | 0.32 | 0.22 | 0.46 | 0.39 | 0.42 | 0.31 |
| 4 | 182696987 | rs12505558 | -0.052 | -0.761 | -1.603 | 0.019 | 0.002 | -0.041 | -0.041 | 1.975 | 0.41 | 0.22 | 0.54 | 0.51 | 0.54 | 0.44 |
| 4 | 182699379 | rs7659598 | 0.744 | -0.966 | -1.055 | -0.095 | -0.01 | -0.021 | -0.021 | 0.95 | 0.06 | 0.18 | 0.02 | 0 | 0.03 | 0.06 |
| 4 | 182704551 | rs6855088 | 0.078 | -0.94 | -0.968 | -0.053 | -0.005 | -0.008 | -0.008 | 0.117 | 0.16 | 0.13 | 0.21 | 0.24 | 0.18 | 0.2 |
| 4 | 182710802 | rs11733364 | 1.377 | -0.646 | -1.1 | -0.216 | -0.079 | 0.07 | 0.07 | 0 | 0.12 | 0.03 | 0.1 | 0 | 0.24 | 0.08 |
| 4 | 182711326 | rs10013118 | -0.194 | -0.909 | -1.058 | -0.215 | -0.004 | -0.003 | -0.003 | 0 | 0.39 | 0.4 | 0.48 | 0.4 | 0.46 | 0.37 |
| 4 | 182712956 | rs4241739 | 0.819 | -1.175 | -1.061 | -0.257 | 0.031 | 0.033 | 0.033 | 0.043 | 0.38 | 0.13 | 0.27 | 0.31 | 0.34 | 0.42 |
| 4 | 182713919 | rs4541554 | 1.558 | -1.158 | -0.834 | -0.183 | 0.003 | 0.032 | 0.032 | 0 | 0.26 | 0.14 | 0.23 | 0.2 | 0.28 | 0.3 |
| 4 | 182714961 | rs17072280 | 1.272 | -0.745 | -0.697 | 0.024 | -0.003 | 0.006 | 0.006 | 4.929 | 0.08 | 0.11 | 0.06 | 0.15 | 0.08 | 0.09 |
| 4 | 182716439 | rs4074866 | -1.247 | -0.674 | -0.751 | -0.02 | 0.007 | -0.003 | -0.003 | 0 | 0.86 | 0.81 | 0.83 | 0.85 | 0.87 | 0.9 |
| 4 | 182720291 | rs6552523 | -0.88 | -0.396 | -1.604 | -0.058 | -0.008 | 0.029 | 0.029 | 1.927 | 0.75 | 0.47 | 0.67 | 0.65 | 0.78 | 0.8 |
| 4 | 182724616 | rs7665942 | 0.902 | -0.41 | -1.963 | -0.172 | 0.024 | 0.033 | 0.033 | 0 | 0.32 | 0.18 | 0.25 | 0.21 | 0.3 | 0.38 |
| 4 | 182724752 | rs17072291 | 0.746 | -0.195 | -2.172 | -0.011 | 0.023 | -0.057 | -0.057 | 0.04 | 0.21 | 0.17 | 0.35 | 0.38 | 0.18 | 0.25 |
| 4 | 182727134 | rs6815030 | -0.638 | 0.48 | -1.895 | -0.714 | 0.037 | 0.079 | 0.079 | 0.209 | 0.55 | 0.3 | 0.53 | 0.3 | 0.6 | 0.57 |
| 4 | 182727482 | rs17283383 | -0.761 | 1.246 | -0.93 | -0.399 | -0.037 | 0.048 | 0.048 | 0.012 | 0.26 | 0.17 | 0.16 | 0.09 | 0.31 | 0.24 |
| 4 | 182733126 | rs4862004 | 1.379 | 0.309 | -0.189 | 0.114 | -0.021 | -0.03 | -0.03 | 0 | 0.88 | 0.74 | 0.89 | 0.93 | 0.92 | 0.82 |
| 4 | 182733805 | rs4333223 | -1.824 | 0.111 | -0.246 | 0.295 | -0.01 | 0 | 0 | 0 | 0.1 | 0.11 | 0.11 | 0.09 | 0.05 | 0.16 |
| 4 | 182740613 | rs10012329 | -0.614 | 0.323 | -0.547 | -0.213 | 0.016 | 0.06 | 0.06 | 0.631 | 0.47 | 0.65 | 0.75 | 0.2 | 0.58 | 0.58 |
| 4 | 182742349 | rs7687804 | 0.765 | -0.157 | -0.114 | 0.578 | -0.052 | 0.113 | 0.113 | 0 | 0.78 | 0.5 | 0.83 | 0.35 | 0.84 | 0.6 |
| 4 | 182744106 | rs11728634 | -0.061 | 0.153 | -0.379 | 0.367 | 0.003 | -0.005 | -0.005 | 0 | 0.38 | 0.4 | 0.45 | 0.3 | 0.38 | 0.26 |
| 4 | 182744904 | rs7681861 | -0.388 | 0.468 | -0.477 | -0.149 | 0.001 | -0.001 | -0.001 | 0.318 | 0.3 | 0.58 | 0.33 | 0.26 | 0.32 | 0.36 |
| 4 | 182747639 | rs7434383 | 0.003 | 0.413 | -0.296 | -0.133 | 0.047 | 0.206 | 0.206 | 0.523 | 0.92 | 0.63 | 0.91 | 0.8 | 0.96 | 0.91 |
| 4 | 182748140 | rs4862009 | 0.611 | -0.085 | -0.385 | 0.045 | -0.028 | 0.174 | 0.174 | 0.797 | 0.85 | 0.54 | 0.88 | 0.28 | 0.91 | 0.82 |
| 4 | 182748677 | rs17072369 | 0.043 | -0.453 | -0.8 | -0.175 | 0.013 | -0.026 | -0.026 | 0 | 0.13 | 0.09 | 0.06 | 0.53 | 0.11 | 0.1 |
| 4 | 182749521 | rs4458490 | 0.682 | -0.447 | -0.788 | -0.139 | 0.007 | 0.067 | 0.067 | 0 | 0.07 | 0.07 | 0.03 | 0.52 | 0.05 | 0.09 |
| 4 | 182749549 | rs4519836 | 0.562 | -0.394 | -0.714 | -0.259 | -0.026 | 0.091 | 0.091 | 1.056 | 0.56 | 0.28 | 0.61 | 0.21 | 0.57 | 0.53 |
| 4 | 182749585 | rs12505472 | 0.941 | -0.557 | -0.772 | -0.374 | 0.009 | 0.011 | 0.011 | 0 | 0.22 | 0.21 | 0.22 | 0.12 | 0.2 | 0.32 |
| 4 | 182754966 | rs749752 | 1.05 | -0.507 | -0.578 | -0.187 | -0.014 | 0.076 | 0.076 | 0.223 | 0.33 | 0.2 | 0.26 | 0.07 | 0.34 | 0.17 |
| 4 | 182756765 | rs10001447 | -0.628 | -0.552 | -0.344 | -0.161 | 0.002 | 0.028 | 0.028 | 0 | 0.59 | 0.28 | 0.64 | 0.74 | 0.61 | 0.69 |
| 4 | 182757120 | rs2048917 | -1.297 | -0.498 | -0.38 | -0.486 | -0.022 | 0.081 | 0.081 | 0.12 | 0.66 | 0.8 | 0.74 | 0.94 | 0.65 | 0.83 |
| 4 | 182770248 | rs12508245 | 0.248 | -0.609 | -0.421 | -0.268 | -0.035 | 0.05 | 0.05 | 0.172 | 0.36 | 0.2 | 0.53 | 0.21 | 0.41 | 0.46 |
| 4 | 182770862 | rs7679999 | -0.328 | -0.335 | -0.192 | -0.06 | 0.006 | 0.087 | 0.087 | 0 | 0.48 | 0.29 | 0.6 | 0.24 | 0.52 | 0.59 |
| 4 | 182775986 | rs11931121 | 0.792 | -0.219 | 0.589 | 1.131 | 0.047 | 0.53 | 0.53 | 0.748 | 0.91 | 0.51 | 0.9 | 0.57 | 0.93 | 0.9 |
| 4 | 182776413 | rs10016994 | 1.332 | -0.177 | 0.075 | 0.587 | -0.017 | 0.078 | 0.078 | 1.994 | 0.24 | 0.08 | 0.22 | 0.05 | 0.24 | 0.17 |
| 4 | 182777782 | rs7655087 | -0.075 | -0.28 | 0.133 | 0.612 | 0.021 | -0.036 | -0.036 | 0.076 | 0.6 | 0.28 | 0.65 | 0.51 | 0.65 | 0.68 |
| 4 | 182780188 | rs6838314 | -0.127 | -0.391 | 0.082 | 0.768 | 0.024 | 0.031 | 0.031 | 1.215 | 0.61 | 0.26 | 0.65 | 0.27 | 0.68 | 0.63 |
| 4 | 182780568 | rs1914369 | -1.423 | -0.384 | 0.152 | 0.753 | 0.002 | 0.05 | 0.05 | 0 | 0.73 | 0.84 | 0.78 | 0.94 | 0.76 | 0.83 |
| 4 | 182781422 | rs2063954 | 0.115 | -0.532 | 0.548 | 1.059 | 0.014 | 0.068 | 0.068 | 0 | 0.42 | 0.3 | 0.36 | 0.15 | 0.41 | 0.46 |
| 4 | 182784916 | rs6823165 | 0.372 | -0.21 | -0.652 | 0.641 | 0.009 | 0.032 | 0.032 | 0 | 0.47 | 0.26 | 0.53 | 0.42 | 0.49 | 0.41 |
| 4 | 182794879 | rs965523 | 0.412 | -0.483 | 0.056 | 0.49 | 0 | -0.003 | -0.003 | 0 | 0.52 | 0.46 | 0.54 | 0.25 | 0.55 | 0.37 |
| 4 | 182798961 | rs1400625 | -0.002 | -0.437 | 0.27 | 1.288 | -0.004 | 0.154 | 0.154 | 0 | 0.9 | 0.8 | 0.89 | 0.44 | 0.9 | 0.83 |

| | | | | | | | | | | | | | | | | |
|---|---|---|---|---|---|---|---|---|---|---|---|---|---|---|---|---|
| 4 | 182805092 | rs11724059 | 1.239 | -0.333 | -0.044 | 0.851 | 0.018 | -0.035 | -0.035 | 0 | 0.38 | 0.43 | 0.43 | 0.19 | 0.36 | 0.26 |
| 4 | 182805286 | rs12644818 | 0.277 | -0.254 | 0.002 | 1.608 | -0.001 | 0.084 | 0.084 | 0 | 0.08 | 0.13 | 0.06 | 0.32 | 0.08 | 0.09 |
| 4 | 182817317 | rs2138743 | -1.274 | -0.476 | 0.363 | 1.019 | -0.001 | 0.004 | 0.004 | 0 | 0.37 | 0.57 | 0.31 | 0.52 | 0.4 | 0.35 |
| 4 | 182817468 | rs1400624 | -0.364 | -0.71 | 0.547 | 1.073 | -0.017 | 0.089 | 0.089 | 0 | 0.11 | 0.25 | 0.11 | 0.27 | 0.09 | 0.13 |
| 4 | 182821968 | rs1517298 | 0.233 | -0.969 | 0.133 | 0.189 | 0.004 | 0.004 | 0.004 | 0.373 | 0.25 | 0.22 | 0.24 | 0.16 | 0.19 | 0.37 |
| 4 | 182825247 | rs2048914 | 0.201 | -0.737 | 0.011 | 0.287 | -0.042 | 0.066 | 0.066 | 0 | 0.18 | 0.03 | 0.13 | 0.06 | 0.22 | 0.13 |
| 4 | 182828390 | rs17072509 | 0.637 | -0.895 | 0.237 | 0.464 | 0.002 | 0.026 | 0.026 | 0 | 0.07 | 0.13 | 0.06 | 0.2 | 0.09 | 0.1 |
| 4 | 182830928 | rs17286513 | 0.097 | -0.952 | 0.112 | 0.217 | 0.005 | 0.042 | 0.042 | 0 | 0.18 | 0.09 | 0.12 | 0.06 | 0.19 | 0.13 |
| 4 | 182834128 | rs6824345 | -0.411 | -0.927 | 0.648 | 0.412 | -0.029 | -0.008 | -0.008 | 0 | 0.82 | 0.71 | 0.78 | 0.84 | 0.88 | 0.67 |
| 4 | 182835130 | rs17072515 | -0.207 | -0.76 | 0.432 | 0.253 | -0.029 | -0.041 | -0.041 | 0 | 0.21 | 0.3 | 0.14 | 0.11 | 0.14 | 0.33 |
| 4 | 182847338 | rs2199118 | 0.441 | -0.429 | 0.274 | 0.468 | 0.004 | 0.02 | 0.02 | 0.006 | 0.13 | 0.18 | 0.1 | 0.22 | 0.16 | 0.09 |
| 4 | 182854996 | rs1458889 | -0.142 | -0.819 | 0.44 | 0.683 | -0.041 | 0.054 | 0.054 | 1.513 | 0.89 | 0.77 | 0.92 | 0.77 | 0.97 | 0.89 |
| 4 | 182859352 | rs6813301 | 0.222 | -0.972 | 0.492 | 0.484 | -0.025 | -0.036 | -0.036 | 0.784 | 0.2 | 0.37 | 0.16 | 0.11 | 0.17 | 0.4 |
| 4 | 182860907 | rs11945595 | 0.669 | -0.951 | 0.216 | 0.493 | 0.004 | 0.005 | 0.005 | 0 | 0.17 | 0.15 | 0.13 | 0.11 | 0.16 | 0.31 |
| 4 | 182863198 | rs2048913 | 1.711 | -0.736 | -0.109 | 0.41 | -0.07 | -0.029 | -0.029 | 0 | 0.4 | 0.15 | 0.53 | 0.37 | 0.43 | 0.34 |
| 4 | 182868840 | rs9993255 | -0.515 | 0.008 | -0.117 | -0.079 | -0.009 | -0.023 | -0.023 | 0.056 | 0.25 | 0.48 | 0.19 | 0.2 | 0.27 | 0.22 |
| 4 | 182879932 | rs4861497 | 0.452 | 0.079 | -0.042 | -0.134 | 0.001 | 0.006 | 0.006 | 0.01 | 0.13 | 0.15 | 0.1 | 0.19 | 0.16 | 0.09 |
| 4 | 182884590 | rs9685589 | -0.007 | 0.11 | -0.543 | -0.042 | 0.008 | -0.016 | -0.016 | 0 | 0.35 | 0.53 | 0.27 | 0.3 | 0.42 | 0.3 |
| 4 | 182886016 | rs13141696 | -0.878 | 0.091 | -0.562 | -0.048 | -0.003 | 0.018 | 0.018 | 0 | 0.52 | 0.68 | 0.41 | 0.42 | 0.57 | 0.62 |
| 4 | 182888219 | rs9992244 | 0.063 | 0.065 | -0.522 | 0.13 | -0.005 | -0.011 | -0.011 | 0 | 0.36 | 0.53 | 0.26 | 0.3 | 0.39 | 0.28 |
| 4 | 182895078 | rs2063956 | -0.31 | 0.508 | -0.736 | 0.406 | -0.01 | 0.014 | 0.014 | 0.023 | 0.53 | 0.71 | 0.39 | 0.42 | 0.56 | 0.56 |
| 4 | 182911139 | rs7676827 | -0.672 | 0.423 | -0.574 | 0.101 | -0.003 | -0.036 | -0.036 | 0.648 | 0.14 | 0.22 | 0.18 | 0.04 | 0.13 | 0.15 |
| 4 | 182913820 | rs17288133 | -0.309 | 0.31 | -0.84 | -0.429 | 0.003 | 0.009 | 0.009 | 0 | 0.2 | 0.15 | 0.14 | 0.13 | 0.21 | 0.25 |
| 4 | 182914650 | rs17072625 | 0.979 | 0.624 | -0.916 | -0.661 | -0.001 | -0.001 | -0.001 | 0 | 0.14 | 0.17 | 0.3 | 0.07 | 0.19 | 0.12 |
| 4 | 182914694 | rs11938148 | 0.506 | 0.441 | -1.395 | -0.901 | -0.016 | 0.111 | 0.111 | 0.104 | 0.37 | 0.12 | 0.29 | 0.16 | 0.33 | 0.33 |
| 4 | 182916588 | rs17222326 | 0.751 | 0.411 | -1.248 | -0.899 | 0.003 | 0.035 | 0.035 | 0 | 0.2 | 0.05 | 0.13 | 0.13 | 0.22 | 0.25 |
| 4 | 182917804 | rs7697411 | 1.378 | 0.311 | -1.216 | -0.881 | 0.005 | 0.047 | 0.047 | 0.095 | 0.26 | 0.11 | 0.12 | 0.12 | 0.21 | 0.11 |
| 4 | 182918564 | rs12503812 | 1.137 | 0.381 | -1.023 | -0.88 | 0.006 | -0.008 | -0.008 | 0.135 | 0.16 | 0.21 | 0.32 | 0.08 | 0.22 | 0.16 |
| 4 | 182919753 | rs11946900 | -1.587 | 0.4 | -0.99 | -0.068 | 0.026 | 0.066 | 0.066 | 0 | 0.8 | 0.64 | 0.77 | 0.68 | 0.77 | 0.78 |
| 4 | 182921858 | rs7667255 | 0.761 | 0.465 | -1.412 | -1.054 | -0.007 | 0.022 | 0.022 | 0 | 0.22 | 0.18 | 0.32 | 0.09 | 0.24 | 0.17 |
| 4 | 182932198 | rs2100695 | -0.16 | 0.468 | -1.541 | -1.272 | -0.014 | 0.112 | 0.112 | 0 | 0.49 | 0.16 | 0.41 | 0.29 | 0.52 | 0.51 |
| 4 | 182932763 | rs12645487 | 1.39 | 0.693 | -1.561 | -1.255 | -0.031 | 0.038 | 0.038 | 0 | 0.25 | 0.07 | 0.12 | 0.13 | 0.23 | 0.11 |
| 4 | 182933601 | rs977361 | 0.086 | 0.622 | -1.074 | -1.034 | 0.013 | 0.055 | 0.055 | 0 | 0.52 | 0.33 | 0.42 | 0.29 | 0.53 | 0.58 |
| 4 | 182937043 | rs9799641 | 0.399 | 1.057 | -1.098 | -0.543 | -0.047 | 0.063 | 0.063 | 0 | 0.39 | 0.12 | 0.24 | 0.25 | 0.42 | 0.31 |
| 4 | 182942414 | rs6816160 | -0.347 | 0.882 | -0.616 | -0.664 | -0.002 | -0.003 | -0.003 | 2.033 | 0.35 | 0.36 | 0.46 | 0.07 | 0.38 | 0.4 |
| 4 | 182951552 | rs9993807 | 0.866 | 0.716 | 0.193 | -0.28 | 0 | -0.076 | -0.076 | 0 | 0.8 | 0.52 | 0.85 | 0.92 | 0.82 | 0.89 |
| 4 | 182959299 | rs10866251 | 1.614 | 1.212 | 0.118 | 0.171 | 0.008 | -0.002 | -0.002 | 0.017 | 0.72 | 0.38 | 0.79 | 0.75 | 0.71 | 0.77 |
| 4 | 182959578 | rs1979296 | 1.545 | 1.302 | 0.139 | 0.173 | 0.017 | 0 | 0 | 0.014 | 0.73 | 0.43 | 0.79 | 0.75 | 0.7 | 0.77 |
| 4 | 182964412 | rs17072678 | -1.397 | 1.175 | 0.971 | -0.106 | 0.002 | -0.044 | -0.044 | 0.823 | 0.21 | 0.3 | 0.15 | 0.09 | 0.22 | 0.18 |
| 4 | 182965163 | rs7693462 | -2.188 | 1.121 | 1.21 | -0.059 | 0.018 | -0.011 | -0.011 | 0 | 0.3 | 0.39 | 0.21 | 0.25 | 0.34 | 0.29 |
| 4 | 182973237 | rs10866253 | 1.762 | 0.887 | 1.621 | -0.078 | -0.002 | -0.022 | -0.022 | 0.003 | 0.8 | 0.3 | 0.82 | 0.91 | 0.81 | 0.83 |
| 4 | 182975220 | rs6812990 | -0.459 | 1.036 | 1.549 | 0.031 | 0.038 | 0.054 | 0.054 | 0 | 0.06 | 0.11 | 0.06 | 0.15 | 0.11 | 0.1 |
| 4 | 182975418 | rs335074 | 1.476 | 0.847 | 1.566 | -0.136 | -0.002 | 0.006 | 0.006 | 0.215 | 0.86 | 0.89 | 0.91 | 0.9 | 0.87 | 0.87 |
| 4 | 182976765 | rs17072702 | -2.327 | 0.718 | 1.973 | -0.3 | 0.003 | -0.132 | -0.132 | 0.236 | 0.19 | 0.53 | 0.15 | 0.01 | 0.16 | 0.12 |
| 4 | 182983862 | rs17072710 | -1.437 | 0.546 | 1.518 | -0.177 | 0.004 | 0.012 | 0.012 | 0 | 0.07 | 0.1 | 0.07 | 0.14 | 0.1 | 0.11 |
| 4 | 182983955 | rs6812461 | -1.572 | 0.282 | 1.458 | -0.287 | 0.001 | -0.016 | -0.016 | 0.962 | 0.13 | 0.12 | 0.06 | 0 | 0.1 | 0.08 |
| 4 | 182988914 | rs2597144 | 2.478 | 0.075 | 1.632 | -0.534 | -0.008 | -0.031 | -0.031 | 0.143 | 0.71 | 0.38 | 0.71 | 0.85 | 0.7 | 0.74 |
| 4 | 182989346 | rs17072729 | -1.245 | 0.038 | 1.573 | -0.531 | 0.007 | -0.005 | -0.005 | 4.251 | 0.13 | 0.08 | 0.13 | 0.14 | 0.11 | 0.15 |
| 4 | 182990553 | rs10520518 | 1.446 | -0.261 | -0.057 | -0.493 | -0.187 | -0.038 | -0.038 | 0 | 0.34 | 0.03 | 0.39 | 0.26 | 0.43 | 0.4 |
| 4 | 182990909 | rs2609554 | 1.482 | -0.441 | 0.546 | -0.687 | -0.045 | -0.053 | -0.053 | 0 | 0.71 | 0.52 | 0.75 | 0.86 | 0.77 | 0.73 |
| 4 | 182995164 | rs17072756 | 0.644 | -0.622 | -0.148 | 0.088 | -0.045 | -0.025 | -0.025 | 0.496 | 0.27 | 0.05 | 0.27 | 0.31 | 0.28 | 0.24 |
| 4 | 182995319 | rs10026007 | -0.33 | -0.476 | 0.442 | 0.067 | -0.013 | -0.027 | -0.027 | 0.057 | 0.09 | 0.18 | 0.07 | 0.02 | 0.07 | 0.1 |
| 4 | 182996793 | rs2597125 | -0.437 | -0.266 | 0.026 | 0.031 | -0.001 | -0.002 | -0.002 | 0 | 0.35 | 0.6 | 0.3 | 0.29 | 0.29 | 0.4 |
| 4 | 182998238 | rs17303659 | 1.492 | -0.299 | -1.092 | -0.04 | -0.036 | -0.011 | -0.011 | 0.28 | 0.12 | 0 | 0.18 | 0.11 | 0.16 | 0.13 |
| 4 | 182998270 | rs1868642 | -0.71 | 0.013 | -1.181 | -0.396 | -0.005 | -0.011 | -0.011 | 0 | 0.62 | 0.62 | 0.51 | 0.27 | 0.65 | 0.49 |
| 4 | 182998326 | rs1868641 | 1.774 | 0.174 | -1.029 | -0.588 | -0.016 | 0.01 | 0.01 | 0 | 0.18 | 0.12 | 0.15 | 0.12 | 0.25 | 0.2 |
| 4 | 183000121 | rs10024819 | 2.075 | 0.193 | -0.944 | -0.033 | 0.004 | 0.003 | 0.003 | 0 | 0.24 | 0.23 | 0.34 | 0.34 | 0.29 | 0.31 |
| 4 | 183000252 | rs17303841 | 0.898 | 0.157 | -0.964 | -0.138 | -0.016 | 0.075 | 0.075 | 0 | 0.22 | 0.13 | 0.12 | 0 | 0.24 | 0.09 |
| 4 | 183000925 | rs13135549 | 1.203 | 0.276 | -0.85 | -0.014 | -0.012 | -0.003 | -0.003 | 0.092 | 0.28 | 0.19 | 0.36 | 0.34 | 0.35 | 0.35 |
| 4 | 183004954 | rs6552541 | 0.216 | -0.137 | -0.942 | -0.068 | 0.024 | 0.027 | 0.027 | 0.017 | 0.44 | 0.62 | 0.46 | 0.37 | 0.35 | 0.43 |
| 4 | 183006470 | rs17072794 | -0.525 | -0.112 | -0.604 | -0.084 | -0.001 | -0.012 | -0.012 | 0 | 0.41 | 0.45 | 0.25 | 0.14 | 0.25 | 0.26 |
| 4 | 183007676 | rs11132111 | -1.169 | 0.397 | -0.743 | 0.368 | -0.038 | -0.01 | -0.01 | 0 | 0.74 | 0.84 | 0.68 | 0.71 | 0.6 | 0.64 |
| 4 | 183007922 | rs2034975 | -2.513 | 0.574 | -0.721 | 0.421 | 0.018 | 0.007 | 0.007 | 0 | 0.86 | 0.84 | 0.81 | 0.82 | 0.77 | 0.77 |
| 4 | 183008572 | rs6839797 | -0.751 | -0.164 | -0.409 | 0.479 | 0.002 | 0.001 | 0.001 | 0 | 0.41 | 0.46 | 0.3 | 0.33 | 0.38 | 0.27 |
| 4 | 183008639 | rs6840333 | 0.283 | -0.1 | -0.592 | 0.602 | 0.001 | -0.001 | -0.001 | 0 | 0.22 | 0.25 | 0.11 | 0.21 | 0.18 | 0.1 |
| 4 | 183011216 | rs10440484 | 1.448 | 0.134 | -0.872 | 0.189 | 0.005 | 0.04 | 0.04 | 0.603 | 0.16 | 0.09 | 0.07 | 0 | 0.16 | 0.04 |
| 4 | 183013816 | rs4862025 | 0.382 | -0.05 | -0.895 | 0.36 | 0.01 | 0.033 | 0.033 | 0.008 | 0.24 | 0.1 | 0.17 | 0.15 | 0.2 | 0.18 |
| 4 | 183014495 | rs335109 | 0.415 | 0.253 | -0.664 | 0.313 | 0.002 | -0.001 | -0.001 | 0.117 | 0.19 | 0.14 | 0.16 | 0.19 | 0.2 | 0.23 |
| 4 | 183018034 | rs749763 | -0.25 | 0.237 | -0.421 | 0.11 | -0.001 | -0.002 | -0.002 | 0.356 | 0.36 | 0.36 | 0.28 | 0.14 | 0.4 | 0.23 |

| | | | | | | | | | | | | | | | | |
|---|---|---|---|---|---|---|---|---|---|---|---|---|---|---|---|---|
| 4 | 183021079 | rs335101 | -0.334 | -0.012 | -0.329 | 0.641 | -0.008 | 0.028 | 0.028 | 0 | 0.24 | 0.28 | 0.23 | 0.48 | 0.2 | 0.35 |
| 4 | 183023753 | rs6858183 | 0.001 | 0.469 | -0.047 | 0.263 | -0.01 | 0.002 | 0.002 | 0.03 | 0.65 | 0.42 | 0.72 | 0.71 | 0.72 | 0.75 |
| 4 | 183023943 | rs10022206 | 0.095 | 0.788 | -0.592 | 0.064 | 0.012 | -0.007 | -0.007 | 1.612 | 0.25 | 0.31 | 0.24 | 0.18 | 0.33 | 0.34 |
| 4 | 183023981 | rs6810793 | 0.952 | 0.822 | -1.175 | 0.14 | 0.02 | -0.045 | -0.045 | 0 | 0.41 | 0.14 | 0.48 | 0.53 | 0.39 | 0.4 |
| 4 | 183026790 | rs1470681 | 1.301 | 1.116 | -0.752 | 0.289 | 0.002 | -0.001 | -0.001 | 1.197 | 0.12 | 0.1 | 0.17 | 0.09 | 0.19 | 0.12 |
| 4 | 183028134 | rs13152542 | 0.821 | 1.069 | -1.058 | 0.258 | -0.008 | 0.003 | 0.003 | 0.23 | 0.09 | 0.07 | 0.04 | 0.05 | 0.12 | 0.17 |
| 4 | 183029476 | rs17304841 | 0.535 | 0.777 | -1.305 | 0.135 | -0.012 | 0.019 | 0.019 | 0 | 0.34 | 0.24 | 0.23 | 0.19 | 0.35 | 0.27 |
| 4 | 183030136 | rs6815885 | 1.02 | 0.29 | -1.478 | 0.327 | 0.046 | -0.059 | -0.059 | 0.194 | 0.33 | 0.16 | 0.4 | 0.53 | 0.25 | 0.39 |
| 4 | 183030731 | rs898131 | 0.924 | 0.353 | -1.208 | 0.343 | 0.002 | -0.003 | -0.003 | 0.015 | 0.14 | 0.12 | 0.18 | 0.1 | 0.19 | 0.12 |
| 4 | 183033029 | rs1458893 | 1.076 | 0.231 | -1.621 | 0.343 | 0.029 | -0.05 | -0.05 | 0 | 0.31 | 0.17 | 0.4 | 0.53 | 0.24 | 0.39 |
| 4 | 183034755 | rs17226385 | 1.109 | 0.683 | -1.832 | 0.05 | -0.077 | 0.001 | 0.001 | 0 | 0.23 | 0.1 | 0.2 | 0.13 | 0.37 | 0.17 |
| 4 | 183035530 | rs11724181 | 1.011 | 0.619 | -1.685 | 0.127 | -0.022 | 0.002 | 0.002 | 0 | 0.12 | 0.03 | 0.17 | 0.1 | 0.18 | 0.12 |
| 4 | 183039092 | rs7661253 | 0.616 | 0.702 | -1.832 | 0.053 | -0.119 | 0.025 | 0.025 | 0.188 | 0.28 | 0.12 | 0.27 | 0.13 | 0.45 | 0.19 |
| 4 | 183039938 | rs1450150 | 0.698 | 0.069 | -1.845 | -0.022 | 0.037 | -0.028 | -0.028 | 0.716 | 0.34 | 0.17 | 0.45 | 0.69 | 0.27 | 0.41 |
| 4 | 183040169 | rs3919765 | 1.177 | 0.335 | -1.862 | -0.448 | -0.05 | 0.047 | 0.047 | 0.383 | 0.08 | 0.01 | 0.05 | 0 | 0.14 | 0.15 |
| 4 | 183040330 | rs6825212 | 0.068 | 0.035 | -0.845 | -0.608 | -0.007 | -0.057 | -0.057 | 0 | 0.23 | 0.38 | 0.18 | 0.09 | 0.21 | 0.2 |
| 4 | 183042695 | rs7662442 | 0.22 | -0.683 | -0.993 | -0.211 | -0.006 | -0.002 | -0.002 | 0 | 0.43 | 0.39 | 0.38 | 0.53 | 0.44 | 0.4 |
| 4 | 183044054 | rs1114531 | 0.326 | -1.197 | -1.272 | -0.141 | 0.06 | -0.059 | -0.059 | 0.196 | 0.18 | 0.06 | 0.11 | 0.37 | 0.11 | 0.24 |
| 4 | 183044215 | rs7669814 | 1.209 | -1.109 | -1.274 | -0.04 | 0.03 | -0.008 | -0.008 | 0 | 0.14 | 0 | 0.1 | 0.12 | 0.09 | 0.17 |
| 4 | 183047903 | rs335068 | 0.256 | -0.993 | -0.777 | -0.176 | -0.013 | 0.006 | 0.006 | 0.148 | 0.41 | 0.47 | 0.51 | 0.46 | 0.35 | 0.39 |
| 4 | 183048696 | rs335067 | -0.51 | -0.824 | -0.898 | -0.501 | 0.023 | 0.069 | 0.069 | 2.305 | 0.53 | 0.69 | 0.67 | 0.62 | 0.43 | 0.54 |
| 4 | 183060694 | rs1450141 | -0.878 | -0.756 | -0.458 | -0.152 | 0.024 | -0.059 | -0.059 | 0.05 | 0.84 | 0.61 | 0.81 | 0.92 | 0.82 | 0.88 |
| 4 | 183061232 | rs1125145 | 0.029 | -0.78 | -1.347 | -0.003 | -0.043 | 0.05 | 0.05 | 0.468 | 0.44 | 0.2 | 0.27 | 0.38 | 0.48 | 0.38 |
| 4 | 183063011 | rs2309626 | -0.479 | 0.383 | -0.552 | 0.155 | 0.025 | -0.046 | -0.046 | 0.184 | 0.67 | 0.51 | 0.72 | 0.84 | 0.63 | 0.84 |
| 4 | 183067139 | rs9312290 | 0.114 | 0.572 | -0.962 | -0.009 | 0.005 | -0.003 | -0.003 | 0 | 0.68 | 0.74 | 0.76 | 0.61 | 0.71 | 0.59 |
| 4 | 183068987 | rs4862031 | 0.337 | 0.637 | -1.006 | -0.029 | 0.003 | 0.001 | 0.001 | 0 | 0.65 | 0.74 | 0.73 | 0.61 | 0.66 | 0.55 |
| 4 | 183072277 | rs7681106 | -0.59 | 1.087 | -0.343 | 0.012 | 0.071 | 0.103 | 0.103 | 0 | 0.11 | 0.4 | 0.29 | 0.21 | 0.12 | 0.09 |
| 4 | 183077514 | rs876836 | -0.3 | 0.935 | -0.672 | -0.26 | 0.007 | -0.03 | -0.03 | 1.152 | 0.25 | 0.43 | 0.4 | 0.23 | 0.24 | 0.12 |
| 4 | 183078161 | rs7695632 | 0.452 | 0.804 | -1.081 | -0.246 | 0.011 | 0.055 | 0.055 | 0.225 | 0.54 | 0.36 | 0.38 | 0.64 | 0.47 | 0.61 |
| 4 | 183081780 | rs12640278 | 0.525 | 0.437 | -1.323 | 0.074 | 0.011 | -0.051 | -0.051 | 0 | 0.23 | 0.13 | 0.19 | 0.38 | 0.19 | 0.26 |
| 4 | 183086701 | rs12504374 | 0.261 | 0.823 | -1.05 | -0.165 | 0.007 | 0.015 | 0.015 | 0 | 0.73 | 0.59 | 0.76 | 0.71 | 0.73 | 0.53 |
| 4 | 183087547 | rs896028 | -0.902 | 0.489 | -1.123 | -0.408 | 0 | -0.001 | -0.001 | 0 | 0.65 | 0.68 | 0.69 | 0.53 | 0.72 | 0.63 |
| 4 | 183091203 | rs13126808 | -0.466 | 0.698 | -1.768 | -0.52 | -0.011 | -0.002 | -0.002 | 0 | 0.55 | 0.53 | 0.44 | 0.41 | 0.65 | 0.55 |
| 4 | 183094353 | rs1450149 | 0.633 | 1.098 | -2.838 | -0.775 | -0.11 | 0.19 | 0.19 | 0.496 | 0.31 | 0.05 | 0.22 | 0.01 | 0.4 | 0.14 |
| 4 | 183111871 | rs1450146 | -1.241 | 0.929 | -2.743 | -0.412 | 0.047 | -0.099 | -0.099 | 0.415 | 0.68 | 0.93 | 0.75 | 0.57 | 0.77 | 0.62 |
| 4 | 183114014 | rs17307191 | 0.96 | 1.454 | -2.582 | -0.251 | -0.054 | -0.024 | -0.024 | 2.094 | 0.1 | 0.01 | 0.16 | 0.12 | 0.15 | 0.22 |
| 4 | 183117920 | rs13115107 | 0.408 | 1.284 | -2.303 | -0.378 | -0.035 | 0.089 | 0.089 | 0.115 | 0.24 | 0.08 | 0.18 | 0.01 | 0.29 | 0.1 |
| 4 | 183121231 | rs2726802 | -0.237 | 0.785 | -1.184 | 0.546 | -0.029 | 0.087 | 0.087 | 0 | 0.84 | 0.69 | 0.79 | 0.56 | 0.92 | 0.79 |
| 4 | 183123850 | rs7349674 | 0.345 | 1.108 | -2.074 | 0.455 | -0.002 | -0.041 | -0.041 | 0 | 0.64 | 0.17 | 0.62 | 0.55 | 0.66 | 0.7 |
| 4 | 183124933 | rs11132117 | -0.151 | 1.926 | -2.405 | 0.311 | -0.128 | -0.002 | -0.002 | 0.213 | 0.25 | 0.04 | 0.36 | 0.18 | 0.41 | 0.27 |
| 4 | 183125630 | rs1450144 | 0.479 | 1.522 | -1.45 | 0.637 | 0.015 | 0.03 | 0.03 | 0 | 0.71 | 0.58 | 0.56 | 0.39 | 0.73 | 0.53 |
| 4 | 183129070 | rs17235541 | 0.572 | 2.031 | -1.567 | 0.123 | -0.026 | 0.028 | 0.028 | 0 | 0.12 | 0.06 | 0.13 | 0.01 | 0.21 | 0.04 |
| 4 | 183132303 | rs1031313 | -0.52 | 1.934 | -0.517 | -0.166 | 0.061 | -0.024 | -0.024 | 0 | 0.12 | 0.35 | 0.11 | 0.07 | 0.16 | 0.11 |
| 4 | 183132543 | rs4861499 | 1.958 | 1.162 | -0.938 | -0.418 | -0.015 | 0.049 | 0.049 | 0.368 | 0.33 | 0.02 | 0.34 | 0.21 | 0.28 | 0.31 |
| 4 | 183136858 | rs17072860 | -0.288 | 1.289 | -0.575 | -0.405 | 0.009 | 0.002 | 0.002 | 0 | 0.08 | 0.29 | 0.06 | 0.06 | 0.06 | 0.07 |
| 4 | 183137216 | rs17308154 | 0.196 | 1.471 | -0.472 | 0.208 | -0.029 | -0.01 | -0.01 | 0 | 0.17 | 0.12 | 0.17 | 0.18 | 0.26 | 0.11 |
| 4 | 183137398 | rs2726807 | 0.748 | 1.282 | -0.002 | 0.121 | 0.001 | 0 | 0 | 0 | 0.59 | 0.42 | 0.61 | 0.4 | 0.64 | 0.55 |
| 4 | 183138649 | rs13152117 | 0.515 | 1.148 | 0.082 | -0.196 | 0.005 | -0.014 | -0.014 | 0 | 0.16 | 0.15 | 0.2 | 0.06 | 0.21 | 0.09 |
| 4 | 183141735 | rs2249112 | 0.211 | 1.343 | -0.134 | -0.178 | -0.012 | -0.005 | -0.005 | 0 | 0.64 | 0.56 | 0.65 | 0.35 | 0.68 | 0.6 |
| 4 | 183143114 | rs896024 | 1.113 | 0.76 | -0.528 | -0.229 | 0.006 | 0.082 | 0.082 | 0.031 | 0.39 | 0.24 | 0.41 | 0.1 | 0.37 | 0.44 |
| 4 | 183157313 | rs4241744 | 0.35 | 0.779 | -0.003 | 0.839 | -0.002 | 0.019 | 0.019 | 0 | 0.93 | 0.83 | 0.96 | 0.88 | 0.95 | 0.93 |
| 4 | 183157458 | rs4241745 | 1.084 | 0.666 | -0.069 | 1.221 | -0.015 | 0.058 | 0.058 | 0.113 | 0.73 | 0.41 | 0.66 | 0.44 | 0.75 | 0.55 |
| 4 | 183157856 | rs17072904 | -0.814 | 0.723 | -0.865 | 0.685 | -0.015 | 0 | 0 | 0 | 0.34 | 0.53 | 0.38 | 0.63 | 0.3 | 0.47 |
| 4 | 183160086 | rs4861500 | 0.813 | 0.602 | -2.066 | 0.165 | -0.011 | 0.107 | 0.107 | 0.273 | 0.38 | 0.03 | 0.38 | 0.16 | 0.31 | 0.35 |
| 4 | 183161287 | rs7663302 | -1.015 | 1.012 | -1.706 | 0.34 | -0.05 | -0.033 | -0.033 | 0 | 0.51 | 0.93 | 0.53 | 0.48 | 0.54 | 0.31 |
| 4 | 183161620 | rs7668736 | -0.999 | 1.268 | -1.019 | -0.625 | -0.03 | 0.095 | 0.095 | 0.004 | 0.35 | 0.8 | 0.38 | 0.12 | 0.41 | 0.21 |
| 4 | 183162284 | rs7694663 | -1.244 | 1.341 | -0.165 | -0.33 | -0.001 | -0.018 | -0.018 | 0.322 | 0.28 | 0.74 | 0.33 | 0.13 | 0.31 | 0.17 |
| 4 | 183166980 | rs7656126 | -1.239 | 1.16 | -0.17 | -0.344 | 0 | -0.019 | -0.019 | 0.059 | 0.27 | 0.73 | 0.32 | 0.13 | 0.3 | 0.16 |
| 4 | 183170004 | rs7687160 | 1.483 | 1.101 | 0.043 | -0.274 | -0.004 | 0.017 | 0.017 | 1.967 | 0.58 | 0.35 | 0.61 | 0.71 | 0.54 | 0.75 |
| 4 | 183172201 | rs11724819 | -0.655 | 1.114 | 0.521 | -0.158 | -0.02 | 0.015 | 0.015 | 0 | 0.15 | 0.12 | 0.21 | 0.12 | 0.25 | 0.1 |
| 4 | 183174982 | rs2726824 | -2.199 | 1.138 | 1.251 | -0.105 | 0.003 | -0.06 | -0.06 | 0.139 | 0.29 | 0.59 | 0.32 | 0.14 | 0.32 | 0.17 |
| 4 | 183176671 | rs12640203 | -1.86 | 1.167 | 0.821 | -0.059 | -0.019 | 0.003 | 0.003 | 0 | 0.07 | 0.16 | 0.13 | 0.11 | 0.05 | 0.04 |
| 4 | 183178219 | rs11736246 | -1.91 | 1.223 | 0.858 | -0.078 | -0.004 | 0.007 | 0.007 | 0.782 | 0.24 | 0.24 | 0.27 | 0.13 | 0.3 | 0.12 |
| 4 | 183182681 | rs11940750 | -2.637 | 0.479 | 1.192 | 0.223 | -0.023 | -0.002 | -0.002 | 0 | 0.1 | 0.25 | 0.16 | 0.12 | 0.07 | 0.1 |
| 4 | 183184352 | rs12648007 | -1.61 | 0.53 | 0.989 | 0.217 | -0.008 | -0.002 | -0.002 | 0 | 0.11 | 0.13 | 0.14 | 0.12 | 0.11 | 0.07 |
| 4 | 183187854 | rs10011107 | -1.391 | 0.994 | 1.457 | 0.305 | 0.024 | 0.006 | 0.006 | 0 | 0.13 | 0.24 | 0.11 | 0.13 | 0.15 | 0.1 |
| 4 | 183193206 | rs6825349 | -1.735 | 0.439 | 1.838 | 0.352 | -0.012 | -0.019 | -0.019 | 0 | 0.2 | 0.33 | 0.17 | 0.1 | 0.19 | 0.09 |
| 4 | 183198752 | rs2063905 | -1.037 | 0.346 | 1.946 | 0.206 | -0.007 | -0.017 | -0.017 | 0.023 | 0.16 | 0.21 | 0.13 | 0.05 | 0.17 | 0.08 |
| 4 | 183206596 | rs11732877 | -1.833 | 0.356 | 2.152 | 0.236 | -0.035 | -0.064 | -0.064 | 3.157 | 0.23 | 0.47 | 0.19 | 0.07 | 0.19 | 0.12 |

| | | | | | | | | | | | | | | | | |
|---|---|---|---|---|---|---|---|---|---|---|---|---|---|---|---|---|
| 4 | 183213736 | rs938526 | -1.785 | 0.331 | 1.57 | 0.225 | -0.034 | -0.058 | -0.058 | 0 | 0.23 | 0.45 | 0.21 | 0.07 | 0.19 | 0.12 |
| 4 | 183229221 | rs6552549 | -1.719 | 0.286 | 0.866 | 0.212 | -0.015 | -0.034 | -0.034 | 0 | 0.36 | 0.42 | 0.16 | 0.17 | 0.28 | 0.17 |
| 4 | 183232270 | rs17073038 | -0.358 | 0.299 | 0.74 | 0.255 | -0.013 | 0.002 | 0.002 | 3.781 | 0.13 | 0.14 | 0.04 | 0.13 | 0.06 | 0.06 |
| 4 | 183237479 | rs2675539 | -2.098 | 0.365 | 0.576 | 0.161 | -0.009 | -0.029 | -0.029 | 0 | 0.37 | 0.37 | 0.15 | 0.15 | 0.3 | 0.18 |
| 4 | 183239761 | rs2675534 | -1.135 | 0.799 | -0.044 | 0.387 | -0.007 | 0.031 | 0.031 | 0 | 0.2 | 0.1 | 0.07 | 0.02 | 0.22 | 0.09 |
| 4 | 183244218 | rs2726822 | 0.864 | 1.018 | -0.364 | 1.414 | -0.042 | -0.101 | -0.101 | 0 | 0.79 | 0.92 | 0.89 | 0.44 | 0.74 | 0.8 |
| 4 | 183247111 | rs2675531 | 1.152 | 1.385 | 0.593 | 1.945 | 0.054 | 0.24 | 0.24 | 1.346 | 0.85 | 0.54 | 0.82 | 0.52 | 0.84 | 0.79 |
| 4 | 183247601 | rs2726805 | -0.688 | 0.978 | 0.207 | 1.276 | 0 | -0.002 | -0.002 | 0.291 | 0.37 | 0.38 | 0.2 | 0.26 | 0.41 | 0.4 |
| 4 | 183248389 | rs957053 | -0.679 | 0.913 | 0.343 | 1.188 | -0.001 | -0.001 | -0.001 | 0 | 0.41 | 0.42 | 0.28 | 0.48 | 0.47 | 0.48 |
| 4 | 183248852 | rs6845440 | -0.871 | 0.657 | 0.436 | 1.045 | -0.003 | -0.001 | -0.001 | 0 | 0.44 | 0.39 | 0.31 | 0.48 | 0.53 | 0.48 |
| 4 | 183254661 | rs955638 | -0.297 | 0.751 | 0.165 | 0.952 | -0.004 | 0.005 | 0.005 | 0 | 0.29 | 0.24 | 0.2 | 0.12 | 0.32 | 0.18 |
| 4 | 183255434 | rs1516537 | 1.334 | 0.234 | 0.056 | 0.764 | -0.007 | -0.014 | -0.014 | 0 | 0.42 | 0.54 | 0.47 | 0.42 | 0.37 | 0.46 |
| 4 | 183256787 | rs7668966 | -0.01 | 0.335 | -0.016 | 0.863 | -0.012 | -0.01 | -0.01 | 0 | 0.83 | 0.8 | 0.75 | 0.89 | 0.89 | 0.93 |
| 4 | 183256975 | rs7669582 | -0.394 | 0.33 | 0.014 | 0.911 | -0.015 | -0.007 | -0.007 | 0 | 0.87 | 0.84 | 0.77 | 0.9 | 0.92 | 0.95 |
| 4 | 183260601 | rs10520528 | 0.961 | 0.551 | -0.131 | 0.859 | -0.002 | -0.001 | -0.001 | 0 | 0.47 | 0.58 | 0.64 | 0.51 | 0.42 | 0.5 |
| 4 | 183260894 | rs1516530 | 0.389 | 0.492 | -0.501 | 1.065 | -0.082 | -0.042 | -0.042 | 0 | 0.18 | 0.01 | 0.12 | 0.12 | 0.24 | 0.12 |
| 4 | 183271349 | rs4861505 | 0.978 | 0.188 | 0.387 | 1.488 | 0.011 | 0.034 | 0.034 | 0 | 0.84 | 0.83 | 0.83 | 0.63 | 0.78 | 0.77 |
| 4 | 183271631 | rs2168682 | -0.943 | 0.024 | 0.367 | 1.525 | 0.006 | 0.033 | 0.033 | 0 | 0.13 | 0.11 | 0.14 | 0.36 | 0.18 | 0.22 |
| 4 | 183272312 | rs13148469 | 1.573 | -0.467 | 0.872 | 1.269 | -0.003 | 0.013 | 0.013 | 0.247 | 0.67 | 0.43 | 0.61 | 0.5 | 0.66 | 0.64 |
| 4 | 183276790 | rs35124573 | -0.321 | -0.141 | 0.785 | 0.725 | 0.028 | -0.035 | -0.035 | 0.981 | 0.08 | 0.09 | 0.11 | 0 | 0.15 | 0.03 |
| 4 | 183285393 | rs6813422 | 1.679 | -0.686 | 1.07 | 0.425 | -0.01 | -0.082 | -0.082 | 1.97 | 0.73 | 0.4 | 0.8 | 0.92 | 0.77 | 0.84 |
| 4 | 183289625 | rs13137861 | -1.87 | -0.764 | 1.171 | 0.423 | -0.014 | -0.089 | -0.089 | 0 | 0.27 | 0.59 | 0.21 | 0.07 | 0.22 | 0.16 |
| 4 | 183291465 | rs13435448 | -0.788 | -0.812 | 1.329 | 0.399 | -0.047 | -0.028 | -0.028 | 0 | 0.11 | 0.37 | 0.06 | 0.07 | 0.03 | 0.1 |
| 4 | 183292629 | rs6552560 | -1.247 | -0.738 | 1.215 | 0.376 | 0.035 | -0.107 | -0.107 | 0 | 0.17 | 0.39 | 0.17 | 0.01 | 0.2 | 0.05 |
| 4 | 183296575 | rs930878 | 1.509 | -1.201 | 0.286 | 0.51 | 0.016 | 0.09 | 0.09 | 0 | 0.46 | 0.2 | 0.49 | 0.25 | 0.54 | 0.49 |
| 4 | 183299086 | rs2309656 | 0.829 | -1.178 | 0.715 | 0.72 | -0.036 | -0.082 | -0.082 | 0 | 0.86 | 0.61 | 0.93 | 0.99 | 0.94 | 0.95 |
| 4 | 183299318 | rs17073211 | -0.986 | -1.094 | 0.518 | 0.767 | -0.052 | -0.093 | -0.093 | 0.423 | 0.19 | 0.41 | 0.09 | 0.03 | 0.08 | 0.13 |
| 4 | 183302059 | rs1878221 | 0.465 | -0.556 | -0.394 | 1.864 | -0.011 | -0.008 | -0.008 | 0 | 0.92 | 0.98 | 0.85 | 0.59 | 0.92 | 0.82 |
| 4 | 183309211 | rs17073223 | -0.457 | -0.444 | -0.333 | 1.548 | -0.006 | -0.007 | -0.007 | 0.301 | 0.07 | 0.05 | 0.26 | 0.18 | 0.1 | 0.17 |
| 4 | 183313353 | rs2045405 | 1.33 | -1.403 | 0.13 | 1.288 | -0.053 | 0.008 | 0.008 | 0.45 | 0.71 | 0.31 | 0.59 | 0.68 | 0.86 | 0.7 |
| 4 | 183314283 | rs11935545 | 0.321 | -0.74 | 0.08 | 1.23 | 0.018 | 0.033 | 0.033 | 1.109 | 0.31 | 0.35 | 0.36 | 0.52 | 0.4 | 0.44 |
| 4 | 183314359 | rs13109717 | -0.125 | -1.103 | -0.666 | 0.832 | 0.026 | -0.054 | -0.054 | 0 | 0.27 | 0.16 | 0.42 | 0.45 | 0.26 | 0.38 |
| 4 | 183315919 | rs10520529 | 0.392 | -0.802 | -0.553 | 0.509 | -0.043 | 0.191 | 0.191 | 0 | 0.37 | 0.13 | 0.18 | 0.03 | 0.34 | 0.19 |
| 4 | 183321916 | rs1516538 | -0.232 | -0.878 | -0.506 | 0.796 | -0.042 | 0.101 | 0.101 | 0.065 | 0.45 | 0.18 | 0.49 | 0.27 | 0.42 | 0.33 |
| 4 | 183324280 | rs17253954 | 0.532 | -0.929 | -0.4 | 0.894 | -0.036 | 0.15 | 0.15 | 0 | 0.36 | 0.18 | 0.26 | 0.08 | 0.35 | 0.19 |
| 4 | 183327907 | rs6830860 | 0.336 | -0.869 | 0.136 | 1.379 | 0 | 0 | 0 | 0 | 0.48 | 0.44 | 0.36 | 0.47 | 0.53 | 0.59 |
| 4 | 183330031 | rs17073255 | 0.227 | -0.692 | -0.241 | 1.296 | -0.016 | -0.042 | -0.042 | 0 | 0.07 | 0 | 0.23 | 0.2 | 0.07 | 0.11 |
| 4 | 183349618 | rs10050018 | -0.776 | -0.735 | -0.185 | 1.022 | -0.015 | 0.103 | 0.103 | 0 | 0.63 | 0.76 | 0.74 | 0.93 | 0.65 | 0.79 |
| 4 | 183361857 | rs6822105 | 0.411 | -0.732 | 0.011 | 1.41 | -0.002 | -0.001 | -0.001 | 0 | 0.46 | 0.41 | 0.39 | 0.47 | 0.53 | 0.59 |
| 4 | 183364714 | rs17073286 | 0.058 | -0.56 | -0.34 | 1.024 | 0.001 | 0.028 | 0.028 | 0 | 0.52 | 0.56 | 0.47 | 0.66 | 0.58 | 0.64 |
| 4 | 183365002 | rs12498833 | -0.421 | -0.15 | -0.698 | 1.424 | -0.013 | -0.043 | -0.043 | 1.873 | 0.07 | 0.01 | 0.19 | 0.21 | 0.06 | 0.11 |
| 4 | 183367131 | rs12505033 | -0.55 | -0.174 | -0.71 | 1.412 | -0.011 | -0.046 | -0.046 | 0 | 0.08 | 0.01 | 0.19 | 0.21 | 0.06 | 0.11 |
| 4 | 183368769 | rs11736941 | -0.269 | 0.111 | -0.77 | 0.568 | -0.004 | 0.013 | 0.013 | 0 | 0.4 | 0.32 | 0.38 | 0.24 | 0.41 | 0.47 |
| 4 | 183373146 | rs727695 | -0.273 | 0.595 | -1.25 | 0.892 | -0.024 | -0.041 | -0.041 | 0 | 0.26 | 0.18 | 0.41 | 0.44 | 0.31 | 0.37 |
| 4 | 183373593 | rs13132492 | -0.65 | 0.378 | -1.319 | 0.008 | 0.006 | 0.078 | 0.078 | 0 | 0.55 | 0.63 | 0.72 | 0.82 | 0.59 | 0.57 |
| 4 | 183376566 | rs2063906 | 0.73 | 0.236 | -1.808 | -0.479 | -0.011 | 0.054 | 0.054 | 0 | 0.27 | 0.16 | 0.19 | 0.17 | 0.25 | 0.25 |
| 4 | 183380952 | rs6830998 | 1.042 | 0.1 | -1.935 | -0.285 | -0.095 | -0.136 | -0.136 | 0.558 | 0.25 | 0.04 | 0.42 | 0.51 | 0.33 | 0.4 |
| 4 | 183381596 | rs10012537 | 1.085 | -0.317 | -1.016 | -0.431 | -0.015 | 0.011 | 0.011 | 0 | 0.23 | 0.21 | 0.15 | 0.28 | 0.19 | 0.23 |
| 4 | 183382591 | rs9312297 | -0.918 | -0.256 | -0.55 | -0.619 | -0.008 | -0.067 | -0.067 | 0 | 0.8 | 0.64 | 0.8 | 0.96 | 0.82 | 0.9 |
| 4 | 183391730 | rs4861512 | 0.224 | -0.165 | -0.349 | -0.575 | -0.003 | 0.117 | 0.117 | 0 | 0.48 | 0.23 | 0.56 | 0.8 | 0.53 | 0.63 |
| 4 | 183396702 | rs13135084 | 1.378 | -0.44 | -0.7 | -0.657 | -0.043 | 0.006 | 0.006 | 0 | 0.17 | 0.1 | 0.17 | 0.17 | 0.24 | 0.12 |
| 4 | 183404353 | rs961533 | 0.564 | -0.688 | -0.529 | -0.769 | 0.002 | 0.029 | 0.029 | 0 | 0.51 | 0.34 | 0.58 | 0.75 | 0.56 | 0.65 |
| 4 | 183404747 | rs2124983 | -1.482 | -0.922 | -0.458 | -0.968 | -0.039 | -0.054 | -0.054 | 0.201 | 0.86 | 0.76 | 0.93 | 1 | 0.94 | 0.97 |
| 4 | 183405200 | rs10461323 | 1.379 | -0.862 | -0.72 | -0.848 | 0.006 | -0.004 | -0.004 | 0 | 0.23 | 0.12 | 0.14 | 0.27 | 0.2 | 0.22 |
| 4 | 183405506 | rs11132133 | 0.43 | -0.636 | -0.443 | -1.084 | 0.001 | -0.021 | -0.021 | 0 | 0.65 | 0.31 | 0.72 | 0.92 | 0.76 | 0.75 |
| 4 | 183405866 | rs4626248 | 1.011 | -0.625 | -0.479 | -1.034 | -0.009 | 0.11 | 0.11 | 0 | 0.48 | 0.21 | 0.56 | 0.76 | 0.52 | 0.62 |
| 4 | 183407652 | rs1350319 | 0.523 | -0.996 | -0.292 | -0.758 | -0.05 | -0.105 | -0.105 | 0 | 0.68 | 0.44 | 0.75 | 0.92 | 0.8 | 0.77 |
| 4 | 183407913 | rs1350318 | -0.476 | -1.313 | -0.314 | -0.361 | -0.051 | -0.064 | -0.064 | 0 | 0.88 | 0.74 | 0.94 | 1 | 0.95 | 0.97 |
| 4 | 183408872 | rs2198923 | -0.083 | -0.664 | -0.965 | 0.201 | -0.029 | -0.017 | -0.017 | 0 | 0.3 | 0.16 | 0.34 | 0.37 | 0.5 | 0.37 |
| 4 | 183409136 | rs13145928 | 0.058 | -1.944 | -0.991 | -0.584 | -0.017 | -0.028 | -0.028 | 0 | 0.72 | 0.73 | 0.9 | 0.95 | 0.89 | 0.81 |
| 4 | 183412942 | rs2309731 | 0.389 | -1.16 | -1.145 | -0.06 | -0.023 | -0.012 | -0.012 | 0 | 0.26 | 0.15 | 0.3 | 0.33 | 0.45 | 0.35 |
| 4 | 183415338 | rs7668237 | 0.383 | -1.353 | -1.117 | 0.172 | -0.006 | -0.005 | -0.005 | 0 | 0.22 | 0.15 | 0.29 | 0.34 | 0.41 | 0.33 |
| 4 | 183417126 | rs6838995 | -1.021 | -1.996 | -1.202 | 0.123 | -0.033 | 0.017 | 0.017 | 0 | 0.58 | 0.78 | 0.54 | 0.63 | 0.56 | 0.48 |
| 4 | 183417526 | rs13107205 | 1.524 | -2.196 | -1.284 | 0.014 | -0.106 | -0.053 | -0.053 | 0.289 | 0.3 | 0.08 | 0.43 | 0.37 | 0.39 | 0.4 |
| 4 | 183417581 | rs978520 | 0.572 | -2.37 | -0.878 | -0.046 | -0.058 | 0.012 | 0.012 | 0 | 0.25 | 0.4 | 0.21 | 0.31 | 0.12 | 0.18 |
| 4 | 183422459 | rs10034371 | -0.938 | -2.18 | -0.8 | 0.156 | 0.004 | 0.006 | 0.006 | 0 | 0.52 | 0.76 | 0.45 | 0.58 | 0.52 | 0.46 |
| 4 | 183422869 | rs13105695 | 0.628 | -2.09 | -0.746 | 0.208 | -0.002 | -0.002 | -0.002 | 0 | 0.26 | 0.21 | 0.28 | 0.3 | 0.4 | 0.3 |
| 4 | 183424808 | rs6856951 | 1.119 | -2.22 | -0.552 | 0.263 | -0.066 | 0.028 | 0.028 | 0 | 0.18 | 0.33 | 0.11 | 0.27 | 0.07 | 0.15 |
| 4 | 183427057 | rs4862073 | 1.088 | -2.169 | -0.799 | 0.249 | -0.08 | 0.031 | 0.031 | 0 | 0.19 | 0.4 | 0.11 | 0.28 | 0.07 | 0.15 |

| | | | | | | | | | | | | | | | | |
|---|---|---|---|---|---|---|---|---|---|---|---|---|---|---|---|---|
| 4 | 183438470 | rs7657508 | 0.061 | -1.822 | -1.23 | 0.384 | 0.002 | 0.023 | 0.023 | 0 | 0.53 | 0.68 | 0.57 | 0.65 | 0.48 | 0.57 |
| 4 | 183439439 | rs11736841 | 1.066 | -1.654 | -1.332 | 0.414 | -0.05 | 0.005 | 0.005 | 0 | 0.3 | 0.18 | 0.33 | 0.29 | 0.34 | 0.39 |
| 4 | 183449806 | rs6552568 | 0.505 | -1.869 | -1.356 | 0.308 | 0.091 | 0.06 | 0.06 | 1.564 | 0.46 | 0.22 | 0.67 | 0.68 | 0.74 | 0.69 |
| 4 | 183450690 | rs7671649 | -0.687 | -1.728 | -1.111 | 0.382 | -0.004 | -0.018 | -0.018 | 0.598 | 0.74 | 0.74 | 0.82 | 0.96 | 0.85 | 0.85 |
| 4 | 183452259 | rs869240 | 1.112 | -1.586 | -1.206 | 0.502 | -0.003 | -0.003 | -0.003 | 0.135 | 0.21 | 0.15 | 0.26 | 0.31 | 0.36 | 0.27 |
| 4 | 183454132 | rs11731599 | 0.2 | -1.582 | -1.187 | 0.511 | 0 | 0 | 0 | 2.551 | 0.43 | 0.55 | 0.39 | 0.59 | 0.46 | 0.45 |
| 4 | 183454812 | rs2309692 | 1.755 | -1.452 | -1.324 | 0.357 | 0.001 | -0.011 | -0.011 | 0 | 0.12 | 0.04 | 0.1 | 0.18 | 0.15 | 0.1 |
| 4 | 183458613 | rs13110757 | -0.509 | -1.548 | -1.356 | 0.178 | 0.011 | 0.005 | 0.005 | 0 | 0.74 | 0.77 | 0.82 | 0.95 | 0.85 | 0.85 |
| 4 | 183460719 | rs6812487 | -1.048 | -1.389 | -1.349 | 0.348 | 0.008 | 0.008 | 0.008 | 0 | 0.82 | 0.85 | 0.87 | 0.96 | 0.91 | 0.86 |
| 4 | 183464677 | rs6840083 | -0.604 | -0.504 | -1.21 | 0.88 | 0.002 | 0.025 | 0.025 | 0 | 0.45 | 0.61 | 0.49 | 0.66 | 0.51 | 0.46 |
| 4 | 183468125 | rs6812970 | -0.218 | 0.4 | -0.919 | 0.852 | 0.006 | 0.006 | 0.006 | 0.449 | 0.23 | 0.2 | 0.24 | 0.3 | 0.36 | 0.3 |
| 4 | 183472582 | rs4428328 | -0.073 | 0.085 | -1.226 | 0.702 | 0.025 | -0.004 | -0.004 | 0 | 0.48 | 0.39 | 0.5 | 0.58 | 0.53 | 0.56 |
| 4 | 183475934 | rs13144500 | -0.052 | 0.509 | -1.69 | 0.943 | -0.047 | -0.062 | -0.062 | 0 | 0.13 | 0.01 | 0.11 | 0.22 | 0.15 | 0.22 |
| 4 | 183476247 | rs7674760 | -0.329 | -0.023 | -1.238 | 0.601 | -0.024 | 0.007 | 0.007 | 0 | 0.28 | 0.56 | 0.41 | 0.24 | 0.24 | 0.33 |
| 4 | 183477329 | rs10013166 | 0.76 | -0.058 | -1.973 | 0.344 | -0.035 | -0.021 | -0.021 | 3.561 | 0.47 | 0.18 | 0.39 | 0.58 | 0.49 | 0.42 |
| 4 | 183479255 | rs17258085 | 0.54 | -0.272 | -1.942 | 0.39 | -0.018 | 0.022 | 0.022 | 0 | 0.21 | 0.09 | 0.24 | 0.1 | 0.25 | 0.31 |
| 4 | 183481315 | rs11937236 | 0.425 | 0.104 | -1.923 | 0.294 | 0.005 | -0.032 | -0.032 | 0 | 0.61 | 0.23 | 0.49 | 0.59 | 0.62 | 0.47 |
| 4 | 183484147 | rs7670152 | 0.577 | -0.151 | -2.154 | 0.374 | -0.03 | -0.028 | -0.028 | 0 | 0.43 | 0.1 | 0.35 | 0.65 | 0.55 | 0.6 |
| 4 | 183486609 | rs17258176 | 1.049 | 0.002 | -2.075 | 0.401 | -0.047 | 0.069 | 0.069 | 0 | 0.13 | 0.01 | 0.1 | 0.01 | 0.13 | 0.05 |
| 4 | 183497776 | rs9996737 | 0.894 | 0.314 | -1.775 | 0.724 | -0.069 | -0.072 | -0.072 | 0 | 0.33 | 0.11 | 0.27 | 0.51 | 0.43 | 0.44 |
| 4 | 183500457 | rs10020119 | -0.969 | 0.222 | -1.806 | 0.579 | 0.009 | 0.026 | 0.026 | 1.924 | 0.76 | 0.79 | 0.78 | 0.92 | 0.77 | 0.85 |
| 4 | 183501781 | rs6552578 | -0.217 | 0.171 | -1.725 | 0.623 | 0.007 | 0.016 | 0.016 | 0 | 0.42 | 0.62 | 0.47 | 0.33 | 0.35 | 0.42 |
| 4 | 183503611 | rs10520536 | 0.307 | 0.876 | -1.004 | 1.149 | 0.029 | 0.025 | 0.025 | 0 | 0.12 | 0.25 | 0.09 | 0.13 | 0.13 | 0.18 |
| 4 | 183504017 | rs13145682 | -0.269 | 0.905 | -1.675 | 1.256 | -0.037 | -0.037 | -0.037 | 0.005 | 0.34 | 0.15 | 0.27 | 0.5 | 0.42 | 0.42 |
| 4 | 183506166 | rs13136589 | -0.318 | 0.922 | -1.666 | 1.102 | -0.036 | -0.048 | -0.048 | 0 | 0.34 | 0.15 | 0.31 | 0.61 | 0.42 | 0.43 |
| 4 | 183516599 | rs10033526 | -0.215 | 0.975 | -1.846 | 0.923 | -0.056 | -0.053 | -0.053 | 0.378 | 0.33 | 0.12 | 0.27 | 0.5 | 0.42 | 0.42 |
| 4 | 183518246 | rs9985535 | -0.092 | 0.614 | -2.267 | 0.45 | 0.03 | 0.021 | 0.021 | 0 | 0.51 | 0.21 | 0.53 | 0.52 | 0.55 | 0.62 |
| 4 | 183521798 | rs6833629 | 0.034 | 0.494 | -2.101 | -0.431 | -0.004 | -0.068 | -0.068 | 0 | 0.72 | 0.42 | 0.86 | 0.93 | 0.77 | 0.82 |
| 4 | 183525697 | rs11132137 | 0.801 | 1.068 | -3.126 | -0.299 | -0.078 | -0.056 | -0.056 | 0 | 0.16 | 0.02 | 0.17 | 0.26 | 0.23 | 0.27 |
| 4 | 183526414 | rs6814485 | 0.323 | 0.436 | -3.537 | -0.417 | 0.033 | -0.129 | -0.129 | 2.05 | 0.18 | 0.01 | 0.32 | 0.54 | 0.11 | 0.24 |
| 4 | 183529154 | rs13109227 | 1.069 | 0.958 | -3.282 | -0.594 | -0.067 | 0.057 | 0.057 | 0 | 0.15 | 0.01 | 0.05 | 0.04 | 0.17 | 0.12 |
| 4 | 183532267 | rs6818350 | 0.687 | 1.03 | -2.702 | -0.617 | 0.029 | -0.092 | -0.092 | 0 | 0.15 | 0.33 | 0.14 | 0.01 | 0.17 | 0.07 |
| 4 | 183533788 | rs17324394 | 0.468 | 0.782 | -3.304 | -0.035 | -0.012 | 0.04 | 0.04 | 0.29 | 0.23 | 0.04 | 0.23 | 0.12 | 0.25 | 0.23 |
| 4 | 183536086 | rs12499296 | 0.37 | 0.797 | -3.324 | -0.048 | -0.014 | 0.046 | 0.046 | 0.065 | 0.24 | 0.04 | 0.28 | 0.12 | 0.25 | 0.23 |
| 4 | 183541307 | rs6843930 | -0.381 | 0.878 | -2.563 | 0.437 | -0.079 | -0.058 | -0.058 | 0 | 0.25 | 0.02 | 0.22 | 0.31 | 0.27 | 0.3 |
| 4 | 183543569 | rs12648727 | -0.281 | 0.385 | -1.97 | 0.604 | 0.004 | -0.006 | -0.006 | 0 | 0.09 | 0.08 | 0.06 | 0.14 | 0.07 | 0.14 |
| 4 | 183543770 | rs11132138 | -0.308 | 0.05 | -2.396 | 0.192 | 0.023 | -0.025 | -0.025 | 0 | 0.57 | 0.25 | 0.47 | 0.52 | 0.65 | 0.66 |
| 4 | 183545589 | rs6552585 | -0.05 | -0.14 | -2.438 | 0.339 | 0.008 | -0.019 | -0.019 | 0.007 | 0.9 | 0.95 | 0.91 | 0.79 | 0.95 | 0.85 |
| 4 | 183546180 | rs12504411 | 0.482 | 0.13 | -2.8 | 0.182 | -0.017 | -0.008 | -0.008 | 0 | 0.73 | 0.9 | 0.75 | 0.71 | 0.76 | 0.71 |
| 4 | 183550589 | rs7693901 | 0.39 | -0.316 | -2.485 | 0.081 | -0.008 | 0.072 | 0.072 | 0.018 | 0.13 | 0.12 | 0.32 | 0.47 | 0.09 | 0.19 |
| 4 | 183550637 | rs1554037 | 1.065 | -0.21 | -2.568 | -0.152 | 0.017 | -0.016 | -0.016 | 0 | 0.59 | 0.73 | 0.64 | 0.57 | 0.64 | 0.55 |
| 4 | 183553052 | rs6841261 | 0.425 | -0.16 | -2.556 | -0.44 | 0.003 | 0.036 | 0.036 | 0.062 | 0.69 | 0.75 | 0.76 | 0.74 | 0.73 | 0.71 |
| 4 | 183553276 | rs12639914 | 0.9 | 0.054 | -2.516 | -0.188 | -0.014 | -0.047 | -0.047 | 0 | 0.08 | 0.01 | 0.07 | 0.13 | 0.08 | 0.14 |
| 4 | 183553837 | rs1013096 | -0.084 | 0.083 | -2.385 | -0.27 | -0.016 | 0.014 | 0.014 | 0.335 | 0.18 | 0.1 | 0.2 | 0.2 | 0.2 | 0.16 |
| 4 | 183556869 | rs17325360 | 1.042 | 0.12 | -2.645 | -0.61 | -0.069 | 0.056 | 0.056 | 0 | 0.27 | 0.07 | 0.17 | 0.14 | 0.37 | 0.31 |
| 4 | 183556999 | rs10520539 | 0.341 | -0.387 | -1.735 | -0.916 | -0.006 | -0.014 | -0.014 | 0.335 | 0.63 | 0.51 | 0.39 | 0.27 | 0.63 | 0.57 |
| 4 | 183559306 | rs7688538 | 0.643 | -0.375 | -1.903 | -0.769 | -0.035 | -0.128 | -0.128 | 0 | 0.2 | 0.03 | 0.4 | 0.51 | 0.22 | 0.26 |
| 4 | 183559425 | rs4507403 | -0.386 | -0.466 | -0.933 | -0.846 | 0.004 | -0.015 | -0.015 | 0 | 0.57 | 0.52 | 0.68 | 0.73 | 0.5 | 0.52 |
| 4 | 183561942 | rs11930604 | 0.569 | -0.507 | -1.054 | -0.894 | -0.003 | 0.006 | 0.006 | 0 | 0.09 | 0.12 | 0.08 | 0.13 | 0.08 | 0.14 |
| 4 | 183565618 | rs11722354 | 0.834 | -0.339 | -1.08 | -1.148 | -0.004 | -0.026 | -0.026 | 0.302 | 0.05 | 0.01 | 0.09 | 0.22 | 0.04 | 0.12 |
| 4 | 183567188 | rs6552587 | -0.999 | -0.227 | -0.946 | -1.626 | -0.052 | 0.067 | 0.067 | 0 | 0.7 | 0.86 | 0.78 | 0.87 | 0.57 | 0.67 |
| 4 | 183572390 | rs6850978 | -0.811 | -0.512 | -0.272 | -1.193 | 0.008 | 0.013 | 0.013 | 0 | 0.42 | 0.41 | 0.62 | 0.87 | 0.37 | 0.55 |
| 4 | 183578598 | rs9312302 | 0.426 | -0.076 | -0.405 | -0.713 | -0.001 | -0.017 | -0.017 | 0 | 0.63 | 0.47 | 0.5 | 0.18 | 0.61 | 0.5 |
| 4 | 183581152 | rs4386638 | 0.038 | 0.055 | -0.301 | -1.076 | -0.001 | -0.052 | -0.052 | 0 | 0.19 | 0.33 | 0.14 | 0 | 0.22 | 0.05 |
| 4 | 183583762 | rs10428431 | -1.133 | 0.126 | -0.559 | -1.146 | -0.009 | 0.027 | 0.027 | 0 | 0.87 | 0.94 | 0.9 | 0.95 | 0.87 | 0.92 |
| 4 | 183584040 | rs17073734 | 1.427 | 0.07 | -0.466 | -0.731 | 0.003 | -0.007 | -0.007 | 0 | 0.26 | 0.19 | 0.4 | 0.39 | 0.22 | 0.25 |
| 4 | 183589823 | rs12648977 | 1.047 | 0.146 | -0.648 | -0.451 | -0.002 | 0.013 | 0.013 | 0 | 0.09 | 0.09 | 0.07 | 0.17 | 0.06 | 0.13 |
| 4 | 183590236 | rs2309774 | 0.333 | 0.542 | -0.502 | -0.867 | 0.003 | -0.07 | -0.07 | 0.031 | 0.66 | 0.53 | 0.74 | 0.89 | 0.63 | 0.83 |
| 4 | 183591707 | rs11132145 | -0.524 | 1.053 | -0.848 | -0.088 | -0.007 | 0.023 | 0.023 | 0 | 0.85 | 0.94 | 0.88 | 0.89 | 0.85 | 0.89 |
| 4 | 183593318 | rs2309759 | -1.115 | 1.288 | -0.697 | -0.194 | 0.007 | -0.039 | -0.039 | 0 | 0.88 | 0.87 | 0.91 | 0.95 | 0.88 | 0.92 |
| 4 | 183594938 | rs4535369 | -0.093 | 1.228 | -0.995 | 0.266 | -0.036 | -0.013 | -0.013 | 0 | 0.27 | 0.17 | 0.25 | 0.35 | 0.34 | 0.44 |
| 4 | 183604892 | rs4618351 | 0.439 | 0.936 | -0.286 | -0.107 | 0.007 | -0.037 | -0.037 | 0.42 | 0.9 | 0.74 | 0.93 | 0.97 | 0.91 | 0.79 |
| 4 | 183605088 | rs11938233 | 0.563 | 0.942 | -0.277 | -0.764 | -0.001 | -0.008 | -0.008 | 0 | 0.57 | 0.45 | 0.37 | 0.19 | 0.58 | 0.39 |
| 4 | 183608189 | rs7697828 | 0.374 | 0.969 | -0.172 | -0.711 | 0.001 | -0.045 | -0.045 | 0 | 0.08 | 0.19 | 0.07 | 0 | 0.07 | 0.04 |
| 4 | 183611553 | rs6845497 | -0.937 | 0.77 | -0.422 | -0.634 | -0.001 | -0.004 | -0.004 | 0.217 | 0.19 | 0.2 | 0.22 | 0.09 | 0.24 | 0.25 |
| 4 | 183612535 | rs7694547 | 1.098 | 0.667 | -0.143 | -0.522 | 0 | 0.007 | 0.007 | 0.255 | 0.18 | 0.19 | 0.38 | 0.7 | 0.17 | 0.32 |
| 4 | 183613431 | rs17073855 | 1.681 | 0.697 | -0.61 | -1.481 | -0.005 | 0.032 | 0.032 | 0 | 0.19 | 0.1 | 0.13 | 0.04 | 0.19 | 0.13 |
| 4 | 183616911 | rs1987079 | -0.495 | 0.535 | -0.28 | -1.641 | -0.003 | -0.026 | -0.026 | 2.325 | 0.55 | 0.53 | 0.46 | 0.19 | 0.61 | 0.47 |
| 4 | 183618777 | rs4241748 | -0.013 | 0.503 | -0.298 | -1.676 | -0.001 | -0.017 | -0.017 | 0 | 0.47 | 0.5 | 0.56 | 0.82 | 0.43 | 0.53 |

| | | | | | | | | | | | | | | | | |
|---|---|---|---|---|---|---|---|---|---|---|---|---|---|---|---|---|
| 4 | 183622901 | rs10434298 | -1.129 | 0.518 | -0.122 | -1.605 | -0.011 | -0.048 | -0.048 | 0 | 0.92 | 0.79 | 0.94 | 1 | 0.94 | 0.96 |
| 4 | 183626332 | rs6858496 | 0.245 | 0.515 | -0.012 | -1.633 | -0.005 | -0.037 | -0.037 | 0 | 0.82 | 0.66 | 0.79 | 0.91 | 0.79 | 0.76 |
| 4 | 183628586 | rs6841510 | 1.274 | 0.388 | -0.437 | -1.963 | -0.044 | 0.1 | 0.1 | 2.679 | 0.31 | 0.13 | 0.17 | 0.09 | 0.32 | 0.21 |
| 4 | 183628872 | rs12498650 | 2.708 | 0.382 | -0.581 | -1.288 | -0.049 | -0.093 | -0.093 | 1.238 | 0.12 | 0.02 | 0.3 | 0.48 | 0.14 | 0.18 |
| 4 | 183632213 | rs17073904 | 0.409 | 0.48 | -0.052 | -2.105 | -0.009 | -0.045 | -0.045 | 0 | 0.34 | 0.36 | 0.23 | 0.1 | 0.3 | 0.23 |
| 4 | 183640777 | rs13101364 | 0.924 | 0.49 | -0.372 | -1.834 | -0.013 | 0.011 | 0.011 | 1.259 | 0.17 | 0.09 | 0.08 | 0.14 | 0.19 | 0.17 |
| 4 | 183647869 | rs11721401 | 1.369 | 0.092 | -0.272 | -1.821 | -0.002 | -0.014 | -0.014 | 0 | 0.08 | 0.08 | 0.03 | 0 | 0.04 | 0.05 |
| 4 | 183650100 | rs13340295 | 0.226 | 0.17 | -0.498 | -1.674 | -0.017 | -0.049 | -0.049 | 0.312 | 0.1 | 0.17 | 0.05 | 0 | 0.06 | 0.04 |
| 4 | 183651765 | rs6829893 | -0.134 | 0.635 | -0.701 | -1.394 | -0.002 | -0.002 | -0.002 | 0.132 | 0.9 | 0.9 | 0.84 | 0.9 | 0.84 | 0.77 |
| 4 | 183654099 | rs13435237 | 0.351 | 0.115 | -0.539 | -1.594 | -0.03 | -0.065 | -0.065 | 0 | 0.19 | 0.32 | 0.11 | 0.05 | 0.08 | 0.14 |
| 4 | 183655636 | rs4241750 | -0.094 | 0.65 | -1.262 | -1.524 | 0.036 | 0.074 | 0.074 | 0.429 | 0.54 | 0.34 | 0.62 | 0.86 | 0.67 | 0.58 |
| 4 | 183655750 | rs4557308 | 0.958 | 1.109 | -1.519 | -1.434 | -0.092 | -0.006 | -0.006 | 0 | 0.13 | 0.01 | 0.1 | 0.13 | 0.26 | 0.16 |
| 4 | 183661413 | rs10021016 | 0.698 | 1.125 | -1.457 | -1.436 | -0.011 | 0.008 | 0.008 | 0.784 | 0.15 | 0.09 | 0.12 | 0.11 | 0.26 | 0.16 |
| 4 | 183661673 | rs12648546 | -0.508 | 1.134 | -1.635 | -1.427 | -0.022 | -0.04 | -0.04 | 0 | 0.58 | 0.51 | 0.64 | 0.81 | 0.7 | 0.56 |
| 4 | 183662527 | rs11734588 | -0.508 | 1.134 | -1.635 | -1.428 | -0.022 | -0.041 | -0.041 | 0 | 0.58 | 0.51 | 0.64 | 0.81 | 0.7 | 0.55 |
| 4 | 183663175 | rs6552603 | -0.926 | 1.117 | -1.654 | -1.458 | -0.027 | -0.069 | -0.069 | 0 | 0.64 | 0.55 | 0.8 | 0.91 | 0.76 | 0.74 |
| 4 | 183664522 | rs7654255 | 0.057 | 1.248 | -1.932 | -1.306 | 0.069 | -0.048 | -0.048 | 0 | 0.2 | 0.35 | 0.14 | 0.14 | 0.3 | 0.17 |
| 4 | 183666740 | rs7690035 | -0.103 | 1.254 | -2.6 | -0.768 | -0.003 | 0.029 | 0.029 | 0 | 0.38 | 0.15 | 0.52 | 0.68 | 0.4 | 0.37 |
| 4 | 183675227 | rs33914616 | 0.611 | 1.592 | -2.518 | -0.216 | -0.029 | -0.097 | -0.097 | 0 | 0.18 | 0.05 | 0.25 | 0.61 | 0.17 | 0.17 |
| 4 | 183677453 | rs11736855 | 0.686 | 1.913 | -2.282 | -0.513 | 0.005 | 0.036 | 0.036 | 0 | 0.26 | 0.17 | 0.2 | 0.13 | 0.26 | 0.23 |
| 4 | 183680961 | rs10028158 | 0.182 | 2.524 | -1.952 | 0.374 | -0.027 | 0.022 | 0.022 | 0 | 0.12 | 0.05 | 0.08 | 0.07 | 0.19 | 0.17 |
| 4 | 183684925 | rs11727669 | -0.377 | 1.833 | -1.855 | 0.446 | -0.083 | 0.044 | 0.044 | 0.125 | 0.38 | 0.14 | 0.5 | 0.76 | 0.47 | 0.49 |
| 4 | 183685402 | rs17074036 | -0.556 | 1.685 | -0.913 | 1.051 | 0 | 0 | 0 | 0.207 | 0.09 | 0.08 | 0.08 | 0.06 | 0.05 | 0.13 |
| 4 | 183686235 | rs17264104 | -1.681 | 2.044 | -0.836 | 1.148 | -0.029 | -0.021 | -0.021 | 0 | 0.05 | 0.01 | 0.18 | 0.09 | 0.12 | 0.14 |
| 4 | 183689629 | rs17074042 | -1.658 | 2.097 | -0.935 | 1.709 | -0.013 | -0.039 | -0.039 | 0.507 | 0.07 | 0.01 | 0.15 | 0.14 | 0.08 | 0.04 |
| 4 | 183689666 | rs17074044 | -2.375 | 1.94 | -0.297 | 1.614 | 0.006 | 0.064 | 0.064 | 0 | 0.13 | 0.24 | 0.34 | 0.26 | 0.18 | 0.18 |
| 4 | 183692079 | rs17330237 | -2.066 | 2.513 | -0.384 | 1.31 | 0.016 | 0.004 | 0.004 | 0 | 0.08 | 0.07 | 0.22 | 0.11 | 0.19 | 0.15 |
| 4 | 183692580 | rs4577620 | 0.933 | 2.814 | -0.367 | 1.22 | 0.029 | -0.004 | -0.004 | 0 | 0.78 | 0.66 | 0.58 | 0.83 | 0.72 | 0.73 |
| 4 | 183692641 | rs4467618 | 1.687 | 2.211 | -0.751 | 0.788 | -0.013 | 0.01 | 0.01 | 0.015 | 0.85 | 0.96 | 0.79 | 0.89 | 0.81 | 0.81 |
| 4 | 183707781 | rs12500335 | -0.538 | 2.63 | -0.884 | 0.894 | -0.049 | -0.037 | -0.037 | 0 | 0.09 | 0.01 | 0.21 | 0.08 | 0.14 | 0.07 |
| 4 | 183707836 | rs12508106 | -0.538 | 2.642 | -0.884 | 0.896 | -0.047 | -0.035 | -0.035 | 0.079 | 0.09 | 0.01 | 0.21 | 0.09 | 0.14 | 0.07 |
| 4 | 183709536 | rs17074066 | -0.484 | 1.861 | 0.385 | 1.094 | -0.012 | 0.045 | 0.045 | 0 | 0.13 | 0.19 | 0.17 | 0.25 | 0.09 | 0.3 |
| 4 | 183709842 | rs3805200 | 0.112 | 1.965 | -0.204 | 0.012 | -0.031 | 0.025 | 0.025 | 0.151 | 0.24 | 0.2 | 0.23 | 0.07 | 0.38 | 0.12 |
| 4 | 183715225 | rs6842610 | -1.182 | 1.687 | 0.127 | 0.666 | 0.01 | 0.046 | 0.046 | 0 | 0.23 | 0.27 | 0.36 | 0.42 | 0.28 | 0.38 |
| 4 | 183716648 | rs2309772 | 0.595 | 1.879 | -0.234 | -0.141 | -0.033 | 0.001 | 0.001 | 0 | 0.08 | 0.01 | 0.13 | 0.04 | 0.1 | 0.05 |
| 4 | 183722743 | rs2309777 | -1.564 | 1.54 | -0.309 | -0.363 | -0.001 | 0.015 | 0.015 | 0.046 | 0.37 | 0.65 | 0.35 | 0.26 | 0.35 | 0.25 |
| 4 | 183730650 | rs17267298 | -1.084 | 1.583 | -0.247 | -0.889 | -0.01 | 0.016 | 0.016 | 0 | 0.22 | 0.14 | 0.2 | 0.04 | 0.26 | 0.1 |
| 4 | 183731737 | rs13124391 | 0.86 | 0.719 | -0.986 | -1.273 | 0.034 | 0.042 | 0.042 | 0.243 | 0.22 | 0.08 | 0.09 | 0.12 | 0.13 | 0.17 |
| 4 | 183735049 | rs6858295 | -1.231 | 0.991 | -0.53 | -1.069 | 0.033 | -0.081 | -0.081 | 0 | 0.26 | 0.35 | 0.24 | 0.07 | 0.31 | 0.11 |
| 4 | 183735309 | rs4862103 | 0.119 | 1.486 | -1.339 | -0.49 | -0.019 | 0.012 | 0.012 | 0 | 0.1 | 0.04 | 0.16 | 0.04 | 0.17 | 0.06 |
| 4 | 183744704 | rs4241754 | 0.069 | 0.576 | -1.079 | -0.497 | 0.003 | -0.053 | -0.053 | 0 | 0.83 | 0.82 | 0.9 | 0.96 | 0.85 | 0.9 |
| 4 | 183750550 | rs13133452 | -0.423 | 0.227 | -1.609 | -0.031 | 0.023 | 0.073 | 0.073 | 0.339 | 0.23 | 0.07 | 0.12 | 0.12 | 0.17 | 0.21 |
| 4 | 183761427 | rs6854616 | -0.742 | 0.35 | -1.164 | 0.526 | 0.003 | -0.001 | -0.001 | 0 | 0.4 | 0.61 | 0.49 | 0.41 | 0.37 | 0.62 |
| 4 | 183763618 | rs4862116 | 0.805 | -0.061 | -0.308 | 0.315 | -0.054 | -0.039 | -0.039 | 0.079 | 0.74 | 0.56 | 0.8 | 0.82 | 0.83 | 0.82 |
| 4 | 183768698 | rs6822595 | 0.649 | 0.528 | -0.827 | 0.168 | -0.018 | 0.046 | 0.046 | 0 | 0.39 | 0.27 | 0.36 | 0.28 | 0.41 | 0.37 |
| 4 | 183771387 | rs7680411 | -0.245 | 0.48 | -0.224 | 0.367 | 0 | -0.016 | -0.016 | 0.021 | 0.14 | 0.14 | 0.12 | 0.01 | 0.13 | 0.06 |
| 4 | 183778572 | rs13118559 | 0.799 | 1.042 | -0.734 | 0.634 | -0.032 | -0.071 | -0.071 | 0.004 | 0.11 | 0.01 | 0.17 | 0.26 | 0.12 | 0.16 |
| 4 | 183778788 | rs11736797 | 0.753 | 1.142 | -0.684 | 0.2 | 0 | -0.012 | -0.012 | 0 | 0.53 | 0.84 | 0.48 | 0.54 | 0.47 | 0.54 |
| 4 | 183782993 | rs4862122 | 0.769 | 1.126 | -0.403 | -0.021 | -0.002 | 0.002 | 0.002 | 1.529 | 0.52 | 0.42 | 0.54 | 0.49 | 0.52 | 0.5 |
| 4 | 183791706 | rs4861534 | -0.353 | 1.483 | -0.273 | -0.573 | 0.005 | -0.076 | -0.076 | 0.564 | 0.12 | 0.32 | 0.13 | 0.02 | 0.08 | 0.03 |
| 4 | 183792885 | rs17074159 | -0.641 | 1.612 | -0.409 | -0.539 | 0.017 | -0.138 | -0.138 | 0 | 0.23 | 0.42 | 0.22 | 0.03 | 0.22 | 0.11 |
| 4 | 183793275 | rs11132155 | -0.548 | 1.669 | -1.136 | 0.241 | -0.018 | -0.058 | -0.058 | 0.057 | 0.24 | 0.12 | 0.42 | 0.35 | 0.25 | 0.3 |
| 4 | 183797355 | rs11132156 | 0.696 | 1.585 | -0.488 | -0.484 | 0.023 | -0.11 | -0.11 | 0 | 0.71 | 0.43 | 0.69 | 0.96 | 0.66 | 0.85 |
| 4 | 183809272 | rs17271644 | -0.687 | 1.081 | -1.287 | 0.1 | 0.009 | 0.007 | 0.007 | 0.042 | 0.46 | 0.75 | 0.43 | 0.39 | 0.43 | 0.54 |
| 4 | 183810141 | rs6829021 | -0.226 | 0.739 | -0.263 | 0.015 | 0.034 | -0.028 | -0.028 | 0 | 0.23 | 0.36 | 0.16 | 0.14 | 0.27 | 0.23 |
| 4 | 183810813 | rs6834938 | -0.413 | 0.66 | -0.122 | 0.014 | -0.041 | 0.021 | 0.021 | 0 | 0.23 | 0.37 | 0.26 | 0.25 | 0.16 | 0.3 |
| 4 | 183810865 | rs6835318 | -1.106 | 0.752 | 0.032 | -0.092 | -0.003 | -0.001 | -0.001 | 0 | 0.3 | 0.62 | 0.32 | 0.25 | 0.23 | 0.33 |
| 4 | 183811487 | rs7278 | 0.18 | 0.876 | 0 | -0.134 | -0.068 | -0.028 | -0.028 | 0 | 0.14 | 0.02 | 0.09 | 0.14 | 0.18 | 0.18 |
| 4 | 183811513 | rs7277 | -1.08 | 0.883 | 0.034 | -0.129 | -0.005 | -0.001 | -0.001 | 0 | 0.3 | 0.62 | 0.32 | 0.25 | 0.23 | 0.33 |
| 4 | 183812287 | rs3190314 | -1.08 | 0.883 | 0.034 | -0.129 | -0.005 | -0.001 | -0.001 | 2.147 | 0.3 | 0.62 | 0.32 | 0.26 | 0.23 | 0.33 |
| 4 | 183812351 | rs9542 | 1.573 | 0.749 | 0.013 | -0.185 | -0.001 | -0.006 | -0.006 | 0.61 | 0.53 | 0.15 | 0.56 | 0.61 | 0.59 | 0.47 |
| 4 | 183813147 | rs2515683 | 1.179 | -0.183 | -0.428 | -0.609 | -0.004 | -0.002 | -0.002 | 0 | 0.56 | 0.46 | 0.59 | 0.61 | 0.61 | 0.49 |
| 4 | 183813425 | rs12507552 | -0.279 | -0.041 | 0.341 | -0.279 | -0.028 | 0.036 | 0.036 | 0 | 0.16 | 0.26 | 0.17 | 0.23 | 0.11 | 0.29 |
| 4 | 183815561 | rs17331744 | -1.04 | 0.261 | 0.062 | -0.177 | -0.024 | -0.006 | -0.006 | 0 | 0.18 | 0.09 | 0.14 | 0.21 | 0.24 | 0.17 |
| 4 | 183817659 | rs6834019 | -1.056 | 0.549 | 0.908 | -0.29 | 0.109 | 0.024 | 0.024 | 0 | 0.07 | 0.39 | 0.05 | 0 | 0.05 | 0.05 |
| 4 | 183819095 | rs6831306 | -1.525 | 0.544 | 1.127 | -0.276 | 0.012 | -0.006 | -0.006 | 0.035 | 0.23 | 0.24 | 0.16 | 0.21 | 0.29 | 0.21 |
| 4 | 183821662 | rs13114435 | -1.364 | 0.559 | 0.963 | -0.269 | -0.038 | 0.006 | 0.006 | 0 | 0.21 | 0.08 | 0.15 | 0.21 | 0.27 | 0.21 |
| 4 | 183823597 | rs13101260 | -1.646 | 0.361 | 0.738 | -0.047 | -0.031 | -0.026 | -0.026 | 0 | 0.25 | 0.08 | 0.15 | 0.31 | 0.3 | 0.24 |
| 4 | 183824984 | rs10016530 | -1.86 | 0.302 | 1.372 | -0.171 | 0.1 | 0.016 | 0.016 | 0.161 | 0.07 | 0.31 | 0.04 | 0.01 | 0.05 | 0.03 |

| | | | | | | | | | | | | | | | | |
|---|---|---|---|---|---|---|---|---|---|---|---|---|---|---|---|---|
| 4 | 183825476 | rs13111117 | -0.744 | 0.37 | 1.034 | -0.133 | 0.002 | -0.019 | -0.019 | 0 | 0.06 | 0.02 | 0.04 | 0.07 | 0.08 | 0.1 |
| 4 | 183833942 | rs6823077 | -2.517 | 0.495 | 1.032 | -0.17 | -0.01 | 0.015 | 0.015 | 0.294 | 0.37 | 0.62 | 0.28 | 0.28 | 0.43 | 0.32 |
| 4 | 183840898 | rs5016499 | -2.029 | 0.552 | 1.116 | -0.125 | 0.002 | 0.024 | 0.024 | 0.694 | 0.25 | 0.49 | 0.21 | 0.34 | 0.28 | 0.26 |
| 4 | 183849072 | rs13117933 | -2.285 | 0.503 | 0.843 | 0.12 | 0.01 | 0.022 | 0.022 | 0.306 | 0.27 | 0.34 | 0.22 | 0.36 | 0.31 | 0.26 |
| 4 | 183849605 | rs10520544 | -0.534 | 0.528 | 0.047 | 0.169 | 0.01 | -0.024 | -0.024 | 0 | 0.05 | 0.02 | 0.02 | 0.08 | 0.06 | 0.06 |
| 4 | 183850395 | rs11737887 | -0.608 | 0.589 | 0.099 | 0.172 | -0.03 | -0.039 | -0.039 | 0 | 0.14 | 0.05 | 0.15 | 0.28 | 0.2 | 0.16 |
| 4 | 183858411 | rs4131396 | 1.368 | 0.573 | -1.439 | 0.167 | -0.089 | 0.005 | 0.005 | 0.343 | 0.27 | 0.02 | 0.25 | 0.18 | 0.33 | 0.16 |
| 4 | 183872630 | rs4607268 | 0.634 | 1.161 | -0.925 | 1.388 | -0.04 | -0.105 | -0.105 | 0 | 0.2 | 0.01 | 0.34 | 0.3 | 0.17 | 0.32 |
| 4 | 183873241 | rs4453974 | -0.439 | 1.488 | -0.049 | 1.35 | -0.06 | -0.073 | -0.073 | 0 | 0.12 | 0.01 | 0.15 | 0.28 | 0.2 | 0.15 |
| 4 | 183880263 | rs2714510 | -2.028 | 1.349 | 0.54 | 1.396 | 0.01 | 0.091 | 0.091 | 0.019 | 0.21 | 0.48 | 0.21 | 0.36 | 0.25 | 0.19 |
| 4 | 183892717 | rs6830570 | -1.95 | 1.54 | 0.494 | 1.265 | 0.03 | 0.036 | 0.036 | 0.037 | 0.22 | 0.49 | 0.22 | 0.29 | 0.23 | 0.17 |
| 4 | 183896828 | rs2709350 | -1.437 | 1.21 | 0.706 | 1.198 | 0.072 | 0.133 | 0.133 | 0 | 0.09 | 0.32 | 0.07 | 0.17 | 0.12 | 0.12 |
| 4 | 183907715 | rs1454371 | 1.221 | 1.23 | 0.538 | 1.152 | 0.095 | 0.15 | 0.15 | 0 | 0.91 | 0.61 | 0.92 | 0.85 | 0.88 | 0.86 |
| 4 | 183916674 | rs4069861 | 1.53 | 1.242 | 0.322 | 1.072 | 0.002 | 0.009 | 0.009 | 0.017 | 0.77 | 0.4 | 0.81 | 0.71 | 0.76 | 0.73 |
| 4 | 183929581 | rs1377943 | 1.005 | 1.377 | 0.29 | 1.243 | 0.041 | 0.083 | 0.083 | 0 | 0.84 | 0.53 | 0.85 | 0.72 | 0.79 | 0.78 |
| 4 | 183931994 | rs9998545 | -0.954 | 1.727 | -0.285 | 1.379 | 0.004 | 0.008 | 0.008 | 0 | 0.1 | 0.08 | 0.14 | 0.18 | 0.17 | 0.08 |
| 4 | 183932844 | rs2278924 | 0.463 | 1.3 | -1.207 | 1.368 | -0.031 | -0.075 | -0.075 | 0.06 | 0.2 | 0.01 | 0.3 | 0.23 | 0.15 | 0.32 |
| 4 | 183934023 | rs2714546 | 0.961 | 1.028 | -0.652 | 0.95 | 0.003 | 0.001 | 0.001 | 1.008 | 0.46 | 0.43 | 0.37 | 0.39 | 0.38 | 0.38 |
| 4 | 183934142 | rs17334067 | 0.573 | 1.45 | -0.925 | 0.752 | -0.025 | 0.002 | 0.002 | 0 | 0.13 | 0.06 | 0.13 | 0.13 | 0.18 | 0.11 |
| 4 | 183935289 | rs2714545 | 0.474 | 0.896 | -0.237 | 0.628 | 0.004 | -0.026 | -0.026 | 0 | 0.06 | 0.24 | 0.04 | 0.01 | 0.03 | 0.05 |
| 4 | 183942308 | rs7660903 | -0.673 | 1.001 | -0.312 | 0.569 | -0.001 | 0.009 | 0.009 | 1.57 | 0.42 | 0.26 | 0.48 | 0.38 | 0.42 | 0.53 |
| 4 | 183944066 | rs17274861 | 0.979 | 1.01 | -0.293 | 0.516 | -0.005 | -0.022 | -0.022 | 0 | 0.06 | 0.08 | 0.03 | 0.01 | 0.03 | 0.05 |
| 4 | 183947384 | rs1902044 | 1.003 | 0.947 | -0.584 | 0.537 | -0.072 | -0.075 | -0.075 | 0 | 0.23 | 0.02 | 0.34 | 0.25 | 0.25 | 0.3 |
| 4 | 183951585 | rs747368 | -0.694 | 0.876 | -0.132 | 0.548 | -0.001 | -0.003 | -0.003 | 0 | 0.38 | 0.38 | 0.44 | 0.32 | 0.38 | 0.5 |
| 4 | 183959226 | rs2016910 | -0.301 | 0.808 | -0.461 | 0.636 | 0.003 | 0.01 | 0.01 | 0.298 | 0.36 | 0.24 | 0.42 | 0.34 | 0.35 | 0.5 |
| 4 | 183963607 | rs4450975 | 0.137 | 0.803 | -0.544 | 0.625 | 0.001 | 0 | 0 | 0.964 | 0.32 | 0.19 | 0.38 | 0.32 | 0.29 | 0.47 |
| 4 | 183989102 | rs6850320 | -0.314 | 0.78 | -0.299 | 0.656 | 0.003 | 0.005 | 0.005 | 0.306 | 0.37 | 0.3 | 0.44 | 0.34 | 0.35 | 0.5 |
| 4 | 183994213 | rs10520548 | 1.489 | 0.566 | -1.056 | 1.05 | -0.011 | -0.078 | -0.078 | 0.605 | 0.13 | 0.01 | 0.11 | 0.25 | 0.11 | 0.1 |
| 4 | 183995589 | rs6825466 | -0.464 | 0.74 | -0.64 | 0.882 | 0.002 | 0.004 | 0.004 | 0.186 | 0.42 | 0.41 | 0.5 | 0.35 | 0.4 | 0.54 |
| 4 | 184060896 | rs6846985 | 0.745 | 1.158 | -1.123 | 1.104 | -0.054 | -0.071 | -0.071 | 0.039 | 0.15 | 0.01 | 0.2 | 0.24 | 0.17 | 0.23 |
| 4 | 184063608 | rs11935093 | 0.686 | 1.011 | -0.878 | 1.252 | -0.002 | 0.011 | 0.011 | 0 | 0.17 | 0.21 | 0.14 | 0.24 | 0.16 | 0.1 |
| 4 | 184070426 | rs10520545 | 1.756 | 1.048 | -0.96 | 1.239 | -0.03 | -0.071 | -0.071 | 0 | 0.12 | 0.01 | 0.1 | 0.23 | 0.12 | 0.07 |
| 4 | 184098334 | rs4469120 | 0.785 | 1.067 | -0.592 | 0.949 | 0.005 | 0.017 | 0.017 | 0 | 0.65 | 0.47 | 0.66 | 0.49 | 0.6 | 0.57 |
| 4 | 184122020 | rs6845251 | -0.667 | 1.028 | -0.287 | 0.946 | 0.023 | 0.034 | 0.034 | 0 | 0.8 | 0.55 | 0.75 | 0.75 | 0.78 | 0.73 |
| 4 | 184127332 | rs6831552 | 0.812 | 1.141 | -0.103 | 0.997 | -0.001 | 0 | 0 | 0.042 | 0.18 | 0.16 | 0.25 | 0.26 | 0.22 | 0.27 |
| 4 | 184132563 | rs1877290 | 0.811 | 1.139 | -0.104 | 0.996 | -0.001 | 0 | 0 | 3.475 | 0.18 | 0.16 | 0.25 | 0.26 | 0.22 | 0.26 |
| 4 | 184139997 | rs12643595 | 0.992 | 1.151 | -0.316 | 1.166 | -0.024 | -0.044 | -0.044 | 0.318 | 0.09 | 0.01 | 0.04 | 0.15 | 0.13 | 0.11 |
| 4 | 184144709 | rs6855711 | -0.518 | 0.685 | -0.712 | 0.465 | -0.005 | -0.002 | -0.002 | 0 | 0.63 | 0.57 | 0.73 | 0.67 | 0.65 | 0.69 |
| 4 | 184155515 | rs7688139 | 1.68 | 0.821 | -0.552 | 0.585 | 0.007 | 0.057 | 0.057 | 0 | 0.1 | 0.17 | 0.11 | 0.24 | 0.1 | 0.06 |
| 4 | 184168117 | rs17290885 | 2.9 | 0.616 | -0.931 | 0.14 | -0.011 | 0.031 | 0.031 | 0 | 0.07 | 0.01 | 0.05 | 0 | 0.06 | 0.02 |
| 4 | 184171187 | rs10020261 | -0.333 | 1.039 | -0.937 | 0.105 | 0.002 | -0.014 | -0.014 | 0 | 0.58 | 0.41 | 0.35 | 0.25 | 0.57 | 0.65 |
| 4 | 184179699 | rs7695317 | 0.007 | 0.797 | -0.928 | 0.388 | 0.004 | 0.009 | 0.009 | 0.524 | 0.38 | 0.37 | 0.34 | 0.3 | 0.37 | 0.5 |
| 4 | 184183588 | rs7438080 | 1.327 | 0.766 | -1.287 | 0.174 | 0.002 | 0.053 | 0.053 | 0 | 0.2 | 0.01 | 0.12 | 0.07 | 0.16 | 0.25 |
| 4 | 184188404 | rs6812460 | -0.562 | 1.073 | -0.672 | 0.42 | 0.006 | 0.001 | 0.001 | 2.621 | 0.59 | 0.5 | 0.45 | 0.54 | 0.58 | 0.67 |
| 4 | 184189952 | rs3926627 | -0.391 | 1.561 | -0.717 | 0.243 | -0.012 | 0.006 | 0.006 | 0 | 0.22 | 0.16 | 0.18 | 0.13 | 0.32 | 0.2 |
| 4 | 184192037 | rs1473070 | 0.071 | 1.045 | -0.093 | 0.422 | 0.002 | 0.065 | 0.065 | 0 | 0.71 | 0.58 | 0.69 | 0.55 | 0.73 | 0.72 |
| 4 | 184196516 | rs9996190 | 1.096 | 0.604 | -0.325 | -0.112 | 0.001 | 0.033 | 0.033 | 0.358 | 0.29 | 0.12 | 0.39 | 0.24 | 0.26 | 0.23 |
| 4 | 184197313 | rs13116042 | 2.042 | 0.814 | -0.392 | -0.149 | -0.04 | 0.036 | 0.036 | 0 | 0.07 | 0 | 0.07 | 0.01 | 0.13 | 0.05 |
| 4 | 184201182 | rs12643425 | 0.007 | 0.702 | 0.128 | -0.157 | -0.009 | 0.039 | 0.039 | 2.842 | 0.44 | 0.38 | 0.52 | 0.28 | 0.46 | 0.35 |
| 4 | 184201909 | rs11132173 | 0.734 | 0.68 | 0.275 | -0.161 | -0.006 | 0.012 | 0.012 | 0.179 | 0.35 | 0.29 | 0.48 | 0.25 | 0.39 | 0.28 |
| 4 | 184206296 | rs17291855 | 1.51 | 0.764 | 0.318 | -0.057 | 0 | 0.008 | 0.008 | 0.019 | 0.07 | 0.02 | 0.04 | 0.02 | 0.05 | 0.06 |
| 4 | 184215675 | rs955748 | 0.18 | 0.773 | 0.444 | -0.051 | -0.01 | 0.059 | 0.059 | 0 | 0.7 | 0.51 | 0.68 | 0.54 | 0.75 | 0.71 |
| 4 | 184215832 | rs2175476 | 1.763 | 0.808 | 0.403 | 0.05 | 0 | 0.002 | 0.002 | 0 | 0.1 | 0.08 | 0.16 | 0.18 | 0.09 | 0.18 |
| 4 | 184221172 | rs6811091 | 0.771 | 0.741 | 0.15 | -1.258 | -0.024 | 0.048 | 0.048 | 1.143 | 0.32 | 0.23 | 0.35 | 0.2 | 0.36 | 0.27 |
| 4 | 184224437 | rs17074589 | 1.923 | 0.754 | 0.546 | -1.344 | -0.002 | -0.006 | -0.006 | 0 | 0.16 | 0.13 | 0.08 | 0.08 | 0.12 | 0.21 |
| 4 | 184225421 | rs11929772 | 0.089 | 0.797 | 0.554 | -1.391 | 0 | 0.014 | 0.014 | 0.056 | 0.31 | 0.24 | 0.2 | 0.12 | 0.3 | 0.34 |
| 4 | 184226187 | rs11730648 | 0.384 | 0.944 | 0.509 | -1.319 | -0.043 | 0.041 | 0.041 | 0.039 | 0.13 | 0.04 | 0.12 | 0.03 | 0.18 | 0.11 |
| 4 | 184227066 | rs2309875 | -0.817 | 0.737 | 0.708 | -1.307 | 0.035 | 0.086 | 0.086 | 0 | 0.76 | 0.59 | 0.68 | 0.56 | 0.72 | 0.72 |
| 4 | 184228248 | rs1397942 | 1.476 | 0.755 | 0.583 | -1.279 | -0.012 | 0.024 | 0.024 | 1.009 | 0.28 | 0.13 | 0.4 | 0.24 | 0.28 | 0.23 |
| 4 | 184230630 | rs10003798 | 1.416 | 0.784 | 0.501 | -1.269 | -0.002 | 0.022 | 0.022 | 1.192 | 0.28 | 0.19 | 0.4 | 0.24 | 0.27 | 0.23 |
| 4 | 184232252 | rs6849819 | 0.39 | 0.661 | 0.394 | -1.183 | -0.009 | 0.024 | 0.024 | 0 | 0.22 | 0.11 | 0.13 | 0.13 | 0.25 | 0.2 |
| 4 | 184234587 | rs907360 | -1.24 | 0.338 | 1.012 | -0.768 | 0.006 | 0.109 | 0.109 | 0 | 0.82 | 0.68 | 0.75 | 0.57 | 0.84 | 0.78 |
| 4 | 184235796 | rs13141994 | -0.532 | 0.301 | 1.031 | -0.789 | 0.002 | 0.078 | 0.078 | 0 | 0.77 | 0.62 | 0.72 | 0.57 | 0.8 | 0.76 |
| 4 | 184236868 | rs4862155 | 1.079 | 0.346 | 0.392 | -0.855 | 0.003 | 0.009 | 0.009 | 0.019 | 0.07 | 0.02 | 0.04 | 0.02 | 0.05 | 0.06 |
| 4 | 184243528 | rs7688467 | -0.61 | 0.435 | 0.49 | -0.816 | 0.004 | 0.073 | 0.073 | 0 | 0.78 | 0.67 | 0.73 | 0.57 | 0.79 | 0.76 |
| 4 | 184249490 | rs2048472 | 0.211 | 0.432 | -0.136 | -0.663 | 0.021 | 0.004 | 0.004 | 0 | 0.23 | 0.18 | 0.13 | 0.19 | 0.18 | 0.29 |
| 4 | 184251370 | rs35432122 | 0.815 | 0.487 | -0.116 | -0.693 | -0.003 | 0.054 | 0.054 | 0 | 0.51 | 0.33 | 0.53 | 0.37 | 0.51 | 0.42 |
| 4 | 184252789 | rs907365 | -0.701 | 0.831 | 0.187 | -0.568 | 0.015 | 0.174 | 0.174 | 0 | 0.81 | 0.49 | 0.73 | 0.57 | 0.83 | 0.77 |
| 4 | 184259614 | rs4616804 | -0.249 | 0.656 | 0.139 | -1.26 | -0.03 | 0.056 | 0.056 | 0.11 | 0.23 | 0.08 | 0.15 | 0.13 | 0.27 | 0.27 |

| | | | | | | | | | | | | | | | | |
|---|---|---|---|---|---|---|---|---|---|---|---|---|---|---|---|---|
| 4 | 184270907 | rs10029290 | -0.267 | 0.672 | 0.954 | -1.177 | 0.003 | 0.002 | 0.002 | 0 | 0.3 | 0.29 | 0.18 | 0.14 | 0.28 | 0.3 |
| 4 | 184271019 | rs7669080 | -1.012 | 0.706 | 0.911 | -1.139 | -0.004 | -0.039 | -0.039 | 5.378 | 0.41 | 0.5 | 0.26 | 0.16 | 0.42 | 0.37 |
| 4 | 184276454 | rs4861543 | 0.94 | 0.865 | 0.556 | -1.097 | 0 | 0.014 | 0.014 | 0 | 0.54 | 0.42 | 0.69 | 0.81 | 0.54 | 0.6 |
| 4 | 184277307 | rs17074631 | 0.178 | 1.161 | 0.394 | -0.97 | 0.022 | -0.027 | -0.027 | 0 | 0.07 | 0.13 | 0.07 | 0.01 | 0.13 | 0.06 |
| 4 | 184278868 | rs907364 | 1.083 | 0.833 | 0.487 | -1.1 | 0.002 | 0.004 | 0.004 | 0 | 0.57 | 0.4 | 0.73 | 0.8 | 0.59 | 0.6 |
| 4 | 184279303 | rs7687627 | 0.482 | 0.702 | 0.473 | -1.218 | -0.016 | -0.033 | -0.033 | 0 | 0.65 | 0.58 | 0.78 | 0.82 | 0.69 | 0.67 |
| 4 | 184279613 | rs4861544 | -0.199 | 0.933 | 0.766 | -0.886 | 0.001 | 0.001 | 0.001 | 0.104 | 0.25 | 0.24 | 0.16 | 0.17 | 0.24 | 0.29 |
| 4 | 184280695 | rs868083 | 1.095 | 0.871 | 0.36 | -0.79 | -0.001 | 0.006 | 0.006 | 0.498 | 0.61 | 0.36 | 0.74 | 0.81 | 0.59 | 0.62 |
| 4 | 184280944 | rs868082 | -0.033 | 1.096 | -0.094 | -0.673 | -0.026 | 0.026 | 0.026 | 0 | 0.07 | 0.04 | 0.06 | 0.01 | 0.13 | 0.06 |
| 4 | 184281811 | rs9992054 | 0.228 | 0.343 | 0.247 | -0.381 | -0.006 | -0.015 | -0.015 | 0 | 0.74 | 0.66 | 0.83 | 0.83 | 0.74 | 0.71 |
| 4 | 184283411 | rs9995832 | -1.157 | 0.299 | 0.291 | -0.31 | -0.008 | -0.057 | -0.057 | 0.62 | 0.11 | 0.31 | 0.06 | 0.01 | 0.09 | 0.07 |
| 4 | 184285017 | rs13151481 | 0.043 | 0.237 | 0.062 | -0.661 | 0.004 | -0.05 | -0.05 | 0.119 | 0.42 | 0.49 | 0.42 | 0.21 | 0.46 | 0.43 |
| 4 | 184285089 | rs6552661 | -0.087 | -0.077 | 0.452 | -0.263 | -0.005 | 0.062 | 0.062 | 0.196 | 0.53 | 0.32 | 0.55 | 0.7 | 0.52 | 0.45 |
| 4 | 184285671 | rs6820687 | -0.241 | -0.615 | 0.551 | -0.769 | 0.006 | -0.004 | -0.004 | 0 | 0.32 | 0.4 | 0.51 | 0.74 | 0.36 | 0.46 |
| 4 | 184289319 | rs1106052 | 1.063 | -1.073 | 0.604 | -0.582 | -0.011 | 0.029 | 0.029 | 0 | 0.37 | 0.29 | 0.37 | 0.22 | 0.43 | 0.4 |
| 4 | 184292528 | rs6821104 | 0.404 | -1.412 | 0.807 | -0.573 | 0.004 | 0.002 | 0.002 | 0 | 0.52 | 0.48 | 0.49 | 0.4 | 0.58 | 0.55 |
| 4 | 184292904 | rs4862171 | 0.733 | -1.456 | 0.761 | -0.484 | -0.057 | -0.103 | -0.103 | 0 | 0.22 | 0.1 | 0.36 | 0.58 | 0.3 | 0.33 |
| 4 | 184293030 | rs17074696 | 0.025 | -1.349 | 0.849 | -0.826 | 0.057 | 0.189 | 0.189 | 0 | 0.06 | 0.23 | 0.03 | 0.13 | 0.06 | 0.13 |
| 4 | 184294510 | rs6834774 | 1.121 | -1.242 | 0.508 | -0.289 | -0.037 | -0.028 | -0.028 | 0 | 0.13 | 0.08 | 0.15 | 0.21 | 0.22 | 0.27 |
| 4 | 184296685 | rs11931607 | -0.354 | -1.348 | 0.798 | -0.98 | -0.052 | -0.022 | -0.022 | 0.409 | 0.29 | 0.36 | 0.15 | 0.15 | 0.18 | 0.24 |
| 4 | 184296883 | rs955229 | 1.097 | -1.196 | 0.156 | -0.971 | -0.041 | 0.026 | 0.026 | 0 | 0.08 | 0 | 0.04 | 0.01 | 0.1 | 0.02 |
| 4 | 184298419 | rs4602536 | -0.722 | -1.257 | 0.221 | -0.995 | -0.058 | 0.088 | 0.088 | 2.263 | 0.4 | 0.17 | 0.53 | 0.79 | 0.44 | 0.51 |
| 4 | 184300125 | rs17074703 | -0.273 | -1.028 | 0.344 | -0.846 | -0.006 | -0.008 | -0.008 | 0.11 | 0.13 | 0.07 | 0.11 | 0.13 | 0.13 | 0.09 |
| 4 | 184301020 | rs1567597 | -0.381 | -1.179 | 0.311 | -0.927 | 0.024 | 0.038 | 0.038 | 0.143 | 0.3 | 0.1 | 0.22 | 0.15 | 0.24 | 0.18 |
| 4 | 184302049 | rs17293543 | 1.857 | -1.342 | 0.459 | -1.012 | 0.015 | -0.043 | -0.043 | 0 | 0.32 | 0.34 | 0.32 | 0.15 | 0.4 | 0.32 |
| 4 | 184302549 | rs17293620 | 0.501 | -1.331 | 0.057 | -0.879 | 0.037 | 0.009 | 0.009 | 0.211 | 0.21 | 0.09 | 0.13 | 0.12 | 0.14 | 0.15 |
| 4 | 184302684 | rs12641554 | 0.288 | -1.175 | -0.094 | -0.782 | -0.049 | -0.067 | -0.067 | 0 | 0.29 | 0.1 | 0.42 | 0.66 | 0.33 | 0.43 |
| 4 | 184306513 | rs1511674 | 0.116 | -1.201 | 0.225 | -0.429 | -0.031 | -0.034 | -0.034 | 0.667 | 0.75 | 0.64 | 0.86 | 0.94 | 0.85 | 0.84 |
| 4 | 184311757 | rs17074720 | -0.303 | -0.738 | 0.086 | -0.132 | -0.006 | 0.071 | 0.071 | 0 | 0.05 | 0.14 | 0.01 | 0.09 | 0.02 | 0.06 |
| 4 | 184311806 | rs11132179 | 0.582 | -0.724 | -0.337 | -0.329 | -0.004 | -0.003 | -0.003 | 0 | 0.61 | 0.4 | 0.74 | 0.81 | 0.73 | 0.75 |
| 4 | 184315862 | rs7699622 | -0.917 | 0.008 | -0.785 | -0.726 | 0.006 | 0.042 | 0.042 | 0 | 0.34 | 0.27 | 0.25 | 0.07 | 0.31 | 0.28 |
| 4 | 184323148 | rs17294373 | -0.669 | 0.421 | -0.71 | -0.622 | -0.023 | 0.022 | 0.022 | 0.137 | 0.12 | 0.05 | 0.09 | 0 | 0.16 | 0.12 |
| 4 | 184323983 | rs17074740 | -1.017 | 0.626 | -0.146 | -0.136 | 0.03 | 0.024 | 0.024 | 0 | 0.17 | 0.33 | 0.22 | 0.23 | 0.22 | 0.17 |
| 4 | 184325258 | rs4862177 | 0.481 | -0.423 | -0.554 | -0.386 | 0 | -0.002 | -0.002 | 0 | 0.68 | 0.73 | 0.87 | 0.77 | 0.79 | 0.7 |
| 4 | 184325450 | rs4862179 | 0.511 | 0.05 | -0.844 | 0.213 | 0 | -0.001 | -0.001 | 0.237 | 0.66 | 0.33 | 0.8 | 0.59 | 0.69 | 0.64 |
| 4 | 184334102 | rs11725216 | 0.4 | 0.393 | -1.74 | 0.262 | -0.052 | -0.085 | -0.085 | 0.379 | 0.36 | 0.14 | 0.46 | 0.4 | 0.41 | 0.3 |
| 4 | 184335306 | rs4415002 | 0.211 | -0.166 | -1.305 | 0.086 | 0.016 | 0.033 | 0.033 | 0.156 | 0.76 | 0.85 | 0.88 | 0.82 | 0.77 | 0.81 |
| 4 | 184338589 | rs6552667 | 0.631 | -0.369 | -0.882 | 0.92 | -0.008 | 0.063 | 0.063 | 0.188 | 0.76 | 0.61 | 0.63 | 0.57 | 0.78 | 0.58 |
| 4 | 184345714 | rs6858319 | 0.912 | -1.387 | -1.807 | 1.045 | 0.075 | -0.005 | -0.005 | 0 | 0.46 | 0.16 | 0.4 | 0.4 | 0.61 | 0.37 |
| 4 | 184353138 | rs4241767 | 0.14 | -2.034 | -0.913 | 0.831 | -0.029 | -0.05 | -0.05 | 0 | 0.72 | 0.67 | 0.7 | 0.89 | 0.83 | 0.8 |
| 4 | 184356197 | rs6842529 | 0.445 | -2.173 | -0.668 | 1.304 | -0.048 | -0.011 | -0.011 | 0.595 | 0.66 | 0.51 | 0.5 | 0.76 | 0.8 | 0.74 |
| 4 | 184356574 | rs13111484 | 1.103 | -2.206 | -1.451 | 0.696 | -0.032 | 0.004 | 0.004 | 0.113 | 0.34 | 0.15 | 0.32 | 0.29 | 0.59 | 0.33 |
| 4 | 184359288 | rs4485840 | -1.339 | -2.705 | -0.707 | 0.829 | -0.003 | 0.012 | 0.012 | 0 | 0.88 | 0.78 | 0.79 | 0.89 | 0.91 | 0.85 |
| 4 | 184362046 | rs4862197 | 0.892 | -2.901 | -0.768 | 0.791 | 0.141 | 0.012 | 0.012 | 0.25 | 0.53 | 0.24 | 0.46 | 0.38 | 0.77 | 0.43 |
| 4 | 184362068 | rs4861551 | 0.996 | -2.873 | -0.753 | 0.73 | 0.164 | 0.09 | 0.09 | 0.222 | 0.5 | 0.22 | 0.43 | 0.29 | 0.73 | 0.41 |
| 4 | 184369040 | rs3187025 | 0.15 | -2.092 | -0.401 | 0.327 | 0.009 | -0.001 | -0.001 | 0 | 0.16 | 0.11 | 0.08 | 0.01 | 0.09 | 0.06 |
| 4 | 184373811 | rs6844630 | 0.47 | -2.146 | -0.573 | 0.355 | 0.022 | 0.069 | 0.069 | 2.28 | 0.14 | 0.03 | 0.07 | 0.01 | 0.08 | 0.06 |
| 4 | 184376851 | rs6816216 | 0.157 | -2.4 | -0.312 | 0.596 | -0.028 | -0.005 | -0.005 | 0 | 0.74 | 0.72 | 0.74 | 0.8 | 0.87 | 0.78 |
| 4 | 184384311 | rs4241775 | 0.322 | -2.068 | -0.382 | 0.526 | -0.015 | -0.006 | -0.006 | 0 | 0.72 | 0.72 | 0.73 | 0.8 | 0.82 | 0.76 |
| 4 | 184389698 | rs4273514 | 0.971 | -2.249 | -0.435 | 0.637 | -0.02 | -0.019 | -0.019 | 3.223 | 0.57 | 0.53 | 0.42 | 0.41 | 0.69 | 0.58 |
| 4 | 184394392 | rs10007707 | 0.78 | -2.203 | -0.436 | 0.71 | -0.016 | 0.043 | 0.043 | 0 | 0.14 | 0.23 | 0.23 | 0.19 | 0.09 | 0.18 |
| 4 | 184399833 | rs4862210 | -0.005 | -2.116 | -0.516 | 0.434 | -0.023 | 0.056 | 0.056 | 0.258 | 0.23 | 0.3 | 0.28 | 0.46 | 0.19 | 0.34 |
| 4 | 184412581 | rs7677714 | -0.25 | -2.396 | -0.437 | 0.481 | -0.048 | -0.013 | -0.013 | 0 | 0.22 | 0.44 | 0.16 | 0.18 | 0.14 | 0.07 |
| 4 | 184415303 | rs6850330 | -1.37 | -2.543 | -0.534 | 0.495 | 0.021 | -0.024 | -0.024 | 0 | 0.89 | 0.94 | 0.78 | 0.8 | 0.94 | 0.82 |
| 4 | 184415338 | rs11723316 | 1.23 | -2.137 | -0.329 | 0.303 | 0.065 | 0.188 | 0.188 | 1.314 | 0.44 | 0.15 | 0.34 | 0.14 | 0.62 | 0.41 |
| 4 | 184416995 | rs11943515 | 0.01 | -1.98 | -0.187 | 0.431 | 0.017 | 0.002 | 0.002 | 3.53 | 0.07 | 0.42 | 0.05 | 0 | 0.02 | 0 |
| 4 | 184426387 | rs8872 | 0.078 | -1.886 | -0.811 | 0.609 | 0.005 | -0.008 | -0.008 | 0.714 | 0.7 | 0.72 | 0.65 | 0.63 | 0.8 | 0.74 |
| 4 | 184429659 | rs6830958 | 0.312 | -1.014 | -0.629 | 0.494 | 0.006 | -0.006 | -0.006 | 2.467 | 0.62 | 0.7 | 0.57 | 0.63 | 0.79 | 0.64 |
| 4 | 184438322 | rs6838723 | 0.511 | 0.196 | -0.596 | 0.325 | -0.031 | 0.043 | 0.043 | 1.27 | 0.13 | 0.23 | 0.15 | 0.17 | 0.06 | 0.13 |
| 4 | 184441694 | rs13141754 | -0.452 | 0.961 | -1.516 | 0.057 | -0.014 | 0.008 | 0.008 | 0.003 | 0.34 | 0.29 | 0.31 | 0.26 | 0.39 | 0.28 |
| 4 | 184441917 | rs11732255 | -0.178 | 0.542 | -1.91 | -0.269 | 0.077 | 0.207 | 0.207 | 2.86 | 0.46 | 0.14 | 0.44 | 0.15 | 0.61 | 0.49 |
| 4 | 184447081 | rs7658620 | -0.343 | 0.784 | -1.041 | 0.526 | -0.001 | -0.002 | -0.002 | 0 | 0.36 | 0.35 | 0.23 | 0.32 | 0.34 | 0.29 |
| 4 | 184454079 | rs4440262 | -0.49 | 1.272 | -1.006 | 0.941 | -0.014 | -0.011 | -0.011 | 0 | 0.24 | 0.26 | 0.41 | 0.31 | 0.34 | 0.36 |
| 4 | 184458946 | rs4524431 | -0.228 | 0.606 | -0.511 | 0.878 | -0.001 | -0.002 | -0.002 | 0 | 0.35 | 0.62 | 0.58 | 0.58 | 0.36 | 0.38 |
| 4 | 184462577 | rs4550975 | -0.124 | 0.61 | 0.135 | 1.05 | 0.022 | 0.042 | 0.042 | 0 | 0.7 | 0.56 | 0.52 | 0.62 | 0.69 | 0.78 |
| 4 | 184463695 | rs7699347 | -1.071 | -0.002 | -0.19 | 0.869 | -0.025 | 0.032 | 0.032 | 0 | 0.55 | 0.74 | 0.68 | 0.62 | 0.53 | 0.37 |
| 4 | 184465856 | rs10017798 | 0.314 | -0.024 | -0.153 | 0.813 | 0.007 | 0.007 | 0.007 | 0 | 0.19 | 0.17 | 0.14 | 0.14 | 0.16 | 0.11 |
| 4 | 184468253 | rs4426827 | -0.832 | 0.022 | -0.145 | 0.854 | 0.014 | 0.045 | 0.045 | 0 | 0.51 | 0.68 | 0.64 | 0.62 | 0.47 | 0.33 |
| 4 | 184471255 | rs11726805 | 0.793 | 0.02 | 0.081 | 0.923 | 0.009 | 0.047 | 0.047 | 0 | 0.26 | 0.34 | 0.45 | 0.38 | 0.25 | 0.18 |

| | | | | | | | | | | | | | | | | |
|---|---|---|---|---|---|---|---|---|---|---|---|---|---|---|---|---|
| 4 | 184472543 | rs6834876 | -0.438 | 0.179 | -0.393 | 0.915 | -0.008 | -0.006 | -0.006 | 0.373 | 0.39 | 0.71 | 0.59 | 0.57 | 0.38 | 0.36 |
| 4 | 184473333 | rs4862223 | 0.96 | 0.146 | -0.594 | 0.863 | -0.08 | 0.057 | 0.057 | 0.097 | 0.39 | 0.09 | 0.28 | 0.28 | 0.47 | 0.48 |
| 4 | 184478353 | rs4292366 | -0.466 | 0.351 | 0.258 | 1.205 | -0.002 | 0.007 | 0.007 | 0 | 0.34 | 0.65 | 0.51 | 0.47 | 0.3 | 0.3 |
| 4 | 184481702 | rs6811805 | -0.968 | 0.17 | 0.695 | 1.08 | 0.029 | 0.072 | 0.072 | 0.476 | 0.07 | 0.24 | 0.09 | 0.08 | 0.07 | 0.06 |
| 4 | 184487521 | rs7694722 | -0.649 | 0.131 | 0.593 | 0.878 | -0.019 | 0 | 0 | 4.39 | 0.18 | 0.34 | 0.15 | 0.19 | 0.14 | 0.17 |
| 4 | 184494270 | rs11723412 | -0.712 | -0.045 | 0.35 | 1.283 | -0.007 | 0.102 | 0.102 | 0 | 0.14 | 0.23 | 0.22 | 0.38 | 0.14 | 0.17 |
| 4 | 184494883 | rs10009455 | -0.621 | -0.328 | 0.49 | 0.945 | -0.043 | -0.031 | -0.031 | 0 | 0.19 | 0.3 | 0.13 | 0.15 | 0.09 | 0.22 |
| 4 | 184501675 | rs7662749 | -0.211 | 0.264 | -0.153 | 0.77 | -0.013 | -0.061 | -0.061 | 0 | 0.7 | 0.51 | 0.74 | 0.77 | 0.71 | 0.7 |
| 4 | 184503980 | rs11132193 | 1.144 | 0.738 | -1.333 | 0.696 | -0.092 | 0.011 | 0.011 | 0.111 | 0.4 | 0.06 | 0.37 | 0.28 | 0.4 | 0.43 |
| 4 | 184504511 | rs13127102 | 0.522 | 0.843 | -0.487 | 0.979 | 0.01 | -0.012 | -0.012 | 0 | 0.11 | 0.01 | 0.09 | 0.12 | 0.07 | 0.06 |
| 4 | 184507283 | rs4861562 | -0.604 | 1.232 | -0.008 | 1.327 | 0.002 | 0.025 | 0.025 | 0 | 0.08 | 0.09 | 0.11 | 0.2 | 0.1 | 0.12 |
| 4 | 184510402 | rs4394039 | 0.734 | 1.324 | 0.237 | 1.001 | 0.027 | 0.053 | 0.053 | 0 | 0.91 | 0.76 | 0.83 | 0.87 | 0.87 | 0.94 |
| 4 | 184517243 | rs7656979 | -0.321 | 1.358 | 0.041 | 0.005 | -0.002 | -0.048 | -0.048 | 0 | 0.38 | 0.39 | 0.28 | 0.2 | 0.31 | 0.21 |
| 4 | 184523965 | rs12641020 | 0.078 | 1.624 | 0.144 | 0.61 | 0.008 | 0.049 | 0.049 | 0 | 0.09 | 0.15 | 0.13 | 0.28 | 0.14 | 0.26 |
| 4 | 184525386 | rs7674701 | -1.229 | 1.51 | 0.309 | 0.114 | 0.01 | 0.001 | 0.001 | 0.525 | 0.26 | 0.36 | 0.3 | 0.29 | 0.34 | 0.4 |
| 4 | 184526440 | rs9884125 | -1.242 | 1.479 | 0.055 | -0.166 | 0.001 | -0.002 | -0.002 | 0 | 0.37 | 0.57 | 0.36 | 0.4 | 0.42 | 0.43 |
| 4 | 184529029 | rs4543168 | 0.262 | 1.803 | 0.058 | 0.157 | 0.009 | 0.051 | 0.051 | 0 | 0.09 | 0.14 | 0.06 | 0.28 | 0.13 | 0.24 |
| 4 | 184529969 | rs7671807 | 0.162 | 1.573 | -0.033 | -0.35 | -0.018 | 0.036 | 0.036 | 0.388 | 0.1 | 0.02 | 0.07 | 0 | 0.15 | 0.01 |
| 4 | 184530109 | rs13118994 | -0.278 | 1.554 | 0.185 | -0.124 | -0.001 | -0.008 | -0.008 | 0 | 0.28 | 0.3 | 0.23 | 0.17 | 0.25 | 0.18 |
| 4 | 184534704 | rs4624699 | 0.094 | 1.539 | 0.365 | 0.116 | 0.016 | -0.041 | -0.041 | 0 | 0.7 | 0.51 | 0.56 | 0.77 | 0.72 | 0.76 |
| 4 | 184535274 | rs13128324 | 0.422 | 1.783 | 0.162 | 0.182 | -0.003 | 0.005 | 0.005 | 0.296 | 0.59 | 0.39 | 0.5 | 0.72 | 0.58 | 0.75 |
| 4 | 184536452 | rs12510912 | 0.619 | 1.924 | 0.307 | 0.498 | 0.011 | -0.01 | -0.01 | 0 | 0.68 | 0.48 | 0.64 | 0.71 | 0.63 | 0.59 |
| 4 | 184537343 | rs12504367 | 0.848 | 1.849 | -0.308 | 0.507 | 0.003 | 0.015 | 0.015 | 0.954 | 0.55 | 0.29 | 0.56 | 0.55 | 0.49 | 0.46 |
| 4 | 184537463 | rs12504410 | 0.546 | 2.004 | -0.03 | 0.593 | 0.005 | 0.004 | 0.004 | 0.569 | 0.67 | 0.43 | 0.68 | 0.64 | 0.68 | 0.6 |
| 4 | 184540054 | rs7656347 | 0.861 | 2 | 0.09 | 0.764 | 0.008 | 0.014 | 0.014 | 0.271 | 0.7 | 0.61 | 0.79 | 0.61 | 0.67 | 0.57 |
| 4 | 184542062 | rs11132199 | 0.893 | 2.052 | 0.144 | 0.742 | 0.006 | -0.005 | -0.005 | 0 | 0.64 | 0.39 | 0.68 | 0.73 | 0.53 | 0.52 |
| 4 | 184546565 | rs13124076 | 1.492 | 1.582 | 0.049 | 0.825 | 0.091 | -0.001 | -0.001 | 0.6 | 0.51 | 0.23 | 0.46 | 0.46 | 0.39 | 0.3 |
| 4 | 184551432 | rs6830164 | -1.521 | 1.527 | 0.064 | 0.71 | -0.001 | 0 | 0 | 0 | 0.44 | 0.59 | 0.48 | 0.41 | 0.49 | 0.57 |
| 4 | 184569516 | rs7679973 | -0.711 | 1.707 | -0.024 | 0.717 | 0.018 | -0.006 | -0.006 | 0 | 0.23 | 0.5 | 0.16 | 0.26 | 0.21 | 0.22 |
| 4 | 184571085 | rs6552695 | -1.149 | 1.378 | -0.776 | 0.405 | 0.009 | -0.054 | -0.054 | 0 | 0.61 | 0.81 | 0.55 | 0.5 | 0.57 | 0.52 |
| 4 | 184572036 | rs4547833 | -0.712 | 1.118 | -0.446 | -0.113 | 0.025 | -0.019 | -0.019 | 0 | 0.18 | 0.22 | 0.27 | 0.13 | 0.26 | 0.23 |
| 4 | 184572951 | rs6832981 | -0.4 | 0.145 | -0.708 | 0.016 | 0.01 | 0.005 | 0.005 | 0 | 0.45 | 0.45 | 0.32 | 0.54 | 0.31 | 0.62 |
| 4 | 184573379 | rs10033002 | 0.073 | -0.311 | -0.86 | -0.139 | 0.037 | 0.019 | 0.019 | 0 | 0.65 | 0.74 | 0.69 | 0.69 | 0.74 | 0.66 |
| 4 | 184575568 | rs9790676 | -0.634 | 0.199 | -0.743 | -0.272 | -0.001 | 0.019 | 0.019 | 1.775 | 0.53 | 0.35 | 0.32 | 0.38 | 0.48 | 0.44 |
| 4 | 184577177 | rs6838230 | 0.027 | 0.56 | -0.777 | -0.124 | 0.015 | -0.02 | -0.02 | 0 | 0.3 | 0.14 | 0.24 | 0.3 | 0.24 | 0.35 |
| 4 | 184577348 | rs5019706 | 1.001 | 0.993 | 0.498 | -0.171 | 0.058 | -0.075 | -0.075 | 0 | 0.88 | 0.53 | 0.91 | 1 | 0.88 | 0.98 |
| 4 | 184586738 | rs4400034 | 0.867 | 0.886 | 0.347 | -0.216 | 0 | -0.08 | -0.08 | 0.049 | 0.86 | 0.78 | 0.92 | 1 | 0.88 | 0.94 |
| 4 | 184588121 | rs4282227 | -0.483 | 1.364 | 0.988 | -0.382 | 0.059 | -0.014 | -0.014 | 0 | 0.17 | 0.41 | 0.21 | 0.16 | 0.23 | 0.18 |
| 4 | 184590150 | rs7684339 | -0.016 | 1.532 | 0.494 | 0.138 | -0.005 | -0.031 | -0.031 | 0 | 0.14 | 0.1 | 0.28 | 0.3 | 0.14 | 0.11 |
| 4 | 184606653 | rs11132206 | -1.69 | 0.983 | 0.629 | 0.34 | -0.003 | -0.042 | -0.042 | 0 | 0.37 | 0.75 | 0.53 | 0.53 | 0.39 | 0.29 |
| 4 | 184609373 | rs10018951 | -0.064 | 0.974 | 0.735 | 0.534 | -0.001 | 0.003 | 0.003 | 1.037 | 0.15 | 0.2 | 0.14 | 0.19 | 0.14 | 0.07 |
| 4 | 184609457 | rs10019182 | -1.726 | 0.889 | 0.647 | 0.448 | -0.003 | -0.007 | -0.007 | 0 | 0.37 | 0.62 | 0.52 | 0.53 | 0.41 | 0.3 |
| 4 | 184610208 | rs13149502 | 0.294 | 0.93 | 0.596 | 0.441 | -0.006 | -0.014 | -0.014 | 0 | 0.13 | 0.1 | 0.13 | 0.19 | 0.14 | 0.07 |
| 4 | 184614874 | rs4862234 | 0.165 | 1.145 | 0.994 | 0.597 | 0.066 | 0.018 | 0.018 | 0 | 0.14 | 0.33 | 0.2 | 0.18 | 0.22 | 0.15 |
| 4 | 184628710 | rs4583804 | -0.271 | 1.108 | 0.918 | 0.578 | 0.055 | 0.002 | 0.002 | 0 | 0.17 | 0.51 | 0.21 | 0.18 | 0.22 | 0.16 |
| 4 | 184629794 | rs9312318 | 0.781 | 0.795 | -0.214 | 0.184 | 0.019 | 0.037 | 0.037 | 0.706 | 0.43 | 0.2 | 0.26 | 0.31 | 0.34 | 0.2 |
| 4 | 184631294 | rs13146475 | 1.038 | 0.815 | -0.653 | 0.133 | -0.044 | 0.047 | 0.047 | 0.49 | 0.39 | 0.09 | 0.23 | 0.26 | 0.32 | 0.19 |
| 4 | 184634741 | rs12642166 | 1.12 | 0.912 | -0.413 | 0.264 | 0.008 | -0.023 | -0.023 | 0 | 0.07 | 0.05 | 0.04 | 0.14 | 0.05 | 0.02 |
| 4 | 184638760 | rs4241780 | -0.185 | 1.024 | -0.281 | 0.019 | -0.001 | -0.006 | -0.006 | 0 | 0.83 | 0.79 | 0.76 | 0.9 | 0.85 | 0.89 |
| 4 | 184639740 | rs4327526 | 1.078 | 0.873 | 0.259 | 0.116 | 0.006 | 0.009 | 0.009 | 1.127 | 0.35 | 0.29 | 0.21 | 0.11 | 0.25 | 0.16 |
| 4 | 184648954 | rs10000388 | -0.313 | 1.514 | -0.158 | 0.855 | -0.018 | 0.015 | 0.015 | 0 | 0.62 | 0.76 | 0.59 | 0.6 | 0.48 | 0.38 |
| 4 | 184651150 | rs12499750 | 0.695 | 1.671 | -0.501 | 0.728 | 0.039 | 0.111 | 0.111 | 0.893 | 0.43 | 0.14 | 0.31 | 0.23 | 0.36 | 0.3 |
| 4 | 184655041 | rs11735908 | 0.147 | 1.655 | -0.422 | 0.688 | 0.052 | 0.026 | 0.026 | 0.215 | 0.51 | 0.25 | 0.5 | 0.41 | 0.43 | 0.37 |
| 4 | 184659447 | rs13132455 | 1.167 | 1.364 | -0.715 | 0.483 | 0.044 | 0.152 | 0.152 | 1.151 | 0.34 | 0.1 | 0.18 | 0.09 | 0.23 | 0.17 |
| 4 | 184660531 | rs4132292 | 0.166 | 1.647 | -0.6 | 0.715 | -0.012 | 0.077 | 0.077 | 0.791 | 0.5 | 0.17 | 0.48 | 0.37 | 0.48 | 0.39 |
| 4 | 184668956 | rs4130368 | 0.161 | 1.657 | -0.637 | 0.633 | -0.016 | 0.098 | 0.098 | 0.793 | 0.5 | 0.16 | 0.49 | 0.37 | 0.49 | 0.38 |
| 4 | 184673268 | rs6552702 | 0.156 | 1.655 | -0.638 | 0.631 | -0.015 | 0.099 | 0.099 | 0.758 | 0.5 | 0.16 | 0.49 | 0.37 | 0.49 | 0.38 |
| 4 | 184686219 | rs4862250 | -0.52 | 2.234 | 0.431 | 0.612 | 0.041 | 0.008 | 0.008 | 0 | 0.18 | 0.31 | 0.25 | 0.17 | 0.25 | 0.23 |
| 4 | 184687644 | rs4862251 | 0.026 | 2.025 | 0.422 | 0.698 | -0.019 | 0.055 | 0.055 | 0 | 0.7 | 0.55 | 0.75 | 0.6 | 0.74 | 0.58 |
| 4 | 184688001 | rs10028767 | -0.156 | 1.998 | 0.166 | 0.618 | 0.014 | -0.039 | -0.039 | 0 | 0.69 | 0.81 | 0.75 | 0.61 | 0.71 | 0.57 |
| 4 | 184689409 | rs4538533 | 0.11 | 2.134 | 0.509 | -0.072 | 0.028 | 0.01 | 0.01 | 0 | 0.92 | 0.83 | 0.75 | 0.89 | 0.86 | 0.89 |
| 4 | 184689417 | rs4568278 | 0.114 | 2.132 | 0.508 | -0.073 | 0.027 | 0.01 | 0.01 | 0 | 0.92 | 0.83 | 0.75 | 0.89 | 0.86 | 0.89 |
| 4 | 184694074 | rs4129955 | -0.548 | 2.047 | 0.51 | -0.114 | 0.006 | -0.013 | -0.013 | 1.023 | 0.42 | 0.48 | 0.41 | 0.67 | 0.48 | 0.58 |
| 4 | 184694959 | rs10018596 | -1.121 | 1.496 | 0.921 | -0.12 | 0.019 | 0.021 | 0.021 | 0 | 0.08 | 0.19 | 0.12 | 0.1 | 0.1 | 0.1 |
| 4 | 184697378 | rs4368651 | -0.019 | 1.623 | 0.512 | -0.312 | -0.01 | 0.004 | 0.004 | 0 | 0.14 | 0.07 | 0.18 | 0.11 | 0.17 | 0.14 |
| 4 | 184697574 | rs6845392 | -1.185 | 1.167 | 0.84 | -0.268 | 0.101 | 0.03 | 0.03 | 1.578 | 0.06 | 0.31 | 0.1 | 0.03 | 0.08 | 0.08 |
| 4 | 184704385 | rs13142457 | -0.441 | 0.962 | 0.661 | -0.241 | -0.017 | 0.002 | 0.002 | 0 | 0.16 | 0.34 | 0.15 | 0.12 | 0.15 | 0.11 |
| 4 | 184712051 | rs4370182 | -0.455 | 1.106 | -0.269 | -0.35 | -0.002 | 0.008 | 0.008 | 0.523 | 0.43 | 0.44 | 0.57 | 0.69 | 0.51 | 0.56 |
| 4 | 184720139 | rs9990869 | -0.209 | 0.893 | -0.445 | -0.534 | 0 | -0.02 | -0.02 | 0 | 0.49 | 0.46 | 0.44 | 0.21 | 0.45 | 0.36 |

| | | | | | | | | | | | | | | | | |
|---|---|---|---|---|---|---|---|---|---|---|---|---|---|---|---|---|
| 4 | 184721809 | rs10019343 | 0.295 | 0.58 | -0.496 | -0.603 | -0.001 | -0.016 | -0.016 | 0 | 0.69 | 0.66 | 0.74 | 0.88 | 0.71 | 0.8 |
| 4 | 184725906 | rs930306 | 0.59 | 0.652 | -0.154 | -0.345 | 0.014 | -0.033 | -0.033 | 0 | 0.83 | 0.37 | 0.79 | 0.86 | 0.85 | 0.92 |
| 4 | 184727641 | rs11722400 | -0.219 | 0.155 | -0.726 | -0.829 | -0.006 | -0.019 | -0.019 | 0 | 0.48 | 0.44 | 0.27 | 0.22 | 0.41 | 0.46 |
| 4 | 184728682 | rs7659026 | -0.35 | 0.309 | -1.013 | -1.121 | 0.016 | 0.043 | 0.043 | 1.221 | 0.52 | 0.55 | 0.29 | 0.19 | 0.43 | 0.47 |
| 4 | 184728730 | rs7659056 | -0.327 | 0.231 | -0.97 | -0.154 | 0.03 | 0.017 | 0.017 | 1.151 | 0.49 | 0.55 | 0.33 | 0.4 | 0.39 | 0.45 |
| 4 | 184730177 | rs7666373 | -0.038 | -0.317 | -0.989 | -0.978 | -0.005 | -0.03 | -0.03 | 0.262 | 0.35 | 0.56 | 0.18 | 0.09 | 0.23 | 0.3 |
| 4 | 184749270 | rs17075058 | 0.155 | 0.407 | -1.326 | -0.565 | -0.022 | 0.025 | 0.025 | 0 | 0.14 | 0.03 | 0.08 | 0.06 | 0.13 | 0.08 |
| 4 | 184757437 | rs7682282 | -0.306 | 0.441 | -0.53 | -0.634 | -0.002 | -0.039 | -0.039 | 2.157 | 0.57 | 0.47 | 0.57 | 0.85 | 0.59 | 0.58 |
| 4 | 184762700 | rs13106255 | 0.878 | 1.044 | 0.157 | 0.416 | 0.033 | 0.047 | 0.047 | 1.008 | 0.87 | 0.74 | 0.88 | 0.8 | 0.87 | 0.89 |
| 4 | 184767521 | rs7695711 | -0.482 | 0.561 | -0.095 | -0.396 | 0.062 | 0.105 | 0.105 | 0.427 | 0.52 | 0.34 | 0.61 | 0.91 | 0.65 | 0.62 |
| 4 | 184770923 | rs6813846 | 0.11 | 0.3 | 0.466 | 0.675 | 0.014 | 0.022 | 0.022 | 0 | 0.26 | 0.33 | 0.38 | 0.43 | 0.35 | 0.25 |
| 4 | 184779187 | rs6552711 | 0.77 | -0.225 | 0.766 | 0.161 | -0.001 | 0.027 | 0.027 | 0 | 0.52 | 0.41 | 0.51 | 0.29 | 0.5 | 0.49 |
| 4 | 184781037 | rs892836 | 0.634 | -0.186 | 0.982 | 0.249 | -0.005 | 0.053 | 0.053 | 0.079 | 0.57 | 0.3 | 0.53 | 0.3 | 0.54 | 0.48 |
| 4 | 184782619 | rs13150058 | 0.981 | 0.133 | 1.727 | 0.549 | 0 | 0.001 | 0.001 | 0 | 0.51 | 0.53 | 0.6 | 0.55 | 0.48 | 0.49 |
| 4 | 184783233 | rs4416516 | 0.066 | -0.184 | 1.674 | 0.672 | -0.005 | 0.008 | 0.008 | 0 | 0.84 | 0.78 | 0.91 | 0.82 | 0.86 | 0.85 |
| 4 | 184784920 | rs6824804 | 0.132 | -0.24 | 1.643 | 0.672 | -0.002 | 0.007 | 0.007 | 0 | 0.83 | 0.77 | 0.91 | 0.82 | 0.86 | 0.85 |
| 4 | 184787159 | rs13435909 | 0.839 | -0.109 | 1.92 | 0.58 | 0.006 | -0.001 | -0.001 | 0 | 0.56 | 0.52 | 0.61 | 0.56 | 0.51 | 0.61 |
| 4 | 184789877 | rs17075108 | -0.181 | -0.258 | 1.893 | 0.517 | -0.013 | -0.028 | -0.028 | 0 | 0.15 | 0.23 | 0.08 | 0.06 | 0.1 | 0.1 |
| 4 | 184790301 | rs11729441 | -0.832 | -0.177 | 1.876 | 0.504 | -0.003 | 0.008 | 0.008 | 0 | 0.4 | 0.36 | 0.36 | 0.31 | 0.43 | 0.34 |
| 4 | 184790373 | rs11737799 | 0.466 | -0.163 | 1.673 | 0.511 | -0.067 | -0.018 | -0.018 | 0 | 0.24 | 0.05 | 0.28 | 0.25 | 0.32 | 0.23 |
| 4 | 184794696 | rs2164303 | -1.17 | -0.224 | 1.996 | 0.503 | -0.002 | -0.009 | -0.009 | 0 | 0.45 | 0.48 | 0.38 | 0.34 | 0.44 | 0.42 |
| 4 | 184796501 | rs7678253 | -0.27 | -0.12 | 2.106 | 0.513 | 0.004 | -0.021 | -0.021 | 0 | 0.7 | 0.62 | 0.67 | 0.76 | 0.65 | 0.68 |
| 4 | 184799985 | rs7688995 | -0.127 | -0.096 | 1.666 | 0.468 | -0.012 | -0.044 | -0.044 | 0.252 | 0.85 | 0.58 | 0.91 | 0.94 | 0.89 | 0.9 |
| 4 | 184805900 | rs7657089 | -0.131 | -0.077 | 1.15 | 0.659 | -0.042 | -0.043 | -0.043 | 1.82 | 0.3 | 0.1 | 0.32 | 0.35 | 0.37 | 0.38 |
| 4 | 184808546 | rs1469809 | 0.952 | 0.596 | 1.644 | 1.03 | 0.089 | 0.297 | 0.297 | 0 | 0.89 | 0.62 | 0.78 | 0.65 | 0.83 | 0.65 |
| 4 | 184811318 | rs2871381 | 0.947 | 0.392 | 1.497 | 0.931 | 0.053 | 0.256 | 0.256 | 1.951 | 0.86 | 0.59 | 0.78 | 0.64 | 0.82 | 0.65 |
| 4 | 184813876 | rs1369445 | 0.33 | 0.171 | 0.718 | 0.329 | -0.01 | -0.029 | -0.029 | 0 | 0.58 | 0.43 | 0.77 | 0.83 | 0.7 | 0.7 |
| 4 | 184825696 | rs17075136 | -0.235 | 0.048 | 0.026 | 0.928 | 0.017 | -0.014 | -0.014 | 0 | 0.37 | 0.29 | 0.27 | 0.42 | 0.27 | 0.23 |
| 4 | 184826026 | rs1560982 | 0.251 | 0.016 | -0.755 | 0.798 | 0.012 | -0.012 | -0.012 | 0.131 | 0.44 | 0.19 | 0.51 | 0.5 | 0.57 | 0.6 |
| 4 | 184836189 | rs1978777 | 0.325 | 0.815 | -1.412 | 0.338 | -0.087 | 0.054 | 0.054 | 0 | 0.24 | 0.07 | 0.33 | 0.11 | 0.35 | 0.29 |
| 4 | 184839829 | rs2309942 | -0.457 | -0.676 | -1.373 | 0.676 | 0.021 | -0.055 | -0.055 | 0 | 0.25 | 0.16 | 0.24 | 0.55 | 0.22 | 0.25 |
| 4 | 184841111 | rs6552719 | -0.735 | -0.762 | -0.87 | 0.684 | -0.002 | 0.034 | 0.034 | 0.475 | 0.22 | 0.27 | 0.34 | 0.4 | 0.26 | 0.2 |
| 4 | 184848691 | rs7696821 | -0.031 | -1.616 | -0.65 | -0.409 | -0.035 | -0.123 | -0.123 | 0.064 | 0.19 | 0.37 | 0.11 | 0 | 0.12 | 0.02 |
| 4 | 184848958 | rs17363081 | -0.535 | -1.805 | -1.31 | -0.469 | 0.015 | 0.019 | 0.019 | 0 | 0.2 | 0.16 | 0.13 | 0.04 | 0.15 | 0.24 |
| 4 | 184849456 | rs892835 | 0.548 | -1.792 | -1.363 | 0.492 | 0.008 | -0.009 | -0.009 | 0 | 0.82 | 0.84 | 0.78 | 0.43 | 0.9 | 0.77 |
| 4 | 184852627 | rs6552722 | -0.787 | -1.594 | -1.466 | 0.193 | 0.017 | -0.014 | -0.014 | 0 | 0.28 | 0.75 | 0.34 | 0.58 | 0.13 | 0.27 |
| 4 | 184858078 | rs2309944 | -0.392 | -0.438 | -1.083 | 0.479 | -0.011 | 0.017 | 0.017 | 0 | 0.35 | 0.4 | 0.37 | 0.48 | 0.27 | 0.34 |
| 4 | 184866656 | rs12502711 | -0.207 | 0.214 | -1.785 | 0.481 | -0.039 | -0.053 | -0.053 | 0 | 0.4 | 0.14 | 0.35 | 0.52 | 0.5 | 0.57 |
| 4 | 184870145 | rs6853748 | -0.269 | 0.233 | -1.43 | 1.147 | -0.001 | 0.009 | 0.009 | 0 | 0.11 | 0.07 | 0.19 | 0.28 | 0.12 | 0.13 |
| 4 | 184875120 | rs2871385 | 0.256 | -0.079 | -1.628 | 0.437 | 0.038 | 0.074 | 0.074 | 0.271 | 0.41 | 0.73 | 0.4 | 0.2 | 0.35 | 0.24 |
| 4 | 184876413 | rs10213203 | -0.109 | 0.984 | -2.016 | 0.253 | -0.007 | 0.096 | 0.096 | 1.485 | 0.54 | 0.74 | 0.61 | 0.24 | 0.56 | 0.35 |
| 4 | 184882782 | rs9999145 | 0.877 | 0.826 | -0.794 | -0.095 | 0.003 | -0.071 | -0.071 | 3.545 | 0.35 | 0.45 | 0.31 | 0.13 | 0.4 | 0.2 |
| 4 | 184886694 | rs4861596 | 0.702 | 1.058 | -0.848 | 0.294 | -0.011 | -0.017 | -0.017 | 0.04 | 0.71 | 0.77 | 0.77 | 0.66 | 0.71 | 0.64 |
| 4 | 184902533 | rs2309949 | 0.386 | 0.657 | -0.752 | 0.443 | 0.005 | 0.019 | 0.019 | 0 | 0.5 | 0.34 | 0.47 | 0.53 | 0.53 | 0.45 |
| 4 | 184905482 | rs4862279 | 1.527 | 0.49 | -0.655 | 0.177 | 0 | 0.086 | 0.086 | 0 | 0.34 | 0.2 | 0.21 | 0.13 | 0.35 | 0.23 |
| 4 | 184905863 | rs12513114 | 0.711 | 0.695 | -0.365 | 0.314 | 0.006 | -0.025 | -0.025 | 0 | 0.28 | 0.1 | 0.28 | 0.29 | 0.27 | 0.24 |
| 4 | 184906674 | rs7685679 | 0.78 | 0.799 | 0.087 | 0.487 | 0.007 | 0.009 | 0.009 | 0.164 | 0.69 | 0.68 | 0.59 | 0.42 | 0.67 | 0.48 |
| 4 | 184911340 | rs4397039 | 0.159 | 1.02 | 0.049 | 0.637 | 0.011 | 0.03 | 0.03 | 0.001 | 0.81 | 0.73 | 0.75 | 0.75 | 0.8 | 0.71 |
| 4 | 184918852 | rs17090673 | 0.643 | 0.911 | -0.676 | 0.694 | -0.039 | -0.041 | -0.041 | 0 | 0.26 | 0.04 | 0.29 | 0.24 | 0.32 | 0.25 |
| 4 | 184921299 | rs12498735 | -0.449 | 0.905 | -0.524 | 0.695 | -0.023 | -0.035 | -0.035 | 0 | 0.39 | 0.18 | 0.46 | 0.58 | 0.44 | 0.47 |
| 4 | 184922669 | rs3749534 | 0.22 | 0.49 | -0.478 | 0.078 | -0.101 | -0.115 | -0.115 | 0.636 | 0.37 | 0.06 | 0.4 | 0.46 | 0.44 | 0.5 |
| 4 | 184924281 | rs10212888 | -0.308 | -0.067 | 0.349 | -0.775 | -0.003 | -0.035 | -0.035 | 1.155 | 0.55 | 0.49 | 0.73 | 0.83 | 0.63 | 0.73 |
| 4 | 184928980 | rs7655396 | 0.88 | 0.072 | 0.484 | -0.243 | -0.039 | -0.055 | -0.055 | 0 | 0.84 | 0.53 | 0.88 | 0.88 | 0.91 | 0.83 |
| 4 | 184931818 | rs4861597 | -0.7 | 0.497 | -0.102 | -0.042 | -0.008 | 0.01 | 0.01 | 0.727 | 0.31 | 0.41 | 0.32 | 0.6 | 0.3 | 0.48 |
| 4 | 184943274 | rs3749531 | -0.536 | 0.146 | -0.33 | -0.913 | -0.025 | -0.114 | -0.114 | 0.205 | 0.61 | 0.5 | 0.68 | 0.89 | 0.69 | 0.83 |
| 4 | 184952005 | rs7690914 | -0.257 | 0.272 | -0.302 | -0.713 | -0.008 | -0.003 | -0.003 | 0 | 0.36 | 0.45 | 0.42 | 0.66 | 0.32 | 0.52 |
| 4 | 184952645 | rs6552733 | -0.2 | 0.57 | -0.077 | -1.105 | 0.017 | -0.002 | -0.002 | 0 | 0.26 | 0.22 | 0.39 | 0.22 | 0.43 | 0.33 |
| 4 | 184958217 | rs4862292 | -0.541 | 0.698 | 0.066 | -1.386 | 0.024 | -0.032 | -0.032 | 1.658 | 0.33 | 0.42 | 0.4 | 0.18 | 0.42 | 0.37 |
| 4 | 184974318 | rs4391081 | -0.394 | 0.324 | 0.097 | -1.485 | -0.008 | -0.011 | -0.011 | 2.594 | 0.68 | 0.62 | 0.55 | 0.34 | 0.67 | 0.52 |
| 4 | 184974967 | rs11725377 | -0.662 | 0.605 | 0.177 | -1.46 | 0.036 | -0.059 | -0.059 | 0 | 0.34 | 0.54 | 0.33 | 0.17 | 0.42 | 0.36 |
| 4 | 184988203 | rs4862298 | -0.543 | 0.913 | 0.115 | -1.485 | -0.029 | 0.044 | 0.044 | 0 | 0.3 | 0.23 | 0.3 | 0.15 | 0.35 | 0.37 |
| 4 | 184989565 | rs12650935 | 0.7 | 0.759 | -0.51 | -1.563 | -0.12 | -0.066 | -0.066 | 0 | 0.26 | 0.02 | 0.42 | 0.69 | 0.36 | 0.46 |
| 4 | 184990090 | rs4862299 | -1.094 | 0.287 | 0.712 | -1.558 | 0.004 | -0.004 | -0.004 | 0 | 0.6 | 0.62 | 0.74 | 0.84 | 0.73 | 0.84 |
| 4 | 184993366 | rs6552744 | 1.159 | 0.238 | 0.762 | -1.429 | 0.026 | 0.036 | 0.036 | 0 | 0.35 | 0.23 | 0.23 | 0.18 | 0.25 | 0.17 |
| 4 | 184995011 | rs11939877 | -1.133 | -0.006 | 1.502 | -1.078 | -0.001 | -0.029 | -0.029 | 0 | 0.07 | 0.27 | 0.07 | 0 | 0.05 | 0.06 |
| 4 | 184996810 | rs4862305 | 0.089 | 0.278 | 1.42 | -0.5 | -0.013 | -0.02 | -0.02 | 1.493 | 0.31 | 0.2 | 0.42 | 0.57 | 0.36 | 0.46 |
| 4 | 185001631 | rs6829678 | 0.434 | 0.287 | 1.448 | -0.469 | 0.003 | 0.004 | 0.004 | 0.632 | 0.59 | 0.34 | 0.49 | 0.42 | 0.59 | 0.48 |
| 4 | 185012398 | rs4479748 | -0.834 | 0.19 | 0.515 | -0.222 | 0.006 | 0.011 | 0.011 | 2.314 | 0.43 | 0.71 | 0.5 | 0.59 | 0.43 | 0.5 |
| 4 | 185016399 | rs6819770 | -0.479 | 0.164 | 0.83 | -0.109 | 0.004 | 0.007 | 0.007 | 0 | 0.41 | 0.34 | 0.46 | 0.63 | 0.4 | 0.57 |

| | | | | | | | | | | | | | | | | |
|---|---|---|---|---|---|---|---|---|---|---|---|---|---|---|---|---|
| 4 | 185019266 | rs7669472 | -0.869 | 0.04 | 1.278 | -0.237 | 0.072 | -0.07 | -0.07 | 0 | 0.13 | 0.34 | 0.17 | 0.02 | 0.15 | 0.2 |
| 4 | 185023528 | rs6810830 | -1.665 | 0.169 | 0.766 | 0.079 | -0.017 | 0.003 | 0.003 | 0.912 | 0.59 | 0.75 | 0.67 | 0.63 | 0.55 | 0.74 |
| 4 | 185023826 | rs12501526 | -0.183 | 0.145 | 0.481 | 0.228 | -0.01 | -0.029 | -0.029 | 0.05 | 0.37 | 0.17 | 0.45 | 0.61 | 0.39 | 0.52 |
| 4 | 185024257 | rs11132222 | -0.252 | 0.146 | 0.47 | 0.229 | -0.001 | -0.013 | -0.013 | 0.149 | 0.37 | 0.26 | 0.46 | 0.61 | 0.39 | 0.52 |
| 4 | 185026377 | rs11132224 | -0.098 | 0.076 | 0.105 | 0.438 | -0.01 | -0.079 | -0.079 | 0.009 | 0.36 | 0.1 | 0.47 | 0.54 | 0.36 | 0.44 |
| 4 | 185027064 | rs10010069 | 1.09 | 0.148 | 0.419 | 0.34 | -0.005 | 0.068 | 0.068 | 0.012 | 0.42 | 0.24 | 0.31 | 0.17 | 0.44 | 0.31 |
| 4 | 185032985 | rs4241784 | 0.336 | -0.527 | -0.005 | 0.441 | 0.025 | -0.011 | -0.011 | 0.007 | 0.56 | 0.58 | 0.52 | 0.47 | 0.64 | 0.52 |
| 4 | 185034183 | rs4862310 | 1.253 | -0.234 | -0.435 | 0.152 | -0.094 | 0.094 | 0.094 | 1.666 | 0.38 | 0.08 | 0.31 | 0.21 | 0.47 | 0.24 |
| 4 | 185042789 | rs4861602 | 0.018 | -0.208 | 0.113 | 0.513 | 0.009 | -0.033 | -0.033 | 0.116 | 0.71 | 0.84 | 0.57 | 0.59 | 0.71 | 0.66 |
| 4 | 185062880 | rs4554119 | -0.132 | -0.238 | 0.41 | 0.544 | -0.006 | 0.007 | 0.007 | 0.97 | 0.7 | 0.65 | 0.55 | 0.59 | 0.71 | 0.71 |
| 4 | 185066950 | rs4421030 | 1.226 | -0.069 | 0.505 | 0.137 | -0.021 | -0.029 | -0.029 | 0 | 0.93 | 0.74 | 0.97 | 1 | 0.99 | 0.97 |
| 4 | 185070555 | rs4862319 | 0.462 | -0.049 | -0.075 | -0.264 | 0.013 | 0.084 | 0.084 | 0.951 | 0.51 | 0.32 | 0.4 | 0.14 | 0.51 | 0.43 |
| 4 | 185071227 | rs4862321 | 0.829 | -0.018 | -0.378 | -0.594 | -0.009 | 0.092 | 0.092 | 0.268 | 0.39 | 0.18 | 0.25 | 0.1 | 0.39 | 0.26 |
| 4 | 185072020 | rs4241790 | 0.387 | -0.275 | -0.376 | -0.518 | -0.019 | -0.012 | -0.012 | 0 | 0.88 | 0.8 | 0.92 | 0.93 | 0.91 | 0.86 |
| 4 | 185074686 | rs13147555 | 1.35 | -0.334 | -0.811 | -0.64 | -0.126 | 0.143 | 0.143 | 1.736 | 0.32 | 0.02 | 0.2 | 0.08 | 0.37 | 0.25 |
| 4 | 185076246 | rs9992036 | 0.066 | -0.272 | 0.111 | 0.662 | -0.013 | -0.004 | -0.004 | 1.081 | 0.36 | 0.22 | 0.48 | 0.4 | 0.43 | 0.26 |
| 4 | 185078076 | rs1921564 | -0.514 | 0.197 | -0.34 | -0.208 | 0.006 | 0.02 | 0.02 | 0.346 | 0.47 | 0.59 | 0.59 | 0.88 | 0.48 | 0.6 |
| 4 | 185086713 | rs2310044 | 0.498 | 0.623 | -0.306 | 0.383 | 0.006 | 0.019 | 0.019 | 0 | 0.76 | 0.71 | 0.56 | 0.66 | 0.74 | 0.72 |
| 4 | 185088648 | rs1921561 | 0.796 | 0.686 | -0.85 | -0.086 | 0.014 | -0.022 | -0.022 | 0 | 0.4 | 0.23 | 0.29 | 0.53 | 0.39 | 0.29 |
| 4 | 185088697 | rs1921560 | 0.474 | 0.58 | -0.464 | -0.141 | 0.01 | -0.001 | -0.001 | 0.949 | 0.44 | 0.37 | 0.31 | 0.53 | 0.4 | 0.38 |
| 4 | 185091025 | rs9637691 | -0.379 | 0.908 | -0.828 | -0.022 | -0.033 | -0.075 | -0.075 | 0 | 0.2 | 0.06 | 0.42 | 0.34 | 0.23 | 0.19 |
| 4 | 185091885 | rs11722344 | 0.442 | 0.145 | -0.791 | -1.121 | -0.006 | -0.049 | -0.049 | 0.009 | 0.7 | 0.33 | 0.86 | 0.93 | 0.83 | 0.71 |
| 4 | 185094473 | rs6810545 | 0.097 | -0.106 | -0.023 | -1.269 | -0.026 | -0.068 | -0.068 | 0 | 0.17 | 0.23 | 0.05 | 0.02 | 0.09 | 0.18 |
| 4 | 185096468 | rs2178570 | -0.203 | 0.038 | 0.665 | -1.126 | 0.025 | -0.067 | -0.067 | 0 | 0.89 | 0.61 | 0.93 | 1 | 0.92 | 0.97 |
| 4 | 185104471 | rs6842767 | 0.174 | 0.073 | 0.649 | -1.109 | 0.012 | -0.056 | -0.056 | 1.306 | 0.1 | 0.27 | 0.06 | 0 | 0.08 | 0.03 |
| 4 | 185109553 | rs6552765 | -0.27 | -0.06 | 0.541 | -1.625 | -0.026 | -0.111 | -0.111 | 0 | 0.22 | 0.37 | 0.14 | 0.03 | 0.18 | 0.25 |
| 4 | 185111580 | rs10520559 | 0.769 | 0.444 | 0.402 | -1.643 | -0.001 | -0.002 | -0.002 | 0 | 0.53 | 0.45 | 0.41 | 0.64 | 0.51 | 0.47 |
| 4 | 185111634 | rs4452474 | 0.185 | 0.067 | 0.582 | -2.277 | -0.024 | -0.096 | -0.096 | 0 | 0.76 | 0.58 | 0.9 | 0.98 | 0.86 | 0.71 |
| 4 | 185116224 | rs11930046 | 0.94 | -0.157 | -0.024 | -2.838 | -0.006 | -0.114 | -0.114 | 0 | 0.39 | 0.45 | 0.24 | 0.03 | 0.35 | 0.29 |
| 4 | 185117077 | rs4862330 | -0.193 | -0.118 | -0.066 | -2.862 | -0.012 | -0.015 | -0.015 | 0 | 0.4 | 0.32 | 0.67 | 0.95 | 0.52 | 0.42 |
| 4 | 185121167 | rs17584314 | 1.316 | -0.251 | -0.485 | -2.661 | -0.084 | 0.242 | 0.242 | 1.089 | 0.34 | 0.02 | 0.2 | 0.03 | 0.34 | 0.3 |
| 4 | 185121619 | rs4862338 | 0.228 | -0.863 | 0.425 | -2.266 | -0.06 | -0.098 | -0.098 | 3.391 | 0.82 | 0.6 | 0.91 | 0.98 | 0.95 | 0.84 |
| 4 | 185123138 | rs12647212 | 0.419 | -0.663 | -0.298 | -1.968 | 0 | 0.098 | 0.098 | 0.016 | 0.48 | 0.27 | 0.49 | 0.75 | 0.52 | 0.38 |
| 4 | 185123650 | rs17075459 | 0.66 | -0.695 | -0.384 | -2.101 | 0 | -0.004 | -0.004 | 0 | 0.1 | 0.1 | 0.04 | 0 | 0.07 | 0.07 |
| 4 | 185123766 | rs1882331 | -0.774 | -0.967 | 0.088 | -2.107 | -0.028 | -0.019 | -0.019 | 0 | 0.29 | 0.45 | 0.39 | 0.78 | 0.23 | 0.44 |
| 4 | 185124123 | rs4862339 | 0.025 | -0.987 | -0.238 | -2.08 | -0.061 | -0.097 | -0.097 | 1.215 | 0.8 | 0.43 | 0.88 | 0.93 | 0.93 | 0.69 |
| 4 | 185127748 | rs10011933 | -0.19 | -0.692 | 0.076 | -1.703 | -0.004 | 0.011 | 0.011 | 0 | 0.57 | 0.61 | 0.56 | 0.8 | 0.52 | 0.63 |
| 4 | 185128095 | rs17679896 | 0.339 | -0.644 | 0.447 | -1.03 | -0.002 | 0.037 | 0.037 | 0.009 | 0.16 | 0.22 | 0.32 | 0.73 | 0.18 | 0.22 |
| 4 | 185137596 | rs6819977 | -1.006 | -0.202 | -0.318 | -0.981 | -0.053 | 0.025 | 0.025 | 0 | 0.72 | 0.87 | 0.68 | 0.83 | 0.57 | 0.73 |
| 4 | 185148372 | rs6853677 | 0.462 | -0.832 | 0.317 | 0.222 | -0.007 | -0.056 | -0.056 | 0 | 0.13 | 0.21 | 0.18 | 0 | 0.13 | 0.08 |
| 4 | 185149599 | rs1217965 | -0.731 | -0.891 | 0.309 | 0.272 | -0.021 | 0.173 | 0.173 | 0 | 0.19 | 0.45 | 0.3 | 0.57 | 0.17 | 0.16 |
| 4 | 185150443 | rs12644741 | -0.569 | -0.804 | -0.203 | 0.776 | 0.007 | -0.009 | -0.009 | 1.002 | 0.1 | 0.06 | 0.09 | 0.19 | 0.06 | 0.3 |
| 4 | 185152834 | rs938016 | -1.242 | -0.781 | -0.23 | 0.198 | -0.025 | 0.046 | 0.046 | 0 | 0.15 | 0.25 | 0.19 | 0.76 | 0.09 | 0.38 |
| 4 | 185160596 | rs17584650 | -0.789 | -0.896 | -0.927 | -0.489 | 0.04 | 0.059 | 0.059 | 0 | 0.22 | 0.09 | 0.1 | 0 | 0.12 | 0.14 |
| 4 | 185162040 | rs1217960 | 0.148 | -0.811 | -0.429 | 0.417 | -0.011 | 0.095 | 0.095 | 0 | 0.13 | 0.19 | 0.19 | 0.51 | 0.11 | 0.21 |
| 4 | 185164636 | rs1217957 | -0.155 | -0.38 | -0.967 | -0.268 | -0.036 | -0.078 | -0.078 | 0 | 0.28 | 0.13 | 0.33 | 0.51 | 0.36 | 0.26 |
| 4 | 185165607 | rs7696184 | 0.302 | -0.596 | -0.131 | -0.039 | -0.015 | 0.077 | 0.077 | 0 | 0.11 | 0.3 | 0.1 | 0.15 | 0.07 | 0.14 |
| 4 | 185170812 | rs1217945 | -1.53 | -1.306 | -0.098 | -0.48 | -0.015 | 0.011 | 0.011 | 0 | 0.62 | 0.72 | 0.57 | 0.7 | 0.55 | 0.5 |
| 4 | 185172565 | rs2248232 | 1.211 | -1.4 | -0.147 | -0.493 | 0.01 | -0.042 | -0.042 | 0 | 0.12 | 0.05 | 0.18 | 0.52 | 0.11 | 0.21 |
| 4 | 185173385 | rs2703731 | -0.506 | -1.511 | 0.057 | -0.61 | -0.035 | 0.041 | 0.041 | 0 | 0.23 | 0.4 | 0.29 | 0.66 | 0.18 | 0.37 |
| 4 | 185173887 | rs13138586 | 0.74 | -1.713 | -0.044 | -0.77 | -0.046 | 0.067 | 0.067 | 0 | 0.25 | 0.11 | 0.23 | 0.04 | 0.35 | 0.11 |
| 4 | 185175421 | rs3946906 | 1.653 | -1.795 | 0.133 | -0.778 | -0.055 | 0.031 | 0.031 | 0 | 0.37 | 0.16 | 0.42 | 0.3 | 0.45 | 0.48 |
| 4 | 185176608 | rs2175292 | 0.829 | -1.622 | -0.104 | -0.902 | 0.009 | 0.003 | 0.003 | 0 | 0.07 | 0.12 | 0.09 | 0.11 | 0.06 | 0.24 |
| 4 | 185176699 | rs938015 | -0.896 | -1.678 | 0.058 | -0.902 | 0 | 0 | 0 | 0.043 | 0.49 | 0.5 | 0.52 | 0.7 | 0.53 | 0.48 |
| 4 | 185177783 | rs13143328 | -0.23 | -1.473 | -0.027 | -0.895 | 0 | -0.034 | -0.034 | 0 | 0.39 | 0.37 | 0.29 | 0.04 | 0.37 | 0.16 |
| 4 | 185177832 | rs13143366 | -0.23 | -1.473 | -0.027 | -0.895 | 0 | -0.034 | -0.034 | 0 | 0.39 | 0.37 | 0.29 | 0.04 | 0.37 | 0.16 |
| 4 | 185179394 | rs12503492 | -0.813 | -1.346 | 0.176 | -0.54 | -0.017 | 0.106 | 0.106 | 0 | 0.86 | 0.8 | 0.79 | 0.48 | 0.89 | 0.79 |
| 4 | 185182695 | rs11727655 | 0.538 | -1.447 | 0.244 | -0.521 | -0.063 | 0.05 | 0.05 | 0.121 | 0.72 | 0.51 | 0.7 | 0.4 | 0.82 | 0.62 |
| 4 | 185183535 | rs6856666 | 0.39 | -0.994 | 0.036 | -0.362 | 0.009 | 0.044 | 0.044 | 0 | 0.06 | 0.11 | 0.07 | 0.16 | 0.06 | 0.27 |
| 4 | 185186535 | rs877378 | 0.401 | -0.883 | -0.102 | -0.718 | 0.005 | 0.039 | 0.039 | 0 | 0.53 | 0.38 | 0.62 | 0.88 | 0.58 | 0.7 |
| 4 | 185191779 | rs4862347 | 0.63 | -1.026 | -0.233 | -0.957 | 0.047 | 0.023 | 0.023 | 2.229 | 0.48 | 0.3 | 0.51 | 0.42 | 0.62 | 0.48 |
| 4 | 185192543 | rs793913 | 0.998 | -1.132 | 0.164 | -0.959 | -0.007 | -0.034 | -0.034 | 8.144 | 0.11 | 0.21 | 0.09 | 0.04 | 0.1 | 0.11 |
| 4 | 185193099 | rs793911 | 0.133 | -1.143 | 0.149 | -0.954 | -0.035 | 0.004 | 0.004 | 0.011 | 0.59 | 0.51 | 0.6 | 0.46 | 0.72 | 0.6 |
| 4 | 185194074 | rs793908 | -0.358 | -0.828 | 0.048 | -1.008 | 0.004 | -0.012 | -0.012 | 0 | 0.33 | 0.37 | 0.3 | 0.24 | 0.38 | 0.41 |
| 4 | 185196778 | rs7670891 | 1.655 | -0.705 | -0.463 | -0.997 | -0.058 | 0.083 | 0.083 | 0 | 0.16 | 0.03 | 0.13 | 0 | 0.22 | 0.05 |
| 4 | 185197061 | rs812666 | -0.4 | -0.813 | -0.115 | -0.963 | -0.005 | 0.004 | 0.004 | 0 | 0.67 | 0.62 | 0.6 | 0.45 | 0.72 | 0.61 |
| 4 | 185203124 | rs17487801 | 1.923 | -0.678 | -0.267 | -0.891 | -0.068 | -0.001 | -0.001 | 0 | 0.25 | 0.08 | 0.29 | 0.22 | 0.34 | 0.18 |
| 4 | 185205186 | rs3796693 | 0.329 | -0.96 | 0.063 | -0.581 | 0.013 | 0.089 | 0.089 | 0 | 0.07 | 0.18 | 0.08 | 0.2 | 0.05 | 0.25 |
| 4 | 185205210 | rs3796692 | 1.198 | -1.165 | 0.077 | -0.594 | -0.002 | -0.013 | -0.013 | 0.237 | 0.09 | 0.14 | 0.1 | 0.04 | 0.08 | 0.1 |

| | | | | | | | | | | | | | | | | |
|---|---|---|---|---|---|---|---|---|---|---|---|---|---|---|---|---|
| 4 | 185209537 | rs1217969 | 0.217 | -1.049 | 0.026 | -0.602 | -0.001 | -0.001 | -0.001 | 0 | 0.33 | 0.34 | 0.4 | 0.54 | 0.29 | 0.39 |
| 4 | 185215134 | rs12650772 | 0.125 | -0.805 | 0.045 | -0.662 | 0.005 | 0.096 | 0.096 | 0 | 0.07 | 0.18 | 0.06 | 0.21 | 0.04 | 0.16 |
| 4 | 185217303 | rs2703737 | 0.862 | -1.034 | -0.145 | -0.914 | 0.002 | 0.006 | 0.006 | 0.033 | 0.42 | 0.39 | 0.44 | 0.2 | 0.56 | 0.24 |
| 4 | 185217356 | rs6552771 | 0.391 | -1.121 | -0.184 | -0.948 | 0.024 | 0.032 | 0.032 | 0 | 0.51 | 0.43 | 0.5 | 0.24 | 0.66 | 0.36 |
| 4 | 185220262 | rs4862356 | 1.785 | -0.711 | -0.546 | -0.964 | -0.076 | 0.019 | 0.019 | 0 | 0.24 | 0.07 | 0.28 | 0.2 | 0.34 | 0.17 |
| 4 | 185221748 | rs6841399 | 1.636 | -0.696 | -0.489 | -0.972 | -0.007 | 0.005 | 0.005 | 0 | 0.25 | 0.27 | 0.29 | 0.2 | 0.34 | 0.18 |
| 4 | 185222701 | rs1467069 | 1.501 | -0.5 | -0.765 | -0.877 | -0.081 | 0.09 | 0.09 | 0.022 | 0.15 | 0.01 | 0.14 | 0 | 0.22 | 0.07 |
| 4 | 185226573 | rs10029721 | 0.565 | -0.595 | -0.162 | -0.417 | 0.002 | 0.079 | 0.079 | 0 | 0.05 | 0.12 | 0.06 | 0.21 | 0.04 | 0.16 |
| 4 | 185227184 | rs10008153 | -0.277 | -0.748 | -0.368 | -0.529 | -0.002 | 0.009 | 0.009 | 0 | 0.47 | 0.6 | 0.44 | 0.44 | 0.48 | 0.46 |
| 4 | 185234505 | rs12506922 | 1.109 | -0.368 | -0.976 | -0.43 | -0.093 | 0.013 | 0.013 | 0 | 0.25 | 0.06 | 0.3 | 0.19 | 0.38 | 0.19 |
| 4 | 185238400 | rs6819481 | -0.024 | -0.637 | -0.866 | -0.333 | -0.009 | 0.05 | 0.05 | 0 | 0.46 | 0.33 | 0.44 | 0.32 | 0.49 | 0.52 |
| 4 | 185241316 | rs12650550 | -0.671 | -0.872 | -0.711 | 0.145 | -0.028 | 0.028 | 0.028 | 0 | 0.16 | 0.16 | 0.13 | 0.31 | 0.04 | 0.39 |
| 4 | 185241610 | rs12641965 | -0.859 | -0.922 | -0.536 | 0.114 | -0.051 | 0.058 | 0.058 | 0 | 0.17 | 0.24 | 0.14 | 0.31 | 0.04 | 0.39 |
| 4 | 185245514 | rs999759 | 0.553 | -0.847 | -0.817 | 0.115 | 0.026 | 0.002 | 0.002 | 0 | 0.11 | 0.05 | 0.05 | 0.08 | 0.03 | 0.26 |
| 4 | 185248266 | rs793890 | -0.157 | -0.877 | -0.862 | 0.15 | -0.01 | 0.131 | 0.131 | 0 | 0.05 | 0.14 | 0.08 | 0.23 | 0.02 | 0.13 |
| 4 | 185249147 | rs793892 | -0.622 | -0.63 | -0.957 | -0.269 | 0 | -0.001 | -0.001 | 0.167 | 0.63 | 0.66 | 0.61 | 0.54 | 0.68 | 0.75 |
| 4 | 185251371 | rs13149699 | 1.136 | -0.196 | -2.074 | -0.21 | -0.142 | -0.145 | -0.145 | 2.122 | 0.29 | 0.03 | 0.38 | 0.41 | 0.39 | 0.32 |
| 4 | 185252818 | rs793899 | 0.568 | -0.274 | -2.007 | -0.218 | -0.049 | -0.064 | -0.064 | 0 | 0.37 | 0.11 | 0.44 | 0.5 | 0.41 | 0.58 |
| 4 | 185256315 | rs793882 | 0.461 | -0.827 | -2.511 | -0.237 | -0.011 | -0.099 | -0.099 | 0 | 0.29 | 0.07 | 0.3 | 0.45 | 0.27 | 0.18 |
| 4 | 185257159 | rs6552774 | -0.218 | -0.29 | -2.183 | 0.08 | -0.007 | -0.033 | -0.033 | 0 | 0.53 | 0.87 | 0.51 | 0.47 | 0.45 | 0.33 |
| 4 | 185259670 | rs4269220 | -0.506 | 0.031 | -1.864 | 0.174 | -0.048 | -0.051 | -0.051 | 0 | 0.62 | 0.83 | 0.55 | 0.56 | 0.48 | 0.58 |
| 4 | 185262740 | rs793878 | -0.68 | -0.078 | -1.222 | 0.181 | 0.002 | 0.019 | 0.019 | 0 | 0.5 | 0.51 | 0.52 | 0.77 | 0.47 | 0.7 |
| 4 | 185263811 | rs793879 | 0.208 | -0.825 | -1.179 | 0.528 | 0.013 | -0.037 | -0.037 | 0 | 0.19 | 0.13 | 0.23 | 0.42 | 0.16 | 0.14 |
| 4 | 185265401 | rs793880 | 0.431 | -0.984 | -1.018 | 0.28 | 0.004 | -0.001 | -0.001 | 0 | 0.62 | 0.62 | 0.64 | 0.43 | 0.68 | 0.5 |
| 4 | 185273483 | rs712032 | 1.023 | -0.952 | -1.19 | 0.182 | 0.001 | 0.002 | 0.002 | 0 | 0.46 | 0.49 | 0.48 | 0.25 | 0.54 | 0.37 |
| 4 | 185278885 | rs4326049 | 0.245 | -0.87 | -0.977 | 0.454 | -0.015 | 0.124 | 0.124 | 0 | 0.83 | 0.66 | 0.79 | 0.67 | 0.85 | 0.84 |
| 4 | 185279284 | rs4590089 | -0.352 | -0.783 | -1.385 | 0.497 | 0.001 | 0.001 | 0.001 | 0 | 0.86 | 0.87 | 0.8 | 0.67 | 0.85 | 0.89 |
| 4 | 185280853 | rs6834535 | 0.763 | -0.986 | -1.785 | 0.098 | -0.068 | -0.097 | -0.097 | 0.459 | 0.4 | 0.09 | 0.43 | 0.54 | 0.53 | 0.46 |
| 4 | 185284684 | rs4861621 | -0.672 | -1.116 | -1.345 | -0.02 | -0.003 | -0.004 | -0.004 | 1.437 | 0.15 | 0.22 | 0.11 | 0.12 | 0.16 | 0.07 |
| 4 | 185286768 | rs17624012 | 0.197 | -1.035 | -1.31 | 0.156 | 0.01 | -0.047 | -0.047 | 0 | 0.26 | 0.13 | 0.23 | 0.42 | 0.19 | 0.34 |
| 4 | 185287351 | rs11727845 | 1.722 | -1.43 | -1.502 | -0.111 | -0.088 | -0.075 | -0.075 | 0 | 0.35 | 0.1 | 0.51 | 0.42 | 0.51 | 0.35 |
| 4 | 185287935 | rs17051405 | 0.288 | -1.384 | -1.407 | -0.165 | 0.055 | 0.089 | 0.089 | 0.387 | 0.61 | 0.21 | 0.73 | 0.85 | 0.71 | 0.71 |
| 4 | 185290163 | rs4642292 | -0.25 | -1 | -0.988 | -0.575 | -0.017 | -0.143 | -0.143 | 0 | 0.7 | 0.62 | 0.79 | 1 | 0.72 | 0.8 |
| 4 | 185299830 | rs12505296 | -0.571 | -0.711 | -0.929 | 0.457 | 0.018 | -0.058 | -0.058 | 0.444 | 0.3 | 0.2 | 0.26 | 0.57 | 0.21 | 0.39 |
| 4 | 185303451 | rs6831978 | 0.278 | -0.496 | -0.361 | -0.136 | -0.025 | -0.066 | -0.066 | 0 | 0.87 | 0.66 | 0.94 | 0.99 | 0.94 | 0.94 |
| 4 | 185304463 | rs1105207 | -0.445 | -0.032 | -1.877 | -0.859 | -0.083 | 0.209 | 0.209 | 0 | 0.57 | 0.87 | 0.41 | 0.14 | 0.48 | 0.4 |
| 4 | 185305366 | rs4629495 | 0.079 | -0.595 | -0.97 | -0.281 | 0.003 | -0.003 | -0.003 | 0 | 0.86 | 0.91 | 0.96 | 0.8 | 0.93 | 0.88 |
| 4 | 185307045 | rs11732927 | 0.317 | -0.188 | -0.795 | -0.646 | -0.002 | -0.028 | -0.028 | 0 | 0.16 | 0.21 | 0.09 | 0.03 | 0.12 | 0.03 |
| 4 | 185308459 | rs6817412 | -0.068 | -0.198 | -0.544 | -0.096 | -0.015 | 0.004 | 0.004 | 1.568 | 0.29 | 0.35 | 0.17 | 0.3 | 0.22 | 0.23 |
| 4 | 185311570 | rs13119453 | 1.273 | 0.247 | -1.288 | -0.378 | -0.054 | -0.086 | -0.086 | 0.5 | 0.28 | 0.03 | 0.36 | 0.32 | 0.26 | 0.46 |
| 4 | 185312355 | rs6856910 | 0.364 | -0.429 | -0.682 | -0.29 | 0.012 | 0.004 | 0.004 | 1.04 | 0.55 | 0.43 | 0.68 | 0.59 | 0.67 | 0.66 |
| 4 | 185312684 | rs793779 | 0.151 | 0.117 | -0.788 | -0.131 | 0.011 | 0.05 | 0.05 | 0 | 0.14 | 0.28 | 0.18 | 0.26 | 0.16 | 0.15 |
| 4 | 185315792 | rs3756089 | 0.783 | 0.26 | -1.016 | 0.032 | 0.006 | -0.008 | -0.008 | 0 | 0.16 | 0.17 | 0.1 | 0.25 | 0.09 | 0.19 |
| 4 | 185316897 | rs965225 | 0.814 | 0.46 | -1.363 | -0.292 | -0.01 | 0.066 | 0.066 | 0.528 | 0.13 | 0.04 | 0.06 | 0 | 0.09 | 0.06 |
| 4 | 185321941 | rs11132242 | 0.002 | 0.631 | -0.968 | -0.114 | 0.001 | 0.004 | 0.004 | 0 | 0.38 | 0.34 | 0.31 | 0.29 | 0.32 | 0.28 |
| 4 | 185323726 | rs724528 | 0.827 | 0.48 | -1.481 | -0.1 | -0.033 | 0.04 | 0.04 | 0 | 0.17 | 0.27 | 0.17 | 0.3 | 0.1 | 0.18 |
| 4 | 185324939 | rs7655371 | -0.061 | 1.448 | -2.246 | 0.479 | -0.002 | 0.038 | 0.038 | 0 | 0.34 | 0.22 | 0.28 | 0.22 | 0.28 | 0.25 |
| 4 | 185325242 | rs6812958 | 0.138 | 1.98 | -1.827 | 1.24 | -0.039 | -0.09 | -0.09 | 0.06 | 0.22 | 0.06 | 0.35 | 0.35 | 0.27 | 0.38 |
| 4 | 185325492 | rs17075783 | -0.821 | 1.687 | -1.278 | 0.803 | 0.002 | 0 | 0 | 0 | 0.07 | 0.07 | 0.04 | 0.12 | 0.1 | 0.14 |
| 4 | 185328817 | rs3775553 | -0.305 | 1.323 | -1.448 | 0.838 | -0.007 | 0.015 | 0.015 | 0.748 | 0.46 | 0.27 | 0.6 | 0.57 | 0.53 | 0.61 |
| 4 | 185329052 | rs2241500 | -0.143 | 1.42 | -1.498 | 0.944 | -0.013 | -0.024 | -0.024 | 0.129 | 0.41 | 0.25 | 0.54 | 0.54 | 0.45 | 0.53 |
| 4 | 185330598 | rs17686080 | -0.679 | 1.218 | -1.639 | 1.473 | -0.023 | -0.052 | -0.052 | 0 | 0.09 | 0.01 | 0.07 | 0.15 | 0.09 | 0.09 |
| 4 | 185330651 | rs6816525 | -0.304 | 1.003 | -1.298 | 1.419 | -0.005 | -0.003 | -0.003 | 0 | 0.27 | 0.4 | 0.18 | 0.31 | 0.26 | 0.25 |
| 4 | 185330669 | rs17624154 | 0.236 | 1.421 | -1.988 | 1.856 | -0.05 | -0.078 | -0.078 | 0 | 0.12 | 0.01 | 0.19 | 0.26 | 0.15 | 0.18 |
| 4 | 185331345 | rs7685555 | 0.13 | 0.448 | -2.283 | 1.533 | -0.027 | -0.024 | -0.024 | 2.595 | 0.4 | 0.24 | 0.57 | 0.45 | 0.51 | 0.63 |
| 4 | 185331977 | rs3775556 | 0.464 | 0.284 | -2.078 | 0.981 | 0 | -0.001 | -0.001 | 0 | 0.3 | 0.31 | 0.37 | 0.22 | 0.26 | 0.29 |
| 4 | 185332571 | rs1425552 | 0.288 | 0.408 | -2.232 | 1.115 | -0.01 | 0.014 | 0.014 | 0 | 0.1 | 0.07 | 0.06 | 0.44 | 0.15 | 0.2 |
| 4 | 185333030 | rs10009261 | 0.485 | 0.155 | -2.341 | 0.592 | -0.136 | 0 | 0 | 0 | 0.23 | 0.03 | 0.35 | 0.2 | 0.39 | 0.35 |
| 4 | 185336262 | rs13115095 | 1.514 | 0.088 | -2.1 | 0.568 | -0.045 | 0.032 | 0.032 | 0 | 0.12 | 0.03 | 0.21 | 0.05 | 0.24 | 0.2 |
| 4 | 185336852 | rs7677486 | -0.045 | -0.794 | -1.67 | 0.177 | 0.036 | 0.095 | 0.095 | 0.243 | 0.48 | 0.29 | 0.46 | 0.21 | 0.57 | 0.4 |
| 4 | 185338921 | rs2288247 | 0.812 | -0.707 | -1.96 | 0.192 | -0.135 | 0.085 | 0.085 | 0 | 0.37 | 0.06 | 0.4 | 0.21 | 0.48 | 0.42 |
| 4 | 185343635 | rs890193 | -0.531 | -0.515 | -1.13 | -0.031 | -0.005 | 0.003 | 0.003 | 3.673 | 0.69 | 0.65 | 0.8 | 0.6 | 0.77 | 0.73 |
| 4 | 185344606 | rs6827018 | 1.587 | -0.37 | -0.7 | 0.141 | -0.002 | 0.006 | 0.006 | 0.016 | 0.11 | 0.08 | 0.08 | 0 | 0.15 | 0.19 |
| 4 | 185344932 | rs6827840 | 1.447 | -0.259 | -1.081 | 0.564 | 0.003 | 0.022 | 0.022 | 0 | 0.06 | 0.07 | 0.04 | 0.24 | 0.12 | 0.11 |
| 4 | 185346090 | rs10011243 | -0.573 | -0.292 | -1.13 | 0.338 | 0.002 | -0.005 | -0.005 | 0 | 0.65 | 0.64 | 0.76 | 0.6 | 0.69 | 0.73 |
| 4 | 185348916 | rs11726275 | -0.171 | 0.116 | -1.338 | -0.022 | 0.002 | -0.009 | -0.009 | 0 | 0.31 | 0.33 | 0.31 | 0.22 | 0.37 | 0.48 |
| 4 | 185350934 | rs3756094 | -0.3 | 0.573 | -1.213 | 0.246 | -0.001 | 0 | 0 | 0 | 0.54 | 0.54 | 0.75 | 0.71 | 0.67 | 0.66 |
| 4 | 185352413 | rs744038 | 0.913 | 1.259 | -1.504 | 0.317 | -0.01 | -0.053 | -0.053 | 0.289 | 0.38 | 0.11 | 0.37 | 0.37 | 0.33 | 0.42 |
| 4 | 185353799 | rs12504466 | 0.521 | 1.805 | -1.339 | 0.479 | -0.019 | -0.003 | -0.003 | 0 | 0.46 | 0.26 | 0.44 | 0.57 | 0.48 | 0.53 |

| | | | | | | | | | | | | | | | | |
|---|---|---|---|---|---|---|---|---|---|---|---|---|---|---|---|---|
| 4 | 185354298 | rs3756095 | 0.152 | 2.012 | -1.212 | 1.122 | -0.021 | 0.02 | 0.02 | 3.086 | 0.28 | 0.31 | 0.31 | 0.41 | 0.15 | 0.15 |
| 4 | 185355556 | rs7685698 | -0.797 | 2.958 | -1.308 | 0.959 | 0.011 | 0.004 | 0.004 | 0 | 0.07 | 0.08 | 0.06 | 0.2 | 0.16 | 0.16 |
| 4 | 185359278 | rs11723606 | -0.483 | 2.72 | -1.287 | 0.911 | -0.03 | 0.028 | 0.028 | 0 | 0.11 | 0.05 | 0.15 | 0.03 | 0.23 | 0.22 |
| 4 | 185359416 | rs13116389 | -0.823 | 1.668 | -1.001 | 1.317 | 0.002 | -0.001 | -0.001 | 0 | 0.08 | 0.08 | 0.08 | 0.21 | 0.18 | 0.12 |
| 4 | 185363037 | rs793794 | -0.681 | -0.179 | -0.804 | 0.064 | -0.015 | -0.075 | -0.075 | 0.064 | 0.3 | 0.35 | 0.15 | 0.01 | 0.17 | 0.08 |
| 4 | 185369452 | rs3775572 | 1.138 | -0.83 | -0.011 | 0.823 | -0.057 | 0.098 | 0.098 | 1.472 | 0.87 | 0.37 | 0.94 | 0.75 | 0.96 | 0.9 |
| 4 | 185370066 | rs3775574 | 0.351 | -0.603 | -1.382 | 0.569 | 0.02 | 0.024 | 0.024 | 4.652 | 0.56 | 0.09 | 0.57 | 0.35 | 0.64 | 0.51 |
| 4 | 185378160 | rs793806 | -0.472 | -0.92 | -0.51 | 0.752 | 0.002 | -0.001 | -0.001 | 0.258 | 0.65 | 0.67 | 0.69 | 0.62 | 0.69 | 0.58 |
| 4 | 185381055 | rs793803 | 0.471 | -1.21 | -0.148 | 0.1 | 0 | -0.028 | -0.028 | 1.922 | 0.28 | 0.25 | 0.13 | 0.04 | 0.17 | 0.07 |
| 4 | 185382061 | rs793798 | 0.263 | -1.172 | 0.262 | 0.262 | -0.019 | 0.008 | 0.008 | 1.402 | 0.41 | 0.51 | 0.4 | 0.49 | 0.33 | 0.52 |
| 4 | 185383746 | rs3775580 | -0.841 | -0.336 | 0.194 | 0.4 | 0.001 | 0.001 | 0.001 | 0 | 0.73 | 0.76 | 0.55 | 0.53 | 0.57 | 0.61 |
| 4 | 185388487 | rs3775582 | -0.014 | -0.567 | -0.136 | 0.365 | -0.001 | -0.001 | -0.001 | 0 | 0.15 | 0.19 | 0.13 | 0.17 | 0.1 | 0.15 |
| 4 | 185395385 | rs1863315 | 0.026 | -0.645 | -0.146 | 0.45 | 0.001 | -0.002 | -0.002 | 0 | 0.15 | 0.19 | 0.13 | 0.22 | 0.1 | 0.15 |
| 4 | 185397811 | rs1863314 | -0.292 | -1.526 | 0.167 | 0.326 | -0.034 | -0.002 | -0.002 | 0 | 0.4 | 0.52 | 0.25 | 0.34 | 0.24 | 0.26 |
| 4 | 185398115 | rs1059492 | 1.05 | -1.3 | 0.054 | 0.411 | 0 | -0.001 | -0.001 | 0.083 | 0.24 | 0.25 | 0.28 | 0.23 | 0.27 | 0.34 |
| 4 | 185400748 | rs7658103 | -0.297 | -1.592 | 0.076 | 0.205 | 0.018 | 0.023 | 0.023 | 0 | 0.5 | 0.61 | 0.3 | 0.24 | 0.34 | 0.24 |
| 4 | 185406285 | rs12172979 | -0.581 | -0.914 | -0.636 | 0.266 | -0.021 | -0.017 | -0.017 | 0.475 | 0.71 | 0.76 | 0.54 | 0.58 | 0.56 | 0.63 |
| 4 | 185406638 | rs7696696 | 1.448 | -0.64 | -0.893 | 0.334 | -0.008 | -0.027 | -0.027 | 0 | 0.21 | 0.03 | 0.23 | 0.24 | 0.22 | 0.34 |
| 4 | 185412322 | rs2014472 | 1.575 | -0.807 | -1.198 | 0.269 | 0.038 | -0.036 | -0.036 | 0.168 | 0.15 | 0.05 | 0.13 | 0.25 | 0.08 | 0.23 |
| 4 | 185424289 | rs17488695 | 1.745 | 0.997 | -1.172 | -0.031 | -0.045 | 0.043 | 0.043 | 0.054 | 0.11 | 0.03 | 0.11 | 0 | 0.21 | 0.04 |
| 4 | 185425574 | rs17585837 | 0.667 | -1.757 | -1.21 | -0.257 | -0.002 | -0.002 | -0.002 | 0 | 0.23 | 0.17 | 0.2 | 0.06 | 0.28 | 0.17 |
| 4 | 185425853 | rs6816265 | -0.414 | -1.22 | -1.396 | 0.471 | -0.055 | -0.047 | -0.047 | 0 | 0.73 | 0.86 | 0.54 | 0.55 | 0.56 | 0.65 |
| 4 | 185434005 | rs1077767 | -0.482 | -2.055 | -0.623 | 0.02 | -0.007 | -0.005 | -0.005 | 0 | 0.45 | 0.52 | 0.39 | 0.66 | 0.38 | 0.5 |
| 4 | 185434615 | rs17585851 | 1.224 | -2.319 | -0.777 | -0.262 | 0.015 | 0.064 | 0.064 | 1.199 | 0.26 | 0.1 | 0.27 | 0.1 | 0.21 | 0.16 |
| 4 | 185436255 | rs6849842 | 0.602 | -2.633 | -0.658 | -0.056 | -0.041 | 0.051 | 0.051 | 0 | 0.27 | 0.33 | 0.24 | 0.4 | 0.21 | 0.34 |
| 4 | 185436378 | rs4862374 | -0.003 | -2.97 | -0.493 | -0.051 | 0.008 | -0.007 | -0.007 | 0 | 0.58 | 0.56 | 0.52 | 0.49 | 0.43 | 0.58 |
| 4 | 185440669 | rs12646629 | 1.712 | -3.043 | -0.746 | 0.368 | 0.033 | -0.032 | -0.032 | 0 | 0.15 | 0.05 | 0.14 | 0.23 | 0.09 | 0.17 |
| 4 | 185449260 | rs6552787 | -0.3 | -2.895 | -0.653 | 0.079 | 0.001 | 0.001 | 0.001 | 0 | 0.43 | 0.55 | 0.39 | 0.62 | 0.38 | 0.48 |
| 4 | 185458566 | rs13111483 | 0.162 | -2.907 | -0.536 | -0.18 | -0.028 | 0.012 | 0.012 | 0 | 0.58 | 0.46 | 0.66 | 0.52 | 0.66 | 0.55 |
| 4 | 185461350 | rs1918633 | 1.369 | -2.464 | -1.407 | -0.046 | -0.12 | -0.108 | -0.108 | 0 | 0.22 | 0.06 | 0.39 | 0.43 | 0.48 | 0.3 |
| 4 | 185462993 | rs6854047 | 1.176 | -2.391 | -1.26 | -0.259 | -0.001 | -0.008 | -0.008 | 0.135 | 0.08 | 0.15 | 0.07 | 0.07 | 0.07 | 0.08 |
| 4 | 185467264 | rs4862378 | 1.388 | -2.632 | -1.347 | -0.363 | 0.048 | 0.043 | 0.043 | 0 | 0.19 | 0.09 | 0.07 | 0.06 | 0.06 | 0.07 |
| 4 | 185467652 | rs10023706 | 1.474 | -2.477 | -1.26 | -0.228 | -0.034 | -0.033 | -0.033 | 0 | 0.23 | 0.15 | 0.4 | 0.44 | 0.48 | 0.32 |
| 4 | 185468885 | rs766908 | 1.076 | -1.721 | -0.943 | -0.519 | 0.028 | -0.028 | -0.028 | 0 | 0.17 | 0.07 | 0.16 | 0.21 | 0.11 | 0.17 |
| 4 | 185469914 | rs2059516 | 1.169 | -0.798 | -0.311 | -0.994 | -0.011 | -0.045 | -0.045 | 0 | 0.54 | 0.47 | 0.75 | 0.83 | 0.69 | 0.7 |
| 4 | 185474558 | rs4862379 | -0.357 | -0.405 | -1.219 | 0.026 | 0.046 | 0.042 | 0.042 | 0 | 0.19 | 0.08 | 0.07 | 0.05 | 0.06 | 0.07 |
| 4 | 185477454 | rs7682529 | -0.784 | 0.969 | -0.752 | 0.548 | 0.001 | -0.01 | -0.01 | 0.083 | 0.08 | 0.16 | 0.07 | 0.09 | 0.08 | 0.07 |
| 4 | 185481019 | rs965938 | -0.972 | 1.839 | -1.114 | 0.929 | -0.03 | -0.034 | -0.034 | 0 | 0.23 | 0.12 | 0.52 | 0.37 | 0.36 | 0.36 |
| 4 | 185482878 | rs6844313 | -0.786 | 2.299 | -0.334 | -0.071 | 0.057 | -0.049 | -0.049 | 0.041 | 0.2 | 0.46 | 0.16 | 0.09 | 0.21 | 0.1 |
| 4 | 185490816 | rs4862385 | -0.252 | 2.199 | -0.898 | -0.548 | -0.035 | 0.051 | 0.051 | 0 | 0.09 | 0.01 | 0.09 | 0 | 0.14 | 0.04 |
| 4 | 185491838 | rs10520563 | 0.903 | 1.312 | -0.693 | -1.274 | -0.017 | -0.068 | -0.068 | 0 | 0.42 | 0.57 | 0.22 | 0.07 | 0.31 | 0.29 |
| 4 | 185510238 | rs6838378 | -0.667 | 1.222 | -0.079 | -1.068 | 0.027 | -0.057 | -0.057 | 2.83 | 0.27 | 0.29 | 0.24 | 0.08 | 0.33 | 0.13 |
| 4 | 185514341 | rs1976955 | -0.92 | 0.835 | -0.718 | -1.257 | -0.012 | 0.049 | 0.049 | 0 | 0.38 | 0.63 | 0.28 | 0.08 | 0.39 | 0.17 |
| 4 | 185520068 | rs3970313 | -0.132 | 0.262 | -0.841 | -0.362 | 0.016 | -0.018 | -0.018 | 0.22 | 0.13 | 0.12 | 0.29 | 0.72 | 0.08 | 0.27 |
| 4 | 185531158 | rs12501768 | -1.427 | 0.755 | -1.559 | -1.14 | -0.03 | 0.183 | 0.183 | 0 | 0.45 | 0.19 | 0.65 | 0.82 | 0.49 | 0.59 |
| 4 | 185532342 | rs2178094 | 1.393 | 0.584 | -0.85 | -1.214 | 0.011 | 0.057 | 0.057 | 0 | 0.51 | 0.52 | 0.33 | 0.18 | 0.52 | 0.41 |
| 4 | 185533691 | rs7675251 | 0.031 | 0.351 | -0.814 | -1.048 | -0.05 | 0.066 | 0.066 | 0.023 | 0.85 | 0.67 | 0.71 | 0.28 | 0.93 | 0.7 |
| 4 | 185543672 | rs4862396 | 0.137 | 0.622 | -0.994 | -1.118 | -0.038 | -0.023 | -0.023 | 0 | 0.1 | 0.03 | 0.21 | 0.1 | 0.17 | 0.13 |
| 4 | 185548951 | rs1049253 | -0.079 | 0.721 | -1.113 | -1.299 | -0.029 | 0.141 | 0.141 | 0 | 0.21 | 0.01 | 0.13 | 0 | 0.19 | 0.11 |
| 4 | 185553098 | rs2705897 | 0.464 | 0.954 | -1.06 | -1.261 | -0.021 | 0 | 0 | 0.073 | 0.77 | 0.85 | 0.49 | 0.2 | 0.73 | 0.58 |
| 4 | 185554300 | rs2720376 | 1.419 | 0.923 | -1.006 | -1.288 | 0.008 | 0.09 | 0.09 | 0 | 0.54 | 0.63 | 0.36 | 0.2 | 0.54 | 0.47 |
| 4 | 185557581 | rs2019978 | -0.57 | 0.577 | -0.785 | -1.142 | 0.003 | -0.032 | -0.032 | 0 | 0.23 | 0.18 | 0.13 | 0 | 0.19 | 0.11 |
| 4 | 185567931 | rs4647610 | -0.663 | 0.547 | -0.931 | -0.84 | 0.019 | -0.061 | -0.061 | 0.148 | 0.17 | 0.09 | 0.32 | 0.69 | 0.12 | 0.3 |
| 4 | 185569730 | rs4647603 | -0.395 | 0.584 | -1.111 | -1.086 | -0.02 | 0.119 | 0.119 | 0 | 0.18 | 0.01 | 0.12 | 0 | 0.15 | 0.07 |
| 4 | 185571557 | rs12108497 | -1.784 | 0.375 | -0.698 | -1.189 | 0.026 | 0.022 | 0.022 | 0 | 0.35 | 0.19 | 0.47 | 0.72 | 0.29 | 0.38 |
| 4 | 185591481 | rs1405940 | -0.18 | 0.489 | -0.002 | -1.131 | 0 | 0.029 | 0.029 | 0.544 | 0.13 | 0.03 | 0.1 | 0 | 0.11 | 0.07 |
| 4 | 185604780 | rs2696051 | 1.526 | 0.35 | 0.151 | -0.539 | 0.004 | -0.013 | -0.013 | 0 | 0.75 | 0.82 | 0.6 | 0.39 | 0.78 | 0.75 |
| 4 | 185615764 | rs14969 | -0.86 | 0.638 | 0.469 | -0.765 | 0.018 | -0.108 | -0.108 | 2.457 | 0.17 | 0.3 | 0.14 | 0 | 0.15 | 0.09 |
| 4 | 185615768 | rs3184982 | -0.86 | 0.638 | 0.469 | -0.765 | 0.018 | -0.108 | -0.108 | 0.003 | 0.17 | 0.3 | 0.14 | 0 | 0.15 | 0.09 |
| 4 | 185616438 | rs13478 | -1.155 | 0.552 | 0.418 | -0.802 | 0.003 | -0.066 | -0.066 | 0 | 0.17 | 0.08 | 0.36 | 0.72 | 0.16 | 0.29 |
| 4 | 185637699 | rs6552804 | -0.804 | 0.573 | 0.488 | -0.819 | 0.017 | -0.105 | -0.105 | 0.03 | 0.17 | 0.3 | 0.14 | 0 | 0.15 | 0.09 |
| 4 | 185639169 | rs2130392 | 2.209 | 0.411 | 0.996 | -0.882 | 0.012 | 0.021 | 0.021 | 0 | 0.62 | 0.28 | 0.48 | 0.27 | 0.69 | 0.56 |
| 4 | 185649551 | rs10025167 | -0.767 | 0.458 | 0.845 | -0.866 | 0.017 | -0.105 | -0.105 | 0 | 0.17 | 0.3 | 0.14 | 0 | 0.15 | 0.09 |
| 4 | 185650154 | rs6818447 | -0.828 | 0.441 | 0.836 | -0.874 | 0.014 | -0.106 | -0.106 | 0 | 0.17 | 0.3 | 0.14 | 0 | 0.15 | 0.09 |
| 4 | 185650821 | rs1401357 | -1.236 | 0.581 | 0.438 | -0.716 | -0.002 | -0.066 | -0.066 | 1.632 | 0.17 | 0.08 | 0.36 | 0.72 | 0.16 | 0.29 |
| 4 | 185654723 | rs2871418 | 1.746 | 1.121 | 0.526 | -0.775 | -0.017 | 0.004 | 0.004 | 0.115 | 0.7 | 0.78 | 0.54 | 0.27 | 0.68 | 0.62 |
| 4 | 185664774 | rs4862411 | 0.961 | 1.804 | 0.578 | -0.813 | 0.005 | 0.016 | 0.016 | 0.692 | 0.79 | 0.67 | 0.63 | 0.28 | 0.78 | 0.61 |
| 4 | 185665118 | rs12513029 | -1.256 | 1.419 | 0.845 | -0.807 | 0.032 | -0.135 | -0.135 | 0.089 | 0.17 | 0.45 | 0.13 | 0 | 0.14 | 0.08 |
| 4 | 185667404 | rs2171838 | -1.278 | 1.437 | 0.802 | -0.791 | 0.046 | -0.129 | -0.129 | 0.686 | 0.18 | 0.42 | 0.14 | 0 | 0.16 | 0.09 |

| | | | | | | | | | | | | | | | | |
|---|---|---|---|---|---|---|---|---|---|---|---|---|---|---|---|---|
| 4 | 185674313 | rs11934173 | 0.374 | 0.546 | -0.335 | -0.606 | 0.008 | 0 | 0 | 0 | 0.29 | 0.22 | 0.32 | 0.25 | 0.23 | 0.19 |
| 4 | 185677363 | rs1056896 | 0.383 | 0.688 | -0.793 | -0.575 | -0.031 | 0.038 | 0.038 | 0 | 0.13 | 0.03 | 0.11 | 0 | 0.16 | 0.11 |
| 4 | 185677421 | rs8086 | 1.475 | 0.711 | -1.38 | -0.661 | -0.091 | 0.131 | 0.131 | 0 | 0.3 | 0.04 | 0.17 | 0.07 | 0.35 | 0.24 |
| 4 | 185678110 | rs2292899 | -0.027 | 0.549 | -0.525 | -0.425 | 0 | 0.037 | 0.037 | 0.152 | 0.22 | 0.32 | 0.37 | 0.68 | 0.25 | 0.39 |
| 4 | 185687233 | rs3749233 | 0.945 | 0.461 | -0.704 | -0.541 | 0.009 | 0.006 | 0.006 | 0.155 | 0.28 | 0.21 | 0.28 | 0.2 | 0.23 | 0.19 |
| 4 | 185687373 | rs3749232 | -0.546 | 0.184 | -0.93 | -0.798 | 0 | 0 | 0 | 0 | 0.29 | 0.31 | 0.43 | 0.71 | 0.29 | 0.42 |
| 4 | 185688754 | rs4862415 | 1.913 | 0.367 | -1.131 | -0.92 | -0.086 | 0.127 | 0.127 | 0 | 0.28 | 0.03 | 0.17 | 0.07 | 0.33 | 0.22 |
| 4 | 185690601 | rs4862417 | -0.06 | 0.337 | -1.032 | -0.884 | -0.001 | 0.002 | 0.002 | 0 | 0.25 | 0.28 | 0.47 | 0.71 | 0.25 | 0.4 |
| 4 | 185694697 | rs7691972 | 1.811 | 0.405 | -1.542 | -0.869 | -0.089 | 0.119 | 0.119 | 0 | 0.28 | 0.03 | 0.17 | 0.07 | 0.33 | 0.22 |
| 4 | 185700776 | rs4069938 | 0.34 | 0.282 | -1.115 | -0.822 | 0.006 | -0.008 | -0.008 | 0 | 0.62 | 0.43 | 0.46 | 0.3 | 0.61 | 0.5 |
| 4 | 185701549 | rs1803898 | 1.849 | 0.406 | -1.745 | -0.837 | -0.091 | 0.135 | 0.135 | 0 | 0.28 | 0.03 | 0.17 | 0.06 | 0.35 | 0.23 |
| 4 | 185705006 | rs2102862 | 0.673 | 0.153 | -1.515 | -0.69 | 0.002 | 0.002 | 0.002 | 0.539 | 0.3 | 0.29 | 0.27 | 0.22 | 0.25 | 0.2 |
| 4 | 185710859 | rs7681334 | -0.467 | 0.228 | -1.354 | -0.845 | 0.001 | 0.02 | 0.02 | 0 | 0.52 | 0.41 | 0.67 | 0.93 | 0.49 | 0.6 |
| 4 | 185718727 | rs11132255 | -0.318 | -0.046 | -1.469 | -0.357 | -0.015 | 0.015 | 0.015 | 0.261 | 0.19 | 0.67 | 0.22 | 0.27 | 0.16 | 0.19 |
| 4 | 185725416 | rs6552828 | -0.265 | 0.178 | -1.75 | -0.242 | 0.025 | 0.029 | 0.029 | 0 | 0.56 | 0.75 | 0.45 | 0.38 | 0.6 | 0.48 |
| 4 | 185728839 | rs902177 | 0.694 | 0.293 | -2.17 | -0.477 | -0.029 | -0.034 | -0.034 | 0.737 | 0.21 | 0.05 | 0.16 | 0.22 | 0.23 | 0.23 |
| 4 | 185731442 | rs13120078 | 1.092 | 0.686 | -1.871 | -0.803 | -0.133 | 0.137 | 0.137 | 0 | 0.31 | 0.04 | 0.2 | 0.08 | 0.39 | 0.25 |
| 4 | 185737843 | rs9997745 | -0.834 | 0.401 | -0.415 | -0.404 | 0.021 | -0.114 | -0.114 | 0.264 | 0.16 | 0.6 | 0.13 | 0 | 0.15 | 0.07 |
| 4 | 185744820 | rs1554336 | 0.062 | -0.007 | -1.049 | 0.021 | -0.002 | -0.119 | -0.119 | 0 | 0.24 | 0.05 | 0.22 | 0.52 | 0.22 | 0.33 |
| 4 | 185746088 | rs12503643 | -1.491 | 0.123 | -0.899 | -0.028 | 0.003 | -0.014 | -0.014 | 0.833 | 0.42 | 0.72 | 0.37 | 0.52 | 0.4 | 0.4 |
| 4 | 185750840 | rs10027540 | -0.694 | -0.077 | -0.272 | -0.398 | 0.004 | 0.02 | 0.02 | 0 | 0.3 | 0.28 | 0.19 | 0.19 | 0.26 | 0.31 |
| 4 | 185753007 | rs10033885 | 0.019 | -0.064 | -0.41 | -0.363 | -0.096 | -0.054 | -0.054 | 0 | 0.23 | 0.59 | 0.13 | 0.18 | 0.08 | 0.23 |
| 4 | 185755328 | rs6827693 | -1.023 | 0.068 | -0.639 | -0.311 | 0.006 | 0.006 | 0.006 | 0 | 0.35 | 0.72 | 0.23 | 0.18 | 0.26 | 0.31 |
| 4 | 185767176 | rs4862428 | -0.166 | -0.2 | -0.384 | 0.132 | -0.043 | -0.005 | -0.005 | 0 | 0.73 | 0.44 | 0.77 | 0.7 | 0.83 | 0.76 |
| 4 | 185771979 | rs4862432 | 0.466 | -0.129 | -0.726 | 0.296 | 0.007 | -0.005 | -0.005 | 0 | 0.58 | 0.4 | 0.66 | 0.48 | 0.69 | 0.58 |
| 4 | 185772925 | rs6852527 | 0.317 | -0.183 | -0.851 | 0.296 | -0.002 | -0.048 | -0.048 | 0.607 | 0.15 | 0.04 | 0.13 | 0.23 | 0.14 | 0.17 |
| 4 | 185773528 | rs6552833 | 0.408 | 0.559 | -0.592 | 0.296 | 0.002 | -0.013 | -0.013 | 0.273 | 0.42 | 0.31 | 0.49 | 0.47 | 0.43 | 0.49 |
| 4 | 185776616 | rs13144344 | -0.209 | 0.699 | -0.906 | 0.221 | -0.031 | -0.003 | -0.003 | 0 | 0.28 | 0.18 | 0.23 | 0.24 | 0.37 | 0.24 |
| 4 | 185776850 | rs12650715 | 0.107 | 0.496 | -0.837 | -0.007 | -0.03 | -0.027 | -0.027 | 0 | 0.38 | 0.56 | 0.44 | 0.29 | 0.29 | 0.29 |
| 4 | 185781300 | rs4862439 | -0.383 | 1.525 | -0.957 | 0.263 | -0.032 | -0.065 | -0.065 | 0 | 0.75 | 0.93 | 0.72 | 0.65 | 0.71 | 0.62 |
| 4 | 185781717 | rs13142179 | -0.976 | 2.186 | -0.216 | -0.216 | 0.034 | -0.028 | -0.028 | 0.213 | 0.09 | 0.11 | 0.08 | 0 | 0.19 | 0.1 |
| 4 | 185788486 | rs4862448 | -0.29 | 1.62 | 0.033 | -0.326 | -0.011 | -0.018 | -0.018 | 0 | 0.28 | 0.33 | 0.17 | 0.14 | 0.24 | 0.14 |
| 4 | 185788762 | rs4862450 | -1.307 | 1.011 | 0.26 | -0.178 | 0 | 0.001 | 0.001 | 0 | 0.49 | 0.54 | 0.37 | 0.32 | 0.54 | 0.42 |
| 4 | 185789995 | rs4862451 | -0.418 | 1.162 | -0.198 | -0.33 | -0.01 | 0.007 | 0.007 | 0 | 0.13 | 0.15 | 0.12 | 0.18 | 0.11 | 0.21 |
| 4 | 185791901 | rs7670324 | -0.87 | 1.182 | -0.274 | -0.671 | 0.013 | 0.081 | 0.081 | 0 | 0.5 | 0.68 | 0.33 | 0.1 | 0.53 | 0.32 |
| 4 | 185801346 | rs6853755 | -1.104 | 1.14 | 0.289 | -0.369 | -0.009 | -0.043 | -0.043 | 0 | 0.2 | 0.28 | 0.07 | 0.05 | 0.11 | 0.08 |
| 4 | 185801567 | rs13434707 | -1.468 | 0.723 | 0.978 | -0.572 | -0.042 | -0.108 | -0.108 | 0.194 | 0.25 | 0.59 | 0.1 | 0.05 | 0.11 | 0.08 |
| 4 | 185802696 | rs11734276 | -1.807 | 1.936 | 0.559 | -0.491 | -0.055 | 0.076 | 0.076 | 0.563 | 0.36 | 0.82 | 0.31 | 0.17 | 0.41 | 0.24 |
| 4 | 185804364 | rs11723505 | -1.351 | 2.035 | 1.316 | -0.074 | 0.04 | 0.043 | 0.043 | 0 | 0.12 | 0.3 | 0.16 | 0.12 | 0.15 | 0.12 |
| 4 | 185804494 | rs11736582 | -0.838 | 2.072 | 1.183 | -0.053 | 0.035 | 0.028 | 0.028 | 0 | 0.09 | 0.19 | 0.1 | 0.1 | 0.15 | 0.12 |
| 4 | 185805191 | rs6832271 | -1.471 | 1.79 | 1.416 | -0.192 | 0.057 | -0.004 | -0.004 | 0 | 0.11 | 0.37 | 0.13 | 0.07 | 0.07 | 0.05 |
| 4 | 185808270 | rs13143841 | -0.375 | 1.391 | 0.711 | -0.277 | -0.003 | -0.004 | -0.004 | 0 | 0.46 | 0.53 | 0.4 | 0.35 | 0.38 | 0.37 |
| 4 | 185808934 | rs6822813 | 0.427 | 1.512 | 0.284 | -1.163 | 0.005 | -0.025 | -0.025 | 0 | 0.31 | 0.29 | 0.18 | 0.01 | 0.26 | 0.21 |
| 4 | 185809272 | rs6849202 | -0.149 | 1.061 | 0.251 | -1.689 | -0.003 | -0.088 | -0.088 | 0.61 | 0.45 | 0.53 | 0.27 | 0.01 | 0.36 | 0.3 |
| 4 | 185809643 | rs6828840 | -0.892 | 1.294 | -0.152 | -1.268 | -0.029 | -0.019 | -0.019 | 0 | 0.77 | 0.87 | 0.57 | 0.32 | 0.64 | 0.57 |
| 4 | 185810760 | rs6835984 | -0.466 | 0.902 | -0.048 | -1.077 | 0.061 | 0.095 | 0.095 | 4.938 | 0.5 | 0.86 | 0.43 | 0.33 | 0.39 | 0.44 |
| 4 | 185828677 | rs6552840 | 0.469 | 1.526 | 0.629 | -1.048 | -0.001 | -0.002 | -0.002 | 2.289 | 0.58 | 0.63 | 0.72 | 0.73 | 0.53 | 0.69 |
| 4 | 185830781 | rs4392546 | -0.058 | 1.695 | 0.624 | -1.199 | 0.017 | -0.032 | -0.032 | 0 | 0.87 | 0.81 | 0.9 | 0.97 | 0.81 | 0.92 |
| 4 | 185835029 | rs4496623 | 0.185 | 1.644 | 0.726 | -1.132 | 0.023 | -0.028 | -0.028 | 0 | 0.86 | 0.74 | 0.89 | 0.95 | 0.81 | 0.92 |
| 4 | 185845735 | rs10005243 | 0.595 | 1.528 | 0.741 | -1.286 | 0.024 | -0.04 | -0.04 | 0 | 0.65 | 0.49 | 0.61 | 0.78 | 0.59 | 0.7 |
| 4 | 185847793 | rs6835458 | 0.392 | 1.659 | 0.72 | -1.275 | -0.003 | 0.001 | 0.001 | 0.022 | 0.21 | 0.21 | 0.28 | 0.19 | 0.23 | 0.22 |
| 4 | 185848004 | rs4318691 | -0.408 | 1.625 | 0.677 | -1.225 | -0.006 | 0.005 | 0.005 | 0 | 0.91 | 0.94 | 0.91 | 0.97 | 0.82 | 0.92 |
| 4 | 185849906 | rs10001154 | 1.655 | 1.48 | 0.469 | -1.054 | 0.01 | 0.028 | 0.028 | 0.739 | 0.45 | 0.26 | 0.45 | 0.58 | 0.39 | 0.6 |
| 4 | 185853636 | rs7682801 | 0.132 | 1.584 | 0.567 | -1.021 | 0.02 | -0.007 | -0.007 | 0.054 | 0.85 | 0.77 | 0.71 | 0.9 | 0.77 | 0.9 |
| 4 | 185859682 | rs10446837 | 0.99 | 1.589 | 0.171 | -1.294 | -0.004 | 0.003 | 0.003 | 4.367 | 0.15 | 0.16 | 0.28 | 0.08 | 0.18 | 0.22 |
| 4 | 185861258 | rs12641377 | 1.164 | 1.478 | 0.19 | -1.252 | 0.005 | 0.006 | 0.006 | 0 | 0.41 | 0.37 | 0.44 | 0.65 | 0.35 | 0.47 |
| 4 | 185864741 | rs4862466 | 0.375 | 1.735 | 0.005 | -1.355 | -0.005 | -0.002 | -0.002 | 0.52 | 0.13 | 0.12 | 0.28 | 0.15 | 0.21 | 0.07 |
| 4 | 185869216 | rs4370185 | -0.3 | 1.503 | 0.029 | -1.456 | -0.001 | -0.012 | -0.012 | 0 | 0.77 | 0.79 | 0.81 | 0.96 | 0.7 | 0.71 |
| 4 | 185870383 | rs6830291 | -0.378 | 1.505 | -0.158 | -1.454 | -0.021 | 0.031 | 0.031 | 1.847 | 0.77 | 0.86 | 0.82 | 0.96 | 0.7 | 0.71 |
| 4 | 185872178 | rs9992806 | 0.312 | 1.401 | -0.001 | -1.402 | 0.002 | -0.034 | -0.034 | 0 | 0.64 | 0.62 | 0.72 | 0.86 | 0.58 | 0.67 |
| 4 | 185874623 | rs12502026 | 1.252 | 0.954 | 0.025 | -0.953 | 0.017 | -0.003 | -0.003 | 0 | 0.33 | 0.23 | 0.37 | 0.64 | 0.27 | 0.38 |
| 4 | 185874675 | rs4861647 | 0.685 | 1.127 | -0.009 | -0.928 | -0.011 | -0.002 | -0.002 | 8.144 | 0.09 | 0.09 | 0.24 | 0.08 | 0.14 | 0.09 |
| 4 | 185875614 | rs11734892 | 0.938 | 1.267 | -0.135 | -0.969 | 0.004 | -0.009 | -0.009 | 1.387 | 0.1 | 0.08 | 0.09 | 0.03 | 0.16 | 0.09 |
| 4 | 185876135 | rs6822322 | 0.203 | 1.071 | -0.16 | -1.008 | -0.002 | -0.005 | -0.005 | 0.074 | 0.47 | 0.48 | 0.48 | 0.67 | 0.43 | 0.47 |
| 4 | 185876473 | rs6822805 | -0.596 | 0.961 | -0.97 | -0.998 | -0.028 | 0.107 | 0.107 | 0.204 | 0.59 | 0.86 | 0.76 | 0.75 | 0.58 | 0.56 |
| 4 | 185877029 | rs6854741 | -0.916 | 0.374 | -0.941 | -0.606 | 0.001 | 0.019 | 0.019 | 0 | 0.88 | 0.94 | 0.91 | 0.91 | 0.87 | 0.96 |
| 4 | 185877443 | rs13143382 | 0.482 | 0.302 | -0.991 | -0.059 | 0.012 | -0.002 | -0.002 | 0 | 0.32 | 0.24 | 0.38 | 0.65 | 0.28 | 0.39 |
| 4 | 185882454 | rs6419946 | 0.196 | -0.136 | -0.671 | 0.136 | -0.033 | 0.018 | 0.018 | 0 | 0.71 | 0.66 | 0.81 | 0.65 | 0.79 | 0.72 |
| 4 | 185884206 | rs10866272 | -0.606 | 0.263 | -0.885 | 0.435 | 0.003 | -0.005 | -0.005 | 0 | 0.81 | 0.83 | 0.85 | 0.63 | 0.85 | 0.75 |

| | | | | | | | | | | | | | | | | |
|---|---|---|---|---|---|---|---|---|---|---|---|---|---|---|---|---|
| 4 | 185885897 | rs4355430 | -0.024 | 0.859 | -1.218 | 0.506 | 0.011 | 0 | 0 | 0.006 | 0.48 | 0.31 | 0.56 | 0.54 | 0.53 | 0.61 |
| 4 | 185894655 | rs9993348 | 0.291 | 0.96 | -1.381 | 0.612 | -0.014 | 0.047 | 0.047 | 0.781 | 0.56 | 0.24 | 0.49 | 0.68 | 0.57 | 0.71 |
| 4 | 185905754 | rs10866273 | 0.225 | 0.768 | -0.618 | 0.313 | -0.005 | -0.059 | -0.059 | 0 | 0.9 | 0.76 | 0.9 | 1 | 0.92 | 0.95 |
| 4 | 185907334 | rs6843954 | -0.12 | 1.648 | -0.952 | 0.627 | -0.044 | -0.041 | -0.041 | 0 | 0.23 | 0.12 | 0.43 | 0.43 | 0.34 | 0.22 |
| 4 | 185907997 | rs4862478 | -0.12 | 1.648 | -0.952 | 0.627 | -0.044 | -0.041 | -0.041 | 1.863 | 0.23 | 0.12 | 0.43 | 0.43 | 0.34 | 0.22 |
| 4 | 185916343 | rs6820983 | -1.115 | 1.243 | -0.618 | 1.005 | -0.022 | -0.012 | -0.012 | 0.362 | 0.37 | 0.66 | 0.61 | 0.59 | 0.44 | 0.28 |
| 4 | 185925922 | rs11132270 | 0.083 | 0.167 | -1.125 | 0.573 | -0.015 | 0.101 | 0.101 | 0.058 | 0.46 | 0.77 | 0.49 | 0.73 | 0.43 | 0.67 |
| 4 | 185928446 | rs11722349 | -0.833 | -0.504 | 0.003 | 1.176 | -0.003 | -0.003 | -0.003 | 0 | 0.3 | 0.34 | 0.23 | 0.21 | 0.22 | 0.2 |
| 4 | 185929997 | rs7664096 | -1.018 | -0.427 | -0.586 | 1.334 | -0.014 | 0.004 | 0.004 | 0.003 | 0.34 | 0.63 | 0.34 | 0.36 | 0.23 | 0.28 |
| 4 | 185930269 | rs16813 | 0.036 | 0.503 | -1.227 | 1.398 | -0.032 | -0.037 | -0.037 | 0 | 0.89 | 0.99 | 0.86 | 0.86 | 0.83 | 0.94 |
| 4 | 185935020 | rs1486664 | -0.149 | 0.449 | -0.932 | 1.309 | -0.001 | 0 | 0 | 1.186 | 0.16 | 0.17 | 0.18 | 0.21 | 0.18 | 0.15 |
| 4 | 185935259 | rs13119549 | 0.643 | 0.117 | -1.609 | 1.237 | -0.006 | 0.044 | 0.044 | 0.885 | 0.48 | 0.14 | 0.42 | 0.45 | 0.53 | 0.6 |
| 4 | 185941196 | rs901434 | 0.104 | -0.263 | -0.337 | 1.767 | -0.008 | 0.068 | 0.068 | 0 | 0.1 | 0.13 | 0.15 | 0.19 | 0.1 | 0.08 |
| 4 | 185943413 | rs11727431 | -0.451 | -0.437 | -0.574 | 1.536 | -0.013 | 0.022 | 0.022 | 0 | 0.23 | 0.13 | 0.22 | 0.14 | 0.25 | 0.26 |
| 4 | 185943856 | rs6854448 | -1.792 | -0.893 | -0.588 | 1.29 | -0.029 | 0.175 | 0.175 | 0 | 0.54 | 0.83 | 0.67 | 0.78 | 0.49 | 0.63 |
| 4 | 185946538 | rs6851788 | -1.723 | -0.882 | 0.601 | 1.627 | -0.001 | -0.002 | -0.002 | 0 | 0.43 | 0.59 | 0.51 | 0.65 | 0.39 | 0.54 |
| 4 | 185948996 | rs10019464 | -0.972 | -0.969 | 0.79 | 1.881 | -0.003 | 0.004 | 0.004 | 0 | 0.31 | 0.36 | 0.37 | 0.35 | 0.29 | 0.27 |
| 4 | 185957817 | rs752632 | -0.062 | -0.955 | 0.156 | 1.855 | -0.022 | 0.025 | 0.025 | 0 | 0.19 | 0.07 | 0.21 | 0.13 | 0.23 | 0.23 |
| 4 | 185965017 | rs1012518 | -0.801 | -0.943 | 0.192 | 1.924 | -0.005 | -0.037 | -0.037 | 3.332 | 0.3 | 0.13 | 0.31 | 0.41 | 0.29 | 0.4 |
| 4 | 185973097 | rs4862495 | -1.672 | -0.988 | 0.024 | 1.658 | 0.039 | 0.051 | 0.051 | 0 | 0.46 | 0.78 | 0.53 | 0.67 | 0.37 | 0.51 |
| 4 | 185977068 | rs728756 | -1.273 | -1.134 | 0.061 | 1.834 | -0.004 | 0.002 | 0.002 | 0.412 | 0.39 | 0.65 | 0.4 | 0.42 | 0.31 | 0.33 |
| 4 | 185977226 | rs4415003 | 0.064 | -1.195 | 0.105 | 1.877 | -0.008 | 0.046 | 0.046 | 0 | 0.09 | 0.15 | 0.12 | 0.25 | 0.06 | 0.16 |
| 4 | 185977354 | rs4577622 | -1.739 | -1.108 | -0.19 | 1.643 | 0.057 | 0.076 | 0.076 | 0.184 | 0.47 | 0.82 | 0.53 | 0.66 | 0.37 | 0.48 |
| 4 | 185979170 | rs6852910 | -1.38 | -1.261 | -0.408 | 1.406 | -0.011 | -0.001 | -0.001 | 0.101 | 0.42 | 0.48 | 0.32 | 0.41 | 0.32 | 0.47 |
| 4 | 185979298 | rs6853461 | -0.841 | -1.129 | -0.193 | 1.317 | -0.024 | 0.02 | 0.02 | 0 | 0.12 | 0.33 | 0.07 | 0.13 | 0.05 | 0.22 |
| 4 | 185981436 | rs6844558 | -0.923 | -1.362 | -1.288 | 1.424 | 0.099 | -0.039 | -0.039 | 0 | 0.31 | 0.82 | 0.29 | 0.35 | 0.15 | 0.46 |
| 4 | 185983220 | rs4861655 | -0.334 | 0.65 | -1.345 | 0.917 | -0.043 | 0 | 0 | 0.14 | 0.3 | 0.19 | 0.42 | 0.69 | 0.39 | 0.45 |
| 4 | 185997791 | rs4862499 | 0.637 | 0.978 | -1.279 | 0.779 | -0.013 | -0.006 | -0.006 | 0 | 0.71 | 0.8 | 0.58 | 0.31 | 0.63 | 0.57 |
| 4 | 185999348 | rs10023344 | 0.407 | 0.941 | -0.918 | 0.91 | -0.014 | -0.014 | -0.014 | 0.226 | 0.74 | 0.85 | 0.6 | 0.37 | 0.64 | 0.59 |
| 4 | 186003688 | rs6819903 | -0.894 | 1.111 | -0.925 | 0.315 | 0 | -0.011 | -0.011 | 0.247 | 0.21 | 0.23 | 0.13 | 0.11 | 0.2 | 0.21 |
| 4 | 186007879 | rs4862502 | -1.293 | 0.341 | -1.181 | -0.489 | 0.003 | 0.004 | 0.004 | 0 | 0.71 | 0.82 | 0.83 | 0.73 | 0.74 | 0.82 |
| 4 | 186008402 | rs4861656 | -1.34 | 0.142 | -0.387 | -0.586 | -0.005 | -0.005 | -0.005 | 0 | 0.69 | 0.65 | 0.82 | 0.73 | 0.74 | 0.82 |
| 4 | 186012601 | rs1545951 | -1.095 | 0.043 | -0.21 | -0.646 | -0.009 | -0.02 | -0.02 | 0.97 | 0.67 | 0.57 | 0.81 | 0.73 | 0.73 | 0.81 |
| 4 | 186013276 | rs11132275 | 0.583 | -0.13 | -0.887 | -0.815 | -0.075 | 0.144 | 0.144 | 0.06 | 0.29 | 0.07 | 0.34 | 0.07 | 0.33 | 0.3 |
| 4 | 186016584 | rs7667259 | 0.787 | 0.152 | -0.445 | -0.367 | 0.004 | 0.016 | 0.016 | 0 | 0.2 | 0.15 | 0.37 | 0.53 | 0.33 | 0.33 |
| 4 | 186019565 | rs13120719 | 0.207 | 0.01 | -0.096 | -0.45 | 0.019 | 0.044 | 0.044 | 0 | 0.25 | 0.31 | 0.41 | 0.56 | 0.36 | 0.34 |
| 4 | 186019962 | rs13121592 | -1.293 | -0.205 | -0.402 | -0.521 | -0.003 | 0.011 | 0.011 | 0 | 0.7 | 0.81 | 0.82 | 0.73 | 0.73 | 0.79 |
| 4 | 186022884 | rs2310156 | -0.682 | -0.347 | -0.168 | -0.712 | -0.001 | -0.001 | -0.001 | 0.003 | 0.45 | 0.42 | 0.51 | 0.7 | 0.54 | 0.48 |
| 4 | 186026298 | rs6828163 | -0.824 | -0.322 | -0.732 | -1.177 | -0.062 | 0.045 | 0.045 | 0 | 0.7 | 0.88 | 0.81 | 0.75 | 0.6 | 0.74 |
| 4 | 186026909 | rs12512707 | 0.448 | -0.67 | -0.951 | -1.287 | 0.036 | 0.033 | 0.033 | 0 | 0.43 | 0.21 | 0.6 | 0.58 | 0.41 | 0.44 |
| 4 | 186027524 | rs11132277 | -0.407 | -0.919 | 0.027 | -0.954 | 0.001 | -0.014 | -0.014 | 0.663 | 0.22 | 0.29 | 0.19 | 0.17 | 0.19 | 0.29 |
| 4 | 186029032 | rs2310155 | -0.407 | -0.961 | 0.027 | -0.951 | 0 | -0.013 | -0.013 | 0.029 | 0.22 | 0.29 | 0.19 | 0.17 | 0.19 | 0.29 |
| 4 | 186031823 | rs6552864 | -1.054 | -0.922 | -0.572 | -0.773 | 0.04 | 0.06 | 0.06 | 1.705 | 0.31 | 0.82 | 0.24 | 0.17 | 0.19 | 0.36 |
| 4 | 186034134 | rs1872719 | -0.362 | -0.955 | -0.235 | -0.78 | -0.08 | -0.105 | -0.105 | 0 | 0.19 | 0.55 | 0.07 | 0.04 | 0.05 | 0.2 |
| 4 | 186037329 | rs3912538 | 0.222 | -0.204 | -0.638 | -0.503 | 0.001 | -0.008 | -0.008 | 0.2 | 0.8 | 0.8 | 0.8 | 0.9 | 0.76 | 0.79 |
| 4 | 186038657 | rs7661225 | -1.034 | -0.899 | -1.043 | 0.067 | 0.022 | -0.032 | -0.032 | 0.43 | 0.85 | 0.92 | 0.82 | 0.8 | 0.87 | 0.8 |
| 4 | 186038842 | rs9884823 | -0.245 | -1.276 | -0.511 | -0.871 | -0.026 | -0.081 | -0.081 | 0 | 0.27 | 0.63 | 0.15 | 0.05 | 0.15 | 0.22 |
| 4 | 186039201 | rs10026207 | 0.109 | -0.564 | -0.971 | -0.556 | 0.009 | -0.004 | -0.004 | 0 | 0.41 | 0.21 | 0.38 | 0.42 | 0.39 | 0.37 |
| 4 | 186044138 | rs6853087 | -0.58 | -0.596 | -0.471 | -0.594 | 0.022 | 0.068 | 0.068 | 0 | 0.51 | 0.68 | 0.37 | 0.34 | 0.41 | 0.42 |
| 4 | 186045532 | rs10002793 | 0.604 | -0.85 | -0.099 | -0.626 | -0.003 | 0.003 | 0.003 | 0 | 0.31 | 0.33 | 0.27 | 0.38 | 0.27 | 0.32 |
| 4 | 186045738 | rs1486668 | 0.882 | -0.763 | -0.58 | -0.693 | -0.025 | -0.005 | -0.005 | 0 | 0.28 | 0.06 | 0.25 | 0.29 | 0.27 | 0.22 |
| 4 | 186046473 | rs6818704 | -0.348 | -0.626 | 0.062 | -0.652 | -0.039 | -0.128 | -0.128 | 1.093 | 0.22 | 0.4 | 0.11 | 0 | 0.13 | 0.11 |
| 4 | 186055546 | rs28678870 | -0.666 | -0.73 | 0.008 | -0.577 | -0.075 | -0.098 | -0.098 | 0 | 0.24 | 0.51 | 0.1 | 0.1 | 0.11 | 0.2 |
| 4 | 186059893 | rs12506141 | 0.166 | -0.414 | -0.997 | -0.581 | -0.095 | -0.015 | -0.015 | 0 | 0.4 | 0.14 | 0.53 | 0.59 | 0.49 | 0.45 |
| 4 | 186063341 | rs10024068 | -0.566 | -0.682 | -0.883 | -0.896 | 0.018 | 0.069 | 0.069 | 0 | 0.59 | 0.79 | 0.68 | 0.76 | 0.62 | 0.75 |
| 4 | 186070432 | rs2046535 | 0.512 | -0.657 | -1.022 | -0.882 | -0.067 | 0.043 | 0.043 | 0 | 0.36 | 0.17 | 0.55 | 0.72 | 0.47 | 0.55 |
| 4 | 186071068 | rs7675110 | 0.778 | -0.686 | -1.039 | -0.895 | -0.075 | 0.023 | 0.023 | 0 | 0.34 | 0.17 | 0.55 | 0.68 | 0.48 | 0.4 |
| 4 | 186071932 | rs11729551 | 0.614 | -0.658 | -1.028 | -0.881 | -0.074 | 0.029 | 0.029 | 0 | 0.35 | 0.17 | 0.55 | 0.68 | 0.48 | 0.45 |
| 4 | 186072110 | rs11729626 | 0.03 | -0.652 | -0.834 | -0.912 | -0.038 | 0.119 | 0.119 | 0 | 0.45 | 0.25 | 0.57 | 0.74 | 0.51 | 0.62 |
| 4 | 186078324 | rs13145878 | 1.289 | -0.96 | -1.065 | -1.087 | 0 | 0.043 | 0.043 | 0 | 0.08 | 0 | 0.03 | 0 | 0.07 | 0.03 |
| 4 | 186086046 | rs6850094 | -0.543 | -0.941 | -0.116 | -0.863 | 0.002 | -0.005 | -0.005 | 0.42 | 0.55 | 0.5 | 0.45 | 0.4 | 0.5 | 0.44 |
| 4 | 186090818 | rs6839413 | -0.551 | -0.944 | -0.118 | -0.865 | 0.002 | -0.005 | -0.005 | 0.41 | 0.55 | 0.5 | 0.45 | 0.4 | 0.5 | 0.44 |
| 4 | 186091191 | rs6814673 | -0.629 | -0.961 | 0.026 | -0.874 | -0.008 | 0.006 | 0.006 | 0 | 0.57 | 0.63 | 0.46 | 0.4 | 0.51 | 0.44 |
| 4 | 186097045 | rs6855305 | -0.663 | -0.963 | 0.056 | -0.876 | -0.012 | 0.008 | 0.008 | 0 | 0.57 | 0.66 | 0.46 | 0.4 | 0.51 | 0.44 |
| 4 | 186111639 | rs1133657 | 0.972 | -1.064 | 0.044 | -0.933 | -0.025 | -0.007 | -0.007 | 0 | 0.36 | 0.25 | 0.52 | 0.58 | 0.48 | 0.41 |
| 4 | 186112111 | rs3108237 | 0.197 | -1.053 | 0.055 | -0.976 | 0.002 | -0.006 | -0.006 | 0 | 0.45 | 0.5 | 0.55 | 0.6 | 0.5 | 0.56 |
| 4 | 186117162 | rs1534140 | 0.291 | -1.035 | 0.044 | -0.966 | -0.008 | 0.006 | 0.006 | 0 | 0.43 | 0.38 | 0.54 | 0.6 | 0.5 | 0.56 |
| 4 | 186125842 | rs3108244 | 0.281 | -1.924 | -0.393 | -1.338 | 0.006 | 0.005 | 0.005 | 0 | 0.55 | 0.41 | 0.61 | 0.61 | 0.63 | 0.59 |
| 4 | 186131934 | rs2177796 | -0.9 | -1.743 | -0.589 | -1.512 | -0.017 | 0.002 | 0.002 | 0 | 0.76 | 0.89 | 0.77 | 0.76 | 0.72 | 0.79 |

| | | | | | | | | | | | | | | | | |
|---|---|---|---|---|---|---|---|---|---|---|---|---|---|---|---|---|
| 4 | 186134204 | rs2310159 | 0.644 | -1.705 | -0.243 | -1.543 | 0 | -0.015 | -0.015 | 0.787 | 0.46 | 0.52 | 0.61 | 0.74 | 0.52 | 0.61 |
| 4 | 186142979 | rs3112898 | -0.064 | -1.769 | -0.329 | -1.572 | -0.008 | -0.032 | -0.032 | 0 | 0.63 | 0.56 | 0.68 | 0.74 | 0.65 | 0.7 |
| 4 | 186150254 | rs3112911 | 0.849 | -1.695 | -0.453 | -1.535 | -0.001 | -0.03 | -0.03 | 0 | 0.16 | 0.18 | 0.09 | 0.01 | 0.13 | 0.08 |
| 4 | 186159445 | rs7675903 | 1.771 | -1.529 | -0.876 | -1.485 | -0.157 | -0.098 | -0.098 | 0 | 0.31 | 0.03 | 0.49 | 0.6 | 0.45 | 0.5 |
| 4 | 186175197 | rs7698980 | 1.412 | -1.513 | -0.85 | -1.404 | -0.082 | -0.054 | -0.054 | 0.307 | 0.37 | 0.1 | 0.45 | 0.56 | 0.44 | 0.49 |
| 4 | 186175566 | rs3112854 | 0.985 | -1.543 | -0.673 | -1.446 | -0.006 | -0.035 | -0.035 | 0 | 0.16 | 0.2 | 0.07 | 0.01 | 0.1 | 0.08 |
| 4 | 186178278 | rs3108227 | 0.954 | -1.571 | -0.653 | -1.437 | -0.014 | 0.03 | 0.03 | 0 | 0.48 | 0.26 | 0.5 | 0.58 | 0.52 | 0.51 |
| 4 | 186185293 | rs2280665 | 1.574 | -1.357 | -0.385 | -1.25 | -0.02 | -0.016 | -0.016 | 0 | 0.33 | 0.28 | 0.44 | 0.58 | 0.42 | 0.51 |
| 4 | 186196453 | rs3108224 | 1.501 | -1.32 | 0.248 | -1.256 | 0.007 | 0.007 | 0.007 | 0 | 0.34 | 0.4 | 0.46 | 0.56 | 0.43 | 0.44 |
| 4 | 186197070 | rs1534146 | -1.613 | -1.28 | -0.372 | -1.377 | -0.07 | -0.003 | -0.003 | 0 | 0.87 | 0.99 | 0.91 | 0.93 | 0.82 | 0.92 |
| 4 | 186204090 | rs2692596 | -0.099 | -1.553 | 0.267 | -1.225 | -0.008 | 0.107 | 0.107 | 0 | 0.53 | 0.71 | 0.6 | 0.75 | 0.55 | 0.69 |
| 4 | 186205791 | rs12641710 | 1.432 | -1.2 | 0.124 | -0.99 | -0.032 | -0.031 | -0.031 | 0.057 | 0.32 | 0.18 | 0.44 | 0.56 | 0.42 | 0.43 |
| 4 | 186209273 | rs11725533 | 1.036 | -1.141 | 0.239 | -0.986 | -0.002 | -0.024 | -0.024 | 0 | 0.05 | 0.08 | 0.03 | 0.01 | 0.05 | 0.03 |
| 4 | 186216136 | rs1288547 | 0.89 | -1.118 | 0.142 | -0.849 | -0.003 | -0.007 | -0.007 | 0 | 0.4 | 0.36 | 0.46 | 0.52 | 0.43 | 0.51 |
| 4 | 186230276 | rs1288569 | 0.769 | -1.119 | 0.131 | -0.847 | -0.005 | -0.004 | -0.004 | 0 | 0.41 | 0.34 | 0.46 | 0.55 | 0.43 | 0.58 |
| 4 | 186242233 | rs10000719 | -1.245 | -2.009 | 1.205 | -0.679 | 0.004 | 0.07 | 0.07 | 0 | 0.15 | 0.35 | 0.15 | 0.17 | 0.11 | 0.06 |
| 4 | 186252138 | rs1288572 | -2.856 | -2.174 | 1.032 | -0.684 | 0.058 | -0.016 | -0.016 | 0 | 0.33 | 0.86 | 0.24 | 0.26 | 0.22 | 0.28 |
| 4 | 186254126 | rs3733644 | -1.481 | -2.007 | 1.194 | -0.609 | -0.023 | 0.066 | 0.066 | 0.103 | 0.16 | 0.38 | 0.14 | 0.18 | 0.12 | 0.07 |
| 4 | 186271154 | rs10030392 | -1.323 | -2.161 | 1.117 | -0.843 | -0.025 | 0.002 | 0.002 | 0.647 | 0.3 | 0.61 | 0.28 | 0.25 | 0.24 | 0.24 |
| 4 | 186272773 | rs3822302 | -2.29 | -2.217 | 1.004 | -0.885 | 0.047 | 0.042 | 0.042 | 0.196 | 0.44 | 0.77 | 0.38 | 0.28 | 0.34 | 0.38 |
| 4 | 186281755 | rs1031128 | -1.907 | -2.242 | 0.987 | -0.856 | 0.03 | 0.017 | 0.017 | 0.445 | 0.4 | 0.74 | 0.33 | 0.26 | 0.29 | 0.38 |
| 4 | 186288846 | rs7664573 | -1.187 | -2.294 | 1.244 | -0.815 | -0.016 | -0.013 | -0.013 | 0 | 0.32 | 0.39 | 0.3 | 0.19 | 0.23 | 0.22 |
| 4 | 186292169 | rs4862522 | 0.224 | -2.239 | 1.037 | -0.782 | 0.011 | -0.028 | -0.028 | 0 | 0.16 | 0.09 | 0.19 | 0.18 | 0.13 | 0.13 |
| 4 | 186292826 | rs3756276 | -0.035 | -2.23 | 1.015 | -0.651 | -0.005 | 0.009 | 0.009 | 0 | 0.06 | 0.09 | 0.02 | 0.07 | 0.05 | 0.1 |
| 4 | 186294198 | rs3797026 | -0.622 | -2.26 | 0.796 | -0.775 | -0.011 | -0.001 | -0.001 | 0.152 | 0.25 | 0.32 | 0.27 | 0.18 | 0.19 | 0.16 |
| 4 | 186299519 | rs1288549 | -2.035 | -2.18 | -0.101 | -0.756 | 0.098 | 0.043 | 0.043 | 1.087 | 0.47 | 0.93 | 0.37 | 0.27 | 0.32 | 0.39 |
| 4 | 186299656 | rs3797032 | -1.56 | -1.939 | 0.247 | -0.492 | 0.052 | 0.016 | 0.016 | 0.183 | 0.4 | 0.75 | 0.33 | 0.28 | 0.27 | 0.38 |
| 4 | 186302406 | rs10018215 | -0.717 | -1.78 | 0.522 | -0.646 | -0.009 | -0.002 | -0.002 | 0 | 0.25 | 0.27 | 0.22 | 0.15 | 0.13 | 0.19 |
| 4 | 186304749 | rs11944575 | 0.961 | -1.916 | 0.991 | -0.593 | -0.03 | -0.002 | -0.002 | 0 | 0.67 | 0.43 | 0.73 | 0.67 | 0.82 | 0.65 |
| 4 | 186309757 | rs6818425 | -0.528 | -2.028 | 0.995 | -1.408 | -0.003 | -0.033 | -0.033 | 0.053 | 0.89 | 0.83 | 0.91 | 0.99 | 0.92 | 0.93 |
| 4 | 186313557 | rs6845326 | 1.735 | -2.337 | 0.93 | -1.712 | 0.126 | 0.207 | 0.207 | 0.164 | 0.53 | 0.22 | 0.73 | 0.8 | 0.7 | 0.56 |
| 4 | 186316633 | rs4862523 | 1.762 | -2.365 | 0.909 | -1.725 | 0.131 | 0.215 | 0.215 | 0.208 | 0.52 | 0.21 | 0.73 | 0.8 | 0.7 | 0.56 |
| 4 | 186316829 | rs4862524 | 1.038 | -2.357 | 1.449 | -1.729 | -0.049 | -0.13 | -0.13 | 0 | 0.66 | 0.46 | 0.81 | 0.95 | 0.8 | 0.72 |
| 4 | 186325429 | rs6815168 | -0.911 | -2.268 | 1.555 | -1.637 | -0.009 | -0.009 | -0.009 | 0 | 0.12 | 0.23 | 0.07 | 0.15 | 0.1 | 0.16 |
| 4 | 186336465 | rs2289722 | 0.682 | -2.257 | 1.181 | -1.633 | 0.003 | 0.008 | 0.008 | 0 | 0.07 | 0.04 | 0.04 | 0.03 | 0.06 | 0.05 |
| 4 | 186336583 | rs10032989 | -0.559 | -2.153 | 1.265 | -1.588 | -0.003 | -0.032 | -0.032 | 0 | 0.88 | 0.84 | 0.91 | 0.99 | 0.92 | 0.93 |
| 4 | 186346239 | rs2028701 | 0.53 | -2.164 | 1.391 | -1.604 | -0.009 | -0.041 | -0.041 | 0 | 0.83 | 0.61 | 0.88 | 0.85 | 0.87 | 0.82 |
| 4 | 186350063 | rs12500107 | 1.373 | -2.23 | 1.022 | -1.636 | 0.077 | 0.118 | 0.118 | 0.198 | 0.57 | 0.28 | 0.75 | 0.81 | 0.71 | 0.62 |
| 4 | 186351612 | rs11934546 | 1.389 | -2.202 | 1.009 | -1.62 | 0.114 | 0.177 | 0.177 | 0.295 | 0.55 | 0.23 | 0.74 | 0.79 | 0.71 | 0.56 |
| 4 | 186353670 | rs7672394 | 0.214 | -1.247 | 1.407 | -0.987 | -0.011 | -0.071 | -0.071 | 0.986 | 0.87 | 0.72 | 0.91 | 0.99 | 0.91 | 0.93 |
| 4 | 186357523 | rs7697970 | 0.551 | -1.231 | 1.256 | -0.979 | -0.019 | -0.088 | -0.088 | 0.029 | 0.85 | 0.63 | 0.9 | 0.99 | 0.91 | 0.93 |
| 4 | 186372826 | rs6838700 | -0.865 | -0.747 | -0.288 | -0.886 | 0.037 | 0.073 | 0.073 | 0 | 0.46 | 0.63 | 0.38 | 0.28 | 0.37 | 0.38 |
| 4 | 186373316 | rs13102235 | -0.468 | -0.217 | -1.012 | -1.025 | -0.018 | 0.148 | 0.148 | 0 | 0.52 | 0.75 | 0.7 | 0.8 | 0.5 | 0.58 |
| 4 | 186375488 | rs10022110 | 0.17 | -0.261 | -0.513 | -0.988 | -0.015 | -0.017 | -0.017 | 0 | 0.32 | 0.36 | 0.29 | 0.28 | 0.25 | 0.34 |
| 4 | 186378790 | rs4279261 | 0.559 | -0.148 | -0.17 | -0.611 | -0.02 | -0.02 | -0.02 | 0 | 0.72 | 0.43 | 0.75 | 0.73 | 0.77 | 0.78 |
| 4 | 186379735 | rs6827370 | 0.532 | -0.147 | -0.111 | -0.582 | -0.01 | 0.044 | 0.044 | 0 | 0.14 | 0.3 | 0.16 | 0.27 | 0.12 | 0.17 |
| 4 | 186380515 | rs11132306 | 0.393 | 0.122 | -1.053 | -0.41 | -0.004 | 0.067 | 0.067 | 0.196 | 0.54 | 0.28 | 0.36 | 0.28 | 0.54 | 0.49 |
| 4 | 186381115 | rs7699687 | 0.369 | 0.155 | -1.043 | -0.415 | -0.004 | 0.064 | 0.064 | 0.238 | 0.54 | 0.28 | 0.36 | 0.28 | 0.54 | 0.49 |
| 4 | 186382206 | rs11132309 | 0.369 | 0.155 | -1.099 | -0.415 | -0.005 | 0.068 | 0.068 | 0.238 | 0.54 | 0.28 | 0.36 | 0.28 | 0.54 | 0.49 |
| 4 | 186383240 | rs4862534 | 0.74 | 0.621 | -1.461 | 0.06 | -0.019 | -0.04 | -0.04 | 0.623 | 0.17 | 0.13 | 0.37 | 0.44 | 0.22 | 0.22 |
| 4 | 186383415 | rs4314326 | 0.097 | 0.416 | -1.682 | -0.508 | -0.053 | 0.074 | 0.074 | 0.112 | 0.4 | 0.15 | 0.32 | 0.28 | 0.45 | 0.31 |
| 4 | 186386587 | rs6849514 | -0.362 | -0.018 | -1.128 | -0.646 | -0.007 | 0.017 | 0.017 | 0 | 0.4 | 0.34 | 0.33 | 0.31 | 0.46 | 0.39 |
| 4 | 186388073 | rs6857096 | -0.477 | -1.527 | -0.802 | -0.451 | 0.041 | 0.075 | 0.075 | 0 | 0.52 | 0.67 | 0.35 | 0.27 | 0.38 | 0.44 |
| 4 | 186390753 | rs9998424 | -0.682 | -1.568 | 0.268 | 0.343 | -0.04 | 0.141 | 0.141 | 0.173 | 0.11 | 0.37 | 0.14 | 0.2 | 0.05 | 0.13 |
| 4 | 186397351 | rs4862535 | 0.542 | -1.621 | -0.365 | 0.474 | 0.008 | -0.005 | -0.005 | 0 | 0.67 | 0.32 | 0.74 | 0.66 | 0.79 | 0.69 |
| 4 | 186400228 | rs4241798 | 0.175 | -1.12 | -0.822 | 0.322 | 0.016 | -0.041 | -0.041 | 0.073 | 0.19 | 0.04 | 0.27 | 0.21 | 0.15 | 0.28 |
| 4 | 186401713 | rs6846185 | -0.837 | -0.646 | 0.153 | 0.196 | -0.021 | -0.035 | -0.035 | 0 | 0.2 | 0.29 | 0.15 | 0.11 | 0.16 | 0.15 |
| 4 | 186414626 | rs1385581 | 0.824 | -0.618 | 0.539 | 0.518 | -0.022 | 0.03 | 0.03 | 0 | 0.91 | 0.66 | 0.9 | 0.94 | 0.98 | 0.87 |
| 4 | 186423637 | rs4635850 | 0.792 | -1.709 | 0.07 | 0.043 | 0.043 | 0.028 | 0.028 | 0.303 | 0.66 | 0.31 | 0.79 | 0.81 | 0.83 | 0.71 |
| 4 | 186423655 | rs12644280 | 0.475 | -0.87 | -0.626 | 0.764 | -0.068 | -0.082 | -0.082 | 0 | 0.12 | 0.02 | 0.15 | 0.31 | 0.16 | 0.1 |
| 4 | 186424667 | rs6552886 | 0.509 | -1.277 | 0.252 | 0.488 | -0.048 | -0.02 | -0.02 | 0 | 0.67 | 0.46 | 0.81 | 0.8 | 0.83 | 0.72 |
| 4 | 186448369 | rs7691287 | 0.565 | -1.648 | 0.528 | 0.508 | -0.067 | -0.024 | -0.024 | 0 | 0.69 | 0.47 | 0.8 | 0.81 | 0.85 | 0.75 |
| 4 | 186449651 | rs3792700 | 0.556 | -1.651 | 0.527 | 0.506 | -0.068 | -0.024 | -0.024 | 0 | 0.69 | 0.47 | 0.8 | 0.81 | 0.85 | 0.75 |
| 4 | 186459026 | rs3762920 | -0.013 | -1.803 | -0.392 | 0.327 | 0.061 | -0.034 | -0.034 | 0.389 | 0.42 | 0.17 | 0.45 | 0.43 | 0.34 | 0.48 |
| 4 | 186467147 | rs2310322 | 0.893 | -2.375 | -0.224 | -0.116 | 0.007 | 0.005 | 0.005 | 0.011 | 0.66 | 0.3 | 0.71 | 0.77 | 0.77 | 0.66 |
| 4 | 186467810 | rs7680845 | 1.08 | -2.456 | -0.494 | -0.074 | 0.022 | -0.037 | -0.037 | 0 | 0.27 | 0.05 | 0.31 | 0.27 | 0.22 | 0.28 |
| 4 | 186468992 | rs11132321 | -0.557 | -2.481 | 0.095 | -0.11 | -0.047 | -0.001 | -0.001 | 0 | 0.19 | 0.32 | 0.17 | 0.16 | 0.13 | 0.21 |
| 4 | 186469493 | rs1158465 | 0.894 | -2.104 | -0.391 | 0.002 | -0.04 | -0.083 | -0.083 | 0.212 | 0.36 | 0.13 | 0.39 | 0.5 | 0.53 | 0.35 |
| 4 | 186470421 | rs12649063 | 0.924 | -2.11 | -0.394 | 0.001 | -0.042 | -0.081 | -0.081 | 1.555 | 0.36 | 0.13 | 0.39 | 0.5 | 0.53 | 0.35 |

| | | | | | | | | | | | | | | | | |
|---|---|---|---|---|---|---|---|---|---|---|---|---|---|---|---|---|
| 4 | 186474700 | rs931305 | 0.436 | -1.091 | -0.277 | -0.037 | 0.003 | 0.001 | 0.001 | 0 | 0.2 | 0.21 | 0.34 | 0.41 | 0.37 | 0.18 |
| 4 | 186483220 | rs11935679 | -0.507 | -1.631 | -0.037 | -0.318 | -0.012 | -0.014 | -0.014 | 0 | 0.27 | 0.34 | 0.15 | 0.16 | 0.21 | 0.29 |
| 4 | 186485293 | rs11132328 | 1.344 | -1.598 | -0.006 | -0.433 | 0.027 | -0.023 | -0.023 | 0 | 0.31 | 0.2 | 0.36 | 0.35 | 0.28 | 0.31 |
| 4 | 186488834 | rs13102445 | 0.805 | -1.567 | -0.263 | -0.523 | -0.019 | 0.071 | 0.071 | 0 | 0.12 | 0.01 | 0.07 | 0 | 0.11 | 0.14 |
| 4 | 186489396 | rs10003413 | -0.823 | -1.93 | 0.15 | -0.387 | -0.013 | -0.056 | -0.056 | 0 | 0.74 | 0.67 | 0.83 | 0.92 | 0.82 | 0.75 |
| 4 | 186499546 | rs10866278 | 0.526 | -1.997 | -0.21 | -0.314 | 0.048 | -0.038 | -0.038 | 0.845 | 0.4 | 0.19 | 0.38 | 0.44 | 0.33 | 0.43 |
| 4 | 186504058 | rs1872445 | 0.373 | -1.919 | -0.45 | -0.388 | -0.109 | -0.093 | -0.093 | 0.002 | 0.29 | 0.08 | 0.42 | 0.47 | 0.45 | 0.3 |
| 4 | 186504441 | rs13123635 | -0.462 | -1.457 | 0.157 | -0.19 | -0.008 | -0.077 | -0.077 | 0 | 0.75 | 0.67 | 0.85 | 0.98 | 0.81 | 0.83 |
| 4 | 186507021 | rs10009018 | -0.235 | -1.817 | -0.013 | 0.23 | 0.008 | -0.025 | -0.025 | 0.005 | 0.14 | 0.08 | 0.08 | 0.19 | 0.11 | 0.14 |
| 4 | 186507187 | rs1046236 | 1.08 | -2.09 | -0.392 | 0.256 | -0.008 | -0.051 | -0.051 | 0 | 0.3 | 0.04 | 0.31 | 0.33 | 0.27 | 0.25 |
| 4 | 186507280 | rs7674108 | -0.619 | -2.119 | 0.77 | 0.183 | -0.036 | -0.041 | -0.041 | 1.579 | 0.8 | 0.65 | 0.9 | 0.91 | 0.89 | 0.85 |
| 4 | 186507409 | rs10031753 | -0.987 | -1.164 | 0.721 | 0.609 | 0.003 | -0.006 | -0.006 | 0 | 0.14 | 0.13 | 0.08 | 0.19 | 0.11 | 0.14 |
| 4 | 186512433 | rs6854874 | -1.904 | -1.131 | 0.96 | 0.543 | -0.041 | 0.028 | 0.028 | 0 | 0.23 | 0.37 | 0.14 | 0.31 | 0.13 | 0.2 |
| 4 | 186518902 | rs9991125 | -2.001 | -0.805 | 0.876 | 0.5 | -0.001 | -0.001 | -0.001 | 0 | 0.19 | 0.24 | 0.15 | 0.24 | 0.14 | 0.24 |
| 4 | 186519992 | rs6823956 | -1.883 | -0.702 | 1.416 | 0.54 | 0.017 | 0.068 | 0.068 | 5.088 | 0.1 | 0.46 | 0.16 | 0.11 | 0.06 | 0.15 |
| 4 | 186522043 | rs10012577 | -1.709 | -0.546 | 1.243 | 0.629 | -0.045 | 0.107 | 0.107 | 0 | 0.18 | 0.36 | 0.16 | 0.31 | 0.1 | 0.22 |
| 4 | 186522953 | rs6818789 | -1.45 | -0.391 | 0.828 | 0.702 | -0.029 | 0.075 | 0.075 | 0.033 | 0.17 | 0.59 | 0.15 | 0.31 | 0.11 | 0.23 |
| 4 | 186524787 | rs4861672 | 0.469 | 0.384 | -0.77 | -0.317 | -0.01 | 0.078 | 0.078 | 1.773 | 0.7 | 0.82 | 0.83 | 0.87 | 0.71 | 0.75 |
| 4 | 186525914 | rs2171046 | 0.099 | 0.519 | -0.909 | -0.928 | 0.011 | 0.039 | 0.039 | 0 | 0.46 | 0.34 | 0.68 | 0.86 | 0.62 | 0.64 |
| 4 | 186528433 | rs13144963 | -0.478 | 0.727 | -1.403 | -0.36 | -0.052 | 0.027 | 0.027 | 0.167 | 0.59 | 0.84 | 0.69 | 0.63 | 0.6 | 0.45 |
| 4 | 186529468 | rs12503636 | 0.265 | 0.516 | -1.136 | 0.115 | -0.004 | 0.058 | 0.058 | 0.096 | 0.14 | 0.16 | 0.28 | 0.45 | 0.15 | 0.13 |
| 4 | 186529560 | rs10008758 | -0.632 | 0.474 | -1.208 | 0.653 | -0.048 | 0.11 | 0.11 | 0.087 | 0.57 | 0.82 | 0.59 | 0.25 | 0.58 | 0.37 |
| 4 | 186531693 | rs9312336 | -0.751 | 0.349 | -0.703 | 0.567 | -0.009 | 0.028 | 0.028 | 0.054 | 0.57 | 0.68 | 0.67 | 0.63 | 0.59 | 0.45 |
| 4 | 186533350 | rs10019065 | -0.901 | 0.19 | -0.296 | 0.425 | -0.001 | -0.007 | -0.007 | 0.011 | 0.35 | 0.63 | 0.42 | 0.21 | 0.35 | 0.18 |
| 4 | 186534984 | rs10002478 | -1.058 | -0.008 | -0.181 | 0.646 | 0 | -0.001 | -0.001 | 0 | 0.33 | 0.68 | 0.46 | 0.56 | 0.28 | 0.22 |
| 4 | 186535247 | rs12502295 | -0.795 | 0.007 | 0.403 | 0.72 | -0.016 | 0.119 | 0.119 | 0.938 | 0.14 | 0.26 | 0.27 | 0.54 | 0.1 | 0.15 |
| 4 | 186537232 | rs13144235 | 0.093 | 0.064 | 0.378 | -0.083 | 0.003 | -0.001 | -0.001 | 0 | 0.1 | 0.09 | 0.06 | 0 | 0.09 | 0.03 |
| 4 | 186540503 | rs10026821 | -1.062 | -0.09 | 0.521 | -0.162 | -0.011 | -0.05 | -0.05 | 2.217 | 0.31 | 0.53 | 0.34 | 0.2 | 0.27 | 0.15 |
| 4 | 186541887 | rs10030246 | -1.176 | -0.077 | 0.527 | -0.155 | -0.009 | -0.049 | -0.049 | 0 | 0.3 | 0.52 | 0.34 | 0.2 | 0.27 | 0.15 |
| 4 | 186542593 | rs13435845 | -0.192 | -0.103 | 0.37 | -0.466 | 0.005 | -0.079 | -0.079 | 0.612 | 0.19 | 0.31 | 0.17 | 0.02 | 0.18 | 0.08 |
| 4 | 186545719 | rs13119106 | -0.336 | 0.348 | 0.023 | -0.353 | -0.063 | 0.078 | 0.078 | 0 | 0.27 | 0.1 | 0.21 | 0.11 | 0.33 | 0.14 |
| 4 | 186550283 | rs10006389 | 0.874 | -0.117 | 0.168 | -0.417 | -0.008 | -0.027 | -0.027 | 0 | 0.4 | 0.49 | 0.3 | 0.29 | 0.35 | 0.55 |
| 4 | 186553480 | rs13138440 | 0.36 | -0.002 | 0.25 | -0.469 | 0.01 | -0.022 | -0.022 | 0 | 0.25 | 0.29 | 0.22 | 0.12 | 0.28 | 0.1 |
| 4 | 186555742 | rs1471795 | -0.884 | -0.165 | 0.081 | -0.508 | -0.001 | -0.003 | -0.003 | 0 | 0.75 | 0.76 | 0.56 | 0.4 | 0.72 | 0.71 |
| 4 | 186558150 | rs13144135 | -0.895 | -0.243 | -0.145 | -0.606 | -0.003 | -0.044 | -0.044 | 0 | 0.4 | 0.52 | 0.29 | 0.12 | 0.37 | 0.17 |
| 4 | 186558860 | rs3816807 | 0.119 | -0.028 | -1.178 | -0.472 | 0.031 | -0.003 | -0.003 | 0 | 0.6 | 0.25 | 0.44 | 0.4 | 0.63 | 0.66 |
| 4 | 186559109 | rs2306707 | -0.058 | 0.619 | -0.92 | -0.061 | -0.012 | 0.022 | 0.022 | 0 | 0.27 | 0.18 | 0.26 | 0.12 | 0.29 | 0.11 |
| 4 | 186563188 | rs4862556 | 0.086 | 0.261 | -0.944 | 0.009 | 0.007 | 0.019 | 0.019 | 0 | 0.51 | 0.47 | 0.33 | 0.29 | 0.45 | 0.63 |
| 4 | 186570521 | rs7663196 | 0.327 | 0.745 | -2.386 | 0.271 | -0.128 | 0.116 | 0.116 | 1.806 | 0.43 | 0.05 | 0.35 | 0.25 | 0.49 | 0.53 |
| 4 | 186571441 | rs6821894 | 0.362 | 1.031 | -1.833 | 0.647 | -0.029 | -0.029 | -0.029 | 0 | 0.67 | 0.82 | 0.72 | 0.63 | 0.6 | 0.65 |
| 4 | 186575288 | rs6848934 | 0.649 | 0.973 | -1.125 | 0.512 | 0.004 | -0.007 | -0.007 | 0 | 0.85 | 0.82 | 0.88 | 0.86 | 0.84 | 0.77 |
| 4 | 186579041 | rs11934819 | 0.087 | 0.786 | -1.616 | 0.247 | -0.042 | 0.1 | 0.1 | 2.075 | 0.42 | 0.17 | 0.3 | 0.25 | 0.44 | 0.36 |
| 4 | 186581859 | rs4241809 | 0.742 | 0.325 | -0.362 | 1.064 | -0.035 | 0.063 | 0.063 | 0 | 0.74 | 0.57 | 0.74 | 0.62 | 0.76 | 0.78 |
| 4 | 186584255 | rs1021217 | -1.265 | 0.821 | -0.676 | 1.36 | -0.003 | 0.016 | 0.016 | 0 | 0.12 | 0.11 | 0.11 | 0.27 | 0.09 | 0.23 |
| 4 | 186589959 | rs9312338 | -0.277 | 1.407 | -0.488 | 1.008 | 0.032 | 0.017 | 0.017 | 0 | 0.31 | 0.38 | 0.47 | 0.38 | 0.4 | 0.27 |
| 4 | 186590237 | rs7686233 | 0.108 | 1.288 | -0.753 | 0.938 | -0.015 | -0.037 | -0.037 | 0 | 0.76 | 0.78 | 0.71 | 0.61 | 0.72 | 0.73 |
| 4 | 186593410 | rs4599460 | 0.748 | 0.885 | -0.043 | 0.243 | 0.003 | -0.022 | -0.022 | 0 | 0.72 | 0.45 | 0.73 | 0.83 | 0.72 | 0.71 |
| 4 | 186593672 | rs4862559 | 0.177 | 0.793 | 0.346 | 0.179 | -0.012 | -0.036 | -0.036 | 0.432 | 0.8 | 0.71 | 0.89 | 0.92 | 0.83 | 0.77 |
| 4 | 186594332 | rs4340826 | -1.006 | 0.992 | 0.292 | 0.363 | 0.095 | 0.019 | 0.019 | 0 | 0.1 | 0.3 | 0.16 | 0.09 | 0.12 | 0.06 |
| 4 | 186597950 | rs13140168 | -1.031 | 0.24 | -0.59 | -0.004 | 0.023 | 0.062 | 0.062 | 0 | 0.13 | 0.02 | 0.05 | 0 | 0.09 | 0.12 |
| 4 | 186602210 | rs4862562 | -0.546 | -0.535 | -0.296 | -0.66 | -0.026 | -0.059 | -0.059 | 0.075 | 0.15 | 0.16 | 0.07 | 0.01 | 0.09 | 0.03 |
| 4 | 186607977 | rs7653904 | -0.326 | -0.79 | -0.363 | -0.302 | -0.012 | 0 | 0 | 0.478 | 0.37 | 0.34 | 0.44 | 0.39 | 0.32 | 0.24 |
| 4 | 186608071 | rs11132333 | -0.234 | -0.602 | -1.293 | -0.308 | 0.001 | -0.001 | -0.001 | 0 | 0.58 | 0.37 | 0.7 | 0.63 | 0.53 | 0.61 |
| 4 | 186610123 | rs13150700 | 1.657 | -0.579 | -1.674 | -0.208 | -0.07 | -0.051 | -0.051 | 0 | 0.16 | 0.05 | 0.14 | 0.24 | 0.28 | 0.22 |
| 4 | 186611663 | rs6823546 | 2.181 | -0.876 | -1.636 | 0.295 | -0.034 | -0.01 | -0.01 | 0.468 | 0.2 | 0.04 | 0.2 | 0.15 | 0.2 | 0.11 |
| 4 | 186615500 | rs10434304 | 0.447 | -1.142 | -0.923 | 0.324 | 0.053 | 0.093 | 0.093 | 0.157 | 0.26 | 0.46 | 0.33 | 0.49 | 0.32 | 0.31 |
| 4 | 186616234 | rs12507995 | 1.048 | -1.189 | -1.1 | 0.363 | -0.026 | -0.043 | -0.043 | 1.448 | 0.25 | 0.03 | 0.27 | 0.25 | 0.22 | 0.39 |
| 4 | 186618175 | rs955264 | 0.736 | -1.284 | -1.17 | 0.301 | -0.087 | -0.053 | -0.053 | 4.381 | 0.47 | 0.05 | 0.48 | 0.41 | 0.45 | 0.51 |
| 4 | 186618304 | rs955263 | 0.375 | -1.19 | -0.93 | 0.272 | -0.025 | 0.039 | 0.039 | 0.07 | 0.3 | 0.43 | 0.47 | 0.5 | 0.23 | 0.49 |
| 4 | 186622179 | rs1547909 | -1.167 | -1.067 | -1.045 | 0.232 | -0.006 | 0.017 | 0.017 | 0.12 | 0.67 | 0.85 | 0.73 | 0.74 | 0.66 | 0.79 |
| 4 | 186630946 | rs904451 | 0.614 | -1.246 | -0.441 | 0.476 | -0.062 | 0.064 | 0.064 | 1.835 | 0.77 | 0.49 | 0.73 | 0.66 | 0.83 | 0.75 |
| 4 | 186636564 | rs6825114 | -0.244 | -1.223 | -0.401 | 0.543 | -0.039 | 0.145 | 0.145 | 3.225 | 0.84 | 0.54 | 0.76 | 0.71 | 0.89 | 0.8 |
| 4 | 186638655 | rs11132340 | 0.253 | -1.662 | -1.034 | 0.376 | -0.015 | 0.001 | 0.001 | 0.418 | 0.57 | 0.55 | 0.41 | 0.4 | 0.61 | 0.4 |
| 4 | 186641002 | rs4862566 | -1.889 | -1.598 | -2.001 | 0.429 | -0.003 | -0.011 | -0.011 | 0.031 | 0.92 | 0.98 | 0.93 | 0.9 | 0.9 | 0.94 |
| 4 | 186641213 | rs13131324 | 0.808 | -1.571 | -1.996 | 0.492 | -0.003 | -0.021 | -0.021 | 0 | 0.26 | 0.07 | 0.36 | 0.3 | 0.27 | 0.4 |
| 4 | 186641563 | rs10017683 | 1.162 | -1.001 | -1.915 | 0.501 | -0.096 | -0.047 | -0.047 | 0 | 0.2 | 0.07 | 0.15 | 0.23 | 0.28 | 0.2 |
| 4 | 186642895 | rs11736488 | 1.548 | -0.955 | -1.555 | 0.477 | -0.026 | 0.03 | 0.03 | 1.392 | 0.24 | 0.11 | 0.21 | 0.14 | 0.24 | 0.12 |
| 4 | 186648646 | rs10011292 | -1.084 | -0.95 | -1.523 | 0.5 | -0.001 | -0.01 | -0.01 | 0 | 0.91 | 0.96 | 0.92 | 0.9 | 0.9 | 0.94 |
| 4 | 186650946 | rs12650192 | 1.315 | -0.902 | -1.393 | 0.532 | -0.026 | 0.024 | 0.024 | 1.45 | 0.24 | 0.11 | 0.21 | 0.15 | 0.24 | 0.12 |

| | | | | | | | | | | | | | | | | |
|---|---|---|---|---|---|---|---|---|---|---|---|---|---|---|---|---|
| 4 | 186652292 | rs3796644 | 0.376 | -0.993 | -1.435 | 0.512 | 0.009 | -0.026 | -0.026 | 0 | 0.38 | 0.22 | 0.47 | 0.45 | 0.39 | 0.32 |
| 4 | 186658094 | rs2276911 | -0.374 | -0.824 | -1.11 | 1.031 | 0.001 | 0.011 | 0.011 | 0 | 0.13 | 0.12 | 0.2 | 0.38 | 0.09 | 0.14 |
| 4 | 186660076 | rs7669883 | 0.298 | -0.289 | -0.901 | 0.846 | -0.003 | 0.002 | 0.002 | 0 | 0.72 | 0.68 | 0.55 | 0.28 | 0.76 | 0.58 |
| 4 | 186661203 | rs1027521 | 0.563 | 0.38 | -1.173 | 0.633 | -0.032 | -0.015 | -0.015 | 0 | 0.14 | 0.06 | 0.1 | 0.11 | 0.18 | 0.23 |
| 4 | 186663336 | rs7667992 | 0.53 | 0.233 | -0.84 | 0.173 | 0.004 | -0.08 | -0.08 | 0.255 | 0.33 | 0.38 | 0.27 | 0.06 | 0.37 | 0.26 |
| 4 | 186663425 | rs904448 | -0.568 | 0.15 | -0.838 | 0.486 | 0.012 | -0.039 | -0.039 | 0 | 0.75 | 0.84 | 0.74 | 0.46 | 0.79 | 0.75 |
| 4 | 186664542 | rs7679506 | -0.577 | 0.349 | -0.343 | 0.551 | 0.017 | 0.154 | 0.154 | 0.753 | 0.06 | 0.28 | 0.19 | 0.17 | 0.03 | 0.17 |
| 4 | 186670089 | rs7678763 | 0.412 | 0.526 | -0.71 | 0.32 | 0.004 | -0.006 | -0.006 | 0 | 0.37 | 0.43 | 0.3 | 0.23 | 0.35 | 0.29 |
| 4 | 186670339 | rs7654366 | 1.528 | 0.525 | -0.768 | 0.45 | -0.003 | 0.009 | 0.009 | 0 | 0.22 | 0.22 | 0.18 | 0.08 | 0.2 | 0.08 |
| 4 | 186671479 | rs2173479 | -0.704 | 0.52 | -0.777 | 0.617 | 0.005 | -0.024 | -0.024 | 0.207 | 0.77 | 0.82 | 0.71 | 0.48 | 0.76 | 0.76 |
| 4 | 186671599 | rs12503317 | 0.989 | 0.662 | -0.892 | 0.697 | -0.011 | -0.037 | -0.037 | 0 | 0.06 | 0.01 | 0.19 | 0.17 | 0.08 | 0.2 |
| 4 | 186674804 | rs10027307 | 1.473 | 0.673 | -0.872 | 0.549 | 0 | 0 | 0 | 0 | 0.26 | 0.27 | 0.19 | 0.08 | 0.21 | 0.08 |
| 4 | 186677253 | rs4543173 | 0.017 | 0.65 | -0.805 | 0.519 | -0.002 | 0.002 | 0.002 | 0 | 0.43 | 0.39 | 0.32 | 0.23 | 0.47 | 0.52 |
| 4 | 186688767 | rs1039237 | 0.762 | 0.868 | -1.235 | 0.192 | 0.002 | -0.008 | -0.008 | 0.018 | 0.06 | 0.02 | 0.19 | 0.2 | 0.08 | 0.2 |
| 4 | 186692629 | rs7655720 | 1.111 | 0.919 | -1.119 | -0.721 | -0.002 | 0.024 | 0.024 | 0.133 | 0.24 | 0.19 | 0.18 | 0.05 | 0.21 | 0.08 |
| 4 | 186694576 | rs7693361 | 0.091 | 1.191 | -0.861 | -0.488 | 0.003 | -0.033 | -0.033 | 0 | 0.28 | 0.3 | 0.22 | 0.07 | 0.33 | 0.3 |
| 4 | 186695208 | rs4487396 | -0.554 | 0.979 | -1.168 | -0.166 | -0.005 | 0.063 | 0.063 | 0 | 0.54 | 0.63 | 0.67 | 0.78 | 0.56 | 0.73 |
| 4 | 186697494 | rs2310357 | 0.27 | 0.553 | -1.102 | 0.368 | 0.005 | 0.011 | 0.011 | 1.748 | 0.65 | 0.35 | 0.8 | 0.74 | 0.69 | 0.75 |
| 4 | 186698737 | rs12640132 | -0.432 | 0.755 | -1.49 | 1.393 | 0.004 | -0.079 | -0.079 | 0 | 0.14 | 0.04 | 0.24 | 0.31 | 0.12 | 0.32 |
| 4 | 186699584 | rs10028428 | -0.426 | 0.76 | -1.487 | 1.396 | 0.004 | -0.078 | -0.078 | 0.058 | 0.14 | 0.04 | 0.24 | 0.3 | 0.11 | 0.32 |
| 4 | 186701046 | rs979922 | -0.517 | 1.006 | -1.223 | 1.184 | 0.002 | 0.002 | 0.002 | 0 | 0.09 | 0.1 | 0.09 | 0.16 | 0.14 | 0.12 |
| 4 | 186707604 | rs6830250 | 0.425 | 0.966 | -1.392 | 1.059 | -0.037 | 0.039 | 0.039 | 0 | 0.68 | 0.8 | 0.75 | 0.81 | 0.6 | 0.74 |
| 4 | 186710312 | rs6858040 | -0.278 | 0.873 | -0.884 | 1.465 | -0.011 | 0.076 | 0.076 | 0.207 | 0.14 | 0.23 | 0.2 | 0.42 | 0.12 | 0.24 |
| 4 | 186714004 | rs4862571 | -1.032 | 1.094 | -1.255 | 1.156 | -0.027 | -0.046 | -0.046 | 0 | 0.3 | 0.19 | 0.33 | 0.45 | 0.38 | 0.44 |
| 4 | 186715008 | rs7684943 | 0.763 | 0.158 | -0.816 | 0.582 | -0.004 | -0.002 | -0.002 | 0 | 0.63 | 0.62 | 0.54 | 0.36 | 0.61 | 0.58 |
| 4 | 186716550 | rs13146524 | -0.632 | -0.799 | -0.923 | -0.461 | 0.025 | 0.012 | 0.012 | 0 | 0.2 | 0.11 | 0.03 | 0.01 | 0.1 | 0.12 |
| 4 | 186716984 | rs10021308 | -0.974 | -0.74 | -0.942 | -1.06 | 0.006 | -0.01 | -0.01 | 0 | 0.6 | 0.53 | 0.48 | 0.33 | 0.52 | 0.53 |
| 4 | 186717043 | rs9998709 | 0.447 | -0.077 | -0.627 | -0.86 | 0.006 | -0.024 | -0.024 | 0.709 | 0.25 | 0.32 | 0.18 | 0.18 | 0.27 | 0.13 |
| 4 | 186717589 | rs13126725 | 0.219 | -0.149 | -0.534 | -0.855 | 0.002 | -0.017 | -0.017 | 0.658 | 0.6 | 0.51 | 0.57 | 0.7 | 0.58 | 0.59 |
| 4 | 186719815 | rs1995193 | -0.346 | 0.3 | -0.423 | -1.694 | 0.003 | 0.011 | 0.011 | 4.018 | 0.5 | 0.45 | 0.56 | 0.15 | 0.5 | 0.4 |
| 4 | 186721885 | rs4861680 | 0.414 | 0.224 | -0.316 | -2.068 | 0.001 | 0.026 | 0.026 | 4.99 | 0.31 | 0.22 | 0.27 | 0.03 | 0.26 | 0.12 |
| 4 | 186722819 | rs11132349 | -0.365 | 0.142 | -0.001 | -2.389 | 0.002 | -0.05 | -0.05 | 0 | 0.43 | 0.43 | 0.36 | 0.03 | 0.36 | 0.19 |
| 4 | 186725449 | rs11132350 | -0.099 | 0.617 | -0.015 | -2.166 | -0.008 | 0.013 | 0.013 | 2.546 | 0.56 | 0.61 | 0.49 | 0.69 | 0.55 | 0.59 |
| 4 | 186730448 | rs878357 | 1.098 | 0.175 | -0.157 | -1.998 | 0.012 | -0.003 | -0.003 | 0 | 0.34 | 0.23 | 0.35 | 0.66 | 0.33 | 0.48 |
| 4 | 186731313 | rs4861681 | 0.446 | 0.722 | -0.056 | -1.93 | 0.005 | 0 | 0 | 1.002 | 0.17 | 0.15 | 0.18 | 0.01 | 0.11 | 0.08 |
| 4 | 186732566 | rs1566347 | -0.195 | 0.892 | 0.37 | -1.738 | 0.053 | 0.025 | 0.025 | 0 | 0.79 | 0.7 | 0.71 | 0.74 | 0.68 | 0.67 |
| 4 | 186734480 | rs6858186 | 0.018 | 0.767 | 0.319 | -1.759 | 0.024 | 0.01 | 0.01 | 0 | 0.22 | 0.22 | 0.33 | 0.28 | 0.33 | 0.36 |
| 4 | 186735780 | rs12502302 | 0.234 | 0.498 | 0.448 | -2.019 | -0.015 | -0.101 | -0.101 | 1.874 | 0.34 | 0.45 | 0.29 | 0.07 | 0.26 | 0.15 |
| 4 | 186738016 | rs7673778 | 0.546 | 0.491 | 0.162 | -2.067 | 0.062 | -0.007 | -0.007 | 1.622 | 0.7 | 0.49 | 0.6 | 0.75 | 0.61 | 0.66 |
| 4 | 186739072 | rs4861682 | -0.331 | 0.198 | -0.166 | -1.44 | 0.003 | 0.003 | 0.003 | 1.33 | 0.54 | 0.54 | 0.6 | 0.29 | 0.58 | 0.48 |
| 4 | 186744774 | rs13132552 | -0.459 | -0.128 | -0.529 | -1.236 | 0.022 | 0.013 | 0.013 | 0 | 0.59 | 0.7 | 0.65 | 0.32 | 0.67 | 0.5 |
| 4 | 186745734 | rs13112696 | 0.942 | 0.166 | -0.482 | -1.5 | 0.001 | 0.007 | 0.007 | 0 | 0.12 | 0.13 | 0.24 | 0.19 | 0.13 | 0.32 |
| 4 | 186746561 | rs11732469 | 1.159 | -0.296 | -0.646 | -1.663 | 0.008 | -0.013 | -0.013 | 0 | 0.25 | 0.17 | 0.21 | 0.63 | 0.21 | 0.46 |
| 4 | 186746956 | rs1039238 | -0.889 | -0.414 | -0.461 | -1.718 | -0.036 | 0.014 | 0.014 | 0.571 | 0.71 | 0.6 | 0.73 | 0.32 | 0.77 | 0.53 |
| 4 | 186747070 | rs11132355 | 1.168 | -0.32 | -1.176 | -1.677 | 0.008 | -0.013 | -0.013 | 0 | 0.25 | 0.17 | 0.21 | 0.63 | 0.21 | 0.46 |
| 4 | 186747694 | rs6814512 | -1.328 | -0.136 | -1.667 | -1.448 | 0.019 | 0.017 | 0.017 | 0 | 0.85 | 0.91 | 0.9 | 1 | 0.89 | 0.97 |
| 4 | 186751133 | rs13111004 | -0.285 | 0.458 | -1.428 | -0.983 | 0.001 | 0.006 | 0.006 | 0 | 0.17 | 0.16 | 0.28 | 0.22 | 0.18 | 0.33 |
| 4 | 186751341 | rs4862577 | 0.502 | 0.881 | -1.873 | -1.168 | -0.133 | 0.157 | 0.157 | 0 | 0.3 | 0.02 | 0.28 | 0.08 | 0.36 | 0.15 |
| 4 | 186758065 | rs9994285 | -0.042 | 1.13 | -0.765 | -0.327 | 0.002 | 0.01 | 0.01 | 0 | 0.12 | 0.14 | 0.24 | 0.18 | 0.13 | 0.32 |
| 4 | 186760232 | rs10028944 | 0.1 | 0.855 | -0.859 | -0.607 | 0.009 | 0.078 | 0.078 | 0 | 0.49 | 0.37 | 0.42 | 0.76 | 0.45 | 0.53 |
| 4 | 186761638 | rs1499010 | -0.113 | 1.189 | 0.275 | -0.64 | 0.017 | -0.009 | -0.009 | 0.448 | 0.73 | 0.67 | 0.59 | 0.77 | 0.69 | 0.65 |
| 4 | 186764476 | rs7694456 | 0.289 | 1.119 | 0.535 | -1.018 | 0.085 | 0.008 | 0.008 | 1.079 | 0.88 | 0.66 | 0.81 | 0.88 | 0.85 | 0.92 |
| 4 | 186764747 | rs11724690 | 0.8 | 1.028 | -0.18 | -1.507 | 0.021 | 0.096 | 0.096 | 0.601 | 0.36 | 0.2 | 0.17 | 0.09 | 0.31 | 0.2 |
| 4 | 186769051 | rs6552928 | -0.048 | 1.041 | -0.035 | -1.47 | 0.001 | 0.042 | 0.042 | 0.147 | 0.52 | 0.43 | 0.64 | 0.78 | 0.54 | 0.74 |
| 4 | 186775572 | rs6848278 | -0.089 | 1.003 | 0.262 | -1.53 | -0.001 | -0.027 | -0.027 | 0 | 0.52 | 0.48 | 0.64 | 0.79 | 0.53 | 0.73 |
| 4 | 186776464 | rs1566355 | 0.647 | 0.852 | 0.078 | -1.639 | -0.024 | 0.004 | 0.004 | 0 | 0.08 | 0.03 | 0.06 | 0.06 | 0.12 | 0.08 |
| 4 | 186782009 | rs10020233 | 0.998 | 1.223 | -0.043 | -1.501 | 0 | 0 | 0 | 0 | 0.25 | 0.25 | 0.29 | 0.54 | 0.27 | 0.38 |
| 4 | 186782291 | rs2648126 | 0.239 | 1.213 | 0.177 | -1.372 | 0.055 | -0.006 | -0.006 | 0 | 0.1 | 0.24 | 0.07 | 0.06 | 0.13 | 0.08 |
| 4 | 186787096 | rs2648117 | -0.842 | 1.015 | -0.286 | -1.308 | -0.009 | -0.012 | -0.012 | 0 | 0.73 | 0.84 | 0.65 | 0.74 | 0.74 | 0.65 |
| 4 | 186791290 | rs2648112 | -0.123 | 0.833 | -0.478 | -1.721 | 0.053 | 0.173 | 0.173 | 0 | 0.48 | 0.77 | 0.37 | 0.2 | 0.42 | 0.32 |
| 4 | 186791524 | rs2648111 | -0.485 | 0.874 | -0.602 | -1.692 | -0.031 | -0.019 | -0.019 | 0.798 | 0.72 | 0.84 | 0.62 | 0.72 | 0.67 | 0.62 |
| 4 | 186793887 | rs4862580 | -0.416 | 0.398 | -0.215 | -1.871 | -0.003 | -0.002 | -0.002 | 0.061 | 0.39 | 0.61 | 0.44 | 0.7 | 0.35 | 0.45 |
| 4 | 186794072 | rs4861688 | 1.247 | 0.388 | -0.201 | -1.867 | -0.018 | -0.105 | -0.105 | 0 | 0.24 | 0.1 | 0.26 | 0.52 | 0.24 | 0.27 |
| 4 | 186796587 | rs7679935 | -0.786 | 0.354 | -0.067 | -1.842 | -0.039 | 0.026 | 0.026 | 0.593 | 0.15 | 0.49 | 0.17 | 0.18 | 0.1 | 0.18 |
| 4 | 186796912 | rs10002028 | -0.252 | 0.072 | -0.546 | -2.184 | 0.004 | 0.035 | 0.035 | 0 | 0.51 | 0.63 | 0.55 | 0.72 | 0.53 | 0.52 |
| 4 | 186799239 | rs11721899 | 0.616 | 0.053 | -0.442 | -2.199 | -0.013 | -0.031 | -0.031 | 0.012 | 0.4 | 0.26 | 0.4 | 0.55 | 0.48 | 0.39 |
| 4 | 186800037 | rs9994907 | -1.445 | -0.08 | -0.498 | -2.087 | -0.009 | -0.017 | -0.017 | 0 | 0.79 | 0.84 | 0.67 | 0.73 | 0.78 | 0.72 |
| 4 | 186801590 | rs4343763 | -0.397 | 0 | -0.508 | -2.124 | -0.009 | 0.052 | 0.052 | 0.065 | 0.68 | 0.51 | 0.52 | 0.56 | 0.73 | 0.52 |
| 4 | 186802492 | rs13132627 | -0.036 | 0.047 | -0.445 | -2.005 | 0.001 | 0.002 | 0.002 | 0 | 0.56 | 0.49 | 0.39 | 0.53 | 0.52 | 0.45 |

| | | | | | | | | | | | | | | | | |
|---|---|---|---|---|---|---|---|---|---|---|---|---|---|---|---|---|
| 4 | 186807834 | rs6552934 | 0.1 | 0.119 | -0.428 | -2.051 | 0.004 | 0.012 | 0.012 | 0 | 0.37 | 0.44 | 0.35 | 0.52 | 0.39 | 0.33 |
| 4 | 186809714 | rs4862585 | -0.469 | -0.442 | -0.406 | -2.057 | 0.005 | -0.009 | -0.009 | 0 | 0.58 | 0.58 | 0.47 | 0.69 | 0.43 | 0.5 |
| 4 | 186809838 | rs4862586 | 1.031 | -0.306 | -0.675 | -1.903 | -0.023 | -0.087 | -0.087 | 0 | 0.25 | 0.12 | 0.27 | 0.52 | 0.26 | 0.25 |
| 4 | 186816480 | rs10031442 | 0.529 | -0.344 | -0.578 | -1.909 | -0.001 | 0.005 | 0.005 | 0 | 0.29 | 0.3 | 0.28 | 0.52 | 0.26 | 0.25 |
| 4 | 186821893 | rs7693781 | -0.898 | -0.313 | -0.291 | -1.622 | 0.01 | 0.032 | 0.032 | 0.282 | 0.8 | 0.72 | 0.69 | 0.72 | 0.77 | 0.66 |
| 4 | 186832817 | rs6819746 | -0.299 | -0.637 | -0.382 | -1.63 | -0.011 | -0.048 | -0.048 | 0 | 0.35 | 0.46 | 0.34 | 0.19 | 0.35 | 0.4 |
| 4 | 186839888 | rs12641600 | 0.084 | -0.468 | -0.519 | -1.533 | 0.002 | -0.003 | -0.003 | 0 | 0.56 | 0.53 | 0.55 | 0.66 | 0.48 | 0.43 |
| 4 | 186840924 | rs7680928 | 1.13 | -0.535 | -0.711 | -1.544 | -0.011 | -0.025 | -0.025 | 0 | 0.07 | 0.13 | 0.03 | 0.01 | 0.05 | 0.06 |
| 4 | 186840968 | rs7656290 | 0.323 | -0.427 | -0.792 | -1.481 | -0.049 | -0.031 | -0.031 | 0.164 | 0.22 | 0.39 | 0.21 | 0.15 | 0.12 | 0.16 |
| 4 | 186840999 | rs7656287 | 0.897 | -0.299 | -0.7 | -1.334 | -0.02 | -0.043 | -0.043 | 0 | 0.18 | 0.2 | 0.09 | 0.01 | 0.09 | 0.1 |
| 4 | 186842789 | rs13140248 | -0.102 | 0.837 | -0.433 | -0.615 | -0.002 | -0.006 | -0.006 | 0 | 0.4 | 0.38 | 0.36 | 0.55 | 0.38 | 0.36 |
| 4 | 186843924 | rs2034787 | -0.36 | 1.464 | -1.509 | -0.392 | 0.008 | 0.023 | 0.023 | 0.156 | 0.81 | 0.8 | 0.75 | 0.72 | 0.77 | 0.67 |
| 4 | 186846999 | rs4862596 | -0.042 | 1.43 | -1.495 | -0.185 | -0.08 | -0.104 | -0.104 | 0 | 0.34 | 0.12 | 0.34 | 0.51 | 0.37 | 0.28 |
| 4 | 186855364 | rs6552938 | 0.025 | 1.321 | -0.641 | 0.286 | 0.017 | 0.061 | 0.061 | 0 | 0.84 | 0.76 | 0.73 | 0.72 | 0.81 | 0.75 |
| 4 | 186861791 | rs10033624 | -0.314 | 0.949 | -1.205 | -0.321 | -0.025 | 0.08 | 0.08 | 0 | 0.15 | 0.03 | 0.07 | 0 | 0.16 | 0.03 |
| 4 | 186866070 | rs10866286 | 0.011 | 0.77 | -0.589 | 0.18 | -0.006 | -0.012 | -0.012 | 0 | 0.89 | 0.89 | 0.91 | 0.95 | 0.93 | 0.8 |
| 4 | 186868062 | rs959585 | 0.339 | 0.725 | -0.251 | 0.098 | -0.019 | -0.057 | -0.057 | 0 | 0.84 | 0.68 | 0.89 | 0.95 | 0.92 | 0.79 |
| 4 | 186872505 | rs1551438 | -0.256 | 1.584 | -0.543 | 1.045 | 0.022 | 0.121 | 0.121 | 0.685 | 0.11 | 0.12 | 0.19 | 0.57 | 0.17 | 0.34 |
| 4 | 186873844 | rs1031505 | -0.466 | 1.475 | -0.724 | 0.991 | -0.001 | 0.044 | 0.044 | 0.806 | 0.12 | 0.09 | 0.2 | 0.57 | 0.18 | 0.35 |
| 4 | 186881436 | rs959621 | -0.614 | 1.256 | -0.938 | -0.264 | 0.002 | -0.071 | -0.071 | 0 | 0.59 | 0.55 | 0.7 | 0.94 | 0.54 | 0.81 |
| 4 | 186881570 | rs959622 | -0.405 | 0.903 | -1.392 | -0.151 | 0.001 | -0.026 | -0.026 | 0 | 0.44 | 0.49 | 0.6 | 0.83 | 0.43 | 0.76 |
| 4 | 186881988 | rs7694098 | -0.104 | 0.8 | -1.229 | -0.114 | 0.003 | 0.031 | 0.031 | 0 | 0.28 | 0.34 | 0.43 | 0.66 | 0.26 | 0.56 |
| 4 | 186882290 | rs10155470 | -0.638 | 0.978 | -1.788 | 0.012 | 0.011 | -0.025 | -0.025 | 0 | 0.33 | 0.17 | 0.29 | 0.28 | 0.32 | 0.26 |
| 4 | 186884336 | rs1454746 | -0.455 | 1.666 | -1.511 | 0.23 | -0.03 | -0.004 | -0.004 | 0 | 0.3 | 0.22 | 0.32 | 0.31 | 0.48 | 0.32 |
| 4 | 186893350 | rs830829 | -0.187 | 1.114 | -1.229 | -0.017 | 0.01 | 0.018 | 0.018 | 0.341 | 0.5 | 0.46 | 0.39 | 0.3 | 0.6 | 0.49 |
| 4 | 186901587 | rs7676342 | 0.047 | 1.088 | -1.43 | -0.328 | 0 | 0.055 | 0.055 | 1.31 | 0.72 | 0.89 | 0.78 | 0.88 | 0.72 | 0.79 |
| 4 | 186902860 | rs4862611 | 0.188 | 1.215 | -0.904 | -0.128 | 0 | 0.011 | 0.011 | 0 | 0.55 | 0.48 | 0.63 | 0.71 | 0.54 | 0.68 |
| 4 | 186913109 | rs830838 | 0.077 | 1.456 | -0.774 | -0.483 | 0.01 | -0.014 | -0.014 | 0 | 0.34 | 0.39 | 0.41 | 0.69 | 0.45 | 0.43 |
| 4 | 186915297 | rs7681323 | 0.387 | 0.868 | -1.203 | -1.428 | 0.012 | 0.133 | 0.133 | 1.088 | 0.35 | 0.11 | 0.25 | 0.03 | 0.34 | 0.16 |
| 4 | 186917039 | rs6852642 | 0.413 | 0.63 | -0.728 | -1.62 | -0.021 | 0.033 | 0.033 | 0.302 | 0.14 | 0.27 | 0.21 | 0.27 | 0.08 | 0.29 |
| 4 | 186917269 | rs830841 | -0.15 | 0.485 | -1.476 | -1.995 | -0.014 | 0.09 | 0.09 | 0.104 | 0.66 | 0.8 | 0.73 | 0.97 | 0.66 | 0.76 |
| 4 | 186917931 | rs830842 | -1.298 | 0.415 | -1.504 | -1.919 | -0.046 | 0.078 | 0.078 | 0 | 0.87 | 1 | 0.9 | 1 | 0.87 | 0.88 |
| 4 | 186921897 | rs13113778 | 0.669 | 0.447 | -1.347 | -1.869 | 0.003 | -0.018 | -0.018 | 0 | 0.16 | 0.11 | 0.1 | 0 | 0.13 | 0.02 |
| 4 | 186923601 | rs2049044 | 1.295 | 0.719 | -1.282 | -1.988 | 0.01 | -0.025 | -0.025 | 0 | 0.34 | 0.59 | 0.41 | 0.69 | 0.41 | 0.43 |
| 4 | 186928529 | rs11132376 | 0.855 | 0.759 | -1.501 | -1.928 | -0.074 | 0.132 | 0.132 | 0 | 0.21 | 0.02 | 0.17 | 0.03 | 0.26 | 0.13 |
| 4 | 186929646 | rs2008461 | -0.169 | 1.219 | -1.515 | -1.933 | -0.063 | 0.192 | 0.192 | 0.93 | 0.37 | 0.08 | 0.29 | 0.03 | 0.45 | 0.15 |
| 4 | 186932780 | rs2603731 | 0.475 | 1.334 | -1.012 | -1.377 | -0.006 | 0.107 | 0.107 | 0.469 | 0.51 | 0.69 | 0.5 | 0.69 | 0.53 | 0.45 |
| 4 | 186935500 | rs7679967 | 0.037 | 1.121 | -0.801 | -0.707 | 0.016 | 0.05 | 0.05 | 0 | 0.06 | 0.08 | 0.16 | 0.25 | 0.12 | 0.24 |
| 4 | 186942380 | rs12643213 | 0.758 | 1.167 | -0.484 | -0.554 | 0.003 | 0.004 | 0.004 | 0.311 | 0.45 | 0.37 | 0.49 | 0.49 | 0.44 | 0.35 |
| 4 | 186943462 | rs1401573 | -0.602 | 0.899 | -0.197 | -0.478 | 0.004 | 0.021 | 0.021 | 0 | 0.19 | 0.28 | 0.21 | 0.33 | 0.18 | 0.32 |
| 4 | 186944931 | rs6815342 | -0.67 | 0.849 | -0.354 | -0.024 | -0.001 | 0.005 | 0.005 | 0 | 0.81 | 0.78 | 0.8 | 0.83 | 0.8 | 0.78 |
| 4 | 186953813 | rs12648771 | 0.355 | 0.813 | -0.594 | -0.214 | -0.01 | 0.008 | 0.008 | 0 | 0.1 | 0.06 | 0.08 | 0.09 | 0.1 | 0.13 |
| 4 | 186957112 | rs11725282 | 0.198 | 0.658 | -0.728 | -0.658 | 0.001 | 0.004 | 0.004 | 0 | 0.14 | 0.11 | 0.09 | 0 | 0.16 | 0.01 |
| 4 | 186959365 | rs6814242 | 0.632 | 0.548 | -0.516 | -0.527 | 0.03 | 0.059 | 0.059 | 1.562 | 0.31 | 0.52 | 0.46 | 0.48 | 0.4 | 0.4 |
| 4 | 186961015 | rs9996894 | -0.247 | 0.177 | -1.049 | -0.473 | 0.031 | 0.06 | 0.06 | 0 | 0.25 | 0.17 | 0.11 | 0.09 | 0.18 | 0.11 |
| 4 | 186961525 | rs7699825 | -0.033 | 0.43 | -1.042 | -0.231 | -0.001 | -0.003 | -0.003 | 0 | 0.46 | 0.5 | 0.56 | 0.54 | 0.53 | 0.52 |
| 4 | 186972741 | rs7691384 | -0.372 | 0.837 | -1.13 | 0.54 | -0.018 | -0.007 | -0.007 | 0 | 0.58 | 0.75 | 0.57 | 0.56 | 0.51 | 0.61 |
| 4 | 186979699 | rs7678294 | 0.112 | 1.058 | -0.97 | 0.784 | -0.001 | -0.001 | -0.001 | 0 | 0.76 | 0.77 | 0.73 | 0.79 | 0.78 | 0.69 |
| 4 | 186983342 | rs956239 | -0.906 | 1.368 | -1.153 | 0.877 | -0.014 | -0.007 | -0.007 | 0 | 0.14 | 0.07 | 0.16 | 0.14 | 0.18 | 0.14 |
| 4 | 186994039 | rs7657186 | 0.486 | 1.201 | -1.315 | 0.958 | 0.01 | 0.011 | 0.011 | 0 | 0.3 | 0.23 | 0.24 | 0.14 | 0.22 | 0.24 |
| 4 | 186994178 | rs13126816 | 0.381 | 1.511 | -1.721 | 0.971 | -0.035 | 0.003 | 0.003 | 0 | 0.23 | 0.1 | 0.28 | 0.21 | 0.25 | 0.27 |
| 4 | 186997767 | rs3775296 | 0.846 | 0.969 | -1.359 | 0.835 | -0.004 | -0.005 | -0.005 | 0 | 0.15 | 0.15 | 0.16 | 0.24 | 0.17 | 0.18 |
| 4 | 187004074 | rs3775291 | 0.293 | 0.992 | -1.69 | 0.786 | -0.124 | -0.078 | -0.078 | 0 | 0.24 | 0.03 | 0.31 | 0.33 | 0.32 | 0.26 |
| 4 | 187006806 | rs10025405 | -0.465 | 0.68 | -1.306 | 0.605 | -0.03 | -0.039 | -0.039 | 0 | 0.33 | 0.26 | 0.38 | 0.52 | 0.41 | 0.47 |
| 4 | 187009972 | rs4862633 | -1.917 | 0.16 | -0.567 | 0.344 | 0.02 | 0.031 | 0.031 | 0 | 0.68 | 0.72 | 0.81 | 0.87 | 0.75 | 0.73 |
| 4 | 187010104 | rs4608848 | -1.309 | 0.304 | -0.597 | 0.486 | -0.004 | 0.003 | 0.003 | 0 | 0.61 | 0.74 | 0.7 | 0.7 | 0.61 | 0.57 |
| 4 | 187011126 | rs6857595 | 0.15 | 0.161 | 0.159 | -0.158 | -0.006 | 0.006 | 0.006 | 0 | 0.25 | 0.33 | 0.27 | 0.33 | 0.25 | 0.29 |
| 4 | 187015089 | rs1519309 | -0.915 | 0.134 | 0.119 | -0.3 | 0.037 | 0.043 | 0.043 | 1.993 | 0.47 | 0.35 | 0.61 | 0.69 | 0.64 | 0.61 |
| 4 | 187016610 | rs6848963 | -2.173 | 0.098 | 0.076 | -0.374 | 0.036 | 0.077 | 0.077 | 0 | 0.7 | 0.86 | 0.83 | 0.88 | 0.76 | 0.73 |
| 4 | 187019433 | rs1983369 | -2.182 | 0.067 | 0.082 | -0.316 | 0.037 | 0.069 | 0.069 | 0 | 0.7 | 0.87 | 0.76 | 0.87 | 0.75 | 0.73 |
| 4 | 187021315 | rs9312342 | -0.441 | 1.004 | 0.147 | -0.387 | 0.024 | -0.016 | -0.016 | 0 | 0.25 | 0.35 | 0.33 | 0.2 | 0.32 | 0.31 |
| 4 | 187029821 | rs1401570 | -1.028 | 0.535 | 0.497 | -0.42 | 0.005 | 0.005 | 0.005 | 0.327 | 0.51 | 0.46 | 0.64 | 0.68 | 0.64 | 0.64 |
| 4 | 187038546 | rs1715052 | -0.732 | 1.087 | 0.196 | -0.332 | 0.027 | -0.023 | -0.023 | 0 | 0.91 | 0.86 | 0.88 | 0.99 | 0.88 | 0.86 |
| 4 | 187038801 | rs1715051 | -1.462 | 0.786 | 0.136 | -0.448 | -0.037 | -0.067 | -0.067 | 5.563 | 0.67 | 0.45 | 0.8 | 0.88 | 0.75 | 0.73 |
| 4 | 187042798 | rs7665411 | 0.211 | 0.812 | -0.118 | -0.444 | 0.014 | 0.002 | 0.002 | 0 | 0.35 | 0.17 | 0.37 | 0.38 | 0.33 | 0.31 |
| 4 | 187043908 | rs10030678 | -0.031 | 0.273 | 0.211 | -0.475 | 0 | -0.006 | -0.006 | 1.114 | 0.54 | 0.52 | 0.49 | 0.61 | 0.51 | 0.53 |
| 4 | 187044161 | rs1706145 | -0.031 | 0.28 | 0.177 | -0.447 | -0.016 | 0.028 | 0.028 | 0 | 0.43 | 0.23 | 0.49 | 0.39 | 0.49 | 0.47 |
| 4 | 187047812 | rs9998530 | 1.337 | 0.521 | -0.634 | -0.345 | -0.003 | -0.009 | -0.009 | 1.572 | 0.3 | 0.32 | 0.18 | 0.12 | 0.25 | 0.26 |
| 4 | 187054136 | rs7656677 | -0.382 | 0.709 | -0.469 | 0.266 | -0.002 | 0.065 | 0.065 | 0 | 0.27 | 0.39 | 0.32 | 0.49 | 0.27 | 0.28 |

| | | | | | | | | | | | | | | | | |
|---|---|---|---|---|---|---|---|---|---|---|---|---|---|---|---|---|
| 4 | 187057812 | rs4862644 | -0.431 | 0.599 | -1.345 | -0.214 | -0.007 | 0.138 | 0.138 | 0 | 0.5 | 0.84 | 0.57 | 0.69 | 0.43 | 0.53 |
| 4 | 187059563 | rs13105400 | -0.255 | 0.086 | -1.093 | -0.366 | -0.015 | -0.006 | -0.006 | 1.899 | 0.42 | 0.37 | 0.34 | 0.52 | 0.44 | 0.5 |
| 4 | 187059719 | rs7660462 | 0.678 | -0.077 | -1.066 | -0.576 | 0.03 | 0.047 | 0.047 | 0 | 0.41 | 0.25 | 0.38 | 0.3 | 0.42 | 0.37 |
| 4 | 187060065 | rs7660915 | 0.631 | -0.079 | -0.976 | -0.577 | 0.03 | 0.044 | 0.044 | 0 | 0.42 | 0.27 | 0.38 | 0.3 | 0.42 | 0.37 |
| 4 | 187061084 | rs7668227 | -0.936 | 0.008 | -0.805 | -0.315 | -0.026 | -0.025 | -0.025 | 0 | 0.9 | 1 | 0.84 | 0.83 | 0.89 | 0.9 |
| 4 | 187072383 | rs7684126 | -0.457 | -0.304 | -0.261 | -0.351 | -0.02 | -0.004 | -0.004 | 0 | 0.85 | 0.76 | 0.84 | 0.82 | 0.89 | 0.88 |
| 4 | 187072932 | rs2276914 | -0.812 | -0.272 | -0.58 | -0.333 | 0.016 | -0.01 | -0.01 | 0 | 0.87 | 0.96 | 0.86 | 0.82 | 0.89 | 0.88 |
| 4 | 187074833 | rs4862650 | -0.508 | -0.326 | -0.549 | -0.359 | 0.02 | -0.002 | -0.002 | 0.101 | 0.86 | 0.93 | 0.86 | 0.82 | 0.89 | 0.88 |
| 4 | 187076793 | rs4862652 | 0.663 | -0.995 | -0.483 | -0.529 | 0.042 | 0.013 | 0.013 | 0.007 | 0.58 | 0.33 | 0.49 | 0.59 | 0.71 | 0.66 |
| 4 | 187077206 | rs4862653 | -1.048 | -0.628 | -0.8 | -0.473 | 0.02 | -0.011 | -0.011 | 0 | 0.87 | 0.95 | 0.86 | 0.82 | 0.89 | 0.88 |
| 4 | 187078785 | rs2276924 | -1.133 | -0.602 | -0.801 | -0.485 | 0.018 | -0.01 | -0.01 | 0 | 0.87 | 0.96 | 0.86 | 0.82 | 0.89 | 0.88 |
| 4 | 187078938 | rs2276920 | 0.853 | -0.425 | -0.383 | -0.844 | -0.012 | -0.043 | -0.043 | 1.374 | 0.23 | 0.32 | 0.15 | 0.1 | 0.2 | 0.23 |
| 4 | 187079017 | rs1877321 | -0.115 | -0.326 | -0.789 | -0.766 | 0.027 | 0.062 | 0.062 | 0.421 | 0.33 | 0.2 | 0.31 | 0.19 | 0.27 | 0.17 |
| 4 | 187079384 | rs13141433 | 0.328 | -0.327 | -0.492 | -0.403 | -0.015 | -0.01 | -0.01 | 0 | 0.21 | 0.29 | 0.26 | 0.19 | 0.17 | 0.2 |
| 4 | 187099043 | rs2036912 | -0.096 | 0.415 | -1.578 | -0.149 | 0.042 | 0.063 | 0.063 | 2.071 | 0.53 | 0.26 | 0.45 | 0.29 | 0.59 | 0.4 |
| 4 | 187099609 | rs4862658 | 0.947 | 0.919 | -2.021 | -0.208 | -0.057 | -0.013 | -0.013 | 0 | 0.21 | 0.02 | 0.14 | 0.16 | 0.25 | 0.1 |
| 4 | 187100384 | rs6816903 | 0.299 | 0.504 | -1.078 | 0.23 | 0.006 | 0.007 | 0.007 | 2.233 | 0.56 | 0.43 | 0.48 | 0.29 | 0.6 | 0.4 |
| 4 | 187107849 | rs10027488 | 0.311 | 0.509 | -1.075 | 0.233 | 0.006 | 0.008 | 0.008 | 2.369 | 0.56 | 0.43 | 0.48 | 0.29 | 0.6 | 0.39 |
| 4 | 187109866 | rs7689823 | 0.311 | 0.509 | -1.075 | 0.233 | 0.006 | 0.008 | 0.008 | 2.369 | 0.56 | 0.43 | 0.48 | 0.29 | 0.6 | 0.39 |
| 4 | 187110782 | rs1877319 | 0.311 | 0.509 | -1.075 | 0.233 | 0.006 | 0.008 | 0.008 | 2.369 | 0.56 | 0.43 | 0.48 | 0.29 | 0.6 | 0.39 |
| 4 | 187114479 | rs10866290 | -0.273 | 0.708 | -1.695 | 0.719 | -0.028 | -0.107 | -0.107 | 0 | 0.31 | 0.09 | 0.36 | 0.5 | 0.33 | 0.46 |
| 4 | 187116227 | rs7684025 | -0.298 | 0.965 | -1.289 | 0.791 | -0.009 | -0.003 | -0.003 | 0 | 0.33 | 0.68 | 0.33 | 0.37 | 0.42 | 0.16 |
| 4 | 187118662 | rs4862662 | 0.18 | 0.989 | -0.205 | 0.857 | 0.033 | 0.016 | 0.016 | 0 | 0.26 | 0.37 | 0.28 | 0.34 | 0.38 | 0.15 |
| 4 | 187120211 | rs13146272 | -0.567 | 0.489 | -0.495 | 0.176 | 0.007 | 0.008 | 0.008 | 0 | 0.54 | 0.6 | 0.47 | 0.41 | 0.65 | 0.63 |
| 4 | 187122009 | rs7687961 | -0.155 | 0.114 | -0.424 | 0.305 | -0.035 | 0.064 | 0.064 | 0.701 | 0.72 | 0.58 | 0.64 | 0.58 | 0.82 | 0.7 |
| 4 | 187122319 | rs3736455 | 0.019 | 0.041 | -0.263 | 0.264 | 0.015 | 0.011 | 0.011 | 0.025 | 0.56 | 0.64 | 0.49 | 0.41 | 0.67 | 0.63 |
| 4 | 187122355 | rs3736456 | 1.122 | -0.132 | -0.275 | 0.474 | 0.021 | -0.031 | -0.031 | 0.615 | 0.14 | 0.07 | 0.19 | 0.35 | 0.07 | 0.17 |
| 4 | 187127817 | rs1473597 | -0.177 | 0.002 | -0.143 | 0.27 | 0.004 | 0.014 | 0.014 | 0.19 | 0.49 | 0.52 | 0.38 | 0.25 | 0.44 | 0.57 |
| 4 | 187131504 | rs2102575 | -1.163 | -0.072 | -0.173 | 0.333 | 0.003 | 0.005 | 0.005 | 0 | 0.9 | 0.91 | 0.88 | 0.94 | 0.92 | 0.89 |
| 4 | 187133576 | rs6842047 | -1.304 | -0.071 | -0.174 | 0.375 | 0.002 | 0.004 | 0.004 | 0 | 0.9 | 0.91 | 0.88 | 0.93 | 0.91 | 0.88 |
| 4 | 187135556 | rs1511800 | -0.12 | -0.096 | -0.023 | 0.379 | -0.007 | -0.016 | -0.016 | 1.327 | 0.86 | 0.56 | 0.86 | 0.93 | 0.91 | 0.88 |
| 4 | 187148071 | rs4253236 | -0.387 | -0.211 | -0.46 | 0.096 | -0.004 | 0.058 | 0.058 | 0.603 | 0.6 | 0.83 | 0.7 | 0.75 | 0.66 | 0.59 |
| 4 | 187148387 | rs4253238 | 0.177 | -0.078 | -0.534 | 0.02 | -0.021 | 0.063 | 0.063 | 0.193 | 0.48 | 0.77 | 0.59 | 0.68 | 0.52 | 0.39 |
| 4 | 187148475 | rs4253239 | 2.143 | -0.148 | -0.629 | -0.048 | -0.055 | -0.039 | -0.039 | 0 | 0.17 | 0.01 | 0.11 | 0.17 | 0.18 | 0.07 |
| 4 | 187149540 | rs1912826 | 0.373 | -0.143 | -0.516 | -0.035 | -0.01 | 0.04 | 0.04 | 0.139 | 0.47 | 0.69 | 0.58 | 0.68 | 0.52 | 0.39 |
| 4 | 187150806 | rs1511802 | 1.956 | -0.229 | -0.021 | 0.08 | 0.005 | 0.003 | 0.003 | 0.011 | 0.27 | 0.33 | 0.33 | 0.36 | 0.38 | 0.14 |
| 4 | 187162048 | rs4253259 | 1.501 | -0.342 | -0.161 | 0.139 | -0.011 | -0.009 | -0.009 | 0.604 | 0.11 | 0.12 | 0.08 | 0.07 | 0.06 | 0.07 |
| 4 | 187168866 | rs4253292 | 2.132 | -0.252 | -0.3 | 0.142 | -0.008 | -0.002 | -0.002 | 0.273 | 0.17 | 0.09 | 0.11 | 0.17 | 0.18 | 0.07 |
| 4 | 187173012 | rs4253301 | 1.095 | -0.259 | -0.44 | 0.23 | -0.002 | 0.013 | 0.013 | 0 | 0.12 | 0.05 | 0.11 | 0.07 | 0.13 | 0.23 |
| 4 | 187174363 | rs4253308 | 1.822 | -0.077 | -0.111 | 0.392 | -0.008 | -0.008 | -0.008 | 0.042 | 0.26 | 0.25 | 0.31 | 0.37 | 0.38 | 0.14 |
| 4 | 187175457 | rs4253315 | 0.008 | -0.632 | 0.094 | 0.297 | -0.037 | 0.073 | 0.073 | 0 | 0.18 | 0.34 | 0.24 | 0.31 | 0.12 | 0.22 |
| 4 | 187177814 | rs3775302 | -0.21 | -0.617 | 0.096 | 0.298 | -0.019 | 0.07 | 0.07 | 0 | 0.19 | 0.34 | 0.26 | 0.32 | 0.14 | 0.22 |
| 4 | 187178473 | rs4253325 | 0.229 | -0.966 | -0.012 | 0.164 | -0.025 | 0.051 | 0.051 | 0 | 0.17 | 0.26 | 0.23 | 0.31 | 0.12 | 0.22 |
| 4 | 187179210 | rs925453 | 0.401 | -0.773 | -0.343 | -0.184 | 0.004 | 0.005 | 0.005 | 1.171 | 0.63 | 0.63 | 0.68 | 0.85 | 0.7 | 0.44 |
| 4 | 187179486 | rs3087505 | -0.922 | -0.659 | -0.124 | 0.402 | -0.001 | -0.002 | -0.002 | 0 | 0.92 | 0.94 | 0.88 | 0.94 | 0.92 | 0.89 |
| 4 | 187186111 | rs3756009 | 0.985 | -0.484 | -0.499 | 0.4 | -0.066 | -0.022 | -0.022 | 0 | 0.3 | 0.12 | 0.32 | 0.32 | 0.39 | 0.32 |
| 4 | 187187005 | rs3822056 | 0.167 | -0.544 | 0.215 | 0.61 | 0.064 | 0.234 | 0.234 | 0 | 0.11 | 0.43 | 0.18 | 0.27 | 0.12 | 0.16 |
| 4 | 187187135 | rs3733403 | 0.297 | -0.545 | 0.311 | 0.56 | 0.067 | 0.243 | 0.243 | 0 | 0.1 | 0.37 | 0.18 | 0.29 | 0.12 | 0.16 |
| 4 | 187187569 | rs925451 | 1.242 | -0.866 | -0.087 | 0.456 | -0.03 | -0.007 | -0.007 | 0 | 0.29 | 0.17 | 0.32 | 0.32 | 0.38 | 0.31 |
| 4 | 187190810 | rs4253405 | -0.166 | -0.891 | -0.44 | -0.114 | 0.041 | 0.129 | 0.129 | 0 | 0.49 | 0.12 | 0.31 | 0.3 | 0.4 | 0.4 |
| 4 | 187192481 | rs2036914 | 0.143 | -0.871 | -0.237 | -0.837 | -0.01 | 0.049 | 0.049 | 0 | 0.45 | 0.67 | 0.54 | 0.77 | 0.53 | 0.48 |
| 4 | 187199888 | rs4253419 | 1.008 | -1.037 | -0.479 | -0.905 | -0.003 | 0.029 | 0.029 | 0.343 | 0.18 | 0.11 | 0.19 | 0.04 | 0.17 | 0.07 |
| 4 | 187201001 | rs2241817 | -0.75 | -1.329 | -0.114 | -1.187 | 0.016 | 0.031 | 0.031 | 0 | 0.5 | 0.57 | 0.4 | 0.16 | 0.38 | 0.34 |
| 4 | 187201211 | rs5974 | 0.229 | -1.497 | -0.116 | -1.319 | -0.003 | -0.03 | -0.03 | 1.187 | 0.21 | 0.26 | 0.14 | 0.04 | 0.16 | 0.2 |
| 4 | 187204937 | rs4253421 | -1.149 | -1.604 | -0.318 | -1.303 | -0.003 | -0.014 | -0.014 | 0 | 0.89 | 0.98 | 0.91 | 0.91 | 0.89 | 0.87 |
| 4 | 187205301 | rs5970 | 0.165 | -1.5 | -0.234 | -1.294 | 0 | -0.023 | -0.023 | 0 | 0.23 | 0.26 | 0.14 | 0.04 | 0.17 | 0.21 |
| 4 | 187207381 | rs2289252 | 1.975 | -1.481 | -0.108 | -1.094 | -0.021 | -0.003 | -0.003 | 0 | 0.3 | 0.21 | 0.34 | 0.35 | 0.39 | 0.34 |
| 4 | 187209559 | rs5966 | 0.898 | -1.686 | -0.11 | -1.875 | -0.02 | -0.007 | -0.007 | 0 | 0.11 | 0.14 | 0.07 | 0.08 | 0.05 | 0.07 |
| 4 | 187209702 | rs5971 | 0.951 | -1.686 | -0.11 | -1.875 | -0.02 | -0.006 | -0.006 | 0 | 0.11 | 0.14 | 0.07 | 0.08 | 0.05 | 0.07 |
| 4 | 187209729 | rs5976 | 1.226 | -1.687 | -0.107 | -1.875 | 0.022 | 0.003 | 0.003 | 0 | 0.1 | 0.03 | 0.06 | 0.08 | 0.05 | 0.07 |
| 4 | 187210033 | rs4253429 | 0.804 | -1.694 | -0.116 | -1.885 | 0 | -0.02 | -0.02 | 0 | 0.18 | 0.17 | 0.2 | 0.04 | 0.17 | 0.07 |
| 4 | 187210064 | rs4253430 | -1.431 | -1.702 | -0.096 | -1.893 | 0.011 | 0.021 | 0.021 | 0 | 0.52 | 0.57 | 0.4 | 0.16 | 0.39 | 0.35 |
| 4 | 187210090 | rs4253865 | 0.884 | -1.701 | -0.099 | -1.892 | -0.023 | -0.008 | -0.008 | 0 | 0.12 | 0.16 | 0.07 | 0.08 | 0.05 | 0.07 |
| 4 | 187210247 | rs1062547 | -1.382 | -1.641 | -0.114 | -1.856 | 0.013 | 0.03 | 0.03 | 0 | 0.52 | 0.58 | 0.4 | 0.16 | 0.39 | 0.35 |
| 4 | 187210620 | rs4253431 | -0.168 | -1.639 | -0.141 | -1.854 | -0.003 | -0.03 | -0.03 | 0 | 0.23 | 0.26 | 0.14 | 0.04 | 0.17 | 0.21 |
| 4 | 187212134 | rs11938564 | -0.253 | -1.598 | -0.108 | -1.822 | -0.013 | -0.037 | -0.037 | 0 | 0.3 | 0.32 | 0.26 | 0.13 | 0.22 | 0.14 |
| 4 | 187213027 | rs6843926 | 1.177 | -1.292 | -0.191 | -1.536 | -0.017 | -0.034 | -0.034 | 0 | 0.06 | 0.2 | 0.04 | 0 | 0.03 | 0.01 |
| 4 | 187215016 | rs13145616 | -0.397 | -1.293 | -0.315 | -1.551 | -0.002 | -0.028 | -0.028 | 0 | 0.23 | 0.26 | 0.14 | 0.04 | 0.17 | 0.21 |
| 4 | 187217008 | rs13136269 | -0.862 | -0.841 | -1.011 | -1.357 | 0.013 | 0.088 | 0.088 | 0 | 0.37 | 0.2 | 0.25 | 0.15 | 0.29 | 0.23 |

| | | | | | | | | | | | | | | | | |
|---|---|---|---|---|---|---|---|---|---|---|---|---|---|---|---|---|
| 4 | 187217373 | rs12507040 | -0.683 | -0.846 | -1.017 | -1.359 | -0.029 | 0.143 | 0.143 | 0 | 0.36 | 0.05 | 0.24 | 0.15 | 0.3 | 0.23 |
| 4 | 187217560 | rs12511874 | -0.846 | -0.808 | -1.031 | -1.341 | 0.014 | 0.089 | 0.089 | 0 | 0.37 | 0.2 | 0.25 | 0.15 | 0.3 | 0.24 |
| 4 | 187223583 | rs4572916 | -0.857 | -0.334 | -0.094 | -0.499 | -0.023 | -0.005 | -0.005 | 0 | 0.22 | 0.35 | 0.26 | 0.16 | 0.17 | 0.18 |
| 4 | 187226673 | rs1008728 | 0.748 | -0.265 | -0.845 | -0.381 | 0.004 | 0.009 | 0.009 | 0 | 0.57 | 0.38 | 0.64 | 0.8 | 0.64 | 0.75 |
| 4 | 187230737 | rs13133050 | 0.257 | 0.462 | -1.156 | -0.158 | 0 | 0.033 | 0.033 | 0.059 | 0.69 | 0.23 | 0.61 | 0.86 | 0.69 | 0.84 |
| 4 | 187233838 | rs12331264 | -0.44 | 0.352 | -1.123 | 0.175 | 0.014 | 0.017 | 0.017 | 1.299 | 0.41 | 0.33 | 0.36 | 0.25 | 0.35 | 0.22 |
| 4 | 187235953 | rs6854650 | -0.58 | 0.119 | -0.865 | 0.407 | -0.009 | 0.006 | 0.006 | 0 | 0.4 | 0.4 | 0.41 | 0.54 | 0.3 | 0.33 |
| 4 | 187235998 | rs10020303 | 0.664 | -0.098 | -1.123 | -0.026 | 0.005 | -0.001 | -0.001 | 0.697 | 0.25 | 0.24 | 0.22 | 0.24 | 0.22 | 0.29 |
| 4 | 187237305 | rs6839415 | 0.147 | -0.111 | -1.123 | -0.033 | -0.003 | 0.021 | 0.021 | 0 | 0.05 | 0.07 | 0.02 | 0.13 | 0.05 | 0.07 |
| 4 | 187238404 | rs6552972 | 0.534 | -0.443 | -1.358 | -0.275 | 0.048 | 0.111 | 0.111 | 0 | 0.29 | 0.08 | 0.13 | 0.05 | 0.19 | 0.13 |
| 4 | 187239747 | rs7687352 | -0.182 | -0.558 | -1.113 | -0.423 | -0.007 | 0.033 | 0.033 | 0 | 0.43 | 0.72 | 0.42 | 0.25 | 0.49 | 0.45 |
| 4 | 187239849 | rs9993749 | 0.268 | -1.07 | -1.033 | -0.076 | 0.008 | -0.055 | -0.055 | 0.94 | 0.27 | 0.16 | 0.36 | 0.49 | 0.26 | 0.27 |
| 4 | 187241497 | rs7700014 | -0.691 | 0.059 | -0.448 | -0.529 | 0 | -0.008 | -0.008 | 0 | 0.62 | 0.47 | 0.63 | 0.71 | 0.62 | 0.59 |
| 4 | 187247089 | rs7349633 | -0.722 | 0.424 | 0.008 | -0.66 | 0.056 | -0.069 | -0.069 | 0.324 | 0.8 | 0.56 | 0.79 | 0.93 | 0.76 | 0.8 |
| 4 | 187251860 | rs11132389 | -0.26 | -0.052 | -0.592 | -0.394 | 0.021 | 0.012 | 0.012 | 0 | 0.45 | 0.36 | 0.46 | 0.64 | 0.38 | 0.39 |
| 4 | 187257004 | rs6829128 | 0.009 | -0.077 | -0.678 | -0.356 | 0.025 | -0.017 | -0.017 | 0 | 0.4 | 0.25 | 0.43 | 0.61 | 0.34 | 0.32 |
| 4 | 187257576 | rs13148903 | 0.87 | 0.613 | -0.537 | -0.742 | -0.03 | 0.027 | 0.027 | 0 | 0.17 | 0.09 | 0.18 | 0.07 | 0.24 | 0.21 |
| 4 | 187262489 | rs10032695 | -1.07 | 0.425 | -0.564 | -0.706 | -0.009 | 0.005 | 0.005 | 0 | 0.7 | 0.77 | 0.75 | 0.72 | 0.63 | 0.74 |
| 4 | 187263675 | rs4861710 | -1.089 | 0.427 | -0.681 | -0.696 | -0.019 | 0.003 | 0.003 | 0.033 | 0.72 | 0.81 | 0.76 | 0.72 | 0.64 | 0.74 |
| 4 | 187271047 | rs7672650 | 0.092 | 0.099 | -0.717 | -0.964 | 0.023 | 0.085 | 0.085 | 0 | 0.4 | 0.23 | 0.27 | 0.14 | 0.3 | 0.27 |
| 4 | 187280138 | rs10020635 | -2.057 | 0.254 | -0.734 | -0.939 | -0.016 | 0.01 | 0.01 | 0.222 | 0.74 | 0.81 | 0.8 | 0.75 | 0.69 | 0.79 |
| 4 | 187283915 | rs11132390 | 0.418 | -0.074 | -0.253 | -1.218 | 0.007 | -0.011 | -0.011 | 0 | 0.21 | 0.15 | 0.3 | 0.61 | 0.15 | 0.47 |
| 4 | 187285215 | rs6816112 | -2.645 | 0.002 | -0.414 | -1.192 | -0.023 | 0.001 | 0.001 | 0.015 | 0.76 | 0.87 | 0.8 | 0.75 | 0.7 | 0.8 |
| 4 | 187286307 | rs1901314 | -1.284 | 0.034 | -0.001 | -1.261 | 0.014 | -0.026 | -0.026 | 0 | 0.62 | 0.5 | 0.61 | 0.75 | 0.47 | 0.74 |
| 4 | 187287475 | rs1868893 | 1.413 | 0.012 | -0.065 | -1.065 | 0.001 | -0.075 | -0.075 | 2.833 | 0.14 | 0.03 | 0.26 | 0.55 | 0.12 | 0.41 |
| 4 | 187289668 | rs4862682 | 2.071 | -0.044 | -0.181 | -1.296 | -0.016 | 0.015 | 0.015 | 0 | 0.13 | 0.05 | 0.17 | 0.09 | 0.16 | 0.07 |
| 4 | 187292020 | rs10014510 | 1.042 | 0.147 | 0.02 | -1.098 | 0.045 | -0.06 | -0.06 | 0.14 | 0.12 | 0.22 | 0.18 | 0 | 0.22 | 0.06 |
| 4 | 187297232 | rs9997314 | -2.271 | -0.113 | -0.286 | -0.758 | -0.007 | 0.004 | 0.004 | 0.74 | 0.75 | 0.79 | 0.81 | 0.75 | 0.71 | 0.8 |
| 4 | 187299297 | rs4862684 | -1.236 | -0.102 | -0.311 | -0.78 | 0.006 | 0.054 | 0.054 | 0.011 | 0.63 | 0.77 | 0.75 | 0.74 | 0.65 | 0.73 |
| 4 | 187301857 | rs4862687 | -1.213 | -0.19 | -0.196 | -0.797 | 0.009 | 0.035 | 0.035 | 0.158 | 0.63 | 0.7 | 0.76 | 0.75 | 0.66 | 0.73 |
| 4 | 187313377 | rs6817039 | 0.223 | -0.11 | -0.194 | -0.8 | -0.032 | 0.059 | 0.059 | 0 | 0.57 | 0.22 | 0.53 | 0.74 | 0.5 | 0.69 |
| 4 | 187314751 | rs12331403 | -0.224 | 0.056 | 0.188 | -0.838 | 0.007 | -0.013 | -0.013 | 0.853 | 0.24 | 0.27 | 0.24 | 0.14 | 0.28 | 0.19 |
| 4 | 187315811 | rs2889188 | 1.024 | 0.055 | 0.212 | -0.837 | 0.02 | -0.061 | -0.061 | 0.056 | 0.22 | 0.49 | 0.25 | 0.11 | 0.25 | 0.13 |
| 4 | 187318091 | rs10018544 | -0.235 | 0.008 | 0.145 | -0.729 | 0.006 | -0.011 | -0.011 | 0.032 | 0.24 | 0.25 | 0.23 | 0.14 | 0.28 | 0.19 |
| 4 | 187329245 | rs2167370 | 0.073 | -0.207 | 0.197 | -0.985 | 0.004 | 0.003 | 0.003 | 0 | 0.6 | 0.61 | 0.6 | 0.32 | 0.73 | 0.36 |
| 4 | 187331659 | rs11945366 | 0.263 | 0.325 | 0.245 | -0.765 | 0.042 | -0.043 | -0.043 | 0 | 0.15 | 0.27 | 0.17 | 0.05 | 0.19 | 0.08 |
| 4 | 187340605 | rs6826848 | 1.753 | 0.424 | -0.187 | -0.7 | -0.007 | 0.009 | 0.009 | 0.28 | 0.23 | 0.19 | 0.19 | 0.18 | 0.29 | 0.18 |
| 4 | 187340688 | rs11931790 | -0.037 | 0.187 | -0.157 | -0.89 | 0 | -0.009 | -0.009 | 0 | 0.28 | 0.23 | 0.2 | 0.08 | 0.24 | 0.22 |
| 4 | 187341533 | rs2060767 | 1.506 | 0.014 | -0.06 | -0.507 | -0.006 | 0.009 | 0.009 | 0.371 | 0.23 | 0.19 | 0.19 | 0.17 | 0.29 | 0.18 |
| 4 | 187347487 | rs7669240 | -0.142 | -0.079 | -0.281 | -0.603 | -0.002 | 0.041 | 0.041 | 0 | 0.52 | 0.67 | 0.57 | 0.82 | 0.53 | 0.68 |
| 4 | 187347655 | rs7691259 | 0.142 | 0.111 | -0.368 | -0.241 | 0.008 | -0.017 | -0.017 | 2.756 | 0.72 | 0.79 | 0.64 | 0.35 | 0.76 | 0.49 |
| 4 | 187358040 | rs4862695 | 0.628 | 0.545 | -0.49 | -0.456 | -0.023 | 0.038 | 0.038 | 0.007 | 0.23 | 0.15 | 0.2 | 0.07 | 0.31 | 0.11 |
| 4 | 187360776 | rs10454254 | -0.918 | -0.207 | 0.257 | -0.406 | 0.004 | 0.01 | 0.01 | 0.292 | 0.43 | 0.36 | 0.48 | 0.76 | 0.38 | 0.74 |
| 4 | 187361302 | rs6844494 | 0.534 | -0.495 | 0.504 | 0.03 | -0.025 | 0.033 | 0.033 | 0.063 | 0.65 | 0.46 | 0.67 | 0.5 | 0.74 | 0.43 |
| 4 | 187361370 | rs10434307 | -0.322 | -0.454 | 0.593 | 0.043 | -0.006 | 0.032 | 0.032 | 0 | 0.15 | 0.22 | 0.2 | 0.25 | 0.14 | 0.17 |
| 4 | 187365460 | rs969372 | 0.126 | -0.568 | 0.506 | 0.06 | -0.002 | -0.001 | -0.001 | 0.186 | 0.79 | 0.74 | 0.77 | 0.79 | 0.81 | 0.73 |
| 4 | 187365581 | rs1812785 | 0.987 | -0.65 | 0.561 | 0.037 | -0.024 | -0.005 | -0.005 | 1.689 | 0.72 | 0.38 | 0.73 | 0.72 | 0.8 | 0.69 |
| 4 | 187365864 | rs2084729 | 0.162 | -0.632 | 0.121 | 0.558 | 0.022 | -0.077 | -0.077 | 0 | 0.13 | 0.02 | 0.06 | 0.45 | 0.08 | 0.28 |
| 4 | 187365960 | rs11132396 | -1.144 | -0.409 | 0.764 | 0.17 | 0.001 | -0.013 | -0.013 | 0.173 | 0.33 | 0.32 | 0.36 | 0.19 | 0.34 | 0.35 |
| 4 | 187366649 | rs9999651 | 1.508 | -0.542 | 0.356 | 0.222 | 0.036 | 0.08 | 0.08 | 0.484 | 0.55 | 0.2 | 0.53 | 0.27 | 0.62 | 0.43 |
| 4 | 187367404 | rs1037354 | 0.55 | -0.495 | 0.366 | 0.274 | -0.014 | -0.019 | -0.019 | 0 | 0.14 | 0.09 | 0.19 | 0.18 | 0.15 | 0.16 |
| 4 | 187367442 | rs7654929 | 1.266 | -0.551 | 0.726 | 0.353 | -0.004 | -0.002 | -0.002 | 0 | 0.61 | 0.54 | 0.57 | 0.39 | 0.63 | 0.48 |
| 4 | 187371043 | rs11723365 | -0.707 | -0.495 | 0.297 | 0.354 | -0.048 | 0.016 | 0.016 | 0.742 | 0.28 | 0.11 | 0.31 | 0.21 | 0.33 | 0.36 |
| 4 | 187372264 | rs2375775 | 0.185 | -0.207 | 0.2 | 0.342 | -0.001 | 0.062 | 0.062 | 0 | 0.09 | 0.07 | 0.1 | 0.4 | 0.07 | 0.17 |
| 4 | 187375370 | rs4241830 | 0.869 | -0.256 | 0.281 | 0.282 | -0.001 | 0 | 0 | 0 | 0.65 | 0.36 | 0.63 | 0.67 | 0.67 | 0.6 |
| 4 | 187376559 | rs1839746 | 0.383 | -0.259 | -0.08 | -0.343 | 0.022 | -0.005 | -0.005 | 0 | 0.68 | 0.26 | 0.65 | 0.31 | 0.79 | 0.57 |
| 4 | 187378007 | rs7670038 | -0.058 | -0.023 | -0.324 | -0.741 | 0.002 | 0.033 | 0.033 | 0.022 | 0.53 | 0.66 | 0.51 | 0.36 | 0.51 | 0.49 |
| 4 | 187378920 | rs10004538 | -0.977 | -0.05 | -0.405 | -0.742 | 0.011 | -0.051 | -0.051 | 0 | 0.8 | 0.89 | 0.72 | 0.42 | 0.83 | 0.71 |
| 4 | 187382294 | rs12506228 | 0.862 | 0.383 | 0.01 | -0.585 | -0.011 | 0.021 | 0.021 | 1.032 | 0.3 | 0.19 | 0.33 | 0.21 | 0.29 | 0.24 |
| 4 | 187382542 | rs6846710 | -0.746 | 0.4 | -0.099 | -0.515 | -0.007 | -0.04 | -0.04 | 0.048 | 0.22 | 0.52 | 0.18 | 0.14 | 0.2 | 0.24 |
| 4 | 187387204 | rs11723770 | 0.335 | 0.673 | -0.417 | -0.647 | -0.029 | 0.041 | 0.041 | 0.237 | 0.32 | 0.14 | 0.34 | 0.22 | 0.32 | 0.25 |
| 4 | 187390680 | rs1901317 | -0.063 | 0.892 | -0.598 | -0.325 | 0.011 | -0.031 | -0.031 | 0 | 0.12 | 0.21 | 0.17 | 0.02 | 0.17 | 0.14 |
| 4 | 187390722 | rs1376535 | 0.076 | 0.647 | -0.603 | -0.297 | 0.001 | -0.031 | -0.031 | 0 | 0.11 | 0.21 | 0.12 | 0.02 | 0.14 | 0.13 |
| 4 | 187390861 | rs1376534 | 0.084 | 0.664 | -1.129 | -0.193 | -0.014 | 0.029 | 0.029 | 1.043 | 0.28 | 0.16 | 0.3 | 0.2 | 0.26 | 0.23 |
| 4 | 187391215 | rs6832915 | -0.313 | 0.027 | -0.99 | -0.319 | -0.017 | -0.02 | -0.02 | 0 | 0.35 | 0.41 | 0.24 | 0.2 | 0.22 | 0.27 |
| 4 | 187403169 | rs6830267 | -0.35 | 1.077 | -1.559 | 0.142 | -0.083 | -0.031 | -0.031 | 0.873 | 0.35 | 0.16 | 0.55 | 0.64 | 0.49 | 0.42 |
| 4 | 187404608 | rs10866292 | 0.262 | 1.271 | -0.622 | -0.253 | 0.037 | -0.058 | -0.058 | 0.505 | 0.84 | 0.71 | 0.85 | 0.94 | 0.82 | 0.8 |
| 4 | 187411297 | rs11723176 | -0.392 | 1.562 | -1.174 | -0.073 | -0.018 | 0.019 | 0.019 | 0 | 0.29 | 0.23 | 0.3 | 0.23 | 0.32 | 0.27 |
| 4 | 187411620 | rs6854525 | -0.138 | 1.369 | -0.724 | -0.248 | 0.009 | -0.064 | -0.064 | 0.329 | 0.16 | 0.39 | 0.12 | 0.05 | 0.17 | 0.2 |

| | | | | | | | | | | | | | | | | |
|---|---|---|---|---|---|---|---|---|---|---|---|---|---|---|---|---|
| 4 | 187411802 | rs921453 | -0.955 | 1.295 | -1.316 | 0.3 | 0.016 | 0.016 | 0.016 | 0.054 | 0.6 | 0.67 | 0.67 | 0.8 | 0.71 | 0.74 |
| 4 | 187414115 | rs2375799 | -0.314 | 1.863 | -1.237 | 1.025 | -0.048 | -0.054 | -0.054 | 0.134 | 0.12 | 0.06 | 0.25 | 0.52 | 0.21 | 0.27 |
| 4 | 187421810 | rs11729609 | -0.339 | 1.458 | -0.934 | 0.04 | -0.001 | -0.019 | -0.019 | 0 | 0.11 | 0.07 | 0.07 | 0 | 0.13 | 0.03 |
| 4 | 187421850 | rs6553000 | 0.4 | 1.232 | -1.081 | 0.235 | 0.002 | -0.024 | -0.024 | 0.009 | 0.73 | 0.78 | 0.69 | 0.39 | 0.75 | 0.46 |
| 4 | 187424564 | rs10022244 | 0.878 | 0.707 | -1.667 | -0.677 | 0 | 0.185 | 0.185 | 0 | 0.38 | 0.1 | 0.25 | 0.07 | 0.34 | 0.24 |
| 4 | 187427709 | rs11736645 | -1.141 | 0.406 | -0.192 | -0.2 | -0.009 | -0.035 | -0.035 | 0 | 0.22 | 0.48 | 0.18 | 0.13 | 0.19 | 0.1 |
| 4 | 187431073 | rs11734081 | 0.413 | 0.42 | -0.225 | -0.206 | -0.001 | -0.012 | -0.012 | 0 | 0.11 | 0.1 | 0.08 | 0 | 0.12 | 0.03 |
| 4 | 187444718 | rs10049809 | 0.354 | 0.315 | -0.113 | -0.26 | -0.004 | -0.002 | -0.002 | 0.51 | 0.89 | 0.86 | 0.9 | 0.93 | 0.92 | 0.9 |
| 4 | 187445552 | rs4862705 | 1.529 | 0.33 | -0.027 | -0.28 | -0.017 | -0.048 | -0.048 | 0 | 0.72 | 0.53 | 0.79 | 0.85 | 0.76 | 0.82 |
| 4 | 187449434 | rs7440284 | 0.524 | 0.323 | -0.006 | -0.319 | -0.017 | -0.102 | -0.102 | 0.401 | 0.8 | 0.64 | 0.88 | 0.99 | 0.83 | 0.89 |
| 4 | 187455946 | rs6553008 | -0.44 | 0.209 | 0.021 | -0.161 | -0.019 | -0.037 | -0.037 | 0.606 | 0.92 | 0.85 | 0.95 | 1 | 0.96 | 0.93 |
| 4 | 187456343 | rs6553010 | -0.475 | 0.285 | 0.053 | 0.087 | 0 | 0.008 | 0.008 | 0 | 0.23 | 0.21 | 0.28 | 0.63 | 0.2 | 0.53 |
| 4 | 187460652 | rs11943499 | 0.091 | 0.238 | 0.147 | 0.161 | 0.001 | 0.03 | 0.03 | 0.026 | 0.19 | 0.16 | 0.25 | 0.57 | 0.16 | 0.5 |
| 4 | 187461706 | rs2165666 | 1.169 | 0.063 | 0.01 | 0.01 | 0.005 | 0.001 | 0.001 | 0.143 | 0.6 | 0.36 | 0.6 | 0.34 | 0.73 | 0.37 |
| 4 | 187465729 | rs13140444 | -0.674 | 0.435 | -0.046 | 0.139 | -0.008 | -0.007 | -0.007 | 0 | 0.37 | 0.32 | 0.44 | 0.63 | 0.4 | 0.6 |
| 4 | 187465991 | rs13113549 | -0.764 | 0.416 | 0.128 | 0.147 | 0 | 0 | 0 | 0 | 0.38 | 0.42 | 0.45 | 0.63 | 0.4 | 0.6 |
| 4 | 187469075 | rs2119883 | -0.43 | 0.222 | 0.486 | 0.218 | -0.028 | -0.045 | -0.045 | 1.134 | 0.92 | 0.81 | 0.97 | 1 | 0.97 | 0.93 |
| 4 | 187482867 | rs7687823 | -1.12 | 0.586 | 0.576 | 0.38 | 0.001 | 0.009 | 0.009 | 0 | 0.38 | 0.54 | 0.42 | 0.63 | 0.37 | 0.58 |
| 4 | 187485933 | rs13131052 | 0.667 | 0.546 | 0.638 | 0.931 | 0.003 | 0.145 | 0.145 | 0 | 0.93 | 0.76 | 0.83 | 0.75 | 0.96 | 0.91 |
| 4 | 187490395 | rs6850080 | -0.34 | 0.818 | -0.013 | 1.079 | -0.006 | -0.021 | -0.021 | 0 | 0.28 | 0.21 | 0.24 | 0.52 | 0.3 | 0.5 |
| 4 | 187491445 | rs1867070 | -0.437 | 0.438 | 0.151 | 0.704 | -0.03 | -0.019 | -0.019 | 0 | 0.36 | 0.53 | 0.39 | 0.74 | 0.25 | 0.53 |
| 4 | 187492881 | rs11930659 | -0.158 | 1.033 | 0.235 | 0.557 | -0.003 | -0.004 | -0.004 | 1.052 | 0.45 | 0.58 | 0.46 | 0.66 | 0.41 | 0.66 |
| 4 | 187508057 | rs10001487 | -0.245 | 0.672 | 0.388 | 0.739 | -0.001 | 0.034 | 0.034 | 0 | 0.51 | 0.3 | 0.46 | 0.4 | 0.46 | 0.29 |
| 4 | 187509321 | rs7675885 | -0.745 | 0.508 | 0.953 | 0.715 | -0.041 | -0.016 | -0.016 | 0.006 | 0.58 | 0.79 | 0.54 | 0.44 | 0.49 | 0.3 |
| 4 | 187513585 | rs3775309 | -0.644 | 0.592 | 1.314 | 1.128 | -0.014 | 0.014 | 0.014 | 0 | 0.57 | 0.69 | 0.61 | 0.58 | 0.5 | 0.31 |
| 4 | 187515047 | rs1280092 | -0.798 | 0.665 | 0.922 | 1.13 | -0.068 | 0.022 | 0.022 | 0 | 0.58 | 0.86 | 0.62 | 0.58 | 0.49 | 0.3 |
| 4 | 187516880 | rs1298865 | -0.884 | 0.01 | 0.855 | 0.816 | 0.021 | 0.026 | 0.026 | 0 | 0.45 | 0.71 | 0.53 | 0.57 | 0.37 | 0.21 |
| 4 | 187518976 | rs2306990 | -0.133 | -0.025 | 0.86 | 0.783 | -0.007 | -0.04 | -0.04 | 0.123 | 0.37 | 0.16 | 0.45 | 0.54 | 0.34 | 0.2 |
| 4 | 187521037 | rs10009030 | -0.077 | -0.027 | 0.948 | 0.859 | 0.008 | 0 | 0 | 0 | 0.37 | 0.24 | 0.36 | 0.38 | 0.3 | 0.19 |
| 4 | 187525020 | rs2637777 | -0.776 | 0.06 | 0.777 | 0.732 | 0.026 | 0.034 | 0.034 | 0 | 0.45 | 0.78 | 0.54 | 0.58 | 0.37 | 0.21 |
| 4 | 187526002 | rs1280103 | -0.202 | 0.031 | 0.568 | 0.477 | -0.02 | -0.003 | -0.003 | 1.676 | 0.41 | 0.5 | 0.48 | 0.56 | 0.29 | 0.16 |
| 4 | 187527472 | rs2584344 | -0.988 | 0.095 | 0.71 | 0.843 | -0.038 | 0.018 | 0.018 | 0 | 0.63 | 0.86 | 0.67 | 0.66 | 0.59 | 0.41 |
| 4 | 187527818 | rs2584345 | -0.05 | 0.009 | 0.771 | 0.763 | 0.003 | 0.015 | 0.015 | 0 | 0.43 | 0.63 | 0.32 | 0.1 | 0.44 | 0.28 |
| 4 | 187531022 | rs28647489 | 0.258 | -0.007 | 0.779 | 0.885 | -0.002 | 0.01 | 0.01 | 0.893 | 0.19 | 0.19 | 0.19 | 0.38 | 0.11 | 0.08 |
| 4 | 187534363 | rs2249916 | -0.909 | -0.012 | 0.811 | 0.855 | -0.014 | 0.026 | 0.026 | 0 | 0.61 | 0.74 | 0.66 | 0.67 | 0.58 | 0.38 |
| 4 | 187534375 | rs2249917 | -0.007 | 0 | 0.772 | 0.841 | 0 | -0.003 | -0.003 | 0 | 0.42 | 0.54 | 0.47 | 0.28 | 0.47 | 0.3 |
| 4 | 187538330 | rs1280099 | 0.006 | -0.012 | 0.779 | 0.848 | 0 | -0.003 | -0.003 | 0.031 | 0.42 | 0.54 | 0.47 | 0.28 | 0.47 | 0.3 |
| 4 | 187550697 | rs10008621 | 0.252 | 0 | 0.369 | 0.878 | 0.004 | -0.009 | -0.009 | 0 | 0.18 | 0.14 | 0.19 | 0.33 | 0.11 | 0.08 |
| 4 | 187556139 | rs189031 | -0.137 | 0.122 | -0.204 | -0.035 | 0.018 | 0.015 | 0.015 | 0 | 0.09 | 0.31 | 0.11 | 0.06 | 0.1 | 0.06 |
| 4 | 187557893 | rs328418 | -0.594 | 0.496 | -1.127 | 0.199 | -0.058 | 0.033 | 0.033 | 0 | 0.63 | 0.87 | 0.65 | 0.68 | 0.56 | 0.39 |
| 4 | 187561798 | rs7663350 | -0.82 | 0.588 | -0.457 | 1.138 | -0.005 | 0.025 | 0.025 | 0.062 | 0.22 | 0.32 | 0.37 | 0.57 | 0.17 | 0.11 |
| 4 | 187573918 | rs328431 | -0.83 | 0.333 | -0.432 | 0.918 | -0.021 | 0.047 | 0.047 | 0.757 | 0.25 | 0.54 | 0.34 | 0.45 | 0.21 | 0.19 |
| 4 | 187574398 | rs1388297 | 0.791 | 0.595 | -1.393 | 0.297 | -0.037 | 0.177 | 0.177 | 0 | 0.39 | 0.14 | 0.27 | 0.06 | 0.4 | 0.26 |
| 4 | 187580706 | rs7672047 | -0.09 | 1.154 | -1.213 | 0.865 | -0.099 | -0.035 | -0.035 | 4.963 | 0.33 | 0.11 | 0.33 | 0.35 | 0.41 | 0.54 |
| 4 | 187585952 | rs167853 | -0.299 | 1.059 | -0.652 | 0.47 | 0.056 | -0.028 | -0.028 | 0 | 0.08 | 0.28 | 0.05 | 0 | 0.09 | 0.08 |
| 4 | 187586210 | rs2130910 | 0.253 | 1.062 | -1.064 | 0.598 | -0.01 | 0.011 | 0.011 | 0 | 0.58 | 0.67 | 0.62 | 0.64 | 0.5 | 0.37 |
| 4 | 187587980 | rs10155467 | -0.4 | 0.947 | -1.699 | 0.665 | -0.027 | -0.011 | -0.011 | 0 | 0.37 | 0.14 | 0.4 | 0.3 | 0.38 | 0.44 |
| 4 | 187588477 | rs2130909 | -0.73 | 0.559 | -0.944 | 0.597 | -0.013 | 0.025 | 0.025 | 0 | 0.35 | 0.56 | 0.33 | 0.43 | 0.29 | 0.47 |
| 4 | 187588695 | rs2375889 | -1.134 | -0.05 | -1.17 | 0.772 | 0.003 | -0.024 | -0.024 | 0 | 0.19 | 0.14 | 0.37 | 0.57 | 0.2 | 0.26 |
| 4 | 187589603 | rs162182 | -0.481 | -0.653 | -1.028 | 0.291 | 0.001 | -0.001 | -0.001 | 0.229 | 0.23 | 0.28 | 0.1 | 0.25 | 0.12 | 0.28 |
| 4 | 187590946 | rs10025804 | -1.207 | -0.285 | -0.692 | 0.305 | 0 | -0.004 | -0.004 | 0 | 0.32 | 0.41 | 0.34 | 0.74 | 0.28 | 0.43 |
| 4 | 187591526 | rs4862723 | -0.4 | -0.141 | -0.771 | 0.496 | 0.022 | 0.128 | 0.128 | 0.52 | 0.09 | 0.14 | 0.24 | 0.5 | 0.16 | 0.16 |
| 4 | 187601962 | rs455600 | 0.716 | -0.576 | -1.251 | 0.195 | -0.041 | -0.09 | -0.09 | 0.444 | 0.84 | 0.92 | 0.64 | 0.37 | 0.79 | 0.79 |
| 4 | 187605476 | rs1877730 | 1.222 | -1.155 | -0.986 | -0.295 | 0.065 | 0.28 | 0.28 | 0.347 | 0.49 | 0.09 | 0.46 | 0.08 | 0.61 | 0.48 |
| 4 | 187614434 | rs10020743 | 1.24 | -1.153 | -0.725 | -0.163 | 0.076 | 0.228 | 0.228 | 0 | 0.5 | 0.14 | 0.49 | 0.14 | 0.61 | 0.48 |
| 4 | 187621281 | rs6818461 | -0.415 | -1.406 | 0.515 | 0.304 | -0.053 | -0.016 | -0.016 | 0 | 0.3 | 0.48 | 0.12 | 0.27 | 0.13 | 0.3 |
| 4 | 187622064 | rs455219 | -0.93 | -1.544 | 0.65 | 0.396 | 0.014 | -0.002 | -0.002 | 0.088 | 0.34 | 0.66 | 0.22 | 0.36 | 0.21 | 0.33 |
| 4 | 187627160 | rs13123522 | 0.309 | -1.303 | 0.728 | 0.393 | 0.009 | 0.085 | 0.085 | 0 | 0.81 | 0.68 | 0.7 | 0.5 | 0.78 | 0.76 |
| 4 | 187627792 | rs11939575 | -0.323 | -1.31 | 0.7 | 0.389 | -0.019 | -0.04 | -0.04 | 0 | 0.84 | 0.91 | 0.72 | 0.5 | 0.78 | 0.77 |
| 4 | 187629140 | rs367863 | 0.096 | -1.282 | 0.634 | 0.414 | 0.005 | 0.055 | 0.055 | 0 | 0.82 | 0.71 | 0.7 | 0.5 | 0.78 | 0.77 |
| 4 | 187629538 | rs3733413 | 1.709 | -1.333 | 0.859 | 0.385 | 0.037 | 0.184 | 0.184 | 0 | 0.49 | 0.16 | 0.48 | 0.14 | 0.58 | 0.44 |
| 4 | 187629770 | rs3733414 | 1.556 | -1.325 | 0.873 | 0.398 | 0.056 | 0.113 | 0.113 | 0.024 | 0.51 | 0.21 | 0.57 | 0.22 | 0.64 | 0.47 |
| 4 | 187630590 | rs3733415 | -0.617 | -1.621 | 1.312 | 0.202 | -0.042 | -0.004 | -0.004 | 0.094 | 0.28 | 0.4 | 0.12 | 0.28 | 0.13 | 0.29 |
| 4 | 187638636 | rs327095 | -1.048 | -1.589 | 0.708 | 0.165 | -0.048 | 0.002 | 0.002 | 0 | 0.35 | 0.51 | 0.22 | 0.36 | 0.2 | 0.32 |
| 4 | 187640836 | rs327102 | -0.845 | -1.678 | 0.487 | 0.213 | -0.031 | 0.003 | 0.003 | 0 | 0.33 | 0.57 | 0.2 | 0.36 | 0.15 | 0.32 |
| 4 | 187652655 | rs7673895 | -1.021 | -1.333 | 0.936 | 0.069 | -0.032 | 0.001 | 0.001 | 0 | 0.14 | 0.49 | 0.16 | 0.08 | 0.11 | 0.07 |
| 4 | 187656456 | rs11132407 | -1.519 | -1.656 | 0.707 | -0.081 | -0.05 | -0.046 | -0.046 | 0 | 0.27 | 0.39 | 0.22 | 0.14 | 0.21 | 0.14 |
| 4 | 187661911 | rs7698262 | -1.018 | -0.894 | 0.693 | 0.07 | 0 | -0.055 | -0.055 | 0 | 0.23 | 0.52 | 0.32 | 0.13 | 0.26 | 0.12 |
| 4 | 187668795 | rs6553019 | 0.49 | -1.069 | -0.108 | 0.197 | 0.008 | 0.015 | 0.015 | 0.22 | 0.51 | 0.45 | 0.71 | 0.83 | 0.58 | 0.62 |

| | | | | | | | | | | | | | | | | |
|---|---|---|---|---|---|---|---|---|---|---|---|---|---|---|---|---|
| 4 | 187668886 | rs6553020 | 0.498 | -1.073 | -0.104 | 0.195 | 0.008 | 0.014 | 0.014 | 0.188 | 0.5 | 0.44 | 0.62 | 0.8 | 0.57 | 0.62 |
| 4 | 187672156 | rs11729956 | 0.198 | -1.193 | -0.565 | 0.686 | 0.005 | -0.001 | -0.001 | 1.94 | 0.55 | 0.42 | 0.64 | 0.59 | 0.63 | 0.65 |
| 4 | 187678866 | rs925642 | 0.336 | -1.039 | -1.035 | 0.623 | 0.054 | 0.036 | 0.036 | 0.9 | 0.51 | 0.27 | 0.5 | 0.44 | 0.6 | 0.56 |
| 4 | 187680241 | rs3733416 | -0.285 | -0.824 | -1.12 | 0.999 | 0.002 | -0.019 | -0.019 | 0.035 | 0.08 | 0.05 | 0.14 | 0.12 | 0.05 | 0.23 |
| 4 | 187688114 | rs7660386 | -0.536 | -1.114 | -1.396 | 0.909 | 0.043 | -0.064 | -0.064 | 0.13 | 0.31 | 0.15 | 0.36 | 0.57 | 0.24 | 0.54 |
| 4 | 187693326 | rs6856002 | 0.544 | -1.238 | -0.868 | 0.115 | 0.008 | -0.008 | -0.008 | 0 | 0.47 | 0.42 | 0.37 | 0.12 | 0.59 | 0.31 |
| 4 | 187696656 | rs161711 | 0.161 | -1.492 | -1.017 | 0.793 | 0.041 | 0.003 | 0.003 | 0 | 0.57 | 0.7 | 0.53 | 0.4 | 0.7 | 0.47 |
| 4 | 187696768 | rs6833702 | 1.114 | -1.441 | -1.27 | 0.4 | -0.036 | 0.09 | 0.09 | 0.218 | 0.36 | 0.19 | 0.32 | 0.13 | 0.56 | 0.24 |
| 4 | 187697015 | rs6834299 | 0.166 | -1.247 | -0.866 | 0.704 | -0.012 | -0.008 | -0.008 | 0 | 0.55 | 0.55 | 0.41 | 0.31 | 0.68 | 0.49 |
| 4 | 187701648 | rs13145229 | -0.47 | -0.58 | -1.169 | 1.618 | 0.031 | -0.003 | -0.003 | 0 | 0.07 | 0 | 0.12 | 0.39 | 0.05 | 0.17 |
| 4 | 187707798 | rs6847347 | 0.543 | -0.174 | -0.734 | 1.038 | 0.028 | 0.017 | 0.017 | 5.444 | 0.75 | 0.58 | 0.75 | 0.76 | 0.73 | 0.74 |
| 4 | 187712375 | rs766685 | 0.548 | -0.427 | -1.894 | 0.557 | 0.052 | 0.11 | 0.11 | 0.015 | 0.54 | 0.14 | 0.45 | 0.27 | 0.62 | 0.52 |
| 4 | 187714691 | rs11733039 | 0.886 | 0.566 | -1.88 | 0.767 | -0.039 | 0.074 | 0.074 | 0 | 0.43 | 0.13 | 0.41 | 0.29 | 0.48 | 0.41 |
| 4 | 187716010 | rs11132408 | 0.887 | 0.562 | -1.882 | 0.799 | -0.039 | 0.101 | 0.101 | 0 | 0.43 | 0.13 | 0.39 | 0.24 | 0.48 | 0.41 |
| 4 | 187718537 | rs12640954 | -0.29 | 0.435 | -2.046 | 0.996 | -0.06 | -0.034 | -0.034 | 0 | 0.37 | 0.04 | 0.31 | 0.31 | 0.34 | 0.4 |
| 4 | 187722479 | rs11132409 | 0.631 | 0.336 | -0.99 | 1.391 | 0 | 0.001 | 0.001 | 0.014 | 0.78 | 0.2 | 0.71 | 0.59 | 0.81 | 0.8 |
| 4 | 187724314 | rs7657648 | -0.991 | 0.985 | -0.085 | 1.144 | 0.052 | 0 | 0 | 0 | 0.17 | 0.36 | 0.21 | 0.15 | 0.26 | 0.14 |
| 4 | 187729614 | rs1019895 | -0.393 | 1.209 | -0.466 | 1.59 | -0.025 | -0.02 | -0.02 | 0 | 0.06 | 0.01 | 0.07 | 0.21 | 0.1 | 0.19 |
| 4 | 187730631 | rs7689213 | -1.294 | 1.141 | 0.504 | 1.346 | 0.144 | 0.113 | 0.113 | 0 | 0.22 | 0.48 | 0.29 | 0.35 | 0.38 | 0.35 |
| 4 | 187745843 | rs9992417 | -0.099 | 0.427 | -0.016 | 0.671 | 0.043 | 0.04 | 0.04 | 0 | 0.15 | 0.07 | 0.06 | 0 | 0.07 | 0.07 |
| 4 | 187747224 | rs10028835 | -0.463 | 0.63 | 0.297 | 0.871 | 0.077 | 0.076 | 0.076 | 0 | 0.18 | 0.33 | 0.17 | 0.34 | 0.28 | 0.32 |
| 4 | 187751154 | rs1583219 | -1.565 | 0.245 | 0.158 | 0.62 | 0 | -0.001 | -0.001 | 0 | 0.4 | 0.39 | 0.27 | 0.32 | 0.4 | 0.45 |
| 4 | 187754871 | rs13105379 | -1.734 | 0.295 | 0.182 | 0.83 | 0.001 | 0.004 | 0.004 | 1.13 | 0.43 | 0.48 | 0.29 | 0.42 | 0.41 | 0.45 |
| 4 | 187756317 | rs4862735 | 1.776 | 0.149 | -0.404 | 0.767 | -0.02 | -0.02 | -0.02 | 0 | 0.5 | 0.12 | 0.56 | 0.58 | 0.48 | 0.51 |
| 4 | 187759082 | rs10033282 | 0.258 | 0.096 | -0.251 | 0.527 | 0.05 | 0.066 | 0.066 | 0 | 0.13 | 0.02 | 0.05 | 0 | 0.07 | 0.07 |
| 4 | 187774488 | rs11132410 | 0.802 | 0.438 | 0.343 | 0.865 | 0.032 | 0.022 | 0.022 | 0 | 0.68 | 0.44 | 0.65 | 0.65 | 0.62 | 0.6 |
| 4 | 187775324 | rs1431929 | -0.246 | 0.513 | 0.706 | 0.82 | 0.055 | 0.075 | 0.075 | 0 | 0.18 | 0.28 | 0.22 | 0.35 | 0.28 | 0.31 |
| 4 | 187779084 | rs443846 | 0.948 | 0.406 | 1.007 | 0.976 | -0.023 | -0.012 | -0.012 | 0 | 0.85 | 0.6 | 0.86 | 0.89 | 0.9 | 0.91 |
| 4 | 187788311 | rs322442 | 0.657 | 0.543 | 1.142 | 0.95 | 0.127 | 0.108 | 0.108 | 0 | 0.77 | 0.53 | 0.74 | 0.65 | 0.65 | 0.67 |
| 4 | 187792665 | rs322447 | 0.643 | 0.452 | 1.13 | 0.941 | -0.002 | -0.017 | -0.017 | 0.057 | 0.85 | 0.74 | 0.84 | 0.89 | 0.85 | 0.9 |
| 4 | 187797022 | rs172818 | 1.224 | 0.566 | 0.935 | 0.786 | 0.014 | 0.017 | 0.017 | 0.008 | 0.66 | 0.39 | 0.62 | 0.54 | 0.58 | 0.59 |
| 4 | 187799603 | rs322453 | 1.571 | 0.14 | 0.442 | 0.708 | -0.042 | -0.038 | -0.038 | 0 | 0.52 | 0.13 | 0.56 | 0.55 | 0.51 | 0.55 |
| 4 | 187802501 | rs4862740 | 0.761 | 0.059 | 0.785 | 0.36 | -0.022 | -0.103 | -0.103 | 0 | 0.76 | 0.7 | 0.81 | 1 | 0.82 | 0.9 |
| 4 | 187804194 | rs170427 | 1.599 | -0.451 | -0.416 | 0.286 | -0.047 | 0.001 | 0.001 | 0.317 | 0.4 | 0.08 | 0.45 | 0.33 | 0.44 | 0.46 |
| 4 | 187813409 | rs736879 | -0.426 | -0.988 | -0.012 | -0.287 | -0.037 | -0.108 | -0.108 | 0.71 | 0.19 | 0.29 | 0.11 | 0 | 0.12 | 0.07 |
| 4 | 187815647 | rs397321 | 0.752 | -0.896 | 0.131 | 0.283 | -0.033 | 0.048 | 0.048 | 0.209 | 0.75 | 0.48 | 0.8 | 0.64 | 0.83 | 0.87 |
| 4 | 187817303 | rs4862742 | 1.329 | -0.847 | -0.286 | 0.336 | -0.066 | 0.029 | 0.029 | 0.083 | 0.43 | 0.17 | 0.52 | 0.36 | 0.51 | 0.52 |
| 4 | 187818100 | rs10030988 | 0.087 | -0.768 | -0.074 | 0 | -0.002 | -0.037 | -0.037 | 0 | 0.17 | 0.18 | 0.09 | 0 | 0.14 | 0.02 |
| 4 | 187818466 | rs9685837 | 0.355 | -0.635 | -0.154 | 0.233 | 0.005 | 0.006 | 0.006 | 0.075 | 0.28 | 0.3 | 0.27 | 0.3 | 0.29 | 0.38 |
| 4 | 187821208 | rs578135 | -0.67 | -0.737 | 0.089 | 0.022 | -0.023 | -0.058 | -0.058 | 0 | 0.89 | 0.57 | 0.95 | 1 | 0.95 | 0.99 |
| 4 | 187825117 | rs9991793 | 1.621 | -0.653 | -0.561 | 0.063 | -0.006 | -0.021 | -0.021 | 0.103 | 0.07 | 0.07 | 0.03 | 0 | 0.04 | 0.01 |
| 4 | 187828093 | rs322398 | 1.321 | -0.599 | -0.994 | 0.201 | -0.056 | -0.067 | -0.067 | 2.625 | 0.36 | 0.1 | 0.44 | 0.42 | 0.41 | 0.46 |
| 4 | 187833759 | rs1895844 | -0.11 | -0.216 | -0.555 | -0.018 | 0.002 | -0.002 | -0.002 | 0 | 0.34 | 0.37 | 0.39 | 0.27 | 0.37 | 0.44 |
| 4 | 187836373 | rs322429 | 0.06 | 0.151 | -0.276 | -0.453 | -0.003 | -0.05 | -0.05 | 0.278 | 0.81 | 0.78 | 0.89 | 1 | 0.84 | 0.98 |
| 4 | 187843726 | rs10018164 | 0.38 | 0.494 | -0.61 | -0.241 | 0.006 | -0.007 | -0.007 | 0 | 0.42 | 0.33 | 0.42 | 0.54 | 0.38 | 0.52 |
| 4 | 187845057 | rs10033110 | -0.442 | 1.069 | -1.006 | -0.455 | -0.015 | 0.047 | 0.047 | 0.029 | 0.75 | 0.89 | 0.71 | 0.87 | 0.73 | 0.78 |
| 4 | 187848075 | rs6829485 | 0.119 | 1.041 | -0.341 | -0.063 | -0.006 | 0.002 | 0.002 | 0 | 0.43 | 0.51 | 0.43 | 0.54 | 0.38 | 0.52 |
| 4 | 187851288 | rs7682123 | -0.112 | 0.926 | -0.578 | -0.309 | -0.002 | -0.002 | -0.002 | 0.352 | 0.48 | 0.54 | 0.48 | 0.56 | 0.41 | 0.55 |
| 4 | 187851352 | rs7657521 | 0.477 | 1.944 | -0.823 | -0.097 | -0.11 | 0.077 | 0.077 | 0.685 | 0.74 | 0.93 | 0.63 | 0.88 | 0.65 | 0.75 |
| 4 | 187854369 | rs999319 | 1.28 | 1.878 | -0.171 | 0.223 | 0.005 | 0.059 | 0.059 | 0.204 | 0.62 | 0.74 | 0.48 | 0.84 | 0.45 | 0.56 |
| 4 | 187857192 | rs1158815 | 0.465 | 2.335 | -0.601 | 0.247 | -0.013 | 0.012 | 0.012 | 0 | 0.81 | 0.85 | 0.79 | 0.93 | 0.72 | 0.77 |
| 4 | 187860363 | rs754548 | -0.601 | 1.673 | -0.463 | 0.848 | 0.007 | 0.147 | 0.147 | 0 | 0.08 | 0.22 | 0.15 | 0.32 | 0.08 | 0.07 |
| 4 | 187864673 | rs13137204 | -0.518 | 1.921 | -0.999 | 0.079 | -0.119 | 0.108 | 0.108 | 0.209 | 0.18 | 0.01 | 0.2 | 0.01 | 0.28 | 0.11 |
| 4 | 187870988 | rs13143485 | -0.818 | 1.226 | -0.307 | 0.576 | 0.021 | -0.001 | -0.001 | 0 | 0.38 | 0.4 | 0.46 | 0.32 | 0.42 | 0.5 |
| 4 | 187871454 | rs9312351 | -0.729 | 1.248 | 0.442 | 0.695 | 0.059 | -0.029 | -0.029 | 0 | 0.23 | 0.46 | 0.4 | 0.2 | 0.3 | 0.17 |
| 4 | 187874691 | rs9998558 | -0.955 | 0.593 | -0.172 | 0.716 | 0.067 | -0.025 | -0.025 | 0 | 0.56 | 0.81 | 0.69 | 0.4 | 0.6 | 0.7 |
| 4 | 187875680 | rs7658351 | -0.741 | 0.54 | 0.47 | 0.711 | 0.036 | 0.022 | 0.022 | 0.091 | 0.52 | 0.38 | 0.66 | 0.35 | 0.59 | 0.69 |
| 4 | 187899660 | rs6836076 | -0.974 | 0.528 | -0.039 | 0.637 | 0.066 | 0.003 | 0.003 | 0.302 | 0.56 | 0.78 | 0.7 | 0.38 | 0.6 | 0.69 |
| 4 | 187903412 | rs6858853 | -0.598 | 0.468 | 0.34 | 0.6 | 0.075 | 0.034 | 0.034 | 0.176 | 0.51 | 0.32 | 0.66 | 0.37 | 0.6 | 0.69 |
| 4 | 187903476 | rs6835681 | -0.987 | 0.459 | 0.063 | 0.609 | 0.073 | 0.008 | 0.008 | 0.489 | 0.56 | 0.78 | 0.7 | 0.37 | 0.6 | 0.7 |
| 4 | 187906366 | rs4862752 | -0.349 | 0.757 | 0.076 | 0.806 | -0.006 | -0.003 | -0.003 | 0.373 | 0.17 | 0.19 | 0.17 | 0.19 | 0.23 | 0.22 |
| 4 | 187912810 | rs13131381 | 0.35 | 0.878 | -0.237 | 0.486 | -0.063 | 0.082 | 0.082 | 0.197 | 0.14 | 0.02 | 0.15 | 0 | 0.18 | 0.05 |
| 4 | 187915635 | rs10471185 | -0.415 | 0.088 | 0.537 | 1.135 | -0.059 | 0.052 | 0.052 | 0 | 0.27 | 0.56 | 0.19 | 0.37 | 0.14 | 0.39 |
| 4 | 187916373 | rs966575 | 0.471 | 0.049 | 0.452 | 1.186 | 0.027 | -0.032 | -0.032 | 0 | 0.19 | 0.1 | 0.14 | 0.31 | 0.1 | 0.37 |
| 4 | 187916415 | rs966576 | -0.651 | 0.417 | 0.462 | 1.009 | 0.011 | 0 | 0 | 0.087 | 0.4 | 0.72 | 0.34 | 0.56 | 0.32 | 0.54 |
| 4 | 187916497 | rs10026367 | -1.147 | 0.343 | 0.747 | 0.983 | 0.05 | 0.057 | 0.057 | 0.726 | 0.06 | 0.26 | 0.02 | 0.06 | 0.04 | 0.01 |
| 4 | 187926758 | rs6829136 | -0.255 | -0.031 | -0.659 | 1.1 | 0.013 | -0.019 | -0.019 | 0 | 0.64 | 0.72 | 0.57 | 0.5 | 0.62 | 0.63 |
| 4 | 187932470 | rs6851529 | -0.364 | 0.045 | -1.15 | 0.839 | -0.039 | 0.098 | 0.098 | 0 | 0.67 | 0.87 | 0.65 | 0.88 | 0.61 | 0.77 |
| 4 | 187932716 | rs6856787 | 0.501 | -0.405 | -1.059 | 0.937 | 0 | 0.036 | 0.036 | 0 | 0.48 | 0.36 | 0.52 | 0.73 | 0.52 | 0.59 |

| | | | | | | | | | | | | | | | | |
|---|---|---|---|---|---|---|---|---|---|---|---|---|---|---|---|---|
| 4 | 187949161 | rs1559779 | 0.722 | -0.39 | -1.312 | 1.194 | -0.123 | -0.058 | -0.058 | 0 | 0.44 | 0.05 | 0.44 | 0.46 | 0.48 | 0.57 |
| 4 | 187954207 | rs715070 | -0.706 | -0.192 | -1.211 | 1.127 | -0.108 | 0.086 | 0.086 | 0 | 0.68 | 0.94 | 0.63 | 0.85 | 0.58 | 0.75 |
| 4 | 187966612 | rs6835306 | -1.02 | -0.587 | 0.015 | 1.063 | -0.075 | 0.033 | 0.033 | 0.224 | 0.19 | 0.57 | 0.15 | 0.23 | 0.06 | 0.15 |
| 4 | 187969016 | rs13126183 | 0.208 | -0.121 | -0.315 | 0.75 | -0.003 | -0.012 | -0.012 | 0.253 | 0.58 | 0.47 | 0.72 | 0.67 | 0.6 | 0.65 |
| 4 | 187969329 | rs6814129 | -0.033 | -0.224 | -0.65 | 0.631 | 0.03 | 0.038 | 0.038 | 0 | 0.51 | 0.68 | 0.63 | 0.42 | 0.51 | 0.62 |
| 4 | 187969452 | rs6814850 | -0.519 | -0.756 | 0.108 | 0.607 | -0.05 | 0.029 | 0.029 | 1.318 | 0.16 | 0.54 | 0.18 | 0.12 | 0.07 | 0.13 |
| 4 | 187978589 | rs4476657 | 1.25 | -0.562 | -0.222 | 0.662 | -0.036 | 0.021 | 0.021 | 0 | 0.26 | 0.16 | 0.39 | 0.2 | 0.26 | 0.35 |
| 4 | 187979306 | rs4333244 | -0.522 | -0.612 | -0.447 | 0.812 | 0.014 | -0.015 | -0.015 | 0 | 0.62 | 0.75 | 0.67 | 0.61 | 0.58 | 0.78 |
| 4 | 187979630 | rs7441526 | 0.285 | -0.937 | -0.016 | 1.204 | -0.001 | 0.009 | 0.009 | 0 | 0.11 | 0.12 | 0.18 | 0.25 | 0.14 | 0.18 |
| 4 | 187979712 | rs4507419 | -0.505 | -0.958 | 0.062 | 1.192 | 0 | 0.005 | 0.005 | 0 | 0.35 | 0.4 | 0.27 | 0.42 | 0.32 | 0.43 |
| 4 | 187980711 | rs3923586 | -0.332 | -0.857 | -0.083 | 1.144 | 0.003 | 0.001 | 0.001 | 0 | 0.33 | 0.38 | 0.22 | 0.38 | 0.31 | 0.42 |
| 4 | 187981262 | rs4862758 | 0.034 | -1.263 | -0.397 | 1.458 | 0.043 | -0.065 | -0.065 | 0.011 | 0.78 | 0.91 | 0.78 | 0.59 | 0.81 | 0.8 |
| 4 | 187999052 | rs13141650 | 0.964 | -0.653 | 0.597 | 1.506 | 0.004 | 0.15 | 0.15 | 0 | 0.84 | 0.71 | 0.81 | 0.57 | 0.85 | 0.88 |
| 4 | 188000355 | rs4861739 | 1.745 | -0.903 | 0.292 | 1.059 | 0.027 | 0.103 | 0.103 | 0.002 | 0.52 | 0.15 | 0.51 | 0.34 | 0.59 | 0.54 |
| 4 | 188000653 | rs13104006 | 1.718 | -0.918 | 0.626 | 1.052 | 0.032 | 0.091 | 0.091 | 0.002 | 0.53 | 0.22 | 0.51 | 0.34 | 0.59 | 0.54 |
| 4 | 188004538 | rs7662795 | -0.849 | -0.912 | 0.975 | 1.065 | 0.003 | 0.085 | 0.085 | 0 | 0.15 | 0.21 | 0.18 | 0.44 | 0.15 | 0.13 |
| 4 | 188005948 | rs2029257 | -2.006 | -1.129 | 1.066 | 0.947 | 0 | -0.001 | -0.001 | 0 | 0.38 | 0.69 | 0.38 | 0.54 | 0.39 | 0.38 |
| 4 | 188007418 | rs1038598 | -0.076 | -1.124 | 1.091 | 0.853 | -0.039 | 0 | 0 | 0 | 0.9 | 0.83 | 0.89 | 0.88 | 0.98 | 0.89 |
| 4 | 188009643 | rs6818214 | -0.556 | -1.054 | 1.194 | 0.793 | -0.007 | -0.05 | -0.05 | 0 | 0.19 | 0.36 | 0.18 | 0.09 | 0.19 | 0.22 |
| 4 | 188013721 | rs7665787 | -1.966 | -1.485 | 1.97 | 1.202 | -0.045 | 0.069 | 0.069 | 0 | 0.26 | 0.49 | 0.28 | 0.53 | 0.17 | 0.24 |
| 4 | 188015375 | rs1487599 | -2.541 | -1.538 | 2.083 | 1.172 | -0.003 | -0.001 | -0.001 | 0.186 | 0.34 | 0.6 | 0.31 | 0.57 | 0.19 | 0.36 |
| 4 | 188015715 | rs6836337 | 1.044 | -1.537 | 2.25 | 1.185 | -0.033 | 0.194 | 0.194 | 0 | 0.84 | 0.66 | 0.84 | 0.57 | 0.9 | 0.81 |
| 4 | 188019647 | rs6553036 | -1.511 | -1.458 | 2.064 | 1.189 | -0.075 | 0.128 | 0.128 | 0 | 0.25 | 0.46 | 0.18 | 0.47 | 0.11 | 0.31 |
| 4 | 188021294 | rs2129654 | -2.478 | -1.49 | 2.157 | 1.167 | 0.018 | -0.007 | -0.007 | 0 | 0.35 | 0.63 | 0.32 | 0.56 | 0.19 | 0.36 |
| 4 | 188021950 | rs7659108 | 1.228 | -1.418 | 2.131 | 1.142 | 0.003 | -0.005 | -0.005 | 0 | 0.87 | 0.8 | 0.84 | 0.9 | 0.88 | 0.92 |
| 4 | 188022194 | rs13129961 | 2.167 | -1.422 | 1.954 | 1.163 | 0.063 | 0.003 | 0.003 | 0 | 0.61 | 0.27 | 0.65 | 0.43 | 0.76 | 0.6 |
| 4 | 188030901 | rs1487590 | 1.658 | -1.552 | 2.111 | 1.492 | 0.026 | 0.078 | 0.078 | 0.792 | 0.91 | 0.67 | 0.88 | 0.89 | 0.94 | 0.94 |
| 4 | 188040526 | rs2376075 | 1.105 | -1.509 | 1.306 | 1.513 | 0.023 | 0.068 | 0.068 | 0.257 | 0.92 | 0.78 | 0.89 | 0.89 | 0.94 | 0.94 |
| 4 | 188041074 | rs4862774 | 1.352 | -1.464 | 0.074 | 1.358 | 0.058 | -0.025 | -0.025 | 0 | 0.69 | 0.27 | 0.73 | 0.43 | 0.87 | 0.65 |
| 4 | 188047143 | rs2029259 | 1.25 | -0.894 | 0.656 | 1.729 | -0.059 | 0.146 | 0.146 | 1.263 | 0.81 | 0.46 | 0.85 | 0.62 | 0.93 | 0.81 |
| 4 | 188050164 | rs10018611 | 1.689 | -0.699 | 1.323 | 1.426 | 0.033 | 0.056 | 0.056 | 4.418 | 0.9 | 0.57 | 0.94 | 0.91 | 0.94 | 0.97 |
| 4 | 188051175 | rs6858203 | 0.839 | -0.722 | 0.469 | 0.86 | -0.008 | 0.091 | 0.091 | 0.317 | 0.46 | 0.33 | 0.5 | 0.29 | 0.52 | 0.55 |
| 4 | 188051690 | rs10024641 | -0.203 | -0.872 | 0.456 | 0.952 | 0 | -0.002 | -0.002 | 1.451 | 0.41 | 0.23 | 0.42 | 0.55 | 0.41 | 0.31 |
| 4 | 188056826 | rs6826891 | 0.983 | -0.98 | 0.615 | 0.937 | -0.029 | 0.07 | 0.07 | 0.864 | 0.9 | 0.64 | 0.94 | 0.85 | 0.98 | 0.86 |
| 4 | 188061467 | rs4611977 | 0.882 | -1.033 | 0.108 | 0.789 | -0.013 | 0.058 | 0.058 | 0.328 | 0.43 | 0.25 | 0.52 | 0.3 | 0.54 | 0.55 |
| 4 | 188071834 | rs13145604 | 0.873 | -1.408 | 0.353 | 0.119 | -0.008 | -0.031 | -0.031 | 0 | 0.6 | 0.43 | 0.74 | 0.85 | 0.73 | 0.67 |
| 4 | 188073672 | rs1487582 | 0.901 | -1.413 | 0.337 | 0.135 | -0.007 | -0.024 | -0.024 | 0 | 0.6 | 0.44 | 0.74 | 0.83 | 0.72 | 0.65 |
| 4 | 188073853 | rs10016164 | -0.149 | -1.473 | 0.4 | 0.187 | -0.006 | 0.004 | 0.004 | 0 | 0.07 | 0.05 | 0.11 | 0.06 | 0.07 | 0.08 |
| 4 | 188077812 | rs882227 | -0.946 | -1.542 | 0.576 | 0.167 | -0.002 | 0.004 | 0.004 | 0.023 | 0.11 | 0.13 | 0.13 | 0.08 | 0.12 | 0.09 |
| 4 | 188079254 | rs11730573 | -0.086 | -1.625 | 0.789 | 0.434 | -0.001 | 0.002 | 0.002 | 0 | 0.22 | 0.19 | 0.34 | 0.59 | 0.24 | 0.15 |
| 4 | 188079777 | rs11132427 | 2.168 | -1.696 | 0.666 | 0.322 | -0.067 | 0.063 | 0.063 | 0.858 | 0.4 | 0.14 | 0.42 | 0.25 | 0.5 | 0.52 |
| 4 | 188080985 | rs10021692 | -1.049 | -1.667 | 0.831 | 0.417 | -0.001 | -0.002 | -0.002 | 0 | 0.12 | 0.14 | 0.13 | 0.08 | 0.12 | 0.09 |
| 4 | 188081872 | rs6815262 | 0.873 | -1.771 | 1.295 | 0.382 | -0.024 | 0.033 | 0.033 | 0 | 0.93 | 0.73 | 0.95 | 0.92 | 0.99 | 0.88 |
| 4 | 188083739 | rs13142468 | 0.373 | -1.656 | 0.671 | 0.354 | 0.03 | 0.045 | 0.045 | 0 | 0.18 | 0.08 | 0.06 | 0 | 0.13 | 0.12 |
| 4 | 188092843 | rs4862782 | -0.169 | -1.421 | 0.573 | 0.692 | -0.01 | -0.017 | -0.017 | 0 | 0.14 | 0.1 | 0.37 | 0.6 | 0.19 | 0.14 |
| 4 | 188095514 | rs10003140 | 0.022 | -1.691 | 0.565 | 0.432 | 0.024 | 0.051 | 0.051 | 0 | 0.22 | 0.16 | 0.09 | 0.02 | 0.16 | 0.12 |
| 4 | 188099394 | rs6826520 | 1.843 | -1.886 | -0.25 | 0.11 | -0.095 | 0.03 | 0.03 | 0 | 0.38 | 0.08 | 0.43 | 0.26 | 0.54 | 0.55 |
| 4 | 188100049 | rs1027058 | 0.459 | -1.958 | 0.841 | 0.243 | -0.014 | 0.02 | 0.02 | 0 | 0.93 | 0.72 | 0.96 | 0.95 | 0.98 | 0.9 |
| 4 | 188103937 | rs10000307 | 1.166 | -2.105 | 1.258 | 0.18 | -0.017 | 0.03 | 0.03 | 0.293 | 0.74 | 0.41 | 0.57 | 0.31 | 0.76 | 0.77 |
| 4 | 188105792 | rs931370 | 1.091 | -2.202 | 1.157 | 0.62 | -0.004 | 0 | 0 | 0 | 0.78 | 0.71 | 0.69 | 0.73 | 0.79 | 0.81 |
| 4 | 188106544 | rs13145216 | -0.302 | -2.28 | 1.227 | 0.647 | -0.017 | 0.209 | 0.209 | 0 | 0.1 | 0.22 | 0.14 | 0.46 | 0.06 | 0.1 |
| 4 | 188108024 | rs1979570 | 2.116 | -2.281 | 1.167 | 0.495 | -0.017 | -0.016 | -0.016 | 0.008 | 0.65 | 0.49 | 0.55 | 0.26 | 0.72 | 0.71 |
| 4 | 188110164 | rs1038599 | 1.462 | -2.219 | 1.398 | 0.748 | -0.009 | 0.063 | 0.063 | 0 | 0.73 | 0.58 | 0.62 | 0.36 | 0.77 | 0.73 |
| 4 | 188114093 | rs4472191 | 0.246 | -2.256 | 1.276 | 0.662 | -0.025 | -0.048 | -0.048 | 1.51 | 0.67 | 0.58 | 0.64 | 0.78 | 0.72 | 0.74 |
| 4 | 188115905 | rs13121701 | 1.52 | -2.026 | 0.8 | 0.893 | 0.022 | 0.006 | 0.006 | 0 | 0.46 | 0.25 | 0.42 | 0.41 | 0.58 | 0.57 |
| 4 | 188116718 | rs1021287 | 1.819 | -1.905 | 0.714 | 0.972 | -0.05 | -0.041 | -0.041 | 0 | 0.38 | 0.13 | 0.37 | 0.41 | 0.53 | 0.56 |
| 4 | 188117428 | rs7684803 | 0.423 | -1.377 | 1.575 | 1.6 | 0.008 | 0.066 | 0.066 | 0 | 0.91 | 0.82 | 0.76 | 0.76 | 0.89 | 0.89 |
| 4 | 188125082 | rs6811993 | 2.169 | -1.54 | 1.268 | 1.558 | 0.04 | -0.017 | -0.017 | 0 | 0.63 | 0.18 | 0.48 | 0.35 | 0.72 | 0.71 |
| 4 | 188127832 | rs7674650 | 0.77 | -1.202 | 1.393 | 1.44 | 0 | 0.05 | 0.05 | 0 | 0.88 | 0.73 | 0.75 | 0.77 | 0.86 | 0.88 |
| 4 | 188131680 | rs6845980 | -1.248 | -1.234 | 1.273 | 1.426 | 0 | 0.002 | 0.002 | 0 | 0.17 | 0.2 | 0.15 | 0.11 | 0.14 | 0.09 |
| 4 | 188136307 | rs1565106 | 0.539 | -1.13 | 0.893 | 1.432 | 0.019 | 0.088 | 0.088 | 0 | 0.88 | 0.73 | 0.71 | 0.72 | 0.85 | 0.88 |
| 4 | 188138856 | rs7699015 | 1.718 | -1.253 | 0.639 | 1.522 | 0.013 | -0.031 | -0.031 | 0.01 | 0.67 | 0.22 | 0.49 | 0.39 | 0.72 | 0.71 |
| 4 | 188157742 | rs7695955 | -1.593 | -1.455 | 0.862 | 1.519 | -0.032 | 0.036 | 0.036 | 0 | 0.31 | 0.53 | 0.44 | 0.63 | 0.23 | 0.2 |
| 4 | 188159641 | rs1532021 | 0.426 | -1.723 | 0.238 | 1.139 | -0.015 | 0.088 | 0.088 | 0 | 0.14 | 0.01 | 0.07 | 0 | 0.13 | 0.09 |
| 4 | 188160927 | rs6816228 | -0.121 | -1.772 | 0.311 | 1.438 | 0.011 | -0.053 | -0.053 | 0.267 | 0.17 | 0.06 | 0.16 | 0.31 | 0.14 | 0.18 |
| 4 | 188166889 | rs6831858 | -0.248 | -1.886 | 0.555 | 0.802 | -0.026 | -0.014 | -0.014 | 0 | 0.17 | 0.33 | 0.1 | 0.13 | 0.11 | 0.04 |
| 4 | 188172182 | rs4321687 | 1.078 | -1.973 | 0.618 | 1.012 | -0.035 | 0.034 | 0.034 | 0 | 0.68 | 0.51 | 0.66 | 0.57 | 0.75 | 0.75 |
| 4 | 188185489 | rs6553050 | 0.491 | -1.631 | -0.118 | 0.686 | 0.006 | 0.053 | 0.053 | 0 | 0.14 | 0.04 | 0.07 | 0 | 0.12 | 0.1 |
| 4 | 188186007 | rs13131356 | 0.296 | -1.629 | -0.128 | 0.65 | 0.014 | 0.029 | 0.029 | 0 | 0.16 | 0.1 | 0.08 | 0 | 0.12 | 0.11 |

| | | | | | | | | | | | | | | | | |
|---|---|---|---|---|---|---|---|---|---|---|---|---|---|---|---|---|
| 4 | 188186955 | rs10461324 | 1.546 | -1.352 | -0.239 | 0.896 | 0.1 | 0.008 | 0.008 | 0.135 | 0.5 | 0.18 | 0.54 | 0.43 | 0.64 | 0.7 |
| 4 | 188191516 | rs4478240 | 0.331 | -0.548 | -0.456 | 0.734 | -0.001 | 0.007 | 0.007 | 0.016 | 0.89 | 0.94 | 0.91 | 0.91 | 0.9 | 0.9 |
| 4 | 188201350 | rs4862792 | 0.8 | -0.453 | -0.379 | 0.82 | 0.001 | 0 | 0 | 0 | 0.85 | 0.86 | 0.89 | 0.83 | 0.88 | 0.91 |
| 4 | 188209801 | rs13126689 | -0.06 | -0.256 | -0.696 | 0.37 | 0.004 | 0.024 | 0.024 | 0 | 0.21 | 0.13 | 0.12 | 0.07 | 0.19 | 0.1 |
| 4 | 188213866 | rs4308418 | -1.337 | -0.499 | -0.086 | 0.131 | -0.008 | -0.054 | -0.054 | 0.612 | 0.39 | 0.37 | 0.24 | 0.09 | 0.34 | 0.19 |
| 4 | 188223037 | rs9993067 | 0.21 | -0.235 | 0.132 | 0.821 | -0.015 | -0.017 | -0.017 | 0 | 0.74 | 0.62 | 0.75 | 0.77 | 0.77 | 0.88 |
| 4 | 188223165 | rs2045725 | 0.376 | 0.473 | -0.493 | 1.161 | -0.005 | -0.004 | -0.004 | 0.088 | 0.39 | 0.32 | 0.47 | 0.43 | 0.43 | 0.55 |
| 4 | 188227951 | rs976000 | -0.697 | 0.562 | -0.839 | 1.03 | -0.01 | -0.01 | -0.01 | 0 | 0.77 | 0.86 | 0.64 | 0.71 | 0.71 | 0.78 |
| 4 | 188230701 | rs6836534 | -0.22 | -0.317 | -0.791 | 1.074 | 0.023 | -0.022 | -0.022 | 0 | 0.41 | 0.26 | 0.36 | 0.53 | 0.33 | 0.37 |
| 4 | 188258978 | rs1319774 | 0.348 | -0.56 | 0.036 | 1.104 | 0.016 | 0.011 | 0.011 | 0.025 | 0.52 | 0.44 | 0.6 | 0.38 | 0.66 | 0.53 |
| 4 | 188265382 | rs1156741 | 0.4 | -0.621 | 0.258 | 0.756 | -0.047 | -0.015 | -0.015 | 0.684 | 0.82 | 0.61 | 0.92 | 0.87 | 0.91 | 0.82 |
| 4 | 188268391 | rs7679761 | 0.138 | -0.839 | 0.378 | 0.85 | 0.012 | -0.01 | -0.01 | 0 | 0.66 | 0.68 | 0.64 | 0.38 | 0.76 | 0.63 |
| 4 | 188280616 | rs1870852 | 0.389 | -0.523 | 0.359 | 1.035 | -0.005 | -0.066 | -0.066 | 0 | 0.26 | 0.16 | 0.27 | 0.51 | 0.24 | 0.31 |
| 4 | 188284813 | rs2376165 | 0.425 | -0.547 | 0.35 | 1.035 | -0.01 | -0.071 | -0.071 | 0.099 | 0.25 | 0.13 | 0.27 | 0.51 | 0.24 | 0.31 |
| 4 | 188289740 | rs13107031 | -0.233 | -0.778 | 0.574 | 0.999 | 0.011 | -0.022 | -0.022 | 0 | 0.33 | 0.25 | 0.29 | 0.46 | 0.26 | 0.32 |
| 4 | 188292015 | rs11132443 | 0.535 | 0.217 | 1.068 | 0.784 | 0.003 | -0.018 | -0.018 | 0 | 0.63 | 0.37 | 0.63 | 0.82 | 0.61 | 0.61 |
| 4 | 188294980 | rs3911752 | 0.224 | 0.28 | 1.393 | 0.741 | 0.016 | -0.039 | -0.039 | 0.829 | 0.69 | 0.58 | 0.61 | 0.86 | 0.65 | 0.68 |
| 4 | 188295836 | rs12505423 | 0.992 | 0.126 | 1.274 | 0.853 | 0.03 | -0.002 | -0.002 | 1.343 | 0.39 | 0.25 | 0.27 | 0.38 | 0.36 | 0.38 |
| 4 | 188311821 | rs10018680 | -0.396 | -0.149 | 1.654 | 1.042 | 0.016 | 0.06 | 0.06 | 0 | 0.71 | 0.59 | 0.66 | 0.51 | 0.7 | 0.69 |
| 4 | 188317381 | rs4862801 | 0.796 | -0.096 | 1.177 | 1.013 | -0.021 | -0.085 | -0.085 | 0 | 0.26 | 0.1 | 0.29 | 0.51 | 0.25 | 0.27 |
| 4 | 188332196 | rs2217486 | 0.734 | -0.042 | 1.148 | 1.028 | -0.027 | -0.084 | -0.084 | 0 | 0.27 | 0.08 | 0.31 | 0.43 | 0.26 | 0.29 |
| 4 | 188345346 | rs11132449 | 0.493 | 0.018 | 1.415 | 0.841 | -0.002 | 0.003 | 0.003 | 0 | 0.71 | 0.71 | 0.65 | 0.76 | 0.71 | 0.69 |
| 4 | 188345368 | rs1431005 | 1.018 | 0.02 | 1.675 | 0.864 | 0.002 | 0.001 | 0.001 | 0.112 | 0.38 | 0.37 | 0.36 | 0.36 | 0.4 | 0.4 |
| 4 | 188351787 | rs11724481 | -0.4 | 0.223 | 1.455 | 0.657 | -0.003 | -0.009 | -0.009 | 0.023 | 0.08 | 0.12 | 0.03 | 0.03 | 0.06 | 0.1 |
| 4 | 188364027 | rs11132451 | 0.103 | 0.21 | 1.521 | 0.693 | 0.001 | 0.003 | 0.003 | 0 | 0.6 | 0.46 | 0.66 | 0.53 | 0.64 | 0.62 |
| 4 | 188364088 | rs12641785 | 0.103 | 0.21 | 1.521 | 0.693 | 0.001 | 0.003 | 0.003 | 0 | 0.6 | 0.46 | 0.66 | 0.53 | 0.63 | 0.62 |
| 4 | 188365395 | rs1864603 | -0.273 | 0.205 | 1.547 | 0.691 | 0.008 | 0.015 | 0.015 | 0 | 0.39 | 0.44 | 0.32 | 0.47 | 0.36 | 0.38 |
| 4 | 188369068 | rs12508101 | -0.145 | 0.202 | 1.606 | 0.684 | -0.002 | 0.01 | 0.01 | 0 | 0.63 | 0.58 | 0.7 | 0.53 | 0.69 | 0.62 |
| 4 | 188376505 | rs12651353 | 1.104 | 0.164 | 1.612 | 0.559 | -0.005 | -0.018 | -0.018 | 0 | 0.66 | 0.37 | 0.62 | 0.77 | 0.7 | 0.72 |
| 4 | 188381449 | rs6553069 | 0.04 | 0.174 | 1.374 | 0.552 | -0.002 | 0.003 | 0.003 | 0 | 0.78 | 0.81 | 0.68 | 0.83 | 0.76 | 0.81 |
| 4 | 188391246 | rs7691606 | 0.697 | 0.343 | 1.678 | 0.607 | 0.002 | -0.003 | -0.003 | 0.336 | 0.72 | 0.64 | 0.65 | 0.78 | 0.71 | 0.73 |
| 4 | 188394768 | rs13141517 | 0.619 | 0.248 | 1.585 | 0.802 | -0.033 | -0.021 | -0.021 | 0 | 0.29 | 0.11 | 0.27 | 0.35 | 0.31 | 0.3 |
| 4 | 188398663 | rs4862809 | 1.486 | 0.254 | 1.745 | 0.686 | 0.053 | 0.018 | 0.018 | 0.306 | 0.94 | 0.73 | 0.94 | 0.97 | 0.94 | 0.99 |
| 4 | 188398695 | rs4862810 | 0.703 | 0.241 | 1.82 | 0.704 | -0.002 | -0.004 | -0.004 | 0 | 0.28 | 0.24 | 0.26 | 0.39 | 0.25 | 0.29 |
| 4 | 188399309 | rs13133599 | 1.513 | 0.343 | 2.776 | 0.543 | 0.02 | -0.013 | -0.013 | 0 | 0.71 | 0.59 | 0.62 | 0.79 | 0.67 | 0.71 |
| 4 | 188406503 | rs7667497 | -1.083 | 0.426 | 2.176 | 0.531 | -0.017 | -0.003 | -0.003 | 0 | 0.24 | 0.13 | 0.33 | 0.21 | 0.27 | 0.28 |
| 4 | 188411479 | rs10027738 | 1.516 | 0.417 | 2.281 | 0.484 | -0.001 | -0.001 | -0.001 | 0 | 0.71 | 0.67 | 0.61 | 0.75 | 0.67 | 0.71 |
| 4 | 188420909 | rs1367555 | 2.069 | 0.753 | 2.277 | 0.677 | 0.024 | -0.026 | -0.026 | 0.001 | 0.95 | 0.79 | 0.94 | 1 | 0.94 | 0.99 |
| 4 | 188423355 | rs10001779 | 2.014 | 0.764 | 2.282 | 0.682 | 0.025 | -0.025 | -0.025 | 0 | 0.95 | 0.79 | 0.94 | 1 | 0.94 | 0.99 |
| 4 | 188426129 | rs4861749 | 1.756 | 0.725 | 1.915 | 0.66 | -0.007 | 0.014 | 0.014 | 0.26 | 0.67 | 0.2 | 0.58 | 0.76 | 0.67 | 0.7 |
| 4 | 188428740 | rs10026177 | 0.669 | 0.926 | 1.489 | 0.734 | -0.004 | -0.006 | -0.006 | 0.364 | 0.78 | 0.79 | 0.66 | 0.69 | 0.74 | 0.65 |
| 4 | 188431411 | rs10032833 | -2.482 | 0.486 | 1.27 | 0.41 | 0.09 | -0.011 | -0.011 | 0 | 0.07 | 0.36 | 0.09 | 0 | 0.08 | 0.01 |
| 4 | 188432233 | rs11929993 | -1.916 | 0.481 | 1.276 | 0.407 | 0.071 | 0.004 | 0.004 | 0.036 | 0.05 | 0.27 | 0.06 | 0 | 0.06 | 0.01 |
| 4 | 188434677 | rs28475332 | -1.746 | 0.537 | 1.113 | 0.441 | 0.09 | -0.011 | -0.011 | 0.076 | 0.07 | 0.36 | 0.09 | 0 | 0.08 | 0.01 |
| 4 | 188469538 | rs6857356 | 0.912 | -0.024 | -0.496 | 0.481 | 0.036 | 0.024 | 0.024 | 0 | 0.74 | 0.16 | 0.82 | 0.8 | 0.85 | 0.76 |
| 4 | 188479688 | rs1532857 | 0.071 | 0.181 | 0.223 | 0.433 | -0.008 | -0.001 | -0.001 | 1.764 | 0.64 | 0.58 | 0.63 | 0.37 | 0.65 | 0.6 |
| 4 | 188480913 | rs1553566 | 0.185 | 0.035 | -0.221 | 0.533 | -0.009 | -0.078 | -0.078 | 0.086 | 0.24 | 0.08 | 0.22 | 0.5 | 0.25 | 0.28 |
| 4 | 188486138 | rs7677258 | 0.394 | 0.427 | -0.157 | -0.197 | -0.023 | 0.001 | 0.001 | 0 | 0.57 | 0.31 | 0.46 | 0.51 | 0.5 | 0.61 |
| 4 | 188493082 | rs6812381 | -0.809 | 0.636 | -0.52 | -0.408 | 0 | 0.016 | 0.016 | 0 | 0.47 | 0.19 | 0.54 | 0.49 | 0.56 | 0.44 |
| 4 | 188498590 | rs11734029 | 0.757 | 0.757 | -0.75 | -0.409 | -0.066 | -0.133 | -0.133 | 0.064 | 0.24 | 0.02 | 0.18 | 0.48 | 0.25 | 0.28 |
| 4 | 188499628 | rs871603 | 0.297 | 0.718 | -0.694 | -0.515 | -0.055 | 0.088 | 0.088 | 0 | 0.16 | 0.05 | 0.29 | 0.01 | 0.24 | 0.11 |
| 4 | 188509004 | rs6826401 | -0.228 | 0.009 | -0.172 | -0.549 | 0.008 | -0.009 | -0.009 | 0 | 0.39 | 0.33 | 0.25 | 0.59 | 0.33 | 0.41 |
| 4 | 188522356 | rs1505505 | -0.079 | -0.052 | -0.267 | -0.587 | -0.026 | -0.049 | -0.049 | 0 | 0.37 | 0.11 | 0.23 | 0.49 | 0.37 | 0.39 |
| 4 | 188526344 | rs13147499 | -0.405 | -0.089 | 0.041 | -0.61 | 0.005 | -0.013 | -0.013 | 0.527 | 0.41 | 0.19 | 0.25 | 0.49 | 0.38 | 0.38 |
| 4 | 188531799 | rs13150936 | -0.342 | -0.065 | 0.018 | -0.619 | 0.008 | -0.022 | -0.022 | 0 | 0.4 | 0.17 | 0.24 | 0.49 | 0.37 | 0.38 |
| 4 | 188536462 | rs1876072 | -0.038 | 0.128 | 0.568 | -0.769 | 0.041 | -0.051 | -0.051 | 4.236 | 0.75 | 0.54 | 0.61 | 0.87 | 0.69 | 0.74 |
| 4 | 188547880 | rs1354634 | -0.138 | 0.126 | 0.592 | -0.77 | 0.03 | -0.036 | -0.036 | 0 | 0.76 | 0.6 | 0.62 | 0.87 | 0.69 | 0.74 |
| 4 | 188554682 | rs2279883 | 0.362 | 0.154 | 0.225 | -0.642 | -0.01 | -0.088 | -0.088 | 0 | 0.32 | 0.08 | 0.21 | 0.49 | 0.32 | 0.34 |
| 4 | 188570203 | rs1505509 | -0.39 | 0.198 | 0.284 | -0.597 | 0.01 | -0.004 | -0.004 | 0 | 0.41 | 0.34 | 0.25 | 0.5 | 0.37 | 0.38 |
| 4 | 188575728 | rs10028895 | -0.953 | 0.179 | 0.358 | -0.859 | -0.019 | -0.059 | -0.059 | 0 | 0.16 | 0.26 | 0.07 | 0.01 | 0.1 | 0.11 |
| 4 | 188576298 | rs10866300 | 0.123 | 0.249 | 0.391 | -1.273 | -0.014 | 0.003 | 0.003 | 0 | 0.73 | 0.79 | 0.54 | 0.97 | 0.64 | 0.78 |
| 4 | 188581325 | rs2292161 | 1.811 | 0.077 | 0.44 | -1.041 | 0.003 | -0.011 | -0.011 | 0 | 0.3 | 0.24 | 0.26 | 0.35 | 0.25 | 0.29 |
| 4 | 188583907 | rs9992899 | -0.073 | 0.406 | 0.52 | -1.134 | 0.008 | -0.053 | -0.053 | 0 | 0.77 | 0.72 | 0.59 | 0.96 | 0.73 | 0.78 |
| 4 | 188586853 | rs11929702 | 0.93 | 0.349 | 0.55 | -1.162 | -0.001 | -0.111 | -0.111 | 0.006 | 0.64 | 0.49 | 0.52 | 0.95 | 0.63 | 0.69 |
| 4 | 188590603 | rs1466956 | -0.566 | 0.434 | 0.444 | -1.134 | 0.002 | -0.043 | -0.043 | 0.08 | 0.82 | 0.8 | 0.7 | 0.99 | 0.78 | 0.89 |
| 4 | 188590745 | rs11930461 | -1.067 | 0.463 | 0.349 | -1.039 | 0.001 | -0.019 | -0.019 | 0 | 0.2 | 0.25 | 0.22 | 0.04 | 0.22 | 0.19 |
| 4 | 188597930 | rs7684752 | 0.731 | 0.574 | 0.264 | -1.061 | 0.007 | -0.035 | -0.035 | 0 | 0.71 | 0.71 | 0.55 | 0.96 | 0.64 | 0.75 |
| 4 | 188608123 | rs6826851 | 1.806 | 0.279 | 0.261 | -1.006 | 0.003 | -0.005 | -0.005 | 0 | 0.31 | 0.25 | 0.33 | 0.34 | 0.28 | 0.3 |
| 4 | 188608857 | rs7687115 | 0.851 | 0.601 | 0.339 | -1.024 | -0.007 | 0.001 | 0.001 | 0.004 | 0.42 | 0.34 | 0.43 | 0.38 | 0.45 | 0.43 |

| | | | | | | | | | | | | | | | | |
|---|---|---|---|---|---|---|---|---|---|---|---|---|---|---|---|---|
| 4 | 188615607 | rs9992966 | -0.996 | 0.719 | 1.144 | -1.204 | -0.063 | 0.027 | 0.027 | 0 | 0.18 | 0.38 | 0.12 | 0.17 | 0.08 | 0.17 |
| 4 | 188616420 | rs9993712 | -1.006 | 0.721 | 1.145 | -1.203 | -0.064 | 0.027 | 0.027 | 0 | 0.18 | 0.38 | 0.11 | 0.17 | 0.08 | 0.17 |
| 4 | 188627935 | rs10026452 | -1.807 | 0.695 | 1.035 | -1.198 | -0.006 | 0.004 | 0.004 | 0.011 | 0.29 | 0.53 | 0.21 | 0.21 | 0.25 | 0.3 |
| 4 | 188628654 | rs12648438 | -0.172 | 0.696 | 1.027 | -1.228 | 0 | -0.043 | -0.043 | 0 | 0.82 | 0.81 | 0.71 | 0.99 | 0.81 | 0.88 |
| 4 | 188632948 | rs4266338 | 1.878 | 0.702 | 0.829 | -1.179 | 0.004 | 0.07 | 0.07 | 0.162 | 0.53 | 0.28 | 0.49 | 0.78 | 0.56 | 0.58 |
| 4 | 188646845 | rs10008174 | -1.368 | 0.932 | 0.653 | -1.314 | 0.058 | -0.047 | -0.047 | 0 | 0.17 | 0.32 | 0.18 | 0.03 | 0.28 | 0.17 |
| 4 | 188648818 | rs11132478 | 2.155 | 0.849 | 0.305 | -1.369 | -0.006 | 0.073 | 0.073 | 0.094 | 0.51 | 0.32 | 0.47 | 0.79 | 0.5 | 0.55 |
| 4 | 188650233 | rs7438821 | 0.259 | 0.971 | -0.693 | -1.428 | -0.035 | 0.082 | 0.082 | 0 | 0.14 | 0.01 | 0.25 | 0.01 | 0.16 | 0.13 |
| 4 | 188652223 | rs11723292 | -0.329 | 0.997 | -0.522 | -1.263 | -0.014 | 0.003 | 0.003 | 0 | 0.09 | 0.05 | 0.12 | 0.05 | 0.17 | 0.13 |
| 4 | 188658741 | rs6553094 | -0.396 | 0.724 | -0.678 | -1.592 | -0.001 | -0.009 | -0.009 | 0.101 | 0.78 | 0.85 | 0.69 | 1 | 0.78 | 0.88 |
| 4 | 188661045 | rs13139954 | -0.068 | 1.09 | -0.353 | -1.073 | 0 | -0.001 | -0.001 | 0 | 0.09 | 0.12 | 0.13 | 0.05 | 0.15 | 0.11 |
| 4 | 188665614 | rs13151215 | -0.281 | 0.93 | -0.349 | -1.055 | -0.003 | -0.002 | -0.002 | 0 | 0.15 | 0.17 | 0.16 | 0.05 | 0.25 | 0.15 |
| 4 | 188669480 | rs13114385 | -0.504 | 0.912 | -0.363 | -0.989 | 0.006 | -0.014 | -0.014 | 0 | 0.19 | 0.26 | 0.18 | 0.08 | 0.28 | 0.22 |
| 4 | 188675224 | rs7658700 | -0.521 | 1.032 | -0.363 | -0.846 | -0.002 | 0.002 | 0.002 | 0.674 | 0.28 | 0.37 | 0.23 | 0.18 | 0.31 | 0.33 |
| 4 | 188680866 | rs9992060 | -0.942 | 1.085 | -0.825 | -0.859 | -0.007 | 0.028 | 0.028 | 0.504 | 0.35 | 0.8 | 0.28 | 0.19 | 0.36 | 0.36 |
| 4 | 188690307 | rs13148546 | 0.719 | 0.946 | -0.501 | -1.032 | 0 | -0.121 | -0.121 | 0 | 0.72 | 0.48 | 0.65 | 0.99 | 0.72 | 0.85 |
| 4 | 188698095 | rs7683009 | -0.675 | 0.938 | -1.223 | -1.441 | -0.039 | 0.158 | 0.158 | 1.565 | 0.43 | 0.22 | 0.49 | 0.11 | 0.5 | 0.53 |
| 4 | 188707091 | rs4861758 | 0.239 | 0.258 | -0.555 | -0.29 | -0.028 | 0.049 | 0.049 | 0 | 0.85 | 0.27 | 0.9 | 0.66 | 0.92 | 0.89 |
| 4 | 188716736 | rs6849009 | 0.29 | 0.869 | -1.27 | -0.712 | -0.081 | 0.08 | 0.08 | 0.507 | 0.21 | 0.02 | 0.23 | 0.08 | 0.26 | 0.23 |
| 4 | 188717853 | rs4318702 | 0.885 | 0.131 | -0.305 | -0.38 | -0.003 | 0.03 | 0.03 | 0.069 | 0.7 | 0.51 | 0.71 | 0.63 | 0.7 | 0.65 |
| 4 | 188721512 | rs4470690 | 1.345 | -0.921 | 0.268 | -0.512 | -0.011 | -0.039 | -0.039 | 0.006 | 0.66 | 0.68 | 0.77 | 0.88 | 0.72 | 0.74 |
| 4 | 188721544 | rs4481281 | 1.732 | -1.143 | 0.696 | 0.448 | -0.004 | 0.004 | 0.004 | 1.086 | 0.73 | 0.31 | 0.7 | 0.67 | 0.83 | 0.79 |
| 4 | 188724806 | rs6845047 | -1.916 | -1.066 | 0.956 | 0.521 | -0.003 | -0.007 | -0.007 | 0 | 0.28 | 0.69 | 0.19 | 0.18 | 0.22 | 0.31 |
| 4 | 188730709 | rs10017989 | 1.3 | -1.65 | 0.854 | 0.444 | -0.011 | 0.017 | 0.017 | 0 | 0.66 | 0.4 | 0.62 | 0.52 | 0.76 | 0.43 |
| 4 | 188731055 | rs10028606 | 2.035 | -1.651 | 0.829 | 0.426 | 0.052 | -0.007 | -0.007 | 0 | 0.51 | 0.27 | 0.53 | 0.52 | 0.64 | 0.39 |
| 4 | 188734615 | rs9996150 | -1.184 | -1.442 | 1.194 | 0.538 | -0.028 | -0.1 | -0.1 | 0.651 | 0.2 | 0.38 | 0.12 | 0.03 | 0.13 | 0.09 |
| 4 | 188735086 | rs6829286 | -1.643 | -1.468 | 1.407 | 0.568 | -0.041 | -0.085 | -0.085 | 0 | 0.23 | 0.57 | 0.13 | 0.13 | 0.14 | 0.24 |
| 4 | 188736913 | rs6816835 | -0.847 | -1.484 | 1.188 | 0.517 | 0.01 | 0.008 | 0.008 | 0.076 | 0.19 | 0.14 | 0.08 | 0.03 | 0.13 | 0.09 |
| 4 | 188744214 | rs4862831 | -0.439 | -1.502 | 1.101 | 0.83 | 0 | 0.027 | 0.027 | 0 | 0.07 | 0.07 | 0.19 | 0.32 | 0.06 | 0.18 |
| 4 | 188744789 | rs13125278 | 1.592 | -1.688 | 1.677 | 0.811 | -0.009 | -0.022 | -0.022 | 0 | 0.69 | 0.43 | 0.83 | 0.9 | 0.8 | 0.77 |
| 4 | 188746046 | rs4566706 | 2.181 | -1.68 | 1.66 | 0.803 | 0.038 | -0.021 | -0.021 | 0 | 0.62 | 0.25 | 0.64 | 0.58 | 0.74 | 0.59 |
| 4 | 188770791 | rs13120072 | 1.179 | -1.668 | 2.024 | 0.708 | -0.052 | -0.141 | -0.141 | 0.502 | 0.76 | 0.58 | 0.88 | 0.97 | 0.85 | 0.86 |
| 4 | 188786838 | rs12648158 | -1.276 | -1.495 | 1.647 | 0.885 | 0.001 | 0.091 | 0.091 | 0 | 0.12 | 0.25 | 0.21 | 0.32 | 0.1 | 0.26 |
| 4 | 188792218 | rs4377642 | 1.656 | -1.399 | 1.684 | 0.814 | 0.004 | 0.047 | 0.047 | 0 | 0.83 | 0.71 | 0.77 | 0.74 | 0.86 | 0.79 |
| 4 | 188801533 | rs4862838 | 1.481 | -1.864 | 1.599 | 0.76 | -0.032 | -0.027 | -0.027 | 0 | 0.73 | 0.47 | 0.85 | 0.81 | 0.83 | 0.69 |
| 4 | 188802381 | rs12644677 | 2.066 | -1.852 | 1.168 | 0.816 | 0.047 | -0.045 | -0.045 | 0 | 0.62 | 0.16 | 0.62 | 0.58 | 0.74 | 0.59 |
| 4 | 188804628 | rs13139939 | 0.157 | -1.884 | 1.037 | 0.691 | 0.014 | 0.038 | 0.038 | 0 | 0.16 | 0.05 | 0.11 | 0.06 | 0.11 | 0.24 |
| 4 | 188805951 | rs4618368 | 1.614 | -1.735 | 1.209 | 0.72 | -0.007 | 0.069 | 0.069 | 0 | 0.91 | 0.79 | 0.94 | 0.81 | 0.95 | 0.75 |
| 4 | 188815970 | rs4315850 | -2.24 | -1.875 | 0.596 | 0.981 | 0.015 | 0.023 | 0.023 | 0 | 0.55 | 0.83 | 0.55 | 0.58 | 0.39 | 0.66 |
| 4 | 188829088 | rs10866303 | -1.508 | -1.914 | 0.731 | 0.801 | 0.012 | 0.014 | 0.014 | 0 | 0.41 | 0.58 | 0.29 | 0.15 | 0.28 | 0.41 |
| 4 | 188834175 | rs11132490 | -1.537 | -1.879 | 0.65 | 0.768 | -0.024 | 0.007 | 0.007 | 0.068 | 0.36 | 0.56 | 0.45 | 0.56 | 0.25 | 0.57 |
| 4 | 188840051 | rs4862840 | -1.517 | -2.017 | 0.46 | 0.709 | 0.004 | 0 | 0 | 0.473 | 0.39 | 0.41 | 0.42 | 0.34 | 0.28 | 0.41 |
| 4 | 188842209 | rs6813823 | -0.121 | -1.321 | 0.238 | 0.948 | -0.037 | 0.121 | 0.121 | 0.192 | 0.16 | 0.24 | 0.28 | 0.34 | 0.1 | 0.3 |
| 4 | 188843305 | rs6553113 | -0.196 | -1.333 | -0.553 | 0.72 | -0.005 | -0.025 | -0.025 | 0.23 | 0.28 | 0.37 | 0.17 | 0.19 | 0.2 | 0.3 |
| 4 | 188844211 | rs10866305 | -1.141 | -0.679 | -1.431 | 0.975 | -0.04 | -0.031 | -0.031 | 0 | 0.59 | 0.89 | 0.58 | 0.57 | 0.41 | 0.67 |
| 4 | 188848924 | rs7667835 | -0.518 | -0.776 | -0.672 | 1.496 | -0.003 | 0.016 | 0.016 | 0.012 | 0.15 | 0.13 | 0.27 | 0.34 | 0.1 | 0.31 |
| 4 | 188851691 | rs6821751 | -1.27 | -1.098 | -0.685 | 1.14 | 0.015 | -0.009 | -0.009 | 0 | 0.24 | 0.21 | 0.19 | 0.35 | 0.15 | 0.51 |
| 4 | 188856373 | rs10018412 | -0.461 | -0.31 | -0.819 | 0.277 | -0.015 | 0.044 | 0.044 | 0.02 | 0.61 | 0.66 | 0.63 | 0.92 | 0.56 | 0.73 |
| 4 | 188856430 | rs11936270 | -0.751 | -0.432 | 0.088 | 0.569 | -0.037 | 0.035 | 0.035 | 0 | 0.13 | 0.28 | 0.11 | 0.2 | 0.05 | 0.12 |
| 4 | 188859908 | rs2131290 | -1.008 | 0.247 | 0.266 | 0.734 | -0.003 | 0.032 | 0.032 | 0 | 0.18 | 0.25 | 0.19 | 0.35 | 0.16 | 0.41 |
| 4 | 188865510 | rs7696985 | 0.015 | 0.807 | 0.099 | 0.224 | -0.003 | -0.011 | -0.011 | 0 | 0.32 | 0.3 | 0.37 | 0.07 | 0.41 | 0.18 |
| 4 | 188887420 | rs12644789 | -1.184 | 1.049 | 0.304 | 1.233 | 0.006 | -0.024 | -0.024 | 2.86 | 0.22 | 0.15 | 0.28 | 0.4 | 0.17 | 0.43 |
| 4 | 188892958 | rs4861762 | -1.25 | 1.183 | 0.552 | 1.118 | -0.002 | 0.045 | 0.045 | 0 | 0.22 | 0.25 | 0.29 | 0.42 | 0.18 | 0.45 |
| 4 | 188898937 | rs6553120 | -0.982 | 1.234 | -0.101 | 0.139 | 0 | 0 | 0 | 0 | 0.18 | 0.22 | 0.1 | 0.18 | 0.15 | 0.29 |
| 4 | 188899298 | rs13146799 | -0.978 | 1.3 | -0.132 | 0.141 | 0.001 | 0 | 0 | 0 | 0.18 | 0.21 | 0.11 | 0.18 | 0.16 | 0.29 |
| 4 | 188904235 | rs7439650 | 0.118 | 1.371 | -0.746 | -0.543 | -0.016 | 0.066 | 0.066 | 1.593 | 0.43 | 0.35 | 0.37 | 0.12 | 0.48 | 0.3 |
| 4 | 188905061 | rs1105763 | -0.251 | 1.265 | -0.474 | -0.281 | 0.004 | 0.017 | 0.017 | 0 | 0.55 | 0.61 | 0.43 | 0.28 | 0.55 | 0.49 |
| 4 | 188909667 | rs6553122 | 0.019 | 0.827 | -0.709 | -0.392 | 0 | 0 | 0 | 0.731 | 0.33 | 0.31 | 0.48 | 0.7 | 0.28 | 0.49 |
| 4 | 188921440 | rs4862848 | -0.51 | 1.277 | -1.452 | -0.786 | -0.059 | 0.14 | 0.14 | 0.545 | 0.68 | 0.91 | 0.76 | 0.96 | 0.64 | 0.8 |
| 4 | 188924979 | rs6815391 | 0.038 | 0.844 | -0.658 | -0.332 | -0.013 | 0.014 | 0.014 | 0 | 0.27 | 0.3 | 0.38 | 0.69 | 0.18 | 0.41 |
| 4 | 188932750 | rs6553126 | 0.114 | 1.29 | -0.739 | -0.843 | -0.012 | 0.012 | 0.012 | 0 | 0.62 | 0.65 | 0.4 | 0.25 | 0.53 | 0.53 |
| 4 | 188933651 | rs6840208 | 0.114 | 1.29 | -0.739 | -0.842 | -0.012 | 0.012 | 0.012 | 0 | 0.62 | 0.64 | 0.4 | 0.24 | 0.53 | 0.53 |
| 4 | 188935662 | rs6553129 | -0.065 | 1.184 | -0.835 | -1.145 | -0.029 | 0.03 | 0.03 | 0.66 | 0.63 | 0.78 | 0.48 | 0.26 | 0.53 | 0.47 |
| 4 | 188936629 | rs11733349 | 0.297 | 1.098 | -1.228 | -2.027 | 0.017 | 0.121 | 0.121 | 0.092 | 0.29 | 0.08 | 0.15 | 0.04 | 0.23 | 0.12 |
| 4 | 188938468 | rs2172340 | 0.545 | 1.103 | -0.928 | -2.186 | 0.001 | -0.033 | -0.033 | 0 | 0.17 | 0.18 | 0.2 | 0 | 0.22 | 0.06 |
| 4 | 188939849 | rs6553132 | -0.189 | 0.992 | -0.575 | -1.944 | 0.001 | -0.038 | -0.038 | 0 | 0.49 | 0.42 | 0.36 | 0.04 | 0.45 | 0.18 |
| 4 | 188943843 | rs2131288 | -0.138 | 1.447 | -0.391 | -1.489 | 0.019 | -0.079 | -0.079 | 0 | 0.15 | 0.25 | 0.09 | 0 | 0.16 | 0.12 |
| 4 | 188943890 | rs2131287 | -0.379 | 1.561 | -0.506 | -0.478 | -0.003 | -0.007 | -0.007 | 0.469 | 0.38 | 0.67 | 0.58 | 0.63 | 0.43 | 0.41 |
| 4 | 188945315 | rs6553136 | 0.592 | 1.49 | -0.63 | -0.765 | 0.008 | -0.008 | -0.008 | 0 | 0.8 | 0.71 | 0.69 | 0.85 | 0.76 | 0.7 |

| | | | | | | | | | | | | | | | | |
|---|---|---|---|---|---|---|---|---|---|---|---|---|---|---|---|---|
| 4 | 188953780 | rs6553138 | 0.014 | 0.648 | -0.749 | -1.34 | -0.008 | -0.047 | -0.047 | 0 | 0.59 | 0.57 | 0.66 | 0.86 | 0.61 | 0.83 |
| 4 | 188956686 | rs6853977 | 0.082 | 0.412 | -0.848 | -2.02 | 0.012 | -0.098 | -0.098 | 0 | 0.4 | 0.42 | 0.33 | 0.09 | 0.48 | 0.25 |
| 4 | 188957800 | rs2889360 | 0.058 | -0.115 | -1.361 | -2.358 | 0.049 | 0.172 | 0.172 | 0.467 | 0.5 | 0.22 | 0.45 | 0.11 | 0.59 | 0.31 |
| 4 | 188959863 | rs2376393 | 0.977 | -0.121 | -1.778 | -1.719 | -0.025 | -0.074 | -0.074 | 0.306 | 0.15 | 0.02 | 0.31 | 0.72 | 0.15 | 0.26 |
| 4 | 188961336 | rs10004698 | -0.166 | -0.187 | -2.216 | -2.246 | 0.038 | 0.138 | 0.138 | 0 | 0.37 | 0.12 | 0.31 | 0.15 | 0.32 | 0.21 |
| 4 | 188962325 | rs7437246 | -0.554 | 0.021 | -2.336 | -2.084 | 0.001 | 0.278 | 0.278 | 0 | 0.49 | 0.08 | 0.61 | 0.81 | 0.46 | 0.36 |
| 4 | 188963197 | rs1586887 | 1.542 | -0.289 | -2.163 | -2.039 | -0.013 | 0.045 | 0.045 | 0 | 0.11 | 0 | 0.15 | 0.01 | 0.12 | 0.04 |
| 4 | 188963851 | rs4440286 | -0.118 | -0.066 | -2.046 | -1.881 | 0.016 | 0.147 | 0.147 | 0 | 0.41 | 0.09 | 0.53 | 0.79 | 0.32 | 0.34 |
| 4 | 188965567 | rs2068445 | 0.501 | -0.285 | -1.899 | -1.767 | 0.035 | 0.109 | 0.109 | 0 | 0.25 | 0.06 | 0.2 | 0.06 | 0.15 | 0.08 |
| 4 | 188965971 | rs6553144 | 0.444 | 0.246 | -1.026 | -1.442 | 0.009 | 0.01 | 0.01 | 0 | 0.08 | 0.16 | 0.05 | 0.11 | 0.07 | 0.35 |
| 4 | 188971489 | rs12501116 | -0.131 | 0.282 | -0.734 | -1.61 | 0.003 | 0.031 | 0.031 | 0 | 0.47 | 0.39 | 0.63 | 0.8 | 0.42 | 0.39 |
| 4 | 188978587 | rs6847145 | -1.311 | 0.399 | -0.728 | -1.581 | 0.007 | -0.073 | -0.073 | 0 | 0.77 | 0.75 | 0.84 | 1 | 0.72 | 0.86 |
| 4 | 188981386 | rs1540663 | 0.178 | 0.281 | -1.151 | -0.604 | -0.035 | -0.021 | -0.021 | 0 | 0.38 | 0.19 | 0.52 | 0.33 | 0.39 | 0.26 |
| 4 | 188985493 | rs11132507 | -0.321 | 0.112 | -0.636 | -0.717 | 0 | -0.076 | -0.076 | 1.554 | 0.7 | 0.56 | 0.65 | 0.88 | 0.68 | 0.49 |
| 4 | 188985500 | rs11132508 | -0.33 | 0.116 | -0.633 | -0.715 | 0 | -0.076 | -0.076 | 0 | 0.7 | 0.56 | 0.65 | 0.88 | 0.68 | 0.49 |
| 4 | 188990434 | rs9997579 | -0.245 | 0.107 | -1.39 | -0.786 | 0.003 | 0.101 | 0.101 | 0 | 0.65 | 0.2 | 0.66 | 0.93 | 0.64 | 0.77 |
| 4 | 188990955 | rs4611976 | 0.76 | -0.011 | -1.287 | 0.686 | 0.021 | -0.025 | -0.025 | 0 | 0.1 | 0.02 | 0.14 | 0.53 | 0.06 | 0.14 |
| 4 | 188998692 | rs12644540 | -0.605 | 0.09 | -1.092 | 0.3 | 0.002 | -0.003 | -0.003 | 0 | 0.25 | 0.24 | 0.23 | 0.64 | 0.17 | 0.54 |
| 4 | 188999227 | rs6853509 | -0.62 | 0.089 | -1.64 | -0.203 | -0.073 | -0.008 | -0.008 | 0 | 0.56 | 0.89 | 0.52 | 0.71 | 0.46 | 0.74 |
| 4 | 189007284 | rs4299646 | 0.938 | -0.016 | -1.365 | -0.316 | 0.006 | 0.017 | 0.017 | 0 | 0.14 | 0.12 | 0.07 | 0.01 | 0.13 | 0.05 |
| 4 | 189011174 | rs12647135 | -1.304 | 0.034 | -1.304 | -0.587 | -0.012 | 0.146 | 0.146 | 0 | 0.55 | 0.81 | 0.57 | 0.91 | 0.46 | 0.73 |
| 4 | 189012809 | rs2279551 | 1.255 | 0.085 | -1.195 | 0.393 | 0.015 | -0.049 | -0.049 | 0.076 | 0.11 | 0.02 | 0.16 | 0.49 | 0.06 | 0.12 |
| 4 | 189018613 | rs10004012 | 0.088 | -0.046 | -0.477 | 0.296 | -0.042 | 0.123 | 0.123 | 0 | 0.17 | 0.33 | 0.2 | 0.55 | 0.1 | 0.18 |
| 4 | 189020348 | rs902910 | 0.424 | 0.659 | -0.386 | 0.39 | 0.006 | 0.006 | 0.006 | 0 | 0.1 | 0.16 | 0.04 | 0.15 | 0.08 | 0.37 |
| 4 | 189022420 | rs4862860 | 0.045 | 0.63 | -0.429 | 0.446 | 0.024 | 0.078 | 0.078 | 0 | 0.1 | 0.18 | 0.23 | 0.19 | 0.14 | 0.09 |
| 4 | 189031028 | rs9684216 | -0.363 | 0.281 | -0.852 | -0.005 | -0.001 | 0.02 | 0.02 | 1.753 | 0.5 | 0.46 | 0.57 | 0.32 | 0.55 | 0.32 |
| 4 | 189034513 | rs10015609 | 0.78 | -0.098 | -1.18 | 0.06 | 0.01 | 0.016 | 0.016 | 0 | 0.15 | 0.11 | 0.06 | 0.05 | 0.13 | 0.08 |
| 4 | 189036322 | rs13116982 | -0.925 | 0.023 | -1.267 | 0.659 | 0.002 | -0.07 | -0.07 | 0 | 0.85 | 0.96 | 0.8 | 0.53 | 0.87 | 0.85 |
| 4 | 189045617 | rs6553155 | -0.545 | 0.035 | -1.326 | 0.809 | 0.001 | -0.058 | -0.058 | 2.009 | 0.82 | 0.91 | 0.72 | 0.44 | 0.82 | 0.8 |
| 4 | 189057415 | rs7376389 | 0.605 | 0.062 | -1.936 | -0.093 | -0.083 | 0.162 | 0.162 | 0.008 | 0.39 | 0.12 | 0.49 | 0.06 | 0.47 | 0.25 |
| 4 | 189059078 | rs7441586 | -1.555 | -0.15 | -1.308 | 0.163 | -0.003 | 0.059 | 0.059 | 0 | 0.86 | 0.81 | 0.78 | 0.6 | 0.86 | 0.84 |
| 4 | 189059174 | rs34645730 | 1.275 | 0.334 | -1.641 | -0.172 | -0.079 | 0.12 | 0.12 | 0.018 | 0.25 | 0.07 | 0.34 | 0 | 0.3 | 0.2 |
| 4 | 189060431 | rs11938607 | -0.139 | -0.247 | -1.795 | -0.303 | 0.063 | -0.014 | -0.014 | 0.138 | 0.63 | 0.19 | 0.68 | 0.27 | 0.74 | 0.43 |
| 4 | 189061106 | rs13131525 | 0.642 | 0.025 | -1.671 | -0.508 | -0.083 | 0.158 | 0.158 | 0.002 | 0.38 | 0.12 | 0.49 | 0.07 | 0.47 | 0.25 |
| 4 | 189062214 | rs6553160 | 0.972 | -0.267 | -0.984 | -0.581 | 0.002 | -0.023 | -0.023 | 0 | 0.26 | 0.24 | 0.14 | 0.01 | 0.22 | 0.11 |
| 4 | 189065934 | rs7694050 | 0.008 | -0.079 | -0.854 | -0.433 | 0.013 | 0.082 | 0.082 | 0 | 0.54 | 0.32 | 0.54 | 0.14 | 0.58 | 0.34 |
| 4 | 189069181 | rs11733655 | -0.865 | -0.105 | -0.548 | -0.332 | -0.011 | 0.036 | 0.036 | 0 | 0.76 | 0.71 | 0.8 | 0.48 | 0.8 | 0.77 |
| 4 | 189070726 | rs11723187 | 0.43 | 0.219 | -1.027 | -1.019 | -0.045 | 0.051 | 0.051 | 0 | 0.37 | 0.23 | 0.61 | 0.16 | 0.48 | 0.25 |
| 4 | 189081654 | rs11132521 | -0.115 | -0.055 | -0.689 | -0.797 | 0.001 | -0.001 | -0.001 | 0 | 0.43 | 0.43 | 0.38 | 0.16 | 0.39 | 0.19 |
| 4 | 189088220 | rs4349672 | 0.334 | 0.088 | -0.55 | -0.576 | -0.003 | -0.017 | -0.017 | 0 | 0.16 | 0.25 | 0.13 | 0.06 | 0.16 | 0.07 |
| 4 | 189092903 | rs4863164 | 0.519 | 0.083 | -0.425 | -0.568 | -0.001 | -0.015 | -0.015 | 0 | 0.15 | 0.27 | 0.13 | 0.07 | 0.16 | 0.06 |
| 4 | 189095240 | rs12501503 | 0.415 | 0.083 | -0.642 | -0.466 | -0.006 | 0.008 | 0.008 | 0 | 0.07 | 0.1 | 0.2 | 0.09 | 0.06 | 0.03 |
| 4 | 189095790 | rs4323178 | -0.201 | 0.082 | -0.414 | -0.395 | 0.001 | 0.096 | 0.096 | 0 | 0.23 | 0.38 | 0.33 | 0.52 | 0.23 | 0.49 |
| 4 | 189098744 | rs6827299 | -1.731 | 0.011 | -1.256 | -1.15 | -0.036 | 0.099 | 0.099 | 0 | 0.83 | 0.99 | 0.92 | 0.99 | 0.82 | 0.93 |
| 4 | 189101482 | rs7375996 | 0.434 | 0.625 | -0.955 | -1.065 | -0.013 | 0.037 | 0.037 | 0 | 0.25 | 0.17 | 0.36 | 0 | 0.27 | 0.16 |
| 4 | 189101773 | rs7665403 | 0.728 | 0.594 | -1.114 | -1.013 | -0.03 | 0.089 | 0.089 | 0 | 0.23 | 0.08 | 0.34 | 0 | 0.25 | 0.15 |
| 4 | 189115231 | rs4492063 | -0.35 | 0.429 | -0.474 | -0.532 | 0.004 | -0.021 | -0.021 | 0 | 0.43 | 0.51 | 0.28 | 0.78 | 0.42 | 0.64 |
| 4 | 189119202 | rs6818741 | -0.076 | 0.893 | -0.004 | -0.562 | 0.038 | 0.067 | 0.067 | 0 | 0.23 | 0.4 | 0.19 | 0.33 | 0.28 | 0.48 |
| 4 | 189120911 | rs6857559 | 0.096 | 0.618 | 0.223 | -1.583 | 0 | -0.044 | -0.044 | 0 | 0.66 | 0.62 | 0.61 | 0.99 | 0.63 | 0.83 |
| 4 | 189120984 | rs4863348 | -0.046 | 0.801 | -0.223 | -1.459 | 0 | -0.036 | -0.036 | 0 | 0.7 | 0.72 | 0.64 | 0.98 | 0.64 | 0.83 |
| 4 | 189133207 | rs7376568 | -0.753 | 0.99 | -0.032 | -1.314 | 0.027 | -0.14 | -0.14 | 0 | 0.24 | 0.55 | 0.21 | 0.05 | 0.22 | 0.35 |
| 4 | 189134108 | rs6813871 | -0.409 | 1.241 | -0.549 | -0.972 | -0.001 | -0.001 | -0.001 | 0.303 | 0.19 | 0.18 | 0.31 | 0.22 | 0.3 | 0.14 |
| 4 | 189138814 | rs4863386 | 0.7 | 0.717 | -0.462 | -1.31 | 0.042 | 0.079 | 0.079 | 0 | 0.52 | 0.34 | 0.47 | 0.68 | 0.56 | 0.47 |
| 4 | 189139258 | rs7677236 | 0.101 | 0.885 | -0.234 | -1.062 | 0.016 | 0.016 | 0.016 | 0.107 | 0.14 | 0.17 | 0.28 | 0.25 | 0.21 | 0.12 |
| 4 | 189139613 | rs7682583 | -0.46 | 0.686 | -0.017 | -1.164 | 0.047 | 0.015 | 0.015 | 0 | 0.19 | 0.32 | 0.32 | 0.25 | 0.28 | 0.12 |
| 4 | 189143475 | rs10001461 | -0.053 | -0.196 | 0.162 | -1.478 | 0.024 | -0.111 | -0.111 | 0.077 | 0.18 | 0.37 | 0.11 | 0.01 | 0.15 | 0.08 |
| 4 | 189144799 | rs3924963 | -1.107 | -0.278 | -0.516 | -1.553 | 0.013 | 0.019 | 0.019 | 0.13 | 0.84 | 0.9 | 0.78 | 0.99 | 0.82 | 0.9 |
| 4 | 189148200 | rs4863391 | -1.215 | -0.278 | -0.693 | -1.552 | -0.002 | 0.081 | 0.081 | 0.196 | 0.84 | 1 | 0.78 | 0.99 | 0.82 | 0.9 |
| 4 | 189150117 | rs12650036 | 0.488 | -0.304 | -0.431 | -1.28 | 0.051 | 0.137 | 0.137 | 0 | 0.58 | 0.24 | 0.57 | 0.88 | 0.65 | 0.57 |
| 4 | 189150656 | rs6553184 | 0.458 | -0.361 | -0.53 | -1.428 | 0.043 | 0.114 | 0.114 | 4.532 | 0.54 | 0.31 | 0.59 | 0.98 | 0.63 | 0.79 |
| 4 | 189151984 | rs4279287 | -1.181 | -0.282 | 0.163 | -0.446 | -0.056 | -0.03 | -0.03 | 8.058 | 0.17 | 0.32 | 0.15 | 0.12 | 0.07 | 0.24 |
| 4 | 189158163 | rs4863421 | -0.106 | 0.531 | -0.23 | 0.294 | 0.002 | -0.037 | -0.037 | 0.049 | 0.62 | 0.76 | 0.6 | 0.45 | 0.62 | 0.36 |
| 4 | 189166087 | rs4863079 | -0.134 | 0.546 | -0.242 | 0.465 | 0.005 | -0.009 | -0.009 | 0.099 | 0.61 | 0.72 | 0.56 | 0.44 | 0.63 | 0.37 |
| 4 | 189171046 | rs4476671 | 0.139 | 0.473 | -0.611 | 0.426 | 0.02 | 0.022 | 0.022 | 0 | 0.42 | 0.37 | 0.35 | 0.61 | 0.35 | 0.58 |
| 4 | 189181266 | rs13145617 | -0.193 | 0.67 | -1.084 | 0.69 | 0.004 | 0 | 0 | 0 | 0.59 | 0.42 | 0.54 | 0.44 | 0.62 | 0.37 |
| 4 | 189188139 | rs12374353 | -0.467 | 0.946 | -1.478 | 0.618 | -0.027 | 0.012 | 0.012 | 0 | 0.26 | 0.08 | 0.24 | 0.18 | 0.27 | 0.42 |
| 4 | 189189170 | rs7375584 | -0.28 | 0.323 | -1.289 | 0.756 | -0.003 | 0.003 | 0.003 | 0 | 0.66 | 0.33 | 0.67 | 0.55 | 0.71 | 0.73 |
| 4 | 189189273 | rs4863124 | 0.656 | 0.163 | -1.568 | 0.774 | 0.014 | -0.023 | -0.023 | 0 | 0.17 | 0.07 | 0.14 | 0.2 | 0.15 | 0.17 |
| 4 | 189194661 | rs7378399 | 0.253 | 0.418 | -1.011 | 0.889 | 0.002 | 0.005 | 0.005 | 0.304 | 0.52 | 0.44 | 0.51 | 0.39 | 0.48 | 0.38 |

| | | | | | | | | | | | | | | | | |
|---|---|---|---|---|---|---|---|---|---|---|---|---|---|---|---|---|
| 4 | 189196692 | rs4077675 | -0.759 | 0.256 | -0.711 | 1.248 | 0.002 | -0.004 | -0.004 | 0 | 0.77 | 0.78 | 0.9 | 0.68 | 0.81 | 0.75 |
| 4 | 189205404 | rs11132538 | -0.399 | 0.43 | -0.165 | 1.201 | 0.004 | 0.104 | 0.104 | 0 | 0.13 | 0.34 | 0.17 | 0.3 | 0.14 | 0.28 |
| 4 | 189213140 | rs7654820 | 0.707 | 0.705 | -0.762 | 0.85 | -0.071 | 0.074 | 0.074 | 0 | 0.12 | 0.01 | 0.29 | 0.01 | 0.19 | 0.07 |
| 4 | 189213925 | rs7661001 | 0.656 | 0.354 | -0.621 | 0.646 | 0.017 | 0.028 | 0.028 | 0 | 0.32 | 0.23 | 0.17 | 0.11 | 0.25 | 0.23 |
| 4 | 189214720 | rs10006245 | 0.504 | 0.352 | -0.464 | 0.916 | -0.019 | -0.007 | -0.007 | 0 | 0.34 | 0.4 | 0.19 | 0.32 | 0.27 | 0.26 |
| 4 | 189216438 | rs4618385 | 0.851 | 0.714 | -0.085 | 1.11 | 0.027 | 0.012 | 0.012 | 0 | 0.73 | 0.48 | 0.52 | 0.69 | 0.67 | 0.66 |
| 4 | 189218156 | rs7666115 | -0.012 | 0.71 | 0.089 | 1.184 | 0.022 | 0.194 | 0.194 | 0 | 0.88 | 0.6 | 0.83 | 0.7 | 0.87 | 0.74 |
| 4 | 189220431 | rs11735401 | 0.404 | 0.831 | -0.355 | 1.357 | -0.075 | -0.047 | -0.047 | 0 | 0.13 | 0.01 | 0.33 | 0.15 | 0.19 | 0.11 |
| 4 | 189221207 | rs6830572 | -0.034 | 0.302 | -0.322 | 0.816 | 0.019 | -0.018 | -0.018 | 0 | 0.75 | 0.42 | 0.75 | 0.84 | 0.73 | 0.71 |
| 4 | 189223524 | rs10017800 | -0.545 | 0.216 | -0.524 | 0.778 | 0.006 | -0.006 | -0.006 | 0.409 | 0.77 | 0.8 | 0.79 | 0.65 | 0.8 | 0.83 |
| 4 | 189227731 | rs4862987 | 1.296 | 0.404 | -0.085 | 1.006 | -0.011 | 0.189 | 0.189 | 0 | 0.06 | 0.1 | 0.09 | 0.43 | 0.03 | 0.23 |
| 4 | 189229216 | rs6553193 | 0.335 | 0.878 | -0.2 | 0.286 | 0.024 | 0.018 | 0.018 | 1.063 | 0.67 | 0.67 | 0.53 | 0.43 | 0.55 | 0.38 |
| 4 | 189229584 | rs7696358 | 0.974 | 0.246 | -0.245 | -0.422 | -0.018 | -0.087 | -0.087 | 0 | 0.2 | 0.27 | 0.08 | 0 | 0.12 | 0.04 |
| 4 | 189230337 | rs11724270 | 0.78 | 0.804 | -0.457 | -0.415 | -0.054 | 0.016 | 0.016 | 0.105 | 0.21 | 0.08 | 0.3 | 0.14 | 0.3 | 0.31 |
| 4 | 189233294 | rs6828177 | -1.733 | 0.444 | -0.647 | -0.831 | -0.015 | 0.059 | 0.059 | 0 | 0.65 | 0.77 | 0.74 | 0.97 | 0.58 | 0.75 |
| 4 | 189233358 | rs7671200 | 0.899 | 0.232 | -0.784 | -0.782 | -0.01 | -0.065 | -0.065 | 0 | 0.21 | 0.26 | 0.08 | 0 | 0.12 | 0.04 |
| 4 | 189236548 | rs12647565 | 1.671 | 0.485 | -1.135 | -0.516 | 0.016 | 0.117 | 0.117 | 0 | 0.29 | 0.12 | 0.25 | 0.03 | 0.31 | 0.17 |
| 4 | 189240218 | rs12639689 | 0.761 | 0.508 | -0.924 | 0.211 | -0.002 | -0.086 | -0.086 | 0 | 0.15 | 0.04 | 0.17 | 0.39 | 0.12 | 0.16 |
| 4 | 189246406 | rs12501166 | -0.857 | 0.667 | -0.164 | -0.206 | 0.006 | -0.016 | -0.016 | 0.035 | 0.39 | 0.42 | 0.38 | 0.22 | 0.42 | 0.36 |
| 4 | 189247480 | rs11935681 | -0.456 | 0.56 | -0.056 | -0.324 | -0.003 | -0.054 | -0.054 | 6.495 | 0.29 | 0.36 | 0.15 | 0.12 | 0.24 | 0.23 |
| 4 | 189247773 | rs11930915 | 0.647 | 0.778 | 0.073 | 0.083 | 0.024 | -0.033 | -0.033 | 0.349 | 0.06 | 0.18 | 0.06 | 0 | 0.08 | 0.03 |
| 4 | 189262246 | rs4863305 | -0.68 | 0.558 | -0.328 | 0.183 | -0.003 | -0.023 | -0.023 | 0 | 0.41 | 0.48 | 0.55 | 0.74 | 0.41 | 0.54 |
| 4 | 189266054 | rs6553197 | -0.367 | 0.477 | 0.089 | 0.649 | 0.001 | 0.101 | 0.101 | 0 | 0.05 | 0.17 | 0.07 | 0.17 | 0.06 | 0.09 |
| 4 | 189271272 | rs952719 | -0.389 | 0.158 | 0.597 | 1.248 | -0.023 | 0.124 | 0.124 | 0 | 0.16 | 0.29 | 0.22 | 0.4 | 0.12 | 0.15 |
| 4 | 189282465 | rs12501958 | 1.059 | 0.39 | -0.31 | 0.512 | -0.025 | 0.013 | 0.013 | 0 | 0.39 | 0.14 | 0.37 | 0.4 | 0.45 | 0.29 |
| 4 | 189285725 | rs11132547 | 1.04 | 0.387 | -0.218 | 0.511 | -0.017 | 0.015 | 0.015 | 0 | 0.39 | 0.16 | 0.38 | 0.4 | 0.46 | 0.38 |
| 4 | 189287271 | rs314086 | 0.342 | 0.366 | -0.196 | -0.016 | -0.023 | 0.027 | 0.027 | 0 | 0.26 | 0.17 | 0.28 | 0.19 | 0.28 | 0.31 |
| 4 | 189288209 | rs11728000 | -1.186 | 0.251 | 0.107 | -0.156 | -0.015 | -0.039 | -0.039 | 0 | 0.38 | 0.45 | 0.46 | 0.79 | 0.33 | 0.48 |
| 4 | 189288591 | rs1507074 | 2.311 | 0.258 | 0.085 | -0.15 | -0.04 | 0.122 | 0.122 | 0 | 0.34 | 0.1 | 0.24 | 0.01 | 0.39 | 0.21 |
| 4 | 189293076 | rs4863321 | -1.023 | 0.458 | 0.462 | 0.203 | -0.03 | -0.041 | -0.041 | 0 | 0.31 | 0.45 | 0.42 | 0.77 | 0.28 | 0.35 |
| 4 | 189295229 | rs6553200 | -1.363 | 0.47 | 0.465 | 0.68 | 0.015 | -0.024 | -0.024 | 6.259 | 0.33 | 0.28 | 0.3 | 0.47 | 0.29 | 0.41 |
| 4 | 189296032 | rs1606234 | -1.291 | 0.696 | 0.578 | 0.496 | -0.001 | -0.001 | -0.001 | 0.741 | 0.47 | 0.54 | 0.5 | 0.57 | 0.42 | 0.49 |
| 4 | 189296076 | rs314078 | -1.022 | 0.564 | 0.783 | 0.506 | 0.006 | -0.002 | -0.002 | 0.558 | 0.38 | 0.49 | 0.52 | 0.4 | 0.34 | 0.4 |
| 4 | 189296513 | rs11726434 | -0.675 | 0.249 | 0.53 | 0.824 | -0.008 | 0.007 | 0.007 | 0 | 0.13 | 0.16 | 0.09 | 0.24 | 0.12 | 0.11 |
| 4 | 189300955 | rs11938549 | 0.107 | 0.282 | 0.556 | 1.142 | -0.002 | 0.02 | 0.02 | 0 | 0.12 | 0.14 | 0.15 | 0.34 | 0.17 | 0.24 |
| 4 | 189300970 | rs2055918 | 1.355 | 0.194 | 0.789 | 1.033 | -0.005 | 0.031 | 0.031 | 0.102 | 0.7 | 0.38 | 0.74 | 0.62 | 0.73 | 0.6 |
| 4 | 189303096 | rs6553201 | -0.143 | 0.158 | 0.814 | 1.04 | 0.008 | -0.005 | -0.005 | 0 | 0.9 | 0.64 | 0.94 | 0.97 | 0.95 | 0.87 |
| 4 | 189303437 | rs7669530 | 0.687 | 0.112 | 0.786 | 0.91 | -0.002 | 0.016 | 0.016 | 0 | 0.18 | 0.16 | 0.18 | 0.06 | 0.18 | 0.2 |
| 4 | 189305026 | rs12505859 | 1.01 | 0.066 | 0.752 | 0.883 | -0.001 | -0.006 | -0.006 | 0.053 | 0.59 | 0.34 | 0.6 | 0.57 | 0.6 | 0.43 |
| 4 | 189306960 | rs314054 | -0.471 | 0.161 | 0.608 | 0.949 | -0.009 | -0.019 | -0.019 | 0 | 0.31 | 0.3 | 0.34 | 0.4 | 0.35 | 0.44 |
| 4 | 189310768 | rs12510602 | 0.658 | 0.178 | 0.32 | 0.829 | -0.002 | 0.086 | 0.086 | 0 | 0.11 | 0.21 | 0.29 | 0.32 | 0.09 | 0.1 |
| 4 | 189313568 | rs2019991 | 0.971 | 0.466 | 0.62 | 1 | -0.002 | -0.002 | -0.002 | 0 | 0.5 | 0.28 | 0.53 | 0.58 | 0.5 | 0.39 |
| 4 | 189313688 | rs2376746 | 1.408 | 0.419 | 0.623 | 1.031 | -0.018 | 0.07 | 0.07 | 0 | 0.45 | 0.21 | 0.45 | 0.41 | 0.47 | 0.31 |
| 4 | 189314208 | rs13104957 | 0.092 | 0.618 | 0.726 | 1.012 | -0.003 | -0.049 | -0.049 | 0.355 | 0.63 | 0.41 | 0.71 | 0.92 | 0.7 | 0.63 |
| 4 | 189315208 | rs10028269 | -0.199 | 0.673 | 0.249 | 1.199 | -0.01 | -0.015 | -0.015 | 0.017 | 0.13 | 0.12 | 0.18 | 0.34 | 0.2 | 0.24 |
| 4 | 189326076 | rs314104 | 0.635 | 0.715 | 0.38 | 0.778 | -0.011 | 0.058 | 0.058 | 0 | 0.26 | 0.15 | 0.22 | 0.05 | 0.23 | 0.23 |
| 4 | 189331063 | rs12648353 | 0.204 | 0.679 | 0.481 | 0.843 | 0.001 | 0.024 | 0.024 | 0.09 | 0.62 | 0.38 | 0.64 | 0.84 | 0.69 | 0.63 |
| 4 | 189333660 | rs10009683 | 1.342 | 0.626 | 0.197 | 0.925 | -0.052 | 0.085 | 0.085 | 0 | 0.42 | 0.22 | 0.31 | 0.29 | 0.51 | 0.4 |
| 4 | 189333901 | rs12512320 | -1.461 | 0.278 | 0.871 | 1.131 | -0.031 | 0.037 | 0.037 | 0 | 0.23 | 0.32 | 0.4 | 0.66 | 0.18 | 0.33 |
| 4 | 189334307 | rs12504808 | 0.117 | 0.33 | 0.789 | 1.338 | -0.031 | -0.008 | -0.008 | 0 | 0.31 | 0.43 | 0.38 | 0.34 | 0.2 | 0.33 |
| 4 | 189334992 | rs314034 | 0.295 | 0.417 | 0.795 | 1.433 | 0 | -0.002 | -0.002 | 1.36 | 0.12 | 0.1 | 0.1 | 0.15 | 0.11 | 0.09 |
| 4 | 189338040 | rs2292432 | 0.4 | 0.4 | 0.796 | 1.523 | 0 | 0.033 | 0.033 | 0 | 0.07 | 0.07 | 0.1 | 0.16 | 0.09 | 0.13 |
| 4 | 189338469 | rs2292433 | -0.928 | 0.388 | 0.94 | 1.476 | 0.003 | 0.197 | 0.197 | 0 | 0.11 | 0.29 | 0.18 | 0.23 | 0.12 | 0.14 |
| 4 | 189338814 | rs10008710 | 1.038 | 0.343 | 0.936 | 1.383 | -0.028 | -0.065 | -0.065 | 0 | 0.23 | 0.34 | 0.15 | 0.05 | 0.14 | 0.17 |
| 4 | 189339341 | rs4362867 | -1.511 | 0.482 | 0.784 | 1.438 | 0.01 | 0.019 | 0.019 | 0 | 0.94 | 0.86 | 0.93 | 0.84 | 0.91 | 0.88 |
| 4 | 189341977 | rs713508 | 1.469 | 0.432 | 0.894 | 1.38 | -0.006 | 0.034 | 0.034 | 0 | 0.46 | 0.38 | 0.49 | 0.3 | 0.5 | 0.35 |
| 4 | 189342143 | rs4395553 | -0.532 | 0.429 | 0.912 | 1.389 | 0.002 | 0.004 | 0.004 | 0 | 0.71 | 0.7 | 0.79 | 0.79 | 0.77 | 0.71 |
| 4 | 189342369 | rs713507 | -0.42 | 0.446 | 0.915 | 1.433 | 0.007 | 0.037 | 0.037 | 0 | 0.16 | 0.27 | 0.14 | 0.26 | 0.16 | 0.21 |
| 4 | 189342799 | rs1857955 | 0.147 | 0.612 | 0.407 | 1.443 | 0.001 | 0.001 | 0.001 | 0 | 0.83 | 0.81 | 0.83 | 0.69 | 0.8 | 0.79 |
| 4 | 189343008 | rs11132551 | 0.504 | 0.85 | 0.184 | 1.516 | 0.05 | 0.08 | 0.08 | 0 | 0.51 | 0.27 | 0.55 | 0.46 | 0.59 | 0.47 |
| 4 | 189344762 | rs13434896 | -1.287 | 1.039 | -0.055 | 1.361 | 0.017 | 0.169 | 0.169 | 0 | 0.17 | 0.39 | 0.25 | 0.39 | 0.21 | 0.26 |
| 4 | 189344863 | rs7690014 | -1.287 | 1.039 | -0.058 | 1.361 | 0.018 | 0.173 | 0.173 | 0 | 0.17 | 0.39 | 0.25 | 0.39 | 0.2 | 0.26 |
| 4 | 189347578 | rs993424 | 0.904 | 1.239 | -0.134 | 0.536 | -0.074 | 0.193 | 0.193 | 0 | 0.37 | 0.11 | 0.25 | 0.01 | 0.44 | 0.19 |
| 4 | 189351268 | rs869570 | -0.899 | 1.295 | 0.548 | 1.618 | 0.039 | 0.097 | 0.097 | 0 | 0.11 | 0.32 | 0.15 | 0.23 | 0.12 | 0.14 |
| 4 | 189355668 | rs12505646 | 0.4 | 1.413 | -0.29 | 1.255 | -0.003 | 0.037 | 0.037 | 0 | 0.38 | 0.22 | 0.4 | 0.27 | 0.38 | 0.31 |
| 4 | 189362292 | rs10026052 | 0.672 | 1.056 | 0.381 | 1.776 | 0.036 | 0.082 | 0.082 | 0 | 0.81 | 0.63 | 0.66 | 0.34 | 0.75 | 0.7 |
| 4 | 189371244 | rs2292434 | -0.097 | 0.969 | 0.476 | 1.716 | -0.012 | 0.011 | 0.011 | 0 | 0.14 | 0.31 | 0.17 | 0.17 | 0.12 | 0.22 |
| 4 | 189377718 | rs425974 | -0.219 | 1.049 | 0.292 | 2.235 | 0.026 | 0.066 | 0.066 | 0 | 0.14 | 0.23 | 0.27 | 0.31 | 0.18 | 0.18 |
| 4 | 189379520 | rs372298 | -1.045 | 1.012 | 0.366 | 1.811 | 0.002 | 0.005 | 0.005 | 0 | 0.29 | 0.31 | 0.44 | 0.64 | 0.33 | 0.51 |

| | | | | | | | | | | | | | | | | |
|---|---|---|---|---|---|---|---|---|---|---|---|---|---|---|---|---|
| 4 | 189384671 | rs10019893 | -0.704 | 0.968 | -0.252 | 1.682 | 0.015 | 0.091 | 0.091 | 0 | 0.28 | 0.5 | 0.43 | 0.49 | 0.3 | 0.41 |
| 4 | 189388320 | rs314065 | -0.847 | 0.756 | -0.214 | 1.612 | 0.015 | 0.041 | 0.041 | 0 | 0.32 | 0.52 | 0.47 | 0.6 | 0.36 | 0.51 |
| 4 | 189390949 | rs13114765 | 0.059 | -0.123 | -0.589 | 1.296 | 0.007 | -0.001 | -0.001 | 0 | 0.33 | 0.3 | 0.22 | 0.26 | 0.13 | 0.33 |
| 4 | 189402063 | rs11734013 | 0.111 | 0.576 | -1.763 | 1.357 | -0.059 | -0.053 | -0.053 | 0.001 | 0.39 | 0.05 | 0.23 | 0.39 | 0.44 | 0.48 |
| 4 | 189402769 | rs2089408 | 1.084 | 0.506 | -1.84 | 1.653 | -0.01 | -0.027 | -0.027 | 5.145 | 0.07 | 0.02 | 0.04 | 0.22 | 0.08 | 0.13 |
| 4 | 189404004 | rs1911504 | 0.555 | 0.596 | -1.631 | 1.033 | 0.025 | -0.01 | -0.01 | 0 | 0.36 | 0.22 | 0.35 | 0.6 | 0.19 | 0.34 |
| 4 | 189404814 | rs11132554 | 0.649 | 0.674 | -1.635 | 0.942 | 0.053 | -0.04 | -0.04 | 0 | 0.34 | 0.14 | 0.34 | 0.43 | 0.17 | 0.29 |
| 4 | 189412980 | rs6855687 | -0.355 | 0.781 | -0.91 | 0.562 | -0.004 | -0.076 | -0.076 | 0 | 0.52 | 0.48 | 0.47 | 0.96 | 0.37 | 0.66 |
| 4 | 189414006 | rs12648691 | 0.304 | 0.707 | -0.957 | 0.751 | 0.061 | -0.002 | -0.002 | 0 | 0.41 | 0.18 | 0.37 | 0.6 | 0.22 | 0.43 |
| 4 | 189414658 | rs12510507 | 0.792 | 0.636 | -0.864 | 0.749 | 0.051 | -0.031 | -0.031 | 0 | 0.34 | 0.18 | 0.35 | 0.43 | 0.18 | 0.3 |
| 4 | 189416683 | rs6833998 | -0.187 | 0.627 | -0.514 | 0.847 | 0.001 | 0.199 | 0.199 | 0 | 0.15 | 0.29 | 0.17 | 0.42 | 0.13 | 0.25 |
| 4 | 189417612 | rs1976536 | 0.113 | 0.948 | -0.53 | 0.776 | 0.004 | -0.004 | -0.004 | 0.035 | 0.45 | 0.31 | 0.32 | 0.45 | 0.43 | 0.5 |
| 4 | 189419282 | rs6812291 | 0.114 | 0.709 | -0.726 | 0.828 | 0.059 | -0.072 | -0.072 | 0 | 0.75 | 0.49 | 0.53 | 0.9 | 0.63 | 0.85 |
| 4 | 189430792 | rs4863330 | 0.961 | 0.736 | -0.809 | 0.737 | 0.059 | -0.04 | -0.04 | 0 | 0.33 | 0.14 | 0.24 | 0.4 | 0.16 | 0.29 |
| 4 | 189433290 | rs1395612 | 0.453 | 0.926 | -0.539 | 0.469 | -0.001 | -0.005 | -0.005 | 0 | 0.39 | 0.32 | 0.29 | 0.27 | 0.39 | 0.38 |
| 4 | 189433525 | rs931887 | 0.387 | 1.134 | -0.5 | 0.546 | 0.041 | 0.049 | 0.049 | 0 | 0.69 | 0.5 | 0.45 | 0.58 | 0.57 | 0.65 |
| 4 | 189433594 | rs931885 | 0.675 | 1.116 | -0.375 | 0.566 | 0.037 | 0.023 | 0.023 | 0 | 0.67 | 0.48 | 0.42 | 0.43 | 0.51 | 0.56 |
| 4 | 189434690 | rs10011727 | 0.929 | 1.482 | -0.159 | -0.122 | -0.008 | -0.005 | -0.005 | 0 | 0.71 | 0.28 | 0.54 | 0.68 | 0.58 | 0.68 |
| 4 | 189436829 | rs10018170 | 0.486 | 1.57 | -0.433 | -0.301 | 0.028 | -0.018 | -0.018 | 0.41 | 0.71 | 0.4 | 0.56 | 0.81 | 0.62 | 0.76 |
| 4 | 189436841 | rs11132556 | 1.154 | 1.418 | -0.516 | -0.493 | 0.042 | -0.022 | -0.022 | 0 | 0.41 | 0.15 | 0.4 | 0.64 | 0.29 | 0.43 |
| 4 | 189437449 | rs7671323 | 0.784 | 1.667 | -0.165 | -0.127 | 0.026 | -0.011 | -0.011 | 0.109 | 0.7 | 0.4 | 0.55 | 0.79 | 0.6 | 0.72 |
| 4 | 189439821 | rs4863333 | 0.986 | 1.694 | -0.135 | -0.091 | 0.023 | 0.012 | 0.012 | 0.507 | 0.7 | 0.39 | 0.53 | 0.69 | 0.58 | 0.66 |
| 4 | 189439872 | rs7677233 | -0.017 | 1.476 | 0.306 | -0.25 | -0.001 | -0.09 | -0.09 | 0 | 0.15 | 0.3 | 0.28 | 0 | 0.15 | 0.08 |
| 4 | 189447377 | rs10033480 | -0.399 | 1.077 | 0.111 | 0.082 | 0 | 0.186 | 0.186 | 0 | 0.07 | 0.24 | 0.06 | 0.19 | 0.03 | 0.11 |
| 4 | 189449669 | rs6553202 | -0.647 | 1.003 | -0.436 | -0.643 | 0 | -0.001 | -0.001 | 0.234 | 0.57 | 0.55 | 0.58 | 0.24 | 0.62 | 0.49 |
| 4 | 189455974 | rs7654439 | -0.542 | 0.93 | -0.422 | -0.51 | 0.068 | 0.063 | 0.063 | 0 | 0.09 | 0.32 | 0.11 | 0.14 | 0.15 | 0.12 |
| 4 | 189456233 | rs7679676 | -0.542 | 0.93 | -0.422 | -0.51 | 0.068 | 0.063 | 0.063 | 0 | 0.09 | 0.32 | 0.11 | 0.14 | 0.15 | 0.12 |
| 4 | 189457782 | rs1511805 | 0.913 | 1.063 | -0.541 | -0.551 | 0.001 | -0.002 | -0.002 | 0.71 | 0.36 | 0.36 | 0.39 | 0.62 | 0.28 | 0.39 |
| 4 | 189463597 | rs4863340 | 1.137 | 1.021 | -0.772 | -0.548 | 0.011 | -0.081 | -0.081 | 0.411 | 0.34 | 0.11 | 0.36 | 0.62 | 0.28 | 0.4 |
| 4 | 189466381 | rs10866310 | 0.915 | 1.181 | -0.785 | -0.721 | -0.01 | 0.019 | 0.019 | 0.456 | 0.54 | 0.12 | 0.43 | 0.66 | 0.41 | 0.59 |
| 4 | 189475684 | rs6826087 | 1.013 | 1.2 | -0.641 | -0.738 | 0.09 | 0.053 | 0.053 | 0.386 | 0.49 | 0.17 | 0.43 | 0.66 | 0.38 | 0.54 |
| 4 | 189478215 | rs2376964 | 0.952 | 1.007 | -0.7 | -0.586 | -0.005 | -0.002 | -0.002 | 0.414 | 0.05 | 0 | 0.03 | 0.23 | 0.03 | 0.14 |
| 4 | 189488132 | rs7667826 | 0.659 | 1.152 | -0.698 | -0.582 | -0.018 | -0.01 | -0.01 | 0 | 0.06 | 0.01 | 0.04 | 0.23 | 0.07 | 0.15 |
| 4 | 189489351 | rs7700176 | 0.031 | 1.36 | -0.415 | -0.994 | 0.019 | -0.158 | -0.158 | 0 | 0.31 | 0.43 | 0.39 | 0.01 | 0.31 | 0.16 |
| 4 | 189492177 | rs6830571 | 0.064 | 1.325 | -0.426 | -1.012 | 0.017 | -0.142 | -0.142 | 0 | 0.31 | 0.41 | 0.39 | 0.01 | 0.31 | 0.16 |
| 4 | 189495641 | rs12645101 | 1.784 | 0.722 | -1.14 | -0.921 | -0.009 | -0.104 | -0.104 | 0 | 0.32 | 0.08 | 0.29 | 0.65 | 0.29 | 0.47 |
| 4 | 189498064 | rs7666129 | 1.039 | 1.135 | -0.705 | -0.458 | -0.092 | 0.07 | 0.07 | 0 | 0.57 | 0.1 | 0.42 | 0.79 | 0.52 | 0.71 |
| 4 | 189499145 | rs12643289 | 1.497 | 1.14 | -0.72 | -0.461 | 0.044 | 0.009 | 0.009 | 0.105 | 0.5 | 0.1 | 0.36 | 0.67 | 0.42 | 0.6 |
| 4 | 189499621 | rs12640778 | 1.041 | 1.14 | -0.713 | -0.464 | -0.092 | 0.07 | 0.07 | 0 | 0.57 | 0.1 | 0.42 | 0.79 | 0.52 | 0.71 |
| 4 | 189499865 | rs12640838 | 1.041 | 1.14 | -0.713 | -0.464 | -0.092 | 0.07 | 0.07 | 0 | 0.57 | 0.1 | 0.42 | 0.79 | 0.52 | 0.71 |
| 4 | 189500458 | rs11132562 | 0.075 | 1.273 | -0.045 | -0.239 | -0.048 | 0.071 | 0.071 | 0 | 0.23 | 0.09 | 0.34 | 0.01 | 0.32 | 0.16 |
| 4 | 189501949 | rs4863049 | 0.647 | 1.122 | 0.306 | -0.038 | 0.011 | 0.032 | 0.032 | 0 | 0.9 | 0.76 | 0.93 | 0.87 | 0.9 | 0.88 |
| 4 | 189502823 | rs7663546 | -0.001 | 1.08 | 0.64 | -0.083 | 0.028 | 0.26 | 0.26 | 0.042 | 0.07 | 0.33 | 0.15 | 0.2 | 0.03 | 0.09 |
| 4 | 189508238 | rs4863054 | 0.685 | 1.242 | -0.194 | -0.052 | 0.023 | 0.026 | 0.026 | 0 | 0.9 | 0.76 | 0.93 | 0.88 | 0.88 | 0.88 |
| 4 | 189510469 | rs12512555 | 1.123 | 1.188 | -0.968 | -0.281 | -0.049 | -0.002 | -0.002 | 0 | 0.55 | 0.1 | 0.53 | 0.68 | 0.46 | 0.63 |
| 4 | 189519649 | rs13128985 | -1.06 | 1.113 | -0.41 | -0.744 | 0.018 | -0.121 | -0.121 | 0 | 0.39 | 0.49 | 0.44 | 0.04 | 0.53 | 0.27 |
| 4 | 189522110 | rs4863354 | -1.367 | 1.094 | -0.42 | -0.583 | -0.003 | -0.019 | -0.019 | 0.962 | 0.45 | 0.48 | 0.49 | 0.15 | 0.61 | 0.4 |
| 4 | 189522393 | rs4549442 | -0.467 | 1.336 | -0.541 | -0.28 | -0.005 | -0.024 | -0.024 | 0 | 0.21 | 0.19 | 0.21 | 0.01 | 0.33 | 0.14 |
| 4 | 189524738 | rs2377343 | 0.445 | 0.202 | -0.318 | 0.584 | -0.001 | -0.001 | -0.001 | 0 | 0.64 | 0.61 | 0.62 | 0.67 | 0.62 | 0.68 |
| 4 | 189525837 | rs12501064 | 0.743 | -0.372 | 0.407 | 0.696 | -0.015 | 0.061 | 0.061 | 0 | 0.92 | 0.69 | 0.96 | 0.92 | 0.97 | 0.84 |
| 4 | 189528831 | rs4241895 | 0.521 | -0.818 | -0.242 | 0.755 | -0.026 | 0 | 0 | 0 | 0.66 | 0.42 | 0.65 | 0.69 | 0.8 | 0.59 |
| 4 | 189541364 | rs6817256 | 1.074 | -0.472 | 0.093 | 0.867 | -0.016 | 0.073 | 0.073 | 0.589 | 0.88 | 0.61 | 0.89 | 0.82 | 0.91 | 0.9 |
| 4 | 189542277 | rs11735347 | -0.207 | 0.059 | -1.246 | 0.076 | -0.066 | 0.102 | 0.102 | 0 | 0.22 | 0.02 | 0.23 | 0.04 | 0.25 | 0.26 |
| 4 | 189544738 | rs12510167 | -0.587 | -0.552 | -1.18 | -0.176 | 0.016 | 0.015 | 0.015 | 0 | 0.23 | 0.17 | 0.19 | 0.07 | 0.14 | 0.11 |
| 4 | 189544807 | rs456469 | 0.333 | -0.749 | -1.058 | 0.693 | 0.011 | -0.008 | -0.008 | 0 | 0.77 | 0.81 | 0.77 | 0.68 | 0.84 | 0.87 |
| 4 | 189545795 | rs10866311 | 0.511 | -0.11 | -0.887 | 0.742 | 0.018 | 0 | 0 | 0 | 0.7 | 0.52 | 0.63 | 0.72 | 0.67 | 0.74 |
| 4 | 189545959 | rs458862 | 0.302 | -0.35 | -1.949 | 0.858 | 0.039 | -0.008 | -0.008 | 0 | 0.52 | 0.12 | 0.52 | 0.4 | 0.62 | 0.51 |
| 4 | 189547648 | rs11729434 | -0.079 | -0.041 | -1.451 | 0.637 | -0.002 | -0.003 | -0.003 | 0 | 0.25 | 0.28 | 0.16 | 0.19 | 0.23 | 0.22 |
| 4 | 189548540 | rs690149 | -0.978 | 0.537 | -2.187 | 1.016 | -0.051 | -0.135 | -0.135 | 2.336 | 0.76 | 0.96 | 0.67 | 0.61 | 0.79 | 0.75 |
| 4 | 189548599 | rs690143 | 0.283 | 0.805 | -1.188 | 0.591 | -0.01 | 0.013 | 0.013 | 0 | 0.14 | 0.11 | 0.13 | 0 | 0.16 | 0.08 |
| 4 | 189554217 | rs464270 | 0.488 | 0.768 | -1.22 | 0.687 | -0.036 | 0.029 | 0.029 | 0 | 0.13 | 0.03 | 0.11 | 0.05 | 0.15 | 0.09 |
| 4 | 189557490 | rs13110503 | 0.027 | 0.747 | -0.708 | 0.593 | -0.006 | 0.031 | 0.031 | 0 | 0.17 | 0.07 | 0.18 | 0.11 | 0.17 | 0.16 |
| 4 | 189558299 | rs4403095 | 0.424 | 0.692 | 0.128 | 1.249 | -0.001 | 0.31 | 0.31 | 0.188 | 0.07 | 0.16 | 0.14 | 0.5 | 0.07 | 0.25 |
| 4 | 189558312 | rs734521 | 1.592 | 0.785 | 0.488 | 0.939 | -0.003 | 0.023 | 0.023 | 0.019 | 0.55 | 0.41 | 0.53 | 0.19 | 0.55 | 0.39 |
| 4 | 189563127 | rs11132565 | 0.232 | 0.571 | 0.537 | 0.995 | 0.008 | 0.155 | 0.155 | 1.218 | 0.09 | 0.17 | 0.28 | 0.37 | 0.12 | 0.31 |
| 4 | 189565170 | rs6834498 | -0.622 | 0.411 | 0.509 | 0.896 | -0.004 | -0.01 | -0.01 | 0 | 0.21 | 0.27 | 0.17 | 0.13 | 0.14 | 0.21 |
| 4 | 189571541 | rs360117 | -1.804 | 0.449 | 0.126 | 0.879 | -0.005 | -0.006 | -0.006 | 0 | 0.43 | 0.72 | 0.59 | 0.5 | 0.44 | 0.61 |
| 4 | 189578979 | rs6553217 | 0.012 | 0.386 | 0 | 0.841 | 0.008 | 0.025 | 0.025 | 0.12 | 0.16 | 0.06 | 0.14 | 0.09 | 0.13 | 0.15 |
| 4 | 189580139 | rs1879985 | -0.074 | 0.502 | 0.687 | 1.052 | 0.025 | 0.221 | 0.221 | 0.298 | 0.11 | 0.25 | 0.3 | 0.37 | 0.14 | 0.31 |

| | | | | | | | | | | | | | | | | |
|---|---|---|---|---|---|---|---|---|---|---|---|---|---|---|---|---|
| 4 | 189584353 | rs7686384 | -0.89 | 0.371 | 0.477 | 1.096 | 0.009 | 0.005 | 0.005 | 0.66 | 0.25 | 0.22 | 0.18 | 0.19 | 0.17 | 0.2 |
| 4 | 189584904 | rs899273 | 1.528 | 0.675 | 0.473 | 0.889 | -0.012 | -0.013 | -0.013 | 0 | 0.59 | 0.31 | 0.4 | 0.45 | 0.54 | 0.42 |
| 4 | 189587079 | rs7659062 | -0.217 | 0.334 | 0.504 | 0.994 | 0.006 | 0.008 | 0.008 | 2.018 | 0.19 | 0.18 | 0.16 | 0.12 | 0.14 | 0.1 |
| 4 | 189589490 | rs1478563 | -1.535 | 0.333 | 0.555 | 1.004 | -0.01 | 0.014 | 0.014 | 0 | 0.34 | 0.61 | 0.46 | 0.54 | 0.28 | 0.46 |
| 4 | 189590859 | rs7662358 | -0.219 | 0.348 | -0.035 | 0.995 | 0.009 | 0.012 | 0.012 | 2.018 | 0.19 | 0.18 | 0.16 | 0.12 | 0.14 | 0.1 |
| 4 | 189592930 | rs4863376 | -1.134 | 0.476 | -0.274 | 0.642 | 0.018 | -0.023 | -0.023 | 0 | 0.35 | 0.71 | 0.48 | 0.55 | 0.29 | 0.46 |
| 4 | 189600713 | rs4863380 | 1.242 | 0.599 | -0.715 | 0.228 | -0.083 | 0.044 | 0.044 | 0 | 0.36 | 0.06 | 0.3 | 0.19 | 0.4 | 0.16 |
| 4 | 189608461 | rs1038095 | -0.14 | 0.606 | 0.331 | 0.164 | 0.002 | -0.041 | -0.041 | 0.068 | 0.09 | 0.15 | 0.06 | 0 | 0.12 | 0.06 |
| 4 | 189616701 | rs10434313 | -1.336 | 0.51 | 0.599 | 0.259 | -0.006 | -0.008 | -0.008 | 0 | 0.35 | 0.53 | 0.31 | 0.32 | 0.3 | 0.33 |
| 4 | 189628999 | rs6815218 | -1.933 | 0.348 | 1.264 | 0.723 | 0.034 | 0.175 | 0.175 | 0.199 | 0.1 | 0.33 | 0.07 | 0.3 | 0.06 | 0.2 |
| 4 | 189633448 | rs6818117 | -1.375 | 0.364 | 1.161 | 0.613 | -0.015 | -0.005 | -0.005 | 0 | 0.28 | 0.55 | 0.25 | 0.3 | 0.19 | 0.29 |
| 4 | 189636244 | rs6837842 | -0.792 | 0.386 | 0.529 | 0.644 | -0.006 | -0.012 | -0.012 | 0.235 | 0.22 | 0.25 | 0.21 | 0.15 | 0.15 | 0.19 |
| 4 | 189636722 | rs13106691 | -0.722 | 0.39 | 0.458 | 0.646 | -0.005 | -0.01 | -0.01 | 0 | 0.22 | 0.24 | 0.2 | 0.15 | 0.14 | 0.19 |
| 4 | 189639545 | rs1461776 | -1.53 | 0.488 | -0.378 | 0.6 | 0.026 | 0.024 | 0.024 | 0 | 0.35 | 0.77 | 0.31 | 0.35 | 0.24 | 0.46 |
| 4 | 189642697 | rs13111649 | -1.38 | 0.961 | -0.288 | 1.255 | -0.005 | 0.034 | 0.034 | 0.083 | 0.07 | 0.05 | 0.17 | 0.38 | 0.1 | 0.29 |
| 4 | 189643752 | rs6832545 | -1.391 | 0.786 | -0.004 | 0.984 | -0.001 | -0.007 | -0.007 | 0 | 0.08 | 0.13 | 0.07 | 0.13 | 0.07 | 0.17 |
| 4 | 189653624 | rs7681177 | 0.481 | 0.954 | 0.172 | 0.919 | 0.015 | 0.088 | 0.088 | 0.272 | 0.73 | 0.57 | 0.66 | 0.47 | 0.71 | 0.65 |
| 4 | 189654390 | rs930995 | 0.681 | 1.164 | 0.068 | 0.732 | 0.028 | -0.017 | -0.017 | 0 | 0.68 | 0.49 | 0.61 | 0.25 | 0.63 | 0.49 |
| 4 | 189660025 | rs6553229 | -0.495 | 0.392 | 0.038 | 0.235 | -0.008 | -0.037 | -0.037 | 0.324 | 0.19 | 0.21 | 0.09 | 0.01 | 0.1 | 0.1 |
| 4 | 189663354 | rs7685306 | -1.359 | 0.457 | 0.137 | 0.493 | -0.025 | -0.012 | -0.012 | 0.875 | 0.32 | 0.49 | 0.25 | 0.32 | 0.22 | 0.44 |
| 4 | 189665842 | rs11132580 | -1.527 | 0.754 | 0.332 | 0.455 | 0.027 | 0.06 | 0.06 | 0 | 0.21 | 0.5 | 0.21 | 0.31 | 0.21 | 0.37 |
| 4 | 189668536 | rs1350865 | 0.826 | 0.985 | 0.71 | -0.038 | 0.012 | -0.069 | -0.069 | 1.186 | 0.85 | 0.76 | 0.74 | 0.99 | 0.84 | 0.79 |
| 4 | 189673054 | rs2377333 | 0.721 | 1.146 | 0.91 | -0.77 | -0.003 | -0.075 | -0.075 | 0 | 0.68 | 0.41 | 0.76 | 0.94 | 0.71 | 0.77 |
| 4 | 189673094 | rs6849122 | 0.622 | 1.612 | 0.509 | -0.248 | -0.02 | 0.006 | 0.006 | 0.554 | 0.11 | 0.08 | 0.24 | 0.53 | 0.2 | 0.2 |
| 4 | 189676850 | rs6827397 | 0.622 | 1.612 | 0.509 | -0.248 | -0.02 | 0.006 | 0.006 | 0.554 | 0.11 | 0.08 | 0.24 | 0.53 | 0.2 | 0.2 |
| 4 | 189680843 | rs12502652 | 0.689 | 1.402 | 0.656 | -0.569 | -0.006 | 0.005 | 0.005 | 0 | 0.05 | 0.05 | 0.18 | 0.01 | 0.11 | 0.15 |
| 4 | 189687397 | rs4863387 | 1.21 | 1.445 | 0.737 | -0.627 | -0.029 | 0.077 | 0.077 | 0 | 0.59 | 0.25 | 0.54 | 0.77 | 0.56 | 0.48 |
| 4 | 189688859 | rs1350864 | 0.533 | 1.464 | 0.699 | -0.749 | -0.012 | 0.01 | 0.01 | 0.275 | 0.06 | 0.06 | 0.22 | 0.01 | 0.14 | 0.18 |
| 4 | 189690332 | rs10000370 | 0.029 | 1.36 | 0.985 | -0.77 | 0.005 | -0.084 | -0.084 | 0 | 0.21 | 0.31 | 0.14 | 0.06 | 0.2 | 0.16 |
| 4 | 189690383 | rs10000272 | -0.979 | 1.355 | 0.972 | -0.764 | -0.005 | -0.052 | -0.052 | 0.111 | 0.91 | 0.77 | 0.94 | 1 | 0.94 | 0.96 |
| 4 | 189701600 | rs11724214 | 0.057 | 1.31 | 1.264 | -0.781 | -0.001 | -0.02 | -0.02 | 1.122 | 0.2 | 0.17 | 0.13 | 0.05 | 0.2 | 0.16 |
| 4 | 189713163 | rs1461780 | 0.522 | 1.319 | 1.178 | -0.783 | -0.004 | -0.079 | -0.079 | 0.225 | 0.7 | 0.42 | 0.79 | 0.95 | 0.74 | 0.8 |
| 4 | 189728235 | rs7660712 | -0.71 | 1.22 | 0.863 | -0.81 | -0.014 | -0.066 | -0.066 | 0.565 | 0.89 | 0.77 | 0.92 | 1 | 0.94 | 0.96 |
| 4 | 189737788 | rs3911069 | 0.305 | 1.337 | 0.834 | -0.816 | 0.02 | -0.083 | -0.083 | 0 | 0.83 | 0.7 | 0.7 | 0.99 | 0.8 | 0.78 |
| 4 | 189742516 | rs1073671 | 1.096 | 1.313 | 0.638 | -0.62 | -0.011 | 0.039 | 0.039 | 0 | 0.61 | 0.36 | 0.55 | 0.82 | 0.59 | 0.61 |
| 4 | 189749745 | rs9999605 | -0.37 | 1.012 | 0.158 | -1.003 | -0.001 | -0.036 | -0.036 | 0 | 0.25 | 0.2 | 0.29 | 0.63 | 0.25 | 0.34 |
| 4 | 189756948 | rs2377459 | 1.407 | 0.939 | -1.135 | -1.851 | -0.087 | 0.052 | 0.052 | 0 | 0.17 | 0.01 | 0.12 | 0.06 | 0.24 | 0.08 |
| 4 | 189757132 | rs2377462 | -0.301 | 0.883 | -0.777 | -1.997 | 0.016 | -0.012 | -0.012 | 0.364 | 0.79 | 0.68 | 0.67 | 0.87 | 0.74 | 0.69 |
| 4 | 189759434 | rs13150491 | 0.627 | 0.513 | -1.218 | -2.04 | 0.012 | -0.03 | -0.03 | 0.382 | 0.38 | 0.42 | 0.25 | 0.21 | 0.45 | 0.26 |
| 4 | 189759591 | rs4359959 | 0.004 | 0.434 | -1.245 | -2.085 | 0 | -0.012 | -0.012 | 0 | 0.54 | 0.49 | 0.36 | 0.23 | 0.49 | 0.34 |
| 4 | 189759697 | rs6553238 | -0.012 | 0.537 | -1.2 | -2.066 | -0.005 | 0 | 0 | 0 | 0.4 | 0.47 | 0.4 | 0.64 | 0.36 | 0.49 |
| 4 | 189762669 | rs1947186 | 0.655 | 0.537 | -1.117 | -2.107 | 0.006 | -0.022 | -0.022 | 0 | 0.38 | 0.36 | 0.25 | 0.19 | 0.45 | 0.26 |
| 4 | 189763385 | rs6840969 | 0.084 | 0.379 | -1.054 | -1.782 | -0.017 | 0.011 | 0.011 | 0.056 | 0.33 | 0.47 | 0.36 | 0.66 | 0.29 | 0.47 |
| 4 | 189765784 | rs6834584 | -0.207 | 0.385 | -0.992 | -1.814 | -0.014 | -0.038 | -0.038 | 0.408 | 0.37 | 0.54 | 0.39 | 0.8 | 0.32 | 0.58 |
| 4 | 189766744 | rs7661693 | 0.233 | 0.672 | -0.755 | -1.757 | -0.001 | -0.017 | -0.017 | 1.7 | 0.45 | 0.42 | 0.3 | 0.18 | 0.52 | 0.28 |
| 4 | 189791103 | rs6815501 | -0.724 | 0.478 | -0.629 | -1.698 | -0.005 | -0.114 | -0.114 | 0 | 0.22 | 0.42 | 0.17 | 0.04 | 0.16 | 0.14 |
| 4 | 189792929 | rs4526017 | 0.171 | 0.558 | -1.604 | -1.94 | -0.055 | 0.102 | 0.102 | 0 | 0.19 | 0.01 | 0.14 | 0.03 | 0.21 | 0.12 |
| 4 | 189792991 | rs6828833 | -0.346 | 0.456 | -1.274 | -1.99 | -0.001 | 0.039 | 0.039 | 0 | 0.26 | 0.17 | 0.17 | 0.01 | 0.21 | 0.21 |
| 4 | 189797616 | rs10434316 | 0.051 | 0.249 | -0.478 | -1.823 | -0.022 | -0.087 | -0.087 | 0.057 | 0.79 | 0.58 | 0.89 | 0.94 | 0.87 | 0.87 |
| 4 | 189797770 | rs10434317 | 1.654 | 0.11 | -1.014 | -2.234 | 0.004 | 0.051 | 0.051 | 4.605 | 0.15 | 0.05 | 0.08 | 0.06 | 0.16 | 0.05 |
| 4 | 189798780 | rs10434319 | 0.037 | -0.151 | -0.868 | -2.611 | 0.013 | 0.016 | 0.016 | 0 | 0.33 | 0.24 | 0.18 | 0.06 | 0.27 | 0.22 |
| 4 | 189800486 | rs2218692 | -0.328 | 0.778 | -1.085 | -2.38 | -0.013 | 0.032 | 0.032 | 0 | 0.39 | 0.29 | 0.48 | 0.18 | 0.48 | 0.32 |
| 4 | 189805789 | rs2376962 | -0.047 | 0.577 | -1.192 | -2.353 | 0.006 | 0.006 | 0.006 | 0 | 0.42 | 0.38 | 0.37 | 0.7 | 0.36 | 0.44 |
| 4 | 189806380 | rs2122061 | -0.524 | 0.148 | -0.897 | -2.435 | -0.003 | -0.044 | -0.044 | 0 | 0.54 | 0.51 | 0.72 | 0.89 | 0.64 | 0.69 |
| 4 | 189812524 | rs12498968 | -0.111 | 0.192 | -1.001 | -2.405 | -0.003 | 0.028 | 0.028 | 0 | 0.48 | 0.36 | 0.64 | 0.7 | 0.55 | 0.52 |
| 4 | 189813964 | rs6553246 | -0.11 | 0.189 | -1.006 | -2.43 | -0.003 | 0.031 | 0.031 | 0 | 0.47 | 0.35 | 0.64 | 0.7 | 0.55 | 0.53 |
| 4 | 189814241 | rs1597230 | -0.036 | 0.793 | -0.751 | -1.936 | -0.031 | -0.023 | -0.023 | 0 | 0.31 | 0.18 | 0.54 | 0.64 | 0.39 | 0.48 |
| 4 | 189817730 | rs10471186 | 1.108 | 0.635 | -0.37 | -1.653 | 0.017 | 0.004 | 0.004 | 0 | 0.05 | 0.14 | 0.06 | 0.09 | 0.08 | 0.15 |
| 4 | 189818663 | rs7669213 | -0.077 | 0.389 | -0.641 | -1.797 | 0.027 | -0.043 | -0.043 | 0 | 0.19 | 0.04 | 0.1 | 0.17 | 0.11 | 0.21 |
| 4 | 189819886 | rs13120352 | 0.79 | 0.441 | -0.404 | -1.883 | -0.003 | -0.007 | -0.007 | 0 | 0.62 | 0.37 | 0.75 | 0.74 | 0.7 | 0.6 |
| 4 | 189824882 | rs1448742 | 0.81 | 0.439 | -0.405 | -1.884 | 0 | -0.001 | -0.001 | 0 | 0.62 | 0.35 | 0.75 | 0.74 | 0.7 | 0.6 |
| 4 | 189827490 | rs11733637 | -0.487 | 0.34 | 0.146 | -1.99 | 0.002 | 0.015 | 0.015 | 0 | 0.9 | 0.67 | 0.94 | 0.94 | 0.95 | 0.92 |
| 4 | 189828192 | rs7682983 | 0.89 | 0.53 | -0.031 | -2.198 | 0.008 | 0.014 | 0.014 | 0 | 0.47 | 0.34 | 0.31 | 0.09 | 0.44 | 0.29 |
| 4 | 189831920 | rs1448726 | 0.98 | 0.528 | -0.116 | -2.199 | 0.016 | 0.115 | 0.115 | 0 | 0.46 | 0.23 | 0.31 | 0.09 | 0.44 | 0.29 |
| 4 | 189842681 | rs13107204 | 1.202 | 0.899 | -0.185 | -2.164 | -0.053 | 0.064 | 0.064 | 0 | 0.15 | 0.04 | 0.09 | 0 | 0.18 | 0.06 |
| 4 | 189844501 | rs7677253 | 0.7 | 0.722 | -0.013 | -2.114 | -0.001 | -0.001 | -0.001 | 0 | 0.08 | 0.13 | 0.05 | 0.06 | 0.05 | 0.08 |
| 4 | 189847976 | rs9312388 | 0.787 | 1.008 | 0.262 | -1.901 | -0.007 | 0.023 | 0.023 | 0.004 | 0.1 | 0.14 | 0.07 | 0.13 | 0.07 | 0.15 |
| 4 | 189851017 | rs3860664 | 0.508 | 0.821 | 0.313 | -2.175 | -0.007 | -0.029 | -0.029 | 0 | 0.54 | 0.51 | 0.34 | 0.16 | 0.41 | 0.33 |
| 4 | 189861178 | rs4863076 | 0.441 | 1.063 | 0.197 | -2.122 | -0.01 | -0.03 | -0.03 | 0 | 0.7 | 0.72 | 0.81 | 1 | 0.69 | 0.77 |

| | | | | | | | | | | | | | | | | |
|---|---|---|---|---|---|---|---|---|---|---|---|---|---|---|---|---|
| 4 | 189861553 | rs6853039 | 0.855 | 1.282 | 0.137 | -1.944 | -0.009 | 0.004 | 0.004 | 0 | 0.16 | 0.09 | 0.1 | 0 | 0.15 | 0.06 |
| 4 | 189868753 | rs3860668 | 0.192 | 0.886 | 0.271 | -1.879 | -0.018 | -0.043 | -0.043 | 0 | 0.35 | 0.56 | 0.21 | 0.12 | 0.26 | 0.15 |
| 4 | 189870956 | rs11937288 | -0.437 | 0.884 | 0.069 | -1.743 | 0 | 0.009 | 0.009 | 0.369 | 0.44 | 0.57 | 0.31 | 0.29 | 0.4 | 0.32 |
| 4 | 189876706 | rs10027878 | -0.408 | 1.064 | -0.049 | -1.6 | 0.066 | 0.033 | 0.033 | 0.656 | 0.14 | 0.3 | 0.14 | 0.17 | 0.18 | 0.18 |
| 4 | 189881357 | rs7658965 | -0.477 | 0.697 | -0.687 | -1.703 | 0.02 | 0.036 | 0.036 | 0 | 0.6 | 0.68 | 0.76 | 0.97 | 0.68 | 0.75 |
| 4 | 189882597 | rs3860674 | 0.48 | 0.718 | -0.515 | -1.561 | -0.008 | -0.002 | -0.002 | 0.117 | 0.34 | 0.4 | 0.22 | 0.29 | 0.29 | 0.18 |
| 4 | 189884512 | rs6840831 | 0.48 | 0.718 | -0.515 | -1.561 | -0.008 | -0.002 | -0.002 | 0.174 | 0.34 | 0.4 | 0.22 | 0.29 | 0.29 | 0.18 |
| 4 | 189885883 | rs12507360 | -0.519 | 1.261 | -0.741 | -0.944 | 0.007 | 0.002 | 0.002 | 3.714 | 0.31 | 0.33 | 0.57 | 0.68 | 0.44 | 0.58 |
| 4 | 189891709 | rs7662770 | -0.638 | 1.343 | -1.056 | -0.696 | -0.043 | 0.015 | 0.015 | 0.627 | 0.12 | 0.02 | 0.1 | 0.07 | 0.16 | 0.19 |
| 4 | 189892200 | rs6820632 | -0.184 | 1.044 | -0.89 | -0.774 | 0.007 | 0.015 | 0.015 | 0.025 | 0.31 | 0.26 | 0.15 | 0.16 | 0.24 | 0.21 |
| 4 | 189896929 | rs11941699 | -1.253 | 1.107 | -0.57 | -0.61 | 0.002 | -0.018 | -0.018 | 0 | 0.38 | 0.48 | 0.26 | 0.25 | 0.35 | 0.42 |
| 4 | 189906617 | rs10012177 | -0.512 | 1.251 | -0.546 | -0.523 | -0.009 | 0.007 | 0.007 | 0 | 0.14 | 0.12 | 0.12 | 0.03 | 0.22 | 0.16 |
| 4 | 189910956 | rs11132593 | -1.398 | 0.988 | -0.612 | -0.603 | 0 | -0.008 | -0.008 | 0 | 0.48 | 0.55 | 0.47 | 0.25 | 0.44 | 0.47 |
| 4 | 189914135 | rs10007514 | 0.343 | 0.297 | -0.131 | -0.32 | -0.024 | -0.114 | -0.114 | 1.552 | 0.82 | 0.6 | 0.93 | 1 | 0.91 | 0.87 |
| 4 | 189924789 | rs4863432 | -1.116 | 0.08 | -0.373 | 0.217 | -0.009 | 0.014 | 0.014 | 0 | 0.14 | 0.15 | 0.23 | 0.24 | 0.06 | 0.07 |
| 4 | 189926941 | rs13117955 | 0.084 | 0.067 | -0.787 | 0.329 | -0.024 | -0.006 | -0.006 | 0 | 0.11 | 0.01 | 0.04 | 0.07 | 0.07 | 0.12 |
| 4 | 189931630 | rs10018868 | 1.198 | -0.351 | -0.958 | -0.713 | 0.007 | 0.022 | 0.022 | 0 | 0.43 | 0.27 | 0.54 | 0.65 | 0.61 | 0.64 |
| 4 | 189939769 | rs1908801 | 0.425 | -0.778 | -0.728 | -0.961 | 0.006 | 0.016 | 0.016 | 0 | 0.19 | 0.13 | 0.15 | 0 | 0.19 | 0.12 |
| 4 | 189942126 | rs11132597 | 0.665 | -0.882 | -0.131 | -0.726 | -0.053 | 0.055 | 0.055 | 0 | 0.78 | 0.48 | 0.73 | 0.72 | 0.86 | 0.85 |
| 4 | 189949002 | rs1390881 | 0.27 | -0.827 | 0.468 | -0.616 | 0.016 | 0.228 | 0.228 | 0 | 0.92 | 0.67 | 0.81 | 0.79 | 0.95 | 0.94 |
| 4 | 189949645 | rs11941513 | -0.464 | -0.827 | 0.509 | -0.616 | 0.011 | 0.212 | 0.212 | 0 | 0.08 | 0.31 | 0.19 | 0.21 | 0.05 | 0.06 |
| 4 | 189950397 | rs10020429 | 0.065 | -0.781 | 0.367 | -0.745 | -0.005 | -0.011 | -0.011 | 0 | 0.89 | 0.83 | 0.94 | 0.94 | 0.93 | 0.95 |
| 4 | 189950840 | rs7656029 | 0.722 | -0.836 | 0.364 | -0.782 | -0.015 | -0.017 | -0.017 | 0 | 0.77 | 0.71 | 0.89 | 0.87 | 0.85 | 0.86 |
| 4 | 189951379 | rs13126066 | 0.309 | -0.759 | 0.377 | -0.726 | -0.016 | -0.008 | -0.008 | 0 | 0.13 | 0.2 | 0.06 | 0.08 | 0.08 | 0.09 |
| 4 | 189952149 | rs10004274 | -0.001 | 0.071 | -0.334 | -0.343 | -0.021 | -0.04 | -0.04 | 0 | 0.36 | 0.19 | 0.46 | 0.47 | 0.4 | 0.46 |
| 4 | 189957977 | rs1463792 | 0.087 | -0.158 | 0.134 | -0.421 | -0.02 | -0.026 | -0.026 | 0.669 | 0.78 | 0.7 | 0.89 | 0.87 | 0.85 | 0.86 |
| 4 | 189958110 | rs6838005 | 0.087 | -0.158 | 0.134 | -0.421 | -0.02 | -0.026 | -0.026 | 0 | 0.78 | 0.7 | 0.89 | 0.87 | 0.85 | 0.86 |
| 4 | 189962722 | rs2029952 | -0.01 | 0.293 | -0.219 | -0.44 | -0.012 | -0.016 | -0.016 | 0 | 0.34 | 0.28 | 0.48 | 0.51 | 0.41 | 0.48 |
| 4 | 189964583 | rs4632707 | 0.67 | -0.347 | -0.51 | -0.797 | 0.036 | 0.056 | 0.056 | 0 | 0.7 | 0.73 | 0.85 | 0.87 | 0.8 | 0.82 |
| 4 | 189967275 | rs1982946 | -0.232 | 0.167 | -0.214 | -0.648 | 0.002 | 0.003 | 0.003 | 0 | 0.64 | 0.4 | 0.48 | 0.45 | 0.56 | 0.48 |
| 4 | 189969670 | rs7688840 | 0.743 | -0.268 | -0.77 | -1.173 | 0 | -0.041 | -0.041 | 0 | 0.22 | 0.19 | 0.14 | 0 | 0.24 | 0.12 |
| 4 | 189982824 | rs6825584 | -0.017 | -0.432 | -1.083 | -1.063 | -0.003 | -0.013 | -0.013 | 2.328 | 0.16 | 0.01 | 0.08 | 0.12 | 0.13 | 0.13 |
| 4 | 189984206 | rs7676159 | 0.597 | -0.449 | -1.097 | -1.077 | -0.002 | -0.027 | -0.027 | 0 | 0.13 | 0.01 | 0.06 | 0.12 | 0.09 | 0.1 |
| 4 | 189987905 | rs4863449 | 0.185 | -0.545 | -0.782 | -1.37 | 0.003 | 0.001 | 0.001 | 0 | 0.59 | 0.41 | 0.73 | 0.81 | 0.68 | 0.75 |
| 4 | 189988249 | rs4863450 | -0.914 | -0.454 | -0.38 | -1.228 | -0.022 | -0.093 | -0.093 | 0 | 0.76 | 0.56 | 0.82 | 0.93 | 0.81 | 0.88 |
| 4 | 189989962 | rs4863452 | 1.28 | -0.268 | -0.459 | -0.659 | 0.005 | -0.076 | -0.076 | 1.804 | 0.21 | 0.05 | 0.23 | 0.26 | 0.16 | 0.21 |
| 4 | 189989989 | rs1353581 | 0.716 | -0.1 | -0.342 | -0.644 | 0.015 | 0.044 | 0.044 | 0 | 0.16 | 0.19 | 0.37 | 0.51 | 0.27 | 0.36 |
| 4 | 189992773 | rs13137544 | -0.031 | -0.261 | -0.293 | -0.98 | -0.006 | 0.009 | 0.009 | 0 | 0.39 | 0.4 | 0.63 | 0.81 | 0.43 | 0.63 |
| 4 | 189996454 | rs10016346 | 0.394 | 0.112 | -0.177 | -0.118 | -0.017 | -0.023 | -0.023 | 0 | 0.22 | 0.16 | 0.51 | 0.6 | 0.31 | 0.4 |
| 4 | 189998904 | rs1948573 | -0.461 | -0.165 | 0.001 | -0.475 | 0.002 | 0.002 | 0.002 | 0 | 0.63 | 0.63 | 0.81 | 0.73 | 0.73 | 0.67 |
| 4 | 190004805 | rs1496937 | 1.232 | -0.149 | -0.12 | -0.643 | -0.034 | 0.122 | 0.122 | 0 | 0.31 | 0.12 | 0.21 | 0 | 0.36 | 0.21 |
| 4 | 190006491 | rs1353582 | -0.033 | -0.054 | 0.267 | -0.104 | 0.002 | 0 | 0 | 0 | 0.1 | 0.23 | 0.07 | 0.09 | 0.07 | 0.11 |
| 4 | 190008231 | rs2889670 | -0.036 | 0.192 | -0.098 | 0.106 | 0.015 | 0.011 | 0.011 | 0 | 0.26 | 0.32 | 0.56 | 0.72 | 0.34 | 0.42 |
| 4 | 190010270 | rs11931092 | -0.213 | -0.149 | -0.28 | 0.31 | 0.008 | 0.08 | 0.08 | 0 | 0.11 | 0.26 | 0.12 | 0.28 | 0.11 | 0.13 |
| 4 | 190033466 | rs6857536 | 0.216 | -0.539 | -1.25 | 0.348 | 0.063 | -0.011 | -0.011 | 0 | 0.25 | 0.1 | 0.1 | 0.24 | 0.13 | 0.25 |
| 4 | 190034291 | rs6841266 | -0.135 | 0.577 | -0.8 | 0.512 | -0.003 | -0.006 | -0.006 | 0 | 0.73 | 0.73 | 0.5 | 0.47 | 0.7 | 0.59 |
| 4 | 190036900 | rs6835666 | 0.199 | 0.72 | -0.591 | -0.074 | 0.006 | -0.027 | -0.027 | 2.676 | 0.6 | 0.55 | 0.48 | 0.24 | 0.6 | 0.39 |
| 4 | 190037254 | rs2889668 | -0.004 | 0.183 | -0.775 | -1.744 | -0.003 | 0.013 | 0.013 | 0 | 0.54 | 0.62 | 0.53 | 0.78 | 0.45 | 0.65 |
| 4 | 190037266 | rs2377560 | 0.073 | 0.313 | -0.144 | -1.702 | 0.021 | 0.016 | 0.016 | 0 | 0.82 | 0.59 | 0.88 | 0.8 | 0.83 | 0.76 |
| 4 | 190037550 | rs2132663 | 0.176 | 0.399 | -1.053 | -2.273 | -0.001 | -0.086 | -0.086 | 0 | 0.34 | 0.3 | 0.31 | 0.01 | 0.35 | 0.2 |
| 4 | 190042584 | rs9631786 | 0.771 | 1.012 | -1.561 | -1.532 | -0.084 | -0.11 | -0.11 | 0 | 0.2 | 0.03 | 0.19 | 0.53 | 0.28 | 0.28 |
| 4 | 190044374 | rs7690324 | 1.528 | 0.734 | -1.297 | -1.769 | 0.001 | 0 | 0 | 0 | 0.1 | 0.12 | 0.07 | 0.05 | 0.08 | 0.1 |
| 4 | 190044666 | rs7695418 | 0.59 | 0.71 | -1.19 | -1.712 | -0.001 | 0 | 0 | 0 | 0.16 | 0.18 | 0.29 | 0.19 | 0.13 | 0.12 |
| 4 | 190050735 | rs4285137 | -0.685 | 1.156 | -1.228 | -1.767 | -0.016 | -0.093 | -0.093 | 1.361 | 0.78 | 0.77 | 0.88 | 1 | 0.81 | 0.93 |
| 4 | 190056070 | rs9993186 | -0.594 | 1.247 | -1.228 | -1.769 | -0.003 | -0.092 | -0.092 | 0.021 | 0.78 | 0.78 | 0.81 | 1 | 0.78 | 0.93 |
| 4 | 190056255 | rs6845602 | -1.033 | 1.495 | -1.584 | -1.445 | -0.02 | 0.04 | 0.04 | 0 | 0.82 | 0.89 | 0.7 | 0.96 | 0.77 | 0.84 |
| 4 | 190064912 | rs13135232 | 0.368 | 1.361 | -1.423 | -0.774 | -0.053 | 0.051 | 0.051 | 0 | 0.09 | 0.01 | 0.08 | 0 | 0.17 | 0.03 |
| 4 | 190065197 | rs6822346 | -0.232 | 0.571 | -1.107 | -0.471 | -0.014 | -0.008 | -0.008 | 0 | 0.36 | 0.47 | 0.36 | 0.35 | 0.29 | 0.44 |
| 4 | 190066616 | rs6834497 | 0.198 | 0.844 | -1.214 | -0.924 | -0.001 | -0.004 | -0.004 | 2.651 | 0.65 | 0.41 | 0.67 | 0.81 | 0.62 | 0.69 |
| 4 | 190069486 | rs4302526 | -0.402 | 0.584 | -1.837 | -0.931 | 0.012 | 0.071 | 0.071 | 0 | 0.28 | 0.16 | 0.18 | 0.17 | 0.26 | 0.26 |
| 4 | 190069501 | rs11944137 | 0.297 | 0.348 | -1.808 | -1.171 | -0.007 | 0.041 | 0.041 | 0.946 | 0.33 | 0.24 | 0.51 | 0.76 | 0.36 | 0.52 |
| 4 | 190070914 | rs4241915 | 0.084 | 0.203 | -1.264 | -1.225 | -0.003 | -0.04 | -0.04 | 0.117 | 0.35 | 0.45 | 0.53 | 0.77 | 0.36 | 0.52 |
| 4 | 190071985 | rs11737011 | 0.2 | 0.2 | -2.6 | -1.7 | -0.131 | 0.149 | 0.149 | 1.249 | 0.38 | 0.07 | 0.33 | 0.19 | 0.45 | 0.23 |
| 4 | 190080067 | rs4863105 | 1.173 | 0.562 | -2.476 | -0.59 | -0.034 | 0.022 | 0.022 | 0 | 0.07 | 0 | 0.23 | 0.46 | 0.1 | 0.19 |
| 4 | 190087087 | rs9996363 | 0.264 | 0.187 | -2.141 | -0.988 | -0.002 | -0.075 | -0.075 | 0 | 0.25 | 0.66 | 0.18 | 0.04 | 0.21 | 0.25 |
| 4 | 190087210 | rs10019172 | -0.201 | 0.346 | -2.487 | -1.007 | -0.002 | 0.082 | 0.082 | 0 | 0.58 | 0.8 | 0.47 | 0.26 | 0.55 | 0.49 |
| 4 | 190087776 | rs7677465 | -0.225 | 0.562 | -1.972 | -0.351 | 0.001 | -0.007 | -0.007 | 0 | 0.77 | 0.8 | 0.71 | 0.51 | 0.81 | 0.73 |
| 4 | 190088167 | rs7699928 | -0.124 | 0.495 | -1.379 | -0.568 | 0.012 | 0.048 | 0.048 | 0.231 | 0.47 | 0.63 | 0.4 | 0.28 | 0.42 | 0.54 |
| 4 | 190093246 | rs6848521 | 0.046 | 0.401 | -1.006 | -0.272 | -0.012 | 0.014 | 0.014 | 0 | 0.4 | 0.41 | 0.41 | 0.73 | 0.51 | 0.47 |

| | | | | | | | | | | | | | | | | |
|---|---|---|---|---|---|---|---|---|---|---|---|---|---|---|---|---|
| 4 | 190098562 | rs4241857 | -0.068 | -0.386 | -0.668 | -0.33 | 0.011 | 0.019 | 0.019 | 0 | 0.71 | 0.81 | 0.69 | 0.87 | 0.77 | 0.76 |
| 4 | 190101262 | rs9312393 | -0.547 | -0.691 | -0.654 | -0.557 | -0.02 | -0.045 | -0.045 | 0.008 | 0.89 | 0.88 | 0.96 | 1 | 0.93 | 0.94 |
| 4 | 190103064 | rs6855006 | 1.241 | -1.27 | -0.853 | -0.124 | 0.08 | -0.015 | -0.015 | 0 | 0.53 | 0.17 | 0.36 | 0.61 | 0.66 | 0.5 |
| 4 | 190111329 | rs12650063 | 0.459 | -1.131 | -0.287 | 0.065 | 0 | -0.003 | -0.003 | 0 | 0.19 | 0.16 | 0.27 | 0.2 | 0.16 | 0.22 |
| 4 | 190117114 | rs11132619 | 0.204 | -1.138 | 0.193 | -0.002 | -0.01 | 0 | 0 | 0.684 | 0.15 | 0.27 | 0.25 | 0.13 | 0.1 | 0.11 |
| 4 | 190122932 | rs6822278 | 1.042 | -1.281 | -0.084 | 0.067 | 0.079 | 0.005 | 0.005 | 0.191 | 0.54 | 0.17 | 0.41 | 0.66 | 0.67 | 0.61 |
| 4 | 190126642 | rs7437007 | -0.564 | -1.526 | 0.585 | 0.073 | -0.039 | -0.038 | -0.038 | 0 | 0.88 | 0.69 | 0.9 | 0.95 | 0.93 | 0.86 |
| 4 | 190129325 | rs11732071 | 1.003 | -1.803 | 0.582 | 0.079 | 0.024 | -0.016 | -0.016 | 0 | 0.63 | 0.31 | 0.46 | 0.52 | 0.74 | 0.51 |
| 4 | 190129958 | rs4863119 | -0.017 | -1.802 | 0.63 | 0.146 | 0.004 | -0.015 | -0.015 | 0 | 0.23 | 0.17 | 0.29 | 0.3 | 0.19 | 0.26 |
| 4 | 190130152 | rs4388153 | 1.599 | -1.801 | 0.652 | 0.107 | 0.08 | -0.004 | -0.004 | 2.786 | 0.49 | 0.17 | 0.35 | 0.52 | 0.64 | 0.47 |
| 4 | 190130349 | rs4863120 | 0.426 | -1.689 | 0.942 | 0.547 | 0 | 0 | 0 | 0 | 0.18 | 0.15 | 0.26 | 0.23 | 0.15 | 0.15 |
| 4 | 190137076 | rs6833202 | 0.511 | -1.296 | 0.946 | 0.301 | -0.036 | -0.033 | -0.033 | 0 | 0.83 | 0.66 | 0.86 | 0.86 | 0.89 | 0.74 |
| 4 | 190146555 | rs10866319 | -0.465 | -1.155 | 0.729 | 0.234 | 0.005 | -0.007 | -0.007 | 0 | 0.91 | 0.93 | 0.97 | 1 | 0.93 | 0.95 |
| 4 | 190147729 | rs11940741 | -0.053 | -1.23 | 0.957 | 0.218 | -0.004 | 0.054 | 0.054 | 0 | 0.07 | 0.2 | 0.18 | 0.12 | 0.06 | 0.11 |
| 4 | 190148795 | rs10446846 | 0.984 | -1.472 | 1.204 | 0.072 | -0.068 | -0.03 | -0.03 | 0 | 0.79 | 0.57 | 0.75 | 0.88 | 0.85 | 0.81 |
| 4 | 190151383 | rs4339259 | -0.709 | -0.966 | 0.455 | 0.339 | -0.002 | -0.004 | -0.004 | 0 | 0.4 | 0.61 | 0.47 | 0.54 | 0.41 | 0.52 |
| 4 | 190158729 | rs4434303 | -0.34 | -0.902 | 0.395 | 0.17 | -0.001 | 0.002 | 0.002 | 0 | 0.11 | 0.12 | 0.1 | 0.05 | 0.11 | 0.07 |
| 4 | 190170631 | rs4249884 | -0.972 | -1.322 | 0.141 | -0.114 | 0.024 | 0.018 | 0.018 | 0 | 0.88 | 0.96 | 0.94 | 1 | 0.92 | 0.95 |
| 4 | 190170679 | rs3922497 | 1.737 | -1.506 | -0.322 | 0.007 | -0.037 | 0.01 | 0.01 | 0 | 0.27 | 0.16 | 0.19 | 0.22 | 0.37 | 0.18 |
| 4 | 190181541 | rs6553297 | -0.259 | -1.629 | 0.208 | 0.181 | 0 | -0.002 | -0.002 | 0 | 0.47 | 0.48 | 0.59 | 0.63 | 0.44 | 0.59 |
| 4 | 190188452 | rs9998868 | -0.092 | -1.562 | -0.343 | 0.157 | 0.008 | 0.011 | 0.011 | 0 | 0.78 | 0.8 | 0.84 | 0.93 | 0.82 | 0.88 |
| 4 | 190189043 | rs11721813 | 1.701 | -1.326 | -0.605 | 0.259 | -0.111 | -0.021 | -0.021 | 0 | 0.27 | 0.05 | 0.18 | 0.22 | 0.37 | 0.18 |
| 4 | 190204029 | rs6553301 | -0.996 | -1.236 | -0.062 | -0.148 | -0.013 | -0.01 | -0.01 | 0 | 0.37 | 0.59 | 0.19 | 0.22 | 0.25 | 0.27 |
| 4 | 190206038 | rs7667607 | 0.025 | -1.118 | 0.329 | -0.083 | -0.001 | -0.001 | -0.001 | 0 | 0.35 | 0.34 | 0.3 | 0.63 | 0.39 | 0.48 |
| 4 | 190209288 | rs4241871 | 1.08 | -1.039 | -0.013 | -0.103 | -0.035 | 0.032 | 0.032 | 0 | 0.41 | 0.22 | 0.52 | 0.31 | 0.45 | 0.32 |
| 4 | 190223523 | rs10006955 | 0.884 | -0.919 | -0.03 | -0.051 | -0.012 | 0.116 | 0.116 | 0.594 | 0.46 | 0.23 | 0.52 | 0.24 | 0.47 | 0.3 |
| 4 | 190243940 | rs6842514 | -1.12 | -0.793 | 0.696 | 0.359 | -0.009 | 0.132 | 0.132 | 1.062 | 0.18 | 0.52 | 0.15 | 0.32 | 0.13 | 0.22 |
| 4 | 190256110 | rs11132639 | -1.008 | -0.877 | 0.794 | 0.352 | -0.01 | 0.159 | 0.159 | 0.091 | 0.15 | 0.41 | 0.15 | 0.3 | 0.12 | 0.16 |
| 4 | 190256300 | rs6841296 | -1.22 | -0.879 | 0.85 | 0.345 | -0.006 | 0.164 | 0.164 | 0.256 | 0.16 | 0.48 | 0.15 | 0.3 | 0.12 | 0.17 |
| 4 | 190261652 | rs4863138 | 1.732 | -0.886 | 0.351 | 0.158 | -0.097 | 0.048 | 0.048 | 0 | 0.31 | 0.07 | 0.44 | 0.21 | 0.38 | 0.26 |
| 4 | 190262811 | rs4608851 | -0.663 | -1 | 0.265 | 0.17 | -0.032 | -0.051 | -0.051 | 0 | 0.73 | 0.89 | 0.76 | 0.61 | 0.68 | 0.59 |
| 4 | 190262923 | rs6828995 | -0.258 | -0.887 | 0.876 | 0.365 | 0.004 | 0.12 | 0.12 | 0.422 | 0.12 | 0.33 | 0.12 | 0.26 | 0.11 | 0.07 |
| 4 | 190264846 | rs13129984 | 0.753 | -0.941 | 0.89 | 0.313 | 0.02 | 0.041 | 0.041 | 0 | 0.2 | 0.12 | 0.14 | 0.07 | 0.15 | 0.11 |
| 4 | 190268759 | rs10032784 | -1.058 | -0.933 | 0.922 | 0.332 | -0.032 | 0.022 | 0.022 | 0 | 0.35 | 0.53 | 0.28 | 0.37 | 0.27 | 0.29 |
| 4 | 190276865 | rs10001776 | 0.481 | -0.891 | 0.865 | 0.345 | 0.02 | 0.023 | 0.023 | 0 | 0.22 | 0.15 | 0.17 | 0.11 | 0.16 | 0.22 |
| 4 | 190278973 | rs11930136 | 0.728 | -0.92 | 0.949 | 0.351 | 0.005 | 0.104 | 0.104 | 0 | 0.09 | 0.19 | 0.08 | 0.26 | 0.09 | 0.06 |
| 4 | 190278985 | rs9991497 | 0.068 | -0.898 | 0.786 | 0.085 | 0.004 | 0.004 | 0.004 | 0 | 0.23 | 0.21 | 0.17 | 0.11 | 0.16 | 0.21 |
| 4 | 190290460 | rs1037644 | 0.424 | -0.877 | 0.686 | -0.151 | -0.023 | -0.06 | -0.06 | 0 | 0.68 | 0.57 | 0.73 | 0.93 | 0.77 | 0.87 |
| 4 | 190292844 | rs11943937 | 0.181 | -0.875 | 0.667 | -0.146 | -0.002 | -0.015 | -0.015 | 0.019 | 0.71 | 0.7 | 0.77 | 0.93 | 0.79 | 0.87 |
| 4 | 190293157 | rs1563455 | 0.399 | -0.865 | 0.657 | -0.144 | -0.023 | -0.06 | -0.06 | 0.875 | 0.68 | 0.57 | 0.73 | 0.93 | 0.77 | 0.87 |
| 4 | 190294537 | rs870190 | -0.173 | -0.442 | 0.749 | 0.243 | -0.002 | -0.009 | -0.009 | 0 | 0.24 | 0.21 | 0.36 | 0.45 | 0.27 | 0.44 |
| 4 | 190296345 | rs4863145 | 0.789 | -0.54 | 0.895 | 0.098 | -0.017 | -0.081 | -0.081 | 2.108 | 0.87 | 0.71 | 0.87 | 0.97 | 0.91 | 0.94 |
| 4 | 190300221 | rs4863155 | 1.032 | -0.518 | 0.679 | 0.187 | 0.002 | -0.003 | -0.003 | 2.492 | 0.62 | 0.41 | 0.5 | 0.52 | 0.64 | 0.5 |
| 4 | 190306460 | rs876623 | 0.118 | -0.371 | 0.566 | -0.019 | -0.011 | -0.068 | -0.068 | 0 | 0.89 | 0.69 | 0.88 | 1 | 0.93 | 0.98 |
| 4 | 190307284 | rs1107680 | 0.138 | -0.226 | 0.441 | 0.176 | -0.014 | -0.036 | -0.036 | 0.005 | 0.23 | 0.13 | 0.35 | 0.42 | 0.3 | 0.41 |
| 4 | 190311936 | rs1563452 | 0.635 | -0.068 | 0.445 | 0.237 | -0.001 | 0.042 | 0.042 | 2.118 | 0.67 | 0.56 | 0.53 | 0.56 | 0.64 | 0.57 |
| 4 | 190317498 | rs11132649 | -0.167 | -0.001 | 0.121 | 0.271 | -0.001 | 0 | 0 | 0.753 | 0.25 | 0.22 | 0.37 | 0.42 | 0.31 | 0.42 |
| 4 | 190328590 | rs6553315 | -0.722 | 0.151 | -0.065 | -0.047 | -0.004 | -0.037 | -0.037 | 0 | 0.93 | 0.88 | 0.92 | 1 | 0.96 | 0.99 |
| 4 | 190333580 | rs4863170 | 0.191 | 0.6 | -0.784 | -0.115 | -0.007 | 0.038 | 0.038 | 0 | 0.34 | 0.27 | 0.33 | 0.2 | 0.34 | 0.35 |
| 4 | 190336803 | rs1350180 | -0.575 | 0.331 | -0.521 | 0.105 | -0.005 | 0.007 | 0.007 | 0.093 | 0.47 | 0.66 | 0.53 | 0.47 | 0.48 | 0.54 |
| 4 | 190338123 | rs10034049 | 0.552 | 0.417 | -0.367 | -0.357 | 0 | 0.009 | 0.009 | 0 | 0.21 | 0.17 | 0.11 | 0 | 0.19 | 0.09 |
| 4 | 190341415 | rs872030 | -1.397 | 0.458 | -0.616 | -0.536 | 0.002 | 0.012 | 0.012 | 1.017 | 0.52 | 0.66 | 0.55 | 0.46 | 0.52 | 0.64 |
| 4 | 190344851 | rs9994370 | 0.614 | 0.565 | -0.04 | -0.265 | 0.006 | 0.03 | 0.03 | 0.308 | 0.18 | 0.32 | 0.34 | 0.31 | 0.21 | 0.35 |
| 4 | 190346910 | rs1869943 | 0.183 | 0.56 | 0.047 | -0.259 | 0 | -0.011 | -0.011 | 0.261 | 0.25 | 0.25 | 0.12 | 0.05 | 0.21 | 0.12 |
| 4 | 190347676 | rs2054178 | -1.036 | 0.496 | -0.015 | -0.259 | -0.001 | -0.002 | -0.002 | 0.001 | 0.42 | 0.6 | 0.46 | 0.37 | 0.41 | 0.47 |
| 4 | 190350300 | rs6823300 | -0.675 | 0.344 | 0.183 | -0.185 | 0.007 | 0.041 | 0.041 | 1.043 | 0.28 | 0.39 | 0.43 | 0.46 | 0.31 | 0.5 |
| 4 | 190351843 | rs4148 | 0.395 | 0.563 | -0.246 | -0.504 | -0.046 | 0.115 | 0.115 | 0 | 0.19 | 0.02 | 0.1 | 0 | 0.19 | 0.09 |
| 4 | 190355589 | rs7680130 | 0.348 | 0.682 | 0.041 | -0.354 | 0.016 | 0.029 | 0.029 | 0 | 0.07 | 0.19 | 0.11 | 0.14 | 0.1 | 0.09 |
| 4 | 190357461 | rs1456354 | -1.459 | 0.414 | 0.073 | -0.182 | -0.004 | 0.074 | 0.074 | 1.589 | 0.9 | 0.92 | 0.95 | 0.6 | 0.92 | 0.87 |
| 4 | 190360400 | rs1125808 | 0.52 | 1.032 | -0.068 | -0.32 | -0.031 | -0.057 | -0.057 | 0 | 0.15 | 0.02 | 0.32 | 0.28 | 0.2 | 0.37 |
| 4 | 190361647 | rs6553316 | -0.681 | 1.004 | -0.085 | -0.491 | -0.007 | 0.021 | 0.021 | 0 | 0.3 | 0.28 | 0.22 | 0.14 | 0.3 | 0.18 |
| 4 | 190371697 | rs10007036 | -0.654 | 1.02 | -0.053 | -0.563 | 0 | -0.022 | -0.022 | 0 | 0.3 | 0.36 | 0.22 | 0.09 | 0.3 | 0.18 |
| 4 | 190374703 | rs867016 | 0.556 | 0.918 | -0.109 | -0.502 | -0.03 | -0.065 | -0.065 | 0 | 0.16 | 0.04 | 0.32 | 0.33 | 0.2 | 0.36 |
| 4 | 190389490 | rs12646102 | -1.692 | 0.902 | 0.076 | -0.519 | 0 | 0.004 | 0.004 | 0.584 | 0.48 | 0.56 | 0.56 | 0.42 | 0.5 | 0.53 |
| 4 | 190394124 | rs1840714 | -0.033 | 1.222 | -0.496 | -0.456 | 0.005 | -0.019 | -0.019 | 0 | 0.08 | 0.17 | 0.07 | 0.04 | 0.11 | 0.08 |
| 4 | 190394821 | rs12504268 | -0.012 | 1.122 | -0.516 | -0.422 | 0.021 | 0.01 | 0.01 | 0.564 | 0.08 | 0.24 | 0.11 | 0.09 | 0.1 | 0.08 |
| 4 | 190407970 | rs4862968 | 0.041 | 1.186 | -0.756 | -0.71 | 0.001 | 0.02 | 0.02 | 0.559 | 0.18 | 0.29 | 0.33 | 0.32 | 0.21 | 0.35 |
| 4 | 190413642 | rs9312400 | -0.143 | 1.216 | -0.893 | -0.793 | 0 | -0.062 | -0.062 | 0.719 | 0.24 | 0.26 | 0.13 | 0 | 0.2 | 0.1 |
| 4 | 190414241 | rs6838957 | -1.175 | 1.23 | -0.925 | -0.785 | 0.002 | -0.024 | -0.024 | 0.183 | 0.41 | 0.53 | 0.46 | 0.32 | 0.42 | 0.43 |

| | | | | | | | | | | | | | | | | |
|---|---|---|---|---|---|---|---|---|---|---|---|---|---|---|---|---|
| 4 | 190414910 | rs11132664 | -0.464 | 1.198 | -1.324 | 0.065 | -0.002 | -0.002 | -0.002 | 0 | 0.22 | 0.26 | 0.39 | 0.35 | 0.31 | 0.41 |
| 4 | 190438606 | rs1551137 | 0.379 | 1.047 | -1.856 | -0.05 | -0.002 | 0.108 | 0.108 | 0 | 0.41 | 0.12 | 0.39 | 0.26 | 0.39 | 0.46 |
| 4 | 190448383 | rs1597619 | 0.787 | 0.989 | -1.694 | -0.415 | -0.023 | -0.071 | -0.071 | 0 | 0.31 | 0.52 | 0.25 | 0.15 | 0.27 | 0.29 |
| 4 | 190450305 | rs1597620 | -0.169 | 1.143 | -1.695 | -0.421 | 0.017 | 0.068 | 0.068 | 0 | 0.43 | 0.68 | 0.4 | 0.17 | 0.37 | 0.35 |
| 4 | 190459134 | rs9985694 | -1.154 | 1.68 | -0.935 | 0.683 | 0 | -0.025 | -0.025 | 0 | 0.12 | 0.16 | 0.15 | 0.02 | 0.1 | 0.06 |
| 4 | 190467123 | rs6826535 | 0.437 | 2.05 | -1.354 | 1.132 | 0.001 | 0.021 | 0.021 | 0 | 0.83 | 0.82 | 0.84 | 0.54 | 0.75 | 0.82 |
| 4 | 190481771 | rs11933070 | -1.172 | 1.106 | -1.459 | 0.24 | 0.003 | 0.037 | 0.037 | 0 | 0.13 | 0.07 | 0.07 | 0 | 0.1 | 0.05 |
| 4 | 190522201 | rs11724261 | -0.744 | 1.575 | -1.296 | 0.755 | -0.025 | -0.008 | -0.008 | 0 | 0.26 | 0.16 | 0.34 | 0.27 | 0.34 | 0.17 |
| 4 | 190543118 | rs4092077 | 0.065 | 0.795 | -0.818 | 0.69 | -0.005 | -0.004 | -0.004 | 0 | 0.55 | 0.48 | 0.49 | 0.54 | 0.39 | 0.32 |
| 4 | 190552585 | rs11729484 | 0.137 | 0.503 | -0.496 | 0.344 | 0.025 | 0.043 | 0.043 | 0 | 0.49 | 0.39 | 0.31 | 0.2 | 0.37 | 0.3 |
| 4 | 190641170 | rs11132733 | 0.611 | 0.094 | -0.645 | 0.223 | 0.006 | 0.002 | 0.002 | 0.414 | 0.76 | 0.27 | 0.73 | 0.75 | 0.8 | 0.8 |
| 4 | 190690945 | rs7684487 | -0.983 | 0.09 | 0.019 | -1.446 | 0.036 | -0.083 | -0.083 | 0.103 | 0.32 | 0.54 | 0.33 | 0.16 | 0.4 | 0.36 |
| 5 | 404804 | rs2672715 | 0.177 | 0.193 | -1.441 | -0.004 | -0.027 | 0.086 | 0.086 | 0 | 0.74 | 0.55 | 0.73 | 0.57 | 0.84 | 0.74 |
| 5 | 407230 | rs732111 | -0.675 | 0.177 | -1.054 | -0.155 | -0.019 | 0.051 | 0.051 | 0.237 | 0.12 | 0.47 | 0.19 | 0.13 | 0.09 | 0.06 |
| 5 | 409760 | rs2671898 | -0.968 | 0.232 | -1.191 | -0.234 | -0.001 | -0.008 | -0.008 | 0 | 0.32 | 0.64 | 0.41 | 0.14 | 0.42 | 0.25 |
| 5 | 412694 | rs4956936 | 1.054 | 0.719 | -1.712 | -0.044 | 0.042 | -0.008 | -0.008 | 0 | 0.61 | 0.23 | 0.55 | 0.42 | 0.72 | 0.67 |
| 5 | 422752 | rs10072668 | -1.145 | 1.201 | -1.97 | -0.289 | -0.116 | 0.114 | 0.114 | 0 | 0.34 | 0.1 | 0.45 | 0.17 | 0.48 | 0.3 |
| 5 | 423385 | rs2672777 | 0.407 | 1.337 | -2.418 | -0.002 | -0.088 | -0.149 | -0.149 | 0.367 | 0.8 | 0.99 | 0.76 | 0.58 | 0.69 | 0.66 |
| 5 | 429989 | rs957792 | 0.038 | 1.174 | -2.316 | -0.145 | 0.001 | -0.021 | -0.021 | 0.019 | 0.08 | 0.02 | 0.14 | 0.13 | 0.07 | 0.06 |
| 5 | 431196 | rs768479 | -0.75 | 1.338 | -2.076 | -0.24 | -0.063 | 0.046 | 0.046 | 0.956 | 0.23 | 0.12 | 0.34 | 0.14 | 0.42 | 0.26 |
| 5 | 433654 | rs2720997 | 0.043 | 0.824 | -1.252 | -0.081 | -0.011 | 0.051 | 0.051 | 0 | 0.83 | 0.82 | 0.79 | 0.58 | 0.86 | 0.75 |
| 5 | 433952 | rs2672724 | 2.007 | 0.496 | -1.264 | -0.186 | 0.033 | 0.029 | 0.029 | 0 | 0.48 | 0.69 | 0.33 | 0.38 | 0.38 | 0.43 |
| 5 | 435055 | rs2672779 | 1.98 | 0.502 | -1.282 | -0.224 | 0.037 | 0.018 | 0.018 | 0 | 0.48 | 0.68 | 0.34 | 0.41 | 0.38 | 0.43 |
| 5 | 436665 | rs349583 | 1.105 | -0.107 | -1.328 | -0.164 | 0.002 | 0.009 | 0.009 | 0 | 0.64 | 0.63 | 0.54 | 0.52 | 0.58 | 0.65 |
| 5 | 438102 | rs10078 | 1.037 | -0.006 | -1.375 | -0.235 | 0.017 | -0.021 | -0.021 | 0.87 | 0.7 | 0.76 | 0.65 | 0.44 | 0.78 | 0.68 |
| 5 | 438564 | rs2241598 | 1.014 | -0.123 | -1.471 | -0.329 | 0.055 | -0.105 | -0.105 | 0 | 0.71 | 0.9 | 0.7 | 0.45 | 0.79 | 0.69 |
| 5 | 442790 | rs10035653 | 0.467 | -0.094 | -1.371 | -0.283 | 0.017 | -0.01 | -0.01 | 0 | 0.12 | 0.03 | 0.15 | 0.13 | 0.07 | 0.08 |
| 5 | 446577 | rs12654521 | 2.182 | -0.178 | -1.435 | -0.373 | 0.028 | 0.01 | 0.01 | 0 | 0.46 | 0.7 | 0.33 | 0.4 | 0.37 | 0.42 |
| 5 | 453567 | rs2672722 | 1.121 | -0.027 | -1.389 | -0.288 | -0.041 | -0.055 | -0.055 | 0 | 0.6 | 0.8 | 0.49 | 0.53 | 0.45 | 0.51 |
| 5 | 456915 | rs2721010 | 1.258 | -0.023 | -1.393 | -0.282 | -0.045 | -0.053 | -0.053 | 0 | 0.6 | 0.8 | 0.49 | 0.53 | 0.45 | 0.51 |
| 5 | 464153 | rs2561667 | 1.814 | 0.15 | -1.387 | -0.217 | 0.036 | 0.016 | 0.016 | 0 | 0.48 | 0.7 | 0.34 | 0.4 | 0.38 | 0.43 |
| 5 | 477968 | rs2278254 | 1.737 | 0.033 | -1.151 | -0.21 | 0.001 | 0.009 | 0.009 | 0 | 0.48 | 0.61 | 0.37 | 0.4 | 0.49 | 0.47 |
| 5 | 480509 | rs2241597 | 0.35 | 0.084 | -1.327 | 0.113 | 0.028 | -0.076 | -0.076 | 0 | 0.83 | 0.93 | 0.81 | 0.57 | 0.87 | 0.75 |
| 5 | 488298 | rs2303736 | 0.382 | 0.035 | -1.235 | 0.322 | 0.027 | -0.039 | -0.039 | 0 | 0.85 | 0.92 | 0.81 | 0.57 | 0.92 | 0.73 |
| 5 | 488862 | rs6869782 | 0.493 | 0.602 | -1.365 | -0.142 | -0.115 | 0.096 | 0.096 | 0 | 0.16 | 0.01 | 0.22 | 0.01 | 0.3 | 0.17 |
| 5 | 492922 | rs10475268 | 1.32 | 0.317 | -0.523 | -0.138 | 0 | 0.004 | 0.004 | 0.044 | 0.51 | 0.46 | 0.39 | 0.55 | 0.52 | 0.47 |
| 5 | 494956 | rs4957043 | 0.755 | -0.008 | -0.095 | -0.307 | -0.007 | 0.078 | 0.078 | 0 | 0.78 | 0.62 | 0.75 | 0.62 | 0.84 | 0.72 |
| 5 | 495377 | rs6868535 | 1.776 | 0.321 | -0.365 | -0.328 | 0.007 | -0.011 | -0.011 | 1.041 | 0.42 | 0.26 | 0.25 | 0.42 | 0.39 | 0.34 |
| 5 | 497277 | rs13359853 | 0.1 | 0.397 | -0.283 | -0.103 | 0 | -0.006 | -0.006 | 0.291 | 0.2 | 0.24 | 0.3 | 0.15 | 0.22 | 0.09 |
| 5 | 499071 | rs4957044 | -0.262 | 0.254 | -0.414 | 0.094 | -0.007 | 0.022 | 0.022 | 0 | 0.16 | 0.17 | 0.18 | 0.35 | 0.1 | 0.31 |
| 5 | 501544 | rs13177896 | -0.97 | 0.57 | -0.75 | -0.097 | -0.003 | 0.013 | 0.013 | 0.157 | 0.39 | 0.57 | 0.52 | 0.51 | 0.35 | 0.42 |
| 5 | 506051 | rs6864158 | 0.652 | 0.716 | -0.908 | 0.136 | -0.002 | -0.015 | -0.015 | 0 | 0.4 | 0.34 | 0.4 | 0.52 | 0.41 | 0.4 |
| 5 | 514785 | rs4956949 | 1.03 | 0.486 | -0.805 | 0.489 | 0.004 | 0.025 | 0.025 | 0 | 0.74 | 0.72 | 0.72 | 0.61 | 0.73 | 0.62 |
| 5 | 516808 | rs2434389 | 0.407 | -0.336 | -1.244 | 0.652 | 0.025 | -0.043 | -0.043 | 0 | 0.82 | 0.9 | 0.86 | 0.64 | 0.9 | 0.71 |
| 5 | 539310 | rs772654 | 0.503 | -0.453 | -1.58 | 0.123 | -0.018 | 0.087 | 0.087 | 0 | 0.52 | 0.19 | 0.54 | 0.66 | 0.54 | 0.66 |
| 5 | 539460 | rs772655 | -0.097 | -0.375 | -1.554 | 0.084 | -0.037 | -0.02 | -0.02 | 0 | 0.66 | 0.9 | 0.64 | 0.67 | 0.62 | 0.74 |
| 5 | 542023 | rs9800013 | 0.876 | -0.233 | -1.602 | -0.683 | -0.077 | 0.089 | 0.089 | 0.041 | 0.15 | 0 | 0.16 | 0.03 | 0.21 | 0.2 |
| 5 | 560476 | rs17563576 | 1.215 | -0.224 | -1.54 | -0.661 | -0.044 | 0.071 | 0.071 | 0 | 0.13 | 0.01 | 0.16 | 0.03 | 0.18 | 0.18 |
| 5 | 560923 | rs6555357 | 0.46 | -0.319 | -0.414 | -0.734 | 0.041 | -0.095 | -0.095 | 0 | 0.17 | 0.29 | 0.18 | 0.04 | 0.2 | 0.19 |
| 5 | 566782 | rs2672027 | -0.811 | -0.411 | -0.925 | -0.345 | -0.05 | -0.139 | -0.139 | 0 | 0.8 | 0.98 | 0.77 | 0.55 | 0.77 | 0.91 |
| 5 | 567246 | rs3853657 | -0.814 | -1.232 | -0.238 | -1.024 | -0.008 | -0.008 | -0.008 | 0 | 0.32 | 0.66 | 0.27 | 0.19 | 0.17 | 0.34 |
| 5 | 572663 | rs16900912 | -0.259 | -1.018 | 0.058 | -0.939 | -0.042 | -0.014 | -0.014 | 0.501 | 0.22 | 0.44 | 0.22 | 0.2 | 0.12 | 0.29 |
| 5 | 573395 | rs3915384 | 1.433 | -0.914 | -0.504 | -0.932 | -0.051 | 0.08 | 0.08 | 0.126 | 0.15 | 0.03 | 0.16 | 0.03 | 0.2 | 0.18 |
| 5 | 581053 | rs4541696 | -1.028 | -0.884 | -0.368 | -0.872 | 0.012 | 0.097 | 0.097 | 0 | 0.44 | 0.74 | 0.43 | 0.24 | 0.36 | 0.48 |
| 5 | 594083 | rs11739663 | -0.145 | -0.907 | -0.106 | -1.035 | -0.009 | -0.179 | -0.179 | 0 | 0.26 | 0.48 | 0.18 | 0.01 | 0.23 | 0.21 |
| 5 | 595870 | rs4045344 | 1.645 | -0.973 | -0.273 | -0.944 | -0.051 | -0.065 | -0.065 | 0 | 0.32 | 0.08 | 0.31 | 0.31 | 0.41 | 0.36 |
| 5 | 598237 | rs9313132 | 0.462 | -0.98 | -0.056 | -0.941 | 0.037 | -0.024 | -0.024 | 0.504 | 0.09 | 0.45 | 0.07 | 0 | 0.04 | 0.04 |
| 5 | 598289 | rs9313133 | 0.462 | -0.98 | -0.075 | -0.941 | 0.035 | -0.025 | -0.025 | 0 | 0.09 | 0.44 | 0.07 | 0 | 0.04 | 0.04 |
| 5 | 606384 | rs16900924 | 0.112 | -0.955 | 0.062 | -0.94 | 0.011 | -0.047 | -0.047 | 0 | 0.1 | 0.34 | 0.06 | 0.01 | 0.06 | 0.05 |
| 5 | 607388 | rs4956954 | 0.313 | -0.982 | -0.003 | -0.958 | -0.004 | -0.002 | -0.002 | 0 | 0.58 | 0.52 | 0.54 | 0.33 | 0.64 | 0.57 |
| 5 | 611753 | rs3810857 | 1.289 | -1.22 | -0.302 | -0.981 | 0.026 | -0.018 | -0.018 | 2.209 | 0.12 | 0.04 | 0.09 | 0.14 | 0.07 | 0.24 |
| 5 | 614963 | rs1697952 | 0.086 | -1.027 | -0.138 | -0.928 | 0.002 | -0.001 | -0.001 | 0 | 0.6 | 0.58 | 0.55 | 0.34 | 0.66 | 0.6 |
| 5 | 618586 | rs7726839 | -0.22 | -0.679 | 0.107 | -0.87 | 0.002 | -0.158 | -0.158 | 0 | 0.27 | 0.51 | 0.23 | 0.03 | 0.25 | 0.22 |
| 5 | 640705 | rs868649 | 0.471 | -0.586 | -0.867 | -0.524 | 0.001 | 0.02 | 0.02 | 0.167 | 0.23 | 0.26 | 0.25 | 0.47 | 0.23 | 0.13 |
| 5 | 652034 | rs12518227 | -0.213 | -0.705 | -0.977 | -0.678 | -0.031 | -0.032 | -0.032 | 1.639 | 0.37 | 0.2 | 0.4 | 0.49 | 0.42 | 0.29 |
| 5 | 653343 | rs2458815 | 0.246 | -0.354 | -0.626 | -0.185 | 0.004 | 0.044 | 0.044 | 0 | 0.22 | 0.31 | 0.24 | 0.47 | 0.23 | 0.14 |
| 5 | 659605 | rs1689671 | 0.457 | -0.118 | -1.452 | -0.06 | -0.024 | -0.105 | -0.105 | 0 | 0.23 | 0.08 | 0.25 | 0.48 | 0.23 | 0.14 |
| 5 | 665280 | rs1697963 | -0.921 | 0.019 | -1.552 | -0.234 | -0.061 | -0.071 | -0.071 | 1.227 | 0.71 | 0.93 | 0.7 | 0.69 | 0.61 | 0.61 |
| 5 | 666776 | rs1709541 | -0.456 | -0.434 | -0.86 | 0.015 | -0.015 | -0.002 | -0.002 | 0.868 | 0.4 | 0.49 | 0.46 | 0.68 | 0.33 | 0.46 |

| | | | | | | | | | | | | | | | | |
|---|---|---|---|---|---|---|---|---|---|---|---|---|---|---|---|---|
| 5 | 670706 | rs3792723 | -0.222 | -0.45 | -1.45 | -0.244 | 0.008 | 0.092 | 0.092 | 0 | 0.32 | 0.16 | 0.37 | 0.19 | 0.3 | 0.45 |
| 5 | 670745 | rs3828569 | 0.301 | -0.552 | -1.496 | -0.296 | 0.014 | 0.015 | 0.015 | 0 | 0.17 | 0.14 | 0.26 | 0.17 | 0.12 | 0.26 |
| 5 | 673408 | rs413666 | -0.055 | -0.013 | -1.122 | -0.185 | 0.005 | 0.001 | 0.001 | 0 | 0.57 | 0.48 | 0.6 | 0.62 | 0.55 | 0.59 |
| 5 | 674921 | rs1709544 | -1.013 | 0.344 | -1.998 | -0.098 | -0.13 | -0.156 | -0.156 | 0.482 | 0.75 | 0.97 | 0.7 | 0.64 | 0.64 | 0.69 |
| 5 | 676806 | rs394465 | -0.999 | 0.423 | -2.004 | -0.135 | -0.145 | -0.15 | -0.15 | 0 | 0.75 | 0.97 | 0.69 | 0.66 | 0.61 | 0.7 |
| 5 | 677718 | rs392088 | -1.009 | 0.363 | -1.984 | -0.131 | -0.123 | -0.142 | -0.142 | 0.024 | 0.75 | 0.96 | 0.7 | 0.66 | 0.64 | 0.71 |
| 5 | 685504 | rs6862648 | -0.192 | -0.165 | -1.542 | 0.017 | -0.036 | -0.057 | -0.057 | 0 | 0.18 | 0.57 | 0.1 | 0.08 | 0.07 | 0.11 |
| 5 | 685748 | rs386057 | -0.038 | 0.022 | -1.232 | 0.307 | 0.014 | -0.008 | -0.008 | 0.047 | 0.44 | 0.4 | 0.5 | 0.59 | 0.4 | 0.41 |
| 5 | 730271 | rs436354 | -0.928 | 0.74 | -0.842 | 0.891 | -0.022 | -0.008 | -0.008 | 0 | 0.19 | 0.14 | 0.28 | 0.24 | 0.23 | 0.37 |
| 5 | 816849 | rs3890125 | -1.679 | -0.476 | -0.143 | 1.253 | -0.042 | 0.089 | 0.089 | 1.307 | 0.2 | 0.62 | 0.3 | 0.4 | 0.09 | 0.19 |
| 5 | 818658 | rs588820 | 0.781 | 0.052 | -0.368 | 1.282 | -0.062 | -0.013 | -0.013 | 0 | 0.1 | 0 | 0.11 | 0.11 | 0.17 | 0.26 |
| 5 | 821726 | rs3747737 | -1.639 | -0.452 | -0.146 | 1.142 | -0.042 | 0.09 | 0.09 | 1.477 | 0.2 | 0.62 | 0.3 | 0.4 | 0.09 | 0.19 |
| 5 | 851046 | rs3747733 | -1.115 | -0.531 | 0.686 | 1.131 | -0.014 | 0.155 | 0.155 | 1.628 | 0.09 | 0.39 | 0.04 | 0.15 | 0.01 | 0.03 |
| 5 | 851582 | rs3810859 | -0.71 | -1.309 | 0.291 | 0.695 | -0.024 | -0.012 | -0.012 | 0.364 | 0.31 | 0.4 | 0.37 | 0.31 | 0.24 | 0.34 |
| 5 | 852207 | rs7702532 | -1.769 | -1.209 | 0.028 | 0.614 | 0.054 | -0.13 | -0.13 | 0.043 | 0.45 | 0.96 | 0.45 | 0.48 | 0.32 | 0.39 |
| 5 | 862682 | rs10475433 | -1.648 | -1.303 | 0.703 | 0.689 | -0.033 | 0.08 | 0.08 | 1.469 | 0.13 | 0.6 | 0.09 | 0.16 | 0.05 | 0.06 |
| 5 | 864053 | rs16877782 | -0.749 | -1.29 | 0.689 | 0.703 | -0.014 | 0.155 | 0.155 | 1.628 | 0.09 | 0.38 | 0.05 | 0.15 | 0.01 | 0.03 |
| 5 | 884251 | rs3747740 | -0.799 | -2.278 | 1.024 | 1.024 | -0.011 | 0.039 | 0.039 | 0 | 0.07 | 0.14 | 0.22 | 0.18 | 0.04 | 0.16 |
| 5 | 884439 | rs10072022 | -1.009 | -2.372 | 1.436 | 0.971 | 0.004 | -0.026 | -0.026 | 0 | 0.06 | 0.16 | 0.03 | 0.01 | 0.04 | 0.03 |
| 5 | 894439 | rs6555568 | -2.511 | -2.473 | 1.951 | 0.923 | -0.017 | -0.055 | -0.055 | 0 | 0.1 | 0.54 | 0.06 | 0.01 | 0.04 | 0.04 |
| 5 | 896983 | rs920980 | -2.96 | -2.747 | 1.719 | 0.907 | 0.079 | -0.053 | -0.053 | 0 | 0.27 | 0.88 | 0.32 | 0.34 | 0.09 | 0.23 |
| 5 | 904374 | rs12518024 | 0.173 | -2.728 | 1.586 | 0.816 | -0.018 | 0.019 | 0.019 | 0 | 0.18 | 0.01 | 0.11 | 0.14 | 0.19 | 0.16 |
| 5 | 917878 | rs14389 | -2.85 | -2.664 | 1.902 | 1.043 | -0.028 | -0.005 | -0.005 | 0 | 0.23 | 0.68 | 0.29 | 0.19 | 0.09 | 0.2 |
| 5 | 918523 | rs7714178 | -0.653 | -2.47 | 2.028 | 1.042 | -0.009 | -0.015 | -0.015 | 0 | 0.05 | 0.18 | 0.02 | 0 | 0 | 0 |
| 5 | 927625 | rs10068712 | -2.583 | -2.494 | 1.827 | 1.147 | -0.036 | 0.064 | 0.064 | 0.924 | 0.13 | 0.61 | 0.09 | 0.17 | 0.06 | 0.07 |
| 5 | 944298 | rs7728035 | -2.844 | -2.71 | 1.365 | 1.049 | 0.067 | -0.063 | -0.063 | 0 | 0.26 | 0.87 | 0.33 | 0.42 | 0.1 | 0.24 |
| 5 | 946418 | rs16903665 | -1.916 | -2.436 | 1.681 | 0.921 | 0.014 | -0.022 | -0.022 | 0 | 0.08 | 0.36 | 0.05 | 0.01 | 0.04 | 0.02 |
| 5 | 956671 | rs4957112 | 1.733 | -2.731 | 1.045 | 0.882 | 0.134 | -0.043 | -0.043 | 0 | 0.55 | 0.13 | 0.56 | 0.5 | 0.7 | 0.58 |
| 5 | 959224 | rs16867926 | -1.592 | -2.661 | 1.223 | 0.962 | -0.031 | 0.015 | 0.015 | 0 | 0.15 | 0.21 | 0.28 | 0.19 | 0.1 | 0.19 |
| 5 | 959731 | rs16868012 | -0.888 | -2.63 | 1.005 | 0.992 | 0.01 | -0.011 | -0.011 | 0 | 0.11 | 0.09 | 0.26 | 0.19 | 0.06 | 0.18 |
| 5 | 964250 | rs16868701 | -1.075 | -2.261 | 1.325 | 1.06 | -0.003 | -0.005 | -0.005 | 0 | 0.05 | 0.3 | 0.03 | 0 | 0 | 0 |
| 5 | 965618 | rs12653180 | 1.759 | -2.417 | 0.722 | 1.105 | 0.035 | -0.011 | -0.011 | 0 | 0.57 | 0.27 | 0.57 | 0.53 | 0.68 | 0.61 |
| 5 | 969285 | rs10071835 | 0.423 | -1.858 | 0.043 | 1.089 | 0.014 | -0.035 | -0.035 | 0.022 | 0.76 | 0.88 | 0.89 | 0.72 | 0.78 | 0.79 |
| 5 | 979148 | rs2289827 | 0.282 | -1.713 | 0.216 | 1.148 | -0.002 | 0.003 | 0.003 | 0 | 0.78 | 0.78 | 0.86 | 0.71 | 0.78 | 0.79 |
| 5 | 981019 | rs10065177 | 1.019 | -1.979 | -0.226 | 1.005 | 0.076 | -0.015 | -0.015 | 0 | 0.64 | 0.11 | 0.6 | 0.65 | 0.74 | 0.58 |
| 5 | 988510 | rs4975533 | 0.603 | -2.072 | 0.174 | 0.834 | -0.049 | -0.016 | -0.016 | 0 | 0.65 | 0.58 | 0.78 | 0.68 | 0.82 | 0.65 |
| 5 | 992509 | rs6882848 | -0.282 | -2.066 | -0.395 | 0.887 | 0.069 | 0.013 | 0.013 | 0 | 0.27 | 0.11 | 0.15 | 0.25 | 0.11 | 0.31 |
| 5 | 994615 | rs13160308 | 0.417 | -2.044 | 0.009 | 0.903 | 0.009 | 0.004 | 0.004 | 0.063 | 0.68 | 0.73 | 0.83 | 0.71 | 0.83 | 0.69 |
| 5 | 996696 | rs7443724 | 0.558 | -1.97 | 0.317 | 1.33 | -0.038 | 0.015 | 0.015 | 0 | 0.64 | 0.59 | 0.79 | 0.58 | 0.79 | 0.67 |
| 5 | 999338 | rs374361 | -0.082 | -1.271 | -0.463 | 1.221 | -0.045 | -0.064 | -0.064 | 0 | 0.17 | 0.01 | 0.28 | 0.27 | 0.2 | 0.18 |
| 5 | 1010083 | rs4078802 | -1.632 | -1.233 | 0.262 | 1.426 | 0.013 | 0.065 | 0.065 | 0 | 0.08 | 0.22 | 0.09 | 0.16 | 0.08 | 0.05 |
| 5 | 1011149 | rs6869901 | -1.632 | -1.233 | 0.262 | 1.426 | 0.013 | 0.065 | 0.065 | 0 | 0.08 | 0.22 | 0.09 | 0.16 | 0.08 | 0.05 |
| 5 | 1012446 | rs12518095 | -0.979 | -1.275 | 0.23 | 1.412 | -0.008 | 0.067 | 0.067 | 1.232 | 0.05 | 0.18 | 0.06 | 0.11 | 0.02 | 0.01 |
| 5 | 1015063 | rs16870629 | 0.37 | -0.961 | -1.166 | 0.716 | -0.119 | -0.158 | -0.158 | 0.555 | 0.34 | 0.05 | 0.38 | 0.46 | 0.42 | 0.44 |
| 5 | 1032032 | rs4975577 | -0.304 | -1.51 | -0.441 | 0.252 | -0.006 | 0.019 | 0.019 | 0 | 0.54 | 0.42 | 0.53 | 0.61 | 0.52 | 0.54 |
| 5 | 1039278 | rs10866472 | 1.409 | -1.515 | 0.036 | 0.542 | -0.018 | 0.058 | 0.058 | 4.807 | 0.88 | 0.29 | 0.83 | 0.84 | 0.94 | 0.94 |
| 5 | 1040657 | rs12517757 | 0.472 | -1.406 | -0.122 | 0.511 | -0.026 | -0.047 | -0.047 | 0.825 | 0.48 | 0.09 | 0.4 | 0.5 | 0.46 | 0.49 |
| 5 | 1041105 | rs4975526 | 0.792 | -1.762 | -0.23 | 0.316 | -0.02 | -0.001 | -0.001 | 0 | 0.79 | 0.28 | 0.8 | 0.82 | 0.91 | 0.81 |
| 5 | 1051687 | rs1058508 | -0.48 | -1.523 | 0.322 | 0.325 | 0.013 | 0.011 | 0.011 | 0 | 0.5 | 0.59 | 0.57 | 0.5 | 0.54 | 0.52 |
| 5 | 1051918 | rs3810853 | -0.106 | -1.521 | 0.503 | 0.304 | 0.023 | 0.144 | 0.144 | 3.843 | 0.05 | 0.31 | 0.11 | 0.09 | 0.01 | 0.01 |
| 5 | 1052360 | rs3810854 | -0.22 | -1.316 | 0.36 | 0.304 | 0.001 | -0.005 | -0.005 | 0 | 0.57 | 0.58 | 0.55 | 0.64 | 0.52 | 0.54 |
| 5 | 1057615 | rs2241606 | -0.187 | -1.333 | -0.409 | 0.131 | -0.008 | -0.02 | -0.02 | 0 | 0.67 | 0.24 | 0.63 | 0.48 | 0.64 | 0.61 |
| 5 | 1063998 | rs7722287 | 0.106 | -0.51 | -0.63 | -0.027 | -0.124 | 0.006 | 0.006 | 0.012 | 0.35 | 0.03 | 0.36 | 0.32 | 0.45 | 0.43 |
| 5 | 1065399 | rs737154 | 0.019 | -0.77 | -0.153 | -0.025 | 0.013 | -0.018 | -0.018 | 0 | 0.58 | 0.51 | 0.57 | 0.63 | 0.53 | 0.55 |
| 5 | 1066615 | rs7737804 | 0.355 | -0.818 | -0.758 | 0.042 | 0.013 | 0.036 | 0.036 | 0 | 0.52 | 0.35 | 0.53 | 0.55 | 0.47 | 0.46 |
| 5 | 1074921 | rs7714914 | 0.383 | -0.018 | -0.965 | -0.132 | -0.008 | 0.027 | 0.027 | 0 | 0.61 | 0.25 | 0.72 | 0.71 | 0.61 | 0.6 |
| 5 | 1076245 | rs11951420 | 0.245 | 0.265 | -1.393 | 0.049 | -0.145 | -0.041 | -0.041 | 0 | 0.26 | 0.02 | 0.26 | 0.24 | 0.39 | 0.28 |
| 5 | 1078642 | rs4975684 | 0.031 | 0.249 | -1.473 | 0.107 | -0.146 | -0.053 | -0.053 | 0 | 0.3 | 0.02 | 0.27 | 0.29 | 0.41 | 0.28 |
| 5 | 1081702 | rs6865765 | -0.588 | 0.012 | -0.41 | 0.015 | 0.005 | -0.007 | -0.007 | 0 | 0.41 | 0.48 | 0.32 | 0.36 | 0.48 | 0.42 |
| 5 | 1081767 | rs4526148 | 0.637 | -0.166 | -0.116 | -0.065 | -0.009 | 0.027 | 0.027 | 0 | 0.56 | 0.33 | 0.65 | 0.67 | 0.52 | 0.59 |
| 5 | 1090534 | rs6864667 | 0.398 | -0.1 | -0.23 | -0.155 | 0.006 | -0.013 | -0.013 | 5.71 | 0.6 | 0.54 | 0.67 | 0.69 | 0.53 | 0.69 |
| 5 | 1097281 | rs4975593 | 0.328 | 0.14 | -0.614 | -0.15 | -0.016 | 0.023 | 0.023 | 0.493 | 0.43 | 0.27 | 0.32 | 0.28 | 0.52 | 0.38 |
| 5 | 1102134 | rs6879835 | -0.372 | 0.283 | -0.899 | 0.394 | 0.025 | 0.001 | 0.001 | 0 | 0.64 | 0.87 | 0.72 | 0.62 | 0.72 | 0.58 |
| 5 | 1103495 | rs6894574 | 0.004 | 0.264 | -0.576 | 0.41 | 0.005 | 0.008 | 0.008 | 0 | 0.6 | 0.67 | 0.68 | 0.63 | 0.66 | 0.57 |
| 5 | 1113244 | rs4580814 | 0.609 | 0.23 | -1.058 | 0.452 | -0.017 | 0.056 | 0.056 | 0.485 | 0.46 | 0.19 | 0.37 | 0.31 | 0.56 | 0.41 |
| 5 | 1113556 | rs4285270 | -0.41 | 0.103 | -1.38 | 0.145 | 0.015 | -0.013 | -0.013 | 4.027 | 0.71 | 0.92 | 0.78 | 0.68 | 0.78 | 0.71 |
| 5 | 1115240 | rs11954838 | 0.225 | 0.535 | -0.625 | 0.122 | 0.012 | -0.033 | -0.033 | 0 | 0.12 | 0.16 | 0.09 | 0.05 | 0.18 | 0.19 |
| 5 | 1120954 | rs13171076 | 0.646 | 0.422 | -0.657 | 0.453 | -0.008 | 0.002 | 0.002 | 0.022 | 0.45 | 0.31 | 0.5 | 0.46 | 0.56 | 0.41 |
| 5 | 1121954 | rs13162116 | 0.055 | 0.646 | -1.015 | 0.152 | -0.009 | 0.009 | 0.009 | 0 | 0.66 | 0.7 | 0.72 | 0.74 | 0.63 | 0.62 |

| | | | | | | | | | | | | | | | | |
|---|---|---|---|---|---|---|---|---|---|---|---|---|---|---|---|---|
| 5 | 1124560 | rs4975614 | 0.335 | -0.036 | -0.925 | 0.285 | 0 | -0.007 | -0.007 | 0 | 0.1 | 0.06 | 0.12 | 0.23 | 0.09 | 0.18 |
| 5 | 1129746 | rs11951225 | -0.169 | -0.469 | -1.16 | -0.048 | 0.012 | 0.014 | 0.014 | 0 | 0.34 | 0.22 | 0.33 | 0.35 | 0.31 | 0.39 |
| 5 | 1132216 | rs4594899 | -0.594 | -0.961 | -1.124 | -0.182 | -0.002 | -0.002 | -0.002 | 0 | 0.53 | 0.51 | 0.56 | 0.45 | 0.44 | 0.54 |
| 5 | 1134166 | rs4076817 | -0.138 | -1.325 | -1.5 | -0.298 | 0.047 | 0.016 | 0.016 | 0 | 0.37 | 0.81 | 0.38 | 0.38 | 0.31 | 0.43 |
| 5 | 1134914 | rs4076818 | 0.294 | -1.138 | -1.036 | -0.303 | -0.015 | -0.029 | -0.029 | 0.547 | 0.26 | 0.6 | 0.22 | 0.12 | 0.21 | 0.19 |
| 5 | 1143298 | rs7379059 | -0.535 | -0.955 | -0.746 | 0.409 | -0.009 | 0.003 | 0.003 | 0 | 0.45 | 0.52 | 0.43 | 0.62 | 0.34 | 0.53 |
| 5 | 1151161 | rs3922757 | -0.06 | -0.677 | -0.745 | 0.529 | -0.001 | -0.013 | -0.013 | 0.008 | 0.2 | 0.16 | 0.22 | 0.29 | 0.21 | 0.23 |
| 5 | 1157313 | rs7726442 | 0.484 | -0.858 | -0.699 | 0.425 | -0.006 | -0.004 | -0.004 | 0 | 0.56 | 0.46 | 0.49 | 0.65 | 0.63 | 0.64 |
| 5 | 1169044 | rs7719062 | 0.116 | -1.012 | -0.806 | 0.354 | 0.038 | 0.021 | 0.021 | 0.199 | 0.2 | 0.11 | 0.26 | 0.09 | 0.09 | 0.11 |
| 5 | 1169112 | rs7702150 | -0.168 | -1.16 | -0.307 | 0.588 | -0.025 | 0.065 | 0.065 | 0 | 0.79 | 0.55 | 0.79 | 0.63 | 0.84 | 0.67 |
| 5 | 1170323 | rs13181306 | 0.596 | -1.406 | -0.747 | 0.514 | 0.129 | -0.013 | -0.013 | 0 | 0.57 | 0.24 | 0.52 | 0.53 | 0.81 | 0.58 |
| 5 | 1176601 | rs4975536 | 0.789 | -1.08 | -0.835 | 0.493 | 0.149 | 0.021 | 0.021 | 0 | 0.52 | 0.16 | 0.46 | 0.55 | 0.72 | 0.52 |
| 5 | 1186086 | rs4246740 | 0.856 | -0.528 | -1.104 | 0.58 | -0.093 | -0.044 | -0.044 | 0 | 0.35 | 0.1 | 0.21 | 0.28 | 0.44 | 0.26 |
| 5 | 1186121 | rs6554634 | 0.569 | -0.777 | -0.263 | 0.665 | -0.077 | -0.016 | -0.016 | 0 | 0.23 | 0.47 | 0.36 | 0.18 | 0.13 | 0.16 |
| 5 | 1186853 | rs4246736 | 0.043 | -0.659 | -0.281 | 0.697 | 0.002 | 0 | 0 | 0 | 0.61 | 0.61 | 0.59 | 0.49 | 0.61 | 0.44 |
| 5 | 1187180 | rs4460173 | -0.584 | -0.731 | -0.186 | 0.693 | -0.015 | 0.05 | 0.05 | 0 | 0.84 | 0.66 | 0.86 | 0.73 | 0.87 | 0.67 |
| 5 | 1188837 | rs4975599 | -0.837 | -0.79 | -0.165 | 0.625 | -0.03 | 0.04 | 0.04 | 0 | 0.85 | 0.66 | 0.88 | 0.77 | 0.91 | 0.7 |
| 5 | 1189347 | rs4975596 | 0.495 | -0.92 | -0.569 | 0.645 | 0.117 | -0.027 | -0.027 | 0 | 0.59 | 0.13 | 0.5 | 0.58 | 0.74 | 0.52 |
| 5 | 1190088 | rs7719875 | -0.128 | -0.622 | -0.198 | 0.603 | 0.003 | 0.013 | 0.013 | 0 | 0.45 | 0.35 | 0.63 | 0.45 | 0.43 | 0.4 |
| 5 | 1190440 | rs6884486 | 0.329 | -0.864 | 0.235 | 0.562 | -0.042 | -0.005 | -0.005 | 0 | 0.66 | 0.59 | 0.56 | 0.68 | 0.79 | 0.56 |
| 5 | 1191061 | rs10060236 | 0.308 | -0.831 | 0.154 | 0.539 | -0.055 | -0.009 | -0.009 | 0.471 | 0.66 | 0.55 | 0.56 | 0.69 | 0.78 | 0.56 |
| 5 | 1200573 | rs11747247 | -0.657 | -0.028 | 0.583 | 0.785 | 0.014 | 0.092 | 0.092 | 0 | 0.89 | 0.69 | 0.9 | 0.78 | 0.92 | 0.8 |
| 5 | 1200882 | rs13183296 | -1.142 | 1.26 | 0.524 | 0.269 | 0.062 | -0.057 | -0.057 | 0.011 | 0.11 | 0.23 | 0.1 | 0.01 | 0.22 | 0.09 |
| 5 | 1203437 | rs13159461 | 0.416 | 0.557 | 0.003 | 0.542 | 0.02 | 0.003 | 0.003 | 0 | 0.33 | 0.36 | 0.26 | 0.35 | 0.44 | 0.32 |
| 5 | 1205636 | rs10072823 | 0.228 | 0.101 | -0.588 | 0.711 | 0.022 | 0.01 | 0.01 | 0 | 0.47 | 0.33 | 0.34 | 0.41 | 0.58 | 0.45 |
| 5 | 1206462 | rs12658538 | -0.199 | 0.307 | -0.652 | 0.916 | 0 | -0.009 | -0.009 | 0.485 | 0.15 | 0.16 | 0.12 | 0.23 | 0.15 | 0.16 |
| 5 | 1211556 | rs7444093 | -0.567 | -0.302 | -0.585 | 0.742 | 0.019 | -0.005 | -0.005 | 0.377 | 0.46 | 0.61 | 0.58 | 0.45 | 0.36 | 0.43 |
| 5 | 1212357 | rs7727745 | -0.207 | -0.115 | 0.118 | 0.683 | -0.019 | 0.03 | 0.03 | 0 | 0.16 | 0.38 | 0.15 | 0.22 | 0.13 | 0.14 |
| 5 | 1213668 | rs7732589 | -0.138 | -0.126 | 0.259 | 0.65 | -0.021 | 0.004 | 0.004 | 0 | 0.16 | 0.32 | 0.14 | 0.18 | 0.13 | 0.14 |
| 5 | 1215474 | rs11133684 | 0.575 | 1.022 | -0.482 | -0.042 | -0.073 | 0.094 | 0.094 | 0 | 0.15 | 0.01 | 0.08 | 0.01 | 0.18 | 0.12 |
| 5 | 1216900 | rs35329108 | 0.172 | 0.79 | 0.036 | -0.016 | 0 | -0.008 | -0.008 | 0.895 | 0.24 | 0.36 | 0.2 | 0.15 | 0.23 | 0.12 |
| 5 | 1217471 | rs6554666 | 0.258 | 0.851 | 0.201 | -0.295 | -0.011 | 0.019 | 0.019 | 0 | 0.35 | 0.27 | 0.43 | 0.2 | 0.42 | 0.29 |
| 5 | 1220032 | rs4975628 | 0.275 | 0.762 | 0.123 | -0.376 | -0.007 | 0.008 | 0.008 | 0 | 0.36 | 0.32 | 0.46 | 0.22 | 0.44 | 0.42 |
| 5 | 1221199 | rs12513763 | -0.802 | 0.682 | -0.082 | 0.124 | 0 | 0.102 | 0.102 | 0 | 0.94 | 0.94 | 0.86 | 0.6 | 0.96 | 0.83 |
| 5 | 1223062 | rs10060827 | 0.765 | 0.613 | 0.088 | 0.054 | 0.029 | -0.005 | -0.005 | 0 | 0.34 | 0.23 | 0.22 | 0.33 | 0.29 | 0.3 |
| 5 | 1225121 | rs9418 | 0.34 | 0.576 | -0.143 | 0.115 | 0.005 | -0.013 | -0.013 | 2.621 | 0.37 | 0.43 | 0.47 | 0.22 | 0.44 | 0.34 |
| 5 | 1225434 | rs7704882 | 0.103 | 0.8 | -0.326 | 0.358 | 0.002 | -0.023 | -0.023 | 1.124 | 0.75 | 0.8 | 0.75 | 0.56 | 0.79 | 0.75 |
| 5 | 1225703 | rs7724858 | 0.452 | 0.945 | 0.198 | 1.121 | 0.001 | 0.157 | 0.157 | 0 | 0.94 | 0.88 | 0.86 | 0.6 | 0.96 | 0.82 |
| 5 | 1236212 | rs7445640 | -0.564 | 1.329 | -0.611 | -0.6 | -0.025 | 0.026 | 0.026 | 0 | 0.33 | 0.23 | 0.37 | 0.23 | 0.38 | 0.46 |
| 5 | 1240848 | rs6554684 | -0.601 | 1.111 | -1.251 | -1.034 | -0.007 | -0.069 | -0.069 | 0.062 | 0.75 | 0.97 | 0.7 | 0.27 | 0.77 | 0.68 |
| 5 | 1241908 | rs10043584 | -0.603 | 1.13 | -1.25 | -1.02 | -0.003 | -0.069 | -0.069 | 0 | 0.75 | 0.97 | 0.7 | 0.28 | 0.78 | 0.68 |
| 5 | 1243427 | rs13361701 | 0.895 | 1.29 | -0.891 | -1.135 | 0 | -0.082 | -0.082 | 0 | 0.42 | 0.51 | 0.32 | 0.04 | 0.42 | 0.27 |
| 5 | 1244425 | rs4073918 | -0.28 | 1.32 | -0.827 | -0.524 | 0.015 | -0.026 | -0.026 | 0.585 | 0.75 | 0.9 | 0.7 | 0.27 | 0.77 | 0.67 |
| 5 | 1253744 | rs2853690 | -0.944 | 1.846 | -0.235 | -0.828 | 0.071 | -0.07 | -0.07 | 0 | 0.11 | 0.33 | 0.11 | 0.01 | 0.17 | 0.08 |
| 5 | 1257621 | rs2736122 | -0.538 | 1.873 | -0.617 | -0.609 | 0.037 | -0.021 | -0.021 | 0 | 0.15 | 0.24 | 0.12 | 0.08 | 0.25 | 0.18 |
| 5 | 1266310 | rs2075786 | 0.038 | 1.089 | -0.722 | -0.528 | -0.001 | -0.012 | -0.012 | 0 | 0.6 | 0.42 | 0.39 | 0.21 | 0.66 | 0.49 |
| 5 | 1279790 | rs10069690 | -0.424 | 0.938 | -0.579 | -0.253 | -0.001 | -0.018 | -0.018 | 0 | 0.29 | 0.66 | 0.22 | 0.17 | 0.28 | 0.27 |
| 5 | 1286516 | rs2736100 | -0.26 | 1.071 | -0.53 | 0.089 | 0.01 | 0.004 | 0.004 | 0 | 0.36 | 0.53 | 0.57 | 0.59 | 0.5 | 0.39 |
| 5 | 1292983 | rs2853672 | -0.386 | 1.387 | -0.089 | 0.28 | 0.086 | 0.086 | 0.086 | 0 | 0.29 | 0.48 | 0.57 | 0.52 | 0.49 | 0.33 |
| 5 | 1294086 | rs2736098 | 0.442 | 0.175 | -1.083 | -0.483 | -0.03 | -0.089 | -0.089 | 6.19 | 0.34 | 0.07 | 0.2 | 0.37 | 0.23 | 0.5 |
| 5 | 1300310 | rs4975612 | -0.053 | 0.675 | -0.211 | 0.055 | -0.023 | 0.149 | 0.149 | 0.226 | 0.21 | 0.38 | 0.34 | 0.52 | 0.2 | 0.31 |
| 5 | 1308552 | rs4635969 | -0.845 | 1.497 | 0.154 | 0.074 | 0.044 | -0.005 | -0.005 | 0.302 | 0.15 | 0.33 | 0.11 | 0.14 | 0.19 | 0.09 |
| 5 | 1315660 | rs4975616 | 1.231 | 1.627 | 0.144 | -0.159 | -0.023 | 0.058 | 0.058 | 0 | 0.65 | 0.25 | 0.71 | 0.81 | 0.58 | 0.81 |
| 5 | 1317102 | rs6554759 | -0.625 | 1.663 | -0.012 | -0.154 | -0.009 | 0.024 | 0.024 | 0 | 0.84 | 0.91 | 0.86 | 0.95 | 0.83 | 0.91 |
| 5 | 1317949 | rs1801075 | -0.561 | 1.672 | 0.12 | -0.15 | -0.001 | 0.001 | 0.001 | 0 | 0.84 | 0.84 | 0.86 | 0.95 | 0.83 | 0.93 |
| 5 | 1320247 | rs380286 | -1.442 | 1.762 | 0.098 | -0.18 | 0 | -0.007 | -0.007 | 0.03 | 0.35 | 0.62 | 0.31 | 0.15 | 0.44 | 0.19 |
| 5 | 1320722 | rs402710 | -0.997 | 1.799 | -0.049 | 0.065 | 0.021 | -0.005 | -0.005 | 0 | 0.28 | 0.48 | 0.35 | 0.31 | 0.33 | 0.16 |
| 5 | 1321873 | rs10073340 | 0.404 | 1.648 | -0.101 | -0.293 | -0.003 | 0.008 | 0.008 | 0 | 0.16 | 0.15 | 0.13 | 0.05 | 0.17 | 0.08 |
| 5 | 1342714 | rs31489 | -0.521 | 1.301 | -0.345 | -0.179 | 0.024 | -0.045 | -0.045 | 0 | 0.32 | 0.49 | 0.28 | 0.19 | 0.41 | 0.17 |
| 5 | 1344458 | rs31490 | -0.859 | 1.312 | -0.327 | -0.171 | 0.018 | -0.045 | -0.045 | 1.148 | 0.35 | 0.5 | 0.29 | 0.19 | 0.44 | 0.19 |
| 5 | 1355058 | rs37006 | -0.711 | 1.419 | -0.469 | -0.058 | 0.01 | -0.021 | -0.021 | 0.614 | 0.34 | 0.57 | 0.31 | 0.19 | 0.44 | 0.18 |
| 5 | 1362793 | rs27061 | 0.093 | 0.674 | -0.93 | 0.265 | 0.031 | 0.012 | 0.012 | 0.044 | 0.5 | 0.35 | 0.56 | 0.55 | 0.53 | 0.62 |
| 5 | 1368997 | rs461193 | -0.03 | 0.119 | -0.578 | 0.471 | -0.005 | -0.004 | -0.004 | 0 | 0.29 | 0.25 | 0.17 | 0.23 | 0.24 | 0.14 |
| 5 | 1370300 | rs10075239 | 0.203 | 1.152 | -0.644 | 0.066 | -0.002 | -0.008 | -0.008 | 0 | 0.18 | 0.19 | 0.15 | 0.02 | 0.27 | 0.11 |
| 5 | 1370832 | rs2963265 | -0.276 | 0.818 | -0.512 | 0.092 | 0.024 | -0.06 | -0.06 | 0.082 | 0.35 | 0.5 | 0.3 | 0.11 | 0.48 | 0.16 |
| 5 | 1376161 | rs246994 | -0.993 | 0.498 | -1.549 | 0.143 | 0.023 | -0.007 | -0.007 | 0 | 0.69 | 0.82 | 0.73 | 0.75 | 0.73 | 0.66 |
| 5 | 1377172 | rs2963259 | 0.644 | 0.295 | -1.459 | -0.134 | -0.001 | -0.029 | -0.029 | 0 | 0.23 | 0.18 | 0.17 | 0.09 | 0.23 | 0.07 |
| 5 | 1378159 | rs2963257 | -0.285 | 0.772 | -2.134 | 0.468 | -0.055 | -0.003 | -0.003 | 0 | 0.56 | 0.84 | 0.52 | 0.41 | 0.51 | 0.46 |
| 5 | 1379762 | rs37017 | -0.325 | 0.835 | -1.556 | 0.529 | -0.047 | -0.015 | -0.015 | 0.008 | 0.86 | 0.99 | 0.84 | 0.87 | 0.86 | 0.8 |

| | | | | | | | | | | | | | | | | |
|---|---|---|---|---|---|---|---|---|---|---|---|---|---|---|---|---|
| 5 | 1388625 | rs4975636 | -0.258 | 0.983 | -1.105 | 0.989 | -0.003 | 0.015 | 0.015 | 0 | 0.07 | 0.06 | 0.2 | 0.21 | 0.08 | 0.13 |
| 5 | 1390349 | rs12516758 | -0.029 | 0.928 | -1.399 | -0.184 | -0.014 | 0.049 | 0.049 | 0 | 0.17 | 0.06 | 0.09 | 0 | 0.18 | 0.14 |
| 5 | 1394522 | rs27072 | 0.164 | 0.611 | -0.774 | 0.096 | 0.007 | -0.014 | -0.014 | 0 | 0.21 | 0.13 | 0.22 | 0.26 | 0.17 | 0.28 |
| 5 | 1394815 | rs1042098 | -0.175 | 0.76 | -0.547 | -0.926 | -0.01 | -0.093 | -0.093 | 0.297 | 0.32 | 0.5 | 0.25 | 0.11 | 0.31 | 0.23 |
| 5 | 1395077 | rs40184 | -0.464 | 0.87 | -0.399 | -0.509 | 0 | 0.002 | 0.002 | 0 | 0.51 | 0.49 | 0.35 | 0.28 | 0.48 | 0.42 |
| 5 | 1407116 | rs3776512 | -0.325 | 0.901 | -1.304 | -0.508 | -0.018 | 0.061 | 0.061 | 0 | 0.19 | 0.06 | 0.15 | 0.09 | 0.23 | 0.12 |
| 5 | 1412645 | rs27048 | -0.257 | 0.986 | -0.972 | -0.534 | -0.015 | 0.047 | 0.047 | 0 | 0.4 | 0.32 | 0.34 | 0.18 | 0.43 | 0.35 |
| 5 | 1416646 | rs2042449 | -0.29 | 0.942 | -1.093 | -0.397 | -0.012 | 0.009 | 0.009 | 0 | 0.19 | 0.11 | 0.24 | 0.13 | 0.22 | 0.28 |
| 5 | 1423905 | rs464049 | -0.581 | 0.688 | -0.821 | -0.455 | 0.009 | 0.071 | 0.071 | 0.024 | 0.5 | 0.67 | 0.55 | 0.64 | 0.47 | 0.68 |
| 5 | 1432825 | rs460000 | 0.577 | 0.112 | -1.211 | -0.81 | -0.013 | 0.044 | 0.044 | 0 | 0.31 | 0.5 | 0.31 | 0.53 | 0.25 | 0.46 |
| 5 | 1438354 | rs403636 | -0.462 | 0.255 | -0.893 | -0.501 | -0.01 | 0.007 | 0.007 | 0.88 | 0.75 | 0.69 | 0.78 | 0.66 | 0.84 | 0.86 |
| 5 | 1443199 | rs6350 | 0.979 | 0.518 | -1.082 | -0.587 | -0.006 | 0.022 | 0.022 | 0 | 0.08 | 0.04 | 0.05 | 0.01 | 0.08 | 0.05 |
| 5 | 1448148 | rs3756450 | 0.355 | 0.137 | -0.332 | -0.13 | -0.006 | 0.208 | 0.208 | 0.099 | 0.17 | 0.47 | 0.28 | 0.44 | 0.13 | 0.38 |
| 5 | 1457554 | rs748209 | -0.8 | 0.109 | -0.765 | -0.684 | 0.004 | -0.034 | -0.034 | 0 | 0.69 | 0.78 | 0.74 | 0.59 | 0.71 | 0.85 |
| 5 | 1461167 | rs7737692 | -0.269 | 0.023 | -0.785 | -0.755 | -0.002 | 0 | 0 | 0 | 0.59 | 0.55 | 0.68 | 0.57 | 0.63 | 0.76 |
| 5 | 1461389 | rs2292023 | 0.643 | 0.05 | -0.456 | -0.591 | -0.005 | 0.149 | 0.149 | 0.02 | 0.17 | 0.4 | 0.3 | 0.43 | 0.15 | 0.38 |
| 5 | 1462098 | rs9728 | 0.665 | 0.054 | -0.362 | -0.59 | -0.004 | 0.147 | 0.147 | 0.014 | 0.17 | 0.36 | 0.29 | 0.44 | 0.15 | 0.38 |
| 5 | 1468287 | rs4975648 | -0.012 | 0.255 | -0.959 | -1.001 | -0.014 | 0.007 | 0.007 | 0 | 0.45 | 0.29 | 0.47 | 0.45 | 0.46 | 0.54 |
| 5 | 1473977 | rs27056 | -1.107 | 0.273 | -1.438 | -0.982 | -0.018 | 0.003 | 0.003 | 0 | 0.81 | 0.96 | 0.78 | 0.87 | 0.76 | 0.74 |
| 5 | 1476785 | rs10043491 | -0.064 | -0.114 | -1.616 | -1.036 | 0.048 | 0.091 | 0.091 | 0 | 0.37 | 0.17 | 0.24 | 0.16 | 0.26 | 0.17 |
| 5 | 1491009 | rs10058172 | -0.35 | 0.221 | -0.978 | -0.905 | 0.01 | -0.118 | -0.118 | 1.024 | 0.28 | 0.37 | 0.18 | 0.06 | 0.28 | 0.26 |
| 5 | 1492559 | rs9800090 | -0.777 | 0.314 | -1.404 | -0.457 | -0.005 | -0.008 | -0.008 | 0 | 0.48 | 0.73 | 0.42 | 0.48 | 0.48 | 0.61 |
| 5 | 1493608 | rs6554859 | 0.674 | 0.351 | -1.135 | 0.136 | 0.024 | 0.126 | 0.126 | 0 | 0.18 | 0.38 | 0.23 | 0.41 | 0.19 | 0.38 |
| 5 | 1494439 | rs40180 | -0.668 | 0.283 | -1.63 | 0.239 | -0.049 | -0.047 | -0.047 | 0 | 0.74 | 0.98 | 0.78 | 0.71 | 0.73 | 0.71 |
| 5 | 1494909 | rs3733796 | 0.843 | 0.674 | -0.927 | 0.623 | 0.024 | 0.109 | 0.109 | 0 | 0.16 | 0.31 | 0.23 | 0.4 | 0.19 | 0.34 |
| 5 | 1497626 | rs36990 | -1.02 | 0.424 | -1.674 | 0.473 | -0.025 | -0.048 | -0.048 | 0.122 | 0.8 | 0.85 | 0.68 | 0.57 | 0.73 | 0.6 |
| 5 | 1498986 | rs36988 | 0.233 | 0.037 | -1.68 | 0.131 | 0 | 0.004 | 0.004 | 0.011 | 0.58 | 0.4 | 0.46 | 0.41 | 0.61 | 0.5 |
| 5 | 1499175 | rs36987 | 0.473 | -0.044 | -1.556 | -0.407 | -0.032 | 0.103 | 0.103 | 1.539 | 0.37 | 0.16 | 0.35 | 0.15 | 0.43 | 0.25 |
| 5 | 1500160 | rs10058381 | 0.072 | -0.158 | -1.456 | -0.191 | -0.044 | 0.012 | 0.012 | 0 | 0.21 | 0.54 | 0.23 | 0.19 | 0.11 | 0.09 |
| 5 | 1501695 | rs17588991 | 0.946 | 0.716 | -1.877 | -0.517 | -0.005 | 0.021 | 0.021 | 0.272 | 0.07 | 0.02 | 0.04 | 0 | 0.06 | 0.04 |
| 5 | 1503205 | rs36982 | -0.211 | 0.855 | -1.834 | -0.401 | -0.035 | 0.115 | 0.115 | 1.547 | 0.37 | 0.14 | 0.35 | 0.15 | 0.43 | 0.25 |
| 5 | 1503610 | rs36979 | 0.054 | 0.849 | -1.984 | -0.259 | -0.047 | 0.099 | 0.099 | 1.686 | 0.29 | 0.07 | 0.3 | 0.12 | 0.33 | 0.13 |
| 5 | 1506303 | rs36971 | -0.022 | 0.785 | -1.939 | -0.286 | -0.035 | 0.099 | 0.099 | 1.872 | 0.3 | 0.09 | 0.29 | 0.11 | 0.33 | 0.15 |
| 5 | 1508270 | rs36966 | -0.443 | 0.49 | -1.814 | -0.006 | 0.017 | 0.079 | 0.079 | 0 | 0.49 | 0.28 | 0.4 | 0.35 | 0.46 | 0.41 |
| 5 | 1508837 | rs2962043 | -0.043 | 0.204 | -1.131 | -0.12 | -0.015 | 0 | 0 | 0 | 0.44 | 0.5 | 0.24 | 0.44 | 0.31 | 0.32 |
| 5 | 1517198 | rs13160153 | -0.367 | 0.979 | -1.032 | 0.167 | -0.023 | -0.023 | -0.023 | 0 | 0.18 | 0.11 | 0.25 | 0.26 | 0.28 | 0.16 |
| 5 | 1517322 | rs2963279 | -0.846 | 0.121 | -1.438 | -0.265 | -0.042 | -0.042 | -0.042 | 0 | 0.57 | 0.85 | 0.45 | 0.48 | 0.49 | 0.46 |
| 5 | 1517642 | rs4371798 | 0.234 | -0.344 | -1.017 | -0.197 | 0.04 | 0.039 | 0.039 | 0.5 | 0.31 | 0.23 | 0.15 | 0.21 | 0.21 | 0.28 |
| 5 | 1519079 | rs4406174 | -0.361 | 0.262 | -1.136 | 0.04 | -0.032 | -0.046 | -0.046 | 0 | 0.08 | 0.01 | 0.15 | 0.18 | 0.11 | 0.05 |
| 5 | 1519693 | rs2963275 | 0.053 | 0.241 | -1.078 | -0.235 | -0.023 | -0.049 | -0.049 | 0.687 | 0.28 | 0.16 | 0.48 | 0.39 | 0.35 | 0.43 |
| 5 | 1521573 | rs7716426 | 0.217 | -0.28 | -0.385 | -0.016 | 0.023 | 0.017 | 0.017 | 1.194 | 0.27 | 0.26 | 0.1 | 0.17 | 0.13 | 0.26 |
| 5 | 1526098 | rs2962057 | 0.785 | 0.298 | -0.752 | 0.039 | -0.046 | -0.049 | -0.049 | 0.039 | 0.78 | 0.94 | 0.73 | 0.64 | 0.67 | 0.8 |
| 5 | 1529358 | rs9312877 | 1.311 | 0.119 | -0.446 | -0.062 | -0.001 | -0.011 | -0.011 | 0 | 0.46 | 0.24 | 0.59 | 0.48 | 0.53 | 0.51 |
| 5 | 1529401 | rs7712396 | 0.955 | 1.672 | -0.094 | 0.448 | -0.025 | -0.028 | -0.028 | 0 | 0.86 | 0.94 | 0.77 | 0.63 | 0.73 | 0.86 |
| 5 | 1536257 | rs4551123 | -0.767 | 1.548 | 0.133 | 0.353 | -0.001 | -0.007 | -0.007 | 0 | 0.09 | 0.1 | 0.05 | 0.01 | 0.06 | 0.08 |
| 5 | 1548764 | rs2962074 | 1.262 | 1.611 | 0.082 | 0.595 | -0.008 | -0.009 | -0.009 | 0 | 0.85 | 0.91 | 0.79 | 0.64 | 0.77 | 0.85 |
| 5 | 1548867 | rs2962073 | 0.821 | 1.64 | 0.042 | 0.604 | -0.007 | 0.002 | 0.002 | 0 | 0.88 | 0.91 | 0.8 | 0.65 | 0.77 | 0.88 |
| 5 | 1549771 | rs6865199 | 1.839 | 2.012 | 0.552 | 0.413 | 0.058 | 0.025 | 0.025 | 0 | 0.7 | 0.55 | 0.67 | 0.64 | 0.58 | 0.74 |
| 5 | 1550734 | rs2963285 | -0.824 | 2.655 | 0.005 | 0.228 | 0.015 | -0.039 | -0.039 | 0 | 0.13 | 0.15 | 0.09 | 0 | 0.19 | 0.11 |
| 5 | 1550942 | rs10079102 | 0.774 | 1.797 | -0.147 | 0.84 | 0.012 | 0.02 | 0.02 | 0 | 0.9 | 0.9 | 0.82 | 0.78 | 0.79 | 0.9 |
| 5 | 1552714 | rs2962066 | -0.501 | 0.561 | 0.039 | 0.069 | -0.04 | -0.057 | -0.057 | 0 | 0.28 | 0.41 | 0.17 | 0.17 | 0.2 | 0.27 |
| 5 | 1553239 | rs2963271 | -0.284 | 1.237 | -0.003 | 0.327 | 0.049 | -0.003 | -0.003 | 0.23 | 0.67 | 0.57 | 0.39 | 0.39 | 0.56 | 0.53 |
| 5 | 1557372 | rs17596501 | -0.219 | 0.962 | -0.568 | -0.637 | -0.011 | 0.135 | 0.135 | 0 | 0.37 | 0.12 | 0.24 | 0.13 | 0.34 | 0.22 |
| 5 | 1557636 | rs4975554 | 0.286 | 1.093 | 0.358 | 0.211 | 0.052 | 0.173 | 0.173 | 0.533 | 0.87 | 0.51 | 0.77 | 0.76 | 0.85 | 0.81 |
| 5 | 1558048 | rs16880246 | 0.022 | 1.205 | 0.352 | -0.225 | 0.004 | -0.028 | -0.028 | 0 | 0.06 | 0.08 | 0.07 | 0.01 | 0.07 | 0.05 |
| 5 | 1558628 | rs6870653 | -0.317 | 1.165 | 0.512 | -0.214 | 0.125 | -0.005 | -0.005 | 0 | 0.08 | 0.46 | 0.09 | 0 | 0.1 | 0.03 |
| 5 | 1564675 | rs4292484 | 0.106 | 0.603 | -0.296 | -0.308 | 0.018 | 0.017 | 0.017 | 0 | 0.48 | 0.36 | 0.59 | 0.39 | 0.59 | 0.51 |
| 5 | 1565371 | rs4975556 | 0.106 | 0.603 | -0.296 | -0.308 | 0.018 | 0.017 | 0.017 | 0 | 0.48 | 0.36 | 0.59 | 0.39 | 0.59 | 0.51 |
| 5 | 1568508 | rs13188808 | 1.027 | 0.902 | -0.032 | -0.212 | -0.009 | -0.003 | -0.003 | 0 | 0.26 | 0.22 | 0.43 | 0.26 | 0.35 | 0.38 |
| 5 | 1594995 | rs4415117 | 0.144 | 0.874 | 0.512 | -0.263 | 0.054 | -0.042 | -0.042 | 1.078 | 0.09 | 0.25 | 0.11 | 0.01 | 0.13 | 0.07 |
| 5 | 1612612 | rs11958097 | 0.162 | 0.875 | 0.516 | -0.262 | 0.056 | -0.041 | -0.041 | 0.083 | 0.08 | 0.25 | 0.1 | 0.01 | 0.13 | 0.07 |
| 5 | 1624351 | rs1968011 | 0.89 | 0.744 | 0.046 | -0.23 | -0.022 | -0.008 | -0.008 | 0 | 0.27 | 0.18 | 0.44 | 0.27 | 0.36 | 0.37 |
| 5 | 1639901 | rs6554915 | 0.02 | 0.24 | -0.081 | -0.906 | 0 | 0.052 | 0.052 | 0.527 | 0.37 | 0.28 | 0.23 | 0.14 | 0.31 | 0.18 |
| 5 | 1641815 | rs6888166 | -0.99 | 0.177 | -0.387 | -0.511 | -0.029 | -0.037 | -0.037 | 0 | 0.76 | 0.85 | 0.5 | 0.46 | 0.65 | 0.52 |
| 5 | 1644315 | rs7705049 | 0.67 | 0.01 | -0.311 | -0.916 | 0.001 | -0.021 | -0.021 | 0 | 0.26 | 0.31 | 0.17 | 0.14 | 0.25 | 0.15 |
| 5 | 1644394 | rs7705207 | 0.747 | 0.075 | 0.113 | -0.901 | 0.072 | -0.026 | -0.026 | 1.827 | 0.08 | 0.33 | 0.1 | 0.01 | 0.11 | 0.07 |
| 5 | 1645242 | rs4551124 | 1.759 | 0.847 | -0.116 | -0.647 | -0.01 | -0.014 | -0.014 | 0 | 0.07 | 0.04 | 0.23 | 0.19 | 0.09 | 0.14 |
| 5 | 1650262 | rs6866927 | -0.591 | 0.246 | -0.261 | -0.871 | -0.002 | 0.026 | 0.026 | 0 | 0.44 | 0.38 | 0.47 | 0.33 | 0.4 | 0.29 |
| 5 | 1653418 | rs6870197 | -1.813 | 0.391 | -0.302 | -0.533 | 0.002 | 0.016 | 0.016 | 0 | 0.82 | 0.79 | 0.7 | 0.63 | 0.73 | 0.64 |

| | | | | | | | | | | | | | | | | |
|---|---|---|---|---|---|---|---|---|---|---|---|---|---|---|---|---|
| 5 | 1659414 | rs1106136 | 0.967 | 0.202 | -0.249 | -0.546 | 0 | 0.012 | 0.012 | 0 | 0.08 | 0.08 | 0.24 | 0.19 | 0.09 | 0.13 |
| 5 | 1660985 | rs2133740 | 1.572 | -0.367 | -0.366 | -0.765 | 0.044 | -0.007 | -0.007 | 0 | 0.26 | 0.18 | 0.12 | 0.29 | 0.18 | 0.24 |
| 5 | 1661397 | rs904734 | -1.77 | 0.134 | -0.233 | -0.614 | -0.004 | 0 | 0 | 0.013 | 0.82 | 0.8 | 0.7 | 0.64 | 0.72 | 0.66 |
| 5 | 1670369 | rs6880499 | -0.168 | 0.062 | -0.233 | -0.917 | 0 | -0.001 | -0.001 | 2.917 | 0.37 | 0.55 | 0.48 | 0.34 | 0.41 | 0.37 |
| 5 | 1671200 | rs931709 | -0.358 | 0.122 | -0.368 | -0.752 | -0.006 | -0.014 | -0.014 | 0.743 | 0.67 | 0.71 | 0.62 | 0.63 | 0.65 | 0.62 |
| 5 | 1676638 | rs10056116 | 0.567 | 0.355 | -0.574 | -0.743 | -0.009 | -0.012 | -0.012 | 0 | 0.24 | 0.17 | 0.35 | 0.33 | 0.26 | 0.26 |
| 5 | 1676963 | rs6897438 | -1.493 | 0.554 | -0.41 | -0.593 | 0.018 | 0.05 | 0.05 | 0 | 0.81 | 0.72 | 0.68 | 0.63 | 0.71 | 0.62 |
| 5 | 1679750 | rs7703975 | 0.195 | 0.156 | -0.297 | -0.845 | -0.01 | -0.026 | -0.026 | 0 | 0.42 | 0.48 | 0.27 | 0.29 | 0.4 | 0.33 |
| 5 | 1683758 | rs10075284 | -0.827 | -0.128 | 0.248 | -0.708 | -0.032 | -0.017 | -0.017 | 1.147 | 0.27 | 0.47 | 0.35 | 0.24 | 0.16 | 0.27 |
| 5 | 1685297 | rs16883078 | 1.125 | -0.085 | 0.275 | -0.672 | 0.018 | 0.059 | 0.059 | 0 | 0.07 | 0.16 | 0.23 | 0.2 | 0.09 | 0.11 |
| 5 | 1685594 | rs10512656 | -0.229 | -0.176 | 0.284 | -0.796 | 0.034 | 0.046 | 0.046 | 0 | 0.21 | 0.13 | 0.11 | 0.04 | 0.1 | 0.15 |
| 5 | 1685655 | rs17597967 | 0.916 | 0.063 | 0.304 | -0.897 | 0.005 | -0.025 | -0.025 | 0 | 0.4 | 0.45 | 0.28 | 0.27 | 0.49 | 0.34 |
| 5 | 1686233 | rs6898158 | 0.369 | 0.083 | 0.123 | -0.881 | 0.014 | 0.019 | 0.019 | 0 | 0.48 | 0.69 | 0.51 | 0.47 | 0.57 | 0.45 |
| 5 | 1687735 | rs13171033 | 1.152 | 0.771 | 0.349 | -0.822 | 0.173 | -0.04 | -0.04 | 0 | 0.08 | 0.26 | 0.13 | 0 | 0.21 | 0.07 |
| 5 | 1687769 | rs10051939 | -0.756 | 0.223 | -0.255 | -0.713 | -0.006 | -0.035 | -0.035 | 0 | 0.67 | 0.82 | 0.62 | 0.51 | 0.65 | 0.6 |
| 5 | 1691268 | rs1875197 | -0.681 | 0.154 | -0.182 | -0.476 | -0.028 | -0.038 | -0.038 | 1.408 | 0.11 | 0.25 | 0.06 | 0.04 | 0.05 | 0.15 |
| 5 | 1693575 | rs9312928 | -0.658 | 0.132 | -0.181 | -0.475 | -0.03 | -0.038 | -0.038 | 0 | 0.11 | 0.24 | 0.05 | 0.04 | 0.05 | 0.15 |
| 5 | 1694562 | rs10071895 | 0.156 | 0.571 | -0.181 | -0.264 | 0.029 | 0.061 | 0.061 | 0.077 | 0.1 | 0.24 | 0.26 | 0.2 | 0.12 | 0.12 |
| 5 | 1698123 | rs6864519 | -2.191 | 0.529 | -0.846 | -0.123 | -0.008 | -0.029 | -0.029 | 0 | 0.87 | 0.93 | 0.74 | 0.65 | 0.85 | 0.72 |
| 5 | 1699151 | rs12519991 | 1.205 | 0.127 | -1.266 | -0.514 | 0.013 | 0.01 | 0.01 | 0.042 | 0.27 | 0.09 | 0.15 | 0.28 | 0.28 | 0.25 |
| 5 | 1705513 | rs16883496 | 0.715 | 0.154 | -1.237 | -0.602 | -0.024 | -0.072 | -0.072 | 0 | 0.34 | 0.13 | 0.43 | 0.49 | 0.38 | 0.38 |
| 5 | 1708983 | rs12523324 | -0.391 | -0.379 | -0.748 | -0.729 | 0.001 | -0.012 | -0.012 | 0 | 0.63 | 0.67 | 0.67 | 0.6 | 0.68 | 0.48 |
| 5 | 1719213 | rs904728 | -0.99 | -0.442 | -0.84 | -0.965 | 0.011 | -0.052 | -0.052 | 0.055 | 0.74 | 0.85 | 0.71 | 0.65 | 0.78 | 0.67 |
| 5 | 1720276 | rs872015 | -0.106 | -0.599 | -0.32 | -1.028 | 0.015 | 0.006 | 0.006 | 0 | 0.5 | 0.48 | 0.59 | 0.52 | 0.64 | 0.56 |
| 5 | 1721003 | rs2133738 | 0.69 | -0.571 | -0.166 | -1.033 | -0.017 | -0.008 | -0.008 | 0 | 0.38 | 0.34 | 0.53 | 0.48 | 0.58 | 0.44 |
| 5 | 1721485 | rs2354124 | -0.153 | -0.85 | 0.336 | -1.504 | -0.003 | -0.016 | -0.016 | 0.383 | 0.57 | 0.45 | 0.37 | 0.3 | 0.66 | 0.44 |
| 5 | 1723063 | rs12515850 | 0.321 | -0.557 | 0.419 | -1.329 | -0.01 | 0.052 | 0.052 | 0 | 0.46 | 0.43 | 0.32 | 0.31 | 0.55 | 0.45 |
| 5 | 1723309 | rs11133883 | -0.613 | -0.357 | -0.142 | -1.429 | -0.006 | -0.01 | -0.01 | 0.023 | 0.64 | 0.82 | 0.42 | 0.44 | 0.67 | 0.57 |
| 5 | 1723360 | rs16883827 | 0.559 | -0.126 | 0.201 | -1.594 | -0.001 | -0.036 | -0.036 | 0.21 | 0.15 | 0.12 | 0.07 | 0 | 0.11 | 0.09 |
| 5 | 1725206 | rs7730084 | -0.18 | -0.001 | 0.102 | -1.527 | -0.012 | -0.079 | -0.079 | 0 | 0.25 | 0.22 | 0.1 | 0 | 0.16 | 0.1 |
| 5 | 1730637 | rs7718539 | 0.577 | 1.239 | 0.357 | -0.87 | 0.068 | -0.075 | -0.075 | 0 | 0.85 | 0.73 | 0.82 | 0.98 | 0.69 | 0.87 |
| 5 | 1734834 | rs11744846 | -0.099 | 0.214 | 0.104 | -1.021 | -0.004 | -0.037 | -0.037 | 0 | 0.22 | 0.19 | 0.09 | 0.02 | 0.11 | 0.11 |
| 5 | 1739266 | rs6554987 | -0.833 | -0.105 | 0.372 | -0.737 | 0.003 | 0.002 | 0.002 | 0 | 0.48 | 0.55 | 0.25 | 0.43 | 0.35 | 0.39 |
| 5 | 1744484 | rs1392466 | -1.395 | 0.315 | 0.362 | -0.95 | -0.015 | 0.008 | 0.008 | 0 | 0.58 | 0.82 | 0.4 | 0.43 | 0.6 | 0.48 |
| 5 | 1744831 | rs1392469 | 0.637 | 0.343 | 0.286 | -0.945 | -0.006 | -0.04 | -0.04 | 4.761 | 0.21 | 0.19 | 0.07 | 0.02 | 0.1 | 0.11 |
| 5 | 1753306 | rs2242411 | -0.49 | 0.39 | 0.493 | -0.935 | 0.007 | 0 | 0 | 0 | 0.36 | 0.47 | 0.31 | 0.41 | 0.49 | 0.37 |
| 5 | 1759000 | rs11741606 | -0.046 | 0.268 | 0.176 | -0.751 | 0.011 | -0.104 | -0.104 | 0 | 0.33 | 0.41 | 0.23 | 0.05 | 0.38 | 0.32 |
| 5 | 1760815 | rs16884541 | 0.868 | 0.224 | -0.545 | -0.722 | -0.013 | -0.107 | -0.107 | 0.644 | 0.16 | 0.01 | 0.09 | 0.38 | 0.16 | 0.14 |
| 5 | 1761033 | rs11750191 | 0.06 | 0.45 | -0.255 | -0.831 | -0.005 | -0.023 | -0.023 | 0 | 0.32 | 0.32 | 0.23 | 0.05 | 0.37 | 0.32 |
| 5 | 1761089 | rs16884543 | -0.634 | 0.014 | -0.208 | 0.33 | -0.021 | 0.042 | 0.042 | 0 | 0.28 | 0.35 | 0.25 | 0.49 | 0.23 | 0.17 |
| 5 | 1764128 | rs1107059 | -0.097 | 0.607 | -0.818 | 0.163 | -0.009 | 0.007 | 0.007 | 0 | 0.54 | 0.34 | 0.45 | 0.52 | 0.46 | 0.39 |
| 5 | 1764532 | rs1107060 | -0.579 | 0.378 | -0.18 | -0.028 | -0.006 | -0.023 | -0.023 | 0 | 0.06 | 0.16 | 0.04 | 0.01 | 0.02 | 0.03 |
| 5 | 1764750 | rs2353591 | 0.046 | 0.467 | -0.802 | -0.331 | 0.001 | -0.011 | -0.011 | 0 | 0.68 | 0.68 | 0.66 | 0.44 | 0.72 | 0.65 |
| 5 | 1766736 | rs7726044 | 0.017 | 0.67 | -0.617 | -0.266 | 0.04 | -0.012 | -0.012 | 0 | 0.06 | 0.13 | 0.08 | 0.03 | 0.1 | 0.12 |
| 5 | 1767774 | rs6884150 | 0.393 | 0.652 | -1.029 | -0.691 | 0.012 | -0.024 | -0.024 | 0 | 0.35 | 0.24 | 0.42 | 0.43 | 0.32 | 0.46 |
| 5 | 1769749 | rs6555008 | 0.382 | 0.512 | -0.822 | -0.728 | -0.011 | -0.006 | -0.006 | 0 | 0.35 | 0.43 | 0.36 | 0.28 | 0.26 | 0.42 |
| 5 | 1770225 | rs13154034 | 0.53 | 0.765 | -1.451 | -1.61 | -0.013 | 0.193 | 0.193 | 0 | 0.3 | 0.04 | 0.16 | 0.01 | 0.28 | 0.13 |
| 5 | 1771032 | rs5000660 | -0.437 | 0.701 | -1.208 | -1.812 | 0.001 | 0.009 | 0.009 | 0 | 0.47 | 0.55 | 0.52 | 0.22 | 0.57 | 0.4 |
| 5 | 1772615 | rs2172532 | -0.781 | 0.98 | -0.747 | -1.599 | 0.051 | -0.019 | -0.019 | 0 | 0.24 | 0.33 | 0.34 | 0.2 | 0.36 | 0.29 |
| 5 | 1776573 | rs924461 | 1.012 | 0.544 | -0.863 | -1.306 | -0.006 | -0.008 | -0.008 | 0 | 0.32 | 0.41 | 0.33 | 0.28 | 0.28 | 0.37 |
| 5 | 1777840 | rs750532 | -1.067 | 0.669 | -0.838 | -1.107 | -0.001 | 0.041 | 0.041 | 0 | 0.86 | 0.82 | 0.87 | 0.5 | 0.9 | 0.77 |
| 5 | 1778342 | rs13186705 | 1.617 | 0.67 | -1.081 | -1.357 | 0.02 | 0.196 | 0.196 | 0.227 | 0.48 | 0.07 | 0.45 | 0.23 | 0.47 | 0.38 |
| 5 | 1778464 | rs11953809 | -0.054 | 0.249 | -0.749 | -1.169 | 0 | 0.001 | 0.001 | 0 | 0.77 | 0.73 | 0.83 | 0.43 | 0.86 | 0.74 |
| 5 | 1778473 | rs11957879 | -0.118 | 0.249 | -0.749 | -1.169 | 0.001 | 0 | 0 | 0 | 0.77 | 0.74 | 0.83 | 0.43 | 0.86 | 0.74 |
| 5 | 1779843 | rs10040940 | -1.06 | 0.277 | -0.819 | -1.111 | 0 | 0.018 | 0.018 | 0 | 0.84 | 0.81 | 0.87 | 0.44 | 0.9 | 0.77 |
| 5 | 1782838 | rs13171074 | 1.301 | 0.791 | -1.04 | -1.251 | -0.018 | 0.149 | 0.149 | 0.063 | 0.56 | 0.09 | 0.61 | 0.23 | 0.68 | 0.45 |
| 5 | 1783237 | rs10866525 | -0.191 | 0.459 | -0.559 | -1.183 | -0.016 | 0.129 | 0.129 | 0.028 | 0.78 | 0.4 | 0.82 | 0.49 | 0.87 | 0.71 |
| 5 | 1783655 | rs12514619 | -0.151 | 0.837 | -0.557 | -1.004 | 0.041 | 0.043 | 0.043 | 0 | 0.18 | 0.31 | 0.3 | 0.25 | 0.21 | 0.22 |
| 5 | 1791180 | rs13186183 | 1.252 | 0.749 | -1.073 | -1.49 | -0.074 | 0.182 | 0.182 | 0.039 | 0.52 | 0.11 | 0.51 | 0.24 | 0.61 | 0.45 |
| 5 | 1794921 | rs4975846 | 1.168 | 1.016 | -0.554 | -1.467 | 0.012 | 0.026 | 0.026 | 0 | 0.41 | 0.32 | 0.39 | 0.28 | 0.34 | 0.46 |
| 5 | 1796564 | rs1546253 | 0.556 | 1.334 | 0.015 | -1.02 | 0.073 | 0.039 | 0.039 | 0 | 0.11 | 0.25 | 0.29 | 0.16 | 0.19 | 0.15 |
| 5 | 1797548 | rs7703762 | 0.593 | 1.336 | 0.131 | -1.019 | 0.009 | 0.003 | 0.003 | 0 | 0.11 | 0.13 | 0.28 | 0.16 | 0.19 | 0.15 |
| 5 | 1797735 | rs2276986 | -0.531 | 1.069 | 0.178 | -0.982 | -0.011 | 0.11 | 0.11 | 0 | 0.86 | 0.77 | 0.9 | 0.45 | 0.91 | 0.79 |
| 5 | 1800436 | rs972890 | 0.469 | 1.085 | -0.011 | -1.033 | -0.012 | 0.07 | 0.07 | 1.801 | 0.8 | 0.36 | 0.85 | 0.44 | 0.88 | 0.71 |
| 5 | 1803901 | rs3776150 | 1.67 | 0.941 | -0.256 | -1.239 | 0.013 | -0.002 | -0.002 | 0 | 0.33 | 0.04 | 0.31 | 0.28 | 0.25 | 0.37 |
| 5 | 1806653 | rs17535721 | -0.391 | 1.661 | 0.984 | -0.122 | 0.026 | 0 | 0 | 0.023 | 0.16 | 0.19 | 0.3 | 0.16 | 0.23 | 0.21 |
| 5 | 1815950 | rs10058270 | -0.024 | 1.29 | 1.259 | 0.089 | 0.09 | 0.032 | 0.032 | 2.389 | 0.05 | 0.34 | 0.04 | 0 | 0.03 | 0.08 |
| 5 | 1818127 | rs4975853 | 0.663 | 1.296 | 1.214 | 0.092 | -0.014 | 0.069 | 0.069 | 1.914 | 0.8 | 0.36 | 0.85 | 0.44 | 0.88 | 0.71 |
| 5 | 1835083 | rs2172530 | -0.821 | 2.155 | -0.089 | -0.224 | -0.005 | -0.001 | -0.001 | 0 | 0.16 | 0.16 | 0.3 | 0.18 | 0.22 | 0.22 |

| | | | | | | | | | | | | | | | | |
|---|---|---|---|---|---|---|---|---|---|---|---|---|---|---|---|---|
| 5 | 1837644 | rs17608534 | -0.941 | 2.435 | -0.095 | -0.239 | -0.002 | 0.003 | 0.003 | 0.02 | 0.17 | 0.15 | 0.32 | 0.1 | 0.32 | 0.21 |
| 5 | 1838826 | rs10061246 | -0.942 | 1.909 | 0.423 | -0.506 | -0.01 | -0.064 | -0.064 | 0 | 0.11 | 0.61 | 0.09 | 0.01 | 0.06 | 0.13 |
| 5 | 1839458 | rs6555016 | -0.85 | 1.955 | 0.979 | -0.484 | 0.045 | -0.03 | -0.03 | 0.262 | 0.09 | 0.48 | 0.06 | 0.01 | 0.05 | 0.13 |
| 5 | 1839498 | rs10512646 | -0.749 | 2.342 | 0.735 | -0.239 | 0.037 | 0.01 | 0.01 | 0 | 0.16 | 0.2 | 0.33 | 0.2 | 0.32 | 0.26 |
| 5 | 1844498 | rs16901305 | -0.743 | 1.555 | 1 | -0.601 | -0.015 | -0.085 | -0.085 | 0.02 | 0.15 | 0.34 | 0.09 | 0.02 | 0.1 | 0.13 |
| 5 | 1847639 | rs12188301 | -0.747 | 1.675 | 0.46 | 0.079 | 0 | 0.045 | 0.045 | 0 | 0.88 | 0.87 | 0.9 | 0.46 | 0.88 | 0.78 |
| 5 | 1850220 | rs10512645 | -0.989 | 2.107 | 0.741 | -0.307 | 0.041 | 0.003 | 0.003 | 0 | 0.18 | 0.23 | 0.34 | 0.2 | 0.33 | 0.23 |
| 5 | 1850563 | rs10512644 | -1.042 | 2.09 | 0.733 | -0.315 | 0.038 | 0.002 | 0.002 | 0 | 0.18 | 0.23 | 0.34 | 0.2 | 0.33 | 0.24 |
| 5 | 1852284 | rs263213 | -0.85 | 1.804 | 0.321 | -0.725 | 0.005 | -0.029 | -0.029 | 0.305 | 0.35 | 0.39 | 0.33 | 0.22 | 0.38 | 0.26 |
| 5 | 1854257 | rs260397 | -0.76 | 1.931 | 0.329 | -0.617 | 0.047 | -0.044 | -0.044 | 0.349 | 0.08 | 0.19 | 0.1 | 0 | 0.18 | 0.08 |
| 5 | 1855301 | rs91315 | -0.661 | 1.772 | 0.038 | -0.846 | -0.001 | -0.006 | -0.006 | 0 | 0.4 | 0.56 | 0.47 | 0.24 | 0.51 | 0.31 |
| 5 | 1856596 | rs260393 | 0.174 | 1.649 | -0.221 | -0.694 | -0.001 | 0.002 | 0.002 | 0 | 0.27 | 0.27 | 0.3 | 0.24 | 0.27 | 0.2 |
| 5 | 1859721 | rs1532272 | 1.291 | 1.343 | -1.15 | -0.342 | -0.023 | -0.074 | -0.074 | 0 | 0.27 | 0.07 | 0.34 | 0.31 | 0.26 | 0.31 |
| 5 | 1860049 | rs4975744 | -0.026 | 0.727 | -0.587 | -0.221 | 0 | 0.003 | 0.003 | 0 | 0.39 | 0.59 | 0.39 | 0.31 | 0.33 | 0.33 |
| 5 | 1860281 | rs1689746 | -0.334 | 1.013 | -0.815 | -0.099 | -0.004 | -0.075 | -0.075 | 0 | 0.73 | 0.89 | 0.83 | 0.63 | 0.74 | 0.72 |
| 5 | 1871880 | rs2353580 | -1.277 | 0.856 | -0.654 | 0.02 | 0.006 | -0.008 | -0.008 | 0.459 | 0.88 | 0.9 | 0.94 | 0.65 | 0.9 | 0.81 |
| 5 | 1872329 | rs883823 | -0.244 | 0.899 | -0.353 | -0.237 | 0 | 0.016 | 0.016 | 0 | 0.51 | 0.54 | 0.5 | 0.33 | 0.49 | 0.34 |
| 5 | 1875037 | rs1632597 | -1.001 | 0.9 | -0.146 | 0.052 | -0.001 | -0.002 | -0.002 | 0.003 | 0.63 | 0.7 | 0.56 | 0.64 | 0.58 | 0.54 |
| 5 | 1877280 | rs4975709 | 0.495 | 0.946 | -0.353 | -0.037 | 0 | 0.007 | 0.007 | 0 | 0.27 | 0.35 | 0.33 | 0.33 | 0.24 | 0.24 |
| 5 | 1880865 | rs4975753 | 0.527 | 0.949 | -0.508 | 0.003 | -0.003 | 0.005 | 0.005 | 0 | 0.27 | 0.34 | 0.32 | 0.33 | 0.23 | 0.26 |
| 5 | 1880891 | rs2232376 | 0.274 | 0.944 | -1.304 | -0.117 | 0.001 | -0.014 | -0.014 | 1.7 | 0.21 | 0.22 | 0.16 | 0.01 | 0.21 | 0.07 |
| 5 | 1881733 | rs260402 | -0.623 | 1.182 | -1.679 | 0.522 | -0.021 | -0.026 | -0.026 | 0 | 0.68 | 0.87 | 0.59 | 0.67 | 0.62 | 0.64 |
| 5 | 1886471 | rs260400 | -0.361 | 1.033 | -1.326 | 0.808 | -0.015 | -0.059 | -0.059 | 0.718 | 0.88 | 0.96 | 0.94 | 0.7 | 0.88 | 0.79 |
| 5 | 1887817 | rs260409 | -0.697 | 1.199 | -1.098 | 0.242 | -0.005 | 0.001 | 0.001 | 0 | 0.17 | 0.16 | 0.16 | 0.12 | 0.21 | 0.17 |
| 5 | 1892759 | rs6861778 | 0.01 | 0.796 | -0.723 | -0.003 | 0.016 | -0.022 | -0.022 | 0.855 | 0.3 | 0.39 | 0.36 | 0.27 | 0.33 | 0.22 |
| 5 | 1895829 | rs12653946 | 0.212 | 0.343 | -0.631 | 0.101 | 0.004 | 0.022 | 0.022 | 0 | 0.47 | 0.4 | 0.5 | 0.35 | 0.45 | 0.53 |
| 5 | 1897640 | rs6861105 | -0.122 | 0.332 | -0.216 | 0.706 | -0.022 | 0.161 | 0.161 | 0 | 0.87 | 0.72 | 0.87 | 0.62 | 0.93 | 0.8 |
| 5 | 1898679 | rs10428597 | 0.58 | 0.677 | -1.18 | 0.519 | -0.003 | -0.057 | -0.057 | 0 | 0.62 | 0.21 | 0.57 | 0.53 | 0.66 | 0.51 |
| 5 | 1901205 | rs17537048 | 0.108 | 1.102 | -1.087 | 0.466 | -0.053 | -0.034 | -0.034 | 0 | 0.17 | 0.06 | 0.23 | 0.22 | 0.25 | 0.22 |
| 5 | 1909022 | rs6894487 | -1.181 | 0.263 | -0.208 | 0.196 | -0.001 | -0.048 | -0.048 | 0 | 0.22 | 0.29 | 0.22 | 0.05 | 0.18 | 0.31 |
| 5 | 1911035 | rs4975760 | -1.343 | 0.242 | -0.166 | 0.272 | -0.01 | -0.051 | -0.051 | 0 | 0.25 | 0.45 | 0.25 | 0.12 | 0.2 | 0.34 |
| 5 | 1911819 | rs4975713 | -0.446 | 0.267 | -0.274 | 0.272 | 0.008 | 0.011 | 0.011 | 0 | 0.15 | 0.1 | 0.11 | 0.08 | 0.1 | 0.21 |
| 5 | 1912921 | rs1994388 | -0.895 | 0.326 | -0.484 | 0.502 | -0.009 | -0.014 | -0.014 | 0 | 0.27 | 0.41 | 0.25 | 0.18 | 0.2 | 0.33 |
| 5 | 1922870 | rs6420033 | 0.171 | 0.199 | -0.846 | 1.168 | 0.013 | -0.035 | -0.035 | 0 | 0.79 | 0.84 | 0.71 | 0.58 | 0.82 | 0.66 |
| 5 | 1927293 | rs10042507 | 1.116 | 0.931 | -0.502 | 0.934 | -0.005 | 0.017 | 0.017 | 0 | 0.56 | 0.37 | 0.45 | 0.28 | 0.5 | 0.34 |
| 5 | 1927412 | rs6555027 | -0.324 | 1.09 | -0.569 | 1 | 0.016 | -0.072 | -0.072 | 0 | 0.79 | 0.89 | 0.71 | 0.58 | 0.81 | 0.66 |
| 5 | 1932463 | rs10058874 | 0.988 | 1.012 | -0.395 | 0.905 | -0.002 | 0.001 | 0.001 | 0 | 0.59 | 0.4 | 0.45 | 0.27 | 0.5 | 0.38 |
| 5 | 1934490 | rs6893465 | 0.212 | 1.369 | -0.087 | 0.72 | 0.117 | 0.066 | 0.066 | 0.027 | 0.79 | 0.53 | 0.75 | 0.69 | 0.67 | 0.69 |
| 5 | 1936116 | rs10065645 | -0.426 | 0.907 | 0.114 | 0.063 | 0.024 | -0.042 | -0.042 | 0 | 0.14 | 0.34 | 0.11 | 0.04 | 0.13 | 0.1 |
| 5 | 1936443 | rs2129470 | 0.202 | 1.439 | -0.268 | 0.103 | -0.047 | 0.001 | 0.001 | 0 | 0.15 | 0.11 | 0.16 | 0.12 | 0.28 | 0.23 |
| 5 | 1937407 | rs11737953 | 0.886 | 0.202 | -0.912 | -1.116 | 0.012 | 0.04 | 0.04 | 0 | 0.28 | 0.14 | 0.24 | 0 | 0.28 | 0.17 |
| 5 | 1938626 | rs901495 | -0.971 | -0.144 | -0.136 | -1.212 | 0.005 | 0.005 | 0.005 | 0.093 | 0.61 | 0.67 | 0.57 | 0.27 | 0.74 | 0.54 |
| 5 | 1939938 | rs6892417 | -0.405 | -0.008 | -0.115 | -1.069 | 0.072 | -0.033 | -0.033 | 0.59 | 0.27 | 0.5 | 0.25 | 0.17 | 0.41 | 0.35 |
| 5 | 1943728 | rs11743544 | 1.433 | -0.508 | -0.754 | -1.209 | 0.034 | 0.055 | 0.055 | 0.285 | 0.12 | 0.03 | 0.06 | 0 | 0.07 | 0.07 |
| 5 | 1944308 | rs10079709 | -1.174 | -0.118 | -0.366 | -1.065 | -0.002 | 0 | 0 | 0.065 | 0.06 | 0.11 | 0.08 | 0.1 | 0.04 | 0.02 |
| 5 | 1947034 | rs11133917 | 0.68 | -0.097 | -0.974 | -1.518 | 0.026 | 0.131 | 0.131 | 0 | 0.34 | 0.12 | 0.23 | 0.08 | 0.29 | 0.22 |
| 5 | 1948303 | rs13183501 | 0.644 | 0.382 | -1.111 | -1.008 | -0.007 | -0.007 | -0.007 | 0 | 0.12 | 0.1 | 0.2 | 0.21 | 0.16 | 0.08 |
| 5 | 1948464 | rs1471886 | 0.668 | 0.411 | -1.08 | -1.021 | 0.007 | -0.001 | -0.001 | 0.081 | 0.15 | 0.2 | 0.14 | 0.14 | 0.26 | 0.22 |
| 5 | 1948519 | rs11746674 | 0.897 | 0.026 | -1.229 | -1.174 | 0.031 | 0.164 | 0.164 | 0 | 0.44 | 0.15 | 0.35 | 0.2 | 0.38 | 0.27 |
| 5 | 1949340 | rs972973 | 0.359 | -0.017 | -1.201 | -1.197 | 0.015 | 0.1 | 0.1 | 0 | 0.54 | 0.29 | 0.41 | 0.22 | 0.6 | 0.46 |
| 5 | 1949562 | rs7736642 | 0.997 | -0.238 | -1.257 | -1.252 | 0.011 | 0.213 | 0.213 | 0 | 0.39 | 0.08 | 0.27 | 0.09 | 0.34 | 0.24 |
| 5 | 1953161 | rs10055976 | -0.415 | -0.075 | -0.897 | -1.061 | -0.012 | 0.002 | 0.002 | 0 | 0.08 | 0.09 | 0.1 | 0.11 | 0.06 | 0.05 |
| 5 | 1956884 | rs13170991 | -0.963 | 0.66 | -1.062 | -1.251 | -0.015 | -0.111 | -0.111 | 0 | 0.81 | 0.93 | 0.73 | 0.48 | 0.8 | 0.6 |
| 5 | 1956973 | rs4246751 | -0.725 | 0.628 | -0.159 | -1.266 | -0.008 | 0.079 | 0.079 | 0 | 0.78 | 0.65 | 0.7 | 0.48 | 0.79 | 0.6 |
| 5 | 1957109 | rs2200559 | 0.09 | 0.854 | -0.232 | -1.423 | -0.054 | -0.013 | -0.013 | 0 | 0.26 | 0.17 | 0.31 | 0.29 | 0.39 | 0.31 |
| 5 | 1962256 | rs12654104 | -0.247 | 0.859 | 0.752 | -1.019 | 0.005 | -0.006 | -0.006 | 0 | 0.11 | 0.3 | 0.11 | 0.08 | 0.08 | 0.05 |
| 5 | 1962436 | rs12654102 | -0.509 | 0.875 | 0.553 | -0.901 | -0.007 | 0.073 | 0.073 | 0 | 0.79 | 0.65 | 0.71 | 0.47 | 0.79 | 0.59 |
| 5 | 1967265 | rs1017923 | 0.176 | 1.753 | 0.527 | -0.781 | -0.019 | 0.006 | 0.006 | 0.099 | 0.16 | 0.09 | 0.1 | 0.14 | 0.27 | 0.26 |
| 5 | 1967849 | rs1023981 | -1.242 | 1.222 | 0.834 | -0.601 | 0.033 | 0.009 | 0.009 | 0.716 | 0.32 | 0.39 | 0.39 | 0.37 | 0.43 | 0.35 |
| 5 | 1976187 | rs1017924 | -0.481 | 0.854 | 0.715 | -0.815 | -0.038 | 0.108 | 0.108 | 1.802 | 0.75 | 0.51 | 0.69 | 0.46 | 0.79 | 0.61 |
| 5 | 1976478 | rs1123391 | -0.932 | 1.147 | 0.594 | -0.636 | -0.026 | -0.025 | -0.025 | 0 | 0.28 | 0.2 | 0.42 | 0.46 | 0.42 | 0.34 |
| 5 | 1983710 | rs13172746 | 0.373 | 1.261 | 0.651 | -0.692 | 0.013 | 0.049 | 0.049 | 0 | 0.12 | 0.12 | 0.26 | 0.32 | 0.17 | 0.08 |
| 5 | 1987633 | rs1601641 | -1.215 | 1.05 | 0.755 | -0.72 | 0.026 | 0.023 | 0.023 | 0 | 0.3 | 0.32 | 0.38 | 0.44 | 0.41 | 0.34 |
| 5 | 1993900 | rs17611544 | 0.256 | 1.237 | 0.787 | -0.589 | -0.041 | 0.011 | 0.011 | 0.041 | 0.14 | 0.05 | 0.07 | 0.12 | 0.21 | 0.25 |
| 5 | 1999840 | rs1872775 | 0.843 | 1.112 | 0.725 | -0.756 | 0.001 | 0.024 | 0.024 | 0.594 | 0.53 | 0.33 | 0.53 | 0.33 | 0.51 | 0.31 |
| 5 | 2001045 | rs13189295 | 0.609 | 1.128 | 0.688 | -0.583 | 0.003 | 0.029 | 0.029 | 0 | 0.11 | 0.11 | 0.29 | 0.35 | 0.19 | 0.1 |
| 5 | 2001784 | rs4493711 | 0.063 | 1.115 | 0.885 | -0.744 | 0.012 | 0.014 | 0.014 | 0.161 | 0.13 | 0.16 | 0.29 | 0.22 | 0.19 | 0.11 |
| 5 | 2002126 | rs10068684 | 1.014 | 1.02 | 1.172 | -0.856 | 0.003 | 0.018 | 0.018 | 1.35 | 0.56 | 0.38 | 0.34 | 0.23 | 0.54 | 0.46 |
| 5 | 2003180 | rs11749073 | -1.809 | 0.812 | 1.147 | -0.941 | 0.031 | -0.063 | -0.063 | 0 | 0.87 | 0.96 | 0.79 | 0.45 | 0.91 | 0.69 |

| | | | | | | | | | | | | | | | | |
|---|---|---|---|---|---|---|---|---|---|---|---|---|---|---|---|---|
| 5 | 2004160 | rs10474733 | -0.34 | 1.054 | 1.718 | -0.748 | -0.019 | 0.018 | 0.018 | 0.264 | 0.69 | 0.59 | 0.64 | 0.45 | 0.73 | 0.55 |
| 5 | 2007007 | rs901508 | -0.179 | 1.256 | 1.608 | -0.784 | 0.017 | -0.014 | -0.014 | 0.313 | 0.1 | 0.22 | 0.05 | 0.05 | 0.1 | 0.08 |
| 5 | 2008571 | rs901498 | 1.037 | 1.208 | 1.253 | -0.754 | 0.002 | -0.001 | -0.001 | 0 | 0.07 | 0.09 | 0.04 | 0.06 | 0.09 | 0.07 |
| 5 | 2008973 | rs7717285 | 0.877 | 1.1 | 1.186 | -0.792 | 0.007 | -0.004 | -0.004 | 0.555 | 0.08 | 0.08 | 0.04 | 0.08 | 0.13 | 0.17 |
| 5 | 2010496 | rs4975792 | -0.649 | 1.048 | 1.187 | -0.852 | -0.003 | -0.012 | -0.012 | 0.081 | 0.4 | 0.54 | 0.43 | 0.73 | 0.42 | 0.6 |
| 5 | 2012474 | rs13190594 | -0.754 | 1.03 | 1.174 | -0.746 | 0.006 | -0.002 | -0.002 | 0 | 0.41 | 0.66 | 0.43 | 0.59 | 0.41 | 0.6 |
| 5 | 2012888 | rs9312971 | -0.795 | 1.03 | 1.174 | -0.748 | 0.007 | 0.019 | 0.019 | 2.153 | 0.41 | 0.66 | 0.44 | 0.73 | 0.42 | 0.6 |
| 5 | 2014366 | rs2220309 | 0.071 | 0.866 | 1.246 | -0.676 | 0.02 | 0.043 | 0.043 | 0.394 | 0.8 | 0.73 | 0.65 | 0.58 | 0.72 | 0.82 |
| 5 | 2014721 | rs4246753 | 0.202 | 0.87 | 1.357 | -0.691 | -0.025 | 0.045 | 0.045 | 0.812 | 0.31 | 0.45 | 0.36 | 0.52 | 0.28 | 0.42 |
| 5 | 2014776 | rs4246754 | 0.364 | 0.893 | 0.52 | -0.696 | 0.023 | -0.061 | -0.061 | 3.008 | 0.28 | 0.17 | 0.35 | 0.52 | 0.27 | 0.43 |
| 5 | 2016422 | rs17612453 | 0.822 | 1.037 | 0.209 | -1.506 | 0.004 | 0.158 | 0.158 | 0.961 | 0.49 | 0.26 | 0.3 | 0.19 | 0.44 | 0.4 |
| 5 | 2032660 | rs4975807 | -1.7 | 2.026 | 0.731 | -0.673 | -0.06 | 0.056 | 0.056 | 1.818 | 0.37 | 0.8 | 0.46 | 0.32 | 0.45 | 0.33 |
| 5 | 2041665 | rs4129455 | -0.092 | 2.473 | 0.895 | -0.379 | 0.005 | -0.003 | -0.003 | 0.106 | 0.08 | 0.11 | 0.05 | 0.05 | 0.1 | 0.08 |
| 5 | 2047463 | rs4975840 | -1.837 | 2.442 | 0.815 | -0.66 | 0.034 | 0.073 | 0.073 | 0.01 | 0.46 | 0.71 | 0.51 | 0.32 | 0.61 | 0.35 |
| 5 | 2050328 | rs10065111 | -0.58 | 2.903 | 1.274 | -0.489 | 0.002 | -0.001 | -0.001 | 0 | 0.16 | 0.24 | 0.3 | 0.19 | 0.2 | 0.07 |
| 5 | 2050483 | rs4975838 | -0.503 | 2.903 | 1.275 | -0.521 | 0.004 | -0.023 | -0.023 | 0 | 0.15 | 0.23 | 0.29 | 0.03 | 0.2 | 0.07 |
| 5 | 2056586 | rs6555046 | -1.239 | 2.916 | 1.145 | -0.517 | 0.007 | -0.007 | -0.007 | 0 | 0.3 | 0.68 | 0.23 | 0.29 | 0.39 | 0.28 |
| 5 | 2058143 | rs4975727 | -1.977 | 2.956 | 1.013 | -0.527 | 0.021 | 0.053 | 0.053 | 0 | 0.44 | 0.71 | 0.51 | 0.31 | 0.6 | 0.35 |
| 5 | 2058183 | rs13163790 | -0.533 | 2.965 | 0.864 | -0.121 | 0 | 0 | 0 | 0 | 0.21 | 0.23 | 0.18 | 0.28 | 0.33 | 0.23 |
| 5 | 2059311 | rs7719270 | -0.587 | 2.305 | 0.658 | 0.093 | 0.024 | -0.048 | -0.048 | 0 | 0.12 | 0.15 | 0.27 | 0.01 | 0.19 | 0.08 |
| 5 | 2075104 | rs10475133 | -1.045 | 2.262 | 0.619 | 0.536 | 0.078 | -0.004 | -0.004 | 4.224 | 0.2 | 0.39 | 0.34 | 0.13 | 0.29 | 0.21 |
| 5 | 2075162 | rs10475134 | -0.947 | 2.275 | 0.885 | 0.482 | 0.079 | -0.006 | -0.006 | 2.898 | 0.2 | 0.49 | 0.35 | 0.13 | 0.29 | 0.22 |
| 5 | 2075565 | rs4361562 | 0.765 | 1.857 | -0.45 | 0.772 | -0.037 | -0.147 | -0.147 | 0.818 | 0.18 | 0 | 0.16 | 0.46 | 0.2 | 0.3 |
| 5 | 2077356 | rs11133927 | -0.433 | 2.502 | -0.006 | 0.517 | -0.008 | 0.005 | 0.005 | 1.336 | 0.35 | 0.24 | 0.3 | 0.61 | 0.39 | 0.5 |
| 5 | 2077642 | rs7704452 | -0.672 | 1.543 | -0.102 | -0.263 | 0.001 | -0.046 | -0.046 | 1.309 | 0.18 | 0.24 | 0.19 | 0.01 | 0.17 | 0.1 |
| 5 | 2083797 | rs4566837 | 0.83 | 1.721 | -0.072 | -0.863 | 0.003 | -0.059 | -0.059 | 0 | 0.41 | 0.41 | 0.35 | 0.01 | 0.34 | 0.36 |
| 5 | 2086448 | rs11743496 | 1.315 | 1.808 | -0.456 | -0.606 | -0.052 | -0.151 | -0.151 | 0 | 0.23 | 0.02 | 0.28 | 0.43 | 0.24 | 0.34 |
| 5 | 2087916 | rs10063808 | 0.961 | 1.94 | -0.123 | -0.76 | -0.008 | -0.092 | -0.092 | 0 | 0.15 | 0.29 | 0.08 | 0 | 0.11 | 0.07 |
| 5 | 2089221 | rs7702195 | 0.009 | 2.019 | -0.139 | -0.545 | 0.001 | 0.074 | 0.074 | 0.052 | 0.58 | 0.73 | 0.57 | 0.23 | 0.58 | 0.46 |
| 5 | 2090652 | rs10060649 | 0.948 | 1.929 | -0.225 | -0.408 | -0.01 | -0.094 | -0.094 | 0.157 | 0.15 | 0.28 | 0.08 | 0 | 0.11 | 0.07 |
| 5 | 2091155 | rs11748879 | 1.575 | 2.026 | -0.422 | -0.311 | -0.047 | -0.15 | -0.15 | 0.402 | 0.2 | 0 | 0.27 | 0.43 | 0.22 | 0.34 |
| 5 | 2094148 | rs10076573 | 0.093 | 2.073 | 0.15 | -0.491 | 0.027 | 0.005 | 0.005 | 0 | 0.18 | 0.25 | 0.18 | 0.23 | 0.23 | 0.1 |
| 5 | 2097011 | rs4568402 | -0.525 | 2.091 | -0.436 | -0.53 | 0.048 | -0.018 | -0.018 | 0 | 0.24 | 0.53 | 0.25 | 0.23 | 0.3 | 0.13 |
| 5 | 2097819 | rs13360716 | 0.233 | 1.455 | -0.385 | -0.557 | -0.022 | -0.142 | -0.142 | 0 | 0.19 | 0.47 | 0.12 | 0 | 0.14 | 0.08 |
| 5 | 2099667 | rs13180333 | -0.015 | 1.705 | -1.17 | 0.008 | -0.014 | 0.004 | 0.004 | 0 | 0.58 | 0.79 | 0.57 | 0.42 | 0.56 | 0.5 |
| 5 | 2103617 | rs4975819 | 0.605 | 2.127 | -0.877 | 0.709 | 0.004 | 0.006 | 0.006 | 0 | 0.6 | 0.49 | 0.52 | 0.51 | 0.48 | 0.58 |
| 5 | 2105709 | rs10051274 | -0.357 | 2.341 | -1.104 | 0.733 | 0.007 | -0.003 | -0.003 | 0 | 0.36 | 0.42 | 0.38 | 0.35 | 0.45 | 0.39 |
| 5 | 2109901 | rs6879627 | 1.084 | 2.323 | -1.188 | 1.526 | -0.027 | 0.014 | 0.014 | 0 | 0.93 | 0.96 | 0.78 | 0.59 | 0.86 | 0.93 |
| 5 | 2113672 | rs11950809 | -0.12 | 1.258 | -0.925 | 0.666 | -0.044 | -0.083 | -0.083 | 0 | 0.17 | 0.41 | 0.24 | 0.06 | 0.1 | 0.21 |
| 5 | 2117328 | rs6861474 | 0.156 | 1.627 | -2.631 | 0.629 | -0.063 | 0.006 | 0.006 | 0.544 | 0.33 | 0.06 | 0.3 | 0.25 | 0.33 | 0.34 |
| 5 | 2123985 | rs4075798 | 0.548 | 1.98 | -2.466 | 1.007 | 0.03 | 0.092 | 0.092 | 0 | 0.38 | 0.22 | 0.28 | 0.13 | 0.3 | 0.32 |
| 5 | 2129233 | rs4077288 | 0.482 | 2.02 | -2.431 | 1.014 | 0.03 | 0.098 | 0.098 | 0 | 0.39 | 0.22 | 0.3 | 0.15 | 0.3 | 0.3 |
| 5 | 2131784 | rs11953155 | -0.217 | 1.729 | -2.477 | 0.825 | -0.114 | -0.041 | -0.041 | 0 | 0.31 | 0.02 | 0.32 | 0.27 | 0.36 | 0.36 |
| 5 | 2133412 | rs6860349 | 0.282 | 1.158 | -1.746 | 0.884 | 0.009 | -0.013 | -0.013 | 0.022 | 0.55 | 0.3 | 0.31 | 0.57 | 0.42 | 0.52 |
| 5 | 2136570 | rs6555064 | -0.19 | 1.03 | -1.429 | 0.846 | 0.002 | -0.003 | -0.003 | 0 | 0.47 | 0.67 | 0.46 | 0.46 | 0.55 | 0.53 |
| 5 | 2137758 | rs7719723 | 0.279 | 0.808 | -0.989 | 0.714 | 0 | -0.001 | -0.001 | 0 | 0.62 | 0.62 | 0.59 | 0.42 | 0.42 | 0.53 |
| 5 | 2140755 | rs11960538 | -0.729 | 0.075 | -0.255 | 0.327 | -0.022 | -0.052 | -0.052 | 0 | 0.17 | 0.36 | 0.12 | 0.06 | 0.1 | 0.05 |
| 5 | 2144662 | rs6893579 | -0.957 | 0.046 | -0.097 | 0.805 | 0.005 | 0.088 | 0.088 | 0 | 0.05 | 0.16 | 0.12 | 0.11 | 0.05 | 0.05 |
| 5 | 2149503 | rs7448302 | -0.177 | -0.655 | -0.982 | 0.777 | 0.047 | -0.043 | -0.043 | 0 | 0.72 | 0.94 | 0.83 | 0.63 | 0.83 | 0.63 |
| 5 | 2150273 | rs7735656 | 0.283 | -0.236 | -0.465 | 0.695 | 0.006 | -0.001 | -0.001 | 0.658 | 0.63 | 0.67 | 0.78 | 0.59 | 0.75 | 0.58 |
| 5 | 2156561 | rs6872691 | -0.487 | -0.232 | -0.486 | 0.528 | 0.012 | 0.018 | 0.018 | 0 | 0.47 | 0.67 | 0.54 | 0.44 | 0.57 | 0.3 |
| 5 | 2156619 | rs17644704 | 0.151 | 0.186 | -0.047 | 0.033 | 0.019 | -0.096 | -0.096 | 1.646 | 0.28 | 0.32 | 0.23 | 0.01 | 0.36 | 0.11 |
| 5 | 2158553 | rs11133930 | -0.075 | 0.049 | 0.1 | 0.192 | -0.003 | -0.002 | -0.002 | 0 | 0.09 | 0.15 | 0.12 | 0.14 | 0.1 | 0.1 |
| 5 | 2159490 | rs10072352 | -1.397 | -0.08 | 0.374 | 0.109 | -0.01 | -0.023 | -0.023 | 0.883 | 0.14 | 0.47 | 0.15 | 0.15 | 0.11 | 0.11 |
| 5 | 2163977 | rs4866721 | 1.694 | -0.696 | 0.379 | -0.246 | 0.032 | 0.024 | 0.024 | 0.02 | 0.45 | 0.26 | 0.43 | 0.55 | 0.38 | 0.67 |
| 5 | 2164702 | rs10076206 | -1.053 | -0.626 | 0.394 | -0.204 | -0.004 | 0.017 | 0.017 | 0 | 0.43 | 0.66 | 0.5 | 0.4 | 0.54 | 0.24 |
| 5 | 2164846 | rs10076280 | -0.776 | -0.609 | 0.425 | -0.195 | -0.009 | -0.019 | -0.019 | 0 | 0.13 | 0.37 | 0.14 | 0.14 | 0.11 | 0.11 |
| 5 | 2166397 | rs13171126 | 0.365 | -0.563 | 0.231 | -0.171 | -0.042 | 0.009 | 0.009 | 0 | 0.29 | 0.16 | 0.34 | 0.25 | 0.42 | 0.13 |
| 5 | 2166433 | rs6863477 | -0.288 | -0.353 | 0.087 | -0.122 | 0 | -0.001 | -0.001 | 0 | 0.89 | 0.87 | 0.87 | 0.86 | 0.88 | 0.9 |
| 5 | 2167740 | rs4591752 | 0.52 | -0.662 | -0.417 | -0.094 | 0.012 | -0.02 | -0.02 | 0 | 0.36 | 0.28 | 0.28 | 0.4 | 0.29 | 0.67 |
| 5 | 2170286 | rs11746413 | 0.051 | -1.277 | -0.694 | -0.375 | 0.012 | 0.009 | 0.009 | 0 | 0.37 | 0.36 | 0.36 | 0.34 | 0.28 | 0.23 |
| 5 | 2173676 | rs17645173 | 1.58 | -0.048 | -0.893 | -0.476 | -0.07 | 0.061 | 0.061 | 0 | 0.13 | 0.02 | 0.11 | 0 | 0.23 | 0.07 |
| 5 | 2174313 | rs6878643 | -0.927 | -0.49 | -0.659 | -0.434 | -0.032 | 0.048 | 0.048 | 0.519 | 0.68 | 0.53 | 0.75 | 0.61 | 0.77 | 0.4 |
| 5 | 2174621 | rs4866475 | 0.431 | -0.518 | -0.816 | -0.446 | 0.021 | 0.038 | 0.038 | 0 | 0.39 | 0.29 | 0.46 | 0.34 | 0.38 | 0.23 |
| 5 | 2175401 | rs6555068 | -0.983 | -0.617 | -0.556 | -0.031 | -0.044 | -0.008 | -0.008 | 0 | 0.74 | 0.69 | 0.9 | 0.8 | 0.87 | 0.64 |
| 5 | 2175605 | rs4262114 | 1.095 | 0.008 | -0.734 | -0.225 | -0.007 | -0.049 | -0.049 | 0 | 0.08 | 0 | 0.16 | 0.26 | 0.1 | 0.05 |
| 5 | 2175760 | rs16903521 | 0.472 | 0.056 | -0.782 | -0.293 | -0.017 | -0.059 | -0.059 | 0.882 | 0.09 | 0.01 | 0.19 | 0.26 | 0.12 | 0.05 |
| 5 | 2178060 | rs12658255 | 1.092 | 0.048 | -0.733 | -0.235 | -0.007 | -0.049 | -0.049 | 0 | 0.08 | 0 | 0.16 | 0.26 | 0.1 | 0.05 |
| 5 | 2178140 | rs10073915 | 0.018 | 0.084 | -0.255 | -0.444 | 0.029 | -0.019 | -0.019 | 0.859 | 0.08 | 0.25 | 0.07 | 0.02 | 0.07 | 0.05 |

| | | | | | | | | | | | | | | | | |
|---|---|---|---|---|---|---|---|---|---|---|---|---|---|---|---|---|
| 5 | 2179628 | rs10057158 | 0.292 | -0.135 | -0.687 | -0.606 | 0.023 | 0.033 | 0.033 | 0 | 0.44 | 0.4 | 0.48 | 0.38 | 0.41 | 0.23 |
| 5 | 2182854 | rs6555069 | 0.293 | -0.072 | -0.837 | -0.806 | 0.021 | 0.054 | 0.054 | 0.001 | 0.43 | 0.36 | 0.49 | 0.32 | 0.43 | 0.24 |
| 5 | 2184365 | rs13172851 | 0.697 | 0.082 | -1.281 | -0.756 | -0.026 | 0.052 | 0.052 | 0.036 | 0.38 | 0.12 | 0.47 | 0.32 | 0.42 | 0.22 |
| 5 | 2191437 | rs11133935 | -0.809 | 0.039 | -1.145 | -0.703 | -0.001 | -0.002 | -0.002 | 0 | 0.6 | 0.64 | 0.74 | 0.61 | 0.62 | 0.35 |
| 5 | 2194596 | rs4866654 | 0.83 | -0.076 | -1.311 | -0.75 | -0.051 | 0.045 | 0.045 | 0.027 | 0.37 | 0.09 | 0.46 | 0.32 | 0.43 | 0.2 |
| 5 | 2194769 | rs16903612 | 1.257 | 0.102 | -1.298 | -0.505 | -0.009 | 0.016 | 0.016 | 0 | 0.08 | 0 | 0.07 | 0.06 | 0.08 | 0.02 |
| 5 | 2196941 | rs16903629 | 0.908 | 0.109 | -1.298 | -0.513 | -0.014 | -0.06 | -0.06 | 0.443 | 0.09 | 0 | 0.2 | 0.29 | 0.12 | 0.05 |
| 5 | 2199605 | rs2405802 | -1.194 | 0.357 | -1.354 | -0.612 | -0.028 | 0.029 | 0.029 | 0 | 0.69 | 0.65 | 0.83 | 0.62 | 0.81 | 0.43 |
| 5 | 2200150 | rs6865374 | -0.131 | 0.009 | -1.243 | -0.567 | 0.007 | 0.041 | 0.041 | 0 | 0.5 | 0.63 | 0.54 | 0.32 | 0.49 | 0.29 |
| 5 | 2202172 | rs12187097 | 0.453 | 0.074 | -1.889 | -0.563 | 0.015 | 0.034 | 0.034 | 0 | 0.17 | 0.11 | 0.13 | 0.13 | 0.15 | 0.1 |
| 5 | 2202971 | rs1542430 | -0.182 | 0.385 | -1.3 | -0.307 | -0.002 | 0 | 0 | 0.257 | 0.44 | 0.45 | 0.6 | 0.49 | 0.45 | 0.24 |
| 5 | 2204833 | rs6555073 | 1.196 | 0.701 | -2.163 | -0.281 | -0.101 | -0.036 | -0.036 | 0 | 0.21 | 0.04 | 0.32 | 0.24 | 0.32 | 0.16 |
| 5 | 2207155 | rs10060730 | -0.369 | 0.455 | -1.831 | -0.245 | -0.013 | -0.012 | -0.012 | 0 | 0.6 | 0.35 | 0.59 | 0.52 | 0.6 | 0.37 |
| 5 | 2208339 | rs11133937 | 0.279 | 0.619 | -1.875 | 0.124 | -0.007 | -0.057 | -0.057 | 0 | 0.09 | 0 | 0.16 | 0.19 | 0.1 | 0.03 |
| 5 | 2214721 | rs10041522 | 0.491 | 1.299 | -1.774 | -0.196 | -0.012 | 0.003 | 0.003 | 0 | 0.09 | 0.05 | 0.1 | 0 | 0.16 | 0.06 |
| 5 | 2215243 | rs10058811 | -0.347 | 1.159 | -2.155 | -0.453 | -0.064 | 0.084 | 0.084 | 0 | 0.23 | 0.06 | 0.21 | 0.11 | 0.32 | 0.2 |
| 5 | 2216296 | rs4263504 | -0.73 | 0.462 | -1.608 | -0.578 | -0.023 | 0.012 | 0.012 | 0.167 | 0.65 | 0.47 | 0.62 | 0.45 | 0.69 | 0.45 |
| 5 | 2217896 | rs10073263 | 0.542 | 0.48 | -1.467 | -0.76 | 0.002 | -0.042 | -0.042 | 0 | 0.18 | 0.18 | 0.14 | 0 | 0.21 | 0.08 |
| 5 | 2218025 | rs11741203 | 1.043 | 0.525 | -1.681 | -0.664 | -0.061 | 0.04 | 0.04 | 0.009 | 0.28 | 0.08 | 0.27 | 0.21 | 0.33 | 0.25 |
| 5 | 2223959 | rs10512689 | -0.07 | 0.096 | -0.763 | 0.03 | 0 | -0.013 | -0.013 | 1.101 | 0.05 | 0.05 | 0.03 | 0.01 | 0.05 | 0.05 |
| 5 | 2229014 | rs1458521 | -0.29 | 0 | -0.713 | 0.28 | -0.023 | -0.003 | -0.003 | 0.495 | 0.37 | 0.53 | 0.54 | 0.58 | 0.31 | 0.56 |
| 5 | 2247093 | rs1458516 | 0.021 | -0.181 | -0.529 | 0.644 | 0.009 | 0 | 0 | 0 | 0.66 | 0.72 | 0.71 | 0.66 | 0.75 | 0.61 |
| 5 | 2251109 | rs6878043 | -0.621 | 0.448 | 0.199 | 0.726 | 0.08 | 0.04 | 0.04 | 0.354 | 0.14 | 0.41 | 0.33 | 0.15 | 0.15 | 0.22 |
| 5 | 2251313 | rs6882404 | -0.26 | 0.473 | 0.621 | 0.725 | 0.043 | 0.042 | 0.042 | 0.22 | 0.1 | 0.24 | 0.29 | 0.15 | 0.12 | 0.21 |
| 5 | 2251369 | rs2018533 | -0.581 | 0.373 | 0.192 | 0.275 | 0.02 | -0.058 | -0.058 | 0 | 0.26 | 0.46 | 0.4 | 0.15 | 0.24 | 0.32 |
| 5 | 2251386 | rs2020430 | -0.673 | 0.16 | 0.08 | 0.29 | 0.011 | 0 | 0 | 0 | 0.66 | 0.74 | 0.71 | 0.65 | 0.74 | 0.6 |
| 5 | 2252304 | rs10061160 | -0.642 | 0.178 | 0.286 | 0.184 | 0.007 | -0.045 | -0.045 | 2.543 | 0.27 | 0.53 | 0.4 | 0.15 | 0.23 | 0.31 |
| 5 | 2252398 | rs10068115 | -0.621 | 0.321 | 0.006 | 0.456 | -0.003 | 0.014 | 0.014 | 0 | 0.56 | 0.68 | 0.6 | 0.42 | 0.41 | 0.64 |
| 5 | 2254124 | rs1458514 | 0.736 | 0.702 | 0.734 | 1.418 | 0.005 | 0.134 | 0.134 | 0 | 0.85 | 0.72 | 0.84 | 0.65 | 0.86 | 0.9 |
| 5 | 2254295 | rs1458513 | 1.228 | 0.566 | 0.569 | 1.257 | -0.002 | -0.004 | -0.004 | 0 | 0.62 | 0.35 | 0.46 | 0.5 | 0.64 | 0.59 |
| 5 | 2256873 | rs10475140 | -1.967 | 0.832 | 0.748 | 1.68 | 0.042 | 0.115 | 0.115 | 0 | 0.19 | 0.49 | 0.34 | 0.34 | 0.23 | 0.19 |
| 5 | 2259217 | rs2004010 | -0.882 | 0.711 | 0.223 | 1.505 | 0.007 | 0.022 | 0.022 | 0 | 0.11 | 0.12 | 0.29 | 0.25 | 0.2 | 0.22 |
| 5 | 2265478 | rs6871126 | -1.853 | 0.213 | 0.363 | 1.51 | -0.005 | 0.075 | 0.075 | 0 | 0.24 | 0.39 | 0.36 | 0.46 | 0.22 | 0.27 |
| 5 | 2266931 | rs6877636 | -1.729 | 0.37 | 0.174 | 1.362 | -0.001 | -0.001 | -0.001 | 0.034 | 0.19 | 0.26 | 0.34 | 0.25 | 0.22 | 0.25 |
| 5 | 2268735 | rs593929 | -1.935 | 0.036 | 0.455 | 1.322 | -0.005 | 0.017 | 0.017 | 0 | 0.19 | 0.3 | 0.34 | 0.26 | 0.17 | 0.26 |
| 5 | 2269218 | rs9765121 | -1.329 | 0.017 | 0.466 | 1.153 | -0.028 | -0.056 | -0.056 | 0.278 | 0.1 | 0.26 | 0.06 | 0 | 0.03 | 0.06 |
| 5 | 2272248 | rs6875273 | -0.091 | 0.008 | -0.054 | 0.716 | -0.003 | 0.002 | 0.002 | 0.245 | 0.4 | 0.45 | 0.47 | 0.36 | 0.52 | 0.3 |
| 5 | 2275109 | rs7732508 | 0.456 | -0.052 | -0.344 | 0.683 | -0.067 | -0.016 | -0.016 | 0 | 0.33 | 0.17 | 0.4 | 0.36 | 0.5 | 0.25 |
| 5 | 2275627 | rs464904 | 0.008 | -0.199 | -0.29 | 0.974 | -0.015 | -0.018 | -0.018 | 0 | 0.08 | 0.03 | 0.27 | 0.27 | 0.09 | 0.17 |
| 5 | 2279208 | rs575414 | -0.391 | -1.106 | 0.106 | 0.272 | -0.035 | -0.062 | -0.062 | 0 | 0.6 | 0.55 | 0.81 | 0.82 | 0.78 | 0.57 |
| 5 | 2279457 | rs2609085 | -0.998 | -1.032 | 0.258 | 0.404 | 0.003 | 0.024 | 0.024 | 0 | 0.29 | 0.64 | 0.44 | 0.47 | 0.29 | 0.34 |
| 5 | 2279813 | rs1870335 | -0.773 | -1.03 | 0.36 | 0.194 | 0.013 | 0.01 | 0.01 | 0 | 0.28 | 0.6 | 0.42 | 0.26 | 0.27 | 0.32 |
| 5 | 2279866 | rs923048 | -0.816 | -1.016 | 0.28 | 0.2 | 0.008 | 0.006 | 0.006 | 0 | 0.28 | 0.61 | 0.43 | 0.26 | 0.28 | 0.32 |
| 5 | 2280105 | rs547706 | -0.212 | -0.736 | 0.171 | 0.402 | 0.004 | 0.048 | 0.048 | 0 | 0.14 | 0.27 | 0.34 | 0.26 | 0.15 | 0.23 |
| 5 | 2282348 | rs635969 | -0.062 | -0.941 | 0.136 | 0.018 | 0.05 | 0.112 | 0.112 | 0 | 0.48 | 0.37 | 0.73 | 0.82 | 0.66 | 0.48 |
| 5 | 2286042 | rs554745 | -0.084 | -1.322 | 0.126 | 0.134 | 0.032 | 0.031 | 0.031 | 0 | 0.53 | 0.62 | 0.66 | 0.66 | 0.73 | 0.52 |
| 5 | 2286126 | rs2652148 | -0.223 | -1.72 | 0.025 | -0.036 | 0.011 | 0.008 | 0.008 | 0 | 0.65 | 0.68 | 0.69 | 0.69 | 0.77 | 0.58 |
| 5 | 2286906 | rs316575 | -0.224 | -1.542 | -0.272 | -0.056 | -0.007 | -0.006 | -0.006 | 0 | 0.63 | 0.6 | 0.67 | 0.69 | 0.75 | 0.58 |
| 5 | 2287730 | rs4866449 | 1.156 | -1.302 | -0.846 | -0.063 | -0.124 | -0.083 | -0.083 | 0 | 0.32 | 0.07 | 0.38 | 0.36 | 0.48 | 0.23 |
| 5 | 2291316 | rs2652147 | -0.576 | -1.64 | -0.304 | 0.237 | 0.007 | -0.008 | -0.008 | 0 | 0.4 | 0.62 | 0.49 | 0.53 | 0.34 | 0.4 |
| 5 | 2294533 | rs316578 | -0.507 | -1.613 | -0.275 | 0.335 | 0.002 | -0.001 | -0.001 | 0 | 0.39 | 0.57 | 0.48 | 0.38 | 0.34 | 0.36 |
| 5 | 2295317 | rs13187359 | 0.523 | -1.78 | -0.081 | 0.295 | -0.015 | -0.012 | -0.012 | 0 | 0.24 | 0.25 | 0.11 | 0.17 | 0.14 | 0.14 |
| 5 | 2301089 | rs882496 | 0.36 | -1.071 | -0.318 | 0.708 | -0.034 | -0.032 | -0.032 | 3.812 | 0.42 | 0.55 | 0.2 | 0.28 | 0.25 | 0.42 |
| 5 | 2301342 | rs882497 | 0.463 | -1.005 | -0.285 | 0.234 | -0.042 | -0.107 | -0.107 | 0 | 0.41 | 0.53 | 0.2 | 0.12 | 0.24 | 0.4 |
| 5 | 2302513 | rs922025 | 0.708 | 0.071 | 0.483 | 1.315 | 0.008 | 0.034 | 0.034 | 0.059 | 0.77 | 0.66 | 0.6 | 0.66 | 0.74 | 0.72 |
| 5 | 2304811 | rs316574 | 0.61 | 0.406 | 0.105 | 1.43 | 0.007 | 0.023 | 0.023 | 0 | 0.79 | 0.73 | 0.56 | 0.7 | 0.74 | 0.7 |
| 5 | 2305235 | rs316573 | 0.109 | 0.467 | -0.034 | 1.498 | 0.004 | 0.018 | 0.018 | 0 | 0.83 | 0.8 | 0.61 | 0.74 | 0.79 | 0.75 |
| 5 | 2305490 | rs7718226 | 0.053 | -0.193 | -0.418 | 1.146 | 0.041 | 0.023 | 0.023 | 0 | 0.5 | 0.37 | 0.31 | 0.44 | 0.35 | 0.48 |
| 5 | 2315086 | rs316582 | -0.361 | -0.117 | -0.354 | 1.552 | 0.001 | -0.003 | -0.003 | 0 | 0.2 | 0.2 | 0.23 | 0.4 | 0.19 | 0.21 |
| 5 | 2317424 | rs7445525 | 0.458 | -0.002 | -1.299 | 1.233 | -0.121 | 0.023 | 0.023 | 0 | 0.31 | 0.06 | 0.3 | 0.26 | 0.43 | 0.35 |
| 5 | 2319964 | rs12189181 | 0.174 | -0.052 | -1.123 | 1.209 | -0.052 | 0.034 | 0.034 | 0 | 0.33 | 0.2 | 0.31 | 0.24 | 0.43 | 0.35 |
| 5 | 2320472 | rs1032112 | 0.473 | -0.002 | -1.164 | 1.227 | 0.001 | -0.002 | -0.002 | 0 | 0.18 | 0.14 | 0.21 | 0.22 | 0.19 | 0.18 |
| 5 | 2322360 | rs160726 | -0.146 | -0.998 | -1.161 | 1.015 | 0.067 | 0.06 | 0.06 | 1.388 | 0.5 | 0.19 | 0.52 | 0.66 | 0.63 | 0.56 |
| 5 | 2323282 | rs320597 | -0.146 | -0.987 | -1.161 | 1.015 | 0.076 | 0.06 | 0.06 | 1.388 | 0.5 | 0.19 | 0.52 | 0.66 | 0.64 | 0.56 |
| 5 | 2329018 | rs458382 | 0.364 | -1.055 | -0.829 | 0.996 | 0.025 | 0.022 | 0.022 | 0.138 | 0.46 | 0.58 | 0.47 | 0.35 | 0.36 | 0.45 |
| 5 | 2330992 | rs898060 | -0.041 | -1.016 | -1.434 | 0.794 | -0.052 | 0.017 | 0.017 | 0.372 | 0.44 | 0.08 | 0.46 | 0.36 | 0.58 | 0.47 |
| 5 | 2333103 | rs16867991 | -0.243 | -1.161 | -0.596 | 0.725 | -0.022 | 0.049 | 0.049 | 0 | 0.1 | 0.31 | 0.06 | 0.15 | 0.06 | 0.06 |
| 5 | 2333303 | rs467939 | 0.552 | -1.577 | -1.204 | 0.234 | -0.022 | 0.021 | 0.021 | 0 | 0.6 | 0.7 | 0.67 | 0.63 | 0.58 | 0.73 |
| 5 | 2333982 | rs2641217 | -0.231 | -2.045 | -0.919 | 0.219 | 0.048 | 0.008 | 0.008 | 0 | 0.38 | 0.82 | 0.41 | 0.56 | 0.27 | 0.44 |

| | | | | | | | | | | | | | | | | |
|---|---|---|---|---|---|---|---|---|---|---|---|---|---|---|---|---|
| 5 | 2335228 | rs1873952 | 0.888 | -1.927 | -0.626 | 0.113 | -0.014 | 0.075 | 0.075 | 0 | 0.39 | 0.22 | 0.33 | 0.17 | 0.42 | 0.41 |
| 5 | 2335313 | rs1873951 | 0.737 | -1.56 | -0.565 | 0.208 | 0.031 | 0.098 | 0.098 | 0 | 0.46 | 0.26 | 0.34 | 0.19 | 0.4 | 0.44 |
| 5 | 2337193 | rs13169221 | 1.072 | -1.752 | -0.392 | 0.54 | -0.005 | 0.064 | 0.064 | 0 | 0.42 | 0.25 | 0.34 | 0.22 | 0.43 | 0.44 |
| 5 | 2338908 | rs92833 | -0.037 | -1.809 | -0.003 | 0.786 | -0.029 | 0.009 | 0.009 | 0.016 | 0.3 | 0.41 | 0.34 | 0.25 | 0.21 | 0.34 |
| 5 | 2339273 | rs462498 | 0.012 | -1.794 | 0.062 | 0.754 | 0.006 | 0.004 | 0.004 | 0 | 0.71 | 0.58 | 0.66 | 0.63 | 0.63 | 0.78 |
| 5 | 2346621 | rs468976 | -0.016 | -1.698 | -0.01 | 0.815 | -0.034 | 0.058 | 0.058 | 0 | 0.31 | 0.43 | 0.34 | 0.4 | 0.22 | 0.35 |
| 5 | 2352620 | rs467723 | -0.006 | -1.693 | -0.008 | 0.818 | -0.034 | 0.058 | 0.058 | 0 | 0.31 | 0.43 | 0.34 | 0.4 | 0.22 | 0.35 |
| 5 | 2352730 | rs10036054 | 0.876 | -1.766 | -0.282 | 0.467 | -0.003 | 0.069 | 0.069 | 0 | 0.42 | 0.22 | 0.32 | 0.22 | 0.42 | 0.43 |
| 5 | 2354731 | rs461425 | -0.541 | -2.78 | 0.663 | 0.956 | -0.054 | 0.057 | 0.057 | 0 | 0.22 | 0.38 | 0.27 | 0.31 | 0.09 | 0.34 |
| 5 | 2355389 | rs462906 | -1.035 | -2.565 | 0.598 | 1.173 | -0.075 | 0.066 | 0.066 | 0.258 | 0.26 | 0.49 | 0.33 | 0.56 | 0.16 | 0.36 |
| 5 | 2355587 | rs458006 | -1.021 | -2.51 | 0.628 | 0.562 | -0.061 | 0.109 | 0.109 | 0.088 | 0.25 | 0.43 | 0.31 | 0.42 | 0.16 | 0.35 |
| 5 | 2358493 | rs7733952 | -0.375 | -1.646 | 0.808 | -0.246 | -0.018 | -0.008 | -0.008 | 0.855 | 0.13 | 0.21 | 0.07 | 0.12 | 0.09 | 0.07 |
| 5 | 2358675 | rs7709574 | -0.441 | -1.618 | 0.432 | -0.233 | -0.039 | -0.022 | -0.022 | 0 | 0.15 | 0.35 | 0.09 | 0.12 | 0.1 | 0.07 |
| 5 | 2360079 | rs10071443 | -0.441 | -1.618 | 0.432 | -0.233 | -0.039 | -0.022 | -0.022 | 0.284 | 0.15 | 0.35 | 0.09 | 0.12 | 0.1 | 0.07 |
| 5 | 2363480 | rs10474742 | -0.441 | -1.618 | 0.432 | -0.233 | -0.039 | -0.022 | -0.022 | 0 | 0.15 | 0.35 | 0.09 | 0.12 | 0.1 | 0.07 |
| 5 | 2366237 | rs6869394 | -0.756 | -0.713 | 0.016 | -0.03 | -0.025 | -0.081 | -0.081 | 0 | 0.19 | 0.33 | 0.08 | 0.02 | 0.11 | 0.12 |
| 5 | 2371873 | rs10036992 | -0.208 | -0.878 | -0.281 | 0.346 | -0.025 | -0.017 | -0.017 | 0 | 0.41 | 0.54 | 0.33 | 0.27 | 0.24 | 0.38 |
| 5 | 2374305 | rs7715102 | -0.316 | -0.468 | -0.111 | 1.03 | 0.01 | 0 | 0 | 0 | 0.43 | 0.62 | 0.36 | 0.4 | 0.27 | 0.38 |
| 5 | 2376332 | rs16899405 | -0.196 | -0.606 | 0.265 | 0.893 | -0.001 | -0.001 | -0.001 | 0 | 0.41 | 0.59 | 0.35 | 0.26 | 0.27 | 0.37 |
| 5 | 2378836 | rs465850 | 0.247 | -1.086 | 0.113 | 1.157 | -0.018 | 0.011 | 0.011 | 0 | 0.66 | 0.58 | 0.7 | 0.6 | 0.77 | 0.63 |
| 5 | 2382270 | rs457979 | 1.084 | -0.796 | -0.005 | 1.536 | -0.007 | 0.051 | 0.051 | 0 | 0.92 | 0.84 | 0.96 | 0.82 | 0.91 | 0.95 |
| 5 | 2383722 | rs462203 | 0.264 | -0.697 | -0.152 | 1.518 | -0.001 | -0.004 | -0.004 | 0 | 0.94 | 0.96 | 0.96 | 0.86 | 0.91 | 0.94 |
| 5 | 2387550 | rs931283 | 0.088 | -0.038 | 0.065 | 0.951 | 0 | 0.002 | 0.002 | 0 | 0.54 | 0.52 | 0.53 | 0.61 | 0.53 | 0.51 |
| 5 | 2389029 | rs6895737 | 0.084 | -0.547 | 0.104 | 0.782 | -0.033 | -0.056 | -0.056 | 0 | 0.14 | 0.25 | 0.06 | 0.05 | 0.08 | 0.03 |
| 5 | 2392831 | rs160730 | 0.131 | -0.157 | -0.141 | 0.995 | -0.004 | 0.012 | 0.012 | 0 | 0.52 | 0.57 | 0.52 | 0.61 | 0.5 | 0.47 |
| 5 | 2397617 | rs6871313 | 0.188 | -0.007 | -0.066 | 0.921 | -0.001 | -0.023 | -0.023 | 0 | 0.4 | 0.45 | 0.39 | 0.11 | 0.34 | 0.45 |
| 5 | 2399225 | rs468085 | 0.089 | 0.671 | -0.449 | 1.21 | 0.01 | 0.007 | 0.007 | 0 | 0.52 | 0.47 | 0.57 | 0.43 | 0.54 | 0.58 |
| 5 | 2400064 | rs518850 | 0.093 | 0.74 | -0.456 | 1.203 | 0.013 | 0.006 | 0.006 | 0 | 0.52 | 0.47 | 0.57 | 0.43 | 0.56 | 0.58 |
| 5 | 2400149 | rs624376 | 0.409 | 1.102 | -0.604 | 1.562 | -0.004 | -0.007 | -0.007 | 0.547 | 0.92 | 0.92 | 0.91 | 0.85 | 0.85 | 0.92 |
| 5 | 2400280 | rs464716 | 0.192 | 0.96 | -0.552 | 1.334 | -0.033 | -0.024 | -0.024 | 1.47 | 0.12 | 0.05 | 0.18 | 0.18 | 0.19 | 0.11 |
| 5 | 2403444 | rs10059366 | -0.552 | -0.152 | -0.468 | 1.147 | 0.001 | 0 | 0 | 0 | 0.43 | 0.42 | 0.29 | 0.44 | 0.36 | 0.43 |
| 5 | 2409988 | rs4866482 | -0.368 | -0.178 | -0.167 | 1.487 | -0.003 | -0.039 | -0.039 | 1.635 | 0.3 | 0.41 | 0.24 | 0.22 | 0.3 | 0.35 |
| 5 | 2417417 | rs17648791 | -0.063 | -0.247 | -0.206 | 1.452 | 0.002 | -0.01 | -0.01 | 0 | 0.27 | 0.29 | 0.23 | 0.22 | 0.3 | 0.38 |
| 5 | 2418025 | rs4866484 | -0.053 | -0.238 | -0.089 | 1.763 | 0.004 | 0.001 | 0.001 | 0 | 0.27 | 0.31 | 0.23 | 0.32 | 0.3 | 0.39 |
| 5 | 2418088 | rs315914 | 1.165 | 0.107 | 0.052 | 1.748 | 0.006 | 0.063 | 0.063 | 0.023 | 0.54 | 0.3 | 0.6 | 0.26 | 0.55 | 0.46 |
| 5 | 2420984 | rs10462757 | 1.449 | -0.019 | -0.308 | 1.437 | 0.026 | -0.009 | -0.009 | 2.286 | 0.05 | 0 | 0.01 | 0.22 | 0.01 | 0.05 |
| 5 | 2424387 | rs13358358 | -0.351 | 0.596 | -0.231 | 0.64 | 0.002 | -0.009 | -0.009 | 0 | 0.27 | 0.29 | 0.23 | 0.22 | 0.3 | 0.38 |
| 5 | 2447447 | rs6875672 | -0.66 | 0.592 | -0.311 | 0.81 | -0.002 | 0.001 | 0.001 | 0 | 0.24 | 0.58 | 0.21 | 0.24 | 0.2 | 0.31 |
| 5 | 2448052 | rs438055 | -0.541 | 0.845 | -1.775 | 0.681 | -0.101 | -0.031 | -0.031 | 0.027 | 0.39 | 0.06 | 0.32 | 0.31 | 0.44 | 0.48 |
| 5 | 2448146 | rs370672 | -0.641 | 0.642 | -1.486 | 0.008 | 0 | -0.009 | -0.009 | 0 | 0.35 | 0.31 | 0.37 | 0.84 | 0.31 | 0.58 |
| 5 | 2448160 | rs6555108 | -0.841 | 0.422 | -1.494 | 0.753 | 0.029 | 0.014 | 0.014 | 0 | 0.5 | 0.75 | 0.45 | 0.53 | 0.41 | 0.59 |
| 5 | 2448538 | rs16869638 | -0.63 | 0.23 | -0.496 | 1.137 | -0.012 | 0.014 | 0.014 | 0 | 0.11 | 0.2 | 0.17 | 0.15 | 0.09 | 0.14 |
| 5 | 2448590 | rs4866650 | -0.54 | 0.199 | -0.524 | 1.195 | -0.014 | 0.016 | 0.016 | 0 | 0.11 | 0.2 | 0.18 | 0.18 | 0.07 | 0.14 |
| 5 | 2453377 | rs1908159 | -1.314 | 0.157 | -0.922 | 1.3 | 0.003 | -0.051 | -0.051 | 0 | 0.45 | 0.82 | 0.47 | 0.53 | 0.41 | 0.61 |
| 5 | 2458254 | rs315893 | -1.162 | 0.277 | -0.766 | 1.296 | 0.037 | -0.061 | -0.061 | 0 | 0.43 | 0.82 | 0.45 | 0.53 | 0.35 | 0.61 |
| 5 | 2462271 | rs11746384 | -0.597 | 0.344 | 0.314 | 1.424 | 0.072 | 0.032 | 0.032 | 0.006 | 0.16 | 0.4 | 0.23 | 0.2 | 0.23 | 0.26 |
| 5 | 2463858 | rs6877519 | -1.036 | 0.057 | 0.559 | 1.761 | 0.003 | 0.028 | 0.028 | 0 | 0.4 | 0.65 | 0.4 | 0.33 | 0.36 | 0.55 |
| 5 | 2465418 | rs412137 | -0.853 | 0.16 | 1.142 | 1.759 | 0.021 | -0.015 | -0.015 | 0 | 0.33 | 0.45 | 0.39 | 0.31 | 0.38 | 0.45 |
| 5 | 2466584 | rs315918 | -1.723 | -0.503 | 1.393 | 2.102 | -0.05 | 0.134 | 0.134 | 0 | 0.17 | 0.6 | 0.18 | 0.41 | 0.1 | 0.17 |
| 5 | 2468375 | rs924134 | -1.681 | -0.359 | 0.538 | 1.77 | 0.018 | -0.044 | -0.044 | 0.285 | 0.45 | 0.87 | 0.43 | 0.55 | 0.38 | 0.57 |
| 5 | 2470082 | rs9312995 | -1.052 | -0.389 | 0.879 | 1.841 | -0.025 | 0.162 | 0.162 | 0 | 0.11 | 0.32 | 0.09 | 0.33 | 0.07 | 0.15 |
| 5 | 2471705 | rs1496447 | 0.153 | -1.077 | -0.471 | 1.044 | 0.059 | 0.045 | 0.045 | 0 | 0.24 | 0.06 | 0.12 | 0.11 | 0.12 | 0.11 |
| 5 | 2474274 | rs3104228 | 1.051 | -0.43 | 0.187 | 1.717 | -0.012 | -0.017 | -0.017 | 0 | 0.05 | 0 | 0.04 | 0.21 | 0.06 | 0.03 |
| 5 | 2479446 | rs10075636 | -0.258 | -0.389 | 0.554 | 1.408 | 0.025 | -0.026 | -0.026 | 0 | 0.24 | 0.38 | 0.24 | 0.18 | 0.27 | 0.32 |
| 5 | 2481088 | rs3112426 | -1.321 | -0.339 | -0.222 | 1.287 | -0.005 | -0.072 | -0.072 | 0 | 0.42 | 0.86 | 0.37 | 0.52 | 0.38 | 0.52 |
| 5 | 2493280 | rs2453814 | -1.288 | -0.288 | -0.03 | 0.866 | -0.069 | -0.011 | -0.011 | 0 | 0.42 | 0.91 | 0.34 | 0.44 | 0.42 | 0.43 |
| 5 | 2494273 | rs895658 | -0.911 | -0.318 | 0.196 | 0.762 | -0.004 | -0.009 | -0.009 | 0 | 0.36 | 0.65 | 0.3 | 0.11 | 0.35 | 0.35 |
| 5 | 2495251 | rs10475158 | -0.516 | -0.365 | 0.488 | 0.697 | 0.03 | -0.132 | -0.132 | 0.078 | 0.3 | 0.51 | 0.27 | 0.08 | 0.34 | 0.31 |
| 5 | 2498479 | rs1493650 | -0.56 | -0.364 | 0.732 | 0.72 | 0.037 | -0.071 | -0.071 | 0 | 0.3 | 0.47 | 0.27 | 0.24 | 0.35 | 0.34 |
| 5 | 2505743 | rs3112415 | -0.996 | -0.375 | 0.549 | 0.72 | -0.03 | 0.079 | 0.079 | 0 | 0.34 | 0.81 | 0.3 | 0.24 | 0.35 | 0.34 |
| 5 | 2509474 | rs16869844 | -1.506 | -0.08 | 1.077 | 0.888 | -0.009 | 0.117 | 0.117 | 0.052 | 0.12 | 0.36 | 0.06 | 0.21 | 0.09 | 0.1 |
| 5 | 2513346 | rs10039326 | -1.316 | -0.337 | 0.771 | 0.584 | 0.004 | -0.004 | -0.004 | 2.898 | 0.25 | 0.32 | 0.19 | 0.22 | 0.26 | 0.23 |
| 5 | 2514880 | rs10462760 | -0.951 | -0.682 | -0.031 | 0.672 | 0 | -0.044 | -0.044 | 0 | 0.34 | 0.16 | 0.28 | 0.4 | 0.35 | 0.45 |
| 5 | 2514931 | rs3112411 | 0.38 | -0.776 | -0.186 | 0.521 | 0.012 | 0.003 | 0.003 | 0 | 0.15 | 0.2 | 0.14 | 0.22 | 0.19 | 0.15 |
| 5 | 2516230 | rs3112409 | 0.488 | -0.736 | -0.268 | 0.468 | 0.002 | -0.003 | -0.003 | 0 | 0.16 | 0.19 | 0.13 | 0.21 | 0.19 | 0.16 |
| 5 | 2516620 | rs17651154 | 0.203 | -0.617 | -0.573 | 0.471 | -0.12 | -0.048 | -0.048 | 0 | 0.22 | 0.02 | 0.27 | 0.2 | 0.34 | 0.35 |
| 5 | 2516934 | rs3112408 | -1.039 | -0.899 | -0.584 | 0.486 | 0.051 | 0.025 | 0.025 | 0.859 | 0.31 | 0.93 | 0.21 | 0.25 | 0.21 | 0.2 |
| 5 | 2518738 | rs378257 | -1.264 | -0.848 | -0.608 | 0.434 | 0.057 | -0.077 | -0.077 | 0.32 | 0.33 | 0.93 | 0.21 | 0.42 | 0.21 | 0.25 |
| 5 | 2518949 | rs7720355 | -0.011 | -0.873 | -0.194 | 0.416 | -0.012 | 0.184 | 0.184 | 2.371 | 0.08 | 0.55 | 0.05 | 0.17 | 0.01 | 0.05 |

| | | | | | | | | | | | | | | | | |
|---|---|---|---|---|---|---|---|---|---|---|---|---|---|---|---|---|
| 5 | 2519149 | rs385263 | -1.461 | -0.829 | -0.241 | 0.383 | 0.001 | 0.009 | 0.009 | 0 | 0.47 | 0.4 | 0.42 | 0.45 | 0.54 | 0.55 |
| 5 | 2522181 | rs13171172 | 1.475 | -0.839 | -0.556 | 0.394 | 0.019 | -0.051 | -0.051 | 0 | 0.12 | 0.03 | 0.12 | 0.19 | 0.1 | 0.21 |
| 5 | 2522212 | rs16869867 | 0.841 | -0.983 | -0.73 | 0.265 | -0.02 | -0.006 | -0.006 | 0.153 | 0.09 | 0.1 | 0.01 | 0.03 | 0.01 | 0.04 |
| 5 | 2522431 | rs374683 | -0.799 | -0.773 | -0.723 | 0.292 | -0.001 | 0 | 0 | 0 | 0.39 | 0.38 | 0.27 | 0.43 | 0.3 | 0.41 |
| 5 | 2523602 | rs419628 | -1.781 | -0.334 | -1.065 | 0.011 | -0.087 | -0.062 | -0.062 | 0 | 0.57 | 0.94 | 0.49 | 0.63 | 0.57 | 0.61 |
| 5 | 2530794 | rs17598228 | 0.643 | 0.349 | -0.55 | -0.07 | 0.034 | -0.027 | -0.027 | 0.311 | 0.11 | 0.15 | 0.16 | 0.04 | 0.21 | 0.14 |
| 5 | 2533617 | rs6875477 | -0.738 | 0.272 | -0.715 | -0.328 | 0.015 | -0.029 | -0.029 | 0 | 0.32 | 0.35 | 0.37 | 0.18 | 0.5 | 0.31 |
| 5 | 2537894 | rs6888076 | 0.522 | 0.385 | -0.645 | -0.271 | 0.045 | -0.036 | -0.036 | 0 | 0.11 | 0.21 | 0.16 | 0.03 | 0.21 | 0.14 |
| 5 | 2538080 | rs12517190 | -0.132 | 0.536 | -1.187 | -0.252 | -0.015 | 0.027 | 0.027 | 0 | 0.23 | 0.15 | 0.21 | 0.14 | 0.29 | 0.17 |
| 5 | 2539161 | rs16869906 | 0.797 | 0.738 | -1.581 | -0.369 | 0.006 | 0.043 | 0.043 | 0.006 | 0.29 | 0.21 | 0.27 | 0.09 | 0.24 | 0.12 |
| 5 | 2539286 | rs6879053 | -0.558 | 0.328 | -1.409 | 0.289 | -0.022 | -0.025 | -0.025 | 0 | 0.55 | 0.77 | 0.49 | 0.55 | 0.53 | 0.61 |
| 5 | 2543192 | rs323689 | -0.191 | -0.683 | -1.2 | 0.229 | 0.051 | -0.004 | -0.004 | 0.351 | 0.46 | 0.7 | 0.43 | 0.59 | 0.39 | 0.56 |
| 5 | 2544880 | rs13436812 | 0.749 | -0.79 | -0.876 | 0.213 | -0.031 | -0.007 | -0.007 | 0 | 0.07 | 0.1 | 0.01 | 0.03 | 0.01 | 0.04 |
| 5 | 2553159 | rs6888228 | -0.019 | -0.831 | -0.601 | 0.22 | -0.019 | 0.015 | 0.015 | 0 | 0.21 | 0.37 | 0.13 | 0.21 | 0.14 | 0.22 |
| 5 | 2557781 | rs323691 | 0.657 | -0.825 | -0.695 | 0.477 | -0.004 | 0.019 | 0.019 | 0.074 | 0.06 | 0.08 | 0.07 | 0.22 | 0.06 | 0.08 |
| 5 | 2563748 | rs16869997 | 1.11 | -1.405 | -1.397 | 0.236 | 0.014 | -0.061 | -0.061 | 0 | 0.18 | 0 | 0.18 | 0.2 | 0.15 | 0.27 |
| 5 | 2564243 | rs323700 | -0.047 | -1.364 | -1.003 | 0.077 | -0.007 | 0 | 0 | 0.093 | 0.18 | 0.5 | 0.17 | 0.22 | 0.19 | 0.11 |
| 5 | 2565830 | rs445296 | -1.028 | -1.802 | -1.836 | -0.169 | -0.079 | -0.121 | -0.121 | 0 | 0.64 | 0.96 | 0.45 | 0.59 | 0.46 | 0.6 |
| 5 | 2569156 | rs437485 | -0.771 | -1.82 | -1.721 | -0.236 | -0.031 | -0.042 | -0.042 | 0.017 | 0.63 | 0.89 | 0.44 | 0.45 | 0.47 | 0.52 |
| 5 | 2571422 | rs323707 | 0.043 | -2.256 | -1.692 | -0.417 | 0.037 | 0.015 | 0.015 | 0 | 0.41 | 0.7 | 0.36 | 0.4 | 0.33 | 0.37 |
| 5 | 2572654 | rs1159055 | 0.795 | -2.193 | -1.639 | -0.403 | 0.001 | -0.008 | -0.008 | 0 | 0.22 | 0.19 | 0.08 | 0.05 | 0.14 | 0.18 |
| 5 | 2574247 | rs11951460 | 0.338 | -2.3 | -1.687 | -0.389 | 0.003 | 0.003 | 0.003 | 0 | 0.25 | 0.21 | 0.1 | 0.19 | 0.15 | 0.24 |
| 5 | 2575174 | rs404016 | 0.111 | -2.313 | -1.446 | -0.336 | -0.015 | 0.002 | 0.002 | 0 | 0.31 | 0.34 | 0.16 | 0.39 | 0.2 | 0.33 |
| 5 | 2575392 | rs380259 | -1.157 | -2.305 | -1.568 | -0.36 | -0.07 | -0.114 | -0.114 | 0 | 0.66 | 0.93 | 0.46 | 0.58 | 0.48 | 0.62 |
| 5 | 2576250 | rs9647565 | 1.662 | -1.521 | -2.001 | -0.689 | -0.14 | -0.139 | -0.139 | 0 | 0.18 | 0.02 | 0.38 | 0.37 | 0.36 | 0.27 |
| 5 | 2577035 | rs10866539 | 1.141 | -1.526 | -2.023 | -0.699 | -0.168 | -0.156 | -0.156 | 0.786 | 0.25 | 0.02 | 0.4 | 0.37 | 0.39 | 0.29 |
| 5 | 2578112 | rs16870061 | 0.74 | -1.751 | -1.939 | -0.742 | -0.017 | -0.022 | -0.022 | 0 | 0.35 | 0.58 | 0.29 | 0.19 | 0.28 | 0.29 |
| 5 | 2578373 | rs403500 | 0.797 | -1.264 | -1.708 | -0.418 | -0.045 | 0.049 | 0.049 | 0.022 | 0.15 | 0.28 | 0.09 | 0.34 | 0.07 | 0.19 |
| 5 | 2579350 | rs6555116 | 0.555 | -1.23 | -1.732 | -0.474 | 0.042 | 0.086 | 0.086 | 0 | 0.52 | 0.68 | 0.38 | 0.24 | 0.43 | 0.45 |
| 5 | 2582828 | rs376205 | 1.243 | -1.223 | -1.685 | -0.501 | 0.004 | -0.003 | -0.003 | 0 | 0.13 | 0.12 | 0.07 | 0.2 | 0.06 | 0.1 |
| 5 | 2586224 | rs7705916 | 0.323 | -1.017 | -1.485 | -0.502 | 0.042 | 0.039 | 0.039 | 0 | 0.52 | 0.66 | 0.38 | 0.37 | 0.42 | 0.53 |
| 5 | 2588264 | rs225752 | -1.263 | -0.372 | -1.063 | -0.551 | -0.034 | 0.022 | 0.022 | 0 | 0.9 | 0.98 | 0.86 | 0.94 | 0.87 | 0.91 |
| 5 | 2588400 | rs7726088 | 0.575 | -0.985 | -0.259 | -0.524 | 0.019 | -0.069 | -0.069 | 0 | 0.11 | 0.3 | 0.09 | 0 | 0.13 | 0.02 |
| 5 | 2599639 | rs569244 | 0.901 | -1.221 | -0.433 | -0.385 | -0.02 | -0.002 | -0.002 | 0.187 | 0.18 | 0.23 | 0.09 | 0.22 | 0.1 | 0.07 |
| 5 | 2601689 | rs390560 | -0.404 | -1.302 | -0.383 | -0.522 | -0.037 | -0.038 | -0.038 | 0 | 0.3 | 0.59 | 0.19 | 0.22 | 0.21 | 0.09 |
| 5 | 2604686 | rs323685 | -1.46 | -0.516 | -1.077 | -0.322 | -0.082 | -0.108 | -0.108 | 0 | 0.79 | 0.95 | 0.63 | 0.65 | 0.74 | 0.73 |
| 5 | 2604903 | rs10512678 | 0.897 | -1.683 | -0.822 | -0.215 | 0.031 | 0.015 | 0.015 | 0 | 0.33 | 0.21 | 0.23 | 0.26 | 0.24 | 0.41 |
| 5 | 2605834 | rs6877761 | 0.623 | -0.595 | -0.837 | -0.138 | -0.003 | -0.064 | -0.064 | 0 | 0.11 | 0.28 | 0.09 | 0 | 0.11 | 0.02 |
| 5 | 2605988 | rs2962561 | -0.12 | -0.03 | -1.576 | 0.304 | -0.044 | -0.057 | -0.057 | 0 | 0.67 | 0.86 | 0.45 | 0.58 | 0.48 | 0.61 |
| 5 | 2610337 | rs395063 | -0.129 | -0.993 | -1.214 | 0.396 | 0.017 | -0.005 | -0.005 | 0.037 | 0.42 | 0.34 | 0.28 | 0.58 | 0.25 | 0.44 |
| 5 | 2610744 | rs434930 | 0.808 | -1.611 | -1.426 | 0.462 | 0.034 | 0.007 | 0.007 | 2.727 | 0.19 | 0.11 | 0.09 | 0.22 | 0.09 | 0.04 |
| 5 | 2611996 | rs10512677 | 0.481 | -1.643 | -1.38 | 0.708 | -0.019 | 0.002 | 0.002 | 0 | 0.21 | 0.23 | 0.16 | 0.24 | 0.14 | 0.32 |
| 5 | 2612747 | rs2004187 | -0.933 | -0.551 | -1.382 | 0.398 | 0.007 | 0.003 | 0.003 | 0 | 0.76 | 0.76 | 0.57 | 0.66 | 0.59 | 0.73 |
| 5 | 2614896 | rs2937764 | 1.099 | -0.674 | -1.298 | 0.325 | 0.043 | 0.004 | 0.004 | 4.528 | 0.21 | 0.11 | 0.09 | 0.24 | 0.09 | 0.04 |
| 5 | 2617235 | rs11742219 | 0.713 | -0.497 | -1.23 | 0.242 | 0.004 | -0.069 | -0.069 | 0.108 | 0.34 | 0.43 | 0.28 | 0.09 | 0.33 | 0.29 |
| 5 | 2617846 | rs17659675 | 1.291 | -0.646 | -1.53 | -0.008 | -0.059 | 0.08 | 0.08 | 0.104 | 0.18 | 0.03 | 0.2 | 0.06 | 0.24 | 0.16 |
| 5 | 2618844 | rs2922062 | 0.172 | -0.522 | -1.108 | 0.509 | -0.028 | -0.001 | -0.001 | 0 | 0.41 | 0.21 | 0.59 | 0.55 | 0.55 | 0.59 |
| 5 | 2620072 | rs13436220 | 0.92 | -0.421 | -1.109 | 0.509 | 0.007 | -0.087 | -0.087 | 5.068 | 0.35 | 0.47 | 0.29 | 0.09 | 0.33 | 0.29 |
| 5 | 2621228 | rs323651 | -1.305 | -0.413 | -1.648 | 0.511 | 0.05 | -0.01 | -0.01 | 0.594 | 0.82 | 0.93 | 0.92 | 0.78 | 0.91 | 0.95 |
| 5 | 2622585 | rs1875539 | 0.536 | -0.257 | -1.388 | 0.108 | -0.006 | 0.048 | 0.048 | 0 | 0.38 | 0.74 | 0.33 | 0.21 | 0.36 | 0.39 |
| 5 | 2622918 | rs172834 | -0.14 | 0.28 | -1.256 | 0.773 | -0.01 | -0.073 | -0.073 | 0 | 0.72 | 0.89 | 0.6 | 0.52 | 0.7 | 0.79 |
| 5 | 2627075 | rs2441949 | -0.021 | 0.504 | -1.344 | 1.079 | -0.007 | -0.07 | -0.07 | 1.524 | 0.14 | 0.02 | 0.13 | 0.26 | 0.12 | 0.3 |
| 5 | 2627545 | rs1868038 | 0.288 | 0.595 | -1.248 | 0.661 | 0.004 | 0.017 | 0.017 | 0 | 0.37 | 0.63 | 0.35 | 0.22 | 0.34 | 0.37 |
| 5 | 2627728 | rs406353 | -0.067 | 0.569 | -1.072 | 0.768 | 0.002 | -0.001 | -0.001 | 0 | 0.37 | 0.47 | 0.25 | 0.32 | 0.39 | 0.44 |
| 5 | 2627745 | rs1868037 | 0.423 | 0.517 | -0.736 | 0.449 | -0.002 | -0.013 | -0.013 | 1.799 | 0.31 | 0.33 | 0.23 | 0.19 | 0.26 | 0.27 |
| 5 | 2627822 | rs437272 | -0.553 | 0.556 | -0.487 | 0.707 | -0.009 | -0.032 | -0.032 | 0 | 0.16 | 0.38 | 0.09 | 0.09 | 0.14 | 0.26 |
| 5 | 2628328 | rs7703939 | -0.303 | 1.107 | -1.384 | 1.268 | -0.02 | -0.065 | -0.065 | 0 | 0.31 | 0.2 | 0.48 | 0.45 | 0.33 | 0.26 |
| 5 | 2630823 | rs323666 | 0.627 | 0.901 | -0.515 | 1.19 | 0 | 0.005 | 0.005 | 2.44 | 0.67 | 0.37 | 0.67 | 0.55 | 0.71 | 0.69 |
| 5 | 2632712 | rs323668 | 0.342 | 0.626 | 0.037 | 0.882 | 0.011 | -0.021 | -0.021 | 0 | 0.8 | 0.54 | 0.75 | 0.84 | 0.86 | 0.78 |
| 5 | 2635891 | rs6555117 | 0.259 | 0.599 | 0.192 | 0.979 | 0.017 | 0.02 | 0.02 | 0 | 0.66 | 0.55 | 0.56 | 0.56 | 0.66 | 0.74 |
| 5 | 2637827 | rs323672 | 0.547 | 0.344 | 0.121 | 1.033 | -0.029 | -0.003 | -0.003 | 0.009 | 0.16 | 0.08 | 0.22 | 0.16 | 0.2 | 0.12 |
| 5 | 2640007 | rs2405037 | 0.907 | -0.428 | 0.947 | 1 | -0.013 | 0.034 | 0.034 | 0 | 0.68 | 0.46 | 0.59 | 0.48 | 0.72 | 0.64 |
| 5 | 2640289 | rs13155678 | -0.228 | -0.365 | 0.752 | 0.634 | -0.011 | -0.011 | -0.011 | 0 | 0.16 | 0.16 | 0.24 | 0 | 0.21 | 0.08 |
| 5 | 2640466 | rs323653 | -0.148 | -0.435 | 1.109 | 0.73 | -0.043 | 0.186 | 0.186 | 0 | 0.9 | 0.68 | 0.86 | 0.72 | 0.98 | 0.87 |
| 5 | 2640771 | rs323654 | 0.888 | -0.47 | 1.099 | 0.822 | -0.081 | 0.184 | 0.184 | 0 | 0.84 | 0.48 | 0.77 | 0.57 | 0.92 | 0.71 |
| 5 | 2641522 | rs905230 | 0.626 | -0.455 | 1.135 | 0.806 | -0.01 | 0.03 | 0.03 | 0 | 0.69 | 0.59 | 0.58 | 0.41 | 0.72 | 0.59 |
| 5 | 2643974 | rs17660394 | 2.014 | -1.068 | 0.548 | 0.632 | -0.028 | -0.011 | -0.011 | 0.038 | 0.35 | 0.07 | 0.2 | 0.27 | 0.39 | 0.36 |
| 5 | 2645159 | rs323658 | 1.356 | -0.948 | 0.546 | 0.644 | -0.038 | -0.011 | -0.011 | 0 | 0.16 | 0.04 | 0.21 | 0.17 | 0.21 | 0.12 |
| 5 | 2646701 | rs921127 | -0.555 | -0.929 | 0.608 | 0.482 | -0.008 | 0.001 | 0.001 | 0.619 | 0.21 | 0.21 | 0.29 | 0.07 | 0.27 | 0.18 |

| | | | | | | | | | | | | | | | | |
|---|---|---|---|---|---|---|---|---|---|---|---|---|---|---|---|---|
| 5 | 2647306 | rs12186830 | 1.16 | -0.992 | 0.627 | 0.605 | 0.032 | 0.029 | 0.029 | 0 | 0.5 | 0.28 | 0.32 | 0.4 | 0.49 | 0.5 |
| 5 | 2650624 | rs10475171 | 0.602 | -1.587 | 0.685 | 0.81 | -0.009 | -0.032 | -0.032 | 0 | 0.07 | 0.12 | 0.15 | 0 | 0.05 | 0.04 |
| 5 | 2652947 | rs6859765 | 0.095 | -1.813 | 0.899 | 0.685 | -0.018 | -0.018 | -0.018 | 0 | 0.19 | 0.27 | 0.16 | 0.1 | 0.17 | 0.16 |
| 5 | 2653554 | rs7705712 | -0.257 | -1.442 | 0.231 | 0.857 | -0.004 | -0.006 | -0.006 | 0 | 0.1 | 0.13 | 0.18 | 0.07 | 0.1 | 0.14 |
| 5 | 2654417 | rs10475173 | 0.482 | -1.492 | 0.245 | 1.008 | -0.003 | -0.015 | -0.015 | 0.41 | 0.71 | 0.76 | 0.57 | 0.49 | 0.68 | 0.67 |
| 5 | 2655116 | rs7713186 | -0.692 | -1.41 | 0.371 | 1.261 | -0.008 | -0.011 | -0.011 | 0.403 | 0.37 | 0.57 | 0.37 | 0.17 | 0.28 | 0.35 |
| 5 | 2655990 | rs12518480 | -0.799 | -1.422 | 0.364 | 1.337 | 0.001 | 0.001 | 0.001 | 0.002 | 0.39 | 0.58 | 0.48 | 0.31 | 0.29 | 0.4 |
| 5 | 2657449 | rs7737578 | 0.683 | -1.31 | -0.243 | 1.207 | 0.087 | -0.031 | -0.031 | 0 | 0.6 | 0.25 | 0.44 | 0.45 | 0.75 | 0.61 |
| 5 | 2663723 | rs2897046 | 0.556 | -0.264 | 0.113 | 1.143 | 0.022 | 0.048 | 0.048 | 0 | 0.77 | 0.63 | 0.61 | 0.58 | 0.73 | 0.79 |
| 5 | 2663888 | rs7732655 | -0.865 | -1.195 | -0.129 | 0.675 | -0.004 | -0.006 | -0.006 | 0.579 | 0.42 | 0.56 | 0.42 | 0.24 | 0.29 | 0.32 |
| 5 | 2664086 | rs2962558 | -0.878 | -1.44 | -0.421 | 0.717 | 0.045 | 0.03 | 0.03 | 0 | 0.5 | 0.69 | 0.49 | 0.4 | 0.42 | 0.49 |
| 5 | 2664214 | rs7356656 | -0.653 | -0.634 | 0.12 | 0.9 | 0.003 | -0.071 | -0.071 | 0 | 0.25 | 0.28 | 0.16 | 0.01 | 0.25 | 0.13 |
| 5 | 2664245 | rs2937761 | -1.2 | -0.266 | -0.226 | 1.512 | -0.008 | 0.054 | 0.054 | 0.603 | 0.11 | 0.2 | 0.25 | 0.31 | 0.08 | 0.12 |
| 5 | 2667632 | rs6555120 | 1.012 | 0.244 | -0.543 | 0.726 | 0.083 | 0.083 | 0.083 | 0 | 0.51 | 0.26 | 0.3 | 0.26 | 0.37 | 0.56 |
| 5 | 2667858 | rs950864 | -0.053 | 1.285 | -0.404 | 1.14 | -0.045 | 0.036 | 0.036 | 0 | 0.12 | 0.06 | 0.15 | 0 | 0.24 | 0.04 |
| 5 | 2668308 | rs7711788 | 0.663 | 0.048 | 0.315 | 1.929 | 0.016 | 0.455 | 0.455 | 0 | 0.94 | 0.75 | 0.76 | 0.42 | 0.96 | 0.85 |
| 5 | 2670222 | rs2937762 | 0.795 | 0.14 | 0.122 | 2.248 | -0.004 | 0.088 | 0.088 | 0 | 0.86 | 0.8 | 0.82 | 0.64 | 0.86 | 0.87 |
| 5 | 2671057 | rs10066976 | -0.485 | 0.413 | 0.311 | 1.773 | 0.003 | -0.003 | -0.003 | 0 | 0.2 | 0.26 | 0.29 | 0.23 | 0.28 | 0.18 |
| 5 | 2674526 | rs10054312 | -0.475 | -1.413 | -0.048 | 1.113 | 0.041 | 0.032 | 0.032 | 0 | 0.49 | 0.56 | 0.36 | 0.33 | 0.36 | 0.37 |
| 5 | 2676993 | rs7721778 | -0.269 | -0.941 | -0.428 | 0.86 | 0.007 | -0.039 | -0.039 | 0 | 0.67 | 0.78 | 0.58 | 0.49 | 0.69 | 0.5 |
| 5 | 2686712 | rs924218 | 1.262 | -0.348 | -0.349 | 0.697 | 0.037 | -0.007 | -0.007 | 0 | 0.19 | 0.27 | 0.21 | 0.16 | 0.29 | 0.13 |
| 5 | 2690668 | rs10071208 | 0.864 | -0.561 | -0.415 | 0.698 | -0.001 | -0.001 | -0.001 | 0.4 | 0.12 | 0.09 | 0.14 | 0.14 | 0.13 | 0.31 |
| 5 | 2692076 | rs2962610 | -0.807 | -0.565 | -0.365 | 1.03 | -0.01 | -0.01 | -0.01 | 0.443 | 0.81 | 0.84 | 0.64 | 0.56 | 0.7 | 0.8 |
| 5 | 2692841 | rs11748873 | 0.61 | -0.796 | -0.293 | 1.087 | -0.004 | 0.023 | 0.023 | 0 | 0.13 | 0.2 | 0.15 | 0.21 | 0.13 | 0.35 |
| 5 | 2694637 | rs10079562 | -0.107 | -1.85 | -0.12 | 0.614 | 0.006 | 0.006 | 0.006 | 0 | 0.51 | 0.59 | 0.55 | 0.39 | 0.59 | 0.58 |
| 5 | 2694684 | rs10079610 | 0.815 | -1.884 | -0.138 | 0.593 | -0.009 | 0.008 | 0.008 | 0 | 0.37 | 0.38 | 0.4 | 0.18 | 0.45 | 0.22 |
| 5 | 2696580 | rs4866662 | 1.141 | -1.854 | -0.116 | 0.638 | -0.035 | -0.015 | -0.015 | 0 | 0.31 | 0.28 | 0.09 | 0.17 | 0.12 | 0.23 |
| 5 | 2696644 | rs16870313 | 0.716 | -1.458 | -0.099 | 0.841 | -0.003 | 0.023 | 0.023 | 0 | 0.13 | 0.2 | 0.15 | 0.21 | 0.14 | 0.35 |
| 5 | 2700386 | rs10066528 | 1.351 | -1.418 | -0.248 | 0.789 | -0.001 | 0 | 0 | 0 | 0.11 | 0.09 | 0.14 | 0.17 | 0.13 | 0.27 |
| 5 | 2701674 | rs11133959 | 0.073 | -2.353 | -0.725 | 0.281 | 0 | -0.003 | -0.003 | 0.283 | 0.46 | 0.38 | 0.38 | 0.51 | 0.48 | 0.6 |
| 5 | 2703282 | rs2860332 | -0.424 | -1.077 | -0.584 | 0.784 | -0.05 | -0.064 | -0.064 | 0 | 0.79 | 0.91 | 0.61 | 0.41 | 0.67 | 0.73 |
| 5 | 2706253 | rs13182525 | 1.226 | -1.199 | -0.363 | 0.73 | -0.052 | 0.072 | 0.072 | 0 | 0.37 | 0.12 | 0.24 | 0.25 | 0.36 | 0.29 |
| 5 | 2706414 | rs10474751 | -0.13 | -0.127 | 0.854 | 1.65 | -0.044 | 0.033 | 0.033 | 0.023 | 0.1 | 0.43 | 0.04 | 0.06 | 0.02 | 0.07 |
| 5 | 2709971 | rs2962587 | 0.896 | 0.024 | 0.326 | 1.498 | 0.004 | 0.003 | 0.003 | 0 | 0.69 | 0.44 | 0.56 | 0.35 | 0.65 | 0.63 |
| 5 | 2716573 | rs2962580 | 0.439 | 1.457 | -0.455 | 1.657 | -0.067 | -0.083 | -0.083 | 0 | 0.81 | 0.89 | 0.58 | 0.52 | 0.66 | 0.78 |
| 5 | 2724362 | rs2173226 | -0.298 | 1.344 | -0.203 | 0.726 | 0 | -0.004 | -0.004 | 0.179 | 0.37 | 0.6 | 0.37 | 0.32 | 0.36 | 0.4 |
| 5 | 2725382 | rs1566163 | 0.473 | 1.732 | 0.141 | 0.763 | 0.002 | 0.012 | 0.012 | 0 | 0.76 | 0.71 | 0.57 | 0.46 | 0.7 | 0.76 |
| 5 | 2726735 | rs6866626 | 0.896 | 1.634 | 0.348 | 0.36 | -0.001 | 0.034 | 0.034 | 0 | 0.54 | 0.32 | 0.4 | 0.33 | 0.49 | 0.66 |
| 5 | 2737502 | rs4866665 | 1.02 | 1.312 | 1.019 | 1.165 | 0.048 | 0.092 | 0.092 | 0 | 0.76 | 0.52 | 0.53 | 0.39 | 0.67 | 0.77 |
| 5 | 2737700 | rs924213 | 0.839 | 1.467 | 0.596 | 1.238 | 0.029 | 0.055 | 0.055 | 0 | 0.78 | 0.66 | 0.55 | 0.39 | 0.67 | 0.77 |
| 5 | 2740131 | rs2962564 | 0.83 | 1.222 | 0.275 | 1.218 | 0.047 | 0.071 | 0.071 | 0 | 0.77 | 0.66 | 0.56 | 0.4 | 0.67 | 0.77 |
| 5 | 2746229 | rs1540709 | 0.393 | 0.213 | -0.138 | 1.295 | -0.007 | 0.237 | 0.237 | 0.214 | 0.81 | 0.54 | 0.69 | 0.43 | 0.84 | 0.82 |
| 5 | 2747127 | rs11432 | 0.8 | -0.136 | -1.104 | 0.819 | 0.025 | 0.011 | 0.011 | 0 | 0.61 | 0.39 | 0.5 | 0.31 | 0.7 | 0.63 |
| 5 | 2757920 | rs1995106 | -0.375 | -0.012 | -1.524 | 0.668 | 0.024 | 0 | 0 | 0 | 0.2 | 0.02 | 0.2 | 0.12 | 0.14 | 0.09 |
| 5 | 2757940 | rs1353630 | -0.877 | 1.016 | -1.237 | 0.606 | -0.005 | 0.005 | 0.005 | 0.744 | 0.12 | 0.1 | 0.19 | 0.05 | 0.2 | 0.22 |
| 5 | 2762286 | rs6555128 | 0.385 | 2.278 | -1.291 | 1.255 | -0.025 | -0.026 | -0.026 | 0 | 0.81 | 0.88 | 0.65 | 0.43 | 0.65 | 0.75 |
| 5 | 2767183 | rs6867840 | 0.145 | 0.626 | -0.891 | 1.053 | 0.012 | 0.001 | 0.001 | 0 | 0.44 | 0.37 | 0.36 | 0.54 | 0.28 | 0.46 |
| 5 | 2767357 | rs6883371 | -0.643 | 1.015 | -0.577 | 0.812 | -0.028 | 0.018 | 0.018 | 0 | 0.64 | 0.76 | 0.63 | 0.67 | 0.62 | 0.68 |
| 5 | 2767527 | rs2935622 | -0.681 | 0.991 | -0.453 | 0.813 | -0.088 | 0.018 | 0.018 | 0 | 0.65 | 0.89 | 0.64 | 0.67 | 0.62 | 0.68 |
| 5 | 2767556 | rs2935623 | -0.27 | 1 | -0.445 | 0.814 | 0.057 | -0.031 | -0.031 | 0 | 0.21 | 0.39 | 0.27 | 0.13 | 0.35 | 0.23 |
| 5 | 2768879 | rs10071246 | 0.44 | 0.979 | -0.077 | 0.723 | 0.03 | 0.008 | 0.008 | 0.059 | 0.47 | 0.59 | 0.39 | 0.54 | 0.3 | 0.45 |
| 5 | 2772332 | rs7712012 | 0.431 | 0.977 | -0.083 | 0.716 | 0.043 | 0.011 | 0.011 | 0.053 | 0.47 | 0.61 | 0.39 | 0.54 | 0.29 | 0.45 |
| 5 | 2774101 | rs13355853 | 0.563 | 0.963 | 0.033 | 0.708 | -0.004 | 0 | 0 | 1.794 | 0.46 | 0.45 | 0.38 | 0.54 | 0.28 | 0.45 |
| 5 | 2775566 | rs12522364 | 0.261 | 0.92 | 0.029 | 0.688 | 0.097 | 0.038 | 0.038 | 0.012 | 0.49 | 0.69 | 0.41 | 0.54 | 0.29 | 0.55 |
| 5 | 2777178 | rs7733720 | -1.671 | 0.56 | 0.608 | 1.078 | 0.008 | 0.47 | 0.47 | 0.16 | 0.09 | 0.42 | 0.14 | 0.43 | 0.04 | 0.19 |
| 5 | 2778782 | rs2068829 | 0.162 | 0.764 | 0.402 | 0.482 | -0.006 | 0.012 | 0.012 | 0 | 0.4 | 0.4 | 0.47 | 0.86 | 0.4 | 0.51 |
| 5 | 2782587 | rs4866498 | -0.013 | 0.877 | 0.002 | 1.382 | -0.038 | -0.043 | -0.043 | 0 | 0.13 | 0.05 | 0.26 | 0.28 | 0.22 | 0.19 |
| 5 | 2783046 | rs2132870 | 0.384 | 0.25 | -0.087 | 1.376 | 0.016 | -0.005 | -0.005 | 0.669 | 0.47 | 0.35 | 0.49 | 0.56 | 0.41 | 0.41 |
| 5 | 2783561 | rs1024048 | 0.989 | 0.081 | -0.166 | 1.394 | 0.019 | -0.005 | -0.005 | 0 | 0.36 | 0.25 | 0.44 | 0.4 | 0.3 | 0.29 |
| 5 | 2783869 | rs6864829 | -0.409 | -0.385 | -0.609 | 1.006 | -0.027 | 0.021 | 0.021 | 0.961 | 0.23 | 0.14 | 0.22 | 0.19 | 0.29 | 0.3 |
| 5 | 2785072 | rs6870714 | 0.573 | -0.368 | -0.595 | 1.048 | 0.019 | -0.005 | -0.005 | 0 | 0.36 | 0.25 | 0.44 | 0.4 | 0.3 | 0.29 |
| 5 | 2788906 | rs7702905 | 1.319 | -0.438 | -0.171 | 0.823 | 0.005 | -0.013 | -0.013 | 0 | 0.29 | 0.33 | 0.17 | 0.18 | 0.3 | 0.19 |
| 5 | 2789155 | rs7727905 | 1.053 | -0.479 | -0.028 | 0.824 | -0.001 | -0.012 | -0.012 | 0 | 0.54 | 0.48 | 0.54 | 0.77 | 0.42 | 0.46 |
| 5 | 2789702 | rs6867209 | 0.406 | -0.46 | -0.096 | 0.827 | -0.005 | 0.004 | 0.004 | 0 | 0.68 | 0.65 | 0.71 | 0.85 | 0.6 | 0.64 |
| 5 | 2789932 | rs1908976 | 0.567 | -0.446 | -0.13 | 0.891 | 0.001 | 0 | 0 | 0 | 0.65 | 0.59 | 0.62 | 0.58 | 0.6 | 0.48 |
| 5 | 2795776 | rs16870453 | 0.53 | -0.363 | -0.646 | 0.871 | 0.002 | -0.007 | -0.007 | 0.043 | 0.13 | 0.17 | 0.17 | 0.08 | 0.18 | 0.18 |
| 5 | 2798478 | rs11959104 | -0.664 | -0.522 | -1.023 | 0.888 | -0.071 | 0.082 | 0.082 | 0 | 0.32 | 0.14 | 0.27 | 0.17 | 0.4 | 0.45 |
| 5 | 2799100 | rs7729458 | 1.653 | -0.576 | -0.975 | 0.957 | 0.045 | -0.036 | -0.036 | 0.203 | 0.22 | 0.1 | 0.27 | 0.34 | 0.12 | 0.11 |
| 5 | 2799759 | rs1497457 | -0.422 | 0.222 | -0.67 | 0.886 | -0.008 | -0.008 | -0.008 | 0.968 | 0.23 | 0.21 | 0.32 | 0.31 | 0.29 | 0.4 |

| | | | | | | | | | | | | | | | | |
|---|---|---|---|---|---|---|---|---|---|---|---|---|---|---|---|---|
| 5 | 2802065 | rs10512672 | -0.038 | -0.129 | -0.132 | 0.755 | 0.002 | -0.002 | -0.002 | 0.092 | 0.12 | 0.15 | 0.06 | 0.1 | 0.12 | 0.12 |
| 5 | 2802282 | rs7710715 | -0.081 | -0.343 | 0.003 | 0.549 | 0 | 0 | 0 | 0.454 | 0.54 | 0.47 | 0.59 | 0.76 | 0.45 | 0.58 |
| 5 | 2809055 | rs10043028 | 0.077 | -0.166 | 0.02 | 0.666 | -0.001 | 0.001 | 0.001 | 0 | 0.15 | 0.19 | 0.33 | 0.2 | 0.14 | 0.18 |
| 5 | 2809713 | rs904117 | 1.427 | -0.588 | -0.435 | 0.369 | -0.096 | 0.05 | 0.05 | 0 | 0.6 | 0.07 | 0.44 | 0.67 | 0.56 | 0.55 |
| 5 | 2811135 | rs4866679 | 0.189 | -0.584 | 0.24 | 0.318 | 0.047 | -0.003 | -0.003 | 0 | 0.86 | 0.55 | 0.84 | 0.88 | 0.84 | 0.83 |
| 5 | 2812230 | rs6878883 | 1.718 | -0.635 | 0.035 | 0.404 | -0.117 | 0.007 | 0.007 | 0 | 0.58 | 0.05 | 0.44 | 0.64 | 0.56 | 0.47 |
| 5 | 2813509 | rs16870505 | -0.193 | -0.683 | 0.166 | 0.388 | 0.004 | -0.019 | -0.019 | 0 | 0.11 | 0.15 | 0.06 | 0.05 | 0.12 | 0.18 |
| 5 | 2814881 | rs4866504 | -0.042 | -0.631 | 0.23 | 0.425 | -0.001 | 0.001 | 0.001 | 0 | 0.18 | 0.21 | 0.34 | 0.24 | 0.17 | 0.27 |
| 5 | 2815488 | rs1122217 | 1.258 | -0.655 | 0.263 | 0.333 | -0.001 | 0.002 | 0.002 | 0 | 0.75 | 0.27 | 0.77 | 0.84 | 0.73 | 0.67 |
| 5 | 2815854 | rs13154311 | 0.097 | -0.653 | 0.277 | 0.339 | 0.006 | -0.021 | -0.021 | 2.324 | 0.1 | 0.12 | 0.05 | 0.03 | 0.11 | 0.1 |
| 5 | 2818217 | rs1540708 | 1.725 | -0.627 | 0.079 | 0.476 | -0.096 | -0.03 | -0.03 | 1.975 | 0.58 | 0.06 | 0.44 | 0.59 | 0.56 | 0.47 |
| 5 | 2818295 | rs7714315 | 0.052 | -0.577 | 0.222 | 0.632 | -0.001 | 0.008 | 0.008 | 0 | 0.18 | 0.21 | 0.34 | 0.29 | 0.16 | 0.27 |
| 5 | 2822959 | rs13170951 | 0.733 | -0.939 | -0.153 | 0.419 | 0.021 | 0.075 | 0.075 | 0 | 0.46 | 0.29 | 0.43 | 0.25 | 0.59 | 0.45 |
| 5 | 2823353 | rs10038374 | 0.053 | -1.052 | -0.054 | 0.514 | 0.053 | 0.016 | 0.016 | 0 | 0.4 | 0.24 | 0.41 | 0.65 | 0.25 | 0.43 |
| 5 | 2828738 | rs7701252 | -0.443 | -1.039 | 0.038 | 0.446 | 0.068 | -0.013 | -0.013 | 0 | 0.87 | 0.6 | 0.85 | 0.9 | 0.84 | 0.85 |
| 5 | 2829191 | rs10038356 | 0.071 | -1.015 | 0.021 | 0.466 | 0.056 | 0 | 0 | 0.942 | 0.39 | 0.2 | 0.4 | 0.62 | 0.25 | 0.35 |
| 5 | 2831070 | rs1018089 | -0.54 | -1.029 | 0.098 | 0.477 | 0.06 | -0.013 | -0.013 | 0 | 0.88 | 0.66 | 0.85 | 0.9 | 0.84 | 0.85 |
| 5 | 2831521 | rs6862810 | -0.286 | -0.961 | -0.417 | 0.465 | 0.037 | -0.004 | -0.004 | 0 | 0.86 | 0.43 | 0.83 | 0.86 | 0.84 | 0.77 |
| 5 | 2832643 | rs13157801 | 0.723 | -0.925 | -0.45 | 0.444 | -0.014 | 0.015 | 0.015 | 0 | 0.09 | 0.03 | 0.05 | 0.03 | 0.1 | 0.08 |
| 5 | 2833623 | rs13163086 | 1.364 | -0.903 | -0.525 | 0.46 | -0.049 | 0.056 | 0.056 | 0 | 0.37 | 0.23 | 0.38 | 0.21 | 0.49 | 0.35 |
| 5 | 2834261 | rs2047379 | -0.027 | -0.867 | -0.698 | 0.54 | 0.057 | 0.002 | 0.002 | 0.009 | 0.39 | 0.2 | 0.4 | 0.62 | 0.25 | 0.34 |
| 5 | 2838420 | rs13178497 | -0.451 | -0.634 | -1.343 | 0.52 | 0.072 | 0.075 | 0.075 | 0 | 0.48 | 0.19 | 0.46 | 0.64 | 0.36 | 0.41 |
| 5 | 2841709 | rs12054687 | 0.368 | -0.44 | -1.466 | 0.629 | 0.072 | 0.085 | 0.085 | 0 | 0.48 | 0.18 | 0.54 | 0.31 | 0.64 | 0.49 |
| 5 | 2843679 | rs10866544 | 0.363 | -0.431 | -1.461 | 0.634 | 0.072 | 0.086 | 0.086 | 0 | 0.48 | 0.18 | 0.54 | 0.31 | 0.64 | 0.49 |
| 5 | 2843763 | rs7442964 | -0.343 | -0.508 | -1.471 | 0.723 | 0.023 | -0.018 | -0.018 | 0.424 | 0.61 | 0.4 | 0.76 | 0.48 | 0.74 | 0.61 |
| 5 | 2844917 | rs17626019 | 0.233 | -0.519 | -1.268 | 1.047 | -0.057 | -0.014 | -0.014 | 0 | 0.13 | 0.01 | 0.17 | 0.1 | 0.18 | 0.14 |
| 5 | 2845113 | rs13160629 | -0.477 | -0.965 | -0.546 | 1.277 | -0.002 | 0.003 | 0.003 | 0 | 0.13 | 0.17 | 0.21 | 0.17 | 0.1 | 0.12 |
| 5 | 2849436 | rs438418 | 0.155 | -1.233 | -0.238 | 1.291 | -0.086 | 0.054 | 0.054 | 0.114 | 0.75 | 0.6 | 0.9 | 0.63 | 0.92 | 0.88 |
| 5 | 2851856 | rs559379 | -0.555 | 0.149 | -0.287 | 1.321 | 0.048 | 0.05 | 0.05 | 0 | 0.33 | 0.47 | 0.43 | 0.57 | 0.44 | 0.28 |
| 5 | 2852127 | rs12655083 | 0.056 | 0.233 | -1.069 | 1.204 | -0.074 | -0.042 | -0.042 | 0.32 | 0.41 | 0.13 | 0.41 | 0.48 | 0.47 | 0.38 |
| 5 | 2853858 | rs903246 | -0.385 | -0.508 | -0.07 | 1.073 | -0.013 | 0.039 | 0.039 | 0 | 0.15 | 0.41 | 0.3 | 0.14 | 0.11 | 0.12 |
| 5 | 2857885 | rs1004953 | 0.016 | -0.831 | -0.252 | 0.748 | -0.047 | -0.041 | -0.041 | 0 | 0.68 | 0.46 | 0.64 | 0.17 | 0.78 | 0.49 |
| 5 | 2858054 | rs1004952 | 0.202 | -1.112 | -0.152 | 1.213 | -0.03 | 0.072 | 0.072 | 0 | 0.21 | 0.36 | 0.16 | 0.46 | 0.12 | 0.34 |
| 5 | 2858513 | rs2974638 | 0.206 | -1.129 | -0.168 | 1.221 | 0.008 | -0.023 | -0.023 | 0 | 0.11 | 0.05 | 0.19 | 0.38 | 0.1 | 0.16 |
| 5 | 2860537 | rs10055154 | 0.605 | -1.139 | -0.122 | 1.191 | -0.025 | -0.021 | -0.021 | 0 | 0.69 | 0.57 | 0.64 | 0.17 | 0.79 | 0.49 |
| 5 | 2861216 | rs16870583 | 0.411 | -1.051 | -0.002 | 1.202 | -0.008 | 0.03 | 0.03 | 0 | 0.19 | 0.22 | 0.15 | 0.46 | 0.12 | 0.34 |
| 5 | 2865272 | rs6884970 | 0.24 | -1.035 | -0.013 | 1.207 | -0.014 | 0.038 | 0.038 | 0 | 0.19 | 0.24 | 0.15 | 0.46 | 0.12 | 0.34 |
| 5 | 2865604 | rs6881327 | 0.223 | -1.035 | -0.01 | 1.207 | -0.022 | 0.059 | 0.059 | 0 | 0.2 | 0.26 | 0.15 | 0.46 | 0.12 | 0.34 |
| 5 | 2866246 | rs9313014 | -0.628 | -0.86 | -0.061 | 0.997 | 0.005 | 0.005 | 0.005 | 0 | 0.3 | 0.34 | 0.36 | 0.83 | 0.24 | 0.51 |
| 5 | 2866346 | rs9313015 | -0.044 | -0.098 | -0.113 | 1.155 | -0.002 | -0.001 | -0.001 | 0 | 0.36 | 0.26 | 0.26 | 0.42 | 0.32 | 0.35 |
| 5 | 2869505 | rs1024029 | -0.432 | 0.024 | -0.258 | 0.984 | 0.008 | 0.011 | 0.011 | 0 | 0.49 | 0.62 | 0.56 | 0.59 | 0.44 | 0.56 |
| 5 | 2870681 | rs6899275 | -0.19 | -0.019 | -0.182 | 0.936 | -0.001 | 0 | 0 | 0 | 0.45 | 0.51 | 0.53 | 0.55 | 0.41 | 0.48 |
| 5 | 2878297 | rs756220 | -0.994 | -0.13 | -0.475 | 0.372 | -0.051 | 0.138 | 0.138 | 0 | 0.61 | 0.79 | 0.77 | 1 | 0.55 | 0.75 |
| 5 | 2878637 | rs2934550 | -1.174 | -0.138 | -0.641 | 0.388 | -0.097 | 0.21 | 0.21 | 0.017 | 0.62 | 0.89 | 0.77 | 1 | 0.55 | 0.75 |
| 5 | 2883677 | rs16870625 | -0.081 | 0.07 | -0.556 | 1.02 | -0.004 | 0.031 | 0.031 | 0 | 0.11 | 0.1 | 0.2 | 0.35 | 0.1 | 0.14 |
| 5 | 2884879 | rs4866693 | -0.27 | 0.24 | -0.349 | 0.841 | -0.001 | -0.001 | -0.001 | 0 | 0.43 | 0.38 | 0.52 | 0.41 | 0.41 | 0.44 |
| 5 | 2886593 | rs4866694 | -0.329 | 0.501 | -0.307 | 0.799 | -0.001 | -0.008 | -0.008 | 0 | 0.46 | 0.49 | 0.52 | 0.36 | 0.47 | 0.53 |
| 5 | 2892699 | rs12654325 | -0.785 | 1.052 | 0.211 | 1.924 | -0.004 | 0.008 | 0.008 | 0 | 0.15 | 0.13 | 0.23 | 0.5 | 0.2 | 0.17 |
| 5 | 2896161 | rs13153727 | -0.188 | 0.902 | -0.526 | 0.712 | 0.004 | -0.004 | -0.004 | 0 | 0.57 | 0.38 | 0.57 | 0.59 | 0.66 | 0.45 |
| 5 | 2897087 | rs11745728 | 0.048 | 0.628 | -0.645 | 0.639 | 0.027 | -0.058 | -0.058 | 0.003 | 0.54 | 0.11 | 0.54 | 0.59 | 0.65 | 0.45 |
| 5 | 2897236 | rs7731319 | 0.176 | 0.543 | -0.143 | 0.935 | 0.055 | 0.027 | 0.027 | 0 | 0.11 | 0.26 | 0.13 | 0.18 | 0.16 | 0.11 |
| 5 | 2900123 | rs7732075 | 0.952 | 0.075 | -0.512 | 0.383 | -0.024 | 0.111 | 0.111 | 2.844 | 0.38 | 0.18 | 0.3 | 0.01 | 0.43 | 0.28 |
| 5 | 2901649 | rs2974657 | -0.568 | -0.081 | -0.691 | 1.596 | -0.026 | -0.061 | -0.061 | 0 | 0.85 | 0.92 | 0.74 | 0.52 | 0.81 | 0.67 |
| 5 | 2902855 | rs1389595 | 0.615 | -0.221 | -0.542 | 1.601 | -0.001 | -0.017 | -0.017 | 0 | 0.38 | 0.39 | 0.3 | 0.15 | 0.34 | 0.32 |
| 5 | 2902999 | rs2860294 | 0.023 | -0.211 | -0.551 | 1.614 | 0.004 | 0.004 | 0.004 | 2.989 | 0.46 | 0.56 | 0.4 | 0.38 | 0.38 | 0.36 |
| 5 | 2907012 | rs956897 | 1.282 | -0.453 | -0.431 | 1.085 | 0.004 | -0.053 | -0.053 | 0.044 | 0.22 | 0.31 | 0.17 | 0.07 | 0.19 | 0.12 |
| 5 | 2908616 | rs1565808 | -0.182 | -0.37 | -0.546 | 1.24 | -0.009 | -0.031 | -0.031 | 0 | 0.4 | 0.47 | 0.31 | 0.3 | 0.34 | 0.35 |
| 5 | 2908851 | rs17673214 | 1.078 | -0.345 | -0.675 | 1.252 | -0.01 | 0.057 | 0.057 | 1.03 | 0.43 | 0.26 | 0.4 | 0.23 | 0.48 | 0.31 |
| 5 | 2910083 | rs2934555 | -0.735 | -0.212 | -0.863 | 1.553 | -0.004 | -0.001 | -0.001 | 2.379 | 0.57 | 0.46 | 0.58 | 0.46 | 0.59 | 0.68 |
| 5 | 2912375 | rs17627501 | 0.933 | 0.058 | -1.116 | 1.458 | -0.014 | -0.008 | -0.008 | 0.783 | 0.41 | 0.24 | 0.4 | 0.44 | 0.43 | 0.3 |
| 5 | 2913619 | rs17678795 | 1.388 | 0.114 | -1.112 | 1.357 | 0 | 0.08 | 0.08 | 0.012 | 0.36 | 0.21 | 0.3 | 0.2 | 0.36 | 0.26 |
| 5 | 2915301 | rs903244 | 0.6 | 0.049 | -1.174 | 1.47 | -0.065 | -0.088 | -0.088 | 0.703 | 0.18 | 0.04 | 0.27 | 0.27 | 0.26 | 0.37 |
| 5 | 2917143 | rs2974690 | -1.258 | -0.042 | -0.778 | 1.167 | -0.001 | -0.002 | -0.002 | 0 | 0.63 | 0.6 | 0.67 | 0.8 | 0.61 | 0.72 |
| 5 | 2922427 | rs13185688 | 1.389 | -0.556 | -0.992 | 1.342 | 0.003 | 0.166 | 0.166 | 0.78 | 0.51 | 0.2 | 0.43 | 0.24 | 0.56 | 0.43 |
| 5 | 2923396 | rs1494064 | 0.342 | -0.67 | -0.979 | 1.533 | -0.06 | -0.058 | -0.058 | 0 | 0.18 | 0.04 | 0.14 | 0.22 | 0.26 | 0.3 |
| 5 | 2929499 | rs11738897 | 1.455 | -0.427 | -1.126 | 1.236 | -0.03 | 0.176 | 0.176 | 0.291 | 0.52 | 0.16 | 0.44 | 0.25 | 0.57 | 0.43 |
| 5 | 2929714 | rs10475186 | -0.239 | -0.4 | -0.808 | 1.356 | 0.025 | 0.018 | 0.018 | 0.013 | 0.8 | 0.72 | 0.84 | 0.8 | 0.74 | 0.72 |
| 5 | 2934319 | rs2404802 | 1.167 | -0.467 | -0.892 | 1.393 | 0.005 | 0.001 | 0.001 | 0.511 | 0.57 | 0.27 | 0.69 | 0.63 | 0.61 | 0.52 |
| 5 | 2940406 | rs2934519 | -0.301 | -0.491 | -0.848 | 1.277 | -0.043 | -0.067 | -0.067 | 0.112 | 0.77 | 0.59 | 0.85 | 0.9 | 0.88 | 0.88 |

| | | | | | | | | | | | | | | | | |
|---|---|---|---|---|---|---|---|---|---|---|---|---|---|---|---|---|
| 5 | 2945834 | rs1389585 | -0.685 | -0.256 | -0.856 | 1.446 | 0.04 | 0.036 | 0.036 | 0.116 | 0.25 | 0.41 | 0.41 | 0.65 | 0.31 | 0.43 |
| 5 | 2947877 | rs2934528 | 0.018 | -1.238 | -1.058 | 1.332 | 0.022 | -0.001 | -0.001 | 0 | 0.39 | 0.45 | 0.53 | 0.58 | 0.52 | 0.49 |
| 5 | 2948632 | rs1389589 | -1.258 | -0.966 | -1.041 | 1.649 | 0.051 | 0.398 | 0.398 | 0 | 0.07 | 0.34 | 0.28 | 0.38 | 0.04 | 0.07 |
| 5 | 2949174 | rs7737453 | 0.667 | -1.404 | -1.193 | 1.506 | 0.003 | 0.009 | 0.009 | 0 | 0.36 | 0.41 | 0.53 | 0.44 | 0.35 | 0.29 |
| 5 | 2950492 | rs6555155 | 0.171 | -1.662 | -1.159 | 1.466 | -0.009 | -0.002 | -0.002 | 0.108 | 0.49 | 0.28 | 0.42 | 0.51 | 0.45 | 0.36 |
| 5 | 2955932 | rs6893622 | 0.529 | -1.33 | -0.907 | 1.524 | 0 | -0.016 | -0.016 | 0 | 0.56 | 0.59 | 0.44 | 0.12 | 0.53 | 0.51 |
| 5 | 2960267 | rs2974658 | 0.281 | -0.706 | -1.842 | 1.16 | -0.145 | -0.062 | -0.062 | 0 | 0.29 | 0.02 | 0.23 | 0.26 | 0.42 | 0.37 |
| 5 | 2965167 | rs1565807 | -1.034 | -2.053 | -0.755 | 1.564 | -0.03 | 0.011 | 0.011 | 0.015 | 0.24 | 0.71 | 0.39 | 0.64 | 0.07 | 0.17 |
| 5 | 2965303 | rs10512669 | 0.527 | -1.553 | -0.695 | 1.433 | 0.007 | -0.003 | -0.003 | 0 | 0.09 | 0.05 | 0.07 | 0.11 | 0.03 | 0.06 |
| 5 | 2967695 | rs10072840 | -0.205 | -1.584 | -0.604 | 1.668 | -0.01 | 0.023 | 0.023 | 0 | 0.12 | 0.13 | 0.08 | 0.42 | 0.04 | 0.09 |
| 5 | 2968466 | rs7724104 | 0.09 | -1.221 | -0.729 | 0.607 | 0.006 | -0.017 | -0.017 | 0 | 0.28 | 0.22 | 0.27 | 0.53 | 0.22 | 0.26 |
| 5 | 2970320 | rs1494066 | -0.203 | -0.837 | -0.285 | 0.329 | -0.032 | -0.042 | -0.042 | 0 | 0.68 | 0.52 | 0.78 | 0.79 | 0.77 | 0.77 |
| 5 | 2972677 | rs13362004 | 0.443 | -0.92 | -0.373 | 0.291 | 0.011 | -0.032 | -0.032 | 0 | 0.31 | 0.2 | 0.28 | 0.51 | 0.24 | 0.29 |
| 5 | 2972851 | rs10454923 | 0.84 | -0.918 | -0.49 | 0.296 | -0.038 | 0.036 | 0.036 | 0 | 0.37 | 0.25 | 0.5 | 0.22 | 0.53 | 0.5 |
| 5 | 2973712 | rs12657444 | 0.268 | -0.597 | -0.795 | 0.467 | 0.002 | -0.006 | -0.006 | 3.084 | 0.31 | 0.26 | 0.28 | 0.51 | 0.24 | 0.29 |
| 5 | 2974992 | rs1494063 | -0.456 | -0.646 | -0.743 | 0.432 | 0.01 | -0.015 | -0.015 | 0 | 0.92 | 0.82 | 0.96 | 0.95 | 0.92 | 0.96 |
| 5 | 2978271 | rs10079415 | 0.001 | -0.365 | -1.033 | 0.687 | 0.008 | -0.021 | -0.021 | 0.658 | 0.32 | 0.25 | 0.36 | 0.5 | 0.27 | 0.31 |
| 5 | 2979710 | rs4242095 | -0.399 | -0.389 | -1.001 | 0.571 | 0.001 | 0.001 | 0.001 | 0 | 0.36 | 0.66 | 0.33 | 0.71 | 0.22 | 0.28 |
| 5 | 2981865 | rs6888766 | -0.017 | 0.471 | -1.169 | 0.389 | 0.006 | 0.005 | 0.005 | 0 | 0.52 | 0.58 | 0.49 | 0.58 | 0.58 | 0.47 |
| 5 | 2984081 | rs6861422 | 0.087 | 0.695 | -1.771 | -0.286 | -0.023 | 0.107 | 0.107 | 0.303 | 0.7 | 0.9 | 0.72 | 0.9 | 0.7 | 0.76 |
| 5 | 2993617 | rs10070337 | 0.163 | 0.686 | -1.145 | -0.081 | 0.002 | 0.01 | 0.01 | 0.126 | 0.47 | 0.58 | 0.52 | 0.61 | 0.48 | 0.46 |
| 5 | 2993645 | rs10075407 | -0.107 | 0.577 | -1.123 | -0.201 | -0.013 | 0.073 | 0.073 | 0.636 | 0.55 | 0.71 | 0.56 | 0.72 | 0.54 | 0.6 |
| 5 | 2999174 | rs6871461 | 0.532 | 0.267 | -0.995 | -0.39 | 0.011 | -0.025 | -0.025 | 0.366 | 0.32 | 0.41 | 0.28 | 0.18 | 0.36 | 0.18 |
| 5 | 2999518 | rs7712222 | -0.045 | 0.292 | -0.759 | -0.292 | -0.001 | -0.004 | -0.004 | 0 | 0.45 | 0.49 | 0.31 | 0.2 | 0.47 | 0.36 |
| 5 | 3001216 | rs10072629 | -0.796 | 0.389 | -1.399 | -0.551 | -0.071 | 0.091 | 0.091 | 0.022 | 0.67 | 0.91 | 0.64 | 0.8 | 0.62 | 0.75 |
| 5 | 3004584 | rs2009378 | 0.743 | -0.43 | -0.093 | 0.029 | -0.048 | 0.094 | 0.094 | 0.147 | 0.82 | 0.61 | 0.67 | 0.4 | 0.89 | 0.65 |
| 5 | 3004742 | rs2009370 | 0.883 | -0.209 | 0.051 | 0.102 | -0.01 | 0.133 | 0.133 | 0.797 | 0.83 | 0.6 | 0.78 | 0.4 | 0.87 | 0.68 |
| 5 | 3007514 | rs2961797 | 0.34 | 0.316 | -0.314 | -0.002 | -0.012 | 0.051 | 0.051 | 0.02 | 0.54 | 0.29 | 0.65 | 0.72 | 0.52 | 0.69 |
| 5 | 3010613 | rs2961793 | 0.266 | -0.571 | 0.396 | -0.104 | -0.023 | -0.072 | -0.072 | 0 | 0.73 | 0.62 | 0.85 | 0.93 | 0.81 | 0.86 |
| 5 | 3011663 | rs2922992 | 0.642 | -0.368 | 0.334 | 0.069 | -0.015 | -0.025 | -0.025 | 0 | 0.65 | 0.4 | 0.77 | 0.79 | 0.75 | 0.8 |
| 5 | 3017791 | rs11952070 | 0.618 | -0.405 | 0.61 | 1.066 | 0.002 | 0.033 | 0.033 | 0 | 0.71 | 0.63 | 0.79 | 0.52 | 0.72 | 0.57 |
| 5 | 3018524 | rs2084547 | 0.507 | -0.402 | 0.725 | 1.231 | 0 | -0.003 | -0.003 | 0.125 | 0.72 | 0.72 | 0.82 | 0.62 | 0.74 | 0.6 |
| 5 | 3019005 | rs12517482 | 1.331 | -0.94 | 0.899 | 0.941 | 0.013 | -0.011 | -0.011 | 0 | 0.61 | 0.33 | 0.75 | 0.52 | 0.7 | 0.45 |
| 5 | 3025452 | rs2404567 | 0.637 | -0.983 | 0.72 | 0.731 | -0.014 | 0.057 | 0.057 | 0 | 0.69 | 0.48 | 0.79 | 0.55 | 0.73 | 0.6 |
| 5 | 3028654 | rs11133974 | 1.15 | -1.247 | 0.736 | 0.664 | -0.019 | 0.008 | 0.008 | 0 | 0.63 | 0.43 | 0.78 | 0.56 | 0.74 | 0.51 |
| 5 | 3029927 | rs10454925 | -1.205 | -1.799 | 0.504 | 0.476 | 0.049 | 0.05 | 0.05 | 2.198 | 0.36 | 0.76 | 0.22 | 0.13 | 0.21 | 0.27 |
| 5 | 3032625 | rs4866516 | -0.689 | -1.762 | 1.022 | 0.485 | -0.032 | -0.036 | -0.036 | 0.814 | 0.29 | 0.47 | 0.17 | 0.13 | 0.2 | 0.28 |
| 5 | 3036596 | rs191168 | 0.449 | -1.845 | 0.915 | 0.924 | -0.012 | 0.11 | 0.11 | 0 | 0.91 | 0.86 | 0.93 | 0.59 | 0.92 | 0.76 |
| 5 | 3042790 | rs11133976 | -0.436 | -1.835 | 1.129 | 0.416 | 0.011 | 0.011 | 0.011 | 0 | 0.26 | 0.24 | 0.15 | 0.13 | 0.2 | 0.29 |
| 5 | 3044895 | rs1037291 | -0.494 | -1.845 | 1.075 | 0.383 | 0.017 | 0.022 | 0.022 | 0.047 | 0.27 | 0.2 | 0.15 | 0.13 | 0.2 | 0.28 |
| 5 | 3045983 | rs10068728 | -1.309 | -1.875 | 1.238 | 0.368 | -0.036 | -0.04 | -0.04 | 0 | 0.34 | 0.59 | 0.2 | 0.13 | 0.21 | 0.29 |
| 5 | 3054500 | rs16870972 | -0.272 | -1.935 | 0.685 | 0.354 | 0.007 | -0.002 | -0.002 | 0 | 0.18 | 0.14 | 0.1 | 0.13 | 0.1 | 0.13 |
| 5 | 3055887 | rs6890167 | -0.98 | -1.773 | 1.35 | 0.18 | -0.022 | -0.122 | -0.122 | 2.663 | 0.18 | 0.63 | 0.12 | 0 | 0.1 | 0.15 |
| 5 | 3057014 | rs7714375 | -0.898 | -1.724 | 1.2 | 0.131 | -0.01 | -0.101 | -0.101 | 1.323 | 0.17 | 0.63 | 0.13 | 0 | 0.11 | 0.15 |
| 5 | 3059961 | rs170225 | -0.553 | -1.919 | 0.775 | 0.783 | -0.002 | -0.028 | -0.028 | 4.236 | 0.92 | 1 | 0.95 | 0.65 | 0.93 | 0.85 |
| 5 | 3060017 | rs167528 | 0.221 | -1.497 | -0.216 | 0.102 | 0 | 0.075 | 0.075 | 0 | 0.49 | 0.71 | 0.51 | 0.31 | 0.53 | 0.55 |
| 5 | 3060489 | rs7706611 | 0.466 | -1.773 | -0.288 | 0.002 | -0.027 | -0.011 | -0.011 | 0.103 | 0.17 | 0.22 | 0.09 | 0.09 | 0.09 | 0.14 |
| 5 | 3060697 | rs6555171 | -0.89 | -1.819 | -0.245 | -0.022 | -0.041 | -0.047 | -0.047 | 0 | 0.33 | 0.5 | 0.19 | 0.16 | 0.19 | 0.29 |
| 5 | 3061140 | rs160874 | 0.773 | -1.135 | -0.853 | -0.356 | -0.021 | -0.005 | -0.005 | 0.068 | 0.34 | 0.21 | 0.29 | 0.36 | 0.4 | 0.49 |
| 5 | 3064150 | rs160884 | 0.245 | -1.175 | -0.773 | -0.319 | -0.004 | 0.031 | 0.031 | 0.122 | 0.78 | 0.75 | 0.84 | 0.49 | 0.83 | 0.65 |
| 5 | 3064862 | rs7728601 | 0.311 | -0.979 | -0.803 | -0.274 | 0.025 | 0.002 | 0.002 | 0 | 0.07 | 0.16 | 0.04 | 0.07 | 0.06 | 0.09 |
| 5 | 3065101 | rs160882 | 1.526 | -1.219 | -1.377 | -0.43 | -0.086 | -0.082 | -0.082 | 0 | 0.24 | 0.02 | 0.24 | 0.29 | 0.33 | 0.28 |
| 5 | 3073199 | rs6891263 | 0.754 | -1.514 | -1.094 | -0.643 | 0.011 | -0.005 | -0.005 | 0 | 0.15 | 0.16 | 0.05 | 0 | 0.04 | 0.15 |
| 5 | 3073671 | rs466896 | -0.24 | -1.392 | -1.477 | -1.005 | 0.03 | 0.045 | 0.045 | 0 | 0.42 | 0.32 | 0.42 | 0.25 | 0.24 | 0.33 |
| 5 | 3074339 | rs461578 | -1.135 | -1.463 | -1.401 | -0.784 | 0.02 | -0.045 | -0.045 | 0 | 0.85 | 0.93 | 0.88 | 0.56 | 0.88 | 0.74 |
| 5 | 3074414 | rs464050 | -0.12 | -1.245 | -1.498 | -1.049 | 0.028 | 0.052 | 0.052 | 0 | 0.43 | 0.32 | 0.44 | 0.24 | 0.28 | 0.36 |
| 5 | 3074790 | rs460017 | -0.133 | -1.187 | -1.496 | -1.057 | 0.027 | 0.056 | 0.056 | 0 | 0.44 | 0.32 | 0.45 | 0.24 | 0.29 | 0.36 |
| 5 | 3076720 | rs10475190 | 0.405 | -0.576 | -1.033 | -1.092 | 0 | -0.039 | -0.039 | 3.731 | 0.43 | 0.61 | 0.28 | 0.08 | 0.43 | 0.25 |
| 5 | 3076956 | rs10475191 | 0.413 | -0.579 | -1.022 | -1.093 | 0 | -0.033 | -0.033 | 1.097 | 0.42 | 0.62 | 0.28 | 0.08 | 0.43 | 0.26 |
| 5 | 3078217 | rs462624 | 0.555 | -0.578 | -0.929 | -1.011 | 0 | -0.005 | -0.005 | 0 | 0.32 | 0.26 | 0.45 | 0.5 | 0.26 | 0.42 |
| 5 | 3079057 | rs464918 | 0.076 | -0.275 | -0.463 | -1.117 | -0.006 | -0.007 | -0.007 | 0.524 | 0.57 | 0.61 | 0.56 | 0.5 | 0.43 | 0.58 |
| 5 | 3079891 | rs16871030 | 1.39 | -0.3 | -0.392 | -1.023 | -0.043 | -0.007 | -0.007 | 0 | 0.26 | 0.43 | 0.15 | 0.25 | 0.18 | 0.25 |
| 5 | 3080654 | rs13172975 | -0.245 | -0.284 | -0.472 | -1.071 | -0.003 | -0.01 | -0.01 | 0.003 | 0.68 | 0.67 | 0.71 | 0.5 | 0.63 | 0.61 |
| 5 | 3080880 | rs3104064 | 0.404 | -0.214 | -0.514 | -1.107 | -0.022 | 0.057 | 0.057 | 0.619 | 0.42 | 0.25 | 0.56 | 0.25 | 0.45 | 0.37 |
| 5 | 3082219 | rs2016714 | 0.842 | 0.006 | -1.251 | -1.08 | -0.001 | 0.087 | 0.087 | 0 | 0.35 | 0.2 | 0.25 | 0 | 0.37 | 0.19 |
| 5 | 3082643 | rs160876 | -0.554 | -0.053 | -1.307 | -0.964 | 0.019 | -0.009 | -0.009 | 0.011 | 0.57 | 0.79 | 0.68 | 0.58 | 0.56 | 0.62 |
| 5 | 3084205 | rs6879233 | 1.38 | 0.323 | -0.992 | -0.765 | -0.003 | 0.002 | 0.002 | 0 | 0.15 | 0.13 | 0.16 | 0.08 | 0.25 | 0.14 |
| 5 | 3086514 | rs11133978 | 1.493 | -0.168 | -0.896 | -0.465 | 0.009 | 0.026 | 0.026 | 0 | 0.13 | 0.15 | 0.2 | 0.25 | 0.21 | 0.21 |
| 5 | 3088781 | rs2442700 | -0.361 | -0.266 | -0.929 | -1.178 | 0.001 | -0.004 | -0.004 | 0 | 0.72 | 0.77 | 0.68 | 0.33 | 0.67 | 0.47 |

| | | | | | | | | | | | | | | | | |
|---|---|---|---|---|---|---|---|---|---|---|---|---|---|---|---|---|
| 5 | 3089964 | rs11744932 | 0.936 | -0.317 | -1.2 | -1.454 | 0.022 | 0.06 | 0.06 | 0 | 0.27 | 0.16 | 0.11 | 0 | 0.2 | 0.15 |
| 5 | 3090084 | rs9313033 | -0.184 | 0.089 | -0.665 | -0.977 | -0.002 | 0.03 | 0.03 | 0 | 0.46 | 0.65 | 0.58 | 0.33 | 0.47 | 0.33 |
| 5 | 3091466 | rs17543060 | 0.437 | 0.361 | -0.524 | -1.256 | 0.004 | -0.05 | -0.05 | 0 | 0.53 | 0.56 | 0.34 | 0 | 0.48 | 0.33 |
| 5 | 3092192 | rs6873793 | 0.346 | 0.744 | -0.622 | -0.943 | -0.007 | 0.005 | 0.005 | 0 | 0.66 | 0.68 | 0.64 | 0.17 | 0.58 | 0.39 |
| 5 | 3092254 | rs6872625 | 0.13 | 1.413 | -0.377 | -0.546 | 0.025 | -0.019 | -0.019 | 1.84 | 0.14 | 0.19 | 0.16 | 0.06 | 0.22 | 0.11 |
| 5 | 3093075 | rs4866523 | -0.172 | 0.963 | 0.173 | -0.142 | 0.033 | 0.057 | 0.057 | 0 | 0.17 | 0.41 | 0.39 | 0.31 | 0.2 | 0.19 |
| 5 | 3094738 | rs6870469 | 0.619 | 0.182 | -0.757 | -0.719 | 0.017 | 0.077 | 0.077 | 2.687 | 0.29 | 0.18 | 0.21 | 0 | 0.3 | 0.09 |
| 5 | 3095124 | rs12109259 | 0.95 | 0.263 | -0.884 | -0.594 | -0.011 | -0.01 | -0.01 | 0.01 | 0.1 | 0.08 | 0.33 | 0.16 | 0.14 | 0.06 |
| 5 | 3095424 | rs11960253 | 0.675 | 0.364 | -0.854 | -0.684 | 0.014 | 0.113 | 0.113 | 2.754 | 0.29 | 0.14 | 0.21 | 0 | 0.3 | 0.09 |
| 5 | 3095939 | rs13187652 | 0.408 | 0.551 | -0.868 | -0.336 | 0 | -0.008 | -0.008 | 0 | 0.16 | 0.11 | 0.34 | 0.33 | 0.16 | 0.27 |
| 5 | 3096001 | rs2442697 | 0.276 | 1.074 | -0.905 | 0.288 | 0.019 | 0.038 | 0.038 | 0 | 0.06 | 0.12 | 0.1 | 0.13 | 0.08 | 0.1 |
| 5 | 3098333 | rs314008 | -0.216 | 0.899 | -1.017 | 0.328 | -0.022 | 0.017 | 0.017 | 0.569 | 0.53 | 0.76 | 0.68 | 0.46 | 0.53 | 0.47 |
| 5 | 3100475 | rs160888 | -0.026 | 0.625 | -1.116 | 0.479 | 0.005 | 0.013 | 0.013 | 0.573 | 0.48 | 0.57 | 0.61 | 0.39 | 0.49 | 0.39 |
| 5 | 3103070 | rs317000 | -0.3 | 0.469 | -1.885 | 0.915 | -0.101 | -0.174 | -0.174 | 0 | 0.72 | 0.98 | 0.76 | 0.55 | 0.63 | 0.63 |
| 5 | 3104201 | rs316998 | -0.167 | -1.002 | -0.832 | 1.043 | -0.052 | 0 | 0 | 0 | 0.7 | 0.57 | 0.75 | 0.75 | 0.82 | 0.82 |
| 5 | 3104267 | rs172700 | -0.587 | -0.151 | -0.805 | 1.325 | -0.024 | -0.006 | -0.006 | 0.045 | 0.18 | 0.14 | 0.2 | 0.16 | 0.23 | 0.24 |
| 5 | 3105242 | rs4866716 | -1.106 | -0.963 | -0.296 | 1.33 | -0.023 | 0.005 | 0.005 | 0 | 0.28 | 0.6 | 0.21 | 0.3 | 0.17 | 0.23 |
| 5 | 3106162 | rs314002 | -1.314 | -0.914 | -0.243 | 0.977 | -0.007 | -0.003 | -0.003 | 0 | 0.4 | 0.51 | 0.31 | 0.33 | 0.29 | 0.42 |
| 5 | 3107008 | rs160577 | -1.473 | -0.833 | -0.254 | 1.066 | 0.012 | -0.021 | -0.021 | 0 | 0.4 | 0.21 | 0.26 | 0.38 | 0.32 | 0.39 |
| 5 | 3107310 | rs171658 | -0.993 | -0.828 | -0.09 | 0.688 | 0.001 | -0.005 | -0.005 | 0.798 | 0.14 | 0.21 | 0.1 | 0.12 | 0.11 | 0.25 |
| 5 | 3107861 | rs160770 | 1.628 | -0.921 | -0.322 | 0.763 | -0.002 | 0.04 | 0.04 | 0 | 0.59 | 0.8 | 0.74 | 0.72 | 0.65 | 0.58 |
| 5 | 3108117 | rs17544008 | 0.174 | -0.956 | -0.405 | 0.726 | 0.003 | 0.004 | 0.004 | 0.574 | 0.12 | 0.07 | 0.07 | 0.08 | 0.09 | 0.08 |
| 5 | 3109410 | rs17621396 | -0.746 | -0.917 | -0.22 | 0.93 | -0.018 | -0.006 | -0.006 | 0 | 0.18 | 0.03 | 0.1 | 0.11 | 0.16 | 0.1 |
| 5 | 3110197 | rs160765 | 1.956 | -0.998 | 0.277 | 0.896 | 0.005 | 0.006 | 0.006 | 0 | 0.64 | 0.74 | 0.75 | 0.71 | 0.7 | 0.6 |
| 5 | 3118894 | rs1389879 | -1.057 | -1.036 | 0.741 | 0.844 | 0.002 | 0.002 | 0.002 | 0 | 0.21 | 0.23 | 0.1 | 0.11 | 0.16 | 0.11 |
| 5 | 3119937 | rs160780 | 0.318 | -0.962 | 0.533 | 0.888 | -0.045 | -0.006 | -0.006 | 0 | 0.9 | 0.99 | 0.91 | 0.94 | 0.9 | 0.74 |
| 5 | 3121671 | rs160776 | 0.442 | -0.884 | 0.554 | 0.909 | -0.049 | -0.004 | -0.004 | 0 | 0.9 | 0.99 | 0.93 | 0.94 | 0.89 | 0.76 |
| 5 | 3121728 | rs16871117 | -1.331 | -1 | 0.646 | 0.891 | 0.021 | 0.007 | 0.007 | 0 | 0.23 | 0.14 | 0.15 | 0.2 | 0.18 | 0.14 |
| 5 | 3121797 | rs160774 | 1.542 | -1.003 | 0.981 | 0.937 | 0.02 | 0.05 | 0.05 | 0.758 | 0.85 | 0.67 | 0.84 | 0.73 | 0.88 | 0.72 |
| 5 | 3123775 | rs12657742 | -0.871 | -1.03 | 0.819 | 0.881 | -0.003 | -0.002 | -0.002 | 0.06 | 0.17 | 0.2 | 0.09 | 0.11 | 0.12 | 0.06 |
| 5 | 3124651 | rs6886617 | -1.072 | -1.101 | 0.854 | 0.828 | -0.005 | -0.004 | -0.004 | 0 | 0.21 | 0.24 | 0.14 | 0.14 | 0.14 | 0.1 |
| 5 | 3124796 | rs11748308 | 1.947 | -1.164 | 0.58 | 0.8 | 0.02 | -0.008 | -0.008 | 0.001 | 0.55 | 0.34 | 0.64 | 0.49 | 0.66 | 0.58 |
| 5 | 3128346 | rs10043332 | 1.422 | -1.008 | 0.505 | 0.683 | 0.01 | 0.049 | 0.049 | 0 | 0.79 | 0.49 | 0.86 | 0.69 | 0.86 | 0.69 |
| 5 | 3128522 | rs10045318 | 2.167 | -0.442 | 0.264 | 0.564 | -0.001 | 0 | 0 | 0 | 0.54 | 0.45 | 0.64 | 0.48 | 0.6 | 0.53 |
| 5 | 3128854 | rs16871149 | 1.821 | -0.676 | 0.428 | 0.523 | -0.014 | -0.002 | -0.002 | 0.066 | 0.71 | 0.45 | 0.77 | 0.64 | 0.77 | 0.59 |
| 5 | 3131064 | rs903397 | 2.297 | -0.86 | 0.314 | 0.407 | 0.021 | 0.003 | 0.003 | 0 | 0.54 | 0.37 | 0.66 | 0.47 | 0.63 | 0.53 |
| 5 | 3131074 | rs17551811 | -1.247 | -0.929 | 0.31 | 0.343 | -0.015 | -0.014 | -0.014 | 0 | 0.14 | 0.09 | 0.14 | 0.19 | 0.18 | 0.26 |
| 5 | 3131474 | rs469280 | 1.69 | -0.879 | 0.412 | 0.358 | -0.014 | -0.014 | -0.014 | 0 | 0.67 | 0.62 | 0.8 | 0.71 | 0.74 | 0.67 |
| 5 | 3132025 | rs467716 | 1.335 | -0.86 | 0.32 | 0.416 | 0 | -0.002 | -0.002 | 2.36 | 0.78 | 0.81 | 0.84 | 0.75 | 0.8 | 0.63 |
| 5 | 3132458 | rs17622688 | 0.386 | -0.84 | 0.252 | 0.412 | -0.043 | -0.022 | -0.022 | 4.002 | 0.07 | 0.01 | 0.07 | 0.07 | 0.12 | 0.23 |
| 5 | 3132559 | rs469508 | -1.064 | -0.906 | 0.416 | 0.435 | -0.039 | 0.013 | 0.013 | 0.59 | 0.22 | 0.43 | 0.16 | 0.29 | 0.14 | 0.09 |
| 5 | 3132692 | rs469402 | 0.787 | -0.883 | 0.37 | 0.449 | -0.001 | -0.001 | -0.001 | 0.541 | 0.86 | 0.9 | 0.92 | 0.82 | 0.92 | 0.85 |
| 5 | 3133471 | rs468334 | 1.08 | -0.83 | 0.373 | 0.411 | 0.001 | 0.004 | 0.004 | 0 | 0.83 | 0.85 | 0.89 | 0.88 | 0.85 | 0.67 |
| 5 | 3134466 | rs160891 | 1.861 | -0.783 | 0.45 | 0.483 | -0.006 | 0.03 | 0.03 | 0 | 0.75 | 0.73 | 0.8 | 0.54 | 0.79 | 0.59 |
| 5 | 3134719 | rs7726783 | 2.497 | -0.854 | 0.469 | 0.433 | 0.016 | -0.007 | -0.007 | 0.106 | 0.55 | 0.37 | 0.68 | 0.45 | 0.66 | 0.53 |
| 5 | 3134879 | rs160890 | -0.275 | -0.757 | 0.365 | 0.501 | 0.008 | 0.151 | 0.151 | 0 | 0.08 | 0.17 | 0.09 | 0.34 | 0.06 | 0.08 |
| 5 | 3138553 | rs167587 | 0.021 | -0.772 | 0.285 | 0.462 | 0 | 0 | 0 | 0 | 0.91 | 0.95 | 0.96 | 0.95 | 0.95 | 0.9 |
| 5 | 3138651 | rs316996 | 0.021 | -0.772 | 0.285 | 0.462 | 0 | 0 | 0 | 0 | 0.91 | 0.95 | 0.96 | 0.95 | 0.95 | 0.9 |
| 5 | 3139158 | rs597099 | -0.959 | -0.782 | 0.262 | 0.474 | 0.008 | 0.093 | 0.093 | 0 | 0.15 | 0.26 | 0.16 | 0.42 | 0.15 | 0.3 |
| 5 | 3140979 | rs467326 | 1.305 | -0.851 | 0.287 | 0.236 | 0.011 | 0.013 | 0.013 | 0.197 | 0.72 | 0.76 | 0.86 | 0.87 | 0.83 | 0.83 |
| 5 | 3141651 | rs160906 | -0.737 | -0.868 | 0.285 | 0.22 | -0.006 | -0.013 | -0.013 | 0.234 | 0.19 | 0.21 | 0.1 | 0.08 | 0.13 | 0.06 |
| 5 | 3142898 | rs828416 | 0.258 | -0.765 | 0.281 | 0.252 | 0.009 | 0.001 | 0.001 | 0 | 0.91 | 0.96 | 0.96 | 0.92 | 0.93 | 0.89 |
| 5 | 3143209 | rs160903 | -0.155 | -0.804 | 0.235 | 0.17 | 0.042 | 0.132 | 0.132 | 0.664 | 0.06 | 0.25 | 0.06 | 0.21 | 0.05 | 0.04 |
| 5 | 3145337 | rs1215429 | 1.632 | -0.844 | 0.175 | 0.029 | -0.027 | 0.015 | 0.015 | 0 | 0.63 | 0.52 | 0.76 | 0.5 | 0.75 | 0.78 |
| 5 | 3146907 | rs181860 | -0.731 | -0.78 | -0.123 | 0.142 | 0.012 | -0.01 | -0.01 | 0.015 | 0.94 | 0.97 | 0.97 | 0.92 | 0.96 | 0.94 |
| 5 | 3150973 | rs1661119 | -1.283 | -0.702 | 0.033 | -0.026 | 0.006 | -0.003 | -0.003 | 0 | 0.37 | 0.3 | 0.26 | 0.45 | 0.3 | 0.38 |
| 5 | 3151352 | rs10069129 | 1.442 | -0.582 | -0.655 | -0.682 | 0.009 | -0.068 | -0.068 | 0 | 0.36 | 0.38 | 0.24 | 0.05 | 0.44 | 0.22 |
| 5 | 3152858 | rs468828 | -0.631 | -0.163 | -0.645 | -0.191 | 0.003 | -0.054 | -0.054 | 0 | 0.75 | 0.84 | 0.52 | 0.5 | 0.73 | 0.57 |
| 5 | 3156312 | rs457099 | 0.641 | -0.396 | -0.651 | -0.523 | 0.001 | 0.016 | 0.016 | 0 | 0.53 | 0.52 | 0.35 | 0.26 | 0.52 | 0.3 |
| 5 | 3156508 | rs461401 | 0.062 | -0.361 | -0.703 | -0.407 | -0.002 | 0.003 | 0.003 | 0 | 0.88 | 0.85 | 0.92 | 0.73 | 0.91 | 0.91 |
| 5 | 3161355 | rs9313034 | 1.443 | -0.579 | -0.65 | -0.58 | -0.004 | -0.006 | -0.006 | 0 | 0.58 | 0.42 | 0.71 | 0.64 | 0.67 | 0.61 |
| 5 | 3162157 | rs1691026 | 0.152 | -0.324 | -1.033 | -0.35 | -0.006 | 0.012 | 0.012 | 0 | 0.88 | 0.83 | 0.91 | 0.74 | 0.91 | 0.91 |
| 5 | 3162549 | rs1661106 | 0.161 | -0.325 | -1.032 | -0.35 | -0.006 | 0.012 | 0.012 | 0 | 0.88 | 0.83 | 0.91 | 0.73 | 0.91 | 0.91 |
| 5 | 3163688 | rs2172766 | 1.417 | -0.286 | -1.008 | -0.388 | -0.003 | -0.053 | -0.053 | 0 | 0.19 | 0.13 | 0.45 | 0.46 | 0.21 | 0.38 |
| 5 | 3166087 | rs1494681 | -0.457 | -0.187 | -0.899 | -0.392 | -0.008 | -0.009 | -0.009 | 0 | 0.92 | 0.86 | 0.96 | 0.94 | 0.94 | 0.95 |
| 5 | 3170573 | rs9313036 | 1.128 | 0.593 | -1.294 | -0.83 | -0.002 | 0.009 | 0.009 | 0 | 0.54 | 0.58 | 0.7 | 0.64 | 0.54 | 0.57 |
| 5 | 3170720 | rs1215656 | -0.318 | 0.615 | -1.05 | -0.579 | 0.001 | -0.005 | -0.005 | 0.039 | 0.23 | 0.28 | 0.18 | 0.21 | 0.22 | 0.29 |
| 5 | 3170959 | rs6892918 | 1.031 | 0.862 | -1.195 | -0.641 | -0.002 | -0.027 | -0.027 | 4.218 | 0.43 | 0.51 | 0.28 | 0.19 | 0.39 | 0.22 |
| 5 | 3174785 | rs12657403 | 0.201 | 0.979 | -1.51 | -0.353 | -0.025 | -0.077 | -0.077 | 0.443 | 0.14 | 0.01 | 0.1 | 0.28 | 0.11 | 0.09 |

| | | | | | | | | | | | | | | | | |
|---|---|---|---|---|---|---|---|---|---|---|---|---|---|---|---|---|
| 5 | 3175819 | rs6861203 | 0.222 | 0.973 | -1.348 | -0.456 | 0.002 | -0.051 | -0.051 | 0 | 0.15 | 0.19 | 0.12 | 0.02 | 0.16 | 0.25 |
| 5 | 3177302 | rs16871346 | -0.201 | 1.118 | -1.465 | -0.074 | -0.048 | -0.078 | -0.078 | 0 | 0.15 | 0.01 | 0.14 | 0.28 | 0.17 | 0.11 |
| 5 | 3177876 | rs251833 | -0.781 | 1.025 | -1.483 | -0.232 | -0.049 | -0.089 | -0.089 | 0 | 0.68 | 0.89 | 0.44 | 0.51 | 0.6 | 0.48 |
| 5 | 3179620 | rs369283 | -0.256 | 0.775 | -1.188 | -0.471 | -0.005 | -0.002 | -0.002 | 0.3 | 0.07 | 0.07 | 0.04 | 0.05 | 0.04 | 0.04 |
| 5 | 3181008 | rs1215665 | 0.558 | 1.076 | -0.961 | -0.559 | 0.003 | 0.029 | 0.029 | 0.02 | 0.41 | 0.67 | 0.25 | 0.2 | 0.34 | 0.22 |
| 5 | 3187567 | rs251830 | -0.287 | 1.051 | -1.158 | -0.394 | -0.001 | 0.135 | 0.135 | 0 | 0.49 | 0.25 | 0.68 | 0.75 | 0.55 | 0.71 |
| 5 | 3190290 | rs392439 | 0.523 | 0.963 | -1.483 | -0.204 | -0.086 | -0.1 | -0.1 | 0 | 0.35 | 0.11 | 0.59 | 0.46 | 0.48 | 0.5 |
| 5 | 3192848 | rs431905 | 0.614 | 0.912 | -1.508 | -0.171 | -0.095 | -0.103 | -0.103 | 0.069 | 0.36 | 0.1 | 0.42 | 0.45 | 0.48 | 0.51 |
| 5 | 3193401 | rs427727 | -0.478 | 0.694 | -0.633 | -0.31 | 0.003 | 0.004 | 0.004 | 0 | 0.55 | 0.4 | 0.73 | 0.65 | 0.62 | 0.72 |
| 5 | 3197490 | rs1215693 | 0.619 | 0.807 | -0.693 | -0.173 | -0.055 | -0.062 | -0.062 | 0.721 | 0.36 | 0.15 | 0.42 | 0.44 | 0.49 | 0.51 |
| 5 | 3200876 | rs875852 | 0.508 | -0.045 | -0.434 | -0.78 | -0.036 | -0.096 | -0.096 | 1.723 | 0.16 | 0.4 | 0.11 | 0.02 | 0.08 | 0.17 |
| 5 | 3200928 | rs875853 | 0.508 | -0.045 | -0.434 | -0.78 | -0.036 | -0.096 | -0.096 | 3.943 | 0.16 | 0.4 | 0.11 | 0.02 | 0.08 | 0.17 |
| 5 | 3202136 | rs13171621 | 0.609 | -0.132 | -0.792 | -1.003 | -0.019 | 0.029 | 0.029 | 0 | 0.27 | 0.1 | 0.17 | 0.24 | 0.26 | 0.18 |
| 5 | 3203444 | rs433274 | -0.206 | -0.408 | -0.575 | -0.949 | -0.013 | 0.011 | 0.011 | 0 | 0.81 | 0.71 | 0.73 | 0.75 | 0.87 | 0.85 |
| 5 | 3204386 | rs10474761 | 0.695 | -0.201 | -0.417 | -0.855 | -0.058 | -0.085 | -0.085 | 0 | 0.14 | 0.4 | 0.1 | 0.02 | 0.05 | 0.08 |
| 5 | 3205018 | rs1310088 | -1.217 | 0.251 | -0.841 | -0.911 | 0.009 | 0.003 | 0.003 | 1.181 | 0.7 | 0.62 | 0.61 | 0.39 | 0.63 | 0.66 |
| 5 | 3205290 | rs639617 | 0.41 | 0.071 | -1.011 | -1.094 | 0.006 | 0.024 | 0.024 | 0 | 0.24 | 0.17 | 0.15 | 0.23 | 0.2 | 0.22 |
| 5 | 3205765 | rs396670 | -0.331 | 0.113 | -0.863 | -0.328 | -0.005 | -0.015 | -0.015 | 3.269 | 0.93 | 0.95 | 0.96 | 0.76 | 0.91 | 0.89 |
| 5 | 3209477 | rs1215666 | 1.73 | 0.155 | -0.769 | -0.411 | -0.004 | -0.005 | -0.005 | 1.104 | 0.23 | 0.18 | 0.34 | 0.4 | 0.31 | 0.24 |
| 5 | 3209668 | rs1215667 | 0.773 | 0.236 | -0.815 | -0.443 | -0.012 | -0.005 | -0.005 | 0 | 0.62 | 0.53 | 0.59 | 0.64 | 0.67 | 0.52 |
| 5 | 3210098 | rs1661066 | -0.389 | 0.78 | -1.032 | 0.164 | -0.004 | -0.005 | -0.005 | 0.058 | 0.94 | 0.95 | 0.96 | 0.77 | 0.93 | 0.83 |
| 5 | 3211052 | rs396319 | -0.031 | 0.805 | -1.041 | 0.16 | -0.014 | -0.018 | -0.018 | 0 | 0.93 | 0.95 | 0.96 | 0.77 | 0.91 | 0.82 |
| 5 | 3211391 | rs421093 | 0.775 | 0.719 | -1.044 | 0.051 | -0.005 | -0.011 | -0.011 | 0 | 0.84 | 0.86 | 0.91 | 0.65 | 0.83 | 0.68 |
| 5 | 3213546 | rs6894463 | 0.254 | 0.535 | -1.005 | 0.071 | -0.003 | -0.012 | -0.012 | 0 | 0.08 | 0.04 | 0.05 | 0.14 | 0.07 | 0.14 |
| 5 | 3215231 | rs1691086 | -0.255 | 0.436 | -0.767 | 0.025 | -0.01 | 0.046 | 0.046 | 0 | 0.14 | 0.39 | 0.09 | 0.14 | 0.07 | 0.14 |
| 5 | 3215661 | rs6555181 | 0.304 | 0.731 | -1.93 | -0.111 | -0.069 | 0.021 | 0.021 | 0 | 0.24 | 0.04 | 0.38 | 0.24 | 0.26 | 0.2 |
| 5 | 3217841 | rs392679 | 0.76 | 0.57 | -1.291 | 0.193 | -0.004 | 0.067 | 0.067 | 0 | 0.53 | 0.15 | 0.68 | 0.65 | 0.57 | 0.43 |
| 5 | 3219694 | rs1632064 | 0.73 | 0.51 | -0.114 | 0.74 | 0.001 | 0.131 | 0.131 | 0 | 0.82 | 0.52 | 0.86 | 0.68 | 0.85 | 0.74 |
| 5 | 3225962 | rs4866534 | -0.503 | 0.551 | -0.114 | 0.293 | 0.004 | -0.031 | -0.031 | 0 | 0.36 | 0.49 | 0.31 | 0.32 | 0.34 | 0.43 |
| 5 | 3226096 | rs4866437 | -0.978 | 0.386 | -0.355 | 0.362 | 0 | 0.004 | 0.004 | 1.404 | 0.54 | 0.58 | 0.5 | 0.45 | 0.53 | 0.68 |
| 5 | 3226865 | rs407190 | 1.833 | 0.339 | -0.533 | 0.628 | -0.028 | -0.046 | -0.046 | 0.017 | 0.31 | 0.11 | 0.22 | 0.36 | 0.37 | 0.24 |
| 5 | 3228264 | rs6876947 | 0.514 | 0.503 | -0.038 | 0.441 | -0.001 | -0.009 | -0.009 | 4.022 | 0.28 | 0.32 | 0.2 | 0.19 | 0.23 | 0.21 |
| 5 | 3231351 | rs4866535 | 0.831 | 0.502 | -0.128 | 0.445 | -0.003 | 0.005 | 0.005 | 0.337 | 0.06 | 0 | 0.05 | 0.06 | 0.03 | 0.04 |
| 5 | 3232706 | rs251810 | 1.248 | 0.223 | 0.399 | 0.369 | -0.025 | 0.021 | 0.021 | 0 | 0.73 | 0.42 | 0.63 | 0.67 | 0.83 | 0.67 |
| 5 | 3234220 | rs173726 | 0.897 | 0.295 | 0.268 | 0.41 | -0.001 | 0.001 | 0.001 | 0 | 0.53 | 0.52 | 0.43 | 0.49 | 0.57 | 0.5 |
| 5 | 3237433 | rs415144 | 1.602 | 0.157 | 0.132 | 0.761 | 0.027 | -0.023 | -0.023 | 0 | 0.52 | 0.12 | 0.4 | 0.52 | 0.61 | 0.42 |
| 5 | 3238376 | rs7717593 | -0.267 | 0.167 | 0.523 | 0.723 | -0.008 | -0.026 | -0.026 | 0.062 | 0.27 | 0.39 | 0.28 | 0.13 | 0.25 | 0.31 |
| 5 | 3238615 | rs441638 | 1.31 | 0.195 | 0.21 | 0.783 | 0.019 | 0.004 | 0.004 | 0 | 0.57 | 0.28 | 0.43 | 0.65 | 0.65 | 0.59 |
| 5 | 3243988 | rs7444147 | 0.826 | 0.205 | -0.172 | 0.745 | -0.002 | -0.001 | -0.001 | 0.301 | 0.18 | 0.21 | 0.21 | 0.12 | 0.19 | 0.23 |
| 5 | 3244550 | rs1691018 | 2.132 | 0.186 | -0.378 | 0.607 | -0.043 | 0 | 0 | 0 | 0.37 | 0.11 | 0.26 | 0.34 | 0.44 | 0.26 |
| 5 | 3245702 | rs427867 | -0.628 | 0.11 | -0.16 | 0.494 | -0.007 | 0.007 | 0.007 | 0 | 0.33 | 0.39 | 0.48 | 0.34 | 0.28 | 0.32 |
| 5 | 3245982 | rs611396 | 1.061 | 0.243 | -0.712 | 0.552 | 0.037 | 0.003 | 0.003 | 0 | 0.57 | 0.17 | 0.43 | 0.66 | 0.66 | 0.6 |
| 5 | 3246919 | rs1691016 | 0.666 | 0.218 | -0.704 | 0.58 | -0.033 | -0.004 | -0.004 | 0 | 0.76 | 0.37 | 0.66 | 0.78 | 0.88 | 0.84 |
| 5 | 3250572 | rs1691101 | 1.224 | 0.64 | -1.261 | 0.622 | -0.051 | -0.049 | -0.049 | 0 | 0.41 | 0.14 | 0.29 | 0.46 | 0.49 | 0.42 |
| 5 | 3253177 | rs1494678 | 0.011 | 0.762 | -1.139 | 0.702 | -0.042 | -0.083 | -0.083 | 0.168 | 0.2 | 0.03 | 0.3 | 0.33 | 0.2 | 0.22 |
| 5 | 3253428 | rs1441540 | -0.83 | 0.753 | -1.218 | 0.661 | -0.045 | -0.118 | -0.118 | 0.146 | 0.29 | 0.05 | 0.36 | 0.46 | 0.27 | 0.4 |
| 5 | 3254137 | rs903405 | 0.909 | 0.604 | -0.427 | -0.092 | -0.003 | -0.034 | -0.034 | 0.021 | 0.67 | 0.42 | 0.68 | 0.84 | 0.69 | 0.57 |
| 5 | 3254413 | rs16871518 | -0.217 | 0.87 | -0.102 | -0.199 | 0.002 | -0.015 | -0.015 | 0 | 0.2 | 0.22 | 0.23 | 0.07 | 0.26 | 0.29 |
| 5 | 3258039 | rs16871519 | -0.808 | 1.252 | -0.09 | -0.179 | -0.008 | 0.022 | 0.022 | 0.65 | 0.3 | 0.2 | 0.27 | 0.22 | 0.33 | 0.43 |
| 5 | 3261155 | rs1441537 | -0.333 | 1.136 | 0.343 | -0.288 | 0.001 | -0.009 | -0.009 | 0 | 0.19 | 0.2 | 0.21 | 0.07 | 0.22 | 0.28 |
| 5 | 3265689 | rs1441534 | -0.883 | 0.913 | 0.028 | -0.432 | -0.001 | -0.022 | -0.022 | 0 | 0.3 | 0.36 | 0.31 | 0.07 | 0.3 | 0.27 |
| 5 | 3265846 | rs1441533 | -1.16 | 0.927 | 0.28 | -0.466 | 0.002 | -0.09 | -0.09 | 0.985 | 0.33 | 0.5 | 0.35 | 0.08 | 0.35 | 0.3 |
| 5 | 3266396 | rs10038264 | -0.836 | 0.851 | 0.122 | -0.396 | -0.002 | -0.054 | -0.054 | 0.31 | 0.31 | 0.43 | 0.32 | 0.07 | 0.3 | 0.28 |
| 5 | 3269778 | rs7706315 | 0.536 | 0.87 | 0.058 | 0.368 | 0.004 | 0.006 | 0.006 | 0.067 | 0.61 | 0.59 | 0.69 | 0.55 | 0.58 | 0.47 |
| 5 | 3269938 | rs16871560 | -0.407 | 0.582 | -0.041 | 0.245 | -0.001 | 0.002 | 0.002 | 0 | 0.29 | 0.31 | 0.22 | 0.4 | 0.25 | 0.35 |
| 5 | 3271982 | rs10063962 | 0.954 | 0.413 | -0.107 | 0.21 | 0.008 | 0.006 | 0.006 | 0.052 | 0.54 | 0.37 | 0.61 | 0.55 | 0.55 | 0.47 |
| 5 | 3272039 | rs17571538 | 0.658 | 0.56 | -0.534 | 0.203 | -0.019 | 0.023 | 0.023 | 0.456 | 0.16 | 0.06 | 0.17 | 0.07 | 0.21 | 0.24 |
| 5 | 3273318 | rs4866539 | -0.52 | 0.507 | -0.373 | 0.421 | -0.005 | -0.028 | -0.028 | 0.717 | 0.79 | 0.89 | 0.85 | 0.74 | 0.79 | 0.73 |
| 5 | 3281517 | rs10041745 | 1.22 | 0.723 | -0.574 | 0.509 | -0.004 | -0.005 | -0.005 | 0 | 0.28 | 0.29 | 0.26 | 0.34 | 0.32 | 0.23 |
| 5 | 3282671 | rs4866542 | -0.954 | 0.408 | -0.696 | 0.548 | 0.001 | 0.002 | 0.002 | 0 | 0.91 | 0.9 | 0.76 | 0.78 | 0.94 | 0.86 |
| 5 | 3285526 | rs4546389 | -0.071 | 0.357 | -0.656 | 0.271 | 0.008 | 0.021 | 0.021 | 0 | 0.45 | 0.59 | 0.35 | 0.37 | 0.4 | 0.41 |
| 5 | 3286292 | rs4866544 | 1.796 | 0.512 | -0.779 | 0.286 | -0.014 | -0.008 | -0.008 | 0 | 0.27 | 0.21 | 0.21 | 0.32 | 0.33 | 0.21 |
| 5 | 3286564 | rs4242087 | -0.454 | 0.496 | -0.009 | 0.35 | -0.005 | -0.021 | -0.021 | 0 | 0.6 | 0.56 | 0.72 | 0.73 | 0.65 | 0.65 |
| 5 | 3286623 | rs4866546 | 0.581 | 0.495 | -0.13 | 0.356 | 0.002 | 0.003 | 0.003 | 0 | 0.47 | 0.34 | 0.4 | 0.43 | 0.55 | 0.46 |
| 5 | 3286715 | rs4866547 | -0.742 | 0.769 | 0.266 | 0.354 | 0.016 | -0.091 | -0.091 | 0 | 0.84 | 0.68 | 0.86 | 1 | 0.85 | 0.9 |
| 5 | 3287958 | rs4476727 | 0.203 | 1.022 | -0.075 | 0.523 | -0.007 | -0.007 | -0.007 | 0 | 0.73 | 0.79 | 0.59 | 0.69 | 0.72 | 0.71 |
| 5 | 3288988 | rs6893182 | 0.528 | 0.734 | -0.801 | 0.387 | -0.012 | -0.039 | -0.039 | 0 | 0.41 | 0.13 | 0.3 | 0.38 | 0.43 | 0.5 |
| 5 | 3292902 | rs10045706 | 0.479 | 0.934 | -0.023 | 0.667 | -0.001 | -0.003 | -0.003 | 0 | 0.28 | 0.26 | 0.22 | 0.33 | 0.28 | 0.25 |
| 5 | 3302781 | rs4505956 | -0.525 | 0.904 | 0.393 | 0.723 | 0.005 | 0.025 | 0.025 | 0 | 0.22 | 0.3 | 0.43 | 0.37 | 0.3 | 0.33 |

| | | | | | | | | | | | | | | | | |
|---|---|---|---|---|---|---|---|---|---|---|---|---|---|---|---|---|
| 5 | 3308312 | rs4866550 | 0.598 | 0.569 | 0.337 | 0.612 | -0.019 | -0.036 | -0.036 | 1.071 | 0.64 | 0.52 | 0.64 | 0.79 | 0.65 | 0.7 |
| 5 | 3310539 | rs7716659 | 0.877 | 0.592 | 0.066 | 0.825 | 0.005 | -0.015 | -0.015 | 0.006 | 0.59 | 0.3 | 0.42 | 0.54 | 0.57 | 0.56 |
| 5 | 3311493 | rs11745750 | 1.904 | 0.156 | 0.939 | 1.184 | -0.057 | -0.079 | -0.079 | 0 | 0.88 | 0.36 | 0.87 | 1 | 0.96 | 0.93 |
| 5 | 3313928 | rs10866553 | -0.769 | 0.51 | 1.022 | 1.149 | -0.001 | 0.006 | 0.006 | 0 | 0.4 | 0.51 | 0.51 | 0.44 | 0.42 | 0.39 |
| 5 | 3316959 | rs11746449 | -0.124 | 0.55 | 1.366 | 1.242 | 0.001 | 0.044 | 0.044 | 0 | 0.33 | 0.2 | 0.34 | 0.18 | 0.36 | 0.25 |
| 5 | 3317260 | rs6896919 | 0.758 | 0.539 | 1.451 | 1.244 | 0.003 | -0.006 | -0.006 | 0.002 | 0.58 | 0.33 | 0.45 | 0.56 | 0.58 | 0.61 |
| 5 | 3318642 | rs9313047 | 0.526 | 0.59 | 1.478 | 1.26 | -0.001 | -0.003 | -0.003 | 0 | 0.61 | 0.62 | 0.47 | 0.56 | 0.58 | 0.62 |
| 5 | 3331005 | rs6861409 | 0.304 | 0.953 | 1.855 | 1.444 | 0 | -0.001 | -0.001 | 0.005 | 0.66 | 0.36 | 0.48 | 0.68 | 0.63 | 0.72 |
| 5 | 3333684 | rs13171721 | 0.334 | 0.992 | 1.911 | 1.42 | -0.002 | 0.01 | 0.01 | 0 | 0.27 | 0.23 | 0.27 | 0.21 | 0.31 | 0.15 |
| 5 | 3333760 | rs7718013 | 2.16 | 1.012 | 1.921 | 1.443 | 0.078 | 0.105 | 0.105 | 0.048 | 0.93 | 0.67 | 0.75 | 0.88 | 0.94 | 0.88 |
| 5 | 3337194 | rs7721657 | -0.036 | 0.995 | 1.872 | 1.446 | -0.003 | -0.018 | -0.018 | 0 | 0.31 | 0.44 | 0.42 | 0.26 | 0.32 | 0.24 |
| 5 | 3337359 | rs7735206 | -0.556 | 0.993 | 1.9 | 1.613 | 0.003 | 0.017 | 0.017 | 0 | 0.37 | 0.53 | 0.52 | 0.46 | 0.38 | 0.32 |
| 5 | 3337491 | rs7711995 | -0.217 | 1.059 | 1.925 | 1.544 | 0.004 | -0.021 | -0.021 | 0 | 0.33 | 0.52 | 0.47 | 0.26 | 0.35 | 0.28 |
| 5 | 3338013 | rs11746659 | 0.408 | 1.064 | 1.594 | 1.478 | -0.002 | 0.009 | 0.009 | 0 | 0.27 | 0.21 | 0.27 | 0.2 | 0.31 | 0.16 |
| 5 | 3338074 | rs10045889 | -0.29 | 1.11 | 1.487 | 1.511 | 0.002 | -0.018 | -0.018 | 0 | 0.71 | 0.65 | 0.68 | 0.8 | 0.67 | 0.83 |
| 5 | 3338456 | rs13360235 | 0.459 | 1.076 | 1.439 | 1.319 | -0.002 | 0.007 | 0.007 | 0.06 | 0.63 | 0.34 | 0.48 | 0.66 | 0.61 | 0.69 |
| 5 | 3339142 | rs7703994 | -0.167 | 0.999 | 1.132 | 1.121 | -0.003 | -0.06 | -0.06 | 0.063 | 0.71 | 0.57 | 0.71 | 0.86 | 0.69 | 0.84 |
| 5 | 3343024 | rs6860525 | -1.851 | 0.927 | 1.267 | 1.531 | 0.013 | 0.073 | 0.073 | 0 | 0.07 | 0.17 | 0.11 | 0.16 | 0.07 | 0.07 |
| 5 | 3347836 | rs10475202 | -0.681 | 1.034 | 0.827 | 1.275 | -0.002 | -0.003 | -0.003 | 0 | 0.37 | 0.52 | 0.41 | 0.34 | 0.38 | 0.36 |
| 5 | 3349583 | rs6869583 | -0.133 | 1.139 | -0.122 | 1.339 | -0.023 | -0.04 | -0.04 | 0.114 | 0.31 | 0.07 | 0.28 | 0.29 | 0.33 | 0.31 |
| 5 | 3356538 | rs4521479 | -0.415 | 1.068 | -0.045 | 1.255 | -0.001 | 0.002 | 0.002 | 0 | 0.36 | 0.38 | 0.39 | 0.4 | 0.38 | 0.35 |
| 5 | 3357818 | rs4602632 | -0.425 | 1.066 | -0.04 | 1.255 | 0.001 | -0.007 | -0.007 | 0 | 0.35 | 0.33 | 0.35 | 0.4 | 0.36 | 0.34 |
| 5 | 3360109 | rs10073886 | -0.311 | 1.069 | -0.506 | 1.148 | -0.001 | -0.008 | -0.008 | 0.523 | 0.36 | 0.3 | 0.39 | 0.38 | 0.38 | 0.35 |
| 5 | 3362336 | rs4397137 | 0.81 | 1.611 | -0.342 | 1.097 | 0.007 | 0.004 | 0.004 | 0.025 | 0.81 | 0.79 | 0.7 | 0.76 | 0.73 | 0.84 |
| 5 | 3364201 | rs10050840 | 1.328 | 1.091 | 0.924 | 0.667 | 0.078 | 0.038 | 0.038 | 4.726 | 0.94 | 0.45 | 0.92 | 1 | 0.96 | 1 |
| 5 | 3365033 | rs4242090 | 1.857 | 1.172 | 0.842 | 0.65 | 0.003 | -0.02 | -0.02 | 0.124 | 0.8 | 0.3 | 0.84 | 0.88 | 0.83 | 0.83 |
| 5 | 3366480 | rs4147089 | 0.336 | 1.237 | 0.188 | 0.374 | -0.015 | 0.015 | 0.015 | 0.003 | 0.39 | 0.33 | 0.52 | 0.26 | 0.52 | 0.41 |
| 5 | 3377309 | rs4866559 | -0.059 | 1.147 | 0.46 | 0.092 | -0.043 | 0.091 | 0.091 | 0.549 | 0.55 | 0.19 | 0.42 | 0.75 | 0.44 | 0.59 |
| 5 | 3380046 | rs251433 | -0.028 | 1.436 | 0.337 | -0.004 | -0.012 | 0.029 | 0.029 | 0 | 0.43 | 0.35 | 0.5 | 0.25 | 0.56 | 0.38 |
| 5 | 3382263 | rs16871658 | 0.487 | 1.141 | -0.617 | 0.059 | -0.034 | -0.013 | -0.013 | 0 | 0.17 | 0.04 | 0.13 | 0.16 | 0.17 | 0.13 |
| 5 | 3383339 | rs426126 | 0.533 | 1.343 | -0.551 | -0.204 | -0.022 | 0.088 | 0.088 | 0 | 0.58 | 0.25 | 0.53 | 0.21 | 0.57 | 0.43 |
| 5 | 3383382 | rs395395 | 0.445 | 1.342 | 0.136 | 0.166 | 0.024 | 0.047 | 0.047 | 0 | 0.8 | 0.69 | 0.81 | 0.71 | 0.76 | 0.66 |
| 5 | 3384836 | rs32689 | 0.336 | 1.302 | 0.471 | -0.184 | 0.066 | -0.025 | -0.025 | 0 | 0.86 | 0.48 | 0.82 | 0.95 | 0.83 | 0.81 |
| 5 | 3386191 | rs32687 | -0.553 | 1.278 | -0.114 | -0.103 | 0.021 | -0.007 | -0.007 | 0 | 0.9 | 0.84 | 0.89 | 0.95 | 0.88 | 0.81 |
| 5 | 3389796 | rs6881435 | -0.205 | 1.013 | 0.046 | -0.036 | -0.005 | -0.007 | -0.007 | 0 | 0.31 | 0.45 | 0.32 | 0.75 | 0.26 | 0.35 |
| 5 | 3393547 | rs924610 | 0.121 | 0.759 | -1.01 | -0.331 | -0.045 | -0.034 | -0.034 | 0.465 | 0.42 | 0.14 | 0.43 | 0.59 | 0.44 | 0.42 |
| 5 | 3395518 | rs32712 | -0.4 | 0.83 | -0.828 | -0.693 | 0.021 | 0.051 | 0.051 | 0 | 0.54 | 0.37 | 0.57 | 0.88 | 0.58 | 0.7 |
| 5 | 3397467 | rs6869954 | 0.605 | 0.956 | -0.779 | -0.66 | -0.02 | -0.03 | -0.03 | 0.173 | 0.33 | 0.2 | 0.37 | 0.61 | 0.32 | 0.39 |
| 5 | 3400160 | rs4866443 | 1.171 | 0.803 | -0.648 | -1.015 | 0.011 | 0.022 | 0.022 | 0 | 0.27 | 0.22 | 0.29 | 0.09 | 0.23 | 0.21 |
| 5 | 3411611 | rs402272 | -0.8 | 0.863 | -0.661 | -1.002 | -0.018 | -0.058 | -0.058 | 0.118 | 0.68 | 0.41 | 0.67 | 0.91 | 0.71 | 0.78 |
| 5 | 3415311 | rs7703631 | 1.432 | 0.827 | -0.664 | -0.952 | -0.002 | 0.047 | 0.047 | 0 | 0.14 | 0.04 | 0.1 | 0.04 | 0.13 | 0.09 |
| 5 | 3415406 | rs6555203 | 0.208 | 0.776 | -0.635 | -0.919 | 0.004 | 0.032 | 0.032 | 0 | 0.47 | 0.23 | 0.51 | 0.65 | 0.48 | 0.46 |
| 5 | 3421142 | rs376855 | 0.671 | 0.692 | -0.673 | -0.761 | -0.004 | -0.01 | -0.01 | 0 | 0.21 | 0.18 | 0.17 | 0.27 | 0.24 | 0.3 |
| 5 | 3428925 | rs17683976 | 0.865 | 0.716 | -0.727 | -0.905 | -0.037 | -0.028 | -0.028 | 1.944 | 0.34 | 0.2 | 0.42 | 0.61 | 0.37 | 0.4 |
| 5 | 3429107 | rs11742901 | 1.396 | 0.593 | -0.42 | -0.954 | 0.001 | -0.053 | -0.053 | 4.403 | 0.14 | 0.22 | 0.11 | 0.04 | 0.13 | 0.09 |
| 5 | 3435141 | rs251444 | -0.85 | 0.591 | -0.487 | -0.822 | -0.017 | -0.032 | -0.032 | 1.003 | 0.7 | 0.62 | 0.7 | 0.88 | 0.74 | 0.79 |
| 5 | 3436677 | rs7732434 | 0.703 | 1.024 | -1.063 | -0.387 | -0.027 | -0.029 | -0.029 | 0.16 | 0.3 | 0.2 | 0.38 | 0.58 | 0.32 | 0.4 |
| 5 | 3445667 | rs863794 | -0.277 | 0.415 | -0.915 | -0.643 | 0.023 | 0.037 | 0.037 | 0 | 0.55 | 0.38 | 0.58 | 0.85 | 0.61 | 0.7 |
| 5 | 3449132 | rs436417 | 0.37 | 1.055 | -1.082 | -0.514 | -0.025 | -0.002 | -0.002 | 0 | 0.26 | 0.15 | 0.23 | 0.26 | 0.32 | 0.39 |
| 5 | 3451906 | rs32723 | -0.547 | 0.679 | -0.71 | -0.752 | 0.002 | 0.002 | 0.002 | 0 | 0.61 | 0.59 | 0.55 | 0.44 | 0.59 | 0.66 |
| 5 | 3452780 | rs876023 | 0.464 | 0.409 | -0.893 | -0.575 | 0.02 | 0.022 | 0.022 | 0 | 0.29 | 0.23 | 0.27 | 0.12 | 0.21 | 0.19 |
| 5 | 3454482 | rs4866447 | 0.564 | 1.345 | -0.926 | 0.37 | -0.04 | -0.053 | -0.053 | 0 | 0.19 | 0.09 | 0.25 | 0.41 | 0.24 | 0.2 |
| 5 | 3462896 | rs32736 | 0.07 | 0.913 | -1.201 | 0.178 | -0.075 | -0.022 | -0.022 | 0 | 0.31 | 0.05 | 0.25 | 0.3 | 0.34 | 0.21 |
| 5 | 3463708 | rs16871863 | -0.057 | 1.55 | -0.494 | 0.296 | 0.039 | 0.022 | 0.022 | 0.038 | 0.07 | 0.16 | 0.06 | 0.15 | 0.1 | 0.19 |
| 5 | 3465678 | rs396398 | 1.065 | 1.599 | 0.798 | 0.179 | 0.131 | 0.038 | 0.038 | 2.088 | 0.88 | 0.49 | 0.88 | 0.85 | 0.83 | 0.81 |
| 5 | 3466243 | rs1847011 | 0.27 | 1.518 | 1.13 | 0.183 | 0.097 | 0.065 | 0.065 | 0.314 | 0.92 | 0.69 | 0.93 | 0.85 | 0.89 | 0.81 |
| 5 | 3483595 | rs11133987 | 0.642 | 1.5 | 1.254 | 0.175 | 0.108 | 0.072 | 0.072 | 0.483 | 0.91 | 0.63 | 0.92 | 0.85 | 0.9 | 0.82 |
| 5 | 3502963 | rs17686103 | 0.664 | 1.584 | 0.67 | -0.054 | -0.117 | 0.116 | 0.116 | 0 | 0.18 | 0.01 | 0.21 | 0 | 0.3 | 0.11 |
| 5 | 3513485 | rs698147 | -0.719 | 0.937 | 0.982 | 0.351 | -0.013 | -0.012 | -0.012 | 1.209 | 0.56 | 0.75 | 0.45 | 0.6 | 0.5 | 0.63 |
| 5 | 3515183 | rs39620 | 0.258 | 1.019 | 0.943 | 0.384 | 0.04 | 0.029 | 0.029 | 0 | 0.08 | 0.15 | 0.06 | 0.16 | 0.1 | 0.18 |
| 5 | 3520064 | rs828544 | -0.884 | 1.079 | 0.539 | 0.477 | -0.032 | -0.006 | -0.006 | 0 | 0.57 | 0.82 | 0.47 | 0.63 | 0.52 | 0.63 |
| 5 | 3524832 | rs11957094 | 0.002 | 1.182 | 0.987 | 0.388 | -0.003 | 0 | 0 | 0 | 0.41 | 0.4 | 0.34 | 0.45 | 0.32 | 0.44 |
| 5 | 3525440 | rs710987 | -0.936 | 1.19 | 0.622 | 0.389 | -0.029 | -0.007 | -0.007 | 0 | 0.52 | 0.82 | 0.45 | 0.6 | 0.49 | 0.62 |
| 5 | 3528233 | rs698144 | -0.77 | 1.099 | 0.635 | 0.402 | -0.014 | 0.005 | 0.005 | 0.093 | 0.5 | 0.81 | 0.42 | 0.6 | 0.42 | 0.62 |
| 5 | 3533532 | rs2115350 | -0.016 | 1.045 | 0.641 | 0.407 | 0.019 | -0.014 | -0.014 | 0 | 0.39 | 0.25 | 0.33 | 0.45 | 0.31 | 0.44 |
| 5 | 3539652 | rs710973 | -1.549 | 0.706 | 0.858 | 0.384 | 0.103 | 0.036 | 0.036 | 2.06 | 0.13 | 0.56 | 0.13 | 0.15 | 0.17 | 0.19 |
| 5 | 3540098 | rs4866451 | 0.426 | -0.714 | 0.3 | 0.358 | 0.049 | -0.012 | -0.012 | 0 | 0.57 | 0.33 | 0.72 | 0.5 | 0.68 | 0.53 |
| 5 | 3542548 | rs1697997 | -2.097 | 0.603 | 1.475 | 0.413 | 0.035 | -0.05 | -0.05 | 0 | 0.1 | 0.59 | 0.06 | 0.01 | 0.08 | 0.03 |
| 5 | 3556260 | rs1664738 | -0.96 | 0.09 | 1.715 | 0.775 | -0.016 | 0.014 | 0.014 | 0 | 0.33 | 0.51 | 0.33 | 0.43 | 0.31 | 0.5 |

| | | | | | | | | | | | | | | | | |
|---|---|---|---|---|---|---|---|---|---|---|---|---|---|---|---|---|
| 5 | 3559242 | rs17687915 | -0.088 | 0.179 | 1.389 | 0.893 | -0.003 | 0 | 0 | 0 | 0.23 | 0.18 | 0.29 | 0.26 | 0.25 | 0.29 |
| 5 | 3559973 | rs1428739 | -0.993 | 0.245 | 1.3 | 0.852 | 0.005 | 0.006 | 0.006 | 0 | 0.32 | 0.34 | 0.35 | 0.42 | 0.36 | 0.51 |
| 5 | 3566024 | rs828311 | -0.924 | 0.039 | 1.107 | 0.934 | -0.002 | 0.006 | 0.006 | 0 | 0.32 | 0.57 | 0.34 | 0.42 | 0.31 | 0.51 |
| 5 | 3569556 | rs2195412 | -1.118 | 0.236 | 1.084 | 1.045 | -0.004 | 0.052 | 0.052 | 0 | 0.06 | 0.16 | 0.04 | 0.15 | 0.05 | 0.22 |
| 5 | 3573395 | rs828302 | -1.734 | 0.138 | 0.955 | 1.158 | -0.012 | 0.199 | 0.199 | 0.144 | 0.1 | 0.4 | 0.05 | 0.27 | 0.06 | 0.25 |
| 5 | 3574619 | rs4866585 | -0.263 | 0.298 | -0.108 | 0.641 | -0.005 | 0.005 | 0.005 | 0.108 | 0.4 | 0.6 | 0.4 | 0.28 | 0.42 | 0.16 |
| 5 | 3575370 | rs12517827 | 1.263 | 0.25 | -0.404 | 0.503 | -0.008 | -0.006 | -0.006 | 0.073 | 0.48 | 0.15 | 0.44 | 0.45 | 0.51 | 0.35 |
| 5 | 3575637 | rs11746591 | 1.264 | 0.253 | -0.431 | 0.504 | -0.011 | -0.008 | -0.008 | 0 | 0.48 | 0.15 | 0.44 | 0.44 | 0.51 | 0.35 |
| 5 | 3579516 | rs10067415 | 1.26 | 0.491 | -0.107 | 0.618 | 0.003 | 0.009 | 0.009 | 0 | 0.49 | 0.18 | 0.45 | 0.45 | 0.51 | 0.35 |
| 5 | 3580400 | rs828316 | -0.776 | 0.401 | -0.184 | 0.845 | 0.02 | -0.04 | -0.04 | 0 | 0.89 | 0.97 | 0.91 | 0.78 | 0.93 | 0.74 |
| 5 | 3581760 | rs10042985 | 1.599 | 0.414 | -0.15 | 0.787 | 0.01 | 0.016 | 0.016 | 0 | 0.5 | 0.21 | 0.45 | 0.45 | 0.51 | 0.35 |
| 5 | 3584368 | rs860439 | -1.32 | 1.281 | 0.435 | 0.271 | 0.009 | -0.008 | -0.008 | 0 | 0.35 | 0.44 | 0.41 | 0.33 | 0.35 | 0.38 |
| 5 | 3591916 | rs710998 | -1.198 | 1.929 | -0.37 | 0.65 | 0.063 | 0.142 | 0.142 | 0.332 | 0.06 | 0.29 | 0.1 | 0.22 | 0.08 | 0.06 |
| 5 | 3597830 | rs2166365 | -0.331 | 2.063 | -0.961 | 0.638 | -0.091 | -0.005 | -0.005 | 0 | 0.17 | 0.01 | 0.12 | 0.1 | 0.21 | 0.17 |
| 5 | 3599320 | rs861185 | -0.746 | 1.504 | -0.102 | 0.741 | 0.019 | 0.008 | 0.008 | 0.093 | 0.16 | 0.25 | 0.35 | 0.23 | 0.18 | 0.21 |
| 5 | 3610658 | rs1373997 | 0.233 | 1.461 | -0.415 | 0.673 | 0 | 0 | 0 | 0 | 0.83 | 0.85 | 0.9 | 0.82 | 0.74 | 0.89 |
| 5 | 3612448 | rs13173543 | -0.557 | 1.537 | 0.075 | 0.264 | 0.025 | -0.039 | -0.039 | 0 | 0.07 | 0.13 | 0.07 | 0 | 0.1 | 0.03 |
| 5 | 3614352 | rs13170992 | -0.545 | 1.603 | 0.096 | 0.258 | 0.014 | -0.034 | -0.034 | 0 | 0.07 | 0.13 | 0.07 | 0 | 0.1 | 0.04 |
| 5 | 3617040 | rs6555208 | -0.613 | 0.923 | -0.357 | 0.395 | 0.042 | -0.001 | -0.001 | 0 | 0.56 | 0.8 | 0.54 | 0.6 | 0.64 | 0.55 |
| 5 | 3618765 | rs7380187 | -0.622 | 0.637 | -0.402 | 0.585 | 0.033 | -0.047 | -0.047 | 0 | 0.55 | 0.87 | 0.54 | 0.49 | 0.64 | 0.52 |
| 5 | 3623182 | rs7722069 | -0.019 | 0.987 | 0.121 | 0.39 | 0.016 | -0.037 | -0.037 | 0 | 0.24 | 0.3 | 0.18 | 0.08 | 0.32 | 0.18 |
| 5 | 3629141 | rs12516914 | 1.349 | 0.427 | -0.715 | 0.352 | -0.048 | -0.087 | -0.087 | 0 | 0.18 | 0.01 | 0.34 | 0.25 | 0.19 | 0.2 |
| 5 | 3636551 | rs4866455 | -0.967 | 0.699 | -0.187 | 0.349 | 0.035 | -0.077 | -0.077 | 0 | 0.55 | 0.9 | 0.53 | 0.58 | 0.63 | 0.51 |
| 5 | 3642021 | rs2166362 | 0.413 | 0.788 | 0.357 | 0.4 | 0.001 | 0.015 | 0.015 | 0 | 0.27 | 0.33 | 0.32 | 0.51 | 0.25 | 0.32 |
| 5 | 3643741 | rs11133999 | -0.783 | 0.596 | 0.419 | 0.351 | 0.049 | -0.007 | -0.007 | 0 | 0.53 | 0.71 | 0.52 | 0.58 | 0.63 | 0.5 |
| 5 | 3651176 | rs6870795 | 0.261 | 0.242 | 0.549 | 0.329 | -0.004 | -0.028 | -0.028 | 0.789 | 0.05 | 0.13 | 0.04 | 0 | 0.03 | 0.03 |
| 5 | 3655804 | rs4866458 | 0.225 | 0.508 | 0.073 | 0.791 | -0.004 | 0 | 0 | 0 | 0.22 | 0.25 | 0.21 | 0.31 | 0.15 | 0.27 |
| 5 | 3657787 | rs10065876 | 0.531 | 0.415 | -0.188 | 0.275 | -0.005 | 0.035 | 0.035 | 0.047 | 0.64 | 0.33 | 0.76 | 0.79 | 0.62 | 0.78 |
| 5 | 3662460 | rs4866595 | 0.154 | 0.28 | 0.57 | 0.403 | 0.051 | -0.033 | -0.033 | 0 | 0.74 | 0.45 | 0.81 | 0.84 | 0.64 | 0.75 |
| 5 | 3663183 | rs1502645 | 0.766 | 0.354 | 0.795 | 0.379 | 0.109 | 0.155 | 0.155 | 1.346 | 0.94 | 0.56 | 0.95 | 0.92 | 0.94 | 0.99 |
| 5 | 3663741 | rs7733481 | 0.766 | 0.354 | 0.79 | 0.379 | 0.112 | 0.159 | 0.159 | 1.346 | 0.94 | 0.56 | 0.95 | 0.92 | 0.93 | 0.99 |
| 5 | 3670973 | rs6883836 | 0.238 | 0.361 | 0.573 | 0.357 | -0.034 | 0.034 | 0.034 | 0 | 0.2 | 0.1 | 0.14 | 0.09 | 0.3 | 0.24 |
| 5 | 3671006 | rs1502644 | 0.987 | -0.019 | 0.877 | 0.485 | 0.11 | 0.156 | 0.156 | 1.193 | 0.94 | 0.56 | 0.95 | 0.92 | 0.94 | 0.99 |
| 5 | 3671350 | rs16872244 | -0.195 | -0.458 | 0.202 | 0.511 | 0.044 | -0.003 | -0.003 | 0 | 0.13 | 0.05 | 0.05 | 0.12 | 0.05 | 0.04 |
| 5 | 3675933 | rs7711236 | 0.597 | -0.47 | -0.315 | 0.711 | 0 | 0.009 | 0.009 | 0 | 0.38 | 0.34 | 0.56 | 0.25 | 0.36 | 0.42 |
| 5 | 3678658 | rs2398622 | -0.042 | 0.068 | -0.884 | 0.786 | 0.012 | -0.026 | -0.026 | 0 | 0.55 | 0.71 | 0.65 | 0.46 | 0.41 | 0.46 |
| 5 | 3684777 | rs6892937 | 0.575 | -0.188 | -0.628 | 0.954 | -0.003 | 0.001 | 0.001 | 0 | 0.74 | 0.76 | 0.52 | 0.32 | 0.82 | 0.57 |
| 5 | 3687018 | rs7728616 | 0.667 | 0.271 | -0.615 | 0.696 | 0.001 | 0.038 | 0.038 | 0 | 0.51 | 0.33 | 0.29 | 0.41 | 0.51 | 0.55 |
| 5 | 3687568 | rs4866462 | -0.724 | 0.273 | -0.367 | 0.878 | -0.009 | 0.005 | 0.005 | 0 | 0.47 | 0.71 | 0.7 | 0.51 | 0.48 | 0.44 |
| 5 | 3689527 | rs10071484 | -0.293 | 0.245 | 0.08 | 0.532 | -0.001 | -0.098 | -0.098 | 0 | 0.25 | 0.52 | 0.21 | 0.05 | 0.23 | 0.11 |
| 5 | 3690564 | rs13173587 | 0.274 | 0.155 | 0.008 | 0.391 | -0.023 | 0.058 | 0.058 | 0 | 0.19 | 0.06 | 0.17 | 0.01 | 0.22 | 0.1 |
| 5 | 3692378 | rs6866062 | -0.094 | 0.158 | 0.476 | 0.394 | 0.001 | -0.083 | -0.083 | 0 | 0.23 | 0.26 | 0.18 | 0.01 | 0.22 | 0.1 |
| 5 | 3695920 | rs4866604 | -1.075 | 0.157 | -0.02 | 0.827 | 0.02 | -0.019 | -0.019 | 0 | 0.42 | 0.71 | 0.44 | 0.44 | 0.37 | 0.37 |
| 5 | 3698654 | rs10042356 | -1.132 | 0.667 | 0.241 | 1.688 | 0.045 | 0.256 | 0.256 | 0 | 0.05 | 0.12 | 0.22 | 0.42 | 0.11 | 0.21 |
| 5 | 3700168 | rs13169861 | -0.474 | 0.73 | -0.486 | 0.796 | -0.008 | 0.065 | 0.065 | 4.92 | 0.17 | 0.06 | 0.13 | 0.01 | 0.17 | 0.1 |
| 5 | 3703076 | rs1039460 | 0.728 | 0.667 | -0.437 | 0.851 | -0.001 | -0.016 | -0.016 | 0 | 0.78 | 0.24 | 0.74 | 0.6 | 0.79 | 0.75 |
| 5 | 3706050 | rs954767 | -0.942 | 0.588 | -0.202 | 0.786 | -0.001 | -0.006 | -0.006 | 0.246 | 0.27 | 0.31 | 0.35 | 0.17 | 0.28 | 0.24 |
| 5 | 3707452 | rs12659333 | 0.869 | 0.576 | -0.654 | 0.898 | -0.04 | -0.003 | -0.003 | 0 | 0.11 | 0 | 0.08 | 0.05 | 0.11 | 0.06 |
| 5 | 3710450 | rs4866607 | -0.356 | 0.722 | -0.192 | 1.102 | 0.053 | 0.234 | 0.234 | 0 | 0.05 | 0.12 | 0.24 | 0.35 | 0.12 | 0.2 |
| 5 | 3710645 | rs10062024 | 1.109 | 0.566 | -0.03 | 0.845 | 0.005 | 0.009 | 0.009 | 4.716 | 0.7 | 0.4 | 0.44 | 0.48 | 0.62 | 0.64 |
| 5 | 3711914 | rs7714769 | 1.344 | 0.795 | -0.432 | 0.55 | -0.012 | 0.005 | 0.005 | 1.476 | 0.6 | 0.39 | 0.36 | 0.4 | 0.51 | 0.47 |
| 5 | 3714522 | rs2398624 | 1.811 | 0.806 | -1.258 | 0.498 | -0.071 | 0.044 | 0.044 | 4.501 | 0.42 | 0.02 | 0.2 | 0.35 | 0.38 | 0.34 |
| 5 | 3715708 | rs6878714 | -0.307 | 1.167 | -0.082 | 0.819 | -0.005 | -0.002 | -0.002 | 0 | 0.14 | 0.12 | 0.09 | 0 | 0.16 | 0.04 |
| 5 | 3717009 | rs2134400 | -1.539 | 1.127 | 0.429 | 1.514 | 0.07 | 0.301 | 0.301 | 0 | 0.08 | 0.3 | 0.26 | 0.35 | 0.12 | 0.2 |
| 5 | 3717737 | rs7729617 | -0.484 | 1.115 | -0.322 | 0.99 | 0.001 | -0.007 | -0.007 | 0 | 0.22 | 0.13 | 0.36 | 0.27 | 0.18 | 0.28 |
| 5 | 3717768 | rs12522688 | 0.547 | 2.087 | -0.762 | 1.413 | -0.054 | -0.048 | -0.048 | 0 | 0.09 | 0.01 | 0.22 | 0.31 | 0.19 | 0.14 |
| 5 | 3718000 | rs1502635 | -1.021 | 1.479 | -0.763 | 0.543 | 0.012 | -0.025 | -0.025 | 0 | 0.43 | 0.75 | 0.51 | 0.51 | 0.39 | 0.44 |
| 5 | 3718709 | rs1604563 | -0.356 | 0.819 | -0.633 | 0.349 | -0.052 | -0.052 | -0.052 | 0 | 0.38 | 0.2 | 0.44 | 0.44 | 0.47 | 0.44 |
| 5 | 3724533 | rs11749388 | 0.322 | 0.551 | -0.071 | 0.147 | 0.021 | -0.024 | -0.024 | 0 | 0.91 | 0.6 | 0.92 | 1 | 0.93 | 0.96 |
| 5 | 3727179 | rs1566691 | 0.004 | 0.548 | 0.106 | 0.146 | 0.027 | -0.015 | -0.015 | 0.058 | 0.92 | 0.68 | 0.93 | 1 | 0.93 | 0.96 |
| 5 | 3727332 | rs10040425 | 0.508 | 0.515 | 0.064 | 0.248 | 0.007 | 0.138 | 0.138 | 0 | 0.12 | 0.4 | 0.22 | 0.3 | 0.11 | 0.19 |
| 5 | 3728440 | rs13174381 | -0.474 | 0.803 | -0.39 | 0.287 | -0.047 | -0.043 | -0.043 | 2.95 | 0.38 | 0.23 | 0.45 | 0.44 | 0.47 | 0.45 |
| 5 | 3730312 | rs13171764 | -0.232 | 0.432 | -0.142 | 0.171 | 0.011 | -0.026 | -0.026 | 0 | 0.08 | 0.27 | 0.07 | 0 | 0.07 | 0.04 |
| 5 | 3735278 | rs16872344 | -0.214 | 0.722 | -0.715 | 0.174 | -0.043 | -0.057 | -0.057 | 0 | 0.32 | 0.22 | 0.38 | 0.39 | 0.38 | 0.4 |
| 5 | 3742035 | rs4866470 | -0.238 | 0.505 | -0.884 | 0.126 | -0.115 | -0.092 | -0.092 | 0 | 0.33 | 0.14 | 0.45 | 0.46 | 0.47 | 0.43 |
| 5 | 3743335 | rs13188410 | -0.576 | 0.698 | -0.767 | 0.309 | -0.007 | -0.005 | -0.005 | 0 | 0.42 | 0.34 | 0.52 | 0.46 | 0.53 | 0.48 |
| 5 | 3752812 | rs7719640 | 0.418 | 0.44 | -1.072 | 0.064 | -0.012 | 0.032 | 0.032 | 0.44 | 0.53 | 0.27 | 0.51 | 0.49 | 0.55 | 0.55 |
| 5 | 3754582 | rs7730021 | 0.264 | 0.582 | -1.319 | -0.923 | -0.007 | 0.042 | 0.042 | 0 | 0.33 | 0.23 | 0.25 | 0.05 | 0.36 | 0.23 |
| 5 | 3757066 | rs13361372 | -0.298 | 0.271 | -0.901 | -0.749 | -0.003 | 0.039 | 0.039 | 0 | 0.49 | 0.66 | 0.52 | 0.42 | 0.5 | 0.41 |

| | | | | | | | | | | | | | | | | |
|---|---|---|---|---|---|---|---|---|---|---|---|---|---|---|---|---|
| 5 | 3760907 | rs905350 | -0.408 | 0.297 | -0.713 | -0.771 | -0.002 | 0.004 | 0.004 | 0 | 0.56 | 0.58 | 0.57 | 0.42 | 0.56 | 0.47 |
| 5 | 3763450 | rs11750498 | 1.298 | 0.179 | -0.644 | -0.793 | -0.004 | -0.035 | -0.035 | 3.693 | 0.15 | 0.17 | 0.08 | 0.01 | 0.15 | 0.06 |
| 5 | 3765085 | rs13436137 | 1.067 | 0.388 | -0.602 | -0.733 | 0.016 | -0.002 | -0.002 | 0 | 0.17 | 0.29 | 0.2 | 0.19 | 0.18 | 0.12 |
| 5 | 3766628 | rs2134404 | 1.359 | 0.396 | -0.621 | -0.809 | 0.019 | -0.005 | -0.005 | 0.811 | 0.13 | 0.26 | 0.14 | 0.07 | 0.15 | 0.03 |
| 5 | 3770600 | rs11741338 | 1.164 | 0.502 | -1.031 | -0.91 | -0.002 | -0.062 | -0.062 | 0 | 0.16 | 0.23 | 0.09 | 0.02 | 0.16 | 0.06 |
| 5 | 3771121 | rs1039462 | -0.364 | 0.683 | -1.718 | -0.592 | -0.01 | 0.036 | 0.036 | 0 | 0.53 | 0.71 | 0.49 | 0.36 | 0.51 | 0.39 |
| 5 | 3773462 | rs13356024 | -0.409 | 0.7 | -2.165 | -0.584 | -0.034 | 0.036 | 0.036 | 0 | 0.54 | 0.8 | 0.5 | 0.36 | 0.51 | 0.39 |
| 5 | 3774574 | rs13156175 | 1.02 | 0.891 | -2.145 | -1.215 | -0.027 | 0.071 | 0.071 | 0 | 0.19 | 0.02 | 0.16 | 0.04 | 0.18 | 0.19 |
| 5 | 3777302 | rs955635 | 2.235 | 0.859 | -1.537 | -0.764 | -0.029 | 0.032 | 0.032 | 0 | 0.07 | 0.01 | 0.09 | 0 | 0.12 | 0.01 |
| 5 | 3779591 | rs6882956 | -1.317 | 0.232 | -1.668 | -0.744 | -0.002 | -0.056 | -0.056 | 0.132 | 0.76 | 0.86 | 0.68 | 0.32 | 0.79 | 0.63 |
| 5 | 3781181 | rs1566689 | -0.323 | -0.116 | -1.329 | -1.295 | 0.004 | 0.006 | 0.006 | 0 | 0.49 | 0.55 | 0.38 | 0.08 | 0.54 | 0.35 |
| 5 | 3781191 | rs2963803 | -1.348 | -0.12 | -1.875 | -1.292 | -0.001 | -0.039 | -0.039 | 0 | 0.72 | 0.87 | 0.61 | 0.29 | 0.73 | 0.55 |
| 5 | 3783340 | rs10059788 | 0.642 | 0.068 | -1.552 | -1.046 | 0.001 | -0.002 | -0.002 | 0 | 0.21 | 0.2 | 0.22 | 0.22 | 0.19 | 0.2 |
| 5 | 3783643 | rs13177139 | 0.403 | 0.37 | -1.597 | -0.949 | -0.088 | 0.13 | 0.13 | 0 | 0.25 | 0.05 | 0.25 | 0.04 | 0.33 | 0.21 |
| 5 | 3787719 | rs2963801 | -0.572 | -0.799 | -1.107 | -0.424 | 0.009 | -0.002 | -0.002 | 0.824 | 0.64 | 0.83 | 0.58 | 0.28 | 0.7 | 0.47 |
| 5 | 3789862 | rs2967831 | -0.089 | -1.205 | 0.864 | 0.178 | -0.037 | 0.011 | 0.011 | 0 | 0.19 | 0.34 | 0.17 | 0.2 | 0.14 | 0.2 |
| 5 | 3790765 | rs2963799 | -0.388 | -1.129 | 0.64 | 0.339 | 0.001 | 0.016 | 0.016 | 0 | 0.92 | 0.86 | 0.9 | 0.88 | 0.93 | 0.84 |
| 5 | 3792631 | rs7726033 | -0.315 | -1.134 | 0.637 | 0.337 | 0.001 | 0.015 | 0.015 | 0 | 0.92 | 0.86 | 0.9 | 0.88 | 0.93 | 0.82 |
| 5 | 3794274 | rs10462777 | 0.628 | -1.206 | 0.799 | 0.266 | -0.027 | 0.016 | 0.016 | 0.058 | 0.72 | 0.43 | 0.73 | 0.68 | 0.79 | 0.65 |
| 5 | 3800097 | rs11948295 | -0.211 | -1.105 | 0.655 | -0.21 | -0.006 | -0.008 | -0.008 | 0.535 | 0.89 | 0.8 | 0.89 | 0.91 | 0.93 | 0.84 |
| 5 | 3801906 | rs10060475 | 1.295 | -1.501 | 0.971 | -0.183 | 0.036 | -0.012 | -0.012 | 0.119 | 0.6 | 0.29 | 0.67 | 0.61 | 0.73 | 0.57 |
| 5 | 3802888 | rs12659399 | 1.983 | -1.211 | 0.757 | -0.392 | -0.029 | -0.029 | -0.029 | 0 | 0.4 | 0.22 | 0.5 | 0.56 | 0.49 | 0.42 |
| 5 | 3805412 | rs2967839 | -1.429 | -1.178 | 0.62 | -0.406 | 0.013 | 0.018 | 0.018 | 0.049 | 0.48 | 0.61 | 0.4 | 0.35 | 0.43 | 0.42 |
| 5 | 3809717 | rs2967840 | 0.386 | -0.863 | 0.712 | -0.152 | -0.01 | 0.036 | 0.036 | 0 | 0.18 | 0.27 | 0.18 | 0.27 | 0.14 | 0.22 |
| 5 | 3815015 | rs7729252 | 0.451 | -0.696 | 0.338 | -0.482 | 0.006 | -0.015 | -0.015 | 0 | 0.19 | 0.11 | 0.17 | 0.21 | 0.17 | 0.22 |
| 5 | 3815297 | rs2174047 | 0.666 | -0.827 | 0.661 | -0.708 | 0.002 | -0.035 | -0.035 | 0.21 | 0.67 | 0.59 | 0.68 | 0.85 | 0.7 | 0.68 |
| 5 | 3815679 | rs2134403 | 1.243 | -0.775 | 0.421 | -0.676 | 0 | -0.001 | -0.001 | 0 | 0.48 | 0.47 | 0.51 | 0.63 | 0.53 | 0.45 |
| 5 | 3822893 | rs12657754 | -0.822 | -0.721 | -0.009 | -0.821 | -0.001 | -0.084 | -0.084 | 0.243 | 0.28 | 0.39 | 0.23 | 0.07 | 0.26 | 0.19 |
| 5 | 3824532 | rs13180945 | 0.322 | -0.699 | -0.223 | -0.811 | -0.051 | 0.066 | 0.066 | 0 | 0.16 | 0.07 | 0.16 | 0.05 | 0.21 | 0.16 |
| 5 | 3833240 | rs12521994 | 0.386 | -0.508 | 0.083 | -0.595 | -0.023 | -0.028 | -0.028 | 0 | 0.1 | 0.15 | 0.06 | 0.02 | 0.05 | 0.03 |
| 5 | 3834405 | rs11750441 | 1.607 | -0.571 | 0.045 | -0.622 | -0.071 | -0.02 | -0.02 | 0 | 0.37 | 0.11 | 0.42 | 0.62 | 0.48 | 0.4 |
| 5 | 3834603 | rs2963794 | -0.27 | -0.261 | 0.02 | -0.444 | -0.009 | 0.014 | 0.014 | 0 | 0.35 | 0.27 | 0.33 | 0.26 | 0.38 | 0.38 |
| 5 | 3840278 | rs2860410 | -0.519 | -0.333 | 0.274 | -0.145 | -0.003 | -0.007 | -0.007 | 0.95 | 0.45 | 0.44 | 0.39 | 0.28 | 0.4 | 0.41 |
| 5 | 3844414 | rs2398647 | 0.635 | -0.318 | -0.829 | 0.271 | 0.039 | 0.027 | 0.027 | 0.358 | 0.47 | 0.15 | 0.56 | 0.59 | 0.61 | 0.45 |
| 5 | 3844541 | rs2963791 | 0.603 | -0.496 | 0.097 | -0.238 | -0.014 | -0.085 | -0.085 | 5.026 | 0.82 | 0.52 | 0.82 | 0.91 | 0.9 | 0.79 |
| 5 | 3845917 | rs1994579 | -0.089 | -0.415 | -0.624 | -0.334 | 0.014 | 0.015 | 0.015 | 0 | 0.24 | 0.18 | 0.1 | 0.02 | 0.18 | 0.16 |
| 5 | 3846180 | rs2967821 | 0.709 | 0.469 | -0.231 | 0.716 | 0.011 | 0.012 | 0.012 | 0 | 0.81 | 0.69 | 0.79 | 0.79 | 0.84 | 0.69 |
| 5 | 3856449 | rs12153260 | -0.2 | 0.127 | -0.718 | 1.282 | 0.007 | -0.033 | -0.033 | 1.092 | 0.15 | 0.08 | 0.16 | 0.34 | 0.13 | 0.22 |
| 5 | 3862889 | rs16872456 | -1.107 | 0.193 | -0.1 | 0.896 | 0.002 | 0.017 | 0.017 | 0 | 0.08 | 0.21 | 0.05 | 0.09 | 0.1 | 0.06 |
| 5 | 3865308 | rs17653429 | 0.633 | 0.404 | -1.421 | 0.293 | -0.092 | 0.105 | 0.105 | 0.648 | 0.41 | 0.09 | 0.36 | 0.25 | 0.43 | 0.36 |
| 5 | 3865417 | rs2895167 | 0.006 | 0.519 | -0.677 | 0.293 | 0.013 | 0.042 | 0.042 | 0.037 | 0.5 | 0.4 | 0.43 | 0.34 | 0.58 | 0.42 |
| 5 | 3873124 | rs583835 | -1.018 | 0.827 | -0.033 | -0.079 | -0.003 | 0.005 | 0.005 | 0 | 0.14 | 0.11 | 0.07 | 0.03 | 0.14 | 0.09 |
| 5 | 3875103 | rs16872476 | -1.401 | 0.554 | 0.225 | 0.13 | -0.003 | 0 | 0 | 0 | 0.16 | 0.16 | 0.14 | 0.19 | 0.19 | 0.09 |
| 5 | 3879123 | rs478794 | 1.989 | 0.492 | 0.586 | -0.445 | 0.001 | 0.025 | 0.025 | 0.587 | 0.59 | 0.3 | 0.65 | 0.76 | 0.56 | 0.57 |
| 5 | 3888310 | rs623918 | 2.136 | 0.444 | 0.67 | -0.073 | 0 | -0.035 | -0.035 | 0 | 0.7 | 0.46 | 0.75 | 0.77 | 0.67 | 0.77 |
| 5 | 3890330 | rs594459 | 1.801 | 0.266 | -0.08 | 0.372 | 0 | -0.01 | -0.01 | 0 | 0.76 | 0.81 | 0.77 | 0.68 | 0.73 | 0.78 |
| 5 | 3891796 | rs550167 | 0.519 | 0.493 | 0.067 | 0.215 | 0.006 | -0.006 | -0.006 | 0 | 0.9 | 0.88 | 0.91 | 0.96 | 0.86 | 0.86 |
| 5 | 3895815 | rs10475241 | -0.91 | -0.083 | -0.015 | 0.386 | 0.009 | -0.008 | -0.008 | 1.626 | 0.1 | 0.07 | 0.05 | 0.17 | 0.08 | 0.04 |
| 5 | 3904292 | rs6886339 | -1.176 | -0.043 | 1.135 | 0.128 | 0.023 | -0.011 | -0.011 | 0 | 0.1 | 0.45 | 0.09 | 0.02 | 0.07 | 0.07 |
| 5 | 3925083 | rs11956711 | -1.162 | -0.02 | 1.447 | 0.109 | 0.018 | -0.021 | -0.021 | 0 | 0.1 | 0.46 | 0.1 | 0.02 | 0.07 | 0.07 |
| 5 | 3926567 | rs11958394 | -1.143 | -0.025 | 1.7 | 0.107 | 0.008 | -0.027 | -0.027 | 0 | 0.1 | 0.38 | 0.08 | 0.02 | 0.07 | 0.07 |
| 5 | 3927833 | rs505800 | 1.506 | 0.001 | 1.384 | 0.209 | 0.006 | 0.03 | 0.03 | 0.485 | 0.65 | 0.23 | 0.59 | 0.75 | 0.68 | 0.61 |
| 5 | 3950622 | rs12186559 | 0.307 | -0.741 | 1.342 | 0.91 | 0.024 | -0.077 | -0.077 | 0 | 0.11 | 0.01 | 0.15 | 0.38 | 0.05 | 0.16 |
| 5 | 3973783 | rs547398 | -0.779 | -0.853 | 2.105 | 0.281 | -0.015 | -0.048 | -0.048 | 0 | 0.08 | 0.18 | 0.05 | 0 | 0.05 | 0.06 |
| 5 | 3974744 | rs1400794 | 1.147 | -0.946 | 2.607 | 0.272 | 0.003 | -0.038 | -0.038 | 0 | 0.68 | 0.55 | 0.62 | 0.76 | 0.67 | 0.63 |
| 5 | 3976564 | rs687595 | 0.4 | -0.972 | 2.468 | 0.225 | -0.004 | 0 | 0 | 0 | 0.78 | 0.82 | 0.7 | 0.76 | 0.74 | 0.7 |
| 5 | 3991294 | rs4702310 | 1.576 | -1.049 | 2.501 | 0.278 | 0.002 | 0.013 | 0.013 | 0 | 0.65 | 0.33 | 0.6 | 0.75 | 0.67 | 0.63 |
| 5 | 3992147 | rs12187878 | 1.517 | -1.095 | 2.498 | 0.229 | 0.003 | 0.018 | 0.018 | 0 | 0.64 | 0.28 | 0.64 | 0.78 | 0.68 | 0.61 |
| 5 | 3993746 | rs680810 | 1.069 | -1.034 | 2.412 | 0.26 | -0.001 | 0.004 | 0.004 | 0 | 0.69 | 0.71 | 0.69 | 0.78 | 0.68 | 0.61 |
| 5 | 3997830 | rs11748824 | -0.44 | -0.979 | 2.212 | 0.273 | 0.002 | -0.005 | -0.005 | 0.287 | 0.14 | 0.06 | 0.06 | 0.15 | 0.12 | 0.19 |
| 5 | 4002470 | rs4455563 | 0.551 | -1.037 | 2.447 | 0.268 | -0.001 | -0.004 | -0.004 | 1.316 | 0.83 | 0.83 | 0.75 | 0.92 | 0.8 | 0.81 |
| 5 | 4003908 | rs10078693 | 1.766 | -1.037 | 2.614 | 0.266 | 0 | -0.007 | -0.007 | 0 | 0.68 | 0.65 | 0.69 | 0.77 | 0.68 | 0.62 |
| 5 | 4012303 | rs3943033 | -0.657 | 0.093 | 1.834 | -0.238 | 0.005 | -0.008 | -0.008 | 1.733 | 0.27 | 0.3 | 0.22 | 0.15 | 0.33 | 0.25 |
| 5 | 4012694 | rs10071096 | -0.657 | 0.093 | 1.834 | -0.238 | 0.005 | -0.008 | -0.008 | 0.003 | 0.27 | 0.3 | 0.22 | 0.15 | 0.33 | 0.25 |
| 5 | 4014659 | rs540937 | 1.724 | -0.087 | 1.748 | -0.251 | 0.007 | 0.021 | 0.021 | 0 | 0.54 | 0.4 | 0.51 | 0.76 | 0.47 | 0.57 |
| 5 | 4020601 | rs11738089 | -0.257 | -0.117 | 1.635 | -0.204 | -0.066 | 0.036 | 0.036 | 0 | 0.25 | 0.07 | 0.2 | 0.15 | 0.33 | 0.25 |
| 5 | 4041154 | rs580641 | 1.583 | 0.071 | 1.66 | 0.047 | -0.041 | 0.041 | 0.041 | 0 | 0.6 | 0.84 | 0.61 | 0.71 | 0.51 | 0.59 |
| 5 | 4048841 | rs16872714 | 0.911 | -0.25 | 1.321 | 0.48 | 0.042 | -0.065 | -0.065 | 0 | 0.13 | 0.03 | 0.23 | 0.32 | 0.06 | 0.16 |
| 5 | 4053766 | rs1201368 | 1.64 | 0.373 | 1.388 | 0.241 | -0.014 | 0.022 | 0.022 | 0 | 0.59 | 0.72 | 0.59 | 0.71 | 0.51 | 0.58 |

| | | | | | | | | | | | | | | | | |
|---|---|---|---|---|---|---|---|---|---|---|---|---|---|---|---|---|
| 5 | 4065252 | rs11134030 | -0.537 | 0.471 | 1.057 | 0.25 | -0.03 | 0.012 | 0.012 | 0.003 | 0.24 | 0.09 | 0.15 | 0.19 | 0.31 | 0.25 |
| 5 | 4071067 | rs1187460 | -0.026 | 0.438 | 1.121 | 0.325 | -0.009 | 0.009 | 0.009 | 0 | 0.85 | 0.89 | 0.75 | 0.9 | 0.82 | 0.83 |
| 5 | 4075178 | rs12659815 | -0.74 | 0.462 | 1.049 | 0.41 | -0.013 | 0.005 | 0.005 | 0 | 0.24 | 0.15 | 0.15 | 0.2 | 0.3 | 0.24 |
| 5 | 4088133 | rs10462783 | 1.691 | 0.544 | 1.192 | 0.467 | 0.015 | -0.009 | -0.009 | 0.476 | 0.6 | 0.56 | 0.64 | 0.69 | 0.54 | 0.58 |
| 5 | 4093896 | rs9313069 | -0.764 | 0.759 | 0.901 | 0.356 | -0.014 | -0.001 | -0.001 | 0 | 0.24 | 0.18 | 0.14 | 0.21 | 0.29 | 0.26 |
| 5 | 4096237 | rs1187477 | 1.02 | 0.433 | 1.09 | 0.389 | -0.006 | -0.023 | -0.023 | 0 | 0.73 | 0.71 | 0.59 | 0.86 | 0.77 | 0.72 |
| 5 | 4096258 | rs1187478 | 1.017 | 0.439 | 1.093 | 0.392 | -0.006 | -0.024 | -0.024 | 0.464 | 0.73 | 0.71 | 0.59 | 0.86 | 0.77 | 0.72 |
| 5 | 4100989 | rs1493470 | -0.774 | 0.647 | 1.084 | 0.56 | 0.038 | 0.008 | 0.008 | 0.037 | 0.18 | 0.28 | 0.16 | 0.21 | 0.28 | 0.14 |
| 5 | 4101370 | rs1187482 | 1.519 | 0.114 | 1.179 | 0.566 | 0.003 | 0.002 | 0.002 | 0 | 0.63 | 0.33 | 0.61 | 0.65 | 0.68 | 0.69 |
| 5 | 4101569 | rs1187483 | 1.53 | 0.098 | 1.204 | 0.572 | 0.005 | 0.003 | 0.003 | 0 | 0.62 | 0.29 | 0.6 | 0.66 | 0.68 | 0.67 |
| 5 | 4104155 | rs1187489 | 0.718 | 0.039 | 1.423 | 0.552 | -0.013 | 0.001 | 0.001 | 0 | 0.85 | 0.83 | 0.92 | 0.86 | 0.92 | 0.88 |
| 5 | 4105915 | rs1187491 | -0.447 | -0.499 | 0.861 | 0.874 | -0.004 | 0.005 | 0.005 | 0 | 0.31 | 0.37 | 0.3 | 0.44 | 0.26 | 0.39 |
| 5 | 4106399 | rs2452883 | 0.824 | -0.556 | 0.965 | 0.913 | -0.054 | 0.018 | 0.018 | 0 | 0.83 | 0.59 | 0.9 | 0.86 | 0.92 | 0.88 |
| 5 | 4110205 | rs10866562 | 1.223 | -0.105 | -0.088 | 0.715 | 0.062 | 0.01 | 0.01 | 0 | 0.53 | 0.24 | 0.61 | 0.41 | 0.67 | 0.51 |
| 5 | 4123457 | rs1911912 | 0.679 | -0.043 | 0.087 | 0.633 | 0 | -0.005 | -0.005 | 0 | 0.54 | 0.49 | 0.47 | 0.38 | 0.57 | 0.49 |
| 5 | 4126106 | rs1911914 | 0.642 | -0.042 | 0.078 | 0.634 | 0 | -0.007 | -0.007 | 2.364 | 0.54 | 0.49 | 0.47 | 0.37 | 0.57 | 0.49 |
| 5 | 4126274 | rs1911915 | 0.695 | -0.048 | 0.148 | 0.631 | -0.001 | 0.007 | 0.007 | 0 | 0.54 | 0.43 | 0.46 | 0.37 | 0.57 | 0.49 |
| 5 | 4136154 | rs6860112 | 0.605 | -0.444 | -0.153 | 0.284 | -0.005 | 0.042 | 0.042 | 0 | 0.51 | 0.28 | 0.4 | 0.37 | 0.55 | 0.51 |
| 5 | 4138095 | rs17665143 | 0.62 | -0.308 | -0.162 | 0.412 | -0.021 | 0.045 | 0.045 | 0.156 | 0.55 | 0.26 | 0.4 | 0.32 | 0.51 | 0.48 |
| 5 | 4140796 | rs10063373 | -0.232 | 0.096 | 0.097 | 0.92 | 0.007 | 0.013 | 0.013 | 0.042 | 0.79 | 0.8 | 0.69 | 0.63 | 0.69 | 0.71 |
| 5 | 4157570 | rs10076711 | -0.796 | -0.559 | -0.107 | 0.265 | -0.04 | -0.057 | -0.057 | 0.053 | 0.12 | 0.39 | 0.06 | 0.03 | 0.04 | 0.12 |
| 5 | 4161404 | rs10077300 | -0.257 | -0.522 | -0.337 | 0.282 | -0.022 | -0.03 | -0.03 | 0 | 0.11 | 0.18 | 0.05 | 0.03 | 0.04 | 0.12 |
| 5 | 4176833 | rs3938039 | 0.716 | -0.373 | -1.889 | -0.012 | -0.02 | 0.065 | 0.065 | 0.007 | 0.33 | 0.04 | 0.34 | 0.2 | 0.29 | 0.25 |
| 5 | 4186632 | rs11953297 | 0.283 | 0.165 | -1.169 | 0.562 | -0.079 | -0.077 | -0.077 | 0.614 | 0.15 | 0.01 | 0.15 | 0.26 | 0.24 | 0.24 |
| 5 | 4193777 | rs10051866 | 0.632 | -0.494 | -0.275 | 0.146 | -0.014 | -0.054 | -0.054 | 0 | 0.91 | 0.74 | 0.96 | 1 | 0.94 | 0.89 |
| 5 | 4198117 | rs6555259 | -0.29 | -0.238 | -0.63 | 0.278 | -0.001 | -0.03 | -0.03 | 0 | 0.94 | 0.87 | 0.97 | 1 | 0.94 | 0.89 |
| 5 | 4205074 | rs7734004 | 0.162 | -0.205 | -1.282 | 0.306 | -0.006 | 0.003 | 0.003 | 0 | 0.61 | 0.78 | 0.68 | 0.67 | 0.59 | 0.69 |
| 5 | 4211418 | rs16872928 | -0.333 | -0.312 | -0.648 | 0.39 | -0.04 | -0.065 | -0.065 | 0 | 0.12 | 0.31 | 0.04 | 0.01 | 0.04 | 0.21 |
| 5 | 4213582 | rs7735751 | -0.824 | -0.391 | -0.634 | 0.353 | -0.062 | -0.091 | -0.091 | 0.001 | 0.15 | 0.39 | 0.07 | 0.01 | 0.05 | 0.21 |
| 5 | 4216069 | rs1035866 | -0.068 | 0.607 | -1.322 | 0.957 | -0.044 | -0.056 | -0.056 | 0 | 0.91 | 0.99 | 0.85 | 0.8 | 0.86 | 0.86 |
| 5 | 4224788 | rs959937 | -0.301 | 0.056 | -1.008 | 0.825 | -0.01 | 0.019 | 0.019 | 1.15 | 0.33 | 0.38 | 0.28 | 0.5 | 0.29 | 0.37 |
| 5 | 4225004 | rs11948368 | -0.126 | -0.121 | -1.859 | 0.139 | -0.03 | 0.056 | 0.056 | 0.114 | 0.35 | 0.09 | 0.3 | 0.21 | 0.35 | 0.37 |
| 5 | 4227497 | rs7706780 | -0.571 | -0.986 | -1.411 | 0.447 | 0.02 | -0.01 | -0.01 | 0 | 0.31 | 0.25 | 0.27 | 0.32 | 0.23 | 0.38 |
| 5 | 4229937 | rs12521129 | -0.468 | -0.451 | -1.325 | 0.239 | 0.002 | 0.017 | 0.017 | 1.644 | 0.24 | 0.1 | 0.24 | 0.17 | 0.23 | 0.25 |
| 5 | 4230268 | rs7723568 | -0.956 | -0.981 | -0.959 | 0.18 | -0.01 | -0.009 | -0.009 | 0 | 0.42 | 0.54 | 0.25 | 0.28 | 0.29 | 0.4 |
| 5 | 4244114 | rs17723858 | -0.348 | 0.01 | -1.414 | 0.522 | -0.008 | -0.002 | -0.002 | 0 | 0.18 | 0.18 | 0.26 | 0.22 | 0.18 | 0.32 |
| 5 | 4244462 | rs265180 | -0.726 | 0.157 | -1.399 | 0.505 | 0.003 | 0.029 | 0.029 | 0 | 0.45 | 0.33 | 0.45 | 0.39 | 0.4 | 0.52 |
| 5 | 4248397 | rs3105127 | 0.487 | 0.215 | -1.33 | 0.787 | 0.002 | -0.007 | -0.007 | 0 | 0.74 | 0.83 | 0.71 | 0.7 | 0.77 | 0.64 |
| 5 | 4256610 | rs263734 | 0.54 | 0.265 | -0.444 | 1.288 | 0.028 | 0.133 | 0.133 | 0 | 0.94 | 0.73 | 0.88 | 0.78 | 0.93 | 0.81 |
| 5 | 4257463 | rs11739066 | -0.522 | -0.15 | -0.869 | 0.646 | 0.003 | 0.003 | 0.003 | 0.296 | 0.28 | 0.28 | 0.17 | 0.14 | 0.2 | 0.21 |
| 5 | 4269490 | rs458649 | 0.518 | -0.066 | -0.757 | 0.689 | -0.001 | 0 | 0 | 0 | 0.76 | 0.75 | 0.77 | 0.79 | 0.81 | 0.82 |
| 5 | 4269803 | rs17543979 | 0.004 | 0.318 | -1.297 | 0.872 | -0.024 | -0.029 | -0.029 | 0.324 | 0.13 | 0.01 | 0.06 | 0.2 | 0.15 | 0.16 |
| 5 | 4275538 | rs462021 | 0.414 | 0.117 | -0.882 | 0.745 | 0.002 | 0 | 0 | 1.816 | 0.77 | 0.78 | 0.77 | 0.77 | 0.81 | 0.79 |
| 5 | 4280113 | rs1532688 | -0.538 | -0.022 | -1.07 | 0.407 | 0.008 | 0.024 | 0.024 | 1.292 | 0.19 | 0.13 | 0.13 | 0.08 | 0.17 | 0.19 |
| 5 | 4281622 | rs11134042 | -0.246 | -0.251 | -1.21 | 0.619 | 0.02 | -0.006 | -0.006 | 0 | 0.16 | 0.07 | 0.12 | 0.11 | 0.12 | 0.12 |
| 5 | 4281719 | rs255675 | 1.155 | -0.001 | -2.109 | 0.338 | -0.04 | 0.014 | 0.014 | 0 | 0.32 | 0.12 | 0.33 | 0.26 | 0.42 | 0.27 |
| 5 | 4282851 | rs12659526 | -0.348 | -0.132 | -1.744 | 0.358 | 0.006 | 0.003 | 0.003 | 0.108 | 0.2 | 0.18 | 0.14 | 0.11 | 0.12 | 0.18 |
| 5 | 4288545 | rs12110273 | -0.119 | -0.083 | -0.861 | -0.013 | -0.002 | 0.004 | 0.004 | 0 | 0.57 | 0.63 | 0.47 | 0.36 | 0.53 | 0.47 |
| 5 | 4289170 | rs155349 | -0.857 | 0.589 | -1.4 | 0.057 | -0.086 | -0.126 | -0.126 | 0.184 | 0.85 | 0.99 | 0.82 | 0.48 | 0.8 | 0.66 |
| 5 | 4293345 | rs11742174 | -0.095 | 0.308 | -1.094 | -0.296 | -0.002 | -0.003 | -0.003 | 0 | 0.22 | 0.21 | 0.17 | 0.1 | 0.17 | 0.22 |
| 5 | 4293859 | rs7702429 | 0.023 | 0.309 | -1.114 | -0.295 | 0.013 | 0.019 | 0.019 | 0 | 0.22 | 0.12 | 0.16 | 0.1 | 0.17 | 0.22 |
| 5 | 4297423 | rs7731506 | 1.805 | 0.379 | -1.063 | -0.61 | -0.011 | -0.018 | -0.018 | 0 | 0.28 | 0.35 | 0.23 | 0.17 | 0.26 | 0.24 |
| 5 | 4303898 | rs1393106 | 0.314 | 0.155 | -0.883 | -0.756 | 0.022 | 0.048 | 0.048 | 0 | 0.55 | 0.7 | 0.41 | 0.28 | 0.44 | 0.47 |
| 5 | 4305161 | rs11134044 | 0.081 | 0.052 | -0.566 | -0.8 | -0.014 | -0.016 | -0.016 | 0 | 0.23 | 0.24 | 0.16 | 0.11 | 0.17 | 0.18 |
| 5 | 4311402 | rs155350 | 0.862 | 0.094 | -1.076 | -1.113 | -0.04 | 0.085 | 0.085 | 0.09 | 0.15 | 0.02 | 0.1 | 0 | 0.16 | 0.06 |
| 5 | 4316307 | rs261133 | -1.941 | 0.12 | -1.043 | -1.166 | 0.001 | 0.001 | 0.001 | 0.244 | 0.67 | 0.37 | 0.73 | 0.79 | 0.72 | 0.7 |
| 5 | 4316470 | rs11749633 | 0.584 | 0.109 | -1.256 | -1.179 | 0.014 | 0.024 | 0.024 | 0.018 | 0.21 | 0.08 | 0.15 | 0.1 | 0.16 | 0.17 |
| 5 | 4321335 | rs155279 | 0.484 | 0.215 | -1.43 | -1.043 | -0.123 | -0.153 | -0.153 | 0 | 0.22 | 0.01 | 0.26 | 0.58 | 0.32 | 0.38 |
| 5 | 4325295 | rs155353 | 0.448 | 0.19 | -1.38 | -1.042 | -0.078 | 0.05 | 0.05 | 0 | 0.19 | 0.02 | 0.3 | 0.09 | 0.24 | 0.09 |
| 5 | 4341114 | rs10069119 | 0.367 | -0.095 | -1.02 | -1.266 | 0.001 | 0.001 | 0.001 | 0 | 0.22 | 0.19 | 0.15 | 0.11 | 0.16 | 0.16 |
| 5 | 4346529 | rs155356 | -2.926 | 0.073 | -1.079 | -1.394 | -0.03 | -0.012 | -0.012 | 0 | 0.75 | 0.62 | 0.79 | 0.79 | 0.8 | 0.74 |
| 5 | 4355708 | rs17727202 | 1.746 | 0.193 | -1.097 | -1.359 | 0 | -0.017 | -0.017 | 0 | 0.07 | 0.08 | 0.04 | 0 | 0.08 | 0.05 |
| 5 | 4356792 | rs261144 | -1.178 | 0.184 | -1.192 | -1.418 | 0.008 | 0.02 | 0.02 | 0 | 0.45 | 0.39 | 0.59 | 0.67 | 0.55 | 0.49 |
| 5 | 4368462 | rs261124 | -2.674 | 0.451 | -0.626 | -1.084 | -0.008 | -0.016 | -0.016 | 0 | 0.71 | 0.64 | 0.58 | 0.79 | 0.73 | 0.75 |
| 5 | 4369887 | rs261125 | -0.863 | 0.426 | -0.79 | -1.053 | -0.017 | 0.008 | 0.008 | 0 | 0.4 | 0.34 | 0.38 | 0.67 | 0.48 | 0.46 |
| 5 | 4372548 | rs1354129 | -0.221 | 0.673 | -0.413 | -0.745 | 0.011 | 0.048 | 0.048 | 0 | 0.27 | 0.1 | 0.18 | 0.11 | 0.24 | 0.24 |
| 5 | 4375021 | rs9313075 | 0.127 | 0.7 | -0.006 | -0.292 | -0.009 | -0.018 | -0.018 | 1.017 | 0.2 | 0.19 | 0.11 | 0.05 | 0.14 | 0.18 |
| 5 | 4375160 | rs261126 | -0.67 | 1.034 | -0.33 | -0.032 | 0.01 | 0.025 | 0.025 | 0 | 0.23 | 0.31 | 0.26 | 0.62 | 0.29 | 0.37 |
| 5 | 4375862 | rs13175033 | -0.745 | 0.777 | -0.834 | -0.059 | -0.069 | -0.089 | -0.089 | 0.124 | 0.35 | 0.09 | 0.33 | 0.55 | 0.37 | 0.41 |

| | | | | | | | | | | | | | | | | |
|---|---|---|---|---|---|---|---|---|---|---|---|---|---|---|---|---|
| 5 | 4376022 | rs261129 | 1.305 | 0.222 | -0.22 | 0.206 | 0.012 | 0.077 | 0.077 | 0.181 | 0.54 | 0.22 | 0.55 | 0.31 | 0.6 | 0.39 |
| 5 | 4380545 | rs12187807 | 0.023 | -0.73 | 0.085 | 0.747 | -0.02 | -0.061 | -0.061 | 0 | 0.1 | 0 | 0.1 | 0.23 | 0.12 | 0.16 |
| 5 | 4380725 | rs4701855 | -1.017 | -0.555 | 0.481 | 0.264 | 0.029 | -0.012 | -0.012 | 1.417 | 0.38 | 0.73 | 0.34 | 0.55 | 0.27 | 0.48 |
| 5 | 4390768 | rs1080648 | 1.18 | -0.601 | -0.168 | 0.063 | 0.058 | -0.01 | -0.01 | 0 | 0.54 | 0.17 | 0.62 | 0.42 | 0.67 | 0.41 |
| 5 | 4391653 | rs12716158 | -0.968 | -1.035 | -0.148 | 0.258 | -0.02 | 0.017 | 0.017 | 0 | 0.35 | 0.43 | 0.32 | 0.47 | 0.26 | 0.43 |
| 5 | 4396901 | rs13168384 | -0.698 | -0.688 | 0.004 | 0.262 | 0.039 | -0.048 | -0.048 | 0 | 0.32 | 0.15 | 0.29 | 0.45 | 0.23 | 0.4 |
| 5 | 4397964 | rs9313078 | -1.238 | 0.407 | -0.073 | -0.144 | -0.018 | -0.043 | -0.043 | 0.003 | 0.19 | 0.36 | 0.24 | 0.12 | 0.12 | 0.18 |
| 5 | 4399737 | rs17677148 | -0.726 | 1.055 | -0.99 | -0.602 | -0.039 | 0.066 | 0.066 | 0 | 0.23 | 0.07 | 0.13 | 0.01 | 0.31 | 0.22 |
| 5 | 4405918 | rs7718259 | -0.801 | 0.358 | -0.364 | 0.617 | -0.015 | 0.133 | 0.133 | 0 | 0.11 | 0.15 | 0.25 | 0.41 | 0.06 | 0.22 |
| 5 | 4409168 | rs4701864 | -0.028 | -0.242 | -0.99 | -0.17 | 0.029 | 0.063 | 0.063 | 0 | 0.22 | 0.09 | 0.19 | 0.1 | 0.2 | 0.19 |
| 5 | 4417450 | rs12518914 | -0.233 | -0.134 | -0.614 | -0.344 | 0.015 | 0.059 | 0.059 | 0 | 0.24 | 0.13 | 0.23 | 0.11 | 0.24 | 0.2 |
| 5 | 4419552 | rs10044501 | 0.026 | -0.185 | 0.522 | -0.319 | 0.028 | 0.058 | 0.058 | 0.535 | 0.93 | 0.64 | 0.93 | 0.94 | 0.95 | 0.93 |
| 5 | 4420716 | rs13186177 | 0.97 | -0.278 | 0.061 | -0.761 | 0.022 | 0.001 | 0.001 | 0 | 0.35 | 0.14 | 0.33 | 0.4 | 0.27 | 0.27 |
| 5 | 4423292 | rs10078927 | -0.338 | -0.29 | 0.204 | -0.752 | 0.011 | 0.038 | 0.038 | 2.211 | 0.92 | 0.65 | 0.93 | 0.94 | 0.95 | 0.93 |
| 5 | 4426699 | rs10076204 | 0.612 | -0.498 | 0.25 | -0.904 | -0.067 | -0.069 | -0.069 | 0.972 | 0.28 | 0.12 | 0.35 | 0.42 | 0.44 | 0.44 |
| 5 | 4428580 | rs11134052 | 1.077 | -0.766 | 0.39 | -0.893 | 0.013 | 0.038 | 0.038 | 0 | 0.3 | 0.14 | 0.12 | 0.08 | 0.21 | 0.11 |
| 5 | 4429187 | rs10054113 | 0.351 | -0.518 | 0.691 | -0.564 | 0.028 | 0.061 | 0.061 | 4.2 | 0.91 | 0.55 | 0.92 | 0.94 | 0.95 | 0.93 |
| 5 | 4436013 | rs4286636 | -0.023 | -0.476 | 0.647 | -0.519 | 0.015 | 0.047 | 0.047 | 1.672 | 0.93 | 0.7 | 0.93 | 0.94 | 0.95 | 0.93 |
| 5 | 4438356 | rs6894740 | 0.767 | -0.762 | 0.41 | -0.703 | -0.051 | -0.045 | -0.045 | 0.72 | 0.83 | 0.64 | 0.93 | 0.94 | 0.93 | 0.89 |
| 5 | 4439260 | rs10064177 | -0.423 | -0.166 | 0.099 | -0.65 | 0.012 | 0.003 | 0.003 | 0 | 0.53 | 0.43 | 0.56 | 0.51 | 0.65 | 0.62 |
| 5 | 4439651 | rs6861699 | 0.648 | 1.234 | 0.575 | -0.784 | 0.095 | -0.015 | -0.015 | 0 | 0.85 | 0.61 | 0.83 | 0.93 | 0.74 | 0.73 |
| 5 | 4439786 | rs6887047 | -0.426 | 0.388 | -0.066 | -0.698 | -0.002 | 0.007 | 0.007 | 0 | 0.24 | 0.27 | 0.25 | 0.13 | 0.3 | 0.22 |
| 5 | 4442390 | rs2043354 | -2.063 | -0.098 | 1.005 | -0.431 | -0.01 | -0.013 | -0.013 | 0 | 0.08 | 0.42 | 0.05 | 0.01 | 0.01 | 0.02 |
| 5 | 4455368 | rs4701877 | -0.031 | 0.172 | 0.257 | -0.607 | -0.065 | 0.074 | 0.074 | 0 | 0.2 | 0.04 | 0.29 | 0.04 | 0.29 | 0.34 |
| 5 | 4456841 | rs1433271 | 1.727 | -0.125 | 1.368 | -0.618 | -0.01 | -0.014 | -0.014 | 0.126 | 0.92 | 0.4 | 0.94 | 0.99 | 0.99 | 0.98 |
| 5 | 4474329 | rs9313082 | 0.028 | -0.244 | 0.927 | -0.865 | -0.012 | -0.05 | -0.05 | 0 | 0.33 | 0.46 | 0.37 | 0.84 | 0.26 | 0.4 |
| 5 | 4474589 | rs1433265 | 0.601 | -0.014 | 0.909 | -1.216 | 0 | -0.104 | -0.104 | 0.088 | 0.68 | 0.41 | 0.65 | 0.96 | 0.68 | 0.58 |
| 5 | 4478547 | rs10066908 | 0.555 | -0.561 | 1.303 | -0.572 | -0.02 | -0.04 | -0.04 | 2.335 | 0.27 | 0.12 | 0.3 | 0.47 | 0.26 | 0.29 |
| 5 | 4488309 | rs10462795 | 0.178 | -0.843 | 1.708 | -1.024 | -0.004 | -0.042 | -0.042 | 0 | 0.73 | 0.69 | 0.69 | 0.97 | 0.76 | 0.62 |
| 5 | 4493944 | rs12516470 | -0.382 | -0.881 | 1.69 | -0.992 | 0.005 | 0.024 | 0.024 | 0 | 0.27 | 0.21 | 0.3 | 0.03 | 0.26 | 0.38 |
| 5 | 4511087 | rs1450827 | 0.09 | -0.795 | 1.668 | -0.931 | -0.023 | -0.099 | -0.099 | 0.005 | 0.74 | 0.59 | 0.69 | 0.97 | 0.77 | 0.65 |
| 5 | 4514532 | rs16873509 | 0.254 | -0.714 | 1.493 | -0.837 | -0.002 | -0.049 | -0.049 | 0 | 0.23 | 0.2 | 0.29 | 0.01 | 0.22 | 0.21 |
| 5 | 4515447 | rs6881826 | 0.155 | -0.764 | 1.66 | -0.849 | -0.005 | -0.091 | -0.091 | 0 | 0.73 | 0.66 | 0.68 | 0.99 | 0.75 | 0.75 |
| 5 | 4516670 | rs11134055 | 0.008 | -0.888 | 1.679 | -0.901 | -0.001 | -0.013 | -0.013 | 0 | 0.32 | 0.31 | 0.4 | 0.86 | 0.3 | 0.41 |
| 5 | 4517509 | rs13187596 | -0.031 | -0.876 | 1.614 | -0.879 | -0.002 | -0.046 | -0.046 | 0 | 0.33 | 0.41 | 0.41 | 0.85 | 0.3 | 0.41 |
| 5 | 4520220 | rs12517115 | 0.256 | -0.551 | 1.381 | -1.044 | -0.002 | -0.045 | -0.045 | 0 | 0.23 | 0.2 | 0.19 | 0.01 | 0.22 | 0.21 |
| 5 | 4520856 | rs1450822 | 0.977 | -0.492 | 1.02 | -0.653 | -0.033 | 0.01 | 0.01 | 0 | 0.44 | 0.24 | 0.47 | 0.61 | 0.51 | 0.33 |
| 5 | 4532227 | rs17690482 | -0.015 | -0.423 | 0.911 | -0.825 | 0.011 | 0.056 | 0.056 | 0 | 0.27 | 0.14 | 0.21 | 0.01 | 0.26 | 0.25 |
| 5 | 4534191 | rs6555299 | -0.477 | -0.27 | 0.924 | -0.556 | 0 | 0.006 | 0.006 | 0 | 0.76 | 0.72 | 0.7 | 0.59 | 0.8 | 0.72 |
| 5 | 4542749 | rs3943533 | 1.041 | -0.829 | 0.975 | -0.974 | 0.007 | -0.023 | -0.023 | 0 | 0.63 | 0.38 | 0.73 | 0.97 | 0.68 | 0.59 |
| 5 | 4547704 | rs10064215 | 0.74 | -0.481 | 0.156 | -0.557 | -0.005 | -0.082 | -0.082 | 0 | 0.66 | 0.63 | 0.74 | 0.95 | 0.68 | 0.6 |
| 5 | 4568974 | rs12515499 | -0.607 | -0.155 | -0.672 | -0.476 | 0.004 | 0.057 | 0.057 | 0 | 0.32 | 0.16 | 0.25 | 0 | 0.33 | 0.26 |
| 5 | 4574945 | rs1030490 | -1.102 | -0.455 | 0.038 | 0.925 | -0.017 | 0.257 | 0.257 | 0 | 0.1 | 0.28 | 0.18 | 0.49 | 0.04 | 0.25 |
| 5 | 4576516 | rs7700360 | -0.541 | -0.489 | -0.607 | -0.029 | 0.001 | 0.048 | 0.048 | 0 | 0.24 | 0.15 | 0.14 | 0 | 0.26 | 0.18 |
| 5 | 4577399 | rs2052887 | -1.009 | -0.657 | -0.046 | 0.284 | -0.005 | 0.006 | 0.006 | 0 | 0.36 | 0.37 | 0.31 | 0.45 | 0.31 | 0.44 |
| 5 | 4577842 | rs12657445 | -1.271 | -1.247 | 0.023 | -0.271 | 0.06 | 0.004 | 0.004 | 0 | 0.5 | 0.75 | 0.39 | 0.48 | 0.38 | 0.53 |
| 5 | 4578875 | rs16873639 | -0.759 | -1.17 | -0.236 | 0.016 | -0.006 | 0.004 | 0.004 | 0 | 0.35 | 0.34 | 0.29 | 0.46 | 0.27 | 0.39 |
| 5 | 4599400 | rs293067 | -0.227 | -1.678 | -0.804 | -0.393 | 0.049 | 0.087 | 0.087 | 0 | 0.57 | 0.81 | 0.62 | 0.31 | 0.32 | 0.59 |
| 5 | 4600961 | rs7730075 | -0.258 | -1.834 | -1.138 | -0.626 | -0.009 | 0.128 | 0.128 | 1.236 | 0.24 | 0.04 | 0.36 | 0 | 0.22 | 0.16 |
| 5 | 4629210 | rs6887582 | 0.415 | -1.125 | -0.665 | -0.081 | -0.017 | -0.047 | -0.047 | 0 | 0.62 | 0.48 | 0.76 | 0.17 | 0.7 | 0.58 |
| 5 | 4632694 | rs293102 | -0.202 | -0.275 | -1.039 | 0.023 | 0.024 | 0.035 | 0.035 | 0 | 0.51 | 0.61 | 0.47 | 0.71 | 0.37 | 0.63 |
| 5 | 4633275 | rs293105 | -0.402 | -0.217 | -1.967 | -0.058 | -0.049 | -0.028 | -0.028 | 0 | 0.46 | 0.82 | 0.61 | 0.49 | 0.47 | 0.31 |
| 5 | 4633984 | rs10512743 | 1.016 | 0.143 | -1.479 | -0.01 | 0.002 | 0.065 | 0.065 | 0 | 0.06 | 0.12 | 0.13 | 0.28 | 0.08 | 0.06 |
| 5 | 4634162 | rs293106 | -0.453 | 0.15 | -1.808 | -0.182 | 0.008 | 0.036 | 0.036 | 0 | 0.43 | 0.78 | 0.4 | 0.38 | 0.4 | 0.28 |
| 5 | 4638885 | rs7713961 | 0.25 | -0.393 | -2.389 | -0.469 | 0.032 | 0.07 | 0.07 | 0 | 0.22 | 0.08 | 0.08 | 0.05 | 0.12 | 0.18 |
| 5 | 4641160 | rs293119 | -1.343 | -0.217 | -2.451 | -0.401 | -0.127 | -0.161 | -0.161 | 0 | 0.7 | 0.98 | 0.7 | 0.47 | 0.59 | 0.47 |
| 5 | 4644270 | rs293123 | -0.422 | -1.106 | -1.636 | -1.219 | 0.023 | 0.061 | 0.061 | 0 | 0.5 | 0.6 | 0.36 | 0.27 | 0.42 | 0.27 |
| 5 | 4644642 | rs11134061 | 1.157 | -1.073 | -1.882 | -1.612 | -0.055 | 0.101 | 0.101 | 0 | 0.17 | 0.04 | 0.15 | 0 | 0.23 | 0.09 |
| 5 | 4645455 | rs2077369 | -0.398 | -0.868 | -1.106 | -1.296 | 0.016 | 0.025 | 0.025 | 0 | 0.08 | 0.18 | 0.29 | 0.14 | 0.08 | 0.13 |
| 5 | 4647436 | rs7713507 | -0.176 | -1.177 | -1.216 | -1.558 | 0.001 | 0.013 | 0.013 | 0 | 0.16 | 0.15 | 0.28 | 0.06 | 0.16 | 0.11 |
| 5 | 4647464 | rs6555302 | -1.168 | -1.51 | -1.056 | -1.878 | 0.009 | 0.011 | 0.011 | 0 | 0.3 | 0.26 | 0.35 | 0.07 | 0.24 | 0.14 |
| 5 | 4647493 | rs7713659 | -1.751 | -1.753 | -0.631 | -1.817 | 0.018 | 0.02 | 0.02 | 0 | 0.4 | 0.68 | 0.5 | 0.35 | 0.29 | 0.3 |
| 5 | 4648002 | rs418752 | -1.389 | -1.656 | -0.72 | -1.69 | 0.035 | 0.014 | 0.014 | 0.382 | 0.3 | 0.74 | 0.24 | 0.28 | 0.14 | 0.19 |
| 5 | 4648027 | rs1428981 | -0.065 | -1.636 | -0.954 | -1.788 | -0.017 | 0.14 | 0.14 | 0.013 | 0.41 | 0.22 | 0.46 | 0.1 | 0.43 | 0.29 |
| 5 | 4648932 | rs6879780 | -0.359 | -1.162 | -0.326 | -1.031 | -0.008 | 0.176 | 0.176 | 0 | 0.11 | 0.4 | 0.15 | 0.28 | 0.05 | 0.16 |
| 5 | 4651632 | rs16873742 | -0.306 | -1.153 | -0.321 | -1.023 | -0.005 | 0.185 | 0.185 | 0 | 0.11 | 0.4 | 0.15 | 0.28 | 0.05 | 0.16 |
| 5 | 4651742 | rs450409 | -0.965 | -1.173 | -0.395 | -1.043 | -0.054 | 0.057 | 0.057 | 0 | 0.18 | 0.43 | 0.19 | 0.28 | 0.09 | 0.17 |
| 5 | 4654214 | rs1366642 | 1.093 | -0.593 | -1.265 | -1.065 | -0.108 | -0.036 | -0.036 | 0 | 0.36 | 0.14 | 0.3 | 0.62 | 0.44 | 0.56 |
| 5 | 4654629 | rs17692869 | 1.494 | -0.586 | -1.271 | -1.107 | -0.081 | -0.045 | -0.045 | 0.011 | 0.32 | 0.14 | 0.27 | 0.62 | 0.38 | 0.55 |

| | | | | | | | | | | | | | | | | |
|---|---|---|---|---|---|---|---|---|---|---|---|---|---|---|---|---|
| 5 | 4657375 | rs1366643 | 0.871 | -0.769 | -0.777 | -1.315 | -0.006 | -0.034 | -0.034 | 0 | 0.13 | 0.23 | 0.13 | 0.03 | 0.12 | 0.11 |
| 5 | 4657945 | rs1346575 | 0.923 | -0.943 | -0.52 | -1.467 | -0.011 | -0.024 | -0.024 | 0 | 0.68 | 0.48 | 0.45 | 0.73 | 0.68 | 0.79 |
| 5 | 4659940 | rs3928432 | 1.343 | -0.935 | -0.445 | -1.407 | 0.001 | 0.002 | 0.002 | 0.312 | 0.67 | 0.34 | 0.43 | 0.73 | 0.66 | 0.78 |
| 5 | 4660902 | rs2115525 | 1.468 | -0.967 | -0.601 | -1.453 | -0.067 | 0.036 | 0.036 | 0 | 0.39 | 0.2 | 0.32 | 0.69 | 0.43 | 0.62 |
| 5 | 4662323 | rs7447371 | -1.199 | -0.876 | -0.53 | -1.332 | -0.014 | 0.112 | 0.112 | 0.217 | 0.94 | 0.78 | 0.91 | 0.79 | 0.98 | 0.96 |
| 5 | 4664704 | rs2162995 | 0.736 | -0.735 | -0.81 | -1.321 | -0.029 | 0.025 | 0.025 | 0.61 | 0.08 | 0.03 | 0.29 | 0.03 | 0.15 | 0.05 |
| 5 | 4664870 | rs11744458 | 0.457 | -0.741 | -0.618 | -1.213 | -0.002 | -0.049 | -0.049 | 0.041 | 0.15 | 0.23 | 0.14 | 0.02 | 0.13 | 0.11 |
| 5 | 4665670 | rs4370242 | 1.449 | -0.85 | -0.709 | -1.303 | -0.085 | 0.036 | 0.036 | 0 | 0.37 | 0.16 | 0.32 | 0.69 | 0.43 | 0.61 |
| 5 | 4665724 | rs2895497 | 0.61 | -0.72 | -0.744 | -1.166 | 0.02 | 0.055 | 0.055 | 0 | 0.51 | 0.39 | 0.46 | 0.71 | 0.56 | 0.72 |
| 5 | 4668289 | rs4407602 | 0.999 | -0.696 | -0.535 | -0.587 | 0.02 | 0 | 0 | 0 | 0.65 | 0.29 | 0.64 | 0.63 | 0.74 | 0.65 |
| 5 | 4668593 | rs7731649 | 0.271 | -0.604 | -0.578 | -0.698 | 0 | -0.008 | -0.008 | 0.362 | 0.23 | 0.16 | 0.16 | 0.05 | 0.25 | 0.15 |
| 5 | 4670864 | rs7724465 | 0.193 | -0.41 | -0.199 | -0.062 | -0.004 | 0.047 | 0.047 | 0 | 0.91 | 0.77 | 0.86 | 0.81 | 0.92 | 0.89 |
| 5 | 4671383 | rs6555306 | 0.498 | -0.368 | -0.209 | -0.063 | -0.006 | 0.026 | 0.026 | 0 | 0.85 | 0.68 | 0.81 | 0.81 | 0.88 | 0.85 |
| 5 | 4673835 | rs7720816 | 0.721 | -0.011 | -1.033 | 0.272 | -0.047 | -0.075 | -0.075 | 0 | 0.41 | 0.1 | 0.32 | 0.47 | 0.4 | 0.48 |
| 5 | 4674868 | rs4412086 | -0.069 | -0.067 | -0.263 | -0.223 | 0.001 | 0.01 | 0.01 | 0 | 0.45 | 0.36 | 0.29 | 0.22 | 0.43 | 0.35 |
| 5 | 4675355 | rs7716432 | 0.761 | -0.094 | -0.393 | 0.015 | -0.024 | 0.033 | 0.033 | 0.015 | 0.41 | 0.19 | 0.44 | 0.62 | 0.44 | 0.53 |
| 5 | 4676429 | rs4568336 | 1.039 | -0.14 | 0.555 | 0.696 | 0.003 | 0.104 | 0.104 | 0 | 0.94 | 0.71 | 0.96 | 0.89 | 0.99 | 0.91 |
| 5 | 4677722 | rs13359697 | -0.913 | 0.084 | 0.711 | 0.364 | 0.02 | -0.01 | -0.01 | 0 | 0.13 | 0.25 | 0.29 | 0.16 | 0.15 | 0.13 |
| 5 | 4678192 | rs7713482 | 1.783 | -0.571 | -0.258 | 0.51 | -0.071 | -0.12 | -0.12 | 0.237 | 0.48 | 0.05 | 0.48 | 0.44 | 0.57 | 0.48 |
| 5 | 4678221 | rs10063789 | -0.468 | -0.445 | -0.014 | 0.584 | 0.016 | -0.013 | -0.013 | 0 | 0.13 | 0.24 | 0.29 | 0.16 | 0.15 | 0.14 |
| 5 | 4678287 | rs7718402 | 0.044 | -0.828 | 1.028 | 0.701 | -0.031 | -0.007 | -0.007 | 0 | 0.85 | 0.74 | 0.95 | 0.89 | 0.92 | 0.88 |
| 5 | 4679559 | rs4235620 | 1.692 | -0.865 | 1.125 | 0.682 | 0.023 | -0.011 | -0.011 | 0.81 | 0.63 | 0.31 | 0.79 | 0.6 | 0.73 | 0.68 |
| 5 | 4680691 | rs4563594 | 0.473 | -0.814 | 0.842 | 0.575 | -0.019 | -0.017 | -0.017 | 0 | 0.82 | 0.74 | 0.91 | 0.88 | 0.86 | 0.87 |
| 5 | 4695085 | rs6885274 | -0.175 | -0.723 | 1.021 | 0.551 | -0.005 | -0.021 | -0.021 | 1.703 | 0.09 | 0.07 | 0.02 | 0.02 | 0.06 | 0.06 |
| 5 | 4700587 | rs4702913 | 1.456 | -0.532 | 1.123 | 0.675 | 0 | 0.04 | 0.04 | 0 | 0.83 | 0.76 | 0.9 | 0.71 | 0.85 | 0.79 |
| 5 | 4705982 | rs12716162 | -0.679 | -0.506 | 1.274 | 0.602 | 0.021 | -0.008 | -0.008 | 0 | 0.11 | 0.26 | 0.28 | 0.14 | 0.12 | 0.14 |
| 5 | 4706519 | rs4532332 | 1.491 | -0.634 | 1.208 | 0.589 | -0.014 | 0.028 | 0.028 | 0 | 0.7 | 0.6 | 0.85 | 0.61 | 0.76 | 0.69 |
| 5 | 4717060 | rs2047728 | 2.218 | -0.608 | 0.382 | 0.559 | -0.044 | -0.092 | -0.092 | 0 | 0.53 | 0.08 | 0.51 | 0.45 | 0.59 | 0.55 |
| 5 | 4724332 | rs7711434 | 1.476 | -0.719 | 0.879 | 1.051 | 0.011 | 0.05 | 0.05 | 0 | 0.87 | 0.82 | 0.74 | 0.64 | 0.84 | 0.84 |
| 5 | 4725405 | rs10035433 | -0.951 | -0.645 | 0.537 | 0.796 | -0.027 | 0 | 0 | 0 | 0.19 | 0.38 | 0.1 | 0.18 | 0.11 | 0.21 |
| 5 | 4730319 | rs6895818 | -1.101 | -0.703 | 0.518 | 0.84 | -0.032 | -0.001 | -0.001 | 0.302 | 0.19 | 0.41 | 0.1 | 0.2 | 0.1 | 0.21 |
| 5 | 4738222 | rs1393120 | 1.021 | -0.265 | -0.026 | 1.288 | -0.009 | 0.03 | 0.03 | 0 | 0.94 | 0.95 | 0.85 | 0.71 | 0.89 | 0.89 |
| 5 | 4755398 | rs7716642 | 1.512 | -0.439 | -0.042 | 1.298 | -0.002 | -0.006 | -0.006 | 0 | 0.83 | 0.85 | 0.75 | 0.63 | 0.83 | 0.8 |
| 5 | 4776191 | rs2088900 | 1.682 | -1.046 | -1.012 | 1.292 | -0.006 | -0.032 | -0.032 | 5.763 | 0.57 | 0.08 | 0.46 | 0.33 | 0.67 | 0.6 |
| 5 | 4776500 | rs4235550 | 1.098 | -0.7 | -0.087 | 1.443 | 0 | 0.016 | 0.016 | 0 | 0.88 | 0.87 | 0.79 | 0.7 | 0.89 | 0.77 |
| 5 | 4777480 | rs10069581 | -1.407 | -0.739 | 0.72 | 0.738 | 0.007 | 0.018 | 0.018 | 0 | 0.11 | 0.37 | 0.06 | 0.14 | 0.07 | 0.16 |
| 5 | 4780629 | rs16874044 | -1.295 | -1.224 | 0.818 | 0.397 | -0.041 | -0.05 | -0.05 | 0 | 0.23 | 0.34 | 0.28 | 0.15 | 0.16 | 0.22 |
| 5 | 4784657 | rs1173229 | 1.157 | -1.349 | 1.117 | 0.78 | -0.025 | 0.023 | 0.023 | 0 | 0.67 | 0.61 | 0.57 | 0.41 | 0.8 | 0.58 |
| 5 | 4785389 | rs1173228 | 1.623 | -1.377 | 1.286 | 0.758 | 0.04 | 0.028 | 0.028 | 0 | 0.61 | 0.29 | 0.51 | 0.31 | 0.73 | 0.57 |
| 5 | 4785594 | rs1173227 | 1.401 | -1.375 | 1.323 | 0.757 | -0.005 | 0.002 | 0.002 | 0 | 0.63 | 0.46 | 0.52 | 0.39 | 0.74 | 0.57 |
| 5 | 4788892 | rs1173218 | 1.524 | -1.423 | 0.622 | 0.659 | 0.065 | 0.037 | 0.037 | 0 | 0.6 | 0.19 | 0.5 | 0.31 | 0.74 | 0.57 |
| 5 | 4795354 | rs1452040 | 1.184 | -1.378 | 0.687 | 0.133 | -0.029 | -0.012 | -0.012 | 2.055 | 0.65 | 0.54 | 0.55 | 0.28 | 0.78 | 0.58 |
| 5 | 4801119 | rs10051649 | -1.826 | -1.511 | -0.065 | -0.139 | 0.081 | 0.133 | 0.133 | 0.053 | 0.44 | 0.84 | 0.49 | 0.8 | 0.28 | 0.44 |
| 5 | 4801996 | rs7728112 | -1.629 | -1.211 | 0.786 | -0.652 | -0.001 | 0.058 | 0.058 | 0.286 | 0.09 | 0.3 | 0.06 | 0.11 | 0.07 | 0.06 |
| 5 | 4803549 | rs1173179 | -0.005 | -0.641 | -0.15 | -1.61 | 0.021 | 0.172 | 0.172 | 0 | 0.5 | 0.28 | 0.49 | 0.13 | 0.5 | 0.51 |
| 5 | 4814945 | rs1173192 | -2.394 | -0.553 | 0.678 | -1.613 | 0.013 | 0.001 | 0.001 | 0 | 0.28 | 0.73 | 0.33 | 0.23 | 0.2 | 0.24 |
| 5 | 4819918 | rs1173201 | -2.147 | -0.654 | 0.711 | -1.613 | -0.008 | 0 | 0 | 0.098 | 0.24 | 0.7 | 0.29 | 0.23 | 0.12 | 0.25 |
| 5 | 4825401 | rs1173213 | -2.641 | -0.661 | 1.688 | -1.674 | 0.05 | 0.007 | 0.007 | 0.058 | 0.31 | 0.83 | 0.36 | 0.24 | 0.21 | 0.27 |
| 5 | 4828015 | rs11134077 | -0.587 | -0.698 | 1.963 | -1.645 | -0.001 | -0.047 | -0.047 | 0 | 0.09 | 0.28 | 0.05 | 0 | 0.03 | 0.02 |
| 5 | 4831601 | rs816475 | -2.101 | -0.7 | 1.841 | -1.646 | -0.038 | -0.031 | -0.031 | 0 | 0.21 | 0.58 | 0.28 | 0.13 | 0.12 | 0.2 |
| 5 | 4831687 | rs2123104 | -0.509 | -0.688 | 1.499 | -1.636 | 0.003 | -0.006 | -0.006 | 0 | 0.08 | 0.07 | 0.21 | 0.12 | 0.06 | 0.12 |
| 5 | 4835205 | rs816480 | -2.864 | -0.783 | 1.582 | -2.106 | 0.046 | 0.038 | 0.038 | 0.759 | 0.28 | 0.81 | 0.31 | 0.13 | 0.14 | 0.18 |
| 5 | 4840284 | rs2652227 | -2.198 | -0.86 | 1.682 | -2.08 | -0.045 | -0.084 | -0.084 | 0 | 0.21 | 0.66 | 0.26 | 0.02 | 0.07 | 0.12 |
| 5 | 4853391 | rs860987 | -2.8 | -0.852 | 1.496 | -2.073 | 0.036 | 0.031 | 0.031 | 1.044 | 0.28 | 0.79 | 0.3 | 0.13 | 0.14 | 0.18 |
| 5 | 4867913 | rs707645 | -1.877 | -0.806 | 1.442 | -2.283 | -0.056 | -0.136 | -0.136 | 0.052 | 0.26 | 0.53 | 0.27 | 0.02 | 0.12 | 0.14 |
| 5 | 4872310 | rs816467 | -1.465 | -0.854 | 1.466 | -2.302 | -0.061 | -0.12 | -0.12 | 1.946 | 0.19 | 0.48 | 0.24 | 0.01 | 0.09 | 0.11 |
| 5 | 4872511 | rs816468 | -2.504 | -0.84 | 0.776 | -2.274 | 0.088 | 0.108 | 0.108 | 0.504 | 0.34 | 0.86 | 0.32 | 0.12 | 0.18 | 0.18 |
| 5 | 4875420 | rs16874171 | -0.628 | -0.65 | 1.182 | -2.167 | -0.018 | -0.053 | -0.053 | 0.056 | 0.1 | 0.22 | 0.03 | 0.01 | 0.03 | 0.03 |
| 5 | 4884392 | rs7728954 | -1.504 | -0.655 | 0.832 | -2.157 | -0.053 | -0.01 | -0.01 | 0 | 0.17 | 0.57 | 0.06 | 0.11 | 0.06 | 0.12 |
| 5 | 4885068 | rs629342 | -1.228 | -0.017 | -0.643 | -2.709 | -0.055 | 0.212 | 0.212 | 0.046 | 0.55 | 0.88 | 0.39 | 0.12 | 0.48 | 0.44 |
| 5 | 4885562 | rs17752880 | 1.142 | -0.54 | -0.817 | -2.72 | -0.081 | 0.085 | 0.085 | 0 | 0.25 | 0.1 | 0.29 | 0.01 | 0.4 | 0.3 |
| 5 | 4887171 | rs10051514 | -1.102 | -0.423 | -0.99 | -2.53 | -0.037 | 0.204 | 0.204 | 0 | 0.57 | 0.91 | 0.44 | 0.12 | 0.58 | 0.46 |
| 5 | 4900062 | rs590199 | -0.884 | -0.51 | 0.118 | -2.292 | -0.042 | -0.076 | -0.076 | 0.403 | 0.18 | 0.48 | 0.1 | 0.07 | 0.08 | 0.06 |
| 5 | 4901816 | rs26875 | -1.448 | -0.416 | -0.999 | -2.3 | 0.04 | 0.074 | 0.074 | 0.252 | 0.31 | 0.77 | 0.17 | 0.09 | 0.16 | 0.11 |
| 5 | 4905892 | rs26880 | -0.36 | -0.138 | -1.392 | -2.397 | -0.053 | 0.26 | 0.26 | 0 | 0.47 | 0.88 | 0.5 | 0.13 | 0.47 | 0.37 |
| 5 | 4907084 | rs6894984 | 0.324 | -0.059 | -1.007 | -2.347 | -0.003 | -0.026 | -0.026 | 0 | 0.09 | 0.1 | 0.05 | 0.01 | 0.07 | 0.02 |
| 5 | 4907778 | rs1392411 | 1.001 | -0.074 | -1.107 | -2.439 | -0.089 | 0.118 | 0.118 | 0 | 0.27 | 0.1 | 0.35 | 0.01 | 0.39 | 0.3 |
| 5 | 4912310 | rs610722 | -0.75 | -0.418 | -1.4 | -2.512 | -0.062 | 0.152 | 0.152 | 0.11 | 0.61 | 0.94 | 0.58 | 0.16 | 0.61 | 0.45 |
| 5 | 4914886 | rs6555321 | 0.837 | -0.26 | -0.445 | -2.563 | -0.13 | 0.187 | 0.187 | 0.034 | 0.33 | 0.07 | 0.4 | 0.01 | 0.49 | 0.37 |

| | | | | | | | | | | | | | | | | |
|---|---|---|---|---|---|---|---|---|---|---|---|---|---|---|---|---|
| 5 | 4917365 | rs484696 | 0.953 | -0.891 | -0.051 | -2.553 | 0.062 | 0.219 | 0.219 | 0.688 | 0.46 | 0.15 | 0.47 | 0.01 | 0.66 | 0.4 |
| 5 | 4924195 | rs10475277 | -0.146 | -0.983 | 0.543 | -2.551 | 0.007 | 0.029 | 0.029 | 2.671 | 0.06 | 0.29 | 0.03 | 0.01 | 0 | 0.02 |
| 5 | 4928318 | rs501031 | 1.001 | -0.927 | 0.119 | -2.567 | 0.058 | 0.277 | 0.277 | 0.664 | 0.46 | 0.12 | 0.47 | 0.01 | 0.66 | 0.4 |
| 5 | 4928520 | rs502854 | 0.704 | -0.944 | 0.416 | -2.542 | 0.039 | 0.057 | 0.057 | 0 | 0.5 | 0.38 | 0.51 | 0.12 | 0.66 | 0.42 |
| 5 | 4930407 | rs30452 | 0.538 | -0.218 | 0.518 | -2.31 | 0.016 | 0.01 | 0.01 | 0.356 | 0.58 | 0.67 | 0.52 | 0.04 | 0.66 | 0.39 |
| 5 | 4936235 | rs654685 | 0.825 | -0.131 | 0.796 | -1.777 | -0.008 | -0.063 | -0.063 | 2.449 | 0.6 | 0.5 | 0.55 | 0.11 | 0.69 | 0.43 |
| 5 | 4938915 | rs6868240 | -1.117 | -0.002 | 0.232 | -1.255 | -0.007 | -0.059 | -0.059 | 0.35 | 0.15 | 0.31 | 0.11 | 0.01 | 0.12 | 0.07 |
| 5 | 4941242 | rs11134086 | 0.613 | 0.195 | -0.015 | -0.588 | 0.005 | 0.048 | 0.048 | 0 | 0.67 | 0.8 | 0.62 | 0.18 | 0.7 | 0.4 |
| 5 | 4946412 | rs30445 | -0.545 | 0.055 | -0.127 | 0.594 | -0.013 | -0.096 | -0.096 | 0.071 | 0.3 | 0.11 | 0.3 | 0.59 | 0.28 | 0.53 |
| 5 | 4948424 | rs643466 | 0.926 | -0.091 | 0.376 | 0.22 | 0.025 | 0.049 | 0.049 | 0.056 | 0.62 | 0.32 | 0.58 | 0.19 | 0.7 | 0.43 |
| 5 | 4948859 | rs466277 | 0.668 | -0.311 | 1.063 | 1.07 | -0.053 | 0.159 | 0.159 | 0.031 | 0.78 | 0.52 | 0.69 | 0.54 | 0.85 | 0.66 |
| 5 | 4950651 | rs458279 | 0.598 | 0.118 | 0.802 | 0.966 | 0.002 | -0.035 | -0.035 | 0 | 0.71 | 0.77 | 0.6 | 0.56 | 0.66 | 0.59 |
| 5 | 4951614 | rs26847 | 1.018 | -0.108 | 1.162 | 0.862 | 0.022 | -0.028 | -0.028 | 0.482 | 0.64 | 0.26 | 0.54 | 0.56 | 0.65 | 0.58 |
| 5 | 4955950 | rs10070917 | 1.007 | -0.075 | 1.298 | 0.888 | 0.032 | -0.035 | -0.035 | 0 | 0.64 | 0.25 | 0.54 | 0.55 | 0.64 | 0.56 |
| 5 | 4959279 | rs1366431 | -1.048 | 0.502 | 1.026 | 0.512 | -0.001 | -0.01 | -0.01 | 0 | 0.37 | 0.59 | 0.42 | 0.24 | 0.38 | 0.42 |
| 5 | 4968029 | rs1452027 | -1.231 | 0.932 | 1.512 | 1.233 | 0.014 | 0.01 | 0.01 | 0.347 | 0.3 | 0.44 | 0.34 | 0.32 | 0.36 | 0.46 |
| 5 | 4977446 | rs42202 | -0.751 | 0.881 | 0.451 | 1.404 | -0.025 | -0.068 | -0.068 | 0 | 0.89 | 1 | 0.85 | 0.6 | 0.91 | 0.87 |
| 5 | 4978270 | rs42401 | -1.331 | 0.982 | 1.058 | 1.287 | -0.006 | -0.014 | -0.014 | 0 | 0.32 | 0.74 | 0.38 | 0.43 | 0.36 | 0.48 |
| 5 | 4987376 | rs30463 | -1.062 | 0.934 | 0.993 | 1.163 | -0.027 | -0.012 | -0.012 | 0 | 0.28 | 0.23 | 0.31 | 0.3 | 0.35 | 0.46 |
| 5 | 4992930 | rs30449 | -1.374 | 0.966 | 0.528 | 1.102 | -0.019 | -0.027 | -0.027 | 0 | 0.33 | 0.84 | 0.38 | 0.4 | 0.36 | 0.47 |
| 5 | 5009457 | rs30485 | -1.344 | 0.791 | 1.633 | 0.47 | -0.006 | -0.074 | -0.074 | 0 | 0.14 | 0.36 | 0.11 | 0.04 | 0.1 | 0.08 |
| 5 | 5023441 | rs458084 | -1.547 | 1.022 | 0.804 | 0.257 | -0.027 | 0.111 | 0.111 | 0 | 0.48 | 0.81 | 0.57 | 0.71 | 0.46 | 0.61 |
| 5 | 5023977 | rs467075 | -0.521 | 1.019 | 0.872 | 0.247 | 0.002 | -0.006 | -0.006 | 0 | 0.13 | 0.1 | 0.07 | 0.04 | 0.09 | 0.05 |
| 5 | 5024621 | rs465871 | -0.943 | 0.98 | 1.011 | 0.238 | 0.008 | -0.014 | -0.014 | 0.003 | 0.39 | 0.52 | 0.5 | 0.67 | 0.41 | 0.58 |
| 5 | 5032533 | rs6878673 | -1.056 | 1.202 | 0.923 | 0.346 | -0.005 | 0.027 | 0.027 | 0.061 | 0.44 | 0.58 | 0.53 | 0.69 | 0.44 | 0.59 |
| 5 | 5038540 | rs446464 | 1.395 | 1.346 | 1.481 | 0.675 | 0.022 | 0.045 | 0.045 | 0.167 | 0.91 | 0.75 | 0.91 | 0.87 | 0.91 | 0.91 |
| 5 | 5051464 | rs272194 | 0.751 | 1.483 | 1.414 | 1.087 | 0.04 | 0.112 | 0.112 | 0 | 0.78 | 0.67 | 0.66 | 0.51 | 0.72 | 0.55 |
| 5 | 5052953 | rs16874638 | -1.108 | 1.282 | 1.989 | 0.978 | 0.002 | -0.027 | -0.027 | 0.706 | 0.07 | 0.27 | 0.04 | 0.01 | 0.04 | 0.02 |
| 5 | 5052998 | rs272195 | 0.606 | 1.321 | 1.79 | 1.013 | 0.044 | 0.118 | 0.118 | 0 | 0.79 | 0.68 | 0.66 | 0.51 | 0.72 | 0.55 |
| 5 | 5063242 | rs156251 | 1.932 | 1.264 | 1.71 | 0.689 | 0.044 | -0.008 | -0.008 | 0 | 0.89 | 0.62 | 0.91 | 0.92 | 0.87 | 0.96 |
| 5 | 5074825 | rs270636 | 1.409 | 1.544 | 1.134 | 0.334 | 0.015 | 0.016 | 0.016 | 0.174 | 0.51 | 0.45 | 0.39 | 0.1 | 0.4 | 0.31 |
| 5 | 5078313 | rs3733787 | -0.624 | 1.831 | 0.822 | 0.558 | -0.02 | -0.013 | -0.013 | 0 | 0.27 | 0.19 | 0.36 | 0.4 | 0.4 | 0.45 |
| 5 | 5080435 | rs6866197 | -0.149 | 1.597 | 0.961 | 0.535 | -0.001 | 0.062 | 0.062 | 1.075 | 0.07 | 0.38 | 0.04 | 0.06 | 0 | 0.01 |
| 5 | 5089229 | rs1382896 | 1.223 | 1.685 | 0.948 | 0.484 | 0.014 | 0.041 | 0.041 | 0.324 | 0.54 | 0.62 | 0.41 | 0.13 | 0.42 | 0.31 |
| 5 | 5093643 | rs1160800 | 1.223 | 1.67 | 0.948 | 0.484 | 0.015 | 0.041 | 0.041 | 0.324 | 0.54 | 0.62 | 0.41 | 0.13 | 0.41 | 0.31 |
| 5 | 5098840 | rs814797 | -1.673 | 1.5 | 0.09 | 0.489 | 0.014 | 0.045 | 0.045 | 0 | 0.54 | 0.63 | 0.62 | 0.82 | 0.59 | 0.7 |
| 5 | 5106180 | rs16874709 | 0.034 | 1.483 | -0.853 | 0.852 | 0.012 | -0.074 | -0.074 | 0 | 0.18 | 0.08 | 0.22 | 0.37 | 0.13 | 0.22 |
| 5 | 5110692 | rs814790 | -0.834 | 2.324 | -0.314 | 0.567 | -0.059 | -0.053 | -0.053 | 0.158 | 0.25 | 0.08 | 0.26 | 0.39 | 0.34 | 0.4 |
| 5 | 5116544 | rs2115010 | -0.282 | 2.157 | 0.623 | 0.006 | 0.111 | -0.019 | -0.019 | 0.505 | 0.07 | 0.22 | 0.21 | 0.02 | 0.13 | 0.07 |
| 5 | 5120745 | rs814783 | 0.464 | 2.18 | 0.169 | -0.193 | -0.017 | -0.006 | -0.006 | 0 | 0.69 | 0.83 | 0.7 | 0.33 | 0.66 | 0.74 |
| 5 | 5120776 | rs861512 | 1.537 | 1.733 | 0.455 | -0.438 | -0.025 | 0.09 | 0.09 | 0.217 | 0.54 | 0.21 | 0.46 | 0.3 | 0.52 | 0.65 |
| 5 | 5121337 | rs1834925 | -0.669 | 1.747 | 0.9 | -0.534 | 0.141 | -0.033 | -0.033 | 0 | 0.1 | 0.4 | 0.22 | 0.02 | 0.13 | 0.08 |
| 5 | 5123214 | rs9313102 | -1.48 | 1.752 | 0.893 | -0.531 | 0.036 | -0.081 | -0.081 | 0 | 0.15 | 0.59 | 0.24 | 0.02 | 0.15 | 0.07 |
| 5 | 5124149 | rs270188 | -0.135 | 1.944 | 0.621 | -0.386 | 0.001 | -0.023 | -0.023 | 0 | 0.83 | 0.9 | 0.82 | 0.43 | 0.84 | 0.89 |
| 5 | 5124263 | rs270187 | 1.075 | 2.064 | 0.362 | -0.355 | 0 | -0.001 | -0.001 | 0 | 0.64 | 0.64 | 0.68 | 0.38 | 0.65 | 0.74 |
| 5 | 5124549 | rs16874791 | -0.625 | 2.253 | 0.482 | -0.657 | -0.008 | 0.017 | 0.017 | 0 | 0.15 | 0.11 | 0.12 | 0.06 | 0.18 | 0.13 |
| 5 | 5125959 | rs10475286 | -0.93 | 2.09 | 0.476 | -0.515 | 0.133 | -0.031 | -0.031 | 0 | 0.1 | 0.37 | 0.22 | 0.02 | 0.14 | 0.09 |
| 5 | 5126400 | rs10042501 | -0.811 | 2.125 | 0.343 | -0.499 | 0.125 | -0.03 | -0.03 | 0.278 | 0.1 | 0.33 | 0.22 | 0.02 | 0.14 | 0.08 |
| 5 | 5129879 | rs12516403 | -0.787 | 1.337 | -0.822 | -0.268 | -0.01 | 0.043 | 0.043 | 0 | 0.17 | 0.07 | 0.13 | 0.09 | 0.2 | 0.2 |
| 5 | 5133444 | rs751905 | -0.092 | 1.084 | -0.484 | -0.467 | 0.004 | -0.095 | -0.095 | 0 | 0.31 | 0.46 | 0.2 | 0.12 | 0.33 | 0.2 |
| 5 | 5136207 | rs7720806 | -0.12 | 0.894 | -0.821 | 0.335 | 0.001 | -0.016 | -0.016 | 0 | 0.33 | 0.31 | 0.45 | 0.47 | 0.3 | 0.29 |
| 5 | 5140970 | rs270206 | 0.134 | 0.56 | -0.702 | -0.3 | -0.003 | -0.019 | -0.019 | 0 | 0.64 | 0.63 | 0.77 | 0.86 | 0.71 | 0.65 |
| 5 | 5141905 | rs270198 | -0.649 | 0.759 | -0.59 | 0.179 | 0 | 0 | 0 | 3.198 | 0.44 | 0.55 | 0.61 | 0.5 | 0.42 | 0.38 |
| 5 | 5142460 | rs2291112 | 0.495 | 0.744 | -0.594 | 0.221 | -0.001 | 0 | 0 | 0 | 0.58 | 0.43 | 0.41 | 0.43 | 0.61 | 0.64 |
| 5 | 5142768 | rs270193 | -0.646 | 0.816 | -0.612 | 0.654 | 0 | -0.003 | -0.003 | 0 | 0.42 | 0.56 | 0.61 | 0.69 | 0.4 | 0.38 |
| 5 | 5145406 | rs1863970 | -0.437 | 0.894 | 0.019 | 1.245 | 0.004 | 0.027 | 0.027 | 0 | 0.79 | 0.77 | 0.51 | 0.59 | 0.75 | 0.88 |
| 5 | 5146395 | rs1863968 | 0.956 | 0.847 | 0.234 | 1.198 | 0.003 | 0 | 0 | 0 | 0.57 | 0.35 | 0.4 | 0.43 | 0.61 | 0.64 |
| 5 | 5147376 | rs2255273 | -1.031 | 1.666 | 0.291 | 1.051 | 0.024 | 0.027 | 0.027 | 0 | 0.83 | 0.94 | 0.94 | 0.9 | 0.91 | 0.81 |
| 5 | 5149621 | rs2560412 | 0.296 | 1.54 | 0.36 | 1.008 | 0.005 | 0.016 | 0.016 | 0 | 0.2 | 0.22 | 0.49 | 0.34 | 0.24 | 0.1 |
| 5 | 5152447 | rs11134095 | 0.318 | 1.5 | 0.374 | 1.005 | -0.006 | 0.034 | 0.034 | 0 | 0.21 | 0.31 | 0.12 | 0.34 | 0.16 | 0.26 |
| 5 | 5153648 | rs978280 | 1.124 | 1.487 | 0.347 | 0.951 | 0.002 | 0.186 | 0.186 | 0.516 | 0.52 | 0.2 | 0.31 | 0.13 | 0.48 | 0.55 |
| 5 | 5153911 | rs16874878 | 0.47 | 1.528 | 0.601 | 0.976 | -0.003 | 0.017 | 0.017 | 0 | 0.21 | 0.27 | 0.12 | 0.34 | 0.16 | 0.26 |
| 5 | 5154215 | rs922542 | -0.162 | 1.535 | 0.544 | 0.954 | 0.042 | 0.051 | 0.051 | 0 | 0.26 | 0.41 | 0.57 | 0.46 | 0.37 | 0.18 |
| 5 | 5154458 | rs6555331 | 1.063 | 1.472 | 0.278 | 0.746 | 0.002 | 0.185 | 0.185 | 0.716 | 0.52 | 0.2 | 0.31 | 0.13 | 0.48 | 0.55 |
| 5 | 5155357 | rs814775 | -0.148 | 1.593 | 0.924 | 0.901 | 0.011 | 0.023 | 0.023 | 0 | 0.23 | 0.32 | 0.53 | 0.45 | 0.31 | 0.17 |
| 5 | 5156655 | rs6899080 | 0.807 | 1.33 | 0.446 | 0.249 | -0.006 | 0.063 | 0.063 | 0 | 0.31 | 0.16 | 0.19 | 0.06 | 0.33 | 0.4 |
| 5 | 5159316 | rs7720897 | -0.32 | 1.289 | 0.362 | 0.435 | -0.003 | 0.014 | 0.014 | 0 | 0.74 | 0.73 | 0.51 | 0.57 | 0.74 | 0.84 |
| 5 | 5162228 | rs992542 | -0.155 | 1.166 | 0.175 | 0.548 | 0.001 | 0.022 | 0.022 | 0 | 0.72 | 0.66 | 0.49 | 0.42 | 0.72 | 0.77 |
| 5 | 5165415 | rs270178 | -0.073 | 1.199 | 0.07 | 0.162 | 0.005 | -0.023 | -0.023 | 0.376 | 0.6 | 0.45 | 0.71 | 0.76 | 0.52 | 0.52 |

| | | | | | | | | | | | | | | | | |
|---|---|---|---|---|---|---|---|---|---|---|---|---|---|---|---|---|
| 5 | 5165870 | rs874428 | 1.207 | 1.13 | -0.385 | -0.128 | 0.006 | 0.041 | 0.041 | 0 | 0.2 | 0.08 | 0.13 | 0 | 0.18 | 0.23 |
| 5 | 5165952 | rs1079295 | 0.291 | 1.246 | 0.177 | 0.359 | 0.013 | 0.049 | 0.049 | 0.306 | 0.7 | 0.55 | 0.44 | 0.43 | 0.64 | 0.77 |
| 5 | 5167587 | rs997427 | 0.473 | 1.473 | 0.164 | 0.401 | 0.026 | 0.007 | 0.007 | 0.704 | 0.68 | 0.43 | 0.4 | 0.28 | 0.58 | 0.7 |
| 5 | 5167648 | rs997428 | -0.769 | 1.139 | 0.186 | 0.422 | 0.038 | 0.073 | 0.073 | 0 | 0.06 | 0.17 | 0.07 | 0.14 | 0.12 | 0.11 |
| 5 | 5176672 | rs4701693 | 0.222 | 0.765 | 0.123 | 0.273 | -0.019 | 0.117 | 0.117 | 0 | 0.17 | 0.26 | 0.45 | 0.4 | 0.14 | 0.16 |
| 5 | 5178624 | rs4702234 | 0.155 | 0.674 | -0.506 | -0.081 | 0.001 | 0 | 0 | 0.121 | 0.41 | 0.35 | 0.6 | 0.41 | 0.4 | 0.41 |
| 5 | 5181634 | rs13177816 | 0.113 | 0.833 | -1.375 | -0.583 | -0.024 | 0.035 | 0.035 | 0 | 0.27 | 0.21 | 0.19 | 0.07 | 0.33 | 0.25 |
| 5 | 5181829 | rs13176907 | 0.324 | 0.88 | -1.904 | -0.53 | -0.088 | 0.099 | 0.099 | 0 | 0.25 | 0.06 | 0.17 | 0.07 | 0.33 | 0.25 |
| 5 | 5181928 | rs7724547 | 0.516 | 0.101 | -1.332 | 0.143 | 0.021 | -0.015 | -0.015 | 0.002 | 0.25 | 0.13 | 0.12 | 0.24 | 0.19 | 0.25 |
| 5 | 5183700 | rs1808529 | -0.219 | -0.12 | -1.017 | 0.087 | 0.049 | -0.026 | -0.026 | 0 | 0.4 | 0.68 | 0.61 | 0.46 | 0.27 | 0.22 |
| 5 | 5183783 | rs10037656 | 0.343 | -0.4 | -0.16 | 0.023 | -0.038 | 0.198 | 0.198 | 0.077 | 0.16 | 0.32 | 0.47 | 0.45 | 0.1 | 0.16 |
| 5 | 5184252 | rs12518196 | 1.431 | 0.379 | -0.876 | -0.232 | -0.069 | 0.03 | 0.03 | 0 | 0.18 | 0.01 | 0.15 | 0.06 | 0.23 | 0.21 |
| 5 | 5187371 | rs4702235 | -0.216 | -0.054 | -0.074 | 0.064 | -0.011 | -0.061 | -0.061 | 0.612 | 0.71 | 0.52 | 0.8 | 0.8 | 0.71 | 0.83 |
| 5 | 5191047 | rs11743270 | 1.596 | -0.195 | -0.028 | 0.409 | -0.017 | 0.143 | 0.143 | 0 | 0.07 | 0.12 | 0.25 | 0.32 | 0.05 | 0.09 |
| 5 | 5191117 | rs12517136 | 1.215 | -0.117 | 0.006 | 0.065 | 0.036 | 0.074 | 0.074 | 0.062 | 0.2 | 0.1 | 0.1 | 0 | 0.15 | 0.07 |
| 5 | 5192434 | rs12520783 | 1.128 | -0.111 | 0.165 | 0.068 | 0.013 | -0.001 | -0.001 | 0.002 | 0.21 | 0.14 | 0.1 | 0 | 0.15 | 0.07 |
| 5 | 5192861 | rs871122 | 0.892 | -0.203 | 0.066 | 0.04 | -0.009 | 0.055 | 0.055 | 0 | 0.15 | 0.19 | 0.46 | 0.4 | 0.13 | 0.15 |
| 5 | 5195069 | rs10039354 | 1.293 | 0.008 | -0.04 | -0.113 | -0.001 | -0.02 | -0.02 | 0 | 0.22 | 0.13 | 0.09 | 0.26 | 0.19 | 0.26 |
| 5 | 5195384 | rs4701694 | -0.253 | 0.206 | 0.068 | -0.123 | -0.017 | -0.049 | -0.049 | 0 | 0.71 | 0.49 | 0.81 | 0.79 | 0.73 | 0.82 |
| 5 | 5195721 | rs16874994 | 1.189 | 0.258 | 0.14 | -0.178 | -0.01 | -0.05 | -0.05 | 0 | 0.21 | 0.23 | 0.1 | 0 | 0.15 | 0.07 |
| 5 | 5196007 | rs2081852 | -0.413 | 0.39 | 0.105 | -0.101 | 0.06 | 0.04 | 0.04 | 0 | 0.93 | 0.82 | 0.91 | 0.84 | 0.86 | 0.9 |
| 5 | 5197553 | rs3943879 | 1.18 | 0.431 | -0.457 | -0.116 | -0.077 | 0.066 | 0.066 | 0.828 | 0.3 | 0.11 | 0.23 | 0.14 | 0.38 | 0.41 |
| 5 | 5198567 | rs16875033 | 1.078 | 0.379 | -0.293 | -0.289 | -0.039 | -0.106 | -0.106 | 0.004 | 0.23 | 0.32 | 0.11 | 0 | 0.15 | 0.07 |
| 5 | 5199350 | rs1346460 | -0.378 | 0.419 | -0.326 | -0.271 | -0.024 | -0.07 | -0.07 | 0 | 0.71 | 0.57 | 0.83 | 0.83 | 0.74 | 0.82 |
| 5 | 5200281 | rs6555335 | -0.384 | 0.455 | -0.317 | -0.239 | -0.025 | -0.054 | -0.054 | 0 | 0.71 | 0.57 | 0.83 | 0.8 | 0.74 | 0.77 |
| 5 | 5200490 | rs17762877 | 1.104 | 0.451 | -0.658 | -0.264 | -0.088 | 0.12 | 0.12 | 0.954 | 0.3 | 0.11 | 0.23 | 0.07 | 0.39 | 0.36 |
| 5 | 5202232 | rs4702241 | 0.852 | 0.167 | -0.784 | -0.031 | -0.007 | 0.057 | 0.057 | 0.013 | 0.17 | 0.22 | 0.49 | 0.44 | 0.15 | 0.16 |
| 5 | 5202753 | rs979708 | 0.213 | 0.399 | -1.114 | 0.273 | -0.012 | -0.042 | -0.042 | 1.35 | 0.7 | 0.63 | 0.83 | 0.82 | 0.72 | 0.78 |
| 5 | 5203046 | rs979710 | -0.647 | 0.536 | -1.667 | 0.337 | 0.023 | 0.011 | 0.011 | 0 | 0.93 | 0.89 | 0.93 | 0.85 | 0.86 | 0.9 |
| 5 | 5203600 | rs1428024 | 0.457 | 0.332 | -1.74 | 0.221 | 0.007 | -0.001 | -0.001 | 0 | 0.22 | 0.16 | 0.1 | 0.03 | 0.15 | 0.11 |
| 5 | 5204149 | rs1366165 | 0.256 | 1.407 | -1.774 | 1.101 | -0.002 | -0.004 | -0.004 | 0.043 | 0.72 | 0.72 | 0.85 | 0.56 | 0.7 | 0.66 |
| 5 | 5208254 | rs6888758 | -0.693 | 1.658 | -2.359 | 1.162 | -0.145 | -0.157 | -0.157 | 0 | 0.29 | 0.02 | 0.21 | 0.47 | 0.38 | 0.23 |
| 5 | 5210050 | rs1560063 | -0.791 | 1.21 | -2.85 | 1.51 | -0.05 | -0.107 | -0.107 | 0.034 | 0.24 | 0.05 | 0.41 | 0.63 | 0.29 | 0.35 |
| 5 | 5210315 | rs7704912 | -0.196 | 0.734 | -3.288 | 0.275 | 0.038 | 0.133 | 0.133 | 0.097 | 0.53 | 0.27 | 0.48 | 0.73 | 0.57 | 0.58 |
| 5 | 5221300 | rs1560062 | -0.138 | 0.774 | -3.037 | 0.214 | 0.002 | 0.008 | 0.008 | 0 | 0.11 | 0.15 | 0.05 | 0.17 | 0.13 | 0.1 |
| 5 | 5222336 | rs4702248 | -0.383 | 0.306 | -2.825 | -0.084 | -0.012 | 0.012 | 0.012 | 0 | 0.46 | 0.32 | 0.32 | 0.38 | 0.43 | 0.43 |
| 5 | 5224299 | rs16875084 | 0.762 | 0.621 | -2.685 | -0.12 | -0.01 | -0.012 | -0.012 | 1.424 | 0.14 | 0.11 | 0.21 | 0.49 | 0.2 | 0.21 |
| 5 | 5225032 | rs2964444 | -0.777 | 0.584 | -2.71 | -0.158 | -0.04 | 0.043 | 0.043 | 0 | 0.67 | 0.92 | 0.74 | 0.8 | 0.71 | 0.68 |
| 5 | 5226211 | rs2913615 | 0.952 | 0.563 | -2.663 | -0.169 | 0.018 | 0.054 | 0.054 | 0.023 | 0.41 | 0.65 | 0.47 | 0.13 | 0.37 | 0.35 |
| 5 | 5227291 | rs12652633 | 0.758 | 0.394 | -2.751 | -0.211 | 0.021 | -0.006 | -0.006 | 0 | 0.12 | 0.05 | 0.15 | 0.13 | 0.1 | 0.09 |
| 5 | 5227314 | rs16875090 | 0.406 | 0.408 | -2.687 | -0.173 | 0.001 | 0.006 | 0.006 | 0 | 0.09 | 0.1 | 0.05 | 0.17 | 0.13 | 0.1 |
| 5 | 5232398 | rs2913624 | -0.231 | 0.353 | -2.73 | -0.218 | -0.02 | 0.048 | 0.048 | 0 | 0.65 | 0.88 | 0.73 | 0.79 | 0.7 | 0.67 |
| 5 | 5234261 | rs10067097 | -0.255 | -0.2 | -1.722 | -0.324 | 0.008 | -0.009 | -0.009 | 0 | 0.4 | 0.5 | 0.23 | 0.34 | 0.4 | 0.38 |
| 5 | 5236356 | rs2913633 | 0.097 | -0.17 | -1.876 | -0.368 | -0.006 | 0.019 | 0.019 | 0 | 0.54 | 0.38 | 0.61 | 0.32 | 0.54 | 0.5 |
| 5 | 5239533 | rs2964431 | 0.291 | -0.247 | -2.241 | -0.637 | -0.006 | 0.003 | 0.003 | 0 | 0.36 | 0.23 | 0.34 | 0.31 | 0.43 | 0.33 |
| 5 | 5239921 | rs11742341 | 1.281 | -0.566 | -2.31 | -0.56 | -0.091 | 0.028 | 0.028 | 0 | 0.22 | 0.01 | 0.11 | 0.08 | 0.19 | 0.22 |
| 5 | 5242683 | rs17713759 | 1.261 | -0.557 | -1.424 | 0.085 | -0.026 | 0.03 | 0.03 | 0 | 0.2 | 0.08 | 0.11 | 0.07 | 0.18 | 0.18 |
| 5 | 5244339 | rs1019746 | 0.632 | -0.818 | -1.508 | -0.019 | 0.016 | -0.011 | -0.011 | 0 | 0.28 | 0.22 | 0.27 | 0.32 | 0.24 | 0.32 |
| 5 | 5245062 | rs2964426 | 1.154 | -0.808 | -1.765 | -0.265 | -0.002 | 0.053 | 0.053 | 0.208 | 0.13 | 0.06 | 0.16 | 0.01 | 0.12 | 0.14 |
| 5 | 5245154 | rs1834928 | 0.399 | -0.949 | -1.313 | -0.365 | 0.016 | 0.021 | 0.021 | 0 | 0.2 | 0.14 | 0.18 | 0.01 | 0.16 | 0.18 |
| 5 | 5246251 | rs934096 | 0.604 | -0.717 | -1.296 | -0.114 | -0.047 | -0.06 | -0.06 | 0.757 | 0.2 | 0.11 | 0.38 | 0.5 | 0.34 | 0.24 |
| 5 | 5246451 | rs10038527 | 0.485 | -1.239 | -0.386 | -0.012 | -0.009 | 0.001 | 0.001 | 0 | 0.31 | 0.34 | 0.27 | 0.32 | 0.25 | 0.31 |
| 5 | 5247857 | rs1808528 | 0.505 | -0.791 | -0.22 | 0.375 | 0.003 | 0.028 | 0.028 | 0 | 0.17 | 0.12 | 0.17 | 0.04 | 0.15 | 0.16 |
| 5 | 5248799 | rs12659315 | 1.173 | -0.784 | -0.381 | 0.843 | -0.09 | 0.079 | 0.079 | 0 | 0.44 | 0.09 | 0.27 | 0.24 | 0.46 | 0.42 |
| 5 | 5249101 | rs10036107 | 1.126 | -0.708 | 0.214 | 0.861 | -0.006 | 0.035 | 0.035 | 0 | 0.46 | 0.36 | 0.3 | 0.24 | 0.46 | 0.42 |
| 5 | 5251790 | rs2913645 | 0.362 | -0.706 | 0.64 | 0.927 | -0.011 | -0.012 | -0.012 | 0 | 0.91 | 0.84 | 0.95 | 0.92 | 0.92 | 0.91 |
| 5 | 5260903 | rs4323203 | 0.678 | -0.881 | 0.994 | 1.404 | -0.077 | 0.181 | 0.181 | 0 | 0.78 | 0.47 | 0.7 | 0.48 | 0.87 | 0.79 |
| 5 | 5261011 | rs2913655 | 0.43 | -1.373 | 0.78 | 1.224 | -0.023 | 0.007 | 0.007 | 0.806 | 0.72 | 0.41 | 0.76 | 0.73 | 0.81 | 0.73 |
| 5 | 5261219 | rs2964399 | -1.251 | -0.894 | 0.592 | 0.872 | 0.015 | -0.021 | -0.021 | 0.053 | 0.36 | 0.73 | 0.41 | 0.61 | 0.32 | 0.34 |
| 5 | 5262547 | rs1428021 | -0.198 | -0.415 | 0.46 | 0.856 | 0.036 | 0.007 | 0.007 | 2.018 | 0.36 | 0.29 | 0.25 | 0.39 | 0.27 | 0.32 |
| 5 | 5263101 | rs2913656 | -0.287 | -0.397 | 0.482 | 0.878 | 0.037 | 0.011 | 0.011 | 1.805 | 0.37 | 0.29 | 0.27 | 0.39 | 0.29 | 0.33 |
| 5 | 5268673 | rs746636 | -0.397 | -0.101 | -0.025 | 0.957 | -0.01 | -0.012 | -0.012 | 0 | 0.85 | 0.87 | 0.78 | 0.77 | 0.8 | 0.82 |
| 5 | 5268710 | rs746635 | 0.596 | 0.116 | 0.106 | 0.499 | -0.005 | 0.02 | 0.02 | 0.302 | 0.46 | 0.28 | 0.48 | 0.34 | 0.49 | 0.47 |
| 5 | 5270436 | rs959693 | 0.64 | 0.134 | -0.069 | 0.49 | -0.011 | 0.029 | 0.029 | 0.302 | 0.46 | 0.24 | 0.48 | 0.33 | 0.49 | 0.47 |
| 5 | 5273945 | rs10044817 | -0.74 | -0.009 | 0.667 | 0.05 | 0.032 | 0.043 | 0.043 | 0 | 0.08 | 0.27 | 0.07 | 0.06 | 0.06 | 0.07 |
| 5 | 5274415 | rs10045494 | -0.132 | 0.3 | 0.184 | -0.261 | -0.016 | -0.016 | -0.016 | 0 | 0.62 | 0.72 | 0.68 | 0.47 | 0.56 | 0.55 |
| 5 | 5277095 | rs7341133 | 1.005 | -0.012 | 0.029 | -0.934 | 0.002 | -0.011 | -0.011 | 0 | 0.18 | 0.16 | 0.12 | 0 | 0.19 | 0.08 |
| 5 | 5277259 | rs13178690 | -0.857 | -0.064 | 0.366 | -0.877 | -0.03 | 0.004 | 0.004 | 0.662 | 0.67 | 0.76 | 0.69 | 0.35 | 0.58 | 0.53 |
| 5 | 5279592 | rs10512778 | -0.337 | -0.372 | 0.256 | -1.107 | -0.007 | 0.024 | 0.024 | 1.013 | 0.56 | 0.63 | 0.63 | 0.28 | 0.5 | 0.4 |

| | | | | | | | | | | | | | | | | |
|---|---|---|---|---|---|---|---|---|---|---|---|---|---|---|---|---|
| 5 | 5281408 | rs10512779 | 1.262 | -0.57 | 0.158 | -1.255 | -0.003 | 0.014 | 0.014 | 1.563 | 0.22 | 0.15 | 0.4 | 0.2 | 0.2 | 0.18 |
| 5 | 5282051 | rs16875221 | 1.337 | -0.623 | 0.178 | -1.384 | 0.008 | 0.009 | 0.009 | 0 | 0.11 | 0.06 | 0.06 | 0.06 | 0.08 | 0.13 |
| 5 | 5283234 | rs11134100 | -0.274 | -0.708 | 0.011 | -1.411 | -0.027 | -0.005 | -0.005 | 0.741 | 0.84 | 0.7 | 0.89 | 0.87 | 0.89 | 0.8 |
| 5 | 5283260 | rs10512780 | 1.572 | -0.646 | 0.029 | -1.374 | 0.017 | -0.008 | -0.008 | 0 | 0.06 | 0.19 | 0.04 | 0 | 0.03 | 0 |
| 5 | 5286424 | rs16875241 | 1.1 | -0.644 | -0.276 | -1.391 | -0.014 | -0.008 | -0.008 | 2.976 | 0.08 | 0.01 | 0.05 | 0.07 | 0.09 | 0.14 |
| 5 | 5287739 | rs16875244 | 0.39 | -0.634 | -0.238 | -1.385 | 0.009 | 0.056 | 0.056 | 0 | 0.27 | 0.2 | 0.18 | 0.07 | 0.28 | 0.22 |
| 5 | 5290195 | rs4076657 | 0.919 | -0.465 | -0.349 | -1.354 | 0.059 | 0.169 | 0.169 | 0 | 0.51 | 0.24 | 0.65 | 0.8 | 0.56 | 0.58 |
| 5 | 5290797 | rs11134106 | 0.738 | -0.488 | -0.494 | -1.341 | -0.078 | -0.029 | -0.029 | 0 | 0.39 | 0.12 | 0.31 | 0.6 | 0.44 | 0.52 |
| 5 | 5292821 | rs6555340 | -0.369 | -0.34 | -0.305 | -1.256 | 0.015 | 0.011 | 0.011 | 0.122 | 0.64 | 0.74 | 0.52 | 0.68 | 0.71 | 0.68 |
| 5 | 5294187 | rs16875287 | 0.454 | -0.166 | -0.559 | -1.158 | -0.01 | -0.013 | -0.013 | 0 | 0.1 | 0.06 | 0.06 | 0.13 | 0.13 | 0.2 |
| 5 | 5298350 | rs7701192 | 0.714 | -0.246 | -0.455 | -1.118 | -0.005 | -0.013 | -0.013 | 1.513 | 0.46 | 0.26 | 0.39 | 0.52 | 0.55 | 0.48 |
| 5 | 5302501 | rs12518997 | -0.967 | -0.262 | -0.369 | -1.144 | -0.007 | 0 | 0 | 0 | 0.75 | 0.67 | 0.58 | 0.75 | 0.81 | 0.81 |
| 5 | 5302512 | rs12518998 | 0.189 | 0.295 | -0.819 | -0.822 | 0.003 | -0.003 | -0.003 | 0.193 | 0.46 | 0.35 | 0.4 | 0.52 | 0.55 | 0.48 |
| 5 | 5303614 | rs11134110 | 0.797 | 0.497 | -1.244 | -0.619 | -0.088 | -0.119 | -0.119 | 0 | 0.28 | 0.08 | 0.26 | 0.52 | 0.36 | 0.39 |
| 5 | 5309697 | rs4702257 | -0.008 | 1.064 | -1.132 | -0.608 | 0.005 | 0.016 | 0.016 | 0.451 | 0.5 | 0.52 | 0.58 | 0.61 | 0.5 | 0.59 |
| 5 | 5309811 | rs4702258 | -0.008 | 1.099 | -1.132 | -0.608 | 0.006 | 0.016 | 0.016 | 0.451 | 0.5 | 0.52 | 0.57 | 0.61 | 0.5 | 0.59 |
| 5 | 5314703 | rs7721079 | 0.221 | 1.461 | -1.002 | -0.202 | 0.012 | 0.029 | 0.029 | 0.548 | 0.32 | 0.32 | 0.46 | 0.5 | 0.37 | 0.43 |
| 5 | 5316866 | rs4128586 | -0.113 | 1.52 | -1.262 | -0.324 | -0.041 | 0.055 | 0.055 | 0.365 | 0.23 | 0.12 | 0.16 | 0.07 | 0.33 | 0.15 |
| 5 | 5317110 | rs2964154 | 0.723 | 1.229 | -0.931 | 0.348 | -0.002 | -0.002 | -0.002 | 0 | 0.87 | 0.83 | 0.9 | 0.82 | 0.86 | 0.76 |
| 5 | 5317882 | rs2964478 | 0.205 | 1.248 | -1.093 | 0.314 | -0.018 | 0.013 | 0.013 | 0 | 0.82 | 0.89 | 0.88 | 0.88 | 0.81 | 0.78 |
| 5 | 5318581 | rs6885895 | -1.236 | 0.522 | -0.338 | 0.492 | -0.01 | 0.028 | 0.028 | 0 | 0.14 | 0.35 | 0.08 | 0.2 | 0.11 | 0.18 |
| 5 | 5323994 | rs11134111 | -0.508 | 0.559 | -0.6 | 0.747 | 0 | 0 | 0 | 0.201 | 0.1 | 0.07 | 0.09 | 0.15 | 0.07 | 0.14 |
| 5 | 5324172 | rs3923919 | -0.431 | 0.265 | -0.898 | 0.899 | 0.018 | -0.006 | -0.006 | 0.072 | 0.52 | 0.61 | 0.43 | 0.5 | 0.45 | 0.49 |
| 5 | 5324373 | rs10512787 | 0.345 | 0.081 | -1.02 | 1.138 | -0.004 | -0.012 | -0.012 | 0.182 | 0.06 | 0.04 | 0.16 | 0.18 | 0.1 | 0.13 |
| 5 | 5324983 | rs16875333 | 0.678 | 0.066 | -1.025 | 1.115 | -0.02 | -0.047 | -0.047 | 0 | 0.09 | 0 | 0.08 | 0.14 | 0.07 | 0.14 |
| 5 | 5325727 | rs10060293 | 0.204 | 0.028 | -0.886 | 1.051 | -0.003 | -0.007 | -0.007 | 0 | 0.06 | 0.04 | 0.16 | 0.2 | 0.1 | 0.13 |
| 5 | 5326138 | rs7725935 | -0.349 | 0.272 | -1.001 | 0.674 | 0.032 | 0.03 | 0.03 | 0.108 | 0.46 | 0.62 | 0.38 | 0.39 | 0.42 | 0.38 |
| 5 | 5326788 | rs11739886 | -0.075 | 0.158 | -0.741 | 0.685 | 0.016 | 0.008 | 0.008 | 0.334 | 0.42 | 0.63 | 0.35 | 0.39 | 0.37 | 0.31 |
| 5 | 5328297 | rs2964474 | -0.551 | -0.203 | -0.91 | 1.491 | 0.008 | -0.008 | -0.008 | 0.161 | 0.89 | 0.91 | 0.74 | 0.81 | 0.93 | 0.84 |
| 5 | 5328681 | rs7731357 | -0.85 | -0.682 | -0.117 | 1.502 | -0.023 | 0.14 | 0.14 | 0 | 0.12 | 0.54 | 0.19 | 0.3 | 0.11 | 0.15 |
| 5 | 5329658 | rs2964153 | 0.953 | 0.163 | -0.613 | 1.065 | 0.002 | -0.011 | -0.011 | 0.589 | 0.64 | 0.33 | 0.47 | 0.51 | 0.63 | 0.63 |
| 5 | 5335483 | rs11747342 | 0.887 | -0.056 | -0.728 | 1.214 | -0.002 | -0.001 | -0.001 | 0 | 0.15 | 0.16 | 0.09 | 0.13 | 0.13 | 0.13 |
| 5 | 5344745 | rs2964149 | -0.511 | 0.239 | -0.603 | 1.365 | 0.015 | -0.016 | -0.016 | 0.903 | 0.91 | 0.93 | 0.74 | 0.83 | 0.95 | 0.83 |
| 5 | 5350456 | rs7735787 | -1.048 | 0.53 | 0.077 | 1.441 | 0.094 | 0.177 | 0.177 | 0.039 | 0.13 | 0.42 | 0.23 | 0.35 | 0.22 | 0.15 |
| 5 | 5354615 | rs7723605 | -0.918 | 0.463 | 0.186 | 1.447 | 0.055 | 0.209 | 0.209 | 0.596 | 0.12 | 0.46 | 0.2 | 0.34 | 0.15 | 0.12 |
| 5 | 5361469 | rs2964459 | 1.476 | 0.389 | 0.106 | 1.055 | -0.021 | 0.07 | 0.07 | 0 | 0.47 | 0.17 | 0.49 | 0.34 | 0.49 | 0.36 |
| 5 | 5361557 | rs16875423 | -0.111 | 0.92 | 0.599 | 0.798 | 0.105 | 0.187 | 0.187 | 0 | 0.07 | 0.31 | 0.19 | 0.21 | 0.13 | 0.08 |
| 5 | 5363136 | rs12655520 | 0.731 | 1.06 | 0.046 | 0.621 | -0.044 | 0 | 0 | 0.099 | 0.13 | 0.01 | 0.07 | 0.07 | 0.09 | 0.18 |
| 5 | 5365240 | rs10474786 | -0.803 | 0.977 | 0.24 | 0.705 | 0.055 | 0.018 | 0.018 | 0 | 0.21 | 0.49 | 0.26 | 0.3 | 0.3 | 0.18 |
| 5 | 5371590 | rs13172631 | 0.35 | 0.93 | 0.235 | 0.687 | 0.003 | -0.009 | -0.009 | 0 | 0.13 | 0.16 | 0.07 | 0.13 | 0.16 | 0.1 |
| 5 | 5373748 | rs17785206 | 0.714 | 0.945 | 0.051 | 0.652 | -0.036 | -0.003 | -0.003 | 0.022 | 0.13 | 0.01 | 0.07 | 0.08 | 0.09 | 0.18 |
| 5 | 5374574 | rs2964453 | 1.242 | 0.945 | 0.614 | 0.778 | 0.027 | 0.004 | 0.004 | 0 | 0.75 | 0.49 | 0.74 | 0.69 | 0.7 | 0.8 |
| 5 | 5376984 | rs6884256 | -0.093 | 1.371 | 0.2 | 0.94 | 0.077 | 0.099 | 0.099 | 0 | 0.06 | 0.23 | 0.18 | 0.18 | 0.13 | 0.08 |
| 5 | 5378905 | rs11748854 | 0.149 | 1.318 | 0.032 | 0.877 | -0.012 | 0.017 | 0.017 | 0 | 0.13 | 0.09 | 0.07 | 0.13 | 0.17 | 0.1 |
| 5 | 5383342 | rs6891687 | -0.351 | 1.379 | 0.249 | 0.976 | 0.072 | 0.09 | 0.09 | 0 | 0.06 | 0.24 | 0.18 | 0.17 | 0.13 | 0.08 |
| 5 | 5388969 | rs11134112 | 0.757 | 1.245 | -0.121 | 0.632 | -0.011 | 0.002 | 0.002 | 0 | 0.09 | 0.07 | 0.05 | 0 | 0.15 | 0.04 |
| 5 | 5396166 | rs2964137 | -0.347 | 1.478 | -0.211 | 0.539 | -0.009 | 0.002 | 0.002 | 0.011 | 0.44 | 0.38 | 0.45 | 0.42 | 0.52 | 0.33 |
| 5 | 5397165 | rs1863943 | -0.072 | 1.846 | -0.467 | 0.601 | -0.044 | -0.042 | -0.042 | 0 | 0.1 | 0.05 | 0.16 | 0.21 | 0.21 | 0.13 |
| 5 | 5397777 | rs2964445 | -0.203 | 1.914 | -0.051 | 0.416 | 0.001 | 0.001 | 0.001 | 0 | 0.35 | 0.38 | 0.41 | 0.42 | 0.41 | 0.3 |
| 5 | 5398279 | rs2964135 | -0.029 | 2.031 | 0.115 | 0.488 | 0.023 | -0.001 | -0.001 | 1.986 | 0.85 | 0.8 | 0.83 | 0.87 | 0.75 | 0.89 |
| 5 | 5398894 | rs2964437 | -0.526 | 1.984 | 0.098 | 0.521 | 0.06 | 0.056 | 0.056 | 0 | 0.94 | 0.84 | 0.87 | 0.87 | 0.87 | 0.92 |
| 5 | 5399472 | rs6868526 | 0.767 | 1.301 | -0.379 | 0.082 | 0.011 | -0.017 | -0.017 | 0.22 | 0.26 | 0.22 | 0.21 | 0.51 | 0.16 | 0.44 |
| 5 | 5401773 | rs2964132 | 0.865 | 1.23 | -0.28 | 0.09 | -0.001 | -0.012 | -0.012 | 0 | 0.58 | 0.44 | 0.59 | 0.28 | 0.56 | 0.43 |
| 5 | 5402163 | rs16875556 | 0.657 | 0.961 | -0.32 | 0.346 | -0.012 | 0.017 | 0.017 | 0.012 | 0.21 | 0.21 | 0.2 | 0.49 | 0.11 | 0.41 |
| 5 | 5404664 | rs4702267 | 0.86 | 1.443 | -0.831 | -0.492 | -0.008 | 0.007 | 0.007 | 0 | 0.28 | 0.21 | 0.33 | 0.06 | 0.32 | 0.23 |
| 5 | 5405699 | rs2913341 | 0.236 | 1.484 | -0.782 | -0.239 | -0.003 | 0.014 | 0.014 | 0.812 | 0.42 | 0.35 | 0.34 | 0.19 | 0.4 | 0.28 |
| 5 | 5406467 | rs1078266 | 0.002 | 1.478 | -0.852 | 0.187 | 0.002 | 0 | 0 | 0 | 0.2 | 0.2 | 0.18 | 0.2 | 0.27 | 0.09 |
| 5 | 5406529 | rs2964383 | 0.259 | 1.216 | -0.809 | 0.335 | -0.001 | 0 | 0 | 0.022 | 0.53 | 0.5 | 0.33 | 0.51 | 0.36 | 0.52 |
| 5 | 5407814 | rs2964475 | 0.192 | 0.496 | -0.362 | 0.951 | 0 | 0 | 0 | 0 | 0.58 | 0.54 | 0.54 | 0.51 | 0.57 | 0.7 |
| 5 | 5415012 | rs2913356 | 0.159 | 0.528 | -0.349 | 0.792 | 0 | 0 | 0 | 0 | 0.58 | 0.54 | 0.54 | 0.53 | 0.57 | 0.7 |
| 5 | 5416398 | rs2913364 | 0.72 | 0.188 | -0.054 | 1.205 | -0.005 | 0.178 | 0.178 | 1.077 | 0.92 | 0.76 | 0.81 | 0.73 | 0.93 | 0.86 |
| 5 | 5418792 | rs2964163 | -0.104 | 0.181 | -0.268 | 1.003 | 0 | -0.001 | -0.001 | 0 | 0.45 | 0.46 | 0.5 | 0.44 | 0.45 | 0.44 |
| 5 | 5419547 | rs2399696 | 0.099 | 0.346 | -0.724 | 0.35 | -0.018 | 0.065 | 0.065 | 0 | 0.41 | 0.23 | 0.25 | 0.21 | 0.43 | 0.34 |
| 5 | 5437380 | rs2578550 | -0.66 | 0.489 | 0.114 | 1.023 | -0.008 | -0.004 | -0.004 | 0 | 0.29 | 0.62 | 0.22 | 0.31 | 0.35 | 0.24 |
| 5 | 5449367 | rs2578553 | -0.884 | 0.589 | 0.052 | 1.58 | -0.002 | 0.007 | 0.007 | 0 | 0.33 | 0.64 | 0.46 | 0.64 | 0.38 | 0.34 |
| 5 | 5460434 | rs16875597 | -0.839 | 0.564 | 0.591 | 1.57 | 0.054 | 0.346 | 0.346 | 0 | 0.06 | 0.34 | 0.28 | 0.33 | 0.03 | 0.11 |
| 5 | 5460569 | rs2578565 | 0.21 | 0.61 | 0.408 | 1.587 | -0.001 | 0.001 | 0.001 | 0 | 0.73 | 0.71 | 0.82 | 0.69 | 0.65 | 0.76 |
| 5 | 5462148 | rs2578500 | 0.158 | 0.657 | 0.41 | 1.61 | 0 | 0.002 | 0.002 | 0 | 0.74 | 0.71 | 0.78 | 0.68 | 0.65 | 0.76 |
| 5 | 5475858 | rs3747730 | -0.57 | 0.402 | 0.066 | 1.307 | -0.003 | -0.031 | -0.031 | 1.156 | 0.09 | 0.06 | 0.18 | 0.3 | 0.08 | 0.2 |

| | | | | | | | | | | | | | | | | |
|---|---|---|---|---|---|---|---|---|---|---|---|---|---|---|---|---|
| 5 | 5495373 | rs252940 | -0.345 | 0.562 | 0.251 | 0.52 | -0.005 | 0.004 | 0.004 | 0 | 0.26 | 0.31 | 0.19 | 0.31 | 0.34 | 0.24 |
| 5 | 5496203 | rs13181223 | 0.469 | 0.585 | 0.096 | 0.249 | -0.009 | 0.005 | 0.005 | 0 | 0.06 | 0.03 | 0.04 | 0 | 0.09 | 0.1 |
| 5 | 5496480 | rs252938 | 0.751 | -0.14 | 0.462 | 0.635 | -0.007 | -0.005 | -0.005 | 0.12 | 0.68 | 0.46 | 0.5 | 0.3 | 0.62 | 0.63 |
| 5 | 5504842 | rs2619855 | 0.845 | -0.046 | 0.358 | 0.504 | 0.002 | 0.001 | 0.001 | 0 | 0.67 | 0.31 | 0.61 | 0.33 | 0.63 | 0.64 |
| 5 | 5509703 | rs252930 | 0.427 | -0.044 | 0.257 | 0.518 | 0.01 | 0.025 | 0.025 | 0 | 0.72 | 0.55 | 0.64 | 0.33 | 0.65 | 0.66 |
| 5 | 5513747 | rs252926 | 0.431 | 0.093 | -0.357 | 0.611 | -0.006 | 0.047 | 0.047 | 0 | 0.1 | 0.16 | 0.07 | 0.25 | 0.11 | 0.13 |
| 5 | 5514517 | rs2652715 | 0.729 | 0.129 | -0.829 | 0.04 | -0.003 | 0 | 0 | 0 | 0.68 | 0.46 | 0.62 | 0.33 | 0.64 | 0.63 |
| 5 | 5536047 | rs16875692 | 0.091 | 0.774 | -1.684 | -0.175 | -0.004 | 0.029 | 0.029 | 0 | 0.1 | 0.18 | 0.08 | 0.21 | 0.1 | 0.1 |
| 5 | 5539266 | rs17770324 | 0.169 | 1.142 | -2.54 | -0.99 | -0.16 | 0.131 | 0.131 | 0 | 0.38 | 0.05 | 0.48 | 0.16 | 0.42 | 0.37 |
| 5 | 5546157 | rs1871473 | -0.52 | 1.392 | -1.442 | -0.987 | -0.013 | 0.015 | 0.015 | 0 | 0.3 | 0.19 | 0.19 | 0.03 | 0.37 | 0.19 |
| 5 | 5560070 | rs252838 | -0.571 | 1.687 | -1.581 | -0.438 | -0.077 | 0.002 | 0.002 | 0 | 0.25 | 0.04 | 0.19 | 0.28 | 0.31 | 0.25 |
| 5 | 5566987 | rs2652701 | -0.297 | 1.589 | -0.985 | -0.518 | -0.057 | -0.011 | -0.011 | 0 | 0.22 | 0.04 | 0.18 | 0.29 | 0.28 | 0.23 |
| 5 | 5570011 | rs252613 | -0.946 | 1.424 | -0.053 | -0.543 | 0.007 | -0.016 | -0.016 | 0.157 | 0.33 | 0.58 | 0.27 | 0.32 | 0.36 | 0.27 |
| 5 | 5577176 | rs11741289 | -1.456 | 1.165 | 0.241 | -0.447 | -0.008 | -0.002 | -0.002 | 3.974 | 0.11 | 0.25 | 0.08 | 0.07 | 0.09 | 0.19 |
| 5 | 5578527 | rs13154779 | -0.743 | 1.351 | 0.368 | -0.907 | 0.024 | -0.09 | -0.09 | 0.427 | 0.15 | 0.4 | 0.14 | 0.02 | 0.17 | 0.13 |
| 5 | 5579270 | rs16875781 | -0.91 | 1.337 | 0.294 | -0.913 | 0.023 | -0.105 | -0.105 | 1.337 | 0.16 | 0.48 | 0.14 | 0.02 | 0.17 | 0.13 |
| 5 | 5580216 | rs246760 | 0.156 | 1.787 | -0.939 | -0.715 | -0.065 | 0.002 | 0.002 | 0.254 | 0.78 | 0.97 | 0.84 | 0.21 | 0.73 | 0.65 |
| 5 | 5581522 | rs246764 | 0.49 | 1.57 | -0.303 | -0.608 | -0.002 | -0.008 | -0.008 | 0 | 0.9 | 0.9 | 0.95 | 0.97 | 0.92 | 0.83 |
| 5 | 5584551 | rs181773 | 0.615 | 1.562 | -0.376 | -0.629 | 0 | 0.003 | 0.003 | 0 | 0.88 | 0.89 | 0.83 | 0.9 | 0.89 | 0.8 |
| 5 | 5585343 | rs153264 | 1.299 | 1.418 | -0.513 | -0.967 | 0.011 | 0.067 | 0.067 | 1.618 | 0.73 | 0.18 | 0.72 | 0.9 | 0.78 | 0.7 |
| 5 | 5597328 | rs16875838 | -1.147 | 2.47 | 0.317 | -0.584 | 0.127 | -0.034 | -0.034 | 0 | 0.07 | 0.24 | 0.07 | 0 | 0.21 | 0.07 |
| 5 | 5598181 | rs11739751 | -0.698 | 1.61 | -0.372 | 0.14 | -0.002 | -0.01 | -0.01 | 0 | 0.18 | 0.14 | 0.13 | 0.24 | 0.17 | 0.18 |
| 5 | 5600683 | rs252596 | -0.49 | 1.415 | -0.245 | 0.314 | 0 | -0.003 | -0.003 | 0 | 0.14 | 0.14 | 0.07 | 0.14 | 0.14 | 0.11 |
| 5 | 5601717 | rs2560286 | -0.619 | 1.507 | -0.277 | 0.461 | -0.007 | -0.034 | -0.034 | 0.014 | 0.16 | 0.05 | 0.18 | 0.22 | 0.17 | 0.13 |
| 5 | 5601744 | rs12522553 | 0.584 | 1.268 | -1.016 | -0.674 | 0 | 0.087 | 0.087 | 0 | 0.27 | 0.12 | 0.2 | 0.02 | 0.25 | 0.22 |
| 5 | 5602308 | rs1318543 | -0.147 | 1.354 | 0.093 | -0.848 | 0.002 | 0.001 | 0.001 | 0 | 0.66 | 0.34 | 0.63 | 0.89 | 0.71 | 0.72 |
| 5 | 5602530 | rs2438479 | -0.256 | 1.359 | 0.124 | -0.852 | -0.002 | -0.066 | -0.066 | 0 | 0.67 | 0.39 | 0.74 | 0.97 | 0.71 | 0.73 |
| 5 | 5602673 | rs2928229 | -0.888 | 1.358 | 0.282 | -0.853 | 0.002 | 0.029 | 0.029 | 0.05 | 0.48 | 0.55 | 0.43 | 0.24 | 0.46 | 0.37 |
| 5 | 5605051 | rs9313114 | 0.113 | 1.339 | 0.317 | -0.853 | -0.006 | -0.152 | -0.152 | 1.47 | 0.33 | 0.51 | 0.26 | 0.02 | 0.29 | 0.24 |
| 5 | 5609695 | rs378932 | -0.497 | 1.407 | 0.575 | -0.814 | 0.001 | -0.069 | -0.069 | 0 | 0.7 | 0.63 | 0.77 | 0.97 | 0.72 | 0.74 |
| 5 | 5611791 | rs2560290 | -0.356 | 1.472 | 0.376 | -0.529 | -0.011 | -0.028 | -0.028 | 0.004 | 0.17 | 0.1 | 0.2 | 0.25 | 0.21 | 0.19 |
| 5 | 5611992 | rs422576 | 0.364 | 1.366 | 0.51 | -0.572 | 0 | 0.015 | 0.015 | 0.086 | 0.82 | 0.75 | 0.79 | 0.76 | 0.79 | 0.79 |
| 5 | 5614515 | rs13185585 | 0.729 | 1.271 | 0.68 | -0.86 | -0.005 | -0.084 | -0.084 | 0 | 0.28 | 0.33 | 0.21 | 0.02 | 0.22 | 0.2 |
| 5 | 5615414 | rs394842 | -0.094 | 1.265 | 0.593 | -0.612 | -0.001 | -0.005 | -0.005 | 0.354 | 0.83 | 0.88 | 0.91 | 0.83 | 0.8 | 0.8 |
| 5 | 5619500 | rs2255659 | 0.058 | 1.485 | 0.084 | 0.081 | -0.007 | -0.008 | -0.008 | 0 | 0.45 | 0.32 | 0.45 | 0.47 | 0.44 | 0.52 |
| 5 | 5621914 | rs7703207 | 0.184 | 1.389 | 0.503 | 0.171 | -0.001 | 0.022 | 0.022 | 0 | 0.51 | 0.28 | 0.39 | 0.42 | 0.51 | 0.42 |
| 5 | 5628443 | rs252609 | -0.198 | 1.612 | 0.03 | 0.273 | -0.005 | -0.004 | -0.004 | 0.112 | 0.26 | 0.21 | 0.26 | 0.34 | 0.34 | 0.3 |
| 5 | 5628903 | rs34123 | 0.031 | 1.712 | -0.193 | 0.45 | -0.029 | -0.014 | -0.014 | 0 | 0.12 | 0.09 | 0.11 | 0.14 | 0.19 | 0.15 |
| 5 | 5629110 | rs10462632 | 0.477 | 1.235 | -0.11 | 0.259 | -0.019 | -0.016 | -0.016 | 0.076 | 0.4 | 0.16 | 0.56 | 0.45 | 0.47 | 0.34 |
| 5 | 5638245 | rs13181152 | 0.755 | 1.294 | 0.589 | -0.373 | 0.001 | 0.02 | 0.02 | 0 | 0.31 | 0.3 | 0.25 | 0.1 | 0.27 | 0.25 |
| 5 | 5641116 | rs6865834 | 0.521 | 1.292 | 0.483 | -0.373 | -0.008 | -0.08 | -0.08 | 2.681 | 0.33 | 0.5 | 0.27 | 0.1 | 0.27 | 0.25 |
| 5 | 5647018 | rs10052145 | 0.52 | 1.267 | 0.459 | -0.361 | -0.01 | -0.077 | -0.077 | 0 | 0.33 | 0.53 | 0.27 | 0.1 | 0.27 | 0.25 |
| 5 | 5649467 | rs13182121 | 0.949 | 1.278 | -0.274 | -0.282 | 0.001 | 0.011 | 0.011 | 0 | 0.28 | 0.22 | 0.22 | 0.1 | 0.24 | 0.21 |
| 5 | 5649753 | rs17791535 | 1.003 | 1.327 | -1.138 | -0.259 | -0.013 | 0.071 | 0.071 | 0.01 | 0.27 | 0.06 | 0.2 | 0.1 | 0.24 | 0.21 |
| 5 | 5670085 | rs256358 | 1.123 | 1.527 | -0.076 | 0.226 | 0.005 | -0.027 | -0.027 | 0.61 | 0.81 | 0.78 | 0.85 | 0.91 | 0.78 | 0.8 |
| 5 | 5672577 | rs256360 | 1.151 | 1.537 | -0.091 | 0.23 | 0.013 | -0.063 | -0.063 | 0 | 0.8 | 0.69 | 0.84 | 0.91 | 0.77 | 0.79 |
| 5 | 5678997 | rs11748708 | 0.071 | 1.43 | -0.632 | 0.684 | -0.06 | -0.029 | -0.029 | 1.17 | 0.16 | 0.01 | 0.11 | 0.16 | 0.23 | 0.12 |
| 5 | 5680337 | rs11959645 | -1.383 | 1.157 | -0.217 | 0.721 | 0.014 | -0.042 | -0.042 | 0 | 0.12 | 0.3 | 0.1 | 0.08 | 0.13 | 0.12 |
| 5 | 5686448 | rs11134122 | -1.597 | 1.152 | -0.153 | 0.719 | 0.014 | -0.05 | -0.05 | 0 | 0.13 | 0.35 | 0.11 | 0.07 | 0.13 | 0.12 |
| 5 | 5689218 | rs10512804 | -0.718 | 1.205 | -0.48 | 0.746 | -0.007 | 0.017 | 0.017 | 0 | 0.09 | 0.07 | 0.08 | 0.07 | 0.11 | 0.12 |
| 5 | 5690605 | rs7736476 | -0.904 | 1.205 | -0.455 | 0.746 | 0 | -0.007 | -0.007 | 0 | 0.1 | 0.14 | 0.08 | 0.07 | 0.11 | 0.12 |
| 5 | 5692072 | rs13157338 | -0.532 | 1.311 | -0.771 | 0.799 | -0.008 | 0.014 | 0.014 | 0 | 0.09 | 0.07 | 0.08 | 0.08 | 0.11 | 0.12 |
| 5 | 5703184 | rs17736004 | -0.633 | 1.364 | -0.56 | 1.053 | -0.036 | 0.013 | 0.013 | 0 | 0.22 | 0.09 | 0.18 | 0.17 | 0.28 | 0.21 |
| 5 | 5711184 | rs12110260 | -1.217 | 1.047 | -1.036 | 1.247 | -0.001 | -0.003 | -0.003 | 0.426 | 0.07 | 0.13 | 0.07 | 0.08 | 0.07 | 0.11 |
| 5 | 5713635 | rs1560073 | -0.744 | 0.93 | -1.236 | 1.86 | -0.022 | -0.075 | -0.075 | 0 | 0.22 | 0.07 | 0.28 | 0.37 | 0.25 | 0.25 |
| 5 | 5713766 | rs7727470 | -0.803 | 0.921 | -1.24 | 1.84 | -0.021 | -0.074 | -0.074 | 0 | 0.23 | 0.07 | 0.29 | 0.37 | 0.25 | 0.25 |
| 5 | 5723290 | rs7725455 | -0.278 | 1.102 | -1.165 | 1.78 | -0.054 | -0.079 | -0.079 | 0 | 0.18 | 0.05 | 0.2 | 0.28 | 0.22 | 0.24 |
| 5 | 5725162 | rs4143541 | -1.692 | 0.427 | -0.095 | 1.999 | -0.044 | 0.08 | 0.08 | 5.684 | 0.12 | 0.76 | 0.11 | 0.28 | 0.02 | 0.09 |
| 5 | 5729063 | rs7725206 | -1.687 | 0.416 | 1.113 | 2.062 | -0.004 | 0.158 | 0.158 | 2.744 | 0.09 | 0.48 | 0.07 | 0.21 | 0.02 | 0.06 |
| 5 | 5737475 | rs2895450 | 0.054 | 0.467 | 0.881 | 1.498 | -0.001 | 0 | 0 | 0 | 0.57 | 0.56 | 0.59 | 0.53 | 0.54 | 0.5 |
| 5 | 5740672 | rs7722440 | 0.011 | 0.46 | 0.984 | 1.495 | -0.001 | -0.015 | -0.015 | 0 | 0.57 | 0.69 | 0.6 | 0.53 | 0.54 | 0.5 |
| 5 | 5743661 | rs406792 | -0.268 | 0.433 | 1.218 | 1.565 | 0.005 | 0.004 | 0.004 | 0.015 | 0.45 | 0.62 | 0.43 | 0.56 | 0.46 | 0.52 |
| 5 | 5750355 | rs449492 | 0.078 | 0.473 | 0.992 | 1.577 | -0.022 | -0.03 | -0.03 | 0.715 | 0.4 | 0.18 | 0.38 | 0.46 | 0.46 | 0.49 |
| 5 | 5750925 | rs434192 | -0.115 | 0.624 | 0.778 | 1.653 | 0.001 | -0.002 | -0.002 | 0 | 0.43 | 0.38 | 0.4 | 0.46 | 0.46 | 0.5 |
| 5 | 5761030 | rs11743797 | -0.11 | 0.683 | 0.337 | 1.266 | 0 | 0 | 0 | 0 | 0.46 | 0.56 | 0.43 | 0.57 | 0.49 | 0.52 |
| 5 | 5768140 | rs34950 | -0.201 | 1.094 | -0.124 | 0.823 | -0.032 | -0.035 | -0.035 | 0 | 0.85 | 0.94 | 0.73 | 0.76 | 0.8 | 0.84 |
| 5 | 5771161 | rs3852153 | -0.688 | 0.614 | 0.44 | 0.379 | -0.006 | -0.042 | -0.042 | 0 | 0.09 | 0.22 | 0.06 | 0 | 0.06 | 0.02 |
| 5 | 5772527 | rs16876243 | -0.723 | 0.601 | 0.45 | 0.372 | -0.006 | -0.043 | -0.043 | 0.147 | 0.09 | 0.22 | 0.07 | 0 | 0.07 | 0.02 |
| 5 | 5773005 | rs16876246 | -0.655 | 0.878 | 0.444 | 0.389 | 0.002 | -0.032 | -0.032 | 0 | 0.09 | 0.18 | 0.06 | 0 | 0.07 | 0.02 |

| | | | | | | | | | | | | | | | | |
|---|---|---|---|---|---|---|---|---|---|---|---|---|---|---|---|---|
| 5 | 5773139 | rs16876248 | -0.626 | 0.596 | 0.448 | 0.394 | -0.003 | -0.031 | -0.031 | 0.006 | 0.09 | 0.16 | 0.05 | 0 | 0.06 | 0.02 |
| 5 | 5779953 | rs166285 | -1.002 | 0.459 | 0.744 | 0.122 | -0.002 | -0.023 | -0.023 | 0.03 | 0.37 | 0.57 | 0.46 | 0.18 | 0.36 | 0.29 |
| 5 | 5786026 | rs6897110 | -0.163 | 0.377 | 1.155 | 0.003 | -0.008 | -0.046 | -0.046 | 0 | 0.09 | 0.15 | 0.05 | 0 | 0.06 | 0.02 |
| 5 | 5786702 | rs187632 | -1.444 | 0.158 | 0.824 | 0.17 | 0.031 | 0.015 | 0.015 | 0 | 0.46 | 0.79 | 0.6 | 0.4 | 0.4 | 0.43 |
| 5 | 5789072 | rs10036598 | -0.541 | 0.577 | 0.947 | -0.431 | -0.004 | -0.074 | -0.074 | 0.482 | 0.12 | 0.44 | 0.08 | 0 | 0.06 | 0.02 |
| 5 | 5790928 | rs42286 | -1.357 | 0.628 | 0.914 | -0.45 | 0 | 0.006 | 0.006 | 0 | 0.4 | 0.64 | 0.5 | 0.18 | 0.39 | 0.3 |
| 5 | 5792507 | rs32566 | -0.525 | 0.591 | 0.244 | 0.063 | -0.003 | 0.005 | 0.005 | 0 | 0.25 | 0.2 | 0.39 | 0.18 | 0.29 | 0.27 |
| 5 | 5792720 | rs28165 | -0.541 | 0.875 | -0.951 | 0.197 | -0.012 | 0.008 | 0.008 | 0 | 0.24 | 0.14 | 0.39 | 0.18 | 0.29 | 0.27 |
| 5 | 5798853 | rs32558 | 0.897 | 0.518 | -0.766 | -0.113 | -0.002 | -0.043 | -0.043 | 0 | 0.82 | 0.82 | 0.88 | 1 | 0.86 | 0.93 |
| 5 | 5800497 | rs4701707 | -1.027 | 0.914 | -1.116 | 0.66 | -0.021 | -0.049 | -0.049 | 0 | 0.19 | 0.12 | 0.22 | 0.31 | 0.29 | 0.13 |
| 5 | 5801024 | rs26806 | -0.601 | 1.025 | -1.107 | 0.46 | -0.001 | 0.001 | 0.001 | 0 | 0.36 | 0.33 | 0.49 | 0.66 | 0.39 | 0.5 |
| 5 | 5802056 | rs3864249 | 0.65 | 0.243 | -2.114 | -0.083 | -0.099 | -0.07 | -0.07 | 0 | 0.26 | 0.03 | 0.36 | 0.69 | 0.37 | 0.36 |
| 5 | 5802680 | rs4702288 | 0.485 | 0.282 | -1.533 | 0.015 | -0.014 | 0.019 | 0.019 | 0 | 0.12 | 0.07 | 0.1 | 0 | 0.18 | 0.05 |
| 5 | 5806153 | rs7711899 | -0.441 | 0.321 | -0.567 | -0.204 | -0.019 | 0.014 | 0.014 | 0.489 | 0.18 | 0.12 | 0.15 | 0 | 0.27 | 0.06 |
| 5 | 5806875 | rs32553 | 0.255 | -0.043 | -0.073 | -0.207 | -0.001 | -0.001 | -0.001 | 0 | 0.46 | 0.55 | 0.45 | 0.49 | 0.45 | 0.38 |
| 5 | 5807278 | rs42285 | -1.242 | -0.053 | -0.168 | -0.113 | 0.027 | -0.002 | -0.002 | 0.064 | 0.71 | 0.84 | 0.8 | 0.72 | 0.8 | 0.76 |
| 5 | 5808963 | rs21757 | 1.175 | -0.035 | -0.341 | -0.115 | -0.013 | 0.031 | 0.031 | 0 | 0.15 | 0.03 | 0.11 | 0.06 | 0.12 | 0.1 |
| 5 | 5809047 | rs32550 | -0.772 | -0.219 | -0.23 | -0.221 | 0.015 | 0.012 | 0.012 | 0 | 0.64 | 0.74 | 0.75 | 0.72 | 0.7 | 0.74 |
| 5 | 5810732 | rs6867853 | -0.441 | 0.005 | 0.082 | -0.19 | -0.005 | 0.012 | 0.012 | 0 | 0.24 | 0.44 | 0.3 | 0.27 | 0.25 | 0.39 |
| 5 | 5811074 | rs2617562 | -0.894 | 0.163 | -0.266 | -0.105 | 0.006 | 0.052 | 0.052 | 0 | 0.48 | 0.7 | 0.5 | 0.33 | 0.47 | 0.5 |
| 5 | 5818164 | rs1864117 | -0.149 | 0.34 | -0.195 | 0.128 | 0.01 | 0.006 | 0.006 | 0 | 0.39 | 0.57 | 0.56 | 0.6 | 0.48 | 0.53 |
| 5 | 5820386 | rs10068484 | 1.297 | 0.342 | 0.047 | 0.441 | 0.022 | -0.015 | -0.015 | 3.375 | 0.23 | 0.27 | 0.16 | 0.23 | 0.3 | 0.23 |
| 5 | 5820530 | rs6892546 | -0.009 | 0.288 | -0.069 | 0.166 | 0.004 | 0.001 | 0.001 | 0.089 | 0.15 | 0.23 | 0.25 | 0.18 | 0.16 | 0.27 |
| 5 | 5822647 | rs3852160 | -0.499 | 0.121 | -0.276 | 0.128 | 0.058 | 0.017 | 0.017 | 0 | 0.51 | 0.7 | 0.66 | 0.6 | 0.65 | 0.58 |
| 5 | 5825953 | rs40516 | -0.605 | 0.283 | -0.995 | 0.473 | 0.022 | 0.036 | 0.036 | 0.078 | 0.73 | 0.66 | 0.6 | 0.56 | 0.67 | 0.73 |
| 5 | 5826272 | rs35741 | -0.06 | -0.158 | -1.621 | 0.509 | 0.031 | -0.04 | -0.04 | 0.33 | 0.63 | 0.92 | 0.77 | 0.67 | 0.72 | 0.66 |
| 5 | 5828648 | rs188684 | -0.372 | 0.624 | -0.823 | 0.601 | 0.004 | 0.042 | 0.042 | 0 | 0.33 | 0.48 | 0.3 | 0.48 | 0.38 | 0.36 |
| 5 | 5828805 | rs309424 | 0.386 | 0.634 | -0.411 | 0.709 | -0.007 | -0.003 | -0.003 | 0 | 0.16 | 0.2 | 0.11 | 0.15 | 0.16 | 0.08 |
| 5 | 5829193 | rs7722388 | 0.362 | 0.619 | -0.203 | 0.691 | -0.002 | 0.002 | 0.002 | 0.018 | 0.16 | 0.11 | 0.16 | 0.33 | 0.22 | 0.28 |
| 5 | 5834614 | rs257073 | -0.386 | 0.86 | -0.604 | 0.473 | 0.007 | 0.012 | 0.012 | 0 | 0.31 | 0.34 | 0.25 | 0.45 | 0.37 | 0.34 |
| 5 | 5836435 | rs3852162 | 1.524 | 0.586 | -0.579 | -0.252 | -0.001 | -0.028 | -0.028 | 0 | 0.25 | 0.24 | 0.33 | 0.03 | 0.22 | 0.31 |
| 5 | 5838642 | rs309421 | -1.004 | 0.544 | -0.296 | -0.271 | -0.006 | 0.031 | 0.031 | 1.002 | 0.77 | 0.67 | 0.73 | 0.61 | 0.83 | 0.76 |
| 5 | 5838682 | rs218317 | 0.939 | 0.396 | -0.242 | -0.515 | 0.002 | 0.002 | 0.002 | 0 | 0.39 | 0.39 | 0.43 | 0.17 | 0.37 | 0.39 |
| 5 | 5839032 | rs182552 | -0.182 | 0.594 | -0.013 | -0.398 | 0.007 | 0.014 | 0.014 | 0 | 0.66 | 0.62 | 0.66 | 0.43 | 0.67 | 0.69 |
| 5 | 5848736 | rs2560461 | -0.6 | 0.764 | -0.033 | -0.477 | 0.025 | 0.014 | 0.014 | 0 | 0.51 | 0.7 | 0.42 | 0.59 | 0.55 | 0.44 |
| 5 | 5851873 | rs2617549 | -0.631 | 0.847 | 0.277 | -0.465 | -0.001 | -0.001 | -0.001 | 0 | 0.49 | 0.57 | 0.38 | 0.53 | 0.5 | 0.42 |
| 5 | 5852963 | rs2560483 | -1.281 | 0.819 | 0.145 | -0.511 | 0.007 | -0.035 | -0.035 | 0 | 0.59 | 0.8 | 0.47 | 0.59 | 0.63 | 0.45 |
| 5 | 5855016 | rs7715747 | 1.694 | 0.681 | 0.168 | -0.582 | 0.006 | 0.004 | 0.004 | 0 | 0.14 | 0.07 | 0.12 | 0.13 | 0.15 | 0.1 |
| 5 | 5855519 | rs2652483 | -1.112 | 0.594 | 0.168 | -0.628 | -0.003 | -0.017 | -0.017 | 0 | 0.57 | 0.71 | 0.44 | 0.54 | 0.58 | 0.44 |
| 5 | 5857554 | rs10512842 | 1.699 | 0.578 | 0.089 | -0.656 | -0.013 | -0.017 | -0.017 | 0 | 0.16 | 0.07 | 0.15 | 0.27 | 0.2 | 0.24 |
| 5 | 5857792 | rs1628345 | 1.817 | 0.577 | 0.006 | -0.658 | -0.027 | -0.014 | -0.014 | 0.547 | 0.13 | 0.05 | 0.07 | 0.12 | 0.14 | 0.08 |
| 5 | 5859422 | rs2561858 | -1.323 | 0.586 | 0.058 | -0.596 | 0.008 | -0.038 | -0.038 | 0.007 | 0.58 | 0.83 | 0.48 | 0.59 | 0.63 | 0.45 |
| 5 | 5860950 | rs2652486 | 0.309 | 0.619 | 0.216 | -0.585 | 0.011 | -0.034 | -0.034 | 0 | 0.72 | 0.5 | 0.81 | 0.8 | 0.71 | 0.88 |
| 5 | 5863733 | rs309016 | -0.016 | 0.611 | 0.296 | -0.566 | -0.002 | -0.025 | -0.025 | 0 | 0.24 | 0.35 | 0.15 | 0.14 | 0.23 | 0.11 |
| 5 | 5865100 | rs12653640 | 1.289 | 0.654 | 0.106 | -0.512 | 0.021 | 0.03 | 0.03 | 0 | 0.17 | 0.24 | 0.17 | 0.26 | 0.19 | 0.24 |
| 5 | 5869722 | rs2246725 | 0.547 | 0.531 | -0.385 | -0.721 | -0.003 | 0.011 | 0.011 | 0.048 | 0.62 | 0.65 | 0.73 | 0.69 | 0.57 | 0.78 |
| 5 | 5879419 | rs16876469 | 0.129 | 0.45 | -0.428 | -0.781 | 0 | -0.001 | -0.001 | 0.053 | 0.21 | 0.24 | 0.13 | 0.12 | 0.22 | 0.1 |
| 5 | 5881234 | rs1913763 | 0.715 | 0.111 | -0.56 | -0.656 | 0.002 | -0.032 | -0.032 | 0.012 | 0.36 | 0.19 | 0.46 | 0.4 | 0.34 | 0.54 |
| 5 | 5890342 | rs1875512 | 1.277 | -0.001 | -0.55 | -0.834 | 0.003 | 0.017 | 0.017 | 0.364 | 0.27 | 0.16 | 0.34 | 0.03 | 0.26 | 0.38 |
| 5 | 5891878 | rs1174930 | -0.262 | 0.004 | -0.69 | -0.844 | -0.031 | -0.014 | -0.014 | 0.006 | 0.59 | 0.27 | 0.62 | 0.54 | 0.56 | 0.66 |
| 5 | 5893255 | rs1174931 | 0.593 | -0.022 | -0.769 | -0.856 | 0.019 | -0.022 | -0.022 | 0.014 | 0.41 | 0.23 | 0.48 | 0.41 | 0.36 | 0.57 |
| 5 | 5894870 | rs2256415 | -0.371 | -0.03 | -0.643 | -0.8 | -0.001 | -0.005 | -0.005 | 0 | 0.8 | 0.79 | 0.85 | 0.87 | 0.79 | 0.9 |
| 5 | 5903890 | rs1032748 | 0.157 | -0.055 | -0.59 | -0.843 | 0 | 0 | 0 | 0 | 0.51 | 0.52 | 0.59 | 0.47 | 0.45 | 0.63 |
| 5 | 5904424 | rs1306085 | 0.948 | 0.186 | -1.008 | -0.579 | -0.007 | -0.085 | -0.085 | 0 | 0.12 | 0.02 | 0.13 | 0.39 | 0.09 | 0.17 |
| 5 | 5910657 | rs2173946 | 0.509 | 0.363 | -0.787 | -0.524 | 0.009 | 0.01 | 0.01 | 0 | 0.3 | 0.31 | 0.27 | 0.4 | 0.35 | 0.27 |
| 5 | 5912054 | rs10454950 | -0.118 | 0.415 | -1.135 | -0.45 | 0.005 | 0.028 | 0.028 | 0.037 | 0.18 | 0.11 | 0.15 | 0.13 | 0.19 | 0.12 |
| 5 | 5916224 | rs13188105 | -0.492 | 0.5 | -1.232 | 0.015 | -0.02 | -0.091 | -0.091 | 0 | 0.21 | 0.06 | 0.17 | 0.53 | 0.17 | 0.21 |
| 5 | 5916337 | rs2455385 | 0.513 | 0.018 | -0.892 | -0.614 | 0.029 | 0.036 | 0.036 | 0 | 0.49 | 0.59 | 0.47 | 0.45 | 0.61 | 0.45 |
| 5 | 5922597 | rs10462810 | 0.099 | 0.029 | -0.794 | -0.434 | 0.003 | -0.021 | -0.021 | 0 | 0.18 | 0.15 | 0.16 | 0.45 | 0.13 | 0.19 |
| 5 | 5922663 | rs2673846 | -1.053 | -0.163 | -0.828 | -0.605 | 0.014 | 0.017 | 0.017 | 0.044 | 0.75 | 0.82 | 0.66 | 0.97 | 0.79 | 0.68 |
| 5 | 5931513 | rs16876615 | 0.386 | 0.442 | -0.16 | -0.785 | 0.162 | 0.014 | 0.014 | 0.747 | 0.08 | 0.28 | 0.17 | 0.05 | 0.16 | 0.11 |
| 5 | 5936810 | rs1174900 | 1.148 | 0.293 | -0.752 | -0.919 | -0.052 | 0.03 | 0.03 | 0 | 0.38 | 0.05 | 0.25 | 0.36 | 0.41 | 0.34 |
| 5 | 5937776 | rs1174901 | 0.033 | 0.321 | -0.648 | -0.877 | 0.028 | -0.001 | -0.001 | 0.393 | 0.56 | 0.64 | 0.5 | 0.53 | 0.65 | 0.49 |
| 5 | 5938780 | rs10068316 | 0.533 | 0.357 | -0.66 | -0.915 | 0 | -0.047 | -0.047 | 0 | 0.17 | 0.13 | 0.16 | 0.48 | 0.13 | 0.19 |
| 5 | 5943861 | rs657865 | -0.928 | 0.383 | -0.476 | -0.788 | -0.059 | 0.232 | 0.232 | 0.05 | 0.49 | 0.92 | 0.6 | 0.18 | 0.46 | 0.49 |
| 5 | 5951796 | rs216578 | -0.123 | 0.548 | -0.456 | -0.722 | -0.011 | 0.04 | 0.04 | 0 | 0.35 | 0.28 | 0.45 | 0.14 | 0.4 | 0.44 |
| 5 | 5956806 | rs160126 | -1.406 | 0.821 | 0.308 | 0.016 | 0.001 | -0.032 | -0.032 | 2.127 | 0.09 | 0.48 | 0.06 | 0 | 0.02 | 0.03 |
| 5 | 5959444 | rs1366154 | -0.031 | 1.056 | -0.707 | 0.299 | -0.035 | -0.002 | -0.002 | 0 | 0.07 | 0.02 | 0.13 | 0.5 | 0.11 | 0.17 |
| 5 | 5966730 | rs160134 | -1.354 | 0.978 | -0.783 | 0.215 | -0.133 | -0.09 | -0.09 | 0 | 0.37 | 0.95 | 0.36 | 0.56 | 0.37 | 0.36 |

| | | | | | | | | | | | | | | | | |
|---|---|---|---|---|---|---|---|---|---|---|---|---|---|---|---|---|
| 5 | 5972299 | rs10474793 | -1.052 | 0.745 | 0.454 | 1.175 | 0.057 | 0.296 | 0.296 | 2.192 | 0.13 | 0.4 | 0.18 | 0.51 | 0.13 | 0.17 |
| 5 | 5973626 | rs302913 | -0.846 | 0.5 | 0.22 | 0.413 | -0.004 | -0.021 | -0.021 | 1.252 | 0.17 | 0.23 | 0.21 | 0.07 | 0.15 | 0.25 |
| 5 | 5975333 | rs6887254 | -0.354 | -0.98 | 0.046 | 0.359 | -0.001 | 0.001 | 0.001 | 0 | 0.16 | 0.14 | 0.05 | 0.19 | 0.05 | 0.17 |
| 5 | 5977222 | rs10213777 | -0.389 | -1.002 | 0.216 | 0.349 | -0.016 | 0 | 0 | 0 | 0.16 | 0.15 | 0.05 | 0.19 | 0.05 | 0.17 |
| 5 | 5981228 | rs306254 | 1.401 | -1.574 | 0.242 | 0.088 | -0.01 | -0.01 | -0.01 | 0.74 | 0.39 | 0.26 | 0.59 | 0.55 | 0.53 | 0.39 |
| 5 | 5989136 | rs159981 | 0.759 | -1.889 | -0.039 | -0.02 | -0.001 | -0.005 | -0.005 | 0 | 0.16 | 0.19 | 0.2 | 0.11 | 0.14 | 0.25 |
| 5 | 5989446 | rs12516740 | 0.596 | -1.894 | 0.293 | -0.045 | -0.005 | -0.002 | -0.002 | 0 | 0.16 | 0.16 | 0.11 | 0.02 | 0.19 | 0.18 |
| 5 | 6012340 | rs458635 | 0.5 | -2.189 | 0.502 | -0.084 | -0.001 | 0.003 | 0.003 | 0 | 0.72 | 0.7 | 0.85 | 0.85 | 0.72 | 0.81 |
| 5 | 6016507 | rs2434744 | 1.555 | -2.42 | 0.304 | -0.188 | 0.008 | 0.085 | 0.085 | 0 | 0.52 | 0.21 | 0.63 | 0.72 | 0.58 | 0.56 |
| 5 | 6021846 | rs419119 | 0.069 | -2.268 | 0.352 | -0.133 | -0.002 | 0.002 | 0.002 | 0 | 0.76 | 0.77 | 0.87 | 0.87 | 0.76 | 0.86 |
| 5 | 6022683 | rs4377680 | -0.776 | -2.285 | 0.264 | -0.134 | 0.017 | 0.058 | 0.058 | 0.001 | 0.34 | 0.2 | 0.18 | 0.22 | 0.27 | 0.31 |
| 5 | 6023131 | rs457990 | 0.851 | -2.22 | 0.772 | -0.157 | 0.012 | -0.078 | -0.078 | 0 | 0.14 | 0.24 | 0.1 | 0.02 | 0.15 | 0.13 |
| 5 | 6023338 | rs3846567 | 0.192 | -2.537 | 0.395 | -0.281 | 0.001 | -0.001 | -0.001 | 0 | 0.22 | 0.21 | 0.16 | 0.1 | 0.15 | 0.28 |
| 5 | 6024988 | rs461240 | 2.059 | -2.374 | 0.006 | -0.089 | -0.094 | -0.099 | -0.099 | 0 | 0.38 | 0.11 | 0.64 | 0.55 | 0.54 | 0.4 |
| 5 | 6028250 | rs154814 | 0.897 | -2.007 | 0.389 | 0.086 | -0.001 | 0.03 | 0.03 | 1.617 | 0.86 | 0.73 | 0.93 | 0.86 | 0.87 | 0.96 |
| 5 | 6034364 | rs173577 | 1.131 | -1.834 | -0.109 | -0.301 | 0.054 | 0.03 | 0.03 | 0.502 | 0.64 | 0.18 | 0.76 | 0.69 | 0.76 | 0.52 |
| 5 | 6034845 | rs40111 | 1.449 | -1.992 | -0.678 | -0.757 | -0.048 | 0.076 | 0.076 | 0 | 0.42 | 0.18 | 0.33 | 0.69 | 0.47 | 0.33 |
| 5 | 6036454 | rs10051928 | -0.059 | -2.164 | -0.887 | -0.824 | 0.004 | 0.034 | 0.034 | 0.455 | 0.29 | 0.11 | 0.29 | 0.22 | 0.3 | 0.33 |
| 5 | 6038449 | rs16876890 | -0.236 | -2.272 | -0.262 | -0.882 | -0.003 | -0.076 | -0.076 | 3.034 | 0.19 | 0.49 | 0.35 | 0.08 | 0.18 | 0.31 |
| 5 | 6038726 | rs1027249 | 1.207 | -2.241 | -0.611 | -0.85 | 0.008 | -0.034 | -0.034 | 0 | 0.09 | 0 | 0.11 | 0.13 | 0.05 | 0.15 |
| 5 | 6041659 | rs28047 | 0.708 | -2.236 | -0.708 | -1.053 | -0.027 | -0.115 | -0.115 | 0.426 | 0.84 | 0.62 | 0.93 | 0.98 | 0.92 | 0.98 |
| 5 | 6043189 | rs17769749 | 1.777 | -2.196 | -0.991 | -1.026 | -0.007 | 0.019 | 0.019 | 0 | 0.06 | 0.03 | 0.03 | 0 | 0.05 | 0.01 |
| 5 | 6043624 | rs13166306 | 0.45 | -2.285 | -0.722 | -1.072 | 0.006 | -0.022 | -0.022 | 0 | 0.15 | 0.23 | 0.33 | 0.08 | 0.17 | 0.3 |
| 5 | 6043644 | rs26647 | 2.268 | -2.3 | -1.203 | -1.089 | -0.169 | -0.019 | -0.019 | 0 | 0.35 | 0.03 | 0.29 | 0.69 | 0.43 | 0.33 |
| 5 | 6044675 | rs10064690 | -1.023 | -1.95 | -0.829 | -0.367 | -0.008 | 0.013 | 0.013 | 0.013 | 0.39 | 0.14 | 0.31 | 0.34 | 0.38 | 0.36 |
| 5 | 6045849 | rs17821004 | 1.054 | -1.966 | -0.86 | -1.033 | 0.009 | -0.029 | -0.029 | 0 | 0.09 | 0 | 0.11 | 0.12 | 0.05 | 0.15 |
| 5 | 6050231 | rs6878493 | -1.163 | -1.718 | -0.484 | -1.029 | -0.003 | -0.019 | -0.019 | 0 | 0.44 | 0.56 | 0.37 | 0.23 | 0.4 | 0.37 |
| 5 | 6054601 | rs11134135 | -0.113 | -1.447 | -1.199 | -0.784 | -0.011 | 0.021 | 0.021 | 0.714 | 0.31 | 0.08 | 0.31 | 0.24 | 0.31 | 0.33 |
| 5 | 6058009 | rs12187733 | -0.242 | -0.894 | -0.74 | -0.579 | -0.012 | 0.017 | 0.017 | 0 | 0.14 | 0.08 | 0.3 | 0.1 | 0.16 | 0.3 |
| 5 | 6075956 | rs16876913 | 0.167 | -0.719 | -0.579 | -0.606 | -0.022 | 0.008 | 0.008 | 0.265 | 0.15 | 0.08 | 0.29 | 0.12 | 0.17 | 0.33 |
| 5 | 6082486 | rs17770545 | -0.479 | -0.602 | -0.42 | -0.671 | -0.023 | 0.019 | 0.019 | 0.131 | 0.25 | 0.17 | 0.37 | 0.17 | 0.34 | 0.39 |
| 5 | 6083408 | rs12515958 | -0.452 | -0.596 | 0.03 | -0.668 | 0.013 | -0.012 | -0.012 | 0 | 0.25 | 0.27 | 0.38 | 0.17 | 0.34 | 0.38 |
| 5 | 6084659 | rs2115365 | 0.69 | -0.141 | -0.401 | -0.823 | 0.007 | 0.003 | 0.003 | 0 | 0.25 | 0.11 | 0.22 | 0.22 | 0.24 | 0.2 |
| 5 | 6087348 | rs307177 | -1.136 | -0.304 | -0.08 | -0.553 | -0.027 | -0.063 | -0.063 | 0 | 0.65 | 0.87 | 0.68 | 0.4 | 0.63 | 0.59 |
| 5 | 6087464 | rs307176 | -0.989 | -0.285 | -0.041 | -0.562 | -0.008 | -0.012 | -0.012 | 1.58 | 0.63 | 0.79 | 0.65 | 0.36 | 0.6 | 0.57 |
| 5 | 6093701 | rs11134138 | -0.194 | 0.052 | 0.071 | -0.37 | 0 | 0.059 | 0.059 | 0.108 | 0.48 | 0.2 | 0.57 | 0.35 | 0.54 | 0.57 |
| 5 | 6104208 | rs409978 | -0.538 | 0.291 | 0.457 | -0.431 | -0.016 | 0.005 | 0.005 | 0 | 0.58 | 0.72 | 0.63 | 0.41 | 0.54 | 0.56 |
| 5 | 6110760 | rs301772 | -0.376 | 0.605 | -1.218 | -0.291 | -0.013 | 0.013 | 0.013 | 0 | 0.56 | 0.75 | 0.62 | 0.41 | 0.53 | 0.56 |
| 5 | 6132166 | rs9313144 | -0.831 | 0.727 | -1.483 | -0.092 | -0.085 | -0.113 | -0.113 | 0 | 0.65 | 0.93 | 0.71 | 0.47 | 0.61 | 0.57 |
| 5 | 6136125 | rs6861277 | -0.596 | 0.635 | -0.461 | 0.675 | -0.005 | 0.043 | 0.043 | 0 | 0.09 | 0.18 | 0.07 | 0.23 | 0.06 | 0.11 |
| 5 | 6141688 | rs4334845 | -0.223 | 0.657 | -0.232 | 0.479 | 0 | -0.006 | -0.006 | 0.09 | 0.15 | 0.14 | 0.12 | 0.11 | 0.17 | 0.15 |
| 5 | 6142003 | rs16876953 | -0.507 | 0.382 | 0.031 | 0.822 | 0.003 | 0.02 | 0.02 | 0 | 0.06 | 0.13 | 0.05 | 0.08 | 0.06 | 0.06 |
| 5 | 6142364 | rs12652811 | -0.507 | 0.382 | 0.031 | 0.822 | 0.003 | 0.02 | 0.02 | 0 | 0.06 | 0.13 | 0.05 | 0.08 | 0.06 | 0.06 |
| 5 | 6148345 | rs12658976 | -1.87 | 0.361 | 0.051 | 0.815 | -0.014 | 0.087 | 0.087 | 0 | 0.16 | 0.34 | 0.12 | 0.31 | 0.13 | 0.17 |
| 5 | 6156355 | rs10077859 | -2.338 | 0.333 | 0.108 | 0.756 | -0.014 | 0.06 | 0.06 | 0.017 | 0.21 | 0.42 | 0.36 | 0.34 | 0.2 | 0.28 |
| 5 | 6158213 | rs12657021 | -2.021 | 0.625 | -0.209 | 0.559 | -0.004 | -0.004 | -0.004 | 0 | 0.13 | 0.24 | 0.29 | 0.11 | 0.13 | 0.17 |
| 5 | 6162445 | rs993046 | -0.556 | 0.855 | -0.808 | 0.507 | 0 | -0.005 | -0.005 | 0.604 | 0.06 | 0.08 | 0.24 | 0.03 | 0.07 | 0.11 |
| 5 | 6164849 | rs11750309 | -1.063 | 1.107 | -0.869 | 0.949 | -0.009 | -0.009 | -0.009 | 0 | 0.11 | 0.11 | 0.09 | 0.26 | 0.15 | 0.14 |
| 5 | 6167165 | rs1560178 | 0.234 | 0.683 | -2.272 | 0.342 | -0.024 | -0.173 | -0.173 | 0 | 0.34 | 0.02 | 0.28 | 0.54 | 0.33 | 0.34 |
| 5 | 6167854 | rs11134144 | -0.328 | 0.427 | -1.621 | 0.151 | 0.007 | 0.158 | 0.158 | 0 | 0.43 | 0.21 | 0.37 | 0.86 | 0.46 | 0.49 |
| 5 | 6172968 | rs10059895 | -0.117 | 0.077 | -1.079 | -0.554 | -0.032 | -0.068 | -0.068 | 0.909 | 0.82 | 0.49 | 0.89 | 0.94 | 0.89 | 0.82 |
| 5 | 6174586 | rs11953803 | 0.716 | 0.081 | -1.077 | -0.56 | -0.034 | 0.072 | 0.072 | 0.019 | 0.39 | 0.27 | 0.51 | 0.08 | 0.43 | 0.33 |
| 5 | 6176473 | rs11745650 | 0.29 | 0.09 | -1.125 | -0.511 | 0.005 | 0.157 | 0.157 | 0 | 0.43 | 0.2 | 0.38 | 0.86 | 0.46 | 0.49 |
| 5 | 6179771 | rs2964722 | -0.24 | 0.011 | -0.081 | -0.414 | -0.013 | -0.033 | -0.033 | 0 | 0.87 | 0.71 | 0.91 | 0.94 | 0.91 | 0.82 |
| 5 | 6197044 | rs2937785 | -0.45 | 0.016 | 0.08 | -0.457 | 0 | -0.023 | -0.023 | 0.014 | 0.88 | 0.75 | 0.91 | 0.95 | 0.91 | 0.83 |
| 5 | 6203128 | rs2964767 | 0.663 | -0.051 | -0.941 | -1.279 | 0.002 | -0.02 | -0.02 | 0 | 0.42 | 0.24 | 0.56 | 0.55 | 0.37 | 0.45 |
| 5 | 6204971 | rs11134145 | 1.109 | -0.036 | -0.886 | -1.255 | -0.004 | 0.025 | 0.025 | 0 | 0.08 | 0.09 | 0.08 | 0.28 | 0.12 | 0.15 |
| 5 | 6205134 | rs11134147 | 0.27 | -0.223 | -0.648 | -1.329 | -0.003 | -0.004 | -0.004 | 0.275 | 0.25 | 0.3 | 0.2 | 0.38 | 0.35 | 0.23 |
| 5 | 6206875 | rs1875542 | 2.025 | -0.528 | -0.909 | -1.497 | 0.022 | -0.15 | -0.15 | 0.374 | 0.18 | 0.02 | 0.16 | 0.48 | 0.13 | 0.24 |
| 5 | 6208293 | rs4143217 | 0.947 | -0.391 | -0.853 | -1.445 | -0.036 | -0.054 | -0.054 | 0 | 0.22 | 0.15 | 0.19 | 0.36 | 0.32 | 0.23 |
| 5 | 6212193 | rs2937791 | -0.959 | -0.343 | -0.7 | -1.442 | -0.001 | -0.049 | -0.049 | 0 | 0.69 | 0.7 | 0.78 | 0.94 | 0.71 | 0.69 |
| 5 | 6212759 | rs2964739 | -1.219 | -0.085 | -0.074 | -1.233 | -0.006 | -0.018 | -0.018 | 0.376 | 0.88 | 0.83 | 0.91 | 0.95 | 0.91 | 0.83 |
| 5 | 6216046 | rs7727548 | 0.986 | -0.317 | -0.386 | -1.358 | -0.074 | 0.067 | 0.067 | 0 | 0.19 | 0.04 | 0.37 | 0.1 | 0.23 | 0.19 |
| 5 | 6218907 | rs7726513 | -0.157 | -0.322 | 0.11 | -1.046 | 0.011 | 0.024 | 0.024 | 0.01 | 0.28 | 0.47 | 0.22 | 0.36 | 0.34 | 0.24 |
| 5 | 6235302 | rs2964755 | 0.625 | -0.193 | -0.567 | -0.935 | -0.003 | -0.006 | -0.006 | 0 | 0.6 | 0.28 | 0.68 | 0.59 | 0.57 | 0.59 |
| 5 | 6247401 | rs1870082 | -0.654 | -0.337 | -0.097 | -0.647 | -0.016 | -0.041 | -0.041 | 1.16 | 0.25 | 0.6 | 0.15 | 0.08 | 0.16 | 0.2 |
| 5 | 6248674 | rs1598734 | -0.212 | -0.263 | -0.217 | -0.2 | 0.004 | -0.007 | -0.007 | 0.457 | 0.61 | 0.62 | 0.63 | 0.49 | 0.65 | 0.58 |
| 5 | 6249440 | rs2124935 | 0.253 | 0.233 | -0.198 | 0.295 | 0.008 | 0 | 0 | 0 | 0.47 | 0.36 | 0.54 | 0.48 | 0.56 | 0.45 |
| 5 | 6251548 | rs2925721 | 0.625 | 0.316 | -0.377 | 0.391 | -0.024 | -0.027 | -0.027 | 3.163 | 0.41 | 0.29 | 0.51 | 0.47 | 0.52 | 0.39 |

| | | | | | | | | | | | | | | | | |
|---|---|---|---|---|---|---|---|---|---|---|---|---|---|---|---|---|
| 5 | 6252929 | rs1457120 | 0.537 | 0.322 | -0.127 | 0.634 | -0.003 | -0.005 | -0.005 | 0 | 0.45 | 0.41 | 0.51 | 0.49 | 0.51 | 0.41 |
| 5 | 6257553 | rs4702325 | 0.007 | 0.442 | -0.359 | 0.588 | -0.004 | -0.019 | -0.019 | 1.448 | 0.08 | 0.03 | 0.1 | 0.12 | 0.06 | 0.09 |
| 5 | 6257996 | rs2914757 | -0.633 | 0.448 | -0.261 | 0.448 | 0.013 | -0.04 | -0.04 | 0.099 | 0.92 | 0.85 | 0.95 | 1 | 0.9 | 0.98 |
| 5 | 6262981 | rs4702326 | -0.191 | 0.051 | -0.41 | 0.766 | 0 | 0.003 | 0.003 | 0 | 0.09 | 0.09 | 0.1 | 0.12 | 0.06 | 0.09 |
| 5 | 6264448 | rs1870083 | 0.765 | 0.203 | -0.659 | 0.412 | -0.005 | 0.079 | 0.079 | 0 | 0.38 | 0.24 | 0.24 | 0.16 | 0.41 | 0.31 |
| 5 | 6266723 | rs3846569 | -0.613 | 0.461 | -0.665 | 0.758 | -0.003 | 0.019 | 0.019 | 0 | 0.89 | 0.86 | 0.88 | 0.87 | 0.92 | 0.91 |
| 5 | 6273004 | rs1457122 | 0.472 | 0.331 | -0.719 | 0.627 | -0.035 | -0.019 | -0.019 | 0 | 0.26 | 0.21 | 0.4 | 0.3 | 0.32 | 0.27 |
| 5 | 6274436 | rs1551572 | -0.072 | 0.21 | -0.711 | 0.689 | 0.009 | -0.023 | -0.023 | 0 | 0.72 | 0.8 | 0.69 | 0.43 | 0.76 | 0.59 |
| 5 | 6276222 | rs4701720 | -0.111 | 0.645 | -0.661 | 0.935 | -0.015 | -0.005 | -0.005 | 0.05 | 0.28 | 0.27 | 0.41 | 0.29 | 0.32 | 0.29 |
| 5 | 6277738 | rs6866699 | -0.218 | 0.535 | -0.631 | 1.169 | -0.002 | 0.001 | 0.001 | 0 | 0.18 | 0.13 | 0.2 | 0.48 | 0.21 | 0.32 |
| 5 | 6279339 | rs2061280 | 0.25 | 0.179 | -0.285 | 0.958 | -0.008 | 0.05 | 0.05 | 0 | 0.77 | 0.67 | 0.78 | 0.55 | 0.79 | 0.67 |
| 5 | 6281648 | rs2914752 | 0.405 | 0.17 | -0.577 | 0.565 | 0.005 | -0.004 | -0.004 | 0 | 0.47 | 0.44 | 0.66 | 0.44 | 0.58 | 0.46 |
| 5 | 6286005 | rs2925703 | 0.157 | 0.748 | -0.667 | 1.255 | -0.003 | 0.073 | 0.073 | 0 | 0.79 | 0.64 | 0.83 | 0.6 | 0.81 | 0.67 |
| 5 | 6286955 | rs2925702 | -0.236 | 1.517 | -1.332 | 1.16 | -0.034 | -0.045 | -0.045 | 0 | 0.2 | 0.15 | 0.38 | 0.3 | 0.29 | 0.23 |
| 5 | 6289200 | rs1379305 | -0.446 | 1.135 | -1.358 | 0.735 | 0.004 | 0.025 | 0.025 | 4.059 | 0.47 | 0.33 | 0.69 | 0.61 | 0.55 | 0.41 |
| 5 | 6296991 | rs2914746 | -0.124 | 0.89 | -1.746 | 0.428 | -0.046 | -0.045 | -0.045 | 0 | 0.31 | 0.15 | 0.55 | 0.38 | 0.38 | 0.27 |
| 5 | 6301470 | rs7703162 | 0.472 | 0.888 | -1.787 | 0.279 | -0.009 | -0.017 | -0.017 | 1.407 | 0.53 | 0.85 | 0.35 | 0.46 | 0.43 | 0.55 |
| 5 | 6302491 | rs12513969 | 0.305 | 0.514 | -1.013 | -0.137 | 0.004 | 0.003 | 0.003 | 0.219 | 0.41 | 0.49 | 0.35 | 0.57 | 0.44 | 0.6 |
| 5 | 6307991 | rs2242132 | -0.31 | 0.208 | -0.336 | -0.208 | 0 | 0.012 | 0.012 | 0.569 | 0.3 | 0.36 | 0.52 | 0.39 | 0.34 | 0.29 |
| 5 | 6310014 | rs10055872 | -1.128 | -0.053 | -0.097 | -0.558 | 0.003 | 0.006 | 0.006 | 0.428 | 0.51 | 0.71 | 0.65 | 0.46 | 0.51 | 0.6 |
| 5 | 6311020 | rs10050369 | -0.213 | -0.008 | -0.158 | -0.024 | 0.069 | 0.061 | 0.061 | 0.343 | 0.05 | 0.32 | 0.05 | 0.03 | 0.05 | 0.01 |
| 5 | 6311100 | rs10059734 | -0.631 | -0.012 | -0.196 | -0.028 | -0.001 | 0.014 | 0.014 | 0.435 | 0.31 | 0.39 | 0.52 | 0.39 | 0.34 | 0.3 |
| 5 | 6311896 | rs17825532 | -0.177 | 0.122 | -0.914 | 0.005 | -0.061 | -0.108 | -0.108 | 0.332 | 0.24 | 0.02 | 0.46 | 0.34 | 0.28 | 0.29 |
| 5 | 6316676 | rs13166082 | -0.389 | -0.438 | -0.737 | -0.12 | 0.03 | -0.022 | -0.022 | 0 | 0.28 | 0.15 | 0.28 | 0.3 | 0.24 | 0.29 |
| 5 | 6317260 | rs7714333 | -0.516 | -0.314 | -0.505 | -0.253 | 0.022 | 0.026 | 0.026 | 0 | 0.51 | 0.73 | 0.42 | 0.4 | 0.44 | 0.56 |
| 5 | 6317306 | rs12716179 | -1.072 | -0.366 | 0.488 | -0.184 | -0.015 | -0.019 | -0.019 | 0 | 0.1 | 0.4 | 0.06 | 0.02 | 0.03 | 0.19 |
| 5 | 6323989 | rs12520404 | -0.646 | 0.004 | 0.106 | -0.231 | 0.007 | -0.043 | -0.043 | 0 | 0.08 | 0.14 | 0.06 | 0 | 0.07 | 0.08 |
| 5 | 6325444 | rs13177370 | 0.144 | 0.186 | -0.43 | -0.86 | -0.036 | 0.047 | 0.047 | 0 | 0.35 | 0.24 | 0.26 | 0.19 | 0.44 | 0.41 |
| 5 | 6325850 | rs4345283 | -0.71 | -0.836 | -0.276 | -0.543 | 0.033 | 0.039 | 0.039 | 0.078 | 0.57 | 0.72 | 0.45 | 0.34 | 0.6 | 0.52 |
| 5 | 6326059 | rs1428986 | 0.289 | -0.847 | 0.038 | -0.581 | -0.001 | -0.003 | -0.003 | 0 | 0.41 | 0.36 | 0.32 | 0.19 | 0.51 | 0.48 |
| 5 | 6328426 | rs6872084 | 0.521 | -0.54 | -0.712 | -0.653 | 0.001 | -0.012 | -0.012 | 0 | 0.1 | 0.12 | 0.03 | 0 | 0.06 | 0.05 |
| 5 | 6335043 | rs7731289 | -0.624 | -1.05 | -1.181 | 0.016 | 0.002 | -0.014 | -0.014 | 0 | 0.75 | 0.77 | 0.54 | 0.61 | 0.74 | 0.85 |
| 5 | 6336397 | rs13189546 | -0.103 | -0.763 | -0.924 | -0.221 | 0.037 | 0.039 | 0.039 | 0.053 | 0.56 | 0.73 | 0.45 | 0.37 | 0.6 | 0.53 |
| 5 | 6340693 | rs271403 | -0.555 | -1.025 | -0.163 | -0.33 | -0.053 | -0.074 | -0.074 | 0 | 0.18 | 0.42 | 0.08 | 0.05 | 0.09 | 0.04 |
| 5 | 6344389 | rs271409 | -0.639 | -1.249 | -1.016 | -0.322 | -0.025 | -0.042 | -0.042 | 0 | 0.79 | 0.99 | 0.86 | 0.78 | 0.79 | 0.7 |
| 5 | 6347695 | rs271413 | -0.22 | -1.179 | -0.383 | 0.15 | -0.042 | 0.079 | 0.079 | 0 | 0.27 | 0.49 | 0.29 | 0.46 | 0.19 | 0.22 |
| 5 | 6353024 | rs271418 | -0.541 | -0.963 | -0.438 | -0.218 | -0.005 | 0.032 | 0.032 | 0.588 | 0.68 | 0.84 | 0.78 | 0.74 | 0.66 | 0.66 |
| 5 | 6353815 | rs10040610 | 0.976 | -1.214 | 0.001 | 0.253 | -0.006 | 0.093 | 0.093 | 0 | 0.13 | 0.24 | 0.16 | 0.33 | 0.11 | 0.19 |
| 5 | 6354132 | rs271421 | 0.593 | -1.228 | -0.329 | 0.31 | -0.042 | 0.022 | 0.022 | 0 | 0.16 | 0.5 | 0.15 | 0.15 | 0.11 | 0.04 |
| 5 | 6355707 | rs271422 | -0.428 | -1.125 | -0.48 | -0.081 | -0.022 | 0.018 | 0.018 | 0.173 | 0.31 | 0.64 | 0.32 | 0.46 | 0.21 | 0.23 |
| 5 | 6358449 | rs7711308 | 1.377 | 0.21 | -0.569 | -0.438 | -0.007 | 0.018 | 0.018 | 0 | 0.23 | 0.15 | 0.19 | 0.16 | 0.29 | 0.13 |
| 5 | 6365689 | rs4701724 | 0.122 | 0.24 | -0.162 | -0.677 | 0.008 | 0.038 | 0.038 | 1.949 | 0.51 | 0.53 | 0.68 | 0.71 | 0.49 | 0.61 |
| 5 | 6372248 | rs4592 | -0.187 | 0.123 | 0.29 | -0.824 | -0.007 | -0.011 | -0.011 | 0 | 0.48 | 0.34 | 0.44 | 0.49 | 0.5 | 0.51 |
| 5 | 6373100 | rs17831845 | 1.883 | 0.023 | 0.348 | -0.908 | -0.001 | 0.033 | 0.033 | 0 | 0.25 | 0.13 | 0.19 | 0.11 | 0.29 | 0.11 |
| 5 | 6374004 | rs2455146 | -0.203 | 0.044 | 0.409 | -0.892 | -0.001 | -0.001 | -0.001 | 0 | 0.49 | 0.55 | 0.45 | 0.49 | 0.5 | 0.51 |
| 5 | 6377973 | rs10052685 | 0.083 | 0.495 | 0.251 | -0.885 | -0.003 | 0.028 | 0.028 | 0 | 0.26 | 0.3 | 0.34 | 0.4 | 0.22 | 0.37 |
| 5 | 6381817 | rs7702608 | 0.585 | 0.482 | 0.298 | -0.977 | 0 | 0.004 | 0.004 | 0 | 0.42 | 0.56 | 0.33 | 0.36 | 0.43 | 0.21 |
| 5 | 6383318 | rs11134158 | 1.667 | 0.461 | 0.291 | -0.976 | -0.006 | -0.058 | -0.058 | 0 | 0.27 | 0.35 | 0.2 | 0.12 | 0.28 | 0.12 |
| 5 | 6386041 | rs4701728 | -1.376 | 0.45 | -0.04 | -1.021 | -0.009 | 0.138 | 0.138 | 0.332 | 0.73 | 0.94 | 0.86 | 0.95 | 0.73 | 0.72 |
| 5 | 6392452 | rs13361971 | 0.943 | 0.526 | -0.063 | -0.94 | 0.003 | 0.006 | 0.006 | 1.547 | 0.16 | 0.21 | 0.12 | 0.24 | 0.15 | 0.1 |
| 5 | 6393771 | rs10050504 | 0.101 | 0.394 | -0.195 | -0.933 | -0.002 | 0.016 | 0.016 | 0 | 0.26 | 0.25 | 0.33 | 0.4 | 0.22 | 0.37 |
| 5 | 6395712 | rs4702345 | 1.453 | 0.335 | -0.322 | -0.936 | -0.007 | -0.079 | -0.079 | 0 | 0.13 | 0.02 | 0.25 | 0.43 | 0.12 | 0.21 |
| 5 | 6398254 | rs12153196 | -0.976 | 0.343 | -0.22 | -0.99 | 0.001 | -0.006 | -0.006 | 0.182 | 0.4 | 0.34 | 0.59 | 0.83 | 0.34 | 0.58 |
| 5 | 6401135 | rs6869702 | -0.491 | 0.323 | -0.362 | -1.004 | -0.013 | -0.126 | -0.126 | 0 | 0.59 | 0.93 | 0.61 | 0.51 | 0.61 | 0.5 |
| 5 | 6401960 | rs10056261 | 0.953 | 0.379 | -0.259 | -0.976 | 0.003 | 0.003 | 0.003 | 1.911 | 0.16 | 0.18 | 0.14 | 0.24 | 0.17 | 0.1 |
| 5 | 6402598 | rs1393067 | 1.081 | 0.563 | -0.658 | -0.941 | -0.018 | 0.042 | 0.042 | 1.559 | 0.11 | 0.16 | 0.2 | 0.16 | 0.04 | 0.27 |
| 5 | 6404896 | rs2398140 | 0.344 | 0.598 | -0.658 | -0.946 | 0.072 | 0.076 | 0.076 | 1.069 | 0.14 | 0.33 | 0.27 | 0.16 | 0.15 | 0.3 |
| 5 | 6405678 | rs17775764 | 1.154 | 0.566 | -1.47 | -0.86 | -0.041 | 0.134 | 0.134 | 0.693 | 0.26 | 0.01 | 0.14 | 0.05 | 0.27 | 0.28 |
| 5 | 6410448 | rs12654055 | 0.103 | 0.662 | -1.202 | -0.719 | -0.025 | -0.024 | -0.024 | 0.752 | 0.48 | 0.83 | 0.53 | 0.49 | 0.48 | 0.47 |
| 5 | 6413449 | rs17775925 | 0.581 | 0.979 | -0.526 | -0.307 | -0.005 | 0.001 | 0.001 | 0.252 | 0.1 | 0.06 | 0.08 | 0.02 | 0.13 | 0.03 |
| 5 | 6415844 | rs16877282 | 0.111 | 0.788 | -0.514 | -0.224 | 0 | -0.001 | -0.001 | 0 | 0.13 | 0.12 | 0.23 | 0.17 | 0.07 | 0.27 |
| 5 | 6418287 | rs16877284 | -0.236 | 0.895 | -0.526 | -0.26 | -0.001 | -0.001 | -0.001 | 0 | 0.14 | 0.14 | 0.29 | 0.16 | 0.18 | 0.29 |
| 5 | 6418689 | rs16877286 | -0.639 | 0.608 | -0.665 | -0.281 | -0.007 | -0.053 | -0.053 | 6.266 | 0.24 | 0.16 | 0.31 | 0.51 | 0.24 | 0.25 |
| 5 | 6418912 | rs16877287 | -0.806 | 0.604 | -0.738 | -0.349 | -0.013 | -0.062 | -0.062 | 0 | 0.28 | 0.17 | 0.33 | 0.5 | 0.29 | 0.26 |
| 5 | 6424822 | rs7737864 | 0.512 | 0.637 | -0.377 | -0.552 | 0.006 | 0.009 | 0.009 | 0 | 0.77 | 0.68 | 0.65 | 0.32 | 0.75 | 0.68 |
| 5 | 6428242 | rs10038921 | 0.397 | 0.697 | -0.47 | -0.598 | 0.006 | 0.017 | 0.017 | 0 | 0.13 | 0.19 | 0.15 | 0.27 | 0.15 | 0.09 |
| 5 | 6428642 | rs6869437 | -0.861 | 0.581 | -0.091 | -0.726 | 0.048 | 0.002 | 0.002 | 0 | 0.23 | 0.44 | 0.22 | 0.27 | 0.28 | 0.13 |
| 5 | 6430604 | rs9687391 | 0.222 | -0.281 | 0.097 | -0.366 | 0.006 | 0.018 | 0.018 | 0 | 0.13 | 0.19 | 0.15 | 0.27 | 0.15 | 0.09 |
| 5 | 6432740 | rs7714347 | 0.591 | -1.072 | 0.222 | -0.019 | -0.002 | 0 | 0 | 0 | 0.74 | 0.72 | 0.54 | 0.31 | 0.79 | 0.45 |

| | | | | | | | | | | | | | | | | |
|---|---|---|---|---|---|---|---|---|---|---|---|---|---|---|---|---|
| 5 | 6439663 | rs824634 | 0.81 | -1.019 | 0.054 | -0.009 | -0.022 | -0.008 | -0.008 | 2.031 | 0.71 | 0.51 | 0.53 | 0.32 | 0.79 | 0.46 |
| 5 | 6450203 | rs169250 | 0.441 | -1.043 | -0.472 | -0.202 | -0.019 | -0.005 | -0.005 | 0 | 0.67 | 0.62 | 0.69 | 0.7 | 0.74 | 0.45 |
| 5 | 6456627 | rs4701734 | 0.233 | -1.025 | -1.016 | -0.567 | 0.038 | 0.093 | 0.093 | 0 | 0.48 | 0.31 | 0.4 | 0.07 | 0.63 | 0.33 |
| 5 | 6457563 | rs2651945 | -1.301 | -1.046 | -0.345 | 0.617 | -0.028 | 0.027 | 0.027 | 1.521 | 0.35 | 0.43 | 0.34 | 0.5 | 0.22 | 0.34 |
| 5 | 6461429 | rs169251 | -0.226 | -0.773 | -0.503 | 0.755 | 0 | 0.002 | 0.002 | 0 | 0.22 | 0.21 | 0.29 | 0.48 | 0.17 | 0.24 |
| 5 | 6464724 | rs272479 | -0.787 | -0.714 | -0.657 | 0.571 | -0.011 | 0.01 | 0.01 | 0.802 | 0.34 | 0.39 | 0.34 | 0.53 | 0.22 | 0.35 |
| 5 | 6469031 | rs272485 | 0.6 | -0.253 | -1.422 | 0.082 | 0.018 | 0.104 | 0.104 | 0.018 | 0.49 | 0.15 | 0.54 | 0.34 | 0.6 | 0.62 |
| 5 | 6471344 | rs507971 | 0.57 | -0.357 | 0.548 | -0.274 | 0.075 | -0.057 | -0.057 | 1.868 | 0.83 | 0.56 | 0.84 | 0.89 | 0.81 | 0.93 |
| 5 | 6473308 | rs12652806 | 1.423 | -0.752 | 0.024 | -0.526 | 0.007 | 0.122 | 0.122 | 0.112 | 0.66 | 0.16 | 0.77 | 0.88 | 0.76 | 0.84 |
| 5 | 6473810 | rs824619 | 0.159 | -0.982 | 0.406 | -0.448 | 0.027 | 0.005 | 0.005 | 0.141 | 0.91 | 0.77 | 0.92 | 0.9 | 0.93 | 0.95 |
| 5 | 6475025 | rs824622 | 1.024 | -1.005 | 0.511 | -0.466 | 0.006 | 0 | 0 | 0 | 0.74 | 0.3 | 0.85 | 0.89 | 0.9 | 0.86 |
| 5 | 6484012 | rs272456 | -1.524 | -0.433 | 0.763 | -0.278 | -0.002 | -0.005 | -0.005 | 0 | 0.36 | 0.69 | 0.57 | 0.43 | 0.38 | 0.5 |
| 5 | 6486876 | rs10071072 | -1.588 | -0.628 | 0.723 | -0.397 | 0.008 | 0.001 | 0.001 | 0.198 | 0.25 | 0.73 | 0.29 | 0.09 | 0.11 | 0.15 |
| 5 | 6488117 | rs272461 | -0.147 | -0.383 | 0.606 | -0.268 | -0.073 | -0.081 | -0.081 | 0 | 0.14 | 0.03 | 0.31 | 0.34 | 0.27 | 0.35 |
| 5 | 6494449 | rs10044719 | -0.49 | -0.977 | 1.161 | -0.625 | -0.001 | -0.008 | -0.008 | 0 | 0.08 | 0.07 | 0.07 | 0.02 | 0.07 | 0.04 |
| 5 | 6495579 | rs883525 | -0.503 | -1.061 | 1.908 | -0.647 | -0.062 | -0.069 | -0.069 | 0.002 | 0.11 | 0.32 | 0.05 | 0.01 | 0.01 | 0.07 |
| 5 | 6504171 | rs9765361 | -1.189 | -0.722 | 1.37 | -0.493 | -0.001 | -0.011 | -0.011 | 0 | 0.09 | 0.16 | 0.09 | 0.08 | 0.07 | 0.04 |
| 5 | 6505467 | rs17834269 | 0.693 | -0.61 | 0.711 | -0.453 | -0.001 | -0.015 | -0.015 | 0.063 | 0.16 | 0.11 | 0.13 | 0.21 | 0.19 | 0.19 |
| 5 | 6511314 | rs573546 | -0.322 | -0.624 | 1.186 | -0.437 | -0.006 | 0.028 | 0.028 | 0 | 0.24 | 0.51 | 0.19 | 0.25 | 0.23 | 0.2 |
| 5 | 6512465 | rs4702357 | 0.766 | -0.571 | 0.679 | -0.949 | 0.01 | 0.001 | 0.001 | 0 | 0.21 | 0.17 | 0.13 | 0.01 | 0.17 | 0.1 |
| 5 | 6517509 | rs566829 | 0.109 | -0.535 | 0.939 | -0.738 | -0.004 | 0.036 | 0.036 | 0 | 0.23 | 0.41 | 0.18 | 0.25 | 0.23 | 0.22 |
| 5 | 6518618 | rs487740 | -0.881 | -0.611 | 0.766 | -0.861 | 0.034 | 0.068 | 0.068 | 0 | 0.51 | 0.8 | 0.51 | 0.31 | 0.44 | 0.42 |
| 5 | 6518928 | rs11748341 | 0.76 | -0.68 | 0.952 | -0.877 | -0.002 | -0.037 | -0.037 | 0.084 | 0.21 | 0.24 | 0.16 | 0.03 | 0.2 | 0.12 |
| 5 | 6521202 | rs4702359 | 0.278 | -0.707 | 1.299 | -0.87 | -0.009 | -0.073 | -0.073 | 0.004 | 0.24 | 0.39 | 0.29 | 0.1 | 0.22 | 0.14 |
| 5 | 6521824 | rs11742039 | 0.943 | -0.69 | 1.271 | -0.867 | -0.002 | -0.059 | -0.059 | 0 | 0.2 | 0.29 | 0.16 | 0.03 | 0.2 | 0.12 |
| 5 | 6521837 | rs563624 | -0.438 | -0.75 | 1.321 | -0.893 | -0.001 | -0.008 | -0.008 | 0.14 | 0.42 | 0.6 | 0.36 | 0.22 | 0.43 | 0.37 |
| 5 | 6522653 | rs4702360 | 0.436 | -0.723 | 1.326 | -0.874 | -0.006 | -0.042 | -0.042 | 0.001 | 0.24 | 0.28 | 0.2 | 0.05 | 0.21 | 0.16 |
| 5 | 6522711 | rs534384 | -0.909 | -0.597 | 1.033 | -0.795 | -0.001 | 0.056 | 0.056 | 0 | 0.53 | 0.76 | 0.52 | 0.3 | 0.49 | 0.44 |
| 5 | 6523738 | rs529524 | -0.405 | -0.567 | 1.067 | -0.795 | -0.001 | -0.004 | -0.004 | 0 | 0.46 | 0.57 | 0.38 | 0.24 | 0.45 | 0.41 |
| 5 | 6528465 | rs540027 | -0.702 | -0.312 | 0.867 | -0.74 | -0.003 | 0.005 | 0.005 | 0 | 0.61 | 0.6 | 0.5 | 0.28 | 0.65 | 0.53 |
| 5 | 6529238 | rs514041 | 0.281 | -0.369 | 0.913 | -0.923 | -0.001 | -0.001 | -0.001 | 0 | 0.32 | 0.28 | 0.26 | 0.1 | 0.31 | 0.26 |
| 5 | 6531185 | rs6881323 | 0.806 | -0.195 | 0.286 | -0.884 | -0.002 | -0.001 | -0.001 | 0.094 | 0.24 | 0.25 | 0.2 | 0.18 | 0.29 | 0.26 |
| 5 | 6532191 | rs884475 | 0.1 | -0.192 | 0.3 | -0.868 | -0.004 | -0.012 | -0.012 | 0 | 0.3 | 0.34 | 0.24 | 0.18 | 0.34 | 0.27 |
| 5 | 6539175 | rs516853 | -1.457 | -0.216 | 0.03 | -0.859 | -0.053 | -0.052 | -0.052 | 0 | 0.67 | 0.87 | 0.65 | 0.34 | 0.67 | 0.55 |
| 5 | 6540593 | rs12189073 | -0.008 | -0.04 | 0.39 | -0.44 | 0 | 0.003 | 0.003 | 0 | 0.28 | 0.22 | 0.24 | 0.18 | 0.33 | 0.27 |
| 5 | 6540747 | rs12189254 | -0.175 | -0.022 | 0.265 | -0.429 | -0.002 | -0.006 | -0.006 | 0 | 0.29 | 0.29 | 0.24 | 0.18 | 0.34 | 0.27 |
| 5 | 6547267 | rs17778905 | 0.54 | 0.087 | -0.411 | -0.195 | -0.019 | -0.026 | -0.026 | 0 | 0.22 | 0.04 | 0.19 | 0.18 | 0.29 | 0.26 |
| 5 | 6547317 | rs13436933 | 0.436 | 0.029 | -0.449 | -0.348 | 0.003 | 0.024 | 0.024 | 0 | 0.09 | 0.04 | 0.07 | 0.01 | 0.08 | 0.04 |
| 5 | 6548889 | rs2617534 | 0.301 | -0.062 | 0.117 | -0.288 | -0.001 | -0.001 | -0.001 | 0 | 0.24 | 0.22 | 0.2 | 0.17 | 0.28 | 0.26 |
| 5 | 6549378 | rs11134165 | 0.455 | -0.116 | 0.149 | -0.314 | 0.002 | 0 | 0 | 0 | 0.23 | 0.15 | 0.19 | 0.16 | 0.28 | 0.26 |
| 5 | 6549456 | rs6887455 | -0.45 | -0.255 | 0.881 | -0.255 | 0.003 | 0.034 | 0.034 | 0.108 | 0.08 | 0.28 | 0.16 | 0.11 | 0.04 | 0.09 |
| 5 | 6550709 | rs11960029 | -1.676 | -0.491 | 1.273 | -0.366 | -0.014 | 0.025 | 0.025 | 1.85 | 0.11 | 0.58 | 0.19 | 0.11 | 0.04 | 0.09 |
| 5 | 6552278 | rs12654436 | -0.227 | -0.459 | 1.158 | -0.359 | -0.018 | 0.003 | 0.003 | 0 | 0.31 | 0.52 | 0.35 | 0.28 | 0.32 | 0.35 |
| 5 | 6552513 | rs524861 | -1.272 | -0.217 | -0.13 | -0.213 | -0.021 | -0.033 | -0.033 | 0.202 | 0.78 | 0.91 | 0.72 | 0.73 | 0.79 | 0.73 |
| 5 | 6554505 | rs487810 | 0.773 | -0.255 | -0.043 | -0.237 | -0.021 | -0.019 | -0.019 | 0.923 | 0.27 | 0.44 | 0.22 | 0.16 | 0.29 | 0.26 |
| 5 | 6556587 | rs474953 | -0.215 | -0.179 | -0.461 | -0.223 | 0.002 | -0.001 | -0.001 | 0 | 0.54 | 0.6 | 0.51 | 0.57 | 0.5 | 0.47 |
| 5 | 6558291 | rs493944 | -0.865 | 0.009 | -1.43 | -0.107 | -0.005 | -0.018 | -0.018 | 0 | 0.74 | 0.84 | 0.66 | 0.7 | 0.75 | 0.65 |
| 5 | 6559304 | rs13165570 | 1.635 | 0.214 | -1.735 | -0.235 | -0.043 | -0.066 | -0.066 | 0 | 0.13 | 0.02 | 0.11 | 0.42 | 0.16 | 0.19 |
| 5 | 6559533 | rs11743719 | 0.59 | 0.17 | -1.596 | -0.551 | -0.027 | 0.108 | 0.108 | 0 | 0.29 | 0.09 | 0.18 | 0.05 | 0.29 | 0.19 |
| 5 | 6560041 | rs567431 | 0.016 | 0.105 | -1.205 | -0.218 | 0.003 | 0.037 | 0.037 | 0.798 | 0.46 | 0.75 | 0.48 | 0.65 | 0.46 | 0.46 |
| 5 | 6562950 | rs477391 | -0.076 | 0.127 | -1.219 | -0.208 | 0.006 | 0.04 | 0.04 | 0.447 | 0.46 | 0.76 | 0.48 | 0.65 | 0.46 | 0.46 |
| 5 | 6568392 | rs532983 | -0.357 | 0.371 | -1.377 | -0.107 | -0.026 | 0.047 | 0.047 | 0.698 | 0.55 | 0.82 | 0.54 | 0.67 | 0.49 | 0.46 |
| 5 | 6572315 | rs16877577 | 1.109 | 0.501 | -1.268 | 0.385 | -0.015 | -0.014 | -0.014 | 0 | 0.1 | 0.04 | 0.11 | 0.42 | 0.13 | 0.16 |
| 5 | 6572542 | rs501592 | 0.623 | 0.036 | -0.471 | -0.086 | -0.014 | -0.061 | -0.061 | 0 | 0.25 | 0.38 | 0.31 | 0.1 | 0.24 | 0.28 |
| 5 | 6572878 | rs522335 | -0.177 | -0.15 | -0.302 | -0.099 | -0.007 | 0.019 | 0.019 | 0.023 | 0.39 | 0.52 | 0.44 | 0.52 | 0.38 | 0.46 |
| 5 | 6573663 | rs532737 | 0.243 | -0.323 | -0.527 | -0.523 | 0.002 | 0.011 | 0.011 | 0 | 0.34 | 0.3 | 0.32 | 0.24 | 0.32 | 0.24 |
| 5 | 6574088 | rs4702370 | 0.533 | -0.133 | -0.915 | -0.569 | -0.011 | -0.033 | -0.033 | 1.131 | 0.37 | 0.24 | 0.31 | 0.46 | 0.42 | 0.45 |
| 5 | 6574609 | rs10055883 | -0.04 | -0.136 | -0.676 | -0.443 | 0.007 | 0.001 | 0.001 | 0.923 | 0.66 | 0.28 | 0.48 | 0.64 | 0.7 | 0.66 |
| 5 | 6576320 | rs10058390 | -0.07 | -0.275 | -0.845 | -0.553 | -0.01 | -0.019 | -0.019 | 0 | 0.57 | 0.45 | 0.75 | 0.83 | 0.67 | 0.59 |
| 5 | 6576489 | rs13152888 | 1.522 | -0.173 | -1.34 | -0.184 | 0.001 | -0.039 | -0.039 | 0 | 0.14 | 0.05 | 0.11 | 0.44 | 0.11 | 0.18 |
| 5 | 6577009 | rs7701143 | 0.209 | 0.142 | -1.125 | -0.717 | 0.004 | -0.03 | -0.03 | 6.593 | 0.42 | 0.52 | 0.43 | 0.17 | 0.47 | 0.24 |
| 5 | 6578382 | rs551574 | 0.508 | 0.4 | -1.21 | -0.028 | -0.006 | -0.01 | -0.01 | 0 | 0.2 | 0.17 | 0.4 | 0.65 | 0.25 | 0.36 |
| 5 | 6580030 | rs498932 | 0.324 | 0.542 | -1.324 | -0.143 | -0.013 | -0.022 | -0.022 | 0 | 0.22 | 0.17 | 0.42 | 0.63 | 0.3 | 0.36 |
| 5 | 6580584 | rs503679 | -0.198 | 0.331 | -1.759 | -0.143 | 0.015 | -0.005 | -0.005 | 0.113 | 0.41 | 0.72 | 0.47 | 0.65 | 0.37 | 0.5 |
| 5 | 6584248 | rs7735244 | 0.378 | 0.522 | -1.533 | 0.503 | -0.007 | -0.042 | -0.042 | 7.792 | 0.11 | 0.02 | 0.09 | 0.19 | 0.1 | 0.06 |
| 5 | 6597622 | rs545493 | -0.527 | 0.374 | -1.901 | 0.313 | -0.029 | -0.015 | -0.015 | 0 | 0.8 | 0.99 | 0.87 | 0.88 | 0.79 | 0.81 |
| 5 | 6597810 | rs7729233 | 0.757 | 0.34 | -1.579 | 0.223 | -0.004 | 0.003 | 0.003 | 0 | 0.36 | 0.36 | 0.21 | 0.44 | 0.27 | 0.4 |
| 5 | 6598553 | rs11743743 | 0.205 | 0.204 | -0.928 | 0.134 | -0.005 | -0.002 | -0.002 | 0 | 0.42 | 0.41 | 0.63 | 0.44 | 0.52 | 0.4 |
| 5 | 6599222 | rs6876835 | 0.091 | 0.646 | -0.969 | 0.101 | -0.001 | -0.003 | -0.003 | 0 | 0.45 | 0.45 | 0.46 | 0.68 | 0.41 | 0.64 |

| | | | | | | | | | | | | | | | | |
|---|---|---|---|---|---|---|---|---|---|---|---|---|---|---|---|---|
| 5 | 6599940 | rs3776448 | 0.435 | 0.771 | -0.999 | 0.217 | -0.016 | 0 | 0 | 0 | 0.05 | 0.03 | 0.24 | 0.24 | 0.12 | 0.18 |
| 5 | 6600150 | rs3822434 | -0.776 | 0.914 | -1.052 | 0.147 | -0.003 | 0.006 | 0.006 | 0 | 0.08 | 0.08 | 0.25 | 0.23 | 0.14 | 0.22 |
| 5 | 6601309 | rs4701741 | -1.119 | 0.664 | -0.91 | 0.134 | 0.008 | 0.014 | 0.014 | 0 | 0.78 | 0.82 | 0.85 | 0.89 | 0.79 | 0.8 |
| 5 | 6602544 | rs2303711 | -0.638 | 0.728 | -0.858 | 0.077 | -0.002 | -0.004 | -0.004 | 0 | 0.75 | 0.74 | 0.62 | 0.67 | 0.68 | 0.64 |
| 5 | 6608361 | rs714713 | 1.989 | 0.464 | -1.221 | -0.092 | -0.035 | -0.117 | -0.117 | 0 | 0.2 | 0.04 | 0.14 | 0.46 | 0.19 | 0.32 |
| 5 | 6610911 | rs11741091 | 0.363 | 0.15 | -0.506 | -0.388 | -0.022 | 0.062 | 0.062 | 0 | 0.39 | 0.26 | 0.42 | 0.21 | 0.44 | 0.2 |
| 5 | 6612464 | rs567289 | -1.124 | 0.072 | -0.472 | -0.545 | -0.026 | -0.043 | -0.043 | 0 | 0.75 | 0.89 | 0.63 | 0.67 | 0.67 | 0.64 |
| 5 | 6613827 | rs4701742 | -0.717 | 0.064 | -0.329 | -0.549 | 0.005 | 0.015 | 0.015 | 0 | 0.73 | 0.63 | 0.61 | 0.67 | 0.67 | 0.63 |
| 5 | 6616277 | rs529439 | 0.025 | -0.022 | -0.488 | -0.55 | -0.004 | 0.003 | 0.003 | 0 | 0.4 | 0.54 | 0.35 | 0.58 | 0.33 | 0.53 |
| 5 | 6620844 | rs12522757 | 0.428 | -0.18 | -0.581 | -0.607 | -0.032 | -0.076 | -0.076 | 0 | 0.15 | 0.26 | 0.05 | 0.01 | 0.07 | 0.1 |
| 5 | 6622017 | rs2303705 | 0.36 | -0.221 | -0.523 | -0.526 | -0.003 | 0.008 | 0.008 | 0 | 0.36 | 0.34 | 0.39 | 0.16 | 0.39 | 0.19 |
| 5 | 6623085 | rs12691276 | -0.021 | -0.173 | -0.462 | -0.476 | 0.002 | 0.003 | 0.003 | 0 | 0.59 | 0.4 | 0.56 | 0.67 | 0.6 | 0.54 |
| 5 | 6629930 | rs518673 | 0.285 | 0.395 | -0.454 | -0.472 | 0.01 | -0.001 | -0.001 | 0 | 0.23 | 0.28 | 0.5 | 0.25 | 0.34 | 0.29 |
| 5 | 6632320 | rs8192120 | -0.024 | -0.038 | -0.586 | -0.487 | 0.005 | 0.02 | 0.02 | 0 | 0.42 | 0.63 | 0.4 | 0.75 | 0.36 | 0.62 |
| 5 | 6635882 | rs477930 | -0.624 | -0.002 | -0.296 | -0.386 | 0.001 | -0.003 | -0.003 | 0 | 0.43 | 0.53 | 0.66 | 0.37 | 0.53 | 0.4 |
| 5 | 6643847 | rs166050 | -1.58 | -0.184 | -0.509 | -0.536 | -0.044 | 0.044 | 0.044 | 0 | 0.77 | 0.99 | 0.85 | 0.91 | 0.78 | 0.82 |
| 5 | 6651970 | rs3822430 | -0.339 | -0.002 | 0.538 | -0.429 | 0 | -0.008 | -0.008 | 0 | 0.35 | 0.33 | 0.39 | 0.16 | 0.39 | 0.19 |
| 5 | 6652252 | rs3733773 | -0.35 | -0.003 | 0.537 | -0.43 | 0 | -0.008 | -0.008 | 0 | 0.35 | 0.33 | 0.39 | 0.16 | 0.39 | 0.19 |
| 5 | 6656161 | rs8192207 | 1.86 | 0.106 | 0.141 | -0.425 | -0.019 | -0.032 | -0.032 | 0 | 0.08 | 0.13 | 0.02 | 0 | 0.02 | 0.1 |
| 5 | 6660400 | rs248800 | 0.368 | 0.239 | -0.011 | -0.318 | 0.014 | 0.053 | 0.053 | 0 | 0.06 | 0.11 | 0.24 | 0.23 | 0.12 | 0.18 |
| 5 | 6661966 | rs6884552 | -0.281 | 0.24 | -0.03 | -0.302 | -0.025 | 0.038 | 0.038 | 0 | 0.33 | 0.19 | 0.38 | 0.22 | 0.39 | 0.21 |
| 5 | 6666922 | rs3797179 | 0.405 | 0.562 | -0.347 | -0.276 | -0.004 | -0.018 | -0.018 | 0 | 0.13 | 0.06 | 0.1 | 0.21 | 0.15 | 0.08 |
| 5 | 6667102 | rs6872996 | 0.425 | 0.404 | -0.39 | -0.422 | -0.026 | 0.044 | 0.044 | 0 | 0.23 | 0.14 | 0.29 | 0.01 | 0.29 | 0.16 |
| 5 | 6669762 | rs39848 | -0.544 | 0.341 | -0.329 | -0.151 | 0.008 | -0.031 | -0.031 | 0 | 0.39 | 0.54 | 0.56 | 0.25 | 0.42 | 0.39 |
| 5 | 6670649 | rs16877779 | 0.555 | 0.437 | -0.677 | 0.192 | -0.006 | -0.036 | -0.036 | 0 | 0.1 | 0.03 | 0.08 | 0.21 | 0.1 | 0.06 |
| 5 | 6671959 | rs39847 | 0.323 | 0.42 | -0.496 | -0.169 | 0 | -0.005 | -0.005 | 0 | 0.66 | 0.71 | 0.77 | 0.79 | 0.68 | 0.78 |
| 5 | 6673322 | rs30434 | -0.385 | 0.453 | -0.28 | 0.259 | -0.027 | -0.038 | -0.038 | 0 | 0.35 | 0.22 | 0.47 | 0.47 | 0.4 | 0.41 |
| 5 | 6676931 | rs1651071 | -1.007 | 0.419 | -0.421 | 0.055 | -0.036 | -0.032 | -0.032 | 0 | 0.46 | 0.83 | 0.56 | 0.49 | 0.49 | 0.46 |
| 5 | 6683000 | rs10512874 | -0.205 | -0.095 | -0.742 | -0.526 | -0.002 | 0.007 | 0.007 | 0 | 0.56 | 0.42 | 0.37 | 0.26 | 0.55 | 0.4 |
| 5 | 6684954 | rs11744535 | 1.216 | 0.075 | -1.504 | -0.47 | -0.069 | 0.009 | 0.009 | 0 | 0.22 | 0.04 | 0.19 | 0.14 | 0.28 | 0.16 |
| 5 | 6687557 | rs371869 | -1.108 | -0.31 | -0.815 | -0.409 | 0.009 | 0.036 | 0.036 | 0.722 | 0.08 | 0.57 | 0.06 | 0.05 | 0.04 | 0.06 |
| 5 | 6688275 | rs1551538 | 0.922 | -0.281 | -0.997 | -0.419 | -0.013 | 0.079 | 0.079 | 0 | 0.26 | 0.06 | 0.13 | 0.1 | 0.24 | 0.19 |
| 5 | 6694692 | rs6555406 | -0.102 | -0.199 | -0.85 | -0.379 | -0.016 | -0.038 | -0.038 | 0 | 0.66 | 0.43 | 0.82 | 0.85 | 0.72 | 0.75 |
| 5 | 6704848 | rs274661 | 0.422 | 0.159 | -1.232 | -0.046 | -0.044 | -0.091 | -0.091 | 0 | 0.31 | 0.08 | 0.3 | 0.39 | 0.31 | 0.36 |
| 5 | 6708483 | rs4701750 | 0.099 | 0.003 | -1.247 | -0.281 | 0.003 | -0.006 | -0.006 | 0.557 | 0.1 | 0.1 | 0.08 | 0.02 | 0.06 | 0.15 |
| 5 | 6712533 | rs274671 | 0.231 | -0.081 | -1.002 | -0.438 | -0.002 | -0.013 | -0.013 | 2.252 | 0.36 | 0.24 | 0.33 | 0.39 | 0.34 | 0.36 |
| 5 | 6723546 | rs274714 | 0.659 | -0.172 | -1.041 | -0.528 | -0.045 | -0.064 | -0.064 | 0.013 | 0.26 | 0.1 | 0.34 | 0.44 | 0.39 | 0.34 |
| 5 | 6724546 | rs16877813 | 1.025 | -0.259 | -1.039 | -0.561 | -0.005 | -0.014 | -0.014 | 0 | 0.07 | 0.07 | 0.06 | 0.02 | 0.03 | 0.12 |
| 5 | 6729695 | rs877238 | 1.896 | -0.204 | -1.051 | -0.53 | -0.029 | 0 | 0 | 0 | 0.18 | 0.06 | 0.16 | 0.16 | 0.25 | 0.12 |
| 5 | 6734625 | rs274692 | -0.25 | -0.321 | -1.012 | -0.49 | 0.015 | 0 | 0 | 1.405 | 0.42 | 0.3 | 0.36 | 0.41 | 0.33 | 0.48 |
| 5 | 6735844 | rs28363340 | 1.435 | -0.068 | -0.895 | -0.329 | 0.012 | 0.032 | 0.032 | 0 | 0.23 | 0.14 | 0.13 | 0.11 | 0.17 | 0.13 |
| 5 | 6739567 | rs416847 | -0.914 | -0.067 | -0.751 | -0.435 | -0.006 | 0.014 | 0.014 | 0.97 | 0.64 | 0.43 | 0.56 | 0.57 | 0.64 | 0.63 |
| 5 | 6742168 | rs187994 | -0.897 | -0.01 | -0.848 | -0.509 | -0.011 | 0.022 | 0.022 | 1.173 | 0.67 | 0.4 | 0.58 | 0.57 | 0.67 | 0.64 |
| 5 | 6747104 | rs2290616 | 1.395 | 0.04 | -1.356 | -0.482 | -0.017 | -0.088 | -0.088 | 4.951 | 0.27 | 0.07 | 0.36 | 0.42 | 0.3 | 0.34 |
| 5 | 6755460 | rs274674 | 0.612 | -0.052 | -1.253 | -0.264 | -0.004 | -0.051 | -0.051 | 0 | 0.32 | 0.14 | 0.29 | 0.39 | 0.3 | 0.36 |
| 5 | 6759714 | rs13163920 | 1.085 | 0.125 | -1.374 | -0.29 | -0.057 | -0.119 | -0.119 | 0 | 0.28 | 0.04 | 0.38 | 0.43 | 0.34 | 0.35 |
| 5 | 6760499 | rs2397977 | -0.297 | -0.2 | -0.838 | -0.333 | -0.007 | 0.018 | 0.018 | 2.954 | 0.63 | 0.43 | 0.67 | 0.54 | 0.69 | 0.63 |
| 5 | 6761511 | rs12188231 | 0.931 | 0.054 | -1.101 | -0.129 | -0.073 | -0.096 | -0.096 | 0 | 0.29 | 0.07 | 0.39 | 0.41 | 0.35 | 0.36 |
| 5 | 6767312 | rs12520016 | 0.387 | 0.277 | -0.478 | -0.075 | 0.002 | -0.026 | -0.026 | 0 | 0.06 | 0.12 | 0.04 | 0 | 0.05 | 0.01 |
| 5 | 6769016 | rs1552914 | -0.188 | 0.266 | -0.342 | -0.136 | 0.001 | 0.014 | 0.014 | 2.876 | 0.64 | 0.4 | 0.65 | 0.56 | 0.64 | 0.65 |
| 5 | 6772593 | rs11947998 | 0.055 | 0.171 | -0.256 | -0.229 | 0.02 | 0.025 | 0.025 | 0.88 | 0.48 | 0.32 | 0.51 | 0.43 | 0.44 | 0.52 |
| 5 | 6777769 | rs429161 | -0.664 | 0.331 | 0.081 | -0.251 | 0.052 | -0.009 | -0.009 | 0 | 0.95 | 0.73 | 0.95 | 1 | 0.95 | 0.98 |
| 5 | 6785435 | rs897712 | 0.299 | 0.217 | -0.222 | -0.299 | 0 | -0.031 | -0.031 | 0.231 | 0.09 | 0.13 | 0.06 | 0 | 0.09 | 0.06 |
| 5 | 6786308 | rs274645 | 0.712 | 0.267 | -0.487 | -0.006 | -0.035 | -0.047 | -0.047 | 0 | 0.09 | 0.01 | 0.17 | 0.34 | 0.13 | 0.19 |
| 5 | 6790678 | rs751594 | -1.041 | 0.206 | -0.035 | -0.137 | 0.048 | 0.002 | 0.002 | 0 | 0.05 | 0.25 | 0.05 | 0 | 0.05 | 0.01 |
| 5 | 6790691 | rs274651 | -0.373 | 0.16 | -0.374 | -0.196 | 0.002 | 0.003 | 0.003 | 0.017 | 0.57 | 0.57 | 0.52 | 0.61 | 0.61 | 0.53 |
| 5 | 6792839 | rs106415 | 0.082 | 0.2 | -0.384 | -0.146 | -0.002 | 0.011 | 0.011 | 1.102 | 0.52 | 0.49 | 0.48 | 0.63 | 0.54 | 0.54 |
| 5 | 6792985 | rs1004758 | 1.152 | 0.195 | -0.476 | -0.241 | -0.017 | -0.029 | -0.029 | 0.044 | 0.32 | 0.12 | 0.43 | 0.37 | 0.32 | 0.38 |
| 5 | 6793959 | rs716976 | 0.677 | 0.327 | -0.338 | -0.154 | -0.002 | 0.006 | 0.006 | 0 | 0.09 | 0.08 | 0.06 | 0 | 0.09 | 0.07 |
| 5 | 6795764 | rs13177725 | 1.23 | 0.501 | -0.258 | 0.117 | -0.001 | -0.003 | -0.003 | 1.527 | 0.21 | 0.21 | 0.13 | 0.17 | 0.18 | 0.15 |
| 5 | 6796405 | rs4702392 | -0.643 | 0.34 | -0.072 | -0.264 | 0.033 | -0.027 | -0.027 | 0.006 | 0.93 | 0.7 | 0.96 | 1 | 0.95 | 0.98 |
| 5 | 6797096 | rs450804 | 0.449 | 0.48 | -1.023 | 0.011 | -0.005 | -0.022 | -0.022 | 0 | 0.28 | 0.21 | 0.3 | 0.5 | 0.31 | 0.33 |
| 5 | 6802179 | rs11956100 | 0.142 | 0.357 | -1.145 | -0.444 | -0.008 | 0.02 | 0.02 | 0.51 | 0.32 | 0.15 | 0.33 | 0.28 | 0.32 | 0.38 |
| 5 | 6803243 | rs12523241 | -0.115 | 0.295 | -0.6 | -0.781 | 0.009 | -0.017 | -0.017 | 0.097 | 0.31 | 0.43 | 0.36 | 0.24 | 0.34 | 0.39 |
| 5 | 6804983 | rs11949128 | 0.495 | 0.22 | -0.542 | -0.862 | -0.026 | 0.004 | 0.004 | 0.212 | 0.24 | 0.14 | 0.31 | 0.24 | 0.31 | 0.35 |
| 5 | 6806216 | rs275460 | 0.136 | -0.053 | -0.352 | -1.144 | -0.004 | 0.016 | 0.016 | 0 | 0.67 | 0.73 | 0.66 | 0.76 | 0.66 | 0.63 |
| 5 | 6810950 | rs7728254 | 0.683 | -0.335 | -0.486 | -1.313 | -0.017 | -0.013 | -0.013 | 0.082 | 0.38 | 0.31 | 0.49 | 0.52 | 0.47 | 0.4 |
| 5 | 6813937 | rs275456 | 1.386 | -0.286 | -0.533 | -1.307 | -0.078 | -0.042 | -0.042 | 0 | 0.23 | 0.01 | 0.29 | 0.25 | 0.28 | 0.35 |
| 5 | 6817709 | rs12109729 | 1.57 | -0.274 | -0.342 | -1.31 | 0.023 | -0.015 | -0.015 | 0 | 0.06 | 0.23 | 0.04 | 0 | 0.03 | 0.02 |

| | | | | | | | | | | | | | | | | |
|---|---|---|---|---|---|---|---|---|---|---|---|---|---|---|---|---|
| 5 | 6818844 | rs2047321 | 1.769 | -0.11 | -0.562 | -1.459 | -0.018 | 0.046 | 0.046 | 0 | 0.23 | 0.12 | 0.19 | 0.07 | 0.27 | 0.18 |
| 5 | 6818876 | rs2047322 | 0.316 | -0.113 | -0.762 | -1.399 | -0.001 | -0.079 | -0.079 | 0 | 0.4 | 0.56 | 0.29 | 0.07 | 0.38 | 0.25 |
| 5 | 6820161 | rs13174522 | 1.369 | -0.094 | -0.594 | -0.797 | -0.001 | -0.001 | -0.001 | 0 | 0.17 | 0.16 | 0.1 | 0.24 | 0.14 | 0.17 |
| 5 | 6821914 | rs1566039 | -0.195 | 0.035 | -0.964 | -0.867 | -0.002 | -0.004 | -0.004 | 0 | 0.35 | 0.38 | 0.41 | 0.68 | 0.34 | 0.39 |
| 5 | 6825016 | rs275447 | -0.278 | 0.023 | -0.557 | -0.427 | 0.018 | 0.177 | 0.177 | 0.079 | 0.82 | 0.64 | 0.71 | 0.57 | 0.83 | 0.77 |
| 5 | 6826269 | rs16878029 | 1.292 | 0.234 | -0.657 | -0.503 | -0.026 | 0.027 | 0.027 | 0 | 0.05 | 0 | 0.03 | 0 | 0.06 | 0.05 |
| 5 | 6827513 | rs275442 | 0.043 | 0.217 | -0.999 | -1.017 | 0 | -0.001 | -0.001 | 0 | 0.54 | 0.49 | 0.55 | 0.53 | 0.58 | 0.63 |
| 5 | 6828995 | rs2453805 | 0.217 | 0.304 | -1.132 | -1.172 | 0.001 | -0.001 | -0.001 | 0 | 0.25 | 0.27 | 0.22 | 0.28 | 0.24 | 0.34 |
| 5 | 6829242 | rs16878054 | 0.671 | 0.309 | -1.36 | -1.181 | 0.002 | 0.013 | 0.013 | 0 | 0.33 | 0.29 | 0.34 | 0.25 | 0.34 | 0.29 |
| 5 | 6829382 | rs275486 | 0.203 | 0.487 | -1.023 | -0.835 | 0 | 0 | 0 | 0 | 0.51 | 0.48 | 0.52 | 0.53 | 0.52 | 0.58 |
| 5 | 6832480 | rs275480 | 0.545 | 0.903 | -1.05 | 0.167 | 0.003 | -0.013 | -0.013 | 0 | 0.63 | 0.34 | 0.57 | 0.5 | 0.7 | 0.7 |
| 5 | 6838445 | rs275468 | -0.066 | 1.525 | -1.002 | 0.408 | -0.033 | -0.028 | -0.028 | 0 | 0.05 | 0.01 | 0.12 | 0.2 | 0.13 | 0.14 |
| 5 | 6838589 | rs275467 | 0.144 | 0.923 | -0.41 | -0.262 | -0.003 | -0.068 | -0.068 | 0.761 | 0.87 | 0.76 | 0.92 | 1 | 0.89 | 0.95 |
| 5 | 6842066 | rs1993450 | -0.306 | 0.605 | -0.627 | -0.026 | 0.018 | 0.062 | 0.062 | 0 | 0.54 | 0.63 | 0.76 | 0.77 | 0.6 | 0.53 |
| 5 | 6842842 | rs6893958 | 0.162 | 0.622 | -0.321 | 0.451 | 0 | -0.014 | -0.014 | 1.005 | 0.47 | 0.44 | 0.4 | 0.32 | 0.53 | 0.38 |
| 5 | 6845035 | rs870347 | 0.45 | 0.679 | -0.81 | 0.562 | -0.002 | 0.127 | 0.127 | 0 | 0.05 | 0.07 | 0.35 | 0.45 | 0.07 | 0.15 |
| 5 | 6845811 | rs11740531 | 0.164 | 0.355 | -1.133 | 0.115 | 0.007 | 0.007 | 0.007 | 0.004 | 0.13 | 0.1 | 0.06 | 0.1 | 0.09 | 0.19 |
| 5 | 6850596 | rs16878070 | 0.825 | 0.29 | -1.289 | 0.211 | 0.016 | -0.013 | -0.013 | 0.226 | 0.08 | 0.04 | 0.03 | 0.1 | 0.03 | 0.15 |
| 5 | 6852600 | rs275490 | -0.722 | 0.562 | -1.109 | 0.228 | 0.006 | 0.026 | 0.026 | 0 | 0.68 | 0.73 | 0.82 | 0.87 | 0.69 | 0.68 |
| 5 | 6854074 | rs275493 | -0.666 | 0.565 | -1.114 | 0.228 | 0.008 | 0.048 | 0.048 | 0 | 0.67 | 0.75 | 0.82 | 0.85 | 0.68 | 0.68 |
| 5 | 6854290 | rs9313172 | -0.112 | 0.822 | -1.316 | 0.256 | -0.024 | 0.04 | 0.04 | 0.005 | 0.41 | 0.2 | 0.33 | 0.34 | 0.43 | 0.44 |
| 5 | 6859895 | rs275510 | 0.264 | 0.707 | -1.031 | 0.444 | -0.002 | 0 | 0 | 0 | 0.52 | 0.56 | 0.64 | 0.45 | 0.52 | 0.58 |
| 5 | 6859946 | rs951476 | -0.034 | 0.019 | -1.594 | 0.163 | 0.041 | -0.004 | -0.004 | 0 | 0.17 | 0.01 | 0.05 | 0.14 | 0.1 | 0.21 |
| 5 | 6864120 | rs12522644 | 0.296 | 0.572 | -1.356 | -0.371 | 0.006 | -0.013 | -0.013 | 0 | 0.17 | 0.23 | 0.21 | 0.06 | 0.21 | 0.17 |
| 5 | 6864573 | rs275499 | 0.726 | 0.596 | -2.219 | -0.723 | -0.037 | 0.021 | 0.021 | 0 | 0.36 | 0.09 | 0.3 | 0.26 | 0.42 | 0.38 |
| 5 | 6864785 | rs721632 | 1.399 | 0.698 | -2.51 | -1.014 | -0.051 | 0.065 | 0.065 | 0 | 0.11 | 0 | 0.23 | 0.01 | 0.12 | 0.11 |
| 5 | 6865801 | rs12513764 | 0.117 | 0.6 | -2.11 | -1.041 | -0.001 | -0.013 | -0.013 | 0.013 | 0.28 | 0.3 | 0.44 | 0.08 | 0.33 | 0.28 |
| 5 | 6866673 | rs275502 | -0.4 | 0.451 | -1.821 | -1.006 | -0.023 | 0.001 | 0.001 | 0.166 | 0.64 | 0.46 | 0.75 | 0.34 | 0.75 | 0.66 |
| 5 | 6867590 | rs275504 | -0.546 | 0.306 | -1.779 | -1.113 | -0.049 | 0.017 | 0.017 | 0.606 | 0.69 | 0.49 | 0.78 | 0.34 | 0.8 | 0.67 |
| 5 | 6869021 | rs997182 | -1.05 | 0.52 | -1.396 | -1.152 | 0.023 | -0.049 | -0.049 | 0 | 0.81 | 0.62 | 0.78 | 0.93 | 0.79 | 0.82 |
| 5 | 6870971 | rs275430 | 0.218 | 0.345 | -1.414 | -1.101 | 0.011 | -0.058 | -0.058 | 0 | 0.65 | 0.56 | 0.52 | 0.92 | 0.62 | 0.7 |
| 5 | 6872924 | rs275425 | 0.982 | 0.383 | -1.849 | -1.087 | -0.029 | 0.026 | 0.026 | 0 | 0.32 | 0.08 | 0.27 | 0.23 | 0.38 | 0.36 |
| 5 | 6873013 | rs275424 | 0.335 | 0.229 | -1.036 | -1.3 | 0.021 | -0.076 | -0.076 | 0.219 | 0.61 | 0.49 | 0.49 | 0.88 | 0.57 | 0.69 |
| 5 | 6873930 | rs2397770 | -0.29 | -0.375 | -1.269 | -1.545 | -0.007 | -0.067 | -0.067 | 0 | 0.29 | 0.32 | 0.33 | 0.06 | 0.24 | 0.17 |
| 5 | 6878651 | rs16878119 | 1.038 | -0.794 | -1.749 | -1.096 | 0.016 | -0.007 | -0.007 | 1.173 | 0.19 | 0.01 | 0.06 | 0.21 | 0.12 | 0.22 |
| 5 | 6879564 | rs7703668 | -0.505 | -0.448 | -1.354 | -1.036 | -0.009 | 0.036 | 0.036 | 0 | 0.24 | 0.29 | 0.23 | 0.49 | 0.24 | 0.15 |
| 5 | 6881857 | rs275438 | -0.098 | -0.179 | -1.626 | -1.054 | 0.072 | 0.044 | 0.044 | 1.555 | 0.53 | 0.23 | 0.72 | 0.57 | 0.61 | 0.67 |
| 5 | 6884986 | rs12514931 | 0.268 | 0.221 | -0.446 | -0.839 | 0.019 | -0.075 | -0.075 | 0 | 0.17 | 0.34 | 0.13 | 0.01 | 0.2 | 0.12 |
| 5 | 6887228 | rs13156798 | 1.228 | 0.232 | -1.057 | -0.776 | 0.046 | 0.066 | 0.066 | 0.333 | 0.13 | 0.03 | 0.03 | 0 | 0.05 | 0.02 |
| 5 | 6892417 | rs10512890 | -0.08 | 0.35 | -0.726 | -0.788 | 0.07 | -0.043 | -0.043 | 0 | 0.14 | 0.54 | 0.13 | 0.06 | 0.18 | 0.15 |
| 5 | 6894060 | rs11959635 | 0.983 | 0.746 | -0.955 | -0.746 | -0.017 | 0.006 | 0.006 | 0 | 0.12 | 0.04 | 0.23 | 0.09 | 0.16 | 0.24 |
| 5 | 6900466 | rs2894967 | 0.044 | 0.159 | -0.786 | -0.283 | -0.033 | 0.061 | 0.061 | 0 | 0.77 | 0.69 | 0.86 | 0.61 | 0.86 | 0.85 |
| 5 | 6902045 | rs1353502 | 0.875 | 0.411 | -0.984 | -0.396 | -0.015 | 0.015 | 0.015 | 0 | 0.24 | 0.15 | 0.46 | 0.1 | 0.29 | 0.37 |
| 5 | 6902924 | rs1353500 | 0.058 | 0.495 | -1.104 | -0.336 | -0.003 | 0.006 | 0.006 | 0 | 0.84 | 0.82 | 0.83 | 0.61 | 0.79 | 0.8 |
| 5 | 6907892 | rs903863 | -0.637 | 0.347 | -1.126 | -0.245 | 0 | 0.114 | 0.114 | 0 | 0.91 | 0.85 | 0.9 | 0.63 | 0.91 | 0.88 |
| 5 | 6913626 | rs12656353 | 0.498 | 0.539 | -1.395 | -0.22 | -0.002 | 0.001 | 0.001 | 0 | 0.53 | 0.51 | 0.4 | 0.52 | 0.57 | 0.48 |
| 5 | 6922386 | rs2397719 | 0.332 | 0.179 | -1.655 | 0.36 | 0.004 | -0.012 | -0.012 | 0 | 0.64 | 0.3 | 0.74 | 0.56 | 0.68 | 0.7 |
| 5 | 6940857 | rs2397718 | -0.274 | -0.12 | -2.277 | -0.11 | 0.014 | 0.011 | 0.011 | 6.692 | 0.21 | 0.03 | 0.18 | 0.16 | 0.14 | 0.26 |
| 5 | 6943022 | rs11134195 | 0.328 | 0.112 | -2.411 | -0.446 | -0.006 | 0.037 | 0.037 | 0 | 0.6 | 0.35 | 0.43 | 0.31 | 0.62 | 0.43 |
| 5 | 6944267 | rs6862164 | -0.732 | 0.371 | -2.306 | -0.01 | 0.016 | 0.062 | 0.062 | 0 | 0.77 | 0.43 | 0.62 | 0.55 | 0.73 | 0.63 |
| 5 | 6950547 | rs6884863 | 0.681 | 0.087 | -3.067 | -0.776 | -0.056 | 0.279 | 0.279 | 0.091 | 0.39 | 0.07 | 0.27 | 0.01 | 0.43 | 0.25 |
| 5 | 6952197 | rs1845852 | -0.186 | -0.048 | -2.9 | -0.778 | -0.038 | 0.051 | 0.051 | 0 | 0.6 | 0.35 | 0.48 | 0.26 | 0.56 | 0.45 |
| 5 | 6959721 | rs7727425 | -0.437 | -0.311 | -3.148 | -1.017 | 0.009 | -0.036 | -0.036 | 0.016 | 0.41 | 0.33 | 0.39 | 0.55 | 0.35 | 0.42 |
| 5 | 6962368 | rs12516580 | 0.322 | -0.311 | -3.144 | -1.043 | 0.009 | 0.172 | 0.172 | 0 | 0.43 | 0.25 | 0.28 | 0.01 | 0.44 | 0.26 |
| 5 | 6963155 | rs7706332 | 1.246 | -0.224 | -3.153 | -0.966 | 0.021 | -0.036 | -0.036 | 0 | 0.18 | 0.1 | 0.2 | 0.24 | 0.12 | 0.19 |
| 5 | 6963922 | rs7729658 | 1.21 | -0.238 | -3.16 | -0.971 | 0.022 | -0.035 | -0.035 | 0 | 0.18 | 0.1 | 0.21 | 0.25 | 0.12 | 0.19 |
| 5 | 6966304 | rs1155297 | 1.016 | -0.196 | -3.153 | -1.002 | -0.056 | 0.238 | 0.238 | 0.361 | 0.34 | 0.05 | 0.22 | 0.01 | 0.37 | 0.19 |
| 5 | 6972638 | rs16878239 | 0.972 | -0.116 | -2.528 | -0.345 | -0.043 | -0.062 | -0.062 | 0 | 0.14 | 0.03 | 0.13 | 0.3 | 0.16 | 0.16 |
| 5 | 6977768 | rs17192980 | 1.301 | -0.149 | -2.541 | -0.31 | 0.011 | 0.011 | 0.011 | 0 | 0.16 | 0.08 | 0.17 | 0.11 | 0.12 | 0.13 |
| 5 | 6978190 | rs1353499 | 0.902 | -0.115 | -2.392 | -0.291 | -0.023 | -0.037 | -0.037 | 0 | 0.14 | 0.07 | 0.13 | 0.3 | 0.16 | 0.16 |
| 5 | 6981399 | rs1353498 | -0.047 | -0.139 | -2.375 | -0.345 | -0.022 | -0.062 | -0.062 | 0 | 0.3 | 0.11 | 0.31 | 0.42 | 0.29 | 0.28 |
| 5 | 6985430 | rs6873965 | 1.023 | -0.019 | -2.56 | -0.639 | 0.001 | 0.001 | 0.001 | 0 | 0.38 | 0.51 | 0.45 | 0.45 | 0.45 | 0.38 |
| 5 | 6993552 | rs13180321 | 0.544 | 0.411 | -2.18 | -0.28 | 0.009 | -0.01 | -0.01 | 0 | 0.64 | 0.68 | 0.65 | 0.58 | 0.7 | 0.71 |
| 5 | 7005949 | rs4446428 | -0.604 | 0.507 | -2.269 | -0.346 | -0.005 | -0.014 | -0.014 | 0.75 | 0.82 | 0.72 | 0.89 | 0.92 | 0.86 | 0.87 |
| 5 | 7006254 | rs4312853 | 0.232 | 0.837 | -2.343 | -0.029 | -0.041 | -0.079 | -0.079 | 0 | 0.17 | 0.03 | 0.24 | 0.32 | 0.17 | 0.16 |
| 5 | 7018060 | rs6555424 | 0.105 | 0.885 | -2.107 | 0.024 | 0.002 | 0.012 | 0.012 | 0 | 0.17 | 0.17 | 0.26 | 0.33 | 0.17 | 0.15 |
| 5 | 7018136 | rs17806359 | 0.647 | 0.542 | -1.962 | 0.004 | 0.011 | 0.027 | 0.027 | 0 | 0.17 | 0.06 | 0.1 | 0.09 | 0.14 | 0.13 |
| 5 | 7019507 | rs10866576 | -0.354 | 1.266 | -1.689 | 0.145 | -0.017 | -0.008 | -0.008 | 0.015 | 0.3 | 0.21 | 0.38 | 0.34 | 0.36 | 0.23 |
| 5 | 7023296 | rs12658529 | 0.108 | 1.298 | -1.474 | 0.257 | -0.025 | -0.023 | -0.023 | 0.141 | 0.65 | 0.66 | 0.57 | 0.45 | 0.54 | 0.64 |

| | | | | | | | | | | | | | | | | |
|---|---|---|---|---|---|---|---|---|---|---|---|---|---|---|---|---|
| 5 | 7025546 | rs4702421 | 0.129 | 0.645 | -1.69 | 0.017 | 0.067 | -0.004 | -0.004 | 0 | 0.46 | 0.65 | 0.43 | 0.45 | 0.3 | 0.57 |
| 5 | 7044847 | rs7705045 | 0.184 | 1.03 | -0.712 | 0.226 | 0.016 | 0.066 | 0.066 | 0 | 0.71 | 0.47 | 0.48 | 0.54 | 0.67 | 0.57 |
| 5 | 7048314 | rs7711636 | 0.773 | 1.007 | -0.978 | 0.131 | 0 | 0.049 | 0.049 | 0.918 | 0.55 | 0.34 | 0.32 | 0.34 | 0.46 | 0.37 |
| 5 | 7053506 | rs6859571 | -0.124 | 0.953 | -0.838 | 0.112 | 0.005 | 0.06 | 0.06 | 0 | 0.72 | 0.44 | 0.5 | 0.54 | 0.69 | 0.63 |
| 5 | 7054372 | rs10462830 | 0.862 | 0.961 | -0.081 | 0.358 | 0.02 | 0.255 | 0.255 | 0 | 0.1 | 0.25 | 0.31 | 0.46 | 0.09 | 0.3 |
| 5 | 7071229 | rs7736268 | 1.173 | 0.561 | -0.463 | -0.365 | -0.003 | 0.041 | 0.041 | 2.996 | 0.29 | 0.18 | 0.19 | 0.12 | 0.29 | 0.15 |
| 5 | 7075590 | rs1874455 | -0.182 | 0.484 | -0.346 | -0.36 | 0.006 | 0.053 | 0.053 | 0.017 | 0.53 | 0.7 | 0.64 | 0.68 | 0.48 | 0.58 |
| 5 | 7083694 | rs13190346 | 1.361 | 0.457 | -0.046 | 0.05 | 0.011 | 0.189 | 0.189 | 0.15 | 0.08 | 0.17 | 0.3 | 0.39 | 0.08 | 0.29 |
| 5 | 7087890 | rs4702423 | 0.929 | 0.552 | -0.454 | 0.025 | -0.02 | -0.015 | -0.015 | 0.068 | 0.18 | 0.12 | 0.18 | 0.2 | 0.22 | 0.27 |
| 5 | 7088994 | rs754335 | 1.481 | 0.324 | -0.34 | -0.104 | 0.009 | 0.003 | 0.003 | 0 | 0.24 | 0.19 | 0.12 | 0.24 | 0.15 | 0.21 |
| 5 | 7091161 | rs2029907 | -0.704 | 0.662 | -0.814 | -0.655 | -0.002 | -0.001 | -0.001 | 0 | 0.8 | 0.79 | 0.8 | 0.8 | 0.77 | 0.73 |
| 5 | 7093653 | rs9654456 | 0.654 | 0.483 | -1.106 | -1.222 | -0.01 | 0.011 | 0.011 | 0 | 0.18 | 0.16 | 0.19 | 0.05 | 0.25 | 0.07 |
| 5 | 7099016 | rs7725101 | 0.089 | -0.006 | -0.996 | -0.789 | -0.012 | 0.05 | 0.05 | 4.145 | 0.28 | 0.15 | 0.17 | 0.12 | 0.29 | 0.16 |
| 5 | 7101299 | rs6861024 | 0.699 | -0.201 | -1.173 | -0.856 | -0.053 | 0.089 | 0.089 | 0 | 0.25 | 0.06 | 0.16 | 0 | 0.33 | 0.11 |
| 5 | 7108672 | rs17194818 | -0.879 | -1.103 | -0.374 | -0.043 | 0.014 | 0.042 | 0.042 | 0 | 0.5 | 0.64 | 0.68 | 0.71 | 0.46 | 0.53 |
| 5 | 7108869 | rs12655214 | -0.082 | -1.455 | -0.132 | 0.323 | 0.001 | 0.003 | 0.003 | 0 | 0.3 | 0.37 | 0.39 | 0.37 | 0.31 | 0.29 |
| 5 | 7111255 | rs16878349 | -0.713 | -1.513 | 0.019 | 0.247 | 0.014 | -0.014 | -0.014 | 0 | 0.25 | 0.21 | 0.31 | 0.34 | 0.2 | 0.28 |
| 5 | 7113315 | rs13190668 | 1.3 | -1.5 | 0.087 | 0.195 | -0.039 | -0.057 | -0.057 | 0.339 | 0.18 | 0.1 | 0.33 | 0.37 | 0.25 | 0.26 |
| 5 | 7116244 | rs2047313 | 0.653 | -1.443 | 0.133 | 0.221 | 0.015 | 0.035 | 0.035 | 0 | 0.21 | 0.3 | 0.35 | 0.37 | 0.25 | 0.27 |
| 5 | 7118376 | rs7709128 | -0.489 | -1.514 | -0.012 | -0.688 | -0.003 | 0.004 | 0.004 | 0 | 0.37 | 0.37 | 0.61 | 0.69 | 0.39 | 0.51 |
| 5 | 7118462 | rs13161773 | 0.862 | -1.44 | -0.074 | -0.563 | -0.021 | -0.038 | -0.038 | 0 | 0.2 | 0.16 | 0.34 | 0.56 | 0.26 | 0.34 |
| 5 | 7121294 | rs6885291 | 0.248 | -1.184 | 0.256 | -0.325 | -0.012 | 0.024 | 0.024 | 0.033 | 0.1 | 0.11 | 0.05 | 0.14 | 0.05 | 0.19 |
| 5 | 7122604 | rs903848 | -1.172 | -1.014 | 0.305 | -0.715 | -0.007 | 0.003 | 0.003 | 0 | 0.6 | 0.64 | 0.73 | 0.83 | 0.53 | 0.74 |
| 5 | 7125022 | rs16878386 | -0.351 | -1.092 | 0.591 | -0.003 | -0.01 | -0.035 | -0.035 | 0.185 | 0.08 | 0.11 | 0.04 | 0 | 0.06 | 0.03 |
| 5 | 7136893 | rs1496361 | -0.652 | -1.073 | 0.279 | 0.013 | 0.004 | 0.002 | 0.002 | 0 | 0.13 | 0.12 | 0.25 | 0.13 | 0.12 | 0.16 |
| 5 | 7138170 | rs11134203 | -1.102 | -0.972 | 0.148 | 0.073 | -0.04 | -0.06 | -0.06 | 0 | 0.24 | 0.46 | 0.31 | 0.13 | 0.18 | 0.18 |
| 5 | 7142618 | rs4702433 | -1.091 | -0.586 | -0.252 | 0.63 | -0.011 | 0.007 | 0.007 | 0 | 0.22 | 0.24 | 0.28 | 0.27 | 0.18 | 0.26 |
| 5 | 7148048 | rs7725195 | 0.46 | -0.466 | -0.078 | 0.821 | -0.005 | 0.039 | 0.039 | 0 | 0.9 | 0.82 | 0.9 | 0.84 | 0.92 | 0.77 |
| 5 | 7148491 | rs7709141 | -1.041 | -0.323 | 0.733 | 0.796 | 0.055 | 0.091 | 0.091 | 0.694 | 0.1 | 0.36 | 0.23 | 0.15 | 0.09 | 0.07 |
| 5 | 7148725 | rs7709479 | 0.278 | -0.181 | -0.341 | 0.682 | -0.004 | -0.01 | -0.01 | 0 | 0.44 | 0.47 | 0.29 | 0.31 | 0.37 | 0.4 |
| 5 | 7149071 | rs16878428 | 0.805 | -0.39 | -0.954 | 0.542 | 0.035 | 0.003 | 0.003 | 0.011 | 0.22 | 0.13 | 0.13 | 0.2 | 0.12 | 0.26 |
| 5 | 7149265 | rs1566035 | -1.056 | -0.532 | -1.035 | 0.357 | -0.029 | -0.027 | -0.027 | 0 | 0.6 | 0.8 | 0.55 | 0.46 | 0.52 | 0.51 |
| 5 | 7155116 | rs10035660 | 0.026 | -0.548 | -0.901 | 0.319 | 0.002 | 0.003 | 0.003 | 0 | 0.46 | 0.57 | 0.29 | 0.31 | 0.37 | 0.41 |
| 5 | 7155612 | rs4702435 | -0.859 | -0.471 | -1.009 | 0.357 | -0.004 | 0.034 | 0.034 | 0 | 0.88 | 0.83 | 0.88 | 0.81 | 0.9 | 0.76 |
| 5 | 7155671 | rs4701770 | -0.341 | -0.477 | -1.015 | 0.353 | 0.015 | 0.01 | 0.01 | 0 | 0.13 | 0.23 | 0.26 | 0.15 | 0.15 | 0.1 |
| 5 | 7155958 | rs12659620 | 1.155 | -0.488 | -0.954 | 0.349 | -0.096 | -0.09 | -0.09 | 0 | 0.34 | 0.09 | 0.53 | 0.51 | 0.47 | 0.32 |
| 5 | 7157361 | rs7444602 | -0.949 | -0.498 | -0.959 | 0.348 | -0.028 | -0.031 | -0.031 | 0 | 0.59 | 0.8 | 0.55 | 0.46 | 0.52 | 0.51 |
| 5 | 7158473 | rs9791081 | 1.709 | -0.457 | -0.973 | 0.387 | -0.096 | -0.062 | -0.062 | 0 | 0.29 | 0.03 | 0.33 | 0.35 | 0.38 | 0.25 |
| 5 | 7164445 | rs11749475 | -0.008 | -0.608 | -1.043 | 0.286 | 0.012 | -0.027 | -0.027 | 0.292 | 0.34 | 0.5 | 0.37 | 0.26 | 0.37 | 0.23 |
| 5 | 7164967 | rs16878438 | 1.293 | -0.207 | -0.826 | 0.548 | 0.014 | -0.016 | -0.016 | 0 | 0.16 | 0.09 | 0.12 | 0.2 | 0.09 | 0.24 |
| 5 | 7169448 | rs7713650 | 0.303 | 0.418 | -0.195 | 0.557 | -0.002 | 0.005 | 0.005 | 0 | 0.83 | 0.82 | 0.84 | 0.81 | 0.84 | 0.73 |
| 5 | 7169851 | rs7727403 | 1.356 | 0.477 | -0.393 | 0.579 | -0.019 | -0.029 | -0.029 | 0 | 0.51 | 0.18 | 0.46 | 0.55 | 0.5 | 0.51 |
| 5 | 7178644 | rs11745983 | 0.226 | 0.974 | -0.177 | 0.812 | 0.005 | 0.015 | 0.015 | 0 | 0.06 | 0.05 | 0.2 | 0.15 | 0.09 | 0.07 |
| 5 | 7179551 | rs13166960 | -0.353 | 0.691 | -0.361 | 0.778 | 0.033 | -0.016 | -0.016 | 0 | 0.3 | 0.5 | 0.36 | 0.26 | 0.35 | 0.22 |
| 5 | 7183381 | rs6420053 | 0.565 | 0.474 | -0.534 | 1.152 | -0.002 | 0.017 | 0.017 | 0 | 0.88 | 0.86 | 0.88 | 0.8 | 0.9 | 0.79 |
| 5 | 7184639 | rs10051593 | 0.438 | 0.939 | -0.693 | 1.47 | 0 | 0.008 | 0.008 | 0 | 0.8 | 0.8 | 0.82 | 0.64 | 0.77 | 0.77 |
| 5 | 7189524 | rs13155334 | -0.77 | 0.918 | -0.592 | 1.015 | -0.003 | -0.009 | -0.009 | 0.637 | 0.13 | 0.19 | 0.06 | 0.05 | 0.12 | 0.08 |
| 5 | 7189925 | rs9313179 | -0.465 | 1.072 | -0.84 | 0.612 | -0.006 | -0.01 | -0.01 | 0 | 0.62 | 0.77 | 0.69 | 0.61 | 0.6 | 0.72 |
| 5 | 7191074 | rs4702440 | 0.683 | 0.915 | -0.511 | 0.362 | 0.01 | -0.037 | -0.037 | 2.532 | 0.24 | 0.34 | 0.32 | 0.16 | 0.26 | 0.18 |
| 5 | 7202086 | rs10058673 | -0.247 | 0.858 | -0.502 | 0.371 | 0.004 | -0.042 | -0.042 | 0 | 0.37 | 0.49 | 0.37 | 0.2 | 0.37 | 0.26 |
| 5 | 7204616 | rs16900420 | 0.616 | 0.848 | -0.49 | 0.407 | -0.002 | 0.006 | 0.006 | 0.343 | 0.27 | 0.31 | 0.32 | 0.4 | 0.24 | 0.47 |
| 5 | 7204935 | rs11740932 | 1.553 | 0.899 | -0.748 | 0.197 | -0.057 | 0.007 | 0.007 | 0.032 | 0.17 | 0.03 | 0.25 | 0.16 | 0.23 | 0.15 |
| 5 | 7209108 | rs10062310 | 0.942 | 0.809 | -0.534 | 0.481 | -0.057 | -0.018 | -0.018 | 0 | 0.21 | 0.56 | 0.16 | 0.19 | 0.09 | 0.27 |
| 5 | 7210079 | rs4569841 | -0.116 | 0.911 | -0.546 | 0.521 | -0.007 | 0.012 | 0.012 | 0 | 0.33 | 0.57 | 0.38 | 0.41 | 0.26 | 0.49 |
| 5 | 7210795 | rs13360543 | 0.033 | 0.899 | -0.548 | 0.534 | -0.019 | 0.021 | 0.021 | 0 | 0.31 | 0.55 | 0.36 | 0.4 | 0.24 | 0.49 |
| 5 | 7216226 | rs13186772 | -1.5 | 0.717 | -0.775 | 0.427 | -0.023 | -0.033 | -0.033 | 0 | 0.64 | 0.86 | 0.69 | 0.6 | 0.61 | 0.72 |
| 5 | 7218137 | rs13155746 | 0.728 | 0.716 | -0.82 | 0.417 | 0.007 | 0.013 | 0.013 | 0 | 0.13 | 0.1 | 0.05 | 0.04 | 0.11 | 0.08 |
| 5 | 7226862 | rs6899057 | -1.349 | 0.739 | -0.872 | 0.46 | -0.013 | -0.023 | -0.023 | 0 | 0.63 | 0.84 | 0.69 | 0.6 | 0.61 | 0.72 |
| 5 | 7230432 | rs4382132 | 1.575 | 0.552 | -0.805 | 0.51 | -0.036 | -0.055 | -0.055 | 6.931 | 0.11 | 0.01 | 0.2 | 0.22 | 0.16 | 0.22 |
| 5 | 7238212 | rs4235576 | 0.386 | 0.552 | -0.737 | 0.482 | 0.043 | -0.007 | -0.007 | 0 | 0.55 | 0.22 | 0.59 | 0.55 | 0.64 | 0.43 |
| 5 | 7244491 | rs4276378 | 0.635 | 0.879 | -0.91 | 0.34 | 0.019 | 0.018 | 0.018 | 0 | 0.51 | 0.28 | 0.35 | 0.57 | 0.41 | 0.52 |
| 5 | 7252066 | rs9986256 | 1.572 | 0.657 | -0.987 | 0.354 | 0.016 | -0.004 | -0.004 | 0 | 0.14 | 0.11 | 0.11 | 0.18 | 0.07 | 0.23 |
| 5 | 7252807 | rs6555443 | -0.155 | 0.782 | -0.689 | 0.323 | 0.008 | 0 | 0 | 0 | 0.64 | 0.54 | 0.47 | 0.63 | 0.58 | 0.63 |
| 5 | 7253812 | rs13360728 | 1.67 | 0.716 | -0.626 | 0.317 | 0.018 | -0.004 | -0.004 | 0 | 0.14 | 0.11 | 0.11 | 0.18 | 0.07 | 0.23 |
| 5 | 7261197 | rs10462834 | -0.027 | 1.059 | -0.62 | 0.183 | 0.009 | -0.003 | -0.003 | 0 | 0.62 | 0.54 | 0.48 | 0.63 | 0.51 | 0.6 |
| 5 | 7270306 | rs6555448 | -0.208 | 1.108 | -0.522 | 0.355 | 0.039 | 0.052 | 0.052 | 0 | 0.83 | 0.6 | 0.73 | 0.81 | 0.81 | 0.73 |
| 5 | 7270417 | rs10462838 | 0.845 | 0.927 | -1.648 | 0.373 | 0.009 | -0.077 | -0.077 | 0.134 | 0.33 | 0.07 | 0.21 | 0.37 | 0.27 | 0.29 |
| 5 | 7271817 | rs16878542 | 0.367 | 1.274 | -1.301 | 0.531 | -0.058 | -0.081 | -0.081 | 0 | 0.14 | 0.01 | 0.21 | 0.23 | 0.18 | 0.27 |
| 5 | 7275068 | rs17810712 | -0.084 | 1.22 | -1.786 | -0.243 | -0.085 | 0.079 | 0.079 | 0 | 0.22 | 0.03 | 0.15 | 0.07 | 0.28 | 0.18 |

| | | | | | | | | | | | | | | | | |
|---|---|---|---|---|---|---|---|---|---|---|---|---|---|---|---|---|
| 5 | 7279325 | rs16878549 | 1.806 | 0.831 | -1.906 | -0.319 | 0.021 | -0.02 | -0.02 | 0 | 0.11 | 0.07 | 0.13 | 0.22 | 0.06 | 0.24 |
| 5 | 7281508 | rs4266373 | -0.101 | 1.218 | -1.693 | -0.432 | -0.086 | 0.078 | 0.078 | 0 | 0.22 | 0.03 | 0.15 | 0.07 | 0.28 | 0.19 |
| 5 | 7281915 | rs10067993 | 0.358 | 1.161 | -1.085 | -0.485 | -0.001 | -0.011 | -0.011 | 0 | 0.74 | 0.76 | 0.82 | 0.93 | 0.69 | 0.81 |
| 5 | 7299845 | rs17842508 | 0.996 | 1.192 | -1.105 | -0.229 | -0.013 | -0.033 | -0.033 | 0 | 0.11 | 0.03 | 0.12 | 0.15 | 0.12 | 0.22 |
| 5 | 7303877 | rs6868691 | 1.259 | 1.041 | -1.146 | -0.081 | -0.002 | -0.009 | -0.009 | 0 | 0.13 | 0.1 | 0.19 | 0.4 | 0.17 | 0.12 |
| 5 | 7317421 | rs10475365 | 1.329 | 0.946 | -1.407 | -0.598 | 0.002 | 0.026 | 0.026 | 0 | 0.14 | 0.08 | 0.07 | 0 | 0.12 | 0.07 |
| 5 | 7317925 | rs4132671 | 0.45 | 1.194 | -1.181 | -0.913 | -0.032 | 0.038 | 0.038 | 0 | 0.24 | 0.16 | 0.18 | 0.02 | 0.32 | 0.18 |
| 5 | 7324451 | rs4702457 | 0.078 | 0.837 | -0.358 | -0.888 | -0.004 | -0.005 | -0.005 | 0.161 | 0.57 | 0.53 | 0.6 | 0.44 | 0.59 | 0.55 |
| 5 | 7326039 | rs6887339 | 1.647 | 0.692 | -0.621 | -0.817 | -0.005 | 0.042 | 0.042 | 0 | 0.13 | 0.04 | 0.06 | 0 | 0.12 | 0.07 |
| 5 | 7336886 | rs1505083 | 0.226 | 0.787 | -0.599 | -0.932 | -0.064 | 0.152 | 0.152 | 0 | 0.33 | 0.13 | 0.16 | 0.01 | 0.37 | 0.23 |
| 5 | 7339482 | rs4492081 | -1.165 | 0.728 | -0.119 | -0.621 | -0.011 | 0.117 | 0.117 | 0 | 0.82 | 0.67 | 0.79 | 0.59 | 0.85 | 0.84 |
| 5 | 7345847 | rs1505080 | -0.302 | 0.825 | -0.178 | -0.668 | -0.011 | -0.011 | -0.011 | 0 | 0.68 | 0.75 | 0.59 | 0.43 | 0.62 | 0.64 |
| 5 | 7347593 | rs3812025 | -0.302 | 0.825 | -0.178 | -0.668 | -0.011 | -0.011 | -0.011 | 0 | 0.68 | 0.75 | 0.59 | 0.43 | 0.62 | 0.64 |
| 5 | 7352602 | rs7710093 | 1.398 | 0.977 | -0.118 | -0.868 | 0 | -0.039 | -0.039 | 0 | 0.17 | 0.13 | 0.08 | 0 | 0.16 | 0.09 |
| 5 | 7356978 | rs884267 | 0.758 | 1.048 | -0.239 | -0.777 | -0.005 | -0.003 | -0.003 | 0 | 0.19 | 0.17 | 0.23 | 0.17 | 0.23 | 0.24 |
| 5 | 7357767 | rs6886365 | 0.61 | 0.923 | -0.469 | -0.826 | 0.003 | 0.004 | 0.004 | 0 | 0.47 | 0.41 | 0.47 | 0.44 | 0.4 | 0.47 |
| 5 | 7359857 | rs17207026 | 0.884 | 1.139 | -0.998 | -0.487 | -0.02 | 0.039 | 0.039 | 0 | 0.18 | 0.11 | 0.1 | 0.01 | 0.22 | 0.16 |
| 5 | 7364187 | rs11134224 | 0.783 | 0.683 | -0.743 | 0.012 | 0.028 | -0.055 | -0.055 | 0.724 | 0.29 | 0.13 | 0.38 | 0.45 | 0.25 | 0.37 |
| 5 | 7367623 | rs10036962 | 0.24 | 1.066 | -0.346 | -0.172 | 0.028 | 0.001 | 0.001 | 0 | 0.19 | 0.28 | 0.25 | 0.17 | 0.29 | 0.26 |
| 5 | 7369839 | rs11949012 | 0.64 | 0.463 | -0.837 | -0.029 | 0.031 | -0.052 | -0.052 | 0.173 | 0.29 | 0.12 | 0.25 | 0.47 | 0.24 | 0.37 |
| 5 | 7375754 | rs10065659 | -0.277 | 0.831 | -0.353 | -0.136 | 0.011 | -0.027 | -0.027 | 0 | 0.27 | 0.37 | 0.26 | 0.16 | 0.29 | 0.25 |
| 5 | 7383873 | rs6875496 | -0.137 | 0.391 | -0.459 | -0.306 | 0.002 | 0 | 0 | 0 | 0.44 | 0.42 | 0.34 | 0.5 | 0.34 | 0.49 |
| 5 | 7391462 | rs12514566 | 0.083 | 0.292 | -0.385 | -0.266 | -0.001 | -0.001 | -0.001 | 0 | 0.4 | 0.37 | 0.32 | 0.45 | 0.32 | 0.49 |
| 5 | 7398546 | rs13153439 | -0.696 | 0.393 | -0.443 | -1.228 | 0.005 | 0.092 | 0.092 | 0 | 0.57 | 0.78 | 0.43 | 0.19 | 0.58 | 0.47 |
| 5 | 7402561 | rs10512920 | -0.267 | 0.452 | -0.084 | -1.356 | -0.001 | -0.119 | -0.119 | 0.143 | 0.33 | 0.43 | 0.13 | 0.02 | 0.3 | 0.21 |
| 5 | 7418630 | rs10061752 | -0.362 | 0.419 | 0.038 | -1.376 | -0.002 | -0.158 | -0.158 | 0.131 | 0.34 | 0.48 | 0.14 | 0.02 | 0.31 | 0.22 |
| 5 | 7422768 | rs6872422 | 0.466 | -0.08 | -0.266 | -1.183 | -0.001 | 0 | 0 | 0 | 0.44 | 0.46 | 0.48 | 0.48 | 0.44 | 0.55 |
| 5 | 7424672 | rs13175846 | 1.03 | -0.125 | 0.025 | -1.218 | -0.017 | -0.044 | -0.044 | 0 | 0.09 | 0.12 | 0.03 | 0 | 0.04 | 0.09 |
| 5 | 7425788 | rs17813652 | 0.648 | -0.057 | -0.283 | -1.274 | -0.06 | 0.117 | 0.117 | 0 | 0.28 | 0.09 | 0.13 | 0 | 0.33 | 0.13 |
| 5 | 7427289 | rs7710510 | -1.043 | 0.261 | -0.237 | -1 | 0.002 | 0.055 | 0.055 | 0 | 0.8 | 0.7 | 0.64 | 0.49 | 0.8 | 0.76 |
| 5 | 7445289 | rs1025291 | 0.496 | 0.224 | -0.687 | -0.627 | 0.007 | -0.025 | -0.025 | 0 | 0.36 | 0.25 | 0.39 | 0.48 | 0.36 | 0.5 |
| 5 | 7447405 | rs902522 | 0.279 | -0.307 | -0.8 | -0.291 | 0.003 | -0.06 | -0.06 | 0.503 | 0.63 | 0.79 | 0.6 | 0.53 | 0.65 | 0.57 |
| 5 | 7447525 | rs13175786 | -0.775 | -0.772 | 0.082 | -0.77 | -0.064 | -0.128 | -0.128 | 0 | 0.2 | 0.37 | 0.07 | 0.01 | 0.08 | 0.18 |
| 5 | 7459043 | rs7712809 | 0.031 | -0.357 | 0.309 | -0.45 | 0.015 | -0.01 | -0.01 | 0 | 0.08 | 0.26 | 0.07 | 0.01 | 0.03 | 0.07 |
| 5 | 7463625 | rs26737 | -0.747 | -0.255 | -1.411 | -0.193 | 0.025 | 0.039 | 0.039 | 0 | 0.45 | 0.76 | 0.54 | 0.65 | 0.35 | 0.4 |
| 5 | 7466176 | rs26739 | 0.742 | 0.035 | -1.775 | 0.258 | -0.041 | -0.139 | -0.139 | 0 | 0.2 | 0.03 | 0.18 | 0.53 | 0.2 | 0.24 |
| 5 | 7466743 | rs10063916 | 1.053 | 0.454 | -2.416 | -0.66 | -0.061 | 0.094 | 0.094 | 0.365 | 0.25 | 0.09 | 0.16 | 0 | 0.36 | 0.09 |
| 5 | 7468560 | rs27378 | 1.044 | 0.559 | -2.341 | -0.075 | -0.037 | -0.13 | -0.13 | 0 | 0.2 | 0.04 | 0.18 | 0.48 | 0.2 | 0.23 |
| 5 | 7478914 | rs252546 | -0.345 | -0.028 | -1.728 | -0.111 | 0.01 | -0.013 | -0.013 | 0 | 0.63 | 0.77 | 0.65 | 0.58 | 0.64 | 0.43 |
| 5 | 7479953 | rs252545 | -0.301 | 0.063 | -1.508 | -0.045 | 0 | -0.004 | -0.004 | 0 | 0.73 | 0.77 | 0.77 | 0.68 | 0.73 | 0.54 |
| 5 | 7493177 | rs11948030 | 0.711 | 0.127 | -1.603 | -0.696 | -0.014 | 0.113 | 0.113 | 0 | 0.19 | 0.03 | 0.09 | 0 | 0.19 | 0.13 |
| 5 | 7497783 | rs4333285 | -0.676 | 0.15 | -1.24 | -0.692 | 0.007 | 0.007 | 0.007 | 0 | 0.58 | 0.65 | 0.48 | 0.33 | 0.66 | 0.41 |
| 5 | 7503123 | rs968119 | 0.835 | 0.443 | -0.726 | -0.876 | 0.013 | -0.07 | -0.07 | 0.37 | 0.16 | 0.21 | 0.1 | 0 | 0.18 | 0.12 |
| 5 | 7506433 | rs7721013 | 0.302 | 0.519 | -0.564 | -0.946 | 0.008 | 0.058 | 0.058 | 0 | 0.14 | 0.19 | 0.21 | 0.33 | 0.16 | 0.12 |
| 5 | 7510019 | rs7734525 | -0.683 | 0.059 | -0.569 | -0.815 | -0.038 | 0.072 | 0.072 | 0 | 0.89 | 0.69 | 0.75 | 0.88 | 0.97 | 0.75 |
| 5 | 7510391 | rs1995276 | -0.046 | 0.492 | -0.406 | -1.181 | 0 | -0.001 | -0.001 | 0 | 0.33 | 0.28 | 0.36 | 0.34 | 0.36 | 0.26 |
| 5 | 7510936 | rs4263480 | -0.438 | 0.427 | -0.366 | -1.115 | 0.03 | 0.025 | 0.025 | 0 | 0.28 | 0.4 | 0.35 | 0.35 | 0.32 | 0.25 |
| 5 | 7515682 | rs17826546 | 0.56 | -0.142 | -0.598 | -1.268 | -0.039 | 0.079 | 0.079 | 0 | 0.19 | 0.07 | 0.09 | 0 | 0.23 | 0.15 |
| 5 | 7516555 | rs17826588 | -0.039 | -0.103 | -0.494 | -1.18 | -0.002 | -0.005 | -0.005 | 0 | 0.4 | 0.32 | 0.2 | 0 | 0.44 | 0.28 |
| 5 | 7517418 | rs10038196 | -0.364 | -0.021 | -0.442 | -0.842 | 0 | 0.007 | 0.007 | 0 | 0.51 | 0.56 | 0.32 | 0.25 | 0.53 | 0.52 |
| 5 | 7518855 | rs10066518 | 0.665 | 0.046 | -0.76 | -0.682 | 0.004 | 0.001 | 0.001 | 4.562 | 0.73 | 0.19 | 0.68 | 0.71 | 0.75 | 0.68 |
| 5 | 7519387 | rs16878741 | 1.74 | 0.083 | -0.344 | -0.699 | 0.042 | -0.053 | -0.053 | 0 | 0.09 | 0.2 | 0.07 | 0 | 0.13 | 0.04 |
| 5 | 7519762 | rs6867567 | 0.527 | 0.138 | -0.178 | -0.838 | -0.007 | 0.03 | 0.03 | 0.013 | 0.78 | 0.52 | 0.73 | 0.7 | 0.81 | 0.71 |
| 5 | 7521083 | rs3828628 | 0.118 | 0.049 | -0.455 | -0.613 | 0.002 | -0.044 | -0.044 | 0.362 | 0.25 | 0.09 | 0.18 | 0.32 | 0.22 | 0.35 |
| 5 | 7524548 | rs7721273 | -0.044 | 0.222 | -0.023 | -0.256 | 0.03 | -0.002 | -0.002 | 2.432 | 0.18 | 0.34 | 0.16 | 0.2 | 0.22 | 0.15 |
| 5 | 7526262 | rs13159560 | 0.496 | 0.189 | -0.667 | -0.191 | 0.002 | -0.034 | -0.034 | 0.208 | 0.14 | 0.07 | 0.14 | 0.28 | 0.12 | 0.11 |
| 5 | 7527521 | rs2055388 | 0.228 | 0.242 | -0.102 | 0.136 | -0.013 | 0.047 | 0.047 | 3.062 | 0.9 | 0.41 | 0.82 | 0.91 | 0.97 | 0.84 |
| 5 | 7528287 | rs7714110 | 0.525 | 0.232 | 0.139 | 0.114 | -0.007 | 0.023 | 0.023 | 0 | 0.06 | 0.15 | 0.14 | 0.09 | 0.02 | 0.16 |
| 5 | 7528558 | rs4702473 | -0.302 | 0.281 | -0.777 | -0.008 | 0 | -0.002 | -0.002 | 0.535 | 0.59 | 0.36 | 0.42 | 0.52 | 0.65 | 0.61 |
| 5 | 7529177 | rs10062110 | 0.912 | 0.334 | -1.315 | 0.018 | -0.039 | -0.068 | -0.068 | 0 | 0.31 | 0.11 | 0.41 | 0.39 | 0.32 | 0.22 |
| 5 | 7529813 | rs6555474 | 0.023 | 0.756 | -1.537 | 0.004 | 0.001 | -0.013 | -0.013 | 0.079 | 0.5 | 0.19 | 0.35 | 0.52 | 0.52 | 0.58 |
| 5 | 7532455 | rs4701786 | 0.225 | 0.551 | -1.426 | 0.131 | 0.004 | 0.044 | 0.044 | 0.022 | 0.43 | 0.16 | 0.28 | 0.32 | 0.43 | 0.47 |
| 5 | 7534221 | rs1027579 | -0.121 | 0.554 | -1.412 | 0.132 | 0.008 | -0.001 | -0.001 | 0 | 0.5 | 0.26 | 0.36 | 0.52 | 0.52 | 0.58 |
| 5 | 7538237 | rs1392481 | 0.234 | 0.568 | -1.679 | 0.164 | 0.001 | -0.011 | -0.011 | 0 | 0.08 | 0.06 | 0.07 | 0.2 | 0.09 | 0.11 |
| 5 | 7539712 | rs10035541 | 0.024 | 0.586 | -1.603 | -0.045 | 0.004 | 0.044 | 0.044 | 0.022 | 0.43 | 0.16 | 0.28 | 0.32 | 0.43 | 0.47 |
| 5 | 7541212 | rs980483 | 0.76 | 1.022 | -1.054 | 0.036 | 0.019 | -0.04 | -0.04 | 0 | 0.09 | 0.12 | 0.05 | 0 | 0.13 | 0.04 |
| 5 | 7543355 | rs16878777 | 0.266 | 0.404 | -1.967 | 0.271 | -0.004 | -0.148 | -0.148 | 0 | 0.24 | 0.01 | 0.38 | 0.39 | 0.2 | 0.25 |
| 5 | 7546015 | rs13436413 | 0.055 | 0.858 | -1.79 | -0.17 | -0.01 | -0.009 | -0.009 | 0 | 0.32 | 0.21 | 0.27 | 0.32 | 0.31 | 0.32 |
| 5 | 7548178 | rs1004784 | 0.009 | 0.836 | -1.757 | -0.855 | -0.06 | 0.113 | 0.113 | 0 | 0.29 | 0.15 | 0.2 | 0.01 | 0.39 | 0.15 |

| | | | | | | | | | | | | | | | | |
|---|---|---|---|---|---|---|---|---|---|---|---|---|---|---|---|---|
| 5 | 7553501 | rs12522444 | 1.186 | 1.203 | -1.773 | -0.872 | -0.044 | 0.084 | 0.084 | 0 | 0.14 | 0.02 | 0.08 | 0 | 0.19 | 0.11 |
| 5 | 7561901 | rs10475378 | -0.507 | 1.012 | -0.706 | -1.057 | 0.009 | -0.108 | -0.108 | 0.197 | 0.32 | 0.42 | 0.22 | 0.01 | 0.39 | 0.15 |
| 5 | 7563591 | rs7717535 | -0.209 | 1.022 | -0.819 | -1.053 | -0.062 | 0.112 | 0.112 | 0 | 0.29 | 0.14 | 0.19 | 0.01 | 0.39 | 0.16 |
| 5 | 7571456 | rs10866584 | -0.939 | 1.05 | -1.019 | -0.931 | 0 | 0 | 0 | 0 | 0.55 | 0.47 | 0.47 | 0.47 | 0.67 | 0.42 |
| 5 | 7575713 | rs6875695 | -0.406 | 1.035 | -0.978 | -0.942 | -0.007 | -0.002 | -0.002 | 0 | 0.43 | 0.33 | 0.4 | 0.48 | 0.53 | 0.39 |
| 5 | 7578342 | rs4541631 | -0.549 | 0.604 | -0.794 | -0.743 | 0.029 | 0.004 | 0.004 | 0 | 0.54 | 0.37 | 0.45 | 0.48 | 0.67 | 0.43 |
| 5 | 7581983 | rs7443591 | -0.734 | 0.609 | -0.83 | -0.741 | -0.014 | 0 | 0 | 0 | 0.56 | 0.53 | 0.48 | 0.48 | 0.67 | 0.42 |
| 5 | 7587939 | rs6555477 | 0.011 | 0.681 | -0.955 | -0.599 | 0.003 | 0.02 | 0.02 | 0 | 0.27 | 0.29 | 0.28 | 0.47 | 0.28 | 0.27 |
| 5 | 7593892 | rs2388875 | 0.69 | 0.931 | -1.227 | -0.563 | -0.001 | 0.018 | 0.018 | 0 | 0.17 | 0.14 | 0.18 | 0.47 | 0.16 | 0.21 |
| 5 | 7595762 | rs13166852 | 0.848 | 1.094 | -1.389 | -0.916 | -0.125 | 0.172 | 0.172 | 0 | 0.25 | 0.03 | 0.15 | 0 | 0.38 | 0.15 |
| 5 | 7609916 | rs16878811 | 0.574 | 1.075 | -1.364 | -0.947 | 0.002 | -0.032 | -0.032 | 0 | 0.15 | 0.14 | 0.12 | 0.03 | 0.17 | 0.17 |
| 5 | 7625795 | rs1428530 | -0.061 | 1.01 | -0.947 | -0.645 | 0.012 | 0.256 | 0.256 | 0 | 0.91 | 0.77 | 0.85 | 0.56 | 0.92 | 0.94 |
| 5 | 7629191 | rs1035905 | 1.267 | 1.012 | -1.109 | -0.729 | -0.105 | 0.155 | 0.155 | 0 | 0.23 | 0.02 | 0.11 | 0 | 0.31 | 0.14 |
| 5 | 7629756 | rs2914290 | 0.677 | 1.084 | -1.142 | -0.792 | -0.018 | -0.01 | -0.01 | 0 | 0.11 | 0.07 | 0.19 | 0.45 | 0.15 | 0.08 |
| 5 | 7632077 | rs7706191 | 0.974 | 1.005 | -1.18 | -0.392 | -0.002 | 0.001 | 0.001 | 0 | 0.14 | 0.14 | 0.1 | 0.17 | 0.13 | 0.17 |
| 5 | 7645474 | rs12520264 | 1.261 | 0.873 | -1.617 | -0.253 | -0.104 | 0.158 | 0.158 | 0 | 0.23 | 0.02 | 0.11 | 0 | 0.31 | 0.14 |
| 5 | 7649860 | rs4702484 | 1.115 | -0.016 | -1.478 | 0.055 | 0.012 | -0.045 | -0.045 | 2.602 | 0.2 | 0.01 | 0.3 | 0.18 | 0.15 | 0.16 |
| 5 | 7657882 | rs2973319 | -0.436 | 0.577 | -0.631 | 0.225 | 0.018 | 0.003 | 0.003 | 0 | 0.54 | 0.6 | 0.47 | 0.62 | 0.65 | 0.42 |
| 5 | 7659279 | rs877886 | 0.096 | 0.665 | -0.852 | -0.601 | -0.003 | -0.003 | -0.003 | 0 | 0.41 | 0.37 | 0.26 | 0.17 | 0.49 | 0.33 |
| 5 | 7663078 | rs727432 | -0.441 | 0.227 | -0.849 | -0.39 | 0 | 0 | 0 | 0 | 0.35 | 0.26 | 0.28 | 0.68 | 0.32 | 0.53 |
| 5 | 7664530 | rs4289537 | -1.069 | 0.01 | -0.748 | -1.255 | -0.015 | 0.024 | 0.024 | 0 | 0.69 | 0.78 | 0.81 | 0.97 | 0.62 | 0.85 |
| 5 | 7666786 | rs7445043 | -0.347 | -0.737 | -0.135 | -1.769 | -0.001 | -0.076 | -0.076 | 0.064 | 0.48 | 0.55 | 0.75 | 0.99 | 0.5 | 0.57 |
| 5 | 7667153 | rs10462643 | -0.505 | -0.657 | 0.186 | -1.447 | -0.01 | 0.02 | 0.02 | 0 | 0.85 | 0.76 | 0.63 | 0.86 | 0.87 | 0.84 |
| 5 | 7668026 | rs9654458 | -0.097 | -0.606 | 0.278 | -1.471 | 0 | 0.035 | 0.035 | 0.057 | 0.83 | 0.62 | 0.6 | 0.84 | 0.84 | 0.83 |
| 5 | 7669054 | rs2914300 | 0.032 | -0.608 | 0.214 | -1.462 | -0.007 | 0.019 | 0.019 | 0.009 | 0.17 | 0.32 | 0.39 | 0.15 | 0.15 | 0.17 |
| 5 | 7681207 | rs6555483 | -0.523 | -0.572 | 0.156 | -1.432 | -0.001 | 0.02 | 0.02 | 0 | 0.85 | 0.77 | 0.61 | 0.86 | 0.86 | 0.84 |
| 5 | 7684243 | rs734986 | 0.545 | -0.905 | -0.083 | -1.848 | -0.008 | 0.09 | 0.09 | 0.012 | 0.44 | 0.25 | 0.56 | 0.16 | 0.54 | 0.33 |
| 5 | 7685079 | rs1025296 | -0.129 | -0.995 | 0.225 | -1.906 | 0.022 | 0.095 | 0.095 | 0 | 0.5 | 0.32 | 0.38 | 0.84 | 0.45 | 0.67 |
| 5 | 7685776 | rs1025298 | 0.025 | -1.2 | 0.1 | -2.026 | 0.007 | 0.043 | 0.043 | 0.404 | 0.43 | 0.32 | 0.36 | 0.84 | 0.4 | 0.62 |
| 5 | 7686874 | rs2914310 | -1.198 | -1.055 | -0.586 | -1.985 | -0.052 | 0.147 | 0.147 | 4.894 | 0.62 | 0.84 | 0.78 | 0.98 | 0.55 | 0.77 |
| 5 | 7690201 | rs7714830 | 1.228 | -0.916 | -0.558 | -1.48 | 0.004 | -0.033 | -0.033 | 0 | 0.14 | 0.02 | 0.09 | 0.18 | 0.12 | 0.11 |
| 5 | 7695845 | rs11134251 | -0.317 | -0.536 | -0.053 | -1.606 | 0.005 | 0.027 | 0.027 | 0 | 0.53 | 0.57 | 0.41 | 0.83 | 0.44 | 0.67 |
| 5 | 7696028 | rs11134252 | -0.325 | -0.535 | -0.053 | -1.606 | 0.005 | 0.026 | 0.026 | 0 | 0.54 | 0.57 | 0.41 | 0.83 | 0.44 | 0.67 |
| 5 | 7704187 | rs4701790 | 0.393 | -0.324 | -0.111 | -1.315 | -0.001 | 0.001 | 0.001 | 0 | 0.14 | 0.12 | 0.37 | 0.15 | 0.14 | 0.15 |
| 5 | 7717396 | rs6878196 | -0.09 | -0.655 | 0.245 | -2.182 | 0.02 | 0.024 | 0.024 | 0 | 0.49 | 0.37 | 0.65 | 0.95 | 0.41 | 0.59 |
| 5 | 7737474 | rs12189243 | 0.907 | -0.657 | 0.346 | -2.292 | -0.115 | 0.234 | 0.234 | 0.474 | 0.37 | 0.08 | 0.22 | 0.03 | 0.45 | 0.29 |
| 5 | 7744625 | rs7730589 | 0.021 | -0.693 | 0.429 | -2.321 | 0.023 | 0.121 | 0.121 | 0.036 | 0.48 | 0.3 | 0.32 | 0.06 | 0.58 | 0.38 |
| 5 | 7747639 | rs4131923 | 0.106 | -0.538 | 0.537 | -2.155 | 0.022 | -0.043 | -0.043 | 0 | 0.88 | 0.62 | 0.89 | 0.97 | 0.88 | 0.91 |
| 5 | 7749240 | rs12186377 | 1.519 | -0.471 | 0.117 | -2.332 | -0.14 | 0.192 | 0.192 | 0 | 0.27 | 0.03 | 0.15 | 0 | 0.37 | 0.16 |
| 5 | 7749260 | rs10060054 | -0.003 | -0.418 | 0.647 | -1.866 | 0.014 | -0.04 | -0.04 | 0 | 0.89 | 0.76 | 0.9 | 0.97 | 0.88 | 0.88 |
| 5 | 7754231 | rs4334848 | -0.092 | -0.968 | 0.209 | -1.616 | -0.011 | -0.011 | -0.011 | 0 | 0.4 | 0.44 | 0.52 | 0.34 | 0.36 | 0.3 |
| 5 | 7756170 | rs11750832 | 0.193 | -0.704 | -0.393 | -1.69 | 0.022 | 0.077 | 0.077 | 0 | 0.53 | 0.27 | 0.39 | 0.66 | 0.56 | 0.67 |
| 5 | 7757812 | rs3935156 | -0.234 | -0.704 | -0.36 | -1.688 | 0.02 | 0.058 | 0.058 | 0 | 0.47 | 0.7 | 0.59 | 0.34 | 0.43 | 0.33 |
| 5 | 7764530 | rs11134261 | 0.687 | -0.508 | -0.581 | -1.382 | -0.004 | -0.042 | -0.042 | 1.287 | 0.16 | 0.1 | 0.18 | 0.63 | 0.14 | 0.38 |
| 5 | 7769905 | rs10051908 | -0.29 | -0.603 | 0.127 | -1.427 | -0.004 | 0.1 | 0.1 | 1.667 | 0.81 | 0.59 | 0.79 | 0.37 | 0.86 | 0.62 |
| 5 | 7772199 | rs7711377 | 0.282 | -1.045 | -0.098 | -1.713 | -0.004 | 0.034 | 0.034 | 0 | 0.4 | 0.27 | 0.49 | 0.32 | 0.44 | 0.31 |
| 5 | 7776646 | rs12520791 | 0.925 | -0.947 | -0.147 | -1.611 | -0.006 | -0.07 | -0.07 | 2.151 | 0.16 | 0.04 | 0.17 | 0.62 | 0.13 | 0.39 |
| 5 | 7782238 | rs4702493 | 0.925 | -0.947 | -0.147 | -1.611 | -0.006 | -0.07 | -0.07 | 0 | 0.16 | 0.04 | 0.17 | 0.62 | 0.13 | 0.38 |
| 5 | 7783259 | rs11748102 | 0.707 | -0.969 | -0.068 | -1.741 | 0.001 | -0.038 | -0.038 | 0 | 0.17 | 0.08 | 0.19 | 0.62 | 0.14 | 0.39 |
| 5 | 7783858 | rs13165346 | 1.249 | -1.237 | -0.072 | -1.894 | -0.061 | 0.143 | 0.143 | 0.026 | 0.35 | 0.14 | 0.21 | 0.06 | 0.41 | 0.28 |
| 5 | 7787698 | rs1563912 | 0.327 | -1.151 | -0.004 | -1.885 | 0.019 | 0.026 | 0.026 | 0 | 0.38 | 0.25 | 0.44 | 0.31 | 0.37 | 0.28 |
| 5 | 7790217 | rs3816620 | 0.252 | -1.274 | -0.049 | -1.697 | 0.021 | 0.028 | 0.028 | 0 | 0.39 | 0.24 | 0.45 | 0.32 | 0.36 | 0.3 |
| 5 | 7791762 | rs326129 | 0.487 | -1.228 | -0.101 | -1.665 | -0.003 | -0.003 | -0.003 | 0.598 | 0.19 | 0.17 | 0.24 | 0.62 | 0.2 | 0.42 |
| 5 | 7791998 | rs326132 | 0.607 | -1.228 | -0.332 | -1.663 | -0.019 | -0.044 | -0.044 | 0.11 | 0.18 | 0.08 | 0.22 | 0.62 | 0.2 | 0.42 |
| 5 | 7792917 | rs16879109 | -0.094 | -1.419 | -0.242 | -1.779 | -0.031 | -0.034 | -0.034 | 0 | 0.08 | 0.34 | 0.1 | 0.01 | 0.01 | 0.05 |
| 5 | 7794326 | rs4629562 | 1.128 | -1.158 | -1.039 | -1.872 | -0.076 | 0.137 | 0.137 | 0.049 | 0.33 | 0.11 | 0.21 | 0.06 | 0.42 | 0.23 |
| 5 | 7794738 | rs1460970 | -0.219 | -0.554 | -0.704 | -1.249 | -0.004 | -0.004 | -0.004 | 0 | 0.82 | 0.87 | 0.78 | 0.38 | 0.8 | 0.58 |
| 5 | 7797291 | rs6864771 | -0.082 | -0.187 | -0.876 | -1.471 | -0.018 | -0.027 | -0.027 | 0 | 0.78 | 0.88 | 0.75 | 0.37 | 0.73 | 0.55 |
| 5 | 7801608 | rs13162758 | 0.692 | 0.167 | -1.225 | -1.971 | -0.094 | 0.215 | 0.215 | 0.294 | 0.32 | 0.04 | 0.19 | 0 | 0.37 | 0.23 |
| 5 | 7802363 | rs2290910 | 0.202 | -0.056 | -0.185 | -0.93 | -0.009 | -0.024 | -0.024 | 0 | 0.23 | 0.12 | 0.42 | 0.28 | 0.26 | 0.18 |
| 5 | 7815809 | rs326153 | -1.818 | -0.228 | 0.999 | -0.82 | 0.066 | 0.044 | 0.044 | 5.371 | 0.06 | 0.46 | 0.05 | 0 | 0.01 | 0 |
| 5 | 7819850 | rs326162 | -1.788 | -0.212 | 0.909 | -0.812 | 0.05 | 0.03 | 0.03 | 5.263 | 0.06 | 0.42 | 0.05 | 0 | 0.01 | 0 |
| 5 | 7822918 | rs989634 | 0.455 | -0.262 | 0.545 | -0.887 | -0.003 | -0.008 | -0.008 | 1.257 | 0.56 | 0.43 | 0.63 | 0.35 | 0.58 | 0.45 |
| 5 | 7826058 | rs326171 | -0.119 | -0.354 | 0.414 | -0.849 | -0.037 | -0.057 | -0.057 | 1.134 | 0.63 | 0.86 | 0.68 | 0.42 | 0.61 | 0.48 |
| 5 | 7828533 | rs326173 | -0.074 | -0.296 | 0.396 | -0.7 | -0.031 | -0.059 | -0.059 | 0.691 | 0.62 | 0.86 | 0.68 | 0.42 | 0.62 | 0.48 |
| 5 | 7831013 | rs2287778 | 0.464 | -0.242 | 0.626 | -0.671 | -0.003 | -0.008 | -0.008 | 0 | 0.55 | 0.44 | 0.64 | 0.35 | 0.58 | 0.45 |
| 5 | 7832387 | rs1993716 | 0.921 | -0.227 | 0.013 | -0.835 | 0 | -0.001 | -0.001 | 0.233 | 0.29 | 0.27 | 0.47 | 0.26 | 0.36 | 0.23 |
| 5 | 7837861 | rs72001 | -0.006 | -0.116 | -0.249 | -0.319 | 0.001 | -0.01 | -0.01 | 0.706 | 0.29 | 0.26 | 0.18 | 0.1 | 0.22 | 0.23 |
| 5 | 7838085 | rs89599 | -0.419 | -0.131 | -0.453 | -0.301 | -0.023 | -0.084 | -0.084 | 0 | 0.77 | 0.88 | 0.77 | 0.44 | 0.77 | 0.55 |

| | | | | | | | | | | | | | | | | |
|---|---|---|---|---|---|---|---|---|---|---|---|---|---|---|---|---|
| 5 | 7838364 | rs729126 | 0.929 | -0.175 | -0.547 | -0.273 | -0.016 | 0.041 | 0.041 | 0 | 0.45 | 0.31 | 0.55 | 0.28 | 0.51 | 0.3 |
| 5 | 7843211 | rs326193 | -0.247 | 0.581 | -1.444 | -0.306 | 0.001 | -0.009 | -0.009 | 0 | 0.29 | 0.25 | 0.18 | 0.1 | 0.22 | 0.23 |
| 5 | 7852282 | rs326205 | 0.329 | 0.687 | -1.777 | -0.017 | -0.021 | 0.008 | 0.008 | 0.505 | 0.53 | 0.78 | 0.61 | 0.39 | 0.53 | 0.32 |
| 5 | 7854289 | rs4702506 | -0.656 | 0.444 | -1.789 | -0.561 | 0.024 | 0.097 | 0.097 | 0.074 | 0.24 | 0.07 | 0.11 | 0 | 0.18 | 0.17 |
| 5 | 7859434 | rs1046014 | -0.505 | 1.422 | -1.253 | -0.796 | -0.007 | -0.016 | -0.016 | 0 | 0.19 | 0.36 | 0.31 | 0.15 | 0.11 | 0.17 |
| 5 | 7859523 | rs1046012 | -0.539 | 0.362 | -1.574 | -1.542 | -0.004 | 0.033 | 0.033 | 0 | 0.28 | 0.23 | 0.19 | 0.05 | 0.32 | 0.14 |
| 5 | 7860404 | rs162024 | -1.533 | 0.305 | -0.933 | -0.659 | -0.038 | 0.023 | 0.023 | 0.17 | 0.52 | 0.86 | 0.66 | 0.41 | 0.5 | 0.34 |
| 5 | 7861638 | rs162027 | -1.147 | 0.129 | -1.774 | -0.231 | 0.002 | -0.043 | -0.043 | 0 | 0.85 | 0.93 | 0.85 | 0.67 | 0.83 | 0.81 |
| 5 | 7861985 | rs2307116 | 1.937 | 0.135 | -1.793 | -0.257 | -0.05 | 0.02 | 0.02 | 0.18 | 0.33 | 0.07 | 0.19 | 0.26 | 0.33 | 0.46 |
| 5 | 7862926 | rs3733784 | 0.614 | 0.073 | -1.775 | -0.245 | 0.01 | 0.023 | 0.023 | 0.191 | 0.61 | 0.31 | 0.38 | 0.31 | 0.65 | 0.6 |
| 5 | 7865891 | rs16879258 | 1.277 | 0.014 | -1.695 | -0.272 | 0.015 | 0.017 | 0.017 | 0 | 0.19 | 0.17 | 0.12 | 0.05 | 0.14 | 0.07 |
| 5 | 7868030 | rs2966952 | -1.063 | 0.132 | -1.71 | -0.191 | 0 | -0.047 | -0.047 | 0 | 0.85 | 0.93 | 0.85 | 0.67 | 0.82 | 0.8 |
| 5 | 7870973 | rs1801394 | 0.914 | 0.154 | -1.403 | -0.29 | -0.006 | 0.063 | 0.063 | 0 | 0.43 | 0.25 | 0.28 | 0.26 | 0.52 | 0.52 |
| 5 | 7874424 | rs13181011 | 0.678 | 0.272 | -1.336 | -0.218 | -0.014 | 0.005 | 0.005 | 0 | 0.09 | 0.06 | 0.07 | 0 | 0.18 | 0.07 |
| 5 | 7875963 | rs7730643 | 0.241 | -0.581 | -1.235 | -0.083 | -0.001 | -0.001 | -0.001 | 0 | 0.22 | 0.25 | 0.18 | 0.23 | 0.17 | 0.4 |
| 5 | 7876288 | rs326121 | -0.518 | -0.963 | -1.607 | -0.142 | 0.01 | -0.006 | -0.006 | 0 | 0.38 | 0.63 | 0.57 | 0.57 | 0.27 | 0.47 |
| 5 | 7876315 | rs3776467 | 0.799 | 0.193 | -0.884 | 0.333 | -0.007 | 0.018 | 0.018 | 0.239 | 0.72 | 0.34 | 0.51 | 0.44 | 0.74 | 0.67 |
| 5 | 7876861 | rs326123 | 1.05 | 0.478 | -0.713 | 0.415 | -0.001 | 0.001 | 0.001 | 0.226 | 0.72 | 0.74 | 0.67 | 0.52 | 0.61 | 0.7 |
| 5 | 7877178 | rs326124 | -0.177 | 0.485 | -0.731 | 0.558 | 0.002 | -0.06 | -0.06 | 0 | 0.83 | 0.94 | 0.85 | 0.67 | 0.81 | 0.8 |
| 5 | 7879950 | rs7703033 | 2.455 | 0.493 | -0.739 | 0.461 | 0.01 | -0.004 | -0.004 | 0 | 0.22 | 0.2 | 0.18 | 0.2 | 0.31 | 0.24 |
| 5 | 7879983 | rs6555501 | 0.885 | 0.42 | -0.679 | 0.424 | 0.002 | -0.001 | -0.001 | 0 | 0.38 | 0.57 | 0.51 | 0.37 | 0.43 | 0.38 |
| 5 | 7880287 | rs162031 | 0.111 | 0.48 | -0.657 | 0.502 | -0.006 | -0.072 | -0.072 | 0.014 | 0.81 | 0.91 | 0.75 | 0.5 | 0.78 | 0.77 |
| 5 | 7884389 | rs161871 | -0.336 | 0.404 | -0.596 | 0.498 | 0.025 | 0.105 | 0.105 | 0.003 | 0.19 | 0.47 | 0.48 | 0.34 | 0.18 | 0.18 |
| 5 | 7885526 | rs16879305 | 0.633 | 0.398 | -0.575 | 0.495 | 0.002 | 0.018 | 0.018 | 0 | 0.16 | 0.38 | 0.33 | 0.17 | 0.11 | 0.14 |
| 5 | 7885959 | rs162036 | 0.237 | 0.402 | -0.601 | 0.498 | 0.015 | 0.014 | 0.014 | 0 | 0.17 | 0.44 | 0.35 | 0.17 | 0.14 | 0.14 |
| 5 | 7887478 | rs162040 | -1.625 | 0.394 | -0.672 | 0.55 | -0.02 | -0.06 | -0.06 | 0 | 0.89 | 0.99 | 0.87 | 0.66 | 0.85 | 0.87 |
| 5 | 7893121 | rs162049 | 0.154 | 0.409 | -0.659 | 0.413 | 0 | -0.066 | -0.066 | 0 | 0.81 | 0.9 | 0.75 | 0.47 | 0.78 | 0.77 |
| 5 | 7896511 | rs3776455 | 1.374 | 0.396 | -0.599 | 0.394 | 0 | -0.009 | -0.009 | 0 | 0.64 | 0.46 | 0.38 | 0.33 | 0.63 | 0.63 |
| 5 | 7897191 | rs10380 | 0.593 | 0.41 | -0.642 | 0.396 | 0.001 | 0 | 0 | 0 | 0.16 | 0.37 | 0.32 | 0.14 | 0.11 | 0.14 |
| 5 | 7897283 | rs12347 | 0.314 | 0.408 | -0.663 | 0.397 | 0.017 | -0.006 | -0.006 | 0 | 0.17 | 0.43 | 0.34 | 0.14 | 0.13 | 0.14 |
| 5 | 7897319 | rs1802059 | 0.617 | 0.42 | -0.724 | 0.337 | 0.035 | 0.102 | 0.102 | 3.526 | 0.41 | 0.26 | 0.21 | 0.13 | 0.34 | 0.39 |
| 5 | 7900712 | rs9332 | 0.225 | 0.612 | -0.714 | 0.419 | 0.017 | -0.006 | -0.006 | 0.035 | 0.17 | 0.44 | 0.35 | 0.14 | 0.14 | 0.14 |
| 5 | 7901230 | rs10520873 | 2.244 | 1.278 | -0.754 | 0.494 | -0.071 | -0.005 | -0.005 | 0 | 0.21 | 0.05 | 0.14 | 0.2 | 0.29 | 0.24 |
| 5 | 7920843 | rs11743287 | -1.318 | 1.044 | -0.835 | 0.647 | -0.02 | -0.059 | -0.059 | 0 | 0.91 | 0.99 | 0.89 | 0.7 | 0.87 | 0.95 |
| 5 | 7923683 | rs327575 | -0.126 | 0.965 | -0.356 | 0.658 | 0.083 | -0.02 | -0.02 | 1.824 | 0.47 | 0.75 | 0.66 | 0.51 | 0.62 | 0.49 |
| 5 | 7928158 | rs16879452 | 1.376 | 1.066 | 0.111 | 0.725 | 0.02 | 0.023 | 0.023 | 1.437 | 0.71 | 0.62 | 0.75 | 0.64 | 0.62 | 0.81 |
| 5 | 7930701 | rs2640660 | 1.065 | 1.077 | 0.138 | 0.734 | 0.006 | 0.012 | 0.012 | 0.264 | 0.73 | 0.7 | 0.77 | 0.64 | 0.65 | 0.82 |
| 5 | 7930866 | rs10440693 | 0.089 | 1.067 | 0.202 | 0.516 | 0.04 | -0.068 | -0.068 | 0 | 0.12 | 0.21 | 0.11 | 0 | 0.19 | 0.08 |
| 5 | 7933629 | rs2640658 | 0.331 | 1.16 | 0.306 | 0.65 | 0.085 | 0.136 | 0.136 | 0 | 0.13 | 0.43 | 0.35 | 0.21 | 0.12 | 0.15 |
| 5 | 7936153 | rs2170498 | -1.234 | 1.154 | 0.102 | 0.642 | 0.013 | -0.002 | -0.002 | 0 | 0.25 | 0.66 | 0.46 | 0.21 | 0.31 | 0.23 |
| 5 | 7936287 | rs10512934 | 0.947 | 0.798 | -0.251 | 0.428 | 0.118 | 0.13 | 0.13 | 3.983 | 0.39 | 0.16 | 0.2 | 0.14 | 0.21 | 0.4 |
| 5 | 7941757 | rs97713 | 0.911 | 1.349 | 0.655 | 1.095 | 0.005 | 0.042 | 0.042 | 0.279 | 0.75 | 0.6 | 0.54 | 0.41 | 0.75 | 0.75 |
| 5 | 7943796 | rs329852 | 1.225 | 1.897 | 0.241 | 1.231 | 0 | 0.001 | 0.001 | 0.006 | 0.08 | 0.05 | 0.11 | 0.36 | 0.12 | 0.05 |
| 5 | 7946262 | rs1471621 | 2.352 | 1.939 | 0.113 | 1.104 | -0.061 | 0.038 | 0.038 | 1.673 | 0.59 | 0.2 | 0.35 | 0.3 | 0.51 | 0.66 |
| 5 | 7950600 | rs1564592 | 2.406 | 1.961 | 0.111 | 1.074 | -0.056 | 0.05 | 0.05 | 1.733 | 0.58 | 0.19 | 0.34 | 0.3 | 0.5 | 0.65 |
| 5 | 7952282 | rs7703389 | 1.578 | 1.839 | 0.253 | 1.079 | 0.006 | 0.004 | 0.004 | 0.919 | 0.67 | 0.37 | 0.47 | 0.43 | 0.59 | 0.73 |
| 5 | 7952536 | rs1542781 | 0.264 | 2.034 | -0.155 | 0.953 | -0.027 | 0.024 | 0.024 | 0 | 0.1 | 0.04 | 0.06 | 0 | 0.17 | 0.07 |
| 5 | 7952808 | rs11744470 | -0.046 | 1.496 | 0.149 | 1.281 | -0.001 | -0.002 | -0.002 | 0.647 | 0.06 | 0.06 | 0.03 | 0.03 | 0.04 | 0.06 |
| 5 | 7953909 | rs329831 | 1.124 | 1.678 | 0.067 | 1.517 | -0.029 | -0.043 | -0.043 | 0 | 0.08 | 0.01 | 0.11 | 0.34 | 0.12 | 0.06 |
| 5 | 7955677 | rs7719963 | -0.777 | 1.676 | 0.775 | 0.905 | 0.082 | 0.115 | 0.115 | 0 | 0.14 | 0.42 | 0.35 | 0.2 | 0.13 | 0.14 |
| 5 | 7958031 | rs331413 | -0.438 | 1.008 | 0.11 | 0.77 | 0.017 | 0 | 0 | 0 | 0.06 | 0.08 | 0.1 | 0.1 | 0.11 | 0.03 |
| 5 | 7967228 | rs9313211 | -1.504 | 0.263 | 0.288 | 0.775 | -0.086 | 0.072 | 0.072 | 0.646 | 0.21 | 0.54 | 0.36 | 0.25 | 0.07 | 0.25 |
| 5 | 7970898 | rs7719216 | -1.455 | 0.459 | 0.232 | 0.743 | -0.045 | 0.029 | 0.029 | 0 | 0.25 | 0.54 | 0.39 | 0.26 | 0.14 | 0.26 |
| 5 | 7980403 | rs11742403 | -0.857 | 0.415 | -0.172 | 0.601 | 0.004 | 0.011 | 0.011 | 0 | 0.08 | 0.05 | 0.05 | 0.03 | 0.06 | 0.11 |
| 5 | 7984232 | rs1482277 | 0.325 | 0.303 | -0.104 | 0.436 | 0.031 | -0.012 | -0.012 | 0 | 0.43 | 0.21 | 0.32 | 0.45 | 0.35 | 0.46 |
| 5 | 7984374 | rs1503322 | 0.36 | 0.28 | 0.079 | 0.249 | -0.002 | -0.008 | -0.008 | 0 | 0.71 | 0.36 | 0.76 | 0.82 | 0.74 | 0.78 |
| 5 | 7985855 | rs2654520 | 0.306 | 0.049 | -0.253 | 0.087 | -0.018 | -0.009 | -0.009 | 0 | 0.65 | 0.51 | 0.76 | 0.72 | 0.74 | 0.54 |
| 5 | 7986155 | rs11134268 | -0.124 | -0.261 | -0.204 | 0.127 | -0.061 | 0.098 | 0.098 | 0 | 0.8 | 0.52 | 0.84 | 0.72 | 0.91 | 0.62 |
| 5 | 7992135 | rs1503317 | 0.802 | -0.006 | -0.834 | 0.031 | 0.003 | -0.04 | -0.04 | 1.468 | 0.32 | 0.08 | 0.27 | 0.35 | 0.29 | 0.18 |
| 5 | 7994914 | rs11134269 | -0.317 | 0.132 | -0.402 | -0.061 | -0.002 | -0.025 | -0.025 | 0 | 0.07 | 0.14 | 0.06 | 0.01 | 0.05 | 0.06 |
| 5 | 7995509 | rs6866169 | -0.237 | -0.004 | -0.401 | 0.04 | -0.007 | 0 | 0 | 1.002 | 0.47 | 0.48 | 0.34 | 0.51 | 0.32 | 0.41 |
| 5 | 7997545 | rs7702491 | -1.228 | 0.242 | -0.519 | -0.037 | 0.001 | 0.023 | 0.023 | 0 | 0.65 | 0.52 | 0.52 | 0.58 | 0.62 | 0.5 |
| 5 | 8004748 | rs4701802 | -0.401 | 0.224 | -0.5 | -0.002 | 0.003 | 0.009 | 0.009 | 0 | 0.55 | 0.38 | 0.45 | 0.38 | 0.56 | 0.35 |
| 5 | 8006656 | rs2966953 | -0.518 | -0.142 | -0.438 | 0.229 | 0.002 | 0 | 0 | 0 | 0.49 | 0.49 | 0.4 | 0.58 | 0.36 | 0.42 |
| 5 | 8013445 | rs2123637 | 0.522 | -0.333 | -0.831 | 0.269 | 0.026 | -0.034 | -0.034 | 0 | 0.3 | 0.1 | 0.24 | 0.37 | 0.25 | 0.21 |
| 5 | 8015429 | rs12520611 | 1.075 | -0.361 | -0.322 | 0.411 | 0.008 | 0.079 | 0.079 | 0 | 0.12 | 0.33 | 0.15 | 0.17 | 0.09 | 0.14 |
| 5 | 8031332 | rs13187219 | 1.139 | -0.285 | -0.454 | 0.123 | -0.014 | -0.063 | -0.063 | 0.716 | 0.31 | 0.11 | 0.37 | 0.39 | 0.33 | 0.49 |
| 5 | 8036385 | rs1377489 | 1.938 | -0.407 | -0.597 | 0.166 | -0.102 | -0.042 | -0.042 | 0 | 0.23 | 0.02 | 0.33 | 0.22 | 0.33 | 0.34 |
| 5 | 8036594 | rs1377490 | 0.367 | -0.364 | -0.438 | -0.137 | -0.004 | 0.031 | 0.031 | 0.05 | 0.63 | 0.29 | 0.68 | 0.83 | 0.66 | 0.75 |

| | | | | | | | | | | | | | | | | |
|---|---|---|---|---|---|---|---|---|---|---|---|---|---|---|---|---|
| 5 | 8047445 | rs6871063 | 0.651 | -0.347 | -0.412 | -0.112 | 0.005 | -0.016 | -0.016 | 0 | 0.33 | 0.22 | 0.32 | 0.45 | 0.32 | 0.26 |
| 5 | 8047543 | rs876712 | 1.359 | -0.375 | -0.199 | -0.135 | -0.014 | 0.05 | 0.05 | 0 | 0.13 | 0.31 | 0.16 | 0.16 | 0.07 | 0.11 |
| 5 | 8049644 | rs6896973 | -1.203 | -0.291 | -0.205 | -0.159 | -0.007 | -0.014 | -0.014 | 1.814 | 0.84 | 0.86 | 0.88 | 1 | 0.77 | 0.92 |
| 5 | 8052369 | rs13167372 | 1.338 | -0.315 | -0.211 | -0.098 | -0.005 | -0.038 | -0.038 | 0 | 0.31 | 0.16 | 0.37 | 0.38 | 0.34 | 0.49 |
| 5 | 8055261 | rs2924471 | -0.784 | 0.006 | 0.081 | -0.44 | 0.019 | -0.012 | -0.012 | 0.036 | 0.1 | 0.26 | 0.13 | 0.09 | 0.1 | 0.11 |
| 5 | 8055319 | rs11959696 | 0.737 | -0.121 | -0.33 | -0.41 | 0.014 | -0.027 | -0.027 | 0.017 | 0.29 | 0.14 | 0.24 | 0.37 | 0.27 | 0.22 |
| 5 | 8055558 | rs3943482 | 0.482 | -0.241 | -0.092 | -0.084 | -0.002 | -0.004 | -0.004 | 0 | 0.45 | 0.49 | 0.54 | 0.54 | 0.44 | 0.6 |
| 5 | 8055690 | rs2938245 | -0.177 | 0.031 | -0.756 | 0.065 | -0.026 | 0.039 | 0.039 | 0 | 0.55 | 0.75 | 0.65 | 0.63 | 0.51 | 0.7 |
| 5 | 8060320 | rs16879792 | -0.112 | -0.292 | -0.558 | 0.515 | 0.001 | 0.037 | 0.037 | 0 | 0.16 | 0.21 | 0.21 | 0.35 | 0.16 | 0.11 |
| 5 | 8062181 | rs964017 | -0.623 | -0.257 | -0.932 | 0.061 | 0.001 | -0.002 | -0.002 | 0 | 0.34 | 0.32 | 0.3 | 0.32 | 0.39 | 0.23 |
| 5 | 8062594 | rs10475403 | 1.057 | -0.69 | -1.172 | 0.114 | -0.03 | 0.065 | 0.065 | 0 | 0.42 | 0.23 | 0.29 | 0.29 | 0.5 | 0.53 |
| 5 | 8062672 | rs6886541 | -1.072 | 0.333 | -0.912 | 0.41 | 0 | -0.001 | -0.001 | 0 | 0.45 | 0.41 | 0.31 | 0.45 | 0.44 | 0.43 |
| 5 | 8062748 | rs6860289 | -0.809 | 0.26 | -0.646 | 0.77 | -0.009 | 0.024 | 0.024 | 0 | 0.2 | 0.24 | 0.16 | 0.32 | 0.17 | 0.18 |
| 5 | 8067973 | rs11744506 | -0.555 | 0.778 | -1.01 | 0.179 | -0.005 | 0.011 | 0.011 | 0.017 | 0.4 | 0.34 | 0.42 | 0.68 | 0.47 | 0.5 |
| 5 | 8073071 | rs2961986 | 0.51 | 0.883 | -1.042 | 0.131 | 0.005 | 0 | 0 | 0.13 | 0.43 | 0.36 | 0.31 | 0.43 | 0.36 | 0.49 |
| 5 | 8073378 | rs1453363 | 0.402 | 1.005 | -1.137 | 0.421 | -0.005 | -0.019 | -0.019 | 0.091 | 0.76 | 0.78 | 0.56 | 0.45 | 0.76 | 0.7 |
| 5 | 8073928 | rs4702523 | -0.952 | 2.002 | -1.115 | -0.453 | -0.01 | -0.009 | -0.009 | 0 | 0.29 | 0.27 | 0.24 | 0.06 | 0.39 | 0.29 |
| 5 | 8074665 | rs10043550 | -0.964 | 2.091 | -1.283 | -0.305 | -0.024 | 0.026 | 0.026 | 0.031 | 0.15 | 0.1 | 0.15 | 0.05 | 0.19 | 0.11 |
| 5 | 8077509 | rs2924466 | -0.219 | 1.433 | -0.975 | -0.144 | 0 | 0 | 0 | 0 | 0.52 | 0.49 | 0.56 | 0.49 | 0.5 | 0.46 |
| 5 | 8083187 | rs11134282 | -0.212 | 1.042 | -0.268 | -0.452 | -0.006 | -0.046 | -0.046 | 0 | 0.64 | 0.44 | 0.67 | 0.9 | 0.63 | 0.69 |
| 5 | 8090525 | rs4702526 | 0.369 | 0.93 | -0.774 | -0.507 | -0.05 | 0.084 | 0.084 | 0 | 0.28 | 0.06 | 0.27 | 0.1 | 0.36 | 0.31 |
| 5 | 8097721 | rs13190226 | -0.066 | 0.815 | -0.592 | -0.429 | -0.012 | 0.037 | 0.037 | 0.341 | 0.41 | 0.26 | 0.27 | 0.25 | 0.39 | 0.34 |
| 5 | 8100893 | rs7716010 | 1.151 | 0.622 | -0.502 | -0.35 | -0.032 | -0.064 | -0.064 | 0 | 0.25 | 0.13 | 0.4 | 0.47 | 0.29 | 0.31 |
| 5 | 8105591 | rs7725153 | -0.394 | 0.417 | 0.174 | -0.407 | 0.004 | 0.034 | 0.034 | 0 | 0.67 | 0.56 | 0.52 | 0.41 | 0.67 | 0.61 |
| 5 | 8113536 | rs6860481 | 0.233 | 0.079 | 0.216 | -0.545 | -0.024 | -0.092 | -0.092 | 0 | 0.27 | 0.45 | 0.22 | 0.08 | 0.26 | 0.26 |
| 5 | 8115307 | rs4702528 | 1.282 | 0.228 | -0.194 | -0.559 | -0.037 | -0.066 | -0.066 | 1.507 | 0.27 | 0.13 | 0.29 | 0.48 | 0.31 | 0.29 |
| 5 | 8121656 | rs1506105 | 0.048 | -0.231 | -0.025 | -0.564 | -0.002 | -0.002 | -0.002 | 0 | 0.53 | 0.49 | 0.62 | 0.58 | 0.57 | 0.54 |
| 5 | 8123555 | rs12521059 | 0.858 | 0.026 | -0.177 | -0.803 | 0.019 | 0.039 | 0.039 | 1.033 | 0.22 | 0.09 | 0.27 | 0.09 | 0.23 | 0.25 |
| 5 | 8123994 | rs10057537 | -0.016 | 0.18 | -0.119 | -0.536 | 0 | -0.005 | -0.005 | 0 | 0.63 | 0.65 | 0.75 | 0.87 | 0.69 | 0.65 |
| 5 | 8130010 | rs13157534 | 0.212 | 0.306 | -0.703 | -0.487 | 0.009 | 0.058 | 0.058 | 0 | 0.31 | 0.16 | 0.19 | 0.13 | 0.24 | 0.33 |
| 5 | 8130362 | rs6555523 | 0.706 | 0.324 | -0.794 | -0.451 | -0.003 | 0.006 | 0.006 | 0 | 0.08 | 0.1 | 0.1 | 0.19 | 0.11 | 0.04 |
| 5 | 8131880 | rs13172069 | 0.018 | 0.042 | -0.807 | -1.059 | 0.009 | 0.062 | 0.062 | 0 | 0.31 | 0.15 | 0.19 | 0.13 | 0.25 | 0.33 |
| 5 | 8135922 | rs6876162 | -0.43 | 0.139 | -0.349 | -1.082 | 0.005 | -0.01 | -0.01 | 0 | 0.49 | 0.5 | 0.45 | 0.14 | 0.43 | 0.47 |
| 5 | 8140979 | rs1506127 | 0.604 | 0.439 | -0.649 | -0.699 | -0.032 | 0.093 | 0.093 | 0.914 | 0.48 | 0.17 | 0.57 | 0.71 | 0.49 | 0.57 |
| 5 | 8145014 | rs13171587 | 0.441 | 0.201 | -0.501 | -0.997 | -0.013 | 0.029 | 0.029 | 0 | 0.21 | 0.15 | 0.24 | 0.11 | 0.25 | 0.16 |
| 5 | 8146371 | rs336138 | 0.469 | 0.148 | -0.437 | -1.088 | 0.004 | 0.045 | 0.045 | 0.096 | 0.19 | 0.08 | 0.09 | 0.08 | 0.16 | 0.2 |
| 5 | 8149540 | rs336164 | -1.157 | 0.099 | 0.504 | -1.101 | -0.002 | -0.02 | -0.02 | 0.022 | 0.45 | 0.55 | 0.37 | 0.19 | 0.43 | 0.37 |
| 5 | 8151217 | rs7723051 | 0.437 | 0.118 | 0.317 | -1.076 | 0.005 | 0.012 | 0.012 | 2.95 | 0.7 | 0.3 | 0.8 | 0.81 | 0.74 | 0.74 |
| 5 | 8151979 | rs10512943 | 0.51 | 0.231 | 0.191 | -0.974 | -0.022 | 0.04 | 0.04 | 0 | 0.2 | 0.11 | 0.23 | 0.1 | 0.24 | 0.16 |
| 5 | 8158804 | rs170936 | 0.346 | -0.097 | -0.375 | -0.875 | 0.018 | 0 | 0 | 0 | 0.69 | 0.22 | 0.68 | 0.74 | 0.72 | 0.79 |
| 5 | 8165111 | rs6555524 | 1.505 | 0.347 | -0.692 | -1.032 | -0.058 | 0.002 | 0.002 | 0 | 0.43 | 0.1 | 0.48 | 0.67 | 0.43 | 0.48 |
| 5 | 8172103 | rs163115 | -0.545 | 0.279 | -0.098 | -1.065 | -0.038 | -0.066 | -0.066 | 0 | 0.77 | 0.54 | 0.74 | 0.89 | 0.82 | 0.82 |
| 5 | 8173436 | rs336178 | 0.567 | 0.362 | -0.543 | -1.079 | -0.001 | 0.071 | 0.071 | 0 | 0.2 | 0.1 | 0.11 | 0.05 | 0.17 | 0.2 |
| 5 | 8174188 | rs13170530 | -0.096 | 0.71 | -0.273 | -0.721 | 0.047 | 0.037 | 0.037 | 0 | 0.14 | 0.26 | 0.14 | 0.17 | 0.23 | 0.14 |
| 5 | 8175049 | rs901893 | 0.183 | 0.553 | -0.122 | -0.776 | 0.02 | 0.041 | 0.041 | 0 | 0.57 | 0.35 | 0.62 | 0.84 | 0.65 | 0.62 |
| 5 | 8175968 | rs730757 | 0.189 | 0.558 | -0.242 | -0.774 | 0.027 | 0.063 | 0.063 | 0 | 0.57 | 0.33 | 0.62 | 0.84 | 0.65 | 0.62 |
| 5 | 8179705 | rs906091 | 0.635 | 0.214 | -1.011 | -0.102 | -0.081 | -0.12 | -0.12 | 2.557 | 0.34 | 0.09 | 0.42 | 0.57 | 0.39 | 0.38 |
| 5 | 8180407 | rs7701145 | 0.371 | 0.17 | -0.494 | -0.247 | -0.004 | -0.039 | -0.039 | 0 | 0.08 | 0.15 | 0.06 | 0 | 0.09 | 0.03 |
| 5 | 8188341 | rs1823809 | 0.608 | 0.256 | -0.75 | 0.2 | -0.025 | -0.057 | -0.057 | 0.007 | 0.36 | 0.16 | 0.42 | 0.43 | 0.35 | 0.35 |
| 5 | 8188395 | rs7702422 | 0.132 | 0.146 | -0.382 | 0.134 | 0.011 | 0.09 | 0.09 | 0 | 0.2 | 0.4 | 0.23 | 0.3 | 0.22 | 0.21 |
| 5 | 8189299 | rs12521485 | 0.989 | 0.116 | -0.373 | 0.135 | -0.028 | -0.071 | -0.071 | 4.23 | 0.35 | 0.12 | 0.41 | 0.43 | 0.34 | 0.34 |
| 5 | 8189392 | rs7707765 | 0.257 | -0.006 | -0.046 | -0.142 | -0.002 | -0.022 | -0.022 | 0 | 0.56 | 0.52 | 0.64 | 0.72 | 0.57 | 0.55 |
| 5 | 8205612 | rs7702825 | -0.78 | -0.364 | 0.777 | -0.158 | -0.021 | -0.076 | -0.076 | 0 | 0.13 | 0.24 | 0.07 | 0 | 0.09 | 0.05 |
| 5 | 8211892 | rs6867552 | -0.209 | -0.377 | 0.569 | 0.56 | -0.002 | 0.039 | 0.039 | 0.04 | 0.83 | 0.84 | 0.71 | 0.51 | 0.89 | 0.64 |
| 5 | 8214282 | rs11738766 | 0.44 | -0.809 | 1.09 | 0.546 | -0.021 | 0.102 | 0.102 | 0 | 0.78 | 0.68 | 0.65 | 0.45 | 0.88 | 0.57 |
| 5 | 8226339 | rs6892570 | 1.434 | -1.12 | 1.638 | 0.238 | 0.012 | -0.013 | -0.013 | 1.411 | 0.64 | 0.39 | 0.56 | 0.45 | 0.77 | 0.51 |
| 5 | 8228590 | rs6875427 | -0.313 | -1.148 | 2.186 | 0.427 | -0.004 | -0.029 | -0.029 | 0 | 0.13 | 0.32 | 0.12 | 0.06 | 0.11 | 0.05 |
| 5 | 8231744 | rs10512947 | -1.122 | -1.171 | 2.186 | 0.422 | -0.031 | -0.015 | -0.015 | 3.625 | 0.17 | 0.38 | 0.16 | 0.12 | 0.12 | 0.11 |
| 5 | 8233351 | rs10512948 | 0.257 | -1.11 | 2.015 | 0.25 | -0.011 | -0.045 | -0.045 | 0 | 0.19 | 0.08 | 0.27 | 0.43 | 0.17 | 0.37 |
| 5 | 8239357 | rs10462854 | 0.395 | -1.114 | 1.873 | 0.248 | -0.046 | -0.099 | -0.099 | 1.22 | 0.18 | 0.02 | 0.27 | 0.44 | 0.17 | 0.38 |
| 5 | 8240153 | rs1857863 | 1.144 | -1.102 | 2.079 | 0.241 | -0.009 | 0.021 | 0.021 | 0 | 0.66 | 0.53 | 0.6 | 0.5 | 0.71 | 0.56 |
| 5 | 8243414 | rs4639181 | -0.121 | -1.002 | 1.884 | 0.157 | -0.005 | -0.074 | -0.074 | 0.115 | 0.12 | 0.26 | 0.08 | 0 | 0.1 | 0.04 |
| 5 | 8244864 | rs1906573 | 1.245 | -1.046 | 1.87 | 0.223 | -0.015 | 0.009 | 0.009 | 0 | 0.63 | 0.52 | 0.59 | 0.5 | 0.72 | 0.51 |
| 5 | 8258881 | rs2171567 | -0.974 | -1.289 | 1.193 | 0.046 | -0.005 | 0.006 | 0.006 | 0 | 0.72 | 0.7 | 0.76 | 0.76 | 0.75 | 0.78 |
| 5 | 8260550 | rs12514960 | 1.097 | -1.847 | 0.555 | -0.263 | -0.126 | 0.112 | 0.112 | 2.1 | 0.35 | 0.09 | 0.29 | 0.16 | 0.54 | 0.3 |
| 5 | 8261891 | rs4235590 | 1.306 | -2.019 | 0.541 | -0.231 | -0.146 | 0.138 | 0.138 | 0.911 | 0.32 | 0.08 | 0.28 | 0.1 | 0.51 | 0.27 |
| 5 | 8265008 | rs7444241 | -0.864 | -2.03 | 0.906 | -0.091 | -0.022 | 0.014 | 0.014 | 0 | 0.68 | 0.58 | 0.71 | 0.69 | 0.76 | 0.72 |
| 5 | 8270536 | rs906087 | 0.935 | -1.944 | 0.932 | -0.128 | -0.005 | -0.038 | -0.038 | 0 | 0.07 | 0.12 | 0.03 | 0 | 0.05 | 0.03 |
| 5 | 8276583 | rs11948974 | 2.25 | -2.087 | 0.781 | 0.004 | -0.029 | 0.054 | 0.054 | 0 | 0.17 | 0.05 | 0.12 | 0.06 | 0.19 | 0.12 |

| | | | | | | | | | | | | | | | | |
|---|---|---|---|---|---|---|---|---|---|---|---|---|---|---|---|---|
| 5 | 8278166 | rs4640759 | -1.18 | -2.069 | 1.113 | 0.115 | -0.002 | 0.002 | 0.002 | 0 | 0.73 | 0.77 | 0.77 | 0.75 | 0.76 | 0.79 |
| 5 | 8278270 | rs4590157 | -0.118 | -2.172 | 1.285 | 0.116 | -0.002 | 0.008 | 0.008 | 0.243 | 0.48 | 0.6 | 0.46 | 0.33 | 0.55 | 0.42 |
| 5 | 8291230 | rs6887466 | 2.617 | -1.669 | 0.63 | -0.218 | -0.091 | 0.062 | 0.062 | 1.447 | 0.16 | 0.01 | 0.13 | 0.04 | 0.27 | 0.13 |
| 5 | 8292668 | rs16880316 | 0.334 | -2.079 | 1.459 | 0.145 | -0.018 | 0.103 | 0.103 | 0.033 | 0.07 | 0.23 | 0.12 | 0.12 | 0.03 | 0.07 |
| 5 | 8295533 | rs10056042 | 0.313 | -2.148 | 1.403 | 0.133 | -0.026 | 0.043 | 0.043 | 0 | 0.33 | 0.43 | 0.34 | 0.37 | 0.27 | 0.28 |
| 5 | 8302197 | rs11134307 | -0.274 | -2.16 | 1.416 | 0.092 | -0.001 | -0.001 | -0.001 | 0.194 | 0.54 | 0.39 | 0.56 | 0.74 | 0.45 | 0.64 |
| 5 | 8303435 | rs6555540 | 0.884 | -2.135 | 1.326 | 0.087 | -0.037 | -0.065 | -0.065 | 0 | 0.24 | 0.05 | 0.3 | 0.43 | 0.21 | 0.36 |
| 5 | 8303911 | rs10512951 | -1.404 | -1.808 | 1.611 | 0.039 | -0.052 | 0.08 | 0.08 | 2.351 | 0.11 | 0.39 | 0.15 | 0.11 | 0.04 | 0.08 |
| 5 | 8319421 | rs2129949 | -0.646 | -1.673 | 1.215 | 0.093 | -0.021 | 0.038 | 0.038 | 0.263 | 0.61 | 0.63 | 0.66 | 0.79 | 0.47 | 0.72 |
| 5 | 8321133 | rs12109465 | 1.322 | -1.639 | 1.226 | 0.128 | -0.01 | 0.053 | 0.053 | 0 | 0.05 | 0.13 | 0.09 | 0.05 | 0.02 | 0.07 |
| 5 | 8322525 | rs17211011 | -0.737 | -1.599 | 0.979 | 0.112 | -0.034 | 0.056 | 0.056 | 0.166 | 0.62 | 0.69 | 0.66 | 0.79 | 0.48 | 0.72 |
| 5 | 8323964 | rs4302555 | -1.034 | -1.54 | 0.796 | 0.115 | -0.06 | 0.092 | 0.092 | 0.137 | 0.65 | 0.78 | 0.7 | 0.85 | 0.48 | 0.73 |
| 5 | 8325714 | rs4492083 | -0.419 | -1.537 | 0.854 | 0.145 | 0 | -0.006 | -0.006 | 0 | 0.57 | 0.57 | 0.6 | 0.8 | 0.45 | 0.66 |
| 5 | 8328783 | rs925203 | -0.157 | -1.29 | 0.716 | 0.436 | 0.006 | -0.011 | -0.011 | 0 | 0.31 | 0.27 | 0.35 | 0.49 | 0.22 | 0.46 |
| 5 | 8330346 | rs7724111 | -0.117 | -1.208 | 0.146 | 0.48 | -0.019 | 0.044 | 0.044 | 0 | 0.07 | 0.22 | 0.1 | 0.05 | 0.03 | 0.07 |
| 5 | 8330492 | rs7710227 | -0.545 | -1.013 | -0.022 | 0.453 | -0.007 | 0.007 | 0.007 | 0 | 0.37 | 0.42 | 0.45 | 0.54 | 0.24 | 0.53 |
| 5 | 8332937 | rs6895861 | 0.646 | -0.801 | -0.303 | 0.447 | -0.012 | 0.015 | 0.015 | 0 | 0.29 | 0.26 | 0.24 | 0.32 | 0.21 | 0.23 |
| 5 | 8337146 | rs2136170 | 0.874 | -0.799 | -0.2 | 0.438 | 0.016 | 0.004 | 0.004 | 0.146 | 0.53 | 0.28 | 0.46 | 0.36 | 0.62 | 0.42 |
| 5 | 8349231 | rs2964129 | -0.133 | -1.122 | -0.369 | 0.459 | -0.096 | -0.069 | -0.069 | 0 | 0.28 | 0.08 | 0.38 | 0.43 | 0.34 | 0.44 |
| 5 | 8349784 | rs2940556 | -1.13 | -1.014 | -0.101 | 0.49 | -0.002 | 0 | 0 | 0 | 0.43 | 0.63 | 0.46 | 0.5 | 0.38 | 0.51 |
| 5 | 8371633 | rs170991 | -0.867 | -0.968 | -0.019 | 0.502 | -0.001 | 0.006 | 0.006 | 0 | 0.4 | 0.61 | 0.45 | 0.5 | 0.37 | 0.5 |
| 5 | 8384247 | rs340677 | -1.45 | -1.028 | 0.173 | 0.757 | 0.003 | 0.003 | 0.003 | 0.464 | 0.46 | 0.62 | 0.48 | 0.66 | 0.38 | 0.58 |
| 5 | 8384936 | rs1448235 | -1.167 | -1.001 | 0.08 | 0.935 | -0.026 | 0.026 | 0.026 | 0.033 | 0.09 | 0.21 | 0.05 | 0.2 | 0.02 | 0.09 |
| 5 | 8385180 | rs340680 | 0.01 | -0.893 | -0.5 | 0.901 | -0.024 | -0.058 | -0.058 | 1.313 | 0.16 | 0.06 | 0.24 | 0.37 | 0.18 | 0.27 |
| 5 | 8393159 | rs12653627 | 1.572 | -0.862 | -0.469 | 0.614 | 0.066 | -0.056 | -0.056 | 0 | 0.6 | 0.15 | 0.58 | 0.42 | 0.72 | 0.58 |
| 5 | 8396885 | rs16880510 | -0.489 | -0.639 | 0.235 | 0.51 | -0.016 | -0.001 | -0.001 | 0 | 0.08 | 0.29 | 0.06 | 0.02 | 0.03 | 0.05 |
| 5 | 8404594 | rs11743533 | 0.593 | -0.444 | -0.493 | 0.345 | -0.054 | -0.009 | -0.009 | 0 | 0.11 | 0.01 | 0.12 | 0.09 | 0.14 | 0.19 |
| 5 | 8404645 | rs11743538 | -0.174 | -0.454 | -0.442 | 0.341 | -0.022 | 0.028 | 0.028 | 0 | 0.16 | 0.07 | 0.15 | 0.09 | 0.16 | 0.22 |
| 5 | 8407305 | rs17279186 | -0.765 | -0.564 | -0.454 | 0.383 | 0.002 | -0.017 | -0.017 | 0 | 0.21 | 0.11 | 0.17 | 0.23 | 0.17 | 0.31 |
| 5 | 8413356 | rs2457108 | -1.295 | -0.17 | 0.111 | -0.065 | -0.002 | 0.044 | 0.044 | 0 | 0.3 | 0.54 | 0.43 | 0.63 | 0.28 | 0.38 |
| 5 | 8417115 | rs448636 | -0.384 | -0.102 | 0.037 | -0.068 | 0.004 | 0.027 | 0.027 | 0 | 0.18 | 0.21 | 0.34 | 0.55 | 0.23 | 0.23 |
| 5 | 8423142 | rs340700 | -0.572 | -0.112 | 0.049 | -0.065 | 0 | 0.01 | 0.01 | 0 | 0.2 | 0.21 | 0.35 | 0.55 | 0.26 | 0.26 |
| 5 | 8424283 | rs4702551 | 1.481 | -0.265 | -0.533 | -0.767 | -0.116 | 0.128 | 0.128 | 1.827 | 0.24 | 0.02 | 0.24 | 0.04 | 0.34 | 0.18 |
| 5 | 8429773 | rs7717779 | -0.954 | -0.755 | 0.815 | 0.007 | -0.074 | -0.068 | -0.068 | 0 | 0.15 | 0.47 | 0.12 | 0.07 | 0.05 | 0.14 |
| 5 | 8436209 | rs340671 | -0.104 | -0.477 | 0.211 | 0.249 | 0.005 | -0.017 | -0.017 | 0.121 | 0.13 | 0.1 | 0.26 | 0.56 | 0.11 | 0.19 |
| 5 | 8438752 | rs10080066 | -0.02 | -0.415 | 0.617 | -0.773 | -0.017 | -0.029 | -0.029 | 0 | 0.05 | 0.15 | 0.03 | 0 | 0.03 | 0.01 |
| 5 | 8439670 | rs12187557 | 1.477 | -0.407 | 0.541 | -0.902 | 0.008 | 0.006 | 0.006 | 0.434 | 0.66 | 0.32 | 0.72 | 0.93 | 0.73 | 0.64 |
| 5 | 8440728 | rs6555545 | -1.481 | -0.293 | 0.405 | -0.832 | -0.004 | -0.055 | -0.055 | 0 | 0.33 | 0.59 | 0.26 | 0.07 | 0.27 | 0.35 |
| 5 | 8442087 | rs16880614 | -0.817 | -0.569 | 0.144 | -0.68 | -0.035 | -0.097 | -0.097 | 0 | 0.23 | 0.38 | 0.18 | 0.07 | 0.14 | 0.32 |
| 5 | 8446336 | rs6555548 | 0.64 | -0.485 | -0.364 | -0.356 | -0.021 | -0.018 | -0.018 | 0 | 0.62 | 0.46 | 0.44 | 0.73 | 0.7 | 0.65 |
| 5 | 8447297 | rs2166870 | 0.278 | -0.548 | -0.5 | -1.066 | -0.009 | 0.008 | 0.008 | 0.602 | 0.37 | 0.35 | 0.32 | 0.74 | 0.44 | 0.39 |
| 5 | 8447327 | rs10475431 | -0.162 | -1.047 | -0.784 | -1.405 | 0.008 | 0 | 0 | 0 | 0.1 | 0.1 | 0.04 | 0.01 | 0.06 | 0.12 |
| 5 | 8451757 | rs6860832 | 0.496 | -1.025 | -0.306 | -1.457 | 0.009 | -0.025 | -0.025 | 0 | 0.41 | 0.53 | 0.4 | 0.75 | 0.5 | 0.39 |
| 5 | 8456731 | rs16880638 | 0.667 | -0.881 | -1.184 | -1.632 | -0.044 | 0.048 | 0.048 | 0.377 | 0.13 | 0.03 | 0.08 | 0.05 | 0.18 | 0.18 |
| 5 | 8459533 | rs9313239 | -0.274 | -1.274 | -0.993 | -1.634 | 0.002 | 0.123 | 0.123 | 1.581 | 0.47 | 0.26 | 0.55 | 0.24 | 0.45 | 0.44 |
| 5 | 8465020 | rs16880661 | 0.767 | -1.16 | -0.983 | -1.423 | -0.011 | 0.012 | 0.012 | 0 | 0.07 | 0.04 | 0.06 | 0.01 | 0.1 | 0.09 |
| 5 | 8466223 | rs11954087 | 0.052 | -2.045 | -1.17 | -1.468 | 0.081 | 0.022 | 0.022 | 1.337 | 0.26 | 0.15 | 0.33 | 0.2 | 0.11 | 0.18 |
| 5 | 8467900 | rs341893 | -0.306 | -1.493 | -0.227 | -0.63 | -0.097 | 0.149 | 0.149 | 0.059 | 0.81 | 0.58 | 0.65 | 0.64 | 0.94 | 0.87 |
| 5 | 8477908 | rs168188 | 0.672 | -1.132 | -1.056 | -0.822 | -0.046 | -0.038 | -0.038 | 0 | 0.3 | 0.18 | 0.32 | 0.59 | 0.38 | 0.32 |
| 5 | 8484044 | rs12657402 | 0.186 | -0.888 | -0.673 | 0.194 | 0.021 | -0.004 | -0.004 | 0.749 | 0.14 | 0.11 | 0.17 | 0.16 | 0.05 | 0.1 |
| 5 | 8493663 | rs13177016 | -0.619 | 0.758 | -1.065 | -0.488 | -0.047 | 0.051 | 0.051 | 0.178 | 0.19 | 0.03 | 0.11 | 0.07 | 0.22 | 0.25 |
| 5 | 8494868 | rs173134 | 0.731 | 0.68 | -0.313 | -0.735 | -0.009 | -0.112 | -0.112 | 0 | 0.76 | 0.44 | 0.81 | 0.94 | 0.78 | 0.82 |
| 5 | 8498184 | rs11750041 | -0.178 | 1.988 | -0.743 | -0.527 | -0.087 | 0.022 | 0.022 | 0 | 0.2 | 0.08 | 0.42 | 0.12 | 0.45 | 0.29 |
| 5 | 8499805 | rs1553550 | -0.765 | 1.562 | -0.57 | 0.066 | -0.031 | -0.024 | -0.024 | 0 | 0.28 | 0.19 | 0.45 | 0.35 | 0.5 | 0.36 |
| 5 | 8504245 | rs16880714 | -0.068 | 1.531 | -0.128 | 0.173 | -0.007 | 0.021 | 0.021 | 0 | 0.08 | 0.12 | 0.03 | 0.14 | 0.04 | 0.07 |
| 5 | 8504567 | rs6892211 | 0.846 | 0.403 | -0.644 | -0.273 | 0.025 | 0.06 | 0.06 | 0 | 0.19 | 0.08 | 0.1 | 0.09 | 0.15 | 0.16 |
| 5 | 8518835 | rs17216061 | 0.523 | 0.528 | -0.556 | -0.226 | -0.067 | -0.015 | -0.015 | 5.613 | 0.2 | 0.11 | 0.42 | 0.21 | 0.45 | 0.3 |
| 5 | 8522021 | rs415234 | -1.231 | 0.095 | 0.167 | -0.439 | 0.07 | -0.005 | -0.005 | 0.013 | 0.57 | 0.76 | 0.63 | 0.44 | 0.69 | 0.54 |
| 5 | 8522181 | rs11948129 | 0.189 | 0.721 | 0.35 | -0.179 | -0.035 | -0.003 | -0.003 | 0 | 0.22 | 0.16 | 0.43 | 0.21 | 0.45 | 0.3 |
| 5 | 8527859 | rs444121 | -0.836 | 1.441 | 0.788 | -0.068 | 0.001 | 0.09 | 0.09 | 0 | 0.19 | 0.37 | 0.29 | 0.34 | 0.15 | 0.28 |
| 5 | 8528431 | rs11949420 | 1.161 | 0.45 | 0.398 | 0.193 | 0.063 | -0.032 | -0.032 | 0 | 0.34 | 0.19 | 0.15 | 0.39 | 0.22 | 0.42 |
| 5 | 8529586 | rs402359 | -0.142 | 0.445 | 1.062 | 0.228 | -0.008 | 0.017 | 0.017 | 0 | 0.12 | 0.25 | 0.06 | 0.14 | 0.08 | 0.08 |
| 5 | 8533359 | rs1613720 | 1.216 | 0.361 | 0.978 | 0.2 | 0.068 | -0.026 | -0.026 | 0.095 | 0.36 | 0.22 | 0.15 | 0.4 | 0.22 | 0.43 |
| 5 | 8538868 | rs1700569 | -0.175 | 0.622 | 1.369 | 0.047 | 0.003 | 0.001 | 0.001 | 0.306 | 0.4 | 0.37 | 0.39 | 0.4 | 0.33 | 0.28 |
| 5 | 8540496 | rs1505016 | -0.351 | 0.721 | 1.363 | 0.051 | 0.024 | 0.001 | 0.001 | 0 | 0.78 | 0.76 | 0.55 | 0.79 | 0.55 | 0.71 |
| 5 | 8540538 | rs1666797 | 1.129 | 0.562 | 1.052 | 0.046 | 0.009 | -0.002 | -0.002 | 0 | 0.38 | 0.37 | 0.16 | 0.4 | 0.22 | 0.43 |
| 5 | 8540995 | rs422977 | 0.498 | 0.637 | 0.552 | 0.128 | 0.035 | 0.019 | 0.019 | 1.554 | 0.28 | 0.14 | 0.32 | 0.25 | 0.25 | 0.2 |
| 5 | 8543442 | rs423297 | 0.6 | 0.391 | 1.115 | 0.105 | -0.011 | 0 | 0 | 0 | 0.44 | 0.56 | 0.24 | 0.41 | 0.26 | 0.48 |
| 5 | 8544489 | rs418262 | 0.761 | 0.322 | 1.129 | 0.554 | -0.019 | -0.011 | -0.011 | 0 | 0.42 | 0.56 | 0.23 | 0.31 | 0.25 | 0.47 |

| | | | | | | | | | | | | | | | | |
|---|---|---|---|---|---|---|---|---|---|---|---|---|---|---|---|---|
| 5 | 8547779 | rs9313241 | 0.686 | 1.257 | 1.732 | 0.675 | 0.095 | 0.017 | 0.017 | 0 | 0.84 | 0.61 | 0.84 | 0.85 | 0.77 | 0.76 |
| 5 | 8549411 | rs9313243 | -0.235 | 1.27 | 1.955 | 0.621 | -0.004 | -0.004 | -0.004 | 0 | 0.47 | 0.33 | 0.59 | 0.55 | 0.54 | 0.3 |
| 5 | 8554971 | rs12653550 | 1.413 | 1.033 | 1.676 | 0.626 | 0.044 | 0.011 | 0.011 | 0.011 | 0.36 | 0.2 | 0.24 | 0.31 | 0.23 | 0.46 |
| 5 | 8558215 | rs6555554 | -0.671 | 1.22 | 1.686 | 0.661 | 0.009 | 0 | 0 | 0 | 0.52 | 0.56 | 0.69 | 0.5 | 0.71 | 0.33 |
| 5 | 8558468 | rs13161601 | 0.876 | 1.273 | 1.153 | 0.647 | 0.043 | -0.047 | -0.047 | 0 | 0.26 | 0.07 | 0.4 | 0.34 | 0.22 | 0.17 |
| 5 | 8562829 | rs756757 | 1.147 | 1.084 | 1.448 | 0.53 | 0.002 | 0.002 | 0.002 | 0.513 | 0.4 | 0.34 | 0.27 | 0.26 | 0.3 | 0.46 |
| 5 | 8565638 | rs1157652 | -1.454 | 1.414 | 1.743 | 0.296 | 0.157 | 0.036 | 0.036 | 0.175 | 0.09 | 0.31 | 0.11 | 0.08 | 0.19 | 0.06 |
| 5 | 8569124 | rs12515836 | -0.048 | 1.363 | 0.607 | 0.366 | -0.026 | -0.038 | -0.038 | 0 | 0.26 | 0.16 | 0.49 | 0.61 | 0.37 | 0.24 |
| 5 | 8572050 | rs11134316 | -0.419 | 1.38 | 0.557 | 0.385 | -0.045 | -0.005 | -0.005 | 0 | 0.3 | 0.16 | 0.58 | 0.7 | 0.55 | 0.31 |
| 5 | 8573076 | rs16880863 | 0.726 | 1.973 | 0.5 | 0.335 | -0.011 | -0.047 | -0.047 | 0 | 0.16 | 0.06 | 0.41 | 0.6 | 0.22 | 0.17 |
| 5 | 8575930 | rs2106316 | -0.227 | 2.254 | 0.499 | 0.002 | -0.014 | -0.011 | -0.011 | 0 | 0.27 | 0.16 | 0.49 | 0.7 | 0.34 | 0.28 |
| 5 | 8578343 | rs6867425 | 2.013 | 1.804 | 0.284 | -0.235 | 0.036 | 0.051 | 0.051 | 0.058 | 0.34 | 0.21 | 0.23 | 0.16 | 0.25 | 0.37 |
| 5 | 8583411 | rs768900 | -0.291 | 1.554 | -0.158 | 0.044 | -0.03 | -0.013 | -0.013 | 0 | 0.28 | 0.16 | 0.51 | 0.69 | 0.4 | 0.28 |
| 5 | 8584086 | rs4702561 | 0.287 | 1.589 | -0.276 | 0.326 | -0.006 | 0.004 | 0.004 | 0 | 0.22 | 0.21 | 0.18 | 0.14 | 0.29 | 0.24 |
| 5 | 8586300 | rs199169 | 1.018 | 1.223 | -0.533 | 0.239 | 0.013 | 0.092 | 0.092 | 0.751 | 0.42 | 0.21 | 0.28 | 0.17 | 0.38 | 0.48 |
| 5 | 8586827 | rs12522315 | 0.725 | 0.689 | -1.041 | 0.25 | -0.001 | -0.009 | -0.009 | 0 | 0.28 | 0.22 | 0.47 | 0.56 | 0.31 | 0.35 |
| 5 | 8588451 | rs1468112 | 0.553 | 0.697 | -1.036 | -0.384 | -0.021 | 0.016 | 0.016 | 0 | 0.25 | 0.18 | 0.21 | 0.25 | 0.29 | 0.3 |
| 5 | 8594764 | rs11952768 | 0.437 | -0.373 | -0.981 | -0.222 | -0.035 | -0.018 | -0.018 | 0 | 0.23 | 0.17 | 0.3 | 0.3 | 0.39 | 0.22 |
| 5 | 8595305 | rs1501340 | -2.059 | -0.668 | -0.278 | -0.263 | 0.009 | -0.005 | -0.005 | 0.432 | 0.66 | 0.72 | 0.6 | 0.63 | 0.77 | 0.7 |
| 5 | 8597846 | rs1604318 | 0.933 | -0.598 | -0.234 | -0.247 | -0.03 | 0.018 | 0.018 | 0 | 0.27 | 0.22 | 0.24 | 0.25 | 0.35 | 0.3 |
| 5 | 8603334 | rs7709563 | -0.55 | -0.713 | 0.009 | -0.175 | -0.013 | -0.015 | -0.015 | 0 | 0.33 | 0.23 | 0.33 | 0.38 | 0.42 | 0.38 |
| 5 | 8613262 | rs1354096 | -0.489 | -0.694 | -0.041 | -0.238 | -0.027 | -0.042 | -0.042 | 0.071 | 0.33 | 0.19 | 0.33 | 0.42 | 0.42 | 0.38 |
| 5 | 8625584 | rs199175 | -1.076 | -0.897 | 0.337 | -0.028 | 0.022 | 0.008 | 0.008 | 0.021 | 0.8 | 0.76 | 0.62 | 0.67 | 0.57 | 0.79 |
| 5 | 8628753 | rs1501347 | -1.361 | -1.336 | 1.015 | -0.591 | 0.021 | 0.05 | 0.05 | 0 | 0.49 | 0.63 | 0.29 | 0.21 | 0.37 | 0.49 |
| 5 | 8634454 | rs10040826 | -0.093 | -1.438 | 1.158 | -0.633 | -0.004 | -0.105 | -0.105 | 0 | 0.22 | 0.34 | 0.15 | 0.01 | 0.21 | 0.1 |
| 5 | 8641457 | rs199990 | 0.656 | -1.414 | 0.54 | -0.582 | 0.025 | 0.02 | 0.02 | 0 | 0.26 | 0.15 | 0.13 | 0.19 | 0.17 | 0.38 |
| 5 | 8645775 | rs200121 | -1.131 | -1.327 | -0.233 | -0.594 | 0.011 | 0.029 | 0.029 | 0 | 0.47 | 0.38 | 0.27 | 0.21 | 0.37 | 0.49 |
| 5 | 8650225 | rs12519783 | -0.567 | -1.02 | -1.288 | -0.332 | 0.004 | 0.008 | 0.008 | 0 | 0.27 | 0.71 | 0.2 | 0.15 | 0.21 | 0.15 |
| 5 | 8650331 | rs200116 | 0.362 | -0.265 | -1.356 | -0.02 | 0.012 | -0.017 | -0.017 | 0 | 0.23 | 0.05 | 0.11 | 0.2 | 0.16 | 0.38 |
| 5 | 8655624 | rs199195 | -0.601 | -0.101 | -0.773 | -0.786 | -0.002 | -0.06 | -0.06 | 0 | 0.22 | 0.22 | 0.17 | 0.01 | 0.22 | 0.12 |
| 5 | 8655966 | rs200104 | -0.618 | -0.824 | -0.267 | -0.416 | -0.025 | 0.012 | 0.012 | 0 | 0.34 | 0.41 | 0.43 | 0.6 | 0.24 | 0.39 |
| 5 | 8657664 | rs9313251 | -0.417 | -0.869 | 0.032 | -0.256 | 0.001 | -0.038 | -0.038 | 0 | 0.07 | 0.15 | 0.04 | 0 | 0.05 | 0.02 |
| 5 | 8661907 | rs200094 | -0.415 | -1.05 | 0.192 | -0.367 | 0.003 | -0.006 | -0.006 | 0.248 | 0.31 | 0.24 | 0.42 | 0.54 | 0.23 | 0.4 |
| 5 | 8663983 | rs200086 | 0.101 | -1.044 | 0.235 | -0.375 | 0.007 | -0.011 | -0.011 | 0 | 0.26 | 0.24 | 0.39 | 0.59 | 0.19 | 0.37 |
| 5 | 8670219 | rs200071 | -0.604 | -0.99 | 0.103 | -0.926 | 0.056 | 0.124 | 0.124 | 0 | 0.52 | 0.71 | 0.77 | 0.8 | 0.64 | 0.62 |
| 5 | 8671855 | rs999419 | -0.069 | -0.966 | 0.373 | -0.899 | 0.031 | -0.01 | -0.01 | 0 | 0.06 | 0.22 | 0.04 | 0 | 0.04 | 0.02 |
| 5 | 8672942 | rs13170506 | 1.341 | -1.607 | 0.271 | -1.177 | 0.02 | 0.096 | 0.096 | 0 | 0.46 | 0.19 | 0.4 | 0.28 | 0.58 | 0.48 |
| 5 | 8676369 | rs199187 | 1.023 | -1.645 | 0.458 | -1.195 | 0.03 | 0.032 | 0.032 | 0 | 0.48 | 0.32 | 0.42 | 0.39 | 0.58 | 0.52 |
| 5 | 8680327 | rs200059 | 0.079 | -1.643 | 0.492 | -1.185 | -0.022 | 0.015 | 0.015 | 0 | 0.32 | 0.54 | 0.45 | 0.59 | 0.23 | 0.39 |
| 5 | 8691906 | rs11134323 | 1.744 | -1.604 | 0.481 | -1.199 | -0.111 | -0.031 | -0.031 | 0 | 0.2 | 0.05 | 0.31 | 0.2 | 0.41 | 0.23 |
| 5 | 8692312 | rs200027 | 1.355 | -1.563 | 0.807 | -0.995 | -0.022 | -0.02 | -0.02 | 0 | 0.73 | 0.38 | 0.81 | 0.82 | 0.8 | 0.85 |
| 5 | 8693465 | rs200016 | -0.338 | -1.377 | 0.495 | -0.897 | 0.002 | -0.007 | -0.007 | 0 | 0.83 | 0.86 | 0.87 | 0.99 | 0.83 | 0.91 |
| 5 | 8694523 | rs200013 | -0.554 | -1.381 | 0.674 | -0.513 | -0.026 | 0.013 | 0.013 | 0 | 0.34 | 0.5 | 0.44 | 0.6 | 0.24 | 0.4 |
| 5 | 8697444 | rs200008 | -0.291 | -1.356 | 0.596 | -0.561 | -0.026 | 0.036 | 0.036 | 0 | 0.33 | 0.4 | 0.44 | 0.54 | 0.23 | 0.39 |
| 5 | 8728874 | rs7727009 | 1.22 | -1.25 | 0.304 | -0.798 | 0.028 | 0.034 | 0.034 | 0 | 0.41 | 0.29 | 0.26 | 0.3 | 0.34 | 0.38 |
| 5 | 8740641 | rs6864374 | 1.22 | -1.25 | 0.301 | -0.774 | 0.029 | 0.034 | 0.034 | 0 | 0.41 | 0.29 | 0.26 | 0.3 | 0.34 | 0.38 |
| 5 | 8743390 | rs156466 | 0.155 | -1.23 | 0.226 | -1.022 | 0 | -0.008 | -0.008 | 0 | 0.82 | 0.87 | 0.89 | 0.99 | 0.83 | 0.92 |
| 5 | 8744557 | rs2388355 | 1.361 | -1.288 | 0.204 | -0.971 | 0.025 | 0.032 | 0.032 | 0 | 0.41 | 0.3 | 0.26 | 0.3 | 0.34 | 0.38 |
| 5 | 8752756 | rs156453 | -0.151 | -1.353 | 0.072 | -0.975 | -0.012 | -0.019 | -0.019 | 0.018 | 0.94 | 0.91 | 0.98 | 1 | 0.98 | 0.99 |
| 5 | 8753733 | rs156452 | 0.429 | -1.209 | -0.023 | -0.988 | -0.11 | -0.059 | -0.059 | 0.029 | 0.34 | 0.12 | 0.56 | 0.58 | 0.47 | 0.49 |
| 5 | 8769738 | rs999428 | 1.337 | -1.442 | 0.196 | -1.125 | -0.014 | -0.053 | -0.053 | 0 | 0.73 | 0.43 | 0.82 | 0.88 | 0.79 | 0.82 |
| 5 | 8770185 | rs999426 | 0.866 | -1.437 | 0.223 | -1.122 | -0.006 | -0.062 | -0.062 | 0 | 0.76 | 0.49 | 0.82 | 0.88 | 0.79 | 0.85 |
| 5 | 8771494 | rs2904956 | 0.607 | -1.39 | 0.129 | -1.101 | -0.097 | -0.046 | -0.046 | 0 | 0.32 | 0.16 | 0.57 | 0.59 | 0.46 | 0.43 |
| 5 | 8777256 | rs1505028 | 0.524 | -1.385 | 0.095 | -1.083 | -0.086 | -0.035 | -0.035 | 0 | 0.33 | 0.15 | 0.57 | 0.59 | 0.47 | 0.43 |
| 5 | 8778454 | rs12655943 | 1.121 | -1.405 | 0.256 | -1.095 | 0.021 | 0.024 | 0.024 | 0 | 0.42 | 0.32 | 0.25 | 0.28 | 0.33 | 0.41 |
| 5 | 8783755 | rs2963391 | 1.555 | -1.09 | 0.38 | -0.776 | -0.012 | -0.044 | -0.044 | 0 | 0.72 | 0.38 | 0.81 | 0.88 | 0.79 | 0.82 |
| 5 | 8786179 | rs877813 | 0.957 | -1.09 | 0.557 | -0.776 | -0.002 | -0.057 | -0.057 | 0 | 0.76 | 0.59 | 0.83 | 0.88 | 0.79 | 0.85 |
| 5 | 8789479 | rs10491224 | 1.518 | -1.253 | -0.139 | -0.916 | -0.017 | 0.126 | 0.126 | 1.704 | 0.26 | 0.01 | 0.1 | 0.07 | 0.2 | 0.24 |
| 5 | 8790996 | rs3111121 | 0.498 | -0.363 | 0.819 | -0.161 | -0.002 | 0.003 | 0.003 | 0 | 0.85 | 0.89 | 0.89 | 0.99 | 0.85 | 0.92 |
| 5 | 8794278 | rs992149 | 1.558 | -0.161 | 0.901 | 0.284 | 0.002 | 0.032 | 0.032 | 0.091 | 0.46 | 0.27 | 0.41 | 0.36 | 0.41 | 0.54 |
| 5 | 8798487 | rs7730627 | 0.13 | -0.156 | 0.931 | 0.312 | -0.011 | -0.012 | -0.012 | 0 | 0.05 | 0.18 | 0.02 | 0 | 0.01 | 0.05 |
| 5 | 8808777 | rs4701822 | 1.504 | -0.079 | 0.525 | 0.312 | 0.001 | 0.034 | 0.034 | 0.204 | 0.46 | 0.25 | 0.41 | 0.36 | 0.41 | 0.54 |
| 5 | 8814924 | rs1505023 | 0.834 | -0.076 | 0.584 | 0.43 | 0 | -0.013 | -0.013 | 0.012 | 0.81 | 0.71 | 0.86 | 0.88 | 0.82 | 0.89 |
| 5 | 8828135 | rs995555 | 1.05 | -0.3 | 0.403 | 0.706 | 0.027 | 0.041 | 0.041 | 0.476 | 0.53 | 0.25 | 0.49 | 0.37 | 0.55 | 0.6 |
| 5 | 8830876 | rs10051161 | 1.073 | -0.59 | 0.478 | 0.694 | 0.015 | 0.053 | 0.053 | 0.539 | 0.53 | 0.3 | 0.49 | 0.35 | 0.55 | 0.6 |
| 5 | 8836536 | rs2963416 | -0.656 | -0.605 | 0.368 | 0.809 | -0.028 | 0.01 | 0.01 | 0 | 0.94 | 0.86 | 0.99 | 0.93 | 0.99 | 0.96 |
| 5 | 8836613 | rs2904930 | 0.133 | -0.536 | 0.161 | 1.323 | -0.011 | 0.02 | 0.02 | 0 | 0.19 | 0.18 | 0.32 | 0.5 | 0.13 | 0.18 |
| 5 | 8837390 | rs1394650 | -0.65 | -0.087 | 0.032 | 0.954 | -0.006 | -0.002 | -0.002 | 0 | 0.35 | 0.3 | 0.47 | 0.68 | 0.43 | 0.33 |
| 5 | 8844967 | rs756666 | -0.215 | -1.403 | 0.303 | 1.513 | 0.015 | -0.017 | -0.017 | 2.241 | 0.25 | 0.21 | 0.34 | 0.65 | 0.17 | 0.18 |

| | | | | | | | | | | | | | | | | |
|---|---|---|---|---|---|---|---|---|---|---|---|---|---|---|---|---|
| 5 | 8857503 | rs4280839 | 0.247 | -1.354 | 0.041 | 1.41 | -0.022 | -0.023 | -0.023 | 1.456 | 0.91 | 0.83 | 0.96 | 0.97 | 0.98 | 0.95 |
| 5 | 8860434 | rs11134335 | -0.295 | -1.319 | -0.327 | 1.507 | -0.045 | 0.027 | 0.027 | 0 | 0.27 | 0.36 | 0.37 | 0.66 | 0.17 | 0.28 |
| 5 | 8862669 | rs16881512 | 0.079 | -0.832 | -1.623 | 1.453 | -0.008 | -0.044 | -0.044 | 0 | 0.1 | 0.04 | 0.07 | 0.13 | 0.06 | 0.17 |
| 5 | 8870267 | rs7702790 | 0.5 | -0.682 | -1.376 | 1.593 | 0.061 | -0.009 | -0.009 | 0.405 | 0.63 | 0.3 | 0.8 | 0.54 | 0.8 | 0.66 |
| 5 | 8871850 | rs10053782 | -1.129 | 0.606 | -1.132 | 1.897 | 0 | 0.093 | 0.093 | 0 | 0.05 | 0.03 | 0.26 | 0.49 | 0.07 | 0.15 |
| 5 | 8873310 | rs17225452 | -1.268 | 1.107 | -1.071 | 0.424 | -0.019 | 0.017 | 0.017 | 0 | 0.08 | 0.02 | 0.08 | 0 | 0.14 | 0.06 |
| 5 | 8891510 | rs11948993 | -0.395 | 0.306 | -1.157 | 0.218 | 0.016 | 0.018 | 0.018 | 0 | 0.38 | 0.65 | 0.31 | 0.25 | 0.3 | 0.43 |
| 5 | 8896731 | rs10447192 | -0.241 | -0.64 | -1.752 | -0.514 | -0.023 | 0.05 | 0.05 | 0.614 | 0.46 | 0.08 | 0.35 | 0.31 | 0.39 | 0.58 |
| 5 | 8901512 | rs16881630 | -0.629 | -0.642 | -0.291 | -0.677 | -0.003 | -0.033 | -0.033 | 0.015 | 0.09 | 0.42 | 0.06 | 0 | 0.04 | 0.05 |
| 5 | 8901696 | rs2962685 | -0.942 | -0.17 | -0.727 | -1.129 | -0.033 | 0.043 | 0.043 | 0 | 0.88 | 0.96 | 0.92 | 0.99 | 0.82 | 0.94 |
| 5 | 8903540 | rs16881639 | 0.299 | -0.445 | -0.924 | -1.337 | 0.008 | 0.008 | 0.008 | 0 | 0.07 | 0.05 | 0.06 | 0.04 | 0.04 | 0.06 |
| 5 | 8903736 | rs2962683 | 0.502 | -0.389 | -0.323 | -1.31 | 0.002 | 0.006 | 0.006 | 0.28 | 0.41 | 0.65 | 0.55 | 0.68 | 0.43 | 0.36 |
| 5 | 8904380 | rs10067616 | 0.23 | -0.393 | -0.287 | -1.308 | -0.007 | 0.06 | 0.06 | 0.571 | 0.46 | 0.12 | 0.35 | 0.31 | 0.39 | 0.57 |
| 5 | 8906784 | rs4702597 | 0.062 | -0.425 | -0.009 | -1.323 | 0.01 | 0.027 | 0.027 | 0.07 | 0.47 | 0.34 | 0.37 | 0.31 | 0.4 | 0.57 |
| 5 | 8908886 | rs2963342 | 0.398 | -0.381 | -0.014 | -1.304 | -0.001 | -0.004 | -0.004 | 0 | 0.4 | 0.6 | 0.55 | 0.69 | 0.44 | 0.36 |
| 5 | 8919975 | rs2962669 | 1.018 | -0.238 | 0.022 | -1.16 | -0.012 | -0.011 | -0.011 | 1.259 | 0.32 | 0.23 | 0.49 | 0.68 | 0.39 | 0.31 |
| 5 | 8923420 | rs2962663 | 0.166 | -0.237 | 0.341 | -1.172 | 0 | 0 | 0 | 0 | 0.46 | 0.47 | 0.59 | 0.69 | 0.57 | 0.38 |
| 5 | 8924770 | rs6555577 | 0.497 | 0.035 | -0.169 | -1.139 | -0.021 | 0.03 | 0.03 | 0 | 0.12 | 0.06 | 0.08 | 0.01 | 0.16 | 0.06 |
| 5 | 8934159 | rs2963327 | -1.16 | -0.292 | -0.026 | -1.461 | 0.003 | -0.072 | -0.072 | 0.389 | 0.35 | 0.43 | 0.25 | 0.13 | 0.32 | 0.22 |
| 5 | 8935632 | rs2963337 | -1.104 | -0.308 | 0.137 | -1.504 | 0.004 | -0.105 | -0.105 | 0 | 0.36 | 0.52 | 0.26 | 0.13 | 0.32 | 0.24 |
| 5 | 8941558 | rs4583857 | 0.314 | -0.356 | -0.132 | -1.5 | -0.031 | 0.052 | 0.052 | 0 | 0.14 | 0.04 | 0.09 | 0.01 | 0.19 | 0.07 |
| 5 | 8955823 | rs6874770 | 1.387 | -0.761 | -0.736 | -0.944 | -0.112 | -0.182 | -0.182 | 0 | 0.26 | 0.02 | 0.41 | 0.57 | 0.37 | 0.29 |
| 5 | 8961561 | rs9313264 | -1.153 | -0.816 | -0.353 | -1.338 | -0.007 | -0.083 | -0.083 | 0 | 0.37 | 0.48 | 0.22 | 0.13 | 0.3 | 0.24 |
| 5 | 8968532 | rs6889183 | 0.344 | -0.844 | -0.324 | -1.401 | -0.001 | -0.022 | -0.022 | 0.015 | 0.14 | 0.14 | 0.09 | 0.02 | 0.09 | 0.13 |
| 5 | 8969172 | rs7728475 | -0.98 | -0.846 | -0.307 | -1.4 | -0.002 | -0.087 | -0.087 | 0 | 0.33 | 0.39 | 0.2 | 0.03 | 0.29 | 0.22 |
| 5 | 8980099 | rs1947584 | 0.257 | -0.813 | -0.298 | -1.179 | 0 | -0.022 | -0.022 | 2.56 | 0.15 | 0.14 | 0.09 | 0.02 | 0.11 | 0.13 |
| 5 | 8983721 | rs11134338 | -0.774 | -1.177 | -0.379 | -1.463 | 0 | -0.014 | -0.014 | 0 | 0.3 | 0.29 | 0.19 | 0.11 | 0.25 | 0.19 |
| 5 | 8997962 | rs1350379 | 0.69 | -1.437 | -0.84 | -1.803 | 0.041 | 0.056 | 0.056 | 0 | 0.21 | 0.11 | 0.09 | 0.05 | 0.14 | 0.11 |
| 5 | 9012115 | rs16881838 | 0.121 | -1.486 | -0.822 | -1.883 | -0.002 | 0.031 | 0.031 | 0 | 0.26 | 0.17 | 0.17 | 0.03 | 0.23 | 0.19 |
| 5 | 9013608 | rs17235037 | 1.092 | -1.463 | -0.868 | -1.959 | -0.008 | 0.058 | 0.058 | 0 | 0.16 | 0.04 | 0.08 | 0.01 | 0.14 | 0.08 |
| 5 | 9013766 | rs985722 | -0.953 | -1.402 | -0.846 | -1.907 | 0.043 | 0.09 | 0.09 | 0 | 0.79 | 0.95 | 0.92 | 0.96 | 0.86 | 0.89 |
| 5 | 9021310 | rs1598826 | 0.999 | -1.501 | -0.428 | -1.893 | 0.012 | 0.041 | 0.041 | 0 | 0.45 | 0.36 | 0.7 | 0.85 | 0.62 | 0.68 |
| 5 | 9026797 | rs13152876 | 1.258 | -1.52 | -0.978 | -1.124 | -0.106 | -0.121 | -0.121 | 0 | 0.28 | 0.01 | 0.35 | 0.38 | 0.36 | 0.44 |
| 5 | 9038043 | rs16881875 | 0.694 | -1.876 | 0.307 | -1.211 | -0.001 | -0.036 | -0.036 | 0.513 | 0.11 | 0.18 | 0.09 | 0.02 | 0.1 | 0.1 |
| 5 | 9041935 | rs786840 | 1.026 | -1.79 | 0.564 | -1.22 | -0.007 | -0.049 | -0.049 | 0 | 0.62 | 0.4 | 0.79 | 0.93 | 0.74 | 0.77 |
| 5 | 9050410 | rs257098 | 0.699 | -1.782 | 0.657 | -1.206 | -0.04 | -0.114 | -0.114 | 0.061 | 0.66 | 0.53 | 0.81 | 0.94 | 0.77 | 0.79 |
| 5 | 9053945 | rs257095 | -0.453 | -1.772 | 0.46 | -1.197 | 0.037 | 0.048 | 0.048 | 0 | 0.78 | 0.9 | 0.92 | 0.96 | 0.86 | 0.89 |
| 5 | 9062943 | rs16881901 | 0.73 | -1.765 | 0.128 | -1.195 | -0.001 | -0.025 | -0.025 | 0.931 | 0.11 | 0.15 | 0.09 | 0.02 | 0.1 | 0.1 |
| 5 | 9066464 | rs786847 | 0.521 | -1.524 | 0.2 | -1.024 | -0.011 | -0.057 | -0.057 | 0.847 | 0.62 | 0.41 | 0.79 | 0.93 | 0.75 | 0.77 |
| 5 | 9066802 | rs786846 | 0.179 | -1.381 | -0.805 | -0.845 | 0.019 | 0.016 | 0.016 | 0 | 0.67 | 0.72 | 0.82 | 0.94 | 0.77 | 0.79 |
| 5 | 9078628 | rs12716205 | 0.081 | -1.296 | -0.953 | -0.804 | 0.02 | 0.023 | 0.023 | 0 | 0.68 | 0.74 | 0.82 | 0.94 | 0.77 | 0.79 |
| 5 | 9087921 | rs1013720 | -0.533 | -1.112 | -0.976 | 0.081 | 0.015 | 0.009 | 0.009 | 0.31 | 0.39 | 0.24 | 0.33 | 0.34 | 0.34 | 0.3 |
| 5 | 9091164 | rs253634 | -0.965 | -1.975 | -0.797 | -0.069 | 0.012 | 0.084 | 0.084 | 0 | 0.42 | 0.64 | 0.31 | 0.11 | 0.37 | 0.33 |
| 5 | 9092642 | rs253635 | -1.889 | -2.453 | -0.965 | -0.046 | -0.048 | -0.098 | -0.098 | 0 | 0.58 | 0.9 | 0.48 | 0.44 | 0.43 | 0.51 |
| 5 | 9100211 | rs17309469 | 0.805 | -2.59 | -0.623 | -0.125 | 0.001 | -0.001 | -0.001 | 0.24 | 0.1 | 0.05 | 0.05 | 0.01 | 0.09 | 0.18 |
| 5 | 9105283 | rs2456237 | -1.968 | -2.517 | -0.659 | -0.095 | -0.054 | -0.101 | -0.101 | 0 | 0.58 | 0.9 | 0.48 | 0.45 | 0.43 | 0.51 |
| 5 | 9107887 | rs7731508 | 0.495 | -2.517 | -0.511 | -0.209 | -0.005 | 0.005 | 0.005 | 9.731 | 0.14 | 0.21 | 0.15 | 0.33 | 0.06 | 0.15 |
| 5 | 9112040 | rs2657031 | -1.898 | -2.643 | -0.73 | 0.381 | -0.045 | 0.058 | 0.058 | 0.065 | 0.6 | 0.86 | 0.67 | 0.7 | 0.46 | 0.58 |
| 5 | 9121472 | rs190423 | -1.509 | -2.635 | -0.349 | 0.479 | -0.017 | -0.002 | -0.002 | 0 | 0.34 | 0.65 | 0.48 | 0.68 | 0.18 | 0.46 |
| 5 | 9121591 | rs3777253 | -0.755 | -2.645 | -0.061 | 0.514 | -0.017 | 0.007 | 0.007 | 3.391 | 0.27 | 0.34 | 0.22 | 0.35 | 0.15 | 0.36 |
| 5 | 9137550 | rs10057636 | -0.861 | -2.516 | 0.864 | 0.66 | -0.013 | 0.252 | 0.252 | 0 | 0.07 | 0.24 | 0.26 | 0.33 | 0.03 | 0.13 |
| 5 | 9140514 | rs6555593 | 0.343 | -2.973 | 0.566 | 0.441 | -0.002 | -0.042 | -0.042 | 0 | 0.31 | 0.33 | 0.21 | 0.1 | 0.31 | 0.23 |
| 5 | 9140848 | rs983856 | -0.314 | -2.776 | 0.544 | 0.606 | -0.014 | 0.213 | 0.213 | 0 | 0.94 | 0.84 | 0.75 | 0.68 | 0.97 | 0.9 |
| 5 | 9142253 | rs13155922 | 0.678 | -2.801 | 0.292 | 0.696 | 0 | -0.001 | -0.001 | 0.043 | 0.14 | 0.19 | 0.16 | 0.32 | 0.06 | 0.16 |
| 5 | 9148278 | rs12189232 | 0.992 | -2.767 | 0.239 | 0.641 | 0.028 | 0.021 | 0.021 | 0 | 0.51 | 0.39 | 0.48 | 0.56 | 0.57 | 0.46 |
| 5 | 9149422 | rs1904454 | 1.938 | -2.676 | -0.167 | 0.409 | -0.121 | 0.033 | 0.033 | 0.096 | 0.35 | 0.05 | 0.31 | 0.24 | 0.51 | 0.3 |
| 5 | 9151237 | rs7723025 | -0.198 | -2.783 | 0.348 | 0.213 | 0.016 | 0.012 | 0.012 | 0 | 0.24 | 0.18 | 0.09 | 0.07 | 0.15 | 0.31 |
| 5 | 9155420 | rs899576 | -0.781 | -2.637 | 0.693 | 0.216 | 0.011 | -0.034 | -0.034 | 0.011 | 0.86 | 0.84 | 0.85 | 0.98 | 0.79 | 0.89 |
| 5 | 9155728 | rs886511 | -0.689 | -2.748 | 0.818 | 1.011 | -0.009 | 0.397 | 0.397 | 0.258 | 0.08 | 0.31 | 0.27 | 0.47 | 0.04 | 0.14 |
| 5 | 9156101 | rs886508 | -0.609 | -2.422 | 0.008 | 0.269 | 0.012 | 0.006 | 0.006 | 0 | 0.24 | 0.19 | 0.1 | 0.07 | 0.15 | 0.31 |
| 5 | 9157454 | rs1806138 | -0.572 | -2.406 | -0.045 | 0.277 | 0.016 | 0.013 | 0.013 | 0 | 0.24 | 0.18 | 0.09 | 0.07 | 0.15 | 0.31 |
| 5 | 9161412 | rs3846574 | 0.923 | -1.935 | 0.132 | 0.968 | -0.001 | 0 | 0 | 0 | 0.61 | 0.59 | 0.51 | 0.44 | 0.65 | 0.44 |
| 5 | 9169940 | rs3777275 | -0.436 | -2.008 | 0.673 | 1.132 | 0.03 | 0.053 | 0.053 | 0 | 0.07 | 0.24 | 0.06 | 0.11 | 0.06 | 0.06 |
| 5 | 9170323 | rs3797928 | 0.062 | -1.98 | 0.259 | 1.341 | 0.02 | -0.038 | -0.038 | 1.661 | 0.15 | 0.03 | 0.04 | 0.16 | 0.09 | 0.13 |
| 5 | 9173880 | rs40718 | 1.027 | -2.191 | 0.139 | 1.255 | 0.002 | 0 | 0 | 4.019 | 0.4 | 0.36 | 0.3 | 0.45 | 0.52 | 0.32 |
| 5 | 9174381 | rs3822773 | 0.811 | -2.337 | 0.271 | 1.338 | -0.032 | 0.06 | 0.06 | 0.818 | 0.07 | 0.23 | 0.16 | 0.14 | 0.03 | 0.09 |
| 5 | 9176431 | rs10044766 | 0.811 | -2.337 | 0.271 | 1.338 | -0.032 | 0.06 | 0.06 | 0.818 | 0.07 | 0.23 | 0.16 | 0.14 | 0.03 | 0.09 |
| 5 | 9177932 | rs3777281 | 0.672 | -2.492 | 0.244 | 1.145 | 0.009 | 0.023 | 0.023 | 0 | 0.23 | 0.18 | 0.16 | 0.09 | 0.2 | 0.24 |
| 5 | 9182376 | rs10513010 | 0.642 | -2.414 | 0.249 | 1.081 | 0.025 | 0.022 | 0.022 | 0 | 0.07 | 0.17 | 0.05 | 0.11 | 0.06 | 0.06 |

| | | | | | | | | | | | | | | | | |
|---|---|---|---|---|---|---|---|---|---|---|---|---|---|---|---|---|
| 5 | 9182717 | rs13171149 | 0.714 | -2.411 | 0.143 | 1.072 | 0.012 | 0.041 | 0.041 | 0 | 0.23 | 0.16 | 0.13 | 0.07 | 0.2 | 0.24 |
| 5 | 9184632 | rs13436248 | 0.624 | -2.279 | 0.134 | 1.205 | -0.032 | 0.089 | 0.089 | 1.136 | 0.08 | 0.27 | 0.17 | 0.15 | 0.03 | 0.09 |
| 5 | 9187035 | rs42353 | 1.411 | -2.284 | -0.071 | 1.249 | 0.006 | 0.016 | 0.016 | 0 | 0.56 | 0.37 | 0.38 | 0.32 | 0.66 | 0.45 |
| 5 | 9189454 | rs40703 | 0.616 | -2.274 | -0.305 | 1.224 | -0.001 | 0.015 | 0.015 | 0 | 0.54 | 0.46 | 0.48 | 0.67 | 0.6 | 0.48 |
| 5 | 9191362 | rs3797937 | -0.735 | -2.281 | -0.933 | 1.335 | -0.035 | 0.034 | 0.034 | 0.4 | 0.3 | 0.43 | 0.29 | 0.55 | 0.2 | 0.35 |
| 5 | 9192881 | rs1806125 | 0.514 | -2.072 | -1.5 | 1.68 | -0.035 | 0.192 | 0.192 | 1.715 | 0.08 | 0.28 | 0.2 | 0.27 | 0.04 | 0.14 |
| 5 | 9194611 | rs40693 | 1.355 | -1.28 | -3.414 | 0.886 | -0.159 | -0.028 | -0.028 | 0.054 | 0.3 | 0.02 | 0.22 | 0.24 | 0.45 | 0.21 |
| 5 | 9195233 | rs40687 | 1.21 | -1.179 | -2.176 | 0.957 | -0.007 | -0.012 | -0.012 | 0.25 | 0.72 | 0.89 | 0.63 | 0.69 | 0.75 | 0.65 |
| 5 | 9197607 | rs1806085 | -0.561 | -1.115 | -2.174 | 0.935 | -0.002 | 0 | 0 | 0 | 0.14 | 0.1 | 0.33 | 0.14 | 0.15 | 0.22 |
| 5 | 9200137 | rs1806112 | -0.704 | -1.689 | -2.063 | 1.034 | 0.021 | -0.015 | -0.015 | 0 | 0.16 | 0.06 | 0.05 | 0.17 | 0.1 | 0.14 |
| 5 | 9203989 | rs40680 | 0.599 | -1.777 | -2.219 | 0.95 | 0.101 | -0.067 | -0.067 | 0 | 0.65 | 0.14 | 0.65 | 0.42 | 0.81 | 0.64 |
| 5 | 9204238 | rs1479651 | -0.345 | -0.732 | -2.079 | 0.437 | 0.041 | 0.035 | 0.035 | 0.044 | 0.3 | 0.12 | 0.14 | 0.22 | 0.24 | 0.34 |
| 5 | 9209831 | rs1805964 | 0.206 | 0.009 | -1.684 | 0.418 | -0.004 | -0.008 | -0.008 | 0 | 0.42 | 0.31 | 0.39 | 0.44 | 0.43 | 0.6 |
| 5 | 9215068 | rs1805962 | -0.179 | -0.599 | -1.07 | -0.03 | -0.01 | -0.004 | -0.004 | 0 | 0.55 | 0.62 | 0.49 | 0.46 | 0.49 | 0.66 |
| 5 | 9218895 | rs10042101 | -0.061 | -0.873 | -0.668 | -0.136 | -0.006 | -0.027 | -0.027 | 0 | 0.1 | 0.1 | 0.07 | 0.01 | 0.05 | 0.05 |
| 5 | 9222811 | rs40669 | -0.516 | -1.358 | -0.354 | 0.124 | -0.003 | 0 | 0 | 0 | 0.27 | 0.29 | 0.34 | 0.33 | 0.18 | 0.21 |
| 5 | 9225088 | rs17254539 | 0.934 | -1.181 | -0.252 | -0.141 | -0.005 | 0.037 | 0.037 | 0.067 | 0.31 | 0.21 | 0.17 | 0.18 | 0.33 | 0.26 |
| 5 | 9228171 | rs40666 | -0.855 | -1.139 | 0.313 | -0.188 | -0.032 | -0.001 | -0.001 | 0 | 0.34 | 0.45 | 0.39 | 0.33 | 0.23 | 0.22 |
| 5 | 9228283 | rs40665 | -0.534 | -1.215 | 0.429 | -0.128 | -0.034 | 0.005 | 0.005 | 0.62 | 0.28 | 0.35 | 0.33 | 0.32 | 0.17 | 0.13 |
| 5 | 9230988 | rs1806018 | 0.127 | -0.674 | 0.385 | -0.171 | -0.026 | 0.014 | 0.014 | 0 | 0.06 | 0.01 | 0.04 | 0.01 | 0.09 | 0.06 |
| 5 | 9231066 | rs42495 | 0.038 | -0.852 | 0.494 | 0.025 | 0.019 | -0.004 | -0.004 | 0.989 | 0.2 | 0.14 | 0.22 | 0.23 | 0.14 | 0.08 |
| 5 | 9234736 | rs1805957 | 0.429 | -0.863 | 0.836 | 0.005 | -0.057 | -0.013 | -0.013 | 0.32 | 0.75 | 0.56 | 0.7 | 0.76 | 0.85 | 0.86 |
| 5 | 9249231 | rs12716206 | 0.507 | -0.875 | 0.921 | -0.1 | 0.001 | 0.042 | 0.042 | 0.034 | 0.46 | 0.39 | 0.32 | 0.26 | 0.46 | 0.44 |
| 5 | 9249806 | rs3912190 | 0.889 | -0.991 | 0.895 | -0.1 | -0.024 | -0.035 | -0.035 | 0 | 0.31 | 0.2 | 0.39 | 0.47 | 0.39 | 0.41 |
| 5 | 9254909 | rs40713 | -0.553 | -0.723 | 0.94 | -0.022 | 0 | 0.019 | 0.019 | 0.008 | 0.54 | 0.59 | 0.63 | 0.73 | 0.54 | 0.5 |
| 5 | 9263933 | rs3797968 | 0.589 | -0.81 | 0.904 | 0.002 | 0.001 | 0.066 | 0.066 | 0.058 | 0.45 | 0.33 | 0.32 | 0.26 | 0.46 | 0.45 |
| 5 | 9264682 | rs40708 | 0.717 | -0.578 | 0.795 | 0.175 | 0.004 | 0.095 | 0.095 | 0 | 0.07 | 0.13 | 0.18 | 0.25 | 0.07 | 0.02 |
| 5 | 9266755 | rs1805939 | 0.863 | -0.521 | 0.442 | 0.123 | 0.002 | 0.122 | 0.122 | 0 | 0.4 | 0.2 | 0.24 | 0.2 | 0.4 | 0.4 |
| 5 | 9268737 | rs1805918 | 0.586 | -0.386 | 0.486 | 0.197 | 0.004 | 0.006 | 0.006 | 0 | 0.31 | 0.3 | 0.44 | 0.48 | 0.4 | 0.47 |
| 5 | 9269658 | rs1034743 | 0.615 | -0.373 | 0.303 | 0.203 | -0.009 | -0.011 | -0.011 | 0 | 0.31 | 0.26 | 0.44 | 0.48 | 0.4 | 0.47 |
| 5 | 9287073 | rs6873909 | 1.171 | -0.479 | 1.015 | 0.748 | -0.032 | 0.016 | 0.016 | 0 | 0.7 | 0.45 | 0.67 | 0.67 | 0.8 | 0.87 |
| 5 | 9292221 | rs4540155 | 1.814 | -0.226 | 1.69 | 0.719 | 0.023 | -0.006 | -0.006 | 0 | 0.92 | 0.62 | 0.92 | 0.97 | 0.96 | 0.97 |
| 5 | 9293213 | rs3822787 | 2.019 | -0.222 | 1.889 | 0.648 | -0.014 | -0.081 | -0.081 | 0 | 0.78 | 0.49 | 0.78 | 0.89 | 0.83 | 0.77 |
| 5 | 9295681 | rs3777306 | -1.332 | -0.21 | 1.898 | 0.615 | 0 | -0.006 | -0.006 | 0 | 0.16 | 0.16 | 0.15 | 0.07 | 0.14 | 0.2 |
| 5 | 9299338 | rs3734152 | 2.151 | -0.283 | 2.097 | 0.87 | -0.001 | 0 | 0 | 0.347 | 0.73 | 0.27 | 0.69 | 0.72 | 0.82 | 0.75 |
| 5 | 9315454 | rs805145 | 2.159 | -0.269 | 2.279 | 0.992 | -0.001 | 0 | 0 | 0 | 0.73 | 0.27 | 0.69 | 0.72 | 0.82 | 0.75 |
| 5 | 9331477 | rs9313273 | -1.919 | -0.207 | 1.721 | 0.992 | -0.016 | -0.037 | -0.037 | 0.12 | 0.19 | 0.27 | 0.14 | 0.07 | 0.12 | 0.19 |
| 5 | 9331490 | rs916990 | -0.342 | 0.083 | 1.286 | 1.286 | -0.033 | -0.048 | -0.048 | 0.006 | 0.08 | 0.01 | 0.1 | 0.14 | 0.1 | 0.15 |
| 5 | 9331848 | rs6896702 | -2.294 | 0.095 | 1.592 | 1.074 | -0.015 | 0.001 | 0.001 | 0 | 0.25 | 0.67 | 0.29 | 0.26 | 0.17 | 0.21 |
| 5 | 9332459 | rs1918779 | -1.09 | 0.233 | 1.012 | 0.999 | 0 | -0.002 | -0.002 | 0 | 0.13 | 0.13 | 0.11 | 0.07 | 0.11 | 0.14 |
| 5 | 9343414 | rs1205726 | 1.772 | 0.155 | 1.15 | 1.122 | -0.01 | -0.013 | -0.013 | 0 | 0.67 | 0.42 | 0.63 | 0.76 | 0.73 | 0.79 |
| 5 | 9343913 | rs1205725 | 1.559 | 0.149 | 1.27 | 1.119 | -0.01 | -0.016 | -0.016 | 0 | 0.69 | 0.54 | 0.64 | 0.76 | 0.73 | 0.79 |
| 5 | 9357007 | rs1024487 | -0.693 | 0.141 | 1.24 | 1.104 | -0.008 | 0.037 | 0.037 | 0 | 0.19 | 0.09 | 0.2 | 0.07 | 0.19 | 0.14 |
| 5 | 9357103 | rs1205718 | 1.638 | 0.312 | 1.294 | 1.247 | -0.008 | -0.009 | -0.009 | 2.945 | 0.69 | 0.57 | 0.64 | 0.75 | 0.73 | 0.79 |
| 5 | 9357942 | rs2463506 | -1.158 | 0.465 | 1.332 | 1.25 | 0.01 | 0.123 | 0.123 | 0 | 0.12 | 0.29 | 0.17 | 0.27 | 0.11 | 0.24 |
| 5 | 9359420 | rs17196621 | -1.038 | 0.521 | 0.322 | 0.871 | -0.009 | 0.036 | 0.036 | 0 | 0.19 | 0.09 | 0.2 | 0.07 | 0.2 | 0.14 |
| 5 | 9371411 | rs40654 | -0.63 | 0.375 | -0.161 | 0.669 | -0.003 | -0.003 | -0.003 | 0 | 0.25 | 0.3 | 0.28 | 0.15 | 0.23 | 0.2 |
| 5 | 9373448 | rs42192 | -0.079 | 0.669 | -0.28 | 0.484 | 0.001 | -0.006 | -0.006 | 0.565 | 0.31 | 0.34 | 0.27 | 0.22 | 0.31 | 0.34 |
| 5 | 9373641 | rs28263 | 0.671 | 0.56 | -0.907 | 0.1 | -0.021 | 0.066 | 0.066 | 0 | 0.25 | 0.06 | 0.19 | 0.14 | 0.28 | 0.15 |
| 5 | 9376118 | rs194261 | 0.597 | 0.535 | -0.81 | 0.045 | -0.007 | 0.06 | 0.06 | 0 | 0.26 | 0.11 | 0.19 | 0.14 | 0.27 | 0.14 |
| 5 | 9380110 | rs1805243 | 1.008 | 0.334 | -1.136 | -0.31 | -0.013 | 0.05 | 0.05 | 0 | 0.09 | 0.02 | 0.05 | 0.02 | 0.08 | 0.12 |
| 5 | 9380932 | rs805153 | -0.185 | 0.534 | -1.221 | -0.405 | -0.001 | 0.001 | 0.001 | 0 | 0.45 | 0.41 | 0.38 | 0.29 | 0.48 | 0.28 |
| 5 | 9382861 | rs1805927 | 0.888 | 0.627 | -0.621 | 0.363 | 0.001 | -0.01 | -0.01 | 0.004 | 0.36 | 0.22 | 0.25 | 0.33 | 0.36 | 0.4 |
| 5 | 9386310 | rs16882600 | 0.35 | 0.695 | -0.541 | 0.652 | 0 | -0.001 | -0.001 | 0 | 0.1 | 0.09 | 0.06 | 0.06 | 0.09 | 0.05 |
| 5 | 9389867 | rs13153629 | 1.594 | 0.866 | -0.703 | 0.591 | -0.021 | 0.011 | 0.011 | 0 | 0.05 | 0 | 0.04 | 0.02 | 0.06 | 0.05 |
| 5 | 9390217 | rs390322 | -0.905 | 0.814 | -0.453 | 0.594 | 0.021 | 0.076 | 0.076 | 0 | 0.3 | 0.51 | 0.48 | 0.53 | 0.29 | 0.42 |
| 5 | 9391753 | rs1877459 | 1.163 | 0.807 | -0.495 | 0.585 | 0.015 | 0.076 | 0.076 | 0 | 0.08 | 0.18 | 0.3 | 0.33 | 0.07 | 0.26 |
| 5 | 9397850 | rs788469 | -0.962 | 0.686 | -0.62 | 0.512 | 0.012 | 0.057 | 0.057 | 0 | 0.32 | 0.53 | 0.48 | 0.55 | 0.32 | 0.43 |
| 5 | 9401606 | rs410741 | -0.823 | 0.481 | -0.679 | 0.453 | 0 | 0 | 0 | 0 | 0.63 | 0.69 | 0.74 | 0.69 | 0.63 | 0.59 |
| 5 | 9403198 | rs10214283 | 0.65 | 0.466 | -0.768 | 0.372 | -0.009 | -0.03 | -0.03 | 0.278 | 0.06 | 0.12 | 0.06 | 0 | 0.05 | 0.01 |
| 5 | 9403456 | rs17323208 | -0.359 | 0.597 | -0.953 | 0.552 | -0.009 | -0.012 | -0.012 | 4.56 | 0.14 | 0.07 | 0.09 | 0.1 | 0.15 | 0.1 |
| 5 | 9409056 | rs2463503 | 0.136 | 0.491 | -0.616 | 0.327 | -0.004 | -0.046 | -0.046 | 0 | 0.87 | 0.83 | 0.94 | 0.97 | 0.9 | 0.9 |
| 5 | 9411249 | rs6879538 | 0.762 | 0.582 | -0.986 | 0.55 | -0.018 | 0.035 | 0.035 | 0 | 0.3 | 0.1 | 0.25 | 0.2 | 0.35 | 0.37 |
| 5 | 9413362 | rs425620 | -0.032 | 0.686 | -0.595 | 1.503 | 0.004 | 0.049 | 0.049 | 0 | 0.07 | 0.12 | 0.32 | 0.29 | 0.07 | 0.2 |
| 5 | 9417622 | rs4701846 | 1.883 | -0.156 | -0.339 | 1.412 | 0.012 | 0.013 | 0.013 | 0 | 0.55 | 0.31 | 0.39 | 0.32 | 0.6 | 0.4 |
| 5 | 9420374 | rs666617 | -0.478 | -0.048 | -0.276 | 1.465 | -0.008 | 0.006 | 0.006 | 0 | 0.23 | 0.26 | 0.21 | 0.32 | 0.24 | 0.3 |
| 5 | 9423834 | rs17258120 | 0.295 | 0.197 | -0.192 | 1.404 | 0.002 | -0.011 | -0.011 | 0 | 0.15 | 0.09 | 0.09 | 0.04 | 0.12 | 0.07 |
| 5 | 9424801 | rs7724626 | 0.758 | 0.546 | -0.021 | 2.317 | 0.003 | 0.054 | 0.054 | 0 | 0.8 | 0.65 | 0.55 | 0.56 | 0.76 | 0.64 |
| 5 | 9426734 | rs268486 | -0.76 | 0.566 | -0.175 | 2.378 | -0.001 | 0.144 | 0.144 | 0.117 | 0.17 | 0.46 | 0.18 | 0.33 | 0.18 | 0.24 |

| | | | | | | | | | | | | | | | | |
|---|---|---|---|---|---|---|---|---|---|---|---|---|---|---|---|---|
| 5 | 9429290 | rs11951666 | -0.21 | 0.951 | -1.294 | 1.908 | -0.001 | -0.003 | -0.003 | 2.606 | 0.13 | 0.17 | 0.11 | 0.14 | 0.18 | 0.19 |
| 5 | 9435478 | rs268482 | -0.034 | 0.942 | -1.254 | 1.805 | -0.018 | -0.01 | -0.01 | 0 | 0.94 | 0.97 | 0.68 | 0.73 | 0.94 | 0.84 |
| 5 | 9438120 | rs3798009 | 0.1 | 1.188 | -1.703 | 0.809 | -0.079 | -0.057 | -0.057 | 0 | 0.2 | 0.01 | 0.12 | 0.24 | 0.26 | 0.15 |
| 5 | 9438454 | rs3822794 | -0.51 | 1.172 | -0.931 | 0.572 | 0.007 | -0.012 | -0.012 | 0 | 0.07 | 0.12 | 0.06 | 0.06 | 0.08 | 0.07 |
| 5 | 9438538 | rs1858428 | 0.086 | 0.795 | -0.697 | 0.178 | 0.005 | 0.017 | 0.017 | 0 | 0.53 | 0.6 | 0.52 | 0.72 | 0.61 | 0.67 |
| 5 | 9438778 | rs268534 | -1.075 | 0.954 | 0.178 | 0.213 | -0.014 | -0.073 | -0.073 | 0 | 0.35 | 0.43 | 0.23 | 0.13 | 0.28 | 0.26 |
| 5 | 9439642 | rs3756725 | 0.303 | 1.17 | 0.313 | 0.602 | 0.036 | 0.075 | 0.075 | 0.111 | 0.06 | 0.13 | 0.1 | 0.2 | 0.12 | 0.06 |
| 5 | 9440094 | rs1512509 | -0.517 | 0.903 | 0.538 | 0.446 | -0.04 | -0.067 | -0.067 | 0 | 0.23 | 0.59 | 0.17 | 0.13 | 0.15 | 0.12 |
| 5 | 9441720 | rs268532 | -1.959 | 1.019 | 0.361 | 0.595 | 0.011 | 0.018 | 0.018 | 0 | 0.47 | 0.79 | 0.4 | 0.51 | 0.42 | 0.47 |
| 5 | 9442007 | rs2135072 | -1.069 | 1.109 | 0.462 | 0.671 | -0.006 | -0.007 | -0.007 | 0 | 0.21 | 0.2 | 0.22 | 0.38 | 0.27 | 0.35 |
| 5 | 9442994 | rs268529 | -1.857 | 0.018 | 0.494 | 0.102 | -0.005 | -0.02 | -0.02 | 0.063 | 0.4 | 0.6 | 0.3 | 0.24 | 0.31 | 0.4 |
| 5 | 9445434 | rs707637 | -1.356 | -0.702 | 0.264 | 0.343 | -0.005 | -0.005 | -0.005 | 0 | 0.32 | 0.65 | 0.2 | 0.23 | 0.16 | 0.25 |
| 5 | 9445461 | rs1512508 | 0.197 | -0.44 | -0.015 | 0.446 | -0.024 | -0.061 | -0.061 | 0 | 0.15 | 0.28 | 0.12 | 0.02 | 0.1 | 0.05 |
| 5 | 9450621 | rs440396 | -1.13 | -0.236 | -0.376 | 0.805 | 0.013 | 0.011 | 0.011 | 1.759 | 0.17 | 0.13 | 0.05 | 0.05 | 0.09 | 0.15 |
| 5 | 9450746 | rs3822799 | -0.301 | 0.338 | -0.583 | 1.056 | 0.022 | 0.042 | 0.042 | 0 | 0.05 | 0.15 | 0.05 | 0.16 | 0.05 | 0.13 |
| 5 | 9453089 | rs436243 | 0.294 | 0.046 | -0.954 | 0.677 | -0.008 | -0.065 | -0.065 | 0 | 0.86 | 0.97 | 0.89 | 0.75 | 0.89 | 0.82 |
| 5 | 9454208 | rs3756727 | 0.259 | 0.124 | -0.56 | 0.479 | -0.02 | 0.001 | 0.001 | 0.68 | 0.64 | 0.54 | 0.74 | 0.63 | 0.75 | 0.7 |
| 5 | 9456166 | rs268542 | -0.062 | 0.823 | -0.786 | 0.387 | -0.016 | -0.012 | -0.012 | 0.246 | 0.32 | 0.28 | 0.36 | 0.57 | 0.47 | 0.55 |
| 5 | 9457243 | rs3798031 | 0.266 | 0.394 | -0.72 | -0.426 | -0.001 | -0.002 | -0.002 | 0.881 | 0.29 | 0.26 | 0.37 | 0.07 | 0.33 | 0.16 |
| 5 | 9463291 | rs397287 | 0.007 | 0.236 | -0.746 | 0.012 | 0.001 | 0 | 0 | 0.212 | 0.47 | 0.54 | 0.39 | 0.55 | 0.37 | 0.55 |
| 5 | 9466215 | rs16882852 | 0.488 | 0.263 | -0.907 | -0.145 | 0.001 | 0.025 | 0.025 | 0 | 0.16 | 0.1 | 0.1 | 0.02 | 0.18 | 0.1 |
| 5 | 9471319 | rs16882862 | 0.662 | 0.091 | -0.795 | -0.112 | -0.007 | 0.011 | 0.011 | 0 | 0.37 | 0.31 | 0.45 | 0.1 | 0.46 | 0.25 |
| 5 | 9473948 | rs171992 | -0.008 | 0.042 | -1.156 | 0.007 | -0.01 | 0.012 | 0.012 | 0 | 0.81 | 0.78 | 0.84 | 0.66 | 0.85 | 0.81 |
| 5 | 9474946 | rs433755 | 0.478 | 0.002 | -0.919 | -0.193 | 0.002 | 0.002 | 0.002 | 0 | 0.48 | 0.53 | 0.53 | 0.13 | 0.56 | 0.43 |
| 5 | 9483209 | rs418446 | -0.194 | 0.041 | -1.181 | -0.121 | 0 | -0.001 | -0.001 | 0 | 0.8 | 0.81 | 0.84 | 0.66 | 0.85 | 0.81 |
| 5 | 9485396 | rs268517 | 0.153 | 0.341 | -1.089 | 0.475 | 0.009 | 0.021 | 0.021 | 0.38 | 0.93 | 0.96 | 0.92 | 0.66 | 0.99 | 0.86 |
| 5 | 9488689 | rs950050 | 0.385 | 0.562 | -1.106 | 0.532 | 0.001 | -0.004 | -0.004 | 0 | 0.07 | 0 | 0.04 | 0.09 | 0.05 | 0.06 |
| 5 | 9495607 | rs12657696 | -0.072 | 0.623 | -0.93 | 0.481 | -0.007 | -0.04 | -0.04 | 0.006 | 0.31 | 0.19 | 0.3 | 0.53 | 0.3 | 0.38 |
| 5 | 9507624 | rs369459 | 0.653 | 0.742 | -0.814 | 0.662 | 0.002 | 0.058 | 0.058 | 1.649 | 0.9 | 0.86 | 0.86 | 0.63 | 0.91 | 0.77 |
| 5 | 9509018 | rs442173 | 0.466 | 0.808 | -0.994 | 0.658 | -0.006 | 0.006 | 0.006 | 1.215 | 0.91 | 0.91 | 0.86 | 0.65 | 0.91 | 0.77 |
| 5 | 9509811 | rs11959773 | -0.244 | 0.336 | -0.882 | 0.613 | -0.003 | -0.018 | -0.018 | 0 | 0.31 | 0.23 | 0.31 | 0.53 | 0.3 | 0.37 |
| 5 | 9519853 | rs268558 | 0.389 | 0.375 | -0.886 | 0.334 | 0.002 | 0.009 | 0.009 | 0 | 0.66 | 0.72 | 0.61 | 0.16 | 0.71 | 0.55 |
| 5 | 9522303 | rs465107 | 0.062 | 1.002 | -1.173 | 1.437 | -0.052 | -0.062 | -0.062 | 0.112 | 0.9 | 0.99 | 0.75 | 0.64 | 0.83 | 0.88 |
| 5 | 9523375 | rs3798068 | 0.502 | 0.678 | -1.11 | 0.397 | 0 | -0.025 | -0.025 | 0.357 | 0.4 | 0.36 | 0.44 | 0.11 | 0.44 | 0.39 |
| 5 | 9529244 | rs465991 | 0.185 | 0.154 | -0.977 | 0.737 | -0.002 | 0.001 | 0.001 | 0 | 0.36 | 0.4 | 0.23 | 0.42 | 0.3 | 0.36 |
| 5 | 9529809 | rs462473 | 0.377 | 0.12 | -0.945 | 0.195 | -0.003 | 0.001 | 0.001 | 0 | 0.31 | 0.29 | 0.19 | 0.13 | 0.29 | 0.26 |
| 5 | 9534206 | rs268551 | 0.421 | 0.104 | -0.982 | 0.127 | -0.003 | -0.002 | -0.002 | 0 | 0.31 | 0.28 | 0.19 | 0.1 | 0.29 | 0.26 |
| 5 | 9534392 | rs3756737 | 0.458 | 0.138 | -1.309 | 0.048 | -0.02 | 0.025 | 0.025 | 0 | 0.37 | 0.19 | 0.45 | 0.25 | 0.43 | 0.41 |
| 5 | 9536983 | rs3798083 | 0.753 | -0.295 | -0.906 | -0.091 | -0.011 | -0.012 | -0.012 | 0.041 | 0.19 | 0.18 | 0.18 | 0.1 | 0.11 | 0.21 |
| 5 | 9539540 | rs6881628 | -0.015 | -0.008 | -0.57 | 0.095 | -0.004 | 0.01 | 0.01 | 0 | 0.84 | 0.77 | 0.82 | 0.76 | 0.84 | 0.83 |
| 5 | 9541289 | rs3798089 | -0.015 | -0.008 | -0.57 | 0.095 | -0.004 | 0.01 | 0.01 | 0 | 0.84 | 0.76 | 0.82 | 0.76 | 0.84 | 0.83 |
| 5 | 9541480 | rs16883082 | 0.433 | 0.936 | -0.55 | -0.006 | 0.028 | -0.004 | -0.004 | 0 | 0.07 | 0.1 | 0.06 | 0.03 | 0.15 | 0.15 |
| 5 | 9542529 | rs3798097 | 0.751 | 1.211 | -1.109 | -0.247 | -0.108 | 0.062 | 0.062 | 0 | 0.25 | 0.03 | 0.17 | 0.09 | 0.3 | 0.26 |
| 5 | 9547129 | rs1651283 | -0.436 | 1.933 | -0.659 | -0.218 | 0.01 | -0.02 | -0.02 | 0 | 0.33 | 0.38 | 0.3 | 0.16 | 0.38 | 0.3 |
| 5 | 9548098 | rs1398337 | 0.593 | 1.794 | -0.614 | -0.215 | 0.007 | 0.023 | 0.023 | 0.349 | 0.22 | 0.17 | 0.2 | 0.13 | 0.18 | 0.15 |
| 5 | 9550642 | rs930076 | 0.341 | 1.503 | -0.376 | 0.34 | 0.016 | 0.001 | 0.001 | 0.524 | 0.19 | 0.24 | 0.14 | 0.18 | 0.29 | 0.34 |
| 5 | 9550770 | rs16883128 | 0.01 | 1.043 | -0.195 | 0.469 | 0.022 | 0.053 | 0.053 | 0 | 0.08 | 0.17 | 0.14 | 0.17 | 0.11 | 0.04 |
| 5 | 9551091 | rs17329702 | -0.042 | 1.006 | -0.193 | 0.471 | 0.032 | 0.049 | 0.049 | 0 | 0.09 | 0.18 | 0.15 | 0.17 | 0.12 | 0.05 |
| 5 | 9551198 | rs16883129 | 1.311 | 0.554 | -0.502 | 0.314 | 0.006 | -0.007 | -0.007 | 0 | 0.3 | 0.22 | 0.36 | 0.41 | 0.18 | 0.3 |
| 5 | 9551296 | rs978896 | -0.36 | 1.053 | -0.502 | 0.246 | 0.004 | -0.01 | -0.01 | 0 | 0.31 | 0.39 | 0.35 | 0.3 | 0.3 | 0.2 |
| 5 | 9558232 | rs218057 | 0.264 | 0.826 | -0.516 | -0.104 | 0.036 | -0.051 | -0.051 | 0 | 0.66 | 0.54 | 0.63 | 0.84 | 0.55 | 0.66 |
| 5 | 9559903 | rs150633 | 0.743 | 0.432 | -0.669 | 0.153 | 0.012 | 0.002 | 0.002 | 0 | 0.46 | 0.41 | 0.43 | 0.57 | 0.31 | 0.35 |
| 5 | 9560249 | rs150631 | 1.074 | 0.221 | -0.796 | 0.062 | -0.01 | 0.007 | 0.007 | 0 | 0.34 | 0.35 | 0.37 | 0.53 | 0.22 | 0.31 |
| 5 | 9562434 | rs1508797 | -1.715 | 0.64 | -0.914 | -0.029 | -0.025 | -0.015 | -0.015 | 0.195 | 0.94 | 0.99 | 0.86 | 0.91 | 0.86 | 0.97 |
| 5 | 9562669 | rs1508796 | 1.514 | 0.44 | -0.627 | -0.295 | -0.035 | -0.082 | -0.082 | 0 | 0.15 | 0.31 | 0.12 | 0 | 0.11 | 0.08 |
| 5 | 9565826 | rs7728549 | -0.493 | 0.861 | -0.677 | 0.033 | 0 | 0.002 | 0.002 | 0 | 0.51 | 0.54 | 0.48 | 0.46 | 0.5 | 0.46 |
| 5 | 9568765 | rs1508801 | 0.63 | 0.957 | -1.045 | 0.125 | -0.016 | -0.028 | -0.028 | 0.669 | 0.17 | 0.09 | 0.19 | 0.31 | 0.23 | 0.28 |
| 5 | 9575041 | rs218041 | 0.553 | 0.565 | -0.945 | 0.111 | 0.003 | 0.011 | 0.011 | 0 | 0.51 | 0.51 | 0.51 | 0.67 | 0.44 | 0.58 |
| 5 | 9578794 | rs218053 | -1.208 | 0.666 | -1.373 | 0.302 | -0.017 | -0.003 | -0.003 | 0.99 | 0.91 | 0.98 | 0.84 | 0.9 | 0.86 | 0.87 |
| 5 | 9580273 | rs218052 | 1.939 | 0.129 | -1.627 | 0.458 | 0.038 | -0.042 | -0.042 | 0.177 | 0.12 | 0.05 | 0.18 | 0.3 | 0.05 | 0.09 |
| 5 | 9592694 | rs41496 | 0.606 | 1.469 | -1.293 | 0.726 | -0.01 | -0.03 | -0.03 | 0 | 0.14 | 0.07 | 0.16 | 0.27 | 0.18 | 0.25 |
| 5 | 9598097 | rs6870771 | 0.957 | 1.181 | -1.085 | 0.563 | 0.022 | 0.002 | 0.002 | 0 | 0.05 | 0.08 | 0.04 | 0.03 | 0.1 | 0.09 |
| 5 | 9602682 | rs218049 | -0.323 | 1.11 | -1.253 | 0.622 | -0.037 | -0.027 | -0.027 | 0.009 | 0.8 | 0.98 | 0.69 | 0.8 | 0.78 | 0.84 |
| 5 | 9604363 | rs390 | 0.957 | 1.064 | -0.918 | 0.434 | -0.002 | 0.003 | 0.003 | 0 | 0.57 | 0.54 | 0.48 | 0.56 | 0.55 | 0.65 |
| 5 | 9606057 | rs400 | -0.535 | 1.354 | -1.001 | 0.202 | 0.022 | 0.045 | 0.045 | 0.043 | 0.11 | 0.23 | 0.13 | 0.2 | 0.12 | 0.09 |
| 5 | 9606166 | rs3846591 | -1.025 | 1.396 | -0.987 | 0.198 | 0.061 | 0.049 | 0.049 | 0.844 | 0.16 | 0.32 | 0.17 | 0.24 | 0.22 | 0.19 |
| 5 | 9606663 | rs3846595 | -0.951 | 1.429 | -0.991 | 0.194 | 0.006 | -0.003 | -0.003 | 0 | 0.28 | 0.33 | 0.23 | 0.24 | 0.32 | 0.23 |
| 5 | 9607318 | rs6897599 | 0.255 | 1.119 | -1.198 | 0.187 | 0.031 | -0.038 | -0.038 | 0 | 0.07 | 0.16 | 0.06 | 0 | 0.1 | 0.07 |
| 5 | 9608903 | rs10474846 | -0.85 | 1.057 | -1.196 | 0.159 | 0.001 | 0.003 | 0.003 | 0 | 0.34 | 0.36 | 0.29 | 0.21 | 0.33 | 0.2 |

| | | | | | | | | | | | | | | | | |
|---|---|---|---|---|---|---|---|---|---|---|---|---|---|---|---|---|
| 5 | 9609808 | rs17265148 | -0.148 | 1.069 | -1.351 | 0.151 | 0.031 | 0.025 | 0.025 | 0 | 0.27 | 0.18 | 0.22 | 0.2 | 0.22 | 0.13 |
| 5 | 9610282 | rs6874254 | -0.505 | 1.044 | -1.4 | 0.291 | 0.004 | 0.016 | 0.016 | 0 | 0.34 | 0.33 | 0.29 | 0.2 | 0.33 | 0.2 |
| 5 | 9616496 | rs218036 | 0.334 | 1.526 | -1.495 | 1.052 | 0.001 | 0.003 | 0.003 | 0 | 0.58 | 0.55 | 0.51 | 0.46 | 0.56 | 0.53 |
| 5 | 9617380 | rs385 | -0.927 | 2.579 | -2.239 | 1.581 | -0.078 | -0.103 | -0.103 | 0 | 0.18 | 0.05 | 0.34 | 0.37 | 0.3 | 0.19 |
| 5 | 9617512 | rs41457 | 0.684 | 3.076 | -2.231 | 0.804 | -0.036 | 0.029 | 0.029 | 0 | 0.79 | 0.87 | 0.76 | 0.81 | 0.62 | 0.73 |
| 5 | 9617945 | rs395 | 0.82 | 2.147 | -2.433 | 1.206 | -0.072 | -0.111 | -0.111 | 0 | 0.78 | 0.94 | 0.77 | 0.6 | 0.69 | 0.77 |
| 5 | 9619905 | rs41461 | -0.232 | 1.667 | -2.775 | 0.361 | -0.122 | 0.056 | 0.056 | 0 | 0.71 | 0.94 | 0.78 | 0.78 | 0.56 | 0.68 |
| 5 | 9621540 | rs41463 | -0.371 | 1.718 | -2.802 | 0.31 | -0.112 | 0.093 | 0.093 | 0.088 | 0.73 | 0.94 | 0.81 | 0.87 | 0.58 | 0.71 |
| 5 | 9629295 | rs2234235 | 0.999 | 0.963 | -3.011 | 0.022 | 0.011 | 0.04 | 0.04 | 0 | 0.08 | 0.01 | 0.02 | 0 | 0.05 | 0.03 |
| 5 | 9629529 | rs2234233 | -0.263 | 0.758 | -3.458 | 0.071 | -0.052 | -0.034 | -0.034 | 0 | 0.12 | 0.03 | 0.13 | 0.11 | 0.15 | 0.19 |
| 5 | 9641628 | rs3213627 | -0.724 | 0.457 | -2.249 | 0.661 | -0.001 | 0.006 | 0.006 | 0 | 0.21 | 0.18 | 0.16 | 0.12 | 0.2 | 0.22 |
| 5 | 9648873 | rs17272325 | -0.799 | 0.116 | -2.356 | 0.478 | 0.001 | 0.022 | 0.022 | 0 | 0.22 | 0.15 | 0.15 | 0.11 | 0.2 | 0.22 |
| 5 | 9659384 | rs27843 | 0.599 | -1.078 | -2.267 | -0.226 | -0.029 | 0.046 | 0.046 | 0.091 | 0.42 | 0.19 | 0.47 | 0.35 | 0.52 | 0.51 |
| 5 | 9661352 | rs1393232 | 0.163 | -0.85 | -1.902 | -0.146 | -0.019 | -0.015 | -0.015 | 1.831 | 0.11 | 0.06 | 0.11 | 0.12 | 0.14 | 0.2 |
| 5 | 9667062 | rs10513029 | 0.051 | -1.969 | -1.845 | -0.684 | 0.008 | 0.003 | 0.003 | 0 | 0.21 | 0.22 | 0.32 | 0.19 | 0.15 | 0.26 |
| 5 | 9677855 | rs1604463 | 0.701 | -1.85 | -1.886 | -0.642 | 0.001 | -0.002 | -0.002 | 0.442 | 0.11 | 0.09 | 0.26 | 0.18 | 0.1 | 0.23 |
| 5 | 9679139 | rs428378 | 0.91 | -2.428 | -1.689 | -1.51 | 0.021 | 0.026 | 0.026 | 0 | 0.51 | 0.42 | 0.39 | 0.22 | 0.67 | 0.44 |
| 5 | 9681798 | rs12109904 | 1.59 | -1.935 | -1.861 | -1.26 | -0.054 | 0.035 | 0.035 | 0 | 0.12 | 0.03 | 0.13 | 0.07 | 0.19 | 0.13 |
| 5 | 9684672 | rs194070 | 0.945 | -2.388 | -1.899 | -1.37 | 0.057 | 0.096 | 0.096 | 0 | 0.46 | 0.18 | 0.34 | 0.27 | 0.59 | 0.52 |
| 5 | 9688551 | rs28738 | 1.604 | -2.129 | -2.186 | -1.578 | -0.124 | 0.109 | 0.109 | 0 | 0.32 | 0.06 | 0.22 | 0.17 | 0.45 | 0.23 |
| 5 | 9689026 | rs10039101 | 1.196 | -2.051 | -1.811 | -1.481 | 0.02 | -0.065 | -0.065 | 0 | 0.17 | 0.3 | 0.14 | 0.06 | 0.18 | 0.12 |
| 5 | 9690464 | rs42372 | 1.411 | -1.363 | -2.259 | -1.134 | -0.08 | 0.073 | 0.073 | 0 | 0.3 | 0.13 | 0.21 | 0.17 | 0.43 | 0.23 |
| 5 | 9692938 | rs734618 | -0.323 | -1.08 | -1.684 | -0.483 | -0.062 | -0.039 | -0.039 | 0 | 0.15 | 0.01 | 0.14 | 0.11 | 0.17 | 0.28 |
| 5 | 9695667 | rs41491 | 0.525 | -1.586 | -1.372 | -0.871 | 0.026 | 0.021 | 0.021 | 0 | 0.64 | 0.36 | 0.47 | 0.26 | 0.75 | 0.62 |
| 5 | 9695843 | rs7713633 | 0.854 | -0.282 | -0.747 | -1.272 | -0.001 | 0 | 0 | 0 | 0.48 | 0.41 | 0.32 | 0.23 | 0.51 | 0.44 |
| 5 | 9698490 | rs960567 | -0.018 | -0.578 | -0.938 | -1.289 | 0.002 | 0.005 | 0.005 | 0 | 0.25 | 0.24 | 0.21 | 0.2 | 0.22 | 0.39 |
| 5 | 9698629 | rs16883397 | -0.085 | -0.553 | -0.902 | -1.286 | 0 | 0 | 0 | 0 | 0.26 | 0.28 | 0.21 | 0.2 | 0.22 | 0.39 |
| 5 | 9699519 | rs41487 | -0.777 | -0.453 | -1.229 | -1.329 | -0.008 | 0.048 | 0.048 | 0 | 0.43 | 0.33 | 0.59 | 0.25 | 0.46 | 0.49 |
| 5 | 9699758 | rs12519432 | 1.502 | 0.201 | -2.16 | -1.123 | -0.067 | -0.027 | -0.027 | 0.674 | 0.21 | 0.07 | 0.14 | 0.25 | 0.28 | 0.21 |
| 5 | 9708576 | rs168788 | 0.142 | -0.142 | -1.822 | -0.819 | 0.054 | 0.053 | 0.053 | 0 | 0.49 | 0.3 | 0.55 | 0.41 | 0.58 | 0.42 |
| 5 | 9709010 | rs194065 | -0.997 | -0.41 | -1.957 | -0.514 | 0.025 | -0.116 | -0.116 | 0.092 | 0.8 | 0.95 | 0.8 | 0.6 | 0.8 | 0.81 |
| 5 | 9714227 | rs10036822 | -0.3 | 0.645 | -1.361 | -0.95 | -0.002 | -0.017 | -0.017 | 0 | 0.15 | 0.16 | 0.07 | 0 | 0.16 | 0.05 |
| 5 | 9715924 | rs118583 | 0.261 | 0.741 | -1.318 | -0.42 | -0.057 | -0.023 | -0.023 | 0 | 0.17 | 0.07 | 0.32 | 0.22 | 0.28 | 0.29 |
| 5 | 9716549 | rs1015855 | 0.537 | 0.727 | -1.258 | -0.451 | -0.01 | 0.018 | 0.018 | 0 | 0.32 | 0.24 | 0.18 | 0.23 | 0.34 | 0.22 |
| 5 | 9716840 | rs4702641 | 0.157 | -0.387 | -1.289 | -0.137 | -0.001 | 0.031 | 0.031 | 0 | 0.1 | 0.05 | 0.22 | 0.04 | 0.09 | 0.08 |
| 5 | 9717016 | rs194060 | 0.21 | -0.43 | -0.652 | 0.265 | -0.008 | 0.002 | 0.002 | 0 | 0.62 | 0.59 | 0.72 | 0.46 | 0.7 | 0.56 |
| 5 | 9721328 | rs1501958 | -0.576 | -0.534 | -0.422 | -0.258 | -0.024 | -0.04 | -0.04 | 0 | 0.28 | 0.34 | 0.16 | 0.19 | 0.16 | 0.19 |
| 5 | 9731900 | rs4701849 | -0.886 | -0.999 | -0.827 | -0.756 | 0.06 | 0.141 | 0.141 | 0.314 | 0.53 | 0.81 | 0.35 | 0.3 | 0.36 | 0.35 |
| 5 | 9732441 | rs10054093 | -0.013 | -0.852 | -0.833 | -0.684 | 0.006 | 0.048 | 0.048 | 0 | 0.13 | 0.02 | 0.09 | 0.03 | 0.13 | 0.11 |
| 5 | 9738382 | rs17274475 | -0.773 | -1.228 | -0.798 | -1.422 | 0.051 | 0.157 | 0.157 | 0.531 | 0.44 | 0.2 | 0.28 | 0.21 | 0.36 | 0.3 |
| 5 | 9739621 | rs3094360 | -0.857 | -0.249 | -1.225 | -1.535 | -0.019 | 0.021 | 0.021 | 0 | 0.84 | 0.9 | 0.87 | 0.93 | 0.78 | 0.83 |
| 5 | 9741016 | rs17334088 | 0.508 | -0.721 | -1.637 | -1.789 | -0.014 | 0.033 | 0.033 | 0.007 | 0.21 | 0.01 | 0.13 | 0.17 | 0.15 | 0.16 |
| 5 | 9758916 | rs3110986 | -0.564 | -0.561 | -0.98 | -1.424 | -0.017 | 0.011 | 0.011 | 0 | 0.82 | 0.88 | 0.84 | 0.91 | 0.75 | 0.81 |
| 5 | 9763915 | rs3094352 | 1.007 | -1.518 | -0.529 | -1.853 | 0.001 | 0 | 0 | 0 | 0.33 | 0.37 | 0.55 | 0.63 | 0.43 | 0.51 |
| 5 | 9764854 | rs3110984 | 1.314 | -1.65 | -0.533 | -0.533 | -0.085 | -0.069 | -0.069 | 0 | 0.3 | 0.08 | 0.51 | 0.37 | 0.42 | 0.4 |
| 5 | 9766668 | rs11134367 | 0.325 | -1.84 | -0.345 | -0.604 | 0.045 | -0.018 | -0.018 | 0.035 | 0.41 | 0.25 | 0.27 | 0.46 | 0.3 | 0.37 |
| 5 | 9770335 | rs3094348 | 1.013 | -1.681 | -0.482 | -0.475 | -0.041 | -0.027 | -0.027 | 1.868 | 0.32 | 0.16 | 0.52 | 0.36 | 0.42 | 0.41 |
| 5 | 9781148 | rs2905184 | 1.212 | -1.628 | -0.652 | -0.453 | -0.089 | -0.065 | -0.065 | 4.391 | 0.3 | 0.07 | 0.51 | 0.35 | 0.42 | 0.4 |
| 5 | 9782994 | rs1476456 | -0.251 | -1.587 | -0.051 | -0.456 | 0.022 | -0.009 | -0.009 | 0 | 0.77 | 0.65 | 0.83 | 0.82 | 0.75 | 0.78 |
| 5 | 9790350 | rs1008624 | -0.943 | -1.55 | -0.135 | -0.38 | -0.009 | 0.001 | 0.001 | 0 | 0.82 | 0.86 | 0.84 | 0.86 | 0.75 | 0.79 |
| 5 | 9793237 | rs1008253 | 0.322 | -1.397 | -0.286 | -0.249 | 0.002 | 0.003 | 0.003 | 0 | 0.37 | 0.4 | 0.54 | 0.41 | 0.43 | 0.43 |
| 5 | 9793538 | rs3110981 | -1.051 | -0.806 | -0.346 | -0.219 | -0.043 | -0.031 | -0.031 | 0 | 0.93 | 1 | 0.95 | 0.9 | 0.86 | 0.93 |
| 5 | 9797374 | rs1006227 | 0.337 | -1.03 | 0.036 | -0.203 | 0.011 | 0.053 | 0.053 | 2.042 | 0.06 | 0.24 | 0.03 | 0.05 | 0 | 0.03 |
| 5 | 9814581 | rs16883565 | 1.102 | -1.279 | -0.157 | -0.316 | 0 | 0 | 0 | 0 | 0.07 | 0.05 | 0.08 | 0.11 | 0.05 | 0.08 |
| 5 | 9815577 | rs2905211 | 0.241 | -1.594 | -0.641 | -0.455 | -0.001 | -0.001 | -0.001 | 0 | 0.38 | 0.35 | 0.55 | 0.41 | 0.43 | 0.42 |
| 5 | 9816942 | rs2027996 | 0.701 | -1.169 | -1.617 | -0.375 | -0.096 | -0.068 | -0.068 | 0 | 0.31 | 0.08 | 0.51 | 0.35 | 0.43 | 0.39 |
| 5 | 9821723 | rs16883572 | 0.517 | -1.069 | -0.827 | -0.17 | 0 | 0 | 0 | 0 | 0.07 | 0.06 | 0.08 | 0.11 | 0.05 | 0.08 |
| 5 | 9827532 | rs2106358 | -0.028 | -1.613 | -0.588 | 0.185 | -0.023 | 0.052 | 0.052 | 0 | 0.16 | 0.27 | 0.26 | 0.27 | 0.11 | 0.22 |
| 5 | 9830381 | rs10491232 | 1.102 | -0.573 | -1.262 | 0.528 | -0.05 | 0.029 | 0.029 | 0.333 | 0.22 | 0.11 | 0.28 | 0.15 | 0.38 | 0.29 |
| 5 | 9831983 | rs1015588 | 0.689 | -0.841 | -1.704 | 0.718 | -0.004 | -0.056 | -0.056 | 3.364 | 0.13 | 0.01 | 0.13 | 0.21 | 0.12 | 0.1 |
| 5 | 9832022 | rs1015587 | -0.751 | -0.914 | -1.52 | 0.677 | 0.015 | -0.02 | -0.02 | 0 | 0.86 | 0.93 | 0.87 | 0.79 | 0.88 | 0.9 |
| 5 | 9834220 | rs10070717 | 1.613 | -1.191 | -1.857 | 0.31 | 0.02 | -0.03 | -0.03 | 0.242 | 0.16 | 0.05 | 0.13 | 0.2 | 0.09 | 0.15 |
| 5 | 9841870 | rs10069487 | 1.112 | -0.82 | -1.597 | 0.365 | 0.023 | 0.015 | 0.015 | 0.264 | 0.25 | 0.28 | 0.43 | 0.38 | 0.38 | 0.4 |
| 5 | 9847262 | rs4701850 | 1.483 | -0.775 | -1.659 | 0.02 | -0.001 | 0.039 | 0.039 | 0.68 | 0.13 | 0.04 | 0.04 | 0 | 0.08 | 0.05 |
| 5 | 9850994 | rs12652744 | 0.972 | -0.609 | -1.333 | 0.489 | 0.021 | -0.022 | -0.022 | 0 | 0.19 | 0.06 | 0.13 | 0.21 | 0.13 | 0.19 |
| 5 | 9852860 | rs2214020 | 0.833 | 0.08 | -0.663 | 0.351 | 0.01 | 0.017 | 0.017 | 0 | 0.2 | 0.34 | 0.16 | 0.17 | 0.22 | 0.21 |
| 5 | 9860402 | rs2963906 | -0.007 | -0.278 | -0.882 | 0.043 | 0.009 | 0.037 | 0.037 | 0 | 0.48 | 0.57 | 0.65 | 0.74 | 0.6 | 0.57 |
| 5 | 9860479 | rs17788970 | 0.179 | -0.863 | -0.787 | -0.375 | 0 | 0.017 | 0.017 | 0 | 0.33 | 0.26 | 0.26 | 0.16 | 0.3 | 0.3 |
| 5 | 9860737 | rs2963905 | -0.285 | -0.505 | -1.417 | -0.17 | 0.016 | -0.014 | -0.014 | 0 | 0.78 | 0.73 | 0.72 | 0.82 | 0.69 | 0.71 |

| | | | | | | | | | | | | | | | | |
|---|---|---|---|---|---|---|---|---|---|---|---|---|---|---|---|---|
| 5 | 9861402 | rs2963904 | -0.377 | -0.425 | -1.165 | 0.19 | -0.012 | 0.013 | 0.013 | 0 | 0.61 | 0.71 | 0.65 | 0.67 | 0.59 | 0.55 |
| 5 | 9868176 | rs11739224 | 0.463 | -0.53 | -1.23 | 0.212 | 0.005 | -0.036 | -0.036 | 0.569 | 0.07 | 0.15 | 0.07 | 0 | 0.07 | 0.02 |
| 5 | 9869692 | rs6898293 | 0.603 | -0.56 | -1.543 | 0.687 | -0.017 | 0.001 | 0.001 | 0.603 | 0.26 | 0.37 | 0.24 | 0.29 | 0.15 | 0.26 |
| 5 | 9870587 | rs2963899 | 0.609 | 0.224 | -2.62 | 0.668 | -0.042 | 0.016 | 0.016 | 0.022 | 0.26 | 0.12 | 0.3 | 0.19 | 0.39 | 0.24 |
| 5 | 9871766 | rs10491227 | -0.413 | -0.771 | -1.841 | 0.825 | 0.042 | -0.02 | -0.02 | 0.353 | 0.43 | 0.72 | 0.5 | 0.51 | 0.28 | 0.35 |
| 5 | 9877169 | rs2963894 | 0.763 | -0.196 | -1.852 | 0.875 | 0.029 | -0.043 | -0.043 | 0.812 | 0.7 | 0.15 | 0.59 | 0.54 | 0.76 | 0.64 |
| 5 | 9885006 | rs17196845 | -0.285 | 0.427 | -1.944 | 0.106 | -0.022 | 0.028 | 0.028 | 0.344 | 0.17 | 0.06 | 0.12 | 0.08 | 0.18 | 0.24 |
| 5 | 9888495 | rs16883772 | -0.647 | 0.005 | -1.208 | 0.434 | -0.007 | -0.005 | -0.005 | 0 | 0.11 | 0.21 | 0.07 | 0.12 | 0.07 | 0.11 |
| 5 | 9894959 | rs17275422 | -0.652 | 0.113 | -1.489 | 0.586 | -0.001 | -0.001 | -0.001 | 0.582 | 0.1 | 0.1 | 0.07 | 0.08 | 0.1 | 0.1 |
| 5 | 9896163 | rs6555620 | 0.205 | -0.376 | -2.231 | 0.454 | 0.082 | 0.013 | 0.013 | 0 | 0.58 | 0.17 | 0.48 | 0.38 | 0.74 | 0.52 |
| 5 | 9896556 | rs10513034 | 0.592 | 0.338 | -2.9 | -0.55 | -0.017 | 0.137 | 0.137 | 0 | 0.27 | 0.07 | 0.16 | 0.04 | 0.28 | 0.14 |
| 5 | 9896602 | rs4701853 | 0.026 | 0.356 | -3.56 | -0.073 | 0.009 | -0.097 | -0.097 | 0 | 0.36 | 0.05 | 0.36 | 0.46 | 0.31 | 0.32 |
| 5 | 9897923 | rs7707030 | -0.713 | 0.15 | -2.469 | -0.329 | -0.023 | -0.085 | -0.085 | 0 | 0.65 | 0.55 | 0.81 | 0.9 | 0.72 | 0.7 |
| 5 | 9898962 | rs10462871 | 0.78 | -0.464 | -2.496 | -0.044 | 0.036 | -0.022 | -0.022 | 0.153 | 0.22 | 0.1 | 0.23 | 0.29 | 0.13 | 0.15 |
| 5 | 9899989 | rs17335228 | 0.372 | 0.474 | -2.423 | -0.081 | -0.089 | -0.058 | -0.058 | 0.028 | 0.17 | 0.01 | 0.19 | 0.21 | 0.23 | 0.2 |
| 5 | 9902403 | rs7732991 | 0.564 | 0.163 | -1.732 | -0.076 | -0.004 | 0 | 0 | 0 | 0.08 | 0.08 | 0.17 | 0.12 | 0.07 | 0.22 |
| 5 | 9905783 | rs10513035 | 1.249 | -0.124 | -2.08 | -0.408 | 0.006 | 0.097 | 0.097 | 0 | 0.17 | 0 | 0.06 | 0.03 | 0.14 | 0.11 |
| 5 | 9907333 | rs16883810 | 0.543 | 0.137 | -1.804 | -0.341 | -0.018 | -0.077 | -0.077 | 1.444 | 0.23 | 0.42 | 0.13 | 0.09 | 0.19 | 0.22 |
| 5 | 9912072 | rs6555624 | 0.226 | 0.332 | -2.092 | -0.097 | -0.021 | -0.023 | -0.023 | 0.052 | 0.09 | 0.02 | 0.2 | 0.1 | 0.11 | 0.23 |
| 5 | 9915399 | rs3925077 | 0.246 | -0.381 | -1.86 | -0.117 | 0.014 | -0.004 | -0.004 | 3.846 | 0.27 | 0.23 | 0.25 | 0.27 | 0.16 | 0.19 |
| 5 | 9917666 | rs6871808 | -0.421 | -0.526 | -0.505 | -0.246 | 0.047 | 0.043 | 0.043 | 1.823 | 0.06 | 0.44 | 0.04 | 0 | 0 | 0 |
| 5 | 9919695 | rs4272105 | 0.523 | -0.158 | -0.955 | -0.447 | -0.001 | -0.001 | -0.001 | 0 | 0.42 | 0.41 | 0.41 | 0.56 | 0.51 | 0.42 |
| 5 | 9921028 | rs10513039 | 1.726 | 0.999 | -1.621 | 0.023 | -0.087 | -0.11 | -0.11 | 0.011 | 0.18 | 0.04 | 0.22 | 0.37 | 0.29 | 0.14 |
| 5 | 9923541 | rs11739710 | 0.637 | 0.296 | -0.911 | -0.377 | -0.002 | -0.035 | -0.035 | 0.015 | 0.23 | 0.24 | 0.14 | 0.06 | 0.24 | 0.21 |
| 5 | 9924597 | rs11949879 | 0.922 | 0.392 | -1.306 | -0.154 | -0.006 | 0.039 | 0.039 | 0 | 0.21 | 0.08 | 0.13 | 0.14 | 0.24 | 0.22 |
| 5 | 9928862 | rs1035961 | 0.189 | 0.56 | -0.843 | 0.217 | -0.003 | -0.006 | -0.006 | 0 | 0.06 | 0.02 | 0.14 | 0.22 | 0.05 | 0.26 |
| 5 | 9933220 | rs13179743 | 0.507 | 0.698 | -1.013 | 0.264 | -0.016 | -0.055 | -0.055 | 1.684 | 0.26 | 0.11 | 0.35 | 0.56 | 0.26 | 0.38 |
| 5 | 9933742 | rs12657074 | 1.34 | 0.689 | -1.026 | 0.31 | -0.018 | -0.072 | -0.072 | 0.133 | 0.2 | 0.08 | 0.21 | 0.34 | 0.21 | 0.12 |
| 5 | 9941041 | rs2914723 | -0.822 | 0.775 | -1.655 | -0.112 | -0.097 | -0.045 | -0.045 | 0.626 | 0.77 | 0.98 | 0.71 | 0.79 | 0.7 | 0.77 |
| 5 | 9941459 | rs1366620 | 0.31 | 0.208 | -1.097 | -0.447 | -0.004 | -0.009 | -0.009 | 0.063 | 0.32 | 0.48 | 0.35 | 0.37 | 0.33 | 0.42 |
| 5 | 9941748 | rs1428945 | -0.081 | 0.086 | -1.166 | -0.499 | -0.045 | 0.078 | 0.078 | 0 | 0.56 | 0.83 | 0.57 | 0.7 | 0.53 | 0.54 |
| 5 | 9941841 | rs1428946 | 0.868 | 0.196 | -0.666 | -0.184 | -0.006 | -0.01 | -0.01 | 0 | 0.08 | 0.03 | 0.19 | 0.22 | 0.12 | 0.26 |
| 5 | 9945590 | rs1428947 | 0.77 | 0.18 | -0.618 | -0.188 | -0.001 | 0 | 0 | 0 | 0.09 | 0.06 | 0.19 | 0.22 | 0.12 | 0.26 |
| 5 | 9946689 | rs2914726 | 0.51 | 0.068 | -0.559 | -0.336 | 0.004 | -0.003 | -0.003 | 0 | 0.44 | 0.41 | 0.35 | 0.41 | 0.36 | 0.35 |
| 5 | 9947222 | rs12152749 | 1.612 | 0.595 | -0.56 | -0.063 | -0.022 | -0.015 | -0.015 | 0 | 0.06 | 0.01 | 0.19 | 0.07 | 0.1 | 0.21 |
| 5 | 9947612 | rs1428949 | 0.104 | 0.365 | -0.513 | -0.142 | -0.012 | -0.007 | -0.007 | 0 | 0.52 | 0.8 | 0.42 | 0.48 | 0.48 | 0.32 |
| 5 | 9947938 | rs16883882 | 1.829 | 0.382 | -0.198 | 0.037 | -0.034 | -0.095 | -0.095 | 0 | 0.2 | 0.04 | 0.22 | 0.34 | 0.22 | 0.13 |
| 5 | 9948603 | rs1366624 | -0.047 | -0.081 | 0.487 | -0.05 | -0.001 | -0.001 | -0.001 | 0.143 | 0.43 | 0.61 | 0.41 | 0.42 | 0.4 | 0.37 |
| 5 | 9948723 | rs1366625 | -0.074 | -0.086 | 0.596 | -0.053 | -0.003 | -0.006 | -0.006 | 1.605 | 0.44 | 0.63 | 0.41 | 0.42 | 0.4 | 0.36 |
| 5 | 9951112 | rs7708238 | 1.602 | -0.182 | 0.527 | -0.023 | 0.012 | 0.04 | 0.04 | 0 | 0.22 | 0.09 | 0.11 | 0.13 | 0.21 | 0.15 |
| 5 | 9956338 | rs16883905 | 1.461 | -0.149 | 0.55 | -0.028 | 0.005 | 0.005 | 0.005 | 0 | 0.1 | 0.07 | 0.11 | 0.07 | 0.09 | 0.12 |
| 5 | 9956601 | rs10065959 | -0.513 | -0.082 | 0.87 | -0.009 | -0.001 | -0.003 | -0.003 | 0 | 0.44 | 0.57 | 0.54 | 0.45 | 0.44 | 0.56 |
| 5 | 9959769 | rs6874073 | 1.843 | 0.05 | 0.545 | 0.029 | 0.006 | 0.02 | 0.02 | 0 | 0.23 | 0.24 | 0.24 | 0.34 | 0.25 | 0.17 |
| 5 | 9963889 | rs1428954 | 0.147 | 0.031 | 0.058 | -0.196 | -0.008 | 0.014 | 0.014 | 0 | 0.6 | 0.71 | 0.49 | 0.56 | 0.57 | 0.45 |
| 5 | 9965794 | rs6867995 | 0.656 | 0.016 | 0.044 | -0.204 | 0 | 0.019 | 0.019 | 0 | 0.49 | 0.65 | 0.38 | 0.48 | 0.48 | 0.33 |
| 5 | 9967137 | rs38091 | 0.611 | -0.107 | -0.023 | -0.223 | -0.069 | -0.014 | -0.014 | 0 | 0.22 | 0.02 | 0.36 | 0.18 | 0.29 | 0.29 |
| 5 | 9971532 | rs11745764 | -0.146 | -0.14 | 0.673 | 0.157 | 0.013 | 0.024 | 0.024 | 0 | 0.38 | 0.67 | 0.27 | 0.2 | 0.32 | 0.29 |
| 5 | 9977436 | rs38083 | -0.109 | -0.223 | 0.813 | 0.297 | -0.018 | -0.014 | -0.014 | 0.728 | 0.62 | 0.45 | 0.68 | 0.74 | 0.7 | 0.72 |
| 5 | 9977993 | rs38081 | -0.097 | -0.071 | 0.581 | 0.505 | 0.008 | 0 | 0 | 0 | 0.33 | 0.41 | 0.49 | 0.42 | 0.41 | 0.53 |
| 5 | 9978292 | rs43224 | 0.122 | 0.014 | 0.765 | 0.118 | -0.001 | -0.001 | -0.001 | 0.328 | 0.53 | 0.48 | 0.53 | 0.62 | 0.59 | 0.6 |
| 5 | 9979072 | rs38076 | -0.578 | 0.044 | 0.893 | -0.011 | 0.006 | 0.006 | 0.006 | 0 | 0.66 | 0.71 | 0.7 | 0.75 | 0.7 | 0.73 |
| 5 | 9979084 | rs553169 | -0.142 | -0.056 | 0.99 | -0.073 | -0.01 | -0.014 | -0.014 | 2.597 | 0.61 | 0.56 | 0.71 | 0.75 | 0.69 | 0.71 |
| 5 | 9979240 | rs38075 | -1.343 | 0.032 | 0.579 | -0.029 | 0.014 | 0.022 | 0.022 | 0 | 0.76 | 0.84 | 0.88 | 0.87 | 0.82 | 0.85 |
| 5 | 9980315 | rs6866016 | 1.09 | 0.276 | 0.126 | 0.261 | -0.001 | -0.001 | -0.001 | 2.456 | 0.1 | 0.11 | 0.18 | 0.12 | 0.11 | 0.12 |
| 5 | 9980435 | rs10052972 | 0.964 | 0.428 | 0.039 | 0.388 | -0.008 | -0.027 | -0.027 | 0 | 0.31 | 0.2 | 0.21 | 0.5 | 0.29 | 0.22 |
| 5 | 9981548 | rs17196873 | 0.328 | 0.567 | -0.023 | 0.362 | -0.001 | 0 | 0 | 0 | 0.11 | 0.14 | 0.16 | 0.12 | 0.11 | 0.14 |
| 5 | 9983519 | rs38068 | -0.66 | 0.618 | 0.289 | 0.121 | 0.01 | -0.066 | -0.066 | 0 | 0.25 | 0.4 | 0.33 | 0.14 | 0.3 | 0.37 |
| 5 | 9984400 | rs38066 | -0.837 | 0.39 | 0.525 | 0.23 | 0.007 | -0.012 | -0.012 | 0.106 | 0.32 | 0.42 | 0.53 | 0.25 | 0.4 | 0.47 |
| 5 | 9984418 | rs38065 | -0.079 | 0.333 | 0.741 | 0.177 | -0.005 | 0.005 | 0.005 | 0 | 0.2 | 0.17 | 0.31 | 0.14 | 0.29 | 0.34 |
| 5 | 9984591 | rs38064 | 0.362 | 0.412 | 0.807 | 0.148 | -0.014 | 0.009 | 0.009 | 1.096 | 0.14 | 0.08 | 0.27 | 0.12 | 0.25 | 0.23 |
| 5 | 9984618 | rs10866598 | 1.018 | 0.045 | 0.491 | 0.201 | -0.021 | -0.047 | -0.047 | 0.153 | 0.38 | 0.19 | 0.3 | 0.44 | 0.38 | 0.33 |
| 5 | 9985893 | rs38061 | -0.666 | -0.196 | 0.633 | 0.17 | 0 | -0.003 | -0.003 | 0 | 0.55 | 0.47 | 0.65 | 0.4 | 0.6 | 0.62 |
| 5 | 9987380 | rs6875901 | 0.991 | -0.685 | 0.915 | 0.222 | 0.009 | 0.013 | 0.013 | 0 | 0.23 | 0.17 | 0.13 | 0.15 | 0.19 | 0.14 |
| 5 | 9987909 | rs10072374 | -0.095 | -0.679 | 1.39 | 0.315 | -0.041 | -0.01 | -0.01 | 0 | 0.12 | 0.4 | 0.23 | 0.06 | 0.08 | 0.09 |
| 5 | 9990529 | rs434147 | 0.585 | -0.538 | 1.276 | 0.349 | 0.002 | 0.001 | 0.001 | 0 | 0.57 | 0.46 | 0.57 | 0.59 | 0.67 | 0.63 |
| 5 | 9997370 | rs16883981 | 0.838 | -0.533 | 0.834 | 0.365 | 0.012 | 0.057 | 0.057 | 0 | 0.26 | 0.07 | 0.13 | 0.15 | 0.22 | 0.15 |
| 5 | 9998256 | rs6890044 | 0.534 | -0.394 | 0.537 | 0.401 | -0.008 | -0.005 | -0.005 | 0 | 0.09 | 0 | 0.18 | 0.06 | 0.1 | 0.12 |
| 5 | 9999146 | rs6891283 | 0.752 | -0.334 | -0.211 | 0.567 | 0 | 0 | 0 | 0 | 0.14 | 0.13 | 0.14 | 0.15 | 0.12 | 0.15 |
| 5 | 10018290 | rs1395093 | 0.326 | -0.5 | 0.132 | 0.391 | 0.017 | 0.025 | 0.025 | 0 | 0.56 | 0.39 | 0.56 | 0.76 | 0.66 | 0.67 |

| | | | | | | | | | | | | | | | | |
|---|---|---|---|---|---|---|---|---|---|---|---|---|---|---|---|---|
| 5 | 10031142 | rs10060537 | -0.318 | -0.289 | -0.023 | 0.973 | -0.004 | 0.004 | 0.004 | 0 | 0.25 | 0.25 | 0.15 | 0.3 | 0.2 | 0.19 |
| 5 | 10031449 | rs314731 | -0.717 | 0.214 | -0.511 | 0.701 | -0.03 | 0.015 | 0.015 | 0 | 0.44 | 0.24 | 0.39 | 0.43 | 0.51 | 0.46 |
| 5 | 10032216 | rs314734 | -0.742 | 0.38 | -0.18 | 0.607 | -0.002 | 0.001 | 0.001 | 0 | 0.47 | 0.6 | 0.44 | 0.48 | 0.48 | 0.51 |
| 5 | 10032708 | rs7735551 | -0.189 | -0.171 | -0.23 | 0.466 | -0.036 | -0.021 | -0.021 | 0.961 | 0.28 | 0.36 | 0.17 | 0.23 | 0.19 | 0.22 |
| 5 | 10034362 | rs10474658 | 0.3 | 0.406 | -0.725 | 0.484 | -0.001 | 0.02 | 0.02 | 0 | 0.52 | 0.41 | 0.55 | 0.59 | 0.51 | 0.47 |
| 5 | 10036432 | rs1476481 | -0.094 | 0.692 | -0.765 | 0.609 | 0 | -0.002 | -0.002 | 0.032 | 0.47 | 0.5 | 0.58 | 0.64 | 0.55 | 0.49 |
| 5 | 10040087 | rs12658915 | -0.354 | 0.397 | -0.999 | 0.698 | 0.009 | -0.001 | -0.001 | 0 | 0.4 | 0.37 | 0.38 | 0.57 | 0.39 | 0.42 |
| 5 | 10042003 | rs314743 | 0.394 | 0.042 | -1.595 | 0.23 | -0.075 | 0.069 | 0.069 | 0 | 0.38 | 0.17 | 0.48 | 0.21 | 0.51 | 0.43 |
| 5 | 10044149 | rs7729241 | 0.083 | -0.457 | -0.771 | 0.787 | 0.003 | 0.05 | 0.05 | 0 | 0.22 | 0.26 | 0.13 | 0.48 | 0.21 | 0.27 |
| 5 | 10055916 | rs1506030 | 0.743 | -0.904 | -0.585 | 0.276 | -0.014 | 0.058 | 0.058 | 0 | 0.43 | 0.3 | 0.49 | 0.19 | 0.54 | 0.44 |
| 5 | 10058975 | rs6554567 | -0.223 | -1.093 | 0.225 | 0.271 | -0.004 | -0.026 | -0.026 | 1.192 | 0.92 | 0.72 | 0.98 | 1 | 0.98 | 0.98 |
| 5 | 10059451 | rs7718749 | 0.767 | -1.448 | -0.267 | 0.332 | 0.02 | 0.026 | 0.026 | 0 | 0.51 | 0.4 | 0.56 | 0.38 | 0.58 | 0.58 |
| 5 | 10060297 | rs16884060 | 0.565 | -1.51 | -0.581 | 0.62 | -0.049 | -0.149 | -0.149 | 0 | 0.21 | 0.01 | 0.14 | 0.48 | 0.23 | 0.3 |
| 5 | 10063991 | rs7703397 | 0.256 | -1.785 | -0.29 | 0.46 | -0.01 | 0.001 | 0.001 | 0 | 0.6 | 0.56 | 0.65 | 0.38 | 0.69 | 0.6 |
| 5 | 10064591 | rs13160983 | 0.922 | -1.904 | -0.437 | 0.307 | 0.01 | 0.016 | 0.016 | 0 | 0.5 | 0.47 | 0.6 | 0.18 | 0.63 | 0.4 |
| 5 | 10067090 | rs7719627 | -0.678 | -1.499 | -0.5 | 0.481 | 0.018 | -0.02 | -0.02 | 0 | 0.21 | 0.18 | 0.26 | 0.3 | 0.15 | 0.25 |
| 5 | 10069158 | rs10066275 | 0.93 | -1.387 | -0.771 | 0.398 | 0.026 | 0.002 | 0.002 | 0 | 0.61 | 0.31 | 0.63 | 0.64 | 0.76 | 0.69 |
| 5 | 10072247 | rs1876154 | -0.558 | -0.527 | -0.62 | 0.475 | 0.001 | 0.007 | 0.007 | 0 | 0.11 | 0.25 | 0.22 | 0.12 | 0.11 | 0.06 |
| 5 | 10073956 | rs11742768 | 0.838 | -0.032 | -1.898 | 0.331 | -0.064 | 0.076 | 0.076 | 0.193 | 0.41 | 0.1 | 0.4 | 0.3 | 0.49 | 0.62 |
| 5 | 10075772 | rs4702667 | -0.078 | -0.021 | -1.392 | 1.037 | -0.024 | -0.1 | -0.1 | 0 | 0.14 | 0.03 | 0.16 | 0.52 | 0.13 | 0.22 |
| 5 | 10080699 | rs975326 | 0.423 | -0.119 | -0.432 | 0.24 | 0.003 | -0.063 | -0.063 | 0 | 0.9 | 0.75 | 0.95 | 1 | 0.93 | 0.98 |
| 5 | 10081435 | rs10052095 | -0.323 | -0.732 | -1.058 | 0.546 | 0.022 | -0.129 | -0.129 | 0 | 0.24 | 0.07 | 0.22 | 0.54 | 0.16 | 0.45 |
| 5 | 10085900 | rs924968 | 0.841 | -0.622 | -0.566 | -0.195 | 0.003 | -0.128 | -0.128 | 0.001 | 0.77 | 0.6 | 0.88 | 0.99 | 0.81 | 0.96 |
| 5 | 10088310 | rs35775016 | 1.184 | -0.546 | -0.548 | -0.538 | 0.002 | -0.043 | -0.043 | 0.596 | 0.13 | 0.16 | 0.09 | 0 | 0.11 | 0.05 |
| 5 | 10098064 | rs11747397 | 1.435 | -0.803 | -0.298 | -0.453 | -0.018 | -0.002 | -0.002 | 0 | 0.43 | 0.28 | 0.58 | 0.41 | 0.58 | 0.45 |
| 5 | 10099927 | rs906067 | -0.351 | -0.686 | -0.438 | 0.011 | 0.009 | -0.111 | -0.111 | 0 | 0.79 | 0.95 | 0.77 | 0.44 | 0.84 | 0.56 |
| 5 | 10105947 | rs1876150 | 0.204 | -0.513 | -0.577 | -0.312 | -0.003 | -0.024 | -0.024 | 0 | 0.16 | 0.18 | 0.09 | 0 | 0.15 | 0.07 |
| 5 | 10107999 | rs11741586 | 0.492 | -0.6 | -0.427 | 0.126 | -0.001 | 0.024 | 0.024 | 0 | 0.59 | 0.67 | 0.57 | 0.24 | 0.64 | 0.46 |
| 5 | 10108500 | rs6886410 | -0.217 | -0.639 | -0.644 | 0.705 | -0.006 | -0.099 | -0.099 | 0.147 | 0.71 | 0.89 | 0.7 | 0.55 | 0.77 | 0.64 |
| 5 | 10109105 | rs906069 | -0.935 | -1.084 | 0.179 | 0.125 | -0.038 | -0.034 | -0.034 | 0 | 0.25 | 0.61 | 0.21 | 0.17 | 0.16 | 0.08 |
| 5 | 10109791 | rs884355 | 1.067 | -0.936 | 0.053 | 0.177 | 0.011 | 0.078 | 0.078 | 0.517 | 0.46 | 0.27 | 0.47 | 0.23 | 0.59 | 0.49 |
| 5 | 10111180 | rs11744434 | -0.428 | -1.088 | 0.482 | 0.146 | 0.001 | -0.015 | -0.015 | 0 | 0.71 | 0.73 | 0.7 | 0.39 | 0.76 | 0.57 |
| 5 | 10111996 | rs10060253 | -0.517 | -1.092 | 0.483 | 0.146 | -0.005 | -0.052 | -0.052 | 2.402 | 0.71 | 0.86 | 0.71 | 0.39 | 0.76 | 0.57 |
| 5 | 10112089 | rs13157877 | 1.766 | -0.883 | 0.335 | -0.068 | -0.13 | 0.082 | 0.082 | 0.817 | 0.37 | 0.04 | 0.37 | 0.16 | 0.43 | 0.39 |
| 5 | 10113245 | rs4702671 | -1.495 | -0.858 | 0.367 | -0.054 | 0.024 | 0.047 | 0.047 | 5.191 | 0.34 | 0.79 | 0.33 | 0.23 | 0.33 | 0.17 |
| 5 | 10113778 | rs1876152 | 0.241 | -0.827 | 0.42 | -0.092 | -0.001 | 0.009 | 0.009 | 0 | 0.6 | 0.66 | 0.6 | 0.34 | 0.58 | 0.5 |
| 5 | 10115230 | rs10058915 | -0.322 | -0.913 | 0.787 | -0.13 | -0.004 | -0.003 | -0.003 | 0 | 0.17 | 0.21 | 0.17 | 0.19 | 0.16 | 0.1 |
| 5 | 10115751 | rs10060606 | -0.457 | -0.918 | 0.742 | -0.127 | 0.011 | 0.077 | 0.077 | 0 | 0.43 | 0.21 | 0.45 | 0.79 | 0.38 | 0.51 |
| 5 | 10116479 | rs4484395 | -1.487 | -0.924 | 0.999 | -0.15 | 0 | -0.011 | -0.011 | 0 | 0.58 | 0.61 | 0.58 | 0.85 | 0.58 | 0.61 |
| 5 | 10117405 | rs17358279 | -0.028 | -0.902 | 0.846 | -0.114 | -0.008 | 0.024 | 0.024 | 0 | 0.11 | 0.06 | 0.1 | 0.05 | 0.18 | 0.09 |
| 5 | 10117618 | rs10073082 | -0.335 | -0.919 | 0.919 | -0.182 | 0 | 0.006 | 0.006 | 0 | 0.39 | 0.29 | 0.4 | 0.66 | 0.41 | 0.51 |
| 5 | 10118153 | rs2662424 | -1.384 | -0.932 | 0.791 | -0.162 | -0.021 | 0.049 | 0.049 | 0 | 0.59 | 0.71 | 0.6 | 0.84 | 0.56 | 0.6 |
| 5 | 10119097 | rs2658094 | -0.485 | -0.997 | 1.124 | 0.021 | 0.053 | 0.049 | 0.049 | 3.764 | 0.05 | 0.34 | 0.04 | 0 | 0 | 0 |
| 5 | 10125673 | rs12654500 | 0.459 | -1.07 | 0.461 | 0.162 | 0.025 | -0.056 | -0.056 | 0 | 0.28 | 0.2 | 0.3 | 0.58 | 0.21 | 0.33 |
| 5 | 10127557 | rs2658110 | 0.153 | -1.138 | 0.669 | 0.117 | 0.002 | -0.004 | -0.004 | 0.299 | 0.32 | 0.32 | 0.29 | 0.58 | 0.19 | 0.37 |
| 5 | 10132963 | rs1395095 | -0.832 | -0.625 | 0.515 | -0.091 | 0.006 | -0.055 | -0.055 | 0 | 0.21 | 0.33 | 0.12 | 0.06 | 0.18 | 0.12 |
| 5 | 10136135 | rs2658084 | -1.067 | -0.693 | -0.229 | -0.189 | -0.078 | 0.142 | 0.142 | 1.119 | 0.58 | 0.91 | 0.48 | 0.85 | 0.44 | 0.53 |
| 5 | 10139143 | rs10050438 | 0.413 | -0.765 | -0.044 | 0.181 | 0.036 | -0.061 | -0.061 | 0.103 | 0.35 | 0.1 | 0.3 | 0.64 | 0.27 | 0.46 |
| 5 | 10140391 | rs2658080 | -0.74 | -0.589 | 0.132 | -0.295 | 0.032 | 0.186 | 0.186 | 0 | 0.53 | 0.81 | 0.48 | 0.88 | 0.45 | 0.58 |
| 5 | 10142175 | rs4409061 | -0.877 | -0.293 | 0.92 | 0.08 | 0.056 | -0.033 | -0.033 | 0.034 | 0.09 | 0.32 | 0.11 | 0.01 | 0.15 | 0.09 |
| 5 | 10146995 | rs7705202 | -1.32 | -0.504 | 0.078 | -0.101 | -0.057 | 0.02 | 0.02 | 0 | 0.62 | 0.77 | 0.53 | 0.71 | 0.55 | 0.61 |
| 5 | 10148156 | rs7443032 | 0.9 | -0.545 | 0.039 | 0.481 | 0.002 | -0.003 | -0.003 | 0.53 | 0.09 | 0.09 | 0.1 | 0.18 | 0.08 | 0.19 |
| 5 | 10148758 | rs955376 | 0.572 | -0.589 | 0.293 | 0.64 | -0.013 | 0.011 | 0.011 | 0 | 0.27 | 0.28 | 0.23 | 0.46 | 0.16 | 0.27 |
| 5 | 10150933 | rs6897624 | -0.556 | -0.337 | -0.053 | 0.385 | 0.017 | 0.036 | 0.036 | 0 | 0.47 | 0.63 | 0.41 | 0.7 | 0.44 | 0.43 |
| 5 | 10152686 | rs6868282 | 0.001 | -0.006 | -0.216 | 0.493 | 0.025 | 0.031 | 0.031 | 0 | 0.09 | 0.17 | 0.07 | 0.16 | 0.12 | 0.07 |
| 5 | 10158297 | rs1911942 | -0.78 | 0.165 | -0.903 | 0.04 | -0.047 | 0.189 | 0.189 | 0 | 0.55 | 0.84 | 0.46 | 0.9 | 0.52 | 0.65 |
| 5 | 10159036 | rs2202815 | -0.698 | 0.65 | -0.007 | 0.323 | 0.008 | 0.058 | 0.058 | 0.067 | 0.21 | 0.4 | 0.12 | 0.33 | 0.2 | 0.27 |
| 5 | 10161876 | rs1508855 | -0.177 | 0.887 | -0.446 | -0.737 | -0.006 | 0.003 | 0.003 | 0 | 0.17 | 0.13 | 0.14 | 0.04 | 0.24 | 0.1 |
| 5 | 10165356 | rs1508854 | -0.199 | 0.814 | -0.05 | -0.957 | 0.004 | -0.057 | -0.057 | 0 | 0.15 | 0.18 | 0.13 | 0.02 | 0.2 | 0.06 |
| 5 | 10166244 | rs13165671 | -0.285 | 0.851 | -0.319 | -1.691 | 0.01 | 0.013 | 0.013 | 0 | 0.55 | 0.41 | 0.6 | 0.12 | 0.65 | 0.46 |
| 5 | 10172430 | rs6861822 | 1.807 | 1.289 | -0.821 | -2.049 | -0.051 | 0.134 | 0.134 | 0.678 | 0.2 | 0.01 | 0.11 | 0 | 0.19 | 0.09 |
| 5 | 10172635 | rs16884244 | 0.326 | 0.783 | -0.217 | -2.204 | -0.014 | -0.041 | -0.041 | 0.702 | 0.1 | 0.28 | 0.08 | 0.03 | 0.08 | 0.11 |
| 5 | 10175162 | rs13185499 | 1.021 | 0.919 | -0.37 | -2.254 | -0.002 | -0.017 | -0.017 | 0 | 0.16 | 0.13 | 0.13 | 0.02 | 0.22 | 0.08 |
| 5 | 10191971 | rs11133603 | 2.053 | 0.701 | -0.71 | -1.94 | 0 | -0.155 | -0.155 | 0 | 0.24 | 0.03 | 0.14 | 0.59 | 0.18 | 0.3 |
| 5 | 10198605 | rs2892069 | -0.966 | 0.83 | -0.053 | -2.593 | 0.018 | 0.087 | 0.087 | 0.282 | 0.95 | 0.82 | 0.98 | 0.85 | 0.96 | 0.89 |
| 5 | 10202891 | rs1508850 | 0.528 | 0.927 | -0.567 | -2.652 | -0.089 | 0.133 | 0.133 | 0 | 0.39 | 0.11 | 0.46 | 0.1 | 0.43 | 0.38 |
| 5 | 10215009 | rs10065995 | -0.453 | 0.838 | -0.02 | -3.227 | -0.016 | -0.154 | -0.154 | 0 | 0.3 | 0.49 | 0.26 | 0.04 | 0.34 | 0.19 |
| 5 | 10217604 | rs17294086 | 0.226 | 0.826 | -0.147 | -3.137 | -0.042 | 0.068 | 0.068 | 1.668 | 0.24 | 0.11 | 0.27 | 0.06 | 0.29 | 0.36 |
| 5 | 10221905 | rs4701865 | -0.044 | 0.58 | 0.149 | -3.019 | 0.011 | 0.088 | 0.088 | 0 | 0.43 | 0.71 | 0.57 | 0.89 | 0.33 | 0.51 |

| | | | | | | | | | | | | | | | | |
|---|---|---|---|---|---|---|---|---|---|---|---|---|---|---|---|---|
| 5 | 10222014 | rs4702678 | 1.575 | 0.53 | 0.042 | -2.999 | 0.039 | 0.003 | 0.003 | 0 | 0.25 | 0.15 | 0.35 | 0.74 | 0.16 | 0.3 |
| 5 | 10223371 | rs4701866 | -0.541 | 0.564 | -0.161 | -3.065 | 0.004 | 0.022 | 0.022 | 0.304 | 0.64 | 0.34 | 0.75 | 0.83 | 0.67 | 0.74 |
| 5 | 10223386 | rs6871733 | 1.321 | 0.526 | -0.361 | -3.109 | -0.089 | 0.103 | 0.103 | 0 | 0.23 | 0.02 | 0.21 | 0.07 | 0.36 | 0.31 |
| 5 | 10224631 | rs12653646 | 1.793 | 0.54 | -0.314 | -3.053 | 0.031 | -0.025 | -0.025 | 0.161 | 0.24 | 0.05 | 0.34 | 0.74 | 0.16 | 0.3 |
| 5 | 10226839 | rs1045369 | 0.912 | 0.681 | -0.207 | -2.988 | -0.004 | -0.012 | -0.012 | 0 | 0.1 | 0.09 | 0.18 | 0 | 0.13 | 0.12 |
| 5 | 10233988 | rs11133604 | 0.442 | 0.414 | -0.352 | -2.977 | -0.034 | 0.041 | 0.041 | 1.568 | 0.34 | 0.25 | 0.4 | 0.07 | 0.49 | 0.43 |
| 5 | 10235490 | rs2292264 | 0.848 | 0.001 | -0.672 | -2.738 | 0.051 | 0.036 | 0.036 | 0 | 0.29 | 0.12 | 0.36 | 0.79 | 0.18 | 0.31 |
| 5 | 10236927 | rs2292263 | 0.236 | 0.195 | -0.236 | -2.594 | -0.005 | 0.017 | 0.017 | 0 | 0.91 | 0.82 | 0.95 | 0.87 | 0.91 | 0.89 |
| 5 | 10239261 | rs2438652 | 0.845 | 0.128 | -0.313 | -2.528 | 0.04 | 0.018 | 0.018 | 0 | 0.26 | 0.12 | 0.34 | 0.79 | 0.16 | 0.3 |
| 5 | 10240549 | rs2607324 | -0.272 | 0.33 | -0.066 | -2.139 | 0.017 | 0.086 | 0.086 | 0.351 | 0.94 | 0.77 | 0.98 | 0.87 | 0.96 | 0.89 |
| 5 | 10241867 | rs2607330 | -0.443 | 0.615 | -1.345 | -2.089 | 0.041 | 0.147 | 0.147 | 0.785 | 0.57 | 0.2 | 0.72 | 0.79 | 0.66 | 0.75 |
| 5 | 10242162 | rs2578628 | -0.305 | 0.529 | -0.651 | -1.027 | -0.049 | 0.014 | 0.014 | 0 | 0.3 | 0.11 | 0.49 | 0.73 | 0.24 | 0.4 |
| 5 | 10245114 | rs2578624 | -0.363 | 0.254 | -0.254 | -1.028 | 0 | 0.005 | 0.005 | 0 | 0.32 | 0.31 | 0.51 | 0.73 | 0.24 | 0.4 |
| 5 | 10248572 | rs3844305 | -1.361 | 0.178 | 0.737 | -1.525 | 0.018 | -0.03 | -0.03 | 0.028 | 0.09 | 0.33 | 0.06 | 0.01 | 0.07 | 0.07 |
| 5 | 10250430 | rs2548546 | 2.027 | -0.053 | 0.78 | -1.87 | -0.014 | -0.04 | -0.04 | 0 | 0.76 | 0.39 | 0.77 | 0.88 | 0.85 | 0.66 |
| 5 | 10250443 | rs2578618 | 2.034 | -0.059 | 0.777 | -1.872 | -0.014 | -0.04 | -0.04 | 0 | 0.76 | 0.39 | 0.77 | 0.88 | 0.85 | 0.66 |
| 5 | 10254117 | rs2259642 | 2.034 | -0.121 | 0.777 | -1.871 | -0.015 | -0.042 | -0.042 | 0 | 0.76 | 0.39 | 0.78 | 0.89 | 0.85 | 0.65 |
| 5 | 10255821 | rs3846599 | 1.803 | -0.663 | -0.085 | -2.388 | -0.038 | 0.287 | 0.287 | 0 | 0.42 | 0.02 | 0.34 | 0.06 | 0.64 | 0.35 |
| 5 | 10260247 | rs10043031 | 0.422 | -0.587 | -0.035 | -2.154 | -0.013 | 0.004 | 0.004 | 1.008 | 0.87 | 0.99 | 0.86 | 0.89 | 0.92 | 0.72 |
| 5 | 10263692 | rs2445872 | 0.206 | -0.667 | 0.177 | -2.13 | 0.038 | 0.07 | 0.07 | 0.076 | 0.3 | 0.12 | 0.39 | 0.82 | 0.21 | 0.31 |
| 5 | 10265277 | rs2578641 | 0.089 | -0.646 | 0.181 | -2.127 | 0.044 | 0.068 | 0.068 | 0.052 | 0.3 | 0.12 | 0.39 | 0.82 | 0.21 | 0.31 |
| 5 | 10281549 | rs3852166 | 0.51 | -0.862 | 0.731 | -2.227 | -0.004 | -0.029 | -0.029 | 0 | 0.6 | 0.6 | 0.57 | 0.08 | 0.78 | 0.55 |
| 5 | 10283530 | rs6871279 | 1.138 | -0.72 | 0.506 | -2.231 | 0.07 | 0.117 | 0.117 | 0 | 0.5 | 0.3 | 0.42 | 0.08 | 0.71 | 0.43 |
| 5 | 10290347 | rs10038054 | -0.117 | -0.446 | 0.776 | -2.036 | 0.033 | 0.055 | 0.055 | 0.363 | 0.7 | 0.82 | 0.62 | 0.19 | 0.8 | 0.7 |
| 5 | 10291867 | rs10060937 | -1.933 | -0.326 | 1.327 | -2.104 | 0.016 | -0.104 | -0.104 | 0 | 0.18 | 0.55 | 0.22 | 0.03 | 0.14 | 0.19 |
| 5 | 10297847 | rs3857348 | 1.081 | -0.126 | 1.661 | -1.927 | 0.004 | -0.039 | -0.039 | 0 | 0.5 | 0.55 | 0.4 | 0.07 | 0.65 | 0.35 |
| 5 | 10301295 | rs13359758 | 0.351 | -0.351 | 1.799 | -1.849 | -0.015 | -0.044 | -0.044 | 0 | 0.6 | 0.47 | 0.46 | 0.2 | 0.74 | 0.56 |
| 5 | 10306644 | rs7702977 | -1.312 | -0.257 | 1.637 | -1.785 | 0.042 | -0.013 | -0.013 | 0.392 | 0.09 | 0.36 | 0.08 | 0.03 | 0.07 | 0.13 |
| 5 | 10307022 | rs11738492 | 0.867 | -0.276 | 1.123 | -1.836 | 0.025 | 0.079 | 0.079 | 0 | 0.51 | 0.35 | 0.39 | 0.17 | 0.65 | 0.44 |
| 5 | 10308240 | rs10072686 | 0.855 | -0.354 | 0.492 | -1.506 | 0.032 | 0.054 | 0.054 | 0 | 0.55 | 0.66 | 0.45 | 0.11 | 0.69 | 0.42 |
| 5 | 10309122 | rs11951923 | -1.079 | -0.046 | 0.621 | -0.498 | -0.016 | 0.002 | 0.002 | 0 | 0.06 | 0.09 | 0.03 | 0.11 | 0.02 | 0.15 |
| 5 | 10315843 | rs1287733 | 1.14 | 0.24 | 0.619 | -0.099 | 0.005 | -0.007 | -0.007 | 0 | 0.85 | 0.84 | 0.67 | 0.33 | 0.91 | 0.69 |
| 5 | 10318455 | rs606490 | -1.235 | 0.436 | 1.217 | -0.59 | 0.018 | -0.08 | -0.08 | 0 | 0.14 | 0.33 | 0.14 | 0.01 | 0.17 | 0.08 |
| 5 | 10343089 | rs4702690 | 1.336 | 0.113 | 1.509 | -0.143 | -0.013 | 0.018 | 0.018 | 0 | 0.85 | 0.79 | 0.67 | 0.29 | 0.92 | 0.69 |
| 5 | 10356885 | rs6887666 | 1.091 | -0.051 | 2.065 | -0.045 | 0.015 | -0.071 | -0.071 | 0 | 0.85 | 0.95 | 0.77 | 0.35 | 0.91 | 0.67 |
| 5 | 10361110 | rs13155132 | 1.395 | -0.031 | 2.043 | -0.202 | -0.004 | 0.002 | 0.002 | 0 | 0.82 | 0.79 | 0.66 | 0.3 | 0.91 | 0.66 |
| 5 | 10413501 | rs2567588 | -1.773 | -0.045 | 2.239 | -0.195 | -0.05 | 0.043 | 0.043 | 0 | 0.2 | 0.42 | 0.36 | 0.7 | 0.09 | 0.34 |
| 5 | 10417589 | rs2967956 | 0.722 | -0.3 | 2.199 | -0.321 | -0.017 | -0.023 | -0.023 | 0.14 | 0.94 | 0.79 | 0.96 | 1 | 0.99 | 0.99 |
| 5 | 10434643 | rs2589668 | -1.453 | -0.266 | 1.914 | -0.303 | -0.006 | 0.004 | 0.004 | 0 | 0.18 | 0.21 | 0.34 | 0.7 | 0.09 | 0.34 |
| 5 | 10448247 | rs1976360 | 2.089 | 0.128 | 2.112 | -0.206 | -0.025 | -0.035 | -0.035 | 0.312 | 0.91 | 0.49 | 0.93 | 1 | 0.99 | 0.99 |
| 5 | 10459942 | rs1814856 | -1.637 | 0.952 | 1.324 | -0.118 | 0 | 0.008 | 0.008 | 0 | 0.26 | 0.34 | 0.43 | 0.67 | 0.24 | 0.39 |
| 5 | 10460700 | rs2567575 | 1.78 | 0.96 | 0.864 | -0.395 | -0.011 | -0.062 | -0.062 | 0 | 0.89 | 0.78 | 0.9 | 1 | 0.91 | 0.98 |
| 5 | 10478341 | rs2589653 | 1.285 | 1.367 | 0.146 | -0.301 | 0.026 | -0.084 | -0.084 | 0 | 0.86 | 0.45 | 0.74 | 0.99 | 0.83 | 0.92 |
| 5 | 10489579 | rs17759853 | -1.138 | 2.458 | -1.066 | -0.321 | -0.119 | 0.069 | 0.069 | 0 | 0.2 | 0.03 | 0.17 | 0.12 | 0.32 | 0.15 |
| 5 | 10492095 | rs2589661 | -0.942 | 2.195 | -0.842 | 1.057 | -0.001 | 0.053 | 0.053 | 0 | 0.11 | 0.14 | 0.27 | 0.47 | 0.13 | 0.12 |
| 5 | 10492322 | rs13180425 | 0.523 | 1.483 | -0.813 | 0.545 | 0.023 | 0.019 | 0.019 | 0.032 | 0.44 | 0.38 | 0.35 | 0.29 | 0.34 | 0.45 |
| 5 | 10497207 | rs7732984 | 0.464 | 1.429 | -0.674 | 0.397 | 0.028 | 0.021 | 0.021 | 0.081 | 0.45 | 0.38 | 0.36 | 0.31 | 0.35 | 0.46 |
| 5 | 10501076 | rs2589658 | 0.141 | 1.239 | -1.169 | 0.471 | -0.012 | 0.002 | 0.002 | 1.325 | 0.62 | 0.71 | 0.56 | 0.39 | 0.52 | 0.55 |
| 5 | 10503977 | rs2919756 | -0.655 | 0.533 | -1.199 | 0.305 | 0.002 | -0.008 | -0.008 | 0.393 | 0.08 | 0.05 | 0.14 | 0.26 | 0.1 | 0.15 |
| 5 | 10505585 | rs17816571 | 0.32 | 0.309 | -1.953 | -0.219 | -0.056 | 0.009 | 0.009 | 0 | 0.36 | 0.09 | 0.36 | 0.26 | 0.34 | 0.21 |
| 5 | 10512330 | rs1971391 | -0.277 | -0.45 | -1.336 | -0.741 | -0.001 | 0.002 | 0.002 | 0 | 0.57 | 0.53 | 0.48 | 0.53 | 0.42 | 0.56 |
| 5 | 10512454 | rs6883094 | -0.031 | -0.349 | -1.259 | -0.938 | 0.007 | -0.042 | -0.042 | 0 | 0.09 | 0.15 | 0.11 | 0.01 | 0.07 | 0.03 |
| 5 | 10512572 | rs4235597 | -0.154 | -0.899 | -1.437 | -0.948 | 0.014 | 0.025 | 0.025 | 0 | 0.54 | 0.64 | 0.39 | 0.27 | 0.36 | 0.4 |
| 5 | 10517908 | rs4702698 | 0.532 | -0.219 | -1.682 | -0.497 | -0.012 | -0.006 | -0.006 | 0.016 | 0.32 | 0.13 | 0.29 | 0.26 | 0.29 | 0.21 |
| 5 | 10523116 | rs2962325 | -0.884 | 0.2 | -1.591 | -0.406 | -0.008 | -0.001 | -0.001 | 0.413 | 0.63 | 0.82 | 0.72 | 0.68 | 0.62 | 0.74 |
| 5 | 10524503 | rs2895545 | 0.31 | -0.031 | -1.41 | -0.428 | -0.016 | -0.015 | -0.015 | 0 | 0.34 | 0.27 | 0.36 | 0.4 | 0.43 | 0.41 |
| 5 | 10525349 | rs2673892 | -1.018 | 0.225 | -1.87 | -0.147 | -0.024 | 0.003 | 0.003 | 0 | 0.62 | 0.83 | 0.72 | 0.67 | 0.59 | 0.72 |
| 5 | 10531620 | rs2936577 | -0.254 | 0.828 | -1.509 | 0.732 | 0.002 | 0.087 | 0.087 | 0 | 0.05 | 0.1 | 0.19 | 0.25 | 0.06 | 0.1 |
| 5 | 10531998 | rs814586 | -0.08 | 1.005 | -1.837 | 0.692 | -0.058 | -0.073 | -0.073 | 0.041 | 0.26 | 0.1 | 0.3 | 0.39 | 0.39 | 0.41 |
| 5 | 10532639 | rs6887463 | -1.079 | 0.43 | -0.846 | 0.576 | -0.002 | 0.026 | 0.026 | 0 | 0.07 | 0.08 | 0.2 | 0.24 | 0.07 | 0.1 |
| 5 | 10537351 | rs2589656 | 0.443 | 0.685 | -1.032 | -0.562 | 0.004 | 0.094 | 0.094 | 0.126 | 0.62 | 0.2 | 0.62 | 0.95 | 0.67 | 0.77 |
| 5 | 10541520 | rs2962322 | -0.969 | 0.663 | -0.011 | 0.233 | 0.019 | 0.005 | 0.005 | 0 | 0.3 | 0.43 | 0.48 | 0.73 | 0.36 | 0.46 |
| 5 | 10541625 | rs814589 | 1.459 | 0.595 | 0.833 | 0.176 | 0.036 | 0.019 | 0.019 | 1.076 | 0.87 | 0.6 | 0.79 | 0.81 | 0.85 | 0.86 |
| 5 | 10547018 | rs2936590 | 0.732 | 0.2 | 0.56 | -0.189 | -0.019 | -0.011 | -0.011 | 0 | 0.62 | 0.58 | 0.66 | 0.69 | 0.71 | 0.47 |
| 5 | 10547032 | rs2673891 | -1.978 | 1.032 | 0.761 | -0.24 | -0.03 | -0.007 | -0.007 | 0 | 0.18 | 0.11 | 0.22 | 0.17 | 0.29 | 0.25 |
| 5 | 10552459 | rs1392006 | 0.406 | 0.806 | 0.748 | -0.371 | -0.004 | -0.001 | -0.001 | 2.047 | 0.9 | 0.93 | 0.93 | 0.98 | 0.89 | 0.9 |
| 5 | 10552667 | rs1399830 | -0.801 | 0.752 | 0.772 | -0.338 | 0.009 | -0.009 | -0.009 | 3.503 | 0.1 | 0.06 | 0.12 | 0.08 | 0.08 | 0.12 |
| 5 | 10553563 | rs2062462 | 1.576 | 0.866 | 1.02 | -0.392 | 0.026 | -0.031 | -0.031 | 0 | 0.79 | 0.68 | 0.82 | 0.9 | 0.71 | 0.81 |
| 5 | 10568417 | rs2399910 | 1.715 | 0.473 | 0.841 | -0.264 | 0.001 | -0.03 | -0.03 | 1.253 | 0.71 | 0.57 | 0.75 | 0.82 | 0.7 | 0.68 |

| | | | | | | | | | | | | | | | | |
|---|---|---|---|---|---|---|---|---|---|---|---|---|---|---|---|---|
| 5 | 10568591 | rs12522627 | -0.68 | 0.733 | -0.325 | -0.136 | -0.006 | 0 | 0 | 1.09 | 0.08 | 0.07 | 0.1 | 0.08 | 0.21 | 0.09 |
| 5 | 10569829 | rs2648802 | -1.085 | 0.562 | -0.441 | -0.109 | -0.026 | 0.011 | 0.011 | 0 | 0.18 | 0.02 | 0.16 | 0.1 | 0.17 | 0.22 |
| 5 | 10574343 | rs959219 | 1.189 | 0.415 | 0.308 | -0.049 | -0.001 | -0.023 | -0.023 | 0 | 0.67 | 0.46 | 0.79 | 0.81 | 0.68 | 0.73 |
| 5 | 10581830 | rs2589663 | 1.611 | 0.589 | 1.133 | 0.056 | 0.108 | 0.044 | 0.044 | 0.373 | 0.93 | 0.69 | 0.92 | 0.94 | 0.9 | 0.88 |
| 5 | 10583537 | rs2648804 | 1.111 | 0.354 | 0.388 | 0.065 | -0.027 | 0.008 | 0.008 | 0 | 0.48 | 0.16 | 0.29 | 0.59 | 0.45 | 0.37 |
| 5 | 10585111 | rs7712927 | -0.897 | 0.665 | 0.877 | 0.145 | 0.128 | 0.112 | 0.112 | 1.038 | 0.05 | 0.22 | 0.05 | 0.1 | 0.06 | 0.04 |
| 5 | 10585287 | rs9790911 | 1.137 | 0.446 | 0.643 | 0.001 | 0.001 | -0.009 | -0.009 | 0 | 0.86 | 0.84 | 0.88 | 0.93 | 0.83 | 0.79 |
| 5 | 10588033 | rs11737977 | 0.53 | 0.51 | 0.698 | 0.03 | 0.036 | 0.003 | 0.003 | 0 | 0.95 | 0.86 | 0.93 | 0.94 | 0.9 | 0.88 |
| 5 | 10589502 | rs6869928 | 0.176 | 0.368 | 0.986 | -0.021 | -0.009 | -0.004 | -0.004 | 0 | 0.29 | 0.36 | 0.52 | 0.24 | 0.31 | 0.35 |
| 5 | 10591672 | rs4702704 | 0.253 | 0.385 | 0.955 | -0.018 | -0.007 | -0.01 | -0.01 | 0 | 0.65 | 0.49 | 0.42 | 0.7 | 0.58 | 0.52 |
| 5 | 10594061 | rs13185308 | -0.506 | 0.323 | 0.635 | -0.003 | 0.009 | 0.011 | 0.011 | 0 | 0.39 | 0.39 | 0.57 | 0.25 | 0.38 | 0.45 |
| 5 | 10604975 | rs12187420 | 1.476 | -0.023 | -0.41 | -0.095 | -0.087 | -0.042 | -0.042 | 0 | 0.44 | 0.08 | 0.29 | 0.59 | 0.45 | 0.38 |
| 5 | 10607797 | rs1384731 | 1.378 | -0.43 | 0.414 | 0.02 | 0.009 | -0.012 | -0.012 | 0 | 0.91 | 0.83 | 0.91 | 0.93 | 0.89 | 0.89 |
| 5 | 10611579 | rs10474877 | -1.1 | -0.625 | 0.508 | -0.05 | -0.052 | -0.116 | -0.116 | 0 | 0.17 | 0.48 | 0.09 | 0.01 | 0.07 | 0.14 |
| 5 | 10612627 | rs10035299 | 2.127 | -0.604 | -0.033 | -0.006 | 0.026 | 0.068 | 0.068 | 0.343 | 0.69 | 0.15 | 0.78 | 0.85 | 0.78 | 0.69 |
| 5 | 10616846 | rs6883387 | 0.805 | -0.872 | 0.218 | 0.203 | 0.012 | -0.002 | -0.002 | 0 | 0.93 | 0.82 | 0.93 | 0.95 | 0.95 | 0.92 |
| 5 | 10617670 | rs6871080 | 1.595 | -0.894 | 0.099 | 0.411 | -0.002 | 0.039 | 0.039 | 0 | 0.91 | 0.86 | 0.93 | 0.82 | 0.94 | 0.86 |
| 5 | 10617984 | rs4701875 | 1.386 | -0.803 | -0.162 | 0.039 | -0.017 | -0.052 | -0.052 | 0.017 | 0.77 | 0.35 | 0.84 | 0.95 | 0.87 | 0.8 |
| 5 | 10619490 | rs11948562 | -1.342 | -0.612 | -0.874 | 0.537 | 0.024 | -0.025 | -0.025 | 0 | 0.1 | 0.02 | 0.14 | 0.1 | 0.05 | 0.12 |
| 5 | 10620085 | rs11748401 | 0.182 | -0.977 | -1.543 | 0.761 | 0.008 | -0.077 | -0.077 | 0 | 0.31 | 0.14 | 0.15 | 0.45 | 0.26 | 0.4 |
| 5 | 10620642 | rs11749155 | -0.281 | -0.843 | -1.634 | 0.2 | 0.005 | 0.019 | 0.019 | 0 | 0.23 | 0.17 | 0.14 | 0.04 | 0.23 | 0.11 |
| 5 | 10621173 | rs1473447 | -0.34 | -0.831 | -1.653 | 0.18 | 0.006 | 0.023 | 0.023 | 0 | 0.23 | 0.17 | 0.15 | 0.04 | 0.24 | 0.12 |
| 5 | 10624887 | rs1531839 | -0.939 | -1.202 | -1.266 | 0.362 | 0.013 | -0.017 | -0.017 | 0 | 0.62 | 0.47 | 0.36 | 0.64 | 0.49 | 0.57 |
| 5 | 10627449 | rs1483300 | 0.529 | -0.982 | -1.609 | 0.276 | 0.027 | -0.112 | -0.112 | 0 | 0.33 | 0.08 | 0.14 | 0.48 | 0.25 | 0.45 |
| 5 | 10631178 | rs6896826 | 0.46 | -1.268 | -1.646 | 0.249 | 0.021 | -0.009 | -0.009 | 0.935 | 0.33 | 0.21 | 0.21 | 0.31 | 0.22 | 0.34 |
| 5 | 10635947 | rs7703985 | -0.586 | -1.343 | -1.48 | 0.302 | 0.009 | 0.004 | 0.004 | 0.299 | 0.54 | 0.34 | 0.35 | 0.34 | 0.43 | 0.44 |
| 5 | 10636151 | rs900531 | 0.307 | -1.394 | -1.528 | 0.259 | 0.001 | 0.002 | 0.002 | 0.239 | 0.37 | 0.3 | 0.27 | 0.29 | 0.35 | 0.36 |
| 5 | 10638180 | rs11745612 | -0.215 | -0.79 | -1.193 | 0.389 | 0.015 | 0.014 | 0.014 | 0.313 | 0.52 | 0.33 | 0.35 | 0.32 | 0.43 | 0.44 |
| 5 | 10652117 | rs6894331 | -0.785 | 0.132 | -0.408 | 1.037 | 0.003 | 0.003 | 0.003 | 0 | 0.28 | 0.31 | 0.29 | 0.26 | 0.31 | 0.21 |
| 5 | 10654769 | rs3822410 | -0.856 | 0.136 | -0.563 | 1.024 | 0.01 | 0.016 | 0.016 | 0 | 0.29 | 0.42 | 0.3 | 0.26 | 0.31 | 0.21 |
| 5 | 10667969 | rs34091808 | -0.796 | -0.643 | -0.471 | 1.169 | -0.021 | 0.017 | 0.017 | 0 | 0.27 | 0.36 | 0.32 | 0.36 | 0.25 | 0.28 |
| 5 | 10671040 | rs4702718 | -0.239 | -1.032 | -0.485 | 1.045 | 0.008 | -0.001 | -0.001 | 0.522 | 0.5 | 0.47 | 0.31 | 0.46 | 0.3 | 0.42 |
| 5 | 10675766 | rs1531842 | -0.495 | -2.23 | -0.646 | 1.335 | -0.01 | -0.004 | -0.004 | 0 | 0.47 | 0.47 | 0.31 | 0.44 | 0.33 | 0.45 |
| 5 | 10676692 | rs13168857 | 0.553 | -2.318 | -0.397 | 1.483 | -0.007 | 0.064 | 0.064 | 0 | 0.06 | 0.14 | 0.11 | 0.18 | 0.05 | 0.06 |
| 5 | 10677339 | rs4702722 | -1.611 | -2.569 | -0.055 | 1.75 | -0.017 | 0.015 | 0.015 | 2.354 | 0.64 | 0.76 | 0.48 | 0.72 | 0.41 | 0.65 |
| 5 | 10678281 | rs5745308 | -0.045 | -2.712 | -0.025 | 1.671 | -0.001 | -0.028 | -0.028 | 0 | 0.07 | 0.08 | 0.06 | 0 | 0.05 | 0.08 |
| 5 | 10679746 | rs7973 | -1.061 | -2.77 | 0.041 | 1.643 | -0.008 | -0.05 | -0.05 | 0 | 0.09 | 0.22 | 0.07 | 0 | 0.05 | 0.08 |
| 5 | 10680487 | rs1049344 | 1.047 | -2.841 | -0.004 | 1.745 | 0.013 | -0.052 | -0.052 | 0 | 0.12 | 0.04 | 0.14 | 0.22 | 0.07 | 0.17 |
| 5 | 10680547 | rs9857 | -0.062 | -2.873 | 0.027 | 1.681 | -0.023 | 0.002 | 0.002 | 2.095 | 0.42 | 0.44 | 0.26 | 0.48 | 0.28 | 0.37 |
| 5 | 10680749 | rs10271 | -0.743 | -2.86 | 0.02 | 1.65 | -0.005 | -0.045 | -0.045 | 0 | 0.08 | 0.2 | 0.07 | 0 | 0.05 | 0.08 |
| 5 | 10682728 | rs1866386 | 0.429 | -2.874 | 0.023 | 1.74 | 0.005 | 0.005 | 0.005 | 3.694 | 0.37 | 0.32 | 0.21 | 0.25 | 0.24 | 0.25 |
| 5 | 10685185 | rs3797111 | -0.647 | -2.829 | 0.017 | 1.749 | 0.001 | -0.002 | -0.002 | 2.174 | 0.22 | 0.25 | 0.22 | 0.26 | 0.14 | 0.26 |
| 5 | 10689562 | rs3776414 | -1.607 | -2.82 | -0.005 | 1.721 | -0.009 | 0.019 | 0.019 | 2.371 | 0.63 | 0.72 | 0.48 | 0.75 | 0.4 | 0.66 |
| 5 | 10689636 | rs3776415 | -1.131 | -3.446 | -0.441 | 1.319 | 0.018 | 0.006 | 0.006 | 0 | 0.33 | 0.25 | 0.26 | 0.26 | 0.22 | 0.3 |
| 5 | 10693687 | rs6891527 | -0.299 | -2.949 | -0.397 | 1.441 | -0.018 | 0.071 | 0.071 | 0 | 0.21 | 0.3 | 0.21 | 0.49 | 0.12 | 0.32 |
| 5 | 10695526 | rs2930047 | -2.08 | -3.095 | -0.441 | 1.287 | -0.016 | 0.028 | 0.028 | 2.831 | 0.64 | 0.76 | 0.48 | 0.75 | 0.4 | 0.66 |
| 5 | 10701514 | rs3756407 | 0.42 | -3.085 | -0.455 | 1.409 | -0.01 | 0.084 | 0.084 | 0 | 0.06 | 0.12 | 0.06 | 0.22 | 0.04 | 0.15 |
| 5 | 10701615 | rs3756408 | 0.189 | -3.132 | -0.395 | 1.471 | -0.013 | 0.094 | 0.094 | 0 | 0.07 | 0.13 | 0.06 | 0.24 | 0.04 | 0.15 |
| 5 | 10702034 | rs2918394 | 0.455 | -3.283 | 0.192 | 1.007 | -0.005 | -0.047 | -0.047 | 0 | 0.11 | 0.13 | 0.05 | 0 | 0.08 | 0.03 |
| 5 | 10702688 | rs3822414 | 0.999 | -3.336 | 0.148 | 1.035 | 0.012 | -0.067 | -0.067 | 0.205 | 0.14 | 0.04 | 0.14 | 0.26 | 0.08 | 0.18 |
| 5 | 10704622 | rs2930051 | -0.394 | -3.317 | 0.256 | 0.89 | -0.008 | -0.006 | -0.006 | 0 | 0.32 | 0.32 | 0.18 | 0.24 | 0.21 | 0.22 |
| 5 | 10707870 | rs3797126 | 0.627 | -3.315 | 0.199 | 0.914 | 0.014 | -0.003 | -0.003 | 0 | 0.22 | 0.17 | 0.14 | 0.24 | 0.14 | 0.19 |
| 5 | 10710716 | rs267988 | -1.261 | -4.146 | -0.032 | 0.566 | 0.007 | 0.005 | 0.005 | 1.484 | 0.33 | 0.69 | 0.2 | 0.13 | 0.18 | 0.25 |
| 5 | 10734647 | rs2918386 | -0.734 | -4.016 | 0.031 | 0.612 | -0.006 | -0.005 | -0.005 | 0.354 | 0.33 | 0.32 | 0.18 | 0.24 | 0.21 | 0.22 |
| 5 | 10736569 | rs1866385 | -0.134 | -4.024 | -0.152 | 0.366 | 0.019 | 0.003 | 0.003 | 3.08 | 0.24 | 0.17 | 0.16 | 0.24 | 0.18 | 0.2 |
| 5 | 10738444 | rs267957 | 0.871 | -4.011 | -0.135 | 0.92 | -0.028 | 0.095 | 0.095 | 0 | 0.84 | 0.68 | 0.86 | 0.65 | 0.9 | 0.74 |
| 5 | 10738938 | rs17831254 | 0.72 | -3.987 | -0.152 | 0.892 | -0.001 | -0.011 | -0.011 | 0 | 0.06 | 0.07 | 0.06 | 0 | 0.05 | 0.08 |
| 5 | 10743929 | rs267949 | 1.392 | -4.037 | -0.152 | 0.888 | 0.053 | 0.023 | 0.023 | 4.305 | 0.51 | 0.35 | 0.67 | 0.41 | 0.7 | 0.52 |
| 5 | 10744450 | rs2930058 | -0.558 | -3.902 | -0.171 | 0.902 | -0.009 | -0.01 | -0.01 | 0 | 0.36 | 0.38 | 0.21 | 0.24 | 0.25 | 0.23 |
| 5 | 10751565 | rs16898820 | 0.364 | -3.914 | -0.28 | 0.925 | 0.007 | -0.013 | -0.013 | 0 | 0.21 | 0.18 | 0.22 | 0.28 | 0.15 | 0.26 |
| 5 | 10752315 | rs267939 | 1.747 | -3.707 | -0.218 | 0.863 | -0.014 | 0.067 | 0.067 | 2.577 | 0.36 | 0.22 | 0.51 | 0.12 | 0.6 | 0.32 |
| 5 | 10753399 | rs2930059 | -1.109 | -3.541 | -0.269 | 0.976 | -0.005 | 0.004 | 0.004 | 3.418 | 0.56 | 0.46 | 0.42 | 0.52 | 0.4 | 0.49 |
| 5 | 10760373 | rs182073 | -0.767 | -3.252 | -0.065 | 0.659 | -0.033 | -0.004 | -0.004 | 0 | 0.29 | 0.37 | 0.14 | 0.31 | 0.19 | 0.24 |
| 5 | 10771228 | rs13153621 | 1.25 | -3.123 | -0.788 | -0.4 | 0.049 | 0.183 | 0.183 | 0 | 0.46 | 0.22 | 0.5 | 0.01 | 0.65 | 0.35 |
| 5 | 10785424 | rs11740309 | 0.041 | -3.311 | -0.639 | 0.473 | 0.011 | 0.106 | 0.106 | 0.258 | 0.09 | 0.17 | 0.13 | 0.32 | 0.1 | 0.17 |
| 5 | 10788833 | rs2964798 | 0.406 | -3.329 | -0.514 | -0.226 | 0.002 | 0.078 | 0.078 | 0 | 0.88 | 0.79 | 0.81 | 0.34 | 0.89 | 0.68 |
| 5 | 10788919 | rs13155209 | 2.116 | -3.521 | -0.739 | -0.42 | -0.028 | 0.244 | 0.244 | 0.025 | 0.41 | 0.1 | 0.46 | 0.01 | 0.61 | 0.31 |
| 5 | 10789570 | rs2964796 | -0.818 | -3.357 | -0.499 | -0.07 | 0.004 | 0.004 | 0.004 | 0.558 | 0.42 | 0.57 | 0.31 | 0.33 | 0.24 | 0.33 |
| 5 | 10795348 | rs2964793 | -0.192 | -2.761 | -0.075 | 0.025 | -0.037 | 0.041 | 0.041 | 3.073 | 0.22 | 0.3 | 0.23 | 0.35 | 0.1 | 0.21 |

| | | | | | | | | | | | | | | | | |
|---|---|---|---|---|---|---|---|---|---|---|---|---|---|---|---|---|
| 5 | 10798221 | rs1866387 | 1.521 | -1.529 | -0.476 | -1.661 | -0.051 | 0.057 | 0.057 | 0 | 0.32 | 0.18 | 0.42 | 0 | 0.47 | 0.25 |
| 5 | 10800150 | rs1470023 | -0.365 | -1.813 | -0.307 | -1.684 | -0.044 | -0.029 | -0.029 | 0.123 | 0.78 | 0.62 | 0.7 | 0.15 | 0.87 | 0.61 |
| 5 | 10800714 | rs7700363 | 0.85 | -2.012 | -0.541 | -1.764 | -0.024 | 0.1 | 0.1 | 0.005 | 0.31 | 0.15 | 0.19 | 0.06 | 0.32 | 0.26 |
| 5 | 10802749 | rs12186813 | 0.151 | -1.949 | -0.488 | -1.627 | 0.005 | 0.021 | 0.021 | 0 | 0.13 | 0.05 | 0.07 | 0.09 | 0.1 | 0.1 |
| 5 | 10810604 | rs7725091 | 0.693 | -1.636 | -0.499 | -1.597 | -0.034 | 0.102 | 0.102 | 0.043 | 0.3 | 0.15 | 0.18 | 0.06 | 0.32 | 0.26 |
| 5 | 10811058 | rs1438715 | 0.823 | -1.849 | -0.448 | -1.759 | 0.061 | 0.075 | 0.075 | 0.399 | 0.62 | 0.29 | 0.6 | 0.07 | 0.79 | 0.51 |
| 5 | 10812811 | rs2083232 | 1.464 | -1.715 | -0.302 | -1.68 | 0.012 | -0.059 | -0.059 | 0 | 0.36 | 0.42 | 0.46 | 0.09 | 0.48 | 0.27 |
| 5 | 10818452 | rs1438712 | 0.805 | -1.741 | 0.135 | -1.694 | -0.007 | 0.017 | 0.017 | 0.022 | 0.31 | 0.23 | 0.2 | 0.06 | 0.32 | 0.26 |
| 5 | 10819285 | rs6554603 | 0.071 | -1.709 | -0.062 | -1.666 | 0.018 | 0.02 | 0.02 | 0.2 | 0.69 | 0.76 | 0.67 | 0.15 | 0.8 | 0.53 |
| 5 | 10827151 | rs4304054 | 0.132 | -1.036 | -0.007 | -1.066 | -0.002 | -0.009 | -0.009 | 0 | 0.32 | 0.3 | 0.21 | 0.06 | 0.32 | 0.27 |
| 5 | 10834014 | rs16901049 | -0.833 | -0.68 | 0.021 | -0.604 | -0.001 | 0 | 0 | 1.262 | 0.18 | 0.21 | 0.16 | 0.18 | 0.14 | 0.1 |
| 5 | 10836485 | rs13157690 | 0.229 | -0.83 | 0.004 | -1.339 | 0.002 | -0.006 | -0.006 | 0 | 0.47 | 0.46 | 0.54 | 0.18 | 0.55 | 0.4 |
| 5 | 10838358 | rs13159908 | 1.562 | -0.8 | -0.482 | -1.462 | -0.086 | 0.095 | 0.095 | 0 | 0.22 | 0.09 | 0.35 | 0.01 | 0.36 | 0.22 |
| 5 | 10847970 | rs4398608 | 0.029 | -0.604 | -0.485 | -1.572 | -0.015 | 0.024 | 0.024 | 0 | 0.36 | 0.31 | 0.35 | 0.04 | 0.41 | 0.28 |
| 5 | 10850197 | rs6554607 | -1.04 | -0.681 | -0.396 | -1.345 | 0 | -0.014 | -0.014 | 0 | 0.6 | 0.5 | 0.5 | 0.27 | 0.55 | 0.46 |
| 5 | 10852652 | rs6895554 | -0.969 | -0.673 | -0.399 | -1.343 | -0.001 | -0.012 | -0.012 | 0 | 0.59 | 0.49 | 0.49 | 0.29 | 0.54 | 0.45 |
| 5 | 10853693 | rs4285191 | 1.162 | 0.126 | -0.069 | -0.934 | 0.012 | 0.018 | 0.018 | 0 | 0.21 | 0.12 | 0.07 | 0.14 | 0.13 | 0.15 |
| 5 | 10855731 | rs12655316 | 0.22 | 0.145 | -0.532 | -0.889 | 0.012 | 0.065 | 0.065 | 0 | 0.26 | 0.11 | 0.12 | 0.14 | 0.2 | 0.2 |
| 5 | 10857683 | rs13185834 | 1.255 | 0.916 | -0.493 | -0.641 | -0.012 | 0.003 | 0.003 | 0 | 0.21 | 0.16 | 0.26 | 0.02 | 0.27 | 0.19 |
| 5 | 10863028 | rs4701885 | 0.446 | 0.412 | -0.617 | 0.029 | 0 | 0.01 | 0.01 | 0 | 0.08 | 0.02 | 0.09 | 0.45 | 0.08 | 0.25 |
| 5 | 10866460 | rs13157531 | 1.479 | 0.517 | -0.57 | -0.204 | -0.058 | 0.055 | 0.055 | 0 | 0.19 | 0.09 | 0.26 | 0.01 | 0.3 | 0.22 |
| 5 | 10876522 | rs6872063 | -0.32 | 0.384 | -0.671 | -0.055 | -0.104 | 0.068 | 0.068 | 0 | 0.63 | 0.87 | 0.68 | 0.73 | 0.57 | 0.67 |
| 5 | 10877143 | rs4361487 | 0.26 | 0.263 | -0.29 | -0.075 | 0.003 | -0.01 | -0.01 | 0 | 0.57 | 0.49 | 0.65 | 0.7 | 0.57 | 0.65 |
| 5 | 10878782 | rs12188457 | 0.428 | 0.256 | -0.472 | -0.034 | 0 | -0.002 | -0.002 | 0.41 | 0.54 | 0.43 | 0.57 | 0.6 | 0.55 | 0.62 |
| 5 | 10883440 | rs6860099 | -0.265 | 0.211 | -0.192 | -0.004 | 0.002 | 0.111 | 0.111 | 0.889 | 0.08 | 0.37 | 0.12 | 0.13 | 0.02 | 0.06 |
| 5 | 10886445 | rs10072111 | -1.506 | 0.406 | -0.444 | -0.087 | -0.067 | 0.036 | 0.036 | 0 | 0.8 | 0.91 | 0.71 | 0.87 | 0.67 | 0.78 |
| 5 | 10886564 | rs10065278 | -1.158 | 0.4 | -0.425 | -0.09 | -0.005 | 0.004 | 0.004 | 0 | 0.78 | 0.79 | 0.71 | 0.87 | 0.67 | 0.78 |
| 5 | 10887902 | rs12522996 | 1.221 | 0.111 | -0.737 | 0.091 | -0.017 | 0.146 | 0.146 | 0.045 | 0.34 | 0.02 | 0.24 | 0.1 | 0.29 | 0.27 |
| 5 | 10891642 | rs17197013 | 0.405 | 0.628 | -0.004 | 0.395 | 0.008 | 0.013 | 0.013 | 0 | 0.2 | 0.16 | 0.07 | 0.14 | 0.13 | 0.13 |
| 5 | 10898059 | rs16901104 | 0.16 | 0.644 | -0.657 | 0.563 | 0.023 | 0.022 | 0.022 | 0 | 0.5 | 0.36 | 0.44 | 0.54 | 0.47 | 0.55 |
| 5 | 10900487 | rs17789137 | 0.155 | 1.092 | -0.552 | 0.754 | 0.002 | 0 | 0 | 0 | 0.68 | 0.65 | 0.52 | 0.69 | 0.56 | 0.68 |
| 5 | 10903781 | rs1995364 | 0.136 | 1.089 | -0.598 | 0.782 | -0.007 | -0.003 | -0.003 | 0 | 0.7 | 0.73 | 0.56 | 0.69 | 0.59 | 0.7 |
| 5 | 10905445 | rs6554614 | -1.022 | 0.173 | 0.251 | 0.766 | -0.029 | 0.058 | 0.058 | 6.085 | 0.12 | 0.4 | 0.13 | 0.13 | 0.05 | 0.06 |
| 5 | 10907870 | rs11133633 | -0.461 | 0.624 | -0.773 | 0.633 | -0.03 | 0.006 | 0.006 | 0 | 0.67 | 0.81 | 0.63 | 0.69 | 0.58 | 0.63 |
| 5 | 10912382 | rs10491237 | -1.267 | 0.026 | -0.39 | 1.113 | -0.048 | 0.135 | 0.135 | 1.654 | 0.19 | 0.55 | 0.21 | 0.37 | 0.11 | 0.25 |
| 5 | 10912945 | rs16901129 | -1.379 | -0.035 | -0.407 | 1.355 | -0.049 | 0.157 | 0.157 | 2.279 | 0.2 | 0.58 | 0.21 | 0.46 | 0.11 | 0.26 |
| 5 | 10913841 | rs10058973 | 0.066 | 0.071 | -0.841 | 0.599 | -0.026 | 0.096 | 0.096 | 0.253 | 0.48 | 0.25 | 0.42 | 0.32 | 0.51 | 0.39 |
| 5 | 10914119 | rs17197055 | 0.643 | 0.303 | -1.375 | 0.47 | -0.062 | 0.07 | 0.07 | 0.07 | 0.38 | 0.07 | 0.36 | 0.23 | 0.4 | 0.35 |
| 5 | 10914151 | rs16898829 | -0.266 | 0.236 | -1.032 | 0.629 | -0.012 | 0.01 | 0.01 | 15.347 | 0.09 | 0.16 | 0.11 | 0.09 | 0.04 | 0.09 |
| 5 | 10914403 | rs17197062 | -0.315 | 0.422 | -1.109 | 0.294 | -0.01 | 0.088 | 0.088 | 0.058 | 0.49 | 0.3 | 0.45 | 0.33 | 0.52 | 0.4 |
| 5 | 10916640 | rs9686531 | 0.515 | 0.565 | -2.04 | -0.255 | -0.034 | 0.057 | 0.057 | 0 | 0.13 | 0.03 | 0.12 | 0.04 | 0.12 | 0.16 |
| 5 | 10918265 | rs12520057 | -0.156 | 0.237 | -2.19 | -0.35 | -0.069 | 0.006 | 0.006 | 0.071 | 0.25 | 0.1 | 0.32 | 0.22 | 0.26 | 0.2 |
| 5 | 10924404 | rs2399984 | 0.682 | -0.575 | -2.188 | -0.441 | -0.004 | 0.097 | 0.097 | 0 | 0.21 | 0.04 | 0.17 | 0 | 0.24 | 0.17 |
| 5 | 10929785 | rs1354134 | -0.631 | -0.411 | -1.642 | 0.157 | -0.026 | 0.029 | 0.029 | 0 | 0.28 | 0.16 | 0.23 | 0.22 | 0.25 | 0.21 |
| 5 | 10935007 | rs13357458 | -0.428 | -1.029 | -0.382 | 0.741 | -0.034 | 0.015 | 0.015 | 0 | 0.07 | 0.21 | 0.03 | 0.08 | 0.02 | 0.02 |
| 5 | 10935557 | rs852556 | -0.059 | -1.128 | -1.122 | 0.789 | -0.045 | -0.014 | -0.014 | 0 | 0.69 | 0.89 | 0.62 | 0.7 | 0.69 | 0.6 |
| 5 | 10935873 | rs16901136 | 0.869 | -1.353 | -0.721 | 0.874 | 0.003 | 0.026 | 0.026 | 0 | 0.08 | 0.12 | 0.03 | 0.14 | 0.08 | 0.12 |
| 5 | 10938199 | rs11133634 | -0.537 | -1.65 | -1.195 | 0.717 | -0.001 | -0.001 | -0.001 | 2.806 | 0.35 | 0.38 | 0.42 | 0.37 | 0.35 | 0.31 |
| 5 | 10938228 | rs10474905 | 0.429 | -1.626 | -1.192 | 0.69 | 0.003 | -0.021 | -0.021 | 0 | 0.22 | 0.28 | 0.22 | 0.15 | 0.23 | 0.14 |
| 5 | 10940352 | rs852562 | 1.321 | -1.826 | -0.982 | 0.599 | 0.001 | 0.003 | 0.003 | 0 | 0.46 | 0.36 | 0.38 | 0.41 | 0.53 | 0.42 |
| 5 | 10941988 | rs852563 | -1.327 | -1.71 | -0.975 | 0.391 | 0.011 | -0.035 | -0.035 | 0.016 | 0.85 | 0.91 | 0.81 | 0.74 | 0.9 | 0.76 |
| 5 | 10942243 | rs852564 | 1.24 | -1.632 | -0.93 | 0.339 | 0 | 0.004 | 0.004 | 0 | 0.46 | 0.35 | 0.37 | 0.45 | 0.53 | 0.42 |
| 5 | 10943255 | rs16901149 | -0.271 | -1.277 | -0.751 | 0.004 | -0.008 | -0.033 | -0.033 | 0 | 0.13 | 0.17 | 0.09 | 0.03 | 0.12 | 0.12 |
| 5 | 10946582 | rs16901155 | -0.276 | -1.262 | -0.776 | 0.011 | -0.011 | -0.044 | -0.044 | 0 | 0.13 | 0.19 | 0.09 | 0.03 | 0.12 | 0.1 |
| 5 | 10951133 | rs970133 | -0.351 | -1.502 | -0.689 | 0.099 | 0 | -0.005 | -0.005 | 0 | 0.41 | 0.54 | 0.29 | 0.25 | 0.41 | 0.3 |
| 5 | 10954186 | rs860434 | -1.124 | -1.439 | -1.037 | -0.046 | 0.031 | -0.006 | -0.006 | 0.06 | 0.79 | 0.85 | 0.86 | 0.81 | 0.88 | 0.75 |
| 5 | 10955488 | rs1501776 | 0.008 | -1.436 | -1 | -0.053 | 0.004 | -0.028 | -0.028 | 0 | 0.36 | 0.52 | 0.28 | 0.19 | 0.38 | 0.28 |
| 5 | 10955552 | rs1501777 | 0.027 | -1.434 | -1.088 | -0.052 | 0.003 | -0.023 | -0.023 | 0 | 0.36 | 0.53 | 0.28 | 0.19 | 0.38 | 0.28 |
| 5 | 10955988 | rs852586 | 1.356 | -1.077 | -1.626 | 0.152 | -0.176 | -0.152 | -0.152 | 0.258 | 0.31 | 0.03 | 0.41 | 0.44 | 0.44 | 0.35 |
| 5 | 10958635 | rs2937509 | -1.237 | 0.151 | -0.48 | -0.21 | 0.01 | 0.064 | 0.064 | 0.382 | 0.47 | 0.77 | 0.44 | 0.38 | 0.44 | 0.4 |
| 5 | 10964557 | rs11133635 | 0.272 | 0.758 | -0.465 | 0.088 | 0.005 | -0.03 | -0.03 | 0 | 0.15 | 0.16 | 0.2 | 0.05 | 0.2 | 0.11 |
| 5 | 10965031 | rs2907122 | -0.537 | 0.627 | -1.031 | 0.383 | 0.001 | 0.02 | 0.02 | 0.222 | 0.41 | 0.66 | 0.42 | 0.39 | 0.39 | 0.35 |
| 5 | 10968872 | rs12188188 | -0.469 | 1.058 | -0.669 | 0.091 | 0 | -0.029 | -0.029 | 2.341 | 0.16 | 0.22 | 0.11 | 0 | 0.15 | 0.04 |
| 5 | 10973727 | rs1566622 | -0.824 | 1.502 | -0.063 | 0.927 | 0.039 | 0.154 | 0.154 | 0 | 0.06 | 0.16 | 0.08 | 0.21 | 0.07 | 0.07 |
| 5 | 10973927 | rs1566624 | -1.451 | 1.407 | 0.46 | 0.866 | 0.034 | 0.031 | 0.031 | 0 | 0.23 | 0.49 | 0.21 | 0.22 | 0.25 | 0.11 |
| 5 | 10976530 | rs7728798 | -1.016 | 1.347 | 0.298 | 1.019 | 0.035 | 0.149 | 0.149 | 0 | 0.07 | 0.16 | 0.09 | 0.21 | 0.07 | 0.07 |
| 5 | 10984610 | rs2062685 | -0.037 | 1.145 | 0.038 | 0.715 | 0.002 | 0.003 | 0.003 | 0 | 0.18 | 0.14 | 0.11 | 0.16 | 0.16 | 0.15 |
| 5 | 10984714 | rs2062688 | -1.496 | 1.621 | 0.486 | 0.815 | 0.008 | 0.002 | 0.002 | 0.278 | 0.39 | 0.57 | 0.31 | 0.37 | 0.4 | 0.24 |
| 5 | 10986947 | rs10434650 | -1.297 | 1.568 | 0.49 | 1.35 | 0.016 | 0.036 | 0.036 | 0.263 | 0.36 | 0.51 | 0.28 | 0.44 | 0.36 | 0.22 |

| | | | | | | | | | | | | | | | | |
|---|---|---|---|---|---|---|---|---|---|---|---|---|---|---|---|---|
| 5 | 10987093 | rs12520815 | 0.26 | 1.579 | 0.492 | 1.363 | 0 | -0.001 | -0.001 | 0 | 0.16 | 0.14 | 0.08 | 0.23 | 0.12 | 0.13 |
| 5 | 10987114 | rs10434651 | -0.599 | 1.56 | 0.6 | 1.258 | 0.031 | 0.02 | 0.02 | 0 | 0.19 | 0.26 | 0.19 | 0.21 | 0.24 | 0.09 |
| 5 | 10998665 | rs706306 | -0.271 | 1.318 | -0.058 | 0.475 | 0.012 | 0.04 | 0.04 | 0 | 0.72 | 0.89 | 0.87 | 0.81 | 0.76 | 0.77 |
| 5 | 11002126 | rs4702781 | -0.136 | 1.462 | -0.015 | -0.016 | 0.006 | -0.037 | -0.037 | 0 | 0.32 | 0.64 | 0.49 | 0.07 | 0.39 | 0.19 |
| 5 | 11005150 | rs6878664 | 0.968 | 1.538 | 0.26 | 0.016 | 0.007 | -0.012 | -0.012 | 0 | 0.16 | 0.18 | 0.36 | 0.08 | 0.23 | 0.18 |
| 5 | 11006477 | rs852614 | 0.371 | 1.506 | 0.008 | -0.022 | 0.019 | 0.083 | 0.083 | 0 | 0.51 | 0.34 | 0.47 | 0.74 | 0.53 | 0.6 |
| 5 | 11011206 | rs852619 | -0.936 | 0.978 | -0.434 | -0.316 | 0.041 | 0.078 | 0.078 | 0 | 0.66 | 0.86 | 0.84 | 0.8 | 0.74 | 0.75 |
| 5 | 11011945 | rs852620 | 0.97 | 0.951 | -0.488 | -0.092 | 0.015 | -0.016 | -0.016 | 2.444 | 0.31 | 0.22 | 0.29 | 0.64 | 0.29 | 0.52 |
| 5 | 11014035 | rs1501772 | -0.11 | 1.044 | -0.911 | -0.155 | -0.035 | -0.028 | -0.028 | 0.319 | 0.08 | 0.01 | 0.1 | 0.11 | 0.14 | 0.09 |
| 5 | 11015971 | rs852623 | 0.891 | 0.722 | -1.144 | 0.16 | 0.015 | 0.023 | 0.023 | 0 | 0.24 | 0.11 | 0.22 | 0.17 | 0.26 | 0.29 |
| 5 | 11016111 | rs852625 | -0.672 | 0.705 | -1.097 | 0.208 | 0.016 | 0.099 | 0.099 | 0 | 0.51 | 0.75 | 0.66 | 0.7 | 0.51 | 0.67 |
| 5 | 11019405 | rs1697901 | 0.408 | 0.647 | -1.099 | 0.213 | -0.02 | -0.003 | -0.003 | 0 | 0.28 | 0.13 | 0.3 | 0.27 | 0.34 | 0.34 |
| 5 | 11020804 | rs1638362 | 0.132 | 0.561 | -1.108 | 0.233 | -0.044 | 0.043 | 0.043 | 0 | 0.39 | 0.22 | 0.41 | 0.27 | 0.51 | 0.37 |
| 5 | 11020918 | rs16901205 | 1.375 | 0.679 | -0.532 | 0.265 | 0.007 | -0.001 | -0.001 | 0 | 0.09 | 0.1 | 0.3 | 0.06 | 0.15 | 0.11 |
| 5 | 11024270 | rs7702184 | -0.252 | 0.031 | -0.039 | 0.733 | -0.065 | 0.237 | 0.237 | 3.377 | 0.17 | 0.6 | 0.13 | 0.54 | 0.07 | 0.24 |
| 5 | 11028344 | rs17179868 | 0.573 | 0.293 | -0.283 | 0.409 | -0.006 | 0.003 | 0.003 | 0 | 0.24 | 0.21 | 0.41 | 0.09 | 0.34 | 0.17 |
| 5 | 11030164 | rs1116698 | -1.344 | 0.029 | -0.898 | 0.098 | 0.012 | 0.073 | 0.073 | 0 | 0.66 | 0.91 | 0.83 | 0.78 | 0.73 | 0.72 |
| 5 | 11032225 | rs1638364 | 1.73 | 0.238 | -1.247 | -0.046 | -0.017 | 0.025 | 0.025 | 0 | 0.17 | 0.05 | 0.18 | 0.08 | 0.21 | 0.2 |
| 5 | 11035367 | rs2400005 | 0.317 | 0.036 | -1.09 | -0.247 | 0 | -0.013 | -0.013 | 0.017 | 0.31 | 0.36 | 0.46 | 0.12 | 0.35 | 0.19 |
| 5 | 11037450 | rs852633 | 0.741 | 0.181 | -1.17 | -0.02 | -0.03 | 0.024 | 0.024 | 0 | 0.25 | 0.14 | 0.29 | 0.12 | 0.36 | 0.33 |
| 5 | 11042858 | rs852635 | 1.417 | 0.139 | -1.287 | 0.011 | -0.031 | 0.015 | 0.015 | 0 | 0.2 | 0.04 | 0.21 | 0.1 | 0.25 | 0.24 |
| 5 | 11045100 | rs852636 | -0.076 | -0.181 | -1.114 | 0.139 | -0.001 | -0.005 | -0.005 | 0 | 0.65 | 0.3 | 0.51 | 0.35 | 0.64 | 0.63 |
| 5 | 11046606 | rs11745793 | 0.933 | -0.216 | -1.217 | 0.02 | -0.005 | -0.007 | -0.007 | 0.021 | 0.07 | 0.1 | 0.06 | 0.04 | 0.04 | 0.04 |
| 5 | 11050264 | rs852640 | -0.97 | -0.191 | -1.013 | -0.026 | 0.006 | -0.023 | -0.023 | 0 | 0.76 | 0.7 | 0.6 | 0.93 | 0.69 | 0.85 |
| 5 | 11054511 | rs1024618 | 1.465 | -0.383 | -1.292 | -0.045 | -0.026 | 0.061 | 0.061 | 0 | 0.31 | 0.03 | 0.15 | 0.18 | 0.24 | 0.26 |
| 5 | 11069269 | rs1860242 | -1.056 | -0.341 | -1.193 | -0.055 | -0.049 | 0.076 | 0.076 | 0 | 0.76 | 0.88 | 0.61 | 0.93 | 0.67 | 0.82 |
| 5 | 11075599 | rs10513065 | 0.312 | -0.435 | -0.996 | 0.036 | -0.016 | -0.022 | -0.022 | 0 | 0.09 | 0.18 | 0.07 | 0.05 | 0.04 | 0.04 |
| 5 | 11075691 | rs2973522 | 0.32 | -0.531 | -0.962 | 0.244 | -0.025 | 0.337 | 0.337 | 1.148 | 0.12 | 0.51 | 0.1 | 0.59 | 0.05 | 0.22 |
| 5 | 11076061 | rs10513066 | 1.47 | -0.291 | -1.366 | -0.01 | -0.04 | 0.008 | 0.008 | 0 | 0.19 | 0.03 | 0.21 | 0.1 | 0.24 | 0.2 |
| 5 | 11086565 | rs2907105 | -0.584 | -0.297 | -1.493 | 0.449 | -0.086 | -0.02 | -0.02 | 0.963 | 0.71 | 0.91 | 0.54 | 0.69 | 0.57 | 0.64 |
| 5 | 11087322 | rs758341 | -1.178 | -0.297 | -1.495 | 0.456 | -0.085 | -0.042 | -0.042 | 0 | 0.76 | 0.97 | 0.61 | 0.71 | 0.67 | 0.75 |
| 5 | 11087563 | rs11133637 | -0.542 | -0.465 | -0.799 | 0.466 | -0.001 | -0.001 | -0.001 | 0 | 0.41 | 0.61 | 0.43 | 0.52 | 0.42 | 0.49 |
| 5 | 11088028 | rs9312754 | 0.099 | -0.481 | -0.524 | 0.62 | -0.03 | 0.164 | 0.164 | 0.936 | 0.16 | 0.4 | 0.14 | 0.41 | 0.09 | 0.19 |
| 5 | 11089822 | rs16901246 | 0.152 | -0.292 | -1.023 | -0.075 | 0.036 | -0.008 | -0.008 | 1.696 | 0.07 | 0.22 | 0.08 | 0.02 | 0.1 | 0.1 |
| 5 | 11091008 | rs2973513 | 0.02 | -0.296 | -1.103 | -0.077 | 0.052 | 0 | 0 | 0.006 | 0.08 | 0.27 | 0.08 | 0.02 | 0.1 | 0.1 |
| 5 | 11095315 | rs2973511 | -0.353 | -0.633 | -1.646 | -0.215 | -0.041 | 0.17 | 0.17 | 0.905 | 0.53 | 0.93 | 0.39 | 0.83 | 0.42 | 0.61 |
| 5 | 11096972 | rs2905997 | 0.267 | -0.324 | -1.392 | -0.47 | -0.002 | -0.017 | -0.017 | 0 | 0.39 | 0.39 | 0.25 | 0.2 | 0.35 | 0.36 |
| 5 | 11104937 | rs2905992 | -0.305 | 0.257 | -1.864 | -0.256 | -0.083 | 0.118 | 0.118 | 0.064 | 0.73 | 0.96 | 0.61 | 0.93 | 0.65 | 0.81 |
| 5 | 11111713 | rs2905986 | 0.409 | -0.338 | -1.607 | -0.223 | 0.018 | 0.121 | 0.121 | 0 | 0.38 | 0.11 | 0.18 | 0.19 | 0.29 | 0.24 |
| 5 | 11111771 | rs6884431 | -0.29 | 0.059 | -0.787 | 0.25 | 0.003 | 0.001 | 0.001 | 0 | 0.31 | 0.37 | 0.45 | 0.63 | 0.39 | 0.51 |
| 5 | 11114201 | rs11954029 | 0.024 | -0.37 | -0.06 | 1.156 | 0.004 | 0.277 | 0.277 | 0 | 0.07 | 0.23 | 0.06 | 0.47 | 0.05 | 0.26 |
| 5 | 11115483 | rs7341140 | -1.045 | -0.45 | 0.315 | 1.128 | -0.008 | 0.359 | 0.359 | 0.273 | 0.11 | 0.42 | 0.1 | 0.47 | 0.06 | 0.27 |
| 5 | 11117583 | rs2285975 | 0.039 | -0.425 | 0.138 | 1.133 | 0.004 | 0.277 | 0.277 | 0 | 0.07 | 0.23 | 0.06 | 0.47 | 0.05 | 0.26 |
| 5 | 11118144 | rs6554619 | -0.946 | -0.749 | -0.143 | 0.92 | -0.024 | -0.01 | -0.01 | 0 | 0.74 | 0.96 | 0.67 | 0.82 | 0.69 | 0.76 |
| 5 | 11124555 | rs7700861 | -1.08 | -0.42 | 0.775 | 1.478 | -0.014 | 0.294 | 0.294 | 0.147 | 0.1 | 0.37 | 0.09 | 0.47 | 0.06 | 0.28 |
| 5 | 11126791 | rs2023923 | -1.138 | -0.42 | 0.664 | 1.478 | -0.005 | 0.377 | 0.377 | 0.543 | 0.11 | 0.45 | 0.1 | 0.47 | 0.06 | 0.27 |
| 5 | 11131114 | rs6865035 | -1.489 | -0.227 | 0.741 | 1.603 | 0.074 | 0.443 | 0.443 | 1.216 | 0.07 | 0.55 | 0.12 | 0.35 | 0.04 | 0.09 |
| 5 | 11134772 | rs10513073 | -0.405 | -0.512 | 0.203 | 1.167 | -0.01 | -0.009 | -0.009 | 0 | 0.56 | 0.6 | 0.69 | 0.72 | 0.63 | 0.58 |
| 5 | 11136893 | rs2214599 | 0.183 | -0.266 | 0.056 | 1.205 | 0.021 | 0.169 | 0.169 | 0.752 | 0.48 | 0.11 | 0.55 | 0.28 | 0.54 | 0.42 |
| 5 | 11145866 | rs12516033 | -0.637 | -0.288 | 0.504 | 1.203 | -0.01 | -0.006 | -0.006 | 0.005 | 0.57 | 0.6 | 0.69 | 0.72 | 0.64 | 0.59 |
| 5 | 11146253 | rs10073056 | -0.675 | -0.279 | 0.463 | 1.207 | -0.004 | -0.002 | -0.002 | 0 | 0.58 | 0.62 | 0.69 | 0.72 | 0.64 | 0.6 |
| 5 | 11147575 | rs12516262 | 0.07 | -0.223 | 0.321 | 1.022 | 0.035 | 0.17 | 0.17 | 1.229 | 0.49 | 0.1 | 0.55 | 0.27 | 0.55 | 0.44 |
| 5 | 11147814 | rs886527 | -0.236 | -0.234 | 0.52 | 1.049 | 0.019 | 0.015 | 0.015 | 1.1 | 0.54 | 0.42 | 0.6 | 0.36 | 0.6 | 0.51 |
| 5 | 11148537 | rs12514212 | 0.148 | 0.089 | 0.431 | 0.898 | -0.005 | 0.013 | 0.013 | 0.342 | 0.43 | 0.42 | 0.52 | 0.27 | 0.44 | 0.39 |
| 5 | 11149808 | rs10055024 | 0.331 | 0.239 | 0.411 | 0.878 | -0.001 | 0.005 | 0.005 | 0.218 | 0.41 | 0.42 | 0.49 | 0.27 | 0.4 | 0.35 |
| 5 | 11154447 | rs13358276 | 0.262 | 0.337 | 0.012 | 0.934 | -0.047 | 0.068 | 0.068 | 0.362 | 0.4 | 0.18 | 0.5 | 0.27 | 0.43 | 0.38 |
| 5 | 11156272 | rs1990004 | -0.454 | 0.031 | -0.02 | 0.511 | 0.021 | 0.012 | 0.012 | 0.564 | 0.54 | 0.31 | 0.58 | 0.43 | 0.54 | 0.46 |
| 5 | 11160043 | rs2277054 | 0.18 | -0.135 | -0.479 | 0.326 | -0.099 | -0.021 | -0.021 | 0 | 0.41 | 0.05 | 0.32 | 0.35 | 0.49 | 0.34 |
| 5 | 11161237 | rs2023916 | -1.582 | -0.011 | -0.03 | 0.432 | 0.015 | 0.033 | 0.033 | 0.804 | 0.64 | 0.72 | 0.72 | 0.86 | 0.66 | 0.65 |
| 5 | 11161625 | rs1978156 | -1.705 | -0.026 | -0.024 | 0.435 | 0.01 | 0.017 | 0.017 | 0.866 | 0.65 | 0.71 | 0.72 | 0.87 | 0.66 | 0.66 |
| 5 | 11162725 | rs2057794 | -0.188 | 0.109 | -0.293 | 0.529 | 0.001 | -0.044 | -0.044 | 0 | 0.14 | 0.19 | 0.09 | 0.03 | 0.11 | 0.05 |
| 5 | 11163984 | rs7703504 | 0.24 | 0.247 | -1.15 | 0.611 | -0.186 | -0.052 | -0.052 | 0 | 0.35 | 0.03 | 0.26 | 0.31 | 0.44 | 0.25 |
| 5 | 11164361 | rs721768 | -0.064 | 0.344 | -0.826 | 0.896 | -0.114 | -0.038 | -0.038 | 0 | 0.41 | 0.07 | 0.48 | 0.41 | 0.44 | 0.41 |
| 5 | 11166948 | rs12716080 | -0.225 | 0.789 | 0.203 | 1.307 | 0.012 | -0.004 | -0.004 | 0 | 0.33 | 0.42 | 0.3 | 0.73 | 0.39 | 0.35 |
| 5 | 11167277 | rs7713461 | 0.115 | 0.876 | -0.35 | 1.304 | -0.013 | -0.005 | -0.005 | 0.537 | 0.28 | 0.25 | 0.21 | 0.31 | 0.32 | 0.19 |
| 5 | 11169601 | rs16901339 | -0.931 | 0.601 | 0.252 | 1.457 | 0.015 | 0.125 | 0.125 | 0.036 | 0.18 | 0.44 | 0.19 | 0.36 | 0.15 | 0.15 |
| 5 | 11172241 | rs13188937 | -1.407 | 0.649 | -0.367 | 1.088 | 0 | 0.006 | 0.006 | 0 | 0.06 | 0.06 | 0.02 | 0.08 | 0.07 | 0.08 |
| 5 | 11172736 | rs6883143 | -0.284 | 0.467 | -0.948 | 0.998 | -0.03 | -0.024 | -0.024 | 1.798 | 0.58 | 0.78 | 0.49 | 0.5 | 0.49 | 0.41 |
| 5 | 11172773 | rs16901347 | -0.623 | -0.151 | 0.016 | 0.914 | -0.053 | 0.014 | 0.014 | 0 | 0.22 | 0.54 | 0.17 | 0.19 | 0.17 | 0.12 |

| | | | | | | | | | | | | | | | | |
|---|---|---|---|---|---|---|---|---|---|---|---|---|---|---|---|---|
| 5 | 11175485 | rs1018965 | 0.559 | 0.135 | 0.486 | 0.663 | -0.002 | -0.004 | -0.004 | 0 | 0.91 | 0.74 | 0.9 | 0.95 | 0.93 | 0.95 |
| 5 | 11176387 | rs1012092 | 0.785 | 0.417 | 0.184 | 0.629 | 0.031 | 0.047 | 0.047 | 0 | 0.56 | 0.25 | 0.58 | 0.73 | 0.6 | 0.61 |
| 5 | 11179193 | rs13179953 | -0.067 | 0.348 | 0.407 | 0.464 | 0.007 | 0.011 | 0.011 | 0.621 | 0.33 | 0.25 | 0.28 | 0.26 | 0.31 | 0.23 |
| 5 | 11179494 | rs10062829 | 0.152 | 0.391 | 0.349 | 0.433 | 0.023 | 0.054 | 0.054 | 1.464 | 0.29 | 0.18 | 0.18 | 0.12 | 0.25 | 0.16 |
| 5 | 11186652 | rs32280 | -1.129 | 0.841 | 0.471 | 0.321 | 0.017 | 0.063 | 0.063 | 0 | 0.51 | 0.67 | 0.47 | 0.27 | 0.45 | 0.38 |
| 5 | 11190998 | rs32274 | -1.751 | 1.181 | -0.065 | 0.576 | -0.138 | -0.163 | -0.163 | 0 | 0.64 | 0.98 | 0.72 | 0.51 | 0.56 | 0.53 |
| 5 | 11194840 | rs153607 | -0.512 | 1.067 | 0.541 | 0.36 | 0.007 | -0.01 | -0.01 | 0 | 0.43 | 0.59 | 0.26 | 0.13 | 0.31 | 0.18 |
| 5 | 11195687 | rs16901390 | -0.783 | 1.038 | 0.804 | 0.344 | -0.008 | -0.034 | -0.034 | 0 | 0.08 | 0.29 | 0.04 | 0 | 0.03 | 0.02 |
| 5 | 11197025 | rs32270 | -0.952 | 1.165 | 0.388 | 0.467 | 0.02 | 0.058 | 0.058 | 0.008 | 0.51 | 0.68 | 0.47 | 0.28 | 0.44 | 0.37 |
| 5 | 11200596 | rs4702794 | -1.093 | 1.327 | 0.259 | 0.866 | -0.018 | -0.02 | -0.02 | 0.338 | 0.08 | 0.05 | 0.17 | 0.13 | 0.12 | 0.17 |
| 5 | 11202649 | rs26461 | -1.994 | 0.886 | 0.562 | 0.591 | -0.015 | -0.069 | -0.069 | 0 | 0.16 | 0.51 | 0.14 | 0.02 | 0.1 | 0.18 |
| 5 | 11203426 | rs1302802 | -0.423 | 1.21 | -0.472 | 0.967 | -0.047 | -0.062 | -0.062 | 0 | 0.12 | 0.01 | 0.24 | 0.27 | 0.14 | 0.12 |
| 5 | 11205980 | rs6879492 | -0.756 | 1.194 | -0.413 | 0.887 | -0.019 | 0.007 | 0.007 | 0 | 0.07 | 0.02 | 0.09 | 0.02 | 0.11 | 0.15 |
| 5 | 11214203 | rs2036319 | -2.121 | 1.372 | 0.064 | 1.008 | 0.03 | 0.04 | 0.04 | 0 | 0.13 | 0.23 | 0.26 | 0.14 | 0.18 | 0.24 |
| 5 | 11218488 | rs27520 | 0.695 | 1.396 | -0.051 | 0.693 | -0.001 | -0.008 | -0.008 | 0 | 0.54 | 0.51 | 0.49 | 0.73 | 0.52 | 0.61 |
| 5 | 11218850 | rs398041 | 0.366 | 1.38 | -0.3 | 0.851 | -0.041 | -0.058 | -0.058 | 0 | 0.1 | 0.01 | 0.23 | 0.27 | 0.12 | 0.12 |
| 5 | 11234229 | rs154717 | -1.328 | 1.329 | -0.053 | 0.55 | 0.002 | -0.008 | -0.008 | 1.106 | 0.09 | 0.14 | 0.16 | 0.03 | 0.12 | 0.18 |
| 5 | 11239180 | rs10474912 | 0.467 | 1.373 | -0.168 | 0.454 | 0.029 | 0.062 | 0.062 | 0 | 0.31 | 0.18 | 0.15 | 0.13 | 0.24 | 0.15 |
| 5 | 11249206 | rs32064 | -0.691 | 1.495 | -0.081 | 0.529 | -0.005 | -0.013 | -0.013 | 0.407 | 0.45 | 0.49 | 0.48 | 0.27 | 0.44 | 0.41 |
| 5 | 11252267 | rs194487 | -0.225 | 1.31 | -0.037 | 0.555 | -0.022 | -0.031 | -0.031 | 0 | 0.38 | 0.46 | 0.34 | 0.23 | 0.32 | 0.24 |
| 5 | 11256115 | rs181889 | -0.339 | 1.448 | -0.042 | 0.512 | -0.012 | -0.025 | -0.025 | 0 | 0.39 | 0.49 | 0.38 | 0.24 | 0.37 | 0.25 |
| 5 | 11269715 | rs26464 | -1.457 | 1.442 | 0.205 | 0.552 | 0.037 | 0.06 | 0.06 | 0 | 0.08 | 0.28 | 0.22 | 0.1 | 0.11 | 0.11 |
| 5 | 11277256 | rs441973 | 0.541 | 1.401 | -0.017 | 0.502 | 0.013 | 0.025 | 0.025 | 0 | 0.3 | 0.22 | 0.15 | 0.11 | 0.22 | 0.11 |
| 5 | 11280739 | rs32100 | -1.757 | 1.555 | -0.312 | 0.984 | -0.058 | -0.037 | -0.037 | 0.213 | 0.6 | 0.94 | 0.76 | 0.61 | 0.59 | 0.46 |
| 5 | 11285548 | rs258634 | -1.502 | 1.465 | -0.253 | 1.085 | -0.094 | -0.034 | -0.034 | 0.106 | 0.57 | 0.94 | 0.72 | 0.59 | 0.54 | 0.41 |
| 5 | 11286173 | rs4702797 | 0.373 | 1.695 | -0.351 | 1.173 | -0.024 | -0.046 | -0.046 | 0 | 0.09 | 0.01 | 0.22 | 0.36 | 0.11 | 0.11 |
| 5 | 11286825 | rs386658 | -0.63 | 1.455 | 0.08 | 0.386 | 0.002 | -0.006 | -0.006 | 0.015 | 0.41 | 0.38 | 0.27 | 0.14 | 0.36 | 0.27 |
| 5 | 11287502 | rs447587 | 0.397 | 1.484 | -0.206 | 0.414 | 0.023 | 0.082 | 0.082 | 0 | 0.29 | 0.14 | 0.14 | 0.11 | 0.22 | 0.12 |
| 5 | 11291539 | rs13160436 | -0.544 | 1.597 | -0.334 | 0.574 | -0.029 | -0.006 | -0.006 | 0 | 0.06 | 0 | 0.08 | 0.03 | 0.1 | 0.15 |
| 5 | 11298223 | rs2530215 | -0.83 | 1.598 | -0.162 | 0.596 | -0.001 | -0.012 | -0.012 | 0.139 | 0.45 | 0.53 | 0.5 | 0.24 | 0.46 | 0.31 |
| 5 | 11299801 | rs40403 | -0.253 | 1.441 | -0.064 | 0.53 | -0.01 | -0.025 | -0.025 | 0 | 0.39 | 0.54 | 0.4 | 0.21 | 0.35 | 0.19 |
| 5 | 11300854 | rs6869903 | -1.667 | 1.814 | 0.053 | 0.903 | 0.05 | 0.016 | 0.016 | 0.052 | 0.18 | 0.36 | 0.36 | 0.19 | 0.26 | 0.16 |
| 5 | 11302022 | rs26000 | 0.251 | 1.311 | -0.383 | 0.787 | 0.021 | -0.01 | -0.01 | 0 | 0.43 | 0.65 | 0.35 | 0.51 | 0.36 | 0.56 |
| 5 | 11302152 | rs28210 | 1.635 | 1.613 | 0.297 | 0.636 | 0.053 | -0.014 | -0.014 | 0.011 | 0.92 | 0.75 | 0.84 | 0.98 | 0.92 | 0.84 |
| 5 | 11303097 | rs6860372 | -1.13 | 1.521 | -0.568 | 0.469 | -0.042 | 0.051 | 0.051 | 0 | 0.14 | 0 | 0.18 | 0.03 | 0.14 | 0.13 |
| 5 | 11303492 | rs27669 | 0.698 | 1.317 | 0.326 | 1.162 | 0.054 | 0.194 | 0.194 | 5.347 | 0.95 | 0.78 | 0.87 | 0.81 | 0.94 | 0.88 |
| 5 | 11303908 | rs26001 | 0.249 | 1.173 | 0.037 | 0.9 | -0.002 | 0 | 0 | 0 | 0.46 | 0.5 | 0.5 | 0.46 | 0.46 | 0.55 |
| 5 | 11304423 | rs26478 | -0.175 | 1.695 | 0.273 | 0.902 | 0.007 | -0.004 | -0.004 | 0 | 0.7 | 0.57 | 0.78 | 0.73 | 0.74 | 0.79 |
| 5 | 11305217 | rs17794637 | 0.774 | 1.72 | -0.005 | 0.512 | -0.011 | 0.062 | 0.062 | 0.087 | 0.2 | 0.07 | 0.14 | 0.09 | 0.2 | 0.08 |
| 5 | 11305991 | rs28863 | 1.664 | 1.696 | -0.301 | 0.089 | 0.035 | 0.158 | 0.158 | 0.055 | 0.42 | 0.13 | 0.29 | 0.12 | 0.37 | 0.41 |
| 5 | 11306627 | rs1015493 | 1.939 | 1.692 | -0.299 | 0.128 | -0.017 | 0.055 | 0.055 | 0 | 0.16 | 0.01 | 0.09 | 0.05 | 0.13 | 0.04 |
| 5 | 11306983 | rs16901472 | 1.217 | 1.648 | 0.394 | 0.111 | -0.005 | -0.033 | -0.033 | 0 | 0.22 | 0.27 | 0.16 | 0.1 | 0.19 | 0.08 |
| 5 | 11310423 | rs406309 | 1.335 | 1.578 | 0.087 | 0.145 | 0.032 | 0.168 | 0.168 | 0 | 0.43 | 0.13 | 0.29 | 0.12 | 0.38 | 0.49 |
| 5 | 11313232 | rs32118 | 0.872 | 1.398 | 0.339 | 1.155 | -0.016 | 0.178 | 0.178 | 5.854 | 0.8 | 0.55 | 0.67 | 0.57 | 0.86 | 0.77 |
| 5 | 11317439 | rs16901481 | 0.695 | 1.564 | -0.507 | 1.234 | -0.014 | -0.036 | -0.036 | 0 | 0.09 | 0.02 | 0.24 | 0.23 | 0.1 | 0.12 |
| 5 | 11320538 | rs1012176 | 0.322 | 1.801 | -0.167 | 1.131 | 0.022 | 0.039 | 0.039 | 0.027 | 0.15 | 0.23 | 0.18 | 0.35 | 0.25 | 0.23 |
| 5 | 11322037 | rs2012187 | 1.02 | 1.437 | -0.449 | 0.81 | 0.01 | 0.021 | 0.021 | 0 | 0.5 | 0.09 | 0.51 | 0.4 | 0.46 | 0.55 |
| 5 | 11324554 | rs4702798 | 0.586 | 0.712 | -0.509 | 0.194 | 0.017 | 0.044 | 0.044 | 0 | 0.59 | 0.32 | 0.66 | 0.76 | 0.67 | 0.73 |
| 5 | 11329757 | rs258839 | -0.276 | 1.035 | -0.72 | 0.378 | -0.001 | 0.009 | 0.009 | 0 | 0.75 | 0.83 | 0.79 | 0.8 | 0.75 | 0.87 |
| 5 | 11335373 | rs32138 | -0.084 | 0.998 | -1.091 | 0.318 | -0.036 | -0.045 | -0.045 | 0.319 | 0.72 | 0.83 | 0.79 | 0.63 | 0.68 | 0.55 |
| 5 | 11336674 | rs32136 | 0.648 | 0.691 | -1.419 | -0.096 | 0.012 | 0.06 | 0.06 | 0.292 | 0.34 | 0.18 | 0.41 | 0.2 | 0.32 | 0.28 |
| 5 | 11337346 | rs6869392 | 0.92 | 0.363 | -0.859 | -0.238 | 0.005 | -0.026 | -0.026 | 0 | 0.28 | 0.25 | 0.19 | 0.11 | 0.34 | 0.1 |
| 5 | 11339984 | rs32135 | 1.309 | 0.334 | -0.57 | -0.117 | 0.003 | -0.004 | -0.004 | 0.83 | 0.29 | 0.28 | 0.47 | 0.36 | 0.25 | 0.35 |
| 5 | 11341218 | rs1423497 | 0.808 | 0.351 | -0.505 | -0.092 | 0.004 | -0.027 | -0.027 | 0 | 0.29 | 0.26 | 0.19 | 0.14 | 0.34 | 0.1 |
| 5 | 11343384 | rs32128 | 0.158 | 0.354 | -0.488 | -0.152 | 0.001 | 0.012 | 0.012 | 1.496 | 0.65 | 0.55 | 0.71 | 0.48 | 0.67 | 0.47 |
| 5 | 11346156 | rs4702799 | -0.059 | 0.237 | -0.319 | -0.259 | -0.001 | -0.018 | -0.018 | 0.507 | 0.39 | 0.38 | 0.25 | 0.22 | 0.43 | 0.16 |
| 5 | 11347020 | rs6881516 | 0.101 | 0.37 | 0.033 | 0.012 | -0.003 | 0 | 0 | 0 | 0.11 | 0.11 | 0.06 | 0.12 | 0.09 | 0.06 |
| 5 | 11349674 | rs40289 | 1.149 | 0.737 | 0.094 | 0.176 | 0.012 | 0.019 | 0.019 | 0.016 | 0.26 | 0.15 | 0.34 | 0.19 | 0.24 | 0.28 |
| 5 | 11361864 | rs1978462 | -0.557 | 0.847 | -0.036 | 0.211 | -0.051 | -0.02 | -0.02 | 0 | 0.61 | 0.86 | 0.73 | 0.64 | 0.6 | 0.57 |
| 5 | 11363645 | rs1423493 | 0.104 | 0.81 | 0.184 | 0.199 | -0.003 | -0.006 | -0.006 | 0.464 | 0.51 | 0.66 | 0.68 | 0.53 | 0.54 | 0.53 |
| 5 | 11365166 | rs2074260 | 0.514 | 0.812 | -0.106 | 0.2 | -0.005 | 0.014 | 0.014 | 0.26 | 0.45 | 0.36 | 0.48 | 0.3 | 0.52 | 0.38 |
| 5 | 11366401 | rs11750088 | 1.26 | 1.021 | -0.042 | -0.278 | -0.086 | -0.075 | -0.075 | 0.501 | 0.18 | 0.01 | 0.23 | 0.26 | 0.27 | 0.13 |
| 5 | 11368864 | rs6875838 | -0.025 | 0.842 | 0.183 | -0.21 | 0.002 | 0.011 | 0.011 | 0.993 | 0.54 | 0.38 | 0.64 | 0.37 | 0.6 | 0.42 |
| 5 | 11372743 | rs1859082 | -0.726 | 1.077 | -0.322 | -0.224 | -0.041 | -0.089 | -0.089 | 0.795 | 0.65 | 0.89 | 0.73 | 0.57 | 0.64 | 0.58 |
| 5 | 11378817 | rs9312768 | -1.174 | 1.18 | -0.492 | -0.159 | -0.031 | -0.09 | -0.09 | 0 | 0.69 | 0.89 | 0.78 | 0.57 | 0.7 | 0.6 |
| 5 | 11380155 | rs2113012 | 0.602 | 1.197 | -0.669 | -1.023 | 0.013 | 0.091 | 0.091 | 0.328 | 0.31 | 0.13 | 0.38 | 0.11 | 0.31 | 0.28 |
| 5 | 11381138 | rs12518049 | 0.586 | 1.327 | 0.064 | -0.054 | -0.005 | 0.022 | 0.022 | 0.492 | 0.46 | 0.38 | 0.49 | 0.24 | 0.54 | 0.36 |
| 5 | 11390193 | rs40485 | 0.704 | 1.967 | -0.463 | 0.495 | 0.021 | 0.021 | 0.021 | 0 | 0.94 | 0.84 | 0.85 | 0.89 | 0.9 | 0.9 |
| 5 | 11392036 | rs154753 | -0.184 | 1.697 | -0.939 | 0.434 | -0.065 | -0.04 | -0.04 | 0 | 0.42 | 0.12 | 0.3 | 0.51 | 0.4 | 0.41 |

| | | | | | | | | | | | | | | | | |
|---|---|---|---|---|---|---|---|---|---|---|---|---|---|---|---|---|
| 5 | 11397573 | rs31837 | -1.474 | 1.268 | 0.898 | 0.387 | 0.031 | 0.071 | 0.071 | 0.046 | 0.11 | 0.27 | 0.08 | 0.13 | 0.08 | 0.17 |
| 5 | 11410718 | rs1469099 | -1.475 | 0.357 | 1.364 | 0.22 | -0.051 | 0.043 | 0.043 | 0.852 | 0.13 | 0.61 | 0.12 | 0.11 | 0.05 | 0.09 |
| 5 | 11423525 | rs10513091 | -1.638 | 1.689 | 1.182 | 0.008 | 0.109 | -0.043 | -0.043 | 0 | 0.08 | 0.19 | 0.1 | 0 | 0.16 | 0.09 |
| 5 | 11423592 | rs31887 | -1.602 | 1.327 | 1.614 | -0.216 | -0.048 | 0.014 | 0.014 | 0 | 0.15 | 0.43 | 0.11 | 0.12 | 0.05 | 0.2 |
| 5 | 11423628 | rs31888 | -2.011 | 1.244 | 0.827 | -0.225 | -0.005 | 0.025 | 0.025 | 0.168 | 0.25 | 0.8 | 0.22 | 0.11 | 0.21 | 0.28 |
| 5 | 11428476 | rs31895 | -0.467 | 0.375 | -0.296 | -0.656 | 0.07 | 0.007 | 0.007 | 0.579 | 0.59 | 0.87 | 0.5 | 0.3 | 0.66 | 0.55 |
| 5 | 11428508 | rs31896 | 0.01 | 0.86 | 0.366 | -1.285 | 0.001 | -0.024 | -0.024 | 0 | 0.32 | 0.4 | 0.23 | 0.12 | 0.27 | 0.29 |
| 5 | 11428523 | rs31897 | -0.175 | 0.811 | 0.499 | -1.359 | 0.012 | -0.034 | -0.034 | 0.367 | 0.36 | 0.47 | 0.27 | 0.15 | 0.36 | 0.35 |
| 5 | 11429782 | rs31899 | -1.016 | 0.428 | 1.03 | -1.217 | -0.019 | -0.011 | -0.011 | 0.016 | 0.2 | 0.74 | 0.11 | 0.09 | 0.06 | 0.15 |
| 5 | 11432888 | rs7710716 | 0.783 | 0.405 | 1.148 | -1.219 | 0.003 | -0.002 | -0.002 | 0 | 0.1 | 0.08 | 0.04 | 0.08 | 0.06 | 0.07 |
| 5 | 11438117 | rs10044905 | -0.821 | 0.412 | 1.914 | -1.253 | -0.042 | -0.016 | -0.016 | 0 | 0.17 | 0.38 | 0.08 | 0.1 | 0.07 | 0.13 |
| 5 | 11440686 | rs31830 | -0.425 | 0.402 | 2.248 | -1.264 | -0.024 | 0.007 | 0.007 | 1.753 | 0.14 | 0.36 | 0.08 | 0.1 | 0.07 | 0.13 |
| 5 | 11441822 | rs249237 | -1.337 | 0.403 | 1.956 | -1.265 | -0.034 | -0.023 | -0.023 | 0 | 0.21 | 0.69 | 0.12 | 0.1 | 0.07 | 0.15 |
| 5 | 11452037 | rs11133654 | 0.692 | 0.557 | 1.552 | -1.178 | 0.008 | -0.008 | -0.008 | 0 | 0.1 | 0.06 | 0.05 | 0.11 | 0.07 | 0.12 |
| 5 | 11461390 | rs31947 | -0.643 | 0.545 | 1.637 | -1.185 | -0.043 | -0.01 | -0.01 | 1.099 | 0.16 | 0.37 | 0.09 | 0.11 | 0.07 | 0.14 |
| 5 | 11461920 | rs10462668 | 0.858 | 0.104 | 0.235 | -1.341 | 0.01 | -0.053 | -0.053 | 0.627 | 0.24 | 0.09 | 0.44 | 0.71 | 0.26 | 0.34 |
| 5 | 11469640 | rs2244828 | -0.006 | 0.506 | 0.011 | -1.889 | -0.008 | 0.107 | 0.107 | 0 | 0.51 | 0.71 | 0.32 | 0.19 | 0.4 | 0.47 |
| 5 | 11479472 | rs249268 | 0.321 | 0.337 | 0.342 | -2.264 | -0.002 | -0.007 | -0.007 | 0.415 | 0.33 | 0.35 | 0.29 | 0.18 | 0.3 | 0.38 |
| 5 | 11481099 | rs249264 | -1.591 | 0.662 | 0.262 | -2.553 | -0.021 | 0.02 | 0.02 | 0.064 | 0.65 | 0.84 | 0.48 | 0.24 | 0.55 | 0.57 |
| 5 | 11482045 | rs10513094 | 0.243 | 0.733 | 0.342 | -2.606 | -0.004 | -0.004 | -0.004 | 0.099 | 0.06 | 0.03 | 0.07 | 0 | 0.12 | 0.04 |
| 5 | 11482412 | rs26147 | -0.598 | 0.659 | 0.536 | -2.638 | 0.002 | -0.011 | -0.011 | 0 | 0.55 | 0.51 | 0.39 | 0.24 | 0.42 | 0.51 |
| 5 | 11484112 | rs17216753 | 0.113 | 0.706 | 0.684 | -2.607 | 0.021 | -0.032 | -0.032 | 0 | 0.06 | 0.09 | 0.07 | 0 | 0.12 | 0.04 |
| 5 | 11487585 | rs26150 | 0.369 | 0.241 | 0.85 | -2.535 | -0.016 | 0.038 | 0.038 | 0 | 0.38 | 0.24 | 0.33 | 0.18 | 0.42 | 0.42 |
| 5 | 11490932 | rs17805650 | -0.203 | 0.571 | 1 | -2.303 | 0.009 | 0.05 | 0.05 | 2.474 | 0.22 | 0.13 | 0.1 | 0.06 | 0.13 | 0.13 |
| 5 | 11502495 | rs249236 | -0.939 | 0.608 | 0.589 | -2.29 | -0.029 | -0.103 | -0.103 | 0.025 | 0.28 | 0.6 | 0.14 | 0.06 | 0.13 | 0.17 |
| 5 | 11507294 | rs26163 | 0.42 | 0.633 | 0.283 | -1.636 | -0.007 | 0.001 | 0.001 | 0 | 0.33 | 0.28 | 0.34 | 0.24 | 0.4 | 0.39 |
| 5 | 11507368 | rs16901689 | -0.932 | 0.487 | 0.415 | -1.498 | -0.027 | -0.116 | -0.116 | 0 | 0.29 | 0.54 | 0.15 | 0.07 | 0.15 | 0.19 |
| 5 | 11512457 | rs10038337 | -0.791 | 0.91 | 0.582 | -1.175 | 0.058 | -0.087 | -0.087 | 0 | 0.14 | 0.44 | 0.09 | 0.03 | 0.1 | 0.03 |
| 5 | 11518138 | rs4702809 | 0.394 | 0.613 | 0.195 | -1.101 | 0.009 | -0.038 | -0.038 | 0.974 | 0.1 | 0.16 | 0.24 | 0.05 | 0.08 | 0.03 |
| 5 | 11518163 | rs32625 | -0.042 | 0.665 | -0.572 | -1.064 | -0.018 | 0.037 | 0.037 | 0 | 0.46 | 0.29 | 0.39 | 0.29 | 0.48 | 0.56 |
| 5 | 11518704 | rs12523633 | 0.986 | 0.611 | -0.905 | -0.531 | 0.007 | 0.017 | 0.017 | 1.257 | 0.11 | 0.06 | 0.04 | 0.05 | 0.06 | 0.14 |
| 5 | 11523516 | rs32434 | -0.582 | 0.735 | -0.867 | -0.301 | -0.005 | 0.001 | 0.001 | 0.059 | 0.61 | 0.79 | 0.48 | 0.32 | 0.55 | 0.58 |
| 5 | 11528719 | rs31780 | 1.099 | 0.818 | 0.074 | 1.084 | -0.008 | 0.054 | 0.054 | 0 | 0.69 | 0.53 | 0.67 | 0.48 | 0.69 | 0.67 |
| 5 | 11529117 | rs13160599 | -0.268 | 1.269 | 0.089 | 1.558 | -0.006 | -0.007 | -0.007 | 0 | 0.16 | 0.14 | 0.34 | 0.24 | 0.19 | 0.15 |
| 5 | 11530088 | rs7731185 | -0.725 | 1.115 | 0.378 | 2.124 | -0.011 | 0.044 | 0.044 | 0 | 0.2 | 0.32 | 0.22 | 0.29 | 0.18 | 0.24 |
| 5 | 11530762 | rs31784 | 1.391 | 1.117 | 0.345 | 2.02 | -0.004 | 0.066 | 0.066 | 0 | 0.7 | 0.53 | 0.66 | 0.5 | 0.69 | 0.67 |
| 5 | 11538361 | rs6871714 | 2.088 | 2.132 | 0.735 | 2.127 | 0.002 | -0.009 | -0.009 | 0 | 0.59 | 0.44 | 0.34 | 0.26 | 0.51 | 0.54 |
| 5 | 11538379 | rs6893111 | 2.128 | 2.155 | 0.93 | 2.135 | -0.006 | 0.006 | 0.006 | 0.173 | 0.59 | 0.39 | 0.35 | 0.33 | 0.53 | 0.55 |
| 5 | 11538504 | rs6554630 | 0.184 | 2.158 | 0.706 | 2.151 | -0.009 | 0.008 | 0.008 | 0 | 0.87 | 0.94 | 0.9 | 0.87 | 0.86 | 0.84 |
| 5 | 11539795 | rs13170650 | -1.035 | 2.17 | 1.054 | 2.275 | 0.025 | 0.074 | 0.074 | 0 | 0.16 | 0.28 | 0.21 | 0.29 | 0.2 | 0.22 |
| 5 | 11543318 | rs2530904 | -1.403 | 2.096 | 1.304 | 2.213 | 0.106 | 0.057 | 0.057 | 0.006 | 0.14 | 0.45 | 0.23 | 0.14 | 0.19 | 0.11 |
| 5 | 11544026 | rs2561621 | 1.851 | 1.816 | 1.291 | 2.432 | -0.001 | 0.006 | 0.006 | 0.738 | 0.71 | 0.63 | 0.56 | 0.63 | 0.75 | 0.67 |
| 5 | 11548050 | rs11750073 | -1.204 | 1.857 | 1.211 | 2.622 | -0.032 | -0.045 | -0.045 | 0 | 0.13 | 0.05 | 0.34 | 0.26 | 0.17 | 0.17 |
| 5 | 11548147 | rs2561596 | -0.975 | 1.878 | 1.342 | 2.681 | 0.068 | 0.034 | 0.034 | 0.203 | 0.12 | 0.43 | 0.21 | 0.11 | 0.15 | 0.1 |
| 5 | 11552229 | rs11740843 | -0.92 | 1.781 | 1.136 | 2.938 | -0.033 | -0.027 | -0.027 | 0.332 | 0.08 | 0.02 | 0.17 | 0.26 | 0.14 | 0.13 |
| 5 | 11552620 | rs13173482 | -0.359 | 1.72 | 1.132 | 2.957 | 0.029 | -0.049 | -0.049 | 0 | 0.1 | 0.03 | 0.18 | 0.24 | 0.05 | 0.14 |
| 5 | 11556824 | rs2530908 | -1.251 | 1.696 | 1.218 | 2.844 | 0.098 | 0.011 | 0.011 | 0 | 0.14 | 0.45 | 0.22 | 0.09 | 0.19 | 0.11 |
| 5 | 11563028 | rs2530910 | -2.16 | 1.698 | 1.122 | 2.881 | 0.069 | 0.118 | 0.118 | 0.064 | 0.23 | 0.52 | 0.38 | 0.4 | 0.3 | 0.23 |
| 5 | 11564995 | rs7719578 | 2.536 | 1.703 | 1.211 | 2.842 | -0.01 | -0.012 | -0.012 | 0 | 0.64 | 0.34 | 0.52 | 0.5 | 0.58 | 0.61 |
| 5 | 11571940 | rs2727588 | 2.351 | 1.869 | 1.493 | 2.402 | 0.002 | 0.051 | 0.051 | 0.071 | 0.67 | 0.73 | 0.5 | 0.17 | 0.72 | 0.44 |
| 5 | 11572318 | rs2190988 | -0.769 | 1.946 | 1.425 | 2.502 | -0.036 | 0.008 | 0.008 | 0 | 0.08 | 0.02 | 0.26 | 0.47 | 0.15 | 0.27 |
| 5 | 11573514 | rs2727590 | 1.085 | 1.92 | 1.423 | 2.423 | 0.013 | -0.03 | -0.03 | 0 | 0.77 | 0.85 | 0.77 | 0.68 | 0.83 | 0.7 |
| 5 | 11577141 | rs2727591 | -2.44 | 2.319 | 2.123 | 2.796 | 0.038 | 0.055 | 0.055 | 0 | 0.27 | 0.51 | 0.47 | 0.6 | 0.32 | 0.41 |
| 5 | 11584449 | rs7731344 | -1.363 | 2.338 | 2.122 | 2.298 | 0.013 | -0.03 | -0.03 | 1.108 | 0.19 | 0.33 | 0.24 | 0.1 | 0.24 | 0.15 |
| 5 | 11585455 | rs2530913 | -0.024 | 2.276 | 1.81 | 2.51 | -0.017 | -0.046 | -0.046 | 0 | 0.1 | 0.02 | 0.14 | 0.22 | 0.1 | 0.12 |
| 5 | 11586703 | rs2429308 | 0.818 | 1.816 | 1.889 | 2.666 | -0.045 | -0.017 | -0.017 | 0.658 | 0.91 | 0.97 | 0.7 | 0.49 | 0.84 | 0.73 |
| 5 | 11588818 | rs364327 | 1.212 | 1.829 | 1.993 | 2.671 | -0.006 | 0.039 | 0.039 | 0.17 | 0.9 | 0.9 | 0.68 | 0.47 | 0.83 | 0.71 |
| 5 | 11593348 | rs7713093 | -1.313 | 1.842 | 2.113 | 2.503 | 0.025 | -0.035 | -0.035 | 0.313 | 0.21 | 0.32 | 0.23 | 0.1 | 0.27 | 0.18 |
| 5 | 11593436 | rs247648 | 1.706 | 1.472 | 2.301 | 2.232 | 0.059 | 0.038 | 0.038 | 2.571 | 0.69 | 0.52 | 0.44 | 0.4 | 0.55 | 0.51 |
| 5 | 11607481 | rs10055710 | 1.401 | 0.899 | 2.215 | 1.411 | -0.004 | 0.004 | 0.004 | 0 | 0.61 | 0.65 | 0.32 | 0.19 | 0.49 | 0.39 |
| 5 | 11623406 | rs2727583 | -2.198 | 0.498 | 2.735 | 2.264 | -0.005 | 0.087 | 0.087 | 0.167 | 0.09 | 0.39 | 0.06 | 0.1 | 0.03 | 0.02 |
| 5 | 11636016 | rs1428514 | -1.959 | 0.995 | 2.189 | 2.649 | 0.006 | 0.037 | 0.037 | 3.885 | 0.33 | 0.59 | 0.44 | 0.46 | 0.34 | 0.38 |
| 5 | 11640694 | rs7721305 | -1.954 | 1.315 | 2.522 | 2.956 | -0.033 | 0.064 | 0.064 | 0 | 0.07 | 0.17 | 0.04 | 0.18 | 0.01 | 0.06 |
| 5 | 11641039 | rs2561598 | -2.208 | 1.561 | 2.113 | 2.692 | 0.027 | 0.044 | 0.044 | 0.008 | 0.25 | 0.69 | 0.34 | 0.51 | 0.31 | 0.28 |
| 5 | 11655830 | rs2727594 | -2.092 | 0.124 | 2.393 | 2.208 | -0.033 | -0.045 | -0.045 | 0 | 0.16 | 0.52 | 0.11 | 0.07 | 0.08 | 0.11 |
| 5 | 11658814 | rs10474928 | -1.675 | 0.145 | 2.593 | 2.265 | -0.008 | -0.007 | -0.007 | 0 | 0.12 | 0.19 | 0.06 | 0.07 | 0.08 | 0.1 |
| 5 | 11677173 | rs10462670 | -1.019 | 0.446 | 2.423 | 2.641 | -0.01 | -0.01 | -0.01 | 0.426 | 0.09 | 0.04 | 0.12 | 0.16 | 0.17 | 0.11 |
| 5 | 11696211 | rs6883905 | -2.694 | 0.118 | 2.472 | 2.727 | 0.009 | -0.087 | -0.087 | 0 | 0.31 | 0.87 | 0.27 | 0.57 | 0.27 | 0.28 |
| 5 | 11710193 | rs10041174 | -2.278 | -0.359 | 3.687 | 2.445 | -0.006 | 0.04 | 0.04 | 0.008 | 0.14 | 0.47 | 0.11 | 0.12 | 0.11 | 0.13 |

| | | | | | | | | | | | | | | | | |
|---|---|---|---|---|---|---|---|---|---|---|---|---|---|---|---|---|
| 5 | 11711684 | rs13186564 | -1.68 | -0.302 | 4.101 | 2.469 | -0.017 | -0.002 | -0.002 | 0 | 0.12 | 0.24 | 0.06 | 0.07 | 0.07 | 0.1 |
| 5 | 11712057 | rs1896703 | -1.915 | -0.356 | 3.961 | 2.619 | -0.01 | 0.04 | 0.04 | 1.204 | 0.1 | 0.19 | 0.08 | 0.21 | 0.07 | 0.19 |
| 5 | 11714657 | rs1025294 | -2.549 | -0.333 | 3.917 | 2.439 | -0.022 | 0.022 | 0.022 | 0.671 | 0.26 | 0.62 | 0.18 | 0.34 | 0.17 | 0.33 |
| 5 | 11720017 | rs6861373 | -2.454 | -0.208 | 3.649 | 2.494 | 0.009 | -0.012 | -0.012 | 0.436 | 0.26 | 0.73 | 0.19 | 0.35 | 0.17 | 0.3 |
| 5 | 11722359 | rs12332227 | -1.994 | -0.149 | 3.704 | 2.522 | -0.028 | 0.073 | 0.073 | 0.098 | 0.2 | 0.42 | 0.14 | 0.29 | 0.14 | 0.28 |
| 5 | 11736524 | rs11952405 | -1.628 | -0.232 | 1.914 | 2.504 | -0.007 | 0.008 | 0.008 | 0 | 0.17 | 0.17 | 0.1 | 0.29 | 0.12 | 0.27 |
| 5 | 11737875 | rs11955159 | -1.537 | -0.082 | 1.978 | 2.599 | -0.009 | 0.018 | 0.018 | 0 | 0.15 | 0.17 | 0.1 | 0.29 | 0.11 | 0.26 |
| 5 | 11746124 | rs10039367 | -2.381 | 0.131 | 3.254 | 2.61 | 0.039 | 0.219 | 0.219 | 0 | 0.05 | 0.28 | 0.04 | 0.13 | 0.01 | 0.01 |
| 5 | 11775395 | rs6867416 | 3.4 | 0.039 | 3.812 | 2.664 | -0.025 | 0.134 | 0.134 | 1.848 | 0.91 | 0.36 | 0.91 | 0.83 | 0.97 | 0.93 |
| 5 | 11801562 | rs6554641 | 3.358 | 0.051 | 3.813 | 2.31 | -0.021 | 0.131 | 0.131 | 2.474 | 0.91 | 0.39 | 0.89 | 0.78 | 0.97 | 0.91 |
| 5 | 11807432 | rs6554644 | -2.504 | 0.105 | 3.343 | 2.307 | 0.044 | 0.201 | 0.201 | 0 | 0.06 | 0.29 | 0.07 | 0.18 | 0.02 | 0.03 |
| 5 | 11822833 | rs3924181 | -2.482 | 0.052 | 2.999 | 2.14 | 0.011 | -0.011 | -0.011 | 0 | 0.07 | 0.28 | 0.05 | 0 | 0.02 | 0 |
| 5 | 11831211 | rs7731468 | -2.567 | 0.036 | 2.639 | 2.272 | -0.002 | 0.238 | 0.238 | 3.505 | 0.09 | 0.52 | 0.1 | 0.25 | 0.03 | 0.07 |
| 5 | 11854197 | rs11951681 | -0.517 | 1.066 | 0.5 | 1.552 | -0.025 | -0.01 | -0.01 | 0 | 0.22 | 0.11 | 0.29 | 0.26 | 0.31 | 0.2 |
| 5 | 11855867 | rs7379525 | 0.616 | 0.987 | 1.613 | 1.49 | 0.028 | 0.005 | 0.005 | 0 | 0.76 | 0.68 | 0.68 | 0.75 | 0.69 | 0.8 |
| 5 | 11859518 | rs13177858 | 0.999 | 0.726 | 2.56 | 1.428 | 0.018 | -0.037 | -0.037 | 0 | 0.72 | 0.48 | 0.68 | 0.82 | 0.69 | 0.8 |
| 5 | 11861884 | rs6897411 | 0.945 | 0.267 | 1.796 | 1.372 | 0.045 | 0.036 | 0.036 | 0 | 0.42 | 0.3 | 0.31 | 0.3 | 0.29 | 0.49 |
| 5 | 11864077 | rs12652699 | 0.049 | 0.484 | 2.233 | 1.263 | -0.01 | 0.018 | 0.018 | 0 | 0.24 | 0.21 | 0.27 | 0.14 | 0.28 | 0.19 |
| 5 | 11865658 | rs4702836 | 0.316 | 0.426 | 2.355 | 1.238 | 0.018 | -0.042 | -0.042 | 0 | 0.72 | 0.49 | 0.68 | 0.82 | 0.69 | 0.8 |
| 5 | 11867393 | rs4379193 | 0.29 | 0.424 | 2.242 | 1.237 | 0.013 | -0.032 | -0.032 | 0.219 | 0.72 | 0.47 | 0.68 | 0.83 | 0.69 | 0.8 |
| 5 | 11870138 | rs6554653 | -0.033 | 0.39 | 2.117 | 1.403 | 0.011 | 0.006 | 0.006 | 0.023 | 0.65 | 0.43 | 0.58 | 0.42 | 0.56 | 0.65 |
| 5 | 11870554 | rs4701924 | 0.262 | 0.235 | 1.727 | 1.335 | -0.024 | -0.003 | -0.003 | 0 | 0.22 | 0.1 | 0.27 | 0.22 | 0.26 | 0.18 |
| 5 | 11874565 | rs1870329 | 0.764 | 0.18 | 1.833 | 1.441 | 0.004 | -0.007 | -0.007 | 0.136 | 0.46 | 0.49 | 0.33 | 0.17 | 0.33 | 0.47 |
| 5 | 11878654 | rs1870330 | -0.415 | 0.166 | 1.717 | 1.398 | 0.02 | 0.016 | 0.016 | 0 | 0.7 | 0.62 | 0.62 | 0.39 | 0.61 | 0.67 |
| 5 | 11892360 | rs2168878 | 0.265 | 0.309 | 1.047 | 0.978 | -0.066 | 0.004 | 0.004 | 1.679 | 0.19 | 0.02 | 0.29 | 0.18 | 0.24 | 0.15 |
| 5 | 11893757 | rs12514516 | -0.157 | 0.229 | 1.507 | 0.988 | 0.001 | -0.002 | -0.002 | 0.345 | 0.23 | 0.2 | 0.32 | 0.26 | 0.24 | 0.16 |
| 5 | 11906448 | rs6554656 | 0.527 | 0.135 | 0.868 | 0.262 | -0.001 | 0.001 | 0.001 | 0.312 | 0.65 | 0.67 | 0.62 | 0.74 | 0.66 | 0.76 |
| 5 | 11908641 | rs2061392 | 0.367 | 0.38 | 0.289 | 0.363 | -0.079 | 0.019 | 0.019 | 1.043 | 0.21 | 0.02 | 0.32 | 0.18 | 0.26 | 0.17 |
| 5 | 11913471 | rs13173187 | -0.396 | -0.141 | 0.475 | 0.809 | -0.013 | 0.085 | 0.085 | 0 | 0.19 | 0.28 | 0.23 | 0.44 | 0.19 | 0.15 |
| 5 | 11918903 | rs7713607 | -0.051 | 0.036 | 0.301 | -0.391 | 0.028 | -0.065 | -0.065 | 0 | 0.68 | 0.59 | 0.61 | 0.84 | 0.61 | 0.66 |
| 5 | 11924237 | rs2125388 | 1.017 | -0.609 | -0.433 | -1.069 | 0.002 | 0.019 | 0.019 | 0.018 | 0.44 | 0.41 | 0.5 | 0.74 | 0.42 | 0.59 |
| 5 | 11926837 | rs10064613 | 0.45 | -0.499 | -0.857 | -1.099 | -0.001 | -0.04 | -0.04 | 0 | 0.17 | 0.14 | 0.08 | 0 | 0.16 | 0.03 |
| 5 | 11929913 | rs6884647 | 0.822 | -0.544 | -0.761 | -1.126 | 0.014 | 0.071 | 0.071 | 0 | 0.48 | 0.68 | 0.53 | 0.83 | 0.43 | 0.6 |
| 5 | 11945135 | rs12651763 | 0.283 | -0.492 | -0.958 | -1.051 | -0.016 | -0.038 | -0.038 | 0.492 | 0.1 | 0.03 | 0.16 | 0.59 | 0.13 | 0.25 |
| 5 | 11945254 | rs9284958 | -0.143 | -0.264 | -0.486 | -1.426 | -0.045 | 0.098 | 0.098 | 0 | 0.66 | 0.86 | 0.61 | 0.84 | 0.58 | 0.63 |
| 5 | 11949787 | rs1379902 | 0.937 | -0.074 | -0.448 | -1.305 | -0.072 | 0.103 | 0.103 | 0 | 0.24 | 0.06 | 0.33 | 0.03 | 0.36 | 0.18 |
| 5 | 11953353 | rs1379903 | 0.758 | -0.165 | -0.339 | -1.225 | 0.021 | 0.116 | 0.116 | 0 | 0.52 | 0.72 | 0.55 | 0.84 | 0.43 | 0.61 |
| 5 | 11965483 | rs6898814 | -0.419 | 0.046 | -1.043 | -1.194 | -0.067 | 0.098 | 0.098 | 0 | 0.7 | 0.89 | 0.63 | 0.85 | 0.6 | 0.65 |
| 5 | 11982921 | rs4642366 | 0.417 | 0.206 | -1.044 | -1.211 | -0.027 | 0.084 | 0.084 | 0 | 0.16 | 0.03 | 0.08 | 0 | 0.16 | 0.04 |
| 5 | 11990887 | rs12517373 | 0.214 | 0.351 | -0.648 | -0.831 | -0.052 | 0.07 | 0.07 | 0 | 0.27 | 0.1 | 0.37 | 0.13 | 0.38 | 0.32 |
| 5 | 12003910 | rs1026474 | -0.763 | 0.035 | -0.117 | -0.624 | -0.002 | 0.006 | 0.006 | 0 | 0.44 | 0.55 | 0.46 | 0.21 | 0.55 | 0.4 |
| 5 | 12017548 | rs4541658 | -0.035 | 0.144 | 0.225 | -0.263 | -0.002 | -0.003 | -0.003 | 0 | 0.24 | 0.21 | 0.35 | 0.14 | 0.37 | 0.29 |
| 5 | 12021831 | rs6886871 | 1.395 | -0.715 | -0.231 | 0.13 | -0.06 | 0.049 | 0.049 | 0 | 0.49 | 0.1 | 0.36 | 0.29 | 0.47 | 0.49 |
| 5 | 12035993 | rs12054798 | 1.401 | -0.76 | 0.172 | 0.186 | -0.027 | 0.035 | 0.035 | 0 | 0.49 | 0.19 | 0.37 | 0.33 | 0.46 | 0.35 |
| 5 | 12040493 | rs11746688 | 0.629 | -0.667 | 0.208 | 0.148 | -0.001 | -0.001 | -0.001 | 0 | 0.21 | 0.19 | 0.32 | 0.16 | 0.35 | 0.18 |
| 5 | 12049381 | rs4318778 | 1.384 | -0.68 | 0.348 | 0.182 | 0.008 | -0.013 | -0.013 | 0 | 0.7 | 0.29 | 0.67 | 0.5 | 0.81 | 0.53 |
| 5 | 12060670 | rs10462685 | 1.044 | -0.691 | 1.062 | 0.187 | -0.033 | 0.093 | 0.093 | 0 | 0.74 | 0.47 | 0.7 | 0.5 | 0.81 | 0.57 |
| 5 | 12061296 | rs7448095 | 0.428 | -0.731 | 1.327 | 0.184 | -0.022 | 0.192 | 0.192 | 0 | 0.82 | 0.63 | 0.75 | 0.5 | 0.86 | 0.78 |
| 5 | 12084583 | rs7446603 | 0.986 | -0.753 | 1.45 | 0.172 | -0.033 | 0.131 | 0.131 | 0 | 0.79 | 0.47 | 0.74 | 0.5 | 0.84 | 0.61 |
| 5 | 12102037 | rs4601043 | 1.052 | -0.78 | 1.594 | 0.144 | -0.014 | 0.047 | 0.047 | 0 | 0.77 | 0.73 | 0.73 | 0.5 | 0.83 | 0.71 |
| 5 | 12109995 | rs6875403 | 1.705 | -0.745 | 1.511 | -0.031 | -0.015 | 0.048 | 0.048 | 0 | 0.49 | 0.22 | 0.37 | 0.3 | 0.45 | 0.35 |
| 5 | 12114723 | rs12513897 | -1.24 | -0.81 | 1.808 | -0.097 | -0.016 | -0.058 | -0.058 | 0 | 0.1 | 0.26 | 0.06 | 0 | 0.03 | 0.04 |
| 5 | 12120335 | rs4702850 | -0.559 | -0.84 | 1.923 | 0.077 | 0.018 | -0.01 | -0.01 | 0 | 0.78 | 0.69 | 0.66 | 0.76 | 0.65 | 0.81 |
| 5 | 12132401 | rs32323 | -2.025 | -0.909 | 1.94 | 0.107 | -0.02 | 0.016 | 0.016 | 0 | 0.29 | 0.6 | 0.3 | 0.43 | 0.19 | 0.43 |
| 5 | 12153033 | rs250780 | -1.139 | -0.92 | 2.182 | 0.108 | -0.025 | 0.144 | 0.144 | 0 | 0.19 | 0.38 | 0.24 | 0.43 | 0.16 | 0.39 |
| 5 | 12163018 | rs2907283 | 2.002 | -0.94 | 2.106 | 0.06 | -0.076 | 0.009 | 0.009 | 0 | 0.48 | 0.1 | 0.36 | 0.32 | 0.45 | 0.38 |
| 5 | 12164978 | rs10491238 | 0.502 | -0.946 | 2.2 | 0.057 | 0.013 | -0.006 | -0.006 | 0 | 0.22 | 0.27 | 0.33 | 0.24 | 0.35 | 0.18 |
| 5 | 12167533 | rs2968125 | -0.486 | -0.951 | 2.271 | 0.05 | -0.014 | 0.097 | 0.097 | 0 | 0.16 | 0.26 | 0.22 | 0.43 | 0.14 | 0.22 |
| 5 | 12174121 | rs17789360 | -0.855 | -0.921 | 2.124 | 0.041 | 0.008 | 0.011 | 0.011 | 0.402 | 0.09 | 0.08 | 0.04 | 0.02 | 0.03 | 0.05 |
| 5 | 12181811 | rs11745148 | 0.68 | -0.909 | 2.146 | 0.039 | -0.023 | 0.004 | 0.004 | 0 | 0.2 | 0.14 | 0.31 | 0.23 | 0.35 | 0.19 |
| 5 | 12187959 | rs2968088 | -2.049 | -1.011 | 2.038 | 0.019 | -0.032 | 0.05 | 0.05 | 0 | 0.27 | 0.54 | 0.3 | 0.41 | 0.19 | 0.43 |
| 5 | 12196488 | rs2968095 | -1.422 | -0.989 | 1.985 | 0.062 | -0.034 | 0.112 | 0.112 | 0 | 0.2 | 0.51 | 0.26 | 0.42 | 0.16 | 0.39 |
| 5 | 12202885 | rs2215256 | -0.855 | -0.989 | 2.021 | 0.048 | 0.009 | -0.047 | -0.047 | 0 | 0.33 | 0.51 | 0.4 | 0.25 | 0.39 | 0.24 |
| 5 | 12206786 | rs2191102 | -1.291 | -0.975 | 2.028 | 0.095 | -0.02 | 0.139 | 0.139 | 0 | 0.19 | 0.39 | 0.24 | 0.42 | 0.16 | 0.39 |
| 5 | 12227134 | rs1112926 | 0.877 | -1.001 | 2.012 | 0.079 | -0.002 | 0.195 | 0.195 | 0 | 0.83 | 0.61 | 0.76 | 0.57 | 0.85 | 0.62 |
| 5 | 12238948 | rs2966901 | -2.019 | -0.986 | 1.788 | 0.024 | -0.076 | 0.001 | 0.001 | 0 | 0.52 | 0.9 | 0.64 | 0.67 | 0.54 | 0.63 |
| 5 | 12243796 | rs4440360 | -0.636 | -0.894 | 1.919 | 0.014 | -0.002 | 0 | 0 | 0 | 0.3 | 0.31 | 0.37 | 0.24 | 0.38 | 0.23 |
| 5 | 12261173 | rs17824844 | 0.421 | -0.856 | 1.847 | 0.067 | 0.024 | -0.01 | -0.01 | 1.118 | 0.22 | 0.28 | 0.34 | 0.22 | 0.36 | 0.18 |
| 5 | 12263385 | rs1113173 | 1.014 | -0.896 | 1.935 | 0.173 | -0.014 | 0.103 | 0.103 | 0 | 0.82 | 0.69 | 0.77 | 0.59 | 0.85 | 0.63 |

| | | | | | | | | | | | | | | | | |
|---|---|---|---|---|---|---|---|---|---|---|---|---|---|---|---|---|
| 5 | 12280849 | rs10043127 | -0.91 | -0.927 | 1.896 | 0.176 | 0.013 | -0.034 | -0.034 | 0 | 0.34 | 0.55 | 0.4 | 0.26 | 0.4 | 0.24 |
| 5 | 12285739 | rs2966898 | 0.616 | -0.952 | 2.068 | 0.188 | -0.009 | 0.085 | 0.085 | 0 | 0.84 | 0.77 | 0.79 | 0.6 | 0.86 | 0.79 |
| 5 | 12310717 | rs11133706 | -0.579 | -0.787 | 1.983 | 0.08 | 0.015 | -0.031 | -0.031 | 0 | 0.31 | 0.41 | 0.39 | 0.22 | 0.4 | 0.23 |
| 5 | 12321300 | rs2215207 | 1.594 | -0.949 | 1.894 | 0.323 | -0.037 | 0.106 | 0.106 | 0.167 | 0.78 | 0.51 | 0.74 | 0.59 | 0.85 | 0.7 |
| 5 | 12338203 | rs6891532 | 1.65 | -0.998 | 2.092 | 0.28 | -0.029 | 0.056 | 0.056 | 0.479 | 0.75 | 0.62 | 0.74 | 0.59 | 0.84 | 0.75 |
| 5 | 12342266 | rs10072442 | 0.709 | -0.864 | 2.069 | 0.193 | -0.015 | 0.002 | 0.002 | 0 | 0.19 | 0.15 | 0.32 | 0.22 | 0.35 | 0.16 |
| 5 | 12358990 | rs2400151 | 2.031 | -0.911 | 2.02 | 0.156 | -0.02 | 0.018 | 0.018 | 0.165 | 0.7 | 0.38 | 0.69 | 0.56 | 0.8 | 0.64 |
| 5 | 12361830 | rs12658793 | 1.953 | -0.94 | 1.958 | 0.151 | -0.056 | 0.008 | 0.008 | 0 | 0.49 | 0.12 | 0.36 | 0.36 | 0.46 | 0.48 |
| 5 | 12378106 | rs2215202 | 1.602 | -0.974 | 2.013 | 0.13 | -0.029 | -0.011 | -0.011 | 0 | 0.56 | 0.18 | 0.4 | 0.36 | 0.49 | 0.52 |
| 5 | 12389594 | rs2052036 | 1.14 | -0.931 | 2.191 | 0.135 | -0.016 | 0.088 | 0.088 | 0 | 0.82 | 0.71 | 0.77 | 0.59 | 0.85 | 0.79 |
| 5 | 12389751 | rs2052035 | 1.14 | -0.931 | 2.191 | 0.135 | -0.016 | 0.088 | 0.088 | 0 | 0.82 | 0.71 | 0.77 | 0.59 | 0.85 | 0.79 |
| 5 | 12392815 | rs6554700 | 0.18 | -0.88 | 2.221 | 0.119 | 0.054 | -0.032 | -0.032 | 0 | 0.23 | 0.43 | 0.36 | 0.2 | 0.35 | 0.26 |
| 5 | 12394645 | rs12519082 | -0.095 | -0.873 | 2.158 | 0.102 | -0.002 | 0.036 | 0.036 | 0 | 0.07 | 0.01 | 0.04 | 0 | 0.03 | 0.04 |
| 5 | 12401130 | rs7708669 | -0.273 | -0.82 | 2.226 | 0.157 | 0.037 | -0.021 | -0.021 | 0 | 0.25 | 0.51 | 0.37 | 0.23 | 0.36 | 0.27 |
| 5 | 12407352 | rs7736493 | 0.49 | -0.831 | 2.178 | 0.171 | 0.025 | -0.012 | -0.012 | 0.367 | 0.21 | 0.29 | 0.33 | 0.2 | 0.34 | 0.16 |
| 5 | 12437894 | rs9312801 | -0.402 | -1.044 | 2.057 | 0.157 | 0.036 | -0.034 | -0.034 | 0.315 | 0.28 | 0.51 | 0.38 | 0.23 | 0.39 | 0.18 |
| 5 | 12448703 | rs6554711 | 0.949 | -1.096 | 1.996 | 0.194 | -0.021 | 0.086 | 0.086 | 0 | 0.84 | 0.76 | 0.78 | 0.59 | 0.87 | 0.79 |
| 5 | 12468013 | rs32248 | -0.919 | -1.242 | 1.97 | 0.019 | 0.014 | -0.034 | -0.034 | 0 | 0.33 | 0.57 | 0.43 | 0.23 | 0.43 | 0.28 |
| 5 | 12474672 | rs6869431 | -1.805 | -1.269 | 2.106 | 0.115 | -0.001 | 0.007 | 0.007 | 0 | 0.48 | 0.57 | 0.63 | 0.64 | 0.52 | 0.42 |
| 5 | 12482460 | rs16902224 | -0.734 | -1.279 | 2.047 | 0.118 | 0.016 | -0.051 | -0.051 | 0 | 0.13 | 0.03 | 0.2 | 0.41 | 0.1 | 0.21 |
| 5 | 12483499 | rs32240 | -1.799 | -1.278 | 2.093 | 0.109 | -0.002 | 0.032 | 0.032 | 0 | 0.5 | 0.66 | 0.64 | 0.64 | 0.53 | 0.49 |
| 5 | 12484279 | rs245682 | -0.439 | -1.208 | 2.146 | 0.108 | 0.002 | -0.01 | -0.01 | 0.035 | 0.08 | 0.06 | 0.04 | 0 | 0.03 | 0.04 |
| 5 | 12496452 | rs245684 | 0.272 | -1.206 | 2.139 | 0.109 | 0.004 | 0.01 | 0.01 | 0 | 0.06 | 0.02 | 0.03 | 0 | 0.03 | 0.04 |
| 5 | 12500383 | rs11133718 | 0.058 | -1.205 | 2.313 | 0.1 | 0.024 | -0.021 | -0.021 | 0 | 0.24 | 0.32 | 0.36 | 0.2 | 0.38 | 0.16 |
| 5 | 12507269 | rs2895925 | -0.631 | -1.198 | 2.288 | 0.112 | 0.013 | -0.033 | -0.033 | 7.09 | 0.32 | 0.38 | 0.4 | 0.2 | 0.41 | 0.2 |
| 5 | 12507398 | rs12653370 | -1.293 | -1.43 | 1.889 | -0.11 | 0.029 | -0.085 | -0.085 | 1.209 | 0.2 | 0.04 | 0.21 | 0.41 | 0.13 | 0.21 |
| 5 | 12510848 | rs32374 | -2.054 | -1.444 | 1.904 | -0.107 | -0.004 | 0.009 | 0.009 | 0 | 0.55 | 0.64 | 0.66 | 0.64 | 0.56 | 0.49 |
| 5 | 12515215 | rs32367 | 2.158 | -1.445 | 1.693 | -0.108 | -0.063 | -0.016 | -0.016 | 0 | 0.44 | 0.12 | 0.33 | 0.35 | 0.44 | 0.5 |
| 5 | 12532357 | rs32358 | 2.17 | -1.457 | 1.681 | -0.09 | -0.085 | -0.029 | -0.029 | 0 | 0.44 | 0.1 | 0.33 | 0.36 | 0.44 | 0.51 |
| 5 | 12543792 | rs7703319 | -1.225 | -1.457 | 1.755 | -0.029 | 0.019 | -0.083 | -0.083 | 1.258 | 0.2 | 0.07 | 0.22 | 0.41 | 0.14 | 0.21 |
| 5 | 12572670 | rs4558988 | 0.736 | -1.464 | 1.893 | 0.2 | -0.017 | 0.081 | 0.081 | 0 | 0.86 | 0.79 | 0.79 | 0.59 | 0.9 | 0.79 |
| 5 | 12577738 | rs4566790 | -1.33 | -1.315 | 2.086 | 0.052 | -0.031 | -0.001 | -0.001 | 0 | 0.12 | 0.41 | 0.08 | 0.07 | 0.04 | 0.07 |
| 5 | 12587264 | rs4247241 | 2.054 | -1.335 | 2.403 | 0.116 | -0.031 | -0.005 | -0.005 | 0.582 | 0.84 | 0.3 | 0.87 | 0.89 | 0.95 | 0.85 |
| 5 | 12589411 | rs10474965 | -1.325 | -1.33 | 2.414 | 0.113 | -0.032 | -0.002 | -0.002 | 0 | 0.12 | 0.41 | 0.08 | 0.07 | 0.04 | 0.07 |
| 5 | 12597678 | rs12187460 | 0.071 | -1.223 | 2.353 | 0.044 | -0.042 | 0.056 | 0.056 | 0 | 0.27 | 0.13 | 0.36 | 0.13 | 0.42 | 0.21 |
| 5 | 12597846 | rs10038940 | -1.068 | -1.228 | 2.319 | 0.044 | 0.002 | -0.022 | -0.022 | 0 | 0.38 | 0.55 | 0.44 | 0.2 | 0.45 | 0.28 |
| 5 | 12599348 | rs7449177 | 1.016 | -1.227 | 2.223 | 0.044 | -0.018 | 0.075 | 0.075 | 0 | 0.59 | 0.24 | 0.53 | 0.76 | 0.54 | 0.71 |
| 5 | 12606091 | rs11749872 | 0.084 | -1.177 | 2.168 | 0.05 | -0.05 | 0.065 | 0.065 | 0 | 0.26 | 0.11 | 0.36 | 0.13 | 0.42 | 0.21 |
| 5 | 12633456 | rs12659350 | 1.562 | -1.426 | 1.576 | -0.066 | -0.013 | -0.004 | -0.004 | 0 | 0.79 | 0.29 | 0.86 | 0.89 | 0.93 | 0.8 |
| 5 | 12634856 | rs7449389 | 1.562 | -1.426 | 1.576 | -0.066 | -0.013 | -0.004 | -0.004 | 0 | 0.79 | 0.29 | 0.86 | 0.89 | 0.93 | 0.8 |
| 5 | 12641719 | rs10068669 | 1.149 | -1.248 | 1.484 | 0.028 | -0.045 | 0.022 | 0.022 | 0.33 | 0.87 | 0.37 | 0.9 | 0.89 | 0.96 | 0.84 |
| 5 | 12642449 | rs7733480 | 1.739 | -1.36 | 0.644 | 0.036 | -0.1 | -0.068 | -0.068 | 0.023 | 0.38 | 0.07 | 0.3 | 0.38 | 0.42 | 0.45 |
| 5 | 12652188 | rs4701945 | -1.811 | -1.572 | 1.708 | 0.503 | -0.033 | 0.059 | 0.059 | 0.281 | 0.24 | 0.35 | 0.26 | 0.42 | 0.13 | 0.3 |
| 5 | 12692686 | rs7726441 | -1.917 | -1.362 | 1.448 | 0.709 | -0.017 | 0.034 | 0.034 | 0 | 0.27 | 0.34 | 0.28 | 0.48 | 0.16 | 0.34 |
| 5 | 12697079 | rs6869725 | 0.867 | -1.344 | 1.348 | 0.763 | -0.001 | 0.061 | 0.061 | 0 | 0.05 | 0.13 | 0.04 | 0.09 | 0.02 | 0.06 |
| 5 | 12698832 | rs13182649 | -1.522 | -1.134 | 1.246 | 0.828 | 0.016 | -0.038 | -0.038 | 0 | 0.21 | 0.14 | 0.23 | 0.4 | 0.14 | 0.29 |
| 5 | 12719263 | rs12658137 | -1.371 | -0.732 | 1.724 | 0.935 | -0.001 | 0.033 | 0.033 | 0 | 0.16 | 0.18 | 0.23 | 0.46 | 0.15 | 0.29 |
| 5 | 12726490 | rs6880752 | -1.633 | -0.76 | 1.702 | 0.968 | -0.003 | 0.176 | 0.176 | 0 | 0.18 | 0.37 | 0.27 | 0.49 | 0.15 | 0.3 |
| 5 | 12733263 | rs12659128 | -1.596 | -0.751 | 1.619 | 0.972 | -0.003 | 0.161 | 0.161 | 0 | 0.18 | 0.35 | 0.27 | 0.49 | 0.15 | 0.3 |
| 5 | 12749695 | rs6862501 | 1.738 | -0.86 | 1.756 | 0.944 | -0.013 | 0.193 | 0.193 | 0 | 0.8 | 0.53 | 0.83 | 0.51 | 0.86 | 0.7 |
| 5 | 12766945 | rs11133726 | 1.352 | -0.796 | 1.426 | 0.988 | -0.004 | 0.093 | 0.093 | 0 | 0.83 | 0.75 | 0.86 | 0.51 | 0.86 | 0.71 |
| 5 | 12772609 | rs13164269 | 1.9 | -1.044 | 1.416 | 0.882 | -0.011 | 0.038 | 0.038 | 0.293 | 0.76 | 0.34 | 0.8 | 0.46 | 0.83 | 0.64 |
| 5 | 12778499 | rs9312811 | 0.933 | -0.784 | 0.653 | 0.954 | 0 | 0.093 | 0.093 | 0 | 0.86 | 0.81 | 0.87 | 0.54 | 0.87 | 0.72 |
| 5 | 12781239 | rs4495184 | 0.614 | -0.456 | -0.926 | 0.047 | 0.004 | 0.255 | 0.255 | 1.125 | 0.48 | 0.11 | 0.42 | 0.13 | 0.38 | 0.48 |
| 5 | 12784265 | rs12186744 | 1.025 | -1.201 | -0.823 | 0.661 | 0.057 | -0.067 | -0.067 | 0.018 | 0.7 | 0.24 | 0.7 | 0.49 | 0.83 | 0.59 |
| 5 | 12785362 | rs13355523 | -1.033 | -1.277 | -0.599 | 0.709 | -0.015 | 0 | 0 | 0 | 0.06 | 0.21 | 0.03 | 0.07 | 0.01 | 0.02 |
| 5 | 12785895 | rs12188566 | 1.732 | -0.727 | -1.272 | 0.396 | -0.071 | 0.257 | 0.257 | 0 | 0.47 | 0.06 | 0.48 | 0.11 | 0.5 | 0.44 |
| 5 | 12786912 | rs7720957 | 0.419 | -0.874 | -0.822 | 0.663 | -0.015 | -0.003 | -0.003 | 0 | 0.94 | 0.79 | 0.97 | 0.94 | 0.99 | 0.98 |
| 5 | 12786980 | rs10073652 | -0.81 | -0.742 | 0.073 | 0.725 | -0.062 | 0.126 | 0.126 | 0.097 | 0.24 | 0.45 | 0.24 | 0.45 | 0.16 | 0.39 |
| 5 | 12975063 | rs2118584 | -0.96 | -0.364 | -0.088 | 0.71 | 0.006 | -0.013 | -0.013 | 1.443 | 0.33 | 0.33 | 0.33 | 0.45 | 0.32 | 0.52 |
| 5 | 12975322 | rs1017454 | 1.965 | -0.448 | -0.211 | 0.479 | -0.015 | 0.086 | 0.086 | 0 | 0.41 | 0.12 | 0.41 | 0.24 | 0.35 | 0.34 |
| 5 | 12978580 | rs16902349 | -1.037 | -0.505 | -0.042 | 0.594 | 0.007 | -0.026 | -0.026 | 0.01 | 0.32 | 0.29 | 0.33 | 0.45 | 0.32 | 0.52 |
| 5 | 12986808 | rs745671 | -1.236 | -0.072 | 0.463 | 0.409 | 0.023 | -0.047 | -0.047 | 0 | 0.28 | 0.5 | 0.27 | 0.19 | 0.32 | 0.38 |
| 5 | 13013974 | rs17239989 | 1.889 | -0.625 | -0.753 | -0.103 | -0.098 | 0.208 | 0.208 | 0 | 0.39 | 0.06 | 0.37 | 0.1 | 0.39 | 0.36 |
| 5 | 13015270 | rs13163690 | 0.446 | -1.053 | -0.186 | -0.18 | -0.004 | -0.006 | -0.006 | 0 | 0.65 | 0.64 | 0.64 | 0.25 | 0.66 | 0.57 |
| 5 | 13015303 | rs17240093 | 2.138 | -1.12 | 0.443 | -0.215 | -0.01 | 0.111 | 0.111 | 0 | 0.45 | 0.25 | 0.42 | 0.13 | 0.41 | 0.38 |
| 5 | 13017177 | rs16902371 | 1.346 | -1.159 | 0.494 | 0.368 | -0.015 | -0.041 | -0.041 | 1.546 | 0.18 | 0.1 | 0.16 | 0.3 | 0.25 | 0.13 |
| 5 | 13019834 | rs17253174 | 1.343 | -1.184 | 0.539 | 0.339 | -0.006 | 0.047 | 0.047 | 0.028 | 0.54 | 0.35 | 0.47 | 0.22 | 0.47 | 0.52 |
| 5 | 13078246 | rs16902392 | -0.406 | -1.165 | 0.397 | 0.285 | 0.022 | -0.026 | -0.026 | 0 | 0.19 | 0.42 | 0.21 | 0.12 | 0.22 | 0.16 |

| | | | | | | | | | | | | | | | | |
|---|---|---|---|---|---|---|---|---|---|---|---|---|---|---|---|---|
| 5 | 13149677 | rs32534 | -0.925 | -1.054 | -0.262 | 0.339 | -0.056 | -0.076 | -0.076 | 0 | 0.82 | 0.96 | 0.83 | 0.7 | 0.74 | 0.87 |
| 5 | 13201512 | rs40406 | 1.689 | -1.43 | -0.639 | 0.01 | -0.123 | 0.107 | 0.107 | 0 | 0.39 | 0.05 | 0.33 | 0.15 | 0.36 | 0.36 |
| 5 | 13207967 | rs12716096 | 0.385 | -1.546 | -0.496 | -0.026 | -0.006 | 0.027 | 0.027 | 0.021 | 0.2 | 0.13 | 0.23 | 0.09 | 0.23 | 0.15 |
| 5 | 13207997 | rs1445728 | -1.435 | -1.697 | 0.103 | 0.114 | -0.01 | 0.017 | 0.017 | 0 | 0.39 | 0.51 | 0.49 | 0.59 | 0.4 | 0.54 |
| 5 | 13209817 | rs16902421 | -0.827 | -1.732 | 0.256 | 0.114 | -0.053 | -0.024 | -0.024 | 1.32 | 0.14 | 0.38 | 0.08 | 0.09 | 0.08 | 0.23 |
| 5 | 13213877 | rs1445730 | 0.371 | -1.707 | 0.255 | 0.141 | -0.001 | 0.086 | 0.086 | 0 | 0.09 | 0.14 | 0.19 | 0.39 | 0.1 | 0.19 |
| 5 | 13213930 | rs32518 | 2.017 | -1.759 | 0.471 | 0.069 | -0.024 | 0.067 | 0.067 | 0 | 0.41 | 0.23 | 0.34 | 0.14 | 0.36 | 0.31 |
| 5 | 13218289 | rs12519301 | -1.191 | -1.579 | 0.341 | 0.203 | -0.032 | 0.1 | 0.1 | 0 | 0.19 | 0.43 | 0.26 | 0.42 | 0.18 | 0.36 |
| 5 | 13221409 | rs10073871 | -0.725 | -1.569 | 0.304 | 0.179 | -0.046 | -0.021 | -0.021 | 0.093 | 0.13 | 0.27 | 0.06 | 0.09 | 0.08 | 0.23 |
| 5 | 13221613 | rs1373950 | -1.147 | -1.426 | 0.215 | 0.166 | -0.001 | 0.001 | 0.001 | 0 | 0.38 | 0.39 | 0.47 | 0.58 | 0.4 | 0.54 |
| 5 | 13235570 | rs962673 | -1.537 | -1.308 | -0.152 | 0.266 | -0.009 | -0.019 | -0.019 | 0 | 0.85 | 0.87 | 0.85 | 0.71 | 0.77 | 0.89 |
| 5 | 13236478 | rs1596773 | -1.666 | -1.305 | -0.275 | 0.268 | -0.048 | -0.086 | -0.086 | 0 | 0.85 | 0.98 | 0.87 | 0.71 | 0.77 | 0.89 |
| 5 | 13237036 | rs11744248 | 1.81 | -1.524 | -0.161 | 0.084 | -0.041 | 0.107 | 0.107 | 0 | 0.41 | 0.17 | 0.34 | 0.14 | 0.36 | 0.35 |
| 5 | 13238542 | rs6554760 | 0.544 | -1.416 | 0.221 | 0.164 | -0.006 | 0.042 | 0.042 | 0 | 0.68 | 0.56 | 0.55 | 0.45 | 0.67 | 0.67 |
| 5 | 13242974 | rs10066447 | 1.147 | -1.075 | 0.173 | 0.269 | 0.017 | 0.074 | 0.074 | 0 | 0.16 | 0.29 | 0.17 | 0.31 | 0.25 | 0.11 |
| 5 | 13243126 | rs4391159 | -0.504 | -1.15 | 0.213 | 0.294 | 0.004 | 0.009 | 0.009 | 0 | 0.35 | 0.39 | 0.39 | 0.47 | 0.47 | 0.29 |
| 5 | 13246979 | rs4522991 | 0.731 | -1.074 | -0.167 | 0.098 | -0.013 | 0.015 | 0.015 | 0.191 | 0.18 | 0.08 | 0.23 | 0.17 | 0.21 | 0.2 |
| 5 | 13249445 | rs13163486 | 1.49 | -1.004 | 0.084 | 0.372 | -0.038 | -0.079 | -0.079 | 0.077 | 0.13 | 0.02 | 0.13 | 0.3 | 0.2 | 0.11 |
| 5 | 13252798 | rs12522258 | 1.335 | -1.28 | -0.038 | 0.24 | 0.02 | 0.064 | 0.064 | 1.235 | 0.59 | 0.19 | 0.55 | 0.28 | 0.58 | 0.51 |
| 5 | 13256587 | rs7710793 | 1.412 | -1.442 | -0.103 | 0.179 | -0.032 | -0.091 | -0.091 | 0.085 | 0.15 | 0.02 | 0.13 | 0.3 | 0.2 | 0.11 |
| 5 | 13256986 | rs13164328 | 1.313 | -1.435 | -0.102 | 0.172 | 0.054 | 0.084 | 0.084 | 0.817 | 0.58 | 0.19 | 0.55 | 0.26 | 0.62 | 0.51 |
| 5 | 13270090 | rs6554770 | 0.629 | -1.433 | 0.422 | 0.284 | -0.051 | 0.085 | 0.085 | 0.555 | 0.78 | 0.57 | 0.72 | 0.6 | 0.82 | 0.64 |
| 5 | 13276651 | rs10462705 | 0.789 | -1.457 | 0.379 | 0.291 | -0.05 | 0.104 | 0.104 | 0.561 | 0.77 | 0.54 | 0.71 | 0.56 | 0.81 | 0.63 |
| 5 | 13278199 | rs13153960 | 1.415 | -1.086 | 0.199 | 0.469 | -0.053 | -0.086 | -0.086 | 0 | 0.14 | 0.02 | 0.13 | 0.3 | 0.24 | 0.11 |
| 5 | 13283522 | rs12691269 | 0.697 | -1.234 | 0.326 | 0.35 | 0.01 | 0.076 | 0.076 | 0 | 0.19 | 0.35 | 0.17 | 0.35 | 0.26 | 0.11 |
| 5 | 13287359 | rs1446046 | 0.639 | -1.218 | 0.303 | 0.411 | 0.004 | 0.049 | 0.049 | 0 | 0.19 | 0.26 | 0.16 | 0.36 | 0.26 | 0.11 |
| 5 | 13297813 | rs10055552 | -0.505 | -1.297 | 0.576 | 0.388 | -0.005 | 0.062 | 0.062 | 0 | 0.89 | 0.82 | 0.8 | 0.65 | 0.91 | 0.8 |
| 5 | 13306035 | rs7700799 | 1.588 | -1.236 | 0.519 | 0.416 | -0.051 | -0.089 | -0.089 | 0 | 0.14 | 0.02 | 0.13 | 0.32 | 0.24 | 0.11 |
| 5 | 13329001 | rs1446056 | -0.033 | -1.281 | 0.595 | 0.408 | 0.013 | 0.209 | 0.209 | 0.007 | 0.87 | 0.58 | 0.77 | 0.6 | 0.88 | 0.81 |
| 5 | 13332729 | rs4702930 | 1.206 | -1.506 | 0.638 | 0.278 | 0.019 | 0.028 | 0.028 | 0.721 | 0.6 | 0.25 | 0.58 | 0.29 | 0.59 | 0.55 |
| 5 | 13340781 | rs1900194 | 0.477 | -1.486 | 0.603 | 0.299 | -0.018 | 0.133 | 0.133 | 0.974 | 0.85 | 0.46 | 0.77 | 0.65 | 0.91 | 0.82 |
| 5 | 13344447 | rs6891686 | 1.519 | -1.484 | 0.489 | 0.298 | -0.006 | -0.009 | -0.009 | 0.163 | 0.16 | 0.1 | 0.14 | 0.32 | 0.24 | 0.11 |
| 5 | 13348985 | rs978845 | 1.032 | -1.492 | 0.579 | 0.271 | 0.036 | 0.054 | 0.054 | 0.875 | 0.59 | 0.18 | 0.58 | 0.29 | 0.6 | 0.55 |
| 5 | 13349497 | rs13162209 | 0.526 | -1.131 | 0.994 | 0.578 | -0.005 | 0.167 | 0.167 | 0.081 | 0.86 | 0.5 | 0.77 | 0.65 | 0.9 | 0.82 |
| 5 | 13353727 | rs7713347 | 1.029 | -0.922 | 0.629 | 0.409 | 0 | -0.002 | -0.002 | 0 | 0.16 | 0.12 | 0.22 | 0.11 | 0.2 | 0.18 |
| 5 | 13367397 | rs11133749 | 0.553 | -0.824 | 0.761 | 0.504 | -0.016 | 0.111 | 0.111 | 1.212 | 0.86 | 0.4 | 0.71 | 0.64 | 0.9 | 0.82 |
| 5 | 13372239 | rs1446049 | 0.608 | -0.829 | 0.727 | 0.503 | -0.029 | 0.095 | 0.095 | 1.293 | 0.85 | 0.41 | 0.77 | 0.65 | 0.9 | 0.82 |
| 5 | 13376546 | rs4574541 | 1.126 | -0.73 | 0.614 | 0.457 | 0.007 | -0.009 | -0.009 | 1.122 | 0.15 | 0.18 | 0.21 | 0.13 | 0.2 | 0.16 |
| 5 | 13380973 | rs13167986 | 1.813 | -0.669 | 0.139 | 0.364 | -0.056 | -0.107 | -0.107 | 0 | 0.21 | 0.02 | 0.13 | 0.29 | 0.26 | 0.11 |
| 5 | 13392571 | rs1867742 | 1.269 | -0.627 | 0.43 | 0.312 | 0.016 | -0.015 | -0.015 | 0 | 0.16 | 0.29 | 0.17 | 0.13 | 0.19 | 0.15 |
| 5 | 13393197 | rs2067587 | 2.013 | -0.399 | -0.172 | 0.095 | -0.021 | 0.063 | 0.063 | 0.826 | 0.11 | 0 | 0.09 | 0 | 0.07 | 0.07 |
| 5 | 13393591 | rs10037689 | 0.179 | -0.479 | 0.466 | 0.662 | 0.007 | 0.141 | 0.141 | 0.304 | 0.86 | 0.64 | 0.72 | 0.58 | 0.85 | 0.66 |
| 5 | 13395707 | rs11956313 | 1.646 | -0.483 | 0.428 | 0.668 | -0.047 | -0.105 | -0.105 | 0 | 0.21 | 0.02 | 0.13 | 0.29 | 0.25 | 0.11 |
| 5 | 13397992 | rs7731918 | -0.021 | -0.394 | 0.608 | 0.674 | -0.004 | 0.107 | 0.107 | 0.011 | 0.87 | 0.67 | 0.77 | 0.64 | 0.89 | 0.69 |
| 5 | 13399475 | rs7721512 | 1.438 | -0.448 | 0.537 | 0.538 | -0.08 | 0.141 | 0.141 | 1.17 | 0.37 | 0.08 | 0.35 | 0.12 | 0.34 | 0.34 |
| 5 | 13405946 | rs2081944 | 0.204 | -0.264 | 0.902 | 0.95 | -0.007 | 0.061 | 0.061 | 0 | 0.87 | 0.78 | 0.77 | 0.64 | 0.89 | 0.68 |
| 5 | 13409868 | rs749224 | 0.664 | -0.183 | 0.38 | 0.54 | 0.007 | 0.001 | 0.001 | 0.021 | 0.17 | 0.2 | 0.21 | 0.19 | 0.23 | 0.19 |
| 5 | 13415984 | rs7724160 | 0.59 | -0.196 | 0.548 | 0.534 | 0.016 | 0.004 | 0.004 | 0.046 | 0.17 | 0.29 | 0.22 | 0.19 | 0.23 | 0.19 |
| 5 | 13424707 | rs10085086 | -0.202 | -0.331 | 0.89 | 0.719 | -0.003 | 0.036 | 0.036 | 0 | 0.11 | 0.15 | 0.21 | 0.28 | 0.11 | 0.33 |
| 5 | 13425919 | rs1864096 | 0.686 | -0.315 | 0.748 | 0.723 | 0.017 | 0.005 | 0.005 | 0.026 | 0.17 | 0.29 | 0.22 | 0.19 | 0.23 | 0.19 |
| 5 | 13430032 | rs11960233 | 0.694 | -0.295 | 0.764 | 0.536 | 0.016 | 0.081 | 0.081 | 0 | 0.17 | 0.29 | 0.22 | 0.41 | 0.23 | 0.26 |
| 5 | 13430225 | rs6895189 | -0.295 | -0.315 | 0.736 | 0.503 | 0.001 | -0.023 | -0.023 | 0 | 0.12 | 0.25 | 0.22 | 0.07 | 0.11 | 0.26 |
| 5 | 13434851 | rs6863880 | -1.53 | -0.513 | 0.001 | 0.449 | 0.029 | -0.052 | -0.052 | 0 | 0.41 | 0.83 | 0.51 | 0.53 | 0.4 | 0.58 |
| 5 | 13441359 | rs1366429 | -1.066 | -0.52 | 0.195 | 0.441 | -0.043 | -0.044 | -0.044 | 0 | 0.24 | 0.63 | 0.28 | 0.12 | 0.17 | 0.32 |
| 5 | 13444174 | rs2042955 | -1.025 | -0.574 | 0.172 | 0.292 | -0.056 | -0.1 | -0.1 | 0 | 0.24 | 0.58 | 0.27 | 0.07 | 0.16 | 0.32 |
| 5 | 13448037 | rs4454051 | -0.529 | -0.334 | -0.377 | 0.425 | 0.019 | 0.003 | 0.003 | 0 | 0.45 | 0.67 | 0.4 | 0.37 | 0.41 | 0.4 |
| 5 | 13453436 | rs956195 | -0.555 | -0.286 | -0.745 | 0.424 | 0.023 | -0.002 | -0.002 | 0 | 0.46 | 0.66 | 0.4 | 0.38 | 0.41 | 0.4 |
| 5 | 13458367 | rs1373992 | 0.5 | 0.462 | -1.014 | 0.976 | 0.002 | 0.012 | 0.012 | 0.032 | 0.17 | 0.15 | 0.21 | 0.38 | 0.23 | 0.28 |
| 5 | 13459048 | rs983830 | 0.259 | 0.318 | -1.237 | 0.934 | 0.001 | -0.015 | -0.015 | 0 | 0.24 | 0.2 | 0.14 | 0.31 | 0.27 | 0.12 |
| 5 | 13464078 | rs10474980 | -0.259 | 0.263 | -1.383 | 0.535 | 0.007 | 0.017 | 0.017 | 0 | 0.1 | 0.03 | 0.16 | 0.07 | 0.08 | 0.21 |
| 5 | 13465905 | rs230076 | -0.418 | 0.361 | -2.307 | 0.658 | -0.034 | -0.041 | -0.041 | 0 | 0.54 | 0.88 | 0.47 | 0.39 | 0.55 | 0.41 |
| 5 | 13471156 | rs972707 | 0.088 | 0.554 | -1.833 | 0.579 | -0.005 | -0.007 | -0.007 | 0 | 0.12 | 0.07 | 0.21 | 0.12 | 0.13 | 0.18 |
| 5 | 13476407 | rs10035331 | 0.088 | 0.554 | -1.83 | 0.591 | -0.004 | -0.007 | -0.007 | 0 | 0.12 | 0.07 | 0.21 | 0.12 | 0.13 | 0.18 |
| 5 | 13481754 | rs3853114 | -0.527 | 0.052 | -0.667 | 1.013 | -0.001 | 0.048 | 0.048 | 0.104 | 0.08 | 0.1 | 0.14 | 0.41 | 0.08 | 0.23 |
| 5 | 13482159 | rs10057809 | -0.63 | 0.187 | -0.992 | 0.563 | -0.001 | 0.005 | 0.005 | 6.369 | 0.34 | 0.4 | 0.43 | 0.53 | 0.32 | 0.48 |
| 5 | 13488878 | rs151151 | -0.733 | 0.201 | -1.367 | 0.011 | -0.057 | 0.077 | 0.077 | 0.048 | 0.54 | 0.9 | 0.6 | 0.69 | 0.57 | 0.55 |
| 5 | 13494184 | rs230681 | 0.87 | 0.152 | -0.264 | 0.084 | -0.001 | 0 | 0 | 0.008 | 0.26 | 0.22 | 0.3 | 0.28 | 0.29 | 0.17 |
| 5 | 13498598 | rs230049 | -0.856 | 0.165 | -0.609 | -0.179 | -0.061 | 0.112 | 0.112 | 0 | 0.51 | 0.9 | 0.58 | 0.69 | 0.53 | 0.52 |
| 5 | 13500023 | rs6881358 | 0.328 | 0.331 | -0.426 | -0.238 | 0 | -0.018 | -0.018 | 1.575 | 0.1 | 0.13 | 0.07 | 0 | 0.1 | 0.1 |

| | | | | | | | | | | | | | | | | |
|---|---|---|---|---|---|---|---|---|---|---|---|---|---|---|---|---|
| 5 | 13501102 | rs230042 | -0.412 | 0.236 | -0.423 | -0.23 | 0.026 | 0.039 | 0.039 | 0 | 0.4 | 0.77 | 0.5 | 0.68 | 0.39 | 0.41 |
| 5 | 13504134 | rs3870147 | -0.205 | 0.493 | -0.111 | -0.763 | 0.023 | -0.053 | -0.053 | 0 | 0.12 | 0.25 | 0.09 | 0 | 0.14 | 0.11 |
| 5 | 13510647 | rs7712828 | -0.537 | -0.417 | 0.614 | -0.545 | 0.001 | -0.001 | -0.001 | 0 | 0.44 | 0.54 | 0.42 | 0.4 | 0.45 | 0.29 |
| 5 | 13510918 | rs1992711 | -0.393 | -0.437 | 0.612 | -0.494 | 0.005 | 0.346 | 0.346 | 0.08 | 0.06 | 0.26 | 0.13 | 0.46 | 0.04 | 0.21 |
| 5 | 13512995 | rs10491319 | -0.659 | -0.423 | 0.711 | -1.309 | 0.015 | -0.045 | -0.045 | 0.146 | 0.1 | 0.16 | 0.07 | 0.01 | 0.14 | 0.1 |
| 5 | 13520973 | rs230068 | -0.944 | -0.4 | 0.546 | -1.279 | -0.043 | 0.306 | 0.306 | 0 | 0.53 | 0.91 | 0.55 | 0.86 | 0.49 | 0.5 |
| 5 | 13525819 | rs230063 | -0.805 | -0.964 | 0.862 | -1.062 | 0.038 | 0.173 | 0.173 | 0 | 0.47 | 0.75 | 0.37 | 0.86 | 0.43 | 0.47 |
| 5 | 13572053 | rs233348 | -0.803 | -0.927 | 0.703 | -0.985 | 0.051 | 0.171 | 0.171 | 0 | 0.47 | 0.76 | 0.38 | 0.84 | 0.41 | 0.48 |
| 5 | 13583651 | rs233364 | 0.408 | -0.356 | 0.984 | -0.692 | 0.026 | -0.012 | -0.012 | 0 | 0.05 | 0.13 | 0.04 | 0.01 | 0.07 | 0.07 |
| 5 | 13588726 | rs233362 | -1.299 | -0.397 | 0.045 | -0.924 | -0.027 | 0.066 | 0.066 | 2.653 | 0.76 | 0.98 | 0.76 | 0.86 | 0.8 | 0.8 |
| 5 | 13589212 | rs4702961 | -0.387 | -0.387 | 0.674 | -0.309 | 0.009 | 0.083 | 0.083 | 0 | 0.33 | 0.19 | 0.28 | 0.81 | 0.33 | 0.37 |
| 5 | 13592971 | rs233339 | -0.961 | -0.199 | 0.118 | -0.317 | 0.019 | 0.071 | 0.071 | 0 | 0.43 | 0.7 | 0.36 | 0.84 | 0.41 | 0.47 |
| 5 | 13601207 | rs1428274 | 0.648 | 0.089 | 0.328 | 0.04 | 0.001 | -0.022 | -0.022 | 0 | 0.67 | 0.36 | 0.68 | 0.12 | 0.68 | 0.6 |
| 5 | 13603883 | rs442392 | 0.155 | -0.241 | -0.223 | 0.4 | -0.036 | -0.061 | -0.061 | 0 | 0.74 | 0.55 | 0.78 | 0.18 | 0.79 | 0.78 |
| 5 | 13604919 | rs424270 | 0.378 | 0.138 | -1.059 | 1.271 | -0.015 | 0.192 | 0.192 | 0 | 0.95 | 0.87 | 0.9 | 0.56 | 0.96 | 0.89 |
| 5 | 13604996 | rs13180663 | 0.251 | 0.418 | -1.752 | 0.44 | -0.027 | 0.112 | 0.112 | 9.225 | 0.34 | 0.06 | 0.27 | 0.19 | 0.32 | 0.27 |
| 5 | 13609659 | rs10035874 | 0.162 | 0.62 | -1.041 | 0.739 | 0.003 | 0.002 | 0.002 | 2.706 | 0.54 | 0.49 | 0.42 | 0.48 | 0.5 | 0.37 |
| 5 | 13610706 | rs443542 | -0.073 | 0.907 | -1.91 | 0.795 | -0.128 | -0.099 | -0.099 | 2.359 | 0.67 | 0.93 | 0.5 | 0.58 | 0.59 | 0.55 |
| 5 | 13611631 | rs408619 | 0.44 | 1.083 | -1.581 | 0.745 | -0.009 | -0.015 | -0.015 | 0 | 0.75 | 0.82 | 0.72 | 0.65 | 0.74 | 0.64 |
| 5 | 13619926 | rs389953 | 0.32 | 1.148 | -1.218 | -0.041 | -0.008 | 0.008 | 0.008 | 0 | 0.64 | 0.7 | 0.62 | 0.67 | 0.63 | 0.5 |
| 5 | 13624894 | rs1373947 | 0.621 | 0.786 | -1.342 | -0.484 | -0.002 | 0 | 0 | 2.947 | 0.37 | 0.23 | 0.45 | 0.57 | 0.39 | 0.45 |
| 5 | 13628062 | rs2676245 | -0.174 | 1.153 | -1.025 | -0.09 | -0.027 | 0.005 | 0.005 | 0 | 0.73 | 0.82 | 0.65 | 0.69 | 0.69 | 0.59 |
| 5 | 13635807 | rs1816896 | 0.299 | 0.561 | -0.187 | -0.693 | -0.001 | -0.006 | -0.006 | 0 | 0.46 | 0.54 | 0.52 | 0.63 | 0.47 | 0.47 |
| 5 | 13643405 | rs1445678 | -0.221 | 0.517 | -0.126 | -0.97 | 0.032 | -0.092 | -0.092 | 0 | 0.15 | 0.32 | 0.27 | 0.06 | 0.21 | 0.05 |
| 5 | 13645988 | rs11746515 | 0.852 | 0.288 | -0.517 | -1.233 | -0.099 | 0.013 | 0.013 | 0.05 | 0.39 | 0.13 | 0.49 | 0.63 | 0.48 | 0.47 |
| 5 | 13655022 | rs1867658 | -0.339 | 0.334 | -0.41 | -1.005 | -0.005 | 0.001 | 0.001 | 0 | 0.61 | 0.88 | 0.79 | 0.65 | 0.68 | 0.53 |
| 5 | 13656808 | rs1445682 | -0.323 | 0.348 | 0.027 | -0.999 | 0.008 | 0.01 | 0.01 | 0 | 0.6 | 0.73 | 0.78 | 0.65 | 0.68 | 0.52 |
| 5 | 13668719 | rs6892801 | 0.666 | 0.374 | -0.093 | -0.797 | -0.021 | -0.028 | -0.028 | 0.084 | 0.28 | 0.1 | 0.35 | 0.36 | 0.31 | 0.29 |
| 5 | 13668821 | rs6871272 | -0.06 | 0.338 | 0.149 | -0.733 | -0.011 | 0.011 | 0.011 | 0 | 0.43 | 0.26 | 0.49 | 0.39 | 0.53 | 0.42 |
| 5 | 13689874 | rs10062092 | -0.035 | 0.209 | 0.586 | -0.64 | -0.008 | -0.014 | -0.014 | 0.033 | 0.06 | 0.23 | 0.03 | 0 | 0.01 | 0.01 |
| 5 | 13693277 | rs6872375 | -0.291 | 0.139 | 0.668 | -0.744 | 0.007 | 0.025 | 0.025 | 0 | 0.49 | 0.56 | 0.52 | 0.45 | 0.55 | 0.47 |
| 5 | 13701525 | rs3734111 | 0.068 | 0.129 | 0.836 | -0.749 | -0.001 | 0.006 | 0.006 | 0 | 0.46 | 0.34 | 0.49 | 0.44 | 0.54 | 0.46 |
| 5 | 13701536 | rs3734110 | -0.329 | 0.127 | 0.875 | -0.75 | 0.004 | 0.018 | 0.018 | 0 | 0.52 | 0.53 | 0.52 | 0.45 | 0.55 | 0.47 |
| 5 | 13703493 | rs10040637 | -0.222 | 0.13 | 0.926 | -0.748 | 0.003 | 0.01 | 0.01 | 0 | 0.51 | 0.48 | 0.51 | 0.45 | 0.54 | 0.47 |
| 5 | 13706625 | rs13169297 | 0.572 | 0.329 | 0.889 | -0.754 | 0 | 0 | 0 | 0 | 0.42 | 0.33 | 0.41 | 0.39 | 0.39 | 0.45 |
| 5 | 13714740 | rs30175 | 0.58 | 0.372 | 0.28 | -0.732 | -0.001 | -0.003 | -0.003 | 0 | 0.42 | 0.35 | 0.41 | 0.44 | 0.39 | 0.47 |
| 5 | 13719022 | rs30169 | 0.556 | 0.388 | -0.092 | -0.725 | -0.001 | -0.005 | -0.005 | 0.034 | 0.42 | 0.39 | 0.41 | 0.44 | 0.4 | 0.47 |
| 5 | 13719434 | rs17262853 | 0.555 | 0.445 | -0.148 | -0.686 | -0.03 | 0.034 | 0.034 | 0 | 0.08 | 0.02 | 0.09 | 0.01 | 0.17 | 0.02 |
| 5 | 13721378 | rs17203442 | 0.555 | 0.445 | -0.148 | -0.686 | -0.03 | 0.034 | 0.034 | 0 | 0.08 | 0.02 | 0.09 | 0.01 | 0.17 | 0.02 |
| 5 | 13723743 | rs2652768 | -0.521 | 0.415 | -0.132 | -0.693 | -0.009 | 0.009 | 0.009 | 0 | 0.57 | 0.7 | 0.55 | 0.45 | 0.58 | 0.51 |
| 5 | 13727212 | rs11133760 | -0.648 | 0.464 | -0.258 | -0.645 | 0 | -0.007 | -0.007 | 0 | 0.58 | 0.62 | 0.59 | 0.61 | 0.6 | 0.54 |
| 5 | 13728151 | rs10513151 | 0.73 | 0.954 | -1.374 | -0.317 | -0.01 | 0.091 | 0.091 | 0.039 | 0.35 | 0.19 | 0.2 | 0.19 | 0.34 | 0.29 |
| 5 | 13735892 | rs3734109 | -0.246 | 1.057 | -1.168 | -0.182 | 0 | -0.001 | -0.001 | 0 | 0.56 | 0.53 | 0.57 | 0.56 | 0.6 | 0.54 |
| 5 | 13737444 | rs17263496 | -0.307 | 1.242 | -1.292 | -0.157 | -0.023 | 0.028 | 0.028 | 0 | 0.07 | 0.02 | 0.09 | 0.05 | 0.15 | 0.04 |
| 5 | 13739720 | rs2896104 | 0.041 | 0.986 | -1.61 | -0.361 | -0.02 | 0.088 | 0.088 | 0 | 0.63 | 0.82 | 0.65 | 0.82 | 0.68 | 0.69 |
| 5 | 13740976 | rs6554809 | -0.567 | 0.311 | -0.853 | -0.54 | -0.029 | -0.048 | -0.048 | 0 | 0.23 | 0.51 | 0.25 | 0.11 | 0.16 | 0.1 |
| 5 | 13741163 | rs6881967 | -0.277 | 0.141 | -0.85 | -1.386 | -0.002 | -0.032 | -0.032 | 0 | 0.33 | 0.38 | 0.25 | 0.1 | 0.33 | 0.28 |
| 5 | 13742858 | rs6554811 | -0.368 | -0.091 | -0.888 | -2.058 | 0.007 | 0.048 | 0.048 | 0.367 | 0.44 | 0.33 | 0.32 | 0.13 | 0.45 | 0.54 |
| 5 | 13755672 | rs1393270 | 0.116 | -0.101 | -0.71 | -1.151 | -0.002 | -0.008 | -0.008 | 0 | 0.12 | 0.13 | 0.08 | 0.22 | 0.12 | 0.07 |
| 5 | 13760548 | rs7733137 | 0.702 | 0.181 | -0.425 | -0.854 | 0.002 | 0 | 0 | 0.112 | 0.21 | 0.15 | 0.3 | 0.23 | 0.19 | 0.2 |
| 5 | 13760721 | rs1354187 | -0.474 | 0.112 | -0.362 | -0.438 | 0.001 | -0.005 | -0.005 | 0.238 | 0.34 | 0.36 | 0.38 | 0.55 | 0.31 | 0.27 |
| 5 | 13761033 | rs1354186 | -1.178 | 0.599 | -0.551 | -0.432 | -0.039 | -0.021 | -0.021 | 0 | 0.62 | 0.83 | 0.66 | 0.65 | 0.61 | 0.75 |
| 5 | 13761068 | rs1354185 | 0.785 | 0.597 | -0.527 | -0.44 | -0.001 | 0.03 | 0.03 | 0 | 0.1 | 0.13 | 0.07 | 0.32 | 0.11 | 0.07 |
| 5 | 13762972 | rs6554812 | 0.052 | 0.544 | -0.483 | -0.717 | 0.001 | -0.084 | -0.084 | 0.22 | 0.27 | 0.37 | 0.27 | 0.11 | 0.3 | 0.48 |
| 5 | 13770000 | rs10513153 | 0.806 | 0.605 | -0.57 | -0.56 | -0.02 | 0 | 0 | 0 | 0.36 | 0.23 | 0.34 | 0.34 | 0.38 | 0.23 |
| 5 | 13781480 | rs1910092 | -0.177 | 0.247 | -0.294 | -0.935 | 0.026 | -0.025 | -0.025 | 0.023 | 0.1 | 0.3 | 0.12 | 0.04 | 0.11 | 0.19 |
| 5 | 13782801 | rs4701984 | -0.188 | 0.755 | -0.349 | -0.851 | 0.006 | -0.012 | -0.012 | 1.334 | 0.61 | 0.57 | 0.54 | 0.3 | 0.57 | 0.61 |
| 5 | 13788282 | rs1502048 | 0.496 | 0.787 | -0.853 | -0.811 | 0.009 | -0.054 | -0.054 | 0 | 0.2 | 0.26 | 0.24 | 0.08 | 0.27 | 0.39 |
| 5 | 13788886 | rs10513155 | 0.75 | 0.55 | -1.284 | -0.591 | 0.032 | 0.077 | 0.077 | 0.384 | 0.37 | 0.13 | 0.26 | 0.22 | 0.27 | 0.21 |
| 5 | 13798559 | rs795544 | -0.645 | 0.757 | -0.564 | -0.279 | 0.001 | -0.012 | -0.012 | 0.743 | 0.84 | 0.82 | 0.83 | 0.89 | 0.78 | 0.62 |
| 5 | 13798819 | rs795543 | 0.925 | 0.879 | -0.666 | -0.265 | 0.015 | 0.056 | 0.056 | 0 | 0.43 | 0.23 | 0.38 | 0.26 | 0.39 | 0.29 |
| 5 | 13800577 | rs795542 | 1.468 | 0.996 | -0.876 | -0.411 | -0.003 | 0.072 | 0.072 | 0.73 | 0.24 | 0.08 | 0.17 | 0.01 | 0.22 | 0.18 |
| 5 | 13811775 | rs10078391 | 0 | 1.307 | -0.107 | 0.7 | 0.024 | 0.09 | 0.09 | 0.25 | 0.13 | 0.19 | 0.18 | 0.56 | 0.21 | 0.17 |
| 5 | 13815288 | rs6876105 | -0.563 | 1.275 | 0.008 | 0.827 | 0.064 | 0.195 | 0.195 | 0.486 | 0.16 | 0.4 | 0.23 | 0.56 | 0.22 | 0.17 |
| 5 | 13816740 | rs16902777 | -0.511 | 1.297 | 0.055 | 0.858 | 0.057 | 0.182 | 0.182 | 0.122 | 0.15 | 0.35 | 0.22 | 0.54 | 0.22 | 0.16 |
| 5 | 13817881 | rs6889720 | -0.933 | 1.2 | -0.051 | 0.731 | 0.007 | 0.003 | 0.003 | 0 | 0.35 | 0.44 | 0.42 | 0.62 | 0.38 | 0.31 |
| 5 | 13829042 | rs1445815 | -1.233 | 1.282 | -0.021 | 0.716 | 0.006 | 0.001 | 0.001 | 0 | 0.37 | 0.53 | 0.45 | 0.63 | 0.38 | 0.32 |
| 5 | 13829140 | rs11738619 | 0.003 | 1.618 | -0.238 | 0.92 | 0.026 | 0.106 | 0.106 | 0.081 | 0.12 | 0.19 | 0.18 | 0.55 | 0.21 | 0.17 |
| 5 | 13834445 | rs4547927 | -0.769 | 1.684 | -1.021 | 0.679 | 0.011 | 0.005 | 0.005 | 0 | 0.35 | 0.53 | 0.43 | 0.62 | 0.37 | 0.31 |

| | | | | | | | | | | | | | | | | |
|---|---|---|---|---|---|---|---|---|---|---|---|---|---|---|---|---|
| 5 | 13836402 | rs1596791 | -0.514 | 0.826 | -0.842 | 0.459 | -0.012 | 0.039 | 0.039 | 0 | 0.32 | 0.53 | 0.34 | 0.44 | 0.25 | 0.26 |
| 5 | 13839167 | rs6868995 | -0.01 | 1.185 | -1.502 | -0.241 | -0.023 | 0.003 | 0.003 | 0 | 0.16 | 0.01 | 0.21 | 0.11 | 0.18 | 0.31 |
| 5 | 13846802 | rs17208191 | -0.345 | 1.344 | -0.889 | -0.112 | 0 | -0.001 | -0.001 | 3.383 | 0.13 | 0.14 | 0.18 | 0.1 | 0.13 | 0.15 |
| 5 | 13850750 | rs6554820 | -0.551 | 1.378 | -1.101 | -0.786 | -0.029 | 0.028 | 0.028 | 0.997 | 0.34 | 0.27 | 0.3 | 0.27 | 0.41 | 0.47 |
| 5 | 13851016 | rs795526 | -0.645 | 1.688 | -0.629 | -0.716 | -0.006 | 0.003 | 0.003 | 1.465 | 0.35 | 0.37 | 0.37 | 0.3 | 0.46 | 0.49 |
| 5 | 13862616 | rs7704417 | -0.534 | 1.643 | -0.303 | -0.86 | 0.002 | -0.005 | -0.005 | 0 | 0.33 | 0.38 | 0.31 | 0.26 | 0.41 | 0.47 |
| 5 | 13864189 | rs795510 | 0.25 | 0.725 | 0.235 | -1.214 | -0.032 | -0.023 | -0.023 | 0 | 0.8 | 0.7 | 0.83 | 0.88 | 0.87 | 0.81 |
| 5 | 13865980 | rs7703349 | -0.372 | 0.783 | 0.326 | -1.37 | -0.005 | 0.006 | 0.006 | 0.19 | 0.41 | 0.6 | 0.34 | 0.33 | 0.47 | 0.53 |
| 5 | 13867077 | rs797055 | -0.22 | 0.774 | 0.245 | -1.377 | 0.015 | 0.038 | 0.038 | 0 | 0.85 | 0.93 | 0.84 | 0.95 | 0.89 | 0.87 |
| 5 | 13867819 | rs1445810 | -0.027 | 0.801 | 0.276 | -1.366 | -0.038 | 0.038 | 0.038 | 0.059 | 0.35 | 0.27 | 0.32 | 0.26 | 0.44 | 0.46 |
| 5 | 13867851 | rs13156044 | 0.954 | 1.406 | 0.237 | -1.523 | 0.019 | 0.001 | 0.001 | 0 | 0.08 | 0.13 | 0.08 | 0.1 | 0.15 | 0.04 |
| 5 | 13872094 | rs13159798 | -0.304 | 1.366 | 0.409 | -1.529 | -0.019 | 0.018 | 0.018 | 0.196 | 0.35 | 0.33 | 0.32 | 0.26 | 0.44 | 0.46 |
| 5 | 13872301 | rs10061803 | 1.913 | 0.835 | 0.093 | -1.774 | 0.022 | 0.01 | 0.01 | 0.646 | 0.4 | 0.17 | 0.41 | 0.58 | 0.35 | 0.31 |
| 5 | 13872584 | rs4701996 | 0.929 | 0.437 | 0.003 | -1.843 | -0.097 | 0.015 | 0.015 | 0.259 | 0.18 | 0.01 | 0.19 | 0.15 | 0.22 | 0.32 |
| 5 | 13876664 | rs1445804 | 0.296 | 0.264 | 0.273 | -1.713 | 0.012 | 0.014 | 0.014 | 0 | 0.82 | 0.86 | 0.83 | 0.95 | 0.88 | 0.87 |
| 5 | 13878843 | rs1445803 | -0.642 | 0.127 | 0.37 | -2.114 | 0.009 | 0.012 | 0.012 | 0 | 0.27 | 0.2 | 0.2 | 0.09 | 0.2 | 0.22 |
| 5 | 13882841 | rs10057007 | -0.897 | 0.131 | 0.576 | -2.109 | -0.013 | -0.041 | -0.041 | 0.055 | 0.28 | 0.33 | 0.28 | 0.12 | 0.26 | 0.24 |
| 5 | 13887631 | rs17275618 | -0.24 | 0.195 | 0.422 | -2.218 | 0.007 | -0.024 | -0.024 | 0 | 0.32 | 0.38 | 0.26 | 0.23 | 0.32 | 0.45 |
| 5 | 13889983 | rs11749049 | -1.114 | 0.019 | 0.591 | -1.992 | 0 | 0.02 | 0.02 | 0 | 0.56 | 0.6 | 0.54 | 0.28 | 0.55 | 0.63 |
| 5 | 13900345 | rs1445823 | 0.482 | 0.078 | 0.713 | -2.116 | 0 | 0.047 | 0.047 | 0 | 0.55 | 0.68 | 0.52 | 0.87 | 0.45 | 0.43 |
| 5 | 13900926 | rs10057950 | 0.251 | 0.119 | 0.919 | -2.227 | 0.058 | -0.065 | -0.065 | 0 | 0.23 | 0.39 | 0.31 | 0.1 | 0.3 | 0.4 |
| 5 | 13902220 | rs1530498 | 1.517 | -0.087 | 0.462 | -2.211 | -0.012 | 0.115 | 0.115 | 0.398 | 0.42 | 0.15 | 0.43 | 0.76 | 0.4 | 0.29 |
| 5 | 13905405 | rs4701999 | -0.517 | -0.093 | 0.351 | -2.224 | -0.003 | -0.01 | -0.01 | 0 | 0.78 | 0.84 | 0.77 | 0.96 | 0.73 | 0.8 |
| 5 | 13909656 | rs6859484 | -0.25 | -0.023 | 0.443 | -2.144 | -0.001 | -0.062 | -0.062 | 3.64 | 0.72 | 0.73 | 0.74 | 0.96 | 0.69 | 0.77 |
| 5 | 13909698 | rs10462713 | 1.112 | -0.084 | -0.214 | -1.907 | -0.011 | 0.098 | 0.098 | 1.106 | 0.39 | 0.14 | 0.42 | 0.75 | 0.37 | 0.27 |
| 5 | 13913885 | rs3213936 | 0.812 | 0.215 | -0.38 | -2.078 | 0.006 | 0.003 | 0.003 | 0.145 | 0.45 | 0.57 | 0.45 | 0.85 | 0.38 | 0.36 |
| 5 | 13914775 | rs13180591 | 0.821 | 0.215 | -0.4 | -2.078 | 0.005 | 0 | 0 | 0.91 | 0.45 | 0.57 | 0.45 | 0.85 | 0.38 | 0.36 |
| 5 | 13914819 | rs13180613 | 0.821 | 0.215 | -0.4 | -2.078 | 0.005 | 0 | 0 | 0 | 0.45 | 0.57 | 0.45 | 0.85 | 0.38 | 0.36 |
| 5 | 13918291 | rs7713700 | 0.727 | 0.405 | -0.529 | -1.97 | 0.025 | 0.065 | 0.065 | 0 | 0.48 | 0.64 | 0.45 | 0.85 | 0.38 | 0.38 |
| 5 | 13926057 | rs35339484 | -0.005 | 0.979 | -0.961 | -1.457 | -0.106 | 0.054 | 0.054 | 0 | 0.21 | 0.03 | 0.22 | 0.09 | 0.28 | 0.37 |
| 5 | 13931340 | rs1530496 | 0.449 | 0.017 | -0.479 | -1.178 | -0.004 | -0.001 | -0.001 | 0 | 0.34 | 0.61 | 0.47 | 0.7 | 0.28 | 0.39 |
| 5 | 13933125 | rs983649 | -0.168 | -0.117 | -0.581 | -1.646 | 0.002 | 0.051 | 0.051 | 0 | 0.57 | 0.79 | 0.72 | 0.72 | 0.57 | 0.56 |
| 5 | 13933416 | rs16902953 | 0.822 | -0.146 | -0.545 | -1.672 | 0.001 | 0 | 0 | 0 | 0.35 | 0.65 | 0.47 | 0.7 | 0.28 | 0.39 |
| 5 | 13934119 | rs1445819 | -0.231 | -0.12 | -0.546 | -1.672 | -0.005 | 0.05 | 0.05 | 0 | 0.58 | 0.83 | 0.73 | 0.72 | 0.58 | 0.56 |
| 5 | 13937476 | rs17278234 | 0.356 | -0.11 | -0.551 | -1.669 | -0.01 | 0.001 | 0.001 | 0 | 0.23 | 0.18 | 0.26 | 0.02 | 0.29 | 0.17 |
| 5 | 13937803 | rs10058864 | 0.889 | -0.147 | -0.521 | -1.586 | -0.004 | -0.001 | -0.001 | 0 | 0.34 | 0.61 | 0.47 | 0.7 | 0.28 | 0.39 |
| 5 | 13939830 | rs10070196 | -0.275 | -0.425 | -0.069 | -1.631 | 0.013 | 0.03 | 0.03 | 0 | 0.55 | 0.57 | 0.59 | 0.22 | 0.67 | 0.58 |
| 5 | 13942973 | rs339447 | 0.749 | -0.203 | 0.421 | -1.472 | 0.007 | -0.019 | -0.019 | 0.096 | 0.28 | 0.32 | 0.27 | 0.15 | 0.36 | 0.38 |
| 5 | 13943499 | rs16902962 | 0.483 | -0.557 | 0.225 | -0.948 | 0.003 | -0.007 | -0.007 | 0 | 0.08 | 0.02 | 0.01 | 0.11 | 0.05 | 0.04 |
| 5 | 13944512 | rs339445 | 0.431 | -0.57 | 0.229 | -0.955 | 0.002 | 0.002 | 0.002 | 0 | 0.86 | 0.87 | 0.96 | 0.89 | 0.93 | 0.94 |
| 5 | 13954951 | rs173090 | 0.355 | -0.381 | 0.069 | -0.908 | -0.004 | -0.01 | -0.01 | 0 | 0.36 | 0.37 | 0.39 | 0.71 | 0.3 | 0.32 |
| 5 | 13958443 | rs10050676 | 0.424 | -0.529 | 0.245 | -1.288 | -0.03 | -0.075 | -0.075 | 0 | 0.72 | 0.4 | 0.81 | 0.88 | 0.83 | 0.77 |
| 5 | 13962264 | rs2017013 | -0.296 | -0.609 | 0.595 | -1.369 | -0.027 | -0.126 | -0.126 | 0 | 0.8 | 0.56 | 0.89 | 1 | 0.86 | 0.89 |
| 5 | 13964588 | rs11133773 | 0.804 | -0.489 | 0.037 | -1.095 | -0.102 | 0.049 | 0.049 | 0 | 0.32 | 0.09 | 0.38 | 0.2 | 0.45 | 0.48 |
| 5 | 13972255 | rs1838697 | 1.037 | -0.693 | 0.321 | -0.866 | 0 | -0.003 | -0.003 | 0 | 0.31 | 0.26 | 0.41 | 0.59 | 0.33 | 0.28 |
| 5 | 13976728 | rs10036753 | 1.08 | -0.723 | 0.274 | -0.872 | -0.008 | -0.013 | -0.013 | 0 | 0.31 | 0.19 | 0.41 | 0.61 | 0.34 | 0.3 |
| 5 | 13979101 | rs339429 | -0.509 | -1.059 | 0.598 | -1.188 | 0.012 | -0.009 | -0.009 | 3.849 | 0.76 | 0.79 | 0.89 | 1 | 0.83 | 0.89 |
| 5 | 13979397 | rs339428 | -0.01 | -1.119 | 0.88 | -1.188 | 0.006 | -0.009 | -0.009 | 0 | 0.44 | 0.49 | 0.49 | 0.39 | 0.49 | 0.59 |
| 5 | 13980278 | rs339425 | 0.958 | -0.877 | 0.729 | -1.363 | -0.021 | 0.013 | 0.013 | 0.007 | 0.29 | 0.21 | 0.36 | 0.21 | 0.38 | 0.47 |
| 5 | 13981968 | rs339424 | 1.302 | -0.47 | 0.714 | -0.359 | 0.003 | 0.005 | 0.005 | 0.534 | 0.3 | 0.24 | 0.23 | 0.14 | 0.23 | 0.22 |
| 5 | 13987103 | rs1031006 | 0.508 | -0.747 | 0.873 | -0.506 | 0 | -0.006 | -0.006 | 0.006 | 0.38 | 0.4 | 0.42 | 0.69 | 0.36 | 0.32 |
| 5 | 13987466 | rs10055489 | 0.314 | -0.748 | 0.882 | -0.849 | -0.002 | -0.023 | -0.023 | 0 | 0.41 | 0.42 | 0.48 | 0.79 | 0.38 | 0.39 |
| 5 | 13988485 | rs193154 | 0.047 | -0.72 | 0.924 | -0.92 | 0.007 | -0.03 | -0.03 | 0.013 | 0.68 | 0.7 | 0.81 | 0.99 | 0.77 | 0.86 |
| 5 | 13991502 | rs10045723 | 1.036 | -0.687 | 0.717 | -0.835 | -0.002 | -0.005 | -0.005 | 0 | 0.31 | 0.23 | 0.38 | 0.62 | 0.31 | 0.29 |
| 5 | 13993530 | rs7727798 | -0.47 | -0.768 | 0.811 | -0.739 | -0.012 | 0.12 | 0.12 | 0 | 0.05 | 0.17 | 0.03 | 0.2 | 0.01 | 0.06 |
| 5 | 13993653 | rs2591543 | 0.138 | -0.865 | 0.639 | -0.878 | 0.004 | -0.023 | -0.023 | 0 | 0.66 | 0.67 | 0.79 | 0.98 | 0.72 | 0.86 |
| 5 | 13995605 | rs2102443 | 0.034 | -0.905 | 0.567 | -0.871 | 0.001 | -0.036 | -0.036 | 0 | 0.65 | 0.64 | 0.79 | 0.97 | 0.73 | 0.87 |
| 5 | 14004657 | rs436457 | 0.453 | -1.014 | 0.552 | -0.827 | 0.003 | 0.003 | 0.003 | 0 | 0.62 | 0.61 | 0.78 | 0.77 | 0.74 | 0.79 |
| 5 | 14007930 | rs444601 | 0.483 | -0.938 | 0.621 | -0.91 | -0.001 | -0.042 | -0.042 | 0 | 0.38 | 0.46 | 0.41 | 0.82 | 0.34 | 0.38 |
| 5 | 14009085 | rs12517769 | 0.618 | -0.953 | 0.67 | -0.917 | 0 | -0.021 | -0.021 | 0 | 0.36 | 0.32 | 0.4 | 0.82 | 0.34 | 0.38 |
| 5 | 14009923 | rs429898 | 0.333 | -0.97 | 0.705 | -0.925 | 0 | -0.008 | -0.008 | 0.499 | 0.39 | 0.54 | 0.42 | 0.82 | 0.34 | 0.38 |
| 5 | 14013407 | rs2582689 | 0.698 | -0.743 | 0.056 | -0.696 | -0.004 | 0 | 0 | 0 | 0.31 | 0.34 | 0.4 | 0.36 | 0.42 | 0.56 |
| 5 | 14024415 | rs10462717 | 0.52 | -0.454 | -0.554 | -0.216 | -0.056 | -0.043 | -0.043 | 3.239 | 0.3 | 0.06 | 0.37 | 0.63 | 0.32 | 0.3 |
| 5 | 14026144 | rs413338 | -0.507 | -0.697 | -0.667 | -0.382 | 0.028 | 0 | 0 | 0 | 0.79 | 0.92 | 0.93 | 0.78 | 0.84 | 0.85 |
| 5 | 14028309 | rs1605743 | -0.03 | 0.188 | -0.711 | 0.337 | -0.053 | -0.05 | -0.05 | 0.124 | 0.3 | 0.07 | 0.37 | 0.63 | 0.33 | 0.3 |
| 5 | 14036509 | rs12186457 | 0.333 | 0.12 | -0.048 | 0.138 | -0.004 | -0.028 | -0.028 | 0 | 0.62 | 0.55 | 0.66 | 0.18 | 0.67 | 0.6 |
| 5 | 14036643 | rs11133776 | 0.075 | 0.09 | 0.29 | 0.123 | -0.003 | 0.032 | 0.032 | 0 | 0.65 | 0.76 | 0.69 | 0.18 | 0.67 | 0.6 |
| 5 | 14039647 | rs6889677 | -0.373 | 0.355 | 0.271 | 0.714 | 0.092 | 0.269 | 0.269 | 0.722 | 0.93 | 0.82 | 0.88 | 0.47 | 0.85 | 0.8 |
| 5 | 14041925 | rs7720329 | 1.188 | -0.064 | 0.366 | 0.269 | -0.007 | 0.027 | 0.027 | 0 | 0.38 | 0.32 | 0.52 | 0.15 | 0.45 | 0.44 |

| | | | | | | | | | | | | | | | | |
|---|---|---|---|---|---|---|---|---|---|---|---|---|---|---|---|---|
| 5 | 14046910 | rs769714 | -0.428 | -0.107 | 0.729 | 0.598 | -0.042 | 0.074 | 0.074 | 0.121 | 0.29 | 0.52 | 0.18 | 0.48 | 0.18 | 0.31 |
| 5 | 14048605 | rs339956 | 0.99 | 0.128 | 0.572 | 0.536 | 0.023 | 0.29 | 0.29 | 0.632 | 0.06 | 0.17 | 0.06 | 0.4 | 0.04 | 0.17 |
| 5 | 14052451 | rs339951 | -0.276 | -0.01 | 0.453 | 0.307 | 0 | -0.015 | -0.015 | 0 | 0.68 | 0.72 | 0.77 | 0.77 | 0.69 | 0.8 |
| 5 | 14055676 | rs10223292 | 0.427 | 0.002 | 0.175 | 0.256 | -0.003 | -0.006 | -0.006 | 0 | 0.57 | 0.29 | 0.66 | 0.36 | 0.63 | 0.6 |
| 5 | 14055801 | rs13181584 | 1.474 | -0.068 | -0.122 | 0.224 | -0.1 | 0.118 | 0.118 | 0.208 | 0.35 | 0.06 | 0.56 | 0.12 | 0.45 | 0.51 |
| 5 | 14056276 | rs13185842 | 1.026 | 0.287 | 0.806 | 0.633 | -0.011 | -0.017 | -0.017 | 0 | 0.57 | 0.21 | 0.65 | 0.35 | 0.63 | 0.6 |
| 5 | 14058036 | rs339959 | -0.403 | 0.594 | 0.38 | 0.66 | -0.053 | 0.026 | 0.026 | 0 | 0.79 | 0.93 | 0.7 | 0.77 | 0.73 | 0.8 |
| 5 | 14060769 | rs11133779 | 0.978 | 0.655 | 0.267 | 0.718 | -0.004 | -0.007 | -0.007 | 0 | 0.6 | 0.4 | 0.55 | 0.37 | 0.61 | 0.61 |
| 5 | 14061184 | rs399712 | -0.388 | 0.391 | -0.04 | 1.028 | -0.014 | 0.101 | 0.101 | 0.202 | 0.14 | 0.23 | 0.07 | 0.39 | 0.08 | 0.18 |
| 5 | 14068444 | rs3849080 | 0.346 | 0.537 | -0.501 | 0.52 | 0 | 0 | 0 | 0 | 0.59 | 0.47 | 0.58 | 0.37 | 0.62 | 0.61 |
| 5 | 14069439 | rs163819 | -0.817 | 0.203 | -0.43 | 0.056 | -0.017 | -0.022 | -0.022 | 0 | 0.3 | 0.57 | 0.2 | 0.14 | 0.21 | 0.15 |
| 5 | 14072057 | rs163818 | -0.782 | 0.07 | -1.444 | -0.997 | 0.036 | 0.216 | 0.216 | 0.563 | 0.45 | 0.89 | 0.31 | 0.1 | 0.36 | 0.3 |
| 5 | 14072453 | rs697550 | -0.307 | -0.39 | -0.335 | -1.388 | -0.031 | -0.097 | -0.097 | 0.113 | 0.19 | 0.49 | 0.15 | 0.05 | 0.14 | 0.2 |
| 5 | 14074555 | rs17230252 | -0.205 | -0.299 | 0.016 | -1.43 | -0.003 | -0.01 | -0.01 | 0 | 0.16 | 0.18 | 0.1 | 0.04 | 0.15 | 0.11 |
| 5 | 14075268 | rs11133781 | -0.309 | -0.14 | 0.041 | -1.292 | 0.002 | -0.056 | -0.056 | 0 | 0.1 | 0.2 | 0.06 | 0 | 0.07 | 0.04 |
| 5 | 14075815 | rs769732 | -0.08 | -0.196 | 0.102 | -1.211 | 0 | -0.019 | -0.019 | 0 | 0.1 | 0.18 | 0.09 | 0.02 | 0.11 | 0.11 |
| 5 | 14076584 | rs7708445 | -0.074 | 0.541 | -0.3 | -1.551 | -0.013 | 0.009 | 0.009 | 0 | 0.11 | 0.07 | 0.13 | 0 | 0.18 | 0.15 |
| 5 | 14077685 | rs392421 | -1.282 | 0.855 | 0.023 | -0.659 | 0.013 | 0.015 | 0.015 | 0.005 | 0.31 | 0.47 | 0.38 | 0.38 | 0.36 | 0.4 |
| 5 | 14083031 | rs384723 | -1.745 | 0.741 | 0.054 | -0.698 | 0.004 | 0.002 | 0.002 | 0.439 | 0.41 | 0.6 | 0.48 | 0.42 | 0.43 | 0.51 |
| 5 | 14083816 | rs11954402 | -0.253 | 0.897 | 0.051 | -0.769 | -0.017 | 0.023 | 0.023 | 0 | 0.14 | 0.11 | 0.24 | 0.01 | 0.2 | 0.14 |
| 5 | 14087481 | rs10513131 | -0.499 | 0.906 | 0.129 | -0.773 | -0.009 | 0.019 | 0.019 | 0 | 0.08 | 0.07 | 0.12 | 0.38 | 0.09 | 0.26 |
| 5 | 14087633 | rs697546 | -2.941 | 0.232 | -0.755 | -0.721 | -0.067 | 0.037 | 0.037 | 0 | 0.63 | 0.94 | 0.63 | 0.67 | 0.59 | 0.7 |
| 5 | 14088834 | rs11950716 | 0.502 | 0.273 | -0.726 | -0.733 | -0.029 | 0.042 | 0.042 | 0 | 0.13 | 0.07 | 0.24 | 0.01 | 0.2 | 0.13 |
| 5 | 14089644 | rs6881582 | -1.102 | 0.219 | -0.562 | -0.697 | 0.002 | 0.044 | 0.044 | 0 | 0.26 | 0.38 | 0.23 | 0.44 | 0.23 | 0.37 |
| 5 | 14090114 | rs9312863 | -1.596 | 0.153 | -0.518 | -0.63 | 0.003 | 0.003 | 0.003 | 0.083 | 0.4 | 0.48 | 0.47 | 0.44 | 0.43 | 0.51 |
| 5 | 14090392 | rs769475 | 0.093 | 0.094 | -0.351 | -0.636 | -0.03 | 0.036 | 0.036 | 0 | 0.21 | 0.3 | 0.14 | 0.24 | 0.16 | 0.19 |
| 5 | 14093389 | rs6863620 | -0.33 | 0.195 | -0.672 | -0.615 | -0.033 | 0.1 | 0.1 | 0 | 0.28 | 0.12 | 0.31 | 0.06 | 0.32 | 0.29 |
| 5 | 14093588 | rs6864072 | -1.024 | 0.167 | -0.302 | -0.656 | -0.002 | 0.001 | 0.001 | 0 | 0.38 | 0.4 | 0.36 | 0.06 | 0.38 | 0.31 |
| 5 | 14099152 | rs706259 | -0.057 | -0.222 | -0.094 | -0.476 | -0.053 | 0.075 | 0.075 | 0 | 0.18 | 0.47 | 0.13 | 0.23 | 0.1 | 0.15 |
| 5 | 14106136 | rs386450 | -1.008 | -0.112 | -0.602 | -0.18 | -0.014 | 0.059 | 0.059 | 0.348 | 0.25 | 0.52 | 0.27 | 0.56 | 0.21 | 0.38 |
| 5 | 14106401 | rs425495 | -2.345 | 0.012 | -0.84 | -0.374 | -0.11 | 0.011 | 0.011 | 0.015 | 0.63 | 0.95 | 0.61 | 0.69 | 0.55 | 0.72 |
| 5 | 14109455 | rs183693 | -1.134 | -0.356 | -0.759 | -0.313 | -0.017 | 0.006 | 0.006 | 0.01 | 0.31 | 0.53 | 0.28 | 0.61 | 0.2 | 0.46 |
| 5 | 14111048 | rs11948304 | -0.311 | -0.153 | -0.929 | -0.461 | -0.007 | 0.074 | 0.074 | 0 | 0.28 | 0.2 | 0.3 | 0.07 | 0.31 | 0.25 |
| 5 | 14116396 | rs10078192 | -1.128 | -0.278 | -0.285 | 0.121 | -0.061 | 0.054 | 0.054 | 0.253 | 0.17 | 0.53 | 0.14 | 0.16 | 0.11 | 0.19 |
| 5 | 14116980 | rs250846 | -1.728 | 0.322 | -1.289 | -0.014 | -0.133 | 0.046 | 0.046 | 1.112 | 0.58 | 0.95 | 0.58 | 0.71 | 0.48 | 0.71 |
| 5 | 14118230 | rs32975 | 1.039 | 0.648 | -1.074 | 0.128 | -0.039 | 0.065 | 0.065 | 0 | 0.54 | 0.15 | 0.44 | 0.36 | 0.52 | 0.35 |
| 5 | 14118242 | rs32976 | 0.298 | 0.581 | -0.445 | 0.128 | 0.004 | 0.09 | 0.09 | 0 | 0.08 | 0.21 | 0.11 | 0.13 | 0.06 | 0.12 |
| 5 | 14118565 | rs252217 | 0.74 | 0.404 | -0.366 | 0.071 | 0.003 | 0.006 | 0.006 | 0 | 0.58 | 0.47 | 0.54 | 0.48 | 0.58 | 0.44 |
| 5 | 14123507 | rs153191 | 0.145 | 0.717 | -0.253 | -0.177 | 0.011 | 0.105 | 0.105 | 0 | 0.15 | 0.24 | 0.23 | 0.53 | 0.17 | 0.33 |
| 5 | 14127900 | rs163823 | -1.083 | 0.493 | -0.181 | -0.487 | 0.004 | 0.019 | 0.019 | 0.457 | 0.51 | 0.4 | 0.56 | 0.59 | 0.5 | 0.64 |
| 5 | 14136404 | rs388154 | 1.683 | 0.362 | -0.209 | -0.551 | -0.024 | 0.017 | 0.017 | 0.428 | 0.39 | 0.13 | 0.4 | 0.38 | 0.43 | 0.31 |
| 5 | 14151799 | rs6554843 | -0.569 | 0.576 | 0.023 | -0.903 | 0.006 | 0.015 | 0.015 | 1.573 | 0.39 | 0.32 | 0.36 | 0.08 | 0.35 | 0.36 |
| 5 | 14157376 | rs17298767 | 0.95 | 0.546 | -0.224 | -0.943 | 0.009 | 0.049 | 0.049 | 1.274 | 0.13 | 0.03 | 0.06 | 0.05 | 0.09 | 0.1 |
| 5 | 14159211 | rs152782 | -0.307 | 0.591 | 0.196 | -0.734 | 0.024 | 0.071 | 0.071 | 0.016 | 0.21 | 0.31 | 0.29 | 0.51 | 0.26 | 0.38 |
| 5 | 14166100 | rs11747048 | 0.315 | 0.54 | -0.011 | -0.865 | -0.003 | -0.008 | -0.008 | 0 | 0.25 | 0.21 | 0.28 | 0.03 | 0.25 | 0.26 |
| 5 | 14166402 | rs26286 | -1.358 | 0.582 | -0.369 | -0.73 | 0.006 | 0.001 | 0.001 | 0 | 0.49 | 0.85 | 0.61 | 0.53 | 0.51 | 0.65 |
| 5 | 14169709 | rs32593 | -1.933 | 0.495 | -0.523 | -0.519 | -0.082 | -0.066 | -0.066 | 0.606 | 0.6 | 0.96 | 0.64 | 0.58 | 0.55 | 0.73 |
| 5 | 14172120 | rs32572 | -0.184 | 0.74 | 0.531 | 0.177 | 0.028 | 0.112 | 0.112 | 0 | 0.14 | 0.21 | 0.24 | 0.48 | 0.18 | 0.33 |
| 5 | 14174279 | rs26116 | 0.443 | 0.526 | 0.527 | -0.036 | 0.003 | 0.002 | 0.002 | 0 | 0.06 | 0.13 | 0.07 | 0.04 | 0.08 | 0.05 |
| 5 | 14177235 | rs12652543 | -0.152 | 0.763 | 0.6 | 0.054 | -0.012 | 0.001 | 0.001 | 0 | 0.14 | 0.08 | 0.15 | 0.47 | 0.16 | 0.33 |
| 5 | 14181939 | rs32340 | 1.031 | 0.449 | 0.937 | -0.287 | -0.005 | -0.02 | -0.02 | 0.046 | 0.66 | 0.7 | 0.63 | 0.53 | 0.7 | 0.47 |
| 5 | 14185484 | rs26135 | 0.701 | 0.19 | 1.201 | -0.458 | -0.048 | -0.118 | -0.118 | 0.057 | 0.72 | 0.51 | 0.7 | 0.95 | 0.79 | 0.76 |
| 5 | 14188019 | rs27665 | -0.051 | 0.188 | 1.166 | -0.456 | -0.035 | -0.101 | -0.101 | 0 | 0.78 | 0.66 | 0.77 | 0.99 | 0.87 | 0.82 |
| 5 | 14188684 | rs26291 | 0.998 | 0.172 | 1.299 | -0.491 | 0.014 | 0.012 | 0.012 | 0.12 | 0.69 | 0.23 | 0.59 | 0.95 | 0.74 | 0.7 |
| 5 | 14192951 | rs26058 | 0.079 | 0.13 | 1.2 | -0.526 | -0.058 | -0.152 | -0.152 | 0 | 0.78 | 0.51 | 0.75 | 0.99 | 0.87 | 0.82 |
| 5 | 14197566 | rs151902 | -0.696 | 0.086 | 1.097 | -0.546 | 0.02 | 0.05 | 0.05 | 0 | 0.82 | 0.92 | 0.79 | 0.99 | 0.87 | 0.82 |
| 5 | 14206537 | rs33005 | 1.682 | 0.169 | 1.033 | -0.491 | -0.025 | -0.049 | -0.049 | 0.955 | 0.55 | 0.12 | 0.44 | 0.51 | 0.58 | 0.35 |
| 5 | 14216900 | rs32388 | -0.611 | 0.078 | 0.897 | -0.514 | 0.015 | 0.016 | 0.016 | 0 | 0.82 | 0.9 | 0.79 | 0.99 | 0.87 | 0.82 |
| 5 | 14222340 | rs149123 | 1.655 | 0.113 | 0.718 | -0.477 | -0.027 | -0.056 | -0.056 | 0 | 0.55 | 0.1 | 0.44 | 0.51 | 0.59 | 0.35 |
| 5 | 14227482 | rs544588 | -0.825 | 0.524 | 1.096 | -0.178 | 0.01 | 0.04 | 0.04 | 0 | 0.2 | 0.23 | 0.22 | 0.48 | 0.24 | 0.4 |
| 5 | 14235406 | rs354286 | -0.649 | 0.181 | 0.75 | -0.588 | 0.001 | 0.004 | 0.004 | 0 | 0.33 | 0.32 | 0.28 | 0.47 | 0.29 | 0.42 |
| 5 | 14237265 | rs354288 | -0.253 | -0.031 | 0.799 | -0.799 | -0.002 | -0.048 | -0.048 | 0.016 | 0.83 | 0.78 | 0.71 | 0.99 | 0.83 | 0.78 |
| 5 | 14239606 | rs11133784 | 1.386 | -0.086 | 0.617 | -0.73 | 0.006 | 0.004 | 0.004 | 0 | 0.51 | 0.35 | 0.42 | 0.5 | 0.54 | 0.36 |
| 5 | 14241854 | rs6876763 | 0.716 | 0.156 | 1.015 | -0.649 | -0.001 | -0.034 | -0.034 | 0.029 | 0.75 | 0.31 | 0.62 | 0.96 | 0.74 | 0.71 |
| 5 | 14245046 | rs12522960 | 1.626 | 0.264 | 0.664 | -0.652 | -0.108 | -0.113 | -0.113 | 1.265 | 0.56 | 0.06 | 0.43 | 0.53 | 0.57 | 0.37 |
| 5 | 14251518 | rs6554850 | -0.962 | 0.397 | 1.242 | -0.556 | 0.028 | 0.077 | 0.077 | 0 | 0.25 | 0.59 | 0.26 | 0.41 | 0.24 | 0.37 |
| 5 | 14253187 | rs16903338 | -0.038 | 0.432 | 1.088 | -0.531 | -0.001 | 0.007 | 0.007 | 0 | 0.18 | 0.19 | 0.17 | 0.38 | 0.16 | 0.34 |
| 5 | 14262535 | rs164861 | -0.347 | 0.383 | 1.353 | -0.608 | 0.008 | -0.091 | -0.091 | 0 | 0.83 | 0.69 | 0.72 | 0.99 | 0.82 | 0.76 |
| 5 | 14263981 | rs10513170 | 0.663 | 0.382 | 1.178 | -0.575 | 0.019 | 0.01 | 0.01 | 0 | 0.06 | 0.22 | 0.08 | 0.03 | 0.08 | 0.03 |

| | | | | | | | | | | | | | | | | |
|---|---|---|---|---|---|---|---|---|---|---|---|---|---|---|---|---|
| 5 | 14266629 | rs165094 | 1.567 | 0.282 | 0.911 | -0.612 | -0.057 | -0.026 | -0.026 | 0.038 | 0.57 | 0.11 | 0.45 | 0.58 | 0.58 | 0.39 |
| 5 | 14275499 | rs154150 | 0.226 | 0.288 | 0.897 | -0.548 | -0.052 | -0.085 | -0.085 | 0 | 0.17 | 0.07 | 0.16 | 0.39 | 0.16 | 0.33 |
| 5 | 14277686 | rs30619 | 0.342 | 0.157 | 1.388 | -0.744 | -0.006 | -0.136 | -0.136 | 0 | 0.8 | 0.48 | 0.69 | 0.99 | 0.82 | 0.76 |
| 5 | 14281126 | rs2591277 | -0.488 | 0.155 | 1.018 | -0.745 | -0.008 | -0.077 | -0.077 | 0 | 0.21 | 0.69 | 0.33 | 0.01 | 0.18 | 0.24 |
| 5 | 14292625 | rs2434605 | 0.176 | 0.037 | 0.361 | -1.062 | 0.001 | 0.057 | 0.057 | 0 | 0.27 | 0.15 | 0.32 | 0.01 | 0.25 | 0.25 |
| 5 | 14298470 | rs30633 | 1.194 | 0.161 | 0.751 | -0.603 | -0.013 | -0.029 | -0.029 | 0 | 0.69 | 0.24 | 0.64 | 0.6 | 0.71 | 0.57 |
| 5 | 14306905 | rs557292 | -0.529 | 0.117 | 1.211 | -0.558 | 0.002 | 0.004 | 0.004 | 0.823 | 0.09 | 0.36 | 0.07 | 0.03 | 0.05 | 0.03 |
| 5 | 14312410 | rs42405 | -0.438 | 0.209 | 0.563 | -0.502 | -0.066 | -0.101 | -0.101 | 0 | 0.84 | 0.98 | 0.85 | 0.63 | 0.84 | 0.69 |
| 5 | 14326046 | rs17303923 | 0.985 | 0.362 | 0.329 | -0.854 | -0.012 | 0.042 | 0.042 | 1.114 | 0.18 | 0.08 | 0.22 | 0.01 | 0.2 | 0.21 |
| 5 | 14329404 | rs923616 | 1.068 | 0.377 | 0.328 | -0.855 | -0.011 | 0.039 | 0.039 | 1.118 | 0.17 | 0.08 | 0.22 | 0.01 | 0.19 | 0.2 |
| 5 | 14332325 | rs30623 | 0.686 | 0.345 | 0.543 | -0.832 | -0.002 | 0 | 0 | 0.05 | 0.59 | 0.36 | 0.51 | 0.64 | 0.61 | 0.42 |
| 5 | 14334735 | rs187432 | 1.185 | 0.366 | -0.008 | -0.83 | -0.057 | 0.006 | 0.006 | 0.193 | 0.53 | 0.12 | 0.43 | 0.6 | 0.57 | 0.39 |
| 5 | 14342262 | rs247138 | 1.781 | 0.202 | -0.415 | -0.803 | -0.018 | -0.115 | -0.115 | 2.13 | 0.28 | 0.07 | 0.31 | 0.55 | 0.36 | 0.23 |
| 5 | 14354827 | rs154159 | 0.529 | 0.274 | 0.067 | -0.694 | 0.016 | 0.006 | 0.006 | 1.189 | 0.76 | 0.35 | 0.71 | 0.73 | 0.79 | 0.62 |
| 5 | 14360436 | rs730184 | 0.88 | -0.046 | -0.696 | -1.077 | -0.015 | 0.153 | 0.153 | 0 | 0.37 | 0.08 | 0.26 | 0.11 | 0.32 | 0.31 |
| 5 | 14363446 | rs13172149 | 1.161 | 0.014 | -0.58 | -0.95 | -0.019 | 0.108 | 0.108 | 0 | 0.32 | 0.08 | 0.23 | 0.1 | 0.28 | 0.28 |
| 5 | 14365895 | rs43171 | 1.051 | -0.003 | -0.638 | -0.958 | -0.033 | -0.016 | -0.016 | 0 | 0.4 | 0.08 | 0.43 | 0.63 | 0.51 | 0.35 |
| 5 | 14368975 | rs13189406 | 1.188 | 0.008 | -0.42 | -0.971 | -0.018 | 0.153 | 0.153 | 0 | 0.32 | 0.08 | 0.23 | 0.06 | 0.28 | 0.28 |
| 5 | 14371180 | rs256417 | -0.286 | 0.183 | -0.212 | -0.84 | -0.019 | -0.02 | -0.02 | 0.064 | 0.83 | 0.88 | 0.81 | 0.73 | 0.84 | 0.7 |
| 5 | 14375908 | rs30765 | 2.064 | 0.266 | -0.435 | -0.696 | -0.031 | -0.16 | -0.16 | 0.018 | 0.27 | 0.03 | 0.29 | 0.56 | 0.34 | 0.21 |
| 5 | 14380324 | rs154152 | -0.121 | 0.326 | -0.104 | -0.905 | -0.003 | -0.002 | -0.002 | 0 | 0.82 | 0.82 | 0.81 | 0.73 | 0.84 | 0.68 |
| 5 | 14381174 | rs247137 | 0.88 | 0.055 | 0.479 | -1.67 | 0.005 | 0.027 | 0.027 | 0 | 0.34 | 0.27 | 0.26 | 0.14 | 0.27 | 0.29 |
| 5 | 14389469 | rs7715916 | 0.225 | 0.156 | 0.919 | -1.687 | 0.026 | -0.024 | -0.024 | 0 | 0.06 | 0.26 | 0.1 | 0 | 0.06 | 0.07 |
| 5 | 14393621 | rs10065203 | 2.26 | 0.021 | 0.783 | -1.427 | -0.02 | -0.049 | -0.049 | 0 | 0.25 | 0.04 | 0.17 | 0.26 | 0.33 | 0.14 |
| 5 | 14397238 | rs30774 | -0.538 | -0.267 | 1.196 | -2.013 | 0.015 | 0.046 | 0.046 | 0 | 0.43 | 0.7 | 0.38 | 0.1 | 0.33 | 0.36 |
| 5 | 14400274 | rs40278 | 0.205 | -0.251 | 1.222 | -1.942 | 0.001 | -0.022 | -0.022 | 0 | 0.37 | 0.39 | 0.27 | 0.1 | 0.27 | 0.29 |
| 5 | 14402871 | rs13356126 | 0.26 | -0.152 | 0.852 | -1.984 | 0.004 | -0.011 | -0.011 | 0 | 0.57 | 0.36 | 0.69 | 0.9 | 0.67 | 0.64 |
| 5 | 14411080 | rs30608 | -0.619 | -0.108 | 0.434 | -1.846 | 0.036 | 0.115 | 0.115 | 0.193 | 0.47 | 0.74 | 0.39 | 0.1 | 0.35 | 0.4 |
| 5 | 14415816 | rs187190 | -0.999 | 0.203 | -0.212 | -1.699 | -0.08 | 0.269 | 0.269 | 0.435 | 0.55 | 0.91 | 0.45 | 0.11 | 0.43 | 0.42 |
| 5 | 14417929 | rs42404 | 0.34 | 0.11 | -0.091 | -1.774 | -0.015 | 0.187 | 0.187 | 0 | 0.41 | 0.15 | 0.29 | 0.11 | 0.35 | 0.32 |
| 5 | 14418419 | rs16903497 | -0.45 | 0.09 | 0.388 | -1.599 | 0.014 | -0.051 | -0.051 | 0 | 0.09 | 0.26 | 0.12 | 0 | 0.08 | 0.1 |
| 5 | 14421026 | rs10056415 | -1.265 | 0.125 | -0.653 | -1.582 | -0.013 | -0.056 | -0.056 | 0 | 0.14 | 0.76 | 0.16 | 0 | 0.08 | 0.1 |
| 5 | 14431980 | rs27761 | -0.51 | 0.359 | -0.82 | -1.623 | 0.008 | 0.217 | 0.217 | 0.618 | 0.51 | 0.82 | 0.44 | 0.1 | 0.44 | 0.39 |
| 5 | 14432660 | rs26188 | -0.84 | 0.176 | -1.156 | -1.561 | 0.013 | 0.324 | 0.324 | 0.799 | 0.5 | 0.89 | 0.44 | 0.1 | 0.39 | 0.38 |
| 5 | 14450135 | rs27111 | 0.728 | 0.127 | -1.179 | -1.351 | -0.008 | 0 | 0 | 0 | 0.21 | 0.09 | 0.11 | 0.17 | 0.19 | 0.19 |
| 5 | 14458747 | rs26106 | 1.299 | 0.219 | -1.074 | -1.443 | 0.001 | 0.01 | 0.01 | 0 | 0.18 | 0.1 | 0.19 | 0.02 | 0.16 | 0.11 |
| 5 | 14462816 | rs27100 | -0.576 | 0.325 | -1.398 | -1.453 | -0.006 | 0.226 | 0.226 | 0.697 | 0.5 | 0.91 | 0.45 | 0.19 | 0.42 | 0.39 |
| 5 | 14465355 | rs17500759 | 0.337 | 0.756 | -1.116 | -0.99 | -0.012 | 0.035 | 0.035 | 0 | 0.12 | 0.03 | 0.08 | 0 | 0.11 | 0.07 |
| 5 | 14472040 | rs706282 | -0.28 | 0.457 | -0.484 | -0.516 | 0 | 0.004 | 0.004 | 1.067 | 0.25 | 0.27 | 0.2 | 0.17 | 0.22 | 0.23 |
| 5 | 14478601 | rs6882373 | -1.092 | 0.927 | -0.626 | -0.048 | -0.003 | 0 | 0 | 0 | 0.09 | 0.1 | 0.05 | 0.05 | 0.09 | 0.07 |
| 5 | 14479938 | rs26173 | -0.08 | 0.582 | -0.527 | 0.09 | -0.006 | -0.004 | -0.004 | 0.495 | 0.84 | 0.8 | 0.76 | 0.85 | 0.87 | 0.88 |
| 5 | 14485386 | rs26098 | 0.108 | 0.477 | -0.739 | 0.109 | 0.011 | 0 | 0 | 0 | 0.14 | 0.05 | 0.21 | 0.15 | 0.1 | 0.12 |
| 5 | 14491969 | rs26110 | -0.473 | 0.345 | -1.372 | -0.254 | -0.102 | 0.039 | 0.039 | 0.091 | 0.57 | 0.93 | 0.48 | 0.31 | 0.43 | 0.48 |
| 5 | 14498727 | rs26107 | 0.485 | 0.223 | -1.216 | -0.062 | 0.009 | 0.003 | 0.003 | 0 | 0.13 | 0.05 | 0.21 | 0.13 | 0.09 | 0.12 |
| 5 | 14503893 | rs3789157 | 0.384 | 0.203 | -0.588 | -0.203 | -0.007 | -0.025 | -0.025 | 0 | 0.39 | 0.38 | 0.2 | 0.16 | 0.29 | 0.31 |
| 5 | 14509966 | rs1058312 | 0.467 | 0.488 | -0.568 | -0.197 | 0.013 | 0.037 | 0.037 | 0 | 0.4 | 0.32 | 0.22 | 0.12 | 0.31 | 0.33 |
| 5 | 14514308 | rs32815 | -0.772 | 1.673 | -0.819 | 1.079 | -0.037 | 0.005 | 0.005 | 0.109 | 0.93 | 0.99 | 0.84 | 0.66 | 0.87 | 0.9 |
| 5 | 14517076 | rs32811 | 0.184 | 1.683 | -0.51 | 0.177 | 0.144 | -0.055 | -0.055 | 2.992 | 0.9 | 0.54 | 0.89 | 1 | 0.84 | 0.94 |
| 5 | 14518208 | rs32808 | 0.03 | 1.224 | -1.48 | 0.63 | -0.053 | -0.078 | -0.078 | 0 | 0.31 | 0.1 | 0.36 | 0.44 | 0.42 | 0.44 |
| 5 | 14530993 | rs26208 | -0.446 | 1.249 | -0.998 | 0.056 | 0.007 | -0.029 | -0.029 | 0 | 0.92 | 0.9 | 0.95 | 1 | 0.93 | 0.95 |
| 5 | 14532534 | rs26219 | -0.185 | 1.551 | -1.5 | -0.05 | -0.068 | 0 | 0 | 0 | 0.24 | 0.05 | 0.26 | 0.24 | 0.3 | 0.22 |
| 5 | 14533529 | rs26218 | -0.341 | 1.111 | -1.341 | 0.013 | 0 | -0.005 | -0.005 | 0 | 0.87 | 0.84 | 0.94 | 0.94 | 0.91 | 0.86 |
| 5 | 14534994 | rs26217 | 0.398 | 0.9 | -0.986 | 0.209 | -0.004 | 0.008 | 0.008 | 0.013 | 0.11 | 0.15 | 0.05 | 0.19 | 0.11 | 0.12 |
| 5 | 14537521 | rs12654768 | 0.445 | 0.867 | -1.127 | 0.107 | -0.009 | 0.022 | 0.022 | 0.175 | 0.6 | 0.7 | 0.66 | 0.69 | 0.59 | 0.6 |
| 5 | 14537840 | rs4702038 | 0.828 | 0.724 | -1.155 | 0.204 | 0.003 | 0.028 | 0.028 | 0.047 | 0.43 | 0.59 | 0.41 | 0.23 | 0.4 | 0.28 |
| 5 | 14539821 | rs6881024 | 0.353 | 0.582 | -0.981 | 0.74 | -0.006 | 0.005 | 0.005 | 0 | 0.12 | 0.13 | 0.19 | 0.18 | 0.05 | 0.1 |
| 5 | 14543201 | rs40297 | 0.019 | 0.884 | -1.146 | 0.911 | -0.034 | 0.053 | 0.053 | 0 | 0.73 | 0.88 | 0.72 | 0.88 | 0.7 | 0.74 |
| 5 | 14545898 | rs3993996 | -0.666 | 0.382 | -0.463 | 1.661 | 0.004 | 0.008 | 0.008 | 0 | 0.14 | 0.14 | 0.21 | 0.46 | 0.14 | 0.3 |
| 5 | 14561262 | rs32631 | -0.943 | 0.919 | -0.466 | 0.757 | 0 | 0 | 0 | 0 | 0.17 | 0.18 | 0.23 | 0.27 | 0.18 | 0.23 |
| 5 | 14563425 | rs17307347 | -0.54 | 0.963 | -0.606 | 0.769 | -0.04 | -0.017 | -0.017 | 0.068 | 0.07 | 0.01 | 0.16 | 0.09 | 0.13 | 0.07 |
| 5 | 14570732 | rs39968 | -0.202 | 1.201 | -1.209 | 0.906 | -0.023 | -0.068 | -0.068 | 0 | 0.72 | 0.86 | 0.66 | 0.59 | 0.67 | 0.6 |
| 5 | 14572712 | rs32635 | -0.6 | 0.772 | 0.022 | 0.735 | -0.03 | 0.002 | 0.002 | 0 | 0.12 | 0.29 | 0.09 | 0.18 | 0.06 | 0.18 |
| 5 | 14575698 | rs152624 | -0.264 | 0.666 | -0.011 | 0.909 | -0.008 | 0.034 | 0.034 | 0 | 0.13 | 0.2 | 0.18 | 0.28 | 0.09 | 0.15 |
| 5 | 14578799 | rs152642 | -0.038 | 0.807 | -0.21 | 0.659 | -0.042 | 0.031 | 0.031 | 0.021 | 0.08 | 0.01 | 0.15 | 0.03 | 0.11 | 0.11 |
| 5 | 14583006 | rs152621 | -0.885 | 0.893 | 0.484 | 0.264 | 0.032 | -0.036 | -0.036 | 1.086 | 0.18 | 0.53 | 0.15 | 0.18 | 0.18 | 0.12 |
| 5 | 14584695 | rs152619 | -1.377 | 0.855 | 0.434 | 0.452 | 0.004 | 0.046 | 0.046 | 0.625 | 0.28 | 0.56 | 0.31 | 0.48 | 0.27 | 0.28 |
| 5 | 14591004 | rs7718540 | 0.529 | 0.881 | 0.452 | 0.447 | -0.019 | -0.039 | -0.039 | 0.069 | 0.1 | 0.04 | 0.18 | 0.32 | 0.14 | 0.16 |
| 5 | 14594377 | rs17248579 | 0.518 | 0.156 | 0.303 | -0.772 | 0.014 | 0.032 | 0.032 | 0 | 0.28 | 0.13 | 0.15 | 0.03 | 0.27 | 0.29 |
| 5 | 14617857 | rs457897 | -0.368 | 0.409 | -0.033 | -0.503 | -0.028 | -0.032 | -0.032 | 0 | 0.95 | 0.99 | 0.92 | 0.77 | 0.92 | 0.91 |

| | | | | | | | | | | | | | | | | |
|---|---|---|---|---|---|---|---|---|---|---|---|---|---|---|---|---|
| 5 | 14621711 | rs6895973 | 0.696 | 0.183 | 0.033 | -0.658 | -0.025 | 0.152 | 0.152 | 0 | 0.49 | 0.15 | 0.53 | 0.66 | 0.51 | 0.51 |
| 5 | 14637272 | rs31868 | 0.039 | 1.095 | 0.461 | -0.146 | -0.035 | -0.037 | -0.037 | 0.291 | 0.94 | 0.99 | 0.94 | 0.79 | 0.89 | 0.87 |
| 5 | 14640921 | rs1896783 | 1.883 | 1.009 | 1.34 | -0.201 | 0.049 | 0.061 | 0.061 | 0.555 | 0.86 | 0.59 | 0.77 | 0.76 | 0.84 | 0.8 |
| 5 | 14648689 | rs25943 | -1.406 | 1.134 | 1.632 | -0.318 | 0.084 | -0.044 | -0.044 | 0.507 | 0.13 | 0.55 | 0.22 | 0.03 | 0.14 | 0.09 |
| 5 | 14655699 | rs456180 | 0.181 | 1.159 | 1.565 | -0.297 | -0.009 | -0.009 | -0.009 | 0 | 0.93 | 0.95 | 0.93 | 0.79 | 0.89 | 0.88 |
| 5 | 14657062 | rs1875513 | 1.684 | 1.121 | 1.658 | -0.321 | 0.042 | -0.001 | -0.001 | 0 | 0.79 | 0.36 | 0.7 | 0.76 | 0.75 | 0.78 |
| 5 | 14667143 | rs17297276 | 0.004 | 0.669 | 1.304 | -0.783 | -0.001 | -0.033 | -0.033 | 0.021 | 0.23 | 0.21 | 0.18 | 0.02 | 0.27 | 0.28 |
| 5 | 14673696 | rs16903631 | 0.889 | 0.727 | 0.954 | -0.145 | -0.004 | 0.032 | 0.032 | 0 | 0.09 | 0.05 | 0.17 | 0.41 | 0.11 | 0.17 |
| 5 | 14676218 | rs6892588 | -0.653 | 0.878 | 1.451 | -0.786 | 0.132 | 0.002 | 0.002 | 0.134 | 0.07 | 0.43 | 0.09 | 0 | 0.09 | 0.01 |
| 5 | 14677305 | rs992737 | 0.564 | 0.853 | 1.334 | -0.837 | -0.049 | 0.1 | 0.1 | 0 | 0.17 | 0.02 | 0.12 | 0.01 | 0.23 | 0.16 |
| 5 | 14689707 | rs152788 | 1.996 | 1.101 | 2.334 | -0.195 | 0.084 | 0.044 | 0.044 | 0.837 | 0.84 | 0.43 | 0.75 | 0.76 | 0.8 | 0.82 |
| 5 | 14697849 | rs6886163 | -0.058 | 0.987 | 2.51 | -0.013 | 0.034 | -0.027 | -0.027 | 0 | 0.05 | 0.12 | 0.05 | 0 | 0.09 | 0.01 |
| 5 | 14700035 | rs10065032 | -0.424 | 0.989 | 2.418 | -0.012 | 0.093 | -0.012 | -0.012 | 2.215 | 0.06 | 0.26 | 0.07 | 0 | 0.09 | 0.01 |
| 5 | 14702375 | rs258290 | 1.781 | 0.963 | 1.886 | 0.005 | 0.068 | 0.036 | 0.036 | 1.224 | 0.83 | 0.39 | 0.75 | 0.76 | 0.8 | 0.82 |
| 5 | 14741603 | rs153932 | -2.016 | 2.138 | 2.355 | 0.283 | 0.094 | 0.007 | 0.007 | 0 | 0.16 | 0.44 | 0.29 | 0.13 | 0.26 | 0.09 |
| 5 | 14741921 | rs187483 | -1.235 | 2.143 | 2.578 | 0.283 | 0.084 | -0.071 | -0.071 | 1.746 | 0.11 | 0.29 | 0.14 | 0 | 0.22 | 0.04 |
| 5 | 14745268 | rs149391 | -2.343 | 2.085 | 2.484 | 0.311 | 0.103 | 0.01 | 0.01 | 0 | 0.17 | 0.54 | 0.32 | 0.15 | 0.28 | 0.09 |
| 5 | 14751400 | rs2291943 | -2.828 | 1.949 | 2.289 | 0.189 | 0.097 | 0.136 | 0.136 | 0.92 | 0.07 | 0.49 | 0.28 | 0.1 | 0.06 | 0.06 |
| 5 | 14752850 | rs17250963 | -1.458 | 2.044 | 2.046 | 0.102 | 0.04 | -0.066 | -0.066 | 0.004 | 0.12 | 0.23 | 0.14 | 0 | 0.21 | 0.03 |
| 5 | 14754571 | rs10036729 | -1.796 | 2.044 | 1.416 | 0.101 | 0.094 | -0.106 | -0.106 | 0.043 | 0.14 | 0.51 | 0.15 | 0 | 0.21 | 0.03 |
| 5 | 14755632 | rs1697124 | -2.347 | 2.008 | 0.8 | 0.127 | 0.042 | -0.038 | -0.038 | 0 | 0.18 | 0.64 | 0.41 | 0.09 | 0.27 | 0.1 |
| 5 | 14756567 | rs152363 | -2.063 | 1.795 | 0.874 | 0.297 | 0.053 | -0.004 | -0.004 | 0 | 0.12 | 0.64 | 0.33 | 0.09 | 0.17 | 0.07 |
| 5 | 14759246 | rs4702049 | -0.14 | 1.671 | -0.012 | 0.347 | 0.003 | 0.001 | 0.001 | 0.273 | 0.3 | 0.33 | 0.5 | 0.34 | 0.4 | 0.34 |
| 5 | 14760607 | rs4701616 | -0.515 | 0.684 | -0.811 | -0.113 | 0.005 | 0.001 | 0.001 | 0.146 | 0.6 | 0.85 | 0.72 | 0.36 | 0.67 | 0.61 |
| 5 | 14763707 | rs744165 | -0.212 | 0.372 | -0.384 | 0.407 | 0 | 0.001 | 0.001 | 0 | 0.32 | 0.31 | 0.27 | 0.39 | 0.35 | 0.29 |
| 5 | 14764934 | rs706293 | 0.3 | 0.607 | 0.296 | 0.294 | 0.03 | 0.048 | 0.048 | 0.175 | 0.28 | 0.47 | 0.48 | 0.4 | 0.4 | 0.3 |
| 5 | 14765662 | rs10513186 | 0.702 | 0.294 | 0.017 | 0.046 | 0.013 | 0.077 | 0.077 | 0 | 0.23 | 0.1 | 0.13 | 0.02 | 0.18 | 0.26 |
| 5 | 14768308 | rs3087948 | -0.702 | 0.133 | 0.255 | 0.251 | -0.01 | 0.079 | 0.079 | 0.123 | 0.19 | 0.28 | 0.32 | 0.47 | 0.19 | 0.3 |
| 5 | 14769103 | rs17251667 | 0.966 | 0.378 | -0.136 | 0.187 | -0.006 | 0.113 | 0.113 | 0 | 0.22 | 0.04 | 0.12 | 0.02 | 0.18 | 0.26 |
| 5 | 14770753 | rs4702052 | 0.398 | 0.325 | -0.014 | 0.331 | 0 | 0 | 0 | 0 | 0.07 | 0.08 | 0.12 | 0.08 | 0.1 | 0.07 |
| 5 | 14772245 | rs17251715 | 0.708 | 0.386 | -0.285 | 0.268 | 0.006 | 0.002 | 0.002 | 0 | 0.27 | 0.33 | 0.36 | 0.32 | 0.33 | 0.27 |
| 5 | 14776585 | rs9790982 | -1.39 | 0.329 | -0.633 | 0.139 | -0.02 | 0.044 | 0.044 | 1.981 | 0.71 | 0.87 | 0.84 | 0.82 | 0.73 | 0.87 |
| 5 | 14776897 | rs17251763 | 0.882 | 0.579 | 0.195 | 1.004 | -0.012 | -0.017 | -0.017 | 0 | 0.22 | 0.13 | 0.31 | 0.32 | 0.29 | 0.23 |
| 5 | 14777799 | rs9312873 | -0.33 | 0.298 | -0.067 | 1.257 | 0.003 | 0.012 | 0.012 | 0 | 0.11 | 0.1 | 0.19 | 0.39 | 0.09 | 0.22 |
| 5 | 14794868 | rs879253 | -1.167 | 0.345 | -0.079 | 1.181 | -0.017 | 0.151 | 0.151 | 1.166 | 0.52 | 0.75 | 0.71 | 0.82 | 0.56 | 0.7 |
| 5 | 14804735 | rs4702054 | 0.039 | 0.992 | -0.672 | 1.564 | -0.001 | -0.002 | -0.002 | 0 | 0.07 | 0.1 | 0.13 | 0.08 | 0.11 | 0.07 |
| 5 | 14810110 | rs12514030 | -1.18 | 0.966 | -0.607 | 1.828 | -0.002 | 0.055 | 0.055 | 0 | 0.13 | 0.14 | 0.2 | 0.42 | 0.1 | 0.31 |
| 5 | 14814329 | rs16903710 | -0.555 | 1.314 | -0.373 | 2.079 | 0.005 | 0.137 | 0.137 | 0 | 0.09 | 0.12 | 0.18 | 0.41 | 0.08 | 0.29 |
| 5 | 14814841 | rs10054602 | -0.443 | 1.328 | -0.511 | 2.086 | -0.004 | 0.03 | 0.03 | 0.034 | 0.08 | 0.05 | 0.17 | 0.41 | 0.08 | 0.29 |
| 5 | 14817669 | rs26304 | -0.821 | 1.301 | -0.448 | 2.005 | 0.009 | -0.018 | -0.018 | 0 | 0.37 | 0.43 | 0.56 | 0.74 | 0.43 | 0.58 |
| 5 | 14835061 | rs10474706 | -0.194 | 1.312 | -0.452 | 2.023 | -0.006 | 0.002 | 0.002 | 0.001 | 0.08 | 0.02 | 0.17 | 0.4 | 0.08 | 0.23 |
| 5 | 14835313 | rs9312874 | -0.363 | 1.299 | -0.443 | 1.905 | 0.001 | 0.096 | 0.096 | 0.049 | 0.09 | 0.1 | 0.18 | 0.42 | 0.08 | 0.29 |
| 5 | 14846372 | rs826358 | -0.402 | 1.137 | -0.733 | 1.018 | 0.003 | -0.009 | -0.009 | 0 | 0.37 | 0.42 | 0.56 | 0.74 | 0.42 | 0.58 |
| 5 | 14863370 | rs16903719 | -1.567 | 1.19 | -0.981 | 2.062 | -0.022 | -0.02 | -0.02 | 2.263 | 0.08 | 0.02 | 0.19 | 0.37 | 0.1 | 0.25 |
| 5 | 14869666 | rs10513191 | -1.19 | 1.106 | -0.978 | 2.228 | -0.007 | 0.005 | 0.005 | 0 | 0.05 | 0.02 | 0.18 | 0.3 | 0.06 | 0.14 |
| 5 | 14869709 | rs3006069 | 0.493 | 1.144 | -0.959 | 1.653 | 0.012 | 0.035 | 0.035 | 0 | 0.53 | 0.42 | 0.45 | 0.3 | 0.45 | 0.38 |
| 5 | 14877474 | rs835153 | -0.708 | 1.007 | -1.079 | 1.565 | 0.04 | 0.107 | 0.107 | 0 | 0.5 | 0.78 | 0.57 | 0.71 | 0.55 | 0.62 |
| 5 | 14884822 | rs835146 | -0.697 | 0.735 | -1.144 | 0.979 | 0 | 0.062 | 0.062 | 0 | 0.63 | 0.91 | 0.69 | 0.74 | 0.64 | 0.58 |
| 5 | 14888508 | rs835141 | -0.56 | 0.799 | -0.78 | 0.491 | 0.014 | 0.037 | 0.037 | 0 | 0.49 | 0.61 | 0.43 | 0.35 | 0.55 | 0.34 |
| 5 | 14890877 | rs863118 | -0.471 | 0.877 | -0.751 | 0.503 | -0.005 | 0.025 | 0.025 | 0 | 0.47 | 0.6 | 0.41 | 0.35 | 0.51 | 0.32 |
| 5 | 14896757 | rs1867676 | 0.088 | 1.5 | -0.346 | 0.471 | -0.003 | 0.005 | 0.005 | 0.235 | 0.22 | 0.16 | 0.14 | 0.09 | 0.22 | 0.17 |
| 5 | 14902038 | rs2441022 | 0.054 | 1.555 | -0.372 | 0.444 | 0.043 | 0.06 | 0.06 | 0.264 | 0.2 | 0.42 | 0.25 | 0.26 | 0.26 | 0.12 |
| 5 | 14902964 | rs3006064 | -0.441 | 1.038 | -0.438 | 0.126 | 0.002 | -0.03 | -0.03 | 0 | 0.23 | 0.27 | 0.16 | 0.1 | 0.23 | 0.2 |
| 5 | 14908578 | rs1550825 | 0.133 | 0.771 | -0.517 | 0.389 | 0 | 0.002 | 0.002 | 0 | 0.63 | 0.38 | 0.63 | 0.61 | 0.63 | 0.36 |
| 5 | 14908793 | rs860666 | -0.663 | 0.159 | -0.511 | 0.063 | -0.009 | -0.011 | -0.011 | 0 | 0.18 | 0.24 | 0.08 | 0.12 | 0.16 | 0.12 |
| 5 | 14913369 | rs844335 | -0.288 | 0.405 | -0.466 | 0.002 | 0.012 | 0.064 | 0.064 | 0 | 0.5 | 0.68 | 0.45 | 0.39 | 0.53 | 0.68 |
| 5 | 14917813 | rs966987 | -0.788 | 0.445 | 0.298 | 0.218 | -0.044 | -0.01 | -0.01 | 0 | 0.2 | 0.48 | 0.1 | 0.18 | 0.13 | 0.12 |
| 5 | 14922430 | rs2592019 | 0.205 | 0.747 | 0.014 | 0.057 | 0.006 | 0.003 | 0.003 | 0 | 0.16 | 0.17 | 0.28 | 0.27 | 0.29 | 0.28 |
| 5 | 14924458 | rs4702062 | -0.529 | 0.408 | 0.085 | -0.124 | 0.003 | -0.008 | -0.008 | 0 | 0.42 | 0.48 | 0.4 | 0.36 | 0.45 | 0.67 |
| 5 | 14926057 | rs835136 | 0.228 | 0.397 | -0.125 | -0.087 | 0.007 | -0.011 | -0.011 | 0 | 0.17 | 0.15 | 0.08 | 0.22 | 0.15 | 0.13 |
| 5 | 14927488 | rs728844 | -1.515 | 0.299 | -0.433 | -0.164 | 0.002 | -0.035 | -0.035 | 0.56 | 0.59 | 0.76 | 0.49 | 0.58 | 0.61 | 0.79 |
| 5 | 14935018 | rs10039962 | 0.752 | 0.238 | -0.425 | -0.389 | 0.012 | 0.037 | 0.037 | 0 | 0.25 | 0.11 | 0.12 | 0.12 | 0.2 | 0.42 |
| 5 | 14943456 | rs2921613 | -1.134 | 0.371 | -0.504 | 0.024 | -0.004 | -0.018 | -0.018 | 0 | 0.61 | 0.73 | 0.5 | 0.64 | 0.63 | 0.84 |
| 5 | 14946949 | rs16903741 | 0.14 | 0.019 | -0.798 | 0.12 | 0.021 | -0.029 | -0.029 | 0.416 | 0.14 | 0.05 | 0.05 | 0.17 | 0.09 | 0.18 |
| 5 | 14949696 | rs835102 | -0.71 | 0.626 | -0.328 | -0.138 | -0.008 | -0.002 | -0.002 | 0 | 0.57 | 0.59 | 0.46 | 0.63 | 0.55 | 0.8 |
| 5 | 14949939 | rs13353954 | -0.176 | 0.514 | -0.59 | -0.583 | 0.015 | 0.058 | 0.058 | 0 | 0.28 | 0.13 | 0.11 | 0.11 | 0.21 | 0.36 |
| 5 | 14952142 | rs4702064 | -0.217 | 0.209 | -0.657 | -0.471 | 0.015 | 0.055 | 0.055 | 0 | 0.28 | 0.15 | 0.12 | 0.11 | 0.21 | 0.36 |
| 5 | 14956699 | rs835118 | 0.373 | 0.246 | -0.241 | -0.152 | -0.004 | -0.03 | -0.03 | 0 | 0.67 | 0.61 | 0.82 | 0.81 | 0.67 | 0.82 |
| 5 | 14960920 | rs706297 | -0.302 | 0.229 | -0.261 | -0.289 | 0.005 | 0.021 | 0.021 | 0 | 0.86 | 0.75 | 0.9 | 0.82 | 0.86 | 0.86 |

| | | | | | | | | | | | | | | | | |
|---|---|---|---|---|---|---|---|---|---|---|---|---|---|---|---|---|
| 5 | 14962587 | rs835110 | -0.62 | 0.219 | -0.318 | -0.29 | -0.005 | -0.006 | -0.006 | 0 | 0.88 | 0.89 | 0.9 | 0.85 | 0.86 | 0.87 |
| 5 | 14970328 | rs835092 | 0.328 | 0.523 | 0.211 | 0.466 | 0.02 | 0.001 | 0.001 | 0 | 0.36 | 0.53 | 0.55 | 0.34 | 0.46 | 0.48 |
| 5 | 14973914 | rs10037304 | 0.386 | 0.532 | -0.305 | 0.484 | 0.056 | 0.012 | 0.012 | 1.241 | 0.49 | 0.22 | 0.35 | 0.47 | 0.4 | 0.38 |
| 5 | 14984116 | rs835069 | 0.507 | 0.493 | 0.466 | 0.636 | 0.007 | 0.027 | 0.027 | 0.525 | 0.83 | 0.59 | 0.88 | 0.82 | 0.85 | 0.84 |
| 5 | 14989823 | rs2402022 | -0.849 | 0.275 | 0.52 | 0.914 | -0.012 | 0.011 | 0.011 | 0 | 0.35 | 0.41 | 0.2 | 0.43 | 0.26 | 0.25 |
| 5 | 15001929 | rs9312876 | -0.623 | -0.096 | 0.234 | 0.913 | 0.021 | 0.009 | 0.009 | 3.547 | 0.27 | 0.18 | 0.14 | 0.2 | 0.2 | 0.14 |
| 5 | 15007802 | rs6867275 | -0.909 | -0.172 | 0.356 | 0.897 | 0.015 | 0.009 | 0.009 | 2.399 | 0.29 | 0.26 | 0.15 | 0.21 | 0.2 | 0.14 |
| 5 | 15022112 | rs7727670 | -0.701 | -0.457 | 0.961 | 1.02 | 0.031 | -0.009 | -0.009 | 0 | 0.27 | 0.19 | 0.14 | 0.25 | 0.18 | 0.15 |
| 5 | 15024160 | rs9688328 | -1.062 | -0.491 | 1.312 | 1.005 | -0.049 | -0.01 | -0.01 | 0 | 0.31 | 0.49 | 0.17 | 0.25 | 0.19 | 0.15 |
| 5 | 15024237 | rs9687159 | -0.945 | -0.484 | 1.347 | 1.008 | 0.006 | 0 | 0 | 3.148 | 0.29 | 0.32 | 0.16 | 0.25 | 0.19 | 0.15 |
| 5 | 15039327 | rs735243 | -0.82 | -0.436 | 1.311 | 0.988 | -0.008 | -0.001 | -0.001 | 0 | 0.28 | 0.31 | 0.15 | 0.23 | 0.17 | 0.1 |
| 5 | 15044842 | rs30539 | -1.022 | -0.253 | 1.316 | 0.942 | -0.017 | -0.01 | -0.01 | 0.502 | 0.31 | 0.42 | 0.17 | 0.23 | 0.21 | 0.17 |
| 5 | 15081127 | rs16903801 | 0.078 | -0.558 | 0.987 | 0.564 | 0.018 | 0.017 | 0.017 | 4.081 | 0.11 | 0.11 | 0.05 | 0 | 0.06 | 0.01 |
| 5 | 15082673 | rs2121183 | -1.099 | -0.58 | 1.104 | 0.66 | -0.023 | -0.014 | -0.014 | 0 | 0.16 | 0.34 | 0.1 | 0.08 | 0.11 | 0.11 |
| 5 | 15087791 | rs32466 | -1.573 | -0.561 | 0.885 | 0.611 | -0.006 | -0.074 | -0.074 | 0 | 0.29 | 0.32 | 0.14 | 0 | 0.18 | 0.11 |
| 5 | 15117438 | rs32447 | -1.121 | -1.027 | -0.711 | 0.494 | -0.08 | -0.044 | -0.044 | 0.215 | 0.66 | 0.9 | 0.37 | 0.66 | 0.44 | 0.66 |
| 5 | 15118799 | rs32443 | 0.268 | -1.061 | -0.612 | 0.47 | 0.008 | 0.006 | 0.006 | 0 | 0.16 | 0.16 | 0.07 | 0.08 | 0.11 | 0.08 |
| 5 | 15123737 | rs1446037 | 0.838 | -1.138 | -0.576 | 0.494 | 0.014 | 0.011 | 0.011 | 3.786 | 0.1 | 0.06 | 0.03 | 0 | 0.04 | 0.01 |
| 5 | 15124021 | rs1446040 | -1.369 | -0.062 | -1.039 | 0.231 | -0.114 | -0.105 | -0.105 | 0.227 | 0.74 | 0.97 | 0.49 | 0.67 | 0.64 | 0.73 |
| 5 | 15130618 | rs251544 | -0.877 | 0.189 | -0.629 | -0.202 | -0.011 | 0.029 | 0.029 | 1.704 | 0.36 | 0.32 | 0.24 | 0.09 | 0.37 | 0.2 |
| 5 | 15143511 | rs2455489 | -0.098 | -0.21 | -0.813 | -0.328 | -0.038 | 0.094 | 0.094 | 0 | 0.27 | 0.14 | 0.17 | 0 | 0.28 | 0.13 |
| 5 | 15151678 | rs2455488 | -0.914 | -0.207 | -0.429 | -0.285 | -0.003 | -0.042 | -0.042 | 0.137 | 0.4 | 0.46 | 0.25 | 0.09 | 0.34 | 0.18 |
| 5 | 15155515 | rs301076 | -0.517 | -0.274 | -0.924 | -0.171 | -0.068 | 0.194 | 0.194 | 0 | 0.35 | 0.11 | 0.25 | 0.07 | 0.39 | 0.17 |
| 5 | 15162646 | rs1500232 | -0.809 | -0.681 | -0.877 | -0.355 | -0.01 | 0.12 | 0.12 | 0 | 0.42 | 0.24 | 0.28 | 0.17 | 0.44 | 0.26 |
| 5 | 15168945 | rs7721868 | -0.462 | -0.505 | -0.779 | -0.202 | -0.029 | 0.089 | 0.089 | 0 | 0.36 | 0.19 | 0.25 | 0.17 | 0.4 | 0.2 |
| 5 | 15194140 | rs7723367 | -0.592 | -0.72 | -0.592 | -0.319 | 0.001 | 0.052 | 0.052 | 0 | 0.42 | 0.34 | 0.29 | 0.17 | 0.43 | 0.21 |
| 5 | 15211107 | rs2166391 | -0.498 | -0.693 | -0.756 | -0.257 | -0.005 | 0.084 | 0.084 | 0 | 0.41 | 0.27 | 0.29 | 0.18 | 0.43 | 0.21 |
| 5 | 15235092 | rs6554873 | -0.583 | -0.771 | -0.607 | -0.294 | 0.001 | 0.07 | 0.07 | 0 | 0.42 | 0.32 | 0.29 | 0.18 | 0.43 | 0.21 |
| 5 | 15251079 | rs434412 | -0.583 | -0.771 | -0.607 | -0.294 | 0.001 | 0.07 | 0.07 | 0 | 0.42 | 0.32 | 0.29 | 0.18 | 0.43 | 0.21 |
| 5 | 15278843 | rs7707492 | -0.405 | -0.804 | -0.488 | -0.308 | -0.011 | 0.089 | 0.089 | 0 | 0.4 | 0.24 | 0.28 | 0.18 | 0.42 | 0.21 |
| 5 | 15298906 | rs17703484 | 0.353 | -1.32 | -0.116 | 0.288 | 0.029 | -0.011 | -0.011 | 0 | 0.28 | 0.15 | 0.11 | 0.33 | 0.15 | 0.36 |
| 5 | 15313022 | rs303345 | -0.344 | -1.136 | 0.068 | 0.228 | 0 | -0.004 | -0.004 | 0 | 0.42 | 0.41 | 0.31 | 0.18 | 0.43 | 0.22 |
| 5 | 15329810 | rs11954098 | 0.744 | -0.877 | 0.109 | 0.638 | -0.022 | -0.011 | -0.011 | 0 | 0.06 | 0.01 | 0.08 | 0.07 | 0.11 | 0.04 |
| 5 | 15364484 | rs1447276 | 0.059 | -1.21 | -0.453 | 0.768 | 0.008 | -0.041 | -0.041 | 0 | 0.26 | 0.13 | 0.17 | 0.44 | 0.22 | 0.34 |
| 5 | 15366156 | rs303352 | 0.048 | -1.197 | -1.121 | 0.204 | -0.015 | 0.179 | 0.179 | 0 | 0.35 | 0.12 | 0.2 | 0.06 | 0.34 | 0.19 |
| 5 | 15384680 | rs10065220 | -0.316 | -0.979 | -1.203 | 0.493 | -0.02 | 0.193 | 0.193 | 0 | 0.42 | 0.13 | 0.22 | 0.16 | 0.37 | 0.4 |
| 5 | 15418957 | rs188592 | -1.121 | -0.902 | -0.377 | 0.356 | -0.026 | -0.028 | -0.028 | 0 | 0.94 | 0.81 | 0.98 | 1 | 0.99 | 0.97 |
| 5 | 15432384 | rs304562 | -1.121 | -0.902 | -0.303 | 0.356 | -0.01 | -0.012 | -0.012 | 0 | 0.94 | 0.76 | 0.97 | 1 | 0.99 | 0.97 |
| 5 | 15447939 | rs304586 | 1.063 | -0.63 | -1.426 | 0.115 | -0.095 | 0.084 | 0.084 | 0.375 | 0.46 | 0.11 | 0.6 | 0.72 | 0.53 | 0.56 |
| 5 | 15530640 | rs10462596 | -0.425 | -0.016 | -0.795 | 0.417 | -0.062 | -0.015 | -0.015 | 1.421 | 0.27 | 0.14 | 0.18 | 0.24 | 0.34 | 0.2 |
| 5 | 15548576 | rs581957 | -0.139 | -0.105 | -0.314 | 0.025 | -0.01 | -0.058 | -0.058 | 0.016 | 0.12 | 0.19 | 0.04 | 0 | 0.07 | 0.05 |
| 5 | 15550687 | rs4144522 | -1.041 | -0.073 | -0.151 | -0.129 | 0 | -0.012 | -0.012 | 1.45 | 0.34 | 0.56 | 0.26 | 0.2 | 0.34 | 0.22 |
| 5 | 15551178 | rs496319 | 0.432 | -0.144 | -0.204 | -0.131 | -0.019 | -0.018 | -0.018 | 0 | 0.08 | 0.13 | 0.08 | 0.05 | 0.07 | 0.06 |
| 5 | 15555486 | rs983889 | -1.473 | -0.187 | -0.823 | -0.143 | -0.015 | 0.029 | 0.029 | 0 | 0.54 | 0.76 | 0.42 | 0.38 | 0.48 | 0.51 |
| 5 | 15567629 | rs969351 | -1.255 | -0.204 | -0.558 | -0.167 | 0.003 | 0.038 | 0.038 | 0 | 0.52 | 0.65 | 0.4 | 0.38 | 0.49 | 0.5 |
| 5 | 15571736 | rs1429036 | 2.23 | 0.01 | -1.198 | -0.338 | -0.06 | 0.154 | 0.154 | 0.019 | 0.33 | 0.08 | 0.39 | 0.09 | 0.35 | 0.36 |
| 5 | 15576756 | rs9312894 | -1.238 | -0.211 | -0.821 | -0.192 | -0.003 | 0.036 | 0.036 | 0 | 0.53 | 0.74 | 0.42 | 0.39 | 0.49 | 0.5 |
| 5 | 15582005 | rs1978633 | -1.342 | -0.275 | -0.754 | -0.295 | -0.021 | 0.068 | 0.068 | 0 | 0.54 | 0.8 | 0.44 | 0.35 | 0.49 | 0.52 |
| 5 | 15586293 | rs7706439 | 0.792 | -0.312 | -0.365 | -0.485 | 0.002 | 0.002 | 0.002 | 0 | 0.17 | 0.19 | 0.11 | 0.18 | 0.21 | 0.15 |
| 5 | 15589171 | rs1429040 | -1.082 | -0.198 | -0.594 | -0.396 | 0.009 | 0.068 | 0.068 | 0 | 0.51 | 0.72 | 0.43 | 0.34 | 0.48 | 0.52 |
| 5 | 15597087 | rs1025327 | 0.446 | 0.356 | -0.005 | -0.048 | 0.015 | 0.049 | 0.049 | 0.561 | 0.74 | 0.72 | 0.8 | 0.57 | 0.71 | 0.82 |
| 5 | 15598629 | rs1019810 | 1.341 | 0.229 | 0.249 | -0.138 | 0.003 | 0.004 | 0.004 | 0.387 | 0.63 | 0.46 | 0.68 | 0.47 | 0.62 | 0.67 |
| 5 | 15601489 | rs6891138 | 0.941 | 0.148 | 0.348 | -0.044 | 0.008 | 0.05 | 0.05 | 0 | 0.68 | 0.52 | 0.64 | 0.49 | 0.67 | 0.69 |
| 5 | 15602644 | rs13170478 | -0.918 | 0.152 | 0.415 | -0.068 | 0.003 | -0.023 | -0.023 | 0 | 0.29 | 0.32 | 0.33 | 0.15 | 0.32 | 0.27 |
| 5 | 15604978 | rs4510586 | -0.1 | 0.173 | 0.245 | 0.092 | 0.003 | 0.16 | 0.16 | 0 | 0.88 | 0.79 | 0.87 | 0.54 | 0.89 | 0.8 |
| 5 | 15613657 | rs16867529 | -0.146 | 0.19 | 0.176 | 0.012 | 0.002 | -0.007 | -0.007 | 0.351 | 0.11 | 0.17 | 0.05 | 0.09 | 0.1 | 0.09 |
| 5 | 15618181 | rs12659622 | 0.994 | 0.217 | 0.069 | 0.048 | -0.015 | -0.015 | -0.015 | 0.008 | 0.08 | 0.04 | 0.1 | 0.09 | 0.1 | 0.16 |
| 5 | 15632099 | rs7704791 | -2.026 | 0.196 | 0.134 | -0.071 | -0.016 | 0.005 | 0.005 | 0.83 | 0.43 | 0.68 | 0.4 | 0.58 | 0.42 | 0.38 |
| 5 | 15633377 | rs17601702 | 0.976 | 0.2 | -0.164 | -0.268 | -0.004 | 0.016 | 0.016 | 0 | 0.1 | 0.06 | 0.06 | 0.03 | 0.13 | 0.05 |
| 5 | 15635087 | rs7720476 | -0.528 | 0.322 | -0.015 | -0.099 | 0.008 | -0.023 | -0.023 | 0 | 0.24 | 0.26 | 0.25 | 0.09 | 0.28 | 0.24 |
| 5 | 15639915 | rs4587075 | 0.029 | 0.387 | 0.014 | -0.412 | 0.006 | -0.034 | -0.034 | 0 | 0.19 | 0.18 | 0.22 | 0.05 | 0.22 | 0.11 |
| 5 | 15646811 | rs6878131 | -1.025 | 0.334 | 0.039 | -0.344 | 0.015 | 0.063 | 0.063 | 0 | 0.3 | 0.47 | 0.34 | 0.49 | 0.32 | 0.3 |
| 5 | 15647084 | rs6878966 | 0.658 | 0.347 | 0.063 | -0.364 | 0.001 | -0.032 | -0.032 | 0 | 0.14 | 0.11 | 0.14 | 0 | 0.19 | 0.08 |
| 5 | 15648296 | rs6898779 | -0.949 | 0.315 | 0.082 | -0.337 | 0.019 | 0.075 | 0.075 | 0 | 0.3 | 0.44 | 0.34 | 0.49 | 0.32 | 0.3 |
| 5 | 15650140 | rs17703964 | -0.316 | 0.276 | -0.133 | -0.307 | 0.019 | 0.111 | 0.111 | 0 | 0.14 | 0.24 | 0.16 | 0.43 | 0.17 | 0.19 |
| 5 | 15654479 | rs1127792 | 0.042 | 0.255 | -0.254 | -0.32 | 0.001 | -0.009 | -0.009 | 0.075 | 0.11 | 0.15 | 0.07 | 0.08 | 0.1 | 0.08 |
| 5 | 15655056 | rs13185548 | -0.688 | 0.26 | -0.359 | -0.284 | 0.011 | 0.051 | 0.051 | 0 | 0.31 | 0.42 | 0.35 | 0.49 | 0.33 | 0.3 |
| 5 | 15659551 | rs17647599 | -0.703 | 0.271 | -0.451 | -0.283 | 0.014 | 0.064 | 0.064 | 0.03 | 0.31 | 0.45 | 0.35 | 0.49 | 0.33 | 0.3 |
| 5 | 15674635 | rs12652447 | -1.342 | 0.13 | -0.984 | -0.339 | -0.001 | 0.012 | 0.012 | 1.286 | 0.46 | 0.61 | 0.41 | 0.58 | 0.43 | 0.39 |

| | | | | | | | | | | | | | | | | |
|---|---|---|---|---|---|---|---|---|---|---|---|---|---|---|---|---|
| 5 | 15678556 | rs34341354 | -1.084 | -0.48 | -1.146 | -0.643 | 0 | -0.001 | -0.001 | 0 | 0.43 | 0.58 | 0.35 | 0.55 | 0.39 | 0.38 |
| 5 | 15685126 | rs12655356 | 1.414 | -0.23 | -1.229 | -0.705 | -0.001 | -0.004 | -0.004 | 0 | 0.52 | 0.44 | 0.53 | 0.66 | 0.55 | 0.55 |
| 5 | 15690455 | rs17704215 | 2.2 | -0.353 | -1.705 | -0.78 | -0.099 | -0.05 | -0.05 | 0 | 0.22 | 0.04 | 0.33 | 0.25 | 0.31 | 0.39 |
| 5 | 15694620 | rs4702100 | -0.15 | -0.472 | -1.532 | -0.826 | -0.016 | -0.028 | -0.028 | 0 | 0.75 | 0.63 | 0.76 | 0.84 | 0.76 | 0.8 |
| 5 | 15696499 | rs17523908 | 1.401 | -0.445 | -1.603 | -0.864 | -0.031 | 0.028 | 0.028 | 0 | 0.12 | 0.02 | 0.1 | 0.06 | 0.11 | 0.17 |
| 5 | 15697062 | rs4701641 | 0.199 | -0.518 | -1.597 | -0.882 | -0.011 | -0.023 | -0.023 | 0 | 0.72 | 0.6 | 0.73 | 0.79 | 0.72 | 0.81 |
| 5 | 15702469 | rs7733676 | 0.574 | -0.427 | -1.598 | -0.797 | -0.002 | 0.027 | 0.027 | 0 | 0.15 | 0.07 | 0.19 | 0.06 | 0.13 | 0.19 |
| 5 | 15707132 | rs17602350 | -0.26 | -0.409 | -1.693 | -0.781 | -0.019 | 0.033 | 0.033 | 0 | 0.22 | 0.06 | 0.21 | 0.15 | 0.19 | 0.26 |
| 5 | 15713072 | rs7704277 | 0.96 | -0.371 | -1.527 | -0.718 | -0.004 | -0.009 | -0.009 | 0 | 0.58 | 0.48 | 0.58 | 0.67 | 0.62 | 0.57 |
| 5 | 15719350 | rs4374757 | 2.267 | -0.187 | -1.679 | -0.635 | -0.093 | -0.047 | -0.047 | 0 | 0.21 | 0.02 | 0.32 | 0.24 | 0.31 | 0.4 |
| 5 | 15723633 | rs11748790 | 0.424 | -0.331 | -1.498 | -0.648 | -0.014 | -0.017 | -0.017 | 0 | 0.67 | 0.52 | 0.71 | 0.73 | 0.68 | 0.74 |
| 5 | 15728229 | rs4133004 | 0.993 | -0.34 | -1.503 | -0.661 | 0.002 | 0.005 | 0.005 | 0 | 0.55 | 0.49 | 0.56 | 0.66 | 0.57 | 0.56 |
| 5 | 15734275 | rs6877517 | 1.288 | -0.315 | -1.507 | -0.636 | -0.025 | 0.03 | 0.03 | 0 | 0.12 | 0.02 | 0.15 | 0.06 | 0.11 | 0.17 |
| 5 | 15742599 | rs4293917 | -0.764 | -0.451 | -1.443 | -0.661 | -0.003 | -0.029 | -0.029 | 0 | 0.79 | 0.58 | 0.77 | 0.88 | 0.78 | 0.83 |
| 5 | 15747814 | rs2937075 | 0.439 | -0.468 | -1.451 | -0.701 | -0.018 | -0.018 | -0.018 | 0 | 0.67 | 0.52 | 0.71 | 0.73 | 0.69 | 0.74 |
| 5 | 15747866 | rs924455 | 1.004 | -0.313 | -1.462 | -0.718 | -0.003 | -0.012 | -0.012 | 0.107 | 0.55 | 0.45 | 0.54 | 0.66 | 0.56 | 0.56 |
| 5 | 15751645 | rs2937074 | 0.439 | -0.34 | -1.537 | -0.709 | -0.028 | -0.026 | -0.026 | 0 | 0.67 | 0.47 | 0.7 | 0.73 | 0.7 | 0.73 |
| 5 | 15752604 | rs2937076 | 0.399 | -0.001 | -2.273 | -0.609 | -0.021 | 0.037 | 0.037 | 0 | 0.39 | 0.21 | 0.53 | 0.3 | 0.51 | 0.58 |
| 5 | 15753722 | rs2133761 | 1.43 | 0.137 | -2.291 | -0.562 | -0.066 | 0.003 | 0.003 | 0 | 0.26 | 0.09 | 0.36 | 0.23 | 0.37 | 0.4 |
| 5 | 15766072 | rs10520823 | 0.171 | 0.137 | -1.413 | -0.277 | -0.002 | -0.022 | -0.022 | 0 | 0.07 | 0.03 | 0.04 | 0.11 | 0.07 | 0.07 |
| 5 | 15770553 | rs2937072 | 2.067 | -0.005 | -1.535 | -0.394 | -0.08 | 0.07 | 0.07 | 0.399 | 0.46 | 0.07 | 0.48 | 0.65 | 0.5 | 0.57 |
| 5 | 15771101 | rs10462723 | -1.196 | 0.124 | -0.806 | 0.074 | -0.007 | 0.009 | 0.009 | 0 | 0.2 | 0.17 | 0.23 | 0.18 | 0.2 | 0.24 |
| 5 | 15777817 | rs2055624 | -0.02 | 0.095 | -0.802 | 0.089 | -0.006 | 0.018 | 0.018 | 0 | 0.12 | 0.09 | 0.16 | 0.07 | 0.12 | 0.17 |
| 5 | 15779815 | rs1505039 | -1.823 | 0.052 | -0.76 | 0.089 | 0.004 | 0.003 | 0.003 | 0.744 | 0.24 | 0.21 | 0.22 | 0.25 | 0.21 | 0.26 |
| 5 | 15781244 | rs11953097 | -1.603 | -0.028 | -0.599 | 0.17 | -0.02 | 0.007 | 0.007 | 0 | 0.13 | 0.18 | 0.07 | 0.18 | 0.1 | 0.09 |
| 5 | 15783500 | rs16867713 | -0.16 | 0.074 | -0.612 | 0.192 | -0.023 | 0.004 | 0.004 | 0 | 0.05 | 0.13 | 0.02 | 0.07 | 0.03 | 0.04 |
| 5 | 15789739 | rs16867730 | -1.275 | 0.225 | -0.904 | 0.134 | -0.007 | 0.015 | 0.015 | 0.191 | 0.28 | 0.17 | 0.25 | 0.27 | 0.29 | 0.27 |
| 5 | 15808138 | rs416952 | 1.434 | -0.351 | -0.416 | -0.111 | -0.007 | -0.009 | -0.009 | 0.314 | 0.54 | 0.46 | 0.38 | 0.61 | 0.61 | 0.59 |
| 5 | 15819269 | rs10050940 | -0.857 | -0.613 | -0.184 | -0.208 | -0.02 | -0.008 | -0.008 | 0 | 0.24 | 0.36 | 0.42 | 0.22 | 0.19 | 0.14 |
| 5 | 15821793 | rs183759 | -0.745 | -0.428 | -0.04 | -0.276 | -0.009 | 0.031 | 0.031 | 0 | 0.89 | 0.98 | 0.86 | 0.9 | 0.89 | 0.76 |
| 5 | 15823365 | rs257752 | 1.791 | -0.446 | 0.103 | -0.237 | 0.02 | 0.03 | 0.03 | 0 | 0.53 | 0.32 | 0.37 | 0.62 | 0.6 | 0.58 |
| 5 | 15832573 | rs13158449 | -1.044 | -0.547 | 0.002 | 0.108 | -0.012 | 0.101 | 0.101 | 0.121 | 0.27 | 0.44 | 0.25 | 0.49 | 0.26 | 0.32 |
| 5 | 15843597 | rs257768 | 1.866 | -0.496 | 0.504 | -0.346 | -0.002 | -0.003 | -0.003 | 0.064 | 0.54 | 0.49 | 0.38 | 0.62 | 0.6 | 0.57 |
| 5 | 15850418 | rs257744 | 1.997 | -0.481 | 0.53 | -0.339 | 0.021 | 0.036 | 0.036 | 0 | 0.53 | 0.31 | 0.37 | 0.62 | 0.6 | 0.57 |
| 5 | 15853939 | rs7723811 | -1.371 | -0.5 | 0.598 | -0.383 | -0.012 | -0.03 | -0.03 | 0 | 0.33 | 0.44 | 0.47 | 0.18 | 0.26 | 0.19 |
| 5 | 15900124 | rs7715223 | -0.774 | -0.291 | 0.511 | -0.318 | 0.002 | -0.01 | -0.01 | 0 | 0.21 | 0.24 | 0.24 | 0.08 | 0.18 | 0.11 |
| 5 | 15904024 | rs6897647 | 0.934 | -0.121 | 0.724 | -0.274 | -0.022 | 0.009 | 0.009 | 0 | 0.83 | 0.52 | 0.82 | 0.81 | 0.86 | 0.76 |
| 5 | 15913112 | rs6878591 | 1.962 | -0.085 | 0.65 | -0.242 | -0.004 | -0.009 | -0.009 | 0 | 0.58 | 0.44 | 0.4 | 0.67 | 0.65 | 0.59 |
| 5 | 15913562 | rs6883363 | -0.651 | -0.012 | 0.158 | -0.308 | -0.001 | -0.006 | -0.006 | 0.112 | 0.19 | 0.24 | 0.13 | 0.1 | 0.16 | 0.1 |
| 5 | 15919799 | rs13153362 | -1.323 | 0.051 | 0.343 | -0.16 | 0.004 | 0.026 | 0.026 | 0 | 0.39 | 0.32 | 0.57 | 0.32 | 0.35 | 0.38 |
| 5 | 15921693 | rs6892388 | -1.358 | 0.066 | 0.283 | -0.153 | 0.004 | 0.023 | 0.023 | 0 | 0.4 | 0.33 | 0.57 | 0.34 | 0.35 | 0.39 |
| 5 | 15924697 | rs3935971 | -0.393 | 0.1 | 0.135 | -0.13 | 0.017 | 0.063 | 0.063 | 0.083 | 0.11 | 0.45 | 0.25 | 0.12 | 0.07 | 0.09 |
| 5 | 15925524 | rs12517809 | -1.7 | 0.099 | 0.147 | -0.143 | 0.017 | 0.041 | 0.041 | 0 | 0.44 | 0.71 | 0.61 | 0.33 | 0.35 | 0.41 |
| 5 | 15925573 | rs13185525 | -1.046 | 0.531 | -0.471 | -0.082 | 0.004 | 0.026 | 0.026 | 0 | 0.32 | 0.26 | 0.27 | 0.21 | 0.28 | 0.32 |
| 5 | 15937910 | rs6554906 | -0.928 | 0.965 | 0.03 | 0.053 | -0.004 | -0.08 | -0.08 | 0 | 0.3 | 0.54 | 0.28 | 0.14 | 0.25 | 0.23 |
| 5 | 15938553 | rs3810847 | -0.991 | 0.992 | 0.138 | 0.066 | 0 | -0.043 | -0.043 | 0 | 0.29 | 0.41 | 0.26 | 0.14 | 0.24 | 0.21 |
| 5 | 15940600 | rs7718315 | -1.031 | 0.74 | 0.372 | 0.216 | -0.008 | -0.041 | -0.041 | 0 | 0.08 | 0.2 | 0.05 | 0 | 0.04 | 0.06 |
| 5 | 15943683 | rs982484 | -1.253 | 0.889 | 0.189 | 0.698 | -0.005 | 0.092 | 0.092 | 0 | 0.26 | 0.5 | 0.38 | 0.56 | 0.28 | 0.26 |
| 5 | 15949793 | rs1354182 | -1.534 | 1.527 | 0.069 | 0.547 | 0.002 | -0.003 | -0.003 | 0.13 | 0.43 | 0.53 | 0.5 | 0.66 | 0.51 | 0.39 |
| 5 | 15960831 | rs1502024 | -0.913 | 1.064 | 0.441 | 0.257 | 0.034 | -0.002 | -0.002 | 0 | 0.07 | 0.31 | 0.06 | 0.01 | 0.04 | 0.09 |
| 5 | 15965000 | rs7735201 | -1.129 | 1.041 | 0.454 | 0.251 | 0.044 | 0.009 | 0.009 | 0.72 | 0.07 | 0.37 | 0.06 | 0.01 | 0.03 | 0.09 |
| 5 | 15968662 | rs7727759 | -0.24 | 1.234 | 0.213 | 0.412 | -0.027 | 0 | 0 | 0 | 0.21 | 0.1 | 0.26 | 0.15 | 0.22 | 0.21 |
| 5 | 15973231 | rs250488 | -1.897 | 1.262 | 0.464 | 0.423 | -0.005 | 0.07 | 0.07 | 0 | 0.47 | 0.65 | 0.66 | 0.8 | 0.49 | 0.51 |
| 5 | 15992984 | rs32826 | 0.167 | 1 | 0.481 | 0.487 | -0.004 | 0.009 | 0.009 | 0 | 0.11 | 0.07 | 0.27 | 0.64 | 0.17 | 0.16 |
| 5 | 15993449 | rs32824 | 0.124 | 1 | 0.481 | 0.429 | -0.004 | 0.002 | 0.002 | 0 | 0.11 | 0.07 | 0.27 | 0.67 | 0.18 | 0.17 |
| 5 | 15998273 | rs1393268 | -1.104 | 0.939 | 0.584 | 0.373 | -0.002 | -0.06 | -0.06 | 0 | 0.1 | 0.35 | 0.08 | 0 | 0.06 | 0.17 |
| 5 | 16000639 | rs250497 | -1.076 | 0.971 | 0.52 | 0.427 | 0.003 | 0.04 | 0.04 | 0 | 0.19 | 0.25 | 0.31 | 0.66 | 0.23 | 0.19 |
| 5 | 16010717 | rs168747 | -1.107 | 0.923 | 0.52 | 0.42 | 0.001 | 0.037 | 0.037 | 0 | 0.19 | 0.25 | 0.31 | 0.67 | 0.23 | 0.19 |
| 5 | 16016736 | rs1502027 | -1.878 | 0.722 | 0.361 | 0.489 | -0.01 | 0.013 | 0.013 | 0.264 | 0.27 | 0.64 | 0.38 | 0.68 | 0.25 | 0.35 |
| 5 | 16017401 | rs10520833 | 0.678 | 0.801 | 0.266 | 0.534 | -0.055 | -0.052 | -0.052 | 2.612 | 0.15 | 0.01 | 0.24 | 0.14 | 0.18 | 0.11 |
| 5 | 16021223 | rs11955061 | 0.472 | 0.863 | 0.285 | 0.541 | -0.032 | -0.027 | -0.027 | 0.255 | 0.17 | 0.04 | 0.25 | 0.14 | 0.18 | 0.12 |
| 5 | 16022694 | rs1393266 | -1.855 | 0.769 | 0.346 | 0.589 | -0.01 | 0.017 | 0.017 | 0.275 | 0.28 | 0.62 | 0.39 | 0.65 | 0.26 | 0.34 |
| 5 | 16024011 | rs16867959 | -0.416 | 0.887 | 0.029 | 0.464 | -0.002 | -0.02 | -0.02 | 0 | 0.07 | 0.09 | 0.04 | 0 | 0.04 | 0.14 |
| 5 | 16027376 | rs1566654 | -0.267 | 0.824 | -0.133 | 0.57 | -0.003 | 0.008 | 0.008 | 0 | 0.27 | 0.09 | 0.28 | 0.17 | 0.23 | 0.25 |
| 5 | 16027427 | rs974847 | -1.525 | 0.795 | -0.042 | 0.751 | -0.001 | -0.007 | -0.007 | 0.221 | 0.46 | 0.53 | 0.63 | 0.73 | 0.45 | 0.46 |
| 5 | 16030905 | rs4701647 | -1.615 | 0.736 | -0.131 | 1.522 | 0 | 0.098 | 0.098 | 0 | 0.19 | 0.3 | 0.32 | 0.47 | 0.23 | 0.21 |
| 5 | 16034119 | rs7707378 | -0.484 | 0.824 | -0.357 | 1.307 | -0.004 | -0.032 | -0.032 | 1.615 | 0.06 | 0.19 | 0.05 | 0 | 0.04 | 0.03 |
| 5 | 16034232 | rs6866525 | -0.845 | 0.963 | -0.999 | 1.723 | -0.008 | -0.018 | -0.018 | 0.002 | 0.13 | 0.07 | 0.28 | 0.44 | 0.19 | 0.17 |
| 5 | 16035369 | rs6876369 | -0.288 | 0.816 | -1.615 | 1.015 | -0.001 | -0.016 | -0.016 | 0.122 | 0.48 | 0.8 | 0.42 | 0.59 | 0.43 | 0.37 |

| | | | | | | | | | | | | | | | | |
|---|---|---|---|---|---|---|---|---|---|---|---|---|---|---|---|---|
| 5 | 16037234 | rs32199 | -0.103 | 0.781 | -1.001 | 0.83 | 0 | -0.001 | -0.001 | 0.484 | 0.55 | 0.55 | 0.58 | 0.56 | 0.48 | 0.46 |
| 5 | 16038102 | rs1039443 | -0.292 | -0.042 | -0.213 | 0.546 | -0.029 | -0.051 | -0.051 | 0 | 0.21 | 0.38 | 0.1 | 0.14 | 0.12 | 0.19 |
| 5 | 16044383 | rs1823075 | 0.182 | -0.013 | -0.779 | 0.599 | -0.012 | -0.069 | -0.069 | 0.001 | 0.3 | 0.15 | 0.54 | 0.58 | 0.35 | 0.51 |
| 5 | 16048639 | rs11133827 | -0.725 | -0.548 | 0.187 | 0.253 | 0.004 | -0.004 | -0.004 | 0 | 0.42 | 0.65 | 0.62 | 0.6 | 0.38 | 0.62 |
| 5 | 16054101 | rs7701823 | -0.487 | -0.47 | 0.589 | 0.017 | -0.001 | -0.029 | -0.029 | 0 | 0.07 | 0.23 | 0.03 | 0 | 0.02 | 0.08 |
| 5 | 16081670 | rs1992719 | 0.138 | -0.554 | 0.42 | 0.293 | 0 | -0.002 | -0.002 | 0 | 0.32 | 0.27 | 0.55 | 0.63 | 0.35 | 0.52 |
| 5 | 16088262 | rs7736562 | 0.372 | -0.673 | 0.297 | 0.514 | -0.003 | -0.05 | -0.05 | 0 | 0.3 | 0.15 | 0.5 | 0.59 | 0.34 | 0.52 |
| 5 | 16088655 | rs9312902 | -1.5 | -0.889 | -0.095 | 0.304 | -0.085 | 0.061 | 0.061 | 1.761 | 0.61 | 0.91 | 0.69 | 0.78 | 0.47 | 0.78 |
| 5 | 16103046 | rs10475072 | -0.164 | -0.616 | 0.644 | 0.583 | -0.017 | 0.029 | 0.029 | 0 | 0.34 | 0.49 | 0.53 | 0.57 | 0.35 | 0.52 |
| 5 | 16110431 | rs6554912 | -0.469 | -0.798 | 0.646 | 0.49 | 0.036 | 0.015 | 0.015 | 0 | 0.3 | 0.09 | 0.28 | 0.28 | 0.23 | 0.32 |
| 5 | 16128234 | rs17649190 | -0.814 | -0.195 | 0.682 | 0.623 | 0.015 | 0.071 | 0.071 | 0 | 0.12 | 0.19 | 0.13 | 0.31 | 0.19 | 0.27 |
| 5 | 16142399 | rs344701 | 1.484 | -0.059 | 0.633 | 0.55 | 0.018 | 0.028 | 0.028 | 0 | 0.51 | 0.32 | 0.54 | 0.4 | 0.56 | 0.38 |
| 5 | 16158739 | rs344734 | 0.537 | -0.208 | 0.753 | 0.074 | -0.01 | 0.013 | 0.013 | 0.476 | 0.78 | 0.62 | 0.71 | 0.72 | 0.8 | 0.69 |
| 5 | 16182201 | rs7707491 | 0.501 | -0.1 | 0.378 | -0.554 | -0.001 | 0.001 | 0.001 | 0 | 0.13 | 0.12 | 0.13 | 0.05 | 0.14 | 0.23 |
| 5 | 16182788 | rs237065 | 1.538 | -0.546 | 0.409 | -0.988 | 0.014 | -0.004 | -0.004 | 0 | 0.56 | 0.36 | 0.72 | 0.91 | 0.7 | 0.54 |
| 5 | 16184180 | rs342559 | 0.994 | -0.462 | 0.718 | -0.348 | 0.021 | 0.025 | 0.025 | 0 | 0.73 | 0.81 | 0.89 | 0.95 | 0.84 | 0.77 |
| 5 | 16195381 | rs344739 | 0.757 | -0.255 | 0.862 | -0.244 | -0.011 | -0.024 | -0.024 | 0 | 0.88 | 0.87 | 0.95 | 0.96 | 0.94 | 0.84 |
| 5 | 16205201 | rs344720 | 0.78 | -0.247 | 0.861 | -0.232 | -0.008 | -0.021 | -0.021 | 0 | 0.88 | 0.87 | 0.94 | 0.96 | 0.92 | 0.83 |
| 5 | 16217387 | rs17611228 | 1.927 | -0.09 | 1.279 | -0.465 | 0.021 | 0.186 | 0.186 | 0 | 0.51 | 0.28 | 0.42 | 0.16 | 0.54 | 0.35 |
| 5 | 16221678 | rs4368728 | 0.564 | 0.009 | 1.304 | -0.333 | -0.004 | -0.021 | -0.021 | 0 | 0.87 | 0.87 | 0.93 | 0.96 | 0.91 | 0.83 |
| 5 | 16224737 | rs995765 | 0.469 | 0.164 | 1.292 | -0.326 | -0.002 | -0.016 | -0.016 | 0 | 0.88 | 0.87 | 0.93 | 0.96 | 0.89 | 0.83 |
| 5 | 16228197 | rs12652713 | 1.995 | -0.282 | 1.055 | -0.396 | 0.02 | 0.24 | 0.24 | 0 | 0.51 | 0.17 | 0.41 | 0.16 | 0.56 | 0.39 |
| 5 | 16237236 | rs4701655 | -0.077 | -0.137 | 1.336 | 0.304 | 0.017 | -0.051 | -0.051 | 0 | 0.21 | 0.1 | 0.3 | 0.59 | 0.16 | 0.24 |
| 5 | 16242706 | rs10038426 | -1.062 | -0.148 | 1.539 | 0.319 | -0.001 | 0.003 | 0.003 | 0 | 0.29 | 0.28 | 0.47 | 0.8 | 0.32 | 0.43 |
| 5 | 16245534 | rs12187958 | 2.269 | -0.263 | 1.39 | 0.215 | 0.019 | 0.245 | 0.245 | 0.047 | 0.5 | 0.11 | 0.41 | 0.16 | 0.57 | 0.37 |
| 5 | 16246505 | rs7706076 | 0.311 | -0.254 | 1.308 | 0.23 | -0.006 | 0.061 | 0.061 | 0 | 0.16 | 0.18 | 0.29 | 0.59 | 0.13 | 0.23 |
| 5 | 16253763 | rs7715586 | -1.121 | -0.197 | 1.288 | 0.099 | 0.009 | -0.032 | -0.032 | 0 | 0.3 | 0.42 | 0.48 | 0.8 | 0.34 | 0.43 |
| 5 | 16254310 | rs11133838 | 1.949 | -0.186 | 0.821 | 0.166 | 0.028 | 0.137 | 0.137 | 0 | 0.51 | 0.24 | 0.44 | 0.16 | 0.58 | 0.37 |
| 5 | 16261338 | rs17705925 | -0.093 | 0.289 | 0.818 | 0.693 | 0.001 | -0.05 | -0.05 | 0.274 | 0.17 | 0.07 | 0.21 | 0.58 | 0.15 | 0.22 |
| 5 | 16270044 | rs11739184 | 1.544 | 0.295 | 1.186 | 0.628 | 0.007 | -0.046 | -0.046 | 0 | 0.66 | 0.5 | 0.44 | 0.19 | 0.64 | 0.59 |
| 5 | 16278279 | rs2967940 | 0.239 | 0.464 | 0.892 | 0.801 | -0.009 | -0.012 | -0.012 | 0.26 | 0.87 | 0.9 | 0.68 | 0.75 | 0.8 | 0.82 |
| 5 | 16286973 | rs831657 | 0.813 | 0.464 | 1.317 | 0.698 | 0.043 | 0.085 | 0.085 | 0 | 0.85 | 0.6 | 0.76 | 0.79 | 0.83 | 0.83 |
| 5 | 16287578 | rs1371974 | -0.773 | 0.096 | 1.515 | 0.519 | -0.007 | -0.023 | -0.023 | 0.103 | 0.09 | 0.19 | 0.11 | 0.05 | 0.08 | 0.09 |
| 5 | 16289065 | rs6874136 | 1.319 | 0.108 | 1.731 | 0.706 | 0.008 | 0.042 | 0.042 | 0 | 0.82 | 0.43 | 0.7 | 0.75 | 0.81 | 0.81 |
| 5 | 16306370 | rs10052415 | -1.975 | 0.049 | 2.173 | 0.663 | 0.085 | 0.069 | 0.069 | 3.374 | 0.05 | 0.46 | 0.05 | 0 | 0.01 | 0.01 |
| 5 | 16346572 | rs12656374 | -1.593 | -0.202 | 1.711 | 0.495 | 0.009 | 0.027 | 0.027 | 0 | 0.37 | 0.68 | 0.39 | 0.83 | 0.29 | 0.43 |
| 5 | 16347660 | rs955205 | 0.13 | -0.126 | 1.091 | 0.661 | 0.015 | -0.06 | -0.06 | 0.458 | 0.18 | 0.06 | 0.09 | 0.58 | 0.1 | 0.24 |
| 5 | 16366460 | rs3849096 | -0.34 | 0.003 | 1.182 | 0.483 | 0 | 0.002 | 0.002 | 0 | 0.13 | 0.11 | 0.25 | 0.24 | 0.18 | 0.18 |
| 5 | 16366540 | rs16868502 | -0.851 | -0.009 | 1.366 | 0.606 | 0.025 | 0.04 | 0.04 | 0 | 0.15 | 0.29 | 0.27 | 0.23 | 0.18 | 0.18 |
| 5 | 16366662 | rs10065052 | -0.127 | -0.174 | 1.106 | 1.069 | -0.005 | 0.03 | 0.03 | 0 | 0.2 | 0.22 | 0.24 | 0.54 | 0.12 | 0.24 |
| 5 | 16367508 | rs16868503 | 0.407 | 0.077 | 0.969 | 0.139 | 0.012 | -0.056 | -0.056 | 0 | 0.1 | 0.19 | 0.06 | 0 | 0.1 | 0.06 |
| 5 | 16374042 | rs29965 | -1.406 | -0.104 | 0.574 | -0.275 | 0.002 | 0.06 | 0.06 | 1.562 | 0.9 | 1 | 0.95 | 1 | 0.93 | 0.97 |
| 5 | 16379041 | rs6885471 | -1.475 | 0.797 | 1.271 | 0.321 | 0.021 | -0.028 | -0.028 | 6.337 | 0.26 | 0.55 | 0.22 | 0.16 | 0.27 | 0.29 |
| 5 | 16379078 | rs17649734 | 0.425 | 0.841 | 0.981 | 0.333 | -0.018 | -0.027 | -0.027 | 0 | 0.09 | 0.05 | 0.08 | 0.16 | 0.13 | 0.19 |
| 5 | 16381553 | rs186758 | 0.031 | 0.736 | 0.859 | 0.572 | -0.016 | 0.044 | 0.044 | 0 | 0.23 | 0.29 | 0.28 | 0.49 | 0.17 | 0.27 |
| 5 | 16383215 | rs11133844 | 0.465 | 0.77 | 0.817 | 0.588 | 0.012 | -0.056 | -0.056 | 0 | 0.1 | 0.19 | 0.06 | 0 | 0.09 | 0.06 |
| 5 | 16384646 | rs831179 | -1.791 | 0.814 | 0.021 | 0.614 | -0.047 | 0.01 | 0.01 | 0 | 0.51 | 0.9 | 0.51 | 0.65 | 0.44 | 0.55 |
| 5 | 16385915 | rs17706248 | 0.302 | 0.777 | 0.045 | 0.681 | -0.006 | -0.006 | -0.006 | 0.252 | 0.1 | 0.11 | 0.1 | 0.15 | 0.15 | 0.19 |
| 5 | 16393546 | rs4702142 | 0.473 | 0.73 | -0.313 | 0.736 | 0.003 | -0.109 | -0.109 | 0.076 | 0.2 | 0.02 | 0.25 | 0.49 | 0.17 | 0.27 |
| 5 | 16408600 | rs32141 | -0.425 | 0.91 | -0.062 | 0.666 | -0.003 | -0.001 | -0.001 | 0.328 | 0.41 | 0.5 | 0.42 | 0.65 | 0.39 | 0.5 |
| 5 | 16410418 | rs12187814 | -1.329 | 1.153 | 0.649 | 0.614 | 0.021 | -0.042 | -0.042 | 0 | 0.09 | 0.34 | 0.08 | 0.01 | 0.04 | 0.04 |
| 5 | 16411810 | rs29948 | 0.805 | 1.607 | 0.114 | 0.974 | 0.035 | 0.02 | 0.02 | 0 | 0.88 | 0.83 | 0.88 | 0.84 | 0.82 | 0.8 |
| 5 | 16413014 | rs29949 | 0.795 | 1.092 | 0.137 | 1.18 | -0.001 | 0.004 | 0.004 | 0.092 | 0.66 | 0.3 | 0.68 | 0.45 | 0.7 | 0.5 |
| 5 | 16418829 | rs258879 | -0.007 | 1.043 | 0.345 | 1.021 | -0.006 | 0.016 | 0.016 | 0.513 | 0.18 | 0.23 | 0.18 | 0.26 | 0.2 | 0.32 |
| 5 | 16419249 | rs168934 | 0.475 | 1.375 | 0.409 | 0.818 | -0.004 | -0.032 | -0.032 | 0.221 | 0.64 | 0.24 | 0.63 | 0.45 | 0.66 | 0.49 |
| 5 | 16427050 | rs258881 | -0.157 | 1.419 | 0.474 | 1.236 | -0.012 | -0.035 | -0.035 | 0 | 0.76 | 0.93 | 0.77 | 0.67 | 0.75 | 0.66 |
| 5 | 16451188 | rs466205 | -0.058 | 0.984 | 0.061 | 0.7 | 0.002 | -0.005 | -0.005 | 0.248 | 0.28 | 0.23 | 0.4 | 0.44 | 0.26 | 0.39 |
| 5 | 16451246 | rs27454 | -0.52 | 0.991 | 0.409 | 0.681 | -0.003 | 0.007 | 0.007 | 0.392 | 0.75 | 0.71 | 0.76 | 0.67 | 0.75 | 0.66 |
| 5 | 16454953 | rs27883 | -0.213 | 0.938 | 0.79 | 0.574 | -0.026 | 0.054 | 0.054 | 0 | 0.3 | 0.41 | 0.43 | 0.44 | 0.26 | 0.39 |
| 5 | 16455126 | rs27452 | 0.224 | 0.904 | 0.893 | 0.543 | 0.003 | 0.025 | 0.025 | 0 | 0.68 | 0.61 | 0.67 | 0.44 | 0.68 | 0.5 |
| 5 | 16456131 | rs10520840 | 1.044 | 0.777 | 0.801 | 0.777 | 0.007 | -0.011 | -0.011 | 0 | 0.11 | 0.07 | 0.21 | 0.16 | 0.06 | 0.05 |
| 5 | 16469415 | rs13155194 | 1.246 | 1.021 | 0.536 | 0.02 | -0.02 | 0.047 | 0.047 | 0 | 0.08 | 0 | 0.05 | 0 | 0.09 | 0.03 |
| 5 | 16469593 | rs149511 | 0.013 | 1.02 | 0.815 | 0.233 | 0.013 | -0.037 | -0.037 | 0 | 0.28 | 0.2 | 0.4 | 0.44 | 0.26 | 0.4 |
| 5 | 16473838 | rs26016 | -0.358 | 0.979 | 1.003 | 0.214 | -0.016 | 0.046 | 0.046 | 0 | 0.32 | 0.47 | 0.44 | 0.44 | 0.28 | 0.39 |
| 5 | 16474644 | rs32147 | 0.086 | 1.017 | 0.813 | 0.232 | 0.009 | -0.045 | -0.045 | 0 | 0.28 | 0.18 | 0.4 | 0.45 | 0.26 | 0.39 |
| 5 | 16478120 | rs162849 | 0.798 | 1.134 | 0.797 | -0.106 | -0.008 | 0.036 | 0.036 | 1.424 | 0.45 | 0.39 | 0.31 | 0.23 | 0.46 | 0.25 |
| 5 | 16487697 | rs4702146 | 0.577 | 1.256 | -0.204 | 0.127 | 0.008 | 0.055 | 0.055 | 0 | 0.59 | 0.29 | 0.44 | 0.3 | 0.61 | 0.43 |
| 5 | 16488673 | rs12519984 | -0.19 | 1.101 | -0.315 | 0.252 | -0.005 | -0.02 | -0.02 | 0.061 | 0.27 | 0.16 | 0.54 | 0.34 | 0.27 | 0.32 |
| 5 | 16491052 | rs12187267 | 0.864 | 1.266 | -0.258 | 0.195 | 0.009 | 0.015 | 0.015 | 0.719 | 0.4 | 0.43 | 0.29 | 0.21 | 0.36 | 0.24 |

| | | | | | | | | | | | | | | | | |
|---|---|---|---|---|---|---|---|---|---|---|---|---|---|---|---|---|
| 5 | 16500790 | rs332807 | 0.52 | 1.467 | 0.054 | 0.275 | 0.016 | -0.002 | -0.002 | 3.204 | 0.61 | 0.57 | 0.64 | 0.41 | 0.56 | 0.32 |
| 5 | 16504763 | rs4702149 | -0.8 | 1.352 | -0.378 | 0.378 | 0.002 | -0.018 | -0.018 | 0.199 | 0.91 | 0.97 | 0.83 | 0.89 | 0.93 | 0.78 |
| 5 | 16508674 | rs332811 | 0.783 | 1.505 | 0.533 | 0.131 | -0.001 | -0.001 | -0.001 | 0.858 | 0.6 | 0.48 | 0.66 | 0.41 | 0.64 | 0.35 |
| 5 | 16508898 | rs33684 | 0.48 | 1.406 | 0.399 | 0.361 | 0.004 | 0.046 | 0.046 | 2.001 | 0.67 | 0.47 | 0.67 | 0.54 | 0.69 | 0.4 |
| 5 | 16509336 | rs33685 | 1.016 | 1.226 | 0.633 | 0.032 | -0.022 | 0.031 | 0.031 | 0.851 | 0.55 | 0.26 | 0.61 | 0.39 | 0.54 | 0.32 |
| 5 | 16512364 | rs334470 | -0.12 | 1.255 | 0.928 | -0.006 | -0.002 | 0.008 | 0.008 | 2.043 | 0.85 | 0.68 | 0.77 | 0.83 | 0.87 | 0.74 |
| 5 | 16514269 | rs4702150 | 2.055 | 0.927 | 0.05 | -0.706 | 0.031 | 0.082 | 0.082 | 0.815 | 0.26 | 0.08 | 0.18 | 0 | 0.23 | 0.07 |
| 5 | 16515716 | rs16868716 | -0.141 | 1.38 | 0.498 | -0.456 | 0.016 | -0.028 | -0.028 | 0 | 0.06 | 0.12 | 0.05 | 0 | 0.1 | 0.03 |
| 5 | 16521575 | rs149263 | 0.103 | 1.332 | 0.546 | -0.376 | -0.006 | -0.017 | -0.017 | 0.678 | 0.29 | 0.2 | 0.43 | 0.41 | 0.31 | 0.26 |
| 5 | 16522125 | rs163282 | 0.135 | 1.725 | 1.028 | 0.056 | 0 | 0.019 | 0.019 | 0 | 0.24 | 0.23 | 0.12 | 0.41 | 0.23 | 0.39 |
| 5 | 16525790 | rs17540822 | 0.405 | 1.56 | 0.879 | -0.161 | 0.002 | -0.024 | -0.024 | 0 | 0.08 | 0.1 | 0.04 | 0 | 0.07 | 0.04 |
| 5 | 16528257 | rs334493 | -0.033 | 1.598 | 0.836 | -0.241 | 0.005 | -0.02 | -0.02 | 0 | 0.1 | 0.16 | 0.07 | 0.02 | 0.12 | 0.07 |
| 5 | 16528603 | rs2028843 | 0.423 | 1.58 | 0.898 | -0.053 | 0.001 | 0.008 | 0.008 | 0 | 0.25 | 0.23 | 0.12 | 0.38 | 0.24 | 0.39 |
| 5 | 16529595 | rs334485 | -0.994 | 1.403 | 0.547 | -0.024 | 0 | -0.009 | -0.009 | 0 | 0.87 | 0.9 | 0.81 | 0.78 | 0.87 | 0.74 |
| 5 | 16529754 | rs12188529 | 1.8 | 1.361 | 0.461 | -0.126 | -0.021 | -0.046 | -0.046 | 0 | 0.39 | 0.1 | 0.55 | 0.38 | 0.43 | 0.24 |
| 5 | 16533551 | rs1551050 | 0.802 | 0.753 | 0.723 | -0.086 | -0.003 | -0.01 | -0.01 | 0 | 0.22 | 0.2 | 0.24 | 0.4 | 0.22 | 0.34 |
| 5 | 16535341 | rs13180554 | 1.932 | 0.815 | 0.582 | -0.175 | -0.108 | -0.04 | -0.04 | 0 | 0.41 | 0.07 | 0.43 | 0.35 | 0.51 | 0.26 |
| 5 | 16537473 | rs6870158 | 0.376 | 0.78 | 1.053 | -0.18 | 0.001 | 0.009 | 0.009 | 0.54 | 0.63 | 0.45 | 0.54 | 0.37 | 0.65 | 0.42 |
| 5 | 16540742 | rs10053126 | 0.173 | 0.893 | 0.838 | -0.089 | -0.014 | -0.029 | -0.029 | 0 | 0.11 | 0.17 | 0.06 | 0.02 | 0.06 | 0.12 |
| 5 | 16540912 | rs10053236 | -0.948 | 0.945 | 0.749 | -0.152 | 0 | -0.007 | -0.007 | 0 | 0.87 | 0.89 | 0.8 | 0.78 | 0.87 | 0.75 |
| 5 | 16547911 | rs12187879 | 0.058 | 0.743 | 0.766 | -0.147 | -0.007 | -0.016 | -0.016 | 2.386 | 0.3 | 0.21 | 0.28 | 0.4 | 0.3 | 0.35 |
| 5 | 16556601 | rs1991471 | -0.561 | 0.988 | 0.603 | -0.023 | -0.003 | -0.004 | -0.004 | 0 | 0.84 | 0.86 | 0.8 | 0.77 | 0.82 | 0.72 |
| 5 | 16559581 | rs1240649 | 1.516 | 0.925 | 0.784 | -0.085 | 0.001 | 0.003 | 0.003 | 0.05 | 0.41 | 0.25 | 0.44 | 0.37 | 0.46 | 0.26 |
| 5 | 16560819 | rs31289 | 1.529 | 0.935 | 0.491 | -0.024 | -0.047 | -0.065 | -0.065 | 0 | 0.4 | 0.09 | 0.43 | 0.38 | 0.46 | 0.26 |
| 5 | 16564901 | rs10035580 | -0.021 | 0.906 | 0.717 | -0.049 | -0.001 | 0.003 | 0.003 | 0.133 | 0.32 | 0.31 | 0.3 | 0.4 | 0.29 | 0.42 |
| 5 | 16566212 | rs2560828 | -0.539 | 0.89 | 0.734 | 0.063 | -0.001 | -0.001 | -0.001 | 3.819 | 0.4 | 0.35 | 0.33 | 0.4 | 0.36 | 0.46 |
| 5 | 16578135 | rs2560826 | 0.479 | 0.975 | 0.266 | 0.395 | 0.001 | 0.017 | 0.017 | 0 | 0.52 | 0.4 | 0.43 | 0.39 | 0.53 | 0.4 |
| 5 | 16580567 | rs154244 | -0.435 | 0.904 | 0.273 | 0.651 | 0.013 | 0.08 | 0.08 | 0.008 | 0.87 | 0.74 | 0.78 | 0.76 | 0.89 | 0.74 |
| 5 | 16584227 | rs2451855 | -0.237 | 0.959 | -0.079 | 0.395 | 0.002 | -0.016 | -0.016 | 0.001 | 0.14 | 0.17 | 0.07 | 0.04 | 0.13 | 0.11 |
| 5 | 16585183 | rs431154 | -0.229 | 0.955 | -0.103 | 0.392 | 0.001 | -0.012 | -0.012 | 0 | 0.14 | 0.17 | 0.07 | 0.04 | 0.13 | 0.11 |
| 5 | 16585512 | rs2637146 | 0.482 | 1.096 | -0.319 | 0.504 | -0.001 | 0 | 0 | 0 | 0.3 | 0.29 | 0.32 | 0.42 | 0.28 | 0.33 |
| 5 | 16586718 | rs6887031 | 0.993 | 0.849 | -0.666 | 0.035 | -0.049 | 0.021 | 0.021 | 0.015 | 0.38 | 0.09 | 0.37 | 0.3 | 0.47 | 0.28 |
| 5 | 16589850 | rs2451851 | 0.204 | 0.879 | -0.317 | 0.443 | 0 | -0.013 | -0.013 | 0 | 0.56 | 0.5 | 0.47 | 0.33 | 0.6 | 0.39 |
| 5 | 16605738 | rs2592037 | 0.123 | 0.616 | -0.728 | 0.524 | 0.004 | -0.004 | -0.004 | 0 | 0.57 | 0.53 | 0.54 | 0.52 | 0.59 | 0.48 |
| 5 | 16607196 | rs12521478 | 0.136 | 0.911 | -1.229 | -0.077 | -0.028 | 0.069 | 0.069 | 2.177 | 0.4 | 0.3 | 0.41 | 0.17 | 0.49 | 0.25 |
| 5 | 16608454 | rs17541315 | 0.702 | 1.327 | -1.594 | -0.283 | -0.104 | 0.039 | 0.039 | 3.132 | 0.28 | 0.03 | 0.31 | 0.16 | 0.34 | 0.17 |
| 5 | 16608493 | rs12516207 | 0.288 | 1.263 | -1.147 | -0.304 | -0.002 | -0.021 | -0.021 | 1.988 | 0.13 | 0.13 | 0.09 | 0 | 0.15 | 0.08 |
| 5 | 16610057 | rs1863997 | -0.033 | 0.483 | -0.94 | 0.336 | 0.007 | -0.003 | -0.003 | 0 | 0.6 | 0.44 | 0.5 | 0.46 | 0.67 | 0.42 |
| 5 | 16616264 | rs14153 | 0.307 | 0.768 | -0.563 | 0.512 | -0.001 | -0.002 | -0.002 | 0 | 0.57 | 0.35 | 0.4 | 0.52 | 0.57 | 0.67 |
| 5 | 16617903 | rs35004 | 0.848 | 0.632 | -0.312 | 0.365 | -0.005 | 0.013 | 0.013 | 0.282 | 0.46 | 0.27 | 0.31 | 0.42 | 0.5 | 0.45 |
| 5 | 16618920 | rs35005 | 0.769 | 0.287 | -0.493 | 0.364 | -0.013 | 0.028 | 0.028 | 0.634 | 0.48 | 0.16 | 0.31 | 0.42 | 0.54 | 0.43 |
| 5 | 16624190 | rs256884 | 0.682 | 0.61 | -0.567 | 0.463 | 0.001 | -0.004 | -0.004 | 1.034 | 0.75 | 0.71 | 0.68 | 0.75 | 0.74 | 0.66 |
| 5 | 16626717 | rs10520842 | -0.006 | 0.696 | -0.619 | 0.456 | -0.013 | -0.007 | -0.007 | 0 | 0.08 | 0.02 | 0.08 | 0.11 | 0.09 | 0.26 |
| 5 | 16627761 | rs6861112 | 0.483 | 0.583 | -0.702 | 0.396 | 0.01 | -0.095 | -0.095 | 0.352 | 0.23 | 0.06 | 0.31 | 0.34 | 0.2 | 0.22 |
| 5 | 16628511 | rs257174 | -1.256 | 1.031 | -0.022 | 0.26 | 0.002 | -0.014 | -0.014 | 0 | 0.25 | 0.34 | 0.33 | 0.25 | 0.26 | 0.34 |
| 5 | 16631653 | rs257172 | 0.022 | 1.191 | 0.106 | 0.38 | 0.007 | -0.009 | -0.009 | 0 | 0.14 | 0.21 | 0.24 | 0.13 | 0.17 | 0.09 |
| 5 | 16631835 | rs257170 | -0.066 | 1.201 | 0.049 | 0.385 | 0.013 | -0.014 | -0.014 | 0 | 0.14 | 0.25 | 0.25 | 0.13 | 0.17 | 0.09 |
| 5 | 16642272 | rs386845 | -0.577 | 1.305 | -0.408 | 0.815 | -0.012 | -0.012 | -0.012 | 0 | 0.81 | 0.87 | 0.89 | 0.77 | 0.74 | 0.73 |
| 5 | 16645335 | rs6892658 | 0.411 | 1.179 | 0.044 | 0.267 | 0.021 | -0.054 | -0.054 | 2.082 | 0.83 | 0.64 | 0.76 | 0.88 | 0.83 | 0.82 |
| 5 | 16654960 | rs379803 | -0.153 | 1.218 | -0.46 | 0.733 | -0.013 | -0.016 | -0.016 | 0 | 0.66 | 0.75 | 0.64 | 0.49 | 0.61 | 0.62 |
| 5 | 16658524 | rs388887 | -0.182 | 1.215 | -0.52 | 0.85 | -0.023 | -0.035 | -0.035 | 0 | 0.67 | 0.78 | 0.64 | 0.56 | 0.61 | 0.62 |
| 5 | 16659864 | rs252369 | 0.702 | 0.966 | -0.396 | 0.639 | 0.004 | 0.002 | 0.002 | 0 | 0.51 | 0.49 | 0.4 | 0.44 | 0.44 | 0.44 |
| 5 | 16660145 | rs257159 | -0.54 | 1.068 | -0.936 | 0.571 | -0.029 | 0.003 | 0.003 | 0 | 0.73 | 0.89 | 0.85 | 0.83 | 0.69 | 0.86 |
| 5 | 16661282 | rs7722385 | -0.124 | 0.452 | -1.109 | 0.071 | 0.005 | 0.018 | 0.018 | 0 | 0.24 | 0.19 | 0.23 | 0.12 | 0.21 | 0.26 |
| 5 | 16661569 | rs153202 | 0.266 | 0.599 | -1.376 | 0.519 | -0.047 | -0.022 | -0.022 | 0.625 | 0.17 | 0.07 | 0.1 | 0.18 | 0.19 | 0.27 |
| 5 | 16662297 | rs13537 | 0.178 | -0.024 | -1.006 | 0.426 | -0.002 | -0.006 | -0.006 | 0.896 | 0.43 | 0.52 | 0.57 | 0.38 | 0.43 | 0.47 |
| 5 | 16664769 | rs26742 | -0.411 | 0.043 | -0.837 | 0.402 | 0.007 | -0.019 | -0.019 | 0 | 0.62 | 0.5 | 0.7 | 0.68 | 0.58 | 0.77 |
| 5 | 16670618 | rs27431 | 0.071 | -0.371 | -0.452 | 0.22 | -0.003 | 0 | 0 | 0 | 0.71 | 0.33 | 0.65 | 0.73 | 0.76 | 0.81 |
| 5 | 16673685 | rs26745 | -0.579 | -0.08 | -0.463 | 0.128 | 0.013 | 0.044 | 0.044 | 0 | 0.37 | 0.29 | 0.31 | 0.2 | 0.31 | 0.39 |
| 5 | 16679966 | rs35580 | -1.209 | 0.012 | -0.285 | 0.18 | -0.063 | -0.063 | -0.063 | 0 | 0.6 | 0.83 | 0.62 | 0.49 | 0.53 | 0.56 |
| 5 | 16680772 | rs2288433 | -0.796 | 0.732 | -0.002 | 0.706 | -0.001 | -0.003 | -0.003 | 0 | 0.3 | 0.31 | 0.26 | 0.21 | 0.27 | 0.39 |
| 5 | 16687522 | rs6881621 | -0.299 | 0.685 | -0.057 | 0.917 | -0.007 | 0.004 | 0.004 | 0 | 0.23 | 0.22 | 0.26 | 0.21 | 0.29 | 0.35 |
| 5 | 16690754 | rs31313 | 0.769 | 0.646 | -0.16 | 0.954 | 0.006 | -0.028 | -0.028 | 0 | 0.18 | 0.08 | 0.27 | 0.27 | 0.2 | 0.17 |
| 5 | 16693918 | rs7733988 | -0.421 | 0.86 | 0.469 | 0.947 | -0.024 | 0.028 | 0.028 | 0 | 0.21 | 0.12 | 0.24 | 0.12 | 0.26 | 0.24 |
| 5 | 16705915 | rs3849106 | -0.319 | 0.585 | 0.8 | 1.043 | -0.023 | -0.045 | -0.045 | 0.027 | 0.2 | 0.14 | 0.3 | 0.3 | 0.25 | 0.2 |
| 5 | 16706530 | rs25912 | 1.155 | 0.708 | 0.978 | 0.728 | -0.004 | 0.007 | 0.007 | 1.158 | 0.55 | 0.56 | 0.49 | 0.38 | 0.57 | 0.29 |
| 5 | 16710763 | rs2288442 | -0.466 | 0.506 | 1.002 | 0.904 | 0.01 | 0.083 | 0.083 | 0 | 0.23 | 0.44 | 0.33 | 0.41 | 0.25 | 0.3 |
| 5 | 16713465 | rs31299 | 0.983 | 0.413 | 1.497 | 0.491 | -0.007 | -0.015 | -0.015 | 0.372 | 0.7 | 0.63 | 0.78 | 0.78 | 0.8 | 0.57 |
| 5 | 16721306 | rs40979 | 0.393 | 0.535 | 1.115 | 0.504 | -0.002 | -0.002 | -0.002 | 0 | 0.76 | 0.79 | 0.8 | 0.83 | 0.87 | 0.61 |

| | | | | | | | | | | | | | | | | |
|---|---|---|---|---|---|---|---|---|---|---|---|---|---|---|---|---|
| 5 | 16724163 | rs152726 | 0.877 | 0.635 | 0.635 | 0.543 | -0.003 | -0.007 | -0.007 | 0 | 0.67 | 0.38 | 0.73 | 0.78 | 0.79 | 0.57 |
| 5 | 16732052 | rs10051970 | -1.949 | 0.988 | 1.044 | 0.798 | 0.023 | 0.025 | 0.025 | 0.04 | 0.14 | 0.38 | 0.26 | 0.11 | 0.15 | 0.11 |
| 5 | 16737513 | rs11133857 | -2.138 | 1.31 | 0.814 | 0.862 | 0.005 | 0.004 | 0.004 | 0 | 0.18 | 0.24 | 0.32 | 0.22 | 0.21 | 0.25 |
| 5 | 16741099 | rs10051930 | -2.195 | 1.409 | 1.539 | 1.075 | 0.069 | 0.117 | 0.117 | 0.038 | 0.11 | 0.59 | 0.34 | 0.22 | 0.11 | 0.15 |
| 5 | 16746784 | rs6896958 | -1.886 | 1.535 | 1.419 | 0.767 | 0.123 | 0.058 | 0.058 | 0 | 0.09 | 0.34 | 0.18 | 0.13 | 0.11 | 0.15 |
| 5 | 16751179 | rs6890128 | -0.618 | 1.551 | 0.451 | 0.512 | -0.011 | 0.009 | 0.009 | 0 | 0.25 | 0.17 | 0.18 | 0.13 | 0.3 | 0.35 |
| 5 | 16755049 | rs17429731 | -1.008 | 1.347 | 0.362 | 0.595 | 0.013 | -0.012 | -0.012 | 0.041 | 0.08 | 0.13 | 0.13 | 0.06 | 0.09 | 0.17 |
| 5 | 16758569 | rs250340 | -0.047 | 1.268 | -0.299 | 0.821 | -0.006 | -0.008 | -0.008 | 0 | 0.21 | 0.13 | 0.17 | 0.22 | 0.28 | 0.17 |
| 5 | 16759034 | rs16869106 | -0.412 | 0.549 | -0.449 | 0.509 | 0 | 0 | 0 | 0 | 0.26 | 0.18 | 0.16 | 0.13 | 0.23 | 0.28 |
| 5 | 16760970 | rs31509 | 0.512 | 0.743 | -0.57 | 1.017 | 0.003 | 0 | 0 | 0 | 0.68 | 0.8 | 0.6 | 0.67 | 0.7 | 0.58 |
| 5 | 16763908 | rs39900 | -1.045 | 0.936 | 0.006 | 1.97 | 0.027 | 0.048 | 0.048 | 0 | 0.1 | 0.16 | 0.1 | 0.15 | 0.14 | 0.17 |
| 5 | 16764334 | rs250341 | -1.03 | 0.995 | -0.276 | 1.187 | -0.007 | -0.011 | -0.011 | 0.881 | 0.58 | 0.49 | 0.49 | 0.31 | 0.59 | 0.66 |
| 5 | 16764644 | rs2561231 | -0.93 | 1.294 | -0.232 | 1.067 | -0.007 | 0.01 | 0.01 | 0 | 0.3 | 0.21 | 0.29 | 0.2 | 0.35 | 0.38 |
| 5 | 16769273 | rs11750538 | 1.37 | 0.97 | 0.37 | 1.274 | -0.003 | -0.034 | -0.034 | 0.583 | 0.65 | 0.34 | 0.4 | 0.54 | 0.6 | 0.55 |
| 5 | 16770452 | rs7703315 | 1.348 | 0.844 | 0.352 | 1.16 | 0 | -0.026 | -0.026 | 0.175 | 0.64 | 0.31 | 0.47 | 0.56 | 0.6 | 0.54 |
| 5 | 16771640 | rs10053767 | -0.24 | 0.939 | 0.763 | 1.201 | 0 | -0.006 | -0.006 | 0 | 0.2 | 0.15 | 0.2 | 0.05 | 0.23 | 0.19 |
| 5 | 16772976 | rs4702171 | 1.436 | 0.929 | 0.684 | 1.207 | -0.003 | -0.028 | -0.028 | 0.174 | 0.65 | 0.33 | 0.47 | 0.56 | 0.6 | 0.56 |
| 5 | 16778415 | rs12520877 | -0.146 | 1.071 | 0.723 | 1.164 | -0.004 | 0.009 | 0.009 | 0 | 0.2 | 0.14 | 0.2 | 0.05 | 0.25 | 0.19 |
| 5 | 16778598 | rs12520918 | -0.265 | 1.072 | 0.724 | 1.16 | -0.004 | 0.008 | 0.008 | 0 | 0.2 | 0.14 | 0.2 | 0.06 | 0.25 | 0.19 |
| 5 | 16783021 | rs25906 | -0.654 | 0.909 | 1.324 | 1.186 | 0.011 | 0.025 | 0.025 | 0 | 0.13 | 0.16 | 0.12 | 0.17 | 0.16 | 0.19 |
| 5 | 16783711 | rs4702172 | 1.857 | 1.147 | 1.02 | 1.814 | -0.022 | -0.031 | -0.031 | 0 | 0.6 | 0.3 | 0.44 | 0.43 | 0.54 | 0.51 |
| 5 | 16783993 | rs11133859 | -0.229 | 0.979 | 0.958 | 1.58 | -0.005 | 0.005 | 0.005 | 0 | 0.2 | 0.15 | 0.22 | 0.05 | 0.28 | 0.19 |
| 5 | 16785604 | rs6877325 | 1.206 | 0.933 | 0.873 | 1.753 | 0.006 | 0.151 | 0.151 | 0 | 0.82 | 0.67 | 0.68 | 0.61 | 0.82 | 0.71 |
| 5 | 16788851 | rs250345 | -0.794 | 0.852 | 0.65 | 1.633 | 0.009 | 0.003 | 0.003 | 1.31 | 0.32 | 0.56 | 0.44 | 0.4 | 0.44 | 0.35 |
| 5 | 16790200 | rs250348 | -0.512 | 0.42 | 0.87 | 1.707 | 0.003 | -0.006 | -0.006 | 0.978 | 0.32 | 0.35 | 0.37 | 0.24 | 0.44 | 0.34 |
| 5 | 16790483 | rs153409 | -0.811 | -0.004 | 0.464 | 1.721 | 0.002 | -0.007 | -0.007 | 1.331 | 0.57 | 0.5 | 0.55 | 0.66 | 0.57 | 0.69 |
| 5 | 16793089 | rs250356 | -0.528 | -0.038 | 0.171 | 1.64 | 0 | 0.002 | 0.002 | 0 | 0.54 | 0.33 | 0.47 | 0.43 | 0.54 | 0.64 |
| 5 | 16793151 | rs6888120 | 1.04 | 0.088 | -0.585 | 1.253 | -0.107 | 0.058 | 0.058 | 0 | 0.36 | 0.03 | 0.29 | 0.18 | 0.4 | 0.25 |
| 5 | 16799341 | rs7717298 | 1.039 | -0.066 | -0.383 | 1.351 | 0.027 | -0.009 | -0.009 | 0 | 0.23 | 0.08 | 0.11 | 0.19 | 0.12 | 0.26 |
| 5 | 16804121 | rs40020 | -0.69 | -0.14 | -0.042 | 1.266 | -0.001 | -0.001 | -0.001 | 0 | 0.61 | 0.63 | 0.51 | 0.57 | 0.57 | 0.7 |
| 5 | 16807786 | rs1872004 | 0.069 | -0.309 | -0.057 | 1.202 | -0.014 | 0.003 | 0.003 | 0 | 0.33 | 0.39 | 0.25 | 0.41 | 0.15 | 0.32 |
| 5 | 16810263 | rs17651095 | -0.255 | -0.279 | -0.114 | 1.194 | -0.007 | 0.065 | 0.065 | 0.271 | 0.06 | 0.21 | 0.12 | 0.21 | 0.03 | 0.04 |
| 5 | 16813881 | rs11745637 | 0.105 | -0.261 | -0.116 | 1.171 | -0.003 | -0.005 | -0.005 | 0 | 0.37 | 0.39 | 0.45 | 0.56 | 0.46 | 0.45 |
| 5 | 16818363 | rs12188582 | -0.375 | 0.119 | -0.449 | 1.009 | -0.029 | 0.005 | 0.005 | 0 | 0.7 | 0.56 | 0.73 | 0.74 | 0.84 | 0.7 |
| 5 | 16819642 | rs890825 | 0.866 | 0.151 | -0.837 | 0.764 | -0.049 | -0.05 | -0.05 | 0 | 0.34 | 0.14 | 0.37 | 0.39 | 0.38 | 0.26 |
| 5 | 16820505 | rs7726872 | 0.16 | -0.132 | -0.104 | 0.397 | 0.087 | 0.071 | 0.071 | 0 | 0.94 | 0.59 | 0.84 | 0.96 | 0.96 | 0.97 |
| 5 | 16820769 | rs17651165 | -0.018 | -0.007 | -1.313 | 0.475 | 0.033 | 0.011 | 0.011 | 0 | 0.27 | 0.15 | 0.12 | 0.2 | 0.14 | 0.28 |
| 5 | 16829442 | rs253324 | 0.183 | 1.034 | -1.2 | -0.114 | -0.011 | 0.01 | 0.01 | 0 | 0.68 | 0.74 | 0.73 | 0.77 | 0.6 | 0.77 |
| 5 | 16834274 | rs31506 | -0.147 | 0.783 | -1.347 | -0.438 | 0.025 | 0.09 | 0.09 | 0 | 0.61 | 0.84 | 0.74 | 0.83 | 0.66 | 0.6 |
| 5 | 16834568 | rs31505 | -0.401 | 0.746 | -1.371 | -0.496 | 0.047 | 0.076 | 0.076 | 0 | 0.64 | 0.83 | 0.76 | 0.83 | 0.71 | 0.6 |
| 5 | 16836005 | rs9312937 | 0.217 | 0.955 | -1.087 | -0.308 | -0.002 | 0.001 | 0.001 | 0.002 | 0.48 | 0.65 | 0.64 | 0.58 | 0.51 | 0.44 |
| 5 | 16840145 | rs17707735 | -0.43 | 0.78 | -1.009 | -0.173 | 0.002 | 0.006 | 0.006 | 0 | 0.17 | 0.2 | 0.12 | 0.29 | 0.19 | 0.15 |
| 5 | 16840641 | rs253336 | -1.336 | 0.626 | -0.997 | -0.261 | 0.011 | 0.012 | 0.012 | 0 | 0.69 | 0.81 | 0.73 | 0.8 | 0.72 | 0.54 |
| 5 | 16841399 | rs2560861 | -1.68 | 0.906 | 0.121 | -0.034 | 0.003 | 0 | 0 | 0 | 0.28 | 0.36 | 0.2 | 0.29 | 0.28 | 0.23 |
| 5 | 16843108 | rs31499 | -1.707 | 0.867 | -0.323 | 0.189 | -0.006 | 0.002 | 0.002 | 0.559 | 0.31 | 0.53 | 0.3 | 0.32 | 0.25 | 0.25 |
| 5 | 16844859 | rs716555 | -0.704 | 0.966 | -0.967 | 0.236 | -0.045 | -0.078 | -0.078 | 0 | 0.12 | 0.02 | 0.18 | 0.28 | 0.13 | 0.14 |
| 5 | 16848200 | rs10213765 | -1.069 | 1.101 | -0.694 | 0.342 | -0.001 | -0.001 | -0.001 | 0.002 | 0.29 | 0.66 | 0.29 | 0.27 | 0.24 | 0.2 |
| 5 | 16850125 | rs9312938 | -0.985 | 1.349 | -1.067 | 0.318 | -0.003 | 0.005 | 0.005 | 0 | 0.36 | 0.77 | 0.4 | 0.33 | 0.32 | 0.32 |
| 5 | 16862419 | rs17614174 | -1.267 | 1.642 | -1.088 | 0.363 | -0.035 | 0.048 | 0.048 | 0 | 0.1 | 0.01 | 0.05 | 0 | 0.11 | 0.08 |
| 5 | 16862998 | rs891163 | -1.158 | 1.504 | -0.382 | 0.85 | 0.023 | 0.031 | 0.031 | 0 | 0.17 | 0.3 | 0.23 | 0.22 | 0.17 | 0.18 |
| 5 | 16864632 | rs17707882 | -0.961 | 1.138 | -0.85 | 0.969 | 0.002 | -0.011 | -0.011 | 0 | 0.05 | 0.03 | 0.03 | 0.11 | 0.04 | 0.19 |
| 5 | 16875690 | rs2625210 | -0.372 | 0.932 | -1.443 | 0.937 | 0.016 | -0.034 | -0.034 | 0 | 0.64 | 0.82 | 0.78 | 0.68 | 0.69 | 0.74 |
| 5 | 16876479 | rs2562338 | -0.586 | 0.979 | -0.273 | 0.719 | 0.033 | -0.022 | -0.022 | 0 | 0.07 | 0.22 | 0.07 | 0.01 | 0.07 | 0.05 |
| 5 | 16877640 | rs7356599 | -0.976 | 0.855 | -0.122 | 1.1 | -0.002 | 0.029 | 0.029 | 0 | 0.1 | 0.22 | 0.05 | 0.16 | 0.06 | 0.21 |
| 5 | 16878237 | rs7711618 | -0.265 | 1.545 | -0.508 | 1.106 | -0.019 | -0.013 | -0.013 | 0 | 0.13 | 0.07 | 0.31 | 0.2 | 0.19 | 0.21 |
| 5 | 16880404 | rs166225 | -0.152 | 0.997 | -0.303 | 1.04 | -0.004 | -0.006 | -0.006 | 0.039 | 0.43 | 0.35 | 0.56 | 0.55 | 0.51 | 0.46 |
| 5 | 16880460 | rs254833 | 0.346 | 0.922 | -0.487 | 1.323 | -0.046 | -0.013 | -0.013 | 0 | 0.33 | 0.12 | 0.52 | 0.36 | 0.45 | 0.4 |
| 5 | 16882242 | rs153706 | -1.019 | 0.729 | 0.262 | 1.084 | 0.003 | 0 | 0 | 0 | 0.54 | 0.67 | 0.61 | 0.58 | 0.6 | 0.55 |
| 5 | 16887383 | rs2560851 | -1.408 | 0.722 | 0.739 | 0.934 | -0.001 | -0.006 | -0.006 | 0 | 0.38 | 0.61 | 0.51 | 0.49 | 0.43 | 0.34 |
| 5 | 16887414 | rs17651616 | -1.394 | 0.587 | 1.132 | 1.011 | -0.017 | 0.141 | 0.141 | 0.183 | 0.13 | 0.32 | 0.06 | 0.3 | 0.06 | 0.09 |
| 5 | 16892404 | rs2562343 | -1.79 | 0.274 | 0.623 | 0.547 | -0.008 | -0.017 | -0.017 | 0 | 0.54 | 0.71 | 0.6 | 0.55 | 0.55 | 0.42 |
| 5 | 16892438 | rs17651643 | -0.925 | 0.164 | 0.561 | 0.514 | -0.007 | 0.004 | 0.004 | 0 | 0.25 | 0.25 | 0.14 | 0.35 | 0.15 | 0.14 |
| 5 | 16895512 | rs173789 | -1.153 | 0.357 | 1.056 | 1.001 | -0.061 | 0.097 | 0.097 | 0 | 0.19 | 0.36 | 0.11 | 0.37 | 0.09 | 0.11 |
| 5 | 16897446 | rs253482 | 0.206 | 0.472 | 0.751 | 1.028 | -0.092 | -0.022 | -0.022 | 0 | 0.19 | 0.02 | 0.43 | 0.18 | 0.34 | 0.24 |
| 5 | 16897597 | rs253483 | -1.888 | 0.431 | 0.871 | 0.994 | -0.003 | -0.013 | -0.013 | 0 | 0.46 | 0.74 | 0.59 | 0.56 | 0.48 | 0.39 |
| 5 | 16897799 | rs2562350 | -1.52 | 0.625 | 0.184 | 1.078 | 0.001 | 0.009 | 0.009 | 0.657 | 0.4 | 0.45 | 0.55 | 0.52 | 0.45 | 0.36 |
| 5 | 16901752 | rs10520846 | -1.682 | 0.022 | 0.629 | 1.302 | -0.015 | 0.024 | 0.024 | 0 | 0.25 | 0.34 | 0.15 | 0.41 | 0.16 | 0.17 |
| 5 | 16903718 | rs10520847 | -1.235 | 0.349 | 0.274 | 1.353 | 0.002 | -0.005 | -0.005 | 0 | 0.22 | 0.21 | 0.22 | 0.37 | 0.17 | 0.2 |
| 5 | 16905100 | rs13176126 | -1.422 | 0.302 | 0.589 | 1.332 | -0.016 | 0.041 | 0.041 | 0 | 0.23 | 0.32 | 0.24 | 0.37 | 0.17 | 0.2 |

| | | | | | | | | | | | | | | | | |
|---|---|---|---|---|---|---|---|---|---|---|---|---|---|---|---|---|
| 5 | 16906913 | rs4702182 | -1.441 | 0.383 | 0.533 | 1.251 | 0.002 | 0 | 0 | 0 | 0.42 | 0.57 | 0.39 | 0.59 | 0.37 | 0.44 |
| 5 | 16907538 | rs29448 | -1.926 | 0.78 | 0.195 | 0.901 | 0.002 | 0.009 | 0.009 | 0 | 0.65 | 0.71 | 0.81 | 0.77 | 0.71 | 0.65 |
| 5 | 16910605 | rs29452 | 0.183 | 0.998 | 0.373 | 0.97 | -0.001 | 0.001 | 0.001 | 0 | 0.23 | 0.21 | 0.25 | 0.18 | 0.31 | 0.19 |
| 5 | 16910789 | rs152342 | -0.783 | 0.919 | 0.072 | 0.947 | 0.006 | -0.009 | -0.009 | 0 | 0.36 | 0.36 | 0.52 | 0.59 | 0.35 | 0.43 |
| 5 | 16912847 | rs16869235 | -0.544 | 0.757 | 0.229 | 0.969 | -0.008 | 0.011 | 0.011 | 0 | 0.09 | 0.19 | 0.13 | 0.1 | 0.07 | 0.08 |
| 5 | 16913578 | rs29454 | -2.181 | 0.758 | 0.277 | 0.874 | -0.005 | -0.034 | -0.034 | 0 | 0.65 | 0.64 | 0.81 | 0.84 | 0.71 | 0.66 |
| 5 | 16913719 | rs29455 | -1.129 | 0.8 | 0.251 | 1.082 | 0.002 | 0.034 | 0.034 | 0 | 0.49 | 0.37 | 0.72 | 0.6 | 0.56 | 0.54 |
| 5 | 16914516 | rs39896 | -1.563 | 0.85 | 0.076 | 1.062 | 0.002 | 0.008 | 0.008 | 0.012 | 0.55 | 0.63 | 0.77 | 0.65 | 0.62 | 0.55 |
| 5 | 16921494 | rs249123 | -0.779 | 1.017 | -0.434 | 1.402 | 0.019 | 0.064 | 0.064 | 1.938 | 0.2 | 0.29 | 0.43 | 0.47 | 0.3 | 0.31 |
| 5 | 16923843 | rs2434960 | 1.159 | 0.863 | -0.635 | 1.133 | -0.008 | 0.003 | 0.003 | 0 | 0.54 | 0.61 | 0.34 | 0.47 | 0.44 | 0.5 |
| 5 | 16924422 | rs17651767 | 0.142 | 1.102 | -0.51 | 1.392 | -0.024 | -0.032 | -0.032 | 0.186 | 0.08 | 0.01 | 0.1 | 0.1 | 0.09 | 0.09 |
| 5 | 16924531 | rs2625177 | 1.339 | 1.063 | -0.498 | 1.107 | -0.006 | 0 | 0 | 0.296 | 0.55 | 0.63 | 0.33 | 0.47 | 0.45 | 0.49 |
| 5 | 16930761 | rs31454 | 2.332 | 1.029 | -0.51 | 1.08 | -0.006 | -0.024 | -0.024 | 0 | 0.4 | 0.36 | 0.2 | 0.19 | 0.34 | 0.35 |
| 5 | 16932148 | rs31457 | 1.441 | 0.934 | -0.53 | 1.107 | 0.002 | 0.004 | 0.004 | 0 | 0.5 | 0.45 | 0.27 | 0.3 | 0.42 | 0.43 |
| 5 | 16940614 | rs3866217 | -0.679 | 0.927 | -0.709 | 1.32 | -0.002 | 0.004 | 0.004 | 1.27 | 0.22 | 0.26 | 0.54 | 0.48 | 0.32 | 0.34 |
| 5 | 16945823 | rs10061139 | 1.827 | 0.877 | -0.987 | 0.813 | -0.004 | -0.026 | -0.026 | 0 | 0.4 | 0.35 | 0.19 | 0.16 | 0.34 | 0.35 |
| 5 | 16948041 | rs2402335 | -0.4 | 1.155 | -1.368 | 0.673 | -0.002 | 0.038 | 0.038 | 1.003 | 0.81 | 0.91 | 0.88 | 0.92 | 0.78 | 0.85 |
| 5 | 16949845 | rs12653955 | 1.463 | 0.979 | -1.286 | 0.793 | 0.004 | 0.022 | 0.022 | 0 | 0.39 | 0.25 | 0.19 | 0.2 | 0.35 | 0.38 |
| 5 | 16952807 | rs249116 | -0.91 | 0.701 | -0.613 | 1.415 | 0.003 | 0.026 | 0.026 | 0 | 0.18 | 0.22 | 0.47 | 0.48 | 0.24 | 0.26 |
| 5 | 16961453 | rs249131 | -0.815 | 0.637 | -1.016 | 0.834 | 0.002 | 0.002 | 0.002 | 0.15 | 0.39 | 0.69 | 0.61 | 0.66 | 0.39 | 0.44 |
| 5 | 16969440 | rs17708295 | -0.767 | 0.59 | -1.3 | 1.044 | -0.008 | -0.055 | -0.055 | 0.557 | 0.14 | 0.03 | 0.2 | 0.26 | 0.11 | 0.1 |
| 5 | 16986171 | rs1549672 | -0.41 | 0.505 | -1.45 | 0.082 | -0.002 | 0.014 | 0.014 | 0 | 0.24 | 0.19 | 0.15 | 0.16 | 0.26 | 0.33 |
| 5 | 16989839 | rs10076041 | -0.234 | 0.321 | -1.18 | -0.392 | 0.018 | 0.029 | 0.029 | 0 | 0.49 | 0.46 | 0.57 | 0.64 | 0.64 | 0.49 |
| 5 | 16991693 | rs6877901 | 1.105 | -0.572 | -0.198 | -0.655 | -0.022 | -0.015 | -0.015 | 0.1 | 0.7 | 0.53 | 0.68 | 0.66 | 0.79 | 0.7 |
| 5 | 16993278 | rs11749259 | 1.449 | -1.06 | -0.19 | -0.928 | -0.004 | -0.005 | -0.005 | 0 | 0.6 | 0.41 | 0.65 | 0.66 | 0.74 | 0.65 |
| 5 | 16993868 | rs2018015 | 0.127 | -1.014 | -0.066 | -0.917 | 0.012 | -0.04 | -0.04 | 0 | 0.1 | 0.21 | 0.08 | 0.03 | 0.11 | 0.05 |
| 5 | 17000364 | rs4129967 | 1.181 | -0.876 | 0.123 | -0.917 | -0.008 | -0.008 | -0.008 | 0 | 0.68 | 0.43 | 0.65 | 0.65 | 0.76 | 0.69 |
| 5 | 17003017 | rs6870542 | 0.188 | -0.822 | 0.171 | -0.856 | 0.019 | 0.087 | 0.087 | 0.53 | 0.94 | 0.77 | 0.95 | 0.84 | 0.97 | 0.86 |
| 5 | 17003085 | rs6870564 | 0.034 | -0.794 | -0.246 | -0.829 | 0.04 | 0.014 | 0.014 | 0.019 | 0.85 | 0.55 | 0.87 | 0.81 | 0.86 | 0.81 |
| 5 | 17005751 | rs4360041 | 1.041 | -0.629 | -0.681 | -0.726 | 0.038 | 0.05 | 0.05 | 3.962 | 0.57 | 0.29 | 0.62 | 0.65 | 0.71 | 0.64 |
| 5 | 17018543 | rs4374758 | 0.378 | -0.085 | -0.419 | -0.284 | -0.002 | 0.002 | 0.002 | 0 | 0.85 | 0.79 | 0.78 | 0.82 | 0.88 | 0.87 |
| 5 | 17023701 | rs6859425 | -0.396 | -0.194 | -0.415 | -0.199 | -0.005 | 0.012 | 0.012 | 0 | 0.11 | 0.16 | 0.18 | 0.19 | 0.06 | 0.12 |
| 5 | 17025067 | rs6866220 | -0.039 | -0.622 | 0.235 | -0.549 | -0.035 | -0.038 | -0.038 | 1.047 | 0.92 | 0.76 | 0.98 | 1 | 0.99 | 0.99 |
| 5 | 17030640 | rs11949558 | 0.344 | -0.336 | -0.019 | -0.397 | 0.013 | 0.018 | 0.018 | 0 | 0.78 | 0.62 | 0.71 | 0.7 | 0.82 | 0.82 |
| 5 | 17049441 | rs4701679 | 0.185 | -0.072 | -1.042 | -0.669 | 0.004 | 0.005 | 0.005 | 0 | 0.43 | 0.33 | 0.36 | 0.53 | 0.43 | 0.33 |
| 5 | 17059369 | rs6862915 | 0.646 | -0.113 | -1.147 | -0.544 | -0.01 | -0.041 | -0.041 | 0 | 0.24 | 0.17 | 0.27 | 0.51 | 0.29 | 0.23 |
| 5 | 17063654 | rs13178476 | 0.551 | 0.002 | -0.897 | -0.307 | -0.017 | 0.016 | 0.016 | 0 | 0.16 | 0.1 | 0.23 | 0.15 | 0.14 | 0.11 |
| 5 | 17069833 | rs17708415 | 0.39 | 0.001 | -0.868 | -0.122 | -0.021 | -0.009 | -0.009 | 1.07 | 0.12 | 0.2 | 0.05 | 0.09 | 0.06 | 0.08 |
| 5 | 17078227 | rs12519524 | -0.726 | 0.833 | -1.077 | -0.028 | 0.002 | -0.001 | -0.001 | 0 | 0.26 | 0.33 | 0.41 | 0.32 | 0.26 | 0.22 |
| 5 | 17085379 | rs12519960 | 0.329 | 0.963 | -1.914 | 0.091 | -0.018 | 0.011 | 0.011 | 0 | 0.17 | 0.09 | 0.23 | 0.17 | 0.14 | 0.12 |
| 5 | 17086303 | rs17652001 | -0.017 | 0.856 | -1.465 | 0.247 | -0.023 | -0.031 | -0.031 | 0 | 0.05 | 0.01 | 0.15 | 0.16 | 0.09 | 0.09 |
| 5 | 17092491 | rs6880043 | -0.272 | 0.687 | -1.436 | -0.246 | -0.014 | 0.044 | 0.044 | 0 | 0.3 | 0.15 | 0.23 | 0.18 | 0.35 | 0.5 |
| 5 | 17095269 | rs2624435 | -0.402 | 0.417 | -1.267 | -0.204 | 0.004 | 0.029 | 0.029 | 0 | 0.26 | 0.15 | 0.23 | 0.16 | 0.26 | 0.17 |
| 5 | 17107465 | rs644358 | -0.51 | 0.887 | -1.398 | 0.827 | -0.008 | 0.009 | 0.009 | 0 | 0.9 | 0.95 | 0.92 | 0.65 | 0.87 | 0.82 |
| 5 | 17114476 | rs297177 | 0.971 | -0.07 | -0.886 | 0.352 | -0.001 | 0 | 0 | 0 | 0.44 | 0.41 | 0.6 | 0.39 | 0.51 | 0.5 |
| 5 | 17116221 | rs1351583 | 1.447 | 0.142 | -0.828 | 0.179 | -0.089 | -0.026 | -0.026 | 0.093 | 0.28 | 0.07 | 0.33 | 0.21 | 0.35 | 0.29 |
| 5 | 17118930 | rs1032763 | -0.157 | -0.565 | -0.18 | 0.397 | 0.001 | -0.003 | -0.003 | 0 | 0.46 | 0.49 | 0.31 | 0.25 | 0.36 | 0.32 |
| 5 | 17124559 | rs7704060 | 0.983 | -0.659 | 0.065 | 0.366 | -0.046 | 0.043 | 0.043 | 0 | 0.4 | 0.22 | 0.54 | 0.25 | 0.49 | 0.41 |
| 5 | 17126149 | rs297205 | -1.571 | -0.63 | 0.031 | 0.448 | -0.008 | 0.009 | 0.009 | 0 | 0.9 | 0.95 | 0.92 | 0.63 | 0.87 | 0.82 |
| 5 | 17126682 | rs297206 | 0.179 | -0.914 | 0.199 | 0.302 | 0.003 | -0.002 | -0.002 | 0 | 0.55 | 0.6 | 0.65 | 0.46 | 0.58 | 0.56 |
| 5 | 17129725 | rs2896245 | 0.505 | -0.908 | 0.167 | 0.308 | -0.012 | 0.102 | 0.102 | 0 | 0.74 | 0.57 | 0.82 | 0.42 | 0.77 | 0.66 |
| 5 | 17132022 | rs2652670 | 1.063 | -0.877 | -0.116 | 0.214 | 0.021 | 0.039 | 0.039 | 0 | 0.23 | 0.11 | 0.11 | 0.08 | 0.15 | 0.12 |
| 5 | 17137175 | rs167214 | 0.023 | -0.48 | -0.293 | 0.456 | 0.004 | 0.004 | 0.004 | 0 | 0.39 | 0.57 | 0.32 | 0.29 | 0.31 | 0.31 |
| 5 | 17142051 | rs9312946 | -1.3 | 0.064 | -0.669 | 0.834 | -0.018 | -0.023 | -0.023 | 0 | 0.9 | 0.97 | 0.92 | 0.64 | 0.87 | 0.79 |
| 5 | 17145198 | rs1875538 | 0.784 | 0.015 | -0.854 | 0.802 | -0.113 | -0.012 | -0.012 | 0 | 0.38 | 0.09 | 0.54 | 0.28 | 0.49 | 0.4 |
| 5 | 17148236 | rs587977 | 0.205 | -0.048 | -0.568 | 0.781 | 0.017 | 0.058 | 0.058 | 0.803 | 0.07 | 0.26 | 0.07 | 0.12 | 0.05 | 0.07 |
| 5 | 17155359 | rs683580 | 0.315 | -0.049 | -0.573 | 0.802 | 0.004 | -0.009 | -0.009 | 0 | 0.46 | 0.36 | 0.61 | 0.48 | 0.54 | 0.45 |
| 5 | 17158041 | rs297182 | -0.607 | 0.153 | -1.15 | 0.86 | -0.024 | -0.049 | -0.049 | 1.934 | 0.86 | 0.95 | 0.89 | 0.67 | 0.84 | 0.76 |
| 5 | 17160258 | rs13175997 | 0.2 | 0.363 | -0.904 | 0.692 | -0.102 | -0.03 | -0.03 | 2.262 | 0.38 | 0.1 | 0.55 | 0.31 | 0.49 | 0.39 |
| 5 | 17172790 | rs2402350 | 0.1 | 0.148 | -0.322 | 0.74 | 0.004 | 0.021 | 0.021 | 0 | 0.41 | 0.18 | 0.38 | 0.25 | 0.43 | 0.37 |
| 5 | 17181964 | rs4602628 | -0.858 | 0.107 | -0.18 | 0.932 | -0.012 | 0.02 | 0.02 | 0.019 | 0.06 | 0.2 | 0.04 | 0.13 | 0.01 | 0.09 |
| 5 | 17183851 | rs638191 | -1.551 | 0.106 | 0.086 | 0.908 | -0.008 | 0.025 | 0.025 | 1.322 | 0.14 | 0.48 | 0.21 | 0.26 | 0.1 | 0.19 |
| 5 | 17183891 | rs177053 | -0.709 | 0.28 | -0.617 | 1.018 | -0.003 | 0.023 | 0.023 | 0.031 | 0.06 | 0.17 | 0.04 | 0.17 | 0.03 | 0.11 |
| 5 | 17185433 | rs297197 | -0.38 | 0.532 | -1.459 | 1.083 | -0.103 | -0.081 | -0.081 | 0.591 | 0.7 | 0.98 | 0.81 | 0.69 | 0.69 | 0.69 |
| 5 | 17193633 | rs585991 | 0.277 | -0.136 | -0.738 | 1.215 | 0.003 | 0.006 | 0.006 | 0.814 | 0.3 | 0.25 | 0.26 | 0.19 | 0.25 | 0.24 |
| 5 | 17193708 | rs586087 | -0.658 | 0.105 | -0.633 | 1.021 | -0.009 | 0.019 | 0.019 | 0.053 | 0.62 | 0.7 | 0.76 | 0.74 | 0.6 | 0.64 |
| 5 | 17193856 | rs586971 | -1.216 | -0.014 | -0.578 | 0.809 | -0.042 | 0.012 | 0.012 | 0 | 0.7 | 0.87 | 0.83 | 0.74 | 0.7 | 0.69 |
| 5 | 17194063 | rs587887 | -0.43 | -0.023 | -0.626 | 0.806 | -0.009 | 0.019 | 0.019 | 0.051 | 0.62 | 0.7 | 0.76 | 0.74 | 0.6 | 0.64 |
| 5 | 17194395 | rs2929711 | -1.205 | 0.023 | -0.996 | 0.843 | -0.045 | 0.01 | 0.01 | 0 | 0.7 | 0.87 | 0.83 | 0.74 | 0.7 | 0.68 |

| | | | | | | | | | | | | | | | | |
|---|---|---|---|---|---|---|---|---|---|---|---|---|---|---|---|---|
| 5 | 17197505 | rs1542598 | -0.172 | -0.061 | -0.779 | 0.877 | -0.001 | -0.001 | -0.001 | 0.463 | 0.53 | 0.44 | 0.59 | 0.41 | 0.54 | 0.47 |
| 5 | 17200074 | rs11952802 | 0.263 | 0.301 | -1.173 | 1.047 | -0.02 | -0.02 | -0.02 | 0 | 0.21 | 0.13 | 0.32 | 0.2 | 0.29 | 0.23 |
| 5 | 17211545 | rs2731792 | 0.862 | -0.012 | -1.508 | 0.913 | -0.015 | -0.041 | -0.041 | 5.296 | 0.27 | 0.02 | 0.09 | 0.2 | 0.21 | 0.14 |
| 5 | 17211649 | rs2929726 | -0.03 | 0.311 | -0.929 | 1.044 | -0.006 | -0.001 | -0.001 | 0 | 0.59 | 0.37 | 0.58 | 0.5 | 0.56 | 0.48 |
| 5 | 17221016 | rs1482344 | 0.474 | 0.32 | -0.271 | 1.186 | -0.024 | 0.115 | 0.115 | 0 | 0.73 | 0.48 | 0.55 | 0.47 | 0.77 | 0.7 |
| 5 | 17225333 | rs2242415 | 0.42 | 1.128 | -0.755 | 0.485 | -0.026 | 0.011 | 0.011 | 0 | 0.23 | 0.2 | 0.22 | 0.19 | 0.38 | 0.29 |
| 5 | 17228908 | rs2291236 | -0.359 | 0.737 | -0.725 | -0.267 | -0.02 | 0.065 | 0.065 | 0.052 | 0.43 | 0.34 | 0.31 | 0.19 | 0.51 | 0.34 |
| 5 | 17234794 | rs12716119 | -0.495 | 0.685 | -0.032 | -0.132 | 0.044 | 0.032 | 0.032 | 0 | 0.07 | 0.24 | 0.17 | 0.06 | 0.04 | 0.18 |
| 5 | 17247021 | rs10053617 | 0.201 | 0.491 | -0.422 | -0.289 | 0.004 | 0.013 | 0.013 | 0 | 0.16 | 0.13 | 0.07 | 0 | 0.11 | 0.05 |
| 5 | 17249420 | rs953431 | 0.101 | 0.211 | -0.382 | 0.362 | -0.001 | 0.003 | 0.003 | 0 | 0.2 | 0.19 | 0.29 | 0.29 | 0.21 | 0.3 |
| 5 | 17255026 | rs11133892 | -0.043 | 0.358 | -0.156 | 0.093 | 0.001 | -0.002 | -0.002 | 0 | 0.53 | 0.5 | 0.5 | 0.22 | 0.59 | 0.45 |
| 5 | 17255803 | rs2937754 | 0.11 | 0.435 | -0.287 | 0.634 | 0.003 | 0.03 | 0.03 | 0 | 0.16 | 0.2 | 0.25 | 0.34 | 0.16 | 0.38 |
| 5 | 17264096 | rs1500508 | 0.692 | 0.465 | -1.108 | 0.519 | -0.048 | 0.056 | 0.056 | 0 | 0.3 | 0.07 | 0.41 | 0.22 | 0.37 | 0.39 |
| 5 | 17264914 | rs298592 | -0.602 | -0.01 | -0.952 | 1.135 | 0.043 | -0.129 | -0.129 | 0.92 | 0.75 | 0.96 | 0.88 | 0.62 | 0.81 | 0.84 |
| 5 | 17276986 | rs298563 | -0.245 | -0.145 | -0.198 | 1.173 | -0.042 | 0.044 | 0.044 | 0 | 0.73 | 0.62 | 0.88 | 0.57 | 0.84 | 0.76 |
| 5 | 17283265 | rs177110 | 0.339 | 0.515 | -0.442 | 0.544 | -0.011 | 0.104 | 0.104 | 0 | 0.53 | 0.75 | 0.55 | 0.25 | 0.49 | 0.43 |
| 5 | 17289965 | rs17615463 | 0.702 | 0.18 | -0.457 | 0.411 | 0.061 | 0.14 | 0.14 | 0.375 | 0.45 | 0.12 | 0.37 | 0.22 | 0.36 | 0.4 |
| 5 | 17296342 | rs12523570 | 0.553 | 0.908 | 0.223 | 0.946 | 0.015 | 0.017 | 0.017 | 0 | 0.64 | 0.44 | 0.46 | 0.47 | 0.54 | 0.64 |
| 5 | 17298852 | rs298564 | 0.06 | 1.273 | 0.177 | 0.901 | 0.004 | 0 | 0 | 0.714 | 0.11 | 0.11 | 0.13 | 0.12 | 0.12 | 0.06 |
| 5 | 17306610 | rs16869688 | -0.291 | 1.051 | 0.59 | 1.01 | 0.056 | 0.062 | 0.062 | 2.653 | 0.08 | 0.31 | 0.09 | 0.11 | 0.1 | 0.1 |
| 5 | 17306790 | rs298512 | 0.51 | 0.848 | 0.036 | 1.078 | 0.009 | 0.002 | 0.002 | 0 | 0.48 | 0.26 | 0.5 | 0.43 | 0.48 | 0.47 |
| 5 | 17316993 | rs298510 | -0.231 | 1.883 | 0.626 | 1.414 | -0.074 | -0.052 | -0.052 | 0 | 0.78 | 0.93 | 0.69 | 0.7 | 0.71 | 0.77 |
| 5 | 17319116 | rs7707380 | -0.58 | 1.671 | 0.724 | 1.345 | -0.049 | -0.04 | -0.04 | 0.023 | 0.81 | 0.9 | 0.77 | 0.68 | 0.73 | 0.79 |
| 5 | 17321707 | rs10475103 | -1.641 | 1.328 | 1.221 | 0.777 | 0.026 | -0.02 | -0.02 | 0 | 0.26 | 0.51 | 0.44 | 0.23 | 0.3 | 0.24 |
| 5 | 17327540 | rs298556 | -0.827 | 1.39 | 0.926 | 0.8 | -0.1 | -0.086 | -0.086 | 0.006 | 0.75 | 0.94 | 0.68 | 0.67 | 0.68 | 0.75 |
| 5 | 17328269 | rs1995289 | -0.726 | 1.408 | 1.028 | 0.795 | -0.017 | 0.015 | 0.015 | 0 | 0.17 | 0.09 | 0.32 | 0.11 | 0.19 | 0.14 |
| 5 | 17333950 | rs298552 | 0.148 | 1.428 | 1.138 | 0.795 | 0.021 | 0.013 | 0.013 | 0 | 0.64 | 0.42 | 0.56 | 0.56 | 0.57 | 0.65 |
| 5 | 17347706 | rs10866522 | -0.279 | 1.474 | 0.919 | 0.698 | -0.014 | 0.02 | 0.02 | 0 | 0.11 | 0.03 | 0.17 | 0 | 0.12 | 0.07 |
| 5 | 17351275 | rs2650944 | -0.281 | 1.485 | 1.323 | 0.818 | 0.016 | 0.023 | 0.023 | 0.148 | 0.08 | 0.19 | 0.09 | 0.12 | 0.09 | 0.1 |
| 5 | 17358730 | rs4702207 | 0.198 | 1.481 | 1.139 | 0.827 | 0.035 | 0.044 | 0.044 | 0 | 0.65 | 0.47 | 0.57 | 0.5 | 0.58 | 0.66 |
| 5 | 17361908 | rs2036851 | 1.111 | 0.975 | 1.15 | 1.002 | 0.027 | 0.011 | 0.011 | 0 | 0.47 | 0.35 | 0.24 | 0.38 | 0.38 | 0.51 |
| 5 | 17366805 | rs177108 | -0.669 | 1.033 | 1.126 | 1.012 | -0.114 | -0.078 | -0.078 | 0 | 0.7 | 0.94 | 0.65 | 0.64 | 0.62 | 0.73 |
| 5 | 17368172 | rs299384 | 0.075 | 1.154 | 1.019 | 1.006 | 0.017 | 0.034 | 0.034 | 0 | 0.07 | 0.14 | 0.09 | 0.16 | 0.1 | 0.1 |
| 5 | 17369216 | rs298529 | 0.389 | 1.157 | 0.927 | 1 | 0.004 | -0.001 | -0.001 | 0 | 0.54 | 0.53 | 0.33 | 0.52 | 0.48 | 0.62 |
| 5 | 17375767 | rs16869742 | -0.164 | 0.378 | 0.356 | 0.677 | 0.001 | -0.001 | -0.001 | 0 | 0.13 | 0.15 | 0.29 | 0.12 | 0.13 | 0.11 |
| 5 | 17380270 | rs6876143 | -0.269 | 0.354 | 0.353 | 0.68 | 0.016 | 0.019 | 0.019 | 0 | 0.6 | 0.46 | 0.53 | 0.47 | 0.52 | 0.64 |
| 5 | 17381321 | rs1877128 | -0.222 | 0.414 | 0.353 | 0.705 | 0.013 | -0.005 | -0.005 | 0 | 0.14 | 0.22 | 0.3 | 0.12 | 0.13 | 0.11 |
| 5 | 17391377 | rs7729265 | 0.431 | -0.046 | -0.144 | 0.66 | 0.062 | 0.034 | 0.034 | 0 | 0.47 | 0.28 | 0.24 | 0.35 | 0.39 | 0.53 |
| 5 | 17400274 | rs1397576 | -0.774 | 0.222 | 0.203 | 0.604 | 0.006 | 0.023 | 0.023 | 0 | 0.21 | 0.14 | 0.32 | 0.11 | 0.16 | 0.15 |
| 5 | 17401187 | rs7712657 | 1.523 | 0.429 | 1.086 | 1.306 | 0.046 | 0.256 | 0.256 | 0.671 | 0.94 | 0.67 | 0.96 | 0.79 | 0.98 | 0.96 |
| 5 | 17403662 | rs2173758 | 1.117 | 0.637 | 0.89 | 0.846 | 0.024 | 0.018 | 0.018 | 0.055 | 0.93 | 0.71 | 0.92 | 0.91 | 0.92 | 0.94 |
| 5 | 17404483 | rs2173759 | -0.752 | 0.55 | 0.517 | 0.599 | -0.001 | -0.012 | -0.012 | 0.224 | 0.17 | 0.22 | 0.13 | 0.09 | 0.16 | 0.27 |
| 5 | 17404549 | rs2036339 | 0.919 | 0.46 | 0.13 | 0.906 | 0 | 0.012 | 0.012 | 0 | 0.91 | 0.88 | 0.96 | 0.8 | 0.92 | 0.88 |
| 5 | 17409558 | rs10475105 | 0.398 | 0.224 | -0.779 | -0.244 | 0.016 | 0.166 | 0.166 | 0 | 0.34 | 0.1 | 0.16 | 0.04 | 0.25 | 0.3 |
| 5 | 17409998 | rs11133899 | -0.399 | 0.207 | 0.083 | -0.385 | 0.021 | 0 | 0 | 0 | 0.13 | 0.27 | 0.11 | 0.13 | 0.15 | 0.11 |
| 5 | 17418256 | rs2457761 | 0.136 | -0.113 | -0.025 | -0.138 | -0.001 | -0.022 | -0.022 | 0 | 0.5 | 0.53 | 0.71 | 0.83 | 0.58 | 0.58 |
| 5 | 17418330 | rs2457762 | 0.272 | -0.116 | -0.021 | -0.133 | -0.001 | -0.023 | -0.023 | 0 | 0.5 | 0.53 | 0.71 | 0.83 | 0.59 | 0.58 |
| 5 | 17421158 | rs2941677 | 0.595 | -0.109 | -0.133 | -0.145 | -0.013 | 0.044 | 0.044 | 0 | 0.78 | 0.54 | 0.83 | 0.67 | 0.77 | 0.83 |
| 5 | 17422915 | rs6868322 | 0.678 | -0.008 | -0.631 | -0.291 | 0.021 | 0.117 | 0.117 | 0 | 0.34 | 0.16 | 0.18 | 0.04 | 0.26 | 0.3 |
| 5 | 17430052 | rs1553847 | 0.494 | 0.637 | -0.832 | 0.497 | -0.058 | -0.075 | -0.075 | 0.786 | 0.29 | 0.15 | 0.53 | 0.43 | 0.39 | 0.33 |
| 5 | 17430238 | rs7718456 | 0.168 | 0.557 | -0.885 | 0.7 | 0.042 | -0.028 | -0.028 | 0 | 0.48 | 0.16 | 0.24 | 0.51 | 0.41 | 0.42 |
| 5 | 17434249 | rs2459791 | -0.022 | 0.696 | -0.647 | 0.196 | 0.022 | 0.065 | 0.065 | 0 | 0.51 | 0.62 | 0.64 | 0.72 | 0.61 | 0.51 |
| 5 | 17442045 | rs2459830 | -0.009 | 0.85 | -0.937 | 0.005 | -0.009 | 0.063 | 0.063 | 0 | 0.48 | 0.35 | 0.38 | 0.29 | 0.5 | 0.44 |
| 5 | 17443654 | rs7706512 | 0.028 | 0.792 | -0.753 | -0.039 | -0.007 | 0.049 | 0.049 | 0 | 0.47 | 0.37 | 0.38 | 0.28 | 0.5 | 0.45 |
| 5 | 17453016 | rs2459826 | -0.324 | 1.325 | -0.85 | -0.225 | -0.016 | 0.015 | 0.015 | 0.055 | 0.87 | 0.91 | 0.85 | 0.95 | 0.78 | 0.84 |
| 5 | 17455566 | rs6863447 | 0.949 | 1.229 | -1.327 | -0.512 | -0.012 | -0.01 | -0.01 | 0 | 0.18 | 0.06 | 0.14 | 0.2 | 0.16 | 0.12 |
| 5 | 17457266 | rs4318775 | 1.003 | 0.656 | -0.973 | -0.553 | 0.029 | -0.02 | -0.02 | 0 | 0.29 | 0.19 | 0.36 | 0.53 | 0.19 | 0.29 |
| 5 | 17477812 | rs2459822 | 0.56 | 0.857 | -0.406 | -0.663 | -0.004 | 0.009 | 0.009 | 0 | 0.34 | 0.33 | 0.31 | 0.24 | 0.41 | 0.42 |
| 5 | 17479389 | rs2459821 | -0.895 | 1.039 | -0.826 | -0.573 | -0.013 | 0.015 | 0.015 | 0 | 0.86 | 0.91 | 0.87 | 0.97 | 0.8 | 0.86 |
| 5 | 17484038 | rs1510960 | -0.787 | 0.595 | -0.367 | -0.156 | -0.001 | 0.004 | 0.004 | 0 | 0.79 | 0.85 | 0.85 | 0.83 | 0.81 | 0.8 |
| 5 | 17489383 | rs11133906 | 0.351 | 0.376 | 0.137 | 0.472 | 0.022 | -0.047 | -0.047 | 0 | 0.3 | 0.19 | 0.38 | 0.55 | 0.23 | 0.28 |
| 5 | 17509888 | rs2402396 | 0.159 | 0.919 | 0.146 | 0.418 | -0.041 | 0.019 | 0.019 | 0 | 0.07 | 0.01 | 0.12 | 0.02 | 0.15 | 0.06 |
| 5 | 17697085 | rs7718648 | 0.223 | 0.1 | 0.697 | 0.873 | 0.005 | -0.01 | -0.01 | 0.132 | 0.26 | 0.23 | 0.42 | 0.52 | 0.21 | 0.26 |
| 5 | 17704161 | rs1514846 | 0.392 | 0.023 | 1.042 | 0.808 | 0.02 | -0.021 | -0.021 | 0 | 0.67 | 0.34 | 0.52 | 0.43 | 0.75 | 0.67 |
| 5 | 17710478 | rs13190307 | 0.274 | 0.017 | 0.626 | 0.899 | 0.017 | -0.042 | -0.042 | 0 | 0.26 | 0.19 | 0.41 | 0.52 | 0.21 | 0.26 |
| 5 | 17719837 | rs6865140 | 0.021 | 0.019 | 0.649 | 0.998 | -0.044 | 0.076 | 0.076 | 0 | 0.69 | 0.55 | 0.55 | 0.47 | 0.77 | 0.68 |
| 5 | 17727088 | rs10805605 | 0.043 | 0.205 | 0.388 | 0.997 | 0.004 | -0.012 | -0.012 | 0 | 0.28 | 0.25 | 0.43 | 0.5 | 0.25 | 0.26 |
| 5 | 17727683 | rs6896861 | 0.571 | 0.191 | 0.069 | 0.627 | 0.059 | -0.001 | -0.001 | 0.38 | 0.51 | 0.26 | 0.43 | 0.5 | 0.61 | 0.56 |
| 5 | 17730304 | rs6450962 | 0.527 | 0.002 | 0.12 | 1.099 | -0.001 | 0 | 0 | 0 | 0.23 | 0.21 | 0.34 | 0.43 | 0.24 | 0.26 |

| | | | | | | | | | | | | | | | | |
|---|---|---|---|---|---|---|---|---|---|---|---|---|---|---|---|---|
| 5 | 17732806 | rs1514866 | 0.433 | 0.089 | -0.162 | 0.815 | -0.007 | -0.001 | -0.001 | 0 | 0.35 | 0.34 | 0.28 | 0.38 | 0.38 | 0.32 |
| 5 | 17734581 | rs6894996 | -0.541 | 0.012 | -0.193 | 0.568 | -0.001 | 0.017 | 0.017 | 0 | 0.7 | 0.82 | 0.69 | 0.81 | 0.69 | 0.64 |
| 5 | 17735395 | rs2379265 | -0.624 | 0.103 | -0.23 | 0.652 | -0.015 | 0.001 | 0.001 | 0 | 0.71 | 0.81 | 0.66 | 0.78 | 0.66 | 0.64 |
| 5 | 17738703 | rs7445769 | 0.453 | -0.018 | 0.183 | 0.682 | -0.005 | 0.002 | 0.002 | 0.004 | 0.36 | 0.38 | 0.37 | 0.43 | 0.31 | 0.33 |
| 5 | 17739449 | rs10461717 | 0.529 | 0.009 | 0.211 | 0.727 | -0.005 | 0.003 | 0.003 | 0 | 0.33 | 0.3 | 0.28 | 0.35 | 0.34 | 0.32 |
| 5 | 17745762 | rs1514855 | -0.476 | 0.159 | 0.162 | 0.58 | 0.006 | -0.003 | -0.003 | 0 | 0.69 | 0.64 | 0.64 | 0.77 | 0.65 | 0.64 |
| 5 | 17746787 | rs1514857 | -0.486 | 0.162 | 0.089 | 0.581 | 0.002 | -0.002 | -0.002 | 0 | 0.69 | 0.69 | 0.64 | 0.77 | 0.65 | 0.64 |
| 5 | 17761225 | rs11959046 | 0.347 | -0.081 | -0.159 | 0.671 | 0.037 | -0.005 | -0.005 | 0.46 | 0.5 | 0.28 | 0.53 | 0.48 | 0.56 | 0.63 |
| 5 | 17762135 | rs6451281 | 1.105 | -0.388 | -0.025 | 0.465 | -0.008 | 0.098 | 0.098 | 0 | 0.26 | 0.47 | 0.35 | 0.46 | 0.25 | 0.27 |
| 5 | 17764015 | rs1913602 | -0.072 | -0.414 | -0.051 | 0.33 | 0.001 | -0.018 | -0.018 | 0 | 0.86 | 0.8 | 0.93 | 0.93 | 0.86 | 0.95 |
| 5 | 17764650 | rs1514872 | 0.969 | -0.656 | 0.225 | 0.378 | -0.019 | -0.023 | -0.023 | 0.068 | 0.13 | 0.19 | 0.05 | 0.02 | 0.05 | 0.06 |
| 5 | 17780459 | rs7737273 | -0.256 | -0.593 | 0.335 | 0.548 | 0.002 | -0.022 | -0.022 | 0 | 0.65 | 0.72 | 0.61 | 0.48 | 0.63 | 0.68 |
| 5 | 17784226 | rs10941533 | 1.394 | -0.655 | 0.376 | 0.836 | -0.002 | -0.042 | -0.042 | 0.164 | 0.23 | 0.12 | 0.22 | 0.38 | 0.24 | 0.27 |
| 5 | 17790562 | rs11960755 | 1.176 | -0.56 | 0.163 | 0.523 | -0.028 | 0.006 | 0.006 | 0 | 0.27 | 0.13 | 0.25 | 0.19 | 0.32 | 0.26 |
| 5 | 17790612 | rs7714225 | -1.685 | -0.269 | 0.248 | 0.672 | 0 | -0.027 | -0.027 | 1.326 | 0.79 | 0.91 | 0.7 | 0.55 | 0.78 | 0.74 |
| 5 | 17791030 | rs11958964 | 0.233 | -0.234 | 0.286 | 0.67 | -0.012 | -0.014 | -0.014 | 0 | 0.12 | 0.16 | 0.06 | 0.07 | 0.1 | 0.05 |
| 5 | 17791844 | rs1822399 | -0.407 | -0.323 | 0.262 | 0.605 | -0.003 | 0.006 | 0.006 | 0 | 0.68 | 0.72 | 0.64 | 0.48 | 0.69 | 0.69 |
| 5 | 17794035 | rs11738744 | 0.607 | 0.203 | -0.079 | 0.83 | 0.006 | -0.004 | -0.004 | 0.88 | 0.28 | 0.36 | 0.33 | 0.27 | 0.32 | 0.37 |
| 5 | 17796513 | rs7713161 | -1.559 | -0.348 | -0.078 | 1.236 | -0.017 | -0.089 | -0.089 | 0.051 | 0.81 | 0.95 | 0.71 | 0.56 | 0.79 | 0.73 |
| 5 | 17802524 | rs10064153 | -0.727 | -0.23 | 0.119 | 1.266 | -0.015 | 0.03 | 0.03 | 0 | 0.73 | 0.66 | 0.65 | 0.59 | 0.75 | 0.7 |
| 5 | 17802864 | rs971716 | -0.728 | 0.086 | -0.02 | 0.754 | -0.036 | 0.001 | 0.001 | 0 | 0.72 | 0.78 | 0.67 | 0.72 | 0.6 | 0.69 |
| 5 | 17803067 | rs971715 | 0.136 | -0.213 | 0.458 | 1.457 | -0.017 | 0.05 | 0.05 | 0 | 0.28 | 0.34 | 0.35 | 0.48 | 0.25 | 0.31 |
| 5 | 17803270 | rs6876885 | -0.056 | -0.201 | -0.047 | 1.451 | -0.012 | 0.048 | 0.048 | 0 | 0.29 | 0.5 | 0.37 | 0.48 | 0.28 | 0.31 |
| 5 | 17804005 | rs9637792 | -0.756 | -0.183 | 0.234 | 1.559 | 0.061 | 0.24 | 0.24 | 0.016 | 0.08 | 0.41 | 0.11 | 0.16 | 0.07 | 0.02 |
| 5 | 17805883 | rs9292873 | -0.468 | -0.124 | -0.199 | 1.332 | -0.003 | 0 | 0 | 0.461 | 0.25 | 0.25 | 0.19 | 0.19 | 0.21 | 0.18 |
| 5 | 17806954 | rs16887560 | -0.983 | 0.486 | -0.016 | 1.27 | 0.08 | 0.172 | 0.172 | 0 | 0.08 | 0.39 | 0.1 | 0.14 | 0.07 | 0.02 |
| 5 | 17807202 | rs16887557 | -0.918 | 0.499 | -0.01 | 1.276 | 0.085 | 0.177 | 0.177 | 0 | 0.08 | 0.39 | 0.1 | 0.14 | 0.07 | 0.02 |
| 5 | 17808002 | rs17710639 | 0.716 | 0.248 | -1.725 | 1.043 | 0.02 | -0.061 | -0.061 | 0.205 | 0.55 | 0.11 | 0.61 | 0.55 | 0.65 | 0.57 |
| 5 | 17809455 | rs13166250 | 0.731 | 0.408 | -2.004 | 0.884 | -0.011 | -0.034 | -0.034 | 0 | 0.49 | 0.1 | 0.53 | 0.51 | 0.53 | 0.54 |
| 5 | 17809984 | rs6881131 | 0.605 | 0.598 | -1.845 | 0.745 | 0.074 | -0.035 | -0.035 | 0 | 0.56 | 0.16 | 0.58 | 0.51 | 0.63 | 0.58 |
| 5 | 17818316 | rs7341145 | 0.341 | 0.318 | -0.523 | 0.978 | -0.003 | -0.002 | -0.002 | 0 | 0.63 | 0.61 | 0.7 | 0.73 | 0.71 | 0.59 |
| 5 | 17819987 | rs6872569 | 0.917 | 0.352 | 0.103 | 0.947 | 0.008 | 0.018 | 0.018 | 0 | 0.55 | 0.36 | 0.62 | 0.72 | 0.61 | 0.54 |
| 5 | 17820745 | rs1513088 | 0.543 | 0.309 | 0.562 | 0.956 | -0.033 | -0.025 | -0.025 | 0 | 0.61 | 0.5 | 0.68 | 0.73 | 0.71 | 0.59 |
| 5 | 17828138 | rs10079374 | 0.125 | -0.061 | 0.538 | 0.87 | -0.004 | 0.025 | 0.025 | 0 | 0.26 | 0.06 | 0.25 | 0.16 | 0.25 | 0.21 |
| 5 | 17836897 | rs6896120 | 1.861 | 0.161 | 1.495 | 0.642 | 0.005 | -0.046 | -0.046 | 0 | 0.77 | 0.34 | 0.84 | 0.88 | 0.8 | 0.73 |
| 5 | 17849527 | rs4866172 | 0.908 | 0.159 | 1.378 | 0.667 | -0.003 | 0.018 | 0.018 | 0.683 | 0.88 | 0.63 | 0.94 | 0.89 | 0.93 | 0.79 |
| 5 | 17861315 | rs1513081 | 0.259 | 0.314 | 0.967 | 0.697 | 0.002 | -0.006 | -0.006 | 0 | 0.06 | 0.06 | 0.04 | 0.01 | 0.1 | 0.03 |
| 5 | 17891126 | rs1542128 | 0.07 | 0.326 | 1.142 | 0.794 | 0.013 | 0.015 | 0.015 | 0.7 | 0.94 | 0.85 | 0.97 | 0.89 | 0.93 | 0.82 |
| 5 | 17893556 | rs17710915 | 0.045 | 0.328 | 1.329 | 1.012 | 0.016 | 0.137 | 0.137 | 0 | 0.05 | 0.13 | 0.05 | 0.2 | 0.08 | 0.01 |
| 5 | 17896146 | rs7726178 | 1.692 | 0.084 | 1.324 | 0.819 | 0.003 | 0.003 | 0.003 | 0 | 0.49 | 0.31 | 0.53 | 0.51 | 0.51 | 0.49 |
| 5 | 17899205 | rs6863677 | 1.544 | 0.075 | 1.817 | 0.778 | -0.001 | 0.003 | 0.003 | 2.232 | 0.87 | 0.64 | 0.92 | 0.89 | 0.91 | 0.77 |
| 5 | 17902917 | rs12655829 | 0.468 | 0.113 | 1.747 | 0.79 | 0.024 | 0.025 | 0.025 | 0.169 | 0.92 | 0.78 | 0.96 | 0.89 | 0.93 | 0.81 |
| 5 | 17918575 | rs16887451 | -0.628 | 0.127 | 1.579 | 0.711 | 0.02 | -0.04 | -0.04 | 0 | 0.09 | 0.13 | 0.06 | 0.01 | 0.1 | 0.04 |
| 5 | 17920818 | rs12514079 | 0.084 | 0.027 | 1.355 | 0.77 | 0.022 | 0.016 | 0.016 | 0 | 0.25 | 0.11 | 0.29 | 0.17 | 0.22 | 0.21 |
| 5 | 17922072 | rs1443402 | -1.231 | 0.093 | 1.408 | 0.746 | -0.015 | 0.014 | 0.014 | 0 | 0.4 | 0.48 | 0.39 | 0.49 | 0.39 | 0.45 |
| 5 | 17929608 | rs1513089 | 0.722 | 0.042 | 0.864 | 0.605 | -0.002 | -0.016 | -0.016 | 0 | 0.89 | 0.89 | 0.92 | 1 | 0.87 | 0.96 |
| 5 | 17930264 | rs6869207 | -1.096 | 0.341 | 0.776 | 0.397 | 0.004 | -0.003 | -0.003 | 0.158 | 0.42 | 0.66 | 0.45 | 0.51 | 0.4 | 0.48 |
| 5 | 17933120 | rs13190514 | 1.967 | 0.399 | 0.087 | 0.417 | -0.063 | -0.075 | -0.075 | 0 | 0.43 | 0.08 | 0.44 | 0.48 | 0.45 | 0.44 |
| 5 | 17933360 | rs4866258 | 0.155 | 0.569 | 0.17 | 0.343 | 0.006 | -0.021 | -0.021 | 0 | 0.91 | 0.9 | 0.92 | 1 | 0.87 | 0.96 |
| 5 | 17953026 | rs782847 | -0.158 | 0.305 | -0.166 | 0.292 | -0.016 | 0.01 | 0.01 | 0 | 0.16 | 0.11 | 0.1 | 0.13 | 0.13 | 0.25 |
| 5 | 17956340 | rs1033148 | 1.904 | 0.173 | -0.463 | 0.455 | -0.067 | -0.071 | -0.071 | 0.677 | 0.37 | 0.1 | 0.44 | 0.45 | 0.43 | 0.43 |
| 5 | 17961096 | rs1813454 | 0.201 | 0.097 | -0.064 | 0.89 | 0.021 | 0.035 | 0.035 | 0 | 0.22 | 0.05 | 0.26 | 0.1 | 0.19 | 0.2 |
| 5 | 17981553 | rs10941929 | -0.64 | 0.277 | 0.514 | 1.368 | 0 | 0.118 | 0.118 | 0 | 0.06 | 0.1 | 0.04 | 0.3 | 0.09 | 0.02 |
| 5 | 17996519 | rs6869139 | -0.051 | 0.188 | 0.526 | 0.626 | 0.009 | 0.011 | 0.011 | 0 | 0.32 | 0.26 | 0.32 | 0.17 | 0.23 | 0.3 |
| 5 | 17999554 | rs2662605 | -0.473 | 0.477 | 0.642 | 0.508 | 0.024 | -0.029 | -0.029 | 0 | 0.09 | 0.14 | 0.07 | 0.02 | 0.13 | 0.04 |
| 5 | 18000871 | rs692332 | -0.198 | -0.249 | 0.81 | 0.783 | -0.006 | 0.01 | 0.01 | 0.458 | 0.92 | 0.87 | 0.95 | 0.88 | 0.94 | 0.88 |
| 5 | 18007227 | rs505682 | 0.396 | -0.295 | 0.84 | 0.715 | -0.01 | -0.008 | -0.008 | 0.58 | 0.61 | 0.6 | 0.63 | 0.72 | 0.71 | 0.58 |
| 5 | 18009498 | rs691027 | 0.349 | 0.004 | 1.065 | 1.102 | 0.038 | 0.194 | 0.194 | 3.384 | 0.91 | 0.63 | 0.8 | 0.74 | 0.92 | 0.83 |
| 5 | 18010430 | rs7733771 | -0.431 | 0.005 | 1.144 | 1.103 | 0.017 | 0.139 | 0.139 | 2.953 | 0.09 | 0.3 | 0.21 | 0.26 | 0.08 | 0.17 |
| 5 | 18011605 | rs704029 | -0.665 | -0.152 | 0.989 | 1.069 | 0.022 | 0.091 | 0.091 | 0.83 | 0.47 | 0.72 | 0.56 | 0.68 | 0.43 | 0.56 |
| 5 | 18017619 | rs1976571 | 1.227 | -0.257 | 0.497 | 0.954 | -0.064 | 0.117 | 0.117 | 0 | 0.43 | 0.13 | 0.39 | 0.28 | 0.53 | 0.38 |
| 5 | 18022107 | rs17839277 | 0.138 | -0.1 | 0.565 | 1.025 | 0 | -0.002 | -0.002 | 0.498 | 0.09 | 0.07 | 0.03 | 0.04 | 0.03 | 0.06 |
| 5 | 18030481 | rs10061385 | -0.389 | 0.227 | 0.882 | 1.415 | 0.031 | 0.18 | 0.18 | 3.12 | 0.09 | 0.35 | 0.21 | 0.26 | 0.08 | 0.17 |
| 5 | 18035736 | rs1390630 | -0.072 | 0.236 | 0.374 | 1.314 | 0.001 | -0.054 | -0.054 | 5.462 | 0.35 | 0.2 | 0.35 | 0.4 | 0.36 | 0.39 |
| 5 | 18048683 | rs13186768 | 0.566 | 0.36 | -0.668 | 0.579 | 0.018 | 0.122 | 0.122 | 0.758 | 0.54 | 0.22 | 0.36 | 0.26 | 0.59 | 0.47 |
| 5 | 18050107 | rs10941964 | -0.369 | 0.541 | -0.714 | 0.733 | 0 | -0.005 | -0.005 | 0 | 0.34 | 0.25 | 0.39 | 0.32 | 0.34 | 0.39 |
| 5 | 18051124 | rs7714990 | -0.367 | 0.283 | -0.705 | -0.165 | 0.001 | -0.008 | -0.008 | 0 | 0.25 | 0.23 | 0.14 | 0.05 | 0.24 | 0.22 |
| 5 | 18051144 | rs2455406 | 0.217 | 0.052 | -0.358 | 0.124 | -0.024 | -0.01 | -0.01 | 0 | 0.72 | 0.5 | 0.61 | 0.25 | 0.83 | 0.49 |
| 5 | 18052634 | rs691970 | -0.328 | 0.018 | -0.526 | -0.32 | 0.006 | 0.018 | 0.018 | 0 | 0.59 | 0.59 | 0.76 | 0.8 | 0.61 | 0.65 |

| | | | | | | | | | | | | | | | | |
|---|---|---|---|---|---|---|---|---|---|---|---|---|---|---|---|---|
| 5 | 18058107 | rs17711516 | -0.134 | -0.076 | -0.822 | -0.467 | 0 | 0.021 | 0.021 | 0.086 | 0.1 | 0.03 | 0.08 | 0.06 | 0.13 | 0.07 |
| 5 | 18066721 | rs645642 | -0.514 | -0.321 | -0.542 | 0.254 | -0.006 | 0.322 | 0.322 | 0 | 0.07 | 0.18 | 0.12 | 0.47 | 0.04 | 0.15 |
| 5 | 18068103 | rs618959 | -0.514 | -0.321 | -0.542 | 0.254 | -0.006 | 0.322 | 0.322 | 0 | 0.07 | 0.18 | 0.12 | 0.47 | 0.04 | 0.15 |
| 5 | 18074375 | rs691360 | -0.194 | -0.323 | 0.196 | 0.087 | 0.001 | 0.002 | 0.002 | 0.383 | 0.08 | 0.27 | 0.16 | 0.07 | 0.06 | 0.09 |
| 5 | 18078815 | rs606186 | -1.286 | -0.287 | 0.053 | 0.198 | -0.014 | 0.087 | 0.087 | 0.33 | 0.17 | 0.21 | 0.2 | 0.53 | 0.17 | 0.21 |
| 5 | 18081055 | rs6452084 | 0.549 | -0.102 | 0.276 | -0.002 | -0.018 | 0.03 | 0.03 | 0.008 | 0.37 | 0.23 | 0.43 | 0.21 | 0.41 | 0.35 |
| 5 | 18081343 | rs2561637 | -0.973 | -0.114 | 0.272 | 0.748 | -0.007 | 0.243 | 0.243 | 0 | 0.07 | 0.18 | 0.12 | 0.57 | 0.04 | 0.14 |
| 5 | 18092224 | rs609071 | -0.882 | -0.108 | -0.253 | 0.504 | 0.008 | -0.024 | -0.024 | 0 | 0.09 | 0.02 | 0.18 | 0.68 | 0.08 | 0.15 |
| 5 | 18095876 | rs10052761 | 0.146 | -0.027 | -0.253 | -0.203 | -0.041 | 0.056 | 0.056 | 0 | 0.37 | 0.27 | 0.43 | 0.2 | 0.41 | 0.44 |
| 5 | 18099766 | rs2561642 | -0.894 | -0.236 | -0.086 | -0.432 | 0.021 | 0.113 | 0.113 | 0 | 0.54 | 0.7 | 0.76 | 0.94 | 0.58 | 0.64 |
| 5 | 18126244 | rs6452106 | 0.139 | 0.117 | -0.415 | -0.007 | -0.047 | 0.007 | 0.007 | 0 | 0.34 | 0.19 | 0.44 | 0.28 | 0.42 | 0.44 |
| 5 | 18129805 | rs437171 | -0.765 | -0.505 | -0.309 | 0.368 | 0.01 | -0.043 | -0.043 | 0 | 0.23 | 0.1 | 0.28 | 0.6 | 0.22 | 0.2 |
| 5 | 18142966 | rs2443546 | -0.298 | -0.268 | 0.115 | 0.052 | -0.015 | -0.005 | -0.005 | 0 | 0.71 | 0.68 | 0.56 | 0.62 | 0.61 | 0.58 |
| 5 | 18146907 | rs10057043 | -0.703 | -0.794 | 0.643 | 0.755 | 0.024 | -0.07 | -0.07 | 0 | 0.21 | 0.07 | 0.22 | 0.55 | 0.18 | 0.19 |
| 5 | 18154453 | rs7727170 | -1.023 | -0.715 | 1.327 | 0.781 | -0.03 | 0.105 | 0.105 | 0 | 0.22 | 0.41 | 0.32 | 0.57 | 0.22 | 0.22 |
| 5 | 18168945 | rs10037793 | -1.131 | -0.65 | 0.849 | 0.8 | -0.012 | 0.018 | 0.018 | 0 | 0.25 | 0.3 | 0.35 | 0.61 | 0.2 | 0.27 |
| 5 | 18173795 | rs1504192 | -0.228 | -0.718 | 0.802 | 0.6 | 0.008 | -0.007 | -0.007 | 0 | 0.54 | 0.61 | 0.43 | 0.47 | 0.46 | 0.53 |
| 5 | 18176583 | rs442958 | -0.809 | -0.698 | 0.717 | 0.557 | -0.084 | 0.047 | 0.047 | 0 | 0.63 | 0.86 | 0.64 | 0.68 | 0.55 | 0.66 |
| 5 | 18194227 | rs4866305 | 0.045 | -0.533 | 0.888 | 0.703 | -0.005 | -0.041 | -0.041 | 0.437 | 0.16 | 0.08 | 0.21 | 0.44 | 0.18 | 0.24 |
| 5 | 18203619 | rs3849678 | -1.313 | -0.517 | 1.135 | 0.625 | -0.003 | 0.018 | 0.018 | 0 | 0.08 | 0.1 | 0.2 | 0.21 | 0.08 | 0.15 |
| 5 | 18215931 | rs17661265 | -0.323 | -0.498 | 0.955 | 0.683 | -0.011 | 0.044 | 0.044 | 0 | 0.05 | 0.09 | 0.12 | 0.18 | 0.02 | 0.11 |
| 5 | 18247297 | rs1587796 | 0.376 | -0.838 | 0.829 | 0.172 | 0.063 | 0.15 | 0.15 | 0.001 | 0.43 | 0.76 | 0.32 | 0.05 | 0.35 | 0.32 |
| 5 | 18264463 | rs1543947 | 0.671 | -0.711 | 0.852 | 0.376 | -0.047 | 0.032 | 0.032 | 0 | 0.37 | 0.2 | 0.37 | 0.32 | 0.44 | 0.34 |
| 5 | 18267949 | rs13179760 | 0.085 | -0.61 | 0.78 | 0.775 | -0.033 | -0.106 | -0.106 | 0 | 0.16 | 0.03 | 0.2 | 0.44 | 0.19 | 0.23 |
| 5 | 18268789 | rs6868510 | -1.679 | -0.5 | 1.499 | 0.716 | 0.043 | 0.05 | 0.05 | 0.671 | 0.08 | 0.37 | 0.11 | 0.04 | 0.07 | 0.04 |
| 5 | 18272737 | rs1354424 | 1.024 | -0.609 | 1.181 | 1.202 | -0.007 | 0.042 | 0.042 | 0 | 0.74 | 0.59 | 0.69 | 0.47 | 0.76 | 0.7 |
| 5 | 18277597 | rs1857707 | 0.142 | -0.583 | 0.663 | 1.356 | -0.033 | -0.087 | -0.087 | 0 | 0.16 | 0.01 | 0.09 | 0.3 | 0.16 | 0.22 |
| 5 | 18279260 | rs4866315 | 0.985 | -0.468 | 0.618 | 0.642 | 0.032 | 0.136 | 0.136 | 0 | 0.39 | 0.12 | 0.34 | 0.12 | 0.36 | 0.4 |
| 5 | 18281760 | rs1032714 | 1 | -0.449 | 1.045 | 0.747 | -0.017 | 0.077 | 0.077 | 0 | 0.72 | 0.46 | 0.67 | 0.48 | 0.76 | 0.7 |
| 5 | 18284077 | rs314020 | -0.153 | -0.144 | 0.593 | 0.854 | -0.04 | -0.123 | -0.123 | 0 | 0.19 | 0.03 | 0.23 | 0.48 | 0.2 | 0.26 |
| 5 | 18284394 | rs2962908 | 0.723 | -0.088 | 0.711 | 0.74 | -0.069 | -0.014 | -0.014 | 0 | 0.32 | 0.11 | 0.3 | 0.35 | 0.38 | 0.3 |
| 5 | 18288995 | rs2938443 | 0.45 | -0.253 | 0.861 | 0.671 | -0.004 | 0.001 | 0.001 | 0 | 0.34 | 0.31 | 0.31 | 0.32 | 0.4 | 0.3 |
| 5 | 18293483 | rs17630210 | 0.946 | 0.075 | -0.138 | 0.525 | -0.147 | -0.015 | -0.015 | 0.216 | 0.28 | 0.01 | 0.19 | 0.24 | 0.35 | 0.24 |
| 5 | 18300739 | rs314029 | -0.471 | -0.201 | 0.414 | 0.259 | 0.021 | 0.057 | 0.057 | 0 | 0.58 | 0.65 | 0.59 | 0.85 | 0.6 | 0.56 |
| 5 | 18304552 | rs375593 | -0.507 | -0.119 | 0.218 | 0.519 | -0.003 | -0.011 | -0.011 | 0 | 0.63 | 0.37 | 0.6 | 0.84 | 0.61 | 0.6 |
| 5 | 18319276 | rs314023 | -0.792 | -0.109 | 0.327 | 1.098 | 0.001 | -0.007 | -0.007 | 1.258 | 0.27 | 0.31 | 0.39 | 0.53 | 0.23 | 0.29 |
| 5 | 18324937 | rs2938469 | 0.814 | -0.212 | 0.524 | 0.863 | 0 | 0.009 | 0.009 | 0 | 0.34 | 0.17 | 0.39 | 0.25 | 0.41 | 0.33 |
| 5 | 18327605 | rs314025 | -1.204 | -0.14 | 0.926 | 1.574 | -0.009 | 0.078 | 0.078 | 0 | 0.1 | 0.17 | 0.18 | 0.39 | 0.07 | 0.08 |
| 5 | 18330038 | rs1500206 | -0.084 | -0.247 | 1.156 | 1.47 | 0 | 0 | 0 | 0.003 | 0.53 | 0.43 | 0.44 | 0.35 | 0.52 | 0.59 |
| 5 | 18335042 | rs2036328 | 1.545 | -0.316 | 1.542 | 1.514 | -0.009 | 0.311 | 0.311 | 0.28 | 0.88 | 0.5 | 0.81 | 0.61 | 0.94 | 0.89 |
| 5 | 18338272 | rs1405707 | 0.954 | -0.229 | 1.408 | 1.442 | -0.021 | 0.061 | 0.061 | 0 | 0.73 | 0.42 | 0.73 | 0.56 | 0.79 | 0.67 |
| 5 | 18351036 | rs2650445 | -0.545 | 0.121 | 1.556 | 1.139 | 0.008 | -0.021 | -0.021 | 0 | 0.62 | 0.48 | 0.75 | 0.71 | 0.65 | 0.62 |
| 5 | 18356268 | rs6895355 | -0.22 | -0.102 | 1.948 | 1.436 | 0.013 | -0.036 | -0.036 | 0 | 0.14 | 0.32 | 0.09 | 0.08 | 0.13 | 0.21 |
| 5 | 18356447 | rs11948468 | 1.163 | -0.031 | 2.268 | 1.955 | -0.004 | 0.002 | 0.002 | 0 | 0.78 | 0.4 | 0.72 | 0.76 | 0.81 | 0.74 |
| 5 | 18356582 | rs6879881 | -0.893 | 0.674 | 1.894 | 1.827 | 0.046 | 0.012 | 0.012 | 0 | 0.29 | 0.44 | 0.27 | 0.37 | 0.37 | 0.51 |
| 5 | 18362647 | rs2950483 | 0.269 | 0.732 | 1.904 | 1.89 | -0.003 | 0.102 | 0.102 | 0 | 0.06 | 0.17 | 0.04 | 0.17 | 0.02 | 0.04 |
| 5 | 18368594 | rs2204613 | 0.143 | 0.647 | 0.762 | 1.613 | -0.035 | -0.015 | -0.015 | 0 | 0.24 | 0.14 | 0.19 | 0.3 | 0.26 | 0.31 |
| 5 | 18379534 | rs10942142 | 1.521 | 0.468 | 0.796 | 1.725 | -0.038 | -0.033 | -0.033 | 0 | 0.51 | 0.15 | 0.52 | 0.44 | 0.55 | 0.39 |
| 5 | 18386675 | rs7720380 | -0.588 | 0.538 | 0.646 | 1.575 | -0.076 | 0.001 | 0.001 | 0.155 | 0.37 | 0.12 | 0.26 | 0.37 | 0.39 | 0.55 |
| 5 | 18393514 | rs1918342 | -2.244 | 0.524 | 1.516 | 2.091 | -0.043 | 0.106 | 0.106 | 1.966 | 0.15 | 0.68 | 0.2 | 0.36 | 0.06 | 0.12 |
| 5 | 18393812 | rs6452273 | 2.457 | 0.846 | 1.686 | 1.943 | -0.014 | 0.054 | 0.054 | 0.171 | 0.84 | 0.38 | 0.63 | 0.7 | 0.85 | 0.87 |
| 5 | 18394610 | rs10461822 | -0.902 | 1.02 | 1.619 | 1.48 | 0.024 | -0.024 | -0.024 | 0 | 0.32 | 0.4 | 0.41 | 0.21 | 0.41 | 0.4 |
| 5 | 18412476 | rs6898878 | 0.245 | 0.648 | 1.347 | 1.734 | 0 | -0.017 | -0.017 | 0 | 0.12 | 0.06 | 0.26 | 0.36 | 0.11 | 0.12 |
| 5 | 18414685 | rs17729677 | -0.948 | 0.583 | 1.713 | 1.546 | 0.002 | 0.012 | 0.012 | 0 | 0.34 | 0.39 | 0.41 | 0.5 | 0.32 | 0.41 |
| 5 | 18418083 | rs2962896 | 0.975 | 0.555 | 1.47 | 1.554 | 0.002 | 0.009 | 0.009 | 0.035 | 0.63 | 0.44 | 0.44 | 0.5 | 0.66 | 0.58 |
| 5 | 18423194 | rs2962954 | -0.597 | 0.584 | 1.429 | 1.617 | 0.004 | 0.042 | 0.042 | 0 | 0.17 | 0.33 | 0.16 | 0.22 | 0.18 | 0.15 |
| 5 | 18424199 | rs2962949 | -0.597 | 0.584 | 1.429 | 1.617 | 0.004 | 0.042 | 0.042 | 0 | 0.17 | 0.33 | 0.16 | 0.23 | 0.18 | 0.15 |
| 5 | 18433482 | rs10065090 | -0.869 | 0.571 | 1.464 | 1.649 | 0.012 | 0.141 | 0.141 | 0.008 | 0.1 | 0.33 | 0.12 | 0.22 | 0.11 | 0.07 |
| 5 | 18435446 | rs1405705 | 2.117 | 0.541 | 0.998 | 1.617 | -0.051 | 0.06 | 0.06 | 0 | 0.46 | 0.09 | 0.28 | 0.28 | 0.48 | 0.44 |
| 5 | 18439405 | rs1358319 | 0.325 | 0.283 | 1.196 | 1.572 | -0.015 | 0.069 | 0.069 | 0 | 0.06 | 0.13 | 0.04 | 0.15 | 0.03 | 0.05 |
| 5 | 18453812 | rs12657680 | 1.984 | 0.378 | 0.941 | 1.504 | 0.013 | 0.065 | 0.065 | 0 | 0.47 | 0.24 | 0.28 | 0.27 | 0.46 | 0.43 |
| 5 | 18455136 | rs17662227 | -1.02 | 0.327 | 1.054 | 1.649 | -0.015 | 0.131 | 0.131 | 0 | 0.2 | 0.34 | 0.46 | 0.51 | 0.17 | 0.16 |
| 5 | 18461047 | rs4866333 | 0.299 | 0.333 | 0.916 | 1.718 | 0.002 | 0.085 | 0.085 | 0 | 0.9 | 0.64 | 0.79 | 0.86 | 0.93 | 0.94 |
| 5 | 18464854 | rs9918121 | 1.145 | 0.322 | 1.039 | 1.73 | -0.017 | 0.192 | 0.192 | 0 | 0.79 | 0.52 | 0.52 | 0.49 | 0.83 | 0.84 |
| 5 | 18467699 | rs10063237 | -0.751 | 0.446 | 0.76 | 1.634 | -0.01 | -0.027 | -0.027 | 0 | 0.32 | 0.27 | 0.4 | 0.52 | 0.31 | 0.41 |
| 5 | 18482995 | rs10042810 | -1.494 | 0.559 | 0.783 | 1.441 | 0 | -0.015 | -0.015 | 0 | 0.93 | 0.93 | 0.96 | 1 | 0.92 | 0.93 |
| 5 | 18490517 | rs1980373 | 0.857 | 0.491 | 1.02 | 1.523 | 0.011 | 0.044 | 0.044 | 0 | 0.8 | 0.58 | 0.69 | 0.77 | 0.77 | 0.84 |
| 5 | 18492554 | rs1527383 | -0.64 | 0.482 | 1.074 | 1.518 | -0.007 | 0.013 | 0.013 | 0 | 0.17 | 0.34 | 0.24 | 0.15 | 0.15 | 0.14 |
| 5 | 18492839 | rs11750643 | 1.54 | 0.512 | 0.573 | 1.536 | -0.009 | 0.009 | 0.009 | 0 | 0.57 | 0.3 | 0.52 | 0.62 | 0.56 | 0.52 |

| | | | | | | | | | | | | | | | | |
|---|---|---|---|---|---|---|---|---|---|---|---|---|---|---|---|---|
| 5 | 18495544 | rs10520858 | 0.419 | 0.355 | 0.511 | 1.683 | -0.04 | 0.008 | 0.008 | 0 | 0.18 | 0.04 | 0.12 | 0.15 | 0.18 | 0.26 |
| 5 | 18495883 | rs1405708 | -1.161 | 0.322 | 0.603 | 1.687 | -0.016 | -0.006 | -0.006 | 0 | 0.33 | 0.52 | 0.35 | 0.31 | 0.25 | 0.38 |
| 5 | 18496264 | rs4334889 | -1.499 | 0.302 | 0.272 | 1.67 | 0 | -0.002 | -0.002 | 0 | 0.37 | 0.69 | 0.44 | 0.38 | 0.36 | 0.41 |
| 5 | 18500953 | rs4348220 | -1.308 | 0.192 | 0.082 | 1.709 | -0.003 | 0.005 | 0.005 | 0 | 0.35 | 0.59 | 0.43 | 0.38 | 0.33 | 0.41 |
| 5 | 18501530 | rs10051365 | -1.611 | 0.164 | -0.022 | 1.494 | 0.002 | 0 | 0 | 0 | 0.42 | 0.61 | 0.6 | 0.65 | 0.35 | 0.48 |
| 5 | 18503663 | rs11954903 | -0.047 | 0.638 | -0.735 | 1.016 | -0.027 | 0.014 | 0.014 | 0 | 0.19 | 0.1 | 0.14 | 0.15 | 0.19 | 0.27 |
| 5 | 18513822 | rs7701896 | -0.679 | -0.261 | -0.724 | 1.003 | 0.005 | 0.021 | 0.021 | 0 | 0.13 | 0.03 | 0.2 | 0.12 | 0.14 | 0.2 |
| 5 | 18520930 | rs9688294 | 1.253 | -0.938 | -0.377 | 1.155 | 0.007 | -0.005 | -0.005 | 0.047 | 0.57 | 0.4 | 0.42 | 0.47 | 0.57 | 0.61 |
| 5 | 18521319 | rs9688315 | -0.581 | -0.736 | -0.511 | 1.333 | -0.003 | -0.001 | -0.001 | 0 | 0.06 | 0.05 | 0.08 | 0.13 | 0.11 | 0.24 |
| 5 | 18532785 | rs13188100 | 0.36 | -1.225 | -0.493 | 1.121 | 0 | 0.004 | 0.004 | 0 | 0.09 | 0.02 | 0.19 | 0.11 | 0.13 | 0.17 |
| 5 | 18544459 | rs11749277 | 0.802 | -1.339 | -0.316 | 1.305 | 0.02 | -0.042 | -0.042 | 1.397 | 0.65 | 0.26 | 0.61 | 0.54 | 0.7 | 0.78 |
| 5 | 18548150 | rs9687246 | -1.036 | -1.415 | -0.29 | 1.272 | 0.009 | -0.023 | -0.023 | 0 | 0.27 | 0.74 | 0.35 | 0.46 | 0.22 | 0.14 |
| 5 | 18561967 | rs7724567 | 0.475 | -1.429 | 0.18 | 1.263 | 0.004 | 0.021 | 0.021 | 0 | 0.83 | 0.45 | 0.89 | 0.83 | 0.84 | 0.93 |
| 5 | 18564331 | rs6891237 | 2.319 | -1.55 | -0.206 | 1.089 | 0.006 | 0.083 | 0.083 | 0.005 | 0.48 | 0.14 | 0.33 | 0.34 | 0.52 | 0.33 |
| 5 | 18565153 | rs4334892 | -1.185 | -1.55 | -0.116 | 1.177 | -0.001 | -0.004 | -0.004 | 0 | 0.22 | 0.25 | 0.39 | 0.5 | 0.26 | 0.36 |
| 5 | 18572605 | rs4866340 | -1.116 | -1.588 | -0.203 | 1.181 | -0.02 | -0.004 | -0.004 | 0 | 0.91 | 0.96 | 0.94 | 0.92 | 0.86 | 0.9 |
| 5 | 18573774 | rs981275 | 1.554 | -1.499 | -0.057 | 1.361 | -0.005 | 0.142 | 0.142 | 0 | 0.07 | 0.13 | 0.21 | 0.34 | 0.04 | 0.08 |
| 5 | 18578054 | rs12654064 | 0.249 | -1.573 | -0.062 | 1.273 | 0.005 | 0.027 | 0.027 | 0 | 0.14 | 0.06 | 0.2 | 0.11 | 0.12 | 0.23 |
| 5 | 18578445 | rs6863200 | -0.417 | -1.58 | -0.091 | 1.286 | -0.001 | 0.084 | 0.084 | 0 | 0.12 | 0.19 | 0.32 | 0.41 | 0.12 | 0.24 |
| 5 | 18579606 | rs12658762 | 0.095 | -1.597 | -0.075 | 0.892 | -0.005 | 0.036 | 0.036 | 0 | 0.14 | 0.03 | 0.2 | 0.11 | 0.12 | 0.23 |
| 5 | 18581119 | rs11954300 | -0.106 | -1.421 | 0.881 | 1.089 | 0.029 | 0.069 | 0.069 | 0 | 0.07 | 0.31 | 0.03 | 0.05 | 0.02 | 0.03 |
| 5 | 18586907 | rs1908699 | 0.985 | -1.897 | 0.806 | 0.769 | 0.003 | -0.001 | -0.001 | 0 | 0.47 | 0.37 | 0.33 | 0.48 | 0.51 | 0.37 |
| 5 | 18599038 | rs1826812 | -1.031 | -1.801 | 0.701 | 0.624 | 0.001 | 0.048 | 0.048 | 0 | 0.07 | 0.25 | 0.1 | 0.07 | 0.07 | 0.18 |
| 5 | 18601469 | rs12153172 | 0.866 | -1.893 | 1.028 | 0.653 | 0.002 | -0.001 | -0.001 | 0.17 | 0.48 | 0.46 | 0.33 | 0.49 | 0.51 | 0.37 |
| 5 | 18605940 | rs7730186 | -0.351 | -1.941 | 1.051 | 0.723 | -0.021 | 0.123 | 0.123 | 0 | 0.14 | 0.29 | 0.29 | 0.32 | 0.11 | 0.26 |
| 5 | 18624481 | rs1390566 | -0.315 | -2.014 | 1.08 | 0.62 | -0.001 | 0.002 | 0.002 | 0 | 0.32 | 0.31 | 0.54 | 0.42 | 0.31 | 0.5 |
| 5 | 18629409 | rs1845830 | -0.606 | -2.104 | 1.058 | 0.566 | -0.009 | 0.004 | 0.004 | 0 | 0.36 | 0.48 | 0.55 | 0.42 | 0.31 | 0.49 |
| 5 | 18637320 | rs7716489 | 0.102 | -2.114 | 1.069 | 0.528 | -0.01 | -0.026 | -0.026 | 0 | 0.75 | 0.65 | 0.84 | 0.88 | 0.8 | 0.83 |
| 5 | 18655253 | rs4529196 | -0.151 | -2.077 | 1.134 | 0.517 | 0.006 | 0.018 | 0.018 | 0 | 0.77 | 0.86 | 0.87 | 0.88 | 0.8 | 0.83 |
| 5 | 18662625 | rs13167620 | -1.007 | -2.085 | 0.857 | 0.528 | -0.014 | 0.037 | 0.037 | 0 | 0.82 | 0.97 | 0.89 | 0.91 | 0.82 | 0.86 |
| 5 | 18678904 | rs7710886 | 0.632 | -2.156 | 0.888 | 0.516 | 0.007 | 0.012 | 0.012 | 0 | 0.54 | 0.59 | 0.53 | 0.78 | 0.6 | 0.54 |
| 5 | 18684530 | rs961606 | -0.019 | -2.075 | 0.487 | 0.708 | 0.011 | -0.012 | -0.012 | 0 | 0.73 | 0.8 | 0.7 | 0.66 | 0.78 | 0.79 |
| 5 | 18685677 | rs10057740 | 0.228 | -2.194 | 0.427 | 0.642 | 0.019 | -0.01 | -0.01 | 3.373 | 0.71 | 0.8 | 0.7 | 0.66 | 0.78 | 0.79 |
| 5 | 18688976 | rs959738 | 1.893 | -2.024 | 0.381 | 0.746 | 0 | -0.033 | -0.033 | 0 | 0.06 | 0 | 0.17 | 0.25 | 0.04 | 0.07 |
| 5 | 18689116 | rs329904 | -0.172 | -2.025 | 0.611 | 0.552 | -0.03 | -0.076 | -0.076 | 0.113 | 0.29 | 0.41 | 0.36 | 0.15 | 0.21 | 0.37 |
| 5 | 18691618 | rs13176281 | 1.156 | -2.018 | 0.415 | 0.552 | -0.029 | 0.048 | 0.048 | 0 | 0.19 | 0.03 | 0.3 | 0.12 | 0.19 | 0.3 |
| 5 | 18692207 | rs13162697 | 0.201 | -2.03 | 0.64 | 0.558 | -0.007 | -0.02 | -0.02 | 0.299 | 0.26 | 0.27 | 0.32 | 0.15 | 0.2 | 0.32 |
| 5 | 18692338 | rs16886725 | -0.025 | -2.085 | 0.698 | 0.604 | 0.005 | 0.062 | 0.062 | 0 | 0.06 | 0.21 | 0.06 | 0.06 | 0.07 | 0.17 |
| 5 | 18693810 | rs10940770 | -0.42 | -2.07 | 0.441 | 0.608 | -0.029 | -0.032 | -0.032 | 0.373 | 0.32 | 0.54 | 0.39 | 0.21 | 0.28 | 0.5 |
| 5 | 18694475 | rs393268 | 1.398 | -1.887 | 0.336 | 0.622 | -0.103 | -0.109 | -0.109 | 0 | 0.38 | 0.07 | 0.28 | 0.45 | 0.5 | 0.25 |
| 5 | 18698028 | rs329891 | -1.854 | -1.68 | 0.937 | 0.71 | -0.05 | -0.006 | -0.006 | 0 | 0.15 | 0.56 | 0.1 | 0.09 | 0.09 | 0.24 |
| 5 | 18726660 | rs10064581 | -1.044 | -1.411 | 0.889 | 0.821 | 0.016 | 0.034 | 0.034 | 0 | 0.12 | 0.39 | 0.1 | 0.09 | 0.13 | 0.22 |
| 5 | 18727016 | rs2102146 | 0.832 | -1.562 | 0.674 | 0.787 | -0.007 | -0.05 | -0.05 | 0.344 | 0.44 | 0.14 | 0.31 | 0.44 | 0.53 | 0.24 |
| 5 | 18738297 | rs16886679 | -0.139 | -0.673 | 0.654 | 0.393 | 0.038 | 0.007 | 0.007 | 0 | 0.07 | 0.22 | 0.05 | 0.03 | 0.06 | 0.03 |
| 5 | 18741673 | rs2937515 | -0.23 | -0.713 | -0.25 | 0.516 | -0.002 | -0.003 | -0.003 | 0.593 | 0.2 | 0.21 | 0.1 | 0.08 | 0.13 | 0.13 |
| 5 | 18759551 | rs4482907 | -0.004 | -0.557 | -0.677 | 0.757 | 0 | 0 | 0 | 0 | 0.43 | 0.59 | 0.62 | 0.5 | 0.57 | 0.34 |
| 5 | 18773727 | rs2962228 | -1.146 | -1.148 | -1.049 | 0.598 | 0.039 | -0.045 | -0.045 | 0 | 0.81 | 0.96 | 0.88 | 0.83 | 0.86 | 0.76 |
| 5 | 18777638 | rs7713991 | 0.974 | -0.404 | -0.929 | 0.442 | 0.005 | -0.064 | -0.064 | 0 | 0.2 | 0.41 | 0.16 | 0.11 | 0.19 | 0.08 |
| 5 | 18777746 | rs17631488 | 0.377 | -0.461 | -1.435 | 0.88 | 0.021 | 0.071 | 0.071 | 0 | 0.23 | 0.07 | 0.17 | 0.15 | 0.18 | 0.29 |
| 5 | 18783485 | rs2962233 | -0.403 | 0.27 | -0.986 | 1.26 | 0.076 | 0.025 | 0.025 | 0 | 0.6 | 0.85 | 0.71 | 0.69 | 0.69 | 0.49 |
| 5 | 18789976 | rs1393137 | -1.108 | 1.962 | 0.054 | 2.058 | 0.032 | 0.075 | 0.075 | 0 | 0.17 | 0.21 | 0.26 | 0.4 | 0.27 | 0.2 |
| 5 | 18792568 | rs12515138 | -0.932 | 1.327 | -0.086 | 1.706 | 0.013 | 0.081 | 0.081 | 0 | 0.09 | 0.14 | 0.1 | 0.21 | 0.11 | 0.05 |
| 5 | 18794692 | rs11741301 | -1.532 | 1.511 | -0.203 | 1.119 | 0.003 | -0.019 | -0.019 | 0 | 0.07 | 0.11 | 0.07 | 0 | 0.08 | 0.1 |
| 5 | 18795292 | rs1021771 | -0.578 | 0.883 | -0.043 | 1.528 | -0.008 | 0.013 | 0.013 | 0.042 | 0.17 | 0.16 | 0.23 | 0.23 | 0.14 | 0.26 |
| 5 | 18797589 | rs10040259 | -0.089 | 1.428 | -0.057 | 1.316 | 0.016 | -0.01 | -0.01 | 0 | 0.72 | 0.69 | 0.7 | 0.76 | 0.63 | 0.81 |
| 5 | 18799045 | rs12518042 | -0.319 | 1.307 | -0.053 | 1.21 | -0.081 | 0.009 | 0.009 | 0 | 0.74 | 0.93 | 0.72 | 0.76 | 0.62 | 0.8 |
| 5 | 18803801 | rs13172937 | 1.592 | 0.524 | 0.356 | 1.493 | 0.016 | 0.05 | 0.05 | 0.009 | 0.41 | 0.28 | 0.25 | 0.25 | 0.39 | 0.18 |
| 5 | 18809420 | rs6450728 | 0.234 | 0.663 | 0.41 | 1.522 | -0.007 | 0.004 | 0.004 | 0 | 0.66 | 0.69 | 0.64 | 0.75 | 0.54 | 0.72 |
| 5 | 18809531 | rs17663114 | 0.795 | 0.66 | 0.359 | 1.635 | 0.012 | -0.022 | -0.022 | 0 | 0.06 | 0 | 0.15 | 0.26 | 0.02 | 0.27 |
| 5 | 18814214 | rs10036680 | -0.161 | 0.625 | 0.548 | 1.453 | 0.019 | -0.02 | -0.02 | 0 | 0.69 | 0.62 | 0.68 | 0.77 | 0.62 | 0.79 |
| 5 | 18816719 | rs7715365 | 1.551 | 0.645 | 0.419 | 1.395 | 0.009 | 0.04 | 0.04 | 0 | 0.42 | 0.29 | 0.26 | 0.28 | 0.39 | 0.18 |
| 5 | 18819946 | rs10472175 | -0.162 | 0.895 | 0.303 | 1.095 | -0.004 | 0.002 | 0.002 | 0.013 | 0.72 | 0.76 | 0.7 | 0.78 | 0.63 | 0.8 |
| 5 | 18823641 | rs10062771 | 1.09 | 0.754 | 0.209 | 1.273 | -0.002 | 0.051 | 0.051 | 0.043 | 0.88 | 0.82 | 0.79 | 0.65 | 0.9 | 0.64 |
| 5 | 18833633 | rs4866126 | 0.162 | 1.358 | 0.212 | 0.806 | 0 | 0.051 | 0.051 | 0 | 0.84 | 0.78 | 0.79 | 0.47 | 0.83 | 0.77 |
| 5 | 18833784 | rs905177 | 0.803 | 1.17 | -0.056 | 0.616 | 0.055 | 0.017 | 0.017 | 0 | 0.52 | 0.21 | 0.45 | 0.47 | 0.42 | 0.54 |
| 5 | 18838134 | rs10057220 | -1.769 | 1.196 | 0.684 | 0.378 | 0.086 | -0.022 | -0.022 | 0 | 0.08 | 0.31 | 0.09 | 0.01 | 0.1 | 0.08 |
| 5 | 18839672 | rs9292384 | -1.403 | 1.261 | 0.511 | 0.427 | 0.09 | -0.015 | -0.015 | 0.31 | 0.07 | 0.25 | 0.07 | 0.01 | 0.1 | 0.08 |
| 5 | 18852401 | rs11747895 | 0.029 | 1.149 | -0.601 | 0.061 | -0.041 | 0.081 | 0.081 | 0 | 0.35 | 0.2 | 0.31 | 0.18 | 0.45 | 0.27 |
| 5 | 18854429 | rs9918266 | 0.066 | 0.595 | -0.214 | 0.425 | -0.015 | 0.004 | 0.004 | 0.006 | 0.25 | 0.32 | 0.21 | 0.27 | 0.21 | 0.33 |

| | | | | | | | | | | | | | | | | |
|---|---|---|---|---|---|---|---|---|---|---|---|---|---|---|---|---|
| 5 | 18863659 | rs13169561 | 0.564 | 0.178 | -0.13 | 0.919 | 0 | 0.029 | 0.029 | 0 | 0.16 | 0.23 | 0.18 | 0.41 | 0.17 | 0.2 |
| 5 | 18863799 | rs13165779 | 0.604 | 0.145 | -0.046 | 1.271 | -0.001 | 0.006 | 0.006 | 0 | 0.15 | 0.22 | 0.17 | 0.21 | 0.15 | 0.16 |
| 5 | 18863867 | rs17269457 | 1.359 | -0.19 | -0.433 | 0.779 | -0.123 | 0.121 | 0.121 | 0 | 0.34 | 0.04 | 0.3 | 0.17 | 0.46 | 0.29 |
| 5 | 18867568 | rs6450785 | -0.59 | -0.301 | -0.012 | 0.826 | 0.009 | 0.055 | 0.055 | 0 | 0.45 | 0.72 | 0.42 | 0.69 | 0.41 | 0.56 |
| 5 | 18868357 | rs6878434 | -0.583 | -0.704 | 0.357 | 0.649 | -0.007 | 0.009 | 0.009 | 0 | 0.29 | 0.57 | 0.33 | 0.69 | 0.24 | 0.51 |
| 5 | 18877165 | rs10065818 | 0.117 | -0.694 | 0.273 | 0.635 | 0.01 | 0.031 | 0.031 | 0.582 | 0.07 | 0.27 | 0.03 | 0.1 | 0.04 | 0.08 |
| 5 | 18879807 | rs10059239 | 0.36 | -0.67 | 0.286 | 0.647 | 0.015 | 0.036 | 0.036 | 0.592 | 0.06 | 0.25 | 0.03 | 0.1 | 0.04 | 0.08 |
| 5 | 18881698 | rs6893620 | -1.382 | -0.743 | -0.135 | 0.612 | 0.009 | 0.068 | 0.068 | 0.462 | 0.85 | 0.98 | 0.92 | 0.95 | 0.89 | 0.92 |
| 5 | 18885119 | rs10055996 | 0.491 | -0.299 | 0.324 | 1.136 | -0.01 | -0.02 | -0.02 | 0 | 0.12 | 0.06 | 0.17 | 0.3 | 0.15 | 0.14 |
| 5 | 18885615 | rs16886522 | 0.566 | -0.34 | 0.526 | 1.171 | -0.011 | 0.064 | 0.064 | 0 | 0.09 | 0.12 | 0.12 | 0.28 | 0.04 | 0.28 |
| 5 | 18890253 | rs10472760 | 0.436 | -0.073 | 0.39 | 0.781 | -0.003 | 0.004 | 0.004 | 0 | 0.12 | 0.09 | 0.17 | 0.3 | 0.16 | 0.14 |
| 5 | 18895407 | rs1566038 | 0.45 | -0.124 | 0.411 | 0.267 | 0 | -0.013 | -0.013 | 0 | 0.17 | 0.14 | 0.1 | 0 | 0.17 | 0.05 |
| 5 | 18908745 | rs11957143 | -0.375 | -0.391 | 0.86 | 0.25 | -0.022 | -0.075 | -0.075 | 0 | 0.12 | 0.36 | 0.11 | 0.01 | 0.08 | 0.12 |
| 5 | 18909675 | rs11958034 | -1.566 | -0.446 | 1.117 | 1.138 | -0.031 | 0.129 | 0.129 | 0.351 | 0.24 | 0.54 | 0.4 | 0.43 | 0.17 | 0.38 |
| 5 | 18913137 | rs12652071 | -1.293 | -0.456 | 1.309 | 1.133 | -0.016 | 0.096 | 0.096 | 0.216 | 0.22 | 0.42 | 0.37 | 0.43 | 0.16 | 0.38 |
| 5 | 18917828 | rs4521476 | 1.664 | -0.433 | 0.28 | 0.607 | -0.071 | 0.178 | 0.178 | 0.926 | 0.43 | 0.04 | 0.29 | 0.2 | 0.41 | 0.31 |
| 5 | 18921988 | rs11951878 | 0.111 | -0.421 | 0.323 | 0.636 | -0.001 | 0.107 | 0.107 | 0 | 0.11 | 0.15 | 0.29 | 0.43 | 0.09 | 0.26 |
| 5 | 18929720 | rs4379197 | 1.728 | -0.454 | 0.201 | 0.683 | -0.076 | 0.125 | 0.125 | 0.47 | 0.42 | 0.04 | 0.29 | 0.25 | 0.41 | 0.32 |
| 5 | 18935805 | rs6450835 | 0.284 | -0.124 | 0.348 | 0.826 | -0.003 | 0.009 | 0.009 | 0.226 | 0.1 | 0.11 | 0.15 | 0.31 | 0.12 | 0.17 |
| 5 | 18940647 | rs11955931 | -0.355 | -0.174 | 0.453 | 0.955 | -0.005 | 0.118 | 0.118 | 0 | 0.13 | 0.17 | 0.3 | 0.45 | 0.08 | 0.26 |
| 5 | 18950552 | rs10940958 | -0.448 | 0.074 | 0.838 | 0.479 | 0.007 | -0.029 | -0.029 | 1.805 | 0.8 | 0.78 | 0.85 | 0.94 | 0.74 | 0.9 |
| 5 | 18954716 | rs4309976 | 0.664 | -0.3 | 1.166 | 1.033 | 0.024 | 0.22 | 0.22 | 0.024 | 0.85 | 0.61 | 0.82 | 0.59 | 0.87 | 0.78 |
| 5 | 18962571 | rs13176582 | -0.317 | -0.309 | 1.003 | 1.12 | -0.004 | 0.007 | 0.007 | 0.082 | 0.12 | 0.12 | 0.16 | 0.35 | 0.13 | 0.21 |
| 5 | 18964842 | rs10040022 | 1.001 | -0.591 | 1.122 | 0.902 | -0.009 | 0.035 | 0.035 | 0 | 0.74 | 0.35 | 0.74 | 0.57 | 0.8 | 0.66 |
| 5 | 18971937 | rs4597970 | 1.099 | -0.625 | 1.083 | 0.886 | 0.001 | -0.004 | -0.004 | 0 | 0.73 | 0.28 | 0.73 | 0.57 | 0.8 | 0.66 |
| 5 | 18977475 | rs12110283 | 1.586 | -0.332 | 0.462 | 0.813 | -0.054 | 0.023 | 0.023 | 0.294 | 0.47 | 0.11 | 0.39 | 0.42 | 0.48 | 0.45 |
| 5 | 18982961 | rs10940978 | 0.872 | -0.313 | 0.798 | 0.842 | 0.008 | 0.008 | 0.008 | 0 | 0.61 | 0.56 | 0.5 | 0.5 | 0.55 | 0.58 |
| 5 | 18985998 | rs10035187 | -0.333 | -0.399 | 0.895 | 0.807 | 0.002 | -0.024 | -0.024 | 0.54 | 0.29 | 0.49 | 0.38 | 0.22 | 0.32 | 0.22 |
| 5 | 18994108 | rs4263496 | 0.47 | -0.366 | 0.823 | 0.724 | -0.004 | 0.065 | 0.065 | 0 | 0.77 | 0.61 | 0.77 | 0.63 | 0.8 | 0.67 |
| 5 | 18999510 | rs4866134 | 0.092 | -0.391 | 0.491 | 0.572 | -0.006 | 0.014 | 0.014 | 0 | 0.26 | 0.19 | 0.33 | 0.14 | 0.32 | 0.18 |
| 5 | 18999592 | rs12515368 | 1.494 | -0.438 | -0.078 | 0.686 | -0.079 | -0.028 | -0.028 | 0 | 0.48 | 0.05 | 0.4 | 0.48 | 0.49 | 0.5 |
| 5 | 19001160 | rs7443398 | 0.051 | -0.353 | 0.272 | 0.696 | 0.006 | -0.046 | -0.046 | 0.72 | 0.72 | 0.61 | 0.65 | 0.86 | 0.68 | 0.82 |
| 5 | 19023177 | rs10036524 | 0.127 | -0.191 | -0.365 | 0.986 | 0.001 | 0.049 | 0.049 | 0.829 | 0.1 | 0.15 | 0.15 | 0.31 | 0.11 | 0.18 |
| 5 | 19027541 | rs4515302 | 1.092 | 0.08 | -0.763 | 0.693 | -0.069 | 0.024 | 0.024 | 0 | 0.49 | 0.07 | 0.4 | 0.45 | 0.47 | 0.53 |
| 5 | 19028438 | rs13178830 | 0.988 | 0.464 | -0.934 | 0.215 | 0.002 | -0.041 | -0.041 | 0 | 0.4 | 0.14 | 0.33 | 0.53 | 0.36 | 0.52 |
| 5 | 19031835 | rs6450894 | 0.266 | 0.536 | -0.733 | -0.178 | -0.019 | 0.115 | 0.115 | 0.379 | 0.55 | 0.28 | 0.66 | 0.92 | 0.51 | 0.79 |
| 5 | 19041768 | rs10941008 | -1.319 | 0.423 | -0.038 | -0.347 | 0.017 | -0.123 | -0.123 | 0.226 | 0.82 | 0.62 | 0.88 | 1 | 0.83 | 0.94 |
| 5 | 19059786 | rs4130533 | 1.297 | 0.371 | -0.209 | -0.471 | -0.012 | -0.056 | -0.056 | 0 | 0.12 | 0.21 | 0.08 | 0 | 0.08 | 0.1 |
| 5 | 19066292 | rs7723526 | 0.067 | 0.327 | -0.302 | -0.503 | 0.009 | 0 | 0 | 0 | 0.69 | 0.24 | 0.77 | 1 | 0.74 | 0.84 |
| 5 | 19067546 | rs13163797 | 0.952 | 0.264 | -0.476 | -0.083 | 0.007 | -0.025 | -0.025 | 0 | 0.11 | 0.06 | 0.12 | 0.26 | 0.11 | 0.21 |
| 5 | 19068967 | rs13181998 | 1.198 | 0.286 | -0.527 | -0.096 | 0.018 | -0.042 | -0.042 | 0 | 0.55 | 0.12 | 0.63 | 0.64 | 0.62 | 0.53 |
| 5 | 19077737 | rs4421107 | 0.58 | 0.434 | -0.196 | -0.152 | -0.017 | 0.096 | 0.096 | 0.047 | 0.56 | 0.29 | 0.65 | 0.92 | 0.51 | 0.79 |
| 5 | 19085981 | rs12054808 | 0.996 | -0.114 | -0.621 | -0.014 | 0.005 | -0.007 | -0.007 | 0 | 0.11 | 0.08 | 0.28 | 0.38 | 0.08 | 0.13 |
| 5 | 19086655 | rs7714384 | -0.677 | -0.147 | -0.473 | -0.115 | 0.033 | -0.107 | -0.107 | 0 | 0.68 | 0.51 | 0.74 | 0.92 | 0.59 | 0.88 |
| 5 | 19087783 | rs4391167 | 0.964 | -0.093 | -0.591 | 0.031 | 0.005 | -0.007 | -0.007 | 0 | 0.11 | 0.08 | 0.28 | 0.38 | 0.08 | 0.13 |
| 5 | 19093749 | rs4327600 | 1.168 | -0.019 | -0.615 | -0.02 | -0.02 | 0.024 | 0.024 | 0 | 0.44 | 0.14 | 0.37 | 0.54 | 0.43 | 0.65 |
| 5 | 19096772 | rs6450929 | -1.168 | -0.159 | 0.173 | -0.065 | 0.012 | -0.121 | -0.121 | 0 | 0.82 | 0.64 | 0.87 | 1 | 0.83 | 0.93 |
| 5 | 19103049 | rs4461634 | -1.083 | -0.192 | 0.293 | -0.065 | 0.006 | -0.119 | -0.119 | 0.552 | 0.82 | 0.63 | 0.87 | 1 | 0.83 | 0.93 |
| 5 | 19121428 | rs11950518 | 0.589 | 0.211 | 0.057 | 0.148 | -0.005 | 0.019 | 0.019 | 0 | 0.09 | 0.07 | 0.28 | 0.39 | 0.09 | 0.2 |
| 5 | 19127527 | rs10805579 | 0.434 | -0.059 | -0.055 | 0.029 | 0.042 | 0.096 | 0.096 | 1.122 | 0.62 | 0.18 | 0.76 | 0.9 | 0.74 | 0.74 |
| 5 | 19138854 | rs4479845 | -1.467 | -0.071 | 0.018 | 0 | 0.012 | -0.109 | -0.109 | 3.856 | 0.83 | 0.67 | 0.87 | 1 | 0.83 | 0.93 |
| 5 | 19148922 | rs11950664 | 0.66 | -0.139 | -0.086 | 0.06 | 0.01 | -0.001 | -0.001 | 0 | 0.18 | 0.13 | 0.09 | 0.12 | 0.11 | 0.2 |
| 5 | 19148952 | rs11740707 | 1.087 | -0.039 | 0.151 | -0.24 | 0.007 | -0.035 | -0.035 | 1.297 | 0.54 | 0.18 | 0.48 | 0.49 | 0.63 | 0.56 |
| 5 | 19151180 | rs4640795 | -0.927 | -0.041 | 0.112 | -0.245 | -0.001 | -0.105 | -0.105 | 0.119 | 0.8 | 0.35 | 0.84 | 1 | 0.83 | 0.93 |
| 5 | 19152050 | rs7720434 | 1.106 | -0.031 | -0.041 | -0.235 | 0.002 | -0.046 | -0.046 | 0.057 | 0.54 | 0.14 | 0.47 | 0.49 | 0.63 | 0.56 |
| 5 | 19155786 | rs7709474 | 1.106 | -0.031 | -0.041 | -0.235 | 0.002 | -0.046 | -0.046 | 0 | 0.54 | 0.14 | 0.47 | 0.49 | 0.63 | 0.56 |
| 5 | 19156521 | rs4552600 | -1.003 | 0.056 | 0.193 | -0.268 | 0.015 | -0.144 | -0.144 | 0 | 0.81 | 0.53 | 0.85 | 1 | 0.83 | 0.93 |
| 5 | 19158050 | rs10941077 | 0.991 | 0.001 | -0.373 | -0.166 | 0.002 | -0.047 | -0.047 | 0 | 0.54 | 0.14 | 0.47 | 0.49 | 0.63 | 0.56 |
| 5 | 19160015 | rs9637779 | -0.329 | -0.044 | -0.149 | -0.236 | 0.004 | -0.028 | -0.028 | 0 | 0.26 | 0.21 | 0.37 | 0.51 | 0.2 | 0.37 |
| 5 | 19160113 | rs9637789 | -1.455 | -0.05 | 0.238 | -0.254 | 0.01 | -0.096 | -0.096 | 0 | 0.83 | 0.7 | 0.87 | 1 | 0.83 | 0.93 |
| 5 | 19171872 | rs12518778 | 0.998 | -0.005 | 0.118 | -0.252 | 0.009 | -0.019 | -0.019 | 0 | 0.54 | 0.27 | 0.48 | 0.49 | 0.63 | 0.57 |
| 5 | 19183884 | rs13153741 | 1.251 | -0.482 | -0.977 | -0.539 | -0.003 | -0.079 | -0.079 | 0 | 0.15 | 0.02 | 0.14 | 0.24 | 0.13 | 0.15 |
| 5 | 19185179 | rs4492113 | -0.153 | -0.424 | -0.714 | -0.546 | 0 | 0 | 0 | 0 | 0.27 | 0.28 | 0.37 | 0.51 | 0.2 | 0.37 |
| 5 | 19188359 | rs4505952 | 0.724 | -0.455 | -1.065 | -0.627 | -0.019 | -0.052 | -0.052 | 0.521 | 0.52 | 0.11 | 0.47 | 0.48 | 0.61 | 0.56 |
| 5 | 19189948 | rs4318784 | -0.006 | -0.494 | -0.868 | -0.572 | -0.022 | -0.078 | -0.078 | 0 | 0.68 | 0.16 | 0.55 | 0.48 | 0.71 | 0.66 |
| 5 | 19195667 | rs10059141 | -1.608 | -0.472 | -0.493 | -0.611 | 0.012 | -0.1 | -0.1 | 0 | 0.83 | 0.67 | 0.88 | 1 | 0.83 | 0.94 |
| 5 | 19196465 | rs10061156 | 0.097 | -0.331 | -0.357 | -0.378 | 0.015 | 0.148 | 0.148 | 0 | 0.12 | 0.22 | 0.29 | 0.52 | 0.12 | 0.28 |
| 5 | 19199387 | rs10061411 | -0.374 | -0.346 | -0.498 | -0.385 | 0.049 | 0.328 | 0.328 | 0 | 0.14 | 0.42 | 0.31 | 0.52 | 0.12 | 0.28 |
| 5 | 19200634 | rs10056848 | -0.113 | -0.285 | -0.982 | -0.887 | 0.053 | 0.065 | 0.065 | 1.519 | 0.28 | 0.51 | 0.43 | 0.58 | 0.38 | 0.36 |

| | | | | | | | | | | | | | | | | |
|---|---|---|---|---|---|---|---|---|---|---|---|---|---|---|---|---|
| 5 | 19201028 | rs11948213 | 0.031 | -0.994 | -1.132 | -0.481 | 0.005 | -0.064 | -0.064 | 0 | 0.22 | 0.03 | 0.21 | 0.24 | 0.19 | 0.16 |
| 5 | 19208299 | rs4403177 | 0.026 | -1.009 | -1.132 | -0.481 | 0.01 | -0.064 | -0.064 | 0 | 0.23 | 0.03 | 0.21 | 0.24 | 0.19 | 0.16 |
| 5 | 19235686 | rs4291014 | 1.113 | -0.608 | -0.95 | -0.731 | -0.008 | 0.089 | 0.089 | 0 | 0.38 | 0.2 | 0.47 | 0.18 | 0.4 | 0.39 |
| 5 | 19240987 | rs6868154 | 0.751 | -0.611 | -0.974 | -0.268 | -0.023 | -0.073 | -0.073 | 0 | 0.14 | 0.03 | 0.1 | 0.22 | 0.14 | 0.12 |
| 5 | 19247098 | rs10941199 | -1.098 | -0.759 | -0.498 | -0.462 | 0 | 0 | 0 | 0 | 0.36 | 0.3 | 0.33 | 0.81 | 0.31 | 0.59 |
| 5 | 19266101 | rs4490578 | -0.104 | -0.804 | -0.402 | -0.045 | -0.022 | 0.088 | 0.088 | 0 | 0.2 | 0.28 | 0.12 | 0.34 | 0.16 | 0.3 |
| 5 | 19271333 | rs6888100 | 0.999 | -0.803 | -0.237 | -0.529 | -0.002 | 0.034 | 0.034 | 0 | 0.46 | 0.45 | 0.56 | 0.21 | 0.52 | 0.35 |
| 5 | 19280885 | rs4535457 | 1.008 | -0.803 | -0.226 | -0.529 | -0.003 | 0.045 | 0.045 | 0 | 0.46 | 0.43 | 0.55 | 0.21 | 0.52 | 0.35 |
| 5 | 19291890 | rs4552597 | -0.662 | -0.84 | 0.133 | -0.749 | 0.004 | 0.053 | 0.053 | 0.547 | 0.49 | 0.35 | 0.47 | 0.75 | 0.46 | 0.49 |
| 5 | 19295220 | rs4543268 | -1.461 | -1.095 | 0.182 | -0.991 | 0.008 | -0.062 | -0.062 | 0 | 0.63 | 0.58 | 0.53 | 0.87 | 0.52 | 0.72 |
| 5 | 19309903 | rs7719998 | 2.28 | -0.917 | 0.001 | -0.945 | -0.007 | 0.003 | 0.003 | 0 | 0.15 | 0.13 | 0.11 | 0 | 0.16 | 0.06 |
| 5 | 19328003 | rs2967017 | 0.517 | -1.041 | 0.342 | -0.899 | 0.001 | 0.008 | 0.008 | 0 | 0.5 | 0.45 | 0.52 | 0.25 | 0.54 | 0.53 |
| 5 | 19331778 | rs10058333 | 1.662 | -0.994 | 0.295 | -0.878 | -0.009 | 0.023 | 0.023 | 0.102 | 0.2 | 0.15 | 0.12 | 0 | 0.2 | 0.1 |
| 5 | 19344220 | rs16886123 | 1.546 | -0.795 | 0.19 | -0.324 | -0.007 | 0.013 | 0.013 | 0.144 | 0.11 | 0.09 | 0.24 | 0.46 | 0.12 | 0.21 |
| 5 | 19346137 | rs1512551 | 1.319 | -0.91 | 0.251 | -0.384 | -0.046 | 0.024 | 0.024 | 0 | 0.15 | 0.23 | 0.06 | 0.12 | 0.06 | 0.22 |
| 5 | 19358684 | rs2942295 | 0.001 | -0.862 | 0.183 | -0.921 | -0.001 | -0.001 | -0.001 | 0.56 | 0.62 | 0.58 | 0.76 | 0.65 | 0.65 | 0.63 |
| 5 | 19364608 | rs877314 | 0.445 | -0.777 | 0.209 | -0.91 | 0 | -0.008 | -0.008 | 0 | 0.52 | 0.46 | 0.52 | 0.19 | 0.54 | 0.55 |
| 5 | 19367728 | rs2942291 | 1.348 | -0.649 | -0.075 | -0.894 | -0.032 | 0.083 | 0.083 | 0.246 | 0.35 | 0.24 | 0.46 | 0.13 | 0.49 | 0.33 |
| 5 | 19376830 | rs925511 | 0.35 | -0.674 | -0.085 | -0.996 | -0.012 | 0.013 | 0.013 | 0 | 0.45 | 0.38 | 0.51 | 0.41 | 0.52 | 0.44 |
| 5 | 19385395 | rs7717664 | 1.028 | -0.625 | -0.141 | -0.865 | 0.002 | -0.03 | -0.03 | 0.461 | 0.14 | 0.08 | 0.08 | 0.29 | 0.12 | 0.12 |
| 5 | 19392440 | rs12189480 | 0.184 | -0.924 | 0.295 | -0.98 | 0.017 | -0.018 | -0.018 | 0.013 | 0.27 | 0.19 | 0.13 | 0.36 | 0.18 | 0.33 |
| 5 | 19393347 | rs1040005 | 0.789 | -0.926 | 0.176 | -1.173 | -0.02 | 0.004 | 0.004 | 0 | 0.4 | 0.3 | 0.64 | 0.62 | 0.5 | 0.56 |
| 5 | 19393903 | rs2942329 | 2.176 | -0.876 | 0.139 | -1.255 | -0.025 | 0.041 | 0.041 | 0 | 0.27 | 0.21 | 0.39 | 0.11 | 0.38 | 0.29 |
| 5 | 19394510 | rs12189093 | 2.102 | -0.853 | -0.029 | -1.218 | -0.006 | -0.01 | -0.01 | 0 | 0.19 | 0.16 | 0.12 | 0 | 0.19 | 0.08 |
| 5 | 19398317 | rs1858432 | 0.519 | -0.969 | -0.068 | -0.783 | -0.001 | -0.012 | -0.012 | 1.052 | 0.69 | 0.71 | 0.64 | 0.19 | 0.71 | 0.61 |
| 5 | 19399602 | rs1512548 | -0.292 | -0.959 | -0.069 | -0.732 | 0.003 | 0.107 | 0.107 | 0 | 0.82 | 0.79 | 0.72 | 0.49 | 0.85 | 0.73 |
| 5 | 19400350 | rs2966994 | 1.133 | -0.839 | 0.132 | -0.48 | 0.003 | -0.015 | -0.015 | 0 | 0.55 | 0.5 | 0.57 | 0.13 | 0.63 | 0.41 |
| 5 | 19412775 | rs10520859 | 0.521 | -0.619 | -0.769 | -0.23 | 0.005 | -0.034 | -0.034 | 0 | 0.14 | 0.08 | 0.09 | 0.31 | 0.14 | 0.13 |
| 5 | 19421313 | rs2942302 | -0.578 | -0.544 | -0.68 | -0.14 | 0.006 | 0.1 | 0.1 | 0.078 | 0.86 | 0.85 | 0.75 | 0.49 | 0.86 | 0.75 |
| 5 | 19425092 | rs10039593 | 1.214 | -0.449 | -0.889 | -0.985 | 0.002 | 0.002 | 0.002 | 0 | 0.13 | 0.13 | 0.07 | 0.06 | 0.07 | 0.19 |
| 5 | 19427619 | rs1021470 | 0.331 | 0.084 | -0.798 | -0.647 | 0 | 0.011 | 0.011 | 0 | 0.7 | 0.74 | 0.64 | 0.18 | 0.7 | 0.62 |
| 5 | 19430825 | rs770140 | 0.098 | 0.033 | -0.647 | -0.694 | 0.018 | 0.039 | 0.039 | 3.123 | 0.56 | 0.38 | 0.76 | 0.87 | 0.65 | 0.64 |
| 5 | 19440168 | rs349475 | -0.14 | 0.36 | -0.607 | -0.039 | 0.005 | 0.002 | 0.002 | 1.408 | 0.69 | 0.65 | 0.55 | 0.62 | 0.64 | 0.71 |
| 5 | 19441735 | rs10520861 | -0.122 | 0.225 | -0.822 | -0.123 | -0.02 | -0.007 | -0.007 | 0 | 0.34 | 0.17 | 0.47 | 0.39 | 0.41 | 0.37 |
| 5 | 19446308 | rs623213 | 0.003 | -0.024 | -0.465 | -0.324 | 0.003 | 0.01 | 0.01 | 0.037 | 0.63 | 0.72 | 0.53 | 0.69 | 0.62 | 0.64 |
| 5 | 19448847 | rs17280924 | 1.112 | -0.088 | -1.14 | -0.372 | -0.091 | -0.058 | -0.058 | 0.009 | 0.26 | 0.02 | 0.27 | 0.27 | 0.29 | 0.29 |
| 5 | 19448955 | rs1707015 | -0.479 | -0.153 | -0.97 | -1.256 | 0.002 | 0.007 | 0.007 | 0.015 | 0.64 | 0.72 | 0.53 | 0.69 | 0.63 | 0.67 |
| 5 | 19450474 | rs17281015 | 1.77 | -0.023 | -1.26 | -1.4 | -0.071 | 0.096 | 0.096 | 0.351 | 0.18 | 0.02 | 0.12 | 0 | 0.22 | 0.03 |
| 5 | 19451589 | rs346784 | 0.919 | -0.832 | -0.502 | -0.775 | -0.046 | 0.017 | 0.017 | 0 | 0.43 | 0.18 | 0.4 | 0.57 | 0.47 | 0.48 |
| 5 | 19459826 | rs349468 | 1.341 | -0.904 | -0.311 | -0.815 | -0.008 | -0.04 | -0.04 | 0.577 | 0.11 | 0.15 | 0.07 | 0 | 0.09 | 0.1 |
| 5 | 19462474 | rs349474 | 0.434 | -0.91 | -0.112 | -0.81 | -0.012 | -0.03 | -0.03 | 0 | 0.16 | 0.26 | 0.11 | 0.08 | 0.13 | 0.16 |
| 5 | 19463410 | rs1707025 | 0.817 | -0.983 | -0.214 | -1.016 | 0.049 | 0.031 | 0.031 | 0 | 0.54 | 0.2 | 0.63 | 0.58 | 0.59 | 0.55 |
| 5 | 19463574 | rs17215554 | 1.276 | -0.334 | -0.656 | -1.298 | -0.035 | 0.06 | 0.06 | 0 | 0.11 | 0.01 | 0.22 | 0.03 | 0.12 | 0.1 |
| 5 | 19467722 | rs347703 | -0.531 | -0.437 | -0.521 | -1.196 | -0.009 | -0.03 | -0.03 | 0 | 0.34 | 0.42 | 0.32 | 0.2 | 0.36 | 0.35 |
| 5 | 19469443 | rs347706 | 1.484 | -0.26 | -0.691 | -1.113 | -0.006 | -0.031 | -0.031 | 0 | 0.1 | 0.12 | 0.06 | 0 | 0.07 | 0.08 |
| 5 | 19472907 | rs770297 | -0.47 | 0.033 | -0.572 | -0.851 | -0.002 | -0.005 | -0.005 | 0 | 0.43 | 0.58 | 0.36 | 0.28 | 0.41 | 0.44 |
| 5 | 19484182 | rs3792727 | -0.961 | 0.58 | -0.17 | -0.286 | 0 | -0.055 | -0.055 | 0 | 0.23 | 0.37 | 0.29 | 0.1 | 0.21 | 0.2 |
| 5 | 19488982 | rs13170473 | -0.799 | 0.582 | -0.085 | -0.409 | 0.001 | -0.113 | -0.113 | 0 | 0.21 | 0.41 | 0.29 | 0.03 | 0.19 | 0.16 |
| 5 | 19490899 | rs6894128 | -0.339 | 0.447 | 0.227 | -0.061 | 0.003 | -0.007 | -0.007 | 0 | 0.36 | 0.43 | 0.42 | 0.31 | 0.3 | 0.46 |
| 5 | 19493537 | rs11951215 | -0.294 | 0.327 | 0.575 | -0.125 | -0.001 | 0 | 0 | 0.807 | 0.33 | 0.32 | 0.4 | 0.31 | 0.29 | 0.42 |
| 5 | 19493684 | rs1486849 | -0.074 | 0.267 | 0.655 | -0.201 | -0.011 | -0.005 | -0.005 | 0.016 | 0.3 | 0.24 | 0.39 | 0.31 | 0.28 | 0.39 |
| 5 | 19498390 | rs392488 | -0.436 | 0.242 | 0.92 | -0.761 | -0.003 | -0.08 | -0.08 | 0 | 0.18 | 0.28 | 0.13 | 0.04 | 0.19 | 0.16 |
| 5 | 19502141 | rs1684848 | -1.017 | 0.098 | 1.316 | -0.753 | -0.024 | 0.077 | 0.077 | 0 | 0.51 | 0.82 | 0.55 | 0.35 | 0.47 | 0.57 |
| 5 | 19513668 | rs347720 | -1.008 | 0.107 | 1.54 | -0.747 | -0.003 | 0.028 | 0.028 | 0 | 0.49 | 0.61 | 0.52 | 0.34 | 0.47 | 0.56 |
| 5 | 19518321 | rs1126069 | -0.793 | -0.282 | 1.141 | -0.084 | 0.012 | -0.017 | -0.017 | 0.018 | 0.92 | 0.97 | 0.97 | 0.72 | 0.97 | 0.92 |
| 5 | 19520236 | rs1000727 | -0.177 | -0.217 | 1.238 | -0.185 | -0.003 | -0.077 | -0.077 | 3.158 | 0.18 | 0.27 | 0.13 | 0.04 | 0.19 | 0.17 |
| 5 | 19520405 | rs1000726 | 1.086 | -0.155 | 1.245 | -0.205 | 0.008 | 0.048 | 0.048 | 0 | 0.11 | 0.03 | 0.07 | 0 | 0.13 | 0.12 |
| 5 | 19520932 | rs441865 | -0.335 | -0.165 | 1.315 | -0.196 | -0.005 | 0.009 | 0.009 | 0 | 0.39 | 0.32 | 0.34 | 0.31 | 0.4 | 0.49 |
| 5 | 19527753 | rs3828573 | 1.057 | -0.117 | 1.093 | -0.19 | -0.062 | -0.057 | -0.057 | 0.112 | 0.23 | 0.03 | 0.22 | 0.27 | 0.22 | 0.36 |
| 5 | 19531296 | rs770298 | -0.491 | -0.088 | 1.365 | -0.142 | -0.005 | -0.021 | -0.021 | 0 | 0.42 | 0.52 | 0.35 | 0.27 | 0.37 | 0.51 |
| 5 | 19531541 | rs7734519 | -0.191 | -0.09 | 1.262 | -0.142 | -0.001 | -0.002 | -0.002 | 0 | 0.06 | 0.15 | 0.16 | 0.03 | 0.05 | 0.05 |
| 5 | 19534805 | rs995491 | -0.962 | -0.06 | 1.055 | -0.097 | 0.015 | 0.066 | 0.066 | 0 | 0.49 | 0.73 | 0.49 | 0.32 | 0.42 | 0.57 |
| 5 | 19537726 | rs770301 | -1.047 | -0.077 | 1.055 | -0.088 | -0.005 | 0.063 | 0.063 | 0 | 0.5 | 0.74 | 0.51 | 0.34 | 0.47 | 0.57 |
| 5 | 19547577 | rs347740 | 1.262 | -0.684 | 1.869 | 0.613 | -0.011 | -0.006 | -0.006 | 0.007 | 0.7 | 0.71 | 0.72 | 0.72 | 0.77 | 0.74 |
| 5 | 19548157 | rs347736 | 1.048 | -0.724 | 1.93 | 0.598 | -0.007 | 0.047 | 0.047 | 0.128 | 0.85 | 0.81 | 0.8 | 0.73 | 0.9 | 0.87 |
| 5 | 19548843 | rs10472287 | 1.982 | -0.666 | 1.978 | 0.559 | 0.025 | -0.004 | -0.004 | 0 | 0.59 | 0.26 | 0.57 | 0.57 | 0.66 | 0.69 |
| 5 | 19551292 | rs1631077 | 0.088 | -0.674 | 1.953 | 0.57 | -0.006 | 0.032 | 0.032 | 0 | 0.08 | 0.22 | 0.11 | 0.09 | 0.06 | 0.02 |
| 5 | 19555266 | rs6882679 | 1.705 | -0.624 | 1.527 | 0.525 | 0.006 | -0.002 | -0.002 | 0 | 0.6 | 0.38 | 0.58 | 0.58 | 0.66 | 0.69 |
| 5 | 19574173 | rs1565004 | 0.847 | -0.899 | 2.105 | 0.432 | -0.007 | 0.052 | 0.052 | 0 | 0.79 | 0.67 | 0.71 | 0.71 | 0.78 | 0.82 |

| | | | | | | | | | | | | | | | | |
|---|---|---|---|---|---|---|---|---|---|---|---|---|---|---|---|---|
| 5 | 19575900 | rs16885756 | 0.139 | -1.013 | 2.011 | 0.243 | -0.002 | -0.021 | -0.021 | 0.993 | 0.09 | 0.13 | 0.06 | 0 | 0.08 | 0.03 |
| 5 | 19578538 | rs7704186 | 0.164 | -1.047 | 1.896 | 0.227 | 0.004 | 0.017 | 0.017 | 0 | 0.09 | 0.05 | 0.05 | 0 | 0.08 | 0.03 |
| 5 | 19583251 | rs9292679 | -0.316 | -1.071 | 1.924 | 0.232 | -0.009 | -0.05 | -0.05 | 0 | 0.11 | 0.18 | 0.07 | 0 | 0.08 | 0.04 |
| 5 | 19585940 | rs6877651 | 0.221 | -1.189 | 1.446 | 0.28 | -0.032 | -0.056 | -0.056 | 0 | 0.21 | 0.11 | 0.29 | 0.41 | 0.21 | 0.38 |
| 5 | 19588949 | rs13167417 | 2.266 | -1.167 | 2.298 | 0.195 | 0.018 | 0.009 | 0.009 | 0 | 0.62 | 0.35 | 0.63 | 0.71 | 0.7 | 0.77 |
| 5 | 19597038 | rs2972819 | 0.077 | -1.189 | 2.298 | 0.178 | 0 | -0.006 | -0.006 | 0.239 | 0.85 | 0.87 | 0.79 | 1 | 0.82 | 0.86 |
| 5 | 19610285 | rs1352199 | 2.582 | -1.182 | 2.11 | 0.186 | 0.055 | 0.024 | 0.024 | 0 | 0.61 | 0.21 | 0.62 | 0.71 | 0.7 | 0.77 |
| 5 | 19611680 | rs164448 | 1.656 | -1.172 | 2.137 | 0.191 | -0.016 | 0.005 | 0.005 | 0 | 0.69 | 0.5 | 0.65 | 0.72 | 0.71 | 0.79 |
| 5 | 19613520 | rs7730733 | 0.471 | -1.115 | 1.492 | 0.214 | -0.002 | 0.033 | 0.033 | 0 | 0.08 | 0.03 | 0.05 | 0 | 0.08 | 0.03 |
| 5 | 19619844 | rs4866038 | 2.17 | -1.059 | 1.51 | 0.267 | 0.058 | 0.027 | 0.027 | 0 | 0.61 | 0.21 | 0.62 | 0.71 | 0.7 | 0.77 |
| 5 | 19647970 | rs2078188 | 2.203 | -1.027 | 1.411 | 0.258 | 0.056 | 0.03 | 0.03 | 0 | 0.6 | 0.16 | 0.62 | 0.71 | 0.69 | 0.77 |
| 5 | 19655277 | rs4866148 | 2.138 | -1.139 | 1.408 | 0.156 | 0.057 | 0.039 | 0.039 | 0 | 0.61 | 0.16 | 0.66 | 0.72 | 0.69 | 0.75 |
| 5 | 19661628 | rs11745155 | 1.928 | -1.178 | 1.774 | 0.134 | 0.012 | 0.007 | 0.007 | 0 | 0.65 | 0.33 | 0.68 | 0.72 | 0.7 | 0.76 |
| 5 | 19668948 | rs1521023 | 0.108 | -1.144 | 1.928 | 0.155 | 0.006 | -0.007 | -0.007 | 0.049 | 0.07 | 0.32 | 0.04 | 0.01 | 0.01 | 0.01 |
| 5 | 19678764 | rs4866039 | 1.254 | -0.981 | 2.212 | 0.243 | 0.003 | 0.223 | 0.223 | 0 | 0.92 | 0.73 | 0.95 | 0.73 | 0.96 | 0.96 |
| 5 | 19679108 | rs6887118 | -1.034 | -0.827 | 2.131 | 0.29 | -0.008 | 0.087 | 0.087 | 0.244 | 0.23 | 0.08 | 0.24 | 0.01 | 0.27 | 0.19 |
| 5 | 19744872 | rs6894851 | 1.33 | -1.058 | 1.785 | 0.671 | -0.031 | 0.034 | 0.034 | 0.278 | 0.79 | 0.4 | 0.78 | 0.73 | 0.85 | 0.89 |
| 5 | 19751140 | rs12109816 | 1.871 | -1.245 | 1.158 | 0.621 | 0.056 | -0.002 | -0.002 | 0 | 0.72 | 0.16 | 0.75 | 0.74 | 0.84 | 0.86 |
| 5 | 19772620 | rs7705221 | -2.005 | -1.283 | 1.326 | 0.614 | -0.018 | -0.002 | -0.002 | 0 | 0.27 | 0.72 | 0.13 | 0.25 | 0.12 | 0.1 |
| 5 | 19800575 | rs10068995 | 1.639 | -1.066 | 1.188 | 0.758 | -0.002 | 0.023 | 0.023 | 0 | 0.49 | 0.36 | 0.46 | 0.33 | 0.52 | 0.38 |
| 5 | 19804889 | rs4392624 | 0.433 | -1.088 | 1.6 | 0.713 | 0.009 | 0.005 | 0.005 | 2.366 | 0.67 | 0.65 | 0.52 | 0.33 | 0.62 | 0.42 |
| 5 | 19826902 | rs10078346 | 1.946 | -1.059 | 1.492 | 0.727 | -0.012 | 0.028 | 0.028 | 0 | 0.47 | 0.28 | 0.44 | 0.33 | 0.52 | 0.38 |
| 5 | 19827322 | rs6451476 | -1.18 | -0.923 | 1.135 | 0.691 | 0.013 | -0.005 | -0.005 | 0.095 | 0.19 | 0.15 | 0.07 | 0.01 | 0.09 | 0.05 |
| 5 | 19839695 | rs10076681 | 0.918 | -0.713 | 1.421 | 1.147 | 0.003 | 0.008 | 0.008 | 0.324 | 0.62 | 0.48 | 0.57 | 0.46 | 0.62 | 0.56 |
| 5 | 19844298 | rs6883590 | 1.122 | -0.904 | 1.6 | 1.16 | -0.069 | 0.031 | 0.031 | 0 | 0.79 | 0.56 | 0.87 | 0.76 | 0.91 | 0.9 |
| 5 | 19848872 | rs10074201 | 0.205 | -0.85 | 1.329 | 1.186 | -0.015 | 0.036 | 0.036 | 0 | 0.87 | 0.83 | 0.89 | 0.76 | 0.91 | 0.92 |
| 5 | 19865904 | rs1911861 | 1.27 | -0.501 | 0.538 | 1.306 | -0.001 | 0 | 0 | 0 | 0.53 | 0.41 | 0.49 | 0.43 | 0.54 | 0.49 |
| 5 | 19868580 | rs17222830 | 0.097 | -0.409 | 0.483 | 1.307 | -0.114 | -0.059 | -0.059 | 0 | 0.25 | 0.07 | 0.36 | 0.33 | 0.35 | 0.4 |
| 5 | 19871092 | rs16888157 | 0.063 | -0.592 | 0.459 | 1.261 | -0.006 | 0 | 0 | 1.82 | 0.08 | 0.07 | 0.11 | 0.09 | 0.07 | 0.18 |
| 5 | 19872234 | rs1876603 | 1.134 | -0.663 | 0.268 | 1.228 | -0.003 | 0.002 | 0.002 | 0 | 0.52 | 0.39 | 0.5 | 0.43 | 0.55 | 0.49 |
| 5 | 19877286 | rs7716704 | 0.234 | -0.595 | 0.048 | 1.239 | -0.089 | -0.042 | -0.042 | 1.32 | 0.26 | 0.07 | 0.36 | 0.33 | 0.35 | 0.42 |
| 5 | 19880618 | rs6869352 | 0.161 | -0.608 | 0.072 | 1.249 | -0.057 | -0.021 | -0.021 | 0 | 0.26 | 0.14 | 0.37 | 0.33 | 0.35 | 0.4 |
| 5 | 19892934 | rs4866159 | 0.483 | -0.576 | -0.019 | 1.132 | -0.068 | 0.018 | 0.018 | 0.003 | 0.78 | 0.54 | 0.87 | 0.76 | 0.91 | 0.9 |
| 5 | 19902953 | rs12658322 | 0.792 | 0.019 | -0.866 | 1.041 | -0.065 | 0.031 | 0.031 | 0.056 | 0.43 | 0.06 | 0.36 | 0.29 | 0.48 | 0.33 |
| 5 | 19920554 | rs7734813 | 0.482 | 0.216 | 0.514 | 1.723 | -0.002 | 0.003 | 0.003 | 0.059 | 0.54 | 0.56 | 0.48 | 0.73 | 0.52 | 0.51 |
| 5 | 19923791 | rs12655266 | 0.851 | 0.007 | 1.152 | 1.693 | 0.006 | 0.042 | 0.042 | 0 | 0.88 | 0.69 | 0.91 | 0.84 | 0.91 | 0.98 |
| 5 | 19926525 | rs2202798 | 2.197 | -0.423 | 1.331 | 1.656 | 0.029 | -0.034 | -0.034 | 0.117 | 0.81 | 0.14 | 0.79 | 0.62 | 0.9 | 0.94 |
| 5 | 19929288 | rs1911845 | -1.442 | -0.387 | 1.623 | 1.648 | 0.05 | 0.275 | 0.275 | 3.587 | 0.06 | 0.49 | 0.12 | 0.22 | 0.01 | 0.02 |
| 5 | 19932686 | rs9986203 | 0.693 | -0.089 | 1.374 | 1.751 | 0 | 0.001 | 0.001 | 0 | 0.58 | 0.38 | 0.44 | 0.6 | 0.6 | 0.49 |
| 5 | 19948930 | rs13177416 | 1.281 | -0.105 | 1.027 | 1.837 | -0.056 | -0.05 | -0.05 | 0 | 0.5 | 0.08 | 0.42 | 0.5 | 0.54 | 0.55 |
| 5 | 19949403 | rs6451573 | 0.336 | -0.169 | 1.419 | 1.822 | -0.027 | 0.006 | 0.006 | 0.048 | 0.63 | 0.72 | 0.49 | 0.71 | 0.56 | 0.59 |
| 5 | 19958804 | rs13175060 | 1.36 | -0.121 | 1.386 | 1.771 | -0.057 | -0.051 | -0.051 | 0 | 0.51 | 0.09 | 0.42 | 0.5 | 0.54 | 0.55 |
| 5 | 19970119 | rs2892429 | 0.585 | -0.044 | 1.42 | 1.789 | 0.039 | 0.099 | 0.099 | 0 | 0.94 | 0.82 | 0.92 | 0.84 | 0.93 | 0.99 |
| 5 | 19985234 | rs1540751 | 0.519 | -0.035 | 1.568 | 1.68 | -0.001 | 0 | 0 | 0 | 0.61 | 0.61 | 0.48 | 0.5 | 0.56 | 0.56 |
| 5 | 19987468 | rs7736515 | -1.82 | -0.231 | 1.958 | 1.534 | -0.022 | -0.054 | -0.054 | 0.033 | 0.1 | 0.45 | 0.05 | 0 | 0.03 | 0.01 |
| 5 | 20012392 | rs7704008 | -0.442 | -0.119 | 2.616 | 1.604 | -0.012 | -0.025 | -0.025 | 0 | 0.05 | 0.18 | 0.03 | 0 | 0.02 | 0.01 |
| 5 | 20016599 | rs3112489 | -1.133 | -0.12 | 2.539 | 1.633 | 0.007 | -0.006 | -0.006 | 0 | 0.07 | 0.39 | 0.04 | 0 | 0.02 | 0.01 |
| 5 | 20028798 | rs4292450 | 2.088 | -0.291 | 2.484 | 1.799 | -0.041 | 0.129 | 0.129 | 0.86 | 0.88 | 0.52 | 0.94 | 0.76 | 0.95 | 0.92 |
| 5 | 20030472 | rs6860771 | -0.858 | -0.37 | 2.394 | 2.274 | -0.005 | 0.073 | 0.073 | 0 | 0.06 | 0.14 | 0.08 | 0.13 | 0.05 | 0.07 |
| 5 | 20030652 | rs2892442 | 1.699 | -0.402 | 2.364 | 2.254 | -0.041 | 0.11 | 0.11 | 0.273 | 0.88 | 0.55 | 0.94 | 0.76 | 0.95 | 0.92 |
| 5 | 20031683 | rs11955587 | 0.11 | -0.24 | 1.855 | 2.124 | -0.051 | 0.054 | 0.054 | 0 | 0.29 | 0.09 | 0.41 | 0.11 | 0.37 | 0.34 |
| 5 | 20053583 | rs1347523 | 1.802 | -0.221 | 1.846 | 2.134 | -0.039 | 0.036 | 0.036 | 2.131 | 0.85 | 0.42 | 0.86 | 0.88 | 0.91 | 0.91 |
| 5 | 20073777 | rs6869124 | 0.12 | -0.237 | 1.186 | 2.026 | -0.132 | 0.04 | 0.04 | 0 | 0.29 | 0.02 | 0.4 | 0.11 | 0.37 | 0.34 |
| 5 | 20075855 | rs10434577 | 0.704 | -0.53 | 0.502 | 1.547 | -0.02 | -0.093 | -0.093 | 0 | 0.17 | 0.01 | 0.11 | 0.32 | 0.14 | 0.27 |
| 5 | 20107527 | rs13170493 | 1.236 | -0.074 | 1.167 | 1.802 | 0.015 | 0.008 | 0.008 | 0.187 | 0.5 | 0.22 | 0.41 | 0.49 | 0.53 | 0.51 |
| 5 | 20109561 | rs11744487 | 1.587 | -0.119 | 0.911 | 1.571 | -0.098 | 0.023 | 0.023 | 0.237 | 0.47 | 0.05 | 0.36 | 0.4 | 0.5 | 0.48 |
| 5 | 20124530 | rs1435972 | 0.169 | -0.173 | 0.842 | 1.217 | -0.099 | 0.082 | 0.082 | 0.025 | 0.28 | 0.04 | 0.4 | 0.1 | 0.36 | 0.33 |
| 5 | 20143940 | rs10060069 | 1.162 | -0.114 | 1.187 | 1.327 | 0.016 | 0.071 | 0.071 | 0.027 | 0.53 | 0.3 | 0.46 | 0.78 | 0.56 | 0.53 |
| 5 | 20174836 | rs9292895 | 0.997 | 0.212 | 1.552 | 1.766 | -0.006 | -0.001 | -0.001 | 2.606 | 0.61 | 0.65 | 0.54 | 0.61 | 0.59 | 0.57 |
| 5 | 20176887 | rs7706987 | -0.502 | 0.183 | 1.459 | 1.822 | -0.01 | 0.014 | 0.014 | 0 | 0.08 | 0.08 | 0.08 | 0.12 | 0.05 | 0.13 |
| 5 | 20194982 | rs925185 | 1.784 | 0.235 | 1.303 | 2.013 | -0.046 | 0.184 | 0.184 | 1.384 | 0.49 | 0.07 | 0.47 | 0.26 | 0.52 | 0.49 |
| 5 | 20207714 | rs11738280 | -1.735 | 0.084 | 1.762 | 1.922 | -0.065 | 0.02 | 0.02 | 0.083 | 0.15 | 0.36 | 0.11 | 0.13 | 0.06 | 0.19 |
| 5 | 20208626 | rs10473344 | 1.674 | 0.12 | 1.791 | 1.886 | 0.002 | 0.06 | 0.06 | 0.785 | 0.52 | 0.3 | 0.46 | 0.32 | 0.58 | 0.5 |
| 5 | 20216940 | rs1508584 | 0.103 | 0.205 | 1.507 | 1.829 | -0.088 | -0.002 | -0.002 | 4.682 | 0.7 | 0.89 | 0.63 | 0.72 | 0.65 | 0.69 |
| 5 | 20225602 | rs6887472 | 0.103 | 0.205 | 1.507 | 1.829 | -0.088 | -0.002 | -0.002 | 0 | 0.7 | 0.89 | 0.63 | 0.72 | 0.65 | 0.69 |
| 5 | 20235071 | rs4866047 | 0.173 | 0.154 | 1.501 | 1.806 | -0.072 | -0.004 | -0.004 | 0 | 0.71 | 0.89 | 0.63 | 0.72 | 0.66 | 0.65 |
| 5 | 20238774 | rs7730653 | 1.825 | 0.122 | 1.45 | 1.826 | -0.066 | 0.207 | 0.207 | 1.152 | 0.5 | 0.06 | 0.42 | 0.25 | 0.52 | 0.44 |
| 5 | 20248382 | rs10071248 | -1.692 | 0.029 | 1.692 | 1.888 | 0 | -0.003 | -0.003 | 1.34 | 0.44 | 0.51 | 0.48 | 0.4 | 0.41 | 0.54 |
| 5 | 20261150 | rs10473372 | -1.201 | 0.05 | 1.517 | 1.898 | -0.005 | -0.002 | -0.002 | 0 | 0.39 | 0.33 | 0.45 | 0.4 | 0.4 | 0.48 |

| | | | | | | | | | | | | | | | | |
|---|---|---|---|---|---|---|---|---|---|---|---|---|---|---|---|---|
| 5 | 20290034 | rs2974591 | -1.568 | 0.208 | 1.675 | 1.995 | 0.002 | 0.012 | 0.012 | 0.345 | 0.46 | 0.58 | 0.53 | 0.68 | 0.41 | 0.59 |
| 5 | 20294072 | rs2063229 | -1.624 | 0.072 | 1.631 | 1.866 | -0.018 | 0.048 | 0.048 | 0 | 0.1 | 0.26 | 0.13 | 0.12 | 0.06 | 0.13 |
| 5 | 20322049 | rs10079950 | -0.41 | 1.132 | 1.646 | 1.89 | 0.01 | -0.016 | -0.016 | 0 | 0.07 | 0.07 | 0.1 | 0 | 0.12 | 0.09 |
| 5 | 20325180 | rs6884961 | 0.021 | 0.944 | 1.232 | 1.837 | 0.003 | -0.029 | -0.029 | 0.796 | 0.79 | 0.92 | 0.74 | 0.7 | 0.79 | 0.75 |
| 5 | 20348425 | rs6897700 | -1.87 | 0.939 | 1.644 | 1.817 | 0.006 | 0.369 | 0.369 | 0.156 | 0.08 | 0.35 | 0.14 | 0.43 | 0.04 | 0.2 |
| 5 | 20355032 | rs6870586 | -0.615 | 1.154 | 1.099 | 1.762 | 0.098 | -0.047 | -0.047 | 0.602 | 0.09 | 0.21 | 0.13 | 0 | 0.17 | 0.1 |
| 5 | 20358169 | rs10078676 | -0.521 | 1.158 | 1.083 | 1.764 | 0.076 | -0.045 | -0.045 | 0 | 0.09 | 0.16 | 0.12 | 0 | 0.17 | 0.1 |
| 5 | 20364007 | rs10473391 | -0.827 | 1.053 | 0.96 | 1.848 | 0.07 | 0.149 | 0.149 | 0 | 0.12 | 0.28 | 0.15 | 0.3 | 0.18 | 0.2 |
| 5 | 20366035 | rs10941700 | -1.118 | 1.054 | 0.718 | 0.694 | -0.003 | 0.037 | 0.037 | 0 | 0.06 | 0.13 | 0.11 | 0.13 | 0.04 | 0.2 |
| 5 | 20366537 | rs11741365 | 1.404 | 0.945 | 0.38 | 0.545 | 0.004 | -0.014 | -0.014 | 0 | 0.61 | 0.54 | 0.58 | 0.34 | 0.59 | 0.56 |
| 5 | 20380859 | rs10073450 | -1.334 | 1.588 | 0.71 | 0.378 | 0.09 | -0.012 | -0.012 | 0 | 0.12 | 0.32 | 0.15 | 0.08 | 0.19 | 0.19 |
| 5 | 20401352 | rs10066903 | 0.459 | 0.979 | -0.183 | 0.264 | 0.014 | -0.023 | -0.023 | 0 | 0.62 | 0.5 | 0.52 | 0.28 | 0.6 | 0.52 |
| 5 | 20419469 | rs12514159 | 0.645 | 0.529 | -1.074 | 0.083 | -0.115 | 0.008 | 0.008 | 0 | 0.39 | 0.07 | 0.45 | 0.34 | 0.51 | 0.38 |
| 5 | 20420209 | rs9687957 | -0.081 | 0.211 | -0.567 | 0.393 | 0 | 0 | 0 | 0 | 0.77 | 0.75 | 0.86 | 0.72 | 0.78 | 0.75 |
| 5 | 20429345 | rs10062513 | 0.008 | 0.194 | -0.541 | 0.381 | -0.013 | -0.012 | -0.012 | 0 | 0.23 | 0.19 | 0.14 | 0.3 | 0.22 | 0.25 |
| 5 | 20439891 | rs13357704 | 0.197 | 0.168 | -0.636 | 0.357 | -0.076 | -0.071 | -0.071 | 0 | 0.21 | 0.05 | 0.12 | 0.28 | 0.22 | 0.19 |
| 5 | 20443700 | rs6877123 | -0.075 | -0.186 | 0.062 | 0.234 | -0.026 | 0.012 | 0.012 | 0 | 0.68 | 0.42 | 0.6 | 0.63 | 0.74 | 0.63 |
| 5 | 20446058 | rs6887953 | 0.131 | -0.248 | 0.651 | 0.183 | 0.024 | 0.017 | 0.017 | 0 | 0.75 | 0.65 | 0.85 | 0.72 | 0.77 | 0.75 |
| 5 | 20446378 | rs4866190 | 0.131 | -0.248 | 0.656 | 0.183 | 0.023 | 0.017 | 0.017 | 0 | 0.75 | 0.65 | 0.85 | 0.72 | 0.77 | 0.75 |
| 5 | 20447801 | rs9292940 | 0.31 | -0.468 | 0.62 | 0.063 | 0.013 | -0.088 | -0.088 | 0 | 0.25 | 0.04 | 0.35 | 0.36 | 0.23 | 0.35 |
| 5 | 20451383 | rs13180247 | 1.533 | -0.477 | 0.578 | 0.056 | -0.08 | 0.044 | 0.044 | 0 | 0.43 | 0.12 | 0.46 | 0.35 | 0.52 | 0.38 |
| 5 | 20452114 | rs1508545 | 0.13 | -0.392 | 0.547 | 0.103 | 0.026 | 0.023 | 0.023 | 0 | 0.74 | 0.62 | 0.85 | 0.7 | 0.77 | 0.74 |
| 5 | 20453495 | rs12659684 | 0.332 | -0.378 | 0.072 | 0.117 | 0 | -0.115 | -0.115 | 0 | 0.25 | 0.03 | 0.35 | 0.36 | 0.23 | 0.35 |
| 5 | 20459638 | rs1472892 | -0.243 | -0.294 | 0.171 | 0.139 | -0.015 | -0.015 | -0.015 | 0.05 | 0.78 | 0.84 | 0.87 | 0.72 | 0.78 | 0.81 |
| 5 | 20462781 | rs1508548 | 1.366 | -0.355 | 0.315 | 0.096 | -0.05 | 0.06 | 0.06 | 0 | 0.44 | 0.18 | 0.47 | 0.34 | 0.52 | 0.38 |
| 5 | 20464464 | rs10067934 | -0.463 | -0.482 | 0.905 | -0.284 | 0.002 | -0.003 | -0.003 | 0.784 | 0.3 | 0.33 | 0.37 | 0.36 | 0.26 | 0.37 |
| 5 | 20474634 | rs12697585 | 0.141 | -0.295 | 0.546 | -0.268 | 0.011 | -0.014 | -0.014 | 0 | 0.69 | 0.78 | 0.66 | 0.59 | 0.78 | 0.6 |
| 5 | 20491707 | rs981587 | 0.484 | -0.192 | 0.882 | -0.119 | 0.041 | 0.034 | 0.034 | 0 | 0.94 | 0.77 | 0.97 | 0.96 | 0.98 | 0.98 |
| 5 | 20492277 | rs992958 | 1.347 | -0.154 | 0.218 | -0.582 | -0.098 | -0.042 | -0.042 | 0 | 0.39 | 0.1 | 0.49 | 0.58 | 0.51 | 0.38 |
| 5 | 20502578 | rs1355057 | 1.189 | -0.229 | 0.821 | -0.636 | -0.005 | 0.002 | 0.002 | 0 | 0.43 | 0.39 | 0.53 | 0.59 | 0.51 | 0.42 |
| 5 | 20518529 | rs12520315 | 0.988 | -0.081 | 0.914 | -0.733 | -0.006 | 0.011 | 0.011 | 0 | 0.45 | 0.43 | 0.52 | 0.6 | 0.51 | 0.47 |
| 5 | 20519284 | rs13190303 | 1.353 | -0.06 | 0.905 | -0.721 | -0.06 | 0.01 | 0.01 | 0 | 0.41 | 0.19 | 0.5 | 0.59 | 0.51 | 0.44 |
| 5 | 20520802 | rs2471138 | 0.203 | -0.143 | 0.99 | -0.879 | 0.003 | -0.109 | -0.109 | 0.699 | 0.68 | 0.6 | 0.81 | 0.97 | 0.69 | 0.82 |
| 5 | 20524918 | rs2457040 | 1.471 | -0.08 | 0.667 | -0.809 | -0.056 | 0.011 | 0.011 | 0 | 0.41 | 0.19 | 0.5 | 0.59 | 0.51 | 0.44 |
| 5 | 20528774 | rs2202790 | 1.471 | -0.08 | 0.66 | -0.809 | -0.056 | 0.011 | 0.011 | 0 | 0.41 | 0.2 | 0.5 | 0.59 | 0.51 | 0.44 |
| 5 | 20538115 | rs1703052 | -0.063 | -0.012 | 0.878 | -0.882 | 0.005 | -0.072 | -0.072 | 0 | 0.88 | 0.74 | 0.93 | 1 | 0.92 | 0.98 |
| 5 | 20539266 | rs1651443 | 1.298 | 0.093 | 1.05 | -0.71 | -0.018 | 0.01 | 0.01 | 0 | 0.42 | 0.32 | 0.51 | 0.6 | 0.49 | 0.47 |
| 5 | 20548944 | rs1703042 | 0.628 | 0.718 | 0.669 | -1.027 | 0.001 | -0.087 | -0.087 | 0 | 0.68 | 0.65 | 0.81 | 0.97 | 0.69 | 0.82 |
| 5 | 20559438 | rs1703049 | 0.694 | 0.768 | 0.165 | -0.95 | 0.001 | -0.079 | -0.079 | 0 | 0.66 | 0.64 | 0.8 | 0.97 | 0.68 | 0.81 |
| 5 | 20563135 | rs17839229 | -1.466 | 1.597 | 0.321 | -0.287 | 0.007 | -0.012 | -0.012 | 0 | 0.08 | 0.11 | 0.18 | 0 | 0.16 | 0.05 |
| 5 | 20564905 | rs2434785 | -2.583 | 1.205 | 1.287 | 0.08 | 0.051 | 0.036 | 0.036 | 0.035 | 0.17 | 0.52 | 0.22 | 0.17 | 0.19 | 0.11 |
| 5 | 20570592 | rs2434781 | -1.374 | 1.118 | 1.514 | 0.131 | -0.013 | 0.077 | 0.077 | 0 | 0.06 | 0.18 | 0.02 | 0.17 | 0.03 | 0.06 |
| 5 | 20570987 | rs986251 | 0.499 | 1.278 | 0.885 | -0.035 | 0 | 0.001 | 0.001 | 0.19 | 0.66 | 0.63 | 0.56 | 0.37 | 0.61 | 0.46 |
| 5 | 20574412 | rs1030479 | 0.116 | 1.45 | 0.714 | -0.028 | -0.013 | -0.047 | -0.047 | 0 | 0.75 | 0.87 | 0.63 | 0.37 | 0.68 | 0.55 |
| 5 | 20591074 | rs2115342 | 0.448 | 1.437 | 1.07 | 0.012 | 0.029 | 0.011 | 0.011 | 0.053 | 0.7 | 0.65 | 0.41 | 0.3 | 0.58 | 0.48 |
| 5 | 20599667 | rs13360869 | -1.539 | 1.232 | 1.171 | -0.142 | -0.024 | -0.002 | -0.002 | 0.056 | 0.19 | 0.47 | 0.13 | 0.12 | 0.16 | 0.14 |
| 5 | 20606559 | rs1007844 | 0.347 | 1.295 | 1.124 | -0.071 | 0.018 | 0.01 | 0.01 | 0.053 | 0.7 | 0.69 | 0.41 | 0.31 | 0.59 | 0.49 |
| 5 | 20606656 | rs2010326 | -1.324 | 1.424 | 1.25 | -0.241 | -0.008 | -0.091 | -0.091 | 0 | 0.16 | 0.39 | 0.12 | 0 | 0.15 | 0.1 |
| 5 | 20608142 | rs919344 | -1.526 | 1.769 | 1.053 | -0.42 | 0.005 | -0.111 | -0.111 | 1.481 | 0.23 | 0.4 | 0.16 | 0 | 0.22 | 0.11 |
| 5 | 20616080 | rs2434772 | 1.701 | 1.999 | 0.619 | 0.183 | 0.034 | 0.047 | 0.047 | 0 | 0.53 | 0.2 | 0.37 | 0.38 | 0.42 | 0.46 |
| 5 | 20621653 | rs919345 | 0.15 | 2.48 | 0.722 | 0.571 | -0.058 | -0.114 | -0.114 | 0 | 0.8 | 0.95 | 0.69 | 0.54 | 0.71 | 0.57 |
| 5 | 20636493 | rs7732005 | -1.657 | 2.591 | 1.72 | 0.433 | 0.021 | 0.126 | 0.126 | 0 | 0.19 | 0.37 | 0.21 | 0.5 | 0.18 | 0.21 |
| 5 | 20647735 | rs6864883 | 0.054 | 2.583 | 1.345 | 0.422 | -0.062 | -0.125 | -0.125 | 0.17 | 0.8 | 0.96 | 0.69 | 0.54 | 0.71 | 0.57 |
| 5 | 20662047 | rs9292952 | 0.022 | 2.575 | 1.477 | 0.418 | -0.04 | -0.092 | -0.092 | 0 | 0.8 | 0.88 | 0.68 | 0.54 | 0.71 | 0.57 |
| 5 | 20685297 | rs6890147 | 1.849 | 2.373 | 1.796 | 0.425 | 0.018 | 0.087 | 0.087 | 0.081 | 0.77 | 0.38 | 0.63 | 0.5 | 0.78 | 0.75 |
| 5 | 20740586 | rs10941853 | -0.33 | 1.54 | 1.661 | 0.461 | -0.067 | -0.14 | -0.14 | 0 | 0.22 | 0.05 | 0.33 | 0.49 | 0.3 | 0.45 |
| 5 | 20786582 | rs10941855 | 2.394 | 1.433 | 2.158 | 0.368 | 0.004 | 0.031 | 0.031 | 5.656 | 0.89 | 0.34 | 0.8 | 0.88 | 0.93 | 0.92 |
| 5 | 20824694 | rs10065829 | -1.282 | 1.259 | 1.999 | 0.201 | -0.006 | -0.074 | -0.074 | 0 | 0.13 | 0.26 | 0.1 | 0 | 0.15 | 0.03 |
| 5 | 20832133 | rs4866066 | -1.794 | 1.87 | 2.11 | 0.417 | 0 | -0.023 | -0.023 | 0 | 0.45 | 0.53 | 0.72 | 0.79 | 0.58 | 0.61 |
| 5 | 20865083 | rs7442892 | 1.004 | 1.445 | 2.032 | 0.807 | -0.005 | 0.161 | 0.161 | 0 | 0.86 | 0.66 | 0.79 | 0.58 | 0.89 | 0.84 |
| 5 | 20878263 | rs10079831 | -0.743 | 1.698 | 1.67 | 1.009 | 0.005 | 0.018 | 0.018 | 0.965 | 0.24 | 0.22 | 0.42 | 0.48 | 0.35 | 0.47 |
| 5 | 20892281 | rs6886032 | -0.868 | 1.739 | 1.428 | 0.965 | 0.022 | 0.077 | 0.077 | 0.653 | 0.27 | 0.39 | 0.45 | 0.49 | 0.34 | 0.49 |
| 5 | 20902807 | rs11956899 | -1.098 | 1.743 | 1.264 | 1.105 | 0.014 | 0.029 | 0.029 | 0.256 | 0.18 | 0.41 | 0.29 | 0.2 | 0.23 | 0.1 |
| 5 | 20924403 | rs2201407 | -0.947 | 1.647 | 0.225 | 0.638 | 0.005 | -0.005 | -0.005 | 0 | 0.38 | 0.51 | 0.67 | 0.67 | 0.49 | 0.61 |
| 5 | 20947356 | rs3913348 | -0.992 | 0.986 | -0.051 | 0.681 | -0.002 | 0 | 0 | 8.191 | 0.21 | 0.23 | 0.1 | 0.25 | 0.17 | 0.16 |
| 5 | 20953429 | rs1979833 | -0.232 | 1.277 | 0.38 | 0.209 | -0.001 | 0 | 0 | 0 | 0.32 | 0.29 | 0.54 | 0.67 | 0.32 | 0.56 |
| 5 | 20964216 | rs17238567 | -0.973 | 1.558 | 0.809 | 0.124 | 0.034 | 0.017 | 0.017 | 0 | 0.07 | 0.15 | 0.07 | 0.08 | 0.1 | 0.08 |
| 5 | 20967185 | rs12659921 | -0.803 | 1.681 | 1.276 | -0.187 | 0.001 | 0.003 | 0.003 | 0.082 | 0.45 | 0.54 | 0.63 | 0.78 | 0.44 | 0.66 |
| 5 | 20967318 | rs12657343 | -0.224 | 1.669 | 1.752 | -0.245 | -0.001 | -0.007 | -0.007 | 1.218 | 0.35 | 0.37 | 0.58 | 0.76 | 0.35 | 0.6 |

| | | | | | | | | | | | | | | | | |
|---|---|---|---|---|---|---|---|---|---|---|---|---|---|---|---|---|
| 5 | 20970069 | rs1499730 | 0.156 | 1.656 | 1.714 | -0.247 | 0.001 | -0.002 | -0.002 | 1.989 | 0.1 | 0.09 | 0.04 | 0 | 0.09 | 0.05 |
| 5 | 20978211 | rs13154017 | -0.843 | 1.623 | 1.984 | -0.273 | -0.002 | -0.031 | -0.031 | 0.082 | 0.45 | 0.51 | 0.63 | 0.77 | 0.44 | 0.65 |
| 5 | 20981688 | rs6878853 | -1.438 | 1.681 | 1.875 | -0.269 | 0.01 | 0.105 | 0.105 | 0.054 | 0.53 | 0.68 | 0.7 | 0.86 | 0.54 | 0.73 |
| 5 | 20982268 | rs4613760 | -1.498 | 1.648 | 1.936 | -0.273 | 0.009 | 0.154 | 0.154 | 0.091 | 0.53 | 0.76 | 0.71 | 0.86 | 0.54 | 0.73 |
| 5 | 20991326 | rs10473457 | -0.995 | 1.659 | 2.103 | -0.168 | 0 | -0.014 | -0.014 | 0.23 | 0.43 | 0.55 | 0.64 | 0.81 | 0.46 | 0.66 |
| 5 | 20991405 | rs10473458 | -1.034 | 1.659 | 2.103 | -0.168 | 0 | -0.013 | -0.013 | 0 | 0.44 | 0.55 | 0.64 | 0.81 | 0.46 | 0.66 |
| 5 | 20995098 | rs2221143 | -0.825 | 1.72 | 2.232 | 0.392 | 0.004 | -0.006 | -0.006 | 0 | 0.42 | 0.44 | 0.62 | 0.62 | 0.46 | 0.65 |
| 5 | 21014949 | rs4476767 | 0.734 | 1.549 | 2.573 | 0.838 | 0.057 | 0.127 | 0.127 | 0 | 0.9 | 0.65 | 0.93 | 0.75 | 0.89 | 0.94 |
| 5 | 21018827 | rs2088664 | -0.307 | 1.583 | 2.525 | 0.762 | 0.003 | 0.122 | 0.122 | 0 | 0.22 | 0.44 | 0.42 | 0.55 | 0.21 | 0.41 |
| 5 | 21041496 | rs10055596 | -0.365 | 1.577 | 2.523 | 0.759 | 0.002 | 0.118 | 0.118 | 0 | 0.22 | 0.42 | 0.42 | 0.55 | 0.21 | 0.42 |
| 5 | 21047987 | rs1027544 | -0.142 | 1.581 | 2.52 | 0.761 | 0.004 | 0.084 | 0.084 | 0 | 0.21 | 0.33 | 0.4 | 0.55 | 0.21 | 0.42 |
| 5 | 21069363 | rs9885417 | -1.157 | 1.678 | 2.609 | 0.715 | 0.044 | -0.028 | -0.028 | 0 | 0.11 | 0.33 | 0.07 | 0.06 | 0.11 | 0.05 |
| 5 | 21071305 | rs13436839 | -0.665 | 1.695 | 2.45 | 0.718 | 0.038 | -0.028 | -0.028 | 0.017 | 0.09 | 0.23 | 0.06 | 0.05 | 0.11 | 0.05 |
| 5 | 21082707 | rs10058965 | -0.529 | 1.707 | 2.384 | 0.718 | 0.029 | -0.028 | -0.028 | 0 | 0.09 | 0.19 | 0.06 | 0.05 | 0.11 | 0.05 |
| 5 | 21094862 | rs16888939 | -0.547 | 2.032 | 2.259 | 0.686 | 0.008 | -0.033 | -0.033 | 0 | 0.09 | 0.12 | 0.04 | 0 | 0.11 | 0.03 |
| 5 | 21117980 | rs7446348 | -0.285 | 1.8 | 2.26 | 0.87 | -0.008 | -0.029 | -0.029 | 2.154 | 0.23 | 0.17 | 0.41 | 0.52 | 0.25 | 0.43 |
| 5 | 21120491 | rs6893407 | 1.301 | 1.89 | 2.32 | 0.865 | 0.029 | 0.06 | 0.06 | 0.23 | 0.85 | 0.36 | 0.87 | 0.73 | 0.84 | 0.93 |
| 5 | 21143240 | rs248189 | -0.534 | 1.85 | 2.238 | 0.92 | 0.011 | 0.096 | 0.096 | 0.006 | 0.26 | 0.46 | 0.43 | 0.52 | 0.25 | 0.43 |
| 5 | 21148212 | rs432786 | -0.378 | 1.924 | 2.468 | 0.922 | 0.007 | 0.039 | 0.039 | 0 | 0.24 | 0.28 | 0.42 | 0.52 | 0.25 | 0.43 |
| 5 | 21168358 | rs16888991 | -0.373 | 1.743 | 2.693 | 0.91 | -0.008 | -0.012 | -0.012 | 1.314 | 0.24 | 0.21 | 0.43 | 0.57 | 0.3 | 0.44 |
| 5 | 21185317 | rs159211 | -0.532 | 1.721 | 2.896 | 0.9 | 0.016 | 0.033 | 0.033 | 0 | 0.24 | 0.38 | 0.46 | 0.57 | 0.3 | 0.44 |
| 5 | 21194251 | rs294114 | -0.975 | 1.166 | 2.824 | 0.955 | 0.017 | -0.022 | -0.022 | 0.343 | 0.07 | 0.01 | 0.03 | 0.1 | 0.05 | 0.11 |
| 5 | 21213478 | rs16889078 | -0.879 | 1.301 | 2.869 | 0.876 | 0.025 | -0.064 | -0.064 | 0 | 0.11 | 0.29 | 0.08 | 0 | 0.11 | 0.04 |
| 5 | 21217860 | rs294713 | 2.93 | 1.15 | 2.853 | 0.847 | -0.047 | 0.042 | 0.042 | 2.171 | 0.85 | 0.29 | 0.9 | 0.68 | 0.94 | 0.86 |
| 5 | 21234972 | rs6451935 | 1.631 | 1.132 | 3.104 | 0.826 | 0.009 | 0.048 | 0.048 | 0 | 0.84 | 0.59 | 0.9 | 0.77 | 0.87 | 0.92 |
| 5 | 21235554 | rs1873609 | 2.6 | 1.137 | 2.783 | 0.829 | -0.007 | 0.003 | 0.003 | 0 | 0.77 | 0.3 | 0.86 | 0.68 | 0.83 | 0.82 |
| 5 | 21238932 | rs1388574 | 2.6 | 1.137 | 2.783 | 0.829 | -0.007 | 0.003 | 0.003 | 0 | 0.77 | 0.3 | 0.86 | 0.68 | 0.83 | 0.82 |
| 5 | 21242810 | rs6884724 | -0.195 | 1.169 | 2.512 | 0.786 | -0.02 | 0.029 | 0.029 | 0 | 0.09 | 0.03 | 0.05 | 0 | 0.11 | 0.04 |
| 5 | 21261450 | rs6862720 | -1.55 | 1.212 | 2.266 | 1.542 | -0.003 | 0.044 | 0.044 | 0 | 0.09 | 0.14 | 0.07 | 0.23 | 0.11 | 0.18 |
| 5 | 21265377 | rs4866272 | 2.795 | 1.299 | 2.202 | 1.722 | 0.001 | -0.009 | -0.009 | 0 | 0.73 | 0.26 | 0.8 | 0.52 | 0.74 | 0.73 |
| 5 | 21267757 | rs7731475 | -0.177 | 1.42 | 1.738 | 1.75 | -0.006 | 0.001 | 0.001 | 0 | 0.09 | 0.08 | 0.06 | 0.01 | 0.11 | 0.05 |
| 5 | 21273843 | rs2883171 | 2.465 | 1.295 | 1.957 | 2.001 | -0.007 | 0.14 | 0.14 | 0 | 0.86 | 0.73 | 0.89 | 0.53 | 0.86 | 0.78 |
| 5 | 21319693 | rs10941887 | 1.331 | 1.305 | 1.672 | 1.643 | -0.006 | 0.016 | 0.016 | 0 | 0.95 | 0.9 | 0.96 | 0.9 | 0.97 | 0.95 |
| 5 | 21391244 | rs10055072 | -0.797 | 1.295 | 1.306 | 1.469 | 0.008 | 0 | 0 | 0 | 0.38 | 0.56 | 0.5 | 0.6 | 0.48 | 0.53 |
| 5 | 21394707 | rs10039330 | -1.084 | 1.194 | 1.304 | 1.549 | -0.016 | 0.045 | 0.045 | 0 | 0.05 | 0.08 | 0.08 | 0.13 | 0.03 | 0.03 |
| 5 | 21421537 | rs11746426 | -0.399 | 1.526 | 1.871 | 1.822 | -0.003 | -0.004 | -0.004 | 0 | 0.32 | 0.3 | 0.43 | 0.44 | 0.4 | 0.39 |
| 5 | 21421712 | rs11746478 | -0.399 | 1.526 | 1.871 | 1.822 | -0.003 | -0.004 | -0.004 | 0.675 | 0.32 | 0.3 | 0.43 | 0.44 | 0.4 | 0.39 |
| 5 | 21432444 | rs4645308 | 0.241 | 1.4 | 2.015 | 1.76 | -0.084 | -0.135 | -0.135 | 0 | 0.25 | 0.06 | 0.39 | 0.44 | 0.34 | 0.38 |
| 5 | 21437412 | rs6451953 | -0.82 | 1.122 | 2.01 | 1.594 | 0.018 | 0.004 | 0.004 | 0 | 0.37 | 0.45 | 0.49 | 0.59 | 0.48 | 0.53 |
| 5 | 21547156 | rs6868224 | 1.391 | 1.125 | 1.291 | 1.657 | -0.003 | 0.008 | 0.008 | 0 | 0.74 | 0.49 | 0.68 | 0.23 | 0.74 | 0.65 |
| 5 | 21551543 | rs1817950 | -0.723 | 1.279 | 0.951 | 2.034 | 0.028 | 0.222 | 0.222 | 0 | 0.06 | 0.14 | 0.12 | 0.41 | 0.09 | 0.15 |
| 5 | 21565058 | rs1496455 | -0.299 | 1.112 | 0.503 | 1.177 | -0.008 | 0.016 | 0.016 | 0.13 | 0.32 | 0.25 | 0.37 | 0.18 | 0.38 | 0.36 |
| 5 | 21574347 | rs7720174 | -1.226 | 1.236 | 0.399 | 0.765 | 0.011 | -0.046 | -0.046 | 0.012 | 0.1 | 0.15 | 0.07 | 0 | 0.09 | 0.04 |
| 5 | 21574610 | rs12519981 | 1.192 | 0.753 | -0.472 | 0.59 | -0.075 | 0.058 | 0.058 | 0 | 0.28 | 0.04 | 0.25 | 0.16 | 0.3 | 0.24 |
| 5 | 21575563 | rs7725903 | 0.123 | 0.139 | 0.646 | 1.187 | -0.003 | 0.006 | 0.006 | 0 | 0.52 | 0.58 | 0.48 | 0.66 | 0.5 | 0.4 |
| 5 | 21585236 | rs515853 | 1.37 | 0.045 | 0.593 | 1.124 | -0.007 | -0.015 | -0.015 | 1.207 | 0.25 | 0.17 | 0.35 | 0.23 | 0.29 | 0.35 |
| 5 | 21585420 | rs2123096 | 0.446 | -0.171 | 0.726 | 1.38 | 0.011 | 0.054 | 0.054 | 0 | 0.52 | 0.2 | 0.47 | 0.44 | 0.53 | 0.42 |
| 5 | 21586180 | rs16888770 | 1.202 | -0.168 | 0.697 | 1.362 | -0.004 | -0.027 | -0.027 | 0 | 0.08 | 0.11 | 0.06 | 0.01 | 0.08 | 0.01 |
| 5 | 21586809 | rs17683709 | 0.152 | -0.052 | 0.644 | 1.544 | -0.003 | 0.018 | 0.018 | 0 | 0.13 | 0.06 | 0.09 | 0.02 | 0.12 | 0.11 |
| 5 | 21588112 | rs13163379 | 0.152 | -0.052 | 0.644 | 1.544 | -0.003 | 0.018 | 0.018 | 0 | 0.13 | 0.06 | 0.09 | 0.02 | 0.12 | 0.11 |
| 5 | 21588931 | rs6896002 | 0.556 | -0.158 | 0.662 | 1.715 | -0.063 | 0.201 | 0.201 | 0 | 0.47 | 0.12 | 0.39 | 0.23 | 0.47 | 0.35 |
| 5 | 21600623 | rs4701222 | -0.666 | -0.205 | 1.148 | 1.434 | -0.003 | 0.008 | 0.008 | 0 | 0.64 | 0.48 | 0.55 | 0.68 | 0.63 | 0.51 |
| 5 | 21606517 | rs1823674 | -2.138 | -0.617 | 1.559 | 1.446 | -0.012 | -0.01 | -0.01 | 0 | 0.27 | 0.4 | 0.18 | 0.24 | 0.23 | 0.24 |
| 5 | 21607097 | rs11745859 | -1.821 | -0.434 | 1.107 | 1.256 | 0 | -0.012 | -0.012 | 0.28 | 0.08 | 0.1 | 0.05 | 0 | 0.07 | 0.04 |
| 5 | 21622035 | rs11956543 | -1.987 | -0.757 | 1.687 | 1.738 | -0.027 | 0.09 | 0.09 | 0 | 0.2 | 0.56 | 0.13 | 0.38 | 0.15 | 0.12 |
| 5 | 21633249 | rs6865279 | -2.491 | -0.765 | 2.71 | 1.717 | 0.037 | 0.089 | 0.089 | 0 | 0.1 | 0.39 | 0.08 | 0.11 | 0.08 | 0.14 |
| 5 | 21634979 | rs16888711 | -2.182 | -0.755 | 2.746 | 1.722 | 0.043 | 0.094 | 0.094 | 0 | 0.08 | 0.28 | 0.06 | 0.11 | 0.07 | 0.11 |
| 5 | 21637415 | rs13161353 | -3.05 | -0.927 | 2.244 | 1.573 | 0.013 | -0.011 | -0.011 | 0 | 0.32 | 0.7 | 0.21 | 0.38 | 0.28 | 0.24 |
| 5 | 21640820 | rs16888684 | -2.623 | -0.918 | 2.484 | 1.555 | 0.032 | 0.101 | 0.101 | 0.081 | 0.1 | 0.47 | 0.07 | 0.13 | 0.07 | 0.11 |
| 5 | 21642121 | rs7707385 | -2.892 | -0.907 | 2.046 | 1.543 | 0.012 | -0.01 | -0.01 | 0.904 | 0.32 | 0.7 | 0.22 | 0.39 | 0.28 | 0.24 |
| 5 | 21656148 | rs10075850 | -2.179 | -1.022 | 2.421 | 1.725 | -0.038 | 0.127 | 0.127 | 0.363 | 0.2 | 0.41 | 0.12 | 0.41 | 0.16 | 0.12 |
| 5 | 21660097 | rs533834 | -2.831 | -1.141 | 1.899 | 1.664 | 0.021 | -0.024 | -0.024 | 0.389 | 0.27 | 0.8 | 0.17 | 0.41 | 0.2 | 0.24 |
| 5 | 21665641 | rs663970 | -2.765 | -1.138 | 1.967 | 1.618 | -0.016 | -0.003 | -0.003 | 0 | 0.2 | 0.72 | 0.11 | 0.16 | 0.14 | 0.17 |
| 5 | 21674640 | rs516235 | -3.065 | -1.158 | 1.88 | 1.679 | 0.032 | -0.02 | -0.02 | 0 | 0.33 | 0.79 | 0.19 | 0.38 | 0.25 | 0.27 |
| 5 | 21676669 | rs586654 | -2.788 | -1.094 | 1.96 | 1.716 | 0.02 | -0.016 | -0.016 | 0 | 0.3 | 0.77 | 0.16 | 0.38 | 0.21 | 0.27 |
| 5 | 21685482 | rs2081963 | -1.83 | -1.161 | 2.253 | 1.777 | 0.019 | -0.029 | -0.029 | 0 | 0.14 | 0.08 | 0.05 | 0.23 | 0.1 | 0.11 |
| 5 | 21687377 | rs608141 | -2.453 | -1.171 | 2.385 | 1.729 | -0.027 | 0.029 | 0.029 | 1.335 | 0.12 | 0.66 | 0.08 | 0.13 | 0.06 | 0.15 |
| 5 | 21689837 | rs1390663 | -3.082 | -0.989 | 2.186 | 2.285 | -0.021 | 0.236 | 0.236 | 0 | 0.18 | 0.46 | 0.14 | 0.54 | 0.13 | 0.29 |
| 5 | 21692989 | rs1428670 | -2.638 | -0.859 | 1.763 | 1.617 | -0.024 | -0.023 | -0.023 | 0 | 0.23 | 0.45 | 0.1 | 0.14 | 0.18 | 0.19 |

| | | | | | | | | | | | | | | | | |
|---|---|---|---|---|---|---|---|---|---|---|---|---|---|---|---|---|
| 5 | 21699126 | rs17259353 | -1.777 | -0.785 | 1.55 | 1.457 | -0.007 | 0.026 | 0.026 | 0 | 0.08 | 0.04 | 0.04 | 0 | 0.09 | 0.02 |
| 5 | 21699135 | rs13354466 | -2.207 | -0.813 | 1.727 | 1.599 | -0.017 | 0.051 | 0.051 | 0 | 0.22 | 0.43 | 0.14 | 0.33 | 0.16 | 0.25 |
| 5 | 21701837 | rs2443768 | 2.051 | -0.78 | 1.66 | 1.146 | -0.028 | -0.057 | -0.057 | 0.285 | 0.81 | 0.45 | 0.9 | 0.92 | 0.87 | 0.8 |
| 5 | 21704695 | rs2967900 | 1.577 | -0.584 | 1.376 | 0.957 | -0.01 | -0.064 | -0.064 | 0.013 | 0.8 | 0.4 | 0.78 | 0.95 | 0.83 | 0.76 |
| 5 | 21718946 | rs16888572 | -0.934 | -1.003 | 1.007 | 1.541 | -0.013 | 0.1 | 0.1 | 0.072 | 0.17 | 0.25 | 0.21 | 0.49 | 0.17 | 0.26 |
| 5 | 21727492 | rs13189619 | 1.796 | -1.539 | 0.395 | 1.274 | 0.07 | -0.078 | -0.078 | 0.769 | 0.63 | 0.1 | 0.6 | 0.45 | 0.73 | 0.58 |
| 5 | 21733331 | rs7703394 | 1.552 | -1.664 | 1.55 | 1.279 | -0.026 | 0.02 | 0.02 | 0.236 | 0.69 | 0.36 | 0.63 | 0.44 | 0.78 | 0.67 |
| 5 | 21742462 | rs4701455 | 0.716 | -1.981 | 0.535 | 0.772 | -0.02 | -0.037 | -0.037 | 0 | 0.2 | 0.05 | 0.1 | 0.22 | 0.15 | 0.21 |
| 5 | 21743174 | rs13162954 | 0.885 | -2.083 | 1.185 | 0.659 | 0 | 0.001 | 0.001 | 0.055 | 0.74 | 0.73 | 0.68 | 0.45 | 0.8 | 0.66 |
| 5 | 21747442 | rs12108917 | -0.891 | -2.064 | 1.243 | 0.646 | 0 | -0.017 | -0.017 | 5.068 | 0.08 | 0.36 | 0.05 | 0.01 | 0.05 | 0.03 |
| 5 | 21752056 | rs6451993 | 1.551 | -2.077 | 1.079 | 0.666 | -0.023 | 0.012 | 0.012 | 0.254 | 0.66 | 0.38 | 0.63 | 0.44 | 0.75 | 0.64 |
| 5 | 21760602 | rs2303748 | 1.609 | -2.136 | 0.851 | 0.694 | 0.04 | -0.012 | -0.012 | 2.498 | 0.65 | 0.25 | 0.63 | 0.41 | 0.74 | 0.64 |
| 5 | 21760731 | rs2303749 | 0.091 | -2.118 | 0.757 | 0.683 | -0.002 | 0.007 | 0.007 | 0.021 | 0.07 | 0.02 | 0.12 | 0.02 | 0.06 | 0.11 |
| 5 | 21772052 | rs13172717 | 1.984 | -2.31 | 0.576 | 0.557 | -0.076 | 0.146 | 0.146 | 0.19 | 0.38 | 0.08 | 0.5 | 0.17 | 0.54 | 0.32 |
| 5 | 21800192 | rs11949505 | 1.947 | -2.068 | 1.923 | 1.603 | -0.005 | 0.004 | 0.004 | 0.044 | 0.7 | 0.34 | 0.69 | 0.59 | 0.78 | 0.68 |
| 5 | 21805266 | rs4485890 | 2.465 | -1.993 | 1.888 | 1.656 | 0.034 | -0.009 | -0.009 | 0.033 | 0.65 | 0.22 | 0.63 | 0.4 | 0.74 | 0.63 |
| 5 | 21810739 | rs13359507 | -1.58 | -2.027 | 2.572 | 1.639 | -0.038 | 0.099 | 0.099 | 0 | 0.24 | 0.37 | 0.25 | 0.41 | 0.16 | 0.29 |
| 5 | 21813303 | rs11952632 | 2.246 | -2.028 | 2.621 | 1.641 | -0.009 | 0.009 | 0.009 | 0.324 | 0.7 | 0.36 | 0.71 | 0.58 | 0.78 | 0.68 |
| 5 | 21813967 | rs6884278 | 2 | -2.039 | 2.568 | 1.654 | -0.052 | 0.093 | 0.093 | 0.05 | 0.73 | 0.53 | 0.73 | 0.59 | 0.8 | 0.69 |
| 5 | 21826642 | rs6887681 | 1.543 | -1.979 | 2.621 | 1.726 | -0.002 | 0.04 | 0.04 | 0 | 0.91 | 0.74 | 0.89 | 0.82 | 0.9 | 0.92 |
| 5 | 21827593 | rs4642327 | 2.457 | -2.025 | 2.604 | 1.704 | -0.015 | 0.006 | 0.006 | 0.012 | 0.68 | 0.35 | 0.64 | 0.4 | 0.74 | 0.64 |
| 5 | 21838360 | rs10059819 | -1.469 | -2.019 | 2.86 | 1.539 | -0.013 | 0.016 | 0.016 | 0.054 | 0.22 | 0.27 | 0.23 | 0.33 | 0.15 | 0.28 |
| 5 | 21838995 | rs10052427 | -1.59 | -2.059 | 3.139 | 1.464 | 0.015 | -0.021 | -0.021 | 1.049 | 0.07 | 0.31 | 0.04 | 0.01 | 0.06 | 0.04 |
| 5 | 21844034 | rs9292995 | -2.906 | -2.148 | 2.995 | 1.648 | 0.062 | -0.06 | -0.06 | 0.98 | 0.35 | 0.85 | 0.38 | 0.51 | 0.26 | 0.42 |
| 5 | 21847419 | rs6452006 | 2.969 | -1.909 | 3.244 | 1.142 | 0.016 | -0.002 | -0.002 | 0 | 0.75 | 0.2 | 0.71 | 0.77 | 0.8 | 0.69 |
| 5 | 21847531 | rs12171538 | -1.798 | -1.904 | 3.382 | 1.153 | -0.008 | 0.025 | 0.025 | 0 | 0.15 | 0.31 | 0.21 | 0.19 | 0.14 | 0.21 |
| 5 | 21853462 | rs10060698 | -1.605 | -1.901 | 3.278 | 1.14 | -0.015 | -0.027 | -0.027 | 0 | 0.13 | 0.29 | 0.15 | 0.03 | 0.09 | 0.16 |
| 5 | 21857517 | rs9292996 | -1.359 | -1.899 | 3.206 | 1.14 | -0.012 | -0.021 | -0.021 | 0 | 0.12 | 0.23 | 0.15 | 0.03 | 0.09 | 0.16 |
| 5 | 21857936 | rs7379407 | 2.317 | -1.888 | 2.921 | 1.147 | 0.016 | -0.002 | -0.002 | 0.736 | 0.75 | 0.19 | 0.71 | 0.77 | 0.8 | 0.69 |
| 5 | 21872292 | rs10045440 | -1.345 | -1.708 | 3.536 | 1.329 | -0.011 | 0.076 | 0.076 | 0 | 0.1 | 0.28 | 0.18 | 0.19 | 0.06 | 0.15 |
| 5 | 21881016 | rs9292998 | 2.299 | -1.696 | 3.491 | 1.32 | -0.024 | 0.067 | 0.067 | 1.664 | 0.85 | 0.33 | 0.76 | 0.78 | 0.93 | 0.79 |
| 5 | 21881514 | rs6874819 | 2.292 | -1.672 | 3.246 | 1.317 | -0.023 | 0.031 | 0.031 | 2.295 | 0.84 | 0.25 | 0.75 | 0.78 | 0.92 | 0.79 |
| 5 | 21890139 | rs4077772 | 1.663 | -1.86 | 3.026 | 1.109 | -0.007 | 0.004 | 0.004 | 0 | 0.94 | 0.78 | 0.97 | 0.99 | 1 | 0.93 |
| 5 | 21893148 | rs7713611 | -1.606 | -1.892 | 3.281 | 1.125 | -0.006 | 0.04 | 0.04 | 0 | 0.14 | 0.27 | 0.22 | 0.21 | 0.12 | 0.19 |
| 5 | 21896184 | rs7442799 | 2.131 | -1.898 | 3.191 | 1.122 | -0.041 | 0.039 | 0.039 | 0 | 0.8 | 0.48 | 0.75 | 0.77 | 0.88 | 0.75 |
| 5 | 21896197 | rs7442801 | 2.131 | -1.898 | 3.191 | 1.122 | -0.041 | 0.039 | 0.039 | 0 | 0.8 | 0.48 | 0.75 | 0.77 | 0.88 | 0.75 |
| 5 | 21899432 | rs6862581 | 2.358 | -1.909 | 3.094 | 1.12 | -0.029 | 0.011 | 0.011 | 0.274 | 0.79 | 0.34 | 0.73 | 0.77 | 0.87 | 0.75 |
| 5 | 21901716 | rs6452027 | 2.992 | -1.983 | 2.733 | 0.889 | 0.022 | 0.007 | 0.007 | 0.231 | 0.71 | 0.21 | 0.68 | 0.77 | 0.82 | 0.71 |
| 5 | 21906026 | rs16888815 | -1.266 | -1.971 | 2.786 | 0.879 | -0.004 | 0.02 | 0.02 | 0.001 | 0.12 | 0.23 | 0.21 | 0.21 | 0.11 | 0.18 |
| 5 | 21906641 | rs7379403 | 3.212 | -2.035 | 2.604 | 1.136 | 0.062 | -0.047 | -0.047 | 0.105 | 0.64 | 0.1 | 0.61 | 0.62 | 0.75 | 0.61 |
| 5 | 21907093 | rs6452029 | 2.145 | -1.944 | 2.779 | 0.853 | -0.027 | 0.019 | 0.019 | 0.041 | 0.79 | 0.36 | 0.73 | 0.77 | 0.87 | 0.75 |
| 5 | 21909369 | rs16888797 | -1.18 | -1.922 | 2.742 | 0.253 | 0 | -0.001 | -0.001 | 0.232 | 0.12 | 0.18 | 0.15 | 0.06 | 0.11 | 0.16 |
| 5 | 21917519 | rs13162273 | 3.037 | -2.08 | 2.718 | 0.146 | -0.008 | -0.024 | -0.024 | 0.423 | 0.7 | 0.3 | 0.74 | 0.93 | 0.81 | 0.71 |
| 5 | 21918864 | rs6452031 | 1.721 | -2.072 | 2.915 | 0.23 | -0.027 | -0.035 | -0.035 | 0.008 | 0.92 | 0.86 | 0.98 | 1 | 0.99 | 0.98 |
| 5 | 21921582 | rs16899347 | -1.844 | -1.542 | 2.917 | 0.03 | -0.01 | -0.019 | -0.019 | 0 | 0.16 | 0.51 | 0.2 | 0.07 | 0.11 | 0.22 |
| 5 | 21925080 | rs6452035 | 2.226 | -1.53 | 2.757 | 0.041 | -0.042 | -0.052 | -0.052 | 0.005 | 0.77 | 0.38 | 0.79 | 0.93 | 0.88 | 0.76 |
| 5 | 21954665 | rs995740 | -1.804 | -1.305 | 2.692 | 0.157 | -0.018 | -0.033 | -0.033 | 0 | 0.17 | 0.29 | 0.17 | 0.05 | 0.12 | 0.14 |
| 5 | 21999692 | rs10056037 | -2.187 | -1.223 | 2.897 | 0.14 | -0.015 | -0.035 | -0.035 | 1.806 | 0.18 | 0.36 | 0.19 | 0.07 | 0.14 | 0.22 |
| 5 | 22009253 | rs4327574 | -2.363 | -0.945 | 2.87 | 0.178 | 0.016 | 0.027 | 0.027 | 0.052 | 0.28 | 0.78 | 0.27 | 0.13 | 0.21 | 0.35 |
| 5 | 22016945 | rs6897914 | -2.292 | -0.943 | 2.9 | 0.179 | 0.007 | 0.011 | 0.011 | 0.037 | 0.28 | 0.72 | 0.27 | 0.13 | 0.21 | 0.35 |
| 5 | 22024387 | rs4337821 | -1.64 | -0.931 | 2.842 | 0.185 | -0.002 | -0.005 | -0.005 | 0.515 | 0.23 | 0.24 | 0.23 | 0.13 | 0.19 | 0.35 |
| 5 | 22033716 | rs6897994 | 2.252 | -0.558 | 2.339 | 0.245 | -0.157 | 0.019 | 0.019 | 0 | 0.41 | 0.03 | 0.44 | 0.24 | 0.48 | 0.32 |
| 5 | 22050002 | rs10052686 | -1.367 | -0.145 | 2.696 | -0.022 | -0.002 | -0.006 | -0.006 | 0 | 0.21 | 0.22 | 0.21 | 0.12 | 0.19 | 0.26 |
| 5 | 22062988 | rs4301199 | -1.589 | 0.146 | 2.36 | 0.141 | -0.003 | -0.025 | -0.025 | 0 | 0.25 | 0.54 | 0.26 | 0.14 | 0.23 | 0.36 |
| 5 | 22067437 | rs7444702 | -2.179 | 0.13 | 2.268 | 0.141 | 0.008 | 0.043 | 0.043 | 0.711 | 0.31 | 0.84 | 0.34 | 0.19 | 0.27 | 0.37 |
| 5 | 22076455 | rs4388179 | -1.596 | 0.265 | 2.268 | 0.174 | 0.002 | -0.023 | -0.023 | 1.089 | 0.24 | 0.51 | 0.27 | 0.14 | 0.23 | 0.35 |
| 5 | 22078584 | rs4371716 | -1.73 | 0.365 | 2.031 | 0.173 | -0.001 | -0.006 | -0.006 | 1.955 | 0.26 | 0.68 | 0.28 | 0.14 | 0.24 | 0.35 |
| 5 | 22081601 | rs10055079 | -1.876 | 0.363 | 1.893 | 0.18 | 0.003 | 0.014 | 0.014 | 0.614 | 0.27 | 0.74 | 0.29 | 0.14 | 0.24 | 0.35 |
| 5 | 22083473 | rs12519679 | -1.466 | 0.51 | 1.53 | 0.222 | 0.008 | -0.025 | -0.025 | 0 | 0.23 | 0.4 | 0.26 | 0.13 | 0.22 | 0.34 |
| 5 | 22101322 | rs7710738 | 0.3 | 0.709 | 0.824 | 0.396 | -0.026 | -0.012 | -0.012 | 0 | 0.6 | 0.77 | 0.57 | 0.43 | 0.58 | 0.49 |
| 5 | 22106756 | rs13186048 | 0.569 | 0.696 | 0.613 | 0.39 | 0.006 | 0.018 | 0.018 | 0.088 | 0.51 | 0.43 | 0.49 | 0.39 | 0.51 | 0.36 |
| 5 | 22109314 | rs6452051 | 0.484 | 0.752 | 0.21 | 0.416 | 0.001 | 0.003 | 0.003 | 0.139 | 0.52 | 0.55 | 0.5 | 0.39 | 0.51 | 0.36 |
| 5 | 22119976 | rs10941934 | 0.497 | 0.777 | 0.05 | 0.428 | -0.001 | -0.003 | -0.003 | 0.163 | 0.52 | 0.52 | 0.5 | 0.38 | 0.51 | 0.36 |
| 5 | 22141555 | rs2963652 | -1.356 | 0.86 | 0.141 | 0.439 | 0.033 | -0.006 | -0.006 | 0 | 0.14 | 0.41 | 0.1 | 0.08 | 0.13 | 0.2 |
| 5 | 22156155 | rs7717005 | -0.232 | 0.95 | -1.097 | 0.271 | -0.049 | 0.055 | 0.055 | 0 | 0.4 | 0.17 | 0.57 | 0.74 | 0.45 | 0.38 |
| 5 | 22193967 | rs6883351 | -0.424 | 0.505 | -1.032 | -0.149 | -0.004 | 0.092 | 0.092 | 0 | 0.46 | 0.24 | 0.57 | 0.74 | 0.5 | 0.37 |
| 5 | 22219503 | rs4518351 | -0.351 | 0.853 | -1.182 | 0.835 | -0.058 | -0.031 | -0.031 | 0.15 | 0.3 | 0.18 | 0.52 | 0.37 | 0.45 | 0.17 |
| 5 | 22219836 | rs4560515 | -0.136 | 0.827 | -1.2 | 0.821 | -0.116 | -0.078 | -0.078 | 0.435 | 0.27 | 0.07 | 0.5 | 0.37 | 0.43 | 0.17 |
| 5 | 22222461 | rs12189051 | -0.872 | 0.47 | -1.316 | 0.445 | -0.033 | -0.011 | -0.011 | 0 | 0.39 | 0.22 | 0.55 | 0.4 | 0.53 | 0.33 |

| | | | | | | | | | | | | | | | | |
|---|---|---|---|---|---|---|---|---|---|---|---|---|---|---|---|---|
| 5 | 22222556 | rs4458548 | 1.078 | -0.713 | -0.965 | 0.146 | 0.009 | -0.002 | -0.002 | 0 | 0.47 | 0.28 | 0.35 | 0.49 | 0.42 | 0.52 |
| 5 | 22225912 | rs4437341 | -0.421 | -0.816 | -0.941 | 0.157 | -0.035 | -0.028 | -0.028 | 0.419 | 0.41 | 0.19 | 0.55 | 0.44 | 0.53 | 0.44 |
| 5 | 22227895 | rs6865522 | -0.63 | -0.907 | 0.319 | 0.142 | -0.027 | 0.052 | 0.052 | 0 | 0.92 | 0.79 | 0.94 | 0.91 | 0.97 | 0.96 |
| 5 | 22243707 | rs4385176 | 1.149 | -0.803 | 0.39 | 0.056 | 0.022 | -0.001 | -0.001 | 0 | 0.47 | 0.28 | 0.32 | 0.49 | 0.39 | 0.51 |
| 5 | 22243986 | rs4593232 | 1.116 | -0.794 | 0.559 | 0.06 | 0.007 | 0.001 | 0.001 | 0 | 0.48 | 0.44 | 0.34 | 0.49 | 0.39 | 0.51 |
| 5 | 22250652 | rs6871980 | 0.439 | -1.117 | 0.609 | -0.068 | 0.002 | -0.002 | -0.002 | 0 | 0.11 | 0.08 | 0.04 | 0.03 | 0.09 | 0.17 |
| 5 | 22273415 | rs4701577 | 0.032 | -0.883 | 0.562 | -0.058 | -0.035 | -0.038 | -0.038 | 0.013 | 0.36 | 0.22 | 0.59 | 0.47 | 0.48 | 0.23 |
| 5 | 22297726 | rs961678 | 1.091 | -0.839 | 0.541 | 0.018 | -0.004 | -0.021 | -0.021 | 0.613 | 0.44 | 0.22 | 0.31 | 0.49 | 0.39 | 0.5 |
| 5 | 22298771 | rs1373374 | 0.387 | -1.038 | 0.434 | 0.178 | -0.035 | 0.032 | 0.032 | 0.345 | 0.07 | 0.16 | 0.06 | 0.1 | 0.02 | 0.05 |
| 5 | 22314300 | rs1444120 | 0.96 | -0.52 | 0.141 | 0.386 | 0.001 | 0.001 | 0.001 | 0 | 0.47 | 0.46 | 0.34 | 0.49 | 0.4 | 0.52 |
| 5 | 22317026 | rs2968384 | -0.75 | -0.601 | 0.183 | 0.355 | 0.001 | 0.001 | 0.001 | 0 | 0.48 | 0.4 | 0.64 | 0.5 | 0.58 | 0.39 |
| 5 | 22318776 | rs13156363 | 0.537 | -0.763 | -0.007 | 0.436 | -0.089 | -0.059 | -0.059 | 0 | 0.24 | 0.1 | 0.51 | 0.34 | 0.44 | 0.16 |
| 5 | 22327274 | rs13179082 | 1.447 | -0.941 | 0.534 | 1.007 | 0.004 | 0.049 | 0.049 | 0 | 0.47 | 0.29 | 0.32 | 0.39 | 0.39 | 0.5 |
| 5 | 22329782 | rs7710321 | 0.932 | -0.976 | 0.753 | 0.985 | -0.001 | 0.024 | 0.024 | 0 | 0.53 | 0.44 | 0.36 | 0.38 | 0.43 | 0.52 |
| 5 | 22344250 | rs16894341 | -1.353 | -1.029 | 0.708 | 1.139 | -0.032 | 0.008 | 0.008 | 0 | 0.11 | 0.19 | 0.04 | 0.12 | 0.04 | 0.12 |
| 5 | 22351139 | rs17265218 | 0.355 | -0.358 | 0.426 | 0.941 | -0.042 | 0.04 | 0.04 | 0.028 | 0.1 | 0.02 | 0.1 | 0.02 | 0.18 | 0.08 |
| 5 | 22355323 | rs2883717 | -1.059 | -0.47 | 0.375 | 0.973 | -0.076 | -0.017 | -0.017 | 0 | 0.36 | 0.15 | 0.56 | 0.37 | 0.53 | 0.36 |
| 5 | 22375786 | rs10473592 | 2.009 | -0.955 | -0.227 | 0.427 | 0 | -0.01 | -0.01 | 0.408 | 0.31 | 0.22 | 0.23 | 0.33 | 0.28 | 0.36 |
| 5 | 22383949 | rs1261368 | 0.127 | -0.773 | -0.282 | 0.467 | -0.061 | -0.038 | -0.038 | 0 | 0.24 | 0.11 | 0.51 | 0.31 | 0.43 | 0.17 |
| 5 | 22391075 | rs13359749 | 0.198 | -0.881 | -0.137 | 0.448 | -0.021 | 0.043 | 0.043 | 0 | 0.05 | 0.13 | 0.02 | 0.13 | 0 | 0.09 |
| 5 | 22397625 | rs1261397 | 0.388 | -0.954 | -0.122 | 0.443 | -0.011 | -0.02 | -0.02 | 0.014 | 0.61 | 0.48 | 0.76 | 0.77 | 0.71 | 0.62 |
| 5 | 22398914 | rs959266 | -0.844 | -0.99 | -0.098 | 0.421 | 0.007 | 0.011 | 0.011 | 0.297 | 0.77 | 0.84 | 0.84 | 0.82 | 0.82 | 0.82 |
| 5 | 22402776 | rs3112473 | 1.19 | -0.597 | -0.301 | 0.563 | 0.002 | 0.009 | 0.009 | 0 | 0.09 | 0.06 | 0.04 | 0.05 | 0.08 | 0.18 |
| 5 | 22403473 | rs1911960 | 0.138 | -0.57 | -0.307 | 0.577 | -0.007 | -0.016 | -0.016 | 0.985 | 0.15 | 0.16 | 0.07 | 0.05 | 0.11 | 0.2 |
| 5 | 22403944 | rs2355597 | 1.112 | -0.034 | -0.28 | 0.891 | -0.073 | -0.081 | -0.081 | 0 | 0.13 | 0.02 | 0.4 | 0.29 | 0.25 | 0.09 |
| 5 | 22405345 | rs1155443 | -0.561 | -0.149 | -0.193 | 0.817 | 0.035 | 0.043 | 0.043 | 0.375 | 0.3 | 0.42 | 0.53 | 0.44 | 0.43 | 0.26 |
| 5 | 22431467 | rs894911 | 1.078 | -0.5 | -0.463 | 1.09 | 0.004 | 0.001 | 0.001 | 0 | 0.33 | 0.37 | 0.19 | 0.31 | 0.31 | 0.34 |
| 5 | 22445794 | rs1911963 | -0.187 | -0.469 | -0.704 | 1.105 | 0 | -0.014 | -0.014 | 0 | 0.81 | 0.88 | 0.9 | 0.75 | 0.8 | 0.71 |
| 5 | 22446214 | rs995178 | -0.455 | -0.655 | -0.54 | 1.036 | 0 | 0 | 0 | 0.038 | 0.47 | 0.45 | 0.7 | 0.47 | 0.48 | 0.37 |
| 5 | 22446353 | rs995179 | 0.589 | -0.785 | -0.655 | 1.075 | 0.004 | 0.012 | 0.012 | 3.115 | 0.23 | 0.12 | 0.13 | 0.18 | 0.17 | 0.16 |
| 5 | 22458871 | rs2010548 | 0.971 | -0.603 | -0.699 | 0.969 | -0.035 | -0.057 | -0.057 | 1.869 | 0.18 | 0.03 | 0.51 | 0.29 | 0.28 | 0.18 |
| 5 | 22472039 | rs2551514 | -1.577 | -1.003 | -0.562 | 1.133 | 0 | -0.001 | -0.001 | 0 | 0.55 | 0.55 | 0.75 | 0.62 | 0.54 | 0.52 |
| 5 | 22474350 | rs10039824 | -0.823 | -1.12 | -0.595 | 1.233 | 0.005 | -0.002 | -0.002 | 0 | 0.45 | 0.39 | 0.67 | 0.52 | 0.41 | 0.37 |
| 5 | 22476100 | rs10473594 | -0.049 | -0.907 | -0.823 | 1.321 | -0.004 | -0.057 | -0.057 | 0 | 0.19 | 0.05 | 0.49 | 0.3 | 0.21 | 0.17 |
| 5 | 22483586 | rs1911956 | -1.416 | -1.009 | -0.714 | 1.015 | 0.009 | -0.014 | -0.014 | 0.714 | 0.91 | 0.95 | 0.97 | 0.85 | 0.95 | 0.87 |
| 5 | 22492196 | rs440188 | -1.416 | -1.009 | -0.714 | 1.016 | 0.009 | -0.014 | -0.014 | 4.847 | 0.91 | 0.95 | 0.97 | 0.86 | 0.95 | 0.87 |
| 5 | 22494033 | rs1393114 | 1.974 | -1.168 | -0.547 | 0.917 | 0.009 | -0.024 | -0.024 | 0 | 0.36 | 0.36 | 0.22 | 0.15 | 0.43 | 0.37 |
| 5 | 22494188 | rs16896252 | 0.504 | -1.153 | -0.744 | 1.031 | -0.008 | -0.093 | -0.093 | 0 | 0.14 | 0.01 | 0.44 | 0.38 | 0.13 | 0.12 |
| 5 | 22507308 | rs10040210 | 1.107 | -1.162 | -0.568 | 0.925 | -0.001 | -0.006 | -0.006 | 0.679 | 0.09 | 0.11 | 0.06 | 0.09 | 0.12 | 0.16 |
| 5 | 22516890 | rs4320220 | 1.045 | -1.157 | -0.575 | 0.922 | 0 | -0.007 | -0.007 | 0 | 0.09 | 0.11 | 0.06 | 0.1 | 0.12 | 0.16 |
| 5 | 22527778 | rs10036620 | 1.07 | -1.091 | -0.452 | 0.985 | 0.004 | 0.005 | 0.005 | 0 | 0.17 | 0.12 | 0.08 | 0.1 | 0.14 | 0.2 |
| 5 | 22532710 | rs10053834 | 0.183 | -1.154 | -0.489 | 1.113 | 0.014 | -0.029 | -0.029 | 0.005 | 0.16 | 0.12 | 0.45 | 0.38 | 0.14 | 0.12 |
| 5 | 22536778 | rs10052996 | 0.321 | -1.126 | -0.502 | 1.12 | 0.015 | -0.038 | -0.038 | 0.374 | 0.16 | 0.08 | 0.43 | 0.38 | 0.13 | 0.12 |
| 5 | 22549703 | rs28474955 | -2.185 | -1.149 | -0.426 | 1.091 | 0.006 | -0.016 | -0.016 | 1.119 | 0.65 | 0.65 | 0.78 | 0.84 | 0.57 | 0.67 |
| 5 | 22550328 | rs2134177 | 0.263 | -1.139 | -0.364 | 0.88 | 0.01 | -0.019 | -0.019 | 0 | 0.16 | 0.13 | 0.44 | 0.39 | 0.13 | 0.12 |
| 5 | 22551757 | rs400155 | -2.083 | -1.129 | -0.388 | 0.914 | 0.005 | -0.002 | -0.002 | 0 | 0.64 | 0.65 | 0.77 | 0.71 | 0.55 | 0.53 |
| 5 | 22557624 | rs13357733 | 0.204 | -1.023 | -0.449 | 1.19 | 0.016 | -0.044 | -0.044 | 0 | 0.16 | 0.08 | 0.43 | 0.38 | 0.13 | 0.12 |
| 5 | 22573330 | rs271090 | 0.777 | -0.7 | -0.244 | 0.9 | -0.002 | -0.005 | -0.005 | 0 | 0.18 | 0.16 | 0.13 | 0.04 | 0.17 | 0.11 |
| 5 | 22591057 | rs269876 | -0.141 | -0.442 | -0.014 | 1.276 | -0.008 | -0.058 | -0.058 | 0 | 0.38 | 0.49 | 0.21 | 0.14 | 0.3 | 0.35 |
| 5 | 22591472 | rs10941965 | 0.291 | -0.452 | -0.008 | 1.516 | 0 | 0.009 | 0.009 | 0.003 | 0.12 | 0.08 | 0.43 | 0.34 | 0.13 | 0.11 |
| 5 | 22604172 | rs833344 | -0.947 | -0.356 | -0.46 | 0.662 | 0.006 | 0.009 | 0.009 | 0 | 0.51 | 0.57 | 0.65 | 0.48 | 0.43 | 0.47 |
| 5 | 22635369 | rs1074463 | -0.77 | -0.378 | -0.318 | 1.04 | -0.019 | 0.05 | 0.05 | 0 | 0.17 | 0.32 | 0.46 | 0.33 | 0.13 | 0.15 |
| 5 | 22652137 | rs16897695 | 0.036 | 0.127 | -0.969 | 1.345 | 0.006 | 0.206 | 0.206 | 0 | 0.07 | 0.13 | 0.1 | 0.44 | 0.1 | 0.18 |
| 5 | 22655865 | rs12521573 | -0.031 | 0.272 | -1.43 | 0.886 | -0.019 | -0.063 | -0.063 | 0.301 | 0.11 | 0.01 | 0.43 | 0.31 | 0.14 | 0.11 |
| 5 | 22656527 | rs268972 | -0.435 | 0.334 | -1.157 | -0.07 | -0.013 | 0.072 | 0.072 | 0 | 0.55 | 0.69 | 0.72 | 0.89 | 0.52 | 0.63 |
| 5 | 22657926 | rs10053320 | -0.778 | -0.182 | -1.25 | 0.563 | 0.013 | -0.015 | -0.015 | 0.017 | 0.16 | 0.15 | 0.28 | 0.2 | 0.11 | 0.13 |
| 5 | 22658043 | rs1487414 | -0.294 | -0.434 | -1.507 | 0.336 | 0.011 | 0.015 | 0.015 | 0 | 0.59 | 0.39 | 0.57 | 0.64 | 0.64 | 0.63 |
| 5 | 22658820 | rs10941967 | -0.235 | -0.952 | -1.071 | 0.069 | -0.004 | 0.02 | 0.02 | 0.745 | 0.58 | 0.62 | 0.71 | 0.67 | 0.56 | 0.58 |
| 5 | 22660564 | rs7708701 | 0.327 | -1.391 | -1.137 | 0.029 | 0.034 | 0.022 | 0.022 | 0 | 0.51 | 0.36 | 0.48 | 0.55 | 0.6 | 0.54 |
| 5 | 22664717 | rs198793 | 0.957 | -1.877 | -1.083 | -0.229 | -0.033 | -0.044 | -0.044 | 6.802 | 0.08 | 0.19 | 0.03 | 0 | 0.04 | 0.01 |
| 5 | 22670144 | rs1038591 | -0.358 | -1.9 | -1.099 | -0.218 | -0.002 | 0.014 | 0.014 | 0.236 | 0.64 | 0.71 | 0.77 | 0.68 | 0.65 | 0.7 |
| 5 | 22683956 | rs9293030 | 0.596 | -2.09 | -0.302 | -0.246 | -0.001 | -0.001 | -0.001 | 0 | 0.49 | 0.53 | 0.49 | 0.52 | 0.57 | 0.53 |
| 5 | 22694384 | rs269017 | 0.535 | -1.912 | -0.319 | -0.275 | -0.02 | -0.033 | -0.033 | 1.918 | 0.09 | 0.35 | 0.04 | 0 | 0.04 | 0.01 |
| 5 | 22694679 | rs269019 | -0.756 | -1.953 | -0.214 | -0.336 | -0.022 | 0.009 | 0.009 | 1.45 | 0.68 | 0.84 | 0.77 | 0.68 | 0.67 | 0.69 |
| 5 | 22699592 | rs6860776 | -0.014 | -1.975 | -0.222 | -0.334 | -0.006 | -0.019 | -0.019 | 0 | 0.59 | 0.49 | 0.73 | 0.68 | 0.63 | 0.69 |
| 5 | 22707203 | rs174873 | -0.771 | -1.789 | -0.391 | -0.214 | -0.076 | -0.027 | -0.027 | 0.076 | 0.7 | 0.93 | 0.8 | 0.68 | 0.67 | 0.69 |
| 5 | 22709981 | rs269024 | -0.833 | -1.782 | -0.349 | -0.214 | -0.009 | 0.003 | 0.003 | 1.703 | 0.7 | 0.76 | 0.78 | 0.7 | 0.67 | 0.69 |
| 5 | 22718422 | rs416802 | 0.51 | -1.76 | -0.058 | -0.524 | -0.036 | -0.05 | -0.05 | 0 | 0.76 | 0.52 | 0.66 | 0.84 | 0.78 | 0.74 |
| 5 | 22730468 | rs9293036 | 0.72 | -1.961 | -0.086 | -0.626 | 0 | -0.002 | -0.002 | 0 | 0.46 | 0.53 | 0.49 | 0.59 | 0.49 | 0.45 |

| | | | | | | | | | | | | | | | | |
|---|---|---|---|---|---|---|---|---|---|---|---|---|---|---|---|---|
| 5 | 22733475 | rs12188885 | 1.033 | -1.985 | -0.202 | -0.613 | -0.064 | 0.009 | 0.009 | 1.645 | 0.26 | 0.09 | 0.18 | 0.25 | 0.29 | 0.29 |
| 5 | 22733747 | rs12519845 | 1.555 | -2.217 | -0.457 | -0.267 | -0.025 | -0.045 | -0.045 | 0 | 0.1 | 0.01 | 0.19 | 0.16 | 0.1 | 0.1 |
| 5 | 22740008 | rs2129612 | 0.483 | -2.053 | -0.163 | -0.32 | -0.022 | 0.027 | 0.027 | 0.828 | 0.32 | 0.21 | 0.28 | 0.27 | 0.35 | 0.33 |
| 5 | 22740091 | rs17358356 | 2.365 | -2.445 | 0.058 | -0.298 | -0.008 | -0.009 | -0.009 | 0.573 | 0.25 | 0.22 | 0.23 | 0.34 | 0.35 | 0.26 |
| 5 | 22741835 | rs6452095 | 1.042 | -2.075 | 0.074 | -0.596 | -0.001 | 0.001 | 0.001 | 0 | 0.46 | 0.56 | 0.49 | 0.59 | 0.49 | 0.44 |
| 5 | 22742635 | rs7705310 | -0.367 | -2.118 | 0.03 | -0.588 | 0.009 | 0.006 | 0.006 | 0 | 0.82 | 0.82 | 0.78 | 0.86 | 0.84 | 0.79 |
| 5 | 22743760 | rs1487416 | 0.044 | -2.117 | 0.028 | -0.565 | -0.012 | -0.006 | -0.006 | 0 | 0.78 | 0.77 | 0.77 | 0.85 | 0.84 | 0.78 |
| 5 | 22748607 | rs1867072 | 0.834 | -2.025 | -0.061 | -0.53 | 0.008 | -0.022 | -0.022 | 0 | 0.06 | 0.09 | 0.11 | 0.02 | 0.06 | 0.05 |
| 5 | 22751348 | rs4701298 | -0.23 | -2.07 | -0.033 | -0.472 | -0.001 | 0 | 0 | 0 | 0.81 | 0.8 | 0.78 | 0.85 | 0.84 | 0.79 |
| 5 | 22755474 | rs2119887 | -0.329 | -2.07 | -0.124 | -0.44 | 0.008 | 0.003 | 0.003 | 0 | 0.82 | 0.83 | 0.78 | 0.84 | 0.84 | 0.79 |
| 5 | 22756398 | rs17358685 | 0.697 | -1.436 | -0.514 | -0.42 | -0.05 | 0.067 | 0.067 | 0.357 | 0.23 | 0.07 | 0.24 | 0.13 | 0.31 | 0.28 |
| 5 | 22759508 | rs958614 | 0.721 | -1.439 | -0.507 | -0.412 | -0.052 | 0.065 | 0.065 | 0.287 | 0.23 | 0.08 | 0.24 | 0.13 | 0.31 | 0.28 |
| 5 | 22761272 | rs16899078 | 0.666 | -1.07 | -0.226 | 0.042 | -0.004 | 0.02 | 0.02 | 0 | 0.09 | 0.16 | 0.05 | 0.12 | 0.05 | 0.06 |
| 5 | 22770519 | rs11953049 | 0.78 | -1.01 | -0.338 | -0.039 | 0.002 | -0.003 | -0.003 | 0 | 0.42 | 0.46 | 0.45 | 0.57 | 0.47 | 0.43 |
| 5 | 22776609 | rs17359069 | 0.433 | -0.934 | -0.699 | -0.164 | -0.005 | 0.022 | 0.022 | 0 | 0.13 | 0 | 0.05 | 0.07 | 0.07 | 0.11 |
| 5 | 22777371 | rs7715585 | -0.665 | -1.297 | 0.087 | -0.137 | -0.041 | 0.008 | 0.008 | 0 | 0.21 | 0.37 | 0.24 | 0.24 | 0.12 | 0.21 |
| 5 | 22780471 | rs2584443 | 1.99 | -0.151 | -0.236 | 0.219 | 0.002 | 0.02 | 0.02 | 0 | 0.22 | 0.22 | 0.18 | 0.38 | 0.22 | 0.27 |
| 5 | 22790036 | rs780188 | -1.121 | -0.56 | -0.435 | -0.713 | 0.021 | 0.031 | 0.031 | 0 | 0.41 | 0.64 | 0.37 | 0.24 | 0.29 | 0.35 |
| 5 | 22791335 | rs16899478 | 1.474 | -0.454 | -0.133 | -0.366 | 0.002 | -0.009 | -0.009 | 0 | 0.05 | 0.04 | 0.16 | 0.17 | 0.05 | 0.11 |
| 5 | 22793334 | rs1707082 | 0.156 | -0.477 | 0.273 | -0.376 | 0.02 | 0.132 | 0.132 | 0.131 | 0.08 | 0.32 | 0.19 | 0.17 | 0.05 | 0.11 |
| 5 | 22794657 | rs1561836 | -0.497 | -0.526 | -0.322 | -0.427 | -0.04 | -0.001 | -0.001 | 0 | 0.23 | 0.49 | 0.26 | 0.24 | 0.12 | 0.22 |
| 5 | 22799308 | rs9293037 | 1.118 | 0.011 | -0.551 | -0.01 | 0.002 | -0.01 | -0.01 | 0.442 | 0.06 | 0.04 | 0.16 | 0.17 | 0.05 | 0.11 |
| 5 | 22812264 | rs780179 | -0.894 | 0.293 | -0.747 | -0.398 | -0.009 | 0.022 | 0.022 | 0 | 0.57 | 0.62 | 0.47 | 0.36 | 0.52 | 0.5 |
| 5 | 22813022 | rs13184821 | 0.166 | 0.439 | -0.458 | -0.615 | 0.001 | -0.056 | -0.056 | 0 | 0.12 | 0.22 | 0.1 | 0 | 0.15 | 0.03 |
| 5 | 22813520 | rs16899832 | -0.297 | 0.454 | -0.785 | -0.518 | -0.015 | -0.014 | -0.014 | 0 | 0.26 | 0.15 | 0.23 | 0.3 | 0.33 | 0.38 |
| 5 | 22817855 | rs1442471 | -0.52 | 0.354 | -0.889 | -0.904 | -0.015 | 0.118 | 0.118 | 0 | 0.43 | 0.23 | 0.51 | 0.18 | 0.48 | 0.33 |
| 5 | 22818921 | rs12189454 | 2.275 | 0.205 | -1.365 | -0.956 | -0.026 | -0.005 | -0.005 | 0 | 0.13 | 0 | 0.05 | 0.13 | 0.08 | 0.09 |
| 5 | 22820390 | rs10472487 | -2.131 | 0.109 | -0.839 | -0.949 | -0.001 | 0.009 | 0.009 | 0 | 0.73 | 0.7 | 0.76 | 0.59 | 0.76 | 0.73 |
| 5 | 22823694 | rs9283772 | -0.118 | 0.147 | -0.695 | -1.203 | 0.001 | -0.046 | -0.046 | 0.114 | 0.4 | 0.45 | 0.51 | 0.19 | 0.43 | 0.31 |
| 5 | 22825527 | rs11743201 | -1.421 | -0.267 | -0.758 | -1.024 | -0.002 | 0.002 | 0.002 | 0 | 0.63 | 0.62 | 0.73 | 0.46 | 0.73 | 0.56 |
| 5 | 22826121 | rs1661975 | 1.687 | -0.319 | -0.885 | -1.004 | 0.009 | 0.006 | 0.006 | 0 | 0.14 | 0.11 | 0.05 | 0.12 | 0.06 | 0.2 |
| 5 | 22832585 | rs16899375 | 2.837 | -0.317 | -1.006 | -0.967 | -0.058 | -0.159 | -0.159 | 0 | 0.21 | 0.02 | 0.21 | 0.42 | 0.21 | 0.24 |
| 5 | 22835892 | rs11957688 | 2.469 | -0.386 | -1 | -0.918 | -0.013 | 0.008 | 0.008 | 0 | 0.08 | 0 | 0.03 | 0.05 | 0.06 | 0.09 |
| 5 | 22842456 | rs780176 | -2.595 | -0.326 | -0.772 | -0.887 | -0.002 | -0.028 | -0.028 | 0 | 0.78 | 0.84 | 0.79 | 0.58 | 0.79 | 0.76 |
| 5 | 22845094 | rs12521133 | 1.983 | -0.229 | -0.951 | -0.991 | -0.015 | 0.034 | 0.034 | 0 | 0.06 | 0.01 | 0.03 | 0.01 | 0.08 | 0.11 |
| 5 | 22850914 | rs13172440 | 0.088 | -0.209 | -0.393 | -1.05 | -0.003 | 0.005 | 0.005 | 0 | 0.39 | 0.33 | 0.51 | 0.16 | 0.45 | 0.3 |
| 5 | 22856741 | rs12652791 | -0.125 | -0.163 | -0.227 | -0.669 | -0.024 | -0.051 | -0.051 | 0 | 0.21 | 0.12 | 0.21 | 0.3 | 0.26 | 0.24 |
| 5 | 22866518 | rs4235521 | -1.769 | -0.223 | 0.001 | -0.576 | -0.007 | 0.028 | 0.028 | 0 | 0.73 | 0.69 | 0.77 | 0.59 | 0.77 | 0.71 |
| 5 | 22869771 | rs1707062 | -2.022 | -0.475 | 0.254 | -0.45 | 0 | -0.009 | -0.009 | 0 | 0.73 | 0.78 | 0.74 | 0.59 | 0.74 | 0.71 |
| 5 | 22875782 | rs4501305 | -0.028 | -1.065 | 0.3 | -0.561 | -0.012 | -0.044 | -0.044 | 1.346 | 0.17 | 0.12 | 0.18 | 0.3 | 0.22 | 0.21 |
| 5 | 22882986 | rs12658777 | -0.291 | -1.059 | 0.462 | -0.461 | 0.001 | 0.034 | 0.034 | 0 | 0.5 | 0.57 | 0.56 | 0.21 | 0.52 | 0.48 |
| 5 | 22884601 | rs1372667 | -0.176 | -1.037 | 0.507 | -0.458 | -0.002 | 0.022 | 0.022 | 0 | 0.47 | 0.44 | 0.55 | 0.21 | 0.52 | 0.37 |
| 5 | 22893819 | rs13171806 | -0.115 | -1.047 | 0.441 | -0.425 | -0.005 | 0.046 | 0.046 | 0 | 0.47 | 0.34 | 0.54 | 0.2 | 0.52 | 0.37 |
| 5 | 22902477 | rs11956790 | -0.024 | -1.059 | 0.51 | -0.324 | -0.001 | 0.001 | 0.001 | 0 | 0.18 | 0.21 | 0.18 | 0.26 | 0.19 | 0.17 |
| 5 | 22923818 | rs1820743 | -0.24 | -1.067 | 0.35 | -0.253 | -0.003 | 0.006 | 0.006 | 0 | 0.19 | 0.26 | 0.18 | 0.27 | 0.18 | 0.17 |
| 5 | 22923887 | rs1366421 | -1.847 | -1.046 | 0.35 | -0.261 | 0 | -0.011 | -0.011 | 0.48 | 0.68 | 0.73 | 0.74 | 0.47 | 0.71 | 0.63 |
| 5 | 22941669 | rs6881479 | -0.332 | -0.671 | -0.101 | -0.423 | -0.004 | 0.007 | 0.007 | 0 | 0.19 | 0.26 | 0.19 | 0.27 | 0.18 | 0.18 |
| 5 | 22946602 | rs11748547 | 0.201 | -0.397 | -0.411 | -0.637 | -0.035 | 0.073 | 0.073 | 0 | 0.22 | 0.08 | 0.26 | 0.01 | 0.3 | 0.12 |
| 5 | 22956970 | rs11959007 | 0.763 | 0.055 | 0.379 | -0.015 | 0.002 | 0.005 | 0.005 | 0 | 0.08 | 0.08 | 0.04 | 0.05 | 0.06 | 0.08 |
| 5 | 22987157 | rs2635633 | -0.44 | 0.101 | 0.666 | 0.26 | 0.004 | -0.014 | -0.014 | 0 | 0.59 | 0.57 | 0.63 | 0.28 | 0.54 | 0.56 |
| 5 | 22989071 | rs1607561 | -0.087 | 0.104 | 0.425 | 0.418 | 0.014 | 0.048 | 0.048 | 0 | 0.11 | 0.21 | 0.15 | 0.2 | 0.15 | 0.1 |
| 5 | 22989124 | rs1607562 | 0.483 | 0.069 | 0.438 | 0.556 | -0.007 | 0.03 | 0.03 | 0 | 0.08 | 0.16 | 0.01 | 0.15 | 0.04 | 0.14 |
| 5 | 22991089 | rs2680768 | -0.495 | 0.142 | 0.393 | 0.509 | -0.001 | -0.002 | -0.002 | 0 | 0.6 | 0.64 | 0.71 | 0.43 | 0.55 | 0.59 |
| 5 | 22991845 | rs1896710 | -1.235 | -0.296 | 0.309 | 0.961 | -0.046 | -0.005 | -0.005 | 0 | 0.61 | 0.84 | 0.71 | 0.63 | 0.54 | 0.65 |
| 5 | 23000050 | rs13154490 | -0.081 | -0.422 | 0.762 | 0.932 | -0.006 | -0.011 | -0.011 | 0 | 0.38 | 0.49 | 0.55 | 0.29 | 0.3 | 0.42 |
| 5 | 23002385 | rs13185192 | -0.071 | -0.406 | 0.727 | 0.919 | -0.006 | -0.011 | -0.011 | 0.31 | 0.37 | 0.5 | 0.55 | 0.3 | 0.3 | 0.42 |
| 5 | 23002989 | rs2561710 | -0.98 | -0.355 | 0.218 | 0.949 | -0.051 | -0.045 | -0.045 | 0.102 | 0.55 | 0.83 | 0.63 | 0.51 | 0.45 | 0.62 |
| 5 | 23004208 | rs1879060 | -1.083 | 0 | 0.841 | 1.497 | 0 | 0.016 | 0.016 | 0 | 0.12 | 0.1 | 0.27 | 0.33 | 0.2 | 0.16 |
| 5 | 23019136 | rs2635636 | -1.837 | -0.24 | 1.613 | 1.624 | 0.026 | 0.149 | 0.149 | 0 | 0.21 | 0.35 | 0.3 | 0.52 | 0.22 | 0.29 |
| 5 | 23036721 | rs2680798 | -1.04 | -0.211 | 1.485 | 1.681 | -0.014 | -0.014 | -0.014 | 0.304 | 0.12 | 0.07 | 0.27 | 0.32 | 0.18 | 0.15 |
| 5 | 23038845 | rs2635642 | -1.251 | -0.212 | 1.53 | 1.643 | 0.029 | 0.109 | 0.109 | 0 | 0.12 | 0.19 | 0.28 | 0.34 | 0.18 | 0.15 |
| 5 | 23045219 | rs1969336 | -1.556 | -0.223 | 1.483 | 1.62 | 0.017 | 0.108 | 0.108 | 0 | 0.14 | 0.21 | 0.29 | 0.5 | 0.19 | 0.15 |
| 5 | 23048840 | rs17299353 | 0.441 | -0.268 | 1.291 | 1.451 | -0.023 | 0.026 | 0.026 | 0 | 0.05 | 0 | 0.03 | 0 | 0.07 | 0.02 |
| 5 | 23056410 | rs6879343 | -1.159 | -0.243 | 1.501 | 1.49 | 0.002 | 0.005 | 0.005 | 0 | 0.14 | 0.17 | 0.07 | 0.15 | 0.15 | 0.11 |
| 5 | 23056837 | rs1980221 | 0.648 | -0.249 | 1.503 | 1.438 | 0.006 | -0.025 | -0.025 | 0 | 0.06 | 0.11 | 0.01 | 0 | 0.02 | 0.04 |
| 5 | 23062359 | rs4701339 | -2.268 | -0.547 | 1.635 | 1.505 | 0.004 | 0.069 | 0.069 | 0 | 0.26 | 0.39 | 0.32 | 0.57 | 0.25 | 0.34 |
| 5 | 23062497 | rs4701340 | -1.915 | -0.542 | 1.64 | 1.526 | 0.027 | 0.12 | 0.12 | 0 | 0.21 | 0.36 | 0.32 | 0.39 | 0.24 | 0.22 |
| 5 | 23065165 | rs1401627 | -0.388 | -0.543 | 1.603 | 1.519 | 0.016 | -0.018 | -0.018 | 0 | 0.08 | 0.18 | 0.03 | 0.04 | 0.05 | 0.09 |
| 5 | 23070854 | rs16889985 | -1.19 | -0.539 | 1.545 | 1.533 | 0.011 | 0.07 | 0.07 | 0 | 0.13 | 0.2 | 0.29 | 0.35 | 0.18 | 0.13 |

| | | | | | | | | | | | | | | | | |
|---|---|---|---|---|---|---|---|---|---|---|---|---|---|---|---|---|
| 5 | 23072759 | rs1836564 | -1.839 | -0.542 | 1.53 | 1.533 | 0.003 | 0.021 | 0.021 | 0 | 0.2 | 0.21 | 0.3 | 0.39 | 0.24 | 0.22 |
| 5 | 23073936 | rs10941977 | 2.338 | -0.782 | 1.429 | 1.331 | -0.011 | 0.026 | 0.026 | 0 | 0.68 | 0.57 | 0.63 | 0.43 | 0.71 | 0.66 |
| 5 | 23077403 | rs7709491 | 2.415 | -0.836 | 1.887 | 1.306 | -0.001 | 0 | 0 | 0 | 0.67 | 0.43 | 0.62 | 0.43 | 0.71 | 0.66 |
| 5 | 23078005 | rs12655902 | -1.55 | -0.63 | 1.719 | 1.386 | 0.013 | 0.072 | 0.072 | 0 | 0.19 | 0.29 | 0.3 | 0.35 | 0.21 | 0.17 |
| 5 | 23085145 | rs1519433 | -1.401 | -0.625 | 1.288 | 1.385 | -0.003 | -0.003 | -0.003 | 0.185 | 0.18 | 0.15 | 0.28 | 0.35 | 0.18 | 0.17 |
| 5 | 23092940 | rs12515326 | -2.136 | -0.688 | 1.423 | 1.295 | 0.008 | 0.084 | 0.084 | 0 | 0.25 | 0.39 | 0.31 | 0.56 | 0.22 | 0.31 |
| 5 | 23094809 | rs12653590 | -0.648 | -0.672 | 0.898 | 1.333 | -0.018 | -0.029 | -0.029 | 0 | 0.12 | 0.06 | 0.27 | 0.33 | 0.16 | 0.11 |
| 5 | 23100596 | rs2078301 | -0.787 | -0.679 | 0.919 | 1.33 | -0.003 | 0.013 | 0.013 | 0 | 0.12 | 0.12 | 0.28 | 0.33 | 0.16 | 0.11 |
| 5 | 23101848 | rs10067560 | 0.849 | -1.108 | 0.484 | 1.014 | -0.002 | -0.008 | -0.008 | 0.989 | 0.11 | 0.1 | 0.03 | 0.02 | 0.05 | 0.13 |
| 5 | 23101921 | rs310906 | 2.063 | -0.613 | 0.728 | 1.398 | 0.002 | -0.005 | -0.005 | 0 | 0.67 | 0.34 | 0.61 | 0.44 | 0.74 | 0.68 |
| 5 | 23103741 | rs10520885 | -1.205 | -0.485 | 0.18 | 1.55 | -0.005 | 0.048 | 0.048 | 0 | 0.07 | 0.04 | 0.22 | 0.37 | 0.07 | 0.15 |
| 5 | 23104936 | rs310904 | -0.484 | -0.839 | 0.099 | 0.989 | 0.003 | -0.004 | -0.004 | 0 | 0.94 | 0.92 | 0.94 | 1 | 0.95 | 1 |
| 5 | 23127912 | rs10053648 | -0.057 | -0.854 | 0.245 | 1.068 | 0.039 | 0.221 | 0.221 | 0 | 0.05 | 0.29 | 0.03 | 0.17 | 0.01 | 0.04 |
| 5 | 23132868 | rs1435231 | -0.994 | -0.814 | 0.195 | 1.009 | -0.005 | 0.117 | 0.117 | 0 | 0.23 | 0.39 | 0.34 | 0.54 | 0.18 | 0.22 |
| 5 | 23138951 | rs1995264 | -1.604 | -0.917 | -0.85 | 0.716 | -0.022 | 0.035 | 0.035 | 0 | 0.52 | 0.93 | 0.65 | 0.63 | 0.4 | 0.5 |
| 5 | 23139208 | rs1435247 | -1.926 | -0.782 | 0.173 | 1.427 | -0.013 | 0.02 | 0.02 | 0 | 0.31 | 0.6 | 0.4 | 0.56 | 0.22 | 0.29 |
| 5 | 23150924 | rs10520887 | -0.792 | -0.336 | 0.019 | 1.617 | -0.003 | -0.027 | -0.027 | 0 | 0.17 | 0.11 | 0.25 | 0.33 | 0.14 | 0.14 |
| 5 | 23153562 | rs11958955 | -1.016 | -0.304 | -0.063 | 0.929 | -0.014 | 0.071 | 0.071 | 0 | 0.25 | 0.5 | 0.35 | 0.59 | 0.18 | 0.29 |
| 5 | 23156895 | rs9293057 | 0.288 | -0.664 | -0.729 | -0.19 | 0.027 | 0.065 | 0.065 | 0 | 0.55 | 0.31 | 0.45 | 0.14 | 0.64 | 0.42 |
| 5 | 23157813 | rs178126 | -0.032 | -0.975 | -0.925 | 0.282 | 0.005 | -0.034 | -0.034 | 0 | 0.87 | 0.91 | 0.8 | 0.33 | 0.87 | 0.72 |
| 5 | 23160613 | rs17301584 | 0.098 | -1.359 | -1.252 | -0.054 | -0.066 | 0.058 | 0.058 | 0.726 | 0.12 | 0.02 | 0.13 | 0.05 | 0.18 | 0.05 |
| 5 | 23160757 | rs17301661 | 1.233 | -1.489 | -2.057 | -0.837 | -0.014 | 0.027 | 0.027 | 0.401 | 0.2 | 0.03 | 0.18 | 0.07 | 0.16 | 0.18 |
| 5 | 23179275 | rs1435230 | 1.21 | -1.167 | -0.257 | -0.462 | -0.009 | 0.006 | 0.006 | 0.012 | 0.67 | 0.66 | 0.5 | 0.19 | 0.62 | 0.58 |
| 5 | 23210180 | rs16892093 | 0.348 | -1.34 | -0.351 | -0.602 | -0.001 | -0.001 | -0.001 | 0 | 0.06 | 0.06 | 0.03 | 0.05 | 0.06 | 0.06 |
| 5 | 23210213 | rs10067536 | 0.392 | -1.352 | -0.363 | -0.697 | -0.002 | 0.003 | 0.003 | 0 | 0.77 | 0.72 | 0.65 | 0.25 | 0.8 | 0.63 |
| 5 | 23212267 | rs10064004 | -0.771 | -1.436 | -0.274 | -0.738 | 0.034 | -0.117 | -0.117 | 0 | 0.2 | 0.42 | 0.19 | 0.06 | 0.2 | 0.04 |
| 5 | 23213699 | rs10050974 | -0.761 | -1.427 | -0.267 | -0.734 | 0.036 | -0.118 | -0.118 | 0.392 | 0.2 | 0.43 | 0.2 | 0.06 | 0.2 | 0.04 |
| 5 | 23216237 | rs10072467 | -0.648 | -1.418 | -0.333 | -0.73 | 0.039 | -0.113 | -0.113 | 0 | 0.19 | 0.4 | 0.19 | 0.06 | 0.21 | 0.04 |
| 5 | 23221917 | rs4701351 | 0.625 | -1.18 | -1.052 | -0.626 | 0.017 | -0.004 | -0.004 | 0 | 0.07 | 0.03 | 0.06 | 0.05 | 0.02 | 0.16 |
| 5 | 23237298 | rs7731943 | -0.4 | -1.124 | -0.976 | -0.469 | -0.011 | -0.021 | -0.021 | 1.445 | 0.86 | 0.89 | 0.77 | 0.33 | 0.83 | 0.74 |
| 5 | 23239197 | rs16892262 | -0.122 | -1.168 | -0.885 | -0.439 | -0.001 | -0.002 | -0.002 | 0 | 0.12 | 0.1 | 0.07 | 0.1 | 0.05 | 0.2 |
| 5 | 23239214 | rs975072 | -0.001 | -0.328 | -0.708 | -0.13 | -0.02 | -0.056 | -0.056 | 0.005 | 0.79 | 0.89 | 0.74 | 0.34 | 0.78 | 0.73 |
| 5 | 23239589 | rs6864495 | 1.003 | 0.047 | -0.761 | -0.348 | -0.004 | 0.023 | 0.023 | 0 | 0.58 | 0.29 | 0.45 | 0.19 | 0.61 | 0.59 |
| 5 | 23241797 | rs17375683 | -0.466 | 1.121 | -0.45 | 0.034 | -0.031 | 0.014 | 0.014 | 0 | 0.06 | 0.01 | 0.12 | 0.05 | 0.14 | 0.03 |
| 5 | 23243079 | rs919332 | 1.427 | 0.919 | -0.046 | 0.507 | 0.048 | 0.055 | 0.055 | 0 | 0.81 | 0.63 | 0.7 | 0.66 | 0.77 | 0.65 |
| 5 | 23243400 | rs7726338 | -1.116 | 1.496 | -0.213 | 0.03 | 0.063 | -0.016 | -0.016 | 0 | 0.06 | 0.12 | 0.15 | 0.03 | 0.18 | 0.06 |
| 5 | 23244025 | rs17376026 | -0.27 | 1.679 | -1.045 | -0.681 | -0.114 | 0.154 | 0.154 | 0 | 0.24 | 0.02 | 0.18 | 0 | 0.33 | 0.15 |
| 5 | 23247790 | rs7715927 | -1.013 | 0.942 | -0.455 | -0.192 | 0.026 | 0.043 | 0.043 | 0.091 | 0.23 | 0.12 | 0.23 | 0.16 | 0.2 | 0.37 |
| 5 | 23251380 | rs992276 | -2.083 | 0.792 | -0.225 | -0.038 | 0.019 | 0.048 | 0.048 | 0.41 | 0.41 | 0.86 | 0.41 | 0.32 | 0.4 | 0.5 |
| 5 | 23251835 | rs12519973 | -0.11 | 0.507 | -0.175 | 0.005 | -0.016 | -0.027 | -0.027 | 0 | 0.06 | 0.01 | 0.16 | 0.14 | 0.07 | 0.26 |
| 5 | 23254969 | rs994675 | -2.197 | 0.566 | 0.332 | 0.016 | 0.001 | 0.002 | 0.002 | 0.72 | 0.24 | 0.74 | 0.32 | 0.32 | 0.26 | 0.39 |
| 5 | 23258050 | rs10074118 | -2.25 | 0.546 | 0.977 | -0.121 | 0.012 | 0.039 | 0.039 | 0.123 | 0.37 | 0.71 | 0.38 | 0.23 | 0.36 | 0.44 |
| 5 | 23260501 | rs1428615 | -2.328 | 0.535 | 1.002 | -0.139 | 0.013 | 0.04 | 0.04 | 0.12 | 0.38 | 0.7 | 0.38 | 0.22 | 0.36 | 0.45 |
| 5 | 23261312 | rs17376271 | -1.191 | 0.677 | 1.6 | -0.049 | 0.01 | -0.032 | -0.032 | 0 | 0.07 | 0.1 | 0.07 | 0 | 0.1 | 0.06 |
| 5 | 23261792 | rs1366450 | -2.112 | 0.652 | 1.813 | -0.028 | 0.072 | -0.019 | -0.019 | 0.02 | 0.15 | 0.48 | 0.15 | 0.09 | 0.19 | 0.11 |
| 5 | 23261842 | rs16892299 | -0.576 | 0.464 | 2.584 | -0.007 | 0.057 | -0.01 | -0.01 | 0.702 | 0.06 | 0.23 | 0.07 | 0 | 0.08 | 0.03 |
| 5 | 23280920 | rs10520889 | -1.405 | 0.395 | 2.467 | -0.031 | -0.002 | -0.001 | -0.001 | 1.979 | 0.22 | 0.26 | 0.23 | 0.21 | 0.16 | 0.42 |
| 5 | 23281444 | rs7727513 | -2.891 | 0.401 | 2.495 | -0.032 | 0.019 | 0.056 | 0.056 | 1.083 | 0.38 | 0.85 | 0.38 | 0.29 | 0.35 | 0.55 |
| 5 | 23283435 | rs17310973 | -1.931 | 0.487 | 2.378 | 0.006 | 0.032 | -0.058 | -0.058 | 0.055 | 0.12 | 0.23 | 0.12 | 0 | 0.17 | 0.1 |
| 5 | 23290393 | rs1428624 | -1.49 | 0.442 | 2.071 | 0.047 | -0.005 | 0 | 0 | 0.158 | 0.22 | 0.27 | 0.23 | 0.27 | 0.17 | 0.42 |
| 5 | 23298902 | rs10941982 | -1.19 | 0.463 | 1.996 | 0.033 | 0.017 | 0.021 | 0.021 | 0.213 | 0.21 | 0.18 | 0.21 | 0.13 | 0.16 | 0.41 |
| 5 | 23304881 | rs16899394 | -2.289 | 0.684 | 2.344 | 0.134 | -0.006 | -0.004 | -0.004 | 0 | 0.28 | 0.35 | 0.29 | 0.27 | 0.23 | 0.5 |
| 5 | 23308068 | rs980924 | -0.495 | 0.6 | 2.113 | 0.156 | -0.02 | -0.031 | -0.031 | 0 | 0.94 | 0.99 | 0.95 | 0.89 | 0.93 | 0.97 |
| 5 | 23313059 | rs6859490 | -1.673 | 0.658 | 2.34 | 0.06 | -0.004 | 0.001 | 0.001 | 0 | 0.08 | 0.09 | 0.07 | 0.06 | 0.1 | 0.08 |
| 5 | 23315549 | rs1896726 | -2.11 | 0.643 | 2.369 | 0.095 | 0.006 | 0.014 | 0.014 | 0 | 0.27 | 0.25 | 0.28 | 0.19 | 0.26 | 0.5 |
| 5 | 23325432 | rs17312807 | -0.413 | 0.633 | 2.366 | 0.1 | 0.02 | 0.023 | 0.023 | 0 | 0.13 | 0.06 | 0.11 | 0.04 | 0.11 | 0.13 |
| 5 | 23331446 | rs10051288 | -2.447 | 0.724 | 2.426 | 0.131 | -0.004 | -0.025 | -0.025 | 0 | 0.35 | 0.65 | 0.37 | 0.2 | 0.34 | 0.55 |
| 5 | 23337424 | rs4701369 | -1.619 | 0.451 | 1.733 | 0.256 | 0.003 | 0.011 | 0.011 | 0 | 0.26 | 0.24 | 0.29 | 0.19 | 0.26 | 0.5 |
| 5 | 23356973 | rs3924616 | -1.678 | 0.365 | 1.608 | 0.204 | -0.006 | -0.027 | -0.027 | 0 | 0.27 | 0.35 | 0.29 | 0.14 | 0.25 | 0.51 |
| 5 | 23371163 | rs10058259 | -2.17 | 0.383 | 2.111 | 0.282 | 0.047 | 0.004 | 0.004 | 0 | 0.1 | 0.29 | 0.09 | 0.06 | 0.1 | 0.08 |
| 5 | 23371640 | rs10050856 | -2.007 | 0.111 | 1.952 | 0.198 | 0.01 | 0.034 | 0.034 | 1.676 | 0.32 | 0.86 | 0.28 | 0.23 | 0.28 | 0.54 |
| 5 | 23389253 | rs6861038 | -2.336 | 0.09 | 2.011 | 0.216 | 0.04 | 0.064 | 0.064 | 0.972 | 0.37 | 0.86 | 0.28 | 0.23 | 0.28 | 0.54 |
| 5 | 23403282 | rs9293071 | -1.393 | 0.069 | 2.439 | 0.25 | -0.064 | 0.007 | 0.007 | 0.224 | 0.23 | 0.5 | 0.16 | 0.23 | 0.17 | 0.45 |
| 5 | 23413553 | rs13175283 | -1.647 | 0.211 | 2.271 | 0.285 | 0.035 | -0.047 | -0.047 | 0 | 0.08 | 0.19 | 0.08 | 0 | 0.09 | 0.07 |
| 5 | 23416474 | rs13166680 | 1.009 | 0.376 | 2.29 | 0.824 | 0.034 | 0.113 | 0.113 | 0 | 0.95 | 0.86 | 0.93 | 0.8 | 0.92 | 0.96 |
| 5 | 23422169 | rs4467669 | 2.23 | 0.275 | 1.698 | 0.65 | -0.017 | -0.044 | -0.044 | 0 | 0.52 | 0.08 | 0.54 | 0.57 | 0.56 | 0.32 |
| 5 | 23428613 | rs10941984 | -1.629 | 0.037 | 2.578 | 0.652 | 0 | -0.004 | -0.004 | 0 | 0.32 | 0.31 | 0.31 | 0.18 | 0.3 | 0.61 |
| 5 | 23439312 | rs13359841 | -1.709 | 0.536 | 1.637 | 0.913 | 0 | 0.037 | 0.037 | 0 | 0.13 | 0.41 | 0.17 | 0.14 | 0.09 | 0.32 |
| 5 | 23446583 | rs2973628 | 0.895 | 0.428 | 0.879 | 1.13 | 0.003 | -0.028 | -0.028 | 0.166 | 0.81 | 0.91 | 0.89 | 0.76 | 0.81 | 0.74 |

| | | | | | | | | | | | | | | | | |
|---|---|---|---|---|---|---|---|---|---|---|---|---|---|---|---|---|
| 5 | 23452924 | rs2914263 | -1.782 | 0.093 | 1.241 | 0.976 | -0.048 | 0.011 | 0.011 | 1.1 | 0.18 | 0.46 | 0.21 | 0.19 | 0.1 | 0.4 |
| 5 | 23454500 | rs2914261 | -2.474 | 0.059 | 0.811 | 1.005 | 0.042 | 0.051 | 0.051 | 0.027 | 0.38 | 0.79 | 0.35 | 0.24 | 0.29 | 0.64 |
| 5 | 23454599 | rs1991495 | -0.258 | 0.432 | 0.2 | 1.236 | -0.014 | -0.028 | -0.028 | 0.05 | 0.05 | 0 | 0.1 | 0.17 | 0.05 | 0.3 |
| 5 | 23456700 | rs17379153 | -1.067 | 0.408 | 0.373 | 1.177 | 0.002 | 0.042 | 0.042 | 0 | 0.18 | 0.09 | 0.11 | 0.05 | 0.19 | 0.24 |
| 5 | 23457592 | rs890965 | 0.981 | 1.051 | 0.392 | 2.502 | -0.039 | -0.006 | -0.006 | 0 | 0.92 | 0.99 | 0.7 | 0.53 | 0.88 | 0.94 |
| 5 | 23484372 | rs2934789 | 1.419 | 1.335 | 0.485 | 2.293 | 0.001 | -0.01 | -0.01 | 0 | 0.86 | 0.89 | 0.62 | 0.63 | 0.82 | 0.86 |
| 5 | 23493956 | rs10075972 | -1.114 | 1.655 | 0.46 | 1.518 | -0.008 | 0.005 | 0.005 | 0.205 | 0.14 | 0.09 | 0.1 | 0.04 | 0.19 | 0.25 |
| 5 | 23501599 | rs12523581 | -1.896 | 1.338 | 1.127 | 2.483 | 0.018 | 0.053 | 0.053 | 0 | 0.13 | 0.23 | 0.37 | 0.38 | 0.19 | 0.24 |
| 5 | 23523348 | rs1874165 | 1.325 | 1.18 | 2.768 | 3.054 | -0.032 | 0.122 | 0.122 | 1.718 | 0.87 | 0.57 | 0.88 | 0.77 | 0.94 | 0.91 |
| 5 | 23528007 | rs3805547 | -1.85 | 1.191 | 2.686 | 3.134 | 0.019 | 0.065 | 0.065 | 0 | 0.14 | 0.24 | 0.37 | 0.38 | 0.19 | 0.25 |
| 5 | 23532643 | rs6889665 | -0.786 | 1.131 | 2.69 | 2.985 | -0.034 | -0.021 | -0.021 | 0 | 0.1 | 0.27 | 0.09 | 0.04 | 0.03 | 0.05 |
| 5 | 23564888 | rs7728406 | -1.87 | 1.114 | 1.989 | 2.797 | 0.008 | 0.182 | 0.182 | 0.161 | 0.13 | 0.35 | 0.32 | 0.35 | 0.12 | 0.2 |
| 5 | 23571543 | rs16892558 | -1.558 | 1.013 | 0.989 | 2.453 | -0.005 | -0.035 | -0.035 | 0 | 0.19 | 0.13 | 0.39 | 0.51 | 0.2 | 0.35 |
| 5 | 23578441 | rs2973620 | 0.306 | 1.327 | 1.301 | 1.942 | 0.037 | 0.02 | 0.02 | 0 | 0.83 | 0.74 | 0.83 | 0.81 | 0.75 | 0.87 |
| 5 | 23584544 | rs16892593 | -0.023 | 1.103 | 1.393 | 1.88 | 0.014 | -0.018 | -0.018 | 0 | 0.05 | 0.14 | 0.08 | 0.02 | 0.06 | 0.03 |
| 5 | 23584606 | rs1494594 | -0.394 | 0.1 | 0.959 | 1.131 | -0.025 | -0.049 | -0.049 | 0 | 0.2 | 0.26 | 0.13 | 0.04 | 0.12 | 0.06 |
| 5 | 23593763 | rs2029778 | -0.697 | 0.254 | 0.77 | 1.606 | 0 | 0.04 | 0.04 | 0 | 0.13 | 0.13 | 0.27 | 0.48 | 0.09 | 0.33 |
| 5 | 23594090 | rs16892616 | 0.214 | 0.366 | 0.403 | 1.163 | -0.023 | -0.022 | -0.022 | 0 | 0.05 | 0 | 0.09 | 0.2 | 0.07 | 0.19 |
| 5 | 23596252 | rs10043677 | 1.347 | -0.219 | 0.32 | 0.751 | 0.003 | 0.071 | 0.071 | 1.807 | 0.54 | 0.33 | 0.41 | 0.3 | 0.58 | 0.46 |
| 5 | 23604239 | rs10077618 | -1.379 | -0.358 | 0.551 | 0.916 | -0.001 | 0 | 0 | 2.003 | 0.46 | 0.47 | 0.46 | 0.61 | 0.48 | 0.43 |
| 5 | 23605043 | rs6868206 | -0.075 | -0.433 | 0.556 | 1.008 | -0.007 | 0.02 | 0.02 | 0 | 0.12 | 0.14 | 0.21 | 0.28 | 0.03 | 0.28 |
| 5 | 23614535 | rs6868212 | 0.684 | -0.394 | 0.495 | 1.104 | -0.024 | -0.016 | -0.016 | 0.323 | 0.05 | 0 | 0.08 | 0.22 | 0.07 | 0.16 |
| 5 | 23616098 | rs9293080 | 1.665 | -0.409 | 0.694 | 1.081 | 0.012 | 0.021 | 0.021 | 0.084 | 0.49 | 0.36 | 0.45 | 0.37 | 0.45 | 0.54 |
| 5 | 23616278 | rs12656827 | 0.63 | -0.406 | 0.658 | 1.083 | -0.024 | -0.017 | -0.017 | 0 | 0.05 | 0 | 0.08 | 0.22 | 0.07 | 0.17 |
| 5 | 23621475 | rs1521009 | 1.218 | -0.422 | 0.7 | 1.045 | -0.005 | 0.01 | 0.01 | 0.446 | 0.54 | 0.39 | 0.54 | 0.59 | 0.52 | 0.7 |
| 5 | 23622036 | rs2030419 | 0.747 | -0.453 | 0.685 | 1.012 | 0.001 | 0 | 0 | 3.417 | 0.63 | 0.4 | 0.6 | 0.59 | 0.61 | 0.72 |
| 5 | 23622185 | rs2030420 | 2.616 | -0.429 | 0.54 | 0.896 | -0.053 | 0.178 | 0.178 | 0.421 | 0.36 | 0.09 | 0.23 | 0.09 | 0.42 | 0.26 |
| 5 | 23622492 | rs1521011 | -1.632 | -0.351 | 0.754 | 0.996 | 0.005 | 0.079 | 0.079 | 1.546 | 0.48 | 0.75 | 0.62 | 0.69 | 0.42 | 0.56 |
| 5 | 23623138 | rs7718389 | -0.162 | -0.65 | 1.023 | 1.418 | -0.035 | 0.137 | 0.137 | 0 | 0.08 | 0.22 | 0.24 | 0.29 | 0.02 | 0.22 |
| 5 | 23623207 | rs7705488 | 1.115 | -0.656 | 1.239 | 1.43 | -0.021 | 0.16 | 0.16 | 0.122 | 0.87 | 0.58 | 0.7 | 0.73 | 0.91 | 0.76 |
| 5 | 23623268 | rs7705523 | 1.167 | -0.87 | 1.062 | 1.37 | -0.004 | -0.001 | -0.001 | 0 | 0.79 | 0.68 | 0.8 | 0.78 | 0.77 | 0.78 |
| 5 | 23623397 | rs7719907 | -1.966 | -0.948 | 0.6 | 1.216 | 0.017 | -0.019 | -0.019 | 0.019 | 0.33 | 0.32 | 0.44 | 0.52 | 0.22 | 0.46 |
| 5 | 23623610 | rs17316188 | 0.275 | -0.8 | 0.16 | 0.953 | 0.012 | 0.027 | 0.027 | 0 | 0.1 | 0.03 | 0.06 | 0 | 0.1 | 0.03 |
| 5 | 23635910 | rs1402410 | 1.749 | -0.892 | 0.745 | 1.295 | -0.036 | 0.058 | 0.058 | 0.261 | 0.74 | 0.62 | 0.65 | 0.57 | 0.81 | 0.58 |
| 5 | 23654181 | rs1464391 | 0.712 | -0.957 | 0.573 | 1.138 | -0.004 | -0.019 | -0.019 | 3.861 | 0.86 | 0.86 | 0.92 | 0.96 | 0.88 | 0.92 |
| 5 | 23684585 | rs10462244 | 1.513 | -0.995 | 1.17 | 1.36 | -0.031 | 0.113 | 0.113 | 5.088 | 0.85 | 0.53 | 0.75 | 0.79 | 0.9 | 0.77 |
| 5 | 23685550 | rs7713783 | -0.038 | -0.99 | 1.165 | 1.368 | -0.028 | 0.118 | 0.118 | 0 | 0.07 | 0.2 | 0.2 | 0.21 | 0.01 | 0.19 |
| 5 | 23687013 | rs6452136 | -2.157 | -1.143 | 1.203 | 1.36 | -0.054 | 0.086 | 0.086 | 0.04 | 0.21 | 0.43 | 0.31 | 0.42 | 0.09 | 0.38 |
| 5 | 23687444 | rs1814984 | -0.504 | -0.969 | 1.096 | 1.462 | 0.004 | -0.033 | -0.033 | 1.8 | 0.93 | 0.87 | 0.96 | 1 | 0.91 | 0.97 |
| 5 | 23689295 | rs7716265 | -0.258 | -1.016 | 1.107 | 1.5 | -0.03 | 0.13 | 0.13 | 0 | 0.07 | 0.22 | 0.21 | 0.22 | 0.01 | 0.19 |
| 5 | 23694438 | rs12516553 | -0.079 | -0.999 | 0.945 | 1.493 | -0.026 | 0.146 | 0.146 | 0.001 | 0.07 | 0.24 | 0.21 | 0.22 | 0.01 | 0.19 |
| 5 | 23696111 | rs11953664 | -0.852 | -0.734 | -0.015 | 1.143 | 0.016 | 0.012 | 0.012 | 0 | 0.38 | 0.45 | 0.38 | 0.43 | 0.39 | 0.29 |
| 5 | 23697300 | rs1521006 | 1.152 | -0.373 | -0.788 | 0.679 | -0.015 | 0.1 | 0.1 | 0.016 | 0.49 | 0.19 | 0.37 | 0.35 | 0.51 | 0.49 |
| 5 | 23710644 | rs6875157 | -0.493 | -0.041 | -0.534 | 0.901 | 0.007 | -0.029 | -0.029 | 7.323 | 0.08 | 0.03 | 0.09 | 0.22 | 0.05 | 0.18 |
| 5 | 23716350 | rs11743379 | 1.421 | 0.19 | -0.45 | 0.422 | -0.019 | 0.184 | 0.184 | 0 | 0.44 | 0.16 | 0.29 | 0.14 | 0.47 | 0.34 |
| 5 | 23730041 | rs1879480 | 0.457 | 0.824 | 0.158 | 1.509 | -0.016 | -0.03 | -0.03 | 0 | 0.7 | 0.81 | 0.68 | 0.6 | 0.73 | 0.74 |
| 5 | 23742976 | rs3932338 | 1.543 | 0.642 | 0.159 | 1.55 | -0.012 | 0.058 | 0.058 | 0 | 0.55 | 0.18 | 0.59 | 0.36 | 0.62 | 0.54 |
| 5 | 23755699 | rs4701393 | 0.29 | 0.625 | 0.372 | 1.612 | -0.012 | -0.025 | -0.025 | 0 | 0.69 | 0.8 | 0.68 | 0.59 | 0.73 | 0.74 |
| 5 | 23758878 | rs1520985 | 0.294 | 0.62 | 0.443 | 1.61 | -0.01 | -0.021 | -0.021 | 0 | 0.69 | 0.78 | 0.68 | 0.59 | 0.73 | 0.74 |
| 5 | 23761700 | rs10060443 | -1.474 | 0.616 | 0.886 | 1.633 | -0.046 | 0.104 | 0.104 | 1.389 | 0.14 | 0.53 | 0.08 | 0.22 | 0.08 | 0.17 |
| 5 | 23778344 | rs13174015 | 1.705 | 0.667 | 0.838 | 1.622 | -0.021 | 0.062 | 0.062 | 0 | 0.54 | 0.16 | 0.58 | 0.36 | 0.62 | 0.54 |
| 5 | 23781623 | rs4569836 | 1.644 | 0.675 | 0.868 | 1.627 | -0.017 | 0.038 | 0.038 | 0 | 0.56 | 0.23 | 0.59 | 0.37 | 0.62 | 0.56 |
| 5 | 23820861 | rs7705043 | -1.878 | 0.645 | 1.004 | 1.638 | -0.056 | 0.068 | 0.068 | 3.022 | 0.16 | 0.59 | 0.09 | 0.23 | 0.09 | 0.19 |
| 5 | 23858021 | rs10942009 | 2.345 | 1.344 | 1.169 | 1.595 | 0.006 | 0.134 | 0.134 | 0 | 0.43 | 0.1 | 0.43 | 0.24 | 0.44 | 0.35 |
| 5 | 23861064 | rs6871540 | 0.239 | 1.32 | 1.3 | 1.775 | -0.029 | -0.037 | -0.037 | 0 | 0.7 | 0.88 | 0.69 | 0.63 | 0.71 | 0.72 |
| 5 | 23872046 | rs4701405 | 2.115 | 1.381 | 1.388 | 1.651 | 0.005 | 0.192 | 0.192 | 0 | 0.47 | 0.09 | 0.38 | 0.19 | 0.49 | 0.36 |
| 5 | 23877218 | rs7714862 | -1.664 | 1.419 | 1.942 | 1.77 | 0.014 | 0.171 | 0.171 | 3.27 | 0.12 | 0.57 | 0.12 | 0.26 | 0.1 | 0.15 |
| 5 | 23878207 | rs966297 | 0.246 | 1.581 | 1.789 | 1.726 | -0.008 | -0.036 | -0.036 | 0 | 0.72 | 0.8 | 0.61 | 0.56 | 0.76 | 0.7 |
| 5 | 23884440 | rs1490758 | 0.371 | 1.599 | 1.737 | 1.734 | -0.002 | -0.018 | -0.018 | 0 | 0.71 | 0.77 | 0.61 | 0.56 | 0.76 | 0.7 |
| 5 | 23909410 | rs2055526 | -1.143 | 1.557 | 2.04 | 1.72 | 0.031 | 0.11 | 0.11 | 0.25 | 0.08 | 0.26 | 0.09 | 0.17 | 0.09 | 0.13 |
| 5 | 23919596 | rs4479808 | -1.091 | 1.559 | 2.045 | 1.721 | 0.04 | 0.125 | 0.125 | 0.047 | 0.07 | 0.27 | 0.09 | 0.17 | 0.09 | 0.13 |
| 5 | 23928233 | rs7448471 | -0.725 | 1.554 | 1.977 | 1.683 | 0.029 | 0.074 | 0.074 | 0 | 0.07 | 0.24 | 0.08 | 0.13 | 0.08 | 0.11 |
| 5 | 23961525 | rs17428710 | 1.85 | 1.616 | 1.476 | 1.478 | 0.026 | 0.076 | 0.076 | 0 | 0.49 | 0.26 | 0.39 | 0.3 | 0.5 | 0.37 |
| 5 | 23980894 | rs11950063 | -0.492 | 1.533 | 0.989 | 1.318 | -0.046 | 0.007 | 0.007 | 0 | 0.11 | 0.03 | 0.1 | 0.07 | 0.16 | 0.19 |
| 5 | 24005118 | rs6861036 | 0.97 | 0.837 | 1.469 | 1.543 | 0.001 | 0.014 | 0.014 | 0 | 0.63 | 0.54 | 0.52 | 0.42 | 0.67 | 0.58 |
| 5 | 24020558 | rs1501949 | 0.604 | 0.402 | 1.75 | 1.525 | 0.005 | 0.028 | 0.028 | 0 | 0.74 | 0.66 | 0.59 | 0.53 | 0.74 | 0.68 |
| 5 | 24030812 | rs1910071 | 0.884 | 0.335 | 2.136 | 1.492 | 0.009 | 0.065 | 0.065 | 0 | 0.72 | 0.48 | 0.57 | 0.53 | 0.74 | 0.68 |
| 5 | 24033839 | rs17429271 | 2.433 | 0.344 | 1.935 | 1.43 | -0.049 | 0.139 | 0.139 | 0 | 0.46 | 0.11 | 0.36 | 0.21 | 0.52 | 0.34 |
| 5 | 24034550 | rs6896352 | -0.431 | 0.336 | 2.077 | 1.425 | -0.005 | -0.017 | -0.017 | 0 | 0.06 | 0.11 | 0.03 | 0.04 | 0.06 | 0.03 |

| | | | | | | | | | | | | | | | | |
|---|---|---|---|---|---|---|---|---|---|---|---|---|---|---|---|---|
| 5 | 24041489 | rs11750284 | 2.515 | 0.338 | 1.905 | 1.407 | -0.07 | 0.134 | 0.134 | 0 | 0.46 | 0.06 | 0.35 | 0.21 | 0.53 | 0.34 |
| 5 | 24044704 | rs17501504 | -0.338 | 0.34 | 1.945 | 1.411 | -0.011 | 0.026 | 0.026 | 0 | 0.14 | 0.02 | 0.1 | 0.07 | 0.1 | 0.2 |
| 5 | 24046739 | rs17501608 | -0.4 | 0.335 | 1.973 | 1.358 | 0.003 | 0.044 | 0.044 | 0 | 0.15 | 0.04 | 0.1 | 0.05 | 0.1 | 0.2 |
| 5 | 24066236 | rs17501795 | 1.89 | 0.365 | 2.349 | 1.372 | 0.001 | -0.006 | -0.006 | 0 | 0.52 | 0.35 | 0.45 | 0.45 | 0.58 | 0.45 |
| 5 | 24072617 | rs16893073 | -0.788 | 0.29 | 2.451 | 1.365 | -0.007 | -0.02 | -0.02 | 0 | 0.07 | 0.15 | 0.03 | 0.04 | 0.06 | 0.03 |
| 5 | 24075658 | rs10078293 | -2.028 | 0.277 | 2.578 | 1.373 | -0.016 | 0.007 | 0.007 | 0 | 0.28 | 0.58 | 0.23 | 0.31 | 0.21 | 0.33 |
| 5 | 24088733 | rs2099055 | 0.746 | 0.22 | 2.549 | 1.405 | 0.017 | 0.025 | 0.025 | 0 | 0.72 | 0.59 | 0.53 | 0.38 | 0.68 | 0.57 |
| 5 | 24092063 | rs13177217 | -0.303 | 0.178 | 2.194 | 1.345 | 0 | -0.01 | -0.01 | 3.145 | 0.15 | 0.13 | 0.12 | 0.04 | 0.1 | 0.19 |
| 5 | 24093347 | rs11743115 | 2.455 | 0.175 | 1.853 | 1.363 | -0.065 | 0.135 | 0.135 | 0 | 0.47 | 0.07 | 0.35 | 0.21 | 0.53 | 0.34 |
| 5 | 24100181 | rs10040797 | -1.684 | 0.138 | 2.087 | 1.463 | 0.011 | 0.106 | 0.106 | 0 | 0.1 | 0.25 | 0.1 | 0.27 | 0.11 | 0.15 |
| 5 | 24108374 | rs10472510 | -1.688 | 0.135 | 2.025 | 1.462 | 0.006 | 0.086 | 0.086 | 0 | 0.1 | 0.22 | 0.1 | 0.27 | 0.11 | 0.15 |
| 5 | 24108770 | rs11750295 | 0.378 | 0.159 | 2.223 | 1.459 | -0.009 | -0.025 | -0.025 | 0.583 | 0.76 | 0.77 | 0.61 | 0.52 | 0.74 | 0.67 |
| 5 | 24109062 | rs11748076 | 0.754 | 0.16 | 1.912 | 1.46 | 0.004 | 0.036 | 0.036 | 0 | 0.72 | 0.42 | 0.57 | 0.52 | 0.74 | 0.67 |
| 5 | 24118301 | rs11958182 | -1.61 | 0.202 | 1.757 | 1.425 | 0 | -0.002 | -0.002 | 0.488 | 0.24 | 0.24 | 0.2 | 0.31 | 0.2 | 0.33 |
| 5 | 24128474 | rs10053948 | -1.61 | 0.202 | 1.757 | 1.425 | 0 | -0.002 | -0.002 | 0 | 0.24 | 0.24 | 0.2 | 0.31 | 0.2 | 0.33 |
| 5 | 24129266 | rs10942028 | -1.61 | 0.202 | 1.757 | 1.425 | 0 | -0.002 | -0.002 | 0 | 0.24 | 0.24 | 0.2 | 0.31 | 0.2 | 0.33 |
| 5 | 24129417 | rs11954471 | -1.61 | 0.202 | 1.757 | 1.425 | 0 | -0.002 | -0.002 | 0 | 0.24 | 0.24 | 0.2 | 0.31 | 0.2 | 0.33 |
| 5 | 24133971 | rs16893146 | -1.277 | 0.188 | 1.768 | 1.421 | 0.001 | 0.002 | 0.002 | 0 | 0.22 | 0.19 | 0.13 | 0.17 | 0.15 | 0.23 |
| 5 | 24137390 | rs1428199 | 0.867 | 0.206 | 1.782 | 1.441 | -0.001 | 0.001 | 0.001 | 0 | 0.71 | 0.32 | 0.56 | 0.52 | 0.74 | 0.66 |
| 5 | 24141118 | rs1428201 | 2.044 | 0.202 | 1.649 | 1.413 | -0.009 | 0.018 | 0.018 | 0 | 0.53 | 0.12 | 0.45 | 0.36 | 0.61 | 0.46 |
| 5 | 24146519 | rs6892387 | -0.14 | 0.227 | 1.635 | 1.439 | -0.014 | -0.015 | -0.015 | 0 | 0.05 | 0.04 | 0.08 | 0.15 | 0.08 | 0.13 |
| 5 | 24150299 | rs7718527 | -1.464 | 0.186 | 1.812 | 1.419 | -0.008 | 0.039 | 0.039 | 0 | 0.09 | 0.28 | 0.05 | 0.13 | 0.06 | 0.05 |
| 5 | 24155061 | rs2115122 | 1.36 | 0.218 | 2.111 | 1.408 | 0.007 | 0.03 | 0.03 | 0 | 0.63 | 0.5 | 0.49 | 0.48 | 0.67 | 0.51 |
| 5 | 24162206 | rs17445670 | -0.215 | 0.247 | 1.921 | 1.431 | 0.006 | 0.04 | 0.04 | 0 | 0.14 | 0.06 | 0.1 | 0.04 | 0.09 | 0.19 |
| 5 | 24163109 | rs6452189 | -2.007 | 0.229 | 2.107 | 1.471 | 0.005 | 0.051 | 0.051 | 0 | 0.14 | 0.25 | 0.12 | 0.27 | 0.14 | 0.17 |
| 5 | 24171380 | rs9687325 | -0.248 | 0.223 | 2.037 | 1.45 | 0.008 | 0.022 | 0.022 | 0 | 0.14 | 0.09 | 0.11 | 0.04 | 0.09 | 0.19 |
| 5 | 24172136 | rs1820694 | 2.856 | 0.011 | 1.961 | 1.364 | -0.064 | 0.118 | 0.118 | 0 | 0.45 | 0.06 | 0.35 | 0.21 | 0.52 | 0.33 |
| 5 | 24172252 | rs9637784 | -0.161 | 0.104 | 2.071 | 1.494 | -0.016 | -0.018 | -0.018 | 1.102 | 0.05 | 0.03 | 0.08 | 0.15 | 0.08 | 0.13 |
| 5 | 24177944 | rs6874374 | 0.361 | 0.074 | 2.495 | 1.71 | 0.018 | 0.048 | 0.048 | 0 | 0.78 | 0.64 | 0.7 | 0.59 | 0.76 | 0.7 |
| 5 | 24182088 | rs1157943 | -0.254 | 0.051 | 2.438 | 1.685 | 0.008 | 0.012 | 0.012 | 0 | 0.14 | 0.1 | 0.2 | 0.12 | 0.09 | 0.19 |
| 5 | 24198232 | rs12654722 | -0.138 | 0.123 | 2.546 | 1.748 | -0.023 | -0.026 | -0.026 | 0 | 0.05 | 0.02 | 0.08 | 0.14 | 0.08 | 0.13 |
| 5 | 24200574 | rs4594816 | -1.441 | 0.099 | 2.637 | 1.729 | -0.01 | 0.031 | 0.031 | 0 | 0.09 | 0.23 | 0.04 | 0.12 | 0.06 | 0.05 |
| 5 | 24206738 | rs7723348 | 1.156 | 0.115 | 2.649 | 1.731 | 0.04 | 0.072 | 0.072 | 0 | 0.71 | 0.49 | 0.59 | 0.44 | 0.67 | 0.57 |
| 5 | 24211218 | rs10054081 | -1.244 | 0.132 | 2.364 | 1.674 | -0.008 | 0.031 | 0.031 | 0 | 0.08 | 0.2 | 0.03 | 0.12 | 0.06 | 0.05 |
| 5 | 24219543 | rs11748884 | 2.584 | 0.098 | 2.202 | 1.477 | -0.075 | 0.125 | 0.125 | 0 | 0.46 | 0.04 | 0.35 | 0.2 | 0.52 | 0.33 |
| 5 | 24230024 | rs4554170 | 2.598 | -0.034 | 2.244 | 1.472 | -0.078 | 0.156 | 0.156 | 0 | 0.45 | 0.05 | 0.33 | 0.18 | 0.52 | 0.32 |
| 5 | 24230888 | rs6868533 | -1.86 | -0.004 | 2.354 | 1.525 | -0.007 | 0.023 | 0.023 | 0 | 0.3 | 0.39 | 0.35 | 0.41 | 0.24 | 0.39 |
| 5 | 24235851 | rs7721732 | -0.656 | -0.348 | 2.183 | 1.426 | -0.009 | -0.008 | -0.008 | 0 | 0.18 | 0.21 | 0.24 | 0.13 | 0.12 | 0.21 |
| 5 | 24237693 | rs4490548 | 0.079 | -0.307 | 1.904 | 1.484 | -0.036 | -0.064 | -0.064 | 0 | 0.81 | 0.91 | 0.72 | 0.59 | 0.76 | 0.71 |
| 5 | 24239623 | rs10040676 | 0.576 | -0.277 | 1.991 | 1.483 | 0.013 | 0.052 | 0.052 | 0 | 0.77 | 0.62 | 0.71 | 0.59 | 0.76 | 0.71 |
| 5 | 24243298 | rs6867094 | 0.241 | -0.212 | 1.442 | 1.514 | -0.009 | -0.018 | -0.018 | 0 | 0.79 | 0.81 | 0.72 | 0.59 | 0.76 | 0.71 |
| 5 | 24247673 | rs11952208 | -0.658 | -0.306 | 1.211 | 1.323 | 0.019 | 0.017 | 0.017 | 0 | 0.17 | 0.1 | 0.22 | 0.14 | 0.12 | 0.21 |
| 5 | 24259694 | rs4277873 | 0.507 | -0.261 | 1.462 | 1.605 | -0.001 | 0.009 | 0.009 | 0 | 0.79 | 0.73 | 0.74 | 0.62 | 0.81 | 0.71 |
| 5 | 24260655 | rs10942035 | 2.501 | -0.21 | 1.23 | 1.566 | -0.086 | 0.115 | 0.115 | 0 | 0.46 | 0.05 | 0.37 | 0.23 | 0.51 | 0.31 |
| 5 | 24261955 | rs12518573 | -1.598 | -0.404 | 2.268 | 1.729 | 0.016 | 0.198 | 0.198 | 0.075 | 0.1 | 0.3 | 0.14 | 0.27 | 0.06 | 0.14 |
| 5 | 24264090 | rs5007912 | 0.992 | -0.38 | 2.34 | 1.698 | 0.001 | 0.064 | 0.064 | 0 | 0.76 | 0.57 | 0.7 | 0.59 | 0.78 | 0.69 |
| 5 | 24269794 | rs7700267 | 0.822 | -0.434 | 2.316 | 1.873 | -0.007 | 0.039 | 0.039 | 0 | 0.77 | 0.56 | 0.72 | 0.64 | 0.81 | 0.71 |
| 5 | 24274235 | rs10805759 | 1.5 | -0.44 | 2.202 | 1.885 | -0.005 | 0.066 | 0.066 | 0 | 0.71 | 0.52 | 0.59 | 0.41 | 0.75 | 0.57 |
| 5 | 24274495 | rs10805760 | 1.593 | -0.44 | 2.193 | 1.885 | -0.009 | 0.061 | 0.061 | 0 | 0.7 | 0.48 | 0.58 | 0.41 | 0.75 | 0.57 |
| 5 | 24279987 | rs10214306 | 0.678 | -0.579 | 1.517 | 1.924 | -0.001 | 0.003 | 0.003 | 0.006 | 0.8 | 0.78 | 0.75 | 0.73 | 0.82 | 0.71 |
| 5 | 24287790 | rs9654386 | -0.625 | -0.579 | 1.662 | 1.979 | 0.003 | 0.01 | 0.01 | 0.09 | 0.16 | 0.2 | 0.24 | 0.23 | 0.16 | 0.26 |
| 5 | 24294511 | rs10070171 | 1.022 | -0.824 | 1.739 | 1.657 | -0.013 | -0.109 | -0.109 | 0 | 0.75 | 0.52 | 0.81 | 0.92 | 0.78 | 0.72 |
| 5 | 24296704 | rs13360911 | -1.123 | -0.805 | 1.742 | 1.985 | 0.007 | 0.202 | 0.202 | 0 | 0.21 | 0.48 | 0.39 | 0.52 | 0.19 | 0.37 |
| 5 | 24298575 | rs6886864 | 0.496 | -0.757 | 1.113 | 1.696 | -0.006 | 0.037 | 0.037 | 0 | 0.81 | 0.9 | 0.85 | 0.92 | 0.79 | 0.73 |
| 5 | 24301428 | rs6452194 | 0.483 | -0.765 | 1.035 | 1.692 | -0.013 | 0.055 | 0.055 | 0 | 0.81 | 0.91 | 0.85 | 0.92 | 0.79 | 0.73 |
| 5 | 24320108 | rs10044803 | -0.808 | -1.17 | 0.981 | 1.679 | 0.001 | -0.002 | -0.002 | 0 | 0.2 | 0.18 | 0.17 | 0.24 | 0.15 | 0.24 |
| 5 | 24320400 | rs10037286 | -1.39 | -1.238 | 1.263 | 1.795 | -0.016 | 0.101 | 0.101 | 0 | 0.27 | 0.46 | 0.32 | 0.51 | 0.21 | 0.35 |
| 5 | 24320469 | rs934599 | 0.087 | -1.363 | 0.351 | 1.746 | -0.02 | 0.042 | 0.042 | 0 | 0.82 | 0.97 | 0.85 | 0.9 | 0.8 | 0.75 |
| 5 | 24322935 | rs1469832 | 1.99 | -1.429 | 0.297 | 1.717 | 0.009 | 0.029 | 0.029 | 0.018 | 0.52 | 0.62 | 0.51 | 0.33 | 0.55 | 0.36 |
| 5 | 24334452 | rs331250 | -0.698 | -1.177 | 0.381 | 1.862 | 0.003 | 0.084 | 0.084 | 0 | 0.21 | 0.28 | 0.3 | 0.52 | 0.2 | 0.35 |
| 5 | 24335447 | rs12519312 | 2.485 | -1.22 | 0.017 | 1.834 | -0.07 | 0.042 | 0.042 | 0.377 | 0.48 | 0.04 | 0.46 | 0.32 | 0.55 | 0.37 |
| 5 | 24337141 | rs4482869 | 0.179 | -0.536 | 0.351 | 2.193 | -0.026 | 0.046 | 0.046 | 0 | 0.83 | 0.97 | 0.85 | 0.9 | 0.8 | 0.76 |
| 5 | 24343572 | rs331303 | 0.796 | -0.513 | 0.367 | 2.206 | -0.009 | 0.032 | 0.032 | 0 | 0.77 | 0.91 | 0.82 | 0.85 | 0.76 | 0.73 |
| 5 | 24355555 | rs972303 | 1.453 | -0.669 | 0.778 | 2.16 | 0.002 | 0.042 | 0.042 | 0 | 0.74 | 0.64 | 0.69 | 0.42 | 0.76 | 0.63 |
| 5 | 24357450 | rs331290 | -0.206 | -0.8 | 0.107 | 1.597 | -0.01 | -0.021 | -0.021 | 0 | 0.06 | 0.01 | 0.02 | 0.06 | 0.05 | 0.03 |
| 5 | 24365309 | rs28605677 | -0.784 | -1.17 | 0.473 | 1.799 | -0.014 | 0.012 | 0.012 | 0 | 0.1 | 0.13 | 0.1 | 0.16 | 0.08 | 0.13 |
| 5 | 24382079 | rs11949863 | 1.749 | -1.95 | 0.683 | 1.866 | 0.054 | -0.027 | -0.027 | 0.351 | 0.67 | 0.2 | 0.72 | 0.38 | 0.79 | 0.67 |
| 5 | 24398596 | rs2319567 | 2.402 | -2.096 | 0.188 | 1.806 | 0.113 | 0.149 | 0.149 | 0.854 | 0.55 | 0.09 | 0.51 | 0.17 | 0.71 | 0.43 |
| 5 | 24404807 | rs331214 | 0.841 | -2.009 | 0.797 | 1.929 | -0.019 | 0.042 | 0.042 | 0 | 0.82 | 0.79 | 0.86 | 0.42 | 0.89 | 0.73 |

| | | | | | | | | | | | | | | | | |
|---|---|---|---|---|---|---|---|---|---|---|---|---|---|---|---|---|
| 5 | 24430251 | rs4701245 | 1.496 | -2.037 | 1.614 | 1.915 | -0.075 | 0.091 | 0.091 | 0 | 0.76 | 0.56 | 0.82 | 0.41 | 0.86 | 0.72 |
| 5 | 24438467 | rs331270 | -1.248 | -2.025 | 1.785 | 1.814 | 0.015 | -0.014 | -0.014 | 0 | 0.11 | 0.39 | 0.11 | 0.04 | 0.06 | 0.05 |
| 5 | 24441601 | rs1978608 | 2.009 | -1.928 | 1.231 | 1.856 | 0.032 | -0.023 | -0.023 | 0 | 0.66 | 0.23 | 0.72 | 0.43 | 0.8 | 0.67 |
| 5 | 24457943 | rs2081922 | -1.652 | -1.038 | 1.317 | 1.401 | -0.001 | -0.032 | -0.032 | 0 | 0.13 | 0.34 | 0.1 | 0.07 | 0.1 | 0.05 |
| 5 | 24463538 | rs1346509 | 0.552 | -0.834 | 0.723 | 1.588 | 0 | 0.005 | 0.005 | 0 | 0.87 | 0.87 | 0.77 | 0.8 | 0.88 | 0.78 |
| 5 | 24469962 | rs13188967 | -1.258 | -0.774 | 1.334 | 1.663 | 0.057 | 0.008 | 0.008 | 0 | 0.1 | 0.33 | 0.12 | 0.07 | 0.1 | 0.1 |
| 5 | 24472315 | rs9283777 | -1.868 | -1.327 | 0.068 | 1.968 | 0.02 | -0.038 | -0.038 | 0.698 | 0.31 | 0.79 | 0.28 | 0.53 | 0.22 | 0.42 |
| 5 | 24482475 | rs1428365 | -0.536 | -1.392 | 0.46 | 2.108 | -0.016 | 0.057 | 0.057 | 0 | 0.12 | 0.14 | 0.12 | 0.37 | 0.09 | 0.21 |
| 5 | 24485005 | rs2874027 | 0.896 | -0.809 | 0.243 | 1.362 | -0.001 | 0.001 | 0.001 | 0 | 0.88 | 0.89 | 0.78 | 0.83 | 0.88 | 0.79 |
| 5 | 24486805 | rs1169561 | 0.918 | -0.813 | 0.226 | 1.325 | -0.001 | 0.001 | 0.001 | 0 | 0.88 | 0.87 | 0.77 | 0.83 | 0.88 | 0.79 |
| 5 | 24496862 | rs6896096 | -1.48 | -0.821 | 0.195 | 1.096 | -0.005 | -0.013 | -0.013 | 0 | 0.06 | 0.06 | 0.02 | 0 | 0.02 | 0.02 |
| 5 | 24498690 | rs3815909 | -2.019 | -0.983 | 0.488 | 1.515 | -0.03 | 0.069 | 0.069 | 0 | 0.24 | 0.4 | 0.25 | 0.44 | 0.16 | 0.39 |
| 5 | 24500645 | rs7714385 | 1.687 | -0.857 | 0.746 | 1.039 | -0.022 | -0.002 | -0.002 | 0 | 0.73 | 0.41 | 0.71 | 0.75 | 0.82 | 0.71 |
| 5 | 24503699 | rs6889987 | 2.367 | -0.679 | -0.629 | 0.664 | -0.165 | -0.099 | -0.099 | 0.363 | 0.38 | 0.03 | 0.33 | 0.59 | 0.5 | 0.45 |
| 5 | 24509947 | rs3797171 | -1.546 | -0.909 | 0.802 | 0.571 | -0.025 | 0.066 | 0.066 | 0 | 0.45 | 0.21 | 0.53 | 0.3 | 0.42 | 0.4 |
| 5 | 24528612 | rs13154399 | -0.559 | -0.833 | 0.63 | 0.462 | -0.024 | 0.044 | 0.044 | 0 | 0.29 | 0.13 | 0.32 | 0.21 | 0.28 | 0.28 |
| 5 | 24528847 | rs4701447 | -0.38 | -0.871 | 0.577 | 0.405 | -0.019 | 0.03 | 0.03 | 0 | 0.26 | 0.14 | 0.31 | 0.21 | 0.28 | 0.26 |
| 5 | 24541668 | rs12652231 | 3.076 | -0.842 | 0.74 | 0.477 | -0.167 | -0.093 | -0.093 | 1.129 | 0.38 | 0.03 | 0.33 | 0.59 | 0.51 | 0.46 |
| 5 | 24552177 | rs952168 | 0.922 | -0.902 | 1.543 | 0.442 | -0.002 | -0.025 | -0.025 | 0 | 0.66 | 0.61 | 0.64 | 0.79 | 0.68 | 0.71 |
| 5 | 24552316 | rs7735817 | -2.383 | -0.828 | 1.605 | 0.486 | -0.025 | -0.021 | -0.021 | 0 | 0.28 | 0.58 | 0.31 | 0.2 | 0.17 | 0.25 |
| 5 | 24556172 | rs10462251 | 0.967 | -0.605 | 1.321 | 0.589 | 0.001 | -0.001 | -0.001 | 0 | 0.86 | 0.87 | 0.78 | 0.91 | 0.86 | 0.88 |
| 5 | 24558533 | rs9293138 | 1.106 | -0.613 | 1.33 | 0.585 | 0.006 | -0.006 | -0.006 | 0 | 0.86 | 0.84 | 0.78 | 0.91 | 0.86 | 0.88 |
| 5 | 24562793 | rs17520214 | 2.757 | -0.624 | 0.908 | 0.62 | -0.119 | -0.05 | -0.05 | 0 | 0.4 | 0.06 | 0.34 | 0.59 | 0.51 | 0.46 |
| 5 | 24566690 | rs12187078 | 2.287 | -0.167 | 1.157 | 0.762 | -0.06 | -0.031 | -0.031 | 0 | 0.47 | 0.11 | 0.37 | 0.55 | 0.53 | 0.54 |
| 5 | 24567660 | rs13174538 | 2.248 | -0.18 | 1.319 | 0.756 | -0.031 | -0.007 | -0.007 | 0 | 0.47 | 0.16 | 0.38 | 0.55 | 0.53 | 0.54 |
| 5 | 24568262 | rs17460736 | -0.502 | -0.138 | 1.135 | 0.765 | -0.041 | -0.038 | -0.038 | 0 | 0.1 | 0.02 | 0.19 | 0.13 | 0.11 | 0.07 |
| 5 | 24572261 | rs12516067 | 2.649 | -0.32 | 0.994 | 0.876 | -0.148 | -0.108 | -0.108 | 0 | 0.44 | 0.03 | 0.37 | 0.53 | 0.54 | 0.48 |
| 5 | 24582271 | rs10805767 | 2.662 | -0.289 | 1.001 | 0.837 | -0.144 | -0.104 | -0.104 | 0 | 0.44 | 0.03 | 0.37 | 0.54 | 0.54 | 0.48 |
| 5 | 24585329 | rs7716158 | -1.761 | 0.229 | 1.618 | 0.916 | 0.006 | 0.028 | 0.028 | 0.021 | 0.14 | 0.22 | 0.21 | 0.2 | 0.11 | 0.17 |
| 5 | 24588189 | rs2174529 | -2.295 | 0.358 | 1.763 | 0.726 | 0.003 | 0.02 | 0.02 | 0 | 0.28 | 0.48 | 0.29 | 0.31 | 0.2 | 0.29 |
| 5 | 24592778 | rs1505877 | 1.131 | 0.312 | 1.63 | 0.976 | 0 | -0.001 | -0.001 | 0 | 0.61 | 0.38 | 0.46 | 0.68 | 0.62 | 0.66 |
| 5 | 24593325 | rs16893586 | -2.18 | 0.483 | 2.383 | 1.762 | 0.009 | 0.055 | 0.055 | 0.483 | 0.06 | 0.26 | 0.04 | 0.07 | 0.02 | 0.1 |
| 5 | 24597533 | rs4701453 | 1.88 | 0.645 | 1.633 | 1.835 | 0.014 | -0.002 | -0.002 | 2.211 | 0.74 | 0.21 | 0.75 | 0.75 | 0.85 | 0.79 |
| 5 | 24600075 | rs16893608 | -0.752 | 0.656 | 1.525 | 1.822 | 0.031 | 0.011 | 0.011 | 0 | 0.1 | 0.05 | 0.21 | 0.07 | 0.05 | 0.09 |
| 5 | 24609447 | rs9293145 | -1.887 | 1.892 | 3.158 | 2.502 | 0.09 | 0.157 | 0.157 | 0 | 0.11 | 0.31 | 0.12 | 0.28 | 0.14 | 0.1 |
| 5 | 24617958 | rs17521324 | -1.244 | 1.728 | 3.393 | 2.236 | -0.001 | -0.019 | -0.019 | 1.639 | 0.14 | 0.19 | 0.09 | 0.07 | 0.12 | 0.22 |
| 5 | 24620819 | rs10067393 | -1.796 | 1.84 | 3.362 | 2.298 | 0.091 | 0.113 | 0.113 | 0 | 0.1 | 0.26 | 0.1 | 0.18 | 0.12 | 0.06 |
| 5 | 24624103 | rs10461813 | 2.357 | 1.754 | 2.883 | 2.261 | -0.056 | -0.029 | -0.029 | 0 | 0.72 | 0.11 | 0.77 | 0.75 | 0.75 | 0.68 |
| 5 | 24639574 | rs1505890 | 2.196 | 1.686 | 3.019 | 2.291 | -0.034 | -0.025 | -0.025 | 0 | 0.73 | 0.18 | 0.78 | 0.73 | 0.75 | 0.7 |
| 5 | 24643756 | rs1395026 | 2.541 | 1.481 | 3.984 | 2.819 | 0.112 | 0.149 | 0.149 | 0.738 | 0.87 | 0.52 | 0.88 | 0.82 | 0.87 | 0.93 |
| 5 | 24645673 | rs332010 | 2.18 | 1.309 | 3.514 | 2.898 | -0.004 | 0.019 | 0.019 | 0 | 0.92 | 0.91 | 0.97 | 0.85 | 0.95 | 0.78 |
| 5 | 24646826 | rs332012 | -2.242 | 1.512 | 3.778 | 3.137 | 0.11 | 0.297 | 0.297 | 0 | 0.11 | 0.37 | 0.29 | 0.44 | 0.15 | 0.2 |
| 5 | 24666440 | rs2082058 | 1.65 | 1.488 | 3.621 | 2.629 | 0.017 | 0.041 | 0.041 | 0 | 0.83 | 0.79 | 0.66 | 0.6 | 0.79 | 0.83 |
| 5 | 24691553 | rs10070341 | 1.876 | 1.49 | 3.983 | 2.6 | 0.051 | 0.106 | 0.106 | 0 | 0.82 | 0.66 | 0.64 | 0.6 | 0.79 | 0.83 |
| 5 | 24722992 | rs1975076 | 1.859 | 1.462 | 3.913 | 2.606 | 0.045 | 0.095 | 0.095 | 0 | 0.82 | 0.69 | 0.64 | 0.61 | 0.79 | 0.83 |
| 5 | 24737095 | rs10039614 | -2.725 | 1.521 | 3.723 | 2.413 | 0.055 | 0.067 | 0.067 | 0.732 | 0.1 | 0.46 | 0.06 | 0.11 | 0.06 | 0.2 |
| 5 | 24738922 | rs10044604 | -1.558 | 1.522 | 3.774 | 2.414 | 0.028 | 0.059 | 0.059 | 0 | 0.07 | 0.24 | 0.03 | 0.11 | 0.04 | 0.19 |
| 5 | 24739100 | rs10038996 | -1.265 | 1.523 | 3.673 | 2.414 | 0.021 | 0.05 | 0.05 | 0 | 0.06 | 0.19 | 0.03 | 0.11 | 0.04 | 0.19 |
| 5 | 24756604 | rs7711384 | 0.967 | 1.549 | 3.042 | 2.559 | -0.024 | -0.024 | -0.024 | 0.047 | 0.87 | 0.95 | 0.68 | 0.61 | 0.81 | 0.84 |
| 5 | 24770890 | rs7700796 | -2.777 | 1.1 | 3.301 | 2.447 | -0.016 | 0.05 | 0.05 | 1.708 | 0.13 | 0.66 | 0.09 | 0.18 | 0.07 | 0.26 |
| 5 | 24773509 | rs10472517 | -2.049 | 1.104 | 3.123 | 2.443 | 0.048 | 0.065 | 0.065 | 1.062 | 0.1 | 0.45 | 0.06 | 0.11 | 0.06 | 0.19 |
| 5 | 24779565 | rs1966590 | -3.019 | 1.071 | 3.033 | 2.435 | -0.008 | 0.003 | 0.003 | 5.547 | 0.15 | 0.81 | 0.1 | 0.18 | 0.07 | 0.25 |
| 5 | 24789752 | rs10070741 | -1.475 | 1.362 | 2.235 | 2.576 | 0.065 | 0.075 | 0.075 | 0.581 | 0.08 | 0.34 | 0.06 | 0.11 | 0.08 | 0.17 |
| 5 | 24791039 | rs1835228 | 0.715 | 1.426 | 1.3 | 2.523 | 0.011 | 0.041 | 0.041 | 0 | 0.87 | 0.86 | 0.69 | 0.62 | 0.83 | 0.85 |
| 5 | 24794619 | rs7737195 | -1.879 | 0.597 | 1.465 | 1.698 | 0.043 | 0.151 | 0.151 | 0.92 | 0.11 | 0.5 | 0.08 | 0.18 | 0.09 | 0.24 |
| 5 | 24805795 | rs1493245 | -1.647 | 0.14 | 1.346 | 1.669 | -0.033 | 0.152 | 0.152 | 2.107 | 0.12 | 0.5 | 0.09 | 0.19 | 0.04 | 0.11 |
| 5 | 24806593 | rs1169654 | -0.219 | -0.306 | 0.299 | 1.629 | 0.001 | -0.038 | -0.038 | 0 | 0.74 | 0.88 | 0.69 | 0.73 | 0.76 | 0.83 |
| 5 | 24806889 | rs1169653 | 1.219 | -0.619 | 0.153 | 1.216 | 0.073 | 0.011 | 0.011 | 0 | 0.52 | 0.17 | 0.53 | 0.5 | 0.65 | 0.64 |
| 5 | 24809481 | rs1183308 | 0.858 | -0.649 | 0.353 | 1.193 | 0.045 | -0.012 | -0.012 | 1.222 | 0.57 | 0.26 | 0.57 | 0.54 | 0.7 | 0.69 |
| 5 | 24812268 | rs1908045 | 0.036 | -0.771 | 0.03 | 0.773 | 0.011 | 0.014 | 0.014 | 0.058 | 0.64 | 0.75 | 0.64 | 0.74 | 0.68 | 0.75 |
| 5 | 24817237 | rs12522615 | -0.063 | -0.814 | 1.234 | 0.782 | -0.019 | 0.11 | 0.11 | 0.292 | 0.09 | 0.28 | 0.07 | 0.19 | 0.04 | 0.1 |
| 5 | 24817630 | rs16894002 | -0.458 | -0.81 | 0.991 | 0.784 | -0.019 | 0.181 | 0.181 | 1.547 | 0.11 | 0.47 | 0.09 | 0.19 | 0.04 | 0.1 |
| 5 | 24822128 | rs1960462 | -1.247 | -0.881 | 0.597 | 0.762 | 0.012 | -0.014 | -0.014 | 0 | 0.54 | 0.89 | 0.48 | 0.49 | 0.39 | 0.38 |
| 5 | 24833154 | rs4421074 | -0.631 | -0.583 | 0.551 | 0.633 | -0.004 | -0.036 | -0.036 | 0 | 0.08 | 0.16 | 0.04 | 0 | 0.06 | 0.05 |
| 5 | 24837899 | rs6881214 | -0.059 | -0.74 | 0.396 | 0.613 | 0.01 | 0.003 | 0.003 | 0.098 | 0.64 | 0.73 | 0.72 | 0.7 | 0.68 | 0.79 |
| 5 | 24846954 | rs10071824 | 0.318 | -0.603 | -0.093 | 0.558 | 0.004 | 0 | 0 | 0 | 0.61 | 0.32 | 0.48 | 0.62 | 0.63 | 0.59 |
| 5 | 24852377 | rs10059367 | 0.426 | -0.835 | -0.138 | 0.743 | -0.03 | 0.063 | 0.063 | 0.143 | 0.77 | 0.38 | 0.75 | 0.44 | 0.82 | 0.62 |
| 5 | 24856119 | rs13162819 | -0.513 | 0.391 | -1.077 | -0.065 | -0.004 | 0.052 | 0.052 | 0 | 0.4 | 0.7 | 0.47 | 0.2 | 0.41 | 0.24 |
| 5 | 24859690 | rs10055390 | 0.353 | -0.17 | -1.161 | 0.344 | 0.086 | 0.006 | 0.006 | 0.846 | 0.45 | 0.18 | 0.37 | 0.63 | 0.28 | 0.48 |

| | | | | | | | | | | | | | | | | |
|---|---|---|---|---|---|---|---|---|---|---|---|---|---|---|---|---|
| 5 | 24868444 | rs962087 | 0.422 | 0.581 | -0.235 | 0.323 | 0.012 | 0.005 | 0.005 | 0.218 | 0.86 | 0.78 | 0.88 | 0.87 | 0.84 | 0.95 |
| 5 | 24871031 | rs1353061 | -1.101 | 0.661 | -0.253 | 0.237 | 0.012 | 0.015 | 0.015 | 0 | 0.5 | 0.63 | 0.44 | 0.41 | 0.45 | 0.51 |
| 5 | 24879885 | rs17400935 | 0.048 | 0.911 | 0.161 | 0.386 | 0.036 | -0.001 | -0.001 | 0 | 0.08 | 0.15 | 0.16 | 0.08 | 0.16 | 0.16 |
| 5 | 24884260 | rs1389221 | -0.581 | 0.334 | 0.328 | 0.222 | 0 | 0 | 0 | 0 | 0.27 | 0.25 | 0.16 | 0.31 | 0.19 | 0.25 |
| 5 | 24902336 | rs10042912 | -0.779 | 0.246 | 0.577 | -0.41 | -0.004 | 0.012 | 0.012 | 0 | 0.27 | 0.26 | 0.26 | 0.1 | 0.34 | 0.28 |
| 5 | 24907439 | rs1500198 | 0.94 | -0.082 | 0.93 | -0.081 | -0.013 | 0.003 | 0.003 | 0.382 | 0.68 | 0.42 | 0.78 | 0.67 | 0.74 | 0.76 |
| 5 | 24913167 | rs9283778 | 0.659 | -0.154 | 1.08 | -0.055 | -0.012 | 0.004 | 0.004 | 0.02 | 0.75 | 0.52 | 0.81 | 0.76 | 0.79 | 0.83 |
| 5 | 24916565 | rs7720640 | -0.738 | -0.093 | 0.658 | -0.039 | 0 | -0.001 | -0.001 | 0 | 0.11 | 0.06 | 0.08 | 0.1 | 0.13 | 0.09 |
| 5 | 24917723 | rs16894041 | -0.126 | -0.127 | 0.775 | 0.015 | -0.003 | 0 | 0 | 0 | 0.07 | 0.08 | 0.03 | 0.09 | 0.05 | 0.07 |
| 5 | 24919141 | rs904746 | 0.053 | -0.146 | 1.08 | 0.046 | -0.007 | 0.011 | 0.011 | 0.391 | 0.83 | 0.81 | 0.89 | 0.76 | 0.87 | 0.85 |
| 5 | 24922595 | rs7711040 | -1.073 | -0.06 | 1.293 | -0.328 | -0.001 | -0.016 | -0.016 | 0 | 0.22 | 0.34 | 0.11 | 0.18 | 0.22 | 0.3 |
| 5 | 24922789 | rs6452234 | -0.295 | -0.174 | 1.311 | -0.344 | 0.021 | -0.046 | -0.046 | 0 | 0.17 | 0.32 | 0.1 | 0.08 | 0.17 | 0.24 |
| 5 | 24924797 | rs2055073 | -2.361 | -0.244 | 1.631 | -0.277 | 0 | 0.003 | 0.003 | 0 | 0.42 | 0.58 | 0.33 | 0.39 | 0.4 | 0.45 |
| 5 | 24926647 | rs17401648 | -0.324 | -0.247 | 1.699 | -0.278 | -0.002 | 0 | 0 | 0.633 | 0.19 | 0.12 | 0.23 | 0.2 | 0.2 | 0.16 |
| 5 | 24930124 | rs6452236 | -2.388 | -0.259 | 1.598 | -0.285 | 0 | 0.003 | 0.003 | 0 | 0.42 | 0.58 | 0.34 | 0.38 | 0.4 | 0.42 |
| 5 | 24930711 | rs10520903 | -0.026 | -0.184 | 1.038 | -0.363 | -0.04 | 0.045 | 0.045 | 0 | 0.15 | 0.06 | 0.08 | 0.08 | 0.17 | 0.21 |
| 5 | 24934186 | rs10805771 | -0.723 | -0.533 | 1.221 | -0.224 | -0.003 | -0.012 | -0.012 | 0 | 0.19 | 0.11 | 0.23 | 0.26 | 0.2 | 0.18 |
| 5 | 24946851 | rs12187422 | 0.005 | -0.464 | 1.158 | -0.181 | -0.003 | -0.022 | -0.022 | 0.509 | 0.1 | 0.01 | 0.06 | 0.13 | 0.09 | 0.07 |
| 5 | 24948884 | rs11738003 | -1.624 | -0.394 | 1.158 | -0.375 | -0.003 | 0.028 | 0.028 | 0 | 0.28 | 0.16 | 0.25 | 0.21 | 0.32 | 0.36 |
| 5 | 24967119 | rs2201296 | -0.39 | -0.278 | 1.11 | -0.321 | 0.002 | -0.001 | -0.001 | 0 | 0.13 | 0.12 | 0.17 | 0.12 | 0.17 | 0.13 |
| 5 | 24967172 | rs2201295 | -2.246 | -0.464 | 0.961 | -0.435 | 0.011 | 0.039 | 0.039 | 0.218 | 0.41 | 0.19 | 0.29 | 0.34 | 0.42 | 0.44 |
| 5 | 24976492 | rs13174080 | -2.279 | -0.455 | 1.193 | -0.194 | 0.01 | 0.018 | 0.018 | 0.438 | 0.41 | 0.26 | 0.3 | 0.36 | 0.42 | 0.44 |
| 5 | 25006350 | rs1493478 | -1.536 | -0.454 | 0.872 | -0.198 | 0.005 | -0.019 | -0.019 | 4.838 | 0.28 | 0.06 | 0.12 | 0.26 | 0.25 | 0.31 |
| 5 | 25011524 | rs9784649 | -0.126 | -0.414 | 0.96 | -0.136 | -0.003 | 0.002 | 0.002 | 0 | 0.12 | 0.08 | 0.16 | 0.1 | 0.17 | 0.13 |
| 5 | 25015565 | rs6869061 | -2.359 | -0.436 | 1.438 | -0.153 | 0.005 | 0.008 | 0.008 | 0.608 | 0.42 | 0.4 | 0.3 | 0.37 | 0.42 | 0.43 |
| 5 | 25027051 | rs6866107 | -1.753 | -0.778 | 1.165 | -0.382 | 0.016 | 0.007 | 0.007 | 0.329 | 0.37 | 0.34 | 0.25 | 0.34 | 0.26 | 0.39 |
| 5 | 25027618 | rs17343615 | -0.791 | -0.812 | 0.791 | -0.34 | -0.007 | 0.004 | 0.004 | 0 | 0.09 | 0.06 | 0.06 | 0 | 0.14 | 0.08 |
| 5 | 25030161 | rs2873911 | 2.692 | -0.461 | 0.275 | -0.768 | -0.106 | -0.039 | -0.039 | 0.639 | 0.36 | 0.05 | 0.5 | 0.61 | 0.45 | 0.34 |
| 5 | 25050413 | rs10053798 | -0.964 | -0.527 | 1.194 | -0.989 | 0.005 | 0.001 | 0.001 | 0.203 | 0.3 | 0.26 | 0.13 | 0.28 | 0.16 | 0.3 |
| 5 | 25053641 | rs11742222 | -0.837 | -0.596 | 0.931 | -1.075 | -0.068 | -0.027 | -0.027 | 0 | 0.24 | 0.05 | 0.21 | 0.22 | 0.29 | 0.34 |
| 5 | 25054004 | rs17403440 | 0.17 | -0.387 | 0.952 | -1.055 | -0.056 | 0.038 | 0.038 | 0 | 0.12 | 0.01 | 0.1 | 0.03 | 0.19 | 0.13 |
| 5 | 25058616 | rs16894132 | -1.95 | -0.416 | 1.204 | -1.052 | 0.009 | 0.058 | 0.058 | 1.563 | 0.5 | 0.34 | 0.32 | 0.33 | 0.43 | 0.58 |
| 5 | 25060782 | rs1160303 | 3.172 | 0.038 | 0.834 | -1.165 | -0.027 | -0.003 | -0.003 | 0.879 | 0.36 | 0.12 | 0.57 | 0.67 | 0.4 | 0.32 |
| 5 | 25065470 | rs9293162 | 1.434 | 0.045 | 1.029 | -1.232 | 0 | -0.008 | -0.008 | 0 | 0.08 | 0.04 | 0.03 | 0.02 | 0.04 | 0.01 |
| 5 | 25067372 | rs10520899 | -0.244 | 0.089 | 1.233 | -1.205 | -0.001 | -0.013 | -0.013 | 0.007 | 0.09 | 0.08 | 0.07 | 0 | 0.17 | 0.1 |
| 5 | 25086440 | rs6872919 | 1.325 | 0.078 | 1.638 | -1.237 | 0.05 | -0.117 | -0.117 | 0 | 0.69 | 0.48 | 0.72 | 0.95 | 0.58 | 0.62 |
| 5 | 25089614 | rs7733733 | -1.055 | 0.065 | 1.649 | -1.139 | 0.059 | -0.067 | -0.067 | 0 | 0.11 | 0.33 | 0.1 | 0 | 0.17 | 0.1 |
| 5 | 25090716 | rs7701392 | -1.284 | -0.07 | 1.263 | -1.112 | 0.025 | 0.044 | 0.044 | 2.158 | 0.39 | 0.16 | 0.19 | 0.31 | 0.31 | 0.38 |
| 5 | 25095178 | rs12521551 | 1.768 | 0.043 | 1.612 | -0.946 | 0.015 | 0.038 | 0.038 | 4.62 | 0.56 | 0.31 | 0.7 | 0.7 | 0.65 | 0.46 |
| 5 | 25096593 | rs4701466 | -0.306 | 0.075 | 1.812 | -0.97 | 0.054 | -0.03 | -0.03 | 1.085 | 0.89 | 0.59 | 0.87 | 0.98 | 0.92 | 0.83 |
| 5 | 25097441 | rs10942089 | -1.075 | 0.037 | 1.835 | -0.609 | 0.015 | -0.01 | -0.01 | 0 | 0.33 | 0.26 | 0.27 | 0.34 | 0.27 | 0.37 |
| 5 | 25097916 | rs11746102 | 1.837 | 0.041 | 1.819 | -0.607 | 0.018 | 0.014 | 0.014 | 4.651 | 0.55 | 0.29 | 0.6 | 0.64 | 0.65 | 0.46 |
| 5 | 25120430 | rs10065343 | -0.169 | 0.089 | 1.833 | -0.585 | 0.055 | -0.035 | -0.035 | 0 | 0.88 | 0.57 | 0.87 | 0.98 | 0.91 | 0.83 |
| 5 | 25128890 | rs11742952 | 2.756 | 0.092 | 1.439 | -0.576 | -0.033 | 0.076 | 0.076 | 6.286 | 0.46 | 0.13 | 0.45 | 0.75 | 0.49 | 0.37 |
| 5 | 25130699 | rs12518815 | -1.734 | 0.231 | 0.986 | -0.549 | 0.007 | 0.007 | 0.007 | 2.788 | 0.4 | 0.32 | 0.32 | 0.35 | 0.39 | 0.46 |
| 5 | 25135494 | rs6859537 | -0.195 | 0.057 | 0.894 | -0.496 | 0.052 | -0.055 | -0.055 | 2.501 | 0.2 | 0.09 | 0.17 | 0.31 | 0.09 | 0.27 |
| 5 | 25143399 | rs13181240 | -1.503 | 0.151 | 0.943 | -0.51 | 0.014 | -0.005 | -0.005 | 0 | 0.35 | 0.27 | 0.29 | 0.35 | 0.3 | 0.37 |
| 5 | 25146653 | rs4701473 | -0.343 | 0.103 | 0.526 | -0.43 | 0.051 | -0.056 | -0.056 | 0 | 0.22 | 0.09 | 0.18 | 0.31 | 0.11 | 0.28 |
| 5 | 25164780 | rs414435 | 0.417 | 0.04 | 0.971 | -0.535 | 0 | -0.045 | -0.045 | 0 | 0.85 | 0.75 | 0.91 | 0.97 | 0.85 | 0.86 |
| 5 | 25180973 | rs6890226 | -0.186 | 0.079 | 0.872 | -0.497 | 0.024 | -0.058 | -0.058 | 0.099 | 0.21 | 0.1 | 0.3 | 0.33 | 0.15 | 0.28 |
| 5 | 25184490 | rs11955750 | 2.71 | 0.138 | 1.196 | -0.628 | -0.01 | 0.019 | 0.019 | 4.758 | 0.46 | 0.21 | 0.46 | 0.62 | 0.48 | 0.35 |
| 5 | 25190282 | rs7732673 | 0.411 | 0.124 | 1.49 | -0.646 | 0.046 | -0.067 | -0.067 | 0 | 0.83 | 0.7 | 0.84 | 0.98 | 0.77 | 0.77 |
| 5 | 25228227 | rs3913960 | 1.533 | 0.136 | 1.309 | -0.613 | 0 | -0.004 | -0.004 | 0 | 0.61 | 0.55 | 0.54 | 0.67 | 0.63 | 0.49 |
| 5 | 25248251 | rs6860994 | -0.22 | 0.248 | 1.059 | -0.504 | 0.02 | -0.031 | -0.031 | 0 | 0.21 | 0.13 | 0.3 | 0.31 | 0.15 | 0.27 |
| 5 | 25278588 | rs3843471 | 0.288 | 0.353 | 0.997 | -0.492 | 0.025 | -0.027 | -0.027 | 0.016 | 0.07 | 0.1 | 0.07 | 0 | 0.16 | 0.09 |
| 5 | 25302983 | rs2063184 | -0.551 | -0.103 | 0.807 | -0.389 | 0.035 | -0.037 | -0.037 | 0 | 0.23 | 0.1 | 0.3 | 0.31 | 0.15 | 0.27 |
| 5 | 25315404 | rs13163964 | 1.63 | 0.068 | 0.83 | -0.409 | 0.028 | 0.031 | 0.031 | 4.767 | 0.54 | 0.25 | 0.51 | 0.64 | 0.64 | 0.44 |
| 5 | 25329172 | rs6890531 | 1.172 | -0.138 | 0.914 | -0.556 | -0.036 | -0.011 | -0.011 | 0 | 0.64 | 0.49 | 0.58 | 0.67 | 0.78 | 0.55 |
| 5 | 25329322 | rs7730066 | 0.091 | 0.538 | 1.179 | -0.431 | 0.011 | -0.04 | -0.04 | 0 | 0.1 | 0.22 | 0.1 | 0.02 | 0.07 | 0.16 |
| 5 | 25333696 | rs17523194 | -0.296 | 0.494 | 0.586 | -0.211 | 0.013 | -0.016 | -0.016 | 0.824 | 0.19 | 0.07 | 0.2 | 0.23 | 0.14 | 0.21 |
| 5 | 25334013 | rs1032987 | 0.959 | 0.561 | 0.7 | -0.269 | 0.01 | -0.093 | -0.093 | 0.711 | 0.76 | 0.37 | 0.8 | 0.94 | 0.78 | 0.72 |
| 5 | 25340617 | rs11738869 | -0.459 | 0.608 | 0.289 | -0.072 | 0.01 | -0.033 | -0.033 | 0.087 | 0.19 | 0.04 | 0.2 | 0.23 | 0.14 | 0.21 |
| 5 | 25351609 | rs4235532 | 1.55 | 1 | 0.228 | -0.141 | -0.014 | -0.002 | -0.002 | 0 | 0.51 | 0.18 | 0.38 | 0.57 | 0.48 | 0.5 |
| 5 | 25355836 | rs4701480 | 0.574 | 0.942 | 0.238 | 0.056 | -0.018 | -0.064 | -0.064 | 0.244 | 0.12 | 0.03 | 0.25 | 0.31 | 0.14 | 0.11 |
| 5 | 25384728 | rs12517406 | 0.002 | 0.962 | 0.601 | -0.706 | 0.006 | -0.029 | -0.029 | 1.261 | 0.9 | 0.85 | 0.91 | 1 | 0.84 | 0.91 |
| 5 | 25386522 | rs10462265 | 2.412 | 1.207 | 0.621 | -0.918 | 0.014 | -0.003 | -0.003 | 0.011 | 0.37 | 0.23 | 0.32 | 0.68 | 0.36 | 0.38 |
| 5 | 25386907 | rs10073102 | -1.726 | 1.204 | 0.615 | -0.912 | 0.001 | 0.017 | 0.017 | 0.298 | 0.53 | 0.62 | 0.59 | 0.32 | 0.47 | 0.54 |
| 5 | 25390490 | rs7349777 | -1.265 | 1.224 | 0.954 | -0.96 | -0.018 | 0.02 | 0.02 | 0.668 | 0.38 | 0.27 | 0.37 | 0.26 | 0.38 | 0.38 |
| 5 | 25392632 | rs10473744 | -0.431 | 1.231 | 0.981 | -0.989 | -0.003 | -0.01 | -0.01 | 0 | 0.9 | 0.87 | 0.92 | 1 | 0.84 | 0.91 |

| | | | | | | | | | | | | | | | | |
|---|---|---|---|---|---|---|---|---|---|---|---|---|---|---|---|---|
| 5 | 25393370 | rs2332017 | 0.152 | 1.246 | 0.833 | -1.032 | -0.021 | 0.048 | 0.048 | 0 | 0.16 | 0.08 | 0.07 | 0.04 | 0.18 | 0.24 |
| 5 | 25396006 | rs7711948 | 0.843 | 1.2 | 0.864 | -1.011 | -0.01 | -0.036 | -0.036 | 0 | 0.11 | 0.16 | 0.04 | 0.03 | 0.04 | 0.04 |
| 5 | 25400941 | rs1169845 | -0.418 | 1.223 | 0.963 | -1.017 | 0.011 | -0.039 | -0.039 | 0 | 0.9 | 0.85 | 0.92 | 1 | 0.84 | 0.91 |
| 5 | 25402825 | rs10076119 | -1.415 | 1.207 | 1.036 | -1.026 | 0.001 | -0.001 | -0.001 | 0 | 0.41 | 0.44 | 0.54 | 0.3 | 0.43 | 0.5 |
| 5 | 25403734 | rs6872370 | -0.892 | 1.197 | 1.156 | -1.035 | 0.01 | 0.003 | 0.003 | 0.788 | 0.29 | 0.39 | 0.36 | 0.24 | 0.33 | 0.36 |
| 5 | 25412743 | rs10062995 | -0.828 | 1.26 | 1.122 | -1.002 | 0.003 | -0.088 | -0.088 | 0.82 | 0.21 | 0.33 | 0.13 | 0.03 | 0.21 | 0.12 |
| 5 | 25416205 | rs2252299 | -0.621 | 1.271 | 1.049 | -0.99 | -0.002 | -0.006 | -0.006 | 0.935 | 0.22 | 0.2 | 0.27 | 0.06 | 0.26 | 0.23 |
| 5 | 25417764 | rs4092595 | 0.156 | 1.274 | 1.047 | -0.97 | 0.014 | -0.042 | -0.042 | 0 | 0.1 | 0.15 | 0.09 | 0 | 0.16 | 0.09 |
| 5 | 25419907 | rs2877553 | -2.18 | 1.269 | 1.271 | -1.028 | -0.004 | 0.127 | 0.127 | 0.049 | 0.49 | 0.68 | 0.4 | 0.14 | 0.48 | 0.5 |
| 5 | 25420089 | rs1904173 | -0.207 | 1.302 | 1.291 | -0.964 | 0.046 | -0.064 | -0.064 | 0 | 0.11 | 0.27 | 0.1 | 0 | 0.17 | 0.09 |
| 5 | 25422667 | rs12188009 | 0.96 | 1.297 | 1.036 | -0.898 | -0.032 | -0.052 | -0.052 | 0 | 0.12 | 0.06 | 0.25 | 0.2 | 0.16 | 0.11 |
| 5 | 25431823 | rs1904177 | -1.687 | 1.275 | 1.171 | -0.973 | 0.01 | -0.079 | -0.079 | 0.02 | 0.39 | 0.46 | 0.36 | 0.11 | 0.43 | 0.46 |
| 5 | 25436319 | rs12517129 | 0.686 | 1.167 | 1.065 | -0.881 | 0 | 0.006 | 0.006 | 0 | 0.13 | 0.16 | 0.27 | 0.18 | 0.16 | 0.12 |
| 5 | 25437715 | rs17524358 | 0.117 | 1.198 | 1.017 | -0.913 | -0.009 | 0.023 | 0.023 | 0 | 0.15 | 0.1 | 0.05 | 0.05 | 0.16 | 0.23 |
| 5 | 25439384 | rs6875620 | -2.698 | 1.203 | 0.791 | -0.874 | 0.007 | -0.02 | -0.02 | 0 | 0.63 | 0.84 | 0.65 | 0.34 | 0.6 | 0.68 |
| 5 | 25446757 | rs11740761 | 0.722 | 1.023 | 0.79 | -0.622 | -0.035 | -0.057 | -0.057 | 0.004 | 0.12 | 0.05 | 0.25 | 0.19 | 0.16 | 0.11 |
| 5 | 25451344 | rs17524924 | 0.696 | 1.017 | 0.901 | -0.903 | -0.004 | 0.022 | 0.022 | 0 | 0.11 | 0.02 | 0.18 | 0.07 | 0.08 | 0.21 |
| 5 | 25451819 | rs10036252 | 0.254 | 1.075 | 1.065 | -0.806 | 0.017 | -0.036 | -0.036 | 0.009 | 0.08 | 0.15 | 0.07 | 0 | 0.13 | 0.1 |
| 5 | 25457358 | rs2126965 | -0.809 | 0.916 | 1.199 | -0.552 | -0.002 | -0.041 | -0.041 | 0 | 0.28 | 0.32 | 0.26 | 0.11 | 0.25 | 0.44 |
| 5 | 25463613 | rs10520922 | -0.915 | 0.997 | 0.855 | -0.691 | -0.017 | -0.079 | -0.079 | 0 | 0.19 | 0.38 | 0.12 | 0.04 | 0.13 | 0.15 |
| 5 | 25468690 | rs10473746 | -1.64 | 1.19 | 0.306 | -0.517 | 0.006 | 0.058 | 0.058 | 0 | 0.42 | 0.7 | 0.4 | 0.2 | 0.37 | 0.44 |
| 5 | 25468811 | rs13359266 | -0.633 | 1.188 | 0.416 | -0.538 | -0.005 | -0.022 | -0.022 | 0 | 0.27 | 0.3 | 0.28 | 0.17 | 0.19 | 0.31 |
| 5 | 25470513 | rs2126967 | -1.048 | 1.191 | 0.441 | -0.54 | -0.004 | -0.067 | -0.067 | 0 | 0.32 | 0.53 | 0.33 | 0.17 | 0.28 | 0.34 |
| 5 | 25471283 | rs10214406 | 0.201 | 1.183 | 0.067 | -0.553 | -0.001 | -0.065 | -0.065 | 0 | 0.11 | 0.28 | 0.08 | 0.01 | 0.08 | 0.03 |
| 5 | 25471581 | rs16894485 | 0.524 | 1.188 | -0.154 | -0.539 | -0.002 | -0.019 | -0.019 | 0 | 0.1 | 0.16 | 0.06 | 0.03 | 0.09 | 0.1 |
| 5 | 25471838 | rs1366277 | 0.661 | 1.227 | -0.827 | -0.472 | -0.002 | 0.008 | 0.008 | 0 | 0.16 | 0.03 | 0.2 | 0.16 | 0.11 | 0.28 |
| 5 | 25471919 | rs12186488 | -0.112 | 1.516 | -0.794 | -0.147 | -0.03 | -0.063 | -0.063 | 0.068 | 0.16 | 0.04 | 0.24 | 0.22 | 0.16 | 0.15 |
| 5 | 25488352 | rs7716893 | -1.519 | 1.592 | -0.754 | -0.136 | -0.01 | -0.005 | -0.005 | 1.298 | 0.56 | 0.85 | 0.65 | 0.42 | 0.53 | 0.59 |
| 5 | 25489061 | rs2086142 | -1.089 | 0.94 | -0.285 | -0.713 | 0.025 | 0.077 | 0.077 | 0 | 0.4 | 0.75 | 0.37 | 0.19 | 0.29 | 0.41 |
| 5 | 25495005 | rs10078091 | -0.604 | 0.968 | -0.339 | -0.086 | -0.004 | -0.061 | -0.061 | 1.21 | 0.31 | 0.47 | 0.32 | 0.17 | 0.29 | 0.42 |
| 5 | 25504896 | rs10058099 | 0.58 | 1.459 | -0.66 | 0.55 | -0.005 | -0.003 | -0.003 | 0 | 0.7 | 0.77 | 0.72 | 0.66 | 0.68 | 0.66 |
| 5 | 25513207 | rs9293168 | -1.004 | 1.721 | -0.566 | 0.127 | -0.005 | -0.001 | -0.001 | 0 | 0.09 | 0.08 | 0.1 | 0.02 | 0.12 | 0.15 |
| 5 | 25515935 | rs7705483 | -0.198 | 1.253 | -0.608 | -0.109 | -0.005 | 0.002 | 0.002 | 0 | 0.61 | 0.66 | 0.48 | 0.33 | 0.52 | 0.61 |
| 5 | 25518458 | rs7720944 | 0.828 | 0.686 | -0.896 | -0.531 | 0.008 | 0.014 | 0.014 | 0 | 0.1 | 0.04 | 0.05 | 0 | 0.06 | 0.03 |
| 5 | 25520961 | rs7712799 | 0.089 | 0.852 | -0.688 | -0.486 | 0.003 | 0.011 | 0.011 | 0 | 0.51 | 0.56 | 0.43 | 0.33 | 0.48 | 0.59 |
| 5 | 25523961 | rs1428238 | 1.124 | -0.159 | -0.859 | -0.917 | 0.01 | 0.012 | 0.012 | 0.435 | 0.28 | 0.21 | 0.13 | 0.13 | 0.22 | 0.2 |
| 5 | 25532432 | rs10052032 | 0.12 | -0.476 | -0.773 | -0.769 | 0.007 | 0.02 | 0.02 | 0 | 0.43 | 0.32 | 0.36 | 0.31 | 0.38 | 0.49 |
| 5 | 25537415 | rs4701487 | 0.544 | -0.467 | -0.859 | -0.746 | 0.003 | -0.011 | -0.011 | 0 | 0.06 | 0.06 | 0.16 | 0.17 | 0.06 | 0.15 |
| 5 | 25538295 | rs13181304 | 0.849 | -0.605 | -0.758 | -0.831 | -0.019 | 0.094 | 0.094 | 0.104 | 0.33 | 0.07 | 0.18 | 0.13 | 0.31 | 0.32 |
| 5 | 25538947 | rs12515108 | 0.587 | -0.585 | -0.755 | -0.813 | 0.003 | -0.011 | -0.011 | 0 | 0.06 | 0.06 | 0.16 | 0.17 | 0.06 | 0.15 |
| 5 | 25540646 | rs13154702 | 0.621 | -0.544 | -0.709 | -0.86 | 0.002 | 0.039 | 0.039 | 0.004 | 0.35 | 0.21 | 0.19 | 0.13 | 0.31 | 0.32 |
| 5 | 25557717 | rs13169085 | -0.989 | -0.548 | -0.435 | -0.881 | 0.01 | -0.013 | -0.013 | 0 | 0.6 | 0.5 | 0.46 | 0.27 | 0.51 | 0.62 |
| 5 | 25558308 | rs13166065 | 0.692 | -0.701 | -0.482 | -0.979 | 0.002 | 0.07 | 0.07 | 0 | 0.36 | 0.19 | 0.19 | 0.08 | 0.32 | 0.33 |
| 5 | 25561508 | rs13161787 | -1.049 | -0.65 | -0.523 | -0.884 | 0 | 0 | 0 | 0.004 | 0.61 | 0.64 | 0.48 | 0.27 | 0.52 | 0.62 |
| 5 | 25572301 | rs6876547 | -1.206 | -0.687 | -0.368 | -0.761 | -0.054 | -0.036 | -0.036 | 0 | 0.25 | 0.6 | 0.3 | 0.2 | 0.2 | 0.3 |
| 5 | 25572996 | rs1542563 | -0.946 | -0.834 | -0.833 | -0.935 | 0.007 | -0.007 | -0.007 | 0 | 0.61 | 0.61 | 0.49 | 0.27 | 0.52 | 0.65 |
| 5 | 25573190 | rs11953292 | 0.319 | -0.97 | -0.928 | -0.757 | -0.018 | 0.002 | 0.002 | 0 | 0.4 | 0.07 | 0.35 | 0.29 | 0.39 | 0.49 |
| 5 | 25573209 | rs11952604 | -0.236 | -0.934 | -0.843 | -0.774 | 0.005 | 0.009 | 0.009 | 0 | 0.48 | 0.45 | 0.42 | 0.3 | 0.44 | 0.55 |
| 5 | 25576239 | rs13158186 | 1.012 | -0.983 | -1.009 | -1.08 | 0.024 | 0.043 | 0.043 | 0 | 0.3 | 0.17 | 0.17 | 0.08 | 0.19 | 0.24 |
| 5 | 25582936 | rs2127392 | -0.245 | -0.505 | -0.793 | -0.85 | -0.007 | -0.038 | -0.038 | 0.338 | 0.14 | 0.15 | 0.09 | 0.01 | 0.1 | 0.08 |
| 5 | 25595051 | rs10056747 | -1.458 | -0.532 | -0.789 | -0.61 | -0.002 | -0.036 | -0.036 | 0 | 0.21 | 0.32 | 0.13 | 0.08 | 0.2 | 0.17 |
| 5 | 25608727 | rs13159537 | -1.245 | -0.324 | -0.773 | -0.592 | 0.009 | -0.02 | -0.02 | 0 | 0.07 | 0.16 | 0.21 | 0.05 | 0.1 | 0.07 |
| 5 | 25611482 | rs36997 | -0.962 | -0.137 | -1.158 | -0.381 | 0 | 0.06 | 0.06 | 0 | 0.73 | 0.62 | 0.71 | 0.52 | 0.76 | 0.68 |
| 5 | 25612336 | rs16894618 | 0.485 | -0.344 | -1.724 | -0.638 | -0.064 | -0.087 | -0.087 | 0 | 0.17 | 0.07 | 0.23 | 0.39 | 0.32 | 0.25 |
| 5 | 25615591 | rs2331930 | -0.575 | -0.379 | -1.649 | -0.768 | 0.008 | -0.022 | -0.022 | 3.12 | 0.08 | 0.18 | 0.21 | 0.05 | 0.1 | 0.07 |
| 5 | 25616251 | rs2619780 | -1.015 | -0.404 | -1.596 | -0.645 | -0.004 | 0.082 | 0.082 | 0 | 0.72 | 0.53 | 0.7 | 0.5 | 0.76 | 0.68 |
| 5 | 25618766 | rs12520396 | 1.801 | -0.446 | -1.926 | -0.849 | 0.019 | 0.096 | 0.096 | 0 | 0.28 | 0.02 | 0.15 | 0.07 | 0.19 | 0.24 |
| 5 | 25623001 | rs10037787 | -0.159 | -0.474 | -1.708 | -0.74 | 0 | -0.021 | -0.021 | 0 | 0.59 | 0.38 | 0.47 | 0.45 | 0.63 | 0.56 |
| 5 | 25632475 | rs10942131 | 0.79 | -0.379 | -1.684 | -0.407 | -0.002 | 0.019 | 0.019 | 0 | 0.08 | 0.12 | 0.06 | 0.25 | 0.07 | 0.18 |
| 5 | 25634038 | rs11956371 | -1.15 | -0.399 | -1.509 | -0.508 | 0 | 0.071 | 0.071 | 0 | 0.73 | 0.61 | 0.7 | 0.52 | 0.76 | 0.67 |
| 5 | 25636829 | rs10462273 | 0.701 | 0.355 | -1.47 | -0.624 | -0.034 | -0.065 | -0.065 | 0 | 0.1 | 0.01 | 0.08 | 0.25 | 0.14 | 0.24 |
| 5 | 25636848 | rs13162889 | 0.166 | -0.179 | -1.713 | -0.919 | -0.003 | 0.062 | 0.062 | 0 | 0.44 | 0.26 | 0.39 | 0.26 | 0.56 | 0.39 |
| 5 | 25643355 | rs7733598 | -1.124 | -0.209 | -1.48 | -0.845 | -0.015 | 0.018 | 0.018 | 0 | 0.64 | 0.5 | 0.51 | 0.51 | 0.72 | 0.67 |
| 5 | 25644220 | rs7734006 | -0.381 | 0.032 | -1.635 | -0.719 | 0.042 | -0.012 | -0.012 | 0 | 0.56 | 0.36 | 0.48 | 0.5 | 0.69 | 0.63 |
| 5 | 25646567 | rs1479581 | 0.363 | 0.128 | -1.716 | -0.584 | -0.03 | -0.043 | -0.043 | 0.062 | 0.22 | 0.11 | 0.17 | 0.25 | 0.24 | 0.27 |
| 5 | 25650030 | rs12187966 | 1.259 | 0.22 | -1.734 | -0.604 | 0.009 | 0.003 | 0.003 | 0 | 0.11 | 0.11 | 0.15 | 0.17 | 0.22 | 0.08 |
| 5 | 25650176 | rs7723935 | 0.317 | 0.086 | -1.514 | -0.646 | -0.028 | -0.047 | -0.047 | 0 | 0.21 | 0.1 | 0.16 | 0.25 | 0.24 | 0.27 |
| 5 | 25658882 | rs10076963 | 1.231 | 0.199 | -1.317 | -0.7 | 0.005 | -0.029 | -0.029 | 0.011 | 0.26 | 0.27 | 0.34 | 0.09 | 0.28 | 0.32 |
| 5 | 25668834 | rs949661 | 0.246 | 0.421 | -0.94 | -0.273 | 0.008 | 0.028 | 0.028 | 0 | 0.5 | 0.43 | 0.52 | 0.36 | 0.56 | 0.6 |

| | | | | | | | | | | | | | | | | |
|---|---|---|---|---|---|---|---|---|---|---|---|---|---|---|---|---|
| 5 | 25673651 | rs248755 | -0.666 | 0.245 | -0.71 | 0.048 | -0.02 | 0.047 | 0.047 | 0 | 0.72 | 0.58 | 0.77 | 0.55 | 0.78 | 0.74 |
| 5 | 25676606 | rs1479571 | 0.688 | 0.38 | -0.698 | -0.142 | 0.068 | 0.087 | 0.087 | 0.178 | 0.45 | 0.24 | 0.24 | 0.18 | 0.29 | 0.41 |
| 5 | 25678072 | rs9293175 | 0.524 | 0.583 | -0.834 | 0.165 | -0.006 | -0.026 | -0.026 | 0 | 0.1 | 0.1 | 0.27 | 0.24 | 0.12 | 0.19 |
| 5 | 25679253 | rs387456 | -0.136 | 0.901 | -0.186 | 0.045 | 0 | 0 | 0 | 0.143 | 0.58 | 0.55 | 0.53 | 0.43 | 0.48 | 0.65 |
| 5 | 25680392 | rs954478 | 0.627 | 0.694 | -0.312 | -0.04 | 0.058 | 0.005 | 0.005 | 0 | 0.44 | 0.27 | 0.46 | 0.37 | 0.34 | 0.49 |
| 5 | 25680417 | rs956628 | 1.681 | 0.521 | -0.452 | -0.17 | 0.043 | 0.063 | 0.063 | 0 | 0.29 | 0.04 | 0.16 | 0.14 | 0.21 | 0.29 |
| 5 | 25686469 | rs1479572 | -0.977 | 1.421 | -0.061 | 0.017 | 0 | -0.004 | -0.004 | 0 | 0.73 | 0.77 | 0.71 | 0.62 | 0.75 | 0.79 |
| 5 | 25686495 | rs10069367 | 0.472 | 1.65 | 0.1 | 0.037 | -0.002 | -0.004 | -0.004 | 0.255 | 0.12 | 0.17 | 0.14 | 0.18 | 0.25 | 0.1 |
| 5 | 25687683 | rs187463 | -0.564 | 0.941 | 0.124 | -0.171 | 0.026 | 0.008 | 0.008 | 0 | 0.69 | 0.57 | 0.68 | 0.62 | 0.66 | 0.68 |
| 5 | 25688567 | rs173585 | 0.282 | 0.949 | -0.369 | -0.117 | -0.003 | 0.021 | 0.021 | 0 | 0.55 | 0.41 | 0.62 | 0.57 | 0.59 | 0.59 |
| 5 | 25695377 | rs10056274 | 0.632 | 0.509 | -0.63 | 0.023 | 0.005 | 0.007 | 0.007 | 0 | 0.06 | 0.13 | 0.05 | 0.09 | 0.07 | 0.04 |
| 5 | 25698030 | rs1678761 | -0.984 | 0.374 | -0.487 | -0.069 | -0.027 | 0.027 | 0.027 | 0 | 0.76 | 0.68 | 0.72 | 0.61 | 0.83 | 0.79 |
| 5 | 25699825 | rs9293177 | 0.895 | 0.334 | -1.002 | -0.013 | -0.009 | -0.025 | -0.025 | 0 | 0.12 | 0.14 | 0.04 | 0.03 | 0.06 | 0.04 |
| 5 | 25700070 | rs10223141 | 0.521 | 0.507 | -1.573 | 0.081 | 0.005 | 0.007 | 0.007 | 0 | 0.07 | 0.14 | 0.05 | 0.09 | 0.07 | 0.04 |
| 5 | 25700355 | rs10520920 | 0.275 | 0.491 | -2.244 | -0.075 | -0.035 | -0.034 | -0.034 | 0 | 0.22 | 0.17 | 0.4 | 0.32 | 0.36 | 0.26 |
| 5 | 25711728 | rs2928179 | 1.38 | -0.167 | -2.098 | -0.007 | 0.033 | 0.035 | 0.035 | 0.052 | 0.28 | 0.06 | 0.18 | 0.15 | 0.22 | 0.29 |
| 5 | 25717110 | rs39595 | 0.758 | -0.537 | -1.37 | 0.095 | 0.05 | -0.016 | -0.016 | 0 | 0.53 | 0.24 | 0.6 | 0.46 | 0.64 | 0.58 |
| 5 | 25717662 | rs2972323 | 0.46 | 0.177 | -0.659 | 0.723 | 0.006 | -0.02 | -0.02 | 0.739 | 0.66 | 0.26 | 0.65 | 0.48 | 0.72 | 0.63 |
| 5 | 25739983 | rs168843 | -0.979 | 0.443 | 0.887 | 0.963 | 0.028 | 0.159 | 0.159 | 0 | 0.2 | 0.41 | 0.37 | 0.5 | 0.24 | 0.3 |
| 5 | 25756561 | rs1904458 | 0.944 | 0.079 | -0.179 | 0.281 | 0.046 | 0.079 | 0.079 | 0 | 0.47 | 0.21 | 0.4 | 0.33 | 0.39 | 0.48 |
| 5 | 25757341 | rs4435838 | 0.213 | -0.706 | -0.385 | -0.114 | -0.054 | 0.061 | 0.061 | 0 | 0.33 | 0.12 | 0.24 | 0.18 | 0.41 | 0.27 |
| 5 | 25759320 | rs2972297 | -0.523 | -0.836 | 0.143 | -0.079 | -0.001 | -0.03 | -0.03 | 0 | 0.24 | 0.2 | 0.35 | 0.5 | 0.25 | 0.35 |
| 5 | 25760679 | rs7727656 | 0.886 | -0.823 | 0.061 | -0.073 | -0.006 | 0.023 | 0.023 | 0 | 0.71 | 0.29 | 0.6 | 0.5 | 0.75 | 0.65 |
| 5 | 25768871 | rs12515469 | 1.378 | -0.679 | -0.114 | 0.186 | -0.023 | 0.057 | 0.057 | 0 | 0.43 | 0.05 | 0.41 | 0.31 | 0.4 | 0.38 |
| 5 | 25770271 | rs969400 | -0.051 | -0.703 | 0.023 | 0.232 | 0.003 | 0.001 | 0.001 | 0 | 0.15 | 0.12 | 0.27 | 0.51 | 0.14 | 0.27 |
| 5 | 25774428 | rs1479674 | 0.134 | -0.628 | 0.612 | 0.174 | 0.011 | -0.051 | -0.051 | 0 | 0.62 | 0.45 | 0.71 | 0.83 | 0.59 | 0.72 |
| 5 | 25777784 | rs7736846 | -1.115 | -0.687 | 1 | 0.254 | -0.013 | 0.139 | 0.139 | 0 | 0.21 | 0.47 | 0.32 | 0.53 | 0.18 | 0.37 |
| 5 | 25781531 | rs7718830 | -0.327 | -0.628 | 0.671 | 0.88 | -0.017 | -0.014 | -0.014 | 1.91 | 0.41 | 0.27 | 0.41 | 0.42 | 0.47 | 0.4 |
| 5 | 25782399 | rs12521681 | 1.468 | -0.492 | -0.127 | 0.909 | -0.022 | 0.08 | 0.08 | 0.095 | 0.43 | 0.06 | 0.38 | 0.31 | 0.41 | 0.37 |
| 5 | 25790587 | rs370277 | 0.931 | -0.19 | 0.204 | 1.031 | 0.008 | -0.004 | -0.004 | 0.29 | 0.51 | 0.26 | 0.53 | 0.57 | 0.48 | 0.5 |
| 5 | 25808399 | rs7703602 | 0.729 | -0.184 | -0.52 | 0.782 | -0.099 | 0.028 | 0.028 | 0 | 0.24 | 0.04 | 0.2 | 0.16 | 0.36 | 0.16 |
| 5 | 25811412 | rs384831 | 1.026 | -0.19 | -0.42 | 0.675 | -0.055 | 0.091 | 0.091 | 0.122 | 0.44 | 0.02 | 0.36 | 0.3 | 0.41 | 0.39 |
| 5 | 25817862 | rs366244 | 1.164 | -0.267 | -0.059 | 0.766 | 0.001 | 0.053 | 0.053 | 0 | 0.9 | 0.81 | 0.72 | 0.71 | 0.87 | 0.83 |
| 5 | 25821685 | rs2217326 | -0.042 | -0.215 | -0.173 | 0.702 | -0.077 | -0.077 | -0.077 | 0 | 0.4 | 0.12 | 0.4 | 0.46 | 0.49 | 0.49 |
| 5 | 25841323 | rs438523 | 1.208 | -0.287 | 0.683 | 0.473 | 0.056 | 0.042 | 0.042 | 0.505 | 0.94 | 0.6 | 0.97 | 1 | 0.96 | 0.95 |
| 5 | 25841613 | rs396045 | 0.937 | -0.33 | 0.015 | 0.446 | 0.001 | 0.159 | 0.159 | 0 | 0.44 | 0.14 | 0.28 | 0.26 | 0.38 | 0.33 |
| 5 | 25847977 | rs7447463 | -0.744 | -0.083 | 0.505 | 0.965 | 0.007 | 0.047 | 0.047 | 0 | 0.11 | 0.22 | 0.18 | 0.26 | 0.15 | 0.17 |
| 5 | 25856000 | rs13158538 | 2.228 | -0.198 | 1.05 | 0.824 | -0.004 | 0.01 | 0.01 | 2.275 | 0.8 | 0.26 | 0.74 | 0.73 | 0.8 | 0.8 |
| 5 | 25857050 | rs1904460 | -0.703 | -0.12 | 1.056 | 0.869 | 0.013 | 0.054 | 0.054 | 0 | 0.11 | 0.23 | 0.18 | 0.26 | 0.15 | 0.18 |
| 5 | 25868547 | rs2195402 | -0.703 | -0.12 | 1.042 | 0.901 | 0.013 | 0.072 | 0.072 | 0 | 0.11 | 0.23 | 0.18 | 0.29 | 0.15 | 0.18 |
| 5 | 25869401 | rs12521157 | 1.501 | -0.1 | 0.142 | 0.279 | -0.108 | 0.189 | 0.189 | 1.45 | 0.41 | 0.02 | 0.29 | 0.16 | 0.41 | 0.32 |
| 5 | 25874921 | rs13164425 | 1.484 | -0.154 | 0.621 | 0.329 | -0.002 | 0.14 | 0.14 | 0.368 | 0.43 | 0.17 | 0.3 | 0.14 | 0.43 | 0.34 |
| 5 | 25875678 | rs1366466 | 1.073 | -0.427 | 0.433 | 0.888 | 0.033 | -0.035 | -0.035 | 0.576 | 0.08 | 0.01 | 0.14 | 0.29 | 0.04 | 0.12 |
| 5 | 25878699 | rs6452297 | -0.079 | -0.211 | 0.799 | 0.387 | -0.016 | -0.057 | -0.057 | 0.712 | 0.37 | 0.15 | 0.43 | 0.55 | 0.36 | 0.45 |
| 5 | 25881744 | rs7443354 | 1.653 | -0.089 | 1.41 | 0.557 | 0.06 | 0 | 0 | 1.295 | 0.93 | 0.67 | 0.94 | 1 | 0.95 | 0.98 |
| 5 | 25883408 | rs10039219 | -1.28 | -0.021 | 1.3 | 0.608 | 0.067 | 0.14 | 0.14 | 0.29 | 0.13 | 0.37 | 0.22 | 0.31 | 0.2 | 0.19 |
| 5 | 25898921 | rs7704909 | 1.422 | -0.381 | 0.39 | 0.016 | -0.107 | 0.139 | 0.139 | 0.169 | 0.37 | 0.02 | 0.28 | 0.14 | 0.42 | 0.32 |
| 5 | 25899020 | rs1896731 | -0.455 | -0.256 | 0.807 | 0 | 0.008 | 0.011 | 0.011 | 0 | 0.43 | 0.4 | 0.39 | 0.6 | 0.34 | 0.48 |
| 5 | 25899414 | rs7705715 | -1.993 | -0.265 | 0.819 | 0.002 | -0.046 | 0.148 | 0.148 | 0.169 | 0.63 | 0.91 | 0.72 | 0.86 | 0.58 | 0.68 |
| 5 | 25899711 | rs16895091 | -1.009 | -0.195 | 0.917 | 0.01 | -0.017 | -0.031 | -0.031 | 0 | 0.06 | 0.2 | 0.03 | 0 | 0.02 | 0.01 |
| 5 | 25902342 | rs10038113 | -0.622 | -0.118 | 0.958 | 0.027 | -0.001 | -0.003 | -0.003 | 0.084 | 0.44 | 0.56 | 0.41 | 0.62 | 0.38 | 0.5 |
| 5 | 25907909 | rs6414928 | 0.573 | -0.09 | 0.989 | 0.045 | 0.026 | 0.024 | 0.024 | 0 | 0.82 | 0.63 | 0.68 | 0.76 | 0.79 | 0.82 |
| 5 | 25910929 | rs10942147 | 1.709 | -0.176 | 0.8 | 0.118 | -0.103 | 0.14 | 0.14 | 0.16 | 0.38 | 0.02 | 0.27 | 0.14 | 0.41 | 0.31 |
| 5 | 25912113 | rs12521388 | 1.568 | -0.111 | 0.922 | 0.127 | -0.042 | 0.167 | 0.167 | 0 | 0.39 | 0.09 | 0.27 | 0.14 | 0.41 | 0.31 |
| 5 | 25913210 | rs7727225 | 0.269 | -0.095 | 1.068 | 0.196 | 0.051 | 0.016 | 0.016 | 0 | 0.07 | 0.21 | 0.05 | 0.08 | 0.1 | 0.12 |
| 5 | 25920401 | rs12659830 | -0.064 | -0.123 | 1.024 | 0.159 | -0.003 | 0.021 | 0.021 | 0 | 0.4 | 0.19 | 0.36 | 0.59 | 0.36 | 0.49 |
| 5 | 25928888 | rs10050387 | -0.055 | -0.084 | 1.263 | 0.168 | 0.066 | 0.02 | 0.02 | 0 | 0.08 | 0.29 | 0.06 | 0.08 | 0.1 | 0.12 |
| 5 | 25931434 | rs12659150 | 0.301 | -0.182 | 0.914 | 0.356 | -0.017 | -0.039 | -0.039 | 1.108 | 0.35 | 0.13 | 0.23 | 0.35 | 0.31 | 0.39 |
| 5 | 25933110 | rs4516819 | 0.004 | -0.168 | 1.025 | 0.342 | 0.09 | 0.036 | 0.036 | 0 | 0.08 | 0.31 | 0.06 | 0.08 | 0.1 | 0.12 |
| 5 | 25935070 | rs12187724 | -0.139 | -0.198 | 1.043 | 0.35 | 0.002 | 0.021 | 0.021 | 0 | 0.41 | 0.27 | 0.36 | 0.59 | 0.36 | 0.48 |
| 5 | 25955765 | rs10040577 | 0.034 | -0.174 | 1.123 | 0.348 | 0.079 | 0.029 | 0.029 | 0 | 0.08 | 0.29 | 0.06 | 0.08 | 0.11 | 0.12 |
| 5 | 25962632 | rs10942150 | 0.301 | -0.208 | 0.938 | 0.336 | -0.02 | -0.044 | -0.044 | 0 | 0.35 | 0.13 | 0.23 | 0.35 | 0.31 | 0.39 |
| 5 | 25969379 | rs6891206 | -1.757 | -0.265 | 0.921 | 0.299 | -0.038 | 0.148 | 0.148 | 0 | 0.59 | 0.84 | 0.69 | 0.84 | 0.55 | 0.66 |
| 5 | 25970666 | rs7732252 | -1.835 | -0.222 | 0.989 | 0.323 | -0.002 | 0.148 | 0.148 | 0.098 | 0.22 | 0.51 | 0.44 | 0.49 | 0.24 | 0.27 |
| 5 | 25972821 | rs4327572 | 1.812 | -0.196 | 0.098 | 0.193 | -0.084 | 0.156 | 0.156 | 0.035 | 0.38 | 0.03 | 0.27 | 0.14 | 0.41 | 0.31 |
| 5 | 25974120 | rs10038851 | 1.869 | 0.303 | 0.286 | 0.246 | 0.004 | -0.003 | -0.003 | 0.002 | 0.77 | 0.34 | 0.62 | 0.74 | 0.74 | 0.79 |
| 5 | 25976052 | rs11954820 | 1.228 | 0.357 | -0.12 | 0.25 | -0.032 | -0.014 | -0.014 | 0.621 | 0.05 | 0 | 0.03 | 0.08 | 0.11 | 0.12 |
| 5 | 25976806 | rs4314370 | 0.801 | 0.25 | -0.179 | 0.271 | -0.044 | -0.025 | -0.025 | 0.465 | 0.52 | 0.2 | 0.65 | 0.55 | 0.55 | 0.48 |
| 5 | 25977806 | rs4277876 | 0.104 | 0.234 | -0.103 | 0.288 | -0.02 | -0.011 | -0.011 | 0 | 0.6 | 0.71 | 0.49 | 0.44 | 0.57 | 0.54 |

| | | | | | | | | | | | | | | | | |
|---|---|---|---|---|---|---|---|---|---|---|---|---|---|---|---|---|
| 5 | 25991167 | rs4537025 | -1.634 | 0.373 | 0.719 | 0.748 | 0.014 | 0.054 | 0.054 | 0.008 | 0.16 | 0.53 | 0.32 | 0.25 | 0.2 | 0.18 |
| 5 | 25991253 | rs4544798 | 0.587 | 0.675 | 0.212 | 0.797 | -0.018 | -0.012 | -0.012 | 0 | 0.06 | 0.02 | 0.03 | 0.09 | 0.12 | 0.13 |
| 5 | 25992457 | rs4594818 | -1.97 | 0.205 | 0.389 | 0.817 | -0.029 | 0.09 | 0.09 | 0.017 | 0.17 | 0.4 | 0.4 | 0.41 | 0.15 | 0.16 |
| 5 | 26003284 | rs7700465 | -1.177 | 0.373 | -0.053 | 0.685 | -0.001 | 0.007 | 0.007 | 0 | 0.25 | 0.36 | 0.49 | 0.43 | 0.28 | 0.19 |
| 5 | 26008899 | rs6888835 | -0.613 | 0.507 | -0.175 | 0.634 | -0.001 | -0.006 | -0.006 | 0 | 0.17 | 0.25 | 0.23 | 0.27 | 0.19 | 0.18 |
| 5 | 26013301 | rs4355485 | -1.635 | 0.245 | 0.627 | -0.099 | 0.028 | 0 | 0 | 0.161 | 0.09 | 0.43 | 0.07 | 0.03 | 0.05 | 0.09 |
| 5 | 26016363 | rs10067113 | -0.471 | 0.258 | -0.301 | -0.486 | -0.031 | -0.007 | -0.007 | 0 | 0.7 | 0.85 | 0.65 | 0.32 | 0.66 | 0.68 |
| 5 | 26028389 | rs4404644 | -0.434 | 0.07 | -0.522 | -0.738 | -0.001 | 0.05 | 0.05 | 0.02 | 0.48 | 0.31 | 0.48 | 0.64 | 0.51 | 0.44 |
| 5 | 26030577 | rs6452312 | 1.267 | 0.109 | -0.621 | -0.826 | -0.022 | 0.122 | 0.122 | 0 | 0.37 | 0.14 | 0.21 | 0.14 | 0.36 | 0.37 |
| 5 | 26048462 | rs1330656 | 0.755 | 0.198 | -0.495 | -0.821 | 0 | 0.005 | 0.005 | 0.252 | 0.18 | 0.15 | 0.12 | 0 | 0.17 | 0.1 |
| 5 | 26057117 | rs12514304 | 0.642 | 0.54 | -0.393 | -0.33 | 0.007 | 0.053 | 0.053 | 0 | 0.1 | 0.11 | 0.22 | 0.59 | 0.15 | 0.15 |
| 5 | 26064803 | rs10072518 | 1.052 | 0.371 | -0.497 | -0.452 | 0 | 0.174 | 0.174 | 0 | 0.53 | 0.23 | 0.33 | 0.12 | 0.54 | 0.56 |
| 5 | 26072458 | rs7707114 | 1.075 | 0.237 | -0.602 | -0.538 | 0 | 0.211 | 0.211 | 0.017 | 0.52 | 0.2 | 0.32 | 0.12 | 0.52 | 0.51 |
| 5 | 26073238 | rs1330642 | 0.33 | 0.295 | -0.744 | -0.472 | -0.021 | -0.03 | -0.03 | 0 | 0.13 | 0.07 | 0.24 | 0.59 | 0.17 | 0.18 |
| 5 | 26088877 | rs7714043 | -0.24 | 0.224 | -0.492 | -0.513 | -0.003 | -0.002 | -0.002 | 0 | 0.33 | 0.64 | 0.44 | 0.29 | 0.31 | 0.23 |
| 5 | 26091605 | rs12651738 | 1.086 | 0.219 | -0.757 | -0.572 | -0.009 | 0.254 | 0.254 | 0.006 | 0.49 | 0.15 | 0.31 | 0.12 | 0.52 | 0.46 |
| 5 | 26112192 | rs9293221 | 0.202 | 0.412 | -0.339 | -0.051 | 0.013 | 0.056 | 0.056 | 0 | 0.09 | 0.23 | 0.25 | 0.14 | 0.11 | 0.06 |
| 5 | 26114496 | rs7721783 | -0.244 | 0.931 | -1.283 | -0.01 | -0.049 | -0.06 | -0.06 | 0 | 0.19 | 0.07 | 0.26 | 0.59 | 0.28 | 0.25 |
| 5 | 26118309 | rs11745199 | 0.164 | 0.327 | -0.835 | -0.425 | 0.008 | 0.012 | 0.012 | 0 | 0.22 | 0.16 | 0.15 | 0.06 | 0.19 | 0.13 |
| 5 | 26125639 | rs1330660 | -0.111 | 1.213 | -1.385 | 0.518 | -0.081 | -0.077 | -0.077 | 0.032 | 0.26 | 0.07 | 0.31 | 0.63 | 0.38 | 0.23 |
| 5 | 26125816 | rs1330659 | 0.045 | 1.501 | -1.369 | 0.908 | -0.054 | -0.088 | -0.088 | 0 | 0.23 | 0.09 | 0.29 | 0.53 | 0.35 | 0.21 |
| 5 | 26132948 | rs4048817 | -0.112 | 0.798 | -0.774 | -0.218 | 0 | -0.037 | -0.037 | 0.168 | 0.64 | 0.62 | 0.69 | 0.91 | 0.69 | 0.72 |
| 5 | 26135884 | rs7710398 | -1.139 | 0.109 | -0.691 | -0.267 | 0.01 | 0.015 | 0.015 | 0 | 0.82 | 0.88 | 0.91 | 0.93 | 0.87 | 0.84 |
| 5 | 26136566 | rs7447064 | -0.401 | -0.233 | -0.478 | 0.117 | -0.013 | -0.004 | -0.004 | 0 | 0.12 | 0.13 | 0.06 | 0.07 | 0.06 | 0.15 |
| 5 | 26147657 | rs4081229 | -0.684 | 0.075 | -0.161 | 0.268 | 0.001 | 0.015 | 0.015 | 0 | 0.13 | 0.27 | 0.19 | 0.15 | 0.14 | 0.16 |
| 5 | 26156304 | rs1411589 | 0.826 | 0.312 | -0.574 | 0.544 | -0.007 | 0.003 | 0.003 | 0 | 0.42 | 0.35 | 0.59 | 0.56 | 0.51 | 0.34 |
| 5 | 26157740 | rs1541077 | -0.557 | 0.552 | -0.854 | -0.373 | -0.002 | -0.014 | -0.014 | 0 | 0.62 | 0.65 | 0.67 | 0.91 | 0.62 | 0.67 |
| 5 | 26172906 | rs13173199 | -0.372 | 0.926 | -0.721 | -0.036 | -0.003 | 0.008 | 0.008 | 0 | 0.54 | 0.6 | 0.65 | 0.87 | 0.58 | 0.56 |
| 5 | 26181001 | rs10062689 | 0.076 | 0.525 | -0.737 | -0.023 | -0.011 | -0.001 | -0.001 | 0 | 0.09 | 0.13 | 0.05 | 0.07 | 0.07 | 0.08 |
| 5 | 26182839 | rs6883139 | 0.213 | 0.91 | -0.877 | -0.184 | 0.004 | 0.005 | 0.005 | 0 | 0.51 | 0.47 | 0.57 | 0.8 | 0.51 | 0.49 |
| 5 | 26193999 | rs6884362 | -0.831 | 1.292 | -0.963 | 0.484 | -0.026 | -0.007 | -0.007 | 0 | 0.09 | 0.02 | 0.16 | 0.08 | 0.17 | 0.04 |
| 5 | 26199587 | rs1330676 | 0.374 | 0.375 | 0.065 | 0.152 | -0.037 | -0.036 | -0.036 | 0 | 0.87 | 0.6 | 0.94 | 0.93 | 0.96 | 0.79 |
| 5 | 26201879 | rs10942164 | 0.009 | 0.323 | 0.283 | 0.127 | -0.006 | -0.005 | -0.005 | 0 | 0.9 | 0.87 | 0.96 | 0.93 | 0.97 | 0.79 |
| 5 | 26207219 | rs11745195 | -0.03 | 0.998 | -0.28 | 0.24 | -0.054 | -0.004 | -0.004 | 0 | 0.27 | 0.1 | 0.22 | 0.26 | 0.38 | 0.2 |
| 5 | 26210563 | rs4701531 | 0.83 | 0.022 | -1.195 | 0.384 | -0.092 | -0.048 | -0.048 | 0.112 | 0.35 | 0.05 | 0.4 | 0.33 | 0.39 | 0.39 |
| 5 | 26219645 | rs10942167 | 0.277 | 0.048 | -0.448 | 0.353 | -0.002 | -0.01 | -0.01 | 0.013 | 0.64 | 0.54 | 0.73 | 0.68 | 0.66 | 0.6 |
| 5 | 26221764 | rs12518513 | 0.573 | 0.024 | -0.182 | 0.369 | -0.022 | -0.007 | -0.007 | 0 | 0.23 | 0.14 | 0.21 | 0.24 | 0.31 | 0.19 |
| 5 | 26223985 | rs6865956 | 0.365 | 0.009 | -0.153 | 0.364 | -0.001 | 0.002 | 0.002 | 0 | 0.65 | 0.67 | 0.75 | 0.68 | 0.66 | 0.61 |
| 5 | 26224041 | rs6870351 | 0.365 | 0.009 | -0.153 | 0.364 | -0.001 | 0.002 | 0.002 | 0 | 0.65 | 0.67 | 0.75 | 0.68 | 0.66 | 0.61 |
| 5 | 26224170 | rs6884536 | 1.827 | -0.325 | -0.132 | 0.295 | 0.037 | -0.031 | -0.031 | 0.012 | 0.24 | 0.11 | 0.17 | 0.3 | 0.16 | 0.19 |
| 5 | 26224692 | rs6885388 | 0.368 | -0.28 | -0.043 | 0.291 | 0.001 | -0.001 | -0.001 | 0 | 0.41 | 0.43 | 0.56 | 0.38 | 0.49 | 0.41 |
| 5 | 26230104 | rs10035723 | -0.57 | 0.042 | -0.278 | 0.145 | -0.001 | -0.004 | -0.004 | 0 | 0.7 | 0.7 | 0.85 | 0.84 | 0.78 | 0.75 |
| 5 | 26230491 | rs13153758 | -0.584 | 0.035 | -0.29 | 0.144 | -0.001 | -0.007 | -0.007 | 1.216 | 0.71 | 0.7 | 0.85 | 0.84 | 0.78 | 0.75 |
| 5 | 26231699 | rs4701532 | 0.412 | 0.091 | -0.075 | 0.432 | 0.021 | 0.012 | 0.012 | 1.701 | 0.2 | 0.3 | 0.29 | 0.23 | 0.27 | 0.25 |
| 5 | 26231934 | rs4604153 | 1.278 | 0.04 | -0.169 | 0.478 | 0.002 | -0.003 | -0.003 | 0 | 0.26 | 0.26 | 0.35 | 0.37 | 0.2 | 0.31 |
| 5 | 26238194 | rs9293229 | -0.4 | 0.299 | 0.058 | 0.812 | 0.025 | 0.096 | 0.096 | 2.301 | 0.25 | 0.51 | 0.53 | 0.43 | 0.33 | 0.42 |
| 5 | 26250248 | rs6892024 | -0.231 | 0.208 | -0.017 | 0.859 | 0 | 0 | 0 | 0.012 | 0.4 | 0.52 | 0.64 | 0.57 | 0.52 | 0.44 |
| 5 | 26263051 | rs10068658 | 0.651 | 0.241 | -0.484 | 0.908 | -0.068 | -0.104 | -0.104 | 0.306 | 0.23 | 0.06 | 0.38 | 0.36 | 0.29 | 0.35 |
| 5 | 26265398 | rs10052385 | -0.684 | 0.245 | -0.34 | 0.859 | 0.002 | 0.039 | 0.039 | 0.47 | 0.46 | 0.65 | 0.71 | 0.68 | 0.54 | 0.62 |
| 5 | 26266929 | rs12153689 | 1.103 | 0.18 | -0.267 | 0.967 | -0.094 | -0.079 | -0.079 | 0.201 | 0.2 | 0.02 | 0.32 | 0.22 | 0.26 | 0.18 |
| 5 | 26267717 | rs10056772 | -0.074 | 0.209 | -0.146 | 0.84 | 0.001 | 0.012 | 0.012 | 0.018 | 0.31 | 0.61 | 0.53 | 0.44 | 0.33 | 0.38 |
| 5 | 26283311 | rs4273570 | -0.382 | 0.138 | -0.557 | 0.454 | -0.004 | 0.037 | 0.037 | 0 | 0.7 | 0.79 | 0.83 | 0.84 | 0.7 | 0.76 |
| 5 | 26292815 | rs4406110 | -0.149 | -0.164 | -0.496 | 0.605 | 0.014 | 0.013 | 0.013 | 0.205 | 0.47 | 0.27 | 0.45 | 0.46 | 0.56 | 0.53 |
| 5 | 26293800 | rs9293230 | -0.557 | -0.828 | 0.415 | 1.498 | 0.021 | 0.356 | 0.356 | 0.82 | 0.08 | 0.44 | 0.12 | 0.22 | 0.03 | 0.07 |
| 5 | 26299629 | rs10434626 | -0.14 | -0.813 | -0.211 | 1.37 | 0.048 | -0.012 | -0.012 | 0.26 | 0.41 | 0.67 | 0.28 | 0.49 | 0.2 | 0.3 |
| 5 | 26322668 | rs6862088 | 0.075 | 0.112 | -0.39 | 1.278 | -0.016 | -0.005 | -0.005 | 0 | 0.1 | 0.14 | 0.03 | 0.07 | 0.03 | 0.21 |
| 5 | 26334156 | rs10040524 | -0.088 | 0.76 | -0.446 | 1.462 | 0.029 | 0.058 | 0.058 | 0.033 | 0.23 | 0.34 | 0.4 | 0.52 | 0.33 | 0.35 |
| 5 | 26336760 | rs6882967 | -0.212 | 0.442 | -1.561 | 0.658 | -0.11 | 0.169 | 0.169 | 0.2 | 0.59 | 0.89 | 0.57 | 0.8 | 0.5 | 0.69 |
| 5 | 26344352 | rs1389888 | -0.265 | 1.149 | -0.596 | 1.703 | -0.004 | -0.005 | -0.005 | 0.316 | 0.19 | 0.17 | 0.31 | 0.28 | 0.24 | 0.23 |
| 5 | 26344613 | rs6882953 | -0.265 | 1.149 | -0.596 | 1.703 | -0.004 | -0.005 | -0.005 | 0 | 0.19 | 0.17 | 0.31 | 0.28 | 0.24 | 0.23 |
| 5 | 26349480 | rs6866505 | -0.228 | 1.167 | -0.587 | 1.671 | -0.003 | -0.004 | -0.004 | 0.146 | 0.19 | 0.17 | 0.31 | 0.27 | 0.24 | 0.23 |
| 5 | 26356507 | rs12188862 | -0.887 | 1.198 | -0.372 | 1.649 | -0.027 | 0.033 | 0.033 | 1.203 | 0.22 | 0.11 | 0.13 | 0.14 | 0.28 | 0.22 |
| 5 | 26357027 | rs12189045 | -0.887 | 1.198 | -0.372 | 1.649 | -0.027 | 0.033 | 0.033 | 1.203 | 0.22 | 0.11 | 0.13 | 0.14 | 0.28 | 0.21 |
| 5 | 26364671 | rs12652998 | -1.652 | 0.595 | 0.153 | 2.713 | -0.003 | 0.026 | 0.026 | 0 | 0.08 | 0.02 | 0.09 | 0.45 | 0.09 | 0.13 |
| 5 | 26378196 | rs12654972 | -1.949 | 0.601 | 0.746 | 2.757 | -0.001 | 0.079 | 0.079 | 0 | 0.09 | 0.07 | 0.1 | 0.45 | 0.09 | 0.13 |
| 5 | 26383317 | rs12516042 | -1.759 | 0.606 | 0.767 | 2.78 | -0.001 | 0.094 | 0.094 | 0 | 0.09 | 0.07 | 0.1 | 0.45 | 0.09 | 0.13 |
| 5 | 26394566 | rs1494699 | -1.411 | 0.822 | 0.082 | 1.74 | -0.001 | -0.015 | -0.015 | 0 | 0.07 | 0.02 | 0.07 | 0.18 | 0.08 | 0.13 |
| 5 | 26406901 | rs7706667 | -1.168 | 2.528 | 1.222 | 1.897 | 0.068 | 0.034 | 0.034 | 0 | 0.12 | 0.28 | 0.13 | 0.21 | 0.21 | 0.12 |
| 5 | 26409352 | rs6897774 | 0.64 | 2.054 | 1.804 | 1.408 | 0.024 | 0.045 | 0.045 | 0 | 0.85 | 0.6 | 0.79 | 0.8 | 0.87 | 0.68 |

| | | | | | | | | | | | | | | | | |
|---|---|---|---|---|---|---|---|---|---|---|---|---|---|---|---|---|
| 5 | 26416388 | rs7706669 | 1.266 | 1.657 | 2.469 | 1.064 | -0.018 | -0.034 | -0.034 | 0 | 0.83 | 0.34 | 0.77 | 0.86 | 0.88 | 0.71 |
| 5 | 26427064 | rs11950170 | 1.014 | 1.622 | 3.331 | 1.046 | -0.016 | -0.061 | -0.061 | 0 | 0.87 | 0.64 | 0.88 | 0.96 | 0.91 | 0.83 |
| 5 | 26430825 | rs13169349 | -0.882 | 1.906 | 2.935 | 1.202 | -0.041 | -0.037 | -0.037 | 0.159 | 0.11 | 0.06 | 0.12 | 0.25 | 0.24 | 0.18 |
| 5 | 26431671 | rs10942177 | 1.192 | 1.138 | 1.944 | -0.127 | 0.051 | -0.016 | -0.016 | 0.816 | 0.35 | 0.11 | 0.41 | 0.57 | 0.28 | 0.24 |
| 5 | 26447676 | rs6894058 | -2.225 | 1.026 | 2.977 | -0.007 | 0.067 | 0.248 | 0.248 | 1.004 | 0.07 | 0.44 | 0.15 | 0.19 | 0.03 | 0.16 |
| 5 | 26482424 | rs6871724 | -1.161 | 0.863 | 3.378 | -0.134 | -0.008 | -0.005 | -0.005 | 0 | 0.16 | 0.45 | 0.2 | 0.14 | 0.11 | 0.22 |
| 5 | 26509802 | rs6877069 | -0.531 | 1.66 | 2.959 | -0.598 | 0.003 | 0.021 | 0.021 | 0 | 0.45 | 0.6 | 0.4 | 0.16 | 0.38 | 0.5 |
| 5 | 26523363 | rs16895743 | -0.582 | 1.61 | 3.121 | -0.604 | -0.002 | -0.004 | -0.004 | 2.135 | 0.14 | 0.26 | 0.19 | 0.13 | 0.11 | 0.24 |
| 5 | 26523488 | rs1366492 | 0.966 | 1.664 | 3.065 | -0.645 | -0.031 | 0.153 | 0.153 | 0 | 0.28 | 0.04 | 0.19 | 0.03 | 0.26 | 0.27 |
| 5 | 26534253 | rs1428718 | -0.152 | 1.57 | 3.379 | -0.569 | 0.017 | 0.023 | 0.023 | 0 | 0.11 | 0.26 | 0.16 | 0.13 | 0.09 | 0.22 |
| 5 | 26537760 | rs1025304 | -0.9 | 1.684 | 3.37 | -0.585 | 0.011 | -0.008 | -0.008 | 0.062 | 0.15 | 0.35 | 0.21 | 0.13 | 0.15 | 0.25 |
| 5 | 26551516 | rs4701544 | 1.279 | 1.93 | 2.779 | 0.017 | 0.017 | -0.093 | -0.093 | 0 | 0.4 | 0.06 | 0.44 | 0.48 | 0.37 | 0.28 |
| 5 | 26555788 | rs6867412 | -0.524 | 2.302 | 2.917 | -0.093 | 0.007 | 0.027 | 0.027 | 0.033 | 0.44 | 0.62 | 0.35 | 0.16 | 0.38 | 0.5 |
| 5 | 26560653 | rs1353954 | -0.856 | 2.137 | 2.714 | -0.233 | 0.022 | 0.172 | 0.172 | 0.029 | 0.5 | 0.77 | 0.46 | 0.22 | 0.42 | 0.58 |
| 5 | 26568243 | rs6452345 | -0.116 | 2.237 | 2.571 | -0.119 | -0.044 | -0.043 | -0.043 | 0.065 | 0.91 | 0.97 | 0.91 | 0.7 | 0.79 | 0.86 |
| 5 | 26570795 | rs10520910 | 0.545 | 1.753 | 2.951 | -0.448 | -0.005 | -0.045 | -0.045 | 1.122 | 0.09 | 0.14 | 0.08 | 0.01 | 0.06 | 0.12 |
| 5 | 26573377 | rs10520911 | -0.841 | 1.795 | 3.077 | -0.455 | 0.001 | -0.006 | -0.006 | 0.156 | 0.14 | 0.24 | 0.18 | 0.14 | 0.13 | 0.23 |
| 5 | 26576539 | rs1353956 | -1.367 | 1.761 | 3.125 | -0.455 | 0.013 | 0.047 | 0.047 | 0.304 | 0.05 | 0.15 | 0.11 | 0.13 | 0.07 | 0.11 |
| 5 | 26577401 | rs16895842 | 0.248 | 1.72 | 2.987 | -0.48 | -0.01 | -0.047 | -0.047 | 0 | 0.1 | 0.15 | 0.08 | 0.01 | 0.06 | 0.12 |
| 5 | 26589040 | rs2319698 | -1.025 | 1.609 | 2.854 | -0.586 | -0.004 | 0.231 | 0.231 | 0.041 | 0.49 | 0.89 | 0.48 | 0.22 | 0.42 | 0.57 |
| 5 | 26602131 | rs10942181 | -1.182 | 1.705 | 3.208 | -0.561 | 0.01 | -0.013 | -0.013 | 0 | 0.16 | 0.48 | 0.22 | 0.14 | 0.15 | 0.23 |
| 5 | 26602765 | rs12654244 | -1.149 | 1.688 | 3.058 | -0.549 | 0.023 | 0.063 | 0.063 | 0.164 | 0.05 | 0.18 | 0.12 | 0.13 | 0.07 | 0.09 |
| 5 | 26602880 | rs1604097 | -0.769 | 1.655 | 2.933 | -0.579 | 0.011 | 0.071 | 0.071 | 0 | 0.47 | 0.63 | 0.44 | 0.22 | 0.41 | 0.56 |
| 5 | 26605393 | rs2874065 | 0.188 | 1.646 | 2.568 | -0.523 | -0.001 | -0.034 | -0.034 | 0 | 0.32 | 0.31 | 0.24 | 0.08 | 0.28 | 0.34 |
| 5 | 26606431 | rs6896582 | 0.096 | 1.638 | 2.452 | -0.52 | 0.001 | 0.006 | 0.006 | 0 | 0.33 | 0.27 | 0.25 | 0.08 | 0.3 | 0.34 |
| 5 | 26637996 | rs9687924 | -0.797 | 1.624 | 1.782 | -0.296 | 0.004 | 0.18 | 0.18 | 0.018 | 0.48 | 0.9 | 0.47 | 0.24 | 0.4 | 0.55 |
| 5 | 26643591 | rs6860010 | -0.73 | 1.614 | 2.071 | -0.482 | 0.009 | 0.126 | 0.126 | 0 | 0.45 | 0.73 | 0.37 | 0.09 | 0.4 | 0.47 |
| 5 | 26647400 | rs6452349 | -1.112 | 1.556 | 2.41 | -0.509 | -0.014 | -0.067 | -0.067 | 0.43 | 0.17 | 0.56 | 0.17 | 0.06 | 0.14 | 0.18 |
| 5 | 26655797 | rs1390638 | -0.257 | 1.759 | 1.567 | -0.32 | -0.058 | -0.075 | -0.075 | 0 | 0.85 | 0.97 | 0.86 | 0.65 | 0.77 | 0.87 |
| 5 | 26662394 | rs6863131 | -0.737 | 1.774 | 1.533 | -0.467 | -0.025 | 0.147 | 0.147 | 0.518 | 0.45 | 0.82 | 0.44 | 0.23 | 0.41 | 0.54 |
| 5 | 26673835 | rs10942193 | 1.418 | 1.699 | 1.473 | -0.438 | -0.008 | -0.114 | -0.114 | 0 | 0.39 | 0.09 | 0.5 | 0.43 | 0.36 | 0.31 |
| 5 | 26688754 | rs13164823 | -1.262 | 1.269 | 1.752 | -0.553 | -0.003 | 0.015 | 0.015 | 0 | 0.18 | 0.58 | 0.18 | 0.18 | 0.16 | 0.29 |
| 5 | 26692369 | rs16895967 | 0.799 | 1.532 | 1.195 | -0.609 | 0.008 | -0.039 | -0.039 | 0.512 | 0.08 | 0.13 | 0.08 | 0.01 | 0.09 | 0.15 |
| 5 | 26699956 | rs2132542 | 0.073 | 1.353 | 0.513 | -0.433 | -0.004 | 0.011 | 0.011 | 0 | 0.29 | 0.24 | 0.2 | 0.16 | 0.28 | 0.33 |
| 5 | 26705812 | rs13171386 | 0.292 | 1.303 | -0.267 | -0.649 | -0.055 | 0.162 | 0.162 | 0 | 0.27 | 0.03 | 0.18 | 0.03 | 0.27 | 0.27 |
| 5 | 26722431 | rs10462305 | 0.075 | 0.509 | -0.475 | -0.274 | 0.002 | -0.088 | -0.088 | 2.707 | 0.22 | 0.07 | 0.41 | 0.57 | 0.2 | 0.22 |
| 5 | 26722785 | rs16895993 | 0.782 | 0.65 | -0.618 | -0.469 | 0 | -0.002 | -0.002 | 0 | 0.09 | 0.11 | 0.11 | 0.07 | 0.11 | 0.18 |
| 5 | 26723311 | rs7737624 | -0.27 | 0.074 | -0.124 | -0.846 | 0.008 | -0.046 | -0.046 | 0 | 0.35 | 0.43 | 0.29 | 0.15 | 0.33 | 0.58 |
| 5 | 26726226 | rs4360003 | -1.126 | -0.437 | -0.067 | -0.674 | -0.114 | -0.057 | -0.057 | 0 | 0.76 | 0.96 | 0.8 | 0.73 | 0.59 | 0.85 |
| 5 | 26727405 | rs10068357 | -0.02 | -0.46 | -0.072 | -0.686 | 0.017 | 0.054 | 0.054 | 0 | 0.14 | 0.42 | 0.13 | 0.13 | 0.11 | 0.29 |
| 5 | 26736006 | rs1015610 | 0.595 | -0.543 | -0.231 | -0.67 | -0.067 | -0.055 | -0.055 | 0 | 0.43 | 0.09 | 0.57 | 0.58 | 0.42 | 0.52 |
| 5 | 26737458 | rs10045386 | -0.137 | -0.794 | 0.923 | -0.695 | 0.006 | 0.03 | 0.03 | 0 | 0.13 | 0.28 | 0.13 | 0.13 | 0.11 | 0.29 |
| 5 | 26751379 | rs17188971 | 1.392 | -0.829 | 0.978 | -0.712 | 0.026 | 0.013 | 0.013 | 0 | 0.16 | 0.1 | 0.06 | 0 | 0.06 | 0.04 |
| 5 | 26752242 | rs1505054 | -0.227 | -0.824 | 1.152 | -0.714 | -0.038 | -0.037 | -0.037 | 0 | 0.28 | 0.41 | 0.19 | 0.13 | 0.17 | 0.33 |
| 5 | 26755279 | rs13356265 | -0.431 | -0.794 | 1.412 | -0.736 | -0.015 | -0.049 | -0.049 | 0.076 | 0.3 | 0.65 | 0.21 | 0.06 | 0.17 | 0.23 |
| 5 | 26758206 | rs1505044 | 0.645 | -0.765 | 1.239 | -0.654 | 0.014 | -0.018 | -0.018 | 0 | 0.1 | 0.2 | 0.11 | 0.06 | 0.1 | 0.18 |
| 5 | 26761648 | rs16899515 | 0.679 | -0.731 | 1.15 | -0.644 | 0.018 | -0.02 | -0.02 | 0 | 0.1 | 0.22 | 0.11 | 0.06 | 0.1 | 0.19 |
| 5 | 26762292 | rs10056950 | -0.52 | -0.648 | 0.986 | -0.644 | -0.008 | -0.011 | -0.011 | 0 | 0.31 | 0.65 | 0.22 | 0.14 | 0.19 | 0.34 |
| 5 | 26765497 | rs11949563 | 0.633 | -0.579 | 0.421 | -0.689 | -0.026 | -0.071 | -0.071 | 0 | 0.18 | 0.25 | 0.1 | 0 | 0.09 | 0.05 |
| 5 | 26770247 | rs10062762 | 0.522 | -0.572 | 0.272 | -0.685 | -0.045 | -0.1 | -0.1 | 0.662 | 0.18 | 0.32 | 0.1 | 0 | 0.09 | 0.05 |
| 5 | 26785851 | rs16896091 | -0.169 | -0.411 | 0.053 | -0.58 | -0.02 | -0.056 | -0.056 | 0 | 0.38 | 0.54 | 0.29 | 0.14 | 0.27 | 0.3 |
| 5 | 26787013 | rs6862526 | 0.736 | -0.584 | 0.32 | -0.727 | -0.032 | -0.026 | -0.026 | 2.434 | 0.21 | 0.32 | 0.14 | 0.12 | 0.14 | 0.08 |
| 5 | 26788638 | rs2036569 | -1.29 | -0.193 | -0.372 | -0.237 | -0.044 | -0.07 | -0.07 | 0 | 0.77 | 0.91 | 0.81 | 0.67 | 0.64 | 0.92 |
| 5 | 26788771 | rs13163090 | 1.48 | -0.268 | 0.007 | -0.271 | -0.017 | -0.024 | -0.024 | 0 | 0.2 | 0.23 | 0.11 | 0.07 | 0.13 | 0.09 |
| 5 | 26789357 | rs16874804 | 0.892 | -0.286 | -0.335 | -0.113 | -0.059 | -0.072 | -0.072 | 1.511 | 0.34 | 0.1 | 0.49 | 0.4 | 0.33 | 0.56 |
| 5 | 26805069 | rs1505053 | 0.001 | -0.289 | -0.002 | -0.312 | 0.032 | 0.1 | 0.1 | 0 | 0.42 | 0.73 | 0.33 | 0.1 | 0.34 | 0.34 |
| 5 | 26808325 | rs7722879 | 0.157 | -0.514 | 0.336 | -0.01 | -0.036 | 0.005 | 0.005 | 0.863 | 0.43 | 0.23 | 0.6 | 0.41 | 0.42 | 0.73 |
| 5 | 26811257 | rs12521749 | -0.518 | -0.481 | 0.772 | 0.036 | 0.002 | -0.001 | -0.001 | 5.41 | 0.49 | 0.6 | 0.64 | 0.53 | 0.44 | 0.79 |
| 5 | 26819303 | rs2248868 | 1.412 | -0.445 | 0.628 | 0.107 | -0.028 | -0.046 | -0.046 | 0.83 | 0.24 | 0.15 | 0.24 | 0.42 | 0.4 | 0.08 |
| 5 | 26826645 | rs966986 | 1.474 | -0.431 | 0.736 | 0.099 | -0.047 | -0.072 | -0.072 | 0.017 | 0.23 | 0.09 | 0.23 | 0.39 | 0.39 | 0.08 |
| 5 | 26828780 | rs10447155 | 1.203 | -0.452 | 0.756 | 0.05 | -0.02 | 0.04 | 0.04 | 0 | 0.13 | 0.05 | 0.15 | 0.03 | 0.13 | 0.21 |
| 5 | 26829452 | rs846946 | 1.421 | -0.41 | 0.67 | 0.08 | -0.027 | -0.041 | -0.041 | 0 | 0.24 | 0.15 | 0.24 | 0.39 | 0.4 | 0.08 |
| 5 | 26832200 | rs3857327 | 1.018 | -0.458 | 0.781 | 0.078 | -0.004 | -0.009 | -0.009 | 1.695 | 0.31 | 0.23 | 0.46 | 0.38 | 0.28 | 0.5 |
| 5 | 26832888 | rs818734 | 0.032 | -0.464 | 0.771 | 0.069 | 0.018 | 0.035 | 0.035 | 0.094 | 0.54 | 0.37 | 0.69 | 0.77 | 0.67 | 0.58 |
| 5 | 26833657 | rs17495847 | 1.215 | -0.453 | 0.563 | -0.082 | 0.019 | 0.022 | 0.022 | 0 | 0.27 | 0.2 | 0.12 | 0.07 | 0.16 | 0.15 |
| 5 | 26834436 | rs818739 | -0.876 | 0.303 | 0.586 | 0.302 | 0.004 | -0.032 | -0.032 | 1.813 | 0.86 | 0.79 | 0.85 | 0.97 | 0.86 | 0.79 |
| 5 | 26837635 | rs257789 | 0.725 | 0.308 | 0.524 | 0.309 | -0.026 | -0.055 | -0.055 | 0 | 0.27 | 0.2 | 0.25 | 0.52 | 0.42 | 0.14 |
| 5 | 26838935 | rs12519694 | 0.36 | 0.318 | 0.524 | 0.302 | 0 | 0 | 0 | 0 | 0.58 | 0.54 | 0.59 | 0.45 | 0.45 | 0.65 |
| 5 | 26840350 | rs190476 | 0.649 | 0.215 | 0.417 | 0.385 | -0.028 | -0.051 | -0.051 | 3.427 | 0.27 | 0.19 | 0.24 | 0.47 | 0.41 | 0.14 |

| | | | | | | | | | | | | | | | | |
|---|---|---|---|---|---|---|---|---|---|---|---|---|---|---|---|---|
| 5 | 26847198 | rs3846559 | -0.038 | 0.149 | 0.286 | 0.412 | 0.007 | 0.006 | 0.006 | 0 | 0.61 | 0.57 | 0.69 | 0.45 | 0.49 | 0.67 |
| 5 | 26862104 | rs13167720 | 0.004 | -0.083 | 0.277 | 0.164 | -0.004 | 0.017 | 0.017 | 0 | 0.4 | 0.31 | 0.59 | 0.61 | 0.35 | 0.62 |
| 5 | 26865301 | rs2035281 | 1.53 | 0.06 | -0.043 | 0.164 | -0.032 | -0.005 | -0.005 | 0 | 0.08 | 0 | 0.17 | 0.05 | 0.08 | 0.21 |
| 5 | 26865656 | rs4701549 | -0.42 | -0.646 | 0.232 | -0.175 | 0.042 | -0.012 | -0.012 | 0 | 0.7 | 0.42 | 0.7 | 0.73 | 0.52 | 0.72 |
| 5 | 26866763 | rs1479683 | -1.079 | -0.645 | 0.247 | -0.178 | 0.021 | -0.013 | -0.013 | 0 | 0.75 | 0.71 | 0.76 | 0.82 | 0.56 | 0.83 |
| 5 | 26872598 | rs13170179 | 1.284 | -0.859 | -0.372 | 0.086 | 0.034 | -0.069 | -0.069 | 0 | 0.17 | 0.09 | 0.31 | 0.49 | 0.12 | 0.24 |
| 5 | 26877871 | rs2170033 | 1.01 | -0.754 | -0.377 | -0.081 | -0.035 | 0.042 | 0.042 | 0 | 0.21 | 0.13 | 0.27 | 0.09 | 0.23 | 0.39 |
| 5 | 26886584 | rs1382932 | 0.206 | -0.891 | -0.42 | -0.213 | 0 | 0.186 | 0.186 | 0.917 | 0.48 | 0.21 | 0.37 | 0.19 | 0.4 | 0.47 |
| 5 | 26892008 | rs6870789 | 0.217 | -0.882 | -0.472 | -0.207 | -0.009 | 0.206 | 0.206 | 1.559 | 0.48 | 0.17 | 0.37 | 0.19 | 0.4 | 0.47 |
| 5 | 26893449 | rs6895743 | 0.453 | -1.071 | -0.49 | 0.027 | 0.114 | -0.024 | -0.024 | 0 | 0.42 | 0.16 | 0.4 | 0.57 | 0.25 | 0.32 |
| 5 | 26900903 | rs7734956 | 1.114 | -0.685 | -0.355 | -0.045 | 0.058 | 0.104 | 0.104 | 0.401 | 0.27 | 0.08 | 0.11 | 0.09 | 0.17 | 0.09 |
| 5 | 26901775 | rs16896353 | -0.939 | 0.15 | 0.484 | 0.512 | 0.039 | 0.067 | 0.067 | 0 | 0.06 | 0.2 | 0.06 | 0.12 | 0.05 | 0.12 |
| 5 | 26917996 | rs6866141 | -0.033 | 0.354 | 0.432 | 0.197 | 0.03 | -0.02 | -0.02 | 0.14 | 0.71 | 0.6 | 0.76 | 0.78 | 0.55 | 0.81 |
| 5 | 26927356 | rs11745103 | 0.468 | 0.791 | -0.219 | 0.305 | -0.077 | 0.039 | 0.039 | 0.257 | 0.23 | 0.09 | 0.2 | 0.12 | 0.42 | 0.17 |
| 5 | 26927503 | rs10072177 | -0.519 | 0.039 | 0.497 | 0.601 | 0.009 | 0.042 | 0.042 | 0 | 0.09 | 0.17 | 0.07 | 0.18 | 0.07 | 0.12 |
| 5 | 26932005 | rs10064442 | -0.519 | 0.066 | 0.389 | 0.614 | 0.009 | 0.042 | 0.042 | 0 | 0.09 | 0.17 | 0.07 | 0.18 | 0.07 | 0.12 |
| 5 | 26937771 | rs10473822 | 0.201 | -0.44 | 0.493 | 0.435 | -0.061 | 0.062 | 0.062 | 0.033 | 0.78 | 0.55 | 0.66 | 0.28 | 0.88 | 0.76 |
| 5 | 26939178 | rs6885387 | -0.465 | 0.258 | 0.314 | -0.154 | 0.004 | 0.005 | 0.005 | 0.132 | 0.52 | 0.58 | 0.35 | 0.34 | 0.57 | 0.31 |
| 5 | 26942758 | rs1007586 | 1.39 | 0.616 | 0.78 | -0.848 | 0.087 | 0.001 | 0.001 | 0.864 | 0.93 | 0.44 | 0.91 | 0.99 | 0.9 | 0.98 |
| 5 | 26944286 | rs10078687 | 1.13 | 0.6 | 0.971 | -0.848 | 0.134 | 0.041 | 0.041 | 0.472 | 0.94 | 0.53 | 0.92 | 0.99 | 0.9 | 0.98 |
| 5 | 26953337 | rs2170039 | 1.25 | 0.834 | -0.205 | -1.186 | -0.086 | 0.057 | 0.057 | 3.376 | 0.16 | 0.02 | 0.11 | 0.02 | 0.26 | 0.07 |
| 5 | 26961133 | rs6871019 | 1.043 | 0.341 | 0.032 | -1.141 | -0.001 | -0.039 | -0.039 | 0 | 0.78 | 0.3 | 0.84 | 0.99 | 0.81 | 0.94 |
| 5 | 26961843 | rs13174843 | 0.165 | 0.374 | -0.012 | -1.111 | 0.031 | 0.036 | 0.036 | 0 | 0.13 | 0.05 | 0.07 | 0 | 0.09 | 0.04 |
| 5 | 26965781 | rs6863714 | 0.772 | 0.401 | -0.019 | -1.123 | -0.019 | -0.076 | -0.076 | 0 | 0.78 | 0.36 | 0.82 | 0.99 | 0.81 | 0.93 |
| 5 | 26970581 | rs10942222 | 0.249 | 0.777 | -0.609 | -1.049 | -0.111 | 0.074 | 0.074 | 0 | 0.33 | 0.07 | 0.19 | 0.15 | 0.42 | 0.2 |
| 5 | 26971951 | rs9293258 | -0.273 | 0.524 | 0.426 | -1.137 | 0.007 | -0.024 | -0.024 | 0 | 0.38 | 0.41 | 0.25 | 0.16 | 0.52 | 0.21 |
| 5 | 26972230 | rs1175232 | 0.397 | 0.431 | 0.638 | -1.18 | -0.024 | 0.079 | 0.079 | 0 | 0.59 | 0.28 | 0.72 | 0.84 | 0.48 | 0.79 |
| 5 | 26977421 | rs1175237 | -0.468 | 0.577 | 0.877 | -0.858 | -0.017 | -0.003 | -0.003 | 0.459 | 0.18 | 0.29 | 0.11 | 0.12 | 0.12 | 0.09 |
| 5 | 26977780 | rs1366462 | -0.165 | 1.24 | 1.087 | -1.299 | -0.071 | 0.103 | 0.103 | 0 | 0.36 | 0.14 | 0.23 | 0.15 | 0.48 | 0.22 |
| 5 | 26978090 | rs7719204 | -0.362 | 0.906 | 1.321 | -1.15 | 0.026 | -0.06 | -0.06 | 0 | 0.37 | 0.49 | 0.25 | 0.15 | 0.47 | 0.19 |
| 5 | 26988328 | rs2288466 | -0.605 | 0.945 | 1.404 | -1.18 | -0.017 | 0.045 | 0.045 | 0.201 | 0.39 | 0.69 | 0.27 | 0.15 | 0.47 | 0.19 |
| 5 | 26988424 | rs2288467 | -0.403 | 0.959 | 1.388 | -1.174 | 0.024 | -0.064 | -0.064 | 0 | 0.37 | 0.49 | 0.25 | 0.15 | 0.47 | 0.19 |
| 5 | 27010686 | rs2033236 | -0.121 | 1.145 | 1.138 | -1.083 | -0.079 | 0.089 | 0.089 | 0.077 | 0.33 | 0.12 | 0.22 | 0.15 | 0.47 | 0.19 |
| 5 | 27012531 | rs1549641 | -0.36 | 1.148 | 1.217 | -1.086 | 0.018 | -0.04 | -0.04 | 0.227 | 0.35 | 0.4 | 0.24 | 0.15 | 0.47 | 0.19 |
| 5 | 27022711 | rs10461828 | 0.505 | 0.713 | 0.848 | -0.955 | -0.014 | -0.036 | -0.036 | 0 | 0.08 | 0.01 | 0.05 | 0.11 | 0.09 | 0.09 |
| 5 | 27058850 | rs6897759 | -0.359 | 0.701 | 0.542 | -0.948 | 0.007 | 0.042 | 0.042 | 0 | 0.45 | 0.33 | 0.34 | 0.17 | 0.58 | 0.33 |
| 5 | 27072469 | rs10942232 | -0.11 | 1.168 | -1.911 | -0.684 | -0.052 | 0.145 | 0.145 | 0 | 0.42 | 0.05 | 0.32 | 0.2 | 0.58 | 0.33 |
| 5 | 27083580 | rs17567635 | -0.311 | 1.483 | -1.026 | -0.424 | 0.006 | -0.015 | -0.015 | 1.803 | 0.06 | 0.06 | 0.04 | 0.01 | 0.11 | 0.03 |
| 5 | 27084863 | rs16896509 | 0.207 | 1.192 | -1.298 | 0.091 | -0.007 | -0.038 | -0.038 | 0 | 0.06 | 0.01 | 0.12 | 0.17 | 0.07 | 0.06 |
| 5 | 27088529 | rs10520933 | 0.982 | 1.468 | -1.059 | -1.534 | 0.005 | -0.026 | -0.026 | 0 | 0.18 | 0.26 | 0.15 | 0.04 | 0.21 | 0.04 |
| 5 | 27099427 | rs10057002 | -0.225 | 1.045 | -0.107 | -1.736 | -0.007 | -0.022 | -0.022 | 0.199 | 0.12 | 0.22 | 0.1 | 0.04 | 0.09 | 0.1 |
| 5 | 27108052 | rs2876847 | -0.253 | 0.591 | -0.11 | -2.024 | 0.005 | -0.002 | -0.002 | 3.133 | 0.76 | 0.78 | 0.56 | 0.22 | 0.81 | 0.55 |
| 5 | 27115148 | rs10071841 | -0.672 | 0.671 | -0.015 | -2.373 | -0.005 | 0.048 | 0.048 | 0.075 | 0.46 | 0.61 | 0.34 | 0.08 | 0.48 | 0.26 |
| 5 | 27125142 | rs4527564 | 0.25 | 0.463 | -0.298 | -2.38 | -0.005 | 0.003 | 0.003 | 0 | 0.1 | 0.06 | 0.07 | 0.01 | 0.13 | 0.07 |
| 5 | 27135351 | rs11952975 | -0.265 | 0.435 | -0.162 | -2.444 | 0.007 | 0.063 | 0.063 | 0.296 | 0.32 | 0.16 | 0.19 | 0.04 | 0.31 | 0.15 |
| 5 | 27146918 | rs10044502 | -0.606 | 0.465 | -0.029 | -2.289 | -0.012 | -0.012 | -0.012 | 0.036 | 0.77 | 0.85 | 0.55 | 0.25 | 0.75 | 0.54 |
| 5 | 27154889 | rs11956267 | -0.525 | 0.386 | 0.611 | -2.193 | -0.016 | -0.068 | -0.068 | 0.251 | 0.32 | 0.56 | 0.18 | 0.08 | 0.24 | 0.15 |
| 5 | 27166660 | rs12109333 | -1.265 | 0.517 | 0.88 | -1.789 | -0.01 | -0.067 | -0.067 | 0 | 0.34 | 0.56 | 0.19 | 0.08 | 0.27 | 0.23 |
| 5 | 27171542 | rs4597930 | -0.829 | 0.571 | 0.385 | -1.814 | 0.002 | 0.032 | 0.032 | 0.013 | 0.3 | 0.2 | 0.16 | 0.03 | 0.27 | 0.23 |
| 5 | 27191708 | rs6880526 | -1.305 | 0.555 | 0.17 | -1.794 | 0.005 | 0.005 | 0.005 | 0 | 0.35 | 0.67 | 0.2 | 0.08 | 0.27 | 0.23 |
| 5 | 27199033 | rs6879518 | 0.579 | 0.54 | -0.215 | -1.887 | -0.023 | 0.057 | 0.057 | 0 | 0.14 | 0.01 | 0.07 | 0.03 | 0.13 | 0.09 |
| 5 | 27208707 | rs11744047 | 1.625 | 0.761 | -0.393 | -1.914 | -0.076 | 0.16 | 0.16 | 0 | 0.4 | 0.07 | 0.29 | 0.16 | 0.44 | 0.22 |
| 5 | 27231407 | rs6887000 | -0.655 | 0.784 | 0.885 | -1.548 | 0.038 | -0.071 | -0.071 | 0.091 | 0.11 | 0.48 | 0.09 | 0 | 0.09 | 0.09 |
| 5 | 27249348 | rs4133220 | -0.786 | 1.166 | 0.488 | -1.105 | -0.015 | 0.035 | 0.035 | 0 | 0.16 | 0.04 | 0.19 | 0.06 | 0.16 | 0.27 |
| 5 | 27252466 | rs4382127 | 0.623 | 1.11 | 0.689 | -1.246 | -0.01 | 0.007 | 0.007 | 0 | 0.68 | 0.8 | 0.46 | 0.29 | 0.65 | 0.38 |
| 5 | 27297644 | rs16896621 | 0.039 | 1.049 | -1.121 | -1.246 | 0 | -0.044 | -0.044 | 0.084 | 0.08 | 0.12 | 0.04 | 0 | 0.05 | 0.09 |
| 5 | 27302423 | rs2199214 | -0.114 | 1.162 | -1.889 | -1.166 | 0.006 | -0.044 | -0.044 | 0.398 | 0.17 | 0.62 | 0.14 | 0.09 | 0.15 | 0.14 |
| 5 | 27308254 | rs961208 | -0.405 | 1.313 | -2.404 | -0.467 | -0.004 | 0.009 | 0.009 | 0.037 | 0.25 | 0.11 | 0.49 | 0.76 | 0.27 | 0.57 |
| 5 | 27308663 | rs13167049 | -0.821 | 1.594 | -1.667 | 0.137 | -0.057 | 0.033 | 0.033 | 0 | 0.14 | 0.01 | 0.08 | 0.06 | 0.15 | 0.03 |
| 5 | 27328631 | rs34505 | -0.851 | 1.734 | -1.792 | 0.454 | -0.079 | 0.008 | 0.008 | 0 | 0.34 | 0.11 | 0.5 | 0.77 | 0.43 | 0.49 |
| 5 | 27352499 | rs12188777 | -0.313 | 1.382 | -1.391 | 0.172 | -0.009 | -0.027 | -0.027 | 0 | 0.05 | 0.13 | 0.03 | 0 | 0.04 | 0.06 |
| 5 | 27373862 | rs325355 | 1.308 | 1.342 | 0.445 | 0.146 | 0.008 | 0.114 | 0.114 | 0 | 0.51 | 0.3 | 0.3 | 0.14 | 0.48 | 0.42 |
| 5 | 27376682 | rs170519 | 0.885 | 1.328 | 0.49 | 0.139 | -0.002 | -0.002 | -0.002 | 0 | 0.57 | 0.45 | 0.33 | 0.14 | 0.52 | 0.49 |
| 5 | 27388778 | rs2434677 | 1.226 | 1.299 | 0.678 | 0.138 | 0.002 | 0.042 | 0.042 | 0 | 0.52 | 0.41 | 0.31 | 0.14 | 0.47 | 0.42 |
| 5 | 27390548 | rs2434678 | -0.156 | 1.21 | 0.248 | 1.448 | 0.017 | -0.017 | -0.017 | 0 | 0.92 | 0.98 | 0.85 | 0.63 | 0.93 | 0.75 |
| 5 | 27401935 | rs6866494 | 0.667 | 1.309 | 0.303 | 1.464 | -0.015 | -0.027 | -0.027 | 0 | 0.67 | 0.85 | 0.53 | 0.29 | 0.68 | 0.54 |
| 5 | 27424981 | rs7732187 | 0.978 | 1.483 | 0.24 | 1.544 | -0.078 | 0.022 | 0.022 | 0 | 0.65 | 0.87 | 0.5 | 0.21 | 0.55 | 0.55 |
| 5 | 27447863 | rs4421076 | 0.998 | 1.435 | 0.613 | 1.507 | -0.04 | 0.028 | 0.028 | 0.001 | 0.65 | 0.78 | 0.5 | 0.2 | 0.55 | 0.55 |
| 5 | 27454710 | rs16896818 | -1.423 | 1.152 | 0.911 | 1.364 | -0.014 | 0.028 | 0.028 | 2.339 | 0.11 | 0.56 | 0.21 | 0.08 | 0.09 | 0.08 |

| | | | | | | | | | | | | | | | | |
|---|---|---|---|---|---|---|---|---|---|---|---|---|---|---|---|---|
| 5 | 27456212 | rs6894886 | 1.653 | 1.132 | 1.842 | 1.261 | 0.001 | 0.029 | 0.029 | 0 | 0.55 | 0.43 | 0.31 | 0.12 | 0.47 | 0.47 |
| 5 | 27457278 | rs16896822 | -0.325 | 1.373 | 1.643 | 1.62 | -0.055 | -0.05 | -0.05 | 1.017 | 0.27 | 0.05 | 0.35 | 0.32 | 0.27 | 0.28 |
| 5 | 27509320 | rs7702167 | -2.345 | 1.184 | 2.565 | 1.423 | 0.057 | 0.053 | 0.053 | 5.73 | 0.06 | 0.45 | 0.04 | 0 | 0 | 0 |
| 5 | 27525292 | rs6866687 | 1.123 | 1.443 | 1.686 | 0.943 | -0.016 | -0.01 | -0.01 | 0.876 | 0.47 | 0.54 | 0.21 | 0.21 | 0.24 | 0.42 |
| 5 | 27532751 | rs10213971 | 0.734 | 2.636 | 1.457 | 0.606 | 0.004 | -0.062 | -0.062 | 0.325 | 0.66 | 0.62 | 0.6 | 0.94 | 0.63 | 0.76 |
| 5 | 27539490 | rs2329628 | -0.125 | 3.179 | 0.816 | 0.624 | -0.016 | -0.015 | -0.015 | 0 | 0.17 | 0.15 | 0.34 | 0.72 | 0.29 | 0.33 |
| 5 | 27539756 | rs4701557 | -0.125 | 3.179 | 0.816 | 0.624 | -0.016 | -0.015 | -0.015 | 0 | 0.17 | 0.15 | 0.34 | 0.72 | 0.29 | 0.33 |
| 5 | 27547030 | rs17526423 | 0.777 | 3.081 | 0.763 | 0.535 | -0.003 | 0.061 | 0.061 | 0 | 0.12 | 0.04 | 0.08 | 0 | 0.12 | 0.09 |
| 5 | 27559659 | rs6870385 | 1.01 | 3.077 | 0.823 | 0.614 | -0.002 | 0.136 | 0.136 | 0.132 | 0.61 | 0.25 | 0.57 | 0.95 | 0.62 | 0.76 |
| 5 | 27570571 | rs2329646 | 0.017 | 3.153 | 0.727 | 0.82 | -0.054 | -0.047 | -0.047 | 0.767 | 0.16 | 0.08 | 0.33 | 0.72 | 0.29 | 0.34 |
| 5 | 27659111 | rs7445307 | 1.306 | 2.783 | 1.42 | 0.653 | -0.002 | -0.003 | -0.003 | 0 | 0.51 | 0.48 | 0.42 | 0.3 | 0.43 | 0.5 |
| 5 | 27660532 | rs7721317 | 1.327 | 2.761 | 1.068 | 0.651 | 0.053 | 0.073 | 0.073 | 0.269 | 0.49 | 0.25 | 0.4 | 0.31 | 0.43 | 0.49 |
| 5 | 27662620 | rs7444723 | 1.677 | 2.712 | 0.478 | 0.529 | 0.061 | 0.059 | 0.059 | 1.058 | 0.44 | 0.08 | 0.23 | 0.27 | 0.35 | 0.39 |
| 5 | 27671869 | rs4867530 | -0.266 | 2.53 | 0.558 | 0.635 | -0.052 | -0.051 | -0.051 | 0.213 | 0.16 | 0.06 | 0.36 | 0.68 | 0.28 | 0.32 |
| 5 | 27684515 | rs6859290 | -1.145 | 1.475 | 0.469 | 0.086 | 0.034 | 0.04 | 0.04 | 0.067 | 0.51 | 0.57 | 0.75 | 0.74 | 0.65 | 0.67 |
| 5 | 27699870 | rs4585457 | 0.492 | 0.709 | -0.138 | -0.294 | 0.007 | -0.002 | -0.002 | 0 | 0.74 | 0.77 | 0.79 | 0.96 | 0.84 | 0.73 |
| 5 | 27708038 | rs4242128 | -1.426 | -0.439 | -0.427 | -0.65 | -0.008 | 0.013 | 0.013 | 0 | 0.45 | 0.63 | 0.55 | 0.65 | 0.46 | 0.41 |
| 5 | 27714105 | rs12651876 | 0 | -0.417 | -0.226 | -0.66 | -0.01 | -0.009 | -0.009 | 0 | 0.24 | 0.23 | 0.46 | 0.64 | 0.36 | 0.32 |
| 5 | 27719260 | rs4510599 | -1.177 | -0.362 | 0.039 | -0.374 | 0.008 | 0.029 | 0.029 | 0.146 | 0.49 | 0.39 | 0.67 | 0.68 | 0.52 | 0.58 |
| 5 | 27720584 | rs4379207 | -1.074 | -0.289 | -0.356 | -0.336 | 0.012 | 0.039 | 0.039 | 0.077 | 0.51 | 0.6 | 0.69 | 0.68 | 0.52 | 0.59 |
| 5 | 27723087 | rs4867154 | 0.195 | 0.142 | -0.184 | 0.243 | -0.02 | -0.038 | -0.038 | 0.863 | 0.23 | 0.18 | 0.4 | 0.58 | 0.32 | 0.31 |
| 5 | 27723520 | rs4867155 | -0.262 | -0.096 | -0.207 | 0.431 | 0 | 0.016 | 0.016 | 0 | 0.08 | 0.06 | 0.31 | 0.19 | 0.06 | 0.2 |
| 5 | 27726745 | rs4382181 | 0.281 | -0.386 | -0.162 | 0.349 | -0.014 | 0.002 | 0.002 | 0 | 0.65 | 0.72 | 0.42 | 0.35 | 0.59 | 0.53 |
| 5 | 27729100 | rs4642381 | 0.295 | -0.619 | 0.359 | 1.254 | -0.005 | 0.074 | 0.074 | 0 | 0.91 | 0.89 | 0.69 | 0.81 | 0.94 | 0.8 |
| 5 | 27743375 | rs7702896 | 0.733 | -0.64 | -0.108 | 1.116 | 0.005 | 0.004 | 0.004 | 0 | 0.46 | 0.37 | 0.42 | 0.5 | 0.55 | 0.43 |
| 5 | 27748053 | rs10454836 | 0.606 | -0.17 | -0.815 | 1.028 | -0.058 | -0.065 | -0.065 | 0 | 0.29 | 0.02 | 0.18 | 0.29 | 0.29 | 0.27 |
| 5 | 27770554 | rs6450563 | 0.277 | -0.138 | -0.183 | 0.443 | -0.041 | -0.113 | -0.113 | 0.006 | 0.84 | 0.45 | 0.89 | 1 | 0.89 | 0.92 |
| 5 | 27773823 | rs4254924 | 0.247 | -0.017 | -0.423 | 0.632 | -0.012 | 0.013 | 0.013 | 0 | 0.59 | 0.51 | 0.57 | 0.53 | 0.64 | 0.65 |
| 5 | 27788565 | rs13166968 | 0.815 | -0.001 | -0.685 | 0.612 | -0.025 | -0.136 | -0.136 | 0 | 0.26 | 0.07 | 0.33 | 0.45 | 0.25 | 0.26 |
| 5 | 27792192 | rs4516860 | -0.229 | 0.407 | -0.418 | 0.986 | 0.005 | -0.004 | -0.004 | 0.33 | 0.58 | 0.45 | 0.56 | 0.55 | 0.65 | 0.66 |
| 5 | 27793344 | rs6868258 | -0.276 | 0.505 | -0.794 | 0.999 | -0.008 | 0.006 | 0.006 | 0 | 0.59 | 0.48 | 0.57 | 0.55 | 0.65 | 0.66 |
| 5 | 27804605 | rs6889876 | 0.107 | 0.725 | -0.755 | 0.717 | 0.001 | 0.003 | 0.003 | 0 | 0.5 | 0.54 | 0.52 | 0.42 | 0.57 | 0.43 |
| 5 | 27817189 | rs4074741 | 0.098 | 1.347 | -0.75 | 1.032 | -0.006 | -0.009 | -0.009 | 0 | 0.28 | 0.25 | 0.29 | 0.39 | 0.38 | 0.24 |
| 5 | 27820172 | rs7710218 | 0.752 | 1.064 | -0.939 | 0.177 | 0.005 | 0.013 | 0.013 | 0 | 0.51 | 0.6 | 0.5 | 0.56 | 0.43 | 0.49 |
| 5 | 27823651 | rs4373290 | 0.705 | 0.431 | -0.545 | 0.318 | -0.016 | 0.047 | 0.047 | 0.616 | 0.17 | 0.36 | 0.1 | 0.25 | 0.1 | 0.05 |
| 5 | 27824645 | rs6862995 | 0.333 | 0.004 | -0.714 | -0.047 | 0.001 | 0.021 | 0.021 | 0 | 0.41 | 0.19 | 0.52 | 0.36 | 0.39 | 0.62 |
| 5 | 27825557 | rs7709151 | -0.799 | -0.097 | -0.472 | -0.224 | 0.008 | -0.031 | -0.031 | 0 | 0.94 | 0.91 | 0.97 | 0.99 | 0.93 | 0.97 |
| 5 | 27827756 | rs11745185 | 0.249 | -0.598 | -0.366 | -0.137 | -0.032 | 0.031 | 0.031 | 1.298 | 0.16 | 0.25 | 0.07 | 0.23 | 0.07 | 0.08 |
| 5 | 27837594 | rs10062061 | -1.147 | -0.393 | -0.609 | -0.54 | 0.008 | 0.018 | 0.018 | 0 | 0.36 | 0.28 | 0.19 | 0.26 | 0.33 | 0.45 |
| 5 | 27837898 | rs13158156 | -0.971 | -0.434 | -0.469 | -0.411 | -0.015 | 0.008 | 0.008 | 0.057 | 0.3 | 0.19 | 0.17 | 0.24 | 0.32 | 0.19 |
| 5 | 27839258 | rs7717247 | -0.757 | -0.581 | 0.018 | -0.377 | -0.015 | 0.003 | 0.003 | 0.056 | 0.26 | 0.41 | 0.14 | 0.24 | 0.18 | 0.13 |
| 5 | 27858188 | rs6884792 | 1.054 | -0.752 | -0.267 | 0.073 | -0.014 | 0.037 | 0.037 | 0 | 0.22 | 0.3 | 0.44 | 0.36 | 0.17 | 0.27 |
| 5 | 27858577 | rs4385219 | 0.015 | -0.768 | -0.566 | -0.167 | -0.04 | 0.053 | 0.053 | 0 | 0.19 | 0.04 | 0.09 | 0.09 | 0.17 | 0.11 |
| 5 | 27862523 | rs7708565 | -0.858 | -1.073 | -0.548 | -0.15 | -0.001 | 0.01 | 0.01 | 0 | 0.56 | 0.42 | 0.6 | 0.68 | 0.44 | 0.48 |
| 5 | 27863084 | rs4371767 | 0.147 | -1.087 | -0.54 | -0.138 | -0.002 | -0.001 | -0.001 | 0.153 | 0.38 | 0.38 | 0.51 | 0.58 | 0.27 | 0.37 |
| 5 | 27863479 | rs10043413 | 0.901 | -0.965 | -0.517 | -0.202 | -0.002 | -0.003 | -0.003 | 0.853 | 0.07 | 0.07 | 0.05 | 0 | 0.14 | 0.07 |
| 5 | 27873179 | rs4867034 | -0.302 | -1.02 | -0.66 | -0.179 | -0.008 | 0 | 0 | 0.494 | 0.62 | 0.61 | 0.48 | 0.42 | 0.73 | 0.63 |
| 5 | 27875181 | rs9292229 | 0.642 | -0.825 | -0.686 | -0.353 | 0 | 0.001 | 0.001 | 0 | 0.43 | 0.55 | 0.39 | 0.32 | 0.56 | 0.53 |
| 5 | 27877131 | rs10069219 | 0.724 | -0.831 | -0.602 | -0.404 | 0.001 | -0.009 | -0.009 | 0 | 0.42 | 0.45 | 0.38 | 0.32 | 0.56 | 0.53 |
| 5 | 27896789 | rs1981979 | 1.233 | -0.851 | -0.584 | -0.315 | 0.002 | -0.057 | -0.057 | 0 | 0.21 | 0.11 | 0.45 | 0.59 | 0.21 | 0.34 |
| 5 | 27897183 | rs7445766 | 1.045 | -0.914 | -0.45 | -0.306 | 0.005 | 0.002 | 0.002 | 0 | 0.38 | 0.41 | 0.34 | 0.41 | 0.43 | 0.48 |
| 5 | 27907713 | rs1991500 | 1.019 | -0.824 | -0.621 | -0.264 | 0.001 | 0 | 0 | 0 | 0.38 | 0.41 | 0.33 | 0.4 | 0.43 | 0.48 |
| 5 | 27914927 | rs1428589 | 1.014 | -0.815 | -0.667 | -0.201 | 0.001 | 0 | 0 | 0 | 0.38 | 0.42 | 0.33 | 0.41 | 0.43 | 0.48 |
| 5 | 27932273 | rs13153124 | -1.297 | -0.818 | -1.153 | -0.222 | 0 | 0.004 | 0.004 | 0 | 0.79 | 0.77 | 0.55 | 0.41 | 0.81 | 0.68 |
| 5 | 27932380 | rs13153654 | -0.634 | -0.802 | -1.306 | -0.223 | -0.009 | 0.044 | 0.044 | 0.158 | 0.4 | 0.23 | 0.2 | 0 | 0.38 | 0.21 |
| 5 | 27935737 | rs1366436 | -1.173 | -0.789 | -2.07 | -0.132 | 0.01 | -0.096 | -0.096 | 0 | 0.78 | 0.94 | 0.56 | 0.41 | 0.81 | 0.67 |
| 5 | 27947512 | rs16897223 | 1.554 | -0.535 | -1.929 | -0.447 | -0.006 | -0.096 | -0.096 | 0 | 0.18 | 0.04 | 0.41 | 0.4 | 0.18 | 0.25 |
| 5 | 27947537 | rs10472596 | -1.112 | -0.63 | -1.942 | -0.518 | -0.003 | -0.003 | -0.003 | 0 | 0.61 | 0.59 | 0.64 | 0.4 | 0.55 | 0.47 |
| 5 | 27947751 | rs10520937 | 1.062 | 0.202 | -2.701 | 0.276 | -0.003 | -0.096 | -0.096 | 0 | 0.18 | 0.04 | 0.41 | 0.4 | 0.17 | 0.25 |
| 5 | 27949271 | rs11739378 | 0.546 | -0.374 | -2.98 | -0.215 | -0.055 | 0.155 | 0.155 | 0.667 | 0.23 | 0.02 | 0.13 | 0 | 0.25 | 0.1 |
| 5 | 27952031 | rs30809 | 0.302 | -0.347 | -1.899 | 0.974 | -0.009 | -0.12 | -0.12 | 0 | 0.2 | 0.04 | 0.43 | 0.42 | 0.22 | 0.28 |
| 5 | 27952573 | rs13159803 | -0.974 | 0.134 | -0.979 | 0.58 | 0.001 | 0.065 | 0.065 | 0 | 0.14 | 0.03 | 0.03 | 0 | 0.07 | 0.09 |
| 5 | 27969709 | rs2876935 | 0.573 | 0.46 | -1.219 | 1.665 | 0.013 | -0.08 | -0.08 | 0 | 0.86 | 0.98 | 0.8 | 0.67 | 0.9 | 0.71 |
| 5 | 27973753 | rs919329 | -0.689 | 0.955 | -0.633 | 1.36 | 0.002 | 0.019 | 0.019 | 0 | 0.13 | 0.18 | 0.08 | 0.18 | 0.12 | 0.13 |
| 5 | 27982697 | rs10075153 | -0.127 | 1.552 | -0.402 | 0.134 | 0 | -0.018 | -0.018 | 0.512 | 0.35 | 0.32 | 0.26 | 0.12 | 0.39 | 0.28 |
| 5 | 27983106 | rs10070047 | -0.147 | 1.808 | 0.182 | 1.125 | 0.036 | 0.084 | 0.084 | 0 | 0.9 | 0.76 | 0.74 | 0.66 | 0.81 | 0.84 |
| 5 | 27987104 | rs4537065 | 0.747 | 1.287 | 0.264 | 1.447 | 0.011 | 0.047 | 0.047 | 0 | 0.72 | 0.49 | 0.62 | 0.35 | 0.7 | 0.68 |
| 5 | 27988819 | rs373087 | 0.467 | 1.117 | 0.158 | 1.356 | -0.015 | -0.077 | -0.077 | 0 | 0.11 | 0.01 | 0.08 | 0.31 | 0.09 | 0.14 |
| 5 | 27992491 | rs2876936 | 1.223 | 0.989 | 0.337 | 1.308 | 0.005 | 0.04 | 0.04 | 0 | 0.7 | 0.4 | 0.61 | 0.35 | 0.69 | 0.69 |

| | | | | | | | | | | | | | | | | |
|---|---|---|---|---|---|---|---|---|---|---|---|---|---|---|---|---|
| 5 | 28006650 | rs4398645 | 0.777 | 0.967 | 0.392 | 1.298 | 0.013 | 0.037 | 0.037 | 0 | 0.75 | 0.63 | 0.63 | 0.35 | 0.69 | 0.68 |
| 5 | 28017229 | rs1494617 | 1.191 | 0.713 | 1.051 | 1.173 | 0.011 | 0.052 | 0.052 | 0 | 0.71 | 0.49 | 0.62 | 0.35 | 0.69 | 0.68 |
| 5 | 28025934 | rs7710236 | -0.041 | 0.719 | 1.033 | 1.322 | 0.03 | 0.068 | 0.068 | 0 | 0.12 | 0.24 | 0.27 | 0.38 | 0.22 | 0.17 |
| 5 | 28028745 | rs1603086 | 0.923 | 0.918 | 1.059 | 1.218 | 0 | 0.012 | 0.012 | 0.661 | 0.33 | 0.26 | 0.24 | 0.11 | 0.35 | 0.27 |
| 5 | 28038878 | rs16897331 | 0.649 | 0.914 | 0.692 | 1.376 | -0.039 | -0.045 | -0.045 | 0 | 0.08 | 0.01 | 0.21 | 0.27 | 0.12 | 0.12 |
| 5 | 28046365 | rs11745759 | 1.062 | 0.875 | 0.713 | 1.324 | -0.008 | 0.039 | 0.039 | 0.206 | 0.37 | 0.16 | 0.36 | 0.23 | 0.34 | 0.41 |
| 5 | 28056225 | rs10940703 | -1.098 | 0.889 | 0.727 | 1.356 | 0.042 | 0.085 | 0.085 | 0 | 0.17 | 0.4 | 0.29 | 0.35 | 0.22 | 0.17 |
| 5 | 28059732 | rs1494608 | -1.011 | 0.69 | -0.154 | 1.545 | 0.011 | -0.03 | -0.03 | 0 | 0.93 | 1 | 0.95 | 0.76 | 0.96 | 0.92 |
| 5 | 28084454 | rs7711451 | 0.81 | 0.784 | -0.122 | 1.339 | -0.036 | 0.057 | 0.057 | 0.11 | 0.39 | 0.13 | 0.38 | 0.22 | 0.41 | 0.41 |
| 5 | 28091661 | rs1494605 | -0.444 | 0.824 | -0.146 | 1.259 | -0.053 | -0.074 | -0.074 | 0 | 0.6 | 0.92 | 0.68 | 0.58 | 0.56 | 0.58 |
| 5 | 28102108 | rs1494613 | 0.472 | 0.399 | 0.049 | 0.939 | -0.016 | 0.037 | 0.037 | 0 | 0.61 | 0.82 | 0.64 | 0.68 | 0.62 | 0.55 |
| 5 | 28112135 | rs10067148 | -0.178 | 0.405 | 0.578 | 0.969 | 0.004 | -0.001 | -0.001 | 0 | 0.69 | 0.84 | 0.68 | 0.69 | 0.7 | 0.59 |
| 5 | 28118124 | rs4293931 | 0.826 | 0.68 | 0.916 | 1.325 | 0.067 | 0.112 | 0.112 | 0 | 0.83 | 0.47 | 0.72 | 0.72 | 0.83 | 0.91 |
| 5 | 28134334 | rs10940704 | 0.707 | 0.677 | 1.133 | 1.323 | 0.058 | 0.095 | 0.095 | 0 | 0.83 | 0.6 | 0.72 | 0.72 | 0.83 | 0.91 |
| 5 | 28142099 | rs11745152 | 1.616 | 0.713 | 0.827 | 1.212 | -0.053 | 0.139 | 0.139 | 0.813 | 0.39 | 0.06 | 0.28 | 0.17 | 0.44 | 0.35 |
| 5 | 28142772 | rs10472603 | 1.423 | 0.685 | 0.878 | 1.199 | 0.003 | 0.039 | 0.039 | 0 | 0.43 | 0.32 | 0.31 | 0.17 | 0.44 | 0.35 |
| 5 | 28145629 | rs12513515 | 0.992 | 0.488 | 0.527 | 1.351 | -0.028 | -0.043 | -0.043 | 0 | 0.23 | 0.03 | 0.28 | 0.22 | 0.24 | 0.35 |
| 5 | 28145684 | rs4867495 | 0.106 | 0.537 | 0.547 | 1.346 | -0.01 | -0.065 | -0.065 | 0 | 0.13 | 0.02 | 0.1 | 0.24 | 0.12 | 0.14 |
| 5 | 28146527 | rs13179093 | 0.603 | 0.54 | 0.555 | 1.338 | 0.007 | -0.01 | -0.01 | 0.128 | 0.27 | 0.12 | 0.3 | 0.31 | 0.27 | 0.41 |
| 5 | 28172825 | rs4305648 | -0.317 | 0.593 | 0.408 | 1.23 | 0.012 | -0.047 | -0.047 | 0.331 | 0.4 | 0.16 | 0.41 | 0.55 | 0.39 | 0.56 |
| 5 | 28183468 | rs6893616 | 1.269 | 0.671 | 0.174 | 1.074 | -0.068 | 0.142 | 0.142 | 0.569 | 0.39 | 0.06 | 0.28 | 0.17 | 0.45 | 0.36 |
| 5 | 28184320 | rs10070878 | 0.833 | 0.726 | 0.669 | 1.618 | 0.063 | 0.287 | 0.287 | 0.27 | 0.89 | 0.67 | 0.75 | 0.6 | 0.89 | 0.9 |
| 5 | 28195096 | rs7702050 | -0.173 | 0.363 | 0.046 | 0.93 | 0.001 | 0.005 | 0.005 | 0 | 0.29 | 0.25 | 0.27 | 0.14 | 0.25 | 0.33 |
| 5 | 28236629 | rs1113342 | -1.371 | 0.13 | 0.229 | 1.461 | 0.012 | -0.019 | -0.019 | 0 | 0.29 | 0.77 | 0.4 | 0.67 | 0.2 | 0.24 |
| 5 | 28239056 | rs10472606 | -0.962 | 0.106 | 0.841 | 1.49 | 0.024 | 0.176 | 0.176 | 1.647 | 0.14 | 0.35 | 0.25 | 0.42 | 0.1 | 0.16 |
| 5 | 28261961 | rs6859620 | -0.432 | 0.055 | 0.933 | 1.415 | 0.012 | 0.101 | 0.101 | 0 | 0.11 | 0.22 | 0.24 | 0.41 | 0.1 | 0.16 |
| 5 | 28264465 | rs1501681 | -1.397 | 0.009 | 0.928 | 1.369 | -0.006 | 0.025 | 0.025 | 0.133 | 0.24 | 0.31 | 0.36 | 0.67 | 0.2 | 0.23 |
| 5 | 28281249 | rs2468483 | -0.953 | -0.268 | 0.161 | 0.945 | -0.001 | -0.016 | -0.016 | 0.519 | 0.78 | 0.97 | 0.76 | 0.86 | 0.8 | 0.69 |
| 5 | 28282431 | rs931739 | -0.478 | -0.197 | 0.389 | 1.111 | 0.015 | 0.124 | 0.124 | 0.049 | 0.12 | 0.27 | 0.24 | 0.41 | 0.1 | 0.2 |
| 5 | 28283502 | rs2468482 | -1.665 | -0.241 | 0.326 | 1.047 | -0.007 | 0.01 | 0.01 | 1.324 | 0.28 | 0.69 | 0.4 | 0.67 | 0.2 | 0.27 |
| 5 | 28286502 | rs1393097 | -0.406 | 0.031 | -0.657 | 0.896 | -0.053 | 0.036 | 0.036 | 0.73 | 0.7 | 0.97 | 0.71 | 0.86 | 0.7 | 0.62 |
| 5 | 28292827 | rs2468477 | -1.557 | 0.032 | 0.165 | 1.496 | -0.01 | 0.074 | 0.074 | 0 | 0.24 | 0.47 | 0.38 | 0.67 | 0.2 | 0.27 |
| 5 | 28295497 | rs2434548 | -0.299 | -0.063 | -0.335 | 1.038 | -0.014 | 0.057 | 0.057 | 0 | 0.11 | 0.2 | 0.13 | 0.27 | 0.09 | 0.07 |
| 5 | 28302858 | rs10056131 | -0.168 | 0.101 | -1.219 | -0.18 | -0.028 | 0.12 | 0.12 | 0 | 0.46 | 0.22 | 0.66 | 0.76 | 0.51 | 0.55 |
| 5 | 28310093 | rs1154991 | -0.553 | -0.827 | -0.719 | -0.471 | -0.017 | -0.066 | -0.066 | 0 | 0.24 | 0.7 | 0.14 | 0.02 | 0.1 | 0.11 |
| 5 | 28319612 | rs10061823 | 0.15 | -0.51 | -0.534 | -0.514 | 0.018 | 0.107 | 0.107 | 0.891 | 0.48 | 0.25 | 0.28 | 0.09 | 0.54 | 0.37 |
| 5 | 28324647 | rs1006069 | 0.032 | -0.492 | -0.129 | -0.534 | 0.013 | 0.059 | 0.059 | 1.054 | 0.48 | 0.29 | 0.29 | 0.09 | 0.54 | 0.37 |
| 5 | 28329667 | rs1582924 | -0.221 | -0.514 | -0.002 | -0.624 | 0 | -0.011 | -0.011 | 0 | 0.49 | 0.58 | 0.66 | 0.82 | 0.37 | 0.51 |
| 5 | 28345493 | rs4867532 | 0.18 | -0.533 | 0.191 | -0.714 | 0.01 | 0.013 | 0.013 | 0.899 | 0.43 | 0.42 | 0.64 | 0.82 | 0.35 | 0.47 |
| 5 | 28380056 | rs9292252 | -0.673 | -0.293 | -0.118 | -0.69 | -0.042 | 0.211 | 0.211 | 0.292 | 0.58 | 0.86 | 0.76 | 0.94 | 0.48 | 0.69 |
| 5 | 28387558 | rs4867536 | -0.079 | -0.688 | 0.109 | -0.832 | -0.013 | -0.037 | -0.037 | 0.506 | 0.19 | 0.26 | 0.16 | 0.12 | 0.17 | 0.18 |
| 5 | 28397668 | rs4260681 | -0.02 | -0.808 | 0.037 | -0.857 | 0.006 | 0.012 | 0.012 | 0.875 | 0.37 | 0.36 | 0.6 | 0.85 | 0.32 | 0.51 |
| 5 | 28402022 | rs7735638 | -0.953 | -1.124 | 0.314 | -0.715 | -0.047 | -0.068 | -0.068 | 0 | 0.35 | 0.44 | 0.53 | 0.86 | 0.2 | 0.34 |
| 5 | 28422193 | rs4383731 | 0.309 | -1.228 | 0.314 | 0.407 | -0.002 | 0.006 | 0.006 | 0 | 0.15 | 0.14 | 0.39 | 0.37 | 0.07 | 0.13 |
| 5 | 28444697 | rs10076533 | 0.276 | -1.284 | 0.512 | 0.477 | -0.012 | 0.02 | 0.02 | 0 | 0.16 | 0.21 | 0.4 | 0.4 | 0.07 | 0.13 |
| 5 | 28464893 | rs13355546 | 0.327 | -1.279 | 0.571 | 0.495 | -0.003 | -0.002 | -0.002 | 0 | 0.15 | 0.14 | 0.19 | 0.25 | 0.07 | 0.12 |
| 5 | 28465895 | rs6889117 | 0.535 | -0.7 | 0.737 | 0.521 | 0.029 | 0.013 | 0.013 | 0 | 0.05 | 0.09 | 0.07 | 0.08 | 0.13 | 0.21 |
| 5 | 28472761 | rs7446953 | 0.436 | -1.259 | 1.049 | 0.415 | -0.029 | 0.128 | 0.128 | 0.261 | 0.82 | 0.57 | 0.66 | 0.37 | 0.9 | 0.79 |
| 5 | 28478206 | rs11953455 | 0.629 | -1.311 | 1.135 | 0.343 | -0.032 | 0.096 | 0.096 | 0.633 | 0.81 | 0.48 | 0.65 | 0.35 | 0.9 | 0.78 |
| 5 | 28489162 | rs6891511 | 1.231 | -1.488 | 1.045 | 0.136 | -0.001 | -0.008 | -0.008 | 0.631 | 0.66 | 0.37 | 0.46 | 0.11 | 0.82 | 0.66 |
| 5 | 28489325 | rs4266409 | -0.032 | -1.387 | 1.496 | 0.35 | 0.029 | 0.073 | 0.073 | 0.043 | 0.06 | 0.25 | 0.04 | 0.13 | 0.04 | 0.17 |
| 5 | 28494582 | rs7730170 | 0.314 | -0.796 | 0.531 | -0.562 | -0.06 | 0.091 | 0.091 | 0.119 | 0.32 | 0.14 | 0.24 | 0.02 | 0.46 | 0.26 |
| 5 | 28494602 | rs7712228 | 0.608 | -1.137 | 0.589 | -0.639 | 0.002 | -0.003 | -0.003 | 0 | 0.64 | 0.36 | 0.45 | 0.12 | 0.81 | 0.64 |
| 5 | 28495829 | rs10940731 | -0.92 | -1.27 | 0.475 | -0.391 | 0.022 | -0.07 | -0.07 | 0 | 0.86 | 0.95 | 0.69 | 0.5 | 0.92 | 0.95 |
| 5 | 28498983 | rs9292261 | 0.424 | -1.291 | 0.628 | -0.448 | -0.015 | -0.016 | -0.016 | 4.444 | 0.68 | 0.66 | 0.48 | 0.15 | 0.81 | 0.66 |
| 5 | 28500714 | rs9986277 | 1.553 | -1.109 | 0.414 | -0.362 | -0.018 | 0.056 | 0.056 | 0 | 0.33 | 0.23 | 0.21 | 0.1 | 0.35 | 0.38 |
| 5 | 28513267 | rs10472638 | 1.465 | -1.114 | 0.502 | -0.36 | -0.004 | 0.01 | 0.01 | 0 | 0.33 | 0.3 | 0.22 | 0.13 | 0.36 | 0.4 |
| 5 | 28549181 | rs4867568 | 0.075 | -1.124 | 0.193 | -0.495 | -0.009 | 0.012 | 0.012 | 0 | 0.58 | 0.65 | 0.59 | 0.63 | 0.49 | 0.5 |
| 5 | 28560977 | rs10045332 | 1.658 | -1.245 | 0.828 | 0.219 | 0.067 | 0.065 | 0.065 | 0 | 0.57 | 0.34 | 0.55 | 0.2 | 0.77 | 0.58 |
| 5 | 28561546 | rs9292272 | 1.557 | -1.246 | 0.858 | 0.223 | -0.005 | -0.002 | -0.002 | 0.247 | 0.58 | 0.45 | 0.56 | 0.22 | 0.77 | 0.6 |
| 5 | 28573939 | rs9292273 | -0.032 | -1.01 | 0.935 | -0.224 | -0.046 | 0.018 | 0.018 | 0 | 0.81 | 0.68 | 0.9 | 0.73 | 0.89 | 0.7 |
| 5 | 28581874 | rs10076497 | -0.86 | -0.957 | 0.537 | -0.215 | -0.052 | -0.009 | -0.009 | 0 | 0.29 | 0.5 | 0.14 | 0.3 | 0.15 | 0.37 |
| 5 | 28621293 | rs6450652 | 1.638 | 0.098 | 0.621 | -0.02 | -0.007 | -0.007 | -0.007 | 0 | 0.56 | 0.49 | 0.52 | 0.34 | 0.62 | 0.56 |
| 5 | 28621394 | rs4389700 | -0.418 | 0.03 | 0.826 | -0.005 | 0.02 | 0.01 | 0.01 | 0 | 0.23 | 0.16 | 0.11 | 0.18 | 0.17 | 0.34 |
| 5 | 28621926 | rs7705800 | 0.139 | 0.207 | 0.781 | -0.004 | 0.019 | -0.091 | -0.091 | 0.124 | 0.85 | 0.65 | 0.87 | 1 | 0.82 | 0.93 |
| 5 | 28627343 | rs10039017 | 0.824 | 0.176 | 0.774 | 0.064 | 0.003 | 0.081 | 0.081 | 0 | 0.79 | 0.65 | 0.63 | 0.52 | 0.78 | 0.89 |
| 5 | 28628417 | rs7710054 | 2.29 | 0.055 | 0.7 | 0.009 | 0.046 | 0.082 | 0.082 | 0 | 0.53 | 0.3 | 0.5 | 0.27 | 0.61 | 0.53 |
| 5 | 28629677 | rs4368739 | 1.259 | 0.286 | 0.912 | 0.16 | 0.001 | 0.116 | 0.116 | 0.259 | 0.77 | 0.46 | 0.61 | 0.46 | 0.77 | 0.87 |
| 5 | 28630178 | rs10940741 | 1.224 | 0.284 | 0.912 | 0.126 | 0 | 0.15 | 0.15 | 0.411 | 0.77 | 0.46 | 0.62 | 0.49 | 0.78 | 0.87 |

| | | | | | | | | | | | | | | | | |
|---|---|---|---|---|---|---|---|---|---|---|---|---|---|---|---|---|
| 5 | 28635825 | rs6865063 | 1.933 | 0.185 | 0.76 | -0.314 | -0.01 | 0.029 | 0.029 | 0 | 0.69 | 0.39 | 0.41 | 0.46 | 0.7 | 0.82 |
| 5 | 28636352 | rs4867149 | -0.473 | 0.607 | 0.526 | -0.28 | -0.017 | 0.028 | 0.028 | 0 | 0.93 | 0.95 | 0.74 | 0.5 | 0.93 | 0.97 |
| 5 | 28640898 | rs9292278 | 0.868 | 0.587 | 0.817 | -0.784 | -0.011 | -0.024 | -0.024 | 0.268 | 0.78 | 0.78 | 0.48 | 0.41 | 0.72 | 0.8 |
| 5 | 28646108 | rs13357735 | -0.073 | 0.635 | 0.595 | -1.018 | 0.004 | -0.012 | -0.012 | 1.182 | 0.23 | 0.12 | 0.12 | 0.23 | 0.23 | 0.31 |
| 5 | 28648576 | rs6450656 | -1.748 | 0.717 | 0.518 | -1.238 | 0 | 0.141 | 0.141 | 0 | 0.48 | 0.69 | 0.68 | 0.86 | 0.51 | 0.54 |
| 5 | 28652860 | rs4527615 | -1.678 | 0.769 | 0.37 | -1.31 | 0.001 | 0.052 | 0.052 | 0 | 0.56 | 0.68 | 0.71 | 0.84 | 0.59 | 0.63 |
| 5 | 28653504 | rs12659954 | -0.436 | 0.778 | 0.378 | -1.015 | 0.001 | 0.05 | 0.05 | 0 | 0.15 | 0.22 | 0.3 | 0.64 | 0.13 | 0.11 |
| 5 | 28662328 | rs2042319 | 0.233 | 0.779 | 0.565 | -1.023 | -0.002 | -0.028 | -0.028 | 0 | 0.07 | 0.19 | 0.05 | 0.02 | 0.05 | 0.04 |
| 5 | 28664932 | rs13166838 | -0.227 | 0.722 | 0.493 | -1.231 | -0.008 | 0.012 | 0.012 | 0.02 | 0.29 | 0.09 | 0.19 | 0.19 | 0.28 | 0.36 |
| 5 | 28665804 | rs11950597 | 2.044 | 0.867 | 0.717 | -1.299 | 0.003 | 0.04 | 0.04 | 0 | 0.46 | 0.35 | 0.3 | 0.15 | 0.43 | 0.39 |
| 5 | 28666832 | rs1593019 | 1.206 | 0.873 | 0.787 | -1.295 | 0.004 | 0.023 | 0.023 | 0 | 0.58 | 0.59 | 0.53 | 0.16 | 0.6 | 0.51 |
| 5 | 28673359 | rs1862558 | 0.985 | 0.883 | 0.73 | -1.32 | 0.001 | 0.067 | 0.067 | 0 | 0.6 | 0.7 | 0.54 | 0.18 | 0.61 | 0.55 |
| 5 | 28678123 | rs1862556 | -0.876 | 1.005 | 0.661 | -1.27 | 0.015 | 0.2 | 0.2 | 0 | 0.94 | 0.87 | 0.77 | 0.36 | 0.93 | 0.91 |
| 5 | 28679153 | rs2161536 | 0.618 | 1.002 | 0.283 | -1.277 | -0.029 | 0.028 | 0.028 | 0 | 0.08 | 0.02 | 0.2 | 0 | 0.17 | 0.06 |
| 5 | 28682246 | rs11746337 | 1.751 | 0.765 | 0.356 | -1.154 | 0.003 | 0.041 | 0.041 | 0 | 0.46 | 0.36 | 0.3 | 0.15 | 0.44 | 0.39 |
| 5 | 28685156 | rs2042321 | 0.858 | 0.855 | 0.496 | -0.344 | 0.023 | 0.284 | 0.284 | 0 | 0.91 | 0.71 | 0.75 | 0.36 | 0.93 | 0.85 |
| 5 | 28690695 | rs12186716 | 0.615 | 1.082 | -0.59 | -1.221 | -0.001 | -0.007 | -0.007 | 0 | 0.05 | 0.07 | 0.04 | 0 | 0.05 | 0.01 |
| 5 | 28701288 | rs6865759 | 0.186 | 1.096 | -0.699 | -1.264 | 0 | 0.003 | 0.003 | 0 | 0.11 | 0.1 | 0.13 | 0 | 0.11 | 0.11 |
| 5 | 28703749 | rs11742999 | 0.133 | 1.007 | -1.098 | -1.495 | 0 | 0.006 | 0.006 | 0 | 0.24 | 0.1 | 0.34 | 0.8 | 0.2 | 0.32 |
| 5 | 28707892 | rs4406145 | -0.183 | 1.265 | -0.464 | -1.11 | 0.01 | -0.047 | -0.047 | 0 | 0.11 | 0.14 | 0.19 | 0 | 0.14 | 0.02 |
| 5 | 28713125 | rs6882327 | -1.096 | 1.185 | -0.559 | -1.109 | 0.006 | -0.035 | -0.035 | 0.421 | 0.4 | 0.49 | 0.43 | 0.17 | 0.43 | 0.42 |
| 5 | 28719681 | rs1423285 | 0.196 | 1.267 | -0.91 | -0.396 | -0.022 | -0.02 | -0.02 | 0 | 0.85 | 0.94 | 0.91 | 0.82 | 0.83 | 0.72 |
| 5 | 28733753 | rs1862559 | 1.071 | 1.559 | -0.301 | -0.507 | 0.026 | -0.019 | -0.019 | 0.119 | 0.69 | 0.44 | 0.64 | 0.74 | 0.62 | 0.6 |
| 5 | 28739139 | rs16898050 | -0.757 | 1.697 | 0.036 | -0.538 | 0.093 | 0.078 | 0.078 | 0 | 0.09 | 0.39 | 0.21 | 0.13 | 0.13 | 0.08 |
| 5 | 28747075 | rs12523586 | 0.681 | 1.408 | -0.447 | -0.886 | 0.003 | -0.054 | -0.054 | 0 | 0.1 | 0.02 | 0.25 | 0.7 | 0.08 | 0.2 |
| 5 | 28754303 | rs1345787 | 1.25 | 1.488 | -0.242 | -0.756 | 0.026 | -0.021 | -0.021 | 0 | 0.68 | 0.44 | 0.64 | 0.74 | 0.62 | 0.6 |
| 5 | 28763652 | rs1160444 | 0.209 | 1.153 | -0.424 | -0.725 | -0.043 | 0.028 | 0.028 | 0 | 0.8 | 0.93 | 0.87 | 0.87 | 0.75 | 0.68 |
| 5 | 28775510 | rs16898120 | 0.44 | 0.892 | -0.958 | -0.497 | -0.074 | -0.147 | -0.147 | 0 | 0.27 | 0.03 | 0.25 | 0.52 | 0.27 | 0.32 |
| 5 | 28786844 | rs1363873 | 1.059 | 0.518 | -0.478 | -0.804 | -0.001 | -0.04 | -0.04 | 0.018 | 0.47 | 0.41 | 0.42 | 0.04 | 0.52 | 0.31 |
| 5 | 28831513 | rs13169827 | -0.307 | 0.358 | -0.371 | -0.936 | -0.106 | 0.087 | 0.087 | 0 | 0.41 | 0.09 | 0.36 | 0.71 | 0.44 | 0.6 |
| 5 | 28849970 | rs1587625 | -0.364 | 0.315 | -0.129 | -0.954 | -0.109 | 0.067 | 0.067 | 0 | 0.4 | 0.08 | 0.35 | 0.69 | 0.44 | 0.58 |
| 5 | 28853116 | rs2652684 | -0.851 | 0.236 | 0.326 | -0.991 | -0.002 | 0.005 | 0.005 | 0 | 0.47 | 0.54 | 0.41 | 0.82 | 0.45 | 0.65 |
| 5 | 28855283 | rs2244541 | -0.97 | 0.244 | 0.317 | -0.983 | -0.01 | 0.05 | 0.05 | 0 | 0.25 | 0.56 | 0.33 | 0.41 | 0.21 | 0.35 |
| 5 | 28860081 | rs17616969 | 0.534 | 0.348 | 0.253 | -0.919 | -0.045 | -0.039 | -0.039 | 0.022 | 0.14 | 0.04 | 0.1 | 0.16 | 0.19 | 0.27 |
| 5 | 28874113 | rs13183423 | -0.203 | 0.254 | 0.402 | -0.983 | -0.113 | 0.065 | 0.065 | 0 | 0.4 | 0.08 | 0.35 | 0.69 | 0.44 | 0.58 |
| 5 | 28899999 | rs2081919 | 1.297 | 0.213 | 0.393 | -1.019 | 0.005 | 0.064 | 0.064 | 0.411 | 0.93 | 0.6 | 0.96 | 0.89 | 0.99 | 0.95 |
| 5 | 28902826 | rs1818121 | -0.253 | 0.342 | 0.136 | -0.987 | -0.021 | 0.054 | 0.054 | 0 | 0.42 | 0.25 | 0.37 | 0.71 | 0.44 | 0.6 |
| 5 | 28921260 | rs397425 | -0.529 | 0.49 | -0.353 | -0.917 | -0.001 | -0.003 | -0.003 | 0 | 0.45 | 0.46 | 0.39 | 0.71 | 0.44 | 0.6 |
| 5 | 28927036 | rs434405 | -0.529 | 0.49 | -0.353 | -0.917 | -0.001 | -0.003 | -0.003 | 0 | 0.45 | 0.46 | 0.39 | 0.71 | 0.44 | 0.6 |
| 5 | 28929082 | rs6869025 | 0.604 | 0.455 | -0.4 | -0.979 | 0 | 0.077 | 0.077 | 0 | 0.51 | 0.35 | 0.59 | 0.18 | 0.55 | 0.35 |
| 5 | 28942952 | rs10077940 | 1.029 | 0.861 | -0.639 | -1.191 | 0.003 | 0.042 | 0.042 | 0 | 0.42 | 0.29 | 0.35 | 0.04 | 0.44 | 0.28 |
| 5 | 28968268 | rs16899662 | 0.743 | 0.965 | -0.871 | -1.048 | -0.051 | -0.141 | -0.141 | 1.848 | 0.25 | 0.03 | 0.25 | 0.53 | 0.24 | 0.31 |
| 5 | 28986950 | rs10039754 | 0.168 | 0.98 | -0.261 | -1.353 | 0 | 0.104 | 0.104 | 0 | 0.51 | 0.67 | 0.45 | 0.04 | 0.55 | 0.33 |
| 5 | 29017460 | rs7710990 | 0.622 | 0.688 | -0.856 | -1.5 | -0.001 | -0.03 | -0.03 | 0 | 0.29 | 0.27 | 0.21 | 0.04 | 0.35 | 0.18 |
| 5 | 29023795 | rs10040459 | 0.164 | 0.747 | -0.87 | -1.602 | -0.01 | 0.006 | 0.006 | 0 | 0.43 | 0.35 | 0.37 | 0.04 | 0.51 | 0.3 |
| 5 | 29028609 | rs12153763 | -0.139 | 0.219 | -0.868 | -1.606 | 0.002 | 0.002 | 0.002 | 0 | 0.19 | 0.2 | 0.31 | 0.86 | 0.15 | 0.31 |
| 5 | 29028901 | rs10051091 | 0.518 | 0.726 | -0.835 | -2.068 | 0 | -0.121 | -0.121 | 0.392 | 0.3 | 0.39 | 0.25 | 0.04 | 0.3 | 0.25 |
| 5 | 29031195 | rs12653139 | -0.752 | 0.158 | -0.531 | -2.51 | 0.004 | -0.026 | -0.026 | 0 | 0.6 | 0.66 | 0.59 | 0.87 | 0.49 | 0.61 |
| 5 | 29044517 | rs1692347 | -0.285 | 0.093 | -0.653 | -2.397 | 0.013 | 0.017 | 0.017 | 0 | 0.29 | 0.31 | 0.34 | 0.82 | 0.19 | 0.33 |
| 5 | 29051880 | rs12153251 | 1.089 | 0.115 | -0.939 | -2.764 | -0.077 | -0.005 | -0.005 | 0 | 0.17 | 0.04 | 0.15 | 0.13 | 0.3 | 0.25 |
| 5 | 29054620 | rs631844 | -0.736 | 0.094 | -0.86 | -2.777 | -0.03 | 0.16 | 0.16 | 0 | 0.45 | 0.31 | 0.49 | 0.96 | 0.49 | 0.58 |
| 5 | 29059043 | rs16898675 | 1.46 | -0.049 | -0.861 | -2.289 | 0.003 | 0.002 | 0.002 | 0 | 0.05 | 0.03 | 0.19 | 0.14 | 0.03 | 0.03 |
| 5 | 29065808 | rs505761 | -0.201 | -0.02 | -0.852 | -2.358 | 0.014 | 0.018 | 0.018 | 0 | 0.29 | 0.31 | 0.34 | 0.82 | 0.19 | 0.33 |
| 5 | 29067266 | rs484786 | -0.201 | -0.02 | -0.852 | -2.358 | 0.014 | 0.018 | 0.018 | 0 | 0.29 | 0.31 | 0.34 | 0.82 | 0.19 | 0.33 |
| 5 | 29070491 | rs160305 | -0.284 | -0.06 | -0.696 | -2.389 | -0.004 | -0.006 | -0.006 | 0 | 0.29 | 0.34 | 0.34 | 0.82 | 0.19 | 0.33 |
| 5 | 29074011 | rs16898699 | 1.111 | 0.021 | -0.716 | -2.377 | -0.077 | -0.005 | -0.005 | 0.549 | 0.17 | 0.04 | 0.15 | 0.13 | 0.3 | 0.25 |
| 5 | 29077070 | rs504531 | -0.03 | -0.116 | -0.666 | -2.369 | 0.006 | 0.008 | 0.008 | 0 | 0.26 | 0.31 | 0.33 | 0.83 | 0.19 | 0.33 |
| 5 | 29077957 | rs824328 | -1.252 | -0.108 | -0.465 | -2.432 | 0.002 | -0.04 | -0.04 | 0 | 0.74 | 0.78 | 0.73 | 1 | 0.79 | 0.85 |
| 5 | 29078099 | rs824329 | -0.12 | -0.117 | -0.467 | -2.357 | -0.004 | -0.007 | -0.007 | 0 | 0.27 | 0.33 | 0.34 | 0.83 | 0.19 | 0.32 |
| 5 | 29086561 | rs6863342 | 1.423 | -0.149 | -0.224 | -2.456 | 0 | -0.015 | -0.015 | 2.295 | 0.29 | 0.22 | 0.23 | 0.04 | 0.3 | 0.29 |
| 5 | 29086909 | rs6867873 | 1.214 | -0.173 | -0.186 | -2.464 | -0.004 | -0.086 | -0.086 | 0.58 | 0.32 | 0.33 | 0.24 | 0.05 | 0.31 | 0.31 |
| 5 | 29091685 | rs679895 | 0.008 | -0.16 | -0.191 | -2.016 | 0 | -0.007 | -0.007 | 0.03 | 0.53 | 0.46 | 0.56 | 0.75 | 0.49 | 0.55 |
| 5 | 29110364 | rs12189263 | 0.352 | -0.202 | -0.654 | -2.047 | -0.114 | 0.094 | 0.094 | 0 | 0.4 | 0.11 | 0.4 | 0.25 | 0.5 | 0.41 |
| 5 | 29129212 | rs160320 | 1.003 | -0.251 | -0.527 | -1.787 | -0.012 | -0.092 | -0.092 | 0 | 0.18 | 0.08 | 0.13 | 0.58 | 0.16 | 0.25 |
| 5 | 29135233 | rs824331 | -0.457 | -0.167 | -0.393 | -2.064 | 0.005 | -0.008 | -0.008 | 0 | 0.76 | 0.8 | 0.68 | 0.29 | 0.81 | 0.73 |
| 5 | 29151261 | rs17623890 | 2.193 | -0.06 | -0.673 | -2.109 | -0.025 | 0.111 | 0.111 | 0 | 0.17 | 0.02 | 0.19 | 0 | 0.15 | 0.09 |
| 5 | 29165247 | rs4555829 | -0.4 | 0.116 | -0.245 | -1.791 | 0.007 | -0.015 | -0.015 | 0 | 0.76 | 0.82 | 0.67 | 0.28 | 0.81 | 0.73 |
| 5 | 29193495 | rs4354055 | 0.332 | 0.113 | 0.301 | -2.1 | 0.004 | 0.127 | 0.127 | 0 | 0.54 | 0.69 | 0.47 | 0.03 | 0.45 | 0.39 |
| 5 | 29193998 | rs10050492 | 0.744 | 0.086 | 0.492 | -2.113 | 0.009 | 0.02 | 0.02 | 0 | 0.48 | 0.34 | 0.43 | 0.03 | 0.44 | 0.37 |

| | | | | | | | | | | | | | | | | |
|---|---|---|---|---|---|---|---|---|---|---|---|---|---|---|---|---|
| 5 | 29204186 | rs10060554 | 0.323 | 0.126 | -0.022 | -2.094 | 0.006 | 0.113 | 0.113 | 0 | 0.54 | 0.66 | 0.46 | 0.03 | 0.45 | 0.39 |
| 5 | 29211937 | rs4398647 | 0.608 | 0.407 | -0.571 | -1.986 | 0.014 | 0.091 | 0.091 | 0 | 0.48 | 0.29 | 0.43 | 0.03 | 0.44 | 0.37 |
| 5 | 29213248 | rs4867169 | 0.298 | 0.976 | -0.915 | -1.056 | -0.06 | -0.004 | -0.004 | 0 | 0.32 | 0.12 | 0.43 | 0.31 | 0.43 | 0.27 |
| 5 | 29214777 | rs13166467 | -0.216 | 1.19 | -0.442 | -1.379 | -0.001 | 0.006 | 0.006 | 0 | 0.43 | 0.56 | 0.42 | 0.26 | 0.51 | 0.41 |
| 5 | 29223692 | rs13357720 | 0.511 | 1.336 | -0.265 | -1.557 | 0.026 | 0.087 | 0.087 | 0.047 | 0.35 | 0.19 | 0.29 | 0.16 | 0.28 | 0.18 |
| 5 | 29226086 | rs7726399 | 1.587 | 1.234 | -0.378 | -1.468 | 0.001 | -0.002 | -0.002 | 0.373 | 0.17 | 0.17 | 0.33 | 0.12 | 0.19 | 0.18 |
| 5 | 29246809 | rs4331913 | 0.114 | 1.112 | -0.098 | -1.551 | 0 | -0.009 | -0.009 | 0.19 | 0.48 | 0.5 | 0.41 | 0.26 | 0.45 | 0.41 |
| 5 | 29249396 | rs7717789 | 0.004 | 1.268 | -0.018 | -1.489 | 0.086 | 0.019 | 0.019 | 0 | 0.24 | 0.43 | 0.33 | 0.26 | 0.32 | 0.33 |
| 5 | 29249544 | rs7717683 | 1.269 | 1.286 | -0.003 | -1.477 | 0.13 | 0.048 | 0.048 | 0 | 0.11 | 0.35 | 0.12 | 0.1 | 0.16 | 0.17 |
| 5 | 29260573 | rs4620035 | -0.174 | 1.129 | -0.109 | -1.496 | 0.103 | 0.011 | 0.011 | 0 | 0.25 | 0.48 | 0.35 | 0.26 | 0.39 | 0.36 |
| 5 | 29266323 | rs6450681 | -0.665 | 0.903 | -0.494 | -1.623 | 0.056 | 0.039 | 0.039 | 0 | 0.51 | 0.69 | 0.71 | 0.39 | 0.63 | 0.65 |
| 5 | 29269294 | rs7712253 | -0.713 | 0.927 | -0.434 | -1.603 | 0.002 | -0.009 | -0.009 | 1.066 | 0.73 | 0.71 | 0.79 | 0.39 | 0.75 | 0.73 |
| 5 | 29277556 | rs7718659 | 0.186 | 1.135 | -0.087 | -1.526 | 0.047 | -0.015 | -0.015 | 0 | 0.3 | 0.47 | 0.45 | 0.24 | 0.4 | 0.36 |
| 5 | 29280364 | rs4444992 | 1.044 | 1.126 | -0.077 | -1.491 | 0.001 | -0.015 | -0.015 | 0.135 | 0.2 | 0.3 | 0.35 | 0.13 | 0.24 | 0.17 |
| 5 | 29286698 | rs10044023 | 1.305 | 1.083 | -0.144 | -1.482 | -0.007 | -0.008 | -0.008 | 4.096 | 0.18 | 0.24 | 0.33 | 0.13 | 0.16 | 0.15 |
| 5 | 29305055 | rs6865945 | 0.757 | 1.099 | -0.069 | -1.518 | -0.002 | 0.009 | 0.009 | 0 | 0.21 | 0.18 | 0.26 | 0.15 | 0.22 | 0.26 |
| 5 | 29320222 | rs10079187 | 1.088 | 1.538 | 0.376 | -1.27 | 0.09 | 0.01 | 0.01 | 0 | 0.12 | 0.23 | 0.12 | 0.11 | 0.23 | 0.2 |
| 5 | 29321754 | rs10073729 | 1.566 | 1.556 | 0.453 | -1.27 | 0.041 | 0.009 | 0.009 | 0 | 0.1 | 0.15 | 0.09 | 0.11 | 0.15 | 0.18 |
| 5 | 29323028 | rs11959737 | -0.229 | 1.179 | 0.152 | -1.475 | -0.001 | 0.001 | 0.001 | 0 | 0.7 | 0.67 | 0.77 | 0.39 | 0.71 | 0.68 |
| 5 | 29327911 | rs2195316 | -0.317 | 1.211 | -0.064 | -1.46 | -0.001 | -0.036 | -0.036 | 0 | 0.71 | 0.79 | 0.79 | 0.39 | 0.71 | 0.68 |
| 5 | 29329716 | rs13362367 | 0.953 | 1.584 | 0.079 | -1.046 | 0.09 | 0.01 | 0.01 | 0 | 0.12 | 0.23 | 0.12 | 0.11 | 0.23 | 0.2 |
| 5 | 29336245 | rs4867173 | 0.691 | 0.806 | -0.338 | -0.938 | -0.011 | -0.001 | -0.001 | 1.855 | 0.15 | 0.24 | 0.32 | 0.13 | 0.13 | 0.13 |
| 5 | 29342140 | rs7706299 | -0.772 | 0.861 | -0.817 | -0.535 | -0.024 | -0.068 | -0.068 | 0 | 0.88 | 0.96 | 0.88 | 0.42 | 0.87 | 0.76 |
| 5 | 29354805 | rs1119468 | -0.586 | 0.687 | -0.445 | -1.294 | 0.001 | -0.002 | -0.002 | 0 | 0.35 | 0.58 | 0.6 | 0.28 | 0.39 | 0.4 |
| 5 | 29358362 | rs2162801 | -0.087 | 0.619 | 0.057 | -1.344 | -0.005 | -0.085 | -0.085 | 0.595 | 0.15 | 0.26 | 0.1 | 0 | 0.16 | 0.09 |
| 5 | 29361330 | rs17430828 | 1.882 | 0.082 | -0.413 | -1.513 | 0.05 | 0.115 | 0.115 | 0.131 | 0.28 | 0.02 | 0.09 | 0.05 | 0.15 | 0.1 |
| 5 | 29367474 | rs7717780 | -0.513 | 0.882 | 0.484 | -0.768 | 0.012 | 0.003 | 0.003 | 0 | 0.28 | 0.56 | 0.54 | 0.28 | 0.28 | 0.34 |
| 5 | 29388968 | rs7733983 | -0.033 | 0.972 | 0.51 | -0.689 | 0.034 | -0.043 | -0.043 | 0 | 0.23 | 0.39 | 0.37 | 0.12 | 0.24 | 0.18 |
| 5 | 29391830 | rs4867175 | 1.076 | 1.036 | 0.527 | -0.843 | 0.018 | 0.066 | 0.066 | 0 | 0.53 | 0.33 | 0.39 | 0.23 | 0.47 | 0.48 |
| 5 | 29394857 | rs12523417 | 0.319 | 1.124 | 0.373 | -0.82 | 0.018 | -0.02 | -0.02 | 0 | 0.22 | 0.28 | 0.35 | 0.12 | 0.24 | 0.18 |
| 5 | 29397044 | rs2877056 | -0.603 | 1.133 | -0.057 | -0.678 | -0.043 | -0.071 | -0.071 | 0.289 | 0.78 | 0.9 | 0.79 | 0.36 | 0.71 | 0.66 |
| 5 | 29413495 | rs12514005 | -0.34 | 1.13 | 0.495 | -0.528 | 0.055 | 0.033 | 0.033 | 0 | 0.24 | 0.42 | 0.52 | 0.29 | 0.28 | 0.33 |
| 5 | 29418298 | rs16898950 | 0.86 | 1.19 | 0.304 | -0.618 | 0.009 | -0.002 | -0.002 | 1.107 | 0.16 | 0.16 | 0.34 | 0.12 | 0.19 | 0.15 |
| 5 | 29425243 | rs2115118 | 0.077 | 1.173 | 0.412 | -0.633 | 0.041 | -0.026 | -0.026 | 1.063 | 0.2 | 0.32 | 0.35 | 0.12 | 0.22 | 0.18 |
| 5 | 29458042 | rs788534 | 0.612 | 1.097 | 0.054 | -0.092 | -0.001 | -0.003 | -0.003 | 0 | 0.7 | 0.69 | 0.77 | 0.43 | 0.68 | 0.62 |
| 5 | 29466550 | rs617354 | -0.454 | 1.218 | -0.142 | -0.014 | -0.034 | -0.071 | -0.071 | 0.507 | 0.78 | 0.93 | 0.81 | 0.42 | 0.75 | 0.68 |
| 5 | 29466930 | rs9790969 | 0.269 | 0.827 | -0.419 | -1.32 | 0 | -0.03 | -0.03 | 0.005 | 0.38 | 0.35 | 0.38 | 0.02 | 0.36 | 0.31 |
| 5 | 29467421 | rs642638 | 0.379 | 0.403 | -0.342 | -0.96 | 0.005 | -0.01 | -0.01 | 0 | 0.39 | 0.33 | 0.21 | 0.05 | 0.37 | 0.3 |
| 5 | 29468652 | rs16899004 | -0.158 | 0.266 | -0.256 | -0.665 | -0.008 | -0.064 | -0.064 | 0 | 0.15 | 0.22 | 0.09 | 0.01 | 0.15 | 0.18 |
| 5 | 29469922 | rs2330081 | 0.657 | 0.362 | -0.893 | -0.093 | -0.009 | -0.026 | -0.026 | 0 | 0.45 | 0.17 | 0.25 | 0.41 | 0.38 | 0.36 |
| 5 | 29470186 | rs905860 | 0.657 | 0.362 | -0.893 | -0.093 | -0.009 | -0.026 | -0.026 | 0 | 0.45 | 0.17 | 0.25 | 0.41 | 0.38 | 0.36 |
| 5 | 29470422 | rs905858 | -0.272 | 0.581 | 0.032 | 0.178 | 0.004 | -0.027 | -0.027 | 0.032 | 0.63 | 0.52 | 0.65 | 0.7 | 0.59 | 0.7 |
| 5 | 29482208 | rs1118711 | 0.369 | 0.751 | 0.295 | 0.931 | 0.032 | 0.09 | 0.09 | 0 | 0.13 | 0.32 | 0.29 | 0.23 | 0.15 | 0.29 |
| 5 | 29485841 | rs669916 | -0.564 | 0.665 | 0.128 | 0.659 | -0.009 | 0.044 | 0.044 | 0 | 0.67 | 0.8 | 0.71 | 0.73 | 0.67 | 0.74 |
| 5 | 29490496 | rs12657240 | 2.137 | 0.194 | -0.473 | 0.172 | 0.054 | 0.092 | 0.092 | 0.75 | 0.23 | 0.06 | 0.08 | 0.03 | 0.12 | 0.13 |
| 5 | 29498695 | rs607373 | -0.345 | 0.614 | 0.103 | 0.24 | 0.003 | -0.033 | -0.033 | 0.436 | 0.91 | 0.96 | 0.95 | 0.76 | 0.94 | 0.86 |
| 5 | 29500305 | rs625812 | 1.137 | 0.658 | 0.139 | 0.264 | -0.003 | 0.016 | 0.016 | 0.661 | 0.45 | 0.17 | 0.25 | 0.51 | 0.38 | 0.38 |
| 5 | 29501462 | rs655219 | 1.035 | 0.689 | 0.287 | 0.235 | -0.008 | 0.032 | 0.032 | 0.332 | 0.72 | 0.41 | 0.52 | 0.54 | 0.73 | 0.52 |
| 5 | 29503058 | rs16899039 | 1.095 | 0.716 | 0.3 | 0.216 | -0.002 | -0.01 | -0.01 | 0 | 0.26 | 0.22 | 0.24 | 0.03 | 0.25 | 0.12 |
| 5 | 29503184 | rs1394674 | 1.097 | 0.846 | 0.474 | 0.248 | 0.007 | -0.053 | -0.053 | 0.286 | 0.1 | 0.16 | 0.06 | 0 | 0.12 | 0.14 |
| 5 | 29503454 | rs182780 | 1.124 | 0.79 | 0.111 | 0.334 | -0.018 | 0.085 | 0.085 | 0.618 | 0.43 | 0.14 | 0.23 | 0.27 | 0.37 | 0.37 |
| 5 | 29503553 | rs191312 | 0.981 | 0.427 | 0.163 | 0.006 | -0.025 | 0.013 | 0.013 | 0.529 | 0.46 | 0.18 | 0.29 | 0.51 | 0.48 | 0.4 |
| 5 | 29507496 | rs629225 | -0.453 | 0.605 | 0.006 | 0.39 | 0.002 | 0.019 | 0.019 | 0 | 0.7 | 0.78 | 0.65 | 0.73 | 0.73 | 0.81 |
| 5 | 29508229 | rs16899054 | -0.662 | 0.905 | 0.093 | 0.648 | 0.025 | 0.106 | 0.106 | 0.009 | 0.09 | 0.26 | 0.18 | 0.22 | 0.09 | 0.23 |
| 5 | 29515854 | rs624343 | -0.182 | 0.898 | -0.596 | -0.096 | -0.019 | 0.093 | 0.093 | 0 | 0.66 | 0.87 | 0.7 | 0.79 | 0.69 | 0.65 |
| 5 | 29516289 | rs588359 | 0.676 | 1.003 | 0.011 | 0.042 | -0.002 | 0.023 | 0.023 | 0 | 0.52 | 0.43 | 0.49 | 0.19 | 0.54 | 0.34 |
| 5 | 29517967 | rs16899061 | 0.587 | 0.905 | -0.209 | -0.358 | -0.018 | 0.037 | 0.037 | 0 | 0.26 | 0.18 | 0.17 | 0.13 | 0.26 | 0.22 |
| 5 | 29518236 | rs17633307 | 1.875 | 1.117 | -0.451 | -0.378 | -0.007 | 0.017 | 0.017 | 0 | 0.05 | 0.01 | 0.05 | 0 | 0.05 | 0.01 |
| 5 | 29520650 | rs9885405 | 0.827 | 1.104 | -0.099 | -0.094 | 0.038 | -0.02 | -0.02 | 0 | 0.07 | 0.24 | 0.06 | 0 | 0.05 | 0.04 |
| 5 | 29521275 | rs605927 | 1.55 | 1.05 | -0.364 | -0.232 | -0.004 | -0.022 | -0.022 | 0.723 | 0.28 | 0.28 | 0.32 | 0.07 | 0.28 | 0.12 |
| 5 | 29526304 | rs625886 | -0.241 | 1.087 | -0.265 | -0.229 | 0 | 0.007 | 0.007 | 0 | 0.61 | 0.55 | 0.71 | 0.57 | 0.61 | 0.6 |
| 5 | 29550404 | rs182778 | 0.788 | 1.191 | -0.34 | -0.177 | -0.017 | -0.083 | -0.083 | 0 | 0.36 | 0.51 | 0.38 | 0.08 | 0.34 | 0.17 |
| 5 | 29552540 | rs13185814 | 0.477 | 1.128 | -0.648 | 0.05 | -0.003 | -0.001 | -0.001 | 0.029 | 0.23 | 0.21 | 0.34 | 0.5 | 0.25 | 0.46 |
| 5 | 29557296 | rs10058895 | 0.852 | 0.967 | -0.637 | -0.201 | 0.001 | -0.002 | -0.002 | 0.157 | 0.2 | 0.19 | 0.13 | 0.11 | 0.22 | 0.22 |
| 5 | 29557328 | rs10073723 | -0.604 | 1.004 | -0.832 | -0.413 | 0.023 | 0.054 | 0.054 | 0 | 0.54 | 0.37 | 0.56 | 0.93 | 0.57 | 0.79 |
| 5 | 29566495 | rs10472163 | 1.445 | 0.991 | -0.762 | 0.01 | 0.015 | -0.027 | -0.027 | 0 | 0.32 | 0.2 | 0.37 | 0.3 | 0.34 | 0.16 |
| 5 | 29567773 | rs10069250 | 0.251 | 1.413 | -1.478 | 0.004 | -0.003 | 0 | 0 | 0 | 0.24 | 0.24 | 0.19 | 0.26 | 0.24 | 0.34 |
| 5 | 29569364 | rs17571587 | 1.338 | 1.317 | -1.412 | 0.099 | 0.001 | 0.014 | 0.014 | 0.183 | 0.15 | 0.08 | 0.09 | 0.06 | 0.14 | 0.13 |
| 5 | 29570198 | rs17634115 | 0.94 | 1.294 | -1.466 | 0.516 | -0.005 | 0.017 | 0.017 | 0 | 0.05 | 0.02 | 0.04 | 0 | 0.05 | 0.02 |

| | | | | | | | | | | | | | | | | |
|---|---|---|---|---|---|---|---|---|---|---|---|---|---|---|---|---|
| 5 | 29581858 | rs6880023 | 0.456 | 1.127 | -1.886 | 0.409 | -0.003 | -0.069 | -0.069 | 0 | 0.2 | 0.07 | 0.12 | 0.23 | 0.16 | 0.1 |
| 5 | 29582697 | rs13185717 | 0.11 | 1.213 | -1.358 | -0.277 | -0.022 | 0.006 | 0.006 | 0.418 | 0.15 | 0.09 | 0.13 | 0.14 | 0.13 | 0.16 |
| 5 | 29584827 | rs4634364 | -0.631 | 1.212 | -1.272 | -0.376 | 0 | -0.008 | -0.008 | 2.968 | 0.46 | 0.56 | 0.5 | 0.2 | 0.46 | 0.31 |
| 5 | 29588169 | rs4587088 | -0.284 | 0.641 | -0.76 | -0.388 | -0.018 | -0.116 | -0.116 | 0.022 | 0.31 | 0.47 | 0.38 | 0.06 | 0.33 | 0.15 |
| 5 | 29601381 | rs11748217 | -0.491 | 1.079 | -1.253 | 0.531 | -0.025 | -0.062 | -0.062 | 0 | 0.26 | 0.11 | 0.29 | 0.43 | 0.32 | 0.18 |
| 5 | 29619266 | rs4867187 | -0.237 | 0.639 | -0.994 | -0.114 | -0.008 | 0.006 | 0.006 | 0 | 0.47 | 0.37 | 0.47 | 0.48 | 0.46 | 0.26 |
| 5 | 29625796 | rs4354058 | 0.35 | 0.252 | -0.387 | -0.263 | -0.001 | 0.005 | 0.005 | 0.322 | 0.07 | 0.23 | 0.04 | 0.05 | 0.04 | 0.06 |
| 5 | 29628568 | rs4409113 | 0.889 | 0.373 | -0.547 | -0.463 | -0.001 | 0 | 0 | 0.051 | 0.16 | 0.14 | 0.15 | 0.12 | 0.17 | 0.15 |
| 5 | 29631637 | rs1350215 | 1.094 | 0.356 | -0.433 | -0.465 | 0.002 | -0.015 | -0.015 | 0 | 0.05 | 0.18 | 0.04 | 0.01 | 0.04 | 0.01 |
| 5 | 29634027 | rs9918231 | 1.289 | 0.413 | -0.646 | -0.86 | -0.005 | 0.134 | 0.134 | 0.738 | 0.29 | 0.07 | 0.2 | 0.05 | 0.24 | 0.13 |
| 5 | 29635164 | rs17573102 | 1.163 | 0.408 | -0.514 | -0.742 | -0.032 | 0.004 | 0.004 | 0 | 0.16 | 0.02 | 0.13 | 0.12 | 0.17 | 0.15 |
| 5 | 29640815 | rs1824111 | 0.222 | 0.292 | -0.257 | -0.605 | 0.022 | 0.029 | 0.029 | 0.147 | 0.07 | 0.36 | 0.06 | 0.05 | 0.04 | 0.06 |
| 5 | 29654854 | rs11948670 | 0.233 | 0.292 | -0.286 | -0.606 | 0.016 | 0.022 | 0.022 | 0.217 | 0.07 | 0.34 | 0.05 | 0.05 | 0.04 | 0.06 |
| 5 | 29655884 | rs1869995 | 0.54 | 0.217 | -0.157 | -0.696 | 0.002 | 0.122 | 0.122 | 0.123 | 0.46 | 0.27 | 0.36 | 0.19 | 0.42 | 0.29 |
| 5 | 29668965 | rs3100987 | -0.045 | 0.201 | -0.316 | -0.663 | -0.022 | 0.042 | 0.042 | 0 | 0.73 | 0.47 | 0.68 | 0.65 | 0.75 | 0.51 |
| 5 | 29672338 | rs1456633 | -0.88 | 0.224 | -0.229 | -0.689 | -0.012 | 0.064 | 0.064 | 0 | 0.81 | 0.76 | 0.72 | 0.66 | 0.8 | 0.55 |
| 5 | 29686820 | rs3100935 | -0.88 | 0.224 | -0.229 | -0.726 | -0.012 | 0.072 | 0.072 | 0 | 0.81 | 0.76 | 0.71 | 0.63 | 0.8 | 0.55 |
| 5 | 29688494 | rs13169659 | 1.636 | 0.251 | -0.504 | -1.01 | -0.075 | 0.198 | 0.198 | 1.11 | 0.32 | 0.04 | 0.22 | 0.05 | 0.29 | 0.12 |
| 5 | 29689156 | rs2330148 | 1.315 | 0.778 | -0.031 | -0.739 | -0.002 | -0.017 | -0.017 | 0.36 | 0.11 | 0.08 | 0.08 | 0 | 0.12 | 0.04 |
| 5 | 29689401 | rs1350214 | -0.037 | 0.704 | 0.301 | -0.207 | 0.017 | -0.022 | -0.022 | 0 | 0.08 | 0.31 | 0.05 | 0.01 | 0.05 | 0.04 |
| 5 | 29689730 | rs7728669 | -0.07 | 0.703 | 0.3 | -0.208 | 0.015 | -0.024 | -0.024 | 0.416 | 0.08 | 0.31 | 0.05 | 0.01 | 0.05 | 0.04 |
| 5 | 29690605 | rs1463423 | 0.557 | 0.792 | 0.062 | -0.096 | 0.003 | -0.005 | -0.005 | 0.105 | 0.15 | 0.09 | 0.11 | 0.18 | 0.13 | 0.2 |
| 5 | 29697380 | rs1456631 | 0.346 | 0.824 | 0.546 | 0.521 | -0.008 | 0.073 | 0.073 | 0.106 | 0.8 | 0.73 | 0.7 | 0.62 | 0.76 | 0.56 |
| 5 | 29697762 | rs10461870 | 0.247 | 0.939 | 0.346 | 0.511 | -0.068 | -0.097 | -0.097 | 0 | 0.15 | 0.03 | 0.26 | 0.42 | 0.22 | 0.15 |
| 5 | 29703260 | rs3101006 | 0.933 | 0.206 | 1.187 | 0.894 | -0.015 | 0.14 | 0.14 | 0.318 | 0.78 | 0.64 | 0.7 | 0.56 | 0.76 | 0.51 |
| 5 | 29704121 | rs3101011 | -1.014 | 0.272 | 0.855 | 0.368 | -0.005 | 0.001 | 0.001 | 1.324 | 0.07 | 0.24 | 0.04 | 0.05 | 0.04 | 0.06 |
| 5 | 29715664 | rs3100963 | 0.749 | 0.836 | 0.624 | 1.05 | 0.02 | 0.083 | 0.083 | 0 | 0.77 | 0.5 | 0.63 | 0.56 | 0.77 | 0.54 |
| 5 | 29721456 | rs6450712 | -2.072 | 0.199 | 1.883 | 1.739 | -0.006 | 0.375 | 0.375 | 2.883 | 0.08 | 0.49 | 0.06 | 0.26 | 0.02 | 0.07 |
| 5 | 29727963 | rs7727296 | -0.596 | 0.492 | 1.425 | 1.34 | 0.009 | 0.078 | 0.078 | 0 | 0.24 | 0.41 | 0.4 | 0.39 | 0.22 | 0.17 |
| 5 | 29729165 | rs6874707 | 1.12 | 0.665 | 1.115 | 1.162 | 0.012 | -0.028 | -0.028 | 0 | 0.6 | 0.55 | 0.37 | 0.27 | 0.55 | 0.4 |
| 5 | 29735598 | rs3114523 | 0.479 | 1.023 | 0.662 | 1.483 | -0.049 | -0.035 | -0.035 | 0.76 | 0.08 | 0.02 | 0.27 | 0.31 | 0.18 | 0.41 |
| 5 | 29748193 | rs16899329 | -0.646 | 0.625 | 1.263 | 1.389 | -0.001 | 0.176 | 0.176 | 0.817 | 0.19 | 0.47 | 0.31 | 0.38 | 0.14 | 0.11 |
| 5 | 29752624 | rs11742951 | -0.8 | 0.295 | 1.013 | 1.214 | -0.035 | -0.053 | -0.053 | 0 | 0.22 | 0.15 | 0.35 | 0.36 | 0.29 | 0.49 |
| 5 | 29780462 | rs17576177 | 1.12 | 0.478 | 0.987 | 1.089 | -0.007 | 0.003 | 0.003 | 0 | 0.05 | 0.06 | 0.06 | 0 | 0.07 | 0.04 |
| 5 | 29780670 | rs12513511 | -0.735 | 0.433 | 0.963 | 1.122 | -0.037 | -0.058 | -0.058 | 0 | 0.22 | 0.14 | 0.35 | 0.36 | 0.28 | 0.47 |
| 5 | 29785725 | rs1463414 | 1.136 | 0.337 | 1.154 | 1.041 | -0.014 | -0.025 | -0.025 | 0 | 0.72 | 0.74 | 0.64 | 0.62 | 0.69 | 0.51 |
| 5 | 29788558 | rs6860417 | 0.874 | -0.521 | 0.966 | 0.841 | -0.021 | 0.013 | 0.013 | 0 | 0.06 | 0.06 | 0.02 | 0.11 | 0.02 | 0.08 |
| 5 | 29794387 | rs4597986 | 2.162 | -0.069 | 1.568 | 1.254 | -0.034 | 0.191 | 0.191 | 0 | 0.48 | 0.11 | 0.29 | 0.24 | 0.52 | 0.34 |
| 5 | 29806888 | rs9292342 | 1.407 | -0.326 | 1.823 | 1.338 | -0.005 | 0.012 | 0.012 | 0 | 0.68 | 0.36 | 0.63 | 0.45 | 0.72 | 0.49 |
| 5 | 29810841 | rs12521126 | -0.525 | -0.257 | 1.829 | 1.346 | 0.008 | 0.032 | 0.032 | 0.056 | 0.21 | 0.31 | 0.34 | 0.36 | 0.24 | 0.46 |
| 5 | 29831967 | rs7709002 | 2.054 | -0.201 | 1.596 | 1.297 | -0.03 | 0.206 | 0.206 | 0 | 0.49 | 0.11 | 0.26 | 0.24 | 0.54 | 0.36 |
| 5 | 29833714 | rs6450724 | 0.933 | -0.356 | 1.831 | 1.489 | -0.029 | 0.127 | 0.127 | 0.061 | 0.89 | 0.61 | 0.96 | 0.82 | 0.96 | 0.94 |
| 5 | 29834582 | rs1428331 | -0.399 | -0.304 | 1.458 | 1.481 | -0.006 | -0.013 | -0.013 | 0.122 | 0.93 | 0.91 | 0.98 | 0.98 | 0.97 | 0.97 |
| 5 | 29839458 | rs11956675 | -0.106 | -0.298 | 1.493 | 1.516 | -0.006 | -0.006 | -0.006 | 0.528 | 0.16 | 0.14 | 0.36 | 0.36 | 0.2 | 0.44 |
| 5 | 29839705 | rs1428318 | 0.56 | -0.288 | 1.431 | 1.509 | 0.001 | -0.001 | -0.001 | 0 | 0.13 | 0.09 | 0.25 | 0.21 | 0.11 | 0.1 |
| 5 | 29840035 | rs12697227 | 0.477 | -0.291 | 1.445 | 1.507 | -0.004 | 0.001 | 0.001 | 0.092 | 0.8 | 0.81 | 0.73 | 0.78 | 0.86 | 0.88 |
| 5 | 29849304 | rs9790891 | 0.535 | -0.286 | 1.419 | 1.507 | 0.001 | -0.002 | -0.002 | 0 | 0.13 | 0.1 | 0.25 | 0.21 | 0.11 | 0.1 |
| 5 | 29851658 | rs7449322 | 0.402 | -0.274 | 1.421 | 1.507 | -0.001 | 0.007 | 0.007 | 0 | 0.79 | 0.81 | 0.73 | 0.61 | 0.85 | 0.84 |
| 5 | 29863566 | rs1428342 | -0.217 | -0.314 | 1.296 | 1.404 | 0.004 | 0.028 | 0.028 | 0 | 0.17 | 0.21 | 0.37 | 0.36 | 0.2 | 0.43 |
| 5 | 29873955 | rs6888279 | 1.862 | -0.422 | 1.358 | 1.283 | 0.021 | 0.174 | 0.174 | 0 | 0.54 | 0.2 | 0.33 | 0.23 | 0.62 | 0.39 |
| 5 | 29879015 | rs4867222 | 1.243 | -0.352 | 1.181 | 1.365 | 0.001 | -0.007 | -0.007 | 0 | 0.64 | 0.68 | 0.38 | 0.41 | 0.65 | 0.45 |
| 5 | 29886553 | rs16899677 | -0.927 | -0.178 | 1.224 | 1.504 | 0.007 | 0.161 | 0.161 | 1.95 | 0.08 | 0.32 | 0.03 | 0.18 | 0.04 | 0.05 |
| 5 | 29887070 | rs16899679 | 0.809 | -0.161 | 0.945 | 1.611 | -0.003 | -0.052 | -0.052 | 1.836 | 0.11 | 0.01 | 0.23 | 0.25 | 0.11 | 0.08 |
| 5 | 29890192 | rs7709732 | -0.487 | -0.018 | 1.189 | 1.7 | 0.009 | 0.039 | 0.039 | 0 | 0.18 | 0.25 | 0.37 | 0.32 | 0.21 | 0.43 |
| 5 | 29890461 | rs1991489 | -1.062 | 0.007 | 0.753 | 1.201 | 0.008 | 0.093 | 0.093 | 0.25 | 0.26 | 0.52 | 0.4 | 0.5 | 0.25 | 0.48 |
| 5 | 29894662 | rs1428345 | -0.317 | 0.107 | -0.389 | 0.547 | -0.007 | 0.024 | 0.024 | 0 | 0.43 | 0.24 | 0.37 | 0.38 | 0.5 | 0.36 |
| 5 | 29896350 | rs16899706 | 1.02 | 0.202 | -0.889 | 0.303 | -0.031 | -0.006 | -0.006 | 0.443 | 0.11 | 0.01 | 0.08 | 0.14 | 0.17 | 0.08 |
| 5 | 29899483 | rs13158751 | -0.144 | 0.023 | -0.513 | -0.059 | -0.001 | 0.045 | 0.045 | 0 | 0.75 | 0.63 | 0.52 | 0.39 | 0.75 | 0.52 |
| 5 | 29902666 | rs1428330 | 0.071 | 0.011 | -0.814 | -0.076 | 0.005 | 0.014 | 0.014 | 0.229 | 0.32 | 0.22 | 0.3 | 0.24 | 0.33 | 0.28 |
| 5 | 29905803 | rs13171652 | 0.787 | 0.558 | -0.747 | 0.007 | -0.04 | 0 | 0 | 0.009 | 0.11 | 0.01 | 0.07 | 0.14 | 0.17 | 0.09 |
| 5 | 29914492 | rs1593906 | -0.871 | 0.247 | -0.399 | -0.066 | -0.023 | -0.02 | -0.02 | 0 | 0.94 | 0.9 | 0.98 | 0.99 | 0.99 | 0.96 |
| 5 | 29918268 | rs7737075 | 1.097 | 0.415 | -0.503 | -0.288 | 0.005 | 0.014 | 0.014 | 0 | 0.42 | 0.33 | 0.22 | 0.04 | 0.35 | 0.25 |
| 5 | 29918701 | rs10940845 | -0.367 | 0.691 | -0.398 | 1.166 | 0 | 0.004 | 0.004 | 0 | 0.2 | 0.2 | 0.33 | 0.34 | 0.24 | 0.42 |
| 5 | 29928351 | rs2914129 | 0.591 | -0.091 | -0.158 | 0.424 | 0 | 0 | 0 | 0 | 0.56 | 0.45 | 0.51 | 0.37 | 0.56 | 0.44 |
| 5 | 29928839 | rs2972797 | 0.115 | -0.189 | -0.592 | -0.405 | 0 | -0.003 | -0.003 | 0 | 0.22 | 0.26 | 0.13 | 0.13 | 0.26 | 0.19 |
| 5 | 29931790 | rs2914127 | -0.972 | -0.428 | -0.122 | -0.68 | -0.018 | -0.043 | -0.043 | 0 | 0.9 | 0.77 | 0.93 | 0.99 | 0.93 | 0.89 |
| 5 | 29935020 | rs2923769 | 0.384 | -0.439 | 0.154 | -0.859 | 0.004 | -0.093 | -0.093 | 0.453 | 0.17 | 0.29 | 0.13 | 0 | 0.21 | 0.15 |
| 5 | 29941681 | rs10940848 | 1.362 | -0.643 | -0.138 | -0.984 | 0.009 | -0.014 | -0.014 | 0.343 | 0.11 | 0.07 | 0.37 | 0.21 | 0.06 | 0.12 |
| 5 | 29949620 | rs1366363 | 1.516 | -0.644 | -0.14 | -0.985 | 0.013 | -0.027 | -0.027 | 0 | 0.11 | 0.03 | 0.37 | 0.21 | 0.06 | 0.12 |

| | | | | | | | | | | | | | | | | |
|---|---|---|---|---|---|---|---|---|---|---|---|---|---|---|---|---|
| 5 | 29951868 | rs7701830 | -1.442 | -0.891 | -0.584 | -1.963 | 0.009 | 0.066 | 0.066 | 0 | 0.53 | 0.64 | 0.69 | 0.99 | 0.53 | 0.71 |
| 5 | 29953444 | rs10940849 | 1.049 | -0.723 | -0.715 | -1.869 | 0 | 0.001 | 0.001 | 0.475 | 0.21 | 0.19 | 0.2 | 0.67 | 0.29 | 0.48 |
| 5 | 29955761 | rs17431444 | 2.04 | -0.515 | -0.791 | -2.126 | -0.07 | 0.247 | 0.247 | 0 | 0.36 | 0.05 | 0.23 | 0 | 0.4 | 0.2 |
| 5 | 29971508 | rs4867023 | -0.267 | -0.241 | -0.263 | -1.677 | -0.001 | -0.009 | -0.009 | 0 | 0.34 | 0.3 | 0.59 | 0.88 | 0.36 | 0.6 |
| 5 | 29978110 | rs10461885 | -0.142 | -0.414 | -0.344 | -1.598 | 0 | 0.036 | 0.036 | 0 | 0.47 | 0.32 | 0.61 | 0.72 | 0.54 | 0.65 |
| 5 | 29980985 | rs1428453 | 0.993 | -0.34 | -0.374 | -1.368 | 0.011 | 0.028 | 0.028 | 0.02 | 0.15 | 0.15 | 0.17 | 0.63 | 0.25 | 0.46 |
| 5 | 29991217 | rs10068240 | -1.439 | -0.498 | -0.392 | -1.45 | 0.011 | 0.006 | 0.006 | 0 | 0.84 | 0.86 | 0.73 | 0.91 | 0.91 | 0.89 |
| 5 | 29996418 | rs17431680 | 1.258 | -0.804 | -0.462 | -1.872 | 0 | -0.061 | -0.061 | 0.683 | 0.28 | 0.27 | 0.18 | 0 | 0.29 | 0.18 |
| 5 | 29999068 | rs17584731 | 2.06 | -0.916 | -0.769 | -1.757 | 0.008 | -0.023 | -0.023 | 2.473 | 0.06 | 0 | 0.02 | 0.07 | 0.04 | 0.02 |
| 5 | 30000585 | rs10061369 | 0.5 | -0.886 | -0.49 | -1.732 | -0.009 | 0.01 | 0.01 | 0 | 0.14 | 0.25 | 0.15 | 0.2 | 0.1 | 0.13 |
| 5 | 30009138 | rs10069092 | -0.067 | -0.859 | -0.447 | -1.817 | 0.018 | 0.033 | 0.033 | 0 | 0.48 | 0.44 | 0.53 | 0.29 | 0.4 | 0.33 |
| 5 | 30009633 | rs11956070 | 1.622 | -0.686 | -0.482 | -1.683 | -0.007 | -0.005 | -0.005 | 0 | 0.2 | 0.12 | 0.15 | 0.01 | 0.25 | 0.1 |
| 5 | 30028447 | rs17431811 | 2.065 | -0.534 | -0.792 | -1.459 | -0.049 | 0.114 | 0.114 | 0 | 0.19 | 0.02 | 0.1 | 0 | 0.2 | 0.14 |
| 5 | 30031390 | rs1428465 | 0.071 | -0.625 | -0.476 | -1.523 | 0.002 | 0.017 | 0.017 | 0 | 0.36 | 0.27 | 0.48 | 0.78 | 0.35 | 0.61 |
| 5 | 30043318 | rs2122899 | -0.848 | -0.663 | -0.453 | -1.583 | 0 | 0.049 | 0.049 | 0 | 0.67 | 0.76 | 0.72 | 0.78 | 0.69 | 0.75 |
| 5 | 30044759 | rs2218892 | -0.04 | -0.618 | -0.267 | -1.599 | -0.004 | -0.02 | -0.02 | 0 | 0.37 | 0.37 | 0.49 | 0.78 | 0.35 | 0.61 |
| 5 | 30046292 | rs2330310 | -0.834 | -0.612 | -0.157 | -1.585 | 0.002 | 0.039 | 0.039 | 0 | 0.67 | 0.72 | 0.73 | 0.78 | 0.69 | 0.75 |
| 5 | 30048332 | rs17510261 | 0.438 | -0.642 | -0.318 | -1.565 | 0.001 | -0.011 | -0.011 | 0.585 | 0.42 | 0.37 | 0.46 | 0.09 | 0.39 | 0.26 |
| 5 | 30059081 | rs12517825 | 1.132 | -0.692 | -0.554 | -1.462 | 0.016 | 0.036 | 0.036 | 0.096 | 0.18 | 0.19 | 0.24 | 0.64 | 0.26 | 0.48 |
| 5 | 30063975 | rs10073717 | 0.316 | -0.732 | -0.551 | -1.498 | -0.01 | 0.066 | 0.066 | 0.096 | 0.47 | 0.26 | 0.56 | 0.71 | 0.5 | 0.7 |
| 5 | 30066630 | rs13186175 | 0.63 | -0.709 | -0.653 | -1.49 | 0.001 | 0 | 0 | 0.297 | 0.36 | 0.33 | 0.34 | 0.64 | 0.45 | 0.59 |
| 5 | 30067131 | rs17581870 | 0.94 | -0.61 | -0.851 | -1.477 | -0.007 | -0.032 | -0.032 | 0 | 0.29 | 0.29 | 0.23 | 0.01 | 0.34 | 0.15 |
| 5 | 30081252 | rs6865875 | 0.902 | -0.517 | -0.917 | -1.433 | 0.013 | 0.106 | 0.106 | 0 | 0.33 | 0.14 | 0.38 | 0.09 | 0.31 | 0.21 |
| 5 | 30084474 | rs13169637 | 1.101 | -0.519 | -0.874 | -1.5 | -0.001 | -0.046 | -0.046 | 0.545 | 0.27 | 0.29 | 0.17 | 0 | 0.27 | 0.16 |
| 5 | 30089045 | rs4147400 | 0.17 | -0.531 | -0.834 | -1.464 | -0.003 | -0.007 | -0.007 | 0 | 0.45 | 0.49 | 0.41 | 0.63 | 0.53 | 0.64 |
| 5 | 30091314 | rs7712255 | 0.509 | -0.514 | -0.809 | -1.411 | 0.01 | 0.096 | 0.096 | 0 | 0.43 | 0.29 | 0.45 | 0.09 | 0.4 | 0.25 |
| 5 | 30092982 | rs4391179 | -0.014 | -0.705 | -0.682 | -0.884 | -0.026 | 0.026 | 0.026 | 0 | 0.78 | 0.64 | 0.79 | 0.72 | 0.84 | 0.85 |
| 5 | 30093584 | rs10940870 | 1.41 | -0.465 | -1.001 | -0.455 | 0.005 | 0.005 | 0.005 | 0 | 0.13 | 0.12 | 0.19 | 0.23 | 0.19 | 0.24 |
| 5 | 30101191 | rs13183446 | 1.358 | -1.215 | -0.957 | -0.689 | 0.04 | -0.018 | -0.018 | 0.52 | 0.12 | 0.01 | 0.22 | 0.08 | 0.05 | 0.11 |
| 5 | 30107943 | rs7707649 | 0.633 | -0.889 | -0.543 | -0.428 | -0.036 | 0.026 | 0.026 | 0 | 0.77 | 0.59 | 0.78 | 0.72 | 0.84 | 0.85 |
| 5 | 30107955 | rs7707658 | 0.228 | -0.513 | -0.532 | -0.388 | -0.017 | 0.019 | 0.019 | 0 | 0.82 | 0.65 | 0.81 | 0.78 | 0.88 | 0.87 |
| 5 | 30108604 | rs9292368 | 0.171 | 0.3 | -0.712 | -0.27 | 0.001 | -0.007 | -0.007 | 0.181 | 0.44 | 0.43 | 0.42 | 0.7 | 0.55 | 0.6 |
| 5 | 30113024 | rs10039248 | -0.474 | 0.403 | -0.703 | 0.362 | 0.008 | -0.052 | -0.052 | 0 | 0.1 | 0.22 | 0.07 | 0.01 | 0.09 | 0.09 |
| 5 | 30116588 | rs9647537 | -0.974 | 0.176 | -1.181 | 1.177 | -0.007 | 0.044 | 0.044 | 0 | 0.07 | 0.02 | 0.07 | 0.49 | 0.08 | 0.3 |
| 5 | 30120629 | rs10050709 | -0.357 | -0.16 | -1.083 | 0.271 | 0.011 | 0.011 | 0.011 | 4.668 | 0.85 | 0.87 | 0.94 | 0.95 | 0.92 | 0.96 |
| 5 | 30121929 | rs4464710 | -0.39 | -0.145 | -1.113 | 0.278 | 0.014 | 0.014 | 0.014 | 4.389 | 0.85 | 0.88 | 0.94 | 0.95 | 0.92 | 0.96 |
| 5 | 30125447 | rs10044555 | -0.082 | 0.917 | -0.986 | 0.531 | 0.023 | 0.002 | 0.002 | 0 | 0.37 | 0.45 | 0.53 | 0.35 | 0.43 | 0.28 |
| 5 | 30138382 | rs6888905 | -0.015 | 0.895 | -1.572 | 0.047 | -0.052 | 0.051 | 0.051 | 0 | 0.67 | 0.87 | 0.58 | 0.22 | 0.6 | 0.43 |
| 5 | 30148789 | rs12188858 | 0.274 | 0.219 | -1.213 | -1.263 | 0.001 | 0.018 | 0.018 | 0.324 | 0.49 | 0.41 | 0.36 | 0.09 | 0.48 | 0.34 |
| 5 | 30148855 | rs13154270 | 0.532 | 0.065 | -1.353 | -1.936 | -0.003 | -0.031 | -0.031 | 0.009 | 0.2 | 0.2 | 0.17 | 0.03 | 0.17 | 0.16 |
| 5 | 30155798 | rs4308495 | 1.056 | -0.077 | -1.36 | -1.999 | -0.058 | 0.128 | 0.128 | 0.75 | 0.26 | 0.04 | 0.16 | 0 | 0.32 | 0.17 |
| 5 | 30164476 | rs10520958 | 0.85 | -0.499 | -0.705 | -1.749 | 0.014 | 0.007 | 0.007 | 1.495 | 0.23 | 0.2 | 0.23 | 0.16 | 0.16 | 0.16 |
| 5 | 30179013 | rs13178362 | 0.58 | -0.728 | -0.491 | -1.765 | 0.007 | 0.007 | 0.007 | 0 | 0.22 | 0.16 | 0.2 | 0.05 | 0.17 | 0.16 |
| 5 | 30180440 | rs11749859 | -1.273 | -0.638 | -0.045 | -1.759 | 0.003 | 0.001 | 0.001 | 0 | 0.77 | 0.77 | 0.64 | 0.27 | 0.66 | 0.5 |
| 5 | 30186011 | rs13157739 | 0.998 | -0.721 | 0.075 | -1.848 | 0.018 | 0.059 | 0.059 | 0 | 0.18 | 0.04 | 0.15 | 0.04 | 0.14 | 0.13 |
| 5 | 30190699 | rs4134393 | -1.314 | -0.723 | 0.078 | -1.833 | 0.003 | 0.001 | 0.001 | 0 | 0.77 | 0.77 | 0.64 | 0.27 | 0.66 | 0.5 |
| 5 | 30218153 | rs4403185 | -0.322 | -0.759 | 0.329 | -1.802 | -0.02 | -0.029 | -0.029 | 0 | 0.43 | 0.49 | 0.42 | 0.2 | 0.3 | 0.29 |
| 5 | 30219838 | rs4282330 | 0.692 | -0.76 | 0.388 | -1.82 | -0.04 | -0.022 | -0.022 | 0.062 | 0.25 | 0.42 | 0.26 | 0.16 | 0.16 | 0.16 |
| 5 | 30222607 | rs13162339 | 0.939 | -0.654 | 0.198 | -1.793 | 0.013 | 0.048 | 0.048 | 0 | 0.18 | 0.05 | 0.15 | 0.04 | 0.14 | 0.13 |
| 5 | 30231951 | rs4128341 | -0.47 | -0.684 | 0.335 | -1.986 | -0.017 | 0.018 | 0.018 | 0 | 0.61 | 0.7 | 0.49 | 0.23 | 0.53 | 0.4 |
| 5 | 30236013 | rs10940878 | 0.621 | -0.779 | 0.548 | -2.019 | 0.025 | 0.014 | 0.014 | 1.127 | 0.24 | 0.18 | 0.11 | 0.01 | 0.17 | 0.15 |
| 5 | 30242594 | rs4295402 | -0.037 | -0.779 | 0.929 | -2.007 | 0.002 | -0.041 | -0.041 | 0 | 0.44 | 0.48 | 0.57 | 0.83 | 0.51 | 0.66 |
| 5 | 30245758 | rs4365850 | -0.985 | -0.195 | 0.805 | -2.165 | -0.011 | 0.017 | 0.017 | 0 | 0.87 | 0.92 | 0.88 | 0.98 | 0.83 | 0.94 |
| 5 | 30257911 | rs4309986 | -0.875 | -0.212 | 0.808 | -2.162 | -0.01 | 0.018 | 0.018 | 0 | 0.86 | 0.92 | 0.89 | 0.98 | 0.83 | 0.94 |
| 5 | 30260291 | rs4867032 | 0.054 | -0.347 | 0.874 | -2.125 | 0.011 | 0.025 | 0.025 | 0.104 | 0.62 | 0.68 | 0.65 | 0.81 | 0.67 | 0.78 |
| 5 | 30284836 | rs6872975 | 0.72 | -0.361 | 0.818 | -2.138 | 0.001 | -0.001 | -0.001 | 2.065 | 0.27 | 0.27 | 0.43 | 0.75 | 0.34 | 0.59 |
| 5 | 30292538 | rs4867263 | 0.67 | -0.332 | 0.802 | -2.125 | 0.004 | -0.001 | -0.001 | 0 | 0.28 | 0.29 | 0.44 | 0.75 | 0.34 | 0.59 |
| 5 | 30317636 | rs4594867 | 0.595 | -0.192 | 0.916 | -1.491 | 0.018 | 0.036 | 0.036 | 0 | 0.62 | 0.72 | 0.66 | 0.77 | 0.67 | 0.78 |
| 5 | 30318510 | rs10472733 | 0.587 | -0.159 | 0.935 | -1.475 | 0.017 | 0.034 | 0.034 | 0 | 0.62 | 0.71 | 0.66 | 0.77 | 0.67 | 0.78 |
| 5 | 30389412 | rs10076325 | -0.916 | 0.419 | 1.57 | -1.476 | 0.051 | -0.061 | -0.061 | 0 | 0.2 | 0.29 | 0.37 | 0.05 | 0.32 | 0.17 |
| 5 | 30390398 | rs6450770 | 0.426 | 0.237 | 1.585 | -1.585 | -0.006 | -0.027 | -0.027 | 0.178 | 0.9 | 0.86 | 0.95 | 1 | 0.93 | 0.94 |
| 5 | 30392067 | rs6871440 | -1.036 | 0.832 | 1.323 | -1.195 | 0.052 | -0.06 | -0.06 | 0.291 | 0.2 | 0.29 | 0.37 | 0.05 | 0.32 | 0.17 |
| 5 | 30392426 | rs4867267 | -1.2 | 0.835 | 1.315 | -0.885 | 0.062 | -0.005 | -0.005 | 0 | 0.22 | 0.35 | 0.39 | 0.24 | 0.35 | 0.28 |
| 5 | 30393124 | rs4362953 | -1.111 | 0.699 | 0.823 | -0.791 | -0.008 | 0.073 | 0.073 | 0 | 0.05 | 0.16 | 0.03 | 0.18 | 0.03 | 0.12 |
| 5 | 30395852 | rs12654311 | -0.774 | 0.735 | 0.693 | -0.98 | -0.008 | 0.042 | 0.042 | 0 | 0.05 | 0.12 | 0.03 | 0.15 | 0.03 | 0.11 |
| 5 | 30401682 | rs12517039 | -0.285 | 1.078 | 0.401 | -1.196 | -0.073 | 0.051 | 0.051 | 0 | 0.16 | 0.05 | 0.34 | 0.07 | 0.31 | 0.17 |
| 5 | 30407686 | rs6878545 | -0.6 | 1.047 | 0.701 | -1.211 | 0.035 | -0.035 | -0.035 | 0.042 | 0.17 | 0.21 | 0.36 | 0.07 | 0.31 | 0.18 |
| 5 | 30411717 | rs12515247 | -0.423 | 1.128 | 0.345 | -1.313 | -0.035 | 0.028 | 0.028 | 0 | 0.17 | 0.11 | 0.36 | 0.04 | 0.33 | 0.18 |
| 5 | 30430435 | rs10066493 | -1.381 | -0.479 | 0.969 | -0.794 | 0.006 | -0.006 | -0.006 | 0.063 | 0.19 | 0.14 | 0.23 | 0.69 | 0.1 | 0.3 |

| | | | | | | | | | | | | | | | | |
|---|---|---|---|---|---|---|---|---|---|---|---|---|---|---|---|---|
| 5 | 30430967 | rs11955656 | 0.93 | -0.268 | 0.777 | -1.18 | -0.045 | 0.046 | 0.046 | 0 | 0.16 | 0.04 | 0.13 | 0 | 0.25 | 0.04 |
| 5 | 30459246 | rs6881036 | -1.398 | -1.18 | 1.406 | -0.321 | -0.008 | -0.025 | -0.025 | 0.912 | 0.15 | 0.11 | 0.05 | 0 | 0.05 | 0.09 |
| 5 | 30478631 | rs10071724 | -1.507 | -1.188 | 1.385 | -0.315 | -0.002 | 0 | 0 | 0.063 | 0.15 | 0.13 | 0.22 | 0.69 | 0.08 | 0.28 |
| 5 | 30483308 | rs6888936 | 2.312 | -1.292 | 1.593 | -0.394 | -0.035 | -0.005 | -0.005 | 0 | 0.73 | 0.36 | 0.69 | 0.28 | 0.86 | 0.67 |
| 5 | 30511767 | rs7442662 | 1.371 | -0.942 | 1.01 | -0.197 | -0.039 | 0.04 | 0.04 | 0 | 0.85 | 0.75 | 0.77 | 0.31 | 0.93 | 0.73 |
| 5 | 30536972 | rs4867276 | 1.494 | -0.484 | 0.717 | 0.04 | -0.044 | 0.056 | 0.056 | 0 | 0.82 | 0.58 | 0.73 | 0.28 | 0.89 | 0.71 |
| 5 | 30552961 | rs4867041 | 0.687 | -0.271 | 0.359 | -0.281 | -0.016 | -0.026 | -0.026 | 0 | 0.69 | 0.49 | 0.64 | 0.29 | 0.72 | 0.67 |
| 5 | 30563394 | rs4262120 | 1.05 | -0.458 | 0.338 | -0.363 | -0.036 | -0.029 | -0.029 | 0.008 | 0.66 | 0.46 | 0.63 | 0.29 | 0.74 | 0.64 |
| 5 | 30571997 | rs9292393 | -2.129 | -0.901 | 1.621 | -0.431 | -0.03 | -0.042 | -0.042 | 0 | 0.13 | 0.38 | 0.06 | 0.07 | 0.07 | 0.08 |
| 5 | 30576056 | rs2330534 | -3.05 | -0.856 | 1.991 | -0.626 | -0.036 | -0.068 | -0.068 | 0 | 0.22 | 0.64 | 0.16 | 0.09 | 0.14 | 0.11 |
| 5 | 30594775 | rs10070502 | -1.732 | -0.843 | 2.567 | -0.618 | -0.004 | -0.013 | -0.013 | 0 | 0.08 | 0.27 | 0.05 | 0.07 | 0.06 | 0.03 |
| 5 | 30602416 | rs1816571 | -1.396 | -0.844 | 2.666 | -0.624 | -0.002 | -0.037 | -0.037 | 0 | 0.08 | 0.16 | 0.08 | 0 | 0.07 | 0.03 |
| 5 | 30602495 | rs1816572 | -3.437 | -0.794 | 2.874 | -0.66 | -0.027 | -0.068 | -0.068 | 0 | 0.22 | 0.63 | 0.18 | 0.1 | 0.17 | 0.11 |
| 5 | 30606548 | rs10940907 | -2.153 | -0.725 | 2.807 | -0.624 | 0.006 | -0.043 | -0.043 | 0.023 | 0.09 | 0.33 | 0.05 | 0 | 0.06 | 0.02 |
| 5 | 30659568 | rs1946758 | 3.273 | -0.708 | 2.934 | -0.675 | 0.029 | 0.003 | 0.003 | 1.258 | 0.93 | 0.67 | 0.94 | 0.97 | 0.94 | 0.98 |
| 5 | 30661077 | rs1347639 | 3.217 | -0.809 | 3.118 | -0.268 | -0.025 | 0.015 | 0.015 | 0 | 0.84 | 0.49 | 0.78 | 0.83 | 0.89 | 0.92 |
| 5 | 30661573 | rs9292394 | 3.649 | -0.611 | 3.086 | -0.419 | -0.015 | -0.031 | -0.031 | 0.904 | 0.75 | 0.35 | 0.67 | 0.83 | 0.78 | 0.87 |
| 5 | 30662356 | rs13190617 | -1.502 | -0.482 | 2.768 | -0.432 | 0.007 | -0.044 | -0.044 | 0 | 0.09 | 0.14 | 0.11 | 0 | 0.11 | 0.06 |
| 5 | 30662729 | rs13153001 | -1.394 | -0.476 | 2.621 | -0.429 | -0.002 | -0.006 | -0.006 | 0 | 0.08 | 0.06 | 0.1 | 0 | 0.11 | 0.06 |
| 5 | 30671988 | rs10472747 | 1.685 | -0.412 | 2.657 | -0.323 | 0.006 | -0.014 | -0.014 | 0 | 0.93 | 0.86 | 0.95 | 0.97 | 0.9 | 0.97 |
| 5 | 30696023 | rs6897245 | 2.894 | -0.399 | 2.981 | -0.402 | 0.034 | -0.093 | -0.093 | 0.703 | 0.84 | 0.51 | 0.83 | 0.97 | 0.8 | 0.92 |
| 5 | 30713080 | rs1370182 | 3.195 | -0.459 | 3.026 | -0.281 | -0.007 | -0.033 | -0.033 | 0 | 0.79 | 0.43 | 0.69 | 0.81 | 0.79 | 0.88 |
| 5 | 30721486 | rs16900173 | -3.038 | -0.47 | 2.873 | -0.284 | -0.007 | -0.013 | -0.013 | 0 | 0.17 | 0.25 | 0.25 | 0.17 | 0.15 | 0.1 |
| 5 | 30724347 | rs11739681 | -2.236 | -0.44 | 2.76 | -0.303 | 0.003 | -0.044 | -0.044 | 0 | 0.12 | 0.17 | 0.11 | 0 | 0.14 | 0.06 |
| 5 | 30738438 | rs11742787 | -1.689 | -0.428 | 2.771 | -0.292 | 0.008 | -0.039 | -0.039 | 0 | 0.07 | 0.15 | 0.1 | 0 | 0.1 | 0.04 |
| 5 | 30746632 | rs10078531 | 2.738 | -0.27 | 2.821 | 0.333 | -0.003 | 0.015 | 0.015 | 0 | 0.68 | 0.4 | 0.47 | 0.46 | 0.68 | 0.61 |
| 5 | 30749170 | rs1104987 | 1.556 | -0.162 | 2.815 | 0.41 | 0.015 | 0.108 | 0.108 | 0 | 0.85 | 0.67 | 0.74 | 0.62 | 0.85 | 0.72 |
| 5 | 30758329 | rs10056075 | -0.198 | -0.434 | 2.138 | 0.167 | -0.011 | -0.018 | -0.018 | 0 | 0.08 | 0.11 | 0.09 | 0.04 | 0.04 | 0.11 |
| 5 | 30771294 | rs10520979 | -0.769 | -0.438 | 2.012 | 0.277 | -0.003 | 0.003 | 0.003 | 0 | 0.11 | 0.08 | 0.22 | 0.35 | 0.11 | 0.26 |
| 5 | 30783732 | rs1392453 | -1.764 | -0.687 | 2.376 | 0.457 | -0.04 | 0.097 | 0.097 | 0 | 0.17 | 0.35 | 0.17 | 0.35 | 0.09 | 0.17 |
| 5 | 30783801 | rs1392452 | 0.744 | -0.464 | 1.947 | 0.163 | -0.009 | 0.006 | 0.006 | 0 | 0.91 | 0.92 | 0.88 | 0.97 | 0.81 | 0.89 |
| 5 | 30787271 | rs9283723 | -1.297 | -0.476 | 2.041 | 0.223 | -0.002 | 0.009 | 0.009 | 0 | 0.16 | 0.18 | 0.16 | 0.35 | 0.09 | 0.18 |
| 5 | 30787747 | rs10073304 | -0.532 | -0.31 | 1.821 | 0.116 | -0.002 | 0.007 | 0.007 | 0 | 0.08 | 0.07 | 0.1 | 0.06 | 0.07 | 0.12 |
| 5 | 30796594 | rs1547530 | 1.657 | -0.233 | 1.784 | -0.02 | -0.017 | 0.024 | 0.024 | 0 | 0.85 | 0.9 | 0.86 | 0.97 | 0.76 | 0.87 |
| 5 | 30799521 | rs6873118 | 1.999 | -0.163 | 1.794 | -0.027 | -0.022 | 0.04 | 0.04 | 0 | 0.77 | 0.84 | 0.76 | 0.91 | 0.69 | 0.75 |
| 5 | 30800973 | rs7723097 | 3.388 | -0.366 | 1.924 | 0.265 | -0.019 | -0.075 | -0.075 | 0.578 | 0.55 | 0.11 | 0.42 | 0.38 | 0.58 | 0.55 |
| 5 | 30805404 | rs1553238 | 1.263 | -0.384 | 2.795 | 0.237 | 0.095 | -0.049 | -0.049 | 0.111 | 0.89 | 0.62 | 0.84 | 0.97 | 0.8 | 0.9 |
| 5 | 30807204 | rs1392447 | -2.047 | -0.367 | 2.591 | 0.28 | -0.036 | -0.093 | -0.093 | 0.426 | 0.15 | 0.38 | 0.07 | 0 | 0.08 | 0.06 |
| 5 | 30814262 | rs9292400 | 1.362 | -0.482 | 2.123 | 0.36 | 0.001 | 0.011 | 0.011 | 0 | 0.81 | 0.79 | 0.75 | 0.63 | 0.75 | 0.73 |
| 5 | 30830655 | rs1500208 | -2.408 | -0.821 | 2.109 | 0.42 | -0.028 | 0.09 | 0.09 | 0.099 | 0.24 | 0.6 | 0.37 | 0.52 | 0.17 | 0.25 |
| 5 | 30831977 | rs1392445 | 0.106 | -0.815 | 1.893 | 0.617 | -0.004 | -0.01 | -0.01 | 0 | 0.08 | 0.05 | 0.15 | 0.3 | 0.08 | 0.2 |
| 5 | 30832311 | rs10044014 | -2.241 | -0.872 | 2.024 | 0.587 | -0.016 | 0.035 | 0.035 | 1.085 | 0.22 | 0.7 | 0.33 | 0.57 | 0.11 | 0.21 |
| 5 | 30839331 | rs2178490 | -2.454 | -0.86 | 2.236 | 0.503 | -0.024 | -0.001 | -0.001 | 0.634 | 0.29 | 0.4 | 0.33 | 0.65 | 0.2 | 0.41 |
| 5 | 30839387 | rs2178489 | -1.906 | -0.863 | 2.249 | 0.504 | -0.024 | 0.017 | 0.017 | 0 | 0.25 | 0.38 | 0.31 | 0.64 | 0.17 | 0.39 |
| 5 | 30841033 | rs6891352 | -2.251 | -0.786 | 2.185 | 0.47 | -0.029 | 0.006 | 0.006 | 0 | 0.28 | 0.59 | 0.35 | 0.64 | 0.18 | 0.4 |
| 5 | 30843437 | rs10805559 | -0.312 | -0.687 | 1.796 | 0.585 | 0.005 | 0.027 | 0.027 | 0 | 0.1 | 0.09 | 0.25 | 0.53 | 0.07 | 0.19 |
| 5 | 30843839 | rs2066930 | -2.04 | -0.722 | 1.946 | 0.286 | -0.006 | 0.021 | 0.021 | 0 | 0.26 | 0.35 | 0.39 | 0.46 | 0.18 | 0.36 |
| 5 | 30849338 | rs9292407 | -0.433 | -0.657 | 1.388 | 0.313 | -0.001 | -0.042 | -0.042 | 0 | 0.12 | 0.08 | 0.15 | 0.29 | 0.09 | 0.23 |
| 5 | 30850795 | rs1882202 | 0.235 | -0.331 | 1.306 | 0.019 | -0.007 | 0.001 | 0.001 | 0 | 0.87 | 0.87 | 0.83 | 0.97 | 0.76 | 0.9 |
| 5 | 30856986 | rs4867286 | 0.3 | -0.659 | 0.239 | 0.046 | -0.005 | -0.097 | -0.097 | 4.206 | 0.14 | 0.01 | 0.14 | 0.42 | 0.12 | 0.18 |
| 5 | 30863015 | rs4502841 | 1.517 | -0.328 | 0.264 | -0.311 | -0.029 | -0.019 | -0.019 | 0 | 0.58 | 0.7 | 0.54 | 0.65 | 0.53 | 0.51 |
| 5 | 30866579 | rs12515842 | -0.471 | -0.327 | 0.093 | -0.322 | -0.01 | 0.037 | 0.037 | 0 | 0.12 | 0.07 | 0.11 | 0.02 | 0.1 | 0.12 |
| 5 | 30869275 | rs16900295 | -0.209 | -0.357 | 0.434 | -0.155 | -0.024 | 0.215 | 0.215 | 0 | 0.19 | 0.41 | 0.35 | 0.57 | 0.14 | 0.19 |
| 5 | 30870858 | rs1921111 | 1.411 | -0.075 | -0.047 | -0.791 | 0 | 0.015 | 0.015 | 0 | 0.39 | 0.29 | 0.19 | 0.09 | 0.39 | 0.32 |
| 5 | 30875225 | rs1921115 | 1.455 | -0.002 | -0.771 | -0.75 | -0.01 | 0.131 | 0.131 | 0 | 0.39 | 0.17 | 0.18 | 0.09 | 0.39 | 0.32 |
| 5 | 30880177 | rs13188689 | 1.198 | -0.057 | -0.949 | -0.715 | -0.036 | 0.206 | 0.206 | 0 | 0.43 | 0.12 | 0.24 | 0.1 | 0.42 | 0.38 |
| 5 | 30886586 | rs10520975 | 0.523 | 0.568 | -0.339 | 0.178 | 0.001 | 0 | 0 | 0 | 0.63 | 0.38 | 0.46 | 0.38 | 0.58 | 0.58 |
| 5 | 30886848 | rs13165986 | 0.157 | 0.689 | 0.114 | 0.445 | 0.001 | 0.036 | 0.036 | 0.017 | 0.69 | 0.58 | 0.62 | 0.46 | 0.65 | 0.62 |
| 5 | 30888137 | rs6878771 | -0.768 | 0.651 | -0.102 | 0.423 | -0.001 | -0.009 | -0.009 | 0 | 0.2 | 0.18 | 0.26 | 0.26 | 0.25 | 0.28 |
| 5 | 30899517 | rs6871087 | 0.486 | 0.825 | 0.18 | 0.221 | 0.036 | -0.017 | -0.017 | 0 | 0.05 | 0.18 | 0.04 | 0 | 0.05 | 0.01 |
| 5 | 30906632 | rs1021714 | -0.191 | 0.727 | -0.288 | 0.442 | 0.002 | 0.005 | 0.005 | 0 | 0.14 | 0.18 | 0.2 | 0.15 | 0.21 | 0.14 |
| 5 | 30914552 | rs1732919 | 0.416 | 0.487 | -0.442 | 0.178 | -0.002 | -0.005 | -0.005 | 0 | 0.65 | 0.68 | 0.55 | 0.36 | 0.61 | 0.49 |
| 5 | 30917209 | rs17603912 | 0.931 | 0.527 | -0.964 | -0.572 | 0.006 | 0.057 | 0.057 | 0 | 0.35 | 0.18 | 0.14 | 0.1 | 0.31 | 0.26 |
| 5 | 30918409 | rs4867289 | 0.931 | 0.562 | -1.201 | -0.555 | -0.026 | 0.173 | 0.173 | 0 | 0.34 | 0.06 | 0.12 | 0.1 | 0.3 | 0.26 |
| 5 | 30919960 | rs13356444 | 0.471 | 0.545 | -1.053 | 0.318 | 0.005 | 0.005 | 0.005 | 0 | 0.57 | 0.27 | 0.38 | 0.43 | 0.58 | 0.52 |
| 5 | 30930971 | rs1276219 | 0.466 | 0.039 | -0.217 | 0.782 | -0.045 | 0.126 | 0.126 | 0 | 0.8 | 0.66 | 0.82 | 0.6 | 0.92 | 0.8 |
| 5 | 30933488 | rs1290062 | 1.212 | 0.242 | 0.273 | 0.536 | -0.009 | 0.022 | 0.022 | 0 | 0.72 | 0.4 | 0.7 | 0.6 | 0.8 | 0.75 |
| 5 | 30933517 | rs10520971 | 1.055 | 0.471 | -0.277 | 0.366 | -0.002 | 0.063 | 0.063 | 0.293 | 0.34 | 0.17 | 0.19 | 0.15 | 0.34 | 0.25 |
| 5 | 30933575 | rs11739505 | -0.197 | 0.487 | -0.229 | 0.379 | 0.06 | -0.039 | -0.039 | 0.107 | 0.08 | 0.31 | 0.11 | 0 | 0.13 | 0.05 |

| | | | | | | | | | | | | | | | | |
|---|---|---|---|---|---|---|---|---|---|---|---|---|---|---|---|---|
| 5 | 30935398 | rs1392421 | 0.785 | 0.319 | -0.817 | 0.755 | -0.01 | 0.05 | 0.05 | 0.205 | 0.35 | 0.17 | 0.28 | 0.24 | 0.36 | 0.26 |
| 5 | 30937514 | rs1276226 | 0.568 | 0.184 | -0.091 | 1.603 | -0.013 | 0.113 | 0.113 | 0 | 0.86 | 0.79 | 0.73 | 0.54 | 0.93 | 0.83 |
| 5 | 30940316 | rs12515837 | -0.513 | 0.131 | -0.703 | 1.559 | 0.009 | 0 | 0 | 0.015 | 0.13 | 0.13 | 0.16 | 0.44 | 0.05 | 0.15 |
| 5 | 30941222 | rs1276165 | -0.185 | 0.435 | -0.976 | 0.534 | -0.04 | 0.078 | 0.078 | 0 | 0.59 | 0.76 | 0.68 | 0.72 | 0.56 | 0.56 |
| 5 | 30944477 | rs1276173 | -0.172 | 0.4 | -1.068 | 0.48 | -0.065 | 0.103 | 0.103 | 0 | 0.59 | 0.8 | 0.67 | 0.72 | 0.56 | 0.55 |
| 5 | 30946644 | rs1276177 | -0.299 | 0.16 | -1.366 | 0.597 | -0.01 | -0.002 | -0.002 | 0 | 0.4 | 0.92 | 0.43 | 0.63 | 0.33 | 0.41 |
| 5 | 30947612 | rs10045797 | -0.165 | 0.17 | -1.251 | 0.602 | 0.006 | 0.011 | 0.011 | 0 | 0.38 | 0.77 | 0.41 | 0.63 | 0.33 | 0.41 |
| 5 | 30959804 | rs10472759 | 0.075 | 0.285 | -0.061 | 0.988 | 0.005 | 0.015 | 0.015 | 0 | 0.35 | 0.51 | 0.48 | 0.54 | 0.33 | 0.34 |
| 5 | 30963876 | rs12513485 | -0.113 | 0.273 | -0.039 | 0.983 | 0 | 0.003 | 0.003 | 0 | 0.37 | 0.58 | 0.48 | 0.54 | 0.33 | 0.34 |
| 5 | 30964809 | rs10052667 | 0.31 | 0.447 | 0.102 | 0.845 | -0.001 | -0.001 | -0.001 | 0 | 0.46 | 0.44 | 0.51 | 0.45 | 0.39 | 0.36 |
| 5 | 30969020 | rs1276153 | -0.08 | 0.393 | 0.449 | 0.806 | -0.009 | 0.033 | 0.033 | 0 | 0.49 | 0.75 | 0.54 | 0.54 | 0.41 | 0.38 |
| 5 | 30975237 | rs4867292 | -0.466 | 0.339 | 0.106 | 0.735 | -0.012 | 0.044 | 0.044 | 0 | 0.86 | 0.71 | 0.89 | 0.72 | 0.92 | 0.73 |
| 5 | 30982959 | rs9292412 | 0.006 | 0.474 | 0.395 | 0.578 | -0.001 | -0.004 | -0.004 | 0 | 0.43 | 0.34 | 0.39 | 0.17 | 0.53 | 0.37 |
| 5 | 30984299 | rs1021711 | 0.558 | 0.489 | 0.373 | 0.631 | -0.001 | -0.001 | -0.001 | 0 | 0.46 | 0.51 | 0.51 | 0.46 | 0.37 | 0.38 |
| 5 | 30986203 | rs6865283 | 0.186 | 0.247 | 0.331 | 0.373 | -0.002 | -0.001 | -0.001 | 0 | 0.63 | 0.41 | 0.7 | 0.58 | 0.67 | 0.6 |
| 5 | 30987828 | rs1909716 | -0.133 | 0.64 | 0.36 | 0.487 | -0.011 | 0.019 | 0.019 | 0.972 | 0.65 | 0.6 | 0.73 | 0.5 | 0.64 | 0.6 |
| 5 | 30998450 | rs10040390 | 0.491 | 0.559 | 0.191 | 0.422 | -0.033 | -0.058 | -0.058 | 0 | 0.27 | 0.54 | 0.2 | 0.14 | 0.25 | 0.16 |
| 5 | 30999730 | rs12519137 | 1.226 | 0.38 | -0.177 | 0.172 | 0 | 0.053 | 0.053 | 0 | 0.32 | 0.22 | 0.23 | 0.05 | 0.27 | 0.26 |
| 5 | 31000116 | rs4867297 | -1.519 | 0.185 | -0.48 | 0.497 | 0.003 | 0.002 | 0.002 | 0 | 0.89 | 0.93 | 0.88 | 0.66 | 0.92 | 0.73 |
| 5 | 31001691 | rs1995266 | -0.251 | 0.073 | -0.264 | 0.463 | -0.022 | 0.018 | 0.018 | 0.197 | 0.67 | 0.63 | 0.7 | 0.5 | 0.71 | 0.61 |
| 5 | 31010683 | rs13187383 | -0.158 | 0.528 | -0.24 | 0.606 | -0.062 | -0.105 | -0.105 | 0 | 0.32 | 0.11 | 0.43 | 0.45 | 0.39 | 0.34 |
| 5 | 31012808 | rs16900540 | 0.825 | 0.23 | -0.286 | 0.542 | -0.022 | -0.026 | -0.026 | 0 | 0.21 | 0.29 | 0.18 | 0.16 | 0.21 | 0.12 |
| 5 | 31014317 | rs7733346 | -0.093 | 0.582 | -0.935 | 0.977 | -0.08 | -0.112 | -0.112 | 0 | 0.25 | 0.03 | 0.27 | 0.36 | 0.31 | 0.25 |
| 5 | 31020501 | rs6869417 | -0.26 | 0.12 | -0.726 | 0.604 | 0 | -0.074 | -0.074 | 0.512 | 0.78 | 0.88 | 0.68 | 0.5 | 0.81 | 0.52 |
| 5 | 31020645 | rs13174447 | -0.049 | 0.28 | -0.901 | 0.444 | -0.009 | -0.004 | -0.004 | 0 | 0.09 | 0.01 | 0.04 | 0.08 | 0.07 | 0.13 |
| 5 | 31021518 | rs9292416 | 0.149 | 0.193 | -0.454 | 0.177 | 0.019 | -0.03 | -0.03 | 1.961 | 0.63 | 0.73 | 0.56 | 0.5 | 0.66 | 0.42 |
| 5 | 31024853 | rs1392428 | 0.467 | 0.76 | -0.604 | 0.052 | -0.06 | 0.037 | 0.037 | 0 | 0.2 | 0.09 | 0.15 | 0.13 | 0.31 | 0.09 |
| 5 | 31025013 | rs1500175 | 0.446 | 0.734 | -0.321 | 0.04 | -0.026 | 0.026 | 0.026 | 0 | 0.21 | 0.15 | 0.15 | 0.13 | 0.31 | 0.09 |
| 5 | 31031638 | rs13167639 | 0.839 | 0.692 | 0.321 | 0.12 | 0.002 | 0.014 | 0.014 | 0 | 0.43 | 0.39 | 0.4 | 0.23 | 0.37 | 0.33 |
| 5 | 31032309 | rs13172112 | 0.868 | 0.692 | 0.517 | 0.12 | 0.003 | 0.034 | 0.034 | 0 | 0.43 | 0.33 | 0.4 | 0.23 | 0.37 | 0.33 |
| 5 | 31033816 | rs12658735 | -0.021 | 0.288 | 0.519 | 0.298 | -0.008 | 0.018 | 0.018 | 0 | 0.66 | 0.63 | 0.57 | 0.48 | 0.68 | 0.43 |
| 5 | 31049958 | rs11749998 | 1.03 | 0.402 | 0.135 | 0.252 | 0 | 0 | 0 | 0 | 0.38 | 0.36 | 0.38 | 0.34 | 0.32 | 0.3 |
| 5 | 31052812 | rs1500165 | -0.032 | 0.486 | -0.086 | 0.327 | -0.01 | 0.024 | 0.024 | 0 | 0.66 | 0.59 | 0.65 | 0.46 | 0.68 | 0.44 |
| 5 | 31056220 | rs16900599 | 0.783 | 0.763 | -0.082 | 0.013 | -0.015 | -0.044 | -0.044 | 0 | 0.16 | 0.21 | 0.13 | 0.02 | 0.14 | 0.09 |
| 5 | 31056670 | rs16900604 | 0.695 | 0.776 | -0.127 | 0.02 | -0.023 | -0.057 | -0.057 | 0 | 0.16 | 0.24 | 0.13 | 0.03 | 0.14 | 0.09 |
| 5 | 31058319 | rs13162331 | -0.908 | 0.851 | -0.264 | 0.014 | -0.001 | -0.001 | -0.001 | 0 | 0.43 | 0.4 | 0.31 | 0.29 | 0.5 | 0.23 |
| 5 | 31059476 | rs13167643 | -0.441 | 1.659 | -0.844 | -0.033 | -0.065 | 0.064 | 0.064 | 0.406 | 0.26 | 0.1 | 0.17 | 0.13 | 0.36 | 0.13 |
| 5 | 31063153 | rs1032713 | -0.995 | 1.516 | 0.221 | 0.386 | 0.117 | 0.18 | 0.18 | 0 | 0.07 | 0.29 | 0.22 | 0.15 | 0.12 | 0.13 |
| 5 | 31068198 | rs16900635 | -0.019 | 1.174 | 0.351 | -0.287 | -0.01 | -0.086 | -0.086 | 0.106 | 0.23 | 0.39 | 0.14 | 0.12 | 0.18 | 0.21 |
| 5 | 31069613 | rs10520964 | -0.46 | 1.17 | -0.171 | -0.866 | 0.015 | -0.002 | -0.002 | 0 | 0.29 | 0.41 | 0.39 | 0.28 | 0.3 | 0.3 |
| 5 | 31070883 | rs17672619 | 0.797 | 0.935 | -0.858 | -0.616 | -0.003 | 0.055 | 0.055 | 0.297 | 0.28 | 0.13 | 0.19 | 0.13 | 0.3 | 0.19 |
| 5 | 31073148 | rs10461694 | 0.782 | 0.934 | -0.883 | -0.599 | -0.003 | 0.056 | 0.056 | 0.219 | 0.28 | 0.12 | 0.19 | 0.13 | 0.3 | 0.19 |
| 5 | 31075755 | rs6872040 | -0.872 | 0.99 | -0.939 | -0.3 | -0.026 | -0.101 | -0.101 | 0 | 0.82 | 0.95 | 0.73 | 0.44 | 0.77 | 0.62 |
| 5 | 31076540 | rs7717530 | -0.446 | 0.659 | -0.236 | -0.686 | 0.005 | 0.005 | 0.005 | 2.741 | 0.66 | 0.64 | 0.6 | 0.6 | 0.63 | 0.6 |
| 5 | 31076878 | rs10059237 | 0.027 | 0.588 | 0.203 | -0.954 | -0.009 | 0.019 | 0.019 | 1.303 | 0.69 | 0.49 | 0.63 | 0.31 | 0.67 | 0.47 |
| 5 | 31077415 | rs7722953 | -0.304 | 0.493 | 0.507 | -1.018 | 0.09 | 0.017 | 0.017 | 0 | 0.06 | 0.32 | 0.05 | 0 | 0.07 | 0.07 |
| 5 | 31078814 | rs13175535 | 0.711 | 0.455 | -0.938 | -1.124 | -0.072 | 0.086 | 0.086 | 0.785 | 0.3 | 0.03 | 0.19 | 0.14 | 0.31 | 0.2 |
| 5 | 31080763 | rs6881843 | 0.132 | 0.391 | -0.37 | -1.234 | -0.003 | -0.03 | -0.03 | 0 | 0.4 | 0.52 | 0.44 | 0.17 | 0.37 | 0.33 |
| 5 | 31082855 | rs4562051 | -0.242 | 1.023 | -0.605 | -1.066 | 0.091 | -0.014 | -0.014 | 0 | 0.17 | 0.44 | 0.16 | 0.14 | 0.26 | 0.2 |
| 5 | 31084270 | rs9292420 | 0.667 | 0.783 | -2 | -1.06 | -0.015 | 0.04 | 0.04 | 0 | 0.31 | 0.22 | 0.4 | 0.15 | 0.33 | 0.27 |
| 5 | 31090094 | rs6885825 | -0.461 | 0.545 | -3.006 | -1.05 | 0.026 | 0.097 | 0.097 | 0 | 0.52 | 0.93 | 0.58 | 0.29 | 0.62 | 0.48 |
| 5 | 31091413 | rs10075290 | 0.157 | 0.575 | -2.518 | -0.677 | -0.038 | 0.063 | 0.063 | 0 | 0.32 | 0.15 | 0.41 | 0.15 | 0.38 | 0.28 |
| 5 | 31093782 | rs10045909 | 0.037 | 0.771 | -2.381 | -0.404 | -0.038 | 0.091 | 0.091 | 0 | 0.38 | 0.15 | 0.43 | 0.18 | 0.4 | 0.31 |
| 5 | 31093849 | rs10043381 | 0.13 | 0.743 | -2.329 | -0.349 | -0.031 | 0.075 | 0.075 | 0 | 0.36 | 0.15 | 0.43 | 0.18 | 0.38 | 0.31 |
| 5 | 31094448 | rs4330472 | -0.325 | 0.454 | -2.029 | 0.692 | 0.015 | -0.114 | -0.114 | 1.651 | 0.19 | 0.05 | 0.28 | 0.57 | 0.17 | 0.36 |
| 5 | 31095369 | rs7724768 | -0.085 | 1.107 | -2.324 | 0.564 | -0.025 | -0.005 | -0.005 | 0 | 0.58 | 0.86 | 0.73 | 0.59 | 0.59 | 0.71 |
| 5 | 31095413 | rs12186729 | -0.517 | 2.094 | -1.41 | -0.333 | 0.009 | -0.022 | -0.022 | 0.012 | 0.24 | 0.28 | 0.26 | 0.09 | 0.34 | 0.44 |
| 5 | 31095521 | rs7724797 | -0.848 | 1.828 | -0.607 | -0.359 | 0.074 | -0.094 | -0.094 | 0.139 | 0.25 | 0.52 | 0.31 | 0.13 | 0.38 | 0.24 |
| 5 | 31096643 | rs6872076 | -0.692 | 0.574 | -0.423 | -0.173 | -0.03 | -0.005 | -0.005 | 0 | 0.33 | 0.52 | 0.3 | 0.28 | 0.23 | 0.19 |
| 5 | 31099404 | rs16900670 | 0.874 | 0.353 | -0.777 | -0.126 | -0.025 | -0.139 | -0.139 | 0 | 0.16 | 0.01 | 0.17 | 0.52 | 0.16 | 0.23 |
| 5 | 31101057 | rs16900672 | 0.013 | 0.145 | -0.508 | -0.307 | 0.011 | 0.05 | 0.05 | 0.247 | 0.13 | 0.2 | 0.23 | 0.28 | 0.13 | 0.1 |
| 5 | 31102396 | rs4577701 | 0.954 | 0.588 | -0.456 | -1.022 | 0.043 | -0.081 | -0.081 | 7.098 | 0.16 | 0.3 | 0.35 | 0.03 | 0.29 | 0.09 |
| 5 | 31106256 | rs17674109 | 0.191 | 0.409 | -0.539 | -1.056 | -0.094 | -0.142 | -0.142 | 0.69 | 0.32 | 0.05 | 0.49 | 0.54 | 0.46 | 0.32 |
| 5 | 31106657 | rs4357048 | -1.455 | -0.109 | -0.484 | -1.326 | -0.016 | 0.026 | 0.026 | 0 | 0.79 | 0.86 | 0.87 | 0.97 | 0.75 | 0.75 |
| 5 | 31107736 | rs4867306 | -0.329 | -0.596 | -0.22 | -1.043 | 0.003 | -0.001 | -0.001 | 0 | 0.48 | 0.54 | 0.47 | 0.81 | 0.38 | 0.42 |
| 5 | 31111978 | rs6870237 | 0.018 | -0.535 | 0.082 | -0.916 | -0.017 | 0.015 | 0.015 | 0.499 | 0.27 | 0.42 | 0.27 | 0.29 | 0.18 | 0.17 |
| 5 | 31112867 | rs13358949 | 0.824 | -1.007 | 0.453 | -0.908 | -0.065 | -0.115 | -0.115 | 0.29 | 0.16 | 0.43 | 0.08 | 0.01 | 0.07 | 0.08 |
| 5 | 31114590 | rs4867308 | -0.376 | -1.249 | -0.141 | -1.03 | 0.087 | 0.011 | 0.011 | 0.017 | 0.42 | 0.74 | 0.37 | 0.65 | 0.25 | 0.35 |
| 5 | 31123143 | rs10079279 | 1.238 | -1.076 | 0.014 | -0.987 | -0.026 | -0.058 | -0.058 | 0.007 | 0.14 | 0.2 | 0.05 | 0 | 0.07 | 0.07 |

| | | | | | | | | | | | | | | | | |
|---|---|---|---|---|---|---|---|---|---|---|---|---|---|---|---|---|
| 5 | 31131409 | rs4867309 | -0.601 | -0.39 | -0.041 | -1.083 | 0.004 | 0.002 | 0.002 | 0 | 0.67 | 0.66 | 0.62 | 0.67 | 0.64 | 0.7 |
| 5 | 31131959 | rs16900709 | 1.579 | -0.241 | -0.287 | -0.938 | 0.006 | -0.076 | -0.076 | 0 | 0.16 | 0.06 | 0.26 | 0.53 | 0.16 | 0.24 |
| 5 | 31136051 | rs4266413 | 0.097 | -0.075 | -0.297 | -0.651 | -0.001 | -0.006 | -0.006 | 0.011 | 0.57 | 0.42 | 0.6 | 0.56 | 0.62 | 0.67 |
| 5 | 31138891 | rs13354347 | -0.346 | 0.341 | -0.441 | -0.563 | 0.001 | -0.002 | -0.002 | 0 | 0.14 | 0.16 | 0.17 | 0.08 | 0.14 | 0.14 |
| 5 | 31143872 | rs10520961 | 0.175 | -0.076 | -0.421 | -0.742 | 0 | -0.013 | -0.013 | 0.127 | 0.09 | 0.08 | 0.04 | 0.01 | 0.07 | 0.25 |
| 5 | 31143956 | rs4867311 | -0.193 | 0.215 | -0.475 | -0.124 | -0.005 | -0.001 | -0.001 | 0.001 | 0.3 | 0.25 | 0.3 | 0.28 | 0.33 | 0.5 |
| 5 | 31144547 | rs4867312 | -1.188 | 0.434 | -0.934 | -0.606 | -0.001 | -0.006 | -0.006 | 0 | 0.76 | 0.78 | 0.85 | 0.95 | 0.74 | 0.91 |
| 5 | 31155754 | rs4457095 | 0.757 | -0.057 | -1.323 | 0.475 | -0.063 | -0.104 | -0.104 | 0 | 0.15 | 0.01 | 0.13 | 0.38 | 0.19 | 0.14 |
| 5 | 31169236 | rs11957918 | -0.406 | -0.392 | -0.936 | -0.019 | 0.001 | -0.018 | -0.018 | 0 | 0.4 | 0.28 | 0.52 | 0.49 | 0.39 | 0.35 |
| 5 | 31175696 | rs4394137 | 0.385 | -0.598 | -0.15 | -0.117 | -0.024 | -0.046 | -0.046 | 0 | 0.65 | 0.51 | 0.73 | 0.76 | 0.74 | 0.74 |
| 5 | 31177469 | rs4352611 | -0.164 | -0.718 | 0.047 | -0.225 | -0.041 | -0.002 | -0.002 | 0 | 0.83 | 0.74 | 0.95 | 0.82 | 0.95 | 0.89 |
| 5 | 31177628 | rs4516871 | 1.18 | -0.683 | 0.056 | -0.342 | 0 | -0.024 | -0.024 | 0 | 0.17 | 0.19 | 0.21 | 0.03 | 0.21 | 0.13 |
| 5 | 31179728 | rs10075707 | 0.958 | -0.155 | -0.157 | -0.026 | -0.001 | -0.001 | -0.001 | 0 | 0.56 | 0.47 | 0.47 | 0.5 | 0.53 | 0.59 |
| 5 | 31184402 | rs6890687 | 0.857 | -0.139 | -0.12 | -0.047 | 0.004 | 0.005 | 0.005 | 0.009 | 0.57 | 0.51 | 0.48 | 0.51 | 0.54 | 0.6 |
| 5 | 31185714 | rs4867315 | 0.811 | 0.085 | -0.047 | 0.172 | 0.011 | 0.027 | 0.027 | 0.165 | 0.09 | 0.08 | 0.19 | 0.26 | 0.2 | 0.15 |
| 5 | 31188433 | rs7713805 | 2.51 | -0.257 | 0.018 | 0.069 | -0.001 | -0.016 | -0.016 | 0.247 | 0.05 | 0.04 | 0.02 | 0.01 | 0.02 | 0.05 |
| 5 | 31191910 | rs3806851 | 0.131 | -0.725 | 0.042 | -0.502 | 0.001 | -0.001 | -0.001 | 0 | 0.61 | 0.6 | 0.52 | 0.69 | 0.49 | 0.61 |
| 5 | 31192038 | rs3822365 | 1.088 | -0.797 | -0.028 | -0.595 | -0.005 | 0.005 | 0.005 | 0 | 0.29 | 0.29 | 0.26 | 0.52 | 0.25 | 0.42 |
| 5 | 31192628 | rs3806850 | 1.414 | 0.141 | 0.353 | -0.111 | -0.003 | 0.039 | 0.039 | 0 | 0.26 | 0.17 | 0.23 | 0.16 | 0.22 | 0.14 |
| 5 | 31200936 | rs7716554 | -0.115 | 0.39 | 0.637 | 0.167 | 0.031 | 0.045 | 0.045 | 0 | 0.83 | 0.74 | 0.7 | 0.73 | 0.74 | 0.83 |
| 5 | 31206701 | rs6875836 | -1.637 | -0.015 | 0.787 | 0.153 | -0.004 | -0.038 | -0.038 | 0.011 | 0.93 | 0.85 | 0.88 | 1 | 0.94 | 0.97 |
| 5 | 31209367 | rs4535467 | -0.497 | -0.177 | 0.746 | 0.487 | -0.012 | 0.091 | 0.091 | 0 | 0.87 | 0.77 | 0.8 | 0.57 | 0.86 | 0.9 |
| 5 | 31215158 | rs7734842 | -0.013 | -0.185 | 0.743 | 0.339 | 0.006 | 0.025 | 0.025 | 0 | 0.17 | 0.19 | 0.19 | 0.29 | 0.17 | 0.22 |
| 5 | 31215770 | rs10038069 | 1.494 | -0.31 | 0.37 | -0.426 | 0.011 | 0.072 | 0.072 | 0 | 0.41 | 0.25 | 0.34 | 0.14 | 0.38 | 0.48 |
| 5 | 31219952 | rs7723322 | 0.711 | 0.012 | 0.002 | -0.094 | -0.012 | 0.03 | 0.03 | 0 | 0.27 | 0.21 | 0.26 | 0.14 | 0.3 | 0.15 |
| 5 | 31223681 | rs4596398 | -1.142 | -0.697 | -0.1 | 0.052 | 0.002 | 0.083 | 0.083 | 0.144 | 0.61 | 0.79 | 0.59 | 0.88 | 0.63 | 0.72 |
| 5 | 31229741 | rs10805563 | -1.135 | -1.401 | 0.888 | 1.008 | -0.02 | 0.084 | 0.084 | 0 | 0.17 | 0.25 | 0.12 | 0.56 | 0.11 | 0.13 |
| 5 | 31235230 | rs4457092 | 1.621 | -1.616 | 0.925 | 0.008 | 0.02 | 0.02 | 0.02 | 0.336 | 0.73 | 0.25 | 0.88 | 0.84 | 0.88 | 0.66 |
| 5 | 31237336 | rs4072916 | -0.75 | -1.506 | 0.723 | 0.075 | -0.016 | -0.051 | -0.051 | 0.311 | 0.14 | 0.2 | 0.06 | 0.03 | 0.09 | 0.29 |
| 5 | 31245334 | rs16900783 | -0.133 | -0.855 | 0.335 | 0.681 | -0.012 | 0.042 | 0.042 | 0 | 0.06 | 0.17 | 0.02 | 0.13 | 0.02 | 0.04 |
| 5 | 31250588 | rs6884544 | -0.39 | -0.793 | -0.023 | 0.637 | -0.005 | 0.004 | 0.004 | 0.259 | 0.43 | 0.4 | 0.45 | 0.37 | 0.55 | 0.42 |
| 5 | 31250964 | rs13186740 | -0.663 | -1.056 | 0.063 | 0.23 | -0.003 | 0.068 | 0.068 | 0.369 | 0.39 | 0.74 | 0.31 | 0.19 | 0.41 | 0.43 |
| 5 | 31257120 | rs9687496 | -1.215 | -1.162 | 0.418 | 0.144 | -0.029 | -0.091 | -0.091 | 0 | 0.16 | 0.58 | 0.08 | 0.04 | 0.06 | 0.36 |
| 5 | 31259129 | rs7705566 | -0.704 | -1.01 | 0.21 | 0.279 | 0.008 | -0.015 | -0.015 | 0 | 0.24 | 0.42 | 0.23 | 0.21 | 0.21 | 0.38 |
| 5 | 31268991 | rs6891192 | 0.667 | -1.076 | 0.564 | 0.353 | -0.005 | 0.021 | 0.021 | 1.194 | 0.53 | 0.38 | 0.53 | 0.6 | 0.46 | 0.37 |
| 5 | 31270555 | rs7736710 | -0.06 | -0.833 | 0.775 | 0.336 | -0.004 | -0.034 | -0.034 | 0 | 0.66 | 0.56 | 0.72 | 0.79 | 0.63 | 0.5 |
| 5 | 31273134 | rs10461695 | 1.394 | -1.547 | 0.33 | 0.432 | -0.021 | -0.014 | -0.014 | 0 | 0.31 | 0.2 | 0.39 | 0.35 | 0.35 | 0.15 |
| 5 | 31273152 | rs10461897 | 1.2 | -1.571 | 0.401 | 0.42 | 0.03 | 0.017 | 0.017 | 0.132 | 0.32 | 0.45 | 0.41 | 0.35 | 0.35 | 0.15 |
| 5 | 31273977 | rs4867324 | 1.063 | -1.69 | -0.39 | 0.597 | 0.012 | -0.057 | -0.057 | 0.045 | 0.26 | 0.05 | 0.12 | 0.26 | 0.2 | 0.26 |
| 5 | 31276167 | rs4867061 | 1.096 | -1.676 | -0.383 | 0.624 | 0.012 | -0.06 | -0.06 | 0 | 0.26 | 0.05 | 0.12 | 0.25 | 0.2 | 0.2 |
| 5 | 31278943 | rs10520983 | 1.762 | -1.623 | -0.392 | 0.674 | -0.101 | -0.074 | -0.074 | 0 | 0.27 | 0.04 | 0.35 | 0.33 | 0.33 | 0.14 |
| 5 | 31281799 | rs13187907 | 1.53 | -1.378 | -0.163 | 0.651 | -0.039 | 0.004 | 0.004 | 0.739 | 0.33 | 0.12 | 0.38 | 0.32 | 0.34 | 0.15 |
| 5 | 31283826 | rs16900827 | 1.076 | -1.41 | -0.244 | 0.614 | -0.001 | -0.087 | -0.087 | 0.51 | 0.25 | 0.05 | 0.13 | 0.28 | 0.22 | 0.19 |
| 5 | 31284709 | rs12657313 | -0.043 | -1.213 | 0.616 | 0.339 | -0.001 | 0.001 | 0.001 | 0 | 0.44 | 0.41 | 0.43 | 0.38 | 0.41 | 0.32 |
| 5 | 31291960 | rs10472770 | -0.618 | -1.362 | 0.404 | 0.368 | -0.001 | -0.011 | -0.011 | 0.323 | 0.1 | 0.14 | 0.04 | 0.05 | 0.06 | 0.15 |
| 5 | 31296213 | rs3828638 | 0.356 | -1.5 | 0.412 | 0.351 | -0.035 | 0.02 | 0.02 | 0 | 0.42 | 0.23 | 0.4 | 0.38 | 0.41 | 0.32 |
| 5 | 31299254 | rs2962799 | 0.244 | -1.548 | 0.224 | 0.313 | -0.003 | 0.021 | 0.021 | 0 | 0.49 | 0.32 | 0.54 | 0.57 | 0.58 | 0.46 |
| 5 | 31300815 | rs2330698 | 1.486 | -1.124 | -0.461 | -0.257 | -0.048 | 0.096 | 0.096 | 0 | 0.27 | 0.13 | 0.18 | 0.05 | 0.27 | 0.15 |
| 5 | 31302288 | rs2287582 | 0.106 | -1.75 | -0.284 | 0.428 | -0.117 | -0.119 | -0.119 | 0 | 0.46 | 0.06 | 0.5 | 0.52 | 0.55 | 0.53 |
| 5 | 31305265 | rs2287581 | -0.506 | -1.558 | -0.152 | 0.365 | -0.006 | -0.003 | -0.003 | 0 | 0.07 | 0.06 | 0.03 | 0.07 | 0.05 | 0.14 |
| 5 | 31315377 | rs2909569 | -0.263 | -1.645 | 0.223 | 0.384 | 0 | 0.002 | 0.002 | 0.059 | 0.49 | 0.38 | 0.53 | 0.49 | 0.56 | 0.47 |
| 5 | 31317494 | rs2302904 | -0.199 | -1.642 | 0.164 | 0.385 | -0.003 | 0 | 0 | 0 | 0.49 | 0.35 | 0.54 | 0.49 | 0.56 | 0.47 |
| 5 | 31317952 | rs2229575 | 0.05 | -1.615 | -0.282 | 0.401 | -0.119 | -0.107 | -0.107 | 0 | 0.46 | 0.06 | 0.5 | 0.49 | 0.56 | 0.47 |
| 5 | 31335274 | rs518451 | -1.9 | -1.623 | 0.917 | 0.704 | 0.035 | 0.044 | 0.044 | 0.002 | 0.09 | 0.32 | 0.05 | 0.1 | 0.06 | 0.14 |
| 5 | 31338589 | rs2193769 | -1.215 | -1.747 | 0.855 | 0.613 | 0.001 | -0.002 | -0.002 | 0 | 0.54 | 0.64 | 0.5 | 0.59 | 0.49 | 0.73 |
| 5 | 31343671 | rs16900905 | -1.285 | -1.351 | 0.732 | 0.743 | 0.018 | 0.009 | 0.009 | 0 | 0.08 | 0.22 | 0.04 | 0.07 | 0.06 | 0.08 |
| 5 | 31351588 | rs471245 | -0.236 | -1.469 | 0.239 | 0.75 | 0.005 | -0.024 | -0.024 | 0 | 0.37 | 0.27 | 0.41 | 0.5 | 0.4 | 0.44 |
| 5 | 31352576 | rs7733250 | -0.598 | -1.533 | 0.504 | 0.659 | -0.01 | -0.029 | -0.029 | 0 | 0.13 | 0.27 | 0.07 | 0.08 | 0.09 | 0.27 |
| 5 | 31353951 | rs593729 | -1.162 | -1.518 | 0.427 | 0.676 | 0.005 | 0.003 | 0.003 | 0 | 0.5 | 0.56 | 0.48 | 0.6 | 0.49 | 0.71 |
| 5 | 31356889 | rs637883 | -0.206 | -1.474 | 0.322 | 0.699 | 0.006 | -0.036 | -0.036 | 0.008 | 0.37 | 0.23 | 0.41 | 0.51 | 0.4 | 0.45 |
| 5 | 31357809 | rs652379 | -0.583 | -1.129 | 0.094 | 0.919 | 0.003 | 0.002 | 0.002 | 0.169 | 0.48 | 0.47 | 0.47 | 0.6 | 0.48 | 0.7 |
| 5 | 31360086 | rs334589 | 0.67 | -0.93 | 0.598 | 1.003 | -0.044 | -0.004 | -0.004 | 1.513 | 0.8 | 0.56 | 0.91 | 0.83 | 0.89 | 0.9 |
| 5 | 31361704 | rs4867326 | -0.266 | -0.218 | 0.237 | 1.192 | 0 | 0 | 0 | 0.035 | 0.16 | 0.16 | 0.28 | 0.17 | 0.27 | 0.19 |
| 5 | 31364138 | rs334582 | -0.302 | -0.212 | -0.041 | 1.707 | -0.004 | -0.05 | -0.05 | 0.249 | 0.82 | 0.9 | 0.74 | 0.6 | 0.84 | 0.8 |
| 5 | 31364999 | rs334581 | 0.158 | -0.68 | 0.489 | 1.327 | -0.021 | 0.006 | 0.006 | 0.915 | 0.21 | 0.3 | 0.11 | 0.27 | 0.16 | 0.22 |
| 5 | 31366169 | rs334579 | -1.089 | -0.832 | 0.974 | 1.45 | 0.006 | -0.01 | -0.01 | 0 | 0.39 | 0.62 | 0.48 | 0.51 | 0.44 | 0.39 |
| 5 | 31370030 | rs334573 | -0.016 | -0.969 | 0.687 | 1.47 | -0.04 | 0.001 | 0.001 | 0.617 | 0.22 | 0.34 | 0.12 | 0.24 | 0.14 | 0.21 |
| 5 | 31371127 | rs458586 | -0.605 | -0.827 | 0.574 | 1.529 | -0.004 | -0.02 | -0.02 | 0 | 0.82 | 0.89 | 0.92 | 0.76 | 0.84 | 0.8 |
| 5 | 31371932 | rs451781 | -0.093 | -0.872 | 0.735 | 1.51 | -0.008 | 0.035 | 0.035 | 0 | 0.79 | 0.57 | 0.89 | 0.72 | 0.84 | 0.78 |

| | | | | | | | | | | | | | | | | |
|---|---|---|---|---|---|---|---|---|---|---|---|---|---|---|---|---|
| 5 | 31373920 | rs13175906 | 1.603 | -0.826 | 0.416 | 1.301 | -0.014 | 0.063 | 0.063 | 0 | 0.41 | 0.26 | 0.33 | 0.25 | 0.38 | 0.42 |
| 5 | 31373941 | rs334287 | 0.225 | -0.886 | 0.371 | 1.35 | -0.029 | 0.006 | 0.006 | 1.199 | 0.22 | 0.29 | 0.11 | 0.25 | 0.14 | 0.21 |
| 5 | 31375376 | rs17403765 | 0.178 | -0.772 | 0.173 | 1.438 | 0.025 | 0.042 | 0.042 | 0 | 0.19 | 0.3 | 0.47 | 0.24 | 0.32 | 0.18 |
| 5 | 31376053 | rs13186629 | 1.421 | -0.582 | -0.076 | 1.299 | -0.083 | 0.106 | 0.106 | 0 | 0.39 | 0.09 | 0.32 | 0.23 | 0.38 | 0.4 |
| 5 | 31384224 | rs672990 | -0.106 | -0.487 | 0.15 | 1.32 | 0.002 | -0.005 | -0.005 | 0.341 | 0.8 | 0.84 | 0.9 | 0.73 | 0.84 | 0.77 |
| 5 | 31385996 | rs550353 | -0.138 | -0.235 | -0.042 | 1.143 | -0.042 | -0.025 | -0.025 | 0.022 | 0.26 | 0.45 | 0.13 | 0.23 | 0.18 | 0.24 |
| 5 | 31386997 | rs16901064 | 0.215 | -0.341 | -1.546 | 1.278 | 0.049 | -0.021 | -0.021 | 0 | 0.23 | 0.15 | 0.19 | 0.29 | 0.14 | 0.24 |
| 5 | 31387526 | rs12188316 | 0.066 | 0.202 | -1.333 | 1.024 | 0.013 | -0.015 | -0.015 | 0 | 0.41 | 0.33 | 0.52 | 0.51 | 0.44 | 0.41 |
| 5 | 31388117 | rs607613 | 0.532 | 0.509 | -1.424 | 1.194 | -0.028 | -0.043 | -0.043 | 0.184 | 0.84 | 0.94 | 0.9 | 0.76 | 0.82 | 0.83 |
| 5 | 31391771 | rs10805564 | -0.083 | 0.312 | -1.469 | 0.929 | 0.027 | 0.018 | 0.018 | 0.648 | 0.46 | 0.23 | 0.56 | 0.46 | 0.57 | 0.52 |
| 5 | 31401003 | rs642321 | -0.056 | -0.618 | -1.242 | 1.39 | 0.017 | -0.021 | -0.021 | 0 | 0.67 | 0.72 | 0.71 | 0.51 | 0.77 | 0.68 |
| 5 | 31401447 | rs10719 | 0.405 | -0.904 | -0.669 | 1.227 | 0.008 | 0.007 | 0.007 | 0 | 0.62 | 0.34 | 0.66 | 0.32 | 0.76 | 0.61 |
| 5 | 31401953 | rs3805525 | 1.187 | -1.337 | -0.958 | 0.949 | -0.015 | 0.081 | 0.081 | 0 | 0.41 | 0.2 | 0.47 | 0.25 | 0.55 | 0.42 |
| 5 | 31404766 | rs6450840 | -0.267 | -1.342 | -0.657 | 0.958 | 0.002 | -0.005 | -0.005 | 0 | 0.09 | 0.08 | 0.1 | 0.17 | 0.06 | 0.08 |
| 5 | 31409252 | rs2241337 | 0.13 | -1.482 | -0.43 | 1.001 | 0.035 | -0.032 | -0.032 | 0 | 0.33 | 0.25 | 0.28 | 0.48 | 0.21 | 0.32 |
| 5 | 31410306 | rs3805521 | 1.247 | -1.463 | -0.43 | 0.912 | -0.024 | 0.03 | 0.03 | 0.262 | 0.16 | 0.07 | 0.18 | 0.01 | 0.21 | 0.16 |
| 5 | 31416248 | rs3792830 | -0.529 | -1.663 | -0.008 | 0.971 | -0.019 | 0.034 | 0.034 | 0 | 0.11 | 0.18 | 0.11 | 0.16 | 0.06 | 0.09 |
| 5 | 31423007 | rs2287584 | -0.582 | -1.404 | 0.659 | 0.911 | -0.043 | -0.028 | -0.028 | 1.014 | 0.39 | 0.56 | 0.32 | 0.67 | 0.25 | 0.38 |
| 5 | 31425788 | rs615344 | -0.593 | -1.425 | 0.7 | 1.195 | 0.002 | 0.001 | 0.001 | 0 | 0.39 | 0.66 | 0.29 | 0.57 | 0.25 | 0.38 |
| 5 | 31428028 | rs2279797 | 1.09 | -1.363 | 0.821 | 1.107 | 0.017 | -0.02 | -0.02 | 1.354 | 0.23 | 0.18 | 0.19 | 0.34 | 0.18 | 0.23 |
| 5 | 31433647 | rs639174 | -0.395 | -1.28 | 1.027 | 0.941 | -0.034 | -0.025 | -0.025 | 0 | 0.37 | 0.57 | 0.33 | 0.69 | 0.27 | 0.34 |
| 5 | 31435627 | rs4867329 | 1.141 | -1.208 | 1.02 | 0.846 | -0.012 | 0.037 | 0.037 | 0 | 0.42 | 0.28 | 0.47 | 0.23 | 0.5 | 0.45 |
| 5 | 31437040 | rs493760 | -0.103 | -1.034 | 0.485 | 0.796 | -0.007 | 0.011 | 0.011 | 0 | 0.75 | 0.82 | 0.72 | 0.8 | 0.71 | 0.78 |
| 5 | 31439859 | rs16901165 | -0.559 | -1.149 | 0.484 | 0.868 | -0.009 | 0.014 | 0.014 | 0 | 0.1 | 0.14 | 0.1 | 0.17 | 0.08 | 0.09 |
| 5 | 31440389 | rs3095825 | -1.525 | -0.942 | 0.303 | 0.766 | -0.011 | 0.017 | 0.017 | 0.59 | 0.85 | 0.92 | 0.83 | 0.97 | 0.8 | 0.87 |
| 5 | 31443564 | rs7712155 | 0.252 | -1.03 | 0.301 | 0.924 | 0.003 | -0.001 | -0.001 | 0 | 0.3 | 0.37 | 0.24 | 0.38 | 0.21 | 0.25 |
| 5 | 31444434 | rs573010 | 0.01 | -0.826 | 0.133 | 0.762 | -0.001 | 0.01 | 0.01 | 0.092 | 0.75 | 0.81 | 0.7 | 0.81 | 0.73 | 0.77 |
| 5 | 31452934 | rs492176 | -0.492 | -0.508 | -0.098 | 0.668 | 0.015 | -0.024 | -0.024 | 0 | 0.62 | 0.72 | 0.68 | 0.41 | 0.67 | 0.75 |
| 5 | 31459743 | rs17408227 | 0.627 | -0.914 | -0.393 | 0.413 | -0.029 | 0.368 | 0.368 | 0.348 | 0.46 | 0.06 | 0.47 | 0.03 | 0.59 | 0.63 |
| 5 | 31465137 | rs524138 | -0.877 | -0.777 | 0.596 | 0.597 | 0.009 | 0.21 | 0.21 | 0.349 | 0.07 | 0.27 | 0.13 | 0.21 | 0.04 | 0.04 |
| 5 | 31469172 | rs6450845 | -1.562 | -0.23 | 0.85 | 0.487 | -0.012 | 0.18 | 0.18 | 1.474 | 0.09 | 0.53 | 0.15 | 0.21 | 0.04 | 0.04 |
| 5 | 31471676 | rs10068052 | 0.145 | -0.603 | 0.53 | 0.228 | 0.061 | 0.178 | 0.178 | 0.033 | 0.48 | 0.21 | 0.53 | 0.2 | 0.61 | 0.62 |
| 5 | 31483596 | rs10052174 | 0.497 | -1.321 | 0.206 | 0.232 | -0.058 | 0.188 | 0.188 | 1.818 | 0.81 | 0.56 | 0.8 | 0.55 | 0.89 | 0.92 |
| 5 | 31485230 | rs7712436 | 0.127 | -1.329 | 0.382 | 0.163 | 0.011 | -0.001 | -0.001 | 0.516 | 0.95 | 0.8 | 0.97 | 1 | 0.98 | 0.97 |
| 5 | 31491121 | rs6884823 | 0.502 | -1.305 | 0.158 | 0.201 | -0.019 | 0.021 | 0.021 | 0.834 | 0.13 | 0.24 | 0.08 | 0.24 | 0.07 | 0.04 |
| 5 | 31504251 | rs7737174 | -0.278 | -1.269 | 0.003 | 0.119 | 0.045 | 0.001 | 0.001 | 0 | 0.45 | 0.31 | 0.33 | 0.59 | 0.34 | 0.34 |
| 5 | 31510758 | rs11958935 | -0.478 | -1.118 | -0.041 | 0.034 | 0.011 | 0.03 | 0.03 | 0 | 0.52 | 0.67 | 0.55 | 0.62 | 0.54 | 0.56 |
| 5 | 31511467 | rs17487575 | 0.793 | -1.041 | -0.288 | -0.021 | -0.187 | 0.025 | 0.025 | 0.097 | 0.32 | 0.03 | 0.36 | 0.19 | 0.45 | 0.48 |
| 5 | 31523281 | rs1382883 | 0.448 | -1.036 | 0.189 | -0.01 | 0.004 | -0.01 | -0.01 | 0 | 0.36 | 0.35 | 0.39 | 0.19 | 0.45 | 0.51 |
| 5 | 31525788 | rs6878195 | 0.268 | -1.062 | 0.085 | -0.069 | 0.027 | 0.01 | 0.01 | 0 | 0.38 | 0.45 | 0.48 | 0.39 | 0.48 | 0.53 |
| 5 | 31528733 | rs17409624 | 0.524 | -0.683 | -1.391 | -0.573 | -0.021 | 0.013 | 0.013 | 0.34 | 0.44 | 0.32 | 0.55 | 0.62 | 0.49 | 0.41 |
| 5 | 31531619 | rs17409803 | 1.552 | -0.296 | -1.706 | -0.688 | -0.093 | 0.086 | 0.086 | 0 | 0.16 | 0.01 | 0.12 | 0.05 | 0.26 | 0.26 |
| 5 | 31533691 | rs13169883 | 1.982 | -0.583 | -1.409 | -0.428 | -0.03 | 0.079 | 0.079 | 1.27 | 0.15 | 0.02 | 0.15 | 0.02 | 0.16 | 0.13 |
| 5 | 31536327 | rs4867349 | -1.212 | -0.501 | -1.19 | -0.063 | 0.012 | -0.036 | -0.036 | 2.9 | 0.74 | 0.87 | 0.74 | 0.65 | 0.75 | 0.75 |
| 5 | 31541142 | rs17410035 | 0.31 | -0.247 | -1.643 | -0.548 | -0.09 | 0.052 | 0.052 | 0 | 0.23 | 0.05 | 0.18 | 0.13 | 0.32 | 0.32 |
| 5 | 31546291 | rs7717331 | -1.636 | -0.574 | -1.52 | -0.364 | 0.004 | -0.025 | -0.025 | 0 | 0.78 | 0.88 | 0.74 | 0.59 | 0.78 | 0.78 |
| 5 | 31550331 | rs12654273 | -1.582 | -0.599 | -1.237 | -0.384 | 0.003 | -0.022 | -0.022 | 0 | 0.77 | 0.89 | 0.75 | 0.59 | 0.78 | 0.78 |
| 5 | 31552301 | rs10042915 | 2.399 | -1.023 | -1.052 | -0.632 | -0.026 | -0.037 | -0.037 | 0 | 0.09 | 0.14 | 0.03 | 0 | 0.02 | 0.03 |
| 5 | 31552606 | rs10520986 | 1.5 | -1.046 | -0.884 | -0.83 | -0.001 | -0.045 | -0.045 | 1.853 | 0.26 | 0.33 | 0.2 | 0.02 | 0.19 | 0.16 |
| 5 | 31553968 | rs12658729 | -0.15 | -0.706 | -0.572 | -0.347 | -0.033 | -0.012 | -0.012 | 1.232 | 0.3 | 0.18 | 0.22 | 0.29 | 0.38 | 0.37 |
| 5 | 31557241 | rs4867070 | -0.481 | -0.673 | -0.546 | -0.334 | -0.033 | -0.042 | -0.042 | 0.19 | 0.34 | 0.18 | 0.29 | 0.47 | 0.39 | 0.45 |
| 5 | 31559692 | rs7701461 | 1.085 | -0.383 | -0.276 | -0.347 | 0.023 | -0.072 | -0.072 | 0 | 0.17 | 0.28 | 0.24 | 0.09 | 0.2 | 0.17 |
| 5 | 31571666 | rs7737527 | -1.15 | -0.644 | -0.64 | -0.38 | 0.004 | -0.006 | -0.006 | 0.169 | 0.44 | 0.66 | 0.49 | 0.42 | 0.55 | 0.55 |
| 5 | 31575084 | rs2330727 | -1.421 | -0.616 | -1.095 | -0.389 | 0.02 | -0.075 | -0.075 | 0 | 0.66 | 0.91 | 0.7 | 0.39 | 0.72 | 0.7 |
| 5 | 31580251 | rs995755 | 1.426 | -0.488 | -0.978 | -0.418 | 0.004 | 0.01 | 0.01 | 0.204 | 0.19 | 0.28 | 0.34 | 0.2 | 0.21 | 0.24 |
| 5 | 31580482 | rs1479570 | 1.287 | -0.362 | -1.661 | -0.483 | 0 | 0.015 | 0.015 | 2.869 | 0.19 | 0.12 | 0.17 | 0.02 | 0.2 | 0.16 |
| 5 | 31582523 | rs10040027 | -0.447 | -0.149 | -1.906 | -0.246 | 0.024 | 0.042 | 0.042 | 1.588 | 0.54 | 0.76 | 0.59 | 0.36 | 0.59 | 0.57 |
| 5 | 31582579 | rs13186496 | 0.089 | 0.017 | -1.514 | 0.045 | -0.007 | -0.001 | -0.001 | 0 | 0.25 | 0.36 | 0.21 | 0.18 | 0.22 | 0.19 |
| 5 | 31588686 | rs13152889 | 1.917 | 0.293 | -1.883 | 0.145 | -0.052 | 0.06 | 0.06 | 1.349 | 0.12 | 0.02 | 0.15 | 0.02 | 0.17 | 0.12 |
| 5 | 31589480 | rs6891910 | -0.127 | 0.333 | -1.28 | 0.692 | 0.004 | 0 | 0 | 0 | 0.24 | 0.33 | 0.14 | 0.17 | 0.29 | 0.26 |
| 5 | 31599706 | rs250839 | -0.63 | -0.074 | -1.584 | 0.888 | -0.005 | -0.095 | -0.095 | 0 | 0.72 | 0.98 | 0.72 | 0.53 | 0.76 | 0.72 |
| 5 | 31601976 | rs250835 | 0.169 | 0.469 | -0.88 | 0.572 | -0.005 | 0.011 | 0.011 | 0 | 0.69 | 0.7 | 0.63 | 0.72 | 0.62 | 0.67 |
| 5 | 31602210 | rs250834 | -0.458 | 0.224 | -0.332 | 0.974 | 0.048 | -0.009 | -0.009 | 0 | 0.55 | 0.33 | 0.62 | 0.55 | 0.72 | 0.52 |
| 5 | 31604451 | rs250832 | 1.181 | 0.593 | 0.601 | 1.094 | 0.019 | -0.025 | -0.025 | 0 | 0.64 | 0.33 | 0.5 | 0.56 | 0.69 | 0.72 |
| 5 | 31609654 | rs6884117 | -0.823 | 0.716 | 0.744 | 0.466 | 0.001 | -0.017 | -0.017 | 0 | 0.28 | 0.3 | 0.24 | 0.04 | 0.28 | 0.19 |
| 5 | 31610171 | rs16901319 | -1.271 | 1.25 | 0.835 | 0.948 | 0.087 | 0.054 | 0.054 | 0 | 0.08 | 0.22 | 0.14 | 0.07 | 0.16 | 0.06 |
| 5 | 31630012 | rs13187570 | -0.362 | 1.203 | 0.358 | 0.484 | -0.013 | 0.013 | 0.013 | 0 | 0.21 | 0.16 | 0.26 | 0.07 | 0.33 | 0.14 |
| 5 | 31632899 | rs29766 | -0.74 | 0.378 | 0.45 | 0.948 | 0.059 | 0.273 | 0.273 | 0 | 0.09 | 0.36 | 0.27 | 0.29 | 0.09 | 0.07 |
| 5 | 31633251 | rs185258 | 0.284 | 0.325 | -0.667 | 0.47 | -0.006 | -0.019 | -0.019 | 0 | 0.9 | 0.84 | 0.91 | 0.95 | 0.92 | 0.9 |

| | | | | | | | | | | | | | | | | |
|---|---|---|---|---|---|---|---|---|---|---|---|---|---|---|---|---|
| 5 | 31642204 | rs1560143 | 0.651 | 0.226 | -1.194 | 0.537 | 0.03 | -0.031 | -0.031 | 0 | 0.57 | 0.2 | 0.43 | 0.59 | 0.62 | 0.71 |
| 5 | 31646411 | rs10940960 | 0.319 | -0.018 | -0.84 | -0.079 | -0.002 | -0.048 | -0.048 | 0 | 0.64 | 0.57 | 0.8 | 0.8 | 0.69 | 0.72 |
| 5 | 31646857 | rs805231 | -0.717 | -0.702 | -1.245 | -0.51 | 0.032 | 0.103 | 0.103 | 0 | 0.36 | 0.13 | 0.24 | 0.22 | 0.37 | 0.28 |
| 5 | 31647977 | rs9292436 | -0.777 | -0.836 | -0.334 | -0.665 | -0.012 | -0.069 | -0.069 | 0 | 0.11 | 0.28 | 0.06 | 0 | 0.05 | 0.05 |
| 5 | 31651015 | rs16899780 | 1.325 | -1.372 | -0.622 | -0.165 | 0.043 | -0.026 | -0.026 | 0 | 0.6 | 0.25 | 0.62 | 0.54 | 0.76 | 0.75 |
| 5 | 31652486 | rs13357590 | -0.025 | -1.334 | -0.409 | -0.154 | -0.025 | -0.034 | -0.034 | 0 | 0.11 | 0.19 | 0.05 | 0.03 | 0.06 | 0.08 |
| 5 | 31652765 | rs7727036 | 1.555 | -1.254 | -0.681 | 0.163 | -0.014 | 0.019 | 0.019 | 0 | 0.47 | 0.18 | 0.52 | 0.57 | 0.5 | 0.64 |
| 5 | 31656390 | rs4867360 | 1.489 | -1.333 | 0.276 | 0.81 | -0.028 | 0.026 | 0.026 | 0 | 0.7 | 0.43 | 0.71 | 0.68 | 0.81 | 0.81 |
| 5 | 31656439 | rs7719598 | 1.586 | -1.333 | 0.086 | 0.849 | -0.02 | 0.021 | 0.021 | 0.013 | 0.65 | 0.39 | 0.63 | 0.55 | 0.76 | 0.75 |
| 5 | 31657098 | rs29746 | 1.127 | -1.79 | 0.177 | 0.348 | -0.061 | -0.023 | -0.023 | 0.672 | 0.75 | 0.58 | 0.83 | 0.84 | 0.85 | 0.83 |
| 5 | 31657655 | rs6885718 | -1.184 | -1.317 | -0.005 | 0.486 | -0.048 | -0.03 | -0.03 | 0 | 0.17 | 0.32 | 0.1 | 0.08 | 0.08 | 0.11 |
| 5 | 31658376 | rs6890960 | 0.072 | -0.206 | -0.891 | 0.886 | 0.002 | -0.007 | -0.007 | 0 | 0.51 | 0.22 | 0.41 | 0.51 | 0.56 | 0.43 |
| 5 | 31666904 | rs16901401 | -1.251 | -0.072 | 0.111 | 1.525 | 0.001 | 0.147 | 0.147 | 0.013 | 0.08 | 0.29 | 0.16 | 0.22 | 0.04 | 0.08 |
| 5 | 31670205 | rs168825 | -0.187 | -0.075 | -0.422 | 0.662 | 0.015 | 0.06 | 0.06 | 0.092 | 0.52 | 0.35 | 0.62 | 0.8 | 0.53 | 0.57 |
| 5 | 31673156 | rs7706138 | 0.223 | 0.028 | -0.452 | 0.951 | 0.014 | -0.029 | -0.029 | 0 | 0.28 | 0.2 | 0.21 | 0.59 | 0.24 | 0.44 |
| 5 | 31674226 | rs1651059 | -0.113 | 0.463 | -0.266 | 0.933 | 0.007 | 0.018 | 0.018 | 0 | 0.07 | 0.09 | 0.14 | 0.15 | 0.09 | 0.09 |
| 5 | 31674905 | rs446069 | -0.267 | 0.458 | -0.138 | 0.93 | 0.014 | 0.028 | 0.028 | 0 | 0.07 | 0.12 | 0.14 | 0.15 | 0.09 | 0.09 |
| 5 | 31675161 | rs400810 | -0.185 | 0.459 | -0.138 | 0.924 | 0.014 | 0.022 | 0.022 | 0 | 0.07 | 0.12 | 0.12 | 0.14 | 0.09 | 0.08 |
| 5 | 31676466 | rs1619571 | -1.502 | 0.395 | 0.664 | 0.794 | 0.059 | 0.079 | 0.079 | 0 | 0.13 | 0.34 | 0.26 | 0.22 | 0.14 | 0.18 |
| 5 | 31676987 | rs6869938 | 0.766 | 0.255 | 0.221 | 0.023 | -0.018 | 0.02 | 0.02 | 0 | 0.31 | 0.22 | 0.3 | 0.16 | 0.43 | 0.21 |
| 5 | 31677085 | rs6891459 | 0.316 | 0.118 | 0.248 | -0.074 | -0.003 | 0.002 | 0.002 | 0 | 0.37 | 0.32 | 0.38 | 0.25 | 0.48 | 0.26 |
| 5 | 31677130 | rs426576 | 0.176 | -0.031 | 0.16 | -0.211 | 0.004 | 0.015 | 0.015 | 0 | 0.39 | 0.35 | 0.34 | 0.74 | 0.34 | 0.57 |
| 5 | 31677384 | rs2009350 | -0.421 | -0.371 | 0.586 | -0.045 | 0.001 | 0.074 | 0.074 | 0 | 0.11 | 0.29 | 0.17 | 0.2 | 0.08 | 0.24 |
| 5 | 31678198 | rs4867365 | 0.353 | -0.516 | 0.318 | -0.253 | 0 | -0.046 | -0.046 | 0.033 | 0.26 | 0.26 | 0.18 | 0.01 | 0.23 | 0.18 |
| 5 | 31678239 | rs6450853 | 1.09 | -0.535 | 0.206 | -0.233 | 0.039 | 0.079 | 0.079 | 0.002 | 0.57 | 0.15 | 0.55 | 0.7 | 0.65 | 0.51 |
| 5 | 31678364 | rs4867366 | 1.066 | -0.396 | 0.334 | 0.023 | 0.005 | 0.094 | 0.094 | 0.464 | 0.83 | 0.46 | 0.75 | 0.71 | 0.88 | 0.69 |
| 5 | 31679344 | rs16901423 | 0.367 | -0.102 | -0.627 | -0.289 | 0.017 | 0.018 | 0.018 | 0.112 | 0.24 | 0.18 | 0.15 | 0.01 | 0.19 | 0.25 |
| 5 | 31679565 | rs437499 | -0.332 | -0.065 | -0.398 | 0.304 | 0.001 | 0.002 | 0.002 | 0.155 | 0.36 | 0.35 | 0.32 | 0.26 | 0.32 | 0.4 |
| 5 | 31680821 | rs29738 | 0.039 | -0.106 | -0.789 | 0.241 | 0.017 | 0.022 | 0.022 | 0.709 | 0.31 | 0.18 | 0.27 | 0.19 | 0.27 | 0.37 |
| 5 | 31682155 | rs1531618 | 0.265 | 0.546 | -0.916 | 0.675 | -0.029 | -0.088 | -0.088 | 0.55 | 0.28 | 0.14 | 0.24 | 0.59 | 0.3 | 0.27 |
| 5 | 31687028 | rs418410 | -0.231 | 0.6 | -0.367 | 0.16 | -0.002 | -0.008 | -0.008 | 0 | 0.52 | 0.54 | 0.65 | 0.67 | 0.52 | 0.61 |
| 5 | 31688681 | rs1678916 | -0.434 | 0.499 | -0.093 | 0.048 | 0.001 | -0.008 | -0.008 | 0 | 0.6 | 0.52 | 0.66 | 0.67 | 0.56 | 0.64 |
| 5 | 31690559 | rs6874845 | -0.041 | 0.563 | -0.008 | 0.335 | 0 | -0.002 | -0.002 | 0 | 0.43 | 0.47 | 0.53 | 0.48 | 0.38 | 0.53 |
| 5 | 31692876 | rs400042 | -0.129 | 0.273 | 0.348 | 0.358 | -0.005 | 0.002 | 0.002 | 0 | 0.22 | 0.39 | 0.16 | 0.19 | 0.19 | 0.12 |
| 5 | 31695322 | rs10472779 | 0.321 | 0.494 | 0.074 | 0.278 | 0.002 | 0.006 | 0.006 | 0 | 0.17 | 0.2 | 0.21 | 0.3 | 0.19 | 0.17 |
| 5 | 31695728 | rs443353 | -0.465 | 0.275 | 0.021 | 0.35 | -0.011 | 0.008 | 0.008 | 0.017 | 0.3 | 0.59 | 0.22 | 0.32 | 0.25 | 0.26 |
| 5 | 31696270 | rs451418 | -1.609 | 0.311 | -0.244 | -0.05 | -0.005 | 0.028 | 0.028 | 0.238 | 0.71 | 0.8 | 0.79 | 0.92 | 0.65 | 0.72 |
| 5 | 31696941 | rs369402 | -0.79 | 0.383 | 0.043 | 0.064 | -0.002 | -0.001 | -0.001 | 3.996 | 0.38 | 0.59 | 0.39 | 0.62 | 0.4 | 0.4 |
| 5 | 31710153 | rs380826 | 0.569 | 0.217 | -0.294 | -0.059 | -0.002 | -0.063 | -0.063 | 0.039 | 0.22 | 0.13 | 0.41 | 0.38 | 0.2 | 0.31 |
| 5 | 31711429 | rs6450855 | -0.614 | 0.329 | 0.103 | -0.338 | -0.002 | -0.003 | -0.003 | 0 | 0.42 | 0.57 | 0.38 | 0.54 | 0.52 | 0.41 |
| 5 | 31711663 | rs6894170 | -0.23 | 0.423 | 0.284 | -0.312 | 0.022 | 0.025 | 0.025 | 0.191 | 0.29 | 0.48 | 0.23 | 0.31 | 0.35 | 0.33 |
| 5 | 31711888 | rs805218 | -1.747 | 0.371 | 0.274 | -0.36 | 0.014 | 0.022 | 0.022 | 1.834 | 0.64 | 0.73 | 0.79 | 0.92 | 0.72 | 0.72 |
| 5 | 31713076 | rs390961 | -0.028 | 0.461 | 0.057 | -0.28 | -0.031 | -0.039 | -0.039 | 0 | 0.35 | 0.21 | 0.56 | 0.61 | 0.38 | 0.4 |
| 5 | 31713446 | rs7725382 | 0.762 | 0.473 | 0.345 | -0.217 | 0.032 | 0.022 | 0.022 | 0 | 0.11 | 0.24 | 0.1 | 0.13 | 0.17 | 0.19 |
| 5 | 31715317 | rs253934 | 1.094 | 0.395 | -0.038 | -0.164 | -0.008 | -0.055 | -0.055 | 0.364 | 0.19 | 0.12 | 0.35 | 0.33 | 0.19 | 0.24 |
| 5 | 31717119 | rs253930 | -1.603 | 0.269 | 0.086 | -0.343 | 0.022 | 0.045 | 0.045 | 1.017 | 0.64 | 0.76 | 0.8 | 0.92 | 0.72 | 0.72 |
| 5 | 31717468 | rs190427 | 0.603 | 0.302 | -0.091 | -0.156 | -0.003 | -0.041 | -0.041 | 0 | 0.23 | 0.16 | 0.42 | 0.38 | 0.23 | 0.32 |
| 5 | 31721408 | rs16901473 | 0.733 | 0.374 | 0.063 | -0.208 | 0.002 | 0.003 | 0.003 | 0.055 | 0.17 | 0.23 | 0.15 | 0.17 | 0.18 | 0.11 |
| 5 | 31724153 | rs253919 | -2.031 | 0.334 | -0.616 | -0.377 | -0.008 | 0.093 | 0.093 | 0.119 | 0.74 | 0.92 | 0.82 | 0.92 | 0.74 | 0.75 |
| 5 | 31726664 | rs17414142 | 1.321 | 0.501 | -0.561 | -0.067 | -0.012 | -0.013 | -0.013 | 0 | 0.12 | 0.07 | 0.13 | 0.24 | 0.16 | 0.1 |
| 5 | 31729863 | rs13181582 | -0.123 | -0.053 | -0.501 | 0.008 | -0.005 | 0.002 | 0.002 | 0 | 0.17 | 0.12 | 0.12 | 0.13 | 0.21 | 0.21 |
| 5 | 31730810 | rs16901494 | 0.695 | -0.05 | -0.523 | 0.403 | -0.001 | 0.047 | 0.047 | 0 | 0.09 | 0.17 | 0.07 | 0.17 | 0.09 | 0.07 |
| 5 | 31732692 | rs253910 | 0.252 | -0.051 | -0.555 | -0.26 | 0.001 | 0.06 | 0.06 | 0 | 0.22 | 0.27 | 0.39 | 0.5 | 0.22 | 0.3 |
| 5 | 31736444 | rs16901499 | 1.176 | -0.192 | -0.741 | -0.769 | -0.007 | -0.023 | -0.023 | 0.155 | 0.09 | 0.1 | 0.02 | 0 | 0.03 | 0.03 |
| 5 | 31736582 | rs375183 | -0.401 | 0.259 | -1.097 | -0.559 | -0.004 | -0.002 | -0.002 | 0 | 0.29 | 0.23 | 0.41 | 0.63 | 0.35 | 0.48 |
| 5 | 31748372 | rs16901514 | 0.398 | -0.459 | -1.587 | -0.816 | 0.027 | 0.008 | 0.008 | 4.256 | 0.21 | 0.12 | 0.05 | 0 | 0.07 | 0.06 |
| 5 | 31748611 | rs6862891 | -0.934 | 1.093 | -1.545 | 0.016 | -0.017 | -0.014 | -0.014 | 0 | 0.39 | 0.3 | 0.36 | 0.47 | 0.43 | 0.41 |
| 5 | 31749374 | rs4867081 | -0.917 | 1.325 | -1.517 | 0.508 | -0.01 | -0.017 | -0.017 | 0 | 0.14 | 0.11 | 0.16 | 0.29 | 0.19 | 0.21 |
| 5 | 31750346 | rs7714189 | -0.807 | 0.791 | -1.756 | -0.422 | 0.028 | 0.086 | 0.086 | 0.603 | 0.31 | 0.89 | 0.28 | 0.13 | 0.27 | 0.2 |
| 5 | 31751372 | rs12516825 | -1.073 | 1.36 | -1.016 | -0.435 | 0.001 | -0.033 | -0.033 | 0 | 0.16 | 0.13 | 0.13 | 0.01 | 0.15 | 0.05 |
| 5 | 31751739 | rs12516878 | -1.02 | 1.662 | -1.078 | -0.425 | -0.017 | 0.01 | 0.01 | 0 | 0.15 | 0.1 | 0.14 | 0.01 | 0.19 | 0.05 |
| 5 | 31753135 | rs493276 | -0.036 | 1.477 | -0.91 | -0.645 | -0.002 | 0.012 | 0.012 | 0 | 0.73 | 0.81 | 0.81 | 0.94 | 0.69 | 0.63 |
| 5 | 31758139 | rs6888920 | 0.348 | 1.516 | -0.442 | 0.418 | 0 | 0.001 | 0.001 | 0 | 0.5 | 0.44 | 0.67 | 0.44 | 0.5 | 0.38 |
| 5 | 31759117 | rs498791 | 0.106 | 1.213 | -0.349 | 0.474 | 0.002 | 0.002 | 0.002 | 0 | 0.51 | 0.37 | 0.66 | 0.83 | 0.55 | 0.46 |
| 5 | 31761519 | rs7707027 | 0.056 | 0.936 | -0.664 | 0.868 | 0 | -0.016 | -0.016 | 0 | 0.42 | 0.22 | 0.6 | 0.5 | 0.61 | 0.43 |
| 5 | 31767461 | rs6870260 | -1.084 | 0.271 | 0.583 | 1.593 | -0.028 | 0.231 | 0.231 | 0 | 0.13 | 0.36 | 0.22 | 0.45 | 0.08 | 0.2 |
| 5 | 31770733 | rs919337 | 0.34 | 0.799 | -0.004 | -0.062 | -0.006 | 0.003 | 0.003 | 0 | 0.44 | 0.4 | 0.38 | 0.06 | 0.44 | 0.22 |
| 5 | 31770811 | rs919336 | 0.068 | 0.396 | -0.307 | -0.111 | -0.001 | -0.012 | -0.012 | 0 | 0.56 | 0.48 | 0.43 | 0.19 | 0.56 | 0.43 |
| 5 | 31775225 | rs7736346 | 0.523 | 0.39 | -0.42 | -0.667 | 0.045 | -0.065 | -0.065 | 2.614 | 0.17 | 0.44 | 0.18 | 0.04 | 0.2 | 0.14 |

| | | | | | | | | | | | | | | | | |
|---|---|---|---|---|---|---|---|---|---|---|---|---|---|---|---|---|
| 5 | 31778968 | rs17506712 | 0.001 | 0.04 | -0.453 | -1.068 | 0.001 | -0.025 | -0.025 | 0 | 0.55 | 0.55 | 0.43 | 0.06 | 0.52 | 0.22 |
| 5 | 31780573 | rs7726009 | 0.165 | -0.099 | -0.437 | -0.716 | 0 | -0.003 | -0.003 | 0 | 0.54 | 0.47 | 0.59 | 0.56 | 0.49 | 0.39 |
| 5 | 31782376 | rs1864121 | -1.305 | -0.08 | -0.409 | -0.642 | 0.01 | 0.011 | 0.011 | 0 | 0.72 | 0.61 | 0.65 | 0.68 | 0.68 | 0.59 |
| 5 | 31787381 | rs6450864 | 0.67 | -0.107 | -0.389 | -0.354 | 0.009 | 0.001 | 0.001 | 0 | 0.12 | 0.08 | 0.05 | 0.13 | 0.12 | 0.2 |
| 5 | 31796741 | rs4867381 | -0.073 | -0.107 | -0.538 | -0.402 | 0.007 | -0.091 | -0.091 | 0.387 | 0.16 | 0.05 | 0.22 | 0.61 | 0.16 | 0.41 |
| 5 | 31796992 | rs16901567 | 1.002 | -0.098 | -0.129 | -0.855 | 0.002 | -0.022 | -0.022 | 0 | 0.1 | 0.17 | 0.1 | 0.03 | 0.11 | 0.07 |
| 5 | 31798252 | rs11738489 | 1.378 | -0.255 | -0.289 | -1.3 | 0.009 | 0.045 | 0.045 | 0 | 0.27 | 0.17 | 0.14 | 0.02 | 0.21 | 0.07 |
| 5 | 31798903 | rs13340334 | -0.537 | -0.013 | 0.471 | -1.219 | 0.054 | -0.045 | -0.045 | 0 | 0.1 | 0.33 | 0.1 | 0.01 | 0.1 | 0.07 |
| 5 | 31800544 | rs2162895 | -1.364 | -0.215 | -0.178 | -0.905 | -0.165 | -0.061 | -0.061 | 0.123 | 0.63 | 0.94 | 0.53 | 0.71 | 0.52 | 0.64 |
| 5 | 31800609 | rs12697260 | 1.706 | -0.117 | 0.061 | -1.015 | 0.005 | 0.029 | 0.029 | 0 | 0.23 | 0.11 | 0.1 | 0 | 0.18 | 0.05 |
| 5 | 31801772 | rs13153009 | 0.24 | -0.065 | 0.548 | -0.965 | 0 | -0.07 | -0.07 | 0 | 0.41 | 0.36 | 0.25 | 0.06 | 0.39 | 0.18 |
| 5 | 31802343 | rs13157213 | -0.779 | -0.154 | 0.05 | -0.936 | -0.065 | -0.056 | -0.056 | 0.567 | 0.55 | 0.85 | 0.51 | 0.57 | 0.48 | 0.4 |
| 5 | 31804216 | rs7713111 | -1.99 | -0.148 | 0.008 | -0.933 | -0.078 | -0.079 | -0.079 | 0 | 0.71 | 0.95 | 0.61 | 0.72 | 0.65 | 0.68 |
| 5 | 31804351 | rs17416314 | 1.053 | -0.072 | 0.077 | -0.921 | -0.021 | 0.023 | 0.023 | 0 | 0.06 | 0.03 | 0.06 | 0 | 0.08 | 0.03 |
| 5 | 31806258 | rs6863125 | -0.384 | -0.119 | 0.284 | -0.94 | 0.001 | -0.015 | -0.015 | 0 | 0.18 | 0.28 | 0.12 | 0.15 | 0.19 | 0.31 |
| 5 | 31806434 | rs2010151 | -0.377 | -0.164 | 0.253 | -0.954 | 0.011 | 0.003 | 0.003 | 0 | 0.49 | 0.64 | 0.48 | 0.56 | 0.43 | 0.34 |
| 5 | 31807004 | rs11743256 | 0.816 | -0.109 | 0.038 | -1.015 | 0 | -0.001 | -0.001 | 0 | 0.11 | 0.14 | 0.04 | 0.15 | 0.11 | 0.28 |
| 5 | 31807557 | rs17507380 | 0.538 | -0.05 | 0.038 | -1.282 | -0.018 | 0.112 | 0.112 | 0 | 0.36 | 0.15 | 0.21 | 0.03 | 0.34 | 0.14 |
| 5 | 31809486 | rs16901583 | -0.332 | 0.247 | 0.241 | -1.033 | 0.005 | -0.034 | -0.034 | 0 | 0.17 | 0.25 | 0.15 | 0.03 | 0.2 | 0.15 |
| 5 | 31809645 | rs10940969 | -1.08 | -0.068 | -0.076 | -0.83 | -0.027 | 0.003 | 0.003 | 0 | 0.63 | 0.83 | 0.55 | 0.72 | 0.56 | 0.65 |
| 5 | 31813213 | rs13357918 | -0.371 | -0.112 | 0.019 | -1.055 | -0.058 | 0.302 | 0.302 | 0 | 0.54 | 0.86 | 0.36 | 0.04 | 0.49 | 0.26 |
| 5 | 31814203 | rs7718233 | -0.566 | -0.233 | 0.612 | -0.943 | 0.021 | 0.009 | 0.009 | 0.15 | 0.05 | 0.26 | 0.03 | 0 | 0.02 | 0.01 |
| 5 | 31818948 | rs4867084 | 0.734 | -0.135 | 0.313 | -0.846 | -0.024 | 0.003 | 0.003 | 0.033 | 0.38 | 0.22 | 0.48 | 0.27 | 0.46 | 0.38 |
| 5 | 31819180 | rs2042959 | 0.046 | -0.076 | -0.089 | -0.764 | -0.009 | -0.075 | -0.075 | 3.114 | 0.82 | 0.92 | 0.75 | 0.29 | 0.81 | 0.55 |
| 5 | 31823644 | rs10038413 | 0.181 | -0.102 | -0.155 | -1.02 | 0.009 | 0.014 | 0.014 | 0 | 0.54 | 0.6 | 0.62 | 0.27 | 0.61 | 0.42 |
| 5 | 31826017 | rs13153000 | 1.067 | -0.09 | -0.18 | -0.911 | -0.003 | -0.061 | -0.061 | 0 | 0.26 | 0.29 | 0.13 | 0.01 | 0.2 | 0.1 |
| 5 | 31826929 | rs10035850 | 0.161 | -0.118 | -0.523 | -0.513 | -0.013 | 0.023 | 0.023 | 0 | 0.44 | 0.36 | 0.52 | 0.26 | 0.49 | 0.33 |
| 5 | 31830261 | rs9800104 | 0.113 | -0.093 | -0.625 | -0.524 | -0.006 | 0.013 | 0.013 | 0 | 0.44 | 0.41 | 0.53 | 0.25 | 0.48 | 0.33 |
| 5 | 31833816 | rs10042146 | -0.185 | -0.246 | -1.314 | 0.649 | -0.004 | 0.093 | 0.093 | 0 | 0.91 | 0.88 | 0.85 | 0.58 | 0.9 | 0.86 |
| 5 | 31833944 | rs11740457 | 0.082 | -0.184 | -1.483 | 0.43 | -0.008 | 0.027 | 0.027 | 0 | 0.6 | 0.7 | 0.44 | 0.32 | 0.54 | 0.55 |
| 5 | 31838491 | rs6874198 | 0.915 | -0.389 | -0.882 | 0.771 | -0.02 | 0.039 | 0.039 | 0 | 0.81 | 0.72 | 0.72 | 0.26 | 0.85 | 0.6 |
| 5 | 31845117 | rs9292445 | -0.304 | 0.052 | -1.496 | 0.637 | 0.036 | 0.016 | 0.016 | 0 | 0.48 | 0.86 | 0.43 | 0.66 | 0.41 | 0.56 |
| 5 | 31853346 | rs764336 | -0.075 | 0.493 | -1.268 | 1.271 | 0 | -0.001 | -0.001 | 1.24 | 0.08 | 0.03 | 0.14 | 0.38 | 0.11 | 0.15 |
| 5 | 31854497 | rs6864968 | 0.103 | 0.177 | -0.968 | 0.321 | 0.028 | 0.055 | 0.055 | 0 | 0.46 | 0.65 | 0.25 | 0.29 | 0.33 | 0.42 |
| 5 | 31862003 | rs6450868 | 0.142 | 0.622 | -0.662 | 0.422 | 0 | -0.011 | -0.011 | 0.057 | 0.6 | 0.47 | 0.48 | 0.33 | 0.57 | 0.47 |
| 5 | 31867203 | rs16901644 | -0.938 | 0.148 | -0.22 | 0.978 | -0.024 | 0.112 | 0.112 | 0 | 0.23 | 0.54 | 0.24 | 0.46 | 0.2 | 0.25 |
| 5 | 31868701 | rs13347335 | -0.125 | 0.222 | -0.742 | 0.578 | -0.003 | -0.017 | -0.017 | 0.942 | 0.12 | 0.04 | 0.11 | 0.21 | 0.1 | 0.21 |
| 5 | 31869442 | rs2330778 | -0.202 | 0.548 | -0.372 | 0.081 | 0.006 | -0.035 | -0.035 | 0 | 0.36 | 0.52 | 0.44 | 0.18 | 0.43 | 0.4 |
| 5 | 31870238 | rs10472790 | -1.441 | 0.656 | 0.104 | 0.071 | 0.004 | -0.014 | -0.014 | 0 | 0.11 | 0.82 | 0.1 | 0.05 | 0.07 | 0.06 |
| 5 | 31871346 | rs905836 | -0.118 | 0.809 | 0.077 | -0.218 | 0.01 | 0.079 | 0.079 | 0.019 | 0.49 | 0.69 | 0.42 | 0.24 | 0.47 | 0.59 |
| 5 | 31874377 | rs9292446 | 0.05 | 0.703 | -0.175 | -0.136 | 0.009 | 0.036 | 0.036 | 0 | 0.47 | 0.56 | 0.41 | 0.24 | 0.46 | 0.6 |
| 5 | 31877708 | rs17508455 | 0.487 | 0.751 | -1.316 | -0.222 | 0.023 | 0.147 | 0.147 | 0 | 0.41 | 0.11 | 0.34 | 0.17 | 0.38 | 0.54 |
| 5 | 31885721 | rs1394647 | -0.01 | 0.82 | -1.462 | 0.035 | 0.021 | 0.134 | 0.134 | 0.035 | 0.49 | 0.84 | 0.44 | 0.24 | 0.43 | 0.57 |
| 5 | 31890186 | rs13157311 | -0.706 | 1.45 | -1.067 | 0.327 | 0.003 | 0.002 | 0.002 | 0 | 0.34 | 0.37 | 0.46 | 0.39 | 0.37 | 0.33 |
| 5 | 31891791 | rs1574436 | -0.533 | 1.529 | -1.138 | 0.185 | -0.01 | -0.008 | -0.008 | 0 | 0.28 | 0.26 | 0.49 | 0.4 | 0.4 | 0.27 |
| 5 | 31891901 | rs1107904 | -1.03 | 1.457 | -1.047 | 0.239 | -0.001 | 0 | 0 | 0 | 0.4 | 0.4 | 0.54 | 0.4 | 0.48 | 0.37 |
| 5 | 31893285 | rs12516203 | -0.69 | 1.554 | -1.411 | 0.263 | 0.018 | 0.008 | 0.008 | 0 | 0.5 | 0.72 | 0.66 | 0.53 | 0.55 | 0.48 |
| 5 | 31893865 | rs10472795 | -0.758 | 1.7 | -1.463 | 0.055 | -0.006 | 0.12 | 0.12 | 0 | 0.51 | 0.81 | 0.7 | 0.78 | 0.54 | 0.58 |
| 5 | 31899994 | rs13182372 | 1.37 | 2.005 | -0.708 | 0.252 | -0.001 | 0.068 | 0.068 | 0.71 | 0.56 | 0.17 | 0.41 | 0.29 | 0.57 | 0.48 |
| 5 | 31900372 | rs7736502 | 1.386 | 2.01 | -0.583 | 0.286 | -0.032 | 0.108 | 0.108 | 0.427 | 0.52 | 0.16 | 0.38 | 0.28 | 0.51 | 0.44 |
| 5 | 31900521 | rs7736535 | -0.859 | 1.804 | 0.33 | 1.471 | 0.001 | 0.005 | 0.005 | 0 | 0.14 | 0.23 | 0.18 | 0.16 | 0.16 | 0.12 |
| 5 | 31907082 | rs2007825 | 0.815 | 1.793 | -0.047 | 1.798 | 0 | 0.062 | 0.062 | 0 | 0.9 | 0.84 | 0.72 | 0.46 | 0.9 | 0.67 |
| 5 | 31914816 | rs12521501 | 1.446 | 2.027 | 0.532 | 0.695 | 0.075 | -0.097 | -0.097 | 0 | 0.85 | 0.43 | 0.84 | 0.98 | 0.76 | 0.92 |
| 5 | 31914966 | rs10067626 | -1.554 | 1.989 | 1.064 | 0.687 | 0.05 | -0.006 | -0.006 | 0 | 0.08 | 0.43 | 0.08 | 0.01 | 0.07 | 0.05 |
| 5 | 31916371 | rs7709630 | -1.985 | 2.5 | 0.136 | 0.772 | 0.004 | -0.025 | -0.025 | 0.604 | 0.09 | 0.09 | 0.11 | 0.01 | 0.17 | 0.08 |
| 5 | 31920340 | rs12110032 | -0.548 | 2.042 | -0.215 | 0.659 | -0.002 | 0.024 | 0.024 | 1.225 | 0.47 | 0.62 | 0.43 | 0.35 | 0.46 | 0.53 |
| 5 | 31920673 | rs4340934 | 0.947 | 2.468 | -0.266 | 0.401 | -0.018 | 0.016 | 0.016 | 0 | 0.94 | 0.98 | 0.91 | 0.99 | 0.79 | 0.96 |
| 5 | 31925190 | rs9292449 | -0.731 | 2.197 | 0.183 | 0.494 | -0.01 | 0.004 | 0.004 | 0.293 | 0.16 | 0.22 | 0.16 | 0.18 | 0.14 | 0.27 |
| 5 | 31926148 | rs13189145 | 0.345 | 2.144 | -0.299 | -0.085 | -0.008 | 0.029 | 0.029 | 0 | 0.25 | 0.16 | 0.2 | 0.12 | 0.27 | 0.16 |
| 5 | 31927317 | rs7719829 | 0.961 | 2.065 | 0.324 | -0.707 | 0.019 | -0.023 | -0.023 | 0.013 | 0.7 | 0.48 | 0.56 | 0.77 | 0.61 | 0.57 |
| 5 | 31928322 | rs4867399 | 0.835 | 1.946 | 0.362 | -0.807 | -0.01 | -0.058 | -0.058 | 0 | 0.81 | 0.59 | 0.76 | 0.94 | 0.78 | 0.85 |
| 5 | 31930080 | rs10940979 | 1.291 | 1.979 | 0.389 | -0.729 | 0.005 | 0.009 | 0.009 | 2.223 | 0.84 | 0.77 | 0.69 | 0.77 | 0.78 | 0.69 |
| 5 | 31930769 | rs16901686 | -0.425 | 1.985 | 0.381 | -0.728 | -0.03 | -0.029 | -0.029 | 0 | 0.1 | 0.03 | 0.17 | 0.16 | 0.15 | 0.26 |
| 5 | 31930842 | rs16901687 | -0.741 | 1.989 | 0.377 | -0.726 | 0.004 | 0.003 | 0.003 | 2.512 | 0.11 | 0.13 | 0.18 | 0.17 | 0.15 | 0.26 |
| 5 | 31936833 | rs1553552 | -1.452 | 1.865 | 0.631 | -0.811 | 0.02 | -0.029 | -0.029 | 0.932 | 0.23 | 0.52 | 0.27 | 0.17 | 0.23 | 0.33 |
| 5 | 31939339 | rs1847610 | 0.624 | 1.667 | 0.635 | -1.068 | 0.004 | 0.17 | 0.17 | 0 | 0.5 | 0.34 | 0.31 | 0.12 | 0.43 | 0.4 |
| 5 | 31940435 | rs1505019 | 0.665 | 1.601 | 0.763 | -1.073 | 0.002 | 0.069 | 0.069 | 0 | 0.51 | 0.4 | 0.31 | 0.12 | 0.44 | 0.4 |
| 5 | 31941331 | rs4867403 | -0.53 | 1.62 | 0.827 | -0.989 | -0.001 | -0.013 | -0.013 | 0 | 0.44 | 0.5 | 0.56 | 0.81 | 0.51 | 0.56 |
| 5 | 31951961 | rs16901697 | -0.19 | 1.111 | 0.571 | -1.168 | -0.002 | -0.011 | -0.011 | 0.044 | 0.45 | 0.46 | 0.48 | 0.67 | 0.44 | 0.35 |

| | | | | | | | | | | | | | | | | |
|---|---|---|---|---|---|---|---|---|---|---|---|---|---|---|---|---|
| 5 | 31952401 | rs10940984 | 1.023 | 1.251 | 0.714 | -1.535 | 0.012 | 0.046 | 0.046 | 0.096 | 0.38 | 0.28 | 0.21 | 0.12 | 0.31 | 0.33 |
| 5 | 31952427 | rs2877238 | -0.693 | 1.325 | 0.562 | -1.404 | 0.005 | 0 | 0 | 0.666 | 0.12 | 0.14 | 0.18 | 0.16 | 0.2 | 0.28 |
| 5 | 31952663 | rs4867095 | 0.202 | 1.273 | 0.491 | -1.194 | -0.01 | -0.009 | -0.009 | 1.791 | 0.05 | 0.03 | 0.11 | 0.16 | 0.11 | 0.22 |
| 5 | 31957550 | rs7702447 | 0.478 | 1.16 | 0.56 | -1.119 | -0.001 | 0 | 0 | 0 | 0.36 | 0.37 | 0.4 | 0.62 | 0.34 | 0.27 |
| 5 | 31966417 | rs10045838 | 0.728 | 0.915 | 0.271 | -1.47 | 0.004 | 0.079 | 0.079 | 0.302 | 0.45 | 0.31 | 0.28 | 0.15 | 0.42 | 0.36 |
| 5 | 31967192 | rs7726919 | 0.659 | 1.054 | -0.111 | -1.345 | -0.002 | -0.046 | -0.046 | 0 | 0.48 | 0.5 | 0.3 | 0.15 | 0.43 | 0.37 |
| 5 | 31969830 | rs12110008 | -0.417 | 1.214 | -0.149 | -1.09 | 0.007 | 0 | 0 | 1.954 | 0.16 | 0.24 | 0.19 | 0.21 | 0.23 | 0.3 |
| 5 | 31970353 | rs13181283 | 0.289 | 1.137 | -0.569 | -0.687 | -0.012 | -0.014 | -0.014 | 0 | 0.31 | 0.15 | 0.38 | 0.59 | 0.29 | 0.29 |
| 5 | 31972609 | rs10059023 | -0.324 | 1.609 | -0.223 | -1 | 0.038 | -0.07 | -0.07 | 0 | 0.14 | 0.28 | 0.14 | 0 | 0.24 | 0.2 |
| 5 | 31976593 | rs17432188 | 0.111 | 1.149 | -0.876 | -1.515 | -0.009 | 0.067 | 0.067 | 0 | 0.3 | 0.14 | 0.18 | 0.13 | 0.35 | 0.15 |
| 5 | 31977625 | rs7706312 | -0.131 | 0.944 | -0.978 | -1.724 | 0.013 | 0.167 | 0.167 | 0 | 0.45 | 0.2 | 0.25 | 0.17 | 0.46 | 0.36 |
| 5 | 31982409 | rs7733163 | -0.117 | 0.467 | -1.238 | -1.341 | -0.011 | 0.045 | 0.045 | 0.114 | 0.33 | 0.25 | 0.25 | 0.1 | 0.34 | 0.25 |
| 5 | 31989348 | rs6863159 | -0.171 | 0.165 | -1.045 | -0.735 | 0.001 | 0.002 | 0.002 | 0 | 0.33 | 0.64 | 0.34 | 0.21 | 0.31 | 0.37 |
| 5 | 31991556 | rs1833813 | 0.3 | 0.5 | -0.694 | -0.175 | -0.008 | 0.015 | 0.015 | 0 | 0.18 | 0.24 | 0.08 | 0.22 | 0.15 | 0.19 |
| 5 | 31992572 | rs6861526 | -0.203 | 1.151 | -0.92 | -0.559 | -0.027 | 0.006 | 0.006 | 0 | 0.13 | 0.1 | 0.17 | 0.11 | 0.27 | 0.16 |
| 5 | 31993222 | rs2052852 | -0.336 | 0.641 | -0.75 | -0.622 | -0.003 | 0 | 0 | 0 | 0.32 | 0.33 | 0.3 | 0.16 | 0.43 | 0.34 |
| 5 | 31993822 | rs6860464 | -0.948 | 0.248 | -0.665 | 0.076 | 0.001 | -0.026 | -0.026 | 0 | 0.1 | 0.11 | 0.08 | 0 | 0.09 | 0.1 |
| 5 | 32000483 | rs10054504 | 0.634 | -1 | -0.605 | 0.321 | -0.01 | -0.001 | -0.001 | 0 | 0.69 | 0.33 | 0.69 | 0.31 | 0.86 | 0.78 |
| 5 | 32002418 | rs890998 | 0.123 | -1.435 | -0.256 | 1.375 | -0.028 | 0.037 | 0.037 | 0.143 | 0.85 | 0.85 | 0.95 | 0.63 | 0.96 | 0.94 |
| 5 | 32007713 | rs4867100 | -1.119 | -1.613 | 0.163 | 1.503 | -0.015 | -0.008 | -0.008 | 0 | 0.17 | 0.76 | 0.26 | 0.11 | 0.07 | 0.08 |
| 5 | 32008979 | rs10940992 | 0.319 | -1.52 | -0.708 | 0.867 | -0.006 | 0.005 | 0.005 | 0 | 0.21 | 0.12 | 0.31 | 0.16 | 0.29 | 0.19 |
| 5 | 32012404 | rs16901740 | -0.353 | -1.632 | 0.473 | 0.823 | -0.037 | -0.03 | -0.03 | 0 | 0.25 | 0.55 | 0.32 | 0.16 | 0.24 | 0.18 |
| 5 | 32015725 | rs12656499 | -0.348 | -1.644 | 0.51 | 0.819 | -0.039 | -0.03 | -0.03 | 0 | 0.25 | 0.52 | 0.31 | 0.16 | 0.24 | 0.18 |
| 5 | 32029638 | rs889216 | -0.948 | -1.67 | 0.183 | 0.916 | 0.038 | -0.036 | -0.036 | 0 | 0.91 | 0.98 | 0.96 | 0.64 | 0.96 | 0.94 |
| 5 | 32031477 | rs16901746 | 1.094 | -1.148 | 0.986 | 0.789 | 0.013 | 0.013 | 0.013 | 0 | 0.51 | 0.48 | 0.54 | 0.34 | 0.62 | 0.64 |
| 5 | 32032742 | rs7707783 | 1.104 | -1.152 | 1.021 | 0.787 | 0.025 | 0.025 | 0.025 | 0 | 0.51 | 0.42 | 0.53 | 0.34 | 0.62 | 0.64 |
| 5 | 32032837 | rs6871123 | -0.12 | -1.374 | 1.093 | 0.668 | -0.02 | 0.003 | 0.003 | 0 | 0.18 | 0.28 | 0.21 | 0.17 | 0.14 | 0.19 |
| 5 | 32033526 | rs10940994 | 1.402 | -1.13 | 0.597 | 0.521 | 0.089 | 0.088 | 0.088 | 0.379 | 0.48 | 0.06 | 0.5 | 0.33 | 0.61 | 0.64 |
| 5 | 32035855 | rs12188679 | 1.205 | -0.901 | 0.535 | 0.542 | 0.067 | 0.073 | 0.073 | 0 | 0.5 | 0.31 | 0.52 | 0.33 | 0.61 | 0.64 |
| 5 | 32042760 | rs10039073 | 1.05 | -0.72 | -0.135 | 0.294 | 0.013 | 0.024 | 0.024 | 0.497 | 0.44 | 0.31 | 0.5 | 0.36 | 0.55 | 0.56 |
| 5 | 32044573 | rs323827 | 1.144 | 0 | -0.773 | 0.179 | -0.04 | 0.02 | 0.02 | 0 | 0.38 | 0.21 | 0.46 | 0.33 | 0.46 | 0.54 |
| 5 | 32049094 | rs438834 | 0.334 | 0.065 | -0.821 | 0.555 | 0.016 | -0.036 | -0.036 | 0 | 0.63 | 0.77 | 0.65 | 0.52 | 0.65 | 0.79 |
| 5 | 32049210 | rs403457 | -0.552 | 0.766 | -1.502 | 0.438 | -0.013 | -0.018 | -0.018 | 0 | 0.75 | 0.87 | 0.74 | 0.75 | 0.73 | 0.62 |
| 5 | 32049787 | rs17433352 | 0.035 | 0.753 | -1.837 | 0.257 | -0.032 | 0.088 | 0.088 | 1.31 | 0.3 | 0.08 | 0.33 | 0.14 | 0.35 | 0.08 |
| 5 | 32051287 | rs328618 | 0.309 | 1.023 | -1.25 | 1.005 | -0.01 | -0.012 | -0.012 | 0 | 0.23 | 0.16 | 0.26 | 0.24 | 0.25 | 0.29 |
| 5 | 32051339 | rs328617 | -0.16 | 0.919 | -1.01 | 0.92 | 0.004 | -0.004 | -0.004 | 0.321 | 0.42 | 0.64 | 0.47 | 0.43 | 0.4 | 0.57 |
| 5 | 32056026 | rs161533 | 0.626 | 1.224 | -0.808 | 1.358 | 0.009 | 0.033 | 0.033 | 0 | 0.83 | 0.76 | 0.71 | 0.67 | 0.76 | 0.83 |
| 5 | 32058850 | rs3909431 | -0.488 | 1.059 | -1.619 | 1.162 | -0.015 | -0.048 | -0.048 | 0 | 0.41 | 0.81 | 0.48 | 0.52 | 0.44 | 0.57 |
| 5 | 32060695 | rs157500 | -1.397 | 0.971 | -0.582 | 1.698 | 0.005 | 0.116 | 0.116 | 0 | 0.12 | 0.27 | 0.19 | 0.34 | 0.1 | 0.18 |
| 5 | 32061570 | rs11955676 | 0.65 | 1.18 | -1.437 | 0.898 | -0.061 | 0.103 | 0.103 | 0 | 0.28 | 0.04 | 0.27 | 0.08 | 0.29 | 0.17 |
| 5 | 32062984 | rs12110295 | -0.644 | 0.692 | -0.546 | 1.303 | -0.002 | 0.072 | 0.072 | 0 | 0.06 | 0.12 | 0.13 | 0.23 | 0.05 | 0.19 |
| 5 | 32064273 | rs157502 | -0.435 | 0.247 | -0.367 | 1.212 | 0.005 | -0.008 | -0.008 | 0.034 | 0.39 | 0.64 | 0.35 | 0.53 | 0.34 | 0.36 |
| 5 | 32067373 | rs12054657 | -0.897 | 0.314 | -0.232 | 1.224 | -0.001 | 0.01 | 0.01 | 0 | 0.1 | 0.13 | 0.13 | 0.23 | 0.11 | 0.2 |
| 5 | 32067398 | rs10065850 | -1.503 | 0.289 | -0.086 | 1.2 | 0.017 | 0.047 | 0.047 | 0.056 | 0.12 | 0.31 | 0.16 | 0.23 | 0.15 | 0.21 |
| 5 | 32067671 | rs283129 | 0.722 | 0.189 | -0.05 | 1.194 | -0.026 | 0.009 | 0.009 | 0 | 0.27 | 0.33 | 0.18 | 0.29 | 0.2 | 0.16 |
| 5 | 32068026 | rs12109819 | -0.435 | 0.181 | -0.099 | 1.033 | 0.001 | -0.001 | -0.001 | 0 | 0.39 | 0.62 | 0.3 | 0.4 | 0.35 | 0.35 |
| 5 | 32068774 | rs438135 | 0.958 | 0.157 | -0.129 | 1.117 | 0.011 | -0.022 | -0.022 | 0 | 0.12 | 0.06 | 0.13 | 0.24 | 0.11 | 0.3 |
| 5 | 32072213 | rs3797064 | -1.028 | 0.296 | -0.088 | 0.995 | 0.003 | 0.02 | 0.02 | 0 | 0.13 | 0.21 | 0.16 | 0.23 | 0.15 | 0.15 |
| 5 | 32073566 | rs32793 | -1.352 | 0.068 | 0.096 | 0.942 | -0.025 | 0.044 | 0.044 | 0 | 0.29 | 0.59 | 0.31 | 0.46 | 0.25 | 0.51 |
| 5 | 32074392 | rs2291114 | 0.903 | -0.147 | -0.244 | 0.848 | -0.004 | -0.005 | -0.005 | 0 | 0.26 | 0.26 | 0.14 | 0.17 | 0.22 | 0.15 |
| 5 | 32075598 | rs10071707 | -0.521 | -0.041 | -0.315 | 0.956 | -0.007 | -0.036 | -0.036 | 0 | 0.08 | 0.16 | 0.03 | 0 | 0.05 | 0.02 |
| 5 | 32076782 | rs4867417 | -0.081 | 0.043 | -0.495 | 1.144 | 0.002 | -0.001 | -0.001 | 0 | 0.06 | 0.03 | 0.12 | 0.23 | 0.05 | 0.11 |
| 5 | 32087228 | rs157494 | -0.697 | -0.228 | -0.61 | 0.873 | 0.037 | 0.026 | 0.026 | 0.059 | 0.5 | 0.4 | 0.38 | 0.65 | 0.37 | 0.63 |
| 5 | 32087253 | rs157495 | 0.421 | -0.27 | -0.613 | 0.904 | -0.001 | -0.001 | -0.001 | 3.755 | 0.37 | 0.34 | 0.23 | 0.41 | 0.28 | 0.51 |
| 5 | 32090294 | rs10066063 | -0.398 | 0.234 | -0.976 | 1.224 | 0.001 | -0.031 | -0.031 | 0 | 0.08 | 0.04 | 0.14 | 0.24 | 0.07 | 0.12 |
| 5 | 32091868 | rs788368 | -1.094 | 0.874 | -1.255 | 1.144 | -0.114 | -0.081 | -0.081 | 3.706 | 0.84 | 0.98 | 0.71 | 0.76 | 0.63 | 0.79 |
| 5 | 32093070 | rs16889442 | -0.556 | -0.138 | -0.631 | 1.181 | 0.002 | -0.023 | -0.023 | 0 | 0.08 | 0.04 | 0.14 | 0.24 | 0.08 | 0.12 |
| 5 | 32093563 | rs283122 | -1.06 | -0.178 | -0.391 | 1.15 | -0.011 | 0.025 | 0.025 | 0 | 0.2 | 0.54 | 0.22 | 0.24 | 0.16 | 0.14 |
| 5 | 32101400 | rs283107 | 0.435 | -0.065 | -1.057 | 1.294 | 0.012 | -0.04 | -0.04 | 3.589 | 0.1 | 0.02 | 0.12 | 0.25 | 0.1 | 0.31 |
| 5 | 32107381 | rs157493 | 0.13 | -0.468 | -1.191 | 1.02 | 0.067 | -0.014 | -0.014 | 0 | 0.61 | 0.81 | 0.69 | 0.57 | 0.72 | 0.41 |
| 5 | 32107764 | rs161522 | -0.163 | -0.741 | -1.154 | 0.915 | 0.01 | 0.01 | 0.01 | 0 | 0.7 | 0.77 | 0.78 | 0.74 | 0.77 | 0.76 |
| 5 | 32109122 | rs32800 | -0.295 | -0.065 | -0.656 | 0.93 | -0.006 | -0.029 | -0.029 | 0 | 0.62 | 0.93 | 0.79 | 0.69 | 0.65 | 0.64 |
| 5 | 32123788 | rs283103 | -0.171 | -0.198 | 1.064 | 0.867 | -0.002 | 0 | 0 | 0 | 0.58 | 0.53 | 0.66 | 0.54 | 0.6 | 0.58 |
| 5 | 32123891 | rs157489 | 0.024 | -0.271 | 1.314 | 0.828 | 0.02 | -0.006 | -0.006 | 0 | 0.55 | 0.34 | 0.64 | 0.54 | 0.6 | 0.58 |
| 5 | 32130401 | rs671879 | -2.155 | -0.166 | 1.369 | 0.464 | 0.037 | 0.127 | 0.127 | 2.383 | 0.09 | 0.58 | 0.15 | 0.15 | 0.04 | 0.06 |
| 5 | 32145957 | rs161524 | -0.446 | -0.194 | 1.057 | 0.556 | 0.031 | -0.061 | -0.061 | 0 | 0.61 | 0.92 | 0.79 | 0.65 | 0.64 | 0.63 |
| 5 | 32151119 | rs367556 | -0.38 | -0.232 | 1.088 | 0.546 | 0.048 | -0.031 | -0.031 | 0 | 0.59 | 0.86 | 0.78 | 0.65 | 0.64 | 0.62 |
| 5 | 32157088 | rs4282332 | -0.412 | -0.188 | 0.282 | 0.567 | 0.029 | -0.061 | -0.061 | 0 | 0.61 | 0.93 | 0.78 | 0.64 | 0.64 | 0.63 |
| 5 | 32159517 | rs683547 | 0.27 | -0.183 | 0.283 | 0.569 | 0.048 | -0.031 | -0.031 | 1.49 | 0.41 | 0.14 | 0.22 | 0.35 | 0.36 | 0.38 |

| | | | | | | | | | | | | | | | | |
|---|---|---|---|---|---|---|---|---|---|---|---|---|---|---|---|---|
| 5 | 32169547 | rs452068 | 0.255 | -0.14 | 0.168 | 0.582 | 0.033 | -0.059 | -0.059 | 0 | 0.39 | 0.08 | 0.22 | 0.35 | 0.36 | 0.38 |
| 5 | 32179888 | rs336517 | -0.06 | 0.53 | -0.093 | 0.348 | 0.028 | 0.045 | 0.045 | 0 | 0.7 | 0.62 | 0.52 | 0.46 | 0.57 | 0.57 |
| 5 | 32189340 | rs4489077 | 0.06 | 0.268 | -0.635 | -0.117 | 0.006 | 0.056 | 0.056 | 1.64 | 0.33 | 0.74 | 0.3 | 0.11 | 0.2 | 0.15 |
| 5 | 32193664 | rs7380110 | 0.025 | 0.308 | -0.717 | -0.15 | 0.007 | 0.068 | 0.068 | 1.917 | 0.33 | 0.78 | 0.31 | 0.11 | 0.2 | 0.15 |
| 5 | 32213670 | rs4867105 | -0.133 | 0.583 | -1.431 | -0.219 | -0.125 | -0.018 | -0.018 | 0 | 0.65 | 0.9 | 0.55 | 0.6 | 0.44 | 0.55 |
| 5 | 32217141 | rs11746458 | -0.041 | 1.005 | 0.211 | -0.216 | 0.024 | -0.078 | -0.078 | 0 | 0.28 | 0.45 | 0.27 | 0.12 | 0.22 | 0.13 |
| 5 | 32221259 | rs11745268 | 0.715 | 1.222 | 0.133 | -0.148 | -0.035 | 0.054 | 0.054 | 0 | 0.21 | 0.12 | 0.16 | 0.05 | 0.17 | 0.13 |
| 5 | 32229677 | rs1873921 | -0.415 | 2.252 | 0.365 | 0.509 | -0.069 | -0.084 | -0.084 | 1.202 | 0.83 | 0.98 | 0.78 | 0.66 | 0.74 | 0.82 |
| 5 | 32231832 | rs11959760 | 1.283 | 1.716 | 1.094 | 0.442 | 0.078 | 0.006 | 0.006 | 0.117 | 0.52 | 0.24 | 0.34 | 0.49 | 0.35 | 0.37 |
| 5 | 32243327 | rs7734652 | 0.141 | 1.663 | 1.623 | 0.408 | -0.03 | -0.018 | -0.018 | 0 | 0.67 | 0.77 | 0.49 | 0.54 | 0.48 | 0.5 |
| 5 | 32252395 | rs1123329 | 1.064 | 1.528 | 1.895 | 0.429 | 0.012 | 0.001 | 0.001 | 0.383 | 0.54 | 0.39 | 0.37 | 0.49 | 0.38 | 0.37 |
| 5 | 32278628 | rs10461909 | 1.562 | 1.489 | 1.847 | 0.375 | 0.085 | 0.011 | 0.011 | 0 | 0.47 | 0.22 | 0.28 | 0.48 | 0.31 | 0.36 |
| 5 | 32286948 | rs11959574 | 0.931 | 1.401 | 1.909 | 0.44 | -0.004 | -0.007 | -0.007 | 0 | 0.56 | 0.5 | 0.46 | 0.58 | 0.39 | 0.41 |
| 5 | 32295574 | rs4867424 | 0.038 | 1.473 | 1.844 | 0.421 | -0.025 | 0.003 | 0.003 | 0 | 0.67 | 0.78 | 0.57 | 0.62 | 0.5 | 0.54 |
| 5 | 32308706 | rs2910884 | 1.185 | 1.138 | 1.33 | 0.316 | 0.054 | 0 | 0 | 0 | 0.49 | 0.32 | 0.29 | 0.44 | 0.33 | 0.37 |
| 5 | 32317439 | rs1019512 | -1.036 | 1.419 | 1.238 | 0.446 | -0.006 | 0.001 | 0.001 | 0.323 | 0.44 | 0.48 | 0.67 | 0.57 | 0.63 | 0.61 |
| 5 | 32326234 | rs10077147 | -0.648 | 1.457 | 1.624 | 0.138 | 0.116 | -0.021 | -0.021 | 4.452 | 0.11 | 0.38 | 0.13 | 0.04 | 0.11 | 0.1 |
| 5 | 32331324 | rs2910909 | -0.36 | 1.457 | 1.144 | 0.246 | -0.007 | -0.018 | -0.018 | 0.421 | 0.95 | 0.97 | 0.88 | 0.85 | 0.96 | 0.88 |
| 5 | 32332180 | rs13436901 | -0.647 | 1.391 | 1.646 | 0.149 | 0.078 | -0.041 | -0.041 | 0.042 | 0.13 | 0.4 | 0.12 | 0.04 | 0.12 | 0.1 |
| 5 | 32358866 | rs4867433 | -1.256 | 1.369 | 1.734 | 0.017 | 0.047 | -0.099 | -0.099 | 0.149 | 0.19 | 0.39 | 0.15 | 0.03 | 0.18 | 0.14 |
| 5 | 32366533 | rs9654389 | 0.389 | 1.616 | 1.593 | 0.258 | 0.01 | 0.026 | 0.026 | 0 | 0.25 | 0.34 | 0.43 | 0.55 | 0.42 | 0.45 |
| 5 | 32369377 | rs11750707 | -0.155 | 1.66 | 1.726 | 0.275 | 0.036 | -0.037 | -0.037 | 0.337 | 0.12 | 0.26 | 0.12 | 0.03 | 0.12 | 0.1 |
| 5 | 32373168 | rs10080085 | -0.279 | 1.657 | 1.649 | 0.274 | 0.072 | -0.049 | -0.049 | 0.138 | 0.13 | 0.36 | 0.12 | 0.03 | 0.12 | 0.09 |
| 5 | 32384427 | rs3828593 | -0.682 | 1.662 | 1.178 | 0.289 | -0.013 | -0.023 | -0.023 | 0.619 | 0.95 | 0.98 | 0.88 | 0.86 | 0.96 | 0.88 |
| 5 | 32402232 | rs10063044 | -0.339 | 1.639 | 1.235 | 0.298 | 0.067 | -0.047 | -0.047 | 0 | 0.13 | 0.35 | 0.12 | 0.03 | 0.12 | 0.1 |
| 5 | 32421850 | rs324433 | -1.129 | 1.647 | 0.958 | 0.266 | 0.001 | 0.008 | 0.008 | 0 | 0.45 | 0.76 | 0.59 | 0.59 | 0.59 | 0.6 |
| 5 | 32427587 | rs450816 | 0.244 | 1.757 | 0.937 | 0.367 | 0.012 | 0.032 | 0.032 | 0 | 0.27 | 0.37 | 0.45 | 0.56 | 0.42 | 0.45 |
| 5 | 32428018 | rs10042099 | -0.77 | 1.658 | 0.982 | 0.218 | 0.07 | -0.049 | -0.049 | 0 | 0.13 | 0.35 | 0.12 | 0.03 | 0.12 | 0.09 |
| 5 | 32443612 | rs368410 | 0.084 | 1.763 | -0.264 | 0.338 | -0.029 | -0.025 | -0.025 | 0 | 0.92 | 0.98 | 0.87 | 0.86 | 0.93 | 0.88 |
| 5 | 32447650 | rs6865992 | -0.764 | 1.54 | 0.113 | 0.283 | 0.071 | -0.031 | -0.031 | 0 | 0.14 | 0.43 | 0.13 | 0.05 | 0.11 | 0.09 |
| 5 | 32451077 | rs191241 | 0.702 | 1.335 | -0.061 | 0.435 | 0.03 | -0.022 | -0.022 | 0 | 0.71 | 0.49 | 0.68 | 0.71 | 0.73 | 0.73 |
| 5 | 32454025 | rs324445 | -0.404 | 1.124 | -0.367 | 0.827 | -0.001 | -0.003 | -0.003 | 0 | 0.31 | 0.33 | 0.45 | 0.57 | 0.43 | 0.41 |
| 5 | 32455790 | rs623435 | -0.571 | 0.698 | -0.245 | 1.104 | -0.013 | 0.062 | 0.062 | 0 | 0.05 | 0.12 | 0.06 | 0.2 | 0.02 | 0.05 |
| 5 | 32457757 | rs637396 | 0.305 | 0.721 | -0.315 | 0.527 | 0.002 | 0.003 | 0.003 | 0.991 | 0.52 | 0.49 | 0.41 | 0.37 | 0.52 | 0.45 |
| 5 | 32461286 | rs586932 | -0.537 | 0.453 | 0.002 | 1.13 | 0.005 | -0.005 | -0.005 | 0 | 0.78 | 0.83 | 0.72 | 0.75 | 0.83 | 0.71 |
| 5 | 32462753 | rs10520999 | 0.369 | 0.465 | 0.415 | 1.026 | 0 | 0.041 | 0.041 | 0 | 0.16 | 0.24 | 0.21 | 0.36 | 0.17 | 0.21 |
| 5 | 32462868 | rs635790 | 0.584 | 0.452 | 0.402 | 1.03 | -0.013 | 0.055 | 0.055 | 0 | 0.05 | 0.12 | 0.06 | 0.17 | 0.02 | 0.05 |
| 5 | 32466436 | rs818600 | 1.545 | 1.011 | -0.171 | 0.333 | 0.002 | 0 | 0 | 0 | 0.32 | 0.32 | 0.24 | 0.3 | 0.25 | 0.3 |
| 5 | 32467984 | rs4867450 | -0.49 | 1.314 | 0.419 | 0.327 | -0.006 | -0.008 | -0.008 | 0 | 0.3 | 0.27 | 0.28 | 0.08 | 0.41 | 0.2 |
| 5 | 32468181 | rs603463 | 0.208 | 1.004 | 0.09 | 0.426 | 0.009 | -0.004 | -0.004 | 0 | 0.07 | 0.2 | 0.15 | 0.06 | 0.05 | 0.05 |
| 5 | 32474006 | rs599932 | -0.866 | 1.141 | 0.024 | 0.391 | -0.002 | -0.001 | -0.001 | 0.088 | 0.41 | 0.42 | 0.52 | 0.47 | 0.49 | 0.42 |
| 5 | 32474203 | rs6879073 | 0.226 | 0.98 | 0.247 | 0.407 | -0.003 | -0.035 | -0.035 | 0 | 0.12 | 0.23 | 0.13 | 0.06 | 0.14 | 0.05 |
| 5 | 32481243 | rs632085 | -1.404 | 1.192 | 0.132 | 0.4 | 0.011 | 0 | 0 | 1.509 | 0.52 | 0.59 | 0.64 | 0.52 | 0.61 | 0.46 |
| 5 | 32482683 | rs11744595 | 0.541 | 1.354 | -0.11 | 0.618 | 0.001 | -0.011 | -0.011 | 0.966 | 0.14 | 0.11 | 0.19 | 0.36 | 0.14 | 0.19 |
| 5 | 32487636 | rs632909 | -0.517 | 1.431 | -0.024 | 0.327 | 0.009 | -0.027 | -0.027 | 0 | 0.27 | 0.31 | 0.32 | 0.1 | 0.32 | 0.21 |
| 5 | 32500158 | rs7725679 | 0.368 | 1.061 | -0.045 | 0.947 | -0.005 | 0.086 | 0.086 | 0 | 0.93 | 0.83 | 0.87 | 0.75 | 0.94 | 0.91 |
| 5 | 32506309 | rs818534 | 0.606 | 0.975 | -0.658 | 0.704 | -0.017 | -0.008 | -0.008 | 0 | 0.64 | 0.75 | 0.66 | 0.62 | 0.58 | 0.67 |
| 5 | 32510951 | rs10520997 | 0.401 | 1.125 | -0.536 | 0.658 | -0.022 | 0.03 | 0.03 | 1.648 | 0.17 | 0.11 | 0.11 | 0.04 | 0.22 | 0.13 |
| 5 | 32511233 | rs2442115 | -0.037 | 0.871 | -0.072 | 1.17 | 0 | 0.179 | 0.179 | 0 | 0.83 | 0.68 | 0.72 | 0.47 | 0.81 | 0.74 |
| 5 | 32511270 | rs2457153 | -0.055 | 0.873 | -0.071 | 1.171 | 0.001 | 0.181 | 0.181 | 0 | 0.83 | 0.68 | 0.72 | 0.47 | 0.81 | 0.74 |
| 5 | 32523748 | rs591985 | 0.431 | 0.7 | 0.095 | 0.874 | -0.003 | 0.046 | 0.046 | 0 | 0.14 | 0.22 | 0.21 | 0.31 | 0.15 | 0.2 |
| 5 | 32525644 | rs1008023 | 0.319 | 0.65 | -0.013 | 0.85 | -0.005 | 0.023 | 0.023 | 0 | 0.15 | 0.18 | 0.21 | 0.29 | 0.16 | 0.2 |
| 5 | 32530797 | rs818545 | -0.312 | 0.753 | -0.353 | 0.798 | -0.006 | 0.007 | 0.007 | 0 | 0.83 | 0.88 | 0.83 | 0.84 | 0.81 | 0.9 |
| 5 | 32530874 | rs17530882 | 0.571 | 0.813 | -0.54 | 0.768 | -0.037 | 0.048 | 0.048 | 0.051 | 0.13 | 0.01 | 0.12 | 0.06 | 0.17 | 0.05 |
| 5 | 32531335 | rs6891927 | -0.492 | 0.812 | -0.38 | 0.917 | -0.015 | -0.001 | -0.001 | 0 | 0.32 | 0.23 | 0.35 | 0.35 | 0.41 | 0.27 |
| 5 | 32535179 | rs671118 | 1.353 | 0.634 | -0.997 | 0.646 | -0.083 | 0.012 | 0.012 | 0.391 | 0.43 | 0.04 | 0.34 | 0.25 | 0.38 | 0.47 |
| 5 | 32537592 | rs648658 | 0.328 | 0.182 | -0.978 | 0.235 | -0.001 | 0.01 | 0.01 | 0.291 | 0.53 | 0.35 | 0.46 | 0.43 | 0.55 | 0.5 |
| 5 | 32538504 | rs617933 | 0.577 | 0.442 | -1.211 | 0.096 | -0.074 | 0.101 | 0.101 | 0 | 0.32 | 0.09 | 0.25 | 0.13 | 0.31 | 0.33 |
| 5 | 32539793 | rs10434584 | 1.043 | 0.463 | -1.383 | 0.075 | -0.029 | -0.083 | -0.083 | 0 | 0.13 | 0.01 | 0.19 | 0.27 | 0.18 | 0.23 |
| 5 | 32541362 | rs818537 | -0.398 | 0.391 | -0.796 | 0.235 | 0.023 | 0.203 | 0.203 | 0.23 | 0.88 | 0.68 | 0.78 | 0.64 | 0.87 | 0.86 |
| 5 | 32545706 | rs4867112 | -0.263 | 0.385 | -0.799 | 0.249 | 0.018 | 0.194 | 0.194 | 0.188 | 0.88 | 0.66 | 0.78 | 0.64 | 0.87 | 0.86 |
| 5 | 32551899 | rs7710792 | 1.467 | 0.28 | -0.909 | 0.006 | -0.031 | -0.038 | -0.038 | 0 | 0.33 | 0.13 | 0.35 | 0.39 | 0.37 | 0.43 |
| 5 | 32562887 | rs4580786 | -0.002 | 0.149 | -0.829 | -0.142 | -0.01 | 0.069 | 0.069 | 0 | 0.46 | 0.37 | 0.4 | 0.22 | 0.49 | 0.41 |
| 5 | 32567732 | rs2012709 | 0.412 | 0.242 | -0.986 | -0.298 | -0.076 | 0.214 | 0.214 | 0 | 0.42 | 0.11 | 0.34 | 0.12 | 0.44 | 0.36 |
| 5 | 32581603 | rs6876035 | 1.18 | 0.427 | -0.822 | -0.106 | -0.03 | -0.017 | -0.017 | 0.55 | 0.34 | 0.15 | 0.37 | 0.38 | 0.4 | 0.45 |
| 5 | 32586844 | rs10057987 | -0.712 | 0.375 | -0.703 | -0.141 | 0.022 | 0.199 | 0.199 | 0.068 | 0.89 | 0.73 | 0.79 | 0.62 | 0.88 | 0.86 |
| 5 | 32593108 | rs6873470 | 1.232 | 0.47 | -1.704 | -0.582 | -0.097 | 0.141 | 0.141 | 0 | 0.31 | 0.02 | 0.24 | 0.11 | 0.35 | 0.28 |
| 5 | 32612917 | rs818115 | 0.085 | 0.295 | -0.934 | -0.343 | -0.006 | -0.049 | -0.049 | 0 | 0.78 | 0.78 | 0.83 | 0.96 | 0.81 | 0.84 |
| 5 | 32614500 | rs1673937 | 1.276 | 0.774 | -0.926 | -0.104 | 0.005 | 0.013 | 0.013 | 0 | 0.32 | 0.38 | 0.38 | 0.53 | 0.34 | 0.35 |

| | | | | | | | | | | | | | | | | |
|---|---|---|---|---|---|---|---|---|---|---|---|---|---|---|---|---|
| 5 | 32616019 | rs1644769 | -0.305 | 0.781 | -0.534 | -0.31 | 0.005 | 0.063 | 0.063 | 0 | 0.82 | 0.6 | 0.72 | 0.71 | 0.81 | 0.69 |
| 5 | 32616255 | rs922000 | 0.01 | 0.816 | -1.073 | -0.829 | -0.025 | 0.198 | 0.198 | 0 | 0.35 | 0.08 | 0.2 | 0.07 | 0.35 | 0.18 |
| 5 | 32625480 | rs864674 | 0.973 | 0.973 | -1.151 | -0.593 | -0.004 | 0.002 | 0.002 | 0 | 0.45 | 0.41 | 0.51 | 0.54 | 0.46 | 0.51 |
| 5 | 32631975 | rs6894230 | -0.498 | 0.609 | -1.358 | -0.505 | -0.001 | -0.001 | -0.001 | 0 | 0.41 | 0.58 | 0.35 | 0.38 | 0.38 | 0.35 |
| 5 | 32635409 | rs2112867 | 0.454 | 0.703 | -1.687 | -1.167 | -0.007 | -0.077 | -0.077 | 0 | 0.44 | 0.52 | 0.61 | 0.89 | 0.31 | 0.62 |
| 5 | 32636097 | rs6886553 | 0.194 | 0.944 | -1.504 | -0.882 | 0 | -0.046 | -0.046 | 0 | 0.54 | 0.49 | 0.64 | 0.89 | 0.37 | 0.65 |
| 5 | 32644326 | rs818130 | 0.173 | 0.867 | -1.911 | 0.327 | -0.044 | -0.07 | -0.07 | 0 | 0.61 | 0.88 | 0.52 | 0.49 | 0.55 | 0.57 |
| 5 | 32649037 | rs7729311 | -0.171 | 0.604 | -1.733 | -0.064 | -0.026 | 0.095 | 0.095 | 1.212 | 0.43 | 0.16 | 0.55 | 0.71 | 0.52 | 0.59 |
| 5 | 32651271 | rs4867116 | -0.262 | 0.38 | -1.214 | -0.169 | 0.014 | 0.041 | 0.041 | 0.574 | 0.44 | 0.32 | 0.56 | 0.66 | 0.55 | 0.57 |
| 5 | 32658838 | rs4867461 | -0.613 | -0.355 | -0.593 | -0.793 | 0.039 | -0.079 | -0.079 | 0 | 0.8 | 0.63 | 0.84 | 0.9 | 0.73 | 0.92 |
| 5 | 32661834 | rs7732831 | 0.545 | -0.511 | -0.396 | -0.828 | 0.021 | 0.019 | 0.019 | 2.971 | 0.36 | 0.32 | 0.29 | 0.25 | 0.2 | 0.33 |
| 5 | 32662689 | rs7733802 | -0.389 | -0.703 | -0.261 | -0.875 | 0.081 | 0.134 | 0.134 | 0.198 | 0.49 | 0.28 | 0.36 | 0.25 | 0.39 | 0.43 |
| 5 | 32663139 | rs6859120 | 0.323 | -0.252 | -0.086 | -1.225 | -0.019 | -0.021 | -0.021 | 0.23 | 0.05 | 0.17 | 0.01 | 0 | 0 | 0.02 |
| 5 | 32663496 | rs11739058 | 0.286 | 0.078 | -0.563 | -1.037 | -0.004 | -0.005 | -0.005 | 2.081 | 0.15 | 0.1 | 0.1 | 0 | 0.2 | 0.07 |
| 5 | 32670800 | rs1862523 | -0.26 | 0.032 | -0.468 | -0.594 | -0.015 | 0.01 | 0.01 | 1.401 | 0.63 | 0.47 | 0.5 | 0.4 | 0.71 | 0.55 |
| 5 | 32670974 | rs12521228 | -0.533 | 0.217 | -0.294 | -1.343 | -0.02 | -0.072 | -0.072 | 0 | 0.32 | 0.35 | 0.15 | 0.12 | 0.24 | 0.19 |
| 5 | 32677299 | rs11747082 | -0.001 | 0.85 | -0.653 | -1.137 | -0.031 | -0.023 | -0.023 | 0 | 0.08 | 0.03 | 0.19 | 0.09 | 0.1 | 0.06 |
| 5 | 32688090 | rs10755252 | -1.052 | 0.496 | -0.384 | -1.647 | -0.005 | -0.038 | -0.038 | 0 | 0.26 | 0.35 | 0.29 | 0.1 | 0.2 | 0.15 |
| 5 | 32688860 | rs4867469 | 0.176 | 0.644 | -0.567 | -1.377 | -0.031 | -0.025 | -0.025 | 0 | 0.08 | 0.03 | 0.19 | 0.09 | 0.1 | 0.06 |
| 5 | 32691144 | rs10941021 | -0.555 | 0.704 | -0.585 | -1.798 | 0.002 | 0.005 | 0.005 | 0 | 0.41 | 0.4 | 0.4 | 0.1 | 0.35 | 0.17 |
| 5 | 32692289 | rs818126 | -1.011 | 1.011 | -0.803 | -1.815 | -0.005 | -0.009 | -0.009 | 1.689 | 0.7 | 0.83 | 0.64 | 0.3 | 0.7 | 0.47 |
| 5 | 32693566 | rs4867472 | -0.505 | 0.873 | -0.701 | -1.636 | 0 | 0.036 | 0.036 | 0 | 0.39 | 0.34 | 0.39 | 0.1 | 0.35 | 0.16 |
| 5 | 32716535 | rs11750438 | -0.064 | 0.997 | -1.072 | -1.273 | -0.006 | 0.003 | 0.003 | 0 | 0.12 | 0.12 | 0.21 | 0.09 | 0.12 | 0.07 |
| 5 | 32716589 | rs10057069 | 0.089 | 0.657 | -0.89 | -1.323 | 0.002 | 0.003 | 0.003 | 0.079 | 0.29 | 0.21 | 0.18 | 0 | 0.27 | 0.09 |
| 5 | 32722499 | rs976576 | -0.188 | 0.472 | -0.741 | -0.477 | 0 | -0.002 | -0.002 | 0 | 0.73 | 0.76 | 0.67 | 0.29 | 0.75 | 0.48 |
| 5 | 32725531 | rs696831 | -0.708 | 0.58 | -0.541 | -0.594 | -0.001 | -0.001 | -0.001 | 0 | 0.24 | 0.21 | 0.21 | 0.2 | 0.29 | 0.3 |
| 5 | 32733418 | rs696835 | -1.487 | 0.261 | 0.132 | -0.666 | 0.013 | 0.024 | 0.024 | 0 | 0.2 | 0.38 | 0.26 | 0.25 | 0.22 | 0.29 |
| 5 | 32734327 | rs1833530 | -0.673 | 0.221 | 0.275 | -0.711 | 0 | 0.012 | 0.012 | 0 | 0.08 | 0.08 | 0.18 | 0.24 | 0.13 | 0.25 |
| 5 | 32734408 | rs696836 | -0.488 | -0.633 | 0.531 | -1.407 | -0.039 | -0.087 | -0.087 | 0.792 | 0.19 | 0.26 | 0.11 | 0 | 0.09 | 0.08 |
| 5 | 32734511 | rs16890187 | -0.569 | -0.36 | 0.578 | -1.211 | 0.009 | 0.039 | 0.039 | 0 | 0.08 | 0.13 | 0.19 | 0.24 | 0.13 | 0.25 |
| 5 | 32735408 | rs1060559 | -1.48 | -0.417 | 0.506 | -1.286 | -0.005 | -0.049 | -0.049 | 0 | 0.14 | 0.23 | 0.08 | 0.01 | 0.1 | 0.06 |
| 5 | 32740361 | rs10074149 | -1.624 | -0.343 | 0.419 | -1.261 | 0.001 | 0.007 | 0.007 | 0.042 | 0.16 | 0.19 | 0.21 | 0.24 | 0.17 | 0.29 |
| 5 | 32741083 | rs11749133 | -1.948 | -0.349 | 0.424 | -1.261 | 0.007 | 0.008 | 0.008 | 0.769 | 0.2 | 0.27 | 0.25 | 0.25 | 0.23 | 0.31 |
| 5 | 32745029 | rs700925 | -1.51 | -0.293 | 0.939 | -1.239 | 0.043 | 0.065 | 0.065 | 0.03 | 0.14 | 0.31 | 0.24 | 0.25 | 0.19 | 0.25 |
| 5 | 32745752 | rs3792758 | -1.825 | -0.856 | 0.877 | -1.696 | -0.02 | -0.03 | -0.03 | 0 | 0.38 | 0.54 | 0.33 | 0.24 | 0.27 | 0.42 |
| 5 | 32746547 | rs3828586 | -1.918 | -0.947 | 0.91 | -1.659 | 0.005 | 0.008 | 0.008 | 0.039 | 0.37 | 0.35 | 0.36 | 0.24 | 0.32 | 0.37 |
| 5 | 32749592 | rs973079 | -1.493 | -0.884 | 1.001 | -1.622 | -0.006 | -0.008 | -0.008 | 0.253 | 0.31 | 0.34 | 0.31 | 0.23 | 0.24 | 0.35 |
| 5 | 32752599 | rs6866384 | -0.037 | -0.886 | 1.127 | -1.636 | -0.022 | -0.065 | -0.065 | 0 | 0.14 | 0.21 | 0.11 | 0 | 0.1 | 0.05 |
| 5 | 32768634 | rs3792752 | -1.586 | -0.881 | 1.241 | -1.605 | -0.016 | -0.071 | -0.071 | 4.354 | 0.28 | 0.46 | 0.3 | 0.17 | 0.25 | 0.28 |
| 5 | 32769181 | rs3811965 | -0.819 | -0.877 | 1.023 | -1.598 | 0.012 | 0.025 | 0.025 | 0 | 0.1 | 0.2 | 0.18 | 0.17 | 0.1 | 0.23 |
| 5 | 32770750 | rs11740452 | -0.08 | -0.854 | 0.672 | -1.587 | -0.016 | -0.024 | -0.024 | 0 | 0.08 | 0.02 | 0.17 | 0.17 | 0.1 | 0.23 |
| 5 | 32771938 | rs1173736 | -1.534 | -0.954 | 1.14 | -1.619 | 0.019 | 0.051 | 0.051 | 0 | 0.31 | 0.75 | 0.32 | 0.17 | 0.25 | 0.28 |
| 5 | 32773314 | rs3792751 | -1.573 | -0.654 | 1.565 | -1.74 | -0.005 | -0.038 | -0.038 | 3.33 | 0.32 | 0.42 | 0.36 | 0.17 | 0.34 | 0.35 |
| 5 | 32781196 | rs751432 | -1.073 | -0.691 | 1.829 | -1.427 | 0.004 | 0.014 | 0.014 | 0.141 | 0.13 | 0.22 | 0.23 | 0.2 | 0.19 | 0.33 |
| 5 | 32781521 | rs10075794 | -0.681 | -0.693 | 1.835 | -1.427 | -0.011 | -0.02 | -0.02 | 0 | 0.12 | 0.07 | 0.22 | 0.2 | 0.19 | 0.33 |
| 5 | 32786389 | rs2270915 | -0.601 | -0.691 | 1.944 | -1.427 | -0.003 | -0.006 | -0.006 | 0.239 | 0.11 | 0.1 | 0.21 | 0.2 | 0.19 | 0.31 |
| 5 | 32788369 | rs1158715 | 0.113 | -0.792 | 1.107 | -0.902 | -0.005 | -0.002 | -0.002 | 0 | 0.4 | 0.19 | 0.41 | 0.66 | 0.41 | 0.54 |
| 5 | 32789852 | rs1173756 | -0.79 | -0.827 | 1.456 | -0.982 | -0.015 | 0.014 | 0.014 | 0 | 0.58 | 0.75 | 0.54 | 0.66 | 0.51 | 0.6 |
| 5 | 32810791 | rs10521001 | -0.006 | -0.558 | 0.593 | -0.742 | -0.003 | -0.004 | -0.004 | 0 | 0.12 | 0.14 | 0.24 | 0.2 | 0.2 | 0.32 |
| 5 | 32811911 | rs12659264 | 0.937 | -0.696 | 0.544 | -0.849 | 0.007 | -0.098 | -0.098 | 0 | 0.29 | 0.09 | 0.2 | 0.46 | 0.24 | 0.23 |
| 5 | 32817576 | rs4867478 | 0.903 | -0.7 | 0.542 | -0.851 | 0.008 | -0.098 | -0.098 | 0 | 0.3 | 0.1 | 0.2 | 0.46 | 0.24 | 0.23 |
| 5 | 32834199 | rs1173726 | -0.011 | -0.785 | 1.111 | -1.061 | -0.053 | -0.14 | -0.14 | 0 | 0.18 | 0.5 | 0.13 | 0 | 0.13 | 0.05 |
| 5 | 32835212 | rs1173723 | -1.08 | -0.493 | 0.749 | -1.197 | -0.11 | -0.097 | -0.097 | 0 | 0.64 | 0.88 | 0.46 | 0.47 | 0.53 | 0.51 |
| 5 | 32837489 | rs1173718 | 0.245 | -0.452 | 1.07 | -1.16 | -0.042 | -0.118 | -0.118 | 0 | 0.17 | 0.41 | 0.12 | 0 | 0.13 | 0.05 |
| 5 | 32837647 | rs11950160 | -0.425 | 0.008 | 0.81 | -0.845 | 0.01 | -0.079 | -0.079 | 0 | 0.16 | 0.25 | 0.12 | 0.01 | 0.14 | 0.21 |
| 5 | 32841934 | rs1818116 | 0.389 | -0.044 | 0.443 | -0.772 | 0.031 | 0.001 | 0.001 | 0 | 0.46 | 0.3 | 0.33 | 0.46 | 0.4 | 0.32 |
| 5 | 32843434 | rs7722828 | -0.775 | -0.055 | 0.776 | -1.011 | 0.014 | -0.049 | -0.049 | 0 | 0.09 | 0.25 | 0.07 | 0.02 | 0.08 | 0.17 |
| 5 | 32843583 | rs1173708 | -0.168 | 0.097 | 0.245 | -0.661 | 0.012 | 0.01 | 0.01 | 0 | 0.5 | 0.63 | 0.61 | 0.62 | 0.55 | 0.66 |
| 5 | 32851259 | rs516451 | 0.377 | 0.089 | -0.278 | -0.603 | -0.066 | -0.047 | -0.047 | 0 | 0.38 | 0.09 | 0.39 | 0.46 | 0.39 | 0.4 |
| 5 | 32852872 | rs1147227 | 0.261 | 0.056 | 0.528 | -0.599 | -0.001 | -0.001 | -0.001 | 0 | 0.38 | 0.33 | 0.4 | 0.46 | 0.39 | 0.39 |
| 5 | 32867260 | rs10044112 | -0.418 | 0.002 | 0.88 | -0.593 | -0.002 | -0.002 | -0.002 | 0 | 0.62 | 0.67 | 0.6 | 0.54 | 0.61 | 0.6 |
| 5 | 32869602 | rs10036130 | -0.146 | -0.068 | 0.995 | -0.623 | 0 | -0.004 | -0.004 | 0.114 | 0.56 | 0.51 | 0.57 | 0.55 | 0.56 | 0.6 |
| 5 | 32872478 | rs1501851 | -0.927 | -0.131 | 1.323 | -0.445 | -0.011 | -0.035 | -0.035 | 0 | 0.17 | 0.45 | 0.2 | 0.12 | 0.12 | 0.13 |
| 5 | 32885528 | rs12522129 | 0.529 | 0.067 | 0.587 | -0.429 | -0.048 | -0.047 | -0.047 | 0.48 | 0.13 | 0.01 | 0.21 | 0.16 | 0.17 | 0.25 |
| 5 | 32887610 | rs1472261 | -0.698 | 0.043 | 0.922 | -0.426 | -0.009 | -0.042 | -0.042 | 0.162 | 0.17 | 0.51 | 0.21 | 0.1 | 0.12 | 0.13 |
| 5 | 32887694 | rs16890456 | -0.698 | 0.043 | 0.922 | -0.426 | -0.009 | -0.042 | -0.042 | 0 | 0.17 | 0.51 | 0.21 | 0.1 | 0.12 | 0.13 |
| 5 | 32888818 | rs10472828 | 0.482 | -0.369 | 0.53 | -0.686 | 0.001 | 0.012 | 0.012 | 0 | 0.46 | 0.2 | 0.49 | 0.52 | 0.43 | 0.53 |
| 5 | 32889284 | rs2331101 | 1.391 | -0.356 | 0.559 | -0.668 | 0.017 | -0.039 | -0.039 | 0 | 0.33 | 0.19 | 0.28 | 0.36 | 0.26 | 0.28 |
| 5 | 32891191 | rs4867479 | -0.678 | -0.294 | 0.511 | -0.636 | -0.013 | 0.01 | 0.01 | 0 | 0.62 | 0.71 | 0.7 | 0.62 | 0.55 | 0.66 |

| | | | | | | | | | | | | | | | | |
|---|---|---|---|---|---|---|---|---|---|---|---|---|---|---|---|---|
| 5 | 32903878 | rs6869594 | -0.646 | -0.272 | 0.456 | -0.682 | -0.008 | 0.007 | 0.007 | 0 | 0.62 | 0.69 | 0.73 | 0.64 | 0.56 | 0.62 |
| 5 | 32903894 | rs6869887 | -0.123 | -0.222 | 0.611 | -0.61 | 0.006 | 0 | 0 | 0.065 | 0.48 | 0.5 | 0.5 | 0.47 | 0.39 | 0.4 |
| 5 | 32907028 | rs4867487 | 0.745 | -0.289 | 0.134 | -0.499 | 0.016 | -0.037 | -0.037 | 0 | 0.34 | 0.2 | 0.31 | 0.37 | 0.27 | 0.29 |
| 5 | 32916978 | rs16890493 | -0.377 | -0.107 | 0.522 | -0.742 | -0.02 | -0.1 | -0.1 | 0 | 0.14 | 0.43 | 0.11 | 0.01 | 0.1 | 0.08 |
| 5 | 32920002 | rs4867490 | -0.208 | -0.171 | 0.277 | -0.794 | 0.012 | -0.004 | -0.004 | 0.118 | 0.53 | 0.66 | 0.45 | 0.46 | 0.41 | 0.38 |
| 5 | 32920484 | rs7730787 | -0.804 | -0.159 | 0.324 | -0.767 | -0.025 | -0.091 | -0.091 | 0 | 0.18 | 0.61 | 0.14 | 0.02 | 0.12 | 0.17 |
| 5 | 32932459 | rs12654257 | -1.808 | 0.196 | -0.412 | -0.764 | -0.076 | -0.092 | -0.092 | 0 | 0.84 | 0.98 | 0.83 | 0.66 | 0.75 | 0.89 |
| 5 | 32935733 | rs9292472 | 0.497 | -0.566 | 0.359 | -0.729 | 0.023 | -0.049 | -0.049 | 0 | 0.38 | 0.21 | 0.34 | 0.45 | 0.32 | 0.3 |
| 5 | 32941161 | rs10067052 | -0.575 | -0.454 | 0.373 | -0.845 | 0.004 | -0.006 | -0.006 | 0 | 0.54 | 0.62 | 0.47 | 0.44 | 0.44 | 0.41 |
| 5 | 32946922 | rs12523390 | 0.728 | -0.391 | -0.068 | -0.804 | 0.012 | -0.046 | -0.046 | 0 | 0.36 | 0.22 | 0.34 | 0.44 | 0.31 | 0.29 |
| 5 | 32947698 | rs10079994 | -0.272 | -0.05 | -0.135 | -0.445 | -0.002 | 0.001 | 0.001 | 0 | 0.53 | 0.55 | 0.46 | 0.45 | 0.43 | 0.4 |
| 5 | 32957681 | rs1494654 | -1.247 | -0.394 | 0.129 | -0.449 | -0.011 | -0.096 | -0.096 | 0.971 | 0.18 | 0.7 | 0.19 | 0 | 0.15 | 0.11 |
| 5 | 32961669 | rs10941035 | 0.091 | -0.858 | -0.532 | -0.456 | 0.057 | -0.078 | -0.078 | 0 | 0.36 | 0.11 | 0.29 | 0.42 | 0.27 | 0.27 |
| 5 | 32968720 | rs2331107 | -0.341 | -0.635 | 0.148 | -0.291 | 0.004 | -0.05 | -0.05 | 0.19 | 0.09 | 0.2 | 0.08 | 0 | 0.09 | 0.08 |
| 5 | 32968807 | rs17546321 | 0.014 | -0.632 | 0.168 | -0.29 | -0.005 | 0.008 | 0.008 | 0 | 0.08 | 0.07 | 0.07 | 0 | 0.09 | 0.08 |
| 5 | 32972458 | rs7715554 | -0.014 | -0.658 | 0.214 | -0.275 | 0.002 | -0.041 | -0.041 | 0 | 0.08 | 0.18 | 0.08 | 0 | 0.09 | 0.07 |
| 5 | 32972552 | rs7715836 | -0.238 | -0.662 | 0.219 | -0.277 | 0.002 | -0.047 | -0.047 | 5.542 | 0.08 | 0.2 | 0.08 | 0 | 0.09 | 0.07 |
| 5 | 32973557 | rs2877294 | -0.884 | -0.994 | 0.185 | -0.41 | 0.033 | -0.018 | -0.018 | 0.063 | 0.42 | 0.72 | 0.37 | 0.41 | 0.34 | 0.27 |
| 5 | 32975490 | rs10079889 | -0.315 | -0.867 | 0.328 | -0.391 | -0.036 | 0.041 | 0.041 | 2.714 | 0.34 | 0.56 | 0.29 | 0.41 | 0.27 | 0.2 |
| 5 | 32977787 | rs7726086 | -0.736 | -0.909 | 0.281 | -0.194 | -0.021 | -0.016 | -0.016 | 0 | 0.28 | 0.6 | 0.26 | 0.16 | 0.22 | 0.15 |
| 5 | 32979106 | rs7735691 | -0.126 | -0.485 | 0.05 | -0.256 | 0.005 | -0.04 | -0.04 | 0 | 0.08 | 0.14 | 0.08 | 0 | 0.08 | 0.07 |
| 5 | 32981090 | rs11954821 | -0.242 | -0.653 | -0.1 | -0.467 | -0.009 | -0.049 | -0.049 | 0 | 0.11 | 0.14 | 0.07 | 0 | 0.08 | 0.07 |
| 5 | 32984015 | rs13355979 | -1.284 | -0.363 | -0.592 | -0.252 | -0.084 | -0.07 | -0.07 | 0 | 0.6 | 0.88 | 0.5 | 0.46 | 0.46 | 0.43 |
| 5 | 32985044 | rs6860419 | -0.306 | -0.529 | -0.41 | -0.267 | 0.088 | -0.053 | -0.053 | 0.022 | 0.43 | 0.17 | 0.33 | 0.43 | 0.3 | 0.28 |
| 5 | 32986782 | rs7708003 | -1.291 | -0.024 | 0.555 | -0.112 | 0.066 | -0.124 | -0.124 | 6.098 | 0.16 | 0.48 | 0.15 | 0.01 | 0.17 | 0.11 |
| 5 | 32988178 | rs1396462 | -1.088 | -0.41 | 0.076 | 0.179 | -0.016 | -0.005 | -0.005 | 1.181 | 0.31 | 0.45 | 0.25 | 0.26 | 0.25 | 0.26 |
| 5 | 32989759 | rs13170607 | 0.644 | -0.297 | -0.815 | 0.377 | 0.002 | 0.001 | 0.001 | 0 | 0.48 | 0.2 | 0.37 | 0.42 | 0.51 | 0.51 |
| 5 | 32991719 | rs16890585 | -0.916 | -0.219 | -0.264 | 0.33 | 0 | -0.002 | -0.002 | 0 | 0.12 | 0.14 | 0.13 | 0.05 | 0.11 | 0.13 |
| 5 | 33005008 | rs11744178 | 1.008 | 0.028 | 0.019 | 0.258 | -0.041 | -0.024 | -0.024 | 0 | 0.42 | 0.14 | 0.3 | 0.35 | 0.4 | 0.48 |
| 5 | 33010844 | rs6867192 | 0.403 | -0.113 | 0.805 | 0.097 | -0.007 | -0.004 | -0.004 | 0 | 0.54 | 0.64 | 0.43 | 0.44 | 0.49 | 0.58 |
| 5 | 33011986 | rs4867131 | 0.023 | -0.148 | 0.605 | 0.457 | 0.001 | -0.001 | -0.001 | 0 | 0.12 | 0.08 | 0.18 | 0.21 | 0.13 | 0.14 |
| 5 | 33017254 | rs1508827 | 1.23 | -0.13 | 0.806 | 0.305 | -0.027 | -0.021 | -0.021 | 0 | 0.43 | 0.15 | 0.3 | 0.37 | 0.4 | 0.48 |
| 5 | 33024072 | rs6881818 | 0.696 | -0.119 | 1.078 | 0.317 | 0 | 0 | 0 | 0 | 0.51 | 0.51 | 0.41 | 0.44 | 0.47 | 0.57 |
| 5 | 33032667 | rs17449766 | -0.981 | 0.041 | 1.28 | 0.228 | 0.001 | -0.023 | -0.023 | 0 | 0.11 | 0.07 | 0.11 | 0.01 | 0.14 | 0.11 |
| 5 | 33033738 | rs4099722 | -2.208 | -0.048 | 1.567 | 0.249 | 0.04 | -0.059 | -0.059 | 0 | 0.19 | 0.4 | 0.2 | 0.09 | 0.21 | 0.19 |
| 5 | 33039387 | rs7736123 | -1.489 | -0.305 | 1.514 | 0.427 | -0.021 | -0.074 | -0.074 | 0.675 | 0.13 | 0.43 | 0.07 | 0.02 | 0.06 | 0.14 |
| 5 | 33041344 | rs1593896 | -1.905 | -0.354 | 1.561 | 0.418 | -0.048 | -0.044 | -0.044 | 0 | 0.15 | 0.57 | 0.19 | 0.05 | 0.07 | 0.21 |
| 5 | 33047414 | rs1428211 | 1.197 | -0.212 | 0.739 | 0.524 | -0.079 | -0.054 | -0.054 | 0 | 0.43 | 0.11 | 0.38 | 0.4 | 0.44 | 0.47 |
| 5 | 33055680 | rs10074451 | -0.885 | -0.271 | 1.027 | 0.395 | 0.003 | -0.091 | -0.091 | 0 | 0.15 | 0.34 | 0.1 | 0.02 | 0.11 | 0.17 |
| 5 | 33056345 | rs1428216 | 0.317 | -0.33 | 0.837 | 0.603 | -0.084 | -0.052 | -0.052 | 0 | 0.35 | 0.1 | 0.35 | 0.41 | 0.43 | 0.27 |
| 5 | 33057266 | rs445454 | 0.874 | -0.355 | 0.983 | 0.851 | 0.015 | 0.005 | 0.005 | 0 | 0.43 | 0.34 | 0.38 | 0.4 | 0.4 | 0.48 |
| 5 | 33065853 | rs433771 | 0.229 | -0.456 | 0.791 | 0.392 | -0.003 | 0.059 | 0.059 | 0 | 0.92 | 0.84 | 0.82 | 0.82 | 0.92 | 0.88 |
| 5 | 33066481 | rs16890687 | -0.185 | -0.446 | 0.442 | 0.53 | 0.001 | 0 | 0 | 0 | 0.48 | 0.39 | 0.44 | 0.57 | 0.51 | 0.4 |
| 5 | 33067875 | rs2619864 | 0.942 | -0.258 | 0.027 | 0.606 | -0.018 | 0.064 | 0.064 | 0 | 0.4 | 0.14 | 0.35 | 0.24 | 0.41 | 0.46 |
| 5 | 33068755 | rs2619862 | 1.188 | -0.097 | 0.699 | 0.813 | -0.006 | 0.102 | 0.102 | 0.078 | 0.89 | 0.66 | 0.8 | 0.82 | 0.92 | 0.88 |
| 5 | 33078406 | rs6870782 | -0.666 | -0.41 | 0.17 | 0.123 | -0.008 | -0.037 | -0.037 | 0 | 0.18 | 0.2 | 0.09 | 0.02 | 0.11 | 0.23 |
| 5 | 33085684 | rs16890717 | 0.214 | -0.302 | -0.381 | 0.179 | -0.065 | -0.092 | -0.092 | 0.005 | 0.33 | 0.11 | 0.37 | 0.54 | 0.4 | 0.2 |
| 5 | 33114638 | rs6894272 | 0.532 | -0.472 | 0.569 | 0.361 | 0.006 | -0.036 | -0.036 | 0.279 | 0.1 | 0.01 | 0.03 | 0.14 | 0.09 | 0.06 |
| 5 | 33119690 | rs13168770 | 0.732 | -0.414 | 0.566 | 0.23 | -0.051 | -0.054 | -0.054 | 0 | 0.23 | 0.11 | 0.35 | 0.41 | 0.31 | 0.14 |
| 5 | 33120048 | rs6450946 | -1.649 | -0.843 | 0.674 | -0.011 | -0.038 | -0.059 | -0.059 | 0 | 0.58 | 0.85 | 0.54 | 0.52 | 0.56 | 0.52 |
| 5 | 33151875 | rs7718826 | -0.614 | -0.973 | 0.774 | -0.206 | 0.036 | 0.082 | 0.082 | 0 | 0.26 | 0.11 | 0.12 | 0.11 | 0.2 | 0.3 |
| 5 | 33169439 | rs10038227 | -0.518 | -0.839 | 1.025 | -0.176 | -0.018 | -0.058 | -0.058 | 0.008 | 0.22 | 0.68 | 0.14 | 0.02 | 0.12 | 0.3 |
| 5 | 33176567 | rs11744729 | 0.538 | -0.468 | 0.937 | 0.136 | 0.004 | -0.003 | -0.003 | 0 | 0.56 | 0.55 | 0.48 | 0.37 | 0.52 | 0.52 |
| 5 | 33177406 | rs10041597 | 0.68 | -0.615 | 0.794 | -0.119 | 0 | 0.02 | 0.02 | 0 | 0.15 | 0.09 | 0.13 | 0.02 | 0.14 | 0.16 |
| 5 | 33177987 | rs7709045 | 1.134 | -0.612 | 0.832 | -0.04 | -0.005 | -0.001 | -0.001 | 0 | 0.42 | 0.48 | 0.4 | 0.4 | 0.38 | 0.47 |
| 5 | 33180039 | rs1479643 | 1.416 | -0.637 | 0.795 | -0.074 | 0 | -0.034 | -0.034 | 0 | 0.39 | 0.19 | 0.37 | 0.41 | 0.38 | 0.47 |
| 5 | 33182745 | rs10041417 | -0.341 | -0.59 | 0.889 | 0.109 | -0.023 | -0.017 | -0.017 | 0 | 0.21 | 0.27 | 0.19 | 0.17 | 0.17 | 0.21 |
| 5 | 33191667 | rs6450953 | -0.784 | -0.538 | 1.005 | 0.11 | -0.009 | 0 | 0 | 0 | 0.44 | 0.59 | 0.49 | 0.58 | 0.48 | 0.36 |
| 5 | 33212459 | rs6870292 | 0.427 | 0.069 | 0.902 | 0.239 | 0.021 | 0.109 | 0.109 | 0.187 | 0.1 | 0.13 | 0.18 | 0.43 | 0.13 | 0.09 |
| 5 | 33216867 | rs10075884 | 0.007 | -0.012 | 1.128 | 0.234 | 0.044 | 0.195 | 0.195 | 0 | 0.12 | 0.25 | 0.18 | 0.43 | 0.13 | 0.1 |
| 5 | 33218955 | rs7719179 | -0.085 | -0.167 | 0.908 | 0.072 | 0.01 | 0.009 | 0.009 | 0 | 0.81 | 0.9 | 0.8 | 0.81 | 0.82 | 0.82 |
| 5 | 33222540 | rs12515068 | 1.589 | -0.063 | 0.798 | -0.201 | -0.007 | -0.055 | -0.055 | 0 | 0.37 | 0.15 | 0.37 | 0.38 | 0.37 | 0.42 |
| 5 | 33223220 | rs6450963 | -0.147 | 0.005 | 0.943 | -0.002 | 0.007 | 0.009 | 0.009 | 0 | 0.82 | 0.9 | 0.8 | 0.81 | 0.82 | 0.82 |
| 5 | 33223956 | rs6450965 | 1.405 | 0.003 | 1.16 | -0.006 | 0.005 | -0.013 | -0.013 | 2.306 | 0.64 | 0.32 | 0.59 | 0.39 | 0.67 | 0.65 |
| 5 | 33227083 | rs10941049 | -0.736 | 0.122 | 1.267 | 0.076 | 0.034 | 0.222 | 0.222 | 0 | 0.16 | 0.53 | 0.2 | 0.43 | 0.14 | 0.17 |
| 5 | 33230034 | rs11745439 | 1.311 | 0.187 | 1.048 | 0.052 | 0.005 | -0.006 | -0.006 | 0 | 0.67 | 0.35 | 0.54 | 0.36 | 0.72 | 0.68 |
| 5 | 33242910 | rs6450973 | 0.084 | 0.165 | 1.247 | 0.094 | -0.01 | -0.008 | -0.008 | 0.19 | 0.8 | 0.75 | 0.71 | 0.79 | 0.83 | 0.83 |
| 5 | 33243808 | rs9791054 | 1.31 | 0.212 | 0.943 | -0.124 | -0.029 | -0.036 | -0.036 | 0.048 | 0.4 | 0.15 | 0.32 | 0.37 | 0.42 | 0.43 |
| 5 | 33251492 | rs6891741 | 0.853 | 0.412 | 1.363 | -0.196 | 0.008 | -0.089 | -0.089 | 0 | 0.28 | 0.31 | 0.22 | 0 | 0.3 | 0.24 |

| | | | | | | | | | | | | | | | | |
|---|---|---|---|---|---|---|---|---|---|---|---|---|---|---|---|---|
| 5 | 33261251 | rs2967087 | -0.573 | 0.394 | 1.288 | -0.038 | 0.008 | 0.146 | 0.146 | 0.533 | 0.91 | 0.87 | 0.84 | 0.59 | 0.9 | 0.92 |
| 5 | 33277773 | rs444815 | 0.116 | 0.21 | 1.199 | -0.218 | -0.003 | -0.001 | -0.001 | 0 | 0.58 | 0.4 | 0.59 | 0.59 | 0.57 | 0.67 |
| 5 | 33285417 | rs419517 | -0.054 | -0.228 | 0.671 | -0.335 | 0.027 | -0.057 | -0.057 | 0 | 0.27 | 0.11 | 0.38 | 0.42 | 0.22 | 0.38 |
| 5 | 33296332 | rs268746 | -0.884 | -0.052 | 0.786 | -0.362 | -0.012 | -0.008 | -0.008 | 0 | 0.63 | 0.71 | 0.65 | 0.42 | 0.57 | 0.7 |
| 5 | 33300314 | rs268739 | -0.34 | 0.333 | 0.786 | -0.324 | 0.002 | -0.005 | -0.005 | 0 | 0.27 | 0.21 | 0.38 | 0.42 | 0.24 | 0.41 |
| 5 | 33302632 | rs1427852 | 0.863 | 0.412 | 0.718 | -0.42 | -0.014 | 0.022 | 0.022 | 0.674 | 0.33 | 0.23 | 0.26 | 0 | 0.33 | 0.26 |
| 5 | 33305739 | rs268763 | -0.493 | 0.215 | 0.922 | -0.143 | 0.003 | 0.118 | 0.118 | 0.442 | 0.9 | 0.82 | 0.85 | 0.6 | 0.9 | 0.91 |
| 5 | 33310214 | rs13182969 | 0.852 | 0.275 | 0.528 | -0.273 | -0.023 | 0.049 | 0.049 | 1.25 | 0.36 | 0.23 | 0.27 | 0 | 0.38 | 0.27 |
| 5 | 33313125 | rs156190 | -0.019 | 0.233 | 0.425 | -0.113 | 0.021 | -0.041 | -0.041 | 0 | 0.22 | 0.11 | 0.36 | 0.41 | 0.16 | 0.35 |
| 5 | 33330807 | rs6896697 | 0.789 | 0.291 | 0.519 | -0.33 | -0.01 | 0.001 | 0.001 | 1.502 | 0.37 | 0.29 | 0.28 | 0.03 | 0.38 | 0.27 |
| 5 | 33338606 | rs10040884 | 1.181 | -0.052 | 0.701 | 0.236 | -0.009 | -0.008 | -0.008 | 0 | 0.66 | 0.42 | 0.48 | 0.25 | 0.74 | 0.52 |
| 5 | 33344107 | rs1876670 | 0.872 | -0.028 | 0.663 | 0.27 | -0.02 | -0.007 | -0.007 | 0 | 0.69 | 0.58 | 0.5 | 0.27 | 0.77 | 0.63 |
| 5 | 33370654 | rs268711 | -0.147 | -0.067 | 0.583 | 0.438 | 0.02 | -0.024 | -0.024 | 0.836 | 0.21 | 0.11 | 0.34 | 0.33 | 0.15 | 0.31 |
| 5 | 33375972 | rs11747502 | 0.948 | -0.014 | 0.595 | 0.402 | -0.021 | -0.013 | -0.013 | 0 | 0.68 | 0.52 | 0.53 | 0.26 | 0.76 | 0.56 |
| 5 | 33399200 | rs156319 | -0.059 | 0.445 | 0.495 | 0.585 | 0.048 | 0.233 | 0.233 | 0.237 | 0.91 | 0.75 | 0.88 | 0.6 | 0.91 | 0.94 |
| 5 | 33401470 | rs272735 | -0.063 | 0.056 | -0.083 | 0.369 | 0.01 | 0.001 | 0.001 | 1.093 | 0.51 | 0.43 | 0.46 | 0.47 | 0.51 | 0.41 |
| 5 | 33405597 | rs268706 | -0.075 | 0.027 | -0.499 | 0.707 | 0.002 | -0.02 | -0.02 | 0 | 0.18 | 0.13 | 0.25 | 0.29 | 0.18 | 0.24 |
| 5 | 33410707 | rs17550908 | 0.221 | -0.28 | -0.357 | 0.281 | 0.001 | 0.007 | 0.007 | 0.635 | 0.35 | 0.3 | 0.27 | 0.17 | 0.4 | 0.31 |
| 5 | 33410770 | rs271787 | 0.004 | 0.073 | -0.222 | 0.808 | 0.002 | 0.041 | 0.041 | 0 | 0.1 | 0.1 | 0.12 | 0.31 | 0.12 | 0.29 |
| 5 | 33430144 | rs7728855 | -1.024 | -0.113 | 0.244 | 0.499 | -0.012 | 0.063 | 0.063 | 0 | 0.19 | 0.41 | 0.21 | 0.26 | 0.17 | 0.44 |
| 5 | 33434002 | rs1438728 | -1.039 | -0.259 | -0.053 | 0.43 | -0.019 | 0.023 | 0.023 | 0.051 | 0.2 | 0.55 | 0.22 | 0.26 | 0.17 | 0.45 |
| 5 | 33440306 | rs3756663 | 0.748 | -0.565 | -0.665 | 0.216 | 0.011 | 0.019 | 0.019 | 0 | 0.24 | 0.17 | 0.15 | 0.14 | 0.25 | 0.14 |
| 5 | 33446539 | rs3777079 | -0.729 | -0.517 | -0.991 | 0.288 | 0.025 | 0.09 | 0.09 | 0.038 | 0.47 | 0.69 | 0.49 | 0.73 | 0.4 | 0.65 |
| 5 | 33450530 | rs6450991 | -1.802 | -0.384 | -1.357 | 0.294 | -0.077 | 0.093 | 0.093 | 0.146 | 0.71 | 0.89 | 0.64 | 0.87 | 0.65 | 0.79 |
| 5 | 33455616 | rs6880623 | 0.857 | -0.422 | -1.208 | 0.291 | 0.015 | -0.033 | -0.033 | 0 | 0.1 | 0.16 | 0.11 | 0.03 | 0.12 | 0.3 |
| 5 | 33457618 | rs1003000 | -1.671 | -0.221 | -2.189 | 0.415 | -0.077 | 0.093 | 0.093 | 0.146 | 0.71 | 0.89 | 0.64 | 0.87 | 0.65 | 0.79 |
| 5 | 33458881 | rs3765147 | 0.686 | -0.158 | -2.134 | 0.984 | -0.017 | -0.103 | -0.103 | 0.094 | 0.26 | 0.1 | 0.27 | 0.47 | 0.23 | 0.2 |
| 5 | 33461355 | rs2270905 | 0.456 | 0.024 | -1.537 | 1.425 | -0.001 | 0.013 | 0.013 | 0 | 0.18 | 0.13 | 0.14 | 0.13 | 0.24 | 0.06 |
| 5 | 33464297 | rs12659798 | -0.428 | -0.155 | -1.233 | 1.668 | 0.001 | 0.014 | 0.014 | 0 | 0.07 | 0.05 | 0.09 | 0.24 | 0.05 | 0.15 |
| 5 | 33493797 | rs4867514 | -0.952 | -0.226 | -1.375 | 1.537 | -0.045 | -0.028 | -0.028 | 0.798 | 0.37 | 0.77 | 0.43 | 0.4 | 0.44 | 0.41 |
| 5 | 33494870 | rs2331238 | 0.141 | 0.434 | -1.824 | 1.061 | -0.104 | -0.002 | -0.002 | 0.009 | 0.21 | 0.07 | 0.3 | 0.18 | 0.36 | 0.12 |
| 5 | 33497319 | rs256600 | 0.515 | -0.36 | -1.129 | 1.322 | 0 | -0.045 | -0.045 | 3.012 | 0.91 | 0.99 | 0.9 | 0.83 | 0.92 | 0.84 |
| 5 | 33501919 | rs16891241 | -0.552 | -0.46 | -0.667 | 1.295 | -0.006 | 0.069 | 0.069 | 0 | 0.18 | 0.26 | 0.23 | 0.48 | 0.14 | 0.41 |
| 5 | 33513441 | rs4867517 | 0.08 | -0.986 | -0.662 | -0.376 | -0.002 | -0.021 | -0.021 | 3.367 | 0.52 | 0.54 | 0.34 | 0.17 | 0.43 | 0.61 |
| 5 | 33518227 | rs4867522 | 0.57 | -1.272 | -0.688 | -0.496 | -0.012 | -0.03 | -0.03 | 0 | 0.37 | 0.39 | 0.28 | 0.14 | 0.31 | 0.51 |
| 5 | 33525321 | rs27405 | -0.721 | -0.705 | -0.644 | -0.487 | 0 | 0.001 | 0.001 | 0 | 0.43 | 0.58 | 0.46 | 0.7 | 0.34 | 0.43 |
| 5 | 33529263 | rs10521004 | 0.595 | -0.812 | -0.302 | -0.656 | -0.011 | -0.069 | -0.069 | 0 | 0.14 | 0.28 | 0.06 | 0.02 | 0.11 | 0.08 |
| 5 | 33531645 | rs256792 | -0.565 | -1.032 | -0.353 | -0.822 | -0.008 | -0.031 | -0.031 | 0 | 0.38 | 0.44 | 0.43 | 0.75 | 0.25 | 0.6 |
| 5 | 33535060 | rs25754 | -0.849 | -1.01 | -0.717 | -0.78 | 0.014 | 0.073 | 0.073 | 0.206 | 0.5 | 0.65 | 0.5 | 0.77 | 0.36 | 0.67 |
| 5 | 33546432 | rs29967 | -1.048 | -0.952 | -0.872 | -0.68 | 0.047 | 0.117 | 0.117 | 0.793 | 0.49 | 0.74 | 0.48 | 0.76 | 0.32 | 0.67 |
| 5 | 33549014 | rs1501724 | 1.021 | -0.514 | -0.957 | -0.353 | -0.07 | 0.247 | 0.247 | 0 | 0.38 | 0.08 | 0.25 | 0.04 | 0.41 | 0.24 |
| 5 | 33556366 | rs256639 | -0.363 | -0.15 | -0.459 | 0.167 | -0.012 | 0.053 | 0.053 | 0.918 | 0.28 | 0.43 | 0.37 | 0.58 | 0.18 | 0.27 |
| 5 | 33558544 | rs194381 | -0.044 | 0.582 | -1.341 | 0.144 | -0.048 | 0.006 | 0.006 | 0 | 0.7 | 0.89 | 0.68 | 0.71 | 0.6 | 0.58 |
| 5 | 33565137 | rs4146150 | -0.369 | 0.609 | -0.874 | 0.555 | 0.007 | 0.008 | 0.008 | 0 | 0.5 | 0.41 | 0.53 | 0.49 | 0.55 | 0.58 |
| 5 | 33578441 | rs10042589 | -0.956 | 1.335 | -0.949 | 0.622 | -0.006 | -0.003 | -0.003 | 0 | 0.19 | 0.17 | 0.26 | 0.21 | 0.28 | 0.29 |
| 5 | 33578638 | rs16891337 | -0.86 | 0.83 | -1.132 | 0.233 | 0.008 | 0.018 | 0.018 | 0 | 0.1 | 0.08 | 0.07 | 0.04 | 0.07 | 0.2 |
| 5 | 33579395 | rs6866077 | 0.139 | 0.507 | -1.236 | -0.358 | 0.024 | 0.05 | 0.05 | 0 | 0.27 | 0.2 | 0.16 | 0.09 | 0.23 | 0.27 |
| 5 | 33580181 | rs384821 | 0.745 | 0.811 | -1.384 | -0.344 | -0.003 | 0.004 | 0.004 | 0 | 0.19 | 0.18 | 0.21 | 0.15 | 0.23 | 0.2 |
| 5 | 33585401 | rs428052 | 0.02 | 1.077 | -0.537 | 0.237 | -0.006 | -0.004 | -0.004 | 0 | 0.29 | 0.28 | 0.5 | 0.62 | 0.41 | 0.3 |
| 5 | 33585754 | rs408758 | 1.032 | 0.941 | -0.351 | 0.225 | 0.046 | 0.109 | 0.109 | 0 | 0.45 | 0.26 | 0.33 | 0.24 | 0.41 | 0.46 |
| 5 | 33588146 | rs818726 | -0.538 | 1.168 | -1.562 | 0.233 | -0.022 | 0.021 | 0.021 | 0 | 0.41 | 0.25 | 0.54 | 0.69 | 0.49 | 0.39 |
| 5 | 33589017 | rs4242083 | -0.63 | 1.012 | -1.393 | 0.471 | 0 | 0.006 | 0.006 | 0 | 0.89 | 0.95 | 0.9 | 0.96 | 0.9 | 0.9 |
| 5 | 33591433 | rs16891371 | -0.168 | 1.232 | -1.345 | 1.061 | -0.045 | -0.083 | -0.083 | 0.275 | 0.2 | 0.06 | 0.27 | 0.38 | 0.26 | 0.24 |
| 5 | 33594566 | rs7730682 | -1.013 | 0.849 | -0.845 | 0.913 | 0.001 | 0 | 0 | 0 | 0.22 | 0.27 | 0.2 | 0.22 | 0.23 | 0.11 |
| 5 | 33594916 | rs12188767 | -0.013 | -0.2 | -1.4 | 1.067 | 0.058 | -0.031 | -0.031 | 0 | 0.35 | 0.14 | 0.33 | 0.57 | 0.24 | 0.37 |
| 5 | 33595803 | rs13153550 | 0.093 | -0.362 | -1.338 | 0.775 | 0.009 | -0.011 | -0.011 | 0 | 0.57 | 0.21 | 0.41 | 0.56 | 0.43 | 0.58 |
| 5 | 33598328 | rs10060471 | -0.356 | -0.122 | -0.702 | 0.783 | -0.01 | -0.007 | -0.007 | 0 | 0.66 | 0.33 | 0.48 | 0.56 | 0.55 | 0.63 |
| 5 | 33600365 | rs13171234 | -0.378 | -0.085 | -0.499 | 0.799 | 0.004 | 0.002 | 0.002 | 0.005 | 0.67 | 0.38 | 0.48 | 0.56 | 0.55 | 0.62 |
| 5 | 33603410 | rs9292501 | -0.524 | -0.01 | -0.731 | 0.832 | 0.041 | 0.03 | 0.03 | 0 | 0.69 | 0.58 | 0.49 | 0.56 | 0.55 | 0.63 |
| 5 | 33612335 | rs10941071 | 0.5 | -0.274 | -1.389 | 0.98 | -0.03 | -0.097 | -0.097 | 0 | 0.26 | 0.01 | 0.23 | 0.31 | 0.23 | 0.28 |
| 5 | 33615552 | rs12520271 | 0.276 | -0.275 | -1.275 | 0.917 | 0.004 | -0.018 | -0.018 | 0 | 0.31 | 0.06 | 0.23 | 0.31 | 0.26 | 0.32 |
| 5 | 33616311 | rs4078398 | -0.207 | -0.411 | -1.086 | 0.468 | -0.012 | 0.059 | 0.059 | 0 | 0.62 | 0.8 | 0.69 | 0.78 | 0.6 | 0.62 |
| 5 | 33617767 | rs4574547 | -0.587 | -1.252 | -0.765 | 1.26 | 0.023 | -0.107 | -0.107 | 0 | 0.15 | 0.01 | 0.16 | 0.35 | 0.09 | 0.32 |
| 5 | 33620887 | rs7703871 | -1.307 | -1.446 | -0.165 | 1.017 | -0.007 | 0.007 | 0.007 | 0 | 0.27 | 0.27 | 0.26 | 0.35 | 0.23 | 0.36 |
| 5 | 33621983 | rs9791008 | -0.286 | -1.291 | -0.044 | 1.309 | 0.047 | 0.166 | 0.166 | 0 | 0.07 | 0.22 | 0.11 | 0.27 | 0.07 | 0.16 |
| 5 | 33622158 | rs9791011 | -1.816 | -1.421 | 0.231 | 0.527 | -0.011 | -0.004 | -0.004 | 0 | 0.4 | 0.55 | 0.37 | 0.63 | 0.31 | 0.52 |
| 5 | 33623760 | rs7715934 | -0.743 | -1.387 | 0.24 | -0.024 | -0.022 | -0.085 | -0.085 | 0 | 0.16 | 0.28 | 0.1 | 0 | 0.14 | 0.04 |
| 5 | 33628171 | rs10068310 | -0.826 | -1.214 | -0.336 | 0.353 | 0.03 | 0.009 | 0.009 | 3.941 | 0.45 | 0.59 | 0.3 | 0.33 | 0.27 | 0.32 |
| 5 | 33631739 | rs4337859 | 0.502 | -1.061 | 0.296 | 0.325 | -0.031 | 0.063 | 0.063 | 0 | 0.84 | 0.54 | 0.87 | 0.75 | 0.9 | 0.69 |

| | | | | | | | | | | | | | | | | |
|---|---|---|---|---|---|---|---|---|---|---|---|---|---|---|---|---|
| 5 | 33636035 | rs13164825 | -0.184 | -1.022 | -0.378 | -0.23 | 0.002 | 0.013 | 0.013 | 0.522 | 0.18 | 0.12 | 0.09 | 0.02 | 0.19 | 0.07 |
| 5 | 33636326 | rs4380665 | 1.446 | -1.435 | -0.345 | -0.477 | 0.055 | 0.078 | 0.078 | 0 | 0.5 | 0.27 | 0.62 | 0.64 | 0.61 | 0.47 |
| 5 | 33637596 | rs4076945 | 0.857 | -1.277 | -0.317 | -0.362 | -0.016 | -0.002 | -0.002 | 0 | 0.68 | 0.37 | 0.7 | 0.66 | 0.8 | 0.54 |
| 5 | 33648317 | rs10042292 | 1.011 | -0.718 | -0.171 | -0.277 | 0.016 | 0.035 | 0.035 | 0 | 0.53 | 0.36 | 0.61 | 0.71 | 0.6 | 0.5 |
| 5 | 33663016 | rs12516835 | 1.031 | -0.644 | -0.525 | -0.368 | 0.043 | 0.101 | 0.101 | 0.09 | 0.49 | 0.28 | 0.68 | 0.73 | 0.58 | 0.54 |
| 5 | 33666554 | rs10472873 | -0.846 | -0.361 | -0.04 | -0.215 | -0.012 | -0.061 | -0.061 | 0 | 0.35 | 0.5 | 0.21 | 0.17 | 0.31 | 0.41 |
| 5 | 33667640 | rs12189209 | -1.035 | -0.545 | -0.604 | -0.164 | 0 | 0.075 | 0.075 | 0 | 0.39 | 0.15 | 0.25 | 0.27 | 0.37 | 0.48 |
| 5 | 33667722 | rs4343844 | 1.074 | -0.732 | -0.266 | -0.184 | 0.017 | 0.056 | 0.056 | 0 | 0.58 | 0.64 | 0.63 | 0.72 | 0.61 | 0.52 |
| 5 | 33667753 | rs9292503 | -0.928 | -0.618 | -0.073 | -0.004 | 0 | 0.015 | 0.015 | 0 | 0.37 | 0.35 | 0.31 | 0.26 | 0.35 | 0.43 |
| 5 | 33667805 | rs4321753 | 0.775 | -0.43 | 0.216 | 0.316 | -0.001 | -0.003 | -0.003 | 0 | 0.85 | 0.77 | 0.84 | 0.86 | 0.86 | 0.64 |
| 5 | 33678161 | rs4437379 | -0.972 | -1.122 | 0.62 | 1.142 | -0.067 | 0.125 | 0.125 | 0.529 | 0.25 | 0.54 | 0.31 | 0.44 | 0.13 | 0.5 |
| 5 | 33678250 | rs4437380 | -1.007 | -1.195 | 1.047 | 1.47 | -0.035 | 0.053 | 0.053 | 1.117 | 0.25 | 0.35 | 0.31 | 0.56 | 0.14 | 0.53 |
| 5 | 33682334 | rs6881360 | -0.067 | -1.258 | 0.946 | 1.855 | -0.019 | 0.067 | 0.067 | 0 | 0.2 | 0.41 | 0.15 | 0.26 | 0.2 | 0.1 |
| 5 | 33685863 | rs10062990 | -0.785 | -1.18 | 0.217 | 1.068 | 0.033 | 0.066 | 0.066 | 0.068 | 0.45 | 0.74 | 0.43 | 0.77 | 0.42 | 0.31 |
| 5 | 33687594 | rs6899224 | 0.358 | -1.479 | 0.831 | 1.39 | -0.001 | 0.011 | 0.011 | 0 | 0.21 | 0.25 | 0.19 | 0.44 | 0.21 | 0.14 |
| 5 | 33687640 | rs7719969 | 0.873 | -1.899 | 0.995 | 0.977 | 0.025 | 0.026 | 0.026 | 0 | 0.55 | 0.35 | 0.56 | 0.33 | 0.66 | 0.38 |
| 5 | 33694998 | rs2032877 | 0.394 | -1.807 | 0.677 | 0.926 | 0.03 | -0.052 | -0.052 | 0 | 0.2 | 0.05 | 0.23 | 0.23 | 0.13 | 0.48 |
| 5 | 33699676 | rs966857 | 1.318 | -1.129 | 0.774 | 0.582 | -0.012 | -0.02 | -0.02 | 0 | 0.34 | 0.25 | 0.31 | 0.01 | 0.37 | 0.21 |
| 5 | 33699777 | rs966855 | -0.923 | -1.449 | 0.999 | 0.653 | -0.008 | 0.013 | 0.013 | 0.075 | 0.84 | 0.85 | 0.81 | 0.78 | 0.88 | 0.58 |
| 5 | 33702830 | rs7700370 | 0.238 | -1.721 | 0.925 | 0.607 | -0.014 | 0.01 | 0.01 | 0 | 0.26 | 0.16 | 0.31 | 0.27 | 0.31 | 0.22 |
| 5 | 33703734 | rs1119629 | 1.991 | -1.772 | 0.932 | 0.549 | -0.098 | 0.174 | 0.174 | 0 | 0.32 | 0.08 | 0.31 | 0.01 | 0.39 | 0.2 |
| 5 | 33712059 | rs10078039 | 0.678 | -1.939 | 1.335 | 1.049 | 0.004 | -0.013 | -0.013 | 0.012 | 0.2 | 0.14 | 0.12 | 0.28 | 0.18 | 0.12 |
| 5 | 33712442 | rs12522957 | 1.013 | -1.922 | 1.412 | 0.791 | -0.003 | -0.038 | -0.038 | 0 | 0.56 | 0.55 | 0.57 | 0.14 | 0.68 | 0.35 |
| 5 | 33713661 | rs4400128 | 1.013 | -1.922 | 1.412 | 0.791 | -0.003 | -0.038 | -0.038 | 1.381 | 0.56 | 0.56 | 0.57 | 0.14 | 0.68 | 0.35 |
| 5 | 33715457 | rs1895440 | 0.631 | -1.909 | 1.298 | 0.815 | 0.006 | 0.006 | 0.006 | 0 | 0.6 | 0.39 | 0.62 | 0.28 | 0.7 | 0.43 |
| 5 | 33721765 | rs4866381 | 0.956 | -1.856 | 1.282 | 0.47 | 0.037 | 0.047 | 0.047 | 0 | 0.54 | 0.35 | 0.56 | 0.14 | 0.68 | 0.35 |
| 5 | 33725681 | rs4377712 | 0.741 | -1.633 | 1.069 | 0.14 | 0.013 | 0.019 | 0.019 | 0 | 0.53 | 0.57 | 0.55 | 0.16 | 0.63 | 0.3 |
| 5 | 33727008 | rs4866342 | 1.29 | -1.852 | 0.589 | 0.144 | -0.022 | 0.039 | 0.039 | 0 | 0.12 | 0.03 | 0.1 | 0.04 | 0.16 | 0.03 |
| 5 | 33731503 | rs11948306 | 1.251 | -1.866 | 0.752 | 0.105 | 0.069 | 0.201 | 0.201 | 0 | 0.5 | 0.23 | 0.54 | 0.16 | 0.63 | 0.3 |
| 5 | 33736909 | rs10941084 | 1.292 | -1.868 | 0.718 | 0.098 | 0.07 | 0.23 | 0.23 | 0 | 0.49 | 0.2 | 0.53 | 0.15 | 0.63 | 0.3 |
| 5 | 33739306 | rs9784618 | 0.083 | -1.776 | 0.529 | 0.587 | 0.022 | -0.063 | -0.063 | 0.76 | 0.19 | 0.13 | 0.17 | 0.54 | 0.1 | 0.29 |
| 5 | 33744247 | rs4866383 | 0.651 | -1.522 | 0.835 | 0.729 | -0.008 | 0.018 | 0.018 | 0 | 0.69 | 0.37 | 0.65 | 0.55 | 0.73 | 0.55 |
| 5 | 33744532 | rs4866384 | 0.409 | -1.482 | 0.83 | 0.748 | -0.001 | 0.002 | 0.002 | 0 | 0.72 | 0.67 | 0.67 | 0.55 | 0.73 | 0.55 |
| 5 | 33745589 | rs1966796 | -0.905 | -1.451 | 0.84 | 0.627 | -0.008 | 0.024 | 0.024 | 0 | 0.86 | 0.83 | 0.83 | 0.75 | 0.89 | 0.61 |
| 5 | 33749812 | rs6895637 | 0.371 | -1.402 | 0.8 | 0.646 | -0.015 | 0.031 | 0.031 | 0 | 0.72 | 0.55 | 0.72 | 0.71 | 0.74 | 0.58 |
| 5 | 33757205 | rs13172462 | 0.856 | -1.122 | 0.308 | 0.576 | -0.015 | 0.023 | 0.023 | 0 | 0.11 | 0.07 | 0.07 | 0.03 | 0.16 | 0.06 |
| 5 | 33758513 | rs7712324 | 0.123 | -1.759 | 0.375 | 0.321 | -0.049 | 0.101 | 0.101 | 0.269 | 0.26 | 0.42 | 0.2 | 0.55 | 0.13 | 0.31 |
| 5 | 33758562 | rs10521007 | 1.264 | -1.727 | 0.187 | 0.335 | -0.015 | 0.023 | 0.023 | 0 | 0.11 | 0.07 | 0.07 | 0.03 | 0.16 | 0.06 |
| 5 | 33759312 | rs9790860 | 0.557 | -1.763 | 0.185 | 0.373 | 0.024 | -0.075 | -0.075 | 0 | 0.22 | 0.13 | 0.17 | 0.55 | 0.13 | 0.31 |
| 5 | 33773500 | rs10070109 | 0.788 | -1.747 | 0.107 | 0.481 | 0.016 | -0.1 | -0.1 | 0.255 | 0.21 | 0.06 | 0.12 | 0.39 | 0.13 | 0.28 |
| 5 | 33774414 | rs4866385 | -0.58 | -2.104 | -0.831 | -0.157 | -0.004 | 0 | 0 | 0.139 | 0.65 | 0.67 | 0.6 | 0.59 | 0.77 | 0.53 |
| 5 | 33775653 | rs9968578 | 0.525 | -1.78 | -0.262 | 0.291 | 0.02 | -0.045 | -0.045 | 1.106 | 0.23 | 0.16 | 0.13 | 0.39 | 0.13 | 0.28 |
| 5 | 33792456 | rs7724739 | -0.382 | -1.721 | -0.269 | 0.19 | -0.012 | 0.007 | 0.007 | 0 | 0.37 | 0.53 | 0.27 | 0.57 | 0.29 | 0.37 |
| 5 | 33795056 | rs6451022 | -0.72 | -1.596 | -0.425 | 0.298 | 0.001 | 0.01 | 0.01 | 0 | 0.44 | 0.61 | 0.34 | 0.59 | 0.45 | 0.39 |
| 5 | 33795167 | rs7712316 | 0.955 | -1.26 | -0.53 | 0.703 | 0.026 | 0.052 | 0.052 | 0 | 0.08 | 0.15 | 0.14 | 0.2 | 0.17 | 0.05 |
| 5 | 33796518 | rs7717852 | -0.201 | -1.627 | -0.824 | 0.438 | -0.001 | -0.002 | -0.002 | 0 | 0.77 | 0.76 | 0.72 | 0.59 | 0.73 | 0.59 |
| 5 | 33799699 | rs2032878 | 0.811 | -1.627 | -1.234 | 0.618 | 0.028 | -0.099 | -0.099 | 0 | 0.22 | 0.05 | 0.11 | 0.34 | 0.11 | 0.28 |
| 5 | 33809758 | rs10071596 | -0.082 | -1.378 | -1.3 | 0.459 | 0.002 | -0.083 | -0.083 | 0 | 0.33 | 0.1 | 0.26 | 0.39 | 0.27 | 0.43 |
| 5 | 33811670 | rs10044427 | -0.988 | -1.596 | -1.096 | 0.396 | 0.014 | -0.009 | -0.009 | 0 | 0.83 | 0.89 | 0.85 | 0.79 | 0.89 | 0.72 |
| 5 | 33812113 | rs10055310 | -0.037 | -1.604 | -1.116 | 0.407 | 0.011 | -0.008 | -0.008 | 0 | 0.73 | 0.85 | 0.7 | 0.74 | 0.73 | 0.67 |
| 5 | 33812568 | rs1445911 | 1.194 | -1.395 | -0.593 | 0.264 | -0.002 | -0.006 | -0.006 | 0 | 0.4 | 0.43 | 0.42 | 0.21 | 0.45 | 0.25 |
| 5 | 33813994 | rs10941092 | 1.154 | -1.395 | -0.494 | 0.264 | 0.002 | -0.02 | -0.02 | 1.58 | 0.4 | 0.46 | 0.42 | 0.21 | 0.45 | 0.25 |
| 5 | 33817339 | rs1867718 | 0.409 | -1.23 | -0.618 | 0.313 | 0.007 | 0.007 | 0.007 | 0.005 | 0.48 | 0.51 | 0.56 | 0.4 | 0.62 | 0.31 |
| 5 | 33817901 | rs2548036 | 0.659 | -1.216 | -0.741 | 0.123 | 0.044 | -0.023 | -0.023 | 0 | 0.33 | 0.48 | 0.37 | 0.22 | 0.52 | 0.17 |
| 5 | 33820006 | rs12514368 | 1.076 | -1.33 | -0.858 | 0.091 | -0.033 | 0.006 | 0.006 | 0.907 | 0.25 | 0.08 | 0.34 | 0.23 | 0.27 | 0.18 |
| 5 | 33820448 | rs2548032 | 0.532 | -0.963 | -0.598 | 0.108 | 0.048 | -0.026 | -0.026 | 0 | 0.33 | 0.49 | 0.38 | 0.22 | 0.52 | 0.17 |
| 5 | 33823475 | rs11952033 | -0.106 | -0.959 | -0.624 | 0.272 | 0.008 | 0.002 | 0.002 | 0 | 0.49 | 0.5 | 0.52 | 0.42 | 0.62 | 0.43 |
| 5 | 33828203 | rs977842 | -0.318 | -0.966 | -0.444 | 0.066 | 0 | -0.008 | -0.008 | 0 | 0.57 | 0.53 | 0.63 | 0.78 | 0.55 | 0.62 |
| 5 | 33831305 | rs1363880 | 0.442 | -1.396 | -0.24 | 0.7 | -0.003 | -0.001 | -0.001 | 0 | 0.53 | 0.49 | 0.73 | 0.46 | 0.77 | 0.34 |
| 5 | 33831355 | rs961252 | 0.134 | -1.541 | -0.355 | 1.37 | 0.009 | -0.01 | -0.01 | 0 | 0.76 | 0.82 | 0.81 | 0.65 | 0.87 | 0.62 |
| 5 | 33831673 | rs2548024 | -0.447 | -0.725 | 0.039 | 1.385 | 0.072 | 0.109 | 0.109 | 0 | 0.92 | 0.83 | 0.84 | 0.84 | 0.82 | 0.95 |
| 5 | 33832015 | rs2591724 | -0.756 | -2.044 | -0.057 | 1.376 | -0.035 | 0.014 | 0.014 | 0.004 | 0.33 | 0.53 | 0.23 | 0.63 | 0.19 | 0.53 |
| 5 | 33835353 | rs1374017 | -0.89 | -3.171 | 0.305 | 1.825 | -0.06 | 0.05 | 0.05 | 1.479 | 0.23 | 0.36 | 0.2 | 0.38 | 0.05 | 0.2 |
| 5 | 33839309 | rs1465436 | 0.063 | -3.139 | 0.298 | 1.961 | 0.02 | -0.032 | -0.032 | 0 | 0.09 | 0.06 | 0.11 | 0.17 | 0.02 | 0.12 |
| 5 | 33842268 | rs2548020 | -0.592 | -3.821 | 0.415 | 1.585 | 0.022 | 0.018 | 0.018 | 0 | 0.56 | 0.7 | 0.58 | 0.64 | 0.38 | 0.47 |
| 5 | 33843258 | rs1867720 | 1.115 | -3.791 | 0.573 | 1.517 | -0.059 | 0.051 | 0.051 | 2.897 | 0.11 | 0.23 | 0.08 | 0.21 | 0.01 | 0.05 |
| 5 | 33847033 | rs1374018 | 1.328 | -3.77 | 0.513 | 1.527 | -0.007 | -0.004 | -0.004 | 0 | 0.1 | 0.07 | 0.06 | 0.21 | 0.01 | 0.05 |
| 5 | 33853082 | rs13187441 | 0.837 | -3.839 | 0.538 | 1.493 | -0.005 | -0.015 | -0.015 | 0 | 0.36 | 0.39 | 0.39 | 0.26 | 0.34 | 0.28 |
| 5 | 33854740 | rs13155761 | 1.198 | -3.889 | 0.416 | 1.504 | -0.007 | 0.098 | 0.098 | 0 | 0.31 | 0.07 | 0.34 | 0.09 | 0.3 | 0.16 |

| | | | | | | | | | | | | | | | | |
|---|---|---|---|---|---|---|---|---|---|---|---|---|---|---|---|---|
| 5 | 33855658 | rs2961909 | 1.401 | -3.855 | 0.439 | 1.551 | 0.012 | -0.016 | -0.016 | 0 | 0.1 | 0.05 | 0.06 | 0.21 | 0.01 | 0.06 |
| 5 | 33878205 | rs13160471 | 1.131 | -3.702 | 0.385 | 1.304 | -0.006 | 0.022 | 0.022 | 0 | 0.31 | 0.07 | 0.39 | 0.22 | 0.3 | 0.17 |
| 5 | 33879792 | rs1364044 | 0.701 | -3.711 | 0.503 | 1.325 | -0.007 | -0.002 | -0.002 | 1.267 | 0.36 | 0.4 | 0.44 | 0.38 | 0.34 | 0.3 |
| 5 | 33883233 | rs4049255 | -0.926 | -3.316 | 0.502 | 1.412 | -0.016 | 0.055 | 0.055 | 0 | 0.64 | 0.82 | 0.64 | 0.82 | 0.42 | 0.59 |
| 5 | 33884127 | rs6451031 | -0.05 | -3.22 | 0.467 | 1.407 | -0.004 | 0 | 0 | 0 | 0.53 | 0.45 | 0.52 | 0.49 | 0.34 | 0.38 |
| 5 | 33895326 | rs10521010 | -0.633 | -2.889 | 0.555 | 1.304 | 0.016 | 0.061 | 0.061 | 0 | 0.08 | 0.3 | 0.06 | 0.15 | 0.06 | 0.14 |
| 5 | 33897522 | rs7446464 | 0.464 | -3.658 | 0.053 | 1.279 | 0.018 | -0.019 | -0.019 | 0 | 0.11 | 0.06 | 0.1 | 0.18 | 0.01 | 0.16 |
| 5 | 33909843 | rs1549283 | 1.115 | -4.84 | -0.557 | 1.106 | 0.146 | -0.079 | -0.079 | 0.802 | 0.25 | 0.03 | 0.26 | 0.26 | 0.05 | 0.16 |
| 5 | 33913600 | rs6862507 | -1.919 | -5.029 | -0.115 | 1.115 | -0.016 | 0.059 | 0.059 | 0.01 | 0.93 | 0.8 | 0.96 | 0.89 | 0.99 | 0.96 |
| 5 | 33922558 | rs6883608 | 0.518 | -5.24 | -0.532 | 1.013 | 0.249 | 0.024 | 0.024 | 0 | 0.49 | 0.19 | 0.54 | 0.43 | 0.86 | 0.5 |
| 5 | 33923606 | rs11955226 | 0.495 | -4.911 | -0.274 | 1.049 | -0.033 | 0.038 | 0.038 | 3.557 | 0.11 | 0.21 | 0.11 | 0.17 | 0.01 | 0.16 |
| 5 | 33928902 | rs7713279 | -0.556 | -5.228 | -1.088 | 0.85 | 0.158 | 0.007 | 0.007 | 0 | 0.56 | 0.79 | 0.5 | 0.57 | 0.17 | 0.52 |
| 5 | 33930441 | rs7737188 | -0.729 | -4.507 | -0.42 | 0.54 | -0.054 | 0.023 | 0.023 | 0 | 0.2 | 0.62 | 0.2 | 0.25 | 0.08 | 0.29 |
| 5 | 33934007 | rs7729545 | -1.01 | -4.494 | -0.919 | 0.508 | 0.01 | -0.007 | -0.007 | 0 | 0.27 | 0.75 | 0.23 | 0.34 | 0.09 | 0.31 |
| 5 | 33938447 | rs9292519 | -0.922 | -3.838 | -1.163 | 0.157 | -0.068 | 0.078 | 0.078 | 0 | 0.75 | 0.92 | 0.66 | 0.93 | 0.45 | 0.69 |
| 5 | 33941356 | rs35401 | 0.379 | -4.014 | -1.071 | 0.349 | 0.065 | 0.046 | 0.046 | 0 | 0.53 | 0.26 | 0.48 | 0.66 | 0.37 | 0.51 |
| 5 | 33942805 | rs35402 | 0.374 | -3.958 | -0.709 | 0.49 | -0.02 | 0.015 | 0.015 | 4.457 | 0.11 | 0.13 | 0.11 | 0.21 | 0.01 | 0.15 |
| 5 | 33943592 | rs35403 | 0.053 | -3.099 | -0.807 | 0.379 | -0.005 | 0.041 | 0.041 | 0 | 0.55 | 0.25 | 0.53 | 0.67 | 0.49 | 0.52 |
| 5 | 33944217 | rs10461928 | 0.228 | -3.259 | -0.717 | 0.332 | 0.018 | 0.009 | 0.009 | 0.538 | 0.66 | 0.48 | 0.74 | 0.63 | 0.56 | 0.76 |
| 5 | 33945758 | rs35405 | 0.447 | -3.953 | -0.997 | -0.408 | -0.036 | 0.031 | 0.031 | 0.198 | 0.25 | 0.17 | 0.23 | 0.1 | 0.46 | 0.19 |
| 5 | 33946143 | rs35406 | 0.939 | -6.623 | -0.838 | 0.454 | 0.123 | -0.146 | -0.146 | 0 | 0.21 | 0.03 | 0.32 | 0.45 | 0.02 | 0.21 |
| 5 | 33946555 | rs7380538 | -0.705 | -7.052 | -0.626 | 0.107 | -0.013 | 0.01 | 0.01 | 0 | 0.75 | 0.8 | 0.9 | 0.82 | 0.99 | 0.58 |
| 5 | 33946571 | rs35407 | 0.563 | -6.486 | -0.705 | 0.027 | 0.037 | 0.057 | 0.057 | 0 | 0.36 | 0.29 | 0.51 | 0.12 | 0.96 | 0.22 |
| 5 | 33951551 | rs2278007 | 0.2 | -5.807 | -0.492 | 0.508 | -0.02 | 0.002 | 0.002 | 0 | 0.15 | 0.15 | 0.05 | 0.27 | 0.01 | 0.15 |
| 5 | 33954511 | rs2287949 | -0.537 | -6.486 | -0.628 | 0.37 | 0.066 | 0.031 | 0.031 | 0 | 0.75 | 0.88 | 0.9 | 0.82 | 0.99 | 0.58 |
| 5 | 33955673 | rs35391 | -0.092 | -6.789 | -0.118 | 0.231 | -0.046 | 0.012 | 0.012 | 0.641 | 0.53 | 0.58 | 0.56 | 0.39 | 0.97 | 0.37 |
| 5 | 33958959 | rs28777 | 0.665 | -6.786 | -0.469 | 0.011 | 0.093 | 0.098 | 0.098 | 0 | 0.35 | 0.2 | 0.49 | 0.12 | 0.96 | 0.22 |
| 5 | 33962770 | rs28117 | -0.01 | -6.945 | -0.246 | 0.243 | -0.076 | 0.005 | 0.005 | 0.304 | 0.49 | 0.52 | 0.56 | 0.39 | 0.96 | 0.38 |
| 5 | 33963870 | rs26722 | 1.045 | -7.105 | -1.133 | 0.419 | 0.084 | -0.093 | -0.093 | 0 | 0.22 | 0.04 | 0.31 | 0.39 | 0.02 | 0.21 |
| 5 | 33964210 | rs183671 | 0.59 | -6.615 | -1.05 | -0.101 | 0.09 | 0.097 | 0.097 | 0 | 0.34 | 0.21 | 0.49 | 0.09 | 0.95 | 0.23 |
| 5 | 33964352 | rs6451049 | 0.014 | -5.047 | -0.855 | 0.445 | 0.075 | 0.023 | 0.023 | 0.231 | 0.73 | 0.87 | 0.89 | 0.78 | 0.99 | 0.58 |
| 5 | 33968322 | rs6451050 | -0.229 | -4.048 | -0.426 | 0.72 | -0.018 | 0 | 0 | 0 | 0.92 | 0.93 | 0.94 | 0.91 | 1 | 0.96 |
| 5 | 33971037 | rs3822466 | -0.928 | -4.331 | 0.513 | 1.11 | -0.05 | 0.126 | 0.126 | 0.306 | 0.12 | 0.43 | 0.08 | 0.19 | 0.04 | 0.06 |
| 5 | 33983044 | rs250412 | -0.303 | -3.45 | -0.016 | 1.628 | -0.007 | -0.011 | -0.011 | 0.047 | 0.29 | 0.25 | 0.43 | 0.45 | 0.59 | 0.2 |
| 5 | 33988054 | rs840380 | -0.033 | -3.485 | -0.467 | 2.122 | 0.018 | -0.051 | -0.051 | 0 | 0.1 | 0 | 0.13 | 0.15 | 0.02 | 0.1 |
| 5 | 33988581 | rs12659370 | -0.233 | -3.117 | -0.034 | 1.604 | 0.022 | -0.022 | -0.022 | 0 | 0.09 | 0.04 | 0.05 | 0.19 | 0.04 | 0.06 |
| 5 | 33990623 | rs250414 | -0.178 | -2.414 | 0.117 | 1.73 | -0.024 | -0.02 | -0.02 | 0.047 | 0.19 | 0.18 | 0.33 | 0.35 | 0.54 | 0.09 |
| 5 | 33991136 | rs3822464 | -0.154 | -2.657 | 0.321 | 1.727 | -0.004 | -0.003 | -0.003 | 1.491 | 0.2 | 0.24 | 0.45 | 0.37 | 0.58 | 0.1 |
| 5 | 33991399 | rs2652130 | -0.236 | -4.129 | 0.26 | 1.089 | -0.015 | 0.003 | 0.003 | 1.685 | 0.28 | 0.27 | 0.1 | 0.31 | 0.13 | 0.36 |
| 5 | 33998768 | rs34677 | 0.121 | -3.865 | 0.158 | 1.182 | 0.022 | 0.031 | 0.031 | 3 | 0.23 | 0.06 | 0.07 | 0.17 | 0.14 | 0.14 |
| 5 | 33998883 | rs2287939 | -1.421 | -3.035 | 0.286 | 0.699 | 0.06 | 0.11 | 0.11 | 0 | 0.6 | 0.82 | 0.71 | 0.85 | 0.69 | 0.46 |
| 5 | 34001444 | rs34682 | 0.005 | -3.122 | 0.689 | 0.97 | -0.021 | 0.241 | 0.241 | 0.049 | 0.06 | 0.18 | 0.16 | 0.33 | 0.01 | 0.13 |
| 5 | 34003361 | rs40507 | -0.542 | -3.139 | 0.856 | 0.672 | -0.049 | -0.061 | -0.061 | 0 | 0.36 | 0.47 | 0.14 | 0.17 | 0.18 | 0.23 |
| 5 | 34008100 | rs3195676 | 1.029 | -2.902 | 0.916 | 0.708 | -0.045 | -0.035 | -0.035 | 0 | 0.19 | 0.14 | 0.41 | 0.35 | 0.51 | 0.1 |
| 5 | 34011356 | rs190418 | -0.282 | -3.067 | 1.083 | 0.614 | 0.043 | 0.051 | 0.051 | 5.9 | 0.33 | 0.2 | 0.12 | 0.17 | 0.18 | 0.23 |
| 5 | 34015174 | rs253202 | -1.093 | -2.983 | 0.867 | 0.468 | 0.033 | -0.014 | -0.014 | 0 | 0.57 | 0.66 | 0.55 | 0.53 | 0.69 | 0.33 |
| 5 | 34020277 | rs840386 | 1.097 | -2.829 | 0.771 | 0.336 | 0.011 | -0.002 | -0.002 | 0 | 0.47 | 0.38 | 0.49 | 0.48 | 0.37 | 0.68 |
| 5 | 34024385 | rs299608 | 0.605 | -3.399 | 0.441 | 0.112 | 0.026 | 0 | 0 | 0 | 0.51 | 0.55 | 0.43 | 0.48 | 0.36 | 0.7 |
| 5 | 34024944 | rs299605 | 0.593 | -3.398 | 0.442 | 0.113 | 0.025 | 0 | 0 | 0 | 0.51 | 0.55 | 0.43 | 0.48 | 0.36 | 0.7 |
| 5 | 34026870 | rs299601 | 0.08 | -3.145 | 0.424 | 0.437 | 0 | 0.094 | 0.094 | 0 | 0.06 | 0.07 | 0.15 | 0.37 | 0.02 | 0.13 |
| 5 | 34032014 | rs299596 | -0.037 | -3.539 | 0.211 | 0.167 | -0.001 | 0.001 | 0.001 | 0 | 0.68 | 0.67 | 0.44 | 0.5 | 0.38 | 0.76 |
| 5 | 34036236 | rs10074541 | 0.169 | -3.178 | 0.512 | 0.386 | 0.073 | 0.075 | 0.075 | 0 | 0.27 | 0.47 | 0.53 | 0.44 | 0.53 | 0.19 |
| 5 | 34038854 | rs301506 | 0.272 | -3.184 | 0.356 | 0.722 | -0.007 | -0.007 | -0.007 | 0 | 0.26 | 0.25 | 0.51 | 0.59 | 0.53 | 0.19 |
| 5 | 34041237 | rs6888550 | -0.029 | -3.222 | 0.361 | 0.695 | 0.038 | 0.039 | 0.039 | 0 | 0.28 | 0.52 | 0.53 | 0.44 | 0.53 | 0.18 |
| 5 | 34041339 | rs6889019 | -2.014 | -3.975 | -0.006 | 0.368 | -0.003 | -0.018 | -0.018 | 0 | 0.63 | 0.6 | 0.75 | 0.86 | 0.69 | 0.5 |
| 5 | 34042097 | rs299615 | -0.794 | -4.054 | 0.069 | 0.301 | 0.08 | 0.02 | 0.02 | 0 | 0.33 | 0.17 | 0.22 | 0.25 | 0.16 | 0.28 |
| 5 | 34042757 | rs16892218 | -0.574 | -4.003 | 0.117 | -0.047 | 0.044 | -0.04 | -0.04 | 0 | 0.3 | 0.24 | 0.22 | 0.4 | 0.15 | 0.24 |
| 5 | 34043649 | rs890948 | -1.645 | -3.828 | 0.076 | -1.111 | 0.02 | 0.03 | 0.03 | 0 | 0.65 | 0.72 | 0.75 | 0.89 | 0.7 | 0.51 |
| 5 | 34044232 | rs890951 | -0.988 | -3.613 | 0.23 | -0.997 | -0.006 | -0.031 | -0.031 | 0 | 0.57 | 0.49 | 0.73 | 0.88 | 0.69 | 0.42 |
| 5 | 34052480 | rs9292524 | -0.768 | -2.826 | -0.001 | -0.709 | 0.045 | 0.093 | 0.093 | 0 | 0.46 | 0.7 | 0.72 | 0.85 | 0.65 | 0.37 |
| 5 | 34053621 | rs7379110 | -0.436 | -2.433 | -0.511 | -0.503 | 0.009 | 0.034 | 0.034 | 0 | 0.39 | 0.3 | 0.68 | 0.85 | 0.64 | 0.32 |
| 5 | 34058555 | rs299622 | 0.335 | -0.574 | -0.171 | 0.14 | -0.005 | -0.002 | -0.002 | 0.963 | 0.64 | 0.75 | 0.6 | 0.39 | 0.62 | 0.64 |
| 5 | 34119847 | rs297619 | 0.926 | -0.54 | -0.264 | 0.2 | 0.019 | -0.002 | -0.002 | 0 | 0.56 | 0.21 | 0.55 | 0.43 | 0.6 | 0.58 |
| 5 | 34123933 | rs297622 | 0.586 | -0.327 | -0.017 | 0.216 | 0.002 | 0.033 | 0.033 | 0 | 0.6 | 0.3 | 0.56 | 0.34 | 0.6 | 0.55 |
| 5 | 34266171 | rs11746806 | 0.774 | -0.475 | 0.42 | 0.639 | -0.05 | 0.058 | 0.058 | 2.721 | 0.9 | 0.37 | 0.93 | 0.88 | 0.96 | 0.94 |
| 5 | 34300233 | rs4703462 | -0.319 | -0.18 | -0.406 | 0.483 | 0.007 | -0.031 | -0.031 | 0 | 0.42 | 0.22 | 0.42 | 0.48 | 0.4 | 0.39 |
| 5 | 34333402 | rs10040978 | 0.448 | -0.055 | -0.504 | 0.486 | -0.006 | 0.052 | 0.052 | 0 | 0.49 | 0.2 | 0.46 | 0.4 | 0.47 | 0.54 |
| 5 | 34383192 | rs7379497 | 0.23 | 0.085 | -0.305 | 0.64 | -0.016 | 0.016 | 0.016 | 0.323 | 0.57 | 0.19 | 0.5 | 0.4 | 0.53 | 0.57 |
| 5 | 34402152 | rs17585602 | -0.132 | 0.101 | -1.239 | 0.735 | 0.054 | -0.027 | -0.027 | 1.213 | 0.48 | 0.13 | 0.46 | 0.44 | 0.59 | 0.4 |

| | | | | | | | | | | | | | | | | |
|---|---|---|---|---|---|---|---|---|---|---|---|---|---|---|---|---|
| 5 | 34410175 | rs7715761 | -0.497 | -0.121 | 0.336 | 0.548 | 0.03 | -0.019 | -0.019 | 0 | 0.08 | 0.35 | 0.05 | 0 | 0.04 | 0.06 |
| 5 | 34410258 | rs1992351 | 1.596 | -0.463 | 1.12 | 0.826 | -0.042 | 0.015 | 0.015 | 0.013 | 0.86 | 0.27 | 0.81 | 0.85 | 0.94 | 0.89 |
| 5 | 34428293 | rs4571447 | 0.785 | -0.394 | 1.174 | 0.875 | 0.021 | 0.195 | 0.195 | 0.707 | 0.95 | 0.67 | 0.86 | 0.85 | 0.98 | 0.95 |
| 5 | 34454322 | rs11743037 | 0.829 | 0.113 | 0.686 | 1.008 | -0.055 | -0.038 | -0.038 | 3.188 | 0.57 | 0.18 | 0.53 | 0.54 | 0.53 | 0.37 |
| 5 | 34455165 | rs6860110 | -0.113 | 0.101 | 1.15 | 1.062 | 0.017 | 0.021 | 0.021 | 1.15 | 0.33 | 0.34 | 0.39 | 0.45 | 0.41 | 0.55 |
| 5 | 34486536 | rs16902290 | -0.231 | -0.145 | 1.433 | 1.094 | -0.012 | -0.035 | -0.035 | 0 | 0.08 | 0.29 | 0.06 | 0 | 0.02 | 0.06 |
| 5 | 34489548 | rs10941189 | 0.19 | 0.202 | 0.703 | 1.106 | -0.043 | -0.027 | -0.027 | 0 | 0.3 | 0.11 | 0.26 | 0.29 | 0.36 | 0.52 |
| 5 | 34491553 | rs6867161 | 0.829 | -0.104 | 1.235 | 1.003 | -0.055 | 0.052 | 0.052 | 0 | 0.87 | 0.33 | 0.8 | 0.82 | 0.96 | 0.88 |
| 5 | 34491949 | rs4703374 | 1.001 | -0.361 | 0.701 | 1.182 | 0.049 | 0.07 | 0.07 | 0.024 | 0.53 | 0.23 | 0.53 | 0.39 | 0.63 | 0.36 |
| 5 | 34493934 | rs10056815 | -0.275 | -0.204 | 0.537 | 1.217 | -0.01 | -0.078 | -0.078 | 0 | 0.34 | 0.1 | 0.27 | 0.44 | 0.33 | 0.53 |
| 5 | 34496464 | rs4703439 | 0.356 | -0.305 | 0.808 | 0.915 | 0 | 0.03 | 0.03 | 0 | 0.63 | 0.49 | 0.68 | 0.58 | 0.63 | 0.47 |
| 5 | 34497252 | rs920479 | -0.991 | -0.566 | 0.712 | 0.649 | -0.002 | -0.003 | -0.003 | 0 | 0.54 | 0.5 | 0.48 | 0.51 | 0.57 | 0.65 |
| 5 | 34503352 | rs1371810 | -0.601 | -0.074 | -0.083 | 0.279 | -0.065 | -0.053 | -0.053 | 0.804 | 0.35 | 0.18 | 0.3 | 0.36 | 0.49 | 0.51 |
| 5 | 34509825 | rs10066990 | -0.761 | 0.56 | -0.005 | 0.254 | -0.118 | -0.064 | -0.064 | 0 | 0.25 | 0.08 | 0.24 | 0.27 | 0.39 | 0.35 |
| 5 | 34511480 | rs16902493 | 0.281 | 0.665 | 0.572 | 0.193 | -0.018 | -0.038 | -0.038 | 0 | 0.07 | 0.21 | 0.07 | 0 | 0.02 | 0.06 |
| 5 | 34525931 | rs10075914 | 0.476 | 0.763 | 0.576 | 0.203 | 0.057 | -0.105 | -0.105 | 0.064 | 0.77 | 0.51 | 0.81 | 0.91 | 0.71 | 0.86 |
| 5 | 34528915 | rs4296778 | 0.846 | 0.73 | 0.448 | 0.062 | 0.018 | -0.071 | -0.071 | 0 | 0.73 | 0.38 | 0.76 | 0.91 | 0.66 | 0.85 |
| 5 | 34530891 | rs10058903 | -0.967 | 0.619 | 0.31 | 0.193 | -0.015 | 0.072 | 0.072 | 0.159 | 0.2 | 0.37 | 0.31 | 0.35 | 0.2 | 0.31 |
| 5 | 34533481 | rs903845 | -0.282 | 0.724 | -0.112 | 0.301 | -0.011 | 0.088 | 0.088 | 0 | 0.16 | 0.24 | 0.26 | 0.35 | 0.16 | 0.3 |
| 5 | 34538056 | rs13187551 | 0.884 | 0.733 | -1.64 | 0.16 | -0.089 | 0.058 | 0.058 | 0 | 0.3 | 0.02 | 0.27 | 0.21 | 0.31 | 0.23 |
| 5 | 34538606 | rs9790988 | 1.061 | 1.433 | -0.459 | 0.298 | -0.002 | -0.001 | -0.001 | 0 | 0.65 | 0.38 | 0.57 | 0.65 | 0.57 | 0.58 |
| 5 | 34541312 | rs12153461 | -0.517 | 0.942 | -0.572 | 0.593 | -0.017 | -0.032 | -0.032 | 0 | 0.11 | 0.07 | 0.14 | 0.15 | 0.17 | 0.25 |
| 5 | 34546533 | rs11738452 | 1.274 | 1.266 | -0.475 | 0.517 | -0.024 | -0.021 | -0.021 | 0 | 0.69 | 0.16 | 0.66 | 0.63 | 0.65 | 0.54 |
| 5 | 34547227 | rs11949228 | 0.135 | 1.387 | -0.016 | 0.492 | 0.05 | -0.025 | -0.025 | 0.149 | 0.06 | 0.22 | 0.09 | 0 | 0.11 | 0.04 |
| 5 | 34547539 | rs1496350 | 0.906 | 1.301 | 0.192 | 0.536 | 0.004 | 0.024 | 0.024 | 0.006 | 0.76 | 0.44 | 0.76 | 0.69 | 0.76 | 0.58 |
| 5 | 34549012 | rs2173009 | 0.077 | 1.28 | 0.256 | 0.512 | -0.019 | -0.029 | -0.029 | 0 | 0.85 | 0.9 | 0.82 | 0.69 | 0.79 | 0.66 |
| 5 | 34551865 | rs6870997 | 1.055 | 1.266 | 0.537 | 0.498 | 0.013 | 0.058 | 0.058 | 0 | 0.76 | 0.52 | 0.76 | 0.65 | 0.76 | 0.57 |
| 5 | 34552389 | rs2388731 | 0.073 | 1.325 | -0.001 | -0.229 | -0.019 | -0.014 | -0.014 | 0 | 0.85 | 0.9 | 0.82 | 0.78 | 0.79 | 0.66 |
| 5 | 34559175 | rs13356779 | -1.248 | 0.875 | -0.463 | 0.015 | -0.006 | -0.009 | -0.009 | 0 | 0.17 | 0.75 | 0.17 | 0.11 | 0.14 | 0.12 |
| 5 | 34564835 | rs4703457 | 0.377 | 0.623 | -0.701 | -0.12 | 0.022 | 0.059 | 0.059 | 0 | 0.51 | 0.34 | 0.62 | 0.63 | 0.54 | 0.49 |
| 5 | 34567310 | rs7736168 | -1.179 | 0.252 | 0.41 | -0.581 | -0.014 | -0.112 | -0.112 | 0 | 0.16 | 0.61 | 0.11 | 0 | 0.11 | 0.12 |
| 5 | 34567357 | rs7735808 | -1.688 | 0.219 | 0.341 | -0.933 | -0.003 | -0.063 | -0.063 | 3.096 | 0.23 | 0.74 | 0.23 | 0 | 0.19 | 0.18 |
| 5 | 34568687 | rs9292552 | -0.358 | -0.152 | 0.755 | -0.788 | 0.007 | -0.01 | -0.01 | 1.82 | 0.18 | 0.27 | 0.25 | 0.16 | 0.21 | 0.41 |
| 5 | 34575747 | rs9687397 | -0.508 | -0.499 | 1.009 | -0.314 | -0.035 | 0.103 | 0.103 | 0 | 0.15 | 0.32 | 0.11 | 0.34 | 0.07 | 0.13 |
| 5 | 34576037 | rs10063005 | -1.148 | -0.638 | 1.198 | -0.431 | -0.022 | 0.045 | 0.045 | 0 | 0.33 | 0.5 | 0.36 | 0.5 | 0.25 | 0.55 |
| 5 | 34578602 | rs6859986 | -0.395 | -0.411 | 1.381 | -0.872 | 0.008 | -0.062 | -0.062 | 0 | 0.17 | 0.25 | 0.08 | 0 | 0.18 | 0.11 |
| 5 | 34580651 | rs2029901 | -0.736 | -0.453 | 1.34 | -0.187 | -0.026 | 0.052 | 0.052 | 0 | 0.14 | 0.27 | 0.09 | 0.25 | 0.07 | 0.12 |
| 5 | 34583138 | rs16903125 | -0.141 | -0.507 | 0.181 | -0.389 | 0.004 | 0.005 | 0.005 | 0 | 0.13 | 0.09 | 0.08 | 0 | 0.11 | 0.05 |
| 5 | 34583665 | rs16903138 | -0.925 | -0.165 | 0.372 | 0.114 | -0.032 | 0.053 | 0.053 | 0 | 0.15 | 0.29 | 0.09 | 0.25 | 0.07 | 0.12 |
| 5 | 34587557 | rs6881093 | -1.244 | -0.048 | 0.042 | -0.204 | -0.002 | -0.002 | -0.002 | 0 | 0.28 | 0.69 | 0.21 | 0.23 | 0.24 | 0.18 |
| 5 | 34587609 | rs5019843 | -0.944 | -0.358 | 0.835 | -0.154 | -0.04 | 0.006 | 0.006 | 0.058 | 0.16 | 0.6 | 0.13 | 0.17 | 0.07 | 0.15 |
| 5 | 34587844 | rs6885856 | -1.47 | -0.188 | 0.618 | -0.392 | -0.003 | -0.009 | -0.009 | 1.995 | 0.37 | 0.75 | 0.43 | 0.43 | 0.36 | 0.6 |
| 5 | 34588975 | rs10074961 | -1.157 | -0.766 | 0.945 | -0.872 | -0.038 | -0.016 | -0.016 | 0 | 0.28 | 0.37 | 0.18 | 0.23 | 0.18 | 0.21 |
| 5 | 34589692 | rs11742309 | 0.122 | -0.755 | 0.764 | -0.79 | 0.016 | -0.024 | -0.024 | 0.241 | 0.15 | 0.07 | 0.14 | 0.23 | 0.12 | 0.15 |
| 5 | 34591473 | rs7704404 | -1.305 | -1.487 | 0.637 | -1.188 | 0.029 | 0.015 | 0.015 | 0 | 0.33 | 0.75 | 0.21 | 0.27 | 0.16 | 0.24 |
| 5 | 34593823 | rs749943 | 0.252 | -1.384 | 0.481 | -1.125 | 0.022 | -0.035 | -0.035 | 0.1 | 0.15 | 0.05 | 0.12 | 0.23 | 0.09 | 0.14 |
| 5 | 34596516 | rs10075334 | -0.234 | -1.376 | 0.554 | -1.355 | 0.013 | -0.036 | -0.036 | 0 | 0.24 | 0.3 | 0.24 | 0.15 | 0.25 | 0.43 |
| 5 | 34600047 | rs16903289 | -0.506 | -1.33 | 1.072 | -1.144 | -0.072 | -0.096 | -0.096 | 0 | 0.17 | 0.44 | 0.06 | 0.04 | 0.04 | 0.05 |
| 5 | 34602269 | rs1445889 | -1.259 | -0.984 | 0.671 | -1.197 | -0.018 | -0.017 | -0.017 | 0.274 | 0.39 | 0.58 | 0.29 | 0.26 | 0.28 | 0.24 |
| 5 | 34602657 | rs256297 | -1.212 | -0.405 | 0.75 | -0.969 | 0.004 | 0.005 | 0.005 | 0 | 0.41 | 0.66 | 0.35 | 0.36 | 0.33 | 0.27 |
| 5 | 34603893 | rs10079047 | 0.48 | 0.341 | 0.855 | -1.43 | 0.006 | -0.05 | -0.05 | 0 | 0.18 | 0.26 | 0.25 | 0.06 | 0.2 | 0.14 |
| 5 | 34604830 | rs715718 | 0.403 | 0.322 | 1.243 | -1.439 | 0.015 | -0.078 | -0.078 | 0 | 0.18 | 0.35 | 0.25 | 0.06 | 0.2 | 0.14 |
| 5 | 34604915 | rs1072516 | -0.856 | 0.097 | 1.388 | -1.411 | -0.006 | -0.016 | -0.016 | 0.05 | 0.22 | 0.72 | 0.14 | 0.13 | 0.15 | 0.1 |
| 5 | 34605325 | rs6451151 | -1.57 | 0.339 | 1.062 | -1.204 | -0.04 | 0.103 | 0.103 | 1.019 | 0.45 | 0.83 | 0.47 | 0.33 | 0.48 | 0.54 |
| 5 | 34609282 | rs9292557 | -0.781 | 0.363 | 1.194 | -1.198 | -0.003 | 0.013 | 0.013 | 3.042 | 0.34 | 0.32 | 0.31 | 0.14 | 0.34 | 0.23 |
| 5 | 34610719 | rs728940 | -0.781 | 0.67 | 0.674 | -0.954 | 0.012 | -0.005 | -0.005 | 0 | 0.26 | 0.32 | 0.21 | 0.27 | 0.28 | 0.42 |
| 5 | 34611483 | rs746456 | 0.28 | 0.708 | 0.896 | -0.645 | 0.048 | 0.004 | 0.004 | 0 | 0.09 | 0.22 | 0.14 | 0.1 | 0.14 | 0.31 |
| 5 | 34612522 | rs256302 | 0.843 | 0.554 | 0.616 | -0.787 | 0.002 | 0.029 | 0.029 | 0.014 | 0.61 | 0.36 | 0.59 | 0.76 | 0.64 | 0.51 |
| 5 | 34621851 | rs962435 | 0.502 | 0.944 | 0.671 | -0.165 | 0.025 | 0.044 | 0.044 | 0 | 0.73 | 0.55 | 0.68 | 0.61 | 0.69 | 0.54 |
| 5 | 34622835 | rs256323 | 1.042 | 1.411 | 0.573 | -0.409 | 0.001 | -0.002 | -0.002 | 0.051 | 0.6 | 0.42 | 0.67 | 0.56 | 0.59 | 0.43 |
| 5 | 34624999 | rs418716 | 0.98 | 1.446 | 0.35 | -0.368 | 0.001 | -0.001 | -0.001 | 0.443 | 0.61 | 0.52 | 0.67 | 0.56 | 0.59 | 0.45 |
| 5 | 34625746 | rs388669 | 0.876 | 1.441 | 0.19 | -0.087 | -0.003 | 0 | 0 | 0 | 0.61 | 0.55 | 0.68 | 0.55 | 0.66 | 0.46 |
| 5 | 34626160 | rs410324 | -0.362 | 1.651 | 0.38 | -0.102 | 0.013 | 0.006 | 0.006 | 0 | 0.91 | 0.8 | 0.91 | 0.88 | 0.9 | 0.97 |
| 5 | 34628393 | rs377504 | 0.256 | 1.769 | 0.034 | -0.55 | -0.009 | 0.012 | 0.012 | 0.283 | 0.67 | 0.64 | 0.7 | 0.86 | 0.64 | 0.84 |
| 5 | 34636740 | rs404462 | -0.91 | 1.74 | -0.071 | -0.499 | 0.012 | 0.015 | 0.015 | 0 | 0.92 | 0.79 | 0.93 | 0.88 | 0.92 | 0.99 |
| 5 | 34639266 | rs12516063 | 0.91 | 1.689 | -0.349 | -0.525 | -0.051 | 0.118 | 0.118 | 0.099 | 0.22 | 0.01 | 0.19 | 0.03 | 0.18 | 0.11 |
| 5 | 34648725 | rs455326 | 1.478 | 1.368 | -0.149 | -0.401 | -0.003 | -0.008 | -0.008 | 0 | 0.36 | 0.33 | 0.46 | 0.51 | 0.42 | 0.32 |
| 5 | 34651789 | rs458247 | -0.105 | 1.081 | -0.515 | -0.319 | -0.007 | 0.022 | 0.022 | 0.438 | 0.35 | 0.27 | 0.29 | 0.08 | 0.33 | 0.17 |
| 5 | 34652533 | rs464917 | 0.892 | 1.332 | -0.427 | -0.341 | -0.082 | 0.091 | 0.091 | 0.089 | 0.22 | 0.01 | 0.22 | 0.04 | 0.22 | 0.12 |

| | | | | | | | | | | | | | | | | |
|---|---|---|---|---|---|---|---|---|---|---|---|---|---|---|---|---|
| 5 | 34654477 | rs464609 | -0.039 | 1.915 | 0.184 | 0.311 | 0.007 | 0.011 | 0.011 | 0 | 0.56 | 0.3 | 0.52 | 0.39 | 0.55 | 0.65 |
| 5 | 34660854 | rs4703469 | 0.635 | 1.915 | 0.136 | 0.304 | -0.01 | 0.018 | 0.018 | 0 | 0.36 | 0.53 | 0.41 | 0.5 | 0.35 | 0.29 |
| 5 | 34663716 | rs365737 | -0.296 | 1.686 | -0.846 | -0.092 | 0 | 0.017 | 0.017 | 0 | 0.57 | 0.33 | 0.52 | 0.23 | 0.54 | 0.7 |
| 5 | 34663915 | rs448182 | -0.083 | 1.494 | -1.879 | 0.066 | -0.04 | 0.153 | 0.153 | 0 | 0.48 | 0.11 | 0.46 | 0.2 | 0.44 | 0.67 |
| 5 | 34676280 | rs256294 | 0.687 | 1.953 | -0.762 | 0.708 | 0.042 | 0.043 | 0.043 | 0 | 0.75 | 0.48 | 0.67 | 0.64 | 0.68 | 0.85 |
| 5 | 34678440 | rs173908 | 0.119 | 2.086 | -1.53 | 0.815 | -0.04 | -0.072 | -0.072 | 0 | 0.79 | 0.93 | 0.8 | 0.7 | 0.77 | 0.92 |
| 5 | 34679959 | rs256296 | 0.077 | 1.946 | -1.652 | 0.587 | -0.01 | -0.012 | -0.012 | 0 | 0.22 | 0.2 | 0.34 | 0.46 | 0.28 | 0.25 |
| 5 | 34682767 | rs3816325 | 0.066 | 1.7 | -1.668 | 0.433 | -0.022 | -0.062 | -0.062 | 1.402 | 0.2 | 0.08 | 0.19 | 0.3 | 0.2 | 0.07 |
| 5 | 34683418 | rs1390 | 1.221 | 1.482 | -1.96 | 0.104 | -0.015 | 0.309 | 0.309 | 0 | 0.52 | 0.12 | 0.43 | 0.12 | 0.52 | 0.64 |
| 5 | 34687911 | rs2279604 | 0.796 | 1.811 | -1.726 | 0.443 | -0.038 | -0.042 | -0.042 | 2.596 | 0.65 | 0.79 | 0.6 | 0.51 | 0.62 | 0.7 |
| 5 | 34691121 | rs7702104 | 0.305 | 1.632 | -1.046 | 1.121 | -0.003 | 0.002 | 0.002 | 0 | 0.79 | 0.79 | 0.68 | 0.51 | 0.76 | 0.75 |
| 5 | 34698693 | rs889319 | 0.117 | 0.98 | -1.097 | 0.543 | -0.007 | 0.004 | 0.004 | 0.988 | 0.59 | 0.67 | 0.46 | 0.38 | 0.51 | 0.34 |
| 5 | 34700704 | rs11740818 | 0.063 | 0.811 | -1.017 | 0.537 | -0.039 | 0.006 | 0.006 | 0.962 | 0.63 | 0.8 | 0.48 | 0.35 | 0.55 | 0.37 |
| 5 | 34704282 | rs13172326 | 0.5 | -0.361 | -0.004 | -0.501 | 0.021 | 0.014 | 0.014 | 1.882 | 0.35 | 0.22 | 0.23 | 0.05 | 0.33 | 0.17 |
| 5 | 34710332 | rs10461944 | 0.518 | -0.793 | -0.101 | -0.589 | 0.05 | 0.098 | 0.098 | 4.798 | 0.3 | 0.11 | 0.2 | 0.15 | 0.28 | 0.16 |
| 5 | 34710929 | rs6451162 | 0.596 | -0.802 | 0.327 | -0.618 | 0 | -0.003 | -0.003 | 0 | 0.23 | 0.51 | 0.22 | 0.19 | 0.21 | 0.19 |
| 5 | 34716270 | rs1867716 | 0.59 | -0.83 | 0.387 | -0.667 | 0.041 | 0.104 | 0.104 | 5.248 | 0.29 | 0.1 | 0.2 | 0.04 | 0.28 | 0.11 |
| 5 | 34732224 | rs6893661 | -0.336 | -0.247 | 0.199 | -0.856 | -0.001 | -0.002 | -0.002 | 0 | 0.61 | 0.58 | 0.4 | 0.21 | 0.52 | 0.44 |
| 5 | 34735350 | rs10052395 | 0.739 | -0.45 | 0.063 | -0.983 | 0 | -0.005 | -0.005 | 0 | 0.21 | 0.19 | 0.12 | 0.04 | 0.16 | 0.22 |
| 5 | 34736570 | rs17593021 | 0.005 | -0.913 | -0.261 | -1.303 | -0.002 | 0.105 | 0.105 | 0 | 0.41 | 0.24 | 0.3 | 0.1 | 0.43 | 0.37 |
| 5 | 34737944 | rs16903850 | -0.123 | -0.814 | -0.081 | -0.976 | -0.002 | 0.006 | 0.006 | 0 | 0.63 | 0.66 | 0.51 | 0.28 | 0.64 | 0.57 |
| 5 | 34738757 | rs11950200 | 1.411 | -0.781 | -0.56 | -0.924 | 0.014 | 0.013 | 0.013 | 4.513 | 0.17 | 0.12 | 0.07 | 0.11 | 0.08 | 0.06 |
| 5 | 34742531 | rs162914 | -0.736 | -0.09 | -0.51 | -0.602 | 0.036 | 0.05 | 0.05 | 1.882 | 0.8 | 0.76 | 0.59 | 0.39 | 0.71 | 0.63 |
| 5 | 34749014 | rs4499805 | -0.044 | -0.119 | -0.794 | -0.483 | 0.004 | 0.011 | 0.011 | 0 | 0.17 | 0.1 | 0.16 | 0.13 | 0.18 | 0.1 |
| 5 | 34749416 | rs9283739 | 0.256 | -0.124 | -0.765 | -0.446 | -0.001 | -0.004 | -0.004 | 0 | 0.14 | 0.16 | 0.1 | 0.09 | 0.13 | 0.21 |
| 5 | 34753340 | rs334912 | -0.431 | -0.008 | -1.043 | -0.245 | -0.011 | -0.025 | -0.025 | 0.597 | 0.54 | 0.8 | 0.56 | 0.46 | 0.55 | 0.48 |
| 5 | 34754248 | rs13176284 | 0.861 | -0.089 | -1.374 | -0.186 | -0.01 | -0.091 | -0.091 | 0 | 0.29 | 0.09 | 0.41 | 0.49 | 0.25 | 0.46 |
| 5 | 34754305 | rs334913 | -0.031 | -0.687 | -1.688 | -0.414 | 0.028 | -0.048 | -0.048 | 1.791 | 0.75 | 0.86 | 0.81 | 0.65 | 0.76 | 0.83 |
| 5 | 34755478 | rs9283740 | 0.969 | -0.303 | -0.613 | -0.863 | 0.022 | -0.082 | -0.082 | 0 | 0.18 | 0.27 | 0.15 | 0.01 | 0.21 | 0.13 |
| 5 | 34755871 | rs10039415 | -0.111 | -0.23 | -0.383 | -0.946 | 0.013 | -0.088 | -0.088 | 1.571 | 0.31 | 0.62 | 0.28 | 0.08 | 0.35 | 0.22 |
| 5 | 34757666 | rs17521570 | 0.892 | 0.133 | -0.806 | -0.618 | -0.017 | -0.023 | -0.023 | 0 | 0.11 | 0.01 | 0.11 | 0.11 | 0.13 | 0.17 |
| 5 | 34759463 | rs164317 | 0.501 | 0.067 | 0.025 | -0.661 | -0.007 | -0.005 | -0.005 | 0 | 0.13 | 0.32 | 0.13 | 0.11 | 0.13 | 0.17 |
| 5 | 34760463 | rs164320 | -0.906 | 0.068 | -0.037 | -0.736 | 0.009 | 0.042 | 0.042 | 0.088 | 0.54 | 0.71 | 0.4 | 0.33 | 0.48 | 0.5 |
| 5 | 34771848 | rs12658204 | -0.649 | -0.113 | -0.192 | -0.963 | 0.001 | -0.029 | -0.029 | 1.192 | 0.31 | 0.26 | 0.2 | 0.06 | 0.37 | 0.21 |
| 5 | 34772786 | rs162909 | -1.55 | -0.062 | -0.19 | -0.976 | 0.005 | 0.02 | 0.02 | 0 | 0.45 | 0.37 | 0.32 | 0.26 | 0.49 | 0.4 |
| 5 | 34778371 | rs11949850 | 0.012 | 0.224 | -0.557 | -0.889 | -0.006 | -0.013 | -0.013 | 0 | 0.19 | 0.16 | 0.17 | 0.02 | 0.32 | 0.11 |
| 5 | 34794128 | rs163121 | -1.217 | 0.022 | -0.844 | -0.643 | -0.046 | 0.006 | 0.006 | 0.367 | 0.56 | 0.89 | 0.49 | 0.37 | 0.6 | 0.51 |
| 5 | 34795305 | rs336462 | -0.23 | -0.321 | 0.234 | -0.397 | -0.029 | 0.007 | 0.007 | 0 | 0.2 | 0.43 | 0.12 | 0.2 | 0.15 | 0.29 |
| 5 | 34795510 | rs6893365 | -1.343 | -1.134 | -0.059 | -0.247 | -0.005 | -0.001 | -0.001 | 0.015 | 0.36 | 0.62 | 0.26 | 0.36 | 0.18 | 0.36 |
| 5 | 34795841 | rs336463 | -0.897 | 0.001 | -0.173 | -0.978 | -0.003 | -0.015 | -0.015 | 0.795 | 0.42 | 0.37 | 0.31 | 0.11 | 0.48 | 0.34 |
| 5 | 34795981 | rs6897719 | -1.388 | -0.565 | 0.233 | -1.111 | -0.024 | -0.111 | -0.111 | 0 | 0.21 | 0.42 | 0.13 | 0.04 | 0.16 | 0.12 |
| 5 | 34798601 | rs4703388 | 1.46 | -0.21 | -0.164 | -0.219 | -0.001 | 0.003 | 0.003 | 1.029 | 0.43 | 0.27 | 0.55 | 0.4 | 0.49 | 0.5 |
| 5 | 34799411 | rs469033 | -0.196 | -0.165 | 0.164 | -0.257 | 0.008 | -0.08 | -0.08 | 0 | 0.16 | 0.31 | 0.11 | 0.01 | 0.16 | 0.12 |
| 5 | 34800351 | rs12659689 | 1.63 | -0.163 | 0.16 | -0.226 | 0.002 | 0.004 | 0.004 | 1.028 | 0.43 | 0.27 | 0.54 | 0.4 | 0.47 | 0.48 |
| 5 | 34800987 | rs462492 | 0.082 | -0.072 | 0 | 0.53 | -0.033 | 0.236 | 0.236 | 0 | 0.92 | 0.76 | 0.87 | 0.59 | 0.98 | 0.9 |
| 5 | 34801172 | rs6886361 | -0.312 | 0.443 | -0.665 | -0.374 | -0.024 | 0.04 | 0.04 | 0.274 | 0.32 | 0.18 | 0.21 | 0.19 | 0.32 | 0.3 |
| 5 | 34810095 | rs10472940 | -0.09 | 0.435 | -0.273 | -0.122 | -0.003 | 0.001 | 0.001 | 0 | 0.59 | 0.56 | 0.69 | 0.44 | 0.62 | 0.61 |
| 5 | 34816140 | rs17522697 | -0.245 | 0.526 | -1.365 | -0.606 | -0.02 | 0.104 | 0.104 | 0 | 0.18 | 0.01 | 0.1 | 0.01 | 0.14 | 0.14 |
| 5 | 34817784 | rs12153278 | 0.976 | 0.618 | -1.163 | -0.474 | -0.048 | -0.043 | -0.043 | 0 | 0.31 | 0.1 | 0.5 | 0.35 | 0.38 | 0.4 |
| 5 | 34819280 | rs3844310 | 1.05 | 0.644 | -1.159 | -0.498 | -0.032 | -0.041 | -0.041 | 0 | 0.3 | 0.1 | 0.49 | 0.35 | 0.34 | 0.39 |
| 5 | 34821825 | rs10461947 | 0.567 | 0.665 | -1.13 | -0.513 | -0.022 | -0.016 | -0.016 | 0.653 | 0.37 | 0.11 | 0.54 | 0.35 | 0.39 | 0.42 |
| 5 | 34824555 | rs1048944 | -0.201 | 0.651 | -0.789 | -0.578 | -0.006 | 0.05 | 0.05 | 0 | 0.5 | 0.29 | 0.59 | 0.37 | 0.49 | 0.55 |
| 5 | 34827731 | rs12189090 | -0.217 | 0.298 | -0.941 | -0.64 | -0.037 | 0.085 | 0.085 | 0 | 0.14 | 0.01 | 0.05 | 0.02 | 0.11 | 0.11 |
| 5 | 34831748 | rs12446 | -1.109 | 0.16 | -0.799 | -0.789 | 0.013 | 0.027 | 0.027 | 0 | 0.25 | 0.17 | 0.12 | 0.02 | 0.19 | 0.16 |
| 5 | 34832058 | rs3817264 | 0.674 | 0.576 | -0.264 | -0.556 | -0.003 | -0.002 | -0.002 | 1.573 | 0.37 | 0.35 | 0.57 | 0.4 | 0.42 | 0.48 |
| 5 | 34833339 | rs10472254 | -0.042 | 0.286 | -0.011 | -0.497 | 0.047 | 0.06 | 0.06 | 0.307 | 0.06 | 0.35 | 0.1 | 0.04 | 0.05 | 0.01 |
| 5 | 34839111 | rs336484 | -0.673 | 0.349 | 0.076 | -0.713 | -0.002 | -0.002 | -0.002 | 0 | 0.61 | 0.67 | 0.7 | 0.42 | 0.58 | 0.62 |
| 5 | 34839428 | rs336485 | 0.212 | 0.494 | 0.016 | -0.868 | 0.003 | -0.036 | -0.036 | 0.014 | 0.13 | 0.13 | 0.05 | 0.02 | 0.13 | 0.11 |
| 5 | 34839548 | rs10078917 | -0.102 | 0.448 | 0.091 | -0.808 | 0.07 | 0.09 | 0.09 | 0.637 | 0.06 | 0.43 | 0.1 | 0.04 | 0.05 | 0.01 |
| 5 | 34840841 | rs6451173 | -0.972 | 0.672 | -0.941 | -0.523 | -0.103 | 0.073 | 0.073 | 0 | 0.59 | 0.9 | 0.41 | 0.26 | 0.55 | 0.42 |
| 5 | 34844054 | rs4703479 | 0.73 | 1.014 | -0.524 | -0.073 | 0.007 | 0.017 | 0.017 | 0 | 0.32 | 0.22 | 0.2 | 0.19 | 0.29 | 0.3 |
| 5 | 34845473 | rs336486 | -0.866 | 1.104 | -0.797 | 0.256 | -0.026 | -0.016 | -0.016 | 0.654 | 0.91 | 0.97 | 0.89 | 0.62 | 0.86 | 0.87 |
| 5 | 34854491 | rs368823 | -0.648 | 0.445 | -0.282 | -0.351 | -0.004 | -0.053 | -0.053 | 3.341 | 0.18 | 0.33 | 0.13 | 0.07 | 0.16 | 0.14 |
| 5 | 34859409 | rs336470 | -0.155 | -0.108 | -0.128 | -0.06 | -0.016 | -0.028 | -0.028 | 0.477 | 0.37 | 0.51 | 0.31 | 0.24 | 0.35 | 0.31 |
| 5 | 34862541 | rs4703481 | -0.355 | 0.259 | -0.473 | 0.134 | 0.006 | 0.018 | 0.018 | 0 | 0.17 | 0.15 | 0.13 | 0.06 | 0.12 | 0.14 |
| 5 | 34863220 | rs3846632 | 1.037 | 0.313 | -1.601 | -0.333 | -0.13 | 0.109 | 0.109 | 0.039 | 0.33 | 0.04 | 0.3 | 0.09 | 0.43 | 0.18 |
| 5 | 34863933 | rs3846633 | 0.186 | -0.011 | -1.149 | -0.456 | -0.013 | 0.069 | 0.069 | 0 | 0.46 | 0.39 | 0.44 | 0.18 | 0.52 | 0.28 |
| 5 | 34868792 | rs10461949 | -0.568 | -0.25 | -1.362 | 0.429 | -0.027 | -0.035 | -0.035 | 0 | 0.33 | 0.16 | 0.34 | 0.39 | 0.34 | 0.48 |
| 5 | 34871161 | rs2388866 | 0.094 | -0.235 | -1.25 | 0.441 | -0.003 | 0.015 | 0.015 | 0 | 0.06 | 0.23 | 0.1 | 0.05 | 0.03 | 0.01 |

| | | | | | | | | | | | | | | | | |
|---|---|---|---|---|---|---|---|---|---|---|---|---|---|---|---|---|
| 5 | 34878150 | rs2892623 | 1.249 | -0.418 | -1.596 | 0.512 | 0.015 | 0.07 | 0.07 | 0.33 | 0.23 | 0.06 | 0.13 | 0.07 | 0.16 | 0.23 |
| 5 | 34882421 | rs17303001 | 1.259 | -0.41 | -1.576 | 0.515 | 0.017 | 0.07 | 0.07 | 0.55 | 0.23 | 0.07 | 0.13 | 0.07 | 0.16 | 0.23 |
| 5 | 34886968 | rs17303064 | 0.383 | -0.179 | -1.257 | 0.711 | 0 | -0.007 | -0.007 | 0 | 0.08 | 0.12 | 0.05 | 0.03 | 0.06 | 0.11 |
| 5 | 34890728 | rs153012 | 0.7 | -0.039 | -1.517 | 0.769 | -0.05 | -0.088 | -0.088 | 0.189 | 0.15 | 0.02 | 0.36 | 0.33 | 0.21 | 0.28 |
| 5 | 34898304 | rs251638 | -0.082 | -0.063 | -1.418 | 0.805 | -0.045 | -0.073 | -0.073 | 0.783 | 0.25 | 0.06 | 0.3 | 0.35 | 0.29 | 0.38 |
| 5 | 34907169 | rs4703484 | 0.108 | -0.162 | -0.196 | 1.278 | 0.072 | 0.523 | 0.523 | 0.173 | 0.95 | 0.64 | 0.89 | 0.6 | 0.99 | 0.9 |
| 5 | 34909596 | rs2069478 | 0.14 | -0.211 | -0.145 | 1.184 | -0.003 | 0.023 | 0.023 | 0 | 0.05 | 0.17 | 0.1 | 0.06 | 0.04 | 0.01 |
| 5 | 34915948 | rs2069465 | 0.927 | -0.243 | -0.142 | 1.227 | -0.012 | 0.007 | 0.007 | 0 | 0.07 | 0 | 0.04 | 0.03 | 0.06 | 0.11 |
| 5 | 34917112 | rs37435 | 0.12 | -0.233 | -0.032 | 1.312 | 0.072 | 0.501 | 0.501 | 0.224 | 0.95 | 0.65 | 0.89 | 0.64 | 0.99 | 0.9 |
| 5 | 34920596 | rs185186 | -0.065 | -0.212 | -0.246 | 1.244 | -0.046 | -0.073 | -0.073 | 0.042 | 0.25 | 0.06 | 0.29 | 0.34 | 0.29 | 0.38 |
| 5 | 34924857 | rs2270934 | 1.007 | -0.389 | -0.127 | 1.232 | -0.014 | 0.005 | 0.005 | 0 | 0.06 | 0 | 0.04 | 0.03 | 0.06 | 0.09 |
| 5 | 34927091 | rs27166 | 0.503 | -0.388 | 0.176 | 1.331 | 0.063 | 0.49 | 0.49 | 0.788 | 0.94 | 0.6 | 0.88 | 0.64 | 0.99 | 0.9 |
| 5 | 34931599 | rs27599 | 0.633 | -0.407 | 0.226 | 1.334 | 0.031 | 0.421 | 0.421 | 0.442 | 0.94 | 0.69 | 0.89 | 0.65 | 0.99 | 0.88 |
| 5 | 34938373 | rs40202 | -0.006 | -0.348 | -0.035 | 1.226 | -0.056 | -0.063 | -0.063 | 1.958 | 0.24 | 0.05 | 0.3 | 0.32 | 0.3 | 0.35 |
| 5 | 34943496 | rs251641 | 0.071 | -0.254 | -0.042 | 1.219 | -0.041 | -0.064 | -0.064 | 0 | 0.24 | 0.05 | 0.28 | 0.32 | 0.26 | 0.35 |
| 5 | 34945122 | rs17304200 | 0.906 | -0.278 | -0.098 | 1.178 | -0.011 | 0.013 | 0.013 | 0 | 0.08 | 0 | 0.04 | 0.04 | 0.07 | 0.11 |
| 5 | 34948532 | rs37440 | -1.635 | -0.389 | -0.258 | 1.031 | -0.127 | -0.116 | -0.116 | 0 | 0.66 | 0.96 | 0.59 | 0.55 | 0.53 | 0.69 |
| 5 | 34951150 | rs37439 | -0.552 | -0.453 | -0.075 | 1.092 | -0.052 | 0.058 | 0.058 | 0.084 | 0.69 | 0.5 | 0.77 | 0.52 | 0.82 | 0.67 |
| 5 | 34955550 | rs17304438 | 1.031 | -0.441 | 0.275 | 1.092 | -0.011 | 0.013 | 0.013 | 0 | 0.08 | 0 | 0.04 | 0.04 | 0.07 | 0.11 |
| 5 | 34956609 | rs17304494 | 1.031 | -0.44 | 0.025 | 1.091 | -0.011 | 0.047 | 0.047 | 0 | 0.08 | 0 | 0.04 | 0.01 | 0.07 | 0.11 |
| 5 | 34957444 | rs251636 | 0.23 | -0.443 | 0.077 | 1.093 | -0.034 | 0.022 | 0.022 | 0 | 0.61 | 0.51 | 0.72 | 0.5 | 0.75 | 0.56 |
| 5 | 34960515 | rs7719119 | 0.604 | -0.364 | 0.473 | 1.188 | 0.033 | 0.405 | 0.405 | 1.673 | 0.94 | 0.71 | 0.89 | 0.65 | 0.99 | 0.88 |
| 5 | 34978170 | rs163588 | -0.721 | -0.349 | 0.08 | 1.085 | 0 | -0.003 | -0.003 | 0 | 0.36 | 0.36 | 0.4 | 0.42 | 0.32 | 0.4 |
| 5 | 34978915 | rs6451178 | 0.769 | -0.326 | -0.172 | 1.096 | 0.018 | 0.025 | 0.025 | 0 | 0.27 | 0.18 | 0.21 | 0.17 | 0.21 | 0.23 |
| 5 | 34986704 | rs163594 | 0.304 | -0.287 | -0.431 | 1.134 | -0.038 | -0.055 | -0.055 | 0 | 0.22 | 0.1 | 0.28 | 0.33 | 0.24 | 0.28 |
| 5 | 34991946 | rs163599 | 0.304 | -0.287 | -0.431 | 1.134 | -0.038 | -0.055 | -0.055 | 0 | 0.22 | 0.06 | 0.27 | 0.32 | 0.24 | 0.28 |
| 5 | 34993030 | rs16870205 | -0.14 | -0.337 | -0.208 | 0.684 | -0.02 | -0.034 | -0.034 | 0 | 0.38 | 0.54 | 0.28 | 0.21 | 0.29 | 0.34 |
| 5 | 34993215 | rs11744796 | 1.511 | -0.322 | -0.98 | 0.676 | -0.138 | 0.113 | 0.113 | 0 | 0.33 | 0.03 | 0.33 | 0.12 | 0.46 | 0.2 |
| 5 | 34997038 | rs17245616 | 0.994 | -0.187 | -0.687 | 0.775 | -0.009 | 0.071 | 0.071 | 0.021 | 0.25 | 0.15 | 0.16 | 0.09 | 0.24 | 0.26 |
| 5 | 34998778 | rs16899972 | -0.965 | -0.188 | -0.675 | 1.002 | -0.024 | -0.031 | -0.031 | 0.181 | 0.56 | 0.3 | 0.46 | 0.55 | 0.53 | 0.64 |
| 5 | 34998877 | rs16899974 | -0.141 | -0.156 | -0.694 | 1.312 | -0.011 | -0.082 | -0.082 | 0.622 | 0.22 | 0.05 | 0.26 | 0.42 | 0.23 | 0.29 |
| 5 | 34998894 | rs17245714 | 0.378 | 0.069 | -0.221 | 1.285 | -0.011 | 0.009 | 0.009 | 0 | 0.08 | 0 | 0.04 | 0.03 | 0.06 | 0.11 |
| 5 | 35000312 | rs9292570 | -0.291 | 0.34 | -0.042 | 0.946 | 0.003 | 0.002 | 0.002 | 0 | 0.48 | 0.5 | 0.47 | 0.56 | 0.41 | 0.51 |
| 5 | 35000795 | rs7717823 | 0.669 | 0.567 | -0.023 | 0.619 | -0.043 | 0.048 | 0.048 | 0 | 0.28 | 0.15 | 0.28 | 0.12 | 0.37 | 0.19 |
| 5 | 35005021 | rs4538595 | 0.405 | 0.197 | 0.325 | 0.667 | 0.02 | 0.058 | 0.058 | 2.14 | 0.5 | 0.4 | 0.32 | 0.24 | 0.44 | 0.5 |
| 5 | 35007085 | rs163841 | -0.069 | 0.056 | -0.074 | 0.684 | -0.002 | 0 | 0 | 0.017 | 0.57 | 0.55 | 0.57 | 0.62 | 0.53 | 0.6 |
| 5 | 35010872 | rs11960801 | 0.911 | 0.169 | -0.261 | 0.59 | -0.028 | 0.044 | 0.044 | 0 | 0.28 | 0.17 | 0.29 | 0.11 | 0.35 | 0.15 |
| 5 | 35014723 | rs17246034 | 1.261 | 0.147 | -0.324 | 0.624 | 0.02 | 0.056 | 0.056 | 2.385 | 0.33 | 0.1 | 0.21 | 0.2 | 0.29 | 0.34 |
| 5 | 35015297 | rs163907 | 0.257 | 0.175 | -0.22 | 0.601 | 0.003 | 0.012 | 0.012 | 2.17 | 0.53 | 0.54 | 0.35 | 0.26 | 0.47 | 0.51 |
| 5 | 35033706 | rs2291702 | 0.497 | 0.326 | 0.029 | 0.156 | -0.019 | 0.061 | 0.061 | 0 | 0.52 | 0.2 | 0.56 | 0.67 | 0.49 | 0.66 |
| 5 | 35034419 | rs13185264 | 0.542 | 0.784 | 0.062 | 0.819 | -0.016 | 0.02 | 0.02 | 0 | 0.25 | 0.18 | 0.23 | 0.11 | 0.33 | 0.09 |
| 5 | 35035579 | rs344156 | -0.369 | 0.869 | -0.138 | 0.743 | 0.017 | 0.017 | 0.017 | 0 | 0.73 | 0.62 | 0.71 | 0.67 | 0.67 | 0.82 |
| 5 | 35039486 | rs37370 | -0.008 | 0.727 | -0.728 | 0.905 | -0.021 | -0.041 | -0.041 | 0 | 0.91 | 0.98 | 0.71 | 0.54 | 0.91 | 0.82 |
| 5 | 35042158 | rs37373 | 0.737 | 0.538 | -0.438 | 0.544 | -0.006 | -0.006 | -0.006 | 0 | 0.37 | 0.35 | 0.29 | 0.31 | 0.33 | 0.49 |
| 5 | 35044888 | rs10038101 | 0.269 | 0.393 | -0.538 | 0.577 | 0 | 0.02 | 0.02 | 0 | 0.23 | 0.2 | 0.16 | 0.12 | 0.23 | 0.21 |
| 5 | 35045745 | rs40200 | 0.345 | -0.012 | -0.217 | 1.12 | -0.005 | 0.151 | 0.151 | 0 | 0.89 | 0.79 | 0.7 | 0.54 | 0.91 | 0.82 |
| 5 | 35047845 | rs40201 | 0.042 | -0.08 | -0.529 | 0.285 | 0.009 | 0.007 | 0.007 | 0.527 | 0.36 | 0.31 | 0.25 | 0.3 | 0.28 | 0.45 |
| 5 | 35052896 | rs10941231 | -0.764 | 0.348 | -0.247 | 0.11 | 0.014 | -0.01 | -0.01 | 0.202 | 0.2 | 0.28 | 0.34 | 0.18 | 0.28 | 0.21 |
| 5 | 35054639 | rs10941232 | 0.695 | -1.258 | -0.153 | -0.59 | -0.01 | -0.028 | -0.028 | 5.297 | 0.57 | 0.54 | 0.56 | 0.2 | 0.72 | 0.48 |
| 5 | 35061731 | rs387032 | -0.762 | -1.441 | 0.511 | 0.243 | -0.049 | 0.075 | 0.075 | 0 | 0.21 | 0.49 | 0.16 | 0.23 | 0.12 | 0.12 |
| 5 | 35063117 | rs1010119 | -0.297 | -2.016 | 0.291 | -0.164 | -0.047 | -0.042 | -0.042 | 0.012 | 0.13 | 0.25 | 0.06 | 0.04 | 0.04 | 0.07 |
| 5 | 35072380 | rs37364 | -1.525 | -2.126 | 0.153 | -0.159 | 0.072 | 0.012 | 0.012 | 0 | 0.38 | 0.75 | 0.22 | 0.27 | 0.2 | 0.26 |
| 5 | 35085180 | rs37389 | -1.265 | -2.079 | 0.811 | -0.023 | -0.055 | 0.042 | 0.042 | 0 | 0.23 | 0.5 | 0.11 | 0.25 | 0.11 | 0.18 |
| 5 | 35092969 | rs37383 | -1.773 | -2.557 | 0.97 | -0.102 | -0.009 | 0.001 | 0.001 | 0.425 | 0.29 | 0.7 | 0.15 | 0.24 | 0.1 | 0.19 |
| 5 | 35101178 | rs10447140 | -0.421 | -2.343 | 1.781 | -0.007 | -0.037 | 0.061 | 0.061 | 5.708 | 0.08 | 0.17 | 0.05 | 0.16 | 0.01 | 0.02 |
| 5 | 35127378 | rs930068 | -1.062 | -2.367 | 1.836 | -0.108 | -0.003 | -0.024 | -0.024 | 0 | 0.05 | 0.08 | 0.02 | 0.01 | 0.04 | 0.01 |
| 5 | 35131805 | rs2914108 | -1.977 | -2.469 | 2.274 | -0.054 | -0.058 | 0.038 | 0.038 | 0.891 | 0.14 | 0.49 | 0.07 | 0.14 | 0.05 | 0.03 |
| 5 | 35132350 | rs2962089 | -2.537 | -2.585 | 2.254 | -0.112 | -0.044 | -0.007 | -0.007 | 0 | 0.23 | 0.66 | 0.12 | 0.17 | 0.08 | 0.11 |
| 5 | 35135741 | rs2962101 | -2.488 | -2.625 | 2.206 | -0.278 | -0.071 | -0.02 | -0.02 | 0.174 | 0.25 | 0.56 | 0.12 | 0.17 | 0.11 | 0.12 |
| 5 | 35145363 | rs9292571 | 1.631 | -2.575 | 2.242 | -0.238 | -0.056 | 0.047 | 0.047 | 0.505 | 0.86 | 0.57 | 0.94 | 0.86 | 0.95 | 0.97 |
| 5 | 35146865 | rs1587605 | 1.084 | -2.527 | 2.075 | -0.215 | -0.032 | 0.018 | 0.018 | 0.607 | 0.88 | 0.81 | 0.95 | 0.86 | 0.95 | 0.97 |
| 5 | 35155671 | rs12109986 | -1.254 | -2.338 | 2.353 | -0.255 | -0.034 | -0.017 | -0.017 | 0 | 0.11 | 0.44 | 0.06 | 0.04 | 0.03 | 0.08 |
| 5 | 35161332 | rs6880595 | -1.131 | -2.095 | 1.869 | -0.698 | 0 | -0.024 | -0.024 | 0 | 0.19 | 0.18 | 0.19 | 0.02 | 0.23 | 0.1 |
| 5 | 35162151 | rs6451185 | 1.106 | -2.784 | 2.175 | -0.235 | -0.065 | 0.129 | 0.129 | 0.35 | 0.85 | 0.54 | 0.88 | 0.77 | 0.95 | 0.96 |
| 5 | 35162762 | rs4703503 | 0.286 | -2.258 | 1.032 | -0.716 | -0.003 | -0.016 | -0.016 | 0.525 | 0.24 | 0.14 | 0.32 | 0.72 | 0.21 | 0.28 |
| 5 | 35164895 | rs16872491 | -0.978 | -1.45 | 1.18 | -0.726 | -0.001 | -0.001 | -0.001 | 0.022 | 0.19 | 0.23 | 0.16 | 0.15 | 0.17 | 0.04 |
| 5 | 35165597 | rs1039428 | -1.706 | -1.388 | 1.385 | -0.813 | -0.001 | -0.031 | -0.031 | 0 | 0.3 | 0.38 | 0.29 | 0.15 | 0.3 | 0.12 |
| 5 | 35168145 | rs11953869 | 0.188 | -1.356 | 1.331 | -0.8 | 0.01 | 0.017 | 0.017 | 0.015 | 0.1 | 0.22 | 0.09 | 0.13 | 0.11 | 0.01 |
| 5 | 35179360 | rs11952661 | -1.491 | -1.366 | 1.34 | -0.802 | -0.001 | -0.029 | -0.029 | 0 | 0.3 | 0.38 | 0.29 | 0.16 | 0.3 | 0.12 |

| | | | | | | | | | | | | | | | |
|---|---|---|---|---|---|---|---|---|---|---|---|---|---|---|---|
| 5 | 35179593 | rs6895193 | -0.596 | -1.473 | 1.189 | -0.979 | 0.009 | 0.003 | 0.003 | 0.684 | 0.74 | 0.67 | 0.59 | 0.29 | 0.7 | 0.66 |
| 5 | 35181833 | rs13154347 | -1.284 | -1.191 | 0.601 | -0.901 | -0.001 | 0.007 | 0.007 | 0 | 0.29 | 0.29 | 0.28 | 0.16 | 0.3 | 0.12 |
| 5 | 35185826 | rs13436213 | 1.513 | -1.266 | -0.268 | -0.827 | 0.002 | 0.099 | 0.099 | 0.928 | 0.4 | 0.14 | 0.28 | 0.18 | 0.39 | 0.45 |
| 5 | 35188476 | rs12514742 | 0.375 | -0.979 | 0.101 | -0.553 | -0.006 | 0.011 | 0.011 | 0 | 0.1 | 0.05 | 0.13 | 0.01 | 0.13 | 0.12 |
| 5 | 35188802 | rs4703508 | 0.252 | -1.023 | 0.072 | -0.565 | -0.003 | 0.009 | 0.009 | 0.591 | 0.08 | 0.05 | 0.09 | 0.03 | 0.06 | 0.03 |
| 5 | 35190327 | rs12521272 | -0.763 | -1.013 | 0.083 | -0.551 | -0.02 | 0.06 | 0.06 | 0 | 0.19 | 0.07 | 0.22 | 0.04 | 0.2 | 0.15 |
| 5 | 35197382 | rs12188786 | -0.331 | -1.199 | 0.73 | -0.31 | 0 | 0.017 | 0.017 | 0.818 | 0.1 | 0.27 | 0.08 | 0.1 | 0.1 | 0.01 |
| 5 | 35199433 | rs9292573 | 1.988 | -1.259 | 0.781 | 0.142 | -0.006 | -0.005 | -0.005 | 0 | 0.6 | 0.4 | 0.52 | 0.29 | 0.66 | 0.66 |
| 5 | 35199657 | rs873456 | 1.934 | -1.235 | 0.901 | 0.154 | -0.001 | -0.001 | -0.001 | 0 | 0.61 | 0.34 | 0.52 | 0.29 | 0.67 | 0.67 |
| 5 | 35200194 | rs4703509 | -1.484 | -1.257 | 1.013 | 0.09 | -0.015 | -0.072 | -0.072 | 0 | 0.27 | 0.52 | 0.27 | 0.12 | 0.26 | 0.15 |
| 5 | 35200744 | rs9687437 | 0.21 | -1.21 | 1.133 | 0.128 | 0.031 | -0.015 | -0.015 | 0 | 0.07 | 0.29 | 0.04 | 0 | 0.04 | 0.01 |
| 5 | 35204799 | rs11959588 | -0.745 | -1.2 | 1.018 | 0.145 | 0 | -0.003 | -0.003 | 0.01 | 0.11 | 0.37 | 0.07 | 0.07 | 0.1 | 0.01 |
| 5 | 35209339 | rs4235652 | 0.55 | -1.185 | 0.73 | 0.039 | -0.014 | 0.05 | 0.05 | 0.315 | 0.79 | 0.65 | 0.72 | 0.38 | 0.83 | 0.8 |
| 5 | 35210629 | rs4703511 | 0.136 | -1.24 | 0.832 | 0.03 | 0.001 | -0.002 | -0.002 | 0 | 0.07 | 0.21 | 0.07 | 0.04 | 0.03 | 0.03 |
| 5 | 35211240 | rs1609500 | 1.527 | -1.19 | 0.541 | -0.015 | -0.019 | -0.006 | -0.006 | 0 | 0.62 | 0.46 | 0.52 | 0.33 | 0.67 | 0.66 |
| 5 | 35220865 | rs7731153 | -0.766 | -0.872 | 0.175 | 0.53 | 0.01 | -0.011 | -0.011 | 0.084 | 0.9 | 0.96 | 0.85 | 0.53 | 0.91 | 0.81 |
| 5 | 35223013 | rs10064495 | 1.521 | -1.07 | 0.744 | 0.461 | 0.006 | 0 | 0 | 0 | 0.63 | 0.39 | 0.56 | 0.34 | 0.69 | 0.68 |
| 5 | 35223842 | rs6878684 | 0.163 | -1.095 | 0.776 | 0.49 | -0.003 | 0.027 | 0.027 | 0 | 0.79 | 0.78 | 0.68 | 0.5 | 0.79 | 0.73 |
| 5 | 35225147 | rs13354826 | 1.97 | -1.148 | -0.134 | -0.119 | -0.085 | 0.124 | 0.124 | 0.168 | 0.34 | 0.06 | 0.3 | 0.12 | 0.38 | 0.48 |
| 5 | 35227015 | rs7735260 | -0.677 | -1.054 | 0.5 | 0.219 | 0.001 | -0.001 | -0.001 | 0 | 0.14 | 0.13 | 0.09 | 0.18 | 0.1 | 0.05 |
| 5 | 35230226 | rs9292579 | 0.931 | -0.907 | 0.452 | 0.212 | -0.003 | -0.016 | -0.016 | 0 | 0.05 | 0.08 | 0.07 | 0.04 | 0.04 | 0.03 |
| 5 | 35231987 | rs10080135 | -0.064 | -0.949 | 0.301 | 0.434 | 0.017 | -0.065 | -0.065 | 0 | 0.82 | 0.92 | 0.66 | 0.5 | 0.84 | 0.74 |
| 5 | 35233786 | rs10059874 | -0.29 | -0.973 | 0.781 | 0.325 | 0.033 | 0.105 | 0.105 | 8.507 | 0.1 | 0.45 | 0.09 | 0.13 | 0.06 | 0.02 |
| 5 | 35234843 | rs13361664 | -0.14 | -0.964 | 0.845 | 0.34 | 0.022 | 0.09 | 0.09 | 8.56 | 0.09 | 0.39 | 0.09 | 0.14 | 0.06 | 0.02 |
| 5 | 35237736 | rs7705676 | 1.602 | -1.025 | 0.157 | 0.088 | -0.047 | 0.151 | 0.151 | 1.147 | 0.42 | 0.16 | 0.31 | 0.15 | 0.46 | 0.52 |
| 5 | 35239886 | rs12189146 | -0.349 | -0.843 | 0.601 | 0.265 | 0.013 | 0.09 | 0.09 | 9.01 | 0.11 | 0.5 | 0.1 | 0.14 | 0.05 | 0.02 |
| 5 | 35244852 | rs6897600 | -0.134 | -0.672 | -0.04 | 0.164 | -0.007 | 0.009 | 0.009 | 0 | 0.29 | 0.25 | 0.23 | 0.21 | 0.32 | 0.2 |
| 5 | 35249422 | rs7720773 | 0.925 | -0.724 | 0.059 | 0.143 | -0.008 | -0.024 | -0.024 | 0 | 0.59 | 0.44 | 0.45 | 0.23 | 0.68 | 0.57 |
| 5 | 35263218 | rs10071999 | -0.768 | -0.414 | -0.659 | 0.619 | -0.017 | -0.048 | -0.048 | 0.205 | 0.9 | 0.98 | 0.8 | 0.54 | 0.89 | 0.91 |
| 5 | 35265240 | rs10512632 | 0.179 | -0.513 | -0.481 | 0.142 | -0.005 | 0.038 | 0.038 | 0.377 | 0.21 | 0.12 | 0.29 | 0.11 | 0.2 | 0.29 |
| 5 | 35270560 | rs7701255 | 0.569 | -0.447 | -0.051 | 0.073 | -0.007 | -0.046 | -0.046 | 0 | 0.61 | 0.46 | 0.42 | 0.2 | 0.66 | 0.6 |
| 5 | 35275087 | rs10063181 | 0.242 | -0.453 | 0.328 | 0.321 | 0.001 | 0.003 | 0.003 | 2.512 | 0.64 | 0.64 | 0.45 | 0.32 | 0.67 | 0.65 |
| 5 | 35305181 | rs6869985 | -0.16 | -0.463 | -0.17 | 0.639 | 0.022 | -0.051 | -0.051 | 0 | 0.85 | 0.89 | 0.81 | 0.46 | 0.9 | 0.92 |
| 5 | 35315860 | rs6451195 | 1.066 | -0.007 | -0.259 | 0.205 | 0.005 | 0.032 | 0.032 | 0.7 | 0.4 | 0.29 | 0.24 | 0.13 | 0.38 | 0.5 |
| 5 | 35325555 | rs10068482 | -0.207 | 0.139 | -0.537 | 0.264 | -0.047 | -0.086 | -0.086 | 0 | 0.73 | 0.87 | 0.67 | 0.45 | 0.66 | 0.88 |
| 5 | 35333673 | rs7728464 | 1.13 | -0.208 | -0.484 | -0.043 | 0.027 | 0.062 | 0.062 | 0.633 | 0.36 | 0.22 | 0.17 | 0.13 | 0.26 | 0.46 |
| 5 | 35345497 | rs1025324 | 0.096 | -0.286 | -0.256 | 0.08 | -0.014 | -0.02 | -0.02 | 0.004 | 0.33 | 0.42 | 0.4 | 0.23 | 0.24 | 0.31 |
| 5 | 35358663 | rs32746 | -0.57 | -0.196 | -0.276 | 0.21 | -0.058 | -0.067 | -0.067 | 0 | 0.75 | 0.82 | 0.62 | 0.48 | 0.54 | 0.86 |
| 5 | 35359054 | rs32747 | -0.58 | -0.267 | -0.297 | 0.191 | -0.105 | -0.112 | -0.112 | 1.079 | 0.75 | 0.9 | 0.63 | 0.48 | 0.53 | 0.86 |
| 5 | 35360450 | rs26191 | 0.054 | -0.738 | 0.576 | 0.196 | -0.001 | -0.002 | -0.002 | 0.69 | 0.41 | 0.55 | 0.21 | 0.26 | 0.28 | 0.55 |
| 5 | 35363909 | rs26193 | -0.003 | -0.748 | 0.496 | 0.176 | 0.013 | 0.017 | 0.017 | 1.865 | 0.41 | 0.58 | 0.21 | 0.17 | 0.28 | 0.55 |
| 5 | 35364432 | rs26196 | -1.341 | -0.706 | 0.564 | 0.351 | 0.054 | 0.199 | 0.199 | 0 | 0.07 | 0.38 | 0.06 | 0.13 | 0.03 | 0.09 |
| 5 | 35366255 | rs244777 | -1.35 | -0.929 | 0.392 | 0.07 | -0.03 | -0.006 | -0.006 | 0 | 0.14 | 0.37 | 0.08 | 0.1 | 0.09 | 0.1 |
| 5 | 35369453 | rs413206 | 0.5 | -0.927 | 0.257 | 0.064 | 0.028 | 0.023 | 0.023 | 0.441 | 0.29 | 0.17 | 0.14 | 0.23 | 0.17 | 0.45 |
| 5 | 35371472 | rs12522594 | -0.557 | -0.571 | 0.457 | 0.234 | 0.013 | -0.036 | -0.036 | 0 | 0.08 | 0.21 | 0.07 | 0 | 0.04 | 0.01 |
| 5 | 35375710 | rs7709680 | 0.44 | -0.785 | 0.416 | 0.771 | 0.006 | 0.001 | 0.001 | 0.577 | 0.62 | 0.35 | 0.66 | 0.57 | 0.71 | 0.44 |
| 5 | 35377821 | rs32775 | 0.608 | -0.771 | -0.014 | 0.792 | -0.008 | 0.013 | 0.013 | 0.668 | 0.26 | 0.12 | 0.15 | 0.23 | 0.2 | 0.46 |
| 5 | 35381373 | rs32782 | -0.16 | -0.8 | -0.019 | 0.802 | 0 | 0.023 | 0.023 | 1.518 | 0.33 | 0.26 | 0.18 | 0.24 | 0.28 | 0.53 |
| 5 | 35385610 | rs7734554 | 0.288 | -0.774 | -0.261 | 0.719 | -0.009 | -0.003 | -0.003 | 0 | 0.64 | 0.49 | 0.68 | 0.59 | 0.72 | 0.44 |
| 5 | 35392886 | rs421834 | 0.501 | -0.415 | -1.17 | 0.423 | -0.014 | 0.053 | 0.053 | 0.656 | 0.25 | 0.11 | 0.13 | 0.18 | 0.2 | 0.45 |
| 5 | 35393350 | rs2194251 | 0.415 | -0.862 | -1.438 | 0.774 | 0.05 | -0.035 | -0.035 | 0 | 0.51 | 0.13 | 0.58 | 0.42 | 0.64 | 0.46 |
| 5 | 35395245 | rs404220 | 0.205 | -0.686 | -0.547 | 1.037 | 0.01 | 0.126 | 0.126 | 0.019 | 0.88 | 0.74 | 0.7 | 0.66 | 0.87 | 0.64 |
| 5 | 35395998 | rs421175 | 0.626 | -0.297 | -1.167 | 0.908 | -0.1 | -0.008 | -0.008 | 0 | 0.19 | 0.04 | 0.32 | 0.14 | 0.39 | 0.11 |
| 5 | 35396172 | rs444265 | -0.358 | -1.269 | -0.595 | 1.127 | -0.001 | 0.047 | 0.047 | 0 | 0.91 | 0.9 | 0.73 | 0.67 | 0.91 | 0.64 |
| 5 | 35407672 | rs12519140 | 2.047 | -1.634 | -0.636 | 0.894 | 0 | -0.005 | -0.005 | 0 | 0.07 | 0.04 | 0.05 | 0 | 0.02 | 0 |
| 5 | 35422561 | rs412904 | 1.065 | -2.032 | 0.144 | 0.742 | -0.045 | 0.028 | 0.028 | 0.458 | 0.76 | 0.37 | 0.63 | 0.66 | 0.87 | 0.61 |
| 5 | 35430751 | rs16902013 | 0.156 | -1.924 | 0.117 | 0.641 | -0.07 | -0.092 | -0.092 | 0 | 0.14 | 0.35 | 0.1 | 0.01 | 0.04 | 0.07 |
| 5 | 35455201 | rs284725 | -1.352 | -2.525 | -0.211 | 0.733 | -0.002 | -0.004 | -0.004 | 0 | 0.37 | 0.59 | 0.45 | 0.57 | 0.19 | 0.45 |
| 5 | 35458475 | rs1605685 | -0.041 | -2.195 | -0.636 | 0.635 | -0.021 | 0.006 | 0.006 | 7.9 | 0.15 | 0.15 | 0.1 | 0.24 | 0.07 | 0.08 |
| 5 | 35458531 | rs1605684 | 0.078 | -2.2 | -0.608 | 0.632 | -0.029 | 0.009 | 0.009 | 7.162 | 0.15 | 0.18 | 0.1 | 0.24 | 0.07 | 0.08 |
| 5 | 35461200 | rs284737 | -0.858 | -1.815 | -1.186 | 0.12 | -0.052 | 0.057 | 0.057 | 0 | 0.6 | 0.89 | 0.58 | 0.79 | 0.41 | 0.73 |
| 5 | 35466081 | rs17252095 | 0.092 | -1.3 | -0.486 | 0.28 | 0 | -0.003 | -0.003 | 0 | 0.09 | 0.1 | 0.04 | 0.05 | 0.08 | 0.02 |
| 5 | 35471886 | rs10053766 | -0.864 | -1.515 | -0.081 | 0.165 | -0.016 | -0.05 | -0.05 | 0 | 0.14 | 0.26 | 0.06 | 0.06 | 0.09 | 0.08 |
| 5 | 35490158 | rs16902178 | -0.149 | -1.251 | 0.153 | 0.255 | 0.045 | -0.006 | -0.006 | 0.996 | 0.08 | 0.27 | 0.07 | 0.05 | 0.09 | 0.02 |
| 5 | 35490745 | rs10472966 | -0.365 | -1.266 | 0.578 | 0.243 | 0.057 | -0.006 | -0.006 | 0 | 0.09 | 0.38 | 0.08 | 0.05 | 0.09 | 0.02 |
| 5 | 35503887 | rs13172117 | 1.032 | -1.658 | -0.182 | 0.035 | -0.104 | 0.02 | 0.02 | 0 | 0.35 | 0.1 | 0.36 | 0.22 | 0.56 | 0.22 |
| 5 | 35516582 | rs10059179 | -1.015 | -1.534 | 0.85 | 0.689 | -0.018 | 0.044 | 0.044 | 1.01 | 0.25 | 0.57 | 0.4 | 0.51 | 0.19 | 0.32 |
| 5 | 35529733 | rs10072334 | 0.388 | -1.527 | 0.664 | 0.726 | -0.038 | 0.138 | 0.138 | 0 | 0.16 | 0.27 | 0.34 | 0.46 | 0.1 | 0.31 |
| 5 | 35535125 | rs286427 | 0.873 | -1.494 | 0.606 | 0.67 | -0.022 | 0.069 | 0.069 | 0.888 | 0.74 | 0.42 | 0.65 | 0.48 | 0.8 | 0.66 |

| | | | | | | | | | | | | | | | | |
|---|---|---|---|---|---|---|---|---|---|---|---|---|---|---|---|---|
| 5 | 35542960 | rs868011 | 0.365 | -1.291 | 0.176 | 0.803 | 0.019 | -0.089 | -0.089 | 0 | 0.15 | 0.03 | 0.34 | 0.46 | 0.12 | 0.35 |
| 5 | 35548874 | rs12653412 | 0.487 | -1.269 | 0.308 | 0.774 | -0.004 | -0.024 | -0.024 | 0.775 | 0.08 | 0.01 | 0.3 | 0.47 | 0.09 | 0.31 |
| 5 | 35551484 | rs12659209 | 0.48 | -1.278 | 0.302 | 0.768 | -0.003 | -0.025 | -0.025 | 0.061 | 0.09 | 0.01 | 0.3 | 0.47 | 0.09 | 0.31 |
| 5 | 35559916 | rs286411 | 0.793 | -1.759 | 0.143 | 0.271 | -0.004 | 0.025 | 0.025 | 0 | 0.44 | 0.24 | 0.4 | 0.33 | 0.56 | 0.32 |
| 5 | 35578461 | rs169587 | -0.379 | -1.861 | 0.137 | 0.113 | 0.035 | 0.096 | 0.096 | 0 | 0.47 | 0.72 | 0.3 | 0.2 | 0.35 | 0.37 |
| 5 | 35585387 | rs10512623 | 0.958 | -1.963 | -0.228 | 0.149 | -0.008 | -0.016 | -0.016 | 0 | 0.18 | 0.03 | 0.13 | 0.14 | 0.14 | 0.2 |
| 5 | 35586717 | rs12651905 | 0.186 | -1.498 | 0.123 | 0.553 | -0.001 | -0.023 | -0.023 | 0 | 0.11 | 0.09 | 0.31 | 0.46 | 0.12 | 0.39 |
| 5 | 35588057 | rs16902314 | -1.028 | -1.596 | 0.403 | 0.413 | -0.024 | 0.062 | 0.062 | 0 | 0.26 | 0.43 | 0.42 | 0.47 | 0.21 | 0.52 |
| 5 | 35608310 | rs12657378 | 0.26 | -1.57 | 0.165 | 0.439 | 0 | -0.003 | -0.003 | 0 | 0.11 | 0.11 | 0.31 | 0.46 | 0.12 | 0.39 |
| 5 | 35616001 | rs10941251 | -1.029 | -1.583 | 0.225 | 0.429 | -0.022 | 0.058 | 0.058 | 0 | 0.26 | 0.43 | 0.42 | 0.47 | 0.21 | 0.52 |
| 5 | 35616805 | rs11951251 | 0.111 | -1.108 | -0.441 | 0.805 | 0 | -0.003 | -0.003 | 0.025 | 0.11 | 0.11 | 0.31 | 0.46 | 0.12 | 0.39 |
| 5 | 35618486 | rs1508650 | -0.003 | -1.14 | -0.446 | 0.799 | 0.002 | -0.009 | -0.009 | 0 | 0.11 | 0.12 | 0.31 | 0.45 | 0.11 | 0.39 |
| 5 | 35621098 | rs286437 | -0.583 | -2.102 | -0.373 | 0.094 | -0.002 | 0.002 | 0.002 | 0.068 | 0.27 | 0.62 | 0.19 | 0.17 | 0.13 | 0.19 |
| 5 | 35632052 | rs455000 | -0.9 | -1.684 | -1.101 | -0.431 | -0.069 | 0.096 | 0.096 | 0.257 | 0.65 | 0.81 | 0.73 | 0.9 | 0.54 | 0.86 |
| 5 | 35633433 | rs452060 | 0.346 | -1.711 | -0.316 | -0.351 | -0.013 | -0.076 | -0.076 | 0 | 0.12 | 0.37 | 0.08 | 0 | 0.06 | 0.1 |
| 5 | 35639275 | rs286445 | -0.567 | -1.839 | -0.976 | -0.3 | 0.009 | 0.068 | 0.068 | 0 | 0.55 | 0.74 | 0.65 | 0.75 | 0.47 | 0.72 |
| 5 | 35641582 | rs6897513 | -0.568 | -1.215 | -1.105 | -0.451 | -0.04 | 0.056 | 0.056 | 0 | 0.57 | 0.72 | 0.69 | 0.74 | 0.53 | 0.71 |
| 5 | 35642939 | rs12513652 | 0.863 | -1.583 | -0.863 | -0.389 | -0.041 | -0.064 | -0.064 | 0.081 | 0.3 | 0.14 | 0.44 | 0.5 | 0.39 | 0.48 |
| 5 | 35644621 | rs7706444 | -1.239 | -1.198 | -0.635 | -0.273 | -0.035 | 0.064 | 0.064 | 0 | 0.69 | 0.78 | 0.77 | 0.89 | 0.62 | 0.86 |
| 5 | 35655248 | rs10472978 | 0.394 | -1.224 | -0.064 | -0.213 | -0.031 | -0.08 | -0.08 | 0 | 0.12 | 0.31 | 0.08 | 0 | 0.07 | 0.09 |
| 5 | 35670741 | rs6891905 | -0.412 | -1.069 | -0.244 | -0.109 | -0.05 | 0.058 | 0.058 | 0 | 0.57 | 0.74 | 0.69 | 0.74 | 0.53 | 0.71 |
| 5 | 35675616 | rs16902391 | 0.758 | -1.202 | 0.008 | 0.813 | 0.059 | 0.293 | 0.293 | 0.257 | 0.05 | 0.21 | 0.17 | 0.39 | 0.06 | 0.17 |
| 5 | 35677385 | rs6872795 | -2.07 | -0.806 | -0.656 | 0.511 | -0.088 | -0.016 | -0.016 | 0.05 | 0.83 | 0.97 | 0.82 | 0.89 | 0.69 | 0.86 |
| 5 | 35682295 | rs1013156 | -0.594 | -0.886 | -0.313 | 0.643 | -0.055 | 0.055 | 0.055 | 0 | 0.57 | 0.82 | 0.68 | 0.75 | 0.51 | 0.7 |
| 5 | 35684832 | rs16902409 | 0.839 | -1.022 | -0.206 | 0.64 | 0.013 | 0.01 | 0.01 | 0 | 0.37 | 0.19 | 0.4 | 0.34 | 0.36 | 0.44 |
| 5 | 35686341 | rs6894157 | -0.855 | -0.793 | 0.091 | 0.821 | -0.017 | 0.049 | 0.049 | 0.336 | 0.19 | 0.62 | 0.26 | 0.41 | 0.13 | 0.25 |
| 5 | 35698426 | rs13328131 | 0.017 | -0.767 | 0.348 | 0.177 | -0.034 | -0.089 | -0.089 | 0 | 0.13 | 0.37 | 0.08 | 0.01 | 0.07 | 0.08 |
| 5 | 35700598 | rs6451206 | -0.386 | -0.781 | 0.176 | 0.194 | -0.003 | -0.018 | -0.018 | 0 | 0.58 | 0.53 | 0.41 | 0.26 | 0.57 | 0.4 |
| 5 | 35707388 | rs7713174 | -0.375 | -0.775 | 0.18 | 0.197 | -0.003 | -0.018 | -0.018 | 0 | 0.58 | 0.53 | 0.41 | 0.26 | 0.57 | 0.4 |
| 5 | 35710330 | rs13356541 | -1.082 | -0.644 | 0.447 | -0.233 | -0.013 | 0.015 | 0.015 | 0 | 0.8 | 0.64 | 0.74 | 0.8 | 0.8 | 0.67 |
| 5 | 35712580 | rs13186771 | 0.187 | -0.747 | 0.317 | -0.323 | -0.008 | -0.007 | -0.007 | 0 | 0.68 | 0.37 | 0.67 | 0.8 | 0.73 | 0.58 |
| 5 | 35718745 | rs10434594 | 0.006 | -0.715 | 0.353 | -0.318 | -0.042 | -0.022 | -0.022 | 0.413 | 0.71 | 0.5 | 0.72 | 0.8 | 0.78 | 0.62 |
| 5 | 35722448 | rs7448303 | 0.4 | -0.641 | 0.369 | -0.12 | 0.066 | 0.079 | 0.079 | 0 | 0.06 | 0.2 | 0.12 | 0.14 | 0.11 | 0.05 |
| 5 | 35723861 | rs10061088 | 0.832 | -0.722 | 0.392 | -0.092 | -0.028 | -0.048 | -0.048 | 0 | 0.09 | 0.12 | 0.02 | 0 | 0.02 | 0.05 |
| 5 | 35724053 | rs7710255 | 0.326 | -0.676 | 0.114 | -0.058 | 0.014 | -0.002 | -0.002 | 0 | 0.65 | 0.3 | 0.6 | 0.66 | 0.68 | 0.57 |
| 5 | 35728440 | rs4130679 | -0.478 | -0.559 | -0.124 | -0.274 | -0.008 | -0.005 | -0.005 | 0.464 | 0.66 | 0.81 | 0.57 | 0.31 | 0.64 | 0.57 |
| 5 | 35730729 | rs11960663 | 0.775 | -0.49 | -0.08 | -0.523 | -0.023 | 0.068 | 0.068 | 0 | 0.35 | 0.24 | 0.23 | 0.1 | 0.37 | 0.16 |
| 5 | 35732551 | rs10053840 | 0.25 | -0.27 | 0.039 | -0.439 | -0.002 | 0.068 | 0.068 | 0 | 0.44 | 0.32 | 0.26 | 0.09 | 0.4 | 0.23 |
| 5 | 35756695 | rs7447258 | 0.196 | -0.046 | 0.233 | -0.003 | 0.072 | 0.165 | 0.165 | 0.638 | 0.92 | 0.61 | 0.9 | 0.85 | 0.93 | 0.84 |
| 5 | 35766304 | rs6893048 | 0.632 | -0.02 | 0.188 | 0.021 | 0.07 | 0.134 | 0.134 | 0.033 | 0.06 | 0.27 | 0.09 | 0.11 | 0.07 | 0.01 |
| 5 | 35776239 | rs11749262 | 0.29 | -0.217 | -0.207 | -0.064 | -0.001 | -0.003 | -0.003 | 0 | 0.27 | 0.35 | 0.34 | 0.17 | 0.28 | 0.29 |
| 5 | 35781435 | rs10941263 | 0.871 | -0.088 | -0.619 | 0.059 | -0.072 | -0.184 | -0.184 | 0 | 0.29 | 0.02 | 0.36 | 0.56 | 0.3 | 0.33 |
| 5 | 35787873 | rs10058955 | -0.834 | 0.343 | -0.092 | -0.227 | -0.074 | 0.048 | 0.048 | 0.571 | 0.78 | 0.96 | 0.75 | 0.88 | 0.68 | 0.82 |
| 5 | 35792990 | rs3822736 | -0.011 | -0.061 | 0.005 | 0.18 | -0.006 | 0.002 | 0.002 | 0 | 0.22 | 0.38 | 0.29 | 0.21 | 0.24 | 0.28 |
| 5 | 35799994 | rs3822735 | -0.108 | 0.344 | -0.869 | -0.559 | 0.002 | 0.045 | 0.045 | 0.342 | 0.64 | 0.71 | 0.72 | 0.9 | 0.68 | 0.87 |
| 5 | 35801125 | rs10050755 | -0.302 | -0.218 | -0.923 | -1.006 | -0.011 | -0.033 | -0.033 | 0.111 | 0.29 | 0.35 | 0.25 | 0.1 | 0.18 | 0.19 |
| 5 | 35801338 | rs11948206 | 0.512 | 0.176 | -0.957 | -0.646 | 0.02 | -0.029 | -0.029 | 0 | 0.31 | 0.5 | 0.23 | 0.2 | 0.31 | 0.16 |
| 5 | 35803577 | rs931555 | 1.506 | 0.492 | -1.403 | -0.019 | -0.08 | -0.071 | -0.071 | 0 | 0.17 | 0.03 | 0.19 | 0.2 | 0.31 | 0.24 |
| 5 | 35804725 | rs10155582 | -0.466 | 0.175 | -1.211 | -0.282 | -0.004 | -0.025 | -0.025 | 0 | 0.15 | 0.35 | 0.19 | 0.1 | 0.12 | 0.17 |
| 5 | 35805553 | rs1389836 | 1.209 | 0.125 | -0.758 | -0.528 | 0.014 | -0.057 | -0.057 | 0.668 | 0.25 | 0.3 | 0.16 | 0.09 | 0.26 | 0.14 |
| 5 | 35807455 | rs7736291 | -0.683 | 0.104 | -0.972 | -0.439 | -0.077 | 0.081 | 0.081 | 0.003 | 0.45 | 0.88 | 0.42 | 0.31 | 0.43 | 0.34 |
| 5 | 35808115 | rs1908369 | -0.451 | 0.05 | -0.679 | -0.352 | -0.006 | -0.025 | -0.025 | 0 | 0.15 | 0.35 | 0.19 | 0.1 | 0.12 | 0.17 |
| 5 | 35816128 | rs6892686 | 0.006 | 0.013 | -0.641 | -0.248 | 0.014 | 0.023 | 0.023 | 1.983 | 0.08 | 0.16 | 0.14 | 0.14 | 0.1 | 0.06 |
| 5 | 35817219 | rs1494569 | 1.109 | -0.423 | -1.123 | -0.346 | -0.017 | -0.17 | -0.17 | 0.517 | 0.23 | 0.02 | 0.37 | 0.53 | 0.2 | 0.42 |
| 5 | 35819147 | rs6869232 | 0.083 | -0.513 | -1.106 | -0.639 | 0.011 | 0.022 | 0.022 | 6.235 | 0.23 | 0.16 | 0.2 | 0.13 | 0.17 | 0.1 |
| 5 | 35831936 | rs6872631 | -0.05 | -0.417 | -0.748 | -0.999 | 0 | -0.027 | -0.027 | 0 | 0.44 | 0.5 | 0.26 | 0.13 | 0.34 | 0.3 |
| 5 | 35833717 | rs11746643 | 0.388 | -0.412 | -0.652 | -0.838 | 0.006 | 0.043 | 0.043 | 0 | 0.36 | 0.29 | 0.27 | 0.22 | 0.38 | 0.31 |
| 5 | 35847722 | rs1494565 | 0.396 | -0.315 | -0.206 | -0.682 | -0.002 | -0.015 | -0.015 | 0 | 0.43 | 0.45 | 0.37 | 0.24 | 0.58 | 0.39 |
| 5 | 35855410 | rs4869492 | 0.35 | -0.479 | -0.166 | -0.483 | 0 | 0 | 0 | 0 | 0.13 | 0.13 | 0.17 | 0.15 | 0.14 | 0.06 |
| 5 | 35856575 | rs11567685 | 1.31 | -0.346 | 0.016 | -0.683 | 0.025 | -0.088 | -0.088 | 1.833 | 0.24 | 0.28 | 0.17 | 0.04 | 0.28 | 0.18 |
| 5 | 35861068 | rs1494558 | -0.561 | -0.228 | 0.179 | -0.38 | 0.064 | -0.001 | -0.001 | 0 | 0.57 | 0.76 | 0.54 | 0.43 | 0.7 | 0.5 |
| 5 | 35862841 | rs7717955 | 2.009 | -0.26 | 0.152 | -0.379 | -0.026 | -0.014 | -0.014 | 0 | 0.18 | 0.14 | 0.19 | 0.19 | 0.27 | 0.21 |
| 5 | 35863244 | rs6451226 | -0.71 | -0.335 | 0.226 | -0.4 | 0.078 | -0.019 | -0.019 | 0.06 | 0.58 | 0.84 | 0.55 | 0.43 | 0.7 | 0.5 |
| 5 | 35864590 | rs10044838 | -0.71 | -0.335 | 0.226 | -0.4 | 0.078 | -0.019 | -0.019 | 0 | 0.58 | 0.84 | 0.55 | 0.43 | 0.7 | 0.5 |
| 5 | 35866322 | rs6891095 | 0.427 | -0.321 | 0.202 | -0.399 | 0 | 0 | 0 | 0 | 0.13 | 0.13 | 0.17 | 0.15 | 0.15 | 0.06 |
| 5 | 35871190 | rs1494555 | -0.892 | -0.41 | 0.137 | -0.367 | 0.044 | -0.137 | -0.137 | 0 | 0.61 | 0.93 | 0.56 | 0.48 | 0.7 | 0.54 |
| 5 | 35874575 | rs6897932 | 2.212 | -0.417 | 0.526 | -0.383 | -0.045 | -0.016 | -0.016 | 0 | 0.17 | 0.07 | 0.18 | 0.17 | 0.27 | 0.21 |
| 5 | 35876274 | rs3194051 | 1.413 | -0.407 | 0.593 | -0.424 | 0.04 | -0.094 | -0.094 | 0 | 0.24 | 0.35 | 0.18 | 0.06 | 0.28 | 0.18 |
| 5 | 35877505 | rs10063294 | 0.301 | -0.422 | 0.27 | -0.405 | 0.003 | 0.109 | 0.109 | 1.242 | 0.45 | 0.81 | 0.39 | 0.27 | 0.56 | 0.44 |

| | | | | | | | | | | | | | | | | |
|---|---|---|---|---|---|---|---|---|---|---|---|---|---|---|---|---|
| 5 | 35878927 | rs6451231 | -0.852 | -0.463 | 0.096 | -0.224 | 0.044 | -0.068 | -0.068 | 2.31 | 0.61 | 0.94 | 0.56 | 0.39 | 0.7 | 0.53 |
| 5 | 35880820 | rs10058453 | -0.628 | -0.452 | 0.109 | -0.257 | 0.072 | -0.053 | -0.053 | 0.004 | 0.59 | 0.91 | 0.56 | 0.43 | 0.7 | 0.5 |
| 5 | 35881376 | rs11742240 | 1.558 | -0.414 | 0.178 | -0.257 | -0.008 | -0.002 | -0.002 | 0 | 0.19 | 0.21 | 0.19 | 0.21 | 0.27 | 0.26 |
| 5 | 35883804 | rs2194225 | -0.084 | -0.539 | 0.194 | -0.284 | -0.023 | 0.04 | 0.04 | 0.022 | 0.39 | 0.66 | 0.36 | 0.21 | 0.43 | 0.24 |
| 5 | 35883986 | rs10214273 | 2.099 | -0.405 | -0.112 | -0.287 | -0.045 | -0.013 | -0.013 | 0 | 0.17 | 0.07 | 0.19 | 0.17 | 0.28 | 0.21 |
| 5 | 35885710 | rs10044883 | -0.482 | -0.25 | 0.017 | -0.097 | 0.045 | -0.138 | -0.138 | 0 | 0.62 | 0.93 | 0.57 | 0.48 | 0.71 | 0.53 |
| 5 | 35887640 | rs1423658 | 1.065 | -0.316 | -0.147 | -0.211 | 0.023 | -0.165 | -0.165 | 0 | 0.2 | 0.01 | 0.33 | 0.52 | 0.14 | 0.42 |
| 5 | 35893746 | rs10059688 | 0.375 | -0.122 | 0.207 | -0.28 | 0.002 | 0.004 | 0.004 | 0 | 0.12 | 0.15 | 0.17 | 0.17 | 0.13 | 0.05 |
| 5 | 35896984 | rs6451234 | 0.878 | -0.235 | 0.34 | -0.267 | -0.02 | 0.038 | 0.038 | 0.828 | 0.81 | 0.61 | 0.81 | 0.73 | 0.86 | 0.86 |
| 5 | 35898876 | rs17376481 | 0.148 | -0.566 | -0.01 | -0.631 | 0.024 | -0.087 | -0.087 | 1.266 | 0.35 | 0.5 | 0.27 | 0.08 | 0.41 | 0.23 |
| 5 | 35903997 | rs2287900 | 1.354 | -0.531 | 0.35 | -0.663 | 0.042 | -0.031 | -0.031 | 0 | 0.16 | 0.22 | 0.15 | 0.07 | 0.27 | 0.19 |
| 5 | 35905211 | rs2361403 | 0.173 | -0.644 | -0.051 | -0.633 | 0.033 | -0.103 | -0.103 | 0.277 | 0.34 | 0.48 | 0.27 | 0.07 | 0.41 | 0.23 |
| 5 | 35906234 | rs12522083 | 1.204 | -0.692 | -0.201 | -0.61 | 0.022 | 0.075 | 0.075 | 0 | 0.47 | 0.11 | 0.57 | 0.66 | 0.45 | 0.63 |
| 5 | 35910419 | rs1445899 | 0.791 | -0.611 | -0.138 | -0.611 | 0.048 | 0.165 | 0.165 | 0 | 0.52 | 0.23 | 0.59 | 0.76 | 0.45 | 0.71 |
| 5 | 35910529 | rs1445898 | 0.791 | -0.611 | -0.138 | -0.611 | 0.048 | 0.165 | 0.165 | 0 | 0.52 | 0.23 | 0.59 | 0.76 | 0.45 | 0.71 |
| 5 | 35921069 | rs6859892 | -0.929 | -0.561 | -0.014 | -0.564 | 0.009 | 0.017 | 0.017 | 0 | 0.88 | 0.81 | 0.87 | 0.83 | 0.87 | 0.95 |
| 5 | 35922462 | rs852251 | 0.837 | -0.507 | -0.132 | -0.522 | 0.032 | -0.068 | -0.068 | 0 | 0.23 | 0.12 | 0.38 | 0.51 | 0.17 | 0.42 |
| 5 | 35923014 | rs852249 | -0.68 | -0.397 | -0.119 | -0.43 | 0.005 | 0 | 0 | 0 | 0.58 | 0.59 | 0.65 | 0.57 | 0.58 | 0.66 |
| 5 | 35925214 | rs852242 | -0.417 | -0.86 | -0.389 | 0.092 | 0.004 | -0.009 | -0.009 | 1.636 | 0.43 | 0.46 | 0.43 | 0.37 | 0.45 | 0.55 |
| 5 | 35926341 | rs11948151 | -0.009 | -0.364 | -0.233 | -0.238 | 0.029 | -0.026 | -0.026 | 0.466 | 0.25 | 0.58 | 0.26 | 0.17 | 0.35 | 0.25 |
| 5 | 35931966 | rs13157282 | 0.379 | 0.16 | -0.292 | -0.334 | 0.032 | -0.017 | -0.017 | 0 | 0.13 | 0.18 | 0.14 | 0.05 | 0.22 | 0.07 |
| 5 | 35933203 | rs700174 | -1.313 | 0.059 | -0.549 | -0.515 | 0.028 | 0.112 | 0.112 | 0 | 0.53 | 0.72 | 0.66 | 0.76 | 0.56 | 0.69 |
| 5 | 35933518 | rs6451241 | 1.24 | 0.084 | -0.496 | -0.505 | 0.03 | -0.026 | -0.026 | 1.808 | 0.13 | 0.42 | 0.11 | 0.08 | 0.12 | 0.19 |
| 5 | 35936237 | rs962029 | -1.83 | 0.043 | -1.036 | -0.536 | -0.003 | 0.052 | 0.052 | 0 | 0.83 | 0.93 | 0.89 | 1 | 0.85 | 0.96 |
| 5 | 35940365 | rs10472267 | -1.267 | -0.043 | -1.061 | -0.518 | 0.01 | 0.033 | 0.033 | 0.036 | 0.8 | 0.86 | 0.89 | 0.94 | 0.85 | 0.91 |
| 5 | 35943042 | rs860413 | 2.396 | -0.066 | -1.098 | -0.494 | -0.008 | 0.036 | 0.036 | 0 | 0.27 | 0.09 | 0.22 | 0.18 | 0.29 | 0.22 |
| 5 | 35946296 | rs852235 | 0.509 | 0.176 | -1.121 | -0.367 | -0.021 | -0.051 | -0.051 | 0 | 0.26 | 0.04 | 0.43 | 0.7 | 0.24 | 0.47 |
| 5 | 35950184 | rs1019504 | -1.101 | 0.072 | -0.698 | -0.478 | 0.006 | -0.052 | -0.052 | 0 | 0.42 | 0.54 | 0.58 | 0.82 | 0.46 | 0.54 |
| 5 | 35953320 | rs2242225 | -0.855 | 0.078 | -0.69 | -0.474 | -0.017 | 0.027 | 0.027 | 0 | 0.39 | 0.34 | 0.53 | 0.76 | 0.46 | 0.53 |
| 5 | 35953697 | rs10045685 | -0.963 | 0.076 | -0.664 | -0.477 | -0.011 | 0.019 | 0.019 | 0 | 0.39 | 0.37 | 0.53 | 0.78 | 0.46 | 0.53 |
| 5 | 35953949 | rs17313508 | 2.44 | 0.002 | -0.851 | -0.561 | -0.031 | 0.111 | 0.111 | 0 | 0.27 | 0.03 | 0.22 | 0.09 | 0.28 | 0.24 |
| 5 | 35954143 | rs700165 | 0.262 | 0.232 | -0.611 | -0.296 | 0.082 | -0.003 | -0.003 | 0 | 0.14 | 0.26 | 0.12 | 0.11 | 0.22 | 0.08 |
| 5 | 35954588 | rs700164 | 0.416 | -0.072 | -0.706 | -0.219 | -0.02 | -0.08 | -0.08 | 2.405 | 0.24 | 0.03 | 0.41 | 0.65 | 0.24 | 0.44 |
| 5 | 35959184 | rs997187 | 1.004 | 0.008 | -0.349 | -0.302 | -0.001 | -0.041 | -0.041 | 0 | 0.47 | 0.42 | 0.33 | 0.12 | 0.42 | 0.33 |
| 5 | 35964286 | rs10472991 | 0.901 | 0.277 | 0.065 | 0.244 | 0.024 | -0.035 | -0.035 | 0 | 0.65 | 0.52 | 0.52 | 0.24 | 0.6 | 0.5 |
| 5 | 35965970 | rs3756669 | 0.799 | -0.156 | -0.243 | 0.964 | 0.006 | -0.108 | -0.108 | 0.111 | 0.16 | 0.02 | 0.22 | 0.38 | 0.14 | 0.37 |
| 5 | 35966681 | rs10040617 | 1.681 | -0.196 | -0.332 | 0.681 | 0.014 | 0.104 | 0.104 | 0 | 0.29 | 0.15 | 0.22 | 0.09 | 0.29 | 0.24 |
| 5 | 35973875 | rs1374011 | 0.406 | 0.003 | 0.162 | 0.72 | 0.044 | 0.065 | 0.065 | 0 | 0.83 | 0.7 | 0.77 | 0.65 | 0.79 | 0.85 |
| 5 | 35988950 | rs745300 | 0.613 | -0.021 | 0.26 | 0.48 | -0.002 | -0.014 | -0.014 | 0 | 0.35 | 0.3 | 0.27 | 0.11 | 0.37 | 0.24 |
| 5 | 35992621 | rs16902679 | 1.074 | -0.237 | 0.713 | 1.228 | 0.031 | 0.114 | 0.114 | 0 | 0.79 | 0.44 | 0.72 | 0.58 | 0.8 | 0.85 |
| 5 | 35997920 | rs4288141 | 0.32 | -0.34 | 0.579 | 1.293 | 0 | -0.01 | -0.01 | 0.325 | 0.55 | 0.33 | 0.54 | 0.51 | 0.58 | 0.68 |
| 5 | 36001766 | rs6890283 | 0.61 | -0.188 | 0.343 | 1.434 | 0.021 | 0.04 | 0.04 | 0 | 0.82 | 0.72 | 0.71 | 0.66 | 0.79 | 0.9 |
| 5 | 36003217 | rs7715372 | 0.124 | -0.482 | 0.201 | 1.381 | -0.019 | 0.01 | 0.01 | 0 | 0.42 | 0.48 | 0.38 | 0.46 | 0.35 | 0.61 |
| 5 | 36014599 | rs2218522 | 0.457 | -0.391 | -0.703 | 1.522 | -0.026 | 0.056 | 0.056 | 0.083 | 0.53 | 0.1 | 0.41 | 0.37 | 0.57 | 0.69 |
| 5 | 36027156 | rs1351142 | 1.566 | -0.704 | -0.855 | 1.151 | -0.002 | 0.034 | 0.034 | 0 | 0.22 | 0.04 | 0.16 | 0.15 | 0.21 | 0.21 |
| 5 | 36028200 | rs2447876 | 0.383 | -0.261 | -0.045 | 1.242 | 0.023 | 0.033 | 0.033 | 0 | 0.87 | 0.73 | 0.83 | 0.81 | 0.86 | 0.93 |
| 5 | 36031513 | rs3860791 | 0.83 | -0.148 | -1.73 | 0.868 | -0.155 | 0.231 | 0.231 | 0.363 | 0.4 | 0.02 | 0.31 | 0.13 | 0.47 | 0.29 |
| 5 | 36031870 | rs2453299 | 0.315 | -0.139 | -0.767 | 1.675 | -0.015 | -0.016 | -0.016 | 0 | 0.89 | 0.93 | 0.86 | 0.8 | 0.86 | 0.93 |
| 5 | 36034020 | rs3912137 | 0.958 | -0.509 | -0.592 | 1.74 | -0.003 | 0.048 | 0.048 | 1.201 | 0.71 | 0.47 | 0.62 | 0.52 | 0.77 | 0.85 |
| 5 | 36034056 | rs10063618 | 0.689 | -0.732 | -1.241 | 1.43 | -0.009 | 0.015 | 0.015 | 0 | 0.24 | 0.04 | 0.21 | 0.2 | 0.23 | 0.19 |
| 5 | 36036278 | rs6451254 | 0.049 | -1.148 | -0.945 | 1.697 | 0.045 | -0.032 | -0.032 | 0 | 0.8 | 0.91 | 0.91 | 0.77 | 0.89 | 0.85 |
| 5 | 36038220 | rs2289877 | -0.525 | -0.841 | -0.707 | 1.476 | 0.026 | -0.039 | -0.039 | 0 | 0.23 | 0.09 | 0.2 | 0.27 | 0.15 | 0.18 |
| 5 | 36038317 | rs2289876 | -0.541 | -0.845 | -0.511 | 1.474 | 0.02 | -0.022 | -0.022 | 0 | 0.23 | 0.14 | 0.2 | 0.27 | 0.15 | 0.18 |
| 5 | 36044279 | rs4869459 | -0.793 | -0.602 | -0.556 | 1.282 | -0.019 | -0.027 | -0.027 | 0 | 0.05 | 0.01 | 0.11 | 0.13 | 0.06 | 0.02 |
| 5 | 36047538 | rs6899037 | 1.294 | -0.801 | -1.042 | 1.087 | -0.038 | 0.042 | 0.042 | 0 | 0.22 | 0.01 | 0.2 | 0.1 | 0.25 | 0.2 |
| 5 | 36050048 | rs1287271 | 0.634 | -1.158 | -0.355 | 1.239 | 0.011 | 0.013 | 0.013 | 0 | 0.45 | 0.44 | 0.48 | 0.42 | 0.56 | 0.37 |
| 5 | 36051611 | rs1287267 | 0.855 | -1.15 | -0.355 | 1.221 | -0.003 | 0.013 | 0.013 | 0 | 0.42 | 0.43 | 0.38 | 0.28 | 0.51 | 0.35 |
| 5 | 36056800 | rs2047176 | 1.05 | -1.633 | -0.016 | 1.31 | 0.087 | 0.02 | 0.02 | 0 | 0.6 | 0.13 | 0.55 | 0.37 | 0.76 | 0.55 |
| 5 | 36058174 | rs921020 | 0.489 | -1.625 | 0.203 | 1.342 | -0.05 | 0.036 | 0.036 | 0 | 0.67 | 0.53 | 0.68 | 0.5 | 0.81 | 0.57 |
| 5 | 36059151 | rs1287260 | 1.084 | -1.621 | -0.118 | 1.183 | 0.09 | 0.015 | 0.015 | 0 | 0.6 | 0.12 | 0.55 | 0.37 | 0.75 | 0.54 |
| 5 | 36064980 | rs17342682 | 1.102 | -1.461 | 0.245 | 0.973 | -0.013 | 0.013 | 0.013 | 0 | 0.31 | 0.23 | 0.27 | 0.25 | 0.41 | 0.27 |
| 5 | 36070109 | rs2197513 | 1.034 | -1.565 | 0.341 | 0.96 | -0.033 | 0.064 | 0.064 | 0 | 0.3 | 0.17 | 0.35 | 0.11 | 0.37 | 0.27 |
| 5 | 36071872 | rs267782 | 0.118 | -1.792 | 0.785 | 1.129 | -0.025 | 0.012 | 0.012 | 0 | 0.73 | 0.73 | 0.75 | 0.48 | 0.86 | 0.59 |
| 5 | 36073812 | rs144694 | 0.417 | -1.739 | 1.023 | 0.996 | 0.001 | -0.003 | -0.003 | 0 | 0.42 | 0.46 | 0.38 | 0.36 | 0.47 | 0.32 |
| 5 | 36074891 | rs267786 | -0.608 | -1.671 | 0.953 | 0.992 | 0.007 | 0.022 | 0.022 | 0.028 | 0.06 | 0.12 | 0.11 | 0.13 | 0.05 | 0.03 |
| 5 | 36088903 | rs155550 | 0.178 | -1.755 | 0.946 | 0.98 | -0.01 | 0.028 | 0.028 | 0 | 0.43 | 0.64 | 0.4 | 0.35 | 0.47 | 0.31 |
| 5 | 36094476 | rs33616 | -0.791 | -1.797 | 0.911 | 1.383 | -0.023 | 0.158 | 0.158 | 0 | 0.94 | 0.87 | 0.84 | 0.73 | 0.96 | 0.97 |
| 5 | 36098822 | rs3799 | 0.973 | -1.56 | 0.618 | 0.528 | -0.053 | 0.084 | 0.084 | 0 | 0.31 | 0.14 | 0.34 | 0.11 | 0.39 | 0.27 |
| 5 | 36103905 | rs17343598 | 0.324 | -0.388 | 0.507 | 0.701 | -0.14 | 0.063 | 0.063 | 0 | 0.33 | 0.02 | 0.25 | 0.22 | 0.4 | 0.28 |

| | | | | | | | | | | | | | | | | |
|---|---|---|---|---|---|---|---|---|---|---|---|---|---|---|---|---|
| 5 | 36142648 | rs12658867 | 0.312 | -0.281 | -1.091 | 0.757 | -0.129 | 0.071 | 0.071 | 0 | 0.33 | 0.03 | 0.25 | 0.22 | 0.41 | 0.31 |
| 5 | 36153847 | rs33674 | -1.344 | -0.75 | 0.769 | 1.346 | 0.066 | 0.156 | 0.156 | 2.188 | 0.07 | 0.33 | 0.13 | 0.16 | 0.06 | 0.03 |
| 5 | 36156654 | rs2362974 | 1.228 | -1.966 | 0.429 | 1.43 | 0.011 | -0.012 | -0.012 | 0 | 0.71 | 0.25 | 0.62 | 0.41 | 0.86 | 0.67 |
| 5 | 36160854 | rs7702272 | -0.235 | -2.453 | 0.293 | 1.222 | -0.039 | -0.015 | -0.015 | 0 | 0.11 | 0.21 | 0.05 | 0.05 | 0.05 | 0.16 |
| 5 | 36160987 | rs155544 | -0.49 | -2.666 | 0.244 | 1.097 | -0.003 | -0.01 | -0.01 | 0 | 0.09 | 0.09 | 0.02 | 0 | 0.02 | 0.05 |
| 5 | 36163470 | rs3815779 | 1.952 | -2.711 | 0.606 | 1.157 | 0.108 | 0.007 | 0.007 | 0.006 | 0.61 | 0.17 | 0.61 | 0.38 | 0.81 | 0.5 |
| 5 | 36171150 | rs16902775 | -0.865 | -2.637 | 0.954 | 1.126 | -0.064 | -0.036 | -0.036 | 0 | 0.15 | 0.57 | 0.08 | 0.05 | 0.05 | 0.16 |
| 5 | 36171420 | rs2287934 | -1.124 | -2.632 | 0.839 | 1.367 | -0.018 | 0.012 | 0.012 | 1.996 | 0.17 | 0.71 | 0.18 | 0.22 | 0.1 | 0.17 |
| 5 | 36173843 | rs37217 | 0.25 | -2.595 | 0.787 | 1.42 | -0.044 | 0.081 | 0.081 | 0 | 0.83 | 0.76 | 0.77 | 0.59 | 0.93 | 0.75 |
| 5 | 36178293 | rs42425 | 1.743 | -2.731 | 0.226 | 1.253 | 0.154 | 0 | 0 | 0.024 | 0.6 | 0.12 | 0.58 | 0.37 | 0.81 | 0.48 |
| 5 | 36179044 | rs3804439 | -1.336 | -2.505 | 0.849 | 1.256 | -0.046 | 0.015 | 0.015 | 0 | 0.22 | 0.5 | 0.18 | 0.22 | 0.12 | 0.25 |
| 5 | 36181627 | rs7731023 | 0.827 | -2.112 | -0.682 | 0.292 | -0.03 | 0.157 | 0.157 | 0 | 0.37 | 0.1 | 0.43 | 0.09 | 0.6 | 0.23 |
| 5 | 36182760 | rs27733 | -0.174 | -2.656 | -0.169 | 1.066 | -0.041 | 0.122 | 0.122 | 0 | 0.91 | 0.83 | 0.84 | 0.7 | 0.97 | 0.96 |
| 5 | 36191407 | rs13160147 | 0.857 | -1.838 | -0.837 | 0.338 | -0.076 | 0.039 | 0.039 | 0 | 0.23 | 0.11 | 0.29 | 0.09 | 0.42 | 0.15 |
| 5 | 36192232 | rs7719686 | -0.229 | -1.903 | -0.653 | 0.709 | 0.033 | 0.045 | 0.045 | 0.041 | 0.53 | 0.7 | 0.39 | 0.39 | 0.37 | 0.72 |
| 5 | 36196428 | rs7716788 | 0.836 | -2.508 | -0.917 | 0.549 | -0.059 | 0.029 | 0.029 | 0 | 0.36 | 0.04 | 0.29 | 0.29 | 0.35 | 0.45 |
| 5 | 36220337 | rs6872093 | 0.832 | -2.618 | -0.255 | 0.518 | -0.038 | 0.013 | 0.013 | 0 | 0.07 | 0.17 | 0.07 | 0.1 | 0.01 | 0.24 |
| 5 | 36224240 | rs1364017 | -0.677 | -2.619 | -0.254 | 0.545 | -0.042 | 0.117 | 0.117 | 0 | 0.91 | 0.83 | 0.84 | 0.7 | 0.97 | 0.96 |
| 5 | 36230339 | rs10941279 | 0.63 | -2.602 | 0.358 | 0.401 | 0.002 | 0.032 | 0.032 | 0 | 0.37 | 0.24 | 0.31 | 0.29 | 0.35 | 0.48 |
| 5 | 36233663 | rs11738984 | 0.587 | -2.736 | 0.004 | 0.215 | -0.001 | 0.062 | 0.062 | 0 | 0.12 | 0.01 | 0.06 | 0.02 | 0.11 | 0.28 |
| 5 | 36238287 | rs10454863 | 1.017 | -2.486 | 0.298 | 0.396 | 0.001 | -0.003 | -0.003 | 0 | 0.41 | 0.38 | 0.39 | 0.09 | 0.59 | 0.21 |
| 5 | 36242354 | rs10075150 | -0.12 | -2.528 | -0.025 | 0.497 | 0.03 | 0.107 | 0.107 | 0 | 0.5 | 0.23 | 0.39 | 0.37 | 0.45 | 0.74 |
| 5 | 36248752 | rs7722251 | 0.149 | -2.736 | 0.895 | 1.044 | -0.043 | 0.209 | 0.209 | 0 | 0.9 | 0.65 | 0.83 | 0.71 | 0.96 | 0.94 |
| 5 | 36253937 | rs4869473 | 0.392 | -2.741 | 0.737 | 1.041 | -0.049 | 0.245 | 0.245 | 0 | 0.89 | 0.57 | 0.82 | 0.71 | 0.96 | 0.94 |
| 5 | 36254647 | rs4869632 | 0.184 | -2.218 | 0.56 | 0.934 | 0.001 | 0.021 | 0.021 | 0 | 0.07 | 0.05 | 0.13 | 0.32 | 0.01 | 0.22 |
| 5 | 36277071 | rs10060125 | 0.661 | -2.091 | -0.089 | 0.591 | -0.005 | -0.001 | -0.001 | 0.431 | 0.06 | 0.05 | 0.06 | 0.09 | 0.01 | 0.18 |
| 5 | 36278890 | rs2133766 | 0.82 | -2.085 | 0.277 | 1.108 | -0.033 | 0.173 | 0.173 | 0.166 | 0.93 | 0.79 | 0.75 | 0.29 | 0.98 | 0.87 |
| 5 | 36281625 | rs11741164 | 0.913 | -1.959 | -0.169 | 0.489 | -0.049 | 0.182 | 0.182 | 0.765 | 0.4 | 0.12 | 0.35 | 0.08 | 0.52 | 0.22 |
| 5 | 36283772 | rs990057 | 1.314 | -1.862 | -0.332 | 0.499 | -0.076 | -0.026 | -0.026 | 0 | 0.23 | 0.03 | 0.25 | 0.22 | 0.26 | 0.28 |
| 5 | 36286088 | rs6451271 | 0.361 | -1.728 | -0.154 | 1.161 | -0.033 | 0.164 | 0.164 | 0.173 | 0.94 | 0.81 | 0.75 | 0.31 | 0.98 | 0.87 |
| 5 | 36301420 | rs884694 | -0.65 | -1.594 | 0.066 | 0.283 | -0.007 | 0.055 | 0.055 | 0 | 0.71 | 0.43 | 0.59 | 0.46 | 0.73 | 0.69 |
| 5 | 36309097 | rs16902947 | -0.273 | -1.732 | -0.15 | -0.189 | -0.013 | -0.062 | -0.062 | 0.248 | 0.26 | 0.3 | 0.12 | 0.03 | 0.18 | 0.33 |
| 5 | 36324582 | rs10473013 | -0.989 | -0.867 | -0.227 | 0.277 | -0.021 | -0.03 | -0.03 | 0 | 0.8 | 0.9 | 0.57 | 0.27 | 0.78 | 0.74 |
| 5 | 36336165 | rs1392457 | -1.426 | -0.97 | -0.155 | 0.613 | -0.006 | -0.064 | -0.064 | 0 | 0.82 | 0.91 | 0.65 | 0.46 | 0.84 | 0.77 |
| 5 | 36336334 | rs12658017 | 0.376 | -0.572 | -0.253 | 0.638 | -0.001 | 0.036 | 0.036 | 0 | 0.08 | 0.06 | 0.12 | 0.33 | 0.06 | 0.16 |
| 5 | 36337293 | rs13177348 | 0.102 | -0.406 | -0.482 | 0.423 | -0.034 | -0.044 | -0.044 | 0 | 0.69 | 0.47 | 0.57 | 0.19 | 0.78 | 0.61 |
| 5 | 36343685 | rs7711733 | 0.045 | -0.172 | -0.766 | 0.108 | 0.016 | 0.023 | 0.023 | 0 | 0.35 | 0.31 | 0.25 | 0.21 | 0.29 | 0.4 |
| 5 | 36346515 | rs4869652 | 1.17 | 0.053 | -0.757 | 0.127 | 0 | -0.02 | -0.02 | 0.485 | 0.38 | 0.34 | 0.34 | 0.1 | 0.49 | 0.22 |
| 5 | 36353334 | rs4869658 | -0.162 | -0.247 | -1.252 | 0.439 | -0.004 | 0.009 | 0.009 | 0 | 0.72 | 0.6 | 0.56 | 0.31 | 0.76 | 0.64 |
| 5 | 36354241 | rs12659453 | -0.105 | 0.158 | -2.07 | 0.291 | 0.009 | -0.042 | -0.042 | 0.158 | 0.07 | 0 | 0.05 | 0.15 | 0.06 | 0.16 |
| 5 | 36356516 | rs13158909 | -0.604 | 0.853 | -1.493 | -0.228 | -0.069 | 0.204 | 0.204 | 0 | 0.38 | 0.07 | 0.28 | 0.09 | 0.4 | 0.38 |
| 5 | 36359459 | rs7730756 | -1.248 | 0.259 | -0.646 | 0.621 | 0.007 | -0.026 | -0.026 | 1.772 | 0.64 | 0.76 | 0.5 | 0.48 | 0.66 | 0.72 |
| 5 | 36359661 | rs7731082 | -1.44 | -0.53 | 0.154 | 0.539 | 0.001 | 0.169 | 0.169 | 0.354 | 0.08 | 0.53 | 0.09 | 0.08 | 0.01 | 0.01 |
| 5 | 36362004 | rs4311444 | -1.44 | -0.608 | 0.693 | 0.384 | -0.031 | 0.03 | 0.03 | 0 | 0.11 | 0.32 | 0.1 | 0.07 | 0.03 | 0.03 |
| 5 | 36363158 | rs4317351 | -1.156 | -0.448 | 0.624 | -0.128 | -0.018 | -0.069 | -0.069 | 0 | 0.38 | 0.47 | 0.26 | 0.1 | 0.31 | 0.26 |
| 5 | 36367987 | rs4351174 | -2.208 | -0.269 | 0.718 | 0.134 | -0.002 | 0.007 | 0.007 | 0 | 0.63 | 0.57 | 0.47 | 0.43 | 0.63 | 0.71 |
| 5 | 36369103 | rs4484458 | -1.847 | -1.453 | 1.222 | 0.297 | 0.014 | 0.009 | 0.009 | 0 | 0.47 | 0.55 | 0.3 | 0.33 | 0.35 | 0.48 |
| 5 | 36369722 | rs17284164 | -0.504 | -1.038 | 1.506 | 0.362 | 0.001 | 0.022 | 0.022 | 0.124 | 0.27 | 0.18 | 0.17 | 0.03 | 0.28 | 0.24 |
| 5 | 36382802 | rs12187339 | -1.191 | -1.292 | 1.48 | 0.46 | 0.01 | 0.011 | 0.011 | 0 | 0.34 | 0.28 | 0.2 | 0.26 | 0.29 | 0.43 |
| 5 | 36383780 | rs6870182 | -1.603 | -1.347 | 1.569 | 0.409 | 0.014 | 0.004 | 0.004 | 0 | 0.41 | 0.64 | 0.27 | 0.3 | 0.29 | 0.32 |
| 5 | 36397629 | rs4582306 | -0.732 | -1.269 | 2.274 | 0.759 | -0.035 | 0.121 | 0.121 | 0 | 0.09 | 0.2 | 0.04 | 0.22 | 0.04 | 0.2 |
| 5 | 36398024 | rs4418129 | -2.216 | -1.28 | 2.159 | 0.633 | -0.016 | -0.01 | -0.01 | 0.143 | 0.37 | 0.59 | 0.21 | 0.27 | 0.26 | 0.37 |
| 5 | 36398377 | rs2967072 | -0.114 | -1.313 | 2.023 | 0.839 | 0.003 | -0.004 | -0.004 | 0 | 0.07 | 0.05 | 0.02 | 0.15 | 0.05 | 0.27 |
| 5 | 36402114 | rs2937532 | -2.412 | -1.303 | 1.621 | 0.853 | 0.014 | -0.004 | -0.004 | 0.01 | 0.41 | 0.65 | 0.23 | 0.43 | 0.3 | 0.63 |
| 5 | 36402905 | rs16902999 | -0.455 | -0.981 | 1.224 | 0.865 | -0.014 | 0.036 | 0.036 | 0 | 0.08 | 0.11 | 0.03 | 0.22 | 0.04 | 0.2 |
| 5 | 36415424 | rs4242264 | -0.515 | -0.858 | 1.103 | 0.423 | 0.017 | -0.031 | -0.031 | 0.173 | 0.94 | 0.99 | 0.98 | 0.87 | 0.96 | 0.79 |
| 5 | 36415695 | rs6451286 | 2.149 | -0.812 | 1.288 | 0.396 | 0.016 | 0.002 | 0.002 | 0.655 | 0.54 | 0.41 | 0.56 | 0.57 | 0.61 | 0.37 |
| 5 | 36421611 | rs16903012 | -1.244 | -1.232 | 1.047 | 0.974 | -0.016 | 0.068 | 0.068 | 0 | 0.2 | 0.07 | 0.13 | 0.1 | 0.2 | 0.16 |
| 5 | 36422893 | rs10512652 | -0.245 | -1.008 | 0.957 | 1.064 | 0.007 | 0.003 | 0.003 | 0 | 0.07 | 0.06 | 0.03 | 0.19 | 0.03 | 0.22 |
| 5 | 36423931 | rs7735940 | 1.523 | -0.913 | 0.934 | 1.014 | -0.013 | 0.005 | 0.005 | 0.08 | 0.62 | 0.44 | 0.78 | 0.58 | 0.71 | 0.41 |
| 5 | 36426411 | rs12515142 | -0.709 | -0.426 | 0.304 | 1.164 | -0.007 | 0.075 | 0.075 | 0 | 0.16 | 0.32 | 0.12 | 0.26 | 0.13 | 0.3 |
| 5 | 36430755 | rs4571496 | 0.987 | -0.674 | -0.427 | 0.914 | 0.032 | -0.014 | -0.014 | 0 | 0.63 | 0.33 | 0.79 | 0.58 | 0.75 | 0.41 |
| 5 | 36431978 | rs4602648 | 0.856 | -0.298 | -0.041 | 0.92 | -0.016 | 0.052 | 0.052 | 0 | 0.74 | 0.64 | 0.84 | 0.61 | 0.74 | 0.49 |
| 5 | 36433782 | rs10941292 | -1.206 | 0.441 | -0.628 | 1.467 | -0.015 | -0.035 | -0.035 | 0 | 0.27 | 0.2 | 0.28 | 0.42 | 0.33 | 0.19 |
| 5 | 36434126 | rs13159489 | -0.908 | 0.491 | -1.132 | 1.187 | 0.024 | -0.008 | -0.008 | 0 | 0.4 | 0.21 | 0.31 | 0.47 | 0.37 | 0.61 |
| 5 | 36435476 | rs4242265 | -0.981 | 0.682 | -0.573 | 1.236 | -0.002 | 0.009 | 0.009 | 0 | 0.4 | 0.51 | 0.4 | 0.54 | 0.36 | 0.4 |
| 5 | 36435494 | rs10941293 | 1.386 | 0.299 | -0.727 | 1.082 | -0.003 | 0.076 | 0.076 | 0 | 0.46 | 0.2 | 0.48 | 0.37 | 0.58 | 0.46 |
| 5 | 36435597 | rs10941294 | 0.52 | 0.34 | -0.818 | 1.069 | -0.002 | 0.004 | 0.004 | 0 | 0.07 | 0.02 | 0.04 | 0.09 | 0.07 | 0.2 |
| 5 | 36437361 | rs4571499 | -0.878 | 0.262 | -0.265 | 1.035 | -0.001 | 0.003 | 0.003 | 0 | 0.38 | 0.37 | 0.39 | 0.53 | 0.35 | 0.41 |

| | | | | | | | | | | | | | | | | |
|---|---|---|---|---|---|---|---|---|---|---|---|---|---|---|---|---|
| 5 | 36438254 | rs10512653 | 0.417 | 0.309 | -0.684 | 1.028 | -0.026 | 0.047 | 0.047 | 0 | 0.16 | 0.03 | 0.1 | 0.07 | 0.14 | 0.1 |
| 5 | 36438503 | rs4425532 | 0.688 | 0.302 | -0.598 | 1.046 | -0.009 | -0.05 | -0.05 | 0 | 0.13 | 0.05 | 0.21 | 0.38 | 0.13 | 0.1 |
| 5 | 36440940 | rs4242267 | 0.973 | 0.071 | -0.369 | 0.879 | -0.012 | -0.018 | -0.018 | 0 | 0.68 | 0.4 | 0.79 | 0.84 | 0.78 | 0.77 |
| 5 | 36441351 | rs4334910 | -0.679 | 0.311 | -0.561 | 0.78 | 0.004 | 0.008 | 0.008 | 0 | 0.33 | 0.39 | 0.36 | 0.56 | 0.34 | 0.33 |
| 5 | 36443160 | rs10061011 | -0.225 | 0.249 | -1.041 | 0.573 | 0.016 | -0.013 | -0.013 | 0 | 0.88 | 0.97 | 0.94 | 0.92 | 0.92 | 0.85 |
| 5 | 36446393 | rs6886206 | 1.506 | 0.441 | -0.888 | 0.858 | 0.034 | 0.006 | 0.006 | 0 | 0.6 | 0.27 | 0.74 | 0.68 | 0.71 | 0.56 |
| 5 | 36449552 | rs2468506 | 0.472 | 0.62 | -0.677 | 0.194 | 0 | 0.036 | 0.036 | 0.086 | 0.76 | 0.85 | 0.85 | 0.88 | 0.79 | 0.76 |
| 5 | 36450612 | rs7709504 | 1.979 | 0.675 | -0.896 | 0.335 | -0.04 | 0.02 | 0.02 | 0 | 0.4 | 0.09 | 0.43 | 0.33 | 0.44 | 0.41 |
| 5 | 36450876 | rs2468509 | -0.898 | 0.676 | -0.912 | 0.31 | -0.008 | -0.008 | -0.008 | 0.883 | 0.9 | 1 | 0.94 | 0.94 | 0.93 | 0.85 |
| 5 | 36453541 | rs17286376 | 2.096 | 0.669 | -0.855 | 0.338 | -0.003 | 0.048 | 0.048 | 0 | 0.4 | 0.2 | 0.45 | 0.33 | 0.44 | 0.44 |
| 5 | 36456056 | rs2455274 | 0.414 | -0.009 | -0.527 | 0.486 | 0.004 | 0.015 | 0.015 | 0 | 0.79 | 0.84 | 0.87 | 0.89 | 0.83 | 0.79 |
| 5 | 36456308 | rs2455275 | 0.173 | 0.058 | -0.917 | 0.561 | -0.004 | -0.007 | -0.007 | 0 | 0.09 | 0.13 | 0.05 | 0.05 | 0.07 | 0.2 |
| 5 | 36459584 | rs2455277 | -0.654 | 0.042 | -0.946 | 0.552 | -0.005 | -0.002 | -0.002 | 0 | 0.9 | 0.98 | 0.94 | 0.94 | 0.92 | 0.86 |
| 5 | 36460107 | rs7730299 | 0.094 | 0.478 | -0.897 | 0.547 | -0.023 | -0.063 | -0.063 | 1.944 | 0.26 | 0.17 | 0.34 | 0.47 | 0.3 | 0.18 |
| 5 | 36460425 | rs2455280 | 0.006 | 0.558 | -0.787 | 0.682 | -0.004 | -0.006 | -0.006 | 0.163 | 0.09 | 0.12 | 0.05 | 0.05 | 0.07 | 0.2 |
| 5 | 36460462 | rs17358533 | 1.099 | 0.641 | -1.064 | 0.579 | -0.055 | 0.02 | 0.02 | 0 | 0.42 | 0.1 | 0.45 | 0.36 | 0.46 | 0.41 |
| 5 | 36461331 | rs2468519 | 0.71 | 0.916 | -0.712 | 0.911 | 0 | 0.013 | 0.013 | 0 | 0.5 | 0.39 | 0.5 | 0.4 | 0.5 | 0.6 |
| 5 | 36473076 | rs11737926 | -0.835 | 1.523 | -0.361 | 1.058 | 0.032 | -0.054 | -0.054 | 0 | 0.18 | 0.57 | 0.21 | 0.07 | 0.19 | 0.1 |
| 5 | 36473128 | rs2937553 | -0.225 | 1.34 | -1.384 | 0.695 | -0.008 | 0.013 | 0.013 | 0 | 0.74 | 0.93 | 0.83 | 0.8 | 0.78 | 0.68 |
| 5 | 36473237 | rs2937550 | 1.645 | 1.078 | -1.856 | 0.375 | -0.017 | 0.044 | 0.044 | 0.157 | 0.42 | 0.13 | 0.52 | 0.65 | 0.46 | 0.4 |
| 5 | 36479022 | rs2468529 | -1.575 | 0.971 | -1.246 | 0.419 | -0.032 | 0.131 | 0.131 | 0 | 0.34 | 0.86 | 0.31 | 0.15 | 0.33 | 0.28 |
| 5 | 36481330 | rs2937545 | -1.78 | 0.947 | -1.27 | 0.522 | -0.08 | -0.044 | -0.044 | 0.431 | 0.38 | 0.91 | 0.44 | 0.4 | 0.39 | 0.31 |
| 5 | 36482830 | rs2468532 | -1.119 | 0.944 | -0.932 | 0.558 | -0.001 | -0.056 | -0.056 | 0 | 0.18 | 0.3 | 0.12 | 0.08 | 0.16 | 0.18 |
| 5 | 36483040 | rs2455253 | -1.186 | 0.741 | -0.901 | 0.549 | -0.034 | -0.068 | -0.068 | 1.159 | 0.14 | 0.3 | 0.08 | 0.02 | 0.06 | 0.15 |
| 5 | 36487409 | rs2455257 | -1.349 | 0.784 | -1.055 | 0.578 | -0.012 | 0.027 | 0.027 | 0.005 | 0.17 | 0.27 | 0.2 | 0.26 | 0.13 | 0.19 |
| 5 | 36494037 | rs2937540 | 1.539 | 0.646 | -1.273 | -0.119 | -0.046 | 0.099 | 0.099 | 0.023 | 0.63 | 0.1 | 0.69 | 0.78 | 0.64 | 0.71 |
| 5 | 36501506 | rs12374568 | -0.204 | 0.739 | -1.312 | -0.139 | -0.035 | 0 | 0 | 0 | 0.06 | 0.01 | 0.05 | 0.07 | 0.09 | 0.08 |
| 5 | 36502818 | rs2956582 | 1.864 | 0.659 | -0.652 | -0.361 | -0.011 | 0.061 | 0.061 | 0 | 0.61 | 0.28 | 0.68 | 0.78 | 0.61 | 0.71 |
| 5 | 36515990 | rs2468526 | 0.887 | 0.597 | 0.173 | -0.395 | 0.061 | -0.018 | -0.018 | 0 | 0.77 | 0.57 | 0.76 | 0.79 | 0.7 | 0.82 |
| 5 | 36519527 | rs2937551 | -0.176 | 0.541 | 0.86 | -0.433 | 0.002 | 0 | 0 | 0 | 0.86 | 0.88 | 0.83 | 0.86 | 0.8 | 0.9 |
| 5 | 36520513 | rs4869483 | 2.365 | 0.484 | 0.614 | -0.472 | -0.04 | 0.127 | 0.127 | 0 | 0.6 | 0.12 | 0.67 | 0.79 | 0.61 | 0.71 |
| 5 | 36520578 | rs2455258 | -1.205 | 0.24 | 0.764 | -0.472 | -0.006 | -0.041 | -0.041 | 0 | 0.14 | 0.16 | 0.06 | 0 | 0.08 | 0.11 |
| 5 | 36521550 | rs4371785 | -2.22 | 0.384 | 0.766 | -0.47 | 0.002 | 0.066 | 0.066 | 0 | 0.37 | 0.73 | 0.29 | 0.19 | 0.34 | 0.27 |
| 5 | 36523915 | rs2731888 | -1.98 | 0.395 | 1.243 | -0.541 | 0.008 | -0.07 | -0.07 | 3.108 | 0.16 | 0.4 | 0.1 | 0.07 | 0.14 | 0.16 |
| 5 | 36526810 | rs2731891 | -0.346 | 0.46 | 1.013 | -0.516 | 0.004 | -0.007 | -0.007 | 2.78 | 0.06 | 0.1 | 0.05 | 0.06 | 0.08 | 0.07 |
| 5 | 36529847 | rs735200 | -0.814 | 0.406 | 1.423 | -0.655 | 0.04 | -0.079 | -0.079 | 0 | 0.17 | 0.39 | 0.19 | 0.05 | 0.19 | 0.09 |
| 5 | 36532804 | rs2115520 | -1.82 | 0.319 | 1.158 | -0.681 | 0.01 | -0.067 | -0.067 | 5.367 | 0.29 | 0.59 | 0.28 | 0.11 | 0.33 | 0.17 |
| 5 | 36538074 | rs1549681 | -2.03 | 0.248 | 0.688 | -0.725 | -0.004 | 0.076 | 0.076 | 1.012 | 0.34 | 0.81 | 0.29 | 0.11 | 0.33 | 0.25 |
| 5 | 36541302 | rs9292633 | -2.044 | 0.165 | 0.767 | -0.622 | 0.001 | 0.01 | 0.01 | 0.156 | 0.28 | 0.72 | 0.26 | 0.12 | 0.3 | 0.17 |
| 5 | 36548461 | rs12518030 | 1.725 | 0.089 | -0.347 | -0.473 | -0.078 | -0.048 | -0.048 | 0.749 | 0.45 | 0.07 | 0.51 | 0.62 | 0.46 | 0.61 |
| 5 | 36570469 | rs1366634 | -1.2 | 0.151 | 0.388 | -0.47 | 0.034 | -0.07 | -0.07 | 0 | 0.25 | 0.56 | 0.26 | 0.13 | 0.33 | 0.07 |
| 5 | 36574931 | rs1428970 | 1.415 | 0.049 | 0.012 | -0.485 | -0.015 | 0.023 | 0.023 | 0.978 | 0.52 | 0.19 | 0.56 | 0.59 | 0.48 | 0.73 |
| 5 | 36575021 | rs1428971 | 0.39 | 0.127 | -0.099 | -0.332 | -0.024 | -0.021 | -0.021 | 0 | 0.71 | 0.75 | 0.69 | 0.68 | 0.65 | 0.75 |
| 5 | 36580791 | rs1549688 | 0.37 | 0.137 | -0.081 | -0.327 | 0.02 | -0.001 | -0.001 | 0 | 0.07 | 0.15 | 0.13 | 0.05 | 0.16 | 0.05 |
| 5 | 36584881 | rs10061650 | -0.412 | -0.225 | -0.246 | -0.229 | 0 | -0.038 | -0.038 | 0 | 0.78 | 0.9 | 0.83 | 0.72 | 0.81 | 0.8 |
| 5 | 36590685 | rs4391195 | -1.612 | -0.265 | -0.027 | -0.547 | 0.004 | -0.002 | -0.002 | 0 | 0.14 | 0.12 | 0.05 | 0.01 | 0.06 | 0.16 |
| 5 | 36597841 | rs2562571 | 0.258 | 0.03 | -0.458 | -0.466 | -0.001 | 0 | 0 | 0 | 0.77 | 0.78 | 0.77 | 0.87 | 0.78 | 0.85 |
| 5 | 36599385 | rs2562573 | 0.432 | -0.488 | -0.842 | -0.782 | -0.002 | 0.076 | 0.076 | 0.108 | 0.48 | 0.66 | 0.57 | 0.73 | 0.49 | 0.71 |
| 5 | 36602563 | rs2731878 | -1.083 | -0.741 | -0.646 | 0.277 | -0.029 | 0.085 | 0.085 | 0 | 0.23 | 0.56 | 0.13 | 0.38 | 0.13 | 0.26 |
| 5 | 36604227 | rs13155110 | 0.761 | -0.761 | -1.161 | -0.153 | -0.032 | 0.096 | 0.096 | 0.535 | 0.27 | 0.1 | 0.22 | 0.09 | 0.33 | 0.1 |
| 5 | 36605360 | rs2562582 | -1.006 | -0.935 | -1.457 | -0.266 | -0.037 | -0.008 | -0.008 | 1.805 | 0.84 | 0.99 | 0.86 | 0.9 | 0.81 | 0.88 |
| 5 | 36606164 | rs1428973 | -0.937 | -0.945 | -1.452 | -0.271 | -0.01 | 0.012 | 0.012 | 0 | 0.84 | 0.93 | 0.86 | 0.9 | 0.81 | 0.88 |
| 5 | 36607962 | rs10078147 | -0.674 | -0.859 | -1.119 | 0.275 | -0.028 | 0.101 | 0.101 | 0 | 0.23 | 0.56 | 0.13 | 0.46 | 0.14 | 0.25 |
| 5 | 36608165 | rs4591778 | -0.574 | -0.834 | -1.019 | 0.272 | -0.03 | 0.153 | 0.153 | 0 | 0.23 | 0.49 | 0.12 | 0.41 | 0.13 | 0.25 |
| 5 | 36613053 | rs7728680 | 1.391 | -0.418 | -1.142 | 0.229 | -0.009 | 0.012 | 0.012 | 0 | 0.33 | 0.27 | 0.51 | 0.35 | 0.34 | 0.54 |
| 5 | 36616406 | rs9292637 | -0.448 | 0.199 | 0.149 | 1.326 | 0.03 | 0.249 | 0.249 | 0 | 0.07 | 0.23 | 0.06 | 0.27 | 0.07 | 0.12 |
| 5 | 36616535 | rs1366638 | -0.309 | 0.613 | -0.191 | 1.241 | -0.062 | -0.01 | -0.01 | 0 | 0.88 | 0.98 | 0.8 | 0.9 | 0.75 | 0.92 |
| 5 | 36618403 | rs6451304 | -0.487 | 0.168 | -0.332 | 1.356 | -0.061 | 0.073 | 0.073 | 0 | 0.16 | 0.4 | 0.07 | 0.22 | 0.06 | 0.19 |
| 5 | 36626746 | rs12188287 | 0.028 | 0.331 | -0.129 | 1.492 | 0.009 | 0.131 | 0.131 | 0 | 0.07 | 0.17 | 0.06 | 0.22 | 0.06 | 0.12 |
| 5 | 36626929 | rs17368211 | 0.701 | 0.341 | -0.383 | 1.25 | -0.027 | 0.026 | 0.026 | 0 | 0.12 | 0.03 | 0.14 | 0.08 | 0.16 | 0.15 |
| 5 | 36627769 | rs1864213 | 1.088 | 0.47 | -0.356 | 1.249 | 0.023 | 0.03 | 0.03 | 0 | 0.71 | 0.54 | 0.72 | 0.68 | 0.68 | 0.72 |
| 5 | 36629258 | rs1030239 | -0.425 | 0.441 | -0.33 | 1.152 | -0.018 | 0.072 | 0.072 | 0 | 0.16 | 0.37 | 0.09 | 0.21 | 0.09 | 0.18 |
| 5 | 36631653 | rs3776565 | 0.718 | 0.334 | -0.463 | 1.087 | -0.033 | 0.025 | 0.025 | 0 | 0.13 | 0.01 | 0.17 | 0.09 | 0.16 | 0.16 |
| 5 | 36631733 | rs3776566 | 0.022 | 0.296 | -0.099 | 1.057 | -0.012 | 0.134 | 0.134 | 0 | 0.13 | 0.38 | 0.08 | 0.22 | 0.06 | 0.17 |
| 5 | 36632403 | rs4869680 | 0.658 | 0.562 | -0.181 | 1.091 | 0.038 | 0.063 | 0.063 | 0.175 | 0.78 | 0.66 | 0.63 | 0.63 | 0.67 | 0.77 |
| 5 | 36634558 | rs1645651 | 0.504 | 0.568 | -0.179 | 1.098 | 0.005 | 0.01 | 0.01 | 0.061 | 0.79 | 0.77 | 0.64 | 0.63 | 0.67 | 0.77 |
| 5 | 36635565 | rs1366632 | 1.468 | 0.426 | -0.393 | 1.081 | -0.024 | 0.012 | 0.012 | 0 | 0.63 | 0.28 | 0.53 | 0.41 | 0.58 | 0.6 |
| 5 | 36635692 | rs442867 | 0.132 | 0.367 | -0.334 | 1.092 | -0.005 | 0.093 | 0.093 | 0 | 0.14 | 0.4 | 0.1 | 0.21 | 0.09 | 0.17 |
| 5 | 36635944 | rs366597 | 1.435 | 0.398 | -0.364 | 1.116 | -0.018 | 0.006 | 0.006 | 0 | 0.63 | 0.28 | 0.53 | 0.43 | 0.59 | 0.6 |

| | | | | | | | | | | | | | | | | |
|---|---|---|---|---|---|---|---|---|---|---|---|---|---|---|---|---|
| 5 | 36636816 | rs432268 | -0.039 | 0.253 | -0.574 | 1.063 | -0.014 | 0.099 | 0.099 | 0 | 0.16 | 0.52 | 0.11 | 0.22 | 0.09 | 0.17 |
| 5 | 36641769 | rs10070340 | -0.722 | 0.752 | -1.878 | 1.097 | 0.026 | 0.063 | 0.063 | 0.359 | 0.2 | 0.27 | 0.34 | 0.36 | 0.28 | 0.21 |
| 5 | 36644300 | rs1645660 | 1.419 | 0.521 | -2.394 | 0.707 | 0.01 | 0.065 | 0.065 | 0.548 | 0.49 | 0.1 | 0.37 | 0.38 | 0.44 | 0.44 |
| 5 | 36645257 | rs3776571 | -0.2 | 0.306 | -2.425 | 0.764 | -0.001 | 0.017 | 0.017 | 0 | 0.11 | 0.07 | 0.17 | 0.24 | 0.1 | 0.11 |
| 5 | 36645367 | rs3822469 | -0.104 | 0.673 | -2.405 | 0.086 | -0.043 | 0.002 | 0.002 | 0.12 | 0.14 | 0.04 | 0.18 | 0.14 | 0.17 | 0.21 |
| 5 | 36646946 | rs1428968 | 0.103 | 0.594 | -2.402 | 0.089 | -0.018 | -0.001 | -0.001 | 0.11 | 0.13 | 0.04 | 0.16 | 0.14 | 0.14 | 0.17 |
| 5 | 36647574 | rs17290638 | 1.132 | 0.252 | -2.595 | -0.08 | -0.06 | -0.018 | -0.018 | 0 | 0.51 | 0.07 | 0.53 | 0.54 | 0.55 | 0.5 |
| 5 | 36648328 | rs753813 | -0.119 | 1.073 | -1.656 | -0.188 | 0.002 | -0.001 | -0.001 | 0.165 | 0.12 | 0.13 | 0.17 | 0.04 | 0.21 | 0.12 |
| 5 | 36651986 | rs3776578 | -0.614 | -0.235 | -1.504 | 0.044 | -0.019 | 0.009 | 0.009 | 2.194 | 0.09 | 0.15 | 0.04 | 0.09 | 0.04 | 0.05 |
| 5 | 36653738 | rs4869681 | -0.115 | -0.653 | -1.17 | 0.106 | -0.022 | 0.131 | 0.131 | 0 | 0.86 | 0.54 | 0.92 | 0.79 | 0.93 | 0.83 |
| 5 | 36654493 | rs3776579 | -0.256 | -0.448 | -1.432 | 0.01 | -0.004 | 0.002 | 0.002 | 1.861 | 0.08 | 0.09 | 0.03 | 0.09 | 0.05 | 0.05 |
| 5 | 36655353 | rs17290929 | 0.306 | 0.302 | -1.98 | -0.654 | -0.091 | 0.091 | 0.091 | 0.306 | 0.18 | 0.05 | 0.22 | 0.02 | 0.3 | 0.22 |
| 5 | 36656718 | rs4869682 | 0.69 | -0.82 | -2.301 | -0.988 | -0.01 | 0.078 | 0.078 | 0 | 0.53 | 0.28 | 0.63 | 0.74 | 0.54 | 0.58 |
| 5 | 36659084 | rs4869685 | 0.447 | -0.97 | -2.192 | -0.502 | 0.15 | 0.122 | 0.122 | 3.443 | 0.51 | 0.24 | 0.72 | 0.71 | 0.71 | 0.69 |
| 5 | 36662361 | rs16903252 | -0.234 | 0.036 | -1.374 | -0.153 | -0.017 | -0.056 | -0.056 | 0 | 0.1 | 0.19 | 0.06 | 0 | 0.08 | 0.03 |
| 5 | 36668022 | rs3776585 | -0.085 | 0.245 | -2.341 | 0.379 | -0.109 | -0.048 | -0.048 | 0 | 0.46 | 0.03 | 0.54 | 0.45 | 0.61 | 0.58 |
| 5 | 36673912 | rs2303715 | -0.502 | 0.468 | -1.176 | 0.775 | -0.016 | -0.016 | -0.016 | 0 | 0.06 | 0.02 | 0.06 | 0.13 | 0.08 | 0.11 |
| 5 | 36676161 | rs1549627 | -0.075 | 0.045 | -0.497 | 0.728 | -0.011 | -0.006 | -0.006 | 0 | 0.56 | 0.36 | 0.59 | 0.52 | 0.54 | 0.43 |
| 5 | 36678275 | rs2301066 | -0.116 | 0.162 | -0.428 | 0.698 | -0.009 | -0.005 | -0.005 | 0.04 | 0.57 | 0.41 | 0.59 | 0.52 | 0.54 | 0.41 |
| 5 | 36678540 | rs2301068 | -0.459 | 0.137 | -0.776 | 0.644 | -0.072 | -0.043 | -0.043 | 0 | 0.65 | 0.86 | 0.71 | 0.59 | 0.55 | 0.5 |
| 5 | 36684550 | rs3776586 | 0.243 | 0.194 | -0.935 | 0.803 | -0.01 | -0.056 | -0.056 | 0 | 0.18 | 0.08 | 0.38 | 0.27 | 0.25 | 0.2 |
| 5 | 36684598 | rs3776587 | -0.029 | -0.184 | -0.599 | 0.533 | -0.007 | -0.02 | -0.02 | 0.064 | 0.13 | 0.15 | 0.07 | 0.09 | 0.12 | 0.12 |
| 5 | 36687856 | rs2269272 | 0.583 | -0.112 | -0.819 | 0.609 | -0.009 | -0.066 | -0.066 | 0.1 | 0.17 | 0.06 | 0.38 | 0.27 | 0.23 | 0.18 |
| 5 | 36689181 | rs1122900 | 1.253 | -0.545 | -1.429 | 0.365 | -0.073 | -0.076 | -0.076 | 0 | 0.48 | 0.03 | 0.31 | 0.44 | 0.56 | 0.55 |
| 5 | 36690994 | rs6863386 | 0.558 | -0.444 | -0.626 | 0.28 | 0.003 | 0.003 | 0.003 | 0 | 0.71 | 0.69 | 0.76 | 0.8 | 0.8 | 0.8 |
| 5 | 36696029 | rs12654391 | 0.621 | -0.371 | -0.934 | 0.387 | -0.004 | -0.055 | -0.055 | 0 | 0.17 | 0.07 | 0.37 | 0.26 | 0.2 | 0.2 |
| 5 | 36702290 | rs1541815 | 0.476 | -0.749 | -0.353 | 0.083 | -0.01 | -0.038 | -0.038 | 0.007 | 0.64 | 0.57 | 0.75 | 0.75 | 0.69 | 0.71 |
| 5 | 36702864 | rs1025288 | 1.372 | 0.116 | -0.257 | -0.099 | 0.002 | -0.029 | -0.029 | 1.034 | 0.39 | 0.29 | 0.22 | 0.48 | 0.35 | 0.5 |
| 5 | 36703653 | rs918642 | 0.311 | 0.309 | -0.303 | -0.179 | -0.019 | -0.019 | -0.019 | 0 | 0.22 | 0.15 | 0.44 | 0.26 | 0.33 | 0.21 |
| 5 | 36709244 | rs1025052 | 0.493 | -0.196 | 0.26 | -0.319 | -0.004 | -0.007 | -0.007 | 0.014 | 0.71 | 0.67 | 0.81 | 0.73 | 0.78 | 0.72 |
| 5 | 36709701 | rs1566444 | 1.087 | -0.306 | 0.532 | -0.405 | 0.03 | 0.073 | 0.073 | 0.001 | 0.64 | 0.17 | 0.67 | 0.73 | 0.73 | 0.7 |
| 5 | 36713033 | rs7709455 | 0.567 | -0.361 | 0.499 | -0.309 | -0.009 | -0.012 | -0.012 | 0 | 0.71 | 0.64 | 0.81 | 0.73 | 0.77 | 0.72 |
| 5 | 36723395 | rs36651 | -0.484 | -0.533 | 0.891 | -0.425 | -0.018 | -0.022 | -0.022 | 0.211 | 0.16 | 0.3 | 0.08 | 0.17 | 0.11 | 0.18 |
| 5 | 36724793 | rs7734294 | 0.792 | -0.507 | 0.603 | -0.437 | -0.014 | -0.017 | -0.017 | 0 | 0.71 | 0.65 | 0.81 | 0.77 | 0.81 | 0.74 |
| 5 | 36727274 | rs10061940 | 0.896 | -0.466 | 0.415 | -0.465 | 0.027 | 0.053 | 0.053 | 0 | 0.67 | 0.8 | 0.77 | 0.76 | 0.72 | 0.71 |
| 5 | 36728340 | rs736917 | 1.151 | -0.207 | -0.222 | -0.146 | -0.024 | -0.067 | -0.067 | 0 | 0.16 | 0.03 | 0.35 | 0.3 | 0.22 | 0.22 |
| 5 | 36731004 | rs245674 | 0.195 | -0.492 | 0.974 | -0.245 | -0.044 | -0.078 | -0.078 | 0 | 0.77 | 0.53 | 0.81 | 0.9 | 0.86 | 0.89 |
| 5 | 36731225 | rs16903304 | 0.026 | -0.449 | 0.949 | -0.202 | 0.008 | 0.008 | 0.008 | 0 | 0.12 | 0.1 | 0.06 | 0.1 | 0.1 | 0.11 |
| 5 | 36735284 | rs42914 | -1.041 | -0.539 | 1.016 | -0.278 | -0.022 | -0.041 | -0.041 | 0 | 0.18 | 0.44 | 0.23 | 0.13 | 0.12 | 0.19 |
| 5 | 36739499 | rs6869867 | 1.23 | -0.078 | 0.049 | -0.825 | -0.039 | -0.078 | -0.078 | 0.587 | 0.21 | 0.09 | 0.29 | 0.45 | 0.3 | 0.42 |
| 5 | 36740846 | rs2886219 | 0.858 | -0.181 | 0.221 | -0.945 | -0.025 | 0.013 | 0.013 | 0 | 0.41 | 0.14 | 0.29 | 0.31 | 0.42 | 0.28 |
| 5 | 36749082 | rs6892066 | 0.947 | -0.18 | 0.416 | -0.926 | -0.005 | 0.004 | 0.004 | 0 | 0.4 | 0.33 | 0.3 | 0.32 | 0.41 | 0.27 |
| 5 | 36759368 | rs2115198 | 0.482 | -0.671 | 0.241 | -1.079 | 0.028 | 0.058 | 0.058 | 0.271 | 0.67 | 0.24 | 0.62 | 0.85 | 0.77 | 0.75 |
| 5 | 36759602 | rs245670 | -0.189 | -0.608 | 0.26 | -1.056 | 0.016 | 0.022 | 0.022 | 0 | 0.73 | 0.78 | 0.67 | 0.9 | 0.81 | 0.83 |
| 5 | 36762502 | rs4869488 | 0.099 | -0.578 | 0.12 | -1.04 | -0.024 | -0.07 | -0.07 | 3.015 | 0.69 | 0.54 | 0.64 | 0.85 | 0.77 | 0.75 |
| 5 | 36781359 | rs6451310 | -0.066 | -0.245 | 0.092 | -0.944 | 0.022 | 0.031 | 0.031 | 0.671 | 0.35 | 0.5 | 0.31 | 0.58 | 0.42 | 0.56 |
| 5 | 36786516 | rs6877724 | -0.556 | -0.796 | -0.013 | -0.917 | 0.053 | 0.065 | 0.065 | 0 | 0.34 | 0.19 | 0.48 | 0.2 | 0.24 | 0.25 |
| 5 | 36793233 | rs13158855 | -0.546 | -0.798 | -0.02 | -0.921 | 0.054 | 0.068 | 0.068 | 0 | 0.34 | 0.18 | 0.48 | 0.2 | 0.24 | 0.25 |
| 5 | 36804225 | rs172384 | 1.225 | -0.536 | 0.031 | -0.706 | -0.014 | -0.021 | -0.021 | 0 | 0.13 | 0.07 | 0.13 | 0.3 | 0.19 | 0.27 |
| 5 | 36808631 | rs11743146 | -0.308 | -0.794 | -0.172 | -0.87 | 0.048 | 0.037 | 0.037 | 0 | 0.2 | 0.09 | 0.21 | 0.13 | 0.12 | 0.14 |
| 5 | 36816661 | rs292170 | -1.115 | -0.855 | 0.505 | -0.939 | -0.014 | 0.011 | 0.011 | 0 | 0.38 | 0.57 | 0.37 | 0.43 | 0.32 | 0.42 |
| 5 | 36833002 | rs12520324 | -0.007 | -0.894 | 0.387 | -0.919 | 0.003 | 0.003 | 0.003 | 0 | 0.15 | 0.18 | 0.2 | 0.13 | 0.09 | 0.12 |
| 5 | 36856801 | rs158411 | 0.945 | -0.355 | 0.293 | -0.999 | -0.033 | -0.035 | -0.035 | 0 | 0.16 | 0.08 | 0.17 | 0.3 | 0.3 | 0.28 |
| 5 | 36886836 | rs13187028 | 1.08 | -1.158 | 0.874 | -0.873 | -0.005 | -0.015 | -0.015 | 0 | 0.15 | 0.14 | 0.28 | 0.08 | 0.12 | 0.11 |
| 5 | 36895270 | rs293746 | 1.495 | -1.27 | 0.68 | -0.909 | -0.022 | -0.07 | -0.07 | 0 | 0.17 | 0.03 | 0.12 | 0.3 | 0.19 | 0.27 |
| 5 | 36900843 | rs293748 | 0.933 | -1.299 | 0.646 | -0.942 | -0.006 | -0.079 | -0.079 | 2 | 0.21 | 0.03 | 0.14 | 0.3 | 0.2 | 0.29 |
| 5 | 36921325 | rs2123010 | 1.739 | -1.307 | 0.699 | -0.913 | -0.008 | -0.096 | -0.096 | 0 | 0.16 | 0.01 | 0.1 | 0.26 | 0.15 | 0.22 |
| 5 | 36926756 | rs923957 | 1.17 | -1.381 | 1.25 | -1.017 | -0.016 | -0.048 | -0.048 | 0 | 0.16 | 0.29 | 0.3 | 0.07 | 0.12 | 0.11 |
| 5 | 36954812 | rs292182 | -1.767 | -1.494 | 1.301 | -1.088 | 0.001 | -0.06 | -0.06 | 0 | 0.57 | 0.83 | 0.65 | 0.51 | 0.43 | 0.52 |
| 5 | 36985303 | rs3822471 | 0.41 | -1.523 | 1.173 | -1.101 | 0.02 | 0.013 | 0.013 | 0 | 0.15 | 0.1 | 0.19 | 0.13 | 0.11 | 0.12 |
| 5 | 36988252 | rs468022 | 1.875 | -1.397 | 1.124 | -1.05 | -0.018 | -0.067 | -0.067 | 1.002 | 0.16 | 0.03 | 0.12 | 0.3 | 0.17 | 0.27 |
| 5 | 37007629 | rs159753 | -1.686 | -1.432 | 1.176 | -1.049 | -0.011 | -0.078 | -0.078 | 0 | 0.57 | 0.89 | 0.67 | 0.51 | 0.44 | 0.53 |
| 5 | 37023898 | rs300063 | -1.574 | -1.44 | 1.179 | -1.053 | -0.002 | -0.069 | -0.069 | 2.239 | 0.56 | 0.89 | 0.66 | 0.51 | 0.43 | 0.53 |
| 5 | 37111565 | rs301904 | 0.88 | -1.213 | 1.334 | -1.033 | -0.011 | -0.029 | -0.029 | 0 | 0.1 | 0.22 | 0.07 | 0.06 | 0.09 | 0.09 |
| 5 | 37128315 | rs301895 | 0.88 | -1.213 | 1.339 | -1.033 | -0.011 | -0.03 | -0.03 | 0.056 | 0.1 | 0.24 | 0.07 | 0.07 | 0.09 | 0.08 |
| 5 | 37158123 | rs10512664 | 1.492 | -1.354 | 1.315 | -1.028 | -0.036 | -0.08 | -0.08 | 0 | 0.18 | 0.04 | 0.2 | 0.35 | 0.23 | 0.29 |
| 5 | 37159877 | rs10072064 | 0.433 | -1.413 | 1.353 | -1.064 | -0.015 | -0.037 | -0.037 | 0 | 0.12 | 0.25 | 0.15 | 0.07 | 0.1 | 0.11 |
| 5 | 37171884 | rs10056250 | -0.144 | -1.573 | 1.257 | -1.18 | 0.033 | 0.05 | 0.05 | 0 | 0.19 | 0.12 | 0.1 | 0.06 | 0.14 | 0.11 |
| 5 | 37173930 | rs10076911 | 0.083 | -1.542 | 1.381 | -1.085 | -0.021 | -0.027 | -0.027 | 0 | 0.13 | 0.35 | 0.16 | 0.09 | 0.1 | 0.11 |

| | | | | | | | | | | | | | | | | |
|---|---|---|---|---|---|---|---|---|---|---|---|---|---|---|---|---|
| 5 | 37197160 | rs16903531 | 1.572 | -1.488 | 0.921 | -1.057 | -0.035 | -0.082 | -0.082 | 0 | 0.18 | 0.04 | 0.2 | 0.36 | 0.23 | 0.29 |
| 5 | 37239240 | rs7735138 | -1.174 | -1.562 | 1.143 | -1.224 | -0.008 | -0.003 | -0.003 | 0 | 0.44 | 0.52 | 0.4 | 0.42 | 0.37 | 0.44 |
| 5 | 37250393 | rs2055129 | 0.67 | -1.453 | 1.139 | -1.192 | -0.003 | -0.004 | -0.004 | 0 | 0.11 | 0.15 | 0.15 | 0.09 | 0.1 | 0.12 |
| 5 | 37251893 | rs28429963 | -0.823 | -1.419 | 1.207 | -1.165 | 0.072 | 0.055 | 0.055 | 0.47 | 0.06 | 0.48 | 0.04 | 0 | 0.01 | 0.01 |
| 5 | 37294473 | rs1045908 | -0.129 | -1.589 | 0.551 | -1.23 | 0.078 | 0.003 | 0.003 | 0 | 0.4 | 0.11 | 0.32 | 0.38 | 0.3 | 0.34 |
| 5 | 37313998 | rs4869498 | 1.792 | -1.608 | 0.532 | -1.233 | 0.029 | 0.036 | 0.036 | 0 | 0.12 | 0.01 | 0.1 | 0.07 | 0.07 | 0.11 |
| 5 | 37318295 | rs217797 | -1.804 | -1.586 | 0.536 | -1.264 | -0.03 | -0.049 | -0.049 | 0.037 | 0.58 | 0.9 | 0.53 | 0.49 | 0.45 | 0.55 |
| 5 | 37333727 | rs150594 | -1.103 | -1.344 | 0.778 | -1.116 | 0.012 | 0.036 | 0.036 | 0.061 | 0.2 | 0.85 | 0.22 | 0.12 | 0.15 | 0.22 |
| 5 | 37349895 | rs13165088 | 1.988 | -1.345 | 0.683 | -1.103 | -0.043 | -0.097 | -0.097 | 3.694 | 0.16 | 0.01 | 0.17 | 0.3 | 0.19 | 0.18 |
| 5 | 37361728 | rs16903601 | 0.156 | -1.4 | 0.85 | -1.11 | 0 | -0.001 | -0.001 | 0.034 | 0.13 | 0.15 | 0.17 | 0.12 | 0.15 | 0.21 |
| 5 | 37372017 | rs12055164 | 1.87 | -1.248 | 0.399 | -0.943 | -0.04 | -0.097 | -0.097 | 0.479 | 0.16 | 0.01 | 0.17 | 0.3 | 0.19 | 0.19 |
| 5 | 37409579 | rs13178300 | 0.549 | -1.155 | 0.357 | -0.804 | 0.02 | -0.095 | -0.095 | 0.516 | 0.27 | 0.04 | 0.21 | 0.39 | 0.23 | 0.22 |
| 5 | 37437848 | rs3776597 | 1.607 | -0.822 | 1.29 | -0.819 | 0.023 | -0.03 | -0.03 | 0 | 0.89 | 0.35 | 0.85 | 0.97 | 0.91 | 0.84 |
| 5 | 37472247 | rs6869205 | 1.166 | -0.814 | 1.372 | -0.794 | 0.044 | 0.029 | 0.029 | 0.145 | 0.94 | 0.46 | 0.96 | 1 | 0.99 | 0.99 |
| 5 | 37489442 | rs12657090 | 0.102 | -1.011 | 0.972 | -0.925 | 0.032 | -0.007 | -0.007 | 1.369 | 0.59 | 0.33 | 0.69 | 0.51 | 0.68 | 0.72 |
| 5 | 37498911 | rs3941788 | 1.436 | -1.196 | 0.77 | -1.017 | -0.063 | -0.124 | -0.124 | 0 | 0.4 | 0.05 | 0.47 | 0.41 | 0.54 | 0.44 |
| 5 | 37505646 | rs1829292 | 1.455 | -1.202 | 0.732 | -1.02 | -0.081 | -0.143 | -0.143 | 0 | 0.4 | 0.03 | 0.47 | 0.41 | 0.54 | 0.44 |
| 5 | 37543121 | rs512562 | -0.613 | -1.316 | 0.576 | -0.985 | 0.031 | -0.087 | -0.087 | 0 | 0.64 | 0.9 | 0.73 | 0.51 | 0.69 | 0.72 |
| 5 | 37552852 | rs7700431 | 0.263 | -1.328 | 0.686 | -1.005 | 0.067 | -0.008 | -0.008 | 0.012 | 0.57 | 0.26 | 0.69 | 0.51 | 0.68 | 0.71 |
| 5 | 37556926 | rs2088077 | 1.262 | -0.972 | 0.381 | -1.021 | -0.05 | -0.101 | -0.101 | 0 | 0.4 | 0.07 | 0.48 | 0.41 | 0.53 | 0.44 |
| 5 | 37567851 | rs10076102 | 0.068 | -1.239 | 0.531 | -1.051 | -0.037 | -0.065 | -0.065 | 0 | 0.12 | 0.23 | 0.05 | 0 | 0.04 | 0.03 |
| 5 | 37585849 | rs665327 | -1.127 | -1.128 | 0.371 | -0.971 | 0.009 | -0.079 | -0.079 | 0 | 0.74 | 0.9 | 0.77 | 0.51 | 0.73 | 0.74 |
| 5 | 37586211 | rs677084 | -1.202 | -1.128 | 0.366 | -0.971 | -0.001 | -0.102 | -0.102 | 0.305 | 0.74 | 0.91 | 0.77 | 0.51 | 0.73 | 0.74 |
| 5 | 37590881 | rs10941348 | 0.464 | -1.16 | 0.592 | -1.03 | 0.078 | -0.011 | -0.011 | 0 | 0.51 | 0.18 | 0.52 | 0.41 | 0.58 | 0.46 |
| 5 | 37642232 | rs6451331 | 0.061 | -1.096 | 0.078 | -0.916 | 0.023 | 0.003 | 0.003 | 0 | 0.55 | 0.62 | 0.56 | 0.37 | 0.58 | 0.48 |
| 5 | 37669632 | rs1400854 | 1.099 | -1.24 | -0.49 | -0.966 | -0.077 | -0.065 | -0.065 | 0 | 0.34 | 0.09 | 0.47 | 0.36 | 0.47 | 0.41 |
| 5 | 37671746 | rs12521063 | 1.708 | -0.726 | -0.892 | -1.278 | -0.018 | -0.025 | -0.025 | 0 | 0.2 | 0.13 | 0.38 | 0.36 | 0.28 | 0.35 |
| 5 | 37681646 | rs4343860 | 0.705 | -0.75 | -0.93 | -1.294 | -0.094 | -0.083 | -0.083 | 1.015 | 0.34 | 0.09 | 0.54 | 0.36 | 0.48 | 0.42 |
| 5 | 37684691 | rs4608942 | 1.239 | -1.098 | -0.327 | -0.756 | -0.027 | -0.042 | -0.042 | 0.19 | 0.2 | 0.12 | 0.45 | 0.36 | 0.28 | 0.35 |
| 5 | 37695743 | rs10941361 | 1.22 | -1.308 | -0.469 | -0.861 | -0.09 | -0.099 | -0.099 | 0.066 | 0.43 | 0.1 | 0.38 | 0.54 | 0.53 | 0.31 |
| 5 | 37696178 | rs6451339 | 1.188 | -1.099 | -0.31 | -0.747 | -0.032 | -0.046 | -0.046 | 0.042 | 0.2 | 0.1 | 0.45 | 0.37 | 0.28 | 0.35 |
| 5 | 37716067 | rs4242252 | -0.318 | -1.177 | -0.598 | -0.854 | -0.016 | -0.027 | -0.027 | 0 | 0.82 | 0.7 | 0.9 | 0.9 | 0.86 | 0.72 |
| 5 | 37737047 | rs2910725 | 0.727 | -1.41 | -1.76 | -0.897 | 0.087 | 0.091 | 0.091 | 0 | 0.67 | 0.21 | 0.86 | 0.93 | 0.88 | 0.84 |
| 5 | 37751672 | rs2973086 | 1.075 | -1.102 | -1.843 | -0.755 | -0.128 | -0.108 | -0.108 | 0.298 | 0.43 | 0.05 | 0.38 | 0.49 | 0.54 | 0.32 |
| 5 | 37751724 | rs2973085 | 0.816 | -1.515 | -0.874 | -0.92 | -0.031 | -0.058 | -0.058 | 0.066 | 0.8 | 0.27 | 0.91 | 1 | 0.95 | 0.95 |
| 5 | 37755337 | rs2910721 | -0.236 | -1.337 | -1.15 | -0.718 | 0.062 | 0.066 | 0.066 | 0.06 | 0.18 | 0.07 | 0.06 | 0.03 | 0.07 | 0.12 |
| 5 | 37772780 | rs10472291 | -0.597 | -0.611 | -0.522 | -0.183 | -0.001 | 0.005 | 0.005 | 0 | 0.38 | 0.57 | 0.57 | 0.42 | 0.35 | 0.58 |
| 5 | 37773877 | rs1019496 | -1.552 | -0.485 | -1.069 | -0.112 | -0.011 | -0.022 | -0.022 | 0 | 0.82 | 0.92 | 0.84 | 0.75 | 0.77 | 0.82 |
| 5 | 37776961 | rs10076330 | -0.096 | -0.271 | -0.206 | -0.049 | 0 | -0.001 | -0.001 | 0 | 0.3 | 0.27 | 0.5 | 0.39 | 0.24 | 0.41 |
| 5 | 37777974 | rs11745243 | 0.879 | 0.401 | -0.225 | 0.392 | -0.013 | -0.014 | -0.014 | 0.039 | 0.12 | 0.07 | 0.42 | 0.39 | 0.13 | 0.29 |
| 5 | 37782403 | rs7737481 | -0.536 | 0.339 | 0.36 | -0.051 | 0.077 | 0.007 | 0.007 | 0.934 | 0.08 | 0.34 | 0.07 | 0.03 | 0.1 | 0.17 |
| 5 | 37788642 | rs2910800 | -0.134 | 0.176 | -0.135 | 0.195 | -0.028 | -0.005 | -0.005 | 0 | 0.68 | 0.78 | 0.63 | 0.65 | 0.55 | 0.56 |
| 5 | 37789409 | rs2972933 | 0.439 | -0.018 | 0.282 | 0.312 | -0.001 | 0 | 0 | 0.006 | 0.55 | 0.52 | 0.3 | 0.33 | 0.45 | 0.28 |
| 5 | 37790061 | rs7730024 | 0.574 | -0.026 | 0.314 | 0.321 | 0 | 0.104 | 0.104 | 0 | 0.13 | 0.27 | 0.32 | 0.32 | 0.1 | 0.27 |
| 5 | 37790089 | rs2910718 | -0.606 | 0.068 | -0.066 | 0.283 | -0.008 | -0.012 | -0.012 | 0.013 | 0.73 | 0.79 | 0.67 | 0.65 | 0.63 | 0.62 |
| 5 | 37791451 | rs9283747 | 1.042 | 0.2 | -0.193 | 0.457 | -0.005 | -0.01 | -0.01 | 0 | 0.11 | 0.06 | 0.3 | 0.32 | 0.1 | 0.27 |
| 5 | 37791733 | rs9292673 | 0.141 | 0.208 | -0.177 | 0.448 | -0.01 | -0.027 | -0.027 | 0 | 0.17 | 0.1 | 0.35 | 0.32 | 0.18 | 0.33 |
| 5 | 37799460 | rs10941368 | 0.244 | -0.187 | -0.577 | 0.179 | 0.006 | -0.029 | -0.029 | 0 | 0.48 | 0.22 | 0.56 | 0.5 | 0.5 | 0.38 |
| 5 | 37807003 | rs13180376 | 0.721 | -0.147 | -0.226 | 0.094 | -0.038 | 0.026 | 0.026 | 0 | 0.37 | 0.2 | 0.26 | 0.3 | 0.42 | 0.18 |
| 5 | 37811864 | rs2910709 | -0.335 | -0.188 | 0.532 | 0.006 | 0.013 | -0.006 | -0.006 | 2.9 | 0.65 | 0.53 | 0.68 | 0.66 | 0.6 | 0.68 |
| 5 | 37813591 | rs17379771 | 1.277 | -0.103 | 0.352 | 0.05 | -0.023 | 0.004 | 0.004 | 0 | 0.33 | 0.17 | 0.22 | 0.31 | 0.35 | 0.14 |
| 5 | 37814102 | rs11111 | 0.071 | -0.339 | 0.839 | -0.139 | 0.009 | 0.006 | 0.006 | 0 | 0.2 | 0.32 | 0.36 | 0.19 | 0.16 | 0.22 |
| 5 | 37814148 | rs3749692 | 0.329 | -0.355 | 0.923 | -0.04 | 0.002 | 0.025 | 0.025 | 0 | 0.53 | 0.39 | 0.65 | 0.63 | 0.54 | 0.57 |
| 5 | 37816845 | rs7731209 | -0.299 | -0.026 | 1.304 | 0.071 | 0.039 | -0.105 | -0.105 | 0 | 0.15 | 0.36 | 0.1 | 0 | 0.16 | 0.07 |
| 5 | 37818139 | rs2973049 | -0.946 | -0.106 | 1.244 | 0.044 | -0.005 | -0.015 | -0.015 | 0 | 0.74 | 0.74 | 0.66 | 0.55 | 0.64 | 0.59 |
| 5 | 37820201 | rs11951867 | 0.196 | -0.113 | 1.289 | 0.006 | 0.017 | -0.09 | -0.09 | 0 | 0.13 | 0.34 | 0.08 | 0 | 0.11 | 0.07 |
| 5 | 37821221 | rs1549250 | -0.3 | -0.141 | 1.255 | 0.003 | -0.002 | 0.001 | 0.001 | 0 | 0.62 | 0.64 | 0.36 | 0.36 | 0.56 | 0.37 |
| 5 | 37824240 | rs884344 | -0.61 | -0.108 | 0.943 | 0.028 | 0.007 | -0.024 | -0.024 | 0 | 0.27 | 0.5 | 0.4 | 0.19 | 0.24 | 0.3 |
| 5 | 37827759 | rs2973042 | -0.477 | -0.078 | 1.178 | 0.046 | 0.036 | 0.034 | 0.034 | 0 | 0.25 | 0.35 | 0.52 | 0.3 | 0.31 | 0.49 |
| 5 | 37828844 | rs2216711 | -2.031 | -0.044 | 0.829 | -0.001 | -0.05 | -0.101 | -0.101 | 0 | 0.81 | 0.99 | 0.83 | 0.68 | 0.76 | 0.82 |
| 5 | 37832759 | rs12518844 | 0.816 | -0.327 | 1.022 | 0.062 | 0.021 | 0.046 | 0.046 | 0 | 0.47 | 0.23 | 0.26 | 0.39 | 0.39 | 0.3 |
| 5 | 37832833 | rs3096140 | -0.587 | -0.291 | 1.262 | 0.076 | 0.003 | 0.003 | 0.003 | 0 | 0.7 | 0.68 | 0.76 | 0.65 | 0.65 | 0.78 |
| 5 | 37833419 | rs10941370 | -0.148 | -0.565 | 1.276 | 0.071 | 0 | -0.001 | -0.001 | 0 | 0.61 | 0.62 | 0.35 | 0.41 | 0.54 | 0.4 |
| 5 | 37839633 | rs2973033 | -1.036 | -0.617 | 1.278 | 0.097 | -0.003 | -0.003 | -0.003 | 0.386 | 0.31 | 0.61 | 0.31 | 0.25 | 0.28 | 0.49 |
| 5 | 37841045 | rs2975100 | 1.221 | -0.484 | 1.392 | 0.232 | -0.006 | 0.023 | 0.023 | 0.606 | 0.12 | 0.16 | 0.1 | 0.2 | 0.08 | 0.24 |
| 5 | 37849688 | rs1862573 | -0.489 | -0.46 | 1.504 | 0.224 | 0.031 | 0.055 | 0.055 | 0.18 | 0.07 | 0.27 | 0.15 | 0.05 | 0.09 | 0.19 |
| 5 | 37860074 | rs2910756 | -0.32 | -0.486 | 1.268 | 0.232 | -0.018 | 0.041 | 0.041 | 1.104 | 0.2 | 0.57 | 0.27 | 0.25 | 0.17 | 0.43 |
| 5 | 37861581 | rs1363927 | 0.992 | -0.27 | 0.891 | 0.299 | -0.006 | 0.053 | 0.053 | 0.235 | 0.13 | 0.28 | 0.12 | 0.2 | 0.08 | 0.2 |
| 5 | 37864666 | rs3112464 | -0.35 | -0.355 | 0.465 | 0.27 | -0.005 | 0.003 | 0.003 | 0 | 0.59 | 0.58 | 0.68 | 0.51 | 0.64 | 0.66 |

| | | | | | | | | | | | | | | | | |
|---|---|---|---|---|---|---|---|---|---|---|---|---|---|---|---|---|
| 5 | 37866601 | rs2910750 | 0.235 | -0.45 | 0.662 | 0.482 | 0.001 | 0 | 0 | 0 | 0.47 | 0.46 | 0.51 | 0.47 | 0.46 | 0.39 |
| 5 | 37866788 | rs16903764 | -0.158 | -0.514 | 0.281 | 0.469 | -0.01 | 0.001 | 0.001 | 0 | 0.11 | 0.1 | 0.04 | 0.15 | 0.03 | 0.13 |
| 5 | 37873327 | rs12519276 | 0.322 | -0.176 | -0.004 | 0.081 | -0.05 | 0.004 | 0.004 | 0 | 0.38 | 0.15 | 0.51 | 0.31 | 0.45 | 0.39 |
| 5 | 37874409 | rs10461985 | 0.845 | -0.235 | 0.352 | 0.015 | -0.004 | 0.005 | 0.005 | 0.075 | 0.12 | 0.16 | 0.1 | 0.13 | 0.09 | 0.21 |
| 5 | 37879544 | rs11956009 | 0.724 | 0.384 | -0.305 | -0.107 | -0.018 | 0.019 | 0.019 | 0 | 0.3 | 0.17 | 0.36 | 0.21 | 0.31 | 0.19 |
| 5 | 37880587 | rs10042812 | 0.774 | 0.383 | -0.376 | -0.107 | -0.044 | 0.019 | 0.019 | 0 | 0.3 | 0.12 | 0.36 | 0.21 | 0.32 | 0.19 |
| 5 | 37882172 | rs889268 | 0.216 | 0.573 | -0.356 | -0.006 | -0.008 | 0.025 | 0.025 | 0 | 0.38 | 0.29 | 0.5 | 0.23 | 0.4 | 0.39 |
| 5 | 37887883 | rs2973102 | -0.102 | 0.345 | 0.4 | 0.4 | -0.022 | -0.031 | -0.031 | 3.224 | 0.06 | 0.11 | 0.01 | 0 | 0.02 | 0.01 |
| 5 | 37888797 | rs2910743 | -0.593 | 0.507 | 0.468 | 0.683 | -0.023 | 0.054 | 0.054 | 0 | 0.17 | 0.38 | 0.15 | 0.2 | 0.1 | 0.32 |
| 5 | 37889231 | rs1423439 | -0.63 | 1.227 | 0.075 | 0.238 | 0.007 | -0.008 | -0.008 | 0.215 | 0.23 | 0.34 | 0.31 | 0.14 | 0.24 | 0.13 |
| 5 | 37889546 | rs11750418 | 0.499 | 1.865 | -0.623 | 0.25 | -0.035 | -0.029 | -0.029 | 0 | 0.06 | 0.01 | 0.04 | 0.08 | 0.09 | 0.03 |
| 5 | 37892725 | rs1363926 | -1.036 | 2.356 | -0.681 | -0.186 | 0.039 | 0.066 | 0.066 | 1.745 | 0.5 | 0.71 | 0.57 | 0.29 | 0.53 | 0.4 |
| 5 | 37892952 | rs1895435 | 0.548 | 2.44 | -1.082 | -0.351 | -0.001 | -0.021 | -0.021 | 0 | 0.1 | 0.08 | 0.05 | 0 | 0.09 | 0.05 |
| 5 | 37894278 | rs2910739 | -0.448 | 2.429 | -1.421 | -0.365 | -0.013 | 0.032 | 0.032 | 0 | 0.31 | 0.15 | 0.39 | 0.17 | 0.3 | 0.22 |
| 5 | 37897449 | rs2973093 | -0.85 | 1.22 | -1.361 | -0.196 | 0.04 | 0.092 | 0.092 | 2.561 | 0.51 | 0.85 | 0.59 | 0.3 | 0.52 | 0.39 |
| 5 | 37898176 | rs2910737 | -0.015 | 1.375 | -0.838 | 0.29 | -0.008 | -0.002 | -0.002 | 0.007 | 0.12 | 0.13 | 0.28 | 0.09 | 0.1 | 0.1 |
| 5 | 37899053 | rs3096143 | -0.439 | 1.513 | -1.135 | 0.372 | 0.008 | 0.005 | 0.005 | 0.063 | 0.22 | 0.23 | 0.19 | 0.29 | 0.23 | 0.38 |
| 5 | 37901942 | rs2973084 | -0.194 | 1.045 | -2.114 | 0.125 | 0.025 | -0.043 | -0.043 | 0 | 0.66 | 0.94 | 0.68 | 0.37 | 0.74 | 0.68 |
| 5 | 37905088 | rs1423435 | 0.017 | 1.468 | -1.625 | -0.204 | -0.018 | 0.085 | 0.085 | 0.105 | 0.41 | 0.75 | 0.54 | 0.14 | 0.48 | 0.39 |
| 5 | 37906865 | rs2973080 | -0.435 | 1.421 | -1.159 | 0.494 | 0.004 | 0 | 0 | 0 | 0.44 | 0.48 | 0.41 | 0.48 | 0.39 | 0.37 |
| 5 | 37909973 | rs2910730 | -0.205 | 2.142 | -1.2 | 0.257 | -0.003 | 0.003 | 0.003 | 0.87 | 0.09 | 0.11 | 0.21 | 0.03 | 0.15 | 0.05 |
| 5 | 37909988 | rs2910729 | 0.027 | 1.124 | -2.218 | 0.284 | 0.039 | -0.068 | -0.068 | 0.029 | 0.69 | 0.93 | 0.73 | 0.33 | 0.8 | 0.61 |
| 5 | 37912995 | rs11959616 | 0.12 | 1.531 | -1.671 | 0.185 | 0.019 | -0.006 | -0.006 | 0.135 | 0.11 | 0.2 | 0.14 | 0.11 | 0.18 | 0.2 |
| 5 | 37913527 | rs7713266 | -0.387 | 1.151 | -1.841 | 0.062 | 0.031 | -0.073 | -0.073 | 0 | 0.78 | 0.9 | 0.75 | 0.35 | 0.86 | 0.77 |
| 5 | 37915720 | rs4574573 | -0.269 | 1.214 | -1.574 | -0.333 | 0.001 | -0.005 | -0.005 | 1.409 | 0.29 | 0.31 | 0.23 | 0.24 | 0.31 | 0.28 |
| 5 | 37916982 | rs7729623 | -0.333 | 0.949 | -1.837 | 0.617 | -0.008 | -0.117 | -0.117 | 2.205 | 0.79 | 0.96 | 0.82 | 0.61 | 0.8 | 0.79 |
| 5 | 37917361 | rs13186624 | 0.68 | 0.976 | -1.183 | -0.038 | 0.005 | 0.014 | 0.014 | 0 | 0.52 | 0.44 | 0.43 | 0.4 | 0.46 | 0.55 |
| 5 | 37918703 | rs2216712 | -0.018 | 1.102 | -1.677 | 0.33 | -0.002 | -0.003 | -0.003 | 0 | 0.15 | 0.14 | 0.2 | 0.06 | 0.11 | 0.09 |
| 5 | 37918856 | rs1035500 | -0.066 | 1.8 | -1.755 | 0.748 | -0.084 | -0.102 | -0.102 | 1.148 | 0.77 | 0.92 | 0.65 | 0.45 | 0.67 | 0.66 |
| 5 | 37921235 | rs958502 | -0.041 | 1.014 | -1.347 | 0.198 | 0.008 | 0.07 | 0.07 | 0 | 0.51 | 0.31 | 0.52 | 0.69 | 0.51 | 0.6 |
| 5 | 37922362 | rs4869558 | 0.304 | 0.408 | -1.058 | -0.382 | -0.007 | -0.022 | -0.022 | 0 | 0.57 | 0.54 | 0.6 | 0.7 | 0.55 | 0.62 |
| 5 | 37922538 | rs4869559 | 0.291 | 0.557 | -0.999 | -0.354 | -0.001 | -0.003 | -0.003 | 0 | 0.58 | 0.49 | 0.6 | 0.62 | 0.54 | 0.66 |
| 5 | 37922890 | rs25448 | 0 | -0.537 | -1.362 | -0.798 | 0 | 0.016 | 0.016 | 0.022 | 0.13 | 0.01 | 0.16 | 0.04 | 0.12 | 0.04 |
| 5 | 37923364 | rs6859065 | -0.719 | -1.218 | -1.011 | -1.329 | -0.007 | -0.012 | -0.012 | 0 | 0.37 | 0.54 | 0.35 | 0.22 | 0.34 | 0.36 |
| 5 | 37925239 | rs2366837 | -1.001 | -1.276 | -1.116 | -1.395 | 0.012 | 0.077 | 0.077 | 0.935 | 0.58 | 0.8 | 0.32 | 0.27 | 0.45 | 0.41 |
| 5 | 37927454 | rs10040845 | -0.237 | -1.605 | -0.499 | -1.526 | -0.024 | -0.035 | -0.035 | 0 | 0.42 | 0.47 | 0.21 | 0.17 | 0.27 | 0.28 |
| 5 | 37929958 | rs1423431 | -1.966 | -0.819 | 0.935 | -1.042 | -0.018 | -0.067 | -0.067 | 0 | 0.25 | 0.53 | 0.31 | 0.16 | 0.26 | 0.18 |
| 5 | 37933172 | rs12519198 | -1.117 | -0.619 | 0.793 | -0.814 | -0.003 | -0.007 | -0.007 | 0 | 0.09 | 0.17 | 0.19 | 0.09 | 0.09 | 0.06 |
| 5 | 37938540 | rs2032873 | -2.466 | -1.051 | 1.3 | -1.127 | -0.016 | -0.032 | -0.032 | 0.12 | 0.37 | 0.52 | 0.36 | 0.22 | 0.33 | 0.25 |
| 5 | 37940532 | rs17389528 | -1.586 | -1.04 | 1.164 | -1.125 | 0.001 | -0.003 | -0.003 | 0 | 0.23 | 0.16 | 0.14 | 0.09 | 0.22 | 0.18 |
| 5 | 37941205 | rs1862582 | -1.602 | -1.046 | 1.265 | -1.128 | -0.001 | -0.007 | -0.007 | 0.135 | 0.22 | 0.2 | 0.14 | 0.09 | 0.22 | 0.18 |
| 5 | 37941603 | rs12521230 | 0.018 | -1.038 | 1.033 | -1.117 | 0 | 0 | 0 | 0 | 0.06 | 0.06 | 0.17 | 0.09 | 0.06 | 0.04 |
| 5 | 37944244 | rs4314423 | -1.551 | -1.062 | 1.02 | -0.986 | 0.005 | 0.005 | 0.005 | 0 | 0.21 | 0.15 | 0.24 | 0.13 | 0.15 | 0.14 |
| 5 | 37947775 | rs13187126 | -1.082 | -1.004 | 0.971 | -0.988 | -0.003 | 0.004 | 0.004 | 0 | 0.08 | 0.07 | 0.07 | 0.04 | 0.15 | 0.08 |
| 5 | 37950691 | rs9292683 | -2.904 | -0.94 | 1.111 | -1.006 | -0.016 | -0.039 | -0.039 | 0 | 0.37 | 0.44 | 0.35 | 0.22 | 0.34 | 0.25 |
| 5 | 37950840 | rs16903810 | -0.754 | -0.897 | 1.057 | -1.001 | 0.013 | 0.016 | 0.016 | 0 | 0.15 | 0.08 | 0.07 | 0.05 | 0.09 | 0.1 |
| 5 | 37953948 | rs3749693 | -0.752 | -0.964 | 1.074 | -1.026 | 0.009 | 0.01 | 0.01 | 0 | 0.15 | 0.09 | 0.07 | 0.05 | 0.09 | 0.1 |
| 5 | 37957085 | rs6898456 | -2.158 | -0.957 | 1.054 | -1.033 | -0.011 | -0.041 | -0.041 | 0 | 0.26 | 0.31 | 0.18 | 0.08 | 0.27 | 0.2 |
| 5 | 37964851 | rs10805630 | -2.161 | -0.841 | 0.831 | -0.967 | -0.001 | 0.012 | 0.012 | 0 | 0.18 | 0.15 | 0.26 | 0.13 | 0.24 | 0.15 |
| 5 | 37966672 | rs2292261 | -0.631 | -0.77 | 0.724 | -0.974 | -0.032 | 0.004 | 0.004 | 0.09 | 0.07 | 0 | 0.06 | 0.04 | 0.14 | 0.08 |
| 5 | 37984257 | rs1508841 | -2.714 | -0.784 | 1.119 | -0.93 | -0.021 | -0.057 | -0.057 | 0.354 | 0.35 | 0.47 | 0.36 | 0.22 | 0.32 | 0.25 |
| 5 | 37986493 | rs1304732 | -1.905 | -0.774 | 1.05 | -0.909 | 0.031 | -0.017 | -0.017 | 0 | 0.09 | 0.25 | 0.08 | 0.04 | 0.13 | 0.07 |
| 5 | 37986588 | rs6869247 | -2.366 | -0.805 | 1.056 | -0.906 | 0.002 | -0.03 | -0.03 | 0 | 0.17 | 0.3 | 0.26 | 0.12 | 0.2 | 0.12 |
| 5 | 37988993 | rs12515304 | -2.006 | -0.955 | 1.01 | -0.938 | 0.033 | 0.038 | 0.038 | 0 | 0.28 | 0.17 | 0.28 | 0.15 | 0.18 | 0.16 |
| 5 | 37989879 | rs7726340 | -2.731 | -0.94 | 1.014 | -0.935 | -0.013 | -0.03 | -0.03 | 0 | 0.37 | 0.41 | 0.36 | 0.19 | 0.31 | 0.23 |
| 5 | 37992218 | rs10805632 | -1.631 | -0.92 | 1.037 | -0.909 | 0.019 | 0.02 | 0.02 | 0 | 0.25 | 0.17 | 0.24 | 0.14 | 0.15 | 0.14 |
| 5 | 37996987 | rs17331676 | -0.88 | -0.905 | 0.681 | -0.903 | 0.006 | -0.005 | -0.005 | 0 | 0.06 | 0.09 | 0.06 | 0.04 | 0.13 | 0.07 |
| 5 | 37999585 | rs12188790 | -1.682 | -0.727 | 0.17 | -0.833 | 0.036 | 0.04 | 0.04 | 0 | 0.25 | 0.1 | 0.26 | 0.15 | 0.15 | 0.14 |
| 5 | 38008042 | rs10050775 | -1.67 | -0.604 | 0.694 | -0.67 | -0.02 | -0.018 | -0.018 | 0 | 0.22 | 0.66 | 0.15 | 0.13 | 0.14 | 0.13 |
| 5 | 38011408 | rs11747656 | -2.741 | -0.839 | 0.546 | -0.803 | 0.038 | 0.053 | 0.053 | 0 | 0.39 | 0.75 | 0.24 | 0.25 | 0.31 | 0.21 |
| 5 | 38014253 | rs1508833 | -1.207 | -0.824 | 0.511 | -0.778 | -0.003 | 0.032 | 0.032 | 0.127 | 0.17 | 0.08 | 0.11 | 0.12 | 0.19 | 0.11 |
| 5 | 38015610 | rs10059028 | -1.586 | -0.839 | 1.086 | -0.773 | -0.065 | -0.048 | -0.048 | 0 | 0.21 | 0.57 | 0.12 | 0.12 | 0.11 | 0.11 |
| 5 | 38018463 | rs10044567 | -2.123 | -0.217 | 1.123 | -0.807 | -0.017 | -0.058 | -0.058 | 0.039 | 0.22 | 0.62 | 0.16 | 0.13 | 0.17 | 0.1 |
| 5 | 38019003 | rs11740374 | -2.507 | -0.266 | 1.301 | -0.845 | -0.027 | -0.049 | -0.049 | 0 | 0.25 | 0.58 | 0.17 | 0.18 | 0.22 | 0.15 |
| 5 | 38022514 | rs270569 | 0.622 | -0.18 | 1.111 | -0.797 | 0.02 | 0.019 | 0.019 | 1.864 | 0.86 | 0.98 | 0.95 | 0.87 | 0.92 | 0.94 |
| 5 | 38022879 | rs270568 | 0.645 | -0.182 | 1.007 | -0.768 | 0.021 | 0.02 | 0.02 | 1.992 | 0.86 | 0.98 | 0.95 | 0.87 | 0.92 | 0.94 |
| 5 | 38033088 | rs270559 | 0.98 | -0.191 | 1.077 | -0.767 | -0.003 | 0.025 | 0.025 | 0.752 | 0.83 | 0.9 | 0.91 | 0.85 | 0.82 | 0.88 |
| 5 | 38035083 | rs6451355 | -1.19 | -0.22 | 1.35 | -0.767 | 0.032 | 0.056 | 0.056 | 3.243 | 0.07 | 0.42 | 0.06 | 0.03 | 0.03 | 0.03 |
| 5 | 38036280 | rs10473079 | -1.19 | -0.22 | 1.35 | -0.742 | 0.032 | 0.061 | 0.061 | 3.243 | 0.07 | 0.42 | 0.06 | 0.03 | 0.03 | 0.03 |

| | | | | | | | | | | | | | | | | |
|---|---|---|---|---|---|---|---|---|---|---|---|---|---|---|---|---|
| 5 | 38038345 | rs270552 | 2.511 | -0.063 | 1.184 | -0.886 | 0.011 | 0.054 | 0.054 | 0 | 0.57 | 0.3 | 0.74 | 0.76 | 0.63 | 0.77 |
| 5 | 38040195 | rs270550 | 1.871 | 0.116 | 0.937 | -0.816 | 0 | 0.014 | 0.014 | 0 | 0.64 | 0.71 | 0.8 | 0.79 | 0.67 | 0.79 |
| 5 | 38040629 | rs172011 | 1.853 | 0.123 | 0.792 | -0.813 | -0.001 | 0.035 | 0.035 | 0 | 0.64 | 0.73 | 0.81 | 0.79 | 0.67 | 0.79 |
| 5 | 38041398 | rs1494640 | 1.043 | 0.237 | 0.438 | -0.558 | -0.006 | 0.021 | 0.021 | 0.799 | 0.83 | 0.92 | 0.91 | 0.85 | 0.82 | 0.88 |
| 5 | 38042262 | rs1845496 | 1.153 | 0.614 | 0.642 | -0.361 | 0.007 | -0.029 | -0.029 | 0 | 0.61 | 0.51 | 0.77 | 0.74 | 0.58 | 0.75 |
| 5 | 38046081 | rs2642682 | -1.619 | 0.598 | 0.884 | 0.317 | -0.007 | -0.012 | -0.012 | 0 | 0.22 | 0.28 | 0.12 | 0.22 | 0.15 | 0.1 |
| 5 | 38049522 | rs7705239 | -0.949 | 1.186 | 0.923 | 0.494 | 0.041 | 0.011 | 0.011 | 0 | 0.12 | 0.22 | 0.1 | 0.08 | 0.17 | 0.11 |
| 5 | 38051989 | rs270544 | -0.515 | 1.16 | 0.902 | 0.483 | -0.004 | 0.016 | 0.016 | 0 | 0.14 | 0.02 | 0.08 | 0.13 | 0.12 | 0.06 |
| 5 | 38053488 | rs270537 | -2.231 | 1.161 | 0.887 | 0.508 | -0.005 | 0.002 | 0.002 | 0 | 0.31 | 0.24 | 0.21 | 0.29 | 0.33 | 0.21 |
| 5 | 38059375 | rs6891116 | -1.62 | 1.177 | 0.704 | 0.51 | -0.006 | -0.003 | -0.003 | 0.343 | 0.16 | 0.12 | 0.12 | 0.15 | 0.21 | 0.15 |
| 5 | 38061840 | rs270593 | -0.64 | 1.145 | 0.453 | 0.458 | 0.001 | 0.032 | 0.032 | 0 | 0.16 | 0.04 | 0.1 | 0.13 | 0.14 | 0.08 |
| 5 | 38074854 | rs270584 | -2.729 | 1.111 | 0.266 | 0.582 | -0.137 | -0.002 | -0.002 | 0 | 0.43 | 0.95 | 0.32 | 0.32 | 0.44 | 0.28 |
| 5 | 38083782 | rs1494645 | -1.04 | 1.001 | 0.889 | 0.736 | -0.005 | -0.011 | -0.011 | 0.368 | 0.18 | 0.18 | 0.11 | 0.13 | 0.14 | 0.08 |
| 5 | 38089107 | rs12653109 | 1.276 | 0.985 | 0.757 | 0.778 | 0.009 | -0.034 | -0.034 | 0.069 | 0.7 | 0.54 | 0.79 | 0.8 | 0.69 | 0.81 |
| 5 | 38099612 | rs2013340 | -1.857 | 0.516 | 0.565 | 1.821 | 0.001 | 0 | 0 | 0 | 0.07 | 0.02 | 0.02 | 0.26 | 0.05 | 0.09 |
| 5 | 38102779 | rs2471058 | 1.881 | 0.587 | 1.472 | 1.935 | 0.01 | 0.137 | 0.137 | 0 | 0.83 | 0.64 | 0.74 | 0.69 | 0.84 | 0.84 |
| 5 | 38103062 | rs2453341 | 2.104 | 0.549 | 1.789 | 1.911 | 0.008 | 0.153 | 0.153 | 0.04 | 0.82 | 0.5 | 0.73 | 0.7 | 0.84 | 0.84 |
| 5 | 38104956 | rs2471068 | 2.174 | 0.59 | 1.591 | 2.038 | 0.029 | 0.197 | 0.197 | 1.349 | 0.93 | 0.73 | 0.96 | 0.79 | 0.96 | 0.97 |
| 5 | 38105437 | rs6874732 | -1.421 | 0.479 | 1.484 | 1.299 | -0.007 | -0.086 | -0.086 | 0.454 | 0.14 | 0.36 | 0.06 | 0.01 | 0.09 | 0.04 |
| 5 | 38106089 | rs1997295 | -0.937 | 0.766 | 0.765 | 1.564 | -0.001 | -0.002 | -0.002 | 0.069 | 0.1 | 0.1 | 0.19 | 0.12 | 0.12 | 0.09 |
| 5 | 38108898 | rs7711670 | -0.888 | 0.672 | 0.848 | 1.675 | -0.013 | -0.001 | -0.001 | 0 | 0.1 | 0.18 | 0.14 | 0.1 | 0.05 | 0.03 |
| 5 | 38110201 | rs9686325 | -1.234 | 0.898 | 0.775 | 2.026 | 0.005 | 0.013 | 0.013 | 0 | 0.14 | 0.16 | 0.12 | 0.24 | 0.22 | 0.12 |
| 5 | 38112559 | rs879184 | -1.738 | 0.807 | 0.692 | 1.807 | 0.004 | 0.016 | 0.016 | 0.429 | 0.35 | 0.54 | 0.38 | 0.39 | 0.38 | 0.23 |
| 5 | 38119094 | rs2471075 | 1.077 | 0.216 | 0.25 | 2.057 | 0.002 | -0.001 | -0.001 | 0.71 | 0.89 | 0.89 | 0.83 | 0.7 | 0.89 | 0.94 |
| 5 | 38119545 | rs1845494 | 1.414 | 0.365 | 0.445 | 1.909 | 0.003 | 0.005 | 0.005 | 0 | 0.65 | 0.58 | 0.63 | 0.49 | 0.62 | 0.7 |
| 5 | 38120428 | rs2453328 | 1.639 | 0.148 | 0.822 | 2.032 | -0.045 | 0.181 | 0.181 | 0.501 | 0.88 | 0.66 | 0.8 | 0.68 | 0.94 | 0.89 |
| 5 | 38129168 | rs13355907 | -0.88 | 0.211 | 0.851 | 1.946 | -0.005 | 0.018 | 0.018 | 0.112 | 0.05 | 0.2 | 0.06 | 0.02 | 0.01 | 0.07 |
| 5 | 38129295 | rs4869565 | 1.334 | 0.248 | 0.694 | 1.961 | 0 | 0.003 | 0.003 | 0 | 0.72 | 0.7 | 0.69 | 0.51 | 0.68 | 0.73 |
| 5 | 38131390 | rs11742872 | 2.134 | -0.267 | 0.232 | 1.909 | -0.002 | 0.065 | 0.065 | 0.159 | 0.57 | 0.08 | 0.6 | 0.33 | 0.61 | 0.65 |
| 5 | 38135718 | rs3812048 | 0.026 | -0.162 | 0.306 | 2.085 | -0.095 | -0.073 | -0.073 | 0 | 0.17 | 0.01 | 0.12 | 0.17 | 0.26 | 0.16 |
| 5 | 38136423 | rs7737797 | -0.606 | -0.146 | 1.05 | 1.58 | -0.016 | 0.093 | 0.093 | 0 | 0.1 | 0.34 | 0.06 | 0.18 | 0.05 | 0.06 |
| 5 | 38137906 | rs12518421 | 1.407 | -0.077 | 0.445 | 1.248 | 0.012 | 0.041 | 0.041 | 0 | 0.61 | 0.22 | 0.62 | 0.32 | 0.63 | 0.66 |
| 5 | 38143698 | rs12658891 | -1.109 | -0.311 | 0.978 | 0.916 | -0.005 | -0.006 | -0.006 | 0 | 0.12 | 0.13 | 0.1 | 0.09 | 0.11 | 0.11 |
| 5 | 38144274 | rs3936162 | 1.24 | -0.703 | 0.9 | 0.101 | 0.022 | 0.086 | 0.086 | 0 | 0.5 | 0.36 | 0.38 | 0.04 | 0.57 | 0.27 |
| 5 | 38144492 | rs12659458 | -0.731 | -0.388 | 0.419 | 0.307 | -0.007 | -0.005 | -0.005 | 0 | 0.12 | 0.13 | 0.1 | 0.12 | 0.11 | 0.12 |
| 5 | 38148794 | rs6897900 | 0.316 | -0.351 | 0.419 | 0.395 | -0.004 | 0.024 | 0.024 | 0.06 | 0.67 | 0.72 | 0.51 | 0.17 | 0.7 | 0.43 |
| 5 | 38151713 | rs4562066 | 0.158 | -0.266 | 0.075 | 0.435 | -0.021 | 0.067 | 0.067 | 0.287 | 0.69 | 0.84 | 0.52 | 0.17 | 0.7 | 0.43 |
| 5 | 38151803 | rs6451369 | 0.17 | -0.267 | 0.075 | 0.434 | -0.021 | 0.067 | 0.067 | 0.321 | 0.69 | 0.84 | 0.52 | 0.17 | 0.7 | 0.43 |
| 5 | 38151950 | rs4583910 | 0.16 | -0.257 | 0.079 | 0.439 | -0.021 | 0.067 | 0.067 | 0.287 | 0.69 | 0.84 | 0.52 | 0.17 | 0.7 | 0.43 |
| 5 | 38152044 | rs4289583 | -0.293 | 0.032 | -0.102 | 0.913 | -0.071 | -0.068 | -0.068 | 0 | 0.77 | 0.96 | 0.55 | 0.35 | 0.73 | 0.47 |
| 5 | 38153756 | rs9292698 | -0.172 | 0.061 | -0.085 | 0.934 | -0.066 | -0.065 | -0.065 | 0 | 0.75 | 0.95 | 0.54 | 0.35 | 0.71 | 0.45 |
| 5 | 38158309 | rs4376295 | -1.11 | -0.491 | 1.016 | 1.291 | -0.005 | 0.032 | 0.032 | 0 | 0.11 | 0.26 | 0.1 | 0.19 | 0.12 | 0.11 |
| 5 | 38166586 | rs12109280 | -1.832 | -0.604 | 1.479 | 1.284 | -0.02 | 0.067 | 0.067 | 0.071 | 0.15 | 0.54 | 0.14 | 0.21 | 0.11 | 0.11 |
| 5 | 38176521 | rs6897859 | -2.062 | -0.889 | 1.153 | 1.666 | 0.028 | -0.04 | -0.04 | 0 | 0.27 | 0.82 | 0.28 | 0.41 | 0.15 | 0.34 |
| 5 | 38190138 | rs13362201 | -1.076 | -0.893 | 2.476 | 1.684 | -0.03 | 0.021 | 0.021 | 0.861 | 0.09 | 0.24 | 0.04 | 0.12 | 0.03 | 0.18 |
| 5 | 38191698 | rs10512682 | -1.778 | -0.904 | 2.946 | 1.845 | 0.025 | 0.173 | 0.173 | 0.267 | 0.13 | 0.43 | 0.21 | 0.26 | 0.1 | 0.24 |
| 5 | 38194034 | rs6451373 | -0.58 | -0.868 | 2.697 | 1.752 | -0.021 | 0.025 | 0.025 | 0.748 | 0.07 | 0.17 | 0.04 | 0.11 | 0.03 | 0.1 |
| 5 | 38203031 | rs6451374 | -1.006 | -0.796 | 1.044 | 1.143 | -0.115 | -0.098 | -0.098 | 0 | 0.6 | 0.95 | 0.56 | 0.43 | 0.52 | 0.54 |
| 5 | 38205477 | rs10036647 | -1.66 | -0.812 | 1.979 | 0.724 | -0.006 | -0.007 | -0.007 | 1.002 | 0.15 | 0.24 | 0.08 | 0.13 | 0.12 | 0.19 |
| 5 | 38209532 | rs1428186 | -0.674 | -0.779 | 2.344 | 0.909 | 0.044 | 0.122 | 0.122 | 0 | 0.06 | 0.23 | 0.17 | 0.13 | 0.06 | 0.06 |
| 5 | 38221660 | rs4869578 | 0.481 | -0.87 | 2.14 | 1.069 | -0.013 | 0.01 | 0.01 | 0.096 | 0.92 | 0.88 | 0.95 | 0.84 | 0.97 | 0.86 |
| 5 | 38221718 | rs16903890 | -1.024 | -0.707 | 2.331 | 0.738 | 0.103 | 0.207 | 0.207 | 0 | 0.07 | 0.36 | 0.19 | 0.13 | 0.06 | 0.06 |
| 5 | 38251005 | rs6867736 | 0.666 | -0.899 | 1.049 | 0.277 | -0.013 | 0.059 | 0.059 | 0 | 0.3 | 0.08 | 0.27 | 0.13 | 0.33 | 0.17 |
| 5 | 38256582 | rs2561098 | -1.58 | -1.243 | 1.885 | 0.092 | -0.044 | -0.082 | -0.082 | 0 | 0.24 | 0.62 | 0.15 | 0.07 | 0.12 | 0.24 |
| 5 | 38258707 | rs2561096 | -0.957 | -1.149 | 2.15 | 0.087 | 0.028 | -0.05 | -0.05 | 0 | 0.1 | 0.36 | 0.11 | 0.01 | 0.08 | 0.13 |
| 5 | 38261117 | rs2589789 | -1.1 | -1.095 | 2.096 | -0.019 | -0.007 | 0.043 | 0.043 | 0 | 0.56 | 0.72 | 0.43 | 0.21 | 0.49 | 0.52 |
| 5 | 38261833 | rs16903908 | -0.431 | -1.083 | 2.077 | -0.188 | -0.052 | -0.048 | -0.048 | 0 | 0.15 | 0.35 | 0.05 | 0.06 | 0.05 | 0.1 |
| 5 | 38262918 | rs6886455 | 0.383 | -0.977 | 1.462 | -0.138 | 0.001 | 0.023 | 0.023 | 0 | 0.33 | 0.19 | 0.4 | 0.25 | 0.36 | 0.3 |
| 5 | 38264735 | rs2006889 | -0.794 | -0.994 | 1.501 | -0.156 | -0.005 | 0.033 | 0.033 | 0 | 0.55 | 0.66 | 0.56 | 0.31 | 0.52 | 0.49 |
| 5 | 38267136 | rs17337455 | 0.192 | 0.018 | 1.448 | 0.071 | 0.028 | 0.018 | 0.018 | 0 | 0.08 | 0.14 | 0.19 | 0.12 | 0.11 | 0.04 |
| 5 | 38270279 | rs2561045 | -1.78 | -0.261 | 1.269 | -0.322 | -0.031 | -0.017 | -0.017 | 0 | 0.66 | 0.75 | 0.64 | 0.39 | 0.6 | 0.51 |
| 5 | 38271336 | rs2561144 | -0.37 | -0.388 | 1.869 | -0.372 | 0.091 | 0.059 | 0.059 | 0 | 0.11 | 0.43 | 0.26 | 0.13 | 0.12 | 0.12 |
| 5 | 38271451 | rs2561143 | -1.906 | -0.388 | 1.861 | -0.375 | -0.055 | -0.036 | -0.036 | 0 | 0.66 | 0.81 | 0.64 | 0.39 | 0.59 | 0.51 |
| 5 | 38271652 | rs10512679 | 0.822 | -0.622 | 1.344 | -0.673 | 0.018 | 0.032 | 0.032 | 0 | 0.26 | 0.18 | 0.2 | 0.13 | 0.2 | 0.14 |
| 5 | 38272492 | rs17422587 | -1.346 | -0.622 | 1.332 | -0.673 | 0.007 | 0.073 | 0.073 | 0.799 | 0.54 | 0.32 | 0.37 | 0.26 | 0.48 | 0.39 |
| 5 | 38273460 | rs1428256 | 0.877 | -0.622 | 1.337 | -0.673 | 0.025 | 0.067 | 0.067 | 0 | 0.25 | 0.11 | 0.18 | 0.08 | 0.19 | 0.11 |
| 5 | 38274312 | rs7724224 | -1.293 | -0.565 | 1.383 | -0.635 | 0.007 | 0.069 | 0.069 | 0.799 | 0.54 | 0.32 | 0.37 | 0.26 | 0.48 | 0.39 |
| 5 | 38281851 | rs1428248 | -0.959 | -0.549 | 1.446 | -0.696 | -0.016 | -0.057 | -0.057 | 0 | 0.31 | 0.38 | 0.19 | 0.11 | 0.28 | 0.24 |
| 5 | 38282246 | rs2561133 | 0.814 | -0.567 | 1.193 | -0.68 | 0.023 | 0.051 | 0.051 | 0 | 0.25 | 0.12 | 0.19 | 0.13 | 0.19 | 0.15 |

| | | | | | | | | | | | | | | | | |
|---|---|---|---|---|---|---|---|---|---|---|---|---|---|---|---|---|
| 5 | 38284663 | rs13182235 | 0.346 | -0.527 | 1.42 | -0.753 | 0.014 | 0.02 | 0.02 | 0 | 0.07 | 0.13 | 0.06 | 0.1 | 0.12 | 0.14 |
| 5 | 38285397 | rs10040077 | 0.269 | -0.494 | 1.497 | -0.745 | 0.072 | 0.047 | 0.047 | 0 | 0.1 | 0.31 | 0.25 | 0.13 | 0.12 | 0.12 |
| 5 | 38286606 | rs2367214 | -2.045 | -0.581 | 1.161 | -0.772 | 0 | 0.088 | 0.088 | 1.254 | 0.42 | 0.75 | 0.44 | 0.25 | 0.4 | 0.37 |
| 5 | 38305547 | rs2451018 | -1.405 | -0.398 | 1.258 | -0.864 | -0.004 | -0.063 | -0.063 | 0.014 | 0.32 | 0.54 | 0.39 | 0.21 | 0.3 | 0.27 |
| 5 | 38307820 | rs17423367 | 0.823 | -0.564 | 1.038 | -0.998 | 0.028 | 0.047 | 0.047 | 0 | 0.25 | 0.14 | 0.19 | 0.06 | 0.16 | 0.1 |
| 5 | 38309923 | rs987452 | -0.968 | -0.363 | 1.861 | -0.663 | -0.05 | -0.159 | -0.159 | 0 | 0.23 | 0.43 | 0.15 | 0.01 | 0.16 | 0.1 |
| 5 | 38313811 | rs13177214 | 2.36 | -0.128 | 1.046 | -0.672 | -0.057 | 0.009 | 0.009 | 1.844 | 0.39 | 0.15 | 0.44 | 0.64 | 0.42 | 0.47 |
| 5 | 38313973 | rs1366281 | 0.183 | 0.007 | 1.334 | -0.613 | -0.001 | -0.004 | -0.004 | 0 | 0.7 | 0.66 | 0.67 | 0.8 | 0.74 | 0.84 |
| 5 | 38314070 | rs1366282 | -0.306 | 0.065 | 1.32 | -0.589 | 0.022 | 0.01 | 0.01 | 0 | 0.9 | 0.84 | 0.82 | 0.86 | 0.88 | 0.92 |
| 5 | 38321141 | rs13356132 | 0.812 | -0.052 | 0.971 | -0.597 | 0.021 | 0.036 | 0.036 | 0 | 0.2 | 0.16 | 0.15 | 0.06 | 0.14 | 0.08 |
| 5 | 38322975 | rs16903936 | -0.473 | 0.132 | 1.05 | -0.609 | 0.014 | 0.007 | 0.007 | 0 | 0.08 | 0.12 | 0.08 | 0.1 | 0.14 | 0.18 |
| 5 | 38326508 | rs2589822 | -0.787 | -0.143 | 0.825 | -0.756 | 0.051 | 0.141 | 0.141 | 0 | 0.34 | 0.8 | 0.29 | 0.05 | 0.25 | 0.17 |
| 5 | 38329510 | rs2589829 | -0.876 | -0.141 | 0.8 | -0.762 | 0.057 | 0.161 | 0.161 | 0.058 | 0.34 | 0.82 | 0.29 | 0.05 | 0.25 | 0.17 |
| 5 | 38332224 | rs2561126 | -0.838 | -0.127 | 0.901 | -0.594 | 0.051 | 0.115 | 0.115 | 0.019 | 0.34 | 0.78 | 0.28 | 0.07 | 0.25 | 0.17 |
| 5 | 38334361 | rs12656306 | 0.587 | -0.043 | 0.784 | -0.571 | 0.006 | 0.029 | 0.029 | 8.556 | 0.09 | 0.01 | 0.02 | 0.04 | 0.04 | 0.03 |
| 5 | 38334997 | rs6885417 | -0.535 | -0.015 | 1.432 | -0.555 | 0.008 | -0.08 | -0.08 | 0.405 | 0.13 | 0.26 | 0.11 | 0.01 | 0.13 | 0.1 |
| 5 | 38343053 | rs2162814 | 1.59 | 0.322 | 1.37 | 0.043 | -0.021 | -0.024 | -0.024 | 0 | 0.55 | 0.28 | 0.66 | 0.53 | 0.57 | 0.57 |
| 5 | 38345522 | rs1428261 | 1.215 | 0.266 | 1.72 | 0.066 | 0 | -0.002 | -0.002 | 0.059 | 0.57 | 0.64 | 0.7 | 0.54 | 0.62 | 0.61 |
| 5 | 38348573 | rs2561115 | 0.524 | 0.226 | 1.872 | 0.097 | 0.007 | 0.006 | 0.006 | 0 | 0.08 | 0.08 | 0.06 | 0.14 | 0.12 | 0.08 |
| 5 | 38348663 | rs2961880 | -1.162 | 0.178 | 1.745 | 0.089 | 0.016 | 0.019 | 0.019 | 0 | 0.83 | 0.98 | 0.87 | 0.89 | 0.88 | 0.87 |
| 5 | 38349573 | rs2561113 | -1.908 | 0.129 | 2.119 | 0.132 | 0.022 | 0.021 | 0.021 | 5.096 | 0.26 | 0.52 | 0.18 | 0.34 | 0.26 | 0.24 |
| 5 | 38350643 | rs2561111 | 0.56 | 0.138 | 2.093 | 0.124 | -0.016 | -0.014 | -0.014 | 0 | 0.08 | 0.03 | 0.06 | 0.14 | 0.12 | 0.08 |
| 5 | 38351102 | rs10045133 | 0.347 | 0.063 | 2.1 | 0.113 | -0.001 | -0.003 | -0.003 | 0 | 0.1 | 0.1 | 0.03 | 0.05 | 0.04 | 0.03 |
| 5 | 38351604 | rs2561110 | -0.951 | 0.072 | 2.064 | 0.118 | 0.068 | 0.036 | 0.036 | 0 | 0.16 | 0.32 | 0.15 | 0.24 | 0.22 | 0.21 |
| 5 | 38352938 | rs1114175 | -2.55 | 0.042 | 1.565 | 0.101 | -0.029 | -0.001 | -0.001 | 0 | 0.35 | 0.75 | 0.23 | 0.35 | 0.35 | 0.34 |
| 5 | 38353480 | rs2589811 | -1.912 | 0.05 | 1.335 | 0.105 | -0.031 | -0.016 | -0.016 | 0.053 | 0.27 | 0.78 | 0.22 | 0.3 | 0.31 | 0.3 |
| 5 | 38355045 | rs2561106 | 1.72 | 0.073 | 1.227 | 0.015 | -0.033 | -0.053 | -0.053 | 0 | 0.55 | 0.17 | 0.65 | 0.54 | 0.58 | 0.57 |
| 5 | 38356769 | rs2589810 | 2.159 | 0.094 | 1.168 | 0.022 | -0.091 | -0.076 | -0.076 | 0 | 0.46 | 0.13 | 0.63 | 0.49 | 0.54 | 0.53 |
| 5 | 38359501 | rs2561099 | 2.214 | 0.335 | 1.207 | -0.765 | -0.085 | 0.024 | 0.024 | 0.87 | 0.46 | 0.18 | 0.63 | 0.59 | 0.53 | 0.54 |
| 5 | 38361483 | rs2561140 | 0.77 | 0.385 | 1.81 | -0.758 | 0 | 0.002 | 0.002 | 0 | 0.64 | 0.42 | 0.71 | 0.69 | 0.65 | 0.65 |
| 5 | 38362043 | rs2255207 | -1.691 | 0.46 | 2.036 | -1.24 | 0.07 | -0.027 | -0.027 | 0 | 0.24 | 0.46 | 0.19 | 0.22 | 0.31 | 0.26 |
| 5 | 38366133 | rs1834967 | 0.937 | 0.49 | 2.07 | -1.287 | 0.002 | -0.029 | -0.029 | 0.189 | 0.19 | 0.14 | 0.14 | 0 | 0.13 | 0.08 |
| 5 | 38366662 | rs10214135 | -0.631 | 0.67 | 2.209 | -1.229 | 0.022 | -0.016 | -0.016 | 0 | 0.16 | 0.16 | 0.11 | 0.13 | 0.21 | 0.13 |
| 5 | 38368719 | rs1366285 | -1.592 | 0.658 | 1.999 | -1.205 | 0.076 | -0.024 | -0.024 | 0 | 0.24 | 0.44 | 0.18 | 0.22 | 0.31 | 0.25 |
| 5 | 38370101 | rs2059835 | -2.396 | 0.636 | 1.686 | -1.301 | -0.094 | 0.095 | 0.095 | 1.398 | 0.57 | 0.86 | 0.41 | 0.3 | 0.48 | 0.44 |
| 5 | 38370256 | rs16903957 | 0.378 | 0.673 | 1.685 | -1.265 | 0.003 | 0.005 | 0.005 | 5.915 | 0.1 | 0 | 0.02 | 0.07 | 0.04 | 0.07 |
| 5 | 38370537 | rs1465567 | -1.813 | 0.69 | 1.868 | -1.278 | -0.01 | -0.055 | -0.055 | 0.31 | 0.26 | 0.44 | 0.14 | 0.18 | 0.2 | 0.24 |
| 5 | 38371063 | rs6879031 | -0.031 | 1.022 | 1.895 | -1.269 | 0.035 | 0.002 | 0.002 | 0 | 0.11 | 0.2 | 0.12 | 0.11 | 0.15 | 0.09 |
| 5 | 38372929 | rs2451003 | -1.252 | 0.957 | 2.097 | -1.212 | 0.004 | -0.006 | -0.006 | 0 | 0.37 | 0.6 | 0.26 | 0.11 | 0.29 | 0.26 |
| 5 | 38373398 | rs2932110 | -1.496 | 0.974 | 2.297 | -0.959 | 0.021 | -0.053 | -0.053 | 0 | 0.16 | 0.38 | 0.1 | 0.08 | 0.16 | 0.17 |
| 5 | 38378483 | rs2366981 | -0.719 | 0.723 | 2.139 | -0.709 | 0.019 | 0.043 | 0.043 | 0 | 0.07 | 0.25 | 0.07 | 0.08 | 0.07 | 0.13 |
| 5 | 38380359 | rs7716864 | -0.072 | 0.868 | 2.399 | -0.71 | 0.025 | 0.011 | 0.011 | 0 | 0.1 | 0.17 | 0.18 | 0.14 | 0.12 | 0.1 |
| 5 | 38381015 | rs3110232 | -0.948 | 0.884 | 2.037 | -0.577 | -0.006 | -0.007 | -0.007 | 0 | 0.15 | 0.26 | 0.09 | 0.13 | 0.11 | 0.19 |
| 5 | 38381445 | rs3099130 | -0.648 | 1.026 | 1.831 | -0.59 | 0.004 | 0.113 | 0.113 | 0.376 | 0.51 | 0.68 | 0.45 | 0.26 | 0.4 | 0.4 |
| 5 | 38389768 | rs6879468 | -2.322 | 1.481 | 2.705 | 0.021 | 0.07 | -0.033 | -0.033 | 0 | 0.19 | 0.51 | 0.19 | 0.13 | 0.24 | 0.22 |
| 5 | 38390778 | rs10213982 | -2.02 | 1.485 | 2.774 | 0.025 | 0.066 | -0.017 | -0.017 | 0 | 0.18 | 0.36 | 0.18 | 0.13 | 0.24 | 0.22 |
| 5 | 38396524 | rs6867586 | -1.618 | 1.546 | 2.264 | 0.02 | 0.069 | -0.051 | -0.051 | 0 | 0.15 | 0.35 | 0.07 | 0.05 | 0.18 | 0.07 |
| 5 | 38399161 | rs3110226 | -2.854 | 1.472 | 2.283 | -0.075 | -0.014 | 0.072 | 0.072 | 3.281 | 0.31 | 0.84 | 0.26 | 0.13 | 0.29 | 0.25 |
| 5 | 38399208 | rs6879807 | -1.831 | 1.703 | 1.751 | -0.032 | 0.068 | -0.053 | -0.053 | 0 | 0.15 | 0.35 | 0.07 | 0.05 | 0.18 | 0.07 |
| 5 | 38400072 | rs17425735 | -1.614 | 1.722 | 1.671 | -0.023 | 0.012 | -0.012 | -0.012 | 0.011 | 0.13 | 0.15 | 0.05 | 0.05 | 0.18 | 0.07 |
| 5 | 38401969 | rs3110992 | -1.804 | 1.763 | 1.419 | -0.004 | 0.018 | -0.029 | -0.029 | 0 | 0.16 | 0.22 | 0.06 | 0.05 | 0.19 | 0.07 |
| 5 | 38412674 | rs16903965 | -1.087 | 1.381 | -0.499 | 0.509 | -0.003 | 0.002 | 0.002 | 0 | 0.1 | 0.11 | 0.05 | 0.24 | 0.14 | 0.07 |
| 5 | 38417839 | rs10512684 | -1.41 | 1.268 | -0.496 | 0.441 | -0.002 | -0.003 | -0.003 | 0.102 | 0.19 | 0.15 | 0.09 | 0.24 | 0.18 | 0.11 |
| 5 | 38422880 | rs4376296 | 0.798 | 1.201 | -1.116 | -0.321 | -0.067 | 0.158 | 0.158 | 3.12 | 0.56 | 0.16 | 0.67 | 0.84 | 0.54 | 0.72 |
| 5 | 38422991 | rs3935014 | -0.978 | 0.752 | 0.424 | -0.422 | -0.028 | -0.122 | -0.122 | 1.109 | 0.19 | 0.49 | 0.12 | 0.02 | 0.15 | 0.13 |
| 5 | 38428087 | rs2956593 | -1.422 | 0.628 | 1.413 | -0.323 | -0.017 | -0.033 | -0.033 | 0 | 0.16 | 0.72 | 0.21 | 0.07 | 0.11 | 0.08 |
| 5 | 38434476 | rs891013 | -1.61 | 0.531 | 2.077 | -0.366 | -0.023 | -0.051 | -0.051 | 0 | 0.17 | 0.71 | 0.21 | 0.06 | 0.11 | 0.08 |
| 5 | 38435130 | rs2956598 | -0.652 | 0.599 | 0.472 | -0.929 | -0.079 | 0.078 | 0.078 | 0.507 | 0.6 | 0.94 | 0.45 | 0.26 | 0.55 | 0.38 |
| 5 | 38440601 | rs2956597 | -1.548 | 0.901 | 1.306 | -0.709 | -0.002 | -0.025 | -0.025 | 0 | 0.17 | 0.76 | 0.21 | 0.04 | 0.14 | 0.09 |
| 5 | 38448700 | rs6892556 | -1.341 | 0.923 | 2.222 | -0.675 | 0.053 | -0.059 | -0.059 | 0 | 0.11 | 0.5 | 0.12 | 0.01 | 0.11 | 0.07 |
| 5 | 38450430 | rs2561805 | -1.896 | 0.863 | 1.993 | -0.066 | -0.003 | 0.005 | 0.005 | 10.256 | 0.15 | 0.73 | 0.25 | 0.11 | 0.14 | 0.08 |
| 5 | 38456766 | rs2731968 | -1.956 | 0.87 | 2.042 | -0.221 | 0.009 | -0.045 | -0.045 | 3.192 | 0.12 | 0.72 | 0.18 | 0.04 | 0.09 | 0.05 |
| 5 | 38468420 | rs10067809 | -1.424 | 1.338 | 2.086 | -0.582 | 0.082 | -0.075 | -0.075 | 0 | 0.15 | 0.47 | 0.42 | 0.03 | 0.18 | 0.17 |
| 5 | 38469842 | rs891012 | -0.446 | 0.768 | 0.283 | -0.563 | -0.025 | -0.087 | -0.087 | 0.085 | 0.72 | 0.96 | 0.86 | 0.7 | 0.76 | 0.68 |
| 5 | 38475507 | rs1046224 | 0.402 | 0.508 | 0.102 | -0.674 | -0.014 | -0.043 | -0.043 | 0.112 | 0.32 | 0.16 | 0.37 | 0.66 | 0.34 | 0.41 |
| 5 | 38481335 | rs1005017 | -1.37 | 0.167 | -0.376 | -0.844 | -0.049 | -0.031 | -0.031 | 0 | 0.58 | 0.93 | 0.77 | 0.69 | 0.58 | 0.63 |
| 5 | 38486065 | rs3110234 | 0.392 | 0.155 | -0.09 | -0.873 | -0.014 | 0.016 | 0.016 | 0.013 | 0.16 | 0.12 | 0.27 | 0.02 | 0.19 | 0.15 |
| 5 | 38496262 | rs2162885 | 0.731 | -0.29 | 0.018 | -0.006 | -0.016 | -0.026 | -0.026 | 0 | 0.31 | 0.14 | 0.35 | 0.4 | 0.34 | 0.39 |
| 5 | 38523740 | rs3729598 | 0.473 | -0.287 | 0.563 | 0.145 | -0.021 | 0.044 | 0.044 | 0 | 0.07 | 0.1 | 0.04 | 0.26 | 0.05 | 0.06 |

| | | | | | | | | | | | | | | | | |
|---|---|---|---|---|---|---|---|---|---|---|---|---|---|---|---|---|
| 5 | 38565639 | rs1025963 | 0.636 | -0.166 | 0.891 | -0.089 | -0.001 | -0.003 | -0.003 | 0 | 0.29 | 0.25 | 0.37 | 0.4 | 0.35 | 0.41 |
| 5 | 38579325 | rs1012760 | 0.324 | -0.179 | 1.012 | -0.033 | -0.003 | 0.001 | 0.001 | 1.288 | 0.56 | 0.59 | 0.35 | 0.55 | 0.48 | 0.44 |
| 5 | 38580741 | rs10040852 | 0.32 | -0.176 | 1.085 | -0.031 | -0.001 | 0 | 0 | 0 | 0.56 | 0.57 | 0.35 | 0.55 | 0.48 | 0.44 |
| 5 | 38590551 | rs1562136 | -0.27 | 0.041 | 1 | 0.108 | -0.008 | 0.006 | 0.006 | 0 | 0.86 | 0.88 | 0.75 | 0.98 | 0.82 | 0.86 |
| 5 | 38593688 | rs13157717 | 0.392 | -0.067 | 0.672 | 0.387 | 0.006 | -0.009 | -0.009 | 0 | 0.36 | 0.16 | 0.4 | 0.38 | 0.37 | 0.45 |
| 5 | 38597478 | rs3812043 | 0.808 | -0.143 | 0.605 | 0.434 | -0.04 | -0.016 | -0.016 | 0 | 0.45 | 0.13 | 0.23 | 0.36 | 0.44 | 0.36 |
| 5 | 38601073 | rs1562134 | -0.43 | 0.687 | 0.556 | -1.148 | 0.041 | -0.056 | -0.056 | 0 | 0.15 | 0.35 | 0.29 | 0.04 | 0.18 | 0.17 |
| 5 | 38608138 | rs3099115 | -0.135 | 0.241 | -0.244 | -0.979 | -0.089 | 0.065 | 0.065 | 0.292 | 0.44 | 0.13 | 0.58 | 0.7 | 0.51 | 0.55 |
| 5 | 38615656 | rs1444884 | -1.765 | 0.366 | 1.413 | -0.938 | -0.008 | -0.016 | -0.016 | 0 | 0.1 | 0.71 | 0.12 | 0.07 | 0.06 | 0.12 |
| 5 | 38649445 | rs13167213 | -0.024 | 0.257 | 1.253 | -0.912 | -0.012 | 0.104 | 0.104 | 0 | 0.56 | 0.79 | 0.37 | 0.15 | 0.53 | 0.31 |
| 5 | 38649818 | rs327295 | -0.078 | 0.279 | 1.085 | -0.901 | -0.027 | 0.122 | 0.122 | 0 | 0.57 | 0.83 | 0.37 | 0.18 | 0.53 | 0.35 |
| 5 | 38690423 | rs10075011 | -0.106 | -0.027 | 0.588 | -1.058 | 0.002 | -0.025 | -0.025 | 0 | 0.57 | 0.46 | 0.46 | 0.09 | 0.64 | 0.49 |
| 5 | 38690509 | rs2471156 | -1.103 | -0.503 | 1.611 | 0.029 | 0.026 | 0.136 | 0.136 | 3.189 | 0.08 | 0.45 | 0.05 | 0.15 | 0.03 | 0.09 |
| 5 | 38698659 | rs1428498 | -0.976 | -0.412 | 0.329 | -0.268 | 0.087 | 0.085 | 0.085 | 1.304 | 0.36 | 0.88 | 0.27 | 0.26 | 0.29 | 0.33 |
| 5 | 38708516 | rs10472308 | 0.452 | 0.276 | 1.516 | -0.282 | -0.05 | -0.085 | -0.085 | 0 | 0.7 | 0.57 | 0.84 | 0.91 | 0.8 | 0.66 |
| 5 | 38708578 | rs9292716 | 0.187 | 0.25 | 1.648 | -0.306 | -0.018 | -0.033 | -0.033 | 0.688 | 0.74 | 0.74 | 0.89 | 0.93 | 0.82 | 0.78 |
| 5 | 38709629 | rs12517759 | 1.113 | 0.238 | 1.526 | -0.427 | 0.038 | 0.098 | 0.098 | 0 | 0.63 | 0.16 | 0.76 | 0.81 | 0.71 | 0.59 |
| 5 | 38712721 | rs10473116 | -0.259 | 0.326 | 1.284 | -0.356 | 0.017 | 0.02 | 0.02 | 0 | 0.26 | 0.15 | 0.1 | 0.07 | 0.18 | 0.21 |
| 5 | 38714694 | rs6862694 | -0.591 | 0.359 | 0.929 | -0.33 | -0.038 | -0.058 | -0.058 | 0 | 0.3 | 0.42 | 0.16 | 0.09 | 0.2 | 0.33 |
| 5 | 38716873 | rs4242256 | 1.517 | 1.451 | 0.581 | 0.412 | 0.075 | 0.052 | 0.052 | 0 | 0.92 | 0.72 | 0.87 | 0.84 | 0.87 | 0.81 |
| 5 | 38722734 | rs4283817 | 1.376 | 2.033 | -0.078 | 0.001 | -0.005 | 0.004 | 0.004 | 0.566 | 0.8 | 0.2 | 0.81 | 0.89 | 0.77 | 0.76 |
| 5 | 38723307 | rs10941402 | 0.652 | 1.778 | -1.571 | -0.179 | -0.012 | 0.122 | 0.122 | 0 | 0.51 | 0.1 | 0.56 | 0.71 | 0.56 | 0.4 |
| 5 | 38723422 | rs6883199 | -0.828 | 1.223 | -0.633 | -0.025 | -0.024 | 0 | 0 | 0 | 0.19 | 0.31 | 0.13 | 0.21 | 0.19 | 0.38 |
| 5 | 38723523 | rs4489086 | 1.029 | 1.62 | -1.802 | -0.539 | -0.095 | 0.075 | 0.075 | 0 | 0.28 | 0.04 | 0.27 | 0.12 | 0.3 | 0.18 |
| 5 | 38724883 | rs4869457 | -0.287 | 1.333 | -1.236 | -0.456 | 0.012 | 0.017 | 0.017 | 0.441 | 0.33 | 0.22 | 0.5 | 0.25 | 0.28 | 0.52 |
| 5 | 38725043 | rs10055279 | 0.079 | 1.043 | -1.022 | 0.104 | -0.001 | 0.022 | 0.022 | 0 | 0.14 | 0.14 | 0.12 | 0.33 | 0.08 | 0.12 |
| 5 | 38725149 | rs7729320 | -1.324 | 1.451 | -0.785 | -0.156 | -0.026 | 0.033 | 0.033 | 0 | 0.58 | 0.73 | 0.7 | 0.66 | 0.48 | 0.71 |
| 5 | 38728218 | rs4337869 | -0.197 | 1.456 | -0.051 | -0.117 | 0.037 | 0.005 | 0.005 | 0 | 0.1 | 0.29 | 0.08 | 0.09 | 0.11 | 0.08 |
| 5 | 38728261 | rs4337870 | -0.93 | 1.411 | -0.015 | -0.08 | -0.001 | 0.07 | 0.07 | 0 | 0.24 | 0.38 | 0.19 | 0.41 | 0.19 | 0.2 |
| 5 | 38728597 | rs16867665 | -1.318 | 1.282 | 0.222 | -0.121 | -0.013 | 0.059 | 0.059 | 0 | 0.31 | 0.53 | 0.25 | 0.51 | 0.25 | 0.34 |
| 5 | 38737718 | rs12520160 | 0.781 | 1.138 | 0.081 | -0.174 | 0.011 | 0.031 | 0.031 | 0.378 | 0.24 | 0.11 | 0.48 | 0.17 | 0.23 | 0.37 |
| 5 | 38737883 | rs6893601 | 1.19 | 1.104 | 0.158 | -0.18 | -0.01 | 0.005 | 0.005 | 0 | 0.07 | 0.15 | 0.06 | 0.09 | 0.05 | 0.12 |
| 5 | 38743568 | rs4869598 | -1.266 | 1.379 | 0.297 | -0.273 | 0.004 | 0.012 | 0.012 | 0 | 0.55 | 0.55 | 0.7 | 0.79 | 0.57 | 0.66 |
| 5 | 38750042 | rs4128777 | 0.667 | 0.912 | 0.304 | -0.169 | 0.006 | 0.013 | 0.013 | 0.145 | 0.25 | 0.17 | 0.45 | 0.16 | 0.23 | 0.37 |
| 5 | 38753979 | rs7723735 | 0.609 | 0.975 | 0.257 | -0.187 | 0 | -0.003 | -0.003 | 0 | 0.53 | 0.47 | 0.37 | 0.21 | 0.62 | 0.36 |
| 5 | 38756243 | rs17457012 | 0.475 | 0.767 | -0.331 | -0.396 | -0.004 | 0.109 | 0.109 | 1.619 | 0.29 | 0.08 | 0.5 | 0.13 | 0.34 | 0.33 |
| 5 | 38756294 | rs10941406 | 1.407 | 0.855 | 0.162 | -0.427 | -0.001 | 0.001 | 0.001 | 0 | 0.4 | 0.41 | 0.24 | 0.22 | 0.36 | 0.34 |
| 5 | 38757335 | rs7726283 | -0.251 | 0.644 | 0.003 | -0.368 | 0.02 | 0.08 | 0.08 | 0.353 | 0.67 | 0.81 | 0.49 | 0.86 | 0.65 | 0.64 |
| 5 | 38761236 | rs4256371 | 2.359 | -0.687 | -0.152 | -0.638 | -0.019 | 0.042 | 0.042 | 0 | 0.31 | 0.2 | 0.18 | 0.12 | 0.32 | 0.21 |
| 5 | 38763356 | rs4400150 | 1.484 | -0.501 | -0.224 | -0.315 | 0.009 | -0.049 | -0.049 | 0 | 0.1 | 0.02 | 0.09 | 0.33 | 0.05 | 0.13 |
| 5 | 38765637 | rs10070428 | 0.295 | -0.279 | 0.087 | -0.246 | -0.005 | -0.006 | -0.006 | 0 | 0.52 | 0.74 | 0.34 | 0.54 | 0.48 | 0.48 |
| 5 | 38770162 | rs11746202 | 0.837 | -0.33 | -0.114 | -0.284 | -0.048 | 0.101 | 0.101 | 0 | 0.4 | 0.22 | 0.23 | 0.12 | 0.44 | 0.25 |
| 5 | 38777383 | rs10941407 | -0.179 | -0.395 | 0.168 | -0.289 | 0.028 | 0.025 | 0.025 | 0 | 0.51 | 0.31 | 0.69 | 0.58 | 0.48 | 0.61 |
| 5 | 38778514 | rs9687972 | 0 | -0.285 | 0.321 | -0.257 | -0.014 | 0.016 | 0.016 | 0 | 0.83 | 0.62 | 0.86 | 0.81 | 0.86 | 0.75 |
| 5 | 38780436 | rs9292720 | 0.971 | 0.108 | 0.311 | -0.335 | -0.005 | -0.004 | -0.004 | 0.134 | 0.08 | 0.06 | 0.04 | 0 | 0.12 | 0.06 |
| 5 | 38780778 | rs10805635 | -1.043 | 0.14 | 0.439 | -0.036 | 0.033 | 0.057 | 0.057 | 0 | 0.93 | 0.75 | 0.94 | 0.91 | 0.93 | 0.88 |
| 5 | 38788671 | rs11950054 | -0.351 | 0.104 | 0.358 | -0.14 | -0.002 | -0.027 | -0.027 | 0 | 0.69 | 0.65 | 0.49 | 0.78 | 0.68 | 0.56 |
| 5 | 38792486 | rs7706715 | 0.139 | 0.155 | -0.153 | -0.161 | -0.001 | -0.007 | -0.007 | 0 | 0.31 | 0.33 | 0.51 | 0.21 | 0.33 | 0.43 |
| 5 | 38793697 | rs11744747 | 0.288 | 0.331 | -1.153 | -0.285 | 0.007 | -0.066 | -0.066 | 1.23 | 0.3 | 0.17 | 0.27 | 0.45 | 0.32 | 0.32 |
| 5 | 38794770 | rs719079 | 1.256 | 0.717 | -1.359 | -0.98 | -0.029 | 0.043 | 0.043 | 0 | 0.3 | 0.23 | 0.21 | 0.11 | 0.34 | 0.19 |
| 5 | 38799793 | rs1864511 | -0.018 | 0.628 | -1.088 | -0.408 | -0.009 | 0.052 | 0.052 | 0.104 | 0.23 | 0.12 | 0.45 | 0.1 | 0.26 | 0.29 |
| 5 | 38802148 | rs6888067 | -0.712 | 0.478 | -0.968 | -0.173 | 0.003 | 0.015 | 0.015 | 0.001 | 0.16 | 0.11 | 0.23 | 0.1 | 0.14 | 0.27 |
| 5 | 38803449 | rs1430421 | 0.787 | 0.541 | -0.552 | -0.362 | -0.001 | -0.026 | -0.026 | 0 | 0.87 | 0.83 | 0.84 | 0.94 | 0.84 | 0.77 |
| 5 | 38804071 | rs7729903 | 0.705 | 0.43 | -1.468 | -0.394 | -0.011 | -0.006 | -0.006 | 0 | 0.3 | 0.09 | 0.21 | 0.25 | 0.21 | 0.28 |
| 5 | 38804215 | rs11749831 | 0.997 | 0.843 | -1.413 | -0.33 | -0.024 | 0.024 | 0.024 | 1.014 | 0.26 | 0.19 | 0.23 | 0.17 | 0.32 | 0.25 |
| 5 | 38807040 | rs2082163 | -0.134 | 0.672 | -0.871 | -0.171 | -0.004 | -0.031 | -0.031 | 0.085 | 0.87 | 0.76 | 0.85 | 0.94 | 0.84 | 0.78 |
| 5 | 38807123 | rs1560422 | 0.12 | 0.688 | -0.342 | -0.016 | 0.002 | 0.014 | 0.014 | 0.426 | 0.49 | 0.41 | 0.49 | 0.22 | 0.57 | 0.41 |
| 5 | 38811521 | rs17457979 | 0.926 | 0.642 | -0.878 | -0.067 | 0.009 | -0.01 | -0.01 | 0.236 | 0.32 | 0.04 | 0.21 | 0.25 | 0.22 | 0.4 |
| 5 | 38818281 | rs386994 | -0.212 | 0.905 | -0.513 | -0.056 | 0.006 | -0.025 | -0.025 | 0 | 0.85 | 0.8 | 0.82 | 0.94 | 0.79 | 0.85 |
| 5 | 38827572 | rs476236 | -0.195 | 0.844 | -0.531 | -0.045 | 0.004 | -0.021 | -0.021 | 0.178 | 0.84 | 0.8 | 0.81 | 0.94 | 0.79 | 0.82 |
| 5 | 38833866 | rs393398 | -0.832 | 0.938 | -0.653 | -0.012 | -0.014 | 0.017 | 0.017 | 0 | 0.73 | 0.76 | 0.79 | 0.81 | 0.71 | 0.78 |
| 5 | 38836704 | rs1501742 | 1.413 | 0.52 | -0.856 | 0.451 | 0.043 | 0.001 | 0.001 | 0 | 0.3 | 0.03 | 0.19 | 0.22 | 0.18 | 0.41 |
| 5 | 38837861 | rs357286 | -0.04 | 0.707 | -0.075 | 0.45 | 0.002 | 0.001 | 0.001 | 0 | 0.52 | 0.54 | 0.52 | 0.77 | 0.44 | 0.63 |
| 5 | 38839713 | rs357284 | -0.776 | 0.765 | -0.137 | 0.374 | -0.028 | 0.031 | 0.031 | 0.115 | 0.72 | 0.78 | 0.78 | 0.85 | 0.68 | 0.78 |
| 5 | 38842959 | rs395967 | -0.063 | 0.677 | -0.204 | 0.922 | 0 | 0 | 0 | 0 | 0.51 | 0.49 | 0.52 | 0.51 | 0.44 | 0.63 |
| 5 | 38848503 | rs2043111 | 0.752 | 0.608 | -0.378 | 0.529 | 0.06 | 0.076 | 0.076 | 0 | 0.38 | 0.19 | 0.26 | 0.17 | 0.26 | 0.43 |
| 5 | 38850990 | rs357287 | -0.796 | 0.893 | 0.158 | 0.494 | 0.001 | -0.001 | -0.001 | 0 | 0.74 | 0.72 | 0.78 | 0.73 | 0.72 | 0.74 |
| 5 | 38852259 | rs9292722 | 0.687 | 0.678 | -0.017 | 0.416 | 0.061 | 0.1 | 0.1 | 0 | 0.38 | 0.19 | 0.27 | 0.15 | 0.27 | 0.46 |
| 5 | 38857366 | rs420444 | 0.171 | 0.866 | 0.293 | 0.469 | 0.025 | -0.03 | -0.03 | 0.15 | 0.14 | 0.23 | 0.19 | 0.06 | 0.2 | 0.14 |

| | | | | | | | | | | | | | | | | |
|---|---|---|---|---|---|---|---|---|---|---|---|---|---|---|---|---|
| 5 | 38867732 | rs395157 | -0.115 | 1.078 | 0.336 | 0.379 | -0.002 | 0.025 | 0.025 | 0 | 0.53 | 0.42 | 0.47 | 0.24 | 0.52 | 0.61 |
| 5 | 38868472 | rs357281 | -0.12 | 1.3 | 0.851 | 0.803 | 0.026 | -0.072 | -0.072 | 0.164 | 0.14 | 0.22 | 0.12 | 0 | 0.21 | 0.1 |
| 5 | 38872383 | rs2367704 | 1.151 | 1.038 | 0.917 | 1.327 | -0.007 | -0.016 | -0.016 | 0.03 | 0.17 | 0.07 | 0.28 | 0.15 | 0.21 | 0.12 |
| 5 | 38877333 | rs357270 | -0.14 | 1.135 | 1.122 | 1.15 | 0.058 | -0.057 | -0.057 | 1.021 | 0.1 | 0.23 | 0.09 | 0 | 0.18 | 0.1 |
| 5 | 38877564 | rs357269 | -0.14 | 1.135 | 1.118 | 1.15 | 0.051 | -0.057 | -0.057 | 1.272 | 0.1 | 0.22 | 0.09 | 0 | 0.18 | 0.1 |
| 5 | 38879218 | rs3792856 | 0.271 | 1.033 | 1.25 | 1.303 | -0.01 | 0.06 | 0.06 | 0 | 0.23 | 0.5 | 0.33 | 0.62 | 0.23 | 0.22 |
| 5 | 38880689 | rs555235 | -0.62 | 1.048 | 1.098 | 1.222 | 0.003 | -0.004 | -0.004 | 0.181 | 0.33 | 0.6 | 0.42 | 0.24 | 0.43 | 0.36 |
| 5 | 38881460 | rs617938 | -0.35 | 1.254 | 0.529 | 1.374 | -0.001 | 0.012 | 0.012 | 0 | 0.95 | 0.9 | 0.97 | 0.89 | 0.95 | 0.96 |
| 5 | 38881628 | rs525735 | -0.789 | 1.308 | 0.844 | 1.364 | 0.077 | 0.248 | 0.248 | 0.021 | 0.15 | 0.39 | 0.13 | 0.38 | 0.22 | 0.2 |
| 5 | 38882247 | rs499225 | -0.326 | 1.079 | 0.248 | 1.104 | 0 | -0.014 | -0.014 | 0.287 | 0.76 | 0.78 | 0.69 | 0.65 | 0.73 | 0.77 |
| 5 | 38886040 | rs663845 | -0.341 | 0.972 | 0.557 | 0.313 | 0.033 | -0.103 | -0.103 | 0 | 0.27 | 0.4 | 0.17 | 0.02 | 0.33 | 0.24 |
| 5 | 38887262 | rs1430420 | -0.22 | 0.009 | 0.522 | 0.411 | -0.047 | 0.003 | 0.003 | 1.339 | 0.14 | 0.32 | 0.07 | 0.08 | 0.09 | 0.18 |
| 5 | 38898447 | rs1897140 | -0.608 | 0.397 | 0.571 | 0.97 | -0.011 | 0.034 | 0.034 | 0.174 | 0.33 | 0.44 | 0.17 | 0.49 | 0.23 | 0.42 |
| 5 | 38905288 | rs10040172 | 0.282 | 0.332 | 0.795 | 0.93 | 0 | -0.008 | -0.008 | 0 | 0.23 | 0.19 | 0.11 | 0.03 | 0.17 | 0.16 |
| 5 | 38905985 | rs1239339 | -1.175 | 0.372 | 0.173 | 0.728 | -0.018 | -0.006 | -0.006 | 0 | 0.63 | 0.86 | 0.54 | 0.62 | 0.63 | 0.66 |
| 5 | 38906339 | rs3805569 | 1.765 | 0.414 | 0.25 | 0.727 | -0.003 | 0.012 | 0.012 | 0.686 | 0.2 | 0.06 | 0.27 | 0.16 | 0.21 | 0.13 |
| 5 | 38912195 | rs9632405 | -0.668 | 0.571 | 0.806 | 0.952 | 0.02 | -0.05 | -0.05 | 0 | 0.17 | 0.34 | 0.11 | 0.06 | 0.23 | 0.24 |
| 5 | 38917574 | rs357260 | -0.86 | 0.63 | 0.49 | 1.004 | -0.006 | 0.003 | 0.003 | 0 | 0.63 | 0.73 | 0.52 | 0.63 | 0.61 | 0.65 |
| 5 | 38919158 | rs10941412 | -0.246 | 0.818 | 0.584 | 1.038 | 0.026 | -0.073 | -0.073 | 0 | 0.13 | 0.27 | 0.09 | 0 | 0.18 | 0.13 |
| 5 | 38923779 | rs357291 | -0.175 | 0.661 | -0.311 | 1.111 | -0.002 | -0.001 | -0.001 | 0 | 0.56 | 0.7 | 0.49 | 0.55 | 0.55 | 0.46 |
| 5 | 38933749 | rs1239344 | 0.039 | 0.69 | -0.247 | 0.825 | -0.001 | -0.001 | -0.001 | 0 | 0.38 | 0.4 | 0.39 | 0.57 | 0.44 | 0.43 |
| 5 | 38938664 | rs10941413 | 0.792 | 0.542 | -0.29 | 1.166 | 0.018 | -0.008 | -0.008 | 0.198 | 0.06 | 0.02 | 0.02 | 0.25 | 0.04 | 0.11 |
| 5 | 38955796 | rs2043112 | 0.209 | 0.514 | -0.114 | 1.295 | -0.006 | -0.028 | -0.028 | 0 | 0.34 | 0.26 | 0.36 | 0.45 | 0.43 | 0.34 |
| 5 | 38974929 | rs16867919 | -0.067 | 0.675 | -0.23 | 1.128 | -0.024 | -0.043 | -0.043 | 0 | 0.14 | 0.07 | 0.09 | 0.25 | 0.22 | 0.21 |
| 5 | 38985269 | rs1239259 | 0.595 | 0.6 | 0.66 | 1.195 | -0.002 | -0.004 | -0.004 | 0 | 0.38 | 0.32 | 0.37 | 0.31 | 0.35 | 0.25 |
| 5 | 39054434 | rs6883671 | 0.595 | 0.356 | 1.017 | 1.082 | -0.001 | 0.021 | 0.021 | 0 | 0.21 | 0.16 | 0.11 | 0.03 | 0.17 | 0.21 |
| 5 | 39062977 | rs9292730 | 0.46 | 0.343 | 0.997 | 1.077 | 0 | -0.022 | -0.022 | 0.595 | 0.22 | 0.24 | 0.12 | 0.03 | 0.17 | 0.22 |
| 5 | 39081475 | rs10064513 | 0.197 | 0.573 | 1.314 | 1.524 | 0 | 0 | 0 | 0 | 0.39 | 0.39 | 0.39 | 0.41 | 0.46 | 0.44 |
| 5 | 39081910 | rs9292732 | 0.885 | 0.446 | 1.344 | 1.558 | -0.001 | 0.002 | 0.002 | 0 | 0.16 | 0.16 | 0.08 | 0.08 | 0.12 | 0.21 |
| 5 | 39084471 | rs6895953 | 0.523 | 0.426 | 1.301 | 1.498 | -0.005 | -0.006 | -0.006 | 0 | 0.35 | 0.33 | 0.36 | 0.4 | 0.44 | 0.35 |
| 5 | 39087074 | rs2367894 | 0.364 | 0.734 | 1.192 | 1.678 | 0.008 | 0.043 | 0.043 | 0 | 0.73 | 0.51 | 0.73 | 0.72 | 0.78 | 0.74 |
| 5 | 39098803 | rs418142 | -0.043 | 0.206 | 1.262 | 1.691 | 0.015 | 0.041 | 0.041 | 0.021 | 0.9 | 0.8 | 0.83 | 0.81 | 0.9 | 0.95 |
| 5 | 39104603 | rs358504 | 0.695 | 0.208 | 1.439 | 1.245 | 0.019 | 0.079 | 0.079 | 0.391 | 0.8 | 0.63 | 0.77 | 0.62 | 0.78 | 0.82 |
| 5 | 39104918 | rs17463165 | 1.304 | 0.192 | 1.151 | 1.124 | -0.026 | 0.042 | 0.042 | 1.031 | 0.28 | 0.19 | 0.35 | 0.14 | 0.37 | 0.22 |
| 5 | 39105594 | rs358501 | 0.353 | 0.608 | 1.457 | 1.517 | 0 | 0.01 | 0.01 | 0.334 | 0.84 | 0.8 | 0.8 | 0.63 | 0.8 | 0.83 |
| 5 | 39113637 | rs10061822 | 0.981 | 0.578 | 1.074 | 1.066 | -0.001 | -0.009 | -0.009 | 0.131 | 0.42 | 0.42 | 0.41 | 0.2 | 0.45 | 0.4 |
| 5 | 39113938 | rs444556 | -0.135 | 0.248 | 0.07 | 1.045 | -0.003 | 0.06 | 0.06 | 0 | 0.37 | 0.1 | 0.35 | 0.26 | 0.33 | 0.4 |
| 5 | 39121508 | rs364640 | 1.114 | -0.051 | 2.231 | 1.638 | -0.017 | 0.002 | 0.002 | 0 | 0.63 | 0.57 | 0.75 | 0.61 | 0.69 | 0.72 |
| 5 | 39132459 | rs6887349 | -1.225 | -0.121 | 2.847 | 1.651 | -0.007 | 0.253 | 0.253 | 1.115 | 0.11 | 0.37 | 0.12 | 0.39 | 0.06 | 0.15 |
| 5 | 39145218 | rs6896856 | -0.691 | -0.194 | 2.932 | 1.889 | -0.023 | -0.016 | -0.016 | 0.176 | 0.27 | 0.39 | 0.19 | 0.2 | 0.24 | 0.14 |
| 5 | 39153251 | rs6861690 | -0.936 | -0.101 | 3.107 | 1.851 | -0.01 | 0.163 | 0.163 | 0.442 | 0.09 | 0.19 | 0.09 | 0.51 | 0.05 | 0.17 |
| 5 | 39156172 | rs4264965 | -0.382 | -0.138 | 3.068 | 1.75 | 0.002 | -0.042 | -0.042 | 0 | 0.25 | 0.25 | 0.14 | 0 | 0.23 | 0.12 |
| 5 | 39157794 | rs1549690 | 1.366 | -0.129 | 3.007 | 1.774 | -0.019 | 0.283 | 0.283 | 2.577 | 0.87 | 0.52 | 0.89 | 0.4 | 0.94 | 0.84 |
| 5 | 39159462 | rs7710762 | 1.409 | -0.112 | 2.982 | 1.782 | -0.007 | -0.001 | -0.001 | 0 | 0.63 | 0.41 | 0.76 | 0.4 | 0.71 | 0.71 |
| 5 | 39172131 | rs11956872 | -0.388 | 0.093 | 1.8 | 1.551 | 0.001 | 0.095 | 0.095 | 0 | 0.24 | 0.08 | 0.13 | 0 | 0.23 | 0.13 |
| 5 | 39185613 | rs834251 | -0.808 | 0.154 | 1.655 | 1.243 | -0.009 | -0.047 | -0.047 | 0.03 | 0.27 | 0.31 | 0.14 | 0.1 | 0.24 | 0.14 |
| 5 | 39185998 | rs834249 | -0.45 | 0.255 | 0.807 | 0.937 | 0.007 | 0.059 | 0.059 | 0 | 0.24 | 0.1 | 0.13 | 0 | 0.23 | 0.13 |
| 5 | 39191294 | rs66782 | 0.848 | 0.465 | 0.76 | 1.295 | -0.011 | 0.014 | 0.014 | 0 | 0.66 | 0.46 | 0.74 | 0.4 | 0.71 | 0.68 |
| 5 | 39197725 | rs700184 | -0.561 | 0.962 | 0.851 | 0.843 | 0.006 | -0.05 | -0.05 | 0 | 0.24 | 0.28 | 0.25 | 0.04 | 0.32 | 0.1 |
| 5 | 39200689 | rs17465965 | -0.072 | 1.06 | 0.131 | 0.557 | -0.037 | 0.024 | 0.024 | 0 | 0.14 | 0.03 | 0.13 | 0.07 | 0.14 | 0.13 |
| 5 | 39201180 | rs260077 | 0.977 | 0.659 | 1.054 | 1.265 | 0.021 | 0.24 | 0.24 | 0.035 | 0.94 | 0.67 | 0.91 | 0.81 | 0.98 | 0.95 |
| 5 | 39205914 | rs13187857 | 1.138 | 0.067 | 1.494 | 1.179 | -0.026 | 0.062 | 0.062 | 0.058 | 0.82 | 0.31 | 0.76 | 0.58 | 0.9 | 0.85 |
| 5 | 39206778 | rs700185 | 0.978 | 0.08 | 1.286 | 1.186 | -0.042 | 0.181 | 0.181 | 0 | 0.83 | 0.45 | 0.76 | 0.58 | 0.9 | 0.85 |
| 5 | 39207939 | rs696743 | 0.964 | 0.27 | 1.331 | 1.126 | 0 | 0 | 0 | 0 | 0.59 | 0.37 | 0.58 | 0.58 | 0.58 | 0.73 |
| 5 | 39211189 | rs696746 | -1.442 | -0.183 | 2.189 | 2.144 | -0.063 | 0.265 | 0.265 | 0.768 | 0.15 | 0.52 | 0.24 | 0.47 | 0.07 | 0.21 |
| 5 | 39213696 | rs696748 | 0.899 | 0.213 | 2.169 | 2.169 | 0.034 | -0.008 | -0.008 | 0 | 0.9 | 0.69 | 0.91 | 0.95 | 0.89 | 0.91 |
| 5 | 39216880 | rs700190 | 1.366 | 0.192 | 2.269 | 2.431 | 0.003 | 0.03 | 0.03 | 0.014 | 0.88 | 0.8 | 0.91 | 0.8 | 0.87 | 0.85 |
| 5 | 39219615 | rs6863066 | 2.157 | 0.099 | 2.683 | 2.477 | -0.01 | 0.051 | 0.051 | 0.363 | 0.78 | 0.31 | 0.7 | 0.58 | 0.84 | 0.71 |
| 5 | 39220055 | rs13188259 | 2.401 | 0.047 | 2.417 | 2.452 | 0.022 | -0.065 | -0.065 | 0 | 0.77 | 0.22 | 0.68 | 0.58 | 0.83 | 0.7 |
| 5 | 39222427 | rs2542714 | 1.411 | 0.129 | 2.513 | 2.404 | 0.01 | 0.021 | 0.021 | 0.133 | 0.87 | 0.74 | 0.89 | 0.87 | 0.88 | 0.85 |
| 5 | 39238087 | rs1645732 | -2.084 | 0.016 | 2.632 | 2.461 | 0.006 | 0.389 | 0.389 | 0.692 | 0.07 | 0.52 | 0.16 | 0.3 | 0.01 | 0.21 |
| 5 | 39253022 | rs7700754 | -2.377 | 0.598 | 2.195 | 1.955 | -0.001 | 0.003 | 0.003 | 0.106 | 0.24 | 0.61 | 0.33 | 0.22 | 0.23 | 0.28 |
| 5 | 39263661 | rs12655171 | -0.245 | 0.775 | 2.098 | 1.92 | 0.007 | -0.002 | -0.002 | 0.65 | 0.07 | 0.06 | 0.06 | 0.04 | 0.11 | 0.08 |
| 5 | 39264004 | rs4957426 | -1.243 | 0.686 | 2.581 | 1.959 | -0.022 | 0.037 | 0.037 | 0 | 0.1 | 0.21 | 0.16 | 0.11 | 0.07 | 0.08 |
| 5 | 39268371 | rs169048 | 0.258 | 0.245 | 1.445 | 1.336 | -0.013 | 0.01 | 0.01 | 0.168 | 0.33 | 0.26 | 0.32 | 0.25 | 0.43 | 0.39 |
| 5 | 39271431 | rs35250 | 1.182 | 0.258 | 1.25 | 1.335 | -0.051 | 0.023 | 0.023 | 0.628 | 0.24 | 0.09 | 0.25 | 0.21 | 0.33 | 0.31 |
| 5 | 39283650 | rs35251 | 1.182 | 0.258 | 1.25 | 1.335 | -0.051 | 0.023 | 0.023 | 0.628 | 0.24 | 0.09 | 0.25 | 0.21 | 0.33 | 0.31 |
| 5 | 39292091 | rs462477 | 0.415 | 0.232 | 1.302 | 1.338 | -0.066 | 0.032 | 0.032 | 0.109 | 0.31 | 0.15 | 0.31 | 0.25 | 0.43 | 0.39 |
| 5 | 39310443 | rs187452 | 0.742 | -0.088 | 1.429 | 1.08 | -0.006 | 0.047 | 0.047 | 0.794 | 0.52 | 0.26 | 0.49 | 0.64 | 0.46 | 0.5 |

| | | | | | | | | | | | | | | | | |
|---|---|---|---|---|---|---|---|---|---|---|---|---|---|---|---|---|
| 5 | 39312722 | rs155375 | 0.574 | -0.463 | 1.05 | 0.704 | -0.012 | 0.012 | 0.012 | 0.949 | 0.52 | 0.2 | 0.44 | 0.57 | 0.46 | 0.46 |
| 5 | 39315177 | rs263275 | 0.364 | -0.431 | 0.852 | 0.594 | -0.069 | 0.036 | 0.036 | 1.394 | 0.35 | 0.16 | 0.32 | 0.27 | 0.47 | 0.42 |
| 5 | 39315745 | rs187875 | 1.316 | -0.394 | 0.741 | 0.589 | -0.065 | 0.051 | 0.051 | 2.013 | 0.27 | 0.1 | 0.26 | 0.2 | 0.37 | 0.32 |
| 5 | 39320143 | rs155378 | -0.538 | -0.391 | 0.956 | 0.638 | -0.006 | 0.037 | 0.037 | 0.335 | 0.47 | 0.72 | 0.5 | 0.36 | 0.54 | 0.5 |
| 5 | 39325342 | rs261753 | -1.189 | -0.365 | 0.245 | 0.74 | 0.005 | -0.103 | -0.103 | 0.378 | 0.7 | 0.96 | 0.74 | 0.63 | 0.82 | 0.66 |
| 5 | 39328506 | rs261752 | -0.423 | -0.214 | 0.762 | 1.036 | -0.005 | -0.019 | -0.019 | 0 | 0.38 | 0.72 | 0.45 | 0.47 | 0.43 | 0.41 |
| 5 | 39331216 | rs696758 | -0.756 | -0.211 | 0.69 | 1.11 | -0.022 | 0.033 | 0.033 | 0.729 | 0.1 | 0.29 | 0.16 | 0.1 | 0.07 | 0.07 |
| 5 | 39338358 | rs3822462 | -0.433 | -0.067 | 0.592 | 1.141 | -0.016 | 0.035 | 0.035 | 0.32 | 0.09 | 0.19 | 0.11 | 0.09 | 0.05 | 0.07 |
| 5 | 39340135 | rs13179150 | 0.832 | 0.092 | 0.016 | 1.117 | -0.077 | -0.067 | -0.067 | 0.075 | 0.25 | 0.04 | 0.25 | 0.38 | 0.36 | 0.34 |
| 5 | 39342493 | rs696764 | 1.178 | 0.069 | 0.768 | 1.179 | -0.018 | 0.104 | 0.104 | 0.105 | 0.82 | 0.41 | 0.83 | 0.65 | 0.85 | 0.84 |
| 5 | 39343163 | rs4957473 | -0.076 | 0.432 | -0.032 | 0.862 | -0.022 | 0.018 | 0.018 | 0 | 0.32 | 0.08 | 0.32 | 0.26 | 0.35 | 0.36 |
| 5 | 39356591 | rs700220 | 0.679 | 1.008 | -0.037 | 0.427 | 0.012 | 0.145 | 0.145 | 2.818 | 0.44 | 0.23 | 0.37 | 0.07 | 0.41 | 0.36 |
| 5 | 39359323 | rs835218 | 0.584 | 0.923 | 0.752 | 0.987 | 0.004 | 0.012 | 0.012 | 1.129 | 0.47 | 0.42 | 0.4 | 0.26 | 0.43 | 0.43 |
| 5 | 39364339 | rs700231 | 0.689 | 0.97 | 0.825 | 0.954 | 0.011 | 0.125 | 0.125 | 1.697 | 0.46 | 0.27 | 0.38 | 0.14 | 0.44 | 0.37 |
| 5 | 39364554 | rs700233 | 0.768 | 1.022 | 0.799 | 0.904 | 0.02 | 0.16 | 0.16 | 2.017 | 0.45 | 0.24 | 0.37 | 0.07 | 0.42 | 0.37 |
| 5 | 39370907 | rs3812042 | -0.227 | 0.98 | 1.238 | 1.068 | -0.033 | 0.035 | 0.035 | 0.101 | 0.43 | 0.2 | 0.43 | 0.3 | 0.47 | 0.41 |
| 5 | 39372150 | rs1046033 | -0.133 | 0.952 | 1.245 | 1.057 | -0.049 | 0.03 | 0.03 | 0.073 | 0.42 | 0.14 | 0.42 | 0.3 | 0.47 | 0.41 |
| 5 | 39373253 | rs4578 | -0.274 | 0.956 | 1.236 | 1.059 | -0.031 | 0.036 | 0.036 | 0.076 | 0.43 | 0.2 | 0.43 | 0.3 | 0.47 | 0.41 |
| 5 | 39375082 | rs2686162 | -0.287 | 0.941 | 1.23 | 1.053 | -0.031 | 0.037 | 0.037 | 0.07 | 0.43 | 0.2 | 0.43 | 0.3 | 0.47 | 0.41 |
| 5 | 39395397 | rs2855510 | 0.217 | 1.123 | 1.251 | 0.952 | 0.013 | 0.01 | 0.01 | 0 | 0.28 | 0.16 | 0.31 | 0.2 | 0.22 | 0.25 |
| 5 | 39417846 | rs12697392 | 0.169 | 1.002 | 1.239 | 0.953 | 0.007 | 0.004 | 0.004 | 0.12 | 0.21 | 0.29 | 0.14 | 0.22 | 0.25 | 0.23 |
| 5 | 39418421 | rs6872083 | -0.153 | 0.993 | 1.199 | 0.912 | 0.014 | 0.007 | 0.007 | 0.157 | 0.24 | 0.29 | 0.17 | 0.26 | 0.29 | 0.25 |
| 5 | 39418887 | rs3849768 | 0.443 | 0.906 | 0.73 | 0.612 | 0.019 | 0.005 | 0.005 | 0 | 0.26 | 0.16 | 0.3 | 0.2 | 0.19 | 0.24 |
| 5 | 39419750 | rs11739328 | 0.109 | 0.751 | 0.696 | -0.184 | -0.096 | 0.096 | 0.096 | 0 | 0.2 | 0.02 | 0.13 | 0.04 | 0.28 | 0.17 |
| 5 | 39423907 | rs10512696 | 0.844 | 0.691 | 1.153 | -0.12 | -0.004 | 0.023 | 0.023 | 1.145 | 0.39 | 0.32 | 0.34 | 0.19 | 0.43 | 0.39 |
| 5 | 39426020 | rs3812040 | -1.342 | 0.428 | 1.503 | 0.447 | -0.009 | -0.011 | -0.011 | 0 | 0.33 | 0.59 | 0.21 | 0.22 | 0.28 | 0.28 |
| 5 | 39427155 | rs9292741 | -1.24 | 0.508 | 1.466 | 0.475 | -0.02 | -0.021 | -0.021 | 0 | 0.32 | 0.52 | 0.21 | 0.22 | 0.27 | 0.26 |
| 5 | 39428050 | rs6894577 | -1.079 | 0.515 | 1.327 | 0.48 | -0.009 | -0.011 | -0.011 | 0.067 | 0.31 | 0.4 | 0.19 | 0.22 | 0.28 | 0.26 |
| 5 | 39434349 | rs7703879 | -0.54 | 0.669 | 0.638 | 0.358 | -0.011 | -0.06 | -0.06 | 0.429 | 0.23 | 0.27 | 0.13 | 0.04 | 0.17 | 0.16 |
| 5 | 39437622 | rs3849774 | -0.844 | 0.802 | -0.06 | 0.725 | 0 | -0.004 | -0.004 | 0 | 0.34 | 0.6 | 0.28 | 0.27 | 0.36 | 0.3 |
| 5 | 39438575 | rs3860796 | -0.786 | 0.097 | -0.069 | 1.147 | -0.035 | 0.033 | 0.033 | 2.971 | 0.28 | 0.54 | 0.24 | 0.41 | 0.23 | 0.43 |
| 5 | 39439306 | rs12516046 | -0.497 | 0.933 | -0.811 | 0.978 | -0.038 | 0.003 | 0.003 | 0 | 0.31 | 0.17 | 0.36 | 0.27 | 0.34 | 0.44 |
| 5 | 39439979 | rs16868886 | -0.819 | 0.679 | -0.548 | 1.229 | -0.001 | 0.002 | 0.002 | 0 | 0.08 | 0.13 | 0.05 | 0.14 | 0.06 | 0.13 |
| 5 | 39444239 | rs12652609 | -0.645 | 0.433 | -0.378 | 1.225 | -0.001 | 0 | 0 | 0.606 | 0.63 | 0.66 | 0.78 | 0.67 | 0.6 | 0.71 |
| 5 | 39447059 | rs4596408 | 0.441 | 0.473 | -0.525 | 1.141 | -0.067 | -0.031 | -0.031 | 1.103 | 0.28 | 0.1 | 0.33 | 0.25 | 0.32 | 0.36 |
| 5 | 39451901 | rs13168791 | 0.378 | 0.335 | 0.246 | 1.385 | 0.002 | 0.002 | 0.002 | 0.002 | 0.34 | 0.31 | 0.27 | 0.32 | 0.27 | 0.3 |
| 5 | 39451982 | rs16868954 | 0.9 | 0.375 | 0.2 | 1.435 | -0.005 | -0.029 | -0.029 | 0.109 | 0.06 | 0 | 0.03 | 0.15 | 0.03 | 0.16 |
| 5 | 39471176 | rs3909475 | -0.832 | 0.296 | 0.494 | 1.386 | -0.001 | 0 | 0 | 0 | 0.63 | 0.67 | 0.78 | 0.67 | 0.61 | 0.7 |
| 5 | 39473358 | rs13167273 | 1.126 | 0.249 | 0.367 | 1.362 | -0.008 | 0.04 | 0.04 | 0.536 | 0.26 | 0.05 | 0.21 | 0.17 | 0.24 | 0.13 |
| 5 | 39481169 | rs7735747 | -0.381 | 0.217 | 0.463 | 1.447 | 0.008 | 0.121 | 0.121 | 0 | 0.11 | 0.28 | 0.11 | 0.31 | 0.08 | 0.21 |
| 5 | 39500920 | rs3849781 | 0.7 | 0.534 | 0.612 | 1.06 | -0.003 | -0.008 | -0.008 | 0.438 | 0.32 | 0.39 | 0.41 | 0.26 | 0.29 | 0.23 |
| 5 | 39502755 | rs4957476 | 0.369 | 0.509 | 0.345 | 0.976 | -0.001 | -0.001 | -0.001 | 3.945 | 0.37 | 0.63 | 0.49 | 0.41 | 0.34 | 0.27 |
| 5 | 39503551 | rs268095 | -0.513 | 0.508 | 0.056 | 0.751 | -0.021 | 0.042 | 0.042 | 0 | 0.76 | 0.87 | 0.72 | 0.88 | 0.72 | 0.69 |
| 5 | 39517220 | rs1546281 | 0.435 | 0.04 | -0.015 | 0.452 | 0.007 | 0.048 | 0.048 | 0 | 0.34 | 0.26 | 0.21 | 0.16 | 0.31 | 0.22 |
| 5 | 39519303 | rs16869017 | 0.612 | -1.078 | -0.535 | 0.75 | 0.078 | -0.044 | -0.044 | 0.019 | 0.21 | 0.06 | 0.05 | 0.27 | 0.07 | 0.11 |
| 5 | 39525184 | rs2910950 | 0.63 | -0.833 | -0.217 | 0.636 | -0.013 | -0.019 | -0.019 | 0.001 | 0.08 | 0.15 | 0.07 | 0.03 | 0.03 | 0.14 |
| 5 | 39526442 | rs989248 | 0.176 | -0.904 | 0.352 | 0.818 | 0.019 | 0.033 | 0.033 | 0 | 0.44 | 0.64 | 0.46 | 0.29 | 0.34 | 0.37 |
| 5 | 39527943 | rs17397864 | 1.794 | -1.092 | -0.07 | 0.467 | 0.007 | 0.12 | 0.12 | 0 | 0.24 | 0.05 | 0.14 | 0.04 | 0.21 | 0.15 |
| 5 | 39533237 | rs12188502 | 1.699 | -1.072 | 0.11 | 0.462 | 0.019 | 0.079 | 0.079 | 0 | 0.25 | 0.12 | 0.14 | 0.04 | 0.21 | 0.15 |
| 5 | 39533992 | rs16869048 | 1.131 | -1.066 | 0.298 | 0.502 | 0.008 | 0.003 | 0.003 | 0.017 | 0.1 | 0.17 | 0.07 | 0.13 | 0.1 | 0.07 |
| 5 | 39534183 | rs11951152 | 1.425 | -1.071 | 0.33 | 0.499 | 0.007 | 0.004 | 0.004 | 0 | 0.09 | 0.12 | 0.07 | 0.13 | 0.1 | 0.07 |
| 5 | 39539188 | rs2937418 | 0.882 | -1.134 | 0.453 | 0.568 | -0.012 | -0.005 | -0.005 | 0.086 | 0.07 | 0.13 | 0.07 | 0.05 | 0.03 | 0.14 |
| 5 | 39542061 | rs2134166 | 0.646 | -1.24 | 0.915 | 0.692 | -0.002 | -0.005 | -0.005 | 0 | 0.36 | 0.4 | 0.38 | 0.26 | 0.32 | 0.22 |
| 5 | 39543680 | rs1501702 | -0.176 | -1.194 | 0.789 | 0.712 | 0.024 | 0.036 | 0.036 | 0 | 0.44 | 0.67 | 0.46 | 0.31 | 0.35 | 0.37 |
| 5 | 39544711 | rs1846975 | -0.031 | -1.143 | 0.787 | 0.714 | -0.011 | -0.018 | -0.018 | 0 | 0.43 | 0.53 | 0.45 | 0.31 | 0.35 | 0.37 |
| 5 | 39556334 | rs13167600 | 1.077 | -0.72 | 0.702 | 0.897 | 0.038 | 0.038 | 0.038 | 0.053 | 0.1 | 0.27 | 0.09 | 0.13 | 0.1 | 0.07 |
| 5 | 39557453 | rs17472933 | 0.387 | -0.678 | 0.02 | 0.75 | -0.078 | 0.06 | 0.06 | 0 | 0.36 | 0.15 | 0.41 | 0.22 | 0.53 | 0.52 |
| 5 | 39562234 | rs17473058 | 1.511 | 0.184 | -0.353 | 0.714 | -0.09 | -0.026 | -0.026 | 0.701 | 0.18 | 0.03 | 0.21 | 0.15 | 0.31 | 0.23 |
| 5 | 39565445 | rs2962490 | 0.894 | -0.122 | 0.027 | 0.83 | -0.012 | -0.004 | -0.004 | 1.447 | 0.07 | 0.15 | 0.08 | 0.05 | 0.03 | 0.14 |
| 5 | 39568609 | rs2962492 | -0.668 | -0.129 | -0.166 | 0.855 | -0.013 | 0.038 | 0.038 | 0.267 | 0.2 | 0.62 | 0.34 | 0.28 | 0.14 | 0.22 |
| 5 | 39569965 | rs2368096 | -0.048 | -0.003 | -1.064 | 0.519 | -0.075 | 0.051 | 0.051 | 0.02 | 0.35 | 0.18 | 0.42 | 0.22 | 0.51 | 0.49 |
| 5 | 39571224 | rs2939337 | -0.316 | -0.891 | -0.553 | 0.601 | -0.006 | 0.018 | 0.018 | 0 | 0.46 | 0.4 | 0.35 | 0.36 | 0.46 | 0.45 |
| 5 | 39572134 | rs6883007 | -0.545 | -1.183 | -0.269 | 0.737 | 0.041 | 0.028 | 0.028 | 0 | 0.5 | 0.6 | 0.56 | 0.35 | 0.65 | 0.66 |
| 5 | 39572571 | rs10052827 | 1.714 | -1.018 | -0.153 | 0.892 | -0.031 | -0.012 | -0.012 | 0.045 | 0.16 | 0.11 | 0.21 | 0.18 | 0.32 | 0.24 |
| 5 | 39573051 | rs2937429 | -0.21 | -1.283 | -0.042 | 0.797 | 0.007 | -0.003 | -0.003 | 0 | 0.29 | 0.27 | 0.14 | 0.31 | 0.2 | 0.31 |
| 5 | 39573464 | rs2939338 | 0.899 | -1.026 | -0.157 | 0.881 | 0.019 | -0.015 | -0.015 | 0.006 | 0.2 | 0.12 | 0.07 | 0.25 | 0.15 | 0.15 |
| 5 | 39576829 | rs10076408 | 0.858 | -1.005 | 0.079 | 1.031 | -0.016 | -0.016 | -0.016 | 0 | 0.22 | 0.17 | 0.45 | 0.38 | 0.36 | 0.25 |
| 5 | 39578660 | rs6451433 | -1.02 | -1.24 | 0.101 | 1.112 | -0.02 | 0.037 | 0.037 | 0 | 0.81 | 0.8 | 0.89 | 0.59 | 0.9 | 0.78 |
| 5 | 39578989 | rs7721839 | -0.325 | -1.288 | 0.112 | 1.072 | -0.054 | 0.082 | 0.082 | 0 | 0.77 | 0.47 | 0.81 | 0.59 | 0.86 | 0.75 |

| | | | | | | | | | | | | | | | | |
|---|---|---|---|---|---|---|---|---|---|---|---|---|---|---|---|---|
| 5 | 39579258 | rs7725868 | 0.142 | -1.126 | -0.107 | 0.671 | 0.004 | -0.01 | -0.01 | 0.002 | 0.29 | 0.31 | 0.29 | 0.17 | 0.3 | 0.4 |
| 5 | 39581163 | rs3947262 | 0.184 | -1.106 | -0.113 | 0.665 | 0.002 | -0.006 | -0.006 | 0 | 0.28 | 0.29 | 0.29 | 0.17 | 0.3 | 0.41 |
| 5 | 39582834 | rs2962497 | -0.203 | -0.967 | 0.05 | 0.655 | -0.009 | 0.015 | 0.015 | 0.664 | 0.41 | 0.3 | 0.28 | 0.35 | 0.4 | 0.51 |
| 5 | 39583375 | rs1260809 | -0.793 | -1.307 | 0.127 | 0.278 | 0.005 | 0.004 | 0.004 | 0.016 | 0.57 | 0.62 | 0.55 | 0.33 | 0.73 | 0.66 |
| 5 | 39588084 | rs7703694 | -0.396 | -0.928 | 0.14 | 0.528 | -0.006 | 0.164 | 0.164 | 0 | 0.79 | 0.64 | 0.73 | 0.54 | 0.8 | 0.67 |
| 5 | 39606335 | rs6869413 | 1.338 | -0.642 | -0.041 | 0.335 | -0.011 | -0.005 | -0.005 | 0 | 0.2 | 0.15 | 0.21 | 0.01 | 0.22 | 0.19 |
| 5 | 39609832 | rs12653467 | 0.624 | -0.539 | -0.086 | 0.593 | -0.038 | -0.001 | -0.001 | 0 | 0.28 | 0.16 | 0.24 | 0.24 | 0.38 | 0.32 |
| 5 | 39612157 | rs10062538 | 0.607 | -0.525 | -0.086 | 0.591 | -0.021 | 0.003 | 0.003 | 0 | 0.28 | 0.21 | 0.25 | 0.24 | 0.38 | 0.32 |
| 5 | 39612629 | rs10462004 | -1.005 | -0.744 | 0.409 | 1.031 | -0.013 | 0.077 | 0.077 | 0 | 0.85 | 0.8 | 0.81 | 0.62 | 0.88 | 0.79 |
| 5 | 39612783 | rs10462005 | -1.005 | -0.744 | 0.409 | 0.773 | -0.013 | 0.103 | 0.103 | 0 | 0.85 | 0.8 | 0.81 | 0.58 | 0.88 | 0.79 |
| 5 | 39615631 | rs11956118 | 0.305 | -0.694 | 0.371 | 0.673 | -0.018 | -0.02 | -0.02 | 0 | 0.07 | 0.23 | 0.04 | 0.05 | 0.02 | 0.05 |
| 5 | 39625080 | rs980793 | -0.35 | -0.751 | -0.002 | 0.608 | -0.02 | 0.092 | 0.092 | 0 | 0.77 | 0.66 | 0.76 | 0.54 | 0.83 | 0.8 |
| 5 | 39632634 | rs977370 | 0.147 | -0.522 | -0.048 | 0.32 | 0.005 | -0.048 | -0.048 | 0 | 0.11 | 0.22 | 0.07 | 0.02 | 0.1 | 0.06 |
| 5 | 39633043 | rs1501707 | -0.453 | -0.539 | -0.488 | 0.198 | -0.003 | 0.008 | 0.008 | 0 | 0.66 | 0.75 | 0.62 | 0.33 | 0.61 | 0.58 |
| 5 | 39637885 | rs13185971 | 0.822 | -0.806 | 0.01 | 0.444 | 0.046 | 0.095 | 0.095 | 0 | 0.33 | 0.14 | 0.13 | 0.1 | 0.24 | 0.34 |
| 5 | 39641154 | rs7732292 | -0.065 | -0.818 | 0.025 | 0.629 | 0.001 | 0.001 | 0.001 | 0 | 0.56 | 0.63 | 0.52 | 0.27 | 0.67 | 0.54 |
| 5 | 39652260 | rs12659111 | -1.341 | -0.424 | 0.257 | 0.952 | -0.001 | 0.286 | 0.286 | 0 | 0.91 | 0.77 | 0.82 | 0.56 | 0.94 | 0.9 |
| 5 | 39662924 | rs6451445 | 0.769 | -0.445 | 0.302 | 0.543 | 0.013 | -0.023 | -0.023 | 0 | 0.06 | 0.27 | 0.08 | 0 | 0.04 | 0.05 |
| 5 | 39666907 | rs7730375 | 0.876 | -0.728 | 0.014 | 0.329 | 0.039 | 0.072 | 0.072 | 0.191 | 0.33 | 0.17 | 0.13 | 0.1 | 0.24 | 0.33 |
| 5 | 39671922 | rs12654371 | -1.41 | -0.607 | 0.42 | 0.944 | 0.006 | 0.339 | 0.339 | 0 | 0.91 | 0.77 | 0.82 | 0.53 | 0.94 | 0.91 |
| 5 | 39679489 | rs6884834 | 1.762 | -0.769 | 0.119 | 0.589 | 0.002 | -0.002 | -0.002 | 0 | 0.22 | 0.21 | 0.23 | 0.28 | 0.15 | 0.22 |
| 5 | 39680721 | rs10473145 | 1.806 | -0.774 | 0.102 | 0.601 | -0.008 | 0.001 | 0.001 | 0.316 | 0.22 | 0.23 | 0.23 | 0.26 | 0.15 | 0.22 |
| 5 | 39702215 | rs10041204 | -1.603 | -0.831 | 0.296 | 0.807 | -0.004 | 0.24 | 0.24 | 0 | 0.92 | 0.81 | 0.82 | 0.44 | 0.93 | 0.9 |
| 5 | 39706933 | rs10077158 | -1.435 | -0.832 | 0.295 | 0.743 | -0.008 | 0.146 | 0.146 | 0 | 0.91 | 0.8 | 0.68 | 0.28 | 0.94 | 0.9 |
| 5 | 39707555 | rs11743326 | -0.389 | -0.87 | 0.388 | 0.599 | 0.023 | 0.028 | 0.028 | 0 | 0.64 | 0.31 | 0.51 | 0.18 | 0.75 | 0.63 |
| 5 | 39722109 | rs6451452 | 0.178 | -0.426 | 0.039 | 0.274 | -0.004 | 0.135 | 0.135 | 0 | 0.51 | 0.28 | 0.32 | 0.18 | 0.55 | 0.51 |
| 5 | 39727753 | rs12656570 | 1.58 | -0.253 | -0.137 | 0.947 | -0.067 | -0.072 | -0.072 | 0 | 0.14 | 0.02 | 0.19 | 0.25 | 0.21 | 0.13 |
| 5 | 39734189 | rs2887979 | 1.008 | -0.256 | -0.063 | 0.68 | -0.003 | -0.004 | -0.004 | 0 | 0.18 | 0.13 | 0.1 | 0.01 | 0.19 | 0.15 |
| 5 | 39735991 | rs12516833 | -0.188 | -0.159 | 0.021 | 1.142 | 0.002 | -0.014 | -0.014 | 0 | 0.57 | 0.55 | 0.53 | 0.35 | 0.6 | 0.57 |
| 5 | 39737356 | rs10941454 | -1.301 | -0.344 | -0.117 | 1.227 | -0.003 | 0.141 | 0.141 | 0 | 0.92 | 0.81 | 0.82 | 0.69 | 0.95 | 0.91 |
| 5 | 39743005 | rs13169069 | 1.286 | -0.08 | -1.051 | 0.289 | 0.008 | 0.044 | 0.044 | 0.009 | 0.24 | 0.08 | 0.13 | 0.13 | 0.23 | 0.24 |
| 5 | 39746640 | rs7733848 | 1.631 | -0.289 | -1.05 | 0.274 | 0.028 | 0.026 | 0.026 | 0.127 | 0.22 | 0.06 | 0.1 | 0.13 | 0.19 | 0.23 |
| 5 | 39748470 | rs1353835 | -0.285 | -0.069 | -0.577 | 0.393 | -0.001 | 0.007 | 0.007 | 0 | 0.55 | 0.49 | 0.51 | 0.32 | 0.54 | 0.5 |
| 5 | 39753535 | rs7704513 | -0.069 | -0.195 | -0.915 | -0.087 | -0.002 | -0.018 | -0.018 | 0 | 0.28 | 0.36 | 0.26 | 0.19 | 0.23 | 0.2 |
| 5 | 39763732 | rs12521441 | -0.955 | -0.191 | -0.714 | -0.075 | 0.047 | 0.119 | 0.119 | 0 | 0.77 | 0.62 | 0.61 | 0.45 | 0.73 | 0.74 |
| 5 | 39770157 | rs11739973 | -0.526 | -0.478 | -0.748 | 0.148 | -0.008 | 0.032 | 0.032 | 0 | 0.43 | 0.36 | 0.32 | 0.17 | 0.44 | 0.39 |
| 5 | 39770956 | rs11960343 | -0.594 | -0.482 | -0.743 | 0.176 | -0.005 | 0.018 | 0.018 | 0.021 | 0.43 | 0.38 | 0.45 | 0.3 | 0.45 | 0.4 |
| 5 | 39780202 | rs11740862 | 0.757 | -0.395 | -0.599 | -0.042 | -0.017 | 0.034 | 0.034 | 0 | 0.18 | 0.11 | 0.11 | 0 | 0.21 | 0.16 |
| 5 | 39786324 | rs7731629 | -1.931 | -0.322 | -0.374 | 0.251 | 0.038 | 0.478 | 0.478 | 0.182 | 0.92 | 0.66 | 0.69 | 0.43 | 0.94 | 0.93 |
| 5 | 39787302 | rs10941455 | -1.583 | -0.322 | -0.411 | 0.228 | 0.055 | 0.523 | 0.523 | 1.701 | 0.92 | 0.57 | 0.8 | 0.56 | 0.95 | 0.93 |
| 5 | 39801230 | rs4957507 | 0.74 | -0.301 | -0.463 | 0.058 | 0.029 | -0.019 | -0.019 | 2.598 | 0.2 | 0.29 | 0.36 | 0.14 | 0.25 | 0.21 |
| 5 | 39801338 | rs1499235 | -1.732 | -0.944 | -0.266 | 0.347 | 0.048 | 0.465 | 0.465 | 0.62 | 0.92 | 0.66 | 0.81 | 0.56 | 0.95 | 0.94 |
| 5 | 39806164 | rs1994882 | -0.608 | -1.212 | -0.634 | 0.147 | -0.014 | -0.012 | -0.012 | 1.113 | 0.63 | 0.44 | 0.56 | 0.23 | 0.73 | 0.66 |
| 5 | 39806217 | rs12658486 | 1.812 | -1.224 | -0.653 | 0.151 | 0.037 | -0.01 | -0.01 | 0 | 0.24 | 0.07 | 0.11 | 0.2 | 0.2 | 0.27 |
| 5 | 39808036 | rs9292751 | -0.448 | -0.934 | -0.362 | 0.438 | -0.018 | 0.301 | 0.301 | 7.041 | 0.87 | 0.52 | 0.79 | 0.56 | 0.94 | 0.94 |
| 5 | 39816835 | rs980306 | 0.833 | -0.984 | -0.976 | -0.154 | -0.019 | 0.019 | 0.019 | 0 | 0.22 | 0.12 | 0.2 | 0.13 | 0.27 | 0.19 |
| 5 | 39817167 | rs10447141 | 0.181 | -0.983 | -0.737 | -0.183 | 0.007 | 0.01 | 0.01 | 0 | 0.27 | 0.41 | 0.36 | 0.27 | 0.3 | 0.22 |
| 5 | 39817764 | rs10941473 | -0.595 | -1.237 | -0.876 | -0.284 | -0.045 | 0.251 | 0.251 | 0.548 | 0.83 | 0.62 | 0.79 | 0.47 | 0.91 | 0.85 |
| 5 | 39820671 | rs4957225 | 1.8 | -0.986 | -0.954 | -0.556 | -0.042 | 0.041 | 0.041 | 3.482 | 0.17 | 0.1 | 0.24 | 0.01 | 0.22 | 0.21 |
| 5 | 39821317 | rs17194068 | 0.857 | -1.119 | -0.609 | -0.345 | -0.03 | 0.02 | 0.02 | 0 | 0.22 | 0.11 | 0.2 | 0.13 | 0.27 | 0.19 |
| 5 | 39825740 | rs6895329 | -0.047 | -1.009 | -0.598 | -0.294 | -0.011 | -0.026 | -0.026 | 0.061 | 0.31 | 0.4 | 0.23 | 0.18 | 0.28 | 0.26 |
| 5 | 39829284 | rs6876969 | -1.051 | -1.011 | -0.543 | -0.235 | -0.004 | 0 | 0 | 0 | 0.64 | 0.6 | 0.66 | 0.34 | 0.68 | 0.62 |
| 5 | 39835837 | rs487149 | 1.243 | -0.991 | -0.364 | -0.059 | 0.025 | 0.231 | 0.231 | 0 | 0.12 | 0.29 | 0.22 | 0.48 | 0.15 | 0.13 |
| 5 | 39852807 | rs1423255 | 1.628 | -0.918 | -0.725 | -0.346 | -0.069 | 0.071 | 0.071 | 4.62 | 0.18 | 0.08 | 0.2 | 0.01 | 0.25 | 0.23 |
| 5 | 39853616 | rs639441 | -0.417 | -0.815 | -0.313 | -0.205 | -0.023 | 0.084 | 0.084 | 2.311 | 0.26 | 0.57 | 0.26 | 0.53 | 0.19 | 0.2 |
| 5 | 39856991 | rs979024 | 1.412 | -1.152 | -0.881 | -0.434 | 0.004 | 0.022 | 0.022 | 0 | 0.27 | 0.05 | 0.14 | 0.18 | 0.25 | 0.3 |
| 5 | 39858621 | rs500241 | -0.345 | -1.051 | -0.545 | -0.414 | 0.013 | 0.038 | 0.038 | 5.519 | 0.54 | 0.68 | 0.41 | 0.71 | 0.44 | 0.5 |
| 5 | 39861826 | rs1816088 | -1.784 | -1.035 | -0.743 | -0.498 | -0.012 | -0.002 | -0.002 | 0 | 0.83 | 0.9 | 0.81 | 0.86 | 0.75 | 0.8 |
| 5 | 39866913 | rs618051 | -1.138 | -0.903 | -0.659 | -0.421 | -0.011 | 0.013 | 0.013 | 0 | 0.78 | 0.88 | 0.79 | 0.86 | 0.71 | 0.75 |
| 5 | 39878219 | rs544283 | -0.526 | -1.067 | -0.59 | -0.34 | 0.003 | 0.142 | 0.142 | 6.66 | 0.56 | 0.8 | 0.58 | 0.86 | 0.45 | 0.52 |
| 5 | 39898872 | rs503621 | 0.2 | -1.257 | -0.369 | -0.272 | 0.027 | 0.002 | 0.002 | 0.469 | 0.4 | 0.32 | 0.34 | 0.37 | 0.3 | 0.37 |
| 5 | 39907157 | rs1113439 | 0.814 | -1.418 | -0.362 | -0.295 | -0.009 | 0.002 | 0.002 | 0 | 0.33 | 0.36 | 0.32 | 0.67 | 0.29 | 0.36 |
| 5 | 39919202 | rs16869416 | 0.876 | -0.835 | -0.011 | 0.078 | 0 | 0.186 | 0.186 | 0.005 | 0.15 | 0.33 | 0.23 | 0.48 | 0.15 | 0.13 |
| 5 | 39924922 | rs10076713 | 1.914 | -0.833 | -0.211 | 0.013 | 0.032 | 0.011 | 0.011 | 0 | 0.25 | 0.08 | 0.15 | 0.19 | 0.21 | 0.29 |
| 5 | 39929770 | rs568179 | 0.121 | -0.695 | -0.136 | -0.052 | 0.003 | 0.003 | 0.003 | 0 | 0.62 | 0.53 | 0.57 | 0.39 | 0.57 | 0.62 |
| 5 | 39932268 | rs542708 | -2.214 | -0.638 | -0.149 | 0.019 | -0.067 | -0.038 | -0.038 | 0.005 | 0.89 | 0.99 | 0.86 | 0.9 | 0.77 | 0.85 |
| 5 | 39946160 | rs506990 | 0.47 | -0.564 | 0.14 | 0.098 | -0.008 | -0.001 | -0.001 | 0.179 | 0.4 | 0.42 | 0.35 | 0.39 | 0.3 | 0.38 |
| 5 | 39960584 | rs10473156 | 1.849 | -0.573 | 0.118 | 0.249 | 0.037 | 0.015 | 0.015 | 0 | 0.25 | 0.08 | 0.11 | 0.19 | 0.2 | 0.29 |
| 5 | 39990379 | rs7711972 | -0.27 | -0.518 | 0.276 | 0.315 | 0.016 | 0.011 | 0.011 | 0 | 0.62 | 0.48 | 0.55 | 0.51 | 0.59 | 0.64 |

| | | | | | | | | | | | | | | | | |
|---|---|---|---|---|---|---|---|---|---|---|---|---|---|---|---|---|
| 5 | 39991855 | rs10068269 | 0.941 | -0.493 | 0.079 | 0.346 | -0.014 | 0.042 | 0.042 | 0 | 0.46 | 0.1 | 0.31 | 0.32 | 0.49 | 0.55 |
| 5 | 39996666 | rs2939382 | 1.278 | -0.408 | 0.322 | 0.433 | 0.013 | 0.191 | 0.191 | 0 | 0.12 | 0.23 | 0.21 | 0.48 | 0.14 | 0.13 |
| 5 | 39996786 | rs2939383 | 0.001 | -0.511 | 0.324 | 0.343 | -0.002 | 0.006 | 0.006 | 0 | 0.65 | 0.69 | 0.74 | 0.82 | 0.62 | 0.66 |
| 5 | 40001006 | rs6899212 | -0.232 | -0.471 | 0.406 | 0.397 | 0.006 | 0.001 | 0.001 | 0.049 | 0.63 | 0.43 | 0.41 | 0.37 | 0.58 | 0.64 |
| 5 | 40002579 | rs1567009 | -0.408 | -0.471 | 0.441 | 0.407 | 0.035 | 0.027 | 0.027 | 0.512 | 0.65 | 0.54 | 0.55 | 0.51 | 0.58 | 0.65 |
| 5 | 40014986 | rs6878420 | -0.344 | -0.466 | 0.414 | 0.397 | 0.012 | 0.004 | 0.004 | 0.253 | 0.64 | 0.48 | 0.42 | 0.37 | 0.58 | 0.64 |
| 5 | 40018284 | rs1506188 | 1.245 | -0.445 | 0.473 | 0.434 | 0.014 | 0.199 | 0.199 | 0 | 0.12 | 0.23 | 0.21 | 0.48 | 0.14 | 0.12 |
| 5 | 40022278 | rs1605202 | 1.636 | -0.486 | 0.237 | 0.304 | 0.033 | 0.024 | 0.024 | 0.086 | 0.27 | 0.1 | 0.11 | 0.2 | 0.21 | 0.29 |
| 5 | 40025084 | rs6865476 | 1.193 | -0.482 | 0.348 | 0.263 | 0.012 | 0.019 | 0.019 | 0 | 0.31 | 0.27 | 0.16 | 0.2 | 0.27 | 0.32 |
| 5 | 40025962 | rs17218399 | 1.277 | -0.502 | 0.263 | 0.22 | -0.016 | -0.041 | -0.041 | 0 | 0.22 | 0.09 | 0.36 | 0.27 | 0.28 | 0.26 |
| 5 | 40027012 | rs12516115 | 1.608 | -0.582 | 0.246 | 0.136 | -0.071 | 0.087 | 0.087 | 1.482 | 0.22 | 0.09 | 0.22 | 0.01 | 0.27 | 0.23 |
| 5 | 40031013 | rs954017 | -0.514 | -0.62 | 0.389 | 0.219 | -0.02 | -0.054 | -0.054 | 0 | 0.17 | 0.35 | 0.08 | 0.05 | 0.09 | 0.08 |
| 5 | 40038458 | rs756318 | 0.435 | -0.559 | 0.397 | 0.304 | 0.008 | 0.006 | 0.006 | 0.374 | 0.34 | 0.33 | 0.43 | 0.49 | 0.42 | 0.35 |
| 5 | 40040278 | rs10039937 | 1.812 | -0.556 | 0.145 | 0.293 | 0.039 | 0.014 | 0.014 | 0 | 0.26 | 0.08 | 0.11 | 0.2 | 0.2 | 0.3 |
| 5 | 40042245 | rs2939378 | 0.398 | -0.372 | 0.444 | 0.401 | -0.001 | -0.011 | -0.011 | 0 | 0.62 | 0.56 | 0.69 | 0.79 | 0.62 | 0.66 |
| 5 | 40049919 | rs4957233 | -0.453 | -0.259 | 0.456 | 0.396 | -0.004 | -0.003 | -0.003 | 0 | 0.53 | 0.52 | 0.64 | 0.36 | 0.6 | 0.54 |
| 5 | 40050301 | rs1567010 | 1.325 | -0.262 | 0.403 | 0.363 | -0.032 | 0.005 | 0.005 | 0 | 0.2 | 0.07 | 0.23 | 0.16 | 0.29 | 0.26 |
| 5 | 40050381 | rs10063169 | 0.28 | -0.256 | 0.503 | 0.35 | 0.003 | -0.007 | -0.007 | 0 | 0.32 | 0.36 | 0.4 | 0.2 | 0.31 | 0.28 |
| 5 | 40055257 | rs4957237 | 1.124 | -0.083 | 0.601 | 0.412 | 0.017 | -0.014 | -0.014 | 0 | 0.24 | 0.31 | 0.38 | 0.15 | 0.27 | 0.23 |
| 5 | 40066219 | rs2036610 | 1.465 | -0.093 | 0.459 | 0.46 | 0.029 | 0.035 | 0.035 | 0.924 | 0.28 | 0.08 | 0.12 | 0.18 | 0.22 | 0.31 |
| 5 | 40069162 | rs6874438 | 1.45 | -0.045 | 0.529 | 0.501 | 0.012 | 0.048 | 0.048 | 0.404 | 0.3 | 0.08 | 0.14 | 0.18 | 0.28 | 0.32 |
| 5 | 40078936 | rs10038914 | -0.214 | 0.087 | 0.686 | 0.537 | 0.002 | 0.004 | 0.004 | 0.021 | 0.51 | 0.53 | 0.61 | 0.36 | 0.52 | 0.53 |
| 5 | 40093632 | rs1857869 | -0.946 | 0.23 | 0.464 | 0.874 | 0.011 | 0.184 | 0.184 | 0 | 0.86 | 0.7 | 0.78 | 0.54 | 0.86 | 0.87 |
| 5 | 40096079 | rs4957247 | 1.285 | 0.24 | 0.263 | 0.808 | -0.044 | 0.063 | 0.063 | 0.306 | 0.21 | 0.09 | 0.24 | 0.01 | 0.28 | 0.24 |
| 5 | 40098605 | rs17220794 | 1.279 | 0.362 | -0.385 | 0.408 | -0.008 | 0.021 | 0.021 | 0 | 0.21 | 0.07 | 0.16 | 0.12 | 0.21 | 0.28 |
| 5 | 40121487 | rs10067281 | 0.583 | 0.308 | -0.243 | 0.513 | 0.004 | 0.015 | 0.015 | 0.071 | 0.41 | 0.36 | 0.22 | 0.23 | 0.36 | 0.36 |
| 5 | 40122187 | rs10076961 | -0.669 | 0.38 | -0.238 | 0.691 | 0.008 | 0.194 | 0.194 | 0.066 | 0.85 | 0.67 | 0.78 | 0.53 | 0.86 | 0.87 |
| 5 | 40140378 | rs919208 | 0.125 | 0.4 | -0.554 | 0.493 | -0.032 | 0.036 | 0.036 | 0.974 | 0.43 | 0.27 | 0.55 | 0.31 | 0.5 | 0.52 |
| 5 | 40140556 | rs1811337 | -0.53 | 0.298 | -0.4 | 0.405 | -0.002 | -0.002 | -0.002 | 0 | 0.55 | 0.5 | 0.61 | 0.36 | 0.59 | 0.59 |
| 5 | 40146458 | rs1427737 | -0.242 | 0.292 | -0.433 | 0.266 | -0.002 | -0.008 | -0.008 | 1.025 | 0.64 | 0.67 | 0.49 | 0.24 | 0.65 | 0.6 |
| 5 | 40146945 | rs4546429 | 0.134 | 0.29 | -0.434 | 0.265 | 0.007 | -0.017 | -0.017 | 0.307 | 0.11 | 0.23 | 0.06 | 0.05 | 0.09 | 0.07 |
| 5 | 40148763 | rs4455605 | -0.01 | 0.29 | -0.459 | 0.291 | 0.012 | -0.068 | -0.068 | 0 | 0.33 | 0.43 | 0.32 | 0.07 | 0.37 | 0.31 |
| 5 | 40154769 | rs1427744 | 1.319 | 0.318 | -1.253 | 0.231 | 0.013 | 0.03 | 0.03 | 0.075 | 0.31 | 0.25 | 0.18 | 0.17 | 0.28 | 0.29 |
| 5 | 40159642 | rs4957254 | 1.1 | 0.456 | -1.456 | 0.413 | -0.006 | -0.061 | -0.061 | 1.551 | 0.22 | 0.07 | 0.29 | 0.29 | 0.21 | 0.28 |
| 5 | 40174446 | rs350089 | 0.207 | 0.798 | -0.765 | 0.332 | -0.011 | 0.062 | 0.062 | 1.459 | 0.78 | 0.49 | 0.61 | 0.31 | 0.81 | 0.83 |
| 5 | 40174663 | rs11952580 | 0.422 | 1.253 | -1.129 | 0.185 | -0.052 | 0.079 | 0.079 | 0.285 | 0.21 | 0.08 | 0.24 | 0.03 | 0.28 | 0.24 |
| 5 | 40183220 | rs690488 | 0.447 | 0.716 | -0.536 | 0.886 | -0.001 | 0.077 | 0.077 | 1.13 | 0.74 | 0.51 | 0.6 | 0.45 | 0.78 | 0.79 |
| 5 | 40183749 | rs350027 | 0.15 | 0.66 | 0.112 | 1.077 | 0.003 | 0.017 | 0.017 | 0 | 0.84 | 0.68 | 0.75 | 0.82 | 0.87 | 0.88 |
| 5 | 40186716 | rs10067548 | 0.613 | 0.382 | 0.213 | 1.165 | -0.001 | -0.001 | -0.001 | 0 | 0.05 | 0.04 | 0.02 | 0 | 0.04 | 0.03 |
| 5 | 40188442 | rs448997 | 0.186 | 1.059 | 0.102 | 1.228 | -0.002 | 0.001 | 0.001 | 0 | 0.24 | 0.31 | 0.28 | 0.2 | 0.36 | 0.23 |
| 5 | 40189966 | rs184782 | -0.928 | 0.265 | 0.047 | 0.673 | 0.005 | -0.032 | -0.032 | 0 | 0.42 | 0.6 | 0.38 | 0.2 | 0.47 | 0.29 |
| 5 | 40192935 | rs350022 | -1.086 | 0.248 | 0.047 | 0.672 | -0.001 | 0 | 0 | 0 | 0.44 | 0.62 | 0.51 | 0.36 | 0.47 | 0.29 |
| 5 | 40193497 | rs10512727 | 1.667 | -0.129 | -0.688 | 0.412 | -0.01 | -0.071 | -0.071 | 0 | 0.21 | 0.03 | 0.16 | 0.29 | 0.21 | 0.29 |
| 5 | 40193851 | rs187040 | -0.944 | -0.06 | -0.638 | -0.118 | -0.003 | 0 | 0 | 0 | 0.65 | 0.66 | 0.67 | 0.66 | 0.68 | 0.58 |
| 5 | 40195443 | rs351634 | 1.124 | 0.104 | -0.658 | -0.105 | -0.014 | 0.008 | 0.008 | 0 | 0.22 | 0.22 | 0.27 | 0.06 | 0.35 | 0.19 |
| 5 | 40203162 | rs353342 | 0.897 | 0.528 | -0.77 | -0.387 | -0.001 | 0 | 0 | 0 | 0.21 | 0.26 | 0.24 | 0.2 | 0.29 | 0.22 |
| 5 | 40203232 | rs4621594 | 1.646 | 0.784 | -0.807 | -0.437 | -0.016 | 0.028 | 0.028 | 0 | 0.16 | 0.1 | 0.19 | 0.06 | 0.23 | 0.18 |
| 5 | 40206804 | rs350042 | 1.749 | 0.609 | -0.748 | -0.452 | 0.001 | -0.018 | -0.018 | 0.719 | 0.05 | 0.04 | 0.03 | 0 | 0.04 | 0.03 |
| 5 | 40207083 | rs10462008 | 1.593 | 0.692 | -0.893 | -0.452 | 0.009 | 0.024 | 0.024 | 0 | 0.26 | 0.11 | 0.14 | 0.16 | 0.23 | 0.34 |
| 5 | 40210933 | rs353325 | -0.146 | 0.895 | -0.694 | -0.357 | 0.04 | -0.13 | -0.13 | 0.192 | 0.3 | 0.51 | 0.3 | 0.06 | 0.37 | 0.25 |
| 5 | 40217324 | rs164764 | -0.548 | 0.638 | -0.413 | -0.209 | 0.006 | 0.005 | 0.005 | 0.01 | 0.56 | 0.62 | 0.63 | 0.65 | 0.65 | 0.56 |
| 5 | 40217572 | rs193433 | -0.681 | 0.635 | 0.057 | -0.18 | 0.016 | 0.008 | 0.008 | 0.581 | 0.13 | 0.35 | 0.08 | 0.13 | 0.08 | 0.1 |
| 5 | 40218050 | rs10472329 | -0.702 | 0.787 | -0.431 | -0.693 | 0.011 | -0.095 | -0.095 | 0 | 0.31 | 0.42 | 0.19 | 0.07 | 0.33 | 0.35 |
| 5 | 40223183 | rs46261 | -0.186 | 0.591 | -0.599 | -0.337 | 0.011 | 0.027 | 0.027 | 0 | 0.53 | 0.72 | 0.35 | 0.39 | 0.58 | 0.61 |
| 5 | 40227586 | rs351651 | 0.097 | 0.629 | -0.347 | -0.503 | 0.088 | -0.009 | -0.009 | 0 | 0.08 | 0.42 | 0.06 | 0 | 0.06 | 0.04 |
| 5 | 40228374 | rs1478903 | 0.866 | 0.477 | -1.71 | -0.674 | -0.04 | 0.123 | 0.123 | 1.971 | 0.28 | 0.07 | 0.26 | 0.05 | 0.28 | 0.14 |
| 5 | 40228613 | rs353365 | -0.924 | 0.577 | -1.713 | -0.345 | 0.022 | -0.095 | -0.095 | 0.114 | 0.83 | 0.96 | 0.62 | 0.44 | 0.86 | 0.76 |
| 5 | 40233337 | rs353397 | -0.401 | 0.655 | -1.864 | -0.673 | -0.005 | 0.251 | 0.251 | 2.475 | 0.53 | 0.83 | 0.43 | 0.11 | 0.53 | 0.42 |
| 5 | 40233662 | rs353396 | -0.045 | 0.654 | -1.755 | -0.639 | -0.022 | 0.187 | 0.187 | 0.657 | 0.46 | 0.73 | 0.4 | 0.11 | 0.47 | 0.38 |
| 5 | 40260929 | rs348609 | 0.257 | 0.898 | -1.791 | -0.56 | -0.014 | 0.04 | 0.04 | 0 | 0.16 | 0.07 | 0.11 | 0.06 | 0.22 | 0.25 |
| 5 | 40267930 | rs2279170 | 0.752 | 0.695 | -1.757 | -0.159 | -0.028 | -0.024 | -0.024 | 0 | 0.3 | 0.14 | 0.2 | 0.33 | 0.34 | 0.35 |
| 5 | 40286497 | rs4957263 | -0.194 | 0.571 | -1.77 | 0.266 | -0.069 | 0.072 | 0.072 | 0 | 0.57 | 0.81 | 0.55 | 0.31 | 0.51 | 0.51 |
| 5 | 40286821 | rs348620 | -0.193 | 0.293 | -0.438 | 0.74 | 0.003 | -0.007 | -0.007 | 0.216 | 0.14 | 0.46 | 0.11 | 0.09 | 0.14 | 0.24 |
| 5 | 40286862 | rs7723858 | -0.177 | 0.652 | -1.277 | 1.205 | -0.022 | -0.057 | -0.057 | 0.021 | 0.88 | 0.94 | 0.75 | 0.66 | 0.86 | 0.83 |
| 5 | 40287885 | rs529750 | -0.916 | 0.372 | -1.263 | 1.166 | 0.009 | -0.054 | -0.054 | 0 | 0.25 | 0.17 | 0.28 | 0.47 | 0.24 | 0.43 |
| 5 | 40288878 | rs4245975 | -1.061 | 0.234 | -0.19 | 1.485 | -0.013 | 0.154 | 0.154 | 3.199 | 0.2 | 0.46 | 0.43 | 0.64 | 0.2 | 0.26 |
| 5 | 40291889 | rs348583 | -0.302 | 0.19 | 0.183 | 1.9 | -0.016 | 0.188 | 0.188 | 0.231 | 0.07 | 0.18 | 0.15 | 0.36 | 0.05 | 0.09 |
| 5 | 40293299 | rs6893565 | -0.993 | -0.319 | -1.057 | 0.791 | 0.021 | -0.082 | -0.082 | 0 | 0.7 | 0.95 | 0.86 | 0.71 | 0.76 | 0.57 |
| 5 | 40302452 | rs6864103 | 1.118 | 0.665 | -0.901 | -0.06 | -0.046 | 0.168 | 0.168 | 0 | 0.38 | 0.13 | 0.31 | 0.07 | 0.4 | 0.3 |

| | | | | | | | | | | | | | | | | |
|---|---|---|---|---|---|---|---|---|---|---|---|---|---|---|---|---|
| 5 | 40306777 | rs4340954 | 0.339 | 0.997 | -0.145 | 0.38 | 0.003 | 0.015 | 0.015 | 3.126 | 0.12 | 0.15 | 0.26 | 0.25 | 0.15 | 0.17 |
| 5 | 40309999 | rs16869807 | 0.524 | 1.114 | 0.103 | 0.112 | 0.034 | -0.034 | -0.034 | 0.047 | 0.08 | 0.14 | 0.08 | 0 | 0.14 | 0.01 |
| 5 | 40320006 | rs348601 | -0.356 | 0.555 | -0.227 | -0.098 | 0.006 | 0.031 | 0.031 | 0 | 0.42 | 0.32 | 0.53 | 0.86 | 0.41 | 0.6 |
| 5 | 40320992 | rs437342 | -0.314 | 0.405 | -0.304 | 0.331 | 0.002 | 0.044 | 0.044 | 3.592 | 0.2 | 0.25 | 0.42 | 0.63 | 0.23 | 0.35 |
| 5 | 40324220 | rs10043093 | 1.23 | 0.31 | -0.201 | -0.106 | -0.036 | 0.172 | 0.172 | 0.01 | 0.44 | 0.19 | 0.32 | 0.06 | 0.44 | 0.25 |
| 5 | 40324532 | rs348593 | -0.127 | 0.47 | -0.168 | 0.016 | -0.001 | 0.031 | 0.031 | 4.331 | 0.2 | 0.24 | 0.42 | 0.63 | 0.22 | 0.35 |
| 5 | 40336546 | rs6897022 | 0.781 | 0.347 | 0.202 | -0.228 | -0.002 | 0.079 | 0.079 | 0 | 0.5 | 0.42 | 0.39 | 0.14 | 0.46 | 0.43 |
| 5 | 40345721 | rs12518245 | 0.704 | 0.438 | -0.002 | 0.059 | -0.004 | -0.016 | -0.016 | 0 | 0.06 | 0.03 | 0.24 | 0.11 | 0.06 | 0.09 |
| 5 | 40355836 | rs7725639 | -0.464 | 0.204 | 0.037 | -0.147 | 0.004 | 0.018 | 0.018 | 0 | 0.75 | 0.63 | 0.83 | 0.77 | 0.74 | 0.8 |
| 5 | 40372223 | rs7725523 | 0.513 | 0.268 | 0.116 | -0.172 | -0.001 | 0.012 | 0.012 | 0.269 | 0.73 | 0.68 | 0.53 | 0.36 | 0.71 | 0.6 |
| 5 | 40386555 | rs1002922 | -0.259 | 0.183 | 0.018 | -0.386 | 0.016 | 0.032 | 0.032 | 0.011 | 0.35 | 0.19 | 0.29 | 0.75 | 0.35 | 0.47 |
| 5 | 40389129 | rs1550761 | 1.898 | 0.164 | 0.009 | -0.154 | 0.02 | -0.012 | -0.012 | 1.628 | 0.14 | 0.02 | 0.07 | 0.16 | 0.12 | 0.12 |
| 5 | 40392728 | rs4613763 | 0.797 | 0.322 | 0.504 | -0.3 | 0.049 | -0.046 | -0.046 | 0 | 0.09 | 0.19 | 0.07 | 0 | 0.13 | 0.01 |
| 5 | 40393605 | rs10512734 | -0.027 | 0.309 | 0.474 | -0.294 | 0.016 | 0.031 | 0.031 | 0.002 | 0.35 | 0.18 | 0.29 | 0.75 | 0.35 | 0.47 |
| 5 | 40394936 | rs2166194 | -0.351 | 0.329 | 0.45 | -0.224 | 0 | -0.001 | -0.001 | 0 | 0.93 | 0.95 | 0.75 | 0.88 | 0.94 | 0.94 |
| 5 | 40401509 | rs17234657 | 0.733 | 0.373 | 0.487 | -0.228 | 0.049 | -0.046 | -0.046 | 0 | 0.09 | 0.19 | 0.07 | 0 | 0.13 | 0.01 |
| 5 | 40404912 | rs7723981 | -0.385 | 0.369 | 0.547 | -0.228 | 0.002 | 0.009 | 0.009 | 0.158 | 0.93 | 0.88 | 0.74 | 0.9 | 0.92 | 0.91 |
| 5 | 40406985 | rs6896243 | 0.737 | 0.399 | 0.518 | -0.232 | 0.049 | -0.046 | -0.046 | 0 | 0.09 | 0.19 | 0.07 | 0 | 0.13 | 0.01 |
| 5 | 40410043 | rs10512737 | 0.737 | 0.45 | 0.526 | -0.232 | 0.057 | -0.046 | -0.046 | 0.607 | 0.09 | 0.19 | 0.07 | 0 | 0.14 | 0.01 |
| 5 | 40410584 | rs11742570 | 0.458 | 0.31 | 0.504 | -0.28 | 0.004 | 0.009 | 0.009 | 0 | 0.59 | 0.61 | 0.44 | 0.14 | 0.59 | 0.45 |
| 5 | 40411291 | rs6451494 | 0.458 | 0.31 | 0.525 | -0.28 | 0.004 | 0.007 | 0.007 | 0 | 0.59 | 0.61 | 0.44 | 0.14 | 0.59 | 0.45 |
| 5 | 40427798 | rs1899980 | 0.853 | 0.399 | 0.415 | -0.212 | -0.002 | -0.002 | -0.002 | 0 | 0.06 | 0.03 | 0.24 | 0.1 | 0.07 | 0.06 |
| 5 | 40429192 | rs6880934 | 0.652 | 0.438 | 0.467 | -0.3 | 0 | -0.01 | -0.01 | 0 | 0.56 | 0.45 | 0.61 | 0.24 | 0.52 | 0.5 |
| 5 | 40431183 | rs1373692 | 0.043 | -0.109 | -0.553 | -0.787 | 0.008 | 0.037 | 0.037 | 0 | 0.59 | 0.7 | 0.45 | 0.14 | 0.59 | 0.45 |
| 5 | 40433552 | rs6878370 | 0.043 | -0.109 | -0.553 | -0.787 | 0.008 | 0.037 | 0.037 | 0 | 0.59 | 0.69 | 0.45 | 0.14 | 0.59 | 0.45 |
| 5 | 40437948 | rs9292777 | 0.043 | -0.109 | -0.553 | -0.787 | 0.008 | 0.037 | 0.037 | 0 | 0.59 | 0.7 | 0.45 | 0.14 | 0.59 | 0.45 |
| 5 | 40445681 | rs11957215 | -0.137 | -0.51 | -0.349 | -0.677 | -0.009 | -0.025 | -0.025 | 0 | 0.4 | 0.44 | 0.39 | 0.78 | 0.37 | 0.56 |
| 5 | 40467272 | rs4532399 | -0.05 | -0.551 | -0.42 | -0.696 | -0.004 | -0.012 | -0.012 | 0.418 | 0.61 | 0.59 | 0.58 | 0.22 | 0.63 | 0.46 |
| 5 | 40481654 | rs6882351 | -1.116 | -0.432 | -0.534 | -0.162 | 0.011 | 0.1 | 0.1 | 0.601 | 0.91 | 0.91 | 0.85 | 0.55 | 0.86 | 0.84 |
| 5 | 40484938 | rs10077544 | 1.008 | -1.157 | -0.889 | -0.452 | 0.039 | -0.084 | -0.084 | 0 | 0.23 | 0.04 | 0.16 | 0.31 | 0.19 | 0.34 |
| 5 | 40486896 | rs7720838 | 1.234 | -1.592 | -0.7 | -0.875 | 0.004 | 0.012 | 0.012 | 0.747 | 0.42 | 0.41 | 0.46 | 0.16 | 0.56 | 0.23 |
| 5 | 40490790 | rs10440635 | 1.091 | -1.554 | -0.678 | -0.842 | 0.003 | 0 | 0 | 0.151 | 0.46 | 0.57 | 0.48 | 0.16 | 0.56 | 0.28 |
| 5 | 40496423 | rs9687958 | 1.367 | -1.153 | -0.443 | -0.559 | -0.008 | -0.007 | -0.007 | 0 | 0.6 | 0.44 | 0.58 | 0.32 | 0.68 | 0.41 |
| 5 | 40497604 | rs4286721 | 1.169 | -1.143 | -0.738 | -0.485 | -0.005 | 0.003 | 0.003 | 0.193 | 0.62 | 0.59 | 0.59 | 0.41 | 0.68 | 0.44 |
| 5 | 40520217 | rs7736920 | 0.479 | -0.766 | -0.498 | -0.116 | 0.003 | -0.005 | -0.005 | 0 | 0.73 | 0.78 | 0.71 | 0.41 | 0.8 | 0.6 |
| 5 | 40521648 | rs1876143 | 0.39 | -0.764 | -0.54 | -0.11 | 0.011 | -0.03 | -0.03 | 0.147 | 0.74 | 0.82 | 0.71 | 0.5 | 0.81 | 0.64 |
| 5 | 40522212 | rs10941516 | 1.691 | -0.368 | -0.541 | -0.317 | -0.093 | 0.079 | 0.079 | 0 | 0.3 | 0.13 | 0.42 | 0.18 | 0.47 | 0.29 |
| 5 | 40530886 | rs10043301 | 1.66 | -0.457 | -0.167 | 0.23 | -0.017 | 0.003 | 0.003 | 0 | 0.65 | 0.52 | 0.61 | 0.41 | 0.7 | 0.57 |
| 5 | 40559387 | rs1024098 | 0.46 | -0.405 | -0.131 | 0.27 | 0.02 | 0.065 | 0.065 | 0 | 0.09 | 0.3 | 0.1 | 0.08 | 0.1 | 0.08 |
| 5 | 40573745 | rs9292780 | -1.288 | -0.399 | -0.369 | 0.352 | 0 | 0.05 | 0.05 | 0 | 0.89 | 0.89 | 0.84 | 0.63 | 0.89 | 0.85 |
| 5 | 40588364 | rs2036597 | 1.728 | -0.14 | -0.399 | 0.214 | -0.006 | -0.012 | -0.012 | 0 | 0.61 | 0.5 | 0.59 | 0.37 | 0.64 | 0.53 |
| 5 | 40594758 | rs9292783 | -0.493 | -0.404 | -0.422 | 0.429 | -0.004 | 0.047 | 0.047 | 0 | 0.84 | 0.85 | 0.81 | 0.54 | 0.87 | 0.79 |
| 5 | 40597618 | rs6869535 | -0.316 | -0.233 | -0.693 | 0.134 | -0.004 | -0.019 | -0.019 | 0 | 0.17 | 0.26 | 0.19 | 0.08 | 0.16 | 0.17 |
| 5 | 40603203 | rs16870155 | 0.865 | -0.153 | -1.144 | 0.228 | 0.012 | 0.008 | 0.008 | 0 | 0.1 | 0.06 | 0.1 | 0.08 | 0.06 | 0.14 |
| 5 | 40607910 | rs13163402 | 0.406 | -0.096 | -0.868 | 0.235 | -0.019 | -0.008 | -0.008 | 0 | 0.64 | 0.5 | 0.61 | 0.38 | 0.7 | 0.56 |
| 5 | 40622940 | rs4957138 | 0.499 | 0.082 | -1.031 | 0.316 | 0.002 | -0.005 | -0.005 | 2.254 | 0.63 | 0.4 | 0.6 | 0.46 | 0.71 | 0.6 |
| 5 | 40627066 | rs11745901 | -0.306 | -0.265 | -1.402 | 0.291 | -0.067 | 0.029 | 0.029 | 0 | 0.31 | 0.17 | 0.51 | 0.2 | 0.45 | 0.39 |
| 5 | 40629347 | rs11739531 | -1.093 | -1.052 | -1.033 | 0.349 | 0.002 | -0.005 | -0.005 | 0 | 0.61 | 0.66 | 0.71 | 0.53 | 0.66 | 0.64 |
| 5 | 40646560 | rs7716982 | -1.099 | -1.056 | -1.039 | 0.495 | 0.002 | -0.001 | -0.001 | 0.028 | 0.61 | 0.66 | 0.71 | 0.6 | 0.66 | 0.64 |
| 5 | 40647216 | rs12651829 | 0.982 | -1.143 | -0.668 | 0.495 | -0.008 | 0.008 | 0.008 | 0 | 0.23 | 0.27 | 0.12 | 0.36 | 0.14 | 0.21 |
| 5 | 40649263 | rs6893430 | 0.164 | -1.13 | -0.609 | 0.501 | -0.036 | 0.016 | 0.016 | 0 | 0.3 | 0.45 | 0.18 | 0.36 | 0.21 | 0.25 |
| 5 | 40668727 | rs6861121 | 0.941 | -1.097 | -0.677 | 0.578 | -0.01 | 0.011 | 0.011 | 0 | 0.23 | 0.26 | 0.12 | 0.36 | 0.14 | 0.21 |
| 5 | 40678049 | rs4957341 | 1.268 | -1.313 | -0.824 | 0.496 | -0.015 | 0.022 | 0.022 | 0 | 0.15 | 0.26 | 0.1 | 0.25 | 0.11 | 0.13 |
| 5 | 40679789 | rs10039983 | 0.249 | -1.314 | -1.083 | 0.353 | -0.063 | 0.016 | 0.016 | 0 | 0.3 | 0.18 | 0.52 | 0.24 | 0.45 | 0.39 |
| 5 | 40681254 | rs2228058 | 1.916 | -1.279 | -0.884 | 0.265 | -0.003 | -0.03 | -0.03 | 0 | 0.07 | 0.11 | 0.06 | 0 | 0.07 | 0.04 |
| 5 | 40689996 | rs10060234 | -1.236 | -1.645 | -1.146 | 0.293 | 0.004 | 0.01 | 0.01 | 0 | 0.63 | 0.68 | 0.77 | 0.8 | 0.66 | 0.73 |
| 5 | 40692940 | rs16870224 | 1.525 | -1.538 | -1.388 | 0.16 | -0.039 | -0.057 | -0.057 | 0.114 | 0.13 | 0.01 | 0.11 | 0.25 | 0.14 | 0.13 |
| 5 | 40716153 | rs10737963 | -2.897 | -1.665 | -1.094 | -0.036 | -0.003 | -0.029 | -0.029 | 0 | 0.78 | 0.79 | 0.84 | 1 | 0.71 | 0.91 |
| 5 | 40726496 | rs434778 | -1.21 | -1.621 | -1.051 | 0.079 | 0.005 | 0.012 | 0.012 | 0 | 0.63 | 0.69 | 0.77 | 0.8 | 0.66 | 0.73 |
| 5 | 40735926 | rs10941525 | 0.476 | -1.693 | -1.172 | 0.151 | -0.056 | 0.024 | 0.024 | 0 | 0.3 | 0.13 | 0.5 | 0.21 | 0.4 | 0.4 |
| 5 | 40753611 | rs12523329 | 0.74 | -1.268 | -1.429 | -0.043 | -0.039 | -0.019 | -0.019 | 0 | 0.25 | 0.06 | 0.43 | 0.2 | 0.24 | 0.36 |
| 5 | 40753877 | rs3792826 | 1.798 | -1.292 | -1.33 | 0.07 | 0.027 | -0.051 | -0.051 | 0.12 | 0.11 | 0.01 | 0.07 | 0.31 | 0.04 | 0.17 |
| 5 | 40765862 | rs6882903 | -1.894 | -1.043 | -1.209 | -0.428 | 0.007 | 0.036 | 0.036 | 0.044 | 0.69 | 0.43 | 0.65 | 0.86 | 0.69 | 0.18 |
| 5 | 40767397 | rs1002424 | 0.198 | -0.962 | -1.131 | -0.379 | -0.005 | 0.018 | 0.018 | 1.879 | 0.31 | 0.35 | 0.25 | 0.55 | 0.29 | 0.33 |
| 5 | 40770087 | rs17239241 | 1.965 | -0.87 | -1.358 | -0.515 | -0.038 | 0.033 | 0.033 | 0 | 0.06 | 0 | 0.03 | 0.01 | 0.1 | 0.06 |
| 5 | 40771216 | rs257009 | -1.483 | -0.89 | -1.253 | -0.482 | 0 | -0.001 | -0.001 | 0 | 0.63 | 0.63 | 0.76 | 0.79 | 0.69 | 0.73 |
| 5 | 40772190 | rs11745121 | 2.23 | -0.842 | -1.307 | -0.492 | -0.028 | 0.028 | 0.028 | 0 | 0.05 | 0.01 | 0.06 | 0.01 | 0.17 | 0.04 |
| 5 | 40781559 | rs12188129 | 2.23 | -0.842 | -1.303 | -0.492 | -0.014 | 0.02 | 0.02 | 0 | 0.05 | 0.02 | 0.06 | 0.01 | 0.16 | 0.04 |
| 5 | 40785811 | rs10053664 | -0.207 | -0.871 | -1.261 | -0.489 | -0.027 | 0.017 | 0.017 | 0 | 0.31 | 0.22 | 0.5 | 0.2 | 0.4 | 0.4 |

| | | | | | | | | | | | | | | | | |
|---|---|---|---|---|---|---|---|---|---|---|---|---|---|---|---|---|
| 5 | 40790551 | rs10074991 | 0.256 | -0.972 | -1.302 | -0.444 | -0.006 | 0.022 | 0.022 | 1.631 | 0.31 | 0.37 | 0.26 | 0.55 | 0.29 | 0.33 |
| 5 | 40798644 | rs10035235 | 0.296 | -0.825 | -1.319 | -0.35 | -0.032 | -0.013 | -0.013 | 0 | 0.25 | 0.06 | 0.43 | 0.2 | 0.24 | 0.36 |
| 5 | 40810426 | rs1001684 | 0.379 | -1.238 | -1.278 | -0.335 | -0.009 | 0.016 | 0.016 | 0.14 | 0.32 | 0.35 | 0.25 | 0.53 | 0.27 | 0.33 |
| 5 | 40821324 | rs462366 | -1.49 | -1.336 | -1.303 | -0.336 | 0.001 | 0.001 | 0.001 | 0 | 0.67 | 0.68 | 0.81 | 0.81 | 0.71 | 0.74 |
| 5 | 40827307 | rs7725810 | -0.215 | -1.362 | -1.338 | -0.344 | -0.033 | 0.012 | 0.012 | 0 | 0.32 | 0.23 | 0.52 | 0.25 | 0.42 | 0.4 |
| 5 | 40829311 | rs6876367 | 0.296 | -1.384 | -1.434 | -0.366 | -0.01 | 0.017 | 0.017 | 0.018 | 0.34 | 0.38 | 0.27 | 0.52 | 0.29 | 0.33 |
| 5 | 40830927 | rs1631660 | -2.652 | -1.014 | -1.739 | -0.226 | 0.024 | -0.041 | -0.041 | 0 | 0.93 | 1 | 1 | 0.9 | 0.98 | 0.87 |
| 5 | 40831808 | rs2291782 | -0.024 | -0.237 | -1.249 | 0.019 | -0.049 | -0.03 | -0.03 | 0 | 0.26 | 0.06 | 0.45 | 0.24 | 0.26 | 0.37 |
| 5 | 40839644 | rs1070444 | 1.416 | -0.149 | -0.887 | 0.017 | -0.007 | 0.005 | 0.005 | 0 | 0.06 | 0.06 | 0.07 | 0.05 | 0.16 | 0.03 |
| 5 | 40841741 | rs10512747 | 1.094 | -0.168 | -0.914 | -0.003 | -0.048 | 0.044 | 0.044 | 0.093 | 0.08 | 0.01 | 0.06 | 0.01 | 0.13 | 0.06 |
| 5 | 40841956 | rs173969 | -0.683 | -1.468 | -0.406 | 0.5 | 0.002 | -0.005 | -0.005 | 0.117 | 0.38 | 0.58 | 0.34 | 0.35 | 0.44 | 0.25 |
| 5 | 40847428 | rs369331 | -0.028 | -1.717 | -0.822 | 0.781 | -0.024 | -0.043 | -0.043 | 0 | 0.16 | 0.08 | 0.27 | 0.37 | 0.19 | 0.27 |
| 5 | 40848803 | rs323565 | -1.357 | -1.855 | -0.673 | 0.768 | 0.011 | 0.01 | 0.01 | 0 | 0.49 | 0.74 | 0.52 | 0.59 | 0.49 | 0.5 |
| 5 | 40853048 | rs16870407 | 1.034 | -1.724 | -0.665 | 0.658 | -0.015 | -0.058 | -0.058 | 0 | 0.09 | 0.03 | 0.2 | 0.33 | 0.08 | 0.24 |
| 5 | 40865863 | rs16870425 | 0.275 | -1.724 | -0.594 | 0.658 | -0.037 | -0.051 | -0.051 | 0.732 | 0.14 | 0.06 | 0.27 | 0.33 | 0.19 | 0.26 |
| 5 | 40871745 | rs323561 | -2.164 | -1.995 | -0.687 | 0.514 | -0.037 | 0.057 | 0.057 | 0 | 0.64 | 0.79 | 0.62 | 0.89 | 0.55 | 0.7 |
| 5 | 40872152 | rs7715597 | -0.348 | -1.701 | -0.397 | 0.603 | -0.015 | -0.023 | -0.023 | 0 | 0.33 | 0.42 | 0.22 | 0.2 | 0.3 | 0.23 |
| 5 | 40879839 | rs11746650 | 0.119 | -1.459 | -0.143 | 0.957 | -0.035 | -0.047 | -0.047 | 0 | 0.13 | 0.06 | 0.27 | 0.33 | 0.19 | 0.25 |
| 5 | 40881478 | rs11744923 | 0.157 | -1.474 | -0.191 | 0.885 | -0.01 | -0.016 | -0.016 | 4.132 | 0.23 | 0.44 | 0.2 | 0.2 | 0.23 | 0.19 |
| 5 | 40885080 | rs619491 | -0.347 | -1.362 | -0.102 | 0.961 | 0.053 | 0.12 | 0.12 | 0 | 0.15 | 0.32 | 0.3 | 0.38 | 0.19 | 0.26 |
| 5 | 40887781 | rs7717457 | -0.347 | -1.362 | -0.102 | 0.921 | 0.053 | 0.117 | 0.117 | 0 | 0.15 | 0.32 | 0.3 | 0.37 | 0.19 | 0.26 |
| 5 | 40904992 | rs11744119 | 0.018 | -1.428 | -0.216 | 0.832 | -0.012 | -0.019 | -0.019 | 0 | 0.25 | 0.44 | 0.2 | 0.2 | 0.24 | 0.21 |
| 5 | 40908901 | rs1376178 | -1.115 | -1.458 | -0.181 | 0.859 | -0.005 | -0.018 | -0.018 | 0.401 | 0.4 | 0.72 | 0.49 | 0.58 | 0.41 | 0.45 |
| 5 | 40911592 | rs1910016 | -1.977 | -1.578 | -0.278 | 0.919 | -0.064 | 0.122 | 0.122 | 0.735 | 0.62 | 0.85 | 0.63 | 0.9 | 0.51 | 0.7 |
| 5 | 40915165 | rs16870514 | 0.698 | -0.625 | 0.048 | 1.551 | 0.042 | -0.067 | -0.067 | 0.482 | 0.12 | 0.01 | 0.03 | 0.32 | 0.05 | 0.21 |
| 5 | 40929543 | rs2455307 | -0.963 | 0.599 | 0.831 | 1.637 | 0.033 | 0.079 | 0.079 | 0 | 0.15 | 0.28 | 0.3 | 0.34 | 0.19 | 0.24 |
| 5 | 40929815 | rs2455305 | -1.025 | 0.624 | 0.542 | 1.171 | -0.01 | -0.008 | -0.008 | 1.087 | 0.26 | 0.63 | 0.22 | 0.24 | 0.23 | 0.21 |
| 5 | 40933240 | rs324056 | -0.09 | 0.678 | 0.283 | 0.966 | 0.001 | 0.006 | 0.006 | 0 | 0.17 | 0.08 | 0.14 | 0.13 | 0.18 | 0.14 |
| 5 | 40934832 | rs10512751 | 0.091 | 0.649 | 0.226 | 0.991 | -0.03 | -0.021 | -0.021 | 0.228 | 0.17 | 0.01 | 0.14 | 0.14 | 0.19 | 0.15 |
| 5 | 40936669 | rs16870528 | 0.032 | 0.697 | 0.347 | 1.027 | 0.005 | 0.008 | 0.008 | 0 | 0.18 | 0.11 | 0.15 | 0.14 | 0.19 | 0.15 |
| 5 | 40941071 | rs2597738 | -1.833 | 0.766 | 0.235 | 1.089 | -0.003 | 0.002 | 0.002 | 0 | 0.39 | 0.63 | 0.46 | 0.38 | 0.42 | 0.46 |
| 5 | 40942420 | rs17175825 | 0.202 | 0.771 | 0.059 | 1.059 | -0.013 | -0.05 | -0.05 | 0.353 | 0.09 | 0.03 | 0.16 | 0.18 | 0.08 | 0.22 |
| 5 | 40943218 | rs16870537 | 0.173 | 0.77 | 0.161 | 1.076 | 0.062 | 0.117 | 0.117 | 0.187 | 0.07 | 0.34 | 0.05 | 0.11 | 0.04 | 0.07 |
| 5 | 40943442 | rs6875226 | -0.355 | 0.954 | -0.906 | 0.803 | -0.001 | -0.002 | -0.002 | 0.322 | 0.19 | 0.2 | 0.16 | 0.14 | 0.19 | 0.17 |
| 5 | 40949776 | rs1450657 | 1.07 | 0.765 | -0.91 | 0.854 | -0.001 | 0 | 0 | 0 | 0.67 | 0.7 | 0.59 | 0.68 | 0.62 | 0.61 |
| 5 | 40949997 | rs1450656 | 1.047 | 0.808 | -1.332 | 0.875 | -0.008 | 0 | 0 | 0 | 0.67 | 0.74 | 0.59 | 0.68 | 0.62 | 0.61 |
| 5 | 40953191 | rs2329431 | -0.823 | 0.57 | -1.331 | 1.138 | -0.013 | -0.045 | -0.045 | 0.029 | 0.09 | 0.03 | 0.15 | 0.17 | 0.08 | 0.22 |
| 5 | 40960278 | rs2675982 | -0.721 | 0.656 | -1.199 | 0.763 | -0.01 | -0.004 | -0.004 | 0 | 0.08 | 0 | 0.1 | 0.06 | 0.07 | 0.04 |
| 5 | 40961164 | rs1901167 | 0.086 | -0.364 | -1.309 | 0.839 | 0.014 | -0.009 | -0.009 | 0 | 0.36 | 0.35 | 0.16 | 0.41 | 0.25 | 0.39 |
| 5 | 40961894 | rs324058 | -1.011 | 1.001 | -0.993 | 0.54 | 0.037 | -0.015 | -0.015 | 0 | 0.87 | 0.81 | 0.77 | 0.92 | 0.74 | 0.95 |
| 5 | 40962547 | rs324059 | -1.011 | 1.001 | -0.993 | 0.54 | 0.037 | -0.015 | -0.015 | 0 | 0.87 | 0.81 | 0.77 | 0.92 | 0.74 | 0.95 |
| 5 | 40962863 | rs10941528 | 1.083 | 1.208 | -1.628 | 0.421 | -0.045 | -0.044 | -0.044 | 0 | 0.22 | 0.02 | 0.21 | 0.21 | 0.22 | 0.36 |
| 5 | 40962987 | rs324060 | 0.108 | 1.314 | -1.184 | 0.438 | 0 | -0.001 | -0.001 | 0 | 0.12 | 0.09 | 0.18 | 0.18 | 0.12 | 0.06 |
| 5 | 40964852 | rs13157656 | 1.1 | 1.308 | -1.305 | 0.428 | -0.043 | -0.042 | -0.042 | 0.352 | 0.22 | 0.03 | 0.21 | 0.21 | 0.22 | 0.36 |
| 5 | 40968030 | rs7712533 | 0.816 | 1.313 | -0.901 | 0.687 | -0.004 | 0.003 | 0.003 | 2.677 | 0.25 | 0.15 | 0.23 | 0.21 | 0.24 | 0.39 |
| 5 | 40974005 | rs3805226 | 0.277 | 1.284 | -0.916 | 0.73 | -0.026 | -0.045 | -0.045 | 0.22 | 0.12 | 0.02 | 0.17 | 0.19 | 0.12 | 0.07 |
| 5 | 40977760 | rs1078375 | -0.053 | 1.167 | -0.674 | 0.646 | 0 | -0.013 | -0.013 | 0 | 0.42 | 0.43 | 0.38 | 0.28 | 0.39 | 0.51 |
| 5 | 40980102 | rs2292727 | 1.2 | 1.132 | -0.678 | 0.738 | -0.007 | 0 | 0 | 0.011 | 0.35 | 0.37 | 0.16 | 0.3 | 0.24 | 0.36 |
| 5 | 40980585 | rs971075 | 0.035 | 1.156 | -0.701 | 0.752 | -0.022 | -0.001 | -0.001 | 0 | 0.77 | 0.82 | 0.59 | 0.74 | 0.63 | 0.88 |
| 5 | 40981103 | rs1450665 | 1.001 | 1.038 | -0.541 | 0.619 | -0.011 | 0 | 0 | 0 | 0.35 | 0.38 | 0.16 | 0.3 | 0.24 | 0.35 |
| 5 | 40982031 | rs10473230 | 1.13 | 1.103 | -0.704 | 0.6 | 0.015 | -0.054 | -0.054 | 4.349 | 0.16 | 0.26 | 0.15 | 0.07 | 0.16 | 0.13 |
| 5 | 40982622 | rs1061443 | 1.194 | 0.869 | -0.711 | 0.51 | 0.035 | -0.076 | -0.076 | 0.068 | 0.32 | 0.06 | 0.13 | 0.3 | 0.24 | 0.36 |
| 5 | 40982780 | rs8264 | 1.135 | 0.853 | -0.532 | 0.395 | 0.016 | -0.053 | -0.053 | 0 | 0.16 | 0.26 | 0.15 | 0.07 | 0.16 | 0.13 |
| 5 | 40985604 | rs4957361 | -0.104 | 0.478 | -0.432 | 0.504 | 0.004 | -0.004 | -0.004 | 0.349 | 0.4 | 0.65 | 0.46 | 0.47 | 0.36 | 0.51 |
| 5 | 40987374 | rs2122564 | -0.019 | 0.878 | -0.147 | 0.58 | 0.055 | -0.051 | -0.051 | 0 | 0.7 | 0.47 | 0.6 | 0.83 | 0.57 | 0.8 |
| 5 | 40988783 | rs12659224 | 0.472 | -0.334 | -0.892 | 0.677 | 0.03 | -0.043 | -0.043 | 0 | 0.23 | 0.11 | 0.1 | 0.27 | 0.18 | 0.29 |
| 5 | 40991344 | rs7736582 | -0.07 | -0.497 | -0.352 | 0.725 | 0.004 | -0.002 | -0.002 | 0 | 0.87 | 0.9 | 0.86 | 0.86 | 0.9 | 0.92 |
| 5 | 40994519 | rs6884830 | 0.591 | -0.693 | -0.641 | 0.496 | 0.011 | 0.005 | 0.005 | 0 | 0.37 | 0.46 | 0.44 | 0.45 | 0.39 | 0.31 |
| 5 | 40995847 | rs768682 | 0.507 | -0.901 | -1.045 | 0.445 | -0.028 | -0.057 | -0.057 | 0 | 0.23 | 0.09 | 0.15 | 0.28 | 0.25 | 0.24 |
| 5 | 40997996 | rs2271707 | 0.48 | -0.909 | -1.041 | 0.441 | -0.025 | -0.054 | -0.054 | 0.097 | 0.23 | 0.09 | 0.15 | 0.28 | 0.25 | 0.24 |
| 5 | 40998196 | rs704040 | -0.447 | -0.992 | -0.901 | 0.381 | 0 | 0.001 | 0.001 | 0 | 0.6 | 0.6 | 0.59 | 0.73 | 0.65 | 0.55 |
| 5 | 40998235 | rs722575 | 1.051 | -1.102 | -0.482 | 0.557 | -0.011 | 0.013 | 0.013 | 0 | 0.34 | 0.3 | 0.36 | 0.3 | 0.39 | 0.3 |
| 5 | 40998851 | rs2329435 | 0.031 | -1.104 | -0.796 | 0.568 | 0.001 | 0.017 | 0.017 | 1.461 | 0.51 | 0.42 | 0.46 | 0.57 | 0.51 | 0.61 |
| 5 | 40999419 | rs896117 | 1.322 | -1.101 | -0.787 | 0.566 | -0.008 | -0.025 | -0.025 | 2.524 | 0.27 | 0.33 | 0.27 | 0.14 | 0.26 | 0.37 |
| 5 | 41000343 | rs1001420 | -0.72 | -1.053 | -0.789 | 0.58 | -0.009 | -0.007 | -0.007 | 0 | 0.65 | 0.6 | 0.63 | 0.71 | 0.68 | 0.56 |
| 5 | 41001492 | rs7724505 | 0.531 | -0.977 | -0.698 | 0.592 | 0.006 | -0.029 | -0.029 | 0.128 | 0.41 | 0.51 | 0.43 | 0.29 | 0.43 | 0.29 |
| 5 | 41002758 | rs16870603 | 0.736 | -0.873 | -0.889 | 0.55 | -0.022 | -0.061 | -0.061 | 0.029 | 0.18 | 0.08 | 0.13 | 0.28 | 0.19 | 0.21 |
| 5 | 41006444 | rs13155300 | -0.995 | -0.888 | -0.744 | 0.555 | -0.004 | -0.019 | -0.019 | 0 | 0.83 | 0.88 | 0.88 | 0.72 | 0.83 | 0.81 |
| 5 | 41008650 | rs2271705 | -0.734 | -0.942 | -0.767 | 0.465 | -0.002 | -0.034 | -0.034 | 0 | 0.81 | 0.86 | 0.83 | 0.58 | 0.81 | 0.79 |

| | | | | | | | | | | | | | | | | |
|---|---|---|---|---|---|---|---|---|---|---|---|---|---|---|---|---|
| 5 | 41008780 | rs2271704 | -0.734 | -0.942 | -0.767 | 0.465 | -0.002 | -0.034 | -0.034 | 0 | 0.81 | 0.87 | 0.83 | 0.59 | 0.81 | 0.79 |
| 5 | 41009282 | rs10061334 | 0.585 | -1.255 | -0.778 | 0.531 | 0.001 | 0.002 | 0.002 | 0.423 | 0.36 | 0.34 | 0.39 | 0.24 | 0.35 | 0.43 |
| 5 | 41010431 | rs7706084 | 0.252 | -0.967 | -0.751 | 0.316 | 0.003 | -0.014 | -0.014 | 0 | 0.45 | 0.51 | 0.44 | 0.34 | 0.46 | 0.36 |
| 5 | 41017006 | rs13361985 | 0.691 | -0.729 | -0.905 | 0.492 | -0.026 | 0.021 | 0.021 | 0 | 0.37 | 0.25 | 0.37 | 0.34 | 0.43 | 0.34 |
| 5 | 41020018 | rs13154853 | 0.54 | -0.67 | -0.644 | 0.797 | -0.001 | 0 | 0 | 0.65 | 0.69 | 0.67 | 0.69 | 0.28 | 0.73 | 0.63 |
| 5 | 41032998 | rs325831 | 0.417 | -0.918 | -0.804 | 0.924 | -0.026 | 0.01 | 0.01 | 0 | 0.73 | 0.67 | 0.77 | 0.67 | 0.81 | 0.76 |
| 5 | 41037603 | rs1156442 | 1.116 | -0.787 | -0.505 | 0.989 | -0.008 | -0.008 | -0.008 | 0 | 0.58 | 0.47 | 0.63 | 0.36 | 0.64 | 0.57 |
| 5 | 41037965 | rs325836 | 0.953 | -0.188 | -0.51 | 1.032 | 0.006 | 0.019 | 0.019 | 0 | 0.14 | 0.17 | 0.13 | 0.28 | 0.17 | 0.19 |
| 5 | 41041963 | rs13157706 | 1.1 | -0.444 | -0.698 | 0.184 | -0.027 | 0.069 | 0.069 | 0 | 0.32 | 0.16 | 0.33 | 0.12 | 0.38 | 0.26 |
| 5 | 41042284 | rs325854 | -0.006 | -1.02 | -0.358 | 0.495 | -0.055 | 0.061 | 0.061 | 0 | 0.77 | 0.57 | 0.78 | 0.67 | 0.84 | 0.79 |
| 5 | 41045087 | rs325858 | -0.697 | -0.802 | -0.026 | 0.622 | -0.016 | 0.165 | 0.165 | 0 | 0.87 | 0.63 | 0.81 | 0.68 | 0.89 | 0.85 |
| 5 | 41045768 | rs325860 | -1.174 | -0.764 | -0.165 | 0.637 | -0.011 | 0.066 | 0.066 | 0 | 0.88 | 0.8 | 0.83 | 0.68 | 0.89 | 0.85 |
| 5 | 41049389 | rs325863 | -1.174 | -0.764 | -0.17 | 0.637 | -0.011 | 0.069 | 0.069 | 0 | 0.88 | 0.8 | 0.83 | 0.68 | 0.89 | 0.85 |
| 5 | 41052152 | rs325871 | -1.36 | -0.756 | -0.162 | 0.67 | -0.011 | 0.039 | 0.039 | 0 | 0.89 | 0.8 | 0.87 | 0.74 | 0.91 | 0.85 |
| 5 | 41054934 | rs325874 | 0.414 | -0.627 | -0.211 | 0.454 | -0.002 | 0 | 0 | 0.633 | 0.23 | 0.21 | 0.16 | 0.22 | 0.23 | 0.24 |
| 5 | 41056295 | rs6876819 | 0.777 | -0.511 | -0.284 | 0.282 | -0.009 | 0.069 | 0.069 | 0 | 0.42 | 0.3 | 0.37 | 0.15 | 0.4 | 0.28 |
| 5 | 41056370 | rs10454864 | 0.656 | -0.666 | -0.217 | 0.489 | 0.011 | -0.003 | -0.003 | 0 | 0.31 | 0.22 | 0.27 | 0.32 | 0.25 | 0.32 |
| 5 | 41057562 | rs618334 | -1.049 | -0.597 | 0.019 | 0.403 | -0.008 | 0.061 | 0.061 | 0.111 | 0.87 | 0.8 | 0.76 | 0.66 | 0.85 | 0.83 |
| 5 | 41061716 | rs865093 | -1.082 | -0.597 | 0.021 | 0.403 | -0.007 | 0.059 | 0.059 | 0.052 | 0.87 | 0.81 | 0.76 | 0.66 | 0.85 | 0.83 |
| 5 | 41061871 | rs587412 | 0.357 | -0.698 | 0.098 | 0.295 | 0.016 | 0.013 | 0.013 | 0 | 0.53 | 0.41 | 0.34 | 0.35 | 0.58 | 0.47 |
| 5 | 41069852 | rs13174484 | 1.509 | -0.725 | -0.002 | 0.24 | -0.025 | 0.034 | 0.034 | 0 | 0.3 | 0.23 | 0.22 | 0.14 | 0.37 | 0.21 |
| 5 | 41070247 | rs13158729 | 1.638 | -0.656 | 0.002 | 0.283 | -0.069 | 0.092 | 0.092 | 0 | 0.3 | 0.09 | 0.2 | 0.12 | 0.38 | 0.22 |
| 5 | 41070840 | rs1423391 | 0.866 | -0.641 | 0.065 | 0.295 | -0.018 | 0.042 | 0.042 | 0 | 0.38 | 0.3 | 0.25 | 0.14 | 0.43 | 0.27 |
| 5 | 41085635 | rs13173552 | 1.52 | -0.681 | 0.008 | 0.33 | 0.008 | -0.024 | -0.024 | 0 | 0.2 | 0.1 | 0.34 | 0.3 | 0.18 | 0.3 |
| 5 | 41085670 | rs178986 | 1.576 | -0.65 | 0.043 | 0.322 | -0.003 | -0.003 | -0.003 | 0 | 0.14 | 0.12 | 0.09 | 0.21 | 0.16 | 0.2 |
| 5 | 41092809 | rs6883572 | 0.84 | -0.67 | 0.105 | 0.224 | 0.004 | -0.031 | -0.031 | 0 | 0.41 | 0.46 | 0.26 | 0.14 | 0.43 | 0.27 |
| 5 | 41094409 | rs1423406 | -0.151 | -0.682 | 0.077 | 0.262 | 0.001 | 0.018 | 0.018 | 0 | 0.72 | 0.69 | 0.67 | 0.45 | 0.69 | 0.62 |
| 5 | 41103175 | rs10059016 | 1.467 | -0.585 | -0.266 | 0.263 | -0.002 | -0.019 | -0.019 | 0 | 0.12 | 0.12 | 0.06 | 0.01 | 0.08 | 0.05 |
| 5 | 41116225 | rs11958837 | 1.467 | -0.575 | -0.571 | 0.35 | -0.024 | -0.065 | -0.065 | 0 | 0.19 | 0.01 | 0.33 | 0.22 | 0.18 | 0.25 |
| 5 | 41120899 | rs325813 | 0.916 | -0.379 | -0.349 | 0.475 | -0.01 | -0.028 | -0.028 | 0 | 0.18 | 0.11 | 0.26 | 0.44 | 0.21 | 0.22 |
| 5 | 41142606 | rs9200 | -0.816 | -0.626 | -0.163 | 0.322 | 0.006 | 0.045 | 0.045 | 0.022 | 0.56 | 0.87 | 0.71 | 0.68 | 0.46 | 0.58 |
| 5 | 41143756 | rs1423414 | 0.24 | -0.455 | -0.013 | 0.374 | -0.023 | -0.012 | -0.012 | 0 | 0.33 | 0.44 | 0.42 | 0.31 | 0.26 | 0.35 |
| 5 | 41154150 | rs4957374 | -1.624 | -0.193 | -0.236 | 0.489 | -0.004 | -0.005 | -0.005 | 0 | 0.86 | 0.89 | 0.91 | 0.78 | 0.84 | 0.83 |
| 5 | 41154528 | rs2301247 | 0.61 | -0.307 | 0.045 | 0.273 | -0.003 | 0.001 | 0.001 | 0 | 0.41 | 0.43 | 0.26 | 0.15 | 0.44 | 0.3 |
| 5 | 41156008 | rs13158601 | 1.229 | -0.085 | 0.02 | 0.245 | -0.03 | 0.054 | 0.054 | 0 | 0.31 | 0.2 | 0.21 | 0.12 | 0.37 | 0.25 |
| 5 | 41157340 | rs10512764 | 0.638 | -0.34 | 0.141 | 0.5 | -0.005 | 0.043 | 0.043 | 0.383 | 0.13 | 0.15 | 0.24 | 0.35 | 0.14 | 0.17 |
| 5 | 41185607 | rs17201075 | 1.226 | 0.006 | -0.126 | 0 | -0.008 | 0.02 | 0.02 | 0 | 0.06 | 0.02 | 0.04 | 0 | 0.06 | 0.03 |
| 5 | 41190399 | rs7727839 | 0.44 | -0.096 | 0.262 | -0.081 | 0 | -0.02 | -0.02 | 0 | 0.42 | 0.44 | 0.26 | 0.12 | 0.47 | 0.29 |
| 5 | 41195700 | rs4245977 | 1.481 | -0.22 | 0.347 | 0.049 | -0.005 | 0.001 | 0.001 | 0 | 0.09 | 0.13 | 0.09 | 0.12 | 0.09 | 0.12 |
| 5 | 41196311 | rs6892389 | 0.773 | -0.44 | 0.405 | 0.073 | -0.009 | 0.005 | 0.005 | 0 | 0.16 | 0.18 | 0.28 | 0.16 | 0.13 | 0.11 |
| 5 | 41197075 | rs13168926 | 0.881 | -0.573 | 0.44 | 0.068 | 0.027 | 0.013 | 0.013 | 0 | 0.51 | 0.4 | 0.5 | 0.44 | 0.59 | 0.54 |
| 5 | 41199959 | rs1801033 | 0.372 | -0.567 | 0.418 | 0.074 | -0.014 | 0.007 | 0.007 | 0 | 0.6 | 0.55 | 0.55 | 0.44 | 0.65 | 0.58 |
| 5 | 41203344 | rs2305060 | 1.561 | -0.623 | 0.401 | 0.015 | -0.005 | 0.001 | 0.001 | 0 | 0.09 | 0.13 | 0.09 | 0.12 | 0.09 | 0.12 |
| 5 | 41215708 | rs16871009 | 1.289 | -0.633 | 0.364 | 0.003 | -0.01 | -0.031 | -0.031 | 0.315 | 0.09 | 0.15 | 0.05 | 0 | 0.06 | 0.04 |
| 5 | 41217415 | rs16871019 | 1.404 | -0.59 | 0.309 | 0.044 | -0.005 | 0.01 | 0.01 | 0 | 0.1 | 0.23 | 0.09 | 0.12 | 0.09 | 0.12 |
| 5 | 41219940 | rs10071904 | 0.683 | -0.368 | 0.23 | -0.028 | -0.004 | 0.003 | 0.003 | 0 | 0.17 | 0.19 | 0.32 | 0.16 | 0.16 | 0.13 |
| 5 | 41220990 | rs10058883 | 1.141 | -0.393 | 0.191 | -0.056 | -0.007 | -0.013 | -0.013 | 0 | 0.07 | 0.04 | 0.26 | 0.16 | 0.12 | 0.09 |
| 5 | 41223861 | rs2962288 | -0.205 | -0.416 | 0.239 | -0.005 | 0 | -0.004 | -0.004 | 0 | 0.69 | 0.71 | 0.6 | 0.59 | 0.71 | 0.7 |
| 5 | 41229277 | rs10064818 | 1.172 | -0.387 | 0.161 | 0.018 | -0.007 | -0.014 | -0.014 | 0 | 0.07 | 0.04 | 0.26 | 0.15 | 0.12 | 0.09 |
| 5 | 41231813 | rs4957381 | 0.453 | -0.376 | 0.186 | 0.006 | 0.028 | -0.013 | -0.013 | 0.107 | 0.58 | 0.27 | 0.54 | 0.56 | 0.65 | 0.66 |
| 5 | 41234403 | rs4957152 | 1.654 | -0.327 | -0.015 | -0.021 | 0.002 | -0.054 | -0.054 | 0.372 | 0.19 | 0.06 | 0.29 | 0.32 | 0.17 | 0.32 |
| 5 | 41234643 | rs10512781 | 0.795 | -0.318 | 0.036 | -0.039 | -0.089 | 0.091 | 0.091 | 0 | 0.39 | 0.11 | 0.25 | 0.23 | 0.48 | 0.33 |
| 5 | 41241610 | rs751138 | 2.197 | 0.194 | -0.09 | -0.213 | -0.001 | 0.001 | 0.001 | 0 | 0.05 | 0.05 | 0.06 | 0 | 0.14 | 0.01 |
| 5 | 41244885 | rs4957383 | 0.87 | -0.902 | 0.129 | 0.025 | -0.086 | 0.095 | 0.095 | 0 | 0.39 | 0.11 | 0.25 | 0.23 | 0.48 | 0.33 |
| 5 | 41247288 | rs2921184 | -2.18 | -0.809 | 0.095 | 0.152 | -0.005 | -0.03 | -0.03 | 0 | 0.86 | 0.93 | 0.92 | 0.75 | 0.87 | 0.82 |
| 5 | 41250433 | rs2004385 | 1.51 | -0.758 | -0.009 | 0.001 | 0.004 | -0.055 | -0.055 | 3.034 | 0.18 | 0.06 | 0.28 | 0.32 | 0.17 | 0.33 |
| 5 | 41253208 | rs2921170 | -0.128 | -0.552 | -0.143 | 0.191 | -0.008 | 0.034 | 0.034 | 0 | 0.68 | 0.59 | 0.6 | 0.55 | 0.71 | 0.7 |
| 5 | 41253853 | rs403624 | 1.561 | -0.494 | -0.728 | 0.179 | -0.004 | -0.034 | -0.034 | 0.208 | 0.08 | 0.01 | 0.08 | 0.13 | 0.09 | 0.12 |
| 5 | 41259942 | rs2910647 | 0.008 | -0.274 | -0.443 | 0.272 | -0.001 | 0.003 | 0.003 | 0 | 0.69 | 0.66 | 0.6 | 0.55 | 0.71 | 0.7 |
| 5 | 41262141 | rs368743 | 0.105 | -0.199 | -0.435 | -0.062 | 0.005 | 0.001 | 0.001 | 0 | 0.57 | 0.4 | 0.57 | 0.39 | 0.64 | 0.47 |
| 5 | 41262690 | rs440781 | -0.408 | -0.073 | -0.593 | -0.182 | 0.014 | 0.019 | 0.019 | 0 | 0.51 | 0.58 | 0.47 | 0.61 | 0.46 | 0.58 |
| 5 | 41263582 | rs399443 | 0.316 | -0.03 | -0.157 | -0.244 | 0.031 | -0.019 | -0.019 | 0 | 0.1 | 0.31 | 0.19 | 0.07 | 0.11 | 0.09 |
| 5 | 41263849 | rs380649 | -0.076 | -0.174 | -0.106 | -0.125 | 0.002 | 0.017 | 0.017 | 0.014 | 0.25 | 0.57 | 0.29 | 0.34 | 0.25 | 0.25 |
| 5 | 41264019 | rs2921178 | 1.382 | -0.149 | -0.351 | -0.113 | -0.001 | -0.009 | -0.009 | 0 | 0.15 | 0.12 | 0.09 | 0.26 | 0.15 | 0.17 |
| 5 | 41264243 | rs391781 | -1.045 | -0.283 | -0.901 | -0.243 | -0.004 | -0.055 | -0.055 | 0 | 0.48 | 0.89 | 0.63 | 0.5 | 0.43 | 0.35 |
| 5 | 41264563 | rs399760 | 0.026 | -0.64 | -0.507 | -0.104 | 0.006 | 0.023 | 0.023 | 0 | 0.25 | 0.44 | 0.28 | 0.33 | 0.25 | 0.25 |
| 5 | 41267218 | rs384267 | -0.576 | -0.421 | -1.323 | 0.098 | 0.023 | -0.087 | -0.087 | 0.146 | 0.73 | 0.92 | 0.78 | 0.63 | 0.76 | 0.56 |
| 5 | 41267328 | rs390035 | -0.516 | -0.229 | -0.544 | -0.203 | 0.004 | 0.006 | 0.006 | 1.609 | 0.35 | 0.71 | 0.34 | 0.34 | 0.3 | 0.28 |
| 5 | 41268036 | rs401035 | -0.076 | 0.363 | -0.503 | -0.071 | 0.039 | -0.029 | -0.029 | 0 | 0.11 | 0.19 | 0.21 | 0.08 | 0.19 | 0.1 |

| | | | | | | | | | | | | | | | | |
|---|---|---|---|---|---|---|---|---|---|---|---|---|---|---|---|---|
| 5 | 41269103 | rs454232 | 0.118 | 0 | -1.266 | 0.259 | 0 | -0.008 | -0.008 | 0 | 0.4 | 0.65 | 0.39 | 0.09 | 0.4 | 0.21 |
| 5 | 41271357 | rs452949 | -0.073 | 0.494 | -1.544 | 0.554 | 0.03 | 0.052 | 0.052 | 0.826 | 0.47 | 0.71 | 0.4 | 0.38 | 0.41 | 0.33 |
| 5 | 41273830 | rs373469 | -0.05 | 0.598 | -1.54 | 0.554 | 0.025 | 0.045 | 0.045 | 0.815 | 0.47 | 0.69 | 0.39 | 0.37 | 0.41 | 0.33 |
| 5 | 41278228 | rs12655804 | -0.447 | 0.731 | -1.922 | 1.071 | -0.047 | -0.108 | -0.108 | 0 | 0.24 | 0.01 | 0.3 | 0.3 | 0.22 | 0.32 |
| 5 | 41278513 | rs973152 | -0.286 | 0.822 | -0.801 | 0.644 | 0.021 | -0.035 | -0.035 | 0 | 0.17 | 0.55 | 0.34 | 0.16 | 0.17 | 0.07 |
| 5 | 41282364 | rs7711831 | -0.856 | 1.213 | -0.71 | 1.186 | -0.003 | -0.005 | -0.005 | 0.134 | 0.25 | 0.71 | 0.36 | 0.31 | 0.29 | 0.27 |
| 5 | 41283011 | rs16871167 | 0.318 | 1.132 | 0.482 | 1.103 | 0.021 | -0.08 | -0.08 | 0.139 | 0.13 | 0.24 | 0.08 | 0 | 0.15 | 0.03 |
| 5 | 41284862 | rs10078083 | -0.889 | 1.102 | 0.826 | 1.217 | -0.022 | 0.16 | 0.16 | 0 | 0.06 | 0.15 | 0.14 | 0.31 | 0.05 | 0.19 |
| 5 | 41288078 | rs11745204 | -1.427 | 1.115 | 0.798 | 1.209 | -0.003 | -0.006 | -0.006 | 0 | 0.25 | 0.71 | 0.36 | 0.32 | 0.29 | 0.24 |
| 5 | 41288697 | rs920908 | 0.454 | 1.129 | 0.275 | 1.319 | -0.026 | -0.059 | -0.059 | 0 | 0.19 | 0.01 | 0.16 | 0.18 | 0.15 | 0.29 |
| 5 | 41293360 | rs959808 | -1.213 | 1.027 | 1.171 | 1.439 | 0.009 | 0.346 | 0.346 | 0.172 | 0.07 | 0.35 | 0.16 | 0.31 | 0.05 | 0.19 |
| 5 | 41293370 | rs768763 | 1.098 | 0.711 | 0.123 | 0.957 | -0.035 | -0.053 | -0.053 | 0.64 | 0.21 | 0.03 | 0.11 | 0.22 | 0.2 | 0.2 |
| 5 | 41298726 | rs11749734 | -0.416 | 0.276 | 0.501 | 0.992 | -0.02 | 0.199 | 0.199 | 0 | 0.07 | 0.19 | 0.14 | 0.3 | 0.05 | 0.18 |
| 5 | 41301008 | rs16871201 | -0.324 | 0.302 | 0.306 | 1.014 | 0.009 | 0.021 | 0.021 | 0 | 0.06 | 0.01 | 0.13 | 0.36 | 0.05 | 0.21 |
| 5 | 41301337 | rs11739338 | -0.208 | 0.293 | 0.609 | 0.749 | -0.008 | 0.062 | 0.062 | 0.387 | 0.2 | 0.59 | 0.23 | 0.38 | 0.18 | 0.19 |
| 5 | 41302815 | rs320614 | -1.174 | 0.288 | 0.059 | 0.333 | -0.081 | -0.059 | -0.059 | 0 | 0.59 | 0.98 | 0.72 | 0.66 | 0.65 | 0.52 |
| 5 | 41310735 | rs7701760 | -0.185 | 0 | 0.925 | 0.876 | -0.004 | 0.1 | 0.1 | 0 | 0.05 | 0.07 | 0.13 | 0.37 | 0.04 | 0.17 |
| 5 | 41310957 | rs1373639 | -1.291 | -0.009 | 1.19 | 0.872 | 0.032 | 0.503 | 0.503 | 0.238 | 0.08 | 0.45 | 0.16 | 0.38 | 0.04 | 0.17 |
| 5 | 41311975 | rs11744499 | -0.674 | -0.006 | 0.763 | 0.873 | -0.019 | 0.15 | 0.15 | 0 | 0.06 | 0.15 | 0.14 | 0.37 | 0.04 | 0.17 |
| 5 | 41312164 | rs11737958 | -0.192 | 0.022 | 0.754 | 0.886 | -0.004 | 0.1 | 0.1 | 0 | 0.05 | 0.07 | 0.13 | 0.37 | 0.04 | 0.17 |
| 5 | 41315243 | rs11740787 | -0.31 | 0 | 0.76 | 0.886 | -0.017 | 0.16 | 0.16 | 0 | 0.06 | 0.15 | 0.14 | 0.37 | 0.04 | 0.17 |
| 5 | 41324783 | rs1904178 | -0.25 | 0.001 | 0.71 | 0.849 | -0.011 | 0.122 | 0.122 | 0 | 0.05 | 0.11 | 0.14 | 0.37 | 0.04 | 0.17 |
| 5 | 41336050 | rs6872806 | -0.504 | -0.04 | 0.798 | 0.776 | -0.008 | 0.034 | 0.034 | 0.481 | 0.21 | 0.66 | 0.24 | 0.38 | 0.19 | 0.2 |
| 5 | 41340045 | rs2329601 | -0.075 | -0.01 | 0.455 | 0.792 | 0.005 | 0.069 | 0.069 | 0 | 0.05 | 0.05 | 0.13 | 0.37 | 0.04 | 0.17 |
| 5 | 41345349 | rs2197422 | -0.322 | 0.025 | 0.533 | 0.803 | -0.001 | 0.119 | 0.119 | 0.03 | 0.2 | 0.49 | 0.22 | 0.38 | 0.19 | 0.2 |
| 5 | 41347002 | rs13179463 | 0.943 | 0.12 | 0.002 | 0.494 | -0.013 | -0.036 | -0.036 | 0.056 | 0.14 | 0 | 0.1 | 0.1 | 0.09 | 0.08 |
| 5 | 41363888 | rs319003 | -0.112 | 0.39 | 0.466 | 0.513 | -0.011 | 0.004 | 0.004 | 0 | 0.1 | 0.08 | 0.07 | 0.01 | 0.17 | 0.07 |
| 5 | 41365183 | rs318838 | 0.182 | 0.581 | 0.656 | 1.048 | 0.015 | 0.062 | 0.062 | 0 | 0.95 | 0.97 | 0.84 | 0.54 | 0.95 | 0.79 |
| 5 | 41373205 | rs318843 | 1.01 | 0.537 | 1.011 | 0.896 | 0.033 | 0.104 | 0.104 | 0 | 0.77 | 0.49 | 0.6 | 0.54 | 0.7 | 0.73 |
| 5 | 41374770 | rs318845 | 1.086 | 0.429 | 0.984 | 0.802 | -0.008 | 0.056 | 0.056 | 0.112 | 0.75 | 0.62 | 0.58 | 0.55 | 0.74 | 0.72 |
| 5 | 41382681 | rs319013 | 1.439 | 0.447 | 0.509 | 0.803 | 0.016 | 0.044 | 0.044 | 0 | 0.71 | 0.46 | 0.53 | 0.52 | 0.65 | 0.69 |
| 5 | 41385427 | rs167628 | -0.073 | 0.44 | 0.485 | 0.8 | 0.014 | 0.064 | 0.064 | 0 | 0.95 | 0.97 | 0.84 | 0.56 | 0.95 | 0.8 |
| 5 | 41386225 | rs1380996 | 1.46 | 0.421 | 0.677 | 0.799 | -0.009 | -0.032 | -0.032 | 0.116 | 0.69 | 0.2 | 0.51 | 0.52 | 0.64 | 0.69 |
| 5 | 41405404 | rs4380707 | 0.143 | 0.631 | 0.151 | 0.903 | 0.006 | 0.012 | 0.012 | 0 | 0.89 | 0.91 | 0.75 | 0.55 | 0.89 | 0.76 |
| 5 | 41408040 | rs672658 | 0.195 | 1.098 | 0.395 | 0.205 | -0.007 | 0.018 | 0.018 | 0 | 0.12 | 0.06 | 0.15 | 0.01 | 0.14 | 0.05 |
| 5 | 41409846 | rs6451574 | -0.832 | 1.11 | 0.395 | 0.213 | 0.054 | -0.115 | -0.115 | 0.346 | 0.18 | 0.39 | 0.22 | 0.03 | 0.25 | 0.07 |
| 5 | 41415656 | rs10512771 | -0.484 | 0.981 | -0.008 | 0.79 | 0.007 | -0.055 | -0.055 | 0.027 | 0.12 | 0.01 | 0.11 | 0.19 | 0.08 | 0.18 |
| 5 | 41417693 | rs897807 | -1.079 | 1.466 | 0.611 | 0.56 | 0.138 | 0.01 | 0.01 | 0 | 0.06 | 0.32 | 0.06 | 0.01 | 0.11 | 0.03 |
| 5 | 41435715 | rs897808 | -0.588 | 1.654 | -0.189 | 0.867 | -0.002 | -0.001 | -0.001 | 0.224 | 0.22 | 0.22 | 0.23 | 0.25 | 0.26 | 0.27 |
| 5 | 41435950 | rs897809 | 0.134 | 1.059 | -0.867 | 0.641 | 0.001 | 0.004 | 0.004 | 0.002 | 0.69 | 0.31 | 0.78 | 0.69 | 0.74 | 0.68 |
| 5 | 41438380 | rs1457128 | 0.134 | 1.059 | -0.89 | 0.642 | 0.003 | 0.01 | 0.01 | 0.002 | 0.69 | 0.29 | 0.78 | 0.69 | 0.74 | 0.68 |
| 5 | 41439239 | rs318060 | 0.099 | 0.917 | -0.164 | 0.695 | -0.009 | 0.052 | 0.052 | 0 | 0.89 | 0.77 | 0.96 | 0.71 | 0.93 | 0.75 |
| 5 | 41448187 | rs318074 | 0.914 | 0.966 | -0.042 | 0.818 | -0.004 | 0.024 | 0.024 | 0 | 0.86 | 0.79 | 0.9 | 0.73 | 0.89 | 0.76 |
| 5 | 41458237 | rs318081 | 0.022 | 0.887 | 0.19 | 0.345 | -0.003 | -0.007 | -0.007 | 4.909 | 0.2 | 0.14 | 0.11 | 0 | 0.22 | 0.11 |
| 5 | 41461984 | rs1379310 | 1.237 | 0.86 | 0.668 | 0.36 | 0.003 | 0.009 | 0.009 | 0 | 0.64 | 0.32 | 0.76 | 0.71 | 0.69 | 0.64 |
| 5 | 41462456 | rs1457132 | 1.229 | 0.851 | 1.042 | 0.36 | -0.004 | -0.013 | -0.013 | 0 | 0.66 | 0.43 | 0.78 | 0.73 | 0.69 | 0.65 |
| 5 | 41474732 | rs318041 | -1.737 | 1.09 | 0.606 | 0.197 | -0.016 | 0.077 | 0.077 | 0.513 | 0.52 | 0.86 | 0.47 | 0.32 | 0.53 | 0.58 |
| 5 | 41476347 | rs7714031 | 0.247 | 1.207 | 0.836 | 0.195 | 0.001 | -0.001 | -0.001 | 0 | 0.08 | 0.09 | 0.02 | 0.08 | 0.05 | 0.2 |
| 5 | 41485573 | rs4588635 | -0.707 | 0.728 | 0.947 | 0.288 | -0.004 | 0.032 | 0.032 | 0 | 0.31 | 0.44 | 0.28 | 0.35 | 0.27 | 0.45 |
| 5 | 41485610 | rs11746987 | 0.513 | 0.314 | 0.555 | -0.001 | -0.008 | 0.049 | 0.049 | 0 | 0.21 | 0.1 | 0.12 | 0 | 0.22 | 0.11 |
| 5 | 41504813 | rs662066 | -0.214 | 0.259 | 0.838 | 0.094 | -0.018 | -0.004 | -0.004 | 0 | 0.11 | 0.27 | 0.04 | 0.09 | 0.06 | 0.2 |
| 5 | 41508421 | rs611647 | -0.856 | -0.459 | 0.249 | -0.002 | 0.011 | -0.019 | -0.019 | 0 | 0.84 | 0.87 | 0.86 | 0.44 | 0.92 | 0.77 |
| 5 | 41509015 | rs624733 | -0.087 | -0.217 | 1.02 | -0.003 | -0.004 | 0.016 | 0.016 | 0 | 0.11 | 0.39 | 0.05 | 0.09 | 0.06 | 0.2 |
| 5 | 41513971 | rs4957155 | 0.747 | -0.251 | 0.858 | -0.133 | -0.004 | 0.041 | 0.041 | 0 | 0.21 | 0.12 | 0.12 | 0 | 0.22 | 0.12 |
| 5 | 41523175 | rs16871449 | 0.849 | 0.118 | 0.551 | -0.187 | -0.005 | 0.009 | 0.009 | 0 | 0.19 | 0.17 | 0.2 | 0.26 | 0.15 | 0.23 |
| 5 | 41529448 | rs2644046 | 0.173 | 0.014 | 0.622 | -0.202 | -0.017 | -0.003 | -0.003 | 0 | 0.11 | 0.25 | 0.04 | 0.09 | 0.06 | 0.2 |
| 5 | 41538044 | rs4957158 | 0.951 | 0.271 | -0.067 | -0.093 | -0.052 | 0.023 | 0.023 | 0 | 0.46 | 0.17 | 0.58 | 0.64 | 0.5 | 0.44 |
| 5 | 41541583 | rs7703079 | 0.202 | 0.348 | -0.189 | 0.279 | 0.001 | 0.004 | 0.004 | 0 | 0.2 | 0.18 | 0.23 | 0.26 | 0.22 | 0.23 |
| 5 | 41546004 | rs17274014 | 0.875 | 0.616 | -0.77 | -0.251 | -0.048 | 0.094 | 0.094 | 0.002 | 0.16 | 0.02 | 0.09 | 0 | 0.17 | 0.07 |
| 5 | 41560079 | rs3913723 | -0.319 | -0.304 | -0.305 | 0.285 | -0.034 | 0.121 | 0.121 | 0 | 0.13 | 0.21 | 0.06 | 0.46 | 0.05 | 0.31 |
| 5 | 41561373 | rs625487 | 0.931 | -0.858 | -1.196 | -0.307 | 0.028 | -0.049 | -0.049 | 0 | 0.45 | 0.2 | 0.66 | 0.47 | 0.64 | 0.49 |
| 5 | 41564562 | rs687174 | -0.16 | -0.926 | -0.335 | -0.503 | 0.013 | 0.012 | 0.012 | 0 | 0.4 | 0.34 | 0.26 | 0.02 | 0.31 | 0.2 |
| 5 | 41572722 | rs589757 | 0.36 | -1.088 | -0.167 | -0.633 | 0.002 | -0.012 | -0.012 | 1.59 | 0.28 | 0.25 | 0.18 | 0.01 | 0.23 | 0.14 |
| 5 | 41574343 | rs669684 | -2.193 | -1.042 | -0.662 | -0.706 | -0.082 | -0.01 | -0.01 | 0 | 0.72 | 0.84 | 0.62 | 0.72 | 0.54 | 0.77 |
| 5 | 41579054 | rs6864343 | 1.622 | -1.063 | -0.461 | -0.697 | 0.013 | 0.048 | 0.048 | 0 | 0.12 | 0.07 | 0.07 | 0 | 0.08 | 0.06 |
| 5 | 41586405 | rs583442 | -0.818 | -1.018 | -0.258 | -0.599 | 0.005 | 0.003 | 0.003 | 0 | 0.53 | 0.58 | 0.31 | 0.48 | 0.36 | 0.51 |
| 5 | 41613672 | rs16871652 | 1.407 | -0.762 | -0.241 | -0.572 | 0.025 | -0.02 | -0.02 | 0.202 | 0.07 | 0.18 | 0.09 | 0.01 | 0.08 | 0.07 |
| 5 | 41616226 | rs4277955 | 2.089 | -0.867 | -0.521 | -0.777 | -0.029 | 0.092 | 0.092 | 0.067 | 0.16 | 0.01 | 0.08 | 0 | 0.15 | 0.06 |
| 5 | 41617256 | rs17276531 | 2.089 | -0.867 | -0.532 | -0.79 | -0.033 | 0.097 | 0.097 | 0.067 | 0.16 | 0.01 | 0.09 | 0 | 0.15 | 0.06 |

| | | | | | | | | | | | | | | | | |
|---|---|---|---|---|---|---|---|---|---|---|---|---|---|---|---|---|
| 5 | 41619040 | rs10512791 | -0.189 | -1.041 | -0.464 | -0.815 | -0.01 | -0.061 | -0.061 | 0 | 0.24 | 0.32 | 0.17 | 0.02 | 0.16 | 0.14 |
| 5 | 41620475 | rs7702075 | 1.654 | -1.055 | -0.486 | -0.82 | 0.013 | 0.051 | 0.051 | 0 | 0.12 | 0.07 | 0.07 | 0.02 | 0.08 | 0.06 |
| 5 | 41623306 | rs3933957 | 0.124 | -0.878 | -0.269 | -0.59 | -0.005 | -0.052 | -0.052 | 0.146 | 0.12 | 0.25 | 0.1 | 0.01 | 0.08 | 0.07 |
| 5 | 41625892 | rs3851448 | 0.176 | -0.897 | -0.415 | -0.228 | -0.018 | 0.056 | 0.056 | 0 | 0.13 | 0.14 | 0.05 | 0.46 | 0.05 | 0.31 |
| 5 | 41629963 | rs316720 | -1.566 | -0.697 | -0.785 | -0.545 | -0.065 | -0.005 | -0.005 | 0 | 0.71 | 0.81 | 0.62 | 0.73 | 0.56 | 0.77 |
| 5 | 41637293 | rs13186086 | 0.581 | -1.128 | -0.898 | -0.96 | 0.02 | 0.154 | 0.154 | 0 | 0.3 | 0.12 | 0.15 | 0 | 0.23 | 0.14 |
| 5 | 41645551 | rs316738 | -0.13 | -1.482 | -0.595 | -1.075 | 0.042 | 0.064 | 0.064 | 0 | 0.4 | 0.75 | 0.36 | 0.03 | 0.29 | 0.27 |
| 5 | 41648335 | rs675785 | 1.514 | -1.336 | -0.061 | -0.889 | 0.018 | -0.025 | -0.025 | 0.315 | 0.06 | 0.21 | 0.05 | 0 | 0.05 | 0.06 |
| 5 | 41669628 | rs316717 | 0.909 | -1.328 | 0.344 | -0.884 | 0.016 | -0.021 | -0.021 | 0 | 0.07 | 0.25 | 0.17 | 0.03 | 0.05 | 0.13 |
| 5 | 41676293 | rs316753 | -1.379 | -0.804 | -0.205 | -0.919 | -0.011 | -0.043 | -0.043 | 0.664 | 0.78 | 0.87 | 0.78 | 0.33 | 0.74 | 0.73 |
| 5 | 41684432 | rs1423240 | 1.198 | -0.958 | -0.514 | -1.204 | -0.143 | -0.072 | -0.072 | 0 | 0.27 | 0.04 | 0.34 | 0.25 | 0.43 | 0.29 |
| 5 | 41685344 | rs624097 | 0.418 | -1.042 | -0.146 | -1.361 | 0.038 | 0.05 | 0.05 | 0 | 0.42 | 0.71 | 0.37 | 0.03 | 0.31 | 0.28 |
| 5 | 41698513 | rs643505 | 1.611 | -0.827 | 0.107 | -1.053 | 0.009 | -0.021 | -0.021 | 0 | 0.07 | 0.22 | 0.17 | 0.04 | 0.05 | 0.13 |
| 5 | 41699025 | rs631069 | 1.611 | -0.827 | 0.107 | -1.063 | 0.009 | -0.026 | -0.026 | 0 | 0.07 | 0.22 | 0.17 | 0.03 | 0.05 | 0.13 |
| 5 | 41719678 | rs316725 | 0.984 | -0.855 | 0.158 | -1.08 | 0.033 | -0.023 | -0.023 | 0 | 0.08 | 0.51 | 0.08 | 0 | 0.05 | 0.06 |
| 5 | 41751702 | rs10512793 | 0.288 | -0.822 | -0.027 | -1.18 | -0.001 | 0.003 | 0.003 | 0 | 0.15 | 0.09 | 0.13 | 0.05 | 0.12 | 0.2 |
| 5 | 41786266 | rs1027348 | 1.767 | -0.836 | 0.181 | -1.075 | 0.071 | 0.011 | 0.011 | 0 | 0.07 | 0.38 | 0.07 | 0 | 0.05 | 0.06 |
| 5 | 41802771 | rs12186512 | 0.744 | -1.048 | -0.14 | -1.307 | 0.002 | 0.204 | 0.204 | 0 | 0.39 | 0.14 | 0.2 | 0.04 | 0.33 | 0.3 |
| 5 | 41842746 | rs13164201 | 1.095 | -1.511 | 0.263 | -0.982 | 0.018 | 0.167 | 0.167 | 0 | 0.31 | 0.12 | 0.16 | 0 | 0.25 | 0.15 |
| 5 | 41851458 | rs6881873 | 0.751 | -1.495 | 0.335 | -0.972 | 0.015 | -0.008 | -0.008 | 0.186 | 0.34 | 0.29 | 0.29 | 0.03 | 0.25 | 0.21 |
| 5 | 41853514 | rs7723992 | 1.114 | -1.152 | 0.484 | -0.611 | 0.056 | -0.003 | -0.003 | 0.021 | 0.07 | 0.38 | 0.08 | 0 | 0.05 | 0.07 |
| 5 | 41895460 | rs7713580 | 1.497 | -1.344 | -0.203 | -1.162 | 0.028 | 0.077 | 0.077 | 0 | 0.16 | 0.06 | 0.07 | 0 | 0.1 | 0.08 |
| 5 | 41906206 | rs10059424 | 0.62 | -1.114 | 0.547 | -0.792 | 0.021 | -0.021 | -0.021 | 0.219 | 0.08 | 0.35 | 0.17 | 0.03 | 0.05 | 0.08 |
| 5 | 41907381 | rs10076689 | 1.529 | -1.116 | 0.603 | -0.793 | 0.003 | -0.034 | -0.034 | 0 | 0.06 | 0.16 | 0.04 | 0 | 0.05 | 0.08 |
| 5 | 41907628 | rs277410 | -1.207 | -1.214 | 0.461 | -0.744 | 0.005 | -0.035 | -0.035 | 0 | 0.83 | 0.88 | 0.77 | 0.35 | 0.83 | 0.75 |
| 5 | 41916688 | rs10062703 | 0.326 | -1.09 | 1.037 | -1.059 | 0.021 | -0.021 | -0.021 | 0.219 | 0.08 | 0.35 | 0.17 | 0.03 | 0.05 | 0.08 |
| 5 | 41918457 | rs10077481 | 0.326 | -1.09 | 1.042 | -1.059 | 0.022 | -0.02 | -0.02 | 0 | 0.08 | 0.35 | 0.17 | 0.03 | 0.05 | 0.08 |
| 5 | 41918947 | rs10064380 | 1.161 | -1.09 | 1.034 | -1.059 | 0.009 | -0.023 | -0.023 | 0 | 0.06 | 0.2 | 0.16 | 0.03 | 0.05 | 0.08 |
| 5 | 41924949 | rs2231910 | 0.381 | -1.086 | 0.966 | -1.057 | 0.021 | -0.021 | -0.021 | 0 | 0.08 | 0.35 | 0.17 | 0.03 | 0.05 | 0.08 |
| 5 | 41927423 | rs277414 | 0.017 | -1.128 | 0.782 | -1.133 | 0.039 | 0.035 | 0.035 | 0.021 | 0.42 | 0.74 | 0.37 | 0.03 | 0.29 | 0.2 |
| 5 | 41943776 | rs10045473 | 0.461 | -0.675 | 0.821 | -0.459 | 0.007 | -0.028 | -0.028 | 0.791 | 0.07 | 0.26 | 0.17 | 0.03 | 0.05 | 0.08 |
| 5 | 42019110 | rs2011105 | 0.362 | -0.685 | 0.783 | -0.5 | 0.003 | -0.056 | -0.056 | 0 | 0.1 | 0.27 | 0.06 | 0 | 0.06 | 0.08 |
| 5 | 42039712 | rs202299 | -0.06 | -1.174 | 0.617 | -0.944 | 0.039 | 0.093 | 0.093 | 0 | 0.43 | 0.73 | 0.4 | 0.12 | 0.3 | 0.21 |
| 5 | 42044673 | rs275036 | 0.949 | -1.252 | 0.894 | -1.001 | 0.024 | 0.103 | 0.103 | 0.567 | 0.32 | 0.17 | 0.18 | 0 | 0.23 | 0.11 |
| 5 | 42046381 | rs17619781 | 0.004 | -1.43 | 0.59 | -0.784 | 0.002 | -0.005 | -0.005 | 0 | 0.23 | 0.18 | 0.2 | 0.32 | 0.21 | 0.29 |
| 5 | 42073476 | rs276269 | -0.214 | -1.194 | 0.613 | -1.118 | 0.04 | 0.109 | 0.109 | 0 | 0.57 | 0.26 | 0.56 | 0.88 | 0.7 | 0.79 |
| 5 | 42084883 | rs687238 | -0.206 | -1.197 | 0.611 | -1.121 | 0.041 | 0.111 | 0.111 | 0 | 0.57 | 0.26 | 0.56 | 0.91 | 0.7 | 0.79 |
| 5 | 42120061 | rs9292840 | -1.235 | -1.17 | 0.861 | -1.115 | 0.006 | -0.039 | -0.039 | 0 | 0.68 | 0.7 | 0.8 | 1 | 0.76 | 0.88 |
| 5 | 42147328 | rs2134101 | -1.291 | -1.116 | 1.248 | -1.008 | 0.007 | 0.066 | 0.066 | 0.177 | 0.65 | 0.78 | 0.78 | 1 | 0.7 | 0.89 |
| 5 | 42148371 | rs1079193 | 0.878 | -1.143 | 0.987 | -0.772 | -0.032 | -0.088 | -0.088 | 0.096 | 0.13 | 0.01 | 0.11 | 0.3 | 0.17 | 0.22 |
| 5 | 42152335 | rs666581 | 1.095 | -0.855 | 1.322 | -0.795 | 0 | -0.046 | -0.046 | 0 | 0.08 | 0.16 | 0.05 | 0 | 0.06 | 0.07 |
| 5 | 42204561 | rs10941565 | -0.105 | -0.846 | 1.379 | -0.823 | 0.034 | 0.164 | 0.164 | 0 | 0.52 | 0.27 | 0.53 | 0.86 | 0.63 | 0.79 |
| 5 | 42216965 | rs6887597 | 0.941 | -0.984 | 1.196 | -0.82 | -0.077 | -0.073 | -0.073 | 0 | 0.32 | 0.16 | 0.38 | 0.58 | 0.42 | 0.52 |
| 5 | 42220979 | rs7710353 | 1.382 | -0.901 | 1.183 | -0.809 | -0.001 | -0.021 | -0.021 | 0 | 0.08 | 0.09 | 0.04 | 0 | 0.06 | 0.07 |
| 5 | 42231896 | rs4245983 | 1.246 | -0.907 | 1.237 | -0.811 | -0.002 | -0.041 | -0.041 | 0 | 0.08 | 0.15 | 0.05 | 0 | 0.06 | 0.07 |
| 5 | 42243481 | rs11959168 | 0.789 | -0.803 | 1.183 | -0.675 | -0.074 | -0.063 | -0.063 | 2.016 | 0.33 | 0.14 | 0.38 | 0.58 | 0.43 | 0.51 |
| 5 | 42278624 | rs13189999 | 0.101 | -0.62 | 1.256 | -0.937 | 0.031 | 0.095 | 0.095 | 0 | 0.5 | 0.31 | 0.53 | 0.81 | 0.63 | 0.77 |
| 5 | 42300305 | rs10041272 | 0.058 | -0.473 | 0.553 | -0.856 | -0.001 | -0.024 | -0.024 | 0.13 | 0.51 | 0.45 | 0.54 | 0.8 | 0.63 | 0.75 |
| 5 | 42307714 | rs10462034 | 0.097 | -0.417 | 0.356 | -0.81 | 0.019 | 0.05 | 0.05 | 0 | 0.51 | 0.37 | 0.53 | 0.8 | 0.63 | 0.75 |
| 5 | 42315700 | rs10064948 | 0.366 | -0.293 | 0.507 | -0.638 | 0 | -0.052 | -0.052 | 0 | 0.09 | 0.22 | 0.06 | 0 | 0.07 | 0.07 |
| 5 | 42381962 | rs13153060 | 0.974 | -0.331 | 0.052 | -1.205 | -0.022 | 0.025 | 0.025 | 0.884 | 0.19 | 0.11 | 0.12 | 0 | 0.22 | 0.04 |
| 5 | 42385855 | rs10079480 | 0.567 | -0.341 | 0.136 | -1.182 | -0.008 | -0.007 | -0.007 | 0 | 0.37 | 0.28 | 0.44 | 0.55 | 0.4 | 0.57 |
| 5 | 42390038 | rs2972394 | -0.246 | -0.3 | 0.427 | -1.217 | 0.009 | 0.07 | 0.07 | 0 | 0.53 | 0.6 | 0.58 | 0.86 | 0.59 | 0.76 |
| 5 | 42396541 | rs7379192 | -1.067 | -0.304 | 0.25 | -1.183 | 0.04 | 0.092 | 0.092 | 0.617 | 0.73 | 0.86 | 0.84 | 1 | 0.81 | 0.86 |
| 5 | 42410060 | rs10473282 | 1.395 | -0.426 | 0.171 | -1.248 | -0.033 | -0.055 | -0.055 | 0 | 0.26 | 0.13 | 0.38 | 0.56 | 0.33 | 0.52 |
| 5 | 42422389 | rs2940927 | 0.756 | -0.454 | 0.345 | -1.131 | -0.021 | 0.08 | 0.08 | 0 | 0.43 | 0.29 | 0.51 | 0.86 | 0.52 | 0.69 |
| 5 | 42422855 | rs1876790 | -0.548 | -0.511 | 0.262 | -1.035 | -0.01 | -0.023 | -0.023 | 0 | 0.72 | 0.63 | 0.69 | 0.86 | 0.81 | 0.8 |
| 5 | 42455956 | rs2972407 | 0.461 | -0.507 | 0.178 | -0.992 | 0.016 | 0.02 | 0.02 | 0.301 | 0.28 | 0.29 | 0.3 | 0.14 | 0.18 | 0.21 |
| 5 | 42467455 | rs2940917 | 0.457 | -0.513 | 0.202 | -0.995 | 0.016 | 0.02 | 0.02 | 0.32 | 0.28 | 0.29 | 0.3 | 0.14 | 0.18 | 0.21 |
| 5 | 42473555 | rs13188386 | -0.117 | -0.415 | 0.091 | -1.855 | 0.015 | -0.121 | -0.121 | 0 | 0.29 | 0.35 | 0.18 | 0 | 0.31 | 0.13 |
| 5 | 42474696 | rs11744988 | 0.861 | -0.43 | 0.017 | -1.424 | -0.003 | -0.047 | -0.047 | 0 | 0.08 | 0.15 | 0.05 | 0 | 0.07 | 0.07 |
| 5 | 42475327 | rs7728337 | 0.166 | -0.322 | -0.575 | -1.174 | 0.02 | 0.005 | 0.005 | 0.583 | 0.27 | 0.24 | 0.16 | 0 | 0.18 | 0.14 |
| 5 | 42483693 | rs2972418 | -0.487 | -0.067 | -0.559 | -1.169 | -0.01 | 0.063 | 0.063 | 0 | 0.56 | 0.68 | 0.5 | 0.15 | 0.52 | 0.34 |
| 5 | 42483877 | rs2972419 | 0.469 | -0.248 | -0.53 | -1.424 | 0.031 | 0.038 | 0.038 | 0.633 | 0.27 | 0.19 | 0.15 | 0 | 0.18 | 0.15 |
| 5 | 42489142 | rs2940944 | -0.638 | -0.355 | -0.14 | -1.488 | 0 | -0.028 | -0.028 | 0.227 | 0.55 | 0.53 | 0.48 | 0.15 | 0.5 | 0.34 |
| 5 | 42490129 | rs2972391 | 0.756 | -0.678 | -0.422 | -1.672 | 0.015 | 0.026 | 0.026 | 0.33 | 0.2 | 0.15 | 0.12 | 0 | 0.14 | 0.12 |
| 5 | 42507333 | rs13153388 | -0.8 | -0.483 | -0.218 | -1.541 | 0.001 | -0.04 | -0.04 | 0.675 | 0.55 | 0.48 | 0.47 | 0.15 | 0.5 | 0.34 |
| 5 | 42523172 | rs4129472 | 0.624 | 0.049 | -0.431 | -1.304 | -0.077 | 0.077 | 0.077 | 3.682 | 0.13 | 0.01 | 0.1 | 0 | 0.22 | 0.06 |
| 5 | 42547255 | rs4365846 | -0.673 | -0.136 | 0.011 | -1.35 | 0.025 | -0.125 | -0.125 | 4.436 | 0.27 | 0.57 | 0.19 | 0.01 | 0.31 | 0.13 |

| | | | | | | | | | | | | | | | | |
|---|---|---|---|---|---|---|---|---|---|---|---|---|---|---|---|---|
| 5 | 42553879 | rs12153009 | 0.994 | 0.197 | -0.778 | -1.06 | -0.084 | -0.072 | -0.072 | 0.141 | 0.21 | 0.01 | 0.24 | 0.27 | 0.29 | 0.27 |
| 5 | 42555931 | rs7703713 | -1.126 | 0.273 | 0.383 | -0.499 | 0.01 | -0.028 | -0.028 | 0 | 0.29 | 0.59 | 0.24 | 0.18 | 0.31 | 0.19 |
| 5 | 42564748 | rs6883523 | 0.25 | 0.317 | 0.573 | -0.599 | 0.01 | -0.039 | -0.039 | 0.005 | 0.15 | 0.17 | 0.11 | 0 | 0.24 | 0.05 |
| 5 | 42569642 | rs4410646 | -0.805 | 0.324 | 0.05 | -0.072 | -0.023 | 0.03 | 0.03 | 0.171 | 0.95 | 0.99 | 0.7 | 0.63 | 0.93 | 0.85 |
| 5 | 42580760 | rs11955075 | -0.773 | 0.061 | 0.503 | -0.988 | 0.049 | -0.156 | -0.156 | 0.795 | 0.25 | 0.46 | 0.17 | 0.01 | 0.3 | 0.12 |
| 5 | 42610409 | rs13354167 | -0.337 | -0.256 | 0.261 | -0.878 | 0.025 | -0.14 | -0.14 | 0.391 | 0.21 | 0.4 | 0.12 | 0 | 0.22 | 0.05 |
| 5 | 42617492 | rs4866941 | 0.982 | -0.28 | -0.027 | -0.794 | -0.003 | -0.024 | -0.024 | 0 | 0.66 | 0.37 | 0.71 | 0.83 | 0.7 | 0.64 |
| 5 | 42619316 | rs4554233 | 0.627 | -0.242 | 0.167 | -0.435 | -0.003 | -0.058 | -0.058 | 0 | 0.75 | 0.51 | 0.82 | 0.84 | 0.77 | 0.87 |
| 5 | 42626206 | rs4590183 | -0.849 | -0.483 | 0.032 | -0.592 | -0.015 | -0.051 | -0.051 | 0 | 0.37 | 0.53 | 0.27 | 0.21 | 0.31 | 0.26 |
| 5 | 42631264 | rs6873545 | -0.791 | -0.505 | 0.193 | -0.603 | -0.013 | -0.052 | -0.052 | 0 | 0.36 | 0.45 | 0.26 | 0.2 | 0.31 | 0.26 |
| 5 | 42640471 | rs11948463 | 0.238 | -0.63 | 0.402 | -0.751 | 0.016 | -0.027 | -0.027 | 0.123 | 0.12 | 0.16 | 0.14 | 0.01 | 0.2 | 0.06 |
| 5 | 42650470 | rs7701605 | 0.978 | -0.977 | 0.471 | -0.906 | -0.004 | -0.09 | -0.09 | 0.235 | 0.68 | 0.43 | 0.75 | 0.94 | 0.7 | 0.87 |
| 5 | 42667871 | rs6878512 | -0.435 | -0.84 | 0.344 | -0.85 | 0.022 | -0.037 | -0.037 | 0 | 0.88 | 0.78 | 0.85 | 1 | 0.81 | 0.94 |
| 5 | 42716326 | rs2972781 | 0.837 | -1.237 | -0.078 | 0.349 | 0.005 | 0.04 | 0.04 | 0.982 | 0.06 | 0 | 0.06 | 0.44 | 0.02 | 0.09 |
| 5 | 42719239 | rs6180 | 1.211 | -0.848 | -0.477 | -0.632 | 0 | 0.001 | 0.001 | 0 | 0.43 | 0.38 | 0.4 | 0.57 | 0.42 | 0.46 |
| 5 | 42719826 | rs2910875 | 1.195 | -0.854 | -0.468 | -0.636 | 0 | 0.001 | 0.001 | 0 | 0.43 | 0.41 | 0.41 | 0.57 | 0.42 | 0.46 |
| 5 | 42724518 | rs1345825 | -0.351 | -0.894 | -0.39 | -0.753 | 0.001 | -0.024 | -0.024 | 0 | 0.3 | 0.18 | 0.42 | 0.42 | 0.3 | 0.42 |
| 5 | 42742798 | rs11950069 | 0.443 | -0.89 | -0.387 | -0.753 | -0.021 | -0.004 | -0.004 | 0 | 0.23 | 0.12 | 0.39 | 0.27 | 0.29 | 0.22 |
| 5 | 42757540 | rs2972995 | 0.794 | -0.893 | -0.377 | -0.738 | -0.005 | -0.004 | -0.004 | 0.085 | 0.07 | 0.09 | 0.04 | 0.15 | 0.01 | 0.21 |
| 5 | 42782492 | rs2973011 | 1.285 | -0.961 | -0.188 | -0.798 | -0.002 | -0.008 | -0.008 | 0 | 0.44 | 0.45 | 0.42 | 0.57 | 0.42 | 0.46 |
| 5 | 42800724 | rs6413428 | 0.139 | -0.917 | -0.178 | -0.83 | 0.005 | -0.086 | -0.086 | 0 | 0.26 | 0.34 | 0.16 | 0 | 0.29 | 0.12 |
| 5 | 42800808 | rs7579 | 0.478 | -0.919 | -0.254 | -0.828 | -0.021 | -0.004 | -0.004 | 0 | 0.23 | 0.12 | 0.39 | 0.27 | 0.29 | 0.22 |
| 5 | 42801268 | rs3877899 | 0.244 | -0.874 | -0.146 | -0.817 | -0.004 | -0.029 | -0.029 | 0 | 0.25 | 0.27 | 0.15 | 0 | 0.29 | 0.12 |
| 5 | 42820125 | rs2972756 | -1.247 | -0.563 | -0.304 | -0.73 | -0.005 | -0.007 | -0.007 | 0.095 | 0.56 | 0.54 | 0.59 | 0.43 | 0.61 | 0.55 |
| 5 | 42823799 | rs1035516 | 1.142 | -0.632 | -0.357 | -0.778 | -0.017 | -0.009 | -0.009 | 0 | 0.18 | 0.14 | 0.37 | 0.24 | 0.26 | 0.24 |
| 5 | 42827221 | rs7703289 | -0.868 | -0.797 | -0.373 | -0.801 | 0.002 | 0.001 | 0.001 | 1.23 | 0.39 | 0.4 | 0.22 | 0.18 | 0.34 | 0.31 |
| 5 | 42828101 | rs13154178 | -0.403 | -0.776 | -0.362 | -0.826 | 0 | -0.07 | -0.07 | 0 | 0.33 | 0.4 | 0.19 | 0.03 | 0.34 | 0.16 |
| 5 | 42839189 | rs16872824 | 1.047 | -0.702 | -0.585 | -0.573 | 0.029 | -0.028 | -0.028 | 0 | 0.06 | 0 | 0.02 | 0.15 | 0.01 | 0.16 |
| 5 | 42843771 | rs12520759 | 0.91 | -0.331 | -0.496 | -0.517 | -0.012 | -0.013 | -0.013 | 3.605 | 0.18 | 0.13 | 0.37 | 0.26 | 0.23 | 0.26 |
| 5 | 42854408 | rs11951328 | 0.619 | -0.333 | -0.583 | -0.704 | -0.027 | 0.024 | 0.024 | 0 | 0.13 | 0.09 | 0.09 | 0 | 0.23 | 0.07 |
| 5 | 42856126 | rs4409101 | -1.256 | -0.478 | -0.217 | -0.554 | -0.002 | 0.002 | 0.002 | 0 | 0.58 | 0.69 | 0.59 | 0.44 | 0.57 | 0.52 |
| 5 | 42856350 | rs1423652 | 0.157 | -0.217 | -0.298 | -0.046 | 0.006 | 0.006 | 0.006 | 0 | 0.19 | 0.22 | 0.38 | 0.26 | 0.23 | 0.26 |
| 5 | 42871462 | rs1105983 | -0.503 | -0.825 | -0.246 | -0.636 | -0.006 | -0.048 | -0.048 | 0.01 | 0.14 | 0.2 | 0.1 | 0 | 0.14 | 0.07 |
| 5 | 42877570 | rs427799 | -0.298 | -1.865 | -0.106 | -1.266 | -0.035 | -0.06 | -0.06 | 0 | 0.17 | 0.23 | 0.08 | 0.01 | 0.07 | 0.04 |
| 5 | 42888925 | rs315259 | -0.02 | -1.767 | -0.092 | -1.227 | -0.007 | -0.026 | -0.026 | 3.916 | 0.12 | 0.12 | 0.07 | 0 | 0.07 | 0.02 |
| 5 | 42896083 | rs10072083 | -0.063 | -1.669 | -0.076 | -1.176 | -0.008 | -0.054 | -0.054 | 1.022 | 0.14 | 0.2 | 0.1 | 0 | 0.13 | 0.07 |
| 5 | 42906572 | rs2253157 | -1.493 | -2.426 | -0.195 | -1.179 | -0.001 | -0.005 | -0.005 | 0.048 | 0.39 | 0.66 | 0.23 | 0.16 | 0.23 | 0.25 |
| 5 | 42908384 | rs10036147 | -0.082 | -2.58 | -0.161 | -1.215 | -0.024 | 0.001 | 0.001 | 0.057 | 0.17 | 0.34 | 0.13 | 0.16 | 0.13 | 0.15 |
| 5 | 42910025 | rs12659760 | -1.426 | -2.464 | -0.375 | -1.159 | -0.039 | -0.043 | -0.043 | 0.056 | 0.39 | 0.58 | 0.22 | 0.16 | 0.23 | 0.25 |
| 5 | 42915470 | rs13177679 | 0.927 | -1.017 | -1.29 | -1.418 | -0.123 | -0.003 | -0.003 | 0 | 0.15 | 0.02 | 0.16 | 0.12 | 0.36 | 0.19 |
| 5 | 42918296 | rs2444853 | 0.14 | -1.827 | -0.927 | -1.381 | 0.004 | -0.014 | -0.014 | 0 | 0.16 | 0.14 | 0.08 | 0 | 0.08 | 0.03 |
| 5 | 42919685 | rs6451656 | -0.535 | -2.02 | -0.804 | -1.019 | -0.077 | -0.043 | -0.043 | 0.145 | 0.23 | 0.52 | 0.18 | 0.14 | 0.15 | 0.23 |
| 5 | 42922816 | rs2548370 | -1.082 | -1.996 | -0.884 | -1.026 | -0.061 | -0.017 | -0.017 | 0 | 0.27 | 0.69 | 0.18 | 0.21 | 0.1 | 0.18 |
| 5 | 42938617 | rs2548379 | -1.054 | -2.474 | -1.476 | -1.367 | -0.003 | 0.005 | 0.005 | 0 | 0.69 | 0.71 | 0.45 | 0.38 | 0.69 | 0.6 |
| 5 | 42951941 | rs3893725 | -0.853 | -2.792 | -1.614 | -1.519 | -0.001 | -0.001 | -0.001 | 0 | 0.63 | 0.71 | 0.42 | 0.37 | 0.61 | 0.53 |
| 5 | 42953802 | rs3849698 | 0.585 | -3.155 | -1.293 | -1.683 | 0.001 | 0.038 | 0.038 | 0 | 0.4 | 0.32 | 0.28 | 0.18 | 0.53 | 0.35 |
| 5 | 42954436 | rs2548334 | -2.277 | -3.022 | -1.479 | -1.642 | 0.012 | 0.05 | 0.05 | 0.593 | 0.78 | 0.76 | 0.49 | 0.39 | 0.74 | 0.61 |
| 5 | 42961249 | rs6883097 | -0.396 | -3.054 | -1.4 | -1.652 | -0.003 | -0.003 | -0.003 | 0.797 | 0.56 | 0.62 | 0.39 | 0.38 | 0.59 | 0.52 |
| 5 | 42967215 | rs2924819 | -0.841 | -2.946 | -1.39 | -1.127 | -0.009 | 0.022 | 0.022 | 0.474 | 0.63 | 0.63 | 0.43 | 0.48 | 0.66 | 0.59 |
| 5 | 42969416 | rs309924 | -2.016 | -2.818 | -1.331 | -1.595 | 0.012 | 0.048 | 0.048 | 0.807 | 0.77 | 0.75 | 0.48 | 0.39 | 0.74 | 0.61 |
| 5 | 42984559 | rs12188691 | 0.823 | -3.009 | -1.579 | -1.77 | -0.131 | 0.087 | 0.087 | 0 | 0.36 | 0.06 | 0.24 | 0.19 | 0.5 | 0.34 |
| 5 | 42984638 | rs186322 | -0.886 | -3.097 | -1.475 | -1.888 | -0.03 | -0.018 | -0.018 | 1.039 | 0.36 | 0.65 | 0.23 | 0.21 | 0.21 | 0.28 |
| 5 | 42985370 | rs309906 | -1.386 | -3.237 | -1.158 | -1.972 | 0.14 | 0.113 | 0.113 | 0.964 | 0.43 | 0.8 | 0.24 | 0.21 | 0.21 | 0.29 |
| 5 | 42992040 | rs2603656 | -1.001 | -3.296 | -1.595 | -1.951 | 0.016 | 0 | 0 | 0 | 0.63 | 0.29 | 0.41 | 0.38 | 0.68 | 0.51 |
| 5 | 42996470 | rs10941605 | -0.986 | -3.304 | -1.639 | -1.955 | 0.019 | -0.003 | -0.003 | 0.008 | 0.63 | 0.24 | 0.41 | 0.38 | 0.68 | 0.52 |
| 5 | 42997599 | rs13174226 | 1.187 | -3.157 | -1.69 | -1.946 | 0.023 | 0.082 | 0.082 | 0.547 | 0.14 | 0.02 | 0.06 | 0.01 | 0.11 | 0.09 |
| 5 | 43002823 | rs13173064 | 0.773 | -2.861 | -1.548 | -1.789 | -0.078 | 0.096 | 0.096 | 0 | 0.34 | 0.11 | 0.23 | 0.16 | 0.5 | 0.34 |
| 5 | 43007609 | rs309914 | -1.543 | -2.049 | -0.976 | -1.279 | 0.01 | 0.026 | 0.026 | 0 | 0.75 | 0.76 | 0.48 | 0.37 | 0.72 | 0.61 |
| 5 | 43009986 | rs12697439 | -1.5 | -2.008 | -0.976 | -1.281 | 0.016 | 0.037 | 0.037 | 0.004 | 0.75 | 0.68 | 0.48 | 0.37 | 0.72 | 0.61 |
| 5 | 43034142 | rs585302 | -0.082 | -2.24 | -0.688 | -1.216 | -0.065 | 0.112 | 0.112 | 0.091 | 0.12 | 0.47 | 0.08 | 0.2 | 0.03 | 0.15 |
| 5 | 43037389 | rs673855 | 0.737 | -2.125 | -0.614 | -1.193 | 0.02 | 0.005 | 0.005 | 2.273 | 0.07 | 0.44 | 0.06 | 0 | 0.03 | 0.08 |
| 5 | 43066774 | rs316408 | -1.822 | -2.15 | -0.906 | -1.399 | 0 | 0.109 | 0.109 | 0 | 0.84 | 0.77 | 0.65 | 0.37 | 0.86 | 0.69 |
| 5 | 43093541 | rs10941609 | 0.621 | -2.147 | -1.205 | -1.812 | 0.004 | 0.053 | 0.053 | 0.025 | 0.2 | 0.1 | 0.28 | 0 | 0.21 | 0.1 |
| 5 | 43116830 | rs1532331 | -0.62 | -1.364 | -1.231 | -1.99 | 0.065 | -0.012 | -0.012 | 0 | 0.8 | 0.67 | 0.82 | 0.84 | 0.67 | 0.83 |
| 5 | 43120938 | rs11748652 | 0.584 | -1.82 | -1.572 | -2.251 | 0.022 | 0.116 | 0.116 | 0 | 0.27 | 0.07 | 0.29 | 0 | 0.22 | 0.12 |
| 5 | 43123623 | rs567884 | 1.665 | -2.143 | -1.563 | -2.284 | -0.005 | 0.036 | 0.036 | 0 | 0.24 | 0.24 | 0.39 | 0.62 | 0.23 | 0.37 |
| 5 | 43124688 | rs782971 | 1.659 | -2.171 | -1.518 | -2.402 | -0.009 | 0.058 | 0.058 | 0 | 0.24 | 0.29 | 0.39 | 0.61 | 0.23 | 0.37 |
| 5 | 43143799 | rs6892422 | 1.745 | -2.16 | -1.5 | -2.393 | -0.004 | 0.024 | 0.024 | 0 | 0.23 | 0.21 | 0.39 | 0.61 | 0.23 | 0.37 |
| 5 | 43150395 | rs12110174 | 0.938 | -1.949 | -1.672 | -2.502 | 0.015 | 0.105 | 0.105 | 0.474 | 0.25 | 0.06 | 0.28 | 0.01 | 0.22 | 0.12 |

| | | | | | | | | | | | | | | | | |
|---|---|---|---|---|---|---|---|---|---|---|---|---|---|---|---|---|
| 5 | 43164060 | rs11744749 | 1.976 | -2.144 | -1.868 | -2.67 | -0.024 | 0.048 | 0.048 | 0 | 0.11 | 0 | 0.04 | 0.01 | 0.08 | 0.14 |
| 5 | 43170654 | rs12514198 | 0.989 | -2.061 | -1.779 | -2.649 | 0.016 | 0.102 | 0.102 | 0.462 | 0.25 | 0.07 | 0.28 | 0.01 | 0.22 | 0.12 |
| 5 | 43175103 | rs35397154 | 1.35 | -1.808 | -1.68 | -2.434 | -0.005 | -0.053 | -0.053 | 0 | 0.13 | 0.05 | 0.08 | 0.21 | 0.12 | 0.14 |
| 5 | 43180535 | rs4866808 | -0.722 | -1.873 | -1.542 | -2.488 | 0.094 | -0.027 | -0.027 | 1.22 | 0.77 | 0.53 | 0.81 | 0.84 | 0.66 | 0.76 |
| 5 | 43191033 | rs7730004 | -0.103 | -1.899 | -1.513 | -2.498 | 0.053 | -0.045 | -0.045 | 0 | 0.73 | 0.41 | 0.81 | 0.84 | 0.65 | 0.76 |
| 5 | 43191823 | rs10074873 | -0.671 | -1.645 | -1.439 | -2.368 | 0.092 | -0.025 | -0.025 | 0 | 0.78 | 0.53 | 0.82 | 0.84 | 0.66 | 0.76 |
| 5 | 43192421 | rs17243515 | 1.004 | -1.475 | -1.281 | -1.973 | -0.003 | -0.049 | -0.049 | 0.439 | 0.13 | 0.05 | 0.08 | 0.21 | 0.12 | 0.14 |
| 5 | 43192453 | rs17302931 | 1.004 | -1.475 | -1.281 | -1.973 | -0.003 | -0.049 | -0.049 | 0.947 | 0.13 | 0.05 | 0.08 | 0.21 | 0.12 | 0.14 |
| 5 | 43199867 | rs6871636 | 0.653 | -1.022 | -0.979 | -2.305 | 0.003 | -0.008 | -0.008 | 0 | 0.29 | 0.29 | 0.22 | 0.14 | 0.37 | 0.24 |
| 5 | 43203125 | rs2444779 | 1.308 | -0.628 | -0.53 | -1.896 | 0.004 | 0.037 | 0.037 | 0 | 0.22 | 0.24 | 0.41 | 0.68 | 0.26 | 0.37 |
| 5 | 43208238 | rs6887169 | 1.426 | -0.634 | -0.563 | -1.843 | -0.019 | -0.028 | -0.028 | 0.704 | 0.06 | 0.14 | 0.03 | 0 | 0.02 | 0.08 |
| 5 | 43212590 | rs10512821 | 1.082 | -0.686 | -0.756 | -1.937 | -0.012 | 0.058 | 0.058 | 0.025 | 0.18 | 0.06 | 0.23 | 0.01 | 0.18 | 0.1 |
| 5 | 43222326 | rs7701546 | 1.446 | -0.65 | -0.604 | -2.015 | -0.019 | -0.028 | -0.028 | 0.022 | 0.06 | 0.19 | 0.03 | 0 | 0.02 | 0.08 |
| 5 | 43232311 | rs11747314 | 0.94 | -0.584 | -1.029 | -2.082 | -0.019 | 0.073 | 0.073 | 0 | 0.19 | 0.05 | 0.23 | 0.01 | 0.19 | 0.11 |
| 5 | 43232637 | rs10056788 | 1.325 | -0.539 | -0.968 | -2.074 | -0.02 | -0.03 | -0.03 | 0 | 0.06 | 0.19 | 0.03 | 0 | 0.02 | 0.08 |
| 5 | 43233265 | rs11748138 | 1.015 | -0.524 | -0.994 | -2.07 | -0.02 | 0.07 | 0.07 | 0 | 0.19 | 0.05 | 0.23 | 0.01 | 0.19 | 0.11 |
| 5 | 43236708 | rs17303482 | 0.536 | -0.385 | -0.368 | -1.286 | -0.016 | 0.045 | 0.045 | 0 | 0.09 | 0 | 0.02 | 0 | 0.07 | 0.07 |
| 5 | 43237188 | rs4866811 | 0.416 | -0.217 | 0.144 | -0.856 | 0.002 | 0.024 | 0.024 | 0 | 0.22 | 0.23 | 0.41 | 0.67 | 0.26 | 0.37 |
| 5 | 43237644 | rs6867229 | -0.274 | -0.14 | 0.036 | -1.214 | -0.024 | -0.034 | -0.034 | 1.445 | 0.06 | 0.2 | 0.03 | 0 | 0.02 | 0.08 |
| 5 | 43243774 | rs10473304 | -0.028 | -0.073 | 0.055 | -1.182 | -0.02 | -0.03 | -0.03 | 0 | 0.06 | 0.19 | 0.02 | 0 | 0.02 | 0.08 |
| 5 | 43256445 | rs10941615 | 0.461 | -0.015 | 0.184 | -1.18 | -0.013 | -0.024 | -0.024 | 0.146 | 0.94 | 0.72 | 0.97 | 1 | 0.99 | 0.95 |
| 5 | 43271547 | rs10805671 | 0.013 | 0.656 | -0.482 | -1.409 | -0.102 | 0.033 | 0.033 | 0 | 0.31 | 0.07 | 0.48 | 0.8 | 0.44 | 0.57 |
| 5 | 43297032 | rs3733768 | -0.035 | 0.691 | 0.792 | -1.536 | 0.012 | -0.014 | -0.014 | 0.746 | 0.68 | 0.64 | 0.51 | 0.2 | 0.55 | 0.45 |
| 5 | 43304402 | rs9292871 | -0.035 | 0.663 | 0.764 | -1.537 | 0.012 | -0.014 | -0.014 | 0.011 | 0.68 | 0.64 | 0.51 | 0.2 | 0.55 | 0.45 |
| 5 | 43347344 | rs7380703 | 0.728 | 0.371 | -0.236 | -1.835 | -0.001 | 0.009 | 0.009 | 0 | 0.26 | 0.18 | 0.2 | 0.12 | 0.27 | 0.17 |
| 5 | 43358759 | rs7704588 | -0.411 | 0.709 | -0.266 | -1.582 | -0.076 | 0.037 | 0.037 | 0 | 0.69 | 0.88 | 0.54 | 0.2 | 0.55 | 0.45 |
| 5 | 43368066 | rs10512827 | 0.057 | 0.702 | -0.19 | -1.62 | 0.001 | -0.011 | -0.011 | 0 | 0.63 | 0.58 | 0.49 | 0.15 | 0.53 | 0.39 |
| 5 | 43370366 | rs922439 | 0.627 | 0.222 | -0.446 | -1.921 | 0.011 | 0.044 | 0.044 | 0 | 0.32 | 0.18 | 0.22 | 0.12 | 0.28 | 0.2 |
| 5 | 43374933 | rs7727047 | 0.477 | 0.854 | -0.415 | -1.186 | -0.001 | -0.009 | -0.009 | 0 | 0.62 | 0.55 | 0.43 | 0.15 | 0.54 | 0.39 |
| 5 | 43394778 | rs7701642 | -0.093 | 0.981 | -0.994 | -0.951 | -0.095 | 0.005 | 0.005 | 0 | 0.71 | 0.95 | 0.5 | 0.22 | 0.63 | 0.47 |
| 5 | 43401980 | rs9292877 | -0.93 | 0.618 | -0.294 | -0.747 | 0.003 | 0.044 | 0.044 | 0 | 0.05 | 0.21 | 0.04 | 0.07 | 0.02 | 0.07 |
| 5 | 43430273 | rs750582 | 0.137 | 0.969 | -0.82 | -1.013 | -0.065 | 0.065 | 0.065 | 0 | 0.66 | 0.82 | 0.43 | 0.2 | 0.57 | 0.4 |
| 5 | 43444233 | rs10054519 | 0.495 | 0.995 | -0.732 | -1.101 | -0.008 | -0.005 | -0.005 | 0 | 0.6 | 0.63 | 0.38 | 0.09 | 0.5 | 0.33 |
| 5 | 43453361 | rs6451696 | 0.178 | 0.993 | -0.804 | -1.053 | -0.057 | 0.127 | 0.127 | 0 | 0.65 | 0.81 | 0.42 | 0.09 | 0.55 | 0.39 |
| 5 | 43462210 | rs10038608 | -0.167 | 0.985 | -0.82 | -1.048 | -0.121 | 0.042 | 0.042 | 0 | 0.68 | 0.94 | 0.44 | 0.2 | 0.57 | 0.4 |
| 5 | 43490142 | rs10036954 | -0.282 | 0.998 | -0.812 | -1.082 | -0.087 | 0.161 | 0.161 | 0 | 0.34 | 0.1 | 0.58 | 0.91 | 0.45 | 0.61 |
| 5 | 43514996 | rs10055801 | 0.495 | 0.983 | -0.732 | -1.086 | -0.008 | -0.004 | -0.004 | 0 | 0.6 | 0.63 | 0.38 | 0.09 | 0.5 | 0.33 |
| 5 | 43525322 | rs4072918 | 0.505 | 0.983 | -0.718 | -1.094 | -0.008 | -0.005 | -0.005 | 0 | 0.6 | 0.63 | 0.38 | 0.09 | 0.5 | 0.33 |
| 5 | 43539868 | rs10058558 | -0.199 | 1 | -0.766 | -1.061 | -0.072 | 0.064 | 0.064 | 0 | 0.34 | 0.12 | 0.57 | 0.8 | 0.44 | 0.59 |
| 5 | 43544592 | rs6890419 | 0.584 | 0.999 | -0.747 | -1.066 | 0.037 | -0.016 | -0.016 | 0.048 | 0.05 | 0.16 | 0.04 | 0 | 0.05 | 0.06 |
| 5 | 43558629 | rs12521249 | -0.751 | 0.984 | -0.749 | -1.073 | -0.018 | 0.03 | 0.03 | 0 | 0.4 | 0.29 | 0.62 | 0.91 | 0.5 | 0.67 |
| 5 | 43561759 | rs10042992 | -0.188 | 0.995 | -0.811 | -1.063 | -0.078 | 0.065 | 0.065 | 0 | 0.34 | 0.1 | 0.57 | 0.8 | 0.44 | 0.6 |
| 5 | 43562931 | rs12652547 | -0.025 | 1.023 | -0.898 | -1.026 | -0.113 | 0.038 | 0.038 | 0 | 0.33 | 0.06 | 0.56 | 0.79 | 0.43 | 0.58 |
| 5 | 43601610 | rs1366410 | 0.47 | 0.93 | -0.766 | -1.129 | -0.002 | -0.029 | -0.029 | 0 | 0.59 | 0.58 | 0.38 | 0.09 | 0.5 | 0.33 |
| 5 | 43604258 | rs10065689 | 0.419 | 0.944 | -0.779 | -1.123 | -0.005 | -0.015 | -0.015 | 2.388 | 0.59 | 0.62 | 0.38 | 0.09 | 0.5 | 0.33 |
| 5 | 43609343 | rs10057103 | -0.355 | 1.171 | -1.078 | -1.06 | -0.106 | 0.035 | 0.035 | 0 | 0.7 | 0.94 | 0.45 | 0.2 | 0.59 | 0.45 |
| 5 | 43648914 | rs16873438 | -1.109 | 0.849 | -0.703 | -0.821 | 0.023 | 0.048 | 0.048 | 0.303 | 0.1 | 0.3 | 0.07 | 0.12 | 0.09 | 0.12 |
| 5 | 43711378 | rs6451722 | -0.608 | 0.102 | -1.342 | -1.262 | 0.069 | 0.06 | 0.06 | 0.72 | 0.35 | 0.88 | 0.21 | 0.22 | 0.21 | 0.22 |
| 5 | 43727542 | rs4327605 | -1.44 | 0.768 | -1.545 | -0.902 | -0.056 | -0.081 | -0.081 | 0.106 | 0.88 | 0.99 | 0.7 | 0.39 | 0.85 | 0.86 |
| 5 | 43731692 | rs4574552 | -0.75 | 0.004 | -1.294 | -1.387 | -0.02 | 0.073 | 0.073 | 0 | 0.58 | 0.81 | 0.39 | 0.28 | 0.41 | 0.36 |
| 5 | 43768939 | rs10075553 | -0.75 | 0.058 | -0.456 | -0.661 | 0.021 | 0.019 | 0.019 | 0.362 | 0.3 | 0.74 | 0.19 | 0.21 | 0.2 | 0.2 |
| 5 | 43780972 | rs6892300 | -1.038 | 0.661 | -0.596 | 0.017 | 0.015 | 0.08 | 0.08 | 0 | 0.08 | 0.19 | 0.05 | 0.18 | 0.08 | 0.11 |
| 5 | 43865891 | rs10045432 | -2.209 | 0.623 | -0.718 | 0.261 | 0.012 | 0.16 | 0.16 | 0 | 0.06 | 0.21 | 0.05 | 0.24 | 0.06 | 0.04 |
| 5 | 43873092 | rs7723677 | -1.402 | 0.474 | -0.667 | 0.02 | 0.002 | -0.045 | -0.045 | 0 | 0.08 | 0.26 | 0.04 | 0 | 0.05 | 0.07 |
| 5 | 43881065 | rs7729400 | -1.91 | 0.469 | -0.671 | 0.05 | 0 | 0.052 | 0.052 | 0 | 0.1 | 0.27 | 0.06 | 0.17 | 0.07 | 0.08 |
| 5 | 43899602 | rs7736219 | -1.437 | 0.62 | -0.768 | 0.206 | 0.032 | 0.109 | 0.109 | 0 | 0.06 | 0.24 | 0.14 | 0.2 | 0.06 | 0.06 |
| 5 | 43899639 | rs7735852 | -1.609 | 0.592 | -0.81 | 0.193 | 0.01 | 0.068 | 0.068 | 0 | 0.07 | 0.19 | 0.13 | 0.2 | 0.06 | 0.03 |
| 5 | 43927216 | rs4866756 | -2.058 | 0.592 | -0.806 | 0.192 | 0.014 | 0.076 | 0.076 | 0 | 0.08 | 0.25 | 0.14 | 0.2 | 0.06 | 0.06 |
| 5 | 43973081 | rs7721923 | -1.156 | 0.122 | -1.173 | -0.348 | -0.006 | -0.01 | -0.01 | 0.185 | 0.28 | 0.36 | 0.21 | 0.23 | 0.19 | 0.2 |
| 5 | 43985957 | rs4354051 | -2.109 | 0.293 | -1.04 | -0.088 | 0.016 | 0.077 | 0.077 | 0 | 0.09 | 0.28 | 0.15 | 0.2 | 0.08 | 0.09 |
| 5 | 43989069 | rs4583891 | -1.736 | 0.263 | -1.053 | -0.11 | 0.013 | -0.04 | -0.04 | 0 | 0.08 | 0.28 | 0.06 | 0 | 0.05 | 0.05 |
| 5 | 44002170 | rs4515309 | 0.258 | -0.21 | -1.808 | -0.419 | 0.015 | 0.041 | 0.041 | 0.408 | 0.16 | 0.01 | 0.14 | 0.06 | 0.15 | 0.12 |
| 5 | 44003444 | rs13159468 | -1.443 | 0.198 | -1.363 | -0.106 | 0.017 | 0.074 | 0.074 | 0 | 0.07 | 0.21 | 0.13 | 0.2 | 0.07 | 0.04 |
| 5 | 44051305 | rs6879107 | 0.893 | -0.286 | -1.624 | -0.097 | 0.007 | -0.001 | -0.001 | 0 | 0.8 | 0.84 | 0.7 | 0.75 | 0.85 | 0.72 |
| 5 | 44059937 | rs4334895 | 0.88 | 0.546 | -2.516 | 0.079 | -0.117 | -0.16 | -0.16 | 0 | 0.19 | 0.02 | 0.3 | 0.43 | 0.3 | 0.29 |
| 5 | 44066093 | rs6859263 | -0.156 | 0.24 | -2.238 | -0.009 | -0.005 | -0.014 | -0.014 | 0 | 0.12 | 0.11 | 0.13 | 0.22 | 0.13 | 0.14 |
| 5 | 44066669 | rs7705250 | -0.342 | 0.268 | -2.25 | 0.013 | -0.003 | -0.038 | -0.038 | 0.598 | 0.86 | 0.98 | 0.72 | 0.75 | 0.87 | 0.74 |
| 5 | 44088038 | rs10076380 | -0.601 | 0.141 | -1.963 | -0.023 | 0.004 | -0.001 | -0.001 | 0 | 0.19 | 0.17 | 0.3 | 0.24 | 0.15 | 0.28 |
| 5 | 44098765 | rs4502833 | -1.579 | -0.057 | -1.484 | -0.104 | -0.01 | 0.016 | 0.016 | 0 | 0.33 | 0.41 | 0.51 | 0.48 | 0.28 | 0.43 |

| | | | | | | | | | | | | | | | | |
|---|---|---|---|---|---|---|---|---|---|---|---|---|---|---|---|---|
| 5 | 44129083 | rs4242107 | -1.27 | 0.052 | -1.406 | -0.039 | -0.005 | 0.031 | 0.031 | 0.015 | 0.27 | 0.36 | 0.49 | 0.46 | 0.24 | 0.3 |
| 5 | 44132277 | rs7730111 | -0.096 | 0.325 | -1.408 | -0.019 | 0.001 | 0.025 | 0.025 | 1.366 | 0.13 | 0.2 | 0.28 | 0.27 | 0.15 | 0.22 |
| 5 | 44150391 | rs4866869 | -1.654 | 0.322 | -1.421 | -0.222 | 0 | 0.008 | 0.008 | 0.073 | 0.31 | 0.37 | 0.51 | 0.47 | 0.32 | 0.39 |
| 5 | 44190962 | rs4242110 | -1.624 | 0.374 | -0.834 | -0.204 | 0.001 | 0.015 | 0.015 | 0 | 0.29 | 0.36 | 0.5 | 0.47 | 0.3 | 0.39 |
| 5 | 44208106 | rs4866772 | -0.002 | 0.365 | -0.827 | -0.139 | 0.001 | 0.028 | 0.028 | 1.144 | 0.13 | 0.2 | 0.27 | 0.3 | 0.14 | 0.22 |
| 5 | 44224808 | rs4643965 | -0.608 | 0.33 | -0.868 | -0.212 | 0.047 | 0.119 | 0.119 | 0.228 | 0.17 | 0.33 | 0.46 | 0.46 | 0.24 | 0.24 |
| 5 | 44228425 | rs10512836 | 1.075 | -0.432 | -2.136 | -1.046 | -0.026 | 0.072 | 0.072 | 3.3 | 0.23 | 0.01 | 0.18 | 0.09 | 0.22 | 0.16 |
| 5 | 44233778 | rs726941 | 1.399 | -0.078 | -1.787 | -1.028 | -0.046 | -0.09 | -0.09 | 0.141 | 0.19 | 0.11 | 0.15 | 0.42 | 0.25 | 0.26 |
| 5 | 44241741 | rs2053783 | 0.245 | -0.323 | -1.484 | -0.875 | 0.029 | 0.054 | 0.054 | 0 | 0.12 | 0.03 | 0.04 | 0 | 0.06 | 0.15 |
| 5 | 44263397 | rs6883600 | -0.985 | -0.361 | -1.075 | -0.863 | -0.001 | -0.001 | -0.001 | 0 | 0.26 | 0.26 | 0.5 | 0.43 | 0.27 | 0.27 |
| 5 | 44265078 | rs10044215 | -0.664 | -0.38 | -1.055 | -0.843 | -0.001 | 0.011 | 0.011 | 0 | 0.22 | 0.18 | 0.28 | 0.12 | 0.21 | 0.14 |
| 5 | 44267175 | rs6883160 | 0.872 | -0.304 | -0.81 | -0.848 | 0.009 | -0.002 | -0.002 | 0 | 0.72 | 0.6 | 0.46 | 0.72 | 0.68 | 0.73 |
| 5 | 44279496 | rs1542219 | 1.296 | -0.452 | -0.606 | -0.275 | 0.011 | -0.057 | -0.057 | 0 | 0.62 | 0.22 | 0.43 | 0.46 | 0.66 | 0.6 |
| 5 | 44289499 | rs10062585 | 0.373 | -0.702 | -0.724 | -0.599 | -0.009 | 0 | 0 | 0 | 0.15 | 0.17 | 0.26 | 0.18 | 0.12 | 0.22 |
| 5 | 44296986 | rs1448044 | -1.191 | -0.706 | -0.637 | -0.512 | 0.005 | -0.019 | -0.019 | 0 | 0.38 | 0.66 | 0.56 | 0.54 | 0.33 | 0.4 |
| 5 | 44303760 | rs1374961 | -1.24 | -0.672 | -0.701 | -0.498 | 0.011 | -0.046 | -0.046 | 0 | 0.39 | 0.74 | 0.57 | 0.54 | 0.33 | 0.4 |
| 5 | 44308252 | rs10473352 | 0.877 | -0.656 | -0.768 | -0.511 | -0.011 | -0.006 | -0.006 | 0.23 | 0.13 | 0.14 | 0.37 | 0.18 | 0.19 | 0.21 |
| 5 | 44313151 | rs10060796 | 0.852 | -0.711 | -0.764 | -0.636 | -0.019 | 0.196 | 0.196 | 0.572 | 0.1 | 0.39 | 0.04 | 0.26 | 0.02 | 0.12 |
| 5 | 44335820 | rs1011814 | 0.311 | -0.234 | -1.642 | -0.884 | 0.007 | -0.05 | -0.05 | 0 | 0.61 | 0.24 | 0.42 | 0.47 | 0.67 | 0.6 |
| 5 | 44362204 | rs17234079 | 1.003 | -0.293 | -2.217 | -1.186 | 0.002 | 0.103 | 0.103 | 0.179 | 0.25 | 0 | 0.15 | 0.07 | 0.19 | 0.21 |
| 5 | 44365545 | rs2121875 | 0.545 | -0.138 | -1.364 | -0.747 | 0.008 | -0.054 | -0.054 | 0.861 | 0.61 | 0.23 | 0.42 | 0.47 | 0.66 | 0.61 |
| 5 | 44373060 | rs11750845 | 0.936 | -0.1 | -1.75 | -0.639 | -0.078 | -0.083 | -0.083 | 0 | 0.5 | 0.06 | 0.3 | 0.47 | 0.51 | 0.57 |
| 5 | 44381698 | rs16901816 | -0.213 | -0.192 | -0.851 | -0.237 | 0.009 | 0 | 0 | 0.069 | 0.15 | 0.22 | 0.33 | 0.18 | 0.19 | 0.24 |
| 5 | 44385415 | rs1482679 | 0.658 | -0.362 | -1.464 | -0.324 | 0.003 | -0.079 | -0.079 | 0.309 | 0.6 | 0.15 | 0.47 | 0.47 | 0.66 | 0.61 |
| 5 | 44396015 | rs723166 | -0.237 | 0.122 | -1.483 | -0.447 | -0.043 | 0.066 | 0.066 | 0.028 | 0.26 | 0.04 | 0.3 | 0.14 | 0.27 | 0.36 |
| 5 | 44398696 | rs10941665 | 0.85 | 0.198 | -1.305 | -0.474 | 0.005 | 0.006 | 0.006 | 0 | 0.26 | 0.26 | 0.36 | 0.45 | 0.38 | 0.24 |
| 5 | 44400659 | rs16901841 | 0.145 | -0.085 | -1.672 | -0.707 | 0.016 | 0.033 | 0.033 | 0 | 0.23 | 0.03 | 0.28 | 0.15 | 0.21 | 0.27 |
| 5 | 44403117 | rs16901843 | 0.132 | -0.09 | -1.674 | -0.709 | 0.017 | 0.034 | 0.034 | 0 | 0.23 | 0.03 | 0.28 | 0.15 | 0.21 | 0.27 |
| 5 | 44419254 | rs1384451 | 1.003 | -0.074 | -1.339 | -0.568 | 0.021 | 0.034 | 0.034 | 0 | 0.27 | 0.32 | 0.42 | 0.52 | 0.39 | 0.25 |
| 5 | 44422633 | rs10068476 | 0.545 | -0.173 | -1.118 | -0.598 | -0.006 | 0.002 | 0.002 | 0 | 0.19 | 0.26 | 0.31 | 0.23 | 0.17 | 0.26 |
| 5 | 44428516 | rs9292907 | -0.392 | -0.106 | -1.35 | -0.764 | 0.005 | 0.014 | 0.014 | 0 | 0.51 | 0.48 | 0.7 | 0.68 | 0.59 | 0.52 |
| 5 | 44430824 | rs9292908 | -0.483 | -0.219 | -0.718 | -0.728 | -0.002 | 0.002 | 0.002 | 0 | 0.7 | 0.73 | 0.84 | 0.68 | 0.77 | 0.63 |
| 5 | 44432885 | rs13357659 | 0.387 | 0.476 | -0.458 | -0.218 | 0.034 | 0.057 | 0.057 | 0 | 0.28 | 0.41 | 0.42 | 0.52 | 0.39 | 0.25 |
| 5 | 44442578 | rs12188017 | 0.33 | 0.185 | -0.503 | -0.451 | -0.003 | -0.031 | -0.031 | 0 | 0.16 | 0.19 | 0.12 | 0.03 | 0.15 | 0.1 |
| 5 | 44447093 | rs13160481 | -0.202 | 0.016 | -0.434 | -0.721 | -0.005 | -0.03 | -0.03 | 0 | 0.07 | 0.13 | 0.06 | 0 | 0.06 | 0.02 |
| 5 | 44452052 | rs922853 | 0.385 | -0.047 | -0.211 | -0.771 | -0.006 | -0.074 | -0.074 | 0 | 0.16 | 0.23 | 0.1 | 0 | 0.15 | 0.1 |
| 5 | 44473605 | rs6879342 | -0.62 | -0.458 | -0.345 | -0.837 | -0.008 | 0.076 | 0.076 | 0 | 0.76 | 0.61 | 0.87 | 0.65 | 0.79 | 0.6 |
| 5 | 44480144 | rs1384450 | 0.869 | -0.546 | -0.425 | -0.556 | 0.03 | -0.028 | -0.028 | 0.02 | 0.51 | 0.18 | 0.65 | 0.53 | 0.58 | 0.48 |
| 5 | 44482784 | rs13171351 | 0.448 | -0.516 | -0.388 | -0.555 | -0.001 | -0.016 | -0.016 | 0.152 | 0.06 | 0.08 | 0.06 | 0 | 0.06 | 0.02 |
| 5 | 44490799 | rs6865330 | -0.25 | -0.527 | -0.366 | -0.563 | -0.014 | -0.037 | -0.037 | 0 | 0.24 | 0.39 | 0.22 | 0.12 | 0.22 | 0.12 |
| 5 | 44496415 | rs2062140 | 0.519 | -0.468 | -0.495 | -0.525 | -0.001 | -0.017 | -0.017 | 0.046 | 0.06 | 0.08 | 0.06 | 0 | 0.06 | 0.01 |
| 5 | 44513809 | rs2128434 | 0.714 | -0.66 | -0.716 | -0.635 | 0.034 | -0.028 | -0.028 | 0 | 0.51 | 0.13 | 0.65 | 0.53 | 0.58 | 0.48 |
| 5 | 44515935 | rs13153459 | -0.093 | -0.838 | -0.564 | -0.729 | -0.013 | -0.035 | -0.035 | 0 | 0.24 | 0.4 | 0.22 | 0.12 | 0.22 | 0.12 |
| 5 | 44517392 | rs922855 | 0.857 | -0.861 | -0.448 | -0.812 | -0.002 | -0.043 | -0.043 | 0 | 0.17 | 0.2 | 0.1 | 0 | 0.15 | 0.1 |
| 5 | 44543851 | rs1351633 | -0.419 | -0.683 | -0.398 | -0.757 | -0.006 | 0.083 | 0.083 | 0 | 0.75 | 0.57 | 0.86 | 0.65 | 0.79 | 0.6 |
| 5 | 44560424 | rs13355205 | -0.332 | -0.87 | -0.03 | -0.52 | -0.015 | -0.06 | -0.06 | 0 | 0.24 | 0.44 | 0.22 | 0.1 | 0.23 | 0.12 |
| 5 | 44569226 | rs7708449 | 0.517 | -0.135 | -0.478 | -0.801 | -0.008 | -0.074 | -0.074 | 0 | 0.4 | 0.11 | 0.45 | 0.55 | 0.35 | 0.43 |
| 5 | 44570040 | rs13169538 | 0.546 | -0.118 | -0.438 | -0.827 | -0.003 | -0.029 | -0.029 | 0.603 | 0.06 | 0.13 | 0.06 | 0 | 0.06 | 0.01 |
| 5 | 44576171 | rs10054521 | -0.729 | -0.173 | -0.16 | -0.845 | -0.001 | -0.001 | -0.001 | 0 | 0.64 | 0.68 | 0.68 | 0.65 | 0.58 | 0.55 |
| 5 | 44598274 | rs13158953 | 0.046 | -0.205 | -0.175 | -0.829 | -0.007 | -0.037 | -0.037 | 0.182 | 0.07 | 0.17 | 0.07 | 0 | 0.06 | 0.01 |
| 5 | 44611650 | rs4573006 | -0.508 | -0.132 | -0.077 | -0.704 | 0.008 | 0.003 | 0.003 | 0 | 0.63 | 0.58 | 0.68 | 0.64 | 0.58 | 0.55 |
| 5 | 44628098 | rs6872069 | 0.19 | -0.396 | 0.667 | 0.138 | -0.01 | -0.003 | -0.003 | 0 | 0.17 | 0.35 | 0.16 | 0.12 | 0.16 | 0.11 |
| 5 | 44632008 | rs5004228 | -0.868 | -0.706 | 0.721 | -0.041 | -0.014 | 0.016 | 0.016 | 0 | 0.09 | 0.31 | 0.15 | 0.06 | 0.04 | 0.07 |
| 5 | 44662515 | rs4415084 | -0.863 | -0.439 | 0.966 | 0.036 | 0.018 | 0 | 0 | 0.124 | 0.46 | 0.65 | 0.56 | 0.55 | 0.41 | 0.48 |
| 5 | 44685698 | rs12522626 | -0.851 | -0.383 | 0.53 | 0.068 | 0.006 | -0.002 | -0.002 | 0.083 | 0.44 | 0.4 | 0.54 | 0.55 | 0.4 | 0.49 |
| 5 | 44687188 | rs4571480 | -0.741 | -0.364 | 0.405 | 0.039 | 0.01 | -0.014 | -0.014 | 1.676 | 0.43 | 0.32 | 0.48 | 0.49 | 0.4 | 0.48 |
| 5 | 44699051 | rs920328 | -0.503 | -0.362 | 0.226 | 0.055 | 0.015 | -0.037 | -0.037 | 0 | 0.41 | 0.23 | 0.46 | 0.49 | 0.38 | 0.48 |
| 5 | 44737710 | rs11747159 | -0.53 | -0.461 | 0.765 | 0.274 | 0.021 | -0.013 | -0.013 | 0.005 | 0.41 | 0.25 | 0.46 | 0.54 | 0.38 | 0.45 |
| 5 | 44777878 | rs11746980 | -0.813 | -0.472 | 0.847 | 0.28 | -0.008 | -0.003 | -0.003 | 0 | 0.45 | 0.56 | 0.49 | 0.55 | 0.4 | 0.45 |
| 5 | 44800665 | rs10462081 | -0.561 | -0.469 | 0.763 | 0.291 | 0.019 | -0.007 | -0.007 | 0 | 0.41 | 0.32 | 0.46 | 0.54 | 0.37 | 0.45 |
| 5 | 44812566 | rs11746506 | -0.527 | -0.473 | 0.824 | 0.288 | 0.022 | -0.016 | -0.016 | 1.229 | 0.41 | 0.25 | 0.46 | 0.54 | 0.37 | 0.45 |
| 5 | 44814004 | rs12188871 | -0.629 | -0.486 | 0.784 | 0.273 | 0.023 | -0.006 | -0.006 | 0.428 | 0.42 | 0.31 | 0.46 | 0.54 | 0.37 | 0.45 |
| 5 | 44816984 | rs7716571 | -0.901 | -0.507 | 0.92 | 0.266 | -0.005 | -0.002 | -0.002 | 0 | 0.46 | 0.57 | 0.49 | 0.55 | 0.4 | 0.45 |
| 5 | 44819646 | rs9637783 | -0.645 | -0.506 | 0.796 | 0.268 | 0.015 | -0.003 | -0.003 | 0 | 0.42 | 0.38 | 0.47 | 0.54 | 0.37 | 0.45 |
| 5 | 44875005 | rs7716600 | 0.032 | -0.691 | 0.259 | 0.235 | 0.017 | -0.084 | -0.084 | 0 | 0.74 | 0.83 | 0.7 | 0.52 | 0.8 | 0.74 |
| 5 | 44901349 | rs10462084 | 0.436 | -0.603 | 0.192 | 0.233 | 0.009 | 0.045 | 0.045 | 0 | 0.17 | 0.05 | 0.15 | 0.05 | 0.17 | 0.19 |
| 5 | 44927365 | rs10044096 | 0.665 | -0.42 | 0.859 | 0.347 | 0.012 | 0.004 | 0.004 | 0.251 | 0.55 | 0.39 | 0.51 | 0.45 | 0.6 | 0.55 |
| 5 | 44941630 | rs6875287 | -0.72 | -0.667 | 0.792 | 0.335 | 0.041 | -0.016 | -0.016 | 0 | 0.43 | 0.24 | 0.45 | 0.52 | 0.37 | 0.47 |
| 5 | 44961444 | rs3935213 | -0.299 | -1.131 | 0.991 | 0.023 | -0.009 | -0.027 | -0.027 | 0 | 0.25 | 0.27 | 0.28 | 0.05 | 0.2 | 0.19 |

| | | | | | | | | | | | | | | | | |
|---|---|---|---|---|---|---|---|---|---|---|---|---|---|---|---|---|
| 5 | 45003293 | rs11742346 | -0.121 | -1.29 | 0.892 | 0.556 | 0.012 | -0.04 | -0.04 | 0.12 | 0.2 | 0.08 | 0.15 | 0.28 | 0.13 | 0.26 |
| 5 | 45038026 | rs7731099 | -0.403 | -1.281 | 1.198 | 0.38 | -0.048 | -0.109 | -0.109 | 0 | 0.27 | 0.44 | 0.29 | 0.07 | 0.21 | 0.19 |
| 5 | 45057320 | rs12054976 | -0.229 | -1.283 | 1.332 | 0.505 | -0.013 | -0.027 | -0.027 | 0 | 0.25 | 0.27 | 0.28 | 0.1 | 0.21 | 0.19 |
| 5 | 45071911 | rs4389695 | 0.142 | -1.293 | 1.059 | 0.5 | 0.013 | 0.057 | 0.057 | 0.007 | 0.22 | 0.03 | 0.26 | 0.08 | 0.21 | 0.19 |
| 5 | 45074633 | rs13183434 | 0.006 | -1.28 | 1.028 | 1.14 | 0.005 | -0.065 | -0.065 | 0 | 0.19 | 0.08 | 0.14 | 0.4 | 0.12 | 0.27 |
| 5 | 45090306 | rs10051592 | -1.43 | -1.451 | 1.323 | 1.263 | 0.077 | 0 | 0 | 0 | 0.48 | 0.74 | 0.45 | 0.52 | 0.34 | 0.46 |
| 5 | 45099778 | rs10941692 | 0.081 | -1.316 | 1.174 | 1.011 | 0.018 | 0.057 | 0.057 | 0.034 | 0.22 | 0.05 | 0.26 | 0.08 | 0.21 | 0.19 |
| 5 | 45170011 | rs12516998 | -0.497 | -1.34 | 1.378 | 1.022 | -0.032 | -0.038 | -0.038 | 0 | 0.25 | 0.34 | 0.28 | 0.12 | 0.18 | 0.19 |
| 5 | 45182131 | rs7711528 | -0.42 | -1.332 | 1.379 | 1.026 | -0.033 | -0.04 | -0.04 | 0 | 0.24 | 0.34 | 0.28 | 0.12 | 0.18 | 0.19 |
| 5 | 45229673 | rs1392970 | 0.182 | -1.332 | 1.024 | 1.031 | 0.029 | 0.032 | 0.032 | 0 | 0.21 | 0.07 | 0.25 | 0.12 | 0.17 | 0.18 |
| 5 | 45265768 | rs13180087 | 0.471 | -1.296 | 0.78 | 1.345 | -0.004 | -0.042 | -0.042 | 0.006 | 0.15 | 0.08 | 0.13 | 0.38 | 0.13 | 0.21 |
| 5 | 45266589 | rs4866929 | 1.464 | -0.799 | -0.173 | 0.874 | -0.154 | -0.082 | -0.082 | 3.001 | 0.29 | 0.03 | 0.43 | 0.32 | 0.47 | 0.21 |
| 5 | 45277112 | rs6451793 | -0.101 | -0.848 | 0.6 | 0.846 | -0.047 | -0.068 | -0.068 | 0.33 | 0.26 | 0.46 | 0.29 | 0.12 | 0.2 | 0.19 |
| 5 | 45285752 | rs16902086 | -0.98 | -0.766 | 0.132 | 0.898 | 0.063 | -0.02 | -0.02 | 0.006 | 0.48 | 0.73 | 0.46 | 0.47 | 0.36 | 0.47 |
| 5 | 45291514 | rs9790873 | -0.241 | -0.623 | 0.864 | 1.268 | -0.015 | 0.087 | 0.087 | 0 | 0.22 | 0.44 | 0.16 | 0.36 | 0.14 | 0.24 |
| 5 | 45301035 | rs9292918 | -0.789 | -0.805 | 1.093 | 1.31 | -0.009 | 0.031 | 0.031 | 0.19 | 0.27 | 0.63 | 0.32 | 0.47 | 0.18 | 0.3 |
| 5 | 45346189 | rs34835217 | 0.095 | -0.811 | 1.369 | 1.277 | -0.004 | 0.024 | 0.024 | 0 | 0.2 | 0.25 | 0.15 | 0.34 | 0.14 | 0.24 |
| 5 | 45378207 | rs1501362 | -1.107 | -0.883 | 1.147 | 1.195 | 0.005 | -0.012 | -0.012 | 0 | 0.29 | 0.69 | 0.33 | 0.46 | 0.19 | 0.31 |
| 5 | 50229851 | rs27246 | -0.172 | 0.234 | -1.194 | -0.237 | -0.02 | -0.086 | -0.086 | 0.255 | 0.23 | 0.04 | 0.35 | 0.62 | 0.19 | 0.43 |
| 5 | 50230371 | rs27628 | -0.936 | 0.3 | -1.125 | -0.41 | 0.004 | 0.064 | 0.064 | 0 | 0.35 | 0.16 | 0.49 | 0.82 | 0.27 | 0.54 |
| 5 | 50269870 | rs250244 | -0.324 | 0.328 | -1.249 | -0.392 | -0.041 | -0.107 | -0.107 | 0 | 0.79 | 0.97 | 0.66 | 0.4 | 0.81 | 0.56 |
| 5 | 50279857 | rs622304 | -0.618 | 0.607 | -0.612 | -0.371 | 0.016 | -0.006 | -0.006 | 0 | 0.23 | 0.49 | 0.22 | 0.23 | 0.21 | 0.23 |
| 5 | 50281358 | rs250216 | 0.133 | 0.2 | -0.796 | -0.257 | -0.013 | 0.019 | 0.019 | 0 | 0.12 | 0.19 | 0.15 | 0.2 | 0.09 | 0.1 |
| 5 | 50289496 | rs250223 | -0.266 | 0.615 | -0.997 | -0.208 | 0.001 | -0.029 | -0.029 | 0 | 0.8 | 0.88 | 0.67 | 0.42 | 0.83 | 0.55 |
| 5 | 50292595 | rs250212 | -1.499 | 0.481 | -1.143 | 0.336 | 0.028 | -0.026 | -0.026 | 0 | 0.92 | 0.99 | 0.87 | 0.56 | 0.97 | 0.74 |
| 5 | 50297517 | rs13175573 | 0.816 | 0.694 | -1.11 | 0.172 | -0.028 | -0.034 | -0.034 | 0 | 0.09 | 0.03 | 0.15 | 0.2 | 0.12 | 0.09 |
| 5 | 50306946 | rs10055437 | 0.184 | 0.323 | -0.942 | 0.077 | 0.03 | 0.054 | 0.054 | 0 | 0.49 | 0.67 | 0.46 | 0.35 | 0.61 | 0.4 |
| 5 | 50318550 | rs7715247 | -0.346 | 0.211 | -0.984 | -0.309 | 0.014 | 0 | 0 | 0.499 | 0.21 | 0.3 | 0.21 | 0.24 | 0.24 | 0.24 |
| 5 | 50326227 | rs10064652 | -0.285 | -0.482 | -1.334 | -0.339 | 0.001 | -0.002 | -0.002 | 0.004 | 0.35 | 0.37 | 0.34 | 0.39 | 0.24 | 0.37 |
| 5 | 50332780 | rs7446150 | 0.141 | -0.079 | -1.292 | -0.917 | -0.001 | -0.004 | -0.004 | 0.16 | 0.56 | 0.51 | 0.48 | 0.32 | 0.5 | 0.41 |
| 5 | 50351642 | rs4429809 | 0.593 | 0.298 | -1.146 | -1.101 | -0.011 | 0.017 | 0.017 | 0 | 0.14 | 0.08 | 0.1 | 0.11 | 0.18 | 0.13 |
| 5 | 50352482 | rs10064446 | 0.21 | 0.349 | -0.964 | -1.325 | 0.002 | -0.03 | -0.03 | 0 | 0.11 | 0.12 | 0.06 | 0 | 0.13 | 0.07 |
| 5 | 50372657 | rs17220752 | 0.119 | 0.197 | -1.08 | -1.346 | -0.002 | 0.01 | 0.01 | 0.079 | 0.2 | 0.23 | 0.29 | 0.21 | 0.19 | 0.2 |
| 5 | 50374894 | rs4446421 | 1.284 | 0.164 | -1.053 | -1.299 | 0.056 | 0.067 | 0.067 | 0 | 0.06 | 0.22 | 0.12 | 0.06 | 0.06 | 0.06 |
| 5 | 50392093 | rs2961807 | 1.94 | 0.019 | -1.677 | -1.305 | -0.084 | -0.132 | -0.132 | 0 | 0.37 | 0.06 | 0.39 | 0.57 | 0.41 | 0.42 |
| 5 | 50400307 | rs6866012 | 0.592 | 0.295 | -0.611 | -1.254 | -0.011 | -0.006 | -0.006 | 0.025 | 0.15 | 0.23 | 0.15 | 0.11 | 0.09 | 0.18 |
| 5 | 50403297 | rs2962255 | 0.713 | 0.877 | -0.54 | -1.136 | -0.001 | -0.011 | -0.011 | 0.465 | 0.75 | 0.77 | 0.84 | 0.88 | 0.68 | 0.81 |
| 5 | 50413703 | rs2961811 | 0.917 | 0.565 | -0.272 | -0.977 | -0.006 | -0.004 | -0.004 | 0 | 0.82 | 0.75 | 0.78 | 0.86 | 0.84 | 0.9 |
| 5 | 50418252 | rs16890389 | 0.064 | 0.929 | -0.118 | -0.653 | -0.003 | 0.092 | 0.092 | 0 | 0.06 | 0.1 | 0.15 | 0.26 | 0.06 | 0.07 |
| 5 | 50428578 | rs6871271 | -0.075 | 1.644 | 0.225 | -0.927 | 0.052 | -0.037 | -0.037 | 0.412 | 0.11 | 0.3 | 0.1 | 0.04 | 0.16 | 0.16 |
| 5 | 50432178 | rs2962271 | 0.592 | 1.842 | 0.166 | -1.097 | -0.007 | 0.004 | 0.004 | 0 | 0.92 | 0.93 | 0.93 | 0.99 | 0.89 | 0.95 |
| 5 | 50434358 | rs2962276 | 1.334 | 1.838 | 0.28 | -1.136 | 0.065 | -0.094 | -0.094 | 0.745 | 0.79 | 0.54 | 0.81 | 0.95 | 0.72 | 0.79 |
| 5 | 50445762 | rs12696972 | -0.296 | 1.583 | 0.527 | -1.218 | -0.008 | 0.127 | 0.127 | 0 | 0.09 | 0.21 | 0.18 | 0.26 | 0.06 | 0.09 |
| 5 | 50451964 | rs2962247 | 1.88 | 1.561 | 0.359 | -1.219 | 0.009 | 0.002 | 0.002 | 0 | 0.71 | 0.33 | 0.64 | 0.69 | 0.67 | 0.7 |
| 5 | 50459798 | rs11740760 | 2.147 | 1.612 | -0.113 | -1.609 | 0.027 | 0.025 | 0.025 | 0 | 0.48 | 0.06 | 0.45 | 0.63 | 0.42 | 0.5 |
| 5 | 50465915 | rs13172994 | 1.165 | 2.351 | 0.371 | -1.584 | 0.051 | 0.019 | 0.019 | 0 | 0.78 | 0.38 | 0.69 | 0.77 | 0.73 | 0.74 |
| 5 | 50473298 | rs11749314 | 1.213 | 2.308 | 0.278 | -1.587 | 0.022 | 0.008 | 0.008 | 0 | 0.77 | 0.3 | 0.68 | 0.76 | 0.73 | 0.74 |
| 5 | 50495353 | rs1014076 | 2.015 | 2.111 | 0.265 | -1.632 | 0.016 | 0.056 | 0.056 | 0 | 0.55 | 0.27 | 0.53 | 0.76 | 0.45 | 0.61 |
| 5 | 50495914 | rs987756 | 1.221 | 2.188 | 0.538 | -1.37 | 0.006 | 0.022 | 0.022 | 0 | 0.83 | 0.69 | 0.75 | 0.79 | 0.83 | 0.86 |
| 5 | 50500628 | rs1039440 | -1.409 | 2.063 | 0.472 | -1.734 | 0.003 | -0.028 | -0.028 | 0.484 | 0.35 | 0.4 | 0.33 | 0.09 | 0.54 | 0.33 |
| 5 | 50523369 | rs7716272 | -2.249 | 1.626 | 0.721 | -1.352 | -0.01 | 0.047 | 0.047 | 0 | 0.43 | 0.73 | 0.4 | 0.28 | 0.54 | 0.4 |
| 5 | 50533202 | rs10080044 | -1.271 | 1.825 | 0.728 | -1.292 | 0.013 | -0.015 | -0.015 | 0.014 | 0.24 | 0.43 | 0.23 | 0.22 | 0.27 | 0.27 |
| 5 | 50560963 | rs1039441 | -0.124 | 1.89 | 0.224 | -1.293 | -0.031 | 0.02 | 0.02 | 0 | 0.13 | 0.05 | 0.1 | 0.06 | 0.19 | 0.11 |
| 5 | 50561259 | rs13181365 | -0.646 | 1.156 | 0.728 | -1.004 | 0.012 | -0.036 | -0.036 | 1.466 | 0.15 | 0.24 | 0.15 | 0.03 | 0.21 | 0.19 |
| 5 | 50563823 | rs1501988 | -1.447 | 1.17 | 0.089 | -0.909 | 0.002 | -0.017 | -0.017 | 0.041 | 0.3 | 0.63 | 0.29 | 0.24 | 0.34 | 0.29 |
| 5 | 50591476 | rs10052407 | -0.297 | 1.697 | 0.622 | -0.92 | 0.038 | -0.014 | -0.014 | 0 | 0.15 | 0.25 | 0.17 | 0.14 | 0.19 | 0.14 |
| 5 | 50596784 | rs2033237 | -1.257 | 1.533 | 1.035 | -1.068 | 0.045 | -0.068 | -0.068 | 1.262 | 0.23 | 0.52 | 0.23 | 0.14 | 0.25 | 0.21 |
| 5 | 50606563 | rs7714854 | -0.62 | 1.642 | 1.02 | -1.08 | 0.004 | -0.034 | -0.034 | 0 | 0.07 | 0.11 | 0.05 | 0 | 0.06 | 0.08 |
| 5 | 50612558 | rs10036382 | -1.193 | 1.636 | 1.244 | -1.084 | 0.056 | -0.071 | -0.071 | 0 | 0.23 | 0.51 | 0.23 | 0.14 | 0.26 | 0.21 |
| 5 | 50623533 | rs4141540 | -0.673 | 1.644 | 1.363 | -1.473 | 0.001 | -0.049 | -0.049 | 0.752 | 0.16 | 0.24 | 0.15 | 0.02 | 0.19 | 0.23 |
| 5 | 50634878 | rs6449586 | -2.178 | 1.109 | 0.809 | -2.028 | -0.033 | 0.183 | 0.183 | 0.006 | 0.53 | 0.91 | 0.45 | 0.23 | 0.55 | 0.49 |
| 5 | 50635643 | rs10037522 | -1.819 | 1.13 | 1.314 | -1.815 | -0.019 | 0.053 | 0.053 | 0.092 | 0.35 | 0.77 | 0.35 | 0.16 | 0.4 | 0.39 |
| 5 | 50636521 | rs9291750 | 0.211 | 1.121 | 1.535 | -1.824 | 0.016 | -0.011 | -0.011 | 0 | 0.15 | 0.18 | 0.15 | 0.14 | 0.19 | 0.14 |
| 5 | 50636650 | rs7702670 | -1.645 | 1.034 | 1.374 | -1.796 | -0.014 | 0.033 | 0.033 | 0.044 | 0.34 | 0.74 | 0.35 | 0.16 | 0.4 | 0.39 |
| 5 | 50659788 | rs4865656 | -1.489 | 1.034 | 1.487 | -1.731 | -0.003 | -0.001 | -0.001 | 0.212 | 0.33 | 0.7 | 0.36 | 0.17 | 0.35 | 0.35 |
| 5 | 50663234 | rs6869844 | -1.022 | 1.066 | 1.878 | -1.745 | -0.021 | -0.092 | -0.092 | 0.157 | 0.17 | 0.42 | 0.1 | 0.03 | 0.13 | 0.2 |
| 5 | 50683655 | rs2288468 | -1.548 | 1.002 | 1.877 | -1.743 | -0.001 | -0.008 | -0.008 | 0 | 0.33 | 0.67 | 0.35 | 0.18 | 0.36 | 0.34 |
| 5 | 50684451 | rs3811911 | 0.274 | 1.009 | 1.925 | -1.733 | 0.019 | -0.012 | -0.012 | 0 | 0.15 | 0.17 | 0.25 | 0.15 | 0.22 | 0.15 |
| 5 | 50695877 | rs6895297 | -1.435 | 0.995 | 1.916 | -1.736 | 0.017 | -0.056 | -0.056 | 0 | 0.32 | 0.61 | 0.35 | 0.18 | 0.36 | 0.34 |

| | | | | | | | | | | | | | | | | |
|---|---|---|---|---|---|---|---|---|---|---|---|---|---|---|---|---|
| 5 | 50704234 | rs6449613 | -0.335 | 0.965 | 1.818 | -1.715 | -0.008 | -0.034 | -0.034 | 0.974 | 0.13 | 0.21 | 0.06 | 0.03 | 0.12 | 0.17 |
| 5 | 50722238 | rs4865664 | -1.534 | 0.987 | 1.778 | -1.732 | -0.012 | 0.026 | 0.026 | 0 | 0.33 | 0.72 | 0.37 | 0.18 | 0.36 | 0.34 |
| 5 | 50734670 | rs16875094 | 0.317 | 1.044 | 1.66 | -1.716 | 0.025 | -0.012 | -0.012 | 0 | 0.14 | 0.18 | 0.25 | 0.15 | 0.22 | 0.14 |
| 5 | 50736314 | rs11951998 | -1.438 | 0.987 | 1.858 | -1.721 | 0.003 | -0.019 | -0.019 | 0 | 0.32 | 0.64 | 0.36 | 0.18 | 0.36 | 0.34 |
| 5 | 50739152 | rs10039723 | -1.194 | 1.012 | 1.804 | -1.709 | 0.033 | -0.068 | -0.068 | 0 | 0.3 | 0.46 | 0.35 | 0.18 | 0.36 | 0.33 |
| 5 | 50743281 | rs7719442 | -0.338 | 1.119 | 1.59 | -1.622 | 0.08 | -0.045 | -0.045 | 0.022 | 0.18 | 0.37 | 0.29 | 0.15 | 0.24 | 0.16 |
| 5 | 50771082 | rs17824668 | 1.356 | 0.478 | 0.746 | -2.145 | 0.006 | 0.066 | 0.066 | 0.317 | 0.11 | 0.01 | 0.03 | 0 | 0.06 | 0.04 |
| 5 | 50771114 | rs1423611 | -1.72 | 0.184 | 0.476 | -2.296 | -0.027 | 0.066 | 0.066 | 0 | 0.59 | 0.84 | 0.49 | 0.3 | 0.54 | 0.48 |
| 5 | 50781062 | rs10053849 | 1.156 | 0.165 | 0.693 | -2.335 | 0.011 | -0.006 | -0.006 | 0.002 | 0.15 | 0.16 | 0.26 | 0.15 | 0.23 | 0.15 |
| 5 | 50781973 | rs1363999 | 0.363 | 0.057 | 0.652 | -2.369 | -0.021 | -0.046 | -0.046 | 0.19 | 0.17 | 0.34 | 0.08 | 0.06 | 0.12 | 0.12 |
| 5 | 50784546 | rs6891554 | -1.943 | 0.064 | 0.25 | -2.343 | -0.049 | 0.064 | 0.064 | 0 | 0.59 | 0.9 | 0.5 | 0.29 | 0.54 | 0.48 |
| 5 | 50788606 | rs9291789 | -0.685 | 0.304 | 0.472 | -2.221 | 0.004 | -0.015 | -0.015 | 0.474 | 0.34 | 0.65 | 0.37 | 0.24 | 0.36 | 0.31 |
| 5 | 50791584 | rs7446508 | 0.581 | 0.534 | 0.715 | -1.93 | 0.063 | -0.033 | -0.033 | 0.924 | 0.19 | 0.38 | 0.29 | 0.18 | 0.25 | 0.19 |
| 5 | 50794221 | rs1559278 | -0.311 | 0.744 | 0.614 | -1.98 | 0.058 | -0.022 | -0.022 | 0 | 0.27 | 0.5 | 0.37 | 0.26 | 0.35 | 0.31 |
| 5 | 50821162 | rs12189442 | 0.594 | 0.675 | 0.74 | -1.723 | -0.001 | -0.001 | -0.001 | 0 | 0.09 | 0.08 | 0.07 | 0.08 | 0.11 | 0.12 |
| 5 | 50828866 | rs6896357 | 0.305 | 0.584 | 0.767 | -1.914 | 0.009 | -0.054 | -0.054 | 0 | 0.34 | 0.56 | 0.35 | 0.19 | 0.32 | 0.26 |
| 5 | 50840436 | rs1946265 | -1.6 | 0.384 | 0.345 | -2.27 | -0.006 | 0.003 | 0.003 | 0 | 0.68 | 0.89 | 0.54 | 0.32 | 0.65 | 0.61 |
| 5 | 50851575 | rs6449708 | -0.935 | 0.354 | 0.222 | -2.246 | -0.007 | 0.101 | 0.101 | 0 | 0.58 | 0.79 | 0.46 | 0.24 | 0.53 | 0.49 |
| 5 | 50853897 | rs10471608 | -0.995 | 0.435 | 0.305 | -2.237 | -0.02 | -0.081 | -0.081 | 0.02 | 0.29 | 0.42 | 0.14 | 0.06 | 0.22 | 0.24 |
| 5 | 50890863 | rs10063114 | 0.682 | 0.647 | 0.159 | -1.826 | 0.011 | -0.042 | -0.042 | 0.111 | 0.3 | 0.35 | 0.32 | 0.15 | 0.31 | 0.24 |
| 5 | 50920764 | rs17236135 | 0.385 | 0.478 | -0.435 | -1.193 | -0.011 | 0.041 | 0.041 | 0 | 0.08 | 0.02 | 0.04 | 0 | 0.09 | 0.07 |
| 5 | 50922671 | rs1423304 | -1.202 | 0.346 | -0.045 | -1.26 | -0.008 | 0 | 0 | 0 | 0.32 | 0.6 | 0.26 | 0.23 | 0.29 | 0.32 |
| 5 | 50928810 | rs4865673 | 1.146 | 0.348 | -0.315 | -1.59 | 0 | -0.036 | -0.036 | 0 | 0.3 | 0.27 | 0.3 | 0.06 | 0.31 | 0.25 |
| 5 | 50928871 | rs244545 | -1.248 | 0.398 | -0.18 | -1.446 | -0.018 | -0.043 | -0.043 | 0 | 0.67 | 0.9 | 0.57 | 0.29 | 0.64 | 0.6 |
| 5 | 50971593 | rs244554 | -0.271 | 0.228 | 0.37 | -1.06 | 0 | 0.032 | 0.032 | 0 | 0.18 | 0.28 | 0.19 | 0.23 | 0.19 | 0.25 |
| 5 | 50984183 | rs4865518 | -0.957 | 0.04 | 0.029 | -0.914 | 0.019 | -0.045 | -0.045 | 0 | 0.88 | 0.96 | 0.91 | 0.8 | 0.9 | 0.84 |
| 5 | 51012870 | rs10056533 | 0.706 | 0.273 | -0.065 | -1.564 | -0.012 | 0.001 | 0.001 | 0 | 0.4 | 0.33 | 0.36 | 0.06 | 0.47 | 0.31 |
| 5 | 51017612 | rs17834513 | 0.22 | 0.289 | -0.575 | -1.519 | -0.011 | -0.042 | -0.042 | 0.049 | 0.39 | 0.11 | 0.43 | 0.64 | 0.35 | 0.43 |
| 5 | 51022880 | rs6862321 | -0.342 | 0.103 | 0.03 | -1.074 | 0.001 | -0.005 | -0.005 | 0 | 0.82 | 0.86 | 0.83 | 0.7 | 0.82 | 0.74 |
| 5 | 51029098 | rs6867918 | -0.089 | 0.258 | -0.223 | -1.091 | -0.005 | 0.041 | 0.041 | 0 | 0.11 | 0.03 | 0.04 | 0 | 0.08 | 0.07 |
| 5 | 51059051 | rs4865684 | 0.652 | 0.074 | 0.237 | -0.964 | -0.018 | 0.022 | 0.022 | 0 | 0.72 | 0.54 | 0.76 | 0.72 | 0.73 | 0.67 |
| 5 | 51070843 | rs4242024 | -0.584 | 0.199 | 0.091 | -0.785 | 0 | 0.013 | 0.013 | 0 | 0.85 | 0.86 | 0.83 | 0.74 | 0.83 | 0.77 |
| 5 | 51108566 | rs6872636 | 0.898 | 0.298 | -0.134 | -0.765 | -0.008 | -0.017 | -0.017 | 0 | 0.28 | 0.22 | 0.4 | 0.63 | 0.36 | 0.38 |
| 5 | 51108839 | rs12188469 | 0.734 | 0.24 | 0.184 | -0.853 | -0.004 | 0.041 | 0.041 | 0.173 | 0.15 | 0.19 | 0.16 | 0.29 | 0.14 | 0.24 |
| 5 | 51113147 | rs4242025 | 1.722 | 0.187 | 0.088 | -1.245 | -0.004 | 0.059 | 0.059 | 0 | 0.31 | 0.19 | 0.31 | 0.08 | 0.3 | 0.22 |
| 5 | 51116418 | rs12659291 | 0.949 | 0.2 | 0.057 | -1.049 | 0.001 | -0.029 | -0.029 | 0 | 0.14 | 0.08 | 0.15 | 0.26 | 0.13 | 0.23 |
| 5 | 51123678 | rs4146568 | 0.983 | 0.294 | 0.127 | -1.071 | -0.004 | -0.01 | -0.01 | 0.419 | 0.28 | 0.25 | 0.4 | 0.63 | 0.35 | 0.39 |
| 5 | 51124949 | rs7713485 | 0.587 | 0.192 | 0.52 | -0.927 | -0.01 | 0.073 | 0.073 | 0.212 | 0.16 | 0.28 | 0.17 | 0.28 | 0.14 | 0.23 |
| 5 | 51130912 | rs16879212 | 0.81 | 0.216 | 0.398 | -0.882 | -0.005 | 0.039 | 0.039 | 0.207 | 0.16 | 0.21 | 0.17 | 0.3 | 0.14 | 0.24 |
| 5 | 51135688 | rs6872018 | 0.18 | -0.11 | 0.291 | -1.025 | -0.004 | -0.002 | -0.002 | 0 | 0.37 | 0.35 | 0.46 | 0.67 | 0.44 | 0.48 |
| 5 | 51146016 | rs3111109 | -0.491 | -0.342 | 0.001 | -1.276 | -0.016 | -0.077 | -0.077 | 1.177 | 0.17 | 0.26 | 0.08 | 0 | 0.13 | 0.08 |
| 5 | 51153019 | rs3111112 | -0.826 | -0.067 | -0.206 | -0.752 | -0.008 | 0.051 | 0.051 | 0 | 0.47 | 0.74 | 0.51 | 0.73 | 0.51 | 0.54 |
| 5 | 51155430 | rs3111113 | -0.023 | 0.148 | -0.257 | -0.821 | -0.002 | -0.024 | -0.024 | 0 | 0.08 | 0.11 | 0.03 | 0 | 0.07 | 0.02 |
| 5 | 51159583 | rs3111066 | -1.256 | 0.217 | -0.557 | -0.782 | -0.018 | 0.116 | 0.116 | 0 | 0.62 | 0.83 | 0.64 | 0.89 | 0.6 | 0.72 |
| 5 | 51163584 | rs4296746 | 0.234 | 0.51 | -0.624 | -0.064 | -0.084 | 0.014 | 0.014 | 0.443 | 0.2 | 0.02 | 0.22 | 0.15 | 0.28 | 0.31 |
| 5 | 51174404 | rs6449873 | 0.167 | 0.448 | 0.134 | -0.001 | 0.004 | 0.01 | 0.01 | 0 | 0.24 | 0.25 | 0.4 | 0.68 | 0.32 | 0.47 |
| 5 | 51176115 | rs16879231 | 0.108 | -0.242 | 0.733 | 0.378 | 0.019 | -0.016 | -0.016 | 1.097 | 0.18 | 0.11 | 0.13 | 0.16 | 0.1 | 0.2 |
| 5 | 51179679 | rs4331855 | 1.003 | 0.555 | 1.354 | 0.101 | 0.013 | -0.04 | -0.04 | 0.079 | 0.91 | 0.84 | 0.94 | 1 | 0.88 | 0.95 |
| 5 | 51182660 | rs10036878 | 2.175 | 0.31 | 1.57 | 0.034 | 0.012 | -0.111 | -0.111 | 0 | 0.81 | 0.51 | 0.87 | 0.95 | 0.81 | 0.91 |
| 5 | 51185421 | rs4502769 | 1.089 | 0.356 | 1.098 | -0.043 | 0.015 | -0.05 | -0.05 | 0.057 | 0.9 | 0.79 | 0.93 | 1 | 0.87 | 0.94 |
| 5 | 51189407 | rs6862940 | -0.684 | 0.378 | 1.125 | -0.027 | -0.007 | -0.043 | -0.043 | 0 | 0.08 | 0.19 | 0.05 | 0 | 0.06 | 0.02 |
| 5 | 51199072 | rs4865694 | 1.229 | 0.554 | 0.718 | 0.304 | 0.014 | -0.044 | -0.044 | 0 | 0.9 | 0.81 | 0.94 | 1 | 0.88 | 0.94 |
| 5 | 51203281 | rs9291901 | -1.78 | 0.182 | 1.086 | 0.272 | -0.004 | -0.037 | -0.037 | 1.953 | 0.08 | 0.23 | 0.05 | 0 | 0.05 | 0.05 |
| 5 | 51231060 | rs17254098 | -1.314 | 0.263 | 0.854 | 0.122 | -0.006 | 0.006 | 0.006 | 0 | 0.08 | 0.05 | 0.04 | 0.01 | 0.09 | 0.12 |
| 5 | 51257127 | rs4296747 | 1.525 | 0.186 | 1.638 | 0.08 | -0.006 | -0.039 | -0.039 | 0 | 0.92 | 0.72 | 0.94 | 1 | 0.94 | 0.97 |
| 5 | 51257336 | rs4618396 | 2.652 | 0.08 | 2.102 | -0.02 | 0.006 | -0.118 | -0.118 | 0.486 | 0.84 | 0.46 | 0.88 | 1 | 0.85 | 0.87 |
| 5 | 51263501 | rs10051973 | 1.614 | 0.122 | 1.998 | 0.684 | 0.006 | -0.031 | -0.031 | 0.06 | 0.92 | 0.65 | 0.93 | 1 | 0.94 | 0.96 |
| 5 | 51267268 | rs11750419 | -2.448 | 0.587 | 2.149 | 1.384 | 0.048 | 0.202 | 0.202 | 0 | 0.19 | 0.44 | 0.26 | 0.54 | 0.22 | 0.35 |
| 5 | 51276536 | rs10461541 | -2.063 | 0.599 | 2.101 | 1.316 | 0.011 | -0.054 | -0.054 | 0 | 0.12 | 0.22 | 0.08 | 0.01 | 0.12 | 0.14 |
| 5 | 51277529 | rs10067022 | -0.232 | 0.665 | 2.051 | 1.447 | 0.04 | 0.296 | 0.296 | 0 | 0.07 | 0.2 | 0.18 | 0.53 | 0.1 | 0.22 |
| 5 | 51282797 | rs4429802 | -0.392 | 0.7 | 2.087 | 0.902 | 0.02 | -0.028 | -0.028 | 0.688 | 0.06 | 0.16 | 0.1 | 0 | 0.1 | 0.07 |
| 5 | 51283559 | rs4627945 | -1.964 | 0.68 | 2.124 | 1.1 | 0.014 | -0.007 | -0.007 | 0.269 | 0.13 | 0.22 | 0.19 | 0.09 | 0.17 | 0.15 |
| 5 | 51303575 | rs4075972 | -1.002 | 0.499 | 1.97 | 1.164 | 0.002 | 0.003 | 0.003 | 0 | 0.07 | 0.09 | 0.09 | 0.09 | 0.07 | 0.08 |
| 5 | 51311778 | rs17255242 | -0.809 | 0.316 | 1.768 | 0.977 | -0.001 | -0.008 | -0.008 | 0 | 0.06 | 0.06 | 0.04 | 0.01 | 0.06 | 0.08 |
| 5 | 51367031 | rs13355028 | -1.035 | -0.237 | 1.331 | 0.694 | -0.026 | -0.042 | -0.042 | 0 | 0.32 | 0.5 | 0.23 | 0.14 | 0.24 | 0.28 |
| 5 | 51367454 | rs4540127 | -1.23 | -0.238 | 1.337 | 0.717 | -0.031 | -0.021 | -0.021 | 0 | 0.34 | 0.51 | 0.27 | 0.21 | 0.25 | 0.28 |
| 5 | 51371819 | rs4131677 | -0.584 | -0.373 | 1.412 | 0.663 | -0.053 | -0.029 | -0.029 | 1.251 | 0.28 | 0.49 | 0.2 | 0.14 | 0.18 | 0.21 |
| 5 | 51388888 | rs4612018 | -0.322 | -0.343 | 1.391 | 0.679 | -0.05 | -0.029 | -0.029 | 0 | 0.26 | 0.43 | 0.21 | 0.13 | 0.17 | 0.2 |
| 5 | 51393342 | rs6861077 | -0.322 | -0.313 | 1.227 | 0.679 | -0.05 | -0.029 | -0.029 | 0.158 | 0.26 | 0.43 | 0.21 | 0.13 | 0.17 | 0.2 |

| | | | | | | | | | | | | | | | | |
|---|---|---|---|---|---|---|---|---|---|---|---|---|---|---|---|---|
| 5 | 51401580 | rs9291954 | 0.225 | -0.109 | 0.265 | 0.066 | -0.001 | 0.013 | 0.013 | 0 | 0.56 | 0.63 | 0.49 | 0.23 | 0.52 | 0.31 |
| 5 | 51405600 | rs12655411 | -0.49 | -0.079 | 0.05 | -0.089 | 0.006 | 0.023 | 0.023 | 0 | 0.64 | 0.78 | 0.54 | 0.23 | 0.61 | 0.43 |
| 5 | 51412949 | rs7733928 | 0.53 | 0.119 | -0.126 | 0.557 | -0.012 | -0.008 | -0.008 | 0 | 0.09 | 0.05 | 0.16 | 0.28 | 0.21 | 0.2 |
| 5 | 51415798 | rs6450047 | 0.471 | -0.183 | 0.113 | 0.106 | 0 | 0.018 | 0.018 | 0 | 0.5 | 0.45 | 0.48 | 0.32 | 0.47 | 0.24 |
| 5 | 51425511 | rs4434347 | 0.907 | -0.514 | 0.108 | -0.155 | -0.011 | -0.001 | -0.001 | 0 | 0.18 | 0.16 | 0.13 | 0.11 | 0.1 | 0.13 |
| 5 | 51427729 | rs12187185 | 0.106 | -0.41 | 0.162 | -0.22 | 0.008 | 0.018 | 0.018 | 0 | 0.55 | 0.39 | 0.46 | 0.22 | 0.55 | 0.33 |
| 5 | 51429429 | rs6864053 | -0.401 | -0.834 | 0.223 | -1.213 | -0.051 | -0.142 | -0.142 | 0.156 | 0.31 | 0.5 | 0.17 | 0.04 | 0.18 | 0.23 |
| 5 | 51431680 | rs10064799 | 0.61 | -0.441 | -0.066 | -1.114 | -0.026 | 0.004 | 0.004 | 0 | 0.63 | 0.71 | 0.53 | 0.37 | 0.52 | 0.46 |
| 5 | 51432669 | rs4507452 | 0.603 | -0.344 | -0.104 | -1.094 | -0.017 | -0.019 | -0.019 | 0 | 0.18 | 0.14 | 0.39 | 0.57 | 0.34 | 0.4 |
| 5 | 51442271 | rs4339299 | 0.191 | -0.612 | 0.06 | -1.302 | 0.004 | 0.027 | 0.027 | 0 | 0.27 | 0.16 | 0.11 | 0.07 | 0.22 | 0.2 |
| 5 | 51448021 | rs7443372 | 1.357 | -0.726 | 0.573 | -1.342 | -0.002 | 0.037 | 0.037 | 0 | 0.4 | 0.29 | 0.42 | 0.31 | 0.39 | 0.33 |
| 5 | 51448042 | rs7445986 | 1.3 | -0.742 | 0.618 | -1.349 | -0.007 | 0.044 | 0.044 | 0 | 0.39 | 0.25 | 0.41 | 0.31 | 0.39 | 0.33 |
| 5 | 51449637 | rs7714590 | 0.015 | -0.66 | 0.764 | -2.046 | -0.006 | -0.051 | -0.051 | 0 | 0.28 | 0.35 | 0.14 | 0.07 | 0.22 | 0.21 |
| 5 | 51464618 | rs7703986 | -0.023 | -0.608 | 0.495 | -2.01 | 0.014 | 0.012 | 0.012 | 0 | 0.72 | 0.64 | 0.55 | 0.37 | 0.61 | 0.55 |
| 5 | 51483914 | rs13165318 | 1.158 | -1.136 | 0.249 | -1.852 | 0.004 | 0.005 | 0.005 | 0.327 | 0.17 | 0.11 | 0.08 | 0.07 | 0.13 | 0.12 |
| 5 | 51491064 | rs4348174 | 0.49 | -0.837 | 0.321 | -1.651 | -0.001 | -0.006 | -0.006 | 0 | 0.43 | 0.46 | 0.44 | 0.3 | 0.4 | 0.33 |
| 5 | 51494293 | rs13180551 | 1.428 | -0.918 | -0.108 | -1.794 | -0.021 | 0.045 | 0.045 | 0 | 0.23 | 0.12 | 0.26 | 0.13 | 0.26 | 0.08 |
| 5 | 51495855 | rs10041461 | 0.634 | -0.512 | -0.022 | -1.537 | -0.003 | -0.015 | -0.015 | 0 | 0.43 | 0.54 | 0.45 | 0.3 | 0.4 | 0.33 |
| 5 | 51508533 | rs7708815 | -0.815 | 0.101 | -0.037 | -1.247 | -0.008 | 0.002 | 0.002 | 0.248 | 0.34 | 0.32 | 0.5 | 0.66 | 0.46 | 0.69 |
| 5 | 51515468 | rs4865529 | 0.676 | -0.306 | 0.197 | -1.434 | -0.02 | -0.024 | -0.024 | 0.406 | 0.91 | 0.54 | 0.94 | 0.99 | 0.98 | 0.96 |
| 5 | 51528556 | rs16879486 | 1.26 | -0.279 | -0.296 | -1.811 | -0.074 | 0.034 | 0.034 | 0 | 0.32 | 0.03 | 0.29 | 0.22 | 0.34 | 0.13 |
| 5 | 51531910 | rs10512950 | 1.586 | 0.749 | 0.695 | -0.896 | -0.02 | -0.073 | -0.073 | 0.334 | 0.76 | 0.43 | 0.89 | 0.92 | 0.84 | 0.73 |
| 5 | 51535809 | rs12187948 | -0.128 | 0.983 | 0.512 | -0.683 | -0.013 | -0.024 | -0.024 | 0 | 0.19 | 0.17 | 0.44 | 0.59 | 0.31 | 0.46 |
| 5 | 51554332 | rs6896294 | -0.835 | 0.905 | 0.903 | -0.716 | 0.005 | -0.002 | -0.002 | 0 | 0.11 | 0.34 | 0.07 | 0.07 | 0.08 | 0.15 |
| 5 | 51560998 | rs10072937 | -0.689 | 0.696 | 0.947 | -0.696 | -0.008 | -0.022 | -0.022 | 0.004 | 0.1 | 0.32 | 0.06 | 0.03 | 0.06 | 0.1 |
| 5 | 51563266 | rs16879548 | -0.49 | 0.783 | 0.956 | -0.75 | 0.001 | -0.018 | -0.018 | 0 | 0.07 | 0.07 | 0.02 | 0 | 0.06 | 0.08 |
| 5 | 51565236 | rs7703314 | -1.686 | 0.643 | 1.065 | -0.561 | 0.006 | -0.08 | -0.08 | 2.863 | 0.17 | 0.44 | 0.09 | 0.03 | 0.12 | 0.18 |
| 5 | 51577564 | rs10447095 | 0.011 | 0.752 | 0.661 | -0.4 | -0.011 | -0.018 | -0.018 | 0 | 0.18 | 0.14 | 0.44 | 0.59 | 0.31 | 0.46 |
| 5 | 51608348 | rs13155284 | -0.53 | 0.716 | 0.678 | -0.556 | -0.006 | -0.018 | -0.018 | 0 | 0.09 | 0.27 | 0.06 | 0.03 | 0.06 | 0.1 |
| 5 | 51625646 | rs10471360 | -0.248 | 0.735 | 0.085 | -0.547 | -0.005 | -0.008 | -0.008 | 2.687 | 0.09 | 0.16 | 0.06 | 0.03 | 0.07 | 0.09 |
| 5 | 51626698 | rs7705145 | 0.104 | 0.922 | -0.247 | -0.371 | -0.013 | -0.031 | -0.031 | 0 | 0.17 | 0.12 | 0.38 | 0.55 | 0.3 | 0.53 |
| 5 | 51643282 | rs10058189 | -1.622 | 0.898 | -0.081 | -0.447 | -0.005 | 0.011 | 0.011 | 0.188 | 0.41 | 0.71 | 0.5 | 0.6 | 0.47 | 0.7 |
| 5 | 51646929 | rs1593280 | 1.089 | 0.972 | -0.027 | -0.496 | 0.008 | 0.005 | 0.005 | 0.033 | 0.65 | 0.41 | 0.53 | 0.43 | 0.56 | 0.38 |
| 5 | 51649222 | rs16879651 | -0.6 | 1.013 | -0.101 | -0.495 | 0.002 | -0.027 | -0.027 | 0 | 0.08 | 0.11 | 0.03 | 0.01 | 0.07 | 0.07 |
| 5 | 51657466 | rs16879682 | -0.715 | 1.058 | -0.117 | -0.503 | 0.007 | -0.033 | -0.033 | 0 | 0.09 | 0.12 | 0.04 | 0.01 | 0.08 | 0.08 |
| 5 | 51660682 | rs1559578 | 0.157 | 0.37 | 0.072 | -0.213 | -0.023 | -0.037 | -0.037 | 0.335 | 0.78 | 0.85 | 0.61 | 0.46 | 0.64 | 0.47 |
| 5 | 51663821 | rs10063953 | -0.37 | -0.068 | 0.28 | -0.149 | -0.009 | -0.039 | -0.039 | 1.321 | 0.13 | 0.43 | 0.07 | 0.03 | 0.08 | 0.09 |
| 5 | 51680179 | rs2961725 | 0.544 | -0.15 | -0.401 | -0.095 | -0.009 | -0.026 | -0.026 | 1.787 | 0.72 | 0.83 | 0.55 | 0.46 | 0.61 | 0.45 |
| 5 | 51688530 | rs10067693 | -0.225 | -0.184 | -0.309 | -0.19 | -0.015 | -0.064 | -0.064 | 0.033 | 0.13 | 0.46 | 0.07 | 0.01 | 0.07 | 0.09 |
| 5 | 51688602 | rs2911762 | 1.306 | -0.071 | -1.133 | -0.09 | 0.037 | 0.056 | 0.056 | 0 | 0.54 | 0.22 | 0.44 | 0.4 | 0.52 | 0.32 |
| 5 | 51690015 | rs1424894 | 0.534 | 1.038 | -1.196 | 0.814 | -0.015 | -0.03 | -0.03 | 2.093 | 0.8 | 0.87 | 0.61 | 0.45 | 0.68 | 0.46 |
| 5 | 51700018 | rs10074792 | -0.908 | 0.758 | -1.429 | 0.574 | 0 | 0.003 | 0.003 | 0.498 | 0.35 | 0.18 | 0.44 | 0.63 | 0.37 | 0.53 |
| 5 | 51704359 | rs11739296 | -1.06 | 0.555 | -0.528 | 1.289 | 0.008 | 0.1 | 0.1 | 0 | 0.08 | 0.25 | 0.1 | 0.2 | 0.06 | 0.18 |
| 5 | 51704435 | rs2961726 | -0.17 | 0.428 | -1.756 | 0.756 | -0.025 | 0.016 | 0.016 | 0 | 0.72 | 0.96 | 0.74 | 0.79 | 0.7 | 0.79 |
| 5 | 51708099 | rs16879721 | -0.543 | 0.576 | -1.558 | 0.592 | 0.012 | 0.058 | 0.058 | 0 | 0.1 | 0.24 | 0.12 | 0.19 | 0.11 | 0.21 |
| 5 | 51709218 | rs6865723 | 0.926 | 0.355 | -1.417 | 0.37 | -0.012 | 0.026 | 0.026 | 0.114 | 0.33 | 0.17 | 0.31 | 0.28 | 0.38 | 0.2 |
| 5 | 51710027 | rs1424908 | 0.402 | -0.446 | -1.18 | -0.024 | -0.005 | 0 | 0 | 0 | 0.44 | 0.57 | 0.55 | 0.44 | 0.48 | 0.51 |
| 5 | 51711739 | rs7717591 | -0.595 | -0.253 | -1.19 | -0.001 | -0.004 | 0.09 | 0.09 | 0 | 0.6 | 0.82 | 0.68 | 0.79 | 0.61 | 0.74 |
| 5 | 51717417 | rs2161956 | 0.632 | -0.334 | -1.074 | 0.026 | -0.005 | 0.029 | 0.029 | 0.013 | 0.54 | 0.38 | 0.42 | 0.29 | 0.5 | 0.35 |
| 5 | 51717496 | rs1345984 | -0.82 | -0.33 | -1.105 | 0.029 | -0.02 | 0.025 | 0.025 | 0 | 0.72 | 0.86 | 0.75 | 0.79 | 0.7 | 0.78 |
| 5 | 51717912 | rs4865534 | 1.468 | -0.263 | -1.059 | 0.14 | -0.002 | 0.048 | 0.048 | 0 | 0.06 | 0.08 | 0.2 | 0.29 | 0.09 | 0.23 |
| 5 | 51720930 | rs9647518 | 1.25 | -0.432 | -1.224 | -0.118 | -0.034 | 0.118 | 0.118 | 0 | 0.43 | 0.1 | 0.37 | 0.28 | 0.45 | 0.23 |
| 5 | 51724318 | rs10074909 | 1.728 | -0.447 | -1.101 | -0.121 | -0.017 | -0.026 | -0.026 | 0 | 0.09 | 0.19 | 0.04 | 0.01 | 0.04 | 0.11 |
| 5 | 51733627 | rs2405853 | 2.379 | -0.296 | -1.175 | -0.112 | -0.002 | -0.005 | -0.005 | 0 | 0.06 | 0.03 | 0.03 | 0 | 0.05 | 0.01 |
| 5 | 51734233 | rs13173579 | 1.163 | -0.174 | -1.196 | 0.08 | -0.03 | 0.147 | 0.147 | 0 | 0.42 | 0.09 | 0.36 | 0.21 | 0.45 | 0.23 |
| 5 | 51736293 | rs7719826 | 0.256 | -0.155 | -1.137 | 0.124 | 0.006 | 0.005 | 0.005 | 0 | 0.53 | 0.34 | 0.48 | 0.47 | 0.56 | 0.44 |
| 5 | 51737413 | rs1113266 | 0.235 | -0.153 | -1.128 | 0.116 | 0.004 | 0.015 | 0.015 | 0 | 0.53 | 0.34 | 0.49 | 0.41 | 0.56 | 0.44 |
| 5 | 51741996 | rs16879748 | 0.003 | -0.168 | -1.046 | 0.099 | 0.007 | 0.047 | 0.047 | 0 | 0.12 | 0.27 | 0.12 | 0.2 | 0.12 | 0.22 |
| 5 | 51743012 | rs1364625 | 0.623 | -0.207 | -1.14 | 0.091 | -0.002 | 0.012 | 0.012 | 0 | 0.5 | 0.22 | 0.57 | 0.57 | 0.54 | 0.46 |
| 5 | 51743427 | rs13172170 | -0.354 | -0.257 | -1.083 | -0.032 | 0.001 | 0.005 | 0.005 | 0 | 0.63 | 0.66 | 0.7 | 0.72 | 0.66 | 0.68 |
| 5 | 51749055 | rs750544 | 0.764 | -0.344 | -0.622 | -0.012 | 0.004 | -0.022 | -0.022 | 0 | 0.05 | 0.17 | 0.03 | 0 | 0.03 | 0.01 |
| 5 | 51749342 | rs17260318 | 1.309 | 0.046 | -1.212 | -0.11 | -0.025 | 0.029 | 0.029 | 0 | 0.37 | 0.08 | 0.27 | 0.28 | 0.4 | 0.23 |
| 5 | 51757715 | rs11948045 | 0.936 | 0.059 | -0.852 | -0.112 | -0.013 | -0.007 | -0.007 | 0 | 0.08 | 0.07 | 0.11 | 0.13 | 0.13 | 0.15 |
| 5 | 51759418 | rs11956028 | 0.806 | 0.046 | -0.254 | -0.094 | -0.006 | 0.033 | 0.033 | 0.457 | 0.46 | 0.24 | 0.47 | 0.59 | 0.5 | 0.5 |
| 5 | 51762532 | rs2406113 | -1.282 | -0.108 | -0.26 | -0.272 | 0.001 | 0.02 | 0.02 | 0 | 0.73 | 0.8 | 0.76 | 0.8 | 0.74 | 0.81 |
| 5 | 51774963 | rs13159991 | 0.157 | -0.09 | -0.203 | -0.271 | 0.008 | -0.004 | -0.004 | 0 | 0.14 | 0.2 | 0.13 | 0.13 | 0.17 | 0.24 |
| 5 | 51779197 | rs6864159 | 1.158 | -0.101 | -0.137 | -0.219 | 0.021 | 0.175 | 0.175 | 0.096 | 0.09 | 0.23 | 0.22 | 0.3 | 0.1 | 0.26 |
| 5 | 51780385 | rs13162834 | 1.323 | -0.138 | -0.205 | -0.256 | -0.034 | 0.019 | 0.019 | 0 | 0.42 | 0.08 | 0.37 | 0.36 | 0.45 | 0.29 |
| 5 | 51783027 | rs6860155 | -1.37 | -0.147 | -0.158 | -0.268 | 0 | 0.02 | 0.02 | 0 | 0.73 | 0.82 | 0.77 | 0.8 | 0.74 | 0.81 |

| | | | | | | | | | | | | | | | | |
|---|---|---|---|---|---|---|---|---|---|---|---|---|---|---|---|---|
| 5 | 51787940 | rs10940253 | 0.627 | -0.082 | -0.33 | -0.214 | 0.003 | 0.055 | 0.055 | 0.588 | 0.51 | 0.29 | 0.59 | 0.66 | 0.55 | 0.54 |
| 5 | 51794898 | rs13176389 | 1.184 | -0.071 | -0.85 | 0.253 | -0.04 | 0.016 | 0.016 | 0 | 0.42 | 0.08 | 0.37 | 0.36 | 0.45 | 0.28 |
| 5 | 51796625 | rs12188733 | 1.896 | 0.244 | -0.878 | 0.01 | -0.029 | 0.079 | 0.079 | 0 | 0.31 | 0.07 | 0.25 | 0.16 | 0.35 | 0.2 |
| 5 | 51797570 | rs13178052 | -0.498 | 0.249 | -0.274 | 0.206 | 0.005 | -0.006 | -0.006 | 0 | 0.15 | 0.2 | 0.13 | 0.13 | 0.17 | 0.25 |
| 5 | 51799156 | rs7713283 | 0.622 | 0.366 | -0.344 | 0.293 | 0.011 | 0.001 | 0.001 | 0 | 0.09 | 0.12 | 0.11 | 0.12 | 0.13 | 0.14 |
| 5 | 51804761 | rs7706977 | -1.328 | -0.013 | -0.612 | 0.196 | 0.003 | -0.008 | -0.008 | 0 | 0.78 | 0.88 | 0.8 | 0.74 | 0.78 | 0.8 |
| 5 | 51805742 | rs7702276 | -1.116 | -0.156 | -0.671 | 0.122 | -0.011 | -0.02 | -0.02 | 1.877 | 0.37 | 0.76 | 0.44 | 0.42 | 0.36 | 0.52 |
| 5 | 51809445 | rs6450064 | -1.603 | -0.098 | -0.577 | 0.253 | 0.006 | 0 | 0 | 0.18 | 0.77 | 0.84 | 0.8 | 0.74 | 0.78 | 0.8 |
| 5 | 51816961 | rs4434348 | -0.789 | -0.2 | -0.195 | 0.226 | 0.018 | 0.101 | 0.101 | 0.391 | 0.27 | 0.49 | 0.36 | 0.45 | 0.29 | 0.48 |
| 5 | 51819014 | rs4296761 | 0.486 | 0.021 | -0.049 | 0.297 | 0.002 | -0.001 | -0.001 | 0 | 0.14 | 0.17 | 0.12 | 0.16 | 0.17 | 0.22 |
| 5 | 51820344 | rs4242034 | -1.644 | -0.025 | -0.008 | 0.264 | 0.006 | 0.01 | 0.01 | 0.067 | 0.77 | 0.81 | 0.8 | 0.76 | 0.78 | 0.8 |
| 5 | 51821771 | rs7721698 | -0.058 | -0.017 | 0.023 | 0.21 | -0.001 | 0.003 | 0.003 | 0.568 | 0.63 | 0.62 | 0.67 | 0.61 | 0.61 | 0.58 |
| 5 | 51825613 | rs4593228 | 1.207 | 0.026 | 0.149 | 0.354 | 0.029 | 0.167 | 0.167 | 0.126 | 0.1 | 0.21 | 0.23 | 0.3 | 0.12 | 0.26 |
| 5 | 51835355 | rs7732397 | -0.316 | -0.035 | 0.077 | 0.278 | -0.001 | 0.024 | 0.024 | 0 | 0.68 | 0.6 | 0.57 | 0.45 | 0.65 | 0.55 |
| 5 | 51840420 | rs10512968 | 0.891 | -0.042 | 0.117 | 0.355 | 0.038 | 0.22 | 0.22 | 0.347 | 0.11 | 0.29 | 0.23 | 0.29 | 0.12 | 0.28 |
| 5 | 51840628 | rs10512969 | 1.476 | -0.096 | -0.105 | 0.159 | -0.032 | 0.016 | 0.016 | 0 | 0.41 | 0.07 | 0.36 | 0.31 | 0.42 | 0.28 |
| 5 | 51841173 | rs1600171 | -0.611 | -0.238 | 0.18 | 0.209 | -0.002 | -0.008 | -0.008 | 0.263 | 0.72 | 0.62 | 0.75 | 0.75 | 0.74 | 0.79 |
| 5 | 51851927 | rs1600177 | 0.519 | -0.208 | -0.164 | 0.154 | 0.007 | -0.003 | -0.003 | 0 | 0.55 | 0.29 | 0.48 | 0.45 | 0.58 | 0.51 |
| 5 | 51859242 | rs1600178 | 0.43 | -1.03 | 0.088 | 0.091 | -0.002 | 0 | 0 | 0 | 0.3 | 0.27 | 0.33 | 0.25 | 0.36 | 0.44 |
| 5 | 51861275 | rs12153325 | 1.104 | -1.166 | -0.047 | 0.17 | -0.034 | 0.03 | 0.03 | 0 | 0.42 | 0.15 | 0.35 | 0.32 | 0.42 | 0.27 |
| 5 | 51867115 | rs10067327 | -1.202 | -1.194 | 0.23 | 0.318 | -0.009 | 0.044 | 0.044 | 0.156 | 0.79 | 0.71 | 0.73 | 0.57 | 0.81 | 0.74 |
| 5 | 51869235 | rs4475224 | -1.202 | -1.194 | 0.23 | 0.306 | -0.009 | 0.046 | 0.046 | 0.156 | 0.79 | 0.71 | 0.73 | 0.56 | 0.81 | 0.74 |
| 5 | 51877090 | rs1303763 | 1.187 | -1.193 | 0.1 | 0.313 | 0.003 | 0.105 | 0.105 | 0 | 0.1 | 0.19 | 0.2 | 0.24 | 0.12 | 0.26 |
| 5 | 51886039 | rs1599728 | -1.761 | -1.151 | -0.219 | 0.374 | 0.002 | -0.003 | -0.003 | 0.367 | 0.83 | 0.83 | 0.83 | 0.7 | 0.86 | 0.84 |
| 5 | 51896454 | rs350436 | 0.871 | -0.934 | -0.643 | 0.288 | -0.075 | 0.002 | 0.002 | 0 | 0.41 | 0.09 | 0.35 | 0.33 | 0.43 | 0.25 |
| 5 | 51911181 | rs17204488 | 1.147 | -1.311 | -0.675 | 0.261 | -0.091 | -0.037 | -0.037 | 0.386 | 0.34 | 0.09 | 0.32 | 0.32 | 0.4 | 0.24 |
| 5 | 51912216 | rs830890 | 0.384 | -1.017 | 0.141 | 0.481 | -0.012 | 0.004 | 0.004 | 0 | 0.1 | 0.14 | 0.13 | 0.13 | 0.07 | 0.11 |
| 5 | 51919136 | rs1450949 | 0.705 | -1.186 | 0.274 | 0.173 | -0.022 | 0.027 | 0.027 | 0 | 0.71 | 0.46 | 0.62 | 0.63 | 0.79 | 0.58 |
| 5 | 51949770 | rs1823538 | 1.109 | -1.227 | -0.223 | 0.079 | -0.014 | -0.04 | -0.04 | 3.908 | 0.47 | 0.17 | 0.45 | 0.55 | 0.52 | 0.39 |
| 5 | 51954991 | rs351956 | 0.334 | -1.402 | 0.045 | -0.017 | -0.048 | -0.091 | -0.091 | 0.357 | 0.14 | 0.36 | 0.09 | 0 | 0.06 | 0.03 |
| 5 | 51956733 | rs889289 | 1.488 | -1.395 | -0.089 | -0.054 | -0.04 | 0.083 | 0.083 | 0 | 0.19 | 0.05 | 0.14 | 0.01 | 0.24 | 0.14 |
| 5 | 51956999 | rs351960 | -0.463 | -1.336 | 0.126 | 0.033 | -0.005 | 0.058 | 0.058 | 0 | 0.81 | 0.73 | 0.69 | 0.62 | 0.83 | 0.57 |
| 5 | 51959557 | rs4865737 | 1.065 | -1.249 | -0.317 | 0.149 | -0.014 | -0.023 | -0.023 | 0 | 0.46 | 0.21 | 0.44 | 0.54 | 0.5 | 0.4 |
| 5 | 51963086 | rs10512982 | 1.055 | -1.231 | -0.336 | 0.002 | -0.008 | 0.036 | 0.036 | 0 | 0.46 | 0.24 | 0.44 | 0.62 | 0.51 | 0.4 |
| 5 | 51963919 | rs7715220 | 0.68 | -1.416 | -0.215 | 0.127 | -0.011 | 0.003 | 0.003 | 0 | 0.67 | 0.38 | 0.59 | 0.62 | 0.75 | 0.55 |
| 5 | 51970107 | rs6893614 | 0.383 | -1.237 | -0.39 | 0.158 | 0.006 | 0.02 | 0.02 | 0 | 0.54 | 0.74 | 0.51 | 0.6 | 0.55 | 0.43 |
| 5 | 51972277 | rs638304 | 0.423 | -1.362 | -0.434 | -0.057 | 0.005 | 0.055 | 0.055 | 0 | 0.27 | 0.15 | 0.18 | 0.01 | 0.27 | 0.13 |
| 5 | 51972326 | rs638715 | -0.075 | -1.444 | -0.13 | 0.074 | -0.017 | -0.066 | -0.066 | 0.137 | 0.33 | 0.56 | 0.22 | 0.07 | 0.28 | 0.16 |
| 5 | 51979514 | rs623193 | -0.29 | -1.641 | 0.069 | 0.553 | 0 | -0.009 | -0.009 | 0 | 0.81 | 0.85 | 0.69 | 0.6 | 0.82 | 0.57 |
| 5 | 51984886 | rs10452448 | 1.019 | -1.821 | -0.174 | 0.538 | 0.001 | -0.002 | -0.002 | 0.003 | 0.2 | 0.18 | 0.15 | 0.29 | 0.12 | 0.16 |
| 5 | 51987238 | rs350450 | 0.37 | -1.78 | -0.237 | 0.493 | 0.002 | 0.001 | 0.001 | 0.102 | 0.6 | 0.62 | 0.53 | 0.31 | 0.71 | 0.39 |
| 5 | 51988075 | rs350445 | 0.736 | -1.806 | -0.346 | 0.287 | -0.002 | 0.001 | 0.001 | 0.046 | 0.68 | 0.31 | 0.61 | 0.55 | 0.78 | 0.51 |
| 5 | 51990269 | rs4865535 | 1.016 | -1.903 | -0.554 | 0.217 | 0.055 | -0.017 | -0.017 | 0.004 | 0.59 | 0.22 | 0.52 | 0.51 | 0.73 | 0.52 |
| 5 | 51991651 | rs164610 | 0.367 | -1.937 | -0.015 | 0.663 | -0.074 | 0.089 | 0.089 | 0.483 | 0.81 | 0.51 | 0.78 | 0.68 | 0.93 | 0.89 |
| 5 | 51996032 | rs651377 | -0.028 | -1.974 | 0.133 | 0.645 | -0.03 | 0.035 | 0.035 | 0 | 0.83 | 0.75 | 0.8 | 0.68 | 0.93 | 0.9 |
| 5 | 52000533 | rs10940264 | 1.161 | -2.231 | -0.178 | 0.36 | 0.005 | 0.232 | 0.232 | 0.766 | 0.43 | 0.1 | 0.39 | 0.1 | 0.62 | 0.36 |
| 5 | 52002578 | rs4865739 | 1.531 | -2.287 | 0.02 | 0.403 | -0.01 | 0.008 | 0.008 | 1.247 | 0.2 | 0.2 | 0.16 | 0.29 | 0.15 | 0.18 |
| 5 | 52003387 | rs351929 | 0.434 | -2.342 | 0.003 | 0.403 | 0.094 | -0.049 | -0.049 | 0 | 0.6 | 0.2 | 0.62 | 0.39 | 0.77 | 0.72 |
| 5 | 52005409 | rs351935 | 0.316 | -2.342 | 0.003 | 0.403 | 0.094 | -0.046 | -0.046 | 0 | 0.6 | 0.2 | 0.61 | 0.39 | 0.77 | 0.72 |
| 5 | 52006802 | rs183013 | -1.454 | -2.326 | 0.097 | 0.441 | -0.017 | 0.092 | 0.092 | 0 | 0.93 | 0.8 | 0.85 | 0.83 | 0.98 | 0.92 |
| 5 | 52007218 | rs722822 | 1.499 | -2.259 | -0.411 | 0.441 | -0.011 | 0.006 | 0.006 | 1.258 | 0.2 | 0.21 | 0.16 | 0.28 | 0.15 | 0.18 |
| 5 | 52012715 | rs351944 | 0.209 | -1.797 | -0.456 | 0.603 | 0.003 | -0.055 | -0.055 | 0 | 0.17 | 0.11 | 0.22 | 0.3 | 0.16 | 0.35 |
| 5 | 52024627 | rs13185633 | 1.398 | -1.428 | -1.004 | -0.029 | -0.055 | 0.121 | 0.121 | 0 | 0.19 | 0.01 | 0.13 | 0.01 | 0.24 | 0.16 |
| 5 | 52031088 | rs13176939 | 0.655 | -1.394 | -0.934 | 0.068 | 0.005 | 0.186 | 0.186 | 0.921 | 0.42 | 0.1 | 0.39 | 0.16 | 0.62 | 0.36 |
| 5 | 52034098 | rs1450952 | 1.353 | -1.327 | -1.281 | 0.124 | -0.1 | 0.051 | 0.051 | 0.322 | 0.25 | 0.06 | 0.27 | 0.16 | 0.38 | 0.25 |
| 5 | 52039850 | rs7713338 | 0.408 | -1.805 | -1.046 | 0.442 | 0.101 | 0.124 | 0.124 | 0 | 0.49 | 0.15 | 0.41 | 0.32 | 0.66 | 0.39 |
| 5 | 52046104 | rs171288 | 0.303 | -1.72 | -0.762 | 0.718 | 0.046 | 0.045 | 0.045 | 0.714 | 0.29 | 0.1 | 0.18 | 0.21 | 0.19 | 0.18 |
| 5 | 52048958 | rs16880156 | 0.558 | -1.697 | -0.757 | 0.748 | 0.005 | -0.001 | -0.001 | 0 | 0.05 | 0.05 | 0.02 | 0.16 | 0.03 | 0.02 |
| 5 | 52051514 | rs351902 | -0.072 | -1.61 | -0.706 | 0.701 | 0.055 | -0.006 | -0.006 | 0.441 | 0.34 | 0.17 | 0.2 | 0.37 | 0.22 | 0.2 |
| 5 | 52064075 | rs244486 | 0.279 | -1.466 | -0.068 | 0.837 | -0.012 | 0.126 | 0.126 | 0 | 0.07 | 0.24 | 0.16 | 0.18 | 0.02 | 0.09 |
| 5 | 52064120 | rs12658846 | 0.308 | -1.744 | -0.638 | 0.738 | 0.052 | 0.045 | 0.045 | 0.908 | 0.29 | 0.1 | 0.18 | 0.2 | 0.18 | 0.18 |
| 5 | 52064349 | rs10072364 | -0.113 | -1.531 | -0.517 | 0.823 | -0.002 | -0.004 | -0.004 | 0.412 | 0.6 | 0.43 | 0.54 | 0.43 | 0.59 | 0.62 |
| 5 | 52065181 | rs173417 | -0.25 | -1.494 | -1.24 | 0.695 | 0.067 | -0.013 | -0.013 | 0.735 | 0.63 | 0.83 | 0.77 | 0.55 | 0.75 | 0.79 |
| 5 | 52069314 | rs17208683 | 0.543 | -1.296 | -1.402 | 0.574 | -0.084 | 0.026 | 0.026 | 0.799 | 0.3 | 0.07 | 0.27 | 0.22 | 0.36 | 0.26 |
| 5 | 52069435 | rs10512992 | 0.718 | -1.123 | -1.392 | 0.958 | -0.101 | -0.075 | -0.075 | 0 | 0.26 | 0.05 | 0.32 | 0.23 | 0.38 | 0.41 |
| 5 | 52074401 | rs2648727 | 0.111 | -1.35 | -0.689 | 0.76 | -0.013 | -0.002 | -0.002 | 0 | 0.29 | 0.27 | 0.25 | 0.3 | 0.24 | 0.25 |
| 5 | 52078773 | rs16880227 | -0.436 | -1.474 | -0.075 | 0.631 | 0.094 | 0.03 | 0.03 | 0.412 | 0.06 | 0.42 | 0.06 | 0 | 0.05 | 0.02 |
| 5 | 52079002 | rs13189973 | 0.018 | -1.482 | -0.231 | 0.662 | 0.001 | 0.001 | 0.001 | 0 | 0.69 | 0.39 | 0.72 | 0.7 | 0.76 | 0.73 |
| 5 | 52081992 | rs10940270 | 0.475 | -1.258 | -0.757 | 0.447 | -0.062 | -0.058 | -0.058 | 0 | 0.45 | 0.12 | 0.5 | 0.48 | 0.51 | 0.56 |

| | | | | | | | | | | | | | | | | |
|---|---|---|---|---|---|---|---|---|---|---|---|---|---|---|---|---|
| 5 | 52084843 | rs6867040 | 1.223 | -1.16 | -0.777 | 0.468 | -0.027 | 0.002 | 0.002 | 0 | 0.22 | 0.17 | 0.21 | 0.22 | 0.25 | 0.17 |
| 5 | 52091647 | rs13188662 | 1.113 | -1.349 | -0.993 | 0.426 | -0.096 | -0.031 | -0.031 | 0 | 0.33 | 0.07 | 0.38 | 0.24 | 0.41 | 0.44 |
| 5 | 52094912 | rs4865747 | 1.252 | -1.522 | -0.933 | 0.289 | 0.013 | 0.018 | 0.018 | 1.147 | 0.21 | 0.13 | 0.15 | 0.15 | 0.17 | 0.17 |
| 5 | 52096889 | rs1499280 | -1.386 | -1.473 | -0.951 | 0.337 | 0.003 | 0.04 | 0.04 | 0.446 | 0.93 | 0.91 | 0.9 | 0.85 | 0.93 | 0.92 |
| 5 | 52097764 | rs11886 | 1.377 | -1.4 | -0.938 | 0.325 | -0.024 | 0.01 | 0.01 | 0 | 0.22 | 0.19 | 0.21 | 0.2 | 0.25 | 0.16 |
| 5 | 52099436 | rs9291995 | -1.348 | -1.396 | -0.94 | 0.342 | 0.002 | 0.034 | 0.034 | 0.362 | 0.93 | 0.91 | 0.9 | 0.85 | 0.93 | 0.92 |
| 5 | 52101169 | rs17275890 | 1.367 | -1.364 | -0.976 | 0.235 | -0.029 | 0.011 | 0.011 | 0 | 0.22 | 0.17 | 0.21 | 0.2 | 0.25 | 0.16 |
| 5 | 52105776 | rs6896291 | 1.304 | -1.357 | -0.989 | 0.278 | -0.013 | 0.002 | 0.002 | 0 | 0.22 | 0.21 | 0.21 | 0.22 | 0.25 | 0.17 |
| 5 | 52110676 | rs889294 | -0.441 | -1.432 | -0.942 | 0.175 | -0.012 | 0.045 | 0.045 | 0.383 | 0.56 | 0.7 | 0.5 | 0.36 | 0.52 | 0.47 |
| 5 | 52111796 | rs17210759 | 1.203 | -1.418 | -0.907 | 0.031 | 0.012 | 0.017 | 0.017 | 0.964 | 0.21 | 0.14 | 0.15 | 0.15 | 0.17 | 0.17 |
| 5 | 52114611 | rs6887205 | -0.059 | -1.349 | -0.993 | 0.072 | -0.04 | -0.075 | -0.075 | 0.366 | 0.34 | 0.5 | 0.29 | 0.16 | 0.27 | 0.3 |
| 5 | 52119356 | rs1820167 | -0.146 | -1.065 | -0.824 | 0.424 | 0.016 | 0.034 | 0.034 | 0 | 0.67 | 0.55 | 0.67 | 0.42 | 0.68 | 0.75 |
| 5 | 52125265 | rs2454585 | 0.028 | -0.871 | -1.111 | 0.257 | 0.004 | 0.002 | 0.002 | 1.044 | 0.36 | 0.62 | 0.46 | 0.52 | 0.42 | 0.46 |
| 5 | 52125378 | rs2454586 | 0.263 | -1.398 | -1.219 | 0.555 | 0.034 | -0.038 | -0.038 | 0 | 0.63 | 0.82 | 0.64 | 0.53 | 0.72 | 0.56 |
| 5 | 52128374 | rs17211415 | 1.07 | -1.326 | -1.136 | 0.437 | 0.028 | 0.046 | 0.046 | 1.455 | 0.19 | 0.01 | 0.11 | 0.12 | 0.12 | 0.23 |
| 5 | 52129172 | rs1391984 | 0.247 | -0.704 | -0.733 | 0.67 | 0.003 | -0.004 | -0.004 | 0 | 0.13 | 0.14 | 0.18 | 0.08 | 0.14 | 0.15 |
| 5 | 52131315 | rs2432143 | 0.403 | -0.4 | -0.748 | 0.768 | 0.007 | 0.035 | 0.035 | 1.916 | 0.08 | 0.1 | 0.11 | 0.18 | 0.09 | 0.12 |
| 5 | 52131591 | rs2938793 | 0.178 | -0.346 | -1.044 | 0.63 | -0.027 | 0.017 | 0.017 | 0 | 0.35 | 0.4 | 0.35 | 0.42 | 0.29 | 0.46 |
| 5 | 52133266 | rs2432144 | 0.543 | -0.118 | -1.267 | 0.591 | 0.009 | -0.012 | -0.012 | 3.373 | 0.74 | 0.78 | 0.72 | 0.66 | 0.82 | 0.88 |
| 5 | 52135543 | rs6867399 | -0.275 | 0.634 | -1.291 | 0.045 | -0.016 | 0.047 | 0.047 | 4.107 | 0.26 | 0.16 | 0.18 | 0.05 | 0.26 | 0.21 |
| 5 | 52139462 | rs16880353 | -0.363 | 0.259 | -0.607 | 0.272 | 0.001 | 0.001 | 0.001 | 1.134 | 0.32 | 0.35 | 0.19 | 0.23 | 0.25 | 0.42 |
| 5 | 52141943 | rs1423570 | -1.283 | 0.372 | -1.085 | 0.361 | -0.069 | -0.03 | -0.03 | 4.034 | 0.62 | 0.85 | 0.52 | 0.64 | 0.45 | 0.73 |
| 5 | 52153066 | rs11750248 | 0.031 | 0.894 | -0.739 | 0.575 | -0.004 | 0.001 | 0.001 | 0.05 | 0.1 | 0.08 | 0.06 | 0.03 | 0.08 | 0.14 |
| 5 | 52153756 | rs10038615 | -0.099 | 0.839 | -0.697 | 0.598 | -0.021 | -0.029 | -0.029 | 0 | 0.34 | 0.47 | 0.2 | 0.24 | 0.25 | 0.44 |
| 5 | 52163091 | rs2456221 | -1.135 | 1.182 | -0.57 | 1.076 | -0.013 | -0.012 | -0.012 | 0 | 0.15 | 0.1 | 0.28 | 0.41 | 0.17 | 0.22 |
| 5 | 52166732 | rs2456220 | -0.436 | 1.031 | -0.648 | 0.19 | 0.006 | -0.04 | -0.04 | 0 | 0.44 | 0.47 | 0.67 | 0.75 | 0.47 | 0.42 |
| 5 | 52168400 | rs2927686 | -0.47 | 0.987 | -0.648 | 0.149 | 0.005 | -0.036 | -0.036 | 0.641 | 0.44 | 0.48 | 0.67 | 0.76 | 0.48 | 0.43 |
| 5 | 52168715 | rs2447858 | -0.921 | 1.062 | -0.59 | 1.036 | -0.013 | -0.01 | -0.01 | 0.585 | 0.06 | 0.04 | 0.22 | 0.23 | 0.11 | 0.1 |
| 5 | 52171623 | rs11948124 | -0.817 | 0.091 | -0.216 | 0.487 | -0.009 | -0.034 | -0.034 | 0 | 0.35 | 0.45 | 0.17 | 0.21 | 0.23 | 0.46 |
| 5 | 52172630 | rs1105757 | -0.771 | -0.007 | -0.39 | 0.34 | -0.009 | -0.041 | -0.041 | 0.556 | 0.34 | 0.44 | 0.17 | 0.16 | 0.22 | 0.43 |
| 5 | 52175899 | rs4865749 | -0.545 | -0.106 | -0.692 | 1.088 | -0.007 | 0.01 | 0.01 | 0.002 | 0.3 | 0.64 | 0.28 | 0.44 | 0.18 | 0.45 |
| 5 | 52179008 | rs13163497 | -0.135 | 0.103 | -1.399 | 0.734 | -0.025 | -0.002 | -0.002 | 0 | 0.16 | 0.01 | 0.14 | 0.09 | 0.13 | 0.05 |
| 5 | 52179298 | rs12520768 | -0.92 | -0.397 | -2.115 | 0.675 | 0.004 | 0.007 | 0.007 | 0.091 | 0.38 | 0.09 | 0.45 | 0.29 | 0.36 | 0.29 |
| 5 | 52179949 | rs2860365 | -1.31 | -0.613 | -1.903 | 0.71 | -0.024 | -0.021 | -0.021 | 0 | 0.63 | 0.7 | 0.63 | 0.56 | 0.56 | 0.62 |
| 5 | 52184268 | rs4865540 | 0.062 | -0.918 | -1.693 | 0.46 | 0.01 | 0.038 | 0.038 | 0 | 0.24 | 0.16 | 0.23 | 0.09 | 0.2 | 0.14 |
| 5 | 52184613 | rs10940277 | -0.739 | -0.93 | -1.256 | 0.578 | 0.023 | 0 | 0 | 0.023 | 0.41 | 0.24 | 0.47 | 0.39 | 0.37 | 0.33 |
| 5 | 52184921 | rs7706201 | -1.518 | -0.873 | -1.256 | 0.573 | -0.028 | -0.028 | -0.028 | 0 | 0.63 | 0.71 | 0.63 | 0.56 | 0.55 | 0.62 |
| 5 | 52185695 | rs16880442 | -0.167 | -1.071 | -1.042 | 0.896 | -0.004 | -0.003 | -0.003 | 0 | 0.09 | 0.08 | 0.16 | 0.06 | 0.07 | 0.06 |
| 5 | 52185966 | rs1904162 | -1.336 | -1.183 | -0.66 | 0.693 | -0.002 | -0.001 | -0.001 | 0 | 0.6 | 0.35 | 0.58 | 0.61 | 0.5 | 0.56 |
| 5 | 52185996 | rs1904163 | 1.029 | -1.288 | -0.845 | 0.813 | -0.028 | -0.078 | -0.078 | 0 | 0.12 | 0.01 | 0.22 | 0.28 | 0.18 | 0.15 |
| 5 | 52186515 | rs6870949 | 0.061 | -1.398 | -0.807 | 0.725 | 0.019 | -0.036 | -0.036 | 0 | 0.29 | 0.11 | 0.33 | 0.38 | 0.26 | 0.2 |
| 5 | 52188028 | rs7731949 | -1.34 | -1.39 | -0.921 | 0.721 | -0.009 | 0.001 | 0.001 | 0.143 | 0.58 | 0.32 | 0.58 | 0.61 | 0.51 | 0.56 |
| 5 | 52189049 | rs1110350 | -0.259 | -1.327 | -1.175 | 0.73 | -0.001 | 0.024 | 0.024 | 0 | 0.31 | 0.19 | 0.25 | 0.23 | 0.25 | 0.37 |
| 5 | 52193125 | rs4074793 | 1.562 | -1.156 | -1.452 | 0.701 | 0.011 | 0.019 | 0.019 | 4.869 | 0.16 | 0.01 | 0.09 | 0.07 | 0.08 | 0.05 |
| 5 | 52193287 | rs1531545 | -1.338 | -0.868 | -1.16 | 0.784 | -0.038 | -0.018 | -0.018 | 0 | 0.63 | 0.73 | 0.62 | 0.62 | 0.51 | 0.65 |
| 5 | 52195033 | rs1551943 | 0.136 | -0.782 | -1.362 | 0.669 | -0.061 | 0.031 | 0.031 | 0 | 0.24 | 0.04 | 0.11 | 0.18 | 0.21 | 0.31 |
| 5 | 52214197 | rs2938789 | -0.591 | -1.402 | -1.232 | 1.788 | 0.072 | -0.023 | -0.023 | 0 | 0.65 | 0.9 | 0.86 | 0.67 | 0.77 | 0.82 |
| 5 | 52215178 | rs2279585 | -0.2 | -1.442 | -0.524 | 1.746 | 0.024 | 0.036 | 0.036 | 0.247 | 0.26 | 0.38 | 0.41 | 0.39 | 0.25 | 0.44 |
| 5 | 52218333 | rs2406370 | 1.782 | -1.186 | -1.305 | 1.252 | -0.14 | 0.009 | 0.009 | 0 | 0.24 | 0.05 | 0.33 | 0.13 | 0.44 | 0.25 |
| 5 | 52220743 | rs2447871 | 0.703 | -1.13 | -0.951 | 1.405 | 0.002 | 0.001 | 0.001 | 0 | 0.41 | 0.48 | 0.47 | 0.44 | 0.52 | 0.37 |
| 5 | 52222508 | rs2934216 | -0.316 | -1.165 | -0.888 | 1.138 | 0.068 | -0.05 | -0.05 | 0 | 0.59 | 0.83 | 0.6 | 0.5 | 0.7 | 0.73 |
| 5 | 52223516 | rs10038773 | 0.187 | -0.791 | -0.823 | 0.516 | 0.03 | -0.012 | -0.012 | 0.031 | 0.22 | 0.39 | 0.14 | 0.23 | 0.18 | 0.39 |
| 5 | 52230527 | rs1275660 | -0.895 | -0.838 | -1.166 | 0.531 | 0.036 | 0.054 | 0.054 | 0.041 | 0.69 | 0.94 | 0.87 | 0.83 | 0.76 | 0.85 |
| 5 | 52230770 | rs16880533 | -0.431 | 0.093 | -0.507 | 0.767 | 0.046 | -0.002 | -0.002 | 0 | 0.23 | 0.42 | 0.17 | 0.25 | 0.21 | 0.41 |
| 5 | 52235225 | rs10513004 | 0.664 | 0.255 | -1.319 | 0.319 | -0.015 | 0.08 | 0.08 | 0.552 | 0.27 | 0.09 | 0.14 | 0.08 | 0.22 | 0.09 |
| 5 | 52235802 | rs1904165 | -0.484 | 0.525 | -1.105 | 0.199 | -0.001 | -0.007 | -0.007 | 0 | 0.86 | 0.84 | 0.92 | 0.89 | 0.84 | 0.77 |
| 5 | 52237991 | rs2406371 | -0.017 | 0.489 | -0.76 | 0.686 | 0 | 0.014 | 0.014 | 0 | 0.28 | 0.34 | 0.44 | 0.51 | 0.29 | 0.3 |
| 5 | 52239555 | rs16880541 | 1.371 | 0.399 | -0.657 | 0.898 | 0.012 | 0.006 | 0.006 | 0 | 0.11 | 0.13 | 0.17 | 0.17 | 0.15 | 0.18 |
| 5 | 52240476 | rs1363189 | -0.265 | 0.505 | -0.389 | 1.306 | 0 | 0.004 | 0.004 | 0 | 0.27 | 0.33 | 0.43 | 0.36 | 0.28 | 0.28 |
| 5 | 52246736 | rs7718758 | 1.101 | 0.5 | -0.627 | 1.399 | -0.003 | -0.001 | -0.001 | 0 | 0.3 | 0.27 | 0.33 | 0.29 | 0.35 | 0.38 |
| 5 | 52247233 | rs2193968 | 1.055 | 0.474 | -0.656 | 1.376 | 0.001 | -0.001 | -0.001 | 0 | 0.58 | 0.38 | 0.47 | 0.37 | 0.57 | 0.5 |
| 5 | 52247276 | rs11748504 | 2.049 | 0.476 | -0.652 | 1.374 | -0.007 | 0.067 | 0.067 | 0.462 | 0.27 | 0.11 | 0.14 | 0.09 | 0.22 | 0.09 |
| 5 | 52248904 | rs6890588 | 1.111 | 0.455 | -0.574 | 1.457 | 0.001 | -0.002 | -0.002 | 0 | 0.58 | 0.38 | 0.47 | 0.38 | 0.57 | 0.5 |
| 5 | 52249226 | rs3088365 | 1.111 | 0.455 | -0.574 | 1.441 | 0.001 | -0.001 | -0.001 | 0 | 0.58 | 0.38 | 0.47 | 0.38 | 0.57 | 0.5 |
| 5 | 52249476 | rs1047481 | 1.046 | 0.445 | -0.578 | 1.437 | 0.001 | -0.001 | -0.001 | 4.95 | 0.58 | 0.38 | 0.47 | 0.38 | 0.57 | 0.5 |
| 5 | 52252191 | rs7035 | 0.609 | 0.355 | -0.623 | 1.405 | -0.003 | 0.004 | 0.004 | 0 | 0.65 | 0.42 | 0.7 | 0.59 | 0.67 | 0.59 |
| 5 | 52257301 | rs2279584 | 1.144 | 0.465 | -0.707 | 1.334 | -0.005 | -0.019 | -0.019 | 0 | 0.07 | 0.03 | 0.22 | 0.19 | 0.1 | 0.09 |
| 5 | 52275103 | rs246409 | -0.754 | 0.383 | -0.61 | 1.339 | -0.003 | 0.01 | 0.01 | 0 | 0.34 | 0.57 | 0.34 | 0.44 | 0.33 | 0.4 |
| 5 | 52281914 | rs33445 | -0.851 | 0.455 | -0.437 | 1.3 | 0.001 | 0 | 0 | 0.052 | 0.58 | 0.53 | 0.58 | 0.57 | 0.62 | 0.71 |

| | | | | | | | | | | | | | | | | |
|---|---|---|---|---|---|---|---|---|---|---|---|---|---|---|---|---|
| 5 | 52284386 | rs26680 | -0.159 | 0.392 | -0.305 | 1.348 | 0.021 | 0.138 | 0.138 | 0.524 | 0.88 | 0.69 | 0.73 | 0.68 | 0.85 | 0.87 |
| 5 | 52288173 | rs27504 | 0.986 | 0.266 | -0.171 | 1.21 | -0.01 | -0.013 | -0.013 | 0.229 | 0.13 | 0.19 | 0.07 | 0.11 | 0.14 | 0.2 |
| 5 | 52291187 | rs4865755 | 1.982 | 0.402 | -0.403 | 1.074 | -0.011 | 0.105 | 0.105 | 0.49 | 0.28 | 0.1 | 0.2 | 0.07 | 0.22 | 0.1 |
| 5 | 52292237 | rs35235 | -1.187 | 0.37 | -0.294 | 1.271 | -0.004 | -0.007 | -0.007 | 1.208 | 0.42 | 0.52 | 0.49 | 0.63 | 0.44 | 0.55 |
| 5 | 52292555 | rs35236 | -0.112 | 0.37 | -0.271 | 1.506 | 0.032 | 0.248 | 0.248 | 0.481 | 0.87 | 0.68 | 0.68 | 0.51 | 0.84 | 0.81 |
| 5 | 52293087 | rs1421942 | 1.488 | 0.291 | -0.357 | 1.385 | 0.001 | 0.125 | 0.125 | 0.113 | 0.53 | 0.19 | 0.46 | 0.23 | 0.52 | 0.4 |
| 5 | 52294551 | rs10042752 | 0.59 | 0.2 | -0.285 | 1.311 | 0 | -0.011 | -0.011 | 3.354 | 0.23 | 0.26 | 0.16 | 0.09 | 0.23 | 0.2 |
| 5 | 52297242 | rs1645761 | -0.584 | 0.146 | -0.228 | 1.535 | 0.028 | 0.161 | 0.161 | 0.598 | 0.88 | 0.69 | 0.73 | 0.67 | 0.85 | 0.84 |
| 5 | 52301716 | rs6859355 | 0.936 | -0.014 | -0.61 | 1.211 | -0.003 | 0.153 | 0.153 | 3.379 | 0.49 | 0.28 | 0.34 | 0.16 | 0.42 | 0.29 |
| 5 | 52304996 | rs153141 | 0.521 | -0.08 | -0.586 | 1.321 | -0.007 | -0.01 | -0.01 | 0.193 | 0.35 | 0.23 | 0.32 | 0.34 | 0.39 | 0.51 |
| 5 | 52306013 | rs1421940 | 1.541 | -0.041 | -0.698 | 1.35 | -0.061 | 0.004 | 0.004 | 0.613 | 0.26 | 0.05 | 0.26 | 0.16 | 0.3 | 0.3 |
| 5 | 52307370 | rs1862639 | 0.887 | -0.019 | -0.525 | 1.359 | -0.002 | -0.021 | -0.021 | 0 | 0.21 | 0.26 | 0.15 | 0.09 | 0.2 | 0.16 |
| 5 | 52309015 | rs152088 | 0.187 | -0.172 | -0.597 | 1.497 | -0.008 | 0.092 | 0.092 | 3.75 | 0.16 | 0.29 | 0.29 | 0.38 | 0.19 | 0.33 |
| 5 | 52312422 | rs152090 | 0.546 | -0.033 | -0.255 | 1.373 | -0.006 | 0.014 | 0.014 | 0 | 0.64 | 0.54 | 0.55 | 0.43 | 0.65 | 0.65 |
| 5 | 52313564 | rs4865756 | 1.402 | -0.033 | -0.291 | 1.341 | 0.024 | 0.084 | 0.084 | 0.141 | 0.54 | 0.26 | 0.49 | 0.24 | 0.56 | 0.4 |
| 5 | 52319076 | rs246406 | 0.553 | -0.017 | -0.294 | 1.399 | -0.006 | 0.016 | 0.016 | 0 | 0.64 | 0.54 | 0.55 | 0.44 | 0.65 | 0.65 |
| 5 | 52320940 | rs43188 | 1.2 | 0.024 | -0.3 | 1.405 | -0.005 | 0.05 | 0.05 | 0.568 | 0.1 | 0.22 | 0.06 | 0.18 | 0.09 | 0.21 |
| 5 | 52322721 | rs26678 | 1.327 | 0.114 | -0.51 | 1.192 | -0.006 | 0.034 | 0.034 | 0 | 0.36 | 0.22 | 0.26 | 0.26 | 0.32 | 0.3 |
| 5 | 52323568 | rs7705328 | 0.504 | -0.032 | -0.347 | 0.954 | -0.009 | 0.114 | 0.114 | 1.288 | 0.79 | 0.39 | 0.82 | 0.51 | 0.82 | 0.69 |
| 5 | 52325775 | rs3212430 | 0.602 | 0.034 | -0.331 | 0.717 | -0.004 | 0.01 | 0.01 | 0.212 | 0.33 | 0.32 | 0.32 | 0.21 | 0.36 | 0.37 |
| 5 | 52327230 | rs3756537 | 0.453 | -0.001 | -0.34 | 0.729 | 0 | 0 | 0 | 0 | 0.26 | 0.25 | 0.35 | 0.22 | 0.29 | 0.27 |
| 5 | 52329341 | rs11955860 | -1.609 | 0.008 | -0.35 | 0.92 | 0 | 0.023 | 0.023 | 0 | 0.64 | 0.59 | 0.73 | 0.51 | 0.69 | 0.7 |
| 5 | 52336982 | rs988557 | -0.964 | -0.245 | -0.83 | 0.537 | 0.015 | 0.037 | 0.037 | 0 | 0.68 | 0.56 | 0.7 | 0.43 | 0.69 | 0.66 |
| 5 | 52337326 | rs3212435 | 1.276 | -0.195 | -0.98 | 0.486 | -0.057 | 0.012 | 0.012 | 0.916 | 0.27 | 0.08 | 0.28 | 0.19 | 0.31 | 0.32 |
| 5 | 52347369 | rs1126643 | -0.101 | -0.228 | -1.224 | 0.24 | 0.024 | 0.038 | 0.038 | 0 | 0.42 | 0.28 | 0.44 | 0.28 | 0.4 | 0.33 |
| 5 | 52351876 | rs3212523 | 2.319 | -0.201 | -1.172 | 0.226 | -0.008 | -0.036 | -0.036 | 1.43 | 0.14 | 0.17 | 0.06 | 0.03 | 0.09 | 0.1 |
| 5 | 52356006 | rs3212537 | 1.969 | -0.168 | -1.227 | 0.259 | -0.075 | -0.057 | -0.057 | 1.031 | 0.16 | 0.04 | 0.25 | 0.19 | 0.24 | 0.28 |
| 5 | 52366138 | rs2112290 | -1.775 | -0.328 | -1.026 | -0.029 | -0.009 | -0.019 | -0.019 | 0.472 | 0.74 | 0.78 | 0.77 | 0.45 | 0.73 | 0.7 |
| 5 | 52367146 | rs3212578 | 1.855 | -0.598 | -1.06 | -0.297 | 0 | -0.013 | -0.013 | 0 | 0.09 | 0.07 | 0.06 | 0 | 0.05 | 0.01 |
| 5 | 52368472 | rs3212583 | 2.109 | -0.497 | -1.009 | -0.237 | -0.013 | -0.051 | -0.051 | 0 | 0.15 | 0.23 | 0.07 | 0.03 | 0.09 | 0.09 |
| 5 | 52369002 | rs3212327 | 2.284 | -0.497 | -1.107 | -0.238 | -0.008 | -0.038 | -0.038 | 0.666 | 0.14 | 0.2 | 0.07 | 0.03 | 0.09 | 0.09 |
| 5 | 52369086 | rs3213805 | 2.416 | -0.495 | -1.106 | -0.241 | -0.008 | -0.045 | -0.045 | 0 | 0.14 | 0.2 | 0.07 | 0.02 | 0.09 | 0.05 |
| 5 | 52369495 | rs3212588 | 2.53 | -0.494 | -1.109 | -0.24 | 0.002 | -0.009 | -0.009 | 0 | 0.14 | 0.12 | 0.05 | 0.02 | 0.09 | 0.05 |
| 5 | 52374088 | rs1445942 | 1.302 | -0.528 | -1.168 | -0.28 | 0.02 | -0.03 | -0.03 | 0 | 0.22 | 0.31 | 0.27 | 0.15 | 0.27 | 0.28 |
| 5 | 52379277 | rs2303122 | -0.164 | -0.583 | -1.224 | -0.178 | 0.017 | 0.017 | 0.017 | 0 | 0.38 | 0.28 | 0.43 | 0.29 | 0.37 | 0.33 |
| 5 | 52387150 | rs1109527 | -2.023 | -0.611 | -1.187 | -0.096 | 0.002 | 0.183 | 0.183 | 0.319 | 0.91 | 0.83 | 0.93 | 0.51 | 0.9 | 0.81 |
| 5 | 52387323 | rs1109526 | -1.922 | -0.601 | -1.164 | -0.287 | -0.013 | -0.023 | -0.023 | 0 | 0.74 | 0.79 | 0.77 | 0.41 | 0.73 | 0.7 |
| 5 | 52387681 | rs1042324 | 2.168 | -0.603 | -1.502 | -0.255 | -0.009 | -0.039 | -0.039 | 0 | 0.15 | 0.2 | 0.07 | 0.03 | 0.09 | 0.09 |
| 5 | 52390415 | rs7725246 | -0.284 | -0.413 | -1.489 | -0.189 | 0.016 | 0.019 | 0.019 | 0 | 0.39 | 0.28 | 0.45 | 0.29 | 0.39 | 0.33 |
| 5 | 52395310 | rs2068247 | -0.568 | -0.326 | -1.551 | -0.558 | 0.007 | 0.086 | 0.086 | 0 | 0.47 | 0.32 | 0.47 | 0.2 | 0.46 | 0.4 |
| 5 | 52395525 | rs3756532 | -0.568 | -0.326 | -1.539 | -0.558 | 0.007 | 0.078 | 0.078 | 0 | 0.47 | 0.32 | 0.46 | 0.2 | 0.46 | 0.4 |
| 5 | 52397586 | rs12515434 | 1.032 | -0.368 | -1.597 | -0.576 | -0.007 | -0.018 | -0.018 | 0 | 0.11 | 0.09 | 0.1 | 0.01 | 0.08 | 0.02 |
| 5 | 52399103 | rs17298431 | 0.153 | -0.23 | -1.608 | -0.553 | -0.007 | 0.065 | 0.065 | 0 | 0.29 | 0.13 | 0.34 | 0.11 | 0.33 | 0.29 |
| 5 | 52408132 | rs10471372 | 0.955 | -0.179 | -1.427 | -0.204 | 0 | 0 | 0 | 0 | 0.18 | 0.19 | 0.12 | 0.1 | 0.13 | 0.11 |
| 5 | 52409245 | rs16880892 | 1.777 | -0.185 | -1.514 | -0.266 | 0.004 | 0.004 | 0.004 | 0 | 0.14 | 0.11 | 0.05 | 0.02 | 0.1 | 0.05 |
| 5 | 52417217 | rs7722066 | 0.01 | 0.165 | -1.97 | -0.525 | -0.062 | 0.076 | 0.076 | 0.373 | 0.34 | 0.19 | 0.29 | 0.17 | 0.44 | 0.49 |
| 5 | 52417299 | rs30101 | -0.452 | -0.318 | -1.475 | 0.053 | 0.026 | 0.001 | 0.001 | 0.43 | 0.51 | 0.37 | 0.48 | 0.54 | 0.61 | 0.68 |
| 5 | 52417741 | rs6868966 | 1.262 | -0.294 | -0.399 | 1.028 | -0.03 | 0.103 | 0.103 | 0 | 0.4 | 0.28 | 0.35 | 0.13 | 0.4 | 0.32 |
| 5 | 52418582 | rs30099 | -0.217 | 0.739 | 0.605 | 2.356 | 0.001 | 0.171 | 0.171 | 0 | 0.09 | 0.17 | 0.06 | 0.43 | 0.09 | 0.18 |
| 5 | 52419944 | rs30727 | -0.187 | 0.725 | 0.598 | 2.152 | 0.001 | 0.145 | 0.145 | 0 | 0.09 | 0.17 | 0.06 | 0.39 | 0.09 | 0.18 |
| 5 | 52420445 | rs4631137 | 1.674 | 0.676 | 0.785 | 2.046 | 0 | -0.032 | -0.032 | 0 | 0.61 | 0.51 | 0.48 | 0.28 | 0.54 | 0.41 |
| 5 | 52420841 | rs25826 | -0.637 | 0.565 | 0.809 | 2.345 | 0.008 | 0.178 | 0.178 | 0 | 0.1 | 0.2 | 0.05 | 0.39 | 0.1 | 0.18 |
| 5 | 52421565 | rs30725 | 1.56 | 0.584 | 0.387 | 2.382 | 0 | -0.03 | -0.03 | 0 | 0.82 | 0.86 | 0.69 | 0.38 | 0.78 | 0.7 |
| 5 | 52423178 | rs30094 | 0.364 | 0.575 | 0.302 | 2.383 | 0.005 | -0.029 | -0.029 | 0.094 | 0.89 | 0.95 | 0.85 | 0.77 | 0.89 | 0.88 |
| 5 | 52436224 | rs251329 | -1.823 | 0.717 | 0.476 | 2.12 | -0.009 | -0.039 | -0.039 | 0.37 | 0.34 | 0.7 | 0.46 | 0.54 | 0.37 | 0.45 |
| 5 | 52436899 | rs11955696 | -0.342 | 0.709 | 0.45 | 2.048 | -0.011 | 0.042 | 0.042 | 0.007 | 0.18 | 0.1 | 0.26 | 0.07 | 0.2 | 0.2 |
| 5 | 52448212 | rs254487 | -0.925 | 0.6 | 0.86 | 2.295 | 0.006 | 0.155 | 0.155 | 0.071 | 0.1 | 0.2 | 0.15 | 0.41 | 0.12 | 0.17 |
| 5 | 52448598 | rs254489 | -0.934 | 0.6 | 0.873 | 2.295 | 0.001 | 0.098 | 0.098 | 0.028 | 0.1 | 0.13 | 0.15 | 0.41 | 0.12 | 0.17 |
| 5 | 52453385 | rs254480 | -0.676 | 0.614 | 0.652 | 2.302 | 0 | 0.064 | 0.064 | 0.245 | 0.09 | 0.11 | 0.14 | 0.41 | 0.12 | 0.17 |
| 5 | 52460354 | rs1846703 | -1.268 | 0.605 | 0.783 | 2.258 | 0.042 | 0.034 | 0.034 | 0.884 | 0.08 | 0.56 | 0.06 | 0.07 | 0.05 | 0.1 |
| 5 | 52460637 | rs1445990 | -0.468 | 0.608 | 0.776 | 2.304 | -0.003 | 0.029 | 0.029 | 0.192 | 0.09 | 0.06 | 0.14 | 0.41 | 0.12 | 0.16 |
| 5 | 52464757 | rs1500367 | -1.31 | 0.463 | 0.82 | 1.929 | 0.042 | 0.034 | 0.034 | 0.884 | 0.08 | 0.56 | 0.06 | 0.07 | 0.05 | 0.1 |
| 5 | 52477590 | rs153683 | 0.277 | 0.557 | 0.147 | 2.21 | 0.004 | -0.031 | -0.031 | 1.317 | 0.9 | 0.96 | 0.86 | 0.78 | 0.91 | 0.89 |
| 5 | 52478715 | rs2112876 | -0.178 | 0.697 | -0.065 | 2.187 | -0.067 | 0.015 | 0.015 | 0.006 | 0.18 | 0.02 | 0.26 | 0.08 | 0.22 | 0.2 |
| 5 | 52487595 | rs251580 | -1.828 | 0.54 | 0.465 | 2.318 | 0.009 | 0.034 | 0.034 | 0.663 | 0.34 | 0.43 | 0.45 | 0.56 | 0.39 | 0.47 |
| 5 | 52491141 | rs16881049 | -0.739 | 0.551 | 0.523 | 2.243 | 0.057 | 0.03 | 0.03 | 0.013 | 0.08 | 0.32 | 0.06 | 0.07 | 0.08 | 0.12 |
| 5 | 52492677 | rs254503 | -2.486 | 0.501 | 0.518 | 2.232 | 0 | 0.059 | 0.059 | 0.79 | 0.49 | 0.61 | 0.6 | 0.84 | 0.5 | 0.7 |
| 5 | 52495682 | rs254481 | -0.906 | 0.5 | 0.399 | 2.41 | -0.012 | 0.138 | 0.138 | 0.041 | 0.11 | 0.17 | 0.14 | 0.47 | 0.1 | 0.25 |
| 5 | 52502644 | rs251513 | -0.321 | 0.595 | 0.151 | 2.154 | 0.005 | -0.029 | -0.029 | 0 | 0.1 | 0.04 | 0.14 | 0.23 | 0.09 | 0.13 |

| | | | | | | | | | | | | | | | | |
|---|---|---|---|---|---|---|---|---|---|---|---|---|---|---|---|---|
| 5 | 52510519 | rs39859 | -0.39 | 0.634 | 0.153 | 2.172 | 0.004 | -0.016 | -0.016 | 0 | 0.1 | 0.05 | 0.14 | 0.22 | 0.09 | 0.12 |
| 5 | 52539486 | rs731383 | -1.017 | 0.665 | 0.298 | 1.934 | 0.009 | -0.03 | -0.03 | 0.022 | 0.27 | 0.36 | 0.31 | 0.15 | 0.3 | 0.32 |
| 5 | 52542897 | rs251504 | -0.522 | 0.596 | 0.324 | 2.274 | -0.001 | 0.042 | 0.042 | 0 | 0.09 | 0.07 | 0.13 | 0.39 | 0.09 | 0.16 |
| 5 | 52544231 | rs1445985 | -1.204 | 0.577 | 0.313 | 2.273 | 0.004 | -0.011 | -0.011 | 0 | 0.29 | 0.36 | 0.31 | 0.22 | 0.3 | 0.42 |
| 5 | 52545852 | rs6866517 | -0.927 | 0.614 | 0.153 | 2.242 | 0.004 | -0.015 | -0.015 | 2.363 | 0.27 | 0.3 | 0.31 | 0.15 | 0.3 | 0.32 |
| 5 | 52548545 | rs254142 | -0.332 | 0.695 | 0.14 | 2.318 | -0.002 | 0.034 | 0.034 | 0 | 0.08 | 0.06 | 0.14 | 0.39 | 0.1 | 0.16 |
| 5 | 52550580 | rs27324 | -1.933 | 0.698 | 0.298 | 2.169 | 0.001 | 0.009 | 0.009 | 0.838 | 0.39 | 0.48 | 0.45 | 0.6 | 0.41 | 0.6 |
| 5 | 52552239 | rs27323 | 1.651 | 0.698 | 0.268 | 2.177 | -0.001 | 0.009 | 0.009 | 0.42 | 0.61 | 0.55 | 0.55 | 0.4 | 0.61 | 0.4 |
| 5 | 52571758 | rs419816 | -1.306 | 0.833 | 0.092 | 2.604 | -0.001 | 0.026 | 0.026 | 0 | 0.16 | 0.13 | 0.27 | 0.45 | 0.17 | 0.22 |
| 5 | 52578073 | rs10059098 | -0.132 | 0.813 | 0.242 | 2.61 | 0.016 | 0.055 | 0.055 | 0.042 | 0.05 | 0.23 | 0.04 | 0.06 | 0.03 | 0.1 |
| 5 | 52586993 | rs28088 | -0.254 | 0.885 | 0.148 | 2.635 | -0.007 | -0.01 | -0.01 | 0.31 | 0.1 | 0.06 | 0.15 | 0.22 | 0.1 | 0.13 |
| 5 | 52595653 | rs40229 | -0.226 | 0.806 | 0.153 | 2.64 | 0 | -0.009 | -0.009 | 0 | 0.1 | 0.06 | 0.15 | 0.22 | 0.09 | 0.13 |
| 5 | 52613555 | rs408393 | 0.692 | 1.033 | -0.029 | 2.574 | -0.075 | -0.043 | -0.043 | 0 | 0.75 | 0.95 | 0.7 | 0.73 | 0.69 | 0.69 |
| 5 | 52614762 | rs16881275 | -0.915 | 0.717 | 0.274 | 2.709 | 0.002 | 0.036 | 0.036 | 0.04 | 0.05 | 0.06 | 0.11 | 0.22 | 0.08 | 0.09 |
| 5 | 52636896 | rs1007984 | -1.501 | 0.804 | 0.215 | 2.501 | 0.013 | 0.001 | 0.001 | 0 | 0.17 | 0.34 | 0.21 | 0.19 | 0.19 | 0.27 |
| 5 | 52637544 | rs40220 | 1.789 | 0.676 | 0.145 | 2.907 | -0.031 | 0.139 | 0.139 | 0 | 0.8 | 0.62 | 0.82 | 0.54 | 0.86 | 0.68 |
| 5 | 52642056 | rs37787 | 1.405 | 0.669 | 0.048 | 2.93 | -0.024 | 0.063 | 0.063 | 0 | 0.83 | 0.72 | 0.88 | 0.64 | 0.88 | 0.69 |
| 5 | 52645739 | rs16881347 | -0.299 | 0.752 | -0.022 | 2.783 | -0.006 | 0.044 | 0.044 | 0 | 0.05 | 0.18 | 0.03 | 0.1 | 0.02 | 0.1 |
| 5 | 52650747 | rs252046 | -0.75 | 0.985 | -0.924 | 2.813 | -0.042 | -0.088 | -0.088 | 0 | 0.13 | 0.01 | 0.18 | 0.23 | 0.17 | 0.17 |
| 5 | 52652777 | rs12188879 | 0.712 | 0.936 | -0.538 | 2.67 | 0.001 | -0.045 | -0.045 | 0 | 0.88 | 0.94 | 0.87 | 0.64 | 0.89 | 0.77 |
| 5 | 52653978 | rs7729080 | 1.142 | 0.99 | -0.407 | 2.002 | -0.001 | 0.007 | 0.007 | 0.018 | 0.72 | 0.69 | 0.7 | 0.54 | 0.71 | 0.52 |
| 5 | 52663996 | rs6450127 | -1.332 | 1.613 | -0.253 | 2.032 | 0.023 | 0.035 | 0.035 | 0.053 | 0.26 | 0.39 | 0.34 | 0.37 | 0.35 | 0.29 |
| 5 | 52664035 | rs6450128 | -1.564 | 1.638 | -0.179 | 2.454 | 0.026 | 0.116 | 0.116 | 0.087 | 0.22 | 0.4 | 0.41 | 0.39 | 0.25 | 0.27 |
| 5 | 52664796 | rs11948445 | -0.854 | 1.891 | -1.105 | 2.235 | -0.1 | -0.045 | -0.045 | 0.165 | 0.29 | 0.1 | 0.5 | 0.34 | 0.46 | 0.26 |
| 5 | 52674671 | rs4621519 | -0.835 | 0.752 | -1.971 | 1.625 | 0.003 | -0.068 | -0.068 | 0 | 0.82 | 0.99 | 0.91 | 0.79 | 0.87 | 0.83 |
| 5 | 52675782 | rs13157909 | -0.267 | 1.27 | -2.071 | 1.586 | -0.034 | -0.036 | -0.036 | 1.537 | 0.07 | 0.03 | 0.16 | 0.2 | 0.11 | 0.13 |
| 5 | 52684170 | rs4865544 | 1.347 | 1.31 | -1.713 | 1.193 | 0.012 | -0.007 | -0.007 | 0 | 0.18 | 0.35 | 0.2 | 0.18 | 0.18 | 0.36 |
| 5 | 52697383 | rs6866562 | -0.376 | 1.22 | -1.523 | 1.232 | -0.012 | 0 | 0 | 0.187 | 0.46 | 0.67 | 0.52 | 0.44 | 0.53 | 0.58 |
| 5 | 52698387 | rs4501299 | -1.01 | 1.148 | -1.568 | 1.259 | 0.05 | 0.057 | 0.057 | 0 | 0.53 | 0.74 | 0.69 | 0.64 | 0.64 | 0.71 |
| 5 | 52698518 | rs4279297 | 1.085 | 1.218 | -1.441 | 1.303 | -0.026 | -0.021 | -0.021 | 0 | 0.05 | 0.01 | 0.15 | 0.2 | 0.09 | 0.13 |
| 5 | 52716489 | rs12653439 | 1.361 | 1.168 | -1.347 | 1.095 | 0.014 | -0.009 | -0.009 | 0 | 0.18 | 0.35 | 0.2 | 0.18 | 0.18 | 0.36 |
| 5 | 52720020 | rs17242960 | 1.373 | 1.016 | -2.094 | 0.935 | 0.005 | 0.086 | 0.086 | 0 | 0.28 | 0.11 | 0.2 | 0.16 | 0.23 | 0.12 |
| 5 | 52725158 | rs10428541 | -0.138 | 0.786 | -2.069 | 1.005 | 0.006 | -0.007 | -0.007 | 0 | 0.65 | 0.34 | 0.56 | 0.39 | 0.67 | 0.37 |
| 5 | 52727875 | rs13161622 | 1.542 | 0.795 | -2.156 | 0.916 | -0.002 | 0.003 | 0.003 | 0 | 0.16 | 0.16 | 0.1 | 0.2 | 0.11 | 0.16 |
| 5 | 52728629 | rs1862606 | 1.044 | 1.01 | -2.253 | 0.768 | -0.107 | 0.127 | 0.127 | 0 | 0.21 | 0.03 | 0.24 | 0.02 | 0.32 | 0.08 |
| 5 | 52746246 | rs745321 | -0.755 | 1.051 | -2.326 | 0.985 | -0.005 | -0.036 | -0.036 | 0.012 | 0.62 | 0.82 | 0.61 | 0.55 | 0.65 | 0.7 |
| 5 | 52751154 | rs3943928 | 1.13 | 1.185 | -2.393 | 0.835 | -0.113 | 0.128 | 0.128 | 0 | 0.21 | 0.03 | 0.25 | 0.03 | 0.33 | 0.08 |
| 5 | 52751356 | rs2042332 | 1.44 | 0.819 | -2.259 | 1.062 | 0.012 | 0.086 | 0.086 | 0 | 0.28 | 0.09 | 0.22 | 0.15 | 0.23 | 0.12 |
| 5 | 52752242 | rs13189156 | 0.749 | 0.777 | -2.142 | 1.217 | 0.046 | 0.018 | 0.018 | 0 | 0.42 | 0.25 | 0.3 | 0.36 | 0.32 | 0.29 |
| 5 | 52765331 | rs6450136 | -0.65 | 0.98 | -2.071 | 1.137 | -0.007 | -0.031 | -0.031 | 0 | 0.66 | 0.88 | 0.65 | 0.64 | 0.71 | 0.71 |
| 5 | 52778210 | rs3756498 | 1.165 | 0.81 | -2.028 | 0.863 | 0.032 | 0.073 | 0.073 | 0.042 | 0.26 | 0.11 | 0.2 | 0.14 | 0.18 | 0.11 |
| 5 | 52783356 | rs6450138 | 0.956 | 0.884 | -1.425 | 0.836 | -0.022 | -0.02 | -0.02 | 0 | 0.22 | 0.41 | 0.19 | 0.21 | 0.18 | 0.41 |
| 5 | 52783408 | rs1469101 | 0.392 | 1.558 | -2.223 | 0.238 | -0.009 | 0.192 | 0.192 | 1.432 | 0.43 | 0.18 | 0.3 | 0.16 | 0.38 | 0.19 |
| 5 | 52787926 | rs11954686 | 0.766 | 1.742 | -1.624 | 0.192 | -0.057 | 0.078 | 0.078 | 0.038 | 0.14 | 0.02 | 0.09 | 0.02 | 0.18 | 0.07 |
| 5 | 52788201 | rs12521153 | 0.127 | 1.159 | -1.187 | 0.627 | -0.005 | -0.008 | -0.008 | 0.262 | 0.29 | 0.28 | 0.31 | 0.32 | 0.35 | 0.18 |
| 5 | 52799618 | rs1363972 | -0.27 | 0.879 | -0.603 | 0.554 | -0.004 | 0.108 | 0.108 | 0.199 | 0.24 | 0.37 | 0.17 | 0.51 | 0.22 | 0.35 |
| 5 | 52805638 | rs7703235 | 0.49 | 0.802 | -0.912 | -0.26 | -0.003 | 0.024 | 0.024 | 0 | 0.39 | 0.41 | 0.51 | 0.26 | 0.41 | 0.26 |
| 5 | 52808229 | rs1593054 | 0.436 | 0.742 | -1.014 | -0.204 | -0.019 | 0.072 | 0.072 | 0.007 | 0.26 | 0.46 | 0.19 | 0.44 | 0.18 | 0.52 |
| 5 | 52824476 | rs10050519 | 0.591 | 0.8 | -0.957 | -0.731 | -0.001 | 0.01 | 0.01 | 0 | 0.4 | 0.46 | 0.52 | 0.26 | 0.41 | 0.26 |
| 5 | 52828383 | rs6896666 | 0.131 | 0.72 | -0.893 | -0.755 | 0.003 | 0.025 | 0.025 | 1.175 | 0.53 | 0.51 | 0.6 | 0.29 | 0.56 | 0.31 |
| 5 | 52833426 | rs12516704 | 0.041 | 0.699 | -0.679 | -0.936 | 0.001 | -0.018 | -0.018 | 0 | 0.34 | 0.37 | 0.44 | 0.14 | 0.35 | 0.28 |
| 5 | 52835755 | rs11743522 | 1.363 | 0.647 | -0.927 | -1.069 | -0.018 | 0.028 | 0.028 | 0 | 0.12 | 0.04 | 0.09 | 0.01 | 0.14 | 0.06 |
| 5 | 52842569 | rs1030228 | -1.856 | 0.43 | -0.982 | -0.877 | -0.008 | -0.016 | -0.016 | 0 | 0.78 | 0.83 | 0.82 | 0.79 | 0.75 | 0.76 |
| 5 | 52847778 | rs6872379 | -0.532 | 0.372 | -0.958 | -1.046 | 0.006 | 0.022 | 0.022 | 0.518 | 0.51 | 0.59 | 0.43 | 0.67 | 0.47 | 0.49 |
| 5 | 52847798 | rs6872395 | 0.585 | 0.208 | -0.943 | -1.115 | -0.011 | 0.054 | 0.054 | 0 | 0.29 | 0.19 | 0.27 | 0.17 | 0.32 | 0.2 |
| 5 | 52847995 | rs9790964 | 1.073 | 0.212 | -0.851 | -0.989 | -0.038 | 0.366 | 0.366 | 0 | 0.11 | 0.33 | 0.1 | 0.48 | 0.04 | 0.25 |
| 5 | 52850807 | rs889291 | -0.81 | 0.423 | -0.425 | -1.032 | 0.005 | -0.023 | -0.023 | 0 | 0.6 | 0.55 | 0.67 | 0.78 | 0.59 | 0.56 |
| 5 | 52856797 | rs1532162 | 0.828 | 0.19 | -0.791 | -1.034 | -0.008 | 0.132 | 0.132 | 0 | 0.22 | 0.38 | 0.19 | 0.49 | 0.17 | 0.3 |
| 5 | 52867330 | rs7728496 | -1.94 | 0.269 | -0.806 | -1.051 | -0.002 | -0.008 | -0.008 | 0.015 | 0.77 | 0.84 | 0.82 | 0.8 | 0.76 | 0.76 |
| 5 | 52872506 | rs7717681 | 0.938 | 0.301 | -0.744 | -1.02 | -0.006 | 0.087 | 0.087 | 0 | 0.22 | 0.3 | 0.17 | 0.49 | 0.18 | 0.3 |
| 5 | 52876423 | rs10036010 | 1.804 | 0.331 | -0.954 | -1.062 | 0 | -0.025 | -0.025 | 0 | 0.12 | 0.15 | 0.08 | 0.02 | 0.11 | 0.04 |
| 5 | 52885100 | rs1994648 | 1.853 | 0.428 | -1.019 | -1.067 | -0.005 | -0.018 | -0.018 | 0.026 | 0.16 | 0.11 | 0.14 | 0.01 | 0.18 | 0.2 |
| 5 | 52912891 | rs381575 | -0.073 | 0.368 | -1.034 | -1.068 | -0.003 | -0.006 | -0.006 | 0 | 0.4 | 0.58 | 0.34 | 0.5 | 0.36 | 0.5 |
| 5 | 52927597 | rs4362905 | 0.043 | 0.374 | -1.066 | -1.049 | -0.009 | 0.046 | 0.046 | 0.245 | 0.43 | 0.3 | 0.55 | 0.28 | 0.47 | 0.42 |
| 5 | 52942197 | rs31303 | -2.006 | 0.326 | -1.201 | -1.084 | -0.003 | -0.011 | -0.011 | 0 | 0.77 | 0.86 | 0.82 | 0.79 | 0.76 | 0.76 |
| 5 | 52942463 | rs31302 | -1.09 | 0.323 | -1.219 | -1.098 | -0.012 | 0.026 | 0.026 | 0.114 | 0.67 | 0.8 | 0.75 | 0.77 | 0.65 | 0.72 |
| 5 | 52945258 | rs1388111 | 0.79 | 0.344 | -1.222 | -1.078 | -0.025 | 0.038 | 0.038 | 0.749 | 0.27 | 0.14 | 0.39 | 0.13 | 0.31 | 0.27 |
| 5 | 52962251 | rs256116 | -2.013 | 0.381 | -1.148 | -0.96 | -0.011 | -0.017 | -0.017 | 0 | 0.78 | 0.88 | 0.82 | 0.8 | 0.76 | 0.74 |
| 5 | 52972924 | rs256094 | 0.201 | 0.365 | -1.051 | -0.962 | -0.004 | 0.04 | 0.04 | 0 | 0.33 | 0.45 | 0.26 | 0.51 | 0.29 | 0.32 |

| | | | | | | | | | | | | | | | | |
|---|---|---|---|---|---|---|---|---|---|---|---|---|---|---|---|---|
| 5 | 52979097 | rs567 | 0.121 | 0.349 | -1.058 | -1.001 | -0.012 | 0.059 | 0.059 | 0.145 | 0.43 | 0.27 | 0.55 | 0.28 | 0.47 | 0.41 |
| 5 | 53002783 | rs379439 | 0.039 | 0.36 | -1.187 | -0.966 | -0.005 | 0.02 | 0.02 | 0 | 0.34 | 0.56 | 0.27 | 0.51 | 0.29 | 0.32 |
| 5 | 53011223 | rs11745611 | -0.059 | 0.382 | -1.215 | -0.988 | -0.006 | 0.038 | 0.038 | 0.009 | 0.44 | 0.31 | 0.55 | 0.28 | 0.47 | 0.41 |
| 5 | 53018423 | rs11957871 | -0.486 | 0.374 | -1.225 | -1.003 | 0.007 | 0.036 | 0.036 | 3.788 | 0.54 | 0.34 | 0.62 | 0.31 | 0.58 | 0.46 |
| 5 | 53020039 | rs6886811 | 0.407 | 0.183 | -1.468 | -0.523 | 0.007 | 0.009 | 0.009 | 0 | 0.27 | 0.2 | 0.41 | 0.27 | 0.29 | 0.23 |
| 5 | 53023460 | rs256090 | -1.495 | 0.269 | -1.379 | -0.329 | -0.002 | -0.008 | -0.008 | 0 | 0.77 | 0.83 | 0.82 | 0.79 | 0.76 | 0.74 |
| 5 | 53025465 | rs155783 | 0.175 | 0.366 | -1.186 | -0.007 | 0.007 | 0.021 | 0.021 | 0 | 0.38 | 0.46 | 0.33 | 0.48 | 0.35 | 0.47 |
| 5 | 53041341 | rs7721921 | 0.519 | 0.301 | -1.354 | -0.104 | 0.005 | 0.008 | 0.008 | 0 | 0.16 | 0.13 | 0.16 | 0.15 | 0.16 | 0.15 |
| 5 | 53056428 | rs17247384 | 0.704 | 0.085 | -1.87 | -0.396 | 0 | 0.103 | 0.103 | 6.472 | 0.27 | 0.03 | 0.18 | 0.08 | 0.21 | 0.18 |
| 5 | 53060893 | rs11956638 | 0.23 | 0.589 | -1.313 | 0.255 | -0.016 | 0 | 0 | 0 | 0.39 | 0.3 | 0.34 | 0.36 | 0.44 | 0.29 |
| 5 | 53062881 | rs1394218 | 0.544 | 0.712 | -1.57 | 0.169 | -0.013 | 0.016 | 0.016 | 0 | 0.1 | 0.05 | 0.12 | 0.08 | 0.13 | 0.13 |
| 5 | 53062988 | rs271210 | -0.493 | 0.3 | -0.772 | 0.693 | -0.007 | 0.282 | 0.282 | 0.128 | 0.94 | 0.75 | 0.94 | 0.67 | 0.99 | 0.77 |
| 5 | 53068791 | rs6892188 | -0.04 | 0.321 | -0.916 | 0.619 | 0.022 | 0.017 | 0.017 | 5.235 | 0.49 | 0.3 | 0.45 | 0.44 | 0.56 | 0.43 |
| 5 | 53075037 | rs271234 | 1.058 | 0.067 | -1.182 | -0.073 | -0.001 | 0.002 | 0.002 | 4.206 | 0.16 | 0.12 | 0.12 | 0.05 | 0.15 | 0.23 |
| 5 | 53077259 | rs13165006 | 1.176 | 0.221 | -1.543 | 0.028 | -0.029 | 0.011 | 0.011 | 0.051 | 0.14 | 0.02 | 0.07 | 0.09 | 0.13 | 0.08 |
| 5 | 53080847 | rs16881743 | -0.006 | 0.127 | -0.424 | 0.534 | -0.001 | -0.014 | -0.014 | 0 | 0.12 | 0.19 | 0.13 | 0.08 | 0.13 | 0.16 |
| 5 | 53083975 | rs11955732 | 0.538 | 0.139 | -0.682 | 0.625 | -0.013 | -0.025 | -0.025 | 0 | 0.15 | 0.03 | 0.26 | 0.19 | 0.17 | 0.1 |
| 5 | 53104869 | rs6881775 | -0.039 | -0.028 | -0.038 | 0.776 | -0.021 | 0.151 | 0.151 | 1.307 | 0.79 | 0.52 | 0.85 | 0.57 | 0.86 | 0.6 |
| 5 | 53106463 | rs1986932 | 1.695 | 0.12 | -0.271 | 0.527 | -0.028 | 0.071 | 0.071 | 6.847 | 0.38 | 0.21 | 0.35 | 0.21 | 0.43 | 0.29 |
| 5 | 53110493 | rs10940341 | -0.403 | 0.142 | -0.237 | 0.438 | -0.004 | -0.008 | -0.008 | 0 | 0.31 | 0.22 | 0.37 | 0.35 | 0.33 | 0.29 |
| 5 | 53111334 | rs16881757 | -0.002 | 0.174 | -0.218 | 0.461 | -0.002 | -0.002 | -0.002 | 0 | 0.15 | 0.14 | 0.14 | 0.21 | 0.17 | 0.19 |
| 5 | 53116153 | rs17248426 | 1.614 | 0.127 | 0.241 | 0.268 | -0.003 | 0.02 | 0.02 | 5.979 | 0.38 | 0.3 | 0.35 | 0.22 | 0.4 | 0.27 |
| 5 | 53124836 | rs2007701 | 1.167 | 0.149 | 0.387 | 0.257 | -0.003 | 0.035 | 0.035 | 0.506 | 0.48 | 0.37 | 0.43 | 0.22 | 0.5 | 0.29 |
| 5 | 53125334 | rs271218 | -0.019 | 0.202 | 0.299 | 0.521 | -0.024 | 0.304 | 0.304 | 0.353 | 0.94 | 0.78 | 0.98 | 0.58 | 0.99 | 0.77 |
| 5 | 53127645 | rs12518752 | 0.541 | 0.172 | 0.091 | 0.39 | -0.001 | -0.004 | -0.004 | 0.057 | 0.14 | 0.12 | 0.27 | 0.15 | 0.16 | 0.08 |
| 5 | 53130488 | rs4865789 | 1.702 | 0.039 | 0.167 | 0.29 | -0.008 | 0.074 | 0.074 | 6.379 | 0.4 | 0.21 | 0.34 | 0.25 | 0.41 | 0.3 |
| 5 | 53154963 | rs271251 | -0.878 | -0.058 | 0.306 | 0.372 | -0.001 | 0.013 | 0.013 | 0.937 | 0.46 | 0.38 | 0.57 | 0.29 | 0.53 | 0.47 |
| 5 | 53156145 | rs271254 | 0.092 | -0.113 | 0.359 | 0.493 | -0.021 | 0.353 | 0.353 | 0.131 | 0.94 | 0.73 | 0.98 | 0.54 | 0.99 | 0.76 |
| 5 | 53156777 | rs6866140 | 1.767 | -0.111 | 0.297 | 0.443 | -0.013 | 0.072 | 0.072 | 5.904 | 0.37 | 0.13 | 0.31 | 0.25 | 0.36 | 0.26 |
| 5 | 53158093 | rs271255 | 0.042 | 0.009 | 0.331 | 0.619 | -0.024 | 0.315 | 0.315 | 0.002 | 0.94 | 0.76 | 0.98 | 0.54 | 0.99 | 0.76 |
| 5 | 53166636 | rs702604 | -1.048 | 0.104 | 0.19 | 0.292 | -0.003 | 0.021 | 0.021 | 0 | 0.46 | 0.52 | 0.58 | 0.23 | 0.53 | 0.35 |
| 5 | 53167408 | rs271257 | -0.386 | 0.179 | 0.243 | 0.523 | -0.02 | 0.334 | 0.334 | 0.069 | 0.95 | 0.75 | 0.98 | 0.54 | 0.99 | 0.77 |
| 5 | 53167426 | rs4543225 | 0.068 | 0.378 | 0.296 | 0.367 | 0.016 | -0.022 | -0.022 | 0 | 0.18 | 0.25 | 0.2 | 0.12 | 0.23 | 0.24 |
| 5 | 53169520 | rs702612 | 1.056 | 0.356 | 0.016 | 0.424 | 0 | 0 | 0 | 0 | 0.08 | 0.06 | 0.23 | 0.11 | 0.08 | 0.06 |
| 5 | 53176402 | rs2407068 | 1.793 | 0.279 | -0.121 | 0.36 | -0.036 | 0.1 | 0.1 | 6.706 | 0.4 | 0.13 | 0.38 | 0.25 | 0.43 | 0.26 |
| 5 | 53178047 | rs37540 | -0.089 | 0.238 | -0.007 | 0.592 | -0.012 | 0.392 | 0.392 | 0.959 | 0.94 | 0.71 | 0.97 | 0.53 | 0.99 | 0.76 |
| 5 | 53181731 | rs271246 | 0.944 | 0.305 | -0.527 | 0.44 | -0.029 | 0.009 | 0.009 | 1.506 | 0.12 | 0.01 | 0.08 | 0.05 | 0.11 | 0.14 |
| 5 | 53183038 | rs271247 | 1.232 | 0.449 | -0.194 | 0.605 | -0.007 | 0.051 | 0.051 | 0.072 | 0.53 | 0.3 | 0.39 | 0.26 | 0.53 | 0.4 |
| 5 | 53184552 | rs702613 | 0.168 | 0.007 | -0.343 | 0.897 | -0.018 | -0.017 | -0.017 | 0 | 0.14 | 0.04 | 0.25 | 0.11 | 0.16 | 0.09 |
| 5 | 53187303 | rs271245 | 0.583 | 0.044 | 0.185 | 1.131 | 0.003 | 0.492 | 0.492 | 1.025 | 0.94 | 0.65 | 0.93 | 0.5 | 1 | 0.76 |
| 5 | 53191143 | rs7729734 | 0.728 | 0.18 | -0.741 | 0.66 | 0 | 0.057 | 0.057 | 9.717 | 0.45 | 0.64 | 0.36 | 0.23 | 0.46 | 0.24 |
| 5 | 53192812 | rs271205 | 0.054 | 0.138 | -1.233 | 0.69 | -0.045 | -0.039 | -0.039 | 0 | 0.29 | 0.05 | 0.41 | 0.28 | 0.3 | 0.3 |
| 5 | 53205703 | rs7713737 | 0.805 | 0.468 | -0.922 | 0.747 | -0.02 | 0.061 | 0.061 | 2.354 | 0.4 | 0.27 | 0.28 | 0.23 | 0.42 | 0.24 |
| 5 | 53210473 | rs3822502 | 0.246 | 0.652 | -0.8 | 1.04 | -0.001 | -0.001 | -0.001 | 3.509 | 0.75 | 0.74 | 0.76 | 0.59 | 0.76 | 0.54 |
| 5 | 53218984 | rs10513024 | 0.079 | 0.422 | -1.86 | 0.82 | -0.07 | 0.027 | 0.027 | 0 | 0.31 | 0.03 | 0.31 | 0.19 | 0.27 | 0.28 |
| 5 | 53219787 | rs6884046 | 0.31 | 0.995 | -2.097 | 1.041 | 0.048 | 0.045 | 0.045 | 0 | 0.49 | 0.74 | 0.38 | 0.43 | 0.41 | 0.48 |
| 5 | 53220783 | rs753124 | 0.144 | 1.034 | -1.557 | 0.798 | 0.032 | 0.005 | 0.005 | 0 | 0.43 | 0.62 | 0.33 | 0.43 | 0.35 | 0.44 |
| 5 | 53226422 | rs702620 | 0.679 | 0.829 | -0.78 | 0.476 | -0.016 | 0.04 | 0.04 | 0 | 0.74 | 0.62 | 0.76 | 0.61 | 0.77 | 0.64 |
| 5 | 53227974 | rs702621 | -0.335 | 0.73 | -0.7 | 0.418 | 0.043 | -0.064 | -0.064 | 0 | 0.85 | 0.95 | 0.92 | 0.61 | 0.91 | 0.71 |
| 5 | 53228675 | rs713209 | 0.858 | 0.59 | -0.627 | 0.33 | -0.015 | -0.054 | -0.054 | 0 | 0.26 | 0.31 | 0.15 | 0.09 | 0.21 | 0.16 |
| 5 | 53228705 | rs713210 | -0.971 | 0.71 | -0.41 | 0.461 | -0.009 | 0.015 | 0.015 | 0 | 0.25 | 0.26 | 0.22 | 0.36 | 0.22 | 0.33 |
| 5 | 53240729 | rs13187145 | 0.178 | 0.71 | -0.619 | -0.221 | -0.018 | -0.069 | -0.069 | 0 | 0.32 | 0.37 | 0.2 | 0.09 | 0.25 | 0.18 |
| 5 | 53246614 | rs788514 | 0.074 | 0.843 | -0.672 | 0.419 | 0 | 0.011 | 0.011 | 1.452 | 0.81 | 0.76 | 0.8 | 0.62 | 0.76 | 0.64 |
| 5 | 53246677 | rs17250527 | 0.46 | 0.915 | -0.743 | 0.468 | -0.015 | -0.023 | -0.023 | 0 | 0.09 | 0.02 | 0.06 | 0.18 | 0.11 | 0.13 |
| 5 | 53248363 | rs809798 | 1.352 | 0.945 | -0.447 | 0.089 | -0.005 | -0.001 | -0.001 | 0.082 | 0.38 | 0.28 | 0.44 | 0.36 | 0.37 | 0.29 |
| 5 | 53250497 | rs16881895 | 0.186 | 0.847 | -0.393 | -0.302 | -0.01 | -0.079 | -0.079 | 0 | 0.31 | 0.39 | 0.26 | 0.08 | 0.26 | 0.2 |
| 5 | 53260679 | rs697112 | 1.039 | 1.035 | 0.305 | 0.243 | 0.002 | 0.001 | 0.001 | 1.406 | 0.51 | 0.44 | 0.59 | 0.43 | 0.48 | 0.4 |
| 5 | 53267387 | rs702639 | 1.258 | 1.221 | -0.255 | 0.147 | 0.004 | -0.033 | -0.033 | 0.039 | 0.46 | 0.14 | 0.47 | 0.42 | 0.38 | 0.36 |
| 5 | 53271420 | rs702634 | -0.066 | 1.449 | 0.161 | 0.067 | 0.005 | -0.008 | -0.008 | 0.261 | 0.78 | 0.74 | 0.79 | 0.83 | 0.69 | 0.74 |
| 5 | 53272664 | rs4865796 | -0.015 | 1.435 | 0.151 | 0.043 | 0.003 | -0.006 | -0.006 | 2.671 | 0.78 | 0.75 | 0.79 | 0.84 | 0.7 | 0.74 |
| 5 | 53279558 | rs1694063 | 1.545 | 1.433 | -0.266 | 0.051 | 0.015 | -0.048 | -0.048 | 0 | 0.46 | 0.09 | 0.45 | 0.41 | 0.36 | 0.36 |
| 5 | 53281711 | rs1664786 | 1.459 | 1.477 | -0.223 | 0.065 | 0 | -0.004 | -0.004 | 1.501 | 0.47 | 0.16 | 0.49 | 0.41 | 0.42 | 0.37 |
| 5 | 53282649 | rs1664789 | 1.497 | 1.481 | -0.301 | 0.067 | -0.011 | -0.016 | -0.016 | 0 | 0.47 | 0.14 | 0.49 | 0.41 | 0.42 | 0.37 |
| 5 | 53282979 | rs878781 | 0.26 | 1.222 | 0.305 | 0.125 | 0.052 | -0.038 | -0.038 | 1.227 | 0.77 | 0.57 | 0.75 | 0.83 | 0.65 | 0.74 |
| 5 | 53298025 | rs6450176 | -0.796 | 0.691 | 0.306 | 0.002 | -0.002 | 0.023 | 0.023 | 0.02 | 0.29 | 0.32 | 0.24 | 0.46 | 0.28 | 0.34 |
| 5 | 53299435 | rs6893522 | 0.178 | 0.528 | -0.135 | -0.048 | 0.02 | 0.053 | 0.053 | 0 | 0.11 | 0.02 | 0.05 | 0 | 0.06 | 0.05 |
| 5 | 53305742 | rs1542167 | 0.198 | 0.655 | 0.09 | 0.094 | -0.003 | 0.019 | 0.019 | 0 | 0.14 | 0.16 | 0.15 | 0.46 | 0.17 | 0.29 |
| 5 | 53314757 | rs31127 | 0.123 | 0.673 | 0.416 | -0.171 | -0.006 | -0.041 | -0.041 | 1.746 | 0.77 | 0.59 | 0.81 | 0.83 | 0.76 | 0.75 |
| 5 | 53316271 | rs697105 | 1.11 | 0.419 | 0.468 | -0.267 | 0.002 | 0.002 | 0.002 | 0.731 | 0.59 | 0.35 | 0.65 | 0.38 | 0.58 | 0.46 |

| | | | | | | | | | | | | | | | |
|---|---|---|---|---|---|---|---|---|---|---|---|---|---|---|---|
| 5 | 53320109 | rs255754 | 0.903 | 0.374 | 0.293 | -0.444 | -0.01 | -0.045 | -0.045 | 0.015 | 0.65 | 0.55 | 0.74 | 0.83 | 0.68 | 0.71 |
| 5 | 53324930 | rs697098 | 1.321 | 0.055 | 0.189 | -0.536 | 0.004 | 0.036 | 0.036 | 0.562 | 0.54 | 0.16 | 0.58 | 0.39 | 0.53 | 0.42 |
| 5 | 53326381 | rs696195 | 2.18 | 0.301 | 0.03 | -0.605 | -0.109 | -0.033 | -0.033 | 0 | 0.41 | 0.04 | 0.51 | 0.38 | 0.42 | 0.4 |
| 5 | 53327571 | rs31226 | 0.912 | 0.244 | 0.461 | -0.65 | 0 | -0.019 | -0.019 | 0 | 0.6 | 0.42 | 0.7 | 0.85 | 0.6 | 0.71 |
| 5 | 53333477 | rs10513033 | 0.32 | 0.188 | 0.49 | -0.59 | -0.005 | 0.129 | 0.129 | 0 | 0.19 | 0.34 | 0.18 | 0.47 | 0.18 | 0.32 |
| 5 | 53336820 | rs274362 | -0.277 | 0.169 | 0.778 | -0.656 | 0.004 | -0.022 | -0.022 | 0 | 0.74 | 0.64 | 0.79 | 0.85 | 0.68 | 0.76 |
| 5 | 53339518 | rs25858 | -1.092 | -0.061 | 0.933 | -0.495 | -0.023 | 0.033 | 0.033 | 0 | 0.33 | 0.52 | 0.25 | 0.47 | 0.26 | 0.4 |
| 5 | 53342469 | rs6884003 | 0.016 | -0.026 | 0.578 | -0.482 | -0.033 | -0.078 | -0.078 | 0.675 | 0.14 | 0.21 | 0.07 | 0 | 0.08 | 0.05 |
| 5 | 53343576 | rs11742692 | 0.328 | 0.223 | 0.277 | -0.322 | 0.002 | -0.041 | -0.041 | 0 | 0.18 | 0.11 | 0.16 | 0.46 | 0.18 | 0.35 |
| 5 | 53360059 | rs3776695 | 0.17 | 0.132 | 0.659 | -0.607 | -0.009 | 0.019 | 0.019 | 0.012 | 0.13 | 0.35 | 0.08 | 0.16 | 0.11 | 0.18 |
| 5 | 53360174 | rs3776694 | -1.144 | -0.05 | 0.773 | -0.814 | -0.019 | -0.028 | -0.028 | 0.295 | 0.28 | 0.64 | 0.21 | 0.16 | 0.21 | 0.26 |
| 5 | 53360984 | rs6867281 | -0.832 | -0.066 | 0.795 | -0.805 | -0.029 | -0.034 | -0.034 | 0 | 0.26 | 0.61 | 0.17 | 0.16 | 0.18 | 0.26 |
| 5 | 53364068 | rs10067187 | -0.765 | -0.04 | 0.924 | -0.799 | -0.043 | -0.055 | -0.055 | 0 | 0.25 | 0.56 | 0.16 | 0.16 | 0.18 | 0.26 |
| 5 | 53364230 | rs156828 | 0.008 | 0.287 | 0.446 | -0.75 | -0.018 | -0.039 | -0.039 | 1.41 | 0.69 | 0.77 | 0.71 | 0.53 | 0.63 | 0.62 |
| 5 | 53368869 | rs31242 | 1.858 | 0.402 | 0.319 | -0.884 | -0.01 | 0.007 | 0.007 | 0 | 0.1 | 0.08 | 0.09 | 0 | 0.14 | 0.03 |
| 5 | 53371864 | rs6863779 | -0.41 | 0.143 | 0.742 | -0.8 | -0.025 | -0.036 | -0.036 | 0 | 0.23 | 0.36 | 0.15 | 0.12 | 0.18 | 0.28 |
| 5 | 53373029 | rs156442 | -0.539 | 0.166 | 0.802 | -0.745 | -0.001 | 0.007 | 0.007 | 0.126 | 0.79 | 0.8 | 0.8 | 0.48 | 0.76 | 0.68 |
| 5 | 53376613 | rs156436 | 1.306 | 0.142 | 0.721 | -0.786 | 0.03 | 0.07 | 0.07 | 0 | 0.54 | 0.22 | 0.61 | 0.36 | 0.56 | 0.39 |
| 5 | 53379302 | rs9686938 | 0.386 | 0.134 | 0.459 | -0.825 | 0.008 | -0.007 | -0.007 | 0 | 0.13 | 0.12 | 0.08 | 0 | 0.07 | 0.11 |
| 5 | 53382156 | rs10060587 | 0.709 | 0.202 | 0.479 | -0.825 | 0.006 | 0.024 | 0.024 | 0 | 0.1 | 0.24 | 0.06 | 0.11 | 0.1 | 0.18 |
| 5 | 53382703 | rs392714 | -0.916 | 0.083 | 0.487 | -0.82 | 0.002 | -0.004 | -0.004 | 0 | 0.35 | 0.34 | 0.28 | 0.53 | 0.31 | 0.45 |
| 5 | 53385529 | rs695922 | -0.315 | 0.299 | 0.056 | -1.357 | 0.018 | -0.003 | -0.003 | 0.012 | 0.88 | 0.75 | 0.9 | 0.89 | 0.84 | 0.78 |
| 5 | 53386267 | rs696193 | 0.726 | 0.248 | -0.036 | -1.381 | -0.005 | -0.02 | -0.02 | 0 | 0.74 | 0.67 | 0.82 | 0.89 | 0.77 | 0.68 |
| 5 | 53393965 | rs156824 | -0.151 | 0.624 | -0.576 | -1.968 | -0.048 | 0.067 | 0.067 | 0 | 0.8 | 0.93 | 0.85 | 0.98 | 0.7 | 0.88 |
| 5 | 53394938 | rs682103 | 1.178 | 0.31 | -1.586 | -2.401 | -0.074 | -0.022 | -0.022 | 0 | 0.3 | 0.04 | 0.33 | 0.29 | 0.31 | 0.28 |
| 5 | 53399217 | rs1703368 | 0.006 | 0.128 | -1.558 | -2.483 | -0.004 | 0.023 | 0.023 | 3.076 | 0.84 | 0.91 | 0.88 | 0.98 | 0.84 | 0.86 |
| 5 | 53408103 | rs17266918 | 2.005 | 0.222 | -1.096 | -2.423 | -0.033 | 0.046 | 0.046 | 0 | 0.1 | 0.02 | 0.08 | 0 | 0.14 | 0.03 |
| 5 | 53412620 | rs279744 | 1.232 | 0.09 | -1.2 | -2.498 | -0.088 | -0.023 | -0.023 | 0 | 0.31 | 0.02 | 0.4 | 0.29 | 0.31 | 0.29 |
| 5 | 53415114 | rs279743 | 0.557 | 0.082 | -0.794 | -2.497 | -0.051 | 0.086 | 0.086 | 0 | 0.42 | 0.18 | 0.48 | 0.29 | 0.46 | 0.31 |
| 5 | 53431670 | rs17404688 | 1.991 | 0.239 | -0.752 | -2.378 | -0.035 | 0.046 | 0.046 | 0 | 0.1 | 0.02 | 0.08 | 0 | 0.15 | 0.03 |
| 5 | 53441597 | rs26608 | 0.648 | -0.003 | -0.658 | -2.624 | -0.004 | -0.013 | -0.013 | 0 | 0.23 | 0.15 | 0.3 | 0.68 | 0.26 | 0.43 |
| 5 | 53444498 | rs2441026 | 0.917 | 0.255 | -0.347 | -2.566 | 0.009 | 0.094 | 0.094 | 0.236 | 0.5 | 0.36 | 0.55 | 0.13 | 0.44 | 0.34 |
| 5 | 53445527 | rs151806 | 0.014 | 0.26 | -0.225 | -2.576 | -0.006 | -0.024 | -0.024 | 0 | 0.28 | 0.35 | 0.29 | 0.85 | 0.26 | 0.54 |
| 5 | 53448014 | rs2454916 | 1.621 | 0.232 | -0.327 | -2.577 | -0.079 | 0.035 | 0.035 | 0 | 0.23 | 0.02 | 0.38 | 0.13 | 0.26 | 0.19 |
| 5 | 53452846 | rs277332 | 0.256 | 0.354 | 0.136 | -2.199 | -0.002 | -0.011 | -0.011 | 0 | 0.77 | 0.78 | 0.88 | 0.81 | 0.79 | 0.78 |
| 5 | 53457867 | rs172139 | -0.644 | 0.319 | 0.135 | -2.086 | 0 | -0.051 | -0.051 | 2.21 | 0.39 | 0.48 | 0.37 | 0.85 | 0.4 | 0.57 |
| 5 | 53463117 | rs154021 | 1.261 | 0.315 | -0.24 | -1.917 | -0.03 | 0.035 | 0.035 | 0 | 0.2 | 0.03 | 0.36 | 0.13 | 0.24 | 0.17 |
| 5 | 53463790 | rs13153747 | -0.117 | 0.664 | 0.03 | -1.849 | 0.003 | -0.021 | -0.021 | 0.119 | 0.24 | 0.31 | 0.15 | 0.17 | 0.24 | 0.12 |
| 5 | 53468276 | rs7718177 | 0.161 | 0.672 | -0.008 | -1.848 | -0.001 | -0.001 | -0.001 | 0 | 0.23 | 0.22 | 0.15 | 0.17 | 0.24 | 0.12 |
| 5 | 53469763 | rs277340 | -1.197 | 0.864 | -0.027 | -1.82 | -0.004 | 0.006 | 0.006 | 0 | 0.89 | 0.92 | 0.95 | 0.99 | 0.89 | 0.9 |
| 5 | 53469829 | rs16882383 | 1.877 | 0.79 | 0.014 | -1.901 | -0.007 | -0.084 | -0.084 | 0 | 0.31 | 0.34 | 0.19 | 0.01 | 0.22 | 0.18 |
| 5 | 53472156 | rs26775 | 1.079 | 0.817 | 0.037 | -1.823 | 0.004 | 0.022 | 0.022 | 0 | 0.16 | 0.21 | 0.26 | 0.69 | 0.19 | 0.43 |
| 5 | 53474256 | rs25860 | -1.15 | 0.826 | -0.026 | -1.875 | 0 | -0.079 | -0.079 | 2.635 | 0.59 | 0.6 | 0.77 | 0.99 | 0.66 | 0.72 |
| 5 | 53474849 | rs28033 | -0.407 | 0.838 | -0.064 | -1.833 | -0.006 | 0.02 | 0.02 | 0 | 0.43 | 0.4 | 0.51 | 0.28 | 0.47 | 0.29 |
| 5 | 53476717 | rs17413044 | 1.746 | 0.89 | -0.356 | -1.912 | -0.048 | 0.061 | 0.061 | 0 | 0.12 | 0.03 | 0.12 | 0 | 0.19 | 0.03 |
| 5 | 53486661 | rs181916 | 0.677 | 0.967 | -0.212 | -1.952 | -0.002 | -0.113 | -0.113 | 0 | 0.44 | 0.5 | 0.27 | 0.01 | 0.4 | 0.3 |
| 5 | 53490737 | rs1345815 | -0.54 | 0.999 | -0.45 | -1.823 | -0.008 | 0.002 | 0.002 | 0 | 0.74 | 0.77 | 0.63 | 0.28 | 0.63 | 0.54 |
| 5 | 53490881 | rs13435967 | 1.472 | 0.687 | -0.38 | -1.848 | -0.007 | -0.1 | -0.1 | 0 | 0.34 | 0.35 | 0.21 | 0.01 | 0.28 | 0.2 |
| 5 | 53492282 | rs16882424 | 1.645 | 0.702 | -0.384 | -1.853 | -0.015 | -0.108 | -0.108 | 2.337 | 0.33 | 0.35 | 0.19 | 0.01 | 0.23 | 0.19 |
| 5 | 53496587 | rs6871446 | 1.562 | 0.674 | -0.341 | -1.812 | -0.012 | -0.122 | -0.122 | 0.954 | 0.33 | 0.36 | 0.2 | 0.01 | 0.26 | 0.19 |
| 5 | 53499591 | rs3797254 | -0.109 | 0.733 | -0.289 | -1.741 | 0.001 | 0.002 | 0.002 | 0 | 0.32 | 0.33 | 0.4 | 0.28 | 0.29 | 0.25 |
| 5 | 53501752 | rs3797251 | 0.579 | 0.703 | -0.317 | -1.787 | -0.019 | 0.007 | 0.007 | 0.168 | 0.13 | 0.25 | 0.04 | 0.17 | 0.05 | 0.1 |
| 5 | 53504705 | rs4865808 | -0.363 | 0.771 | -0.743 | -1.817 | -0.002 | 0 | 0 | 0 | 0.71 | 0.75 | 0.62 | 0.28 | 0.58 | 0.52 |
| 5 | 53504764 | rs4865809 | -1.251 | 0.796 | -0.801 | -1.834 | 0.009 | 0.009 | 0.009 | 0 | 0.83 | 0.79 | 0.74 | 0.28 | 0.78 | 0.56 |
| 5 | 53507238 | rs16882451 | -0.259 | -0.344 | -0.513 | -1.439 | -0.012 | -0.014 | -0.014 | 0 | 0.3 | 0.42 | 0.39 | 0.27 | 0.23 | 0.24 |
| 5 | 53508218 | rs3776658 | 0.35 | -0.16 | -1.099 | -0.894 | 0.015 | 0.033 | 0.033 | 0 | 0.42 | 0.27 | 0.29 | 0.27 | 0.35 | 0.37 |
| 5 | 53509186 | rs3797248 | 0.768 | -0.554 | -1.048 | -1.987 | 0.001 | 0.027 | 0.027 | 0 | 0.17 | 0.11 | 0.34 | 0.1 | 0.18 | 0.15 |
| 5 | 53510482 | rs3797247 | 0.076 | -0.072 | -1.11 | -1.737 | 0.021 | 0.024 | 0.024 | 0 | 0.39 | 0.26 | 0.3 | 0.04 | 0.3 | 0.27 |
| 5 | 53510501 | rs3797246 | -0.213 | 0.26 | -0.668 | -0.375 | -0.011 | 0.037 | 0.037 | 0 | 0.08 | 0.02 | 0.12 | 0.04 | 0.09 | 0.08 |
| 5 | 53513531 | rs13164469 | -0.859 | 0.238 | -0.199 | 0.032 | -0.017 | -0.039 | -0.039 | 0.514 | 0.3 | 0.53 | 0.29 | 0.21 | 0.23 | 0.24 |
| 5 | 53517015 | rs27636 | -1.414 | 0.421 | -0.634 | 0.756 | 0.003 | -0.009 | -0.009 | 0 | 0.11 | 0.1 | 0.16 | 0.21 | 0.08 | 0.15 |
| 5 | 53532777 | rs6879661 | -1.047 | -0.488 | -0.273 | 0.52 | -0.024 | -0.006 | -0.006 | 0 | 0.19 | 0.29 | 0.09 | 0.14 | 0.1 | 0.13 |
| 5 | 53544688 | rs10461408 | 1.408 | -0.488 | -0.908 | -0.735 | 0.039 | 0.224 | 0.224 | 0.234 | 0.48 | 0.15 | 0.29 | 0 | 0.58 | 0.45 |
| 5 | 53547960 | rs3797235 | -1.33 | -0.444 | 0.009 | 0.472 | 0.045 | 0.272 | 0.272 | 1.281 | 0.14 | 0.47 | 0.2 | 0.42 | 0.16 | 0.21 |
| 5 | 53553740 | rs26887 | 0.472 | -0.617 | 0.641 | 0.349 | -0.01 | -0.028 | -0.028 | 0 | 0.06 | 0.12 | 0.05 | 0.02 | 0.04 | 0.02 |
| 5 | 53553830 | rs6450182 | -0.788 | -0.571 | 0.824 | 0.365 | 0.029 | -0.024 | -0.024 | 0.011 | 0.08 | 0.37 | 0.08 | 0 | 0.07 | 0.07 |
| 5 | 53555349 | rs3797232 | 0.417 | -0.501 | 0.573 | 0.418 | 0.01 | 0.144 | 0.144 | 0.119 | 0.06 | 0.1 | 0.13 | 0.42 | 0.09 | 0.15 |
| 5 | 53566211 | rs35920 | 0.08 | -0.699 | 0.38 | 0.503 | 0.011 | -0.059 | -0.059 | 0 | 0.9 | 0.99 | 0.65 | 0.62 | 0.93 | 0.81 |
| 5 | 53572117 | rs10940369 | 2.007 | -0.74 | 0.422 | 0.403 | 0.063 | 0.023 | 0.023 | 0 | 0.54 | 0.32 | 0.42 | 0.42 | 0.66 | 0.59 |

| | | | | | | | | | | | | | | | | |
|---|---|---|---|---|---|---|---|---|---|---|---|---|---|---|---|---|
| 5 | 53575073 | rs13183117 | 1.79 | -0.62 | 0.53 | 0.533 | -0.011 | -0.001 | -0.001 | 1.318 | 0.59 | 0.48 | 0.48 | 0.42 | 0.73 | 0.66 |
| 5 | 53581765 | rs3776644 | 0.457 | -0.585 | 0.556 | 0.543 | 0.02 | 0.181 | 0.181 | 0 | 0.06 | 0.12 | 0.12 | 0.42 | 0.08 | 0.15 |
| 5 | 53582605 | rs250382 | -1.25 | -0.617 | 0.526 | 0.526 | -0.033 | -0.028 | -0.028 | 0.022 | 0.31 | 0.51 | 0.17 | 0.2 | 0.2 | 0.15 |
| 5 | 53587702 | rs7704911 | 1.894 | 0.45 | 0.969 | 0.675 | -0.021 | 0.011 | 0.011 | 0 | 0.64 | 0.53 | 0.48 | 0.42 | 0.72 | 0.65 |
| 5 | 53587857 | rs3797228 | 0.287 | 0.365 | 1.039 | 0.711 | 0.006 | 0.123 | 0.123 | 0.119 | 0.06 | 0.08 | 0.12 | 0.42 | 0.09 | 0.15 |
| 5 | 53588584 | rs35955 | -1.354 | 0.35 | 1.092 | 0.64 | -0.025 | -0.014 | -0.014 | 0.632 | 0.25 | 0.45 | 0.16 | 0.2 | 0.18 | 0.16 |
| 5 | 53590220 | rs35954 | 1.262 | 0.373 | 0.709 | 0.815 | 0.001 | -0.006 | -0.006 | 0 | 0.84 | 0.87 | 0.52 | 0.2 | 0.84 | 0.66 |
| 5 | 53592163 | rs35952 | 1.079 | 0.315 | 0.757 | 0.754 | -0.064 | 0.083 | 0.083 | 0 | 0.73 | 0.51 | 0.54 | 0.45 | 0.85 | 0.71 |
| 5 | 53594702 | rs7704507 | 0.644 | 0.3 | 0.438 | 0.814 | 0.002 | -0.003 | -0.003 | 0 | 0.77 | 0.77 | 0.64 | 0.28 | 0.83 | 0.6 |
| 5 | 53597197 | rs11949800 | -1.083 | 1.034 | 0.603 | 0.605 | 0.019 | 0.017 | 0.017 | 1.567 | 0.27 | 0.38 | 0.31 | 0.29 | 0.28 | 0.2 |
| 5 | 53600443 | rs35947 | -0.567 | 1.729 | 0.561 | 0.528 | 0.008 | -0.006 | -0.006 | 0 | 0.31 | 0.35 | 0.22 | 0.25 | 0.36 | 0.28 |
| 5 | 53606295 | rs35941 | -0.511 | 1.233 | -0.261 | 0.802 | 0.027 | -0.054 | -0.054 | 0 | 0.79 | 0.94 | 0.8 | 0.72 | 0.84 | 0.79 |
| 5 | 53607091 | rs35940 | 0.596 | 1.519 | -0.239 | 0.737 | -0.015 | -0.018 | -0.018 | 0 | 0.2 | 0.13 | 0.15 | 0.26 | 0.2 | 0.23 |
| 5 | 53607261 | rs35938 | -0.595 | 1.476 | -0.3 | 0.689 | 0.026 | -0.043 | -0.043 | 0 | 0.79 | 0.91 | 0.79 | 0.69 | 0.84 | 0.81 |
| 5 | 53614482 | rs1900165 | 1.738 | 1.776 | 0.051 | 0.445 | 0.017 | 0.074 | 0.074 | 1.482 | 0.4 | 0.23 | 0.34 | 0 | 0.41 | 0.3 |
| 5 | 53622417 | rs250380 | 0.606 | 1.678 | 0.067 | 0.521 | 0.024 | -0.035 | -0.035 | 0.429 | 0.2 | 0.1 | 0.21 | 0.31 | 0.16 | 0.18 |
| 5 | 53624687 | rs7704345 | -0.331 | 1.696 | 0.399 | 0.538 | 0.031 | 0.216 | 0.216 | 1.706 | 0.16 | 0.29 | 0.23 | 0.47 | 0.17 | 0.27 |
| 5 | 53628179 | rs1445829 | 1.752 | 1.701 | 0.312 | 0.519 | 0.013 | 0.134 | 0.134 | 1.52 | 0.4 | 0.18 | 0.34 | 0 | 0.42 | 0.3 |
| 5 | 53631815 | rs10063411 | -0.627 | 1.726 | 0.335 | 0.519 | 0.076 | 0.196 | 0.196 | 1.645 | 0.18 | 0.33 | 0.29 | 0.47 | 0.24 | 0.3 |
| 5 | 53637054 | rs2407508 | 0.36 | 1.62 | 0.24 | 0.728 | 0.011 | -0.019 | -0.019 | 3.974 | 0.59 | 0.65 | 0.64 | 0.48 | 0.66 | 0.6 |
| 5 | 53643036 | rs2407509 | -0.587 | 1.774 | 0.104 | 0.653 | -0.006 | -0.003 | -0.003 | 0 | 0.78 | 0.79 | 0.83 | 0.79 | 0.79 | 0.77 |
| 5 | 53656177 | rs1544780 | 0.343 | 1.038 | -0.14 | 0.878 | 0.002 | -0.004 | -0.004 | 0 | 0.21 | 0.23 | 0.21 | 0.31 | 0.16 | 0.18 |
| 5 | 53659481 | rs12515078 | -0.039 | 1.329 | -0.181 | 0.962 | -0.001 | 0.077 | 0.077 | 0 | 0.53 | 0.19 | 0.55 | 0.31 | 0.59 | 0.44 |
| 5 | 53684474 | rs6873666 | -1.147 | 1.863 | -0.116 | 1.779 | 0.01 | 0.028 | 0.028 | 1.37 | 0.13 | 0.18 | 0.25 | 0.28 | 0.19 | 0.14 |
| 5 | 53686888 | rs3111643 | -0.438 | 1.464 | -0.425 | 1.39 | -0.004 | -0.025 | -0.025 | 0 | 0.26 | 0.17 | 0.35 | 0.62 | 0.28 | 0.42 |
| 5 | 53693147 | rs3096162 | 1.042 | 2.28 | 0.62 | 1.062 | 0.168 | 0.08 | 0.08 | 0 | 0.94 | 0.71 | 0.89 | 0.94 | 0.9 | 0.93 |
| 5 | 53694111 | rs6877553 | -0.31 | 1.922 | -0.187 | 0.966 | 0.011 | -0.002 | -0.002 | 1.585 | 0.59 | 0.73 | 0.55 | 0.57 | 0.61 | 0.62 |
| 5 | 53694171 | rs6882064 | -0.296 | 1.863 | -0.207 | 0.962 | 0.016 | -0.002 | -0.002 | 0.001 | 0.59 | 0.73 | 0.55 | 0.57 | 0.62 | 0.62 |
| 5 | 53694941 | rs755188 | 0.361 | 1.812 | -0.509 | 0.9 | 0.014 | -0.024 | -0.024 | 0 | 0.39 | 0.27 | 0.26 | 0.42 | 0.36 | 0.41 |
| 5 | 53698535 | rs1981845 | -0.006 | 0.956 | -0.848 | 0.959 | 0.043 | 0.013 | 0.013 | 0.511 | 0.42 | 0.19 | 0.43 | 0.37 | 0.35 | 0.36 |
| 5 | 53707175 | rs7734716 | -0.013 | 0.598 | -0.232 | 0.444 | 0.02 | 0.038 | 0.038 | 1.349 | 0.7 | 0.8 | 0.82 | 0.93 | 0.77 | 0.8 |
| 5 | 53711122 | rs7731258 | -0.764 | 0.736 | 0.223 | 1.424 | -0.009 | 0.155 | 0.155 | 0.166 | 0.13 | 0.25 | 0.24 | 0.38 | 0.12 | 0.24 |
| 5 | 53720950 | rs10513043 | -0.253 | 1.546 | -0.643 | 1.432 | -0.063 | -0.064 | -0.064 | 1.953 | 0.16 | 0.04 | 0.28 | 0.24 | 0.27 | 0.26 |
| 5 | 53721698 | rs10471903 | -1.118 | 1.33 | 0.195 | 1.072 | 0.045 | 0.027 | 0.027 | 0 | 0.23 | 0.4 | 0.34 | 0.24 | 0.33 | 0.27 |
| 5 | 53723601 | rs6896176 | 0.025 | 1.877 | 0.185 | 1.465 | 0.005 | 0.029 | 0.029 | 0.26 | 0.11 | 0.13 | 0.19 | 0.3 | 0.14 | 0.28 |
| 5 | 53726016 | rs4272091 | 0.312 | 2.072 | 0.112 | 1.509 | -0.062 | -0.053 | -0.053 | 0 | 0.11 | 0.01 | 0.23 | 0.14 | 0.19 | 0.2 |
| 5 | 53727092 | rs3923330 | -1.026 | 1.591 | -0.488 | 0.93 | -0.018 | -0.047 | -0.047 | 5.299 | 0.69 | 0.59 | 0.8 | 0.93 | 0.78 | 0.83 |
| 5 | 53729982 | rs4597914 | 0.164 | 1.549 | -0.206 | 0.917 | 0.034 | 0.04 | 0.04 | 0 | 0.52 | 0.3 | 0.58 | 0.63 | 0.61 | 0.53 |
| 5 | 53730029 | rs4547886 | 0.741 | 1.537 | -0.273 | 0.914 | -0.062 | -0.055 | -0.055 | 0 | 0.11 | 0.01 | 0.23 | 0.14 | 0.19 | 0.2 |
| 5 | 53732977 | rs4865826 | -0.697 | 1.647 | -0.188 | 0.848 | -0.016 | -0.052 | -0.052 | 0 | 0.65 | 0.43 | 0.68 | 0.93 | 0.74 | 0.8 |
| 5 | 53733216 | rs4865827 | -1.502 | 1.674 | -0.167 | 0.933 | 0.024 | 0.028 | 0.028 | 0.043 | 0.32 | 0.39 | 0.41 | 0.54 | 0.43 | 0.54 |
| 5 | 53735108 | rs7714196 | -0.994 | 1.467 | 0.132 | 1.024 | 0.007 | 0.103 | 0.103 | 0 | 0.2 | 0.44 | 0.26 | 0.36 | 0.2 | 0.31 |
| 5 | 53735299 | rs4533844 | 1.428 | 1.206 | -0.391 | 0.962 | -0.035 | -0.038 | -0.038 | 0 | 0.36 | 0.05 | 0.28 | 0.38 | 0.31 | 0.26 |
| 5 | 53738058 | rs12654643 | 0.45 | 1.422 | -0.429 | 1.26 | 0.003 | 0.023 | 0.023 | 0.334 | 0.12 | 0.13 | 0.19 | 0.3 | 0.14 | 0.3 |
| 5 | 53739403 | rs4865829 | 0.07 | 1.205 | -0.24 | 0.83 | 0.033 | 0.066 | 0.066 | 0.041 | 0.58 | 0.22 | 0.61 | 0.81 | 0.65 | 0.71 |
| 5 | 53739530 | rs7722951 | -0.237 | 1.339 | -0.537 | 0.747 | 0.005 | 0.013 | 0.013 | 0.147 | 0.63 | 0.64 | 0.67 | 0.85 | 0.67 | 0.73 |
| 5 | 53744849 | rs17342535 | 0.139 | 1.582 | -0.494 | 1.112 | -0.044 | -0.039 | -0.039 | 0.082 | 0.09 | 0.02 | 0.12 | 0.12 | 0.14 | 0.16 |
| 5 | 53750530 | rs12517462 | 0.451 | 1.086 | -0.14 | 1.085 | 0.011 | 0.034 | 0.034 | 1.054 | 0.69 | 0.46 | 0.46 | 0.37 | 0.63 | 0.55 |
| 5 | 53751988 | rs7823 | 0.391 | 1.157 | -0.2 | 1.102 | -0.006 | -0.013 | -0.013 | 1.252 | 0.71 | 0.76 | 0.49 | 0.38 | 0.64 | 0.55 |
| 5 | 53761426 | rs7708321 | 0.071 | 0.77 | -0.392 | 0.827 | 0.003 | -0.016 | -0.016 | 0 | 0.06 | 0.03 | 0.05 | 0.09 | 0.04 | 0.06 |
| 5 | 53768556 | rs6891530 | -0.739 | 0.96 | -0.061 | 0.981 | 0.017 | 0.073 | 0.073 | 0.003 | 0.3 | 0.4 | 0.59 | 0.58 | 0.32 | 0.38 |
| 5 | 53770217 | rs4865572 | -0.175 | 0.249 | -0.723 | -0.08 | -0.004 | 0.003 | 0.003 | 0 | 0.6 | 0.52 | 0.74 | 0.58 | 0.67 | 0.59 |
| 5 | 53770434 | rs10940400 | 0.758 | 0.003 | -0.917 | -0.068 | 0.011 | 0.001 | 0.001 | 0 | 0.3 | 0.08 | 0.14 | 0.22 | 0.23 | 0.24 |
| 5 | 53770581 | rs10223063 | 0.192 | 0.023 | -0.69 | -0.315 | -0.001 | 0.01 | 0.01 | 0 | 0.48 | 0.53 | 0.39 | 0.2 | 0.51 | 0.39 |
| 5 | 53770686 | rs6889172 | -0.324 | 0.063 | -0.64 | -0.149 | 0.041 | 0.203 | 0.203 | 1.399 | 0.91 | 0.55 | 0.87 | 0.8 | 0.91 | 0.83 |
| 5 | 53772796 | rs13167369 | 1.281 | 0.026 | -0.905 | -0.202 | 0.001 | 0.108 | 0.108 | 0 | 0.24 | 0.05 | 0.1 | 0.06 | 0.19 | 0.18 |
| 5 | 53775596 | rs10066174 | 0.889 | 0.023 | -0.483 | -0.163 | 0.008 | 0.104 | 0.104 | 0.654 | 0.08 | 0.25 | 0.11 | 0.2 | 0.08 | 0.16 |
| 5 | 53776281 | rs10513048 | 1.862 | -0.302 | -0.658 | -0.434 | -0.067 | 0.134 | 0.134 | 0 | 0.27 | 0.08 | 0.18 | 0.01 | 0.34 | 0.19 |
| 5 | 53785234 | rs12520518 | -0.532 | -0.332 | -0.511 | -0.407 | -0.001 | -0.011 | -0.011 | 0 | 0.39 | 0.44 | 0.61 | 0.72 | 0.38 | 0.45 |
| 5 | 53788472 | rs10075503 | -1.22 | -0.41 | -0.428 | -0.314 | 0.019 | 0.056 | 0.056 | 0 | 0.74 | 0.82 | 0.9 | 0.93 | 0.8 | 0.81 |
| 5 | 53791374 | rs880944 | -0.413 | -0.406 | -0.42 | -0.217 | -0.009 | -0.004 | -0.004 | 0.053 | 0.66 | 0.52 | 0.79 | 0.71 | 0.71 | 0.65 |
| 5 | 53793487 | rs7448373 | 1.474 | -0.757 | -0.748 | -0.467 | 0.028 | -0.062 | -0.062 | 0 | 0.12 | 0.01 | 0.08 | 0.16 | 0.07 | 0.16 |
| 5 | 53810279 | rs4364350 | -1.44 | -0.639 | -0.665 | -0.371 | 0.017 | 0.064 | 0.064 | 0 | 0.74 | 0.84 | 0.9 | 0.93 | 0.8 | 0.81 |
| 5 | 53812952 | rs2548617 | 0.915 | -0.648 | -0.571 | -0.031 | 0.015 | 0.183 | 0.183 | 1.125 | 0.08 | 0.3 | 0.2 | 0.28 | 0.08 | 0.16 |
| 5 | 53815495 | rs2548612 | -0.158 | -0.457 | -0.539 | -0.494 | 0.009 | -0.004 | -0.004 | 0 | 0.52 | 0.45 | 0.53 | 0.45 | 0.57 | 0.46 |
| 5 | 53815560 | rs13162502 | 1.59 | -0.446 | -0.608 | -0.492 | -0.001 | 0.094 | 0.094 | 0 | 0.24 | 0.06 | 0.09 | 0.07 | 0.19 | 0.19 |
| 5 | 53818895 | rs2548611 | -1.447 | -0.43 | -0.597 | -0.484 | 0.017 | 0.079 | 0.079 | 0 | 0.74 | 0.86 | 0.9 | 0.93 | 0.8 | 0.8 |
| 5 | 53823285 | rs2565004 | -0.054 | -0.41 | -0.535 | -0.486 | -0.004 | 0.012 | 0.012 | 0 | 0.29 | 0.37 | 0.36 | 0.44 | 0.25 | 0.36 |
| 5 | 53824374 | rs2012870 | 1.097 | -0.195 | -0.318 | -0.33 | 0.016 | 0.116 | 0.116 | 0.129 | 0.2 | 0.32 | 0.37 | 0.48 | 0.23 | 0.35 |

| | | | | | | | | | | | | | | | | |
|---|---|---|---|---|---|---|---|---|---|---|---|---|---|---|---|---|
| 5 | 53828095 | rs2241403 | 0.324 | -0.258 | -0.446 | -0.3 | 0.001 | 0.001 | 0.001 | 0 | 0.45 | 0.48 | 0.54 | 0.5 | 0.55 | 0.45 |
| 5 | 53833598 | rs2564998 | -0.725 | -0.179 | -0.284 | -0.172 | 0.003 | 0.061 | 0.061 | 0.186 | 0.78 | 0.75 | 0.63 | 0.54 | 0.75 | 0.62 |
| 5 | 53838471 | rs1003371 | 1.559 | -0.065 | -0.93 | -0.467 | -0.085 | 0.154 | 0.154 | 0 | 0.25 | 0.03 | 0.18 | 0.01 | 0.31 | 0.11 |
| 5 | 53840532 | rs11741770 | 1.379 | -0.187 | -0.869 | -0.306 | -0.036 | 0.077 | 0.077 | 0 | 0.22 | 0.03 | 0.08 | 0.06 | 0.2 | 0.18 |
| 5 | 53843527 | rs2548644 | 0.386 | -0.732 | -0.218 | 0.108 | -0.004 | 0.027 | 0.027 | 0 | 0.19 | 0.27 | 0.25 | 0.46 | 0.14 | 0.32 |
| 5 | 53848745 | rs2548642 | -0.599 | -0.361 | 0.083 | -0.117 | 0.01 | -0.021 | -0.021 | 2.456 | 0.93 | 0.89 | 0.92 | 0.93 | 0.89 | 0.89 |
| 5 | 53850293 | rs10042600 | 1.453 | -0.398 | -0.555 | -0.547 | -0.071 | 0.14 | 0.14 | 0 | 0.21 | 0.02 | 0.19 | 0.01 | 0.2 | 0.19 |
| 5 | 53850772 | rs6869920 | 1.252 | -0.628 | -0.688 | -0.343 | -0.037 | -0.029 | -0.029 | 0 | 0.23 | 0.13 | 0.24 | 0.38 | 0.37 | 0.32 |
| 5 | 53851203 | rs875988 | -0.647 | -0.883 | -0.274 | -0.431 | -0.009 | 0.002 | 0.002 | 0 | 0.65 | 0.61 | 0.69 | 0.56 | 0.8 | 0.66 |
| 5 | 53854288 | rs7711608 | 0.54 | -0.765 | -0.102 | -0.493 | 0.008 | -0.075 | -0.075 | 5.214 | 0.31 | 0.34 | 0.35 | 0.08 | 0.33 | 0.3 |
| 5 | 53854712 | rs10079383 | -0.411 | -0.924 | -0.294 | -0.211 | 0.015 | -0.013 | -0.013 | 0.055 | 0.65 | 0.67 | 0.69 | 0.55 | 0.8 | 0.66 |
| 5 | 53861604 | rs10073644 | 0.563 | -0.487 | -0.381 | -0.731 | -0.026 | 0.062 | 0.062 | 0 | 0.3 | 0.14 | 0.28 | 0.01 | 0.3 | 0.22 |
| 5 | 53867201 | rs16883031 | 0.483 | -1.469 | 0.117 | -0.258 | 0.02 | 0.021 | 0.021 | 1.48 | 0.22 | 0.15 | 0.16 | 0.13 | 0.14 | 0.1 |
| 5 | 53869576 | rs2548638 | -0.686 | -1.402 | 0.718 | 0.493 | -0.002 | 0.004 | 0.004 | 0.086 | 0.8 | 0.81 | 0.77 | 0.72 | 0.84 | 0.68 |
| 5 | 53874176 | rs10067633 | 0.116 | -1.403 | 0.893 | 0.389 | -0.024 | 0.079 | 0.079 | 0.405 | 0.23 | 0.39 | 0.26 | 0.38 | 0.17 | 0.33 |
| 5 | 53874272 | rs1007139 | 0.871 | -1.429 | 0.485 | 0.446 | 0.046 | 0.022 | 0.022 | 0.259 | 0.47 | 0.25 | 0.5 | 0.43 | 0.6 | 0.53 |
| 5 | 53875982 | rs1122401 | -0.324 | -0.796 | 0.62 | 0.698 | -0.003 | 0.054 | 0.054 | 0.006 | 0.26 | 0.42 | 0.32 | 0.38 | 0.24 | 0.35 |
| 5 | 53878831 | rs1006885 | 0.09 | -0.787 | 0.182 | 0.754 | -0.008 | 0.045 | 0.045 | 0 | 0.2 | 0.25 | 0.27 | 0.37 | 0.18 | 0.31 |
| 5 | 53879281 | rs2194249 | 0.123 | -0.943 | -0.121 | 0.385 | 0.007 | 0.003 | 0.003 | 1.011 | 0.7 | 0.74 | 0.74 | 0.72 | 0.75 | 0.83 |
| 5 | 53881925 | rs2548628 | -0.212 | -0.932 | -0.51 | 0.389 | 0.032 | 0.02 | 0.02 | 0 | 0.75 | 0.94 | 0.83 | 0.77 | 0.83 | 0.85 |
| 5 | 53886610 | rs2565009 | 0.577 | -0.504 | -0.882 | 0.418 | -0.117 | -0.094 | -0.094 | 0 | 0.36 | 0.08 | 0.4 | 0.41 | 0.44 | 0.35 |
| 5 | 53887529 | rs10491381 | 0.502 | -0.907 | -0.828 | -0.043 | 0.03 | 0.024 | 0.024 | 0.002 | 0.26 | 0.09 | 0.15 | 0.22 | 0.21 | 0.16 |
| 5 | 53888168 | rs2548627 | 1.498 | -0.837 | -1.055 | -0.108 | -0.107 | -0.081 | -0.081 | 0 | 0.21 | 0.07 | 0.37 | 0.35 | 0.36 | 0.19 |
| 5 | 53889908 | rs2910810 | -0.388 | -1.145 | 0.034 | -0.142 | -0.043 | 0.005 | 0.005 | 0 | 0.7 | 0.49 | 0.8 | 0.68 | 0.78 | 0.83 |
| 5 | 53892279 | rs2249654 | 1.714 | -1.062 | -0.139 | -0.266 | -0.02 | 0.154 | 0.154 | 0 | 0.23 | 0.02 | 0.18 | 0.01 | 0.19 | 0.22 |
| 5 | 53897218 | rs7719215 | 0.477 | -1.025 | 0.065 | -0.218 | 0.016 | 0.015 | 0.015 | 0 | 0.45 | 0.37 | 0.36 | 0.33 | 0.35 | 0.63 |
| 5 | 53898950 | rs889304 | -0.677 | -0.961 | 0.109 | -0.359 | 0.073 | 0.064 | 0.064 | 0 | 0.71 | 0.53 | 0.51 | 0.55 | 0.55 | 0.79 |
| 5 | 53899535 | rs2548623 | -0.08 | -0.997 | 0.161 | -0.343 | 0.012 | -0.01 | -0.01 | 0.561 | 0.47 | 0.27 | 0.53 | 0.54 | 0.56 | 0.34 |
| 5 | 53900858 | rs10471395 | 1.182 | -0.954 | 0.106 | -0.378 | -0.017 | 0.116 | 0.116 | 0 | 0.27 | 0.05 | 0.25 | 0.01 | 0.26 | 0.24 |
| 5 | 53902940 | rs2548620 | 0.79 | -0.75 | 0.227 | -0.253 | -0.031 | -0.016 | -0.016 | 0 | 0.27 | 0.23 | 0.44 | 0.39 | 0.43 | 0.3 |
| 5 | 53906687 | rs6866215 | 0.599 | -0.791 | 0.114 | -0.385 | -0.002 | -0.043 | -0.043 | 0 | 0.31 | 0.39 | 0.29 | 0.15 | 0.3 | 0.33 |
| 5 | 53912105 | rs2059217 | -1.679 | -0.776 | -0.284 | -0.348 | -0.074 | -0.068 | -0.068 | 0.019 | 0.79 | 0.9 | 0.63 | 0.63 | 0.67 | 0.81 |
| 5 | 53914384 | rs10471919 | 1.218 | -1.071 | -0.373 | -0.546 | 0.028 | 0.021 | 0.021 | 0.811 | 0.2 | 0.16 | 0.1 | 0.01 | 0.12 | 0.06 |
| 5 | 53914475 | rs17626937 | 1.217 | -0.574 | -0.217 | -0.25 | -0.005 | 0.024 | 0.024 | 0 | 0.21 | 0.23 | 0.18 | 0.3 | 0.17 | 0.3 |
| 5 | 53920334 | rs16883146 | 0.235 | -0.477 | -0.151 | -0.346 | 0.003 | 0.004 | 0.004 | 0 | 0.13 | 0.17 | 0.12 | 0.21 | 0.15 | 0.22 |
| 5 | 53921870 | rs10461584 | 0.277 | -0.769 | -0.205 | -0.347 | 0 | -0.019 | -0.019 | 0 | 0.46 | 0.54 | 0.35 | 0.11 | 0.35 | 0.29 |
| 5 | 53925541 | rs9292067 | 0.232 | -0.629 | -0.482 | -0.153 | -0.005 | -0.01 | -0.01 | 3.483 | 0.53 | 0.25 | 0.42 | 0.44 | 0.45 | 0.64 |
| 5 | 53926318 | rs11738700 | 0.333 | -0.853 | 0.255 | 0.533 | 0.011 | 0.007 | 0.007 | 0 | 0.15 | 0.15 | 0.1 | 0.01 | 0.12 | 0.06 |
| 5 | 53927780 | rs12108894 | -0.384 | -0.853 | 0.34 | 0.553 | -0.042 | -0.024 | -0.024 | 1.286 | 0.19 | 0.43 | 0.13 | 0.11 | 0.14 | 0.07 |
| 5 | 53929224 | rs10045090 | 0.366 | -0.584 | -0.513 | 0.37 | 0.019 | 0.022 | 0.022 | 0 | 0.14 | 0.11 | 0.06 | 0.01 | 0.09 | 0.07 |
| 5 | 53931756 | rs16883196 | 0.41 | 0.272 | -0.084 | 1.095 | -0.009 | -0.021 | -0.021 | 0 | 0.06 | 0.02 | 0.05 | 0.15 | 0.08 | 0.11 |
| 5 | 53934304 | rs2409101 | -0.238 | 0.133 | 0.271 | 1.111 | -0.011 | 0.053 | 0.053 | 0 | 0.2 | 0.32 | 0.22 | 0.29 | 0.16 | 0.28 |
| 5 | 53934608 | rs13162512 | -0.087 | 0.213 | -0.87 | 1.149 | 0.004 | -0.058 | -0.058 | 0 | 0.19 | 0.08 | 0.21 | 0.29 | 0.16 | 0.28 |
| 5 | 53935602 | rs2197502 | 0.618 | 0.857 | -0.257 | 1.244 | 0.029 | 0.083 | 0.083 | 0 | 0.82 | 0.67 | 0.72 | 0.66 | 0.76 | 0.89 |
| 5 | 53937626 | rs6450228 | 0.427 | -0.042 | -0.878 | 0.606 | 0.034 | 0.034 | 0.034 | 0 | 0.83 | 0.91 | 0.92 | 0.93 | 0.9 | 0.88 |
| 5 | 53939221 | rs7444603 | -0.576 | 0.9 | -0.784 | 1.323 | -0.002 | 0.021 | 0.021 | 0 | 0.15 | 0.14 | 0.28 | 0.6 | 0.23 | 0.36 |
| 5 | 53941552 | rs895327 | 0.644 | -0.188 | -0.942 | 0.736 | 0.06 | 0.037 | 0.037 | 0 | 0.57 | 0.71 | 0.65 | 0.72 | 0.68 | 0.64 |
| 5 | 53943096 | rs7713171 | -0.267 | -0.425 | -0.254 | 0.691 | -0.041 | -0.015 | -0.015 | 0.009 | 0.83 | 0.78 | 0.89 | 0.89 | 0.91 | 0.87 |
| 5 | 53948513 | rs972837 | 0.296 | -0.466 | -0.091 | 0.732 | 0.01 | 0.01 | 0.01 | 0 | 0.27 | 0.2 | 0.26 | 0.17 | 0.23 | 0.2 |
| 5 | 53949598 | rs1445836 | -0.346 | -0.947 | 0.214 | 0.165 | -0.037 | -0.014 | -0.014 | 0 | 0.85 | 0.81 | 0.91 | 0.9 | 0.93 | 0.86 |
| 5 | 53950691 | rs13179508 | -0.346 | -0.947 | 0.214 | 0.165 | -0.037 | -0.014 | -0.014 | 4.751 | 0.85 | 0.81 | 0.91 | 0.9 | 0.93 | 0.86 |
| 5 | 53951585 | rs9790977 | -0.087 | -0.825 | 0.059 | 0.547 | -0.001 | 0.029 | 0.029 | 0 | 0.07 | 0.14 | 0.03 | 0.24 | 0.04 | 0.23 |
| 5 | 53954201 | rs6865208 | -0.827 | -0.928 | -0.119 | 0.465 | 0.031 | 0.002 | 0.002 | 0 | 0.88 | 0.94 | 0.94 | 0.91 | 0.96 | 0.91 |
| 5 | 53959926 | rs7708176 | 1.762 | -0.474 | -0.206 | 0.438 | -0.12 | 0.219 | 0.219 | 0 | 0.39 | 0.08 | 0.32 | 0.06 | 0.48 | 0.3 |
| 5 | 53962772 | rs4865857 | 0.792 | -0.503 | 0.028 | 0.396 | -0.037 | 0.015 | 0.015 | 0 | 0.79 | 0.68 | 0.87 | 0.66 | 0.87 | 0.68 |
| 5 | 53966360 | rs12513800 | -1.085 | -0.574 | -0.264 | 0.235 | -0.032 | 0.282 | 0.282 | 0 | 0.5 | 0.85 | 0.62 | 0.89 | 0.48 | 0.58 |
| 5 | 53969715 | rs4605745 | -0.22 | -0.68 | 0.1 | 0.444 | 0.023 | 0.148 | 0.148 | 1.023 | 0.07 | 0.54 | 0.06 | 0.07 | 0.03 | 0.03 |
| 5 | 53969773 | rs4865859 | -0.126 | -0.665 | -0.072 | 0.481 | -0.018 | -0.029 | -0.029 | 0 | 0.34 | 0.21 | 0.5 | 0.54 | 0.36 | 0.32 |
| 5 | 53970118 | rs4293864 | -0.978 | -0.52 | -0.071 | 0.431 | 0.022 | -0.001 | -0.001 | 0 | 0.89 | 0.95 | 0.94 | 0.9 | 0.96 | 0.91 |
| 5 | 53975846 | rs1373989 | 0.545 | -0.479 | 0.243 | 0.454 | 0.044 | 0.174 | 0.174 | 0 | 0.05 | 0.31 | 0.04 | 0.07 | 0.03 | 0.03 |
| 5 | 53975974 | rs6450233 | 0.366 | -0.58 | 0.324 | 0.437 | -0.101 | 0.053 | 0.053 | 0 | 0.82 | 0.56 | 0.82 | 0.82 | 0.93 | 0.88 |
| 5 | 53979790 | rs4565166 | 0.349 | -0.58 | 0.329 | 0.437 | -0.099 | 0.052 | 0.052 | 0 | 0.82 | 0.57 | 0.82 | 0.82 | 0.93 | 0.88 |
| 5 | 53980001 | rs4549482 | 0.694 | -0.552 | 0.175 | 0.416 | -0.075 | 0.019 | 0.019 | 0 | 0.78 | 0.48 | 0.78 | 0.82 | 0.88 | 0.83 |
| 5 | 53983251 | rs11748572 | -0.46 | -0.396 | -0.589 | 0.151 | -0.011 | 0 | 0 | 1.762 | 0.87 | 0.87 | 0.86 | 0.9 | 0.96 | 0.91 |
| 5 | 53983759 | rs7724650 | 0.783 | 0.136 | -0.908 | -0.059 | -0.024 | 0.193 | 0.193 | 0 | 0.45 | 0.17 | 0.28 | 0.2 | 0.53 | 0.42 |
| 5 | 53987533 | rs6877210 | -0.565 | 0.038 | -0.465 | 0.086 | -0.001 | -0.003 | -0.003 | 1.99 | 0.46 | 0.47 | 0.37 | 0.35 | 0.4 | 0.42 |
| 5 | 53989831 | rs4395600 | 0.317 | -0.299 | -0.171 | -0.172 | 0.01 | 0.012 | 0.012 | 0 | 0.15 | 0.14 | 0.09 | 0.07 | 0.1 | 0.15 |
| 5 | 53991267 | rs2034237 | 0.095 | -0.601 | 0.156 | 0.095 | -0.014 | 0.018 | 0.018 | 0 | 0.24 | 0.31 | 0.15 | 0.4 | 0.14 | 0.14 |
| 5 | 53993648 | rs4865582 | -1.313 | -1.022 | 0.321 | -0.28 | -0.016 | 0.006 | 0.006 | 0 | 0.75 | 0.65 | 0.73 | 0.72 | 0.78 | 0.71 |

| | | | | | | | | | | | | | | | | |
|---|---|---|---|---|---|---|---|---|---|---|---|---|---|---|---|---|
| 5 | 53994294 | rs6894873 | -1.362 | -1.028 | 0.288 | -0.283 | 0.001 | -0.001 | -0.001 | 0.338 | 0.75 | 0.78 | 0.74 | 0.72 | 0.79 | 0.71 |
| 5 | 53995324 | rs6870244 | 1.023 | -0.947 | 0.612 | -0.363 | -0.036 | 0.022 | 0.022 | 0 | 0.36 | 0.24 | 0.46 | 0.24 | 0.51 | 0.38 |
| 5 | 53997352 | rs1823053 | 1.254 | -0.889 | 0.579 | -0.137 | -0.004 | 0.022 | 0.022 | 0 | 0.16 | 0.18 | 0.12 | 0.4 | 0.11 | 0.1 |
| 5 | 54004735 | rs1562231 | -0.42 | -0.895 | 0.58 | -0.19 | -0.001 | 0 | 0 | 0.889 | 0.59 | 0.6 | 0.61 | 0.31 | 0.68 | 0.61 |
| 5 | 54010478 | rs16883342 | 0.496 | -0.868 | 0.535 | -0.241 | -0.009 | -0.029 | -0.029 | 0 | 0.06 | 0.15 | 0.04 | 0.01 | 0.04 | 0.06 |
| 5 | 54011945 | rs4282276 | -2.134 | -0.868 | 0.312 | -0.181 | 0.001 | -0.002 | -0.002 | 2.069 | 0.81 | 0.83 | 0.76 | 0.73 | 0.82 | 0.79 |
| 5 | 54014442 | rs6869164 | 1.758 | -0.505 | -0.101 | -0.343 | -0.037 | -0.009 | -0.009 | 0 | 0.25 | 0.09 | 0.29 | 0.17 | 0.31 | 0.23 |
| 5 | 54016510 | rs4460095 | 1.124 | -0.287 | 0.159 | -0.444 | -0.005 | 0.003 | 0.003 | 0 | 0.17 | 0.14 | 0.19 | 0.09 | 0.24 | 0.2 |
| 5 | 54018232 | rs3886214 | 1.11 | -0.291 | 0.273 | -0.438 | 0.032 | -0.032 | -0.032 | 1.566 | 0.17 | 0.22 | 0.19 | 0.09 | 0.23 | 0.2 |
| 5 | 54019772 | rs11749269 | 0.483 | -0.521 | 0.323 | -0.331 | -0.011 | -0.007 | -0.007 | 0 | 0.55 | 0.49 | 0.59 | 0.33 | 0.64 | 0.58 |
| 5 | 54021166 | rs4624748 | -0.782 | -0.39 | 0.221 | -0.447 | -0.026 | 0.017 | 0.017 | 0 | 0.74 | 0.57 | 0.72 | 0.71 | 0.79 | 0.7 |
| 5 | 54021527 | rs4562000 | -0.087 | -0.342 | 0.065 | -0.42 | -0.032 | -0.006 | -0.006 | 1.853 | 0.68 | 0.47 | 0.66 | 0.7 | 0.75 | 0.64 |
| 5 | 54026811 | rs4485879 | -0.457 | -0.279 | -0.122 | -0.366 | -0.027 | 0.013 | 0.013 | 0 | 0.73 | 0.56 | 0.69 | 0.7 | 0.78 | 0.67 |
| 5 | 54031153 | rs12516870 | -0.681 | -0.271 | -0.086 | -0.448 | -0.025 | 0.045 | 0.045 | 0 | 0.75 | 0.57 | 0.71 | 0.63 | 0.8 | 0.78 |
| 5 | 54031705 | rs4482860 | -0.369 | 0.039 | -0.144 | -0.512 | 0.009 | 0.012 | 0.012 | 0 | 0.52 | 0.54 | 0.47 | 0.61 | 0.42 | 0.6 |
| 5 | 54032140 | rs973959 | -0.234 | 0.222 | -0.055 | -0.774 | -0.001 | -0.002 | -0.002 | 0 | 0.48 | 0.43 | 0.46 | 0.67 | 0.42 | 0.46 |
| 5 | 54035726 | rs10077272 | 0.707 | -0.806 | -0.17 | -0.717 | -0.009 | -0.013 | -0.013 | 0 | 0.31 | 0.22 | 0.37 | 0.34 | 0.34 | 0.42 |
| 5 | 54035929 | rs12520102 | -0.293 | -0.339 | -0.413 | -0.643 | 0.005 | 0.08 | 0.08 | 0 | 0.54 | 0.71 | 0.54 | 0.7 | 0.51 | 0.63 |
| 5 | 54038676 | rs10940422 | 0.125 | -0.015 | -0.721 | -0.411 | 0 | 0.001 | 0.001 | 0 | 0.2 | 0.2 | 0.15 | 0.35 | 0.16 | 0.25 |
| 5 | 54043954 | rs13188578 | 0.074 | 0.045 | -0.82 | -0.491 | -0.005 | 0.008 | 0.008 | 0 | 0.64 | 0.59 | 0.61 | 0.37 | 0.67 | 0.7 |
| 5 | 54047137 | rs10066710 | 0.177 | -0.007 | 0.012 | 0.065 | 0 | 0.003 | 0.003 | 0 | 0.4 | 0.47 | 0.42 | 0.37 | 0.37 | 0.54 |
| 5 | 54047476 | rs13180905 | -0.321 | 0.306 | -0.385 | -0.478 | -0.006 | 0.03 | 0.03 | 0.017 | 0.6 | 0.71 | 0.56 | 0.71 | 0.53 | 0.68 |
| 5 | 54048853 | rs1445837 | -0.254 | 0.215 | -0.874 | -0.404 | -0.021 | 0.058 | 0.058 | 0.044 | 0.59 | 0.79 | 0.56 | 0.71 | 0.51 | 0.64 |
| 5 | 54055062 | rs6867643 | -0.121 | 0.03 | 0.01 | -0.8 | 0.001 | -0.085 | -0.085 | 1.366 | 0.3 | 0.32 | 0.22 | 0 | 0.32 | 0.18 |
| 5 | 54059096 | rs1867685 | -0.974 | -0.476 | -0.047 | -1.047 | -0.091 | 0.256 | 0.256 | 0.163 | 0.54 | 0.89 | 0.38 | 0.17 | 0.48 | 0.6 |
| 5 | 54083387 | rs4865870 | 0.816 | -1.149 | 0.523 | -1.012 | 0.073 | 0.196 | 0.196 | 0 | 0.48 | 0.19 | 0.67 | 0.83 | 0.65 | 0.45 |
| 5 | 54087671 | rs4472223 | 0.208 | -1.505 | 1.066 | -1.126 | 0.005 | -0.101 | -0.101 | 0 | 0.29 | 0.34 | 0.22 | 0 | 0.31 | 0.19 |
| 5 | 54104560 | rs10058571 | 0.316 | -1.539 | 1.11 | -1.098 | -0.084 | -0.056 | -0.056 | 1.329 | 0.77 | 0.42 | 0.88 | 0.84 | 0.92 | 0.63 |
| 5 | 54126400 | rs12515327 | 0.793 | -1.532 | 1.014 | -1.124 | 0.045 | 0.155 | 0.155 | 0 | 0.5 | 0.26 | 0.68 | 0.84 | 0.61 | 0.43 |
| 5 | 54128691 | rs4582235 | 0.047 | -1.499 | 1.219 | -1.09 | -0.119 | -0.054 | -0.054 | 0.144 | 0.79 | 0.48 | 0.92 | 0.84 | 0.94 | 0.63 |
| 5 | 54140334 | rs3935813 | 0.867 | -1.341 | 1.005 | -0.704 | 0.01 | 0.189 | 0.189 | 0 | 0.49 | 0.11 | 0.67 | 0.71 | 0.6 | 0.45 |
| 5 | 54140573 | rs3935814 | 0.396 | -1.38 | 1.083 | -0.772 | -0.011 | 0.008 | 0.008 | 0 | 0.28 | 0.16 | 0.2 | 0 | 0.32 | 0.17 |
| 5 | 54153803 | rs10040872 | 0.623 | -1.618 | 0.54 | -0.283 | -0.101 | 0.066 | 0.066 | 0 | 0.75 | 0.47 | 0.91 | 0.62 | 0.93 | 0.64 |
| 5 | 54164371 | rs10940431 | 1.357 | -1.579 | 0.483 | -0.285 | -0.052 | -0.026 | -0.026 | 0 | 0.42 | 0.1 | 0.66 | 0.62 | 0.59 | 0.44 |
| 5 | 54165910 | rs13181749 | 0.809 | -1.665 | 0.458 | -0.269 | -0.072 | 0.052 | 0.052 | 0 | 0.73 | 0.46 | 0.88 | 0.62 | 0.91 | 0.62 |
| 5 | 54170032 | rs1991498 | -0.203 | -1.673 | 0.273 | -0.266 | -0.092 | 0.053 | 0.053 | 0.138 | 0.82 | 0.58 | 0.92 | 0.72 | 0.94 | 0.64 |
| 5 | 54170865 | rs1560167 | 0.348 | -1.684 | 0.267 | -0.273 | -0.109 | 0.125 | 0.125 | 0.071 | 0.78 | 0.54 | 0.91 | 0.62 | 0.93 | 0.64 |
| 5 | 54174285 | rs36000196 | 0.431 | -1.481 | -0.956 | -0.194 | -0.039 | 0.134 | 0.134 | 0 | 0.29 | 0.08 | 0.18 | 0 | 0.31 | 0.17 |
| 5 | 54174906 | rs10940434 | 0.704 | -0.7 | -0.889 | 0.247 | -0.011 | 0 | 0 | 0 | 0.43 | 0.17 | 0.67 | 0.62 | 0.6 | 0.44 |
| 5 | 54184663 | rs2075401 | 0.141 | -0.738 | -1.376 | 0.259 | 0.012 | -0.011 | -0.011 | 0 | 0.06 | 0.04 | 0.01 | 0.1 | 0.01 | 0.01 |
| 5 | 54186769 | rs2897758 | 0.165 | -0.756 | -1.513 | 0.114 | 0.048 | 0.189 | 0.189 | 0 | 0.49 | 0.11 | 0.66 | 0.72 | 0.62 | 0.46 |
| 5 | 54187813 | rs10805470 | 0.469 | -0.272 | -1.435 | 0.371 | -0.048 | -0.066 | -0.066 | 0 | 0.42 | 0.07 | 0.65 | 0.61 | 0.61 | 0.45 |
| 5 | 54191031 | rs17627141 | -0.768 | -0.433 | 0.054 | 0.78 | -0.035 | 0.106 | 0.106 | 1.408 | 0.15 | 0.4 | 0.25 | 0.32 | 0.07 | 0.31 |
| 5 | 54193725 | rs17527251 | 0.172 | -0.623 | -1.134 | -0.515 | -0.02 | 0.141 | 0.141 | 0.024 | 0.26 | 0.05 | 0.17 | 0.01 | 0.29 | 0.15 |
| 5 | 54197973 | rs2407759 | -0.681 | -0.247 | -0.946 | -0.577 | -0.001 | -0.001 | -0.001 | 0 | 0.24 | 0.69 | 0.3 | 0.23 | 0.24 | 0.29 |
| 5 | 54203474 | rs10491367 | -0.398 | -0.341 | -0.391 | -0.626 | 0 | 0.005 | 0.005 | 0.997 | 0.13 | 0.16 | 0.2 | 0.23 | 0.15 | 0.28 |
| 5 | 54209053 | rs11954161 | -0.657 | -0.611 | -0.317 | -1.029 | 0.002 | 0.001 | 0.001 | 1.286 | 0.22 | 0.16 | 0.24 | 0.23 | 0.2 | 0.35 |
| 5 | 54210950 | rs12055221 | -0.657 | -0.611 | -0.317 | -1.019 | 0.002 | 0.004 | 0.004 | 1.286 | 0.22 | 0.16 | 0.24 | 0.17 | 0.2 | 0.35 |
| 5 | 54225580 | rs7707725 | 1.081 | -0.673 | -0.417 | -1.176 | -0.044 | 0.122 | 0.122 | 0.239 | 0.26 | 0.07 | 0.2 | 0.01 | 0.35 | 0.15 |
| 5 | 54225704 | rs9292075 | -1.161 | -0.713 | 0.599 | -1.223 | -0.035 | -0.044 | -0.044 | 1.139 | 0.28 | 0.4 | 0.26 | 0.18 | 0.2 | 0.35 |
| 5 | 54228585 | rs4865890 | -0.559 | 0.051 | -0.052 | -0.449 | -0.003 | -0.007 | -0.007 | 0 | 0.54 | 0.47 | 0.49 | 0.68 | 0.41 | 0.66 |
| 5 | 54229297 | rs4422488 | -0.461 | 0.081 | 0.06 | -0.413 | 0.037 | 0.057 | 0.057 | 0 | 0.81 | 0.65 | 0.72 | 0.69 | 0.76 | 0.82 |
| 5 | 54229845 | rs10471404 | -0.105 | 0.049 | -0.45 | -0.361 | 0.005 | 0.009 | 0.009 | 1.497 | 0.21 | 0.12 | 0.23 | 0.18 | 0.2 | 0.34 |
| 5 | 54232022 | rs17589045 | 1.151 | 0.045 | -0.496 | -0.356 | 0.005 | -0.141 | -0.141 | 0.15 | 0.27 | 0.07 | 0.24 | 0.5 | 0.21 | 0.32 |
| 5 | 54232372 | rs13180157 | 0.194 | 0.182 | -0.445 | -0.384 | -0.009 | 0.113 | 0.113 | 6.528 | 0.51 | 0.25 | 0.45 | 0.27 | 0.57 | 0.52 |
| 5 | 54232607 | rs6450262 | 1.047 | 0.458 | -0.951 | 0.16 | -0.029 | 0.08 | 0.08 | 0.052 | 0.55 | 0.72 | 0.46 | 0.26 | 0.51 | 0.43 |
| 5 | 54233268 | rs1991497 | 0.239 | 0.221 | -1.257 | 0.305 | -0.05 | 0.021 | 0.021 | 3.1 | 0.57 | 0.84 | 0.53 | 0.35 | 0.57 | 0.44 |
| 5 | 54245799 | rs10491368 | -0.064 | 0.505 | -1.453 | 0.401 | -0.084 | -0.095 | -0.095 | 0.002 | 0.35 | 0.05 | 0.38 | 0.66 | 0.3 | 0.54 |
| 5 | 54247654 | rs2195394 | 0.141 | 0.422 | -0.972 | 0.835 | -0.029 | -0.051 | -0.051 | 0 | 0.09 | 0.01 | 0.19 | 0.42 | 0.07 | 0.19 |
| 5 | 54248010 | rs13357261 | 0.459 | 0.508 | -0.276 | -0.086 | 0 | -0.031 | -0.031 | 1.222 | 0.37 | 0.59 | 0.29 | 0.05 | 0.38 | 0.14 |
| 5 | 54250627 | rs17443195 | 0.693 | 0.751 | -1.033 | -0.075 | -0.029 | 0.143 | 0.143 | 0 | 0.33 | 0.1 | 0.22 | 0 | 0.37 | 0.13 |
| 5 | 54251169 | rs1811345 | -1.414 | 0.607 | 0.103 | 1.346 | -0.034 | 0.157 | 0.157 | 0.484 | 0.13 | 0.59 | 0.09 | 0.28 | 0.08 | 0.17 |
| 5 | 54251194 | rs9292076 | -1.498 | 1.522 | -0.069 | 1.015 | 0.024 | 0.005 | 0.005 | 0 | 0.1 | 0.18 | 0.08 | 0.08 | 0.15 | 0.07 |
| 5 | 54251533 | rs6450270 | -1.061 | 1.445 | -0.67 | 1.073 | -0.008 | -0.011 | -0.011 | 0 | 0.33 | 0.69 | 0.47 | 0.42 | 0.34 | 0.25 |
| 5 | 54252825 | rs3815916 | 1.175 | 1.272 | -0.345 | 0.568 | 0.008 | -0.022 | -0.022 | 0 | 0.6 | 0.51 | 0.54 | 0.81 | 0.57 | 0.5 |
| 5 | 54253124 | rs3815919 | 0.443 | 1.525 | -0.393 | 0.442 | -0.021 | -0.024 | -0.024 | 0 | 0.11 | 0.05 | 0.19 | 0.17 | 0.15 | 0.32 |
| 5 | 54253615 | rs11954366 | -0.948 | 1.32 | -0.125 | 0.362 | -0.002 | -0.038 | -0.038 | 0 | 0.26 | 0.25 | 0.26 | 0.02 | 0.29 | 0.18 |
| 5 | 54256550 | rs1820132 | -0.712 | 1.235 | -0.681 | 0.431 | -0.079 | -0.046 | -0.046 | 0.044 | 0.28 | 0.09 | 0.35 | 0.26 | 0.34 | 0.44 |
| 5 | 54258180 | rs1579319 | -0.177 | 1.14 | -0.242 | 0.306 | 0.015 | -0.101 | -0.101 | 0 | 0.16 | 0.36 | 0.14 | 0.01 | 0.17 | 0.1 |

| | | | | | | | | | | | | | | | | |
|---|---|---|---|---|---|---|---|---|---|---|---|---|---|---|---|---|
| 5 | 54259343 | rs11748356 | -1.224 | 1.046 | 0.041 | 0.377 | 0.008 | -0.146 | -0.146 | 0.392 | 0.28 | 0.5 | 0.26 | 0.05 | 0.28 | 0.18 |
| 5 | 54259795 | rs10940446 | 0.745 | 1.095 | -0.363 | 0.463 | -0.04 | -0.058 | -0.058 | 0.061 | 0.17 | 0.06 | 0.23 | 0.26 | 0.22 | 0.38 |
| 5 | 54261589 | rs1363864 | -1.832 | 0.915 | 0.168 | 0.463 | -0.005 | 0.087 | 0.087 | 0 | 0.49 | 0.77 | 0.52 | 0.3 | 0.51 | 0.56 |
| 5 | 54264447 | rs6890825 | -0.471 | 0.696 | -0.002 | 0.636 | 0.025 | 0.038 | 0.038 | 0.169 | 0.26 | 0.4 | 0.28 | 0.32 | 0.3 | 0.4 |
| 5 | 54265532 | rs1023674 | -1.87 | 0.12 | -0.749 | 0.939 | -0.092 | 0.039 | 0.039 | 0 | 0.54 | 0.94 | 0.73 | 0.65 | 0.55 | 0.69 |
| 5 | 54268487 | rs2112935 | -1.429 | 0.034 | -0.499 | 0.924 | -0.034 | 0.073 | 0.073 | 0.007 | 0.19 | 0.65 | 0.36 | 0.36 | 0.12 | 0.17 |
| 5 | 54272446 | rs1423249 | -1.032 | 0.048 | -0.494 | 0.932 | -0.025 | 0.113 | 0.113 | 0.129 | 0.16 | 0.62 | 0.35 | 0.36 | 0.12 | 0.17 |
| 5 | 54274151 | rs11740219 | 0.725 | 0.128 | -1.18 | 1.002 | -0.055 | -0.081 | -0.081 | 0 | 0.19 | 0.02 | 0.22 | 0.29 | 0.22 | 0.39 |
| 5 | 54274874 | rs3822313 | -1.655 | -0.035 | -1.096 | 0.747 | -0.088 | 0.042 | 0.042 | 0 | 0.54 | 0.93 | 0.72 | 0.65 | 0.55 | 0.69 |
| 5 | 54286886 | rs13159954 | -0.533 | 0.146 | -0.517 | 0.759 | -0.003 | -0.013 | -0.013 | 0 | 0.19 | 0.31 | 0.14 | 0.13 | 0.21 | 0.13 |
| 5 | 54289155 | rs4865901 | 0.098 | 0.254 | -1.197 | 0.902 | -0.043 | -0.05 | -0.05 | 0.285 | 0.21 | 0.07 | 0.26 | 0.28 | 0.25 | 0.42 |
| 5 | 54307421 | rs2407827 | -0.819 | 0.138 | -1.8 | 0.348 | -0.065 | -0.022 | -0.022 | 0.261 | 0.67 | 0.91 | 0.76 | 0.65 | 0.57 | 0.64 |
| 5 | 54314853 | rs4865907 | 0.074 | -0.007 | -1.766 | 0.391 | 0.062 | -0.027 | -0.027 | 0 | 0.46 | 0.84 | 0.5 | 0.49 | 0.33 | 0.26 |
| 5 | 54316028 | rs4865908 | 0.751 | 0.105 | -1.262 | 0.453 | 0.063 | 0.265 | 0.265 | 0.595 | 0.16 | 0.49 | 0.39 | 0.48 | 0.19 | 0.17 |
| 5 | 54317900 | rs6891823 | -0.748 | -0.02 | -1.982 | 0.064 | -0.109 | -0.04 | -0.04 | 0.019 | 0.66 | 0.95 | 0.76 | 0.65 | 0.54 | 0.65 |
| 5 | 54323955 | rs1396500 | 0.529 | -0.78 | -1.034 | -0.205 | 0.066 | 0.065 | 0.065 | 0.146 | 0.21 | 0.06 | 0.06 | 0.01 | 0.09 | 0.1 |
| 5 | 54324622 | rs6875661 | -0.591 | -0.235 | -0.99 | 0.152 | 0.063 | 0.042 | 0.042 | 0.619 | 0.43 | 0.79 | 0.48 | 0.35 | 0.3 | 0.27 |
| 5 | 54327955 | rs2112938 | -0.377 | 1.399 | 0.162 | 1.005 | 0.093 | 0.13 | 0.13 | 0.884 | 0.16 | 0.52 | 0.22 | 0.31 | 0.21 | 0.16 |
| 5 | 54329913 | rs17681401 | -0.876 | 1.886 | -0.354 | 0.361 | -0.049 | 0.052 | 0.052 | 0 | 0.12 | 0.02 | 0.11 | 0 | 0.2 | 0.11 |
| 5 | 54334424 | rs1876680 | -0.857 | 1.121 | -0.028 | 0.55 | -0.003 | 0.001 | 0.001 | 0 | 0.08 | 0.1 | 0.21 | 0.07 | 0.07 | 0.13 |
| 5 | 54370089 | rs10037065 | -0.466 | 0.831 | 1.769 | 1.141 | -0.005 | -0.022 | -0.022 | 0.005 | 0.11 | 0.1 | 0.09 | 0.28 | 0.13 | 0.16 |
| 5 | 54378192 | rs3111191 | -1.341 | 0.892 | 2.124 | 1.079 | 0.022 | 0.157 | 0.157 | 0 | 0.16 | 0.49 | 0.13 | 0.28 | 0.16 | 0.16 |
| 5 | 54392373 | rs3105807 | -2.434 | 1.037 | 1.676 | 1.023 | 0.005 | 0.001 | 0.001 | 0 | 0.44 | 0.71 | 0.49 | 0.54 | 0.44 | 0.58 |
| 5 | 54398049 | rs2069187 | 0.782 | 1.042 | 1.904 | 1.054 | 0.014 | 0.111 | 0.111 | 0 | 0.84 | 0.65 | 0.88 | 0.73 | 0.84 | 0.84 |
| 5 | 54402970 | rs8192914 | -0.074 | 1.076 | 1.655 | 1.071 | -0.005 | -0.025 | -0.025 | 0 | 0.11 | 0.1 | 0.09 | 0.27 | 0.13 | 0.15 |
| 5 | 54409587 | rs389075 | -0.8 | 1.124 | 1.72 | 0.939 | -0.001 | -0.005 | -0.005 | 0.054 | 0.19 | 0.2 | 0.16 | 0.19 | 0.21 | 0.3 |
| 5 | 54410099 | rs444527 | -0.694 | 1.125 | 1.595 | 0.94 | -0.03 | -0.006 | -0.006 | 0 | 0.18 | 0.04 | 0.15 | 0.19 | 0.21 | 0.3 |
| 5 | 54417181 | rs6883587 | -1.027 | 1.075 | 1.858 | 0.919 | -0.004 | -0.016 | -0.016 | 0 | 0.21 | 0.26 | 0.17 | 0.19 | 0.22 | 0.3 |
| 5 | 54420799 | rs382402 | -0.687 | 1.098 | 1.685 | 0.989 | -0.027 | -0.013 | -0.013 | 0 | 0.18 | 0.04 | 0.15 | 0.19 | 0.21 | 0.3 |
| 5 | 54426248 | rs336106 | 2.426 | 1.02 | 1.838 | 1.012 | 0.004 | 0 | 0 | 0 | 0.55 | 0.26 | 0.51 | 0.47 | 0.56 | 0.41 |
| 5 | 54428310 | rs10043664 | -0.306 | 1.204 | 1.584 | 1.092 | -0.015 | -0.051 | -0.051 | 1.462 | 0.12 | 0.06 | 0.08 | 0.27 | 0.15 | 0.16 |
| 5 | 54429974 | rs336102 | -0.766 | 1.202 | 1.56 | 1.084 | -0.027 | -0.003 | -0.003 | 0 | 0.18 | 0.04 | 0.15 | 0.19 | 0.21 | 0.3 |
| 5 | 54439466 | rs1021580 | 0.973 | 1.318 | 1.876 | 1.201 | 0.042 | 0.175 | 0.175 | 0 | 0.84 | 0.56 | 0.87 | 0.73 | 0.83 | 0.84 |
| 5 | 54439892 | rs336076 | 1.783 | 1.291 | 1.832 | 1.165 | 0.01 | 0.062 | 0.062 | 0 | 0.68 | 0.54 | 0.75 | 0.54 | 0.64 | 0.58 |
| 5 | 54446050 | rs1335673 | -1.058 | 1.25 | 1.695 | 1.194 | 0 | 0.053 | 0.053 | 0 | 0.15 | 0.25 | 0.1 | 0.27 | 0.15 | 0.16 |
| 5 | 54455302 | rs17682812 | -0.728 | 1.157 | 1.738 | 1.15 | -0.001 | 0.054 | 0.054 | 0 | 0.13 | 0.24 | 0.1 | 0.26 | 0.13 | 0.16 |
| 5 | 54455439 | rs336082 | 0.689 | 1.158 | 1.873 | 1.117 | -0.003 | 0.003 | 0.003 | 0 | 0.83 | 0.85 | 0.86 | 0.83 | 0.79 | 0.75 |
| 5 | 54458763 | rs436282 | 1.956 | 1.203 | 2.05 | 1.118 | 0 | 0.032 | 0.032 | 0 | 0.68 | 0.5 | 0.75 | 0.56 | 0.65 | 0.59 |
| 5 | 54459961 | rs381852 | 0.607 | 1.137 | 1.713 | 1.164 | -0.007 | 0.006 | 0.006 | 0 | 0.83 | 0.91 | 0.86 | 0.83 | 0.79 | 0.75 |
| 5 | 54461990 | rs3747725 | -0.108 | 1.152 | 1.69 | 1.166 | 0 | -0.045 | -0.045 | 0 | 0.1 | 0.05 | 0.06 | 0.26 | 0.1 | 0.14 |
| 5 | 54468921 | rs336115 | 1.76 | 1.133 | 1.735 | 1.141 | -0.002 | 0.007 | 0.007 | 0 | 0.7 | 0.68 | 0.78 | 0.56 | 0.69 | 0.63 |
| 5 | 54470075 | rs336116 | 1.591 | 1.385 | 1.862 | 1.247 | 0.005 | 0.031 | 0.031 | 0 | 0.75 | 0.68 | 0.79 | 0.56 | 0.7 | 0.68 |
| 5 | 54474427 | rs336123 | 1.636 | 1.372 | 1.958 | 1.241 | 0.006 | 0.038 | 0.038 | 0 | 0.75 | 0.66 | 0.78 | 0.56 | 0.7 | 0.68 |
| 5 | 54477561 | rs1109967 | -0.771 | 1.357 | 1.95 | 1.274 | -0.009 | 0.12 | 0.12 | 0 | 0.12 | 0.33 | 0.09 | 0.26 | 0.1 | 0.12 |
| 5 | 54478686 | rs10041502 | -0.354 | 1.381 | 1.679 | 1.285 | 0.001 | 0.009 | 0.009 | 0 | 0.11 | 0.14 | 0.07 | 0.26 | 0.1 | 0.12 |
| 5 | 54492136 | rs375631 | 1.011 | 1.59 | 1.775 | 1.175 | 0.043 | 0.007 | 0.007 | 0 | 0.86 | 0.74 | 0.85 | 0.83 | 0.79 | 0.83 |
| 5 | 54518687 | rs334877 | -0.685 | 1.38 | 1.608 | 1.353 | -0.004 | 0.055 | 0.055 | 0 | 0.12 | 0.23 | 0.08 | 0.28 | 0.11 | 0.08 |
| 5 | 54523420 | rs2783478 | -2.094 | 1.665 | 1.879 | 1.33 | 0.01 | 0.043 | 0.043 | 0.358 | 0.27 | 0.43 | 0.26 | 0.45 | 0.31 | 0.26 |
| 5 | 54523778 | rs334885 | -0.807 | 1.554 | 1.835 | 1.355 | -0.004 | -0.003 | -0.003 | 0.004 | 0.12 | 0.11 | 0.17 | 0.17 | 0.19 | 0.16 |
| 5 | 54524616 | rs334888 | -1.051 | 1.555 | 1.83 | 1.355 | 0.012 | 0.004 | 0.004 | 0.246 | 0.12 | 0.19 | 0.17 | 0.17 | 0.19 | 0.17 |
| 5 | 54532144 | rs151237 | 0.83 | 1.465 | 1.792 | 1.343 | -0.007 | 0.028 | 0.028 | 0 | 0.87 | 0.8 | 0.92 | 0.72 | 0.9 | 0.92 |
| 5 | 54534752 | rs151240 | 0.83 | 1.465 | 1.823 | 1.343 | -0.007 | 0.03 | 0.03 | 0 | 0.87 | 0.8 | 0.92 | 0.72 | 0.9 | 0.92 |
| 5 | 54548830 | rs163098 | 1.438 | 1.396 | 2.396 | 1.413 | -0.035 | 0.125 | 0.125 | 0 | 0.84 | 0.61 | 0.77 | 0.66 | 0.9 | 0.83 |
| 5 | 54561643 | rs170779 | 0.68 | 1.393 | 2.412 | 1.422 | -0.014 | 0.062 | 0.062 | 0 | 0.88 | 0.74 | 0.91 | 0.71 | 0.9 | 0.92 |
| 5 | 54563480 | rs17647768 | -0.84 | 1.591 | 2.113 | 1.061 | -0.016 | 0.02 | 0.02 | 0 | 0.09 | 0.04 | 0.09 | 0 | 0.09 | 0.05 |
| 5 | 54596867 | rs7730157 | -1.151 | 1.488 | 2.319 | 1.165 | -0.022 | 0.076 | 0.076 | 0 | 0.14 | 0.33 | 0.1 | 0.29 | 0.1 | 0.08 |
| 5 | 54601452 | rs17281809 | -0.8 | 1.437 | 2.084 | 0.984 | -0.007 | 0.001 | 0.001 | 0.301 | 0.15 | 0.13 | 0.19 | 0.16 | 0.22 | 0.2 |
| 5 | 54603822 | rs2290595 | -2.219 | 1.429 | 2.147 | 0.986 | 0 | 0.041 | 0.041 | 1.098 | 0.32 | 0.62 | 0.43 | 0.5 | 0.33 | 0.38 |
| 5 | 54676222 | rs10041616 | -2.832 | 1.415 | 2.198 | 1.118 | -0.002 | -0.008 | -0.008 | 0.967 | 0.39 | 0.68 | 0.51 | 0.5 | 0.41 | 0.43 |
| 5 | 54702185 | rs12522703 | -0.887 | 1.38 | 1.943 | 1.174 | -0.008 | 0.002 | 0.002 | 0.28 | 0.16 | 0.13 | 0.19 | 0.16 | 0.22 | 0.2 |
| 5 | 54718415 | rs230754 | 0.429 | 1.386 | 1.998 | 1.195 | -0.017 | 0.077 | 0.077 | 0 | 0.88 | 0.72 | 0.91 | 0.71 | 0.9 | 0.92 |
| 5 | 54737008 | rs11741874 | 2.022 | 1.391 | 1.891 | 1.157 | 0.001 | 0.048 | 0.048 | 0.949 | 0.69 | 0.4 | 0.57 | 0.5 | 0.66 | 0.62 |
| 5 | 54774169 | rs10066754 | -0.743 | 1.056 | 1.72 | 1.212 | -0.001 | -0.001 | -0.001 | 0 | 0.12 | 0.13 | 0.15 | 0.1 | 0.13 | 0.09 |
| 5 | 54790559 | rs10079783 | 1.133 | 1.115 | 1.854 | 1.329 | -0.017 | 0.181 | 0.181 | 0 | 0.85 | 0.56 | 0.76 | 0.62 | 0.88 | 0.82 |
| 5 | 54807076 | rs10069504 | 0.571 | 1.111 | 1.968 | 1.354 | -0.011 | 0.052 | 0.052 | 0 | 0.88 | 0.81 | 0.9 | 0.68 | 0.89 | 0.92 |
| 5 | 54814447 | rs16884281 | -0.731 | 1.203 | 1.875 | 1.272 | -0.008 | 0.008 | 0.008 | 0 | 0.07 | 0.04 | 0.07 | 0 | 0.07 | 0.05 |
| 5 | 54834537 | rs10041669 | 1.814 | 1 | 1.936 | 1.425 | -0.024 | 0.144 | 0.144 | 0 | 0.82 | 0.53 | 0.74 | 0.63 | 0.85 | 0.82 |
| 5 | 54843080 | rs11747258 | 0.814 | 1.123 | 1.448 | 1.404 | -0.012 | 0.063 | 0.063 | 0 | 0.88 | 0.77 | 0.9 | 0.69 | 0.89 | 0.92 |
| 5 | 54843488 | rs12521812 | 0.814 | 1.123 | 1.448 | 1.404 | -0.012 | 0.063 | 0.063 | 0 | 0.88 | 0.77 | 0.9 | 0.69 | 0.89 | 0.92 |

| | | | | | | | | | | | | | | | | |
|---|---|---|---|---|---|---|---|---|---|---|---|---|---|---|---|---|
| 5 | 54850717 | rs6450316 | 0.713 | 1.137 | 1.307 | 1.636 | 0.016 | 0.128 | 0.128 | 0 | 0.93 | 0.82 | 0.88 | 0.72 | 0.91 | 0.93 |
| 5 | 54873285 | rs10940473 | 1.388 | 0.936 | 1.094 | 1.319 | 0 | 0.005 | 0.005 | 0.01 | 0.72 | 0.68 | 0.68 | 0.67 | 0.69 | 0.81 |
| 5 | 54880701 | rs10940474 | 2.044 | 0.703 | 1.301 | 1.263 | 0.001 | -0.002 | -0.002 | 0 | 0.62 | 0.38 | 0.57 | 0.57 | 0.66 | 0.67 |
| 5 | 54904720 | rs11749518 | 1.604 | -0.159 | 0.898 | 1.053 | 0.007 | -0.008 | -0.008 | 0 | 0.61 | 0.4 | 0.58 | 0.53 | 0.68 | 0.68 |
| 5 | 54906367 | rs3846501 | 1.267 | -0.289 | 1.145 | 1.07 | -0.017 | 0.049 | 0.049 | 0 | 0.67 | 0.54 | 0.6 | 0.57 | 0.69 | 0.73 |
| 5 | 54915930 | rs6866871 | -1.48 | -0.447 | 1.349 | 0.883 | -0.019 | -0.047 | -0.047 | 0.233 | 0.11 | 0.22 | 0.04 | 0.04 | 0.04 | 0.08 |
| 5 | 54931106 | rs2161504 | 2.105 | 0.472 | 1.185 | 1.132 | 0.021 | -0.001 | -0.001 | 0 | 0.56 | 0.28 | 0.53 | 0.43 | 0.6 | 0.66 |
| 5 | 54938579 | rs2897834 | 0.487 | -0.128 | 0.766 | 1.016 | -0.023 | -0.065 | -0.065 | 0 | 0.86 | 0.99 | 0.89 | 0.8 | 0.87 | 0.93 |
| 5 | 54945675 | rs2014458 | 2.063 | 0.008 | 0.922 | 1.065 | 0.003 | -0.003 | -0.003 | 0 | 0.61 | 0.43 | 0.58 | 0.46 | 0.66 | 0.71 |
| 5 | 54954478 | rs10056287 | -0.172 | -0.029 | 1.054 | 1.103 | 0.001 | -0.01 | -0.01 | 0 | 0.16 | 0.22 | 0.17 | 0.09 | 0.19 | 0.1 |
| 5 | 54955071 | rs7716219 | 0.951 | -0.032 | 1.128 | 1.093 | 0 | 0.005 | 0.005 | 0 | 0.73 | 0.72 | 0.72 | 0.72 | 0.7 | 0.84 |
| 5 | 54960609 | rs4865614 | 1.823 | 0.01 | 0.921 | 0.771 | 0 | -0.001 | -0.001 | 1.978 | 0.62 | 0.43 | 0.59 | 0.46 | 0.66 | 0.71 |
| 5 | 54972296 | rs13165328 | 1.823 | 0.01 | 0.921 | 0.771 | 0 | -0.001 | -0.001 | 7.284 | 0.62 | 0.43 | 0.59 | 0.46 | 0.66 | 0.71 |
| 5 | 54985062 | rs3846504 | 1.72 | 0.012 | 0.924 | 0.774 | -0.001 | 0 | 0 | 0 | 0.63 | 0.44 | 0.59 | 0.46 | 0.66 | 0.71 |
| 5 | 55002265 | rs11749532 | 1.979 | 0.146 | -0.633 | 0.159 | -0.099 | -0.115 | -0.115 | 0.143 | 0.27 | 0.02 | 0.36 | 0.36 | 0.3 | 0.44 |
| 5 | 55005408 | rs11741159 | 1.905 | -0.087 | -0.037 | 0.606 | 0.043 | 0.006 | 0.006 | 0 | 0.56 | 0.29 | 0.56 | 0.42 | 0.65 | 0.65 |
| 5 | 55010963 | rs7707639 | -1.681 | -0.076 | 0.275 | 0.272 | -0.037 | -0.069 | -0.069 | 0 | 0.15 | 0.27 | 0.04 | 0.04 | 0.04 | 0.09 |
| 5 | 55016916 | rs4440309 | 1.224 | -0.026 | 0.179 | 0.337 | -0.02 | -0.047 | -0.047 | 0 | 0.69 | 0.48 | 0.79 | 0.88 | 0.77 | 0.81 |
| 5 | 55034088 | rs3761772 | -1.49 | 0.841 | 0.779 | 0.95 | 0.028 | 0.171 | 0.171 | 0 | 0.12 | 0.31 | 0.37 | 0.3 | 0.11 | 0.15 |
| 5 | 55038487 | rs17699426 | -0.585 | 0.872 | 0.616 | 0.901 | 0 | 0.004 | 0.004 | 0 | 0.15 | 0.07 | 0.14 | 0.16 | 0.14 | 0.1 |
| 5 | 55046489 | rs6863874 | -1.347 | 0.881 | 0.653 | 0.918 | 0.03 | 0.174 | 0.174 | 0 | 0.12 | 0.31 | 0.37 | 0.3 | 0.11 | 0.15 |
| 5 | 55054678 | rs3843458 | -2.379 | 0.891 | 0.638 | 0.878 | -0.001 | -0.003 | -0.003 | 0.004 | 0.31 | 0.62 | 0.54 | 0.47 | 0.3 | 0.3 |
| 5 | 55058992 | rs3761776 | -2.472 | 0.894 | 0.565 | 0.865 | 0 | -0.001 | -0.001 | 0 | 0.32 | 0.62 | 0.33 | 0.48 | 0.26 | 0.28 |
| 5 | 55060656 | rs7730742 | -2.716 | 0.825 | 0.522 | 0.792 | 0.001 | -0.015 | -0.015 | 0.691 | 0.34 | 0.63 | 0.55 | 0.5 | 0.31 | 0.3 |
| 5 | 55062523 | rs10035707 | -2.716 | 0.825 | 0.522 | 0.792 | 0.001 | -0.015 | -0.015 | 0 | 0.34 | 0.63 | 0.55 | 0.5 | 0.31 | 0.3 |
| 5 | 55078065 | rs3843460 | -2.185 | 0.806 | 0.583 | 0.78 | 0.014 | 0.109 | 0.109 | 0 | 0.26 | 0.47 | 0.52 | 0.5 | 0.27 | 0.26 |
| 5 | 55081694 | rs2305123 | 0.618 | 0.818 | 0.368 | 0.794 | -0.02 | -0.021 | -0.021 | 0 | 0.1 | 0.03 | 0.12 | 0.16 | 0.13 | 0.09 |
| 5 | 55082474 | rs957459 | -2.742 | 0.763 | 0.381 | 0.808 | 0.001 | -0.014 | -0.014 | 0.106 | 0.34 | 0.63 | 0.55 | 0.5 | 0.31 | 0.3 |
| 5 | 55122436 | rs162088 | -0.536 | 0.83 | 0.166 | 0.687 | -0.001 | -0.012 | -0.012 | 0 | 0.95 | 0.95 | 0.97 | 1 | 0.96 | 0.99 |
| 5 | 55122700 | rs3846516 | 0.293 | 0.884 | 0.1 | 1.036 | -0.021 | -0.023 | -0.023 | 1.294 | 0.84 | 0.94 | 0.85 | 0.78 | 0.82 | 0.89 |
| 5 | 55135283 | rs10076398 | 0.896 | 0.787 | 0.119 | 1.079 | -0.022 | -0.048 | -0.048 | 0.117 | 0.79 | 0.84 | 0.53 | 0.45 | 0.76 | 0.77 |
| 5 | 55139267 | rs6871296 | 2.38 | 0.717 | 0.35 | 0.98 | 0 | 0.001 | 0.001 | 0 | 0.66 | 0.41 | 0.45 | 0.42 | 0.68 | 0.68 |
| 5 | 55145935 | rs6876491 | 2.306 | 0.593 | 0.311 | 0.915 | -0.001 | -0.005 | -0.005 | 0.995 | 0.66 | 0.39 | 0.45 | 0.42 | 0.69 | 0.68 |
| 5 | 55149235 | rs10940486 | -1.242 | 0.702 | 0.408 | 0.928 | 0.005 | -0.067 | -0.067 | 2.859 | 0.11 | 0.39 | 0.07 | 0 | 0.06 | 0.04 |
| 5 | 55155402 | rs1009639 | 1.372 | 0.8 | 0.222 | 1.05 | 0.02 | 0.065 | 0.065 | 0 | 0.76 | 0.65 | 0.49 | 0.45 | 0.73 | 0.77 |
| 5 | 55161102 | rs10055201 | 0.808 | 0.842 | 0.095 | 1.08 | -0.014 | -0.022 | -0.022 | 0 | 0.81 | 0.83 | 0.53 | 0.45 | 0.75 | 0.77 |
| 5 | 55167146 | rs9292103 | 1.03 | 0.951 | -0.483 | 0.407 | -0.008 | 0.009 | 0.009 | 0 | 0.56 | 0.65 | 0.37 | 0.39 | 0.53 | 0.65 |
| 5 | 55178747 | rs6873519 | 0.773 | 1.169 | -0.717 | 0.595 | -0.028 | -0.088 | -0.088 | 0 | 0.22 | 0.09 | 0.29 | 0.39 | 0.25 | 0.28 |
| 5 | 55180909 | rs327240 | 0.714 | 1.41 | -0.483 | 0.397 | 0.01 | 0.002 | 0.002 | 0 | 0.78 | 0.7 | 0.66 | 0.75 | 0.77 | 0.84 |
| 5 | 55181941 | rs918626 | 0.405 | 1.291 | -0.484 | 0.185 | -0.085 | -0.022 | -0.022 | 0 | 0.61 | 0.86 | 0.41 | 0.39 | 0.56 | 0.64 |
| 5 | 55186121 | rs166233 | 1.494 | 1.385 | -0.142 | 0.139 | 0 | 0.022 | 0.022 | 0 | 0.47 | 0.39 | 0.33 | 0.35 | 0.48 | 0.57 |
| 5 | 55189983 | rs16884632 | 0.251 | 1.177 | 0.008 | -0.02 | 0.012 | -0.029 | -0.029 | 0.438 | 0.07 | 0.22 | 0.04 | 0 | 0.04 | 0.04 |
| 5 | 55193329 | rs254987 | 1.283 | 1.544 | -0.393 | 0.088 | -0.003 | 0.044 | 0.044 | 0 | 0.47 | 0.36 | 0.28 | 0.25 | 0.47 | 0.52 |
| 5 | 55202551 | rs324454 | 0.671 | 0.992 | -1.068 | -0.68 | -0.007 | 0.13 | 0.13 | 1.036 | 0.57 | 0.82 | 0.54 | 0.8 | 0.57 | 0.69 |
| 5 | 55204187 | rs9632389 | 1.199 | 1.075 | -1.257 | -0.618 | 0.015 | 0.078 | 0.078 | 0.115 | 0.17 | 0.04 | 0.1 | 0.01 | 0.16 | 0.11 |
| 5 | 55206444 | rs161704 | 1.693 | 1.101 | -1.1 | -0.4 | 0.006 | 0.009 | 0.009 | 0 | 0.26 | 0.33 | 0.22 | 0.33 | 0.3 | 0.34 |
| 5 | 55210440 | rs161702 | 0.092 | 1.125 | -1.115 | -0.138 | 0.002 | -0.013 | -0.013 | 0 | 0.76 | 0.73 | 0.66 | 0.81 | 0.78 | 0.83 |
| 5 | 55213631 | rs10052798 | 0.407 | 0.973 | -1.343 | 0.325 | 0 | -0.094 | -0.094 | 0 | 0.24 | 0.07 | 0.28 | 0.48 | 0.25 | 0.35 |
| 5 | 55219716 | rs162085 | 0.139 | 1.38 | -0.979 | -0.219 | -0.008 | -0.025 | -0.025 | 0.75 | 0.7 | 0.29 | 0.69 | 0.49 | 0.65 | 0.55 |
| 5 | 55220344 | rs7728232 | -0.634 | 1.744 | -0.263 | -1.309 | 0.031 | -0.045 | -0.045 | 0 | 0.22 | 0.33 | 0.4 | 0.17 | 0.32 | 0.2 |
| 5 | 55220819 | rs7730168 | -0.259 | 1.56 | -0.345 | -1.102 | 0.001 | -0.01 | -0.01 | 0.139 | 0.26 | 0.25 | 0.4 | 0.19 | 0.31 | 0.23 |
| 5 | 55223929 | rs13357543 | -0.245 | 1.533 | -0.79 | -1.079 | -0.052 | 0.061 | 0.061 | 0 | 0.26 | 0.1 | 0.41 | 0.2 | 0.34 | 0.24 |
| 5 | 55236374 | rs10471960 | 0.162 | 0.927 | 0.653 | -1.219 | -0.027 | -0.092 | -0.092 | 0 | 0.16 | 0.51 | 0.12 | 0.03 | 0.11 | 0.08 |
| 5 | 55262660 | rs10940495 | 0.212 | 1.024 | 0.117 | -1.623 | -0.038 | 0.087 | 0.087 | 0 | 0.26 | 0.07 | 0.17 | 0.08 | 0.29 | 0.29 |
| 5 | 55267814 | rs11741953 | 0.898 | 0.985 | 0.336 | -1.631 | -0.009 | -0.035 | -0.035 | 0 | 0.13 | 0.17 | 0.09 | 0.03 | 0.11 | 0.08 |
| 5 | 55271621 | rs6861772 | 0.276 | 1.076 | -0.077 | -1.644 | -0.018 | -0.085 | -0.085 | 0 | 0.17 | 0.62 | 0.13 | 0.03 | 0.13 | 0.09 |
| 5 | 55292036 | rs2112979 | 0.296 | 1.103 | -0.361 | -1.653 | -0.077 | 0.089 | 0.089 | 0 | 0.25 | 0.06 | 0.17 | 0.08 | 0.29 | 0.28 |
| 5 | 55294328 | rs6870870 | -0.798 | 1.17 | -0.266 | -1.527 | -0.001 | 0.058 | 0.058 | 0 | 0.43 | 0.7 | 0.31 | 0.11 | 0.42 | 0.37 |
| 5 | 55322895 | rs17348299 | 0.505 | 1.184 | -1.186 | -0.99 | -0.019 | 0.045 | 0.045 | 0 | 0.16 | 0.05 | 0.08 | 0.05 | 0.15 | 0.24 |
| 5 | 55324494 | rs11949857 | 0.942 | 1.145 | -1.481 | -1.426 | -0.008 | 0.064 | 0.064 | 0.607 | 0.22 | 0.11 | 0.16 | 0.03 | 0.21 | 0.15 |
| 5 | 55333925 | rs10045084 | -0.582 | 0.992 | -1.298 | -1.112 | 0.022 | 0.061 | 0.061 | 2.131 | 0.62 | 0.82 | 0.65 | 0.25 | 0.66 | 0.6 |
| 5 | 55345347 | rs10069751 | -1.313 | 1.851 | -0.469 | -0.809 | 0.019 | -0.053 | -0.053 | 0 | 0.27 | 0.38 | 0.24 | 0.07 | 0.35 | 0.29 |
| 5 | 55349049 | rs11742921 | 0.117 | 1.732 | -1.11 | 0.135 | -0.055 | -0.103 | -0.103 | 0 | 0.16 | 0.03 | 0.15 | 0.38 | 0.23 | 0.25 |
| 5 | 55349691 | rs11957145 | -1.904 | 1.548 | -0.732 | 0.057 | -0.007 | -0.012 | -0.012 | 1.934 | 0.45 | 0.71 | 0.38 | 0.47 | 0.55 | 0.5 |
| 5 | 55352562 | rs16884734 | -0.349 | 1.264 | -0.808 | -0.5 | -0.031 | -0.012 | -0.012 | 0.118 | 0.15 | 0.03 | 0.09 | 0.09 | 0.16 | 0.2 |
| 5 | 55352770 | rs10042443 | -1.166 | 1.587 | -0.348 | -0.209 | 0.002 | 0 | 0 | 0 | 0.29 | 0.54 | 0.27 | 0.31 | 0.35 | 0.28 |
| 5 | 55355008 | rs10036423 | -0.434 | 1.65 | -0.433 | -0.18 | -0.014 | -0.017 | -0.017 | 0 | 0.2 | 0.13 | 0.21 | 0.33 | 0.27 | 0.27 |
| 5 | 55355145 | rs10059243 | -1.832 | 1.448 | -0.877 | -0.391 | 0.026 | 0.024 | 0.024 | 0 | 0.52 | 0.75 | 0.41 | 0.39 | 0.58 | 0.49 |
| 5 | 55364571 | rs10471966 | -0.625 | 1.926 | 0.094 | -0.383 | 0.011 | 0.011 | 0.011 | 0.435 | 0.19 | 0.21 | 0.21 | 0.29 | 0.27 | 0.27 |

| | | | | | | | | | | | | | | | |
|---|---|---|---|---|---|---|---|---|---|---|---|---|---|---|---|
| 5 | 55371201 | rs10063964 | -1.079 | 1.739 | -0.464 | -0.36 | 0.003 | 0.001 | 0.001 | 0 | 0.27 | 0.3 | 0.29 | 0.29 | 0.35 | 0.27 |
| 5 | 55371234 | rs7700857 | -1.237 | 1.653 | -0.662 | -0.195 | -0.046 | 0.052 | 0.052 | 0 | 0.41 | 0.74 | 0.36 | 0.29 | 0.48 | 0.3 |
| 5 | 55381015 | rs10066150 | -1.072 | 0.9 | -0.525 | 0.599 | 0.002 | 0.011 | 0.011 | 0 | 0.18 | 0.21 | 0.2 | 0.29 | 0.21 | 0.25 |
| 5 | 55382424 | rs16884781 | -0.233 | 0.631 | -0.866 | 0.797 | -0.027 | -0.048 | -0.048 | 0 | 0.09 | 0.01 | 0.11 | 0.33 | 0.11 | 0.19 |
| 5 | 55384909 | rs9647530 | 0.102 | 0.633 | -1.313 | 0.305 | -0.172 | 0.063 | 0.063 | 0.016 | 0.33 | 0.06 | 0.44 | 0.21 | 0.46 | 0.27 |
| 5 | 55385145 | rs322021 | -0.969 | 0.092 | -0.888 | 0.122 | 0.004 | -0.026 | -0.026 | 0 | 0.7 | 0.75 | 0.68 | 0.5 | 0.74 | 0.62 |
| 5 | 55389477 | rs689379 | -0.232 | -0.498 | 0.408 | -0.161 | -0.032 | -0.028 | -0.028 | 0.244 | 0.36 | 0.45 | 0.15 | 0.21 | 0.29 | 0.29 |
| 5 | 55395112 | rs186451 | -0.559 | -0.499 | 0.172 | -0.131 | 0.082 | 0.065 | 0.065 | 0 | 0.4 | 0.74 | 0.23 | 0.23 | 0.29 | 0.39 |
| 5 | 55406952 | rs321775 | -0.061 | -0.388 | -0.442 | -0.063 | 0.045 | -0.028 | -0.028 | 0 | 0.33 | 0.09 | 0.13 | 0.31 | 0.28 | 0.3 |
| 5 | 55407542 | rs321776 | -0.04 | -0.37 | -0.457 | -0.2 | 0.042 | -0.018 | -0.018 | 0 | 0.32 | 0.07 | 0.13 | 0.29 | 0.28 | 0.3 |
| 5 | 55409421 | rs160919 | -0.36 | -0.274 | 0.045 | -0.331 | -0.033 | -0.023 | -0.023 | 0.11 | 0.37 | 0.5 | 0.27 | 0.26 | 0.29 | 0.38 |
| 5 | 55410585 | rs318802 | 0.06 | 0.546 | -0.314 | -0.016 | 0.005 | 0.008 | 0.008 | 0 | 0.47 | 0.4 | 0.29 | 0.2 | 0.43 | 0.33 |
| 5 | 55414956 | rs415407 | -0.162 | 0.726 | -1.503 | 0.122 | 0.004 | 0.085 | 0.085 | 0 | 0.54 | 0.7 | 0.43 | 0.27 | 0.48 | 0.48 |
| 5 | 55422681 | rs321768 | -0.197 | 1.062 | -2.185 | 0.503 | -0.091 | -0.09 | -0.09 | 0 | 0.59 | 0.86 | 0.63 | 0.49 | 0.59 | 0.71 |
| 5 | 55424472 | rs318791 | -0.152 | 1.091 | -2.123 | 0.143 | 0.019 | 0.042 | 0.042 | 0 | 0.4 | 0.72 | 0.63 | 0.72 | 0.37 | 0.52 |
| 5 | 55424740 | rs17645980 | 0.637 | 1.62 | -2.219 | 0.276 | -0.02 | 0.031 | 0.031 | 0 | 0.17 | 0.06 | 0.13 | 0.09 | 0.24 | 0.21 |
| 5 | 55428838 | rs2431873 | -0.043 | 1.242 | -2.032 | 0.894 | -0.001 | 0.029 | 0.029 | 0 | 0.08 | 0.05 | 0.21 | 0.37 | 0.09 | 0.16 |
| 5 | 55434432 | rs6876431 | 0.217 | 1.288 | -1.365 | -0.284 | 0.003 | 0.009 | 0.009 | 0 | 0.48 | 0.54 | 0.25 | 0.21 | 0.37 | 0.22 |
| 5 | 55435220 | rs10040327 | 0.34 | 1.156 | -1.341 | -0.194 | 0.007 | -0.028 | -0.028 | 0 | 0.08 | 0.12 | 0.07 | 0 | 0.11 | 0.02 |
| 5 | 55438851 | rs10065637 | -0.049 | 0.887 | -1.869 | -0.437 | -0.04 | 0.146 | 0.146 | 0 | 0.22 | 0.02 | 0.13 | 0.02 | 0.22 | 0.1 |
| 5 | 55444040 | rs10214316 | 0.31 | 0.835 | -1.296 | 0.368 | 0.001 | -0.004 | -0.004 | 0 | 0.7 | 0.72 | 0.56 | 0.44 | 0.73 | 0.58 |
| 5 | 55445044 | rs10057763 | 0.047 | 0.883 | -0.967 | 0.134 | -0.002 | -0.01 | -0.01 | 0 | 0.29 | 0.29 | 0.25 | 0.22 | 0.26 | 0.21 |
| 5 | 55445129 | rs309622 | 0.215 | 0.993 | -0.662 | 0.566 | 0 | 0 | 0 | 0 | 0.58 | 0.57 | 0.64 | 0.53 | 0.66 | 0.59 |
| 5 | 55448425 | rs4700266 | 0.117 | 1.013 | -0.725 | 0.575 | 0.003 | 0.027 | 0.027 | 0 | 0.31 | 0.26 | 0.23 | 0.19 | 0.27 | 0.21 |
| 5 | 55452505 | rs182572 | -0.007 | 1.067 | -0.623 | 0.757 | -0.001 | -0.001 | -0.001 | 0 | 0.74 | 0.77 | 0.79 | 0.73 | 0.75 | 0.78 |
| 5 | 55458219 | rs159546 | 0.343 | 0.748 | -0.735 | 0.886 | 0.001 | 0.032 | 0.032 | 0 | 0.53 | 0.33 | 0.53 | 0.35 | 0.51 | 0.6 |
| 5 | 55466213 | rs1471655 | 0.087 | 0.491 | -0.415 | 0.998 | -0.006 | -0.005 | -0.005 | 0 | 0.06 | 0.14 | 0.09 | 0.05 | 0.03 | 0.02 |
| 5 | 55473321 | rs6450371 | -1.229 | 0.283 | 0.209 | 1.565 | 0.022 | 0.199 | 0.199 | 0 | 0.1 | 0.5 | 0.08 | 0.21 | 0.06 | 0.12 |
| 5 | 55475759 | rs3846522 | -0.756 | 0.12 | 0.375 | 2.199 | -0.005 | 0.175 | 0.175 | 0 | 0.09 | 0.36 | 0.1 | 0.25 | 0.04 | 0.08 |
| 5 | 55477779 | rs7706204 | -1.071 | 0.148 | 0.391 | 2.145 | 0.041 | 0.233 | 0.233 | 0 | 0.09 | 0.45 | 0.07 | 0.21 | 0.06 | 0.12 |
| 5 | 55481812 | rs160935 | -1.22 | 0.168 | -0.251 | 1.752 | 0.018 | -0.1 | -0.1 | 0 | 0.31 | 0.83 | 0.28 | 0.49 | 0.3 | 0.49 |
| 5 | 55482313 | rs7732186 | 0.179 | 0.185 | -0.102 | 1.595 | -0.008 | -0.004 | -0.004 | 0 | 0.07 | 0.22 | 0.1 | 0.05 | 0.03 | 0.02 |
| 5 | 55488928 | rs36747 | 0.531 | 0.064 | 0.066 | 2.261 | 0.001 | -0.027 | -0.027 | 0 | 0.16 | 0.14 | 0.13 | 0.38 | 0.21 | 0.42 |
| 5 | 55491180 | rs161222 | -0.428 | 0.271 | -0.593 | 1.686 | -0.062 | -0.048 | -0.048 | 0 | 0.77 | 0.93 | 0.64 | 0.66 | 0.72 | 0.78 |
| 5 | 55492679 | rs160011 | 0.645 | -0.281 | 0.026 | 2.247 | 0 | -0.043 | -0.043 | 0 | 0.16 | 0.11 | 0.12 | 0.38 | 0.21 | 0.4 |
| 5 | 55493266 | rs3843464 | -0.688 | -0.271 | 0.384 | 2.11 | 0.073 | 0.286 | 0.286 | 0 | 0.08 | 0.4 | 0.05 | 0.21 | 0.05 | 0.1 |
| 5 | 55507046 | rs159572 | -1.073 | -0.405 | 0.092 | 1.638 | -0.002 | 0 | 0 | 0.076 | 0.33 | 0.6 | 0.33 | 0.65 | 0.28 | 0.47 |
| 5 | 55509554 | rs159566 | -0.663 | -0.403 | 0.115 | 1.658 | -0.012 | 0.035 | 0.035 | 0 | 0.27 | 0.59 | 0.24 | 0.46 | 0.23 | 0.45 |
| 5 | 55509576 | rs16885032 | 0.227 | -0.275 | 0.046 | 1.03 | -0.018 | 0.006 | 0.006 | 0 | 0.07 | 0.19 | 0.09 | 0.07 | 0.03 | 0.01 |
| 5 | 55511394 | rs12652011 | -0.04 | -0.289 | 0.007 | 1.023 | -0.022 | 0.007 | 0.007 | 0 | 0.08 | 0.26 | 0.1 | 0.07 | 0.03 | 0.01 |
| 5 | 55511421 | rs12654690 | -0.743 | -0.299 | 0.026 | 1.002 | -0.021 | -0.007 | -0.007 | 3.863 | 0.11 | 0.32 | 0.11 | 0.07 | 0.06 | 0.04 |
| 5 | 55511683 | rs12654754 | -0.092 | -0.279 | -0.171 | 1.023 | -0.023 | 0.007 | 0.007 | 0 | 0.08 | 0.27 | 0.1 | 0.07 | 0.03 | 0.01 |
| 5 | 55516988 | rs149140 | -0.748 | -0.323 | -0.075 | 0.998 | -0.003 | -0.013 | -0.013 | 0.019 | 0.36 | 0.45 | 0.32 | 0.84 | 0.29 | 0.55 |
| 5 | 55527213 | rs149138 | -0.027 | -0.338 | -0.903 | 1.031 | 0.004 | 0.008 | 0.008 | 0 | 0.27 | 0.17 | 0.22 | 0.77 | 0.25 | 0.53 |
| 5 | 55548797 | rs32498 | -0.713 | -0.119 | -1.192 | 0.818 | 0.055 | 0.049 | 0.049 | 0.334 | 0.42 | 0.86 | 0.37 | 0.68 | 0.31 | 0.45 |
| 5 | 55560577 | rs152343 | -0.629 | -0.067 | -1.123 | 0.844 | 0.048 | 0.043 | 0.043 | 0.185 | 0.41 | 0.82 | 0.37 | 0.68 | 0.32 | 0.46 |
| 5 | 55565275 | rs375808 | 0.511 | 0.308 | -0.825 | 0.584 | -0.003 | 0.004 | 0.004 | 0 | 0.42 | 0.37 | 0.39 | 0.39 | 0.45 | 0.35 |
| 5 | 55566149 | rs463677 | 0.63 | 0.204 | -0.765 | 0.535 | -0.002 | 0.001 | 0.001 | 0 | 0.41 | 0.42 | 0.39 | 0.39 | 0.45 | 0.35 |
| 5 | 55566600 | rs27008 | 0.085 | -0.004 | -1.19 | 0.64 | 0.012 | -0.003 | -0.003 | 0 | 0.8 | 0.88 | 0.76 | 0.77 | 0.87 | 0.75 |
| 5 | 55567087 | rs13328207 | 1.332 | 0.332 | -1.157 | 0.647 | -0.029 | 0.041 | 0.041 | 0 | 0.08 | 0.02 | 0.05 | 0 | 0.12 | 0.02 |
| 5 | 55567581 | rs27006 | 0.34 | 0.127 | -0.638 | 0.812 | -0.01 | -0.03 | -0.03 | 0.006 | 0.25 | 0.2 | 0.28 | 0.38 | 0.29 | 0.37 |
| 5 | 55567749 | rs27573 | 0.433 | -0.438 | -0.708 | 0.401 | -0.042 | 0.019 | 0.019 | 0 | 0.86 | 0.74 | 0.82 | 0.81 | 0.95 | 0.87 |
| 5 | 55567861 | rs637033 | 0.2 | 0.4 | -1.326 | 0.35 | -0.011 | 0.027 | 0.027 | 0 | 0.47 | 0.7 | 0.42 | 0.39 | 0.49 | 0.38 |
| 5 | 55568321 | rs32510 | 0.722 | 1.822 | -1.326 | 0.376 | -0.019 | 0.002 | 0.002 | 0.195 | 0.83 | 0.87 | 0.85 | 0.86 | 0.7 | 0.82 |
| 5 | 55568487 | rs32508 | -0.07 | 1.379 | -1.225 | 0.106 | -0.005 | 0.034 | 0.034 | 0 | 0.55 | 0.7 | 0.65 | 0.72 | 0.52 | 0.57 |
| 5 | 55574191 | rs2342888 | 0.028 | 1.686 | -0.784 | 0.842 | 0.006 | 0.064 | 0.064 | 0 | 0.11 | 0.11 | 0.25 | 0.61 | 0.14 | 0.41 |
| 5 | 55575558 | rs152318 | -0.337 | 1.42 | -0.673 | 0.055 | 0.002 | 0.005 | 0.005 | 0.221 | 0.3 | 0.34 | 0.42 | 0.64 | 0.36 | 0.46 |
| 5 | 55576026 | rs158487 | -0.545 | 1.159 | -0.877 | -1.063 | 0 | 0.054 | 0.054 | 5.213 | 0.7 | 0.8 | 0.75 | 0.94 | 0.7 | 0.85 |
| 5 | 55579789 | rs160078 | 1.29 | 1.085 | -0.715 | -1.027 | -0.002 | -0.05 | -0.05 | 0 | 0.17 | 0.22 | 0.15 | 0 | 0.21 | 0.04 |
| 5 | 55593724 | rs158666 | -0.684 | 1.078 | -0.817 | -1.121 | -0.006 | 0.084 | 0.084 | 4.025 | 0.69 | 0.82 | 0.73 | 0.95 | 0.67 | 0.85 |
| 5 | 55593735 | rs158667 | -0.41 | 0.9 | -0.092 | -1.202 | -0.006 | -0.112 | -0.112 | 0 | 0.66 | 0.52 | 0.71 | 0.95 | 0.67 | 0.85 |
| 5 | 55595142 | rs154828 | 1.21 | 1.13 | -0.244 | -0.455 | 0.005 | 0.076 | 0.076 | 0 | 0.1 | 0.11 | 0.09 | 0.44 | 0.1 | 0.33 |
| 5 | 55595606 | rs11954699 | 1.349 | 1.201 | -0.383 | -0.66 | 0.002 | -0.014 | -0.014 | 0 | 0.12 | 0.11 | 0.13 | 0.03 | 0.13 | 0.06 |
| 5 | 55597064 | rs4699912 | 0.649 | 0.956 | -0.06 | -0.611 | -0.014 | -0.05 | -0.05 | 4.698 | 0.38 | 0.15 | 0.46 | 0.5 | 0.37 | 0.46 |
| 5 | 55598424 | rs32489 | -0.23 | 0.47 | 0.413 | -0.225 | 0.005 | 0.006 | 0.006 | 0.108 | 0.36 | 0.65 | 0.41 | 0.25 | 0.3 | 0.21 |
| 5 | 55601387 | rs1990977 | 0.836 | 0.478 | 0.482 | -0.218 | -0.007 | 0.011 | 0.011 | 0 | 0.17 | 0.19 | 0.25 | 0.24 | 0.14 | 0.17 |
| 5 | 55601889 | rs32493 | -0.816 | 0.461 | 0.369 | -0.227 | 0.03 | 0.041 | 0.041 | 0 | 0.47 | 0.75 | 0.5 | 0.69 | 0.39 | 0.53 |
| 5 | 55603036 | rs152332 | 0.905 | 0.533 | 0.198 | -0.208 | 0.003 | 0.04 | 0.04 | 0 | 0.12 | 0.11 | 0.26 | 0.44 | 0.13 | 0.33 |
| 5 | 55608881 | rs158642 | 1.276 | 0.458 | 0.341 | -0.232 | -0.004 | -0.042 | -0.042 | 0 | 0.17 | 0.23 | 0.14 | 0 | 0.16 | 0.04 |

| | | | | | | | | | | | | | | | | |
|---|---|---|---|---|---|---|---|---|---|---|---|---|---|---|---|---|
| 5 | 55611438 | rs32516 | 1.065 | 0.461 | 0.384 | -0.231 | -0.028 | -0.112 | -0.112 | 0 | 0.18 | 0.42 | 0.15 | 0 | 0.16 | 0.04 |
| 5 | 55611694 | rs158640 | -0.766 | 0.475 | 0.163 | -0.203 | 0.03 | 0.042 | 0.042 | 0 | 0.47 | 0.78 | 0.5 | 0.69 | 0.39 | 0.53 |
| 5 | 55611710 | rs158639 | -0.309 | 0.679 | 0.213 | -0.062 | -0.015 | 0.049 | 0.049 | 0.263 | 0.29 | 0.52 | 0.24 | 0.44 | 0.26 | 0.36 |
| 5 | 55622233 | rs158346 | 0.321 | 0.594 | -0.14 | -0.538 | -0.022 | -0.099 | -0.099 | 3.112 | 0.18 | 0.32 | 0.14 | 0 | 0.16 | 0.04 |
| 5 | 55625333 | rs158342 | 0.386 | 0.708 | -0.653 | -0.523 | -0.006 | -0.061 | -0.061 | 0.053 | 0.19 | 0.24 | 0.14 | 0 | 0.18 | 0.05 |
| 5 | 55627679 | rs158340 | -0.749 | 0.764 | -1.478 | -0.555 | -0.001 | -0.032 | -0.032 | 0 | 0.62 | 0.82 | 0.61 | 0.42 | 0.57 | 0.42 |
| 5 | 55641055 | rs32497 | 0.4 | 0.753 | -1.357 | -0.478 | 0.005 | 0.016 | 0.016 | 1.65 | 0.18 | 0.13 | 0.14 | 0 | 0.18 | 0.06 |
| 5 | 55641683 | rs1423115 | -0.266 | 1.015 | -1.071 | 0.243 | -0.023 | -0.01 | -0.01 | 0.003 | 0.23 | 0.13 | 0.17 | 0.22 | 0.25 | 0.19 |
| 5 | 55643774 | rs32494 | 0.339 | 0.669 | -0.974 | 0.267 | -0.014 | -0.009 | -0.009 | 0 | 0.41 | 0.35 | 0.62 | 0.54 | 0.46 | 0.54 |
| 5 | 55645120 | rs158355 | -0.824 | 0.738 | -1.364 | 0.055 | -0.058 | 0.036 | 0.036 | 0 | 0.9 | 0.98 | 0.92 | 0.96 | 0.81 | 0.91 |
| 5 | 55646812 | rs289782 | -0.7 | -0.181 | -0.803 | 0.759 | 0.005 | 0.027 | 0.027 | 0 | 0.56 | 0.73 | 0.44 | 0.59 | 0.45 | 0.51 |
| 5 | 55647984 | rs11950693 | 0.406 | -0.433 | -1.105 | 0.818 | -0.042 | -0.063 | -0.063 | 0 | 0.12 | 0.02 | 0.12 | 0.17 | 0.14 | 0.19 |
| 5 | 55648104 | rs7703496 | 0.236 | -0.688 | -0.704 | 0.875 | 0.035 | 0.082 | 0.082 | 0 | 0.33 | 0.11 | 0.15 | 0.19 | 0.27 | 0.2 |
| 5 | 55652830 | rs290756 | -0.23 | -0.702 | -0.016 | 0.885 | 0.003 | 0.286 | 0.286 | 0 | 0.09 | 0.41 | 0.15 | 0.23 | 0.04 | 0.12 |
| 5 | 55653889 | rs6892034 | 0.163 | -0.73 | -0.138 | 0.859 | 0.028 | 0.051 | 0.051 | 0 | 0.34 | 0.17 | 0.16 | 0.19 | 0.27 | 0.2 |
| 5 | 55654145 | rs12517669 | 0.617 | -0.894 | -0.441 | 0.731 | 0.013 | 0.021 | 0.021 | 0 | 0.27 | 0.02 | 0.1 | 0.19 | 0.19 | 0.18 |
| 5 | 55655396 | rs1278057 | -0.902 | -0.772 | -0.287 | 0.811 | -0.019 | 0.094 | 0.094 | 0 | 0.23 | 0.61 | 0.28 | 0.4 | 0.18 | 0.31 |
| 5 | 55660509 | rs10471421 | -0.459 | -0.591 | 0.274 | 0.862 | 0.015 | 0.017 | 0.017 | 0 | 0.47 | 0.38 | 0.3 | 0.36 | 0.41 | 0.39 |
| 5 | 55660690 | rs1508900 | -0.809 | -0.297 | -0.533 | 0.412 | 0.008 | 0.03 | 0.03 | 0 | 0.56 | 0.72 | 0.44 | 0.59 | 0.44 | 0.51 |
| 5 | 55671741 | rs7736704 | -0.83 | -0.431 | -0.493 | 0.559 | 0.032 | 0.013 | 0.013 | 0 | 0.47 | 0.62 | 0.39 | 0.57 | 0.34 | 0.43 |
| 5 | 55672099 | rs10940503 | -0.26 | -0.277 | -0.883 | 0.176 | 0.014 | 0.023 | 0.023 | 0 | 0.23 | 0.04 | 0.09 | 0.2 | 0.17 | 0.16 |
| 5 | 55678605 | rs7703778 | -0.63 | -0.472 | -0.525 | 0.17 | -0.016 | -0.015 | -0.015 | 0 | 0.27 | 0.29 | 0.12 | 0.2 | 0.17 | 0.14 |
| 5 | 55695116 | rs6863774 | -0.304 | -0.106 | -0.888 | 0.426 | 0.025 | 0.006 | 0.006 | 0.276 | 0.22 | 0.13 | 0.09 | 0.22 | 0.16 | 0.15 |
| 5 | 55699358 | rs10940504 | -0.46 | 0.978 | -0.739 | 0.631 | 0.008 | 0.002 | 0.002 | 0 | 0.23 | 0.26 | 0.41 | 0.28 | 0.33 | 0.25 |
| 5 | 55699403 | rs158082 | -1.187 | 0.321 | -1.228 | 0.441 | -0.059 | -0.06 | -0.06 | 0.482 | 0.62 | 0.87 | 0.61 | 0.58 | 0.55 | 0.54 |
| 5 | 55702460 | rs286012 | 0.265 | 0.21 | -0.316 | 0.227 | 0.008 | -0.021 | -0.021 | 0 | 0.32 | 0.25 | 0.15 | 0.01 | 0.25 | 0.17 |
| 5 | 55704176 | rs286007 | -0.319 | 0.437 | -0.446 | 0.17 | 0.004 | 0.004 | 0.004 | 0 | 0.4 | 0.61 | 0.22 | 0.3 | 0.31 | 0.29 |
| 5 | 55706081 | rs286004 | -1.303 | 0.461 | -0.637 | 0.171 | -0.048 | -0.06 | -0.06 | 0.038 | 0.62 | 0.88 | 0.62 | 0.58 | 0.58 | 0.55 |
| 5 | 55711291 | rs158214 | 0.636 | 0.589 | 0.158 | 0.58 | 0 | 0.004 | 0.004 | 0 | 0.69 | 0.31 | 0.52 | 0.44 | 0.66 | 0.62 |
| 5 | 55720796 | rs158085 | 0.461 | 0.586 | 0.825 | 0.706 | 0.011 | 0.082 | 0.082 | 0 | 0.93 | 0.73 | 0.95 | 0.78 | 0.94 | 0.89 |
| 5 | 55727176 | rs590575 | 0.269 | 0.68 | 0.704 | 0.529 | 0.041 | 0.035 | 0.035 | 0 | 0.74 | 0.52 | 0.56 | 0.65 | 0.66 | 0.72 |
| 5 | 55728632 | rs285150 | -0.15 | 0.501 | 0.644 | 0.415 | 0.028 | 0.025 | 0.025 | 0 | 0.23 | 0.44 | 0.41 | 0.3 | 0.28 | 0.26 |
| 5 | 55730588 | rs285149 | -0.272 | 0.797 | 0.093 | 0.484 | -0.001 | -0.002 | -0.002 | 0 | 0.06 | 0.14 | 0.04 | 0.17 | 0.06 | 0.05 |
| 5 | 55730683 | rs30356 | 0.394 | 0.401 | -0.718 | -0.632 | 0.014 | 0.11 | 0.11 | 0 | 0.36 | 0.12 | 0.25 | 0.04 | 0.34 | 0.29 |
| 5 | 55730742 | rs30357 | 0.398 | 0.393 | -0.722 | -0.636 | 0.013 | 0.109 | 0.109 | 0 | 0.36 | 0.12 | 0.25 | 0.04 | 0.34 | 0.29 |
| 5 | 55734556 | rs168666 | -2.007 | 0.461 | -1.025 | 0.012 | -0.054 | -0.066 | -0.066 | 0 | 0.87 | 0.98 | 0.86 | 0.76 | 0.84 | 0.84 |
| 5 | 55735500 | rs285146 | 0.759 | 0.411 | -0.869 | 0.038 | -0.001 | 0 | 0 | 0 | 0.06 | 0.14 | 0.04 | 0.17 | 0.06 | 0.06 |
| 5 | 55737637 | rs158210 | 0.929 | 0.157 | -1.061 | 0.092 | 0.016 | -0.051 | -0.051 | 0 | 0.12 | 0.12 | 0.07 | 0.27 | 0.08 | 0.14 |
| 5 | 55738764 | rs245806 | -0.375 | 0.003 | -0.963 | -0.134 | 0.005 | 0.011 | 0.011 | 0 | 0.61 | 0.38 | 0.71 | 0.63 | 0.67 | 0.67 |
| 5 | 55740340 | rs285160 | -0.342 | 0.085 | -0.764 | 0.264 | -0.025 | -0.038 | -0.038 | 0.2 | 0.25 | 0.25 | 0.47 | 0.6 | 0.32 | 0.39 |
| 5 | 55741171 | rs12522646 | 0.467 | 0.478 | -0.328 | 0.212 | -0.01 | -0.006 | -0.006 | 0 | 0.13 | 0.13 | 0.39 | 0.32 | 0.24 | 0.26 |
| 5 | 55742395 | rs13355587 | 0.703 | 0.105 | 0.174 | -0.177 | 0.001 | -0.084 | -0.084 | 0 | 0.14 | 0.34 | 0.08 | 0.01 | 0.12 | 0.08 |
| 5 | 55742710 | rs3844303 | 0.122 | 0.134 | 0.025 | -0.313 | 0 | -0.093 | -0.093 | 0 | 0.19 | 0.31 | 0.12 | 0.01 | 0.19 | 0.1 |
| 5 | 55746342 | rs17656329 | 0.145 | -0.082 | -0.378 | -0.36 | 0.019 | 0.037 | 0.037 | 0 | 0.11 | 0.04 | 0.06 | 0 | 0.06 | 0.03 |
| 5 | 55747268 | rs6450392 | 0.668 | -0.191 | -0.349 | -0.367 | 0.003 | 0.006 | 0.006 | 0.007 | 0.16 | 0.1 | 0.08 | 0.13 | 0.12 | 0.15 |
| 5 | 55748742 | rs17707459 | 0.879 | -0.128 | -0.404 | -0.543 | -0.003 | -0.011 | -0.011 | 0 | 0.13 | 0.11 | 0.11 | 0.04 | 0.08 | 0.23 |
| 5 | 55750231 | rs17656407 | -0.88 | -0.052 | -0.39 | -0.656 | 0.012 | -0.007 | -0.007 | 0.929 | 0.33 | 0.26 | 0.25 | 0.04 | 0.24 | 0.3 |
| 5 | 55750383 | rs10471977 | 1.88 | 0.036 | -0.992 | -0.602 | -0.002 | -0.019 | -0.019 | 2.511 | 0.11 | 0.1 | 0.06 | 0.02 | 0.08 | 0.07 |
| 5 | 55750805 | rs3902134 | 1.141 | 0.585 | -1.186 | -0.15 | -0.002 | 0.017 | 0.017 | 0 | 0.11 | 0.11 | 0.28 | 0.33 | 0.17 | 0.2 |
| 5 | 55750914 | rs3852139 | -1.915 | 0.56 | -1.037 | -0.097 | -0.041 | -0.091 | -0.091 | 0 | 0.61 | 0.79 | 0.65 | 0.5 | 0.59 | 0.65 |
| 5 | 55756394 | rs3852141 | 1.096 | 1.058 | -0.745 | 0.281 | -0.002 | 0.013 | 0.013 | 0 | 0.11 | 0.11 | 0.23 | 0.33 | 0.15 | 0.17 |
| 5 | 55757723 | rs17656738 | 0.285 | 1.024 | -0.737 | 0.257 | -0.008 | 0.008 | 0.008 | 0 | 0.12 | 0.08 | 0.06 | 0.13 | 0.13 | 0.11 |
| 5 | 55758400 | rs12697147 | 0.963 | 0.854 | -0.695 | 0.345 | -0.032 | -0.078 | -0.078 | 0.157 | 0.18 | 0.31 | 0.1 | 0.02 | 0.11 | 0.09 |
| 5 | 55758956 | rs10471978 | -0.024 | 1.389 | -1.52 | 0.823 | -0.017 | -0.021 | -0.021 | 0.087 | 0.15 | 0.11 | 0.33 | 0.33 | 0.24 | 0.22 |
| 5 | 55761177 | rs3911664 | -0.473 | 0.636 | -1.438 | 0.811 | -0.035 | -0.012 | -0.012 | 0 | 0.55 | 0.77 | 0.52 | 0.58 | 0.48 | 0.58 |
| 5 | 55764861 | rs16885510 | -0.088 | 0.195 | -0.243 | 1.397 | 0.015 | 0.2 | 0.2 | 0 | 0.08 | 0.23 | 0.07 | 0.21 | 0.05 | 0.05 |
| 5 | 55765448 | rs284019 | 0.284 | 0.846 | -0.565 | 1.018 | -0.01 | 0.008 | 0.008 | 0 | 0.19 | 0.2 | 0.43 | 0.1 | 0.37 | 0.29 |
| 5 | 55766005 | rs16885511 | 0.098 | -0.062 | -1.223 | 1.006 | 0.003 | -0.032 | -0.032 | 0 | 0.18 | 0.06 | 0.12 | 0.22 | 0.15 | 0.08 |
| 5 | 55766056 | rs4515248 | -0.286 | -0.243 | -1.216 | 1.239 | 0.015 | -0.065 | -0.065 | 1.756 | 0.21 | 0.09 | 0.13 | 0.38 | 0.17 | 0.11 |
| 5 | 55767865 | rs11959048 | 0.451 | -0.238 | -1.271 | 0.95 | -0.008 | -0.043 | -0.043 | 0.244 | 0.16 | 0.05 | 0.12 | 0.21 | 0.15 | 0.06 |
| 5 | 55767937 | rs176118 | -0.903 | -1.26 | -0.726 | 0.583 | 0.046 | 0.069 | 0.069 | 0 | 0.56 | 0.73 | 0.7 | 0.69 | 0.66 | 0.48 |
| 5 | 55771827 | rs157848 | -0.679 | -1.137 | -0.923 | 0.609 | 0.043 | 0.114 | 0.114 | 0 | 0.5 | 0.73 | 0.65 | 0.69 | 0.58 | 0.45 |
| 5 | 55776404 | rs593979 | -0.351 | -0.997 | -0.849 | 0.29 | -0.026 | 0.115 | 0.115 | 0.053 | 0.63 | 0.84 | 0.44 | 0.89 | 0.44 | 0.51 |
| 5 | 55778051 | rs4374718 | -0.726 | -1.594 | 0.241 | 1.359 | -0.059 | 0.1 | 0.1 | 0 | 0.2 | 0.32 | 0.14 | 0.43 | 0.12 | 0.18 |
| 5 | 55780101 | rs7712360 | -1.445 | -1.499 | 0.225 | 1.406 | 0 | -0.001 | -0.001 | 0 | 0.28 | 0.69 | 0.22 | 0.67 | 0.19 | 0.22 |
| 5 | 55780331 | rs7712385 | -1.112 | -1.363 | 0.365 | 1.519 | 0.005 | 0.176 | 0.176 | 0 | 0.16 | 0.37 | 0.13 | 0.35 | 0.14 | 0.08 |
| 5 | 55782130 | rs12187346 | -0.57 | -1.592 | 0.449 | 1.705 | -0.027 | 0.019 | 0.019 | 0 | 0.17 | 0.2 | 0.11 | 0.28 | 0.09 | 0.15 |
| 5 | 55786198 | rs7730486 | -0.612 | -1.502 | 0.603 | 1.764 | 0.025 | 0.296 | 0.296 | 0 | 0.1 | 0.34 | 0.09 | 0.35 | 0.1 | 0.06 |
| 5 | 55788355 | rs2408361 | -1.068 | -1.591 | 0.241 | 1.412 | -0.076 | 0.114 | 0.114 | 0 | 0.24 | 0.45 | 0.16 | 0.52 | 0.13 | 0.2 |

| | | | | | | | | | | | | | | | | |
|---|---|---|---|---|---|---|---|---|---|---|---|---|---|---|---|---|
| 5 | 55797585 | rs245109 | 0.331 | -1.731 | -0.248 | 1.077 | 0.006 | -0.007 | -0.007 | 0 | 0.21 | 0.19 | 0.14 | 0.3 | 0.15 | 0.17 |
| 5 | 55797753 | rs10940505 | 0.138 | -2.443 | -0.459 | 0.913 | 0.036 | 0.025 | 0.025 | 0 | 0.15 | 0.09 | 0.07 | 0.13 | 0.07 | 0.07 |
| 5 | 55805388 | rs29994 | 0.751 | -2.699 | 0.069 | 1.04 | -0.015 | 0.265 | 0.265 | 0.188 | 0.87 | 0.61 | 0.85 | 0.66 | 0.91 | 0.86 |
| 5 | 55811092 | rs456867 | 0.152 | -2.681 | -0.625 | 0.837 | 0.025 | -0.019 | -0.019 | 0 | 0.27 | 0.21 | 0.17 | 0.34 | 0.2 | 0.27 |
| 5 | 55816081 | rs458036 | 0.707 | -2.249 | -0.893 | 0.221 | 0.146 | 0.188 | 0.188 | 1.474 | 0.51 | 0.17 | 0.64 | 0.22 | 0.72 | 0.62 |
| 5 | 55821031 | rs157521 | 0.297 | -1.619 | -1 | -0.203 | -0.027 | 0.061 | 0.061 | 0 | 0.36 | 0.25 | 0.54 | 0.12 | 0.51 | 0.26 |
| 5 | 55824104 | rs29996 | 0.039 | -1.95 | -0.894 | -0.086 | 0.001 | 0.004 | 0.004 | 0 | 0.39 | 0.59 | 0.38 | 0.76 | 0.36 | 0.57 |
| 5 | 55825904 | rs29995 | 0.98 | -2.211 | -0.307 | 0.719 | -0.027 | 0.1 | 0.1 | 0 | 0.82 | 0.73 | 0.88 | 0.63 | 0.88 | 0.87 |
| 5 | 55827152 | rs10461607 | -0.211 | -2.08 | -0.545 | 0.345 | 0.018 | 0.012 | 0.012 | 0.584 | 0.11 | 0.06 | 0.03 | 0.06 | 0.07 | 0.08 |
| 5 | 55828578 | rs40566 | 0.61 | -2.776 | -0.325 | 0.402 | -0.056 | -0.031 | -0.031 | 0 | 0.63 | 0.5 | 0.86 | 0.8 | 0.81 | 0.81 |
| 5 | 55828785 | rs30365 | 0.679 | -1.363 | -0.595 | 0.83 | 0.02 | 0.108 | 0.108 | 0 | 0.07 | 0.12 | 0.18 | 0.22 | 0.12 | 0.09 |
| 5 | 55829376 | rs30364 | 0.028 | -0.45 | -1.164 | 0.426 | -0.021 | 0.029 | 0.029 | 0 | 0.45 | 0.28 | 0.53 | 0.44 | 0.46 | 0.43 |
| 5 | 55830865 | rs39837 | 0.279 | -0.257 | -1.248 | 0.129 | -0.001 | -0.063 | -0.063 | 0 | 0.28 | 0.16 | 0.29 | 0.37 | 0.29 | 0.36 |
| 5 | 55832217 | rs11949724 | -0.025 | -0.174 | -0.726 | -0.049 | 0.014 | -0.041 | -0.041 | 0 | 0.32 | 0.44 | 0.37 | 0.19 | 0.39 | 0.24 |
| 5 | 55833519 | rs30360 | -0.321 | -0.327 | -0.666 | 0.153 | -0.002 | 0.002 | 0.002 | 0 | 0.65 | 0.65 | 0.71 | 0.61 | 0.68 | 0.66 |
| 5 | 55834230 | rs28130 | 0.012 | -0.261 | -1.004 | 0.09 | 0.011 | -0.002 | -0.002 | 0 | 0.52 | 0.66 | 0.62 | 0.49 | 0.54 | 0.51 |
| 5 | 55841462 | rs30335 | -0.216 | -0.222 | -1.256 | 0.032 | -0.002 | -0.043 | -0.043 | 2.781 | 0.56 | 0.77 | 0.64 | 0.49 | 0.55 | 0.49 |
| 5 | 55847019 | rs2245134 | 0.397 | 0.249 | -1.6 | -0.213 | -0.094 | 0.009 | 0.009 | 0.119 | 0.38 | 0.13 | 0.5 | 0.31 | 0.41 | 0.41 |
| 5 | 55853721 | rs621127 | 0.097 | -0.004 | -0.741 | -0.176 | 0 | 0.001 | 0.001 | 0 | 0.53 | 0.45 | 0.48 | 0.44 | 0.52 | 0.5 |
| 5 | 55854153 | rs6867983 | 0.018 | -0.029 | -0.153 | -0.25 | -0.003 | -0.054 | -0.054 | 0 | 0.15 | 0.24 | 0.25 | 0.07 | 0.15 | 0.15 |
| 5 | 55861359 | rs13173241 | -0.678 | -0.289 | -0.63 | -0.42 | 0.001 | -0.004 | -0.004 | 0 | 0.25 | 0.23 | 0.28 | 0.14 | 0.19 | 0.17 |
| 5 | 55862596 | rs3852144 | -0.083 | -0.082 | -0.871 | -0.377 | -0.006 | 0.008 | 0.008 | 0 | 0.38 | 0.32 | 0.47 | 0.36 | 0.41 | 0.43 |
| 5 | 55868097 | rs13179413 | -0.496 | 1.104 | -1.158 | -0.276 | -0.01 | 0.013 | 0.013 | 0 | 0.24 | 0.22 | 0.27 | 0.13 | 0.29 | 0.24 |
| 5 | 55869009 | rs4700377 | -0.266 | 0.801 | -1.755 | 0.044 | -0.028 | -0.047 | -0.047 | 0 | 0.68 | 0.78 | 0.64 | 0.47 | 0.59 | 0.73 |
| 5 | 55870452 | rs1585795 | 0.226 | 1.404 | -1.988 | 0.154 | -0.072 | -0.053 | -0.053 | 0.439 | 0.72 | 0.88 | 0.7 | 0.59 | 0.57 | 0.81 |
| 5 | 55874173 | rs4406105 | 0.465 | 1.142 | -2.007 | 0.033 | -0.034 | 0.051 | 0.051 | 0 | 0.12 | 0.03 | 0.06 | 0.04 | 0.12 | 0.06 |
| 5 | 55874809 | rs16885750 | 0.75 | 0.841 | -1.728 | 0.077 | -0.005 | -0.002 | -0.002 | 0 | 0.15 | 0.24 | 0.12 | 0.16 | 0.1 | 0.1 |
| 5 | 55875713 | rs6450394 | 1.135 | 0.667 | -1.819 | -0.118 | 0.002 | -0.003 | -0.003 | 0 | 0.35 | 0.25 | 0.33 | 0.35 | 0.31 | 0.52 |
| 5 | 55876283 | rs11948950 | 0.867 | 1.014 | -2.188 | 0.161 | -0.027 | -0.012 | -0.012 | 0 | 0.1 | 0.01 | 0.18 | 0.09 | 0.11 | 0.11 |
| 5 | 55882179 | rs6895773 | 0.118 | 0.804 | -1.797 | 0.18 | -0.012 | -0.024 | -0.024 | 0 | 0.18 | 0.3 | 0.1 | 0.1 | 0.11 | 0.09 |
| 5 | 55882435 | rs4700382 | -0.005 | 1.73 | -2.229 | 0.712 | -0.038 | -0.046 | -0.046 | 0.382 | 0.83 | 0.9 | 0.77 | 0.67 | 0.74 | 0.87 |
| 5 | 55883187 | rs13155854 | -0.172 | 1.23 | -2.67 | 0.219 | -0.095 | 0.001 | 0.001 | 0.036 | 0.42 | 0.11 | 0.52 | 0.36 | 0.5 | 0.47 |
| 5 | 55889598 | rs6891726 | -0.211 | 0.694 | -2.38 | -0.271 | 0.015 | 0.173 | 0.173 | 0 | 0.53 | 0.76 | 0.34 | 0.16 | 0.52 | 0.46 |
| 5 | 55892384 | rs7712875 | 0.13 | 0.939 | -2.437 | -0.729 | 0 | 0.062 | 0.062 | 0 | 0.36 | 0.7 | 0.21 | 0.03 | 0.31 | 0.27 |
| 5 | 55892458 | rs7727229 | 1.163 | 0.885 | -1.526 | -0.936 | 0.038 | -0.125 | -0.125 | 2.804 | 0.2 | 0.33 | 0.14 | 0.01 | 0.21 | 0.07 |
| 5 | 55896630 | rs10040902 | 0.72 | 0.996 | -1.489 | -0.783 | 0 | -0.015 | -0.015 | 0 | 0.09 | 0.12 | 0.04 | 0 | 0.08 | 0.05 |
| 5 | 55897451 | rs13178221 | 1.07 | 1.009 | -1.438 | -0.731 | -0.033 | 0.093 | 0.093 | 0 | 0.23 | 0.09 | 0.13 | 0.03 | 0.23 | 0.21 |
| 5 | 55903787 | rs16885817 | 1.099 | 0.675 | -1.62 | -0.54 | -0.011 | 0.008 | 0.008 | 0 | 0.12 | 0.01 | 0.25 | 0.07 | 0.12 | 0.16 |
| 5 | 55904947 | rs17716231 | 0.552 | 0.778 | -1.529 | -0.429 | -0.056 | -0.017 | -0.017 | 0 | 0.17 | 0.03 | 0.13 | 0.13 | 0.2 | 0.21 |
| 5 | 55910300 | rs2134206 | 0.427 | 0.569 | -1.002 | 0.45 | 0.004 | 0.053 | 0.053 | 0.073 | 0.71 | 0.64 | 0.82 | 0.53 | 0.7 | 0.71 |
| 5 | 55913197 | rs1290217 | 0.455 | 0.859 | -1.039 | 0.719 | 0.013 | 0.024 | 0.024 | 0 | 0.06 | 0.08 | 0.15 | 0.15 | 0.09 | 0.06 |
| 5 | 55913552 | rs1296959 | -0.968 | 0.636 | -1.153 | 0.566 | -0.039 | 0.053 | 0.053 | 0.155 | 0.71 | 0.55 | 0.72 | 0.39 | 0.74 | 0.7 |
| 5 | 55917956 | rs2448425 | 0.351 | 0.794 | -1.116 | -0.07 | -0.006 | 0.016 | 0.016 | 0 | 0.44 | 0.38 | 0.55 | 0.25 | 0.45 | 0.43 |
| 5 | 55918201 | rs2448424 | 0.324 | 0.518 | -1.074 | -0.062 | 0.006 | 0.03 | 0.03 | 0 | 0.46 | 0.37 | 0.56 | 0.25 | 0.53 | 0.45 |
| 5 | 55918787 | rs1910023 | 0.546 | 0.489 | -1.123 | 0.317 | -0.004 | -0.012 | -0.012 | 0 | 0.07 | 0.1 | 0.02 | 0 | 0.02 | 0.03 |
| 5 | 55920691 | rs2455377 | -0.461 | 0.844 | -1.217 | 0.416 | -0.001 | 0.025 | 0.025 | 0.049 | 0.43 | 0.67 | 0.59 | 0.35 | 0.42 | 0.45 |
| 5 | 55927125 | rs10043543 | 1.049 | -0.12 | -1.819 | 0.341 | -0.001 | -0.046 | -0.046 | 0.004 | 0.16 | 0.01 | 0.12 | 0.18 | 0.14 | 0.19 |
| 5 | 55927765 | rs1479607 | -0.341 | 0.309 | -1.414 | 0.072 | 0 | 0.001 | 0.001 | 0.01 | 0.47 | 0.5 | 0.55 | 0.23 | 0.48 | 0.52 |
| 5 | 55929388 | rs10940509 | -0.224 | 0.329 | -1.639 | 0.51 | -0.067 | -0.013 | -0.013 | 0.009 | 0.17 | 0.02 | 0.17 | 0.11 | 0.21 | 0.25 |
| 5 | 55930580 | rs1279627 | 0.299 | 0.257 | -0.989 | 0.664 | -0.01 | -0.004 | -0.004 | 0 | 0.56 | 0.46 | 0.63 | 0.36 | 0.64 | 0.51 |
| 5 | 55943160 | rs6874062 | 0.402 | 0.877 | -0.679 | 0.866 | 0.021 | 0.016 | 0.016 | 0 | 0.64 | 0.49 | 0.6 | 0.57 | 0.59 | 0.59 |
| 5 | 55948898 | rs1465447 | 0.262 | 0.642 | -0.49 | 0.816 | -0.069 | -0.065 | -0.065 | 0 | 0.68 | 0.84 | 0.63 | 0.56 | 0.58 | 0.6 |
| 5 | 55951138 | rs13167060 | 0.768 | 0.152 | -0.873 | 0.468 | -0.002 | 0.003 | 0.003 | 0 | 0.34 | 0.32 | 0.29 | 0.34 | 0.34 | 0.33 |
| 5 | 55955224 | rs2408543 | 0.768 | 0.152 | -0.866 | 0.468 | -0.002 | 0.004 | 0.004 | 0 | 0.34 | 0.31 | 0.29 | 0.35 | 0.34 | 0.33 |
| 5 | 55962593 | rs11742315 | 0.811 | 0.029 | -0.747 | 0.361 | 0.039 | 0.052 | 0.052 | 0 | 0.27 | 0.06 | 0.28 | 0.14 | 0.18 | 0.24 |
| 5 | 55965167 | rs889310 | -0.544 | 0.963 | -0.437 | 0.671 | 0.003 | 0.004 | 0.004 | 0 | 0.61 | 0.61 | 0.56 | 0.61 | 0.58 | 0.69 |
| 5 | 55968018 | rs1991007 | -0.311 | 1.543 | -0.553 | 0.711 | -0.01 | 0.004 | 0.004 | 0 | 0.91 | 0.92 | 0.81 | 0.93 | 0.84 | 0.92 |
| 5 | 55970422 | rs10428696 | 0.443 | 1.606 | -0.544 | 0.863 | 0.008 | 0.04 | 0.04 | 0 | 0.21 | 0.31 | 0.22 | 0.44 | 0.25 | 0.34 |
| 5 | 55970973 | rs1292144 | -0.474 | 1.251 | -0.275 | 0.602 | 0.004 | 0.066 | 0.066 | 0.535 | 0.52 | 0.7 | 0.57 | 0.28 | 0.48 | 0.47 |
| 5 | 55977624 | rs7711726 | -0.087 | 0.941 | -0.066 | 0.547 | 0.023 | 0.022 | 0.022 | 0.148 | 0.43 | 0.58 | 0.37 | 0.26 | 0.3 | 0.39 |
| 5 | 55978051 | rs16886018 | 1.058 | 1.014 | -0.112 | 0.574 | -0.011 | 0.017 | 0.017 | 0 | 0.06 | 0.02 | 0.03 | 0.01 | 0.07 | 0.09 |
| 5 | 55978114 | rs10054678 | 1.672 | 1.041 | -0.339 | 1.046 | -0.068 | -0.115 | -0.115 | 4.193 | 0.24 | 0.02 | 0.18 | 0.34 | 0.25 | 0.26 |
| 5 | 55980650 | rs10060816 | -0.33 | 0.529 | 0.254 | 1.033 | 0.007 | 0.007 | 0.007 | 0.12 | 0.42 | 0.56 | 0.38 | 0.26 | 0.3 | 0.4 |
| 5 | 55981589 | rs13185691 | 0.064 | 0.559 | 0.009 | 1.08 | 0.056 | 0.051 | 0.051 | 0.607 | 0.34 | 0.19 | 0.31 | 0.17 | 0.23 | 0.26 |
| 5 | 55982643 | rs1292150 | -1.119 | 1.321 | 0.897 | 1.767 | 0.055 | 0.037 | 0.037 | 0 | 0.15 | 0.32 | 0.25 | 0.18 | 0.21 | 0.11 |
| 5 | 55990143 | rs13163065 | 0.615 | 0.56 | 0.109 | 1.417 | -0.001 | 0.004 | 0.004 | 0.261 | 0.59 | 0.41 | 0.61 | 0.34 | 0.57 | 0.48 |
| 5 | 55990436 | rs10461611 | 0.725 | 0.333 | 0.238 | 1.381 | 0 | 0.001 | 0.001 | 0.197 | 0.62 | 0.42 | 0.63 | 0.32 | 0.65 | 0.5 |
| 5 | 55994947 | rs6897035 | -0.239 | 0.5 | 0.816 | 1.928 | 0.034 | 0.173 | 0.173 | 0 | 0.05 | 0.18 | 0.04 | 0.23 | 0.04 | 0.22 |
| 5 | 55995035 | rs16886113 | -0.129 | 0.367 | 0.785 | 1.922 | -0.005 | 0.015 | 0.015 | 0 | 0.06 | 0.12 | 0.07 | 0.14 | 0.06 | 0.09 |

| | | | | | | | | | | | | | | | | |
|---|---|---|---|---|---|---|---|---|---|---|---|---|---|---|---|---|
| 5 | 55996012 | rs1318111 | 0.658 | 0.057 | 0.906 | 1.713 | -0.002 | -0.003 | -0.003 | 0 | 0.59 | 0.61 | 0.63 | 0.32 | 0.63 | 0.48 |
| 5 | 55998242 | rs6450398 | 0.656 | 0.09 | 1.138 | 1.715 | 0.004 | 0.012 | 0.012 | 0.29 | 0.59 | 0.68 | 0.63 | 0.32 | 0.62 | 0.47 |
| 5 | 56001363 | rs6450399 | 0.808 | -0.044 | 1.098 | 1.717 | 0.002 | 0.075 | 0.075 | 0 | 0.7 | 0.49 | 0.6 | 0.42 | 0.66 | 0.73 |
| 5 | 56009012 | rs6862199 | 1.624 | -0.073 | 1.152 | 1.717 | 0.011 | 0.105 | 0.105 | 0.241 | 0.52 | 0.3 | 0.53 | 0.2 | 0.56 | 0.42 |
| 5 | 56009256 | rs11747611 | -0.066 | -0.102 | 1.142 | 1.956 | -0.002 | 0.008 | 0.008 | 0.87 | 0.09 | 0.12 | 0.14 | 0.26 | 0.15 | 0.18 |
| 5 | 56009384 | rs10051671 | -0.243 | -0.209 | 1.131 | 1.794 | -0.007 | 0.003 | 0.003 | 0 | 0.85 | 0.8 | 0.81 | 0.81 | 0.86 | 0.94 |
| 5 | 56022907 | rs16886164 | -0.364 | -0.283 | 1.296 | 1.742 | 0.056 | 0.101 | 0.101 | 0.206 | 0.07 | 0.35 | 0.09 | 0.12 | 0.05 | 0.03 |
| 5 | 56023083 | rs16886165 | -0.626 | -0.355 | 1.184 | 1.799 | -0.019 | 0.033 | 0.033 | 0.558 | 0.23 | 0.36 | 0.15 | 0.34 | 0.15 | 0.35 |
| 5 | 56026843 | rs12656399 | 0.472 | -0.353 | 1.097 | 1.815 | -0.012 | -0.076 | -0.076 | 0.056 | 0.17 | 0 | 0.06 | 0.23 | 0.1 | 0.31 |
| 5 | 56031884 | rs889312 | 1.273 | -0.389 | 1.096 | 1.778 | 0.009 | -0.015 | -0.015 | 0 | 0.62 | 0.66 | 0.59 | 0.47 | 0.72 | 0.61 |
| 5 | 56032940 | rs1862626 | 1.283 | -0.392 | 1.107 | 1.771 | 0.01 | -0.017 | -0.017 | 0 | 0.62 | 0.66 | 0.59 | 0.46 | 0.71 | 0.61 |
| 5 | 56034207 | rs4700485 | 1.049 | -0.391 | 0.976 | 1.772 | 0.02 | -0.038 | -0.038 | 0 | 0.63 | 0.76 | 0.6 | 0.46 | 0.72 | 0.61 |
| 5 | 56040949 | rs961848 | 0.669 | -0.455 | 0.835 | 1.89 | 0.019 | -0.033 | -0.033 | 0.205 | 0.68 | 0.84 | 0.66 | 0.57 | 0.76 | 0.64 |
| 5 | 56041064 | rs961847 | 1.375 | -0.459 | 1.013 | 1.865 | -0.007 | 0.007 | 0.007 | 0 | 0.61 | 0.47 | 0.57 | 0.46 | 0.71 | 0.61 |
| 5 | 56042288 | rs4312834 | -0.389 | -0.454 | 1.007 | 1.871 | -0.003 | 0 | 0 | 0.517 | 0.85 | 0.86 | 0.82 | 0.81 | 0.87 | 0.94 |
| 5 | 56044186 | rs4700008 | 1.772 | -0.374 | 1.036 | 1.635 | 0.011 | 0.044 | 0.044 | 0.83 | 0.52 | 0.38 | 0.44 | 0.2 | 0.57 | 0.41 |
| 5 | 56044635 | rs7708148 | 1.277 | -0.419 | 0.948 | 1.653 | 0.011 | 0.033 | 0.033 | 0.024 | 0.59 | 0.76 | 0.53 | 0.31 | 0.62 | 0.44 |
| 5 | 56046352 | rs2897972 | 0.641 | -0.467 | 0.908 | 1.672 | -0.004 | -0.079 | -0.079 | 0 | 0.16 | 0 | 0.14 | 0.23 | 0.09 | 0.3 |
| 5 | 56047451 | rs7709874 | 0.642 | -0.433 | 0.982 | 1.653 | 0 | -0.001 | -0.001 | 0 | 0.76 | 0.77 | 0.68 | 0.54 | 0.72 | 0.74 |
| 5 | 56061564 | rs16886256 | -0.045 | -0.536 | 1.069 | 1.594 | -0.042 | 0.058 | 0.058 | 0.039 | 0.2 | 0.48 | 0.18 | 0.27 | 0.1 | 0.3 |
| 5 | 56062821 | rs16886257 | 0.671 | -0.533 | 0.904 | 1.564 | 0.002 | -0.088 | -0.088 | 0 | 0.16 | 0 | 0.15 | 0.27 | 0.09 | 0.3 |
| 5 | 56064133 | rs12657615 | 0.671 | -0.533 | 0.904 | 1.564 | 0.002 | -0.088 | -0.088 | 0 | 0.16 | 0 | 0.14 | 0.27 | 0.08 | 0.3 |
| 5 | 56069600 | rs6874733 | 0.629 | -0.516 | 1.003 | 1.455 | 0 | -0.007 | -0.007 | 0 | 0.76 | 0.79 | 0.61 | 0.58 | 0.71 | 0.77 |
| 5 | 56100006 | rs6893754 | 1.403 | -0.635 | 0.996 | 1.454 | 0.059 | 0.073 | 0.073 | 0 | 0.59 | 0.25 | 0.47 | 0.25 | 0.71 | 0.49 |
| 5 | 56100996 | rs1423620 | 0.894 | -0.634 | 1.259 | 1.451 | -0.03 | 0.061 | 0.061 | 0.402 | 0.75 | 0.43 | 0.63 | 0.57 | 0.81 | 0.78 |
| 5 | 56109154 | rs11960484 | 1.68 | -0.61 | 0.995 | 1.298 | 0.06 | 0.206 | 0.206 | 0.074 | 0.54 | 0.21 | 0.4 | 0.14 | 0.64 | 0.43 |
| 5 | 56119190 | rs252906 | 1.745 | -0.73 | 0.958 | 1.243 | 0.067 | 0.195 | 0.195 | 0.09 | 0.53 | 0.24 | 0.41 | 0.13 | 0.64 | 0.43 |
| 5 | 56127866 | rs726501 | 0.856 | -0.755 | 1.021 | 1.332 | -0.016 | -0.097 | -0.097 | 0 | 0.16 | 0 | 0.14 | 0.3 | 0.09 | 0.3 |
| 5 | 56142999 | rs16886420 | 0.699 | -0.751 | 1.051 | 1.346 | -0.003 | -0.007 | -0.007 | 0.107 | 0.05 | 0.03 | 0.06 | 0.11 | 0.07 | 0.06 |
| 5 | 56149429 | rs860580 | 0.087 | -1.185 | 0.673 | 0.725 | -0.032 | 0.091 | 0.091 | 0 | 0.77 | 0.65 | 0.65 | 0.57 | 0.81 | 0.78 |
| 5 | 56159578 | rs832573 | 0.86 | -1.183 | 0.728 | 0.715 | -0.013 | -0.016 | -0.016 | 1.239 | 0.59 | 0.49 | 0.5 | 0.25 | 0.71 | 0.48 |
| 5 | 56161787 | rs832575 | -0.933 | -1.228 | 0.79 | 1.048 | -0.015 | 0.026 | 0.026 | 0 | 0.84 | 0.77 | 0.72 | 0.72 | 0.87 | 0.94 |
| 5 | 56163787 | rs832577 | 1.534 | -1.242 | 0.816 | 1.038 | 0.068 | 0.22 | 0.22 | 0.175 | 0.52 | 0.21 | 0.4 | 0.13 | 0.64 | 0.43 |
| 5 | 56166028 | rs16886445 | 0.614 | -1.244 | 0.821 | 1.055 | 0.007 | -0.017 | -0.017 | 3.126 | 0.07 | 0.04 | 0.07 | 0.15 | 0.06 | 0.15 |
| 5 | 56177443 | rs702689 | 0.957 | -1.223 | 0.831 | 0.946 | 0 | 0 | 0 | 0 | 0.59 | 0.46 | 0.49 | 0.24 | 0.71 | 0.48 |
| 5 | 56177743 | rs832582 | 0.121 | -1.213 | 0.776 | 0.952 | -0.013 | 0.032 | 0.032 | 0 | 0.77 | 0.71 | 0.65 | 0.57 | 0.81 | 0.78 |
| 5 | 56178111 | rs3822625 | 0.606 | -1.213 | 0.79 | 0.952 | 0 | 0.009 | 0.009 | 0 | 0.06 | 0.12 | 0.08 | 0.11 | 0.08 | 0.06 |
| 5 | 56178217 | rs832583 | 0.146 | -1.18 | 0.78 | 0.945 | -0.013 | 0.032 | 0.032 | 0 | 0.77 | 0.71 | 0.65 | 0.57 | 0.81 | 0.78 |
| 5 | 56180480 | rs3736430 | 1.097 | -1.001 | 0.721 | 1.045 | 0.003 | -0.009 | -0.009 | 0 | 0.16 | 0.07 | 0.15 | 0.33 | 0.09 | 0.3 |
| 5 | 56196604 | rs96844 | 0.71 | -1.204 | -0.073 | 0.76 | 0.039 | 0.034 | 0.034 | 0 | 0.64 | 0.3 | 0.5 | 0.27 | 0.74 | 0.53 |
| 5 | 56199202 | rs832540 | 1.476 | -1.199 | -0.216 | 0.718 | 0.063 | 0.258 | 0.258 | 0.178 | 0.52 | 0.16 | 0.39 | 0.14 | 0.63 | 0.42 |
| 5 | 56203773 | rs252925 | 1.411 | -1.198 | -0.211 | 0.719 | 0.061 | 0.238 | 0.238 | 0.104 | 0.52 | 0.21 | 0.39 | 0.14 | 0.63 | 0.42 |
| 5 | 56205357 | rs185220 | -0.183 | -1.205 | 0.065 | 0.939 | -0.021 | 0.144 | 0.144 | 0.187 | 0.83 | 0.7 | 0.68 | 0.6 | 0.84 | 0.8 |
| 5 | 56206529 | rs33320 | 0.657 | -0.975 | 0.18 | 0.874 | 0.003 | 0.003 | 0.003 | 0.041 | 0.64 | 0.42 | 0.5 | 0.26 | 0.74 | 0.53 |
| 5 | 56209735 | rs40497 | 2.291 | -0.905 | -0.074 | 0.69 | -0.111 | 0.203 | 0.203 | 0 | 0.3 | 0.08 | 0.18 | 0.03 | 0.34 | 0.12 |
| 5 | 56215753 | rs12654125 | 0.317 | -0.901 | 0.299 | 1.062 | -0.001 | 0 | 0 | 0 | 0.11 | 0.09 | 0.09 | 0.13 | 0.11 | 0.1 |
| 5 | 56217529 | rs16886496 | 0.423 | -0.943 | 0.31 | 1.073 | -0.001 | 0 | 0 | 0 | 0.11 | 0.09 | 0.09 | 0.13 | 0.11 | 0.1 |
| 5 | 56220999 | rs12657064 | 0.395 | -0.945 | 0.309 | 1.072 | -0.001 | 0 | 0 | 0 | 0.11 | 0.09 | 0.09 | 0.13 | 0.11 | 0.11 |
| 5 | 56222120 | rs11957276 | 1.53 | -1.042 | 0.172 | 0.988 | -0.067 | 0.051 | 0.051 | 0 | 0.14 | 0.03 | 0.16 | 0.05 | 0.22 | 0.2 |
| 5 | 56222739 | rs6882657 | -0.049 | -1.773 | 0.915 | 1.673 | -0.001 | 0.001 | 0.001 | 0 | 0.11 | 0.12 | 0.09 | 0.13 | 0.11 | 0.1 |
| 5 | 56224124 | rs252895 | -0.671 | -1.805 | 1.045 | 1.726 | -0.011 | 0.098 | 0.098 | 0 | 0.88 | 0.74 | 0.75 | 0.72 | 0.9 | 0.93 |
| 5 | 56224339 | rs252894 | 2.006 | -1.821 | 0.854 | 1.58 | 0.056 | 0.289 | 0.289 | 0.057 | 0.5 | 0.12 | 0.39 | 0.14 | 0.63 | 0.42 |
| 5 | 56226211 | rs12655082 | 0.487 | -1.777 | 0.898 | 1.636 | -0.026 | 0.052 | 0.052 | 0 | 0.18 | 0.29 | 0.17 | 0.33 | 0.08 | 0.22 |
| 5 | 56233541 | rs2591968 | -1.559 | -1.528 | 0.897 | 1.718 | -0.025 | 0.068 | 0.068 | 0 | 0.28 | 0.39 | 0.41 | 0.53 | 0.27 | 0.36 |
| 5 | 56251660 | rs331497 | -2.048 | -1.493 | 0.553 | 1.314 | 0.045 | 0.124 | 0.124 | 0.077 | 0.48 | 0.65 | 0.59 | 0.86 | 0.38 | 0.59 |
| 5 | 56254485 | rs2662027 | 0.243 | -1.406 | 0.684 | 1.619 | -0.004 | 0.095 | 0.095 | 0 | 0.12 | 0.24 | 0.25 | 0.28 | 0.11 | 0.07 |
| 5 | 56259853 | rs978405 | 0.169 | -1.126 | -0.014 | 1.561 | 0.006 | 0.005 | 0.005 | 0 | 0.92 | 0.85 | 0.94 | 0.9 | 0.9 | 0.96 |
| 5 | 56260606 | rs6890270 | -0.775 | -1.407 | -0.434 | 1.633 | -0.006 | -0.147 | -0.147 | 0 | 0.28 | 0.03 | 0.24 | 0.45 | 0.2 | 0.31 |
| 5 | 56260998 | rs6895442 | 1.84 | -1.422 | -0.187 | 1.544 | 0.032 | 0.038 | 0.038 | 0.533 | 0.58 | 0.28 | 0.5 | 0.28 | 0.67 | 0.42 |
| 5 | 56265168 | rs832402 | -1.615 | -0.998 | -0.141 | 1.78 | -0.011 | 0.042 | 0.042 | 0.44 | 0.23 | 0.63 | 0.34 | 0.42 | 0.21 | 0.33 |
| 5 | 56265992 | rs832403 | -0.523 | -0.985 | 0.022 | 1.786 | -0.001 | 0.108 | 0.108 | 0 | 0.15 | 0.26 | 0.25 | 0.37 | 0.17 | 0.29 |
| 5 | 56267431 | rs832404 | -0.676 | -0.967 | -0.005 | 1.71 | -0.037 | -0.084 | -0.084 | 0 | 0.84 | 0.99 | 0.85 | 0.7 | 0.88 | 0.74 |
| 5 | 56268205 | rs1445998 | 1.1 | -0.852 | 0.157 | 1.089 | 0.007 | 0.007 | 0.007 | 0.608 | 0.62 | 0.36 | 0.51 | 0.28 | 0.67 | 0.42 |
| 5 | 56271822 | rs7703149 | 0.393 | -0.385 | -0.177 | 0.988 | 0.009 | 0.062 | 0.062 | 0 | 0.9 | 0.77 | 0.81 | 0.8 | 0.9 | 0.93 |
| 5 | 56279030 | rs6870905 | -0.761 | -0.758 | -0.736 | 0.311 | 0.017 | 0.011 | 0.011 | 1.06 | 0.16 | 0.09 | 0.09 | 0 | 0.11 | 0.02 |
| 5 | 56280245 | rs1876588 | 0.177 | -0.493 | -0.859 | 0.419 | -0.017 | -0.005 | -0.005 | 0 | 0.63 | 0.67 | 0.44 | 0.68 | 0.52 | 0.7 |
| 5 | 56282546 | rs2408654 | -0.373 | -1.181 | -0.681 | 0.235 | -0.032 | 0.034 | 0.034 | 0.169 | 0.35 | 0.45 | 0.2 | 0.5 | 0.24 | 0.59 |
| 5 | 56282580 | rs4624740 | -0.477 | -1.45 | -0.759 | 0.66 | -0.066 | 0.046 | 0.046 | 0.63 | 0.2 | 0.45 | 0.12 | 0.22 | 0.1 | 0.33 |
| 5 | 56283551 | rs16886587 | 1.164 | -1.381 | -0.948 | 0.784 | 0.012 | -0.02 | -0.02 | 0 | 0.05 | 0.03 | 0.02 | 0.16 | 0.03 | 0.21 |

| | | | | | | | | | | | | | | | | |
|---|---|---|---|---|---|---|---|---|---|---|---|---|---|---|---|---|
| 5 | 56284925 | rs6890927 | 0.732 | -1.268 | -0.794 | 0.655 | 0.002 | -0.015 | -0.015 | 0 | 0.12 | 0.09 | 0.05 | 0 | 0.07 | 0.01 |
| 5 | 56285684 | rs9292128 | 0.245 | -1.79 | -1.008 | 0.713 | 0.022 | -0.029 | -0.029 | 0 | 0.2 | 0.13 | 0.15 | 0.32 | 0.13 | 0.31 |
| 5 | 56285987 | rs4505906 | 1.725 | -1.696 | -1.11 | 0.81 | -0.022 | -0.068 | -0.068 | 0 | 0.12 | 0.01 | 0.07 | 0.27 | 0.11 | 0.2 |
| 5 | 56286209 | rs4309923 | -1.142 | -1.706 | -1.138 | 0.722 | -0.092 | -0.011 | -0.011 | 0 | 0.78 | 0.93 | 0.66 | 0.86 | 0.61 | 0.8 |
| 5 | 56288810 | rs10940522 | 1.348 | -1.629 | -0.085 | 0.682 | -0.005 | 0.042 | 0.042 | 0 | 0.13 | 0.25 | 0.09 | 0.27 | 0.11 | 0.2 |
| 5 | 56289473 | rs6863639 | 0.662 | -1.707 | 0.04 | 0.686 | 0 | -0.001 | -0.001 | 0 | 0.44 | 0.59 | 0.44 | 0.53 | 0.41 | 0.48 |
| 5 | 56296304 | rs12518339 | 0.513 | -1.702 | 0.018 | 0.671 | 0.004 | 0.006 | 0.006 | 0.042 | 0.51 | 0.57 | 0.47 | 0.51 | 0.46 | 0.5 |
| 5 | 56301089 | rs17732457 | 1.801 | -1.822 | -0.16 | 0.381 | -0.056 | 0.127 | 0.127 | 0.236 | 0.26 | 0.04 | 0.17 | 0.05 | 0.28 | 0.14 |
| 5 | 56301267 | rs17732494 | 0.537 | -1.241 | 0.376 | 1.046 | -0.005 | -0.009 | -0.009 | 0.352 | 0.56 | 0.68 | 0.55 | 0.57 | 0.46 | 0.5 |
| 5 | 56304373 | rs4700657 | 0.789 | -1.067 | 0.708 | 1.376 | -0.008 | 0.066 | 0.066 | 0 | 0.12 | 0.25 | 0.09 | 0.26 | 0.1 | 0.2 |
| 5 | 56307959 | rs16886638 | 0.873 | -1.06 | 0.189 | 1.379 | 0 | 0 | 0 | 0 | 0.12 | 0.12 | 0.09 | 0.26 | 0.1 | 0.2 |
| 5 | 56320577 | rs2063224 | 0.055 | -0.834 | -0.017 | 1.136 | 0.004 | -0.002 | -0.002 | 2.088 | 0.2 | 0.22 | 0.12 | 0.26 | 0.18 | 0.3 |
| 5 | 56324374 | rs941110 | -1.101 | -0.416 | -0.395 | 1.181 | 0.017 | 0.011 | 0.011 | 0 | 0.46 | 0.66 | 0.42 | 0.58 | 0.4 | 0.66 |
| 5 | 56324912 | rs941109 | -0.786 | -0.203 | -0.412 | 1.269 | -0.004 | 0.013 | 0.013 | 0.361 | 0.32 | 0.48 | 0.33 | 0.35 | 0.3 | 0.49 |
| 5 | 56326423 | rs6898062 | 0.126 | -0.201 | -0.322 | 1.082 | -0.025 | -0.02 | -0.02 | 0.192 | 0.21 | 0.45 | 0.18 | 0.21 | 0.16 | 0.4 |
| 5 | 56327919 | rs16886650 | 0.343 | -0.176 | -0.36 | 1.07 | 0.003 | 0.002 | 0.002 | 1.033 | 0.19 | 0.19 | 0.17 | 0.21 | 0.16 | 0.4 |
| 5 | 56330143 | rs10070167 | -0.408 | -0.049 | -0.683 | 1.093 | -0.013 | -0.006 | -0.006 | 0.568 | 0.29 | 0.48 | 0.23 | 0.28 | 0.24 | 0.45 |
| 5 | 56333946 | rs12659958 | -0.318 | 0.579 | -1.622 | 0.764 | 0.007 | 0.082 | 0.082 | 0 | 0.55 | 0.8 | 0.45 | 0.73 | 0.41 | 0.69 |
| 5 | 56334695 | rs6878762 | 0.624 | 1.441 | -0.875 | 1.316 | 0.025 | 0.042 | 0.042 | 0.021 | 0.71 | 0.46 | 0.48 | 0.61 | 0.59 | 0.7 |
| 5 | 56336311 | rs831774 | -0.799 | 1.943 | -1.452 | 1.033 | -0.022 | 0.004 | 0.004 | 0.229 | 0.2 | 0.11 | 0.29 | 0.22 | 0.26 | 0.29 |
| 5 | 56338395 | rs6450414 | 0.3 | 2.016 | -2.239 | 1.252 | -0.139 | -0.083 | -0.083 | 0.352 | 0.73 | 0.95 | 0.71 | 0.74 | 0.58 | 0.71 |
| 5 | 56338538 | rs10461619 | -0.234 | 0.629 | -1.79 | 1.627 | 0 | 0.051 | 0.051 | 0 | 0.1 | 0.05 | 0.26 | 0.47 | 0.08 | 0.14 |
| 5 | 56343247 | rs16886678 | -0.861 | 0.213 | -1.271 | 0.748 | -0.024 | 0.075 | 0.075 | 0 | 0.3 | 0.49 | 0.38 | 0.48 | 0.23 | 0.36 |
| 5 | 56343864 | rs331035 | -0.333 | 0.121 | -1.69 | -0.524 | 0.025 | 0.044 | 0.044 | 0 | 0.2 | 0.1 | 0.19 | 0.09 | 0.17 | 0.16 |
| 5 | 56344981 | rs4358486 | -0.172 | 0.21 | -1.824 | -0.428 | 0.012 | 0.045 | 0.045 | 0 | 0.17 | 0.09 | 0.15 | 0.03 | 0.15 | 0.14 |
| 5 | 56346338 | rs331031 | -0.586 | 0.354 | -1.395 | -0.151 | 0.022 | 0.034 | 0.034 | 0 | 0.2 | 0.12 | 0.09 | 0.09 | 0.16 | 0.17 |
| 5 | 56347075 | rs831776 | -0.713 | 1.062 | -1.312 | 0.073 | -0.03 | -0.005 | -0.005 | 0 | 0.06 | 0.01 | 0.12 | 0.07 | 0.1 | 0.12 |
| 5 | 56351240 | rs831796 | -0.057 | 0.902 | -1.135 | -0.741 | 0.018 | 0.074 | 0.074 | 0 | 0.5 | 0.4 | 0.5 | 0.18 | 0.55 | 0.5 |
| 5 | 56368573 | rs10059300 | 0.561 | 0.4 | -1.004 | -1.167 | -0.001 | -0.002 | -0.002 | 0 | 0.25 | 0.29 | 0.13 | 0.11 | 0.24 | 0.21 |
| 5 | 56371406 | rs331040 | -0.406 | 0.362 | -0.914 | -1.167 | -0.036 | 0.013 | 0.013 | 0 | 0.75 | 0.63 | 0.64 | 0.3 | 0.8 | 0.71 |
| 5 | 56373363 | rs7711479 | 0.98 | 0.519 | -0.964 | -1.328 | -0.01 | 0.041 | 0.041 | 0 | 0.42 | 0.33 | 0.39 | 0.1 | 0.45 | 0.34 |
| 5 | 56375477 | rs831824 | -0.512 | 0.403 | -0.623 | -1.166 | -0.017 | 0.005 | 0.005 | 0 | 0.75 | 0.74 | 0.57 | 0.3 | 0.8 | 0.71 |
| 5 | 56376026 | rs6450416 | 0.713 | 0.487 | -0.416 | -1.231 | -0.025 | 0.144 | 0.144 | 0 | 0.45 | 0.23 | 0.39 | 0.11 | 0.48 | 0.4 |
| 5 | 56377971 | rs11748741 | 0.35 | 0.412 | -0.216 | -0.52 | -0.014 | -0.033 | -0.033 | 0 | 0.7 | 0.62 | 0.53 | 0.21 | 0.73 | 0.61 |
| 5 | 56382328 | rs331047 | 0.081 | 0.351 | -0.429 | -0.225 | -0.005 | 0.003 | 0.003 | 0 | 0.75 | 0.76 | 0.63 | 0.29 | 0.79 | 0.69 |
| 5 | 56390346 | rs831831 | 0.08 | 0.372 | -0.45 | -0.332 | -0.008 | 0 | 0 | 0 | 0.76 | 0.75 | 0.64 | 0.27 | 0.79 | 0.71 |
| 5 | 56409289 | rs2174999 | 0.211 | 0.806 | -1.132 | -0.963 | -0.036 | 0.077 | 0.077 | 0 | 0.23 | 0.12 | 0.18 | 0 | 0.26 | 0.12 |
| 5 | 56412370 | rs831847 | -0.06 | 0.548 | -0.639 | -1.106 | -0.025 | -0.025 | -0.025 | 1.142 | 0.72 | 0.6 | 0.55 | 0.22 | 0.76 | 0.58 |
| 5 | 56415918 | rs831852 | 0.806 | 0.511 | -0.589 | -1.06 | -0.002 | 0.026 | 0.026 | 0 | 0.47 | 0.42 | 0.36 | 0.11 | 0.49 | 0.45 |
| 5 | 56457598 | rs7712889 | 0.749 | 0.1 | -0.249 | -1.073 | -0.003 | 0.067 | 0.067 | 0 | 0.44 | 0.34 | 0.32 | 0.09 | 0.45 | 0.44 |
| 5 | 56472530 | rs831648 | 1.251 | 0.219 | -0.706 | -1.593 | -0.058 | 0.203 | 0.203 | 0.225 | 0.3 | 0.03 | 0.17 | 0 | 0.34 | 0.23 |
| 5 | 56481011 | rs7718423 | 1.374 | 0.201 | -0.464 | -1.624 | 0.001 | -0.028 | -0.028 | 0 | 0.1 | 0.14 | 0.06 | 0.01 | 0.1 | 0.13 |
| 5 | 56482501 | rs864963 | -0.107 | 0.13 | -0.337 | -1.635 | 0.002 | 0.002 | 0.002 | 0 | 0.49 | 0.46 | 0.43 | 0.14 | 0.46 | 0.51 |
| 5 | 56488165 | rs2545012 | 1.344 | 0.112 | -0.351 | -1.664 | -0.058 | 0.204 | 0.204 | 0.181 | 0.31 | 0.03 | 0.17 | 0 | 0.35 | 0.23 |
| 5 | 56493746 | rs831649 | 0.547 | 0.113 | -0.34 | -1.664 | -0.027 | 0.167 | 0.167 | 0 | 0.41 | 0.17 | 0.22 | 0.01 | 0.45 | 0.36 |
| 5 | 56499686 | rs16886794 | 1.159 | 0.119 | -0.312 | -1.925 | -0.024 | 0.02 | 0.02 | 0 | 0.26 | 0.2 | 0.2 | 0 | 0.31 | 0.13 |
| 5 | 56500426 | rs831652 | -0.129 | 0.131 | 0.134 | -1.738 | -0.011 | -0.016 | -0.016 | 0 | 0.7 | 0.66 | 0.52 | 0.23 | 0.75 | 0.58 |
| 5 | 56514016 | rs10472000 | 1.434 | 0.14 | 0.132 | -1.729 | 0 | -0.027 | -0.027 | 0 | 0.11 | 0.14 | 0.06 | 0.01 | 0.1 | 0.13 |
| 5 | 56526054 | rs11955116 | 1.247 | 0.083 | 0.17 | -1.656 | -0.019 | 0.016 | 0.016 | 0 | 0.25 | 0.19 | 0.19 | 0 | 0.29 | 0.12 |
| 5 | 56527661 | rs12659589 | 1.229 | 0.088 | 0.146 | -1.651 | 0.002 | -0.008 | -0.008 | 0 | 0.25 | 0.28 | 0.2 | 0.13 | 0.3 | 0.14 |
| 5 | 56574452 | rs4700181 | -0.686 | 0.087 | 0.154 | -1.622 | 0 | 0.058 | 0.058 | 0 | 0.56 | 0.66 | 0.68 | 0.91 | 0.55 | 0.56 |
| 5 | 56587609 | rs12153607 | -0.532 | -0.138 | 0.23 | -1.233 | -0.017 | -0.017 | -0.017 | 0 | 0.3 | 0.35 | 0.47 | 0.77 | 0.24 | 0.42 |
| 5 | 56590794 | rs2042175 | 0.118 | -0.152 | 0.293 | -0.726 | -0.002 | 0.006 | 0.006 | 0.064 | 0.26 | 0.25 | 0.37 | 0.64 | 0.24 | 0.35 |
| 5 | 56595262 | rs2161139 | -0.029 | -0.3 | 0.174 | 0.465 | 0 | 0.105 | 0.105 | 0 | 0.23 | 0.35 | 0.46 | 0.47 | 0.22 | 0.25 |
| 5 | 56603577 | rs32870 | 0.441 | -0.361 | 0.023 | -0.138 | -0.005 | -0.022 | -0.022 | 0 | 0.7 | 0.63 | 0.5 | 0.18 | 0.71 | 0.55 |
| 5 | 56603810 | rs6866482 | 1.262 | -0.493 | 0.094 | -0.235 | -0.027 | -0.003 | -0.003 | 1.657 | 0.11 | 0.15 | 0.04 | 0.09 | 0.06 | 0.16 |
| 5 | 56607445 | rs168848 | 0.441 | -0.058 | 0.221 | 0.015 | -0.006 | -0.028 | -0.028 | 0 | 0.7 | 0.61 | 0.5 | 0.18 | 0.7 | 0.55 |
| 5 | 56623761 | rs1363162 | -0.207 | -0.017 | -0.377 | 0.164 | -0.001 | 0 | 0 | 0.039 | 0.58 | 0.59 | 0.6 | 0.43 | 0.64 | 0.58 |
| 5 | 56631872 | rs10805495 | -0.256 | -0.534 | -0.774 | 0.158 | -0.014 | 0.01 | 0.01 | 0.013 | 0.72 | 0.71 | 0.72 | 0.57 | 0.83 | 0.66 |
| 5 | 56640572 | rs10940535 | 0.707 | 0.21 | -1.37 | -0.303 | -0.021 | -0.05 | -0.05 | 0 | 0.08 | 0.02 | 0.09 | 0.2 | 0.12 | 0.13 |
| 5 | 56643050 | rs11954324 | -0.049 | 0.7 | -1.99 | -0.891 | -0.168 | -0.078 | -0.078 | 0 | 0.34 | 0.06 | 0.42 | 0.31 | 0.44 | 0.29 |
| 5 | 56644757 | rs255231 | -1.208 | -0.076 | -1.884 | -1.044 | -0.074 | 0.124 | 0.124 | 0 | 0.75 | 0.54 | 0.72 | 0.45 | 0.86 | 0.67 |
| 5 | 56653871 | rs10071932 | -0.114 | 0.715 | -1.576 | -0.933 | -0.021 | 0.004 | 0.004 | 0 | 0.37 | 0.31 | 0.46 | 0.32 | 0.47 | 0.3 |
| 5 | 56655375 | rs877454 | -0.189 | 0.724 | -1.564 | -0.929 | -0.001 | 0 | 0 | 0 | 0.38 | 0.41 | 0.47 | 0.31 | 0.47 | 0.3 |
| 5 | 56666500 | rs10052004 | -0.108 | 0.935 | -1.63 | -1.02 | -0.068 | -0.028 | -0.028 | 0 | 0.19 | 0.08 | 0.3 | 0.24 | 0.31 | 0.17 |
| 5 | 56671715 | rs2408759 | -0.254 | 0.769 | -1.461 | -0.889 | -0.002 | 0 | 0 | 0 | 0.38 | 0.43 | 0.47 | 0.31 | 0.46 | 0.31 |
| 5 | 56681929 | rs7720225 | -0.471 | -0.328 | -1.691 | -0.777 | 0.122 | 0.047 | 0.047 | 0.349 | 0.54 | 0.25 | 0.64 | 0.31 | 0.72 | 0.49 |
| 5 | 56688452 | rs13357131 | 1.392 | -0.602 | -1.809 | -0.836 | 0.021 | 0.029 | 0.029 | 0 | 0.19 | 0.07 | 0.08 | 0.15 | 0.14 | 0.18 |
| 5 | 56690868 | rs10940547 | -0.537 | 0.929 | -1.545 | -0.79 | -0.096 | 0.059 | 0.059 | 0.064 | 0.36 | 0.1 | 0.47 | 0.24 | 0.56 | 0.35 |

| | | | | | | | | | | | | | | | | |
|---|---|---|---|---|---|---|---|---|---|---|---|---|---|---|---|---|
| 5 | 56695644 | rs2408760 | 0.897 | 1.27 | -1.429 | -0.602 | -0.039 | -0.047 | -0.047 | 0.002 | 0.08 | 0.01 | 0.1 | 0.23 | 0.14 | 0.13 |
| 5 | 56704132 | rs960702 | -1.03 | 1.2 | -1.283 | -0.684 | 0.15 | 0.154 | 0.154 | 0.481 | 0.5 | 0.17 | 0.55 | 0.26 | 0.7 | 0.43 |
| 5 | 56707316 | rs11741561 | -0.745 | 1.125 | -1.195 | -0.786 | 0.044 | 0.081 | 0.081 | 0.575 | 0.43 | 0.23 | 0.53 | 0.26 | 0.64 | 0.39 |
| 5 | 56711877 | rs11744731 | -1.26 | 1.132 | -1.18 | -0.838 | 0.029 | 0.014 | 0.014 | 0 | 0.53 | 0.43 | 0.63 | 0.33 | 0.7 | 0.47 |
| 5 | 56718875 | rs2172771 | 0.374 | 1.455 | -1.378 | -0.881 | -0.144 | 0.102 | 0.102 | 0.999 | 0.22 | 0.02 | 0.34 | 0.04 | 0.38 | 0.22 |
| 5 | 56727379 | rs10805497 | 0.329 | 0.538 | -1.306 | -0.401 | -0.018 | -0.066 | -0.066 | 0 | 0.13 | 0.02 | 0.16 | 0.29 | 0.15 | 0.17 |
| 5 | 56727958 | rs10054252 | -0.374 | 0.092 | -0.83 | -0.216 | 0.011 | 0.008 | 0.008 | 0.004 | 0.52 | 0.48 | 0.64 | 0.33 | 0.7 | 0.47 |
| 5 | 56730383 | rs10472003 | -0.479 | 1.28 | -0.42 | -0.493 | 0.045 | -0.07 | -0.07 | 0.252 | 0.28 | 0.5 | 0.28 | 0.14 | 0.31 | 0.22 |
| 5 | 56730670 | rs10471436 | -0.815 | 2.054 | -0.651 | -1.093 | -0.098 | 0.108 | 0.108 | 0 | 0.3 | 0.14 | 0.35 | 0.01 | 0.43 | 0.24 |
| 5 | 56730691 | rs10472004 | 0.42 | 2.367 | -0.708 | -1.051 | -0.054 | 0.052 | 0.052 | 0.017 | 0.1 | 0 | 0.17 | 0.01 | 0.15 | 0.14 |
| 5 | 56738101 | rs7729777 | 0.023 | 1.529 | -0.385 | -1.337 | -0.032 | -0.076 | -0.076 | 0.33 | 0.12 | 0.32 | 0.07 | 0 | 0.07 | 0.03 |
| 5 | 56738129 | rs7733945 | -0.869 | 1.663 | -0.303 | -1.565 | 0.024 | -0.148 | -0.148 | 0 | 0.25 | 0.43 | 0.25 | 0.01 | 0.27 | 0.18 |
| 5 | 56739381 | rs6878837 | 0.528 | 1.694 | -0.371 | -1.545 | -0.071 | 0.066 | 0.066 | 0 | 0.13 | 0.02 | 0.17 | 0.01 | 0.19 | 0.15 |
| 5 | 56741903 | rs16886951 | -0.092 | 0.951 | -0.485 | -2.021 | 0.027 | 0.017 | 0.017 | 0.352 | 0.18 | 0.12 | 0.07 | 0.01 | 0.1 | 0.07 |
| 5 | 56742262 | rs11950872 | 1.438 | 0.955 | -0.532 | -2.063 | -0.002 | -0.016 | -0.016 | 0.442 | 0.12 | 0.09 | 0.07 | 0 | 0.12 | 0.05 |
| 5 | 56749587 | rs6450425 | -0.977 | 0.859 | -0.351 | -1.802 | -0.001 | -0.089 | -0.089 | 0.147 | 0.45 | 0.5 | 0.33 | 0.03 | 0.42 | 0.39 |
| 5 | 56751453 | rs11741009 | 1.042 | 0.741 | -1.21 | -1.296 | -0.029 | -0.043 | -0.043 | 0 | 0.18 | 0.02 | 0.15 | 0.24 | 0.19 | 0.12 |
| 5 | 56762458 | rs903410 | -0.609 | 0.503 | -0.377 | -1.759 | 0 | -0.047 | -0.047 | 0 | 0.48 | 0.5 | 0.33 | 0.02 | 0.43 | 0.39 |
| 5 | 56765375 | rs952277 | 0.517 | 0.078 | -0.697 | -1.304 | -0.012 | 0.002 | 0.002 | 0.293 | 0.34 | 0.3 | 0.44 | 0.68 | 0.45 | 0.38 |
| 5 | 56775578 | rs1979725 | -0.522 | 0.2 | -0.84 | -1.683 | 0.006 | 0.053 | 0.053 | 0.904 | 0.49 | 0.41 | 0.33 | 0.03 | 0.43 | 0.4 |
| 5 | 56777677 | rs16886992 | 0.968 | 0.104 | -0.893 | -1.693 | -0.011 | -0.032 | -0.032 | 0 | 0.12 | 0.17 | 0.02 | 0 | 0.04 | 0.06 |
| 5 | 56782006 | rs4699905 | -0.434 | 0.42 | -0.859 | -1.582 | 0.003 | 0.051 | 0.051 | 0.766 | 0.49 | 0.44 | 0.34 | 0.03 | 0.45 | 0.41 |
| 5 | 56789327 | rs2968230 | -0.455 | 0.292 | -0.938 | -1.561 | 0.002 | -0.013 | -0.013 | 1.324 | 0.5 | 0.48 | 0.35 | 0.03 | 0.44 | 0.4 |
| 5 | 56792127 | rs2972351 | 0.843 | 1.061 | -1.472 | -1.004 | -0.06 | 0.061 | 0.061 | 0 | 0.11 | 0.03 | 0.16 | 0.01 | 0.18 | 0.14 |
| 5 | 56794319 | rs12658182 | 0.467 | 0.181 | -1.356 | -0.345 | 0.006 | -0.024 | -0.024 | 0 | 0.12 | 0.06 | 0.15 | 0.19 | 0.14 | 0.13 |
| 5 | 56795050 | rs4145581 | 0.485 | 0.181 | -1.356 | -0.342 | 0.006 | -0.023 | -0.023 | 0.458 | 0.12 | 0.06 | 0.15 | 0.19 | 0.14 | 0.13 |
| 5 | 56797293 | rs16887016 | -0.357 | -0.195 | -1.578 | -0.643 | 0.033 | -0.003 | -0.003 | 0 | 0.25 | 0.11 | 0.2 | 0.21 | 0.22 | 0.3 |
| 5 | 56799445 | rs10471439 | 1.043 | 0.063 | -1.174 | -0.807 | 0.023 | -0.028 | -0.028 | 0 | 0.06 | 0.13 | 0.05 | 0 | 0.07 | 0.03 |
| 5 | 56801671 | rs17687301 | 1.002 | 0.658 | -1.602 | -1.152 | -0.059 | 0.06 | 0.06 | 0 | 0.13 | 0.04 | 0.14 | 0 | 0.17 | 0.09 |
| 5 | 56803101 | rs10057549 | 0.746 | -0.055 | -1.886 | -1.757 | 0.019 | 0.024 | 0.024 | 0 | 0.23 | 0.15 | 0.11 | 0 | 0.16 | 0.1 |
| 5 | 56803300 | rs16887026 | 1.244 | -0.101 | -1.656 | -0.97 | 0.048 | 0.142 | 0.142 | 0 | 0.21 | 0.43 | 0.27 | 0.45 | 0.29 | 0.31 |
| 5 | 56806662 | rs1158044 | -2.1 | -0.478 | -2.065 | -0.889 | 0.023 | -0.073 | -0.073 | 0 | 0.83 | 0.94 | 0.69 | 0.62 | 0.86 | 0.77 |
| 5 | 56808015 | rs2968217 | 0.182 | -0.333 | -1.977 | -1.319 | 0.006 | 0.019 | 0.019 | 0 | 0.29 | 0.2 | 0.18 | 0.03 | 0.26 | 0.22 |
| 5 | 56808503 | rs969627 | -0.078 | -0.211 | -2.113 | -1.392 | -0.065 | 0.144 | 0.144 | 0.484 | 0.38 | 0.18 | 0.28 | 0.03 | 0.39 | 0.29 |
| 5 | 56809189 | rs10051391 | 2.21 | -0.681 | -1.913 | -1.11 | -0.016 | 0.076 | 0.076 | 0 | 0.16 | 0.04 | 0.09 | 0 | 0.15 | 0.07 |
| 5 | 56810798 | rs2968225 | 2.21 | -0.681 | -1.925 | -1.11 | -0.025 | 0.09 | 0.09 | 0 | 0.16 | 0.03 | 0.09 | 0 | 0.15 | 0.07 |
| 5 | 56811700 | rs6898854 | 2.016 | -0.992 | -1.791 | -0.955 | -0.002 | -0.001 | -0.001 | 0 | 0.07 | 0.09 | 0.11 | 0.11 | 0.04 | 0.03 |
| 5 | 56811719 | rs2408818 | -0.563 | -1.219 | -1.85 | -0.989 | 0.001 | -0.002 | -0.002 | 0.396 | 0.73 | 0.77 | 0.67 | 0.59 | 0.73 | 0.66 |
| 5 | 56819519 | rs7349782 | 0.062 | -1.308 | -2.023 | -1.034 | 0.032 | 0.016 | 0.016 | 0 | 0.48 | 0.74 | 0.58 | 0.57 | 0.55 | 0.48 |
| 5 | 56830603 | rs9292139 | -0.818 | -1.345 | -1.819 | -0.659 | 0.013 | -0.083 | -0.083 | 0 | 0.77 | 0.93 | 0.72 | 0.59 | 0.76 | 0.67 |
| 5 | 56836190 | rs6881897 | 1.608 | -1.329 | -1.731 | -0.553 | -0.006 | 0.002 | 0.002 | 0.502 | 0.07 | 0.09 | 0.11 | 0.13 | 0.03 | 0.03 |
| 5 | 56837920 | rs7727791 | 0.021 | -1.624 | -1.599 | -0.64 | 0.015 | -0.018 | -0.018 | 2.066 | 0.66 | 0.73 | 0.66 | 0.57 | 0.69 | 0.54 |
| 5 | 56849212 | rs1423141 | -0.214 | -2.312 | -1.201 | -0.451 | -0.001 | -0.003 | -0.003 | 0 | 0.57 | 0.59 | 0.39 | 0.26 | 0.53 | 0.41 |
| 5 | 56850241 | rs1423142 | 0.881 | -2.665 | -1.144 | -0.449 | -0.094 | -0.041 | -0.041 | 0 | 0.2 | 0.4 | 0.06 | 0.14 | 0.03 | 0.17 |
| 5 | 56850710 | rs13357378 | 0.683 | -1.303 | -1.311 | -0.034 | -0.003 | 0.025 | 0.025 | 0 | 0.09 | 0.16 | 0.07 | 0.11 | 0.1 | 0.07 |
| 5 | 56853136 | rs1423143 | -1.266 | -1.141 | -1.799 | 0.273 | 0.008 | -0.028 | -0.028 | 0 | 0.87 | 0.94 | 0.79 | 0.72 | 0.91 | 0.82 |
| 5 | 56860922 | rs12521025 | 1.181 | -0.576 | -1.817 | 0.246 | -0.083 | 0.063 | 0.063 | 0 | 0.12 | 0.02 | 0.15 | 0.01 | 0.22 | 0.1 |
| 5 | 56863759 | rs2910479 | -1.365 | -1.047 | -1.502 | 0.756 | -0.003 | -0.053 | -0.053 | 0 | 0.9 | 0.99 | 0.82 | 0.75 | 0.93 | 0.86 |
| 5 | 56865572 | rs2968214 | -0.324 | -1.061 | -1.231 | 0.559 | -0.002 | 0.024 | 0.024 | 0.254 | 0.81 | 0.75 | 0.72 | 0.63 | 0.81 | 0.77 |
| 5 | 56865689 | rs16887106 | 0.121 | -1.57 | -1.125 | 0.351 | -0.029 | -0.009 | -0.009 | 0 | 0.14 | 0.23 | 0.06 | 0.14 | 0.03 | 0.17 |
| 5 | 56870298 | rs2910498 | 1.18 | -1.274 | -1.475 | 0.107 | -0.032 | 0.104 | 0.104 | 0 | 0.17 | 0.01 | 0.11 | 0 | 0.17 | 0.07 |
| 5 | 56871273 | rs7714264 | 0.824 | -0.722 | -0.97 | 0.395 | 0.007 | -0.025 | -0.025 | 0.419 | 0.06 | 0.11 | 0.06 | 0 | 0.07 | 0.08 |
| 5 | 56879903 | rs985363 | 0.219 | -0.339 | -1.203 | 0.656 | -0.01 | -0.002 | -0.002 | 0 | 0.64 | 0.7 | 0.51 | 0.61 | 0.5 | 0.53 |
| 5 | 56881357 | rs12518745 | 0.842 | 0.148 | -1.127 | 0.152 | 0.04 | -0.01 | -0.01 | 0.192 | 0.13 | 0.12 | 0.19 | 0.07 | 0.28 | 0.2 |
| 5 | 56884904 | rs16887120 | 0.461 | -0.507 | -1.084 | 0.427 | -0.035 | -0.012 | -0.012 | 0 | 0.11 | 0.24 | 0.05 | 0.1 | 0.03 | 0.02 |
| 5 | 56886931 | rs16887124 | 0.43 | -0.066 | -1.756 | 0.707 | 0.008 | -0.027 | -0.027 | 0.183 | 0.21 | 0.18 | 0.3 | 0.39 | 0.18 | 0.21 |
| 5 | 56887318 | rs13172186 | 0.031 | 0.254 | -1.565 | 0.089 | 0 | 0.001 | 0.001 | 0.021 | 0.28 | 0.23 | 0.23 | 0.22 | 0.28 | 0.25 |
| 5 | 56891257 | rs929809 | 0.371 | 0.257 | -1.58 | 0.098 | 0.003 | 0.019 | 0.019 | 0.23 | 0.64 | 0.76 | 0.73 | 0.72 | 0.62 | 0.63 |
| 5 | 56891952 | rs10454814 | -0.193 | 0.487 | -2.147 | 0.185 | -0.053 | 0.01 | 0.01 | 0.595 | 0.3 | 0.02 | 0.23 | 0.22 | 0.31 | 0.26 |
| 5 | 56896963 | rs230841 | 0.246 | 1.311 | -1.657 | 0.377 | -0.012 | -0.004 | -0.004 | 0 | 0.73 | 0.81 | 0.73 | 0.62 | 0.69 | 0.56 |
| 5 | 56901907 | rs108609 | 0.667 | 0.931 | -1.606 | 0.246 | 0.001 | -0.005 | -0.005 | 1.105 | 0.27 | 0.25 | 0.14 | 0.33 | 0.23 | 0.26 |
| 5 | 56903081 | rs17315098 | 0.301 | 1.315 | -1.929 | -0.038 | -0.038 | 0.021 | 0.021 | 0 | 0.07 | 0.01 | 0.06 | 0.01 | 0.13 | 0.07 |
| 5 | 56908982 | rs10059916 | 0.27 | 0.725 | -1.634 | -0.601 | 0.004 | -0.016 | -0.016 | 2.083 | 0.26 | 0.28 | 0.14 | 0.02 | 0.22 | 0.13 |
| 5 | 56914261 | rs4699908 | 0.261 | 0.87 | -1.434 | -0.592 | -0.001 | 0.014 | 0.014 | 0 | 0.39 | 0.3 | 0.37 | 0.31 | 0.36 | 0.54 |
| 5 | 56915745 | rs230832 | -1.309 | 0.848 | -1.316 | -0.593 | 0.015 | 0.054 | 0.054 | 0 | 0.93 | 0.92 | 0.66 | 0.65 | 0.81 | 0.93 |
| 5 | 56919630 | rs2910495 | -0.05 | 0.789 | -1.28 | -0.64 | -0.01 | 0.023 | 0.023 | 0 | 0.43 | 0.35 | 0.43 | 0.31 | 0.46 | 0.6 |
| 5 | 56922085 | rs2910490 | -1.002 | 0.803 | -1.257 | -0.608 | 0.013 | 0.032 | 0.032 | 0 | 0.53 | 0.64 | 0.42 | 0.32 | 0.42 | 0.54 |
| 5 | 56922315 | rs230844 | 1.474 | 0.822 | -1.188 | -0.572 | -0.009 | -0.007 | -0.007 | 0.009 | 0.38 | 0.27 | 0.25 | 0.34 | 0.39 | 0.39 |
| 5 | 56923759 | rs10513124 | 1.204 | 0.417 | -0.974 | -0.41 | 0.025 | 0.043 | 0.043 | 0.171 | 0.12 | 0.03 | 0.1 | 0 | 0.07 | 0.09 |

| | | | | | | | | | | | | | | | | |
|---|---|---|---|---|---|---|---|---|---|---|---|---|---|---|---|---|
| 5 | 56926094 | rs1363788 | 1.083 | 0.414 | -0.97 | -0.417 | -0.001 | 0.051 | 0.051 | 0 | 0.16 | 0.29 | 0.17 | 0.3 | 0.14 | 0.32 |
| 5 | 56928624 | rs12523405 | 1.206 | 0.387 | -0.977 | -0.421 | 0.025 | 0.043 | 0.043 | 0.171 | 0.12 | 0.03 | 0.1 | 0 | 0.07 | 0.09 |
| 5 | 56929080 | rs230880 | 1.077 | 0.398 | -0.97 | -0.406 | 0 | 0.008 | 0.008 | 0 | 0.46 | 0.34 | 0.31 | 0.34 | 0.51 | 0.45 |
| 5 | 56930116 | rs7718757 | 0.728 | 0.375 | -0.91 | -0.366 | -0.032 | -0.068 | -0.068 | 0 | 0.18 | 0.27 | 0.08 | 0.01 | 0.09 | 0.07 |
| 5 | 56930563 | rs110872 | 0.366 | 0.446 | -0.927 | -0.349 | -0.005 | -0.012 | -0.012 | 0 | 0.8 | 0.86 | 0.57 | 0.65 | 0.76 | 0.84 |
| 5 | 56933347 | rs230874 | 0.4 | 0.872 | -0.823 | -0.119 | 0.003 | 0.002 | 0.002 | 0.372 | 0.62 | 0.61 | 0.49 | 0.65 | 0.67 | 0.78 |
| 5 | 56937591 | rs179507 | 0.221 | 2.134 | -0.494 | 0.951 | 0.024 | 0.071 | 0.071 | 0 | 0.93 | 0.92 | 0.68 | 0.65 | 0.82 | 0.9 |
| 5 | 56939184 | rs11959890 | -1.215 | 2.09 | -0.077 | 0.319 | 0.067 | -0.05 | -0.05 | 0 | 0.12 | 0.3 | 0.12 | 0.02 | 0.22 | 0.08 |
| 5 | 56939590 | rs2534842 | 0.65 | 1.652 | 0.527 | 1.443 | 0.009 | 0.253 | 0.253 | 0 | 0.93 | 0.76 | 0.69 | 0.59 | 0.94 | 0.85 |
| 5 | 56944239 | rs2109479 | -0.717 | 1.975 | 0.967 | 1.413 | 0.046 | 0.156 | 0.156 | 0.143 | 0.17 | 0.45 | 0.2 | 0.3 | 0.2 | 0.14 |
| 5 | 56947141 | rs702697 | 1.096 | 1.534 | -0.001 | 0.706 | 0.053 | 0.144 | 0.144 | 0 | 0.33 | 0.07 | 0.14 | 0.1 | 0.24 | 0.2 |
| 5 | 56949522 | rs854140 | -0.858 | 2.071 | -0.015 | 0.635 | 0.005 | -0.036 | -0.036 | 0 | 0.66 | 0.72 | 0.62 | 0.48 | 0.68 | 0.64 |
| 5 | 56950505 | rs13161897 | -1.613 | 2.451 | 0.711 | 0.929 | 0.076 | 0.046 | 0.046 | 0 | 0.25 | 0.41 | 0.3 | 0.33 | 0.38 | 0.2 |
| 5 | 56951744 | rs2191450 | -0.147 | 2.482 | -0.075 | 0.98 | 0.002 | 0.022 | 0.022 | 0 | 0.12 | 0.14 | 0.16 | 0.3 | 0.19 | 0.13 |
| 5 | 56952654 | rs170741 | -0.33 | 2.017 | -0.113 | 0.469 | -0.048 | 0.041 | 0.041 | 0 | 0.1 | 0.03 | 0.12 | 0.03 | 0.2 | 0.07 |
| 5 | 56974635 | rs11742166 | 1.701 | 1.292 | 0.09 | 0.673 | 0 | -0.003 | -0.003 | 0 | 0.75 | 0.29 | 0.7 | 0.69 | 0.77 | 0.84 |
| 5 | 56986451 | rs229960 | -0.574 | 1.575 | -0.003 | 0.202 | -0.015 | 0.011 | 0.011 | 0 | 0.95 | 0.97 | 0.89 | 0.98 | 0.83 | 0.97 |
| 5 | 56988506 | rs12654351 | -0.368 | 1.304 | 0.753 | 0.264 | -0.028 | 0.006 | 0.006 | 0.275 | 0.09 | 0.14 | 0.04 | 0.13 | 0.03 | 0.02 |
| 5 | 57004423 | rs2041402 | 2.108 | 1.585 | 0.717 | -0.124 | 0.002 | -0.016 | -0.016 | 0 | 0.67 | 0.4 | 0.67 | 0.82 | 0.63 | 0.81 |
| 5 | 57019784 | rs12654963 | -0.162 | 1.529 | 0.756 | 0.263 | -0.021 | 0.005 | 0.005 | 0.285 | 0.09 | 0.09 | 0.04 | 0.14 | 0.03 | 0.02 |
| 5 | 57042657 | rs6876911 | -0.776 | 1.416 | 1.231 | 0.204 | -0.034 | 0.009 | 0.009 | 1.27 | 0.11 | 0.27 | 0.05 | 0.11 | 0.03 | 0.05 |
| 5 | 57043739 | rs1861052 | 0.09 | 1.824 | 0.981 | 0.186 | -0.04 | -0.005 | -0.005 | 0.079 | 0.91 | 0.96 | 0.86 | 0.93 | 0.8 | 0.96 |
| 5 | 57044866 | rs741182 | 2.043 | 1.864 | 0.937 | 0.262 | 0.004 | -0.001 | -0.001 | 0 | 0.66 | 0.33 | 0.62 | 0.74 | 0.54 | 0.58 |
| 5 | 57044981 | rs741183 | 0.39 | 1.915 | 1.02 | 0.272 | -0.002 | 0 | 0 | 0 | 0.11 | 0.1 | 0.09 | 0.19 | 0.12 | 0.17 |
| 5 | 57050359 | rs12652479 | 0.921 | 1.864 | 1.016 | 0.247 | -0.008 | 0.007 | 0.007 | 0.269 | 0.05 | 0.06 | 0.03 | 0.11 | 0.02 | 0.02 |
| 5 | 57064861 | rs6875712 | -0.354 | 1.855 | 0.846 | 0.201 | -0.047 | -0.004 | -0.004 | 0.017 | 0.9 | 0.96 | 0.86 | 0.93 | 0.79 | 0.95 |
| 5 | 57065117 | rs1544621 | 0.245 | 1.775 | 1.256 | 0.158 | 0.01 | -0.047 | -0.047 | 0.983 | 0.11 | 0.18 | 0.14 | 0 | 0.13 | 0.16 |
| 5 | 57070495 | rs1023571 | 2.279 | 1.687 | 1.319 | 0.246 | 0.008 | 0.007 | 0.007 | 0 | 0.67 | 0.44 | 0.71 | 0.62 | 0.64 | 0.72 |
| 5 | 57073155 | rs7713500 | -0.592 | 1.349 | 1.382 | 0.304 | -0.03 | 0.007 | 0.007 | 0.651 | 0.1 | 0.23 | 0.05 | 0.12 | 0.03 | 0.06 |
| 5 | 57077047 | rs2215813 | 2.757 | 1.752 | 1.223 | 0.275 | -0.023 | -0.001 | -0.001 | 0 | 0.56 | 0.25 | 0.5 | 0.58 | 0.53 | 0.56 |
| 5 | 57078397 | rs10063478 | -0.107 | 1.726 | 1.344 | 0.272 | -0.033 | -0.003 | -0.003 | 0 | 0.09 | 0.11 | 0.12 | 0.12 | 0.02 | 0.02 |
| 5 | 57080169 | rs1861059 | 0.001 | 1.494 | 1.469 | 0.794 | -0.039 | 0 | 0 | 1.175 | 0.9 | 0.97 | 0.87 | 0.91 | 0.81 | 0.95 |
| 5 | 57089056 | rs854032 | 2.754 | 1.497 | 1.525 | 0.782 | -0.003 | -0.005 | -0.005 | 0.669 | 0.67 | 0.37 | 0.62 | 0.58 | 0.66 | 0.72 |
| 5 | 57091157 | rs17746968 | -0.142 | 1.339 | 1.406 | 0.739 | -0.005 | -0.005 | -0.005 | 0 | 0.1 | 0.08 | 0.12 | 0 | 0.13 | 0.16 |
| 5 | 57098603 | rs854045 | -0.344 | 1.481 | 1.296 | 0.71 | -0.04 | 0.031 | 0.031 | 0 | 0.9 | 0.97 | 0.87 | 0.95 | 0.8 | 0.95 |
| 5 | 57105894 | rs10487616 | 0.03 | 1.46 | 1.441 | 0.729 | 0.008 | 0.01 | 0.01 | 0.082 | 0.08 | 0.1 | 0.08 | 0.12 | 0.11 | 0.13 |
| 5 | 57106067 | rs854050 | 0.145 | 1.474 | 1.394 | 0.742 | -0.054 | 0.007 | 0.007 | 0 | 0.88 | 0.97 | 0.85 | 0.91 | 0.78 | 0.94 |
| 5 | 57107989 | rs981883 | 2.854 | 1.43 | 1.542 | 0.829 | 0.013 | -0.018 | -0.018 | 3.102 | 0.64 | 0.53 | 0.58 | 0.68 | 0.6 | 0.68 |
| 5 | 57108088 | rs854051 | -0.417 | 1.446 | 1.53 | 0.948 | -0.035 | 0.007 | 0.007 | 1.083 | 0.09 | 0.12 | 0.12 | 0.15 | 0.02 | 0.02 |
| 5 | 57109644 | rs854054 | 1.007 | 1.71 | 1.391 | 1.046 | -0.075 | -0.045 | -0.045 | 0.561 | 0.85 | 0.96 | 0.81 | 0.82 | 0.73 | 0.88 |
| 5 | 57114080 | rs7724301 | 1.052 | 1.616 | 1.567 | 1.041 | 0.008 | 0.002 | 0.002 | 0 | 0.84 | 0.8 | 0.82 | 0.83 | 0.74 | 0.87 |
| 5 | 57114951 | rs4700229 | 2.119 | 1.609 | 1.589 | 1.026 | 0.019 | 0.018 | 0.018 | 0 | 0.65 | 0.48 | 0.54 | 0.55 | 0.6 | 0.7 |
| 5 | 57122376 | rs13156567 | 0.245 | 1.645 | 1.422 | 1.04 | 0.001 | 0 | 0 | 0.062 | 0.1 | 0.11 | 0.15 | 0.12 | 0.12 | 0.15 |
| 5 | 57133250 | rs1835874 | -1.683 | 1.414 | 1.575 | 1.04 | -0.008 | 0.052 | 0.052 | 0 | 0.24 | 0.54 | 0.3 | 0.37 | 0.17 | 0.35 |
| 5 | 57138990 | rs1968422 | -1.191 | 1.36 | 1.826 | 1.012 | 0.01 | 0.051 | 0.051 | 0 | 0.19 | 0.38 | 0.27 | 0.28 | 0.14 | 0.26 |
| 5 | 57139075 | rs4285175 | 0.137 | 1.371 | 1.303 | 1.026 | -0.051 | -0.009 | -0.009 | 0 | 0.89 | 0.97 | 0.84 | 0.9 | 0.76 | 0.96 |
| 5 | 57141306 | rs16887464 | 3.173 | 1.238 | 1.215 | 1.034 | -0.015 | -0.03 | -0.03 | 0.035 | 0.51 | 0.13 | 0.3 | 0.51 | 0.5 | 0.48 |
| 5 | 57145774 | rs4618400 | 2.675 | 1.659 | 1.492 | 1.219 | 0.003 | 0.002 | 0.002 | 0 | 0.67 | 0.37 | 0.45 | 0.63 | 0.62 | 0.72 |
| 5 | 57150532 | rs7705954 | -0.355 | 1.663 | 1.257 | 1.16 | 0.002 | -0.045 | -0.045 | 0.395 | 0.1 | 0.2 | 0.07 | 0 | 0.1 | 0.12 |
| 5 | 57156325 | rs4699913 | -0.182 | 1.691 | 0.83 | 1.206 | -0.041 | 0.018 | 0.018 | 0 | 0.9 | 0.97 | 0.86 | 0.95 | 0.78 | 0.96 |
| 5 | 57167434 | rs4460105 | -0.758 | 1.497 | 0.94 | 1.246 | -0.022 | 0.13 | 0.13 | 0.226 | 0.1 | 0.26 | 0.32 | 0.26 | 0.04 | 0.11 |
| 5 | 57172078 | rs4432842 | -1.954 | 1.519 | 0.713 | 1.236 | 0.003 | -0.029 | -0.029 | 0.274 | 0.33 | 0.81 | 0.54 | 0.39 | 0.27 | 0.47 |
| 5 | 57173905 | rs7734050 | 1.791 | 1.398 | 0.777 | 1.485 | 0.019 | 0.058 | 0.058 | 0 | 0.77 | 0.66 | 0.52 | 0.63 | 0.72 | 0.85 |
| 5 | 57179504 | rs1020661 | 2.188 | 1.432 | 0.608 | 1.507 | 0.01 | 0.012 | 0.012 | 0 | 0.69 | 0.61 | 0.48 | 0.63 | 0.63 | 0.72 |
| 5 | 57180012 | rs35310736 | -1.965 | 0.943 | 0.885 | 1.538 | -0.029 | 0.035 | 0.035 | 0 | 0.2 | 0.36 | 0.37 | 0.27 | 0.13 | 0.23 |
| 5 | 57183685 | rs1454911 | 1.48 | 1.364 | 0.306 | 1.232 | 0.004 | -0.003 | -0.003 | 0 | 0.82 | 0.82 | 0.81 | 0.88 | 0.74 | 0.87 |
| 5 | 57189995 | rs2898074 | -0.078 | 1.198 | -0.023 | 1.368 | 0.018 | -0.013 | -0.013 | 0.001 | 0.09 | 0.01 | 0.02 | 0.08 | 0.04 | 0.02 |
| 5 | 57194537 | rs1563263 | 0.241 | 1.37 | 0.103 | 1.281 | -0.017 | -0.006 | -0.006 | 0 | 0.91 | 0.95 | 0.87 | 0.88 | 0.82 | 0.97 |
| 5 | 57199577 | rs6872726 | -1.07 | 1.181 | 0.221 | 1.18 | 0.002 | -0.008 | -0.008 | 0.186 | 0.08 | 0.07 | 0.05 | 0 | 0.07 | 0.08 |
| 5 | 57199879 | rs921770 | 0.288 | 1.367 | 0.379 | 1.246 | 0.024 | 0.002 | 0.002 | 0 | 0.9 | 0.87 | 0.86 | 0.88 | 0.82 | 0.97 |
| 5 | 57214817 | rs10041935 | -1.356 | 0.799 | 0.654 | 2.109 | -0.037 | -0.056 | -0.056 | 0 | 0.17 | 0.08 | 0.23 | 0.29 | 0.29 | 0.23 |
| 5 | 57226653 | rs1026371 | 1.584 | 0.802 | 0.772 | 2.113 | 0.001 | -0.003 | -0.003 | 0 | 0.86 | 0.85 | 0.87 | 0.83 | 0.87 | 0.69 |
| 5 | 57231934 | rs12658525 | 0.095 | 0.567 | 0.717 | 1.957 | 0.014 | 0.003 | 0.003 | 3.002 | 0.11 | 0.07 | 0.03 | 0.09 | 0.02 | 0.02 |
| 5 | 57232003 | rs13175618 | 2.079 | 0.537 | 0.907 | 2.176 | -0.002 | 0.003 | 0.003 | 0.441 | 0.65 | 0.59 | 0.68 | 0.44 | 0.64 | 0.53 |
| 5 | 57235886 | rs4147094 | 1.527 | 0.565 | 0.79 | 2.095 | 0.007 | 0 | 0 | 0 | 0.76 | 0.78 | 0.85 | 0.76 | 0.84 | 0.67 |
| 5 | 57244070 | rs1378244 | 2.024 | 0.72 | 0.911 | 1.986 | 0.001 | 0.013 | 0.013 | 0.559 | 0.67 | 0.59 | 0.66 | 0.47 | 0.64 | 0.57 |
| 5 | 57244979 | rs1948275 | 1.553 | 0.876 | 0.751 | 1.92 | 0.036 | 0.065 | 0.065 | 0 | 0.77 | 0.6 | 0.72 | 0.65 | 0.65 | 0.68 |
| 5 | 57246446 | rs1809660 | 0.985 | 0.994 | 0.423 | 2.044 | -0.002 | -0.004 | -0.004 | 0 | 0.89 | 0.88 | 0.9 | 0.81 | 0.88 | 0.76 |
| 5 | 57252033 | rs1027163 | 0.738 | 0.997 | 0.334 | 2.047 | -0.008 | 0.009 | 0.009 | 0.513 | 0.89 | 0.91 | 0.9 | 0.95 | 0.88 | 0.78 |

| | | | | | | | | | | | | | | | | |
|---|---|---|---|---|---|---|---|---|---|---|---|---|---|---|---|---|
| 5 | 57254667 | rs10472028 | 0.529 | 0.77 | -0.055 | 1.773 | 0.002 | -0.063 | -0.063 | 0.356 | 0.21 | 0 | 0.09 | 0.25 | 0.14 | 0.14 |
| 5 | 57274320 | rs13189262 | -0.219 | 1.36 | 1.078 | 2.456 | -0.019 | 0.137 | 0.137 | 2.516 | 0.11 | 0.3 | 0.29 | 0.27 | 0.05 | 0.07 |
| 5 | 57277305 | rs10054634 | 1.924 | 1.998 | 0.54 | 1.742 | 0.003 | 0.03 | 0.03 | 0.372 | 0.88 | 0.77 | 0.89 | 0.81 | 0.88 | 0.76 |
| 5 | 57277522 | rs9292147 | 1.678 | 2.001 | 0.474 | 1.707 | 0.002 | -0.007 | -0.007 | 1.623 | 0.89 | 0.82 | 0.9 | 0.95 | 0.88 | 0.78 |
| 5 | 57278956 | rs10077767 | -0.722 | 2.001 | 0.446 | 1.706 | -0.002 | 0.004 | 0.004 | 0.028 | 0.05 | 0.05 | 0.06 | 0.01 | 0.07 | 0.07 |
| 5 | 57291294 | rs7715831 | 0.529 | 1.972 | 0.442 | 1.7 | -0.017 | 0.022 | 0.022 | 0.705 | 0.9 | 0.94 | 0.91 | 0.96 | 0.88 | 0.77 |
| 5 | 57309999 | rs11741819 | -0.943 | 1.878 | 0.187 | 1.22 | 0.02 | 0.014 | 0.014 | 0 | 0.1 | 0.11 | 0.16 | 0.15 | 0.23 | 0.08 |
| 5 | 57320343 | rs984926 | 0.739 | 1.84 | -0.839 | 1.235 | -0.013 | -0.012 | -0.012 | 0 | 0.9 | 0.92 | 0.85 | 0.84 | 0.78 | 0.92 |
| 5 | 57321413 | rs1352609 | 0.739 | 1.84 | -0.839 | 1.235 | -0.013 | -0.012 | -0.012 | 0 | 0.9 | 0.92 | 0.85 | 0.84 | 0.78 | 0.92 |
| 5 | 57334825 | rs1454924 | 0.739 | 1.84 | -0.839 | 1.235 | -0.013 | -0.012 | -0.012 | 0 | 0.9 | 0.92 | 0.85 | 0.84 | 0.78 | 0.92 |
| 5 | 57336494 | rs1454925 | 0.739 | 1.84 | -0.839 | 1.26 | -0.013 | -0.012 | -0.012 | 1.721 | 0.9 | 0.92 | 0.85 | 0.84 | 0.77 | 0.92 |
| 5 | 57337744 | rs1826106 | 0.739 | 1.84 | -0.839 | 1.187 | -0.013 | -0.012 | -0.012 | 0 | 0.9 | 0.92 | 0.85 | 0.85 | 0.78 | 0.92 |
| 5 | 57340073 | rs1470990 | 0.274 | 1.233 | -0.98 | 0.151 | 0.019 | 0.075 | 0.075 | 0.137 | 0.52 | 0.64 | 0.41 | 0.19 | 0.43 | 0.64 |
| 5 | 57340902 | rs7735334 | -0.713 | 0.907 | -0.908 | 0.673 | -0.003 | 0.013 | 0.013 | 0 | 0.09 | 0.12 | 0.27 | 0.17 | 0.1 | 0.14 |
| 5 | 57341758 | rs10940592 | -0.373 | 0.661 | -0.666 | 0.241 | -0.009 | -0.015 | -0.015 | 0 | 0.17 | 0.26 | 0.23 | 0.13 | 0.13 | 0.39 |
| 5 | 57342308 | rs1985870 | -0.41 | 1.357 | -0.176 | 0.619 | 0.006 | 0.004 | 0.004 | 0 | 0.13 | 0.18 | 0.11 | 0.15 | 0.14 | 0.11 |
| 5 | 57343495 | rs10940593 | -0.771 | 1.178 | -0.265 | 0.978 | -0.001 | 0.017 | 0.017 | 0 | 0.14 | 0.19 | 0.11 | 0.24 | 0.15 | 0.11 |
| 5 | 57343578 | rs1352615 | -0.615 | 1.036 | -0.648 | 0.665 | -0.001 | 0.002 | 0.002 | 0 | 0.44 | 0.58 | 0.56 | 0.42 | 0.43 | 0.57 |
| 5 | 57343925 | rs10057214 | 0.565 | 0.897 | -0.705 | 0.556 | -0.006 | -0.02 | -0.02 | 3.209 | 0.14 | 0.07 | 0.11 | 0.17 | 0.13 | 0.11 |
| 5 | 57346196 | rs17750724 | 0.464 | 0.941 | -0.654 | 0.533 | 0.006 | 0 | 0 | 0 | 0.13 | 0.18 | 0.11 | 0.15 | 0.15 | 0.11 |
| 5 | 57351996 | rs10461642 | 0.513 | 0.91 | -0.653 | 0.535 | 0.004 | 0.001 | 0.001 | 0 | 0.13 | 0.18 | 0.11 | 0.15 | 0.14 | 0.11 |
| 5 | 57353359 | rs6899128 | 1.048 | 0.865 | -0.518 | 0.485 | 0 | 0.016 | 0.016 | 0 | 0.08 | 0.05 | 0.04 | 0.01 | 0.06 | 0.03 |
| 5 | 57355531 | rs6859725 | -1.957 | 0.855 | -0.446 | 0.512 | -0.018 | -0.03 | -0.03 | 0 | 0.58 | 0.83 | 0.68 | 0.65 | 0.58 | 0.68 |
| 5 | 57359024 | rs7716601 | -1.579 | 0.831 | -0.475 | 0.426 | -0.024 | 0.014 | 0.014 | 0 | 0.52 | 0.78 | 0.64 | 0.64 | 0.5 | 0.65 |
| 5 | 57359476 | rs17750857 | 1.55 | 0.651 | -0.547 | 0.453 | -0.01 | -0.011 | -0.011 | 0 | 0.06 | 0 | 0.03 | 0.06 | 0.06 | 0.05 |
| 5 | 57359818 | rs11741803 | 1.124 | 0.672 | -0.514 | 0.459 | 0.025 | 0.009 | 0.009 | 0 | 0.08 | 0.18 | 0.08 | 0.08 | 0.09 | 0.06 |
| 5 | 57360670 | rs1387638 | 0.978 | 0.655 | -0.287 | 0.479 | -0.004 | 0.046 | 0.046 | 0.041 | 0.07 | 0.13 | 0.07 | 0.18 | 0.05 | 0.08 |
| 5 | 57361321 | rs6892519 | -0.481 | 0.661 | -0.212 | 0.434 | -0.011 | -0.006 | -0.006 | 0.002 | 0.31 | 0.41 | 0.34 | 0.35 | 0.28 | 0.49 |
| 5 | 57367770 | rs11954292 | 0.856 | 0.647 | -0.172 | 0.431 | -0.004 | -0.009 | -0.009 | 0.23 | 0.08 | 0.14 | 0.17 | 0.08 | 0.06 | 0.07 |
| 5 | 57369279 | rs6867541 | 0.117 | 0.632 | -0.186 | 0.426 | -0.001 | -0.001 | -0.001 | 0.008 | 0.23 | 0.23 | 0.25 | 0.26 | 0.19 | 0.43 |
| 5 | 57369327 | rs6867569 | 1.198 | 0.633 | -0.178 | 0.417 | -0.006 | -0.016 | -0.016 | 0 | 0.16 | 0.22 | 0.22 | 0.12 | 0.13 | 0.37 |
| 5 | 57369919 | rs16887769 | 0.849 | 0.657 | -0.038 | 0.441 | -0.004 | -0.008 | -0.008 | 0 | 0.08 | 0.11 | 0.16 | 0.08 | 0.06 | 0.07 |
| 5 | 57372783 | rs2171825 | -1.227 | 0.529 | 0.012 | 0.455 | 0.002 | -0.004 | -0.004 | 0 | 0.39 | 0.61 | 0.42 | 0.52 | 0.33 | 0.56 |
| 5 | 57372815 | rs6885426 | 1.06 | 0.546 | -0.02 | 0.418 | -0.008 | 0.023 | 0.023 | 0 | 0.07 | 0.03 | 0.05 | 0.01 | 0.08 | 0.03 |
| 5 | 57373820 | rs7731570 | -0.131 | 0.346 | 0.154 | 0.546 | 0.006 | 0.039 | 0.039 | 0 | 0.14 | 0.23 | 0.15 | 0.27 | 0.13 | 0.14 |
| 5 | 57374342 | rs1125324 | 0.62 | 0.387 | 0.14 | 0.535 | -0.01 | -0.036 | -0.036 | 0.526 | 0.07 | 0.01 | 0.03 | 0.14 | 0.07 | 0.05 |
| 5 | 57377708 | rs4700254 | 0.799 | 0.396 | 0.167 | 0.48 | -0.015 | -0.033 | -0.033 | 0 | 0.17 | 0.36 | 0.24 | 0.12 | 0.13 | 0.37 |
| 5 | 57378468 | rs1472034 | -0.617 | 0.411 | 0.051 | 0.495 | -0.005 | 0.002 | 0.002 | 0.05 | 0.32 | 0.6 | 0.38 | 0.39 | 0.25 | 0.52 |
| 5 | 57387770 | rs7735526 | -0.227 | 0.345 | 0.016 | 0.341 | -0.005 | -0.003 | -0.003 | 0 | 0.09 | 0.11 | 0.04 | 0.08 | 0.07 | 0.07 |
| 5 | 57391846 | rs6894496 | -1.389 | 0.278 | 0.48 | 0.282 | 0.003 | 0.017 | 0.017 | 0 | 0.29 | 0.33 | 0.36 | 0.42 | 0.3 | 0.24 |
| 5 | 57394960 | rs4700256 | -1.68 | 0.149 | 0.593 | 0.213 | -0.009 | -0.003 | -0.003 | 0 | 0.32 | 0.6 | 0.25 | 0.35 | 0.24 | 0.24 |
| 5 | 57395499 | rs12654580 | -0.521 | 0.158 | 0.592 | 0.221 | -0.005 | 0.013 | 0.013 | 0 | 0.21 | 0.3 | 0.18 | 0.31 | 0.18 | 0.18 |
| 5 | 57401646 | rs12055201 | 0.25 | 0.165 | 0.583 | 0.226 | 0.009 | 0.02 | 0.02 | 0 | 0.09 | 0.11 | 0.19 | 0.15 | 0.12 | 0.06 |
| 5 | 57403925 | rs16887819 | 1.057 | 0.141 | 0.954 | 0.222 | -0.005 | 0.024 | 0.024 | 0.202 | 0.07 | 0.11 | 0.07 | 0.19 | 0.04 | 0.07 |
| 5 | 57405259 | rs13185082 | 0.137 | 0.174 | 0.983 | 0.224 | 0.001 | 0.002 | 0.002 | 0.343 | 0.08 | 0.07 | 0.04 | 0.04 | 0.06 | 0.07 |
| 5 | 57405510 | rs16887824 | -0.462 | 0.191 | 0.897 | 0.232 | -0.004 | 0.012 | 0.012 | 0 | 0.21 | 0.3 | 0.18 | 0.31 | 0.18 | 0.17 |
| 5 | 57411570 | rs9283718 | -2.123 | 0.117 | 0.633 | 0.151 | -0.09 | -0.081 | -0.081 | 0 | 0.56 | 0.88 | 0.52 | 0.54 | 0.49 | 0.65 |
| 5 | 57413380 | rs1981840 | -0.744 | 0.145 | 0.653 | 0.161 | -0.002 | -0.006 | -0.006 | 0 | 0.17 | 0.12 | 0.22 | 0.19 | 0.19 | 0.13 |
| 5 | 57419661 | rs11958057 | 1.051 | 0.13 | 0.7 | 0.102 | -0.005 | -0.042 | -0.042 | 0 | 0.16 | 0.19 | 0.09 | 0.04 | 0.14 | 0.36 |
| 5 | 57419679 | rs9942337 | -0.155 | 0.124 | 0.472 | 0.161 | 0 | 0.004 | 0.004 | 0.793 | 0.12 | 0.11 | 0.19 | 0.15 | 0.12 | 0.06 |
| 5 | 57429105 | rs995476 | -2.234 | 0.112 | 0.33 | 0.106 | -0.105 | -0.088 | -0.088 | 0 | 0.57 | 0.91 | 0.52 | 0.53 | 0.49 | 0.65 |
| 5 | 57429577 | rs6867740 | -1.227 | 0.079 | 0.647 | 0.103 | 0.02 | 0.036 | 0.036 | 0 | 0.4 | 0.74 | 0.3 | 0.35 | 0.31 | 0.53 |
| 5 | 57430832 | rs723852 | -0.131 | 0.003 | 0.871 | 0.113 | -0.029 | 0.008 | 0.008 | 0.066 | 0.15 | 0.32 | 0.11 | 0.19 | 0.08 | 0.07 |
| 5 | 57431053 | rs733250 | -0.512 | 0.188 | 0.628 | 0.184 | 0.034 | 0.046 | 0.046 | 0 | 0.16 | 0.25 | 0.26 | 0.26 | 0.2 | 0.12 |
| 5 | 57435640 | rs2194191 | -0.549 | 0.102 | 0.37 | 0.239 | -0.006 | -0.01 | -0.01 | 0.019 | 0.36 | 0.37 | 0.21 | 0.28 | 0.28 | 0.52 |
| 5 | 57435801 | rs4389640 | -0.436 | 1.123 | -0.673 | 0.17 | -0.06 | -0.025 | -0.025 | 0 | 0.23 | 0.11 | 0.36 | 0.26 | 0.38 | 0.27 |
| 5 | 57440256 | rs10053377 | -1.013 | 0.344 | -0.863 | 0.122 | -0.032 | 0.023 | 0.023 | 0.492 | 0.63 | 0.74 | 0.5 | 0.27 | 0.58 | 0.61 |
| 5 | 57442823 | rs6882495 | 0.396 | -0.172 | -0.316 | -0.516 | -0.006 | -0.131 | -0.131 | 0.437 | 0.23 | 0.39 | 0.21 | 0 | 0.27 | 0.17 |
| 5 | 57442961 | rs6868757 | -1.218 | -0.31 | -0.363 | -0.393 | -0.013 | 0.011 | 0.011 | 0 | 0.57 | 0.67 | 0.51 | 0.36 | 0.54 | 0.62 |
| 5 | 57443601 | rs989579 | 1.546 | -0.244 | -0.465 | -0.415 | 0.001 | 0.004 | 0.004 | 4.294 | 0.08 | 0.03 | 0.03 | 0.06 | 0.08 | 0.07 |
| 5 | 57443632 | rs989578 | -1.85 | -0.195 | -0.501 | -0.355 | -0.022 | -0.022 | -0.022 | 0 | 0.65 | 0.74 | 0.54 | 0.42 | 0.61 | 0.69 |
| 5 | 57444953 | rs10043517 | 0.647 | -0.259 | -0.54 | -0.103 | 0.01 | 0.034 | 0.034 | 0 | 0.13 | 0.15 | 0.25 | 0.29 | 0.19 | 0.18 |
| 5 | 57445023 | rs16887903 | 1.091 | -0.391 | -0.67 | -0.218 | 0.033 | 0.048 | 0.048 | 0 | 0.19 | 0.11 | 0.05 | 0.07 | 0.08 | 0.27 |
| 5 | 57454970 | rs764630 | 0.758 | -0.466 | -0.221 | 0.033 | -0.006 | 0.007 | 0.007 | 0 | 0.21 | 0.19 | 0.18 | 0.34 | 0.1 | 0.35 |
| 5 | 57473240 | rs16887917 | 2.119 | -0.392 | -0.563 | 0.327 | -0.002 | -0.027 | -0.027 | 0.004 | 0.16 | 0.05 | 0.18 | 0.25 | 0.16 | 0.18 |
| 5 | 57475024 | rs2962009 | -0.989 | -0.492 | -0.271 | 0.091 | 0 | 0.002 | 0.002 | 4.567 | 0.89 | 0.95 | 0.97 | 0.93 | 0.91 | 0.91 |
| 5 | 57477898 | rs13173077 | 2.029 | -0.412 | -0.118 | -0.068 | -0.022 | -0.02 | -0.02 | 0 | 0.34 | 0.23 | 0.46 | 0.59 | 0.39 | 0.32 |
| 5 | 57478318 | rs2964185 | 0.239 | -0.503 | 0.033 | -0.236 | -0.006 | -0.07 | -0.07 | 0.172 | 0.21 | 0.41 | 0.19 | 0.01 | 0.22 | 0.13 |
| 5 | 57482900 | rs2964188 | 0.467 | -0.324 | 0.081 | -0.124 | 0.012 | -0.006 | -0.006 | 0 | 0.28 | 0.22 | 0.22 | 0.34 | 0.15 | 0.42 |

| | | | | | | | | | | | | | | | | |
|---|---|---|---|---|---|---|---|---|---|---|---|---|---|---|---|---|
| 5 | 57482938 | rs2964189 | -1.283 | -0.325 | -0.219 | -0.146 | -0.008 | 0.02 | 0.02 | 0 | 0.55 | 0.72 | 0.52 | 0.35 | 0.52 | 0.59 |
| 5 | 57487610 | rs2910597 | -1.239 | -0.316 | -0.225 | -0.127 | -0.007 | 0.005 | 0.005 | 0 | 0.55 | 0.71 | 0.52 | 0.41 | 0.52 | 0.6 |
| 5 | 57491648 | rs17438760 | 1.705 | -0.252 | -0.458 | -0.091 | -0.017 | 0.022 | 0.022 | 0 | 0.14 | 0.07 | 0.16 | 0 | 0.21 | 0.1 |
| 5 | 57494261 | rs2194194 | 1.811 | -0.289 | -0.505 | 0.097 | -0.019 | -0.028 | -0.028 | 0 | 0.34 | 0.23 | 0.43 | 0.52 | 0.39 | 0.32 |
| 5 | 57495341 | rs2964206 | 0.701 | -0.456 | -0.58 | 0.098 | 0.061 | 0.049 | 0.049 | 0.017 | 0.26 | 0.11 | 0.11 | 0.16 | 0.13 | 0.33 |
| 5 | 57499494 | rs2194193 | 0.977 | 0.475 | 0.273 | 0.499 | -0.002 | 0.001 | 0.001 | 0 | 0.62 | 0.4 | 0.57 | 0.69 | 0.52 | 0.65 |
| 5 | 57503024 | rs2964209 | -1.513 | 0.237 | -0.061 | 0.459 | -0.009 | 0.016 | 0.016 | 5.841 | 0.93 | 1 | 0.97 | 0.95 | 0.94 | 0.93 |
| 5 | 57506467 | rs7701350 | -0.505 | 0.344 | 0.362 | 0.379 | 0 | -0.069 | -0.069 | 0 | 0.25 | 0.43 | 0.34 | 0.05 | 0.31 | 0.15 |
| 5 | 57506770 | rs2964173 | -0.136 | 0.283 | 0.462 | 0.399 | -0.016 | -0.031 | -0.031 | 0.296 | 0.9 | 0.76 | 0.95 | 0.94 | 0.94 | 0.93 |
| 5 | 57508816 | rs2962004 | 1.345 | 0.306 | 0.424 | 0.519 | -0.005 | -0.027 | -0.027 | 3.589 | 0.63 | 0.44 | 0.61 | 0.81 | 0.6 | 0.76 |
| 5 | 57517906 | rs12658992 | 1.122 | 0.348 | 0.249 | 0.522 | -0.01 | 0.005 | 0.005 | 0.018 | 0.1 | 0.03 | 0.17 | 0.05 | 0.13 | 0.05 |
| 5 | 57523164 | rs2910587 | 1.715 | 0.465 | 0.34 | 1.096 | 0 | 0.003 | 0.003 | 0 | 0.6 | 0.43 | 0.45 | 0.53 | 0.55 | 0.59 |
| 5 | 57525196 | rs2910584 | 1.426 | 0.386 | 0.431 | 0.898 | 0.01 | 0.04 | 0.04 | 0.055 | 0.48 | 0.38 | 0.55 | 0.71 | 0.52 | 0.54 |
| 5 | 57526780 | rs1035473 | 0.634 | 0.541 | 0.353 | 1.02 | 0.008 | -0.014 | -0.014 | 0 | 0.12 | 0.22 | 0.19 | 0.05 | 0.14 | 0.05 |
| 5 | 57530181 | rs11742503 | -0.085 | 0.556 | 0.349 | 1.109 | 0.019 | 0.11 | 0.11 | 0 | 0.15 | 0.24 | 0.34 | 0.32 | 0.18 | 0.22 |
| 5 | 57532246 | rs17695268 | 0.333 | 0.41 | 0.229 | 0.746 | -0.012 | -0.032 | -0.032 | 0 | 0.27 | 0.35 | 0.22 | 0.17 | 0.25 | 0.3 |
| 5 | 57540013 | rs11745219 | 0.953 | 0.629 | -0.085 | 0.781 | -0.007 | 0.004 | 0.004 | 0.006 | 0.11 | 0.07 | 0.17 | 0.05 | 0.13 | 0.05 |
| 5 | 57566073 | rs2434584 | -0.291 | 0.296 | -0.022 | 0.745 | 0.011 | 0.023 | 0.023 | 3.924 | 0.88 | 0.97 | 0.97 | 0.92 | 0.92 | 0.89 |
| 5 | 57578037 | rs43217 | 0.4 | 0.187 | -0.056 | 0.683 | 0.016 | 0.082 | 0.082 | 2.586 | 0.65 | 0.87 | 0.78 | 0.77 | 0.69 | 0.59 |
| 5 | 57583686 | rs37512 | 0.794 | 0.112 | 0.437 | 0.647 | -0.02 | -0.059 | -0.059 | 0.072 | 0.62 | 0.54 | 0.75 | 0.76 | 0.69 | 0.59 |
| 5 | 57594172 | rs17707535 | 0.873 | 0.28 | -0.028 | 0.651 | 0.017 | 0.047 | 0.047 | 0 | 0.21 | 0.09 | 0.15 | 0 | 0.17 | 0.12 |
| 5 | 57607955 | rs151913 | 0.87 | 0.534 | 0.049 | 0.879 | -0.001 | 0.005 | 0.005 | 0 | 0.61 | 0.69 | 0.71 | 0.7 | 0.61 | 0.52 |
| 5 | 57617905 | rs42964 | 1.021 | 0.549 | 0.068 | 1.148 | 0.008 | 0.075 | 0.075 | 0.368 | 0.83 | 0.78 | 0.75 | 0.5 | 0.81 | 0.62 |
| 5 | 57618101 | rs173425 | -1.149 | 0.566 | 0.091 | 1.163 | 0.042 | 0.228 | 0.228 | 0 | 0.08 | 0.19 | 0.22 | 0.41 | 0.11 | 0.26 |
| 5 | 57619964 | rs40210 | -0.683 | 0.58 | -0.004 | 1.166 | 0.005 | 0.016 | 0.016 | 0.533 | 0.9 | 0.97 | 0.97 | 0.9 | 0.92 | 0.88 |
| 5 | 57620409 | rs152012 | 1.031 | 0.597 | 0.08 | 1.077 | -0.001 | 0.003 | 0.003 | 0 | 0.59 | 0.6 | 0.56 | 0.51 | 0.6 | 0.47 |
| 5 | 57620534 | rs151868 | -0.98 | 0.634 | -0.11 | 1.033 | 0.004 | 0.01 | 0.01 | 0.565 | 0.92 | 0.97 | 0.97 | 0.92 | 0.93 | 0.9 |
| 5 | 57625508 | rs10513141 | 1.307 | 1.055 | -0.215 | 1.005 | -0.008 | 0.002 | 0.002 | 0.095 | 0.09 | 0.03 | 0.15 | 0.06 | 0.11 | 0.05 |
| 5 | 57630221 | rs185425 | 0.492 | 0.929 | -0.102 | 1.11 | 0 | -0.001 | -0.001 | 0.703 | 0.66 | 0.71 | 0.66 | 0.74 | 0.71 | 0.65 |
| 5 | 57635488 | rs245493 | 0.181 | 0.817 | -0.174 | 1.164 | -0.008 | -0.022 | -0.022 | 0.351 | 0.88 | 0.96 | 0.92 | 0.76 | 0.86 | 0.76 |
| 5 | 57635544 | rs152010 | 0.181 | 0.817 | -0.174 | 1.164 | -0.008 | -0.022 | -0.022 | 0 | 0.88 | 0.96 | 0.92 | 0.76 | 0.86 | 0.76 |
| 5 | 57636114 | rs6450473 | 0.493 | 0.849 | -0.153 | 1.156 | 0 | -0.002 | -0.002 | 0 | 0.25 | 0.25 | 0.32 | 0.18 | 0.25 | 0.24 |
| 5 | 57636145 | rs245494 | 0.671 | 0.857 | -0.149 | 1.161 | -0.003 | -0.005 | -0.005 | 0.608 | 0.63 | 0.71 | 0.6 | 0.57 | 0.61 | 0.53 |
| 5 | 57647762 | rs36644 | -0.017 | 0.831 | 0.01 | 1.221 | 0.002 | 0.001 | 0.001 | 0 | 0.28 | 0.29 | 0.38 | 0.3 | 0.31 | 0.34 |
| 5 | 57647874 | rs37533 | -0.388 | 0.796 | -0.129 | 1.145 | -0.004 | 0.027 | 0.027 | 0 | 0.88 | 0.97 | 0.93 | 0.92 | 0.87 | 0.88 |
| 5 | 57652016 | rs27210 | -0.388 | 0.774 | -0.129 | 1.365 | -0.002 | 0.012 | 0.012 | 0 | 0.88 | 0.97 | 0.93 | 0.89 | 0.87 | 0.87 |
| 5 | 57665935 | rs151915 | 1.201 | 1.06 | 0.013 | 0.961 | -0.016 | 0.004 | 0.004 | 0 | 0.4 | 0.5 | 0.31 | 0.45 | 0.29 | 0.4 |
| 5 | 57696464 | rs889303 | -0.783 | 1.238 | -0.134 | 0.889 | 0 | 0.002 | 0.002 | 0 | 0.34 | 0.38 | 0.46 | 0.43 | 0.39 | 0.37 |
| 5 | 57697254 | rs11739229 | 0.5 | 1.187 | -0.221 | 0.835 | -0.002 | 0.014 | 0.014 | 0 | 0.12 | 0.03 | 0.07 | 0.1 | 0.13 | 0.12 |
| 5 | 57698294 | rs905199 | -0.939 | 1.131 | -0.09 | 0.935 | -0.018 | 0.073 | 0.073 | 0 | 0.18 | 0.32 | 0.23 | 0.36 | 0.13 | 0.28 |
| 5 | 57700852 | rs10472060 | 1.287 | 1.163 | -0.176 | 0.908 | -0.003 | -0.014 | -0.014 | 0 | 0.23 | 0.16 | 0.19 | 0 | 0.24 | 0.14 |
| 5 | 57703549 | rs2548599 | 1.315 | 1.159 | -0.108 | 0.968 | 0.011 | -0.007 | -0.007 | 0 | 0.39 | 0.35 | 0.3 | 0.45 | 0.29 | 0.41 |
| 5 | 57710739 | rs719428 | 0.258 | 1.344 | -0.372 | 0.226 | 0 | 0.049 | 0.049 | 0 | 0.57 | 0.68 | 0.53 | 0.81 | 0.42 | 0.68 |
| 5 | 57713579 | rs17766068 | 1.491 | 1.489 | -0.691 | 0.388 | -0.04 | 0.036 | 0.036 | 0 | 0.16 | 0.06 | 0.23 | 0.07 | 0.26 | 0.09 |
| 5 | 57718996 | rs154475 | 0.207 | 1.454 | -0.691 | 0.392 | -0.043 | 0.059 | 0.059 | 0 | 0.72 | 0.91 | 0.71 | 0.83 | 0.61 | 0.79 |
| 5 | 57722165 | rs184968 | 0.605 | 1.482 | -0.276 | 0.499 | -0.021 | 0.004 | 0.004 | 0 | 0.19 | 0.15 | 0.17 | 0.18 | 0.34 | 0.17 |
| 5 | 57722451 | rs154477 | 0.515 | 1.067 | -0.442 | 0.217 | 0.019 | 0.2 | 0.2 | 0 | 0.51 | 0.8 | 0.59 | 0.84 | 0.4 | 0.64 |
| 5 | 57724792 | rs158281 | -0.241 | 1.016 | -0.336 | -0.178 | -0.014 | -0.001 | -0.001 | 0 | 0.86 | 0.92 | 0.89 | 0.84 | 0.84 | 0.7 |
| 5 | 57725558 | rs158282 | 0.533 | 0.25 | -0.066 | -0.1 | -0.004 | 0 | 0 | 0.207 | 0.64 | 0.62 | 0.54 | 0.64 | 0.46 | 0.56 |
| 5 | 57728017 | rs27856 | 1.245 | 0.297 | 0.014 | -0.207 | 0.052 | -0.004 | -0.004 | 0 | 0.45 | 0.18 | 0.43 | 0.53 | 0.35 | 0.46 |
| 5 | 57736111 | rs158150 | 0.291 | 0.192 | -0.027 | -0.246 | -0.019 | 0.023 | 0.023 | 0 | 0.66 | 0.7 | 0.55 | 0.82 | 0.47 | 0.56 |
| 5 | 57743302 | rs32617 | 0.917 | 0.286 | 0.086 | -0.186 | 0.024 | 0.049 | 0.049 | 0 | 0.48 | 0.34 | 0.47 | 0.69 | 0.37 | 0.46 |
| 5 | 57749769 | rs1848511 | 2.047 | 0.238 | 0.049 | -0.099 | 0.011 | 0.012 | 0.012 | 0 | 0.14 | 0.08 | 0.05 | 0.12 | 0.1 | 0.1 |
| 5 | 57750267 | rs32613 | -1.93 | 0.307 | -0.054 | -0.112 | -0.017 | -0.01 | -0.01 | 0 | 0.95 | 1 | 0.95 | 0.92 | 0.94 | 0.85 |
| 5 | 57751443 | rs1042994 | 0.613 | 0.338 | 0.128 | -0.106 | 0.027 | -0.048 | -0.048 | 0 | 0.09 | 0.21 | 0.21 | 0 | 0.13 | 0.07 |
| 5 | 57753116 | rs17710502 | 0.677 | 0.345 | 0.15 | -0.103 | 0.032 | -0.048 | -0.048 | 0 | 0.09 | 0.2 | 0.2 | 0 | 0.13 | 0.07 |
| 5 | 57753149 | rs17767072 | 0.677 | 0.345 | 0.14 | -0.103 | 0.029 | -0.047 | -0.047 | 0 | 0.09 | 0.2 | 0.2 | 0 | 0.13 | 0.06 |
| 5 | 57754851 | rs702722 | 1.534 | 0.211 | 0.249 | 0.017 | -0.021 | 0.032 | 0.032 | 0 | 0.16 | 0.23 | 0.07 | 0.3 | 0.1 | 0.1 |
| 5 | 57763091 | rs6890785 | 0.416 | 0.408 | -0.001 | -0.147 | 0.026 | -0.059 | -0.059 | 0 | 0.11 | 0.22 | 0.22 | 0 | 0.16 | 0.07 |
| 5 | 57765914 | rs11948619 | 0.223 | 0.212 | -0.009 | -0.249 | -0.001 | 0 | 0 | 0.357 | 0.53 | 0.53 | 0.68 | 0.69 | 0.49 | 0.51 |
| 5 | 57773368 | rs10471452 | 0.284 | -0.695 | -0.551 | -0.367 | 0.026 | 0.009 | 0.009 | 0.011 | 0.38 | 0.26 | 0.4 | 0.63 | 0.24 | 0.47 |
| 5 | 57774538 | rs16888206 | 1.317 | -1.054 | -0.873 | -0.087 | 0.009 | -0.027 | -0.027 | 0 | 0.17 | 0.07 | 0.12 | 0.23 | 0.11 | 0.12 |
| 5 | 57776880 | rs2662619 | -0.448 | -0.59 | -0.565 | -0.93 | 0.017 | 0.001 | 0.001 | 0 | 0.67 | 0.78 | 0.6 | 0.35 | 0.74 | 0.56 |
| 5 | 57782141 | rs10074748 | 0.847 | -0.215 | 0.207 | -1.137 | 0.007 | -0.019 | -0.019 | 0 | 0.28 | 0.37 | 0.22 | 0.17 | 0.36 | 0.12 |
| 5 | 57786110 | rs10070522 | -0.706 | -0.331 | -0.096 | -1.21 | 0.055 | 0.121 | 0.121 | 1.253 | 0.5 | 0.74 | 0.53 | 0.27 | 0.64 | 0.45 |
| 5 | 57787376 | rs2307117 | 0.348 | -0.249 | 0.053 | -1.239 | 0.025 | -0.058 | -0.058 | 0.097 | 0.36 | 0.53 | 0.42 | 0.17 | 0.49 | 0.17 |
| 5 | 57790337 | rs7704785 | -0.645 | -0.349 | -0.104 | -1.352 | 0.058 | 0.202 | 0.202 | 1.11 | 0.5 | 0.74 | 0.54 | 0.11 | 0.63 | 0.44 |
| 5 | 57791311 | rs1389308 | -0.671 | -0.353 | -0.106 | -1.354 | 0.056 | 0.199 | 0.199 | 0.975 | 0.51 | 0.74 | 0.54 | 0.11 | 0.63 | 0.44 |
| 5 | 57794296 | rs1858086 | -0.166 | -0.433 | -0.068 | -1.365 | -0.01 | 0.015 | 0.015 | 0.226 | 0.42 | 0.58 | 0.33 | 0.12 | 0.5 | 0.39 |

| | | | | | | | | | | | | | | | | |
|---|---|---|---|---|---|---|---|---|---|---|---|---|---|---|---|---|
| 5 | 57795822 | rs10057028 | -0.685 | -0.429 | -0.186 | -1.373 | 0.051 | 0.236 | 0.236 | 1.069 | 0.51 | 0.81 | 0.54 | 0.11 | 0.62 | 0.44 |
| 5 | 57803076 | rs7717166 | -0.043 | -0.334 | -0.104 | -1.309 | 0.005 | -0.047 | -0.047 | 0 | 0.41 | 0.45 | 0.32 | 0.11 | 0.5 | 0.39 |
| 5 | 57830080 | rs13178332 | -0.721 | -0.308 | -0.463 | -1.256 | 0.026 | 0.075 | 0.075 | 1.375 | 0.49 | 0.67 | 0.52 | 0.11 | 0.62 | 0.43 |
| 5 | 57830713 | rs12153332 | 1.489 | -0.07 | -0.364 | -1.022 | -0.02 | 0.044 | 0.044 | 0 | 0.08 | 0.01 | 0.05 | 0 | 0.1 | 0.13 |
| 5 | 57839048 | rs2136222 | 1.433 | -0.441 | -0.451 | -0.899 | -0.017 | 0.021 | 0.021 | 0 | 0.27 | 0.12 | 0.21 | 0.21 | 0.23 | 0.27 |
| 5 | 57845244 | rs10070704 | -1.128 | 0.522 | -0.308 | -1.275 | 0.026 | 0.04 | 0.04 | 0 | 0.57 | 0.74 | 0.57 | 0.28 | 0.7 | 0.47 |
| 5 | 57847893 | rs10514853 | 0.283 | 0.686 | -0.724 | -1.186 | -0.029 | 0.056 | 0.056 | 0 | 0.34 | 0.2 | 0.25 | 0.18 | 0.43 | 0.17 |
| 5 | 57862640 | rs6450480 | 0.735 | 0.811 | -0.902 | -0.892 | 0.026 | -0.038 | -0.038 | 0 | 0.07 | 0.16 | 0.19 | 0 | 0.11 | 0.05 |
| 5 | 57868045 | rs6450482 | -0.344 | 0.745 | -0.881 | -0.896 | 0.041 | -0.065 | -0.065 | 0 | 0.11 | 0.36 | 0.22 | 0 | 0.11 | 0.05 |
| 5 | 57878863 | rs1876691 | 0.161 | 1.148 | -1.416 | -0.681 | 0.038 | -0.047 | -0.047 | 0 | 0.08 | 0.24 | 0.2 | 0 | 0.11 | 0.05 |
| 5 | 57889306 | rs6896790 | -0.046 | 0.919 | -2.046 | -0.65 | -0.032 | 0.076 | 0.076 | 0 | 0.41 | 0.29 | 0.44 | 0.18 | 0.54 | 0.21 |
| 5 | 57894922 | rs2591919 | -0.493 | 1.208 | -1.892 | -0.146 | -0.045 | 0.006 | 0.006 | 0 | 0.12 | 0.01 | 0.1 | 0.09 | 0.16 | 0.25 |
| 5 | 57905474 | rs167132 | -0.521 | 1.293 | -1.761 | -0.021 | -0.047 | 0.026 | 0.026 | 0 | 0.84 | 0.99 | 0.89 | 0.95 | 0.82 | 0.87 |
| 5 | 57909545 | rs292966 | 0.019 | 1.041 | -1.475 | 0.156 | 0.002 | -0.004 | -0.004 | 0 | 0.32 | 0.31 | 0.36 | 0.57 | 0.23 | 0.36 |
| 5 | 57910859 | rs984664 | -0.131 | 1.427 | -1.346 | -0.047 | 0.008 | -0.023 | -0.023 | 0.001 | 0.08 | 0.1 | 0.18 | 0 | 0.14 | 0.11 |
| 5 | 57911848 | rs292970 | -0.286 | 1.14 | -1.513 | 0.417 | -0.006 | -0.005 | -0.005 | 0.012 | 0.13 | 0.09 | 0.33 | 0.18 | 0.18 | 0.27 |
| 5 | 57915271 | rs433751 | -1.14 | 1.311 | -1.841 | 0.207 | -0.043 | 0.042 | 0.042 | 0 | 0.84 | 0.99 | 0.87 | 0.95 | 0.81 | 0.87 |
| 5 | 57915926 | rs430763 | -0.73 | 1.204 | -1.407 | 0.09 | 0 | -0.005 | -0.005 | 0 | 0.18 | 0.24 | 0.37 | 0.18 | 0.2 | 0.29 |
| 5 | 57916579 | rs386161 | -0.849 | 1.16 | -1.354 | -0.087 | 0.018 | 0.082 | 0.082 | 1.468 | 0.56 | 0.84 | 0.72 | 0.72 | 0.57 | 0.66 |
| 5 | 57917792 | rs425203 | -0.844 | 1.605 | -1.147 | -0.18 | 0.005 | 0 | 0 | 0 | 0.28 | 0.32 | 0.47 | 0.34 | 0.37 | 0.4 |
| 5 | 57919019 | rs425872 | -1.366 | 1.355 | -1.452 | -0.357 | -0.034 | 0.194 | 0.194 | 1.32 | 0.62 | 0.96 | 0.81 | 0.93 | 0.65 | 0.68 |
| 5 | 57931376 | rs292985 | -0.585 | 1.457 | -1.335 | -0.265 | -0.004 | -0.001 | -0.001 | 0 | 0.86 | 0.9 | 0.66 | 0.82 | 0.81 | 0.72 |
| 5 | 57933998 | rs292988 | 0.218 | 1.465 | -1.348 | -0.278 | -0.013 | 0.002 | 0.002 | 1.189 | 0.55 | 0.16 | 0.39 | 0.44 | 0.59 | 0.45 |
| 5 | 57934280 | rs292990 | 0.776 | 1.459 | -1.353 | -0.309 | 0.021 | 0.091 | 0.091 | 1.424 | 0.44 | 0.14 | 0.3 | 0.28 | 0.43 | 0.34 |
| 5 | 57935865 | rs158966 | 1.815 | 1.498 | -1.358 | -0.152 | -0.025 | 0.011 | 0.011 | 0.005 | 0.16 | 0.02 | 0.12 | 0.09 | 0.17 | 0.14 |
| 5 | 57937600 | rs13354637 | 1.119 | 1.455 | -0.792 | -0.142 | 0.001 | 0.064 | 0.064 | 0 | 0.27 | 0.44 | 0.23 | 0.35 | 0.21 | 0.25 |
| 5 | 57939227 | rs4997647 | 1.515 | 1.386 | -0.858 | -0.188 | 0.004 | 0.075 | 0.075 | 0 | 0.05 | 0.11 | 0.1 | 0.21 | 0.07 | 0.02 |
| 5 | 57940720 | rs13155394 | 2.267 | 1.357 | -0.926 | -0.253 | 0.012 | 0.125 | 0.125 | 3.371 | 0.22 | 0.03 | 0.05 | 0.02 | 0.15 | 0.19 |
| 5 | 57944829 | rs965795 | -0.452 | 1.413 | -0.914 | -0.002 | -0.042 | 0.089 | 0.089 | 0.82 | 0.46 | 0.82 | 0.46 | 0.67 | 0.46 | 0.39 |
| 5 | 57945250 | rs12054892 | 1.127 | 1.435 | -0.973 | -0.037 | 0.001 | 0.049 | 0.049 | 0 | 0.27 | 0.44 | 0.23 | 0.31 | 0.21 | 0.25 |
| 5 | 57948000 | rs9292170 | 0.167 | 1.424 | -0.962 | -0.038 | 0.011 | 0.018 | 0.018 | 0 | 0.38 | 0.46 | 0.33 | 0.47 | 0.38 | 0.37 |
| 5 | 57955360 | rs158955 | -0.193 | 1.691 | -0.898 | 0.129 | 0.002 | 0.017 | 0.017 | 0 | 0.81 | 0.8 | 0.56 | 0.58 | 0.73 | 0.7 |
| 5 | 57964018 | rs10078211 | 0.377 | 1.663 | -0.787 | 0.081 | 0.003 | -0.001 | -0.001 | 1.476 | 0.14 | 0.21 | 0.11 | 0.15 | 0.17 | 0.12 |
| 5 | 57964175 | rs293031 | 0.819 | 1.607 | -0.77 | 0.035 | -0.011 | -0.024 | -0.024 | 0 | 0.4 | 0.22 | 0.35 | 0.39 | 0.39 | 0.39 |
| 5 | 57967004 | rs17281110 | 1.816 | 1.558 | -0.794 | 0.312 | -0.027 | 0.01 | 0.01 | 0 | 0.13 | 0.04 | 0.16 | 0.06 | 0.13 | 0.17 |
| 5 | 57972853 | rs158990 | 0.045 | 1.889 | -0.395 | 0.395 | 0.006 | 0.031 | 0.031 | 0 | 0.78 | 0.71 | 0.52 | 0.57 | 0.7 | 0.7 |
| 5 | 57976523 | rs10060872 | -0.229 | 2.251 | 0.055 | 0.846 | 0.022 | -0.003 | -0.003 | 0 | 0.14 | 0.29 | 0.12 | 0.15 | 0.17 | 0.12 |
| 5 | 57981160 | rs158999 | -0.1 | 1.458 | 0.09 | 0.801 | 0.032 | 0.032 | 0.032 | 0.037 | 0.51 | 0.25 | 0.42 | 0.53 | 0.54 | 0.52 |
| 5 | 58009228 | rs293020 | 1.237 | 1.441 | -0.312 | 0.792 | -0.051 | -0.029 | -0.029 | 0.534 | 0.11 | 0.01 | 0.13 | 0.09 | 0.17 | 0.15 |
| 5 | 58013991 | rs293025 | 0.063 | 0.816 | 0.208 | 0.666 | 0.02 | 0.013 | 0.013 | 0.283 | 0.53 | 0.32 | 0.4 | 0.54 | 0.58 | 0.53 |
| 5 | 58028655 | rs293012 | -0.559 | 0.829 | -0.317 | 0.403 | 0.018 | 0.034 | 0.034 | 1.168 | 0.62 | 0.71 | 0.73 | 0.73 | 0.65 | 0.62 |
| 5 | 58031027 | rs10472071 | -0.348 | 0.843 | -0.282 | 0.412 | 0.004 | 0.022 | 0.022 | 0 | 0.07 | 0.24 | 0.17 | 0.13 | 0.07 | 0.07 |
| 5 | 58034966 | rs293002 | -0.159 | 0.917 | -0.498 | 0.389 | -0.002 | -0.004 | -0.004 | 0.022 | 0.55 | 0.47 | 0.56 | 0.6 | 0.58 | 0.55 |
| 5 | 58039641 | rs720084 | -0.366 | 1.555 | -0.444 | 1.278 | -0.046 | -0.057 | -0.057 | 0 | 0.11 | 0.01 | 0.08 | 0.21 | 0.15 | 0.15 |
| 5 | 58039977 | rs7720815 | -1.743 | 1.419 | -0.065 | 1.234 | -0.003 | -0.015 | -0.015 | 0 | 0.31 | 0.69 | 0.4 | 0.54 | 0.34 | 0.25 |
| 5 | 58040838 | rs4296764 | -1.067 | 1.581 | 0.282 | 1.334 | 0.023 | 0.122 | 0.122 | 0 | 0.16 | 0.49 | 0.29 | 0.34 | 0.18 | 0.1 |
| 5 | 58041996 | rs11953345 | -0.507 | 1.626 | 0.665 | 1.233 | 0.009 | 0.047 | 0.047 | 0 | 0.11 | 0.16 | 0.27 | 0.3 | 0.16 | 0.09 |
| 5 | 58043722 | rs4700305 | -0.482 | 1.635 | 0.625 | 1.238 | 0.009 | 0.046 | 0.046 | 0 | 0.11 | 0.16 | 0.26 | 0.3 | 0.16 | 0.09 |
| 5 | 58044070 | rs7717684 | -1.841 | 1.672 | 0.581 | 1.275 | 0.026 | 0.061 | 0.061 | 0 | 0.26 | 0.5 | 0.37 | 0.48 | 0.33 | 0.22 |
| 5 | 58045259 | rs292958 | -0.709 | 1.911 | -0.227 | 1.388 | 0.013 | 0.046 | 0.046 | 0 | 0.12 | 0.16 | 0.28 | 0.31 | 0.18 | 0.1 |
| 5 | 58049332 | rs10514874 | -0.5 | 2.078 | -0.677 | 1.084 | -0.062 | -0.048 | -0.048 | 0.645 | 0.11 | 0.01 | 0.09 | 0.2 | 0.18 | 0.13 |
| 5 | 58049391 | rs292957 | 0.596 | 1.788 | -1.169 | 0.517 | -0.043 | 0.074 | 0.074 | 0.017 | 0.39 | 0.23 | 0.26 | 0.19 | 0.43 | 0.39 |
| 5 | 58050530 | rs895377 | 0.09 | 1.966 | -0.621 | 0.491 | 0.003 | 0.009 | 0.009 | 0 | 0.48 | 0.44 | 0.26 | 0.28 | 0.42 | 0.47 |
| 5 | 58051351 | rs10940623 | -0.353 | 1.785 | -0.448 | 0.858 | 0.002 | -0.002 | -0.002 | 0.184 | 0.19 | 0.18 | 0.14 | 0.19 | 0.19 | 0.21 |
| 5 | 58056208 | rs292950 | -0.521 | 2.025 | -0.563 | 1.149 | -0.019 | -0.025 | -0.025 | 0 | 0.3 | 0.25 | 0.38 | 0.48 | 0.4 | 0.35 |
| 5 | 58059327 | rs1031199 | 1.7 | 1.538 | -0.614 | 0.941 | 0.045 | 0.136 | 0.136 | 0.056 | 0.42 | 0.15 | 0.29 | 0.15 | 0.34 | 0.45 |
| 5 | 58061855 | rs10073306 | 0.153 | 1.353 | -0.157 | 1.516 | -0.014 | -0.012 | -0.012 | 0 | 0.1 | 0.07 | 0.1 | 0.17 | 0.14 | 0.13 |
| 5 | 58061888 | rs292960 | -0.423 | 0.95 | -0.085 | 1.51 | -0.003 | -0.003 | -0.003 | 0.016 | 0.19 | 0.18 | 0.27 | 0.31 | 0.24 | 0.22 |
| 5 | 58063494 | rs1422228 | 0.153 | 0.807 | -0.13 | 1.419 | -0.005 | -0.019 | -0.019 | 0 | 0.79 | 0.82 | 0.75 | 0.62 | 0.77 | 0.82 |
| 5 | 58063797 | rs12522690 | 2.044 | 0.67 | -0.214 | 1.309 | 0.014 | 0.095 | 0.095 | 2.939 | 0.52 | 0.23 | 0.54 | 0.27 | 0.57 | 0.61 |
| 5 | 58070983 | rs10051270 | 0.264 | 0.989 | -0.46 | 1.481 | -0.035 | 0.001 | 0.001 | 0 | 0.1 | 0.05 | 0.09 | 0.09 | 0.13 | 0.14 |
| 5 | 58071799 | rs963443 | 1.789 | 0.924 | 0.037 | 1.472 | 0.001 | -0.008 | -0.008 | 0 | 0.6 | 0.44 | 0.57 | 0.34 | 0.59 | 0.65 |
| 5 | 58073010 | rs6898086 | -0.401 | 1.063 | -0.404 | 1.943 | -0.053 | -0.055 | -0.055 | 1.311 | 0.91 | 0.99 | 0.8 | 0.8 | 0.84 | 0.92 |
| 5 | 58076467 | rs1010757 | -0.243 | 1.028 | -0.289 | 1.417 | -0.015 | -0.05 | -0.05 | 0 | 0.79 | 0.85 | 0.76 | 0.63 | 0.77 | 0.83 |
| 5 | 58077903 | rs372279 | 0.052 | 0.99 | -0.116 | 1.348 | -0.026 | 0.115 | 0.115 | 0 | 0.08 | 0.26 | 0.08 | 0.2 | 0.04 | 0.02 |
| 5 | 58079865 | rs380346 | 0.673 | 1.019 | -0.183 | 1.339 | -0.044 | -0.074 | -0.074 | 0.343 | 0.69 | 0.78 | 0.66 | 0.53 | 0.64 | 0.68 |
| 5 | 58081681 | rs406992 | -0.738 | 0.877 | -0.207 | 1.263 | -0.002 | 0.034 | 0.034 | 0 | 0.18 | 0.32 | 0.17 | 0.28 | 0.16 | 0.16 |
| 5 | 58093288 | rs2675380 | 0.674 | 0.937 | -0.915 | 0.396 | 0.001 | 0.068 | 0.068 | 0.755 | 0.56 | 0.24 | 0.56 | 0.27 | 0.58 | 0.58 |
| 5 | 58095222 | rs4700309 | -0.522 | 0.926 | -0.471 | 0.014 | 0.003 | -0.007 | -0.007 | 0.811 | 0.36 | 0.49 | 0.4 | 0.64 | 0.36 | 0.32 |

| | | | | | | | | | | | | | | | | |
|---|---|---|---|---|---|---|---|---|---|---|---|---|---|---|---|---|
| 5 | 58096085 | rs1400226 | 0.638 | 1.28 | 0.049 | -0.305 | 0.025 | -0.031 | -0.031 | 0 | 0.66 | 0.5 | 0.55 | 0.23 | 0.6 | 0.59 |
| 5 | 58100626 | rs1423377 | 0.849 | 0.978 | 0.746 | -0.616 | 0.002 | -0.006 | -0.006 | 0 | 0.62 | 0.41 | 0.45 | 0.28 | 0.55 | 0.48 |
| 5 | 58106295 | rs7728528 | -1.55 | 0.904 | 1.192 | -0.53 | 0.153 | 0.212 | 0.212 | 0.015 | 0.07 | 0.45 | 0.09 | 0.19 | 0.07 | 0.16 |
| 5 | 58110324 | rs10060763 | -0.499 | 1.387 | 1.092 | -0.61 | 0.044 | -0.034 | -0.034 | 0 | 0.21 | 0.3 | 0.21 | 0.18 | 0.26 | 0.28 |
| 5 | 58115106 | rs1423368 | 1.374 | 1.254 | 0.863 | -0.785 | -0.002 | -0.001 | -0.001 | 0.101 | 0.6 | 0.46 | 0.46 | 0.22 | 0.55 | 0.5 |
| 5 | 58115159 | rs1423369 | 1.158 | 0.944 | 0.099 | -1.592 | 0.032 | 0.075 | 0.075 | 0 | 0.32 | 0.2 | 0.28 | 0.07 | 0.22 | 0.29 |
| 5 | 58119439 | rs2548241 | 0.072 | 0.905 | 0.159 | -1.571 | -0.03 | -0.008 | -0.008 | 0 | 0.69 | 0.87 | 0.55 | 0.36 | 0.64 | 0.64 |
| 5 | 58121059 | rs3749734 | 1.174 | 0.907 | 0.156 | -1.582 | 0.034 | 0.075 | 0.075 | 0 | 0.32 | 0.2 | 0.28 | 0.07 | 0.21 | 0.29 |
| 5 | 58122015 | rs13354482 | 0.931 | 0.979 | 0.095 | -1.53 | 0.041 | 0.035 | 0.035 | 2.125 | 0.07 | 0.21 | 0.07 | 0.13 | 0.08 | 0.14 |
| 5 | 58122819 | rs2548242 | 0.695 | 0.997 | 0.011 | -1.534 | 0.004 | -0.009 | -0.009 | 0 | 0.62 | 0.67 | 0.48 | 0.22 | 0.55 | 0.5 |
| 5 | 58126248 | rs299919 | 1.438 | 0.989 | -0.047 | -1.565 | -0.004 | 0.024 | 0.024 | 1.344 | 0.27 | 0.23 | 0.17 | 0.15 | 0.31 | 0.2 |
| 5 | 58128623 | rs9292179 | 0.431 | 1.736 | 0.105 | -0.72 | -0.01 | 0.011 | 0.011 | 0 | 0.17 | 0.09 | 0.15 | 0.16 | 0.2 | 0.16 |
| 5 | 58128813 | rs1516430 | 0.676 | 1.867 | -0.015 | -0.59 | 0.011 | 0.002 | 0.002 | 0.52 | 0.67 | 0.64 | 0.53 | 0.38 | 0.63 | 0.64 |
| 5 | 58135616 | rs1914023 | 0.695 | 1.877 | 0.022 | -0.746 | -0.001 | -0.001 | -0.001 | 0 | 0.64 | 0.65 | 0.51 | 0.34 | 0.61 | 0.57 |
| 5 | 58138459 | rs1423359 | 0.9 | 1.923 | -0.221 | -0.728 | 0.003 | -0.02 | -0.02 | 1.176 | 0.57 | 0.59 | 0.46 | 0.2 | 0.52 | 0.44 |
| 5 | 58140131 | rs1423363 | -0.565 | 1.889 | -0.264 | -0.731 | -0.016 | -0.021 | -0.021 | 0 | 0.88 | 0.95 | 0.7 | 0.5 | 0.85 | 0.76 |
| 5 | 58147456 | rs295669 | 0.54 | 1.152 | 0.051 | -0.803 | 0.003 | -0.02 | -0.02 | 0 | 0.57 | 0.59 | 0.47 | 0.2 | 0.52 | 0.44 |
| 5 | 58152601 | rs297039 | 0.59 | 1.273 | -0.615 | -0.75 | -0.01 | 0.055 | 0.055 | 3.008 | 0.54 | 0.39 | 0.44 | 0.2 | 0.51 | 0.43 |
| 5 | 58159443 | rs190915 | 0.436 | 1.586 | -0.514 | -0.196 | 0.01 | -0.006 | -0.006 | 0 | 0.73 | 0.72 | 0.79 | 0.71 | 0.64 | 0.74 |
| 5 | 58167380 | rs159384 | 0.776 | 1.605 | -0.333 | -0.3 | 0.023 | -0.028 | -0.028 | 0 | 0.69 | 0.58 | 0.75 | 0.71 | 0.6 | 0.73 |
| 5 | 58170563 | rs152692 | 1.424 | 1.501 | -0.479 | -0.315 | 0 | -0.009 | -0.009 | 0 | 0.54 | 0.44 | 0.45 | 0.21 | 0.46 | 0.54 |
| 5 | 58170733 | rs152693 | 1.424 | 1.497 | -0.479 | -0.312 | 0 | -0.01 | -0.01 | 0 | 0.54 | 0.44 | 0.45 | 0.2 | 0.46 | 0.54 |
| 5 | 58171166 | rs1478433 | -0.244 | 1.538 | -0.457 | -0.295 | 0.003 | -0.017 | -0.017 | 0 | 0.26 | 0.29 | 0.19 | 0.2 | 0.3 | 0.24 |
| 5 | 58171932 | rs159381 | 0.425 | 1.668 | -0.456 | -0.109 | 0.007 | 0.017 | 0.017 | 0 | 0.8 | 0.71 | 0.64 | 0.41 | 0.76 | 0.78 |
| 5 | 58172093 | rs753701 | 0.256 | 1.741 | -0.522 | -0.063 | -0.008 | -0.028 | -0.028 | 0 | 0.14 | 0.1 | 0.28 | 0.5 | 0.14 | 0.19 |
| 5 | 58172462 | rs159382 | -0.032 | 1.762 | -0.37 | -0.311 | 0.036 | 0.015 | 0.015 | 0 | 0.95 | 0.85 | 0.94 | 0.91 | 0.9 | 0.97 |
| 5 | 58172533 | rs6870230 | -0.271 | 1.511 | -0.58 | -0.488 | 0.002 | -0.007 | -0.007 | 0 | 0.23 | 0.23 | 0.19 | 0.18 | 0.27 | 0.24 |
| 5 | 58172884 | rs925824 | 0.517 | 1.177 | -0.922 | -0.239 | -0.011 | 0.018 | 0.018 | 0 | 0.68 | 0.74 | 0.45 | 0.28 | 0.64 | 0.63 |
| 5 | 58175030 | rs482491 | -0.669 | 1.069 | -0.529 | -0.461 | 0 | -0.028 | -0.028 | 0 | 0.23 | 0.28 | 0.25 | 0.08 | 0.24 | 0.21 |
| 5 | 58178353 | rs12655408 | -0.396 | 0.876 | 0.094 | 0.229 | -0.016 | 0.248 | 0.248 | 0 | 0.16 | 0.42 | 0.23 | 0.51 | 0.15 | 0.32 |
| 5 | 58178433 | rs1423213 | 1.5 | 1.114 | -0.953 | -0.481 | -0.077 | 0.048 | 0.048 | 0 | 0.28 | 0.03 | 0.19 | 0.12 | 0.31 | 0.2 |
| 5 | 58178530 | rs1423212 | -0.943 | 0.996 | -0.727 | -0.619 | -0.009 | 0.113 | 0.113 | 0 | 0.24 | 0.56 | 0.3 | 0.51 | 0.24 | 0.36 |
| 5 | 58181908 | rs922438 | 0.369 | 1.276 | -0.958 | -0.728 | -0.005 | 0.031 | 0.031 | 0 | 0.32 | 0.2 | 0.3 | 0.25 | 0.3 | 0.21 |
| 5 | 58183187 | rs538776 | -1.021 | 1.168 | -0.856 | -0.554 | -0.001 | 0.003 | 0.003 | 0 | 0.1 | 0.09 | 0.07 | 0.07 | 0.1 | 0.1 |
| 5 | 58184061 | rs10472076 | 0.282 | 1.331 | -0.735 | -0.581 | -0.012 | 0.01 | 0.01 | 0 | 0.33 | 0.25 | 0.33 | 0.3 | 0.36 | 0.26 |
| 5 | 58186109 | rs467708 | -0.523 | 1.2 | -0.668 | -0.475 | -0.003 | -0.007 | -0.007 | 0 | 0.08 | 0.06 | 0.06 | 0 | 0.09 | 0.04 |
| 5 | 58189880 | rs7707901 | -0.418 | 1.126 | -0.73 | -0.513 | 0.011 | -0.011 | -0.011 | 0 | 0.34 | 0.72 | 0.43 | 0.62 | 0.3 | 0.51 |
| 5 | 58196646 | rs6897261 | 0.037 | 1.065 | -0.337 | -0.434 | -0.027 | 0.081 | 0.081 | 0 | 0.29 | 0.44 | 0.39 | 0.52 | 0.23 | 0.4 |
| 5 | 58197893 | rs299905 | -0.899 | 1.152 | -0.376 | -0.443 | 0.019 | 0.005 | 0.005 | 0 | 0.41 | 0.78 | 0.5 | 0.63 | 0.37 | 0.57 |
| 5 | 58201950 | rs299902 | -0.853 | 1.186 | -0.389 | -0.459 | 0.016 | 0 | 0 | 0 | 0.4 | 0.74 | 0.5 | 0.62 | 0.37 | 0.57 |
| 5 | 58208218 | rs159497 | -1.103 | 1.667 | 0.124 | -0.175 | 0.001 | -0.01 | -0.01 | 0 | 0.44 | 0.53 | 0.41 | 0.36 | 0.49 | 0.38 |
| 5 | 58223798 | rs2968007 | 0.099 | 1.817 | -0.592 | -0.088 | -0.025 | 0.027 | 0.027 | 0 | 0.67 | 0.77 | 0.68 | 0.74 | 0.64 | 0.76 |
| 5 | 58237883 | rs159722 | 0.333 | 1.364 | -0.485 | 0.098 | -0.017 | -0.003 | -0.003 | 0 | 0.7 | 0.76 | 0.69 | 0.69 | 0.67 | 0.71 |
| 5 | 58248718 | rs702519 | -0.588 | 1.344 | -0.555 | 0.115 | -0.006 | 0.016 | 0.016 | 0.001 | 0.8 | 0.92 | 0.89 | 0.88 | 0.83 | 0.83 |
| 5 | 58259569 | rs702524 | 1.109 | 1.378 | -0.277 | 0.09 | 0.015 | -0.003 | -0.003 | 0.204 | 0.63 | 0.5 | 0.63 | 0.63 | 0.61 | 0.62 |
| 5 | 58265231 | rs11950492 | -0.781 | 1.418 | -0.456 | 0.116 | 0.019 | 0.013 | 0.013 | 0 | 0.1 | 0.15 | 0.2 | 0.18 | 0.16 | 0.13 |
| 5 | 58265373 | rs11959349 | -0.781 | 1.418 | -0.456 | 0.116 | 0.019 | 0.013 | 0.013 | 0 | 0.1 | 0.15 | 0.2 | 0.18 | 0.16 | 0.13 |
| 5 | 58266452 | rs10071088 | -0.781 | 1.418 | -0.456 | 0.116 | 0.019 | 0.013 | 0.013 | 0 | 0.1 | 0.15 | 0.2 | 0.18 | 0.16 | 0.13 |
| 5 | 58267723 | rs702531 | 0.732 | 1.417 | -0.444 | 0.114 | 0.005 | 0.004 | 0.004 | 0 | 0.68 | 0.6 | 0.68 | 0.64 | 0.67 | 0.67 |
| 5 | 58271696 | rs10051847 | -0.753 | 1.177 | -0.349 | 0.211 | 0.016 | 0.011 | 0.011 | 0 | 0.1 | 0.15 | 0.2 | 0.18 | 0.16 | 0.14 |
| 5 | 58284208 | rs3805557 | 0.04 | 0.729 | -0.786 | 0.278 | -0.002 | -0.002 | -0.002 | 0 | 0.77 | 0.74 | 0.88 | 0.82 | 0.83 | 0.82 |
| 5 | 58285055 | rs4700314 | -0.319 | 0.751 | -0.771 | 0.291 | -0.003 | 0.019 | 0.019 | 0.007 | 0.79 | 0.92 | 0.89 | 0.88 | 0.83 | 0.85 |
| 5 | 58286625 | rs1553114 | -0.09 | 0.807 | -0.811 | 0.296 | -0.002 | -0.002 | -0.002 | 0 | 0.77 | 0.74 | 0.88 | 0.82 | 0.83 | 0.82 |
| 5 | 58291462 | rs6876057 | 0.297 | 0.659 | -0.572 | 0.715 | -0.019 | 0.121 | 0.121 | 0 | 0.1 | 0.14 | 0.09 | 0.47 | 0.06 | 0.12 |
| 5 | 58295014 | rs7714708 | 1.022 | 0.703 | -0.698 | 0.45 | -0.005 | -0.008 | -0.008 | 0 | 0.61 | 0.58 | 0.69 | 0.3 | 0.65 | 0.58 |
| 5 | 58296165 | rs10472084 | 0.301 | 0.778 | -0.775 | 0.52 | -0.034 | -0.006 | -0.006 | 0.002 | 0.06 | 0.01 | 0.11 | 0.06 | 0.13 | 0.13 |
| 5 | 58301611 | rs7727206 | -0.884 | 0.801 | -0.768 | 0.528 | 0 | 0.017 | 0.017 | 0 | 0.17 | 0.17 | 0.19 | 0.52 | 0.19 | 0.25 |
| 5 | 58313560 | rs1874858 | 0.372 | 0.751 | -0.738 | 0.518 | -0.019 | 0.119 | 0.119 | 0 | 0.1 | 0.14 | 0.09 | 0.47 | 0.06 | 0.11 |
| 5 | 58317865 | rs1078368 | 1.303 | 0.987 | -0.754 | 0.341 | -0.001 | -0.013 | -0.013 | 0 | 0.56 | 0.52 | 0.63 | 0.29 | 0.57 | 0.49 |
| 5 | 58318034 | rs2968006 | -0.514 | 0.713 | -0.553 | 0.401 | -0.003 | -0.005 | -0.005 | 1.837 | 0.8 | 0.8 | 0.9 | 0.87 | 0.84 | 0.85 |
| 5 | 58318052 | rs2968005 | 0.361 | 0.764 | -0.672 | 0.179 | -0.006 | 0.003 | 0.003 | 0 | 0.64 | 0.6 | 0.76 | 0.4 | 0.7 | 0.65 |
| 5 | 58334645 | rs2279737 | 0.638 | 0.783 | -1.582 | -0.341 | -0.018 | 0.137 | 0.137 | 0 | 0.34 | 0.09 | 0.24 | 0.09 | 0.35 | 0.19 |
| 5 | 58337549 | rs1498606 | -0.322 | 0.722 | -0.855 | 0.207 | -0.005 | 0.088 | 0.088 | 0.281 | 0.14 | 0.23 | 0.23 | 0.69 | 0.13 | 0.24 |
| 5 | 58344518 | rs1498609 | 0.868 | 0.699 | -1.222 | -0.175 | -0.014 | 0.147 | 0.147 | 0 | 0.34 | 0.06 | 0.24 | 0.09 | 0.33 | 0.19 |
| 5 | 58351607 | rs2968009 | -1.022 | 1.194 | -1.6 | -0.183 | -0.011 | -0.015 | -0.015 | 2.486 | 0.26 | 0.2 | 0.4 | 0.72 | 0.31 | 0.45 |
| 5 | 58355032 | rs1498599 | -0.479 | 0.853 | -0.672 | -0.461 | 0 | 0.002 | 0.002 | 0 | 0.64 | 0.67 | 0.69 | 0.87 | 0.66 | 0.66 |
| 5 | 58357931 | rs17720639 | 1.343 | 0.644 | 0.131 | -0.532 | -0.004 | 0.011 | 0.011 | 0 | 0.35 | 0.31 | 0.4 | 0.2 | 0.39 | 0.29 |
| 5 | 58362303 | rs17777280 | 1.496 | 0.622 | 0.026 | -0.516 | -0.034 | 0.177 | 0.177 | 0 | 0.34 | 0.06 | 0.23 | 0.06 | 0.34 | 0.17 |
| 5 | 58365249 | rs6892860 | 0.437 | 0.801 | 0.172 | -0.492 | 0 | 0.008 | 0.008 | 0 | 0.16 | 0.15 | 0.17 | 0.61 | 0.18 | 0.22 |

| | | | | | | | | | | | | | | | | |
|---|---|---|---|---|---|---|---|---|---|---|---|---|---|---|---|---|
| 5 | 58372937 | rs11739293 | -0.834 | 0.66 | 0.193 | -0.031 | -0.021 | -0.027 | -0.027 | 0 | 0.27 | 0.17 | 0.41 | 0.7 | 0.32 | 0.46 |
| 5 | 58376455 | rs1391651 | -0.125 | 0.681 | 0.217 | -0.139 | -0.004 | -0.018 | -0.018 | 0 | 0.58 | 0.59 | 0.67 | 0.81 | 0.63 | 0.61 |
| 5 | 58388213 | rs12109774 | 1.417 | 0.767 | 0.127 | -0.071 | -0.002 | -0.02 | -0.02 | 1.39 | 0.33 | 0.35 | 0.26 | 0.11 | 0.32 | 0.19 |
| 5 | 58396248 | rs2910641 | -1.369 | 0.674 | -0.225 | -0.566 | -0.042 | 0.001 | 0.001 | 0.033 | 0.83 | 0.99 | 0.88 | 0.88 | 0.79 | 0.85 |
| 5 | 58399201 | rs6865647 | -0.114 | 0.493 | 0.021 | -0.678 | -0.016 | -0.035 | -0.035 | 0 | 0.16 | 0.31 | 0.18 | 0.07 | 0.13 | 0.12 |
| 5 | 58406998 | rs17721878 | 1.574 | 0.654 | -1.243 | -0.97 | -0.035 | 0.157 | 0.157 | 0 | 0.3 | 0.04 | 0.21 | 0.06 | 0.36 | 0.19 |
| 5 | 58407771 | rs10052657 | 0.206 | 1.197 | -0.93 | -0.325 | -0.023 | 0.001 | 0.001 | 0.013 | 0.15 | 0.11 | 0.19 | 0.13 | 0.18 | 0.15 |
| 5 | 58409273 | rs10071163 | 0.664 | 0.962 | -0.83 | -0.412 | -0.003 | 0.001 | 0.001 | 0 | 0.65 | 0.67 | 0.51 | 0.26 | 0.65 | 0.49 |
| 5 | 58418387 | rs13174180 | 0.272 | 0.707 | -0.271 | -0.375 | -0.006 | -0.003 | -0.003 | 0 | 0.15 | 0.18 | 0.07 | 0.12 | 0.12 | 0.15 |
| 5 | 58420346 | rs10068809 | -0.477 | 0.711 | -0.01 | -0.612 | -0.014 | -0.06 | -0.06 | 0.159 | 0.16 | 0.34 | 0.2 | 0.03 | 0.13 | 0.13 |
| 5 | 58423010 | rs10035950 | 0.264 | 1.281 | -0.448 | -0.665 | -0.03 | 0.036 | 0.036 | 0 | 0.12 | 0.04 | 0.24 | 0 | 0.16 | 0.19 |
| 5 | 58424440 | rs10044993 | 0.137 | 1.107 | 0.215 | -0.511 | 0 | -0.013 | -0.013 | 0 | 0.84 | 0.62 | 0.87 | 0.84 | 0.88 | 0.84 |
| 5 | 58428033 | rs2059191 | 0.502 | 1.239 | 0.265 | -0.617 | 0.003 | -0.036 | -0.036 | 0 | 0.16 | 0.2 | 0.1 | 0.06 | 0.16 | 0.14 |
| 5 | 58428799 | rs13176475 | -0.854 | 1.021 | 0.144 | -0.102 | -0.003 | 0.008 | 0.008 | 1.856 | 0.31 | 0.3 | 0.3 | 0.36 | 0.29 | 0.43 |
| 5 | 58429642 | rs997421 | -0.231 | 1.169 | 0.171 | -0.063 | 0.002 | -0.07 | -0.07 | 4.497 | 0.14 | 0.02 | 0.2 | 0.3 | 0.13 | 0.3 |
| 5 | 58430196 | rs17778950 | -1.147 | 1.282 | 0.331 | -0.254 | 0.006 | -0.034 | -0.034 | 0 | 0.35 | 0.6 | 0.37 | 0.16 | 0.37 | 0.33 |
| 5 | 58430239 | rs16889248 | -0.14 | 1.204 | 0.255 | -0.233 | 0.018 | -0.048 | -0.048 | 0 | 0.19 | 0.34 | 0.28 | 0.06 | 0.21 | 0.17 |
| 5 | 58430399 | rs7734070 | 1.608 | 1.261 | 0.161 | -0.322 | -0.018 | 0.017 | 0.017 | 0 | 0.11 | 0.06 | 0.12 | 0 | 0.13 | 0.12 |
| 5 | 58431386 | rs1551939 | 0.151 | 1.262 | 0.273 | -0.163 | 0.009 | -0.025 | -0.025 | 0 | 0.86 | 0.91 | 0.79 | 0.61 | 0.87 | 0.7 |
| 5 | 58436920 | rs6869149 | -0.117 | 1.333 | 0.377 | -0.224 | 0.01 | -0.036 | -0.036 | 0 | 0.18 | 0.29 | 0.27 | 0.05 | 0.2 | 0.17 |
| 5 | 58440258 | rs1824788 | 1.016 | 1.083 | 0.264 | -0.35 | -0.001 | 0.004 | 0.004 | 0.076 | 0.5 | 0.47 | 0.52 | 0.45 | 0.51 | 0.34 |
| 5 | 58444322 | rs2409627 | 1.258 | 1.004 | 0.424 | -0.37 | 0 | -0.012 | -0.012 | 0 | 0.14 | 0.12 | 0.08 | 0.05 | 0.16 | 0.12 |
| 5 | 58445393 | rs1961 | -0.347 | 0.954 | 0.488 | -0.432 | 0.009 | -0.048 | -0.048 | 0 | 0.34 | 0.45 | 0.42 | 0.11 | 0.34 | 0.33 |
| 5 | 58446582 | rs16889280 | 2.148 | 0.911 | 0.343 | -0.325 | 0.005 | -0.018 | -0.018 | 0 | 0.31 | 0.23 | 0.25 | 0.4 | 0.31 | 0.17 |
| 5 | 58448733 | rs1105577 | 2.26 | 0.935 | 0.25 | -0.334 | 0.007 | -0.039 | -0.039 | 0 | 0.31 | 0.16 | 0.25 | 0.4 | 0.31 | 0.17 |
| 5 | 58453822 | rs1345793 | 1.555 | 0.888 | 1.081 | 0.543 | 0.003 | 0.014 | 0.014 | 0 | 0.62 | 0.51 | 0.5 | 0.52 | 0.61 | 0.44 |
| 5 | 58454031 | rs1345792 | 1.555 | 0.888 | 1.081 | 0.543 | 0.003 | 0.014 | 0.014 | 0 | 0.62 | 0.5 | 0.5 | 0.52 | 0.61 | 0.45 |
| 5 | 58455403 | rs1559251 | 0.791 | 0.772 | 0.842 | 0.611 | 0.001 | -0.035 | -0.035 | 0.161 | 0.71 | 0.85 | 0.72 | 0.56 | 0.71 | 0.56 |
| 5 | 58460599 | rs1971940 | 1.431 | 0.78 | 0.614 | 0.6 | -0.01 | 0.021 | 0.021 | 2.216 | 0.63 | 0.52 | 0.65 | 0.51 | 0.64 | 0.48 |
| 5 | 58476117 | rs9292195 | 0.962 | 0.477 | 0.487 | 0.599 | 0.013 | -0.026 | -0.026 | 0 | 0.8 | 0.84 | 0.75 | 0.52 | 0.82 | 0.59 |
| 5 | 58481760 | rs8180396 | 1.02 | 0.627 | 0.394 | 0.545 | -0.003 | 0.023 | 0.023 | 0 | 0.7 | 0.61 | 0.66 | 0.54 | 0.72 | 0.53 |
| 5 | 58484508 | rs10051191 | 1.67 | 0.319 | 0.202 | 0.448 | 0.013 | 0.044 | 0.044 | 0.514 | 0.56 | 0.32 | 0.44 | 0.31 | 0.56 | 0.37 |
| 5 | 58487764 | rs10055769 | 1.514 | 0.537 | 0.419 | 0.774 | 0.001 | 0.028 | 0.028 | 0.063 | 0.63 | 0.28 | 0.63 | 0.29 | 0.65 | 0.45 |
| 5 | 58491406 | rs2112957 | 1.373 | 0.438 | 0.32 | 1.212 | -0.003 | 0.048 | 0.048 | 0 | 0.88 | 0.85 | 0.74 | 0.39 | 0.88 | 0.64 |
| 5 | 58491538 | rs7444432 | -0.207 | 0.614 | 0.372 | 0.929 | 0.017 | 0.033 | 0.033 | 0 | 0.07 | 0.13 | 0.23 | 0.13 | 0.11 | 0.09 |
| 5 | 58500382 | rs27727 | -1.163 | 0.64 | 0.499 | 0.895 | -0.003 | 0.001 | 0.001 | 0 | 0.39 | 0.36 | 0.25 | 0.53 | 0.32 | 0.58 |
| 5 | 58500896 | rs154222 | -1.541 | 0.698 | 0.727 | 0.813 | 0.015 | 0.032 | 0.032 | 0 | 0.47 | 0.67 | 0.32 | 0.57 | 0.4 | 0.62 |
| 5 | 58502533 | rs10051456 | -0.892 | 0.71 | 0.969 | 0.997 | -0.001 | -0.013 | -0.013 | 0 | 0.33 | 0.3 | 0.21 | 0.13 | 0.28 | 0.29 |
| 5 | 58502651 | rs10067048 | -1.575 | 0.721 | 0.455 | 1.012 | 0.01 | 0.017 | 0.017 | 0 | 0.43 | 0.67 | 0.32 | 0.56 | 0.36 | 0.59 |
| 5 | 58506279 | rs1867724 | -0.881 | 0.958 | 0.623 | 1.096 | 0.021 | 0.086 | 0.086 | 0 | 0.2 | 0.31 | 0.31 | 0.54 | 0.2 | 0.42 |
| 5 | 58508712 | rs6867591 | 0.234 | 0.869 | -0.056 | 0.732 | 0.06 | 0.034 | 0.034 | 0.104 | 0.06 | 0.24 | 0.06 | 0.02 | 0.07 | 0.04 |
| 5 | 58510326 | rs425384 | -0.052 | 0.884 | -1.164 | 0.701 | -0.015 | 0.02 | 0.02 | 0 | 0.14 | 0.02 | 0.11 | 0.08 | 0.16 | 0.13 |
| 5 | 58511501 | rs6450512 | -1.125 | 0.653 | -1.261 | 0.57 | -0.03 | 0.052 | 0.052 | 0.129 | 0.44 | 0.85 | 0.48 | 0.57 | 0.37 | 0.63 |
| 5 | 58525186 | rs4700325 | -0.813 | 0.316 | -0.457 | 0.187 | -0.012 | -0.03 | -0.03 | 0.281 | 0.23 | 0.65 | 0.32 | 0.08 | 0.14 | 0.17 |
| 5 | 58529335 | rs27957 | -0.2 | 0.948 | -0.468 | -0.095 | 0.002 | -0.003 | -0.003 | 0.049 | 0.21 | 0.23 | 0.2 | 0.1 | 0.27 | 0.2 |
| 5 | 58529636 | rs10440630 | 0.788 | 0.992 | -0.558 | -0.549 | 0 | -0.003 | -0.003 | 0 | 0.12 | 0.09 | 0.09 | 0.07 | 0.13 | 0.15 |
| 5 | 58531652 | rs28054 | -0.522 | 0.848 | -0.578 | -0.628 | -0.049 | 0.043 | 0.043 | 0 | 0.29 | 0.18 | 0.26 | 0.16 | 0.36 | 0.32 |
| 5 | 58534107 | rs26705 | -1.034 | 0.292 | -1.215 | -0.904 | 0.053 | 0.125 | 0.125 | 0 | 0.6 | 0.9 | 0.75 | 0.74 | 0.68 | 0.73 |
| 5 | 58534348 | rs10055446 | 1.957 | 0.369 | -1.071 | -0.785 | -0.017 | 0.006 | 0.006 | 0 | 0.08 | 0.01 | 0.06 | 0.06 | 0.1 | 0.12 |
| 5 | 58545163 | rs6858946 | 1.061 | 0.163 | 0.685 | -0.86 | -0.007 | 0.028 | 0.028 | 0 | 0.22 | 0.19 | 0.42 | 0.06 | 0.26 | 0.21 |
| 5 | 58548231 | rs35305 | -0.174 | 0.138 | 0.665 | -0.909 | -0.001 | -0.035 | -0.035 | 4.585 | 0.44 | 0.56 | 0.63 | 0.16 | 0.53 | 0.4 |
| 5 | 58549317 | rs26700 | 1.183 | 0.196 | 0.546 | -0.895 | -0.029 | 0.023 | 0.023 | 0 | 0.2 | 0.14 | 0.19 | 0.09 | 0.27 | 0.19 |
| 5 | 58550539 | rs6450514 | -0.131 | 0.048 | 0.534 | -0.977 | 0.001 | 0.002 | 0.002 | 0 | 0.37 | 0.65 | 0.53 | 0.63 | 0.36 | 0.53 |
| 5 | 58550823 | rs27949 | -0.735 | 0.044 | 0.399 | -0.942 | 0.041 | 0.1 | 0.1 | 0 | 0.57 | 0.81 | 0.73 | 0.72 | 0.63 | 0.72 |
| 5 | 58554904 | rs27547 | -1.22 | -0.125 | 0.069 | -1.043 | 0.007 | 0.012 | 0.012 | 0 | 0.68 | 0.78 | 0.79 | 0.71 | 0.68 | 0.74 |
| 5 | 58558388 | rs16889495 | 0.753 | -0.053 | 0.069 | -1.237 | -0.003 | -0.029 | -0.029 | 0 | 0.16 | 0.16 | 0.12 | 0.02 | 0.11 | 0.12 |
| 5 | 58558595 | rs27549 | 1.398 | -0.009 | -0.044 | -1.219 | -0.004 | 0.003 | 0.003 | 0 | 0.19 | 0.19 | 0.2 | 0.1 | 0.27 | 0.19 |
| 5 | 58564232 | rs27723 | -0.973 | -0.175 | -0.093 | -1.071 | 0.005 | 0.005 | 0.005 | 0 | 0.67 | 0.73 | 0.78 | 0.21 | 0.7 | 0.57 |
| 5 | 58569284 | rs2112959 | 1.208 | -0.127 | -0.378 | -0.896 | -0.034 | 0.017 | 0.017 | 0.043 | 0.11 | 0.01 | 0.07 | 0.07 | 0.11 | 0.13 |
| 5 | 58572727 | rs1947090 | 1.207 | -0.175 | -0.111 | -0.985 | 0 | 0 | 0 | 0 | 0.19 | 0.22 | 0.39 | 0.05 | 0.23 | 0.14 |
| 5 | 58575323 | rs27184 | -0.416 | -0.655 | -0.549 | -1.269 | 0.023 | 0.1 | 0.1 | 1.146 | 0.44 | 0.76 | 0.61 | 0.75 | 0.44 | 0.47 |
| 5 | 58575831 | rs1374028 | 1.4 | -0.728 | -0.263 | -0.324 | -0.001 | 0.001 | 0.001 | 0 | 0.14 | 0.13 | 0.09 | 0.22 | 0.11 | 0.12 |
| 5 | 58583213 | rs2028836 | 1.747 | -0.653 | -0.241 | -0.384 | -0.001 | 0.023 | 0.023 | 0 | 0.19 | 0.16 | 0.38 | 0.05 | 0.23 | 0.13 |
| 5 | 58588269 | rs28044 | -0.45 | -0.833 | -0.182 | -0.387 | -0.011 | 0.003 | 0.003 | 0.017 | 0.28 | 0.59 | 0.25 | 0.69 | 0.23 | 0.34 |
| 5 | 58588580 | rs10472102 | 0.814 | -0.844 | 0.045 | -0.283 | -0.017 | 0.013 | 0.013 | 0 | 0.18 | 0.23 | 0.12 | 0.24 | 0.14 | 0.15 |
| 5 | 58588629 | rs27183 | -1.034 | -0.934 | -0.14 | -0.531 | -0.122 | 0.094 | 0.094 | 0 | 0.59 | 0.92 | 0.69 | 0.75 | 0.55 | 0.63 |
| 5 | 58589499 | rs12655617 | 0.862 | -1.145 | 0.014 | -0.369 | -0.016 | 0.014 | 0.014 | 0 | 0.18 | 0.22 | 0.12 | 0.24 | 0.14 | 0.15 |
| 5 | 58589786 | rs17780175 | 0.734 | -1.173 | -0.228 | -0.6 | 0.008 | 0.046 | 0.046 | 0 | 0.3 | 0.22 | 0.46 | 0.05 | 0.29 | 0.21 |
| 5 | 58594684 | rs12519611 | 1.883 | -1.13 | -0.256 | -0.566 | 0.001 | 0.026 | 0.026 | 0 | 0.19 | 0.13 | 0.37 | 0.06 | 0.23 | 0.14 |

| | | | | | | | | | | | | | | | | |
|---|---|---|---|---|---|---|---|---|---|---|---|---|---|---|---|---|
| 5 | 58595606 | rs7703022 | -0.005 | -1.194 | -0.26 | -0.641 | 0.013 | 0.043 | 0.043 | 0 | 0.43 | 0.36 | 0.53 | 0.06 | 0.39 | 0.38 |
| 5 | 58598034 | rs154221 | 0.388 | -1.234 | 0.081 | -0.36 | -0.012 | 0.165 | 0.165 | 0 | 0.18 | 0.34 | 0.21 | 0.47 | 0.12 | 0.29 |
| 5 | 58598181 | rs4700335 | 1.296 | -1.117 | -0.23 | -0.337 | -0.002 | 0 | 0 | 0 | 0.15 | 0.12 | 0.09 | 0.22 | 0.1 | 0.12 |
| 5 | 58605684 | rs255647 | -0.655 | -1.201 | -0.563 | -0.505 | -0.047 | 0.043 | 0.043 | 0 | 0.61 | 0.84 | 0.66 | 0.7 | 0.52 | 0.57 |
| 5 | 58609468 | rs153981 | -0.28 | -1.368 | -0.38 | 0.151 | 0.04 | 0.047 | 0.047 | 0 | 0.47 | 0.74 | 0.61 | 0.35 | 0.4 | 0.55 |
| 5 | 58612016 | rs37353 | -0.825 | -1.316 | -0.373 | 0.149 | -0.016 | 0.021 | 0.021 | 0 | 0.62 | 0.69 | 0.69 | 0.69 | 0.49 | 0.69 |
| 5 | 58612733 | rs2013979 | 0.838 | -1.398 | -0.331 | 0.121 | -0.004 | -0.052 | -0.052 | 1.117 | 0.29 | 0.32 | 0.42 | 0.07 | 0.31 | 0.29 |
| 5 | 58613132 | rs27168 | 1.06 | -1.269 | -0.851 | 0.238 | 0.008 | -0.018 | -0.018 | 0 | 0.15 | 0.1 | 0.14 | 0.27 | 0.08 | 0.28 |
| 5 | 58618506 | rs27170 | -0.692 | -1.078 | -1.022 | -0.119 | -0.029 | 0.029 | 0.029 | 0 | 0.62 | 0.78 | 0.65 | 0.68 | 0.48 | 0.67 |
| 5 | 58618889 | rs27171 | -0.67 | -1.161 | -0.583 | 0.133 | 0.018 | 0.009 | 0.009 | 0 | 0.43 | 0.58 | 0.32 | 0.63 | 0.25 | 0.54 |
| 5 | 58623098 | rs17780860 | 0.286 | -0.716 | -1.016 | -0.282 | 0.004 | 0.118 | 0.118 | 0 | 0.22 | 0.03 | 0.14 | 0.01 | 0.15 | 0.24 |
| 5 | 58629497 | rs16889615 | 0.419 | -0.99 | -0.34 | 0.36 | -0.028 | 0.032 | 0.032 | 0.006 | 0.2 | 0.3 | 0.08 | 0.31 | 0.1 | 0.14 |
| 5 | 58632660 | rs10472105 | 0.429 | -0.991 | -0.34 | 0.36 | -0.028 | 0.032 | 0.032 | 0.015 | 0.2 | 0.3 | 0.08 | 0.32 | 0.1 | 0.14 |
| 5 | 58633472 | rs40328 | 0.571 | -0.462 | -1.187 | 0.114 | -0.088 | 0.047 | 0.047 | 0 | 0.4 | 0.13 | 0.33 | 0.33 | 0.44 | 0.48 |
| 5 | 58637853 | rs244574 | 0.632 | -0.484 | -1.198 | 0.144 | -0.09 | 0.079 | 0.079 | 0 | 0.39 | 0.13 | 0.26 | 0.28 | 0.44 | 0.48 |
| 5 | 58641024 | rs17781354 | 1.659 | -0.411 | -1.027 | -0.377 | -0.001 | -0.013 | -0.013 | 0.798 | 0.09 | 0.1 | 0.08 | 0.01 | 0.13 | 0.06 |
| 5 | 58642372 | rs13176940 | -0.649 | -0.566 | -0.737 | -0.543 | 0.013 | 0.041 | 0.041 | 0 | 0.53 | 0.36 | 0.33 | 0.26 | 0.38 | 0.5 |
| 5 | 58643239 | rs13153653 | 1.926 | -0.643 | -0.817 | -0.585 | 0.025 | -0.04 | -0.04 | 0 | 0.18 | 0.06 | 0.06 | 0.22 | 0.1 | 0.14 |
| 5 | 58643390 | rs35267 | 1.642 | -0.486 | -0.794 | -0.472 | -0.077 | -0.05 | -0.05 | 0 | 0.28 | 0.12 | 0.25 | 0.38 | 0.38 | 0.27 |
| 5 | 58646112 | rs40125 | -1.357 | -0.466 | -0.662 | -0.498 | 0 | 0.135 | 0.135 | 0 | 0.8 | 0.46 | 0.53 | 0.62 | 0.76 | 0.78 |
| 5 | 58655740 | rs35281 | -1.063 | -0.45 | -0.564 | -0.568 | -0.004 | -0.006 | -0.006 | 2.057 | 0.76 | 0.81 | 0.55 | 0.69 | 0.64 | 0.8 |
| 5 | 58656328 | rs35284 | 1.655 | -0.403 | -0.813 | -0.629 | -0.077 | -0.053 | -0.053 | 0.944 | 0.28 | 0.12 | 0.24 | 0.39 | 0.38 | 0.27 |
| 5 | 58663831 | rs17782374 | 0.778 | -0.501 | -0.444 | -0.891 | 0 | -0.02 | -0.02 | 0.2 | 0.21 | 0.15 | 0.14 | 0.02 | 0.2 | 0.26 |
| 5 | 58666122 | rs1457145 | 1.351 | -0.456 | -0.63 | -0.708 | 0.006 | 0.038 | 0.038 | 0 | 0.13 | 0.05 | 0.13 | 0.01 | 0.09 | 0.1 |
| 5 | 58676328 | rs2702365 | 1.322 | 0.115 | -0.553 | -0.591 | 0.001 | -0.001 | -0.001 | 0 | 0.21 | 0.14 | 0.1 | 0.23 | 0.16 | 0.15 |
| 5 | 58677233 | rs181736 | 0.635 | 0.224 | -0.323 | -0.424 | 0.006 | 0.004 | 0.004 | 0 | 0.34 | 0.4 | 0.3 | 0.43 | 0.43 | 0.3 |
| 5 | 58677499 | rs2591748 | 0.105 | 0.024 | -0.259 | -0.368 | -0.011 | -0.007 | -0.007 | 0 | 0.38 | 0.42 | 0.18 | 0.31 | 0.24 | 0.43 |
| 5 | 58681656 | rs433161 | 0.831 | 0.082 | -0.224 | -0.287 | 0.008 | 0.053 | 0.053 | 0 | 0.28 | 0.64 | 0.26 | 0.47 | 0.3 | 0.3 |
| 5 | 58681857 | rs1445918 | 0.746 | -0.109 | -0.221 | -0.388 | -0.01 | 0.004 | 0.004 | 0.321 | 0.27 | 0.61 | 0.14 | 0.28 | 0.18 | 0.18 |
| 5 | 58682026 | rs444552 | 0.8 | -0.008 | -0.177 | -0.294 | 0.008 | 0.051 | 0.051 | 0 | 0.29 | 0.64 | 0.26 | 0.47 | 0.3 | 0.3 |
| 5 | 58689985 | rs454702 | 1.541 | 0.076 | -0.439 | -0.263 | -0.032 | -0.051 | -0.051 | 0 | 0.23 | 0.14 | 0.2 | 0.4 | 0.29 | 0.26 |
| 5 | 58690251 | rs525099 | -1.207 | -0.062 | -0.556 | -0.484 | -0.013 | -0.011 | -0.011 | 0 | 0.77 | 0.84 | 0.52 | 0.72 | 0.62 | 0.78 |
| 5 | 58698145 | rs258110 | 0.502 | -0.276 | -0.389 | -0.59 | 0 | -0.005 | -0.005 | 0.017 | 0.41 | 0.44 | 0.39 | 0.48 | 0.39 | 0.39 |
| 5 | 58698524 | rs258111 | 0.266 | -0.14 | -0.604 | -0.52 | 0.002 | -0.006 | -0.006 | 0 | 0.43 | 0.66 | 0.4 | 0.49 | 0.39 | 0.4 |
| 5 | 58704650 | rs258121 | 0.028 | -0.03 | -0.539 | -0.428 | 0.001 | -0.004 | -0.004 | 0 | 0.6 | 0.49 | 0.35 | 0.65 | 0.53 | 0.66 |
| 5 | 58707111 | rs1348709 | 0.682 | -0.029 | -0.501 | -0.403 | -0.02 | -0.022 | -0.022 | 0 | 0.18 | 0.29 | 0.19 | 0.08 | 0.1 | 0.13 |
| 5 | 58707299 | rs7735572 | 0.667 | -0.027 | -0.517 | -0.402 | -0.027 | -0.029 | -0.029 | 0 | 0.18 | 0.35 | 0.19 | 0.09 | 0.1 | 0.13 |
| 5 | 58708427 | rs258125 | 0.91 | 0.185 | -0.637 | -0.508 | 0.004 | -0.001 | -0.001 | 0 | 0.44 | 0.48 | 0.29 | 0.59 | 0.43 | 0.39 |
| 5 | 58710841 | rs7735228 | 0.284 | 0.549 | -0.184 | -0.223 | -0.027 | -0.03 | -0.03 | 1.753 | 0.18 | 0.35 | 0.19 | 0.09 | 0.1 | 0.13 |
| 5 | 58713680 | rs2547917 | 0.9 | 0.825 | -0.082 | 0.143 | -0.006 | 0.001 | 0.001 | 0.036 | 0.21 | 0.11 | 0.1 | 0.22 | 0.18 | 0.15 |
| 5 | 58715782 | rs244568 | 0.205 | 0.639 | 0.041 | 0.337 | -0.001 | -0.004 | -0.004 | 0.15 | 0.55 | 0.66 | 0.43 | 0.51 | 0.44 | 0.64 |
| 5 | 58715833 | rs13165581 | -0.12 | 0.534 | -0.165 | 0.451 | -0.025 | 0.001 | 0.001 | 0 | 0.12 | 0.29 | 0.1 | 0.08 | 0.05 | 0.07 |
| 5 | 58719893 | rs2702373 | -0.322 | 1.196 | -0.816 | 0.591 | 0.006 | 0.029 | 0.029 | 0 | 0.88 | 0.86 | 0.78 | 0.73 | 0.82 | 0.89 |
| 5 | 58721609 | rs2607347 | -0.163 | 0.972 | -1.233 | -0.227 | -0.026 | 0.113 | 0.113 | 0.451 | 0.42 | 0.16 | 0.42 | 0.22 | 0.43 | 0.29 |
| 5 | 58722249 | rs13160457 | -0.032 | 0.744 | -1.007 | 0.034 | 0.011 | -0.023 | -0.023 | 7.667 | 0.31 | 0.09 | 0.22 | 0.33 | 0.25 | 0.3 |
| 5 | 58730285 | rs2081092 | -0.902 | 0.916 | 0.215 | 0.077 | -0.001 | -0.008 | -0.008 | 0 | 0.13 | 0.62 | 0.11 | 0.07 | 0.11 | 0.1 |
| 5 | 58753267 | rs6870632 | -1.378 | 1.181 | 0.849 | -0.071 | -0.005 | -0.002 | -0.002 | 1.56 | 0.3 | 0.78 | 0.35 | 0.27 | 0.32 | 0.23 |
| 5 | 58754510 | rs17796040 | 0.342 | 1.185 | 1.184 | -0.072 | 0.001 | -0.035 | -0.035 | 0 | 0.08 | 0.14 | 0.07 | 0.01 | 0.1 | 0.06 |
| 5 | 58755580 | rs16889869 | -1.156 | 1.183 | 0.982 | -0.082 | -0.008 | 0.014 | 0.014 | 0.89 | 0.15 | 0.48 | 0.11 | 0.12 | 0.13 | 0.15 |
| 5 | 58757820 | rs992726 | 0.404 | 1.216 | 0.844 | -0.059 | -0.006 | -0.004 | -0.004 | 0.04 | 0.13 | 0.13 | 0.22 | 0.15 | 0.19 | 0.07 |
| 5 | 58776204 | rs7711639 | -1.271 | 1.21 | 0.83 | -0.164 | 0.008 | -0.016 | -0.016 | 0.63 | 0.17 | 0.42 | 0.13 | 0.12 | 0.19 | 0.16 |
| 5 | 58782840 | rs16889904 | -0.466 | 1.391 | 0.522 | -0.525 | 0.031 | -0.114 | -0.114 | 0 | 0.23 | 0.34 | 0.33 | 0.01 | 0.32 | 0.13 |
| 5 | 58783343 | rs17796734 | 0.097 | 0.679 | -0.105 | -0.673 | 0.013 | 0.035 | 0.035 | 0 | 0.12 | 0.04 | 0.07 | 0.02 | 0.09 | 0.07 |
| 5 | 58785305 | rs16889907 | -1.238 | 1.166 | 0.41 | -0.699 | 0.017 | -0.075 | -0.075 | 0.392 | 0.18 | 0.51 | 0.19 | 0.07 | 0.19 | 0.13 |
| 5 | 58797572 | rs10514862 | -1.04 | 1.126 | 0.419 | -0.697 | 0.027 | -0.045 | -0.045 | 0 | 0.14 | 0.5 | 0.17 | 0.07 | 0.16 | 0.12 |
| 5 | 58801291 | rs4551023 | 0.523 | 1.509 | 0.375 | -1.013 | 0.004 | -0.05 | -0.05 | 1.269 | 0.2 | 0.15 | 0.19 | 0.01 | 0.27 | 0.13 |
| 5 | 58804379 | rs12186707 | 0.12 | 1.711 | 0.152 | -1.214 | -0.038 | 0.04 | 0.04 | 0 | 0.07 | 0.01 | 0.07 | 0 | 0.15 | 0.04 |
| 5 | 58810366 | rs6450521 | -1.593 | 1.592 | 0.239 | -1.143 | -0.009 | 0.019 | 0.019 | 0.134 | 0.4 | 0.66 | 0.48 | 0.09 | 0.49 | 0.31 |
| 5 | 58813031 | rs16889921 | -0.89 | 1.6 | 0.401 | -0.906 | 0.021 | -0.049 | -0.049 | 0.934 | 0.14 | 0.48 | 0.17 | 0.07 | 0.16 | 0.12 |
| 5 | 58814518 | rs6896121 | 0.147 | 1.332 | 0.171 | -0.868 | -0.018 | -0.026 | -0.026 | 0.378 | 0.1 | 0.17 | 0.07 | 0.02 | 0.07 | 0.11 |
| 5 | 58818722 | rs7728286 | -0.585 | 1.498 | -0.055 | -0.291 | 0 | 0 | 0 | 0 | 0.33 | 0.67 | 0.3 | 0.3 | 0.34 | 0.28 |
| 5 | 58829777 | rs1876209 | 0.969 | 0.79 | -0.917 | -0.045 | -0.044 | -0.087 | -0.087 | 0 | 0.33 | 0.03 | 0.3 | 0.38 | 0.27 | 0.4 |
| 5 | 58830145 | rs37576 | 0.776 | 0.822 | -0.299 | -1.165 | 0.001 | 0.026 | 0.026 | 0 | 0.38 | 0.32 | 0.41 | 0.8 | 0.31 | 0.61 |
| 5 | 58833192 | rs167161 | -0.817 | 0.811 | -0.418 | -1.522 | -0.01 | -0.111 | -0.111 | 0.106 | 0.59 | 0.49 | 0.57 | 0.05 | 0.69 | 0.38 |
| 5 | 58834033 | rs159195 | -0.521 | 0.651 | -0.297 | -0.985 | -0.003 | -0.052 | -0.052 | 0 | 0.24 | 0.3 | 0.16 | 0.01 | 0.23 | 0.14 |
| 5 | 58836950 | rs40216 | 0.841 | 0.822 | -0.566 | -0.847 | 0.01 | 0.009 | 0.009 | 1.545 | 0.62 | 0.28 | 0.72 | 0.64 | 0.72 | 0.66 |
| 5 | 58838138 | rs7733296 | -0.335 | 0.671 | 0.117 | -0.92 | 0.021 | -0.046 | -0.046 | 0.233 | 0.17 | 0.45 | 0.17 | 0.11 | 0.13 | 0.23 |
| 5 | 58842774 | rs702543 | 1.697 | 1.082 | -0.418 | -1.186 | -0.125 | 0.009 | 0.009 | 0.583 | 0.54 | 0.06 | 0.54 | 0.63 | 0.57 | 0.62 |
| 5 | 58845915 | rs294482 | 0.62 | 0.864 | 0.191 | -1.057 | -0.001 | -0.008 | -0.008 | 0 | 0.79 | 0.74 | 0.86 | 0.83 | 0.81 | 0.92 |

| | | | | | | | | | | | | | | | | |
|---|---|---|---|---|---|---|---|---|---|---|---|---|---|---|---|---|
| 5 | 58849394 | rs294478 | 0.205 | 0.87 | 0.215 | -1.046 | -0.009 | -0.003 | -0.003 | 0 | 0.82 | 0.74 | 0.91 | 0.83 | 0.88 | 0.93 |
| 5 | 58851811 | rs697076 | 1.27 | 0.87 | 0.386 | -1.068 | -0.01 | -0.004 | -0.004 | 0.114 | 0.72 | 0.53 | 0.75 | 0.75 | 0.77 | 0.85 |
| 5 | 58862842 | rs1395337 | 1.281 | 0.933 | 0.475 | -1.04 | 0.006 | 0.024 | 0.024 | 0 | 0.75 | 0.59 | 0.75 | 0.71 | 0.77 | 0.82 |
| 5 | 58862944 | rs1395336 | 1.405 | 0.951 | 0.386 | -1.044 | 0.016 | 0.008 | 0.008 | 0 | 0.73 | 0.63 | 0.72 | 0.71 | 0.7 | 0.8 |
| 5 | 58882102 | rs1035321 | -0.722 | 0.778 | 0.518 | -1.257 | 0.021 | -0.01 | -0.01 | 0 | 0.05 | 0.13 | 0.13 | 0.04 | 0.08 | 0.07 |
| 5 | 58886873 | rs295943 | 0.457 | 0.671 | 0.454 | -1.223 | -0.006 | 0.026 | 0.026 | 0.046 | 0.83 | 0.64 | 0.9 | 0.79 | 0.89 | 0.92 |
| 5 | 58902233 | rs296410 | 1.453 | 0.622 | -1.104 | -1.805 | -0.12 | 0.102 | 0.102 | 0 | 0.46 | 0.05 | 0.52 | 0.72 | 0.53 | 0.52 |
| 5 | 58935530 | rs12152938 | 0.176 | 0.74 | -0.869 | -1.873 | -0.023 | 0.082 | 0.082 | 0 | 0.24 | 0.09 | 0.19 | 0.07 | 0.24 | 0.28 |
| 5 | 58936943 | rs298034 | -0.029 | 0.547 | -0.47 | -1.695 | -0.001 | 0.005 | 0.005 | 0 | 0.75 | 0.79 | 0.83 | 0.83 | 0.77 | 0.92 |
| 5 | 58939840 | rs298028 | 0.038 | 0.401 | -0.11 | -1.701 | 0.001 | -0.037 | -0.037 | 0.17 | 0.72 | 0.59 | 0.81 | 0.83 | 0.76 | 0.92 |
| 5 | 58940034 | rs6863462 | -0.546 | 0.98 | -0.194 | -1.715 | -0.005 | 0.009 | 0.009 | 0 | 0.32 | 0.32 | 0.34 | 0.1 | 0.35 | 0.39 |
| 5 | 58954992 | rs297975 | 0.545 | 0.49 | -0.107 | -1.596 | 0.007 | 0.044 | 0.044 | 0 | 0.59 | 0.68 | 0.75 | 0.81 | 0.66 | 0.73 |
| 5 | 58957641 | rs1370230 | 1.546 | 0.395 | 0.126 | -1.729 | -0.004 | 0.015 | 0.015 | 0 | 0.41 | 0.3 | 0.56 | 0.68 | 0.43 | 0.46 |
| 5 | 58959977 | rs7729853 | 0.182 | 0.396 | 0.578 | -1.642 | 0.02 | 0.017 | 0.017 | 1.706 | 0.06 | 0.28 | 0.03 | 0 | 0 | 0 |
| 5 | 58960124 | rs7716837 | 0.11 | 0.39 | 0.568 | -1.645 | 0.036 | 0.032 | 0.032 | 1.945 | 0.06 | 0.34 | 0.03 | 0 | 0 | 0 |
| 5 | 58979946 | rs17382202 | 0.805 | 0.569 | 0.097 | -1.597 | -0.059 | -0.042 | -0.042 | 0 | 0.11 | 0.03 | 0.16 | 0.14 | 0.23 | 0.24 |
| 5 | 59001077 | rs4700340 | 0.771 | 0.567 | 0.222 | -1.598 | -0.008 | -0.006 | -0.006 | 0 | 0.11 | 0.07 | 0.16 | 0.14 | 0.23 | 0.24 |
| 5 | 59020054 | rs1992112 | -1.713 | 0.352 | 0.276 | -1.698 | -0.002 | 0.008 | 0.008 | 0 | 0.6 | 0.65 | 0.45 | 0.33 | 0.57 | 0.53 |
| 5 | 59027953 | rs13179321 | -1.034 | 0.119 | -0.158 | -1.931 | 0.02 | 0.147 | 0.147 | 0 | 0.53 | 0.8 | 0.33 | 0.19 | 0.39 | 0.31 |
| 5 | 59032937 | rs4700341 | -1.061 | 0.094 | -0.303 | -2.15 | -0.013 | 0.059 | 0.059 | 0 | 0.45 | 0.59 | 0.39 | 0.15 | 0.51 | 0.39 |
| 5 | 59035558 | rs10472114 | 0.344 | 0.083 | -0.757 | -2.143 | -0.053 | 0.158 | 0.158 | 0 | 0.26 | 0.06 | 0.19 | 0.01 | 0.27 | 0.15 |
| 5 | 59042993 | rs7732908 | 0.435 | 0.094 | -0.812 | -2.188 | -0.002 | 0.081 | 0.081 | 0.813 | 0.16 | 0.06 | 0.11 | 0.01 | 0.15 | 0.12 |
| 5 | 59061282 | rs4235477 | -2.188 | -0.098 | -0.725 | -2.074 | -0.048 | -0.036 | -0.036 | 0.184 | 0.63 | 0.89 | 0.48 | 0.32 | 0.58 | 0.54 |
| 5 | 59073475 | rs4699942 | -1.15 | 0 | -0.417 | -2.092 | -0.014 | 0.09 | 0.09 | 0.012 | 0.44 | 0.65 | 0.39 | 0.14 | 0.47 | 0.38 |
| 5 | 59109014 | rs2963820 | 0.084 | -0.245 | -0.229 | -2.17 | -0.001 | 0.052 | 0.052 | 0 | 0.25 | 0.12 | 0.15 | 0 | 0.22 | 0.07 |
| 5 | 59111349 | rs2938784 | -2.246 | -0.24 | -0.445 | -2.176 | -0.042 | -0.057 | -0.057 | 0 | 0.64 | 0.92 | 0.65 | 0.32 | 0.61 | 0.57 |
| 5 | 59114805 | rs2963818 | -0.175 | -0.223 | 0.039 | -2.117 | -0.077 | 0.008 | 0.008 | 1.553 | 0.3 | 0.12 | 0.48 | 0.18 | 0.46 | 0.3 |
| 5 | 59121757 | rs6859121 | -0.702 | -0.35 | 0.189 | -2.131 | -0.01 | -0.014 | -0.014 | 3.121 | 0.23 | 0.3 | 0.1 | 0.13 | 0.11 | 0.15 |
| 5 | 59138161 | rs16890276 | 0.105 | -0.383 | 0.409 | -2.155 | 0.006 | 0.006 | 0.006 | 0 | 0.17 | 0.17 | 0.06 | 0.13 | 0.06 | 0.14 |
| 5 | 59140876 | rs256350 | -1.583 | -0.571 | 0.372 | -2.383 | 0.061 | 0.171 | 0.171 | 0.014 | 0.46 | 0.77 | 0.27 | 0.14 | 0.28 | 0.28 |
| 5 | 59141187 | rs11951422 | 1.315 | -0.406 | 0.351 | -2.296 | -0.058 | -0.058 | -0.058 | 0.441 | 0.12 | 0.03 | 0.35 | 0.18 | 0.27 | 0.23 |
| 5 | 59143129 | rs16890284 | 0.158 | -1.14 | 0.722 | -2.498 | 0.026 | 0.023 | 0.023 | 2.286 | 0.06 | 0.34 | 0.03 | 0 | 0 | 0 |
| 5 | 59152509 | rs10514888 | 0.457 | -1.096 | 0.13 | -2.479 | 0.014 | 0.036 | 0.036 | 0 | 0.15 | 0.1 | 0.05 | 0.01 | 0.06 | 0.14 |
| 5 | 59165423 | rs6889984 | -0.995 | -0.978 | -0.148 | -2.398 | -0.055 | -0.051 | -0.051 | 0.022 | 0.27 | 0.58 | 0.12 | 0.15 | 0.08 | 0.2 |
| 5 | 59168351 | rs256348 | -1.304 | -0.662 | -0.54 | -2.266 | 0.046 | 0.282 | 0.282 | 0.069 | 0.48 | 0.9 | 0.29 | 0.15 | 0.33 | 0.28 |
| 5 | 59174103 | rs159608 | -1.28 | -0.651 | -0.455 | -2.261 | 0.058 | 0.216 | 0.216 | 0.031 | 0.48 | 0.84 | 0.29 | 0.15 | 0.33 | 0.28 |
| 5 | 59201471 | rs7710463 | -0.383 | -0.422 | -0.136 | -2.021 | 0.005 | -0.053 | -0.053 | 0 | 0.14 | 0.24 | 0.07 | 0.02 | 0.11 | 0.13 |
| 5 | 59209183 | rs13362124 | 0.052 | -0.56 | -1.215 | -2.136 | -0.15 | 0.053 | 0.053 | 0.552 | 0.66 | 0.97 | 0.5 | 0.86 | 0.52 | 0.67 |
| 5 | 59220129 | rs2164240 | 1.049 | -0.571 | -0.558 | -1.975 | 0.01 | 0.116 | 0.116 | 0 | 0.44 | 0.17 | 0.39 | 0.83 | 0.43 | 0.45 |
| 5 | 59225762 | rs1017410 | 1.172 | -0.587 | -0.702 | -1.698 | -0.029 | 0.066 | 0.066 | 0 | 0.43 | 0.08 | 0.39 | 0.75 | 0.42 | 0.45 |
| 5 | 59230059 | rs1836542 | -0.403 | -0.728 | -0.051 | -1.631 | 0.021 | -0.006 | -0.006 | 0 | 0.29 | 0.59 | 0.43 | 0.2 | 0.39 | 0.29 |
| 5 | 59230179 | rs1529842 | -0.312 | -0.965 | -0.404 | -1.837 | -0.007 | 0.159 | 0.159 | 0 | 0.74 | 0.97 | 0.89 | 0.98 | 0.83 | 0.79 |
| 5 | 59230407 | rs1529843 | 1.15 | -0.963 | -0.293 | -1.824 | 0.008 | 0.076 | 0.076 | 0 | 0.45 | 0.25 | 0.45 | 0.79 | 0.44 | 0.5 |
| 5 | 59233347 | rs7379657 | -0.107 | -0.716 | 0.416 | -1.613 | 0.036 | -0.031 | -0.031 | 0 | 0.27 | 0.41 | 0.42 | 0.13 | 0.38 | 0.29 |
| 5 | 59233819 | rs10064439 | 0.337 | -0.856 | 0.26 | -1.66 | -0.048 | -0.052 | -0.052 | 3.094 | 0.18 | 0.55 | 0.11 | 0.08 | 0.1 | 0.1 |
| 5 | 59249174 | rs10940650 | 1.161 | -0.815 | -0.307 | -1.525 | -0.013 | 0.069 | 0.069 | 0.028 | 0.44 | 0.13 | 0.4 | 0.74 | 0.42 | 0.49 |
| 5 | 59254778 | rs7712937 | 0.747 | -0.478 | 0.323 | -1.389 | 0.004 | -0.019 | -0.019 | 0.003 | 0.6 | 0.45 | 0.53 | 0.87 | 0.52 | 0.58 |
| 5 | 59259372 | rs714291 | 0.51 | -0.291 | 0.232 | -1.373 | 0.039 | 0.003 | 0.003 | 0 | 0.13 | 0.17 | 0.34 | 0.1 | 0.3 | 0.22 |
| 5 | 59261429 | rs6896912 | -0.217 | -0.384 | 0.394 | -1.455 | 0.009 | 0.052 | 0.052 | 0 | 0.75 | 0.87 | 0.88 | 0.98 | 0.83 | 0.8 |
| 5 | 59272806 | rs6889660 | 1.27 | -0.665 | -0.033 | -1.614 | -0.026 | 0.035 | 0.035 | 0.006 | 0.38 | 0.13 | 0.38 | 0.75 | 0.41 | 0.5 |
| 5 | 59280611 | rs1369285 | 0.695 | -1.052 | 0.047 | -1.852 | 0.064 | 0.115 | 0.115 | 0.395 | 0.51 | 0.3 | 0.73 | 0.85 | 0.71 | 0.72 |
| 5 | 59288732 | rs16890418 | 0.38 | -0.964 | 0.005 | -1.448 | 0.004 | -0.007 | -0.007 | 0.204 | 0.23 | 0.2 | 0.13 | 0.05 | 0.11 | 0.08 |
| 5 | 59304532 | rs2409679 | -0.126 | -1.096 | -0.341 | -1.568 | -0.028 | -0.053 | -0.053 | 0 | 0.63 | 0.63 | 0.78 | 0.9 | 0.74 | 0.75 |
| 5 | 59307813 | rs1533019 | -0.097 | -1.076 | -0.351 | -1.567 | -0.025 | -0.048 | -0.048 | 0 | 0.62 | 0.63 | 0.78 | 0.89 | 0.74 | 0.75 |
| 5 | 59311463 | rs1876671 | 0.229 | -0.998 | -0.183 | -1.706 | 0.003 | 0.006 | 0.006 | 0 | 0.23 | 0.16 | 0.13 | 0.04 | 0.17 | 0.26 |
| 5 | 59322616 | rs7703240 | -0.058 | -0.604 | -0.312 | -1.649 | 0.002 | 0.021 | 0.021 | 0 | 0.51 | 0.6 | 0.45 | 0.82 | 0.44 | 0.59 |
| 5 | 59330489 | rs7703245 | 1.296 | -0.656 | -0.262 | -1.613 | -0.027 | -0.01 | -0.01 | 0 | 0.27 | 0.19 | 0.31 | 0.75 | 0.27 | 0.33 |
| 5 | 59336110 | rs16890459 | 0.337 | -0.669 | -0.21 | -1.627 | 0.004 | 0.053 | 0.053 | 0 | 0.23 | 0.11 | 0.13 | 0.04 | 0.17 | 0.26 |
| 5 | 59360341 | rs10940659 | -0.105 | -0.723 | -0.594 | -1.586 | 0.007 | 0.127 | 0.127 | 0 | 0.49 | 0.31 | 0.43 | 0.79 | 0.44 | 0.59 |
| 5 | 59366269 | rs7732249 | -1.897 | -0.987 | -0.758 | -1.729 | 0.003 | -0.034 | -0.034 | 0 | 0.89 | 0.99 | 0.93 | 0.89 | 0.88 | 0.88 |
| 5 | 59369794 | rs1588265 | 0.964 | -0.352 | -0.239 | -1.296 | -0.026 | -0.006 | -0.006 | 0.923 | 0.28 | 0.21 | 0.31 | 0.75 | 0.28 | 0.33 |
| 5 | 59371938 | rs12153039 | -0.024 | -0.283 | -0.221 | -1.256 | 0.005 | 0.018 | 0.018 | 0 | 0.23 | 0.18 | 0.12 | 0.07 | 0.17 | 0.26 |
| 5 | 59398304 | rs17376761 | 0.154 | -0.334 | -0.235 | -1.262 | 0.005 | 0.06 | 0.06 | 0 | 0.22 | 0.11 | 0.12 | 0.04 | 0.17 | 0.25 |
| 5 | 59420684 | rs11949645 | 0.439 | -0.349 | -0.168 | -1.255 | 0.004 | 0.029 | 0.029 | 0 | 0.5 | 0.49 | 0.56 | 0.22 | 0.56 | 0.42 |
| 5 | 59427742 | rs4700365 | 0.343 | -0.334 | -0.167 | -1.249 | 0.001 | 0.009 | 0.009 | 0 | 0.49 | 0.47 | 0.55 | 0.17 | 0.55 | 0.41 |
| 5 | 59439082 | rs1544791 | -0.95 | -0.482 | -0.175 | -1.185 | 0.008 | -0.005 | -0.005 | 0.019 | 0.72 | 0.63 | 0.68 | 0.23 | 0.73 | 0.68 |
| 5 | 59445137 | rs983280 | -0.952 | -0.473 | -0.266 | -1.185 | -0.005 | -0.002 | -0.002 | 0 | 0.72 | 0.69 | 0.68 | 0.26 | 0.73 | 0.67 |
| 5 | 59467203 | rs2910830 | -0.099 | -0.554 | -0.425 | -1.21 | -0.001 | 0.003 | 0.003 | 0 | 0.6 | 0.53 | 0.64 | 0.24 | 0.67 | 0.61 |
| 5 | 59469899 | rs2910829 | 0.612 | -0.535 | -0.436 | -1.262 | -0.001 | 0.006 | 0.006 | 0 | 0.48 | 0.47 | 0.52 | 0.18 | 0.56 | 0.41 |

| | | | | | | | | | | | | | | | | |
|---|---|---|---|---|---|---|---|---|---|---|---|---|---|---|---|---|
| 5 | 59474215 | rs13158277 | 0.653 | -0.599 | -0.41 | -0.865 | 0.008 | 0.008 | 0.008 | 0.576 | 0.11 | 0.07 | 0.04 | 0.04 | 0.07 | 0.06 |
| 5 | 59481878 | rs17450114 | -0.366 | -0.391 | -0.443 | -0.835 | -0.001 | 0.01 | 0.01 | 0 | 0.12 | 0.09 | 0.11 | 0.04 | 0.1 | 0.2 |
| 5 | 59486768 | rs7717864 | -0.821 | -0.523 | -0.248 | -0.788 | -0.038 | -0.027 | -0.027 | 0.289 | 0.26 | 0.5 | 0.16 | 0.16 | 0.16 | 0.24 |
| 5 | 59502520 | rs966221 | 0.285 | -0.14 | -0.477 | -0.703 | 0.004 | 0.013 | 0.013 | 0 | 0.54 | 0.53 | 0.59 | 0.17 | 0.62 | 0.46 |
| 5 | 59515334 | rs2910827 | 1.067 | -0.256 | -1.186 | -1.184 | 0.058 | 0.153 | 0.153 | 0 | 0.35 | 0.11 | 0.18 | 0.09 | 0.25 | 0.23 |
| 5 | 59517876 | rs10059334 | -0.097 | -0.011 | -0.376 | -1.173 | 0.115 | -0.105 | -0.105 | 0.224 | 0.22 | 0.43 | 0.44 | 0.05 | 0.39 | 0.23 |
| 5 | 59520445 | rs7442640 | 0.297 | 1.036 | -0.099 | -1.281 | 0.012 | -0.037 | -0.037 | 0 | 0.12 | 0.12 | 0.18 | 0 | 0.18 | 0.11 |
| 5 | 59522505 | rs4302546 | -1.276 | 0.259 | -0.671 | -1.092 | 0.067 | 0.109 | 0.109 | 0 | 0.62 | 0.84 | 0.79 | 0.85 | 0.74 | 0.66 |
| 5 | 59522613 | rs4513649 | 0.495 | 0.453 | -0.823 | -1.138 | -0.005 | 0.047 | 0.047 | 0.012 | 0.14 | 0.06 | 0.07 | 0 | 0.12 | 0.1 |
| 5 | 59526714 | rs13157125 | 0.271 | 0.443 | -0.803 | -1.143 | -0.001 | -0.027 | -0.027 | 0 | 0.15 | 0.19 | 0.09 | 0 | 0.13 | 0.1 |
| 5 | 59544653 | rs4235479 | -0.887 | 0.468 | -1.326 | -0.79 | -0.01 | 0.014 | 0.014 | 0 | 0.38 | 0.66 | 0.55 | 0.18 | 0.52 | 0.33 |
| 5 | 59548973 | rs12188020 | 0.847 | 0.755 | -1.688 | -0.52 | -0.013 | 0.04 | 0.04 | 0.895 | 0.1 | 0.03 | 0.06 | 0 | 0.11 | 0.07 |
| 5 | 59553742 | rs11739760 | 1.069 | 0.581 | -1.631 | 0.713 | -0.05 | -0.113 | -0.113 | 0.038 | 0.21 | 0.01 | 0.14 | 0.41 | 0.2 | 0.22 |
| 5 | 59556461 | rs10434525 | 0.927 | 0.547 | -1.643 | 0.087 | -0.064 | -0.134 | -0.134 | 0.542 | 0.23 | 0.02 | 0.24 | 0.66 | 0.22 | 0.33 |
| 5 | 59569008 | rs7714765 | -0.829 | 0.998 | -0.715 | -0.432 | -0.017 | 0.005 | 0.005 | 0 | 0.32 | 0.68 | 0.51 | 0.06 | 0.48 | 0.31 |
| 5 | 59586116 | rs10939840 | -0.617 | 0.445 | -0.592 | -0.111 | 0.002 | -0.001 | -0.001 | 0.201 | 0.09 | 0.07 | 0.06 | 0 | 0.09 | 0.06 |
| 5 | 59602340 | rs16874969 | -0.601 | -0.311 | -0.305 | 0.57 | -0.054 | 0.054 | 0.054 | 0 | 0.29 | 0.51 | 0.26 | 0.36 | 0.15 | 0.45 |
| 5 | 59618379 | rs13178509 | -0.088 | -0.604 | 0.042 | 0.742 | 0.015 | -0.012 | -0.012 | 0 | 0.18 | 0.14 | 0.16 | 0.2 | 0.1 | 0.34 |
| 5 | 59627555 | rs4394088 | -1.248 | -0.658 | 0.319 | 0.687 | 0.002 | -0.003 | -0.003 | 0 | 0.32 | 0.32 | 0.28 | 0.48 | 0.18 | 0.51 |
| 5 | 59637451 | rs6865762 | -1.144 | -0.721 | 0.614 | 0.665 | -0.008 | 0.021 | 0.021 | 0.001 | 0.17 | 0.57 | 0.15 | 0.15 | 0.13 | 0.12 |
| 5 | 59637749 | rs4547892 | -1.341 | -0.809 | 0.526 | 0.604 | -0.017 | 0.071 | 0.071 | 0 | 0.18 | 0.59 | 0.19 | 0.27 | 0.14 | 0.13 |
| 5 | 59644785 | rs12522161 | 0.914 | -0.202 | -0.392 | -0.131 | -0.109 | 0.116 | 0.116 | 0.002 | 0.21 | 0.03 | 0.23 | 0.05 | 0.37 | 0.21 |
| 5 | 59655640 | rs16877926 | 0.613 | -0.714 | -0.113 | 0.284 | -0.009 | -0.069 | -0.069 | 1.993 | 0.13 | 0.02 | 0.13 | 0.19 | 0.09 | 0.32 |
| 5 | 59657695 | rs10514893 | -0.302 | -0.72 | 0.428 | -0.009 | 0.059 | 0.15 | 0.15 | 0 | 0.05 | 0.24 | 0.06 | 0.12 | 0.04 | 0.06 |
| 5 | 59661456 | rs1423246 | -1.973 | -0.821 | 0.399 | -0.083 | 0.045 | -0.011 | -0.011 | 0 | 0.44 | 0.72 | 0.36 | 0.47 | 0.28 | 0.56 |
| 5 | 59665111 | rs10939844 | -0.279 | -0.852 | 0.309 | -0.316 | 0 | -0.016 | -0.016 | 0 | 0.14 | 0.17 | 0.07 | 0.02 | 0.09 | 0.08 |
| 5 | 59665684 | rs16877953 | 0.289 | -0.885 | 0.335 | -0.066 | 0.011 | -0.028 | -0.028 | 0 | 0.14 | 0.07 | 0.16 | 0.4 | 0.1 | 0.34 |
| 5 | 59673366 | rs1345782 | -0.257 | -0.962 | 0.76 | -0.147 | -0.013 | 0.011 | 0.011 | 0 | 0.14 | 0.25 | 0.09 | 0.12 | 0.07 | 0.14 |
| 5 | 59690935 | rs16877986 | 0.407 | -0.985 | 0.58 | -0.233 | -0.027 | -0.049 | -0.049 | 0 | 0.12 | 0.18 | 0.03 | 0 | 0.04 | 0.08 |
| 5 | 59697303 | rs918592 | -1.462 | -1.04 | 0.605 | -0.191 | 0.011 | -0.012 | -0.012 | 0.021 | 0.32 | 0.69 | 0.29 | 0.54 | 0.21 | 0.52 |
| 5 | 59697808 | rs918590 | -1.416 | -1.003 | 0.013 | -0.174 | -0.034 | 0.048 | 0.048 | 0.097 | 0.3 | 0.51 | 0.29 | 0.54 | 0.22 | 0.52 |
| 5 | 59706800 | rs6887545 | -1.393 | -0.981 | 0.016 | -0.317 | -0.032 | -0.035 | -0.035 | 0.416 | 0.27 | 0.56 | 0.16 | 0.15 | 0.16 | 0.22 |
| 5 | 59732614 | rs10045433 | -0.385 | -0.732 | -0.109 | -0.37 | -0.001 | -0.046 | -0.046 | 0 | 0.14 | 0.18 | 0.1 | 0.01 | 0.15 | 0.1 |
| 5 | 59734769 | rs789396 | -0.972 | -0.929 | -0.591 | -0.336 | -0.004 | 0 | 0 | 0 | 0.31 | 0.33 | 0.24 | 0.58 | 0.17 | 0.54 |
| 5 | 59735799 | rs404202 | -1.644 | -0.852 | -0.777 | -0.308 | -0.009 | -0.003 | -0.003 | 0.144 | 0.45 | 0.49 | 0.34 | 0.59 | 0.32 | 0.63 |
| 5 | 59735865 | rs364917 | -1.415 | -0.897 | -0.803 | -0.332 | -0.024 | 0.014 | 0.014 | 0.508 | 0.35 | 0.49 | 0.32 | 0.59 | 0.29 | 0.55 |
| 5 | 59736177 | rs371424 | -1.625 | -0.847 | -0.79 | -0.309 | 0 | 0 | 0 | 0.003 | 0.45 | 0.46 | 0.34 | 0.59 | 0.32 | 0.63 |
| 5 | 59742759 | rs702554 | 0.54 | -0.763 | -1.067 | -0.421 | -0.001 | -0.088 | -0.088 | 0 | 0.13 | 0.03 | 0.15 | 0.41 | 0.1 | 0.35 |
| 5 | 59746920 | rs6449471 | -0.489 | -0.854 | -1.014 | -0.389 | -0.036 | -0.004 | -0.004 | 0 | 0.22 | 0.33 | 0.12 | 0.18 | 0.08 | 0.19 |
| 5 | 59752640 | rs27223 | -2.197 | -0.635 | -1.849 | -0.8 | -0.061 | -0.083 | -0.083 | 0 | 0.7 | 0.92 | 0.64 | 0.61 | 0.65 | 0.83 |
| 5 | 59756031 | rs4699956 | 0.872 | -0.053 | -1.655 | -0.381 | -0.107 | 0.121 | 0.121 | 0.385 | 0.22 | 0.02 | 0.25 | 0.02 | 0.3 | 0.2 |
| 5 | 59759657 | rs26951 | 0.18 | -0.296 | -0.923 | 0.284 | -0.002 | -0.087 | -0.087 | 0 | 0.13 | 0.03 | 0.16 | 0.41 | 0.1 | 0.35 |
| 5 | 59777491 | rs27224 | -0.156 | -0.299 | -0.923 | 0.274 | 0.008 | -0.091 | -0.091 | 0.252 | 0.14 | 0.04 | 0.16 | 0.4 | 0.1 | 0.35 |
| 5 | 59777561 | rs1423471 | -0.543 | -0.063 | -0.382 | 0.18 | -0.012 | 0.093 | 0.093 | 0.294 | 0.11 | 0.35 | 0.1 | 0.18 | 0.05 | 0.12 |
| 5 | 59783317 | rs12188950 | -0.681 | 0.277 | -1.243 | -0.117 | -0.002 | -0.014 | -0.014 | 0 | 0.11 | 0.11 | 0.09 | 0 | 0.16 | 0.08 |
| 5 | 59790985 | rs26956 | -0.194 | 0.134 | -1.889 | 0.001 | -0.055 | -0.123 | -0.123 | 0 | 0.26 | 0.04 | 0.27 | 0.41 | 0.27 | 0.46 |
| 5 | 59791348 | rs10516154 | -0.521 | -0.134 | -1.805 | -0.324 | -0.001 | -0.021 | -0.021 | 0.074 | 0.13 | 0.13 | 0.08 | 0 | 0.15 | 0.11 |
| 5 | 59793535 | rs26955 | -0.517 | -0.766 | -1.997 | -1.03 | -0.014 | -0.005 | -0.005 | 0 | 0.6 | 0.73 | 0.5 | 0.44 | 0.46 | 0.65 |
| 5 | 59794824 | rs16878165 | 0.315 | -1.188 | -1.022 | -1.155 | -0.073 | -0.108 | -0.108 | 0.186 | 0.19 | 0.31 | 0.07 | 0 | 0.06 | 0.11 |
| 5 | 59802783 | rs35386 | -1.433 | -0.596 | -1.193 | -0.844 | -0.046 | -0.043 | -0.043 | 0 | 0.73 | 0.87 | 0.59 | 0.45 | 0.61 | 0.76 |
| 5 | 59805467 | rs40512 | -1.123 | -0.375 | -1.551 | -0.995 | -0.117 | -0.085 | -0.085 | 0 | 0.71 | 0.91 | 0.58 | 0.47 | 0.6 | 0.75 |
| 5 | 59814314 | rs26953 | -0.512 | -1.391 | -2.118 | -1.379 | -0.026 | -0.104 | -0.104 | 0 | 0.69 | 0.93 | 0.7 | 0.55 | 0.7 | 0.46 |
| 5 | 59817795 | rs11960912 | 0.063 | -1.758 | -1.907 | -1.657 | 0.006 | -0.006 | -0.006 | 0 | 0.15 | 0.12 | 0.09 | 0 | 0.13 | 0.07 |
| 5 | 59818246 | rs4133470 | -1.323 | -1.871 | -1.905 | -1.706 | 0.04 | 0.212 | 0.212 | 0 | 0.38 | 0.85 | 0.21 | 0.03 | 0.28 | 0.2 |
| 5 | 59818675 | rs3857234 | 0.518 | -1.893 | -0.829 | -1.729 | 0.048 | 0.039 | 0.039 | 0.557 | 0.05 | 0.32 | 0.03 | 0 | 0.01 | 0.05 |
| 5 | 59818778 | rs7723306 | 0.568 | -1.845 | -0.868 | -1.717 | 0.048 | -0.061 | -0.061 | 0 | 0.12 | 0.38 | 0.08 | 0.01 | 0.11 | 0.05 |
| 5 | 59821543 | rs35382 | 0.58 | -1.969 | -1.078 | -1.707 | 0.006 | 0.005 | 0.005 | 0.006 | 0.48 | 0.48 | 0.59 | 0.55 | 0.55 | 0.34 |
| 5 | 59821598 | rs10065640 | -0.278 | -1.974 | -1.079 | -1.724 | 0.018 | -0.104 | -0.104 | 0.285 | 0.17 | 0.42 | 0.09 | 0.01 | 0.14 | 0.08 |
| 5 | 59823198 | rs7727343 | -0.075 | -1.958 | -1.129 | -1.717 | 0.018 | -0.089 | -0.089 | 0.317 | 0.17 | 0.38 | 0.09 | 0.02 | 0.14 | 0.08 |
| 5 | 59825300 | rs26949 | 1.802 | -1.923 | -1.038 | -1.625 | -0.12 | -0.105 | -0.105 | 0 | 0.32 | 0.09 | 0.5 | 0.54 | 0.42 | 0.26 |
| 5 | 59829465 | rs10056275 | -0.616 | -1.882 | -0.75 | -1.519 | 0.005 | -0.024 | -0.024 | 0 | 0.09 | 0.38 | 0.05 | 0 | 0.03 | 0.03 |
| 5 | 59837476 | rs8176070 | 0.268 | -2.103 | -1.608 | -1.307 | -0.063 | 0.015 | 0.015 | 0.09 | 0.32 | 0.04 | 0.28 | 0.21 | 0.33 | 0.44 |
| 5 | 59837591 | rs27565 | 0.844 | -2.595 | -1.25 | -1.226 | 0.014 | -0.006 | -0.006 | 0 | 0.45 | 0.72 | 0.59 | 0.51 | 0.53 | 0.39 |
| 5 | 59841361 | rs10939851 | -0.894 | -2.629 | -0.833 | -1.479 | -0.056 | -0.134 | -0.134 | 0 | 0.23 | 0.4 | 0.13 | 0 | 0.16 | 0.18 |
| 5 | 59877538 | rs1562235 | 1.219 | -2.534 | -1.085 | -1.352 | -0.032 | 0.004 | 0.004 | 0.599 | 0.42 | 0.24 | 0.58 | 0.57 | 0.54 | 0.28 |
| 5 | 59890278 | rs4326096 | -0.474 | -2.624 | -1.242 | -1.28 | -0.041 | -0.049 | -0.049 | 0 | 0.65 | 0.97 | 0.69 | 0.66 | 0.67 | 0.55 |
| 5 | 59891782 | rs6449480 | -0.314 | -2.558 | -1.211 | -1.092 | -0.029 | -0.131 | -0.131 | 0 | 0.63 | 0.96 | 0.68 | 0.53 | 0.67 | 0.48 |
| 5 | 59935739 | rs286157 | 0.089 | -2.271 | -0.703 | -1.175 | -0.01 | -0.021 | -0.021 | 0 | 0.07 | 0.24 | 0.03 | 0 | 0.02 | 0.13 |
| 5 | 59939718 | rs7732792 | 0.424 | -2.42 | -0.775 | -1.324 | 0.006 | -0.011 | -0.011 | 0 | 0.15 | 0.15 | 0.1 | 0 | 0.13 | 0.04 |

| | | | | | | | | | | | | | | | | |
|---|---|---|---|---|---|---|---|---|---|---|---|---|---|---|---|---|
| 5 | 59977058 | rs11741754 | 2.261 | -2.425 | -0.939 | -1.324 | -0.127 | -0.092 | -0.092 | 0 | 0.28 | 0.08 | 0.48 | 0.57 | 0.41 | 0.24 |
| 5 | 59988547 | rs7719611 | 1.17 | -2.498 | -1.011 | -1.451 | -0.015 | 0.04 | 0.04 | 0 | 0.42 | 0.35 | 0.57 | 0.78 | 0.46 | 0.45 |
| 5 | 59995393 | rs6449483 | 0.415 | -2.236 | -0.87 | -1.228 | -0.001 | -0.011 | -0.011 | 0 | 0.57 | 0.49 | 0.67 | 0.78 | 0.58 | 0.49 |
| 5 | 59995742 | rs1379112 | 0.47 | -1.953 | -0.469 | -1.042 | 0.08 | -0.012 | -0.012 | 0.022 | 0.09 | 0.37 | 0.05 | 0.01 | 0.08 | 0.05 |
| 5 | 60012235 | rs4298206 | -0.544 | -2.306 | -0.668 | -1.248 | 0.003 | 0.01 | 0.01 | 0 | 0.46 | 0.53 | 0.34 | 0.23 | 0.41 | 0.55 |
| 5 | 60013913 | rs4307050 | 0.212 | -2.381 | -0.639 | -1.294 | 0.008 | 0.062 | 0.062 | 0 | 0.32 | 0.1 | 0.28 | 0.21 | 0.31 | 0.45 |
| 5 | 60020520 | rs1107233 | 0.068 | -2.502 | -0.478 | -1.086 | -0.049 | 0.075 | 0.075 | 0.002 | 0.1 | 0.26 | 0.06 | 0.21 | 0.02 | 0.16 |
| 5 | 60048028 | rs16878354 | 0.494 | -2.486 | -0.709 | -1.11 | -0.039 | 0.038 | 0.038 | 0 | 0.09 | 0.15 | 0.06 | 0.21 | 0.02 | 0.16 |
| 5 | 60057836 | rs7733069 | 0.524 | -2.485 | -0.708 | -1.109 | -0.039 | 0.039 | 0.039 | 0 | 0.08 | 0.15 | 0.06 | 0.21 | 0.02 | 0.16 |
| 5 | 60063511 | rs1563517 | -1.338 | -2.611 | -1.118 | -1.249 | 0.007 | 0.005 | 0.005 | 0 | 0.27 | 0.7 | 0.15 | 0.22 | 0.15 | 0.27 |
| 5 | 60063576 | rs1460958 | -2.265 | -2.662 | -1.712 | -1.292 | -0.136 | -0.097 | -0.097 | 0 | 0.71 | 0.92 | 0.52 | 0.43 | 0.59 | 0.76 |
| 5 | 60065519 | rs16878397 | 0.74 | -2.549 | -1.864 | -1.984 | -0.008 | -0.011 | -0.011 | 0.007 | 0.17 | 0.01 | 0.14 | 0.15 | 0.15 | 0.21 |
| 5 | 60074000 | rs6874699 | -1.862 | -2.378 | -1.38 | -1.945 | -0.038 | 0.056 | 0.056 | 0 | 0.65 | 0.77 | 0.45 | 0.15 | 0.57 | 0.65 |
| 5 | 60081318 | rs1870683 | 0.271 | -2.571 | -1.568 | -2.149 | -0.036 | 0.118 | 0.118 | 0 | 0.24 | 0.09 | 0.17 | 0 | 0.29 | 0.19 |
| 5 | 60088148 | rs6872863 | 1.171 | -2.882 | -1.255 | -2.404 | 0.016 | 0.091 | 0.091 | 0 | 0.5 | 0.69 | 0.56 | 0.9 | 0.53 | 0.44 |
| 5 | 60089436 | rs921898 | 0.212 | -2.885 | -1.027 | -2.366 | -0.005 | -0.04 | -0.04 | 0 | 0.15 | 0.46 | 0.11 | 0.04 | 0.11 | 0.09 |
| 5 | 60094266 | rs4700392 | -0.599 | -2.851 | -1.191 | -2.339 | -0.003 | -0.016 | -0.016 | 0.003 | 0.8 | 0.85 | 0.75 | 0.9 | 0.84 | 0.7 |
| 5 | 60103452 | rs10062839 | 0.116 | -2.806 | -1.189 | -2.341 | -0.002 | -0.021 | -0.021 | 0.323 | 0.74 | 0.79 | 0.73 | 0.9 | 0.81 | 0.64 |
| 5 | 60119737 | rs1445293 | 1.209 | -2.786 | -1.225 | -2.371 | 0.016 | 0.098 | 0.098 | 0 | 0.5 | 0.69 | 0.56 | 0.91 | 0.54 | 0.44 |
| 5 | 60127724 | rs1807017 | 2.076 | -2.788 | -1.067 | -2.276 | -0.034 | 0.061 | 0.061 | 0 | 0.35 | 0.21 | 0.45 | 0.85 | 0.43 | 0.35 |
| 5 | 60135985 | rs4700398 | 0.027 | -1.95 | -0.455 | -2.429 | -0.027 | -0.07 | -0.07 | 0 | 0.74 | 0.63 | 0.72 | 0.86 | 0.81 | 0.67 |
| 5 | 60169151 | rs4647167 | 0.814 | -1.892 | -0.422 | -2.513 | -0.043 | 0.126 | 0.126 | 0 | 0.23 | 0.07 | 0.17 | 0 | 0.27 | 0.21 |
| 5 | 60170337 | rs3117 | -1.38 | -2.017 | -0.195 | -2.594 | -0.003 | 0.044 | 0.044 | 0 | 0.43 | 0.39 | 0.3 | 0.05 | 0.43 | 0.33 |
| 5 | 60217651 | rs976080 | -0.952 | -2.02 | -0.317 | -2.542 | -0.008 | -0.022 | -0.022 | 0 | 0.86 | 0.73 | 0.89 | 0.95 | 0.89 | 0.89 |
| 5 | 60238590 | rs158933 | 1.4 | -1.183 | -0.438 | -2.317 | 0.002 | 0.005 | 0.005 | 0 | 0.54 | 0.35 | 0.68 | 0.95 | 0.57 | 0.67 |
| 5 | 60241826 | rs4647028 | 0.881 | -1.147 | -0.499 | -2.316 | -0.04 | 0.12 | 0.12 | 0 | 0.23 | 0.08 | 0.17 | 0 | 0.27 | 0.21 |
| 5 | 60261889 | rs1609041 | 1.333 | -1.277 | -0.616 | -2.362 | -0.017 | 0.01 | 0.01 | 0 | 0.17 | 0.02 | 0.23 | 0.13 | 0.16 | 0.28 |
| 5 | 60298360 | rs290506 | 1.058 | -1.259 | -0.564 | -2.35 | -0.003 | 0.04 | 0.04 | 0 | 0.57 | 0.61 | 0.7 | 0.95 | 0.57 | 0.67 |
| 5 | 60317433 | rs248688 | 0.131 | -1.215 | -0.522 | -2.304 | -0.002 | 0.015 | 0.015 | 0 | 0.64 | 0.65 | 0.72 | 0.95 | 0.62 | 0.69 |
| 5 | 60318015 | rs7709522 | 1.804 | -1.114 | -0.412 | -2.171 | 0.002 | -0.03 | -0.03 | 0 | 0.4 | 0.52 | 0.54 | 0.82 | 0.41 | 0.41 |
| 5 | 60323753 | rs7723349 | 1.215 | -0.591 | -0.924 | -2.502 | -0.001 | -0.012 | -0.012 | 0 | 0.25 | 0.29 | 0.43 | 0.81 | 0.25 | 0.37 |
| 5 | 60329624 | rs34592 | 0.038 | -0.915 | -0.599 | -2.16 | 0 | -0.044 | -0.044 | 0 | 0.65 | 0.61 | 0.72 | 0.95 | 0.63 | 0.69 |
| 5 | 60336004 | rs2545493 | -1.497 | -0.793 | -0.998 | -2.182 | -0.005 | 0.009 | 0.009 | 3.056 | 0.89 | 0.93 | 0.9 | 0.95 | 0.87 | 0.93 |
| 5 | 60344399 | rs12520484 | 2.158 | -0.675 | -1.125 | -2.035 | -0.052 | 0.061 | 0.061 | 0.359 | 0.11 | 0.01 | 0.08 | 0 | 0.15 | 0.17 |
| 5 | 60357869 | rs248684 | -1.968 | -0.779 | -1.066 | -2.308 | 0.015 | 0.043 | 0.043 | 0.745 | 0.88 | 0.94 | 0.92 | 1 | 0.91 | 0.97 |
| 5 | 60401606 | rs295557 | 0.845 | -0.752 | -1.047 | -2.248 | -0.011 | 0.017 | 0.017 | 0 | 0.17 | 0.02 | 0.15 | 0.13 | 0.16 | 0.26 |
| 5 | 60402210 | rs295556 | 0.691 | -0.771 | -0.95 | -2.257 | 0.014 | 0.031 | 0.031 | 0 | 0.18 | 0.09 | 0.15 | 0.13 | 0.16 | 0.26 |
| 5 | 60410899 | rs11744226 | 1.145 | -0.964 | -1.013 | -2.372 | -0.001 | 0.025 | 0.025 | 0.009 | 0.11 | 0.03 | 0.09 | 0.1 | 0.13 | 0.22 |
| 5 | 60411832 | rs1460961 | 2.107 | -0.881 | -0.859 | -2.49 | -0.016 | 0.036 | 0.036 | 0 | 0.35 | 0.27 | 0.53 | 0.81 | 0.41 | 0.36 |
| 5 | 60437381 | rs1563908 | 3.133 | -1.093 | -1.146 | -2.349 | -0.122 | 0.095 | 0.095 | 0.032 | 0.26 | 0.04 | 0.42 | 0.09 | 0.34 | 0.11 |
| 5 | 60444276 | rs159371 | 0.092 | -0.922 | -0.456 | -2.17 | -0.001 | -0.103 | -0.103 | 0.205 | 0.63 | 0.47 | 0.71 | 0.95 | 0.62 | 0.69 |
| 5 | 60459040 | rs2270822 | 0.743 | -0.855 | -0.82 | -2.137 | 0.005 | -0.017 | -0.017 | 0.042 | 0.12 | 0.23 | 0.11 | 0.05 | 0.13 | 0.11 |
| 5 | 60468683 | rs124899 | -1.062 | -0.865 | -1.519 | -2.101 | -0.038 | 0.123 | 0.123 | 0 | 0.77 | 0.93 | 0.84 | 1 | 0.75 | 0.8 |
| 5 | 60471080 | rs10076568 | 0.539 | -1.32 | -1.334 | -1.843 | 0.021 | 0.043 | 0.043 | 0.1 | 0.16 | 0.02 | 0.09 | 0.12 | 0.13 | 0.22 |
| 5 | 60485778 | rs159542 | 0.661 | -1.05 | -1.29 | -1.876 | -0.001 | 0.155 | 0.155 | 0 | 0.56 | 0.79 | 0.5 | 0.83 | 0.56 | 0.46 |
| 5 | 60486515 | rs159543 | 2.137 | -1.146 | -1.143 | -2.013 | -0.011 | 0.031 | 0.031 | 0 | 0.35 | 0.27 | 0.34 | 0.78 | 0.39 | 0.33 |
| 5 | 60488167 | rs295544 | -0.852 | -1.132 | -0.828 | -1.931 | -0.012 | -0.066 | -0.066 | 0.198 | 0.25 | 0.47 | 0.22 | 0.08 | 0.2 | 0.25 |
| 5 | 60511401 | rs764067 | -0.124 | -0.995 | -0.382 | -1.773 | 0.073 | 0.047 | 0.047 | 0.417 | 0.05 | 0.39 | 0.05 | 0 | 0.03 | 0.02 |
| 5 | 60513501 | rs34635 | -0.633 | -1.069 | -0.521 | -1.894 | -0.006 | 0.138 | 0.138 | 0 | 0.4 | 0.17 | 0.42 | 0.13 | 0.4 | 0.42 |
| 5 | 60521741 | rs6867285 | 0.655 | -1.083 | -0.538 | -1.937 | -0.032 | -0.1 | -0.1 | 0 | 0.73 | 0.54 | 0.85 | 0.87 | 0.8 | 0.75 |
| 5 | 60522414 | rs159364 | 0.265 | -0.996 | -0.483 | -1.632 | 0.001 | 0.073 | 0.073 | 0.007 | 0.9 | 0.58 | 0.88 | 0.91 | 0.94 | 0.86 |
| 5 | 60533515 | rs7446842 | 0.042 | -1.16 | -0.494 | -1.666 | 0.013 | -0.154 | -0.154 | 0.166 | 0.71 | 0.51 | 0.59 | 1 | 0.68 | 0.76 |
| 5 | 60549110 | rs12516321 | 0.605 | -1.006 | -0.768 | -1.63 | -0.043 | 0.107 | 0.107 | 0 | 0.24 | 0.09 | 0.33 | 0 | 0.27 | 0.21 |
| 5 | 60550041 | rs6868457 | 1.32 | -0.953 | -0.626 | -1.52 | -0.004 | 0.012 | 0.012 | 0.31 | 0.45 | 0.4 | 0.48 | 0.79 | 0.5 | 0.43 |
| 5 | 60555922 | rs4540126 | 1.22 | -0.659 | -0.162 | -1.157 | 0.004 | 0.003 | 0.003 | 1.348 | 0.71 | 0.36 | 0.86 | 0.86 | 0.82 | 0.68 |
| 5 | 60558282 | rs12696982 | 0.602 | -0.692 | -0.037 | -1.187 | -0.026 | -0.076 | -0.076 | 0 | 0.79 | 0.44 | 0.87 | 0.91 | 0.87 | 0.87 |
| 5 | 60562640 | rs10044874 | -0.775 | -0.743 | 0.086 | -1.129 | -0.034 | -0.004 | -0.004 | 2.065 | 0.11 | 0.24 | 0.05 | 0.06 | 0.07 | 0.19 |
| 5 | 60562651 | rs4315878 | 0.364 | -0.589 | 0.054 | -1.065 | -0.015 | -0.077 | -0.077 | 0.066 | 0.81 | 0.6 | 0.88 | 0.91 | 0.87 | 0.87 |
| 5 | 60564322 | rs10939892 | -0.736 | -0.408 | -0.281 | -0.969 | -0.015 | -0.012 | -0.012 | 0 | 0.2 | 0.22 | 0.11 | 0.15 | 0.17 | 0.29 |
| 5 | 60570104 | rs7721603 | 0.684 | -0.07 | -0.339 | -0.97 | 0.003 | 0.015 | 0.015 | 0 | 0.11 | 0.03 | 0.06 | 0.09 | 0.1 | 0.09 |
| 5 | 60571540 | rs6870703 | 0.973 | 0.118 | -0.022 | -0.844 | 0.028 | -0.036 | -0.036 | 0 | 0.91 | 0.5 | 0.95 | 1 | 0.97 | 0.97 |
| 5 | 60572119 | rs10070347 | 0.567 | 0.172 | -0.13 | -0.808 | 0.051 | -0.01 | -0.01 | 0 | 0.92 | 0.57 | 0.95 | 1 | 0.97 | 0.97 |
| 5 | 60586625 | rs4352559 | -1.194 | -0.152 | -1.899 | -0.254 | -0.16 | -0.086 | -0.086 | 0 | 0.58 | 0.96 | 0.6 | 0.47 | 0.49 | 0.62 |
| 5 | 60595439 | rs12188621 | 0.012 | -0.632 | -0.753 | -0.639 | -0.027 | 0.01 | 0.01 | 1.046 | 0.09 | 0.38 | 0.06 | 0.03 | 0.02 | 0.12 |
| 5 | 60598543 | rs4391122 | -1.105 | -0.946 | -0.873 | -0.747 | -0.075 | 0.227 | 0.227 | 3.572 | 0.51 | 0.9 | 0.54 | 0.22 | 0.47 | 0.51 |
| 5 | 60600524 | rs10939897 | 0.754 | -0.85 | -0.622 | -0.773 | -0.008 | -0.001 | -0.001 | 0 | 0.14 | 0.07 | 0.1 | 0.19 | 0.15 | 0.18 |
| 5 | 60603462 | rs10471513 | 0.585 | -0.99 | -0.824 | -0.868 | -0.026 | -0.011 | -0.011 | 0.551 | 0.14 | 0.01 | 0.06 | 0.07 | 0.17 | 0.19 |
| 5 | 60604261 | rs10044518 | 0.387 | -0.794 | -0.514 | -0.755 | -0.027 | 0.01 | 0.01 | 1.046 | 0.09 | 0.39 | 0.06 | 0.03 | 0.02 | 0.12 |
| 5 | 60664356 | rs7700289 | -0.152 | -0.81 | -0.075 | -0.866 | -0.008 | -0.011 | -0.011 | 0 | 0.22 | 0.3 | 0.15 | 0.21 | 0.17 | 0.31 |

| | | | | | | | | | | | | | | | | |
|---|---|---|---|---|---|---|---|---|---|---|---|---|---|---|---|---|
| 5 | 60685757 | rs4490539 | 0.338 | -0.799 | -0.219 | -1.155 | 0.026 | -0.137 | -0.137 | 0 | 0.26 | 0.34 | 0.35 | 0 | 0.29 | 0.18 |
| 5 | 60705710 | rs6861729 | 1.581 | -0.632 | -0.127 | -1.171 | 0.089 | -0.035 | -0.035 | 0 | 0.08 | 0.22 | 0.21 | 0 | 0.11 | 0.11 |
| 5 | 60721611 | rs17422192 | 0.924 | -0.685 | -0.297 | -1.214 | 0 | -0.004 | -0.004 | 1.215 | 0.19 | 0.16 | 0.15 | 0 | 0.18 | 0.07 |
| 5 | 60732552 | rs7714712 | -0.678 | -0.72 | -0.319 | -1.227 | -0.002 | -0.018 | -0.018 | 0 | 0.4 | 0.45 | 0.3 | 0.21 | 0.35 | 0.38 |
| 5 | 60736544 | rs13171757 | -0.886 | -0.795 | -0.453 | -1.278 | 0.005 | 0.017 | 0.017 | 0 | 0.43 | 0.42 | 0.3 | 0.22 | 0.39 | 0.44 |
| 5 | 60754419 | rs1986252 | -1.631 | -0.908 | -0.798 | -1.317 | -0.092 | 0.209 | 0.209 | 0.468 | 0.55 | 0.92 | 0.56 | 0.21 | 0.52 | 0.57 |
| 5 | 60759524 | rs10514908 | 1.007 | -0.82 | -1.186 | -1.26 | -0.015 | 0.045 | 0.045 | 0 | 0.15 | 0.08 | 0.11 | 0.07 | 0.17 | 0.17 |
| 5 | 60800336 | rs10223052 | -2.772 | -1.017 | -1.66 | -0.741 | -0.117 | -0.167 | -0.167 | 0.004 | 0.73 | 0.96 | 0.66 | 0.49 | 0.66 | 0.82 |
| 5 | 60806765 | rs16891983 | 1.297 | -1.364 | -1.231 | -0.482 | 0.023 | -0.036 | -0.036 | 0.203 | 0.06 | 0 | 0.02 | 0.14 | 0.02 | 0.2 |
| 5 | 60839907 | rs16892374 | 1.287 | -1.364 | -1.231 | -0.489 | 0.023 | -0.035 | -0.035 | 0 | 0.06 | 0 | 0.02 | 0.14 | 0.02 | 0.2 |
| 5 | 60842210 | rs10053166 | 0.42 | -1.208 | -1.151 | -0.562 | -0.016 | -0.025 | -0.025 | 0.021 | 0.13 | 0.01 | 0.06 | 0.08 | 0.14 | 0.15 |
| 5 | 60847019 | rs1550816 | -1.118 | -1.111 | -0.839 | -0.371 | 0.001 | -0.001 | -0.001 | 1.718 | 0.52 | 0.49 | 0.51 | 0.5 | 0.45 | 0.62 |
| 5 | 60847272 | rs6890611 | -2.659 | -1.037 | -1.369 | -0.472 | -0.116 | -0.142 | -0.142 | 0 | 0.73 | 0.94 | 0.64 | 0.56 | 0.64 | 0.82 |
| 5 | 60848153 | rs16892406 | 0.021 | -1.193 | -1.32 | -0.576 | -0.002 | 0.002 | 0.002 | 0 | 0.26 | 0.26 | 0.29 | 0.3 | 0.22 | 0.44 |
| 5 | 60848185 | rs16892408 | 0.794 | -1.138 | -0.952 | -0.948 | -0.017 | -0.042 | -0.042 | 0.123 | 0.07 | 0 | 0.08 | 0.19 | 0.08 | 0.12 |
| 5 | 60848204 | rs13164150 | 0.864 | -1.076 | -0.659 | -0.906 | -0.002 | -0.025 | -0.025 | 0 | 0.14 | 0.16 | 0.09 | 0.07 | 0.16 | 0.18 |
| 5 | 60848409 | rs16892410 | 0.508 | -1.106 | -0.671 | -0.895 | -0.011 | 0.009 | 0.009 | 0 | 0.22 | 0.07 | 0.21 | 0.2 | 0.23 | 0.17 |
| 5 | 60848745 | rs4582229 | -0.515 | -0.995 | 0.031 | -0.759 | -0.001 | -0.017 | -0.017 | 0 | 0.19 | 0.21 | 0.14 | 0.07 | 0.21 | 0.2 |
| 5 | 60850467 | rs2084130 | -1.499 | -0.821 | 0.133 | -0.3 | 0.001 | -0.062 | -0.062 | 0 | 0.11 | 0.54 | 0.06 | 0 | 0.04 | 0.04 |
| 5 | 60858429 | rs16892456 | 0.009 | -0.525 | -0.674 | -0.046 | 0.015 | -0.01 | -0.01 | 0 | 0.1 | 0.02 | 0.04 | 0.07 | 0.1 | 0.12 |
| 5 | 60858543 | rs10035484 | -0.313 | -0.253 | -0.208 | -0.053 | 0.027 | -0.023 | -0.023 | 0 | 0.06 | 0.16 | 0.04 | 0 | 0.04 | 0.03 |
| 5 | 60859416 | rs10053423 | -0.523 | -0.698 | -0.128 | 0.164 | -0.038 | 0.051 | 0.051 | 1.248 | 0.22 | 0.41 | 0.25 | 0.27 | 0.2 | 0.15 |
| 5 | 60861517 | rs1833831 | -0.304 | -0.479 | -0.214 | -0.55 | 0.014 | -0.089 | -0.089 | 0.624 | 0.18 | 0.25 | 0.18 | 0 | 0.16 | 0.22 |
| 5 | 60861551 | rs11741170 | 1.532 | -0.052 | -0.801 | -0.724 | -0.029 | 0.075 | 0.075 | 0 | 0.17 | 0.05 | 0.15 | 0.04 | 0.25 | 0.12 |
| 5 | 60861742 | rs1582545 | -0.115 | -0.352 | -0.743 | -0.424 | -0.051 | 0.055 | 0.055 | 1.157 | 0.23 | 0.42 | 0.27 | 0.31 | 0.19 | 0.14 |
| 5 | 60862917 | rs933685 | -0.718 | 0.036 | -1.055 | 0.213 | 0.002 | 0.072 | 0.072 | 0 | 0.94 | 0.96 | 0.89 | 0.59 | 0.97 | 0.75 |
| 5 | 60863461 | rs1971513 | -0.691 | 0.033 | -1.057 | 0.211 | 0.002 | 0.069 | 0.069 | 0 | 0.94 | 0.95 | 0.89 | 0.59 | 0.97 | 0.75 |
| 5 | 60867365 | rs10067201 | 0.704 | 0.373 | -1.152 | -0.791 | -0.045 | 0.103 | 0.103 | 0 | 0.21 | 0.04 | 0.18 | 0.05 | 0.3 | 0.15 |
| 5 | 60871157 | rs11954764 | -0.348 | 0.583 | -0.883 | -0.834 | -0.007 | -0.018 | -0.018 | 0 | 0.3 | 0.23 | 0.31 | 0.05 | 0.4 | 0.29 |
| 5 | 60879560 | rs10939908 | -0.131 | 0.305 | -0.905 | -0.198 | 0.004 | 0.023 | 0.023 | 0.252 | 0.43 | 0.4 | 0.31 | 0.16 | 0.46 | 0.32 |
| 5 | 60883190 | rs10058219 | 0.121 | 0.189 | -1.063 | 0.195 | -0.014 | 0.005 | 0.005 | 0.804 | 0.41 | 0.33 | 0.47 | 0.39 | 0.44 | 0.43 |
| 5 | 60883593 | rs863088 | 0.183 | 0.151 | -1.399 | -0.756 | -0.035 | 0.057 | 0.057 | 0.01 | 0.39 | 0.21 | 0.33 | 0.24 | 0.39 | 0.4 |
| 5 | 60886299 | rs420518 | 0.138 | -0.191 | -1.227 | -0.151 | 0.008 | 0.017 | 0.017 | 0 | 0.91 | 0.95 | 0.79 | 0.51 | 0.95 | 0.84 |
| 5 | 60890409 | rs7713549 | 0.309 | -0.171 | -1.044 | -0.462 | 0 | -0.001 | -0.001 | 0 | 0.58 | 0.59 | 0.54 | 0.27 | 0.58 | 0.63 |
| 5 | 60892205 | rs381034 | 0.855 | -0.184 | -1.263 | -0.377 | -0.006 | 0.005 | 0.005 | 0.098 | 0.26 | 0.21 | 0.21 | 0.23 | 0.3 | 0.18 |
| 5 | 60898157 | rs696888 | -0.393 | -0.052 | -1.262 | -0.257 | -0.003 | 0.079 | 0.079 | 0 | 0.93 | 0.94 | 0.71 | 0.51 | 0.93 | 0.85 |
| 5 | 60903201 | rs173780 | 0.042 | -0.443 | -1.058 | -1.265 | 0.009 | 0.009 | 0.009 | 2.079 | 0.26 | 0.18 | 0.2 | 0.18 | 0.17 | 0.2 |
| 5 | 60904429 | rs36084 | 1.603 | -0.014 | -1.134 | -0.692 | -0.087 | -0.032 | -0.032 | 0 | 0.16 | 0.01 | 0.16 | 0.18 | 0.22 | 0.11 |
| 5 | 60908030 | rs895382 | 0.427 | -0.281 | -1.105 | -0.714 | -0.003 | 0.007 | 0.007 | 2.763 | 0.59 | 0.37 | 0.69 | 0.76 | 0.57 | 0.73 |
| 5 | 60910078 | rs4699977 | 0.664 | 0.089 | -1.708 | -0.572 | -0.101 | 0.009 | 0.009 | 0.04 | 0.44 | 0.09 | 0.6 | 0.58 | 0.48 | 0.67 |
| 5 | 60910890 | rs253184 | 0.019 | 0.994 | -1.148 | -0.437 | 0.004 | -0.002 | -0.002 | 0.102 | 0.74 | 0.71 | 0.78 | 0.76 | 0.69 | 0.81 |
| 5 | 60918763 | rs409912 | -0.301 | 0.312 | -1.306 | -0.388 | 0.002 | 0.006 | 0.006 | 0.055 | 0.12 | 0.1 | 0.07 | 0.06 | 0.1 | 0.1 |
| 5 | 60923475 | rs36091 | -0.31 | 0.594 | -1.469 | -0.749 | 0.006 | -0.029 | -0.029 | 0.179 | 0.39 | 0.47 | 0.29 | 0.15 | 0.45 | 0.28 |
| 5 | 60928329 | rs7727255 | 0.264 | 1.143 | -2.39 | -0.651 | -0.067 | 0.096 | 0.096 | 0 | 0.28 | 0.14 | 0.26 | 0.1 | 0.35 | 0.21 |
| 5 | 60928771 | rs26914 | -0.405 | 0.91 | -2.584 | -0.456 | -0.045 | -0.024 | -0.024 | 0 | 0.7 | 0.9 | 0.59 | 0.74 | 0.63 | 0.62 |
| 5 | 60933255 | rs1354138 | -0.189 | 1.35 | -2.004 | -0.712 | -0.079 | 0.109 | 0.109 | 0 | 0.27 | 0.12 | 0.19 | 0.09 | 0.34 | 0.19 |
| 5 | 60936095 | rs26912 | -0.465 | 0.777 | -1.722 | -0.74 | 0.007 | 0.034 | 0.034 | 0 | 0.3 | 0.24 | 0.3 | 0.08 | 0.28 | 0.33 |
| 5 | 60936296 | rs7715669 | -0.685 | 0.456 | -2.041 | -0.914 | 0.01 | 0.03 | 0.03 | 0 | 0.26 | 0.16 | 0.16 | 0.21 | 0.24 | 0.28 |
| 5 | 60936935 | rs1867732 | -0.105 | -0.002 | -1.806 | -1.224 | 0.017 | 0.029 | 0.029 | 0 | 0.45 | 0.37 | 0.54 | 0.7 | 0.4 | 0.47 |
| 5 | 60937964 | rs16893212 | -1.235 | 1.146 | -0.942 | -1.445 | 0.164 | -0.041 | -0.041 | 0 | 0.1 | 0.39 | 0.09 | 0 | 0.13 | 0.05 |
| 5 | 60938249 | rs13174785 | 0.124 | 1.132 | -0.55 | -1.721 | 0.012 | -0.002 | -0.002 | 0 | 0.3 | 0.45 | 0.25 | 0.33 | 0.28 | 0.27 |
| 5 | 60941930 | rs379212 | -0.535 | 0.927 | -0.402 | -1.501 | -0.007 | -0.002 | -0.002 | 0 | 0.1 | 0.12 | 0.14 | 0.13 | 0.17 | 0.08 |
| 5 | 60944856 | rs12658207 | 0.761 | 0.418 | 0.048 | -1.022 | -0.004 | 0.03 | 0.03 | 0 | 0.33 | 0.41 | 0.41 | 0.46 | 0.35 | 0.4 |
| 5 | 60945612 | rs253177 | -0.102 | -0.143 | -0.109 | -1.315 | 0.048 | -0.025 | -0.025 | 0 | 0.32 | 0.5 | 0.31 | 0.29 | 0.34 | 0.31 |
| 5 | 60947483 | rs1501841 | 0.085 | -0.087 | -0.244 | -0.681 | -0.064 | -0.003 | -0.003 | 0 | 0.56 | 0.91 | 0.6 | 0.63 | 0.53 | 0.63 |
| 5 | 60950211 | rs11738335 | 1.096 | 0.083 | 0.395 | -0.457 | 0.011 | -0.022 | -0.022 | 0.114 | 0.2 | 0.33 | 0.17 | 0.17 | 0.18 | 0.22 |
| 5 | 60951124 | rs424876 | -0.528 | 0.062 | 0.046 | -0.507 | 0.034 | 0.079 | 0.079 | 0 | 0.6 | 0.87 | 0.67 | 0.78 | 0.61 | 0.64 |
| 5 | 60960727 | rs36033 | 0.534 | -0.018 | -0.085 | -0.4 | -0.059 | -0.034 | -0.034 | 0.134 | 0.34 | 0.14 | 0.48 | 0.61 | 0.43 | 0.38 |
| 5 | 60960764 | rs4700423 | 1.84 | -0.016 | -0.097 | -0.178 | -0.025 | -0.113 | -0.113 | 0 | 0.24 | 0.06 | 0.35 | 0.44 | 0.27 | 0.31 |
| 5 | 60962986 | rs6883938 | 0.756 | 0.054 | 0.492 | -0.697 | -0.003 | 0.01 | 0.01 | 0 | 0.21 | 0.23 | 0.16 | 0.17 | 0.18 | 0.23 |
| 5 | 60971778 | rs410850 | -0.019 | 0.114 | 0.276 | -0.627 | 0.019 | 0.005 | 0.005 | 0 | 0.3 | 0.39 | 0.29 | 0.35 | 0.35 | 0.28 |
| 5 | 60974357 | rs6880870 | 0.478 | -0.048 | 0.029 | -0.824 | -0.008 | 0.021 | 0.021 | 0 | 0.43 | 0.36 | 0.53 | 0.72 | 0.44 | 0.52 |
| 5 | 60976610 | rs1501845 | 0.783 | -0.064 | 0.056 | -0.816 | 0.001 | -0.018 | -0.018 | 0 | 0.24 | 0.29 | 0.18 | 0.17 | 0.2 | 0.2 |
| 5 | 60981186 | rs16893676 | 0.706 | -0.433 | 0.126 | -1.128 | -0.008 | 0.003 | 0.003 | 0 | 0.13 | 0.22 | 0.13 | 0.09 | 0.08 | 0.25 |
| 5 | 60982841 | rs16893687 | 0.769 | -0.423 | 0.105 | -1.144 | -0.003 | -0.009 | -0.009 | 0 | 0.35 | 0.45 | 0.49 | 0.65 | 0.34 | 0.57 |
| 5 | 60987340 | rs11743540 | -0.277 | -0.365 | -0.237 | -1.184 | -0.007 | 0.085 | 0.085 | 0 | 0.59 | 0.74 | 0.68 | 0.81 | 0.54 | 0.78 |
| 5 | 60994164 | rs2950220 | -2.118 | -0.215 | -0.089 | -1.321 | -0.006 | -0.019 | -0.019 | 0.059 | 0.79 | 0.81 | 0.64 | 0.36 | 0.76 | 0.69 |
| 5 | 60995261 | rs2974452 | -2.145 | -0.209 | -0.199 | -1.354 | -0.012 | -0.04 | -0.04 | 0.017 | 0.8 | 0.85 | 0.64 | 0.36 | 0.76 | 0.69 |
| 5 | 60995351 | rs253180 | 1.327 | 0.138 | -0.192 | -1.434 | -0.033 | -0.031 | -0.031 | 5.892 | 0.05 | 0.01 | 0.09 | 0.1 | 0.13 | 0.05 |

| | | | | | | | | | | | | | | | | |
|---|---|---|---|---|---|---|---|---|---|---|---|---|---|---|---|---|
| 5 | 60996988 | rs253531 | 0.714 | -0.262 | -0.25 | -1.745 | -0.013 | 0.07 | 0.07 | 0 | 0.28 | 0.12 | 0.2 | 0 | 0.33 | 0.17 |
| 5 | 60997309 | rs4700430 | 1.08 | -0.378 | -0.275 | -1.752 | 0.001 | -0.016 | -0.016 | 0.004 | 0.23 | 0.25 | 0.18 | 0.15 | 0.2 | 0.21 |
| 5 | 60999768 | rs436696 | 0.528 | -0.332 | -0.233 | -1.747 | -0.002 | -0.087 | -0.087 | 0 | 0.3 | 0.34 | 0.21 | 0 | 0.33 | 0.17 |
| 5 | 61002359 | rs153378 | 0.438 | -0.189 | -0.133 | -1.013 | -0.028 | -0.013 | -0.013 | 0 | 0.15 | 0.05 | 0.14 | 0.13 | 0.17 | 0.06 |
| 5 | 61003342 | rs4699979 | 0.349 | -0.271 | 0.24 | -0.835 | 0.002 | -0.015 | -0.015 | 1.925 | 0.23 | 0.25 | 0.18 | 0.16 | 0.21 | 0.21 |
| 5 | 61011200 | rs12514378 | 0.446 | -0.188 | 0.415 | -0.74 | 0.018 | -0.037 | -0.037 | 0 | 0.22 | 0.4 | 0.18 | 0.16 | 0.19 | 0.21 |
| 5 | 61016374 | rs436985 | -0.289 | 0.264 | 0.417 | -0.916 | 0.017 | -0.011 | -0.011 | 0 | 0.33 | 0.42 | 0.33 | 0.27 | 0.36 | 0.3 |
| 5 | 61021402 | rs432999 | -0.064 | -0.064 | 0.497 | -1.036 | -0.017 | 0.009 | 0.009 | 0 | 0.19 | 0.13 | 0.17 | 0.13 | 0.21 | 0.14 |
| 5 | 61021484 | rs446297 | 0.486 | -0.4 | 0.499 | -1.515 | -0.022 | -0.115 | -0.115 | 13.022 | 0.25 | 0.34 | 0.14 | 0 | 0.22 | 0.09 |
| 5 | 61025371 | rs153383 | 0.848 | -0.429 | 0.646 | -1.534 | 0.011 | 0.01 | 0.01 | 0 | 0.23 | 0.16 | 0.13 | 0 | 0.22 | 0.09 |
| 5 | 61029525 | rs6873285 | 1.166 | -0.408 | 0.875 | -1.509 | 0.003 | -0.026 | -0.026 | 1.843 | 0.19 | 0.3 | 0.21 | 0.14 | 0.17 | 0.16 |
| 5 | 61030954 | rs2974454 | -0.695 | -0.728 | 0.426 | -1.538 | 0.004 | 0.027 | 0.027 | 0.001 | 0.44 | 0.41 | 0.31 | 0.13 | 0.43 | 0.24 |
| 5 | 61036436 | rs16894458 | 0.896 | -0.65 | 0.445 | -0.994 | -0.025 | 0.021 | 0.021 | 0.036 | 0.08 | 0.15 | 0.08 | 0.08 | 0.02 | 0.17 |
| 5 | 61038510 | rs13177369 | 1.017 | -0.572 | 0.487 | -1.058 | -0.002 | 0.068 | 0.068 | 0.758 | 0.3 | 0.18 | 0.26 | 0.16 | 0.26 | 0.22 |
| 5 | 61042352 | rs1037106 | -0.071 | -0.557 | 0.196 | -0.878 | -0.012 | 0.035 | 0.035 | 0 | 0.32 | 0.21 | 0.25 | 0.17 | 0.36 | 0.21 |
| 5 | 61044083 | rs16894569 | 0.649 | 0.107 | 0.763 | -0.405 | -0.025 | 0.022 | 0.022 | 0.036 | 0.08 | 0.15 | 0.08 | 0.08 | 0.02 | 0.17 |
| 5 | 61045934 | rs187574 | -1.03 | 0.146 | 0.695 | -0.426 | -0.005 | -0.003 | -0.003 | 0 | 0.53 | 0.79 | 0.51 | 0.44 | 0.49 | 0.59 |
| 5 | 61047818 | rs1446001 | 0.111 | 0.183 | 0.645 | -0.454 | -0.011 | -0.069 | -0.069 | 0 | 0.34 | 0.48 | 0.3 | 0.15 | 0.27 | 0.22 |
| 5 | 61049167 | rs2030889 | -0.446 | 0.35 | 0.721 | 0.392 | 0.003 | 0.001 | 0.001 | 0 | 0.1 | 0.11 | 0.12 | 0.17 | 0.15 | 0.12 |
| 5 | 61053123 | rs2077786 | -1.672 | 0.102 | 0.654 | 0.308 | -0.011 | 0.012 | 0.012 | 0.766 | 0.33 | 0.75 | 0.3 | 0.25 | 0.38 | 0.26 |
| 5 | 61055557 | rs1501847 | -1.395 | 0.256 | 0.765 | 0.336 | 0.013 | -0.008 | -0.008 | 0 | 0.3 | 0.67 | 0.28 | 0.24 | 0.37 | 0.23 |
| 5 | 61061963 | rs414568 | -0.546 | 0.333 | 0.73 | 0.243 | -0.004 | -0.118 | -0.118 | 0 | 0.18 | 0.38 | 0.14 | 0 | 0.18 | 0.08 |
| 5 | 61071912 | rs10471519 | -1.098 | 0.116 | -0.468 | 0.519 | -0.019 | 0.005 | 0.005 | 0.268 | 0.39 | 0.81 | 0.37 | 0.34 | 0.39 | 0.44 |
| 5 | 61080390 | rs6882661 | -1.061 | 0.176 | -0.641 | 0.52 | -0.028 | 0 | 0 | 0 | 0.4 | 0.85 | 0.38 | 0.35 | 0.4 | 0.46 |
| 5 | 61085552 | rs6449558 | -0.774 | -0.012 | -0.241 | 0.515 | 0 | 0.003 | 0.003 | 0.106 | 0.32 | 0.65 | 0.28 | 0.26 | 0.29 | 0.39 |
| 5 | 61092625 | rs12516216 | -1.422 | -0.569 | 0.699 | 0.682 | 0 | 0.044 | 0.044 | 0.58 | 0.07 | 0.31 | 0.13 | 0.05 | 0.02 | 0.02 |
| 5 | 61098274 | rs10514922 | -0.978 | -1.214 | 0.516 | 0.839 | 0.004 | -0.002 | -0.002 | 0.452 | 0.2 | 0.16 | 0.13 | 0.18 | 0.14 | 0.12 |
| 5 | 61101470 | rs6861457 | -1.438 | -1.471 | 0.927 | 1.144 | -0.017 | -0.004 | -0.004 | 0.006 | 0.25 | 0.27 | 0.14 | 0.2 | 0.15 | 0.3 |
| 5 | 61106070 | rs6895749 | -1.959 | -1.954 | 0.744 | 0.86 | 0.037 | 0.013 | 0.013 | 0.378 | 0.37 | 0.7 | 0.36 | 0.28 | 0.25 | 0.23 |
| 5 | 61106657 | rs16895276 | -0.505 | -1.952 | 0.697 | 0.896 | 0.005 | -0.006 | -0.006 | 0 | 0.09 | 0.06 | 0.09 | 0.11 | 0.1 | 0.09 |
| 5 | 61107460 | rs6449559 | -2.377 | -2.029 | 0.771 | 1.06 | 0.109 | 0.028 | 0.028 | 0.024 | 0.43 | 0.81 | 0.39 | 0.32 | 0.27 | 0.41 |
| 5 | 61111747 | rs12658967 | -1.756 | -2.181 | 0.941 | 1.11 | -0.037 | -0.008 | -0.008 | 0 | 0.24 | 0.33 | 0.15 | 0.25 | 0.13 | 0.3 |
| 5 | 61117272 | rs4700442 | -2.729 | -2.35 | 0.881 | 1.078 | 0.104 | 0.029 | 0.029 | 0 | 0.44 | 0.77 | 0.4 | 0.37 | 0.29 | 0.42 |
| 5 | 61122159 | rs16895538 | -2.01 | -2.307 | 1.24 | 1.154 | -0.016 | 0.049 | 0.049 | 0 | 0.14 | 0.23 | 0.22 | 0.27 | 0.14 | 0.13 |
| 5 | 61125018 | rs11743431 | -1.558 | -2.205 | 1.362 | 1.132 | -0.019 | 0.052 | 0.052 | 0 | 0.12 | 0.22 | 0.11 | 0.24 | 0.11 | 0.12 |
| 5 | 61137914 | rs975134 | -1.339 | -2.004 | 1.303 | 1.245 | 0.002 | -0.008 | -0.008 | 0 | 0.1 | 0.06 | 0.09 | 0.25 | 0.1 | 0.1 |
| 5 | 61145102 | rs7718109 | -1.936 | -2.309 | 1.28 | 1.139 | -0.022 | 0.007 | 0.007 | 0.145 | 0.25 | 0.32 | 0.18 | 0.33 | 0.18 | 0.23 |
| 5 | 61150440 | rs4699986 | -2.883 | -2.381 | 1.257 | 1.048 | 0.098 | -0.015 | -0.015 | 0 | 0.4 | 0.88 | 0.41 | 0.38 | 0.3 | 0.41 |
| 5 | 61152975 | rs2055380 | -1.106 | -2.188 | 1.69 | 0.872 | -0.001 | -0.003 | -0.003 | 0 | 0.08 | 0.18 | 0.18 | 0.05 | 0.11 | 0.16 |
| 5 | 61165626 | rs1374002 | -1.821 | -2.187 | 1.605 | 0.87 | -0.022 | -0.019 | -0.019 | 0 | 0.25 | 0.65 | 0.27 | 0.12 | 0.15 | 0.25 |
| 5 | 61165960 | rs1995274 | -1.829 | -2.186 | 1.543 | 0.87 | -0.01 | -0.007 | -0.007 | 0.121 | 0.26 | 0.68 | 0.27 | 0.12 | 0.15 | 0.25 |
| 5 | 61172759 | rs10075885 | -1.835 | -2.176 | 1.087 | 0.904 | 0.041 | 0.035 | 0.035 | 1.41 | 0.26 | 0.78 | 0.27 | 0.13 | 0.15 | 0.24 |
| 5 | 61190572 | rs12517264 | 2.255 | -2.102 | 1.065 | 0.975 | 0.065 | -0.022 | -0.022 | 0 | 0.57 | 0.27 | 0.51 | 0.55 | 0.64 | 0.58 |
| 5 | 61191394 | rs1157898 | 0.989 | -1.594 | 1.463 | 1.114 | -0.002 | 0.026 | 0.026 | 0.239 | 0.86 | 0.81 | 0.71 | 0.79 | 0.83 | 0.86 |
| 5 | 61192012 | rs980496 | 1.097 | -1.596 | 1.534 | 1.111 | -0.001 | 0.048 | 0.048 | 0.457 | 0.85 | 0.73 | 0.7 | 0.79 | 0.83 | 0.85 |
| 5 | 61194553 | rs12054936 | -1.372 | -1.536 | 0.84 | 1.144 | -0.003 | -0.009 | -0.009 | 0 | 0.08 | 0.04 | 0.1 | 0.16 | 0.12 | 0.15 |
| 5 | 61196185 | rs4700450 | 1.145 | -1.591 | 0.987 | 1.063 | -0.003 | -0.012 | -0.012 | 0 | 0.82 | 0.88 | 0.65 | 0.63 | 0.74 | 0.71 |
| 5 | 61197403 | rs7717058 | 1.111 | -1.587 | 1.104 | 0.486 | -0.006 | -0.007 | -0.007 | 0.004 | 0.64 | 0.5 | 0.74 | 0.66 | 0.69 | 0.75 |
| 5 | 61199286 | rs1445880 | 0.166 | -0.976 | 0.928 | 0.549 | -0.085 | -0.097 | -0.097 | 0 | 0.78 | 0.94 | 0.82 | 0.65 | 0.72 | 0.76 |
| 5 | 61200407 | rs1445881 | 1.664 | -1.088 | 1.41 | 0.433 | -0.002 | -0.001 | -0.001 | 0 | 0.65 | 0.38 | 0.74 | 0.62 | 0.67 | 0.63 |
| 5 | 61200460 | rs1445882 | 0.197 | -0.833 | 1.282 | 0.615 | -0.077 | -0.087 | -0.087 | 0 | 0.78 | 0.93 | 0.82 | 0.66 | 0.72 | 0.76 |
| 5 | 61216333 | rs10471525 | 0.212 | -0.838 | 1.421 | 0.615 | -0.01 | -0.012 | -0.012 | 0 | 0.78 | 0.81 | 0.8 | 0.66 | 0.72 | 0.78 |
| 5 | 61242239 | rs6449570 | -0.082 | -0.691 | -0.673 | 0.309 | 0.039 | -0.018 | -0.018 | 0 | 0.45 | 0.77 | 0.48 | 0.48 | 0.36 | 0.45 |
| 5 | 61248873 | rs263663 | -1.134 | -0.854 | -0.884 | 0.163 | -0.001 | -0.009 | -0.009 | 0.422 | 0.44 | 0.84 | 0.33 | 0.45 | 0.35 | 0.45 |
| 5 | 61265820 | rs12654555 | -0.521 | -0.824 | -0.843 | 0.138 | 0 | -0.034 | -0.034 | 0 | 0.08 | 0.01 | 0.07 | 0.19 | 0.05 | 0.09 |
| 5 | 61265883 | rs1374005 | 0.748 | -0.592 | -0.048 | 0.24 | 0.001 | -0.003 | -0.003 | 0 | 0.54 | 0.36 | 0.57 | 0.41 | 0.58 | 0.67 |
| 5 | 61279000 | rs1363937 | 1.476 | -0.448 | -0.379 | 0.075 | 0 | -0.013 | -0.013 | 0 | 0.08 | 0.07 | 0.03 | 0 | 0.08 | 0.1 |
| 5 | 61280487 | rs4699994 | -0.24 | -0.552 | -0.094 | 0.096 | -0.035 | -0.022 | -0.022 | 0.33 | 0.65 | 0.82 | 0.67 | 0.6 | 0.63 | 0.76 |
| 5 | 61288869 | rs7734412 | -0.203 | -0.552 | 0.061 | 0.041 | -0.023 | -0.05 | -0.05 | 0 | 0.64 | 0.75 | 0.66 | 0.47 | 0.63 | 0.78 |
| 5 | 61304705 | rs918616 | 0.007 | -0.611 | 0.234 | 0.038 | 0.006 | 0.021 | 0.021 | 0 | 0.62 | 0.47 | 0.64 | 0.46 | 0.63 | 0.78 |
| 5 | 61311715 | rs13177362 | -0.092 | -0.588 | 0.18 | 0.049 | -0.002 | -0.007 | -0.007 | 0.188 | 0.63 | 0.57 | 0.65 | 0.46 | 0.63 | 0.78 |
| 5 | 61312387 | rs2342989 | 0.784 | -0.687 | -0.019 | -0.048 | -0.026 | -0.032 | -0.032 | 0 | 0.48 | 0.15 | 0.59 | 0.46 | 0.55 | 0.63 |
| 5 | 61338462 | rs1972651 | -0.172 | -0.356 | 0.088 | 0.128 | -0.008 | -0.026 | -0.026 | 0 | 0.64 | 0.68 | 0.67 | 0.46 | 0.66 | 0.79 |
| 5 | 61342784 | rs4395592 | -0.377 | -0.251 | -0.355 | -0.01 | -0.044 | -0.041 | -0.041 | 0 | 0.68 | 0.82 | 0.69 | 0.6 | 0.67 | 0.81 |
| 5 | 61348807 | rs4700465 | -0.451 | 0.071 | -0.396 | -0.033 | -0.002 | -0.037 | -0.037 | 0 | 0.69 | 0.8 | 0.72 | 0.6 | 0.73 | 0.8 |
| 5 | 61359597 | rs4388169 | 0.383 | -0.379 | -0.874 | -0.155 | -0.026 | 0.064 | 0.064 | 0 | 0.51 | 0.15 | 0.58 | 0.57 | 0.56 | 0.59 |
| 5 | 61366182 | rs10060036 | -0.564 | -0.449 | -0.671 | -0.21 | 0.032 | 0.036 | 0.036 | 0 | 0.71 | 0.46 | 0.64 | 0.56 | 0.69 | 0.8 |
| 5 | 61368432 | rs1423129 | 0.508 | -0.5 | -0.712 | -0.326 | 0 | 0.049 | 0.049 | 0 | 0.51 | 0.27 | 0.59 | 0.58 | 0.56 | 0.6 |
| 5 | 61369333 | rs13359031 | 0.389 | -0.444 | -1.402 | -0.307 | 0.006 | -0.012 | -0.012 | 0 | 0.19 | 0.17 | 0.1 | 0.02 | 0.13 | 0.22 |

| | | | | | | | | | | | | | | | | |
|---|---|---|---|---|---|---|---|---|---|---|---|---|---|---|---|---|
| 5 | 61376320 | rs1895394 | 0.232 | -0.477 | -1.339 | -0.269 | -0.023 | 0.033 | 0.033 | 0 | 0.55 | 0.74 | 0.62 | 0.58 | 0.56 | 0.6 |
| 5 | 61389786 | rs1423125 | 0.822 | -0.533 | -2.015 | -0.739 | 0.003 | 0.063 | 0.063 | 0 | 0.13 | 0.02 | 0.06 | 0 | 0.12 | 0.12 |
| 5 | 61391723 | rs16898731 | 1.362 | -0.461 | -2.261 | -0.423 | -0.059 | -0.038 | -0.038 | 0.299 | 0.18 | 0 | 0.15 | 0.14 | 0.2 | 0.23 |
| 5 | 61394933 | rs9291747 | 0.45 | -0.379 | -1.754 | -0.455 | 0.012 | 0.021 | 0.021 | 0 | 0.2 | 0.14 | 0.11 | 0.02 | 0.14 | 0.22 |
| 5 | 61396019 | rs7704011 | -0.831 | 0.177 | -1.511 | 0.236 | -0.065 | -0.071 | -0.071 | 0 | 0.74 | 0.91 | 0.73 | 0.6 | 0.7 | 0.82 |
| 5 | 61404673 | rs2910468 | -0.76 | 0.21 | -0.69 | 0.532 | 0.022 | 0.112 | 0.112 | 0.149 | 0.1 | 0.24 | 0.23 | 0.31 | 0.11 | 0.1 |
| 5 | 61428137 | rs1423127 | 0.574 | 0.025 | -1.457 | 0.437 | -0.062 | 0.03 | 0.03 | 0 | 0.38 | 0.07 | 0.41 | 0.26 | 0.4 | 0.42 |
| 5 | 61428613 | rs1363785 | -0.332 | -0.042 | -0.997 | 0.428 | -0.014 | -0.014 | -0.014 | 0.001 | 0.24 | 0.27 | 0.18 | 0.17 | 0.21 | 0.33 |
| 5 | 61433325 | rs9291749 | 1.578 | 0.094 | -1.482 | 0.391 | -0.039 | 0.013 | 0.013 | 1.62 | 0.13 | 0.01 | 0.12 | 0.07 | 0.17 | 0.15 |
| 5 | 61436505 | rs2962027 | -1.136 | -0.109 | -0.965 | 0.792 | -0.096 | -0.072 | -0.072 | 0 | 0.73 | 0.98 | 0.86 | 0.73 | 0.72 | 0.85 |
| 5 | 61440450 | rs1423131 | -0.093 | -0.163 | -0.826 | 0.849 | 0.01 | 0.065 | 0.065 | 0 | 0.08 | 0.11 | 0.22 | 0.3 | 0.11 | 0.1 |
| 5 | 61443773 | rs3846462 | 1.218 | -0.217 | -1.11 | 0.789 | -0.043 | 0.039 | 0.039 | 0 | 0.38 | 0.11 | 0.43 | 0.26 | 0.4 | 0.42 |
| 5 | 61446385 | rs2910475 | -1.588 | -0.321 | -0.841 | 0.882 | 0.025 | -0.065 | -0.065 | 0 | 0.35 | 0.87 | 0.43 | 0.47 | 0.32 | 0.44 |
| 5 | 61456769 | rs10057846 | 0.046 | -0.228 | -0.755 | 0.852 | 0.009 | 0.033 | 0.033 | 0 | 0.12 | 0.06 | 0.1 | 0.04 | 0.12 | 0.22 |
| 5 | 61461609 | rs3846465 | 0.79 | 0.295 | -0.392 | 1.302 | -0.002 | -0.03 | -0.03 | 0 | 0.05 | 0.01 | 0.05 | 0.13 | 0.07 | 0.07 |
| 5 | 61472904 | rs12655910 | 1.496 | 0.245 | -0.332 | 1.246 | 0.012 | -0.018 | -0.018 | 0 | 0.73 | 0.17 | 0.62 | 0.66 | 0.77 | 0.71 |
| 5 | 61474164 | rs1862526 | 1.58 | 0.266 | 0.078 | 1.28 | -0.004 | 0.035 | 0.035 | 0 | 0.72 | 0.62 | 0.61 | 0.53 | 0.71 | 0.67 |
| 5 | 61476901 | rs10939931 | 0.534 | 0.328 | -0.375 | 1.017 | -0.073 | -0.12 | -0.12 | 0 | 0.31 | 0.04 | 0.19 | 0.4 | 0.33 | 0.22 |
| 5 | 61479184 | rs10038739 | -0.757 | 0.262 | -0.145 | 0.949 | -0.019 | -0.049 | -0.049 | 0.819 | 0.14 | 0.2 | 0.09 | 0.04 | 0.1 | 0.16 |
| 5 | 61480289 | rs3909436 | -0.299 | 0.289 | -0.238 | 1.074 | -0.081 | -0.088 | -0.088 | 0 | 0.64 | 0.9 | 0.58 | 0.47 | 0.6 | 0.71 |
| 5 | 61496655 | rs1423119 | -0.128 | 0.397 | -0.344 | 1.029 | -0.031 | -0.044 | -0.044 | 2.512 | 0.92 | 0.98 | 0.74 | 0.74 | 0.87 | 0.9 |
| 5 | 61497716 | rs10069830 | 1.385 | 0.244 | -0.26 | 0.726 | -0.054 | 0.027 | 0.027 | 0 | 0.41 | 0.1 | 0.4 | 0.31 | 0.43 | 0.49 |
| 5 | 61499198 | rs10471540 | 0.497 | 0.454 | 0.069 | 1.044 | -0.01 | 0.013 | 0.013 | 0 | 0.57 | 0.35 | 0.49 | 0.34 | 0.53 | 0.65 |
| 5 | 61552812 | rs1025032 | -0.062 | 0.283 | 0.56 | 1.33 | 0 | 0.025 | 0.025 | 0 | 0.93 | 0.91 | 0.79 | 0.72 | 0.93 | 0.87 |
| 5 | 61553091 | rs7706828 | -0.497 | 0.264 | 0.835 | 1.24 | 0.034 | 0.013 | 0.013 | 0 | 0.06 | 0.18 | 0.06 | 0.04 | 0.04 | 0.12 |
| 5 | 61553881 | rs10062049 | -1.421 | 0.187 | 0.699 | 1.336 | -0.015 | 0.006 | 0.006 | 0.249 | 0.16 | 0.25 | 0.11 | 0.17 | 0.13 | 0.19 |
| 5 | 61556730 | rs1108159 | -0.985 | 0.182 | 0.666 | 1.296 | -0.018 | -0.048 | -0.048 | 0 | 0.12 | 0.21 | 0.06 | 0.04 | 0.06 | 0.15 |
| 5 | 61564841 | rs6882514 | 0.51 | 0.159 | 0.432 | 1.28 | -0.001 | -0.001 | -0.001 | 0 | 0.28 | 0.25 | 0.15 | 0.24 | 0.26 | 0.17 |
| 5 | 61566896 | rs10067200 | 1.197 | 0.238 | 0.191 | 1.246 | -0.069 | 0.087 | 0.087 | 0 | 0.47 | 0.12 | 0.49 | 0.31 | 0.52 | 0.5 |
| 5 | 61585240 | rs6449593 | 0.936 | 0.274 | 0.236 | 1.244 | -0.008 | 0.003 | 0.003 | 0.04 | 0.51 | 0.17 | 0.53 | 0.45 | 0.6 | 0.55 |
| 5 | 61593277 | rs4700005 | 1.302 | 0.061 | 0.576 | 1.294 | -0.031 | 0.024 | 0.024 | 2.186 | 0.78 | 0.42 | 0.7 | 0.68 | 0.87 | 0.71 |
| 5 | 61605567 | rs2344396 | -0.018 | 0.216 | 1.013 | 1.31 | 0.004 | 0.015 | 0.015 | 0 | 0.64 | 0.6 | 0.61 | 0.49 | 0.66 | 0.69 |
| 5 | 61633997 | rs7704066 | 0.64 | 0.221 | 1.15 | 1.356 | -0.005 | -0.002 | -0.002 | 0 | 0.27 | 0.19 | 0.16 | 0.22 | 0.27 | 0.18 |
| 5 | 61642831 | rs16890656 | -0.17 | 0.258 | 1.014 | 1.349 | 0 | 0.02 | 0.02 | 0 | 0.08 | 0.11 | 0.22 | 0.29 | 0.08 | 0.13 |
| 5 | 61650843 | rs2289883 | -0.739 | 0.222 | 1.186 | 1.279 | -0.003 | 0.009 | 0.009 | 0 | 0.43 | 0.63 | 0.46 | 0.64 | 0.42 | 0.35 |
| 5 | 61657228 | rs247265 | -0.721 | 0.237 | 1.194 | 1.286 | -0.004 | 0.013 | 0.013 | 0 | 0.42 | 0.66 | 0.46 | 0.64 | 0.42 | 0.35 |
| 5 | 61665955 | rs1309156 | -0.901 | 0.022 | 1.096 | 1.199 | -0.001 | 0.018 | 0.018 | 0 | 0.45 | 0.64 | 0.46 | 0.65 | 0.42 | 0.35 |
| 5 | 61672733 | rs784901 | 0.584 | 0.284 | 1.079 | 1.361 | -0.001 | 0 | 0 | 0 | 0.28 | 0.24 | 0.16 | 0.22 | 0.27 | 0.18 |
| 5 | 61676911 | rs247254 | -1.456 | 0.263 | 0.806 | 1.332 | -0.053 | 0.081 | 0.081 | 0.028 | 0.53 | 0.86 | 0.52 | 0.68 | 0.48 | 0.49 |
| 5 | 61684030 | rs37032 | -0.261 | 0.459 | 0.775 | 1.492 | 0 | 0.019 | 0.019 | 0 | 0.09 | 0.11 | 0.22 | 0.29 | 0.08 | 0.13 |
| 5 | 61686647 | rs2257389 | -1.008 | 0.469 | 0.765 | 1.403 | 0.005 | 0.078 | 0.078 | 0 | 0.13 | 0.22 | 0.28 | 0.42 | 0.15 | 0.17 |
| 5 | 61690703 | rs247256 | 0.607 | 0.602 | 0.529 | 1.249 | -0.009 | -0.005 | -0.005 | 0 | 0.27 | 0.17 | 0.15 | 0.22 | 0.26 | 0.18 |
| 5 | 61691963 | rs185048 | 0.583 | 0.604 | 0.529 | 1.249 | -0.009 | -0.004 | -0.004 | 0 | 0.27 | 0.17 | 0.15 | 0.22 | 0.26 | 0.18 |
| 5 | 61728012 | rs247240 | -0.569 | 0.498 | 0.883 | 1.478 | -0.004 | 0.002 | 0.002 | 0 | 0.39 | 0.4 | 0.42 | 0.62 | 0.42 | 0.31 |
| 5 | 61733348 | rs153857 | -0.11 | 0.463 | 1.005 | 1.599 | -0.013 | -0.041 | -0.041 | 0 | 0.67 | 0.79 | 0.64 | 0.52 | 0.66 | 0.73 |
| 5 | 61735733 | rs784916 | 1.225 | 0.436 | 1.129 | 1.4 | 0 | 0 | 0 | 0 | 0.48 | 0.41 | 0.49 | 0.32 | 0.49 | 0.53 |
| 5 | 61738245 | rs152184 | 0.333 | 0.491 | 1.124 | 1.401 | -0.005 | 0.002 | 0.002 | 0 | 0.61 | 0.6 | 0.58 | 0.38 | 0.58 | 0.69 |
| 5 | 61744277 | rs247236 | -1.059 | 0.433 | 1.082 | 1.46 | -0.087 | -0.046 | -0.046 | 0.008 | 0.75 | 0.98 | 0.86 | 0.81 | 0.73 | 0.83 |
| 5 | 61746142 | rs247237 | 1.548 | 0.361 | 1.269 | 1.414 | -0.041 | 0.065 | 0.065 | 0.235 | 0.45 | 0.12 | 0.48 | 0.32 | 0.48 | 0.5 |
| 5 | 61747500 | rs2441115 | 0.955 | 0.546 | 1.268 | 1.508 | -0.001 | 0.002 | 0.002 | 0 | 0.05 | 0.1 | 0.06 | 0.13 | 0.08 | 0.04 |
| 5 | 61749048 | rs247229 | -0.186 | 0.542 | 1.34 | 1.505 | -0.032 | -0.082 | -0.082 | 0 | 0.67 | 0.87 | 0.66 | 0.52 | 0.66 | 0.73 |
| 5 | 61750087 | rs4700493 | 0.465 | 0.542 | 1.319 | 1.506 | -0.008 | -0.025 | -0.025 | 0 | 0.07 | 0.02 | 0.2 | 0.3 | 0.07 | 0.09 |
| 5 | 61750357 | rs17382964 | -0.853 | 0.549 | 1.338 | 1.501 | 0.003 | 0.017 | 0.017 | 0 | 0.12 | 0.1 | 0.09 | 0.06 | 0.09 | 0.16 |
| 5 | 61769985 | rs26633 | -0.997 | 0.544 | 1.32 | 1.498 | -0.088 | -0.047 | -0.047 | 0.743 | 0.75 | 0.98 | 0.86 | 0.81 | 0.73 | 0.83 |
| 5 | 61772535 | rs26635 | 1.103 | 0.551 | 1.448 | 1.18 | 0 | 0 | 0 | 0 | 0.48 | 0.41 | 0.49 | 0.32 | 0.49 | 0.53 |
| 5 | 61822907 | rs32185 | 0.854 | 0.564 | 1.065 | 1.208 | -0.001 | 0.002 | 0.002 | 0 | 0.05 | 0.1 | 0.06 | 0.13 | 0.08 | 0.04 |
| 5 | 61854650 | rs32165 | 0.694 | 0.555 | 1.068 | 1.203 | 0 | 0.015 | 0.015 | 0 | 0.05 | 0.14 | 0.06 | 0.13 | 0.08 | 0.04 |
| 5 | 61857577 | rs32163 | 0.157 | 0.546 | 1.07 | 1.188 | -0.015 | -0.017 | -0.017 | 2.918 | 0.62 | 0.73 | 0.58 | 0.46 | 0.59 | 0.65 |
| 5 | 61866559 | rs1562244 | 0.227 | 0.56 | 1.083 | 1.205 | -0.007 | 0.002 | 0.002 | 0 | 0.61 | 0.63 | 0.59 | 0.38 | 0.58 | 0.69 |
| 5 | 61867920 | rs32153 | 0.46 | 0.567 | 0.841 | 1.208 | 0.006 | 0.051 | 0.051 | 0 | 0.06 | 0.2 | 0.07 | 0.13 | 0.08 | 0.04 |
| 5 | 61880915 | rs181912 | -0.3 | 0.605 | 0.646 | 1.189 | -0.015 | -0.043 | -0.043 | 0 | 0.67 | 0.79 | 0.64 | 0.52 | 0.66 | 0.73 |
| 5 | 61941043 | rs7719851 | 0.908 | 0.589 | 0.235 | 1.004 | 0.003 | 0.037 | 0.037 | 0 | 0.48 | 0.38 | 0.44 | 0.21 | 0.48 | 0.52 |
| 5 | 61951416 | rs2216707 | -1.336 | 0.537 | -0.659 | 1.317 | -0.006 | -0.013 | -0.013 | 0.148 | 0.17 | 0.1 | 0.13 | 0.19 | 0.17 | 0.21 |
| 5 | 61967808 | rs2112981 | -0.988 | 0.295 | -1.048 | 1.735 | -0.015 | -0.116 | -0.116 | 0 | 0.19 | 0.04 | 0.16 | 0.39 | 0.16 | 0.12 |
| 5 | 61977629 | rs11952678 | 0.403 | 1.247 | -1.556 | 1.528 | -0.061 | -0.028 | -0.028 | 0.321 | 0.19 | 0.09 | 0.18 | 0.19 | 0.31 | 0.25 |
| 5 | 61978989 | rs4700505 | -1.641 | 0.745 | -0.47 | 1.542 | 0 | 0.004 | 0.004 | 0 | 0.52 | 0.58 | 0.39 | 0.62 | 0.51 | 0.51 |
| 5 | 61983848 | rs1469095 | -0.612 | 0.681 | -0.383 | 1.442 | -0.018 | -0.007 | -0.007 | 0 | 0.29 | 0.22 | 0.21 | 0.28 | 0.34 | 0.37 |
| 5 | 61996644 | rs4146184 | -0.658 | 0.483 | -0.366 | 1.323 | 0.005 | -0.017 | -0.017 | 1.447 | 0.22 | 0.14 | 0.17 | 0.37 | 0.16 | 0.14 |
| 5 | 62007687 | rs903421 | -1.713 | 0.197 | -0.32 | 1.222 | 0.007 | 0.018 | 0.018 | 0 | 0.46 | 0.73 | 0.46 | 0.58 | 0.41 | 0.43 |

| | | | | | | | | | | | | | | | | |
|---|---|---|---|---|---|---|---|---|---|---|---|---|---|---|---|---|
| 5 | 62010640 | rs10434537 | -0.353 | 0.201 | -0.138 | 1.204 | -0.002 | 0 | 0 | 0.372 | 0.22 | 0.22 | 0.16 | 0.26 | 0.16 | 0.13 |
| 5 | 62010847 | rs7706346 | 1.128 | -0.144 | -0.968 | 0.415 | -0.006 | 0.081 | 0.081 | 0 | 0.13 | 0.01 | 0.09 | 0 | 0.12 | 0.03 |
| 5 | 62018297 | rs4700506 | -0.622 | -0.055 | -0.471 | 0.663 | 0.012 | 0.033 | 0.033 | 0 | 0.25 | 0.59 | 0.3 | 0.32 | 0.25 | 0.31 |
| 5 | 62020357 | rs10514950 | 1.191 | -0.001 | -0.656 | 0.607 | 0.015 | -0.034 | -0.034 | 0.021 | 0.06 | 0.1 | 0.08 | 0 | 0.07 | 0.03 |
| 5 | 62021806 | rs13171600 | 0.491 | -0.015 | -0.488 | 0.657 | 0.007 | 0.011 | 0.011 | 0 | 0.15 | 0.21 | 0.19 | 0.23 | 0.22 | 0.2 |
| 5 | 62022336 | rs10461474 | 0.979 | -0.002 | -0.619 | 0.663 | -0.059 | -0.063 | -0.063 | 0 | 0.12 | 0.06 | 0.17 | 0.23 | 0.22 | 0.19 |
| 5 | 62024479 | rs1121882 | 0.136 | -0.134 | -0.299 | 0.607 | -0.01 | -0.002 | -0.002 | 0 | 0.07 | 0.11 | 0.08 | 0.09 | 0.03 | 0.11 |
| 5 | 62025023 | rs11740546 | -2.18 | -0.462 | -0.771 | 0.352 | 0 | 0.053 | 0.053 | 0 | 0.49 | 0.81 | 0.46 | 0.59 | 0.43 | 0.46 |
| 5 | 62027073 | rs251170 | -0.881 | -0.533 | -0.602 | 0.391 | -0.069 | 0.02 | 0.02 | 0.043 | 0.28 | 0.49 | 0.19 | 0.36 | 0.14 | 0.23 |
| 5 | 62029339 | rs251172 | -1.892 | -0.48 | -0.584 | 0.379 | 0.013 | 0.01 | 0.01 | 0 | 0.43 | 0.69 | 0.39 | 0.59 | 0.37 | 0.43 |
| 5 | 62029827 | rs16890992 | 0.583 | -0.486 | -0.331 | 0.413 | -0.051 | 0.068 | 0.068 | 0 | 0.17 | 0.33 | 0.1 | 0.27 | 0.09 | 0.11 |
| 5 | 62031426 | rs7723751 | 2.515 | -0.686 | -1.18 | -0.038 | -0.016 | 0.012 | 0.012 | 0 | 0.19 | 0.11 | 0.26 | 0.14 | 0.21 | 0.23 |
| 5 | 62035429 | rs4487447 | -1.546 | -0.715 | -1.162 | -0.123 | 0.004 | 0.025 | 0.025 | 0 | 0.68 | 0.82 | 0.71 | 0.74 | 0.65 | 0.7 |
| 5 | 62037885 | rs1005051 | -0.821 | -0.738 | -1.074 | -0.109 | 0 | 0.021 | 0.021 | 0 | 0.62 | 0.7 | 0.63 | 0.73 | 0.58 | 0.66 |
| 5 | 62039960 | rs16891019 | 0.79 | -0.197 | -0.578 | 0.059 | -0.015 | 0.059 | 0.059 | 0 | 0.19 | 0.37 | 0.16 | 0.28 | 0.16 | 0.12 |
| 5 | 62041610 | rs1900180 | 0.164 | -0.104 | -0.856 | -0.027 | 0.008 | 0.006 | 0.006 | 0 | 0.1 | 0.09 | 0.08 | 0.09 | 0.05 | 0.12 |
| 5 | 62048009 | rs2100036 | 1.283 | -0.181 | -0.8 | 0.13 | -0.025 | -0.031 | -0.031 | 0 | 0.4 | 0.16 | 0.39 | 0.4 | 0.4 | 0.32 |
| 5 | 62055300 | rs9784594 | -0.124 | -0.175 | -0.646 | 0.253 | 0.001 | 0.006 | 0.006 | 0 | 0.2 | 0.23 | 0.19 | 0.22 | 0.18 | 0.33 |
| 5 | 62060350 | rs1979727 | -0.097 | -0.51 | -0.49 | -0.363 | 0.031 | 0.035 | 0.035 | 0 | 0.6 | 0.81 | 0.68 | 0.63 | 0.62 | 0.61 |
| 5 | 62064028 | rs1540532 | 0.783 | -0.286 | -0.02 | -0.377 | 0.005 | 0.051 | 0.051 | 0 | 0.47 | 0.32 | 0.53 | 0.62 | 0.53 | 0.48 |
| 5 | 62067238 | rs1826675 | 0.203 | -0.272 | 0.056 | -0.235 | 0.005 | 0.006 | 0.006 | 0 | 0.56 | 0.61 | 0.66 | 0.62 | 0.62 | 0.51 |
| 5 | 62067375 | rs1826676 | 0.182 | -0.253 | -0.197 | -0.226 | 0.023 | 0.024 | 0.024 | 0 | 0.56 | 0.66 | 0.66 | 0.62 | 0.62 | 0.51 |
| 5 | 62067885 | rs10939944 | 0.44 | -0.659 | 0.07 | 0.209 | 0 | -0.001 | -0.001 | 0 | 0.66 | 0.65 | 0.73 | 0.59 | 0.78 | 0.55 |
| 5 | 62069552 | rs2591786 | 1.124 | 0.025 | 0.04 | 0.274 | -0.006 | 0.069 | 0.069 | 0 | 0.36 | 0.13 | 0.26 | 0.21 | 0.39 | 0.27 |
| 5 | 62069673 | rs2642776 | 1.019 | 0.012 | 0.299 | 0.268 | 0.005 | 0.037 | 0.037 | 0 | 0.37 | 0.24 | 0.26 | 0.21 | 0.39 | 0.27 |
| 5 | 62073834 | rs2642780 | -0.126 | 0.198 | 0.319 | 0.505 | -0.001 | 0 | 0 | 0 | 0.78 | 0.8 | 0.7 | 0.71 | 0.83 | 0.72 |
| 5 | 62078323 | rs6449611 | -0.192 | 0.346 | 0.584 | 0.593 | 0.049 | 0.061 | 0.061 | 0.455 | 0.17 | 0.53 | 0.22 | 0.25 | 0.16 | 0.23 |
| 5 | 62084196 | rs185170 | 0.926 | 0.17 | 0.585 | 0.518 | 0.012 | 0.018 | 0.018 | 0 | 0.43 | 0.41 | 0.35 | 0.32 | 0.42 | 0.48 |
| 5 | 62084506 | rs903420 | 1.082 | 0.216 | 0.621 | 0.534 | 0.063 | 0.037 | 0.037 | 0 | 0.1 | 0.2 | 0.14 | 0.14 | 0.14 | 0.15 |
| 5 | 62085087 | rs251166 | -0.267 | 0.187 | 0.619 | 0.541 | -0.001 | 0 | 0 | 0 | 0.78 | 0.78 | 0.7 | 0.73 | 0.83 | 0.72 |
| 5 | 62091271 | rs12189374 | 0.99 | 0.263 | 0.544 | 0.569 | 0.066 | 0.036 | 0.036 | 0 | 0.1 | 0.2 | 0.14 | 0.14 | 0.15 | 0.15 |
| 5 | 62091796 | rs16891097 | 0.524 | 0.074 | 0.591 | 0.541 | -0.019 | 0.042 | 0.042 | 1.103 | 0.07 | 0.34 | 0.08 | 0.1 | 0.02 | 0.08 |
| 5 | 62091992 | rs2132020 | -0.256 | 0.207 | 0.6 | 0.564 | -0.001 | 0 | 0 | 0 | 0.78 | 0.78 | 0.7 | 0.73 | 0.83 | 0.72 |
| 5 | 62094034 | rs7708428 | 0.301 | 0.204 | 0.513 | 0.559 | 0.015 | 0.013 | 0.013 | 0.076 | 0.34 | 0.37 | 0.35 | 0.41 | 0.41 | 0.24 |
| 5 | 62094462 | rs10078232 | 1.371 | 0.204 | 0.477 | 0.56 | -0.003 | -0.009 | -0.009 | 0 | 0.24 | 0.17 | 0.21 | 0.27 | 0.27 | 0.09 |
| 5 | 62101475 | rs2642749 | 0.827 | 0.501 | 0.585 | 0.746 | 0.025 | 0.03 | 0.03 | 0 | 0.68 | 0.58 | 0.56 | 0.59 | 0.69 | 0.57 |
| 5 | 62107278 | rs1874183 | -0.984 | 0.586 | 0.3 | 0.817 | 0.006 | 0.039 | 0.039 | 0.003 | 0.91 | 0.87 | 0.73 | 0.75 | 0.89 | 0.82 |
| 5 | 62110910 | rs6864017 | 0.904 | 0.731 | 0.044 | 0.782 | 0.015 | 0.007 | 0.007 | 0 | 0.09 | 0.11 | 0.13 | 0.14 | 0.14 | 0.15 |
| 5 | 62124231 | rs2642761 | 0.216 | 0.258 | -0.154 | 0.628 | 0.002 | 0.004 | 0.004 | 0.454 | 0.5 | 0.51 | 0.47 | 0.46 | 0.53 | 0.63 |
| 5 | 62131163 | rs11740952 | 0.87 | -0.032 | -0.986 | 0.748 | 0 | -0.001 | -0.001 | 0 | 0.28 | 0.24 | 0.23 | 0.28 | 0.3 | 0.09 |
| 5 | 62166774 | rs3107556 | 0.607 | -0.086 | -1.299 | 0.783 | -0.004 | 0.002 | 0.002 | 0 | 0.4 | 0.27 | 0.39 | 0.39 | 0.54 | 0.51 |
| 5 | 62169769 | rs3107558 | 0.653 | -0.233 | -0.333 | 0.897 | -0.046 | 0.004 | 0.004 | 0.706 | 0.69 | 0.46 | 0.63 | 0.66 | 0.83 | 0.6 |
| 5 | 62172239 | rs3111611 | 0.818 | 0.474 | -0.895 | 0.794 | 0.001 | -0.018 | -0.018 | 0 | 0.64 | 0.28 | 0.54 | 0.64 | 0.7 | 0.51 |
| 5 | 62175815 | rs1402912 | 0.57 | 1.345 | -0.753 | 1.21 | 0.001 | 0.011 | 0.011 | 0 | 0.85 | 0.85 | 0.65 | 0.64 | 0.8 | 0.72 |
| 5 | 62184884 | rs17474789 | 0.334 | 0.78 | -1.044 | 0.212 | 0.002 | 0.024 | 0.024 | 0 | 0.16 | 0.06 | 0.1 | 0 | 0.17 | 0.09 |
| 5 | 62193486 | rs1402915 | -0.484 | 0.568 | -0.827 | 0.117 | -0.002 | 0.007 | 0.007 | 0 | 0.53 | 0.57 | 0.38 | 0.38 | 0.47 | 0.33 |
| 5 | 62194602 | rs13359556 | 0.928 | 0.232 | -0.701 | -0.304 | -0.001 | -0.026 | -0.026 | 0 | 0.19 | 0.2 | 0.1 | 0 | 0.12 | 0.11 |
| 5 | 62204534 | rs16891251 | 1.872 | 0.405 | -0.937 | -0.126 | 0.017 | -0.037 | -0.037 | 0 | 0.25 | 0.14 | 0.24 | 0.33 | 0.27 | 0.27 |
| 5 | 62205619 | rs4700537 | -0.278 | 0.388 | -0.676 | -0.043 | 0 | 0.009 | 0.009 | 0 | 0.51 | 0.54 | 0.38 | 0.24 | 0.46 | 0.24 |
| 5 | 62232886 | rs13179171 | -0.195 | 0.468 | -0.665 | -0.055 | 0.001 | -0.024 | -0.024 | 0 | 0.38 | 0.53 | 0.29 | 0.3 | 0.3 | 0.24 |
| 5 | 62237088 | rs939783 | -0.185 | 0.469 | -0.654 | -0.054 | 0.001 | -0.025 | -0.025 | 0 | 0.37 | 0.53 | 0.29 | 0.3 | 0.3 | 0.24 |
| 5 | 62239608 | rs6889201 | 1.121 | 0.576 | -0.38 | -0.011 | 0.013 | 0.017 | 0.017 | 0 | 0.1 | 0.15 | 0.16 | 0.17 | 0.15 | 0.05 |
| 5 | 62255744 | rs939782 | -1.287 | 0.827 | -0.551 | 0.161 | -0.032 | -0.032 | -0.032 | 0 | 0.93 | 0.99 | 0.8 | 0.71 | 0.89 | 0.83 |
| 5 | 62262952 | rs10073117 | 0.294 | 0.768 | -0.431 | 0.102 | 0.007 | -0.02 | -0.02 | 0 | 0.3 | 0.41 | 0.26 | 0.17 | 0.27 | 0.15 |
| 5 | 62271454 | rs1521984 | -1.242 | 0.277 | -0.474 | 0.179 | -0.032 | -0.033 | -0.033 | 0 | 0.93 | 0.99 | 0.8 | 0.72 | 0.89 | 0.83 |
| 5 | 62277081 | rs1402919 | -0.258 | 0.141 | -0.364 | -0.034 | 0 | -0.022 | -0.022 | 0 | 0.47 | 0.53 | 0.36 | 0.17 | 0.44 | 0.24 |
| 5 | 62309979 | rs1033184 | -0.873 | 0.224 | -0.041 | 0.085 | -0.007 | -0.016 | -0.016 | 0 | 0.86 | 0.92 | 0.73 | 0.7 | 0.83 | 0.7 |
| 5 | 62312150 | rs6866618 | 1.205 | 0.139 | -0.283 | -0.057 | 0.008 | 0.021 | 0.021 | 0 | 0.18 | 0.12 | 0.1 | 0.13 | 0.18 | 0.17 |
| 5 | 62320063 | rs2885927 | 0.5 | 0.091 | -0.397 | -0.215 | -0.001 | -0.001 | -0.001 | 0 | 0.14 | 0.14 | 0.17 | 0.17 | 0.19 | 0.08 |
| 5 | 62325295 | rs6865447 | 1.101 | 0.126 | -0.481 | -0.211 | -0.001 | -0.003 | -0.003 | 0.46 | 0.19 | 0.2 | 0.11 | 0.13 | 0.18 | 0.17 |
| 5 | 62332053 | rs16891411 | 2.25 | 0.045 | -0.649 | -0.157 | 0.007 | -0.055 | -0.055 | 0 | 0.24 | 0.05 | 0.23 | 0.29 | 0.24 | 0.24 |
| 5 | 62345572 | rs10454821 | 0.517 | 0.355 | 0.078 | -0.661 | -0.005 | -0.01 | -0.01 | 0 | 0.2 | 0.21 | 0.11 | 0.13 | 0.18 | 0.16 |
| 5 | 62354249 | rs10056765 | -1.064 | 0.394 | 0.061 | -0.66 | 0 | 0.016 | 0.016 | 0 | 0.54 | 0.62 | 0.39 | 0.31 | 0.5 | 0.35 |
| 5 | 62355496 | rs7735880 | -0.243 | 0.345 | -0.011 | -0.695 | 0.012 | -0.029 | -0.029 | 0 | 0.32 | 0.42 | 0.27 | 0.17 | 0.31 | 0.18 |
| 5 | 62357866 | rs1514577 | -0.893 | 0.33 | -0.002 | -0.662 | -0.001 | -0.005 | -0.005 | 0 | 0.52 | 0.54 | 0.38 | 0.31 | 0.5 | 0.35 |
| 5 | 62363212 | rs9291774 | -0.315 | 0.536 | 0.587 | -0.005 | 0.022 | 0.01 | 0.01 | 0 | 0.15 | 0.28 | 0.19 | 0.17 | 0.19 | 0.08 |
| 5 | 62368902 | rs10066030 | -0.834 | 0.425 | 0.545 | 0.113 | -0.001 | -0.018 | -0.018 | 2.794 | 0.37 | 0.51 | 0.32 | 0.3 | 0.38 | 0.26 |
| 5 | 62369102 | rs4323191 | -0.288 | 0.498 | 0.346 | 0.048 | 0.002 | -0.01 | -0.01 | 0 | 0.31 | 0.35 | 0.29 | 0.17 | 0.36 | 0.17 |
| 5 | 62372522 | rs4700548 | 0.491 | 0.583 | 0.099 | 0.277 | -0.017 | -0.02 | -0.02 | 0 | 0.05 | 0.02 | 0.1 | 0.08 | 0.08 | 0.1 |

| | | | | | | | | | | | | | | | | |
|---|---|---|---|---|---|---|---|---|---|---|---|---|---|---|---|---|
| 5 | 62372947 | rs970963 | -0.787 | 0.542 | 0.349 | 0.189 | 0.013 | 0.006 | 0.006 | 0 | 0.37 | 0.48 | 0.38 | 0.4 | 0.42 | 0.35 |
| 5 | 62377385 | rs6881303 | -0.893 | 0.598 | 0.202 | 0.193 | 0.002 | -0.001 | -0.001 | 0.027 | 0.38 | 0.53 | 0.39 | 0.39 | 0.42 | 0.35 |
| 5 | 62392211 | rs4700554 | -0.105 | 0.731 | 0.106 | 0.094 | 0.006 | -0.01 | -0.01 | 0 | 0.26 | 0.3 | 0.26 | 0.17 | 0.32 | 0.14 |
| 5 | 62394367 | rs1160346 | -1.062 | 0.265 | 0.036 | -0.597 | 0.001 | 0.053 | 0.053 | 0.198 | 0.39 | 0.69 | 0.35 | 0.22 | 0.37 | 0.29 |
| 5 | 62397076 | rs6889630 | -0.98 | 0.285 | 0.05 | -0.662 | -0.002 | 0.047 | 0.047 | 0.05 | 0.38 | 0.67 | 0.35 | 0.2 | 0.37 | 0.29 |
| 5 | 62407243 | rs9688293 | -0.887 | 0.019 | 0.456 | -0.658 | 0.015 | 0.141 | 0.141 | 0.326 | 0.09 | 0.52 | 0.12 | 0.16 | 0.05 | 0.11 |
| 5 | 62409385 | rs7731343 | -1.335 | -0.024 | 0.439 | -0.683 | -0.024 | 0.071 | 0.071 | 0.209 | 0.13 | 0.54 | 0.15 | 0.18 | 0.06 | 0.19 |
| 5 | 62414243 | rs1514585 | 0.507 | 0.041 | 0.241 | -0.853 | -0.049 | 0.085 | 0.085 | 0 | 0.14 | 0.02 | 0.1 | 0.01 | 0.14 | 0.03 |
| 5 | 62416806 | rs10066621 | -0.828 | -0.008 | 0.466 | -0.789 | 0 | -0.049 | -0.049 | 0 | 0.29 | 0.62 | 0.23 | 0.03 | 0.28 | 0.09 |
| 5 | 62416834 | rs10066628 | 0.412 | -0.291 | -0.291 | -1.542 | -0.012 | 0.039 | 0.039 | 0 | 0.29 | 0.21 | 0.12 | 0 | 0.24 | 0.33 |
| 5 | 62427908 | rs168416 | 1.057 | -0.261 | -0.494 | -1.058 | -0.001 | -0.026 | -0.026 | 0 | 0.58 | 0.29 | 0.47 | 0.53 | 0.58 | 0.55 |
| 5 | 62429113 | rs350302 | 0.041 | -0.125 | 0.018 | -0.909 | -0.005 | 0.001 | 0.001 | 0 | 0.72 | 0.77 | 0.61 | 0.71 | 0.65 | 0.8 |
| 5 | 62430533 | rs171227 | 0.188 | -0.104 | 0.06 | -0.821 | 0.004 | 0.002 | 0.002 | 0 | 0.7 | 0.69 | 0.6 | 0.68 | 0.65 | 0.8 |
| 5 | 62440012 | rs664540 | 1.319 | -0.28 | -0.203 | -0.589 | -0.033 | -0.15 | -0.15 | 0.541 | 0.29 | 0.07 | 0.35 | 0.52 | 0.35 | 0.28 |
| 5 | 62463639 | rs1355963 | 0.082 | -0.224 | -0.201 | -1.153 | 0 | -0.082 | -0.082 | 3.336 | 0.48 | 0.54 | 0.31 | 0.03 | 0.46 | 0.24 |
| 5 | 62463774 | rs601429 | 1.412 | -0.229 | -1.417 | -0.952 | -0.028 | -0.103 | -0.103 | 0.064 | 0.29 | 0.07 | 0.36 | 0.59 | 0.34 | 0.24 |
| 5 | 62465904 | rs1514582 | 0.823 | -0.217 | -1.386 | -1.018 | 0.005 | 0.051 | 0.051 | 0 | 0.1 | 0.01 | 0.04 | 0.02 | 0.07 | 0.22 |
| 5 | 62472635 | rs691311 | -0.18 | -0.191 | -1.358 | -0.957 | 0.002 | -0.026 | -0.026 | 0.048 | 0.85 | 0.93 | 0.77 | 0.73 | 0.85 | 0.6 |
| 5 | 62480129 | rs2199466 | -0.607 | -0.193 | -1.123 | -1.964 | 0.012 | 0.128 | 0.128 | 0 | 0.55 | 0.8 | 0.41 | 0.17 | 0.51 | 0.37 |
| 5 | 62484878 | rs1824640 | 0.155 | 0.02 | -1.458 | -2.112 | -0.018 | 0.066 | 0.066 | 0 | 0.39 | 0.3 | 0.24 | 0.04 | 0.38 | 0.21 |
| 5 | 62510356 | rs1466736 | -0.051 | 0.328 | -2.204 | -1.82 | 0.005 | 0.138 | 0.138 | 0 | 0.49 | 0.71 | 0.37 | 0.16 | 0.46 | 0.36 |
| 5 | 62523339 | rs347668 | -1.155 | 0.362 | -2.411 | -1.881 | -0.044 | 0.07 | 0.07 | 0 | 0.68 | 0.92 | 0.47 | 0.17 | 0.59 | 0.6 |
| 5 | 62529212 | rs346420 | -0.453 | 0.042 | -1.583 | -0.835 | 0.003 | 0.094 | 0.094 | 0 | 0.17 | 0.04 | 0.08 | 0.01 | 0.12 | 0.25 |
| 5 | 62529686 | rs10514958 | -0.668 | 0.465 | -0.774 | -0.59 | 0 | -0.036 | -0.036 | 0 | 0.12 | 0.15 | 0.09 | 0 | 0.14 | 0.08 |
| 5 | 62537479 | rs346410 | -0.669 | 0.419 | -0.814 | -0.398 | -0.001 | -0.02 | -0.02 | 0.39 | 0.15 | 0.24 | 0.08 | 0.11 | 0.14 | 0.19 |
| 5 | 62541894 | rs346416 | -0.366 | -0.005 | -2.05 | -1.007 | -0.016 | 0.154 | 0.154 | 0 | 0.56 | 0.88 | 0.35 | 0.25 | 0.45 | 0.41 |
| 5 | 62549338 | rs347662 | -0.446 | 0.519 | -2.341 | -1.125 | -0.173 | -0.03 | -0.03 | 0.026 | 0.78 | 0.97 | 0.45 | 0.25 | 0.58 | 0.47 |
| 5 | 62557698 | rs349488 | -0.044 | 0.149 | -2.408 | -0.936 | -0.026 | 0.004 | 0.004 | 0 | 0.7 | 0.83 | 0.64 | 0.71 | 0.66 | 0.62 |
| 5 | 62559522 | rs462972 | -0.093 | 0.011 | -2.496 | -0.861 | -0.078 | -0.045 | -0.045 | 0 | 0.72 | 0.93 | 0.65 | 0.66 | 0.68 | 0.66 |
| 5 | 62560237 | rs58736 | 0.024 | -0.48 | -2.57 | -1.31 | -0.015 | -0.052 | -0.052 | 0.667 | 0.27 | 0.16 | 0.35 | 0.53 | 0.34 | 0.47 |
| 5 | 62566536 | rs418226 | -0.247 | -0.419 | -2.41 | -1.125 | -0.069 | -0.021 | -0.021 | 0 | 0.69 | 0.91 | 0.65 | 0.67 | 0.66 | 0.61 |
| 5 | 62567045 | rs427421 | -0.247 | -0.419 | -2.421 | -1.125 | -0.075 | -0.024 | -0.024 | 0 | 0.69 | 0.92 | 0.65 | 0.67 | 0.66 | 0.61 |
| 5 | 62582619 | rs379152 | -0.434 | -0.511 | -2.448 | -1.147 | -0.067 | -0.05 | -0.05 | 2.71 | 0.82 | 0.93 | 0.71 | 0.69 | 0.71 | 0.65 |
| 5 | 62585230 | rs1389843 | 1.296 | -0.596 | -2.47 | -1.026 | -0.046 | -0.075 | -0.075 | 0.004 | 0.11 | 0.02 | 0.25 | 0.42 | 0.18 | 0.27 |
| 5 | 62598453 | rs2169520 | 0.65 | -1.723 | -1.861 | -1.721 | -0.05 | -0.069 | -0.069 | 0 | 0.36 | 0.45 | 0.22 | 0.12 | 0.19 | 0.09 |
| 5 | 62602454 | rs346437 | 0.539 | -1.922 | -1.521 | -1.935 | 0.033 | 0.066 | 0.066 | 3.122 | 0.33 | 0.19 | 0.19 | 0.14 | 0.25 | 0.21 |
| 5 | 62603647 | rs10514959 | 1.91 | -1.907 | -1.525 | -1.913 | 0.031 | 0.122 | 0.122 | 4.017 | 0.19 | 0.02 | 0.1 | 0 | 0.12 | 0.05 |
| 5 | 62613641 | rs1462460 | 1.266 | -1.86 | -0.705 | -2.021 | -0.038 | -0.087 | -0.087 | 0.005 | 0.23 | 0.35 | 0.13 | 0 | 0.12 | 0.06 |
| 5 | 62641998 | rs4260622 | 0.192 | -1.736 | -0.794 | -2.139 | -0.019 | -0.041 | -0.041 | 0.174 | 0.25 | 0.32 | 0.16 | 0.12 | 0.19 | 0.1 |
| 5 | 62650476 | rs4259134 | 1.276 | -1.112 | -1.043 | -1.811 | -0.099 | -0.088 | -0.088 | 0 | 0.14 | 0.02 | 0.33 | 0.32 | 0.31 | 0.33 |
| 5 | 62670313 | rs4585392 | 0.336 | -1.411 | -0.785 | -2.018 | 0.012 | 0.003 | 0.003 | 2.474 | 0.46 | 0.56 | 0.26 | 0.14 | 0.3 | 0.13 |
| 5 | 62680095 | rs6449656 | -0.207 | -1.205 | -0.626 | -1.963 | 0.01 | 0.012 | 0.012 | 5.779 | 0.46 | 0.39 | 0.7 | 0.86 | 0.62 | 0.77 |
| 5 | 62688370 | rs4406100 | -0.916 | -1.318 | -0.791 | -1.992 | 0.043 | 0.016 | 0.016 | 5.901 | 0.6 | 0.64 | 0.79 | 1 | 0.76 | 0.83 |
| 5 | 62689646 | rs13165578 | 1.29 | -0.824 | -0.818 | -1.623 | -0.001 | 0.003 | 0.003 | 0.085 | 0.06 | 0.04 | 0.06 | 0.13 | 0.06 | 0.06 |
| 5 | 62702322 | rs4605741 | 0.293 | -0.729 | -0.912 | -1.652 | -0.112 | 0.003 | 0.003 | 3.177 | 0.27 | 0.09 | 0.55 | 0.73 | 0.46 | 0.62 |
| 5 | 62705129 | rs9885204 | 1.02 | -1.187 | -0.766 | -1.834 | 0.012 | -0.016 | -0.016 | 0.006 | 0.23 | 0.21 | 0.12 | 0 | 0.13 | 0.04 |
| 5 | 62716281 | rs10078498 | 1.129 | -0.841 | -0.436 | -1.505 | 0.03 | 0.055 | 0.055 | 0.799 | 0.14 | 0.15 | 0.34 | 0.46 | 0.3 | 0.3 |
| 5 | 62725286 | rs7716031 | 0.264 | -1.196 | -0.266 | -1.661 | -0.01 | 0.012 | 0.012 | 0.074 | 0.35 | 0.47 | 0.33 | 0.39 | 0.33 | 0.41 |
| 5 | 62734168 | rs4538568 | 0.751 | -1.226 | -0.39 | -1.679 | -0.031 | -0.041 | -0.041 | 0 | 0.32 | 0.19 | 0.43 | 0.46 | 0.39 | 0.33 |
| 5 | 62735476 | rs4700581 | 0.751 | -1.226 | -0.39 | -1.679 | -0.031 | -0.041 | -0.041 | 0.272 | 0.32 | 0.19 | 0.43 | 0.46 | 0.39 | 0.33 |
| 5 | 62739123 | rs12522826 | 1.261 | -0.928 | -0.193 | -1.489 | 0.017 | 0.037 | 0.037 | 0.804 | 0.14 | 0.13 | 0.34 | 0.46 | 0.3 | 0.3 |
| 5 | 62742726 | rs10056084 | -0.579 | -0.966 | -0.296 | -1.559 | 0.004 | 0.006 | 0.006 | 0 | 0.71 | 0.77 | 0.78 | 0.86 | 0.75 | 0.75 |
| 5 | 62749746 | rs16891858 | 1.492 | -0.64 | -0.348 | -1.341 | -0.023 | -0.022 | -0.022 | 0 | 0.13 | 0.06 | 0.31 | 0.33 | 0.29 | 0.3 |
| 5 | 62761734 | rs2112508 | 0.356 | -0.991 | 0.023 | -1.549 | 0.101 | 0.132 | 0.132 | 2.556 | 0.21 | 0.4 | 0.38 | 0.48 | 0.36 | 0.32 |
| 5 | 62763172 | rs585879 | -0.513 | -1.234 | -0.41 | -1.677 | 0.022 | 0.04 | 0.04 | 0.103 | 0.7 | 0.8 | 0.78 | 0.84 | 0.76 | 0.75 |
| 5 | 62766998 | rs644101 | -0.513 | -1.234 | -0.41 | -1.677 | 0.022 | 0.04 | 0.04 | 0.103 | 0.7 | 0.8 | 0.78 | 0.84 | 0.76 | 0.76 |
| 5 | 62769216 | rs595456 | -0.513 | -1.234 | -0.41 | -1.677 | 0.022 | 0.04 | 0.04 | 0.103 | 0.7 | 0.8 | 0.78 | 0.84 | 0.76 | 0.75 |
| 5 | 62811418 | rs665579 | 0.397 | -1.233 | -0.496 | -1.719 | 0.023 | 0.047 | 0.047 | 0.079 | 0.3 | 0.2 | 0.23 | 0.13 | 0.23 | 0.24 |
| 5 | 62820508 | rs669245 | -0.689 | -1.409 | -0.39 | -1.783 | 0.004 | 0.007 | 0.007 | 0 | 0.64 | 0.59 | 0.55 | 0.46 | 0.6 | 0.64 |
| 5 | 62823933 | rs9885552 | 1.138 | -1.48 | -0.351 | -1.916 | -0.003 | -0.013 | -0.013 | 0.505 | 0.19 | 0.16 | 0.24 | 0.33 | 0.23 | 0.3 |
| 5 | 62824362 | rs30903 | -0.76 | -1.607 | -0.441 | -2.065 | 0.035 | 0.006 | 0.006 | 0.571 | 0.66 | 0.7 | 0.82 | 1 | 0.82 | 0.87 |
| 5 | 62825112 | rs40647 | 0.925 | -1.594 | -0.575 | -2.056 | 0.028 | 0.062 | 0.062 | 0.055 | 0.27 | 0.15 | 0.21 | 0.12 | 0.2 | 0.23 |
| 5 | 62832795 | rs659217 | -0.689 | -1.652 | -0.457 | -2.071 | -0.032 | -0.121 | -0.121 | 0 | 0.66 | 0.59 | 0.81 | 1 | 0.82 | 0.87 |
| 5 | 62838363 | rs255402 | -1.297 | -1.183 | -0.227 | -1.684 | -0.015 | -0.055 | -0.055 | 0 | 0.65 | 0.95 | 0.7 | 0.67 | 0.68 | 0.65 |
| 5 | 62842815 | rs653244 | -0.059 | -0.86 | -0.009 | -1.524 | 0.021 | 0.017 | 0.017 | 0 | 0.4 | 0.3 | 0.27 | 0.12 | 0.26 | 0.3 |
| 5 | 62844222 | rs626392 | -0.675 | -0.79 | -0.158 | -1.516 | -0.001 | -0.003 | -0.003 | 0 | 0.58 | 0.54 | 0.62 | 0.67 | 0.6 | 0.63 |
| 5 | 62853232 | rs166244 | 1.293 | -0.921 | -0.353 | -1.612 | 0.02 | 0.144 | 0.144 | 1.458 | 0.22 | 0.03 | 0.1 | 0 | 0.15 | 0.15 |
| 5 | 62855619 | rs16891970 | 0.69 | -0.838 | 0.067 | -1.479 | -0.034 | -0.065 | -0.065 | 0 | 0.15 | 0.25 | 0.08 | 0 | 0.07 | 0.08 |
| 5 | 62858801 | rs1867735 | 0.623 | -0.754 | 0.028 | -1.465 | 0.13 | 0.059 | 0.059 | 3.069 | 0.05 | 0.4 | 0.06 | 0 | 0.03 | 0.01 |

| | | | | | | | | | | | | | | | | |
|---|---|---|---|---|---|---|---|---|---|---|---|---|---|---|---|---|
| 5 | 62867025 | rs722851 | 0.429 | -0.633 | 0.039 | -1.318 | -0.036 | -0.138 | -0.138 | 0.396 | 0.17 | 0.47 | 0.11 | 0 | 0.1 | 0.09 |
| 5 | 62867084 | rs673743 | -1.143 | -0.592 | -0.662 | -1.233 | -0.019 | -0.062 | -0.062 | 0 | 0.67 | 0.95 | 0.7 | 0.67 | 0.69 | 0.65 |
| 5 | 62874332 | rs1559090 | -1.14 | -0.164 | -0.418 | -1.097 | 0.003 | -0.031 | -0.031 | 0 | 0.63 | 0.95 | 0.67 | 0.66 | 0.69 | 0.6 |
| 5 | 62886869 | rs11959809 | 1.414 | 0.201 | -0.399 | -0.991 | -0.083 | -0.086 | -0.086 | 0 | 0.14 | 0.07 | 0.31 | 0.53 | 0.32 | 0.3 |
| 5 | 62912745 | rs10471588 | -0.729 | -0.091 | -0.204 | -1.167 | 0.047 | 0.073 | 0.073 | 0.395 | 0.71 | 0.88 | 0.8 | 0.88 | 0.79 | 0.8 |
| 5 | 62921104 | rs7715514 | 0.812 | 0.278 | 0.442 | -1.067 | -0.004 | -0.072 | -0.072 | 0 | 0.12 | 0.31 | 0.08 | 0 | 0.08 | 0.05 |
| 5 | 62921438 | rs6449668 | -0.213 | 0.278 | 0.277 | -1.07 | 0.039 | -0.123 | -0.123 | 1.999 | 0.16 | 0.5 | 0.12 | 0 | 0.16 | 0.06 |
| 5 | 62931346 | rs16892078 | 0.543 | 0.261 | 0.154 | -1.045 | -0.005 | -0.083 | -0.083 | 0.159 | 0.13 | 0.38 | 0.09 | 0 | 0.08 | 0.05 |
| 5 | 62942257 | rs10036059 | 0.979 | 0.231 | -0.138 | -1.007 | 0.019 | -0.029 | -0.029 | 0 | 0.37 | 0.08 | 0.33 | 0.37 | 0.31 | 0.4 |
| 5 | 62946149 | rs1566780 | -0.208 | 0.159 | 0.008 | -0.865 | 0.087 | 0.096 | 0.096 | 1.886 | 0.27 | 0.53 | 0.38 | 0.51 | 0.37 | 0.38 |
| 5 | 62964798 | rs2365873 | 0.235 | 0.019 | -0.465 | -0.951 | -0.005 | -0.037 | -0.037 | 2.66 | 0.55 | 0.54 | 0.7 | 0.88 | 0.68 | 0.75 |
| 5 | 62980393 | rs6885945 | 0.154 | 0.238 | -0.602 | -0.571 | 0 | 0.001 | 0.001 | 0 | 0.52 | 0.53 | 0.46 | 0.36 | 0.47 | 0.46 |
| 5 | 62982117 | rs6893075 | -1.213 | -0.152 | -1.23 | -0.879 | 0.033 | 0.121 | 0.121 | 1.2 | 0.81 | 0.98 | 0.92 | 1 | 0.89 | 0.9 |
| 5 | 62983524 | rs4700056 | -0.583 | -0.153 | -1.248 | -0.552 | 0.042 | 0.039 | 0.039 | 2.377 | 0.76 | 0.93 | 0.76 | 0.86 | 0.84 | 0.83 |
| 5 | 62983877 | rs13157106 | 1.089 | 0.422 | -1.523 | -0.246 | -0.084 | -0.109 | -0.109 | 0.791 | 0.17 | 0.03 | 0.3 | 0.48 | 0.3 | 0.38 |
| 5 | 62986318 | rs11742891 | 0.519 | 0.283 | -0.783 | -0.641 | 0.006 | 0.008 | 0.008 | 0 | 0.48 | 0.51 | 0.36 | 0.37 | 0.42 | 0.41 |
| 5 | 62999183 | rs7728227 | 0.426 | -0.233 | -0.678 | -1.393 | 0.002 | -0.016 | -0.016 | 0.446 | 0.36 | 0.62 | 0.24 | 0.02 | 0.33 | 0.13 |
| 5 | 63007060 | rs10035747 | 0.992 | -0.233 | -0.205 | -1.359 | -0.007 | -0.063 | -0.063 | 0 | 0.29 | 0.26 | 0.16 | 0.02 | 0.22 | 0.11 |
| 5 | 63017492 | rs7734344 | 0.002 | -0.27 | -0.258 | -1.371 | -0.025 | 0.059 | 0.059 | 0.719 | 0.39 | 0.29 | 0.51 | 0.85 | 0.51 | 0.7 |
| 5 | 63019264 | rs7726949 | 0.641 | -0.617 | -0.513 | -1.614 | -0.013 | -0.083 | -0.083 | 0.714 | 0.14 | 0.25 | 0.07 | 0.01 | 0.11 | 0.06 |
| 5 | 63020706 | rs1503526 | 0.078 | -0.422 | -0.936 | -1.46 | -0.074 | 0.133 | 0.133 | 0.401 | 0.39 | 0.13 | 0.49 | 0.85 | 0.51 | 0.71 |
| 5 | 63031226 | rs7701186 | 0.097 | -0.384 | -0.926 | -1.442 | -0.018 | 0.05 | 0.05 | 0.392 | 0.4 | 0.34 | 0.51 | 0.83 | 0.51 | 0.7 |
| 5 | 63035923 | rs10054295 | 0.513 | -0.296 | -0.523 | -1.194 | 0.22 | 0.011 | 0.011 | 2.153 | 0.07 | 0.41 | 0.1 | 0 | 0.14 | 0.02 |
| 5 | 63037537 | rs13173277 | -0.388 | -0.544 | -1.145 | -1.241 | -0.002 | -0.018 | -0.018 | 1.892 | 0.63 | 0.42 | 0.62 | 0.86 | 0.68 | 0.78 |
| 5 | 63039074 | rs12517026 | 0.307 | -0.285 | -1.417 | -0.46 | 0.013 | -0.035 | -0.035 | 1.707 | 0.22 | 0.11 | 0.2 | 0.36 | 0.21 | 0.33 |
| 5 | 63043437 | rs260995 | -0.456 | 0.447 | -1.971 | 0.293 | -0.084 | -0.132 | -0.132 | 0 | 0.72 | 0.93 | 0.7 | 0.51 | 0.62 | 0.55 |
| 5 | 63047375 | rs260990 | -0.135 | -0.494 | -1.845 | -0.009 | 0.013 | 0.021 | 0.021 | 0 | 0.78 | 0.85 | 0.86 | 0.85 | 0.81 | 0.88 |
| 5 | 63049105 | rs155020 | 0.351 | -0.565 | -1.988 | 0.217 | 0.001 | -0.016 | -0.016 | 5.241 | 0.26 | 0.17 | 0.25 | 0.33 | 0.22 | 0.38 |
| 5 | 63057033 | rs12374530 | 0.458 | -0.784 | -1.528 | 0.234 | -0.002 | 0 | 0 | 0.236 | 0.37 | 0.39 | 0.4 | 0.36 | 0.45 | 0.35 |
| 5 | 63060649 | rs378268 | 1.682 | -0.874 | -1.58 | 0.237 | 0 | 0.005 | 0.005 | 0.059 | 0.16 | 0.17 | 0.21 | 0.33 | 0.13 | 0.35 |
| 5 | 63066733 | rs1503519 | 1.058 | -0.8 | -1.487 | 0.101 | -0.016 | -0.085 | -0.085 | 7.442 | 0.13 | 0.34 | 0.07 | 0 | 0.1 | 0.03 |
| 5 | 63068130 | rs429752 | 1.969 | -0.577 | -2.119 | 0.336 | -0.019 | -0.037 | -0.037 | 0 | 0.07 | 0 | 0.07 | 0.15 | 0.08 | 0.07 |
| 5 | 63069566 | rs11959818 | 1.622 | -0.491 | -1.95 | 0.224 | -0.139 | -0.112 | -0.112 | 0 | 0.23 | 0.02 | 0.33 | 0.35 | 0.35 | 0.32 |
| 5 | 63074648 | rs1978467 | 0.575 | -1.065 | -1.862 | -0.292 | 0.003 | 0.192 | 0.192 | 0 | 0.5 | 0.17 | 0.52 | 0.27 | 0.56 | 0.4 |
| 5 | 63075583 | rs9291798 | 2.685 | -0.923 | -1.593 | -0.372 | 0.008 | 0.052 | 0.052 | 0 | 0.13 | 0.03 | 0.15 | 0.02 | 0.14 | 0.07 |
| 5 | 63077057 | rs1478497 | -0.359 | -1.23 | -1.081 | -0.236 | -0.007 | 0.012 | 0.012 | 0 | 0.6 | 0.45 | 0.62 | 0.5 | 0.69 | 0.51 |
| 5 | 63077992 | rs10061598 | -0.501 | -1.249 | -0.877 | -0.245 | -0.002 | 0.003 | 0.003 | 0 | 0.62 | 0.58 | 0.65 | 0.5 | 0.69 | 0.51 |
| 5 | 63079104 | rs1955277 | 0.075 | -1.126 | -0.925 | -0.17 | 0.012 | 0 | 0 | 0 | 0.5 | 0.38 | 0.58 | 0.49 | 0.59 | 0.47 |
| 5 | 63082296 | rs6895545 | 0.163 | -0.59 | -0.219 | 0.149 | -0.011 | 0.015 | 0.015 | 0.812 | 0.15 | 0.27 | 0.15 | 0.12 | 0.13 | 0.13 |
| 5 | 63084896 | rs165538 | 1.378 | -0.273 | -0.622 | 0.082 | -0.01 | -0.033 | -0.033 | 0 | 0.07 | 0 | 0.04 | 0.16 | 0.05 | 0.06 |
| 5 | 63096804 | rs1382381 | -0.298 | -0.133 | -0.376 | -0.108 | 0.019 | -0.014 | -0.014 | 0.519 | 0.38 | 0.51 | 0.38 | 0.34 | 0.42 | 0.32 |
| 5 | 63098552 | rs10075443 | -0.354 | -0.164 | -0.548 | -0.109 | 0.005 | -0.006 | -0.006 | 0.065 | 0.39 | 0.56 | 0.38 | 0.34 | 0.42 | 0.32 |
| 5 | 63098804 | rs12658738 | 0.588 | 0.516 | -1.191 | 0.452 | -0.097 | -0.066 | -0.066 | 0.088 | 0.23 | 0.04 | 0.31 | 0.34 | 0.34 | 0.3 |
| 5 | 63114051 | rs1478491 | -0.374 | -0.297 | -0.599 | 0.294 | -0.05 | -0.013 | -0.013 | 0 | 0.22 | 0.49 | 0.15 | 0.13 | 0.13 | 0.09 |
| 5 | 63116471 | rs13179925 | 0.872 | 0.407 | -1.229 | 0.411 | -0.093 | -0.042 | -0.042 | 0 | 0.2 | 0.02 | 0.18 | 0.26 | 0.29 | 0.32 |
| 5 | 63117788 | rs358824 | 0.815 | 0.265 | -1.058 | 0.359 | -0.018 | -0.027 | -0.027 | 0.01 | 0.07 | 0.01 | 0.11 | 0.17 | 0.12 | 0.11 |
| 5 | 63131024 | rs2216721 | -0.194 | -0.361 | -0.87 | -0.427 | 0.034 | 0.046 | 0.046 | 2.137 | 0.39 | 0.69 | 0.32 | 0.13 | 0.28 | 0.17 |
| 5 | 63147022 | rs17181344 | 0.265 | -0.203 | -0.582 | -0.366 | -0.007 | 0.009 | 0.009 | 0 | 0.16 | 0.1 | 0.15 | 0.08 | 0.18 | 0.1 |
| 5 | 63156739 | rs10060813 | 0.924 | 0.184 | -1.046 | -0.185 | -0.14 | -0.082 | -0.082 | 0.474 | 0.24 | 0.02 | 0.29 | 0.3 | 0.34 | 0.35 |
| 5 | 63173373 | rs2059204 | 1.212 | -0.109 | -0.962 | -0.234 | -0.101 | -0.075 | -0.075 | 0 | 0.22 | 0.02 | 0.26 | 0.23 | 0.3 | 0.32 |
| 5 | 63185329 | rs16892364 | -0.047 | -0.019 | -0.288 | -0.095 | 0.006 | -0.019 | -0.019 | 0 | 0.18 | 0.24 | 0.16 | 0.09 | 0.19 | 0.1 |
| 5 | 63193047 | rs1580312 | -0.785 | -0.354 | -1.407 | -0.281 | -0.067 | -0.096 | -0.096 | 1.534 | 0.68 | 0.91 | 0.63 | 0.38 | 0.61 | 0.53 |
| 5 | 63207016 | rs7721270 | -0.637 | -0.71 | -0.407 | -0.399 | 0.008 | -0.035 | -0.035 | 0.652 | 0.2 | 0.51 | 0.19 | 0.09 | 0.2 | 0.1 |
| 5 | 63216232 | rs17484586 | 1.112 | -0.677 | -1.007 | -0.318 | 0.077 | 0.06 | 0.06 | 6.113 | 0.18 | 0.01 | 0.13 | 0.06 | 0.09 | 0.08 |
| 5 | 63216847 | rs16892399 | -0.994 | 0.492 | -0.274 | 0.031 | 0.003 | -0.01 | -0.01 | 0 | 0.24 | 0.71 | 0.22 | 0.13 | 0.23 | 0.13 |
| 5 | 63217612 | rs358523 | -1.273 | 0.728 | 0.416 | 0.434 | 0.079 | 0.103 | 0.103 | 1.637 | 0.15 | 0.46 | 0.18 | 0.14 | 0.2 | 0.05 |
| 5 | 63221842 | rs7735151 | 0.216 | 0.316 | 0.578 | -0.277 | -0.019 | -0.097 | -0.097 | 0 | 0.31 | 0.46 | 0.27 | 0.1 | 0.23 | 0.17 |
| 5 | 63226159 | rs358528 | -0.59 | 0.373 | 0.323 | -0.182 | -0.032 | 0.024 | 0.024 | 6.134 | 0.64 | 0.76 | 0.55 | 0.82 | 0.46 | 0.57 |
| 5 | 63229897 | rs358532 | -0.17 | 0.166 | 1.113 | 0.049 | -0.001 | 0 | 0 | 0 | 0.59 | 0.43 | 0.48 | 0.75 | 0.45 | 0.52 |
| 5 | 63249424 | rs4521432 | -0.662 | 0.141 | 1.07 | 0.039 | -0.077 | 0.045 | 0.045 | 6.328 | 0.65 | 0.81 | 0.55 | 0.82 | 0.46 | 0.57 |
| 5 | 63250851 | rs1364043 | 0.375 | 0.095 | 1.114 | 0.026 | 0.02 | -0.036 | -0.036 | 0 | 0.31 | 0.2 | 0.24 | 0.67 | 0.21 | 0.35 |
| 5 | 63256018 | rs6449693 | -0.678 | 0.158 | 1.195 | 0.11 | -0.035 | 0.03 | 0.03 | 6.852 | 0.65 | 0.77 | 0.55 | 0.83 | 0.46 | 0.57 |
| 5 | 63261329 | rs10042486 | -0.678 | 0.153 | 1.251 | 0.125 | -0.032 | 0.027 | 0.027 | 6.852 | 0.65 | 0.76 | 0.55 | 0.82 | 0.46 | 0.57 |
| 5 | 63265891 | rs13361335 | 0.753 | -0.215 | 0.43 | 0.016 | 0.002 | 0.01 | 0.01 | 0 | 0.15 | 0.12 | 0.12 | 0.66 | 0.09 | 0.2 |
| 5 | 63288619 | rs1364041 | 0.739 | -0.141 | 0.419 | 0.178 | 0.008 | -0.003 | -0.003 | 0.056 | 0.15 | 0.1 | 0.2 | 0.64 | 0.09 | 0.18 |
| 5 | 63294697 | rs382098 | -0.931 | -0.047 | 0.62 | -0.065 | -0.024 | 0.022 | 0.022 | 5.395 | 0.65 | 0.74 | 0.64 | 0.83 | 0.47 | 0.55 |
| 5 | 63299929 | rs440776 | -0.66 | -0.131 | 1.012 | -0.143 | -0.032 | -0.006 | -0.006 | 0 | 0.34 | 0.44 | 0.36 | 0.72 | 0.22 | 0.38 |
| 5 | 63300089 | rs421321 | -0.913 | -0.03 | 0.957 | -0.176 | -0.022 | 0.02 | 0.02 | 5.143 | 0.65 | 0.73 | 0.65 | 0.83 | 0.48 | 0.55 |
| 5 | 63305263 | rs10077379 | 1.716 | -0.036 | 0.877 | -0.363 | 0 | 0.007 | 0.007 | 8.025 | 0.26 | 0.26 | 0.23 | 0.11 | 0.19 | 0.12 |

| | | | | | | | | | | | | | | | | |
|---|---|---|---|---|---|---|---|---|---|---|---|---|---|---|---|---|
| 5 | 63309892 | rs10071371 | -0.169 | -0.133 | 0.865 | -0.373 | 0.007 | 0.027 | 0.027 | 2.148 | 0.49 | 0.59 | 0.43 | 0.15 | 0.38 | 0.33 |
| 5 | 63331109 | rs356562 | -0.11 | 0.534 | 0.862 | -0.462 | 0.001 | -0.005 | -0.005 | 0.554 | 0.48 | 0.54 | 0.42 | 0.15 | 0.38 | 0.33 |
| 5 | 63358227 | rs356611 | 0.791 | 0.448 | 0.989 | -0.488 | -0.001 | -0.012 | -0.012 | 0 | 0.14 | 0.2 | 0.11 | 0.04 | 0.11 | 0.15 |
| 5 | 63369409 | rs441080 | 0.068 | 0.439 | 0.987 | -0.486 | 0.001 | -0.005 | -0.005 | 1.144 | 0.52 | 0.46 | 0.58 | 0.85 | 0.62 | 0.67 |
| 5 | 63394447 | rs1494629 | 0.095 | 0.539 | 0.984 | -0.447 | -0.001 | -0.015 | -0.015 | 0 | 0.46 | 0.52 | 0.43 | 0.16 | 0.37 | 0.34 |
| 5 | 63441140 | rs12520115 | 1.123 | 0.557 | 0.982 | -0.441 | -0.003 | 0.009 | 0.009 | 1.045 | 0.12 | 0.09 | 0.1 | 0 | 0.11 | 0.15 |
| 5 | 63452939 | rs10036420 | 1.836 | 0.668 | 0.952 | -0.504 | 0.001 | -0.002 | -0.002 | 0.5 | 0.25 | 0.28 | 0.21 | 0.11 | 0.16 | 0.1 |
| 5 | 63454211 | rs356640 | -0.222 | 0.72 | 0.895 | -0.494 | 0.014 | 0.041 | 0.041 | 1.681 | 0.48 | 0.67 | 0.44 | 0.2 | 0.37 | 0.35 |
| 5 | 63493203 | rs13355554 | 1.781 | 0.73 | 0.645 | -0.448 | -0.005 | -0.011 | -0.011 | 0 | 0.25 | 0.29 | 0.21 | 0.1 | 0.16 | 0.1 |
| 5 | 63535701 | rs6449703 | 0.896 | 0.454 | 0.736 | -0.354 | -0.001 | -0.004 | -0.004 | 0 | 0.33 | 0.38 | 0.31 | 0.16 | 0.26 | 0.2 |
| 5 | 63569990 | rs13174761 | 1.025 | 0.768 | 0.238 | -0.69 | -0.042 | 0.053 | 0.053 | 0 | 0.09 | 0.01 | 0.07 | 0 | 0.13 | 0.01 |
| 5 | 63587011 | rs6859427 | 1.402 | 0.534 | 0.473 | -0.294 | 0.003 | 0.009 | 0.009 | 5.95 | 0.3 | 0.29 | 0.28 | 0.11 | 0.25 | 0.13 |
| 5 | 63624426 | rs2162804 | 1.097 | 0.52 | -0.014 | 0.232 | -0.01 | -0.048 | -0.048 | 0.548 | 0.11 | 0 | 0.12 | 0.22 | 0.09 | 0.08 |
| 5 | 63640062 | rs7446357 | 1.368 | 0.598 | 0.019 | -0.484 | -0.028 | 0.047 | 0.047 | 0.787 | 0.1 | 0.02 | 0.09 | 0 | 0.1 | 0.14 |
| 5 | 63650110 | rs6449720 | -0.778 | 0.606 | 0.127 | -0.385 | -0.029 | 0.032 | 0.032 | 1.751 | 0.63 | 0.78 | 0.66 | 0.81 | 0.5 | 0.54 |
| 5 | 63665861 | rs10064715 | 0.599 | 0.809 | 0.206 | -0.493 | -0.019 | -0.026 | -0.026 | 1.197 | 0.35 | 0.57 | 0.33 | 0.18 | 0.26 | 0.2 |
| 5 | 63671056 | rs10038839 | 0.648 | 0.76 | 0.128 | -0.483 | -0.023 | -0.03 | -0.03 | 0 | 0.34 | 0.55 | 0.32 | 0.18 | 0.26 | 0.2 |
| 5 | 63676204 | rs7724701 | 0.479 | 0.779 | -0.128 | -0.56 | 0 | -0.026 | -0.026 | 2.397 | 0.13 | 0.13 | 0.11 | 0 | 0.11 | 0.15 |
| 5 | 63708388 | rs6859806 | 1.004 | 0.791 | -0.196 | -0.582 | 0.001 | 0.012 | 0.012 | 2.36 | 0.32 | 0.31 | 0.29 | 0.1 | 0.27 | 0.18 |
| 5 | 63731862 | rs10045003 | -1.266 | 0.8 | -0.157 | -0.471 | 0.002 | 0.009 | 0.009 | 1.371 | 0.69 | 0.69 | 0.71 | 0.88 | 0.74 | 0.83 |
| 5 | 63732305 | rs4557363 | -0.083 | 0.77 | -0.131 | -0.44 | 0.02 | 0.064 | 0.064 | 0.213 | 0.52 | 0.3 | 0.56 | 0.8 | 0.63 | 0.63 |
| 5 | 63741058 | rs271699 | 0.457 | 0.401 | -0.185 | -0.546 | -0.025 | 0.056 | 0.056 | 0 | 0.21 | 0.33 | 0.21 | 0.72 | 0.18 | 0.22 |
| 5 | 63744056 | rs271710 | 0.654 | 0.691 | -0.76 | -0.809 | -0.1 | 0.03 | 0.03 | 3.749 | 0.31 | 0.09 | 0.39 | 0.18 | 0.47 | 0.45 |
| 5 | 63745097 | rs2012899 | -1.551 | 0.627 | -0.645 | -0.898 | 0.002 | -0.009 | -0.009 | 0 | 0.67 | 0.71 | 0.79 | 0.92 | 0.76 | 0.83 |
| 5 | 63753398 | rs34403320 | 0.435 | 0.604 | -0.241 | -0.529 | -0.008 | 0.032 | 0.032 | 0.919 | 0.12 | 0.09 | 0.17 | 0 | 0.11 | 0.13 |
| 5 | 63769508 | rs1579841 | -0.579 | 0.681 | -0.165 | -0.356 | -0.005 | -0.003 | -0.003 | 0 | 0.37 | 0.61 | 0.4 | 0.75 | 0.3 | 0.38 |
| 5 | 63790221 | rs13183954 | -0.433 | 1.536 | -0.585 | -0.236 | -0.001 | -0.005 | -0.005 | 0 | 0.06 | 0.07 | 0.07 | 0.01 | 0.1 | 0.06 |
| 5 | 63793107 | rs17798594 | 1.014 | 1.737 | -0.599 | -0.352 | -0.002 | -0.008 | -0.008 | 1.384 | 0.06 | 0.05 | 0.15 | 0 | 0.07 | 0.08 |
| 5 | 63793417 | rs17190940 | 0.466 | 1.733 | -1.079 | -0.529 | -0.067 | -0.031 | -0.031 | 0 | 0.26 | 0.1 | 0.35 | 0.22 | 0.39 | 0.37 |
| 5 | 63795467 | rs13186658 | -0.512 | 1.668 | -0.98 | -0.497 | 0.003 | -0.013 | -0.013 | 12.235 | 0.06 | 0.11 | 0.07 | 0.01 | 0.1 | 0.06 |
| 5 | 63805283 | rs16892778 | 0.428 | 1.158 | -0.743 | -0.74 | -0.041 | 0.022 | 0.022 | 0 | 0.29 | 0.4 | 0.23 | 0.72 | 0.21 | 0.26 |
| 5 | 63807078 | rs272629 | -0.293 | 1.382 | -0.778 | -0.648 | -0.015 | 0.009 | 0.009 | 0.021 | 0.34 | 0.34 | 0.54 | 0.23 | 0.47 | 0.46 |
| 5 | 63810722 | rs272627 | 0.174 | 1.174 | -0.578 | -0.374 | -0.012 | -0.002 | -0.002 | 0 | 0.28 | 0.24 | 0.38 | 0.23 | 0.39 | 0.38 |
| 5 | 63815993 | rs272620 | 0.331 | 0.862 | -0.661 | -0.331 | -0.053 | -0.016 | -0.016 | 0 | 0.26 | 0.08 | 0.28 | 0.2 | 0.32 | 0.38 |
| 5 | 63827509 | rs272637 | -1.382 | 0.768 | -0.685 | -0.435 | -0.048 | 0.142 | 0.142 | 0 | 0.64 | 0.85 | 0.69 | 0.96 | 0.63 | 0.72 |
| 5 | 63831964 | rs6870654 | -0.365 | 0.38 | -0.044 | 0.486 | -0.042 | 0.024 | 0.024 | 0 | 0.32 | 0.42 | 0.32 | 0.69 | 0.23 | 0.29 |
| 5 | 63841538 | rs4700636 | 0.134 | 0.413 | -0.655 | 0.9 | 0.027 | -0.111 | -0.111 | 0 | 0.72 | 0.92 | 0.79 | 0.46 | 0.78 | 0.72 |
| 5 | 63850157 | rs4404633 | 0.261 | 0.147 | -0.846 | 1 | 0.052 | -0.088 | -0.088 | 1.473 | 0.7 | 0.92 | 0.79 | 0.6 | 0.77 | 0.76 |
| 5 | 63851332 | rs7734083 | 1.265 | 0.67 | -0.542 | 0.704 | -0.013 | 0.058 | 0.058 | 0 | 0.45 | 0.35 | 0.36 | 0.23 | 0.41 | 0.38 |
| 5 | 63855700 | rs16892838 | 0.383 | 1.142 | -0.278 | 1.094 | -0.063 | -0.09 | -0.09 | 0 | 0.16 | 0.04 | 0.33 | 0.31 | 0.27 | 0.35 |
| 5 | 63860337 | rs3797049 | -0.918 | 1.065 | 0.247 | 1.006 | 0.146 | 0.147 | 0.147 | 0.334 | 0.05 | 0.37 | 0.08 | 0.05 | 0.1 | 0.03 |
| 5 | 63861495 | rs10038212 | -0.191 | 1.144 | 0.162 | 1.074 | 0.066 | 0.124 | 0.124 | 0.693 | 0.21 | 0.41 | 0.4 | 0.42 | 0.35 | 0.37 |
| 5 | 63863233 | rs16892850 | -0.563 | 0.935 | 0.363 | 0.7 | -0.008 | 0.035 | 0.035 | 0 | 0.27 | 0.41 | 0.36 | 0.36 | 0.3 | 0.41 |
| 5 | 63863622 | rs16892853 | -0.445 | 0.94 | 0.445 | 0.665 | 0.002 | 0.025 | 0.025 | 0 | 0.26 | 0.34 | 0.37 | 0.34 | 0.34 | 0.38 |
| 5 | 63863679 | rs10036915 | -0.063 | 1.713 | 0.121 | 0.301 | 0.014 | -0.018 | -0.018 | 0 | 0.09 | 0.11 | 0.13 | 0.01 | 0.19 | 0.14 |
| 5 | 63865046 | rs1814198 | -0.268 | 1.33 | -0.192 | 0.058 | 0.015 | -0.035 | -0.035 | 0 | 0.76 | 0.85 | 0.86 | 0.61 | 0.83 | 0.78 |
| 5 | 63867550 | rs10805387 | -1.112 | 1.252 | -0.043 | -0.523 | 0.03 | 0.021 | 0.021 | 2.844 | 0.49 | 0.66 | 0.7 | 0.57 | 0.66 | 0.65 |
| 5 | 63872962 | rs16892876 | 0.845 | 1.253 | 0.141 | -0.493 | 0.002 | 0.008 | 0.008 | 1.824 | 0.13 | 0.19 | 0.2 | 0.2 | 0.14 | 0.12 |
| 5 | 63877325 | rs7730085 | 1.22 | 1.113 | 0.145 | -0.457 | 0.011 | 0.085 | 0.085 | 0 | 0.42 | 0.24 | 0.34 | 0.23 | 0.33 | 0.25 |
| 5 | 63880395 | rs7708672 | -0.247 | 1.042 | 0.419 | -0.404 | -0.026 | 0.071 | 0.071 | 0 | 0.76 | 0.59 | 0.82 | 0.61 | 0.84 | 0.79 |
| 5 | 63881011 | rs3914039 | 0.334 | 1.093 | 0.457 | -0.424 | 0.005 | 0.018 | 0.018 | 0 | 0.7 | 0.62 | 0.71 | 0.6 | 0.68 | 0.66 |
| 5 | 63883738 | rs1509134 | -0.148 | 0.935 | 0.428 | -0.402 | -0.007 | 0.026 | 0.026 | 0 | 0.29 | 0.42 | 0.38 | 0.34 | 0.34 | 0.4 |
| 5 | 63888627 | rs10044070 | -0.205 | 0.92 | 0.298 | -0.409 | -0.004 | 0.003 | 0.003 | 0 | 0.3 | 0.57 | 0.4 | 0.34 | 0.34 | 0.4 |
| 5 | 63888835 | rs10805388 | 0.209 | 0.956 | 0.184 | -0.392 | 0 | 0.006 | 0.006 | 0 | 0.26 | 0.23 | 0.36 | 0.34 | 0.34 | 0.4 |
| 5 | 63891146 | rs993706 | 0.268 | 0.991 | -0.117 | -0.346 | -0.075 | -0.071 | -0.071 | 0 | 0.26 | 0.07 | 0.36 | 0.35 | 0.39 | 0.42 |
| 5 | 63892333 | rs16892941 | -0.046 | 0.359 | 0.08 | -0.7 | -0.044 | -0.1 | -0.1 | 0 | 0.15 | 0.4 | 0.08 | 0.02 | 0.07 | 0.09 |
| 5 | 63893302 | rs4432837 | -0.303 | 0.435 | 0.227 | -0.559 | -0.026 | -0.044 | -0.044 | 0 | 0.17 | 0.32 | 0.11 | 0.07 | 0.11 | 0.08 |
| 5 | 63901304 | rs10060705 | 1.48 | 0.553 | -0.215 | -0.504 | 0.005 | 0.053 | 0.053 | 0.936 | 0.53 | 0.21 | 0.48 | 0.36 | 0.53 | 0.54 |
| 5 | 63905068 | rs889248 | 0.007 | 0.557 | 0.092 | -0.056 | 0.002 | 0.054 | 0.054 | 0 | 0.87 | 0.84 | 0.85 | 0.61 | 0.84 | 0.78 |
| 5 | 63906901 | rs1559261 | 0.49 | 0.624 | -0.104 | -0.191 | 0 | 0 | 0 | 0 | 0.24 | 0.2 | 0.34 | 0.34 | 0.34 | 0.44 |
| 5 | 63911811 | rs7445190 | 1.149 | 0.033 | 0.311 | -0.08 | -0.008 | 0.023 | 0.023 | 0 | 0.71 | 0.66 | 0.76 | 0.54 | 0.72 | 0.7 |
| 5 | 63913687 | rs918604 | 0.908 | 0.351 | 0.275 | 0.21 | 0.007 | 0.194 | 0.194 | 0.238 | 0.84 | 0.43 | 0.81 | 0.61 | 0.84 | 0.77 |
| 5 | 63917733 | rs3857267 | 0.484 | 0.372 | -0.284 | 0.223 | -0.004 | 0.052 | 0.052 | 0 | 0.87 | 0.81 | 0.93 | 0.63 | 0.91 | 0.87 |
| 5 | 63919584 | rs876354 | 2.221 | -0.112 | -0.876 | -0.491 | 0.025 | 0.065 | 0.065 | 0 | 0.31 | 0.16 | 0.21 | 0.03 | 0.23 | 0.12 |
| 5 | 63923514 | rs2366901 | 0.042 | 0.271 | -0.881 | -0.417 | 0.003 | 0 | 0 | 0 | 0.34 | 0.56 | 0.48 | 0.34 | 0.42 | 0.56 |
| 5 | 63926199 | rs2548253 | -0.114 | 0.334 | -0.964 | -0.389 | -0.008 | -0.021 | -0.021 | 0 | 0.17 | 0.08 | 0.2 | 0.21 | 0.18 | 0.13 |
| 5 | 63926883 | rs10050523 | -1.181 | 0.249 | -0.512 | 0.044 | 0.001 | -0.002 | -0.002 | 0 | 0.42 | 0.55 | 0.54 | 0.37 | 0.52 | 0.63 |
| 5 | 63928254 | rs6886740 | 0.113 | 0.149 | -0.127 | 0.215 | 0.04 | 0.032 | 0.032 | 0 | 0.27 | 0.5 | 0.4 | 0.31 | 0.34 | 0.49 |
| 5 | 63932450 | rs2366778 | -0.288 | 0.117 | -0.458 | 0.257 | -0.001 | -0.046 | -0.046 | 0 | 0.85 | 0.92 | 0.86 | 0.63 | 0.84 | 0.82 |

| | | | | | | | | | | | | | | | | |
|---|---|---|---|---|---|---|---|---|---|---|---|---|---|---|---|---|
| 5 | 63942398 | rs10461497 | 1.455 | 0.128 | -0.46 | 0.186 | -0.008 | 0.072 | 0.072 | 0 | 0.53 | 0.28 | 0.49 | 0.3 | 0.49 | 0.31 |
| 5 | 63946919 | rs10074802 | 0.79 | 0.145 | -0.683 | 0.243 | -0.105 | -0.088 | -0.088 | 0 | 0.24 | 0.06 | 0.31 | 0.32 | 0.35 | 0.51 |
| 5 | 63952264 | rs6877406 | -0.596 | 0.474 | -0.349 | 0.331 | 0.011 | -0.068 | -0.068 | 0 | 0.83 | 0.91 | 0.85 | 0.63 | 0.84 | 0.81 |
| 5 | 63954744 | rs4700646 | 1.628 | 0.405 | -0.413 | 0.238 | -0.019 | 0.08 | 0.08 | 0 | 0.44 | 0.25 | 0.45 | 0.23 | 0.44 | 0.25 |
| 5 | 63959834 | rs10491469 | -0.334 | 0.665 | -0.593 | 0.084 | 0.006 | 0.006 | 0.006 | 0 | 0.37 | 0.53 | 0.39 | 0.39 | 0.4 | 0.54 |
| 5 | 63960187 | rs10940001 | -0.334 | 0.665 | -0.593 | 0.05 | 0.006 | 0.004 | 0.004 | 0 | 0.37 | 0.53 | 0.39 | 0.37 | 0.4 | 0.54 |
| 5 | 63972658 | rs6891856 | -0.897 | 0.322 | -1.045 | -1.375 | -0.011 | 0.118 | 0.118 | 0 | 0.58 | 0.77 | 0.6 | 0.91 | 0.54 | 0.74 |
| 5 | 63978255 | rs2610245 | -0.488 | 0.597 | -0.757 | -1.203 | 0.015 | 0.115 | 0.115 | 0 | 0.5 | 0.35 | 0.55 | 0.9 | 0.52 | 0.73 |
| 5 | 63982806 | rs275828 | -0.972 | 0.586 | -0.346 | -0.899 | 0.009 | -0.052 | -0.052 | 0 | 0.49 | 0.91 | 0.59 | 0.53 | 0.53 | 0.53 |
| 5 | 63995050 | rs7716441 | -0.76 | 0.561 | 0.385 | -0.926 | 0.059 | 0.06 | 0.06 | 4.034 | 0.06 | 0.51 | 0.06 | 0.02 | 0.01 | 0 |
| 5 | 64000612 | rs10461498 | 0.218 | 0.999 | 0.339 | -1.042 | 0 | 0.016 | 0.016 | 0 | 0.31 | 0.35 | 0.41 | 0.5 | 0.28 | 0.46 |
| 5 | 64018803 | rs1459196 | 0.225 | 1.066 | 0.701 | -1.009 | 0 | 0.018 | 0.018 | 0 | 0.31 | 0.34 | 0.4 | 0.5 | 0.28 | 0.45 |
| 5 | 64019841 | rs10075967 | -1.254 | 1.089 | 0.62 | -1.02 | -0.018 | -0.069 | -0.069 | 0 | 0.49 | 0.93 | 0.59 | 0.53 | 0.53 | 0.55 |
| 5 | 64039871 | rs13165926 | 0.637 | 1.172 | 0.731 | -0.911 | 0.001 | -0.032 | -0.032 | 0 | 0.27 | 0.08 | 0.35 | 0.33 | 0.23 | 0.42 |
| 5 | 64041741 | rs188027 | -1.27 | 1.146 | 0.671 | -0.917 | 0.009 | -0.043 | -0.043 | 0 | 0.49 | 0.91 | 0.59 | 0.53 | 0.53 | 0.55 |
| 5 | 64048451 | rs10471639 | 0.043 | 0.846 | 0.682 | -0.745 | 0.001 | 0.019 | 0.019 | 0 | 0.31 | 0.35 | 0.4 | 0.5 | 0.28 | 0.45 |
| 5 | 64064943 | rs1363953 | -0.866 | 0.952 | 0.475 | -0.951 | 0.02 | -0.086 | -0.086 | 0 | 0.27 | 0.85 | 0.2 | 0.54 | 0.18 | 0.34 |
| 5 | 64067752 | rs2042329 | -0.72 | 0.709 | 0.805 | -1.318 | -0.001 | -0.01 | -0.01 | 0 | 0.59 | 0.44 | 0.59 | 0.9 | 0.59 | 0.83 |
| 5 | 64072172 | rs10434542 | -0.73 | 0.657 | 1.726 | -0.59 | 0.001 | 0.072 | 0.072 | 0 | 0.51 | 0.45 | 0.51 | 0.85 | 0.53 | 0.77 |
| 5 | 64076999 | rs1037768 | -0.821 | 0.659 | 1.712 | -0.589 | -0.001 | 0.033 | 0.033 | 0 | 0.52 | 0.53 | 0.51 | 0.85 | 0.53 | 0.77 |
| 5 | 64093658 | rs6868760 | 0.024 | 0.688 | 1.15 | -0.469 | 0.04 | 0.089 | 0.089 | 0 | 0.28 | 0.54 | 0.22 | 0.51 | 0.34 | 0.41 |
| 5 | 64096170 | rs6449760 | 0.105 | 0.685 | 1.198 | -0.47 | 0.029 | 0.065 | 0.065 | 0 | 0.28 | 0.45 | 0.21 | 0.51 | 0.34 | 0.41 |
| 5 | 64098899 | rs4518332 | -0.161 | 0.746 | 1.114 | -0.497 | 0.016 | -0.035 | -0.035 | 0 | 0.91 | 0.88 | 0.9 | 1 | 0.89 | 0.95 |
| 5 | 64099769 | rs725127 | -0.104 | 0.739 | 1.143 | -0.503 | 0.015 | 0.034 | 0.034 | 0 | 0.3 | 0.64 | 0.22 | 0.52 | 0.34 | 0.42 |
| 5 | 64099813 | rs725128 | 0.538 | 0.808 | 1.146 | -0.5 | 0.012 | 0.115 | 0.115 | 0 | 0.24 | 0.5 | 0.17 | 0.49 | 0.24 | 0.34 |
| 5 | 64106918 | rs10059206 | 0.318 | -0.049 | 1.332 | -0.426 | 0.039 | -0.037 | -0.037 | 0 | 0.23 | 0.08 | 0.29 | 0.34 | 0.16 | 0.36 |
| 5 | 64118974 | rs10043634 | -1.086 | -0.046 | 1.325 | -0.425 | 0.013 | -0.016 | -0.016 | 0.188 | 0.94 | 0.92 | 0.92 | 1 | 0.93 | 0.96 |
| 5 | 64119275 | rs12521901 | 1.126 | -0.041 | 1.141 | -0.4 | -0.004 | 0.101 | 0.101 | 0 | 0.39 | 0.15 | 0.38 | 0.13 | 0.39 | 0.18 |
| 5 | 64119615 | rs16893170 | 0.668 | -0.042 | 1.103 | -0.347 | 0.014 | 0.031 | 0.031 | 0 | 0.27 | 0.36 | 0.21 | 0.51 | 0.34 | 0.4 |
| 5 | 64156668 | rs11749214 | 0.73 | -0.04 | 0.716 | -0.346 | -0.008 | -0.012 | -0.012 | 0.944 | 0.26 | 0.23 | 0.19 | 0.51 | 0.34 | 0.4 |
| 5 | 64166882 | rs7733752 | 0.477 | -0.036 | 0.69 | -0.344 | 0.032 | 0.071 | 0.071 | 0 | 0.28 | 0.49 | 0.21 | 0.51 | 0.34 | 0.4 |
| 5 | 64167550 | rs10050395 | 1.046 | -0.034 | 0.548 | -0.345 | 0.026 | -0.086 | -0.086 | 0 | 0.17 | 0.02 | 0.27 | 0.34 | 0.14 | 0.31 |
| 5 | 64179886 | rs6897941 | 0.216 | 0.297 | 0.72 | -0.14 | 0.027 | 0.061 | 0.061 | 0 | 0.3 | 0.6 | 0.23 | 0.53 | 0.35 | 0.41 |
| 5 | 64188526 | rs12657887 | 0.146 | 0.301 | 0.61 | -0.151 | 0.032 | 0.08 | 0.08 | 0 | 0.3 | 0.55 | 0.23 | 0.52 | 0.34 | 0.4 |
| 5 | 64214987 | rs10471642 | 0.286 | 0.047 | 0.565 | 0.051 | 0.033 | -0.03 | -0.03 | 0 | 0.22 | 0.09 | 0.29 | 0.33 | 0.15 | 0.36 |
| 5 | 64225533 | rs13179963 | 0.055 | 0.536 | -0.98 | 0.446 | 0.004 | 0.003 | 0.003 | 0 | 0.62 | 0.35 | 0.55 | 0.65 | 0.72 | 0.57 |
| 5 | 64238731 | rs11738728 | -0.424 | 1.488 | -0.814 | 0.871 | 0.01 | 0.014 | 0.014 | 0 | 0.27 | 0.29 | 0.19 | 0.45 | 0.37 | 0.4 |
| 5 | 64239523 | rs1347246 | 0.308 | 0.499 | -0.506 | 0.191 | -0.039 | -0.039 | -0.039 | 0 | 0.36 | 0.45 | 0.29 | 0.26 | 0.23 | 0.43 |
| 5 | 64260661 | rs1433611 | 0.164 | 1.147 | -0.245 | -0.258 | -0.006 | 0.05 | 0.05 | 0 | 0.42 | 0.26 | 0.39 | 0.3 | 0.41 | 0.45 |
| 5 | 64267595 | rs2278351 | 0.756 | 1.524 | -0.203 | 0.115 | -0.008 | 0.073 | 0.073 | 0 | 0.27 | 0.41 | 0.18 | 0.47 | 0.24 | 0.33 |
| 5 | 64272107 | rs1311245 | 0.384 | 1.569 | -0.255 | 0.132 | -0.001 | -0.01 | -0.01 | 0 | 0.46 | 0.47 | 0.44 | 0.34 | 0.43 | 0.49 |
| 5 | 64273394 | rs1309580 | 1.085 | 1.602 | -0.309 | 0.026 | 0.002 | 0 | 0 | 0 | 0.11 | 0.12 | 0.12 | 0.06 | 0.17 | 0.05 |
| 5 | 64278005 | rs1309572 | 0.536 | 1.589 | -0.247 | 0.02 | -0.013 | 0.061 | 0.061 | 0 | 0.42 | 0.2 | 0.38 | 0.3 | 0.41 | 0.45 |
| 5 | 64280127 | rs4632749 | 1.25 | 1.56 | -0.28 | -0.088 | 0.032 | 0.07 | 0.07 | 0 | 0.31 | 0.08 | 0.27 | 0.24 | 0.24 | 0.4 |
| 5 | 64290779 | rs1309544 | -0.334 | 1.615 | -0.354 | -0.132 | -0.06 | 0.022 | 0.022 | 0 | 0.73 | 0.92 | 0.62 | 0.8 | 0.66 | 0.81 |
| 5 | 64293570 | rs1309540 | 0.829 | 1.744 | -0.574 | -0.304 | -0.007 | 0.001 | 0.001 | 0 | 0.16 | 0.03 | 0.28 | 0.11 | 0.16 | 0.11 |
| 5 | 64306758 | rs2968210 | -0.701 | 1.805 | -0.399 | -0.315 | -0.022 | 0.008 | 0.008 | 0 | 0.89 | 0.94 | 0.89 | 0.9 | 0.8 | 0.93 |
| 5 | 64311542 | rs266580 | 0.869 | 1.621 | -0.096 | -0.375 | -0.002 | 0.033 | 0.033 | 0 | 0.42 | 0.29 | 0.38 | 0.29 | 0.4 | 0.43 |
| 5 | 64313887 | rs166475 | 1.07 | 1.629 | -0.186 | -0.377 | 0.011 | 0.004 | 0.004 | 0 | 0.16 | 0.04 | 0.27 | 0.11 | 0.14 | 0.1 |
| 5 | 64314865 | rs2915439 | -0.692 | 1.626 | -0.187 | -0.383 | -0.016 | 0.008 | 0.008 | 0 | 0.89 | 0.93 | 0.89 | 0.91 | 0.8 | 0.93 |
| 5 | 64317015 | rs66360 | -0.974 | 1.451 | -0.046 | -0.284 | 0.001 | 0.008 | 0.008 | 0 | 0.84 | 0.96 | 0.72 | 0.89 | 0.85 | 0.89 |
| 5 | 64322549 | rs11960938 | 0.289 | 1.629 | 0.429 | -0.019 | -0.018 | 0.042 | 0.042 | 0 | 0.31 | 0.48 | 0.22 | 0.51 | 0.26 | 0.39 |
| 5 | 64328615 | rs266585 | 0.862 | 1.765 | 0.191 | 0.017 | 0.02 | -0.006 | -0.006 | 0 | 0.11 | 0.16 | 0.12 | 0.05 | 0.17 | 0.05 |
| 5 | 64340029 | rs12656778 | 0.249 | 1.281 | 0.274 | 0.103 | -0.006 | 0.025 | 0.025 | 0 | 0.29 | 0.37 | 0.18 | 0.5 | 0.25 | 0.39 |
| 5 | 64340353 | rs166473 | -0.411 | 1.835 | 0.354 | -0.08 | 0.048 | 0.004 | 0.004 | 0 | 0.2 | 0.29 | 0.21 | 0.21 | 0.31 | 0.11 |
| 5 | 64347621 | rs266590 | -0.064 | 1.135 | 0.112 | -0.937 | -0.014 | -0.028 | -0.028 | 0 | 0.63 | 0.51 | 0.53 | 0.27 | 0.65 | 0.52 |
| 5 | 64353312 | rs434882 | -0.86 | 1.319 | -0.103 | -1.356 | 0.01 | -0.054 | -0.054 | 0 | 0.22 | 0.54 | 0.18 | 0.08 | 0.23 | 0.08 |
| 5 | 64356942 | rs439327 | 0.149 | 1.681 | -0.359 | -1.563 | 0 | -0.015 | -0.015 | 0 | 0.48 | 0.44 | 0.4 | 0.26 | 0.46 | 0.45 |
| 5 | 64357890 | rs1159105 | 1.714 | 1.744 | -0.657 | -1.511 | -0.06 | 0.059 | 0.059 | 0.807 | 0.28 | 0.03 | 0.24 | 0.19 | 0.24 | 0.38 |
| 5 | 64368149 | rs264742 | 0.127 | 1.749 | -0.59 | -1.507 | -0.019 | 0.019 | 0.019 | 0 | 0.15 | 0.08 | 0.13 | 0.08 | 0.22 | 0.07 |
| 5 | 64371037 | rs10055645 | 1.686 | 1.673 | -0.661 | -1.491 | -0.061 | 0.057 | 0.057 | 0.678 | 0.28 | 0.02 | 0.24 | 0.19 | 0.24 | 0.38 |
| 5 | 64372897 | rs16893364 | 0.505 | 1.62 | -0.18 | -1.462 | 0.002 | -0.063 | -0.063 | 0 | 0.19 | 0.27 | 0.15 | 0 | 0.2 | 0.07 |
| 5 | 64373264 | rs264750 | -0.61 | 1.666 | -0.042 | -1.239 | 0.04 | 0.035 | 0.035 | 0 | 0.2 | 0.37 | 0.3 | 0.22 | 0.27 | 0.1 |
| 5 | 64376348 | rs10434530 | 0.933 | 0.919 | 0.039 | -1.226 | -0.003 | 0.006 | 0.006 | 0 | 0.31 | 0.35 | 0.42 | 0.36 | 0.27 | 0.43 |
| 5 | 64379812 | rs10042857 | 2.632 | 0.901 | -0.332 | -1.283 | -0.025 | 0.01 | 0.01 | 0.014 | 0.2 | 0.04 | 0.23 | 0.19 | 0.19 | 0.37 |
| 5 | 64387048 | rs155338 | -1.26 | 0.808 | -0.558 | -1.104 | -0.042 | -0.149 | -0.149 | 0.07 | 0.72 | 0.92 | 0.73 | 0.49 | 0.7 | 0.6 |
| 5 | 64389333 | rs264768 | -0.19 | 0.791 | -0.308 | -1.148 | -0.007 | 0.034 | 0.034 | 0 | 0.35 | 0.29 | 0.25 | 0.11 | 0.38 | 0.15 |
| 5 | 64391283 | rs972653 | 0.809 | 0.895 | -0.486 | -0.868 | -0.001 | -0.014 | -0.014 | 0 | 0.08 | 0.04 | 0.15 | 0.17 | 0.07 | 0.06 |
| 5 | 64392588 | rs4700661 | 0.809 | 0.895 | -0.496 | -0.874 | -0.001 | -0.017 | -0.017 | 0 | 0.08 | 0.04 | 0.15 | 0.17 | 0.07 | 0.06 |

| | | | | | | | | | | | | | | | | |
|---|---|---|---|---|---|---|---|---|---|---|---|---|---|---|---|---|
| 5 | 64400653 | rs6872582 | -1.053 | 0.909 | -0.699 | -0.962 | -0.032 | -0.136 | -0.136 | 0 | 0.7 | 0.91 | 0.72 | 0.47 | 0.68 | 0.6 |
| 5 | 64403371 | rs11742800 | 0.149 | 0.747 | -0.458 | -1.071 | 0.014 | -0.02 | -0.02 | 0.685 | 0.07 | 0.23 | 0.06 | 0.01 | 0.05 | 0.02 |
| 5 | 64417806 | rs7710076 | -0.26 | 0.724 | -0.996 | -1.158 | -0.024 | 0.039 | 0.039 | 0 | 0.55 | 0.68 | 0.61 | 0.36 | 0.48 | 0.51 |
| 5 | 64422082 | rs6892539 | -1.484 | 0.603 | -1.698 | -0.876 | -0.031 | -0.155 | -0.155 | 0.085 | 0.7 | 0.92 | 0.73 | 0.47 | 0.69 | 0.6 |
| 5 | 64422902 | rs17205854 | 0.953 | 0.622 | -1.022 | -0.999 | 0.002 | 0.039 | 0.039 | 0 | 0.17 | 0.04 | 0.12 | 0 | 0.16 | 0.07 |
| 5 | 64433400 | rs17206232 | -0.448 | 0.56 | -0.939 | -0.581 | -0.013 | -0.003 | -0.003 | 1.346 | 0.1 | 0.02 | 0.09 | 0.09 | 0.1 | 0.06 |
| 5 | 64440243 | rs1363902 | 1.478 | 0.445 | -1.867 | -0.839 | -0.105 | 0.018 | 0.018 | 3.269 | 0.33 | 0.03 | 0.33 | 0.24 | 0.33 | 0.36 |
| 5 | 64444922 | rs16893435 | 0.974 | 0.357 | -1.795 | -0.836 | -0.026 | -0.013 | -0.013 | 0 | 0.07 | 0 | 0.06 | 0.06 | 0.07 | 0.03 |
| 5 | 64445708 | rs17812189 | 1.364 | 0.044 | -2.027 | -1.067 | 0.022 | 0.084 | 0.084 | 0 | 0.14 | 0 | 0.06 | 0 | 0.09 | 0.06 |
| 5 | 64447660 | rs6879867 | -0.524 | -0.2 | -1.053 | -1.077 | 0.033 | 0.03 | 0.03 | 7.79 | 0.05 | 0.58 | 0.05 | 0 | 0 | 0 |
| 5 | 64447777 | rs17206779 | 1.212 | -0.023 | -1.344 | -0.947 | -0.036 | 0.003 | 0.003 | 3.237 | 0.43 | 0.17 | 0.52 | 0.39 | 0.49 | 0.43 |
| 5 | 64449497 | rs10940016 | -1.428 | -0.109 | -1.103 | -0.766 | -0.038 | -0.121 | -0.121 | 0.117 | 0.74 | 0.89 | 0.73 | 0.46 | 0.68 | 0.57 |
| 5 | 64450308 | rs1423352 | 0.476 | -0.035 | -0.458 | -0.759 | 0.007 | -0.012 | -0.012 | 0.499 | 0.61 | 0.47 | 0.62 | 0.37 | 0.52 | 0.5 |
| 5 | 64450343 | rs17812416 | 0.448 | 0.169 | -0.532 | -0.56 | 0.013 | 0.008 | 0.008 | 0 | 0.1 | 0.14 | 0.09 | 0.09 | 0.16 | 0.07 |
| 5 | 64459691 | rs2367239 | 0.748 | -0.002 | -0.592 | -0.827 | -0.006 | 0.045 | 0.045 | 0 | 0.48 | 0.35 | 0.55 | 0.39 | 0.44 | 0.43 |
| 5 | 64467770 | rs16893468 | 0.758 | -0.317 | -0.099 | -0.816 | -0.026 | -0.065 | -0.065 | 0 | 0.16 | 0.24 | 0.09 | 0 | 0.09 | 0.06 |
| 5 | 64483759 | rs16893475 | -0.561 | -0.385 | 0.094 | -0.836 | -0.003 | -0.009 | -0.009 | 0 | 0.26 | 0.27 | 0.16 | 0.11 | 0.18 | 0.13 |
| 5 | 64484802 | rs1862571 | -1.177 | -0.323 | -0.296 | -0.713 | 0 | -0.097 | -0.097 | 0 | 0.72 | 0.87 | 0.75 | 0.53 | 0.71 | 0.52 |
| 5 | 64485563 | rs10805394 | 1.697 | -0.138 | -0.103 | -0.792 | -0.045 | 0.025 | 0.025 | 3.102 | 0.34 | 0.2 | 0.48 | 0.31 | 0.37 | 0.33 |
| 5 | 64491123 | rs10044007 | 1.508 | -0.154 | 0.348 | -0.81 | 0 | -0.002 | -0.002 | 0 | 0.37 | 0.37 | 0.49 | 0.32 | 0.37 | 0.33 |
| 5 | 64495058 | rs1895431 | 1.486 | 0.008 | 0.542 | -0.76 | -0.012 | 0 | 0 | 3.064 | 0.29 | 0.19 | 0.42 | 0.32 | 0.26 | 0.37 |
| 5 | 64496326 | rs10940018 | 0.27 | 0.039 | 0.755 | -0.75 | 0.039 | 0.018 | 0.018 | 0 | 0.09 | 0.15 | 0.1 | 0.1 | 0.19 | 0.07 |
| 5 | 64499782 | rs6864345 | -1.151 | -0.253 | 0.751 | -0.815 | -0.024 | -0.067 | -0.067 | 0.257 | 0.6 | 0.82 | 0.67 | 0.52 | 0.58 | 0.55 |
| 5 | 64505977 | rs7728957 | 0.424 | -0.24 | 0.751 | -0.808 | 0.029 | 0.02 | 0.02 | 0 | 0.09 | 0.14 | 0.09 | 0.1 | 0.17 | 0.07 |
| 5 | 64506320 | rs16893539 | -0.533 | -0.265 | 0.843 | -0.82 | -0.014 | -0.025 | -0.025 | 0 | 0.19 | 0.25 | 0.15 | 0.1 | 0.15 | 0.11 |
| 5 | 64509707 | rs6871603 | 1.432 | -0.213 | 0.904 | -0.772 | 0 | -0.001 | -0.001 | 2.32 | 0.31 | 0.33 | 0.43 | 0.32 | 0.26 | 0.37 |
| 5 | 64512523 | rs6449775 | -1.307 | -0.464 | 0.857 | -0.814 | -0.004 | -0.026 | -0.026 | 0 | 0.28 | 0.4 | 0.24 | 0.2 | 0.32 | 0.18 |
| 5 | 64522124 | rs11746271 | 0.384 | -0.023 | 1.189 | -0.641 | 0.072 | 0.033 | 0.033 | 0 | 0.06 | 0.16 | 0.08 | 0.06 | 0.16 | 0.06 |
| 5 | 64525521 | rs11747988 | 1.191 | 0.024 | 1.14 | -0.652 | -0.001 | 0.002 | 0.002 | 2.452 | 0.29 | 0.21 | 0.41 | 0.31 | 0.25 | 0.37 |
| 5 | 64529878 | rs1593038 | -0.405 | -0.027 | 1.069 | -0.549 | -0.049 | -0.015 | -0.015 | 0 | 0.5 | 0.85 | 0.6 | 0.48 | 0.48 | 0.49 |
| 5 | 64531723 | rs6449777 | -0.37 | -0.07 | 1.251 | -0.546 | 0.122 | 0.064 | 0.064 | 0.247 | 0.08 | 0.33 | 0.09 | 0.06 | 0.16 | 0.06 |
| 5 | 64541034 | rs9291832 | -0.889 | -0.154 | 1.17 | -0.549 | -0.086 | -0.057 | -0.057 | 0.049 | 0.51 | 0.91 | 0.6 | 0.48 | 0.48 | 0.49 |
| 5 | 64542430 | rs7719056 | -0.868 | -0.092 | 1.49 | -0.518 | 0.113 | 0.059 | 0.059 | 1.433 | 0.09 | 0.47 | 0.1 | 0.06 | 0.15 | 0.06 |
| 5 | 64545633 | rs11749052 | -0.318 | -0.031 | 1.227 | -0.49 | 0.117 | 0.065 | 0.065 | 0.146 | 0.08 | 0.3 | 0.1 | 0.06 | 0.15 | 0.06 |
| 5 | 64547588 | rs16893582 | 0.185 | -0.032 | 1.008 | -0.483 | 0.006 | 0 | 0 | 0 | 0.1 | 0.08 | 0.08 | 0.11 | 0.07 | 0.06 |
| 5 | 64548634 | rs12514074 | 0.603 | -0.034 | 1.066 | -0.515 | 0.014 | 0.002 | 0.002 | 0.456 | 0.36 | 0.42 | 0.49 | 0.37 | 0.41 | 0.42 |
| 5 | 64551463 | rs16893612 | -0.546 | 0.001 | 0.988 | -0.47 | 0.019 | 0.008 | 0.008 | 0.246 | 0.17 | 0.27 | 0.17 | 0.16 | 0.23 | 0.12 |
| 5 | 64565261 | rs10057851 | -0.075 | -0.034 | 0.863 | -0.497 | 0.001 | 0 | 0 | 0 | 0.46 | 0.53 | 0.58 | 0.47 | 0.48 | 0.49 |
| 5 | 64582052 | rs2287904 | -0.323 | 0.001 | 0.369 | -0.509 | -0.014 | 0.007 | 0.007 | 0 | 0.48 | 0.77 | 0.6 | 0.46 | 0.48 | 0.47 |
| 5 | 64585621 | rs1547454 | -1.097 | 0.381 | 0.48 | -0.443 | 0.015 | 0.006 | 0.006 | 0.544 | 0.17 | 0.25 | 0.17 | 0.16 | 0.23 | 0.12 |
| 5 | 64625512 | rs2307121 | 0.48 | 0.235 | -0.27 | -0.68 | -0.046 | 0.087 | 0.087 | 0 | 0.3 | 0.08 | 0.31 | 0.17 | 0.3 | 0.39 |
| 5 | 64631395 | rs7713110 | -1.136 | 0.085 | 0.302 | -0.703 | 0.013 | 0.095 | 0.095 | 0.274 | 0.5 | 0.79 | 0.48 | 0.39 | 0.48 | 0.53 |
| 5 | 64640161 | rs10066284 | -0.937 | 0.09 | 0.284 | -0.7 | 0.011 | 0.099 | 0.099 | 0.214 | 0.5 | 0.81 | 0.47 | 0.39 | 0.48 | 0.54 |
| 5 | 64640912 | rs2640717 | 1.514 | 0.061 | -0.042 | -0.606 | -0.074 | -0.122 | -0.122 | 0 | 0.38 | 0.04 | 0.46 | 0.53 | 0.39 | 0.36 |
| 5 | 64654109 | rs11739829 | -1.365 | 0.26 | 0.921 | -0.611 | 0.009 | 0.002 | 0.002 | 3.838 | 0.19 | 0.65 | 0.17 | 0.23 | 0.17 | 0.15 |
| 5 | 64659076 | rs7729401 | -0.395 | 0.47 | 0.948 | -0.598 | 0.049 | -0.023 | -0.023 | 0 | 0.07 | 0.19 | 0.07 | 0.02 | 0.12 | 0.09 |
| 5 | 64659167 | rs4700674 | 0.069 | 0.46 | 1.094 | -0.596 | 0 | 0 | 0 | 0.232 | 0.07 | 0.07 | 0.06 | 0.12 | 0.05 | 0.03 |
| 5 | 64665569 | rs6899042 | -0.426 | 0.524 | 1.023 | -0.578 | 0.052 | -0.022 | -0.022 | 0 | 0.07 | 0.19 | 0.07 | 0.02 | 0.12 | 0.09 |
| 5 | 64679283 | rs6867518 | -1.113 | 0.425 | 0.525 | -0.526 | 0.009 | 0.001 | 0.001 | 0.033 | 0.09 | 0.14 | 0.07 | 0.1 | 0.12 | 0.11 |
| 5 | 64680454 | rs11950228 | -0.778 | 0.44 | 0.517 | -0.535 | 0.01 | -0.016 | -0.016 | 0.198 | 0.08 | 0.13 | 0.06 | 0.02 | 0.12 | 0.09 |
| 5 | 64688741 | rs7708972 | -0.803 | 0.39 | 0.457 | -0.718 | 0.014 | 0.108 | 0.108 | 0.929 | 0.49 | 0.76 | 0.45 | 0.28 | 0.48 | 0.53 |
| 5 | 64691405 | rs6449783 | -0.474 | 0.45 | -0.053 | -0.527 | 0.008 | -0.01 | -0.01 | 0 | 0.08 | 0.12 | 0.06 | 0.03 | 0.12 | 0.09 |
| 5 | 64697866 | rs10042191 | -0.619 | 0.341 | -0.23 | -0.565 | 0.02 | 0.138 | 0.138 | 0.189 | 0.51 | 0.84 | 0.49 | 0.29 | 0.51 | 0.54 |
| 5 | 64709775 | rs6886919 | -0.9 | 0.274 | 0.485 | -0.681 | 0.082 | 0.038 | 0.038 | 1.838 | 0.11 | 0.46 | 0.1 | 0.11 | 0.12 | 0.12 |
| 5 | 64715661 | rs786703 | 1.573 | 0.007 | 0.091 | -0.75 | -0.068 | -0.092 | -0.092 | 5.396 | 0.35 | 0.03 | 0.43 | 0.62 | 0.35 | 0.37 |
| 5 | 64719601 | rs9291838 | 0.123 | 0.111 | 0.385 | -0.757 | -0.006 | 0.034 | 0.034 | 0.357 | 0.4 | 0.37 | 0.39 | 0.18 | 0.39 | 0.41 |
| 5 | 64725524 | rs10072030 | -0.729 | 0.115 | 0.313 | -0.755 | 0.015 | 0.151 | 0.151 | 0.609 | 0.51 | 0.83 | 0.49 | 0.29 | 0.51 | 0.52 |
| 5 | 64729075 | rs10940034 | 0.707 | 0.061 | 0.179 | -0.886 | -0.046 | 0.122 | 0.122 | 0.155 | 0.31 | 0.12 | 0.31 | 0.08 | 0.34 | 0.38 |
| 5 | 64734702 | rs7728131 | -1.059 | 0.06 | 0.183 | -0.87 | 0.001 | 0.141 | 0.141 | 0.879 | 0.52 | 0.89 | 0.5 | 0.29 | 0.51 | 0.52 |
| 5 | 64745925 | rs2131587 | -1.049 | 0.056 | 0.195 | -0.88 | 0.001 | 0.146 | 0.146 | 0.875 | 0.52 | 0.89 | 0.5 | 0.29 | 0.51 | 0.52 |
| 5 | 64746627 | rs10471661 | 0.563 | 0.03 | 0.169 | -0.919 | -0.044 | 0.121 | 0.121 | 0.128 | 0.31 | 0.12 | 0.31 | 0.08 | 0.34 | 0.38 |
| 5 | 64753321 | rs7708044 | -0.238 | 0.114 | 0.182 | -0.866 | -0.011 | 0.071 | 0.071 | 0.276 | 0.41 | 0.33 | 0.38 | 0.18 | 0.39 | 0.41 |
| 5 | 64759531 | rs397400 | 0.315 | 0.184 | 0.026 | -0.693 | 0 | -0.001 | -0.001 | 0 | 0.07 | 0.09 | 0.07 | 0.1 | 0.05 | 0.03 |
| 5 | 64759675 | rs386188 | -0.15 | 0.231 | 0.08 | -0.802 | -0.02 | 0.078 | 0.078 | 0.373 | 0.39 | 0.26 | 0.39 | 0.19 | 0.39 | 0.4 |
| 5 | 64761316 | rs245525 | -0.455 | 0.07 | 0.408 | -0.775 | 0.009 | 0.016 | 0.016 | 0 | 0.5 | 0.55 | 0.47 | 0.22 | 0.52 | 0.5 |
| 5 | 64762465 | rs245526 | -1.38 | 0.142 | 0.962 | -0.653 | 0.071 | 0.061 | 0.061 | 0 | 0.08 | 0.37 | 0.06 | 0.07 | 0.05 | 0.05 |
| 5 | 64763225 | rs245528 | -0.786 | 0.055 | -0.359 | -0.868 | -0.013 | 0.086 | 0.086 | 1.3 | 0.45 | 0.79 | 0.43 | 0.33 | 0.43 | 0.51 |
| 5 | 64772879 | rs11740656 | -0.344 | 0.178 | -0.016 | -0.454 | -0.025 | 0.027 | 0.027 | 0.014 | 0.05 | 0.01 | 0.05 | 0 | 0.08 | 0.03 |
| 5 | 64773576 | rs13155773 | -0.566 | -0.096 | 0.33 | -0.31 | -0.031 | 0.034 | 0.034 | 1.506 | 0.07 | 0.09 | 0.05 | 0.13 | 0.03 | 0.05 |

| | | | | | | | | | | | | | | | |
|---|---|---|---|---|---|---|---|---|---|---|---|---|---|---|---|
| 5 | 64778944 | rs33394 | -0.549 | -0.109 | 0.404 | -0.625 | -0.002 | -0.001 | -0.001 | 0.866 | 0.56 | 0.49 | 0.47 | 0.33 | 0.59 | 0.58 |
| 5 | 64814175 | rs11952932 | -0.151 | -0.439 | 0.632 | -0.74 | -0.04 | 0.003 | 0.003 | 0 | 0.08 | 0.13 | 0.04 | 0.09 | 0.02 | 0.11 |
| 5 | 64842156 | rs11746096 | 0.039 | -0.162 | 0.716 | -1.203 | 0.024 | -0.037 | -0.037 | 0 | 0.68 | 0.45 | 0.75 | 0.81 | 0.66 | 0.75 |
| 5 | 64842825 | rs6895641 | 0.877 | 0.031 | 0.544 | -1.267 | -0.005 | -0.002 | -0.002 | 0 | 0.41 | 0.39 | 0.47 | 0.68 | 0.35 | 0.37 |
| 5 | 64850432 | rs17828812 | 0.954 | -0.154 | 0.869 | -1.284 | 0.002 | 0.004 | 0.004 | 1.276 | 0.06 | 0.08 | 0.07 | 0.08 | 0.06 | 0.04 |
| 5 | 64851737 | rs149514 | -0.411 | -0.157 | 0.896 | -1.29 | 0.004 | 0.005 | 0.005 | 0.221 | 0.51 | 0.51 | 0.41 | 0.24 | 0.54 | 0.54 |
| 5 | 64858315 | rs16893963 | 0.411 | -0.15 | 0.9 | -1.247 | -0.032 | 0.001 | 0.001 | 0 | 0.08 | 0.11 | 0.04 | 0.09 | 0.02 | 0.11 |
| 5 | 64858380 | rs10514984 | 0.926 | -0.085 | 0.83 | -1.228 | 0.003 | 0.004 | 0.004 | 1.197 | 0.06 | 0.08 | 0.07 | 0.08 | 0.06 | 0.04 |
| 5 | 64863688 | rs467362 | -1.618 | -0.157 | 0.941 | -1.272 | -0.004 | 0.004 | 0.004 | 1.716 | 0.64 | 0.63 | 0.51 | 0.41 | 0.62 | 0.68 |
| 5 | 64870687 | rs16894005 | 0.936 | -0.164 | 0.85 | -1.33 | -0.069 | 0.054 | 0.054 | 0 | 0.25 | 0.02 | 0.23 | 0.14 | 0.26 | 0.31 |
| 5 | 64875105 | rs37334 | 0.277 | -0.192 | 1.183 | -1.343 | 0.018 | -0.032 | -0.032 | 0 | 0.25 | 0.4 | 0.17 | 0.11 | 0.28 | 0.22 |
| 5 | 64881578 | rs27140 | -1.697 | -0.197 | 1.173 | -1.335 | -0.004 | 0.004 | 0.004 | 1.813 | 0.64 | 0.63 | 0.51 | 0.41 | 0.62 | 0.68 |
| 5 | 64881936 | rs27141 | -1.688 | -0.201 | 1.171 | -1.336 | -0.004 | 0.004 | 0.004 | 1.687 | 0.64 | 0.63 | 0.51 | 0.41 | 0.62 | 0.68 |
| 5 | 64909487 | rs16894099 | 0.304 | -0.06 | 1.165 | -1.349 | -0.034 | 0.047 | 0.047 | 0.475 | 0.31 | 0.1 | 0.3 | 0.22 | 0.32 | 0.35 |
| 5 | 64919618 | rs468308 | -1.761 | -0.056 | 1.376 | -1.345 | 0.003 | -0.03 | -0.03 | 0.001 | 0.66 | 0.8 | 0.52 | 0.41 | 0.62 | 0.68 |
| 5 | 64920803 | rs37254 | -1.692 | -0.038 | 1.262 | -1.337 | -0.004 | 0.005 | 0.005 | 1.813 | 0.64 | 0.63 | 0.51 | 0.41 | 0.62 | 0.68 |
| 5 | 64936304 | rs10057241 | 1.148 | -0.002 | 0.951 | -1.368 | -0.078 | 0.02 | 0.02 | 0 | 0.17 | 0.02 | 0.09 | 0.05 | 0.2 | 0.17 |
| 5 | 64954159 | rs28084 | -0.949 | -0.298 | 1.363 | -1.304 | 0.004 | -0.016 | -0.016 | 0 | 0.37 | 0.54 | 0.28 | 0.29 | 0.36 | 0.37 |
| 5 | 64956428 | rs37349 | 0.726 | -0.285 | 1.408 | -1.299 | -0.025 | 0.023 | 0.023 | 0 | 0.23 | 0.2 | 0.15 | 0.11 | 0.28 | 0.22 |
| 5 | 64965217 | rs13189159 | 0.449 | -0.294 | 1.169 | -1.283 | -0.005 | -0.001 | -0.001 | 0.02 | 0.08 | 0.05 | 0.03 | 0.09 | 0.02 | 0.11 |
| 5 | 64965757 | rs27394 | -1.496 | -0.245 | 1.104 | -1.283 | -0.004 | 0.002 | 0.002 | 1.573 | 0.63 | 0.61 | 0.5 | 0.42 | 0.62 | 0.68 |
| 5 | 64965900 | rs1549192 | 0.806 | -0.15 | 0.393 | -1.232 | -0.069 | 0.068 | 0.068 | 0 | 0.25 | 0.02 | 0.23 | 0.12 | 0.26 | 0.31 |
| 5 | 64968615 | rs28342 | -0.401 | -0.151 | 0.802 | -1.178 | 0 | -0.012 | -0.012 | 0 | 0.32 | 0.41 | 0.19 | 0.21 | 0.31 | 0.32 |
| 5 | 64976763 | rs1978468 | -0.205 | -0.33 | 0.626 | -1.22 | 0.018 | 0.047 | 0.047 | 0 | 0.34 | 0.16 | 0.27 | 0.23 | 0.28 | 0.44 |
| 5 | 64982321 | rs112314 | -1.541 | -0.152 | 0.657 | -1.115 | 0 | -0.002 | -0.002 | 0.023 | 0.65 | 0.7 | 0.52 | 0.42 | 0.62 | 0.68 |
| 5 | 65006234 | rs16894235 | 0.773 | -0.11 | 0.181 | -1.123 | -0.065 | 0.046 | 0.046 | 0 | 0.25 | 0.02 | 0.23 | 0.14 | 0.25 | 0.33 |
| 5 | 65010764 | rs972501 | -1.189 | -0.032 | 0.583 | -1.054 | -0.006 | 0.004 | 0.004 | 0.396 | 0.63 | 0.47 | 0.5 | 0.4 | 0.62 | 0.68 |
| 5 | 65013703 | rs10067229 | -0.179 | -0.045 | 0.228 | -0.763 | 0.006 | 0.015 | 0.015 | 0.014 | 0.36 | 0.22 | 0.25 | 0.24 | 0.34 | 0.47 |
| 5 | 65037451 | rs34989 | 0.016 | -0.026 | 0.551 | -0.79 | 0 | 0.009 | 0.009 | 0.063 | 0.33 | 0.25 | 0.25 | 0.17 | 0.33 | 0.43 |
| 5 | 65038375 | rs2247193 | 1.071 | 0.043 | 0.589 | -0.818 | 0 | -0.013 | -0.013 | 0 | 0.27 | 0.3 | 0.14 | 0.11 | 0.23 | 0.16 |
| 5 | 65051755 | rs40105 | 0.085 | -0.101 | 0.506 | -0.79 | 0.009 | 0.04 | 0.04 | 0.068 | 0.54 | 0.37 | 0.35 | 0.27 | 0.54 | 0.53 |
| 5 | 65052015 | rs1734262 | 0.119 | -0.109 | 0.446 | -0.793 | 0.012 | 0.068 | 0.068 | 0.125 | 0.54 | 0.28 | 0.33 | 0.27 | 0.54 | 0.53 |
| 5 | 65061915 | rs252642 | -0.621 | -0.144 | 0.195 | -0.779 | -0.001 | 0 | 0 | 0.165 | 0.63 | 0.61 | 0.4 | 0.34 | 0.58 | 0.62 |
| 5 | 65067041 | rs1301451 | 1.528 | -0.005 | -0.84 | -0.649 | -0.036 | -0.133 | -0.133 | 0.111 | 0.25 | 0.07 | 0.47 | 0.56 | 0.32 | 0.31 |
| 5 | 65073224 | rs34979 | -0.628 | -0.01 | -0.226 | -0.186 | -0.025 | -0.03 | -0.03 | 0 | 0.93 | 1 | 0.92 | 0.91 | 0.91 | 0.93 |
| 5 | 65077910 | rs1374060 | -0.615 | -0.223 | 0.153 | -0.264 | -0.009 | 0.012 | 0.012 | 0 | 0.1 | 0.26 | 0.06 | 0.07 | 0.05 | 0.09 |
| 5 | 65080091 | rs1309823 | -0.007 | 0.104 | -0.807 | -0.115 | -0.055 | -0.002 | -0.002 | 0 | 0.73 | 0.94 | 0.79 | 0.84 | 0.68 | 0.63 |
| 5 | 65083796 | rs6862886 | -1.637 | 0.036 | 0.243 | -0.044 | -0.01 | -0.021 | -0.021 | 0 | 0.18 | 0.35 | 0.17 | 0.13 | 0.16 | 0.3 |
| 5 | 65083865 | rs6862580 | -1.466 | 0.051 | -0.073 | -0.392 | -0.001 | -0.02 | -0.02 | 0.719 | 0.23 | 0.25 | 0.2 | 0.09 | 0.2 | 0.49 |
| 5 | 65086627 | rs7711077 | -0.515 | 0.072 | 0.878 | -0.165 | 0.038 | 0.07 | 0.07 | 0.056 | 0.09 | 0.21 | 0.13 | 0.13 | 0.11 | 0.21 |
| 5 | 65087420 | rs2441130 | 0.311 | -0.573 | 0.397 | -0.172 | -0.003 | -0.004 | -0.004 | 0 | 0.56 | 0.61 | 0.46 | 0.41 | 0.55 | 0.44 |
| 5 | 65088175 | rs6896574 | 0.027 | -1.019 | -0.242 | -0.592 | -0.038 | -0.059 | -0.059 | 2.427 | 0.25 | 0.57 | 0.12 | 0.21 | 0.14 | 0.19 |
| 5 | 65090545 | rs2591933 | 0.205 | -0.731 | -0.126 | -0.731 | 0 | 0.001 | 0.001 | 0 | 0.47 | 0.5 | 0.28 | 0.3 | 0.47 | 0.36 |
| 5 | 65093845 | rs10069699 | 0.736 | -0.852 | -0.047 | -0.768 | -0.01 | -0.028 | -0.028 | 0.439 | 0.18 | 0.34 | 0.09 | 0.14 | 0.11 | 0.14 |
| 5 | 65099626 | rs6893191 | -0.516 | -0.988 | -0.186 | -0.864 | -0.005 | -0.008 | -0.008 | 0 | 0.29 | 0.37 | 0.28 | 0.26 | 0.2 | 0.24 |
| 5 | 65100962 | rs2262994 | 2.21 | -0.973 | -0.291 | -0.881 | -0.074 | 0.036 | 0.036 | 2.741 | 0.24 | 0.07 | 0.18 | 0.17 | 0.35 | 0.2 |
| 5 | 65105622 | rs2561196 | -1.048 | -1.057 | -0.081 | -0.611 | 0.021 | -0.048 | -0.048 | 0 | 0.67 | 0.81 | 0.6 | 0.57 | 0.68 | 0.68 |
| 5 | 65110789 | rs895379 | 0.214 | -1.042 | -0.084 | -0.635 | 0.032 | 0.066 | 0.066 | 0.194 | 0.51 | 0.38 | 0.43 | 0.35 | 0.54 | 0.41 |
| 5 | 65115003 | rs2250861 | -0.141 | -1.052 | -0.171 | -0.572 | 0.036 | 0.011 | 0.011 | 0 | 0.56 | 0.75 | 0.4 | 0.44 | 0.58 | 0.57 |
| 5 | 65115288 | rs1542168 | -1.706 | -1.345 | -0.544 | -0.738 | -0.054 | -0.183 | -0.183 | 0.072 | 0.76 | 0.98 | 0.64 | 0.56 | 0.71 | 0.75 |
| 5 | 65118738 | rs6860508 | 0.225 | -1.227 | 0.206 | -0.586 | -0.035 | -0.054 | -0.054 | 0.204 | 0.09 | 0.21 | 0.05 | 0 | 0.03 | 0.06 |
| 5 | 65120027 | rs9291848 | -0.508 | -1.257 | 0.445 | -0.572 | 0.035 | 0.088 | 0.088 | 0.012 | 0.13 | 0.36 | 0.16 | 0.22 | 0.13 | 0.27 |
| 5 | 65120054 | rs1550818 | -0.106 | -1.18 | 0.479 | -0.299 | 0.042 | 0.129 | 0.129 | 0.019 | 0.11 | 0.28 | 0.16 | 0.25 | 0.12 | 0.25 |
| 5 | 65120144 | rs10096 | -2.372 | -1.367 | 0.389 | -0.512 | 0.051 | 0.016 | 0.016 | 2.847 | 0.37 | 0.77 | 0.4 | 0.34 | 0.26 | 0.44 |
| 5 | 65120598 | rs11740088 | 0.95 | -1.349 | 0.26 | -0.399 | -0.011 | -0.036 | -0.036 | 0 | 0.08 | 0 | 0.16 | 0.13 | 0.08 | 0.08 |
| 5 | 65121368 | rs3733657 | -1.748 | -1.352 | 0.539 | -0.394 | -0.067 | -0.043 | -0.043 | 0 | 0.27 | 0.42 | 0.25 | 0.21 | 0.14 | 0.22 |
| 5 | 65122967 | rs7723430 | -2.255 | -1.313 | 0.2 | -0.352 | 0.012 | 0.001 | 0.001 | 1.913 | 0.35 | 0.61 | 0.38 | 0.35 | 0.25 | 0.44 |
| 5 | 65125226 | rs1550819 | -0.796 | -1.295 | 0.292 | -0.371 | -0.007 | -0.006 | -0.006 | 0 | 0.19 | 0.18 | 0.21 | 0.13 | 0.12 | 0.17 |
| 5 | 65126029 | rs7711329 | -1.551 | -1.123 | 0.363 | -0.334 | -0.011 | -0.006 | -0.006 | 1.04 | 0.26 | 0.6 | 0.22 | 0.22 | 0.18 | 0.34 |
| 5 | 65127087 | rs11957292 | 0.52 | -1.149 | 0.073 | -0.448 | -0.004 | -0.001 | -0.001 | 0 | 0.37 | 0.33 | 0.34 | 0.36 | 0.47 | 0.45 |
| 5 | 65127219 | rs16894413 | -0.86 | -1.513 | 0.551 | -0.255 | -0.033 | -0.067 | -0.067 | 0 | 0.11 | 0.26 | 0.05 | 0 | 0.05 | 0.07 |
| 5 | 65128863 | rs7727072 | 0.578 | -1.401 | 0.466 | -0.2 | 0.016 | 0.029 | 0.029 | 0.009 | 0.1 | 0.19 | 0.13 | 0.14 | 0.13 | 0.22 |
| 5 | 65129485 | rs16894422 | 0.578 | -1.401 | 0.567 | -0.2 | 0.021 | 0.039 | 0.039 | 0.009 | 0.1 | 0.22 | 0.14 | 0.14 | 0.13 | 0.22 |
| 5 | 65130420 | rs16894427 | -0.89 | -1.439 | 0.444 | -0.286 | -0.019 | -0.026 | -0.026 | 0.112 | 0.22 | 0.32 | 0.13 | 0.22 | 0.14 | 0.26 |
| 5 | 65132797 | rs10805402 | 1.003 | -1.673 | -0.175 | -0.433 | -0.027 | 0.106 | 0.106 | 0.151 | 0.42 | 0.19 | 0.31 | 0.22 | 0.42 | 0.24 |
| 5 | 65132930 | rs16894435 | -0.108 | -1 | 0.473 | 0.005 | 0.031 | 0.068 | 0.068 | 0 | 0.15 | 0.33 | 0.16 | 0.22 | 0.16 | 0.3 |
| 5 | 65138341 | rs6866449 | -0.787 | -1.41 | 0.235 | 0.02 | -0.005 | -0.009 | -0.009 | 0.7 | 0.17 | 0.18 | 0.09 | 0.13 | 0.11 | 0.15 |
| 5 | 65142159 | rs2367844 | -1.046 | -1.278 | 0.353 | 0.185 | -0.024 | -0.027 | -0.027 | 0 | 0.18 | 0.28 | 0.11 | 0.14 | 0.11 | 0.16 |
| 5 | 65160065 | rs11948927 | -0.525 | -1.533 | -0.041 | 0.227 | 0.023 | -0.003 | -0.003 | 0.097 | 0.55 | 0.69 | 0.38 | 0.44 | 0.57 | 0.58 |

| | | | | | | | | | | | | | | | | |
|---|---|---|---|---|---|---|---|---|---|---|---|---|---|---|---|---|
| 5 | 65162023 | rs16894473 | 0.101 | -1.236 | 0.056 | -0.054 | 0.004 | 0.011 | 0.011 | 0 | 0.08 | 0.17 | 0.05 | 0.13 | 0.07 | 0.09 |
| 5 | 65163769 | rs16894475 | -1.021 | -1.234 | 0.004 | -0.054 | -0.019 | -0.028 | -0.028 | 0 | 0.18 | 0.26 | 0.11 | 0.13 | 0.11 | 0.17 |
| 5 | 65169440 | rs7712590 | -1.323 | -1.378 | -0.835 | 0.027 | 0.027 | 0.009 | 0.009 | 0 | 0.28 | 0.2 | 0.29 | 0.33 | 0.21 | 0.27 |
| 5 | 65171507 | rs784463 | -0.722 | -1.436 | -0.818 | -0.002 | 0.013 | 0.007 | 0.007 | 0 | 0.2 | 0.2 | 0.23 | 0.16 | 0.12 | 0.17 |
| 5 | 65177085 | rs10066424 | 0.442 | -1.37 | -1.132 | 0.041 | -0.023 | -0.031 | -0.031 | 0 | 0.11 | 0.03 | 0.12 | 0.14 | 0.12 | 0.25 |
| 5 | 65177890 | rs237953 | -1.446 | -1.529 | -0.998 | -0.051 | -0.013 | 0.033 | 0.033 | 1.273 | 0.64 | 0.55 | 0.59 | 0.56 | 0.65 | 0.7 |
| 5 | 65179001 | rs346091 | -0.722 | -1.125 | -0.724 | -0.091 | -0.002 | -0.005 | -0.005 | 0.031 | 0.53 | 0.51 | 0.48 | 0.42 | 0.52 | 0.45 |
| 5 | 65180258 | rs173193 | 0.068 | -0.939 | -0.996 | 0.006 | 0.006 | -0.011 | -0.011 | 0.242 | 0.11 | 0.1 | 0.2 | 0.16 | 0.09 | 0.11 |
| 5 | 65180299 | rs784576 | 0.795 | -0.888 | -0.964 | -0.159 | -0.06 | 0.041 | 0.041 | 0.359 | 0.32 | 0.12 | 0.23 | 0.26 | 0.41 | 0.29 |
| 5 | 65183502 | rs251315 | 0.886 | -0.926 | -0.968 | -0.214 | -0.059 | 0.044 | 0.044 | 0 | 0.31 | 0.11 | 0.23 | 0.25 | 0.4 | 0.29 |
| 5 | 65185421 | rs2113030 | 0.13 | -0.897 | -0.75 | -0.003 | 0.003 | 0.007 | 0.007 | 0 | 0.11 | 0.07 | 0.05 | 0.11 | 0.09 | 0.1 |
| 5 | 65185999 | rs251312 | -0.335 | -0.699 | -0.668 | 0.145 | 0 | 0.026 | 0.026 | 0.557 | 0.54 | 0.6 | 0.33 | 0.29 | 0.5 | 0.37 |
| 5 | 65191876 | rs251288 | 1.384 | -0.491 | -0.966 | 0.206 | -0.02 | 0.02 | 0.02 | 0.415 | 0.3 | 0.21 | 0.22 | 0.18 | 0.37 | 0.21 |
| 5 | 65194123 | rs251287 | 0.336 | 0.23 | -1.022 | 0.83 | 0 | 0.018 | 0.018 | 0.765 | 0.54 | 0.53 | 0.34 | 0.3 | 0.51 | 0.39 |
| 5 | 65195360 | rs784577 | -0.685 | 0.12 | -1.006 | 0.763 | -0.037 | -0.189 | -0.189 | 2.467 | 0.71 | 0.97 | 0.64 | 0.55 | 0.67 | 0.71 |
| 5 | 65203347 | rs4418055 | 0.035 | 0.156 | -0.525 | 0.794 | 0 | 0 | 0 | 0 | 0.43 | 0.35 | 0.54 | 0.53 | 0.48 | 0.59 |
| 5 | 65203799 | rs9291850 | -0.057 | 0.216 | -0.811 | 0.822 | 0 | 0 | 0 | 0.27 | 0.44 | 0.49 | 0.54 | 0.53 | 0.48 | 0.59 |
| 5 | 65206531 | rs2548469 | 0.767 | -0.255 | -0.794 | 1.35 | -0.004 | 0.017 | 0.017 | 0 | 0.83 | 0.8 | 0.7 | 0.53 | 0.9 | 0.9 |
| 5 | 65213108 | rs152999 | 0.897 | -0.379 | -0.004 | 1.288 | -0.036 | 0.103 | 0.103 | 0 | 0.83 | 0.61 | 0.68 | 0.53 | 0.91 | 0.9 |
| 5 | 65247893 | rs251608 | 1.531 | -0.795 | 0.874 | 0.902 | -0.045 | 0.076 | 0.076 | 0.02 | 0.77 | 0.41 | 0.66 | 0.54 | 0.86 | 0.84 |
| 5 | 65264090 | rs251295 | 0.376 | -0.174 | -0.725 | -0.151 | -0.125 | 0.043 | 0.043 | 0 | 0.29 | 0.06 | 0.28 | 0.17 | 0.41 | 0.18 |
| 5 | 65268670 | rs16894638 | 1.815 | -0.701 | -0.93 | -0.179 | 0.006 | -0.102 | -0.102 | 0 | 0.14 | 0 | 0.08 | 0.27 | 0.11 | 0.15 |
| 5 | 65288791 | rs27132 | 0.861 | 0.082 | 0.208 | 0.082 | -0.052 | 0.152 | 0.152 | 0 | 0.8 | 0.48 | 0.67 | 0.54 | 0.88 | 0.86 |
| 5 | 65321311 | rs3213837 | 0.483 | 0.325 | 0.031 | -0.036 | -0.057 | -0.008 | -0.008 | 0.264 | 0.09 | 0.01 | 0.17 | 0.09 | 0.15 | 0.23 |
| 5 | 65350374 | rs36303 | -0.471 | -0.164 | 0.236 | 0.228 | 0.008 | -0.045 | -0.045 | 0 | 0.14 | 0.09 | 0.3 | 0.46 | 0.12 | 0.14 |
| 5 | 65382500 | rs796728 | -1.207 | -0.057 | 0.033 | 0.325 | -0.041 | 0.15 | 0.15 | 0 | 0.2 | 0.52 | 0.34 | 0.46 | 0.14 | 0.15 |
| 5 | 65422236 | rs153387 | -0.022 | 0.125 | -0.671 | 0.458 | -0.003 | -0.032 | -0.032 | 0 | 0.08 | 0.04 | 0.07 | 0.33 | 0.09 | 0.07 |
| 5 | 65450196 | rs27076 | -1.847 | 0.024 | -0.802 | 0.203 | 0.035 | -0.033 | -0.033 | 0 | 0.44 | 0.87 | 0.52 | 0.59 | 0.38 | 0.57 |
| 5 | 65459360 | rs27588 | -1.801 | -0.404 | 0.579 | 0.917 | -0.042 | 0.084 | 0.084 | 0 | 0.26 | 0.57 | 0.35 | 0.46 | 0.19 | 0.19 |
| 5 | 65459773 | rs3763061 | -0.725 | -0.413 | 0.568 | 0.983 | 0.005 | -0.012 | -0.012 | 0 | 0.13 | 0.11 | 0.24 | 0.2 | 0.12 | 0.17 |
| 5 | 65464936 | rs36921 | -1.666 | -0.467 | 0.792 | 0.547 | -0.028 | 0.091 | 0.091 | 0 | 0.14 | 0.36 | 0.11 | 0.26 | 0.08 | 0.02 |
| 5 | 65469463 | rs27655 | -0.392 | -0.317 | 1.006 | 0.718 | 0.002 | 0.006 | 0.006 | 0 | 0.08 | 0.05 | 0.07 | 0.26 | 0.07 | 0.02 |
| 5 | 65499401 | rs253214 | -1.994 | -0.42 | 0.813 | 0.622 | -0.05 | 0.052 | 0.052 | 0 | 0.26 | 0.5 | 0.23 | 0.4 | 0.2 | 0.22 |
| 5 | 65503947 | rs253211 | -2.168 | -0.384 | 0.816 | 0.723 | -0.024 | 0.017 | 0.017 | 0 | 0.33 | 0.59 | 0.29 | 0.46 | 0.24 | 0.29 |
| 5 | 65519426 | rs253229 | 2.314 | -0.056 | 0.827 | 0.724 | -0.004 | 0 | 0 | 0 | 0.6 | 0.43 | 0.54 | 0.59 | 0.63 | 0.51 |
| 5 | 65522888 | rs103219 | 2.189 | -0.018 | 0.615 | 0.727 | 0.001 | -0.001 | -0.001 | 0.053 | 0.6 | 0.37 | 0.54 | 0.59 | 0.62 | 0.51 |
| 5 | 65525422 | rs253221 | -0.072 | 0.185 | 0.316 | 0.857 | -0.001 | 0.012 | 0.012 | 0 | 0.07 | 0.06 | 0.06 | 0.12 | 0.06 | 0.02 |
| 5 | 65526035 | rs253223 | -1.257 | 0.201 | 0.222 | 0.809 | 0.042 | 0.032 | 0.032 | 0 | 0.2 | 0.46 | 0.25 | 0.22 | 0.21 | 0.3 |
| 5 | 65527911 | rs4700098 | 0.902 | -0.131 | -0.593 | 1.192 | -0.005 | -0.039 | -0.039 | 0 | 0.83 | 0.97 | 0.81 | 0.8 | 0.84 | 0.81 |
| 5 | 65532167 | rs10940057 | -1.609 | -0.062 | -0.147 | 1.356 | -0.006 | 0.011 | 0.011 | 0 | 0.16 | 0.16 | 0.12 | 0.19 | 0.13 | 0.27 |
| 5 | 65533655 | rs29462 | 1.723 | -0.016 | -0.171 | 1.418 | 0.002 | -0.01 | -0.01 | 0 | 0.72 | 0.8 | 0.71 | 0.57 | 0.71 | 0.54 |
| 5 | 65533865 | rs29461 | 0.797 | 0.267 | -0.386 | 1.298 | 0.001 | -0.015 | -0.015 | 0 | 0.84 | 0.94 | 0.82 | 0.79 | 0.84 | 0.83 |
| 5 | 65534403 | rs29460 | 1.068 | 0.11 | 0.05 | 1.153 | -0.021 | 0.032 | 0.032 | 0 | 0.71 | 0.58 | 0.72 | 0.52 | 0.76 | 0.67 |
| 5 | 65537197 | rs11739182 | -0.974 | 1.154 | 0.246 | 0.709 | 0.04 | -0.009 | -0.009 | 0 | 0.12 | 0.18 | 0.13 | 0.08 | 0.18 | 0.24 |
| 5 | 65537947 | rs16895034 | -1.05 | 0.177 | 0.614 | 0.312 | -0.006 | -0.029 | -0.029 | 0 | 0.19 | 0.33 | 0.17 | 0.13 | 0.19 | 0.2 |
| 5 | 65538791 | rs187569 | 1.728 | -0.495 | 0.312 | -0.112 | -0.016 | 0.123 | 0.123 | 0 | 0.42 | 0.22 | 0.39 | 0.15 | 0.54 | 0.3 |
| 5 | 65539847 | rs29456 | 0.367 | -0.907 | 1.138 | -0.31 | -0.012 | -0.008 | -0.008 | 0 | 0.68 | 0.73 | 0.6 | 0.28 | 0.75 | 0.51 |
| 5 | 65540329 | rs735129 | -1.19 | -0.922 | 1.016 | -0.391 | -0.043 | -0.044 | -0.044 | 0 | 0.19 | 0.5 | 0.14 | 0.13 | 0.1 | 0.17 |
| 5 | 65541414 | rs7732890 | 0.943 | -1.148 | 1.267 | 0.283 | -0.002 | 0.006 | 0.006 | 0 | 0.78 | 0.73 | 0.76 | 0.57 | 0.85 | 0.66 |
| 5 | 65549315 | rs29437 | -0.292 | -1.128 | 1.176 | 0.301 | 0.002 | -0.002 | -0.002 | 0.018 | 0.88 | 0.9 | 0.85 | 0.68 | 0.92 | 0.82 |
| 5 | 65551304 | rs10461510 | -0.075 | -1.11 | 1.208 | 0.243 | -0.005 | 0.023 | 0.023 | 0.373 | 0.2 | 0.17 | 0.25 | 0.41 | 0.17 | 0.33 |
| 5 | 65567281 | rs4699861 | -0.122 | -1.112 | 1.207 | 0.261 | -0.004 | 0.021 | 0.021 | 0.657 | 0.2 | 0.17 | 0.25 | 0.41 | 0.17 | 0.33 |
| 5 | 65568906 | rs10072213 | 2.643 | -1.131 | 1.243 | 0.209 | -0.004 | 0.071 | 0.071 | 0 | 0.42 | 0.27 | 0.39 | 0.14 | 0.54 | 0.29 |
| 5 | 65572385 | rs7700366 | 0.103 | -0.844 | 1.208 | 0.423 | -0.014 | -0.044 | -0.044 | 0.144 | 0.08 | 0.24 | 0.05 | 0 | 0.02 | 0 |
| 5 | 65573132 | rs10471675 | -0.024 | -0.912 | 1.185 | 0.419 | -0.004 | -0.027 | -0.027 | 0 | 0.12 | 0.14 | 0.06 | 0.05 | 0.12 | 0.18 |
| 5 | 65574416 | rs4699862 | -2.023 | -0.921 | 1.084 | 0.428 | -0.013 | 0.012 | 0.012 | 0 | 0.39 | 0.44 | 0.41 | 0.41 | 0.35 | 0.43 |
| 5 | 65580717 | rs12655575 | -0.292 | -0.713 | 0.577 | 0.551 | 0.017 | -0.043 | -0.043 | 0.002 | 0.12 | 0.05 | 0.23 | 0.4 | 0.11 | 0.19 |
| 5 | 65583098 | rs12522102 | -1.65 | -0.77 | 0.889 | 0.321 | -0.008 | 0.023 | 0.023 | 0 | 0.33 | 0.38 | 0.38 | 0.42 | 0.35 | 0.45 |
| 5 | 65589342 | rs10515008 | 0.455 | -0.383 | 0.637 | 0.054 | 0.004 | -0.026 | -0.026 | 0 | 0.08 | 0.12 | 0.08 | 0 | 0.11 | 0.04 |
| 5 | 65590269 | rs1864037 | -0.657 | -0.553 | 0.544 | 0.343 | 0.001 | 0.014 | 0.014 | 0 | 0.15 | 0.09 | 0.1 | 0.04 | 0.18 | 0.26 |
| 5 | 65594860 | rs1428236 | -0.963 | -0.637 | 0.805 | 0.356 | -0.024 | -0.065 | -0.065 | 0 | 0.17 | 0.31 | 0.09 | 0.04 | 0.14 | 0.22 |
| 5 | 65596821 | rs16895073 | -0.403 | -0.394 | 0.208 | 0.627 | 0.001 | 0.028 | 0.028 | 0 | 0.05 | 0.12 | 0.09 | 0.05 | 0.03 | 0.08 |
| 5 | 65598729 | rs17251088 | -0.61 | -0.237 | -0.475 | 0.667 | 0.005 | 0.009 | 0.009 | 1.133 | 0.15 | 0.1 | 0.06 | 0.04 | 0.14 | 0.22 |
| 5 | 65599040 | rs7717523 | 0.217 | 0.567 | -0.612 | 1.161 | 0.001 | 0.003 | 0.003 | 1.238 | 0.9 | 0.91 | 0.85 | 0.73 | 0.86 | 0.83 |
| 5 | 65599360 | rs1366275 | 0.803 | -0.077 | -0.821 | 1.22 | 0.026 | -0.058 | -0.058 | 0 | 0.81 | 0.85 | 0.73 | 0.58 | 0.86 | 0.73 |
| 5 | 65600341 | rs2914030 | 0.329 | 0.22 | -0.813 | 0.522 | -0.003 | -0.003 | -0.003 | 0.67 | 0.58 | 0.56 | 0.64 | 0.63 | 0.64 | 0.52 |
| 5 | 65601617 | rs2968393 | -0.217 | 1.078 | -1.209 | 0.303 | -0.034 | -0.013 | -0.013 | 0.177 | 0.93 | 0.98 | 0.81 | 0.92 | 0.86 | 0.97 |
| 5 | 65603639 | rs10515005 | 0.01 | 1.024 | -1.69 | -0.154 | -0.005 | 0.043 | 0.043 | 0 | 0.11 | 0.04 | 0.05 | 0.04 | 0.1 | 0.16 |
| 5 | 65604718 | rs2801640 | 0.82 | 0.861 | -2.388 | -1.289 | -0.023 | 0.126 | 0.126 | 0 | 0.35 | 0.17 | 0.26 | 0.07 | 0.36 | 0.33 |

| | | | | | | | | | | | | | | | | |
|---|---|---|---|---|---|---|---|---|---|---|---|---|---|---|---|---|
| 5 | 65614238 | rs6893041 | 1.277 | 0.549 | -2.548 | -1.619 | 0.013 | 0.063 | 0.063 | 0 | 0.18 | 0.07 | 0.07 | 0.05 | 0.13 | 0.09 |
| 5 | 65614321 | rs2914029 | -1.548 | 0.706 | -2.5 | -1.488 | -0.038 | -0.02 | -0.02 | 0.369 | 0.93 | 0.98 | 0.8 | 0.91 | 0.85 | 0.95 |
| 5 | 65624559 | rs2561078 | 1.444 | 0.347 | -2.711 | -1.816 | -0.056 | 0.149 | 0.149 | 0 | 0.3 | 0.05 | 0.23 | 0.06 | 0.34 | 0.36 |
| 5 | 65629780 | rs981885 | -0.128 | -0.078 | -1.735 | -1.199 | 0.001 | 0.002 | 0.002 | 0 | 0.4 | 0.38 | 0.35 | 0.28 | 0.36 | 0.37 |
| 5 | 65630552 | rs10471680 | 0.09 | -0.136 | -1.676 | -1.243 | -0.002 | 0.052 | 0.052 | 0 | 0.37 | 0.16 | 0.32 | 0.28 | 0.35 | 0.35 |
| 5 | 65631005 | rs16895099 | 1.069 | -0.313 | -1.741 | -1.169 | -0.012 | -0.029 | -0.029 | 1.253 | 0.17 | 0.05 | 0.23 | 0.23 | 0.19 | 0.26 |
| 5 | 65635913 | rs1560132 | -0.292 | -0.178 | -1.235 | -1.062 | 0.012 | 0.022 | 0.022 | 0 | 0.44 | 0.39 | 0.39 | 0.3 | 0.37 | 0.37 |
| 5 | 65638377 | rs1465566 | 1.764 | -0.171 | -1.263 | -1.052 | -0.004 | -0.026 | -0.026 | 0.002 | 0.07 | 0.1 | 0.05 | 0 | 0.05 | 0 |
| 5 | 65638862 | rs16895111 | 1.325 | -0.313 | -1.465 | -1.171 | -0.001 | -0.012 | -0.012 | 0 | 0.11 | 0.09 | 0.03 | 0.03 | 0.07 | 0.12 |
| 5 | 65640456 | rs2801646 | 1.688 | -0.154 | -1.635 | -1.152 | -0.12 | 0.156 | 0.156 | 0.336 | 0.3 | 0.03 | 0.27 | 0.06 | 0.38 | 0.37 |
| 5 | 65641170 | rs7727166 | 0.309 | -0.48 | -1.394 | -0.895 | 0.015 | 0.037 | 0.037 | 0 | 0.36 | 0.13 | 0.3 | 0.3 | 0.32 | 0.34 |
| 5 | 65642855 | rs1428233 | -0.997 | -0.152 | -0.256 | -0.711 | 0.008 | -0.023 | -0.023 | 0 | 0.63 | 0.64 | 0.58 | 0.85 | 0.55 | 0.66 |
| 5 | 65643187 | rs1781654 | 0 | -0.049 | 0.212 | 0.141 | 0.038 | 0.491 | 0.491 | 0 | 0.06 | 0.34 | 0.11 | 0.51 | 0.03 | 0.09 |
| 5 | 65644187 | rs1651025 | 0.106 | -0.035 | 0.186 | 0.148 | 0.037 | 0.484 | 0.484 | 0 | 0.06 | 0.33 | 0.11 | 0.51 | 0.03 | 0.08 |
| 5 | 65651509 | rs2561072 | -0.413 | -0.039 | -0.186 | -0.521 | 0.008 | 0.014 | 0.014 | 0.018 | 0.45 | 0.62 | 0.49 | 0.83 | 0.39 | 0.48 |
| 5 | 65652904 | rs1004433 | -0.649 | 0.097 | 0.36 | -0.13 | -0.027 | 0.011 | 0.011 | 0 | 0.17 | 0.41 | 0.21 | 0.23 | 0.14 | 0.26 |
| 5 | 65653071 | rs2115128 | -0.404 | 0.341 | -0.191 | 0.565 | -0.003 | -0.014 | -0.014 | 0 | 0.14 | 0.09 | 0.21 | 0.26 | 0.17 | 0.26 |
| 5 | 65653941 | rs2801644 | -0.052 | 0.782 | -0.77 | 0.837 | -0.039 | -0.04 | -0.04 | 0.103 | 0.16 | 0.06 | 0.27 | 0.26 | 0.23 | 0.07 |
| 5 | 65654992 | rs10045766 | -0.437 | 0.348 | -0.845 | 0.853 | 0.008 | -0.004 | -0.004 | 0 | 0.22 | 0.15 | 0.25 | 0.23 | 0.22 | 0.46 |
| 5 | 65655217 | rs2997127 | -0.174 | 0.154 | -1.059 | 0.678 | 0.032 | -0.033 | -0.033 | 0.115 | 0.61 | 0.28 | 0.71 | 0.53 | 0.7 | 0.7 |
| 5 | 65655412 | rs2997125 | -0.465 | 0.119 | -0.718 | 1.071 | -0.013 | -0.001 | -0.001 | 2.359 | 0.65 | 0.46 | 0.79 | 0.7 | 0.74 | 0.76 |
| 5 | 65656048 | rs17265296 | 0.859 | 0.212 | -0.807 | 0.837 | -0.001 | -0.007 | -0.007 | 0 | 0.11 | 0.07 | 0.14 | 0.16 | 0.11 | 0.23 |
| 5 | 65658783 | rs16895146 | -0.612 | 0.21 | -0.593 | 0.69 | -0.007 | 0.029 | 0.029 | 0 | 0.25 | 0.37 | 0.33 | 0.4 | 0.25 | 0.51 |
| 5 | 65659121 | rs1469419 | 0.823 | 0.316 | -1.011 | -0.119 | -0.081 | 0.055 | 0.055 | 0 | 0.4 | 0.09 | 0.45 | 0.3 | 0.48 | 0.24 |
| 5 | 65662133 | rs3121685 | 0.879 | 0.23 | -0.851 | -0.103 | -0.036 | 0.027 | 0.027 | 0 | 0.4 | 0.16 | 0.45 | 0.36 | 0.48 | 0.25 |
| 5 | 65662570 | rs1366273 | -0.238 | 0.445 | -0.888 | -0.17 | -0.002 | -0.001 | -0.001 | 0 | 0.22 | 0.24 | 0.33 | 0.33 | 0.29 | 0.25 |
| 5 | 65665913 | rs3121688 | -1.751 | 0.045 | -0.506 | -0.546 | -0.026 | -0.081 | -0.081 | 0.054 | 0.81 | 0.76 | 0.89 | 1 | 0.9 | 0.94 |
| 5 | 65669141 | rs979031 | 1.277 | 0.748 | -0.888 | 0.075 | -0.02 | 0.063 | 0.063 | 0 | 0.3 | 0.15 | 0.22 | 0.11 | 0.3 | 0.2 |
| 5 | 65673986 | rs2968194 | -0.393 | 0.747 | -0.868 | 0.097 | 0.019 | 0.01 | 0.01 | 0 | 0.53 | 0.42 | 0.54 | 0.45 | 0.59 | 0.47 |
| 5 | 65675641 | rs6884736 | 0.246 | 0.925 | -0.464 | 0.57 | 0 | 0.005 | 0.005 | 0 | 0.12 | 0.13 | 0.08 | 0.09 | 0.11 | 0.23 |
| 5 | 65688151 | rs2371230 | -1.022 | 0.932 | -0.472 | 0.574 | -0.002 | -0.002 | -0.002 | 0 | 0.23 | 0.24 | 0.32 | 0.34 | 0.29 | 0.28 |
| 5 | 65689041 | rs4235460 | 0.474 | 0.995 | -0.466 | 0.633 | 0.003 | 0.006 | 0.006 | 0 | 0.11 | 0.09 | 0.24 | 0.25 | 0.18 | 0.05 |
| 5 | 65695268 | rs10805411 | 0.607 | 1.004 | -0.613 | 0.666 | -0.039 | -0.046 | -0.046 | 0 | 0.11 | 0.04 | 0.26 | 0.27 | 0.18 | 0.05 |
| 5 | 65703259 | rs2371382 | -0.745 | 0.93 | -0.527 | 0.484 | -0.001 | -0.001 | -0.001 | 0 | 0.25 | 0.27 | 0.32 | 0.34 | 0.3 | 0.29 |
| 5 | 65708646 | rs16895169 | 0.963 | 0.819 | -0.756 | 0.559 | -0.029 | -0.033 | -0.033 | 0 | 0.06 | 0.03 | 0.19 | 0.22 | 0.08 | 0.03 |
| 5 | 65709993 | rs3121692 | 1.025 | 1.12 | -1.089 | 0.191 | -0.015 | 0.029 | 0.029 | 0 | 0.21 | 0.11 | 0.16 | 0.1 | 0.24 | 0.16 |
| 5 | 65714450 | rs3121690 | -0.434 | 0.446 | -0.994 | 0.185 | 0.013 | -0.004 | -0.004 | 0.309 | 0.55 | 0.4 | 0.54 | 0.52 | 0.6 | 0.54 |
| 5 | 65724966 | rs13360861 | 0.456 | 0.963 | -0.708 | 0.31 | 0.008 | 0.027 | 0.027 | 0 | 0.07 | 0.15 | 0.21 | 0.22 | 0.09 | 0.03 |
| 5 | 65730895 | rs3111632 | 0.061 | 0.891 | -0.625 | -0.259 | 0.012 | -0.019 | -0.019 | 0 | 0.65 | 0.39 | 0.57 | 0.71 | 0.6 | 0.68 |
| 5 | 65746468 | rs1317632 | -0.611 | 0.805 | -0.368 | 0.055 | 0.004 | 0.025 | 0.025 | 0 | 0.33 | 0.45 | 0.39 | 0.42 | 0.33 | 0.38 |
| 5 | 65746540 | rs13180603 | 0.012 | 0.75 | -0.98 | -0.15 | -0.057 | -0.038 | -0.038 | 0 | 0.24 | 0.01 | 0.13 | 0.2 | 0.23 | 0.31 |
| 5 | 65758455 | rs1937740 | 0.047 | 0.359 | -0.927 | -0.602 | -0.014 | -0.003 | -0.003 | 0 | 0.3 | 0.15 | 0.49 | 0.69 | 0.35 | 0.42 |
| 5 | 65768007 | rs1962397 | -0.686 | 0.02 | 0.131 | -0.936 | -0.002 | -0.005 | -0.005 | 0.514 | 0.44 | 0.47 | 0.56 | 0.69 | 0.42 | 0.54 |
| 5 | 65773712 | rs16895233 | 1.741 | 0.134 | -0.08 | -0.642 | -0.026 | -0.036 | -0.036 | 0 | 0.07 | 0 | 0.14 | 0.1 | 0.14 | 0.09 |
| 5 | 65778398 | rs6859445 | 0.98 | 0.091 | -0.001 | -0.772 | -0.031 | -0.079 | -0.079 | 0 | 0.14 | 0.32 | 0.07 | 0 | 0.07 | 0.12 |
| 5 | 65783709 | rs10069397 | -1.555 | 0.072 | -0.024 | -0.945 | -0.044 | 0.191 | 0.191 | 0 | 0.55 | 0.8 | 0.67 | 0.91 | 0.52 | 0.7 |
| 5 | 65789364 | rs4129014 | -0.56 | 0.105 | 0.104 | -0.914 | -0.003 | 0.011 | 0.011 | 0 | 0.41 | 0.56 | 0.59 | 0.89 | 0.45 | 0.57 |
| 5 | 65796062 | rs6897379 | -1.373 | 0.186 | -0.031 | -0.86 | -0.005 | 0.021 | 0.021 | 0 | 0.53 | 0.52 | 0.63 | 0.89 | 0.52 | 0.66 |
| 5 | 65802754 | rs6896201 | 0.021 | 0.14 | 0.11 | -0.463 | -0.025 | -0.012 | -0.012 | 0 | 0.23 | 0.36 | 0.13 | 0.2 | 0.17 | 0.24 |
| 5 | 65828056 | rs13181346 | 0.331 | 0.267 | -0.804 | -0.542 | -0.053 | -0.099 | -0.099 | 2.303 | 0.27 | 0.06 | 0.46 | 0.6 | 0.28 | 0.36 |
| 5 | 65831439 | rs13181758 | 1.65 | 0.255 | -0.679 | -0.264 | -0.009 | -0.076 | -0.076 | 3.485 | 0.14 | 0.03 | 0.2 | 0.39 | 0.12 | 0.26 |
| 5 | 65834747 | rs7707868 | 1.091 | 0.049 | -0.22 | -0.6 | -0.031 | -0.085 | -0.085 | 0 | 0.15 | 0.37 | 0.08 | 0.02 | 0.08 | 0.13 |
| 5 | 65835876 | rs4547878 | -2.93 | 0.293 | -1.224 | -0.877 | -0.078 | 0.015 | 0.015 | 0 | 0.88 | 0.99 | 0.88 | 0.91 | 0.76 | 0.84 |
| 5 | 65837538 | rs6865758 | -1.258 | -0.457 | -0.706 | -0.586 | -0.031 | 0.147 | 0.147 | 0.001 | 0.54 | 0.79 | 0.62 | 0.81 | 0.46 | 0.65 |
| 5 | 65849279 | rs4235463 | -1.146 | -0.415 | -0.699 | -0.61 | 0.018 | 0.182 | 0.182 | 0.004 | 0.5 | 0.78 | 0.6 | 0.81 | 0.42 | 0.62 |
| 5 | 65853564 | rs10940065 | 1.176 | -0.374 | -0.495 | -0.246 | 0.002 | -0.016 | -0.016 | 0 | 0.14 | 0.09 | 0.2 | 0.4 | 0.12 | 0.26 |
| 5 | 65855856 | rs10940066 | 0.252 | -0.764 | -0.865 | -0.487 | 0.029 | -0.046 | -0.046 | 0 | 0.32 | 0.14 | 0.29 | 0.4 | 0.24 | 0.35 |
| 5 | 65856589 | rs7713842 | -0.604 | -0.827 | -0.658 | -0.52 | 0.04 | 0.027 | 0.027 | 0 | 0.46 | 0.71 | 0.42 | 0.6 | 0.34 | 0.55 |
| 5 | 65859130 | rs10060367 | 0.563 | -0.805 | -0.711 | -0.456 | 0.03 | -0.052 | -0.052 | 2.941 | 0.28 | 0.12 | 0.26 | 0.4 | 0.2 | 0.32 |
| 5 | 65860092 | rs11953252 | 1.884 | -0.771 | -0.646 | -0.428 | 0.029 | 0.051 | 0.051 | 1.98 | 0.14 | 0.04 | 0.07 | 0 | 0.08 | 0.06 |
| 5 | 65860228 | rs13183635 | -0.288 | -0.568 | -0.369 | -0.197 | 0.013 | 0.039 | 0.039 | 0.65 | 0.52 | 0.28 | 0.58 | 0.63 | 0.42 | 0.49 |
| 5 | 65861226 | rs12657904 | -0.48 | -0.578 | -0.02 | -0.201 | 0 | 0 | 0 | 0 | 0.54 | 0.56 | 0.6 | 0.61 | 0.41 | 0.49 |
| 5 | 65861849 | rs6449823 | -0.115 | -0.405 | 0.1 | -0.121 | -0.001 | 0.004 | 0.004 | 0 | 0.21 | 0.35 | 0.3 | 0.23 | 0.18 | 0.14 |
| 5 | 65864189 | rs10055880 | 0.382 | -0.003 | 0.398 | 0.314 | 0.04 | 0.094 | 0.094 | 0.017 | 0.1 | 0.29 | 0.25 | 0.21 | 0.11 | 0.07 |
| 5 | 65872475 | rs6875246 | 0.495 | -0.025 | -0.089 | 0.358 | 0.018 | 0.038 | 0.038 | 0 | 0.1 | 0.19 | 0.26 | 0.21 | 0.13 | 0.08 |
| 5 | 65873081 | rs12655442 | 0.025 | -0.089 | -0.488 | -0.004 | 0.005 | 0.012 | 0.012 | 0.485 | 0.54 | 0.61 | 0.6 | 0.63 | 0.41 | 0.52 |
| 5 | 65875938 | rs4274939 | -0.402 | 0.276 | -0.873 | -0.187 | -0.065 | 0.078 | 0.078 | 0 | 0.65 | 0.85 | 0.68 | 0.81 | 0.51 | 0.68 |
| 5 | 65880241 | rs4699877 | -0.3 | 0.201 | -0.338 | 0.293 | -0.007 | -0.003 | -0.003 | 0 | 0.24 | 0.17 | 0.42 | 0.28 | 0.37 | 0.22 |
| 5 | 65882092 | rs7715969 | -1.159 | -0.403 | 0.22 | 0.315 | 0.001 | 0.001 | 0.001 | 0 | 0.35 | 0.66 | 0.51 | 0.41 | 0.41 | 0.53 |

| | | | | | | | | | | | | | | | | |
|---|---|---|---|---|---|---|---|---|---|---|---|---|---|---|---|---|
| 5 | 65885692 | rs6868283 | -0.185 | -0.398 | 0.354 | 0.34 | -0.031 | 0.016 | 0.016 | 0.893 | 0.12 | 0.6 | 0.11 | 0.12 | 0.06 | 0.32 |
| 5 | 65890467 | rs6877176 | -0.453 | -0.28 | 0.643 | 0.379 | 0 | 0.045 | 0.045 | 0 | 0.17 | 0.33 | 0.24 | 0.32 | 0.17 | 0.39 |
| 5 | 65891203 | rs4700135 | -1.209 | -0.707 | 0.312 | 0.266 | 0.001 | 0.034 | 0.034 | 0 | 0.45 | 0.66 | 0.62 | 0.67 | 0.47 | 0.56 |
| 5 | 65895193 | rs4587034 | -0.893 | -0.66 | 0.362 | 0.37 | 0.001 | -0.001 | -0.001 | 0 | 0.32 | 0.68 | 0.38 | 0.64 | 0.23 | 0.55 |
| 5 | 65903601 | rs16895292 | 0.781 | -0.178 | -0.368 | 0.652 | -0.021 | -0.026 | -0.026 | 0 | 0.1 | 0.04 | 0.2 | 0.25 | 0.15 | 0.16 |
| 5 | 65904324 | rs4699881 | -1.053 | -0.631 | -0.448 | 0.183 | -0.094 | -0.04 | -0.04 | 0.087 | 0.69 | 0.9 | 0.72 | 0.68 | 0.58 | 0.73 |
| 5 | 65905174 | rs10471689 | -0.9 | -0.405 | -0.246 | 0.414 | -0.011 | 0.007 | 0.007 | 0.598 | 0.61 | 0.7 | 0.68 | 0.71 | 0.53 | 0.67 |
| 5 | 65907571 | rs16895297 | -0.104 | 0.321 | 0.263 | 1.222 | 0.033 | 0.074 | 0.074 | 0.448 | 0.06 | 0.28 | 0.07 | 0.09 | 0.05 | 0.2 |
| 5 | 65907879 | rs12173252 | 0.822 | 0.281 | -0.259 | 1.281 | 0.005 | -0.003 | -0.003 | 0 | 0.11 | 0.06 | 0.11 | 0.29 | 0.06 | 0.12 |
| 5 | 65908187 | rs4128347 | -0.764 | 0.521 | -0.179 | 1.277 | 0.004 | 0.051 | 0.051 | 0.094 | 0.27 | 0.53 | 0.28 | 0.4 | 0.26 | 0.41 |
| 5 | 65908385 | rs16895303 | 0.387 | 0.661 | -0.508 | 1.37 | -0.012 | -0.027 | -0.027 | 0 | 0.17 | 0.07 | 0.18 | 0.29 | 0.2 | 0.13 |
| 5 | 65908576 | rs4699882 | 0.379 | 0.665 | -0.852 | 1.121 | -0.002 | -0.01 | -0.01 | 0 | 0.28 | 0.2 | 0.43 | 0.33 | 0.33 | 0.3 |
| 5 | 65909231 | rs10065304 | -0.149 | 0.745 | -0.716 | 1.134 | 0.012 | 0.067 | 0.067 | 0 | 0.25 | 0.46 | 0.27 | 0.38 | 0.26 | 0.32 |
| 5 | 65910316 | rs6890124 | -0.224 | 0.657 | -1.054 | 0.625 | 0.003 | 0.044 | 0.044 | 0 | 0.54 | 0.66 | 0.71 | 0.73 | 0.6 | 0.64 |
| 5 | 65913392 | rs4613646 | -0.308 | 0.38 | -1.657 | 0.812 | -0.004 | 0.036 | 0.036 | 0.086 | 0.31 | 0.2 | 0.22 | 0.18 | 0.31 | 0.21 |
| 5 | 65913576 | rs17268691 | 0.903 | 0.67 | -2.128 | 0.568 | -0.025 | 0.016 | 0.016 | 0 | 0.13 | 0.03 | 0.09 | 0.08 | 0.16 | 0.11 |
| 5 | 65914287 | rs4455504 | -0.469 | 0.847 | -1.846 | 0.623 | -0.001 | 0 | 0 | 0 | 0.8 | 0.81 | 0.64 | 0.68 | 0.77 | 0.82 |
| 5 | 65916383 | rs10805414 | -0.617 | 0.259 | -1.378 | 0.151 | 0.019 | 0.021 | 0.021 | 0.397 | 0.71 | 0.82 | 0.65 | 0.78 | 0.75 | 0.56 |
| 5 | 65916784 | rs10940072 | 0.95 | 0.157 | -1.038 | 0.097 | 0.003 | -0.005 | -0.005 | 0 | 0.4 | 0.27 | 0.32 | 0.58 | 0.38 | 0.34 |
| 5 | 65916978 | rs17209451 | -0.128 | 0.188 | -0.766 | 0.109 | 0.008 | 0.035 | 0.035 | 1.365 | 0.58 | 0.73 | 0.56 | 0.7 | 0.59 | 0.45 |
| 5 | 65917185 | rs10063997 | 0.976 | 0.119 | -0.707 | 0.027 | 0.01 | -0.007 | -0.007 | 0.113 | 0.15 | 0.18 | 0.21 | 0.12 | 0.21 | 0.11 |
| 5 | 65921192 | rs4235471 | -0.541 | -0.31 | -0.699 | 0.101 | 0.018 | 0.027 | 0.027 | 0.325 | 0.72 | 0.82 | 0.65 | 0.8 | 0.75 | 0.56 |
| 5 | 65922787 | rs4532319 | -0.291 | -0.487 | -0.97 | -0.223 | 0.049 | 0.089 | 0.089 | 0 | 0.56 | 0.74 | 0.57 | 0.82 | 0.66 | 0.49 |
| 5 | 65924603 | rs10940073 | 0.551 | -0.525 | -0.742 | 0.103 | -0.009 | -0.008 | -0.008 | 0 | 0.22 | 0.2 | 0.19 | 0.3 | 0.26 | 0.31 |
| 5 | 65929212 | rs4272083 | -1.331 | -1.014 | -0.595 | -0.346 | -0.051 | 0.055 | 0.055 | 0 | 0.69 | 0.81 | 0.75 | 0.88 | 0.55 | 0.81 |
| 5 | 65931332 | rs16895360 | 0.257 | -1.542 | -0.356 | -0.536 | -0.022 | -0.049 | -0.049 | 0.01 | 0.09 | 0.16 | 0.09 | 0 | 0.05 | 0.03 |
| 5 | 65932156 | rs7728510 | 1.375 | -1.659 | -0.229 | -0.421 | -0.016 | 0.151 | 0.151 | 0 | 0.11 | 0.22 | 0.15 | 0.43 | 0.07 | 0.09 |
| 5 | 65933932 | rs7721030 | -1.35 | -1.589 | -0.21 | -0.497 | -0.054 | -0.047 | -0.047 | 0 | 0.66 | 0.82 | 0.68 | 0.44 | 0.58 | 0.74 |
| 5 | 65936799 | rs13190448 | 0.172 | -1.747 | -0.014 | -0.632 | 0.013 | 0.017 | 0.017 | 0 | 0.31 | 0.23 | 0.38 | 0.15 | 0.26 | 0.38 |
| 5 | 65940604 | rs10515016 | 0.276 | -1.721 | -0.236 | -0.619 | 0.038 | 0.065 | 0.065 | 0 | 0.31 | 0.13 | 0.37 | 0.15 | 0.26 | 0.38 |
| 5 | 65945752 | rs4700144 | 0.562 | -1.248 | -0.129 | -0.687 | 0.002 | -0.002 | -0.002 | 0 | 0.08 | 0.04 | 0.1 | 0 | 0.07 | 0.04 |
| 5 | 65951378 | rs4613647 | 1.096 | -1.325 | -0.037 | -0.636 | -0.008 | -0.028 | -0.028 | 0 | 0.06 | 0.1 | 0.09 | 0 | 0.04 | 0.03 |
| 5 | 65961205 | rs6896102 | -0.327 | -1.782 | 0.023 | -0.205 | -0.03 | 0.021 | 0.021 | 0 | 0.28 | 0.41 | 0.4 | 0.3 | 0.24 | 0.4 |
| 5 | 65961733 | rs6881653 | -0.736 | -1.823 | -0.045 | -0.257 | 0.002 | 0.028 | 0.028 | 0 | 0.5 | 0.61 | 0.6 | 0.6 | 0.51 | 0.7 |
| 5 | 65961802 | rs12656033 | 1.156 | -1.745 | -0.115 | -0.203 | -0.009 | -0.008 | -0.008 | 0 | 0.22 | 0.2 | 0.19 | 0.3 | 0.26 | 0.3 |
| 5 | 65963421 | rs16895434 | 1.076 | -1.776 | -0.19 | -0.121 | -0.005 | 0.018 | 0.018 | 0 | 0.17 | 0.18 | 0.33 | 0.31 | 0.15 | 0.35 |
| 5 | 65967123 | rs4499783 | -0.628 | -1.752 | -0.155 | -0.232 | 0 | -0.011 | -0.011 | 0 | 0.39 | 0.6 | 0.35 | 0.3 | 0.39 | 0.38 |
| 5 | 65968172 | rs9687339 | 0.462 | -1.739 | -0.142 | -0.22 | 0.008 | 0.007 | 0.007 | 0 | 0.27 | 0.42 | 0.22 | 0.3 | 0.26 | 0.33 |
| 5 | 65978497 | rs4700148 | 1.015 | -1.709 | -0.218 | -0.203 | 0.003 | 0.008 | 0.008 | 0 | 0.11 | 0.06 | 0.13 | 0 | 0.13 | 0.04 |
| 5 | 65982329 | rs10057708 | -0.523 | -1.868 | 0.042 | -0.157 | -0.025 | 0.008 | 0.008 | 0 | 0.32 | 0.36 | 0.45 | 0.32 | 0.28 | 0.37 |
| 5 | 65983977 | rs16895458 | 1.518 | -2.1 | -0.108 | -0.236 | -0.033 | 0.013 | 0.013 | 0.714 | 0.19 | 0.28 | 0.15 | 0.26 | 0.1 | 0.1 |
| 5 | 65985348 | rs4273563 | 1.291 | -2.022 | -0.157 | -0.182 | -0.007 | 0.022 | 0.022 | 0 | 0.17 | 0.16 | 0.32 | 0.32 | 0.16 | 0.27 |
| 5 | 65986885 | rs7380830 | -1 | -2.014 | -0.461 | -0.184 | 0.08 | 0.028 | 0.028 | 0 | 0.5 | 0.63 | 0.61 | 0.58 | 0.38 | 0.48 |
| 5 | 65988164 | rs10072488 | 0.461 | -1.656 | -0.66 | -0.019 | -0.029 | 0.052 | 0.052 | 0 | 0.21 | 0.28 | 0.33 | 0.32 | 0.16 | 0.33 |
| 5 | 65993985 | rs11749980 | -2.018 | -1.513 | -0.704 | -0.086 | -0.049 | 0.04 | 0.04 | 0 | 0.75 | 0.84 | 0.81 | 0.88 | 0.64 | 0.84 |
| 5 | 66000724 | rs10940080 | -1.524 | -1.46 | -1.025 | -0.048 | 0.032 | -0.054 | -0.054 | 6.332 | 0.69 | 0.59 | 0.8 | 0.84 | 0.63 | 0.66 |
| 5 | 66002170 | rs16895509 | 1.04 | -1.339 | -1.594 | 0.297 | 0.003 | -0.081 | -0.081 | 0 | 0.15 | 0.03 | 0.31 | 0.28 | 0.14 | 0.15 |
| 5 | 66009984 | rs4282272 | -0.082 | -1.344 | -1.459 | 0.333 | 0.047 | -0.038 | -0.038 | 0 | 0.35 | 0.16 | 0.47 | 0.56 | 0.26 | 0.26 |
| 5 | 66011174 | rs13161696 | 1.145 | -0.768 | -0.866 | 0.591 | 0 | 0.036 | 0.036 | 0.05 | 0.1 | 0.11 | 0.13 | 0.27 | 0.09 | 0.1 |
| 5 | 66012846 | rs17278159 | 0.641 | -0.46 | -1.331 | 0.538 | -0.083 | -0.028 | -0.028 | 0 | 0.16 | 0.01 | 0.11 | 0.14 | 0.21 | 0.19 |
| 5 | 66017952 | rs6878085 | -0.604 | -0.738 | -0.649 | 0.443 | 0.002 | -0.014 | -0.014 | 0 | 0.33 | 0.38 | 0.3 | 0.24 | 0.37 | 0.34 |
| 5 | 66018196 | rs9791090 | -0.125 | -1.206 | -0.225 | 0.855 | 0.027 | 0.143 | 0.143 | 0 | 0.14 | 0.41 | 0.14 | 0.34 | 0.11 | 0.31 |
| 5 | 66034905 | rs3111626 | -2.027 | -1.108 | -1.43 | 0.384 | -0.133 | -0.013 | -0.013 | 0 | 0.74 | 0.98 | 0.82 | 0.85 | 0.65 | 0.83 |
| 5 | 66038209 | rs4699891 | 0.061 | -1.477 | -1.434 | 0.117 | -0.029 | 0.037 | 0.037 | 0.019 | 0.24 | 0.17 | 0.32 | 0.13 | 0.36 | 0.14 |
| 5 | 66050568 | rs258275 | 0.826 | -1.659 | -1.286 | 0.597 | 0.008 | -0.041 | -0.041 | 0 | 0.16 | 0.07 | 0.11 | 0.28 | 0.11 | 0.18 |
| 5 | 66059828 | rs258277 | 0.203 | -1.723 | -0.594 | 0.834 | -0.02 | 0.09 | 0.09 | 0 | 0.21 | 0.33 | 0.16 | 0.46 | 0.13 | 0.31 |
| 5 | 66064026 | rs33723 | 0.44 | -1.721 | -0.672 | 0.539 | 0.005 | -0.006 | -0.006 | 0 | 0.19 | 0.16 | 0.14 | 0.47 | 0.12 | 0.25 |
| 5 | 66078314 | rs33696 | -0.575 | -1.866 | -0.322 | 0.4 | -0.031 | -0.021 | -0.021 | 0 | 0.73 | 0.61 | 0.69 | 0.8 | 0.84 | 0.75 |
| 5 | 66089535 | rs6887024 | 1.885 | -1.754 | -0.435 | 0.364 | -0.011 | 0.011 | 0.011 | 0.863 | 0.31 | 0.21 | 0.22 | 0.22 | 0.36 | 0.3 |
| 5 | 66095512 | rs33719 | 0.73 | -1.876 | -0.353 | 0.37 | -0.001 | -0.002 | -0.002 | 2.901 | 0.52 | 0.45 | 0.37 | 0.66 | 0.5 | 0.61 |
| 5 | 66099356 | rs258267 | -0.926 | -1.999 | 0.034 | 0.304 | 0.014 | 0.009 | 0.009 | 0 | 0.75 | 0.79 | 0.71 | 0.8 | 0.84 | 0.76 |
| 5 | 66104174 | rs7736147 | 0.204 | -1.885 | 0.316 | 0.294 | 0.024 | -0.033 | -0.033 | 1.261 | 0.23 | 0.3 | 0.34 | 0.14 | 0.35 | 0.15 |
| 5 | 66115982 | rs7725568 | 0.372 | -2.066 | 0 | 0.239 | 0.002 | 0.001 | 0.001 | 0 | 0.58 | 0.57 | 0.56 | 0.36 | 0.71 | 0.45 |
| 5 | 66127646 | rs13175464 | 0.323 | -1.854 | -0.062 | 0.198 | -0.039 | 0.058 | 0.058 | 0 | 0.23 | 0.11 | 0.31 | 0.11 | 0.31 | 0.14 |
| 5 | 66129815 | rs9918275 | 2.742 | -1.832 | -0.181 | 0.192 | -0.079 | 0.032 | 0.032 | 0.113 | 0.26 | 0.06 | 0.17 | 0.12 | 0.34 | 0.16 |
| 5 | 66131056 | rs258081 | -0.866 | -1.759 | -0.027 | 0.324 | 0.003 | 0.002 | 0.002 | 0 | 0.77 | 0.75 | 0.71 | 0.8 | 0.84 | 0.76 |
| 5 | 66133356 | rs190480 | 1.236 | -1.732 | 0.065 | 0.369 | 0.01 | 0.026 | 0.026 | 0 | 0.48 | 0.33 | 0.34 | 0.57 | 0.51 | 0.47 |
| 5 | 66134711 | rs257699 | 0.74 | -1.576 | 0.012 | 0.436 | 0.001 | 0.038 | 0.038 | 0 | 0.17 | 0.18 | 0.15 | 0.45 | 0.15 | 0.23 |
| 5 | 66139238 | rs2441109 | -0.368 | -1.686 | -0.184 | 0.417 | 0.015 | 0.009 | 0.009 | 0 | 0.75 | 0.78 | 0.71 | 0.8 | 0.84 | 0.76 |

| | | | | | | | | | | | | | | | | |
|---|---|---|---|---|---|---|---|---|---|---|---|---|---|---|---|---|
| 5 | 66147348 | rs17281085 | 1.097 | -1.145 | -0.315 | 0.48 | -0.017 | 0.033 | 0.033 | 0 | 0.09 | 0.03 | 0.13 | 0.01 | 0.15 | 0.05 |
| 5 | 66148984 | rs1007500 | 0.738 | -1.158 | -0.335 | 0.46 | -0.014 | 0.041 | 0.041 | 0 | 0.11 | 0.03 | 0.13 | 0.01 | 0.15 | 0.05 |
| 5 | 66149113 | rs28636 | 0.745 | -1.132 | -0.177 | 0.741 | 0.011 | 0.032 | 0.032 | 0 | 0.2 | 0.26 | 0.2 | 0.34 | 0.21 | 0.19 |
| 5 | 66151551 | rs30732 | -1.392 | -1.588 | -0.301 | 0.544 | 0.014 | -0.011 | -0.011 | 0 | 0.85 | 0.92 | 0.81 | 0.82 | 0.89 | 0.86 |
| 5 | 66152110 | rs13163792 | 0.562 | -1.14 | -0.057 | 0.287 | 0 | -0.018 | -0.018 | 0.236 | 0.11 | 0.1 | 0.14 | 0.01 | 0.15 | 0.05 |
| 5 | 66155743 | rs11745025 | 0.282 | -0.909 | -0.108 | 0.395 | 0.002 | -0.006 | -0.006 | 0 | 0.13 | 0.11 | 0.15 | 0.23 | 0.1 | 0.14 |
| 5 | 66156139 | rs16895744 | 0.297 | -0.9 | -0.143 | 0.332 | 0.003 | -0.007 | -0.007 | 0 | 0.13 | 0.08 | 0.14 | 0.22 | 0.1 | 0.14 |
| 5 | 66157391 | rs27422 | -0.382 | -0.936 | -0.175 | 0.021 | 0.003 | 0.027 | 0.027 | 0 | 0.38 | 0.44 | 0.44 | 0.52 | 0.36 | 0.41 |
| 5 | 66162044 | rs39700 | -0.062 | -0.837 | -0.517 | -0.779 | 0.009 | 0.08 | 0.08 | 0 | 0.57 | 0.26 | 0.48 | 0.26 | 0.64 | 0.49 |
| 5 | 66163119 | rs4700168 | 0.955 | -0.837 | -0.797 | -0.875 | -0.105 | 0.173 | 0.173 | 0.072 | 0.42 | 0.09 | 0.34 | 0.16 | 0.52 | 0.38 |
| 5 | 66166614 | rs36138 | 1.258 | -0.649 | -0.566 | -0.666 | -0.023 | -0.02 | -0.02 | 0 | 0.08 | 0.03 | 0.1 | 0.09 | 0.11 | 0.05 |
| 5 | 66170504 | rs26923 | -0.854 | -1.076 | 0.249 | -0.409 | -0.034 | 0.01 | 0.01 | 0.012 | 0.81 | 0.66 | 0.8 | 0.78 | 0.9 | 0.82 |
| 5 | 66170578 | rs190482 | 0.518 | -1.068 | 0.299 | -0.481 | 0.012 | 0.053 | 0.053 | 0 | 0.23 | 0.28 | 0.3 | 0.53 | 0.26 | 0.34 |
| 5 | 66172672 | rs26921 | 1.583 | -0.894 | -0.011 | -0.679 | -0.124 | 0.131 | 0.131 | 0.013 | 0.33 | 0.08 | 0.27 | 0.11 | 0.43 | 0.2 |
| 5 | 66181023 | rs27876 | -0.521 | -0.911 | 0.132 | -0.687 | -0.001 | 0.009 | 0.009 | 0.353 | 0.43 | 0.36 | 0.47 | 0.67 | 0.46 | 0.57 |
| 5 | 66189792 | rs461536 | 0.516 | -0.89 | 0.142 | -0.719 | 0 | 0.007 | 0.007 | 0.774 | 0.54 | 0.59 | 0.52 | 0.32 | 0.54 | 0.43 |
| 5 | 66192423 | rs10515028 | 0.484 | -0.92 | 0.341 | -0.687 | 0.001 | -0.004 | -0.004 | 0 | 0.16 | 0.14 | 0.12 | 0.02 | 0.17 | 0.17 |
| 5 | 66195774 | rs7711116 | 1.044 | -0.724 | 0.434 | -0.45 | -0.007 | 0.001 | 0.001 | 0.017 | 0.19 | 0.21 | 0.19 | 0.22 | 0.1 | 0.17 |
| 5 | 66197511 | rs465962 | 1.465 | -0.723 | 0.56 | -0.454 | -0.03 | 0.048 | 0.048 | 0.067 | 0.33 | 0.22 | 0.27 | 0.11 | 0.43 | 0.21 |
| 5 | 66199501 | rs456775 | 0.279 | -0.729 | 0.312 | -0.447 | 0 | 0.009 | 0.009 | 0.486 | 0.56 | 0.62 | 0.52 | 0.32 | 0.54 | 0.43 |
| 5 | 66200248 | rs154620 | 1.515 | -0.727 | 0.1 | -0.484 | -0.124 | 0.127 | 0.127 | 0.01 | 0.33 | 0.07 | 0.26 | 0.11 | 0.43 | 0.21 |
| 5 | 66200489 | rs17283091 | 0.491 | -0.712 | 0.097 | -0.479 | 0 | 0 | 0 | 0 | 0.22 | 0.22 | 0.23 | 0.21 | 0.11 | 0.22 |
| 5 | 66203615 | rs10056426 | 1.057 | -0.754 | 0.058 | -0.469 | -0.007 | 0.001 | 0.001 | 0.049 | 0.19 | 0.21 | 0.19 | 0.22 | 0.09 | 0.17 |
| 5 | 66204207 | rs152407 | -0.887 | -0.557 | -0.614 | -0.383 | -0.001 | -0.002 | -0.002 | 0 | 0.78 | 0.84 | 0.73 | 0.47 | 0.77 | 0.64 |
| 5 | 66208840 | rs27219 | -0.534 | -0.353 | -0.214 | -0.488 | -0.004 | 0.01 | 0.01 | 0.298 | 0.57 | 0.68 | 0.53 | 0.36 | 0.54 | 0.43 |
| 5 | 66216892 | rs152448 | 0.315 | 0.534 | -1.379 | -0.244 | -0.108 | 0.086 | 0.086 | 0.001 | 0.33 | 0.08 | 0.26 | 0.15 | 0.43 | 0.21 |
| 5 | 66221262 | rs734828 | -0.32 | 0.081 | -0.141 | 0.056 | -0.03 | 0.009 | 0.009 | 0.169 | 0.19 | 0.29 | 0.21 | 0.21 | 0.1 | 0.17 |
| 5 | 66226798 | rs4090993 | -1.159 | 0.381 | 0.322 | -0.343 | -0.003 | -0.045 | -0.045 | 0 | 0.08 | 0.19 | 0.04 | 0 | 0.07 | 0.07 |
| 5 | 66230254 | rs11956688 | -1.549 | 0.245 | 1.047 | -0.412 | 0.016 | -0.038 | -0.038 | 0 | 0.08 | 0.31 | 0.05 | 0 | 0.07 | 0.07 |
| 5 | 66230903 | rs4102 | -0.66 | 0.261 | 0.893 | -0.404 | 0 | -0.035 | -0.035 | 0 | 0.06 | 0.12 | 0.03 | 0 | 0.07 | 0.07 |
| 5 | 66241285 | rs1159576 | -1.491 | 1.001 | 0.973 | -0.499 | 0.017 | -0.055 | -0.055 | 0.111 | 0.1 | 0.38 | 0.07 | 0 | 0.09 | 0.08 |
| 5 | 66256426 | rs1423463 | -1.545 | 0.937 | 1.102 | -0.3 | 0.012 | -0.062 | -0.062 | 0 | 0.11 | 0.43 | 0.07 | 0 | 0.09 | 0.11 |
| 5 | 66260128 | rs16896062 | 0.09 | 1.35 | 0.974 | -0.94 | 0 | -0.01 | -0.01 | 0 | 0.23 | 0.25 | 0.19 | 0.11 | 0.25 | 0.14 |
| 5 | 66265041 | rs16896069 | -1.461 | 1.463 | 0.799 | -0.512 | 0.008 | -0.066 | -0.066 | 0 | 0.11 | 0.43 | 0.07 | 0 | 0.08 | 0.11 |
| 5 | 66275052 | rs1946260 | -1.493 | 1.463 | 0.482 | -0.512 | 0.012 | -0.067 | -0.067 | 0.476 | 0.11 | 0.45 | 0.07 | 0 | 0.08 | 0.11 |
| 5 | 66276513 | rs42325 | -0.235 | 1.704 | -0.849 | -0.337 | 0.005 | -0.034 | -0.034 | 0 | 0.4 | 0.54 | 0.54 | 0.81 | 0.39 | 0.49 |
| 5 | 66276998 | rs40206 | -1.111 | 2.479 | -1.307 | 0.934 | -0.025 | -0.028 | -0.028 | 0.22 | 0.15 | 0.09 | 0.25 | 0.36 | 0.27 | 0.3 |
| 5 | 66277117 | rs706701 | -0.845 | 2.087 | -1.42 | 1.03 | -0.031 | -0.045 | -0.045 | 0 | 0.17 | 0.12 | 0.2 | 0.37 | 0.26 | 0.24 |
| 5 | 66278913 | rs1083523 | 0.352 | 1.155 | -1.85 | 0.384 | -0.031 | 0.035 | 0.035 | 0.364 | 0.46 | 0.19 | 0.49 | 0.63 | 0.51 | 0.53 |
| 5 | 66280426 | rs17218390 | 0.025 | 0.783 | -1.944 | 0.05 | -0.023 | 0.001 | 0.001 | 0 | 0.22 | 0.09 | 0.23 | 0.2 | 0.26 | 0.18 |
| 5 | 66287598 | rs39664 | -0.19 | 0.465 | -1.133 | -0.183 | -0.001 | -0.002 | -0.002 | 0 | 0.8 | 0.72 | 0.86 | 0.86 | 0.83 | 0.81 |
| 5 | 66293644 | rs10068079 | 0.064 | 0.477 | -0.899 | -0.265 | -0.001 | 0 | 0 | 0 | 0.2 | 0.14 | 0.26 | 0.19 | 0.23 | 0.19 |
| 5 | 66302605 | rs10044452 | 0.097 | 0.402 | -0.937 | -0.313 | -0.025 | 0.001 | 0.001 | 0 | 0.19 | 0.08 | 0.25 | 0.18 | 0.24 | 0.19 |
| 5 | 66324295 | rs463204 | -1.263 | -0.002 | 0.374 | -0.918 | 0.003 | 0.012 | 0.012 | 0.094 | 0.47 | 0.56 | 0.39 | 0.14 | 0.43 | 0.4 |
| 5 | 66326646 | rs37483 | 0.082 | -0.107 | 0.22 | -0.945 | 0.033 | 0.033 | 0.033 | 0 | 0.22 | 0.08 | 0.12 | 0.11 | 0.16 | 0.21 |
| 5 | 66328309 | rs1991492 | 0.039 | -0.045 | 0.33 | -0.963 | 0.02 | -0.116 | -0.116 | 1.424 | 0.22 | 0.33 | 0.18 | 0 | 0.24 | 0.13 |
| 5 | 66331444 | rs152634 | -1.464 | -0.109 | 0.553 | -0.995 | 0.023 | 0.186 | 0.186 | 0.365 | 0.5 | 0.74 | 0.39 | 0.14 | 0.43 | 0.4 |
| 5 | 66334631 | rs12523505 | 0.024 | -0.045 | 0.263 | -0.913 | 0.009 | -0.084 | -0.084 | 0 | 0.22 | 0.29 | 0.18 | 0 | 0.24 | 0.13 |
| 5 | 66348322 | rs462483 | -0.061 | -0.125 | 0.208 | -0.958 | 0.015 | 0.016 | 0.016 | 0 | 0.22 | 0.16 | 0.13 | 0.12 | 0.17 | 0.2 |
| 5 | 66348604 | rs455788 | -1.193 | -0.102 | 0.086 | -1.002 | 0 | -0.013 | -0.013 | 2.157 | 0.44 | 0.41 | 0.3 | 0.11 | 0.39 | 0.33 |
| 5 | 66355956 | rs467655 | -0.598 | -0.287 | -0.311 | -0.56 | 0.038 | 0.038 | 0.038 | 0 | 0.25 | 0.15 | 0.19 | 0.15 | 0.18 | 0.25 |
| 5 | 66357828 | rs2548585 | -0.192 | -0.781 | -0.263 | -1.394 | -0.01 | -0.062 | -0.062 | 6.136 | 0.35 | 0.39 | 0.13 | 0.08 | 0.21 | 0.22 |
| 5 | 66362459 | rs2548588 | -0.656 | -0.549 | 0.271 | -1.374 | 0.002 | -0.017 | -0.017 | 0 | 0.41 | 0.59 | 0.22 | 0.17 | 0.26 | 0.29 |
| 5 | 66364865 | rs1030171 | -0.7 | -0.522 | 0.269 | -1.375 | 0.004 | -0.015 | -0.015 | 0 | 0.42 | 0.58 | 0.22 | 0.16 | 0.28 | 0.29 |
| 5 | 66370975 | rs1295186 | 1.49 | 0.674 | -0.558 | -1.583 | -0.071 | -0.011 | -0.011 | 0 | 0.11 | 0.01 | 0.14 | 0.05 | 0.22 | 0.06 |
| 5 | 66383979 | rs2545385 | -0.096 | -0.055 | -0.067 | -1.548 | 0.034 | 0.142 | 0.142 | 2.024 | 0.46 | 0.27 | 0.26 | 0.23 | 0.37 | 0.37 |
| 5 | 66384463 | rs1896686 | 1.165 | -0.185 | -0.068 | -1.64 | -0.133 | 0.042 | 0.042 | 1.181 | 0.38 | 0.07 | 0.55 | 0.64 | 0.49 | 0.49 |
| 5 | 66403500 | rs2432154 | -0.756 | -0.61 | 0.516 | -1.472 | 0.069 | 0.078 | 0.078 | 0.099 | 0.44 | 0.74 | 0.29 | 0.35 | 0.29 | 0.37 |
| 5 | 66406143 | rs2545394 | 0.391 | -0.626 | 0.448 | -1.481 | 0.045 | 0.043 | 0.043 | 0.108 | 0.32 | 0.26 | 0.16 | 0.23 | 0.18 | 0.27 |
| 5 | 66419434 | rs17221226 | 0.03 | -0.627 | 0.485 | -1.486 | 0.049 | 0.089 | 0.089 | 0 | 0.35 | 0.24 | 0.18 | 0.13 | 0.23 | 0.24 |
| 5 | 66431290 | rs6878808 | -0.607 | -0.599 | 0.597 | -1.457 | 0.015 | 0.02 | 0.02 | 0 | 0.07 | 0.16 | 0.09 | 0.11 | 0.07 | 0.08 |
| 5 | 66437122 | rs13361529 | 0.668 | -0.629 | 0.34 | -1.458 | 0.078 | 0.086 | 0.086 | 0.32 | 0.55 | 0.24 | 0.71 | 0.65 | 0.72 | 0.63 |
| 5 | 66442385 | rs12516375 | -0.314 | -0.624 | 0.161 | -1.392 | 0.043 | 0.04 | 0.04 | 0.123 | 0.67 | 0.74 | 0.83 | 0.75 | 0.8 | 0.74 |
| 5 | 66462069 | rs4699896 | -1.635 | -0.478 | 1.277 | -1.038 | 0.044 | 0.256 | 0.256 | 0.324 | 0.06 | 0.38 | 0.2 | 0.21 | 0.01 | 0.1 |
| 5 | 66465433 | rs938050 | 0.617 | -0.127 | 0.852 | -1.272 | -0.003 | 0.001 | 0.001 | 0 | 0.67 | 0.39 | 0.58 | 0.66 | 0.66 | 0.71 |
| 5 | 66473092 | rs1560159 | -1.425 | -0.454 | -0.085 | -1.372 | -0.152 | -0.088 | -0.088 | 0 | 0.69 | 0.94 | 0.58 | 0.41 | 0.54 | 0.49 |
| 5 | 66473137 | rs1560160 | -1.229 | -0.35 | 0.84 | -1.113 | -0.015 | 0.021 | 0.021 | 0.051 | 0.39 | 0.75 | 0.45 | 0.36 | 0.37 | 0.28 |
| 5 | 66478626 | rs2230525 | -1.77 | -0.198 | 1.767 | -0.988 | 0.005 | 0.067 | 0.067 | 4.469 | 0.13 | 0.6 | 0.25 | 0.24 | 0.09 | 0.14 |
| 5 | 66481069 | rs1697143 | 0.482 | -0.137 | 1.328 | -1.005 | -0.092 | 0.013 | 0.013 | 0 | 0.2 | 0.01 | 0.17 | 0.11 | 0.25 | 0.12 |

| | | | | | | | | | | | | | | | | |
|---|---|---|---|---|---|---|---|---|---|---|---|---|---|---|---|---|
| 5 | 66490093 | rs1428469 | -2.255 | -0.725 | 1.649 | -0.747 | -0.012 | 0.018 | 0.018 | 0.983 | 0.17 | 0.6 | 0.21 | 0.24 | 0.12 | 0.21 |
| 5 | 66490818 | rs5744469 | -0.944 | -0.666 | 1.121 | -0.743 | 0.002 | -0.029 | -0.029 | 0.666 | 0.09 | 0.1 | 0.06 | 0 | 0.08 | 0.08 |
| 5 | 66493080 | rs3756561 | -1.406 | -1.111 | 0.999 | -0.791 | -0.06 | 0.035 | 0.035 | 0.271 | 0.16 | 0.51 | 0.18 | 0.2 | 0.09 | 0.17 |
| 5 | 66495847 | rs1697142 | -1.215 | -0.853 | -0.141 | -0.874 | -0.034 | 0.024 | 0.024 | 1.009 | 0.52 | 0.93 | 0.44 | 0.38 | 0.38 | 0.44 |
| 5 | 66498483 | rs10058233 | -0.517 | -1.125 | 0.353 | -0.774 | 0.025 | 0.033 | 0.033 | 0 | 0.41 | 0.26 | 0.31 | 0.33 | 0.33 | 0.4 |
| 5 | 66499478 | rs6864699 | -0.043 | -0.448 | 0.221 | -0.904 | -0.005 | 0.027 | 0.027 | 0 | 0.27 | 0.17 | 0.19 | 0.15 | 0.27 | 0.18 |
| 5 | 66500025 | rs7706539 | -0.84 | -0.41 | 0.15 | -0.89 | 0.009 | 0.046 | 0.046 | 0.467 | 0.36 | 0.68 | 0.29 | 0.21 | 0.3 | 0.25 |
| 5 | 66500556 | rs6891720 | -0.905 | -0.405 | -0.185 | -0.888 | 0.013 | 0.077 | 0.077 | 0.65 | 0.37 | 0.75 | 0.3 | 0.21 | 0.31 | 0.25 |
| 5 | 66502865 | rs10940094 | -0.313 | 0.046 | -0.708 | -0.622 | -0.008 | 0.04 | 0.04 | 0 | 0.28 | 0.15 | 0.19 | 0.15 | 0.27 | 0.2 |
| 5 | 66513326 | rs1627990 | 0.1 | 0.652 | -0.352 | -0.574 | -0.006 | 0.031 | 0.031 | 0 | 0.55 | 0.54 | 0.39 | 0.18 | 0.5 | 0.42 |
| 5 | 66515817 | rs10434544 | -0.775 | 0.396 | 0.041 | 0.012 | 0.019 | 0.14 | 0.14 | 0.449 | 0.14 | 0.43 | 0.17 | 0.31 | 0.12 | 0.24 |
| 5 | 66526209 | rs9291879 | -1.735 | 0.135 | 0.22 | -0.172 | 0.002 | -0.003 | -0.003 | 0 | 0.23 | 0.75 | 0.24 | 0.43 | 0.15 | 0.28 |
| 5 | 66526556 | rs6887604 | 0.2 | -0.03 | 0.675 | -0.125 | 0.015 | 0.008 | 0.008 | 0.868 | 0.06 | 0.2 | 0.03 | 0.03 | 0.03 | 0 |
| 5 | 66529020 | rs6859520 | 0.972 | 0.047 | 0.242 | -0.236 | 0.027 | 0.192 | 0.192 | 0.058 | 0.46 | 0.18 | 0.31 | 0.07 | 0.43 | 0.34 |
| 5 | 66532920 | rs7721781 | -1.398 | 0.042 | 0.619 | -0.165 | -0.04 | 0.054 | 0.054 | 3.416 | 0.21 | 0.66 | 0.2 | 0.42 | 0.11 | 0.26 |
| 5 | 66538016 | rs1428477 | -0.344 | 0.352 | 0.052 | -0.071 | -0.013 | 0.046 | 0.046 | 0.216 | 0.13 | 0.27 | 0.11 | 0.27 | 0.08 | 0.19 |
| 5 | 66544879 | rs10070739 | -0.372 | 0.484 | 0.004 | 0.005 | -0.006 | 0.069 | 0.069 | 0.149 | 0.13 | 0.29 | 0.11 | 0.3 | 0.09 | 0.19 |
| 5 | 66545138 | rs4700177 | 0.031 | 0.83 | -0.273 | 0.057 | 0.013 | 0.007 | 0.007 | 0 | 0.61 | 0.62 | 0.46 | 0.48 | 0.52 | 0.59 |
| 5 | 66553983 | rs17224116 | 0.067 | 1.162 | -1.11 | -0.675 | -0.087 | 0.078 | 0.078 | 0.112 | 0.2 | 0.02 | 0.16 | 0.06 | 0.3 | 0.06 |
| 5 | 66562959 | rs7715561 | -1.477 | 0.973 | 0.616 | -0.548 | 0.055 | 0.022 | 0.022 | 3.623 | 0.09 | 0.46 | 0.09 | 0.01 | 0.07 | 0.06 |
| 5 | 66567056 | rs9291881 | -0.221 | 0.735 | -0.022 | -0.524 | 0.01 | -0.026 | -0.026 | 0 | 0.14 | 0.03 | 0.18 | 0.21 | 0.09 | 0.23 |
| 5 | 66568384 | rs919308 | -0.392 | 0.928 | 0.171 | -0.428 | -0.015 | -0.032 | -0.032 | 0 | 0.68 | 0.84 | 0.61 | 0.44 | 0.66 | 0.73 |
| 5 | 66578495 | rs7723521 | -1.814 | 0.603 | 0.692 | -0.329 | 0.011 | -0.016 | -0.016 | 0.502 | 0.1 | 0.6 | 0.11 | 0.02 | 0.07 | 0.04 |
| 5 | 66583829 | rs1835019 | -0.461 | 0.897 | 0.025 | -0.395 | -0.111 | -0.108 | -0.108 | 1.052 | 0.69 | 0.98 | 0.62 | 0.43 | 0.63 | 0.65 |
| 5 | 66585307 | rs7732830 | -2.733 | 0.912 | 1.731 | -0.245 | 0.021 | -0.035 | -0.035 | 0 | 0.12 | 0.65 | 0.15 | 0.02 | 0.11 | 0.03 |
| 5 | 66596800 | rs10940099 | -2.335 | 0.914 | 2.106 | -0.242 | 0.106 | 0.025 | 0.025 | 5.205 | 0.07 | 0.5 | 0.1 | 0 | 0.06 | 0.01 |
| 5 | 66610910 | rs2059849 | -2.813 | 0.952 | 1.926 | -0.242 | 0.046 | -0.026 | -0.026 | 0 | 0.11 | 0.62 | 0.15 | 0.01 | 0.12 | 0.03 |
| 5 | 66639898 | rs6449857 | -2.86 | 0.76 | 2.276 | -0.429 | 0.044 | -0.029 | -0.029 | 0 | 0.09 | 0.45 | 0.08 | 0 | 0.05 | 0.01 |
| 5 | 66642457 | rs6878607 | -3.046 | 0.755 | 2.027 | -0.434 | 0.008 | -0.092 | -0.092 | 0 | 0.14 | 0.58 | 0.13 | 0 | 0.11 | 0.02 |
| 5 | 66650273 | rs7714757 | -2.709 | 0.525 | 2.249 | -0.417 | 0.025 | -0.067 | -0.067 | 0 | 0.11 | 0.4 | 0.09 | 0 | 0.08 | 0.02 |
| 5 | 66655247 | rs7717511 | -2.911 | 0.36 | 2.224 | -0.509 | -0.022 | -0.098 | -0.098 | 0 | 0.15 | 0.65 | 0.14 | 0.01 | 0.08 | 0.07 |
| 5 | 66663691 | rs6449859 | -2.42 | 0.347 | 2.815 | -0.469 | -0.013 | -0.027 | -0.027 | 0.184 | 0.08 | 0.39 | 0.06 | 0.01 | 0.01 | 0.02 |
| 5 | 66670452 | rs160844 | -0.392 | 0.311 | 1.763 | -0.239 | -0.034 | -0.041 | -0.041 | 2.17 | 0.59 | 0.79 | 0.53 | 0.57 | 0.52 | 0.59 |
| 5 | 66672954 | rs1346533 | 0.798 | -0.134 | 1.414 | -0.495 | 0.002 | -0.028 | -0.028 | 0.01 | 0.38 | 0.27 | 0.44 | 0.56 | 0.44 | 0.5 |
| 5 | 66680616 | rs1428609 | -0.182 | 0.335 | 1.232 | -0.16 | 0.002 | -0.002 | -0.002 | 0.999 | 0.54 | 0.61 | 0.5 | 0.57 | 0.47 | 0.56 |
| 5 | 66686570 | rs317969 | 0.439 | 0.472 | 1.049 | -0.228 | -0.013 | -0.003 | -0.003 | 0 | 0.37 | 0.29 | 0.47 | 0.51 | 0.44 | 0.45 |
| 5 | 66689271 | rs319247 | -1.804 | 0.425 | 1.138 | -0.268 | 0.051 | -0.04 | -0.04 | 0.362 | 0.09 | 0.37 | 0.09 | 0.01 | 0.08 | 0.07 |
| 5 | 66705097 | rs317976 | -1.314 | 0.307 | 0.917 | -0.349 | -0.015 | -0.089 | -0.089 | 0.104 | 0.15 | 0.32 | 0.09 | 0.01 | 0.1 | 0.14 |
| 5 | 66714141 | rs186460 | -1.433 | 0.244 | 1.069 | -0.388 | -0.011 | -0.094 | -0.094 | 0.286 | 0.15 | 0.35 | 0.09 | 0.01 | 0.1 | 0.14 |
| 5 | 66716144 | rs160978 | -0.348 | 0.219 | 0.699 | -0.292 | 0.024 | 0.023 | 0.023 | 0 | 0.49 | 0.66 | 0.58 | 0.52 | 0.52 | 0.52 |
| 5 | 66727284 | rs160846 | -1.038 | 0.115 | -0.136 | -0.397 | -0.002 | -0.034 | -0.034 | 0 | 0.14 | 0.2 | 0.1 | 0 | 0.08 | 0.08 |
| 5 | 66735551 | rs32035 | -0.12 | 0.201 | -0.768 | -0.702 | 0.047 | 0.146 | 0.146 | 0 | 0.59 | 0.73 | 0.75 | 0.84 | 0.65 | 0.58 |
| 5 | 66739174 | rs32031 | 0.895 | 0.599 | -0.879 | -0.226 | -0.063 | -0.049 | -0.049 | 0 | 0.38 | 0.13 | 0.5 | 0.44 | 0.47 | 0.42 |
| 5 | 66739338 | rs32030 | -0.081 | 0.276 | -0.725 | -0.367 | 0.057 | 0.116 | 0.116 | 0 | 0.55 | 0.62 | 0.74 | 0.84 | 0.66 | 0.58 |
| 5 | 66740702 | rs3906345 | -0.985 | 0.181 | -0.833 | -0.413 | 0.056 | 0.111 | 0.111 | 0 | 0.67 | 0.81 | 0.83 | 0.83 | 0.75 | 0.71 |
| 5 | 66751430 | rs1274867 | -1.314 | 0.353 | -0.822 | -0.378 | 0.04 | 0.096 | 0.096 | 0 | 0.69 | 0.84 | 0.83 | 0.83 | 0.74 | 0.66 |
| 5 | 66758065 | rs10041096 | 1.099 | 1.034 | -1.025 | -0.358 | -0.009 | -0.005 | -0.005 | 0 | 0.64 | 0.34 | 0.49 | 0.58 | 0.6 | 0.53 |
| 5 | 66763696 | rs13362272 | 1.436 | 0.985 | -1.059 | -0.397 | 0.043 | 0.109 | 0.109 | 0 | 0.31 | 0.14 | 0.15 | 0.17 | 0.26 | 0.34 |
| 5 | 66763977 | rs6877544 | 1.035 | 0.983 | -1.04 | -0.183 | -0.011 | -0.034 | -0.034 | 0 | 0.33 | 0.22 | 0.28 | 0.38 | 0.35 | 0.24 |
| 5 | 66764060 | rs6896610 | -1.053 | 1.482 | -0.361 | 0.314 | 0.016 | 0.045 | 0.045 | 0.068 | 0.26 | 0.56 | 0.47 | 0.45 | 0.3 | 0.35 |
| 5 | 66768553 | rs1387630 | -0.524 | 1.529 | -0.426 | 0.299 | -0.002 | 0.003 | 0.003 | 0 | 0.9 | 0.91 | 0.9 | 1 | 0.9 | 0.92 |
| 5 | 66770416 | rs937029 | -1.139 | 1.409 | -0.245 | 0.488 | 0.04 | 0.077 | 0.077 | 0.035 | 0.25 | 0.46 | 0.41 | 0.43 | 0.3 | 0.35 |
| 5 | 66773955 | rs751768 | -0.619 | 1.41 | -0.169 | 0.653 | 0.082 | 0.231 | 0.231 | 0 | 0.19 | 0.4 | 0.36 | 0.45 | 0.25 | 0.17 |
| 5 | 66776884 | rs6869587 | -0.572 | 1.637 | -0.425 | 0.7 | -0.003 | 0.004 | 0.004 | 0 | 0.08 | 0.1 | 0.2 | 0.06 | 0.08 | 0.16 |
| 5 | 66780590 | rs13183818 | -1.311 | 1.47 | -0.537 | 0.589 | -0.001 | -0.004 | -0.004 | 0 | 0.34 | 0.67 | 0.53 | 0.49 | 0.35 | 0.47 |
| 5 | 66792177 | rs747919 | 0.781 | 1.952 | -0.377 | 0.822 | 0.066 | 0.115 | 0.115 | 1.54 | 0.76 | 0.5 | 0.55 | 0.55 | 0.64 | 0.75 |
| 5 | 66792556 | rs10061750 | 0.281 | 2.092 | -1.637 | 1.138 | -0.045 | -0.099 | -0.099 | 3.848 | 0.21 | 0.03 | 0.24 | 0.32 | 0.26 | 0.14 |
| 5 | 66792794 | rs2200863 | -0.136 | 1.674 | -1.926 | 0.808 | -0.032 | -0.08 | -0.08 | 0.178 | 0.15 | 0.01 | 0.11 | 0.24 | 0.15 | 0.19 |
| 5 | 66794632 | rs6860822 | -0.687 | 1.975 | -1.486 | 0.393 | 0.028 | -0.029 | -0.029 | 0 | 0.95 | 0.87 | 0.93 | 1 | 0.89 | 0.95 |
| 5 | 66797033 | rs11948764 | -0.809 | 2.19 | -1.784 | -0.176 | -0.001 | -0.002 | -0.002 | 0 | 0.37 | 0.62 | 0.54 | 0.77 | 0.44 | 0.29 |
| 5 | 66797307 | rs1532121 | -0.407 | 2.337 | -1.275 | -0.047 | 0.012 | 0.018 | 0.018 | 0 | 0.28 | 0.41 | 0.42 | 0.69 | 0.32 | 0.19 |
| 5 | 66797723 | rs1532122 | -0.003 | 2.523 | -1.319 | -0.031 | -0.013 | 0.043 | 0.043 | 0 | 0.25 | 0.39 | 0.38 | 0.68 | 0.24 | 0.18 |
| 5 | 66798668 | rs13357663 | 2.029 | 2.202 | -2.093 | -0.853 | 0.006 | 0.186 | 0.186 | 0.989 | 0.38 | 0.1 | 0.26 | 0.12 | 0.31 | 0.37 |
| 5 | 66799242 | rs6449870 | -0.951 | 2.087 | -1.884 | -0.637 | -0.001 | -0.011 | -0.011 | 0 | 0.36 | 0.62 | 0.56 | 0.81 | 0.38 | 0.31 |
| 5 | 66799452 | rs17308588 | -0.608 | 1.501 | -1.83 | -0.081 | 0.005 | 0.003 | 0.003 | 0 | 0.06 | 0.05 | 0.1 | 0.11 | 0.11 | 0.17 |
| 5 | 66800982 | rs4976077 | 0.502 | 0.82 | -2.032 | 0.086 | 0.004 | -0.05 | -0.05 | 0.279 | 0.29 | 0.15 | 0.35 | 0.53 | 0.27 | 0.22 |
| 5 | 66810839 | rs13183882 | -1.533 | 1.069 | -0.645 | 0.536 | -0.001 | 0.047 | 0.047 | 0 | 0.23 | 0.49 | 0.34 | 0.71 | 0.2 | 0.22 |
| 5 | 66810900 | rs10940112 | -2.749 | 1.043 | -0.873 | 0.39 | -0.03 | 0.28 | 0.28 | 1.804 | 0.51 | 0.89 | 0.61 | 0.86 | 0.47 | 0.53 |
| 5 | 66818485 | rs2171811 | 0.148 | 1.114 | -0.79 | 0.171 | 0.049 | 0.095 | 0.095 | 0 | 0.15 | 0.04 | 0.04 | 0 | 0.08 | 0.1 |

| | | | | | | | | | | | | | | | | |
|---|---|---|---|---|---|---|---|---|---|---|---|---|---|---|---|---|
| 5 | 66822255 | rs11958641 | -1.426 | 1.29 | -0.651 | 0.266 | 0.028 | 0.017 | 0.017 | 1.585 | 0.15 | 0.27 | 0.22 | 0.16 | 0.21 | 0.27 |
| 5 | 66827114 | rs6864405 | -0.371 | 1.111 | -0.783 | 0.192 | 0.026 | 0.046 | 0.046 | 0 | 0.18 | 0.07 | 0.07 | 0 | 0.12 | 0.11 |
| 5 | 66833257 | rs7717572 | -0.435 | 1.17 | -0.605 | 0.551 | -0.003 | 0.143 | 0.143 | 0 | 0.06 | 0.1 | 0.12 | 0.42 | 0.05 | 0.09 |
| 5 | 66845236 | rs4976084 | -2.084 | 1.261 | -0.822 | 0.377 | 0.026 | 0.023 | 0.023 | 0.019 | 0.39 | 0.71 | 0.4 | 0.74 | 0.28 | 0.31 |
| 5 | 66872361 | rs4565172 | -2.594 | 1.315 | -0.858 | 0.381 | 0.025 | 0.135 | 0.135 | 0.519 | 0.45 | 0.82 | 0.5 | 0.79 | 0.35 | 0.43 |
| 5 | 66875825 | rs4305607 | -2.471 | 1.328 | -0.812 | 0.391 | 0.016 | 0.056 | 0.056 | 0.099 | 0.44 | 0.71 | 0.5 | 0.79 | 0.35 | 0.43 |
| 5 | 66876159 | rs6862758 | -2.185 | 1.305 | -0.751 | 0.417 | 0.006 | 0.01 | 0.01 | 0.137 | 0.39 | 0.63 | 0.48 | 0.78 | 0.3 | 0.38 |
| 5 | 66878487 | rs4267824 | -1.838 | 1.287 | -0.752 | 0.42 | -0.012 | -0.003 | -0.003 | 0.332 | 0.35 | 0.61 | 0.38 | 0.71 | 0.23 | 0.26 |
| 5 | 66886475 | rs4308439 | -2.052 | 1.037 | -0.817 | 0.369 | 0.039 | 0.079 | 0.079 | 0.58 | 0.4 | 0.75 | 0.48 | 0.79 | 0.3 | 0.39 |
| 5 | 66908386 | rs10057614 | -2.048 | -0.301 | -0.616 | 0.538 | 0.025 | 0.047 | 0.047 | 0.189 | 0.39 | 0.68 | 0.46 | 0.79 | 0.3 | 0.39 |
| 5 | 66910080 | rs4128309 | -1.281 | -0.224 | -0.59 | 0.524 | 0.041 | 0.011 | 0.011 | 0.039 | 0.13 | 0.31 | 0.15 | 0.08 | 0.17 | 0.15 |
| 5 | 66912233 | rs4976096 | -1.543 | -0.295 | -0.663 | 0.678 | -0.003 | 0.014 | 0.014 | 0.632 | 0.17 | 0.15 | 0.34 | 0.45 | 0.26 | 0.25 |
| 5 | 66917073 | rs12657621 | -2.752 | -0.385 | -0.75 | 0.543 | -0.057 | 0.316 | 0.316 | 2.191 | 0.52 | 0.93 | 0.62 | 0.87 | 0.46 | 0.5 |
| 5 | 66933755 | rs13172995 | -1.31 | -0.248 | -0.332 | 0.917 | -0.007 | 0.001 | 0.001 | 0.207 | 0.3 | 0.62 | 0.24 | 0.42 | 0.16 | 0.21 |
| 5 | 66934212 | rs4451012 | -0.42 | -0.161 | -0.323 | 0.967 | 0.003 | 0.013 | 0.013 | 0.117 | 0.08 | 0.11 | 0.13 | 0.08 | 0.12 | 0.07 |
| 5 | 66936150 | rs4078050 | -0.908 | -0.225 | -0.209 | 0.963 | -0.017 | 0.356 | 0.356 | 0.249 | 0.12 | 0.46 | 0.17 | 0.42 | 0.05 | 0.1 |
| 5 | 66936418 | rs12521113 | -1.094 | -0.279 | -0.232 | 0.995 | 0.009 | 0.017 | 0.017 | 0.802 | 0.11 | 0.15 | 0.19 | 0.15 | 0.17 | 0.17 |
| 5 | 66939093 | rs6891500 | -0.982 | -0.337 | -0.294 | 0.93 | 0.011 | -0.018 | -0.018 | 0.038 | 0.23 | 0.15 | 0.25 | 0.3 | 0.22 | 0.18 |
| 5 | 66969658 | rs6896755 | -3.184 | -0.269 | -0.277 | 0.923 | -0.017 | 0.282 | 0.282 | 2.333 | 0.48 | 0.85 | 0.62 | 0.87 | 0.46 | 0.51 |
| 5 | 66975032 | rs11952144 | -0.228 | -0.365 | -0.165 | 1.041 | 0.025 | 0.161 | 0.161 | 0 | 0.06 | 0.1 | 0.15 | 0.3 | 0.1 | 0.07 |
| 5 | 66979905 | rs7723092 | -1.991 | -0.373 | -0.155 | 1.037 | 0.016 | 0.007 | 0.007 | 0 | 0.36 | 0.63 | 0.42 | 0.72 | 0.28 | 0.28 |
| 5 | 66982204 | rs10063787 | -1.324 | -0.133 | -0.133 | 1.134 | 0.007 | 0.288 | 0.288 | 0.26 | 0.14 | 0.46 | 0.21 | 0.42 | 0.1 | 0.11 |
| 5 | 66985073 | rs12657185 | 0.165 | -0.048 | -0.416 | 1.167 | -0.008 | 0.038 | 0.038 | 0 | 0.05 | 0.02 | 0.15 | 0.3 | 0.1 | 0.07 |
| 5 | 66991015 | rs11950551 | 0.005 | -0.066 | -0.38 | 1.132 | -0.013 | -0.014 | -0.014 | 0 | 0.07 | 0.01 | 0.13 | 0.32 | 0.1 | 0.07 |
| 5 | 66992708 | rs4246770 | -1.592 | -0.22 | -0.322 | 1.043 | -0.006 | 0.003 | 0.003 | 1.377 | 0.32 | 0.67 | 0.29 | 0.44 | 0.22 | 0.26 |
| 5 | 66993222 | rs6887552 | -1.942 | -0.083 | -0.237 | 1.089 | 0.013 | -0.026 | -0.026 | 0.947 | 0.36 | 0.75 | 0.42 | 0.51 | 0.34 | 0.29 |
| 5 | 66993280 | rs6891461 | -1.28 | -0.05 | -0.283 | 0.968 | 0.012 | 0.055 | 0.055 | 0 | 0.14 | 0.17 | 0.25 | 0.38 | 0.17 | 0.2 |
| 5 | 66993865 | rs10077665 | -1.509 | -0.037 | -0.197 | 0.97 | -0.035 | 0.045 | 0.045 | 1.035 | 0.29 | 0.56 | 0.28 | 0.45 | 0.2 | 0.22 |
| 5 | 66994075 | rs6893277 | -0.359 | -0.014 | -0.31 | 0.924 | 0.036 | 0.033 | 0.033 | 0 | 0.08 | 0.2 | 0.13 | 0.09 | 0.14 | 0.08 |
| 5 | 66996304 | rs4976023 | -2.295 | -0.031 | -0.307 | 0.931 | 0.015 | 0.09 | 0.09 | 0.786 | 0.43 | 0.72 | 0.52 | 0.81 | 0.36 | 0.42 |
| 5 | 67010538 | rs4607305 | 0.124 | -0.097 | 0.204 | 1.472 | 0.006 | 0.015 | 0.015 | 0 | 0.06 | 0.07 | 0.11 | 0.08 | 0.12 | 0.04 |
| 5 | 67012900 | rs7701312 | -0.392 | -0.45 | 0.284 | 1.494 | 0.03 | 0.016 | 0.016 | 0 | 0.05 | 0.08 | 0.1 | 0.05 | 0.07 | 0.13 |
| 5 | 67022879 | rs4470720 | 1.428 | -0.605 | 0.332 | 1.603 | -0.038 | 0.267 | 0.267 | 2.118 | 0.85 | 0.53 | 0.81 | 0.56 | 0.93 | 0.87 |
| 5 | 67027882 | rs4131890 | 1.954 | -0.344 | -0.319 | 1.741 | -0.001 | 0.06 | 0.06 | 0 | 0.89 | 0.82 | 0.83 | 0.56 | 0.92 | 0.85 |
| 5 | 67031977 | rs7708333 | -1.062 | -0.825 | -0.452 | 1.681 | -0.005 | 0.02 | 0.02 | 0 | 0.32 | 0.36 | 0.28 | 0.47 | 0.29 | 0.31 |
| 5 | 67038975 | rs9291891 | -1.399 | -0.769 | -0.41 | 1.618 | 0.002 | 0.001 | 0.001 | 0 | 0.39 | 0.37 | 0.47 | 0.82 | 0.34 | 0.39 |
| 5 | 67047907 | rs7711033 | -1.391 | -0.792 | -0.406 | 1.622 | 0.002 | 0.001 | 0.001 | 0 | 0.39 | 0.37 | 0.47 | 0.81 | 0.34 | 0.41 |
| 5 | 67060541 | rs4976024 | -1.294 | -0.216 | -0.9 | 1.935 | 0.017 | 0.013 | 0.013 | 0 | 0.06 | 0.01 | 0.07 | 0.39 | 0.02 | 0.09 |
| 5 | 67084526 | rs6868781 | -1.272 | -0.136 | -1.042 | 0.866 | 0.02 | 0.323 | 0.323 | 0 | 0.48 | 0.91 | 0.56 | 0.89 | 0.37 | 0.38 |
| 5 | 67087499 | rs6896115 | -0.767 | -0.258 | -0.21 | 1.474 | 0.003 | 0.009 | 0.009 | 0 | 0.34 | 0.38 | 0.43 | 0.41 | 0.35 | 0.32 |
| 5 | 67092939 | rs11958666 | -0.015 | 0.073 | -1.051 | 1.596 | -0.007 | -0.035 | -0.035 | 0 | 0.09 | 0.01 | 0.23 | 0.31 | 0.08 | 0.05 |
| 5 | 67093326 | rs6889533 | -1.224 | 0.115 | -1.068 | 0.45 | 0.01 | 0.208 | 0.208 | 0 | 0.46 | 0.91 | 0.57 | 0.81 | 0.38 | 0.39 |
| 5 | 67095390 | rs10054692 | -0.454 | 0.194 | -0.893 | 0.522 | -0.004 | 0.014 | 0.014 | 0.001 | 0.13 | 0.17 | 0.33 | 0.71 | 0.09 | 0.1 |
| 5 | 67096077 | rs3813668 | -0.683 | 0.661 | -0.475 | 0.221 | -0.007 | -0.106 | -0.106 | 2.882 | 0.32 | 0.51 | 0.22 | 0.09 | 0.29 | 0.28 |
| 5 | 67098541 | rs17316988 | -0.96 | 1.986 | -1.158 | 0.816 | -0.077 | -0.074 | -0.074 | 8.233 | 0.13 | 0.02 | 0.13 | 0.25 | 0.24 | 0.2 |
| 5 | 67098598 | rs7728135 | -0.045 | 1.772 | -1.885 | -0.032 | -0.008 | 0.054 | 0.054 | 0 | 0.27 | 0.17 | 0.2 | 0.13 | 0.25 | 0.31 |
| 5 | 67099642 | rs6414812 | 0.476 | 1.521 | -1.698 | -0.423 | -0.039 | -0.009 | -0.009 | 0 | 0.74 | 0.8 | 0.76 | 0.75 | 0.61 | 0.64 |
| 5 | 67099715 | rs4976132 | 0.613 | 1.334 | -0.89 | -0.527 | 0.019 | -0.002 | -0.002 | 0 | 0.08 | 0.16 | 0.13 | 0.09 | 0.11 | 0.12 |
| 5 | 67101321 | rs1490822 | 0.01 | 0.457 | -0.533 | -0.926 | -0.026 | -0.049 | -0.049 | 0 | 0.2 | 0.45 | 0.18 | 0.13 | 0.17 | 0.18 |
| 5 | 67104334 | rs7378773 | -0.028 | 1.434 | -0.676 | -0.685 | -0.021 | -0.012 | -0.012 | 0.351 | 0.82 | 0.85 | 0.82 | 0.78 | 0.74 | 0.72 |
| 5 | 67110964 | rs16897054 | -0.597 | 1.162 | -0.389 | -1.404 | 0.013 | -0.059 | -0.059 | 0.745 | 0.24 | 0.49 | 0.35 | 0.13 | 0.26 | 0.29 |
| 5 | 67112960 | rs11744310 | -0.739 | 1.741 | -0.573 | -1.404 | -0.016 | 0.011 | 0.011 | 0.481 | 0.28 | 0.28 | 0.39 | 0.09 | 0.4 | 0.38 |
| 5 | 67121201 | rs10042680 | -0.439 | 1.025 | -0.786 | -1.268 | -0.05 | 0.039 | 0.039 | 0 | 0.2 | 0.1 | 0.28 | 0.11 | 0.29 | 0.2 |
| 5 | 67125958 | rs10461533 | 1.243 | 0.422 | -1.356 | -1.531 | 0.047 | 0.009 | 0.009 | 0 | 0.17 | 0.02 | 0.07 | 0.12 | 0.11 | 0.16 |
| 5 | 67126863 | rs12520934 | 1.011 | 0.563 | -1.653 | -1.445 | -0.101 | -0.078 | -0.078 | 0 | 0.15 | 0.02 | 0.17 | 0.25 | 0.27 | 0.35 |
| 5 | 67127316 | rs2372179 | 0.47 | 0.44 | -0.989 | -1.463 | 0.002 | -0.025 | -0.025 | 0 | 0.14 | 0.29 | 0.22 | 0.09 | 0.15 | 0.1 |
| 5 | 67128187 | rs7723542 | 1.42 | 0.519 | -1.029 | -1.424 | 0.047 | 0.009 | 0.009 | 0 | 0.17 | 0.02 | 0.07 | 0.12 | 0.11 | 0.16 |
| 5 | 67130616 | rs10040640 | -0.641 | 0.279 | -0.864 | -1.665 | 0.016 | 0.007 | 0.007 | 0 | 0.54 | 0.58 | 0.47 | 0.43 | 0.61 | 0.62 |
| 5 | 67131310 | rs10052518 | 0.223 | 0.276 | -0.586 | -1.687 | -0.001 | 0.111 | 0.111 | 0 | 0.37 | 0.18 | 0.27 | 0.17 | 0.33 | 0.28 |
| 5 | 67137260 | rs10051387 | 0.644 | 0.313 | -0.316 | -1.687 | -0.003 | -0.007 | -0.007 | 1.708 | 0.31 | 0.26 | 0.15 | 0.18 | 0.2 | 0.19 |
| 5 | 67137684 | rs13174062 | 0.647 | 0.275 | -0.31 | -1.697 | 0.046 | 0.078 | 0.078 | 0 | 0.3 | 0.09 | 0.14 | 0.18 | 0.2 | 0.19 |
| 5 | 67139113 | rs1490813 | -0.22 | 0.402 | -0.336 | -1.42 | -0.003 | 0.008 | 0.008 | 0 | 0.47 | 0.53 | 0.35 | 0.43 | 0.48 | 0.54 |
| 5 | 67141138 | rs16897098 | 1.151 | 0.427 | -0.497 | -1.408 | 0.057 | 0.034 | 0.034 | 0.393 | 0.16 | 0.27 | 0.19 | 0.25 | 0.27 | 0.34 |
| 5 | 67150765 | rs10038754 | -1.599 | 0.352 | -0.563 | -1.46 | 0.037 | -0.058 | -0.058 | 0.615 | 0.69 | 0.84 | 0.68 | 0.52 | 0.75 | 0.72 |
| 5 | 67152355 | rs28647834 | 0.864 | -0.917 | -0.393 | -1.293 | -0.002 | 0.022 | 0.022 | 0 | 0.14 | 0.07 | 0.2 | 0.09 | 0.14 | 0.1 |
| 5 | 67158729 | rs10065943 | 1.695 | -0.953 | -0.423 | -1.313 | -0.038 | 0.044 | 0.044 | 0 | 0.12 | 0.01 | 0.09 | 0.06 | 0.13 | 0.03 |
| 5 | 67158880 | rs11738149 | 0.861 | -0.854 | -0.343 | -1.327 | -0.002 | 0.021 | 0.021 | 0 | 0.14 | 0.07 | 0.2 | 0.09 | 0.14 | 0.1 |
| 5 | 67166547 | rs1490800 | 0.442 | -0.699 | 0.023 | -1.103 | 0.03 | 0.001 | 0.001 | 0.426 | 0.24 | 0.29 | 0.31 | 0.25 | 0.4 | 0.43 |
| 5 | 67179335 | rs1490791 | -0.63 | -1.753 | -0.299 | -0.448 | 0.036 | -0.077 | -0.077 | 2.496 | 0.7 | 0.87 | 0.8 | 0.53 | 0.76 | 0.81 |

| | | | | | | | | | | | | | | | | |
|---|---|---|---|---|---|---|---|---|---|---|---|---|---|---|---|---|
| 5 | 67179407 | rs6859671 | 0.248 | -0.898 | 0.112 | -0.446 | 0.01 | 0.009 | 0.009 | 0 | 0.18 | 0.14 | 0.05 | 0.12 | 0.08 | 0.16 |
| 5 | 67194469 | rs10940138 | 0.529 | -0.222 | -0.804 | -0.046 | 0.051 | 0.054 | 0.054 | 0 | 0.32 | 0.19 | 0.25 | 0.15 | 0.22 | 0.14 |
| 5 | 67197955 | rs2372183 | -0.907 | -0.037 | -0.56 | -0.567 | -0.035 | -0.023 | -0.023 | 0 | 0.79 | 0.78 | 0.89 | 0.89 | 0.89 | 0.74 |
| 5 | 67209979 | rs13172284 | 0.619 | 0.044 | -0.336 | -0.905 | -0.003 | -0.017 | -0.017 | 0 | 0.19 | 0.19 | 0.23 | 0.07 | 0.17 | 0.17 |
| 5 | 67210430 | rs7723716 | -0.452 | 0.715 | -0.497 | -0.975 | -0.003 | 0.02 | 0.02 | 0 | 0.67 | 0.78 | 0.77 | 0.87 | 0.68 | 0.66 |
| 5 | 67220196 | rs16897180 | 1.613 | 0.758 | -0.646 | -0.835 | 0.05 | 0.044 | 0.044 | 0 | 0.28 | 0.17 | 0.25 | 0.18 | 0.2 | 0.13 |
| 5 | 67231863 | rs4976139 | 0.367 | 0.879 | -0.179 | -0.333 | 0.003 | -0.039 | -0.039 | 0.545 | 0.26 | 0.26 | 0.39 | 0.07 | 0.34 | 0.26 |
| 5 | 67235057 | rs7720751 | 0.852 | 0.505 | -0.357 | -0.236 | 0.005 | 0.036 | 0.036 | 0 | 0.36 | 0.3 | 0.39 | 0.79 | 0.34 | 0.41 |
| 5 | 67237504 | rs10072910 | -0.861 | 0.369 | -0.23 | -0.305 | -0.061 | -0.054 | -0.054 | 0.59 | 0.75 | 0.63 | 0.88 | 0.89 | 0.86 | 0.72 |
| 5 | 67238423 | rs2372189 | 1.225 | 0.47 | -0.623 | -0.108 | -0.045 | -0.025 | -0.025 | 0 | 0.07 | 0.02 | 0.12 | 0.61 | 0.14 | 0.27 |
| 5 | 67239774 | rs6872249 | -0.703 | 0.355 | -0.547 | -0.336 | 0.052 | 0.048 | 0.048 | 0.058 | 0.47 | 0.36 | 0.6 | 0.71 | 0.66 | 0.58 |
| 5 | 67240236 | rs9291908 | -0.001 | 0.536 | -0.608 | -0.401 | -0.044 | 0.001 | 0.001 | 0.62 | 0.3 | 0.21 | 0.49 | 0.68 | 0.44 | 0.49 |
| 5 | 67241677 | rs10515059 | 0.764 | 0.626 | -0.661 | -0.406 | -0.016 | 0.007 | 0.007 | 0 | 0.17 | 0.12 | 0.11 | 0.03 | 0.22 | 0.09 |
| 5 | 67246171 | rs991123 | 1.716 | 0.522 | -0.715 | -0.406 | 0.036 | 0.043 | 0.043 | 0 | 0.27 | 0.19 | 0.29 | 0.18 | 0.21 | 0.14 |
| 5 | 67253065 | rs1948330 | 0.781 | 0.731 | -0.285 | -0.214 | -0.07 | 0.078 | 0.078 | 0 | 0.15 | 0.03 | 0.11 | 0.03 | 0.23 | 0.09 |
| 5 | 67255464 | rs10471333 | -0.413 | 0.458 | -0.151 | -0.223 | -0.096 | -0.073 | -0.073 | 2.202 | 0.74 | 0.6 | 0.89 | 0.88 | 0.86 | 0.73 |
| 5 | 67255635 | rs12656263 | 0.468 | 1.137 | 0.019 | 0.367 | 0.004 | 0.006 | 0.006 | 0 | 0.49 | 0.39 | 0.65 | 0.48 | 0.49 | 0.37 |
| 5 | 67256569 | rs1366315 | 0.023 | 0.302 | 0.183 | -0.574 | -0.012 | -0.067 | -0.067 | 0.095 | 0.59 | 0.59 | 0.78 | 0.86 | 0.65 | 0.64 |
| 5 | 67267840 | rs1593911 | 0.191 | 0.412 | -0.398 | -0.494 | 0.017 | 0.043 | 0.043 | 0 | 0.57 | 0.43 | 0.77 | 0.86 | 0.65 | 0.63 |
| 5 | 67288272 | rs281455 | 1.061 | 0.711 | -0.037 | 0.524 | 0.021 | 0.146 | 0.146 | 0 | 0.93 | 0.81 | 0.92 | 0.73 | 0.92 | 0.75 |
| 5 | 67290972 | rs10940145 | 0.423 | 0.465 | 0.205 | -0.098 | -0.009 | 0.021 | 0.021 | 0 | 0.28 | 0.37 | 0.31 | 0.65 | 0.25 | 0.15 |
| 5 | 67293409 | rs157414 | 0.842 | 0.592 | 0.157 | 0.154 | 0.025 | 0.169 | 0.169 | 0 | 0.93 | 0.78 | 0.91 | 0.73 | 0.92 | 0.75 |
| 5 | 67294907 | rs157411 | 0.805 | 1.011 | -0.03 | -0.349 | 0.048 | 0.052 | 0.052 | 0 | 0.79 | 0.64 | 0.78 | 0.69 | 0.72 | 0.56 |
| 5 | 67295522 | rs281448 | 0.413 | 0.385 | -1.1 | -1.094 | -0.094 | 0.105 | 0.105 | 0 | 0.26 | 0.06 | 0.37 | 0.06 | 0.35 | 0.32 |
| 5 | 67297484 | rs4976029 | -0.073 | -0.128 | -1.209 | -1.186 | -0.003 | 0.097 | 0.097 | 0 | 0.6 | 0.84 | 0.74 | 0.78 | 0.63 | 0.61 |
| 5 | 67300449 | rs9291919 | 1.169 | -0.049 | -0.795 | -1.193 | -0.008 | 0.015 | 0.015 | 0 | 0.3 | 0.39 | 0.32 | 0.67 | 0.26 | 0.15 |
| 5 | 67301383 | rs11744801 | 0.58 | 0.229 | -1.302 | -1.146 | -0.062 | 0.023 | 0.023 | 0 | 0.12 | 0.02 | 0.11 | 0.03 | 0.19 | 0.15 |
| 5 | 67301948 | rs13354141 | 0.013 | 0.181 | -1.311 | -1.181 | 0.006 | 0.175 | 0.175 | 0 | 0.34 | 0.09 | 0.18 | 0.03 | 0.3 | 0.26 |
| 5 | 67302176 | rs13354232 | 1.085 | -0.167 | -1.123 | -1.03 | 0.07 | 0.079 | 0.079 | 0.176 | 0.22 | 0.07 | 0.07 | 0 | 0.11 | 0.11 |
| 5 | 67307462 | rs10067755 | 0.979 | 1.037 | -0.286 | -0.174 | 0.005 | 0.002 | 0.002 | 0.523 | 0.53 | 0.42 | 0.39 | 0.63 | 0.38 | 0.27 |
| 5 | 67308471 | rs382190 | 0.337 | 0.929 | -0.281 | 0.047 | -0.003 | 0.045 | 0.045 | 0 | 0.89 | 0.84 | 0.9 | 0.78 | 0.91 | 0.61 |
| 5 | 67308533 | rs382480 | 0.064 | 1.231 | -0.561 | -0.007 | -0.055 | 0.054 | 0.054 | 0.303 | 0.22 | 0.11 | 0.36 | 0.1 | 0.35 | 0.19 |
| 5 | 67308629 | rs9986158 | 0.458 | 1.107 | -0.472 | 0.089 | 0.014 | -0.006 | -0.006 | 0.021 | 0.65 | 0.55 | 0.53 | 0.68 | 0.56 | 0.42 |
| 5 | 67310302 | rs282292 | -0.264 | 1.116 | -0.631 | 0.34 | -0.001 | 0 | 0 | 0 | 0.45 | 0.54 | 0.6 | 0.35 | 0.53 | 0.55 |
| 5 | 67323811 | rs383542 | -0.186 | 1.378 | -0.843 | 0.096 | -0.02 | 0.037 | 0.037 | 0 | 0.08 | 0.02 | 0.06 | 0 | 0.11 | 0.07 |
| 5 | 67324941 | rs16897333 | -0.376 | 1.955 | -0.665 | 0.101 | 0.05 | 0.015 | 0.015 | 0 | 0.07 | 0.16 | 0.06 | 0.07 | 0.11 | 0.08 |
| 5 | 67326258 | rs714458 | 0.059 | 2.062 | -1.126 | 0.857 | -0.009 | -0.045 | -0.045 | 0 | 0.88 | 0.97 | 0.91 | 0.77 | 0.89 | 0.83 |
| 5 | 67332561 | rs281463 | 1.085 | 2.055 | -0.921 | 1.033 | 0 | 0.003 | 0.003 | 1.436 | 0.82 | 0.84 | 0.84 | 0.71 | 0.78 | 0.73 |
| 5 | 67335548 | rs12697058 | 1.035 | 2.135 | -0.524 | 0.776 | 0.01 | 0.012 | 0.012 | 0.504 | 0.66 | 0.52 | 0.56 | 0.38 | 0.6 | 0.63 |
| 5 | 67335658 | rs281460 | 0.076 | 2.019 | -0.559 | 0.741 | -0.048 | -0.092 | -0.092 | 0 | 0.87 | 0.98 | 0.82 | 0.42 | 0.82 | 0.74 |
| 5 | 67336674 | rs10078053 | 0.582 | 1.268 | -0.307 | -0.043 | 0 | -0.027 | -0.027 | 0 | 0.41 | 0.41 | 0.18 | 0.05 | 0.27 | 0.25 |
| 5 | 67342328 | rs13158718 | -0.905 | 1.25 | -0.407 | 0 | -0.022 | -0.062 | -0.062 | 0.023 | 0.89 | 0.99 | 0.91 | 0.78 | 0.89 | 0.82 |
| 5 | 67344993 | rs6877998 | 0.269 | 1.1 | -0.062 | -0.362 | 0.009 | -0.009 | -0.009 | 0 | 0.4 | 0.36 | 0.16 | 0.05 | 0.24 | 0.24 |
| 5 | 67348542 | rs11953134 | -0.161 | 1.408 | 0.064 | 0.185 | 0.041 | -0.004 | -0.004 | 0 | 0.07 | 0.14 | 0.09 | 0.06 | 0.14 | 0.12 |
| 5 | 67352080 | rs718291 | 0.66 | 0.563 | 0.134 | 0.437 | -0.01 | 0.101 | 0.101 | 0 | 0.11 | 0.17 | 0.17 | 0.34 | 0.07 | 0.2 |
| 5 | 67353323 | rs964279 | 0.125 | 0.24 | -0.181 | -0.286 | -0.012 | -0.081 | -0.081 | 0 | 0.4 | 0.51 | 0.2 | 0.11 | 0.32 | 0.24 |
| 5 | 67375691 | rs6874150 | -0.166 | 0.253 | -0.195 | -0.183 | 0.023 | -0.001 | -0.001 | 0.015 | 0.51 | 0.61 | 0.37 | 0.45 | 0.4 | 0.43 |
| 5 | 67384201 | rs7737981 | 0.258 | 0.32 | -0.097 | -0.272 | -0.005 | -0.045 | -0.045 | 0 | 0.35 | 0.54 | 0.18 | 0.11 | 0.29 | 0.23 |
| 5 | 67386576 | rs176003 | 0.939 | 0.93 | -0.848 | 0.021 | -0.07 | 0.062 | 0.062 | 0 | 0.23 | 0.02 | 0.16 | 0.11 | 0.28 | 0.23 |
| 5 | 67387273 | rs1864049 | -0.18 | 0.835 | 0.278 | 0.622 | -0.003 | -0.114 | -0.114 | 0.471 | 0.27 | 0.46 | 0.16 | 0.08 | 0.23 | 0.15 |
| 5 | 67391776 | rs17309175 | 0.218 | 0.976 | -0.397 | 0.679 | -0.001 | -0.015 | -0.015 | 0 | 0.24 | 0.21 | 0.13 | 0.06 | 0.23 | 0.15 |
| 5 | 67393410 | rs7723962 | 0.274 | 0.996 | -0.704 | 0.689 | -0.005 | 0.022 | 0.022 | 0.267 | 0.23 | 0.15 | 0.13 | 0.06 | 0.23 | 0.15 |
| 5 | 67397704 | rs465531 | -0.227 | 1.343 | -1.056 | 1.087 | 0.008 | -0.008 | -0.008 | 0 | 0.4 | 0.68 | 0.33 | 0.41 | 0.31 | 0.32 |
| 5 | 67405115 | rs282280 | -0.241 | 1.219 | -0.758 | 1.41 | 0 | 0.012 | 0.012 | 1.867 | 0.48 | 0.68 | 0.41 | 0.4 | 0.43 | 0.31 |
| 5 | 67409441 | rs7714988 | -1.291 | 2.4 | -0.393 | 2.327 | 0.05 | 0.113 | 0.113 | 0 | 0.15 | 0.25 | 0.3 | 0.51 | 0.26 | 0.24 |
| 5 | 67412203 | rs17237251 | -1.247 | 2.412 | -0.393 | 2.291 | 0.053 | 0.129 | 0.129 | 0 | 0.14 | 0.25 | 0.29 | 0.49 | 0.26 | 0.2 |
| 5 | 67450499 | rs10515068 | -1.324 | 2.989 | -0.471 | 2.162 | 0.006 | 0.026 | 0.026 | 0 | 0.1 | 0.12 | 0.26 | 0.27 | 0.24 | 0.25 |
| 5 | 67457160 | rs4524465 | -0.627 | 2.485 | -1.74 | 0.985 | -0.022 | 0.07 | 0.07 | 0 | 0.68 | 0.94 | 0.81 | 0.81 | 0.73 | 0.68 |
| 5 | 67463231 | rs10805422 | -0.636 | 1.847 | -2.001 | 0.497 | -0.019 | 0.159 | 0.159 | 0 | 0.75 | 0.97 | 0.88 | 0.94 | 0.82 | 0.86 |
| 5 | 67475952 | rs4277860 | -0.703 | 2.02 | -0.969 | 1.351 | 0.013 | 0.115 | 0.115 | 0 | 0.09 | 0.14 | 0.15 | 0.42 | 0.12 | 0.14 |
| 5 | 67487328 | rs6886884 | -1.443 | 1.879 | -0.333 | 1.212 | 0.036 | 0.185 | 0.185 | 0 | 0.18 | 0.38 | 0.24 | 0.44 | 0.23 | 0.2 |
| 5 | 67493961 | rs12654191 | 0.039 | 1.931 | -0.303 | 1.258 | 0.007 | 0.066 | 0.066 | 0.244 | 0.06 | 0.09 | 0.19 | 0.25 | 0.13 | 0.21 |
| 5 | 67494991 | rs16897466 | -0.842 | 1.729 | 0.156 | 1.174 | 0.004 | 0.178 | 0.178 | 0 | 0.11 | 0.21 | 0.18 | 0.43 | 0.12 | 0.11 |
| 5 | 67514509 | rs7701498 | -1.028 | 1.303 | 0.609 | -0.127 | 0.03 | -0.127 | -0.127 | 0.47 | 0.18 | 0.47 | 0.17 | 0.02 | 0.19 | 0.1 |
| 5 | 67523267 | rs13173003 | -0.417 | 1.262 | 0.212 | -0.006 | 0.019 | 0.025 | 0.025 | 0 | 0.18 | 0.33 | 0.33 | 0.67 | 0.19 | 0.35 |
| 5 | 67527326 | rs7713645 | -0.962 | 0.957 | -0.991 | -0.187 | -0.017 | 0.196 | 0.196 | 0 | 0.47 | 0.82 | 0.56 | 0.72 | 0.49 | 0.5 |
| 5 | 67534039 | rs7709243 | -0.896 | 1.006 | -0.907 | -0.21 | -0.01 | 0.13 | 0.13 | 0 | 0.47 | 0.73 | 0.57 | 0.74 | 0.5 | 0.52 |
| 5 | 67534057 | rs12652661 | -0.452 | 1.19 | -0.531 | 0.446 | 0.021 | 0.033 | 0.033 | 0.019 | 0.26 | 0.52 | 0.3 | 0.58 | 0.28 | 0.42 |
| 5 | 67536974 | rs173704 | -0.193 | 1.037 | -1.643 | 0.277 | -0.047 | -0.026 | -0.026 | 0 | 0.84 | 0.99 | 0.77 | 0.86 | 0.83 | 0.91 |

| | | | | | | | | | | | | | | | | |
|---|---|---|---|---|---|---|---|---|---|---|---|---|---|---|---|---|
| 5 | 67539967 | rs173702 | 0.314 | 0.832 | -2.616 | 0.391 | -0.122 | -0.08 | -0.08 | 0 | 0.35 | 0.05 | 0.46 | 0.38 | 0.4 | 0.4 |
| 5 | 67545624 | rs4122269 | 0.428 | 0.344 | -2.132 | -0.02 | -0.011 | -0.04 | -0.04 | 0 | 0.36 | 0.51 | 0.25 | 0.25 | 0.36 | 0.28 |
| 5 | 67546406 | rs1823023 | -0.559 | 1.153 | -2.602 | -0.205 | -0.024 | -0.019 | -0.019 | 0.055 | 0.63 | 0.85 | 0.43 | 0.44 | 0.64 | 0.55 |
| 5 | 67547276 | rs10940159 | 1.962 | 1.037 | -2.562 | -0.275 | 0.002 | -0.003 | -0.003 | 0 | 0.08 | 0.1 | 0.06 | 0.16 | 0.09 | 0.19 |
| 5 | 67547326 | rs173703 | 1.62 | 0.98 | -2.657 | 0.085 | -0.059 | -0.092 | -0.092 | 1.28 | 0.2 | 0.02 | 0.31 | 0.36 | 0.21 | 0.35 |
| 5 | 67553771 | rs6893676 | 0.849 | 0.769 | -2.476 | 0.027 | -0.002 | -0.001 | -0.001 | 3.219 | 0.18 | 0.19 | 0.14 | 0.17 | 0.21 | 0.25 |
| 5 | 67553827 | rs34303 | 0.171 | 0.588 | -2.575 | -0.057 | -0.004 | -0.022 | -0.022 | 0 | 0.62 | 0.72 | 0.61 | 0.61 | 0.62 | 0.64 |
| 5 | 67554023 | rs863818 | 0.799 | 0.641 | -2.961 | -0.349 | -0.137 | -0.049 | -0.049 | 0.02 | 0.34 | 0.03 | 0.46 | 0.36 | 0.42 | 0.38 |
| 5 | 67560716 | rs16897561 | 1.35 | 0.502 | -2.762 | -0.354 | -0.032 | -0.009 | -0.009 | 0 | 0.26 | 0.56 | 0.13 | 0.25 | 0.19 | 0.26 |
| 5 | 67562930 | rs831122 | -1.972 | 0.501 | -2.825 | -0.357 | -0.041 | -0.088 | -0.088 | 0.51 | 0.84 | 0.99 | 0.77 | 0.78 | 0.85 | 0.9 |
| 5 | 67564383 | rs34309 | 1.236 | 0.564 | -2.878 | -0.402 | -0.115 | -0.074 | -0.074 | 0.052 | 0.28 | 0.03 | 0.42 | 0.36 | 0.37 | 0.38 |
| 5 | 67569479 | rs2302975 | -0.27 | 0.488 | -2.578 | -0.439 | 0.027 | 0.035 | 0.035 | 0.076 | 0.51 | 0.84 | 0.32 | 0.42 | 0.46 | 0.52 |
| 5 | 67569602 | rs2302976 | 0.856 | 0.386 | -2.412 | -0.285 | -0.001 | -0.001 | -0.001 | 0 | 0.19 | 0.2 | 0.14 | 0.18 | 0.21 | 0.25 |
| 5 | 67570138 | rs3730082 | 1.149 | 0.383 | -2.404 | -0.284 | -0.031 | -0.01 | -0.01 | 0.004 | 0.26 | 0.56 | 0.13 | 0.24 | 0.19 | 0.26 |
| 5 | 67571413 | rs6876003 | -0.502 | 0.426 | -2.604 | -0.273 | 0.045 | 0.033 | 0.033 | 0.1 | 0.49 | 0.84 | 0.3 | 0.42 | 0.42 | 0.52 |
| 5 | 67571771 | rs7716675 | 0.93 | 0.494 | -2.489 | -0.139 | -0.021 | -0.009 | -0.009 | 0 | 0.27 | 0.62 | 0.14 | 0.24 | 0.19 | 0.26 |
| 5 | 67573042 | rs12656176 | 0.511 | 0.965 | -2.329 | 0.102 | -0.003 | -0.003 | -0.003 | 5.218 | 0.17 | 0.17 | 0.14 | 0.18 | 0.21 | 0.25 |
| 5 | 67575642 | rs3815701 | 0.467 | 1.057 | -2.259 | 0.078 | -0.001 | 0.001 | 0.001 | 0.025 | 0.18 | 0.21 | 0.25 | 0.17 | 0.24 | 0.27 |
| 5 | 67578745 | rs34306 | -1.106 | 1.165 | -2.715 | 0.266 | -0.05 | -0.094 | -0.094 | 0.99 | 0.82 | 0.98 | 0.77 | 0.78 | 0.83 | 0.88 |
| 5 | 67583807 | rs1550805 | 1.374 | 1.219 | -2.803 | 0.044 | -0.032 | 0.006 | 0.006 | 0 | 0.13 | 0 | 0.04 | 0.1 | 0.09 | 0.13 |
| 5 | 67587434 | rs831125 | 1.308 | 1.428 | -2.726 | 0.172 | -0.032 | -0.015 | -0.015 | 0.277 | 0.1 | 0.01 | 0.1 | 0.09 | 0.11 | 0.08 |
| 5 | 67588148 | rs3730089 | 0.406 | 1.327 | -1.221 | 0.333 | 0.057 | 0.051 | 0.051 | 0 | 0.14 | 0.42 | 0.21 | 0.17 | 0.17 | 0.07 |
| 5 | 67594324 | rs1043526 | 1.977 | 1.389 | -1.76 | 0.091 | -0.029 | 0.053 | 0.053 | 0 | 0.09 | 0.01 | 0.08 | 0.01 | 0.13 | 0.04 |
| 5 | 67596088 | rs3756668 | -1.282 | 1.037 | -2.267 | 0.22 | -0.088 | -0.073 | -0.073 | 0.758 | 0.44 | 0.96 | 0.55 | 0.48 | 0.45 | 0.51 |
| 5 | 67605884 | rs411751 | -1.695 | 1.06 | -1.757 | 0.418 | 0 | -0.001 | -0.001 | 0 | 0.9 | 0.88 | 0.83 | 0.88 | 0.87 | 0.77 |
| 5 | 67608679 | rs9291928 | 0.197 | 1.001 | -1.522 | 0.333 | 0.013 | 0.034 | 0.034 | 0 | 0.24 | 0.45 | 0.31 | 0.35 | 0.25 | 0.26 |
| 5 | 67611486 | rs184886 | -0.526 | 1.174 | -1.004 | 0.446 | -0.002 | 0.012 | 0.012 | 0 | 0.78 | 0.8 | 0.73 | 0.86 | 0.73 | 0.71 |
| 5 | 67613643 | rs192282 | 0.026 | 1.137 | -0.902 | 0.425 | -0.007 | -0.087 | -0.087 | 0 | 0.73 | 0.57 | 0.7 | 0.86 | 0.71 | 0.71 |
| 5 | 67613801 | rs10515077 | 1.873 | 1.494 | -1.653 | 0.806 | -0.011 | -0.014 | -0.014 | 0 | 0.07 | 0.04 | 0.16 | 0.16 | 0.09 | 0.05 |
| 5 | 67615731 | rs625637 | -0.603 | 1.044 | -1.424 | 0.525 | -0.002 | 0.011 | 0.011 | 0 | 0.8 | 0.81 | 0.74 | 0.86 | 0.73 | 0.71 |
| 5 | 67625070 | rs12519613 | 0.625 | 1.011 | -1.583 | 1.155 | -0.027 | -0.069 | -0.069 | 0.319 | 0.12 | 0.03 | 0.18 | 0.23 | 0.16 | 0.2 |
| 5 | 67625955 | rs34289 | -0.253 | 0.743 | -1.813 | 0.612 | 0.009 | -0.015 | -0.015 | 0 | 0.43 | 0.15 | 0.31 | 0.48 | 0.36 | 0.43 |
| 5 | 67628886 | rs6859287 | -0.28 | 0.327 | -0.824 | 0.459 | 0.027 | 0.028 | 0.028 | 0 | 0.23 | 0.45 | 0.27 | 0.22 | 0.28 | 0.19 |
| 5 | 67638051 | rs6871452 | -0.259 | -0.323 | -0.223 | -0.489 | 0.039 | -0.087 | -0.087 | 0 | 0.14 | 0.33 | 0.11 | 0 | 0.18 | 0.05 |
| 5 | 67639915 | rs34299 | -1.578 | -0.554 | -0.36 | -0.653 | -0.004 | -0.007 | -0.007 | 0 | 0.68 | 0.78 | 0.49 | 0.4 | 0.66 | 0.66 |
| 5 | 67642788 | rs34297 | -1.417 | -0.513 | 0.251 | -0.013 | 0.004 | -0.121 | -0.121 | 0 | 0.39 | 0.53 | 0.35 | 0.07 | 0.44 | 0.27 |
| 5 | 67646565 | rs34296 | -0.009 | -0.487 | 0.053 | 0.079 | -0.006 | 0.05 | 0.05 | 0 | 0.22 | 0.08 | 0.22 | 0.1 | 0.23 | 0.22 |
| 5 | 67649611 | rs16897721 | -0.482 | -0.504 | 0.378 | 0.028 | -0.02 | 0.019 | 0.019 | 0.28 | 0.09 | 0.17 | 0.03 | 0.18 | 0.03 | 0.28 |
| 5 | 67651015 | rs12108976 | -0.991 | -0.472 | 0.532 | 0.028 | -0.023 | -0.002 | -0.002 | 1.892 | 0.16 | 0.37 | 0.07 | 0.18 | 0.09 | 0.29 |
| 5 | 67653782 | rs34294 | -0.75 | -0.469 | 0.44 | 0.016 | -0.008 | 0.022 | 0.022 | 0 | 0.34 | 0.33 | 0.33 | 0.19 | 0.41 | 0.28 |
| 5 | 67655495 | rs256497 | -1.778 | -0.453 | 0.272 | 0.041 | -0.002 | 0.048 | 0.048 | 0 | 0.51 | 0.71 | 0.4 | 0.38 | 0.5 | 0.58 |
| 5 | 67660888 | rs256499 | -1.688 | -0.551 | -0.12 | 0.031 | -0.001 | -0.002 | -0.002 | 0.037 | 0.48 | 0.54 | 0.39 | 0.36 | 0.5 | 0.59 |
| 5 | 67660897 | rs256500 | -1.104 | -0.06 | -0.562 | 0.047 | -0.014 | 0.038 | 0.038 | 0 | 0.34 | 0.28 | 0.32 | 0.17 | 0.41 | 0.28 |
| 5 | 67665310 | rs256504 | -0.576 | -0.487 | -0.823 | 0.423 | -0.025 | 0.001 | 0.001 | 0.015 | 0.22 | 0.03 | 0.21 | 0.16 | 0.22 | 0.24 |
| 5 | 67676341 | rs437204 | -0.446 | -1.075 | -0.67 | 0.18 | 0.019 | 0.035 | 0.035 | 0 | 0.14 | 0.07 | 0.12 | 0.01 | 0.11 | 0.06 |
| 5 | 67678506 | rs7703560 | -1.936 | -1.516 | -0.443 | 0.529 | 0.051 | -0.034 | -0.034 | 0 | 0.41 | 0.74 | 0.4 | 0.47 | 0.3 | 0.53 |
| 5 | 67681235 | rs16897779 | -0.957 | -1.603 | 0.067 | 0.687 | -0.017 | -0.008 | -0.008 | 0 | 0.18 | 0.2 | 0.08 | 0.19 | 0.08 | 0.43 |
| 5 | 67681425 | rs7719763 | -2.135 | -1.595 | 0.108 | 0.694 | 0.063 | -0.04 | -0.04 | 0 | 0.41 | 0.77 | 0.4 | 0.47 | 0.3 | 0.53 |
| 5 | 67682512 | rs7725083 | -0.822 | -1.212 | 0.053 | 0.581 | -0.025 | -0.005 | -0.005 | 0 | 0.14 | 0.18 | 0.06 | 0.18 | 0.05 | 0.28 |
| 5 | 67699045 | rs10213829 | -0.303 | -0.659 | -0.111 | 0.494 | 0 | 0.058 | 0.058 | 0 | 0.13 | 0.03 | 0.15 | 0.01 | 0.13 | 0.05 |
| 5 | 67699693 | rs7721613 | -2.05 | -0.79 | 0.154 | 0.595 | -0.002 | 0.001 | 0.001 | 0 | 0.29 | 0.66 | 0.37 | 0.28 | 0.25 | 0.25 |
| 5 | 67708511 | rs10066761 | -0.127 | -0.814 | -0.049 | 0.516 | 0.009 | 0.052 | 0.052 | 0 | 0.13 | 0.04 | 0.14 | 0.01 | 0.12 | 0.05 |
| 5 | 67712110 | rs6880711 | -1.227 | -0.905 | 0.378 | 0.565 | -0.001 | 0.002 | 0.002 | 0 | 0.22 | 0.24 | 0.23 | 0.35 | 0.16 | 0.1 |
| 5 | 67712993 | rs10075380 | -0.848 | -0.892 | 0.542 | 0.534 | 0 | 0.006 | 0.006 | 0 | 0.17 | 0.16 | 0.19 | 0.01 | 0.19 | 0.07 |
| 5 | 67714246 | rs4976033 | -1.968 | -0.882 | 0.445 | 0.623 | 0.033 | -0.005 | -0.005 | 0 | 0.45 | 0.71 | 0.44 | 0.52 | 0.41 | 0.31 |
| 5 | 67716636 | rs16897820 | -0.425 | -0.605 | 0.812 | 0.847 | -0.01 | 0.124 | 0.124 | 0.366 | 0.09 | 0.18 | 0.09 | 0.34 | 0.05 | 0.06 |
| 5 | 67728508 | rs1428080 | -0.715 | -0.564 | 0.44 | 0.813 | -0.039 | 0.163 | 0.163 | 0.622 | 0.16 | 0.39 | 0.13 | 0.33 | 0.06 | 0.1 |
| 5 | 67729637 | rs6862818 | -0.832 | -0.465 | -0.21 | 0.864 | -0.019 | 0.075 | 0.075 | 1.324 | 0.21 | 0.54 | 0.19 | 0.34 | 0.14 | 0.12 |
| 5 | 67732517 | rs830474 | 0.228 | -0.521 | -1.033 | 0.199 | 0.032 | 0.038 | 0.038 | 0.739 | 0.83 | 0.9 | 0.9 | 0.96 | 0.89 | 0.92 |
| 5 | 67738430 | rs853812 | -0.237 | 0.325 | -1.242 | 0.086 | -0.004 | -0.013 | -0.013 | 0 | 0.53 | 0.27 | 0.51 | 0.51 | 0.54 | 0.36 |
| 5 | 67740062 | rs256456 | 0.379 | 0.503 | -2.39 | -0.71 | -0.03 | 0.211 | 0.211 | 0 | 0.5 | 0.12 | 0.54 | 0.77 | 0.5 | 0.46 |
| 5 | 67741066 | rs11745891 | 1.053 | 0.623 | -1.819 | -0.785 | 0.002 | -0.028 | -0.028 | 0.158 | 0.07 | 0.12 | 0.08 | 0 | 0.08 | 0.03 |
| 5 | 67751221 | rs42905 | -0.019 | 0.611 | -1.397 | -0.871 | 0.004 | -0.033 | -0.033 | 0 | 0.59 | 0.48 | 0.62 | 0.75 | 0.57 | 0.45 |
| 5 | 67751350 | rs853808 | 1.436 | 0.723 | -1.783 | -0.775 | -0.054 | -0.094 | -0.094 | 0 | 0.16 | 0.02 | 0.13 | 0.32 | 0.18 | 0.21 |
| 5 | 67751381 | rs36297 | -1.556 | 0.847 | -1.355 | -0.847 | -0.043 | -0.049 | -0.049 | 0.101 | 0.88 | 0.99 | 0.85 | 0.83 | 0.84 | 0.59 |
| 5 | 67752638 | rs853807 | 0.328 | 0.721 | -1.243 | -0.891 | 0 | -0.005 | -0.005 | 0 | 0.38 | 0.35 | 0.42 | 0.48 | 0.38 | 0.29 |
| 5 | 67755091 | rs853806 | -0.314 | 0.846 | -0.944 | -0.953 | 0.002 | -0.036 | -0.036 | 0 | 0.59 | 0.51 | 0.67 | 0.79 | 0.63 | 0.46 |
| 5 | 67755706 | rs853805 | -0.077 | 0.564 | -0.746 | -0.701 | 0 | 0.005 | 0.005 | 0 | 0.38 | 0.44 | 0.49 | 0.46 | 0.43 | 0.29 |
| 5 | 67758584 | rs4635898 | -0.805 | 0.747 | -0.394 | -0.786 | 0.016 | -0.045 | -0.045 | 0 | 0.12 | 0.28 | 0.12 | 0.03 | 0.15 | 0.03 |

| | | | | | | | | | | | | | | | | |
|---|---|---|---|---|---|---|---|---|---|---|---|---|---|---|---|---|
| 5 | 67762103 | rs853803 | 0.882 | 0.744 | -0.18 | -0.847 | 0.005 | -0.02 | -0.02 | 0 | 0.3 | 0.3 | 0.45 | 0.45 | 0.28 | 0.33 |
| 5 | 67762172 | rs706723 | -0.05 | 0.618 | 0.184 | -1.121 | 0 | -0.016 | -0.016 | 2.177 | 0.5 | 0.58 | 0.61 | 0.8 | 0.46 | 0.42 |
| 5 | 67763676 | rs10056743 | 1.38 | 0.773 | 0.319 | -1.246 | -0.003 | -0.002 | -0.002 | 0 | 0.24 | 0.15 | 0.14 | 0.06 | 0.22 | 0.16 |
| 5 | 67767428 | rs853799 | 1.272 | 0.501 | 0.323 | -1.188 | 0.004 | -0.022 | -0.022 | 0.043 | 0.29 | 0.28 | 0.45 | 0.44 | 0.28 | 0.28 |
| 5 | 67768278 | rs16897859 | 0.073 | 0.576 | 0.422 | -1.187 | -0.008 | -0.037 | -0.037 | 0 | 0.08 | 0.17 | 0.05 | 0 | 0.07 | 0.01 |
| 5 | 67768380 | rs1863982 | 0.793 | 0.784 | 0.502 | -1.054 | -0.008 | 0.029 | 0.029 | 2.046 | 0.19 | 0.25 | 0.22 | 0.36 | 0.18 | 0.1 |
| 5 | 67770083 | rs853795 | 1.29 | 1.98 | 1.02 | -1.69 | 0.051 | 0.096 | 0.096 | 0.072 | 0.44 | 0.31 | 0.46 | 0.77 | 0.35 | 0.37 |
| 5 | 67772740 | rs11957631 | -0.844 | 1.81 | 1.01 | -1.563 | 0.063 | -0.015 | -0.015 | 0 | 0.11 | 0.39 | 0.16 | 0.05 | 0.16 | 0.03 |
| 5 | 67773706 | rs706725 | 0.268 | 1.869 | 0.95 | -1.595 | -0.015 | 0.086 | 0.086 | 1.447 | 0.58 | 0.79 | 0.65 | 0.81 | 0.52 | 0.45 |
| 5 | 67776535 | rs17256961 | 0 | 1.677 | 1.075 | -1.39 | 0.004 | -0.018 | -0.018 | 0 | 0.08 | 0.13 | 0.12 | 0.03 | 0.11 | 0.02 |
| 5 | 67779809 | rs6449962 | -1.094 | 1.77 | 0.854 | -1.404 | 0.004 | 0 | 0 | 0.495 | 0.49 | 0.53 | 0.47 | 0.23 | 0.56 | 0.6 |
| 5 | 67784239 | rs13182495 | -0.703 | 1.226 | 0.621 | -1.03 | 0.044 | 0.114 | 0.114 | 0.463 | 0.52 | 0.67 | 0.5 | 0.24 | 0.6 | 0.58 |
| 5 | 67813663 | rs6870517 | -0.644 | 1.026 | 0.826 | -0.97 | 0.009 | 0.06 | 0.06 | 0.077 | 0.48 | 0.58 | 0.44 | 0.24 | 0.49 | 0.56 |
| 5 | 67819584 | rs6893396 | -0.048 | 1.015 | 0.51 | -0.688 | -0.022 | 0.051 | 0.051 | 2.047 | 0.38 | 0.19 | 0.3 | 0.2 | 0.38 | 0.54 |
| 5 | 67823586 | rs10940168 | -0.297 | 1.201 | -0.093 | -0.542 | 0.01 | -0.05 | -0.05 | 0 | 0.39 | 0.47 | 0.32 | 0.2 | 0.39 | 0.51 |
| 5 | 67829847 | rs13356158 | 0.114 | 1.314 | -0.74 | -0.461 | -0.095 | 0.098 | 0.098 | 1.731 | 0.35 | 0.08 | 0.26 | 0.2 | 0.36 | 0.51 |
| 5 | 67845698 | rs1344367 | -0.069 | 1.177 | -0.449 | -0.395 | 0.004 | -0.025 | -0.025 | 0 | 0.4 | 0.39 | 0.35 | 0.2 | 0.41 | 0.5 |
| 5 | 67852968 | rs176065 | -0.472 | 0.747 | -0.262 | 0.259 | 0.002 | -0.005 | -0.005 | 0 | 0.45 | 0.44 | 0.66 | 0.53 | 0.57 | 0.4 |
| 5 | 67853689 | rs697614 | -1.55 | 0.529 | 0.902 | -0.213 | -0.001 | -0.118 | -0.118 | 0.846 | 0.22 | 0.59 | 0.18 | 0.03 | 0.22 | 0.13 |
| 5 | 67873190 | rs853506 | -1.848 | 0.57 | 0.935 | -0.098 | -0.001 | -0.064 | -0.064 | 0 | 0.24 | 0.65 | 0.21 | 0.13 | 0.24 | 0.14 |
| 5 | 67883394 | rs706733 | -1.043 | 0.727 | 0.986 | -0.049 | 0.002 | -0.005 | -0.005 | 0.116 | 0.11 | 0.17 | 0.11 | 0.11 | 0.15 | 0.12 |
| 5 | 67887939 | rs853645 | -1.085 | 0.576 | 1.04 | -0.168 | 0.002 | -0.036 | -0.036 | 0 | 0.11 | 0.18 | 0.1 | 0 | 0.13 | 0.12 |
| 5 | 67888580 | rs16897939 | -1.325 | 0.46 | 1.367 | -0.085 | 0.075 | 0.034 | 0.034 | 2.609 | 0.06 | 0.31 | 0.06 | 0.03 | 0.07 | 0.01 |
| 5 | 67900727 | rs6883966 | -0.665 | 0.173 | 0.973 | -0.097 | -0.004 | -0.013 | -0.013 | 0 | 0.06 | 0.07 | 0.02 | 0 | 0.03 | 0 |
| 5 | 67901292 | rs706753 | -2.419 | 0.284 | 1.556 | -0.313 | 0.004 | -0.131 | -0.131 | 0.019 | 0.22 | 0.54 | 0.17 | 0.02 | 0.22 | 0.13 |
| 5 | 67903602 | rs6449969 | -1.583 | 0.25 | 1.26 | -0.288 | -0.016 | -0.033 | -0.033 | 1.798 | 0.09 | 0.15 | 0.04 | 0.02 | 0.05 | 0.01 |
| 5 | 67915129 | rs2547985 | -2.044 | 0.186 | 1.287 | -0.27 | 0 | -0.027 | -0.027 | 1.332 | 0.2 | 0.25 | 0.14 | 0.1 | 0.2 | 0.13 |
| 5 | 67917283 | rs2607391 | -0.966 | -0.273 | 0.243 | -0.108 | -0.04 | -0.112 | -0.112 | 0 | 0.73 | 0.98 | 0.71 | 0.43 | 0.76 | 0.76 |
| 5 | 67928146 | rs2112931 | -2.369 | -0.18 | 1.428 | 0.22 | -0.001 | -0.121 | -0.121 | 0 | 0.24 | 0.5 | 0.18 | 0.11 | 0.23 | 0.14 |
| 5 | 67930745 | rs1990991 | -1.522 | -0.41 | 1.547 | 0.52 | 0.011 | -0.008 | -0.008 | 0 | 0.07 | 0.34 | 0.06 | 0.02 | 0.03 | 0.03 |
| 5 | 67938981 | rs282422 | 1.764 | -0.415 | 1.459 | 1.321 | -0.052 | 0.282 | 0.282 | 1.407 | 0.87 | 0.43 | 0.86 | 0.6 | 0.96 | 0.91 |
| 5 | 67950831 | rs33353 | 0.894 | -0.365 | 0.992 | 1.255 | -0.006 | 0.09 | 0.09 | 1.522 | 0.92 | 0.91 | 0.91 | 0.53 | 0.97 | 0.94 |
| 5 | 67950910 | rs33352 | 2.076 | -0.09 | 0.943 | 1.192 | -0.007 | 0.069 | 0.069 | 0.486 | 0.8 | 0.33 | 0.72 | 0.5 | 0.83 | 0.83 |
| 5 | 67954756 | rs255258 | 1.232 | -0.268 | 1.182 | 1.346 | 0.047 | 0.595 | 0.595 | 2.588 | 0.93 | 0.51 | 0.9 | 0.56 | 0.99 | 0.94 |
| 5 | 67959925 | rs2607374 | 1.85 | -0.203 | 0.846 | 1.292 | -0.023 | 0.095 | 0.095 | 0.394 | 0.8 | 0.39 | 0.73 | 0.49 | 0.86 | 0.84 |
| 5 | 67964088 | rs255257 | 1.173 | -0.957 | 0.426 | 0.913 | 0.039 | -0.027 | -0.027 | 0 | 0.65 | 0.3 | 0.56 | 0.48 | 0.8 | 0.68 |
| 5 | 67965602 | rs16898051 | -0.232 | -0.481 | 0.133 | 1.198 | 0.024 | 0.027 | 0.027 | 0.588 | 0.22 | 0.07 | 0.31 | 0.11 | 0.18 | 0.27 |
| 5 | 67966660 | rs255249 | -0.251 | -0.415 | 0.133 | 1.301 | 0.017 | -0.022 | -0.022 | 0 | 0.22 | 0.08 | 0.39 | 0.22 | 0.19 | 0.28 |
| 5 | 67986294 | rs10070379 | 0.104 | -0.017 | 0.405 | 0.828 | -0.048 | 0.088 | 0.088 | 0 | 0.74 | 0.48 | 0.81 | 0.56 | 0.82 | 0.83 |
| 5 | 68000986 | rs10471341 | -0.073 | 0.187 | 0.11 | -0.371 | -0.018 | -0.105 | -0.105 | 0 | 0.23 | 0.37 | 0.16 | 0 | 0.19 | 0.12 |
| 5 | 68003457 | rs12658051 | -0.52 | 1.675 | -0.376 | 0.251 | 0.047 | -0.015 | -0.015 | 7.754 | 0.16 | 0.19 | 0.19 | 0.14 | 0.31 | 0.41 |
| 5 | 68005609 | rs3844301 | -0.713 | 0.996 | -0.814 | 0.285 | -0.011 | -0.005 | -0.005 | 0 | 0.08 | 0.01 | 0.16 | 0.07 | 0.07 | 0.16 |
| 5 | 68006543 | rs246976 | -0.87 | 1.2 | -0.799 | -0.535 | -0.005 | 0.024 | 0.024 | 0 | 0.39 | 0.6 | 0.44 | 0.19 | 0.44 | 0.53 |
| 5 | 68009024 | rs6898192 | -0.998 | 1.204 | -0.03 | -0.747 | -0.015 | -0.073 | -0.073 | 0.004 | 0.18 | 0.4 | 0.27 | 0.07 | 0.15 | 0.17 |
| 5 | 68010984 | rs35247 | 0.26 | 1.316 | -0.038 | -0.756 | 0.008 | -0.017 | -0.017 | 0 | 0.21 | 0.21 | 0.16 | 0.12 | 0.28 | 0.36 |
| 5 | 68016360 | rs3852133 | -0.703 | 1.204 | 0.214 | -0.792 | 0 | 0.031 | 0.031 | 0 | 0.15 | 0.08 | 0.23 | 0.07 | 0.14 | 0.15 |
| 5 | 68018262 | rs35243 | 0.151 | 1.426 | 0.658 | -0.808 | 0.016 | -0.039 | -0.039 | 0 | 0.21 | 0.3 | 0.17 | 0.12 | 0.26 | 0.36 |
| 5 | 68019774 | rs35240 | -0.228 | 0.634 | 0.014 | -1.887 | -0.004 | -0.036 | -0.036 | 0 | 0.43 | 0.39 | 0.23 | 0.13 | 0.37 | 0.42 |
| 5 | 68029078 | rs35239 | -2.116 | 1.032 | -0.382 | -1.077 | -0.008 | -0.015 | -0.015 | 1.929 | 0.84 | 0.86 | 0.85 | 0.74 | 0.8 | 0.8 |
| 5 | 68029602 | rs12653535 | -0.156 | 1.249 | 0.266 | -0.538 | -0.007 | -0.018 | -0.018 | 0 | 0.17 | 0.35 | 0.27 | 0.14 | 0.14 | 0.15 |
| 5 | 68030007 | rs33278 | 1.925 | 1.448 | -0.229 | -0.434 | -0.05 | -0.118 | -0.118 | 0 | 0.15 | 0.01 | 0.31 | 0.45 | 0.18 | 0.2 |
| 5 | 68033010 | rs26685 | -0.071 | 1.61 | -0.004 | -0.626 | 0.001 | -0.004 | -0.004 | 0 | 0.37 | 0.36 | 0.42 | 0.26 | 0.46 | 0.48 |
| 5 | 68038077 | rs12516844 | 0.975 | 1.136 | -0.481 | -0.303 | -0.006 | -0.025 | -0.025 | 0.369 | 0.16 | 0.06 | 0.16 | 0.25 | 0.12 | 0.13 |
| 5 | 68042974 | rs17327213 | 0.513 | 0.901 | -0.238 | -0.097 | 0.001 | 0.036 | 0.036 | 0 | 0.41 | 0.32 | 0.23 | 0.11 | 0.4 | 0.45 |
| 5 | 68044120 | rs6898649 | 0.335 | 0.896 | -0.238 | -0.091 | -0.001 | 0.018 | 0.018 | 0 | 0.43 | 0.33 | 0.34 | 0.35 | 0.42 | 0.46 |
| 5 | 68044854 | rs13177007 | 1.084 | 0.869 | -0.398 | -0.505 | 0 | -0.01 | -0.01 | 0 | 0.07 | 0.13 | 0.06 | 0.04 | 0.08 | 0.04 |
| 5 | 68046285 | rs9291933 | -0.296 | 0.884 | -0.594 | -0.494 | -0.062 | 0.245 | 0.245 | 0.501 | 0.52 | 0.92 | 0.34 | 0.15 | 0.45 | 0.48 |
| 5 | 68054574 | rs26337 | 0.093 | 1.219 | 0.196 | 0.187 | 0.002 | 0.006 | 0.006 | 0 | 0.61 | 0.5 | 0.57 | 0.56 | 0.63 | 0.64 |
| 5 | 68058660 | rs13175238 | 0.46 | 1.178 | -0.446 | 0.241 | -0.001 | -0.002 | -0.002 | 0 | 0.07 | 0.09 | 0.05 | 0.02 | 0.08 | 0.04 |
| 5 | 68060559 | rs28289 | 0.764 | 1.15 | -0.431 | 0.174 | 0 | 0 | 0 | 0 | 0.5 | 0.56 | 0.5 | 0.52 | 0.54 | 0.55 |
| 5 | 68060579 | rs35248 | -0.156 | 1.202 | -0.423 | 0.384 | -0.054 | 0.013 | 0.013 | 0 | 0.13 | 0.01 | 0.2 | 0.1 | 0.13 | 0.14 |
| 5 | 68071181 | rs33295 | -0.739 | 1.224 | -1.203 | 0.754 | -0.022 | -0.075 | -0.075 | 0.025 | 0.74 | 0.96 | 0.87 | 0.71 | 0.8 | 0.8 |
| 5 | 68072719 | rs412501 | 0.096 | 1.141 | -0.803 | 0.579 | -0.026 | -0.019 | -0.019 | 0 | 0.57 | 0.71 | 0.53 | 0.51 | 0.57 | 0.6 |
| 5 | 68080712 | rs33307 | 0.08 | 1.108 | -0.93 | 0.582 | -0.031 | -0.021 | -0.021 | 0 | 0.57 | 0.72 | 0.53 | 0.51 | 0.57 | 0.61 |
| 5 | 68099139 | rs2913276 | -0.531 | 1.21 | -1.569 | 0.864 | -0.068 | -0.031 | -0.031 | 0 | 0.58 | 0.95 | 0.59 | 0.57 | 0.52 | 0.7 |
| 5 | 68103205 | rs2913277 | -0.388 | 1.244 | -1.53 | 0.863 | -0.082 | -0.084 | -0.084 | 0 | 0.54 | 0.93 | 0.47 | 0.49 | 0.42 | 0.63 |
| 5 | 68110727 | rs1582897 | 0.254 | 1.185 | -0.732 | 0.792 | 0.01 | -0.009 | -0.009 | 0.057 | 0.4 | 0.59 | 0.35 | 0.42 | 0.29 | 0.32 |
| 5 | 68119704 | rs10940176 | 0.191 | 1.296 | -0.587 | 0.745 | 0.002 | -0.004 | -0.004 | 0 | 0.11 | 0.15 | 0.12 | 0.08 | 0.13 | 0.3 |
| 5 | 68143193 | rs10039512 | -0.157 | 0.899 | -0.006 | 0.777 | 0.004 | -0.01 | -0.01 | 0 | 0.59 | 0.33 | 0.66 | 0.53 | 0.69 | 0.62 |

| | | | | | | | | | | | | | | | | |
|---|---|---|---|---|---|---|---|---|---|---|---|---|---|---|---|---|
| 5 | 68158818 | rs12153013 | 0.052 | 0.769 | 1.101 | 0.727 | 0.006 | -0.053 | -0.053 | 2.356 | 0.57 | 0.13 | 0.64 | 0.54 | 0.69 | 0.61 |
| 5 | 68179333 | rs7737876 | -0.426 | 0.911 | 0.942 | 0.925 | -0.004 | 0.001 | 0.001 | 0 | 0.64 | 0.74 | 0.8 | 0.67 | 0.7 | 0.69 |
| 5 | 68186380 | rs7709393 | -0.391 | 1.048 | 0.709 | 0.935 | -0.026 | 0.022 | 0.022 | 0 | 0.83 | 0.96 | 0.89 | 0.86 | 0.86 | 0.86 |
| 5 | 68199678 | rs7713600 | 0.827 | 0.804 | 0.775 | 0.884 | 0 | 0 | 0 | 0 | 0.25 | 0.26 | 0.11 | 0.16 | 0.18 | 0.22 |
| 5 | 68200642 | rs7718291 | -1.047 | 0.796 | 1.021 | 0.868 | 0.024 | 0.044 | 0.044 | 0 | 0.16 | 0.44 | 0.28 | 0.2 | 0.17 | 0.38 |
| 5 | 68208104 | rs10461543 | -0.22 | 0.653 | 0.26 | 0.927 | -0.003 | 0.017 | 0.017 | 0 | 0.11 | 0.2 | 0.13 | 0.19 | 0.1 | 0.28 |
| 5 | 68209329 | rs10940184 | -0.034 | 0.646 | 0.16 | 0.768 | 0 | -0.002 | -0.002 | 0.076 | 0.1 | 0.09 | 0.13 | 0.15 | 0.1 | 0.28 |
| 5 | 68209687 | rs11748007 | -0.751 | 0.814 | -0.054 | 0.656 | 0.01 | 0.022 | 0.022 | 0 | 0.46 | 0.78 | 0.32 | 0.34 | 0.41 | 0.58 |
| 5 | 68214816 | rs6888082 | 0.987 | 0.808 | -0.2 | 0.629 | -0.006 | -0.001 | -0.001 | 0 | 0.25 | 0.29 | 0.11 | 0.19 | 0.18 | 0.22 |
| 5 | 68219714 | rs7725683 | -0.955 | 1.105 | -0.864 | 0.475 | 0.046 | -0.048 | -0.048 | 1.937 | 0.18 | 0.44 | 0.21 | 0.11 | 0.22 | 0.38 |
| 5 | 68220909 | rs11743274 | -0.253 | 1.285 | -2.268 | 0.594 | -0.008 | 0.005 | 0.005 | 0 | 0.13 | 0.1 | 0.13 | 0.1 | 0.17 | 0.3 |
| 5 | 68225003 | rs4976054 | 0.937 | 1.181 | -2.742 | 0.924 | -0.037 | -0.042 | -0.042 | 0.542 | 0.27 | 0.04 | 0.26 | 0.28 | 0.3 | 0.18 |
| 5 | 68231686 | rs10044075 | -0.202 | 0.995 | -2.107 | 0.627 | -0.046 | 0.057 | 0.057 | 0 | 0.46 | 0.91 | 0.33 | 0.31 | 0.41 | 0.58 |
| 5 | 68234769 | rs6449986 | -0.295 | 1.077 | -2.085 | 0.665 | -0.053 | 0.063 | 0.063 | 0 | 0.48 | 0.91 | 0.35 | 0.31 | 0.43 | 0.58 |
| 5 | 68242147 | rs6883472 | 0.739 | 1.447 | -1.543 | 1.163 | -0.021 | -0.044 | -0.044 | 0 | 0.73 | 0.8 | 0.79 | 0.56 | 0.72 | 0.59 |
| 5 | 68243246 | rs6870478 | -1.06 | 0.856 | -1.113 | 1.493 | -0.001 | 0.007 | 0.007 | 0 | 0.18 | 0.16 | 0.18 | 0.42 | 0.14 | 0.26 |
| 5 | 68243972 | rs10063661 | 0.808 | 0.769 | -1.864 | 0.572 | -0.03 | 0.104 | 0.104 | 0.025 | 0.34 | 0.11 | 0.39 | 0.16 | 0.39 | 0.18 |
| 5 | 68245290 | rs10056163 | 0.822 | 0.717 | -1.442 | 0.539 | -0.004 | 0.07 | 0.07 | 0 | 0.36 | 0.22 | 0.4 | 0.14 | 0.38 | 0.17 |
| 5 | 68245459 | rs10064315 | -0.334 | -0.124 | -1.485 | 0.26 | -0.078 | -0.077 | -0.077 | 1.197 | 0.57 | 0.9 | 0.63 | 0.57 | 0.52 | 0.44 |
| 5 | 68245920 | rs10071585 | 1.178 | 0.12 | -1.409 | 0.172 | -0.028 | 0.103 | 0.103 | 0.005 | 0.34 | 0.12 | 0.39 | 0.14 | 0.39 | 0.17 |
| 5 | 68263602 | rs6414814 | 1.22 | 0.087 | -1.341 | 0.134 | -0.015 | 0.113 | 0.113 | 0.121 | 0.35 | 0.14 | 0.4 | 0.11 | 0.39 | 0.19 |
| 5 | 68287747 | rs10940190 | 0.881 | 0.228 | -0.218 | 0.149 | -0.019 | 0.015 | 0.015 | 0 | 0.68 | 0.38 | 0.8 | 0.6 | 0.77 | 0.67 |
| 5 | 68294662 | rs4976056 | 0.536 | 0.162 | 0.299 | 0.124 | -0.017 | 0.033 | 0.033 | 0 | 0.73 | 0.65 | 0.83 | 0.6 | 0.79 | 0.71 |
| 5 | 68296704 | rs7733454 | 0.603 | 0.209 | 0.308 | 0.113 | -0.02 | 0.038 | 0.038 | 0 | 0.72 | 0.63 | 0.83 | 0.61 | 0.78 | 0.71 |
| 5 | 68304118 | rs10042916 | 0.963 | 0.09 | 0.133 | 0.124 | -0.046 | 0.029 | 0.029 | 0 | 0.69 | 0.43 | 0.81 | 0.61 | 0.81 | 0.69 |
| 5 | 68322358 | rs10940194 | 1.534 | 0.255 | -0.288 | -0.185 | -0.07 | 0.159 | 0.159 | 2.578 | 0.37 | 0.08 | 0.42 | 0.13 | 0.41 | 0.19 |
| 5 | 68322642 | rs6861079 | 0.262 | 0.111 | 0.06 | -0.01 | 0.01 | -0.014 | -0.014 | 0 | 0.76 | 0.82 | 0.87 | 0.63 | 0.83 | 0.71 |
| 5 | 68337426 | rs164796 | -1.345 | 0.155 | 0.192 | -0.063 | -0.028 | 0.14 | 0.14 | 1.193 | 0.6 | 0.82 | 0.57 | 0.87 | 0.58 | 0.81 |
| 5 | 68342235 | rs4976178 | 1.365 | 0.213 | 0.036 | -0.371 | -0.002 | 0.068 | 0.068 | 0.105 | 0.36 | 0.23 | 0.38 | 0.12 | 0.34 | 0.18 |
| 5 | 68345648 | rs149292 | 0.062 | 0.113 | -0.009 | -0.391 | 0.011 | -0.057 | -0.057 | 0 | 0.77 | 0.9 | 0.83 | 0.62 | 0.79 | 0.73 |
| 5 | 68349917 | rs6897246 | 1.223 | 0.112 | 0.141 | -0.42 | -0.002 | -0.053 | -0.053 | 0 | 0.37 | 0.46 | 0.4 | 0.12 | 0.35 | 0.17 |
| 5 | 68364871 | rs2434343 | 0.181 | 0.202 | 0.189 | -0.328 | 0.006 | 0.011 | 0.011 | 0 | 0.4 | 0.49 | 0.44 | 0.5 | 0.44 | 0.55 |
| 5 | 68371896 | rs2434349 | 0.225 | 0.172 | 0.124 | -0.325 | -0.018 | -0.016 | -0.016 | 0 | 0.39 | 0.3 | 0.43 | 0.5 | 0.46 | 0.55 |
| 5 | 68372674 | rs6450018 | 1.024 | -0.048 | -0.122 | -0.514 | 0.002 | 0.008 | 0.008 | 0.081 | 0.34 | 0.3 | 0.37 | 0.12 | 0.32 | 0.18 |
| 5 | 68379836 | rs163192 | -0.222 | 0.016 | -0.236 | -0.197 | -0.002 | -0.001 | -0.001 | 0 | 0.57 | 0.56 | 0.49 | 0.54 | 0.48 | 0.51 |
| 5 | 68396580 | rs12522805 | 0.23 | -0.508 | -0.645 | -0.638 | -0.004 | -0.025 | -0.025 | 1.672 | 0.36 | 0.35 | 0.25 | 0.16 | 0.28 | 0.3 |
| 5 | 68401941 | rs4976217 | 0.83 | -0.363 | -0.991 | -0.587 | -0.049 | 0.056 | 0.056 | 0 | 0.27 | 0.01 | 0.18 | 0.16 | 0.21 | 0.24 |
| 5 | 68417643 | rs164578 | -0.086 | -0.121 | -0.949 | -0.356 | 0.009 | 0.006 | 0.006 | 0 | 0.48 | 0.33 | 0.42 | 0.54 | 0.42 | 0.44 |
| 5 | 68417755 | rs164577 | -0.188 | -0.12 | -0.946 | -0.356 | 0.011 | 0.008 | 0.008 | 0 | 0.49 | 0.33 | 0.42 | 0.54 | 0.43 | 0.44 |
| 5 | 68423069 | rs337253 | -0.104 | -0.001 | -1.123 | -0.292 | 0.011 | 0.007 | 0.007 | 0 | 0.48 | 0.27 | 0.41 | 0.54 | 0.42 | 0.45 |
| 5 | 70980721 | rs13153562 | -1.387 | -1.001 | -0.178 | -0.179 | 0.002 | -0.008 | -0.008 | 0.78 | 0.54 | 0.76 | 0.55 | 0.51 | 0.6 | 0.62 |
| 5 | 70986022 | rs281121 | 0.179 | -1.061 | 0.131 | -0.119 | -0.001 | 0.002 | 0.002 | 0 | 0.05 | 0.13 | 0.04 | 0.05 | 0.05 | 0.04 |
| 5 | 70986475 | rs2115031 | -0.781 | -1.148 | 0.375 | -0.111 | 0.009 | -0.021 | -0.021 | 0 | 0.36 | 0.62 | 0.43 | 0.26 | 0.43 | 0.4 |
| 5 | 70990736 | rs1549604 | -0.054 | -1.115 | 0.75 | -0.039 | 0.033 | 0.026 | 0.026 | 0.674 | 0.11 | 0.31 | 0.23 | 0.13 | 0.12 | 0.2 |
| 5 | 70992769 | rs1028161 | -0.199 | -1.384 | 0.736 | -0.113 | -0.015 | -0.025 | -0.025 | 0 | 0.22 | 0.41 | 0.3 | 0.15 | 0.21 | 0.26 |
| 5 | 70993914 | rs2560665 | -1.732 | -1.447 | 0.806 | -0.128 | -0.007 | -0.014 | -0.014 | 0 | 0.55 | 0.8 | 0.58 | 0.48 | 0.6 | 0.6 |
| 5 | 70997465 | rs11741424 | -0.492 | -1.232 | 0.581 | -0.184 | -0.002 | 0.019 | 0.019 | 0 | 0.38 | 0.24 | 0.31 | 0.29 | 0.43 | 0.36 |
| 5 | 70998053 | rs6860511 | -1.491 | -1.337 | 0.511 | -0.195 | -0.037 | 0.021 | 0.021 | 0 | 0.52 | 0.88 | 0.55 | 0.43 | 0.56 | 0.55 |
| 5 | 71006645 | rs6866864 | -0.983 | -1.205 | 0.567 | -0.137 | -0.001 | -0.002 | -0.002 | 1.261 | 0.43 | 0.42 | 0.42 | 0.44 | 0.46 | 0.46 |
| 5 | 71006707 | rs4704168 | -0.483 | -1.202 | 0.56 | -0.141 | 0 | 0.029 | 0.029 | 0 | 0.38 | 0.2 | 0.37 | 0.32 | 0.43 | 0.36 |
| 5 | 71010664 | rs10515116 | 0.363 | -1.097 | 0.663 | -0.138 | 0.075 | -0.021 | -0.021 | 0 | 0.07 | 0.28 | 0.2 | 0.01 | 0.09 | 0.1 |
| 5 | 71013466 | rs6859438 | 0.533 | -1.131 | 0.477 | -0.069 | 0.01 | 0.072 | 0.072 | 0.025 | 0.06 | 0.22 | 0.04 | 0.13 | 0.03 | 0.1 |
| 5 | 71015503 | rs2239670 | -0.661 | -2.212 | -0.317 | 0.587 | 0.047 | -0.026 | -0.026 | 0 | 0.78 | 0.88 | 0.87 | 0.73 | 0.86 | 0.81 |
| 5 | 71017951 | rs16871468 | -0.699 | -0.624 | 0.04 | 0.432 | 0.027 | -0.024 | -0.024 | 0 | 0.13 | 0.24 | 0.25 | 0.11 | 0.18 | 0.26 |
| 5 | 71020547 | rs16871475 | -0.775 | -1.656 | -0.113 | 0.739 | -0.076 | 0.014 | 0.014 | 0 | 0.27 | 0.34 | 0.14 | 0.29 | 0.15 | 0.28 |
| 5 | 71022741 | rs6897188 | -0.905 | -1.022 | -0.548 | 0.48 | 0.007 | 0.013 | 0.013 | 0 | 0.19 | 0.1 | 0.13 | 0.18 | 0.16 | 0.24 |
| 5 | 71024072 | rs7707837 | -1.183 | -1.282 | 0.752 | 0.353 | -0.029 | -0.044 | -0.044 | 0.01 | 0.22 | 0.49 | 0.17 | 0.17 | 0.16 | 0.24 |
| 5 | 71029743 | rs4057258 | -0.705 | -1.798 | 0.628 | 0.327 | 0 | 0 | 0 | 0 | 0.11 | 0.12 | 0.11 | 0.08 | 0.07 | 0.1 |
| 5 | 71033592 | rs13362349 | -1.783 | -2.025 | 0.555 | 0.235 | -0.019 | -0.016 | -0.016 | 0 | 0.36 | 0.61 | 0.32 | 0.26 | 0.28 | 0.39 |
| 5 | 71036658 | rs4361494 | -1.86 | -2.247 | 0.706 | 0.116 | 0.035 | 0.032 | 0.032 | 0.132 | 0.38 | 0.74 | 0.31 | 0.26 | 0.27 | 0.39 |
| 5 | 71048614 | rs4583865 | -0.506 | -2.134 | 0.838 | 0.369 | -0.002 | -0.001 | -0.001 | 0 | 0.2 | 0.21 | 0.15 | 0.21 | 0.15 | 0.22 |
| 5 | 71053835 | rs13178975 | 0.829 | -1.692 | -0.151 | 0.284 | -0.122 | 0.006 | 0.006 | 0 | 0.31 | 0.06 | 0.29 | 0.23 | 0.48 | 0.22 |
| 5 | 71054304 | rs6891713 | -0.53 | -1.454 | 0.56 | 0.331 | -0.005 | 0.034 | 0.034 | 0 | 0.09 | 0.31 | 0.12 | 0.08 | 0.04 | 0.12 |
| 5 | 71064766 | rs13186462 | 0.512 | -1.385 | -0.591 | 0.262 | -0.07 | 0.056 | 0.056 | 0.118 | 0.36 | 0.11 | 0.32 | 0.22 | 0.52 | 0.2 |
| 5 | 71073881 | rs16872236 | -0.259 | -2.039 | -0.172 | 0.357 | -0.005 | 0 | 0 | 0.039 | 0.18 | 0.3 | 0.25 | 0.21 | 0.18 | 0.24 |
| 5 | 71074550 | rs12517242 | 0.746 | -2.167 | 0.486 | 0.287 | 0.003 | -0.015 | -0.015 | 0.124 | 0.38 | 0.39 | 0.34 | 0.22 | 0.52 | 0.21 |
| 5 | 71077534 | rs7706490 | -0.12 | -2.142 | 0.421 | 0.457 | 0.017 | 0.002 | 0.002 | 0 | 0.57 | 0.69 | 0.59 | 0.42 | 0.71 | 0.45 |
| 5 | 71079676 | rs10942750 | -0.146 | -2.137 | 0.311 | 0.458 | 0.051 | 0.005 | 0.005 | 0 | 0.57 | 0.77 | 0.6 | 0.42 | 0.71 | 0.44 |
| 5 | 71105662 | rs1798575 | -0.087 | -2.174 | 0.09 | 0.56 | -0.002 | 0 | 0 | 0 | 0.66 | 0.36 | 0.61 | 0.65 | 0.76 | 0.67 |

| | | | | | | | | | | | | | | | | |
|---|---|---|---|---|---|---|---|---|---|---|---|---|---|---|---|---|
| 5 | 71111779 | rs2037202 | -0.121 | -2.261 | -0.576 | 0.297 | 0.07 | -0.018 | -0.018 | 0.573 | 0.45 | 0.24 | 0.46 | 0.51 | 0.31 | 0.54 |
| 5 | 71114678 | rs2721750 | 0.039 | -2.173 | -0.489 | 0.327 | 0.057 | -0.016 | -0.016 | 0 | 0.44 | 0.27 | 0.44 | 0.51 | 0.26 | 0.53 |
| 5 | 71132351 | rs2721775 | -0.501 | -1.725 | -0.788 | 0.175 | -0.007 | 0.005 | 0.005 | 0 | 0.57 | 0.52 | 0.51 | 0.53 | 0.34 | 0.59 |
| 5 | 71133485 | rs16872519 | -0.46 | -1.08 | -1.043 | 0.073 | -0.003 | 0.023 | 0.023 | 0.27 | 0.14 | 0.11 | 0.05 | 0.03 | 0.1 | 0.05 |
| 5 | 71135107 | rs4354016 | -0.123 | -0.217 | -1.385 | 0.156 | -0.056 | -0.009 | -0.009 | 0 | 0.86 | 0.99 | 0.82 | 0.82 | 0.83 | 0.71 |
| 5 | 71135257 | rs10052794 | -0.052 | -0.03 | -0.96 | 0.142 | 0 | 0 | 0 | 0 | 0.61 | 0.31 | 0.52 | 0.58 | 0.43 | 0.53 |
| 5 | 71137775 | rs12659559 | 0.14 | -1.373 | -0.387 | 0.42 | -0.057 | 0.187 | 0.187 | 0 | 0.19 | 0.39 | 0.13 | 0.48 | 0.11 | 0.26 |
| 5 | 71138138 | rs10942785 | 0.443 | -1.504 | -0.355 | 0.325 | 0.002 | 0.024 | 0.024 | 0.077 | 0.73 | 0.64 | 0.67 | 0.4 | 0.69 | 0.44 |
| 5 | 71140325 | rs1911936 | 0.041 | -1.834 | -0.534 | 0.193 | -0.011 | -0.042 | -0.042 | 0 | 0.76 | 0.86 | 0.73 | 0.49 | 0.73 | 0.46 |
| 5 | 71155607 | rs1911923 | 0.457 | -1.794 | -0.5 | 0.196 | -0.001 | -0.006 | -0.006 | 0 | 0.7 | 0.7 | 0.67 | 0.41 | 0.67 | 0.43 |
| 5 | 71156117 | rs11747407 | 2.059 | -2.207 | -0.475 | -0.252 | -0.044 | -0.035 | -0.035 | 0.005 | 0.23 | 0.21 | 0.32 | 0.33 | 0.37 | 0.2 |
| 5 | 71156895 | rs12055280 | 1.109 | -2.839 | -0.238 | -0.616 | 0.025 | 0.009 | 0.009 | 0 | 0.49 | 0.42 | 0.58 | 0.38 | 0.59 | 0.36 |
| 5 | 71158621 | rs3857391 | 0.462 | -2.891 | -0.199 | -0.644 | 0.001 | -0.006 | -0.006 | 0 | 0.66 | 0.66 | 0.65 | 0.4 | 0.66 | 0.43 |
| 5 | 71162904 | rs16872637 | 0.378 | -3.117 | -0.363 | -0.662 | -0.06 | -0.082 | -0.082 | 0 | 0.21 | 0.28 | 0.09 | 0.03 | 0.08 | 0.07 |
| 5 | 71163187 | rs16872647 | 1.048 | -2.911 | -0.774 | -0.565 | 0.039 | 0.01 | 0.01 | 0 | 0.48 | 0.3 | 0.57 | 0.38 | 0.57 | 0.35 |
| 5 | 71169505 | rs6890936 | 2.267 | -2.336 | -0.722 | -0.333 | -0.063 | -0.051 | -0.051 | 0 | 0.22 | 0.19 | 0.31 | 0.33 | 0.37 | 0.19 |
| 5 | 71174721 | rs3846669 | -0.373 | -2.445 | -0.481 | -0.377 | -0.032 | 0.027 | 0.027 | 0 | 0.32 | 0.51 | 0.3 | 0.48 | 0.23 | 0.34 |
| 5 | 71176041 | rs11950306 | 1.57 | -2.584 | -1.195 | -0.649 | 0.012 | 0.029 | 0.029 | 0.201 | 0.17 | 0.09 | 0.22 | 0.02 | 0.15 | 0.11 |
| 5 | 71179121 | rs1911933 | -0.931 | -2.23 | -0.917 | -0.238 | -0.07 | 0.015 | 0.015 | 0 | 0.84 | 0.99 | 0.88 | 0.86 | 0.8 | 0.67 |
| 5 | 71180449 | rs4704363 | 1.391 | -2.28 | -0.761 | -0.216 | 0.009 | 0.116 | 0.116 | 0 | 0.09 | 0.13 | 0.18 | 0.38 | 0.11 | 0.24 |
| 5 | 71184918 | rs6453274 | 0.576 | -2.262 | -0.925 | -0.667 | 0.011 | 0.033 | 0.033 | 0 | 0.67 | 0.81 | 0.82 | 0.83 | 0.73 | 0.6 |
| 5 | 71188232 | rs7736888 | 0.266 | -2.015 | -0.431 | -0.454 | -0.036 | -0.062 | -0.062 | 0 | 0.16 | 0.17 | 0.06 | 0.03 | 0.07 | 0.07 |
| 5 | 71199056 | rs4072550 | 0.755 | -1.72 | -0.631 | -0.136 | 0.009 | 0.011 | 0.011 | 0 | 0.51 | 0.48 | 0.61 | 0.59 | 0.62 | 0.6 |
| 5 | 71199839 | rs1574299 | 0.797 | -1.832 | -0.707 | -0.242 | 0.002 | -0.001 | -0.001 | 0 | 0.47 | 0.46 | 0.44 | 0.48 | 0.55 | 0.35 |
| 5 | 71200670 | rs4602602 | -0.962 | -1.962 | -1.003 | -0.141 | -0.038 | -0.058 | -0.058 | 0.585 | 0.88 | 0.97 | 0.82 | 0.64 | 0.83 | 0.63 |
| 5 | 71201644 | rs16873144 | 1.631 | -2.26 | -1.012 | -0.371 | 0.011 | -0.019 | -0.019 | 0.921 | 0.08 | 0 | 0.03 | 0.05 | 0.03 | 0.05 |
| 5 | 71204261 | rs2879027 | 2.112 | -1.854 | -1.056 | -0.284 | -0.054 | -0.029 | -0.029 | 0 | 0.24 | 0.17 | 0.27 | 0.3 | 0.38 | 0.17 |
| 5 | 71208228 | rs3857394 | -0.273 | -2.004 | -0.433 | -0.294 | -0.091 | -0.047 | -0.047 | 0 | 0.29 | 0.39 | 0.14 | 0.16 | 0.13 | 0.15 |
| 5 | 71209789 | rs16873192 | 1.318 | -1.914 | -0.392 | -0.259 | 0.025 | 0.018 | 0.018 | 0.892 | 0.17 | 0.08 | 0.05 | 0.09 | 0.07 | 0.1 |
| 5 | 71213762 | rs12653952 | 0.452 | -1.656 | -0.301 | -0.146 | -0.045 | 0.002 | 0.002 | 0 | 0.11 | 0.31 | 0.07 | 0.07 | 0.06 | 0.04 |
| 5 | 71223173 | rs1459213 | -0.086 | -1.695 | -0.308 | -0.245 | -0.003 | -0.013 | -0.013 | 0 | 0.22 | 0.23 | 0.3 | 0.08 | 0.2 | 0.18 |
| 5 | 71223201 | rs1026507 | 1.051 | -1.672 | -0.345 | -0.24 | 0.004 | 0.011 | 0.011 | 1.787 | 0.14 | 0.07 | 0.23 | 0.01 | 0.14 | 0.14 |
| 5 | 71226221 | rs2610425 | -0.507 | -1.696 | -0.397 | -0.244 | -0.06 | -0.107 | -0.107 | 0.074 | 0.25 | 0.42 | 0.32 | 0.08 | 0.2 | 0.18 |
| 5 | 71226289 | rs9293717 | 1.015 | -1.664 | -1.043 | -0.153 | 0.014 | -0.018 | -0.018 | 0.022 | 0.63 | 0.32 | 0.48 | 0.55 | 0.64 | 0.47 |
| 5 | 71230022 | rs10071437 | 0.868 | -1.553 | -1.68 | -0.29 | 0.01 | -0.001 | -0.001 | 0.37 | 0.49 | 0.44 | 0.46 | 0.49 | 0.61 | 0.39 |
| 5 | 71230743 | rs1692420 | 1.198 | -1.415 | -1.667 | -0.261 | 0.009 | 0.06 | 0.06 | 1.765 | 0.13 | 0.02 | 0.2 | 0.01 | 0.11 | 0.12 |
| 5 | 71233603 | rs3857397 | 0.905 | -1.274 | -1.639 | -0.163 | 0.034 | -0.014 | -0.014 | 0.039 | 0.47 | 0.25 | 0.45 | 0.46 | 0.59 | 0.38 |
| 5 | 71235359 | rs898216 | -0.16 | -1.071 | -1.826 | -0.095 | 0.009 | -0.022 | -0.022 | 2.874 | 0.69 | 0.35 | 0.71 | 0.56 | 0.77 | 0.56 |
| 5 | 71239769 | rs754101 | -0.474 | -1.172 | -1.808 | -0.694 | -0.013 | -0.015 | -0.015 | 6.585 | 0.45 | 0.55 | 0.42 | 0.25 | 0.29 | 0.3 |
| 5 | 71240138 | rs898219 | -0.403 | 0.388 | -2.292 | -0.513 | -0.053 | 0.16 | 0.16 | 0.792 | 0.32 | 0.09 | 0.37 | 0.07 | 0.35 | 0.39 |
| 5 | 71241023 | rs1217741 | -0.804 | 0.459 | -1.501 | 0.218 | -0.001 | 0 | 0 | 0.66 | 0.17 | 0.21 | 0.12 | 0.19 | 0.14 | 0.13 |
| 5 | 71241377 | rs9293720 | 0.207 | 0.885 | -2.34 | -0.331 | -0.079 | 0.051 | 0.051 | 0.029 | 0.23 | 0.01 | 0.1 | 0.08 | 0.2 | 0.15 |
| 5 | 71241513 | rs2125627 | -0.037 | 0.624 | -1.243 | -0.4 | -0.005 | -0.006 | -0.006 | 0 | 0.82 | 0.59 | 0.74 | 0.88 | 0.85 | 0.79 |
| 5 | 71245810 | rs1217744 | 0.1 | 0.902 | -1.678 | -0.288 | -0.037 | 0.039 | 0.039 | 0 | 0.43 | 0.24 | 0.41 | 0.68 | 0.49 | 0.55 |
| 5 | 71245964 | rs1217745 | 0.235 | 1.184 | -1.503 | 0.624 | 0.001 | 0.024 | 0.024 | 0 | 0.13 | 0.17 | 0.29 | 0.37 | 0.19 | 0.34 |
| 5 | 71249440 | rs1217750 | 0.409 | 0.882 | -1.24 | 0.096 | -0.002 | -0.015 | -0.015 | 0 | 0.26 | 0.32 | 0.18 | 0.17 | 0.24 | 0.15 |
| 5 | 71249940 | rs2662334 | -0.544 | 0.65 | -0.993 | -0.166 | 0.003 | 0.033 | 0.033 | 1.884 | 0.72 | 0.83 | 0.62 | 0.86 | 0.73 | 0.7 |
| 5 | 71256659 | rs2610424 | -0.505 | 0.464 | -0.25 | -0.298 | -0.002 | 0.001 | 0.001 | 0 | 0.47 | 0.5 | 0.5 | 0.55 | 0.48 | 0.62 |
| 5 | 71258005 | rs2250816 | 0.561 | 0.485 | -0.363 | -0.318 | 0.001 | 0 | 0 | 0 | 0.43 | 0.57 | 0.21 | 0.36 | 0.36 | 0.33 |
| 5 | 71258017 | rs2721761 | -1.331 | 0.498 | -0.385 | -0.327 | -0.029 | 0.041 | 0.041 | 0.102 | 0.81 | 0.95 | 0.67 | 0.88 | 0.78 | 0.83 |
| 5 | 71258063 | rs6896246 | 0.608 | 0.493 | -0.193 | -0.213 | 0.026 | 0.137 | 0.137 | 0 | 0.13 | 0.35 | 0.31 | 0.36 | 0.19 | 0.35 |
| 5 | 71258187 | rs1692384 | -0.363 | 0.51 | -0.533 | -0.26 | -0.087 | 0.081 | 0.081 | 0 | 0.56 | 0.94 | 0.52 | 0.72 | 0.55 | 0.68 |
| 5 | 71260828 | rs1217825 | -0.675 | 0.749 | 0.946 | 0.596 | -0.003 | 0.035 | 0.035 | 0.086 | 0.22 | 0.61 | 0.37 | 0.35 | 0.26 | 0.41 |
| 5 | 71266713 | rs2662317 | -0.458 | 0.906 | 0.946 | 0.536 | 0.017 | 0.12 | 0.12 | 0 | 0.17 | 0.33 | 0.13 | 0.36 | 0.21 | 0.25 |
| 5 | 71295157 | rs13187253 | -0.899 | 0.884 | 1.168 | 0.526 | 0.001 | 0.097 | 0.097 | 0.781 | 0.2 | 0.64 | 0.16 | 0.36 | 0.21 | 0.25 |
| 5 | 71295640 | rs7709845 | -1.092 | 0.877 | 1.012 | 0.523 | 0.001 | -0.005 | -0.005 | 0 | 0.22 | 0.79 | 0.17 | 0.36 | 0.21 | 0.25 |
| 5 | 71303308 | rs1010376 | -0.801 | 1.078 | 0.812 | 0.586 | 0.021 | 0.03 | 0.03 | 0 | 0.25 | 0.34 | 0.2 | 0.33 | 0.31 | 0.28 |
| 5 | 71309869 | rs1217769 | -0.481 | 0.935 | 0.553 | 0.62 | 0.002 | 0.009 | 0.009 | 0.011 | 0.21 | 0.24 | 0.35 | 0.36 | 0.28 | 0.42 |
| 5 | 71319262 | rs1692423 | -0.446 | 0.959 | 0.417 | 0.812 | -0.005 | -0.015 | -0.015 | 0 | 0.15 | 0.13 | 0.29 | 0.27 | 0.18 | 0.41 |
| 5 | 71319752 | rs1692421 | -0.399 | 0.955 | 0.387 | 0.81 | -0.008 | -0.021 | -0.021 | 0.643 | 0.15 | 0.11 | 0.29 | 0.27 | 0.18 | 0.41 |
| 5 | 71331087 | rs7711863 | -0.233 | 0.974 | 1.382 | 1.043 | 0.11 | 0.265 | 0.265 | 0.068 | 0.11 | 0.51 | 0.09 | 0.24 | 0.15 | 0.11 |
| 5 | 71345282 | rs1692357 | -0.23 | 0.842 | 1.525 | 1.32 | -0.004 | 0.099 | 0.099 | 0 | 0.14 | 0.26 | 0.28 | 0.31 | 0.14 | 0.33 |
| 5 | 71359777 | rs1692404 | -1.193 | 0.733 | 1.8 | 1.373 | -0.018 | 0.152 | 0.152 | 0 | 0.19 | 0.5 | 0.34 | 0.38 | 0.17 | 0.39 |
| 5 | 71367006 | rs1217780 | -0.698 | 0.591 | 1.588 | 1.425 | -0.004 | 0.03 | 0.03 | 0 | 0.16 | 0.2 | 0.3 | 0.33 | 0.14 | 0.31 |
| 5 | 71368837 | rs1217785 | -0.165 | 0.566 | 1.055 | 0.714 | -0.007 | 0.008 | 0.008 | 0 | 0.63 | 0.71 | 0.63 | 0.81 | 0.53 | 0.69 |
| 5 | 71369445 | rs16874468 | 1.138 | 0.756 | 0.868 | 0.795 | 0.007 | 0.044 | 0.044 | 0 | 0.08 | 0.1 | 0.06 | 0.25 | 0.14 | 0.17 |
| 5 | 71372881 | rs1217793 | 0.744 | 0.676 | 0.747 | 0.68 | -0.001 | 0.018 | 0.018 | 0 | 0.1 | 0.11 | 0.28 | 0.3 | 0.12 | 0.25 |
| 5 | 71375965 | rs17300302 | 1.915 | 0.551 | 0.7 | 0.392 | 0.04 | 0.093 | 0.093 | 0.23 | 0.34 | 0.17 | 0.14 | 0.01 | 0.22 | 0.15 |
| 5 | 71377132 | rs1217800 | -0.665 | 0.885 | 0.445 | 0.439 | -0.026 | 0.059 | 0.059 | 0 | 0.7 | 0.83 | 0.7 | 0.88 | 0.66 | 0.78 |

| | | | | | | | | | | | | | | | | |
|---|---|---|---|---|---|---|---|---|---|---|---|---|---|---|---|---|
| 5 | 71380914 | rs6897294 | 1.61 | 0.729 | 0.42 | 0.652 | 0.015 | 0.021 | 0.021 | 0 | 0.42 | 0.4 | 0.28 | 0.21 | 0.27 | 0.2 |
| 5 | 71383923 | rs1217809 | 0.416 | 0.932 | 0.513 | 1.016 | 0.003 | 0.062 | 0.062 | 0 | 0.1 | 0.18 | 0.28 | 0.3 | 0.12 | 0.26 |
| 5 | 71388338 | rs1217816 | 0.146 | 1.141 | 0.404 | 0.806 | -0.011 | 0.024 | 0.024 | 0 | 0.66 | 0.76 | 0.68 | 0.86 | 0.62 | 0.7 |
| 5 | 71394115 | rs1217823 | -2.347 | 1.465 | 0.771 | 1.75 | 0 | 0 | 0 | 0 | 0.33 | 0.66 | 0.44 | 0.63 | 0.4 | 0.43 |
| 5 | 71394869 | rs3112400 | -0.97 | 1.556 | 0.853 | 1.79 | 0.004 | -0.023 | -0.023 | 0 | 0.22 | 0.16 | 0.23 | 0.32 | 0.2 | 0.36 |
| 5 | 71398049 | rs6880685 | -1.493 | 1.142 | 1.206 | 1.663 | 0.119 | 0.032 | 0.032 | 2.689 | 0.07 | 0.29 | 0.09 | 0.08 | 0.19 | 0.02 |
| 5 | 71402339 | rs3806907 | -1.465 | 1.143 | 1.103 | 1.664 | 0.126 | 0.036 | 0.036 | 2.482 | 0.07 | 0.32 | 0.09 | 0.08 | 0.19 | 0.02 |
| 5 | 71408867 | rs2337390 | 0.56 | 1.451 | 0.48 | 1.791 | -0.027 | -0.049 | -0.049 | 0 | 0.79 | 0.85 | 0.73 | 0.57 | 0.73 | 0.67 |
| 5 | 71409383 | rs7732739 | -0.478 | 1.575 | 0.413 | 1.423 | 0.024 | -0.018 | -0.018 | 0 | 0.13 | 0.17 | 0.12 | 0.08 | 0.26 | 0.1 |
| 5 | 71411718 | rs1531312 | 1.352 | 1.111 | 0.469 | 1.732 | 0.009 | 0.028 | 0.028 | 0 | 0.72 | 0.52 | 0.59 | 0.33 | 0.72 | 0.62 |
| 5 | 71413628 | rs1466344 | -0.616 | 0.932 | -0.036 | 1.705 | -0.002 | 0.032 | 0.032 | 0 | 0.11 | 0.15 | 0.18 | 0.31 | 0.1 | 0.25 |
| 5 | 71422008 | rs10474585 | -0.098 | 0.702 | -0.199 | 1.309 | -0.009 | 0.006 | 0.006 | 0 | 0.38 | 0.6 | 0.45 | 0.45 | 0.39 | 0.35 |
| 5 | 71430895 | rs4704553 | -0.077 | 0.794 | -0.22 | 1.259 | -0.005 | 0.002 | 0.002 | 0 | 0.38 | 0.58 | 0.44 | 0.43 | 0.4 | 0.38 |
| 5 | 71437262 | rs4703774 | 0.636 | 0.861 | -0.35 | 1.209 | 0.036 | 0.066 | 0.066 | 0 | 0.71 | 0.5 | 0.68 | 0.57 | 0.66 | 0.53 |
| 5 | 71441589 | rs4569852 | 0.076 | 0.621 | -0.19 | 0.844 | 0 | 0 | 0 | 0 | 0.43 | 0.57 | 0.34 | 0.29 | 0.33 | 0.3 |
| 5 | 71450015 | rs10061898 | -0.17 | 0.715 | 0.101 | 0.914 | -0.009 | -0.016 | -0.016 | 0.576 | 0.07 | 0.14 | 0.05 | 0.04 | 0.07 | 0.02 |
| 5 | 71450955 | rs12655013 | 0.052 | 0.925 | 0.053 | 0.928 | 0.004 | -0.006 | -0.006 | 0 | 0.09 | 0.13 | 0.08 | 0.04 | 0.13 | 0.05 |
| 5 | 71451160 | rs6882965 | -0.018 | 1.395 | 0.152 | 1.072 | 0.035 | 0.066 | 0.066 | 0 | 0.85 | 0.76 | 0.71 | 0.43 | 0.77 | 0.64 |
| 5 | 71451175 | rs12655064 | -0.345 | 1.026 | 0.261 | 0.889 | 0.011 | -0.023 | -0.023 | 0 | 0.11 | 0.23 | 0.11 | 0.04 | 0.13 | 0.06 |
| 5 | 71454175 | rs6872830 | 0.986 | 1.014 | -0.051 | 1.359 | 0.019 | 0.169 | 0.169 | 0.215 | 0.85 | 0.54 | 0.83 | 0.66 | 0.84 | 0.93 |
| 5 | 71455618 | rs2337692 | 1.566 | 1.014 | -0.109 | 1.349 | 0.005 | 0.031 | 0.031 | 0.001 | 0.75 | 0.38 | 0.73 | 0.61 | 0.71 | 0.88 |
| 5 | 71457008 | rs1545170 | 0.974 | 0.669 | -0.436 | 1.467 | 0.016 | 0.122 | 0.122 | 0.354 | 0.87 | 0.62 | 0.84 | 0.71 | 0.86 | 0.96 |
| 5 | 71457028 | rs1079383 | 0.662 | 0.995 | -1.524 | 1.701 | -0.011 | -0.011 | -0.011 | 0 | 0.07 | 0.07 | 0.09 | 0.14 | 0.12 | 0.19 |
| 5 | 71462632 | rs3098382 | 0.394 | 0.488 | -1.587 | 1.391 | 0.007 | 0.054 | 0.054 | 0 | 0.91 | 0.87 | 0.88 | 0.71 | 0.87 | 0.97 |
| 5 | 71467670 | rs3098378 | 0.828 | 0.002 | -2.444 | 0.28 | -0.031 | 0.115 | 0.115 | 0 | 0.36 | 0.15 | 0.34 | 0.14 | 0.39 | 0.26 |
| 5 | 71471713 | rs12186804 | -0.068 | -0.913 | -2.09 | 0.61 | 0.051 | -0.025 | -0.025 | 0 | 0.29 | 0.08 | 0.21 | 0.25 | 0.18 | 0.28 |
| 5 | 71472020 | rs9293794 | -0.33 | -0.772 | -2.174 | 0.666 | 0.017 | -0.033 | -0.033 | 4.039 | 0.34 | 0.03 | 0.22 | 0.26 | 0.25 | 0.29 |
| 5 | 71474274 | rs10055619 | -0.828 | -0.619 | -1.018 | 0.731 | -0.034 | 0.004 | 0.004 | 0 | 0.22 | 0.32 | 0.1 | 0.18 | 0.14 | 0.14 |
| 5 | 71476905 | rs2337694 | -0.597 | -0.676 | -1.942 | 0.409 | -0.003 | 0.006 | 0.006 | 0 | 0.21 | 0.03 | 0.09 | 0.17 | 0.17 | 0.32 |
| 5 | 71477115 | rs6874186 | 0.633 | -1.336 | -2.074 | -0.179 | 0.024 | 0.044 | 0.044 | 0.395 | 0.61 | 0.7 | 0.64 | 0.19 | 0.65 | 0.4 |
| 5 | 71481887 | rs3792806 | -0.284 | -1.039 | -2.598 | -0.633 | -0.031 | 0.039 | 0.039 | 0 | 0.86 | 0.99 | 0.69 | 0.96 | 0.83 | 0.92 |
| 5 | 71484112 | rs9885364 | 0.155 | -1.699 | -2.168 | -0.408 | -0.029 | 0.014 | 0.014 | 0 | 0.26 | 0.35 | 0.21 | 0.31 | 0.18 | 0.17 |
| 5 | 71486785 | rs1561398 | -0.255 | -2.557 | -2.131 | -1.027 | 0.078 | 0.019 | 0.019 | 0.177 | 0.41 | 0.79 | 0.28 | 0.34 | 0.26 | 0.26 |
| 5 | 71488904 | rs7736611 | 1.635 | -2.616 | -2.492 | -1.329 | -0.092 | 0.085 | 0.085 | 0 | 0.24 | 0.08 | 0.17 | 0.08 | 0.36 | 0.31 |
| 5 | 71489305 | rs12516429 | 1.293 | -2.822 | -2.529 | -1.325 | -0.026 | 0.071 | 0.071 | 0 | 0.18 | 0.34 | 0.19 | 0.31 | 0.13 | 0.16 |
| 5 | 71490962 | rs1866374 | -1.635 | -2.517 | -2.68 | -1.254 | -0.025 | 0.048 | 0.048 | 0 | 0.86 | 0.99 | 0.7 | 0.96 | 0.84 | 0.93 |
| 5 | 71492734 | rs7716079 | 1.362 | -2.513 | -1.745 | -1.191 | 0.006 | 0.007 | 0.007 | 0 | 0.12 | 0.12 | 0.04 | 0.03 | 0.08 | 0.09 |
| 5 | 71495041 | rs3805452 | 0.362 | -2.353 | -1.874 | -1.007 | 0.007 | -0.082 | -0.082 | 0 | 0.25 | 0.11 | 0.27 | 0.55 | 0.26 | 0.37 |
| 5 | 71496419 | rs3828616 | 1.127 | -0.664 | -1.576 | -1.143 | -0.023 | 0.021 | 0.021 | 0 | 0.24 | 0.14 | 0.16 | 0.09 | 0.33 | 0.28 |
| 5 | 71506434 | rs10039706 | 0.129 | -1.003 | -2.009 | -1.058 | -0.044 | -0.067 | -0.067 | 0 | 0.66 | 0.88 | 0.45 | 0.39 | 0.59 | 0.56 |
| 5 | 71509807 | rs6453517 | 0.52 | -0.862 | -0.933 | -1.077 | 0.004 | -0.001 | -0.001 | 2.147 | 0.39 | 0.36 | 0.25 | 0.37 | 0.24 | 0.24 |
| 5 | 71512802 | rs17375040 | 1.506 | -0.613 | -0.875 | -1.17 | -0.035 | 0.011 | 0.011 | 0 | 0.24 | 0.11 | 0.17 | 0.02 | 0.37 | 0.31 |
| 5 | 71515616 | rs6831 | -0.359 | -0.82 | -0.476 | -1.123 | 0.002 | 0.02 | 0.02 | 0.689 | 0.71 | 0.6 | 0.46 | 0.39 | 0.66 | 0.59 |
| 5 | 71517497 | rs17303157 | 0.852 | -0.785 | -0.208 | -1.232 | 0.028 | -0.099 | -0.099 | 0 | 0.3 | 0.29 | 0.2 | 0.02 | 0.41 | 0.34 |
| 5 | 71521551 | rs6889892 | -1.105 | -0.79 | -0.784 | -0.663 | 0.008 | -0.072 | -0.072 | 0 | 0.81 | 0.91 | 0.61 | 0.53 | 0.78 | 0.65 |
| 5 | 71521682 | rs2289281 | 1.936 | -0.821 | -0.795 | -0.55 | 0.01 | -0.05 | -0.05 | 0.964 | 0.12 | 0.06 | 0.12 | 0.43 | 0.12 | 0.33 |
| 5 | 71526092 | rs1982193 | 0.601 | -1.03 | -0.743 | -0.603 | 0.061 | -0.034 | -0.034 | 0.029 | 0.38 | 0.23 | 0.26 | 0.39 | 0.26 | 0.24 |
| 5 | 71532182 | rs10073728 | 2.168 | -1.544 | -1.056 | -0.766 | 0.003 | 0.056 | 0.056 | 0.019 | 0.16 | 0 | 0.04 | 0.03 | 0.12 | 0.07 |
| 5 | 71534854 | rs11956209 | -2.209 | -1.33 | -0.818 | -0.66 | 0 | 0.005 | 0.005 | 0 | 0.92 | 0.97 | 0.72 | 0.94 | 0.91 | 0.98 |
| 5 | 71542203 | rs2289279 | 1.905 | -1.313 | -1.008 | -0.607 | -0.022 | -0.1 | -0.1 | 0 | 0.15 | 0.02 | 0.13 | 0.37 | 0.15 | 0.34 |
| 5 | 71543014 | rs10454918 | -1.23 | -1.221 | -0.894 | -0.726 | -0.002 | -0.089 | -0.089 | 3.377 | 0.77 | 0.92 | 0.59 | 0.56 | 0.76 | 0.63 |
| 5 | 71543142 | rs10454919 | 1.49 | -1.29 | -0.466 | -0.817 | 0.016 | -0.013 | -0.013 | 0.065 | 0.06 | 0.18 | 0.02 | 0 | 0.04 | 0.05 |
| 5 | 71544320 | rs11949953 | -0.606 | -1.12 | -0.521 | -0.865 | -0.032 | -0.093 | -0.093 | 0.011 | 0.72 | 0.88 | 0.49 | 0.41 | 0.66 | 0.58 |
| 5 | 71549048 | rs7710169 | -0.619 | -1.143 | -0.045 | -0.681 | -0.031 | -0.06 | -0.06 | 0 | 0.1 | 0.61 | 0.05 | 0 | 0.04 | 0.07 |
| 5 | 71598904 | rs961468 | 0.007 | -1.119 | 0.265 | -0.67 | 0.039 | -0.001 | -0.001 | 0 | 0.08 | 0.34 | 0.03 | 0 | 0.04 | 0.07 |
| 5 | 71621428 | rs9293273 | 0.598 | -1.149 | -1.201 | -0.58 | 0.033 | -0.018 | -0.018 | 2.093 | 0.29 | 0.16 | 0.17 | 0.31 | 0.17 | 0.18 |
| 5 | 71625561 | rs12186595 | 0.933 | -0.365 | -0.885 | -0.598 | 0.023 | -0.007 | -0.007 | 0 | 0.05 | 0.18 | 0.02 | 0 | 0.04 | 0.02 |
| 5 | 71654758 | rs7734699 | 0.858 | -0.569 | -0.806 | -0.807 | 0.021 | -0.008 | -0.008 | 0 | 0.05 | 0.18 | 0.02 | 0 | 0.04 | 0.02 |
| 5 | 71654937 | rs7718007 | 0.812 | -0.572 | -0.807 | -0.808 | 0.019 | -0.01 | -0.01 | 0 | 0.05 | 0.18 | 0.02 | 0 | 0.04 | 0.02 |
| 5 | 71664031 | rs17377355 | 1.296 | 0.269 | -1.685 | -0.923 | -0.16 | 0.164 | 0.164 | 0 | 0.24 | 0.02 | 0.18 | 0.01 | 0.35 | 0.29 |
| 5 | 71667678 | rs9293282 | 0.932 | -0.17 | -1.294 | -0.839 | -0.002 | -0.015 | -0.015 | 3.39 | 0.19 | 0.18 | 0.06 | 0.02 | 0.1 | 0.07 |
| 5 | 71674855 | rs7709552 | 0.383 | -0.023 | -1.196 | -0.471 | 0.001 | 0.019 | 0.019 | 0.067 | 0.46 | 0.63 | 0.37 | 0.37 | 0.46 | 0.48 |
| 5 | 71683885 | rs9293289 | 0.532 | 0.589 | -1.465 | -0.178 | -0.078 | -0.133 | -0.133 | 0 | 0.24 | 0.06 | 0.26 | 0.51 | 0.33 | 0.35 |
| 5 | 71688223 | rs13158670 | 0.542 | 0.262 | -1.205 | -0.487 | -0.012 | -0.002 | -0.002 | 0 | 0.44 | 0.28 | 0.35 | 0.43 | 0.44 | 0.52 |
| 5 | 71693463 | rs4703854 | -0.632 | 0.039 | -0.684 | -1.104 | -0.002 | -0.025 | -0.025 | 0 | 0.73 | 0.69 | 0.65 | 0.91 | 0.77 | 0.88 |
| 5 | 71693899 | rs4703855 | 0.506 | 0.303 | -0.764 | -0.704 | -0.105 | -0.113 | -0.113 | 1.794 | 0.26 | 0.04 | 0.27 | 0.57 | 0.34 | 0.36 |
| 5 | 71695880 | rs10040192 | -0.207 | 0.289 | -0.899 | -1.077 | -0.043 | 0.204 | 0.204 | 0.14 | 0.42 | 0.15 | 0.48 | 0.88 | 0.52 | 0.56 |
| 5 | 71696942 | rs9293292 | 1.167 | 0.326 | -0.933 | -1.018 | -0.008 | -0.121 | -0.121 | 0 | 0.22 | 0.03 | 0.21 | 0.55 | 0.2 | 0.29 |
| 5 | 71697977 | rs7445013 | -0.329 | 0.442 | -0.345 | -1.349 | -0.001 | -0.029 | -0.029 | 0 | 0.45 | 0.47 | 0.49 | 0.88 | 0.48 | 0.54 |

| | | | | | | | | | | | | | | | | |
|---|---|---|---|---|---|---|---|---|---|---|---|---|---|---|---|---|
| 5 | 71717605 | rs7723734 | 0.247 | 0.481 | -0.786 | -1.269 | 0.01 | 0.148 | 0.148 | 0.283 | 0.37 | 0.14 | 0.4 | 0.88 | 0.35 | 0.5 |
| 5 | 71729221 | rs12054750 | -0.076 | 0.412 | -0.915 | -1.171 | -0.093 | 0.197 | 0.197 | 0.059 | 0.42 | 0.1 | 0.46 | 0.84 | 0.48 | 0.54 |
| 5 | 71741094 | rs6452473 | -1.581 | 0.117 | -0.493 | -1.371 | -0.002 | -0.022 | -0.022 | 0.431 | 0.89 | 0.81 | 0.7 | 0.96 | 0.87 | 0.97 |
| 5 | 71741790 | rs7710292 | -0.048 | -0.291 | -0.449 | -1.787 | 0.016 | 0.135 | 0.135 | 0.146 | 0.55 | 0.19 | 0.52 | 0.87 | 0.58 | 0.65 |
| 5 | 71742156 | rs17378840 | 2.302 | -0.488 | -0.301 | -1.947 | -0.018 | 0.038 | 0.038 | 0 | 0.21 | 0.1 | 0.12 | 0.02 | 0.26 | 0.25 |
| 5 | 71743322 | rs10043368 | -0.048 | -0.163 | -0.556 | -1.742 | 0.017 | 0.139 | 0.139 | 0.241 | 0.55 | 0.19 | 0.52 | 0.87 | 0.58 | 0.65 |
| 5 | 71743954 | rs13177417 | 0.376 | -0.248 | -0.724 | -1.496 | -0.024 | -0.008 | -0.008 | 0 | 0.34 | 0.18 | 0.31 | 0.38 | 0.38 | 0.34 |
| 5 | 71744199 | rs10068109 | -1.245 | -0.701 | -0.359 | -1.162 | -0.008 | -0.045 | -0.045 | 0.603 | 0.87 | 0.71 | 0.68 | 0.96 | 0.87 | 0.97 |
| 5 | 71744664 | rs10070001 | -0.121 | -0.678 | -0.332 | -1.192 | -0.003 | 0.001 | 0.001 | 0 | 0.67 | 0.57 | 0.48 | 0.47 | 0.69 | 0.67 |
| 5 | 71746801 | rs13189566 | 1.727 | -0.637 | -0.554 | -0.95 | 0 | 0.027 | 0.027 | 0.443 | 0.15 | 0.13 | 0.17 | 0.38 | 0.13 | 0.21 |
| 5 | 71747448 | rs16878635 | 0.865 | -0.611 | -0.85 | -0.703 | 0.024 | -0.095 | -0.095 | 2.056 | 0.23 | 0.1 | 0.22 | 0.49 | 0.2 | 0.31 |
| 5 | 71750904 | rs3112483 | -1.565 | -0.315 | -0.57 | -0.726 | -0.006 | -0.012 | -0.012 | 0.14 | 0.92 | 0.85 | 0.73 | 0.96 | 0.93 | 0.98 |
| 5 | 71754962 | rs6862221 | 0.381 | -0.381 | -0.362 | -0.325 | -0.007 | 0.029 | 0.029 | 0 | 0.23 | 0.3 | 0.22 | 0.46 | 0.2 | 0.29 |
| 5 | 71756102 | rs7721922 | -0.366 | -0.347 | -0.684 | -0.516 | -0.011 | -0.072 | -0.072 | 1.298 | 0.39 | 0.5 | 0.41 | 0.86 | 0.34 | 0.5 |
| 5 | 71756670 | rs2278600 | 0.465 | 0.01 | -0.471 | -0.211 | -0.007 | 0.097 | 0.097 | 0.413 | 0.18 | 0.38 | 0.2 | 0.41 | 0.13 | 0.2 |
| 5 | 71757894 | rs10447170 | 0.43 | 0.482 | -1.037 | -0.463 | -0.002 | -0.031 | -0.031 | 3.285 | 0.18 | 0.18 | 0.13 | 0 | 0.18 | 0.13 |
| 5 | 71763154 | rs1019457 | -0.199 | -0.527 | -1.448 | 0.089 | 0.03 | -0.09 | -0.09 | 0.106 | 0.29 | 0.13 | 0.24 | 0.48 | 0.26 | 0.4 |
| 5 | 71763715 | rs6874676 | 0.342 | -0.434 | -1.055 | 0.23 | -0.013 | 0.022 | 0.022 | 0 | 0.35 | 0.45 | 0.24 | 0.56 | 0.24 | 0.36 |
| 5 | 71781839 | rs10043659 | 0.132 | -0.713 | -0.963 | 0.223 | 0.016 | -0.037 | -0.037 | 0 | 0.24 | 0.15 | 0.2 | 0.43 | 0.21 | 0.39 |
| 5 | 71789021 | rs10042132 | 0.7 | -0.85 | -1.06 | 0.196 | 0.002 | -0.05 | -0.05 | 0.18 | 0.18 | 0.11 | 0.18 | 0.41 | 0.21 | 0.33 |
| 5 | 71803446 | rs13358894 | 0.478 | -0.98 | -1.241 | 0.261 | 0.005 | -0.04 | -0.04 | 0 | 0.19 | 0.1 | 0.18 | 0.41 | 0.2 | 0.33 |
| 5 | 71809046 | rs6895716 | 0.858 | -1.354 | -1.875 | -0.024 | -0.024 | -0.093 | -0.093 | 0 | 0.2 | 0.02 | 0.18 | 0.32 | 0.13 | 0.17 |
| 5 | 71809247 | rs246565 | 0.916 | -1.178 | -1.448 | 0.019 | -0.002 | -0.053 | -0.053 | 0 | 0.17 | 0.07 | 0.17 | 0.41 | 0.2 | 0.33 |
| 5 | 71814128 | rs891004 | 1.155 | -0.702 | -1.611 | -0.423 | -0.009 | 0.061 | 0.061 | 0 | 0.25 | 0.15 | 0.15 | 0.05 | 0.27 | 0.31 |
| 5 | 71846721 | rs33419 | -0.519 | -1.08 | -1.224 | -0.147 | -0.034 | 0.01 | 0.01 | 0 | 0.38 | 0.52 | 0.27 | 0.51 | 0.25 | 0.4 |
| 5 | 71863336 | rs33415 | 0.337 | -0.475 | -1.215 | -0.585 | 0.051 | 0.121 | 0.121 | 0 | 0.56 | 0.25 | 0.77 | 0.85 | 0.66 | 0.59 |
| 5 | 71892912 | rs33425 | -0.428 | 0.146 | -1.45 | -0.487 | -0.102 | 0.1 | 0.1 | 0 | 0.68 | 0.92 | 0.8 | 0.89 | 0.59 | 0.68 |
| 5 | 71908620 | rs17590866 | 0.258 | 0.114 | -0.583 | -0.021 | -0.011 | 0.005 | 0.005 | 0 | 0.15 | 0.13 | 0.18 | 0.12 | 0.19 | 0.23 |
| 5 | 71914988 | rs17628947 | 0.044 | 0.055 | -0.089 | -0.041 | 0.006 | -0.007 | -0.007 | 2.059 | 0.17 | 0.2 | 0.19 | 0.12 | 0.19 | 0.23 |
| 5 | 71924577 | rs9765670 | -1.031 | -0.247 | 0.771 | -0.053 | -0.004 | -0.005 | -0.005 | 1.779 | 0.25 | 0.7 | 0.24 | 0.17 | 0.2 | 0.3 |
| 5 | 71927013 | rs34909640 | 0.254 | 0.146 | 0.612 | -0.059 | -0.01 | 0.006 | 0.006 | 0 | 0.16 | 0.13 | 0.18 | 0.12 | 0.19 | 0.23 |
| 5 | 71954583 | rs889234 | 0.077 | 0.173 | 0.585 | -0.066 | 0 | -0.001 | -0.001 | 0 | 0.18 | 0.2 | 0.19 | 0.12 | 0.19 | 0.26 |
| 5 | 71988303 | rs2925684 | -0.055 | 0.158 | 0.568 | -0.124 | 0.002 | -0.008 | -0.008 | 0 | 0.18 | 0.2 | 0.19 | 0.11 | 0.2 | 0.26 |
| 5 | 71999107 | rs2943789 | -0.158 | 0.148 | 0.687 | -0.129 | 0.007 | -0.023 | -0.023 | 0 | 0.19 | 0.26 | 0.19 | 0.11 | 0.2 | 0.26 |
| 5 | 72013573 | rs3010278 | -0.777 | 0.117 | 0.59 | -0.172 | -0.004 | -0.028 | -0.028 | 0 | 0.24 | 0.65 | 0.22 | 0.11 | 0.2 | 0.23 |
| 5 | 72023208 | rs10050720 | 0.022 | 0.157 | 0.228 | -0.224 | -0.001 | -0.01 | -0.01 | 1.157 | 0.1 | 0.08 | 0.16 | 0.04 | 0.13 | 0.06 |
| 5 | 72032166 | rs13156244 | -0.308 | 0.16 | 0.774 | -0.115 | -0.021 | -0.02 | -0.02 | 0 | 0.23 | 0.32 | 0.16 | 0.15 | 0.13 | 0.26 |
| 5 | 72053706 | rs6882763 | 1.027 | 0.344 | 0.752 | -0.274 | 0.006 | -0.023 | -0.023 | 0 | 0.18 | 0.33 | 0.09 | 0.09 | 0.16 | 0.16 |
| 5 | 72061540 | rs6897924 | -0.579 | 0.45 | 0.643 | 0.409 | 0.004 | 0.006 | 0.006 | 0 | 0.41 | 0.61 | 0.25 | 0.24 | 0.29 | 0.41 |
| 5 | 72080422 | rs7709903 | 0.657 | 0.407 | 0.811 | 0.386 | 0.002 | -0.043 | -0.043 | 1.396 | 0.21 | 0.51 | 0.1 | 0.09 | 0.16 | 0.16 |
| 5 | 72084557 | rs10077575 | 0.181 | 0.461 | 0.756 | 0.424 | -0.003 | -0.003 | -0.003 | 0 | 0.22 | 0.24 | 0.16 | 0.15 | 0.13 | 0.26 |
| 5 | 72117128 | rs155426 | 0.427 | 0.463 | 0.721 | 0.376 | -0.002 | -0.003 | -0.003 | 0 | 0.21 | 0.22 | 0.15 | 0.14 | 0.13 | 0.26 |
| 5 | 72121124 | rs155430 | 0.003 | 0.587 | 0.696 | 0.433 | 0.044 | 0.105 | 0.105 | 0.01 | 0.95 | 0.92 | 0.9 | 0.68 | 0.83 | 0.95 |
| 5 | 72129604 | rs13157620 | 1.004 | 0.452 | 0.759 | 0.536 | -0.003 | 0.055 | 0.055 | 0 | 0.51 | 0.22 | 0.64 | 0.37 | 0.54 | 0.53 |
| 5 | 72140173 | rs155434 | -0.562 | 0.491 | 0.847 | 0.57 | 0.103 | 0.241 | 0.241 | 0.211 | 0.06 | 0.17 | 0.11 | 0.39 | 0.17 | 0.05 |
| 5 | 72144209 | rs34652 | -0.203 | 0.498 | 0.859 | 0.569 | 0.091 | 0.225 | 0.225 | 0.01 | 0.05 | 0.13 | 0.1 | 0.39 | 0.16 | 0.05 |
| 5 | 72162264 | rs34669 | 0.505 | 0.477 | 0.854 | 0.513 | -0.001 | -0.004 | -0.004 | 0 | 0.57 | 0.4 | 0.75 | 0.76 | 0.71 | 0.58 |
| 5 | 72168857 | rs3797358 | 0.947 | 0.476 | 0.761 | 0.503 | -0.004 | 0.02 | 0.02 | 0 | 0.52 | 0.24 | 0.65 | 0.43 | 0.55 | 0.54 |
| 5 | 72180377 | rs266443 | -0.48 | 0.475 | 0.815 | 0.03 | -0.007 | -0.01 | -0.01 | 0 | 0.78 | 0.72 | 0.84 | 0.86 | 0.87 | 0.74 |
| 5 | 72206862 | rs155625 | -0.347 | 0.444 | 0.665 | -0.081 | 0.037 | 0.077 | 0.077 | 0 | 0.29 | 0.49 | 0.2 | 0.48 | 0.33 | 0.2 |
| 5 | 72213483 | rs4443401 | -0.504 | 0.551 | 0.576 | 0.064 | -0.008 | -0.006 | -0.006 | 1.194 | 0.78 | 0.74 | 0.85 | 0.83 | 0.87 | 0.74 |
| 5 | 72236312 | rs268793 | 0.976 | 0.59 | -0.13 | 0.082 | 0.001 | 0.056 | 0.056 | 0 | 0.51 | 0.26 | 0.65 | 0.35 | 0.55 | 0.53 |
| 5 | 72247868 | rs2253215 | -0.547 | 0.615 | -0.102 | 0.052 | -0.025 | -0.021 | -0.021 | 0 | 0.78 | 0.59 | 0.84 | 0.83 | 0.87 | 0.73 |
| 5 | 72250961 | rs11746344 | 1.475 | 0.641 | -0.306 | 0.053 | -0.015 | 0.014 | 0.014 | 0.305 | 0.17 | 0.05 | 0.07 | 0.09 | 0.15 | 0.16 |
| 5 | 72252475 | rs7713398 | 0.977 | 0.668 | -0.289 | 0.08 | -0.004 | 0.064 | 0.064 | 0 | 0.51 | 0.25 | 0.65 | 0.35 | 0.56 | 0.52 |
| 5 | 72257180 | rs2548331 | 1.364 | 0.64 | -0.312 | 0.071 | -0.029 | 0.046 | 0.046 | 0 | 0.45 | 0.1 | 0.49 | 0.3 | 0.48 | 0.35 |
| 5 | 72289521 | rs10462470 | -0.086 | 0.683 | -0.272 | 0.142 | 0.063 | 0.164 | 0.164 | 0.087 | 0.05 | 0.09 | 0.1 | 0.39 | 0.16 | 0.05 |
| 5 | 72289671 | rs187731 | -0.511 | 0.624 | -0.282 | 0.122 | -0.031 | -0.027 | -0.027 | 0 | 0.77 | 0.54 | 0.83 | 0.83 | 0.87 | 0.72 |
| 5 | 72294458 | rs26575 | -0.98 | 0.668 | -0.304 | 0.122 | 0.037 | 0.057 | 0.057 | 0 | 0.48 | 0.74 | 0.32 | 0.65 | 0.33 | 0.47 |
| 5 | 72295577 | rs10942525 | 0.038 | 0.691 | -0.285 | 0.145 | -0.004 | 0.039 | 0.039 | 0.699 | 0.25 | 0.28 | 0.15 | 0.48 | 0.2 | 0.2 |
| 5 | 72307142 | rs7731235 | 0.129 | 0.48 | -0.285 | 0.166 | 0.082 | 0.191 | 0.191 | 0.171 | 0.06 | 0.12 | 0.1 | 0.39 | 0.17 | 0.05 |
| 5 | 72315858 | rs16878881 | 0.75 | 0.418 | -0.3 | 0.146 | 0.004 | 0.017 | 0.017 | 3.252 | 0.2 | 0.14 | 0.09 | 0.09 | 0.15 | 0.16 |
| 5 | 72347223 | rs7712838 | -0.137 | 0.455 | -0.368 | 0.16 | 0.012 | 0.029 | 0.029 | 0.274 | 0.27 | 0.29 | 0.19 | 0.48 | 0.32 | 0.2 |
| 5 | 72348272 | rs185435 | 0.385 | 0.403 | -0.38 | 0.118 | -0.028 | -0.026 | -0.026 | 0 | 0.23 | 0.46 | 0.17 | 0.17 | 0.13 | 0.27 |
| 5 | 72354897 | rs11749704 | 1.362 | 0.554 | -0.461 | 0.173 | -0.033 | 0.006 | 0.006 | 0.358 | 0.17 | 0.03 | 0.08 | 0.09 | 0.15 | 0.16 |
| 5 | 72385914 | rs6892973 | -0.309 | 0.709 | -0.205 | 0.369 | 0.007 | 0.019 | 0.019 | 1.428 | 0.28 | 0.29 | 0.2 | 0.48 | 0.33 | 0.21 |
| 5 | 72386139 | rs1046064 | -0.051 | 0.917 | -0.563 | 0.471 | 0.081 | 0.195 | 0.195 | 0.117 | 0.06 | 0.12 | 0.1 | 0.39 | 0.17 | 0.05 |
| 5 | 72387486 | rs10942547 | 1 | 0.568 | -1.113 | 0.236 | -0.03 | 0.015 | 0.015 | 1.555 | 0.18 | 0.03 | 0.09 | 0.09 | 0.16 | 0.16 |
| 5 | 72400190 | rs683633 | 0.503 | 0.388 | -0.794 | 0.267 | -0.031 | -0.016 | -0.016 | 0 | 0.23 | 0.52 | 0.17 | 0.18 | 0.13 | 0.24 |

| | | | | | | | | | | | | | | | | |
|---|---|---|---|---|---|---|---|---|---|---|---|---|---|---|---|---|
| 5 | 72407538 | rs608104 | 0.027 | 0.636 | -1.719 | 0.134 | 0.02 | 0.01 | 0.01 | 0.449 | 0.41 | 0.36 | 0.29 | 0.38 | 0.31 | 0.51 |
| 5 | 72408589 | rs623297 | 0.514 | 0.892 | -2.365 | -0.157 | -0.029 | 0.05 | 0.05 | 0 | 0.26 | 0.07 | 0.33 | 0.1 | 0.29 | 0.11 |
| 5 | 72419041 | rs7448965 | -0.121 | 1.257 | -1.989 | 0.333 | -0.056 | -0.04 | -0.04 | 0 | 0.82 | 0.99 | 0.82 | 0.8 | 0.76 | 0.79 |
| 5 | 72419617 | rs636926 | -0.351 | 1.137 | -1.06 | 0.659 | -0.001 | 0.025 | 0.025 | 0.091 | 0.31 | 0.57 | 0.33 | 0.59 | 0.27 | 0.41 |
| 5 | 72425661 | rs703871 | 0.645 | 1.534 | -1.52 | 0.125 | -0.064 | 0.083 | 0.083 | 0.261 | 0.86 | 0.97 | 0.76 | 0.99 | 0.76 | 0.92 |
| 5 | 72426775 | rs1083428 | 0.579 | 1.534 | -1.516 | 0.129 | -0.056 | 0.079 | 0.079 | 0.379 | 0.86 | 0.97 | 0.78 | 0.99 | 0.79 | 0.92 |
| 5 | 72429346 | rs488174 | -0.662 | 0.918 | -0.553 | 1.081 | -0.008 | 0.218 | 0.218 | 0 | 0.09 | 0.29 | 0.16 | 0.32 | 0.03 | 0.19 |
| 5 | 72431457 | rs549579 | -0.201 | 0.849 | -1.171 | 0.719 | -0.001 | -0.009 | -0.009 | 4.664 | 0.55 | 0.45 | 0.51 | 0.59 | 0.44 | 0.63 |
| 5 | 72434224 | rs13160226 | 0.832 | 0.785 | -1.795 | 0.388 | -0.072 | -0.108 | -0.108 | 0 | 0.18 | 0.02 | 0.22 | 0.24 | 0.21 | 0.27 |
| 5 | 72434301 | rs471507 | 1.017 | 1.049 | -1.73 | 0.248 | 0.004 | -0.014 | -0.014 | 0 | 0.39 | 0.52 | 0.43 | 0.4 | 0.43 | 0.35 |
| 5 | 72439645 | rs635444 | 0.942 | 1.022 | -1.471 | 0.273 | 0.002 | -0.006 | -0.006 | 0.907 | 0.39 | 0.52 | 0.51 | 0.5 | 0.43 | 0.37 |
| 5 | 72451309 | rs6881806 | 0.816 | 1.153 | -1.424 | 0.173 | 0.031 | 0.07 | 0.07 | 1.069 | 0.28 | 0.12 | 0.14 | 0.03 | 0.22 | 0.16 |
| 5 | 72451721 | rs71542 | 0.045 | 1.19 | -1.083 | 0.324 | 0.003 | 0.02 | 0.02 | 0 | 0.11 | 0.11 | 0.1 | 0.25 | 0.13 | 0.19 |
| 5 | 72452547 | rs1916642 | 1.367 | 1.019 | -0.883 | 0.267 | 0 | 0 | 0 | 1.11 | 0.33 | 0.38 | 0.33 | 0.47 | 0.33 | 0.33 |
| 5 | 72456171 | rs9293527 | 1.525 | 0.735 | -1.22 | -0.144 | -0.013 | 0.099 | 0.099 | 0.244 | 0.16 | 0.02 | 0.06 | 0 | 0.14 | 0.06 |
| 5 | 72456421 | rs17731153 | 0.869 | 0.738 | -1.322 | -0.148 | -0.06 | -0.099 | -0.099 | 0.119 | 0.19 | 0.02 | 0.21 | 0.24 | 0.19 | 0.28 |
| 5 | 72459066 | rs1403728 | 1.002 | 0.784 | -0.411 | 0.075 | 0.001 | -0.011 | -0.011 | 0 | 0.65 | 0.49 | 0.64 | 0.74 | 0.61 | 0.72 |
| 5 | 72460126 | rs575415 | -0.645 | 1.369 | 0.277 | 0.564 | 0.081 | 0.259 | 0.259 | 0 | 0.09 | 0.32 | 0.1 | 0.24 | 0.1 | 0.18 |
| 5 | 72460195 | rs576243 | 0.433 | 1.33 | 0.12 | 0.472 | -0.003 | -0.007 | -0.007 | 0 | 0.91 | 0.89 | 0.8 | 1 | 0.85 | 0.97 |
| 5 | 72460429 | rs648422 | 0.528 | 1.336 | 0.113 | 0.475 | 0.005 | -0.027 | -0.027 | 0 | 0.9 | 0.86 | 0.8 | 1 | 0.85 | 0.97 |
| 5 | 72463436 | rs7719668 | 1.302 | 1.424 | 0.154 | 0.659 | 0.057 | 0.08 | 0.08 | 0 | 0.82 | 0.51 | 0.7 | 0.75 | 0.75 | 0.79 |
| 5 | 72466319 | rs4703595 | 0.884 | 0.684 | 0.813 | 0.387 | 0.029 | -0.026 | -0.026 | 2.881 | 0.89 | 0.65 | 0.91 | 0.98 | 0.89 | 0.96 |
| 5 | 72467952 | rs4235664 | -0.148 | 0.69 | 0.79 | 0.448 | 0.005 | -0.005 | -0.005 | 0 | 0.93 | 0.9 | 0.93 | 0.97 | 0.9 | 0.99 |
| 5 | 72470655 | rs1104891 | -0.373 | 0.881 | 0.521 | 0.397 | -0.045 | -0.017 | -0.017 | 4.503 | 0.33 | 0.21 | 0.48 | 0.27 | 0.42 | 0.32 |
| 5 | 72470715 | rs4302565 | 0.676 | 0.929 | 0.383 | 0.401 | 0 | 0.031 | 0.031 | 0.037 | 0.75 | 0.43 | 0.81 | 0.71 | 0.75 | 0.72 |
| 5 | 72470853 | rs998243 | -0.104 | 0.874 | 0.275 | 0.37 | -0.122 | -0.07 | -0.07 | 0 | 0.32 | 0.11 | 0.48 | 0.27 | 0.4 | 0.33 |
| 5 | 72472764 | rs3733859 | 1.812 | 0.833 | 0.383 | 0.417 | 0.022 | 0.026 | 0.026 | 7.12 | 0.41 | 0.07 | 0.31 | 0.44 | 0.33 | 0.39 |
| 5 | 72473814 | rs4235665 | 0.667 | 0.86 | 0.539 | 0.449 | 0 | 0.002 | 0.002 | 0.273 | 0.75 | 0.34 | 0.8 | 0.71 | 0.75 | 0.72 |
| 5 | 72473899 | rs6893265 | 0.718 | 0.929 | 0.006 | 0.527 | 0.004 | 0.034 | 0.034 | 0 | 0.13 | 0.17 | 0.1 | 0.26 | 0.15 | 0.25 |
| 5 | 72493266 | rs7702108 | 0.878 | 1.047 | -0.02 | 0.344 | -0.022 | -0.001 | -0.001 | 0 | 0.58 | 0.39 | 0.42 | 0.71 | 0.49 | 0.66 |
| 5 | 72496179 | rs4703597 | -0.393 | 1.467 | 0.19 | 0.6 | -0.009 | 0.004 | 0.004 | 3.495 | 0.93 | 0.97 | 0.93 | 0.97 | 0.9 | 0.99 |
| 5 | 72501096 | rs3935430 | 1.059 | 1.588 | 0.209 | 0.7 | -0.017 | 0.004 | 0.004 | 1.802 | 0.61 | 0.32 | 0.57 | 0.51 | 0.55 | 0.49 |
| 5 | 72502029 | rs12651896 | 0.388 | 1.549 | 0.512 | 0.473 | 0.036 | 0.035 | 0.035 | 0.76 | 0.78 | 0.66 | 0.68 | 0.81 | 0.71 | 0.75 |
| 5 | 72521202 | rs4262084 | -0.638 | 1.401 | 0.355 | 0.271 | 0.057 | 0.011 | 0.011 | 0.585 | 0.22 | 0.43 | 0.33 | 0.16 | 0.33 | 0.26 |
| 5 | 72541096 | rs6893178 | 0.098 | 1.336 | 0.183 | 0.204 | 0.068 | 0.013 | 0.013 | 0.006 | 0.17 | 0.34 | 0.21 | 0.12 | 0.26 | 0.27 |
| 5 | 72546478 | rs4540159 | -0.073 | 1.336 | -0.16 | 0.204 | 0.057 | 0.005 | 0.005 | 0.088 | 0.18 | 0.46 | 0.23 | 0.12 | 0.26 | 0.26 |
| 5 | 72551134 | rs7702331 | -0.734 | 1.2 | -0.932 | 0.064 | -0.002 | 0.005 | 0.005 | 0 | 0.33 | 0.54 | 0.31 | 0.13 | 0.39 | 0.29 |
| 5 | 72555166 | rs9293551 | -0.885 | 1.415 | -1.956 | 0.212 | -0.084 | -0.062 | -0.062 | 0 | 0.5 | 0.89 | 0.43 | 0.43 | 0.53 | 0.53 |
| 5 | 72565488 | rs4704066 | -0.179 | 1.123 | -1.002 | -0.05 | -0.191 | -0.043 | -0.043 | 0 | 0.27 | 0.03 | 0.39 | 0.18 | 0.42 | 0.4 |
| 5 | 72569945 | rs7727448 | -0.707 | 0.772 | -0.448 | 0.174 | 0.003 | 0.01 | 0.01 | 0 | 0.44 | 0.57 | 0.53 | 0.19 | 0.55 | 0.42 |
| 5 | 72573021 | rs4429830 | -0.757 | 0.754 | -0.605 | 0.192 | 0.007 | 0.017 | 0.017 | 0 | 0.45 | 0.6 | 0.53 | 0.2 | 0.57 | 0.42 |
| 5 | 72578131 | rs4533870 | -1.221 | 0.915 | -0.547 | 0.31 | 0.008 | -0.017 | -0.017 | 0 | 0.22 | 0.32 | 0.33 | 0.17 | 0.29 | 0.16 |
| 5 | 72578446 | rs4131513 | -0.853 | 0.739 | -0.88 | 0.477 | -0.001 | 0.003 | 0.003 | 0 | 0.1 | 0.16 | 0.25 | 0.11 | 0.11 | 0.11 |
| 5 | 72580411 | rs4582253 | -0.099 | 0.466 | -1.04 | 0.312 | 0.015 | 0.006 | 0.006 | 0.056 | 0.9 | 0.94 | 0.93 | 0.9 | 0.92 | 0.94 |
| 5 | 72580837 | rs4704068 | 0.75 | 0.688 | -1.009 | 0.451 | -0.003 | -0.002 | -0.002 | 0 | 0.62 | 0.59 | 0.76 | 0.62 | 0.66 | 0.74 |
| 5 | 72587455 | rs4422515 | 0.548 | 1.037 | -1.487 | 0.186 | -0.016 | 0.005 | 0.005 | 0.046 | 0.57 | 0.19 | 0.53 | 0.4 | 0.56 | 0.62 |
| 5 | 72591905 | rs10079621 | 0.595 | 1.31 | -0.906 | -0.289 | 0.019 | -0.112 | -0.112 | 0.118 | 0.77 | 0.4 | 0.79 | 0.97 | 0.69 | 0.93 |
| 5 | 72596647 | rs4704077 | 0.607 | 0.711 | -1.366 | 0.031 | 0.009 | -0.047 | -0.047 | 0.004 | 0.37 | 0.18 | 0.43 | 0.43 | 0.35 | 0.57 |
| 5 | 72601498 | rs9293560 | 1.077 | 1.063 | -1.282 | -0.009 | -0.034 | 0.083 | 0.083 | 0.251 | 0.5 | 0.12 | 0.63 | 0.64 | 0.52 | 0.79 |
| 5 | 72610530 | rs4389670 | 0.593 | 1.039 | -0.14 | -0.308 | -0.002 | -0.077 | -0.077 | 0 | 0.64 | 0.62 | 0.78 | 1 | 0.65 | 0.85 |
| 5 | 72618120 | rs7725251 | -0.919 | 0.964 | 0.323 | -0.398 | -0.011 | -0.086 | -0.086 | 0 | 0.25 | 0.44 | 0.15 | 0.07 | 0.21 | 0.09 |
| 5 | 72618265 | rs11948073 | 2.021 | 0.94 | -0.209 | -0.36 | -0.024 | 0.082 | 0.082 | 0.247 | 0.51 | 0.1 | 0.63 | 0.65 | 0.53 | 0.8 |
| 5 | 72619502 | rs3922417 | -1.065 | 0.524 | -0.045 | -0.13 | 0.021 | -0.052 | -0.052 | 0 | 0.1 | 0.23 | 0.08 | 0 | 0.11 | 0.05 |
| 5 | 72622740 | rs2973930 | -1.804 | 0.413 | -0.339 | 0.04 | 0.008 | -0.04 | -0.04 | 0 | 0.31 | 0.94 | 0.24 | 0.31 | 0.2 | 0.08 |
| 5 | 72628898 | rs6892889 | -0.265 | 0.329 | 0.312 | -0.285 | -0.025 | -0.051 | -0.051 | 0.045 | 0.13 | 0.2 | 0.06 | 0 | 0.05 | 0.01 |
| 5 | 72639746 | rs2972192 | -1.236 | 0.357 | 0.378 | -0.445 | 0.002 | -0.024 | -0.024 | 0 | 0.23 | 0.67 | 0.12 | 0.03 | 0.12 | 0.03 |
| 5 | 72640187 | rs2973916 | -1.221 | 0.343 | 1.418 | -0.452 | 0.105 | 0.056 | 0.056 | 0 | 0.06 | 0.39 | 0.05 | 0 | 0.02 | 0 |
| 5 | 72650185 | rs4704082 | 0.663 | 1.127 | 1.286 | -0.871 | 0.062 | -0.108 | -0.108 | 0 | 0.79 | 0.66 | 0.77 | 0.98 | 0.65 | 0.89 |
| 5 | 72654304 | rs4443403 | -0.442 | 0.988 | 1.262 | -0.739 | -0.028 | -0.089 | -0.089 | 0 | 0.17 | 0.32 | 0.07 | 0.03 | 0.1 | 0.03 |
| 5 | 72658437 | rs2972210 | 0.786 | 1.026 | 1.525 | -0.791 | 0.082 | -0.098 | -0.098 | 0.507 | 0.79 | 0.61 | 0.77 | 0.96 | 0.61 | 0.9 |
| 5 | 72660785 | rs2972212 | 1.802 | 0.862 | 0.63 | 0.126 | -0.033 | 0.014 | 0.014 | 2.363 | 0.58 | 0.13 | 0.62 | 0.64 | 0.54 | 0.85 |
| 5 | 72671738 | rs12659148 | -1.363 | 0.594 | 0.56 | 0.523 | 0.045 | -0.123 | -0.123 | 0.116 | 0.25 | 0.88 | 0.19 | 0.42 | 0.14 | 0.07 |
| 5 | 72672861 | rs6452976 | -1.464 | 0.598 | 0.646 | 0.525 | 0.039 | -0.07 | -0.07 | 0 | 0.25 | 0.83 | 0.19 | 0.43 | 0.14 | 0.07 |
| 5 | 72690460 | rs4398617 | -1.445 | 0.533 | 0.455 | 0.611 | 0.052 | -0.139 | -0.139 | 0 | 0.26 | 0.85 | 0.2 | 0.48 | 0.13 | 0.11 |
| 5 | 72709904 | rs1449274 | -0.046 | 0.265 | 0.251 | 0.684 | 0.025 | -0.053 | -0.053 | 0 | 0.14 | 0.07 | 0.12 | 0.44 | 0.07 | 0.09 |
| 5 | 72722802 | rs958583 | 0.137 | 0.738 | -0.012 | 0.215 | -0.083 | 0.085 | 0.085 | 0 | 0.85 | 0.98 | 0.79 | 1 | 0.67 | 0.9 |
| 5 | 72734410 | rs7718668 | -1.254 | 0.289 | 0.249 | 0.348 | 0.012 | -0.019 | -0.019 | 0 | 0.24 | 0.71 | 0.16 | 0.47 | 0.09 | 0.13 |
| 5 | 72735544 | rs1375502 | 0.208 | 0.257 | -0.293 | 0.135 | -0.093 | 0.105 | 0.105 | 0 | 0.83 | 0.98 | 0.8 | 1 | 0.65 | 0.88 |
| 5 | 72737895 | rs874973 | -1.237 | -0.037 | -0.137 | 0.256 | 0.069 | -0.078 | -0.078 | 1.756 | 0.26 | 0.84 | 0.18 | 0.47 | 0.09 | 0.13 |

| | | | | | | | | | | | | | | | | |
|---|---|---|---|---|---|---|---|---|---|---|---|---|---|---|---|---|
| 5 | 72741402 | rs6888459 | -1.327 | -0.058 | 0.212 | 0.226 | -0.021 | -0.025 | -0.025 | 0 | 0.08 | 0.62 | 0.06 | 0 | 0 | 0.01 |
| 5 | 72750163 | rs1449272 | -1.093 | 0.012 | -0.188 | 0.292 | 0.054 | -0.06 | -0.06 | 0 | 0.26 | 0.83 | 0.18 | 0.46 | 0.09 | 0.13 |
| 5 | 72756457 | rs2922716 | 1.541 | 0.341 | -0.253 | 0.183 | 0.033 | -0.065 | -0.065 | 0 | 0.6 | 0.21 | 0.68 | 0.49 | 0.58 | 0.76 |
| 5 | 72768166 | rs968660 | -1.057 | 0.374 | 0.329 | 0.227 | 0.012 | 0.465 | 0.465 | 0.786 | 0.11 | 0.44 | 0.12 | 0.5 | 0.07 | 0.13 |
| 5 | 72772062 | rs347251 | 0.796 | 0.352 | 0.317 | 0.032 | 0 | -0.047 | -0.047 | 0 | 0.72 | 0.65 | 0.79 | 0.99 | 0.66 | 0.88 |
| 5 | 72779156 | rs2081989 | -1.335 | 0.281 | 0.356 | 0.274 | -0.016 | 0.32 | 0.32 | 1.384 | 0.12 | 0.54 | 0.12 | 0.54 | 0.07 | 0.13 |
| 5 | 72790693 | rs347243 | -0.109 | 0.386 | -0.182 | 0.24 | -0.044 | 0.06 | 0.06 | 0 | 0.87 | 0.98 | 0.85 | 1 | 0.71 | 0.9 |
| 5 | 72791309 | rs17663872 | 0.689 | 0.409 | -0.765 | 0.273 | -0.026 | -0.04 | -0.04 | 0 | 0.07 | 0.01 | 0.24 | 0.18 | 0.1 | 0.11 |
| 5 | 72793051 | rs347245 | 1.687 | 0.182 | -0.555 | 0.372 | 0.028 | -0.036 | -0.036 | 0 | 0.61 | 0.18 | 0.67 | 0.45 | 0.59 | 0.75 |
| 5 | 72804412 | rs347232 | -0.417 | 0.24 | -0.595 | 0.383 | -0.068 | 0.077 | 0.077 | 0 | 0.87 | 0.99 | 0.85 | 1 | 0.71 | 0.9 |
| 5 | 72809831 | rs2914544 | 1.499 | 0.202 | -1.072 | -0.024 | -0.042 | 0.114 | 0.114 | 0.025 | 0.44 | 0.2 | 0.55 | 0.26 | 0.45 | 0.41 |
| 5 | 72811157 | rs10462345 | 0.459 | 0.864 | -0.811 | 0.471 | -0.011 | -0.023 | -0.023 | 0 | 0.18 | 0.07 | 0.12 | 0.19 | 0.14 | 0.34 |
| 5 | 72815787 | rs6453008 | 0.596 | 0.744 | -0.797 | 0.299 | -0.006 | 0.027 | 0.027 | 0 | 0.58 | 0.43 | 0.61 | 0.26 | 0.51 | 0.43 |
| 5 | 72818878 | rs7722653 | -0.426 | 0.664 | -1.103 | 0.994 | -0.011 | -0.048 | -0.048 | 0 | 0.14 | 0.05 | 0.09 | 0.57 | 0.14 | 0.21 |
| 5 | 72819206 | rs819561 | -0.436 | -0.202 | -0.444 | 0.091 | -0.048 | -0.09 | -0.09 | 0 | 0.23 | 0.44 | 0.17 | 0.04 | 0.18 | 0.09 |
| 5 | 72822466 | rs865340 | -0.868 | 0.245 | -0.299 | 0.072 | 0.005 | -0.01 | -0.01 | 1.145 | 0.09 | 0.13 | 0.1 | 0.04 | 0.14 | 0.17 |
| 5 | 72822630 | rs12651911 | -0.207 | 0.268 | -0.653 | 0.107 | -0.002 | 0.018 | 0.018 | 0 | 0.29 | 0.61 | 0.34 | 0.62 | 0.26 | 0.39 |
| 5 | 72823819 | rs819554 | -1.198 | 0.479 | 0.043 | 0.408 | 0.01 | 0.01 | 0.01 | 0 | 0.31 | 0.49 | 0.43 | 0.65 | 0.39 | 0.38 |
| 5 | 72828213 | rs819571 | -1.36 | -0.256 | 0.243 | 0.759 | -0.019 | 0.218 | 0.218 | 0 | 0.17 | 0.54 | 0.36 | 0.47 | 0.14 | 0.23 |
| 5 | 72830030 | rs17663979 | 0.136 | -0.352 | 0.23 | 0.833 | -0.008 | 0.167 | 0.167 | 0 | 0.08 | 0.12 | 0.24 | 0.48 | 0.06 | 0.21 |
| 5 | 72832193 | rs819569 | -1.56 | -0.265 | -0.079 | 0.354 | 0.016 | 0.014 | 0.014 | 0.497 | 0.39 | 0.76 | 0.51 | 0.65 | 0.4 | 0.52 |
| 5 | 72837582 | rs819600 | -0.861 | -0.286 | -0.06 | 0.015 | -0.008 | 0.025 | 0.025 | 0 | 0.59 | 0.75 | 0.74 | 0.64 | 0.63 | 0.64 |
| 5 | 72840063 | rs13173874 | 1.39 | -0.32 | 0.873 | 0.807 | -0.037 | 0.051 | 0.051 | 0.095 | 0.52 | 0.12 | 0.56 | 0.67 | 0.44 | 0.58 |
| 5 | 72843780 | rs819591 | 0.455 | -0.301 | 0.923 | 0.791 | -0.029 | 0.022 | 0.022 | 0 | 0.11 | 0.06 | 0.15 | 0.06 | 0.22 | 0.14 |
| 5 | 72847601 | rs343101 | 0.84 | -0.365 | 1.18 | 0.813 | 0.009 | 0.016 | 0.016 | 0.032 | 0.63 | 0.2 | 0.71 | 0.72 | 0.66 | 0.71 |
| 5 | 72850282 | rs343104 | 0.529 | -0.418 | 2.065 | 0.802 | -0.003 | -0.063 | -0.063 | 0 | 0.67 | 0.43 | 0.74 | 0.85 | 0.71 | 0.8 |
| 5 | 72861309 | rs343108 | 0.449 | -1.201 | 2.539 | 0.884 | -0.004 | 0.003 | 0.003 | 0 | 0.11 | 0.08 | 0.16 | 0.06 | 0.22 | 0.14 |
| 5 | 72861366 | rs343109 | -1.104 | -1.211 | 2.602 | 0.899 | 0.021 | 0.034 | 0.034 | 0.23 | 0.18 | 0.61 | 0.22 | 0.18 | 0.27 | 0.22 |
| 5 | 72865307 | rs347234 | -1.178 | -1.221 | 2.624 | 0.839 | 0.017 | 0.023 | 0.023 | 0.163 | 0.18 | 0.61 | 0.22 | 0.18 | 0.27 | 0.22 |
| 5 | 72874907 | rs343122 | 0.363 | -1.196 | 2.433 | 0.847 | -0.004 | 0.003 | 0.003 | 0 | 0.11 | 0.1 | 0.15 | 0.06 | 0.22 | 0.14 |
| 5 | 72876035 | rs343120 | -1.058 | -1.213 | 2.515 | 0.843 | 0.028 | 0.036 | 0.036 | 0.048 | 0.18 | 0.56 | 0.21 | 0.18 | 0.27 | 0.22 |
| 5 | 72876119 | rs343119 | -0.617 | -1.191 | 2.212 | 0.854 | 0.031 | 0.037 | 0.037 | 0 | 0.16 | 0.3 | 0.18 | 0.18 | 0.27 | 0.22 |
| 5 | 72876776 | rs343118 | -0.617 | -1.191 | 2.212 | 0.854 | 0.031 | 0.037 | 0.037 | 0 | 0.16 | 0.3 | 0.18 | 0.18 | 0.27 | 0.22 |
| 5 | 72880699 | rs343112 | 0.336 | -1.189 | 2.195 | 0.85 | -0.004 | 0.003 | 0.003 | 0 | 0.11 | 0.1 | 0.15 | 0.06 | 0.22 | 0.14 |
| 5 | 72891758 | rs980631 | 1.677 | -1.128 | 1.305 | 0.759 | -0.021 | 0.061 | 0.061 | 0.346 | 0.51 | 0.09 | 0.59 | 0.66 | 0.46 | 0.59 |
| 5 | 72895239 | rs3108888 | 0.214 | -0.659 | 1.295 | 0.511 | -0.005 | 0.004 | 0.004 | 0 | 0.11 | 0.09 | 0.15 | 0.06 | 0.22 | 0.14 |
| 5 | 72896224 | rs2544652 | -0.719 | -0.708 | 1.221 | 0.582 | -0.014 | -0.04 | -0.04 | 0 | 0.91 | 0.82 | 0.94 | 1 | 0.94 | 0.93 |
| 5 | 72920029 | rs1981809 | 0.732 | -0.661 | 1.423 | 0.969 | -0.002 | 0.009 | 0.009 | 0 | 0.56 | 0.59 | 0.64 | 0.63 | 0.47 | 0.57 |
| 5 | 72928247 | rs17633441 | 0.99 | -0.764 | 1.183 | 1.012 | -0.027 | 0 | 0 | 0.015 | 0.49 | 0.18 | 0.6 | 0.43 | 0.44 | 0.56 |
| 5 | 72950536 | rs4703608 | 0.124 | -0.834 | 1.043 | 0.096 | 0.024 | 0.065 | 0.065 | 0 | 0.66 | 0.76 | 0.82 | 0.83 | 0.71 | 0.76 |
| 5 | 72979253 | rs750345 | 1.39 | -0.871 | 1.169 | 0.244 | -0.031 | 0.06 | 0.06 | 0 | 0.44 | 0.22 | 0.6 | 0.67 | 0.44 | 0.51 |
| 5 | 72980837 | rs16870681 | -0.254 | -0.798 | 1.413 | 0.233 | 0.002 | -0.015 | -0.015 | 0 | 0.17 | 0.22 | 0.16 | 0.04 | 0.23 | 0.17 |
| 5 | 72985444 | rs17633770 | -0.051 | -0.785 | 1.111 | 0.239 | -0.022 | 0.035 | 0.035 | 0 | 0.16 | 0.11 | 0.15 | 0.04 | 0.23 | 0.17 |
| 5 | 72987410 | rs10942685 | 1.236 | -0.986 | 1.068 | 0.273 | -0.066 | 0.074 | 0.074 | 0.036 | 0.46 | 0.11 | 0.6 | 0.66 | 0.43 | 0.5 |
| 5 | 72991879 | rs6888810 | 1.765 | -1.015 | 0.714 | 0.558 | -0.138 | -0.162 | -0.162 | 0 | 0.36 | 0.03 | 0.5 | 0.52 | 0.4 | 0.36 |
| 5 | 72995398 | rs17552357 | 1.291 | -0.851 | 1.085 | -0.137 | -0.052 | 0.07 | 0.07 | 0 | 0.45 | 0.15 | 0.6 | 0.65 | 0.43 | 0.51 |
| 5 | 73016810 | rs16870743 | -1.832 | -0.67 | 2.23 | 0.134 | 0.002 | -0.02 | -0.02 | 0 | 0.23 | 0.66 | 0.25 | 0.16 | 0.28 | 0.22 |
| 5 | 73021455 | rs16870745 | -1.119 | -0.565 | 2.401 | 0.107 | 0.007 | -0.042 | -0.042 | 0.004 | 0.17 | 0.25 | 0.15 | 0.04 | 0.21 | 0.17 |
| 5 | 73037073 | rs11746513 | -1.935 | -0.25 | 2.119 | 0.225 | 0.025 | -0.043 | -0.043 | 0 | 0.25 | 0.45 | 0.23 | 0.16 | 0.33 | 0.22 |
| 5 | 73045506 | rs17552626 | -1.22 | -0.392 | 1.567 | 0.278 | -0.019 | 0.003 | 0.003 | 0 | 0.18 | 0.11 | 0.19 | 0.16 | 0.27 | 0.2 |
| 5 | 73046715 | rs905159 | -2.048 | -0.538 | 2.182 | 0.205 | -0.003 | -0.038 | -0.038 | 0 | 0.26 | 0.54 | 0.24 | 0.14 | 0.3 | 0.2 |
| 5 | 73052741 | rs6897882 | -1.997 | -0.517 | 1.897 | 0.216 | 0.004 | -0.034 | -0.034 | 0.743 | 0.25 | 0.39 | 0.23 | 0.15 | 0.3 | 0.2 |
| 5 | 73061688 | rs10064449 | -1.017 | 0.011 | 0.125 | 0.523 | -0.026 | -0.006 | -0.006 | 2.009 | 0.16 | 0.05 | 0.17 | 0.15 | 0.26 | 0.19 |
| 5 | 73068888 | rs10062885 | -1.093 | -0.014 | 0.12 | 0.337 | -0.016 | 0.027 | 0.027 | 0.053 | 0.16 | 0.06 | 0.16 | 0.05 | 0.25 | 0.18 |
| 5 | 73069226 | rs13164180 | 0.718 | -0.373 | 0.264 | 0.568 | -0.018 | 0.003 | 0.003 | 0 | 0.62 | 0.44 | 0.59 | 0.57 | 0.71 | 0.62 |
| 5 | 73070032 | rs10065074 | -0.748 | -0.666 | 0.012 | 0.3 | 0.006 | 0.018 | 0.018 | 0 | 0.11 | 0.06 | 0.06 | 0.07 | 0.09 | 0.05 |
| 5 | 73072354 | rs7714670 | -0.563 | -0.992 | 0.328 | 0.114 | -0.011 | 0.016 | 0.016 | 0 | 0.38 | 0.34 | 0.37 | 0.32 | 0.46 | 0.38 |
| 5 | 73076457 | rs11949860 | -0.534 | -1.017 | 0.638 | 0.104 | -0.011 | 0.016 | 0.016 | 0 | 0.37 | 0.33 | 0.36 | 0.32 | 0.46 | 0.38 |
| 5 | 73076511 | rs6453022 | -1.412 | -1.394 | 0.781 | 0.639 | -0.01 | -0.013 | -0.013 | 4.227 | 0.52 | 0.73 | 0.53 | 0.51 | 0.5 | 0.56 |
| 5 | 73082942 | rs4599517 | -0.293 | -1.461 | 0.321 | 0.639 | -0.086 | 0.042 | 0.042 | 0 | 0.35 | 0.07 | 0.34 | 0.28 | 0.44 | 0.37 |
| 5 | 73088068 | rs4597943 | -1.362 | -1.81 | 0.817 | 0.661 | 0.015 | -0.01 | -0.01 | 0 | 0.51 | 0.79 | 0.52 | 0.47 | 0.48 | 0.56 |
| 5 | 73090261 | rs7716253 | -1.432 | -1.67 | 0.71 | 0.618 | -0.029 | -0.016 | -0.016 | 2.039 | 0.52 | 0.82 | 0.53 | 0.53 | 0.51 | 0.58 |
| 5 | 73092269 | rs6883409 | -1.665 | -1.724 | 1.256 | 0.642 | -0.035 | 0.177 | 0.177 | 1.643 | 0.13 | 0.53 | 0.17 | 0.25 | 0.05 | 0.2 |
| 5 | 73095635 | rs1565005 | -0.716 | -1.551 | 1.124 | 0.483 | -0.006 | 0.012 | 0.012 | 0 | 0.4 | 0.37 | 0.37 | 0.28 | 0.46 | 0.38 |
| 5 | 73099158 | rs2129403 | -0.509 | -1.599 | 1.084 | 0.515 | -0.014 | 0.009 | 0.009 | 0 | 0.37 | 0.26 | 0.35 | 0.32 | 0.42 | 0.36 |
| 5 | 73104710 | rs2931417 | 1.697 | -1.849 | 0.718 | 0.572 | -0.073 | -0.054 | -0.054 | 0.005 | 0.4 | 0.1 | 0.42 | 0.42 | 0.46 | 0.37 |
| 5 | 73105275 | rs1009200 | 1.903 | -2.286 | 0.454 | 0.303 | -0.081 | -0.079 | -0.079 | 1.077 | 0.32 | 0.1 | 0.4 | 0.43 | 0.42 | 0.38 |
| 5 | 73108573 | rs4703613 | -0.376 | -3.009 | 0.839 | -0.104 | -0.02 | 0.016 | 0.016 | 0 | 0.27 | 0.39 | 0.15 | 0.3 | 0.21 | 0.29 |
| 5 | 73110265 | rs4455542 | 0.272 | -3.044 | 0.834 | -0.105 | 0.03 | -0.045 | -0.045 | 0 | 0.22 | 0.06 | 0.11 | 0.25 | 0.17 | 0.22 |

| | | | | | | | | | | | | | | | | |
|---|---|---|---|---|---|---|---|---|---|---|---|---|---|---|---|---|
| 5 | 73111815 | rs13359267 | 2.433 | -2.845 | 0.551 | -0.245 | -0.107 | -0.113 | -0.113 | 0.187 | 0.3 | 0.09 | 0.39 | 0.49 | 0.41 | 0.37 |
| 5 | 73111953 | rs11745883 | 1.955 | -2.856 | 0.563 | -0.25 | -0.014 | -0.014 | -0.014 | 0.303 | 0.35 | 0.29 | 0.43 | 0.53 | 0.45 | 0.44 |
| 5 | 73112348 | rs6868665 | 0.567 | -2.825 | 0.874 | -0.319 | -0.005 | -0.018 | -0.018 | 0.893 | 0.62 | 0.58 | 0.57 | 0.79 | 0.66 | 0.66 |
| 5 | 73113732 | rs1038559 | -1.07 | -2.907 | 1.029 | -0.251 | -0.013 | 0 | 0 | 0 | 0.4 | 0.52 | 0.31 | 0.38 | 0.29 | 0.43 |
| 5 | 73114769 | rs6860016 | -0.446 | -2.922 | 1.333 | -0.268 | 0 | 0.011 | 0.011 | 0 | 0.16 | 0.3 | 0.18 | 0.17 | 0.13 | 0.24 |
| 5 | 73115001 | rs7705309 | 2.626 | -3.029 | 1.17 | -0.343 | -0.069 | -0.074 | -0.074 | 0 | 0.31 | 0.18 | 0.4 | 0.54 | 0.41 | 0.37 |
| 5 | 73115275 | rs4704098 | 0.555 | -3.085 | 1.237 | -0.351 | 0.006 | -0.003 | -0.003 | 0 | 0.2 | 0.16 | 0.09 | 0.2 | 0.12 | 0.19 |
| 5 | 73115831 | rs4329007 | 0.538 | -3.088 | 1.24 | -0.352 | 0.002 | -0.002 | -0.002 | 0 | 0.2 | 0.16 | 0.09 | 0.2 | 0.12 | 0.19 |
| 5 | 73125146 | rs2931431 | -0.634 | -3.063 | 0.967 | -0.495 | -0.019 | 0.056 | 0.056 | 0 | 0.78 | 0.91 | 0.74 | 0.93 | 0.74 | 0.83 |
| 5 | 73129027 | rs10515164 | -0.535 | -2.979 | 1.221 | -0.371 | -0.028 | -0.048 | -0.048 | 0.009 | 0.11 | 0.26 | 0.06 | 0.03 | 0.08 | 0.03 |
| 5 | 73135485 | rs10040601 | 1.518 | -2.916 | 0.878 | -0.296 | -0.027 | -0.022 | -0.022 | 0 | 0.38 | 0.31 | 0.43 | 0.55 | 0.45 | 0.39 |
| 5 | 73141894 | rs2973572 | -1.411 | -2.895 | 0.786 | -0.344 | 0.029 | 0.031 | 0.031 | 0 | 0.5 | 0.61 | 0.41 | 0.37 | 0.44 | 0.42 |
| 5 | 73142296 | rs2973571 | 0.325 | -2.898 | 0.88 | -0.387 | -0.015 | 0.055 | 0.055 | 0 | 0.23 | 0.12 | 0.25 | 0.08 | 0.26 | 0.2 |
| 5 | 73142380 | rs13161779 | 0.758 | -2.948 | 1.08 | -0.372 | -0.012 | 0.023 | 0.023 | 0 | 0.13 | 0.14 | 0.07 | 0.24 | 0.08 | 0.18 |
| 5 | 73144010 | rs6897754 | 0.68 | -3.118 | 0.451 | -0.344 | -0.033 | -0.014 | -0.014 | 0.073 | 0.15 | 0.22 | 0.16 | 0.11 | 0.08 | 0.19 |
| 5 | 73144845 | rs2973568 | -2.499 | -3.144 | 0.317 | -0.384 | -0.181 | -0.183 | -0.183 | 2.338 | 0.66 | 0.95 | 0.59 | 0.51 | 0.56 | 0.65 |
| 5 | 73146026 | rs2973567 | -1.952 | -3.145 | 0.358 | -0.383 | -0.043 | -0.073 | -0.073 | 0 | 0.6 | 0.84 | 0.54 | 0.49 | 0.48 | 0.58 |
| 5 | 73148294 | rs9637820 | 0.521 | -2.956 | 0.458 | -0.428 | -0.038 | 0.002 | 0.002 | 0 | 0.15 | 0.26 | 0.16 | 0.17 | 0.08 | 0.19 |
| 5 | 73148481 | rs2973566 | 0.391 | -2.812 | 0.476 | -0.456 | -0.024 | 0.063 | 0.063 | 0 | 0.23 | 0.09 | 0.25 | 0.08 | 0.27 | 0.2 |
| 5 | 73148810 | rs16870974 | -0.206 | -2.802 | 0.493 | -0.449 | -0.016 | -0.009 | -0.009 | 0 | 0.2 | 0.34 | 0.22 | 0.2 | 0.15 | 0.26 |
| 5 | 73151033 | rs2973564 | 0.29 | -2.516 | 0.145 | -0.568 | -0.021 | 0.056 | 0.056 | 0 | 0.23 | 0.09 | 0.25 | 0.08 | 0.26 | 0.2 |
| 5 | 73153754 | rs2973563 | -2.032 | -2.77 | 0.024 | -0.372 | -0.036 | -0.024 | -0.024 | 0 | 0.62 | 0.77 | 0.55 | 0.44 | 0.54 | 0.51 |
| 5 | 73159098 | rs2931434 | 0.341 | -2.805 | 0.048 | -0.388 | -0.035 | 0.068 | 0.068 | 0 | 0.24 | 0.11 | 0.35 | 0.08 | 0.3 | 0.19 |
| 5 | 73159601 | rs6869486 | -1.124 | -3.024 | 0.11 | -0.47 | -0.003 | 0 | 0 | 0 | 0.37 | 0.6 | 0.2 | 0.37 | 0.22 | 0.34 |
| 5 | 73163831 | rs2931423 | 0.28 | -2.963 | 0.149 | -0.41 | -0.016 | 0.037 | 0.037 | 0 | 0.26 | 0.18 | 0.36 | 0.14 | 0.3 | 0.19 |
| 5 | 73164206 | rs2973557 | -2.066 | -2.632 | -0.039 | -0.271 | -0.034 | -0.041 | -0.041 | 0 | 0.66 | 0.77 | 0.59 | 0.51 | 0.58 | 0.53 |
| 5 | 73165380 | rs2973556 | -0.885 | -2.631 | 0.166 | -0.194 | 0.007 | -0.006 | -0.006 | 0 | 0.44 | 0.62 | 0.47 | 0.49 | 0.39 | 0.42 |
| 5 | 73167334 | rs1352200 | -0.461 | -1.845 | 0.133 | -0.228 | 0.006 | -0.014 | -0.014 | 0 | 0.19 | 0.16 | 0.1 | 0 | 0.17 | 0.06 |
| 5 | 73179672 | rs3749645 | -0.235 | -1.336 | 0.882 | 0.54 | -0.019 | 0.03 | 0.03 | 0 | 0.17 | 0.17 | 0.1 | 0.26 | 0.11 | 0.19 |
| 5 | 73185504 | rs729272 | -1.562 | -1.428 | 0.788 | 0.491 | -0.02 | 0.008 | 0.008 | 0 | 0.31 | 0.59 | 0.21 | 0.36 | 0.2 | 0.26 |
| 5 | 73187660 | rs6453031 | -0.977 | -1.245 | 0.365 | 0.55 | -0.054 | 0.045 | 0.045 | 0 | 0.24 | 0.38 | 0.15 | 0.29 | 0.14 | 0.26 |
| 5 | 73196609 | rs3828583 | -1.063 | -0.642 | -0.468 | 0.511 | -0.002 | 0 | 0 | 0 | 0.38 | 0.63 | 0.46 | 0.38 | 0.39 | 0.47 |
| 5 | 73196810 | rs3749640 | 0.097 | -0.58 | -0.682 | 0.243 | -0.056 | 0.054 | 0.054 | 0 | 0.19 | 0.06 | 0.32 | 0.08 | 0.28 | 0.19 |
| 5 | 73198364 | rs283622 | -1.032 | -0.487 | -0.67 | 0.495 | -0.008 | -0.003 | -0.003 | 3.439 | 0.41 | 0.64 | 0.49 | 0.44 | 0.45 | 0.48 |
| 5 | 73201991 | rs2434215 | -1.128 | -0.671 | -0.94 | 0.698 | 0.024 | -0.026 | -0.026 | 0 | 0.3 | 0.77 | 0.22 | 0.39 | 0.21 | 0.46 |
| 5 | 73202480 | rs2931425 | -0.878 | -0.958 | -0.948 | -0.04 | 0.002 | -0.021 | -0.021 | 0.05 | 0.41 | 0.41 | 0.38 | 0.24 | 0.45 | 0.37 |
| 5 | 73203410 | rs283619 | -0.503 | -0.579 | -1.497 | -0.132 | -0.002 | -0.026 | -0.026 | 0 | 0.32 | 0.36 | 0.34 | 0.19 | 0.35 | 0.24 |
| 5 | 73207372 | rs1478453 | -0.275 | 0.054 | -1.523 | 0.2 | 0.013 | -0.019 | -0.019 | 0.091 | 0.29 | 0.38 | 0.22 | 0.22 | 0.42 | 0.22 |
| 5 | 73209251 | rs2931426 | -0.54 | 0.379 | -2.559 | 0.793 | -0.108 | -0.045 | -0.045 | 0 | 0.54 | 0.95 | 0.5 | 0.65 | 0.56 | 0.57 |
| 5 | 73211316 | rs411585 | -0.77 | 1.423 | -1.728 | 1.304 | -0.058 | -0.113 | -0.113 | 0 | 0.2 | 0.06 | 0.5 | 0.54 | 0.29 | 0.2 |
| 5 | 73214781 | rs434008 | -0.899 | 1.581 | -1.276 | 0.716 | -0.011 | 0.009 | 0.009 | 0.616 | 0.07 | 0.05 | 0.18 | 0.07 | 0.11 | 0.22 |
| 5 | 73215282 | rs17634951 | -0.179 | 2.029 | -1.701 | 0.405 | -0.056 | 0.041 | 0.041 | 0.494 | 0.15 | 0.01 | 0.09 | 0.04 | 0.17 | 0.3 |
| 5 | 73215462 | rs403031 | -1.219 | 1.443 | -1.162 | 0.746 | -0.01 | 0.011 | 0.011 | 1.069 | 0.08 | 0.07 | 0.18 | 0.07 | 0.11 | 0.22 |
| 5 | 73218861 | rs16871035 | -0.983 | 1.612 | -0.207 | 0.776 | 0.066 | 0.078 | 0.078 | 0.624 | 0.08 | 0.43 | 0.16 | 0.06 | 0.05 | 0.11 |
| 5 | 73218954 | rs423333 | -1.461 | 1.502 | -0.176 | 0.743 | 0.028 | -0.026 | -0.026 | 0 | 0.17 | 0.56 | 0.36 | 0.13 | 0.17 | 0.34 |
| 5 | 73221801 | rs366733 | 0.691 | 1.264 | -0.613 | 1.12 | -0.006 | 0.009 | 0.009 | 0 | 0.62 | 0.45 | 0.53 | 0.55 | 0.58 | 0.54 |
| 5 | 73222133 | rs448000 | -1.033 | 1.377 | -0.431 | 1.806 | 0.001 | -0.015 | -0.015 | 0.083 | 0.23 | 0.2 | 0.45 | 0.45 | 0.22 | 0.27 |
| 5 | 73224678 | rs2931428 | 0.787 | 1.873 | 0.454 | 1.301 | 0.018 | -0.053 | -0.053 | 0.591 | 0.9 | 0.76 | 0.93 | 0.98 | 0.91 | 0.91 |
| 5 | 73224689 | rs2973534 | 0.787 | 1.873 | 0.454 | 1.301 | 0.018 | -0.053 | -0.053 | 0 | 0.9 | 0.76 | 0.93 | 0.98 | 0.91 | 0.91 |
| 5 | 73225890 | rs283628 | -0.998 | 2.084 | 0.171 | 1.409 | 0.009 | -0.002 | -0.002 | 0 | 0.46 | 0.7 | 0.51 | 0.6 | 0.37 | 0.55 |
| 5 | 73229361 | rs12652554 | -1.119 | 2.124 | -0.042 | 1.678 | -0.006 | 0.013 | 0.013 | 0.633 | 0.1 | 0.05 | 0.29 | 0.45 | 0.14 | 0.23 |
| 5 | 73232058 | rs190742 | 0.814 | 2.49 | 0.334 | 1.197 | 0.027 | 0.001 | 0.001 | 0 | 0.85 | 0.8 | 0.86 | 0.83 | 0.77 | 0.82 |
| 5 | 73234583 | rs157557 | -0.865 | 2.016 | 0.603 | 1.611 | 0.007 | -0.036 | -0.036 | 0 | 0.36 | 0.23 | 0.38 | 0.42 | 0.32 | 0.42 |
| 5 | 73236235 | rs282413 | -1.392 | 2.159 | 0.805 | 1.546 | -0.01 | 0.009 | 0.009 | 0.005 | 0.43 | 0.31 | 0.42 | 0.58 | 0.41 | 0.5 |
| 5 | 73248512 | rs283610 | -1.436 | 2.057 | 1.032 | 1.518 | -0.003 | -0.002 | -0.002 | 0.076 | 0.43 | 0.37 | 0.42 | 0.41 | 0.41 | 0.49 |
| 5 | 73250492 | rs283612 | -1.063 | 2.092 | 0.959 | 1.659 | -0.013 | 0.052 | 0.052 | 0 | 0.37 | 0.22 | 0.34 | 0.14 | 0.36 | 0.31 |
| 5 | 73254540 | rs2450438 | -2.427 | 2.129 | 1.322 | 1.643 | 0.028 | 0.182 | 0.182 | 0 | 0.2 | 0.51 | 0.19 | 0.45 | 0.19 | 0.36 |
| 5 | 73258210 | rs17733544 | -1.461 | 2.103 | 1.121 | 1.706 | 0.012 | 0.051 | 0.051 | 0 | 0.14 | 0.2 | 0.15 | 0.29 | 0.16 | 0.31 |
| 5 | 73260230 | rs373729 | -1.715 | 2.112 | 1.455 | 1.649 | 0.003 | -0.001 | -0.001 | 0 | 0.46 | 0.53 | 0.44 | 0.42 | 0.43 | 0.53 |
| 5 | 73262113 | rs416286 | -0.44 | 2.078 | 1.355 | 1.529 | -0.043 | 0.06 | 0.06 | 0 | 0.29 | 0.06 | 0.28 | 0.14 | 0.26 | 0.19 |
| 5 | 73263957 | rs453036 | -1.425 | 2.079 | 1.411 | 1.49 | 0.001 | -0.041 | -0.041 | 0.093 | 0.4 | 0.45 | 0.35 | 0.14 | 0.37 | 0.32 |
| 5 | 73265522 | rs519689 | -1.581 | 2.038 | 1.42 | 1.659 | -0.001 | 0.176 | 0.176 | 0 | 0.09 | 0.21 | 0.12 | 0.44 | 0.07 | 0.24 |
| 5 | 73267367 | rs9986187 | -0.45 | 1.939 | 1.305 | 1.539 | 0.004 | -0.029 | -0.029 | 0 | 0.08 | 0.1 | 0.05 | 0.01 | 0.09 | 0.1 |
| 5 | 73268665 | rs414915 | -0.985 | 2.035 | 1.318 | 1.604 | -0.006 | -0.015 | -0.015 | 0 | 0.37 | 0.28 | 0.37 | 0.38 | 0.32 | 0.43 |
| 5 | 73272981 | rs3909800 | -0.292 | 2.161 | 1.124 | 1.624 | -0.027 | -0.049 | -0.049 | 0 | 0.17 | 0.08 | 0.14 | 0.3 | 0.16 | 0.3 |
| 5 | 73273696 | rs384834 | -2.137 | 2.128 | 1.397 | 1.458 | 0.001 | -0.004 | -0.004 | 0 | 0.45 | 0.53 | 0.42 | 0.39 | 0.41 | 0.52 |
| 5 | 73277373 | rs282381 | -2.48 | 2.031 | 1.257 | 1.211 | 0.004 | 0.06 | 0.06 | 0.018 | 0.55 | 0.7 | 0.72 | 0.88 | 0.5 | 0.67 |
| 5 | 73278287 | rs282384 | -2.246 | 2.067 | 1.269 | 1.209 | 0.007 | 0.025 | 0.025 | 0 | 0.53 | 0.62 | 0.69 | 0.88 | 0.49 | 0.65 |
| 5 | 73280358 | rs2973536 | -2.122 | 2.09 | 1.274 | 1.214 | 0.013 | 0.107 | 0.107 | 0.1 | 0.49 | 0.32 | 0.65 | 0.86 | 0.48 | 0.64 |

| | | | | | | | | | | | | | | | | |
|---|---|---|---|---|---|---|---|---|---|---|---|---|---|---|---|---|
| 5 | 73283692 | rs1904170 | -2.017 | 2.081 | 1.327 | 1.296 | 0.003 | -0.002 | -0.002 | 0.534 | 0.45 | 0.45 | 0.43 | 0.57 | 0.43 | 0.55 |
| 5 | 73287080 | rs10058557 | 1.351 | 2.138 | 1.164 | 1.335 | -0.021 | -0.011 | -0.011 | 0 | 0.59 | 0.75 | 0.61 | 0.44 | 0.6 | 0.5 |
| 5 | 73287411 | rs255592 | 0.743 | 2.178 | 1.152 | 1.336 | -0.008 | -0.015 | -0.015 | 0 | 0.74 | 0.79 | 0.82 | 0.53 | 0.73 | 0.59 |
| 5 | 73293204 | rs2434214 | -0.616 | 2.159 | 1.112 | 1.284 | -0.004 | 0.027 | 0.027 | 0 | 0.22 | 0.13 | 0.26 | 0.1 | 0.23 | 0.19 |
| 5 | 73293417 | rs13158300 | -0.118 | 2.186 | 1.091 | 1.385 | -0.029 | -0.046 | -0.046 | 0 | 0.18 | 0.09 | 0.13 | 0.28 | 0.18 | 0.3 |
| 5 | 73295327 | rs17666430 | -0.72 | 2.185 | 1.099 | 1.415 | 0 | 0.029 | 0.029 | 0.142 | 0.1 | 0.12 | 0.29 | 0.31 | 0.08 | 0.16 |
| 5 | 73307367 | rs1677458 | -2.321 | 2.122 | 0.902 | 1.193 | 0.006 | 0.051 | 0.051 | 0.189 | 0.55 | 0.69 | 0.71 | 0.7 | 0.49 | 0.67 |
| 5 | 73308343 | rs194274 | 0.319 | 1.818 | 0.605 | 1.192 | 0.004 | 0.024 | 0.024 | 0 | 0.16 | 0.04 | 0.21 | 0.09 | 0.13 | 0.1 |
| 5 | 73308764 | rs12153044 | -1.283 | 1.842 | 0.735 | 1.194 | 0.004 | 0.01 | 0.01 | 0.706 | 0.29 | 0.37 | 0.42 | 0.59 | 0.25 | 0.46 |
| 5 | 73308816 | rs11954800 | 1.388 | 1.584 | 0.532 | 1.199 | -0.002 | -0.001 | -0.001 | 0.594 | 0.58 | 0.63 | 0.38 | 0.15 | 0.63 | 0.46 |
| 5 | 73309105 | rs11952583 | 0.185 | 0.75 | 0.113 | 1.763 | -0.038 | 0.027 | 0.027 | 0 | 0.82 | 0.71 | 0.77 | 0.74 | 0.9 | 0.89 |
| 5 | 73309201 | rs255600 | 1.209 | 1.629 | -0.383 | 1.82 | 0 | 0.02 | 0.02 | 0 | 0.82 | 0.83 | 0.68 | 0.51 | 0.84 | 0.77 |
| 5 | 73312298 | rs193543 | 0.76 | 1.426 | -0.491 | 1.526 | -0.001 | 0 | 0 | 0.223 | 0.65 | 0.64 | 0.46 | 0.44 | 0.61 | 0.47 |
| 5 | 73313974 | rs10078294 | -0.397 | 1.473 | -0.828 | 1.833 | -0.099 | -0.074 | -0.074 | 0 | 0.24 | 0.05 | 0.24 | 0.32 | 0.3 | 0.47 |
| 5 | 73317388 | rs9293615 | -0.566 | 1.439 | -0.871 | 1.829 | -0.118 | -0.069 | -0.069 | 0 | 0.25 | 0.03 | 0.26 | 0.3 | 0.34 | 0.49 |
| 5 | 73326518 | rs376049 | 0.776 | 0.437 | -0.306 | 2.01 | -0.067 | 0.115 | 0.115 | 0.158 | 0.85 | 0.67 | 0.67 | 0.73 | 0.95 | 0.94 |
| 5 | 73330159 | rs410390 | -0.698 | 0.817 | -0.367 | 1.792 | 0.023 | 0.048 | 0.048 | 0.136 | 0.3 | 0.54 | 0.36 | 0.57 | 0.33 | 0.54 |
| 5 | 73330562 | rs3097137 | 0.097 | 0.258 | -1.254 | 0.702 | -0.042 | 0.006 | 0.006 | 0 | 0.34 | 0.25 | 0.27 | 0.3 | 0.39 | 0.35 |
| 5 | 73343946 | rs255585 | -0.83 | 0.309 | -0.344 | 1.272 | 0.003 | 0.001 | 0.001 | 0 | 0.36 | 0.67 | 0.5 | 0.63 | 0.35 | 0.56 |
| 5 | 73364042 | rs12514574 | -1.515 | 0.472 | 0.685 | 1.239 | 0.082 | 0.224 | 0.224 | 1.793 | 0.08 | 0.57 | 0.16 | 0.16 | 0.05 | 0.08 |
| 5 | 73364934 | rs1351143 | -1.75 | 0.206 | 0.7 | 0.961 | 0.051 | 0.125 | 0.125 | 3.289 | 0.08 | 0.53 | 0.13 | 0.1 | 0.07 | 0.08 |
| 5 | 73366695 | rs286645 | 0.757 | 0.317 | 0.519 | 1.177 | -0.002 | 0.012 | 0.012 | 0 | 0.69 | 0.36 | 0.63 | 0.47 | 0.68 | 0.52 |
| 5 | 73366839 | rs286644 | 0.113 | 0.175 | 0.833 | 1.145 | -0.007 | 0.04 | 0.04 | 0.049 | 0.81 | 0.75 | 0.75 | 0.61 | 0.82 | 0.55 |
| 5 | 73368238 | rs286641 | 0.311 | -0.268 | 0.477 | 0.295 | -0.026 | -0.066 | -0.066 | 0.017 | 0.29 | 0.45 | 0.32 | 0.16 | 0.25 | 0.13 |
| 5 | 73371678 | rs152044 | -0.315 | 0.15 | -0.581 | 0.033 | -0.051 | -0.038 | -0.038 | 0.077 | 0.69 | 0.89 | 0.69 | 0.59 | 0.61 | 0.47 |
| 5 | 73375454 | rs10942704 | 0.211 | 0.343 | -0.308 | 0.039 | -0.015 | -0.039 | -0.039 | 2.854 | 0.32 | 0.25 | 0.43 | 0.45 | 0.38 | 0.32 |
| 5 | 73376219 | rs1818123 | 0.105 | 0.324 | -0.405 | 0.001 | -0.044 | -0.057 | -0.057 | 0.786 | 0.33 | 0.19 | 0.43 | 0.45 | 0.41 | 0.37 |
| 5 | 73381607 | rs17636369 | 0.759 | 0.631 | -0.255 | 0.305 | -0.023 | -0.007 | -0.007 | 0 | 0.21 | 0.15 | 0.25 | 0.19 | 0.26 | 0.24 |
| 5 | 73383115 | rs17636389 | 1.054 | 0.244 | -0.556 | 0.048 | -0.011 | 0.049 | 0.049 | 0.827 | 0.12 | 0 | 0.12 | 0.01 | 0.08 | 0.15 |
| 5 | 73383170 | rs1501973 | -0.127 | -0.142 | 0.199 | -0.14 | 0.021 | -0.002 | -0.002 | 0 | 0.65 | 0.46 | 0.61 | 0.6 | 0.58 | 0.58 |
| 5 | 73389757 | rs1501975 | -0.066 | -0.164 | 0.323 | -0.145 | -0.023 | 0.025 | 0.025 | 0 | 0.34 | 0.19 | 0.36 | 0.21 | 0.34 | 0.39 |
| 5 | 73390020 | rs1160582 | -0.238 | -0.216 | 0.639 | -0.155 | 0.008 | -0.001 | -0.001 | 0 | 0.66 | 0.55 | 0.62 | 0.61 | 0.58 | 0.58 |
| 5 | 73390580 | rs12516798 | 1.697 | -0.261 | 0.688 | -0.145 | -0.004 | 0.001 | 0.001 | 0 | 0.31 | 0.25 | 0.24 | 0.4 | 0.24 | 0.19 |
| 5 | 73392369 | rs2840108 | 0.605 | -0.208 | 0.482 | -0.16 | -0.001 | -0.001 | -0.001 | 0 | 0.53 | 0.5 | 0.51 | 0.59 | 0.51 | 0.43 |
| 5 | 73393077 | rs17636626 | 1.199 | -0.221 | 0.335 | -0.167 | 0.008 | 0.041 | 0.041 | 0.584 | 0.13 | 0.04 | 0.11 | 0.01 | 0.08 | 0.15 |
| 5 | 73393578 | rs2120727 | -0.558 | -0.323 | 0.665 | -0.172 | -0.007 | -0.004 | -0.004 | 0 | 0.69 | 0.72 | 0.66 | 0.6 | 0.62 | 0.59 |
| 5 | 73394674 | rs12655370 | 1.706 | -0.469 | 0.435 | -0.133 | 0.05 | -0.057 | -0.057 | 0 | 0.3 | 0.11 | 0.23 | 0.39 | 0.24 | 0.19 |
| 5 | 73403215 | rs288855 | -0.984 | -0.382 | 0.516 | -0.101 | 0.102 | -0.019 | -0.019 | 0 | 0.58 | 0.84 | 0.67 | 0.59 | 0.72 | 0.71 |
| 5 | 73409309 | rs464313 | -0.962 | -0.384 | 0.551 | -0.102 | 0.099 | -0.013 | -0.013 | 0 | 0.57 | 0.83 | 0.67 | 0.59 | 0.73 | 0.71 |
| 5 | 73420143 | rs10072489 | 1.051 | -0.303 | 0.6 | -0.126 | -0.035 | -0.004 | -0.004 | 0.121 | 0.21 | 0.14 | 0.26 | 0.19 | 0.31 | 0.27 |
| 5 | 73420538 | rs208118 | -0.023 | -0.293 | 0.684 | -0.146 | 0.014 | -0.002 | -0.002 | 0.006 | 0.34 | 0.46 | 0.38 | 0.4 | 0.41 | 0.45 |
| 5 | 73422196 | rs287955 | -0.924 | -0.456 | 0.199 | 0.094 | 0.103 | 0.033 | 0.033 | 0 | 0.58 | 0.83 | 0.67 | 0.68 | 0.73 | 0.72 |
| 5 | 73424221 | rs158338 | -1.762 | -0.298 | 0.065 | -0.001 | 0.044 | -0.072 | -0.072 | 0.02 | 0.69 | 0.91 | 0.77 | 0.62 | 0.75 | 0.8 |
| 5 | 73425233 | rs167024 | -0.566 | -0.192 | 0.38 | -0.068 | 0.01 | 0.036 | 0.036 | 0 | 0.47 | 0.68 | 0.44 | 0.43 | 0.44 | 0.51 |
| 5 | 73428043 | rs288864 | -0.139 | -0.098 | 0.319 | -0.257 | 0.001 | 0.004 | 0.004 | 0 | 0.37 | 0.33 | 0.38 | 0.4 | 0.53 | 0.48 |
| 5 | 73428190 | rs288865 | -2.107 | -0.203 | -0.186 | -0.279 | 0.044 | -0.084 | -0.084 | 0 | 0.7 | 0.93 | 0.77 | 0.62 | 0.75 | 0.81 |
| 5 | 73428789 | rs288866 | 0.038 | 0.227 | 0.014 | -0.196 | -0.022 | 0.004 | 0.004 | 0 | 0.35 | 0.25 | 0.35 | 0.4 | 0.49 | 0.47 |
| 5 | 73430161 | rs4703622 | 0.165 | -0.427 | -0.462 | -0.914 | 0.037 | -0.021 | -0.021 | 3.777 | 0.31 | 0.08 | 0.33 | 0.22 | 0.28 | 0.33 |
| 5 | 73432001 | rs12517914 | 0.005 | -0.49 | 0.458 | -0.712 | -0.03 | -0.004 | -0.004 | 0 | 0.29 | 0.37 | 0.36 | 0.21 | 0.21 | 0.26 |
| 5 | 73433727 | rs12187319 | 0.308 | 0.386 | 0.438 | -0.854 | 0.076 | -0.075 | -0.075 | 0 | 0.13 | 0.26 | 0.1 | 0.01 | 0.19 | 0.04 |
| 5 | 73434500 | rs158580 | -2.046 | 0.321 | 0.14 | -0.671 | 0.027 | 0.002 | 0.002 | 0 | 0.69 | 0.9 | 0.81 | 0.71 | 0.73 | 0.74 |
| 5 | 73435250 | rs288869 | -2.026 | 0.318 | 0.138 | -0.673 | 0.028 | 0.003 | 0.003 | 0 | 0.69 | 0.9 | 0.81 | 0.71 | 0.73 | 0.74 |
| 5 | 73437503 | rs4704110 | 1.192 | 0.366 | 0.44 | -0.694 | -0.014 | 0.03 | 0.03 | 0 | 0.17 | 0.31 | 0.32 | 0.2 | 0.18 | 0.21 |
| 5 | 73442438 | rs2120729 | 0.258 | 0.34 | -0.174 | -0.779 | -0.017 | -0.038 | -0.038 | 8.507 | 0.27 | 0.19 | 0.33 | 0.49 | 0.33 | 0.45 |
| 5 | 73445424 | rs2937773 | 0.748 | 0.392 | -0.228 | -0.869 | -0.023 | -0.07 | -0.07 | 0 | 0.24 | 0.12 | 0.3 | 0.51 | 0.3 | 0.44 |
| 5 | 73445817 | rs191899 | -0.639 | 0.303 | -0.233 | -0.975 | 0.106 | 0.131 | 0.131 | 0 | 0.49 | 0.2 | 0.56 | 0.7 | 0.65 | 0.66 |
| 5 | 73446042 | rs290716 | -0.696 | 0.273 | -0.179 | -0.971 | 0.093 | 0.089 | 0.089 | 0 | 0.52 | 0.28 | 0.6 | 0.7 | 0.68 | 0.68 |
| 5 | 73447378 | rs10515174 | 1.585 | 0.231 | -0.287 | -0.977 | -0.027 | -0.031 | -0.031 | 0.091 | 0.12 | 0.16 | 0.06 | 0.02 | 0.04 | 0.07 |
| 5 | 73452427 | rs12188320 | 0.052 | 0.413 | -0.456 | -0.933 | -0.076 | -0.002 | -0.002 | 0 | 0.25 | 0.08 | 0.26 | 0.19 | 0.36 | 0.23 |
| 5 | 73452467 | rs17735123 | 1.517 | 0.352 | -0.435 | -0.9 | 0.007 | -0.029 | -0.029 | 2.184 | 0.16 | 0.06 | 0.18 | 0.19 | 0.17 | 0.21 |
| 5 | 73455955 | rs6891167 | -0.843 | 0.415 | -0.218 | -0.936 | 0.052 | -0.024 | -0.024 | 0 | 0.09 | 0.43 | 0.07 | 0 | 0.05 | 0.04 |
| 5 | 73459885 | rs290726 | -1.837 | 0.312 | -0.889 | -0.805 | 0.036 | 0.035 | 0.035 | 0 | 0.68 | 0.89 | 0.71 | 0.74 | 0.72 | 0.75 |
| 5 | 73460035 | rs975949 | 1.355 | 1.199 | -0.77 | -0.453 | -0.011 | 0.007 | 0.007 | 0 | 0.08 | 0.04 | 0.07 | 0.01 | 0.16 | 0.02 |
| 5 | 73460615 | rs2313877 | -0.34 | 1.155 | -0.785 | -0.456 | -0.069 | -0.066 | -0.066 | 2.808 | 0.32 | 0.15 | 0.37 | 0.53 | 0.46 | 0.46 |
| 5 | 73463987 | rs288837 | 0.408 | 0.641 | -0.663 | -0.203 | -0.031 | -0.088 | -0.088 | 0.077 | 0.24 | 0.1 | 0.3 | 0.51 | 0.29 | 0.44 |
| 5 | 73470103 | rs184084 | 0.359 | 0.654 | -0.677 | -0.248 | -0.042 | -0.1 | -0.1 | 0 | 0.26 | 0.11 | 0.31 | 0.5 | 0.31 | 0.39 |
| 5 | 73470740 | rs1444565 | -0.566 | 0.863 | -0.569 | -0.301 | 0.021 | 0.032 | 0.032 | 0 | 0.34 | 0.57 | 0.4 | 0.53 | 0.38 | 0.42 |
| 5 | 73470877 | rs288849 | 0.089 | 0.854 | -0.796 | -0.196 | 0.012 | 0.072 | 0.072 | 0 | 0.29 | 0.43 | 0.35 | 0.52 | 0.31 | 0.39 |
| 5 | 73471677 | rs10071742 | 0.631 | 1.005 | -1.725 | -0.079 | -0.063 | -0.026 | -0.026 | 0 | 0.25 | 0.07 | 0.25 | 0.21 | 0.32 | 0.25 |

| | | | | | | | | | | | | | | | | |
|---|---|---|---|---|---|---|---|---|---|---|---|---|---|---|---|---|
| 5 | 73478536 | rs1444562 | -0.447 | 0.518 | -1.407 | 0.252 | 0.008 | -0.015 | -0.015 | 0 | 0.72 | 0.95 | 0.72 | 0.73 | 0.74 | 0.73 |
| 5 | 73478693 | rs288846 | 0.121 | 1.214 | -1.501 | 0.407 | -0.067 | -0.113 | -0.113 | 0 | 0.18 | 0.04 | 0.34 | 0.54 | 0.27 | 0.36 |
| 5 | 73483215 | rs4704111 | 0.414 | 0.832 | -1.031 | 0.132 | -0.01 | 0.101 | 0.101 | 0 | 0.56 | 0.77 | 0.39 | 0.22 | 0.51 | 0.37 |
| 5 | 73486650 | rs158805 | -0.616 | 1.151 | -1.02 | 0.134 | 0.014 | -0.001 | -0.001 | 0.049 | 0.73 | 0.84 | 0.72 | 0.73 | 0.74 | 0.72 |
| 5 | 73489192 | rs13169946 | 0.979 | 0.923 | -0.333 | -0.031 | 0.001 | 0.005 | 0.005 | 0 | 0.47 | 0.45 | 0.32 | 0.21 | 0.44 | 0.32 |
| 5 | 73490274 | rs158812 | -0.645 | 1.053 | -0.268 | 0.042 | 0.023 | 0.053 | 0.053 | 0.502 | 0.26 | 0.38 | 0.41 | 0.5 | 0.3 | 0.4 |
| 5 | 73490426 | rs158466 | 0.472 | 0.855 | -0.611 | 0.121 | -0.059 | -0.115 | -0.115 | 0 | 0.17 | 0.03 | 0.34 | 0.5 | 0.23 | 0.35 |
| 5 | 73490564 | rs1444560 | 0.901 | 0.81 | -0.572 | -0.224 | -0.003 | 0.03 | 0.03 | 0 | 0.42 | 0.35 | 0.3 | 0.2 | 0.42 | 0.23 |
| 5 | 73497997 | rs158815 | 0.026 | 0.923 | -0.805 | -0.183 | 0.018 | 0.089 | 0.089 | 0.015 | 0.25 | 0.41 | 0.4 | 0.51 | 0.27 | 0.36 |
| 5 | 73503184 | rs10061061 | 0.343 | 0.864 | -1.323 | -0.244 | -0.01 | 0.057 | 0.057 | 0 | 0.42 | 0.29 | 0.29 | 0.2 | 0.43 | 0.22 |
| 5 | 73507192 | rs167111 | 0.236 | 0.923 | -1.465 | 0.49 | -0.015 | -0.023 | -0.023 | 0 | 0.68 | 0.74 | 0.66 | 0.57 | 0.6 | 0.64 |
| 5 | 73509349 | rs12519034 | 0.237 | 0.537 | -1.694 | 0.356 | -0.027 | -0.137 | -0.137 | 0 | 0.26 | 0.05 | 0.4 | 0.54 | 0.25 | 0.39 |
| 5 | 73509753 | rs209770 | 0.456 | 0.974 | -1.743 | -0.306 | -0.072 | 0.088 | 0.088 | 3.117 | 0.18 | 0.05 | 0.14 | 0 | 0.27 | 0.04 |
| 5 | 73510652 | rs2004500 | 0.264 | 0.855 | -1.358 | -0.274 | 0.006 | 0.021 | 0.021 | 1.803 | 0.51 | 0.42 | 0.62 | 0.77 | 0.47 | 0.68 |
| 5 | 73511687 | rs680500 | -0.05 | 0.848 | -1.381 | 0.641 | -0.001 | 0 | 0 | 4.878 | 0.72 | 0.77 | 0.78 | 0.79 | 0.72 | 0.72 |
| 5 | 73516925 | rs7728448 | -0.47 | 0.8 | -1.644 | 1.164 | -0.018 | -0.102 | -0.102 | 2.679 | 0.25 | 0.07 | 0.31 | 0.53 | 0.25 | 0.39 |
| 5 | 73519510 | rs16871495 | -1.45 | 0.46 | -0.98 | 1.055 | -0.011 | 0.013 | 0.013 | 0 | 0.09 | 0.22 | 0.05 | 0.1 | 0.07 | 0.17 |
| 5 | 73522355 | rs6866246 | -1.213 | -0.604 | -0.789 | 1.593 | 0.001 | -0.002 | -0.002 | 3.02 | 0.27 | 0.74 | 0.46 | 0.43 | 0.21 | 0.39 |
| 5 | 73523745 | rs1460811 | -1.427 | -0.751 | 0.517 | 1.599 | -0.002 | 0.031 | 0.031 | 0.47 | 0.11 | 0.47 | 0.08 | 0.09 | 0.07 | 0.14 |
| 5 | 73527328 | rs4479825 | -0.154 | -0.785 | 0.982 | 1.582 | -0.012 | 0.048 | 0.048 | 0 | 0.14 | 0.16 | 0.35 | 0.35 | 0.14 | 0.23 |
| 5 | 73527824 | rs10515165 | -1.227 | -0.11 | -0.049 | 0.848 | -0.039 | 0.021 | 0.021 | 0 | 0.51 | 0.87 | 0.64 | 0.54 | 0.52 | 0.59 |
| 5 | 73528671 | rs16871513 | -0.463 | -0.031 | 0.193 | 0.851 | 0.017 | 0.04 | 0.04 | 0.031 | 0.1 | 0.4 | 0.08 | 0.08 | 0.07 | 0.14 |
| 5 | 73535978 | rs290708 | -0.496 | -0.093 | -0.615 | 0.059 | 0.038 | -0.005 | -0.005 | 1.563 | 0.46 | 0.71 | 0.59 | 0.5 | 0.42 | 0.55 |
| 5 | 73536750 | rs1824495 | 1.206 | -0.04 | -0.674 | -0.013 | -0.055 | 0.053 | 0.053 | 0.585 | 0.25 | 0.08 | 0.19 | 0.1 | 0.32 | 0.21 |
| 5 | 73539736 | rs627834 | 1.508 | -0.075 | -0.846 | -0.153 | -0.006 | 0.086 | 0.086 | 0.54 | 0.14 | 0.02 | 0.06 | 0 | 0.14 | 0.09 |
| 5 | 73542940 | rs17735564 | 0.065 | -0.079 | -0.368 | -0.128 | 0.023 | 0.028 | 0.028 | 0 | 0.14 | 0.24 | 0.13 | 0.16 | 0.15 | 0.21 |
| 5 | 73543591 | rs292213 | -1.198 | -0.103 | -0.508 | -0.177 | -0.021 | -0.038 | -0.038 | 0.062 | 0.72 | 0.87 | 0.79 | 0.61 | 0.74 | 0.76 |
| 5 | 73544647 | rs292215 | -0.437 | -0.131 | -0.255 | 0.099 | -0.01 | 0.042 | 0.042 | 0.267 | 0.31 | 0.38 | 0.49 | 0.48 | 0.3 | 0.5 |
| 5 | 73547069 | rs654937 | 0.217 | 0.806 | -0.448 | 0.204 | 0.016 | 0.029 | 0.029 | 0.783 | 0.23 | 0.29 | 0.48 | 0.39 | 0.27 | 0.38 |
| 5 | 73547476 | rs10069262 | 0.1 | 1.031 | -0.696 | -0.144 | -0.001 | -0.01 | -0.01 | 0 | 0.45 | 0.49 | 0.32 | 0.24 | 0.43 | 0.29 |
| 5 | 73548044 | rs7723844 | 0.437 | 0.952 | -1.141 | -0.188 | -0.006 | 0 | 0 | 0 | 0.13 | 0.2 | 0.1 | 0.1 | 0.09 | 0.15 |
| 5 | 73548271 | rs2962622 | -0.936 | 0.916 | -1.784 | -0.087 | -0.024 | -0.136 | -0.136 | 0 | 0.7 | 0.96 | 0.83 | 0.55 | 0.7 | 0.68 |
| 5 | 73548364 | rs10074415 | 0.909 | 0.944 | -1.516 | -0.093 | -0.012 | 0.084 | 0.084 | 2.974 | 0.3 | 0.12 | 0.21 | 0.13 | 0.33 | 0.14 |
| 5 | 73554344 | rs292209 | 0.107 | 0.884 | -0.868 | 0.279 | -0.002 | 0.012 | 0.012 | 0 | 0.29 | 0.37 | 0.35 | 0.31 | 0.27 | 0.38 |
| 5 | 73554618 | rs16871563 | 0.099 | 0.811 | -0.861 | 0.089 | 0.031 | 0.015 | 0.015 | 0.031 | 0.15 | 0.37 | 0.15 | 0.13 | 0.13 | 0.17 |
| 5 | 73555448 | rs176764 | -0.592 | 0.779 | -2.032 | -0.002 | -0.005 | -0.003 | -0.003 | 1.293 | 0.44 | 0.72 | 0.5 | 0.38 | 0.38 | 0.55 |
| 5 | 73557669 | rs1460806 | -0.236 | 0.816 | -1.718 | -0.534 | -0.016 | 0.067 | 0.067 | 3.452 | 0.42 | 0.32 | 0.36 | 0.16 | 0.54 | 0.31 |
| 5 | 73559582 | rs11738478 | -0.118 | -0.082 | -2.328 | 0.017 | 0.033 | -0.124 | -0.124 | 0 | 0.73 | 0.98 | 0.69 | 0.34 | 0.78 | 0.56 |
| 5 | 73561298 | rs6887081 | -0.349 | 0.596 | -1.873 | -0.722 | -0.001 | 0 | 0 | 0 | 0.33 | 0.34 | 0.43 | 0.21 | 0.5 | 0.27 |
| 5 | 73562278 | rs1460812 | -0.529 | 0.343 | -2.066 | -0.925 | 0.005 | 0.044 | 0.044 | 5.77 | 0.41 | 0.31 | 0.47 | 0.24 | 0.56 | 0.42 |
| 5 | 73564549 | rs2126039 | -1.167 | 0.016 | -2.37 | -0.815 | 0.047 | -0.093 | -0.093 | 0 | 0.76 | 0.96 | 0.71 | 0.33 | 0.82 | 0.54 |
| 5 | 73565675 | rs17643294 | 1.618 | -0.124 | -2.655 | -1.195 | -0.071 | 0.038 | 0.038 | 2.6 | 0.27 | 0.01 | 0.17 | 0.1 | 0.27 | 0.12 |
| 5 | 73566814 | rs505085 | -0.847 | 1.135 | -1.133 | -1.353 | 0.001 | -0.084 | -0.084 | 0 | 0.22 | 0.64 | 0.19 | 0.05 | 0.18 | 0.18 |
| 5 | 73569263 | rs569120 | -0.608 | 0.851 | -0.758 | -1.265 | -0.027 | 0.03 | 0.03 | 3.369 | 0.35 | 0.25 | 0.42 | 0.23 | 0.52 | 0.28 |
| 5 | 73575358 | rs611708 | -0.557 | 1.185 | -0.594 | -1.149 | -0.011 | 0.006 | 0.006 | 3.084 | 0.3 | 0.28 | 0.4 | 0.23 | 0.47 | 0.22 |
| 5 | 73576479 | rs1870658 | 0.54 | 0.802 | -0.254 | -0.874 | -0.001 | -0.056 | -0.056 | 0 | 0.11 | 0.2 | 0.07 | 0.01 | 0.05 | 0.16 |
| 5 | 73578406 | rs624943 | 0.537 | 0.817 | -0.349 | -0.867 | -0.001 | -0.057 | -0.057 | 0 | 0.11 | 0.18 | 0.07 | 0.01 | 0.05 | 0.15 |
| 5 | 73579258 | rs630098 | -0.981 | 1.116 | -0.404 | -0.992 | 0.006 | -0.083 | -0.083 | 0 | 0.21 | 0.3 | 0.16 | 0 | 0.22 | 0.08 |
| 5 | 73579842 | rs6889925 | -0.935 | 0.659 | -1.168 | -0.259 | 0.017 | -0.006 | -0.006 | 0 | 0.44 | 0.8 | 0.43 | 0.35 | 0.35 | 0.39 |
| 5 | 73580484 | rs10515167 | -0.522 | 0.239 | -0.301 | 0.068 | 0.015 | 0.027 | 0.027 | 0 | 0.27 | 0.4 | 0.31 | 0.29 | 0.27 | 0.27 |
| 5 | 73581167 | rs12518802 | 0.209 | 0.26 | -0.258 | 0.094 | -0.008 | -0.01 | -0.01 | 0 | 0.19 | 0.17 | 0.29 | 0.29 | 0.24 | 0.24 |
| 5 | 73582315 | rs292202 | -0.181 | 0.063 | -0.061 | 0.243 | -0.015 | 0.023 | 0.023 | 0.68 | 0.31 | 0.2 | 0.3 | 0.24 | 0.35 | 0.46 |
| 5 | 73589129 | rs922202 | 1.879 | 0.143 | -0.32 | 0.34 | -0.032 | 0.002 | 0.002 | 0.613 | 0.45 | 0.18 | 0.46 | 0.6 | 0.49 | 0.35 |
| 5 | 73591094 | rs292201 | -0.938 | 0.114 | 0.08 | 0.441 | -0.007 | 0.011 | 0.011 | 0 | 0.81 | 0.48 | 0.77 | 0.85 | 0.87 | 0.85 |
| 5 | 73593736 | rs9293622 | 0.689 | 0.004 | -0.392 | 0.496 | -0.012 | 0.035 | 0.035 | 0.064 | 0.5 | 0.2 | 0.47 | 0.6 | 0.51 | 0.39 |
| 5 | 73596750 | rs1114755 | -0.42 | 0.236 | -0.148 | 0.705 | -0.031 | -0.032 | -0.032 | 0 | 0.76 | 0.8 | 0.7 | 0.65 | 0.64 | 0.54 |
| 5 | 73597979 | rs16871632 | 2.287 | 0.042 | 0.601 | 0.685 | -0.011 | 0.026 | 0.026 | 1.04 | 0.08 | 0.17 | 0.14 | 0.06 | 0.05 | 0.02 |
| 5 | 73598221 | rs9293623 | -0.447 | 0.042 | 0.772 | 0.684 | 0.002 | 0.002 | 0.002 | 8.497 | 0.75 | 0.7 | 0.7 | 0.65 | 0.64 | 0.54 |
| 5 | 73599644 | rs4704126 | 1.159 | -0.316 | 0.616 | 0.487 | -0.026 | -0.016 | -0.016 | 6.86 | 0.48 | 0.19 | 0.46 | 0.51 | 0.52 | 0.39 |
| 5 | 73618819 | rs12521529 | 0.348 | -0.207 | 0.889 | 0.536 | 0.001 | 0.002 | 0.002 | 0.853 | 0.65 | 0.39 | 0.54 | 0.6 | 0.58 | 0.51 |
| 5 | 73626450 | rs898601 | 0.523 | 0.141 | 0.635 | 0.538 | 0.006 | 0.012 | 0.012 | 0 | 0.14 | 0.11 | 0.08 | 0.09 | 0.08 | 0.13 |
| 5 | 73630325 | rs6859957 | -0.392 | 0.15 | 0.617 | 0.613 | -0.016 | -0.016 | -0.016 | 0 | 0.76 | 0.75 | 0.7 | 0.65 | 0.63 | 0.54 |
| 5 | 73633836 | rs4704134 | 0.63 | -0.028 | 0.873 | 0.548 | 0.005 | 0.002 | 0.002 | 6.559 | 0.6 | 0.39 | 0.61 | 0.58 | 0.57 | 0.42 |
| 5 | 73636857 | rs11956917 | -0.563 | -0.446 | 0.573 | 0.53 | 0.013 | 0.002 | 0.002 | 0 | 0.25 | 0.18 | 0.38 | 0.23 | 0.17 | 0.27 |
| 5 | 73653987 | rs376518 | -0.295 | -0.435 | 0.564 | 0.549 | 0.003 | -0.006 | -0.006 | 0.54 | 0.86 | 0.88 | 0.69 | 0.77 | 0.9 | 0.82 |
| 5 | 73656720 | rs1873363 | 1.127 | -0.452 | 0.482 | 0.522 | 0.008 | 0.024 | 0.024 | 0 | 0.11 | 0.05 | 0.07 | 0.01 | 0.07 | 0.09 |
| 5 | 73657760 | rs409773 | 0.357 | -0.432 | 0.882 | 0.543 | -0.012 | -0.005 | -0.005 | 0 | 0.73 | 0.73 | 0.62 | 0.77 | 0.83 | 0.74 |
| 5 | 73670471 | rs10805886 | 1.121 | 0.098 | 0.634 | 0.634 | 0.006 | 0.016 | 0.016 | 2.312 | 0.46 | 0.3 | 0.32 | 0.34 | 0.45 | 0.25 |
| 5 | 73670507 | rs192679 | -0.034 | 0.304 | 0.649 | 0.234 | 0.032 | 0 | 0 | 9.36 | 0.27 | 0.45 | 0.31 | 0.25 | 0.36 | 0.48 |

| | | | | | | | | | | | | | | | | |
|---|---|---|---|---|---|---|---|---|---|---|---|---|---|---|---|---|
| 5 | 73671678 | rs17644127 | 0.62 | 0.645 | 0.169 | -0.025 | -0.048 | -0.015 | -0.015 | 0 | 0.15 | 0.05 | 0.13 | 0.14 | 0.23 | 0.28 |
| 5 | 73672540 | rs10805887 | 0.596 | 0.183 | 0.067 | -0.218 | -0.016 | -0.037 | -0.037 | 0 | 0.37 | 0.45 | 0.27 | 0.22 | 0.33 | 0.22 |
| 5 | 73674157 | rs10515178 | 0.585 | 0.191 | 0.02 | -0.215 | -0.008 | -0.019 | -0.019 | 0 | 0.37 | 0.49 | 0.27 | 0.22 | 0.33 | 0.22 |
| 5 | 73676124 | rs172438 | 0.708 | 0.308 | -0.159 | -0.177 | -0.002 | 0.002 | 0.002 | 0 | 0.18 | 0.21 | 0.29 | 0.22 | 0.17 | 0.27 |
| 5 | 73676439 | rs188404 | -0.318 | 0.556 | -0.246 | -0.193 | -0.02 | 0.018 | 0.018 | 1.288 | 0.39 | 0.35 | 0.47 | 0.37 | 0.45 | 0.56 |
| 5 | 73676565 | rs176930 | -0.657 | 0.577 | 0.1 | -0.064 | -0.011 | 0.064 | 0.064 | 0 | 0.74 | 0.7 | 0.72 | 0.53 | 0.79 | 0.77 |
| 5 | 73677709 | rs1687349 | 0.16 | 1.603 | 0.502 | 0.436 | 0.022 | -0.022 | -0.022 | 0 | 0.23 | 0.35 | 0.22 | 0.17 | 0.29 | 0.45 |
| 5 | 73678186 | rs1393085 | 1.459 | 1.588 | 0.337 | 0.451 | -0.001 | -0.041 | -0.041 | 0 | 0.08 | 0.02 | 0.2 | 0.21 | 0.07 | 0.1 |
| 5 | 73679925 | rs9293625 | 1.601 | 1.518 | 0.262 | 0.262 | 0.032 | 0.091 | 0.091 | 0.663 | 0.41 | 0.21 | 0.28 | 0.15 | 0.42 | 0.15 |
| 5 | 73680093 | rs412467 | -0.371 | 1.464 | 0.289 | 0.429 | -0.011 | -0.045 | -0.045 | 0.017 | 0.8 | 0.89 | 0.63 | 0.53 | 0.79 | 0.76 |
| 5 | 73683256 | rs17644264 | 0.919 | 1.364 | 0.423 | 0.223 | -0.001 | -0.012 | -0.012 | 0 | 0.53 | 0.49 | 0.34 | 0.16 | 0.49 | 0.22 |
| 5 | 73685103 | rs10454913 | 0.639 | 1.355 | 0.444 | 0.206 | 0.014 | -0.005 | -0.005 | 0 | 0.61 | 0.51 | 0.55 | 0.36 | 0.56 | 0.32 |
| 5 | 73693614 | rs295698 | -0.468 | 1.45 | 0.533 | 0.154 | -0.004 | 0.003 | 0.003 | 5.914 | 0.78 | 0.75 | 0.79 | 0.91 | 0.73 | 0.56 |
| 5 | 73694418 | rs10474415 | 0.148 | 1.423 | 0.537 | 0.153 | -0.01 | 0.007 | 0.007 | 0.505 | 0.68 | 0.69 | 0.57 | 0.71 | 0.66 | 0.45 |
| 5 | 73704926 | rs295692 | 1.792 | 1.392 | 0.201 | 0.04 | -0.002 | -0.024 | -0.024 | 0.039 | 0.1 | 0.2 | 0.09 | 0.03 | 0.08 | 0.17 |
| 5 | 73705802 | rs464581 | -1.172 | 1.504 | 0.238 | 0.055 | -0.005 | 0.002 | 0.002 | 1.819 | 0.87 | 0.89 | 0.87 | 0.93 | 0.81 | 0.74 |
| 5 | 73710311 | rs1501651 | 1.638 | 1.393 | 0.278 | 0.188 | 0.009 | -0.02 | -0.02 | 0 | 0.09 | 0.04 | 0.21 | 0.23 | 0.07 | 0.1 |
| 5 | 73710424 | rs413430 | 1.593 | 1.399 | 0.35 | 0.016 | -0.002 | -0.044 | -0.044 | 0 | 0.11 | 0.27 | 0.09 | 0.03 | 0.08 | 0.18 |
| 5 | 73712303 | rs583195 | 0.531 | 1.397 | 0.344 | 0.022 | -0.011 | -0.003 | -0.003 | 0 | 0.2 | 0.31 | 0.31 | 0.25 | 0.15 | 0.27 |
| 5 | 73713209 | rs435958 | -1.06 | 1.497 | 0.311 | -0.038 | -0.001 | 0.061 | 0.061 | 0 | 0.48 | 0.68 | 0.62 | 0.71 | 0.41 | 0.58 |
| 5 | 73715484 | rs10515181 | 1.641 | 1.29 | 0.268 | -0.049 | -0.028 | 0.053 | 0.053 | 0 | 0.1 | 0.01 | 0.05 | 0.01 | 0.08 | 0.08 |
| 5 | 73716118 | rs295688 | -0.132 | 1.288 | 0.395 | -0.041 | -0.012 | -0.036 | -0.036 | 0 | 0.31 | 0.46 | 0.38 | 0.25 | 0.24 | 0.35 |
| 5 | 73716608 | rs294985 | -0.196 | 1.161 | 0.104 | 0.061 | 0.003 | 0.091 | 0.091 | 0.014 | 0.27 | 0.48 | 0.31 | 0.49 | 0.24 | 0.39 |
| 5 | 73717645 | rs6863488 | 0.573 | 1.595 | -0.373 | 0.347 | -0.005 | 0.008 | 0.008 | 0 | 0.14 | 0.1 | 0.22 | 0.4 | 0.16 | 0.21 |
| 5 | 73719660 | rs460738 | 0.965 | 1.637 | -0.396 | 0.229 | 0.021 | 0.078 | 0.078 | 0.414 | 0.43 | 0.26 | 0.28 | 0.23 | 0.43 | 0.16 |
| 5 | 73720455 | rs463328 | -1.443 | 1.591 | -0.35 | 0.393 | -0.02 | 0.001 | 0.001 | 1.853 | 0.89 | 0.95 | 0.9 | 0.93 | 0.84 | 0.77 |
| 5 | 73721544 | rs417421 | 1.373 | 1.379 | -0.296 | 0.255 | -0.007 | -0.001 | -0.001 | 0 | 0.2 | 0.29 | 0.31 | 0.25 | 0.14 | 0.28 |
| 5 | 73722209 | rs294983 | 0.071 | 1.416 | -0.287 | 0.252 | -0.001 | -0.001 | -0.001 | 0 | 0.34 | 0.4 | 0.52 | 0.65 | 0.3 | 0.49 |
| 5 | 73735096 | rs2134152 | 0.73 | 2.533 | -0.008 | 1.016 | -0.033 | -0.048 | -0.048 | 0 | 0.1 | 0.04 | 0.09 | 0.3 | 0.14 | 0.19 |
| 5 | 73736889 | rs376759 | -0.35 | 2.669 | -0.092 | 0.536 | -0.023 | 0.045 | 0.045 | 0 | 0.26 | 0.14 | 0.17 | 0.02 | 0.31 | 0.15 |
| 5 | 73737608 | rs457664 | -0.709 | 2.864 | -0.004 | 0.638 | 0.028 | 0.026 | 0.026 | 1.824 | 0.89 | 0.84 | 0.88 | 0.87 | 0.83 | 0.77 |
| 5 | 73743061 | rs16871870 | 0.593 | 1.959 | 0.033 | 0.038 | 0.001 | -0.026 | -0.026 | 0 | 0.11 | 0.1 | 0.08 | 0 | 0.15 | 0.02 |
| 5 | 73744566 | rs445653 | -0.516 | 1.985 | -0.18 | 0.082 | -0.018 | -0.025 | -0.025 | 0.088 | 0.78 | 0.84 | 0.8 | 0.64 | 0.69 | 0.57 |
| 5 | 73745060 | rs294971 | 0.781 | 1.083 | -0.451 | 0.466 | 0.053 | 0.029 | 0.029 | 0.957 | 0.46 | 0.41 | 0.47 | 0.47 | 0.34 | 0.37 |
| 5 | 73746587 | rs296941 | 0.06 | 1.458 | -1.757 | 0.707 | -0.056 | 0.081 | 0.081 | 0 | 0.15 | 0.01 | 0.09 | 0.02 | 0.16 | 0.13 |
| 5 | 73747281 | rs294975 | 0.327 | 1.736 | -0.89 | 1.516 | 0.036 | 0.004 | 0.004 | 0.109 | 0.64 | 0.57 | 0.58 | 0.5 | 0.51 | 0.5 |
| 5 | 73762307 | rs2927625 | 0.583 | 1.535 | -1.565 | 0.855 | -0.046 | 0.014 | 0.014 | 0.801 | 0.82 | 0.94 | 0.75 | 0.86 | 0.75 | 0.82 |
| 5 | 73764923 | rs11948697 | -0.196 | 1.422 | -1.356 | 1.132 | 0 | 0.013 | 0.013 | 0.38 | 0.19 | 0.27 | 0.15 | 0.31 | 0.17 | 0.19 |
| 5 | 73771666 | rs6453061 | 1.299 | 1.446 | -2.042 | 1.641 | 0.013 | 0.075 | 0.075 | 0.027 | 0.51 | 0.66 | 0.35 | 0.24 | 0.43 | 0.38 |
| 5 | 73773153 | rs16871881 | 0.875 | 0.997 | -1.735 | 0.882 | -0.008 | 0.022 | 0.022 | 0.91 | 0.16 | 0.23 | 0.11 | 0.19 | 0.14 | 0.21 |
| 5 | 73775945 | rs2927630 | 0.409 | 1.111 | -1.573 | 0.837 | -0.007 | 0.006 | 0.006 | 0 | 0.6 | 0.62 | 0.53 | 0.3 | 0.49 | 0.44 |
| 5 | 73777378 | rs12153302 | 0.791 | 1.19 | -1.66 | 0.837 | -0.024 | -0.073 | -0.073 | 0 | 0.11 | 0.01 | 0.23 | 0.3 | 0.15 | 0.26 |
| 5 | 73778656 | rs2927632 | 1.15 | 1.123 | -1.034 | 0.355 | 0.003 | 0.018 | 0.018 | 0.066 | 0.33 | 0.22 | 0.23 | 0 | 0.28 | 0.15 |
| 5 | 73778774 | rs2962189 | -0.988 | 1.327 | -0.911 | 0.899 | -0.007 | -0.001 | -0.001 | 0 | 0.91 | 0.93 | 0.91 | 0.91 | 0.81 | 0.9 |
| 5 | 73780784 | rs2927633 | 0.155 | 0.749 | -0.813 | 0.632 | 0.011 | 0.002 | 0.002 | 0.411 | 0.5 | 0.56 | 0.51 | 0.55 | 0.45 | 0.66 |
| 5 | 73784037 | rs2115281 | 0.149 | 0.573 | -0.564 | 0.43 | -0.007 | -0.015 | -0.015 | 0 | 0.3 | 0.39 | 0.34 | 0.24 | 0.27 | 0.35 |
| 5 | 73786880 | rs6874566 | 0.106 | 0.561 | -0.708 | 0.425 | -0.016 | -0.032 | -0.032 | 0 | 0.31 | 0.48 | 0.36 | 0.25 | 0.27 | 0.35 |
| 5 | 73788266 | rs890995 | -0.059 | 0.949 | -0.486 | 0.586 | 0.018 | 0.028 | 0.028 | 1.563 | 0.64 | 0.58 | 0.72 | 0.52 | 0.59 | 0.61 |
| 5 | 73791092 | rs11741747 | -0.824 | 1.531 | 0.109 | 0.326 | 0.069 | -0.068 | -0.068 | 0 | 0.21 | 0.41 | 0.29 | 0.07 | 0.28 | 0.18 |
| 5 | 73794009 | rs298380 | -1.016 | 1.725 | -0.222 | 0.671 | -0.022 | -0.025 | -0.025 | 0 | 0.2 | 0.2 | 0.26 | 0.32 | 0.21 | 0.33 |
| 5 | 73795111 | rs298379 | -2.018 | 0.592 | -0.368 | 0.084 | -0.006 | 0.057 | 0.057 | 0.195 | 0.49 | 0.76 | 0.49 | 0.39 | 0.49 | 0.55 |
| 5 | 73800321 | rs298372 | -0.274 | 0.604 | -0.383 | 0.132 | -0.051 | -0.058 | -0.058 | 0 | 0.19 | 0.07 | 0.25 | 0.32 | 0.21 | 0.33 |
| 5 | 73800618 | rs17645426 | -0.489 | 0.367 | 0.096 | 0.406 | 0.01 | -0.009 | -0.009 | 0.012 | 0.16 | 0.22 | 0.15 | 0.12 | 0.22 | 0.11 |
| 5 | 73802976 | rs1001114 | 2.485 | 0.246 | -0.281 | 0.368 | -0.061 | 0.173 | 0.173 | 0.236 | 0.38 | 0.04 | 0.22 | 0.13 | 0.38 | 0.31 |
| 5 | 73805765 | rs16871933 | 0.634 | 0.171 | 0.332 | 0.398 | 0 | 0 | 0 | 0.083 | 0.09 | 0.11 | 0.08 | 0.02 | 0.07 | 0.11 |
| 5 | 73808746 | rs16871943 | 0.138 | 0.148 | 0.534 | 0.387 | -0.024 | -0.025 | -0.025 | 0.04 | 0.1 | 0.18 | 0.08 | 0.02 | 0.07 | 0.11 |
| 5 | 73810478 | rs212925 | -2.085 | 0.107 | 1.016 | 0.404 | 0.002 | -0.001 | -0.001 | 4.572 | 0.51 | 0.67 | 0.52 | 0.51 | 0.51 | 0.57 |
| 5 | 73811250 | rs11745925 | -1.316 | 0.129 | 0.799 | 0.39 | -0.01 | -0.017 | -0.017 | 2.68 | 0.26 | 0.43 | 0.24 | 0.19 | 0.29 | 0.21 |
| 5 | 73821242 | rs298393 | 2.111 | 0.162 | 0.569 | 0.422 | -0.038 | -0.041 | -0.041 | 0 | 0.49 | 0.13 | 0.47 | 0.49 | 0.49 | 0.43 |
| 5 | 73825889 | rs17645605 | -0.453 | 0.389 | 0.813 | 0.523 | 0.01 | 0.006 | 0.006 | 0 | 0.15 | 0.22 | 0.14 | 0.17 | 0.21 | 0.11 |
| 5 | 73833952 | rs1651349 | 1.286 | 0.276 | 0.723 | 0.411 | -0.004 | 0.004 | 0.004 | 0.033 | 0.65 | 0.48 | 0.62 | 0.6 | 0.68 | 0.56 |
| 5 | 73834112 | rs1510936 | 0.552 | 0.256 | 0.656 | 0.313 | 0.012 | 0.011 | 0.011 | 0 | 0.75 | 0.64 | 0.71 | 0.68 | 0.77 | 0.67 |
| 5 | 73837896 | rs10515182 | -0.429 | 0.121 | 0.146 | 0.453 | 0.008 | 0.008 | 0.008 | 0 | 0.1 | 0.11 | 0.1 | 0.02 | 0.09 | 0.11 |
| 5 | 73839750 | rs4357015 | -0.629 | 0.048 | 0.268 | 0.857 | -0.029 | 0.068 | 0.068 | 0 | 0.21 | 0.32 | 0.32 | 0.36 | 0.16 | 0.17 |
| 5 | 73840051 | rs515549 | 0.553 | 0.217 | 0.021 | 0.512 | 0.05 | 0.01 | 0.01 | 0.254 | 0.71 | 0.54 | 0.66 | 0.63 | 0.62 | 0.62 |
| 5 | 73841827 | rs435624 | 1.238 | 0.083 | 0.006 | 0.489 | 0 | -0.001 | -0.001 | 0 | 0.53 | 0.43 | 0.37 | 0.3 | 0.47 | 0.46 |
| 5 | 73841936 | rs13189660 | 1.692 | -0.107 | -0.38 | 0.184 | -0.004 | -0.009 | -0.009 | 0 | 0.34 | 0.36 | 0.25 | 0.28 | 0.32 | 0.35 |
| 5 | 73843948 | rs452166 | 0.383 | 0.261 | -0.405 | 0.255 | 0 | -0.001 | -0.001 | 0.476 | 0.61 | 0.6 | 0.69 | 0.74 | 0.65 | 0.6 |
| 5 | 73847044 | rs2173671 | -0.525 | 0.511 | -0.803 | 0.555 | -0.011 | -0.026 | -0.026 | 0 | 0.25 | 0.21 | 0.44 | 0.46 | 0.33 | 0.24 |

| | | | | | | | | | | | | | | | | |
|---|---|---|---|---|---|---|---|---|---|---|---|---|---|---|---|---|
| 5 | 73850292 | rs17645885 | 0.943 | 0.147 | -0.789 | 0.302 | 0 | -0.016 | -0.016 | 0 | 0.08 | 0.08 | 0.06 | 0.01 | 0.1 | 0.06 |
| 5 | 73851778 | rs1428558 | -1.011 | -0.241 | -1.378 | 0.55 | 0.042 | -0.065 | -0.065 | 1.011 | 0.54 | 0.92 | 0.53 | 0.44 | 0.4 | 0.35 |
| 5 | 73854583 | rs298386 | 0.855 | 0.087 | -0.842 | 0.31 | 0.013 | 0.002 | 0.002 | 0 | 0.35 | 0.43 | 0.36 | 0.33 | 0.39 | 0.41 |
| 5 | 73855164 | rs298385 | 1.212 | -0.177 | -0.899 | 0.286 | -0.044 | 0.032 | 0.032 | 0.268 | 0.4 | 0.1 | 0.29 | 0.31 | 0.42 | 0.37 |
| 5 | 73855858 | rs17560029 | -0.53 | 0.349 | -0.257 | 0.573 | 0.007 | 0.005 | 0.005 | 0 | 0.11 | 0.15 | 0.18 | 0.11 | 0.14 | 0.1 |
| 5 | 73856207 | rs1428556 | -0.551 | 0.347 | -0.258 | 0.572 | 0.007 | 0.005 | 0.005 | 0 | 0.11 | 0.15 | 0.18 | 0.11 | 0.14 | 0.09 |
| 5 | 73861278 | rs4704147 | -0.024 | 0.106 | 0.014 | 0.779 | 0 | 0.002 | 0.002 | 0 | 0.17 | 0.13 | 0.32 | 0.27 | 0.24 | 0.11 |
| 5 | 73879195 | rs300248 | 0.7 | -0.454 | 0.423 | 0.587 | -0.017 | 0.015 | 0.015 | 0 | 0.67 | 0.37 | 0.57 | 0.43 | 0.78 | 0.58 |
| 5 | 73883887 | rs300252 | 0.838 | -0.387 | 0.686 | 0.422 | 0.004 | -0.004 | -0.004 | 0 | 0.59 | 0.59 | 0.51 | 0.46 | 0.65 | 0.54 |
| 5 | 73886691 | rs300245 | 1.645 | -0.321 | 0.656 | 0.181 | 0.001 | -0.002 | -0.002 | 1.303 | 0.46 | 0.19 | 0.43 | 0.43 | 0.54 | 0.48 |
| 5 | 73899431 | rs6895845 | 0.497 | -0.165 | 1.125 | 0.37 | 0.007 | -0.056 | -0.056 | 0 | 0.11 | 0.2 | 0.07 | 0.01 | 0.11 | 0.07 |
| 5 | 73904734 | rs17670147 | 0.429 | -0.077 | 0.596 | 0.414 | -0.028 | -0.01 | -0.01 | 0 | 0.11 | 0.03 | 0.17 | 0.09 | 0.13 | 0.11 |
| 5 | 73905694 | rs300263 | 0.635 | -0.524 | 1.189 | 0.634 | -0.004 | 0.005 | 0.005 | 0 | 0.75 | 0.72 | 0.8 | 0.61 | 0.81 | 0.64 |
| 5 | 73918616 | rs10079609 | -0.177 | 0.637 | 1.581 | 0.618 | 0 | -0.024 | -0.024 | 0.509 | 0.11 | 0.11 | 0.06 | 0.01 | 0.12 | 0.07 |
| 5 | 73927752 | rs1039320 | -1.38 | 0.74 | 1.971 | 0.759 | 0.017 | -0.087 | -0.087 | 0 | 0.13 | 0.6 | 0.13 | 0.01 | 0.12 | 0.1 |
| 5 | 73934274 | rs10474420 | -0.339 | 0.93 | 0.791 | 0.855 | 0.005 | -0.045 | -0.045 | 0.969 | 0.1 | 0.18 | 0.06 | 0.01 | 0.12 | 0.07 |
| 5 | 73937506 | rs1039321 | 0.568 | 1.028 | 0.421 | 1.038 | -0.068 | -0.079 | -0.079 | 0 | 0.85 | 0.98 | 0.83 | 0.68 | 0.81 | 0.66 |
| 5 | 73944109 | rs7704467 | 1.568 | 0.954 | 0.662 | 1.014 | 0.025 | 0.006 | 0.006 | 0.304 | 0.73 | 0.5 | 0.75 | 0.66 | 0.7 | 0.56 |
| 5 | 73944444 | rs6864196 | -0.839 | 1.118 | 0.684 | 0.445 | 0.029 | -0.069 | -0.069 | 0.544 | 0.11 | 0.36 | 0.08 | 0.01 | 0.12 | 0.13 |
| 5 | 73947591 | rs13173835 | 0.97 | 1.089 | -0.139 | 0.841 | 0.001 | 0.005 | 0.005 | 1.061 | 0.7 | 0.69 | 0.57 | 0.5 | 0.66 | 0.54 |
| 5 | 73949009 | rs10058135 | 1.276 | 0.766 | 0.056 | 0.196 | -0.008 | -0.049 | -0.049 | 0 | 0.9 | 0.8 | 0.94 | 1 | 0.92 | 0.89 |
| 5 | 73950760 | rs904714 | -0.042 | 0.433 | 0.04 | -0.355 | -0.002 | -0.008 | -0.008 | 0 | 0.32 | 0.29 | 0.44 | 0.16 | 0.22 | 0.27 |
| 5 | 73951733 | rs4703638 | 0.891 | 1.18 | 0.21 | 0.059 | 0.004 | 0.003 | 0.003 | 0 | 0.87 | 0.87 | 0.85 | 0.78 | 0.8 | 0.77 |
| 5 | 73954619 | rs12152861 | 0.951 | 1.071 | 0.415 | -0.085 | -0.002 | 0.086 | 0.086 | 0.016 | 0.51 | 0.18 | 0.39 | 0.61 | 0.55 | 0.5 |
| 5 | 73955280 | rs16872154 | 0.253 | 1.145 | 0.531 | -0.042 | -0.004 | -0.011 | -0.011 | 0 | 0.31 | 0.31 | 0.42 | 0.16 | 0.21 | 0.27 |
| 5 | 73958245 | rs4703640 | 1.323 | 1.215 | -0.133 | -0.057 | -0.002 | 0 | 0 | 3.823 | 0.81 | 0.83 | 0.83 | 0.78 | 0.76 | 0.75 |
| 5 | 73959632 | rs2454847 | -0.234 | 1.284 | -0.366 | -0.08 | 0.03 | 0.011 | 0.011 | 0 | 0.13 | 0.16 | 0.14 | 0.2 | 0.23 | 0.17 |
| 5 | 73960725 | rs820858 | -0.513 | 1.297 | -0.358 | -0.123 | 0.001 | -0.001 | -0.001 | 0 | 0.19 | 0.19 | 0.18 | 0.18 | 0.29 | 0.26 |
| 5 | 73962506 | rs820853 | -0.248 | 1.063 | -0.508 | 0.227 | -0.001 | 0.006 | 0.006 | 0 | 0.58 | 0.63 | 0.64 | 0.4 | 0.57 | 0.57 |
| 5 | 73966557 | rs6453071 | -0.271 | 0.93 | -0.699 | -0.09 | 0.005 | 0.025 | 0.025 | 0 | 0.47 | 0.59 | 0.45 | 0.31 | 0.42 | 0.43 |
| 5 | 73966901 | rs6453072 | 0.291 | 0.836 | -1.241 | 0.067 | -0.038 | -0.008 | -0.008 | 0 | 0.62 | 0.76 | 0.55 | 0.6 | 0.55 | 0.7 |
| 5 | 73969058 | rs16872173 | 0.114 | 0.447 | -1.337 | -0.318 | 0.002 | 0.003 | 0.003 | 0.064 | 0.16 | 0.18 | 0.1 | 0.05 | 0.13 | 0.1 |
| 5 | 73970597 | rs820841 | -0.333 | 0.6 | -2.541 | 0.046 | -0.034 | -0.017 | -0.017 | 0 | 0.48 | 0.2 | 0.33 | 0.39 | 0.54 | 0.48 |
| 5 | 73971107 | rs820840 | 0.129 | 0.557 | -1.943 | 0.031 | 0.018 | 0.039 | 0.039 | 1.224 | 0.51 | 0.31 | 0.48 | 0.63 | 0.58 | 0.5 |
| 5 | 73971777 | rs11749103 | 0.951 | 0.706 | -1.817 | 0.184 | -0.038 | -0.053 | -0.053 | 0 | 0.14 | 0.01 | 0.19 | 0.25 | 0.11 | 0.07 |
| 5 | 73973651 | rs3894712 | 0.344 | 0.621 | -1.577 | 0.151 | -0.007 | -0.011 | -0.011 | 0 | 0.18 | 0.06 | 0.22 | 0.27 | 0.16 | 0.11 |
| 5 | 73975094 | rs12233980 | 0.55 | 0.692 | -1.549 | 0.143 | -0.006 | -0.008 | -0.008 | 0 | 0.18 | 0.11 | 0.22 | 0.27 | 0.16 | 0.11 |
| 5 | 73980065 | rs820869 | 0.167 | 0.2 | -0.74 | -0.046 | 0.005 | 0.004 | 0.004 | 0.035 | 0.57 | 0.49 | 0.37 | 0.43 | 0.57 | 0.55 |
| 5 | 73982965 | rs820884 | -0.821 | 0.466 | -0.835 | -0.109 | 0.004 | 0.003 | 0.003 | 0 | 0.73 | 0.64 | 0.59 | 0.69 | 0.73 | 0.63 |
| 5 | 73985215 | rs11556045 | 1.313 | 0.539 | -0.621 | 0.295 | 0.012 | 0.013 | 0.013 | 0 | 0.16 | 0.27 | 0.24 | 0.17 | 0.18 | 0.25 |
| 5 | 73985534 | rs820889 | -1.553 | 0.506 | -0.691 | 0.282 | -0.002 | -0.004 | -0.004 | 0 | 0.89 | 0.88 | 0.83 | 0.86 | 0.91 | 0.87 |
| 5 | 73987775 | rs3756444 | -0.348 | 0.571 | -0.399 | 0.157 | 0.007 | 0.008 | 0.008 | 0.532 | 0.34 | 0.59 | 0.47 | 0.43 | 0.33 | 0.33 |
| 5 | 73992406 | rs1696979 | 0.407 | 0.293 | -1.017 | -0.023 | 0.034 | 0.056 | 0.056 | 0 | 0.5 | 0.29 | 0.48 | 0.52 | 0.6 | 0.64 |
| 5 | 73999205 | rs820834 | -0.847 | 0.038 | -1.094 | -0.599 | -0.021 | -0.019 | -0.019 | 0 | 0.91 | 0.98 | 0.84 | 0.84 | 0.91 | 0.86 |
| 5 | 74013611 | rs6453074 | -1.422 | -0.7 | -1.59 | -0.541 | -0.001 | -0.043 | -0.043 | 0 | 0.83 | 0.91 | 0.76 | 0.61 | 0.81 | 0.7 |
| 5 | 74015350 | rs6867972 | -0.083 | -0.889 | -1.745 | -0.763 | 0.006 | 0.086 | 0.086 | 0 | 0.55 | 0.78 | 0.44 | 0.27 | 0.43 | 0.37 |
| 5 | 74017499 | rs1048167 | 0.906 | -0.492 | -0.162 | -0.391 | -0.002 | -0.02 | -0.02 | 0 | 0.2 | 0.23 | 0.11 | 0.02 | 0.14 | 0.08 |
| 5 | 74030338 | rs13164140 | 1.099 | -0.266 | -0.232 | -0.249 | -0.031 | -0.022 | -0.022 | 0 | 0.13 | 0.05 | 0.16 | 0.16 | 0.25 | 0.18 |
| 5 | 74031525 | rs6877188 | 0.862 | -0.412 | 0.048 | -0.301 | -0.03 | -0.066 | -0.066 | 0 | 0.21 | 0.29 | 0.12 | 0.02 | 0.14 | 0.08 |
| 5 | 74033944 | rs6453077 | -0.621 | -0.575 | 0.128 | -0.314 | 0.043 | 0.058 | 0.058 | 0.256 | 0.36 | 0.75 | 0.33 | 0.21 | 0.24 | 0.24 |
| 5 | 74054595 | rs10515191 | -0.621 | -0.562 | 0.134 | -0.309 | 0.044 | 0.058 | 0.058 | 0.31 | 0.36 | 0.75 | 0.34 | 0.21 | 0.24 | 0.24 |
| 5 | 74055209 | rs957680 | 0.852 | -0.565 | 0.293 | -0.31 | -0.046 | -0.092 | -0.092 | 0 | 0.22 | 0.34 | 0.12 | 0.02 | 0.14 | 0.08 |
| 5 | 74058398 | rs10942719 | -1.727 | -0.556 | 0.016 | -0.288 | -0.006 | -0.066 | -0.066 | 0 | 0.61 | 0.85 | 0.62 | 0.51 | 0.6 | 0.51 |
| 5 | 74059091 | rs10515193 | 1.08 | -1.661 | 0.743 | -0.023 | -0.026 | -0.025 | -0.025 | 0 | 0.12 | 0.05 | 0.13 | 0.14 | 0.11 | 0.08 |
| 5 | 74069863 | rs6874609 | -1.167 | -1.597 | 1.085 | -0.18 | 0.044 | 0.059 | 0.059 | 0.382 | 0.36 | 0.75 | 0.34 | 0.21 | 0.24 | 0.24 |
| 5 | 74073542 | rs15694 | -1.169 | -1.363 | 1.2 | -0.068 | 0.039 | 0.053 | 0.053 | 0.052 | 0.35 | 0.75 | 0.34 | 0.21 | 0.24 | 0.24 |
| 5 | 74076210 | rs3733794 | -0.628 | -1.362 | 1.169 | -0.064 | -0.002 | -0.007 | -0.007 | 0 | 0.29 | 0.29 | 0.27 | 0.19 | 0.22 | 0.22 |
| 5 | 74077474 | rs16872269 | -0.555 | -1.241 | 1.387 | 0.003 | 0.026 | 0.038 | 0.038 | 2.137 | 0.05 | 0.32 | 0.05 | 0.01 | 0.02 | 0.02 |
| 5 | 74092800 | rs6883061 | -0.602 | -1.241 | 1.333 | -0.007 | -0.003 | -0.008 | -0.008 | 0.011 | 0.28 | 0.29 | 0.27 | 0.19 | 0.22 | 0.22 |
| 5 | 74107614 | rs6453086 | -0.749 | -1.254 | 1.461 | -0.013 | -0.021 | -0.041 | -0.041 | 0 | 0.31 | 0.52 | 0.29 | 0.19 | 0.22 | 0.22 |
| 5 | 74142865 | rs1164624 | 1.602 | -1.049 | 0.289 | 0.114 | 0.017 | 0.083 | 0.083 | 0 | 0.48 | 0.18 | 0.37 | 0.26 | 0.58 | 0.44 |
| 5 | 74149478 | rs16872345 | -0.19 | -1.158 | 0.855 | -0.748 | -0.005 | -0.025 | -0.025 | 0.107 | 0.2 | 0.23 | 0.11 | 0.02 | 0.13 | 0.08 |
| 5 | 74154745 | rs17671140 | -0.376 | -0.841 | 0.111 | -0.584 | 0.001 | 0.001 | 0.001 | 0 | 0.09 | 0.06 | 0.16 | 0.17 | 0.09 | 0.14 |
| 5 | 74163159 | rs6453087 | -1.118 | -0.834 | 0.013 | -0.573 | 0.052 | 0.07 | 0.07 | 0.633 | 0.37 | 0.76 | 0.34 | 0.21 | 0.24 | 0.26 |
| 5 | 74169045 | rs10062244 | -0.599 | 0.077 | 0.521 | 0.133 | 0 | -0.001 | -0.001 | 0 | 0.11 | 0.06 | 0.16 | 0.17 | 0.08 | 0.13 |
| 5 | 74172468 | rs11739301 | -0.766 | 0.452 | 1.178 | 0.334 | 0.036 | 0.012 | 0.012 | 0 | 0.11 | 0.29 | 0.22 | 0.17 | 0.15 | 0.12 |
| 5 | 74184144 | rs10942722 | -1.611 | 0.364 | 1.102 | 0.359 | 0 | -0.004 | -0.004 | 0 | 0.38 | 0.5 | 0.43 | 0.35 | 0.37 | 0.38 |
| 5 | 74197953 | rs12519932 | -1.782 | 0.336 | 1.253 | 0.796 | 0.001 | -0.003 | -0.003 | 0.144 | 0.31 | 0.78 | 0.4 | 0.34 | 0.27 | 0.32 |
| 5 | 74198611 | rs11745686 | -1.511 | 0.481 | 0.941 | 0.865 | -0.002 | -0.002 | -0.002 | 0 | 0.27 | 0.3 | 0.36 | 0.34 | 0.27 | 0.32 |

| | | | | | | | | | | | | | | | | |
|---|---|---|---|---|---|---|---|---|---|---|---|---|---|---|---|---|
| 5 | 74212755 | rs6888163 | -0.895 | 0.297 | 0.274 | 0.893 | -0.006 | 0.028 | 0.028 | 0.027 | 0.17 | 0.52 | 0.18 | 0.15 | 0.12 | 0.19 |
| 5 | 74239376 | rs7728102 | 0.618 | 0.147 | -0.366 | 1.253 | -0.012 | 0.036 | 0.036 | 0.117 | 0.73 | 0.35 | 0.61 | 0.38 | 0.76 | 0.72 |
| 5 | 74239784 | rs10069954 | 0.412 | 0.364 | -1.733 | 0.256 | -0.085 | 0.239 | 0.239 | 0.772 | 0.43 | 0.04 | 0.36 | 0.12 | 0.51 | 0.46 |
| 5 | 74239946 | rs9293636 | 0.489 | 0.67 | -0.72 | 1.323 | 0.004 | 0.007 | 0.007 | 0 | 0.91 | 0.89 | 0.94 | 0.85 | 0.87 | 0.94 |
| 5 | 74243120 | rs7356745 | -1.005 | 0.682 | -0.783 | 1.916 | -0.003 | 0.029 | 0.029 | 0.778 | 0.1 | 0.09 | 0.21 | 0.41 | 0.11 | 0.3 |
| 5 | 74253941 | rs6884528 | -1.453 | -0.195 | -0.958 | 1.299 | 0.029 | -0.028 | -0.028 | 0 | 0.37 | 0.75 | 0.37 | 0.44 | 0.28 | 0.48 |
| 5 | 74254502 | rs10060240 | -0.337 | -0.091 | -0.881 | 1.352 | -0.001 | 0.04 | 0.04 | 0.668 | 0.09 | 0.1 | 0.21 | 0.4 | 0.11 | 0.3 |
| 5 | 74258458 | rs10805892 | -1.611 | 0.33 | -0.906 | 1.456 | -0.012 | 0.007 | 0.007 | 0.239 | 0.38 | 0.33 | 0.56 | 0.74 | 0.47 | 0.61 |
| 5 | 74259550 | rs6453099 | -1.122 | -0.241 | -1.554 | 0.844 | -0.013 | 0.007 | 0.007 | 0 | 0.35 | 0.32 | 0.52 | 0.76 | 0.41 | 0.56 |
| 5 | 74262162 | rs4703642 | -1.529 | -0.376 | -1.398 | 0.786 | 0.021 | 0.05 | 0.05 | 0.252 | 0.51 | 0.38 | 0.59 | 0.77 | 0.57 | 0.72 |
| 5 | 74262650 | rs7719121 | -0.281 | -0.174 | -1.443 | 0.847 | -0.002 | -0.015 | -0.015 | 0.69 | 0.25 | 0.21 | 0.29 | 0.36 | 0.27 | 0.25 |
| 5 | 74262980 | rs4392601 | -0.458 | -0.145 | -1.535 | 0.84 | -0.043 | -0.055 | -0.055 | 1.032 | 0.25 | 0.11 | 0.3 | 0.4 | 0.3 | 0.47 |
| 5 | 74263455 | rs10069744 | 0.995 | -0.124 | -1.495 | 0.869 | 0.014 | 0.063 | 0.063 | 0 | 0.13 | 0.03 | 0.04 | 0.03 | 0.1 | 0.12 |
| 5 | 74266343 | rs10043960 | -0.144 | -0.107 | -1.405 | 0.947 | -0.035 | -0.069 | -0.069 | 0 | 0.22 | 0.08 | 0.26 | 0.4 | 0.24 | 0.43 |
| 5 | 74269724 | rs13182265 | -0.002 | -0.148 | -0.109 | 0.892 | 0.044 | 0.104 | 0.104 | 0 | 0.13 | 0.38 | 0.27 | 0.3 | 0.17 | 0.12 |
| 5 | 74274675 | rs4382144 | -1.64 | -0.259 | 0.045 | 0.833 | 0.01 | 0.039 | 0.039 | 0.476 | 0.55 | 0.73 | 0.65 | 0.75 | 0.57 | 0.72 |
| 5 | 74276566 | rs6870053 | -1.259 | -0.246 | 0.147 | 0.799 | -0.002 | -0.003 | -0.003 | 0 | 0.47 | 0.48 | 0.39 | 0.41 | 0.41 | 0.6 |
| 5 | 74281643 | rs6888013 | -0.114 | 0.084 | -0.275 | 1.052 | 0 | 0.025 | 0.025 | 0.663 | 0.11 | 0.1 | 0.28 | 0.39 | 0.15 | 0.31 |
| 5 | 74289069 | rs6453104 | -1.267 | -0.748 | -0.186 | 0.434 | 0.059 | 0.103 | 0.103 | 0 | 0.51 | 0.8 | 0.42 | 0.38 | 0.43 | 0.38 |
| 5 | 74289267 | rs6887521 | -0.5 | -0.856 | 0.325 | 0.316 | -0.032 | -0.07 | -0.07 | 0.018 | 0.19 | 0.31 | 0.08 | 0.04 | 0.11 | 0.1 |
| 5 | 74292694 | rs10067703 | -0.912 | -0.58 | -0.254 | 0.602 | 0.021 | 0.021 | 0.021 | 0 | 0.38 | 0.68 | 0.36 | 0.33 | 0.31 | 0.28 |
| 5 | 74300599 | rs6895821 | -1.164 | -0.497 | -0.18 | 0.491 | 0.049 | 0.069 | 0.069 | 0 | 0.44 | 0.68 | 0.36 | 0.27 | 0.29 | 0.36 |
| 5 | 74301852 | rs7721887 | 0.485 | -0.686 | 0.831 | 0.326 | 0.006 | -0.064 | -0.064 | 0 | 0.11 | 0.23 | 0.07 | 0 | 0.1 | 0.09 |
| 5 | 74302658 | rs4555772 | -1.761 | -0.032 | 0.528 | 0.809 | 0.015 | 0.034 | 0.034 | 0 | 0.5 | 0.73 | 0.45 | 0.41 | 0.4 | 0.6 |
| 5 | 74314884 | rs7706858 | -1.083 | -0.344 | 0.519 | 0.759 | 0.065 | 0.086 | 0.086 | 0 | 0.45 | 0.7 | 0.36 | 0.26 | 0.29 | 0.36 |
| 5 | 74324437 | rs3811986 | -1.24 | -0.475 | 0.433 | 0.703 | 0.101 | 0.122 | 0.122 | 0 | 0.47 | 0.75 | 0.37 | 0.29 | 0.3 | 0.35 |
| 5 | 74324548 | rs3811987 | -1.24 | -0.475 | 0.433 | 0.703 | 0.101 | 0.122 | 0.122 | 0 | 0.47 | 0.75 | 0.37 | 0.29 | 0.3 | 0.35 |
| 5 | 74327436 | rs11957690 | -1.24 | -0.475 | 0.433 | 0.703 | 0.101 | 0.122 | 0.122 | 0 | 0.47 | 0.75 | 0.37 | 0.29 | 0.3 | 0.35 |
| 5 | 74331528 | rs12657184 | -0.912 | -0.483 | 0.752 | 0.755 | 0 | -0.004 | -0.004 | 0.081 | 0.11 | 0.11 | 0.21 | 0.22 | 0.08 | 0.13 |
| 5 | 74342313 | rs10052034 | -1.159 | -0.584 | 1.576 | 1.016 | 0.156 | 0.238 | 0.238 | 0 | 0.08 | 0.31 | 0.06 | 0.16 | 0.11 | 0.25 |
| 5 | 74344465 | rs6861101 | -0.494 | -1.116 | 1.554 | 0.593 | -0.033 | -0.089 | -0.089 | 0 | 0.27 | 0.37 | 0.12 | 0.06 | 0.2 | 0.2 |
| 5 | 74350535 | rs11951174 | -1.249 | -1.424 | 1.492 | 0.813 | -0.037 | -0.029 | -0.029 | 0 | 0.27 | 0.37 | 0.25 | 0.15 | 0.16 | 0.2 |
| 5 | 74352180 | rs3935470 | -2.399 | -1.395 | 0.976 | 1.056 | 0.023 | 0.004 | 0.004 | 0 | 0.54 | 0.86 | 0.47 | 0.58 | 0.37 | 0.61 |
| 5 | 74353702 | rs7718341 | -2.209 | -1.759 | 0.891 | 0.992 | 0.103 | 0.019 | 0.019 | 0 | 0.46 | 0.81 | 0.31 | 0.32 | 0.29 | 0.47 |
| 5 | 74371635 | rs7720035 | 0.977 | -1.54 | 1.482 | 1.04 | -0.033 | -0.039 | -0.039 | 7.056 | 0.07 | 0.17 | 0.03 | 0 | 0.01 | 0 |
| 5 | 74376571 | rs4345300 | -2.718 | -1.744 | 1.323 | 1.085 | 0.024 | -0.068 | -0.068 | 0 | 0.52 | 0.9 | 0.42 | 0.48 | 0.39 | 0.6 |
| 5 | 74382081 | rs4298239 | -1.916 | -1.833 | 1.663 | 1.135 | 0.032 | -0.005 | -0.005 | 0.547 | 0.25 | 0.78 | 0.2 | 0.28 | 0.14 | 0.17 |
| 5 | 74384115 | rs10462511 | -3.039 | -1.727 | 1.372 | 1.105 | -0.042 | -0.011 | -0.011 | 0 | 0.55 | 0.9 | 0.38 | 0.37 | 0.44 | 0.62 |
| 5 | 74408979 | rs4301212 | -0.819 | -1.702 | 2.007 | 1.161 | 0.012 | 0.001 | 0.001 | 0 | 0.27 | 0.2 | 0.21 | 0.19 | 0.18 | 0.42 |
| 5 | 74425217 | rs10053125 | -1.469 | -1.786 | 2.021 | 1.407 | -0.006 | -0.001 | -0.001 | 0.294 | 0.35 | 0.35 | 0.25 | 0.27 | 0.26 | 0.43 |
| 5 | 74436574 | rs1600074 | -2.802 | -1.789 | 1.992 | 1.414 | 0.028 | -0.063 | -0.063 | 0 | 0.52 | 0.89 | 0.42 | 0.48 | 0.38 | 0.6 |
| 5 | 74442964 | rs6893216 | -0.178 | -1.798 | 2.424 | 1.386 | -0.003 | 0.012 | 0.012 | 0 | 0.12 | 0.25 | 0.14 | 0.18 | 0.11 | 0.14 |
| 5 | 74443132 | rs1422698 | -2.848 | -1.798 | 2.423 | 1.386 | 0.026 | -0.069 | -0.069 | 0 | 0.52 | 0.9 | 0.42 | 0.48 | 0.39 | 0.6 |
| 5 | 74446804 | rs10054860 | -0.811 | -1.813 | 2.427 | 1.383 | -0.004 | -0.001 | -0.001 | 0 | 0.27 | 0.11 | 0.22 | 0.19 | 0.25 | 0.42 |
| 5 | 74453027 | rs10462354 | -2.831 | -1.79 | 2.36 | 1.397 | 0.026 | -0.065 | -0.065 | 0 | 0.52 | 0.9 | 0.42 | 0.48 | 0.39 | 0.6 |
| 5 | 74456477 | rs6872586 | -1.231 | -1.79 | 2.371 | 1.373 | -0.006 | 0.03 | 0.03 | 0.07 | 0.16 | 0.51 | 0.16 | 0.21 | 0.13 | 0.17 |
| 5 | 74463185 | rs1363577 | -2.786 | -1.812 | 2.265 | 1.481 | 0.026 | -0.069 | -0.069 | 0 | 0.52 | 0.9 | 0.42 | 0.48 | 0.39 | 0.6 |
| 5 | 74486778 | rs6898116 | -2.781 | -1.797 | 1.452 | 1.479 | 0.027 | -0.068 | -0.068 | 0 | 0.52 | 0.9 | 0.42 | 0.48 | 0.39 | 0.6 |
| 5 | 74490547 | rs6890143 | -0.173 | -1.798 | 1.453 | 1.479 | -0.003 | 0.013 | 0.013 | 0 | 0.12 | 0.25 | 0.14 | 0.18 | 0.11 | 0.14 |
| 5 | 74497842 | rs6885887 | -2.656 | -1.729 | 0.633 | 1.454 | 0.031 | -0.068 | -0.068 | 1.057 | 0.52 | 0.9 | 0.42 | 0.47 | 0.38 | 0.6 |
| 5 | 74506658 | rs2219745 | -1.157 | -1.521 | 0.712 | 1.377 | 0.004 | 0.005 | 0.005 | 0 | 0.29 | 0.21 | 0.23 | 0.19 | 0.25 | 0.42 |
| 5 | 74510023 | rs1422700 | -1.01 | -1.512 | 0.649 | 1.382 | 0.004 | 0.006 | 0.006 | 0 | 0.28 | 0.17 | 0.22 | 0.19 | 0.25 | 0.42 |
| 5 | 74512987 | rs2216638 | -2.66 | -1.734 | 0.562 | 1.461 | 0.026 | -0.069 | -0.069 | 0.674 | 0.52 | 0.9 | 0.42 | 0.48 | 0.39 | 0.6 |
| 5 | 74517954 | rs2161370 | -1.029 | -1.736 | 0.614 | 1.456 | 0.002 | 0.006 | 0.006 | 0 | 0.29 | 0.2 | 0.23 | 0.19 | 0.25 | 0.44 |
| 5 | 74524837 | rs1477937 | -2.608 | -1.722 | 0.599 | 1.444 | 0.028 | -0.067 | -0.067 | 0 | 0.52 | 0.89 | 0.42 | 0.48 | 0.39 | 0.6 |
| 5 | 74537751 | rs4704197 | -2.649 | -1.754 | 0.587 | 1.428 | 0.026 | -0.068 | -0.068 | 0 | 0.52 | 0.9 | 0.42 | 0.48 | 0.4 | 0.61 |
| 5 | 74540397 | rs7711235 | -1.081 | -1.768 | 0.61 | 1.437 | 0.004 | 0.002 | 0.002 | 0 | 0.3 | 0.22 | 0.23 | 0.22 | 0.26 | 0.46 |
| 5 | 74558893 | rs7717415 | -2.382 | -1.633 | 0.42 | 1.38 | 0.036 | -0.041 | -0.041 | 0 | 0.52 | 0.88 | 0.42 | 0.48 | 0.4 | 0.61 |
| 5 | 74565153 | rs10062361 | -1.356 | -1.495 | 0.749 | 1.246 | -0.011 | -0.022 | -0.022 | 0 | 0.19 | 0.16 | 0.23 | 0.3 | 0.22 | 0.38 |
| 5 | 74569856 | rs4704200 | -2.854 | -2.325 | 0.897 | 0.75 | 0.01 | -0.013 | -0.013 | 0.061 | 0.55 | 0.76 | 0.51 | 0.54 | 0.43 | 0.6 |
| 5 | 74570531 | rs1551894 | -0.914 | -2.305 | 0.616 | 0.758 | -0.037 | -0.063 | -0.063 | 2.111 | 0.19 | 0.08 | 0.21 | 0.3 | 0.22 | 0.38 |
| 5 | 74574984 | rs2126736 | -2.691 | -2.212 | 0.718 | 0.788 | 0.006 | -0.006 | -0.006 | 0 | 0.55 | 0.67 | 0.5 | 0.54 | 0.43 | 0.61 |
| 5 | 74605220 | rs10056811 | -1.853 | -2.152 | 0.509 | 0.564 | 0.017 | 0.031 | 0.031 | 0.058 | 0.42 | 0.31 | 0.32 | 0.31 | 0.36 | 0.57 |
| 5 | 74608732 | rs11957260 | -3.159 | -2.245 | 0.352 | 0.598 | -0.102 | -0.159 | -0.159 | 0 | 0.57 | 0.97 | 0.54 | 0.51 | 0.44 | 0.6 |
| 5 | 74616843 | rs10474433 | -2.059 | -2.11 | 0.608 | 0.584 | 0.008 | 0.014 | 0.014 | 0.111 | 0.44 | 0.39 | 0.36 | 0.31 | 0.37 | 0.57 |
| 5 | 74619132 | rs13354746 | -0.895 | -2.079 | 0.543 | 0.705 | -0.041 | -0.017 | -0.017 | 0 | 0.19 | 0.08 | 0.22 | 0.25 | 0.24 | 0.37 |
| 5 | 74620912 | rs6878576 | -0.461 | -1.962 | 0.634 | 0.648 | -0.001 | -0.001 | -0.001 | 0 | 0.22 | 0.21 | 0.1 | 0.06 | 0.12 | 0.2 |
| 5 | 74631139 | rs3761738 | -0.264 | -1.73 | 0.48 | 0.796 | -0.021 | 0.023 | 0.023 | 0 | 0.08 | 0.03 | 0.1 | 0.03 | 0.1 | 0.13 |
| 5 | 74631501 | rs3761739 | -1.262 | -1.727 | 0.557 | 0.845 | 0.014 | 0.01 | 0.01 | 0 | 0.1 | 0.12 | 0.21 | 0.18 | 0.15 | 0.16 |

| | | | | | | | | | | | | | | | | |
|---|---|---|---|---|---|---|---|---|---|---|---|---|---|---|---|---|
| 5 | 74632133 | rs3761740 | -0.263 | -1.73 | 0.521 | 0.807 | -0.021 | 0.023 | 0.023 | 0 | 0.08 | 0.03 | 0.1 | 0.03 | 0.1 | 0.12 |
| 5 | 74641560 | rs10515198 | -0.263 | -1.732 | 0.483 | 0.805 | -0.021 | 0.023 | 0.023 | 0 | 0.08 | 0.03 | 0.1 | 0.03 | 0.1 | 0.13 |
| 5 | 74646878 | rs11742194 | -0.263 | -1.732 | 0.483 | 0.805 | -0.021 | 0.023 | 0.023 | 0 | 0.08 | 0.03 | 0.1 | 0.03 | 0.1 | 0.13 |
| 5 | 74651084 | rs3846662 | -2.982 | -1.88 | 0.301 | 0.805 | -0.042 | -0.092 | -0.092 | 0 | 0.54 | 0.94 | 0.51 | 0.51 | 0.43 | 0.6 |
| 5 | 74655498 | rs17238540 | 1.625 | -1.852 | 0.565 | 0.796 | -0.016 | -0.026 | -0.026 | 0 | 0.07 | 0.11 | 0.02 | 0 | 0.02 | 0 |
| 5 | 74656175 | rs5909 | -0.156 | -1.778 | 0.461 | 0.834 | -0.021 | 0.023 | 0.023 | 0 | 0.08 | 0.03 | 0.1 | 0.03 | 0.1 | 0.13 |
| 5 | 74656539 | rs12916 | -2.025 | -1.85 | 0.571 | 0.842 | -0.008 | -0.041 | -0.041 | 0 | 0.4 | 0.23 | 0.43 | 0.51 | 0.41 | 0.56 |
| 5 | 74663223 | rs16872521 | -0.425 | -1.579 | 0.78 | 0.977 | 0.033 | 0.008 | 0.008 | 0 | 0.15 | 0.07 | 0.06 | 0.06 | 0.09 | 0.16 |
| 5 | 74675717 | rs16872526 | -0.466 | -1.564 | 0.788 | 0.976 | 0.034 | -0.01 | -0.01 | 0 | 0.15 | 0.08 | 0.06 | 0.1 | 0.09 | 0.16 |
| 5 | 74676823 | rs7706933 | 0.636 | -1.662 | 1.497 | 0.907 | -0.032 | -0.048 | -0.048 | 0.082 | 0.09 | 0.27 | 0.04 | 0 | 0.02 | 0 |
| 5 | 74678937 | rs3761742 | -1.753 | -1.695 | 1.783 | 0.959 | 0.005 | -0.009 | -0.009 | 0.606 | 0.27 | 0.27 | 0.17 | 0.3 | 0.22 | 0.43 |
| 5 | 74681773 | rs698912 | -1.763 | -1.701 | 1.78 | 0.956 | 0.005 | -0.009 | -0.009 | 0.641 | 0.27 | 0.27 | 0.17 | 0.33 | 0.22 | 0.44 |
| 5 | 74683646 | rs7356637 | -2.759 | -1.733 | 2.164 | 0.953 | -0.001 | 0 | 0 | 0 | 0.38 | 0.67 | 0.23 | 0.3 | 0.24 | 0.44 |
| 5 | 74685520 | rs3761743 | -2.631 | -1.734 | 2.181 | 0.953 | -0.029 | -0.002 | -0.002 | 0 | 0.37 | 0.6 | 0.22 | 0.33 | 0.24 | 0.44 |
| 5 | 74692776 | rs6453133 | -3.051 | -1.722 | 2.186 | 1.011 | 0.071 | -0.074 | -0.074 | 0 | 0.41 | 0.89 | 0.36 | 0.48 | 0.29 | 0.46 |
| 5 | 74700095 | rs6866607 | 1.099 | -1.723 | 2.184 | 0.995 | -0.029 | -0.042 | -0.042 | 4.512 | 0.08 | 0.21 | 0.03 | 0 | 0.02 | 0 |
| 5 | 74717653 | rs10038723 | -1.743 | -1.7 | 1.997 | 1.018 | 0.005 | -0.009 | -0.009 | 0 | 0.27 | 0.26 | 0.17 | 0.33 | 0.22 | 0.44 |
| 5 | 74734362 | rs4385188 | -2.566 | -1.694 | 2.051 | 1.025 | -0.03 | -0.001 | -0.001 | 0.386 | 0.37 | 0.59 | 0.22 | 0.33 | 0.24 | 0.44 |
| 5 | 74745228 | rs10070207 | -2.472 | -1.659 | 1.999 | 1.034 | -0.047 | -0.006 | -0.006 | 0 | 0.36 | 0.56 | 0.22 | 0.27 | 0.21 | 0.4 |
| 5 | 74757556 | rs4704220 | -3.403 | -1.621 | 1.642 | 0.981 | 0.008 | -0.07 | -0.07 | 1.902 | 0.49 | 0.94 | 0.46 | 0.51 | 0.39 | 0.59 |
| 5 | 74757758 | rs12659791 | -1.163 | -1.618 | 1.65 | 0.981 | -0.01 | -0.008 | -0.008 | 0 | 0.09 | 0.05 | 0.21 | 0.18 | 0.15 | 0.16 |
| 5 | 74764347 | rs4440336 | -2.882 | -1.711 | 1.853 | 0.63 | 0.077 | 0.009 | 0.009 | 0.028 | 0.4 | 0.81 | 0.24 | 0.33 | 0.24 | 0.44 |
| 5 | 74771387 | rs10061689 | 0.068 | -1.572 | 1.645 | 0.669 | -0.004 | -0.009 | -0.009 | 0 | 0.11 | 0.1 | 0.11 | 0.23 | 0.13 | 0.25 |
| 5 | 74798156 | rs10055011 | 0.17 | -1.565 | 1.651 | 0.672 | -0.004 | -0.008 | -0.008 | 0 | 0.11 | 0.09 | 0.11 | 0.23 | 0.13 | 0.24 |
| 5 | 74808417 | rs5744545 | 0.324 | -1.546 | 1.41 | 0.683 | -0.025 | -0.045 | -0.045 | 0 | 0.1 | 0.04 | 0.1 | 0.19 | 0.11 | 0.22 |
| 5 | 74810384 | rs10059022 | 0.219 | -1.547 | 1.413 | 0.673 | -0.004 | -0.007 | -0.007 | 1.504 | 0.11 | 0.09 | 0.11 | 0.2 | 0.13 | 0.24 |
| 5 | 74835023 | rs16872590 | 0.43 | -1.567 | 1.455 | 0.607 | -0.04 | -0.054 | -0.054 | 0 | 0.09 | 0.23 | 0.04 | 0 | 0.02 | 0 |
| 5 | 74840745 | rs5744578 | -0.627 | -1.448 | 0.692 | 0.705 | 0.033 | 0.01 | 0.01 | 0 | 0.15 | 0.06 | 0.05 | 0.06 | 0.09 | 0.16 |
| 5 | 74846613 | rs5744598 | -1.878 | -1.439 | 0.823 | 0.745 | 0.019 | -0.026 | -0.026 | 0 | 0.28 | 0.21 | 0.18 | 0.33 | 0.22 | 0.4 |
| 5 | 74877266 | rs3213801 | -1.857 | -1.425 | 0.736 | 0.766 | 0.019 | -0.026 | -0.026 | 0 | 0.27 | 0.21 | 0.18 | 0.33 | 0.22 | 0.4 |
| 5 | 74890618 | rs5744707 | -0.614 | -1.359 | 0.548 | 0.709 | -0.02 | 0.022 | 0.022 | 0 | 0.08 | 0.03 | 0.1 | 0.03 | 0.1 | 0.15 |
| 5 | 74898752 | rs2287711 | -1.833 | -2.195 | 0.619 | 0.806 | 0.018 | -0.026 | -0.026 | 0 | 0.27 | 0.21 | 0.18 | 0.33 | 0.22 | 0.4 |
| 5 | 74925162 | rs9293656 | -1.587 | -2.183 | 0.566 | 0.799 | 0.018 | -0.025 | -0.025 | 0 | 0.26 | 0.16 | 0.17 | 0.3 | 0.22 | 0.4 |
| 5 | 74926778 | rs253407 | 2.184 | -2.311 | 0.46 | 0.698 | 0.045 | -0.017 | -0.017 | 0 | 0.63 | 0.14 | 0.73 | 0.66 | 0.73 | 0.56 |
| 5 | 74929312 | rs16872670 | -0.844 | -2.318 | 0.892 | 0.702 | -0.024 | 0.012 | 0.012 | 0 | 0.11 | 0.19 | 0.04 | 0.08 | 0.05 | 0.2 |
| 5 | 74937913 | rs1422441 | -0.514 | -2.282 | 1.206 | 0.635 | -0.061 | -0.035 | -0.035 | 0.069 | 0.12 | 0.54 | 0.08 | 0.04 | 0.02 | 0 |
| 5 | 74946455 | rs16872693 | 0.349 | -2.286 | 1.555 | 0.63 | -0.043 | -0.059 | -0.059 | 0 | 0.1 | 0.3 | 0.05 | 0 | 0.02 | 0 |
| 5 | 74947435 | rs16872698 | -0.085 | -2.283 | 1.531 | 0.634 | -0.049 | -0.022 | -0.022 | 0 | 0.11 | 0.44 | 0.07 | 0.04 | 0.02 | 0 |
| 5 | 74951901 | rs7717355 | -1.381 | -2.09 | 1.044 | 1.35 | 0.007 | -0.022 | -0.022 | 0.709 | 0.13 | 0.07 | 0.18 | 0.27 | 0.12 | 0.17 |
| 5 | 74956517 | rs253414 | 2.578 | -2.178 | 1.156 | 1.361 | 0.004 | 0.002 | 0.002 | 0.503 | 0.61 | 0.36 | 0.6 | 0.4 | 0.66 | 0.42 |
| 5 | 74957819 | rs2287709 | -0.576 | -2.226 | 1.263 | 1.394 | -0.029 | 0.016 | 0.016 | 0 | 0.1 | 0.12 | 0.03 | 0.1 | 0.03 | 0.16 |
| 5 | 74959970 | rs34355 | -0.355 | -2.171 | 0.863 | 1.432 | -0.052 | -0.047 | -0.047 | 0 | 0.87 | 0.96 | 0.85 | 0.79 | 0.81 | 0.78 |
| 5 | 74962768 | rs6453139 | 0.687 | -2.177 | 0.9 | 1.498 | 0 | 0.031 | 0.031 | 0 | 0.06 | 0.04 | 0.12 | 0.25 | 0.06 | 0.05 |
| 5 | 74963600 | rs11750595 | -0.492 | -2.195 | 0.847 | 1.368 | 0.005 | 0.026 | 0.026 | 0 | 0.08 | 0.04 | 0.08 | 0.03 | 0.07 | 0.13 |
| 5 | 74964582 | rs40059 | 1.486 | -2.218 | 1.007 | 1.393 | 0.001 | 0.001 | 0.001 | 1.203 | 0.77 | 0.7 | 0.75 | 0.71 | 0.74 | 0.63 |
| 5 | 74965122 | rs34358 | 2.663 | -2.258 | 1.105 | 1.393 | -0.013 | -0.003 | -0.003 | 0 | 0.62 | 0.51 | 0.6 | 0.38 | 0.65 | 0.42 |
| 5 | 74967386 | rs40060 | 2.638 | -2.262 | 1.102 | 1.382 | -0.013 | -0.003 | -0.003 | 0.015 | 0.62 | 0.51 | 0.6 | 0.38 | 0.65 | 0.42 |
| 5 | 74969415 | rs10056022 | -1.532 | -2.22 | 1.224 | 0.875 | -0.012 | 0.009 | 0.009 | 0 | 0.11 | 0.27 | 0.1 | 0.09 | 0.09 | 0.16 |
| 5 | 74970850 | rs888786 | -2.551 | -2.24 | 1.134 | 0.875 | -0.066 | 0.12 | 0.12 | 0 | 0.25 | 0.45 | 0.25 | 0.4 | 0.16 | 0.34 |
| 5 | 74971167 | rs17563512 | -0.307 | -2.21 | 1.018 | 0.854 | 0.005 | 0.026 | 0.026 | 0 | 0.08 | 0.04 | 0.08 | 0.03 | 0.07 | 0.13 |
| 5 | 74972877 | rs2216572 | -2.001 | -2.223 | 1.21 | 0.907 | -0.054 | 0.185 | 0.185 | 0 | 0.18 | 0.5 | 0.18 | 0.37 | 0.1 | 0.24 |
| 5 | 74976105 | rs16872766 | -0.139 | -2.189 | 1.188 | 0.902 | -0.019 | 0.011 | 0.011 | 0 | 0.05 | 0.07 | 0.02 | 0.05 | 0.02 | 0.1 |
| 5 | 74987108 | rs10462356 | -1.194 | -1.999 | 1.121 | 0.78 | -0.014 | 0.043 | 0.043 | 0.116 | 0.22 | 0.15 | 0.24 | 0.15 | 0.26 | 0.34 |
| 5 | 74988336 | rs888788 | 0.369 | -2.052 | 1.083 | 0.749 | -0.022 | 0.026 | 0.026 | 0 | 0.13 | 0.07 | 0.16 | 0.12 | 0.19 | 0.21 |
| 5 | 74997756 | rs10057967 | -2.798 | -2.094 | 1.23 | 0.799 | -0.009 | 0.015 | 0.015 | 1.635 | 0.38 | 0.51 | 0.41 | 0.51 | 0.36 | 0.58 |
| 5 | 75001432 | rs4704234 | -1.489 | -2.099 | 1.307 | 0.817 | 0.007 | -0.039 | -0.039 | 0.436 | 0.23 | 0.37 | 0.26 | 0.19 | 0.26 | 0.37 |
| 5 | 75001582 | rs17672542 | -1.393 | -2.095 | 1.309 | 0.824 | -0.014 | 0.041 | 0.041 | 0 | 0.15 | 0.15 | 0.15 | 0.32 | 0.09 | 0.21 |
| 5 | 75002895 | rs3797581 | 0.173 | -2.089 | 1.302 | 0.824 | 0.034 | -0.011 | -0.011 | 0 | 0.13 | 0.29 | 0.18 | 0.16 | 0.19 | 0.21 |
| 5 | 75003678 | rs2307111 | -3.251 | -2.406 | 1.293 | 0.616 | 0.066 | -0.053 | -0.053 | 0 | 0.5 | 0.92 | 0.48 | 0.53 | 0.4 | 0.6 |
| 5 | 75008193 | rs2047059 | -1.507 | -2.139 | 1.376 | 0.769 | 0.01 | -0.018 | -0.018 | 0 | 0.23 | 0.37 | 0.26 | 0.22 | 0.28 | 0.35 |
| 5 | 75014916 | rs12234025 | -1.492 | -2.054 | 1.234 | 0.77 | -0.014 | 0.043 | 0.043 | 0 | 0.14 | 0.15 | 0.15 | 0.32 | 0.09 | 0.23 |
| 5 | 75020023 | rs17564079 | -0.397 | -1.492 | 0.457 | 0.354 | 0.001 | -0.021 | -0.021 | 0.7 | 0.09 | 0.14 | 0.09 | 0.03 | 0.09 | 0.13 |
| 5 | 75024380 | rs4704235 | -0.181 | -1.586 | 0.064 | 0.302 | -0.014 | 0.028 | 0.028 | 0 | 0.16 | 0.14 | 0.17 | 0.1 | 0.21 | 0.21 |
| 5 | 75034387 | rs7715806 | -1.797 | -1.758 | -0.04 | 0.392 | 0.054 | 0.011 | 0.011 | 1.031 | 0.45 | 0.87 | 0.47 | 0.47 | 0.38 | 0.44 |
| 5 | 75040180 | rs1001935 | -1.106 | -1.808 | 0.948 | -0.026 | -0.051 | -0.076 | -0.076 | 0.006 | 0.17 | 0.38 | 0.17 | 0.05 | 0.08 | 0.08 |
| 5 | 75042913 | rs258496 | 1.813 | -2.057 | 0.186 | -0.117 | -0.093 | -0.03 | -0.03 | 0.396 | 0.37 | 0.06 | 0.45 | 0.35 | 0.5 | 0.4 |
| 5 | 75048465 | rs39656 | 1.391 | -2.456 | 1.893 | 0.163 | 0.008 | 0.001 | 0.001 | 0.442 | 0.5 | 0.54 | 0.56 | 0.47 | 0.57 | 0.56 |
| 5 | 75096559 | rs7725316 | -0.292 | -2.371 | 2.099 | 0.022 | 0 | -0.039 | -0.039 | 0 | 0.33 | 0.27 | 0.24 | 0.53 | 0.36 | 0.34 |
| 5 | 75097385 | rs1525774 | -1.548 | -2.432 | 2.156 | 0.002 | 0.054 | 0.015 | 0.015 | 0.027 | 0.49 | 0.86 | 0.51 | 0.59 | 0.44 | 0.47 |

| | | | | | | | | | | | | | | | | |
|---|---|---|---|---|---|---|---|---|---|---|---|---|---|---|---|---|
| 5 | 75099610 | rs7733891 | 0.657 | -3.076 | 2.193 | 0.003 | 0 | 0 | 0 | 0 | 0.08 | 0.06 | 0.09 | 0.05 | 0.07 | 0.05 |
| 5 | 75103809 | rs1464919 | -0.985 | -2.919 | 2.434 | 0.042 | 0.026 | -0.02 | -0.02 | 0 | 0.39 | 0.74 | 0.27 | 0.53 | 0.32 | 0.38 |
| 5 | 75104101 | rs1997096 | -0.929 | -2.908 | 2.33 | 0.047 | -0.001 | -0.001 | -0.001 | 0 | 0.39 | 0.64 | 0.27 | 0.53 | 0.32 | 0.38 |
| 5 | 75114764 | rs4317299 | -0.666 | -2.705 | 2.069 | 0.15 | -0.026 | 0.05 | 0.05 | 0 | 0.33 | 0.51 | 0.25 | 0.53 | 0.3 | 0.37 |
| 5 | 75122815 | rs2049495 | -0.286 | -2.631 | 1.24 | 0.196 | 0.003 | -0.008 | -0.008 | 0 | 0.29 | 0.34 | 0.21 | 0.52 | 0.29 | 0.33 |
| 5 | 75123529 | rs6453166 | -0.917 | -2.3 | 0.894 | 0.346 | 0.109 | 0.041 | 0.041 | 0 | 0.49 | 0.86 | 0.48 | 0.64 | 0.39 | 0.46 |
| 5 | 75127897 | rs1880648 | -0.974 | -2.028 | 1.636 | 0.45 | -0.031 | 0.001 | 0.001 | 0 | 0.11 | 0.14 | 0.05 | 0.09 | 0.02 | 0.12 |
| 5 | 75132546 | rs7727150 | -1.351 | -2.078 | 1.278 | 0.427 | -0.054 | 0.02 | 0.02 | 0 | 0.6 | 0.92 | 0.52 | 0.69 | 0.46 | 0.59 |
| 5 | 75138443 | rs6891632 | -1.031 | -2.047 | 1.665 | 0.418 | 0.009 | 0.018 | 0.018 | 0 | 0.55 | 0.57 | 0.38 | 0.66 | 0.45 | 0.57 |
| 5 | 75138486 | rs6887352 | -0.29 | -1.899 | 1.005 | 0.503 | 0.01 | -0.003 | -0.003 | 0 | 0.43 | 0.34 | 0.33 | 0.59 | 0.43 | 0.45 |
| 5 | 75160466 | rs6891276 | -1.345 | -1.882 | 0.876 | 0.41 | 0.023 | 0.009 | 0.009 | 4.511 | 0.27 | 0.72 | 0.17 | 0.2 | 0.14 | 0.15 |
| 5 | 75164644 | rs16872966 | -0.927 | -1.906 | 1.649 | 0.395 | -0.06 | -0.021 | -0.021 | 0 | 0.22 | 0.51 | 0.13 | 0.16 | 0.14 | 0.15 |
| 5 | 75174598 | rs6859261 | -1.69 | -1.601 | 1.716 | 0.436 | 0.003 | 0.003 | 0.003 | 0.058 | 0.1 | 0.25 | 0.17 | 0.07 | 0.08 | 0.06 |
| 5 | 75175282 | rs10062196 | -1.614 | -1.749 | 1.364 | 0.486 | -0.036 | 0.027 | 0.027 | 0 | 0.58 | 0.85 | 0.53 | 0.65 | 0.48 | 0.58 |
| 5 | 75181228 | rs6868456 | -1.124 | -1.799 | 2.04 | 0.509 | 0.024 | -0.021 | -0.021 | 0 | 0.11 | 0.36 | 0.18 | 0.03 | 0.08 | 0.08 |
| 5 | 75182449 | rs16872989 | -1.118 | -1.803 | 2.365 | 0.506 | 0.012 | -0.027 | -0.027 | 0.445 | 0.11 | 0.26 | 0.17 | 0.03 | 0.08 | 0.08 |
| 5 | 75191450 | rs4704262 | 0.412 | -1.836 | 1.679 | 0.494 | 0.01 | -0.064 | -0.064 | 0 | 0.29 | 0.1 | 0.24 | 0.43 | 0.28 | 0.39 |
| 5 | 75200884 | rs1525763 | -1.9 | -1.682 | 1.983 | 0.319 | -0.055 | -0.07 | -0.07 | 1.08 | 0.12 | 0.64 | 0.08 | 0 | 0.01 | 0.03 |
| 5 | 75215970 | rs17564993 | -0.737 | -1.516 | 1.481 | 0.433 | -0.001 | 0 | 0 | 0.999 | 0.21 | 0.24 | 0.24 | 0.18 | 0.21 | 0.16 |
| 5 | 75216411 | rs10942747 | -1.164 | -1.55 | 1.079 | 0.463 | 0.041 | 0.044 | 0.044 | 0 | 0.54 | 0.81 | 0.39 | 0.6 | 0.42 | 0.52 |
| 5 | 75220355 | rs10061050 | -1.316 | -1.468 | 1.187 | 0.516 | -0.001 | 0.005 | 0.005 | 0 | 0.57 | 0.63 | 0.53 | 0.64 | 0.5 | 0.57 |
| 5 | 75220933 | rs1917759 | -0.728 | -1.566 | 0.568 | 0.614 | 0.006 | -0.009 | -0.009 | 0 | 0.22 | 0.22 | 0.25 | 0.23 | 0.21 | 0.17 |
| 5 | 75226673 | rs10045727 | -0.257 | -1.47 | 0.317 | 0.425 | 0.017 | 0.004 | 0.004 | 0 | 0.23 | 0.13 | 0.12 | 0.19 | 0.19 | 0.19 |
| 5 | 75229614 | rs4487476 | -0.987 | -1.233 | 0.026 | 0.495 | -0.008 | 0.036 | 0.036 | 0.061 | 0.55 | 0.78 | 0.52 | 0.6 | 0.48 | 0.55 |
| 5 | 75230527 | rs7729037 | 0.495 | -1.131 | -0.07 | 0.445 | -0.037 | -0.122 | -0.122 | 0.197 | 0.28 | 0.06 | 0.35 | 0.41 | 0.29 | 0.35 |
| 5 | 75235544 | rs4704272 | -1.108 | -1.049 | -0.069 | 0.445 | -0.042 | 0.063 | 0.063 | 0 | 0.58 | 0.86 | 0.55 | 0.66 | 0.49 | 0.55 |
| 5 | 75239130 | rs6895132 | -0.14 | -0.509 | -0.336 | 0.534 | -0.034 | -0.021 | -0.021 | 0 | 0.25 | 0.44 | 0.22 | 0.17 | 0.21 | 0.18 |
| 5 | 75240469 | rs1525783 | -0.358 | 0.284 | -1.416 | 0.939 | -0.024 | -0.101 | -0.101 | 0.006 | 0.27 | 0.07 | 0.31 | 0.41 | 0.29 | 0.35 |
| 5 | 75244914 | rs10051972 | -0.079 | 0.483 | -1.015 | 0.257 | -0.002 | 0.003 | 0.003 | 0 | 0.4 | 0.39 | 0.43 | 0.3 | 0.42 | 0.34 |
| 5 | 75246226 | rs10805896 | -0.063 | 0.071 | -0.8 | -0.779 | 0.001 | 0.043 | 0.043 | 0 | 0.51 | 0.59 | 0.32 | 0.09 | 0.46 | 0.3 |
| 5 | 75249163 | rs10058719 | 0.114 | -0.17 | -1.004 | -1.143 | 0.002 | -0.059 | -0.059 | 0 | 0.38 | 0.52 | 0.24 | 0.07 | 0.35 | 0.11 |
| 5 | 75249499 | rs10079572 | 0.598 | -0.571 | -1.004 | -1.34 | -0.094 | 0.151 | 0.151 | 0 | 0.29 | 0.09 | 0.18 | 0 | 0.34 | 0.09 |
| 5 | 75254943 | rs4704273 | -0.991 | -0.983 | -1.257 | -1.753 | -0.027 | 0.048 | 0.048 | 0 | 0.69 | 0.82 | 0.5 | 0.15 | 0.65 | 0.41 |
| 5 | 75260512 | rs11747018 | 0.348 | -0.833 | -1.161 | -1.601 | 0.003 | -0.071 | -0.071 | 0 | 0.38 | 0.51 | 0.24 | 0.07 | 0.35 | 0.11 |
| 5 | 75269747 | rs4704275 | 0.558 | -0.827 | -0.488 | -1.016 | 0.009 | 0.024 | 0.024 | 0 | 0.18 | 0.14 | 0.18 | 0.08 | 0.2 | 0.12 |
| 5 | 75273274 | rs7731991 | -0.475 | -0.502 | -0.121 | -0.633 | -0.022 | 0.049 | 0.049 | 0.05 | 0.09 | 0.44 | 0.06 | 0.07 | 0.01 | 0.02 |
| 5 | 75281297 | rs12055083 | -0.473 | -0.079 | -0.135 | 0.117 | 0.012 | 0.13 | 0.13 | 0 | 0.13 | 0.28 | 0.32 | 0.59 | 0.14 | 0.19 |
| 5 | 75281554 | rs6867123 | 0.163 | -0.067 | -0.145 | -0.212 | 0.002 | 0.004 | 0.004 | 0 | 0.12 | 0.11 | 0.07 | 0.05 | 0.12 | 0.09 |
| 5 | 75283032 | rs6414964 | -0.538 | 0.491 | 0.201 | -0.376 | 0.034 | 0.023 | 0.023 | 0.979 | 0.81 | 0.72 | 0.81 | 0.72 | 0.79 | 0.67 |
| 5 | 75291817 | rs12657643 | -0.554 | 0.404 | 0.068 | -0.281 | 0.003 | 0.042 | 0.042 | 0.101 | 0.18 | 0.26 | 0.33 | 0.62 | 0.17 | 0.28 |
| 5 | 75294346 | rs4425488 | 1.161 | 0.114 | 0.001 | -0.688 | 0.004 | 0.046 | 0.046 | 0.162 | 0.52 | 0.38 | 0.38 | 0.11 | 0.53 | 0.24 |
| 5 | 75294410 | rs10214163 | -1.084 | 0.69 | -0.453 | -0.433 | -0.015 | -0.028 | -0.028 | 0 | 0.85 | 0.9 | 0.81 | 0.74 | 0.83 | 0.69 |
| 5 | 75297146 | rs13180297 | -0.517 | 0.422 | -0.152 | 0.066 | 0.001 | 0.014 | 0.014 | 0 | 0.16 | 0.19 | 0.33 | 0.64 | 0.16 | 0.27 |
| 5 | 75302629 | rs13165217 | -0.59 | -0.063 | -0.194 | 0.482 | -0.008 | 0.03 | 0.03 | 0 | 0.35 | 0.44 | 0.38 | 0.45 | 0.37 | 0.41 |
| 5 | 75304106 | rs7735728 | -0.723 | -0.194 | -0.063 | 0.74 | -0.023 | -0.002 | -0.002 | 0.23 | 0.05 | 0.11 | 0.02 | 0.04 | 0.02 | 0.04 |
| 5 | 75305574 | rs4235691 | 0.292 | 0.259 | -0.11 | 0.586 | 0 | -0.003 | -0.003 | 0 | 0.63 | 0.57 | 0.5 | 0.34 | 0.6 | 0.38 |
| 5 | 75306047 | rs4490566 | -0.061 | 0.236 | -0.389 | 1.154 | 0.011 | -0.041 | -0.041 | 0 | 0.22 | 0.19 | 0.34 | 0.45 | 0.17 | 0.22 |
| 5 | 75315327 | rs4374745 | 0.881 | 0.199 | -0.364 | 1.092 | -0.017 | 0.023 | 0.023 | 0 | 0.63 | 0.74 | 0.52 | 0.31 | 0.6 | 0.38 |
| 5 | 75320034 | rs9293664 | -0.829 | 0.267 | -0.544 | 1.231 | -0.07 | -0.065 | -0.065 | 0 | 0.87 | 0.99 | 0.86 | 0.81 | 0.78 | 0.63 |
| 5 | 75322493 | rs1501903 | -0.868 | -0.055 | 0.316 | 1.238 | -0.024 | 0.039 | 0.039 | 0.888 | 0.31 | 0.45 | 0.37 | 0.49 | 0.23 | 0.29 |
| 5 | 75337594 | rs10044546 | -0.029 | -0.094 | 0.336 | 1.26 | 0.017 | -0.064 | -0.064 | 0.037 | 0.24 | 0.14 | 0.31 | 0.44 | 0.19 | 0.25 |
| 5 | 75365799 | rs258612 | -1.475 | -0.212 | 0.779 | 1.123 | -0.001 | 0.048 | 0.048 | 0 | 0.43 | 0.32 | 0.52 | 0.7 | 0.45 | 0.67 |
| 5 | 75378769 | rs30198 | -1.048 | -0.219 | 0.731 | 1.088 | -0.026 | 0.043 | 0.043 | 0.013 | 0.38 | 0.22 | 0.49 | 0.7 | 0.42 | 0.62 |
| 5 | 75381803 | rs28621 | -1.585 | -0.33 | 1.201 | 1.022 | 0.008 | 0.064 | 0.064 | 1.811 | 0.46 | 0.7 | 0.54 | 0.7 | 0.45 | 0.66 |
| 5 | 75395063 | rs1423102 | 0.127 | -0.39 | 0.987 | 1.116 | -0.068 | -0.064 | -0.064 | 0 | 0.18 | 0.02 | 0.17 | 0.23 | 0.25 | 0.44 |
| 5 | 75395725 | rs7702044 | 0.019 | -0.366 | 1.038 | 1.139 | 0.018 | -0.058 | -0.058 | 0 | 0.24 | 0.15 | 0.34 | 0.45 | 0.2 | 0.24 |
| 5 | 75396236 | rs2731735 | -1.596 | -0.368 | 1.093 | 1.119 | 0.009 | 0.048 | 0.048 | 2.113 | 0.46 | 0.67 | 0.55 | 0.72 | 0.45 | 0.69 |
| 5 | 75404137 | rs17673786 | -0.517 | -0.402 | 1.107 | 1.12 | -0.013 | 0.019 | 0.019 | 0.489 | 0.28 | 0.67 | 0.39 | 0.45 | 0.2 | 0.24 |
| 5 | 75404185 | rs187727 | 0.233 | -0.355 | 0.679 | 1.062 | -0.069 | -0.053 | -0.053 | 0.005 | 0.17 | 0.02 | 0.15 | 0.21 | 0.25 | 0.44 |
| 5 | 75407667 | rs10063621 | -0.455 | -0.443 | 1.034 | 1.216 | -0.029 | 0.072 | 0.072 | 0 | 0.28 | 0.6 | 0.37 | 0.49 | 0.2 | 0.25 |
| 5 | 75408908 | rs17650787 | 1.05 | -0.03 | -0.325 | 0.525 | -0.017 | 0.047 | 0.047 | 1.442 | 0.13 | 0.06 | 0.08 | 0 | 0.15 | 0.04 |
| 5 | 75409553 | rs13160383 | -0.219 | -0.273 | -0.237 | 0.767 | 0.03 | -0.046 | -0.046 | 0 | 0.29 | 0.18 | 0.32 | 0.44 | 0.23 | 0.31 |
| 5 | 75410631 | rs183766 | -1.069 | -0.218 | -0.379 | 0.665 | -0.102 | -0.048 | -0.048 | 0.037 | 0.68 | 0.98 | 0.68 | 0.71 | 0.67 | 0.72 |
| 5 | 75422106 | rs10434645 | -0.213 | -0.444 | 0.144 | 1.097 | 0 | 0.012 | 0.012 | 0 | 0.17 | 0.2 | 0.27 | 0.46 | 0.18 | 0.21 |
| 5 | 75427518 | rs1423099 | 0.929 | -0.93 | -0.302 | 0.798 | 0.062 | 0.011 | 0.011 | 0.063 | 0.63 | 0.17 | 0.43 | 0.32 | 0.74 | 0.64 |
| 5 | 75427935 | rs10070440 | 0.821 | -0.874 | -0.019 | 0.795 | 0.032 | 0.006 | 0.006 | 0.106 | 0.64 | 0.26 | 0.45 | 0.32 | 0.74 | 0.66 |
| 5 | 75429603 | rs2112865 | 0.462 | -0.749 | 0.618 | 0.666 | -0.004 | 0.003 | 0.003 | 0 | 0.13 | 0.15 | 0.24 | 0.2 | 0.1 | 0.11 |
| 5 | 75429982 | rs7706479 | -0.425 | -0.788 | 0.729 | 0.65 | -0.042 | 0.003 | 0.003 | 0 | 0.19 | 0.57 | 0.3 | 0.2 | 0.1 | 0.11 |
| 5 | 75431699 | rs12522597 | -0.16 | -0.098 | -0.188 | 0.042 | -0.005 | 0.052 | 0.052 | 1.506 | 0.54 | 0.65 | 0.55 | 0.67 | 0.48 | 0.59 |

| | | | | | | | | | | | | | | | | |
|---|---|---|---|---|---|---|---|---|---|---|---|---|---|---|---|---|
| 5 | 75441876 | rs10051982 | -0.941 | -0.406 | 0.247 | 0.6 | -0.01 | 0.005 | 0.005 | 0 | 0.87 | 0.83 | 0.81 | 0.85 | 0.93 | 0.93 |
| 5 | 75443228 | rs6873987 | 0.288 | -0.373 | -0.29 | -0.035 | -0.031 | -0.001 | -0.001 | 0.071 | 0.43 | 0.21 | 0.43 | 0.43 | 0.49 | 0.42 |
| 5 | 75445925 | rs1895391 | 0.187 | -0.431 | -0.194 | -0.051 | -0.024 | 0.005 | 0.005 | 0.906 | 0.44 | 0.24 | 0.45 | 0.43 | 0.5 | 0.43 |
| 5 | 75446430 | rs6866404 | 0.532 | -0.345 | -0.203 | -0.125 | -0.077 | 0.214 | 0.214 | 0.166 | 0.4 | 0.09 | 0.26 | 0.03 | 0.47 | 0.33 |
| 5 | 75447123 | rs12514991 | 0.175 | -0.272 | -0.166 | -0.065 | -0.025 | 0.007 | 0.007 | 0.92 | 0.44 | 0.24 | 0.45 | 0.43 | 0.5 | 0.43 |
| 5 | 75450127 | rs6879305 | -0.037 | -0.19 | -0.318 | -0.67 | 0 | 0 | 0 | 0 | 0.46 | 0.5 | 0.4 | 0.42 | 0.47 | 0.53 |
| 5 | 75450373 | rs6879785 | 0.769 | -0.189 | -0.279 | -0.654 | 0.004 | 0.015 | 0.015 | 0 | 0.31 | 0.41 | 0.32 | 0.42 | 0.35 | 0.45 |
| 5 | 75458401 | rs4610441 | -0.667 | -0.097 | -0.365 | -0.759 | -0.032 | 0.103 | 0.103 | 0 | 0.61 | 0.82 | 0.76 | 0.97 | 0.56 | 0.67 |
| 5 | 75462736 | rs10072459 | 0.207 | -0.158 | -0.472 | -0.6 | -0.001 | -0.002 | -0.002 | 0 | 0.45 | 0.45 | 0.39 | 0.45 | 0.47 | 0.51 |
| 5 | 75464430 | rs6453208 | -0.236 | 0.295 | -1.043 | -1.294 | -0.024 | 0.101 | 0.101 | 0 | 0.61 | 0.82 | 0.76 | 0.91 | 0.57 | 0.67 |
| 5 | 75471717 | rs1553325 | -0.334 | 0.342 | -1.408 | -1.31 | -0.051 | 0.173 | 0.173 | 0 | 0.62 | 0.88 | 0.78 | 0.97 | 0.58 | 0.67 |
| 5 | 75479786 | rs2358531 | -0.358 | 0.674 | -1.335 | -0.999 | -0.026 | 0.063 | 0.063 | 0.29 | 0.76 | 0.94 | 0.87 | 0.9 | 0.72 | 0.69 |
| 5 | 75482902 | rs1002541 | 0.356 | 0.663 | -1.41 | -1.145 | -0.026 | 0.161 | 0.161 | 0.757 | 0.61 | 0.88 | 0.79 | 0.9 | 0.59 | 0.62 |
| 5 | 75485855 | rs12153396 | 0.356 | 0.453 | -1.037 | -0.552 | -0.006 | 0.021 | 0.021 | 0 | 0.14 | 0.05 | 0.07 | 0.06 | 0.13 | 0.08 |
| 5 | 75492702 | rs12655684 | -0.201 | 0.436 | -0.582 | -0.667 | -0.003 | -0.015 | -0.015 | 0.05 | 0.19 | 0.24 | 0.12 | 0.07 | 0.17 | 0.15 |
| 5 | 75495349 | rs7445050 | -0.473 | 0.657 | -0.633 | -0.866 | -0.037 | 0.037 | 0.037 | 0.489 | 0.67 | 0.78 | 0.7 | 0.91 | 0.52 | 0.71 |
| 5 | 75496098 | rs6859341 | 0.704 | 0.496 | -0.632 | -0.915 | 0.027 | 0.008 | 0.008 | 5.728 | 0.39 | 0.24 | 0.37 | 0.6 | 0.31 | 0.41 |
| 5 | 75504243 | rs17566351 | 0.434 | 0.325 | -0.864 | -0.784 | 0.003 | -0.004 | -0.004 | 0 | 0.2 | 0.15 | 0.12 | 0 | 0.18 | 0.14 |
| 5 | 75509233 | rs12522470 | 0.217 | 0.077 | -0.758 | -0.378 | 0.001 | 0.001 | 0.001 | 4.93 | 0.41 | 0.41 | 0.38 | 0.6 | 0.29 | 0.41 |
| 5 | 75514986 | rs11960832 | -0.407 | 0.001 | -1.009 | -0.151 | 0.052 | 0.033 | 0.033 | 1.563 | 0.42 | 0.75 | 0.46 | 0.31 | 0.29 | 0.31 |
| 5 | 75515187 | rs13171899 | -0.256 | 0.831 | -1.267 | 0.347 | -0.051 | -0.045 | -0.045 | 0 | 0.12 | 0.04 | 0.25 | 0.22 | 0.26 | 0.12 |
| 5 | 75518746 | rs12514992 | -0.822 | 0.253 | -0.539 | 0.214 | -0.001 | 0.015 | 0.015 | 0 | 0.27 | 0.58 | 0.36 | 0.33 | 0.28 | 0.27 |
| 5 | 75520928 | rs981113 | -0.523 | 0.2 | -0.418 | 0.21 | 0.014 | 0.004 | 0.004 | 2.636 | 0.51 | 0.76 | 0.69 | 0.53 | 0.58 | 0.36 |
| 5 | 75521980 | rs10068332 | -0.615 | 0.174 | -0.145 | 0.24 | 0.004 | 0.001 | 0.001 | 1.192 | 0.45 | 0.53 | 0.61 | 0.53 | 0.54 | 0.35 |
| 5 | 75523779 | rs7448883 | -0.754 | 0.08 | -0.116 | 0.164 | 0.012 | 0.02 | 0.02 | 0 | 0.53 | 0.7 | 0.43 | 0.37 | 0.48 | 0.43 |
| 5 | 75524778 | rs7443699 | -0.32 | -0.096 | 0.287 | 0.062 | -0.071 | -0.095 | -0.095 | 0.178 | 0.22 | 0.47 | 0.1 | 0.07 | 0.11 | 0.22 |
| 5 | 75525141 | rs1553322 | -0.709 | -0.009 | 0.423 | 0.133 | 0.024 | 0.009 | 0.009 | 1.522 | 0.07 | 0.41 | 0.05 | 0.01 | 0.01 | 0.05 |
| 5 | 75525786 | rs6885213 | 0.353 | -0.036 | 0.239 | 0.139 | -0.016 | -0.021 | -0.021 | 0 | 0.3 | 0.18 | 0.32 | 0.3 | 0.37 | 0.21 |
| 5 | 75526922 | rs11750299 | 0.359 | -0.029 | 0.242 | 0.17 | -0.016 | -0.025 | -0.025 | 0 | 0.3 | 0.18 | 0.33 | 0.31 | 0.37 | 0.21 |
| 5 | 75534570 | rs1532699 | -0.353 | -0.008 | 0.166 | 0.175 | -0.001 | -0.001 | -0.001 | 0 | 0.38 | 0.63 | 0.38 | 0.33 | 0.38 | 0.26 |
| 5 | 75540841 | rs11750504 | 0.28 | 0.082 | -0.349 | 0.213 | -0.036 | -0.013 | -0.013 | 0 | 0.29 | 0.14 | 0.32 | 0.25 | 0.37 | 0.21 |
| 5 | 75545035 | rs6879020 | -0.539 | 0.026 | 0.028 | 0.131 | 0.009 | 0.014 | 0.014 | 0 | 0.48 | 0.39 | 0.38 | 0.37 | 0.47 | 0.38 |
| 5 | 75549507 | rs6891999 | -0.656 | 0.588 | -0.778 | 0.462 | -0.052 | -0.049 | -0.049 | 0 | 0.91 | 0.97 | 0.69 | 0.77 | 0.79 | 0.9 |
| 5 | 75550325 | rs12654150 | 1.271 | 0.365 | -0.576 | 0.403 | 0.018 | -0.016 | -0.016 | 0 | 0.37 | 0.14 | 0.25 | 0.4 | 0.32 | 0.47 |
| 5 | 75552105 | rs10462538 | -0.015 | 0.477 | -0.812 | 0.739 | -0.016 | -0.005 | -0.005 | 1.837 | 0.27 | 0.14 | 0.28 | 0.24 | 0.3 | 0.2 |
| 5 | 75552501 | rs17651538 | -0.174 | 0.832 | -0.274 | 1.062 | 0.004 | 0.008 | 0.008 | 0.049 | 0.92 | 0.91 | 0.7 | 0.76 | 0.8 | 0.92 |
| 5 | 75562798 | rs10064140 | -0.71 | 0.494 | -0.257 | 1.027 | -0.006 | 0.002 | 0.002 | 0 | 0.4 | 0.57 | 0.33 | 0.39 | 0.31 | 0.33 |
| 5 | 75563413 | rs6453212 | 0.111 | 0.425 | -0.332 | 1.17 | -0.012 | -0.001 | -0.001 | 0.018 | 0.19 | 0.24 | 0.09 | 0.12 | 0.11 | 0.17 |
| 5 | 75563830 | rs6882321 | 0.126 | 0.433 | -0.328 | 1.158 | -0.012 | -0.002 | -0.002 | 0 | 0.19 | 0.24 | 0.09 | 0.13 | 0.11 | 0.17 |
| 5 | 75564181 | rs2937720 | 0.525 | 0.608 | -0.387 | 1.152 | -0.002 | -0.007 | -0.007 | 0.007 | 0.72 | 0.73 | 0.62 | 0.65 | 0.7 | 0.72 |
| 5 | 75564437 | rs2937719 | 1.044 | 0.114 | -0.266 | 0.675 | -0.002 | 0.026 | 0.026 | 0 | 0.51 | 0.37 | 0.38 | 0.38 | 0.5 | 0.56 |
| 5 | 75566506 | rs2972831 | -0.571 | -0.433 | -0.296 | 0.513 | -0.012 | -0.008 | -0.008 | 0 | 0.21 | 0.25 | 0.09 | 0.11 | 0.12 | 0.19 |
| 5 | 75568053 | rs2937715 | 0.139 | -0.225 | -0.576 | 0.539 | 0.028 | 0.004 | 0.004 | 0 | 0.48 | 0.47 | 0.37 | 0.6 | 0.39 | 0.54 |
| 5 | 75568179 | rs16873292 | -0.476 | -0.115 | -0.326 | 0.244 | -0.005 | -0.031 | -0.031 | 0.271 | 0.11 | 0.11 | 0.04 | 0 | 0.07 | 0.06 |
| 5 | 75572032 | rs2937748 | -0.946 | -0.106 | 0.238 | 0.495 | 0.005 | 0.055 | 0.055 | 0.814 | 0.09 | 0.32 | 0.15 | 0.09 | 0.09 | 0.03 |
| 5 | 75574855 | rs6874435 | -1.763 | -0.286 | 0.432 | 0.282 | -0.02 | -0.018 | -0.018 | 0.052 | 0.17 | 0.28 | 0.17 | 0.09 | 0.13 | 0.12 |
| 5 | 75580687 | rs2358712 | -0.048 | -0.281 | 0.418 | 0.3 | 0.014 | 0.055 | 0.055 | 0 | 0.06 | 0.16 | 0.14 | 0.09 | 0.08 | 0.02 |
| 5 | 75586936 | rs17566680 | 0.572 | 0.128 | -0.219 | -0.364 | -0.05 | 0.08 | 0.08 | 0 | 0.13 | 0.02 | 0.1 | 0 | 0.21 | 0.05 |
| 5 | 75589021 | rs17747572 | 0.837 | 0 | -0.621 | 0.486 | -0.027 | -0.058 | -0.058 | 0 | 0.09 | 0.01 | 0.05 | 0.28 | 0.07 | 0.28 |
| 5 | 75594360 | rs31243 | -1.654 | -0.291 | -0.121 | 0.603 | -0.031 | 0.013 | 0.013 | 0.035 | 0.17 | 0.3 | 0.16 | 0.17 | 0.11 | 0.13 |
| 5 | 75594743 | rs31244 | 0.497 | -0.078 | -0.238 | 0.642 | 0.02 | 0.058 | 0.058 | 0 | 0.05 | 0.16 | 0.14 | 0.09 | 0.08 | 0.02 |
| 5 | 75602703 | rs2937746 | -0.115 | -0.165 | -0.442 | 0.728 | -0.001 | 0.017 | 0.017 | 0 | 0.07 | 0.11 | 0.15 | 0.11 | 0.08 | 0.03 |
| 5 | 75604851 | rs2972844 | -0.245 | 0.004 | -0.986 | 0.368 | -0.042 | 0.018 | 0.018 | 0 | 0.56 | 0.83 | 0.63 | 0.65 | 0.46 | 0.42 |
| 5 | 75606708 | rs4703705 | 1.153 | -0.008 | -0.851 | 0.212 | -0.003 | -0.022 | -0.022 | 0 | 0.35 | 0.19 | 0.4 | 0.46 | 0.31 | 0.29 |
| 5 | 75608993 | rs2913261 | 0.499 | 0.01 | -0.845 | 0.207 | 0 | -0.01 | -0.01 | 0.886 | 0.42 | 0.3 | 0.5 | 0.58 | 0.41 | 0.32 |
| 5 | 75615107 | rs2012412 | 0.898 | 0.222 | -0.897 | -0.037 | 0.003 | -0.018 | -0.018 | 3.706 | 0.3 | 0.22 | 0.31 | 0.47 | 0.26 | 0.28 |
| 5 | 75622814 | rs3733860 | 1.098 | 0.411 | -1.247 | -0.083 | -0.028 | -0.058 | -0.058 | 0 | 0.1 | 0.01 | 0.23 | 0.46 | 0.11 | 0.13 |
| 5 | 75623689 | rs2913251 | 0.14 | 0.492 | -0.962 | -0.508 | -0.005 | 0.007 | 0.007 | 0.601 | 0.6 | 0.54 | 0.62 | 0.41 | 0.64 | 0.64 |
| 5 | 75623775 | rs6892721 | -0.789 | 0.476 | -1.022 | -0.583 | -0.017 | -0.05 | -0.05 | 0.676 | 0.73 | 0.84 | 0.7 | 0.53 | 0.74 | 0.7 |
| 5 | 75624514 | rs17651829 | 0.782 | 0.759 | -1.527 | -1.136 | -0.089 | 0.053 | 0.053 | 0 | 0.23 | 0.02 | 0.23 | 0.11 | 0.27 | 0.25 |
| 5 | 75625414 | rs31270 | -0.041 | 0.564 | -0.787 | -1.223 | 0.003 | 0.035 | 0.035 | 0.259 | 0.54 | 0.41 | 0.48 | 0.14 | 0.59 | 0.45 |
| 5 | 75625952 | rs31269 | 0.453 | 0.783 | -1.312 | -1.138 | -0.078 | 0.214 | 0.214 | 0.019 | 0.41 | 0.16 | 0.44 | 0.09 | 0.52 | 0.27 |
| 5 | 75632745 | rs2913248 | 0.15 | 0.548 | -1.581 | -1.034 | 0.025 | 0.024 | 0.024 | 0.151 | 0.5 | 0.18 | 0.56 | 0.39 | 0.59 | 0.5 |
| 5 | 75637562 | rs246819 | 0.519 | 0.457 | -1.447 | -2.038 | -0.014 | 0.206 | 0.206 | 0.772 | 0.48 | 0.15 | 0.55 | 0.84 | 0.57 | 0.6 |
| 5 | 75642486 | rs246804 | -0.656 | 0.445 | -1.451 | -2.043 | -0.022 | 0.196 | 0.196 | 0.905 | 0.53 | 0.85 | 0.44 | 0.16 | 0.43 | 0.42 |
| 5 | 75647630 | rs246797 | 0.161 | 0.344 | -1.076 | -2.358 | 0.016 | 0.086 | 0.086 | 0 | 0.38 | 0.25 | 0.35 | 0.05 | 0.31 | 0.26 |
| 5 | 75648199 | rs246794 | -0.86 | 0.332 | -0.739 | -2.328 | 0.009 | 0.072 | 0.072 | 0 | 0.49 | 0.64 | 0.46 | 0.13 | 0.42 | 0.47 |
| 5 | 75651786 | rs11748035 | -0.006 | -0.083 | -0.43 | -1.985 | 0.052 | 0.046 | 0.046 | 1.755 | 0.12 | 0.44 | 0.13 | 0.09 | 0.1 | 0.2 |
| 5 | 75659679 | rs405030 | 0.647 | -0.31 | -1.3 | -2.025 | 0.019 | 0.026 | 0.026 | 0 | 0.24 | 0.18 | 0.14 | 0.02 | 0.18 | 0.17 |

| | | | | | | | | | | | | | | | | |
|---|---|---|---|---|---|---|---|---|---|---|---|---|---|---|---|---|
| 5 | 75669542 | rs246783 | -0.593 | 0.315 | -1.887 | -1.986 | -0.041 | 0.063 | 0.063 | 0 | 0.66 | 0.87 | 0.53 | 0.16 | 0.61 | 0.48 |
| 5 | 75675319 | rs10514065 | -0.468 | 0.053 | -1.621 | -1.878 | 0.003 | -0.057 | -0.057 | 2.09 | 0.28 | 0.32 | 0.26 | 0.02 | 0.33 | 0.23 |
| 5 | 75676658 | rs1811340 | 0.036 | 0.523 | -2.458 | -1.317 | -0.068 | 0.158 | 0.158 | 0 | 0.28 | 0.02 | 0.14 | 0.04 | 0.26 | 0.15 |
| 5 | 75684160 | rs890708 | 0 | 0.565 | -1.789 | -0.589 | -0.025 | 0.057 | 0.057 | 0.578 | 0.65 | 0.87 | 0.56 | 0.2 | 0.58 | 0.57 |
| 5 | 75692976 | rs875543 | -0.545 | 0.346 | -1.158 | 0.447 | 0.003 | 0.083 | 0.083 | 0 | 0.14 | 0.26 | 0.16 | 0.48 | 0.12 | 0.19 |
| 5 | 75698291 | rs1984636 | -0.342 | 0.352 | -1.122 | 0.443 | 0.005 | 0.091 | 0.091 | 0 | 0.13 | 0.22 | 0.16 | 0.48 | 0.12 | 0.19 |
| 5 | 75700310 | rs9293683 | -0.575 | 0.208 | -2.108 | -0.511 | -0.018 | 0.133 | 0.133 | 0 | 0.73 | 0.89 | 0.8 | 0.99 | 0.67 | 0.81 |
| 5 | 75710783 | rs4296785 | -0.458 | -0.663 | -2.432 | -0.573 | 0.079 | -0.071 | -0.071 | 1.28 | 0.36 | 0.92 | 0.38 | 0.55 | 0.17 | 0.4 |
| 5 | 75718104 | rs11747835 | 0.852 | -0.35 | -2.298 | -0.946 | -0.04 | 0.077 | 0.077 | 0 | 0.13 | 0.01 | 0.1 | 0 | 0.17 | 0.05 |
| 5 | 75722520 | rs4563602 | 0.18 | -0.886 | -1.246 | -0.554 | -0.082 | -0.075 | -0.075 | 0.776 | 0.21 | 0.56 | 0.2 | 0.06 | 0.05 | 0.23 |
| 5 | 75738858 | rs11743541 | -0.036 | 0.253 | -1.44 | 0.243 | -0.092 | -0.088 | -0.088 | 0.044 | 0.4 | 0.06 | 0.43 | 0.55 | 0.5 | 0.38 |
| 5 | 75746813 | rs6880993 | 0.651 | 0.095 | 0.006 | 0.731 | -0.021 | 0.037 | 0.037 | 0 | 0.87 | 0.78 | 0.88 | 0.77 | 0.92 | 0.75 |
| 5 | 75747296 | rs10067617 | -0.882 | 0.851 | -0.589 | 0.051 | -0.039 | 0.034 | 0.034 | 1.727 | 0.28 | 0.83 | 0.31 | 0.23 | 0.32 | 0.4 |
| 5 | 75751461 | rs6898214 | 0.07 | 0.679 | -0.615 | -0.222 | 0 | 0.042 | 0.042 | 2.19 | 0.56 | 0.67 | 0.33 | 0.25 | 0.45 | 0.24 |
| 5 | 75753324 | rs10474476 | 0.515 | 0.625 | 0.299 | 0.114 | -0.011 | -0.004 | -0.004 | 0.802 | 0.26 | 0.29 | 0.15 | 0.23 | 0.19 | 0.15 |
| 5 | 75757556 | rs10045155 | 0.512 | 0.751 | -0.039 | 0.258 | -0.005 | -0.023 | -0.023 | 1.158 | 0.18 | 0.11 | 0.37 | 0.49 | 0.23 | 0.25 |
| 5 | 75759072 | rs4529172 | 0.7 | 0.707 | 0.238 | 0.131 | -0.012 | -0.004 | -0.004 | 0 | 0.27 | 0.31 | 0.16 | 0.24 | 0.19 | 0.15 |
| 5 | 75760174 | rs10035095 | 0.908 | 0.761 | -0.213 | 0.157 | 0.004 | -0.03 | -0.03 | 0 | 0.25 | 0.06 | 0.13 | 0.24 | 0.19 | 0.15 |
| 5 | 75760204 | rs4452539 | 0.198 | 0.976 | 0.309 | -0.378 | -0.006 | 0.073 | 0.073 | 0 | 0.44 | 0.28 | 0.3 | 0.03 | 0.47 | 0.21 |
| 5 | 75768739 | rs13168171 | -0.225 | 0.839 | -0.085 | -0.164 | -0.001 | -0.049 | -0.049 | 1.323 | 0.53 | 0.54 | 0.36 | 0.08 | 0.57 | 0.39 |
| 5 | 75779471 | rs2068434 | -0.139 | 0.954 | -0.827 | 0.437 | 0 | -0.017 | -0.017 | 0 | 0.59 | 0.52 | 0.63 | 0.72 | 0.58 | 0.63 |
| 5 | 75783886 | rs4703710 | -0.4 | 1.337 | -1.228 | 0.913 | 0.007 | 0.006 | 0.006 | 0 | 0.36 | 0.37 | 0.4 | 0.59 | 0.41 | 0.58 |
| 5 | 75795407 | rs7727095 | -0.314 | 1.263 | -2.229 | 0.59 | -0.088 | -0.058 | -0.058 | 0 | 0.43 | 0.13 | 0.61 | 0.54 | 0.48 | 0.43 |
| 5 | 75801337 | rs7710225 | 0.432 | 1.475 | -2.184 | 0.146 | -0.054 | -0.003 | -0.003 | 0 | 0.64 | 0.83 | 0.39 | 0.41 | 0.53 | 0.42 |
| 5 | 75801583 | rs10054825 | 0.008 | 0.82 | -1.209 | -0.143 | -0.008 | -0.076 | -0.076 | 0 | 0.15 | 0.32 | 0.12 | 0.01 | 0.11 | 0.07 |
| 5 | 75801845 | rs7711045 | 0.22 | 1.18 | -1.507 | 0.18 | 0 | -0.008 | -0.008 | 0.119 | 0.81 | 0.83 | 0.7 | 0.37 | 0.77 | 0.61 |
| 5 | 75805105 | rs6453227 | 0.022 | 1.19 | -1.529 | 0.097 | 0.001 | -0.006 | -0.006 | 0.492 | 0.83 | 0.86 | 0.76 | 0.46 | 0.8 | 0.63 |
| 5 | 75809208 | rs10079855 | 0.062 | 0.543 | -1.208 | -0.981 | -0.027 | -0.055 | -0.055 | 0 | 0.2 | 0.39 | 0.14 | 0.05 | 0.13 | 0.16 |
| 5 | 75810243 | rs4516825 | -0.937 | 1.117 | -1.635 | -0.799 | -0.001 | -0.003 | -0.003 | 0 | 0.25 | 0.24 | 0.3 | 0.89 | 0.25 | 0.51 |
| 5 | 75820998 | rs6453230 | -0.312 | 1.411 | -2.282 | -0.827 | -0.107 | 0.084 | 0.084 | 0 | 0.81 | 0.98 | 0.62 | 0.96 | 0.7 | 0.86 |
| 5 | 75822474 | rs4704322 | -0.566 | 0.443 | -1.459 | -0.002 | -0.002 | 0.011 | 0.011 | 0 | 0.16 | 0.08 | 0.23 | 0.88 | 0.19 | 0.38 |
| 5 | 75826154 | rs10474479 | -0.88 | 0.521 | -1.381 | 0.153 | -0.001 | 0.012 | 0.012 | 0.165 | 0.2 | 0.09 | 0.23 | 0.86 | 0.2 | 0.44 |
| 5 | 75827508 | rs4704325 | -0.795 | 0.468 | -1.354 | 0.153 | 0.001 | 0.01 | 0.01 | 0.238 | 0.19 | 0.09 | 0.23 | 0.87 | 0.2 | 0.44 |
| 5 | 75828617 | rs6886903 | 0.367 | -0.438 | -1.128 | 0.816 | 0.017 | -0.021 | -0.021 | 0 | 0.71 | 0.77 | 0.56 | 0.52 | 0.81 | 0.78 |
| 5 | 75831037 | rs7707801 | -0.848 | -0.311 | -0.477 | 0.738 | 0.024 | -0.017 | -0.017 | 0 | 0.64 | 0.55 | 0.63 | 0.72 | 0.48 | 0.79 |
| 5 | 75839165 | rs9293688 | -0.229 | -0.633 | -0.246 | 0.705 | -0.004 | -0.003 | -0.003 | 0 | 0.47 | 0.49 | 0.3 | 0.62 | 0.33 | 0.64 |
| 5 | 75839350 | rs4703712 | 0.705 | -0.677 | 0.175 | 0.671 | -0.002 | -0.039 | -0.039 | 0 | 0.17 | 0.25 | 0.35 | 0.09 | 0.13 | 0.12 |
| 5 | 75841888 | rs10065424 | -1.134 | -0.638 | 0.166 | 0.68 | -0.019 | 0.016 | 0.016 | 0 | 0.64 | 0.73 | 0.65 | 0.69 | 0.47 | 0.77 |
| 5 | 75856354 | rs3913475 | 0.564 | -0.691 | 0.632 | 0.634 | -0.005 | -0.072 | -0.072 | 2.681 | 0.17 | 0.33 | 0.35 | 0.05 | 0.13 | 0.11 |
| 5 | 75856455 | rs6453231 | -1.297 | -0.509 | 0.533 | 0.627 | -0.063 | 0.03 | 0.03 | 0 | 0.66 | 0.83 | 0.65 | 0.71 | 0.48 | 0.78 |
| 5 | 75860695 | rs6453232 | -1.278 | -0.509 | 0.536 | 0.614 | -0.057 | 0.038 | 0.038 | 0 | 0.66 | 0.82 | 0.66 | 0.74 | 0.48 | 0.78 |
| 5 | 75869228 | rs7735600 | -0.791 | -0.52 | 0.695 | 0.902 | -0.001 | 0 | 0 | 0 | 0.6 | 0.64 | 0.52 | 0.33 | 0.44 | 0.68 |
| 5 | 75871279 | rs3756514 | 0.491 | -0.454 | 0.542 | 0.843 | -0.001 | 0.078 | 0.078 | 0 | 0.05 | 0.03 | 0.12 | 0.4 | 0.04 | 0.11 |
| 5 | 75872448 | rs3797373 | 0.389 | -1.124 | 0.753 | 0.5 | 0.027 | 0.026 | 0.026 | 0 | 0.32 | 0.26 | 0.12 | 0.02 | 0.22 | 0.43 |
| 5 | 75874655 | rs17680811 | 0.431 | -0.997 | 0.674 | 0.669 | -0.002 | 0.068 | 0.068 | 0 | 0.05 | 0.03 | 0.12 | 0.41 | 0.05 | 0.11 |
| 5 | 75875629 | rs17748322 | 0.348 | -0.986 | 0.737 | 0.65 | -0.003 | -0.019 | -0.019 | 0.049 | 0.2 | 0.2 | 0.47 | 0.46 | 0.18 | 0.23 |
| 5 | 75888468 | rs3736395 | 0.883 | -0.969 | 0.714 | 0.479 | -0.001 | -0.003 | -0.003 | 0.093 | 0.17 | 0.18 | 0.35 | 0.05 | 0.14 | 0.13 |
| 5 | 75888788 | rs3736394 | -1.108 | -0.835 | 0.982 | 0.523 | -0.003 | -0.002 | -0.002 | 0 | 0.57 | 0.66 | 0.64 | 0.49 | 0.45 | 0.75 |
| 5 | 75890020 | rs1501788 | -0.394 | -0.817 | 0.845 | 0.542 | -0.009 | 0.001 | 0.001 | 0 | 0.4 | 0.49 | 0.29 | 0.44 | 0.3 | 0.62 |
| 5 | 75891003 | rs9293690 | 0.224 | -0.813 | 0.756 | 0.492 | 0.003 | -0.038 | -0.038 | 0 | 0.33 | 0.38 | 0.16 | 0.02 | 0.25 | 0.51 |
| 5 | 75895138 | rs11948805 | -1.308 | -0.792 | 0.698 | 0.498 | -0.029 | -0.021 | -0.021 | 0.01 | 0.59 | 0.78 | 0.64 | 0.5 | 0.45 | 0.76 |
| 5 | 75897600 | rs10056943 | 0.161 | -0.685 | 0.345 | 0.302 | 0.014 | 0.01 | 0.01 | 0 | 0.33 | 0.29 | 0.15 | 0.02 | 0.25 | 0.51 |
| 5 | 75907065 | rs3797390 | -0.703 | -0.516 | 0.724 | 0.578 | -0.005 | 0.032 | 0.032 | 0 | 0.31 | 0.41 | 0.53 | 0.5 | 0.25 | 0.34 |
| 5 | 75908982 | rs4704346 | 0.683 | -0.281 | 1.068 | 0.96 | 0.011 | 0.173 | 0.173 | 0.25 | 0.84 | 0.67 | 0.81 | 0.52 | 0.83 | 0.92 |
| 5 | 75911399 | rs1047530 | 0.881 | -0.641 | 0.745 | 1.065 | -0.084 | 0.085 | 0.085 | 0 | 0.38 | 0.13 | 0.29 | 0.26 | 0.49 | 0.31 |
| 5 | 75911464 | rs1047529 | 0.882 | -0.642 | 0.747 | 1.064 | -0.09 | 0.086 | 0.086 | 0 | 0.38 | 0.12 | 0.29 | 0.26 | 0.49 | 0.32 |
| 5 | 75911715 | rs2069693 | 0.891 | -0.644 | 0.746 | 1.063 | -0.085 | 0.085 | 0.085 | 0 | 0.38 | 0.13 | 0.28 | 0.26 | 0.48 | 0.31 |
| 5 | 75911987 | rs1047494 | 0.683 | -0.665 | 0.92 | 1.068 | -0.028 | 0.039 | 0.039 | 0 | 0.42 | 0.24 | 0.34 | 0.34 | 0.52 | 0.41 |
| 5 | 75912401 | rs2069689 | 0.673 | -0.665 | 0.92 | 1.068 | -0.028 | 0.04 | 0.04 | 0 | 0.42 | 0.24 | 0.34 | 0.34 | 0.52 | 0.41 |
| 5 | 75912843 | rs961536 | 0.702 | -0.664 | 0.919 | 1.124 | -0.03 | 0.02 | 0.02 | 0 | 0.41 | 0.23 | 0.34 | 0.4 | 0.52 | 0.41 |
| 5 | 75913305 | rs2069685 | 0.008 | -0.658 | 1.029 | 0.984 | 0.013 | 0.033 | 0.033 | 0 | 0.47 | 0.43 | 0.52 | 0.18 | 0.39 | 0.54 |
| 5 | 75914752 | rs2069681 | -0.766 | -0.354 | 0.115 | 0.628 | -0.021 | -0.023 | -0.023 | 0 | 0.06 | 0.21 | 0.27 | 0.01 | 0.02 | 0.07 |
| 5 | 75916434 | rs2069664 | 0.817 | 0.099 | -0.498 | 0.73 | -0.056 | 0.095 | 0.095 | 0 | 0.34 | 0.17 | 0.26 | 0.14 | 0.44 | 0.28 |
| 5 | 75916887 | rs2069698 | -0.613 | 0.059 | -0.144 | 1.591 | -0.005 | -0.036 | -0.036 | 0 | 0.07 | 0.01 | 0.04 | 0.19 | 0.05 | 0.12 |
| 5 | 75917633 | rs2069656 | 1.452 | 0.009 | 0.472 | 1.452 | -0.015 | -0.012 | -0.012 | 0.464 | 0.71 | 0.51 | 0.53 | 0.3 | 0.74 | 0.62 |
| 5 | 75917720 | rs2069655 | -1.608 | 0.272 | 0.524 | 1.992 | 0.006 | 0.059 | 0.059 | 0 | 0.14 | 0.15 | 0.13 | 0.42 | 0.17 | 0.12 |
| 5 | 75922504 | rs1697845 | 0.348 | 0.731 | 0.453 | 1.972 | -0.001 | 0.01 | 0.01 | 0 | 0.67 | 0.69 | 0.53 | 0.48 | 0.67 | 0.68 |
| 5 | 75923842 | rs10077302 | 0.87 | 0.63 | 0.288 | 2.071 | -0.035 | -0.043 | -0.043 | 0 | 0.17 | 0.04 | 0.11 | 0.21 | 0.23 | 0.12 |
| 5 | 75926863 | rs457233 | 1.162 | 0.429 | 0.448 | 1.734 | -0.002 | 0.002 | 0.002 | 0 | 0.6 | 0.55 | 0.5 | 0.47 | 0.61 | 0.64 |

| | | | | | | | | | | | | | | | | |
|---|---|---|---|---|---|---|---|---|---|---|---|---|---|---|---|---|
| 5 | 75932717 | rs2455229 | 1.613 | 0.354 | 0.333 | 1.272 | -0.014 | 0.007 | 0.007 | 0 | 0.32 | 0.25 | 0.27 | 0.25 | 0.39 | 0.39 |
| 5 | 75932869 | rs2910819 | 0.578 | 0.21 | 0.261 | 1.206 | -0.003 | 0.003 | 0.003 | 0 | 0.58 | 0.54 | 0.49 | 0.47 | 0.61 | 0.62 |
| 5 | 75934785 | rs10474483 | 0.155 | 0.402 | 0.236 | 1.131 | 0.039 | 0.101 | 0.101 | 0 | 0.35 | 0.16 | 0.25 | 0.23 | 0.24 | 0.37 |
| 5 | 75935088 | rs2455232 | 1.337 | 0.404 | 0.355 | 1.174 | -0.011 | -0.014 | -0.014 | 0 | 0.35 | 0.3 | 0.31 | 0.41 | 0.42 | 0.42 |
| 5 | 75936944 | rs457821 | -0.091 | 0.332 | 0.218 | 1.191 | 0.015 | 0.043 | 0.043 | 0 | 0.71 | 0.56 | 0.56 | 0.63 | 0.67 | 0.79 |
| 5 | 75942936 | rs458059 | 0.283 | 0.288 | -0.051 | 1.216 | 0 | 0.014 | 0.014 | 0 | 0.51 | 0.44 | 0.44 | 0.63 | 0.51 | 0.72 |
| 5 | 75943156 | rs2059222 | -0.55 | 0.456 | -0.079 | 1.063 | 0.035 | 0.053 | 0.053 | 0 | 0.44 | 0.33 | 0.29 | 0.24 | 0.28 | 0.42 |
| 5 | 75950707 | rs465731 | 0.016 | 1.006 | 0.548 | 1.721 | 0.035 | 0.067 | 0.067 | 0.046 | 0.74 | 0.62 | 0.57 | 0.44 | 0.66 | 0.83 |
| 5 | 75954096 | rs2287930 | 1.033 | 0.833 | -0.031 | 1.238 | 0.005 | 0.107 | 0.107 | 0.307 | 0.51 | 0.37 | 0.39 | 0.13 | 0.51 | 0.41 |
| 5 | 75957624 | rs1038920 | -0.124 | 1.387 | -0.73 | 1.282 | -0.001 | 0.003 | 0.003 | 0.408 | 0.22 | 0.19 | 0.15 | 0.38 | 0.31 | 0.16 |
| 5 | 75957922 | rs3797443 | -0.159 | 1.475 | -0.784 | 1.284 | -0.004 | -0.003 | -0.003 | 0 | 0.22 | 0.2 | 0.16 | 0.39 | 0.31 | 0.17 |
| 5 | 75959063 | rs3797446 | -0.283 | 1.366 | -1.471 | 0.316 | -0.054 | 0.164 | 0.164 | 0 | 0.44 | 0.16 | 0.27 | 0.23 | 0.47 | 0.48 |
| 5 | 75959492 | rs6869765 | -0.586 | 1.311 | -0.687 | 0.392 | -0.005 | -0.005 | -0.005 | 0 | 0.59 | 0.58 | 0.43 | 0.36 | 0.65 | 0.62 |
| 5 | 75959705 | rs6890835 | 0.298 | 1.373 | -0.293 | 0.371 | 0.027 | 0.073 | 0.073 | 0 | 0.1 | 0.32 | 0.11 | 0.11 | 0.1 | 0.09 |
| 5 | 75960121 | rs7734540 | 0.153 | 1.468 | -0.573 | 0.244 | 0.036 | 0.174 | 0.174 | 0 | 0.5 | 0.26 | 0.32 | 0.25 | 0.56 | 0.53 |
| 5 | 75962386 | rs4704350 | 0.125 | 1.27 | 0.024 | 0.458 | 0.014 | 0.046 | 0.046 | 0 | 0.1 | 0.25 | 0.11 | 0.1 | 0.1 | 0.09 |
| 5 | 75963210 | rs17568154 | 0.585 | 1.084 | -0.223 | -0.227 | 0.003 | -0.012 | -0.012 | 1.214 | 0.2 | 0.12 | 0.15 | 0 | 0.15 | 0.1 |
| 5 | 75964702 | rs3764935 | -0.019 | 0.712 | -1.042 | -1.077 | 0.032 | 0.112 | 0.112 | 0 | 0.43 | 0.24 | 0.26 | 0.08 | 0.33 | 0.47 |
| 5 | 75967232 | rs7734335 | 0.552 | 0.889 | -1.023 | -1.094 | 0.014 | -0.008 | -0.008 | 0 | 0.32 | 0.39 | 0.28 | 0.25 | 0.42 | 0.21 |
| 5 | 75968099 | rs17652917 | 1.041 | 0.846 | -1.037 | -1.145 | -0.038 | 0.043 | 0.043 | 0 | 0.25 | 0.13 | 0.16 | 0.14 | 0.34 | 0.14 |
| 5 | 75968518 | rs4704352 | -1.229 | 0.707 | -0.824 | -1.153 | -0.002 | -0.011 | -0.011 | 0 | 0.8 | 0.83 | 0.57 | 0.33 | 0.78 | 0.69 |
| 5 | 75972951 | rs2303162 | 1.22 | 0.444 | -0.986 | -1.424 | -0.021 | 0.062 | 0.062 | 0 | 0.21 | 0.07 | 0.14 | 0.08 | 0.21 | 0.37 |
| 5 | 75974119 | rs11741978 | 1.249 | 0.208 | -0.791 | -1.198 | 0.031 | 0.035 | 0.035 | 0.154 | 0.22 | 0.08 | 0.12 | 0 | 0.13 | 0.1 |
| 5 | 75982670 | rs10805907 | -0.675 | -0.551 | -1.268 | -1.208 | 0.024 | -0.046 | -0.046 | 0 | 0.69 | 0.85 | 0.61 | 0.41 | 0.74 | 0.64 |
| 5 | 75986581 | rs7708326 | 0.572 | -0.456 | -0.64 | -1.391 | -0.021 | -0.139 | -0.139 | 0 | 0.27 | 0.41 | 0.2 | 0 | 0.22 | 0.06 |
| 5 | 75990144 | rs1472215 | -0.324 | -0.574 | -0.736 | -1.324 | 0.017 | 0.002 | 0.002 | 0 | 0.61 | 0.66 | 0.53 | 0.37 | 0.71 | 0.64 |
| 5 | 75994211 | rs10514071 | 1.338 | 0.138 | -1.314 | -1.591 | -0.073 | 0.097 | 0.097 | 0 | 0.21 | 0.02 | 0.12 | 0.03 | 0.2 | 0.39 |
| 5 | 75996837 | rs10942789 | 0.595 | 0.48 | -0.73 | -1.509 | 0.029 | 0.016 | 0.016 | 0 | 0.21 | 0.31 | 0.27 | 0.28 | 0.36 | 0.19 |
| 5 | 75996865 | rs468648 | 0.192 | 0.483 | -0.671 | -0.145 | 0.012 | 0.154 | 0.154 | 0 | 0.06 | 0.13 | 0.1 | 0.41 | 0.08 | 0.05 |
| 5 | 76003258 | rs464494 | -0.478 | 0.159 | -0.858 | -0.36 | 0.006 | 0.001 | 0.001 | 0.342 | 0.53 | 0.65 | 0.41 | 0.46 | 0.44 | 0.52 |
| 5 | 76003482 | rs12735 | 0.901 | -0.124 | -0.637 | -0.616 | -0.005 | 0.012 | 0.012 | 0 | 0.27 | 0.23 | 0.17 | 0.03 | 0.26 | 0.41 |
| 5 | 76004014 | rs7723344 | -0.42 | -0.024 | -0.843 | -0.516 | 0.044 | 0.096 | 0.096 | 0.699 | 0.5 | 0.62 | 0.32 | 0.07 | 0.38 | 0.47 |
| 5 | 76004933 | rs7728560 | 1.956 | -0.145 | -0.821 | -0.566 | 0.028 | 0.046 | 0.046 | 1.091 | 0.17 | 0.06 | 0.09 | 0 | 0.14 | 0.06 |
| 5 | 76006172 | rs27903 | -0.137 | -0.035 | -0.643 | -0.286 | 0.001 | -0.009 | -0.009 | 0.053 | 0.37 | 0.62 | 0.33 | 0.46 | 0.31 | 0.46 |
| 5 | 76008682 | rs12517193 | 0.402 | 0.272 | -0.762 | -0.301 | -0.026 | 0.023 | 0.023 | 0.016 | 0.35 | 0.23 | 0.32 | 0.28 | 0.48 | 0.22 |
| 5 | 76010349 | rs2227744 | 0.276 | 0.26 | -0.754 | -0.192 | -0.027 | 0.03 | 0.03 | 0.017 | 0.37 | 0.23 | 0.32 | 0.28 | 0.49 | 0.22 |
| 5 | 76012843 | rs27135 | 0.186 | 0.06 | -0.689 | -0.594 | -0.041 | 0.023 | 0.023 | 0 | 0.4 | 0.23 | 0.39 | 0.68 | 0.55 | 0.28 |
| 5 | 76014000 | rs253061 | -0.984 | -0.16 | -0.245 | -0.367 | 0.001 | 0 | 0 | 0 | 0.72 | 0.77 | 0.61 | 0.73 | 0.79 | 0.7 |
| 5 | 76016946 | rs37248 | -0.265 | 0.002 | -0.497 | -0.392 | 0.011 | 0.008 | 0.008 | 0 | 0.59 | 0.65 | 0.52 | 0.7 | 0.72 | 0.61 |
| 5 | 76024819 | rs253072 | 0.152 | 0.876 | -0.502 | -0.184 | 0.001 | 0.031 | 0.031 | 2.62 | 0.35 | 0.45 | 0.25 | 0.54 | 0.39 | 0.22 |
| 5 | 76027823 | rs153311 | -1.614 | 0.618 | -0.769 | -0.23 | 0.007 | -0.015 | -0.015 | 0 | 0.73 | 0.93 | 0.66 | 0.75 | 0.79 | 0.69 |
| 5 | 76029372 | rs2227800 | 2.438 | 0.493 | -0.177 | -0.005 | -0.041 | -0.042 | -0.042 | 0 | 0.08 | 0.16 | 0.02 | 0 | 0.01 | 0.01 |
| 5 | 76034355 | rs250736 | 1.289 | 1.091 | -0.413 | 0.31 | 0 | 0.03 | 0.03 | 0 | 0.15 | 0.16 | 0.1 | 0.47 | 0.18 | 0.1 |
| 5 | 76041910 | rs250744 | 0.375 | 0.998 | -0.373 | 0.4 | -0.003 | -0.007 | -0.007 | 6.493 | 0.28 | 0.26 | 0.25 | 0.52 | 0.38 | 0.2 |
| 5 | 76048623 | rs2054192 | 0.561 | 0.794 | -0.405 | 0.54 | 0.014 | 0.025 | 0.025 | 0.002 | 0.08 | 0.11 | 0.14 | 0.21 | 0.12 | 0.08 |
| 5 | 76049832 | rs193943 | -0.47 | 0.015 | -0.362 | 0.066 | -0.001 | -0.001 | -0.001 | 0 | 0.51 | 0.57 | 0.45 | 0.53 | 0.54 | 0.41 |
| 5 | 76056054 | rs250723 | 0.499 | 0.227 | -0.845 | 0.255 | -0.059 | -0.091 | -0.091 | 0 | 0.29 | 0.18 | 0.3 | 0.5 | 0.44 | 0.22 |
| 5 | 76059094 | rs250728 | -1.138 | -0.162 | -1.076 | -0.096 | -0.057 | 0.039 | 0.039 | 0 | 0.62 | 0.89 | 0.63 | 0.77 | 0.62 | 0.47 |
| 5 | 76061256 | rs250724 | 0.37 | -0.326 | -0.261 | 0.021 | 0.016 | 0.028 | 0.028 | 0 | 0.34 | 0.49 | 0.38 | 0.54 | 0.46 | 0.23 |
| 5 | 76067624 | rs250760 | -1.07 | -0.201 | -0.714 | 0.01 | -0.054 | 0.048 | 0.048 | 0.044 | 0.6 | 0.89 | 0.62 | 0.77 | 0.61 | 0.47 |
| 5 | 76086597 | rs250763 | 1.502 | -0.516 | -0.706 | -0.084 | 0.002 | -0.016 | -0.016 | 0 | 0.23 | 0.1 | 0.28 | 0.21 | 0.21 | 0.21 |
| 5 | 76086795 | rs17682153 | 0.036 | -0.624 | -0.433 | -0.279 | 0.006 | 0.004 | 0.004 | 0 | 0.25 | 0.19 | 0.23 | 0.23 | 0.16 | 0.27 |
| 5 | 76092225 | rs463466 | -0.352 | -0.544 | -0.189 | -0.236 | 0.016 | 0.008 | 0.008 | 0 | 0.53 | 0.56 | 0.55 | 0.39 | 0.38 | 0.49 |
| 5 | 76092687 | rs2607351 | -0.879 | -0.402 | 0.066 | -0.443 | 0.013 | 0.028 | 0.028 | 0 | 0.68 | 0.57 | 0.63 | 0.39 | 0.58 | 0.79 |
| 5 | 76093207 | rs615986 | 0.037 | -0.397 | -0.033 | -0.426 | 0.017 | 0.01 | 0.01 | 0 | 0.5 | 0.39 | 0.52 | 0.39 | 0.38 | 0.49 |
| 5 | 76097709 | rs7712660 | 0.534 | -0.418 | -0.756 | -0.527 | -0.009 | 0.056 | 0.056 | 0 | 0.43 | 0.28 | 0.39 | 0.18 | 0.41 | 0.58 |
| 5 | 76107152 | rs6453251 | -0.667 | -0.394 | -0.678 | -0.43 | -0.015 | 0.005 | 0.005 | 0 | 0.64 | 0.7 | 0.59 | 0.22 | 0.51 | 0.71 |
| 5 | 76114859 | rs2242991 | 1.634 | -0.227 | -1.096 | -0.483 | -0.007 | 0.027 | 0.027 | 0 | 0.09 | 0.03 | 0.11 | 0.05 | 0.15 | 0.07 |
| 5 | 76114963 | rs1529505 | -1.194 | -0.474 | -0.697 | -0.283 | -0.058 | -0.005 | -0.005 | 0 | 0.67 | 0.83 | 0.6 | 0.22 | 0.51 | 0.71 |
| 5 | 76115521 | rs2242993 | -0.45 | -0.268 | -0.912 | -0.237 | -0.008 | -0.091 | -0.091 | 0 | 0.38 | 0.43 | 0.27 | 0.05 | 0.3 | 0.43 |
| 5 | 76119105 | rs2243022 | -0.631 | -0.133 | -0.67 | 0.031 | 0.014 | -0.011 | -0.011 | 0 | 0.62 | 0.52 | 0.56 | 0.22 | 0.51 | 0.72 |
| 5 | 76127135 | rs2243057 | -0.534 | -0.173 | -0.893 | 0.042 | 0.008 | -0.01 | -0.01 | 0.629 | 0.61 | 0.45 | 0.55 | 0.23 | 0.52 | 0.72 |
| 5 | 76128367 | rs2243060 | 0.949 | -0.103 | -1.586 | 0.536 | -0.067 | 0.078 | 0.078 | 0 | 0.15 | 0.01 | 0.08 | 0.01 | 0.18 | 0.32 |
| 5 | 76137222 | rs4704366 | 0.57 | -1.349 | -1.325 | -0.35 | -0.021 | -0.065 | -0.065 | 2.206 | 0.23 | 0.01 | 0.31 | 0.16 | 0.19 | 0.32 |
| 5 | 76138696 | rs7712957 | -0.952 | -2.106 | -0.917 | -1.06 | 0.03 | 0.03 | 0.03 | 0.226 | 0.28 | 0.74 | 0.23 | 0.04 | 0.13 | 0.13 |
| 5 | 76140337 | rs6883291 | -0.485 | -1.981 | -0.479 | -0.939 | 0.043 | 0.057 | 0.057 | 0 | 0.52 | 0.59 | 0.51 | 0.18 | 0.31 | 0.44 |
| 5 | 76142020 | rs7729082 | 0.037 | -2.242 | -0.272 | -1.02 | 0.047 | 0.05 | 0.05 | 0 | 0.38 | 0.24 | 0.37 | 0.15 | 0.23 | 0.42 |
| 5 | 76145235 | rs6885207 | -0.322 | -1.852 | 0.422 | -1.023 | 0.009 | 0.001 | 0.001 | 0 | 0.44 | 0.49 | 0.41 | 0.11 | 0.26 | 0.38 |
| 5 | 76147593 | rs6897460 | -1.059 | -1.047 | 1.302 | -0.368 | -0.007 | -0.031 | -0.031 | 0 | 0.14 | 0.49 | 0.13 | 0.05 | 0.08 | 0.02 |

| | | | | | | | | | | | | | | | | |
|---|---|---|---|---|---|---|---|---|---|---|---|---|---|---|---|---|
| 5 | 76152824 | rs12697859 | -0.014 | -0.756 | 0.991 | -0.661 | 0.011 | -0.054 | -0.054 | 0.356 | 0.63 | 0.53 | 0.37 | 0.13 | 0.43 | 0.68 |
| 5 | 76155670 | rs11956840 | 0.292 | -0.735 | 1.084 | -0.639 | -0.005 | -0.055 | -0.055 | 0.98 | 0.09 | 0.32 | 0.06 | 0 | 0.05 | 0.01 |
| 5 | 76156381 | rs2460508 | 0.026 | -0.448 | 0.695 | -0.4 | 0.006 | -0.008 | -0.008 | 0 | 0.67 | 0.62 | 0.43 | 0.18 | 0.46 | 0.69 |
| 5 | 76166018 | rs6453259 | -0.113 | -0.056 | 0.553 | 0.363 | 0.023 | 0.013 | 0.013 | 0 | 0.36 | 0.55 | 0.46 | 0.66 | 0.4 | 0.3 |
| 5 | 76171063 | rs1320309 | 0.086 | -0.833 | 0.379 | 0.832 | 0.035 | -0.004 | -0.004 | 0 | 0.52 | 0.42 | 0.61 | 0.55 | 0.67 | 0.27 |
| 5 | 76174519 | rs1320306 | 0.189 | -0.932 | 0.111 | 0.797 | 0.081 | 0.016 | 0.016 | 0.231 | 0.5 | 0.3 | 0.66 | 0.58 | 0.69 | 0.35 |
| 5 | 76187308 | rs7722773 | 0.278 | -0.87 | -0.286 | 0.788 | 0.089 | 0 | 0 | 0 | 0.48 | 0.15 | 0.59 | 0.48 | 0.68 | 0.28 |
| 5 | 76219181 | rs1320021 | 1.151 | -1.006 | 0.265 | 0.746 | -0.056 | 0.042 | 0.042 | 5.869 | 0.9 | 0.29 | 0.88 | 0.89 | 0.98 | 0.92 |
| 5 | 76219972 | rs1700654 | 0.195 | -0.839 | -0.216 | 0.737 | 0.018 | 0.006 | 0.006 | 0 | 0.53 | 0.58 | 0.66 | 0.47 | 0.68 | 0.3 |
| 5 | 76221198 | rs987951 | -0.09 | -0.473 | -1.075 | 0.406 | 0.024 | 0.026 | 0.026 | 2.334 | 0.39 | 0.13 | 0.27 | 0.34 | 0.28 | 0.61 |
| 5 | 76224925 | rs874228 | 0.324 | -0.503 | -0.831 | -0.385 | 0.013 | 0 | 0 | 0.34 | 0.28 | 0.24 | 0.17 | 0 | 0.23 | 0.07 |
| 5 | 76225153 | rs1715762 | 0.126 | -0.556 | -1.191 | -0.716 | -0.008 | 0.044 | 0.044 | 0 | 0.18 | 0.08 | 0.12 | 0.06 | 0.18 | 0.1 |
| 5 | 76226055 | rs1613519 | 0.662 | -0.518 | -1.469 | -0.056 | -0.1 | -0.127 | -0.127 | 0 | 0.3 | 0.09 | 0.51 | 0.5 | 0.4 | 0.68 |
| 5 | 76231193 | rs11747190 | 0.558 | -0.653 | -0.739 | 0.096 | 0.001 | 0 | 0 | 0 | 0.15 | 0.16 | 0.08 | 0.06 | 0.11 | 0.07 |
| 5 | 76232047 | rs10942804 | 0.681 | -0.895 | -0.72 | 0.113 | -0.001 | -0.004 | -0.004 | 0.037 | 0.14 | 0.16 | 0.07 | 0.06 | 0.08 | 0.05 |
| 5 | 76232103 | rs10942805 | 1.001 | -0.836 | -0.955 | 0.141 | 0.007 | 0.023 | 0.023 | 0 | 0.12 | 0.07 | 0.06 | 0.06 | 0.08 | 0.05 |
| 5 | 76232818 | rs1715760 | 0.657 | -0.673 | -0.947 | 0.282 | -0.123 | -0.116 | -0.116 | 0 | 0.31 | 0.1 | 0.51 | 0.49 | 0.44 | 0.7 |
| 5 | 76234482 | rs2055628 | 0.158 | -0.426 | 0.042 | 0.507 | 0.012 | 0.029 | 0.029 | 0 | 0.1 | 0.14 | 0.1 | 0.18 | 0.12 | 0.13 |
| 5 | 76238478 | rs1700680 | -0.234 | -0.376 | -0.006 | 0.565 | -0.002 | -0.006 | -0.006 | 0 | 0.6 | 0.62 | 0.7 | 0.72 | 0.64 | 0.78 |
| 5 | 76239704 | rs1715751 | -0.32 | -0.355 | -0.155 | 0.575 | -0.008 | -0.02 | -0.02 | 0 | 0.6 | 0.55 | 0.7 | 0.72 | 0.64 | 0.78 |
| 5 | 76247784 | rs3792738 | -0.245 | -0.268 | -0.242 | 0.987 | 0.001 | -0.001 | -0.001 | 0 | 0.1 | 0.1 | 0.08 | 0.16 | 0.09 | 0.03 |
| 5 | 76250378 | rs7721799 | -0.18 | -0.244 | -0.379 | 1.003 | 0.01 | -0.024 | -0.024 | 0 | 0.09 | 0.04 | 0.08 | 0.17 | 0.09 | 0.04 |
| 5 | 76250972 | rs32897 | -0.951 | -0.169 | 0.274 | 0.758 | 0.01 | 0.016 | 0.016 | 0.019 | 0.18 | 0.32 | 0.14 | 0.24 | 0.16 | 0.16 |
| 5 | 76258048 | rs7718461 | -1.099 | -0.287 | -0.343 | 0.631 | -0.003 | -0.019 | -0.019 | 0.005 | 0.48 | 0.81 | 0.38 | 0.53 | 0.39 | 0.46 |
| 5 | 76264354 | rs10062367 | -0.956 | -0.393 | 0.485 | 0.586 | 0.007 | -0.012 | -0.012 | 0 | 0.2 | 0.39 | 0.16 | 0.18 | 0.2 | 0.26 |
| 5 | 76264982 | rs1875999 | -1.061 | -0.517 | 0.797 | 0.622 | 0.001 | 0.013 | 0.013 | 0.924 | 0.4 | 0.45 | 0.33 | 0.54 | 0.34 | 0.43 |
| 5 | 76265035 | rs1053989 | -1.558 | -0.5 | 0.527 | 0.635 | 0.005 | -0.006 | -0.006 | 0 | 0.47 | 0.79 | 0.37 | 0.53 | 0.39 | 0.46 |
| 5 | 76266447 | rs10514082 | -0.281 | -0.279 | 0.359 | 0.799 | -0.011 | -0.01 | -0.01 | 0 | 0.12 | 0.07 | 0.12 | 0.19 | 0.15 | 0.24 |
| 5 | 76270853 | rs10474485 | -0.24 | -0.241 | 0.731 | 0.995 | -0.012 | 0.05 | 0.05 | 0.625 | 0.27 | 0.44 | 0.21 | 0.36 | 0.19 | 0.2 |
| 5 | 76274537 | rs1715747 | -0.847 | -0.354 | -0.138 | 0.663 | 0.024 | -0.007 | -0.007 | 0 | 0.45 | 0.74 | 0.37 | 0.55 | 0.34 | 0.41 |
| 5 | 76292643 | rs1651094 | -0.874 | 0.262 | 0.37 | 0.56 | 0.01 | -0.024 | -0.024 | 0.023 | 0.26 | 0.49 | 0.3 | 0.2 | 0.26 | 0.36 |
| 5 | 76293603 | rs247751 | -0.013 | -0.273 | 0.437 | 0.601 | -0.007 | 0.033 | 0.033 | 0 | 0.73 | 0.68 | 0.67 | 0.51 | 0.8 | 0.84 |
| 5 | 76296410 | rs10066702 | 1.007 | -0.187 | -0.022 | 0.22 | 0.016 | 0.125 | 0.125 | 3.782 | 0.42 | 0.2 | 0.29 | 0.14 | 0.42 | 0.23 |
| 5 | 76298772 | rs247746 | 0.072 | -0.179 | 0.309 | 0.264 | 0.006 | 0.008 | 0.008 | 0.115 | 0.26 | 0.33 | 0.33 | 0.38 | 0.35 | 0.57 |
| 5 | 76303398 | rs6898403 | -0.206 | -0.17 | -0.208 | 0.263 | 0.024 | 0.044 | 0.044 | 0.967 | 0.3 | 0.54 | 0.51 | 0.43 | 0.35 | 0.61 |
| 5 | 76305011 | rs11745330 | 1.436 | -0.122 | -1.01 | -0.203 | 0.004 | 0.087 | 0.087 | 4.173 | 0.33 | 0.14 | 0.23 | 0.11 | 0.34 | 0.16 |
| 5 | 76305056 | rs10473985 | 0.355 | -0.044 | -0.924 | -0.038 | -0.006 | 0.031 | 0.031 | 0 | 0.1 | 0.02 | 0.08 | 0.03 | 0.09 | 0.09 |
| 5 | 76310780 | rs12653511 | 1.249 | -0.042 | -0.875 | -0.007 | 0.001 | 0.073 | 0.073 | 4.374 | 0.32 | 0.14 | 0.23 | 0.12 | 0.35 | 0.16 |
| 5 | 76311491 | rs13178007 | -0.096 | 0.19 | -0.979 | -0.023 | -0.037 | 0.032 | 0.032 | 0.998 | 0.26 | 0.17 | 0.31 | 0.2 | 0.39 | 0.53 |
| 5 | 76311992 | rs11955295 | -0.096 | 0.351 | -1.162 | 0.006 | -0.098 | 0.014 | 0.014 | 2.01 | 0.25 | 0.05 | 0.28 | 0.24 | 0.38 | 0.54 |
| 5 | 76312118 | rs6865782 | -0.116 | -0.369 | -0.857 | -0.301 | -0.012 | -0.043 | -0.043 | 0 | 0.66 | 0.6 | 0.76 | 0.87 | 0.75 | 0.77 |
| 5 | 76312265 | rs17750118 | 1.714 | -0.734 | -1.165 | 0.2 | -0.021 | -0.003 | -0.003 | 0.969 | 0.25 | 0.07 | 0.17 | 0.22 | 0.3 | 0.13 |
| 5 | 76316299 | rs4484404 | -0.497 | -0.893 | -0.692 | -0.08 | -0.039 | -0.044 | -0.044 | 0 | 0.79 | 0.67 | 0.89 | 0.9 | 0.89 | 0.88 |
| 5 | 76319344 | rs9293701 | 0.932 | 0.002 | -1.314 | -0.008 | -0.078 | 0.055 | 0.055 | 0 | 0.15 | 0.02 | 0.19 | 0.02 | 0.21 | 0.14 |
| 5 | 76319894 | rs247753 | 0.957 | -0.026 | -1.4 | -0.13 | -0.037 | -0.021 | -0.021 | 0.049 | 0.16 | 0.01 | 0.2 | 0.16 | 0.2 | 0.23 |
| 5 | 76323206 | rs382543 | 1.66 | -0.076 | -1.586 | 0.116 | -0.009 | 0.008 | 0.008 | 1.353 | 0.28 | 0.08 | 0.32 | 0.27 | 0.29 | 0.14 |
| 5 | 76341228 | rs7737897 | -0.224 | 0.039 | -0.992 | 0.28 | 0 | -0.004 | -0.004 | 0 | 0.82 | 0.83 | 0.87 | 0.73 | 0.83 | 0.63 |
| 5 | 76342829 | rs995743 | 0.674 | -0.152 | -0.44 | 0.184 | 0.004 | -0.005 | -0.005 | 2.239 | 0.62 | 0.39 | 0.62 | 0.57 | 0.63 | 0.4 |
| 5 | 76343999 | rs13155212 | 0.577 | -0.085 | -0.35 | 0.116 | 0.011 | -0.01 | -0.01 | 0.26 | 0.19 | 0.24 | 0.23 | 0.16 | 0.24 | 0.24 |
| 5 | 76363075 | rs13162491 | -0.437 | -0.422 | -0.178 | 0.379 | -0.034 | 0.032 | 0.032 | 0.006 | 0.83 | 0.67 | 0.87 | 0.75 | 0.9 | 0.65 |
| 5 | 76368663 | rs10052297 | -0.684 | -0.261 | -0.596 | 0.422 | 0.002 | -0.003 | -0.003 | 0 | 0.82 | 0.84 | 0.89 | 0.74 | 0.87 | 0.64 |
| 5 | 76369337 | rs1048834 | 0.834 | -0.366 | -0.704 | 0.516 | 0.004 | -0.035 | -0.035 | 0 | 0.15 | 0.13 | 0.09 | 0.27 | 0.11 | 0.08 |
| 5 | 76370827 | rs1529460 | 1.392 | -0.55 | -0.516 | 0.678 | 0.006 | 0.009 | 0.009 | 1.157 | 0.28 | 0.17 | 0.33 | 0.28 | 0.3 | 0.13 |
| 5 | 76371119 | rs1001960 | 0.121 | -0.583 | -0.026 | 1.192 | 0.001 | -0.002 | -0.002 | 0.2 | 0.82 | 0.84 | 0.89 | 0.74 | 0.87 | 0.65 |
| 5 | 76378192 | rs2914141 | -1.036 | -0.19 | 0.267 | 1.106 | -0.022 | -0.028 | -0.028 | 0 | 0.21 | 0.37 | 0.23 | 0.16 | 0.19 | 0.25 |
| 5 | 76380000 | rs2914143 | -0.875 | 0.24 | -0.002 | 1.615 | 0 | -0.002 | -0.002 | 0.522 | 0.18 | 0.12 | 0.2 | 0.33 | 0.21 | 0.25 |
| 5 | 76383933 | rs2914146 | -1.411 | 0.006 | 0.547 | 0.973 | 0.004 | 0.045 | 0.045 | 0.15 | 0.1 | 0.4 | 0.08 | 0.09 | 0.06 | 0.07 |
| 5 | 76386843 | rs13154434 | -0.401 | 0.687 | -0.55 | 0.874 | -0.071 | -0.029 | -0.029 | 0 | 0.19 | 0.02 | 0.4 | 0.2 | 0.27 | 0.16 |
| 5 | 76390859 | rs7709295 | -0.545 | 0.29 | 0.07 | 0.716 | 0.024 | 0.016 | 0.016 | 2.172 | 0.05 | 0.29 | 0.04 | 0 | 0.01 | 0 |
| 5 | 76395184 | rs4418080 | -0.596 | 0.572 | -0.8 | 0.859 | -0.013 | 0.002 | 0.002 | 0 | 0.22 | 0.11 | 0.4 | 0.21 | 0.27 | 0.16 |
| 5 | 76395815 | rs13170206 | -0.485 | 0.657 | -0.858 | 0.984 | -0.023 | -0.019 | -0.019 | 1.498 | 0.21 | 0.1 | 0.4 | 0.24 | 0.27 | 0.16 |
| 5 | 76407577 | rs4703727 | 0.322 | 0.621 | -1.152 | 0.822 | -0.052 | -0.005 | -0.005 | 0.236 | 0.75 | 0.96 | 0.6 | 0.8 | 0.72 | 0.84 |
| 5 | 76418218 | rs12519589 | -0.256 | 1.585 | -0.916 | 1.077 | -0.015 | -0.01 | -0.01 | 2.202 | 0.08 | 0.05 | 0.14 | 0.14 | 0.13 | 0.15 |
| 5 | 76419094 | rs10070420 | -0.395 | 0.976 | -0.599 | 1.171 | -0.002 | 0.084 | 0.084 | 0 | 0.14 | 0.22 | 0.13 | 0.38 | 0.1 | 0.25 |
| 5 | 76423967 | rs4128741 | 0.534 | 0.584 | -1.37 | -0.154 | -0.006 | 0.089 | 0.089 | 0 | 0.35 | 0.2 | 0.27 | 0.15 | 0.35 | 0.19 |
| 5 | 76425867 | rs7708285 | -0.561 | 0.44 | -1.588 | -0.593 | -0.022 | 0.085 | 0.085 | 0 | 0.73 | 0.89 | 0.69 | 0.95 | 0.68 | 0.84 |
| 5 | 76426987 | rs12521058 | 1.21 | 0.277 | -1.318 | -0.958 | -0.001 | 0.05 | 0.05 | 0 | 0.36 | 0.26 | 0.32 | 0.13 | 0.36 | 0.29 |
| 5 | 76427000 | rs6859104 | 0.25 | 0.339 | -1.229 | -0.233 | 0.011 | 0.143 | 0.143 | 0 | 0.13 | 0.23 | 0.12 | 0.49 | 0.13 | 0.26 |
| 5 | 76435346 | rs7732628 | 0.323 | 0.145 | -1.091 | -0.219 | -0.002 | 0.001 | 0.001 | 0 | 0.56 | 0.58 | 0.54 | 0.47 | 0.54 | 0.54 |

| | | | | | | | | | | | | | | | | |
|---|---|---|---|---|---|---|---|---|---|---|---|---|---|---|---|---|
| 5 | 76439250 | rs6885904 | 0.243 | 0.149 | -1.542 | -0.191 | -0.036 | 0.008 | 0.008 | 0 | 0.56 | 0.79 | 0.55 | 0.42 | 0.51 | 0.59 |
| 5 | 76439667 | rs9293707 | 1.147 | 0.046 | -1.933 | -0.982 | -0.099 | 0.061 | 0.061 | 0 | 0.35 | 0.03 | 0.34 | 0.17 | 0.39 | 0.37 |
| 5 | 76444754 | rs10474489 | 0.913 | 0.323 | -1.134 | -1.071 | -0.013 | 0.101 | 0.101 | 0 | 0.61 | 0.24 | 0.56 | 0.87 | 0.58 | 0.62 |
| 5 | 76448284 | rs7708818 | -0.194 | 0.526 | -0.51 | -0.891 | 0.026 | 0.009 | 0.009 | 0.601 | 0.07 | 0.27 | 0.04 | 0.06 | 0.06 | 0.08 |
| 5 | 76451520 | rs6866080 | 0.377 | 0.602 | -0.675 | -0.875 | -0.008 | 0.007 | 0.007 | 0 | 0.71 | 0.76 | 0.64 | 0.94 | 0.64 | 0.71 |
| 5 | 76454559 | rs6862414 | -0.435 | 0.759 | -0.712 | -0.651 | -0.015 | 0.021 | 0.021 | 0.666 | 0.8 | 0.87 | 0.68 | 0.92 | 0.74 | 0.87 |
| 5 | 76454612 | rs10035405 | 0.708 | 0.3 | -0.502 | -1.016 | -0.008 | -0.047 | -0.047 | 0.009 | 0.39 | 0.5 | 0.36 | 0.15 | 0.38 | 0.38 |
| 5 | 76455453 | rs10044684 | -0.58 | 0.007 | -0.864 | -0.831 | -0.004 | 0.003 | 0.003 | 0.19 | 0.29 | 0.14 | 0.25 | 0.78 | 0.26 | 0.44 |
| 5 | 76460134 | rs12171500 | 0.078 | -0.115 | -1.049 | -0.645 | -0.018 | -0.084 | -0.084 | 0 | 0.2 | 0.04 | 0.19 | 0.76 | 0.16 | 0.28 |
| 5 | 76460800 | rs4426900 | 0.974 | -0.21 | -1.168 | -0.957 | -0.065 | 0.125 | 0.125 | 0 | 0.37 | 0.05 | 0.29 | 0.14 | 0.37 | 0.35 |
| 5 | 76463560 | rs7721952 | 0.206 | -0.166 | -0.077 | 0.122 | 0.024 | 0.226 | 0.226 | 0 | 0.09 | 0.29 | 0.13 | 0.39 | 0.08 | 0.09 |
| 5 | 76464610 | rs12514758 | 1.139 | -0.364 | -0.667 | -0.233 | -0.006 | 0.045 | 0.045 | 0 | 0.35 | 0.25 | 0.31 | 0.13 | 0.38 | 0.34 |
| 5 | 76467984 | rs1479561 | -0.793 | -0.296 | -0.629 | -0.141 | -0.012 | 0.095 | 0.095 | 1.425 | 0.19 | 0.6 | 0.2 | 0.44 | 0.16 | 0.17 |
| 5 | 76468843 | rs2972341 | -0.209 | -0.239 | -1.47 | -0.21 | -0.156 | 0.177 | 0.177 | 3.41 | 0.65 | 0.95 | 0.59 | 0.94 | 0.56 | 0.72 |
| 5 | 76470423 | rs963913 | 0.199 | 0.206 | -0.89 | -0.003 | 0 | -0.023 | -0.023 | 0 | 0.46 | 0.76 | 0.45 | 0.52 | 0.46 | 0.44 |
| 5 | 76480761 | rs6453292 | 0.104 | 0.288 | -0.457 | 0.058 | -0.002 | 0 | 0 | 0 | 0.53 | 0.5 | 0.44 | 0.52 | 0.57 | 0.57 |
| 5 | 76482340 | rs6889575 | 0.627 | 0.758 | -0.006 | -0.673 | -0.007 | 0.037 | 0.037 | 0.597 | 0.35 | 0.23 | 0.29 | 0.09 | 0.38 | 0.32 |
| 5 | 76486226 | rs2928164 | -0.819 | 0.138 | -0.458 | -0.978 | -0.017 | 0.052 | 0.052 | 0.7 | 0.72 | 0.9 | 0.59 | 0.97 | 0.64 | 0.78 |
| 5 | 76488613 | rs1119208 | 0.839 | 0.281 | -0.51 | -0.818 | -0.001 | -0.088 | -0.088 | 0 | 0.36 | 0.43 | 0.29 | 0.09 | 0.36 | 0.32 |
| 5 | 76496152 | rs10462550 | -0.143 | 0.548 | -1.238 | 0.249 | -0.036 | -0.078 | -0.078 | 1.047 | 0.12 | 0.01 | 0.1 | 0.38 | 0.1 | 0.16 |
| 5 | 76499542 | rs1982658 | -0.703 | 0.25 | -0.896 | -0.824 | 0.003 | -0.001 | -0.001 | 0 | 0.38 | 0.65 | 0.31 | 0.87 | 0.29 | 0.49 |
| 5 | 76503432 | rs11739698 | 0.467 | 0.563 | -1.956 | -1.213 | -0.061 | 0.201 | 0.201 | 0 | 0.34 | 0.02 | 0.23 | 0.09 | 0.32 | 0.33 |
| 5 | 76514019 | rs11953611 | -0.382 | 0.739 | -1.145 | -1.026 | -0.001 | -0.011 | -0.011 | 0 | 0.31 | 0.33 | 0.3 | 0.82 | 0.29 | 0.47 |
| 5 | 76515824 | rs9687206 | -1.41 | 0.913 | -1.053 | -1.459 | -0.013 | 0.164 | 0.164 | 0.92 | 0.47 | 0.77 | 0.6 | 0.84 | 0.44 | 0.62 |
| 5 | 76517982 | rs13158164 | 1.326 | 1.142 | -1.206 | -1.212 | -0.035 | 0.029 | 0.029 | 0 | 0.06 | 0.01 | 0.23 | 0 | 0.13 | 0.08 |
| 5 | 76520512 | rs12520862 | 0.455 | 0.771 | -0.841 | -0.157 | 0 | 0.03 | 0.03 | 2.081 | 0.14 | 0.17 | 0.14 | 0.33 | 0.15 | 0.26 |
| 5 | 76521421 | rs12516568 | 0.695 | 0.781 | -1.068 | -0.055 | -0.033 | -0.082 | -0.082 | 0 | 0.13 | 0.01 | 0.12 | 0.32 | 0.15 | 0.25 |
| 5 | 76539972 | rs12515498 | -0.681 | 0.705 | -0.632 | -0.535 | -0.002 | -0.03 | -0.03 | 0 | 0.29 | 0.41 | 0.3 | 0.83 | 0.28 | 0.47 |
| 5 | 76544942 | rs1351283 | -0.639 | 0.607 | -0.293 | -0.547 | 0.002 | -0.014 | -0.014 | 0 | 0.36 | 0.44 | 0.53 | 0.75 | 0.38 | 0.54 |
| 5 | 76550329 | rs832785 | -0.957 | 0.313 | -0.731 | -1.254 | -0.01 | 0.027 | 0.027 | 0 | 0.69 | 0.8 | 0.76 | 0.96 | 0.66 | 0.84 |
| 5 | 76551010 | rs4704400 | -0.2 | 0.245 | -0.896 | -0.753 | -0.001 | -0.002 | -0.002 | 0 | 0.55 | 0.56 | 0.63 | 0.54 | 0.55 | 0.65 |
| 5 | 76554807 | rs7702192 | 0.067 | 0.167 | -1.013 | -0.725 | 0.001 | 0.008 | 0.008 | 0 | 0.48 | 0.44 | 0.58 | 0.45 | 0.51 | 0.57 |
| 5 | 76555245 | rs4704401 | 1.492 | 0.102 | -0.37 | -0.354 | 0.006 | 0.02 | 0.02 | 0 | 0.07 | 0.1 | 0.05 | 0.08 | 0.05 | 0.07 |
| 5 | 76562245 | rs7714529 | -0.441 | 0.068 | -0.987 | -0.639 | 0 | -0.002 | -0.002 | 0 | 0.34 | 0.65 | 0.24 | 0.12 | 0.26 | 0.21 |
| 5 | 76569611 | rs6885813 | 0.591 | 0.257 | -1.511 | -0.202 | -0.075 | -0.157 | -0.157 | 0 | 0.23 | 0.03 | 0.36 | 0.42 | 0.25 | 0.49 |
| 5 | 76572671 | rs251419 | 1.131 | 0.444 | -1.128 | -0.283 | -0.012 | -0.004 | -0.004 | 3.035 | 0.17 | 0.12 | 0.16 | 0.13 | 0.2 | 0.2 |
| 5 | 76572970 | rs251421 | 1.227 | 0.443 | -1.153 | -0.28 | -0.021 | -0.01 | -0.01 | 0 | 0.17 | 0.07 | 0.15 | 0.13 | 0.2 | 0.2 |
| 5 | 76574841 | rs13167306 | 0.391 | 0.01 | -0.525 | -0.51 | -0.002 | 0.006 | 0.006 | 0 | 0.36 | 0.43 | 0.43 | 0.42 | 0.32 | 0.52 |
| 5 | 76576011 | rs251426 | -1.358 | 0.024 | -1.274 | -0.708 | -0.04 | 0.125 | 0.125 | 3.415 | 0.69 | 0.98 | 0.79 | 0.9 | 0.71 | 0.82 |
| 5 | 76591698 | rs12514694 | 1.858 | 0.36 | -0.619 | -0.306 | -0.006 | 0.015 | 0.015 | 0 | 0.08 | 0.06 | 0.13 | 0.33 | 0.15 | 0.08 |
| 5 | 76594229 | rs832787 | 1.3 | 0.357 | -0.2 | -0.587 | -0.006 | 0.009 | 0.009 | 0.32 | 0.17 | 0.13 | 0.16 | 0.03 | 0.19 | 0.14 |
| 5 | 76611264 | rs7731628 | 1.065 | 0.145 | 0.016 | -0.657 | 0 | -0.013 | -0.013 | 0 | 0.28 | 0.25 | 0.2 | 0.07 | 0.27 | 0.19 |
| 5 | 76614059 | rs10045589 | 0.586 | -0.114 | 0.382 | -0.626 | 0 | 0 | 0 | 0 | 0.34 | 0.37 | 0.23 | 0.22 | 0.3 | 0.33 |
| 5 | 76615691 | rs2359797 | -2.2 | -0.053 | 0.488 | -0.702 | -0.023 | 0.035 | 0.035 | 0.397 | 0.85 | 0.93 | 0.84 | 0.98 | 0.79 | 0.89 |
| 5 | 76616738 | rs6863296 | 0.189 | -0.088 | 0.849 | -0.69 | 0.018 | 0.129 | 0.129 | 0 | 0.1 | 0.43 | 0.09 | 0.15 | 0.07 | 0.15 |
| 5 | 76618857 | rs1531615 | 0.122 | -0.121 | 0.798 | -0.69 | 0 | 0.008 | 0.008 | 0 | 0.48 | 0.45 | 0.57 | 0.76 | 0.46 | 0.55 |
| 5 | 76620582 | rs6873924 | 0.978 | -0.113 | 0.769 | -0.554 | -0.02 | 0.097 | 0.097 | 0 | 0.07 | 0.27 | 0.05 | 0.15 | 0.03 | 0.15 |
| 5 | 76622250 | rs16874184 | 1.301 | -0.091 | 0.46 | -0.544 | -0.019 | 0.056 | 0.056 | 0 | 0.07 | 0.17 | 0.04 | 0.15 | 0.03 | 0.15 |
| 5 | 76623421 | rs9686502 | -0.622 | -0.074 | 0.458 | -0.695 | -0.048 | 0.178 | 0.178 | 0 | 0.56 | 0.85 | 0.62 | 0.9 | 0.49 | 0.7 |
| 5 | 76623830 | rs9686527 | 0.327 | -0.221 | 0.669 | -0.695 | -0.015 | 0.02 | 0.02 | 0.169 | 0.25 | 0.54 | 0.22 | 0.33 | 0.2 | 0.15 |
| 5 | 76627436 | rs9687030 | -0.088 | -0.203 | 0.578 | -0.778 | 0.002 | 0.008 | 0.008 | 0 | 0.52 | 0.56 | 0.4 | 0.4 | 0.47 | 0.33 |
| 5 | 76629834 | rs12523314 | 1.651 | -0.214 | 0.615 | -0.721 | -0.01 | -0.041 | -0.041 | 0 | 0.24 | 0.13 | 0.36 | 0.42 | 0.26 | 0.4 |
| 5 | 76630561 | rs11951639 | 0.322 | -0.195 | 0.677 | -0.758 | -0.013 | 0.027 | 0.027 | 0.085 | 0.25 | 0.48 | 0.21 | 0.33 | 0.2 | 0.15 |
| 5 | 76632104 | rs6893108 | 1.894 | -0.203 | 0.794 | -0.769 | -0.002 | 0 | 0 | 0 | 0.13 | 0.11 | 0.14 | 0.33 | 0.15 | 0.11 |
| 5 | 76638632 | rs10064824 | 0.781 | -0.219 | 0.603 | -0.765 | 0.001 | -0.006 | -0.006 | 0 | 0.3 | 0.29 | 0.4 | 0.58 | 0.29 | 0.56 |
| 5 | 76642311 | rs922540 | -0.285 | -0.172 | 0.571 | -0.78 | -0.001 | -0.002 | -0.002 | 1.711 | 0.63 | 0.62 | 0.62 | 0.5 | 0.58 | 0.63 |
| 5 | 76644510 | rs11949597 | 0.639 | -0.203 | 0.704 | -0.903 | -0.013 | -0.08 | -0.08 | 0.028 | 0.12 | 0.39 | 0.08 | 0 | 0.05 | 0.04 |
| 5 | 76644962 | rs2046046 | -0.747 | -0.129 | 0.346 | -0.846 | 0.003 | -0.003 | -0.003 | 0.109 | 0.75 | 0.69 | 0.76 | 0.82 | 0.73 | 0.74 |
| 5 | 76653752 | rs17683162 | -0.633 | -0.131 | 0.221 | -0.845 | 0.016 | 0.027 | 0.027 | 0.065 | 0.46 | 0.84 | 0.51 | 0.59 | 0.38 | 0.61 |
| 5 | 76656678 | rs10942819 | 0.728 | -0.141 | 0.428 | -0.728 | -0.016 | -0.075 | -0.075 | 0.009 | 0.39 | 0.46 | 0.27 | 0.11 | 0.32 | 0.22 |
| 5 | 76662518 | rs4704416 | 1.483 | -0.031 | 0.184 | -0.557 | -0.016 | -0.057 | -0.057 | 0.334 | 0.24 | 0.12 | 0.35 | 0.42 | 0.26 | 0.41 |
| 5 | 76670628 | rs6453304 | 1.194 | 0.037 | 0.142 | -0.365 | -0.001 | -0.001 | -0.001 | 0 | 0.16 | 0.16 | 0.18 | 0.29 | 0.19 | 0.11 |
| 5 | 76671544 | rs7732963 | 1.398 | 0.006 | 0.191 | -0.398 | -0.017 | 0.046 | 0.046 | 0 | 0.06 | 0.17 | 0.04 | 0.1 | 0.03 | 0.14 |
| 5 | 76672186 | rs11750710 | 0.721 | 0.125 | -0.13 | -0.667 | -0.001 | 0.049 | 0.049 | 0.064 | 0.34 | 0.25 | 0.25 | 0.11 | 0.33 | 0.19 |
| 5 | 76676958 | rs13178853 | -1.943 | 0.278 | -0.334 | -0.506 | -0.046 | 0.04 | 0.04 | 0.288 | 0.84 | 0.96 | 0.83 | 0.94 | 0.79 | 0.89 |
| 5 | 76694631 | rs335598 | -0.743 | 0.223 | -0.253 | -0.402 | -0.032 | -0.035 | -0.035 | 0.642 | 0.72 | 0.9 | 0.7 | 0.62 | 0.65 | 0.79 |
| 5 | 76695327 | rs4704417 | -0.559 | 0.19 | 0.213 | -0.324 | 0.042 | 0.038 | 0.038 | 0 | 0.48 | 0.67 | 0.54 | 0.69 | 0.37 | 0.53 |
| 5 | 76695540 | rs2359875 | 0.883 | 0.14 | 0.675 | -0.492 | 0.007 | -0.007 | -0.007 | 0.591 | 0.3 | 0.28 | 0.37 | 0.37 | 0.23 | 0.41 |
| 5 | 76696334 | rs3733952 | 1.276 | 0.095 | 0.384 | -0.495 | 0.031 | -0.056 | -0.056 | 1.479 | 0.26 | 0.11 | 0.33 | 0.37 | 0.19 | 0.41 |

| | | | | | | | | | | | | | | | | |
|---|---|---|---|---|---|---|---|---|---|---|---|---|---|---|---|---|
| 5 | 76701739 | rs6874493 | -0.789 | 0.261 | 0.589 | -0.939 | 0.006 | -0.015 | -0.015 | 0 | 0.64 | 0.56 | 0.65 | 0.82 | 0.55 | 0.69 |
| 5 | 76703374 | rs3101866 | 0.115 | 0.359 | 0.952 | -0.813 | 0 | 0.001 | 0.001 | 0 | 0.47 | 0.42 | 0.42 | 0.47 | 0.53 | 0.31 |
| 5 | 76704251 | rs11956168 | 2.328 | 0.243 | 0.403 | -0.895 | -0.008 | 0.006 | 0.006 | 0 | 0.11 | 0.06 | 0.05 | 0 | 0.1 | 0.05 |
| 5 | 76708987 | rs335614 | -0.975 | 0.343 | 0.857 | -0.494 | -0.007 | 0.004 | 0.004 | 0.048 | 0.76 | 0.82 | 0.73 | 0.82 | 0.67 | 0.76 |
| 5 | 76709426 | rs2279095 | 1.752 | -0.068 | 0.689 | -0.486 | -0.024 | 0.063 | 0.063 | 0 | 0.16 | 0.21 | 0.12 | 0.36 | 0.09 | 0.31 |
| 5 | 76712162 | rs12521494 | 0.226 | -0.144 | 0.728 | -0.562 | -0.02 | -0.002 | -0.002 | 0 | 0.31 | 0.34 | 0.27 | 0.7 | 0.24 | 0.41 |
| 5 | 76712575 | rs890717 | 1.204 | 0.031 | 0.523 | -0.499 | -0.028 | 0.081 | 0.081 | 0 | 0.18 | 0.34 | 0.17 | 0.35 | 0.15 | 0.11 |
| 5 | 76714380 | rs2242159 | 0.071 | 0.238 | -0.264 | -0.547 | 0.002 | 0.005 | 0.005 | 0 | 0.52 | 0.46 | 0.53 | 0.59 | 0.43 | 0.75 |
| 5 | 76715131 | rs2242160 | 1.026 | 0.298 | -0.808 | -0.589 | -0.011 | 0.041 | 0.041 | 0 | 0.35 | 0.17 | 0.4 | 0.24 | 0.34 | 0.44 |
| 5 | 76715163 | rs2242161 | 1.598 | 0.349 | -0.843 | -0.543 | -0.027 | -0.025 | -0.025 | 0 | 0.07 | 0.02 | 0.1 | 0.12 | 0.12 | 0.13 |
| 5 | 76716913 | rs335644 | -0.341 | 0.2 | -0.513 | -0.457 | 0.028 | 0.085 | 0.085 | 0 | 0.5 | 0.64 | 0.47 | 0.75 | 0.46 | 0.52 |
| 5 | 76718070 | rs335643 | 1.915 | 0.269 | -0.78 | -0.132 | -0.043 | 0.027 | 0.027 | 0.248 | 0.14 | 0.04 | 0.16 | 0.06 | 0.21 | 0.11 |
| 5 | 76718187 | rs1124742 | 0.593 | 0.072 | -0.656 | 0.008 | -0.007 | 0.045 | 0.045 | 0 | 0.36 | 0.18 | 0.4 | 0.24 | 0.34 | 0.45 |
| 5 | 76718626 | rs6860674 | 0.25 | -0.584 | -0.307 | 0.585 | -0.043 | 0.076 | 0.076 | 0 | 0.17 | 0.29 | 0.13 | 0.35 | 0.08 | 0.29 |
| 5 | 76720244 | rs335640 | -0.208 | -0.001 | -0.884 | 0.301 | -0.032 | -0.019 | -0.019 | 0 | 0.75 | 0.88 | 0.77 | 0.67 | 0.67 | 0.87 |
| 5 | 76720537 | rs2359877 | -1.221 | -0.116 | -0.086 | -0.283 | 0.038 | -0.036 | -0.036 | 0.401 | 0.08 | 0.29 | 0.1 | 0 | 0.08 | 0.02 |
| 5 | 76721089 | rs90682 | 0.392 | -0.226 | -0.443 | -0.577 | -0.016 | 0.027 | 0.027 | 0 | 0.26 | 0.16 | 0.27 | 0.09 | 0.33 | 0.12 |
| 5 | 76722443 | rs40594 | -0.433 | -1.267 | -0.349 | -0.554 | -0.058 | -0.029 | -0.029 | 0 | 0.64 | 0.85 | 0.71 | 0.57 | 0.49 | 0.65 |
| 5 | 76722895 | rs459366 | 0.43 | -1.679 | -0.195 | -0.681 | -0.08 | 0.048 | 0.048 | 0.319 | 0.32 | 0.53 | 0.21 | 0.43 | 0.12 | 0.37 |
| 5 | 76724599 | rs335636 | -0.71 | -1.568 | -0.223 | -0.634 | -0.087 | -0.046 | -0.046 | 0.373 | 0.65 | 0.9 | 0.71 | 0.58 | 0.49 | 0.66 |
| 5 | 76725582 | rs335635 | -0.589 | -1.615 | -0.26 | -0.666 | -0.094 | -0.056 | -0.056 | 0.014 | 0.64 | 0.91 | 0.71 | 0.57 | 0.49 | 0.66 |
| 5 | 76726202 | rs2114950 | 1.941 | -1.406 | -0.393 | -0.613 | -0.045 | -0.041 | -0.041 | 0.595 | 0.09 | 0.01 | 0.11 | 0.15 | 0.17 | 0.18 |
| 5 | 76728039 | rs335633 | 1.176 | -1.67 | -0.454 | -0.468 | 0.003 | -0.001 | -0.001 | 0 | 0.08 | 0.09 | 0.21 | 0.16 | 0.07 | 0.27 |
| 5 | 76729378 | rs335630 | -0.775 | -1.645 | -0.614 | -0.732 | -0.049 | -0.012 | -0.012 | 0 | 0.74 | 0.92 | 0.8 | 0.72 | 0.66 | 0.83 |
| 5 | 76730567 | rs335625 | 2.325 | -1.678 | -0.551 | -0.821 | -0.034 | 0.026 | 0.026 | 0 | 0.18 | 0.09 | 0.24 | 0.09 | 0.3 | 0.11 |
| 5 | 76730936 | rs335624 | 0.378 | -1.351 | -0.114 | -0.373 | -0.011 | -0.009 | -0.009 | 0.573 | 0.57 | 0.68 | 0.63 | 0.47 | 0.47 | 0.53 |
| 5 | 76731495 | rs335622 | 0.61 | -1.527 | -0.06 | -0.471 | -0.001 | -0.001 | -0.001 | 0 | 0.42 | 0.42 | 0.58 | 0.47 | 0.42 | 0.51 |
| 5 | 76733379 | rs919224 | 1.15 | -1.602 | -0.126 | -0.457 | -0.002 | -0.001 | -0.001 | 0 | 0.34 | 0.33 | 0.38 | 0.35 | 0.35 | 0.25 |
| 5 | 76734084 | rs33204 | 0.31 | -1.593 | -0.086 | -0.449 | 0.002 | -0.002 | -0.002 | 0.347 | 0.45 | 0.64 | 0.61 | 0.46 | 0.42 | 0.51 |
| 5 | 76734260 | rs33205 | 0.538 | -1.633 | -0.085 | -0.45 | -0.021 | 0.023 | 0.023 | 0 | 0.1 | 0.31 | 0.24 | 0.12 | 0.07 | 0.25 |
| 5 | 76738927 | rs448293 | 0.11 | -1.609 | -0.113 | -0.447 | -0.015 | -0.015 | -0.015 | 0 | 0.58 | 0.74 | 0.65 | 0.47 | 0.47 | 0.52 |
| 5 | 76744977 | rs17751031 | 2.261 | -1.486 | 0.02 | -0.287 | -0.049 | -0.066 | -0.066 | 0.013 | 0.1 | 0.01 | 0.09 | 0.28 | 0.14 | 0.08 |
| 5 | 76749435 | rs179815 | -0.703 | -1.902 | 0.556 | 0.215 | 0.007 | 0.022 | 0.022 | 0 | 0.53 | 0.86 | 0.69 | 0.57 | 0.45 | 0.64 |
| 5 | 76759233 | rs690280 | 0.052 | -1.932 | 0.564 | 0.204 | 0.003 | -0.008 | -0.008 | 0 | 0.41 | 0.71 | 0.61 | 0.46 | 0.4 | 0.47 |
| 5 | 76780042 | rs163035 | -1.211 | -1.884 | 0.366 | 0.131 | -0.079 | -0.044 | -0.044 | 0 | 0.65 | 0.89 | 0.71 | 0.57 | 0.49 | 0.66 |
| 5 | 76780753 | rs163032 | -0.703 | -1.876 | 0.38 | 0.136 | 0.01 | 0.024 | 0.024 | 0 | 0.52 | 0.85 | 0.69 | 0.57 | 0.45 | 0.64 |
| 5 | 76789265 | rs33198 | -0.632 | -1.952 | 0.352 | 0.136 | -0.021 | -0.007 | -0.007 | 0 | 0.58 | 0.82 | 0.67 | 0.57 | 0.45 | 0.63 |
| 5 | 76803523 | rs4273585 | -0.407 | -1.892 | 0.248 | 0.189 | 0.005 | -0.002 | -0.002 | 0 | 0.54 | 0.58 | 0.48 | 0.46 | 0.42 | 0.4 |
| 5 | 76805269 | rs2359879 | 0.029 | -1.914 | 0.206 | 0.192 | -0.026 | -0.022 | -0.022 | 0 | 0.25 | 0.26 | 0.12 | 0.11 | 0.1 | 0.18 |
| 5 | 76807827 | rs11742115 | 0.964 | -1.742 | 0.173 | 0.312 | -0.002 | -0.002 | -0.002 | 0.031 | 0.28 | 0.25 | 0.34 | 0.36 | 0.32 | 0.22 |
| 5 | 76808135 | rs10514104 | 1.3 | -1.834 | -0.617 | 0.306 | -0.05 | -0.043 | -0.043 | 0 | 0.1 | 0.01 | 0.1 | 0.15 | 0.16 | 0.18 |
| 5 | 76825975 | rs2360040 | -0.954 | -1.831 | -0.46 | 0.306 | 0.004 | 0 | 0 | 0 | 0.66 | 0.63 | 0.74 | 0.64 | 0.61 | 0.7 |
| 5 | 76828957 | rs1123168 | 1.056 | -1.41 | -0.238 | 0.213 | -0.001 | -0.001 | -0.001 | 0 | 0.32 | 0.33 | 0.33 | 0.21 | 0.32 | 0.25 |
| 5 | 76832956 | rs6872745 | 0.056 | -1.522 | -0.213 | 0.217 | 0.055 | -0.008 | -0.008 | 0 | 0.51 | 0.71 | 0.47 | 0.45 | 0.37 | 0.36 |
| 5 | 76833452 | rs10042636 | 0.235 | -1.511 | -0.207 | 0.223 | 0.011 | -0.002 | -0.002 | 0 | 0.12 | 0.07 | 0.25 | 0.12 | 0.1 | 0.3 |
| 5 | 76835304 | rs3846672 | -0.005 | -1.524 | -0.158 | 0.22 | -0.006 | -0.008 | -0.008 | 0 | 0.45 | 0.51 | 0.59 | 0.32 | 0.42 | 0.54 |
| 5 | 76835373 | rs3846673 | -1.025 | -1.514 | -0.225 | 0.231 | -0.065 | -0.033 | -0.033 | 0.135 | 0.64 | 0.9 | 0.73 | 0.57 | 0.47 | 0.65 |
| 5 | 76835928 | rs7729096 | 0.605 | -1.443 | -0.152 | 0.216 | 0.003 | 0.245 | 0.245 | 0.01 | 0.07 | 0.3 | 0.11 | 0.24 | 0.01 | 0.09 |
| 5 | 76836055 | rs10070168 | 1.083 | -1.37 | -0.23 | 0.202 | -0.001 | -0.008 | -0.008 | 0 | 0.31 | 0.33 | 0.33 | 0.21 | 0.31 | 0.25 |
| 5 | 76854563 | rs9688203 | -0.712 | -1.366 | 0.069 | 0.53 | -0.006 | -0.003 | -0.003 | 0.123 | 0.14 | 0.23 | 0.26 | 0.11 | 0.1 | 0.3 |
| 5 | 76857870 | rs10052921 | -1.462 | -1.378 | 0.042 | 0.595 | 0.005 | 0 | 0 | 0.718 | 0.37 | 0.69 | 0.44 | 0.36 | 0.19 | 0.42 |
| 5 | 76862566 | rs10063453 | -0.745 | -1.32 | 0.04 | 0.694 | -0.048 | 0.111 | 0.111 | 0 | 0.16 | 0.33 | 0.2 | 0.36 | 0.08 | 0.33 |
| 5 | 76864478 | rs1903262 | -0.054 | -0.696 | -0.216 | 0.702 | 0.023 | 0.097 | 0.097 | 0 | 0.22 | 0.39 | 0.39 | 0.42 | 0.23 | 0.32 |
| 5 | 76868950 | rs6453323 | -0.995 | -0.641 | -1.006 | 0.722 | 0.032 | -0.005 | -0.005 | 0 | 0.47 | 0.69 | 0.55 | 0.53 | 0.35 | 0.59 |
| 5 | 76871345 | rs6453330 | -0.886 | 0.608 | -0.293 | 1.385 | 0.012 | 0.085 | 0.085 | 0.02 | 0.23 | 0.56 | 0.4 | 0.42 | 0.24 | 0.32 |
| 5 | 76890568 | rs4704434 | -0.242 | 0.519 | -0.938 | 1.01 | -0.001 | 0.016 | 0.016 | 0 | 0.46 | 0.67 | 0.48 | 0.38 | 0.46 | 0.45 |
| 5 | 76891021 | rs7703420 | -0.835 | -0.561 | -0.567 | 1.49 | -0.027 | 0.019 | 0.019 | 0 | 0.21 | 0.22 | 0.19 | 0.31 | 0.12 | 0.35 |
| 5 | 76891359 | rs4590167 | -0.768 | -0.323 | -0.188 | 1.396 | -0.02 | -0.002 | -0.002 | 0 | 0.24 | 0.41 | 0.17 | 0.23 | 0.22 | 0.13 |
| 5 | 76903692 | rs3933483 | -0.398 | 0.473 | -0.474 | 1.728 | -0.002 | 0.01 | 0.01 | 0 | 0.16 | 0.16 | 0.14 | 0.19 | 0.12 | 0.11 |
| 5 | 76904032 | rs2124937 | 0.082 | 0.989 | -0.987 | 1.104 | -0.034 | 0.017 | 0.017 | 0.663 | 0.71 | 0.84 | 0.8 | 0.73 | 0.67 | 0.65 |
| 5 | 76919925 | rs6868555 | 0.373 | 0.601 | -0.701 | 0.81 | -0.005 | 0.028 | 0.028 | 0 | 0.54 | 0.67 | 0.47 | 0.33 | 0.51 | 0.48 |
| 5 | 76924534 | rs10055404 | 1.383 | 0.719 | -1.534 | 0.131 | -0.051 | 0.228 | 0.228 | 0.146 | 0.32 | 0.01 | 0.26 | 0.02 | 0.35 | 0.14 |
| 5 | 76928202 | rs6864981 | 0.467 | 0.919 | -0.971 | 0.399 | 0.002 | 0.008 | 0.008 | 0 | 0.49 | 0.55 | 0.44 | 0.33 | 0.47 | 0.42 |
| 5 | 76932225 | rs2241367 | 0.937 | 0.808 | -1.673 | 0.121 | 0.019 | 0.094 | 0.094 | 0.006 | 0.39 | 0.24 | 0.33 | 0.13 | 0.38 | 0.35 |
| 5 | 76933506 | rs11746588 | 0.886 | 1.032 | -1.865 | 0.17 | -0.05 | -0.048 | -0.048 | 0 | 0.14 | 0.02 | 0.15 | 0.22 | 0.2 | 0.18 |
| 5 | 76938952 | rs1000586 | 0.994 | 0.692 | -1.491 | -0.365 | 0.021 | 0.11 | 0.11 | 0.067 | 0.39 | 0.21 | 0.34 | 0.13 | 0.38 | 0.35 |
| 5 | 76942489 | rs1895254 | 0.075 | 0.88 | -0.684 | 0.203 | 0.043 | 0.154 | 0.154 | 1.815 | 0.13 | 0.44 | 0.15 | 0.24 | 0.09 | 0.12 |
| 5 | 76943223 | rs1024985 | 0 | 1.037 | -1.3 | 0.194 | -0.003 | -0.004 | -0.004 | 0 | 0.25 | 0.26 | 0.44 | 0.69 | 0.32 | 0.31 |
| 5 | 76943827 | rs4541640 | -0.084 | 0.785 | -1.475 | -0.032 | -0.012 | 0.079 | 0.079 | 0.035 | 0.84 | 0.97 | 0.93 | 0.98 | 0.89 | 0.83 |

| | | | | | | | | | | | | | | | | |
|---|---|---|---|---|---|---|---|---|---|---|---|---|---|---|---|---|
| 5 | 76949758 | rs4266383 | -0.022 | 0.81 | -1.076 | 0.123 | -0.001 | -0.001 | -0.001 | 0 | 0.73 | 0.72 | 0.65 | 0.79 | 0.66 | 0.77 |
| 5 | 76961468 | rs7341016 | 0.03 | 0.316 | -1.35 | 0.358 | -0.002 | -0.014 | -0.014 | 0 | 0.88 | 0.97 | 0.93 | 0.82 | 0.91 | 0.82 |
| 5 | 76961781 | rs7701985 | 0.581 | 0.144 | -0.863 | 1.067 | -0.006 | 0.059 | 0.059 | 0 | 0.87 | 0.8 | 0.89 | 0.62 | 0.91 | 0.79 |
| 5 | 76967147 | rs10045408 | 0.352 | 0.047 | -0.713 | 0.56 | 0.002 | -0.033 | -0.033 | 0 | 0.45 | 0.48 | 0.54 | 0.26 | 0.47 | 0.47 |
| 5 | 76967446 | rs1363443 | -0.555 | 0.008 | -0.677 | 0.626 | 0.004 | -0.003 | -0.003 | 0 | 0.88 | 0.92 | 0.93 | 0.83 | 0.92 | 0.83 |
| 5 | 76973108 | rs385375 | 0.641 | -0.271 | -0.81 | 0.842 | -0.039 | 0.164 | 0.164 | 1.286 | 0.12 | 0.29 | 0.09 | 0.29 | 0.04 | 0.12 |
| 5 | 76973352 | rs441492 | 0.938 | -0.265 | -0.895 | 0.75 | -0.088 | -0.029 | -0.029 | 0.424 | 0.3 | 0.08 | 0.29 | 0.25 | 0.39 | 0.24 |
| 5 | 76976039 | rs351871 | 0.421 | -0.443 | -1.01 | 0.266 | 0.005 | -0.034 | -0.034 | 0.469 | 0.36 | 0.45 | 0.5 | 0.25 | 0.38 | 0.37 |
| 5 | 76979394 | rs13156814 | 0.303 | -0.733 | -1.029 | 0.209 | -0.005 | -0.028 | -0.028 | 0.874 | 0.19 | 0.31 | 0.1 | 0.06 | 0.16 | 0.28 |
| 5 | 76979432 | rs181270 | -0.484 | -0.815 | -0.97 | 0.272 | -0.008 | 0.005 | 0.005 | 0.677 | 0.32 | 0.62 | 0.19 | 0.36 | 0.2 | 0.39 |
| 5 | 76981809 | rs16874482 | 0.096 | -0.775 | -0.915 | 0.102 | -0.008 | -0.03 | -0.03 | 0 | 0.22 | 0.31 | 0.11 | 0.07 | 0.17 | 0.28 |
| 5 | 76982066 | rs7701420 | 0.426 | -0.776 | -0.86 | 0.124 | -0.009 | 0.008 | 0.008 | 0 | 0.09 | 0.2 | 0.03 | 0.07 | 0.04 | 0.16 |
| 5 | 76983089 | rs422564 | 0.712 | -0.636 | -0.951 | 0.128 | -0.002 | 0.006 | 0.006 | 0 | 0.43 | 0.31 | 0.34 | 0.4 | 0.44 | 0.4 |
| 5 | 76999353 | rs17789838 | 0.661 | -0.789 | -1.098 | 0.126 | -0.008 | -0.051 | -0.051 | 0 | 0.14 | 0.01 | 0.08 | 0.18 | 0.09 | 0.19 |
| 5 | 77008253 | rs2652210 | -0.526 | -0.817 | -1.08 | -0.011 | 0.008 | 0.026 | 0.026 | 0 | 0.66 | 0.62 | 0.45 | 0.47 | 0.61 | 0.68 |
| 5 | 77011066 | rs9791058 | 0.204 | -0.863 | -1.164 | 0.006 | -0.016 | 0.041 | 0.041 | 0 | 0.18 | 0.21 | 0.1 | 0.36 | 0.09 | 0.22 |
| 5 | 77019909 | rs384109 | 1.728 | -0.741 | -1.295 | -0.201 | -0.103 | 0.146 | 0.146 | 0.368 | 0.27 | 0.01 | 0.24 | 0.04 | 0.35 | 0.19 |
| 5 | 77026166 | rs11954532 | 1.468 | -0.651 | -1.093 | -0.056 | 0.001 | 0.008 | 0.008 | 0 | 0.14 | 0.1 | 0.07 | 0 | 0.13 | 0.12 |
| 5 | 77079857 | rs254411 | 0.415 | -0.651 | -0.774 | -0.03 | -0.001 | -0.016 | -0.016 | 0 | 0.47 | 0.46 | 0.36 | 0.11 | 0.51 | 0.43 |
| 5 | 77083651 | rs1542891 | 0.524 | -1.616 | -0.64 | 0.163 | 0.004 | 0 | 0 | 0 | 0.17 | 0.14 | 0.09 | 0.18 | 0.1 | 0.23 |
| 5 | 77094403 | rs6453351 | 0.463 | -1.36 | -0.761 | 0.188 | -0.012 | -0.049 | -0.049 | 0.006 | 0.3 | 0.4 | 0.28 | 0.21 | 0.26 | 0.15 |
| 5 | 77097396 | rs353948 | -0.102 | -1.186 | -0.666 | 0.373 | -0.012 | 0.157 | 0.157 | 0 | 0.11 | 0.25 | 0.08 | 0.39 | 0.06 | 0.27 |
| 5 | 77113617 | rs353958 | -0.725 | -1.445 | -0.738 | 0.34 | -0.008 | 0.004 | 0.004 | 0 | 0.67 | 0.79 | 0.71 | 0.7 | 0.63 | 0.74 |
| 5 | 77114598 | rs1949026 | 1.708 | -1.479 | -0.855 | 0.294 | -0.117 | 0.143 | 0.143 | 0 | 0.29 | 0.08 | 0.49 | 0.04 | 0.47 | 0.24 |
| 5 | 77116391 | rs16874730 | 0.945 | -1.499 | -0.787 | 0.349 | -0.008 | 0.004 | 0.004 | 0 | 0.14 | 0.14 | 0.09 | 0.26 | 0.06 | 0.19 |
| 5 | 77117657 | rs353961 | 0.651 | -1.437 | -1.569 | 0.378 | -0.01 | -0.016 | -0.016 | 0 | 0.42 | 0.35 | 0.58 | 0.43 | 0.55 | 0.54 |
| 5 | 77120649 | rs10942832 | -1.177 | -1.659 | -1.402 | 0.942 | 0.068 | 0.071 | 0.071 | 0.132 | 0.39 | 0.74 | 0.23 | 0.84 | 0.16 | 0.53 |
| 5 | 77121824 | rs1396612 | -1.18 | -1.65 | -1.434 | 0.946 | 0.073 | 0.077 | 0.077 | 0.269 | 0.39 | 0.77 | 0.23 | 0.84 | 0.16 | 0.53 |
| 5 | 77122856 | rs354613 | -0.177 | -0.851 | -0.825 | 1.546 | -0.051 | 0.208 | 0.208 | 1.551 | 0.17 | 0.58 | 0.13 | 0.52 | 0.06 | 0.19 |
| 5 | 77126210 | rs6884279 | -0.075 | -0.1 | -1.611 | 0.627 | 0.082 | 0.08 | 0.08 | 0 | 0.45 | 0.73 | 0.29 | 0.76 | 0.26 | 0.48 |
| 5 | 77128654 | rs10514122 | -0.38 | 0.716 | -1.625 | 1.085 | -0.066 | -0.008 | -0.008 | 0 | 0.2 | 0.02 | 0.2 | 0.14 | 0.25 | 0.17 |
| 5 | 77131099 | rs353975 | -1.145 | 0.436 | -0.672 | 2.13 | 0.015 | 0.325 | 0.325 | 0 | 0.05 | 0.18 | 0.13 | 0.38 | 0.04 | 0.18 |
| 5 | 77134397 | rs354599 | -0.621 | 0.868 | -0.876 | 1.428 | -0.006 | -0.004 | -0.004 | 0 | 0.13 | 0.08 | 0.3 | 0.15 | 0.15 | 0.18 |
| 5 | 77134589 | rs354598 | -0.517 | 0.251 | -0.81 | 0.724 | -0.003 | -0.006 | -0.006 | 0 | 0.31 | 0.31 | 0.19 | 0.25 | 0.23 | 0.26 |
| 5 | 77142481 | rs7728693 | -0.351 | 0.147 | -1.15 | 0.404 | -0.056 | 0.107 | 0.107 | 0 | 0.62 | 0.87 | 0.42 | 0.2 | 0.55 | 0.52 |
| 5 | 77144066 | rs6881736 | -0.382 | 0.07 | -1.235 | 0.36 | -0.074 | 0.082 | 0.082 | 0 | 0.62 | 0.91 | 0.47 | 0.23 | 0.55 | 0.57 |
| 5 | 77152160 | rs2361312 | -0.227 | 0.048 | -0.391 | 0.24 | -0.003 | 0 | 0 | 0 | 0.57 | 0.6 | 0.52 | 0.6 | 0.54 | 0.63 |
| 5 | 77156734 | rs1443096 | -0.119 | -0.411 | -0.877 | -0.331 | 0.002 | -0.004 | -0.004 | 0 | 0.3 | 0.7 | 0.35 | 0.43 | 0.21 | 0.42 |
| 5 | 77156883 | rs1443097 | -1.034 | -0.418 | -1.063 | -0.359 | -0.018 | -0.021 | -0.021 | 0.015 | 0.62 | 0.81 | 0.56 | 0.6 | 0.59 | 0.67 |
| 5 | 77159396 | rs10514124 | 0.476 | -0.347 | -1.587 | -0.604 | -0.06 | 0.068 | 0.068 | 0 | 0.31 | 0.07 | 0.21 | 0.17 | 0.37 | 0.25 |
| 5 | 77159956 | rs7730738 | 0.649 | -0.586 | -1.156 | -0.523 | -0.01 | -0.055 | -0.055 | 0.051 | 0.2 | 0.25 | 0.14 | 0.03 | 0.13 | 0.19 |
| 5 | 77161489 | rs12518218 | 0.266 | -0.445 | -1.823 | -0.557 | -0.083 | 0.114 | 0.114 | 0.676 | 0.36 | 0.07 | 0.24 | 0.17 | 0.42 | 0.25 |
| 5 | 77165430 | rs6860985 | 0.316 | -0.734 | -1.991 | -0.525 | -0.032 | 0.085 | 0.085 | 0 | 0.33 | 0.13 | 0.21 | 0.16 | 0.39 | 0.25 |
| 5 | 77168090 | rs4490567 | -0.25 | -0.423 | -2.117 | -0.278 | -0.024 | 0.197 | 0.197 | 0.896 | 0.52 | 0.9 | 0.41 | 0.23 | 0.45 | 0.31 |
| 5 | 77170420 | rs9687161 | -1.087 | -0.214 | -2.041 | 0.021 | -0.131 | -0.133 | -0.133 | 0 | 0.71 | 0.97 | 0.6 | 0.62 | 0.61 | 0.62 |
| 5 | 77170584 | rs2028595 | -0.832 | -0.329 | -1.314 | -0.486 | 0.021 | 0.151 | 0.151 | 2.58 | 0.45 | 0.85 | 0.39 | 0.22 | 0.4 | 0.26 |
| 5 | 77173522 | rs7734027 | -0.301 | 0.008 | -0.782 | -0.074 | -0.001 | -0.01 | -0.01 | 0 | 0.31 | 0.65 | 0.27 | 0.18 | 0.32 | 0.24 |
| 5 | 77177468 | rs9293725 | -0.729 | 0.218 | -0.877 | 0.646 | -0.009 | -0.027 | -0.027 | 1.669 | 0.24 | 0.19 | 0.35 | 0.44 | 0.25 | 0.3 |
| 5 | 77182655 | rs10514128 | -0.637 | 0.087 | -0.516 | 0.941 | 0 | 0.083 | 0.083 | 0 | 0.18 | 0.31 | 0.21 | 0.49 | 0.17 | 0.21 |
| 5 | 77183087 | rs10440685 | -1.063 | -0.411 | -0.029 | 0.308 | -0.02 | -0.104 | -0.104 | 0 | 0.19 | 0.42 | 0.28 | 0.03 | 0.1 | 0.13 |
| 5 | 77201838 | rs2099891 | -0.523 | -0.187 | -1.141 | 0.153 | 0.005 | 0.026 | 0.026 | 0 | 0.15 | 0.09 | 0.25 | 0.03 | 0.1 | 0.15 |
| 5 | 77202560 | rs10069818 | 0.638 | 0.048 | -1.765 | 0.058 | -0.077 | -0.007 | -0.007 | 1.605 | 0.36 | 0.12 | 0.35 | 0.37 | 0.46 | 0.33 |
| 5 | 77208149 | rs2289599 | 0.054 | -0.106 | -1.962 | -0.262 | -0.064 | 0.052 | 0.052 | 0.145 | 0.8 | 0.98 | 0.83 | 0.91 | 0.7 | 0.71 |
| 5 | 77209661 | rs5014235 | 0.073 | -0.257 | -2.031 | -0.329 | -0.057 | 0.077 | 0.077 | 0.007 | 0.78 | 0.98 | 0.83 | 0.91 | 0.7 | 0.71 |
| 5 | 77210168 | rs4704458 | 0.582 | -0.408 | -1.762 | -0.551 | -0.012 | 0.079 | 0.079 | 0 | 0.73 | 0.88 | 0.8 | 0.91 | 0.67 | 0.69 |
| 5 | 77210880 | rs2120101 | 1.357 | -0.887 | -1.517 | -0.752 | 0.019 | 0.074 | 0.074 | 0 | 0.51 | 0.37 | 0.59 | 0.75 | 0.5 | 0.55 |
| 5 | 77211670 | rs355425 | -0.644 | -0.87 | -1.507 | -0.695 | 0.005 | 0.004 | 0.004 | 0 | 0.21 | 0.1 | 0.17 | 0.15 | 0.18 | 0.16 |
| 5 | 77216237 | rs902523 | -0.579 | -0.867 | -1.565 | -0.572 | 0.011 | -0.001 | -0.001 | 0 | 0.18 | 0.07 | 0.11 | 0.1 | 0.15 | 0.13 |
| 5 | 77218488 | rs6453360 | 0.076 | -0.72 | -0.926 | -0.59 | 0.001 | 0.02 | 0.02 | 0 | 0.69 | 0.75 | 0.78 | 0.96 | 0.63 | 0.71 |
| 5 | 77219131 | rs4704462 | 1.03 | -0.542 | -0.707 | 0.109 | 0.01 | -0.002 | -0.002 | 1.371 | 0.51 | 0.44 | 0.56 | 0.56 | 0.51 | 0.5 |
| 5 | 77222803 | rs10942835 | -0.977 | -0.523 | -0.79 | 0.058 | 0.002 | 0.009 | 0.009 | 0 | 0.52 | 0.63 | 0.4 | 0.5 | 0.5 | 0.5 |
| 5 | 77222819 | rs4704463 | -1.015 | -0.83 | -0.206 | 0.927 | -0.024 | 0.275 | 0.275 | 0 | 0.12 | 0.42 | 0.1 | 0.36 | 0.05 | 0.11 |
| 5 | 77224214 | rs355438 | -1.246 | -1.194 | -0.402 | 0.718 | -0.076 | 0.082 | 0.082 | 0 | 0.26 | 0.5 | 0.22 | 0.52 | 0.1 | 0.19 |
| 5 | 77224666 | rs7712357 | -0.332 | -0.077 | -1.523 | 0.314 | -0.047 | -0.009 | -0.009 | 0.769 | 0.52 | 0.89 | 0.7 | 0.5 | 0.51 | 0.43 |
| 5 | 77225805 | rs355443 | -0.051 | -0.578 | -1.385 | 0.669 | 0.068 | -0.027 | -0.027 | 0 | 0.17 | 0.02 | 0.03 | 0.14 | 0.07 | 0.09 |
| 5 | 77231482 | rs4549507 | -0.694 | -0.417 | -1.323 | 0.735 | -0.026 | -0.065 | -0.065 | 0.443 | 0.8 | 0.95 | 0.87 | 0.63 | 0.83 | 0.74 |
| 5 | 77239920 | rs837036 | 1.177 | -0.114 | -0.632 | 0.217 | 0.026 | -0.003 | -0.003 | 0 | 0.28 | 0.42 | 0.48 | 0.27 | 0.38 | 0.26 |
| 5 | 77244796 | rs7725100 | 0.811 | -0.038 | -0.31 | 0.053 | 0.001 | 0.033 | 0.033 | 0.019 | 0.35 | 0.26 | 0.24 | 0.14 | 0.37 | 0.33 |
| 5 | 77245012 | rs4484406 | -0.766 | -0.005 | -0.365 | 0.236 | 0.002 | 0.023 | 0.023 | 0 | 0.79 | 0.77 | 0.75 | 0.54 | 0.8 | 0.68 |

| | | | | | | | | | | | | | | | | |
|---|---|---|---|---|---|---|---|---|---|---|---|---|---|---|---|---|
| 5 | 77247878 | rs10044946 | 1.141 | 0.592 | -0.192 | 0.269 | 0.016 | 0.019 | 0.019 | 1.103 | 0.27 | 0.42 | 0.46 | 0.35 | 0.33 | 0.25 |
| 5 | 77249424 | rs12521066 | 0.392 | 0.549 | -0.35 | -0.222 | 0.008 | 0.045 | 0.045 | 0.141 | 0.33 | 0.21 | 0.22 | 0.14 | 0.34 | 0.34 |
| 5 | 77259149 | rs7709783 | -0.835 | 0.699 | 0.188 | 0.074 | 0.155 | 0.053 | 0.053 | 0 | 0.05 | 0.33 | 0.06 | 0 | 0.08 | 0 |
| 5 | 77262282 | rs355421 | 0.042 | 0.17 | -0.473 | 0.074 | -0.006 | -0.042 | -0.042 | 0 | 0.11 | 0 | 0.04 | 0.13 | 0.08 | 0.07 |
| 5 | 77262582 | rs13154492 | -0.186 | 0.362 | -0.223 | 0.239 | 0.111 | 0.219 | 0.219 | 0 | 0.05 | 0.36 | 0.14 | 0.07 | 0.04 | 0.07 |
| 5 | 77272926 | rs355416 | 0.441 | 0.467 | -1.59 | -0.799 | -0.006 | 0.031 | 0.031 | 0 | 0.32 | 0.23 | 0.47 | 0.01 | 0.37 | 0.18 |
| 5 | 77273254 | rs355415 | -0.792 | 0.045 | -1.539 | 0.09 | 0.018 | -0.019 | -0.019 | 0 | 0.19 | 0.01 | 0.11 | 0.15 | 0.11 | 0.16 |
| 5 | 77277101 | rs6874954 | 0.057 | 0.536 | -1.177 | 0.266 | -0.002 | -0.003 | -0.003 | 0.191 | 0.55 | 0.5 | 0.43 | 0.48 | 0.55 | 0.49 |
| 5 | 77278872 | rs10064609 | -0.323 | -0.033 | -1.246 | 0.274 | -0.007 | 0 | 0 | 0 | 0.48 | 0.51 | 0.39 | 0.42 | 0.37 | 0.38 |
| 5 | 77280135 | rs355412 | 0.08 | 0.64 | -2.102 | -0.313 | -0.13 | 0.188 | 0.188 | 0.063 | 0.65 | 0.98 | 0.56 | 0.92 | 0.54 | 0.74 |
| 5 | 77282521 | rs355410 | -0.357 | -0.03 | -0.797 | 0.667 | -0.023 | 0.065 | 0.065 | 0 | 0.3 | 0.49 | 0.31 | 0.46 | 0.27 | 0.3 |
| 5 | 77284789 | rs7708242 | 0.875 | 0.539 | 0.07 | 0.568 | -0.006 | 0.048 | 0.048 | 0 | 0.76 | 0.58 | 0.72 | 0.67 | 0.76 | 0.81 |
| 5 | 77289748 | rs1873425 | -0.826 | 0.827 | -0.46 | 0.218 | -0.01 | -0.033 | -0.033 | 0 | 0.57 | 0.77 | 0.51 | 0.5 | 0.61 | 0.45 |
| 5 | 77290512 | rs17189696 | 0.193 | 0.981 | -0.543 | 0.423 | -0.045 | -0.028 | -0.028 | 0.837 | 0.1 | 0.03 | 0.1 | 0.12 | 0.17 | 0.12 |
| 5 | 77290902 | rs10078715 | -0.519 | 0.505 | 0.235 | 0.47 | -0.012 | 0.09 | 0.09 | 0 | 0.11 | 0.22 | 0.16 | 0.21 | 0.07 | 0.06 |
| 5 | 77292714 | rs6887042 | 0.135 | 0.56 | 0.272 | -0.253 | -0.003 | -0.038 | -0.038 | 0.375 | 0.32 | 0.39 | 0.2 | 0.17 | 0.32 | 0.21 |
| 5 | 77296281 | rs16874991 | -1.228 | 0.793 | 0.948 | 0.15 | 0.03 | 0.122 | 0.122 | 0.032 | 0.16 | 0.43 | 0.45 | 0.28 | 0.2 | 0.16 |
| 5 | 77297771 | rs13168272 | -0.695 | 0.85 | 1.017 | 0.169 | 0.043 | 0.018 | 0.018 | 0 | 0.12 | 0.26 | 0.33 | 0.12 | 0.21 | 0.12 |
| 5 | 77306390 | rs3776924 | -1.63 | 0.827 | 1.127 | 0.198 | 0.04 | 0.137 | 0.137 | 0 | 0.17 | 0.45 | 0.47 | 0.29 | 0.22 | 0.17 |
| 5 | 77314586 | rs4704474 | -0.068 | 0.828 | 0.999 | -0.063 | -0.053 | -0.044 | -0.044 | 0 | 0.09 | 0.02 | 0.31 | 0.12 | 0.21 | 0.12 |
| 5 | 77336576 | rs380913 | -1.654 | 0.686 | 1.156 | -0.005 | 0.02 | 0.076 | 0.076 | 0.211 | 0.16 | 0.42 | 0.45 | 0.22 | 0.19 | 0.16 |
| 5 | 77339846 | rs11746090 | 0.348 | 0.936 | 0.529 | -0.577 | -0.113 | 0.121 | 0.121 | 0.239 | 0.19 | 0.02 | 0.16 | 0.03 | 0.29 | 0.3 |
| 5 | 77343939 | rs444908 | -1.497 | 0.801 | 0.932 | -0.263 | 0.023 | 0.044 | 0.044 | 0.021 | 0.16 | 0.39 | 0.45 | 0.19 | 0.19 | 0.17 |
| 5 | 77346011 | rs252800 | 0.41 | 0.829 | 0.7 | -0.259 | -0.047 | -0.011 | -0.011 | 0 | 0.08 | 0.01 | 0.29 | 0.07 | 0.17 | 0.11 |
| 5 | 77352850 | rs252782 | -1.272 | 0.827 | 0.785 | -0.251 | 0.024 | 0.046 | 0.046 | 0.02 | 0.16 | 0.38 | 0.45 | 0.19 | 0.19 | 0.17 |
| 5 | 77359088 | rs34436 | -1.625 | 0.311 | 0.465 | -0.395 | 0.063 | -0.1 | -0.1 | 0.165 | 0.78 | 0.96 | 0.88 | 0.46 | 0.89 | 0.76 |
| 5 | 77389973 | rs252749 | -1.151 | 1.427 | 1.019 | -0.401 | 0.061 | -0.033 | -0.033 | 2.566 | 0.14 | 0.42 | 0.35 | 0.08 | 0.21 | 0.12 |
| 5 | 77412011 | rs42360 | -0.561 | 1.442 | 1.068 | -0.394 | 0.029 | -0.017 | -0.017 | 2.3 | 0.12 | 0.24 | 0.33 | 0.08 | 0.21 | 0.12 |
| 5 | 77458150 | rs4704484 | -1.722 | 1.399 | 0.92 | -0.337 | 0.023 | 0.018 | 0.018 | 0.112 | 0.19 | 0.4 | 0.43 | 0.19 | 0.22 | 0.17 |
| 5 | 77461566 | rs6873309 | -1.448 | 1.417 | 1.026 | -0.32 | -0.009 | 0.077 | 0.077 | 0 | 0.09 | 0.3 | 0.12 | 0.12 | 0.02 | 0.06 |
| 5 | 77464544 | rs4388193 | -0.661 | 1.437 | 0.964 | -0.311 | -0.012 | 0.051 | 0.051 | 0 | 0.07 | 0.16 | 0.1 | 0.12 | 0.02 | 0.06 |
| 5 | 77473165 | rs4532349 | 0.06 | 1.483 | 0.851 | -0.31 | -0.001 | 0 | 0 | 0 | 0.1 | 0.1 | 0.3 | 0.07 | 0.2 | 0.11 |
| 5 | 77493085 | rs10039112 | -0.18 | 1.438 | 0.954 | -0.553 | -0.002 | -0.014 | -0.014 | 0 | 0.39 | 0.51 | 0.33 | 0.32 | 0.36 | 0.28 |
| 5 | 77498846 | rs9293729 | -0.192 | 1.439 | 0.955 | -0.566 | -0.003 | -0.017 | -0.017 | 0.006 | 0.39 | 0.51 | 0.33 | 0.32 | 0.36 | 0.27 |
| 5 | 77499566 | rs11742673 | -0.881 | 1.476 | 0.894 | -0.573 | 0.008 | 0.031 | 0.031 | 0 | 0.49 | 0.61 | 0.63 | 0.39 | 0.55 | 0.38 |
| 5 | 77500665 | rs7733511 | 0.486 | 1.471 | 0.989 | -0.575 | 0.002 | -0.016 | -0.016 | 0 | 0.32 | 0.34 | 0.22 | 0.2 | 0.34 | 0.22 |
| 5 | 77501557 | rs4505927 | -0.824 | 1.464 | 0.78 | -0.559 | 0.007 | 0.03 | 0.03 | 0 | 0.49 | 0.61 | 0.63 | 0.39 | 0.55 | 0.38 |
| 5 | 77515737 | rs7721185 | 0.534 | 1.171 | 0.7 | -0.534 | -0.011 | 0.037 | 0.037 | 0 | 0.31 | 0.22 | 0.2 | 0.1 | 0.34 | 0.21 |
| 5 | 77537371 | rs6865361 | -0.238 | 1.131 | 0.605 | -0.505 | -0.002 | -0.018 | -0.018 | 3.231 | 0.39 | 0.48 | 0.32 | 0.32 | 0.36 | 0.27 |
| 5 | 77546796 | rs7726585 | -0.839 | 1.805 | 0.485 | -0.586 | 0.018 | 0.021 | 0.021 | 0.15 | 0.52 | 0.56 | 0.62 | 0.39 | 0.59 | 0.44 |
| 5 | 77572439 | rs10059285 | 0.428 | 1.673 | -0.131 | -0.635 | -0.048 | 0.053 | 0.053 | 0 | 0.3 | 0.07 | 0.19 | 0.2 | 0.31 | 0.23 |
| 5 | 77579205 | rs10062813 | -0.791 | 1.71 | 0.42 | -0.624 | 0.006 | 0.013 | 0.013 | 0.096 | 0.5 | 0.55 | 0.52 | 0.4 | 0.56 | 0.43 |
| 5 | 77582571 | rs4593248 | -0.953 | 1.702 | 0.45 | -0.628 | 0 | 0.001 | 0.001 | 0.135 | 0.5 | 0.52 | 0.51 | 0.4 | 0.56 | 0.43 |
| 5 | 77592924 | rs10805919 | -1.002 | 1.795 | 0.573 | -0.575 | 0.052 | 0.002 | 0.002 | 0.365 | 0.19 | 0.36 | 0.32 | 0.19 | 0.25 | 0.2 |
| 5 | 77598854 | rs9790855 | -0.131 | 1.767 | 0.354 | -0.666 | 0.001 | 0.002 | 0.002 | 0 | 0.39 | 0.4 | 0.25 | 0.33 | 0.34 | 0.32 |
| 5 | 77606215 | rs4572979 | -0.756 | 1.58 | 0.059 | -0.616 | 0.002 | 0.003 | 0.003 | 0.568 | 0.53 | 0.57 | 0.52 | 0.38 | 0.58 | 0.47 |
| 5 | 77616951 | rs7727630 | -0.506 | 1.601 | -0.653 | -1.266 | 0.016 | 0.159 | 0.159 | 0.791 | 0.49 | 0.26 | 0.47 | 0.09 | 0.56 | 0.44 |
| 5 | 77618179 | rs10474543 | 0.083 | 1.165 | -0.81 | -1.103 | 0.016 | 0.121 | 0.121 | 0 | 0.33 | 0.14 | 0.18 | 0.05 | 0.29 | 0.26 |
| 5 | 77624910 | rs4704495 | 0.357 | 1.032 | -0.578 | -0.76 | 0.014 | 0.031 | 0.031 | 0.357 | 0.48 | 0.62 | 0.59 | 0.63 | 0.37 | 0.54 |
| 5 | 77624948 | rs4704496 | 0.266 | 0.966 | -1.226 | -0.919 | -0.069 | 0.011 | 0.011 | 0.146 | 0.13 | 0.01 | 0.43 | 0.06 | 0.19 | 0.16 |
| 5 | 77630854 | rs6881634 | -0.078 | 0.639 | -0.894 | -1.222 | 0.03 | 0.215 | 0.215 | 0.847 | 0.49 | 0.26 | 0.4 | 0.1 | 0.63 | 0.38 |
| 5 | 77631634 | rs4588572 | 0.091 | 0.837 | -0.677 | -0.927 | -0.037 | 0.02 | 0.02 | 0 | 0.15 | 0.06 | 0.43 | 0.07 | 0.22 | 0.18 |
| 5 | 77640190 | rs11749979 | 0.807 | 1.021 | -0.294 | -1.136 | 0.002 | 0 | 0 | 0 | 0.57 | 0.63 | 0.28 | 0.54 | 0.4 | 0.45 |
| 5 | 77645784 | rs7722126 | 0.588 | 0.975 | 0.007 | -1.011 | 0.007 | 0.022 | 0.022 | 0 | 0.78 | 0.75 | 0.51 | 0.6 | 0.75 | 0.69 |
| 5 | 77647764 | rs4267850 | 0.978 | 1.078 | 0.163 | -1.183 | 0.004 | 0.001 | 0.001 | 0.135 | 0.61 | 0.59 | 0.36 | 0.54 | 0.53 | 0.51 |
| 5 | 77656112 | rs4530741 | 1.335 | 0.374 | 0.401 | -1.102 | 0.023 | -0.021 | -0.021 | 0 | 0.36 | 0.24 | 0.16 | 0.53 | 0.26 | 0.41 |
| 5 | 77669593 | rs4235707 | -0.845 | 0.369 | 0.318 | -1.038 | 0 | 0.048 | 0.048 | 0 | 0.95 | 0.95 | 0.96 | 0.7 | 0.97 | 0.91 |
| 5 | 77670793 | rs10078289 | 0.412 | 0.538 | 0.418 | -1.119 | -0.016 | 0.008 | 0.008 | 0 | 0.14 | 0.08 | 0.44 | 0.08 | 0.21 | 0.16 |
| 5 | 77670996 | rs9293746 | -0.646 | 0.519 | 0.433 | -1.107 | -0.001 | 0.052 | 0.052 | 0 | 0.95 | 0.93 | 0.96 | 0.7 | 0.98 | 0.91 |
| 5 | 77673641 | rs10054586 | 0.103 | 0.505 | 0.7 | -1.264 | 0.003 | -0.078 | -0.078 | 0 | 0.42 | 0.42 | 0.34 | 0.08 | 0.51 | 0.34 |
| 5 | 77683490 | rs4703756 | 0.408 | 0.572 | 0.486 | -1.168 | -0.016 | 0.008 | 0.008 | 0 | 0.14 | 0.08 | 0.44 | 0.08 | 0.21 | 0.16 |
| 5 | 77684726 | rs3922654 | 0.4 | 0.512 | 0.513 | -1.167 | -0.014 | 0.007 | 0.007 | 0 | 0.14 | 0.08 | 0.44 | 0.08 | 0.21 | 0.16 |
| 5 | 77716621 | rs7701154 | -0.648 | 0.483 | 0.502 | -1.167 | -0.015 | 0.008 | 0.008 | 0 | 0.86 | 0.92 | 0.56 | 0.92 | 0.79 | 0.84 |
| 5 | 77724891 | rs13159112 | 0.241 | 0.405 | 0.311 | -1.289 | -0.101 | 0.242 | 0.242 | 0.555 | 0.39 | 0.1 | 0.31 | 0.09 | 0.51 | 0.34 |
| 5 | 77739350 | rs6860842 | 1.298 | 0.378 | 0.405 | -1.207 | 0.009 | -0.005 | -0.005 | 0 | 0.37 | 0.32 | 0.17 | 0.54 | 0.26 | 0.41 |
| 5 | 77759362 | rs10063742 | -0.698 | 0.421 | 0.343 | -1.194 | -0.019 | 0.011 | 0.011 | 0 | 0.86 | 0.93 | 0.56 | 0.92 | 0.79 | 0.84 |
| 5 | 77772702 | rs6899253 | 0.779 | 0.398 | 0.295 | -1.116 | 0 | 0.048 | 0.048 | 0 | 0.05 | 0.05 | 0.04 | 0.3 | 0.02 | 0.09 |
| 5 | 77773152 | rs1046819 | 1.292 | 0.397 | 0.266 | -1.154 | 0.011 | -0.007 | -0.007 | 0.031 | 0.37 | 0.28 | 0.16 | 0.54 | 0.26 | 0.4 |
| 5 | 77784542 | rs11740697 | 0.655 | 0.045 | -0.276 | -1.464 | -0.149 | 0.222 | 0.222 | 0.199 | 0.35 | 0.05 | 0.29 | 0.09 | 0.51 | 0.27 |

| | | | | | | | | | | | | | | | | |
|---|---|---|---|---|---|---|---|---|---|---|---|---|---|---|---|---|
| 5 | 77788791 | rs16875506 | 0.525 | -0.099 | 0.388 | -0.957 | 0.055 | 0.086 | 0.086 | 0.431 | 0.45 | 0.74 | 0.23 | 0.74 | 0.28 | 0.51 |
| 5 | 77796711 | rs11737914 | 1.236 | -0.077 | 0.236 | -0.709 | 0.022 | -0.05 | -0.05 | 0 | 0.33 | 0.17 | 0.16 | 0.47 | 0.24 | 0.4 |
| 5 | 77799836 | rs6453397 | 0.777 | -0.194 | 0.621 | -0.341 | -0.005 | -0.006 | -0.006 | 0 | 0.38 | 0.39 | 0.2 | 0.69 | 0.27 | 0.5 |
| 5 | 77801359 | rs876605 | 0.99 | 0.148 | 0.541 | -0.665 | 0.002 | 0.008 | 0.008 | 0.062 | 0.73 | 0.34 | 0.47 | 0.66 | 0.74 | 0.7 |
| 5 | 77802188 | rs751659 | -0.433 | 0.184 | 0.879 | -0.736 | -0.02 | -0.122 | -0.122 | 0 | 0.16 | 0.49 | 0.25 | 0 | 0.09 | 0.05 |
| 5 | 77803267 | rs10942858 | 1.159 | 0.186 | 0.666 | -0.743 | -0.011 | -0.05 | -0.05 | 0 | 0.76 | 0.38 | 0.49 | 0.93 | 0.79 | 0.84 |
| 5 | 77809989 | rs11749358 | 1.254 | 0.149 | 0.374 | -0.693 | 0.002 | -0.005 | -0.005 | 0.181 | 0.33 | 0.28 | 0.17 | 0.49 | 0.25 | 0.38 |
| 5 | 77812321 | rs2362834 | -0.528 | 0.2 | 0.232 | -0.716 | -0.013 | -0.013 | -0.013 | 0 | 0.93 | 0.96 | 0.75 | 0.93 | 0.88 | 0.89 |
| 5 | 77812753 | rs6875879 | -0.027 | -0.334 | 0.127 | -1.127 | -0.011 | 0.016 | 0.016 | 0 | 0.33 | 0.55 | 0.36 | 0.48 | 0.28 | 0.41 |
| 5 | 77812808 | rs6861942 | 0.001 | -0.287 | 0.201 | -1.037 | 0.007 | 0.008 | 0.008 | 0 | 0.44 | 0.56 | 0.4 | 0.59 | 0.34 | 0.44 |
| 5 | 77813139 | rs890744 | -0.992 | -0.112 | -0.132 | -0.982 | 0.005 | -0.023 | -0.023 | 2.292 | 0.68 | 0.75 | 0.81 | 0.52 | 0.74 | 0.62 |
| 5 | 77815056 | rs17804233 | 1.099 | -0.155 | -0.317 | -1.122 | -0.133 | 0.193 | 0.193 | 0.235 | 0.41 | 0.08 | 0.29 | 0.21 | 0.52 | 0.36 |
| 5 | 77816007 | rs10474556 | 0.089 | -0.208 | 0.184 | -0.846 | -0.003 | 0.002 | 0.002 | 0.408 | 0.44 | 0.4 | 0.39 | 0.43 | 0.35 | 0.43 |
| 5 | 77816427 | rs12189541 | 0.621 | 0.019 | -0.12 | -1.106 | 0.015 | 0.043 | 0.043 | 0 | 0.53 | 0.38 | 0.35 | 0.41 | 0.53 | 0.43 |
| 5 | 77820081 | rs4704514 | 0.3 | 0.468 | 0.154 | -0.773 | 0.002 | 0 | 0 | 0.118 | 0.17 | 0.16 | 0.29 | 0.2 | 0.28 | 0.18 |
| 5 | 77826346 | rs17215893 | 0.247 | 0.057 | 0.107 | -0.825 | -0.008 | 0.089 | 0.089 | 0 | 0.41 | 0.22 | 0.29 | 0.25 | 0.49 | 0.32 |
| 5 | 77837789 | rs10057188 | 0.274 | 0.144 | -0.117 | -0.814 | -0.105 | 0.163 | 0.163 | 0.05 | 0.39 | 0.04 | 0.28 | 0.17 | 0.48 | 0.31 |
| 5 | 77848586 | rs344652 | 1.109 | -0.004 | -0.025 | -0.774 | 0.066 | -0.058 | -0.058 | 0 | 0.33 | 0.16 | 0.14 | 0.47 | 0.16 | 0.4 |
| 5 | 77852998 | rs344651 | 1.093 | -0.012 | -0.009 | -0.758 | 0.07 | -0.058 | -0.058 | 0 | 0.33 | 0.16 | 0.14 | 0.49 | 0.14 | 0.4 |
| 5 | 77866228 | rs1561735 | -0.221 | 0.102 | 0.253 | -0.893 | 0.003 | 0.024 | 0.024 | 0.008 | 0.45 | 0.6 | 0.33 | 0.26 | 0.5 | 0.31 |
| 5 | 77874205 | rs10060400 | -0.082 | 0.139 | -0.219 | -0.901 | -0.003 | 0.057 | 0.057 | 0 | 0.42 | 0.29 | 0.3 | 0.25 | 0.5 | 0.31 |
| 5 | 77881710 | rs890746 | -0.08 | 0.136 | -0.251 | -0.902 | -0.009 | 0.095 | 0.095 | 0 | 0.42 | 0.27 | 0.3 | 0.25 | 0.5 | 0.31 |
| 5 | 77894770 | rs17455697 | 0.034 | 0.076 | -0.14 | -1.744 | -0.009 | 0.075 | 0.075 | 0 | 0.4 | 0.26 | 0.3 | 0.25 | 0.5 | 0.31 |
| 5 | 77901235 | rs12186748 | -0.01 | 0.084 | -0.305 | -1.74 | -0.038 | 0.107 | 0.107 | 0 | 0.39 | 0.15 | 0.29 | 0.25 | 0.5 | 0.31 |
| 5 | 77903262 | rs10052968 | 1.381 | 0.473 | -0.524 | -1.709 | -0.062 | 0.099 | 0.099 | 0 | 0.16 | 0.01 | 0.1 | 0.02 | 0.21 | 0.13 |
| 5 | 77909939 | rs1372319 | -0.812 | -0.363 | -0.128 | -1.533 | -0.006 | -0.002 | -0.002 | 0 | 0.54 | 0.55 | 0.58 | 0.5 | 0.73 | 0.46 |
| 5 | 77916642 | rs10491474 | -0.476 | -0.537 | 0.176 | -1.13 | -0.006 | 0.017 | 0.017 | 0 | 0.37 | 0.33 | 0.28 | 0.24 | 0.47 | 0.26 |
| 5 | 77922539 | rs12651937 | -0.143 | 0.365 | -0.133 | -0.844 | -0.018 | 0.018 | 0.018 | 0 | 0.26 | 0.28 | 0.27 | 0.19 | 0.37 | 0.22 |
| 5 | 77924020 | rs9293757 | -0.518 | 0.48 | -0.776 | -0.788 | 0 | -0.006 | -0.006 | 0 | 0.3 | 0.69 | 0.31 | 0.19 | 0.37 | 0.22 |
| 5 | 77926281 | rs7722214 | -0.318 | 0.276 | -0.66 | -0.826 | 0.002 | -0.016 | -0.016 | 0 | 0.28 | 0.69 | 0.22 | 0.17 | 0.35 | 0.21 |
| 5 | 77933091 | rs341903 | -0.257 | -1.07 | -0.732 | -0.501 | -0.007 | 0 | 0 | 1.302 | 0.36 | 0.38 | 0.38 | 0.41 | 0.23 | 0.38 |
| 5 | 77936965 | rs168190 | 0.543 | 0.135 | -0.882 | -0.951 | 0.013 | 0.124 | 0.124 | 0 | 0.51 | 0.25 | 0.46 | 0.29 | 0.48 | 0.54 |
| 5 | 77940652 | rs163633 | 0.085 | 0.609 | -0.569 | -0.837 | 0.019 | -0.033 | -0.033 | 0 | 0.64 | 0.52 | 0.54 | 0.28 | 0.58 | 0.6 |
| 5 | 77949565 | rs341917 | 0.257 | 0.577 | -1.031 | -0.669 | -0.017 | 0.021 | 0.021 | 0 | 0.61 | 0.68 | 0.51 | 0.28 | 0.52 | 0.58 |
| 5 | 77961474 | rs184652 | -0.003 | 0.683 | -1.747 | -1.348 | -0.013 | 0.169 | 0.169 | 0 | 0.34 | 0.1 | 0.42 | 0.01 | 0.3 | 0.22 |
| 5 | 77962439 | rs6868351 | -0.314 | 0.987 | -1.164 | -0.624 | 0.028 | 0.129 | 0.129 | 0 | 0.26 | 0.49 | 0.22 | 0.41 | 0.29 | 0.37 |
| 5 | 77964717 | rs341908 | -0.306 | 0.984 | -0.809 | -0.646 | 0.021 | 0.076 | 0.076 | 0 | 0.24 | 0.35 | 0.21 | 0.41 | 0.28 | 0.37 |
| 5 | 77966625 | rs341911 | -0.133 | 1.12 | -0.862 | -0.66 | 0.019 | 0.191 | 0.191 | 0 | 0.15 | 0.31 | 0.17 | 0.41 | 0.17 | 0.3 |
| 5 | 77998786 | rs340085 | 1.048 | 0.963 | -1.275 | -0.807 | 0.002 | 0.02 | 0.02 | 0 | 0.5 | 0.5 | 0.67 | 0.23 | 0.55 | 0.34 |
| 5 | 77999276 | rs920581 | 0.567 | 1.285 | -1.737 | -0.536 | -0.025 | 0.056 | 0.056 | 0.004 | 0.11 | 0.01 | 0.06 | 0 | 0.12 | 0.11 |
| 5 | 78012193 | rs6893444 | -0.822 | 1.159 | -1.467 | 0.181 | -0.001 | 0.031 | 0.031 | 1.069 | 0.84 | 0.86 | 0.86 | 0.57 | 0.87 | 0.82 |
| 5 | 78016860 | rs745981 | -0.658 | 1.383 | -1.841 | -0.016 | -0.08 | -0.087 | -0.087 | 0 | 0.19 | 0.01 | 0.11 | 0.34 | 0.21 | 0.33 |
| 5 | 78020166 | rs539990 | 1.061 | 1.238 | -1.388 | -0.352 | 0.006 | 0.128 | 0.128 | 0 | 0.45 | 0.26 | 0.52 | 0.14 | 0.43 | 0.41 |
| 5 | 78021218 | rs7714373 | 0.065 | 1.103 | -0.856 | 0.868 | -0.009 | 0.111 | 0.111 | 0.902 | 0.82 | 0.69 | 0.75 | 0.54 | 0.84 | 0.79 |
| 5 | 78024366 | rs569523 | 0.599 | 1.363 | -0.989 | 0.871 | -0.012 | -0.096 | -0.096 | 0 | 0.76 | 0.88 | 0.68 | 0.42 | 0.71 | 0.63 |
| 5 | 78024759 | rs565802 | 0.04 | 1.055 | -0.84 | 0.672 | -0.023 | -0.016 | -0.016 | 2.93 | 0.06 | 0.01 | 0.04 | 0.08 | 0.07 | 0.15 |
| 5 | 78029909 | rs252242 | -0.968 | 0.491 | -0.245 | 0.968 | -0.003 | 0.008 | 0.008 | 0 | 0.26 | 0.26 | 0.34 | 0.49 | 0.2 | 0.36 |
| 5 | 78034521 | rs6897874 | 1.035 | 0.55 | -0.212 | 0.382 | -0.017 | 0.087 | 0.087 | 0 | 0.44 | 0.31 | 0.39 | 0.1 | 0.5 | 0.47 |
| 5 | 78034805 | rs6878810 | 0.577 | 0.672 | 0.08 | 1.484 | -0.003 | 0.062 | 0.062 | 0 | 0.88 | 0.87 | 0.74 | 0.54 | 0.9 | 0.73 |
| 5 | 78035233 | rs10514145 | -0.901 | 0.458 | 0.126 | 1.491 | 0 | -0.007 | -0.007 | 1.704 | 0.24 | 0.23 | 0.24 | 0.42 | 0.24 | 0.16 |
| 5 | 78037138 | rs2453766 | -0.394 | 0.442 | 0.3 | 1.526 | 0.002 | 0.045 | 0.045 | 0 | 0.1 | 0.09 | 0.18 | 0.36 | 0.19 | 0.06 |
| 5 | 78041976 | rs13164947 | 1.402 | -0.08 | 0.31 | 1.248 | 0.017 | 0.087 | 0.087 | 0 | 0.55 | 0.18 | 0.44 | 0.28 | 0.57 | 0.55 |
| 5 | 78042215 | rs16875859 | -1.44 | -0.613 | 1.142 | 1.302 | -0.039 | -0.074 | -0.074 | 0 | 0.13 | 0.38 | 0.07 | 0.01 | 0.05 | 0.04 |
| 5 | 78044557 | rs550550 | 1.058 | -0.574 | 0.944 | 1.337 | -0.023 | 0.044 | 0.044 | 0.007 | 0.84 | 0.75 | 0.72 | 0.69 | 0.9 | 0.71 |
| 5 | 78044921 | rs660567 | 1.105 | -0.572 | 0.935 | 1.34 | -0.025 | 0.042 | 0.042 | 0.034 | 0.83 | 0.74 | 0.72 | 0.69 | 0.9 | 0.71 |
| 5 | 78051223 | rs2173013 | 1.27 | -0.436 | 0.476 | 1.38 | -0.06 | 0.102 | 0.102 | 0.032 | 0.81 | 0.47 | 0.72 | 0.68 | 0.9 | 0.7 |
| 5 | 78061140 | rs252235 | 1.025 | -0.29 | 0.086 | 1.475 | -0.043 | 0.122 | 0.122 | 0 | 0.82 | 0.63 | 0.71 | 0.59 | 0.89 | 0.65 |
| 5 | 78066631 | rs4704521 | 0.191 | 0.079 | -0.99 | 0.848 | 0 | -0.004 | -0.004 | 0 | 0.57 | 0.71 | 0.68 | 0.51 | 0.61 | 0.42 |
| 5 | 78067984 | rs2132537 | 0.149 | -0.054 | -1.423 | 0.791 | -0.042 | -0.042 | -0.042 | 0 | 0.58 | 0.88 | 0.68 | 0.53 | 0.61 | 0.42 |
| 5 | 78072760 | rs7732938 | -0.453 | 0.176 | -2.357 | 0.064 | -0.051 | -0.138 | -0.138 | 0.008 | 0.74 | 0.97 | 0.76 | 0.58 | 0.75 | 0.59 |
| 5 | 78073025 | rs16875887 | 0.669 | -0.35 | -1.697 | -0.398 | -0.02 | -0.036 | -0.036 | 0 | 0.11 | 0.13 | 0.05 | 0 | 0.05 | 0.03 |
| 5 | 78074198 | rs7704939 | -0.418 | 0.067 | -1.138 | 0.096 | 0.002 | 0.004 | 0.004 | 0.105 | 0.4 | 0.53 | 0.43 | 0.46 | 0.4 | 0.37 |
| 5 | 78074918 | rs754566 | -0.122 | 0.194 | -1.016 | -0.19 | -0.007 | -0.002 | -0.002 | 0.005 | 0.55 | 0.57 | 0.54 | 0.44 | 0.53 | 0.43 |
| 5 | 78074963 | rs754567 | 1.085 | 0.373 | -0.502 | -1.287 | -0.005 | -0.035 | -0.035 | 0 | 0.25 | 0.21 | 0.22 | 0 | 0.32 | 0.11 |
| 5 | 78079883 | rs6871810 | -0.901 | -0.24 | 0.291 | -0.586 | -0.042 | -0.057 | -0.057 | 0 | 0.26 | 0.55 | 0.25 | 0.13 | 0.17 | 0.15 |
| 5 | 78084051 | rs16875913 | 0.18 | -0.248 | 0.245 | -0.64 | 0.014 | 0.011 | 0.011 | 0 | 0.17 | 0.11 | 0.06 | 0.01 | 0.11 | 0.05 |
| 5 | 78089433 | rs3846677 | -1.173 | -0.161 | 0.187 | -0.605 | -0.018 | -0.038 | -0.038 | 0.236 | 0.3 | 0.58 | 0.29 | 0.13 | 0.21 | 0.18 |
| 5 | 78094383 | rs16875919 | -0.077 | -0.088 | 0.312 | -0.53 | 0 | -0.042 | -0.042 | 0 | 0.2 | 0.21 | 0.09 | 0 | 0.15 | 0.08 |
| 5 | 78097220 | rs11960862 | -1.333 | -0.182 | 0.318 | -0.237 | 0.01 | 0.013 | 0.013 | 0 | 0.3 | 0.7 | 0.23 | 0.22 | 0.23 | 0.19 |

| | | | | | | | | | | | | | | | | |
|---|---|---|---|---|---|---|---|---|---|---|---|---|---|---|---|---|
| 5 | 78101191 | rs16875951 | -0.134 | -0.125 | 0.442 | -0.194 | -0.014 | -0.083 | -0.083 | 0 | 0.2 | 0.3 | 0.1 | 0.02 | 0.13 | 0.07 |
| 5 | 78103605 | rs16876279 | -0.681 | -0.09 | -0.418 | -0.166 | -0.021 | -0.047 | -0.047 | 1.611 | 0.24 | 0.55 | 0.13 | 0.11 | 0.16 | 0.11 |
| 5 | 78105347 | rs10514108 | -1.56 | 0.015 | 0.107 | 0.5 | 0.012 | 0.13 | 0.13 | 0.016 | 0.11 | 0.57 | 0.17 | 0.21 | 0.1 | 0.12 |
| 5 | 78106062 | rs10474567 | -0.033 | 0.05 | -1.343 | 0.209 | -0.017 | -0.005 | -0.005 | 0 | 0.18 | 0 | 0.22 | 0.11 | 0.15 | 0.2 |
| 5 | 78108455 | rs4704524 | -0.598 | 0.174 | -0.853 | 0.286 | 0.003 | 0.029 | 0.029 | 0.347 | 0.07 | 0.16 | 0.05 | 0.17 | 0.08 | 0.05 |
| 5 | 78110117 | rs13152961 | -0.395 | -0.117 | -1.713 | -0.221 | -0.011 | 0.139 | 0.139 | 0 | 0.24 | 0.01 | 0.28 | 0.04 | 0.19 | 0.22 |
| 5 | 78110532 | rs6453417 | -0.651 | 0.353 | -1.465 | 0.251 | -0.008 | 0.012 | 0.012 | 0 | 0.56 | 0.77 | 0.51 | 0.38 | 0.52 | 0.64 |
| 5 | 78115511 | rs16875961 | -1.021 | 0.257 | -0.463 | 0.262 | -0.01 | 0.009 | 0.009 | 0 | 0.14 | 0.31 | 0.07 | 0.09 | 0.11 | 0.12 |
| 5 | 78117584 | rs3857405 | -0.757 | 0.542 | -1.292 | 0.327 | -0.005 | -0.015 | -0.015 | 0.67 | 0.6 | 0.8 | 0.56 | 0.51 | 0.66 | 0.6 |
| 5 | 78123484 | rs4704525 | -0.524 | 0.54 | -1.17 | 0.732 | 0 | 0.002 | 0.002 | 0 | 0.26 | 0.3 | 0.18 | 0.35 | 0.29 | 0.37 |
| 5 | 78124029 | rs12655291 | 0.541 | 0.611 | -1.079 | 0.304 | 0.003 | -0.002 | -0.002 | 0 | 0.16 | 0.1 | 0.1 | 0.15 | 0.18 | 0.17 |
| 5 | 78126375 | rs10058261 | 0.21 | 0.649 | -0.959 | 0.147 | 0.001 | 0.009 | 0.009 | 0 | 0.06 | 0.05 | 0.04 | 0 | 0.06 | 0.03 |
| 5 | 78131336 | rs10454897 | 0.271 | 0.598 | -1.511 | -0.064 | -0.005 | 0.036 | 0.036 | 0 | 0.4 | 0.17 | 0.34 | 0.25 | 0.42 | 0.3 |
| 5 | 78131547 | rs784589 | 0.085 | 0.6 | -0.684 | 0.383 | 0.015 | 0.134 | 0.134 | 0.759 | 0.79 | 0.64 | 0.76 | 0.48 | 0.8 | 0.67 |
| 5 | 78135201 | rs25413 | 0.467 | 0.29 | -0.853 | -0.386 | -0.04 | -0.135 | -0.135 | 0.08 | 0.29 | 0.56 | 0.15 | 0.05 | 0.18 | 0.22 |
| 5 | 78152192 | rs3098681 | 0.237 | 0.802 | -1.205 | -0.395 | 0.02 | 0.007 | 0.007 | 0.071 | 0.69 | 0.65 | 0.67 | 0.3 | 0.61 | 0.53 |
| 5 | 78153325 | rs12186484 | 1.123 | -0.039 | -1.9 | -1.24 | -0.065 | 0.008 | 0.008 | 0 | 0.4 | 0.12 | 0.43 | 0.28 | 0.46 | 0.3 |
| 5 | 78179604 | rs4704531 | 0.99 | -0.19 | -1.783 | -1.304 | 0.011 | 0.1 | 0.1 | 1.501 | 0.19 | 0.05 | 0.2 | 0.01 | 0.15 | 0.19 |
| 5 | 78180482 | rs337835 | 0.11 | -0.245 | -1.573 | -1.296 | 0.04 | 0.048 | 0.048 | 4.234 | 0.5 | 0.35 | 0.64 | 0.93 | 0.67 | 0.59 |
| 5 | 78180736 | rs337836 | 0.799 | -0.176 | -1.691 | -1.019 | 0.014 | 0.101 | 0.101 | 0.957 | 0.2 | 0.31 | 0.21 | 0.55 | 0.2 | 0.35 |
| 5 | 78181477 | rs1065757 | 1.128 | -0.172 | -1.702 | -1.02 | -0.14 | -0.104 | -0.104 | 0.604 | 0.31 | 0.04 | 0.43 | 0.37 | 0.47 | 0.24 |
| 5 | 78181850 | rs180049 | -0.424 | -0.184 | -1.635 | -0.991 | -0.006 | -0.003 | -0.003 | 0 | 0.41 | 0.43 | 0.43 | 0.63 | 0.35 | 0.59 |
| 5 | 78182431 | rs17220863 | 0.713 | -0.092 | -1.643 | -0.846 | 0.021 | 0.044 | 0.044 | 1.427 | 0.21 | 0.12 | 0.23 | 0.07 | 0.15 | 0.25 |
| 5 | 78184403 | rs6893857 | -0.439 | -0.273 | -1.198 | -0.632 | -0.017 | -0.025 | -0.025 | 0 | 0.29 | 0.65 | 0.29 | 0.16 | 0.18 | 0.39 |
| 5 | 78186558 | rs11741352 | 0.776 | 0.023 | -1.174 | -0.442 | -0.049 | -0.031 | -0.031 | 1.035 | 0.32 | 0.16 | 0.45 | 0.37 | 0.47 | 0.25 |
| 5 | 78190471 | rs16876062 | -0.742 | -0.029 | -0.382 | -0.075 | 0.002 | 0.043 | 0.043 | 2.436 | 0.08 | 0.51 | 0.06 | 0.04 | 0.01 | 0.07 |
| 5 | 78202826 | rs337853 | 0.398 | 0.078 | -0.162 | -0.228 | -0.001 | -0.001 | -0.001 | 0 | 0.41 | 0.39 | 0.32 | 0.55 | 0.39 | 0.47 |
| 5 | 78202968 | rs11747224 | -0.128 | -0.246 | 0.244 | -0.469 | -0.037 | -0.03 | -0.03 | 0 | 0.14 | 0.35 | 0.09 | 0.1 | 0.07 | 0.22 |
| 5 | 78204390 | rs337851 | -0.811 | -0.073 | 0.258 | -0.426 | -0.008 | -0.009 | -0.009 | 0.191 | 0.69 | 0.52 | 0.6 | 0.78 | 0.69 | 0.78 |
| 5 | 78206783 | rs13184587 | 1.754 | -0.068 | 0.057 | -0.437 | -0.029 | -0.036 | -0.036 | 0 | 0.22 | 0.06 | 0.35 | 0.21 | 0.29 | 0.18 |
| 5 | 78206969 | rs337849 | -0.881 | -0.077 | 0.226 | -0.439 | -0.001 | -0.001 | -0.001 | 0 | 0.71 | 0.68 | 0.62 | 0.78 | 0.69 | 0.78 |
| 5 | 78210725 | rs9293758 | -0.285 | -0.048 | 0.045 | -0.471 | -0.023 | 0.044 | 0.044 | 2.852 | 0.32 | 0.16 | 0.32 | 0.29 | 0.35 | 0.37 |
| 5 | 78220029 | rs10053925 | 1.301 | 0.228 | -0.198 | -0.425 | -0.062 | -0.043 | -0.043 | 0 | 0.11 | 0.01 | 0.09 | 0.19 | 0.19 | 0.07 |
| 5 | 78220709 | rs4704537 | 0.328 | 0.032 | -0.091 | -0.425 | -0.055 | 0.051 | 0.051 | 1.428 | 0.27 | 0.1 | 0.29 | 0.23 | 0.33 | 0.24 |
| 5 | 78220996 | rs10452492 | 1.808 | 0.042 | 0.066 | -0.392 | -0.01 | -0.015 | -0.015 | 0 | 0.22 | 0.11 | 0.36 | 0.22 | 0.29 | 0.18 |
| 5 | 78221946 | rs337895 | 0.537 | -0.011 | 0.292 | -0.381 | 0 | 0 | 0 | 0 | 0.19 | 0.19 | 0.15 | 0.4 | 0.19 | 0.24 |
| 5 | 78228199 | rs337888 | -1.064 | -0.116 | 0.555 | -0.519 | 0.008 | 0.063 | 0.063 | 0 | 0.53 | 0.75 | 0.49 | 0.7 | 0.53 | 0.61 |
| 5 | 78232453 | rs4421085 | 0.318 | -0.036 | 0.754 | -0.288 | 0.002 | 0.116 | 0.116 | 2.349 | 0.19 | 0.31 | 0.18 | 0.45 | 0.19 | 0.25 |
| 5 | 78234241 | rs337841 | -1.005 | 0.086 | 0.735 | -0.427 | 0.006 | 0.026 | 0.026 | 0 | 0.54 | 0.66 | 0.52 | 0.66 | 0.56 | 0.5 |
| 5 | 78249078 | rs449839 | 1.175 | 0.163 | 0.616 | -0.444 | -0.001 | 0.01 | 0.01 | 0.003 | 0.25 | 0.32 | 0.36 | 0.26 | 0.28 | 0.3 |
| 5 | 78250133 | rs447998 | 1.721 | 0.257 | 0.173 | -0.391 | -0.007 | -0.02 | -0.02 | 0.021 | 0.22 | 0.1 | 0.34 | 0.22 | 0.27 | 0.19 |
| 5 | 78253122 | rs234687 | 0.657 | 0.275 | 0.336 | -0.397 | 0.003 | 0.006 | 0.006 | 0 | 0.49 | 0.52 | 0.51 | 0.35 | 0.45 | 0.51 |
| 5 | 78259645 | rs7725928 | 0.904 | 0.36 | -0.152 | -0.495 | -0.037 | -0.115 | -0.115 | 1.454 | 0.29 | 0.53 | 0.19 | 0.13 | 0.18 | 0.32 |
| 5 | 78259888 | rs337847 | -0.733 | 0.895 | -0.644 | 0.369 | -0.002 | 0.017 | 0.017 | 0.371 | 0.91 | 0.92 | 0.9 | 0.71 | 0.92 | 0.85 |
| 5 | 78265041 | rs3733895 | -0.564 | 0.888 | -0.688 | -0.188 | -0.039 | 0.059 | 0.059 | 0.051 | 0.31 | 0.13 | 0.3 | 0.22 | 0.37 | 0.25 |
| 5 | 78267797 | rs6871197 | -0.774 | 0.382 | -0.469 | -0.591 | 0.002 | -0.027 | -0.027 | 0 | 0.32 | 0.34 | 0.25 | 0.08 | 0.24 | 0.32 |
| 5 | 78268128 | rs13178105 | 0.292 | 0.307 | -0.57 | -0.698 | -0.012 | 0.078 | 0.078 | 0 | 0.17 | 0.07 | 0.2 | 0.03 | 0.15 | 0.16 |
| 5 | 78268889 | rs6453423 | -0.736 | 0.388 | -0.476 | -0.59 | 0.003 | -0.024 | -0.024 | 0 | 0.32 | 0.33 | 0.25 | 0.08 | 0.24 | 0.32 |
| 5 | 78272540 | rs4704542 | -0.147 | 0.795 | -0.552 | 0.126 | 0.003 | 0.002 | 0.002 | 0 | 0.12 | 0.13 | 0.31 | 0.14 | 0.15 | 0.12 |
| 5 | 78276148 | rs1841991 | 0.066 | 0.502 | -0.505 | 0.243 | 0.002 | -0.015 | -0.015 | 0 | 0.37 | 0.46 | 0.48 | 0.33 | 0.42 | 0.28 |
| 5 | 78276474 | rs1717570 | 0.071 | 0.502 | -0.505 | 0.243 | 0.003 | -0.015 | -0.015 | 0.116 | 0.37 | 0.46 | 0.48 | 0.33 | 0.41 | 0.27 |
| 5 | 78278229 | rs672413 | -0.103 | 0.093 | -0.511 | 0.499 | 0.001 | 0 | 0 | 0 | 0.56 | 0.48 | 0.73 | 0.42 | 0.68 | 0.44 |
| 5 | 78279820 | rs163129 | 0.213 | -0.195 | -0.858 | 0.85 | -0.004 | 0.003 | 0.003 | 2.259 | 0.21 | 0.22 | 0.11 | 0.32 | 0.13 | 0.27 |
| 5 | 78282249 | rs163126 | -0.931 | -0.168 | -1.324 | 0.837 | -0.015 | -0.045 | -0.045 | 0.131 | 0.81 | 0.92 | 0.89 | 0.76 | 0.82 | 0.68 |
| 5 | 78283003 | rs163124 | 0.035 | -0.475 | -0.708 | 0.669 | -0.022 | 0.01 | 0.01 | 0.704 | 0.61 | 0.5 | 0.81 | 0.6 | 0.72 | 0.56 |
| 5 | 78290215 | rs9293761 | 0.495 | 0.124 | -2.136 | 0.418 | -0.125 | -0.071 | -0.071 | 1.227 | 0.33 | 0.03 | 0.48 | 0.33 | 0.44 | 0.28 |
| 5 | 78291960 | rs705415 | 1.574 | 0.801 | -1.767 | 0.31 | -0.036 | -0.024 | -0.024 | 0.136 | 0.06 | 0.01 | 0.09 | 0.08 | 0.12 | 0.12 |
| 5 | 78295854 | rs28326 | 0.709 | 0.538 | -1.747 | 0.072 | -0.015 | -0.003 | -0.003 | 0 | 0.16 | 0.07 | 0.22 | 0.21 | 0.17 | 0.17 |
| 5 | 78297271 | rs7710824 | 0.387 | 0.339 | -1.863 | -0.376 | -0.034 | 0.081 | 0.081 | 0 | 0.23 | 0.08 | 0.13 | 0.03 | 0.28 | 0.23 |
| 5 | 78302196 | rs250513 | 0.365 | 0.312 | -1.74 | -0.374 | -0.011 | 0.017 | 0.017 | 0 | 0.23 | 0.14 | 0.24 | 0.21 | 0.21 | 0.19 |
| 5 | 78302859 | rs479405 | 0.246 | 0.193 | -1.989 | -0.082 | -0.043 | 0.034 | 0.034 | 0.142 | 0.59 | 0.86 | 0.66 | 0.68 | 0.57 | 0.67 |
| 5 | 78303487 | rs10514151 | 0.243 | -0.704 | -1.296 | 0.084 | -0.062 | 0.096 | 0.096 | 0 | 0.17 | 0.52 | 0.07 | 0.29 | 0.06 | 0.29 |
| 5 | 78307309 | rs570207 | 0.187 | -0.68 | -1.375 | -0.174 | 0.002 | 0.01 | 0.01 | 0 | 0.65 | 0.69 | 0.7 | 0.72 | 0.7 | 0.71 |
| 5 | 78308731 | rs653275 | -0.987 | -0.445 | -1.473 | -0.038 | -0.056 | -0.039 | -0.039 | 0.054 | 0.78 | 0.94 | 0.77 | 0.73 | 0.77 | 0.83 |
| 5 | 78311315 | rs16876302 | -0.552 | -0.654 | -0.762 | -0.551 | -0.013 | -0.066 | -0.066 | 0.313 | 0.33 | 0.45 | 0.19 | 0.14 | 0.3 | 0.3 |
| 5 | 78315964 | rs2034900 | -0.721 | -0.705 | -0.252 | -0.586 | -0.013 | -0.063 | -0.063 | 0 | 0.34 | 0.52 | 0.2 | 0.14 | 0.3 | 0.3 |
| 5 | 78317413 | rs731124 | -0.704 | -0.712 | -0.14 | -0.589 | -0.016 | -0.076 | -0.076 | 0.091 | 0.34 | 0.49 | 0.2 | 0.14 | 0.3 | 0.3 |
| 5 | 78320085 | rs2303128 | 1.396 | -1.253 | 0.154 | -0.684 | -0.045 | 0.029 | 0.029 | 0.148 | 0.12 | 0.2 | 0.01 | 0.15 | 0.02 | 0.12 |
| 5 | 78321109 | rs2431332 | 1.227 | -1.255 | -0.283 | -1.071 | -0.083 | 0.08 | 0.08 | 0.892 | 0.21 | 0.03 | 0.2 | 0.08 | 0.24 | 0.16 |

| | | | | | | | | | | | | | | | | |
|---|---|---|---|---|---|---|---|---|---|---|---|---|---|---|---|---|
| 5 | 78323434 | rs10514154 | 0.635 | -0.966 | 0.408 | -0.733 | -0.001 | -0.003 | -0.003 | 1.473 | 0.23 | 0.23 | 0.13 | 0.14 | 0.2 | 0.25 |
| 5 | 78324075 | rs933684 | -0.677 | -0.987 | 0.59 | -0.729 | -0.014 | -0.065 | -0.065 | 0 | 0.33 | 0.5 | 0.2 | 0.14 | 0.3 | 0.3 |
| 5 | 78324352 | rs1805074 | -0.677 | -0.987 | 0.587 | -0.729 | -0.016 | -0.074 | -0.074 | 0 | 0.33 | 0.49 | 0.2 | 0.14 | 0.3 | 0.3 |
| 5 | 78328560 | rs1805072 | 0.588 | -0.932 | 0.302 | -0.649 | -0.006 | -0.009 | -0.009 | 0 | 0.23 | 0.24 | 0.13 | 0.14 | 0.2 | 0.25 |
| 5 | 78339755 | rs248383 | 0.629 | -0.79 | 0.33 | -0.654 | -0.008 | 0.023 | 0.023 | 0 | 0.57 | 0.62 | 0.59 | 0.78 | 0.51 | 0.56 |
| 5 | 78350390 | rs631151 | 1.178 | -0.927 | 0.366 | -0.621 | -0.016 | 0.033 | 0.033 | 0.014 | 0.45 | 0.25 | 0.56 | 0.63 | 0.51 | 0.45 |
| 5 | 78351512 | rs6873326 | 0.948 | -1 | 0.218 | -0.714 | -0.033 | 0.057 | 0.057 | 0.552 | 0.1 | 0.01 | 0.05 | 0 | 0.1 | 0.06 |
| 5 | 78351636 | rs2253262 | -0.096 | -1.047 | 0.556 | -0.7 | 0.007 | -0.022 | -0.022 | 2.694 | 0.67 | 0.59 | 0.65 | 0.78 | 0.63 | 0.62 |
| 5 | 78353947 | rs7705204 | 1.44 | -1.056 | 0.49 | -0.668 | -0.053 | 0.032 | 0.032 | 0.376 | 0.12 | 0.21 | 0.02 | 0.15 | 0.02 | 0.11 |
| 5 | 78354446 | rs476620 | 0.803 | -1.079 | 0.55 | -0.7 | 0.023 | 0.042 | 0.042 | 0 | 0.54 | 0.23 | 0.62 | 0.63 | 0.61 | 0.51 |
| 5 | 78356436 | rs1984685 | 1.411 | -1.075 | 0.334 | -0.619 | -0.053 | 0.032 | 0.032 | 0.376 | 0.12 | 0.21 | 0.02 | 0.15 | 0.02 | 0.11 |
| 5 | 78359036 | rs1644004 | 1.492 | -0.978 | -0.132 | -0.66 | -0.103 | -0.008 | -0.008 | 2.555 | 0.42 | 0.11 | 0.56 | 0.63 | 0.51 | 0.45 |
| 5 | 78364687 | rs642431 | 0.055 | -0.817 | 0.358 | -0.627 | -0.001 | -0.011 | -0.011 | 4.517 | 0.73 | 0.48 | 0.76 | 0.78 | 0.79 | 0.75 |
| 5 | 78364844 | rs642934 | -0.197 | -0.782 | 0.527 | -0.636 | 0.015 | 0.002 | 0.002 | 0 | 0.76 | 0.55 | 0.76 | 0.78 | 0.79 | 0.76 |
| 5 | 78366705 | rs671960 | 0.902 | -0.838 | -0.154 | -0.665 | 0.028 | 0.066 | 0.066 | 0 | 0.51 | 0.24 | 0.6 | 0.64 | 0.59 | 0.51 |
| 5 | 78373431 | rs682985 | 0.91 | -0.856 | -0.144 | -0.591 | 0.029 | 0.048 | 0.048 | 0 | 0.51 | 0.24 | 0.6 | 0.6 | 0.6 | 0.51 |
| 5 | 78374865 | rs1422086 | 0.835 | -0.87 | -0.076 | -0.597 | 0.024 | 0.041 | 0.041 | 0 | 0.52 | 0.24 | 0.6 | 0.6 | 0.6 | 0.51 |
| 5 | 78389912 | rs575425 | 0.047 | -0.892 | 0.434 | -0.548 | 0.009 | -0.001 | -0.001 | 1.171 | 0.66 | 0.55 | 0.65 | 0.68 | 0.63 | 0.6 |
| 5 | 78397980 | rs586199 | 1.838 | -0.943 | 0.151 | -0.674 | -0.084 | -0.04 | -0.04 | 0 | 0.4 | 0.15 | 0.55 | 0.6 | 0.51 | 0.45 |
| 5 | 78403378 | rs631305 | -2.416 | -0.871 | 0.097 | -0.685 | -0.051 | 0.002 | 0.002 | 0 | 0.9 | 0.98 | 0.83 | 0.92 | 0.83 | 0.87 |
| 5 | 78405673 | rs600473 | -0.09 | -0.875 | 0.246 | -0.749 | 0.01 | -0.006 | -0.006 | 0 | 0.64 | 0.52 | 0.63 | 0.68 | 0.6 | 0.58 |
| 5 | 78406032 | rs3756548 | 1.293 | -0.912 | 0.38 | -0.757 | -0.045 | -0.03 | -0.03 | 1.995 | 0.13 | 0.29 | 0.04 | 0.08 | 0.02 | 0.09 |
| 5 | 78406434 | rs562487 | 1.962 | -0.909 | 0.014 | -0.804 | -0.145 | -0.108 | -0.108 | 1.267 | 0.38 | 0.04 | 0.52 | 0.6 | 0.48 | 0.43 |
| 5 | 78407261 | rs16876512 | 0.484 | -0.684 | 0.184 | -0.706 | 0 | -0.037 | -0.037 | 0 | 0.12 | 0.12 | 0.06 | 0 | 0.1 | 0.06 |
| 5 | 78409987 | rs492842 | 0.185 | -0.542 | 0.154 | -0.543 | 0.006 | -0.005 | -0.005 | 1.375 | 0.63 | 0.49 | 0.62 | 0.68 | 0.61 | 0.58 |
| 5 | 78411324 | rs7700970 | -0.624 | -0.637 | 0.09 | -0.762 | -0.012 | -0.102 | -0.102 | 0.187 | 0.36 | 0.5 | 0.2 | 0.1 | 0.32 | 0.3 |
| 5 | 78413504 | rs3797546 | 0.648 | -1.403 | -0.532 | 0.161 | 0.052 | -0.035 | -0.035 | 0 | 0.2 | 0.1 | 0.06 | 0.28 | 0.06 | 0.13 |
| 5 | 78414337 | rs506500 | -0.182 | -0.227 | -0.35 | -0.068 | 0.019 | -0.012 | -0.012 | 0 | 0.81 | 0.79 | 0.82 | 0.89 | 0.69 | 0.85 |
| 5 | 78421780 | rs694290 | 0.259 | -0.374 | 0.117 | 0.127 | 0.073 | 0.012 | 0.012 | 0 | 0.77 | 0.52 | 0.8 | 0.78 | 0.69 | 0.82 |
| 5 | 78421959 | rs3733890 | 0.506 | -0.146 | -0.465 | 0.218 | -0.002 | -0.005 | -0.005 | 0 | 0.28 | 0.2 | 0.38 | 0.32 | 0.32 | 0.29 |
| 5 | 78436211 | rs1915706 | 0.282 | -0.285 | -0.55 | 0.405 | 0.019 | 0.062 | 0.062 | 0.101 | 0.55 | 0.7 | 0.48 | 0.75 | 0.38 | 0.6 |
| 5 | 78453899 | rs2364594 | 1.584 | -0.467 | -0.733 | 0.3 | -0.039 | -0.101 | -0.101 | 0 | 0.34 | 0.04 | 0.41 | 0.39 | 0.33 | 0.44 |
| 5 | 78463395 | rs13158309 | 0.442 | -0.533 | -0.682 | 0.304 | 0.029 | 0.018 | 0.018 | 0 | 0.52 | 0.67 | 0.46 | 0.56 | 0.37 | 0.57 |
| 5 | 78468004 | rs2121107 | 1.625 | -0.548 | -0.717 | 0.293 | -0.04 | -0.101 | -0.101 | 10.406 | 0.34 | 0.04 | 0.41 | 0.38 | 0.33 | 0.44 |
| 5 | 78477017 | rs8180502 | 0.532 | -0.649 | -1.03 | -0.021 | 0.028 | -0.006 | -0.006 | 0 | 0.42 | 0.65 | 0.39 | 0.53 | 0.3 | 0.49 |
| 5 | 78517143 | rs12189248 | 0.94 | -0.501 | -0.064 | 0.178 | 0.032 | -0.014 | -0.014 | 0 | 0.46 | 0.26 | 0.43 | 0.47 | 0.36 | 0.52 |
| 5 | 78522895 | rs10078815 | 1.659 | -0.452 | -0.311 | 0.212 | -0.046 | -0.126 | -0.126 | 0 | 0.34 | 0.03 | 0.41 | 0.41 | 0.34 | 0.44 |
| 5 | 78532658 | rs3733893 | 1.538 | -0.425 | -0.228 | 0.091 | 0.005 | -0.031 | -0.031 | 0 | 0.36 | 0.15 | 0.42 | 0.37 | 0.34 | 0.45 |
| 5 | 78546293 | rs2591392 | 1.053 | -0.428 | -0.215 | 0.09 | 0.031 | -0.011 | -0.011 | 0 | 0.45 | 0.2 | 0.41 | 0.43 | 0.36 | 0.52 |
| 5 | 78558425 | rs2591387 | 1.646 | -0.439 | -0.214 | 0.085 | -0.04 | -0.095 | -0.095 | 0 | 0.34 | 0.06 | 0.4 | 0.37 | 0.34 | 0.45 |
| 5 | 78576210 | rs6866405 | 1.94 | -0.399 | -0.164 | 0.096 | -0.034 | -0.027 | -0.027 | 0.151 | 0.11 | 0.17 | 0.02 | 0.06 | 0.02 | 0.07 |
| 5 | 78587297 | rs16876619 | 0.161 | -0.013 | -0.262 | 0.042 | 0.041 | 0.046 | 0.046 | 0.703 | 0.21 | 0.36 | 0.39 | 0.34 | 0.3 | 0.21 |
| 5 | 78593204 | rs1870057 | -0.143 | -0.43 | 0.016 | -0.092 | 0.005 | 0 | 0 | 0 | 0.45 | 0.36 | 0.54 | 0.49 | 0.58 | 0.39 |
| 5 | 78596044 | rs10514159 | -1.22 | -0.384 | -0.43 | -0.076 | -0.05 | -0.073 | -0.073 | 0 | 0.62 | 0.94 | 0.59 | 0.63 | 0.62 | 0.51 |
| 5 | 78599519 | rs7711939 | -1.561 | -0.281 | -0.628 | -0.022 | -0.039 | -0.089 | -0.089 | 0 | 0.65 | 0.94 | 0.6 | 0.63 | 0.66 | 0.55 |
| 5 | 78607147 | rs4704556 | -0.706 | -0.058 | -0.474 | 0.067 | 0.021 | -0.017 | -0.017 | 0 | 0.52 | 0.86 | 0.58 | 0.57 | 0.6 | 0.44 |
| 5 | 78617534 | rs1062326 | -0.989 | -0.039 | -0.612 | 0.073 | 0.033 | -0.006 | -0.006 | 0.294 | 0.55 | 0.81 | 0.59 | 0.57 | 0.64 | 0.48 |
| 5 | 78629346 | rs9293769 | -0.502 | 0.117 | -0.617 | 0.283 | 0.001 | 0.001 | 0.001 | 0 | 0.48 | 0.51 | 0.61 | 0.48 | 0.6 | 0.39 |
| 5 | 78630605 | rs10462567 | -0.818 | -0.128 | -0.859 | 0.666 | 0.019 | -0.014 | -0.014 | 0 | 0.45 | 0.85 | 0.47 | 0.6 | 0.38 | 0.4 |
| 5 | 78640296 | rs9293772 | 0.101 | -0.156 | -0.968 | 0.375 | 0.004 | 0.143 | 0.143 | 0.167 | 0.3 | 0.08 | 0.17 | 0.06 | 0.25 | 0.23 |
| 5 | 78641327 | rs7719464 | -0.498 | -0.025 | 0.001 | 0.936 | 0.049 | 0.261 | 0.261 | 0.104 | 0.11 | 0.32 | 0.32 | 0.47 | 0.13 | 0.14 |
| 5 | 78660354 | rs7702071 | -0.629 | -0.103 | 0.24 | 1.088 | 0.059 | 0.282 | 0.282 | 0.113 | 0.11 | 0.36 | 0.31 | 0.38 | 0.12 | 0.13 |
| 5 | 78663158 | rs6879014 | -1.05 | -0.506 | -0.957 | 0.805 | 0.018 | -0.097 | -0.097 | 0 | 0.45 | 0.88 | 0.5 | 0.54 | 0.35 | 0.39 |
| 5 | 78664650 | rs10075154 | 0.171 | -0.484 | -1.039 | 0.436 | 0.008 | 0.139 | 0.139 | 0.059 | 0.3 | 0.09 | 0.16 | 0.05 | 0.25 | 0.22 |
| 5 | 78668358 | rs2290638 | 0.247 | -0.438 | -1.057 | 0.49 | 0.009 | 0.127 | 0.127 | 0.015 | 0.31 | 0.1 | 0.17 | 0.1 | 0.25 | 0.32 |
| 5 | 78668550 | rs2290637 | 1.757 | -0.52 | -1.228 | 0.462 | 0.025 | 0.02 | 0.02 | 0 | 0.1 | 0.02 | 0.01 | 0.05 | 0.03 | 0.07 |
| 5 | 78670802 | rs3822568 | -0.734 | 0.006 | -0.947 | 0.948 | -0.044 | -0.049 | -0.049 | 0.011 | 0.49 | 0.97 | 0.53 | 0.65 | 0.4 | 0.5 |
| 5 | 78673236 | rs12187625 | 0.586 | -0.214 | -1.087 | 0.239 | -0.197 | 0.253 | 0.253 | 0 | 0.35 | 0.02 | 0.26 | 0.01 | 0.53 | 0.28 |
| 5 | 78685117 | rs13360983 | 0.446 | -0.518 | 0.226 | 0.764 | -0.005 | 0.005 | 0.005 | 0.073 | 0.42 | 0.53 | 0.47 | 0.55 | 0.35 | 0.49 |
| 5 | 78691033 | rs6892318 | -0.078 | -0.566 | 0.397 | 0.743 | 0.034 | -0.032 | -0.032 | 0 | 0.47 | 0.83 | 0.5 | 0.55 | 0.35 | 0.49 |
| 5 | 78691697 | rs6868393 | -0.152 | -0.57 | 0.4 | 0.742 | 0.04 | -0.035 | -0.035 | 0 | 0.47 | 0.86 | 0.5 | 0.55 | 0.35 | 0.49 |
| 5 | 78694007 | rs10038376 | 1.271 | -0.588 | 0.717 | 0.717 | 0.023 | 0.099 | 0.099 | 0.105 | 0.33 | 0.17 | 0.18 | 0.12 | 0.23 | 0.34 |
| 5 | 78710606 | rs4455546 | -0.741 | -0.527 | 1.026 | 0.801 | 0.078 | 0.368 | 0.368 | 0 | 0.09 | 0.37 | 0.3 | 0.42 | 0.11 | 0.15 |
| 5 | 78719016 | rs7708478 | -0.205 | -0.562 | 0.874 | 0.76 | 0.028 | -0.053 | -0.053 | 0 | 0.47 | 0.88 | 0.5 | 0.54 | 0.35 | 0.49 |
| 5 | 78721444 | rs7705514 | -0.371 | -0.399 | 1.156 | 0.72 | 0.007 | -0.028 | -0.028 | 0 | 0.61 | 0.6 | 0.7 | 0.87 | 0.47 | 0.74 |
| 5 | 78728923 | rs7719054 | 0.69 | -0.409 | 1.475 | 0.738 | -0.013 | -0.007 | -0.007 | 0 | 0.23 | 0.28 | 0.48 | 0.75 | 0.19 | 0.36 |
| 5 | 78738510 | rs6453450 | 0.376 | -0.43 | 1.252 | 1.251 | -0.011 | -0.001 | -0.001 | 0 | 0.43 | 0.51 | 0.3 | 0.6 | 0.34 | 0.44 |
| 5 | 78740930 | rs10039490 | 1.315 | -0.425 | 1.268 | 1.377 | 0.017 | 0.146 | 0.146 | 0.19 | 0.33 | 0.09 | 0.17 | 0.15 | 0.24 | 0.36 |
| 5 | 78744395 | rs12514775 | 0.288 | -0.414 | 1.349 | 1.426 | -0.022 | 0.058 | 0.058 | 2.181 | 0.25 | 0.34 | 0.44 | 0.4 | 0.17 | 0.33 |

| | | | | | | | | | | | | | | | | |
|---|---|---|---|---|---|---|---|---|---|---|---|---|---|---|---|---|
| 5 | 78750724 | rs6874510 | 0.552 | -0.388 | 1.146 | 1.52 | -0.007 | 0.024 | 0.024 | 2.361 | 0.24 | 0.23 | 0.43 | 0.4 | 0.17 | 0.33 |
| 5 | 78753539 | rs4637531 | -0.996 | -0.324 | 0.351 | 1.02 | -0.117 | 0.17 | 0.17 | 0 | 0.67 | 0.9 | 0.73 | 0.99 | 0.48 | 0.8 |
| 5 | 78759672 | rs9293782 | 0.905 | -0.369 | 0.448 | 1.103 | 0.016 | 0.073 | 0.073 | 0.373 | 0.37 | 0.26 | 0.21 | 0.16 | 0.29 | 0.4 |
| 5 | 78761776 | rs12522716 | 1.42 | -0.36 | -0.12 | 1.2 | -0.004 | -0.031 | -0.031 | 0.178 | 0.18 | 0.01 | 0.19 | 0.14 | 0.11 | 0.23 |
| 5 | 78763345 | rs4323213 | -0.458 | 0.082 | 0.506 | 1.348 | -0.028 | 0.007 | 0.007 | 0 | 0.64 | 0.71 | 0.68 | 0.72 | 0.46 | 0.75 |
| 5 | 78769369 | rs9293785 | 0.728 | -0.106 | 0.686 | 1.233 | 0.015 | -0.002 | -0.002 | 0 | 0.39 | 0.29 | 0.26 | 0.43 | 0.3 | 0.5 |
| 5 | 78772071 | rs10474588 | -0.859 | 0.022 | 0.22 | 1.057 | -0.161 | 0.228 | 0.228 | 0 | 0.67 | 0.95 | 0.71 | 0.99 | 0.47 | 0.79 |
| 5 | 78775086 | rs10070879 | -0.375 | 0.227 | 1.586 | 2.1 | 0.001 | -0.001 | -0.001 | 0.705 | 0.22 | 0.17 | 0.38 | 0.33 | 0.14 | 0.26 |
| 5 | 78812909 | rs4704559 | -1.549 | 1.144 | 2.597 | 2.194 | 0.027 | 0.017 | 0.017 | 0 | 0.08 | 0.31 | 0.24 | 0.08 | 0.09 | 0.1 |
| 5 | 78823934 | rs12520582 | -2.183 | 1.181 | 2.616 | 2.338 | 0.003 | 0.008 | 0.008 | 0 | 0.06 | 0.07 | 0.14 | 0.11 | 0.06 | 0.07 |
| 5 | 78829154 | rs11746854 | -1.276 | 1.529 | 2.794 | 2.499 | 0.043 | 0.078 | 0.078 | 0 | 0.16 | 0.27 | 0.16 | 0.24 | 0.21 | 0.16 |
| 5 | 78831504 | rs7704425 | -3.163 | 1.526 | 2.749 | 1.724 | 0.027 | 0.151 | 0.151 | 0.014 | 0.17 | 0.58 | 0.43 | 0.34 | 0.17 | 0.2 |
| 5 | 78831995 | rs10942894 | -2.785 | 1.609 | 2.433 | 0.94 | 0.035 | 0.047 | 0.047 | 0 | 0.14 | 0.43 | 0.38 | 0.19 | 0.16 | 0.18 |
| 5 | 78840610 | rs7707252 | -2.123 | 1.541 | 1.706 | 0.938 | 0.034 | -0.029 | -0.029 | 0 | 0.22 | 0.3 | 0.27 | 0.13 | 0.26 | 0.21 |
| 5 | 78849505 | rs6865469 | -2.083 | 1.447 | 1.434 | 0.931 | 0.021 | -0.02 | -0.02 | 0 | 0.21 | 0.27 | 0.27 | 0.12 | 0.27 | 0.2 |
| 5 | 78849697 | rs17237492 | -2.125 | 1.671 | 1.044 | 1.322 | -0.015 | -0.006 | -0.006 | 0 | 0.11 | 0.08 | 0.21 | 0.13 | 0.21 | 0.17 |
| 5 | 78852686 | rs17237618 | -1.599 | 1.333 | 0.937 | 1.297 | -0.016 | -0.016 | -0.016 | 0 | 0.06 | 0.02 | 0.14 | 0.08 | 0.07 | 0.1 |
| 5 | 78860792 | rs16876942 | -2.281 | 1.195 | 1.282 | 1.294 | 0.006 | 0.013 | 0.013 | 0 | 0.08 | 0.09 | 0.15 | 0.09 | 0.07 | 0.12 |
| 5 | 78863936 | rs7720424 | -2.329 | 1.068 | 0.955 | 1.506 | -0.029 | -0.05 | -0.05 | 0 | 0.35 | 0.84 | 0.52 | 0.51 | 0.36 | 0.39 |
| 5 | 78863953 | rs7702211 | -2.345 | 1.065 | 0.953 | 1.505 | -0.028 | -0.05 | -0.05 | 0 | 0.35 | 0.84 | 0.52 | 0.51 | 0.36 | 0.39 |
| 5 | 78867249 | rs7737048 | -0.964 | 1.116 | 1.925 | 1.682 | -0.029 | 0.002 | 0.002 | 0 | 0.18 | 0.36 | 0.26 | 0.17 | 0.11 | 0.12 |
| 5 | 78876335 | rs7704901 | -1.981 | 0.889 | 2.104 | 1.564 | 0.007 | 0.028 | 0.028 | 0 | 0.32 | 0.57 | 0.49 | 0.45 | 0.33 | 0.28 |
| 5 | 78881844 | rs4479827 | -2.283 | 0.991 | 2.198 | 1.583 | 0.021 | 0.028 | 0.028 | 0.566 | 0.33 | 0.53 | 0.52 | 0.54 | 0.36 | 0.4 |
| 5 | 78899835 | rs2161176 | -2.309 | 0.797 | 2.694 | 1.884 | 0.091 | 0.229 | 0.229 | 1.94 | 0.09 | 0.39 | 0.23 | 0.3 | 0.12 | 0.1 |
| 5 | 78909324 | rs3829808 | -1.712 | 0.635 | 2.651 | 1.713 | -0.043 | -0.014 | -0.014 | 0 | 0.18 | 0.48 | 0.23 | 0.13 | 0.1 | 0.13 |
| 5 | 78938315 | rs6453468 | -3.174 | 0.642 | 2.587 | 1.701 | -0.029 | -0.12 | -0.12 | 0 | 0.35 | 0.88 | 0.48 | 0.51 | 0.28 | 0.37 |
| 5 | 78940783 | rs16876991 | 0.092 | 0.724 | 2.587 | 1.735 | 0 | -0.045 | -0.045 | 0.88 | 0.09 | 0.17 | 0.04 | 0.01 | 0.07 | 0.06 |
| 5 | 78945171 | rs7712162 | -2.642 | 0.729 | 2.261 | 1.746 | -0.004 | -0.002 | -0.002 | 0 | 0.27 | 0.69 | 0.29 | 0.23 | 0.18 | 0.28 |
| 5 | 78945736 | rs7736467 | -2.426 | 0.73 | 2.255 | 1.75 | 0.004 | 0.078 | 0.078 | 1.731 | 0.23 | 0.58 | 0.42 | 0.43 | 0.19 | 0.22 |
| 5 | 78945869 | rs16876996 | -0.349 | 0.758 | 1.527 | 1.764 | -0.034 | 0.054 | 0.054 | 0 | 0.07 | 0.17 | 0.18 | 0.13 | 0.02 | 0.07 |
| 5 | 78951714 | rs6876607 | -2.48 | 0.831 | 0.58 | 1.799 | -0.008 | -0.06 | -0.06 | 1.666 | 0.33 | 0.81 | 0.47 | 0.52 | 0.27 | 0.37 |
| 5 | 78961093 | rs11738224 | -0.889 | 0.84 | 0.584 | 1.769 | 0.024 | 0.083 | 0.083 | 0.016 | 0.06 | 0.1 | 0.19 | 0.29 | 0.09 | 0.08 |
| 5 | 78963323 | rs12657392 | -0.34 | 0.849 | 0.454 | 1.748 | -0.033 | -0.031 | -0.031 | 0.003 | 0.07 | 0 | 0.04 | 0.09 | 0.08 | 0.14 |
| 5 | 78963999 | rs6877794 | -1.983 | 0.818 | 0.508 | 1.662 | -0.008 | -0.062 | -0.062 | 0.601 | 0.33 | 0.81 | 0.47 | 0.51 | 0.27 | 0.36 |
| 5 | 78981550 | rs3087813 | -1.64 | 0.407 | 0.368 | 0.989 | -0.004 | -0.002 | -0.002 | 0 | 0.27 | 0.69 | 0.29 | 0.23 | 0.18 | 0.28 |
| 5 | 78984173 | rs34590047 | -0.707 | 0.541 | -0.121 | 1.161 | 0.008 | 0.067 | 0.067 | 0.316 | 0.23 | 0.37 | 0.25 | 0.38 | 0.23 | 0.16 |
| 5 | 78989486 | rs4704576 | -1.402 | 0.807 | -0.889 | 0.844 | -0.048 | 0.037 | 0.037 | 1.163 | 0.53 | 0.95 | 0.7 | 0.65 | 0.5 | 0.52 |
| 5 | 78994362 | rs9784710 | 0.501 | 0.965 | -0.591 | 0.779 | 0.006 | -0.043 | -0.043 | 2.85 | 0.16 | 0.22 | 0.09 | 0.03 | 0.14 | 0.15 |
| 5 | 78996050 | rs6870619 | -0.527 | 1.037 | -0.5 | 0.863 | 0.007 | 0.086 | 0.086 | 0.233 | 0.29 | 0.51 | 0.56 | 0.46 | 0.27 | 0.24 |
| 5 | 78996244 | rs4441859 | -1.731 | 0.458 | -0.614 | 0.521 | 0.005 | 0.026 | 0.026 | 0.013 | 0.4 | 0.78 | 0.45 | 0.29 | 0.29 | 0.37 |
| 5 | 78998903 | rs16877062 | -0.834 | 1.468 | -0.746 | 1.078 | -0.009 | -0.016 | -0.016 | 0 | 0.09 | 0.03 | 0.04 | 0.16 | 0.09 | 0.15 |
| 5 | 79003360 | rs12651804 | -0.794 | 1.404 | -0.503 | 0.804 | 0.008 | -0.003 | -0.003 | 0 | 0.2 | 0.23 | 0.36 | 0.13 | 0.2 | 0.23 |
| 5 | 79012338 | rs13170837 | -0.263 | 0.852 | -0.868 | 0.678 | 0.002 | 0.029 | 0.029 | 0.621 | 0.45 | 0.64 | 0.38 | 0.58 | 0.4 | 0.37 |
| 5 | 79013201 | rs13171893 | -0.313 | 0.218 | -0.211 | 1.216 | 0.001 | 0.005 | 0.005 | 0.039 | 0.4 | 0.36 | 0.33 | 0.22 | 0.34 | 0.26 |
| 5 | 79016671 | rs4704583 | -1.243 | 0.019 | 0.312 | 1.785 | 0.049 | 0.145 | 0.145 | 1.218 | 0.12 | 0.47 | 0.08 | 0.22 | 0.08 | 0.02 |
| 5 | 79017377 | rs12518982 | -0.467 | 0.134 | 0.216 | 1.745 | -0.004 | -0.042 | -0.042 | 0 | 0.24 | 0.19 | 0.28 | 0.38 | 0.24 | 0.33 |
| 5 | 79024734 | rs1541813 | -1.665 | -0.151 | 0.02 | 1.043 | -0.002 | 0.075 | 0.075 | 1.947 | 0.46 | 0.81 | 0.39 | 0.6 | 0.38 | 0.43 |
| 5 | 79024779 | rs16877109 | -1.585 | -0.245 | 0.333 | 1.074 | -0.016 | 0.158 | 0.158 | 5.382 | 0.16 | 0.6 | 0.09 | 0.36 | 0.08 | 0.1 |
| 5 | 79025317 | rs1366270 | -1.585 | -0.245 | 0.333 | 1.074 | -0.016 | 0.158 | 0.158 | 5.382 | 0.16 | 0.6 | 0.09 | 0.36 | 0.08 | 0.1 |
| 5 | 79025634 | rs1366271 | -1.311 | -0.252 | 0.523 | 1.071 | 0.023 | 0.299 | 0.299 | 1.279 | 0.13 | 0.44 | 0.07 | 0.36 | 0.08 | 0.1 |
| 5 | 79027605 | rs6893869 | -1.484 | -0.25 | 0.475 | 1.072 | -0.016 | 0.16 | 0.16 | 5.242 | 0.16 | 0.58 | 0.09 | 0.36 | 0.08 | 0.1 |
| 5 | 79028472 | rs4704585 | -1.793 | -0.201 | 0.506 | 1.261 | -0.009 | 0.095 | 0.095 | 1.674 | 0.45 | 0.78 | 0.41 | 0.64 | 0.39 | 0.43 |
| 5 | 79028513 | rs16877133 | -0.788 | -0.258 | 0.75 | 1.208 | 0.034 | 0.303 | 0.303 | 2.716 | 0.08 | 0.27 | 0.05 | 0.36 | 0.06 | 0.09 |
| 5 | 79028586 | rs16877135 | -0.788 | -0.258 | 0.75 | 1.208 | 0.034 | 0.303 | 0.303 | 2.716 | 0.08 | 0.27 | 0.05 | 0.36 | 0.06 | 0.09 |
| 5 | 79028726 | rs13158477 | -0.228 | -0.224 | 0.408 | 1.192 | -0.001 | -0.006 | -0.006 | 0 | 0.12 | 0.12 | 0.13 | 0.02 | 0.16 | 0.13 |
| 5 | 79029383 | rs1428224 | -1.397 | -0.241 | 0.476 | 1.313 | -0.016 | 0.16 | 0.16 | 5.242 | 0.16 | 0.59 | 0.09 | 0.36 | 0.08 | 0.1 |
| 5 | 79030346 | rs16877151 | -1.431 | -0.228 | 0.398 | 1.323 | -0.016 | 0.158 | 0.158 | 5.382 | 0.16 | 0.59 | 0.09 | 0.36 | 0.08 | 0.1 |
| 5 | 79033306 | rs2278239 | -1.429 | -0.223 | 0.536 | 1.325 | -0.014 | 0.178 | 0.178 | 5.013 | 0.16 | 0.56 | 0.08 | 0.36 | 0.08 | 0.1 |
| 5 | 79033391 | rs2278240 | 0.246 | 0.021 | 0.067 | 1.423 | 0.002 | -0.003 | -0.003 | 0.858 | 0.07 | 0.04 | 0.01 | 0.13 | 0.02 | 0.07 |
| 5 | 79041057 | rs12514461 | 0.326 | 0.163 | 0.03 | 1.38 | -0.025 | -0.002 | -0.002 | 0.039 | 0.11 | 0.04 | 0.17 | 0.08 | 0.13 | 0.12 |
| 5 | 79045101 | rs7709188 | -1.415 | -0.015 | 0.713 | 1.588 | 0.018 | 0.148 | 0.148 | 0.646 | 0.21 | 0.45 | 0.18 | 0.38 | 0.22 | 0.22 |
| 5 | 79046415 | rs1428226 | -0.101 | 0.185 | 0.554 | 1.411 | 0.007 | -0.012 | -0.012 | 0 | 0.07 | 0.11 | 0.1 | 0.02 | 0.12 | 0.07 |
| 5 | 79047407 | rs13159668 | -0.015 | 0.329 | -0.036 | 1.391 | -0.025 | 0.007 | 0.007 | 1.4 | 0.34 | 0.25 | 0.49 | 0.24 | 0.45 | 0.38 |
| 5 | 79048562 | rs1366272 | -1.253 | 0.513 | 0.231 | 2.351 | 0.06 | 0.198 | 0.198 | 0.872 | 0.11 | 0.24 | 0.19 | 0.42 | 0.1 | 0.18 |
| 5 | 79049271 | rs6453481 | -1.504 | 0.14 | 0.082 | 2.354 | -0.005 | 0.085 | 0.085 | 0.653 | 0.21 | 0.6 | 0.16 | 0.47 | 0.16 | 0.18 |
| 5 | 79058817 | rs259128 | -0.228 | 0.476 | -0.815 | 2.104 | -0.038 | 0.002 | 0.002 | 0 | 0.11 | 0.03 | 0.08 | 0.04 | 0.13 | 0.16 |
| 5 | 79059417 | rs259129 | -0.536 | 0.341 | -0.65 | 1.999 | 0.021 | 0.053 | 0.053 | 0.051 | 0.17 | 0.28 | 0.14 | 0.3 | 0.19 | 0.16 |
| 5 | 79067033 | rs6453482 | -0.205 | 0.509 | -1.698 | 2.258 | -0.043 | -0.063 | -0.063 | 0 | 0.13 | 0.02 | 0.19 | 0.33 | 0.13 | 0.16 |
| 5 | 79068089 | rs109894 | -0.89 | 0.05 | -1.764 | 1.781 | -0.046 | -0.028 | -0.028 | 0 | 0.4 | 0.83 | 0.39 | 0.63 | 0.35 | 0.35 |

| | | | | | | | | | | | | | | | | |
|---|---|---|---|---|---|---|---|---|---|---|---|---|---|---|---|---|
| 5 | 79071781 | rs11739525 | -1.069 | -0.175 | -0.905 | 2.147 | 0.009 | 0.015 | 0.015 | 0 | 0.08 | 0.06 | 0.07 | 0.4 | 0.04 | 0.14 |
| 5 | 79081713 | rs6453484 | -0.379 | 0.067 | -0.523 | 1.489 | -0.01 | -0.006 | -0.006 | 0.026 | 0.29 | 0.5 | 0.18 | 0.28 | 0.2 | 0.22 |
| 5 | 79083467 | rs402955 | -0.251 | -0.034 | -0.693 | 2.246 | -0.001 | -0.016 | -0.016 | 0 | 0.05 | 0.01 | 0.05 | 0.29 | 0.04 | 0.09 |
| 5 | 79083679 | rs4703789 | 0.084 | -0.508 | -0.751 | 1.926 | -0.008 | 0.01 | 0.01 | 0.047 | 0.17 | 0.24 | 0.37 | 0.2 | 0.1 | 0.16 |
| 5 | 79085726 | rs12657828 | -0.078 | -0.657 | -0.294 | 1.38 | -0.011 | 0.01 | 0.01 | 0 | 0.18 | 0.29 | 0.37 | 0.2 | 0.11 | 0.16 |
| 5 | 79086883 | rs1129770 | 0.868 | -0.318 | -1.045 | 0.468 | 0.002 | 0.031 | 0.031 | 0 | 0.18 | 0.07 | 0.11 | 0.09 | 0.2 | 0.13 |
| 5 | 79086903 | rs10052867 | 0.463 | -0.332 | -0.493 | 0.327 | 0.002 | -0.01 | -0.01 | 0 | 0.25 | 0.28 | 0.15 | 0.02 | 0.17 | 0.2 |
| 5 | 79088995 | rs2287673 | -0.632 | 0.003 | -0.167 | 0.811 | 0.01 | 0.012 | 0.012 | 0 | 0.08 | 0.13 | 0.11 | 0.11 | 0.08 | 0.07 |
| 5 | 79092097 | rs259100 | -0.878 | -0.426 | -0.04 | 1.13 | -0.006 | 0.019 | 0.019 | 0.301 | 0.31 | 0.37 | 0.39 | 0.56 | 0.24 | 0.25 |
| 5 | 79092370 | rs259102 | -0.363 | -0.494 | -0.038 | 1.133 | -0.001 | 0.002 | 0.002 | 0.864 | 0.26 | 0.26 | 0.35 | 0.44 | 0.17 | 0.18 |
| 5 | 79094154 | rs10491460 | 0.011 | -0.548 | -0.126 | 0.787 | -0.001 | -0.013 | -0.013 | 0 | 0.07 | 0.06 | 0.04 | 0.01 | 0.07 | 0.07 |
| 5 | 79094685 | rs259103 | -1.022 | -0.171 | 0.167 | 0.651 | 0.006 | 0.041 | 0.041 | 1.356 | 0.47 | 0.67 | 0.53 | 0.66 | 0.44 | 0.38 |
| 5 | 79095721 | rs7343 | 1.454 | 0.272 | -0.204 | 0.57 | -0.007 | 0.007 | 0.007 | 0 | 0.13 | 0.08 | 0.09 | 0.1 | 0.18 | 0.06 |
| 5 | 79098998 | rs259067 | -0.017 | 0.075 | -0.305 | 0.535 | 0.002 | -0.005 | -0.005 | 0 | 0.27 | 0.25 | 0.37 | 0.44 | 0.19 | 0.2 |
| 5 | 79100717 | rs259065 | -0.768 | -0.128 | -0.299 | 0.615 | 0.01 | -0.004 | -0.004 | 0 | 0.39 | 0.67 | 0.21 | 0.39 | 0.33 | 0.24 |
| 5 | 79102932 | rs16877221 | 0.37 | 0.048 | -0.21 | 0.484 | -0.008 | -0.026 | -0.026 | 0 | 0.06 | 0.11 | 0.02 | 0.01 | 0.05 | 0.05 |
| 5 | 79105021 | rs410079 | -0.553 | -0.166 | -0.477 | 0.68 | -0.024 | -0.02 | -0.02 | 0 | 0.2 | 0.6 | 0.11 | 0.16 | 0.14 | 0.16 |
| 5 | 79105154 | rs393062 | 0.02 | 0.064 | -1.071 | 0.317 | 0.031 | -0.029 | -0.029 | 0 | 0.56 | 0.76 | 0.51 | 0.47 | 0.67 | 0.51 |
| 5 | 79106802 | rs11957025 | -0.395 | -0.006 | -1.542 | 0.325 | 0.026 | 0.067 | 0.067 | 0.688 | 0.63 | 0.94 | 0.58 | 0.76 | 0.74 | 0.64 |
| 5 | 79108476 | rs13185886 | 0.084 | 0.346 | -0.885 | 0.291 | 0.013 | -0.02 | -0.02 | 0 | 0.25 | 0.43 | 0.16 | 0.2 | 0.29 | 0.13 |
| 5 | 79111060 | rs259069 | -0.252 | 0.036 | -1.416 | 0.424 | 0.022 | -0.004 | -0.004 | 0.184 | 0.73 | 0.84 | 0.82 | 0.75 | 0.83 | 0.61 |
| 5 | 79115662 | rs9293789 | -0.276 | 0.386 | -1.202 | -0.05 | 0.013 | 0.073 | 0.073 | 0 | 0.5 | 0.35 | 0.53 | 0.29 | 0.45 | 0.43 |
| 5 | 79115827 | rs9293790 | -0.093 | 0.42 | -1.242 | -0.125 | 0.001 | 0.007 | 0.007 | 0.02 | 0.37 | 0.34 | 0.24 | 0.14 | 0.35 | 0.38 |
| 5 | 79120482 | rs1864035 | 0.061 | -0.153 | -1.7 | -0.432 | -0.014 | 0.037 | 0.037 | 0 | 0.31 | 0.11 | 0.36 | 0.21 | 0.3 | 0.34 |
| 5 | 79120797 | rs7707140 | -0.464 | -0.128 | -0.455 | -0.689 | 0.058 | 0.016 | 0.016 | 0.123 | 0.07 | 0.37 | 0.05 | 0 | 0.02 | 0.04 |
| 5 | 79129532 | rs1529457 | 1.115 | 0.266 | -1.16 | -0.96 | -0.048 | 0.075 | 0.075 | 0.01 | 0.27 | 0.09 | 0.25 | 0.03 | 0.35 | 0.24 |
| 5 | 79129655 | rs16877240 | -0.31 | -0.476 | -0.704 | -0.013 | -0.015 | -0.001 | -0.001 | 0 | 0.16 | 0.22 | 0.1 | 0.16 | 0.11 | 0.09 |
| 5 | 79132520 | rs259072 | 0.337 | -0.156 | -0.679 | 0.391 | 0.001 | 0.007 | 0.007 | 0 | 0.47 | 0.38 | 0.62 | 0.39 | 0.56 | 0.38 |
| 5 | 79132571 | rs259073 | 0.655 | -0.186 | -1.015 | 0.299 | 0.003 | -0.007 | -0.007 | 0.703 | 0.15 | 0.12 | 0.21 | 0.22 | 0.15 | 0.15 |
| 5 | 79132881 | rs4704594 | 0.984 | -0.116 | -0.853 | 0.175 | -0.029 | 0.043 | 0.043 | 0 | 0.32 | 0.2 | 0.39 | 0.15 | 0.42 | 0.23 |
| 5 | 79132907 | rs259074 | 0.195 | -0.144 | -0.834 | 0.431 | 0.017 | 0.025 | 0.025 | 0 | 0.5 | 0.34 | 0.62 | 0.42 | 0.59 | 0.38 |
| 5 | 79133770 | rs1560130 | -0.782 | -0.434 | -1.082 | 0.201 | 0.01 | 0.009 | 0.009 | 0 | 0.85 | 0.87 | 0.91 | 0.9 | 0.9 | 0.82 |
| 5 | 79133987 | rs17472293 | 1.143 | -0.275 | -1.18 | 0.091 | -0.069 | 0.094 | 0.094 | 0 | 0.31 | 0.09 | 0.38 | 0.14 | 0.41 | 0.23 |
| 5 | 79135153 | rs13354102 | 0.194 | -0.057 | -1.055 | 0.651 | -0.004 | -0.003 | -0.003 | 0 | 0.27 | 0.16 | 0.21 | 0.24 | 0.25 | 0.35 |
| 5 | 79135217 | rs2404666 | -0.466 | -0.283 | -0.998 | 0.93 | 0.006 | -0.001 | -0.001 | 0 | 0.73 | 0.79 | 0.69 | 0.74 | 0.71 | 0.75 |
| 5 | 79136757 | rs16877250 | -0.133 | -0.051 | -1 | 1.031 | -0.014 | -0.024 | -0.024 | 0 | 0.31 | 0.16 | 0.25 | 0.54 | 0.31 | 0.44 |
| 5 | 79137556 | rs265009 | -0.032 | -0.404 | -1.05 | 0.639 | 0.014 | 0.01 | 0.01 | 0.995 | 0.59 | 0.39 | 0.65 | 0.65 | 0.66 | 0.65 |
| 5 | 79137792 | rs9790975 | 0.456 | -0.161 | -1.216 | 1.243 | -0.074 | -0.081 | -0.081 | 0 | 0.21 | 0.03 | 0.13 | 0.24 | 0.23 | 0.24 |
| 5 | 79139377 | rs16877256 | -0.5 | -0.679 | -1.165 | 0.964 | 0.035 | -0.042 | -0.042 | 0.124 | 0.17 | 0.11 | 0.12 | 0.27 | 0.07 | 0.23 |
| 5 | 79142191 | rs10056632 | -1.167 | -0.284 | -0.565 | 1.011 | -0.001 | -0.006 | -0.006 | 0.588 | 0.36 | 0.44 | 0.27 | 0.72 | 0.32 | 0.52 |
| 5 | 79142384 | rs10062050 | 0.74 | -0.29 | -0.512 | 1.25 | -0.009 | -0.003 | -0.003 | 0 | 0.14 | 0.12 | 0.11 | 0.43 | 0.2 | 0.26 |
| 5 | 79143194 | rs10079417 | -0.63 | -0.287 | -0.484 | 0.736 | -0.001 | 0.004 | 0.004 | 1.198 | 0.25 | 0.34 | 0.2 | 0.66 | 0.23 | 0.43 |
| 5 | 79145803 | rs264982 | -0.338 | -0.513 | -0.817 | -0.041 | 0.008 | 0.012 | 0.012 | 0 | 0.68 | 0.75 | 0.63 | 0.75 | 0.7 | 0.7 |
| 5 | 79152076 | rs10491456 | -0.352 | -0.44 | -0.678 | 0.391 | -0.003 | -0.019 | -0.019 | 0 | 0.22 | 0.16 | 0.18 | 0.69 | 0.22 | 0.42 |
| 5 | 79153359 | rs7705021 | -0.338 | -0.467 | -0.447 | 0.405 | -0.001 | -0.008 | -0.008 | 0 | 0.22 | 0.2 | 0.17 | 0.69 | 0.22 | 0.4 |
| 5 | 79155579 | rs149767 | -0.084 | -0.536 | -0.427 | -0.229 | 0.009 | 0.01 | 0.01 | 0.011 | 0.59 | 0.4 | 0.63 | 0.86 | 0.69 | 0.85 |
| 5 | 79158432 | rs10068385 | 0.813 | -0.25 | 0.088 | 0.57 | 0.038 | 0.132 | 0.132 | 0.635 | 0.16 | 0.33 | 0.13 | 0.39 | 0.19 | 0.32 |
| 5 | 79159152 | rs265002 | 0.713 | -0.382 | 0.131 | 0.223 | 0 | 0.003 | 0.003 | 0 | 0.62 | 0.44 | 0.62 | 0.45 | 0.58 | 0.48 |
| 5 | 79160486 | rs10942906 | 0.11 | -0.33 | 0.403 | 0.006 | -0.015 | -0.034 | -0.034 | 0.395 | 0.31 | 0.59 | 0.17 | 0.08 | 0.23 | 0.1 |
| 5 | 79161503 | rs10942907 | 0.27 | -0.302 | 0.235 | 0.004 | -0.022 | -0.054 | -0.054 | 0 | 0.3 | 0.4 | 0.15 | 0.07 | 0.23 | 0.11 |
| 5 | 79163127 | rs16877298 | 0.195 | -0.326 | 0.242 | 0.024 | -0.026 | -0.048 | -0.048 | 3.943 | 0.3 | 0.39 | 0.14 | 0.07 | 0.21 | 0.1 |
| 5 | 79166824 | rs264990 | 0.678 | -0.221 | -0.028 | 0.094 | 0.027 | 0.057 | 0.057 | 0 | 0.5 | 0.39 | 0.7 | 0.77 | 0.55 | 0.64 |
| 5 | 79170601 | rs264985 | 0.44 | 0.408 | 0.439 | 0.427 | 0.009 | -0.014 | -0.014 | 0 | 0.73 | 0.62 | 0.76 | 0.8 | 0.69 | 0.68 |
| 5 | 79170694 | rs264984 | 0.138 | 0.419 | 0.303 | 0.42 | -0.014 | 0.02 | 0.02 | 0 | 0.76 | 0.88 | 0.82 | 0.84 | 0.7 | 0.74 |
| 5 | 79171387 | rs182042 | 0.224 | 0.742 | 0.305 | 0.488 | -0.022 | 0.015 | 0.015 | 0 | 0.75 | 0.86 | 0.78 | 0.82 | 0.65 | 0.7 |
| 5 | 79172136 | rs265005 | 1.121 | 0.758 | 0.234 | 0.739 | 0.009 | 0.046 | 0.046 | 0 | 0.33 | 0.16 | 0.39 | 0.25 | 0.24 | 0.31 |
| 5 | 79173521 | rs10474594 | 0.085 | 0.741 | 0.727 | 0.827 | 0 | -0.002 | -0.002 | 0 | 0.4 | 0.53 | 0.34 | 0.59 | 0.39 | 0.42 |
| 5 | 79174137 | rs10491455 | 0.915 | 0.792 | 0.64 | 0.874 | 0.008 | 0.018 | 0.018 | 1.271 | 0.27 | 0.18 | 0.18 | 0.14 | 0.24 | 0.17 |
| 5 | 79175246 | rs6888630 | -0.161 | 0.899 | 0.572 | 0.959 | -0.004 | -0.006 | -0.006 | 0 | 0.39 | 0.61 | 0.27 | 0.25 | 0.35 | 0.19 |
| 5 | 79181467 | rs265008 | 0.846 | 0.989 | 0.475 | 0.916 | 0.001 | 0 | 0 | 0.342 | 0.61 | 0.45 | 0.52 | 0.4 | 0.47 | 0.44 |
| 5 | 79186809 | rs10514162 | 0.52 | 1.023 | 0.392 | 0.857 | -0.003 | 0.003 | 0.003 | 0 | 0.17 | 0.14 | 0.19 | 0.1 | 0.26 | 0.23 |
| 5 | 79187428 | rs7706921 | -0.762 | 0.912 | 0.446 | 0.866 | 0.002 | 0.004 | 0.004 | 0 | 0.57 | 0.62 | 0.53 | 0.65 | 0.65 | 0.66 |
| 5 | 79189682 | rs12656872 | 0.566 | 1.244 | 0.838 | 1.119 | 0.016 | -0.015 | -0.015 | 0 | 0.09 | 0.21 | 0.07 | 0.05 | 0.11 | 0.03 |
| 5 | 79191702 | rs4704604 | -0.271 | 1.306 | 0.591 | 1.121 | 0.042 | 0.15 | 0.15 | 0.894 | 0.21 | 0.41 | 0.29 | 0.47 | 0.26 | 0.34 |
| 5 | 79207070 | rs13170399 | -0.642 | 1.361 | 0.207 | 1.196 | 0.016 | 0.032 | 0.032 | 0 | 0.29 | 0.31 | 0.39 | 0.53 | 0.41 | 0.54 |
| 5 | 79215861 | rs10474600 | 0.518 | 1.161 | 0.01 | 1.345 | 0.011 | 0.078 | 0.078 | 0.45 | 0.13 | 0.18 | 0.22 | 0.43 | 0.16 | 0.31 |
| 5 | 79216901 | rs1565513 | -0.325 | 1.152 | -0.144 | 1.148 | 0 | 0.001 | 0.001 | 0 | 0.49 | 0.43 | 0.34 | 0.57 | 0.48 | 0.44 |
| 5 | 79217017 | rs4703796 | 1.38 | 1.096 | -0.404 | 1.291 | 0.013 | 0.08 | 0.08 | 0.365 | 0.11 | 0.11 | 0.17 | 0.4 | 0.16 | 0.26 |
| 5 | 79218998 | rs13171934 | -0.038 | 0.659 | -0.459 | 0.814 | 0.002 | -0.02 | -0.02 | 0 | 0.19 | 0.26 | 0.18 | 0.08 | 0.24 | 0.25 |

| | | | | | | | | | | | | | | | | |
|---|---|---|---|---|---|---|---|---|---|---|---|---|---|---|---|---|
| 5 | 79228270 | rs6865146 | 0.213 | 0.611 | -0.232 | 1.178 | -0.004 | 0.04 | 0.04 | 0.026 | 0.48 | 0.38 | 0.49 | 0.38 | 0.49 | 0.5 |
| 5 | 79228519 | rs12658005 | -0.037 | 0.748 | -1.13 | 0.582 | 0.006 | -0.035 | -0.035 | 2.967 | 0.12 | 0.21 | 0.08 | 0.05 | 0.12 | 0.03 |
| 5 | 79228649 | rs6453492 | -0.184 | 0.907 | -1.885 | 1.142 | -0.037 | -0.051 | -0.051 | 0 | 0.62 | 0.86 | 0.67 | 0.6 | 0.62 | 0.58 |
| 5 | 79230312 | rs6876520 | -0.215 | 0.909 | -1.866 | 0.954 | -0.034 | -0.018 | -0.018 | 0.11 | 0.62 | 0.85 | 0.67 | 0.65 | 0.62 | 0.59 |
| 5 | 79233056 | rs7707589 | -1.2 | 0.978 | -1.481 | 0.811 | -0.006 | 0.013 | 0.013 | 0.165 | 0.1 | 0.03 | 0.12 | 0.08 | 0.14 | 0.07 |
| 5 | 79237487 | rs7712567 | -0.522 | 1.192 | -1.491 | 0.988 | -0.031 | -0.021 | -0.021 | 0 | 0.62 | 0.77 | 0.62 | 0.52 | 0.55 | 0.59 |
| 5 | 79242663 | rs4535437 | 0.094 | 0.764 | -1.523 | 0.687 | -0.008 | 0.044 | 0.044 | 3.329 | 0.29 | 0.16 | 0.2 | 0.15 | 0.26 | 0.25 |
| 5 | 79251504 | rs2471104 | -0.759 | 0.941 | -0.997 | 0.881 | -0.005 | -0.001 | -0.001 | 0.789 | 0.61 | 0.65 | 0.57 | 0.54 | 0.56 | 0.63 |
| 5 | 79254118 | rs2471106 | -1.459 | 1.001 | -0.941 | 1.024 | -0.021 | -0.001 | -0.001 | 0.035 | 0.69 | 0.78 | 0.66 | 0.68 | 0.62 | 0.67 |
| 5 | 79256562 | rs2404710 | -0.12 | 0.796 | -0.103 | 1.232 | -0.006 | 0.003 | 0.003 | 0.109 | 0.41 | 0.41 | 0.33 | 0.34 | 0.42 | 0.31 |
| 5 | 79260025 | rs4576130 | -0.118 | 1.019 | 0.026 | 1.28 | -0.015 | -0.002 | -0.002 | 0 | 0.41 | 0.37 | 0.36 | 0.37 | 0.45 | 0.34 |
| 5 | 79261984 | rs11749003 | 0.73 | 0.923 | 0.342 | 1.262 | 0.004 | -0.002 | -0.002 | 0 | 0.19 | 0.15 | 0.28 | 0.19 | 0.18 | 0.32 |
| 5 | 79262938 | rs3926218 | 0.91 | 0.897 | 0.335 | 0.885 | -0.025 | 0.065 | 0.065 | 5.28 | 0.24 | 0.12 | 0.14 | 0.05 | 0.25 | 0.13 |
| 5 | 79263211 | rs1017998 | -1.346 | 0.8 | 0.364 | 0.928 | 0 | 0 | 0 | 3.917 | 0.62 | 0.68 | 0.65 | 0.56 | 0.64 | 0.67 |
| 5 | 79273042 | rs7446065 | -1.308 | 0.793 | 0.417 | 0.937 | 0 | 0 | 0 | 0 | 0.62 | 0.66 | 0.65 | 0.56 | 0.64 | 0.66 |
| 5 | 79277215 | rs10064975 | -1.994 | 0.766 | 0.331 | 0.903 | -0.016 | 0.009 | 0.009 | 0.706 | 0.69 | 0.79 | 0.7 | 0.72 | 0.66 | 0.7 |
| 5 | 79277817 | rs6453495 | -1.206 | 0.813 | 0.288 | 0.893 | -0.003 | -0.003 | -0.003 | 0.042 | 0.61 | 0.52 | 0.64 | 0.58 | 0.64 | 0.66 |
| 5 | 79282212 | rs17196895 | 1.594 | 0.761 | 0.072 | 0.856 | -0.014 | 0.009 | 0.009 | 0 | 0.06 | 0 | 0.05 | 0.03 | 0.09 | 0.09 |
| 5 | 79282798 | rs9293796 | 1.694 | 0.763 | -0.018 | 0.734 | -0.037 | 0.082 | 0.082 | 2.255 | 0.15 | 0.02 | 0.08 | 0.01 | 0.14 | 0.11 |
| 5 | 79288995 | rs1438739 | -1.221 | 0.79 | 0.366 | 1 | -0.003 | -0.003 | -0.003 | 0 | 0.61 | 0.57 | 0.65 | 0.58 | 0.64 | 0.66 |
| 5 | 79292899 | rs7707036 | -0.783 | 0.723 | 0.414 | 0.999 | 0 | 0.001 | 0.001 | 0.201 | 0.54 | 0.48 | 0.59 | 0.55 | 0.55 | 0.57 |
| 5 | 79298854 | rs383122 | 0.442 | 0.918 | 0.112 | 1.2 | -0.036 | -0.041 | -0.041 | 1.072 | 0.11 | 0.03 | 0.12 | 0.1 | 0.12 | 0.16 |
| 5 | 79303633 | rs2451932 | -1.389 | 0.767 | 0.333 | 0.807 | -0.03 | -0.01 | -0.01 | 1.183 | 0.63 | 0.75 | 0.61 | 0.63 | 0.58 | 0.6 |
| 5 | 79303948 | rs2434316 | -0.27 | 0.731 | 0.572 | 0.735 | -0.004 | -0.004 | -0.004 | 0.822 | 0.3 | 0.36 | 0.37 | 0.23 | 0.31 | 0.37 |
| 5 | 79305981 | rs12332358 | 1.074 | 0.803 | 0.646 | 0.737 | -0.015 | 0.056 | 0.056 | 2.923 | 0.23 | 0.11 | 0.14 | 0.05 | 0.23 | 0.16 |
| 5 | 79308315 | rs2028269 | -0.441 | 0.795 | 0.763 | 0.71 | 0.001 | 0.008 | 0.008 | 1.761 | 0.48 | 0.56 | 0.44 | 0.25 | 0.42 | 0.46 |
| 5 | 79309458 | rs2438612 | 0.163 | 0.784 | 0.768 | 0.709 | -0.025 | -0.007 | -0.007 | 0 | 0.25 | 0.44 | 0.3 | 0.2 | 0.19 | 0.3 |
| 5 | 79316331 | rs11743110 | 1.023 | 0.825 | 0.398 | 0.725 | -0.013 | 0.051 | 0.051 | 2.961 | 0.23 | 0.12 | 0.14 | 0.05 | 0.23 | 0.16 |
| 5 | 79317779 | rs6875852 | 1.333 | 0.828 | 0.386 | 0.654 | -0.016 | -0.049 | -0.049 | 0 | 0.17 | 0.02 | 0.24 | 0.18 | 0.15 | 0.27 |
| 5 | 79330559 | rs3813667 | -0.062 | 0.95 | 0.554 | 0.713 | -0.001 | 0 | 0 | 0 | 0.33 | 0.29 | 0.24 | 0.29 | 0.35 | 0.24 |
| 5 | 79337307 | rs2434311 | 0.331 | 0.78 | 0.628 | 0.763 | -0.015 | 0.002 | 0.002 | 0 | 0.24 | 0.38 | 0.3 | 0.21 | 0.19 | 0.3 |
| 5 | 79342519 | rs256440 | 0.344 | 0.784 | 0.63 | 0.764 | -0.014 | -0.001 | -0.001 | 0 | 0.24 | 0.38 | 0.3 | 0.21 | 0.19 | 0.3 |
| 5 | 79345549 | rs9293799 | -0.043 | 0.884 | 0.677 | 0.749 | -0.001 | 0.007 | 0.007 | 1.896 | 0.34 | 0.25 | 0.24 | 0.29 | 0.36 | 0.25 |
| 5 | 79348429 | rs445471 | -0.99 | 0.904 | 0.643 | 0.764 | -0.014 | -0.017 | -0.017 | 0 | 0.58 | 0.68 | 0.55 | 0.51 | 0.55 | 0.55 |
| 5 | 79348759 | rs404375 | -0.99 | 0.904 | 0.643 | 0.764 | -0.014 | -0.017 | -0.017 | 0 | 0.58 | 0.68 | 0.55 | 0.51 | 0.55 | 0.55 |
| 5 | 79349157 | rs13174295 | 1.073 | 0.898 | 0.494 | 0.722 | -0.006 | 0.018 | 0.018 | 0 | 0.07 | 0.01 | 0.07 | 0.03 | 0.12 | 0.07 |
| 5 | 79351735 | rs423906 | 0.35 | 0.847 | 0.547 | 0.784 | -0.015 | 0.004 | 0.004 | 0 | 0.24 | 0.37 | 0.31 | 0.22 | 0.2 | 0.3 |
| 5 | 79363273 | rs9293800 | -0.012 | 0.944 | 0.506 | 0.758 | -0.003 | 0.017 | 0.017 | 2.16 | 0.33 | 0.2 | 0.23 | 0.27 | 0.34 | 0.25 |
| 5 | 79366080 | rs256439 | 0.592 | 0.857 | 0.679 | 0.834 | 0.007 | -0.003 | -0.003 | 0 | 0.28 | 0.28 | 0.21 | 0.24 | 0.29 | 0.2 |
| 5 | 79368392 | rs12110039 | 0.247 | 0.819 | 0.55 | 0.769 | -0.048 | 0.05 | 0.05 | 0 | 0.26 | 0.39 | 0.33 | 0.41 | 0.18 | 0.31 |
| 5 | 79370247 | rs13167730 | 1.249 | 0.868 | 0.025 | 0.737 | -0.005 | 0.035 | 0.035 | 0 | 0.07 | 0.01 | 0.07 | 0.01 | 0.1 | 0.07 |
| 5 | 79373803 | rs2288394 | -0.443 | 0.851 | 0.067 | 0.752 | -0.042 | 0.011 | 0.011 | 0 | 0.34 | 0.51 | 0.4 | 0.4 | 0.28 | 0.38 |
| 5 | 79376231 | rs10514175 | 1.732 | 1.139 | -0.431 | 0.648 | -0.074 | 0.152 | 0.152 | 2.51 | 0.26 | 0.02 | 0.14 | 0.01 | 0.31 | 0.23 |
| 5 | 79377900 | rs2913544 | -0.941 | 0.47 | -0.156 | 0.823 | -0.025 | 0.001 | 0.001 | 1.274 | 0.61 | 0.72 | 0.6 | 0.63 | 0.55 | 0.58 |
| 5 | 79381796 | rs2247450 | -0.935 | 0.403 | -0.264 | 0.827 | -0.033 | 0.003 | 0.003 | 1.454 | 0.61 | 0.73 | 0.57 | 0.64 | 0.5 | 0.55 |
| 5 | 79383276 | rs2118732 | 0.055 | 0.052 | -0.141 | 0.351 | -0.047 | 0.007 | 0.007 | 0 | 0.25 | 0.57 | 0.29 | 0.25 | 0.16 | 0.25 |
| 5 | 79392193 | rs2438632 | -0.362 | 0.976 | -0.512 | -0.096 | -0.001 | -0.001 | -0.001 | 0.096 | 0.37 | 0.37 | 0.3 | 0.23 | 0.41 | 0.27 |
| 5 | 79393144 | rs2438631 | -0.042 | 1.015 | -0.344 | -0.22 | 0.007 | -0.037 | -0.037 | 0.141 | 0.59 | 0.52 | 0.62 | 0.76 | 0.53 | 0.64 |
| 5 | 79395455 | rs11738550 | -1.336 | 0.88 | 0.393 | -0.326 | 0.042 | -0.067 | -0.067 | 2.445 | 0.12 | 0.52 | 0.18 | 0.03 | 0.19 | 0.14 |
| 5 | 79395829 | rs425491 | -0.246 | 1.009 | -0.774 | -0.257 | -0.043 | -0.075 | -0.075 | 1.505 | 0.68 | 0.88 | 0.5 | 0.48 | 0.68 | 0.77 |
| 5 | 79396743 | rs1465852 | 0.306 | 0.384 | -1.075 | -0.639 | 0.02 | 0.073 | 0.073 | 0 | 0.53 | 0.64 | 0.43 | 0.32 | 0.62 | 0.61 |
| 5 | 79398567 | rs13158441 | 0.805 | 0.336 | -0.248 | -0.688 | 0 | -0.011 | -0.011 | 0 | 0.45 | 0.44 | 0.29 | 0.29 | 0.45 | 0.51 |
| 5 | 79398709 | rs6865426 | -0.158 | 0.45 | -0.151 | -0.704 | 0.048 | -0.047 | -0.047 | 0 | 0.08 | 0.25 | 0.15 | 0.01 | 0.18 | 0.11 |
| 5 | 79400778 | rs2015459 | -1.419 | 0.058 | -0.207 | -0.947 | -0.021 | -0.063 | -0.063 | 1.542 | 0.3 | 0.53 | 0.27 | 0.19 | 0.29 | 0.27 |
| 5 | 79401417 | rs421837 | -0.918 | 0.202 | -0.663 | -0.716 | -0.082 | -0.163 | -0.163 | 0.036 | 0.74 | 0.98 | 0.56 | 0.48 | 0.75 | 0.75 |
| 5 | 79401691 | rs6863009 | -0.207 | 0.194 | 0.216 | -0.909 | 0.008 | -0.024 | -0.024 | 0 | 0.07 | 0.2 | 0.08 | 0.03 | 0.08 | 0.03 |
| 5 | 79402236 | rs4704608 | -0.5 | 0.134 | 0.134 | -0.742 | 0.009 | 0 | 0 | 3.788 | 0.13 | 0.11 | 0.05 | 0.16 | 0.06 | 0.13 |
| 5 | 79402443 | rs4704609 | -1.096 | 0.292 | 0.405 | -0.623 | -0.014 | -0.002 | -0.002 | 0.41 | 0.17 | 0.34 | 0.16 | 0.19 | 0.17 | 0.19 |
| 5 | 79406018 | rs256436 | -0.093 | 1.181 | -0.113 | -1.244 | -0.019 | 0.016 | 0.016 | 0 | 0.14 | 0.07 | 0.11 | 0.01 | 0.2 | 0.24 |
| 5 | 79411248 | rs4704610 | 0.51 | 1.241 | -0.738 | -1.192 | -0.005 | 0.007 | 0.007 | 1.105 | 0.19 | 0.11 | 0.22 | 0.12 | 0.17 | 0.11 |
| 5 | 79411443 | rs10474010 | 0.419 | 1.878 | -0.527 | -1.257 | 0.057 | -0.039 | -0.039 | 0.009 | 0.07 | 0.14 | 0.24 | 0 | 0.19 | 0.06 |
| 5 | 79422993 | rs11950944 | -0.494 | 1.082 | -1.038 | -1.675 | -0.001 | -0.009 | -0.009 | 0 | 0.6 | 0.62 | 0.62 | 0.21 | 0.61 | 0.54 |
| 5 | 79423347 | rs10057542 | 0.705 | 0.865 | -1.77 | -1.665 | -0.001 | 0.03 | 0.03 | 0 | 0.12 | 0.03 | 0.05 | 0.08 | 0.09 | 0.14 |
| 5 | 79427346 | rs2164993 | 0.379 | 1.165 | -1.359 | -1.587 | -0.043 | 0.052 | 0.052 | 0.813 | 0.27 | 0.09 | 0.4 | 0.11 | 0.29 | 0.12 |
| 5 | 79435683 | rs10072008 | 0.057 | 1.019 | -1.599 | -1.293 | -0.022 | -0.057 | -0.057 | 0 | 0.14 | 0.03 | 0.11 | 0.26 | 0.12 | 0.15 |
| 5 | 79437329 | rs4704618 | -0.917 | 0.917 | -1.306 | -1.22 | 0.009 | 0.023 | 0.023 | 0 | 0.56 | 0.68 | 0.61 | 0.33 | 0.56 | 0.46 |
| 5 | 79438705 | rs4703803 | 0.645 | 0.913 | -1.364 | -1.292 | -0.064 | 0.109 | 0.109 | 0 | 0.25 | 0.1 | 0.4 | 0.07 | 0.29 | 0.12 |
| 5 | 79438926 | rs7707754 | -0.878 | 0.505 | -0.512 | -0.779 | -0.002 | -0.003 | -0.003 | 0 | 0.28 | 0.57 | 0.2 | 0.27 | 0.26 | 0.33 |
| 5 | 79461110 | rs1132801 | -0.718 | 0.562 | -0.18 | -1.101 | -0.007 | -0.131 | -0.131 | 0 | 0.21 | 0.47 | 0.12 | 0.03 | 0.17 | 0.28 |

| | | | | | | | | | | | | | | | | |
|---|---|---|---|---|---|---|---|---|---|---|---|---|---|---|---|---|
| 5 | 79466203 | rs7721370 | -1.079 | 0.848 | -0.188 | -0.924 | 0.027 | -0.07 | -0.07 | 0.04 | 0.24 | 0.56 | 0.18 | 0.09 | 0.24 | 0.34 |
| 5 | 79469529 | rs11959429 | -0.919 | 0.673 | -0.064 | -1.08 | 0.014 | -0.097 | -0.097 | 0.279 | 0.24 | 0.43 | 0.14 | 0.04 | 0.22 | 0.34 |
| 5 | 79470031 | rs13186313 | 0.606 | 1.194 | -1.318 | -1.652 | -0.063 | 0.095 | 0.095 | 0 | 0.19 | 0.05 | 0.28 | 0 | 0.24 | 0.13 |
| 5 | 79471383 | rs10050980 | -0.631 | 0.271 | -1.927 | -1.802 | -0.023 | 0.028 | 0.028 | 0.058 | 0.73 | 0.93 | 0.83 | 0.82 | 0.68 | 0.74 |
| 5 | 79472714 | rs4447967 | -0.09 | -0.016 | -1.367 | -0.678 | 0.023 | -0.056 | -0.056 | 0 | 0.37 | 0.76 | 0.34 | 0.51 | 0.29 | 0.42 |
| 5 | 79473286 | rs6890275 | -0.249 | 0.141 | -1.101 | -1.122 | 0.001 | -0.068 | -0.068 | 0 | 0.22 | 0.67 | 0.16 | 0.01 | 0.18 | 0.24 |
| 5 | 79480620 | rs4704624 | -0.497 | -0.272 | -0.578 | -0.963 | 0.006 | -0.035 | -0.035 | 0.052 | 0.59 | 0.58 | 0.7 | 0.81 | 0.52 | 0.51 |
| 5 | 79488191 | rs4704627 | -0.167 | -0.58 | -0.561 | -0.913 | 0.006 | 0.004 | 0.004 | 0.165 | 0.43 | 0.41 | 0.42 | 0.81 | 0.29 | 0.44 |
| 5 | 79495456 | rs6876176 | -0.574 | -0.292 | -0.409 | -0.682 | 0.004 | -0.014 | -0.014 | 0 | 0.29 | 0.32 | 0.33 | 0.19 | 0.3 | 0.16 |
| 5 | 79496939 | rs4704633 | 1.868 | -0.771 | -0.923 | -0.124 | 0.009 | -0.116 | -0.116 | 0 | 0.14 | 0.01 | 0.24 | 0.46 | 0.1 | 0.22 |
| 5 | 79497061 | rs4704634 | 0.673 | -0.932 | -1.13 | -0.316 | 0.003 | -0.189 | -0.189 | 0.005 | 0.27 | 0.01 | 0.31 | 0.42 | 0.23 | 0.35 |
| 5 | 79497156 | rs10055033 | 0.238 | -0.709 | -0.943 | -0.293 | 0 | 0.005 | 0.005 | 7.546 | 0.16 | 0.21 | 0.1 | 0.18 | 0.13 | 0.1 |
| 5 | 79497415 | rs10075871 | -1.598 | -0.855 | -0.489 | -0.431 | -0.036 | 0.021 | 0.021 | 0 | 0.63 | 0.75 | 0.68 | 0.65 | 0.54 | 0.52 |
| 5 | 79498398 | rs7727507 | 0.089 | -0.573 | -0.537 | -0.334 | 0 | 0.005 | 0.005 | 7.546 | 0.16 | 0.21 | 0.1 | 0.19 | 0.13 | 0.1 |
| 5 | 79499495 | rs6868932 | 0.458 | -0.591 | -0.5 | -0.358 | -0.002 | -0.004 | -0.004 | 0 | 0.12 | 0.11 | 0.08 | 0.19 | 0.12 | 0.1 |
| 5 | 79503869 | rs10058567 | 1.811 | -0.571 | -0.477 | -0.388 | -0.005 | -0.028 | -0.028 | 0 | 0.05 | 0.11 | 0.02 | 0 | 0.02 | 0.02 |
| 5 | 79505996 | rs10514178 | 0.593 | -0.575 | -0.584 | -0.287 | -0.007 | -0.01 | -0.01 | 0 | 0.11 | 0.11 | 0.1 | 0.32 | 0.12 | 0.09 |
| 5 | 79507194 | rs6872447 | 0.109 | -0.404 | -0.671 | -0.47 | -0.017 | 0.013 | 0.013 | 0 | 0.24 | 0.13 | 0.17 | 0.19 | 0.26 | 0.22 |
| 5 | 79509295 | rs10068511 | 1.479 | -0.458 | -0.662 | -0.54 | -0.006 | -0.011 | -0.011 | 0 | 0.13 | 0.09 | 0.23 | 0 | 0.18 | 0.06 |
| 5 | 79509974 | rs10036776 | 0.373 | -0.52 | -0.507 | -0.267 | 0.001 | -0.021 | -0.021 | 0.022 | 0.31 | 0.25 | 0.51 | 0.49 | 0.33 | 0.25 |
| 5 | 79514022 | rs11960254 | -0.711 | -0.611 | -0.157 | -0.357 | -0.009 | -0.04 | -0.04 | 0 | 0.33 | 0.47 | 0.22 | 0.19 | 0.29 | 0.25 |
| 5 | 79514675 | rs6895353 | -1.696 | -0.583 | -0.291 | -0.339 | -0.007 | 0.014 | 0.014 | 0 | 0.64 | 0.73 | 0.73 | 0.67 | 0.62 | 0.5 |
| 5 | 79516926 | rs7711197 | 0.424 | -0.605 | 0.009 | -0.482 | 0.02 | 0.033 | 0.033 | 0 | 0.13 | 0.32 | 0.09 | 0.16 | 0.11 | 0.09 |
| 5 | 79518278 | rs13355369 | -1.436 | -0.529 | -0.75 | -0.041 | -0.008 | -0.011 | -0.011 | 0 | 0.59 | 0.68 | 0.64 | 0.52 | 0.57 | 0.47 |
| 5 | 79521704 | rs7726099 | -0.957 | -0.175 | -0.412 | 0.344 | -0.001 | -0.004 | -0.004 | 0 | 0.46 | 0.62 | 0.55 | 0.51 | 0.44 | 0.33 |
| 5 | 79525305 | rs10041917 | -0.919 | -0.136 | -1.063 | 0.438 | -0.003 | -0.01 | -0.01 | 0 | 0.48 | 0.79 | 0.57 | 0.52 | 0.45 | 0.33 |
| 5 | 79526346 | rs12522787 | 1.022 | -0.248 | -0.827 | 0.743 | 0.005 | -0.014 | -0.014 | 0.255 | 0.16 | 0.11 | 0.22 | 0.35 | 0.14 | 0.17 |
| 5 | 79530306 | rs16877647 | 0.387 | -0.63 | -0.696 | 0.33 | -0.02 | 0.004 | 0.004 | 0.006 | 0.21 | 0.26 | 0.18 | 0.25 | 0.13 | 0.2 |
| 5 | 79532244 | rs17200856 | 0.413 | 0.103 | -0.993 | -0.185 | -0.002 | -0.017 | -0.017 | 0 | 0.12 | 0.1 | 0.24 | 0 | 0.18 | 0.05 |
| 5 | 79543124 | rs1010809 | 0.772 | -0.468 | -1.384 | -0.339 | 0.025 | 0.064 | 0.064 | 0 | 0.17 | 0.04 | 0.08 | 0 | 0.15 | 0.14 |
| 5 | 79548274 | rs16877662 | -0.005 | 0.177 | -1.524 | 0.076 | -0.016 | -0.071 | -0.071 | 0 | 0.3 | 0.2 | 0.34 | 0.55 | 0.34 | 0.51 |
| 5 | 79549684 | rs881708 | 0.082 | 0.179 | -1.156 | 0.074 | 0.001 | 0.014 | 0.014 | 3.649 | 0.31 | 0.32 | 0.35 | 0.55 | 0.34 | 0.51 |
| 5 | 79564658 | rs9293807 | 0.468 | -0.248 | -1.711 | -0.434 | 0.014 | 0.014 | 0.014 | 0.175 | 0.31 | 0.11 | 0.18 | 0.23 | 0.27 | 0.27 |
| 5 | 79564796 | rs9293808 | 0.687 | -0.265 | -1.771 | -0.489 | -0.004 | -0.005 | -0.005 | 0 | 0.09 | 0.07 | 0.1 | 0.23 | 0.13 | 0.15 |
| 5 | 79565071 | rs9293809 | 0.238 | -0.009 | -1.428 | -0.453 | -0.027 | 0.039 | 0.039 | 0 | 0.35 | 0.12 | 0.21 | 0.22 | 0.36 | 0.3 |
| 5 | 79578661 | rs6453512 | 0.493 | 0.014 | -0.977 | -0.377 | 0.033 | 0.095 | 0.095 | 2.699 | 0.42 | 0.17 | 0.23 | 0.16 | 0.36 | 0.27 |
| 5 | 79586085 | rs9293811 | 0.644 | -0.115 | -1.01 | -0.339 | -0.005 | -0.035 | -0.035 | 0.577 | 0.1 | 0.05 | 0.08 | 0.16 | 0.11 | 0.12 |
| 5 | 79590674 | rs6886552 | 0.278 | -0.243 | -0.613 | -0.165 | -0.004 | 0.035 | 0.035 | 0 | 0.15 | 0.18 | 0.23 | 0.44 | 0.13 | 0.26 |
| 5 | 79595396 | rs4704659 | 0.288 | -0.486 | -0.134 | -0.365 | 0 | 0 | 0 | 0.047 | 0.49 | 0.49 | 0.4 | 0.44 | 0.38 | 0.41 |
| 5 | 79596021 | rs11738780 | 2.024 | -0.596 | -0.123 | -0.393 | 0.043 | 0.053 | 0.053 | 5.437 | 0.27 | 0.1 | 0.12 | 0 | 0.16 | 0.12 |
| 5 | 79598589 | rs11738136 | 0.715 | -0.242 | -0.017 | -0.147 | 0.001 | 0.001 | 0.001 | 1.197 | 0.45 | 0.44 | 0.36 | 0.42 | 0.3 | 0.38 |
| 5 | 79600508 | rs10054927 | 0.067 | 0.052 | 0.048 | -0.516 | -0.002 | -0.011 | -0.011 | 0.176 | 0.15 | 0.12 | 0.11 | 0.16 | 0.15 | 0.19 |
| 5 | 79600617 | rs4440342 | -0.754 | 0.632 | -0.253 | -0.795 | -0.032 | 0.01 | 0.01 | 0.414 | 0.82 | 0.87 | 0.81 | 1 | 0.62 | 0.86 |
| 5 | 79600693 | rs4998063 | 0.688 | -0.108 | 0.117 | -0.66 | -0.003 | -0.025 | -0.025 | 0 | 0.59 | 0.52 | 0.52 | 0.79 | 0.42 | 0.58 |
| 5 | 79603708 | rs10805934 | 0.067 | -0.302 | 0.062 | -0.97 | -0.025 | 0.042 | 0.042 | 2.393 | 0.63 | 0.75 | 0.55 | 0.82 | 0.49 | 0.67 |
| 5 | 79606605 | rs6866850 | 0.521 | -0.449 | -0.143 | -0.872 | -0.012 | -0.051 | -0.051 | 0 | 0.6 | 0.81 | 0.56 | 0.58 | 0.4 | 0.54 |
| 5 | 79615048 | rs7718110 | -0.306 | -0.412 | -0.078 | -0.885 | -0.054 | 0.052 | 0.052 | 0.829 | 0.73 | 0.82 | 0.64 | 0.94 | 0.54 | 0.76 |
| 5 | 79615589 | rs11744364 | 1.75 | -0.535 | 0.271 | -0.879 | 0.057 | -0.014 | -0.014 | 0 | 0.37 | 0.23 | 0.35 | 0.46 | 0.21 | 0.33 |
| 5 | 79617357 | rs16876315 | 0.494 | -0.225 | 0.091 | -0.661 | -0.025 | -0.076 | -0.076 | 0.399 | 0.13 | 0.04 | 0.09 | 0.36 | 0.15 | 0.23 |
| 5 | 79617616 | rs1500139 | 0.784 | -0.236 | 0.319 | -0.659 | -0.002 | -0.002 | -0.002 | 0 | 0.15 | 0.18 | 0.13 | 0.12 | 0.14 | 0.13 |
| 5 | 79618649 | rs10060273 | 0.534 | -0.251 | 0.42 | -0.662 | -0.003 | -0.003 | -0.003 | 0 | 0.16 | 0.29 | 0.15 | 0.12 | 0.15 | 0.13 |
| 5 | 79619086 | rs11749321 | 0.486 | -0.253 | -0.108 | -0.716 | -0.009 | -0.053 | -0.053 | 0 | 0.15 | 0.06 | 0.12 | 0.38 | 0.18 | 0.23 |
| 5 | 79621016 | rs879738 | -0.821 | -0.93 | 0.314 | -0.602 | -0.011 | 0.03 | 0.03 | 5.038 | 0.3 | 0.62 | 0.41 | 0.57 | 0.22 | 0.39 |
| 5 | 79627154 | rs1990858 | -0.388 | -0.838 | 0.228 | -0.648 | -0.045 | -0.047 | -0.047 | 0.48 | 0.63 | 0.81 | 0.56 | 0.57 | 0.46 | 0.57 |
| 5 | 79630844 | rs17202384 | 0.596 | -0.871 | -0.142 | -0.8 | 0.001 | -0.001 | -0.001 | 0 | 0.14 | 0.15 | 0.14 | 0.11 | 0.15 | 0.13 |
| 5 | 79632803 | rs13168595 | 0.875 | -1.199 | -0.749 | -0.849 | 0.042 | 0.051 | 0.051 | 1.37 | 0.34 | 0.17 | 0.14 | 0 | 0.24 | 0.18 |
| 5 | 79660190 | rs12652103 | -0.642 | -0.617 | -0.193 | 0.077 | -0.011 | 0.071 | 0.071 | 0 | 0.16 | 0.29 | 0.3 | 0.46 | 0.15 | 0.23 |
| 5 | 79663875 | rs12654489 | -0.524 | -0.603 | -0.509 | 0.093 | -0.001 | 0.026 | 0.026 | 0 | 0.16 | 0.24 | 0.29 | 0.45 | 0.14 | 0.23 |
| 5 | 79664768 | rs13179066 | 1.107 | -0.556 | -1.074 | -0.297 | 0.066 | -0.007 | -0.007 | 0 | 0.41 | 0.15 | 0.19 | 0.39 | 0.29 | 0.4 |
| 5 | 79665635 | rs10062770 | 0.705 | 0.639 | -1.027 | -0.936 | -0.021 | 0.067 | 0.067 | 0.114 | 0.66 | 0.77 | 0.57 | 0.94 | 0.45 | 0.71 |
| 5 | 79666031 | rs7715187 | 0.071 | 0.83 | -1.546 | -0.988 | -0.001 | -0.007 | -0.007 | 0 | 0.06 | 0.03 | 0.03 | 0 | 0.09 | 0.04 |
| 5 | 79669697 | rs4467688 | 0.058 | 0.872 | -1.557 | -0.779 | -0.163 | 0.167 | 0.167 | 0 | 0.75 | 0.95 | 0.62 | 1 | 0.54 | 0.82 |
| 5 | 79681727 | rs6861026 | 0.288 | -0.132 | -0.936 | -0.558 | 0.029 | 0.036 | 0.036 | 0.441 | 0.54 | 0.61 | 0.3 | 0.21 | 0.34 | 0.44 |
| 5 | 79686227 | rs28496723 | -0.163 | -1.228 | -0.334 | -0.92 | -0.075 | -0.037 | -0.037 | 0.757 | 0.17 | 0.36 | 0.07 | 0.07 | 0.07 | 0.14 |
| 5 | 79695370 | rs259035 | -0.951 | -1.61 | -0.066 | -1.101 | -0.077 | -0.023 | -0.023 | 1.003 | 0.23 | 0.57 | 0.16 | 0.17 | 0.11 | 0.25 |
| 5 | 79700854 | rs7730069 | 1.676 | -1.799 | -0.304 | -1.468 | 0.049 | 0.09 | 0.09 | 3.126 | 0.26 | 0.06 | 0.12 | 0 | 0.16 | 0.14 |
| 5 | 79701839 | rs13188088 | 1.313 | -1.776 | -0.317 | -1.455 | 0.04 | 0.11 | 0.11 | 2.644 | 0.29 | 0.06 | 0.13 | 0.06 | 0.21 | 0.23 |
| 5 | 79703254 | rs2544589 | -0.206 | -1.759 | -0.123 | -1.402 | -0.083 | -0.039 | -0.039 | 0 | 0.83 | 0.6 | 0.92 | 0.92 | 0.93 | 0.87 |
| 5 | 79704614 | rs259053 | -0.23 | -1.77 | -0.124 | -1.412 | -0.077 | -0.039 | -0.039 | 0 | 0.83 | 0.64 | 0.93 | 0.92 | 0.93 | 0.87 |

| | | | | | | | | | | | | | | | | |
|---|---|---|---|---|---|---|---|---|---|---|---|---|---|---|---|---|
| 5 | 79721720 | rs192313 | 0.342 | -1.799 | -0.086 | -1.386 | -0.076 | -0.062 | -0.062 | 0.091 | 0.79 | 0.45 | 0.88 | 0.92 | 0.89 | 0.82 |
| 5 | 79733079 | rs2544600 | 0.399 | -1.794 | -0.088 | -1.387 | -0.075 | -0.059 | -0.059 | 0 | 0.78 | 0.44 | 0.88 | 0.92 | 0.89 | 0.82 |
| 5 | 79734297 | rs259028 | -0.361 | -1.816 | 0.15 | -1.396 | -0.063 | -0.026 | -0.026 | 4.861 | 0.84 | 0.69 | 0.93 | 0.92 | 0.93 | 0.87 |
| 5 | 79734409 | rs259029 | -0.981 | -1.924 | 0.33 | -1.448 | -0.051 | -0.002 | -0.002 | 0 | 0.89 | 0.75 | 0.94 | 0.92 | 0.94 | 0.87 |
| 5 | 79735858 | rs168939 | 0.849 | -1.977 | 0.646 | -1.472 | -0.057 | -0.047 | -0.047 | 0.364 | 0.77 | 0.38 | 0.87 | 0.92 | 0.89 | 0.82 |
| 5 | 79739685 | rs259033 | 0.628 | -1.961 | 0.665 | -1.465 | -0.065 | -0.054 | -0.054 | 0.248 | 0.78 | 0.42 | 0.87 | 0.92 | 0.89 | 0.81 |
| 5 | 79745469 | rs249038 | -0.176 | -1.972 | 0.667 | -1.47 | -0.063 | -0.026 | -0.026 | 0.913 | 0.83 | 0.69 | 0.93 | 0.92 | 0.93 | 0.86 |
| 5 | 79757298 | rs249017 | 0.242 | -2.076 | 0.865 | -1.533 | 0.105 | 0.153 | 0.153 | 1.454 | 0.48 | 0.25 | 0.73 | 0.86 | 0.68 | 0.64 |
| 5 | 79774171 | rs187479 | 0.736 | -1.919 | 0.998 | -1.338 | -0.066 | -0.055 | -0.055 | 0 | 0.78 | 0.42 | 0.87 | 0.91 | 0.89 | 0.82 |
| 5 | 79774906 | rs10433 | 0.717 | -1.942 | 0.941 | -1.348 | -0.075 | -0.062 | -0.062 | 0.281 | 0.78 | 0.46 | 0.87 | 0.92 | 0.89 | 0.82 |
| 5 | 79782530 | rs171596 | -0.043 | -1.861 | 0.729 | -1.298 | -0.062 | -0.025 | -0.025 | 0 | 0.84 | 0.7 | 0.93 | 0.92 | 0.93 | 0.86 |
| 5 | 79801520 | rs248998 | -0.785 | -1.489 | 0.5 | -0.558 | -0.02 | 0.01 | 0.01 | 0 | 0.36 | 0.55 | 0.33 | 0.44 | 0.3 | 0.43 |
| 5 | 79812343 | rs896389 | -0.147 | -0.921 | 0.141 | -0.82 | 0.002 | -0.045 | -0.045 | 0 | 0.09 | 0.16 | 0.07 | 0 | 0.06 | 0.05 |
| 5 | 79815658 | rs369998 | -0.488 | -1.133 | 0.228 | -0.819 | -0.001 | -0.01 | -0.01 | 0 | 0.81 | 0.83 | 0.7 | 0.61 | 0.79 | 0.71 |
| 5 | 79818229 | rs1008902 | -0.243 | -1.162 | 0.277 | -0.852 | 0.001 | -0.048 | -0.048 | 3.228 | 0.09 | 0.17 | 0.07 | 0 | 0.06 | 0.05 |
| 5 | 79820724 | rs171598 | -0.52 | -1.152 | 0.181 | -0.853 | 0 | -0.012 | -0.012 | 0 | 0.81 | 0.83 | 0.7 | 0.61 | 0.79 | 0.71 |
| 5 | 79821353 | rs187261 | -0.655 | -1.151 | 0.182 | -0.845 | -0.002 | -0.011 | -0.011 | 0.018 | 0.81 | 0.84 | 0.7 | 0.6 | 0.79 | 0.71 |
| 5 | 79822916 | rs10474629 | 0.314 | -1.277 | 0.666 | -0.83 | -0.048 | -0.056 | -0.056 | 0.017 | 0.15 | 0.26 | 0.06 | 0.01 | 0.06 | 0.1 |
| 5 | 79833821 | rs2443549 | -0.222 | -1.142 | 0.721 | -0.77 | 0.006 | -0.05 | -0.05 | 0 | 0.09 | 0.19 | 0.07 | 0 | 0.06 | 0.05 |
| 5 | 79836881 | rs17664442 | 0.702 | 0.139 | 0.414 | -1.045 | -0.006 | 0.002 | 0.002 | 0 | 0.18 | 0.16 | 0.35 | 0.52 | 0.3 | 0.21 |
| 5 | 79837249 | rs42416 | -0.19 | -0.302 | 0.363 | -0.937 | 0.006 | -0.05 | -0.05 | 0 | 0.09 | 0.19 | 0.07 | 0 | 0.06 | 0.05 |
| 5 | 79841683 | rs9687094 | -0.332 | -0.155 | -0.264 | -0.299 | -0.06 | -0.094 | -0.094 | 0 | 0.15 | 0.04 | 0.3 | 0.43 | 0.25 | 0.25 |
| 5 | 79844769 | rs10071064 | -0.098 | -0.866 | 0.088 | -1.091 | -0.05 | -0.052 | -0.052 | 0 | 0.17 | 0.25 | 0.06 | 0.02 | 0.06 | 0.15 |
| 5 | 79846926 | rs10076737 | -0.144 | -0.869 | 0.066 | -1.092 | -0.052 | -0.054 | -0.054 | 4.439 | 0.17 | 0.25 | 0.05 | 0.02 | 0.06 | 0.15 |
| 5 | 79852478 | rs6453521 | -0.917 | -1.27 | 0.047 | -1.071 | -0.092 | -0.032 | -0.032 | 0 | 0.17 | 0.39 | 0.09 | 0.12 | 0.05 | 0.19 |
| 5 | 79856771 | rs11949327 | -0.032 | -0.077 | -0.476 | -1.253 | 0.002 | -0.006 | -0.006 | 0 | 0.11 | 0.12 | 0.1 | 0.05 | 0.15 | 0.19 |
| 5 | 79858271 | rs249219 | -0.31 | -0.624 | -0.895 | -1.74 | 0.007 | 0.065 | 0.065 | 0.042 | 0.41 | 0.28 | 0.66 | 0.88 | 0.59 | 0.5 |
| 5 | 79867209 | rs249209 | -0.77 | -0.777 | -1.092 | -1.84 | 0.027 | 0.108 | 0.108 | 1.601 | 0.45 | 0.28 | 0.68 | 0.96 | 0.6 | 0.57 |
| 5 | 79870334 | rs6865790 | -0.001 | -0.884 | -1.175 | -1.901 | -0.027 | -0.019 | -0.019 | 0 | 0.12 | 0.22 | 0.07 | 0.08 | 0.06 | 0.17 |
| 5 | 79870727 | rs185105 | 0.086 | -0.479 | -1.564 | -1.749 | -0.101 | 0.12 | 0.12 | 0.739 | 0.31 | 0.08 | 0.61 | 0.88 | 0.55 | 0.4 |
| 5 | 79883154 | rs393061 | 0.495 | -0.362 | -1.157 | -1.479 | -0.003 | 0.009 | 0.009 | 0 | 0.07 | 0.04 | 0.05 | 0.03 | 0.09 | 0.13 |
| 5 | 79888450 | rs2405867 | 0.705 | 1.055 | -1.397 | -0.359 | -0.058 | -0.065 | -0.065 | 0 | 0.11 | 0.01 | 0.37 | 0.62 | 0.18 | 0.22 |
| 5 | 79889117 | rs13164793 | 2.565 | -0.558 | -1.277 | -0.991 | 0.023 | 0.007 | 0.007 | 2.102 | 0.22 | 0.12 | 0.11 | 0 | 0.13 | 0.14 |
| 5 | 79893203 | rs1072841 | 0.92 | -0.892 | -0.86 | -1.193 | 0.006 | -0.037 | -0.037 | 0.086 | 0.41 | 0.37 | 0.22 | 0.04 | 0.29 | 0.34 |
| 5 | 79912044 | rs836788 | -0.119 | -0.016 | -0.614 | -0.764 | -0.001 | 0.028 | 0.028 | 0.008 | 0.35 | 0.51 | 0.36 | 0.38 | 0.38 | 0.43 |
| 5 | 79917517 | rs1222809 | 1.363 | -0.242 | -1.067 | -1.079 | 0.03 | 0.009 | 0.009 | 3.056 | 0.38 | 0.27 | 0.19 | 0.02 | 0.24 | 0.29 |
| 5 | 79920073 | rs12517451 | 1.089 | -0.065 | -1.437 | -1.103 | -0.018 | -0.11 | -0.11 | 0.379 | 0.24 | 0.06 | 0.25 | 0.34 | 0.25 | 0.33 |
| 5 | 79922030 | rs1650723 | 2.789 | -0.299 | -1.297 | -1.273 | 0.035 | 0.099 | 0.099 | 1.106 | 0.22 | 0.04 | 0.12 | 0 | 0.15 | 0.15 |
| 5 | 79939471 | rs11951910 | 1.084 | -0.276 | -1.055 | -1.219 | -0.022 | -0.055 | -0.055 | 1.453 | 0.16 | 0.21 | 0.07 | 0.02 | 0.09 | 0.14 |
| 5 | 79945140 | rs10072026 | -0.366 | 0.093 | -0.746 | -1.234 | 0.037 | -0.069 | -0.069 | 0 | 0.12 | 0.53 | 0.11 | 0.02 | 0.13 | 0.07 |
| 5 | 79948005 | rs863215 | 1.536 | -0.053 | -0.898 | -1.376 | 0.032 | 0.018 | 0.018 | 2.839 | 0.4 | 0.29 | 0.21 | 0.05 | 0.25 | 0.33 |
| 5 | 79949575 | rs1478834 | 1.536 | -0.053 | -0.898 | -1.376 | 0.032 | 0.018 | 0.018 | 2.839 | 0.4 | 0.29 | 0.21 | 0.05 | 0.25 | 0.33 |
| 5 | 79950781 | rs1650697 | -1.36 | 0.107 | -1.072 | -1.244 | -0.02 | -0.11 | -0.11 | 1.09 | 0.77 | 0.93 | 0.76 | 0.66 | 0.76 | 0.68 |
| 5 | 79951032 | rs6151599 | 1.206 | 0.091 | -0.373 | -1.238 | 0.063 | -0.026 | -0.026 | 0 | 0.07 | 0.21 | 0.08 | 0.02 | 0.12 | 0.06 |
| 5 | 79952034 | rs380691 | 1.306 | 0.297 | -0.598 | -1.253 | -0.054 | -0.062 | -0.062 | 0 | 0.17 | 0.1 | 0.4 | 0.58 | 0.32 | 0.23 |
| 5 | 79952154 | rs1382539 | 1.677 | 0.022 | -0.646 | -1.266 | 0.031 | 0.02 | 0.02 | 2.769 | 0.39 | 0.28 | 0.21 | 0.06 | 0.25 | 0.34 |
| 5 | 79957737 | rs1677703 | 1.793 | 0.024 | -0.592 | -1.264 | 0.016 | -0.011 | -0.011 | 0.072 | 0.4 | 0.32 | 0.21 | 0.06 | 0.26 | 0.34 |
| 5 | 79957835 | rs7712332 | 0.421 | 0.289 | -0.488 | -0.84 | 0.085 | 0.008 | 0.008 | 0 | 0.22 | 0.35 | 0.24 | 0.26 | 0.39 | 0.2 |
| 5 | 79962439 | rs1650666 | 1.918 | 0.243 | -0.399 | -0.871 | -0.001 | -0.043 | -0.043 | 0 | 0.4 | 0.34 | 0.21 | 0.06 | 0.26 | 0.34 |
| 5 | 79966029 | rs1805355 | 0.76 | 0.397 | -0.471 | -0.549 | 0.013 | 0.197 | 0.197 | 0.022 | 0.06 | 0.2 | 0.27 | 0.34 | 0.05 | 0.08 |
| 5 | 79966178 | rs6151630 | 2.777 | 0.458 | -0.71 | -0.586 | -0.016 | 0.028 | 0.028 | 0.031 | 0.06 | 0 | 0.03 | 0 | 0.05 | 0.04 |
| 5 | 79970423 | rs1650647 | -0.719 | 0.457 | -0.441 | -0.484 | 0.004 | -0.046 | -0.046 | 0 | 0.71 | 0.85 | 0.71 | 0.64 | 0.71 | 0.63 |
| 5 | 79971612 | rs6151653 | 1.241 | 0.735 | -0.195 | -0.489 | 0.025 | 0.006 | 0.006 | 0 | 0.15 | 0.17 | 0.16 | 0.24 | 0.27 | 0.14 |
| 5 | 79992418 | rs836817 | -0.741 | 0.554 | -0.308 | -0.503 | 0.01 | -0.043 | -0.043 | 0 | 0.69 | 0.84 | 0.71 | 0.65 | 0.7 | 0.62 |
| 5 | 79996632 | rs2035256 | -0.885 | 0.452 | -0.076 | -0.687 | 0.084 | 0.072 | 0.072 | 0 | 0.29 | 0.5 | 0.51 | 0.62 | 0.45 | 0.33 |
| 5 | 80000152 | rs865644 | -0.815 | 0.447 | -0.052 | -0.69 | 0.009 | -0.029 | -0.029 | 0 | 0.68 | 0.8 | 0.7 | 0.65 | 0.7 | 0.62 |
| 5 | 80001170 | rs7709909 | -0.153 | 0.533 | 0.116 | -0.611 | 0.042 | 0 | 0 | 0 | 0.24 | 0.31 | 0.25 | 0.28 | 0.4 | 0.26 |
| 5 | 80008704 | rs1428030 | 1.144 | 0.558 | 0.137 | -0.548 | 0.034 | 0.258 | 0.258 | 0 | 0.05 | 0.21 | 0.27 | 0.34 | 0.05 | 0.07 |
| 5 | 80012251 | rs836794 | 1.749 | 0.461 | 0.104 | -0.654 | 0.05 | 0.049 | 0.049 | 2.47 | 0.39 | 0.25 | 0.18 | 0.01 | 0.25 | 0.29 |
| 5 | 80016874 | rs181747 | 1.49 | 0.471 | -0.109 | -0.632 | -0.015 | -0.009 | -0.009 | 0 | 0.43 | 0.46 | 0.45 | 0.36 | 0.3 | 0.36 |
| 5 | 80021168 | rs32951 | 1.435 | 0.89 | -0.254 | -0.517 | 0.024 | 0.006 | 0.006 | 0 | 0.15 | 0.17 | 0.16 | 0.24 | 0.27 | 0.15 |
| 5 | 80025471 | rs6151735 | 1.396 | 0.787 | -0.496 | -0.483 | 0.004 | -0.007 | -0.007 | 0.069 | 0.06 | 0.05 | 0.07 | 0.02 | 0.12 | 0.06 |
| 5 | 80049892 | rs26784 | -0.087 | 0.482 | -0.576 | -0.437 | -0.023 | -0.041 | -0.041 | 0 | 0.37 | 0.17 | 0.36 | 0.37 | 0.42 | 0.44 |
| 5 | 80056961 | rs6151792 | 1.032 | 0.698 | -0.564 | 0.462 | -0.018 | -0.022 | -0.022 | 0 | 0.09 | 0.05 | 0.13 | 0.3 | 0.12 | 0.07 |
| 5 | 80060016 | rs33013 | 0.498 | 0.109 | -0.213 | 0.11 | 0.067 | 0.114 | 0.114 | 0 | 0.22 | 0.34 | 0.42 | 0.57 | 0.33 | 0.21 |
| 5 | 80073634 | rs6151816 | 0.993 | -0.149 | -0.491 | 0.264 | 0 | 0.026 | 0.026 | 0 | 0.13 | 0.12 | 0.18 | 0.28 | 0.14 | 0.1 |
| 5 | 80082865 | rs6151838 | 0.839 | -0.787 | -0.68 | -0.537 | 0.001 | -0.011 | -0.011 | 0 | 0.17 | 0.2 | 0.06 | 0.03 | 0.1 | 0.14 |
| 5 | 80101773 | rs245397 | -2.314 | -0.752 | -0.828 | -0.665 | 0.019 | 0.03 | 0.03 | 0.734 | 0.77 | 0.88 | 0.86 | 1 | 0.83 | 0.87 |
| 5 | 80116090 | rs32989 | -1.345 | -1.107 | -0.541 | -0.312 | -0.008 | -0.001 | -0.001 | 0 | 0.66 | 0.65 | 0.69 | 0.76 | 0.72 | 0.79 |

| | | | | | | | | | | | | | | | | |
|---|---|---|---|---|---|---|---|---|---|---|---|---|---|---|---|---|
| 5 | 80117918 | rs32985 | 1.724 | -1.009 | -0.462 | -0.363 | 0.009 | 0.002 | 0.002 | 0 | 0.16 | 0.16 | 0.21 | 0.3 | 0.3 | 0.14 |
| 5 | 80132177 | rs10075024 | 1.03 | -1.406 | -0.457 | -0.545 | 0.018 | 0.018 | 0.018 | 1.273 | 0.17 | 0.16 | 0.06 | 0.03 | 0.1 | 0.15 |
| 5 | 80137400 | rs245369 | -2.944 | -1.424 | -0.412 | -0.606 | 0.027 | 0.019 | 0.019 | 1.328 | 0.78 | 0.88 | 0.87 | 1 | 0.86 | 0.87 |
| 5 | 80148234 | rs12515548 | 1.558 | -1.437 | -0.377 | -0.573 | 0.013 | 0.112 | 0.112 | 0 | 0.12 | 0.23 | 0.18 | 0.24 | 0.14 | 0.08 |
| 5 | 80149981 | rs184967 | -3.022 | -1.416 | -0.432 | -0.593 | 0.03 | 0.032 | 0.032 | 1.037 | 0.78 | 0.9 | 0.87 | 1 | 0.86 | 0.87 |
| 5 | 80157824 | rs245343 | -0.114 | -1.445 | -0.485 | -0.596 | -0.001 | 0.025 | 0.025 | 1.856 | 0.41 | 0.26 | 0.36 | 0.38 | 0.56 | 0.53 |
| 5 | 80164943 | rs249634 | 0.933 | -1.346 | -0.524 | -0.621 | -0.003 | 0.072 | 0.072 | 0.129 | 0.25 | 0.11 | 0.14 | 0.08 | 0.26 | 0.43 |
| 5 | 80168005 | rs16878204 | 0.593 | -1.575 | -0.286 | -0.389 | 0.027 | 0.23 | 0.23 | 0 | 0.06 | 0.21 | 0.28 | 0.36 | 0.06 | 0.11 |
| 5 | 80168937 | rs26279 | -1.552 | -1.564 | -0.403 | -0.382 | -0.01 | -0.002 | -0.002 | 4.485 | 0.65 | 0.66 | 0.69 | 0.75 | 0.72 | 0.79 |
| 5 | 80171134 | rs33003 | -0.714 | -1.452 | -0.485 | -0.381 | -0.009 | -0.004 | -0.004 | 0 | 0.54 | 0.45 | 0.54 | 0.62 | 0.71 | 0.63 |
| 5 | 80187929 | rs12522063 | 2.018 | -1.226 | -0.745 | -0.256 | 0.026 | -0.02 | -0.02 | 0 | 0.25 | 0.13 | 0.35 | 0.33 | 0.18 | 0.18 |
| 5 | 80199400 | rs10942932 | 0.889 | -1.849 | -0.694 | -0.536 | 0.014 | -0.016 | -0.016 | 4.077 | 0.23 | 0.16 | 0.3 | 0.35 | 0.17 | 0.21 |
| 5 | 80200144 | rs4704689 | -0.951 | -2.028 | -1.026 | -0.848 | -0.004 | 0.091 | 0.091 | 1.602 | 0.73 | 0.92 | 0.83 | 0.89 | 0.73 | 0.61 |
| 5 | 80201140 | rs13178429 | 1.288 | -2.505 | -0.41 | -0.44 | -0.001 | 0.058 | 0.058 | 0 | 0.14 | 0.18 | 0.18 | 0.25 | 0.14 | 0.08 |
| 5 | 80201284 | rs6864713 | 2.585 | -2.468 | -0.414 | -0.482 | -0.001 | 0.002 | 0.002 | 0 | 0.15 | 0.17 | 0.3 | 0.33 | 0.14 | 0.14 |
| 5 | 80203678 | rs7737445 | 0.941 | -2.547 | -0.398 | -0.61 | -0.005 | 0.006 | 0.006 | 0 | 0.27 | 0.34 | 0.27 | 0.25 | 0.24 | 0.17 |
| 5 | 80206473 | rs958575 | -0.304 | -2.438 | -0.385 | -0.756 | 0.011 | 0.013 | 0.013 | 1.028 | 0.51 | 0.55 | 0.57 | 0.6 | 0.41 | 0.37 |
| 5 | 80207733 | rs7700440 | 2.326 | -2.168 | -0.676 | -0.694 | 0.001 | -0.023 | -0.023 | 3.01 | 0.14 | 0.15 | 0.09 | 0 | 0.11 | 0.08 |
| 5 | 80211574 | rs10067194 | -0.396 | -2.14 | -0.621 | -0.574 | 0.018 | 0.031 | 0.031 | 1.588 | 0.55 | 0.67 | 0.59 | 0.63 | 0.41 | 0.42 |
| 5 | 80212435 | rs16878218 | 0.675 | -2.24 | -0.442 | -0.861 | -0.024 | 0.023 | 0.023 | 0.28 | 0.23 | 0.28 | 0.3 | 0.35 | 0.14 | 0.26 |
| 5 | 80212643 | rs11949801 | 0.726 | -2.507 | -0.906 | -1.332 | -0.011 | -0.01 | -0.01 | 0 | 0.32 | 0.26 | 0.45 | 0.56 | 0.51 | 0.31 |
| 5 | 80213816 | rs4703820 | 1.156 | -2.538 | -0.524 | -1.183 | -0.027 | 0.024 | 0.024 | 1.449 | 0.22 | 0.31 | 0.3 | 0.31 | 0.14 | 0.16 |
| 5 | 80215220 | rs1373968 | 0.13 | -3.025 | -0.981 | -1.556 | 0.08 | 0.059 | 0.059 | 4.086 | 0.49 | 0.33 | 0.52 | 0.68 | 0.69 | 0.65 |
| 5 | 80215251 | rs1823024 | 0.51 | -3.039 | -1.037 | -1.552 | 0.061 | 0.035 | 0.035 | 2.88 | 0.46 | 0.3 | 0.51 | 0.65 | 0.68 | 0.59 |
| 5 | 80217130 | rs10079494 | 1.093 | -2.988 | -1.07 | -1.427 | -0.002 | 0 | 0 | 0 | 0.23 | 0.19 | 0.2 | 0.27 | 0.28 | 0.36 |
| 5 | 80219978 | rs571653 | -1.348 | -2.71 | -1.235 | -1.472 | 0.047 | 0.076 | 0.076 | 1.231 | 0.86 | 0.97 | 0.9 | 1 | 0.91 | 0.89 |
| 5 | 80220759 | rs6871067 | 1.732 | -2.075 | -0.926 | -1.093 | -0.086 | -0.071 | -0.071 | 0.118 | 0.22 | 0.09 | 0.28 | 0.37 | 0.4 | 0.23 |
| 5 | 80220997 | rs555470 | 0.379 | -2.394 | -0.247 | -1.477 | -0.047 | -0.134 | -0.134 | 0.179 | 0.26 | 0.47 | 0.12 | 0.05 | 0.15 | 0.23 |
| 5 | 80223536 | rs511038 | 2.153 | -1.479 | -0.865 | -1.064 | 0 | -0.03 | -0.03 | 0.856 | 0.13 | 0.12 | 0.07 | 0 | 0.08 | 0.08 |
| 5 | 80228327 | rs4704691 | -0.512 | -2.252 | -0.715 | -0.834 | 0.011 | 0.003 | 0.003 | 0 | 0.42 | 0.63 | 0.42 | 0.39 | 0.32 | 0.54 |
| 5 | 80230698 | rs715406 | 1.892 | -2.004 | -0.987 | -0.665 | -0.004 | -0.073 | -0.073 | 0 | 0.11 | 0 | 0.25 | 0.29 | 0.08 | 0.06 |
| 5 | 80231174 | rs10072255 | 0.457 | -2.407 | -0.97 | -1.293 | 0.004 | 0.052 | 0.052 | 0 | 0.2 | 0.1 | 0.13 | 0.07 | 0.21 | 0.29 |
| 5 | 80231585 | rs12513906 | 0.101 | -3.056 | -0.69 | -1.668 | -0.082 | -0.112 | -0.112 | 0.103 | 0.23 | 0.37 | 0.11 | 0.02 | 0.12 | 0.18 |
| 5 | 80232068 | rs6863791 | -0.767 | -2.804 | -1.08 | -1.544 | 0.121 | 0.179 | 0.179 | 0.007 | 0.46 | 0.77 | 0.26 | 0.1 | 0.34 | 0.48 |
| 5 | 80244913 | rs960990 | 0.846 | -1.222 | -0.62 | -1.382 | -0.002 | -0.07 | -0.07 | 0 | 0.23 | 0.3 | 0.19 | 0 | 0.24 | 0.1 |
| 5 | 80252534 | rs10491488 | -0.33 | -1.508 | -0.875 | -1.135 | 0.037 | 0.027 | 0.027 | 0.02 | 0.52 | 0.72 | 0.48 | 0.4 | 0.4 | 0.41 |
| 5 | 80256909 | rs888821 | 1.162 | -1.329 | -0.786 | -1.164 | -0.001 | -0.007 | -0.007 | 0 | 0.22 | 0.2 | 0.18 | 0 | 0.24 | 0.1 |
| 5 | 80257798 | rs888817 | -0.787 | -0.813 | -0.707 | -0.732 | -0.023 | -0.02 | -0.02 | 0 | 0.7 | 0.82 | 0.63 | 0.48 | 0.63 | 0.71 |
| 5 | 80263865 | rs6874731 | 0.095 | -0.422 | -0.621 | -0.978 | -0.006 | 0.062 | 0.062 | 0 | 0.41 | 0.29 | 0.33 | 0.09 | 0.48 | 0.4 |
| 5 | 80267066 | rs34992 | -0.58 | 0.447 | -0.35 | 0.179 | 0.014 | 0.098 | 0.098 | 0 | 0.11 | 0.19 | 0.16 | 0.41 | 0.16 | 0.2 |
| 5 | 80270992 | rs34995 | -0.556 | -0.021 | -0.311 | -0.602 | 0.001 | 0.002 | 0.002 | 0 | 0.36 | 0.36 | 0.49 | 0.76 | 0.34 | 0.5 |
| 5 | 80277045 | rs34999 | -0.391 | 0.01 | -0.176 | -0.789 | 0.006 | -0.013 | -0.013 | 0 | 0.64 | 0.65 | 0.71 | 0.74 | 0.64 | 0.66 |
| 5 | 80278212 | rs857671 | -0.415 | -0.005 | -0.157 | -0.853 | -0.006 | 0.008 | 0.008 | 0 | 0.66 | 0.77 | 0.71 | 0.76 | 0.65 | 0.66 |
| 5 | 80278998 | rs7717830 | -0.954 | 0.182 | -0.096 | -0.847 | -0.009 | -0.042 | -0.042 | 0 | 0.08 | 0.26 | 0.05 | 0 | 0.04 | 0.03 |
| 5 | 80282584 | rs2052482 | 1.37 | 0.255 | -0.827 | -1.049 | -0.069 | 0.174 | 0.174 | 0 | 0.26 | 0.04 | 0.17 | 0 | 0.28 | 0.14 |
| 5 | 80282815 | rs2052481 | 1.37 | 0.255 | -0.827 | -1.049 | -0.069 | 0.174 | 0.174 | 0 | 0.26 | 0.04 | 0.17 | 0 | 0.28 | 0.14 |
| 5 | 80283127 | rs401758 | -0.564 | 0.281 | -0.431 | -0.78 | 0.033 | 0.036 | 0.036 | 0 | 0.64 | 0.74 | 0.69 | 0.76 | 0.75 | 0.57 |
| 5 | 80287475 | rs7706448 | 0.181 | 0.745 | -0.595 | -0.757 | -0.013 | 0.12 | 0.12 | 0 | 0.44 | 0.26 | 0.27 | 0.07 | 0.52 | 0.37 |
| 5 | 80291972 | rs503367 | -0.02 | 0.728 | -0.403 | -0.2 | 0.017 | 0.012 | 0.012 | 0 | 0.49 | 0.59 | 0.57 | 0.45 | 0.36 | 0.5 |
| 5 | 80296969 | rs483970 | 0.09 | 1.18 | -0.122 | 0.457 | 0.005 | 0.111 | 0.111 | 0 | 0.93 | 0.87 | 0.85 | 0.51 | 0.86 | 0.88 |
| 5 | 80298768 | rs12153097 | -0.341 | 1.272 | 0.064 | 0.037 | 0.014 | -0.016 | -0.016 | 0 | 0.15 | 0.19 | 0.1 | 0.06 | 0.22 | 0.22 |
| 5 | 80300491 | rs499090 | -0.122 | 1.243 | 0.329 | 0.291 | 0.021 | 0.154 | 0.154 | 0 | 0.92 | 0.87 | 0.85 | 0.51 | 0.82 | 0.86 |
| 5 | 80300524 | rs6879497 | -0.167 | 0.707 | 0.386 | -0.272 | -0.01 | -0.013 | -0.013 | 3.516 | 0.31 | 0.61 | 0.31 | 0.19 | 0.23 | 0.32 |
| 5 | 80300607 | rs12153055 | -0.068 | 0.177 | -0.022 | -0.777 | 0.049 | 0.097 | 0.097 | 0 | 0.42 | 0.23 | 0.36 | 0.17 | 0.36 | 0.42 |
| 5 | 80301687 | rs9293827 | 0.049 | 0.305 | 0.233 | -0.79 | 0.002 | 0.025 | 0.025 | 0 | 0.28 | 0.23 | 0.18 | 0.04 | 0.32 | 0.33 |
| 5 | 80302654 | rs13182907 | 0.959 | 0.283 | 0.219 | -0.749 | 0.003 | 0.004 | 0.004 | 0.032 | 0.24 | 0.22 | 0.25 | 0.15 | 0.19 | 0.21 |
| 5 | 80307193 | rs565476 | 0.405 | 0.341 | 0.144 | -0.689 | -0.006 | -0.009 | -0.009 | 0 | 0.64 | 0.65 | 0.67 | 0.48 | 0.5 | 0.51 |
| 5 | 80308016 | rs503363 | 1.773 | 0.191 | 0.04 | -0.982 | -0.06 | 0.016 | 0.016 | 0.236 | 0.33 | 0.06 | 0.36 | 0.29 | 0.28 | 0.21 |
| 5 | 80311835 | rs508636 | 1.686 | 0.181 | 0.065 | -0.889 | -0.05 | 0.118 | 0.118 | 0.02 | 0.34 | 0.08 | 0.36 | 0.2 | 0.28 | 0.21 |
| 5 | 80312586 | rs9293828 | -0.156 | 0.255 | 0.461 | -0.878 | 0.049 | -0.028 | -0.028 | 0 | 0.14 | 0.36 | 0.11 | 0.06 | 0.18 | 0.21 |
| 5 | 80313117 | rs6872530 | 1.243 | 0.194 | 0.389 | -0.879 | 0.028 | 0.066 | 0.066 | 0 | 0.18 | 0.07 | 0.11 | 0.02 | 0.17 | 0.21 |
| 5 | 80315929 | rs9293829 | 0.225 | 0.033 | -0.042 | -1.049 | 0 | -0.004 | -0.004 | 0 | 0.6 | 0.58 | 0.51 | 0.31 | 0.5 | 0.41 |
| 5 | 80316686 | rs476036 | 0.82 | -0.141 | -0.521 | -0.996 | 0.014 | 0.042 | 0.042 | 0 | 0.5 | 0.4 | 0.44 | 0.29 | 0.36 | 0.3 |
| 5 | 80316844 | rs13183890 | 0.794 | -0.099 | -0.611 | -1.096 | -0.02 | -0.05 | -0.05 | 1.661 | 0.14 | 0.21 | 0.07 | 0 | 0.08 | 0.06 |
| 5 | 80319619 | rs551463 | 1.49 | 0.084 | -1.021 | -1.134 | -0.059 | 0.258 | 0.258 | 0.876 | 0.39 | 0.06 | 0.39 | 0.1 | 0.33 | 0.24 |
| 5 | 80319719 | rs7717573 | -0.028 | -0.375 | -0.696 | -0.998 | 0.043 | 0.046 | 0.046 | 0.104 | 0.32 | 0.27 | 0.17 | 0.02 | 0.23 | 0.27 |
| 5 | 80322231 | rs440583 | -0.085 | 0.517 | -0.363 | 0.106 | 0.013 | 0.023 | 0.023 | 0 | 0.14 | 0.19 | 0.36 | 0.21 | 0.2 | 0.15 |
| 5 | 80322712 | rs13187484 | 1.372 | 0.059 | -0.505 | -0.663 | -0.001 | -0.069 | -0.069 | 0.251 | 0.36 | 0.36 | 0.22 | 0 | 0.33 | 0.15 |
| 5 | 80336741 | rs17475899 | 1.201 | -0.035 | -0.784 | -0.396 | 0.017 | 0.088 | 0.088 | 0.038 | 0.17 | 0.01 | 0.09 | 0.01 | 0.15 | 0.17 |

| | | | | | | | | | | | | | | | | |
|---|---|---|---|---|---|---|---|---|---|---|---|---|---|---|---|---|
| 5 | 80337765 | rs477016 | 1.048 | 0.03 | -0.631 | -0.268 | 0 | -0.025 | -0.025 | 0 | 0.53 | 0.46 | 0.3 | 0.14 | 0.41 | 0.3 |
| 5 | 80337995 | rs367647 | -0.091 | 0.555 | -1.1 | 0.15 | -0.005 | 0.038 | 0.038 | 0 | 0.89 | 0.86 | 0.77 | 0.44 | 0.81 | 0.77 |
| 5 | 80338581 | rs6453538 | 0.715 | 0.741 | -1.105 | 0.074 | -0.002 | 0.057 | 0.057 | 0.01 | 0.05 | 0.06 | 0.06 | 0.4 | 0.15 | 0.16 |
| 5 | 80360012 | rs13359431 | 1.746 | 0.386 | -1.192 | -0.529 | -0.006 | 0.026 | 0.026 | 0 | 0.25 | 0.2 | 0.15 | 0 | 0.26 | 0.09 |
| 5 | 80362119 | rs26903 | -0.604 | 0.443 | -0.804 | -0.082 | -0.028 | 0.092 | 0.092 | 0.697 | 0.72 | 0.58 | 0.73 | 0.51 | 0.85 | 0.65 |
| 5 | 80362905 | rs26904 | -0.582 | 0.546 | -0.771 | -0.076 | -0.013 | 0.064 | 0.064 | 1.165 | 0.69 | 0.59 | 0.72 | 0.51 | 0.81 | 0.64 |
| 5 | 80364540 | rs26905 | 0.077 | 0.772 | -0.642 | 0.013 | 0 | -0.001 | -0.001 | 1.135 | 0.53 | 0.58 | 0.63 | 0.5 | 0.66 | 0.47 |
| 5 | 80365054 | rs7736964 | -0.449 | 1.548 | -0.6 | 1.127 | 0.002 | 0.035 | 0.035 | 0 | 0.06 | 0.08 | 0.09 | 0.38 | 0.18 | 0.2 |
| 5 | 80365315 | rs26907 | 0.04 | 1.499 | -0.553 | 0.808 | 0.01 | 0.014 | 0.014 | 0 | 0.08 | 0.14 | 0.31 | 0.11 | 0.15 | 0.12 |
| 5 | 80368312 | rs153227 | -0.223 | 1.689 | -0.516 | 0.975 | -0.007 | -0.006 | -0.006 | 0.637 | 0.11 | 0.11 | 0.34 | 0.12 | 0.22 | 0.14 |
| 5 | 80370915 | rs153226 | -1.381 | 0.982 | -0.562 | 0.562 | 0.006 | 0.011 | 0.011 | 0 | 0.34 | 0.35 | 0.49 | 0.14 | 0.37 | 0.37 |
| 5 | 80371321 | rs190409 | -1.64 | 0.977 | -0.537 | 0.571 | -0.001 | -0.001 | -0.001 | 0 | 0.38 | 0.44 | 0.57 | 0.56 | 0.54 | 0.56 |
| 5 | 80372278 | rs252585 | 0.195 | 1.04 | -0.476 | 0.593 | 0.012 | 0.01 | 0.01 | 0 | 0.1 | 0.12 | 0.34 | 0.12 | 0.22 | 0.14 |
| 5 | 80373479 | rs10514203 | -0.189 | 0.844 | -0.53 | 0.403 | -0.014 | -0.036 | -0.036 | 0 | 0.11 | 0.2 | 0.09 | 0 | 0.07 | 0.08 |
| 5 | 80376142 | rs27650 | -1.318 | 0.738 | -0.792 | 0.358 | 0 | -0.003 | -0.003 | 0 | 0.43 | 0.43 | 0.57 | 0.56 | 0.52 | 0.56 |
| 5 | 80384221 | rs458300 | -0.243 | 0.545 | -1.243 | 0.265 | -0.016 | 0.017 | 0.017 | 0 | 0.27 | 0.08 | 0.4 | 0.13 | 0.32 | 0.29 |
| 5 | 80384331 | rs696879 | -0.48 | 0.65 | -1.677 | 0.134 | -0.069 | -0.114 | -0.114 | 0 | 0.74 | 0.89 | 0.83 | 0.42 | 0.7 | 0.62 |
| 5 | 80390225 | rs505748 | 1.26 | 0.406 | -0.555 | 0.421 | -0.016 | 0.026 | 0.026 | 0 | 0.56 | 0.25 | 0.44 | 0.42 | 0.46 | 0.43 |
| 5 | 80393802 | rs460641 | -0.505 | 0.497 | -0.248 | 0.267 | 0.004 | 0.01 | 0.01 | 0 | 0.29 | 0.28 | 0.42 | 0.12 | 0.33 | 0.32 |
| 5 | 80393949 | rs455461 | -0.283 | 0.576 | -0.45 | 0.306 | 0.008 | -0.005 | -0.005 | 1.093 | 0.19 | 0.3 | 0.36 | 0.11 | 0.26 | 0.17 |
| 5 | 80399926 | rs252581 | -0.035 | 0.672 | -1.066 | 0.459 | -0.035 | -0.042 | -0.042 | 2.409 | 0.88 | 0.92 | 0.87 | 0.55 | 0.78 | 0.76 |
| 5 | 80405616 | rs17212829 | 0.706 | 0.156 | -0.955 | -0.008 | 0.021 | 0.024 | 0.024 | 0 | 0.11 | 0.07 | 0.07 | 0.01 | 0.07 | 0.12 |
| 5 | 80405669 | rs34114 | 0.423 | 0.785 | -0.984 | 0.366 | 0.002 | 0.021 | 0.021 | 0 | 0.81 | 0.81 | 0.71 | 0.51 | 0.71 | 0.7 |
| 5 | 80406899 | rs153225 | 0.054 | 0.963 | -1.39 | 0.538 | -0.034 | -0.054 | -0.054 | 0 | 0.14 | 0.05 | 0.35 | 0.25 | 0.23 | 0.14 |
| 5 | 80411221 | rs4703822 | 0.803 | 0.706 | -1.828 | -0.245 | 0.032 | 0.112 | 0.112 | 0 | 0.43 | 0.15 | 0.26 | 0.27 | 0.31 | 0.24 |
| 5 | 80416697 | rs26898 | 0.341 | 1.189 | -1.871 | -0.301 | 0.033 | 0.103 | 0.103 | 0 | 0.59 | 0.79 | 0.33 | 0.16 | 0.3 | 0.37 |
| 5 | 80421700 | rs26889 | 0.428 | 1.152 | -1.783 | -0.315 | 0.025 | 0.061 | 0.061 | 0 | 0.58 | 0.7 | 0.33 | 0.17 | 0.3 | 0.38 |
| 5 | 80425860 | rs153239 | 0.444 | 1.174 | -1.914 | -0.281 | 0.043 | 0.086 | 0.086 | 0 | 0.58 | 0.73 | 0.33 | 0.16 | 0.29 | 0.36 |
| 5 | 80433460 | rs10223133 | 0.326 | 2.273 | -2.428 | 0.578 | -0.053 | -0.05 | -0.05 | 0 | 0.08 | 0.02 | 0.34 | 0.19 | 0.24 | 0.13 |
| 5 | 80436528 | rs17213885 | -0.252 | 1.759 | -2.482 | -0.037 | -0.031 | 0.14 | 0.14 | 0 | 0.41 | 0.18 | 0.27 | 0.17 | 0.36 | 0.29 |
| 5 | 80437241 | rs17213955 | 0.244 | 2.079 | -1.876 | 0.295 | -0.04 | 0.094 | 0.094 | 0 | 0.21 | 0.06 | 0.14 | 0 | 0.25 | 0.13 |
| 5 | 80442724 | rs42430 | -0.772 | 2.039 | -1.179 | 1.51 | -0.024 | -0.001 | -0.001 | 0 | 0.09 | 0.05 | 0.11 | 0.42 | 0.17 | 0.25 |
| 5 | 80449911 | rs2114972 | -1.125 | 1.427 | -0.722 | 0.916 | 0.005 | 0.027 | 0.027 | 0 | 0.14 | 0.2 | 0.27 | 0.35 | 0.21 | 0.1 |
| 5 | 80459446 | rs7703816 | -1.276 | 0.852 | 0.085 | -0.228 | 0 | -0.045 | -0.045 | 0 | 0.17 | 0.32 | 0.13 | 0.11 | 0.13 | 0.22 |
| 5 | 80460782 | rs13361589 | -1.223 | 0.394 | 0.826 | -0.444 | -0.001 | -0.045 | -0.045 | 0 | 0.18 | 0.57 | 0.13 | 0.11 | 0.13 | 0.26 |
| 5 | 80462439 | rs12697916 | 1.282 | -0.003 | 0.053 | -0.823 | -0.018 | -0.032 | -0.032 | 0 | 0.62 | 0.45 | 0.77 | 0.77 | 0.69 | 0.53 |
| 5 | 80462935 | rs10942943 | 1.283 | 0.005 | 0.056 | -0.82 | -0.018 | -0.032 | -0.032 | 1.603 | 0.62 | 0.45 | 0.77 | 0.77 | 0.69 | 0.53 |
| 5 | 80463635 | rs4704705 | -0.035 | 0.102 | -0.309 | -0.749 | -0.007 | -0.004 | -0.004 | 0 | 0.81 | 0.82 | 0.88 | 0.87 | 0.84 | 0.79 |
| 5 | 80464097 | rs10474653 | -0.656 | 0.043 | -0.398 | -0.798 | -0.017 | 0.005 | 0.005 | 0 | 0.88 | 0.86 | 0.93 | 0.87 | 0.88 | 0.84 |
| 5 | 80467798 | rs4144527 | 0.032 | 0.084 | -0.659 | -0.779 | 0.017 | 0.01 | 0.01 | 0 | 0.82 | 0.94 | 0.89 | 0.87 | 0.83 | 0.79 |
| 5 | 80468463 | rs1379270 | 0.957 | 0.201 | -0.756 | -0.722 | 0.044 | 0.081 | 0.081 | 0.768 | 0.59 | 0.23 | 0.73 | 0.77 | 0.67 | 0.51 |
| 5 | 80468617 | rs1379271 | 0.423 | 0.591 | -1.214 | -0.924 | -0.037 | 0.128 | 0.128 | 0 | 0.24 | 0.05 | 0.14 | 0.01 | 0.22 | 0.09 |
| 5 | 80472887 | rs1020721 | 0.361 | 0.678 | -0.326 | -0.099 | -0.001 | -0.004 | -0.004 | 0 | 0.86 | 0.78 | 0.73 | 0.9 | 0.84 | 0.82 |
| 5 | 80479164 | rs7701594 | -0.125 | 0.749 | -0.589 | 0.207 | -0.001 | 0.001 | 0.001 | 0 | 0.86 | 0.9 | 0.63 | 0.77 | 0.81 | 0.81 |
| 5 | 80479910 | rs4703825 | 0.227 | 0.789 | -0.601 | -0.271 | -0.002 | 0.013 | 0.013 | 0 | 0.52 | 0.58 | 0.41 | 0.63 | 0.5 | 0.44 |
| 5 | 80482514 | rs10514211 | 0.168 | 0.017 | -0.806 | -0.949 | 0.008 | -0.019 | -0.019 | 0.301 | 0.2 | 0.15 | 0.08 | 0.02 | 0.1 | 0.07 |
| 5 | 80482875 | rs7718438 | -0.385 | 0.645 | -0.721 | -1.214 | 0.017 | -0.021 | -0.021 | 0.081 | 0.35 | 0.45 | 0.3 | 0.26 | 0.39 | 0.3 |
| 5 | 80484109 | rs12521784 | -0.396 | -0.669 | -0.72 | -1.662 | 0.006 | 0.037 | 0.037 | 0 | 0.52 | 0.41 | 0.34 | 0.23 | 0.41 | 0.4 |
| 5 | 80486098 | rs10079031 | 0.01 | -0.895 | -0.859 | -1.957 | 0.005 | 0.006 | 0.006 | 0 | 0.4 | 0.36 | 0.3 | 0.21 | 0.27 | 0.33 |
| 5 | 80486198 | rs10055900 | -1.358 | -0.939 | -0.671 | -2.017 | 0.015 | -0.014 | -0.014 | 0 | 0.66 | 0.63 | 0.58 | 0.29 | 0.57 | 0.52 |
| 5 | 80486819 | rs13188677 | -0.257 | -1.081 | -0.582 | -1.851 | -0.003 | 0.008 | 0.008 | 0 | 0.15 | 0.19 | 0.21 | 0.2 | 0.18 | 0.18 |
| 5 | 80492967 | rs7710039 | -1.032 | -1.354 | -0.526 | -1.91 | 0.024 | 0.088 | 0.088 | 0 | 0.34 | 0.18 | 0.25 | 0.08 | 0.3 | 0.26 |
| 5 | 80496533 | rs13179530 | -1.08 | -1.236 | -0.409 | -1.795 | 0.022 | 0.075 | 0.075 | 0 | 0.29 | 0.2 | 0.24 | 0.14 | 0.28 | 0.26 |
| 5 | 80498379 | rs6883756 | -1.368 | -1.32 | -0.3 | -1.865 | 0.003 | -0.041 | -0.041 | 0 | 0.57 | 0.59 | 0.49 | 0.22 | 0.48 | 0.48 |
| 5 | 80498443 | rs9968730 | 0.92 | -1.24 | -0.25 | -1.964 | -0.004 | 0.012 | 0.012 | 0 | 0.24 | 0.16 | 0.16 | 0.01 | 0.19 | 0.16 |
| 5 | 80499626 | rs4704706 | -0.574 | -1.246 | -0.183 | -1.778 | -0.003 | 0.001 | 0.001 | 0 | 0.21 | 0.22 | 0.17 | 0.28 | 0.15 | 0.24 |
| 5 | 80501981 | rs6890619 | -0.785 | -1.164 | -0.106 | -1.673 | 0.001 | 0.001 | 0.001 | 0 | 0.48 | 0.55 | 0.35 | 0.29 | 0.35 | 0.42 |
| 5 | 80503528 | rs7732064 | -1.517 | -1.25 | -0.041 | -1.406 | -0.018 | 0.02 | 0.02 | 0 | 0.58 | 0.76 | 0.49 | 0.38 | 0.49 | 0.56 |
| 5 | 80505879 | rs16878538 | 0.361 | -1.134 | 0.077 | -1.14 | -0.001 | -0.001 | -0.001 | 0.15 | 0.08 | 0.08 | 0.13 | 0.08 | 0.11 | 0.1 |
| 5 | 80508670 | rs11958318 | -1.188 | -1.185 | 0.147 | -1.147 | 0.002 | -0.015 | -0.015 | 0 | 0.55 | 0.57 | 0.64 | 0.3 | 0.47 | 0.5 |
| 5 | 80509137 | rs11949363 | -0.508 | -1.237 | 0.193 | -1.168 | -0.011 | 0.036 | 0.036 | 0 | 0.1 | 0.26 | 0.12 | 0.18 | 0.06 | 0.11 |
| 5 | 80509936 | rs6893150 | -1.347 | -1.358 | 0.484 | -0.967 | 0.019 | 0.037 | 0.037 | 0.077 | 0.24 | 0.18 | 0.2 | 0.11 | 0.22 | 0.22 |
| 5 | 80512974 | rs6868762 | -1.256 | -1.354 | 0.497 | -0.976 | 0.011 | 0.042 | 0.042 | 0.779 | 0.48 | 0.42 | 0.55 | 0.13 | 0.42 | 0.4 |
| 5 | 80513068 | rs6870105 | -1.389 | -1.392 | 0.479 | -0.971 | 0.015 | 0.021 | 0.021 | 0 | 0.25 | 0.22 | 0.21 | 0.11 | 0.22 | 0.2 |
| 5 | 80516195 | rs2124888 | -1.154 | -1.24 | 0.531 | -0.9 | 0.006 | 0.027 | 0.027 | 0.63 | 0.48 | 0.42 | 0.53 | 0.13 | 0.42 | 0.4 |
| 5 | 80521485 | rs7725743 | 0.881 | -1.114 | 0.506 | -0.645 | 0.001 | 0.001 | 0.001 | 0 | 0.06 | 0.06 | 0.09 | 0.05 | 0.08 | 0.09 |
| 5 | 80526113 | rs6898834 | -1.269 | -1.145 | 0.536 | -0.563 | -0.014 | 0.075 | 0.075 | 0 | 0.1 | 0.17 | 0.13 | 0.32 | 0.05 | 0.16 |
| 5 | 80528135 | rs4703827 | 0.687 | -1.116 | 0.507 | -0.772 | -0.002 | -0.019 | -0.019 | 0 | 0.24 | 0.19 | 0.31 | 0.01 | 0.19 | 0.16 |
| 5 | 80528457 | rs1870061 | -0.624 | -1.139 | 0.54 | -0.729 | -0.012 | -0.016 | -0.016 | 0 | 0.12 | 0.19 | 0.11 | 0.07 | 0.1 | 0.13 |

| | | | | | | | | | | | | | | | | |
|---|---|---|---|---|---|---|---|---|---|---|---|---|---|---|---|---|
| 5 | 80535187 | rs10942945 | 0.779 | -1.278 | 0.521 | -0.741 | -0.006 | 0.014 | 0.014 | 0 | 0.24 | 0.16 | 0.31 | 0.01 | 0.19 | 0.16 |
| 5 | 80546499 | rs3756636 | -0.749 | -1.349 | 0.706 | -0.712 | 0 | 0.001 | 0.001 | 0 | 0.41 | 0.42 | 0.5 | 0.35 | 0.33 | 0.36 |
| 5 | 80551423 | rs4704717 | -0.581 | -1.037 | 0.157 | -0.611 | -0.007 | -0.015 | -0.015 | 0 | 0.37 | 0.45 | 0.5 | 0.25 | 0.29 | 0.35 |
| 5 | 80562120 | rs545 | -0.457 | -1.087 | 0.168 | -0.614 | 0.002 | -0.012 | -0.012 | 0 | 0.35 | 0.35 | 0.42 | 0.08 | 0.28 | 0.28 |
| 5 | 80580063 | rs11745275 | -0.024 | -1.062 | 0.146 | -0.621 | 0 | 0.032 | 0.032 | 0 | 0.12 | 0.06 | 0.07 | 0.01 | 0.11 | 0.08 |
| 5 | 80582099 | rs11953113 | -0.264 | -1.062 | 0.152 | -0.618 | 0.002 | 0.014 | 0.014 | 0 | 0.13 | 0.1 | 0.1 | 0.04 | 0.11 | 0.07 |
| 5 | 80593392 | rs12516414 | -0.498 | -1.074 | 0.164 | -0.621 | -0.008 | -0.017 | -0.017 | 0.003 | 0.37 | 0.45 | 0.5 | 0.25 | 0.29 | 0.33 |
| 5 | 80594346 | rs10061143 | -1.529 | -1.074 | 0.169 | -0.607 | 0.009 | 0.017 | 0.017 | 1.253 | 0.53 | 0.73 | 0.62 | 0.42 | 0.41 | 0.45 |
| 5 | 80596222 | rs16878586 | 0.362 | -1.006 | -0.58 | -1.231 | 0.002 | 0.002 | 0.002 | 0 | 0.05 | 0.05 | 0.09 | 0.18 | 0.03 | 0.12 |
| 5 | 80598005 | rs10052929 | -1.545 | -0.988 | -0.645 | -1.273 | 0.007 | 0.017 | 0.017 | 1.056 | 0.5 | 0.61 | 0.61 | 0.29 | 0.4 | 0.41 |
| 5 | 80599578 | rs7709149 | -0.466 | -0.83 | -1.215 | -1.235 | -0.003 | 0.059 | 0.059 | 0 | 0.35 | 0.25 | 0.38 | 0.02 | 0.3 | 0.23 |
| 5 | 80599928 | rs11740916 | 0.641 | -0.757 | -1.428 | -1.159 | -0.01 | 0.03 | 0.03 | 0 | 0.08 | 0.02 | 0.04 | 0.01 | 0.08 | 0.06 |
| 5 | 80608545 | rs2270825 | 0.528 | -0.682 | -1.362 | -1.123 | -0.006 | -0.004 | -0.004 | 0 | 0.07 | 0.11 | 0.09 | 0.05 | 0.08 | 0.07 |
| 5 | 80609768 | rs1039326 | 0.004 | -0.191 | -1.526 | -0.873 | -0.009 | 0.024 | 0.024 | 0 | 0.06 | 0.09 | 0.12 | 0.21 | 0.03 | 0.12 |
| 5 | 80616365 | rs10053706 | 0.226 | 0.344 | -1.613 | -1.039 | -0.011 | 0.051 | 0.051 | 0 | 0.22 | 0.1 | 0.3 | 0.01 | 0.18 | 0.15 |
| 5 | 80619046 | rs10056565 | -0.113 | 0.414 | -1.315 | -0.744 | -0.015 | 0.105 | 0.105 | 0 | 0.34 | 0.16 | 0.38 | 0.02 | 0.3 | 0.24 |
| 5 | 80625946 | rs9546 | -0.088 | 0.82 | -0.802 | -0.039 | 0.019 | -0.009 | -0.009 | 0 | 0.64 | 0.49 | 0.69 | 0.32 | 0.53 | 0.52 |
| 5 | 80629372 | rs6881927 | -0.726 | 0.941 | -0.718 | -0.251 | 0 | -0.001 | -0.001 | 0 | 0.27 | 0.33 | 0.2 | 0.17 | 0.18 | 0.19 |
| 5 | 80631511 | rs6452399 | -0.957 | 1.401 | -0.607 | -0.872 | 0.001 | -0.07 | -0.07 | 0 | 0.42 | 0.52 | 0.33 | 0.06 | 0.4 | 0.27 |
| 5 | 80648695 | rs7729671 | -0.968 | 1.402 | -0.371 | -1.089 | -0.004 | 0.058 | 0.058 | 2.648 | 0.46 | 0.69 | 0.41 | 0.13 | 0.47 | 0.3 |
| 5 | 80650462 | rs13154467 | -1.027 | 1.299 | 0.002 | -1.099 | -0.001 | 0 | 0 | 2.878 | 0.45 | 0.47 | 0.39 | 0.14 | 0.46 | 0.3 |
| 5 | 80650890 | rs7706593 | -1.033 | 1.316 | -0.176 | -1.091 | -0.004 | 0.058 | 0.058 | 2.841 | 0.46 | 0.69 | 0.41 | 0.14 | 0.47 | 0.3 |
| 5 | 80651829 | rs10053334 | -0.222 | 1.457 | -0.119 | -1.252 | -0.007 | 0.009 | 0.009 | 0 | 0.24 | 0.21 | 0.26 | 0.14 | 0.27 | 0.18 |
| 5 | 80653460 | rs7717804 | 0.112 | 1.51 | 0.256 | -1.269 | 0.005 | -0.111 | -0.111 | 0 | 0.19 | 0.36 | 0.12 | 0 | 0.18 | 0.1 |
| 5 | 80654966 | rs4703516 | -1.093 | 1.753 | -0.841 | -0.473 | -0.039 | -0.007 | -0.007 | 0 | 0.92 | 0.99 | 0.79 | 0.51 | 0.89 | 0.86 |
| 5 | 80655266 | rs4703517 | -1.078 | 1.319 | -0.629 | -1.302 | -0.002 | 0.098 | 0.098 | 2.707 | 0.47 | 0.72 | 0.42 | 0.14 | 0.46 | 0.3 |
| 5 | 80655709 | rs13163821 | 0.454 | 1.41 | -0.809 | -0.818 | -0.062 | 0.015 | 0.015 | 0 | 0.15 | 0.02 | 0.17 | 0.06 | 0.24 | 0.13 |
| 5 | 80657729 | rs13174473 | 0.029 | 1.566 | -0.91 | -0.929 | -0.089 | 0.032 | 0.032 | 0 | 0.18 | 0.02 | 0.17 | 0.06 | 0.27 | 0.17 |
| 5 | 80660945 | rs7713774 | -0.32 | 0.259 | -0.654 | -1.58 | -0.014 | -0.063 | -0.063 | 3.74 | 0.37 | 0.53 | 0.28 | 0.11 | 0.32 | 0.32 |
| 5 | 80661696 | rs10078088 | -1.34 | -0.074 | -1.541 | -2.196 | -0.126 | 0.051 | 0.051 | 0 | 0.7 | 0.93 | 0.59 | 0.2 | 0.55 | 0.49 |
| 5 | 80665700 | rs10042996 | -0.21 | -1.051 | -0.876 | -2.627 | -0.01 | -0.048 | -0.048 | 0 | 0.35 | 0.37 | 0.25 | 0.09 | 0.23 | 0.25 |
| 5 | 80666733 | rs4703520 | -0.551 | -0.651 | -0.99 | -2.633 | -0.008 | 0.101 | 0.101 | 0.951 | 0.61 | 0.8 | 0.47 | 0.08 | 0.46 | 0.42 |
| 5 | 80675496 | rs10942262 | -0.566 | -0.645 | -0.987 | -2.539 | -0.008 | 0.097 | 0.097 | 0.962 | 0.61 | 0.8 | 0.48 | 0.11 | 0.46 | 0.44 |
| 5 | 80680149 | rs13154060 | 0.452 | -0.414 | -1.053 | -2.379 | -0.025 | -0.003 | -0.003 | 1.401 | 0.08 | 0.02 | 0.1 | 0.07 | 0.09 | 0.04 |
| 5 | 80682715 | rs6872459 | -0.52 | -1.365 | -0.544 | -2.864 | -0.041 | -0.071 | -0.071 | 0 | 0.23 | 0.4 | 0.19 | 0.06 | 0.12 | 0.19 |
| 5 | 80684302 | rs17230425 | 1.787 | -1.242 | -1.027 | -2.764 | -0.038 | 0.088 | 0.088 | 0.139 | 0.19 | 0.01 | 0.19 | 0.03 | 0.24 | 0.13 |
| 5 | 80700023 | rs10070789 | -0.473 | -1.323 | -0.289 | -2.826 | -0.001 | -0.017 | -0.017 | 0 | 0.5 | 0.55 | 0.44 | 0.17 | 0.46 | 0.38 |
| 5 | 80704269 | rs1013009 | 1.128 | -1.027 | -0.765 | -2.636 | -0.011 | -0.004 | -0.004 | 0 | 0.07 | 0.05 | 0.1 | 0.08 | 0.09 | 0.03 |
| 5 | 80708693 | rs3763142 | -1.153 | -1.325 | -0.79 | -2.865 | -0.04 | 0.084 | 0.084 | 0 | 0.62 | 0.87 | 0.48 | 0.18 | 0.5 | 0.43 |
| 5 | 80715913 | rs9641 | 1.176 | -1.392 | -0.274 | -2.862 | -0.039 | -0.049 | -0.049 | 0.045 | 0.09 | 0.23 | 0.03 | 0.01 | 0.02 | 0.03 |
| 5 | 80716163 | rs10746 | 1.511 | -1.477 | -0.42 | -3.026 | -0.021 | 0.102 | 0.102 | 0.528 | 0.22 | 0.04 | 0.18 | 0.04 | 0.26 | 0.22 |
| 5 | 80719125 | rs7732320 | 0.529 | -1.631 | 0.127 | -2.772 | -0.055 | -0.077 | -0.077 | 0 | 0.22 | 0.5 | 0.15 | 0.07 | 0.11 | 0.1 |
| 5 | 80723458 | rs11740719 | -0.821 | -1.677 | -0.559 | -2.896 | 0.024 | 0.06 | 0.06 | 0 | 0.41 | 0.67 | 0.35 | 0.15 | 0.28 | 0.19 |
| 5 | 80737212 | rs3792923 | -1.026 | -0.438 | -0.052 | -2.444 | 0.017 | -0.076 | -0.076 | 0 | 0.26 | 0.4 | 0.28 | 0.09 | 0.28 | 0.16 |
| 5 | 80737420 | rs13177962 | 0.672 | -0.478 | -0.229 | -2.396 | -0.059 | -0.007 | -0.007 | 0 | 0.11 | 0.02 | 0.16 | 0.09 | 0.19 | 0.11 |
| 5 | 80748607 | rs6887937 | -1.829 | -0.868 | -0.264 | -2.555 | 0.045 | 0.154 | 0.154 | 0.648 | 0.42 | 0.89 | 0.38 | 0.18 | 0.31 | 0.33 |
| 5 | 80754434 | rs6452414 | -1.162 | -0.36 | -0.258 | -2.505 | -0.135 | -0.021 | -0.021 | 0 | 0.75 | 0.97 | 0.58 | 0.2 | 0.64 | 0.55 |
| 5 | 80761967 | rs16898954 | 0.019 | -0.412 | -0.031 | -2.493 | -0.051 | -0.006 | -0.006 | 0 | 0.13 | 0.42 | 0.06 | 0.08 | 0.04 | 0.04 |
| 5 | 80779904 | rs9293275 | -1.134 | -0.163 | -0.846 | -2.417 | -0.138 | -0.022 | -0.022 | 0 | 0.75 | 0.97 | 0.58 | 0.2 | 0.64 | 0.54 |
| 5 | 80819553 | rs13355366 | -0.629 | -0.468 | -0.558 | -2.48 | 0.005 | -0.003 | -0.003 | 0 | 0.19 | 0.17 | 0.12 | 0.04 | 0.1 | 0.07 |
| 5 | 80825738 | rs2973349 | 1.985 | -0.411 | -1.17 | -2.624 | -0.045 | 0.242 | 0.242 | 0 | 0.35 | 0.02 | 0.23 | 0.03 | 0.34 | 0.26 |
| 5 | 80841360 | rs11740689 | -1.008 | -0.35 | -0.448 | -2.504 | -0.007 | -0.061 | -0.061 | 0 | 0.32 | 0.47 | 0.34 | 0.16 | 0.28 | 0.28 |
| 5 | 80841753 | rs2973342 | 1.273 | 0.509 | -0.995 | -1.688 | -0.049 | -0.049 | -0.049 | 0.458 | 0.08 | 0.01 | 0.13 | 0.17 | 0.23 | 0.17 |
| 5 | 80845428 | rs6880200 | -1.787 | 0.216 | -0.985 | -2.123 | 0.026 | 0.193 | 0.193 | 0.547 | 0.45 | 0.95 | 0.39 | 0.17 | 0.31 | 0.32 |
| 5 | 80853708 | rs10056236 | 0.184 | 0.156 | -0.676 | -1.972 | -0.008 | -0.034 | -0.034 | 0 | 0.16 | 0.15 | 0.1 | 0.01 | 0.09 | 0.05 |
| 5 | 80870584 | rs4529168 | -0.98 | 0.295 | -0.706 | -1.947 | 0.012 | 0.033 | 0.033 | 1.619 | 0.29 | 0.79 | 0.29 | 0.16 | 0.22 | 0.26 |
| 5 | 80873851 | rs7709916 | -1.175 | 0.366 | -0.418 | -1.595 | -0.008 | -0.03 | -0.03 | 0 | 0.34 | 0.48 | 0.33 | 0.24 | 0.28 | 0.31 |
| 5 | 80876359 | rs2973347 | 0.584 | 0.678 | -0.613 | -1.636 | -0.102 | -0.075 | -0.075 | 0 | 0.19 | 0.03 | 0.42 | 0.72 | 0.33 | 0.43 |
| 5 | 80895033 | rs10942273 | 0.016 | 0.299 | 0.392 | -1.346 | 0.024 | 0.097 | 0.097 | 0 | 0.05 | 0.26 | 0.04 | 0.09 | 0.02 | 0.11 |
| 5 | 80898725 | rs10514220 | 0.318 | 0.524 | 0.127 | -1.352 | 0 | -0.02 | -0.02 | 0 | 0.1 | 0.09 | 0.12 | 0 | 0.11 | 0.03 |
| 5 | 80900598 | rs10942275 | -0.918 | 0.807 | 0.513 | -1.148 | -0.001 | 0.006 | 0.006 | 0 | 0.26 | 0.23 | 0.25 | 0.12 | 0.25 | 0.17 |
| 5 | 80902402 | rs13173845 | 0.197 | 0.93 | 0.513 | -1.034 | -0.026 | 0 | 0 | 0 | 0.16 | 0.07 | 0.2 | 0.12 | 0.19 | 0.14 |
| 5 | 80904142 | rs11743304 | 0.069 | 0.947 | 0.505 | -1.027 | -0.011 | 0.004 | 0.004 | 0 | 0.16 | 0.11 | 0.2 | 0.12 | 0.19 | 0.14 |
| 5 | 80904309 | rs1561150 | -1.383 | 1.013 | 0.532 | -0.53 | 0 | -0.007 | -0.007 | 0 | 0.35 | 0.32 | 0.62 | 0.84 | 0.48 | 0.54 |
| 5 | 80904933 | rs2973335 | 0.048 | 0.79 | 0.385 | 0.149 | -0.072 | -0.076 | -0.076 | 0 | 0.15 | 0.03 | 0.37 | 0.72 | 0.25 | 0.39 |
| 5 | 80938489 | rs6897943 | -1.154 | 0.985 | 1.131 | 0.439 | -0.019 | -0.036 | -0.036 | 0 | 0.13 | 0.38 | 0.08 | 0.08 | 0.1 | 0.04 |
| 5 | 80940436 | rs10069094 | -0.299 | 1.009 | 0.941 | 0.451 | -0.01 | -0.015 | -0.015 | 0 | 0.11 | 0.15 | 0.05 | 0.08 | 0.1 | 0.04 |
| 5 | 80972255 | rs13360536 | -0.183 | 0.91 | 0.864 | 0.38 | -0.033 | -0.042 | -0.042 | 0 | 0.1 | 0.37 | 0.04 | 0 | 0.02 | 0.02 |
| 5 | 80990358 | rs433366 | 0.629 | 1.772 | -0.238 | 0.945 | -0.067 | -0.065 | -0.065 | 0 | 0.9 | 0.99 | 0.81 | 0.29 | 0.72 | 0.65 |

| | | | | | | | | | | | | | | | | |
|---|---|---|---|---|---|---|---|---|---|---|---|---|---|---|---|---|
| 5 | 81009067 | rs6860990 | -0.592 | 0.302 | -0.047 | 0.722 | -0.007 | -0.001 | -0.001 | 0 | 0.3 | 0.25 | 0.19 | 0.39 | 0.15 | 0.26 |
| 5 | 81012281 | rs649872 | -0.42 | 0.722 | 0.217 | 0.952 | 0.03 | 0.078 | 0.078 | 0 | 0.91 | 0.86 | 0.89 | 0.71 | 0.83 | 0.78 |
| 5 | 81012751 | rs396231 | 2.121 | 1.096 | -0.267 | 0.528 | -0.077 | 0.243 | 0.243 | 0.236 | 0.4 | 0.03 | 0.39 | 0.05 | 0.39 | 0.35 |
| 5 | 81014958 | rs411840 | 1.482 | 0.336 | 0.486 | 0.713 | 0.004 | 0.017 | 0.017 | 0 | 0.51 | 0.5 | 0.51 | 0.21 | 0.53 | 0.43 |
| 5 | 81018408 | rs34044243 | 0.682 | 0.345 | 0.59 | 0.829 | 0.008 | 0.006 | 0.006 | 0 | 0.09 | 0.1 | 0.19 | 0.11 | 0.13 | 0.07 |
| 5 | 81022494 | rs10035917 | -0.457 | 0.135 | 0.641 | 0.757 | -0.024 | 0.181 | 0.181 | 0 | 0.13 | 0.26 | 0.11 | 0.52 | 0.09 | 0.29 |
| 5 | 81029595 | rs16899116 | 0.436 | 0.164 | 0.483 | 0.82 | -0.035 | 0.124 | 0.124 | 0 | 0.1 | 0.18 | 0.07 | 0.39 | 0.05 | 0.21 |
| 5 | 81036805 | rs417590 | 0.557 | 0.883 | 0.574 | 1.007 | 0.033 | 0.064 | 0.064 | 0 | 0.82 | 0.76 | 0.7 | 0.6 | 0.68 | 0.71 |
| 5 | 81049187 | rs393766 | 1.182 | 0.819 | 0.539 | 0.419 | -0.005 | 0.178 | 0.178 | 0 | 0.49 | 0.3 | 0.48 | 0.09 | 0.49 | 0.37 |
| 5 | 81053445 | rs1550802 | 0.127 | 1.471 | 0.38 | 1.027 | 0.033 | 0.095 | 0.095 | 0 | 0.9 | 0.84 | 0.89 | 0.68 | 0.81 | 0.79 |
| 5 | 81056637 | rs4703842 | -0.514 | 0.92 | -0.042 | 0.973 | 0.006 | 0.085 | 0.085 | 0 | 0.13 | 0.17 | 0.24 | 0.43 | 0.16 | 0.26 |
| 5 | 81061504 | rs435740 | 0.218 | 0.534 | -0.678 | 0.625 | 0.024 | -0.021 | -0.021 | 0 | 0.59 | 0.3 | 0.7 | 0.52 | 0.67 | 0.64 |
| 5 | 81062082 | rs394961 | 0.741 | 1.009 | -1.543 | 0.026 | -0.112 | 0.26 | 0.26 | 0 | 0.45 | 0.07 | 0.44 | 0.12 | 0.47 | 0.38 |
| 5 | 81062529 | rs4703845 | 0.096 | 0.596 | -1.123 | -0.313 | 0.001 | 0.003 | 0.003 | 0 | 0.49 | 0.46 | 0.65 | 0.61 | 0.5 | 0.55 |
| 5 | 81063858 | rs4703847 | 0.277 | -0.251 | -0.805 | -0.277 | -0.008 | 0.002 | 0.002 | 0.015 | 0.37 | 0.61 | 0.33 | 0.48 | 0.23 | 0.28 |
| 5 | 81065317 | rs7728918 | 1.102 | -0.439 | -0.724 | -1.042 | -0.002 | -0.05 | -0.05 | 0 | 0.27 | 0.26 | 0.15 | 0 | 0.16 | 0.09 |
| 5 | 81067435 | rs377843 | 0.19 | -0.45 | -1.104 | -0.888 | 0.037 | 0.051 | 0.051 | 0 | 0.26 | 0.14 | 0.11 | 0.13 | 0.12 | 0.22 |
| 5 | 81068347 | rs401654 | 0.314 | -0.337 | -1.338 | -0.861 | 0.045 | 0.07 | 0.07 | 0 | 0.23 | 0.06 | 0.1 | 0.13 | 0.11 | 0.22 |
| 5 | 81068474 | rs445410 | -0.462 | 0.199 | -1.173 | -0.588 | -0.027 | 0.056 | 0.056 | 0 | 0.44 | 0.25 | 0.37 | 0.33 | 0.47 | 0.44 |
| 5 | 81072979 | rs9791117 | -0.41 | 0.12 | -0.724 | -0.047 | 0.001 | -0.021 | -0.021 | 0.171 | 0.25 | 0.19 | 0.39 | 0.47 | 0.24 | 0.4 |
| 5 | 81074655 | rs457008 | -0.017 | 0.068 | -1.297 | -0.415 | -0.159 | -0.155 | -0.155 | 0 | 0.38 | 0.04 | 0.38 | 0.52 | 0.47 | 0.46 |
| 5 | 81075969 | rs465059 | -2.032 | -0.072 | -0.265 | -0.788 | -0.008 | -0.107 | -0.107 | 0 | 0.67 | 0.65 | 0.82 | 0.99 | 0.72 | 0.86 |
| 5 | 81080090 | rs7714413 | 0.424 | 0.347 | -0.134 | -0.667 | 0.019 | -0.062 | -0.062 | 0 | 0.11 | 0.23 | 0.12 | 0 | 0.11 | 0.04 |
| 5 | 81080808 | rs462647 | -1.251 | 0.132 | -0.207 | -0.65 | 0.003 | 0.006 | 0.006 | 0 | 0.47 | 0.42 | 0.39 | 0.45 | 0.46 | 0.48 |
| 5 | 81081641 | rs11948076 | 0.205 | 0.057 | -0.36 | -0.776 | -0.01 | -0.014 | -0.014 | 0 | 0.08 | 0.1 | 0.02 | 0 | 0.01 | 0.05 |
| 5 | 81083050 | rs6869736 | -0.672 | 0.114 | -0.341 | -0.804 | -0.017 | -0.096 | -0.096 | 1.884 | 0.19 | 0.32 | 0.15 | 0 | 0.13 | 0.09 |
| 5 | 81087004 | rs384075 | 0.608 | 0.19 | -0.665 | -0.497 | -0.081 | -0.143 | -0.143 | 0 | 0.25 | 0.02 | 0.26 | 0.45 | 0.31 | 0.28 |
| 5 | 81090395 | rs462151 | -1.831 | 0.402 | -0.813 | -0.973 | 0.011 | 0.073 | 0.073 | 0 | 0.71 | 0.85 | 0.87 | 0.99 | 0.76 | 0.86 |
| 5 | 81091203 | rs6885263 | 0.176 | 1.073 | -0.317 | -0.134 | -0.005 | -0.004 | -0.004 | 0 | 0.09 | 0.07 | 0.1 | 0 | 0.12 | 0.04 |
| 5 | 81095080 | rs10514224 | 0.042 | 1.055 | -0.251 | -0.142 | 0.005 | -0.035 | -0.035 | 0 | 0.1 | 0.15 | 0.11 | 0 | 0.12 | 0.04 |
| 5 | 81096255 | rs6895201 | -1.422 | 0.944 | -0.906 | 0.207 | -0.062 | 0.033 | 0.033 | 0 | 0.52 | 0.89 | 0.61 | 0.59 | 0.49 | 0.44 |
| 5 | 81098921 | rs957222 | -1.325 | 0.86 | -0.5 | 0.239 | -0.025 | 0.042 | 0.042 | 0.037 | 0.51 | 0.83 | 0.59 | 0.58 | 0.48 | 0.44 |
| 5 | 81106956 | rs4703849 | -1.034 | 0.53 | 0.277 | 0.152 | -0.002 | 0.001 | 0.001 | 0 | 0.42 | 0.36 | 0.54 | 0.58 | 0.43 | 0.44 |
| 5 | 81107024 | rs6452427 | 1.606 | 0.284 | -0.011 | -0.412 | 0.004 | -0.057 | -0.057 | 0 | 0.41 | 0.41 | 0.2 | 0.01 | 0.34 | 0.16 |
| 5 | 81109049 | rs6899114 | -1.18 | -0.33 | -0.131 | 0.529 | -0.005 | -0.001 | -0.001 | 0.929 | 0.3 | 0.66 | 0.36 | 0.17 | 0.17 | 0.19 |
| 5 | 81111682 | rs11949897 | -1.177 | -0.331 | -0.101 | 0.586 | -0.019 | -0.008 | -0.008 | 0.201 | 0.3 | 0.63 | 0.35 | 0.19 | 0.17 | 0.17 |
| 5 | 81117472 | rs6452429 | -2.086 | -0.166 | -0.907 | 0.844 | 0.026 | -0.034 | -0.034 | 0 | 0.49 | 0.86 | 0.57 | 0.48 | 0.43 | 0.46 |
| 5 | 81129663 | rs12653216 | 0.997 | 0.411 | -1.022 | 0.926 | -0.026 | -0.034 | -0.034 | 0 | 0.06 | 0 | 0.11 | 0.14 | 0.2 | 0.1 |
| 5 | 81130158 | rs11747644 | 0.679 | 0.415 | -0.961 | 0.669 | -0.099 | -0.104 | -0.104 | 0.9 | 0.18 | 0.01 | 0.17 | 0.3 | 0.26 | 0.26 |
| 5 | 81134423 | rs12518812 | 0.71 | 0.196 | -0.953 | 0.646 | -0.089 | -0.107 | -0.107 | 0 | 0.18 | 0.01 | 0.16 | 0.31 | 0.25 | 0.26 |
| 5 | 81144874 | rs1472611 | -1.417 | 0.153 | -0.695 | 0.547 | 0.029 | -0.02 | -0.02 | 0 | 0.43 | 0.78 | 0.53 | 0.44 | 0.38 | 0.42 |
| 5 | 81146156 | rs17241969 | 0.338 | 0.094 | -0.553 | 0.418 | -0.002 | 0.014 | 0.014 | 0.112 | 0.08 | 0.05 | 0.06 | 0 | 0.09 | 0.04 |
| 5 | 81154906 | rs17242130 | 0.095 | 0.078 | -0.504 | 0.404 | -0.001 | 0.001 | 0.001 | 0 | 0.08 | 0.05 | 0.06 | 0 | 0.09 | 0.04 |
| 5 | 81157622 | rs6891774 | 0.863 | -0.131 | -0.317 | 0.691 | 0.009 | -0.004 | -0.004 | 0 | 0.64 | 0.36 | 0.5 | 0.39 | 0.69 | 0.61 |
| 5 | 81160103 | rs6877846 | 0.257 | -0.316 | 0.091 | 0.92 | -0.051 | 0.157 | 0.157 | 0.887 | 0.86 | 0.5 | 0.68 | 0.7 | 0.94 | 0.86 |
| 5 | 81167030 | rs7717239 | 1.251 | -0.473 | 0.274 | 0.858 | -0.024 | 0.066 | 0.066 | 0 | 0.87 | 0.66 | 0.69 | 0.78 | 0.9 | 0.86 |
| 5 | 81168651 | rs12523001 | -1.382 | -0.441 | 0.064 | 0.853 | 0.011 | -0.001 | -0.001 | 0 | 0.39 | 0.62 | 0.29 | 0.39 | 0.33 | 0.31 |
| 5 | 81169660 | rs6859406 | 1.933 | -0.46 | 0.06 | 0.846 | 0.06 | 0.032 | 0.032 | 1.013 | 0.52 | 0.15 | 0.42 | 0.39 | 0.59 | 0.55 |
| 5 | 81170704 | rs1501791 | 1.712 | -0.494 | 0.169 | 0.755 | 0.023 | 0.001 | 0.001 | 0.955 | 0.7 | 0.27 | 0.59 | 0.7 | 0.81 | 0.79 |
| 5 | 81172913 | rs12652012 | -1.221 | -0.22 | 0.122 | 0.764 | 0.01 | -0.004 | -0.004 | 0 | 0.38 | 0.62 | 0.3 | 0.41 | 0.33 | 0.31 |
| 5 | 81174174 | rs2897524 | -0.139 | -0.041 | 0.434 | 0.634 | -0.036 | 0.034 | 0.034 | 1.84 | 0.09 | 0.3 | 0.05 | 0.08 | 0.01 | 0.03 |
| 5 | 81176570 | rs12186353 | -0.249 | 1.005 | -0.78 | 0.522 | -0.055 | -0.048 | -0.048 | 0.393 | 0.14 | 0.01 | 0.24 | 0.19 | 0.22 | 0.31 |
| 5 | 81179751 | rs6886425 | -1.564 | 1.333 | 0.073 | -0.222 | 0.042 | -0.109 | -0.109 | 2.275 | 0.19 | 0.43 | 0.13 | 0.04 | 0.18 | 0.19 |
| 5 | 81185254 | rs6880017 | -1.436 | 1.324 | -0.406 | -0.265 | -0.012 | -0.024 | -0.024 | 7.302 | 0.58 | 0.78 | 0.68 | 0.58 | 0.6 | 0.75 |
| 5 | 81188142 | rs11745641 | -0.828 | 1.306 | 0.006 | -0.267 | 0.044 | -0.091 | -0.091 | 0 | 0.15 | 0.57 | 0.11 | 0.01 | 0.15 | 0.12 |
| 5 | 81203203 | rs6881638 | -0.181 | 1.032 | -0.194 | -0.386 | -0.022 | -0.078 | -0.078 | 0 | 0.41 | 0.11 | 0.56 | 0.57 | 0.46 | 0.62 |
| 5 | 81203695 | rs12658320 | -0.821 | 1.161 | 0.412 | -0.62 | 0.002 | 0.001 | 0.001 | 0 | 0.43 | 0.45 | 0.57 | 0.58 | 0.47 | 0.65 |
| 5 | 81203903 | rs1001797 | 0.008 | 1.213 | 0.224 | -0.586 | -0.009 | -0.094 | -0.094 | 0 | 0.35 | 0.11 | 0.51 | 0.57 | 0.38 | 0.58 |
| 5 | 81207272 | rs1566629 | -1.017 | 1.152 | 0.403 | -0.616 | 0.02 | -0.002 | -0.002 | 0 | 0.47 | 0.27 | 0.59 | 0.58 | 0.49 | 0.67 |
| 5 | 81214775 | rs611805 | -0.877 | 1.085 | 0.26 | -0.627 | 0.01 | -0.021 | -0.021 | 0 | 0.45 | 0.19 | 0.59 | 0.57 | 0.49 | 0.67 |
| 5 | 81217666 | rs655680 | -0.039 | 1.148 | 0.259 | -0.635 | 0.009 | -0.025 | -0.025 | 0 | 0.05 | 0.15 | 0.03 | 0 | 0.04 | 0.05 |
| 5 | 81219261 | rs9293284 | 0.121 | 1.138 | 0.249 | -0.635 | -0.055 | -0.17 | -0.17 | 0.304 | 0.34 | 0.03 | 0.5 | 0.57 | 0.38 | 0.58 |
| 5 | 81220187 | rs687543 | -0.693 | 1.147 | 0.31 | -0.604 | -0.001 | -0.053 | -0.053 | 0 | 0.12 | 0.16 | 0.09 | 0 | 0.12 | 0.09 |
| 5 | 81221223 | rs542449 | -1.664 | 1.107 | 0.32 | -0.598 | -0.02 | -0.019 | -0.019 | 0.057 | 0.57 | 0.77 | 0.66 | 0.58 | 0.56 | 0.7 |
| 5 | 81232844 | rs4410626 | 0.287 | 1.061 | 0.541 | -0.637 | -0.006 | -0.053 | -0.053 | 0.464 | 0.9 | 0.83 | 0.93 | 1 | 0.92 | 0.95 |
| 5 | 81233027 | rs13184971 | 0.009 | 1.112 | 0.904 | -0.676 | 0.135 | -0.021 | -0.021 | 0 | 0.08 | 0.38 | 0.06 | 0.01 | 0.1 | 0.07 |
| 5 | 81234145 | rs4703857 | 1.543 | 1.106 | 0.66 | -0.664 | 0.044 | -0.126 | -0.126 | 0 | 0.81 | 0.46 | 0.87 | 1 | 0.82 | 0.88 |
| 5 | 81234311 | rs4703859 | 1.543 | 1.106 | 0.66 | -0.664 | 0.044 | -0.126 | -0.126 | 0.397 | 0.81 | 0.46 | 0.87 | 1 | 0.82 | 0.88 |
| 5 | 81247381 | rs1542596 | 1.585 | 1.042 | 0.527 | -0.554 | -0.018 | -0.025 | -0.025 | 0 | 0.42 | 0.26 | 0.34 | 0.42 | 0.43 | 0.31 |

| | | | | | | | | | | | | | | | | |
|---|---|---|---|---|---|---|---|---|---|---|---|---|---|---|---|---|
| 5 | 81253212 | rs2101146 | 1.289 | 0.796 | 0.971 | -0.669 | -0.021 | -0.086 | -0.086 | 0 | 0.87 | 0.58 | 0.91 | 1 | 0.92 | 0.97 |
| 5 | 81255228 | rs17243501 | 0.706 | 0.761 | -0.026 | -0.669 | -0.026 | 0.03 | 0.03 | 0 | 0.18 | 0 | 0.13 | 0.11 | 0.19 | 0.22 |
| 5 | 81260121 | rs2406905 | 1.687 | 0.69 | -0.105 | -0.612 | 0.044 | 0.138 | 0.138 | 0 | 0.54 | 0.23 | 0.68 | 0.75 | 0.6 | 0.5 |
| 5 | 81264006 | rs6898784 | -0.865 | 0.624 | 0.561 | -0.608 | -0.024 | -0.039 | -0.039 | 0 | 0.09 | 0.18 | 0.04 | 0.01 | 0.05 | 0.05 |
| 5 | 81297077 | rs6875409 | 3.099 | 1.277 | -0.305 | -1.275 | 0.005 | 0.123 | 0.123 | 0 | 0.29 | 0.11 | 0.14 | 0.01 | 0.23 | 0.16 |
| 5 | 81307412 | rs10514231 | -1.573 | 1.291 | 0.073 | -1.414 | 0.038 | 0.101 | 0.101 | 0 | 0.54 | 0.3 | 0.71 | 0.9 | 0.65 | 0.74 |
| 5 | 81312256 | rs2406911 | -1.551 | 1.241 | 0.16 | -1.478 | 0.077 | 0.157 | 0.157 | 0.323 | 0.53 | 0.24 | 0.75 | 0.99 | 0.67 | 0.77 |
| 5 | 81324365 | rs6892261 | 0.277 | 1.444 | 0.459 | -1.457 | -0.013 | -0.008 | -0.008 | 0 | 0.22 | 0.35 | 0.18 | 0.22 | 0.2 | 0.35 |
| 5 | 81327042 | rs1485587 | 1.408 | 1.379 | 0.357 | -1.484 | -0.005 | 0.008 | 0.008 | 0 | 0.61 | 0.64 | 0.37 | 0.23 | 0.48 | 0.5 |
| 5 | 81354389 | rs3734114 | 0.655 | 1.398 | 0.047 | -1.329 | -0.001 | -0.007 | -0.007 | 0 | 0.2 | 0.06 | 0.16 | 0.22 | 0.2 | 0.35 |
| 5 | 81362480 | rs11748868 | 1.022 | 1.533 | 0.012 | -1.295 | -0.034 | 0.015 | 0.015 | 0.271 | 0.06 | 0 | 0.04 | 0.02 | 0.09 | 0.07 |
| 5 | 81398182 | rs901135 | -1.182 | 1.485 | 0.101 | -1.345 | -0.008 | -0.031 | -0.031 | 0.025 | 0.94 | 0.9 | 0.96 | 1 | 0.95 | 0.97 |
| 5 | 81407916 | rs1835112 | 0.565 | 1.577 | -0.128 | -0.965 | -0.057 | -0.123 | -0.123 | 0 | 0.17 | 0.02 | 0.32 | 0.38 | 0.2 | 0.23 |
| 5 | 81416305 | rs178957 | 3.341 | 1.419 | -0.192 | -1.628 | 0.014 | 0.107 | 0.107 | 0 | 0.27 | 0.1 | 0.11 | 0.01 | 0.2 | 0.15 |
| 5 | 81480371 | rs4703537 | -1.339 | 1.852 | 0.083 | -1.967 | 0.036 | 0.116 | 0.116 | 0 | 0.52 | 0.29 | 0.65 | 0.9 | 0.59 | 0.73 |
| 5 | 81490277 | rs1109846 | -2.077 | 1.826 | 0.259 | -1.888 | -0.007 | -0.043 | -0.043 | 0 | 0.18 | 0.52 | 0.18 | 0.1 | 0.18 | 0.19 |
| 5 | 81511286 | rs16899425 | -0.756 | 1.814 | 0.273 | -1.868 | 0.048 | 0.06 | 0.06 | 0 | 0.09 | 0.41 | 0.11 | 0.09 | 0.08 | 0.13 |
| 5 | 81515340 | rs9293293 | 0.446 | 1.955 | -0.136 | -1.976 | -0.055 | 0.163 | 0.163 | 0 | 0.59 | 0.83 | 0.43 | 0.12 | 0.54 | 0.37 |
| 5 | 81515446 | rs4703871 | -0.968 | 1.973 | -0.122 | -1.965 | 0.019 | 0.032 | 0.032 | 0 | 0.8 | 0.91 | 0.89 | 0.86 | 0.84 | 0.71 |
| 5 | 81527262 | rs17245874 | 0.905 | 1.959 | -0.124 | -2.004 | -0.048 | -0.097 | -0.097 | 0 | 0.14 | 0.02 | 0.28 | 0.34 | 0.16 | 0.21 |
| 5 | 81533735 | rs6888977 | -3.222 | 1.704 | 0.145 | -2.598 | 0.012 | 0.119 | 0.119 | 0 | 0.72 | 0.89 | 0.85 | 1 | 0.77 | 0.92 |
| 5 | 81537176 | rs6870406 | -2.008 | 1.713 | 0.439 | -2.579 | -0.013 | -0.056 | -0.056 | 0.624 | 0.19 | 0.55 | 0.19 | 0.1 | 0.17 | 0.19 |
| 5 | 81549216 | rs1864183 | -0.329 | 1.74 | 0.234 | -2.599 | -0.054 | 0.174 | 0.174 | 0 | 0.41 | 0.17 | 0.57 | 0.89 | 0.46 | 0.63 |
| 5 | 81549240 | rs1864182 | -0.611 | 1.753 | 0.216 | -2.738 | 0.007 | 0.232 | 0.232 | 0 | 0.46 | 0.18 | 0.61 | 0.91 | 0.55 | 0.7 |
| 5 | 81550043 | rs6884232 | -3.218 | 1.739 | 0.222 | -2.753 | 0.013 | 0.128 | 0.128 | 0 | 0.71 | 0.9 | 0.86 | 0.99 | 0.77 | 0.85 |
| 5 | 81550741 | rs4703878 | -0.334 | 1.809 | 0.136 | -2.732 | -0.054 | 0.183 | 0.183 | 0 | 0.41 | 0.17 | 0.57 | 0.9 | 0.47 | 0.63 |
| 5 | 81555751 | rs10070719 | -1.975 | 1.498 | 0.295 | -2.677 | -0.012 | -0.056 | -0.056 | 0 | 0.19 | 0.55 | 0.19 | 0.09 | 0.17 | 0.12 |
| 5 | 81559412 | rs12652358 | -0.608 | 1.369 | -0.431 | -2.574 | 0.018 | 0.232 | 0.232 | 0 | 0.46 | 0.18 | 0.63 | 0.91 | 0.57 | 0.7 |
| 5 | 81571669 | rs226202 | -2.184 | 1.182 | -0.481 | -2.355 | -0.006 | -0.051 | -0.051 | 0 | 0.67 | 0.64 | 0.79 | 0.9 | 0.7 | 0.78 |
| 5 | 81574182 | rs6880209 | 2.065 | 1.182 | -0.482 | -2.355 | -0.006 | -0.051 | -0.051 | 0 | 0.33 | 0.36 | 0.21 | 0.09 | 0.3 | 0.14 |
| 5 | 81588011 | rs862241 | -2.185 | 1.199 | -0.501 | -2.347 | -0.008 | -0.065 | -0.065 | 1.402 | 0.67 | 0.61 | 0.78 | 0.91 | 0.7 | 0.86 |
| 5 | 81593600 | rs701405 | 0.391 | 1.487 | -0.795 | -2.232 | -0.047 | 0.001 | 0.001 | 0 | 0.22 | 0.01 | 0.16 | 0.16 | 0.22 | 0.29 |
| 5 | 81595084 | rs2215128 | -0.273 | 1.363 | -0.968 | -2.33 | -0.066 | 0.011 | 0.011 | 0 | 0.29 | 0.11 | 0.51 | 0.74 | 0.35 | 0.4 |
| 5 | 81595570 | rs862239 | 1.284 | 1.272 | -0.827 | -2.299 | -0.007 | 0.007 | 0.007 | 0 | 0.16 | 0.06 | 0.12 | 0.14 | 0.18 | 0.32 |
| 5 | 81601575 | rs224821 | 0.539 | 0.877 | -0.44 | -2.437 | 0.022 | 0.286 | 0.286 | 0 | 0.49 | 0.85 | 0.32 | 0.1 | 0.43 | 0.32 |
| 5 | 81605353 | rs224823 | -0.559 | 0.878 | -0.472 | -2.437 | 0.015 | 0.31 | 0.31 | 0 | 0.5 | 0.13 | 0.66 | 0.9 | 0.57 | 0.68 |
| 5 | 81612479 | rs658912 | 0.916 | 0.815 | -0.042 | -2.461 | -0.004 | -0.11 | -0.11 | 0.269 | 0.43 | 0.55 | 0.25 | 0.02 | 0.36 | 0.26 |
| 5 | 81615350 | rs10514237 | 0.066 | 1.016 | -0.307 | -2.148 | 0.027 | -0.024 | -0.024 | 0 | 0.05 | 0.09 | 0.05 | 0.02 | 0.08 | 0.15 |
| 5 | 81619915 | rs1071181 | -0.618 | 0.871 | -0.791 | -2.394 | -0.004 | 0.306 | 0.306 | 0 | 0.49 | 0.12 | 0.67 | 0.9 | 0.56 | 0.7 |
| 5 | 81630324 | rs6867982 | 1.45 | 0.802 | -0.324 | -2.433 | 0.016 | 0.011 | 0.011 | 0 | 0.35 | 0.31 | 0.2 | 0.01 | 0.29 | 0.13 |
| 5 | 81632928 | rs224900 | -0.463 | 0.747 | -0.756 | -2.437 | -0.012 | 0.299 | 0.299 | 0 | 0.48 | 0.11 | 0.66 | 0.9 | 0.55 | 0.7 |
| 5 | 81634599 | rs9293301 | 0.844 | 0.775 | -0.839 | -2.493 | -0.013 | 0.034 | 0.034 | 0 | 0.08 | 0.01 | 0.05 | 0.03 | 0.07 | 0.06 |
| 5 | 81637621 | rs1354123 | 0.534 | 0.817 | -0.353 | -2.281 | 0.023 | 0.135 | 0.135 | 0 | 0.46 | 0.76 | 0.31 | 0.1 | 0.4 | 0.28 |
| 5 | 81652773 | rs17247678 | 2.126 | 0.509 | -0.95 | -2.313 | 0.03 | 0.099 | 0.099 | 0 | 0.27 | 0.09 | 0.16 | 0 | 0.23 | 0.08 |
| 5 | 81653481 | rs224912 | 0.656 | 0.823 | -1.231 | -2.03 | -0.005 | -0.045 | -0.045 | 0 | 0.18 | 0.08 | 0.35 | 0.36 | 0.17 | 0.28 |
| 5 | 81669092 | rs224836 | 0.298 | 0.416 | -0.598 | -1.892 | -0.037 | 0.291 | 0.291 | 0 | 0.48 | 0.11 | 0.65 | 0.9 | 0.53 | 0.69 |
| 5 | 81674497 | rs7717296 | -1.477 | 0.542 | -0.504 | -1.797 | -0.003 | 0.007 | 0.007 | 2.547 | 0.21 | 0.78 | 0.15 | 0.1 | 0.19 | 0.2 |
| 5 | 81686797 | rs224808 | -0.823 | -0.892 | -0.333 | -2.803 | 0.032 | 0.031 | 0.031 | 0 | 0.65 | 0.74 | 0.8 | 1 | 0.72 | 0.89 |
| 5 | 81696698 | rs12109149 | -0.795 | 0.408 | -0.707 | -1.695 | -0.004 | -0.001 | -0.001 | 0.161 | 0.07 | 0.08 | 0.04 | 0.08 | 0.12 | 0.06 |
| 5 | 81698717 | rs224961 | -0.305 | -0.127 | -0.913 | -1.634 | -0.05 | 0.218 | 0.218 | 0 | 0.43 | 0.16 | 0.66 | 0.91 | 0.52 | 0.74 |
| 5 | 81710048 | rs224948 | 0.006 | 0.482 | -1.292 | -1.212 | -0.069 | 0.213 | 0.213 | 0.343 | 0.4 | 0.14 | 0.66 | 0.9 | 0.52 | 0.7 |
| 5 | 81714463 | rs224945 | 0.756 | 0.209 | 0.333 | -0.341 | -0.019 | -0.08 | -0.08 | 0.039 | 0.88 | 0.59 | 0.93 | 1 | 0.94 | 0.97 |
| 5 | 81714913 | rs10491245 | -1.1 | 0.745 | 0.272 | -0.164 | 0.026 | 0.005 | 0.005 | 0 | 0.08 | 0.2 | 0.04 | 0.1 | 0.08 | 0.09 |
| 5 | 81716027 | rs10491244 | -0.206 | 0.195 | -0.194 | -0.103 | 0.01 | 0.009 | 0.009 | 0.258 | 0.31 | 0.19 | 0.32 | 0.27 | 0.21 | 0.29 |
| 5 | 81717102 | rs224939 | 0.367 | 0.647 | -1.058 | 0.276 | -0.097 | -0.054 | -0.054 | 0 | 0.28 | 0.05 | 0.25 | 0.32 | 0.4 | 0.53 |
| 5 | 81717176 | rs224938 | -0.174 | 0.472 | 0.029 | 0.647 | -0.003 | 0.001 | 0.001 | 1.578 | 0.37 | 0.36 | 0.54 | 0.66 | 0.49 | 0.74 |
| 5 | 81717561 | rs1008570 | -0.562 | 0.451 | 0.115 | 0.747 | 0.011 | 0.233 | 0.233 | 0.624 | 0.1 | 0.31 | 0.29 | 0.35 | 0.09 | 0.21 |
| 5 | 81736795 | rs2429239 | -0.56 | -0.101 | 0.378 | 0.648 | -0.012 | -0.009 | -0.009 | 0 | 0.7 | 0.44 | 0.73 | 0.72 | 0.72 | 0.87 |
| 5 | 81738908 | rs224868 | -1.054 | -0.203 | 1.037 | 0.61 | 0 | 0.001 | 0.001 | 0 | 0.75 | 0.77 | 0.75 | 0.74 | 0.72 | 0.88 |
| 5 | 81739666 | rs917385 | 0.928 | -0.131 | 0.975 | 0.583 | 0.008 | 0.064 | 0.064 | 0 | 0.06 | 0.12 | 0.13 | 0.15 | 0.08 | 0.03 |
| 5 | 81743961 | rs224869 | 0.5 | 0.294 | 0.748 | 0.389 | -0.079 | -0.004 | -0.004 | 0.057 | 0.35 | 0.09 | 0.29 | 0.32 | 0.44 | 0.54 |
| 5 | 81764535 | rs523566 | 0.194 | 0.552 | 0.505 | 0.272 | 0.01 | -0.044 | -0.044 | 2.2 | 0.29 | 0.18 | 0.31 | 0.42 | 0.32 | 0.43 |
| 5 | 81771251 | rs224857 | -0.685 | 0.478 | 0.673 | 0.276 | -0.006 | -0.007 | -0.007 | 0.013 | 0.43 | 0.4 | 0.43 | 0.46 | 0.52 | 0.57 |
| 5 | 81776926 | rs224846 | -0.681 | 0.544 | 0.732 | 0.355 | -0.011 | 0.052 | 0.052 | 0.375 | 0.71 | 0.47 | 0.58 | 0.52 | 0.72 | 0.69 |
| 5 | 81788024 | rs6894206 | 1.108 | 0.524 | 0.872 | 0.35 | -0.007 | -0.01 | -0.01 | 0.004 | 0.25 | 0.33 | 0.19 | 0.22 | 0.24 | 0.12 |
| 5 | 81807074 | rs6872917 | 0.217 | 0.639 | 0.651 | 0.144 | 0 | -0.001 | -0.001 | 0 | 0.63 | 0.62 | 0.45 | 0.32 | 0.6 | 0.36 |
| 5 | 81809458 | rs6452462 | -1.091 | 0.621 | 0.483 | 0.239 | 0.001 | -0.053 | -0.053 | 0.002 | 0.77 | 0.95 | 0.82 | 0.71 | 0.76 | 0.59 |
| 5 | 81826276 | rs17249159 | 1.593 | 0.695 | 0.363 | 0.083 | -0.036 | 0.137 | 0.137 | 1.945 | 0.28 | 0.02 | 0.14 | 0.05 | 0.2 | 0.11 |
| 5 | 81829065 | rs13189857 | 0.933 | 0.812 | 0.523 | 0.165 | -0.038 | 0.024 | 0.024 | 0 | 0.1 | 0.03 | 0.12 | 0.05 | 0.2 | 0.15 |

| | | | | | | | | | | | | | | | | |
|---|---|---|---|---|---|---|---|---|---|---|---|---|---|---|---|---|
| 5 | 81832369 | rs6452463 | 0.794 | 0.807 | 0.557 | 0.162 | -0.008 | 0.008 | 0.008 | 0 | 0.11 | 0.07 | 0.12 | 0.05 | 0.2 | 0.15 |
| 5 | 81837045 | rs17249504 | 0.615 | 0.829 | 0.593 | 0.162 | 0.02 | -0.011 | -0.011 | 0.158 | 0.11 | 0.17 | 0.12 | 0.05 | 0.2 | 0.14 |
| 5 | 81840727 | rs7713277 | -0.333 | 0.77 | 0.489 | 0.507 | -0.005 | -0.066 | -0.066 | 0 | 0.7 | 0.9 | 0.69 | 0.61 | 0.65 | 0.56 |
| 5 | 81843333 | rs10462380 | 1.039 | 0.661 | 0.753 | 0.467 | -0.005 | -0.01 | -0.01 | 0.195 | 0.24 | 0.34 | 0.19 | 0.22 | 0.24 | 0.12 |
| 5 | 81847115 | rs17309089 | 0.921 | 0.73 | 0.458 | 0.469 | -0.04 | 0.025 | 0.025 | 0 | 0.11 | 0.04 | 0.12 | 0.05 | 0.2 | 0.14 |
| 5 | 81848595 | rs10474053 | 0.733 | 0.624 | 0.626 | 0.484 | -0.016 | -0.022 | -0.022 | 0 | 0.26 | 0.47 | 0.21 | 0.22 | 0.24 | 0.12 |
| 5 | 81849746 | rs9293305 | -0.884 | 0.536 | -0.115 | 0.529 | -0.013 | -0.054 | -0.054 | 7.62 | 0.76 | 0.98 | 0.82 | 0.75 | 0.76 | 0.57 |
| 5 | 81849888 | rs2891895 | -0.291 | 0.718 | 0.135 | 0.564 | -0.005 | -0.001 | -0.001 | 7.935 | 0.15 | 0.14 | 0.32 | 0.34 | 0.21 | 0.31 |
| 5 | 81855572 | rs4703894 | -0.568 | 0.527 | 0.365 | 0.18 | -0.014 | 0.01 | 0.01 | 0 | 0.6 | 0.62 | 0.73 | 0.93 | 0.6 | 0.74 |
| 5 | 81856537 | rs905221 | -0.388 | 0.444 | 0.398 | 0.525 | -0.006 | 0.005 | 0.005 | 0 | 0.57 | 0.59 | 0.53 | 0.6 | 0.57 | 0.57 |
| 5 | 81857794 | rs1121960 | 0.216 | 0.255 | -0.031 | 0.596 | 0.014 | -0.05 | -0.05 | 0 | 0.3 | 0.13 | 0.32 | 0.39 | 0.33 | 0.46 |
| 5 | 81870084 | rs7729839 | 1.399 | 0.292 | -0.027 | 0.613 | -0.001 | 0.016 | 0.016 | 0 | 0.05 | 0.07 | 0.13 | 0.12 | 0.08 | 0.04 |
| 5 | 81873473 | rs1847051 | 1.417 | 0.31 | -0.09 | 0.376 | 0.006 | 0.122 | 0.122 | 1.861 | 0.28 | 0.09 | 0.14 | 0.03 | 0.2 | 0.11 |
| 5 | 81882182 | rs905219 | -0.519 | 0.296 | -0.213 | 0.726 | 0.01 | -0.054 | -0.054 | 0 | 0.73 | 0.89 | 0.74 | 0.64 | 0.72 | 0.62 |
| 5 | 81883470 | rs1847052 | 1.664 | 0.215 | 0.053 | 0.613 | 0 | -0.012 | -0.012 | 0 | 0.17 | 0.11 | 0.3 | 0.31 | 0.19 | 0.23 |
| 5 | 81892187 | rs17309918 | 0.999 | 0.008 | 0.198 | 0.271 | 0.031 | 0.112 | 0.112 | 1.036 | 0.34 | 0.16 | 0.16 | 0.04 | 0.23 | 0.14 |
| 5 | 81893325 | rs7720861 | 1.044 | 0.422 | 0.447 | 0.43 | 0.004 | 0 | 0 | 0 | 0.1 | 0.11 | 0.15 | 0.07 | 0.19 | 0.12 |
| 5 | 81896198 | rs17250714 | 1.519 | 0.047 | 0.37 | 0.209 | -0.036 | 0.148 | 0.148 | 1.325 | 0.29 | 0.01 | 0.14 | 0.04 | 0.2 | 0.12 |
| 5 | 81896956 | rs1847035 | -0.391 | 0.034 | 0.679 | 0.57 | 0.015 | -0.008 | -0.008 | 0 | 0.72 | 0.84 | 0.74 | 0.7 | 0.72 | 0.62 |
| 5 | 81898168 | rs10072866 | 0.373 | -0.018 | 0.362 | 0.434 | -0.001 | -0.004 | -0.004 | 0.787 | 0.54 | 0.47 | 0.5 | 0.62 | 0.43 | 0.45 |
| 5 | 81917419 | rs10039290 | 1.043 | -0.101 | -0.272 | 0.478 | -0.001 | 0.028 | 0.028 | 0 | 0.22 | 0.26 | 0.35 | 0.33 | 0.23 | 0.3 |
| 5 | 81920484 | rs6876647 | 0.041 | -0.789 | -0.673 | -0.126 | 0.05 | 0.036 | 0.036 | 0.009 | 0.47 | 0.72 | 0.54 | 0.66 | 0.35 | 0.46 |
| 5 | 81921045 | rs11744977 | 1.808 | -0.67 | -1.241 | -0.132 | -0.024 | 0.046 | 0.046 | 0 | 0.16 | 0.01 | 0.08 | 0.05 | 0.13 | 0.08 |
| 5 | 81929609 | rs6452470 | -0.266 | -0.46 | -0.96 | -0.131 | 0.004 | 0.021 | 0.021 | 0 | 0.61 | 0.66 | 0.69 | 0.73 | 0.61 | 0.55 |
| 5 | 81937093 | rs17283841 | 2.257 | -0.442 | -1.051 | 0.065 | -0.001 | -0.037 | -0.037 | 0.325 | 0.2 | 0.09 | 0.33 | 0.31 | 0.21 | 0.31 |
| 5 | 81938017 | rs4703905 | 0.253 | -0.3 | -0.624 | 0.216 | -0.062 | 0.042 | 0.042 | 0.793 | 0.23 | 0.11 | 0.26 | 0.16 | 0.34 | 0.19 |
| 5 | 81938623 | rs1847040 | -1.12 | -0.217 | -0.393 | 0.593 | -0.01 | -0.028 | -0.028 | 0.348 | 0.82 | 0.94 | 0.83 | 0.77 | 0.79 | 0.69 |
| 5 | 81939318 | rs1032757 | 0.431 | 0.116 | 0.403 | 0.464 | -0.039 | 0.091 | 0.091 | 1.648 | 0.16 | 0.38 | 0.1 | 0.26 | 0.06 | 0.07 |
| 5 | 81944343 | rs7712336 | -0.598 | 0.996 | -0.082 | 0.364 | 0.043 | 0.002 | 0.002 | 0 | 0.29 | 0.55 | 0.3 | 0.31 | 0.37 | 0.25 |
| 5 | 81946255 | rs12658345 | 0.392 | 0.363 | -0.379 | 0.34 | -0.001 | -0.005 | -0.005 | 0.152 | 0.31 | 0.3 | 0.43 | 0.51 | 0.34 | 0.35 |
| 5 | 81951284 | rs16899808 | -0.347 | 0.039 | -0.052 | 0.137 | -0.036 | -0.041 | -0.041 | 0 | 0.25 | 0.36 | 0.18 | 0.13 | 0.17 | 0.13 |
| 5 | 81953043 | rs924619 | 1.517 | 0.137 | 0.041 | 0.284 | 0 | 0.004 | 0.004 | 0 | 0.21 | 0.19 | 0.33 | 0.41 | 0.22 | 0.29 |
| 5 | 81953183 | rs2015915 | -1.611 | 0.751 | -0.593 | -0.169 | -0.007 | 0.025 | 0.025 | 0 | 0.67 | 0.86 | 0.76 | 0.69 | 0.66 | 0.65 |
| 5 | 81955591 | rs924615 | 0.46 | 0.692 | -0.072 | -0.205 | -0.019 | 0.017 | 0.017 | 0 | 0.36 | 0.49 | 0.38 | 0.42 | 0.28 | 0.32 |
| 5 | 81957124 | rs17283890 | 1.086 | 1.238 | -0.481 | 0.008 | -0.042 | -0.02 | -0.02 | 0 | 0.07 | 0.02 | 0.16 | 0.09 | 0.18 | 0.14 |
| 5 | 81958134 | rs10805810 | 0.025 | 0.58 | -0.123 | -0.226 | -0.001 | -0.001 | -0.001 | 0 | 0.4 | 0.43 | 0.6 | 0.59 | 0.51 | 0.47 |
| 5 | 81958653 | rs11749713 | 1.354 | 0.602 | -0.731 | -0.102 | -0.007 | -0.023 | -0.023 | 1.84 | 0.06 | 0 | 0.03 | 0.06 | 0.04 | 0.06 |
| 5 | 81960140 | rs4703545 | 0.248 | 0.601 | -0.625 | -0.141 | -0.068 | -0.026 | -0.026 | 0 | 0.36 | 0.19 | 0.51 | 0.57 | 0.44 | 0.49 |
| 5 | 81962506 | rs11740383 | -0.173 | -0.43 | 0.005 | -0.694 | -0.06 | -0.069 | -0.069 | 0 | 0.23 | 0.39 | 0.1 | 0.06 | 0.08 | 0.07 |
| 5 | 81962842 | rs2055437 | -0.617 | -0.236 | -0.065 | -1.013 | -0.021 | -0.071 | -0.071 | 1.29 | 0.71 | 0.63 | 0.81 | 0.93 | 0.75 | 0.78 |
| 5 | 81966957 | rs745648 | -1.363 | 0.216 | -0.242 | -1.074 | -0.016 | 0.12 | 0.12 | 0 | 0.58 | 0.77 | 0.63 | 0.91 | 0.47 | 0.6 |
| 5 | 81968863 | rs9791094 | -1.632 | 0.41 | -0.215 | -1.136 | -0.015 | 0.06 | 0.06 | 0 | 0.62 | 0.73 | 0.69 | 0.91 | 0.57 | 0.64 |
| 5 | 81971113 | rs4703909 | 0.118 | 0.427 | -0.214 | -1.001 | -0.006 | 0 | 0 | 0 | 0.08 | 0.06 | 0.11 | 0.08 | 0.1 | 0.09 |
| 5 | 81975108 | rs1827396 | -1.012 | 0.204 | -0.036 | -0.747 | 0.015 | 0.087 | 0.087 | 0 | 0.53 | 0.67 | 0.53 | 0.83 | 0.46 | 0.6 |
| 5 | 81979166 | rs1847046 | -0.997 | 0.173 | 0.43 | -0.401 | -0.012 | -0.004 | -0.004 | 2.525 | 0.33 | 0.55 | 0.49 | 0.71 | 0.29 | 0.31 |
| 5 | 81983638 | rs11959945 | -0.367 | -0.468 | 0.418 | -0.009 | -0.026 | 0.035 | 0.035 | 0 | 0.21 | 0.32 | 0.39 | 0.71 | 0.16 | 0.29 |
| 5 | 81988068 | rs1971334 | -0.776 | -0.105 | -0.17 | 0.143 | 0 | -0.001 | -0.001 | 0 | 0.18 | 0.18 | 0.15 | 0.17 | 0.16 | 0.17 |
| 5 | 81989527 | rs7449153 | -1.34 | -0.106 | -0.213 | -0.278 | 0.028 | 0.062 | 0.062 | 0 | 0.45 | 0.65 | 0.55 | 0.87 | 0.34 | 0.47 |
| 5 | 81996913 | rs7722067 | 0.29 | -0.164 | 0.612 | 0.862 | -0.037 | 0.039 | 0.039 | 0.567 | 0.13 | 0.31 | 0.21 | 0.2 | 0.06 | 0.03 |
| 5 | 81998898 | rs16899857 | -0.276 | 0.544 | 0.961 | 1.802 | 0.026 | 0.116 | 0.116 | 0.025 | 0.09 | 0.17 | 0.17 | 0.29 | 0.13 | 0.15 |
| 5 | 82002186 | rs7701376 | -2.03 | 0.493 | 1.053 | 1.704 | -0.001 | 0.06 | 0.06 | 0.024 | 0.32 | 0.49 | 0.33 | 0.47 | 0.32 | 0.44 |
| 5 | 82002782 | rs1345808 | -2.06 | 0.487 | 1.094 | 1.701 | -0.002 | 0.042 | 0.042 | 0.095 | 0.33 | 0.57 | 0.33 | 0.47 | 0.32 | 0.42 |
| 5 | 82007207 | rs355289 | 0.097 | 0.657 | 0.945 | 1.726 | 0.035 | 0.073 | 0.073 | 0 | 0.91 | 0.8 | 0.87 | 0.76 | 0.88 | 0.87 |
| 5 | 82012854 | rs4703911 | 0.336 | 0.613 | 0.977 | 1.703 | 0.043 | 0.072 | 0.072 | 0 | 0.89 | 0.72 | 0.85 | 0.75 | 0.85 | 0.87 |
| 5 | 82020118 | rs6452479 | -1.051 | -0.003 | 1.029 | 1.544 | -0.021 | -0.05 | -0.05 | 0 | 0.13 | 0.26 | 0.06 | 0.02 | 0.08 | 0.07 |
| 5 | 82020796 | rs12054791 | -1.481 | -0.018 | 1.084 | 1.723 | -0.022 | 0.049 | 0.049 | 0 | 0.14 | 0.32 | 0.23 | 0.22 | 0.11 | 0.09 |
| 5 | 82020844 | rs12054792 | -1.418 | -0.002 | 1.091 | 1.73 | -0.021 | 0.052 | 0.052 | 0 | 0.14 | 0.32 | 0.23 | 0.22 | 0.11 | 0.09 |
| 5 | 82024079 | rs12188665 | -1.208 | -0.017 | 0.978 | 1.533 | -0.024 | 0.012 | 0.012 | 0.242 | 0.14 | 0.37 | 0.1 | 0.12 | 0.08 | 0.2 |
| 5 | 82024353 | rs11960525 | -0.555 | 0.162 | 0.671 | 1.828 | 0.034 | 0.185 | 0.185 | 0 | 0.09 | 0.22 | 0.17 | 0.42 | 0.13 | 0.15 |
| 5 | 82025104 | rs7703068 | -1.164 | 0.05 | 0.667 | 1.712 | 0.022 | 0.114 | 0.114 | 0.473 | 0.16 | 0.27 | 0.19 | 0.43 | 0.19 | 0.24 |
| 5 | 82028130 | rs9293311 | 0.063 | -0.848 | -0.512 | 0.448 | 0.043 | 0.005 | 0.005 | 0 | 0.49 | 0.74 | 0.68 | 0.58 | 0.59 | 0.37 |
| 5 | 82031196 | rs2113024 | 0.877 | -0.724 | -0.833 | 0.168 | -0.061 | 0.037 | 0.037 | 0 | 0.34 | 0.16 | 0.28 | 0.24 | 0.43 | 0.25 |
| 5 | 82032840 | rs1465444 | 0.928 | -0.387 | -0.643 | 0.205 | 0.058 | -0.057 | -0.057 | 0 | 0.11 | 0.24 | 0.28 | 0 | 0.16 | 0.09 |
| 5 | 82044105 | rs10514243 | 0.956 | -0.755 | -1.131 | -0.155 | 0 | 0.01 | 0.01 | 0 | 0.14 | 0.12 | 0.07 | 0.01 | 0.16 | 0.08 |
| 5 | 82044554 | rs2113025 | -0.596 | -1.014 | -1.755 | -0.611 | -0.033 | 0.068 | 0.068 | 0 | 0.57 | 0.79 | 0.4 | 0.29 | 0.48 | 0.5 |
| 5 | 82047695 | rs17205439 | 1.462 | -1.041 | -1.724 | -0.615 | 0.039 | 0.043 | 0.043 | 0 | 0.22 | 0.12 | 0.12 | 0.06 | 0.1 | 0.15 |
| 5 | 82052937 | rs12517454 | 1.775 | -0.82 | -1.852 | -0.483 | 0.021 | 0.062 | 0.062 | 0 | 0.11 | 0.16 | 0.37 | 0.36 | 0.17 | 0.11 |
| 5 | 82053933 | rs2386072 | 0.343 | -1.055 | -1.91 | -0.624 | -0.013 | 0.016 | 0.016 | 0 | 0.66 | 0.49 | 0.78 | 0.61 | 0.72 | 0.5 |
| 5 | 82054744 | rs10942308 | 0.668 | -1.09 | -2.231 | -0.675 | 0.003 | 0.051 | 0.051 | 0 | 0.43 | 0.21 | 0.58 | 0.37 | 0.6 | 0.28 |

| | | | | | | | | | | | | | | | | |
|---|---|---|---|---|---|---|---|---|---|---|---|---|---|---|---|---|
| 5 | 82055864 | rs6863549 | 1.455 | -1.259 | -2.007 | -0.411 | 0.037 | 0.043 | 0.043 | 0 | 0.21 | 0.12 | 0.12 | 0.06 | 0.1 | 0.15 |
| 5 | 82057045 | rs13165887 | 1.986 | 0.041 | -1.514 | 0.132 | -0.093 | 0.116 | 0.116 | 0 | 0.19 | 0.02 | 0.14 | 0 | 0.28 | 0.09 |
| 5 | 82058500 | rs2113027 | 0.493 | -0.008 | -1.409 | 0.237 | -0.015 | -0.015 | -0.015 | 0.676 | 0.15 | 0.09 | 0.14 | 0.18 | 0.18 | 0.15 |
| 5 | 82061539 | rs12658884 | 0.333 | 0.452 | -1.169 | 0.299 | -0.047 | -0.012 | -0.012 | 0 | 0.3 | 0.18 | 0.51 | 0.37 | 0.44 | 0.2 |
| 5 | 82067145 | rs1363948 | -0.649 | 0.028 | -0.624 | 0.177 | 0.009 | 0.001 | 0.001 | 1.3 | 0.9 | 0.95 | 0.92 | 0.54 | 0.92 | 0.74 |
| 5 | 82068106 | rs9283781 | 0.998 | 0.037 | -0.821 | -0.018 | -0.042 | 0.058 | 0.058 | 0.434 | 0.37 | 0.22 | 0.31 | 0.19 | 0.43 | 0.2 |
| 5 | 82070143 | rs10514244 | 0.066 | -0.325 | -0.723 | 0.138 | 0.027 | 0.019 | 0.019 | 0.819 | 0.4 | 0.27 | 0.34 | 0.33 | 0.34 | 0.46 |
| 5 | 82070913 | rs6892289 | 0.783 | -0.559 | -0.918 | -0.579 | 0.07 | 0.078 | 0.078 | 7.19 | 0.26 | 0.16 | 0.2 | 0.03 | 0.12 | 0.16 |
| 5 | 82071017 | rs6873579 | 0.781 | -0.583 | -0.914 | -0.576 | 0.072 | 0.078 | 0.078 | 7.228 | 0.26 | 0.16 | 0.2 | 0.03 | 0.11 | 0.16 |
| 5 | 82072989 | rs10060008 | 0.835 | 0.094 | -0.53 | -0.003 | -0.009 | -0.013 | -0.013 | 0 | 0.33 | 0.29 | 0.31 | 0.39 | 0.37 | 0.21 |
| 5 | 82073110 | rs6885068 | 0.174 | 0.728 | -1.067 | -1.059 | 0.021 | 0.135 | 0.135 | 1.208 | 0.4 | 0.16 | 0.29 | 0.14 | 0.34 | 0.44 |
| 5 | 82080216 | rs1549262 | -0.997 | 1.243 | -0.756 | -0.541 | 0.014 | -0.039 | -0.039 | 0.869 | 0.14 | 0.16 | 0.29 | 0.01 | 0.21 | 0.1 |
| 5 | 82082648 | rs7725235 | -0.194 | 0.479 | -1.219 | -0.414 | -0.023 | 0.159 | 0.159 | 0 | 0.44 | 0.15 | 0.32 | 0.14 | 0.44 | 0.44 |
| 5 | 82083323 | rs7712500 | 0.584 | -0.376 | 0.16 | -0.039 | -0.043 | -0.039 | -0.039 | 0.472 | 0.77 | 0.41 | 0.69 | 0.85 | 0.87 | 0.85 |
| 5 | 82092653 | rs992452 | -0.73 | 0.65 | 0.6 | 0.625 | 0.003 | 0.015 | 0.015 | 1.255 | 0.36 | 0.53 | 0.47 | 0.49 | 0.41 | 0.44 |
| 5 | 82094671 | rs1862598 | -0.701 | 0.515 | 0.564 | 0.601 | -0.015 | -0.038 | -0.038 | 0.011 | 0.34 | 0.22 | 0.45 | 0.49 | 0.4 | 0.45 |
| 5 | 82111569 | rs6892129 | -0.924 | 0.335 | -0.311 | 0.687 | 0.013 | -0.047 | -0.047 | 1.508 | 0.38 | 0.78 | 0.48 | 0.44 | 0.37 | 0.38 |
| 5 | 82120504 | rs16899961 | -0.735 | -0.059 | -0.47 | 0.929 | 0.019 | -0.019 | -0.019 | 0.051 | 0.24 | 0.12 | 0.35 | 0.32 | 0.17 | 0.36 |
| 5 | 82124904 | rs4605761 | 0.373 | 0.402 | -0.853 | 0.88 | 0.003 | -0.004 | -0.004 | 1.239 | 0.64 | 0.84 | 0.59 | 0.42 | 0.69 | 0.52 |
| 5 | 82127629 | rs1423510 | 0.651 | 0.336 | -0.771 | 0.246 | 0.001 | -0.002 | -0.002 | 0.109 | 0.32 | 0.32 | 0.21 | 0.15 | 0.23 | 0.33 |
| 5 | 82134620 | rs10073948 | 0.65 | 0.152 | -0.237 | 0.219 | 0.013 | -0.009 | -0.009 | 0 | 0.29 | 0.33 | 0.36 | 0.24 | 0.44 | 0.19 |
| 5 | 82147653 | rs10044294 | -0.251 | 0.126 | -0.332 | 0.766 | -0.004 | 0.001 | 0.001 | 0.491 | 0.61 | 0.42 | 0.65 | 0.47 | 0.7 | 0.54 |
| 5 | 82149437 | rs7707735 | -0.582 | 0.232 | -0.834 | 0.156 | 0.002 | -0.01 | -0.01 | 0.002 | 0.66 | 0.67 | 0.8 | 0.84 | 0.78 | 0.67 |
| 5 | 82152745 | rs13176760 | 0.1 | 0.729 | -0.499 | 0.589 | 0.024 | -0.015 | -0.015 | 0 | 0.3 | 0.5 | 0.38 | 0.25 | 0.44 | 0.18 |
| 5 | 82154979 | rs10050614 | 0.269 | 0.204 | -1.19 | 0.74 | 0.022 | -0.007 | -0.007 | 0 | 0.56 | 0.36 | 0.47 | 0.56 | 0.59 | 0.56 |
| 5 | 82158758 | rs12522257 | -0.615 | -0.238 | -1.659 | 1.077 | 0.026 | -0.055 | -0.055 | 0.83 | 0.28 | 0.13 | 0.23 | 0.35 | 0.22 | 0.32 |
| 5 | 82160801 | rs6859331 | 0.722 | -0.709 | -1.276 | 0.818 | 0.004 | 0.005 | 0.005 | 0.443 | 0.48 | 0.4 | 0.53 | 0.36 | 0.61 | 0.43 |
| 5 | 82162360 | rs10054357 | 0.575 | -0.112 | -1.498 | 0.231 | -0.043 | 0.048 | 0.048 | 0 | 0.31 | 0.19 | 0.33 | 0.22 | 0.42 | 0.21 |
| 5 | 82168645 | rs13436921 | -0.594 | -1.199 | -1.223 | 0.799 | 0.012 | 0.095 | 0.095 | 1.396 | 0.78 | 0.61 | 0.68 | 0.49 | 0.77 | 0.73 |
| 5 | 82169620 | rs6880701 | 0.566 | -1.402 | -1.249 | 1.127 | 0.005 | 0.001 | 0.001 | 0 | 0.51 | 0.49 | 0.58 | 0.43 | 0.67 | 0.45 |
| 5 | 82170556 | rs4074603 | -1.079 | -1.415 | -1.248 | 1.111 | 0.002 | 0.032 | 0.032 | 0.772 | 0.8 | 0.67 | 0.75 | 0.7 | 0.81 | 0.78 |
| 5 | 82171190 | rs9293320 | 1.448 | -1.143 | -1.549 | 0.774 | 0.035 | -0.069 | -0.069 | 0 | 0.3 | 0.32 | 0.37 | 0.05 | 0.46 | 0.26 |
| 5 | 82171406 | rs10077352 | 0.272 | -1.076 | -1.497 | 0.828 | 0.012 | 0.027 | 0.027 | 0 | 0.5 | 0.39 | 0.51 | 0.22 | 0.63 | 0.4 |
| 5 | 82177857 | rs4383715 | 0.492 | -1.11 | -1.373 | 0.82 | 0.041 | 0.019 | 0.019 | 0 | 0.28 | 0.17 | 0.15 | 0.26 | 0.14 | 0.34 |
| 5 | 82180306 | rs4518370 | -0.481 | -1.123 | -1.407 | 0.81 | -0.002 | -0.001 | -0.001 | 0.027 | 0.67 | 0.6 | 0.66 | 0.68 | 0.7 | 0.73 |
| 5 | 82182959 | rs4285197 | 0.31 | -0.767 | -1.078 | 0.819 | -0.034 | -0.03 | -0.03 | 0 | 0.4 | 0.68 | 0.52 | 0.43 | 0.56 | 0.4 |
| 5 | 82184403 | rs9293321 | 0.297 | -0.653 | -1.16 | 0.873 | 0.02 | 0.018 | 0.018 | 0 | 0.39 | 0.43 | 0.5 | 0.43 | 0.56 | 0.39 |
| 5 | 82188060 | rs6868050 | 0.354 | -0.637 | -1.386 | 0.648 | -0.001 | -0.061 | -0.061 | 0.25 | 0.44 | 0.39 | 0.44 | 0.04 | 0.57 | 0.31 |
| 5 | 82189299 | rs10462391 | 0.873 | -0.603 | -1.341 | 0.903 | 0.03 | -0.016 | -0.016 | 0 | 0.32 | 0.32 | 0.43 | 0.22 | 0.5 | 0.3 |
| 5 | 82192905 | rs6875265 | -0.16 | -0.733 | -1.34 | 0.887 | 0.003 | 0.001 | 0.001 | 0 | 0.58 | 0.55 | 0.59 | 0.43 | 0.67 | 0.48 |
| 5 | 82201161 | rs1504998 | -0.114 | -0.475 | -1.376 | 1.336 | -0.001 | -0.003 | -0.003 | 0 | 0.44 | 0.43 | 0.59 | 0.58 | 0.58 | 0.46 |
| 5 | 82202898 | rs10051358 | 0.135 | -0.306 | -1.315 | 1.206 | 0.012 | -0.022 | -0.022 | 0 | 0.41 | 0.42 | 0.47 | 0.25 | 0.49 | 0.46 |
| 5 | 82212572 | rs376310 | 0.027 | -0.905 | -1.806 | 0.818 | -0.026 | -0.008 | -0.008 | 0 | 0.21 | 0.03 | 0.15 | 0.14 | 0.18 | 0.26 |
| 5 | 82216170 | rs6452498 | -0.645 | -1.42 | -0.803 | 0.885 | -0.043 | 0.034 | 0.034 | 0 | 0.3 | 0.41 | 0.26 | 0.58 | 0.2 | 0.33 |
| 5 | 82216495 | rs6878182 | -0.645 | -1.42 | -0.803 | 0.885 | -0.043 | 0.034 | 0.034 | 0 | 0.3 | 0.41 | 0.25 | 0.59 | 0.2 | 0.33 |
| 5 | 82221080 | rs423704 | -0.064 | -1.135 | -1.014 | 0.321 | 0.014 | 0.014 | 0.014 | 0 | 0.13 | 0.13 | 0.1 | 0.07 | 0.1 | 0.13 |
| 5 | 82222004 | rs40533 | 0.656 | -1.171 | -1.032 | 0.285 | 0.011 | 0.005 | 0.005 | 0.297 | 0.77 | 0.79 | 0.79 | 0.82 | 0.82 | 0.79 |
| 5 | 82223633 | rs37554 | 1.238 | -1.554 | 0.061 | 0.419 | -0.061 | 0.01 | 0.01 | 0 | 0.75 | 0.42 | 0.77 | 0.74 | 0.84 | 0.7 |
| 5 | 82227023 | rs37561 | 0.886 | -1.499 | 0.205 | 0.289 | -0.05 | -0.057 | -0.057 | 0.508 | 0.78 | 0.53 | 0.79 | 0.9 | 0.87 | 0.73 |
| 5 | 82234988 | rs4703557 | 0.349 | -1.449 | -0.254 | 0.069 | -0.047 | 0.002 | 0.002 | 0.018 | 0.1 | 0.01 | 0.14 | 0.05 | 0.14 | 0.09 |
| 5 | 82248664 | rs12187956 | 0.742 | -1.449 | 0.061 | 0.149 | -0.068 | -0.024 | -0.024 | 0 | 0.78 | 0.5 | 0.73 | 0.85 | 0.88 | 0.73 |
| 5 | 82251766 | rs2089197 | -0.43 | -1.395 | -0.812 | -0.124 | -0.057 | 0.062 | 0.062 | 0.05 | 0.25 | 0.02 | 0.22 | 0.12 | 0.23 | 0.2 |
| 5 | 82257823 | rs423723 | -0.377 | -1.391 | -0.02 | 0.109 | -0.011 | 0.052 | 0.052 | 0 | 0.19 | 0.28 | 0.33 | 0.6 | 0.19 | 0.24 |
| 5 | 82257991 | rs1505074 | -0.325 | -1.845 | -0.218 | -0.268 | 0.011 | 0.02 | 0.02 | 0 | 0.31 | 0.18 | 0.2 | 0.23 | 0.25 | 0.2 |
| 5 | 82259908 | rs166278 | -1.019 | -1.548 | -0.056 | -0.576 | 0.023 | 0.099 | 0.099 | 0 | 0.52 | 0.66 | 0.55 | 0.83 | 0.44 | 0.43 |
| 5 | 82260048 | rs13188099 | 1.415 | -1.665 | -0.183 | -0.676 | -0.016 | 0.192 | 0.192 | 0.022 | 0.42 | 0.2 | 0.41 | 0.05 | 0.53 | 0.44 |
| 5 | 82261054 | rs256809 | 0.728 | -1.425 | 0.39 | 0.065 | -0.018 | 0.023 | 0.023 | 0 | 0.95 | 0.88 | 0.96 | 0.91 | 0.97 | 0.88 |
| 5 | 82275028 | rs7711883 | -0.916 | -0.914 | 0.815 | -0.02 | 0.027 | 0.017 | 0.017 | 0.279 | 0.16 | 0.54 | 0.21 | 0.15 | 0.16 | 0.19 |
| 5 | 82280978 | rs13168088 | 0.162 | -1.063 | 0.706 | 0.002 | -0.027 | -0.008 | -0.008 | 0.161 | 0.1 | 0.05 | 0.17 | 0.07 | 0.16 | 0.19 |
| 5 | 82281043 | rs1366566 | -1.718 | -1.509 | 0.755 | -0.301 | 0.006 | 0.007 | 0.007 | 1.308 | 0.32 | 0.67 | 0.29 | 0.21 | 0.27 | 0.28 |
| 5 | 82284721 | rs905849 | 0.192 | -1.454 | 0.534 | -0.35 | -0.013 | 0 | 0 | 0 | 0.11 | 0.09 | 0.17 | 0.07 | 0.16 | 0.21 |
| 5 | 82285028 | rs16900082 | -0.466 | -1.651 | 0.833 | -0.358 | 0.008 | 0.008 | 0.008 | 0 | 0.16 | 0.1 | 0.07 | 0.06 | 0.1 | 0.08 |
| 5 | 82297362 | rs256795 | -0.944 | -1.567 | 0.075 | -0.604 | -0.003 | 0.064 | 0.064 | 0 | 0.55 | 0.65 | 0.7 | 0.85 | 0.5 | 0.47 |
| 5 | 82299073 | rs376689 | -0.266 | -1.705 | 0.426 | -0.722 | 0.001 | -0.019 | -0.019 | 0.059 | 0.36 | 0.4 | 0.64 | 0.73 | 0.39 | 0.39 |
| 5 | 82303829 | rs702505 | 0.778 | -1.926 | 0.2 | -0.912 | 0.008 | 0.024 | 0.024 | 0.006 | 0.55 | 0.45 | 0.32 | 0.1 | 0.55 | 0.39 |
| 5 | 82319583 | rs40213 | 0.961 | -1.532 | 0.431 | -0.491 | -0.028 | 0.009 | 0.009 | 0 | 0.92 | 0.82 | 0.94 | 0.92 | 0.93 | 0.79 |
| 5 | 82326672 | rs2731853 | -0.464 | -1.382 | 0.221 | -0.451 | 0.001 | -0.002 | -0.002 | 0 | 0.06 | 0.05 | 0.03 | 0.08 | 0.07 | 0.22 |
| 5 | 82334790 | rs42754 | -0.161 | -1.599 | -0.012 | -0.617 | 0 | 0.041 | 0.041 | 0 | 0.14 | 0.01 | 0.04 | 0.06 | 0.13 | 0.07 |
| 5 | 82350810 | rs255561 | 0.918 | -1.744 | 0.663 | -0.831 | -0.002 | 0.015 | 0.015 | 0 | 0.45 | 0.62 | 0.31 | 0.21 | 0.46 | 0.59 |

| | | | | | | | | | | | | | | | | |
|---|---|---|---|---|---|---|---|---|---|---|---|---|---|---|---|---|
| 5 | 82372665 | rs2075685 | 0.92 | -1.746 | 0.679 | -0.832 | -0.002 | 0.005 | 0.005 | 0 | 0.45 | 0.58 | 0.3 | 0.21 | 0.46 | 0.59 |
| 5 | 82374163 | rs6864054 | -0.379 | -2.102 | 0.337 | -0.784 | -0.013 | 0.003 | 0.003 | 0 | 0.06 | 0.08 | 0.03 | 0.07 | 0.06 | 0.21 |
| 5 | 82376926 | rs7720588 | 1.506 | -2.133 | 0.554 | -0.904 | -0.036 | 0.067 | 0.067 | 0 | 0.37 | 0.27 | 0.23 | 0.12 | 0.39 | 0.37 |
| 5 | 82377004 | rs1478486 | 0.945 | -2.21 | 0.79 | -0.981 | -0.004 | 0.004 | 0.004 | 0 | 0.42 | 0.43 | 0.26 | 0.19 | 0.43 | 0.6 |
| 5 | 82377566 | rs1479569 | 1.706 | -2.601 | 0.538 | -0.906 | -0.179 | 0.239 | 0.239 | 0.452 | 0.37 | 0.02 | 0.22 | 0.03 | 0.41 | 0.39 |
| 5 | 82381400 | rs11746977 | 1.71 | -2.537 | 1.042 | -0.871 | -0.085 | 0.159 | 0.159 | 0.155 | 0.36 | 0.16 | 0.21 | 0.03 | 0.39 | 0.39 |
| 5 | 82393549 | rs1011980 | 1.084 | -2.795 | 0.748 | -0.844 | -0.085 | 0.028 | 0.028 | 0 | 0.16 | 0.02 | 0.24 | 0.05 | 0.26 | 0.19 |
| 5 | 82394383 | rs10057303 | 2.045 | -2.819 | 0.807 | -0.856 | -0.059 | 0.111 | 0.111 | 0.079 | 0.37 | 0.21 | 0.21 | 0.03 | 0.39 | 0.38 |
| 5 | 82395874 | rs13180356 | 1.038 | -2.759 | 0.765 | -0.823 | -0.076 | 0.035 | 0.035 | 0 | 0.17 | 0.02 | 0.24 | 0.05 | 0.26 | 0.18 |
| 5 | 82406338 | rs3901654 | 0.539 | -2.276 | 1.513 | 0.128 | 0.008 | 0.049 | 0.049 | 0 | 0.51 | 0.72 | 0.47 | 0.68 | 0.49 | 0.6 |
| 5 | 82422643 | rs1120476 | 0.897 | -2.234 | 1.639 | 0.138 | -0.002 | -0.011 | -0.011 | 0 | 0.44 | 0.62 | 0.27 | 0.14 | 0.44 | 0.53 |
| 5 | 82436811 | rs6452526 | 0.734 | -2.238 | 1.597 | 0.19 | -0.006 | 0.03 | 0.03 | 0 | 0.45 | 0.66 | 0.28 | 0.22 | 0.45 | 0.56 |
| 5 | 82451212 | rs1382367 | 0.023 | -2.377 | 1.469 | 0.141 | 0.001 | 0.018 | 0.018 | 0 | 0.4 | 0.24 | 0.49 | 0.26 | 0.4 | 0.32 |
| 5 | 82481380 | rs2974445 | 0.962 | -2.245 | 1.799 | 0.196 | 0.002 | -0.016 | -0.016 | 1.487 | 0.43 | 0.51 | 0.27 | 0.22 | 0.46 | 0.57 |
| 5 | 82484885 | rs2662242 | 0.902 | -2.243 | 1.721 | 0.197 | 0.003 | -0.021 | -0.021 | 0 | 0.44 | 0.58 | 0.28 | 0.22 | 0.46 | 0.57 |
| 5 | 82500295 | rs3777041 | -1.421 | -2.278 | 1.497 | 0.237 | -0.041 | -0.008 | -0.008 | 0.633 | 0.1 | 0.28 | 0.05 | 0.11 | 0.04 | 0.15 |
| 5 | 82501893 | rs7729841 | -0.539 | -2.252 | 1.224 | 0.245 | -0.002 | 0 | 0 | 0 | 0.07 | 0.07 | 0.04 | 0.11 | 0.04 | 0.15 |
| 5 | 82504856 | rs10514249 | -1.271 | -2.182 | 1.2 | 0.255 | -0.035 | -0.033 | -0.033 | 0.689 | 0.6 | 0.71 | 0.59 | 0.49 | 0.57 | 0.57 |
| 5 | 82521655 | rs10054951 | -1.037 | -2.763 | 1.087 | 0.29 | 0.001 | -0.002 | -0.002 | 0 | 0.09 | 0.11 | 0.04 | 0.1 | 0.06 | 0.18 |
| 5 | 82543086 | rs1193695 | -0.592 | -2.806 | 1.314 | 0.244 | 0.027 | 0.081 | 0.081 | 0 | 0.49 | 0.26 | 0.43 | 0.27 | 0.45 | 0.37 |
| 5 | 82547566 | rs2731866 | -0.647 | -2.727 | 1.902 | 1.364 | 0.013 | -0.017 | -0.017 | 0.166 | 0.08 | 0.03 | 0.05 | 0.11 | 0.05 | 0.02 |
| 5 | 82586489 | rs40123 | -1.599 | -2.264 | 2.307 | 1.658 | -0.068 | 0.001 | 0.001 | 0 | 0.2 | 0.51 | 0.13 | 0.23 | 0.15 | 0.25 |
| 5 | 82588496 | rs35268 | -1.359 | -2.232 | 1.814 | 1.577 | -0.02 | -0.013 | -0.013 | 0 | 0.18 | 0.28 | 0.11 | 0.14 | 0.17 | 0.25 |
| 5 | 82606235 | rs301289 | 1.496 | -2.284 | 1.498 | 1.213 | 0.008 | -0.006 | -0.006 | 0 | 0.72 | 0.26 | 0.67 | 0.3 | 0.81 | 0.73 |
| 5 | 82624287 | rs7706470 | -2.364 | -1.594 | 1.427 | 1.713 | -0.037 | 0.114 | 0.114 | 1.958 | 0.11 | 0.39 | 0.09 | 0.24 | 0.02 | 0.02 |
| 5 | 82626919 | rs3777018 | -0.635 | -1.639 | 1.073 | 2.107 | -0.001 | 0.02 | 0.02 | 0 | 0.07 | 0.02 | 0.18 | 0.46 | 0.08 | 0.14 |
| 5 | 82633951 | rs10514254 | -1.432 | -1.639 | 1.233 | 2.089 | -0.01 | 0.017 | 0.017 | 0.208 | 0.06 | 0.12 | 0.07 | 0.16 | 0.02 | 0.02 |
| 5 | 82636062 | rs16900340 | -2.765 | -1.656 | 1.316 | 2.058 | -0.057 | 0.095 | 0.095 | 0.236 | 0.18 | 0.4 | 0.28 | 0.71 | 0.1 | 0.16 |
| 5 | 82640477 | rs7735781 | -1.586 | -1.57 | 1.091 | 2.118 | -0.028 | 0.165 | 0.165 | 0 | 0.12 | 0.22 | 0.24 | 0.47 | 0.08 | 0.14 |
| 5 | 82642137 | rs16900343 | -1.057 | -1.497 | 1.037 | 1.972 | 0.015 | 0.051 | 0.051 | 0 | 0.06 | 0.17 | 0.09 | 0.15 | 0.06 | 0.03 |
| 5 | 82648883 | rs3777015 | -0.662 | -1.666 | 0.733 | 1.71 | 0.029 | -0.033 | -0.033 | 1.016 | 0.12 | 0.01 | 0.01 | 0.08 | 0.05 | 0.12 |
| 5 | 82648943 | rs1805377 | -2.317 | -0.884 | 1.336 | 2.107 | -0.035 | 0.105 | 0.105 | 0.275 | 0.19 | 0.48 | 0.33 | 0.7 | 0.14 | 0.17 |
| 5 | 82660901 | rs11960003 | -1.563 | -0.651 | 0.734 | 1.335 | -0.011 | 0.048 | 0.048 | 0.478 | 0.12 | 0.39 | 0.11 | 0.21 | 0.06 | 0.05 |
| 5 | 82673517 | rs16900376 | -1.01 | -0.77 | 0.06 | 1.426 | 0.005 | -0.036 | -0.036 | 0 | 0.07 | 0.01 | 0.16 | 0.14 | 0.05 | 0.13 |
| 5 | 82680138 | rs10074082 | -2.28 | -1.072 | 1.068 | 1.854 | -0.052 | 0.091 | 0.091 | 0 | 0.15 | 0.31 | 0.2 | 0.33 | 0.06 | 0.2 |
| 5 | 82683240 | rs12657920 | -2.15 | -1.117 | 1.169 | 1.835 | -0.057 | 0.089 | 0.089 | 0 | 0.15 | 0.35 | 0.2 | 0.32 | 0.06 | 0.18 |
| 5 | 82684424 | rs7706361 | -1.299 | -0.977 | 1.512 | 1.667 | 0.02 | 0.207 | 0.207 | 1.474 | 0.07 | 0.47 | 0.05 | 0.19 | 0.01 | 0.07 |
| 5 | 82685260 | rs2441010 | -0.631 | -1.11 | -0.08 | 1.426 | -0.003 | -0.023 | -0.023 | 0 | 0.1 | 0 | 0.06 | 0.07 | 0.08 | 0.11 |
| 5 | 82709977 | rs1445780 | -0.756 | -0.486 | -1.17 | 0.251 | -0.138 | 0.203 | 0.203 | 0 | 0.64 | 0.94 | 0.7 | 0.93 | 0.54 | 0.67 |
| 5 | 82715152 | rs16900394 | -1.265 | -1.278 | -0.787 | 1.516 | -0.012 | 0.052 | 0.052 | 0 | 0.13 | 0.14 | 0.27 | 0.5 | 0.08 | 0.24 |
| 5 | 82717101 | rs2591455 | -1.046 | -2.576 | -0.254 | 0.582 | -0.072 | -0.091 | -0.091 | 0 | 0.25 | 0.36 | 0.08 | 0.05 | 0.08 | 0.23 |
| 5 | 82717664 | rs2591454 | -0.939 | -3.296 | -0.828 | 0.284 | 0.05 | 0.066 | 0.066 | 0.807 | 0.21 | 0.04 | 0.11 | 0.1 | 0.1 | 0.22 |
| 5 | 82717885 | rs2591453 | -0.79 | -2.572 | -0.885 | -0.025 | 0.04 | 0.009 | 0.009 | 0 | 0.7 | 0.56 | 0.73 | 0.7 | 0.6 | 0.73 |
| 5 | 82720112 | rs1354122 | -0.562 | -2.535 | -1.208 | -0.109 | 0.005 | 0.004 | 0.004 | 0 | 0.5 | 0.5 | 0.5 | 0.64 | 0.3 | 0.6 |
| 5 | 82729928 | rs185356 | -0.472 | -1.765 | -1.082 | -0.266 | -0.011 | 0.008 | 0.008 | 0 | 0.62 | 0.53 | 0.53 | 0.57 | 0.64 | 0.5 |
| 5 | 82731955 | rs40028 | 1.14 | -1.914 | -0.903 | -0.311 | 0.004 | 0.007 | 0.007 | 0.176 | 0.14 | 0.1 | 0.1 | 0.1 | 0.11 | 0.17 |
| 5 | 82737843 | rs10942328 | 1.983 | -2.004 | -0.813 | 0.144 | -0.008 | 0.025 | 0.025 | 0 | 0.17 | 0.2 | 0.11 | 0.3 | 0.15 | 0.15 |
| 5 | 82739803 | rs10474101 | 1.115 | -1.685 | -1.035 | 0.015 | -0.054 | 0.036 | 0.036 | 0 | 0.29 | 0.21 | 0.34 | 0.18 | 0.41 | 0.19 |
| 5 | 82742118 | rs12188947 | 1.433 | -1.696 | -1.006 | 0.019 | -0.115 | 0.084 | 0.084 | 0 | 0.27 | 0.09 | 0.27 | 0.1 | 0.4 | 0.19 |
| 5 | 82744472 | rs924584 | -1.881 | -1.717 | -0.919 | 0.063 | 0.008 | 0.024 | 0.024 | 0 | 0.75 | 0.67 | 0.68 | 0.63 | 0.71 | 0.73 |
| 5 | 82747765 | rs11746897 | 0.618 | -1.434 | -1.459 | -0.323 | -0.056 | 0.059 | 0.059 | 0 | 0.33 | 0.21 | 0.34 | 0.17 | 0.41 | 0.19 |
| 5 | 82748948 | rs871759 | 0.154 | -1.519 | -1.155 | 0.005 | 0.028 | 0.03 | 0.03 | 0.704 | 0.23 | 0.12 | 0.22 | 0.16 | 0.15 | 0.39 |
| 5 | 82753478 | rs831927 | 0.154 | -1.519 | -1.244 | -0.033 | 0.033 | 0.041 | 0.041 | 0.704 | 0.23 | 0.11 | 0.22 | 0.14 | 0.15 | 0.39 |
| 5 | 82770558 | rs11746859 | 0.491 | -0.769 | -1.272 | 0.229 | -0.017 | 0.035 | 0.035 | 0.347 | 0.44 | 0.27 | 0.36 | 0.35 | 0.48 | 0.33 |
| 5 | 82775634 | rs33598 | 0.09 | -0.98 | -0.904 | 0.375 | 0.033 | 0.002 | 0.002 | 0.022 | 0.47 | 0.34 | 0.44 | 0.52 | 0.37 | 0.54 |
| 5 | 82779013 | rs1867667 | -1.398 | -0.444 | -1.036 | 0.52 | 0.031 | 0.105 | 0.105 | 0 | 0.79 | 0.61 | 0.72 | 0.61 | 0.75 | 0.73 |
| 5 | 82782034 | rs10045126 | 0.706 | -0.519 | -0.996 | 0.456 | 0.028 | -0.038 | -0.038 | 1.841 | 0.32 | 0.5 | 0.38 | 0.2 | 0.39 | 0.2 |
| 5 | 82786549 | rs4558964 | 0.956 | -0.27 | -0.905 | 0.476 | 0.028 | 0.106 | 0.106 | 0 | 0.23 | 0.43 | 0.35 | 0.47 | 0.26 | 0.27 |
| 5 | 82788892 | rs6452540 | -0.813 | 0.058 | -1.235 | 0.137 | -0.008 | 0.036 | 0.036 | 0.026 | 0.87 | 0.96 | 0.88 | 0.92 | 0.85 | 0.82 |
| 5 | 82789647 | rs4470745 | 0.147 | 0.113 | -0.843 | 0.459 | -0.017 | 0.018 | 0.018 | 0 | 0.3 | 0.23 | 0.28 | 0.09 | 0.41 | 0.19 |
| 5 | 82792625 | rs2292013 | -1.07 | -0.414 | -0.455 | 1.574 | -0.009 | 0.131 | 0.131 | 0 | 0.06 | 0.12 | 0.13 | 0.6 | 0.05 | 0.23 |
| 5 | 82794036 | rs2652106 | -0.858 | -0.514 | -0.59 | 1.139 | -0.013 | 0.014 | 0.014 | 1.032 | 0.36 | 0.58 | 0.3 | 0.48 | 0.31 | 0.39 |
| 5 | 82805424 | rs251124 | -0.577 | -0.958 | -0.408 | 1.371 | 0.011 | -0.011 | -0.011 | 0.005 | 0.24 | 0.18 | 0.2 | 0.37 | 0.21 | 0.31 |
| 5 | 82810714 | rs160380 | 0.972 | -1.31 | -0.464 | 0.799 | -0.005 | 0 | 0 | 0 | 0.45 | 0.43 | 0.47 | 0.54 | 0.33 | 0.43 |
| 5 | 82811500 | rs173686 | -0.595 | -2.212 | -0.728 | 0.41 | 0.036 | 0.15 | 0.15 | 1.388 | 0.55 | 0.81 | 0.55 | 0.87 | 0.45 | 0.54 |
| 5 | 82812442 | rs308359 | 1.538 | -2.219 | -0.487 | 0.726 | 0.004 | -0.006 | -0.006 | 0 | 0.27 | 0.33 | 0.34 | 0.37 | 0.22 | 0.2 |
| 5 | 82815408 | rs2287926 | 1.231 | -2.081 | -0.367 | 0.578 | -0.083 | 0.015 | 0.015 | 0 | 0.22 | 0.33 | 0.14 | 0.18 | 0.12 | 0.24 |
| 5 | 82819185 | rs2541312 | -1.111 | -1.993 | -0.738 | 0.304 | -0.145 | 0.087 | 0.087 | 0.755 | 0.73 | 0.98 | 0.68 | 0.91 | 0.52 | 0.75 |
| 5 | 82823791 | rs308362 | -0.842 | -2.074 | -0.169 | 0.547 | -0.087 | 0.106 | 0.106 | 0 | 0.75 | 0.61 | 0.86 | 0.59 | 0.86 | 0.66 |

| | | | | | | | | | | | | | | | | |
|---|---|---|---|---|---|---|---|---|---|---|---|---|---|---|---|---|
| 5 | 82826254 | rs16900511 | -0.064 | -1.327 | -0.324 | 0.364 | -0.014 | -0.042 | -0.042 | 0 | 0.21 | 0.24 | 0.15 | 0.07 | 0.21 | 0.18 |
| 5 | 82826483 | rs17348094 | 0.441 | -1.295 | -0.452 | 0.455 | 0.009 | 0.036 | 0.036 | 0 | 0.16 | 0.07 | 0.12 | 0.08 | 0.17 | 0.18 |
| 5 | 82828001 | rs1193355 | -0.638 | -0.569 | -0.935 | 0.761 | -0.112 | -0.046 | -0.046 | 0 | 0.66 | 0.97 | 0.66 | 0.74 | 0.65 | 0.74 |
| 5 | 82830308 | rs309561 | -0.099 | -0.598 | -1.021 | 0.515 | 0.012 | -0.032 | -0.032 | 0 | 0.19 | 0.01 | 0.09 | 0.21 | 0.13 | 0.33 |
| 5 | 82830442 | rs1193354 | -0.783 | -0.709 | -0.72 | 0.656 | -0.022 | -0.039 | -0.039 | 0 | 0.66 | 0.79 | 0.63 | 0.63 | 0.66 | 0.73 |
| 5 | 82831964 | rs3096169 | -1.699 | -0.735 | -0.835 | 0.65 | 0.002 | -0.003 | -0.003 | 0 | 0.82 | 0.84 | 0.92 | 0.79 | 0.88 | 0.83 |
| 5 | 82833369 | rs309559 | 0.709 | -0.835 | -0.244 | 0.633 | 0.002 | -0.01 | -0.01 | 0 | 0.45 | 0.51 | 0.53 | 0.39 | 0.53 | 0.4 |
| 5 | 82834630 | rs309557 | 0.699 | -0.848 | -0.25 | 0.627 | 0.001 | -0.01 | -0.01 | 0 | 0.46 | 0.5 | 0.53 | 0.39 | 0.53 | 0.4 |
| 5 | 82837631 | rs160277 | 1.157 | -0.695 | -0.096 | 0.544 | 0.005 | 0.022 | 0.022 | 0 | 0.38 | 0.36 | 0.46 | 0.32 | 0.43 | 0.27 |
| 5 | 82838532 | rs308363 | -0.811 | -0.662 | -0.418 | 0.594 | -0.041 | -0.064 | -0.064 | 0.104 | 0.67 | 0.85 | 0.65 | 0.63 | 0.66 | 0.73 |
| 5 | 82841476 | rs695103 | 0.028 | -0.676 | -0.354 | 0.642 | 0.024 | -0.028 | -0.028 | 0 | 0.19 | 0.05 | 0.09 | 0.24 | 0.13 | 0.33 |
| 5 | 82844728 | rs309596 | -0.595 | -1.114 | -0.062 | 0.651 | 0.046 | -0.073 | -0.073 | 3.109 | 0.73 | 0.88 | 0.88 | 0.59 | 0.79 | 0.56 |
| 5 | 82848324 | rs17205951 | -0.125 | -0.92 | 0.092 | 0.547 | 0.017 | -0.005 | -0.005 | 0 | 0.06 | 0.11 | 0.05 | 0.07 | 0.08 | 0.1 |
| 5 | 82850587 | rs2290672 | 0.184 | -0.917 | -0.291 | 0.567 | -0.004 | 0.017 | 0.017 | 0.011 | 0.45 | 0.47 | 0.51 | 0.39 | 0.51 | 0.39 |
| 5 | 82856519 | rs309587 | 0.185 | -0.784 | -0.486 | 0.962 | 0.004 | -0.034 | -0.034 | 0.116 | 0.67 | 0.75 | 0.83 | 0.48 | 0.67 | 0.46 |
| 5 | 82858828 | rs12653308 | -0.291 | -0.433 | -0.019 | 1.298 | -0.008 | -0.008 | -0.008 | 0 | 0.12 | 0.09 | 0.11 | 0.31 | 0.22 | 0.45 |
| 5 | 82859485 | rs160283 | 0.516 | -0.906 | -0.447 | 1.128 | 0.049 | -0.017 | -0.017 | 0 | 0.3 | 0.11 | 0.36 | 0.3 | 0.26 | 0.21 |
| 5 | 82861400 | rs13164785 | -0.075 | -0.792 | 0.267 | 1.285 | 0.025 | 0.043 | 0.043 | 0 | 0.13 | 0.17 | 0.11 | 0.29 | 0.2 | 0.44 |
| 5 | 82862227 | rs160187 | -1.135 | -1.095 | -0.041 | 0.917 | -0.004 | -0.004 | -0.004 | 0 | 0.76 | 0.91 | 0.87 | 0.82 | 0.77 | 0.82 |
| 5 | 82871546 | rs309581 | 0.437 | -1.008 | -0.042 | 0.972 | -0.019 | -0.033 | -0.033 | 0.425 | 0.32 | 0.14 | 0.42 | 0.36 | 0.37 | 0.41 |
| 5 | 82874589 | rs831588 | -1.134 | -0.634 | 0.136 | 0.638 | -0.009 | -0.019 | -0.019 | 0.019 | 0.78 | 0.9 | 0.87 | 0.79 | 0.77 | 0.81 |
| 5 | 82879340 | rs10514259 | 1.04 | -0.623 | 0.529 | 0.72 | -0.001 | 0 | 0 | 0 | 0.22 | 0.23 | 0.16 | 0.22 | 0.22 | 0.19 |
| 5 | 82881455 | rs309599 | 0.076 | -0.835 | 0.874 | 0.765 | 0 | 0.001 | 0.001 | 0.371 | 0.56 | 0.46 | 0.65 | 0.6 | 0.59 | 0.61 |
| 5 | 82889910 | rs310501 | -0.51 | -0.766 | 1.077 | 0.756 | 0.027 | -0.001 | -0.001 | 0 | 0.72 | 0.58 | 0.73 | 0.75 | 0.72 | 0.88 |
| 5 | 82897008 | rs10074448 | -0.278 | -0.936 | 1.248 | 0.437 | 0.003 | 0.005 | 0.005 | 0 | 0.44 | 0.59 | 0.26 | 0.24 | 0.37 | 0.36 |
| 5 | 82897465 | rs309564 | 0.351 | -1.024 | 1.508 | 0.433 | 0 | -0.001 | -0.001 | 0 | 0.53 | 0.5 | 0.62 | 0.53 | 0.57 | 0.52 |
| 5 | 82897718 | rs309568 | -1.064 | -0.926 | 0.925 | 0.489 | -0.034 | -0.041 | -0.041 | 1.128 | 0.76 | 0.89 | 0.75 | 0.64 | 0.72 | 0.76 |
| 5 | 82898246 | rs309570 | 0.771 | -0.933 | 0.757 | 0.49 | -0.007 | -0.027 | -0.027 | 0.32 | 0.31 | 0.18 | 0.48 | 0.4 | 0.35 | 0.43 |
| 5 | 82899585 | rs6866362 | -0.243 | -0.921 | 0.595 | 0.471 | 0.003 | 0.005 | 0.005 | 0 | 0.44 | 0.61 | 0.26 | 0.24 | 0.37 | 0.33 |
| 5 | 82903716 | rs566008 | -0.848 | -0.703 | 0.486 | 0.571 | 0.011 | 0.014 | 0.014 | 0 | 0.75 | 0.74 | 0.74 | 0.64 | 0.72 | 0.76 |
| 5 | 82907181 | rs4532345 | 0.185 | -0.632 | 0.426 | 0.54 | -0.014 | -0.022 | -0.022 | 0 | 0.2 | 0.29 | 0.11 | 0.12 | 0.15 | 0.23 |
| 5 | 82908474 | rs12697730 | -0.984 | -0.424 | -0.048 | 0.659 | 0.006 | 0.008 | 0.008 | 0 | 0.77 | 0.78 | 0.75 | 0.64 | 0.74 | 0.75 |
| 5 | 82909863 | rs518706 | -0.988 | -0.42 | -0.077 | 0.661 | 0.005 | 0.006 | 0.006 | 0 | 0.77 | 0.79 | 0.75 | 0.64 | 0.74 | 0.75 |
| 5 | 82911585 | rs2607148 | -0.988 | -0.42 | -0.077 | 0.661 | 0.005 | 0.006 | 0.006 | 0 | 0.77 | 0.79 | 0.75 | 0.64 | 0.74 | 0.75 |
| 5 | 82914623 | rs336944 | -0.415 | -0.417 | -0.103 | 0.653 | -0.014 | -0.016 | -0.016 | 0.312 | 0.57 | 0.76 | 0.64 | 0.52 | 0.52 | 0.64 |
| 5 | 82929501 | rs12657861 | -0.95 | -0.084 | 0.091 | 0.734 | 0 | 0.004 | 0.004 | 0 | 0.79 | 0.8 | 0.75 | 0.58 | 0.72 | 0.72 |
| 5 | 82931384 | rs17206055 | 0.009 | -0.343 | -0.259 | 0.581 | 0.02 | 0.027 | 0.027 | 0 | 0.2 | 0.13 | 0.1 | 0.13 | 0.16 | 0.23 |
| 5 | 82934554 | rs7706851 | -0.862 | 0.136 | 0.031 | 0.635 | -0.029 | -0.035 | -0.035 | 0 | 0.79 | 0.86 | 0.75 | 0.6 | 0.72 | 0.73 |
| 5 | 82948216 | rs2045380 | 0.11 | -0.003 | -0.842 | 0.543 | 0.003 | -0.004 | -0.004 | 0 | 0.15 | 0.14 | 0.08 | 0.13 | 0.13 | 0.19 |
| 5 | 82953592 | rs17284608 | 0.002 | 0.358 | -0.897 | 0.608 | -0.001 | -0.001 | -0.001 | 0.585 | 0.29 | 0.26 | 0.31 | 0.22 | 0.26 | 0.39 |
| 5 | 82955664 | rs3777048 | 0.004 | 1.09 | -1.081 | 1.231 | -0.046 | -0.061 | -0.061 | 0 | 0.77 | 0.89 | 0.76 | 0.57 | 0.7 | 0.73 |
| 5 | 82958596 | rs336948 | -0.403 | 0.238 | -0.613 | 1.038 | -0.009 | 0.001 | 0.001 | 0 | 0.25 | 0.24 | 0.28 | 0.29 | 0.21 | 0.41 |
| 5 | 82960734 | rs3846635 | -0.798 | 0.693 | 0.183 | 1.176 | 0.036 | 0.103 | 0.103 | 0 | 0.07 | 0.22 | 0.2 | 0.17 | 0.07 | 0.04 |
| 5 | 82964746 | rs336956 | -0.341 | 0.883 | 0.066 | 0.986 | 0.002 | 0.024 | 0.024 | 0 | 0.34 | 0.4 | 0.4 | 0.55 | 0.35 | 0.44 |
| 5 | 82966191 | rs170880 | -0.287 | 0.917 | 0.039 | 0.986 | 0.001 | 0.006 | 0.006 | 0 | 0.33 | 0.33 | 0.4 | 0.55 | 0.35 | 0.44 |
| 5 | 82968585 | rs336965 | -0.667 | 0.846 | -0.595 | 0.572 | -0.005 | 0.014 | 0.014 | 0 | 0.44 | 0.61 | 0.62 | 0.71 | 0.48 | 0.49 |
| 5 | 82971512 | rs13168731 | 0.196 | 0.836 | -0.571 | 0.545 | -0.009 | 0.01 | 0.01 | 0.002 | 0.58 | 0.66 | 0.43 | 0.6 | 0.49 | 0.56 |
| 5 | 82971921 | rs10514260 | -0.549 | 0.554 | -0.115 | 0.411 | 0.009 | 0.03 | 0.03 | 0 | 0.47 | 0.55 | 0.63 | 0.8 | 0.56 | 0.55 |
| 5 | 82973210 | rs6886442 | -0.746 | 0.121 | 0.025 | 0.443 | 0 | -0.001 | -0.001 | 0 | 0.59 | 0.6 | 0.74 | 0.55 | 0.75 | 0.51 |
| 5 | 82977764 | rs1154891 | -0.481 | 0.83 | -0.024 | 0.321 | 0.023 | -0.067 | -0.067 | 0 | 0.33 | 0.53 | 0.32 | 0.14 | 0.39 | 0.24 |
| 5 | 82977843 | rs1420655 | 0.734 | 0.814 | -0.078 | 0.329 | -0.052 | -0.072 | -0.072 | 0.745 | 0.27 | 0.1 | 0.42 | 0.42 | 0.36 | 0.27 |
| 5 | 82982945 | rs6894022 | -1.072 | 0.728 | 0.368 | 0.439 | 0.001 | -0.003 | -0.003 | 1.617 | 0.59 | 0.45 | 0.67 | 0.83 | 0.7 | 0.63 |
| 5 | 82985576 | rs4466136 | -1.527 | 0.76 | -0.225 | 0.47 | 0.043 | 0.059 | 0.059 | 0.287 | 0.64 | 0.82 | 0.76 | 0.83 | 0.76 | 0.63 |
| 5 | 82986149 | rs16898 | -0.701 | 0.876 | -0.511 | 0.543 | 0.071 | -0.015 | -0.015 | 0 | 0.54 | 0.27 | 0.65 | 0.56 | 0.68 | 0.49 |
| 5 | 82996199 | rs4703961 | -0.286 | 0.97 | -0.65 | 0.704 | -0.044 | 0.057 | 0.057 | 0.088 | 0.27 | 0.12 | 0.23 | 0.13 | 0.33 | 0.22 |
| 5 | 83020138 | rs975563 | -0.824 | 0.926 | -0.596 | 0.634 | 0.002 | 0.002 | 0.002 | 0.012 | 0.33 | 0.33 | 0.27 | 0.39 | 0.34 | 0.33 |
| 5 | 83021939 | rs1563552 | 0.203 | 0.777 | -1.401 | 0.117 | -0.018 | 0.055 | 0.055 | 0 | 0.2 | 0.08 | 0.18 | 0.07 | 0.23 | 0.12 |
| 5 | 83024710 | rs1563553 | -0.709 | 0.547 | -1.198 | 0.305 | 0.026 | 0.031 | 0.031 | 0.135 | 0.64 | 0.76 | 0.76 | 0.82 | 0.75 | 0.63 |
| 5 | 83026432 | rs10514261 | -0.407 | 1.41 | -1.377 | 0.898 | -0.036 | -0.043 | -0.043 | 0 | 0.09 | 0.01 | 0.2 | 0.2 | 0.1 | 0.12 |
| 5 | 83030867 | rs10058970 | -0.33 | 1.054 | -1.408 | 0.349 | 0 | 0.038 | 0.038 | 0.657 | 0.22 | 0.16 | 0.21 | 0.11 | 0.21 | 0.13 |
| 5 | 83034211 | rs1457075 | 0.307 | 1.314 | -0.908 | 0.794 | 0.025 | 0.042 | 0.042 | 0 | 0.72 | 0.65 | 0.65 | 0.53 | 0.69 | 0.62 |
| 5 | 83036792 | rs6897626 | 0.163 | 1.45 | -2.44 | 0.206 | -0.124 | -0.036 | -0.036 | 0 | 0.66 | 0.96 | 0.73 | 0.7 | 0.61 | 0.62 |
| 5 | 83047195 | rs1457076 | -0.125 | 0.566 | -2.842 | -0.488 | 0.03 | 0.102 | 0.102 | 0 | 0.62 | 0.89 | 0.74 | 0.79 | 0.67 | 0.57 |
| 5 | 83069019 | rs10051286 | 0.498 | 0.467 | -1.798 | -0.679 | -0.031 | 0.03 | 0.03 | 0 | 0.39 | 0.21 | 0.56 | 0.65 | 0.51 | 0.42 |
| 5 | 83088178 | rs11746129 | 0.384 | 0.637 | -1.6 | -0.659 | 0.009 | 0.047 | 0.047 | 0.018 | 0.21 | 0.09 | 0.15 | 0.07 | 0.19 | 0.27 |
| 5 | 83107104 | rs4610439 | 0.905 | 0.635 | -1.654 | -0.659 | -0.04 | 0.012 | 0.012 | 0.312 | 0.17 | 0.01 | 0.14 | 0.07 | 0.18 | 0.27 |
| 5 | 83109006 | rs11741548 | 0.839 | 0.665 | -1.379 | -0.574 | 0.002 | 0.028 | 0.028 | 0.375 | 0.18 | 0.05 | 0.12 | 0.14 | 0.15 | 0.15 |
| 5 | 83120842 | rs7732939 | 0.07 | 0.823 | -0.686 | -0.611 | -0.016 | -0.073 | -0.073 | 0 | 0.25 | 0.49 | 0.2 | 0.14 | 0.19 | 0.16 |
| 5 | 83129532 | rs7714410 | 0.05 | 1.493 | -0.744 | -0.271 | 0.126 | 0.021 | 0.021 | 0 | 0.15 | 0.38 | 0.18 | 0.15 | 0.28 | 0.16 |

| | | | | | | | | | | | | | | | | |
|---|---|---|---|---|---|---|---|---|---|---|---|---|---|---|---|---|
| 5 | 83139409 | rs6882023 | -0.714 | 1.485 | -1.52 | -0.272 | -0.026 | 0.03 | 0.03 | 0.067 | 0.39 | 0.7 | 0.33 | 0.3 | 0.47 | 0.31 |
| 5 | 83145042 | rs10063526 | -1.001 | 0.991 | -1.588 | -0.056 | 0.017 | 0.019 | 0.019 | 0 | 0.58 | 0.8 | 0.47 | 0.36 | 0.59 | 0.56 |
| 5 | 83147033 | rs11745452 | 0.268 | 0.888 | -1.562 | 0.138 | -0.009 | 0.041 | 0.041 | 0.22 | 0.2 | 0.04 | 0.14 | 0.14 | 0.18 | 0.15 |
| 5 | 83154723 | rs10056609 | -0.661 | 1.199 | -0.946 | 0.865 | 0.005 | 0.003 | 0.003 | 0 | 0.48 | 0.49 | 0.47 | 0.55 | 0.56 | 0.49 |
| 5 | 83156500 | rs10044480 | 0.027 | 1.025 | -0.934 | 0.903 | 0.005 | 0.005 | 0.005 | 0 | 0.3 | 0.25 | 0.21 | 0.24 | 0.27 | 0.31 |
| 5 | 83163554 | rs6897010 | -0.306 | 1.46 | -0.839 | 1.21 | 0.007 | 0.051 | 0.051 | 0 | 0.13 | 0.22 | 0.16 | 0.28 | 0.16 | 0.14 |
| 5 | 83165955 | rs10474118 | 0.191 | 1.503 | -1.642 | 0.896 | -0.083 | -0.046 | -0.046 | 0 | 0.26 | 0.04 | 0.24 | 0.26 | 0.3 | 0.31 |
| 5 | 83166175 | rs13355405 | 0.087 | 1.447 | -1.675 | 0.84 | -0.074 | -0.042 | -0.042 | 0 | 0.27 | 0.04 | 0.24 | 0.26 | 0.31 | 0.32 |
| 5 | 83173593 | rs4552569 | -0.546 | 0.88 | -1.759 | 0.428 | 0.017 | 0 | 0 | 0 | 0.7 | 0.78 | 0.72 | 0.75 | 0.73 | 0.81 |
| 5 | 83173714 | rs10059533 | 0.865 | 0.842 | -1.439 | 0.39 | -0.021 | -0.025 | -0.025 | 0 | 0.26 | 0.1 | 0.24 | 0.29 | 0.3 | 0.34 |
| 5 | 83175976 | rs11737928 | 0.233 | 0.753 | -0.96 | 0.332 | -0.005 | 0.008 | 0.008 | 0.069 | 0.12 | 0.13 | 0.11 | 0.1 | 0.15 | 0.11 |
| 5 | 83191479 | rs13177784 | -0.177 | 0.838 | -1.075 | 0.32 | 0.001 | 0.003 | 0.003 | 0.347 | 0.41 | 0.68 | 0.44 | 0.46 | 0.35 | 0.44 |
| 5 | 83196320 | rs4455541 | -0.113 | 1.017 | -1.05 | 0.179 | 0.022 | -0.002 | -0.002 | 0 | 0.41 | 0.74 | 0.43 | 0.47 | 0.3 | 0.45 |
| 5 | 83215930 | rs4703974 | -0.415 | 0.669 | -0.32 | 0.137 | 0.004 | 0.02 | 0.02 | 1.01 | 0.45 | 0.81 | 0.46 | 0.45 | 0.37 | 0.46 |
| 5 | 83217091 | rs12656419 | 0.734 | 0.526 | -0.189 | 0.137 | -0.054 | 0.014 | 0.014 | 0 | 0.3 | 0.43 | 0.3 | 0.35 | 0.15 | 0.35 |
| 5 | 83219474 | rs13160212 | -0.039 | 0.576 | -0.227 | 0.174 | -0.007 | -0.002 | -0.002 | 0.979 | 0.41 | 0.53 | 0.4 | 0.46 | 0.3 | 0.45 |
| 5 | 83221280 | rs10462406 | 0.031 | 0.264 | -0.213 | -0.15 | 0.049 | 0.083 | 0.083 | 0.286 | 0.53 | 0.29 | 0.56 | 0.63 | 0.67 | 0.65 |
| 5 | 83222316 | rs7703659 | -0.22 | 0.312 | 0.169 | -0.236 | 0.05 | -0.027 | -0.027 | 0 | 0.12 | 0.35 | 0.12 | 0.09 | 0.15 | 0.13 |
| 5 | 83222559 | rs4557383 | -0.04 | 0.167 | 0.096 | -0.275 | 0.029 | 0.081 | 0.081 | 0 | 0.54 | 0.25 | 0.56 | 0.63 | 0.68 | 0.63 |
| 5 | 83224173 | rs4364367 | 0.507 | 0.128 | 0.054 | -0.272 | 0 | -0.004 | -0.004 | 0 | 0.09 | 0.18 | 0.08 | 0.1 | 0.07 | 0.13 |
| 5 | 83224184 | rs4364368 | -1.202 | 0.013 | 0.251 | -0.289 | -0.027 | -0.023 | -0.023 | 0 | 0.69 | 0.66 | 0.69 | 0.74 | 0.83 | 0.76 |
| 5 | 83225289 | rs10474120 | 0.291 | 0.713 | -0.345 | -0.442 | -0.017 | 0.077 | 0.077 | 0.03 | 0.35 | 0.27 | 0.33 | 0.16 | 0.39 | 0.45 |
| 5 | 83226533 | rs7725359 | 0.441 | 0.806 | -0.426 | -0.131 | -0.052 | -0.051 | -0.051 | 0 | 0.08 | 0.01 | 0.09 | 0.2 | 0.17 | 0.06 |
| 5 | 83228107 | rs6865941 | 0.106 | 0.706 | -0.291 | -0.381 | -0.016 | 0.026 | 0.026 | 0.095 | 0.38 | 0.38 | 0.36 | 0.16 | 0.46 | 0.46 |
| 5 | 83232813 | rs4368707 | -0.792 | 0.599 | -0.263 | -0.219 | -0.006 | -0.015 | -0.015 | 0.174 | 0.63 | 0.48 | 0.64 | 0.74 | 0.75 | 0.74 |
| 5 | 83234200 | rs4533866 | 0.204 | 0.561 | -0.446 | -0.239 | -0.064 | 0.139 | 0.139 | 0.026 | 0.36 | 0.12 | 0.35 | 0.07 | 0.48 | 0.42 |
| 5 | 83239788 | rs13153252 | 1.114 | 0.148 | 0.219 | 0.061 | 0.004 | 0.02 | 0.02 | 0.875 | 0.32 | 0.21 | 0.36 | 0.32 | 0.28 | 0.21 |
| 5 | 83243314 | rs4703977 | 1.096 | 0.133 | 0.164 | 0.436 | 0.005 | -0.038 | -0.038 | 0 | 0.11 | 0.04 | 0.15 | 0.27 | 0.12 | 0.1 |
| 5 | 83249950 | rs9293361 | -0.059 | 0.161 | 0.713 | 0.368 | -0.007 | 0.003 | 0.003 | 0 | 0.56 | 0.6 | 0.57 | 0.57 | 0.56 | 0.4 |
| 5 | 83250507 | rs4330447 | -0.106 | 0.149 | 0.853 | 0.429 | 0.01 | -0.008 | -0.008 | 0 | 0.56 | 0.64 | 0.57 | 0.58 | 0.56 | 0.4 |
| 5 | 83250826 | rs4557384 | -2.316 | 0.334 | 0.84 | 0.375 | -0.032 | 0.025 | 0.025 | 1.428 | 0.92 | 0.85 | 0.96 | 0.91 | 0.96 | 0.89 |
| 5 | 83258483 | rs13160099 | 0.478 | 0.644 | 0.639 | 0.344 | 0.057 | 0.014 | 0.014 | 0 | 0.12 | 0.21 | 0.15 | 0.14 | 0.23 | 0.06 |
| 5 | 83260938 | rs2301070 | -0.099 | 0.428 | 0.276 | 0.278 | -0.009 | 0.001 | 0.001 | 0 | 0.54 | 0.56 | 0.55 | 0.57 | 0.56 | 0.4 |
| 5 | 83269151 | rs2301071 | 0.823 | 1.214 | 0.43 | 0.164 | 0.007 | -0.013 | -0.013 | 0 | 0.42 | 0.19 | 0.5 | 0.61 | 0.39 | 0.32 |
| 5 | 83284724 | rs4282296 | 0.497 | 0.839 | 0.386 | 0.52 | 0.002 | 0 | 0 | 0 | 0.83 | 0.84 | 0.87 | 0.85 | 0.81 | 0.82 |
| 5 | 83292653 | rs6452558 | -1.241 | 0.624 | 0.617 | 0.485 | 0.054 | 0.049 | 0.049 | 2.855 | 0.07 | 0.43 | 0.04 | 0.03 | 0.03 | 0.17 |
| 5 | 83300717 | rs2216561 | -0.394 | 1.247 | -0.621 | 0.839 | -0.032 | -0.052 | -0.052 | 0.466 | 0.26 | 0.13 | 0.32 | 0.45 | 0.33 | 0.24 |
| 5 | 83306192 | rs2301074 | 0.176 | 1.081 | -0.24 | 0.608 | -0.001 | -0.003 | -0.003 | 0 | 0.4 | 0.38 | 0.41 | 0.45 | 0.39 | 0.33 |
| 5 | 83307574 | rs7725025 | -0.815 | 1.502 | 0.756 | 0.384 | 0.119 | -0.027 | -0.027 | 0 | 0.08 | 0.32 | 0.07 | 0 | 0.11 | 0.06 |
| 5 | 83307587 | rs958643 | 0.913 | 0.804 | 1.161 | 0.558 | -0.023 | 0.011 | 0.011 | 0 | 0.81 | 0.47 | 0.78 | 0.77 | 0.84 | 0.88 |
| 5 | 83307766 | rs720267 | 1.223 | 0.931 | 1.209 | 0.596 | 0.002 | -0.003 | -0.003 | 0.383 | 0.88 | 0.74 | 0.9 | 0.83 | 0.9 | 0.89 |
| 5 | 83316361 | rs194124 | 0.663 | 1.064 | 0.606 | 0.669 | 0 | 0.009 | 0.009 | 1.759 | 0.64 | 0.26 | 0.56 | 0.67 | 0.66 | 0.55 |
| 5 | 83318042 | rs393110 | 0.68 | 1.012 | 0.442 | 0.612 | 0.012 | -0.008 | -0.008 | 0 | 0.89 | 0.62 | 0.9 | 0.87 | 0.91 | 0.83 |
| 5 | 83319613 | rs2301079 | -0.072 | 1.187 | 0.146 | 0.794 | -0.041 | 0.047 | 0.047 | 0.603 | 0.23 | 0.07 | 0.27 | 0.11 | 0.25 | 0.28 |
| 5 | 83329010 | rs1428925 | -0.527 | 1.267 | 0.313 | 0.75 | 0.024 | 0.017 | 0.017 | 4.458 | 0.08 | 0.28 | 0.07 | 0.13 | 0.06 | 0.13 |
| 5 | 83333988 | rs17629239 | 0.457 | 1.304 | -0.281 | 0.936 | -0.054 | -0.084 | -0.084 | 0 | 0.16 | 0.03 | 0.19 | 0.28 | 0.19 | 0.1 |
| 5 | 83340536 | rs1428926 | -1.086 | 1.229 | 0.378 | 0.971 | 0.004 | -0.009 | -0.009 | 0 | 0.28 | 0.36 | 0.33 | 0.23 | 0.31 | 0.36 |
| 5 | 83360520 | rs2269287 | -1.187 | 0.929 | 0.596 | 0.816 | 0.015 | 0.006 | 0.006 | 4.152 | 0.08 | 0.27 | 0.07 | 0.13 | 0.06 | 0.13 |
| 5 | 83372000 | rs4703982 | -0.808 | 1.164 | 0.551 | 1.103 | 0.028 | 0.135 | 0.135 | 0 | 0.2 | 0.41 | 0.31 | 0.42 | 0.21 | 0.31 |
| 5 | 83376382 | rs1559064 | 0.286 | 1.003 | 0.333 | 0.477 | -0.001 | -0.013 | -0.013 | 0 | 0.28 | 0.31 | 0.19 | 0.15 | 0.29 | 0.22 |
| 5 | 83377948 | rs10462408 | -0.614 | 0.922 | -0.129 | 0.14 | 0.007 | 0.027 | 0.027 | 0 | 0.29 | 0.37 | 0.37 | 0.46 | 0.3 | 0.42 |
| 5 | 83378524 | rs6878457 | -1.219 | 0.864 | -0.114 | 0.305 | 0.003 | 0.051 | 0.051 | 0.092 | 0.59 | 0.83 | 0.63 | 0.71 | 0.59 | 0.63 |
| 5 | 83378766 | rs4703574 | -0.583 | 0.8 | 0.17 | 0.455 | 0.006 | 0.02 | 0.02 | 0 | 0.28 | 0.31 | 0.36 | 0.44 | 0.3 | 0.39 |
| 5 | 83382640 | rs10073918 | -0.095 | 0.202 | -0.163 | 0.452 | -0.005 | 0.001 | 0.001 | 0 | 0.41 | 0.42 | 0.46 | 0.38 | 0.37 | 0.35 |
| 5 | 83385090 | rs2269289 | -0.993 | 0.324 | -0.205 | 0.609 | 0.001 | -0.007 | -0.007 | 0 | 0.74 | 0.75 | 0.72 | 0.6 | 0.75 | 0.71 |
| 5 | 83385150 | rs2269290 | 0.878 | 0.44 | -0.814 | 0.605 | 0.014 | 0.002 | 0.002 | 0.172 | 0.29 | 0.18 | 0.32 | 0.23 | 0.28 | 0.27 |
| 5 | 83406044 | rs350477 | -0.73 | 0.887 | -0.149 | 0.431 | -0.007 | 0.016 | 0.016 | 0 | 0.42 | 0.36 | 0.32 | 0.24 | 0.47 | 0.42 |
| 5 | 83408465 | rs2301098 | 0.088 | 1.08 | -0.964 | 0.676 | 0 | -0.01 | -0.01 | 0 | 0.11 | 0.07 | 0.07 | 0.03 | 0.08 | 0.08 |
| 5 | 83410784 | rs348894 | -0.223 | 1.317 | -1.061 | 1.022 | -0.049 | 0.024 | 0.024 | 0 | 0.29 | 0.17 | 0.25 | 0.22 | 0.36 | 0.31 |
| 5 | 83437836 | rs39966 | -0.24 | 0.972 | -1.434 | 0.84 | -0.067 | -0.058 | -0.058 | 0.097 | 0.22 | 0.01 | 0.17 | 0.16 | 0.24 | 0.18 |
| 5 | 83439277 | rs164437 | -0.74 | 0.588 | -0.555 | 0.648 | -0.001 | -0.01 | -0.01 | 0 | 0.37 | 0.38 | 0.5 | 0.75 | 0.37 | 0.43 |
| 5 | 83443132 | rs32598 | -0.578 | 0.561 | -0.356 | 0.843 | -0.003 | 0.036 | 0.036 | 0 | 0.35 | 0.19 | 0.44 | 0.7 | 0.37 | 0.42 |
| 5 | 83450922 | rs12153170 | 0.442 | 0.293 | -1.192 | 0.24 | -0.022 | 0.024 | 0.024 | 0.396 | 0.14 | 0.01 | 0.09 | 0.09 | 0.15 | 0.15 |
| 5 | 83459109 | rs347347 | -0.312 | 0.395 | -0.6 | -0.023 | -0.002 | -0.011 | -0.011 | 0 | 0.42 | 0.47 | 0.47 | 0.7 | 0.4 | 0.44 |
| 5 | 83461429 | rs11747890 | 0.527 | 0.828 | -0.94 | -0.724 | 0.014 | 0.173 | 0.173 | 0 | 0.41 | 0.18 | 0.31 | 0.13 | 0.38 | 0.32 |
| 5 | 83469915 | rs153248 | -0.306 | 0.631 | -0.406 | -0.438 | 0.024 | -0.032 | -0.032 | 0.24 | 0.63 | 0.47 | 0.6 | 0.7 | 0.53 | 0.58 |
| 5 | 83470986 | rs342422 | -0.195 | 0.649 | -0.395 | -0.42 | 0.011 | -0.019 | -0.019 | 1.356 | 0.61 | 0.44 | 0.6 | 0.7 | 0.52 | 0.58 |
| 5 | 83477213 | rs163716 | -0.292 | 0.915 | -0.419 | -0.23 | 0.002 | 0.025 | 0.025 | 1.261 | 0.41 | 0.25 | 0.43 | 0.24 | 0.42 | 0.48 |
| 5 | 83482308 | rs342406 | -0.418 | 1.042 | -0.501 | -0.105 | -0.008 | 0.015 | 0.015 | 0 | 0.86 | 0.92 | 0.76 | 0.93 | 0.85 | 0.73 |

| | | | | | | | | | | | | | | | | |
|---|---|---|---|---|---|---|---|---|---|---|---|---|---|---|---|---|
| 5 | 83495204 | rs13155727 | 0.541 | 1.792 | -0.235 | -0.019 | 0.054 | -0.01 | -0.01 | 0 | 0.05 | 0.08 | 0.09 | 0.02 | 0.15 | 0.03 |
| 5 | 83499643 | rs347344 | 0.648 | 1.432 | -0.114 | -0.132 | -0.002 | 0 | 0 | 0.937 | 0.62 | 0.67 | 0.53 | 0.71 | 0.57 | 0.53 |
| 5 | 83508667 | rs342421 | 0.406 | 0.786 | -0.245 | -0.022 | 0.005 | 0.004 | 0.004 | 2.018 | 0.67 | 0.69 | 0.59 | 0.74 | 0.72 | 0.59 |
| 5 | 83511806 | rs342418 | -0.411 | 0.914 | 0.01 | -0.143 | 0.012 | -0.021 | -0.021 | 3.988 | 0.88 | 0.7 | 0.79 | 0.91 | 0.88 | 0.81 |
| 5 | 83512129 | rs2053677 | 0.003 | 0.561 | -0.774 | -0.112 | -0.023 | -0.013 | -0.013 | 0 | 0.19 | 0.01 | 0.12 | 0.14 | 0.15 | 0.17 |
| 5 | 83513967 | rs342415 | 0.825 | 0.485 | -0.917 | -0.278 | 0.005 | 0.014 | 0.014 | 0.13 | 0.55 | 0.62 | 0.54 | 0.7 | 0.56 | 0.48 |
| 5 | 83514459 | rs342414 | 0.01 | 0.488 | -0.853 | -0.311 | 0 | -0.005 | -0.005 | 0.048 | 0.75 | 0.75 | 0.66 | 0.86 | 0.71 | 0.65 |
| 5 | 83516148 | rs13166899 | 0.333 | 0.598 | -1.102 | 0.207 | -0.002 | -0.017 | -0.017 | 0.06 | 0.2 | 0.15 | 0.14 | 0.33 | 0.19 | 0.14 |
| 5 | 83518105 | rs17206271 | 0.733 | 0.505 | -1.253 | 0.128 | -0.01 | -0.012 | -0.012 | 0.802 | 0.18 | 0.03 | 0.11 | 0.13 | 0.17 | 0.16 |
| 5 | 83518387 | rs10056071 | 1.23 | 0.893 | -1.052 | -0.074 | 0.015 | 0.04 | 0.04 | 0 | 0.27 | 0.42 | 0.35 | 0.38 | 0.27 | 0.32 |
| 5 | 83523495 | rs2121105 | 1.103 | 0.923 | -1.022 | -0.024 | 0.015 | 0.019 | 0.019 | 0 | 0.27 | 0.41 | 0.36 | 0.36 | 0.27 | 0.32 |
| 5 | 83524072 | rs16900992 | 1.333 | 0.761 | -1.133 | -0.102 | -0.019 | -0.033 | -0.033 | 0 | 0.12 | 0.02 | 0.09 | 0.13 | 0.13 | 0.1 |
| 5 | 83524163 | rs10474123 | 1.182 | 0.744 | -0.856 | -0.004 | 0.016 | 0.024 | 0.024 | 0 | 0.26 | 0.41 | 0.36 | 0.34 | 0.26 | 0.32 |
| 5 | 83527862 | rs1373962 | 0.005 | 0.88 | -1.257 | -0.638 | -0.01 | 0.01 | 0.01 | 0 | 0.31 | 0.22 | 0.49 | 0.8 | 0.4 | 0.45 |
| 5 | 83529051 | rs16900997 | 0.836 | 0.358 | -1.488 | -1.049 | 0.027 | 0.052 | 0.052 | 0 | 0.27 | 0.12 | 0.17 | 0.14 | 0.19 | 0.23 |
| 5 | 83530071 | rs3797655 | 1.607 | 0.217 | -1.559 | -1.07 | 0.019 | -0.021 | -0.021 | 0 | 0.12 | 0.03 | 0.08 | 0.14 | 0.09 | 0.18 |
| 5 | 83531299 | rs2218504 | -1.454 | 0.281 | -1.417 | -1.274 | -0.015 | 0.074 | 0.074 | 0 | 0.79 | 0.85 | 0.83 | 0.95 | 0.76 | 0.74 |
| 5 | 83536587 | rs7707528 | -1.344 | 0.274 | -1.203 | -1.276 | -0.002 | 0.023 | 0.023 | 0 | 0.78 | 0.77 | 0.83 | 0.95 | 0.76 | 0.74 |
| 5 | 83564529 | rs17284887 | 1.815 | 0.477 | -1.382 | -1.12 | -0.034 | -0.035 | -0.035 | 0 | 0.1 | 0 | 0.09 | 0.12 | 0.13 | 0.13 |
| 5 | 83570979 | rs3846638 | 1.79 | 0.248 | -1.277 | -1.31 | 0.005 | 0.028 | 0.028 | 0.152 | 0.14 | 0.08 | 0.09 | 0 | 0.1 | 0.05 |
| 5 | 83575190 | rs11745906 | 0.708 | 0.459 | -0.848 | -0.503 | 0 | 0.006 | 0.006 | 0 | 0.23 | 0.2 | 0.2 | 0.43 | 0.29 | 0.23 |
| 5 | 83579008 | rs16901039 | 2.257 | 0.329 | -0.776 | -0.728 | 0 | -0.023 | -0.023 | 0 | 0.18 | 0.16 | 0.13 | 0.01 | 0.15 | 0.05 |
| 5 | 83594712 | rs2173698 | 0.907 | 0.27 | -0.759 | -0.709 | 0.002 | 0.042 | 0.042 | 0 | 0.37 | 0.34 | 0.26 | 0.05 | 0.34 | 0.3 |
| 5 | 83623880 | rs4301211 | 0.209 | 0.269 | -0.842 | -0.597 | 0.019 | 0.107 | 0.107 | 0 | 0.49 | 0.37 | 0.35 | 0.2 | 0.44 | 0.49 |
| 5 | 83625620 | rs1500227 | 1.319 | 0.226 | -1.011 | -0.608 | -0.039 | 0.204 | 0.204 | 0 | 0.35 | 0.1 | 0.24 | 0.05 | 0.34 | 0.3 |
| 5 | 83626673 | rs10072019 | 0.283 | 0.279 | -0.943 | -0.543 | 0.02 | 0.145 | 0.145 | 0 | 0.49 | 0.27 | 0.35 | 0.2 | 0.45 | 0.49 |
| 5 | 83632691 | rs7720807 | 1.34 | 0.593 | -1.058 | 0.311 | -0.01 | 0.016 | 0.016 | 0 | 0.22 | 0.14 | 0.25 | 0.17 | 0.23 | 0.24 |
| 5 | 83638031 | rs12153600 | 1.181 | 0.658 | -1.007 | 0.314 | 0.007 | -0.02 | -0.02 | 3.312 | 0.13 | 0.03 | 0.09 | 0.15 | 0.11 | 0.2 |
| 5 | 83662645 | rs10073440 | -0.08 | 0.688 | -0.841 | 0.24 | 0.007 | 0.004 | 0.004 | 0 | 0.37 | 0.37 | 0.39 | 0.57 | 0.49 | 0.47 |
| 5 | 83681037 | rs305661 | 0.486 | 0.613 | -0.58 | 0.183 | -0.001 | -0.025 | -0.025 | 9.796 | 0.25 | 0.33 | 0.21 | 0.16 | 0.23 | 0.25 |
| 5 | 83686405 | rs2941228 | -1.239 | 0.648 | -0.846 | 0.198 | 0.031 | 0.077 | 0.077 | 0 | 0.56 | 0.78 | 0.5 | 0.71 | 0.62 | 0.66 |
| 5 | 83699343 | rs10061976 | -0.485 | 0.946 | -0.522 | 0.74 | 0.014 | 0.029 | 0.029 | 0 | 0.31 | 0.38 | 0.3 | 0.55 | 0.38 | 0.42 |
| 5 | 83704241 | rs13170840 | 0.335 | 0.894 | -1.168 | 0.617 | -0.052 | -0.006 | -0.006 | 0.64 | 0.15 | 0.01 | 0.1 | 0.09 | 0.19 | 0.15 |
| 5 | 83740880 | rs13172559 | 0.107 | 0.027 | -0.302 | 0.541 | -0.046 | -0.017 | -0.017 | 0 | 0.8 | 0.52 | 0.85 | 0.86 | 0.87 | 0.79 |
| 5 | 83756654 | rs10037310 | 0.899 | 0.515 | -0.893 | 0.321 | -0.037 | 0.061 | 0.061 | 0.109 | 0.11 | 0.02 | 0.06 | 0.04 | 0.14 | 0.13 |
| 5 | 83787691 | rs6881082 | -0.894 | 0.534 | -1.248 | 0.599 | -0.061 | -0.098 | -0.098 | 0 | 0.82 | 0.99 | 0.77 | 0.75 | 0.78 | 0.8 |
| 5 | 83806010 | rs6886927 | -1.024 | 0.473 | -0.488 | 0.231 | 0.03 | 0.017 | 0.017 | 0.098 | 0.49 | 0.72 | 0.58 | 0.46 | 0.4 | 0.46 |
| 5 | 83842687 | rs10067300 | 1.547 | 0.307 | -0.441 | 0.333 | -0.033 | 0.035 | 0.035 | 0 | 0.08 | 0.03 | 0.04 | 0.04 | 0.11 | 0.08 |
| 5 | 83855826 | rs10041822 | -1.402 | 0.356 | -0.063 | 0.37 | 0.005 | 0.02 | 0.02 | 0.289 | 0.82 | 0.8 | 0.62 | 0.72 | 0.81 | 0.89 |
| 5 | 83856400 | rs13165786 | 0.566 | 0.229 | 0.073 | 0.249 | 0.005 | 0.011 | 0.011 | 0 | 0.63 | 0.42 | 0.46 | 0.56 | 0.61 | 0.73 |
| 5 | 83860927 | rs16901159 | -0.125 | 0.175 | 0.302 | 0.143 | -0.017 | -0.033 | -0.033 | 0.011 | 0.15 | 0.16 | 0.06 | 0.03 | 0.05 | 0.17 |
| 5 | 83867744 | rs7706657 | -0.805 | 0.638 | 0.22 | 0.467 | 0.011 | -0.015 | -0.015 | 0.17 | 0.16 | 0.34 | 0.13 | 0.15 | 0.17 | 0.11 |
| 5 | 83870263 | rs13359765 | 0.188 | 0.772 | -0.075 | 0.5 | -0.039 | 0.008 | 0.008 | 0.193 | 0.11 | 0.02 | 0.08 | 0.09 | 0.15 | 0.15 |
| 5 | 83883999 | rs10065928 | -0.417 | 0.72 | 0.369 | 0.536 | 0.036 | 0.101 | 0.101 | 2.283 | 0.06 | 0.3 | 0.07 | 0.1 | 0.05 | 0.02 |
| 5 | 83884321 | rs10059091 | -0.626 | 0.714 | 0.41 | 0.533 | 0.045 | 0.113 | 0.113 | 2.904 | 0.06 | 0.34 | 0.08 | 0.1 | 0.05 | 0.02 |
| 5 | 83890606 | rs10462410 | -0.734 | 0.687 | 0.375 | 0.527 | 0.016 | 0.109 | 0.109 | 1.084 | 0.07 | 0.31 | 0.17 | 0.12 | 0.06 | 0.03 |
| 5 | 83897084 | rs10052280 | -0.587 | 0.718 | 0.351 | 0.542 | 0.011 | 0.099 | 0.099 | 2.657 | 0.07 | 0.27 | 0.17 | 0.12 | 0.06 | 0.03 |
| 5 | 83907654 | rs10077624 | -0.629 | 0.744 | -0.075 | 0.347 | 0.001 | 0 | 0 | 0 | 0.23 | 0.2 | 0.14 | 0.11 | 0.19 | 0.29 |
| 5 | 83915898 | rs9293378 | -1.222 | 0.884 | -0.417 | 0.317 | -0.01 | -0.022 | -0.022 | 0 | 0.28 | 0.62 | 0.36 | 0.25 | 0.27 | 0.28 |
| 5 | 83916464 | rs13436770 | 0.089 | 0.976 | -0.169 | 0.47 | 0.002 | -0.039 | -0.039 | 0 | 0.24 | 0.3 | 0.16 | 0.12 | 0.23 | 0.25 |
| 5 | 83917033 | rs4454038 | -0.116 | 1.129 | -0.784 | 0.522 | -0.002 | -0.029 | -0.029 | 1.389 | 0.31 | 0.6 | 0.33 | 0.24 | 0.3 | 0.29 |
| 5 | 83926167 | rs16901186 | 0.768 | 1.1 | -1.153 | 0.711 | -0.031 | -0.05 | -0.05 | 0 | 0.13 | 0 | 0.09 | 0.17 | 0.15 | 0.11 |
| 5 | 83932189 | rs12153218 | 0.057 | 1.17 | -1.17 | 0.748 | -0.001 | -0.033 | -0.033 | 0.968 | 0.3 | 0.55 | 0.32 | 0.25 | 0.29 | 0.28 |
| 5 | 83932510 | rs10071420 | 0.082 | 1.12 | -0.896 | 0.833 | -0.001 | 0.002 | 0.002 | 0 | 0.43 | 0.59 | 0.42 | 0.42 | 0.44 | 0.4 |
| 5 | 83963326 | rs10514267 | -0.711 | 1.555 | -1.774 | 1.34 | -0.054 | -0.078 | -0.078 | 0 | 0.12 | 0.01 | 0.17 | 0.22 | 0.16 | 0.19 |
| 5 | 83974401 | rs7713225 | -0.75 | 1.081 | -1.978 | 0.48 | -0.048 | 0.074 | 0.074 | 0.466 | 0.26 | 0.1 | 0.21 | 0.11 | 0.29 | 0.39 |
| 5 | 83978496 | rs6866089 | 0.228 | 0.674 | -1.821 | 0.552 | 0.002 | 0.001 | 0.001 | 0 | 0.54 | 0.56 | 0.43 | 0.59 | 0.38 | 0.52 |
| 5 | 83978954 | rs7710912 | -0.554 | 0.674 | -1.662 | 0.347 | 0.043 | -0.004 | -0.004 | 0 | 0.94 | 0.73 | 0.93 | 1 | 0.92 | 0.98 |
| 5 | 83981463 | rs11958291 | 0.485 | 0.66 | -1.516 | 0.356 | 0.001 | 0.001 | 0.001 | 0 | 0.54 | 0.56 | 0.43 | 0.59 | 0.38 | 0.52 |
| 5 | 83981802 | rs6452576 | 0.854 | -0.054 | -1.925 | 0.763 | -0.013 | -0.024 | -0.024 | 0 | 0.41 | 0.51 | 0.35 | 0.3 | 0.32 | 0.27 |
| 5 | 83983783 | rs12523086 | 0.805 | 0.32 | -2.149 | 0.711 | -0.051 | -0.013 | -0.013 | 0 | 0.12 | 0.03 | 0.12 | 0.11 | 0.17 | 0.21 |
| 5 | 83984548 | rs10050693 | 0.062 | 0.248 | -2.234 | 0.616 | -0.071 | -0.048 | -0.048 | 0 | 0.37 | 0.14 | 0.49 | 0.41 | 0.53 | 0.44 |
| 5 | 83986193 | rs4416606 | 0.88 | 0.262 | -2.142 | 0.622 | -0.051 | -0.013 | -0.013 | 3.494 | 0.12 | 0.03 | 0.12 | 0.11 | 0.17 | 0.21 |
| 5 | 83988022 | rs6892080 | 0.844 | 0.078 | -2.01 | 0.384 | -0.088 | -0.069 | -0.069 | 0 | 0.25 | 0.05 | 0.37 | 0.3 | 0.36 | 0.23 |
| 5 | 84003689 | rs10076429 | 0.39 | -0.424 | -1.663 | 0.594 | 0.001 | -0.034 | -0.034 | 0 | 0.76 | 0.81 | 0.79 | 0.62 | 0.75 | 0.6 |
| 5 | 84004601 | rs10462415 | 0.079 | -0.879 | -1.06 | 0.389 | -0.007 | -0.005 | -0.005 | 0 | 0.41 | 0.53 | 0.39 | 0.39 | 0.32 | 0.29 |
| 5 | 84006457 | rs4282316 | -0.473 | -0.8 | -1.377 | 0.393 | 0.059 | -0.008 | -0.008 | 0 | 0.48 | 0.85 | 0.37 | 0.47 | 0.34 | 0.31 |
| 5 | 84018132 | rs10051998 | -1.572 | -0.861 | -1.231 | 0.256 | -0.023 | -0.046 | -0.046 | 0 | 0.22 | 0.32 | 0.13 | 0.07 | 0.16 | 0.24 |
| 5 | 84021236 | rs11748013 | -1.393 | -0.865 | -1.231 | 0.257 | -0.041 | -0.046 | -0.046 | 0.008 | 0.19 | 0.31 | 0.08 | 0.07 | 0.08 | 0.21 |

| | | | | | | | | | | | | | | | | |
|---|---|---|---|---|---|---|---|---|---|---|---|---|---|---|---|---|
| 5 | 84025813 | rs10052448 | -0.542 | -0.681 | -1.214 | 0.328 | 0.038 | 0.029 | 0.029 | 0.343 | 0.49 | 0.75 | 0.41 | 0.44 | 0.43 | 0.33 |
| 5 | 84033417 | rs10462421 | -0.151 | -0.474 | -1.416 | 0.389 | 0.043 | -0.017 | -0.017 | 1.874 | 0.44 | 0.73 | 0.32 | 0.5 | 0.32 | 0.25 |
| 5 | 84046428 | rs7722606 | -0.494 | -0.143 | -1.52 | 0.616 | -0.022 | -0.043 | -0.043 | 0 | 0.55 | 0.81 | 0.44 | 0.5 | 0.45 | 0.4 |
| 5 | 84054083 | rs4605785 | -0.878 | 0.24 | -0.279 | 0.9 | 0.018 | 0.045 | 0.045 | 1.851 | 0.12 | 0.62 | 0.07 | 0.12 | 0.1 | 0.04 |
| 5 | 84054709 | rs4461632 | -1.107 | -0.215 | -0.061 | 0.747 | -0.03 | 0.014 | 0.014 | 0 | 0.2 | 0.52 | 0.19 | 0.26 | 0.13 | 0.22 |
| 5 | 84055287 | rs6414935 | -1.795 | 0.031 | -0.384 | 0.791 | -0.007 | -0.077 | -0.077 | 0 | 0.39 | 0.88 | 0.36 | 0.44 | 0.32 | 0.38 |
| 5 | 84061023 | rs6888165 | -0.502 | -0.063 | 0.062 | 0.943 | 0.017 | 0.054 | 0.054 | 0 | 0.18 | 0.38 | 0.12 | 0.23 | 0.17 | 0.15 |
| 5 | 84062058 | rs4133436 | -1.779 | 0.105 | -0.509 | -0.019 | -0.024 | 0.097 | 0.097 | 0.053 | 0.4 | 0.9 | 0.35 | 0.25 | 0.37 | 0.41 |
| 5 | 84065374 | rs6871305 | -0.379 | 0.143 | -0.305 | 0.002 | 0.02 | -0.057 | -0.057 | 0 | 0.18 | 0.39 | 0.1 | 0.07 | 0.17 | 0.14 |
| 5 | 84071853 | rs7448787 | 0.133 | 0.045 | -1.113 | 0.283 | 0.008 | 0.063 | 0.063 | 0 | 0.06 | 0.17 | 0.12 | 0.16 | 0.06 | 0.14 |
| 5 | 84076305 | rs1158242 | -0.669 | 0.319 | -1.707 | 0.217 | -0.021 | 0.08 | 0.08 | 0 | 0.42 | 0.9 | 0.36 | 0.23 | 0.38 | 0.42 |
| 5 | 84091283 | rs7707956 | 0.069 | 0.291 | -0.938 | -0.086 | -0.012 | -0.037 | -0.037 | 0 | 0.76 | 0.69 | 0.69 | 0.83 | 0.81 | 0.81 |
| 5 | 84097825 | rs10942359 | -0.189 | 0.86 | -1.436 | -0.267 | -0.04 | -0.004 | -0.004 | 0.947 | 0.1 | 0 | 0.08 | 0.07 | 0.1 | 0.18 |
| 5 | 84098894 | rs4565212 | -0.418 | 0.807 | -1.701 | -1.053 | 0.009 | 0.128 | 0.128 | 0 | 0.48 | 0.27 | 0.61 | 0.82 | 0.55 | 0.6 |
| 5 | 84099716 | rs4345320 | -0.367 | 0.734 | -1.739 | -1.053 | -0.018 | 0.206 | 0.206 | 0 | 0.47 | 0.16 | 0.59 | 0.82 | 0.55 | 0.6 |
| 5 | 84111810 | rs4235660 | -0.379 | 0.109 | -1.153 | -1.715 | -0.046 | -0.104 | -0.104 | 1.476 | 0.75 | 0.59 | 0.83 | 0.93 | 0.85 | 0.76 |
| 5 | 84112660 | rs7737976 | 0.983 | 0.337 | -1.383 | -1.453 | -0.052 | 0.069 | 0.069 | 0.018 | 0.25 | 0.06 | 0.2 | 0.12 | 0.29 | 0.17 |
| 5 | 84120872 | rs4432885 | 0.087 | 0.806 | -1.225 | -0.565 | -0.014 | -0.035 | -0.035 | 0.077 | 0.32 | 0.24 | 0.49 | 0.49 | 0.33 | 0.45 |
| 5 | 84122916 | rs2036347 | 1.068 | 0.925 | -1.68 | -0.762 | -0.053 | 0.02 | 0.02 | 0 | 0.13 | 0.03 | 0.11 | 0.08 | 0.2 | 0.09 |
| 5 | 84124824 | rs34280106 | 0.724 | 0.781 | -1.63 | -1.034 | -0.067 | 0.161 | 0.161 | 0 | 0.44 | 0.14 | 0.32 | 0.21 | 0.48 | 0.31 |
| 5 | 84127329 | rs981144 | -0.602 | 0.71 | -0.753 | -0.96 | -0.002 | 0.067 | 0.067 | 0 | 0.68 | 0.46 | 0.5 | 0.52 | 0.73 | 0.57 |
| 5 | 84136777 | rs4451048 | 0.779 | 0.713 | -0.535 | -0.877 | 0.001 | 0.069 | 0.069 | 0 | 0.14 | 0.23 | 0.09 | 0.25 | 0.17 | 0.03 |
| 5 | 84138061 | rs16901251 | 1.681 | 0.699 | -0.898 | -0.801 | -0.017 | -0.053 | -0.053 | 1.053 | 0.09 | 0.01 | 0.08 | 0.25 | 0.14 | 0.1 |
| 5 | 84138612 | rs12655657 | 0.619 | 0.802 | -0.846 | -0.878 | -0.009 | -0.026 | -0.026 | 0 | 0.3 | 0.25 | 0.48 | 0.48 | 0.31 | 0.44 |
| 5 | 84138750 | rs4282315 | -1.357 | 0.705 | -0.945 | -0.932 | 0.012 | 0.074 | 0.074 | 0 | 0.68 | 0.82 | 0.77 | 0.88 | 0.72 | 0.77 |
| 5 | 84155207 | rs12654219 | 0.782 | 0.764 | -1.129 | 0.138 | -0.009 | -0.051 | -0.051 | 0.329 | 0.2 | 0.07 | 0.29 | 0.29 | 0.17 | 0.23 |
| 5 | 84157168 | rs13165966 | 0.6 | 0.505 | -0.911 | -0.022 | 0.008 | -0.015 | -0.015 | 4.098 | 0.44 | 0.3 | 0.39 | 0.48 | 0.46 | 0.38 |
| 5 | 84157521 | rs7713643 | -0.014 | 0.113 | -0.515 | -0.051 | 0.002 | 0 | 0 | 0 | 0.55 | 0.43 | 0.52 | 0.56 | 0.61 | 0.55 |
| 5 | 84160703 | rs6860783 | 0.95 | 0.228 | -0.414 | 0.968 | 0.001 | 0 | 0 | 0 | 0.09 | 0.09 | 0.08 | 0.25 | 0.08 | 0.07 |
| 5 | 84160853 | rs1500619 | -0.584 | -0.041 | -0.285 | 1.066 | -0.014 | 0.04 | 0.04 | 0 | 0.3 | 0.43 | 0.31 | 0.48 | 0.3 | 0.32 |
| 5 | 84161671 | rs16901263 | 1.987 | -0.141 | -0.513 | 0.873 | -0.001 | -0.017 | -0.017 | 0.337 | 0.16 | 0.08 | 0.27 | 0.2 | 0.14 | 0.21 |
| 5 | 84163069 | rs11950801 | 0.872 | -0.175 | -0.423 | 0.942 | -0.005 | -0.011 | -0.011 | 0 | 0.54 | 0.48 | 0.5 | 0.34 | 0.56 | 0.46 |
| 5 | 84163749 | rs10942360 | 2.138 | -0.1 | -0.597 | 0.966 | -0.037 | -0.063 | -0.063 | 0 | 0.16 | 0.01 | 0.26 | 0.2 | 0.14 | 0.21 |
| 5 | 84166481 | rs12520666 | 1.904 | -0.07 | -0.55 | 0.983 | -0.001 | -0.019 | -0.019 | 0 | 0.16 | 0.08 | 0.27 | 0.2 | 0.14 | 0.21 |
| 5 | 84167042 | rs4466170 | 1.34 | 0.081 | -0.63 | 0.688 | -0.002 | 0.005 | 0.005 | 3.457 | 0.24 | 0.18 | 0.26 | 0.23 | 0.25 | 0.26 |
| 5 | 84167325 | rs7720603 | 1.46 | -0.013 | -0.533 | 0.598 | -0.021 | 0.044 | 0.044 | 0.902 | 0.28 | 0.17 | 0.22 | 0.11 | 0.31 | 0.2 |
| 5 | 84167824 | rs4235662 | 1.311 | 0.041 | -0.581 | 0.667 | 0 | 0.005 | 0.005 | 3.225 | 0.25 | 0.17 | 0.27 | 0.23 | 0.25 | 0.26 |
| 5 | 84168406 | rs6896935 | -1.839 | -0.001 | -0.525 | 0.848 | -0.001 | 0.008 | 0.008 | 0 | 0.91 | 0.89 | 0.92 | 0.76 | 0.91 | 0.92 |
| 5 | 84170267 | rs10068633 | -0.643 | -0.014 | -0.619 | 0.844 | -0.006 | 0.023 | 0.023 | 1.549 | 0.73 | 0.68 | 0.64 | 0.56 | 0.77 | 0.71 |
| 5 | 84170604 | rs10060952 | -0.643 | -0.014 | -0.632 | 0.844 | -0.006 | 0.021 | 0.021 | 1.549 | 0.73 | 0.68 | 0.64 | 0.56 | 0.77 | 0.71 |
| 5 | 84177145 | rs12522014 | 2.325 | 0.095 | -0.84 | 0.819 | -0.033 | -0.061 | -0.061 | 0.518 | 0.15 | 0.01 | 0.26 | 0.19 | 0.14 | 0.21 |
| 5 | 84177643 | rs11958924 | 0.099 | 0.089 | -0.714 | 0.739 | 0.001 | 0.005 | 0.005 | 0.225 | 0.49 | 0.55 | 0.38 | 0.33 | 0.52 | 0.45 |
| 5 | 84177660 | rs7380362 | 1.029 | 0.009 | -0.656 | 0.965 | -0.001 | 0.004 | 0.004 | 2.802 | 0.25 | 0.18 | 0.26 | 0.24 | 0.26 | 0.27 |
| 5 | 84178732 | rs10474144 | 0.367 | 0.031 | -0.73 | 0.772 | -0.004 | 0.019 | 0.019 | 0.378 | 0.21 | 0.3 | 0.15 | 0.21 | 0.22 | 0.24 |
| 5 | 84192194 | rs6870856 | -0.005 | -0.216 | -0.907 | 0.668 | -0.004 | 0 | 0 | 5.52 | 0.57 | 0.46 | 0.62 | 0.5 | 0.63 | 0.57 |
| 5 | 84195407 | rs4441875 | 0.175 | 0.151 | -1.694 | 0.586 | -0.003 | -0.004 | -0.004 | 1.592 | 0.42 | 0.21 | 0.56 | 0.41 | 0.57 | 0.46 |
| 5 | 84198417 | rs1500630 | 1.079 | 0.418 | -1.718 | 0.49 | -0.041 | -0.062 | -0.062 | 0.219 | 0.16 | 0.01 | 0.27 | 0.19 | 0.16 | 0.26 |
| 5 | 84216086 | rs13171180 | 0.745 | 0.141 | -1.728 | 0.217 | -0.019 | 0.045 | 0.045 | 0 | 0.1 | 0.01 | 0.07 | 0 | 0.09 | 0.03 |
| 5 | 84219553 | rs10036067 | 1.012 | 0.118 | -1.874 | 0.364 | -0.046 | -0.042 | -0.042 | 0 | 0.21 | 0.01 | 0.25 | 0.2 | 0.21 | 0.3 |
| 5 | 84230881 | rs4507487 | 0.013 | -0.055 | -1.509 | 0.365 | 0.031 | 0.027 | 0.027 | 1.237 | 0.54 | 0.69 | 0.65 | 0.56 | 0.62 | 0.44 |
| 5 | 84234232 | rs3913424 | -0.882 | -0.21 | -1.744 | 0.295 | 0.006 | -0.032 | -0.032 | 0 | 0.78 | 0.97 | 0.89 | 0.75 | 0.84 | 0.75 |
| 5 | 84236586 | rs1912463 | 0.331 | 0.234 | -0.563 | 1.051 | -0.017 | 0.008 | 0.008 | 2.249 | 0.26 | 0.45 | 0.24 | 0.27 | 0.22 | 0.31 |
| 5 | 84240377 | rs10085144 | 1.073 | 0.264 | -0.791 | 0.971 | -0.05 | -0.022 | -0.022 | 2.214 | 0.21 | 0.01 | 0.22 | 0.2 | 0.21 | 0.31 |
| 5 | 84242662 | rs2048343 | -1.528 | 0.198 | -0.802 | 0.866 | 0.004 | -0.015 | -0.015 | 0 | 0.93 | 0.95 | 0.96 | 0.78 | 0.94 | 0.9 |
| 5 | 84248966 | rs9293392 | 1.19 | 0.196 | -0.871 | 0.847 | -0.056 | 0.066 | 0.066 | 3.049 | 0.21 | 0.01 | 0.18 | 0.11 | 0.21 | 0.31 |
| 5 | 84263712 | rs4704018 | 1.001 | 0.515 | -0.564 | 1.069 | 0.017 | 0.02 | 0.02 | 1.382 | 0.42 | 0.27 | 0.39 | 0.33 | 0.37 | 0.41 |
| 5 | 84270117 | rs4704022 | 1.327 | 0.628 | -0.608 | 1.02 | -0.023 | -0.024 | -0.024 | 0.104 | 0.16 | 0.12 | 0.29 | 0.27 | 0.28 | 0.15 |
| 5 | 84281179 | rs2063362 | -0.525 | 0.518 | -0.396 | 1.001 | -0.001 | -0.001 | -0.001 | 0 | 0.7 | 0.66 | 0.76 | 0.66 | 0.73 | 0.59 |
| 5 | 84322004 | rs6861810 | 1.22 | 0.709 | -0.452 | 1.198 | -0.021 | -0.019 | -0.019 | 0 | 0.15 | 0.12 | 0.29 | 0.24 | 0.29 | 0.14 |
| 5 | 84322960 | rs1510400 | 0.093 | 0.518 | -0.367 | 1.2 | -0.005 | 0.001 | 0.001 | 3.568 | 0.6 | 0.42 | 0.67 | 0.52 | 0.69 | 0.51 |
| 5 | 84326295 | rs2136724 | -0.34 | 0.446 | -0.332 | 1.208 | -0.003 | -0.004 | -0.004 | 0 | 0.64 | 0.61 | 0.72 | 0.62 | 0.69 | 0.51 |
| 5 | 84339722 | rs6452630 | -1.128 | 0.46 | -0.76 | 0.855 | 0.004 | 0.005 | 0.005 | 1.666 | 0.64 | 0.7 | 0.67 | 0.71 | 0.66 | 0.66 |
| 5 | 84352281 | rs6452631 | -1.008 | 0.813 | -0.923 | 0.948 | -0.011 | -0.005 | -0.005 | 0 | 0.63 | 0.45 | 0.63 | 0.69 | 0.68 | 0.66 |
| 5 | 84361544 | rs10051690 | -0.182 | 0.809 | -0.882 | 1.004 | -0.002 | 0 | 0 | 1.596 | 0.39 | 0.39 | 0.28 | 0.4 | 0.32 | 0.49 |
| 5 | 84363639 | rs13167091 | 0.788 | 0.945 | -1.355 | 0.923 | -0.021 | 0.048 | 0.048 | 0 | 0.09 | 0.01 | 0.07 | 0 | 0.1 | 0.04 |
| 5 | 84372008 | rs7445269 | -0.241 | 0.622 | -1.05 | 1.035 | 0.005 | -0.002 | -0.002 | 0 | 0.38 | 0.33 | 0.28 | 0.39 | 0.32 | 0.49 |
| 5 | 84383845 | rs4337884 | -0.646 | 0.452 | -1.367 | 0.588 | -0.061 | 0.125 | 0.125 | 1.06 | 0.77 | 0.99 | 0.85 | 0.96 | 0.78 | 0.76 |
| 5 | 84386904 | rs10068915 | -0.503 | 0.568 | -1.22 | 0.731 | 0.003 | 0.02 | 0.02 | 0.192 | 0.75 | 0.83 | 0.81 | 0.86 | 0.78 | 0.7 |
| 5 | 84387741 | rs11956130 | -0.053 | 0.565 | -1.036 | 0.803 | -0.009 | -0.005 | -0.005 | 0 | 0.34 | 0.33 | 0.33 | 0.3 | 0.28 | 0.33 |

| | | | | | | | | | | | | | | | | |
|---|---|---|---|---|---|---|---|---|---|---|---|---|---|---|---|---|
| 5 | 84389994 | rs11960412 | -0.039 | 0.534 | -1.03 | 0.597 | -0.009 | -0.012 | -0.012 | 0 | 0.33 | 0.33 | 0.33 | 0.24 | 0.28 | 0.33 |
| 5 | 84399941 | rs12055053 | -0.401 | 1.332 | -1.697 | 0.906 | -0.109 | -0.142 | -0.142 | 0 | 0.25 | 0.05 | 0.37 | 0.6 | 0.38 | 0.32 |
| 5 | 84400376 | rs11958311 | 0.563 | 1.203 | -1.075 | 0.685 | 0.007 | 0.005 | 0.005 | 0 | 0.58 | 0.49 | 0.62 | 0.42 | 0.51 | 0.57 |
| 5 | 84402001 | rs9968655 | -0.309 | 0.344 | -1.037 | 0.051 | -0.005 | 0.007 | 0.007 | 0 | 0.56 | 0.47 | 0.64 | 0.69 | 0.53 | 0.52 |
| 5 | 84405876 | rs4920688 | 0.946 | -0.288 | -1.226 | -0.351 | -0.03 | 0 | 0 | 0 | 0.11 | 0.18 | 0.17 | 0.11 | 0.04 | 0.14 |
| 5 | 84407859 | rs12521624 | 0.671 | -0.004 | -1.263 | -0.376 | -0.046 | -0.107 | -0.107 | 0 | 0.26 | 0.1 | 0.32 | 0.51 | 0.34 | 0.31 |
| 5 | 84407912 | rs12514722 | -1.955 | -0.052 | -0.949 | -0.484 | -0.002 | 0.006 | 0.006 | 0 | 0.86 | 0.91 | 0.91 | 0.93 | 0.86 | 0.89 |
| 5 | 84409231 | rs10942370 | 1.208 | -0.097 | -0.882 | -0.498 | 0.018 | 0.016 | 0.016 | 0 | 0.3 | 0.48 | 0.3 | 0.3 | 0.33 | 0.37 |
| 5 | 84410666 | rs10474157 | 0.145 | 0.12 | -0.709 | -0.402 | 0.012 | 0.019 | 0.019 | 0 | 0.28 | 0.25 | 0.28 | 0.11 | 0.19 | 0.21 |
| 5 | 84411394 | rs6452635 | -1.864 | 0.17 | -0.791 | -0.404 | -0.007 | 0.018 | 0.018 | 0 | 0.86 | 0.95 | 0.91 | 0.93 | 0.86 | 0.89 |
| 5 | 84411618 | rs7727192 | 0.303 | 0.145 | -0.712 | -0.293 | 0 | -0.003 | -0.003 | 0 | 0.57 | 0.56 | 0.61 | 0.78 | 0.67 | 0.68 |
| 5 | 84417685 | rs4373307 | -0.601 | 0.081 | -0.723 | -0.163 | -0.001 | 0 | 0 | 0 | 0.52 | 0.53 | 0.58 | 0.63 | 0.5 | 0.55 |
| 5 | 84436652 | rs12187560 | 1.219 | -0.014 | -0.943 | -0.289 | 0.015 | 0.029 | 0.029 | 0 | 0.13 | 0.06 | 0.09 | 0 | 0.1 | 0.06 |
| 5 | 84437520 | rs7734159 | 0.791 | -0.015 | -1.026 | -0.24 | -0.035 | -0.063 | -0.063 | 0 | 0.21 | 0.11 | 0.29 | 0.5 | 0.3 | 0.33 |
| 5 | 84439094 | rs10064396 | 0.44 | -0.247 | -0.411 | -0.444 | -0.041 | -0.075 | -0.075 | 0.998 | 0.15 | 0.27 | 0.15 | 0.02 | 0.06 | 0.14 |
| 5 | 84444642 | rs13354392 | -0.094 | -0.058 | -0.45 | -0.327 | -0.027 | -0.028 | -0.028 | 0 | 0.37 | 0.23 | 0.42 | 0.62 | 0.43 | 0.4 |
| 5 | 84465785 | rs10059648 | 1.313 | 0.194 | -0.649 | -0.631 | -0.07 | 0.082 | 0.082 | 0 | 0.13 | 0.02 | 0.1 | 0.01 | 0.19 | 0.12 |
| 5 | 84468609 | rs4516895 | -1.046 | 0.151 | -0.37 | -0.385 | 0 | -0.002 | -0.002 | 0.423 | 0.53 | 0.55 | 0.6 | 0.63 | 0.5 | 0.49 |
| 5 | 84470679 | rs7703927 | -2.191 | 0.156 | -0.44 | -0.542 | -0.008 | 0.014 | 0.014 | 0 | 0.86 | 0.91 | 0.9 | 0.92 | 0.85 | 0.87 |
| 5 | 84473732 | rs9293425 | 0.526 | -0.094 | -0.544 | -0.536 | -0.057 | -0.053 | -0.053 | 1.192 | 0.17 | 0.37 | 0.17 | 0.07 | 0.06 | 0.15 |
| 5 | 84476702 | rs4920709 | -1.884 | -0.088 | -0.502 | -0.346 | 0.017 | 0.027 | 0.027 | 0 | 0.83 | 0.75 | 0.85 | 0.79 | 0.81 | 0.86 |
| 5 | 84477462 | rs12521350 | 1.12 | -0.056 | -0.616 | -0.472 | -0.008 | -0.042 | -0.042 | 0.766 | 0.12 | 0.07 | 0.23 | 0.41 | 0.14 | 0.17 |
| 5 | 84479893 | rs10474180 | 0.067 | -0.083 | -0.465 | -0.379 | -0.054 | -0.022 | -0.022 | 0 | 0.35 | 0.15 | 0.4 | 0.43 | 0.41 | 0.33 |
| 5 | 84480308 | rs10068805 | 0.502 | -0.08 | -0.447 | -0.389 | -0.057 | -0.053 | -0.053 | 0 | 0.17 | 0.36 | 0.16 | 0.07 | 0.06 | 0.15 |
| 5 | 84482974 | rs6884538 | 1.321 | -0.056 | -0.472 | -0.373 | -0.002 | -0.004 | -0.004 | 0.227 | 0.31 | 0.24 | 0.29 | 0.29 | 0.34 | 0.38 |
| 5 | 84485714 | rs4339387 | 1.814 | 0.226 | -0.773 | -0.55 | -0.041 | -0.037 | -0.037 | 0.027 | 0.07 | 0.01 | 0.15 | 0.2 | 0.17 | 0.22 |
| 5 | 84498196 | rs10054183 | 0.516 | -0.154 | -0.443 | -0.589 | -0.038 | -0.035 | -0.035 | 0 | 0.17 | 0.3 | 0.16 | 0.07 | 0.06 | 0.13 |
| 5 | 84506432 | rs13188039 | -0.777 | -0.265 | -0.405 | -0.623 | 0.003 | 0.014 | 0.014 | 0 | 0.52 | 0.44 | 0.55 | 0.49 | 0.46 | 0.48 |
| 5 | 84506668 | rs4432930 | 0.05 | -0.234 | -0.434 | -0.608 | -0.016 | 0.006 | 0.006 | 0 | 0.36 | 0.23 | 0.4 | 0.42 | 0.4 | 0.35 |
| 5 | 84513478 | rs13170861 | -0.786 | -0.265 | -0.414 | -0.608 | -0.003 | 0.006 | 0.006 | 0 | 0.53 | 0.46 | 0.56 | 0.5 | 0.47 | 0.48 |
| 5 | 84516414 | rs9293429 | -0.006 | -0.196 | -0.536 | -0.573 | -0.029 | 0.005 | 0.005 | 0 | 0.37 | 0.23 | 0.4 | 0.42 | 0.41 | 0.34 |
| 5 | 84521675 | rs7444934 | -1.752 | -0.244 | -0.467 | -0.54 | 0.008 | 0.054 | 0.054 | 0.141 | 0.83 | 0.68 | 0.85 | 0.78 | 0.84 | 0.87 |
| 5 | 84529595 | rs12658911 | 0.321 | -0.304 | -0.407 | -0.711 | 0.01 | 0.016 | 0.016 | 0.537 | 0.44 | 0.29 | 0.43 | 0.35 | 0.42 | 0.5 |
| 5 | 84533754 | rs10054679 | 0.458 | -0.316 | -0.437 | -0.766 | -0.014 | 0.03 | 0.03 | 0 | 0.29 | 0.2 | 0.21 | 0.02 | 0.31 | 0.19 |
| 5 | 84534417 | rs11957117 | 1.253 | -0.391 | -0.501 | -0.748 | -0.013 | -0.008 | -0.008 | 0.333 | 0.29 | 0.18 | 0.28 | 0.28 | 0.36 | 0.37 |
| 5 | 84538268 | rs11952876 | 1.67 | -0.341 | -0.717 | -0.787 | 0.004 | -0.011 | -0.011 | 0 | 0.14 | 0.13 | 0.1 | 0 | 0.11 | 0.06 |
| 5 | 84543864 | rs6897958 | 0.238 | -0.333 | -0.785 | -0.655 | 0.01 | 0.015 | 0.015 | 0.537 | 0.44 | 0.3 | 0.46 | 0.35 | 0.42 | 0.5 |
| 5 | 84556276 | rs4602659 | 0.719 | -0.405 | -0.815 | -0.674 | 0.034 | 0.036 | 0.036 | 2.138 | 0.15 | 0.04 | 0.14 | 0.07 | 0.06 | 0.13 |
| 5 | 84562600 | rs7705669 | 0.995 | -0.086 | -0.715 | -0.589 | -0.003 | -0.002 | -0.002 | 0 | 0.3 | 0.25 | 0.29 | 0.28 | 0.37 | 0.37 |
| 5 | 84579502 | rs12518735 | 1.134 | 0.111 | -0.794 | -0.475 | -0.07 | 0.089 | 0.089 | 0 | 0.14 | 0.02 | 0.11 | 0.02 | 0.19 | 0.12 |
| 5 | 84592493 | rs10514269 | 1.166 | 0.092 | -0.765 | -0.308 | 0.009 | -0.004 | -0.004 | 0 | 0.14 | 0.11 | 0.09 | 0 | 0.08 | 0.05 |
| 5 | 84598317 | rs4292478 | -1.053 | 0.427 | -0.733 | 0.204 | -0.004 | 0.007 | 0.007 | 0.621 | 0.92 | 0.94 | 0.8 | 0.59 | 0.9 | 0.82 |
| 5 | 84599662 | rs1486281 | -0.295 | 0.318 | -0.669 | -0.242 | -0.004 | 0.006 | 0.006 | 0.192 | 0.58 | 0.63 | 0.48 | 0.3 | 0.51 | 0.44 |
| 5 | 84605447 | rs416319 | -0.026 | 0.024 | -1.148 | -0.674 | 0.016 | 0.17 | 0.17 | 0.288 | 0.39 | 0.17 | 0.24 | 0.06 | 0.31 | 0.3 |
| 5 | 84609984 | rs10071789 | -0.452 | 0.371 | -0.462 | -0.037 | 0.014 | 0.07 | 0.07 | 0.894 | 0.88 | 0.78 | 0.76 | 0.59 | 0.84 | 0.81 |
| 5 | 84617476 | rs778233 | 0.332 | -0.088 | -0.22 | -0.193 | 0.001 | 0.005 | 0.005 | 0.944 | 0.54 | 0.31 | 0.5 | 0.36 | 0.55 | 0.6 |
| 5 | 84618941 | rs552947 | -0.772 | 0.288 | -0.027 | 0.091 | 0.018 | 0.066 | 0.066 | 0.303 | 0.88 | 0.8 | 0.75 | 0.58 | 0.82 | 0.81 |
| 5 | 84628466 | rs454887 | 1.231 | 0.18 | 0.187 | -0.048 | 0.001 | 0.011 | 0.011 | 0 | 0.16 | 0.17 | 0.15 | 0.22 | 0.18 | 0.14 |
| 5 | 84629509 | rs427755 | 0.162 | 0.14 | 0.509 | -0.011 | 0.026 | 0.068 | 0.068 | 1.931 | 0.7 | 0.51 | 0.59 | 0.36 | 0.64 | 0.7 |
| 5 | 84629958 | rs452618 | 0.058 | 0.146 | 0.497 | -0.008 | 0.023 | 0.056 | 0.056 | 1.534 | 0.71 | 0.56 | 0.59 | 0.36 | 0.64 | 0.7 |
| 5 | 84635697 | rs305734 | -0.648 | 0.182 | 0.386 | 0.036 | 0.027 | 0.082 | 0.082 | 0.218 | 0.88 | 0.73 | 0.73 | 0.58 | 0.82 | 0.84 |
| 5 | 84656911 | rs395729 | -0.07 | 0.3 | 0.361 | -0.427 | 0.003 | 0.009 | 0.009 | 0 | 0.42 | 0.63 | 0.31 | 0.22 | 0.36 | 0.26 |
| 5 | 84664400 | rs305766 | -1.306 | 0.327 | 0.33 | -0.408 | -0.009 | -0.005 | -0.005 | 0 | 0.89 | 0.91 | 0.76 | 0.58 | 0.82 | 0.84 |
| 5 | 84666906 | rs446660 | 0.065 | 0.337 | 0.314 | -0.393 | -0.005 | -0.023 | -0.023 | 0 | 0.41 | 0.45 | 0.29 | 0.22 | 0.35 | 0.26 |
| 5 | 84679869 | rs247722 | 1.913 | 0.335 | 0.12 | -0.422 | -0.008 | -0.022 | -0.022 | 0 | 0.15 | 0.06 | 0.14 | 0.22 | 0.18 | 0.14 |
| 5 | 84680011 | rs11951861 | 1.056 | 0.415 | 0.091 | -0.389 | 0.005 | 0.015 | 0.015 | 0 | 0.22 | 0.16 | 0.13 | 0 | 0.18 | 0.13 |
| 5 | 84696828 | rs307306 | 0.485 | 0.115 | -0.344 | -0.497 | -0.009 | -0.017 | -0.017 | 0 | 0.34 | 0.25 | 0.42 | 0.36 | 0.42 | 0.5 |
| 5 | 84699439 | rs2644818 | -1.266 | 0.039 | -0.572 | -0.312 | -0.028 | -0.044 | -0.044 | 1.052 | 0.88 | 0.95 | 0.76 | 0.57 | 0.82 | 0.84 |
| 5 | 84701663 | rs177710 | 0.967 | 0.076 | -0.34 | -0.739 | 0.004 | -0.039 | -0.039 | 0 | 0.21 | 0.24 | 0.17 | 0.05 | 0.23 | 0.2 |
| 5 | 84703081 | rs187442 | 1.263 | -0.121 | -0.744 | -0.875 | 0.016 | 0.029 | 0.029 | 0 | 0.24 | 0.24 | 0.41 | 0.48 | 0.38 | 0.36 |
| 5 | 84712893 | rs11959646 | 1.105 | -0.524 | -0.961 | -1.23 | 0.042 | 0.086 | 0.086 | 0.077 | 0.31 | 0.15 | 0.15 | 0 | 0.18 | 0.18 |
| 5 | 84724313 | rs6883974 | 0.684 | -0.564 | -0.988 | -1.009 | 0.052 | 0.083 | 0.083 | 1.704 | 0.21 | 0.05 | 0.05 | 0.04 | 0.08 | 0.15 |
| 5 | 84741107 | rs247659 | 0.184 | -0.204 | -0.65 | -0.572 | 0.001 | -0.022 | -0.022 | 0.547 | 0.23 | 0.21 | 0.26 | 0.44 | 0.21 | 0.24 |
| 5 | 84748969 | rs485309 | 1.33 | -0.137 | -0.689 | -0.521 | -0.026 | -0.041 | -0.041 | 0.416 | 0.11 | 0.05 | 0.22 | 0.44 | 0.17 | 0.18 |
| 5 | 84786749 | rs641102 | -2.082 | -0.043 | -0.402 | -0.312 | 0.008 | 0.07 | 0.07 | 0 | 0.84 | 0.75 | 0.84 | 0.7 | 0.82 | 0.85 |
| 5 | 84802685 | rs4457116 | -0.383 | -0.095 | -0.579 | -0.473 | 0.008 | 0.025 | 0.025 | 0 | 0.52 | 0.38 | 0.65 | 0.64 | 0.55 | 0.56 |
| 5 | 84803511 | rs10942387 | 0.915 | 0.085 | -0.809 | -0.341 | -0.032 | 0.011 | 0.011 | 0 | 0.28 | 0.14 | 0.38 | 0.21 | 0.34 | 0.32 |
| 5 | 84810275 | rs10942389 | 1.23 | 0.171 | -0.587 | -0.08 | -0.024 | -0.037 | -0.037 | 1.613 | 0.11 | 0.05 | 0.22 | 0.43 | 0.17 | 0.18 |
| 5 | 84818250 | rs16901822 | 1.392 | 0.155 | -0.661 | -0.246 | -0.01 | -0.005 | -0.005 | 0 | 0.08 | 0.07 | 0.28 | 0.21 | 0.21 | 0.21 |

| | | | | | | | | | | | | | | | | |
|---|---|---|---|---|---|---|---|---|---|---|---|---|---|---|---|---|
| 5 | 84822809 | rs414692 | -0.41 | -0.089 | -0.51 | -0.357 | 0.013 | 0.002 | 0.002 | 0.381 | 0.7 | 0.56 | 0.62 | 0.26 | 0.65 | 0.67 |
| 5 | 84860966 | rs1827347 | 1.656 | -0.347 | -1.197 | -0.575 | 0.031 | 0.072 | 0.072 | 0 | 0.18 | 0.04 | 0.07 | 0 | 0.09 | 0.1 |
| 5 | 84868447 | rs16901859 | 0.932 | -0.179 | -1.318 | -0.35 | -0.027 | -0.048 | -0.048 | 0.893 | 0.13 | 0.05 | 0.24 | 0.43 | 0.2 | 0.2 |
| 5 | 84872810 | rs451838 | 1.255 | -0.438 | -1.625 | -0.787 | -0.081 | 0.156 | 0.156 | 3.367 | 0.29 | 0.03 | 0.19 | 0.06 | 0.31 | 0.29 |
| 5 | 84873823 | rs1428901 | 0.346 | -0.629 | -1.197 | -0.811 | 0.006 | 0.022 | 0.022 | 0 | 0.33 | 0.19 | 0.37 | 0.2 | 0.32 | 0.34 |
| 5 | 84874542 | rs300643 | -2.347 | -0.704 | -0.816 | -0.746 | 0.008 | 0.106 | 0.106 | 0 | 0.84 | 0.64 | 0.84 | 0.71 | 0.85 | 0.87 |
| 5 | 84876628 | rs10514271 | 0.776 | -0.717 | -0.886 | -0.766 | -0.01 | 0.009 | 0.009 | 0.001 | 0.29 | 0.09 | 0.34 | 0.2 | 0.28 | 0.31 |
| 5 | 84886843 | rs1366589 | -0.33 | -0.681 | -0.667 | -0.774 | -0.001 | 0.001 | 0.001 | 0 | 0.44 | 0.43 | 0.62 | 0.65 | 0.49 | 0.51 |
| 5 | 84899446 | rs414536 | -2.728 | -0.598 | -0.414 | -0.547 | 0.001 | -0.003 | -0.003 | 0.192 | 0.86 | 0.81 | 0.9 | 0.89 | 0.86 | 0.89 |
| 5 | 84899961 | rs11746552 | 1.465 | -0.593 | -0.562 | -0.709 | -0.015 | -0.018 | -0.018 | 0 | 0.11 | 0.09 | 0.27 | 0.2 | 0.21 | 0.21 |
| 5 | 84948547 | rs10060598 | -0.862 | -0.429 | -0.241 | -0.312 | 0.022 | 0.014 | 0.014 | 0.295 | 0.74 | 0.59 | 0.59 | 0.26 | 0.68 | 0.7 |
| 5 | 84983412 | rs10474191 | 0.985 | -0.469 | -0.324 | -0.423 | -0.026 | -0.061 | -0.061 | 0.946 | 0.13 | 0.21 | 0.06 | 0 | 0.05 | 0.08 |
| 5 | 84984994 | rs721245 | 1.58 | -0.175 | -0.21 | 0.016 | 0.006 | 0.058 | 0.058 | 0 | 0.1 | 0.13 | 0.27 | 0.45 | 0.17 | 0.17 |
| 5 | 84995031 | rs11749950 | 0.502 | -0.268 | -0.35 | -0.634 | -0.004 | -0.01 | -0.01 | 0.219 | 0.33 | 0.35 | 0.36 | 0.2 | 0.32 | 0.33 |
| 5 | 85038975 | rs6862921 | -0.057 | 0.322 | -0.335 | 0.195 | 0.001 | -0.006 | -0.006 | 0 | 0.66 | 0.62 | 0.57 | 0.25 | 0.64 | 0.63 |
| 5 | 85047893 | rs4920833 | -0.382 | 0.545 | -0.529 | 0.762 | -0.039 | -0.05 | -0.05 | 0 | 0.8 | 0.92 | 0.71 | 0.66 | 0.76 | 0.7 |
| 5 | 85052886 | rs10063186 | 0.732 | 0.662 | 0.562 | 0.75 | 0.046 | 0.064 | 0.064 | 0.72 | 0.68 | 0.49 | 0.57 | 0.51 | 0.63 | 0.6 |
| 5 | 85052916 | rs9293439 | 0.091 | 0.625 | 0.646 | 0.764 | 0.022 | 0.066 | 0.066 | 0.022 | 0.76 | 0.64 | 0.63 | 0.54 | 0.72 | 0.67 |
| 5 | 85056209 | rs4920697 | -0.663 | 0.627 | 0.489 | 0.663 | -0.039 | -0.05 | -0.05 | 0 | 0.8 | 0.92 | 0.71 | 0.66 | 0.76 | 0.7 |
| 5 | 85056929 | rs12520719 | 1.49 | 0.046 | -0.078 | -0.409 | 0.042 | 0.154 | 0.154 | 1.613 | 0.3 | 0.07 | 0.15 | 0 | 0.2 | 0.15 |
| 5 | 85084936 | rs1500148 | 0.812 | -0.062 | -0.2 | -0.012 | -0.085 | -0.08 | -0.08 | 0 | 0.3 | 0.06 | 0.38 | 0.38 | 0.4 | 0.4 |
| 5 | 85093571 | rs7701036 | 0.803 | -0.257 | -0.062 | -0.67 | 0.024 | 0.076 | 0.076 | 0 | 0.39 | 0.2 | 0.21 | 0.02 | 0.28 | 0.27 |
| 5 | 85095480 | rs1500147 | 0.612 | -0.22 | -0.157 | -0.649 | -0.004 | -0.106 | -0.106 | 0 | 0.41 | 0.45 | 0.25 | 0.04 | 0.33 | 0.29 |
| 5 | 85102498 | rs1039324 | 0.626 | -0.196 | -0.051 | -0.25 | -0.014 | -0.015 | -0.015 | 0.912 | 0.62 | 0.31 | 0.58 | 0.52 | 0.61 | 0.63 |
| 5 | 85115583 | rs6877523 | 0.073 | -0.301 | 0.199 | -0.373 | -0.008 | -0.079 | -0.079 | 0 | 0.16 | 0.28 | 0.08 | 0.02 | 0.11 | 0.08 |
| 5 | 85119481 | rs12659004 | 0.802 | -0.35 | -0.645 | -0.298 | -0.045 | -0.076 | -0.076 | 0 | 0.31 | 0.12 | 0.37 | 0.44 | 0.42 | 0.4 |
| 5 | 85127596 | rs10474197 | -1.191 | -0.119 | -0.711 | -0.239 | -0.045 | -0.07 | -0.07 | 0.076 | 0.82 | 0.93 | 0.69 | 0.54 | 0.76 | 0.73 |
| 5 | 85132635 | rs12520429 | 1.297 | -0.357 | -0.199 | -0.437 | 0.042 | 0.053 | 0.053 | 0 | 0.31 | 0.17 | 0.16 | 0 | 0.2 | 0.22 |
| 5 | 85137318 | rs10051236 | 0.383 | -0.004 | 0.021 | 0.38 | -0.044 | -0.048 | -0.048 | 1.139 | 0.27 | 0.17 | 0.38 | 0.42 | 0.41 | 0.3 |
| 5 | 85167780 | rs7721041 | 0.026 | 0.046 | 0.352 | 0.154 | -0.002 | -0.066 | -0.066 | 0.568 | 0.12 | 0.32 | 0.06 | 0.03 | 0.1 | 0.04 |
| 5 | 85168399 | rs6877270 | -1.012 | -0.214 | -0.352 | 0.271 | -0.044 | -0.022 | -0.022 | 0 | 0.49 | 0.92 | 0.54 | 0.54 | 0.55 | 0.46 |
| 5 | 85192537 | rs10070703 | 0.85 | -0.254 | -0.27 | 0.271 | -0.042 | -0.034 | -0.034 | 0.081 | 0.52 | 0.12 | 0.47 | 0.47 | 0.45 | 0.54 |
| 5 | 85194453 | rs1595560 | -0.938 | -0.31 | 0.558 | -0.213 | -0.02 | -0.07 | -0.07 | 0 | 0.86 | 0.75 | 0.95 | 1 | 0.94 | 0.96 |
| 5 | 85194802 | rs12188123 | -0.303 | -0.477 | 1.066 | 0.241 | 0.038 | 0.053 | 0.053 | 0.095 | 0.26 | 0.38 | 0.38 | 0.47 | 0.44 | 0.34 |
| 5 | 85195714 | rs10069395 | 1.004 | -0.519 | 0.86 | 0.226 | -0.016 | -0.036 | -0.036 | 0 | 0.1 | 0.15 | 0.04 | 0 | 0.06 | 0.07 |
| 5 | 85196393 | rs6874335 | 0.857 | -0.54 | 0.767 | 0.274 | -0.03 | 0.035 | 0.035 | 0 | 0.11 | 0.23 | 0.09 | 0.12 | 0.07 | 0.12 |
| 5 | 85198190 | rs2918429 | 1.489 | -0.478 | 0.078 | 0.189 | -0.038 | 0.038 | 0.038 | 0 | 0.49 | 0.11 | 0.47 | 0.41 | 0.43 | 0.51 |
| 5 | 85200195 | rs6888053 | 0.13 | -0.185 | 0.339 | 0.353 | 0 | 0.014 | 0.014 | 0 | 0.11 | 0.13 | 0.06 | 0.17 | 0.12 | 0.12 |
| 5 | 85217869 | rs1583686 | 0.245 | -0.253 | 0.499 | 0.294 | -0.006 | -0.007 | -0.007 | 0 | 0.77 | 0.73 | 0.9 | 0.83 | 0.82 | 0.84 |
| 5 | 85227089 | rs2974062 | 0.815 | -0.163 | 0.329 | 0.355 | -0.004 | -0.004 | -0.004 | 0.645 | 0.67 | 0.3 | 0.52 | 0.58 | 0.56 | 0.62 |
| 5 | 85227531 | rs6892464 | 0.485 | -0.532 | 0.136 | 0.071 | 0.007 | -0.001 | -0.001 | 1.437 | 0.18 | 0.11 | 0.09 | 0.17 | 0.15 | 0.18 |
| 5 | 85231006 | rs13176279 | 0.195 | -0.435 | -0.527 | -0.152 | -0.01 | 0.025 | 0.025 | 0.004 | 0.57 | 0.36 | 0.47 | 0.3 | 0.48 | 0.44 |
| 5 | 85231447 | rs12521803 | -0.082 | -0.478 | -0.555 | -0.194 | 0.014 | 0.018 | 0.018 | 0 | 0.28 | 0.21 | 0.13 | 0.17 | 0.21 | 0.22 |
| 5 | 85237840 | rs7701633 | 0.183 | -0.425 | -0.553 | -0.167 | -0.01 | 0.025 | 0.025 | 0.004 | 0.57 | 0.35 | 0.47 | 0.3 | 0.48 | 0.44 |
| 5 | 85249821 | rs1438743 | 1.834 | -0.481 | -0.575 | -0.17 | -0.008 | 0.067 | 0.067 | 0 | 0.29 | 0.16 | 0.34 | 0.13 | 0.27 | 0.22 |
| 5 | 85260207 | rs6859574 | -0.335 | -0.249 | -0.141 | 0.05 | -0.02 | 0.048 | 0.048 | 0.453 | 0.7 | 0.59 | 0.85 | 0.62 | 0.78 | 0.66 |
| 5 | 85267381 | rs7718298 | -0.066 | -0.124 | -0.43 | 0.049 | -0.008 | 0.028 | 0.028 | 0.548 | 0.67 | 0.45 | 0.81 | 0.62 | 0.72 | 0.65 |
| 5 | 85273284 | rs9293453 | 1.025 | -0.079 | -0.594 | 0.013 | 0.008 | -0.009 | -0.009 | 0.079 | 0.17 | 0.08 | 0.09 | 0.17 | 0.16 | 0.19 |
| 5 | 85299269 | rs12518569 | -0.083 | 0.267 | -0.436 | 0.28 | -0.009 | 0.051 | 0.051 | 0.532 | 0.68 | 0.52 | 0.82 | 0.62 | 0.73 | 0.66 |
| 5 | 85307476 | rs4289589 | 0.95 | 0.939 | -0.456 | 0.357 | 0.027 | 0.029 | 0.029 | 0.148 | 0.12 | 0.15 | 0.37 | 0.32 | 0.29 | 0.21 |
| 5 | 85309578 | rs13172392 | -0.103 | -0.186 | -0.533 | -0.025 | 0 | 0.057 | 0.057 | 0 | 0.92 | 0.81 | 0.95 | 0.9 | 0.94 | 0.88 |
| 5 | 85314047 | rs13160406 | 0.576 | -0.105 | -1.239 | -0.285 | 0.026 | 0.095 | 0.095 | 0 | 0.3 | 0.12 | 0.3 | 0.13 | 0.22 | 0.21 |
| 5 | 85316605 | rs7719439 | -0.585 | 0.183 | -0.642 | 0.535 | -0.007 | 0.006 | 0.006 | 0 | 0.29 | 0.32 | 0.15 | 0.6 | 0.2 | 0.35 |
| 5 | 85321761 | rs7735802 | 0.331 | 0.729 | -0.759 | -0.232 | 0.006 | 0.002 | 0.002 | 0 | 0.68 | 0.68 | 0.39 | 0.6 | 0.54 | 0.62 |
| 5 | 85330526 | rs12518455 | 0.367 | 0.701 | -0.856 | -0.385 | -0.018 | -0.065 | -0.065 | 0 | 0.22 | 0.09 | 0.12 | 0.59 | 0.22 | 0.33 |
| 5 | 85335661 | rs17206928 | 1.032 | 0.829 | -0.714 | -0.279 | 0 | 0 | 0 | 0 | 0.09 | 0.08 | 0.33 | 0.17 | 0.12 | 0.08 |
| 5 | 85337276 | rs2974101 | -0.702 | 0.738 | -0.327 | 0.56 | -0.044 | -0.058 | -0.058 | 0 | 0.81 | 0.89 | 0.73 | 0.61 | 0.67 | 0.69 |
| 5 | 85339189 | rs6898018 | 0.516 | 0.375 | -0.089 | 0.194 | -0.002 | -0.036 | -0.036 | 0.918 | 0.09 | 0.39 | 0.05 | 0 | 0.02 | 0.03 |
| 5 | 85363033 | rs6874607 | -0.275 | 0.809 | -0.243 | 0.847 | 0.013 | -0.02 | -0.02 | 2.647 | 0.23 | 0.15 | 0.11 | 0.34 | 0.19 | 0.31 |
| 5 | 85372505 | rs10071511 | 0.438 | 0.978 | -0.155 | 0.821 | 0.023 | 0.008 | 0.008 | 0.949 | 0.18 | 0.24 | 0.27 | 0.22 | 0.34 | 0.25 |
| 5 | 85385967 | rs12516577 | 0.659 | 1.027 | -0.335 | 1.006 | 0.006 | 0.002 | 0.002 | 0 | 0.18 | 0.06 | 0.09 | 0.18 | 0.15 | 0.26 |
| 5 | 85387177 | rs2216577 | -0.271 | 0.956 | -0.11 | 0.806 | 0.039 | 0.078 | 0.078 | 0.536 | 0.23 | 0.33 | 0.31 | 0.48 | 0.35 | 0.32 |
| 5 | 85388750 | rs1422127 | -1.711 | 0.673 | -0.208 | 0.397 | -0.005 | -0.106 | -0.106 | 0 | 0.56 | 0.5 | 0.75 | 0.99 | 0.65 | 0.74 |
| 5 | 85389354 | rs9293456 | -0.05 | 1.281 | -0.192 | 0.887 | 0.011 | 0.169 | 0.169 | 0 | 0.11 | 0.27 | 0.09 | 0.29 | 0.09 | 0.16 |
| 5 | 85389380 | rs2161208 | 0.102 | 1.375 | -0.677 | 0.824 | -0.036 | -0.027 | -0.027 | 0.663 | 0.14 | 0.06 | 0.22 | 0.19 | 0.26 | 0.2 |
| 5 | 85392336 | rs10061309 | -0.907 | 1.365 | -0.558 | 0.797 | -0.025 | -0.033 | -0.033 | 0.051 | 0.36 | 0.22 | 0.32 | 0.54 | 0.45 | 0.5 |
| 5 | 85396128 | rs2042183 | 0.058 | 0.974 | -1.103 | 0.548 | -0.048 | -0.091 | -0.091 | 0 | 0.19 | 0.01 | 0.12 | 0.31 | 0.19 | 0.2 |
| 5 | 85411181 | rs888804 | 0.081 | 1.561 | -0.576 | 0.602 | 0.013 | 0.031 | 0.031 | 0 | 0.82 | 0.76 | 0.72 | 0.62 | 0.7 | 0.77 |
| 5 | 85413146 | rs13171331 | 2.712 | 1.259 | -1.153 | -0.132 | 0.005 | 0.214 | 0.214 | 0 | 0.31 | 0.03 | 0.17 | 0.01 | 0.24 | 0.24 |

| | | | | | | | | | | | | | | | | |
|---|---|---|---|---|---|---|---|---|---|---|---|---|---|---|---|---|
| 5 | 85417242 | rs10038346 | 1.312 | 1.48 | -0.864 | -0.101 | 0.009 | -0.014 | -0.014 | 0.006 | 0.07 | 0.08 | 0.13 | 0.03 | 0.13 | 0.06 |
| 5 | 85429684 | rs7710893 | 1.416 | 1.454 | -0.539 | -0.005 | -0.017 | 0.043 | 0.043 | 1.335 | 0.57 | 0.23 | 0.41 | 0.34 | 0.53 | 0.53 |
| 5 | 85430133 | rs6874864 | 0.264 | 1.272 | -0.377 | 0.279 | 0.014 | 0.048 | 0.048 | 0.717 | 0.07 | 0.22 | 0.04 | 0.09 | 0.05 | 0.08 |
| 5 | 85432273 | rs6867783 | 0.264 | 1.272 | -0.377 | 0.279 | 0.014 | 0.048 | 0.048 | 0.717 | 0.07 | 0.22 | 0.04 | 0.09 | 0.05 | 0.08 |
| 5 | 85434559 | rs10514275 | 1.156 | 1.059 | 0.035 | 0.284 | -0.032 | 0.13 | 0.13 | 0 | 0.07 | 0.16 | 0.24 | 0.26 | 0.03 | 0.14 |
| 5 | 85435376 | rs1363262 | 0.487 | 1.204 | -0.159 | 0.144 | -0.058 | -0.095 | -0.095 | 2.993 | 0.19 | 0.01 | 0.12 | 0.3 | 0.2 | 0.23 |
| 5 | 85439065 | rs16902074 | -0.079 | 1.193 | 0.459 | -0.189 | 0.039 | -0.047 | -0.047 | 0 | 0.09 | 0.28 | 0.05 | 0 | 0.07 | 0.02 |
| 5 | 85441996 | rs6872709 | -0.722 | 1.132 | 0.501 | -0.142 | 0.104 | 0.021 | 0.021 | 2.207 | 0.05 | 0.32 | 0.04 | 0 | 0.04 | 0.01 |
| 5 | 85448604 | rs987280 | 0.888 | 1.237 | 0.173 | -0.188 | 0.003 | 0.001 | 0.001 | 0.197 | 0.64 | 0.59 | 0.66 | 0.61 | 0.56 | 0.63 |
| 5 | 85450966 | rs1363265 | 1.628 | 1.21 | 0.163 | -0.116 | 0 | 0 | 0 | 0 | 0.57 | 0.43 | 0.42 | 0.45 | 0.52 | 0.49 |
| 5 | 85451901 | rs7700540 | 0.31 | 0.709 | 0.291 | -0.367 | 0.087 | -0.043 | -0.043 | 0.097 | 0.09 | 0.3 | 0.15 | 0.03 | 0.15 | 0.06 |
| 5 | 85451916 | rs12187227 | 2.039 | 0.598 | -0.032 | -0.346 | -0.067 | -0.034 | -0.034 | 0.221 | 0.48 | 0.06 | 0.3 | 0.41 | 0.43 | 0.44 |
| 5 | 85457842 | rs1862231 | 0.993 | 0.541 | 0.238 | -0.284 | -0.021 | 0.131 | 0.131 | 0 | 0.08 | 0.31 | 0.25 | 0.16 | 0.03 | 0.13 |
| 5 | 85458883 | rs4920714 | 1.224 | 0.464 | 0.259 | -0.358 | -0.001 | 0 | 0 | 0.184 | 0.57 | 0.55 | 0.55 | 0.58 | 0.46 | 0.58 |
| 5 | 85466140 | rs1422110 | -0.129 | 0.45 | 0.08 | -0.346 | -0.047 | -0.048 | -0.048 | 0.026 | 0.81 | 0.94 | 0.75 | 0.71 | 0.69 | 0.76 |
| 5 | 85467970 | rs13181836 | 0.815 | 0.403 | -0.195 | -0.467 | -0.057 | -0.128 | -0.128 | 0 | 0.17 | 0.01 | 0.11 | 0.41 | 0.17 | 0.19 |
| 5 | 85469695 | rs1363268 | -0.931 | 0.327 | -0.112 | -0.477 | -0.077 | -0.082 | -0.082 | 0 | 0.86 | 0.98 | 0.77 | 0.72 | 0.72 | 0.84 |
| 5 | 85475137 | rs1363269 | 0.901 | -0.346 | 0.351 | -0.315 | -0.003 | 0.176 | 0.176 | 0 | 0.08 | 0.36 | 0.25 | 0.16 | 0.03 | 0.13 |
| 5 | 85477788 | rs7703008 | 0.945 | -0.212 | 0.312 | -0.388 | -0.016 | 0.001 | 0.001 | 0 | 0.54 | 0.24 | 0.36 | 0.42 | 0.5 | 0.46 |
| 5 | 85479926 | rs13180071 | 2.455 | -0.736 | -0.001 | -0.466 | -0.024 | 0.206 | 0.206 | 0 | 0.31 | 0.04 | 0.18 | 0.01 | 0.25 | 0.25 |
| 5 | 85481442 | rs1006163 | 2.447 | -0.732 | 0.002 | -0.463 | -0.023 | 0.207 | 0.207 | 0 | 0.31 | 0.04 | 0.18 | 0.01 | 0.25 | 0.25 |
| 5 | 85487635 | rs2112386 | 1.132 | -0.711 | 0.36 | -0.408 | -0.032 | 0.097 | 0.097 | 0 | 0.07 | 0.23 | 0.24 | 0.16 | 0.03 | 0.14 |
| 5 | 85488131 | rs17207039 | 2.379 | -0.718 | 0.098 | -0.476 | -0.019 | 0.213 | 0.213 | 0 | 0.31 | 0.04 | 0.18 | 0.01 | 0.25 | 0.25 |
| 5 | 85496114 | rs16902129 | 0.993 | -0.339 | 0.567 | 0.05 | -0.04 | -0.059 | -0.059 | 0 | 0.1 | 0.01 | 0.07 | 0.41 | 0.12 | 0.19 |
| 5 | 85498043 | rs7729431 | 0.781 | -0.344 | 0.623 | 0.056 | -0.011 | 0.001 | 0.001 | 0 | 0.62 | 0.66 | 0.58 | 0.58 | 0.48 | 0.6 |
| 5 | 85500531 | rs1422116 | 1.377 | -0.315 | 0.042 | 0.063 | -0.075 | -0.05 | -0.05 | 0.002 | 0.52 | 0.05 | 0.31 | 0.42 | 0.45 | 0.46 |
| 5 | 85502656 | rs13186437 | 0.818 | -0.034 | 0.197 | 0.267 | -0.039 | -0.061 | -0.061 | 1.558 | 0.1 | 0.01 | 0.07 | 0.41 | 0.12 | 0.19 |
| 5 | 85508164 | rs1422119 | 0.618 | -0.191 | 0.433 | 0.149 | 0.014 | 0.002 | 0.002 | 0.01 | 0.65 | 0.43 | 0.58 | 0.58 | 0.51 | 0.64 |
| 5 | 85512273 | rs2909866 | 1.484 | -0.306 | 0.49 | 0.133 | 0.071 | 0.146 | 0.146 | 0 | 0.49 | 0.28 | 0.48 | 0.16 | 0.37 | 0.44 |
| 5 | 85514313 | rs2909867 | 0.967 | -0.444 | 0.476 | 0.174 | 0.009 | 0 | 0 | 0 | 0.61 | 0.46 | 0.56 | 0.58 | 0.5 | 0.6 |
| 5 | 85519655 | rs11748175 | 0.668 | -0.505 | 0.472 | 0.3 | -0.031 | -0.052 | -0.052 | 2.73 | 0.11 | 0.02 | 0.08 | 0.41 | 0.13 | 0.19 |
| 5 | 85524063 | rs10062043 | -0.832 | -0.613 | 0.536 | 0.134 | 0.023 | 0.01 | 0.01 | 0 | 0.33 | 0.59 | 0.44 | 0.41 | 0.52 | 0.35 |
| 5 | 85547343 | rs11741479 | 0.692 | -0.628 | 0.599 | 0.116 | -0.001 | -0.034 | -0.034 | 0.036 | 0.1 | 0.02 | 0.05 | 0.42 | 0.07 | 0.18 |
| 5 | 85549524 | rs12654068 | -0.852 | -0.739 | 0.666 | 0.071 | -0.03 | 0.094 | 0.094 | 0 | 0.92 | 0.77 | 0.76 | 0.84 | 0.97 | 0.88 |
| 5 | 85554715 | rs6870276 | 1.055 | -0.708 | 0.515 | 0.003 | -0.021 | -0.02 | -0.02 | 0 | 0.61 | 0.32 | 0.35 | 0.45 | 0.51 | 0.59 |
| 5 | 85587649 | rs3888301 | -0.848 | -0.893 | 0.543 | 0.1 | -0.03 | -0.043 | -0.043 | 0 | 0.12 | 0.18 | 0.05 | 0.01 | 0.06 | 0.09 |
| 5 | 85588335 | rs7443049 | -0.254 | -0.871 | 0.556 | 0.115 | -0.029 | -0.03 | -0.03 | 0 | 0.08 | 0.14 | 0.03 | 0.01 | 0.03 | 0.08 |
| 5 | 85593102 | rs10069895 | 1.418 | -0.949 | 0.625 | 0.129 | -0.082 | 0.044 | 0.044 | 0 | 0.7 | 0.5 | 0.66 | 0.41 | 0.84 | 0.59 |
| 5 | 85594695 | rs4551110 | -0.271 | -0.788 | 0.653 | -0.041 | -0.028 | -0.03 | -0.03 | 0 | 0.08 | 0.14 | 0.03 | 0.01 | 0.03 | 0.08 |
| 5 | 85599474 | rs4479876 | -0.66 | -0.545 | 0.249 | 0.104 | -0.03 | 0.108 | 0.108 | 0 | 0.93 | 0.74 | 0.77 | 0.84 | 0.97 | 0.88 |
| 5 | 85606682 | rs4457115 | -1.309 | -0.564 | 0.042 | -0.109 | 0.014 | 0.056 | 0.056 | 0 | 0.53 | 0.63 | 0.55 | 0.83 | 0.59 | 0.59 |
| 5 | 85611175 | rs10051630 | 0.204 | -0.623 | 0.16 | 0.269 | -0.041 | 0.096 | 0.096 | 0 | 0.79 | 0.67 | 0.73 | 0.46 | 0.88 | 0.67 |
| 5 | 85611603 | rs16902179 | 0.068 | -0.433 | 0.471 | 0.136 | 0.095 | 0.03 | 0.03 | 0.249 | 0.06 | 0.37 | 0.04 | 0.01 | 0.05 | 0.05 |
| 5 | 85624417 | rs2964887 | 0.463 | -0.536 | 0.353 | 0.037 | 0.004 | 0.006 | 0.006 | 0 | 0.63 | 0.57 | 0.36 | 0.42 | 0.52 | 0.62 |
| 5 | 85625985 | rs1445701 | -1.155 | -0.293 | 0.631 | 0.331 | -0.016 | 0.147 | 0.147 | 0 | 0.21 | 0.45 | 0.11 | 0.4 | 0.14 | 0.32 |
| 5 | 85637934 | rs7724903 | -0.657 | -0.113 | 0.183 | 0.322 | 0.111 | -0.006 | -0.006 | 0.013 | 0.1 | 0.38 | 0.08 | 0.04 | 0.13 | 0.1 |
| 5 | 85643624 | rs16876651 | 0.782 | -0.104 | 0.09 | 0.474 | 0 | -0.051 | -0.051 | 1.634 | 0.09 | 0.01 | 0.08 | 0.38 | 0.05 | 0.2 |
| 5 | 85646055 | rs16902210 | -0.375 | -0.099 | 0.189 | 0.401 | 0.102 | -0.002 | -0.002 | 0 | 0.09 | 0.29 | 0.08 | 0.03 | 0.13 | 0.09 |
| 5 | 85647100 | rs7735255 | 0.989 | -0.512 | -0.163 | 0.089 | 0.002 | -0.075 | -0.075 | 1.929 | 0.45 | 0.5 | 0.28 | 0.03 | 0.42 | 0.35 |
| 5 | 85650406 | rs6883744 | 1.067 | -0.519 | -0.19 | 0.122 | 0.021 | 0.017 | 0.017 | 0.458 | 0.44 | 0.24 | 0.28 | 0.38 | 0.34 | 0.46 |
| 5 | 85651034 | rs7728021 | 0.712 | -0.572 | -0.071 | 0.21 | -0.002 | -0.001 | -0.001 | 0 | 0.55 | 0.6 | 0.5 | 0.54 | 0.4 | 0.58 |
| 5 | 85654774 | rs4473803 | 0.566 | -0.591 | -0.076 | 0.206 | -0.014 | 0.077 | 0.077 | 0 | 0.11 | 0.21 | 0.1 | 0.37 | 0.05 | 0.2 |
| 5 | 85655306 | rs12657213 | 1.207 | -0.5 | -0.193 | -0.115 | 0.009 | 0.017 | 0.017 | 0 | 0.44 | 0.39 | 0.4 | 0.17 | 0.34 | 0.37 |
| 5 | 85662432 | rs4634387 | -1.091 | 0.464 | -0.231 | 0.417 | -0.034 | -0.026 | -0.026 | 1.1 | 0.92 | 0.98 | 0.83 | 0.9 | 0.85 | 0.91 |
| 5 | 85684428 | rs3930380 | 0.029 | -0.206 | 0.218 | 0.119 | -0.074 | 0.055 | 0.055 | 0 | 0.14 | 0.5 | 0.23 | 0.18 | 0.06 | 0.14 |
| 5 | 85687037 | rs10059045 | -0.513 | -0.002 | -0.075 | 0.235 | -0.077 | -0.115 | -0.115 | 4.238 | 0.71 | 0.9 | 0.6 | 0.51 | 0.55 | 0.75 |
| 5 | 85693190 | rs6867920 | -0.442 | 0.188 | 0.074 | 0.184 | 0.093 | -0.044 | -0.044 | 0.398 | 0.1 | 0.28 | 0.08 | 0.01 | 0.13 | 0.11 |
| 5 | 85727108 | rs16902229 | 0.284 | 0.361 | 0.064 | 0.36 | 0.101 | -0.02 | -0.02 | 0 | 0.07 | 0.21 | 0.07 | 0.01 | 0.11 | 0.08 |
| 5 | 85746871 | rs1500258 | 0.724 | -0.388 | -0.324 | 0.812 | -0.083 | 0.131 | 0.131 | 0 | 0.77 | 0.58 | 0.71 | 0.53 | 0.9 | 0.66 |
| 5 | 85758452 | rs6897770 | -0.066 | 0.071 | -0.857 | 1.161 | -0.047 | 0.025 | 0.025 | 0 | 0.12 | 0.25 | 0.21 | 0.18 | 0.06 | 0.14 |
| 5 | 85764555 | rs1553241 | -0.491 | 1.219 | -1.525 | 1.371 | -0.058 | -0.09 | -0.09 | 0.312 | 0.28 | 0.15 | 0.45 | 0.51 | 0.46 | 0.25 |
| 5 | 85767390 | rs2937096 | 0.636 | 0.898 | -1.099 | 1.782 | -0.003 | 0.013 | 0.013 | 0 | 0.92 | 0.93 | 0.91 | 0.7 | 0.91 | 0.81 |
| 5 | 85767989 | rs3862058 | 0.535 | 0.262 | -1.362 | 1.427 | 0.018 | -0.024 | -0.024 | 0 | 0.64 | 0.73 | 0.77 | 0.53 | 0.73 | 0.63 |
| 5 | 85771107 | rs10035239 | 0.836 | -0.273 | -2.185 | 0.721 | -0.067 | 0.135 | 0.135 | 0 | 0.42 | 0.09 | 0.36 | 0.18 | 0.56 | 0.29 |
| 5 | 85776114 | rs7710314 | 0.558 | -0.583 | -1.014 | 0.933 | 0.018 | 0.037 | 0.037 | 0 | 0.5 | 0.42 | 0.43 | 0.25 | 0.6 | 0.39 |
| 5 | 85824066 | rs10474229 | 0.889 | -0.945 | -0.406 | 0.69 | 0.066 | 0.182 | 0.182 | 0.196 | 0.51 | 0.21 | 0.39 | 0.15 | 0.63 | 0.36 |
| 5 | 85829600 | rs7702789 | 0.318 | -0.969 | 0.302 | 0.804 | -0.027 | 0.085 | 0.085 | 0.118 | 0.09 | 0.35 | 0.07 | 0.09 | 0.04 | 0.11 |
| 5 | 85906658 | rs4605828 | -1.112 | -0.835 | -0.283 | 0.475 | 0.006 | 0.154 | 0.154 | 0.059 | 0.54 | 0.84 | 0.63 | 0.79 | 0.44 | 0.62 |
| 5 | 85910124 | rs17476297 | 1.067 | -1.09 | -0.419 | -0.152 | -0.047 | 0.056 | 0.056 | 0.41 | 0.1 | 0.01 | 0.1 | 0.01 | 0.15 | 0.05 |

| | | | | | | | | | | | | | | | | |
|---|---|---|---|---|---|---|---|---|---|---|---|---|---|---|---|---|
| 5 | 85911654 | rs10805837 | -0.931 | -1.359 | -0.382 | 0.617 | 0.03 | -0.006 | -0.006 | 0 | 0.42 | 0.82 | 0.31 | 0.41 | 0.34 | 0.47 |
| 5 | 85923167 | rs17476583 | 1.149 | -0.29 | -0.692 | 0.101 | -0.087 | 0.149 | 0.149 | 0 | 0.22 | 0.01 | 0.13 | 0 | 0.29 | 0.18 |
| 5 | 85933901 | rs16902295 | -0.632 | -0.257 | 0.184 | 0.654 | 0.012 | 0.052 | 0.052 | 2.234 | 0.06 | 0.31 | 0.08 | 0.04 | 0.03 | 0.07 |
| 5 | 85943936 | rs7733888 | -0.837 | -0.154 | 0.432 | 0.442 | 0.024 | 0.034 | 0.034 | 1.316 | 0.07 | 0.33 | 0.1 | 0.05 | 0.06 | 0.05 |
| 5 | 85946757 | rs13178674 | -0.704 | -0.253 | -0.124 | 0.274 | -0.034 | -0.032 | -0.032 | 0 | 0.73 | 0.88 | 0.49 | 0.39 | 0.68 | 0.64 |
| 5 | 85961838 | rs2410719 | -0.508 | -0.778 | 0.48 | 0.291 | -0.009 | 0.133 | 0.133 | 0 | 0.92 | 0.73 | 0.96 | 0.72 | 0.95 | 0.92 |
| 5 | 85962245 | rs13168605 | -0.018 | -1.103 | 0.335 | -0.176 | -0.003 | -0.109 | -0.109 | 0 | 0.32 | 0.41 | 0.22 | 0.04 | 0.3 | 0.28 |
| 5 | 85964365 | rs6868572 | 0.454 | -1.049 | 0.148 | -0.141 | -0.002 | -0.006 | -0.006 | 0 | 0.6 | 0.45 | 0.74 | 0.67 | 0.65 | 0.64 |
| 5 | 85986822 | rs10491416 | 0.821 | -0.958 | -0.444 | -0.006 | -0.026 | -0.038 | -0.038 | 0 | 0.13 | 0.01 | 0.11 | 0.13 | 0.11 | 0.09 |
| 5 | 85993967 | rs254749 | -0.328 | -0.788 | 0.332 | 0.065 | 0.005 | 0.189 | 0.189 | 0.712 | 0.9 | 0.56 | 0.95 | 0.72 | 0.95 | 0.92 |
| 5 | 85994747 | rs17477971 | 0.046 | -0.866 | 0.27 | -0.206 | 0.004 | 0.034 | 0.034 | 0 | 0.42 | 0.29 | 0.3 | 0.18 | 0.38 | 0.37 |
| 5 | 85998509 | rs254755 | -0.425 | -0.67 | 0.568 | 0.148 | -0.008 | 0.121 | 0.121 | 0 | 0.91 | 0.73 | 0.95 | 0.72 | 0.95 | 0.92 |
| 5 | 86004170 | rs254738 | 0.363 | -0.269 | 0.044 | 0.162 | -0.021 | 0.172 | 0.172 | 4.283 | 0.87 | 0.42 | 0.89 | 0.68 | 0.93 | 0.89 |
| 5 | 86012253 | rs254741 | 0.87 | 0.375 | -0.943 | 0.079 | -0.053 | -0.076 | -0.076 | 0.375 | 0.17 | 0.07 | 0.4 | 0.49 | 0.28 | 0.32 |
| 5 | 86037030 | rs10051486 | -0.255 | -0.093 | -0.862 | -0.215 | 0.009 | -0.012 | -0.012 | 0 | 0.53 | 0.7 | 0.39 | 0.51 | 0.43 | 0.49 |
| 5 | 86042743 | rs2287693 | 0.157 | -0.045 | -0.809 | -0.612 | 0.007 | -0.024 | -0.024 | 0 | 0.65 | 0.64 | 0.4 | 0.09 | 0.53 | 0.59 |
| 5 | 86045601 | rs17285752 | 0.789 | -0.635 | -1.049 | -0.201 | 0 | 0.026 | 0.026 | 0 | 0.16 | 0.1 | 0.09 | 0.03 | 0.16 | 0.1 |
| 5 | 86046005 | rs12332473 | -0.219 | -0.588 | -1.025 | -0.035 | -0.019 | -0.049 | -0.049 | 4.292 | 0.3 | 0.2 | 0.22 | 0.43 | 0.3 | 0.4 |
| 5 | 86059399 | rs10040369 | -0.414 | -0.859 | -1.491 | -0.78 | -0.09 | 0.15 | 0.15 | 0 | 0.24 | 0.04 | 0.18 | 0.01 | 0.27 | 0.31 |
| 5 | 86061783 | rs10474240 | -1.151 | -1.554 | -0.858 | -1.081 | -0.002 | -0.047 | -0.047 | 0.399 | 0.46 | 0.48 | 0.32 | 0.1 | 0.42 | 0.5 |
| 5 | 86062754 | rs1422241 | -1.552 | -1.816 | -0.996 | -1.257 | -0.004 | 0.036 | 0.036 | 0 | 0.55 | 0.6 | 0.35 | 0.1 | 0.45 | 0.52 |
| 5 | 86078660 | rs13158931 | 1.35 | -1.872 | -0.759 | -1.229 | -0.003 | -0.023 | -0.023 | 0.587 | 0.14 | 0.16 | 0.09 | 0.04 | 0.13 | 0.14 |
| 5 | 86081015 | rs970748 | -1.419 | -1.716 | -0.813 | -1.203 | -0.002 | 0.017 | 0.017 | 0 | 0.55 | 0.59 | 0.35 | 0.09 | 0.46 | 0.53 |
| 5 | 86084306 | rs1422248 | -0.564 | -1.841 | -0.823 | -1.284 | -0.004 | -0.122 | -0.122 | 0.148 | 0.3 | 0.47 | 0.22 | 0.05 | 0.26 | 0.37 |
| 5 | 86087631 | rs2112457 | 0.411 | -1.705 | -0.947 | -0.903 | 0.021 | 0.031 | 0.031 | 0 | 0.24 | 0.11 | 0.13 | 0.11 | 0.2 | 0.17 |
| 5 | 86095152 | rs2032817 | 0.49 | -1.694 | -0.981 | -0.9 | 0.02 | 0.07 | 0.07 | 0 | 0.24 | 0.09 | 0.13 | 0.04 | 0.2 | 0.17 |
| 5 | 86097023 | rs16902412 | 0.227 | -1.753 | -1.031 | -0.543 | -0.085 | 0.039 | 0.039 | 0 | 0.21 | 0.03 | 0.29 | 0.12 | 0.24 | 0.33 |
| 5 | 86099406 | rs13175994 | 0.486 | -1.747 | -1.019 | -0.934 | 0.02 | 0.026 | 0.026 | 0 | 0.24 | 0.09 | 0.13 | 0.12 | 0.2 | 0.17 |
| 5 | 86100132 | rs952841 | 0.166 | -1.553 | -0.96 | -0.894 | -0.085 | 0.132 | 0.132 | 0 | 0.21 | 0.03 | 0.16 | 0.01 | 0.24 | 0.32 |
| 5 | 86113699 | rs2898543 | -1.653 | -1.439 | -0.781 | -0.498 | -0.004 | -0.003 | -0.003 | 0 | 0.65 | 0.63 | 0.43 | 0.44 | 0.52 | 0.69 |
| 5 | 86114035 | rs10514286 | -0.206 | -1.633 | -0.773 | -0.105 | 0.033 | 0 | 0 | 0 | 0.29 | 0.11 | 0.15 | 0.31 | 0.23 | 0.3 |
| 5 | 86128792 | rs11746597 | -0.165 | -1.618 | -0.823 | -0.097 | 0.034 | -0.004 | -0.004 | 0 | 0.29 | 0.1 | 0.15 | 0.31 | 0.23 | 0.3 |
| 5 | 86129665 | rs12518171 | -0.468 | -1.439 | -0.729 | -0.167 | -0.038 | 0.01 | 0.01 | 0 | 0.35 | 0.39 | 0.19 | 0.38 | 0.24 | 0.33 |
| 5 | 86131920 | rs6871722 | -0.7 | -0.812 | -1.196 | -0.744 | 0.037 | 0.192 | 0.192 | 0.228 | 0.47 | 0.78 | 0.32 | 0.11 | 0.4 | 0.41 |
| 5 | 86132101 | rs10055138 | 0.399 | 0.436 | -0.885 | -0.876 | -0.004 | -0.005 | -0.005 | 1.63 | 0.22 | 0.18 | 0.19 | 0.03 | 0.26 | 0.16 |
| 5 | 86134848 | rs981308 | -0.642 | 0.107 | -0.551 | -0.663 | -0.004 | 0.004 | 0.004 | 0 | 0.4 | 0.62 | 0.34 | 0.16 | 0.49 | 0.33 |
| 5 | 86136760 | rs2161249 | -0.684 | 0.091 | -0.587 | -0.436 | -0.005 | 0.005 | 0.005 | 0 | 0.41 | 0.62 | 0.38 | 0.19 | 0.49 | 0.47 |
| 5 | 86136965 | rs7701313 | -0.715 | 0.102 | -0.582 | -0.502 | -0.005 | 0.008 | 0.008 | 0 | 0.41 | 0.62 | 0.38 | 0.26 | 0.49 | 0.47 |
| 5 | 86151620 | rs16902429 | 0.288 | 0.317 | -0.897 | -0.377 | -0.002 | 0.009 | 0.009 | 0 | 0.24 | 0.19 | 0.19 | 0.05 | 0.29 | 0.19 |
| 5 | 86155113 | rs4133446 | -1.614 | -0.011 | -1.288 | -0.399 | -0.054 | -0.043 | -0.043 | 0 | 0.61 | 0.86 | 0.49 | 0.37 | 0.58 | 0.66 |
| 5 | 86158455 | rs16902440 | 0.254 | 0.387 | -0.913 | -0.4 | -0.005 | 0.039 | 0.039 | 0 | 0.24 | 0.15 | 0.19 | 0.06 | 0.29 | 0.18 |
| 5 | 86162971 | rs6877480 | -0.68 | 0.465 | -0.799 | 0.016 | 0.023 | 0.021 | 0.021 | 0 | 0.51 | 0.62 | 0.45 | 0.33 | 0.64 | 0.58 |
| 5 | 86163751 | rs4244386 | 0.719 | 0.406 | -0.241 | 0.567 | 0.011 | 0.022 | 0.022 | 0 | 0.83 | 0.7 | 0.81 | 0.78 | 0.8 | 0.9 |
| 5 | 86164803 | rs10942475 | 0.677 | 0.164 | -0.827 | 0.252 | -0.043 | -0.065 | -0.065 | 0 | 0.33 | 0.07 | 0.27 | 0.38 | 0.35 | 0.33 |
| 5 | 86165391 | rs4509057 | 0.696 | -0.056 | -0.265 | -0.06 | -0.005 | 0.045 | 0.045 | 0 | 0.51 | 0.26 | 0.56 | 0.66 | 0.52 | 0.62 |
| 5 | 86170063 | rs10514284 | -0.582 | -0.027 | 0.518 | -0.193 | 0.04 | -0.032 | -0.032 | 0 | 0.13 | 0.34 | 0.17 | 0.03 | 0.16 | 0.1 |
| 5 | 86174147 | rs1493354 | 0.976 | -0.123 | 0.598 | -0.23 | -0.003 | 0.033 | 0.033 | 0 | 0.51 | 0.31 | 0.56 | 0.66 | 0.52 | 0.61 |
| 5 | 86177773 | rs2471042 | -0.098 | -0.203 | 0.958 | -0.183 | 0.001 | -0.015 | -0.015 | 0.505 | 0.65 | 0.75 | 0.66 | 0.66 | 0.68 | 0.71 |
| 5 | 86179008 | rs13161175 | 0.256 | -0.177 | 1.115 | -0.156 | 0.006 | -0.002 | -0.002 | 0 | 0.61 | 0.36 | 0.63 | 0.66 | 0.67 | 0.68 |
| 5 | 86187316 | rs2452761 | -0.561 | -0.275 | 0.859 | -0.229 | -0.012 | -0.015 | -0.015 | 0 | 0.85 | 0.89 | 0.8 | 0.75 | 0.79 | 0.79 |
| 5 | 86195649 | rs2471039 | -0.304 | -0.086 | 1.024 | -0.388 | 0.001 | -0.003 | -0.003 | 0 | 0.31 | 0.31 | 0.23 | 0.08 | 0.26 | 0.19 |
| 5 | 86198387 | rs1908084 | -0.421 | -0.183 | 1.083 | -0.34 | -0.016 | 0.019 | 0.019 | 0.306 | 0.94 | 0.76 | 0.96 | 0.95 | 0.98 | 0.96 |
| 5 | 86204395 | rs983809 | -0.051 | -0.189 | 1.199 | -0.359 | -0.073 | -0.073 | -0.073 | 1.498 | 0.26 | 0.5 | 0.16 | 0.08 | 0.11 | 0.14 |
| 5 | 86243827 | rs10052248 | 1.645 | 0.009 | 0.305 | -0.196 | -0.07 | -0.032 | -0.032 | 0.016 | 0.27 | 0.07 | 0.3 | 0.3 | 0.31 | 0.29 |
| 5 | 86271618 | rs1073933 | 0.444 | 0.078 | 1.064 | -0.386 | 0.004 | 0.001 | 0.001 | 0 | 0.8 | 0.79 | 0.69 | 0.72 | 0.69 | 0.77 |
| 5 | 86278748 | rs7702765 | 0.114 | 0.186 | 1.162 | -0.372 | 0.009 | -0.021 | -0.021 | 0 | 0.09 | 0.09 | 0.06 | 0 | 0.14 | 0.02 |
| 5 | 86282429 | rs10514289 | -0.29 | 0.007 | 1.267 | -0.492 | -0.002 | -0.069 | -0.069 | 0 | 0.29 | 0.28 | 0.11 | 0 | 0.22 | 0.14 |
| 5 | 86286403 | rs11745726 | 0.845 | -0.043 | 1.147 | -0.476 | 0.018 | 0.003 | 0.003 | 0.343 | 0.2 | 0.15 | 0.04 | 0 | 0.08 | 0.08 |
| 5 | 86286426 | rs10462457 | 0.765 | 0.002 | 0.925 | -0.369 | 0.005 | -0.009 | -0.009 | 0 | 0.57 | 0.55 | 0.63 | 0.72 | 0.59 | 0.65 |
| 5 | 86299070 | rs4920782 | 1.091 | 0.015 | 0.405 | -0.338 | -0.006 | 0.056 | 0.056 | 0 | 0.5 | 0.26 | 0.55 | 0.68 | 0.56 | 0.61 |
| 5 | 86319195 | rs10942487 | 0.935 | -0.66 | 0.506 | -0.66 | 0.04 | 0.097 | 0.097 | 0 | 0.62 | 0.25 | 0.8 | 0.92 | 0.74 | 0.83 |
| 5 | 86319339 | rs2452749 | 0.099 | -0.719 | 0.667 | -0.691 | 0 | 0.086 | 0.086 | 0.425 | 0.72 | 0.89 | 0.87 | 0.96 | 0.77 | 0.9 |
| 5 | 86350878 | rs7715840 | 0.733 | -0.709 | 0.868 | -0.737 | 0.016 | 0.015 | 0.015 | 0 | 0.64 | 0.33 | 0.85 | 0.96 | 0.75 | 0.86 |
| 5 | 86378336 | rs2544688 | 0.467 | -0.364 | 0.481 | -0.477 | 0.004 | -0.012 | -0.012 | 0.093 | 0.89 | 0.9 | 0.93 | 0.96 | 0.84 | 0.94 |
| 5 | 86386565 | rs2624189 | 0.386 | -0.614 | 0.336 | -0.668 | 0.014 | 0.07 | 0.07 | 0 | 0.69 | 0.86 | 0.88 | 0.96 | 0.75 | 0.86 |
| 5 | 86393186 | rs2544679 | 1.036 | -0.362 | 0.662 | -0.452 | 0.046 | -0.026 | -0.026 | 0 | 0.87 | 0.36 | 0.86 | 0.96 | 0.85 | 0.94 |
| 5 | 86406135 | rs2544671 | 0.89 | -0.316 | 0.38 | -0.429 | 0.039 | -0.01 | -0.01 | 0.565 | 0.86 | 0.62 | 0.88 | 0.93 | 0.84 | 0.94 |
| 5 | 86409436 | rs2112166 | 0.359 | -0.387 | 0.28 | -0.474 | -0.012 | -0.046 | -0.046 | 0 | 0.63 | 0.58 | 0.83 | 0.92 | 0.75 | 0.83 |
| 5 | 86418751 | rs2624171 | 1.354 | 0.487 | -0.422 | 0.252 | -0.072 | -0.13 | -0.13 | 0.727 | 0.19 | 0.02 | 0.25 | 0.38 | 0.24 | 0.26 |

| | | | | | | | | | | | | | | | | |
|---|---|---|---|---|---|---|---|---|---|---|---|---|---|---|---|---|
| 5 | 86419364 | rs13157029 | 1.351 | 0.525 | 0.253 | -0.06 | -0.029 | 0.059 | 0.059 | 0 | 0.45 | 0.24 | 0.59 | 0.7 | 0.54 | 0.6 |
| 5 | 86424676 | rs4141503 | 0.209 | 0.459 | 0.175 | 0.104 | 0.031 | 0.069 | 0.069 | 0 | 0.15 | 0.3 | 0.23 | 0.26 | 0.19 | 0.2 |
| 5 | 86447940 | rs2544702 | -0.451 | 0.226 | -0.217 | -0.25 | 0.01 | 0.019 | 0.019 | 0 | 0.34 | 0.24 | 0.13 | 0.07 | 0.24 | 0.15 |
| 5 | 86460047 | rs2624164 | -0.536 | 1.602 | 0.076 | 0.942 | -0.025 | 0 | 0 | 0 | 0.07 | 0.03 | 0.05 | 0.07 | 0.13 | 0.03 |
| 5 | 86460269 | rs2624165 | -0.536 | 1.602 | 0.076 | 0.917 | -0.025 | 0.003 | 0.003 | 0 | 0.07 | 0.03 | 0.05 | 0.06 | 0.13 | 0.03 |
| 5 | 86460539 | rs30421 | -0.741 | 0.653 | 0.434 | 1.054 | 0.056 | 0.058 | 0.058 | 0 | 0.14 | 0.34 | 0.21 | 0.17 | 0.18 | 0.16 |
| 5 | 86468013 | rs6870700 | -1.122 | -0.527 | 0.33 | 1.969 | -0.01 | 0.162 | 0.162 | 0 | 0.08 | 0.2 | 0.06 | 0.36 | 0.04 | 0.05 |
| 5 | 86471345 | rs171711 | -1.902 | -1.271 | 0.412 | 1.971 | -0.07 | 0.031 | 0.031 | 0 | 0.28 | 0.49 | 0.38 | 0.61 | 0.12 | 0.25 |
| 5 | 86482632 | rs10942497 | -0.784 | -1.436 | 0.581 | 2.156 | 0.021 | -0.028 | -0.028 | 0 | 0.15 | 0.1 | 0.24 | 0.2 | 0.08 | 0.2 |
| 5 | 86483920 | rs30400 | -1.763 | -1.418 | 0.56 | 1.668 | -0.08 | 0.074 | 0.074 | 0 | 0.22 | 0.47 | 0.35 | 0.62 | 0.09 | 0.23 |
| 5 | 86493926 | rs12523600 | -1.281 | -0.987 | 0.634 | 0.577 | -0.021 | -0.144 | -0.144 | 0.279 | 0.23 | 0.51 | 0.14 | 0.01 | 0.19 | 0.07 |
| 5 | 86518243 | rs11953892 | -1.684 | -1.351 | 0.442 | 1.06 | -0.025 | 0.049 | 0.049 | 0 | 0.32 | 0.46 | 0.33 | 0.5 | 0.27 | 0.23 |
| 5 | 86524315 | rs6890495 | -1.419 | -1.37 | 0.512 | 0.851 | -0.037 | -0.046 | -0.046 | 0 | 0.1 | 0.36 | 0.04 | 0.01 | 0.02 | 0.03 |
| 5 | 86539266 | rs16902601 | -0.817 | -1.289 | 0.187 | 1.203 | -0.017 | -0.017 | -0.017 | 0 | 0.2 | 0.05 | 0.17 | 0.18 | 0.25 | 0.16 |
| 5 | 86540308 | rs10942498 | -1.764 | -1.264 | 0.343 | 1.212 | -0.029 | -0.02 | -0.02 | 0 | 0.3 | 0.41 | 0.31 | 0.19 | 0.25 | 0.19 |
| 5 | 86605943 | rs6452750 | -1.786 | -1.488 | -0.712 | 0.22 | -0.121 | -0.087 | -0.087 | 1.375 | 0.58 | 0.94 | 0.52 | 0.6 | 0.49 | 0.49 |
| 5 | 86607633 | rs10058449 | -0.677 | -1.526 | -0.246 | 0.148 | 0.004 | -0.009 | -0.009 | 0.567 | 0.41 | 0.21 | 0.28 | 0.47 | 0.39 | 0.44 |
| 5 | 86610989 | rs35148638 | -0.321 | -1.542 | -0.266 | 0.139 | -0.018 | -0.077 | -0.077 | 0 | 0.21 | 0.06 | 0.19 | 0.44 | 0.25 | 0.18 |
| 5 | 86628891 | rs2271235 | -1.076 | -1.328 | -0.266 | 0.109 | 0.001 | 0 | 0 | 0 | 0.46 | 0.46 | 0.41 | 0.46 | 0.39 | 0.44 |
| 5 | 86635942 | rs10942502 | -0.952 | -1.297 | -0.258 | 0.116 | 0.019 | -0.008 | -0.008 | 0.166 | 0.44 | 0.23 | 0.39 | 0.48 | 0.4 | 0.45 |
| 5 | 86652624 | rs10045850 | -1.043 | -1.423 | -0.289 | 0.019 | 0.034 | 0.007 | 0.007 | 0 | 0.47 | 0.27 | 0.39 | 0.49 | 0.41 | 0.47 |
| 5 | 86656964 | rs4421140 | -0.102 | -1.406 | -0.31 | 0.032 | -0.003 | -0.077 | -0.077 | 0 | 0.21 | 0.06 | 0.18 | 0.45 | 0.23 | 0.17 |
| 5 | 86670970 | rs10067098 | -1.375 | -1.65 | -0.207 | 0.074 | 0.002 | 0 | 0 | 0 | 0.52 | 0.47 | 0.43 | 0.49 | 0.45 | 0.49 |
| 5 | 86714840 | rs10065575 | -1.527 | -1.708 | -0.042 | 0.101 | 0.003 | 0.001 | 0.001 | 0 | 0.53 | 0.61 | 0.43 | 0.48 | 0.43 | 0.47 |
| 5 | 86727027 | rs7702514 | 0.865 | -1.805 | -0.675 | -0.014 | -0.018 | 0.106 | 0.106 | 0 | 0.2 | 0.01 | 0.09 | 0.03 | 0.16 | 0.27 |
| 5 | 86727165 | rs7702564 | -1.021 | -1.666 | -0.522 | 0.063 | 0.043 | 0.009 | 0.009 | 0 | 0.47 | 0.27 | 0.37 | 0.48 | 0.4 | 0.46 |
| 5 | 86752616 | rs323781 | -1.833 | -1.578 | -0.761 | 0.047 | -0.001 | -0.03 | -0.03 | 0.196 | 0.07 | 0.24 | 0.15 | 0.01 | 0.03 | 0.02 |
| 5 | 86785578 | rs326071 | -1.944 | -1.665 | -0.967 | 0.009 | -0.11 | -0.108 | -0.108 | 0 | 0.61 | 0.95 | 0.51 | 0.58 | 0.47 | 0.51 |
| 5 | 86797499 | rs161520 | -1.371 | -1.392 | -0.714 | 0.1 | 0.011 | 0.007 | 0.007 | 0 | 0.52 | 0.47 | 0.42 | 0.46 | 0.41 | 0.47 |
| 5 | 86826750 | rs1561734 | -0.081 | -1.31 | -0.614 | 0.18 | 0 | -0.002 | -0.002 | 0 | 0.2 | 0.19 | 0.16 | 0.06 | 0.22 | 0.19 |
| 5 | 86835416 | rs35814902 | -0.134 | -1.325 | -0.833 | 0.305 | 0.024 | -0.031 | -0.031 | 0 | 0.25 | 0.18 | 0.21 | 0.4 | 0.16 | 0.29 |
| 5 | 86842168 | rs13177543 | -0.124 | -1.325 | -0.833 | 0.305 | 0.024 | -0.031 | -0.031 | 0.042 | 0.25 | 0.18 | 0.21 | 0.4 | 0.16 | 0.29 |
| 5 | 86895675 | rs1121757 | -0.928 | -1.073 | -1.118 | 0.363 | 0.002 | 0.02 | 0.02 | 0 | 0.26 | 0.36 | 0.32 | 0.41 | 0.3 | 0.25 |
| 5 | 86905677 | rs4976236 | -1.699 | -0.977 | -0.849 | 0.784 | 0.021 | 0.195 | 0.195 | 0 | 0.06 | 0.23 | 0.16 | 0.35 | 0.05 | 0.06 |
| 5 | 86909526 | rs12652533 | -1.836 | -0.931 | -0.838 | 1.032 | 0.032 | 0.311 | 0.311 | 0 | 0.05 | 0.19 | 0.2 | 0.49 | 0.07 | 0.09 |
| 5 | 86910439 | rs3924956 | 0.12 | -1.099 | -0.998 | 0.901 | 0.032 | 0.1 | 0.1 | 0 | 0.22 | 0.01 | 0.07 | 0.04 | 0.11 | 0.23 |
| 5 | 86919884 | rs7737385 | -2.251 | -0.481 | -0.479 | 1.293 | 0.035 | 0.284 | 0.284 | 0 | 0.08 | 0.24 | 0.22 | 0.5 | 0.1 | 0.09 |
| 5 | 86997712 | rs840828 | -0.899 | -0.561 | -0.671 | 1.043 | 0.025 | 0.204 | 0.204 | 0 | 0.05 | 0.18 | 0.14 | 0.35 | 0.05 | 0.05 |
| 5 | 87022986 | rs7733912 | -0.218 | -1.134 | -0.891 | 0.655 | 0.005 | -0.003 | -0.003 | 0 | 0.21 | 0.22 | 0.19 | 0.34 | 0.12 | 0.11 |
| 5 | 87024072 | rs710394 | -0.645 | -1.302 | -1.087 | 0.458 | -0.001 | -0.001 | -0.001 | 0 | 0.3 | 0.31 | 0.32 | 0.22 | 0.32 | 0.21 |
| 5 | 87025144 | rs710392 | -0.122 | -1.389 | -1.167 | 0.401 | -0.017 | -0.008 | -0.008 | 1.795 | 0.21 | 0.28 | 0.19 | 0.14 | 0.19 | 0.07 |
| 5 | 87032597 | rs710386 | -1.184 | -1.443 | -1.133 | 0.474 | 0.003 | 0.001 | 0.001 | 0.01 | 0.51 | 0.52 | 0.5 | 0.56 | 0.43 | 0.32 |
| 5 | 87038565 | rs710384 | -0.447 | -1.445 | -1.165 | 0.487 | 0 | 0.001 | 0.001 | 0 | 0.31 | 0.31 | 0.32 | 0.22 | 0.32 | 0.21 |
| 5 | 87041871 | rs710378 | -1.866 | -1.726 | -1.49 | -0.06 | -0.137 | -0.098 | -0.098 | 0 | 0.62 | 0.96 | 0.57 | 0.59 | 0.49 | 0.51 |
| 5 | 87042023 | rs710377 | -0.457 | -1.436 | -1.16 | 0.08 | -0.001 | 0.022 | 0.022 | 0 | 0.29 | 0.25 | 0.27 | 0.07 | 0.31 | 0.19 |
| 5 | 87050201 | rs723111 | -0.066 | -1.005 | -1.412 | 0.341 | -0.015 | 0.029 | 0.029 | 0 | 0.19 | 0.26 | 0.23 | 0.46 | 0.11 | 0.13 |
| 5 | 87061763 | rs11739923 | -0.564 | -0.555 | -1.807 | -0.292 | -0.007 | 0.06 | 0.06 | 0 | 0.28 | 0.14 | 0.25 | 0.08 | 0.3 | 0.19 |
| 5 | 87068177 | rs17285947 | 0.215 | -1.062 | -1.934 | -0.402 | 0.045 | 0.091 | 0.091 | 0 | 0.16 | 0.02 | 0.08 | 0 | 0.09 | 0.06 |
| 5 | 87068432 | rs17285954 | -0.286 | -0.091 | -2.708 | -0.547 | -0.115 | 0.031 | 0.031 | 0 | 0.33 | 0.06 | 0.33 | 0.24 | 0.56 | 0.16 |
| 5 | 87079042 | rs16902804 | -0.174 | -0.88 | -1.567 | -0.868 | -0.055 | -0.114 | -0.114 | 0 | 0.27 | 0.44 | 0.16 | 0.03 | 0.15 | 0.1 |
| 5 | 87081822 | rs16902806 | 1.415 | -1.044 | -1.949 | -0.426 | 0.047 | -0.064 | -0.064 | 1.365 | 0.37 | 0.08 | 0.35 | 0.38 | 0.27 | 0.47 |
| 5 | 87082276 | rs710375 | 0.831 | -1.041 | -1.605 | -0.227 | -0.005 | -0.02 | -0.02 | 0 | 0.47 | 0.48 | 0.55 | 0.78 | 0.36 | 0.73 |
| 5 | 87086376 | rs698060 | -0.506 | -1.072 | -1.391 | -0.271 | -0.113 | -0.004 | -0.004 | 3.979 | 0.74 | 0.93 | 0.71 | 0.72 | 0.51 | 0.84 |
| 5 | 87089786 | rs16902819 | 1.443 | -1.215 | -0.877 | -0.233 | 0.046 | -0.065 | -0.065 | 0.556 | 0.37 | 0.08 | 0.35 | 0.38 | 0.27 | 0.46 |
| 5 | 87092264 | rs1423633 | -0.297 | -1.243 | -0.569 | -0.555 | -0.054 | -0.11 | -0.11 | 0 | 0.3 | 0.54 | 0.18 | 0.06 | 0.18 | 0.26 |
| 5 | 87092393 | rs2898554 | 0.717 | -1.16 | -0.758 | -0.535 | 0.039 | 0.084 | 0.084 | 0 | 0.21 | 0.08 | 0.1 | 0 | 0.14 | 0.07 |
| 5 | 87096105 | rs1423634 | -0.719 | -0.687 | -0.547 | -0.49 | 0 | -0.005 | -0.005 | 1.512 | 0.38 | 0.63 | 0.35 | 0.15 | 0.27 | 0.31 |
| 5 | 87098260 | rs10942510 | 0.426 | -0.256 | -0.521 | -0.677 | 0.053 | 0.083 | 0.083 | 0 | 0.5 | 0.35 | 0.66 | 0.78 | 0.65 | 0.56 |
| 5 | 87099119 | rs6885160 | -0.257 | -0.361 | -0.278 | -1.016 | -0.006 | -0.067 | -0.067 | 0.439 | 0.45 | 0.5 | 0.26 | 0.04 | 0.3 | 0.35 |
| 5 | 87100609 | rs12520006 | -0.507 | -0.369 | -0.251 | -1.037 | 0 | -0.007 | -0.007 | 0 | 0.11 | 0.09 | 0.05 | 0.01 | 0.09 | 0.07 |
| 5 | 87101610 | rs6876161 | 1.174 | -0.361 | -0.19 | -1.031 | 0.002 | -0.013 | -0.013 | 0 | 0.86 | 0.85 | 0.92 | 0.96 | 0.87 | 0.91 |
| 5 | 87104604 | rs10063127 | 0.562 | -1.036 | -0.076 | -1.194 | 0.029 | 0.004 | 0.004 | 0 | 0.33 | 0.2 | 0.16 | 0.01 | 0.2 | 0.26 |
| 5 | 87146327 | rs1164548 | -0.704 | -1.104 | 0.087 | -1.196 | 0.119 | 0.178 | 0.178 | 0 | 0.46 | 0.8 | 0.27 | 0.02 | 0.27 | 0.33 |
| 5 | 87183433 | rs16902864 | 0.801 | -1.112 | -0.074 | -1.153 | 0.016 | -0.015 | -0.015 | 0.048 | 0.31 | 0.21 | 0.16 | 0.01 | 0.2 | 0.26 |
| 5 | 87190084 | rs10514296 | 0.455 | -1.175 | 0.231 | -1.184 | -0.008 | -0.068 | -0.068 | 0 | 0.33 | 0.26 | 0.16 | 0.01 | 0.2 | 0.27 |
| 5 | 87195373 | rs1458198 | -0.935 | -1.229 | 0.068 | -0.869 | 0.129 | 0.164 | 0.164 | 0 | 0.48 | 0.86 | 0.33 | 0.17 | 0.29 | 0.4 |
| 5 | 87197535 | rs13173741 | 1.604 | -0.395 | -0.545 | -0.46 | -0.128 | -0.057 | -0.057 | 0.263 | 0.27 | 0.04 | 0.44 | 0.38 | 0.4 | 0.29 |
| 5 | 87247068 | rs16902900 | 0.329 | -1.216 | 0.213 | -1.057 | 0.006 | -0.054 | -0.054 | 0 | 0.37 | 0.33 | 0.18 | 0.01 | 0.22 | 0.24 |
| 5 | 87248647 | rs13356439 | 0.554 | -1.209 | 0.213 | -1.054 | 0.009 | -0.045 | -0.045 | 0 | 0.36 | 0.29 | 0.17 | 0.01 | 0.22 | 0.23 |

| | | | | | | | | | | | | | | | | |
|---|---|---|---|---|---|---|---|---|---|---|---|---|---|---|---|---|
| 5 | 87248847 | rs13356445 | 0.554 | -1.209 | 0.213 | -1.054 | 0.009 | -0.045 | -0.045 | 0 | 0.36 | 0.29 | 0.17 | 0.01 | 0.22 | 0.24 |
| 5 | 87249313 | rs2178270 | 1.047 | -1.165 | 0.27 | -1.007 | 0.002 | -0.047 | -0.047 | 0.068 | 0.31 | 0.23 | 0.16 | 0.01 | 0.19 | 0.22 |
| 5 | 87269005 | rs11948159 | 0.922 | -1.157 | 0.141 | -1.004 | 0.005 | -0.044 | -0.044 | 0.035 | 0.32 | 0.27 | 0.16 | 0.01 | 0.2 | 0.23 |
| 5 | 87279077 | rs3902731 | 0.006 | -1.084 | 0.352 | -0.893 | 0.062 | 0.03 | 0.03 | 1.515 | 0.94 | 0.57 | 0.92 | 1 | 0.97 | 0.95 |
| 5 | 87311560 | rs13164055 | -0.167 | -1.217 | -0.108 | -1.017 | 0.065 | 0.065 | 0.065 | 0 | 0.6 | 0.22 | 0.76 | 0.98 | 0.77 | 0.73 |
| 5 | 87315252 | rs965566 | 0.562 | -1.263 | -0.304 | -0.822 | 0.134 | 0.15 | 0.15 | 0.086 | 0.49 | 0.14 | 0.66 | 0.8 | 0.69 | 0.59 |
| 5 | 87318551 | rs1429605 | 0.507 | -0.608 | 0.283 | -0.528 | 0.036 | 0.003 | 0.003 | 2.177 | 0.92 | 0.51 | 0.92 | 1 | 0.97 | 0.95 |
| 5 | 87342183 | rs11740691 | 0.653 | 0.222 | -0.155 | -0.352 | -0.082 | -0.037 | -0.037 | 0 | 0.12 | 0.01 | 0.17 | 0.15 | 0.28 | 0.1 |
| 5 | 87345211 | rs4916879 | 0.959 | -1.443 | 0.367 | -0.361 | 0.051 | -0.015 | -0.015 | 0 | 0.91 | 0.51 | 0.9 | 0.99 | 0.94 | 0.95 |
| 5 | 87387572 | rs1427923 | 1.49 | -1.754 | 0.126 | -0.207 | -0.033 | 0.02 | 0.02 | 3.879 | 0.79 | 0.34 | 0.82 | 0.81 | 0.88 | 0.83 |
| 5 | 87413633 | rs1834910 | -0.207 | -2.404 | -0.286 | -0.715 | -0.016 | -0.011 | -0.011 | 0 | 0.44 | 0.55 | 0.24 | 0.16 | 0.22 | 0.35 |
| 5 | 87422770 | rs1427920 | -0.267 | -2.376 | -0.363 | -0.718 | 0.042 | 0.039 | 0.039 | 0 | 0.44 | 0.65 | 0.25 | 0.16 | 0.22 | 0.35 |
| 5 | 87440497 | rs11948659 | -0.045 | -2.608 | -0.396 | -0.931 | -0.022 | -0.113 | -0.113 | 0 | 0.45 | 0.49 | 0.22 | 0 | 0.28 | 0.34 |
| 5 | 87445490 | rs12522676 | 0.776 | -2.827 | -0.58 | -0.95 | 0.113 | 0.126 | 0.126 | 0.122 | 0.45 | 0.17 | 0.67 | 0.85 | 0.68 | 0.55 |
| 5 | 87470240 | rs6452775 | 1.355 | -2.702 | -0.623 | -0.978 | -0.004 | 0.032 | 0.032 | 0.2 | 0.88 | 0.83 | 0.86 | 0.79 | 0.9 | 0.82 |
| 5 | 87490908 | rs33700 | 0.429 | -2.865 | -0.68 | -1.064 | -0.046 | -0.144 | -0.144 | 0 | 0.4 | 0.48 | 0.2 | 0 | 0.25 | 0.33 |
| 5 | 87491545 | rs33699 | 1.239 | -2.841 | -0.747 | -1.075 | 0.022 | 0.058 | 0.058 | 0.284 | 0.4 | 0.14 | 0.61 | 0.79 | 0.59 | 0.47 |
| 5 | 87495504 | rs358288 | 1.328 | -2.796 | -0.743 | -1.054 | -0.008 | 0.057 | 0.057 | 0.3 | 0.86 | 0.74 | 0.84 | 0.79 | 0.87 | 0.82 |
| 5 | 87498849 | rs3097146 | 1.428 | -2.684 | -1.089 | -0.985 | 0.001 | 0.041 | 0.041 | 0.24 | 0.37 | 0.1 | 0.61 | 0.79 | 0.59 | 0.47 |
| 5 | 87513775 | rs357513 | 1.566 | -2.274 | -0.942 | -0.488 | 0.007 | 0.026 | 0.026 | 0.397 | 0.76 | 0.66 | 0.61 | 0.61 | 0.72 | 0.76 |
| 5 | 87514004 | rs371708 | 0.989 | -2.257 | -1.054 | -0.51 | -0.007 | 0.003 | 0.003 | 0 | 0.8 | 0.8 | 0.81 | 0.77 | 0.73 | 0.84 |
| 5 | 87530253 | rs247916 | 0.376 | -2.7 | -1.129 | -0.77 | -0.037 | -0.114 | -0.114 | 0 | 0.39 | 0.52 | 0.19 | 0 | 0.23 | 0.34 |
| 5 | 87540210 | rs780401 | 1.431 | -2.558 | -1.194 | -0.569 | -0.006 | -0.02 | -0.02 | 0.047 | 0.77 | 0.77 | 0.66 | 0.61 | 0.74 | 0.76 |
| 5 | 87568710 | rs768705 | -1.626 | -2.513 | -1.153 | -0.573 | 0.007 | 0.031 | 0.031 | 0 | 0.22 | 0.33 | 0.35 | 0.37 | 0.26 | 0.24 |
| 5 | 87572026 | rs12163987 | -1.025 | -2.478 | -1.205 | -0.649 | -0.007 | 0.003 | 0.003 | 0 | 0.18 | 0.18 | 0.14 | 0.22 | 0.25 | 0.14 |
| 5 | 87663610 | rs10514299 | -1.002 | -2.483 | -1.247 | -0.652 | -0.007 | 0.003 | 0.003 | 0 | 0.18 | 0.18 | 0.14 | 0.22 | 0.24 | 0.15 |
| 5 | 87675962 | rs11743103 | -1.238 | -2.492 | -1.238 | -0.545 | 0.005 | 0.023 | 0.023 | 0 | 0.2 | 0.26 | 0.33 | 0.38 | 0.26 | 0.24 |
| 5 | 87679052 | rs4916908 | -1.664 | -2.541 | -1.301 | -0.542 | -0.045 | -0.049 | -0.049 | 1.395 | 0.63 | 0.88 | 0.53 | 0.39 | 0.49 | 0.58 |
| 5 | 87712913 | rs6452788 | 0.157 | -2.241 | -1.158 | -0.414 | -0.017 | -0.01 | -0.01 | 0 | 0.41 | 0.54 | 0.4 | 0.18 | 0.27 | 0.43 |
| 5 | 87714704 | rs10474277 | 1.708 | -2.687 | -1.643 | -0.689 | 0.025 | 0.137 | 0.137 | 0 | 0.21 | 0.01 | 0.09 | 0 | 0.13 | 0.2 |
| 5 | 87747781 | rs12659818 | -0.792 | -2.036 | -1.26 | -0.32 | -0.001 | 0 | 0 | 0.023 | 0.16 | 0.16 | 0.11 | 0.23 | 0.22 | 0.16 |
| 5 | 87756918 | rs4571506 | -1.153 | -2.115 | -1.349 | -0.313 | -0.009 | -0.013 | -0.013 | 0.796 | 0.57 | 0.83 | 0.52 | 0.4 | 0.48 | 0.58 |
| 5 | 87776730 | rs16903178 | -1.227 | -1.858 | -1.099 | -0.041 | 0.016 | 0.1 | 0.1 | 0.168 | 0.17 | 0.3 | 0.31 | 0.45 | 0.21 | 0.24 |
| 5 | 87780432 | rs17419291 | 1.867 | -1.639 | -1.667 | -0.673 | -0.017 | 0.058 | 0.058 | 0 | 0.1 | 0 | 0.07 | 0 | 0.09 | 0.14 |
| 5 | 87781168 | rs10044618 | -0.109 | -1.218 | -1.216 | -0.311 | 0.031 | 0.112 | 0.112 | 0 | 0.52 | 0.74 | 0.31 | 0.29 | 0.45 | 0.47 |
| 5 | 87785624 | rs17478824 | -0.216 | -0.887 | -1.169 | -0.206 | 0 | 0.002 | 0.002 | 0.002 | 0.1 | 0.07 | 0.07 | 0.03 | 0.09 | 0.09 |
| 5 | 87792844 | rs6452792 | -1.106 | -0.905 | -1.678 | 0.221 | -0.125 | -0.031 | -0.031 | 0.574 | 0.67 | 0.94 | 0.45 | 0.62 | 0.53 | 0.69 |
| 5 | 87800881 | rs6860708 | -0.176 | -1.219 | -1.628 | 0.4 | 0.089 | -0.048 | -0.048 | 0 | 0.54 | 0.83 | 0.31 | 0.45 | 0.35 | 0.54 |
| 5 | 87801273 | rs2194026 | -0.578 | -0.936 | -0.906 | 0.68 | 0.005 | 0.246 | 0.246 | 0.009 | 0.12 | 0.24 | 0.11 | 0.44 | 0.1 | 0.2 |
| 5 | 87809261 | rs2032805 | -0.102 | -0.92 | -1.16 | 0.591 | -0.025 | -0.025 | -0.025 | 0.006 | 0.12 | 0.04 | 0.12 | 0.18 | 0.17 | 0.15 |
| 5 | 87831734 | rs16903232 | -0.498 | -0.936 | -0.984 | 0.776 | -0.003 | 0.185 | 0.185 | 0.26 | 0.12 | 0.21 | 0.1 | 0.44 | 0.09 | 0.2 |
| 5 | 87854395 | rs4916723 | 1.067 | -0.427 | -1.3 | 0.558 | -0.083 | -0.022 | -0.022 | 0 | 0.31 | 0.11 | 0.54 | 0.36 | 0.41 | 0.28 |
| 5 | 87854985 | rs10491451 | 0.239 | -0.724 | -1.2 | 0.686 | 0.006 | 0.064 | 0.064 | 0 | 0.07 | 0.03 | 0.03 | 0.38 | 0.01 | 0.15 |
| 5 | 87855403 | rs16903246 | 0.648 | -0.72 | -1.189 | 0.688 | 0 | 0.092 | 0.092 | 0.006 | 0.06 | 0.04 | 0.03 | 0.38 | 0.01 | 0.15 |
| 5 | 87855496 | rs918427 | 0.688 | -1.207 | -1.428 | -0.133 | 0.037 | 0.2 | 0.2 | 0.043 | 0.39 | 0.1 | 0.21 | 0.04 | 0.3 | 0.35 |
| 5 | 87870166 | rs10063429 | 0.262 | -1.285 | -0.876 | -0.103 | 0.024 | -0.008 | -0.008 | 0 | 0.08 | 0.6 | 0.07 | 0 | 0.06 | 0.05 |
| 5 | 87870213 | rs10063583 | 0.275 | -1.011 | -1.109 | -0.103 | -0.081 | 0.086 | 0.086 | 0 | 0.42 | 0.88 | 0.62 | 0.26 | 0.5 | 0.37 |
| 5 | 87877474 | rs10078467 | 0.295 | -1.027 | -0.783 | 0.168 | 0.035 | 0.001 | 0.001 | 1.715 | 0.08 | 0.57 | 0.07 | 0 | 0.06 | 0.05 |
| 5 | 87879468 | rs2438466 | 0.065 | -0.969 | -1.359 | 0.225 | -0.039 | -0.024 | -0.024 | 0 | 0.85 | 0.94 | 0.84 | 0.78 | 0.77 | 0.8 |
| 5 | 87894646 | rs2112424 | 0.579 | -0.903 | -1.309 | -0.168 | -0.017 | -0.04 | -0.04 | 0 | 0.78 | 0.91 | 0.8 | 0.29 | 0.75 | 0.65 |
| 5 | 87901163 | rs6859545 | 1.125 | -0.538 | -0.868 | 0.154 | -0.058 | -0.004 | -0.004 | 0 | 0.26 | 0.1 | 0.51 | 0.26 | 0.37 | 0.23 |
| 5 | 87906959 | rs192015 | 0.329 | -0.49 | -1.067 | 0.251 | -0.031 | 0.037 | 0.037 | 0.387 | 0.36 | 0.79 | 0.59 | 0.26 | 0.43 | 0.28 |
| 5 | 87942868 | rs383883 | -0.181 | -1.422 | -0.586 | 0.885 | 0.002 | -0.019 | -0.019 | 0.182 | 0.11 | 0.07 | 0.35 | 0.24 | 0.12 | 0.13 |
| 5 | 87950290 | rs26582 | -1.783 | -1.84 | -0.061 | 0.667 | -0.013 | -0.025 | -0.025 | 0 | 0.36 | 0.54 | 0.53 | 0.75 | 0.29 | 0.42 |
| 5 | 87950543 | rs16903275 | -1.163 | -1.66 | 0.488 | 0.757 | -0.016 | 0.064 | 0.064 | 0 | 0.24 | 0.34 | 0.18 | 0.51 | 0.17 | 0.29 |
| 5 | 87959538 | rs13174131 | -0.443 | -1.599 | 0.453 | 0.782 | -0.004 | 0.077 | 0.077 | 0 | 0.17 | 0.25 | 0.15 | 0.51 | 0.14 | 0.21 |
| 5 | 87963761 | rs1501672 | -0.359 | -1.572 | 0.363 | 0.795 | -0.002 | 0.044 | 0.044 | 0 | 0.16 | 0.21 | 0.15 | 0.51 | 0.14 | 0.21 |
| 5 | 87968864 | rs6882046 | -0.887 | -1.867 | 0.089 | 0.499 | 0.07 | 0.012 | 0.012 | 0 | 0.47 | 0.31 | 0.27 | 0.54 | 0.29 | 0.49 |
| 5 | 87981030 | rs153685 | 1.004 | -1.352 | -0.02 | 0.076 | 0.054 | -0.002 | -0.002 | 0.314 | 0.48 | 0.26 | 0.44 | 0.56 | 0.59 | 0.57 |
| 5 | 87988733 | rs2304607 | -0.744 | -1.148 | -0.069 | 0.143 | -0.004 | 0.024 | 0.024 | 0.001 | 0.19 | 0.25 | 0.16 | 0.32 | 0.16 | 0.2 |
| 5 | 87993365 | rs34338 | 1.244 | -1.027 | -0.156 | 0.008 | 0.026 | -0.008 | -0.008 | 0 | 0.46 | 0.23 | 0.41 | 0.58 | 0.56 | 0.55 |
| 5 | 88001798 | rs2161228 | -0.066 | -1.037 | -0.266 | 0.032 | 0.007 | -0.013 | -0.013 | 0 | 0.17 | 0.19 | 0.15 | 0.33 | 0.13 | 0.13 |
| 5 | 88009593 | rs254778 | 0.211 | -1.012 | -0.053 | -0.128 | -0.022 | -0.115 | -0.115 | 0 | 0.65 | 0.51 | 0.56 | 0.94 | 0.69 | 0.69 |
| 5 | 88021527 | rs34320 | 1.281 | -1.178 | -0.129 | -0.032 | 0.033 | 0.007 | 0.007 | 0 | 0.47 | 0.15 | 0.4 | 0.61 | 0.56 | 0.59 |
| 5 | 88027217 | rs10514302 | -0.627 | -1.184 | -0.183 | -0.015 | 0.024 | 0.167 | 0.167 | 0 | 0.05 | 0.17 | 0.11 | 0.28 | 0.07 | 0.1 |
| 5 | 88029627 | rs412458 | -0.247 | -1.183 | -0.24 | -0.065 | -0.064 | -0.129 | -0.129 | 0 | 0.71 | 0.54 | 0.61 | 0.93 | 0.78 | 0.75 |
| 5 | 88042019 | rs1422189 | 0.302 | -1.14 | -0.911 | -0.085 | -0.005 | 0.06 | 0.06 | 0 | 0.15 | 0.03 | 0.09 | 0.05 | 0.15 | 0.09 |
| 5 | 88055761 | rs664366 | 0.719 | -0.932 | -0.449 | 0.281 | 0.027 | -0.011 | -0.011 | 0 | 0.63 | 0.28 | 0.5 | 0.65 | 0.71 | 0.64 |
| 5 | 88065130 | rs17485836 | 0.607 | -0.895 | -0.922 | -0.208 | 0.036 | 0.118 | 0.118 | 0 | 0.18 | 0.01 | 0.09 | 0 | 0.11 | 0.15 |

| | | | | | | | | | | | | | | | | |
|---|---|---|---|---|---|---|---|---|---|---|---|---|---|---|---|---|
| 5 | 88079010 | rs674747 | -0.023 | -0.831 | -0.165 | -0.069 | -0.021 | -0.032 | -0.032 | 1.728 | 0.72 | 0.67 | 0.66 | 0.92 | 0.83 | 0.76 |
| 5 | 88095785 | rs244755 | 0.689 | -0.59 | -0.044 | 0.445 | 0.018 | -0.012 | -0.012 | 0 | 0.61 | 0.33 | 0.44 | 0.58 | 0.7 | 0.64 |
| 5 | 88102281 | rs17558396 | 1.346 | -0.541 | -0.54 | -0.089 | -0.089 | -0.085 | -0.085 | 0 | 0.38 | 0.11 | 0.29 | 0.56 | 0.46 | 0.49 |
| 5 | 88103213 | rs11959216 | 0.734 | -0.607 | -0.425 | -0.091 | -0.005 | 0.009 | 0.009 | 0 | 0.07 | 0.05 | 0.08 | 0 | 0.1 | 0.06 |
| 5 | 88114476 | rs160044 | -0.718 | -0.788 | -0.451 | -0.037 | 0.056 | 0.019 | 0.019 | 0.345 | 0.44 | 0.71 | 0.52 | 0.33 | 0.31 | 0.38 |
| 5 | 88140962 | rs244749 | -0.047 | -0.894 | -0.376 | -0.027 | -0.007 | 0 | 0 | 0.169 | 0.36 | 0.64 | 0.44 | 0.33 | 0.21 | 0.31 |
| 5 | 88143749 | rs302484 | -0.087 | -0.723 | 0.412 | 0.244 | 0.033 | 0.215 | 0.215 | 4.205 | 0.06 | 0.33 | 0.08 | 0.17 | 0.02 | 0.03 |
| 5 | 88144091 | rs302483 | -1.104 | -0.727 | 0.404 | 0.236 | 0.043 | 0.309 | 0.309 | 7.996 | 0.08 | 0.43 | 0.11 | 0.25 | 0.04 | 0.07 |
| 5 | 88157552 | rs4518438 | 0.911 | -0.73 | 0.037 | 0.283 | -0.009 | -0.009 | -0.009 | 0 | 0.39 | 0.31 | 0.36 | 0.5 | 0.52 | 0.45 |
| 5 | 88181109 | rs3850651 | -0.622 | -0.662 | -0.62 | -0.005 | 0.007 | 0.007 | 0.007 | 0.138 | 0.51 | 0.56 | 0.47 | 0.29 | 0.4 | 0.46 |
| 5 | 88184003 | rs17560451 | 0.045 | -0.631 | -1.244 | -0.221 | 0.003 | -0.022 | -0.022 | 0 | 0.25 | 0.07 | 0.13 | 0.2 | 0.22 | 0.2 |
| 5 | 88194950 | rs770463 | -1.395 | -0.424 | -2.241 | -0.146 | -0.076 | -0.047 | -0.047 | 0 | 0.63 | 0.88 | 0.61 | 0.37 | 0.51 | 0.49 |
| 5 | 88196003 | rs4446500 | -0.679 | -0.068 | -1.962 | -0.001 | -0.004 | 0.009 | 0.009 | 0.312 | 0.28 | 0.09 | 0.17 | 0.19 | 0.27 | 0.22 |
| 5 | 88215594 | rs304132 | 0.098 | 0.324 | -1.864 | 0.124 | -0.02 | 0.1 | 0.1 | 0.518 | 0.57 | 0.87 | 0.58 | 0.76 | 0.6 | 0.6 |
| 5 | 88223420 | rs190982 | -0.039 | 0.288 | -2.019 | -0.042 | -0.049 | 0.169 | 0.169 | 2.533 | 0.59 | 0.94 | 0.81 | 0.87 | 0.63 | 0.62 |
| 5 | 88226422 | rs16903409 | 0.166 | -0.181 | -2.491 | 0.176 | 0.005 | -0.027 | -0.027 | 0.39 | 0.21 | 0.02 | 0.12 | 0.24 | 0.18 | 0.13 |
| 5 | 88233654 | rs301720 | -0.185 | -0.607 | -2.05 | -0.099 | -0.03 | -0.031 | -0.031 | 0 | 0.12 | 0.73 | 0.15 | 0.02 | 0.04 | 0.03 |
| 5 | 88236942 | rs7700950 | 1.027 | -0.593 | -0.859 | -0.093 | 0.035 | 0.032 | 0.032 | 0 | 0.06 | 0.28 | 0.1 | 0.02 | 0.03 | 0.03 |
| 5 | 88242706 | rs301706 | -0.761 | -0.464 | -1.508 | 0.079 | -0.044 | 0.033 | 0.033 | 0.812 | 0.6 | 0.93 | 0.76 | 0.75 | 0.64 | 0.65 |
| 5 | 88253916 | rs214134 | -0.518 | -0.299 | -1.745 | -0.043 | -0.063 | 0.065 | 0.065 | 1.004 | 0.58 | 0.93 | 0.73 | 0.76 | 0.59 | 0.6 |
| 5 | 88270057 | rs188515 | 0.177 | -0.397 | -1.231 | 0.086 | -0.011 | 0.001 | 0.001 | 0.468 | 0.42 | 0.4 | 0.48 | 0.65 | 0.52 | 0.58 |
| 5 | 88332144 | rs4916744 | 0.116 | -0.414 | -0.568 | 0.25 | 0.019 | 0.003 | 0.003 | 1.303 | 0.5 | 0.37 | 0.46 | 0.48 | 0.38 | 0.37 |
| 5 | 88338789 | rs1158464 | 0.898 | -0.694 | -0.883 | 0.112 | -0.02 | -0.027 | -0.027 | 0 | 0.34 | 0.06 | 0.27 | 0.37 | 0.29 | 0.23 |
| 5 | 88347729 | rs6885600 | -0.202 | -0.598 | -0.761 | 0.16 | 0.005 | 0.001 | 0.001 | 1.078 | 0.52 | 0.51 | 0.48 | 0.52 | 0.38 | 0.37 |
| 5 | 88354675 | rs10037512 | -1.111 | -0.579 | -1.48 | 0.182 | -0.111 | -0.043 | -0.043 | 0 | 0.64 | 0.86 | 0.56 | 0.6 | 0.48 | 0.41 |
| 5 | 88392641 | rs10069604 | -0.237 | -0.364 | -1.37 | 0.226 | 0.011 | 0.002 | 0.002 | 0.859 | 0.53 | 0.57 | 0.49 | 0.52 | 0.38 | 0.37 |
| 5 | 88396620 | rs10942546 | 0.928 | -0.289 | -1.412 | -0.087 | 0.006 | -0.002 | -0.002 | 0.023 | 0.08 | 0.04 | 0.05 | 0.08 | 0.08 | 0.04 |
| 5 | 88411214 | rs7728690 | -0.47 | -0.119 | -1.211 | 0.304 | 0.007 | 0.012 | 0.012 | 0.93 | 0.52 | 0.54 | 0.48 | 0.59 | 0.38 | 0.37 |
| 5 | 88412388 | rs10223241 | 0.963 | -0.227 | -1.449 | 0.236 | -0.025 | 0.117 | 0.117 | 1.443 | 0.29 | 0.02 | 0.11 | 0.15 | 0.22 | 0.15 |
| 5 | 88421363 | rs1864180 | -0.267 | 0.464 | -0.973 | 0.456 | 0 | -0.006 | -0.006 | 0 | 0.54 | 0.5 | 0.54 | 0.59 | 0.49 | 0.39 |
| 5 | 88427942 | rs16903468 | 0.569 | 0.159 | -1.884 | 0.482 | 0 | -0.014 | -0.014 | 0 | 0.28 | 0.03 | 0.18 | 0.26 | 0.2 | 0.25 |
| 5 | 88429024 | rs4244201 | 0.003 | 0.442 | -1.538 | 0.595 | 0.003 | 0.045 | 0.045 | 0 | 0.3 | 0.53 | 0.45 | 0.35 | 0.3 | 0.25 |
| 5 | 88429077 | rs4244202 | -1.455 | 0.332 | -1.564 | 0.483 | -0.017 | 0.012 | 0.012 | 0.956 | 0.69 | 0.73 | 0.7 | 0.7 | 0.6 | 0.57 |
| 5 | 88453376 | rs765178 | 1.095 | -0.088 | -1.076 | 0.257 | -0.003 | -0.001 | -0.001 | 0 | 0.22 | 0.19 | 0.14 | 0.16 | 0.13 | 0.1 |
| 5 | 88468185 | rs11746572 | 2.184 | 0.518 | -1.187 | 0.468 | -0.016 | -0.013 | -0.013 | 1.97 | 0.29 | 0.08 | 0.43 | 0.33 | 0.33 | 0.47 |
| 5 | 88472173 | rs12659570 | 1.769 | 0.89 | -1.038 | 0.856 | -0.056 | -0.057 | -0.057 | 1.023 | 0.09 | 0.01 | 0.08 | 0.2 | 0.18 | 0.13 |
| 5 | 88472849 | rs6895825 | 0.863 | 0.273 | -0.555 | 0.838 | -0.06 | -0.039 | -0.039 | 0.011 | 0.26 | 0.54 | 0.17 | 0.16 | 0.13 | 0.1 |
| 5 | 88473328 | rs10514304 | 1.094 | 0.282 | -0.725 | 0.84 | -0.029 | -0.018 | -0.018 | 0 | 0.24 | 0.31 | 0.15 | 0.16 | 0.13 | 0.1 |
| 5 | 88487434 | rs10474293 | 2.639 | 0.329 | -1.641 | 0.826 | -0.04 | -0.009 | -0.009 | 0 | 0.12 | 0.01 | 0.06 | 0.15 | 0.12 | 0.1 |
| 5 | 88499181 | rs6898888 | 0.75 | -0.196 | -1.007 | 0.916 | -0.044 | -0.028 | -0.028 | 0 | 0.26 | 0.56 | 0.17 | 0.16 | 0.13 | 0.1 |
| 5 | 88551836 | rs13355558 | 0.031 | -0.655 | -1.115 | 1.011 | -0.033 | 0.005 | 0.005 | 0.407 | 0.28 | 0.61 | 0.17 | 0.29 | 0.12 | 0.1 |
| 5 | 88588302 | rs4916749 | 1.514 | 0.553 | -1.837 | 1.226 | -0.047 | -0.065 | -0.065 | 0 | 0.28 | 0.04 | 0.4 | 0.33 | 0.31 | 0.54 |
| 5 | 88615707 | rs16867539 | -0.761 | 1.211 | -0.448 | 1.689 | 0.077 | 0.127 | 0.127 | 0 | 0.13 | 0.3 | 0.21 | 0.23 | 0.21 | 0.16 |
| 5 | 88633688 | rs10514305 | 0.844 | 0.543 | -0.665 | 1.42 | 0.03 | -0.021 | -0.021 | 2.499 | 0.19 | 0.08 | 0.07 | 0.2 | 0.1 | 0.06 |
| 5 | 88640130 | rs379717 | 1.036 | 0.382 | -1.045 | 1.284 | -0.043 | 0.074 | 0.074 | 0.052 | 0.25 | 0.08 | 0.14 | 0.16 | 0.25 | 0.18 |
| 5 | 88641366 | rs702849 | -2.188 | 0.245 | -1.204 | 1.281 | -0.066 | -0.1 | -0.1 | 0.16 | 0.66 | 0.95 | 0.47 | 0.58 | 0.59 | 0.4 |
| 5 | 88657310 | rs702847 | -0.461 | 0.218 | -1.111 | 1.07 | 0.006 | 0.031 | 0.031 | 1.395 | 0.53 | 0.62 | 0.26 | 0.36 | 0.38 | 0.24 |
| 5 | 88660918 | rs10514306 | 0.625 | 0.049 | -0.739 | 1.069 | -0.008 | 0 | 0 | 0.009 | 0.21 | 0.2 | 0.09 | 0.2 | 0.1 | 0.06 |
| 5 | 88668305 | rs810029 | -1.133 | 0.274 | -1.458 | 1.448 | -0.071 | -0.076 | -0.076 | 0.256 | 0.66 | 0.95 | 0.4 | 0.65 | 0.59 | 0.42 |
| 5 | 88677293 | rs247587 | -1.388 | 0.185 | -1.437 | 1.114 | -0.068 | 0.029 | 0.029 | 0.368 | 0.68 | 0.97 | 0.57 | 0.78 | 0.62 | 0.47 |
| 5 | 88710519 | rs193689 | -0.238 | 0.029 | -0.953 | 1.26 | 0.004 | 0.005 | 0.005 | 0 | 0.46 | 0.62 | 0.48 | 0.57 | 0.51 | 0.41 |
| 5 | 88726337 | rs7736534 | 0.606 | 0.233 | -0.804 | 1.202 | 0.006 | -0.003 | -0.003 | 0 | 0.2 | 0.13 | 0.07 | 0.2 | 0.11 | 0.06 |
| 5 | 88746405 | rs6892330 | -1.228 | 0.339 | -1.011 | 1.09 | 0 | 0.027 | 0.027 | 0.013 | 0.68 | 0.85 | 0.57 | 0.74 | 0.65 | 0.46 |
| 5 | 88747297 | rs9293516 | 0.197 | 0.33 | -1.032 | 1.169 | -0.004 | -0.01 | -0.01 | 0 | 0.07 | 0.08 | 0.07 | 0.18 | 0.1 | 0.06 |
| 5 | 88747981 | rs2365377 | -0.657 | 0.161 | -0.067 | 1.042 | 0.01 | -0.004 | -0.004 | 0.048 | 0.61 | 0.76 | 0.55 | 0.61 | 0.6 | 0.42 |
| 5 | 88752584 | rs10056443 | 0.381 | 0.217 | 0.196 | 0.144 | 0.021 | -0.067 | -0.067 | 0 | 0.27 | 0.43 | 0.23 | 0.16 | 0.3 | 0.23 |
| 5 | 88752744 | rs7445369 | -1.684 | 0.198 | -0.44 | 0.238 | -0.08 | -0.114 | -0.114 | 0.521 | 0.69 | 0.98 | 0.56 | 0.64 | 0.63 | 0.51 |
| 5 | 88762402 | rs7722399 | 0.018 | -0.312 | 0.132 | 0.506 | -0.042 | -0.024 | -0.024 | 0 | 0.28 | 0.06 | 0.25 | 0.26 | 0.28 | 0.24 |
| 5 | 88774328 | rs4618461 | -1.605 | -0.288 | 0.123 | 0.601 | -0.152 | -0.17 | -0.17 | 0 | 0.63 | 0.97 | 0.46 | 0.55 | 0.51 | 0.47 |
| 5 | 88777537 | rs16867639 | -0.17 | 0.022 | 0.591 | 0.445 | -0.011 | 0.027 | 0.027 | 0 | 0.05 | 0.14 | 0.02 | 0.1 | 0.02 | 0.03 |
| 5 | 88780774 | rs1830362 | 0.938 | 0.015 | 0.362 | 0.305 | -0.013 | 0.114 | 0.114 | 0 | 0.47 | 0.21 | 0.63 | 0.67 | 0.56 | 0.61 |
| 5 | 88780868 | rs10514308 | 1.591 | 0.032 | 0.221 | 0.238 | -0.047 | 0.013 | 0.013 | 0 | 0.15 | 0.02 | 0.08 | 0.14 | 0.16 | 0.1 |
| 5 | 88783422 | rs4916752 | 0.738 | 0.535 | 0.635 | 0.61 | -0.122 | 0.066 | 0.066 | 0.605 | 0.44 | 0.06 | 0.62 | 0.67 | 0.55 | 0.61 |
| 5 | 88785384 | rs6452824 | 1.46 | -0.088 | 1.402 | 0.779 | -0.043 | 0.005 | 0.005 | 3.122 | 0.84 | 0.52 | 0.86 | 0.85 | 0.92 | 0.89 |
| 5 | 88808594 | rs12652212 | -0.187 | 0.12 | 1.316 | 0.911 | -0.001 | -0.001 | -0.001 | 0 | 0.47 | 0.47 | 0.37 | 0.46 | 0.43 | 0.37 |
| 5 | 88829875 | rs11746032 | -0.331 | 0.124 | 1.29 | 0.797 | 0.005 | 0.005 | 0.005 | 0 | 0.49 | 0.55 | 0.37 | 0.46 | 0.43 | 0.37 |
| 5 | 88830987 | rs7732661 | -0.719 | 0.147 | 1.026 | 0.817 | -0.053 | -0.033 | -0.033 | 0 | 0.54 | 0.85 | 0.42 | 0.46 | 0.45 | 0.37 |
| 5 | 88836993 | rs4916755 | 1.182 | 0.176 | 0.933 | 0.777 | -0.173 | -0.06 | -0.06 | 0 | 0.37 | 0.02 | 0.48 | 0.32 | 0.51 | 0.44 |
| 5 | 88847011 | rs12653314 | -0.264 | 0.171 | 1.315 | 0.065 | -0.004 | -0.073 | -0.073 | 0.004 | 0.43 | 0.44 | 0.25 | 0.14 | 0.39 | 0.23 |

| | | | | | | | | | | | | | | | | |
|---|---|---|---|---|---|---|---|---|---|---|---|---|---|---|---|---|
| 5 | 88849142 | rs16867748 | -0.547 | 0.384 | 0.667 | 0.548 | -0.036 | -0.02 | -0.02 | 1.026 | 0.1 | 0.26 | 0.03 | 0.05 | 0.03 | 0.09 |
| 5 | 88853071 | rs6878336 | -0.599 | 0.384 | 0.667 | 0.548 | -0.038 | -0.021 | -0.021 | 1.439 | 0.1 | 0.26 | 0.03 | 0.05 | 0.03 | 0.09 |
| 5 | 88867274 | rs7727304 | -0.523 | 0.482 | 0.288 | 0.534 | -0.012 | -0.018 | -0.018 | 0 | 0.12 | 0.2 | 0.05 | 0.05 | 0.09 | 0.11 |
| 5 | 88869137 | rs10942553 | -0.748 | 1.138 | -0.044 | 0.873 | -0.066 | -0.075 | -0.075 | 0.22 | 0.58 | 0.84 | 0.45 | 0.52 | 0.54 | 0.42 |
| 5 | 88872383 | rs10075489 | 0.276 | 0.789 | 0.017 | -0.093 | 0 | -0.019 | -0.019 | 0 | 0.39 | 0.41 | 0.21 | 0.14 | 0.34 | 0.17 |
| 5 | 88872506 | rs4639270 | -0.351 | 0.763 | -0.14 | 0.2 | -0.012 | 0.079 | 0.079 | 0 | 0.54 | 0.66 | 0.34 | 0.19 | 0.51 | 0.35 |
| 5 | 88880147 | rs7702020 | -0.166 | 0.92 | -0.792 | 0.021 | -0.001 | -0.013 | -0.013 | 0 | 0.52 | 0.5 | 0.32 | 0.16 | 0.51 | 0.3 |
| 5 | 88889955 | rs7727198 | -0.138 | 0.677 | -1.411 | 0.697 | 0.033 | -0.015 | -0.015 | 0 | 0.45 | 0.82 | 0.37 | 0.44 | 0.37 | 0.3 |
| 5 | 88892838 | rs12513521 | 0.437 | 0.325 | -0.912 | 0.736 | -0.005 | -0.001 | -0.001 | 0 | 0.59 | 0.5 | 0.58 | 0.59 | 0.65 | 0.75 |
| 5 | 88893232 | rs4131497 | 0.144 | 0.134 | -1.141 | 0.744 | 0 | -0.018 | -0.018 | 0 | 0.92 | 0.99 | 0.89 | 0.92 | 0.95 | 0.94 |
| 5 | 88893455 | rs4131498 | -0.32 | 0.924 | -0.236 | 1.008 | 0.077 | 0.077 | 0.077 | 0 | 0.15 | 0.38 | 0.25 | 0.15 | 0.22 | 0.18 |
| 5 | 88912685 | rs7380266 | 0.8 | 0.77 | 0.001 | 0.995 | 0.032 | -0.009 | -0.009 | 0.11 | 0.55 | 0.22 | 0.6 | 0.51 | 0.64 | 0.76 |
| 5 | 88958521 | rs4342357 | 0.824 | 0.742 | 0.026 | 0.971 | 0.034 | -0.015 | -0.015 | 0.02 | 0.55 | 0.19 | 0.6 | 0.52 | 0.64 | 0.76 |
| 5 | 88971728 | rs6452845 | 0.167 | 0.476 | 0.369 | 0.717 | 0.005 | -0.03 | -0.03 | 0 | 0.17 | 0.21 | 0.21 | 0.05 | 0.18 | 0.1 |
| 5 | 88980640 | rs2365763 | -0.706 | 0.324 | 0.751 | 0.621 | -0.012 | -0.059 | -0.059 | 0 | 0.16 | 0.28 | 0.07 | 0.04 | 0.1 | 0.09 |
| 5 | 88984265 | rs5002722 | -0.508 | 0.274 | 0.723 | 0.643 | 0 | -0.009 | -0.009 | 0 | 0.27 | 0.23 | 0.25 | 0.1 | 0.25 | 0.19 |
| 5 | 88994840 | rs10069451 | 0.753 | 0.334 | 0.28 | 0.703 | -0.022 | 0.088 | 0.088 | 0 | 0.84 | 0.69 | 0.85 | 0.63 | 0.88 | 0.94 |
| 5 | 88995090 | rs10055858 | -0.548 | 0.331 | 0.378 | 0.701 | 0.018 | -0.025 | -0.025 | 0.17 | 0.95 | 0.86 | 0.97 | 1 | 0.96 | 0.99 |
| 5 | 88995637 | rs16867828 | 0.169 | 0.286 | -0.045 | 0.718 | 0.016 | 0.026 | 0.026 | 0 | 0.06 | 0.01 | 0.05 | 0.36 | 0.03 | 0.04 |
| 5 | 88999206 | rs1449227 | 2.199 | 0.42 | -0.141 | 0.719 | -0.085 | -0.008 | -0.008 | 0 | 0.42 | 0.04 | 0.31 | 0.38 | 0.42 | 0.6 |
| 5 | 89000151 | rs6894060 | 1.636 | -0.066 | 0.314 | 0.535 | 0.04 | -0.011 | -0.011 | 1.005 | 0.55 | 0.17 | 0.59 | 0.55 | 0.64 | 0.76 |
| 5 | 89001243 | rs7718215 | -1.337 | 0.014 | 0.629 | 1.058 | -0.011 | 0.053 | 0.053 | 0 | 0.11 | 0.17 | 0.12 | 0.36 | 0.08 | 0.05 |
| 5 | 89005227 | rs6452846 | -1.681 | 0.278 | 0.753 | 0.884 | 0.001 | 0.043 | 0.043 | 0.202 | 0.24 | 0.44 | 0.23 | 0.33 | 0.22 | 0.11 |
| 5 | 89005564 | rs4443451 | 1.534 | 0.051 | 1.037 | 0.538 | -0.012 | -0.03 | -0.03 | 0 | 0.79 | 0.54 | 0.8 | 0.79 | 0.83 | 0.9 |
| 5 | 89010968 | rs7716630 | 1.66 | 0.164 | 1.016 | 0.673 | -0.017 | 0.003 | 0.003 | 0.339 | 0.82 | 0.39 | 0.77 | 0.8 | 0.88 | 0.89 |
| 5 | 89021578 | rs10514310 | -1.294 | 0.14 | 0.891 | 0.777 | -0.002 | -0.01 | -0.01 | 1.551 | 0.12 | 0.04 | 0.17 | 0.16 | 0.1 | 0.06 |
| 5 | 89032037 | rs7734759 | -2.39 | 0.42 | 1.278 | 0.462 | 0 | -0.026 | -0.026 | 0 | 0.26 | 0.46 | 0.26 | 0.2 | 0.21 | 0.13 |
| 5 | 89041960 | rs10074028 | -2.137 | 0.448 | 1.259 | 0.388 | -0.001 | 0.003 | 0.003 | 0 | 0.23 | 0.17 | 0.22 | 0.14 | 0.19 | 0.1 |
| 5 | 89053605 | rs1449228 | -0.681 | 0.537 | 1.142 | 0.426 | 0.005 | -0.045 | -0.045 | 0 | 0.1 | 0.12 | 0.04 | 0 | 0.09 | 0.03 |
| 5 | 89057750 | rs819344 | 0.349 | -0.253 | 0.15 | 0.121 | 0.069 | 0.053 | 0.053 | 0 | 0.51 | 0.29 | 0.38 | 0.35 | 0.37 | 0.23 |
| 5 | 89067547 | rs9293526 | 0.43 | -1.03 | -0.489 | -0.274 | -0.005 | -0.022 | -0.022 | 0.449 | 0.18 | 0.01 | 0.17 | 0.16 | 0.12 | 0.08 |
| 5 | 89084473 | rs819355 | 0.527 | 0.478 | -0.292 | 0.089 | -0.049 | -0.044 | -0.044 | 0 | 0.32 | 0.2 | 0.42 | 0.44 | 0.37 | 0.47 |
| 5 | 89092732 | rs6896806 | -0.66 | 0.423 | 0.199 | 0.057 | -0.029 | -0.026 | -0.026 | 0 | 0.69 | 0.82 | 0.59 | 0.56 | 0.63 | 0.53 |
| 5 | 89103155 | rs13162065 | 0.377 | 0.29 | 0.339 | -0.014 | 0.023 | 0.017 | 0.017 | 0 | 0.15 | 0.23 | 0.18 | 0.22 | 0.26 | 0.31 |
| 5 | 89110315 | rs16867918 | 1.586 | 0.601 | -0.242 | 0.421 | -0.101 | -0.106 | -0.106 | 0 | 0.2 | 0.01 | 0.35 | 0.39 | 0.28 | 0.41 |
| 5 | 89112538 | rs10514316 | 0.6 | -0.331 | -0.158 | 0.042 | -0.033 | -0.035 | -0.035 | 0 | 0.13 | 0.32 | 0.09 | 0.07 | 0.08 | 0.1 |
| 5 | 89115932 | rs10035607 | -0.92 | -0.637 | -0.366 | -0.172 | 0.036 | 0.02 | 0.02 | 0 | 0.5 | 0.81 | 0.45 | 0.5 | 0.44 | 0.47 |
| 5 | 89124369 | rs13171700 | -0.068 | -0.457 | -0.386 | -0.008 | 0.002 | 0.011 | 0.011 | 0 | 0.23 | 0.22 | 0.22 | 0.35 | 0.31 | 0.39 |
| 5 | 89126824 | rs6872347 | -0.351 | -0.459 | -0.298 | -0.009 | 0.019 | 0.084 | 0.084 | 0 | 0.26 | 0.45 | 0.24 | 0.35 | 0.31 | 0.39 |
| 5 | 89139924 | rs10474308 | 1.124 | -0.609 | -1.101 | -0.017 | 0.029 | 0.02 | 0.02 | 0 | 0.21 | 0.04 | 0.17 | 0.16 | 0.13 | 0.09 |
| 5 | 89153327 | rs6873061 | 1.069 | -0.608 | -1.084 | -0.017 | 0.032 | 0.024 | 0.024 | 0 | 0.22 | 0.06 | 0.18 | 0.16 | 0.13 | 0.09 |
| 5 | 89167360 | rs13157033 | -0.303 | 0.242 | -0.914 | 0.038 | 0.006 | 0.019 | 0.019 | 0 | 0.24 | 0.23 | 0.23 | 0.35 | 0.32 | 0.39 |
| 5 | 89179240 | rs2365904 | 0.97 | -0.079 | -1.258 | -0.315 | 0.001 | 0.002 | 0.002 | 0 | 0.1 | 0.08 | 0.05 | 0.07 | 0.08 | 0.1 |
| 5 | 89189041 | rs277054 | -1.434 | -0.267 | -1.129 | -0.076 | -0.006 | -0.007 | -0.007 | 0.027 | 0.81 | 0.99 | 0.85 | 0.88 | 0.82 | 0.93 |
| 5 | 89209636 | rs4327622 | 0.23 | -0.37 | -0.779 | -0.425 | 0.012 | 0.019 | 0.019 | 0 | 0.24 | 0.15 | 0.23 | 0.07 | 0.15 | 0.17 |
| 5 | 89228470 | rs6870051 | -0.774 | -0.471 | -0.556 | -0.302 | 0.053 | 0.056 | 0.056 | 0.112 | 0.46 | 0.81 | 0.39 | 0.33 | 0.35 | 0.37 |
| 5 | 89230643 | rs16868036 | 1.455 | -0.14 | -1.383 | -0.228 | -0.049 | -0.13 | -0.13 | 0 | 0.25 | 0.02 | 0.35 | 0.4 | 0.26 | 0.39 |
| 5 | 89233833 | rs7720113 | 0.56 | -0.421 | -0.607 | -0.345 | 0.021 | 0.087 | 0.087 | 0 | 0.23 | 0.04 | 0.22 | 0.07 | 0.15 | 0.17 |
| 5 | 89241660 | rs4374783 | 0.56 | -0.381 | -1.049 | -0.327 | 0.021 | 0.087 | 0.087 | 0 | 0.23 | 0.04 | 0.22 | 0.07 | 0.15 | 0.17 |
| 5 | 89248920 | rs9968746 | -0.629 | -0.071 | -0.123 | 0.157 | -0.013 | 0.029 | 0.029 | 1.658 | 0.22 | 0.64 | 0.17 | 0.26 | 0.2 | 0.2 |
| 5 | 89260057 | rs6861634 | -0.51 | -0.075 | -0.317 | -0.074 | 0.004 | 0.014 | 0.014 | 0 | 0.49 | 0.54 | 0.42 | 0.34 | 0.44 | 0.51 |
| 5 | 89261270 | rs10514312 | 0.026 | -0.097 | -0.136 | -0.215 | 0.004 | 0.005 | 0.005 | 0 | 0.2 | 0.22 | 0.18 | 0.23 | 0.26 | 0.28 |
| 5 | 89276486 | rs10474312 | 0.014 | -0.415 | -0.284 | -0.374 | 0.042 | 0.05 | 0.05 | 0.808 | 0.34 | 0.73 | 0.25 | 0.11 | 0.18 | 0.31 |
| 5 | 89287150 | rs749896 | 0.524 | -0.436 | -0.158 | -0.385 | -0.03 | -0.07 | -0.07 | 0.73 | 0.3 | 0.37 | 0.22 | 0.09 | 0.18 | 0.31 |
| 5 | 89288873 | rs1072961 | 1.493 | -0.11 | -1.216 | -0.259 | -0.108 | -0.039 | -0.039 | 0 | 0.19 | 0.02 | 0.17 | 0.16 | 0.31 | 0.17 |
| 5 | 89290604 | rs256426 | 1.188 | -0.084 | -1.174 | -0.223 | -0.003 | -0.004 | -0.004 | 0 | 0.15 | 0.12 | 0.15 | 0.22 | 0.17 | 0.08 |
| 5 | 89306388 | rs256427 | -2.043 | -0.318 | -1.406 | 0.018 | -0.003 | -0.057 | -0.057 | 0 | 0.89 | 0.98 | 0.86 | 0.83 | 0.86 | 0.82 |
| 5 | 89311330 | rs6871792 | 1.541 | -0.257 | -1.126 | -0.53 | 0.041 | 0.092 | 0.092 | 0 | 0.26 | 0.09 | 0.2 | 0.08 | 0.16 | 0.23 |
| 5 | 89314467 | rs2059201 | -0.69 | -0.202 | -1.196 | -0.489 | -0.01 | -0.052 | -0.052 | 0 | 0.73 | 0.85 | 0.7 | 0.62 | 0.66 | 0.73 |
| 5 | 89324079 | rs2366195 | 1.907 | -0.094 | -1.754 | -0.484 | -0.026 | -0.065 | -0.065 | 0 | 0.19 | 0.03 | 0.27 | 0.34 | 0.18 | 0.26 |
| 5 | 89334359 | rs157570 | -0.002 | -0.663 | -1.079 | -0.413 | 0.055 | 0.073 | 0.073 | 0.558 | 0.34 | 0.76 | 0.26 | 0.12 | 0.18 | 0.31 |
| 5 | 89364877 | rs279763 | -0.214 | -0.546 | -1.047 | -0.353 | -0.009 | -0.055 | -0.055 | 0.015 | 0.72 | 0.85 | 0.7 | 0.62 | 0.66 | 0.73 |
| 5 | 89367209 | rs279761 | -0.487 | -0.542 | -1.049 | -0.346 | -0.007 | -0.053 | -0.053 | 0.002 | 0.73 | 0.86 | 0.72 | 0.62 | 0.68 | 0.73 |
| 5 | 89368792 | rs9293534 | 0.274 | -0.478 | -0.138 | -0.542 | 0.028 | 0.044 | 0.044 | 0 | 0.27 | 0.18 | 0.2 | 0.08 | 0.16 | 0.23 |
| 5 | 89399330 | rs1698135 | -0.394 | -0.163 | -1.36 | -0.471 | 0.046 | 0.107 | 0.107 | 0.22 | 0.36 | 0.78 | 0.29 | 0.08 | 0.24 | 0.32 |
| 5 | 89407917 | rs1645412 | -0.317 | -0.597 | -1.276 | -0.399 | 0.058 | 0.082 | 0.082 | 0.179 | 0.33 | 0.77 | 0.26 | 0.11 | 0.19 | 0.31 |
| 5 | 89413164 | rs2652735 | 0.72 | -0.039 | -2.056 | 0.034 | -0.117 | -0.085 | -0.085 | 0.067 | 0.44 | 0.05 | 0.49 | 0.52 | 0.5 | 0.48 |
| 5 | 89418270 | rs2973360 | 0.428 | -0.167 | -1.227 | 0.163 | 0.009 | 0.016 | 0.016 | 0 | 0.5 | 0.31 | 0.6 | 0.61 | 0.53 | 0.51 |
| 5 | 89419677 | rs2920507 | -0.362 | -0.241 | -1.078 | 0.072 | -0.016 | 0.073 | 0.073 | 0 | 0.69 | 0.82 | 0.75 | 0.88 | 0.68 | 0.71 |

| | | | | | | | | | | | | | | | | |
|---|---|---|---|---|---|---|---|---|---|---|---|---|---|---|---|---|
| 5 | 89419754 | rs16868216 | 0.675 | -0.123 | -0.318 | 0.229 | -0.006 | 0.127 | 0.127 | 0 | 0.15 | 0.3 | 0.14 | 0.26 | 0.15 | 0.21 |
| 5 | 89422501 | rs2950026 | -0.197 | 0.133 | -0.714 | -0.025 | -0.035 | 0.063 | 0.063 | 0.02 | 0.75 | 0.93 | 0.8 | 0.87 | 0.72 | 0.75 |
| 5 | 89434524 | rs7721557 | -0.453 | 0.309 | -0.244 | 0.537 | -0.004 | -0.002 | -0.002 | 1.014 | 0.41 | 0.42 | 0.37 | 0.35 | 0.44 | 0.29 |
| 5 | 89434751 | rs2950031 | 0.687 | 0.112 | -0.823 | 0.429 | -0.056 | 0.031 | 0.031 | 0 | 0.5 | 0.09 | 0.52 | 0.51 | 0.47 | 0.63 |
| 5 | 89437963 | rs2366398 | 0.206 | 0.101 | 0.622 | -0.01 | -0.008 | -0.05 | -0.05 | 0.196 | 0.74 | 0.57 | 0.77 | 0.88 | 0.77 | 0.86 |
| 5 | 89441320 | rs10045007 | -0.354 | -0.084 | 1.291 | -0.168 | -0.005 | -0.027 | -0.027 | 0 | 0.26 | 0.35 | 0.22 | 0.12 | 0.23 | 0.13 |
| 5 | 89442298 | rs9293535 | -0.479 | -0.1 | 1.146 | -0.176 | -0.012 | -0.055 | -0.055 | 0 | 0.27 | 0.53 | 0.24 | 0.12 | 0.23 | 0.13 |
| 5 | 89449392 | rs7707262 | -0.473 | -0.104 | 1.233 | -0.178 | -0.011 | -0.055 | -0.055 | 0 | 0.27 | 0.53 | 0.24 | 0.12 | 0.23 | 0.13 |
| 5 | 89457057 | rs1159114 | -0.353 | -0.011 | 0.764 | -0.106 | -0.012 | -0.064 | -0.064 | 0 | 0.81 | 0.86 | 0.67 | 0.45 | 0.81 | 0.7 |
| 5 | 89460396 | rs2973403 | 1.007 | 0.585 | 0.908 | -0.749 | -0.027 | 0.129 | 0.129 | 1.479 | 0.64 | 0.11 | 0.67 | 0.88 | 0.64 | 0.76 |
| 5 | 89462944 | rs930544 | -0.807 | 0.485 | 0.998 | -0.693 | 0.001 | -0.042 | -0.042 | 0 | 0.31 | 0.59 | 0.28 | 0.13 | 0.33 | 0.18 |
| 5 | 89464679 | rs1526168 | -1.531 | 0.417 | 0.309 | -0.674 | -0.011 | 0 | 0 | 0.097 | 0.15 | 0.62 | 0.18 | 0.12 | 0.14 | 0.14 |
| 5 | 89468151 | rs1443888 | 1.697 | 0.397 | -0.204 | -0.723 | -0.093 | 0.127 | 0.127 | 0.878 | 0.6 | 0.05 | 0.65 | 0.8 | 0.59 | 0.71 |
| 5 | 89469753 | rs6899159 | -1.1 | -0.073 | 0.136 | -0.493 | -0.042 | 0.015 | 0.015 | 4.249 | 0.12 | 0.68 | 0.14 | 0.12 | 0.04 | 0.1 |
| 5 | 89476946 | rs4916786 | -0.004 | 0.571 | -0.213 | -0.655 | -0.019 | 0.063 | 0.063 | 0.364 | 0.19 | 0.08 | 0.14 | 0.01 | 0.22 | 0.09 |
| 5 | 89517833 | rs10059737 | -1.44 | -0.101 | 1.255 | -0.252 | -0.022 | -0.065 | -0.065 | 0.99 | 0.11 | 0.41 | 0.07 | 0.01 | 0.05 | 0.15 |
| 5 | 89523227 | rs11949043 | -1.472 | 0.13 | 1.261 | -0.274 | -0.024 | -0.069 | -0.069 | 0 | 0.11 | 0.44 | 0.07 | 0.01 | 0.05 | 0.15 |
| 5 | 89532463 | rs2935505 | -1.853 | 0.203 | 0.742 | -0.409 | 0.003 | -0.112 | -0.112 | 0 | 0.21 | 0.61 | 0.15 | 0.06 | 0.21 | 0.22 |
| 5 | 89546109 | rs12518099 | -0.105 | -0.15 | 0.3 | 0.084 | 0.001 | -0.009 | -0.009 | 0 | 0.26 | 0.22 | 0.27 | 0.39 | 0.26 | 0.33 |
| 5 | 89550034 | rs10054249 | -0.83 | -0.152 | 0.364 | 0.256 | -0.031 | -0.042 | -0.042 | 0.03 | 0.08 | 0.26 | 0.03 | 0 | 0.02 | 0.04 |
| 5 | 89573928 | rs4916794 | -0.349 | 0.45 | -0.901 | 0.39 | 0.005 | 0.004 | 0.004 | 0 | 0.47 | 0.45 | 0.41 | 0.46 | 0.46 | 0.46 |
| 5 | 89587531 | rs10942590 | -0.559 | 0.52 | -1.542 | 0.947 | -0.034 | -0.112 | -0.112 | 0.216 | 0.2 | 0.02 | 0.19 | 0.27 | 0.21 | 0.38 |
| 5 | 89588140 | rs994090 | 0.027 | -0.141 | -2.198 | 0.901 | 0.013 | -0.09 | -0.09 | 0 | 0.23 | 0.05 | 0.34 | 0.43 | 0.23 | 0.17 |
| 5 | 89590812 | rs2935499 | -0.002 | -0.124 | -1.818 | 0.367 | 0.041 | 0.028 | 0.028 | 0.086 | 0.48 | 0.24 | 0.54 | 0.62 | 0.46 | 0.32 |
| 5 | 89622668 | rs1434599 | -1.091 | 0.717 | -0.576 | -0.023 | -0.059 | 0.015 | 0.015 | 0 | 0.79 | 0.92 | 0.86 | 0.86 | 0.78 | 0.76 |
| 5 | 89627079 | rs7707104 | 0.009 | 0.783 | 0.169 | -0.037 | 0 | -0.005 | -0.005 | 0.349 | 0.64 | 0.33 | 0.72 | 0.82 | 0.6 | 0.59 |
| 5 | 89638047 | rs6868587 | 0.674 | 0.237 | -0.163 | -0.031 | 0.009 | -0.056 | -0.056 | 0 | 0.41 | 0.14 | 0.52 | 0.55 | 0.44 | 0.31 |
| 5 | 89642239 | rs4916796 | 0.188 | 0.305 | 0.046 | -0.114 | 0.023 | 0.017 | 0.017 | 0 | 0.5 | 0.3 | 0.62 | 0.57 | 0.61 | 0.39 |
| 5 | 89644476 | rs10041227 | 0.477 | 0.462 | -0.111 | 0.033 | -0.043 | 0.004 | 0.004 | 0 | 0.07 | 0.01 | 0.13 | 0.04 | 0.17 | 0.06 |
| 5 | 89647947 | rs13166637 | 0.678 | 0.132 | -0.169 | 0.051 | 0.012 | -0.056 | -0.056 | 0 | 0.41 | 0.14 | 0.35 | 0.51 | 0.42 | 0.31 |
| 5 | 89649118 | rs1897833 | 0.877 | -0.1 | 0.382 | 0.009 | -0.026 | 0 | 0 | 0.092 | 0.19 | 0.26 | 0.17 | 0.16 | 0.13 | 0.28 |
| 5 | 89651599 | rs7349816 | 0.235 | -0.127 | 0.372 | -0.008 | 0.016 | 0.025 | 0.025 | 0.311 | 0.52 | 0.25 | 0.5 | 0.58 | 0.6 | 0.44 |
| 5 | 89651929 | rs1897832 | -0.691 | 0.351 | 0.79 | -0.164 | 0.012 | -0.02 | -0.02 | 0.795 | 0.73 | 0.59 | 0.69 | 0.82 | 0.75 | 0.72 |
| 5 | 89653677 | rs4916798 | 0.291 | 0.223 | 0.491 | -0.114 | -0.01 | 0.015 | 0.015 | 0 | 0.59 | 0.29 | 0.52 | 0.68 | 0.55 | 0.58 |
| 5 | 89662284 | rs3112537 | 0.632 | -0.084 | -0.137 | -0.332 | 0.026 | 0.023 | 0.023 | 0 | 0.2 | 0.04 | 0.12 | 0.13 | 0.13 | 0.14 |
| 5 | 89668829 | rs2935515 | 1.028 | 0.488 | 0.018 | -0.209 | -0.078 | 0.01 | 0.01 | 0 | 0.2 | 0.05 | 0.15 | 0.15 | 0.25 | 0.25 |
| 5 | 89669924 | rs2935514 | -0.393 | 0.555 | 0.445 | -0.238 | -0.001 | 0 | 0 | 0.497 | 0.34 | 0.69 | 0.45 | 0.22 | 0.41 | 0.35 |
| 5 | 89675329 | rs2935508 | -0.411 | 0.61 | 0.451 | -0.245 | -0.002 | 0 | 0 | 0 | 0.34 | 0.68 | 0.45 | 0.27 | 0.42 | 0.35 |
| 5 | 89679361 | rs2972972 | -0.299 | 0.662 | 0.416 | -0.22 | 0.038 | -0.005 | -0.005 | 0 | 0.34 | 0.54 | 0.44 | 0.27 | 0.41 | 0.33 |
| 5 | 89683864 | rs3794746 | 0.231 | 0.768 | 0.228 | -0.173 | -0.002 | 0.003 | 0.003 | 0 | 0.07 | 0.07 | 0.13 | 0.04 | 0.14 | 0.06 |
| 5 | 89685631 | rs2972964 | -0.128 | 0.276 | 0.374 | -0.179 | 0.044 | -0.005 | -0.005 | 0 | 0.34 | 0.47 | 0.43 | 0.27 | 0.41 | 0.33 |
| 5 | 89686946 | rs6884579 | -0.261 | 0.391 | 0.043 | -0.237 | 0.005 | -0.022 | -0.022 | 0 | 0.07 | 0.07 | 0.07 | 0.01 | 0.14 | 0.05 |
| 5 | 89687104 | rs9293541 | 0.426 | -0.017 | -0.089 | -0.397 | 0.026 | -0.012 | -0.012 | 0 | 0.37 | 0.18 | 0.39 | 0.58 | 0.32 | 0.44 |
| 5 | 89687267 | rs3112539 | -1.074 | -0.535 | 0.246 | -0.485 | -0.003 | -0.005 | -0.005 | 0 | 0.83 | 0.85 | 0.9 | 0.91 | 0.91 | 0.85 |
| 5 | 89731013 | rs4343874 | -0.377 | -0.541 | 1.24 | -0.353 | 0.046 | 0.081 | 0.081 | 0.017 | 0.51 | 0.74 | 0.57 | 0.68 | 0.48 | 0.53 |
| 5 | 89737187 | rs10473934 | 0.244 | -0.57 | 1.31 | -0.323 | -0.005 | 0 | 0 | 0 | 0.39 | 0.39 | 0.38 | 0.59 | 0.33 | 0.45 |
| 5 | 89748142 | rs10214167 | 0.486 | -0.519 | 1.257 | -0.126 | 0.024 | -0.026 | -0.026 | 0.545 | 0.37 | 0.19 | 0.36 | 0.52 | 0.33 | 0.45 |
| 5 | 89751450 | rs7729481 | 0.234 | -0.499 | 1.256 | -0.116 | -0.022 | 0.012 | 0.012 | 0 | 0.39 | 0.46 | 0.38 | 0.52 | 0.33 | 0.45 |
| 5 | 89769647 | rs3763072 | 0.37 | -0.501 | 0.992 | -0.131 | 0.016 | -0.011 | -0.011 | 0 | 0.38 | 0.24 | 0.36 | 0.53 | 0.33 | 0.45 |
| 5 | 89769999 | rs2115501 | 0.322 | -0.406 | 0.481 | -0.122 | 0.005 | -0.024 | -0.024 | 1.776 | 0.07 | 0.08 | 0.06 | 0.01 | 0.14 | 0.05 |
| 5 | 89770768 | rs2303801 | 0.074 | -0.719 | 0.659 | 0.005 | -0.004 | 0 | 0 | 0.004 | 0.4 | 0.54 | 0.39 | 0.53 | 0.33 | 0.45 |
| 5 | 89777895 | rs7712322 | 0.028 | -0.83 | 0.375 | 0.299 | -0.005 | -0.001 | -0.001 | 0 | 0.68 | 0.68 | 0.8 | 0.64 | 0.77 | 0.67 |
| 5 | 89780945 | rs2562520 | 0.272 | -0.86 | 0.555 | 0.285 | -0.027 | -0.003 | -0.003 | 0 | 0.67 | 0.57 | 0.8 | 0.64 | 0.76 | 0.67 |
| 5 | 89786043 | rs6877613 | -0.164 | -0.566 | 0.788 | 0.236 | 0.014 | -0.011 | -0.011 | 1.686 | 0.09 | 0.2 | 0.13 | 0.08 | 0.13 | 0.08 |
| 5 | 89786766 | rs2656953 | 0.195 | -0.837 | 0.974 | 0.16 | -0.009 | 0.008 | 0.008 | 0 | 0.23 | 0.29 | 0.16 | 0.35 | 0.17 | 0.22 |
| 5 | 89791258 | rs2255938 | -0.949 | -0.405 | 0.443 | 0.124 | -0.003 | -0.024 | -0.024 | 0 | 0.64 | 0.84 | 0.55 | 0.62 | 0.66 | 0.67 |
| 5 | 89798873 | rs11743030 | 0.425 | -0.349 | 0.399 | -0.022 | -0.02 | 0.03 | 0.03 | 1.9 | 0.41 | 0.32 | 0.38 | 0.28 | 0.49 | 0.46 |
| 5 | 89807711 | rs6897910 | 0.858 | -0.542 | 0.395 | -0.031 | -0.005 | 0.007 | 0.007 | 1.041 | 0.33 | 0.31 | 0.27 | 0.19 | 0.4 | 0.37 |
| 5 | 89809782 | rs37480 | 0.424 | -0.677 | 0.688 | 0.157 | -0.017 | 0.024 | 0.024 | 0.036 | 0.22 | 0.34 | 0.15 | 0.41 | 0.14 | 0.24 |
| 5 | 89813445 | rs27601 | 0.498 | -0.665 | 0.605 | 0.163 | -0.005 | 0.007 | 0.007 | 0.023 | 0.21 | 0.29 | 0.15 | 0.41 | 0.14 | 0.24 |
| 5 | 89820984 | rs10069050 | -0.339 | -0.695 | 0.705 | 0.143 | -0.01 | -0.007 | -0.007 | 0 | 0.57 | 0.77 | 0.43 | 0.61 | 0.54 | 0.61 |
| 5 | 89824976 | rs37238 | -0.249 | -0.671 | 0.469 | 0.154 | -0.002 | -0.001 | -0.001 | 0.028 | 0.56 | 0.65 | 0.42 | 0.61 | 0.54 | 0.61 |
| 5 | 89825035 | rs603015 | 0.831 | -0.626 | 0.311 | 0.148 | 0.002 | -0.007 | -0.007 | 0 | 0.35 | 0.36 | 0.27 | 0.2 | 0.4 | 0.37 |
| 5 | 89829975 | rs2247158 | 0.61 | -0.59 | 0.057 | 0.431 | 0.007 | -0.016 | -0.016 | 0 | 0.18 | 0.17 | 0.14 | 0.33 | 0.13 | 0.23 |
| 5 | 89841346 | rs37482 | 0.584 | -0.601 | 0.068 | 0.49 | 0.005 | -0.011 | -0.011 | 0 | 0.18 | 0.19 | 0.16 | 0.34 | 0.13 | 0.24 |
| 5 | 89843069 | rs27656 | -1.354 | -0.834 | -0.48 | 0.473 | -0.003 | -0.095 | -0.095 | 0 | 0.8 | 0.97 | 0.64 | 0.7 | 0.85 | 0.74 |
| 5 | 89870822 | rs10057209 | -1.001 | -0.815 | 0.825 | 0.112 | 0.028 | 0.019 | 0.019 | 0.293 | 0.06 | 0.34 | 0.03 | 0 | 0.01 | 0.02 |
| 5 | 89881344 | rs250367 | 0.665 | -0.811 | 0.86 | 0.415 | -0.009 | 0.049 | 0.049 | 0.103 | 0.19 | 0.28 | 0.17 | 0.41 | 0.16 | 0.22 |
| 5 | 89881445 | rs498491 | 2.094 | -0.788 | 0.326 | 0.141 | -0.063 | 0.014 | 0.014 | 0 | 0.25 | 0.07 | 0.2 | 0.16 | 0.31 | 0.25 |

| | | | | | | | | | | | | | | | | |
|---|---|---|---|---|---|---|---|---|---|---|---|---|---|---|---|---|
| 5 | 89882327 | rs645652 | 2.145 | -0.786 | 0.319 | 0.141 | -0.073 | 0.009 | 0.009 | 0 | 0.24 | 0.04 | 0.19 | 0.16 | 0.31 | 0.25 |
| 5 | 89883820 | rs154564 | 0.631 | -0.896 | 1.004 | 0.174 | -0.021 | 0.129 | 0.129 | 0.135 | 0.2 | 0.54 | 0.2 | 0.41 | 0.16 | 0.23 |
| 5 | 89897229 | rs154557 | 0.656 | -0.888 | 1.043 | 0.164 | -0.012 | 0.08 | 0.08 | 0.209 | 0.19 | 0.34 | 0.16 | 0.41 | 0.17 | 0.23 |
| 5 | 89907016 | rs490812 | 0.983 | -0.93 | 0.9 | -0.076 | 0.012 | -0.021 | -0.021 | 2.732 | 0.34 | 0.4 | 0.26 | 0.17 | 0.44 | 0.29 |
| 5 | 89918334 | rs16868864 | 1.47 | -1.049 | 0.789 | -0.204 | 0.004 | -0.002 | -0.002 | 0.006 | 0.1 | 0.05 | 0.05 | 0.01 | 0.07 | 0.13 |
| 5 | 89924976 | rs2366771 | -2.037 | -0.923 | 0.787 | -0.118 | -0.014 | 0.013 | 0.013 | 0 | 0.74 | 0.84 | 0.81 | 0.83 | 0.69 | 0.76 |
| 5 | 89938434 | rs1344030 | 0.857 | -0.827 | 0.231 | 0.117 | -0.001 | 0.009 | 0.009 | 0.092 | 0.19 | 0.22 | 0.15 | 0.41 | 0.16 | 0.23 |
| 5 | 89943433 | rs950692 | 0.89 | -1.083 | 0.156 | 0.157 | -0.001 | 0.005 | 0.005 | 0 | 0.17 | 0.2 | 0.15 | 0.38 | 0.14 | 0.22 |
| 5 | 89943571 | rs2366777 | -2.119 | -1.089 | -0.129 | 0.011 | -0.087 | 0.016 | 0.016 | 0 | 0.75 | 0.95 | 0.81 | 0.83 | 0.67 | 0.74 |
| 5 | 89943699 | rs12186867 | 0.134 | -1.189 | 0.179 | -0.108 | 0.002 | -0.013 | -0.013 | 0 | 0.44 | 0.28 | 0.58 | 0.45 | 0.4 | 0.44 |
| 5 | 89944514 | rs6898209 | -1.989 | -1.011 | 0.218 | -0.105 | -0.048 | 0.025 | 0.025 | 0 | 0.74 | 0.9 | 0.8 | 0.83 | 0.67 | 0.74 |
| 5 | 89951112 | rs10060427 | -0.241 | -1.086 | 0.191 | -0.164 | 0.006 | 0.036 | 0.036 | 0 | 0.52 | 0.48 | 0.68 | 0.82 | 0.47 | 0.53 |
| 5 | 89954184 | rs11958484 | 0.638 | -1.194 | 0.26 | -0.124 | 0.003 | 0.042 | 0.042 | 0 | 0.32 | 0.47 | 0.53 | 0.44 | 0.31 | 0.32 |
| 5 | 89973621 | rs1033290 | -2.151 | -1.163 | -0.298 | -0.262 | -0.073 | 0.023 | 0.023 | 0 | 0.74 | 0.93 | 0.8 | 0.83 | 0.67 | 0.74 |
| 5 | 89979518 | rs17544552 | 1.413 | -1.3 | -0.295 | -0.304 | 0.006 | 0.025 | 0.025 | 0 | 0.11 | 0.06 | 0.05 | 0.02 | 0.08 | 0.14 |
| 5 | 89979589 | rs4916684 | -2.221 | -1.294 | -0.312 | -0.309 | -0.073 | 0.023 | 0.023 | 0 | 0.74 | 0.93 | 0.8 | 0.83 | 0.67 | 0.74 |
| 5 | 89979698 | rs4916685 | 0.873 | -1.415 | -0.196 | -0.197 | -0.01 | -0.026 | -0.026 | 0 | 0.3 | 0.26 | 0.51 | 0.38 | 0.32 | 0.3 |
| 5 | 89988412 | rs10040165 | 0.93 | -1.409 | -0.249 | -0.196 | -0.016 | -0.039 | -0.039 | 0 | 0.3 | 0.21 | 0.5 | 0.38 | 0.32 | 0.3 |
| 5 | 89988504 | rs2366926 | 0.922 | -1.421 | -0.254 | -0.201 | -0.016 | -0.039 | -0.039 | 0 | 0.3 | 0.21 | 0.5 | 0.38 | 0.32 | 0.3 |
| 5 | 89989779 | rs16876822 | 0.931 | -1.414 | -0.251 | -0.198 | -0.016 | -0.039 | -0.039 | 0.28 | 0.3 | 0.21 | 0.5 | 0.38 | 0.32 | 0.3 |
| 5 | 89990324 | rs1878878 | -2.227 | -1.4 | -0.283 | -0.092 | -0.086 | 0.019 | 0.019 | 0 | 0.74 | 0.95 | 0.8 | 0.83 | 0.67 | 0.74 |
| 5 | 89994983 | rs12187771 | 2.079 | -1.39 | -0.454 | -0.003 | -0.044 | -0.099 | -0.099 | 0 | 0.19 | 0.02 | 0.37 | 0.28 | 0.16 | 0.24 |
| 5 | 89999735 | rs11959270 | 1.108 | -1.452 | -0.056 | 0.012 | -0.001 | 0.005 | 0.005 | 0 | 0.18 | 0.22 | 0.15 | 0.43 | 0.14 | 0.21 |
| 5 | 89999954 | rs10942605 | -1.249 | -1.512 | -0.023 | -0.041 | -0.041 | 0.059 | 0.059 | 0 | 0.63 | 0.83 | 0.75 | 0.81 | 0.59 | 0.6 |
| 5 | 89999968 | rs10942606 | -2.039 | -1.338 | 0.245 | 0.246 | -0.068 | 0.023 | 0.023 | 0 | 0.75 | 0.93 | 0.81 | 0.83 | 0.67 | 0.75 |
| 5 | 90000210 | rs16869016 | 1.392 | -1.474 | 0.193 | 0.539 | -0.011 | -0.032 | -0.032 | 0 | 0.05 | 0 | 0.03 | 0.13 | 0.07 | 0.11 |
| 5 | 90001368 | rs6880570 | 0.586 | -1.38 | 0.463 | 0.385 | -0.031 | 0.057 | 0.057 | 0 | 0.19 | 0.36 | 0.17 | 0.43 | 0.14 | 0.21 |
| 5 | 90007196 | rs7732169 | -0.894 | -1.215 | -0.285 | 0.24 | 0 | -0.005 | -0.005 | 2.656 | 0.64 | 0.46 | 0.75 | 0.82 | 0.65 | 0.73 |
| 5 | 90009941 | rs6891645 | 0.235 | -1.279 | -0.19 | 0.214 | -0.024 | -0.022 | -0.022 | 0 | 0.1 | 0.44 | 0.05 | 0.01 | 0.02 | 0.02 |
| 5 | 90016871 | rs16869032 | 0.79 | -1.13 | -0.756 | 0.407 | 0.01 | -0.015 | -0.015 | 0 | 0.18 | 0.17 | 0.15 | 0.43 | 0.14 | 0.21 |
| 5 | 90019080 | rs10036980 | -1.786 | -0.82 | -0.73 | 0.025 | -0.057 | 0.023 | 0.023 | 0 | 0.74 | 0.91 | 0.8 | 0.83 | 0.67 | 0.75 |
| 5 | 90022584 | rs7730204 | 0.182 | -0.776 | -0.869 | -0.028 | 0.015 | -0.005 | -0.005 | 0.782 | 0.23 | 0.18 | 0.18 | 0.3 | 0.23 | 0.25 |
| 5 | 90022780 | rs4916688 | -0.479 | -1.086 | -0.709 | -0.181 | 0.012 | 0.005 | 0.005 | 0 | 0.29 | 0.28 | 0.19 | 0.31 | 0.23 | 0.29 |
| 5 | 90022918 | rs4916689 | 0.365 | -0.784 | -0.944 | -0.177 | 0.006 | -0.017 | -0.017 | 0 | 0.22 | 0.11 | 0.17 | 0.32 | 0.23 | 0.26 |
| 5 | 90024735 | rs2366928 | -2.126 | -0.657 | -1.212 | -0.06 | -0.084 | -0.027 | -0.027 | 0.951 | 0.76 | 0.94 | 0.83 | 0.82 | 0.7 | 0.83 |
| 5 | 90026142 | rs4916816 | 1.325 | -1.055 | -1.697 | -0.151 | -0.026 | -0.094 | -0.094 | 0 | 0.13 | 0.01 | 0.36 | 0.39 | 0.12 | 0.26 |
| 5 | 90036089 | rs2222244 | -1.78 | -0.949 | -1.231 | -0.428 | -0.031 | 0.005 | 0.005 | 0 | 0.72 | 0.85 | 0.79 | 0.81 | 0.66 | 0.7 |
| 5 | 90041161 | rs2438356 | -0.563 | -1.215 | -0.946 | -0.442 | 0.019 | -0.001 | -0.001 | 0 | 0.5 | 0.66 | 0.62 | 0.49 | 0.45 | 0.45 |
| 5 | 90046893 | rs1996549 | 0.07 | -0.95 | -1.185 | -0.304 | -0.032 | -0.07 | -0.07 | 0 | 0.39 | 0.18 | 0.57 | 0.48 | 0.43 | 0.38 |
| 5 | 90053056 | rs2443065 | 0.07 | -0.938 | -1.187 | -0.344 | -0.031 | -0.068 | -0.068 | 5.088 | 0.39 | 0.18 | 0.57 | 0.49 | 0.43 | 0.38 |
| 5 | 90058109 | rs2438351 | -1.056 | -0.495 | -1.35 | -0.257 | -0.051 | 0.086 | 0.086 | 0.055 | 0.63 | 0.8 | 0.75 | 0.83 | 0.55 | 0.7 |
| 5 | 90060782 | rs10514335 | 0.723 | -0.929 | -0.741 | 0.538 | -0.002 | 0.053 | 0.053 | 0 | 0.08 | 0.08 | 0.06 | 0.33 | 0.06 | 0.12 |
| 5 | 90079820 | rs17554631 | 0.989 | -0.044 | -0.347 | -0.24 | -0.071 | 0.064 | 0.064 | 0 | 0.11 | 0.01 | 0.14 | 0.01 | 0.25 | 0.1 |
| 5 | 90089314 | rs17554723 | 1.816 | -0.341 | -0.832 | -0.866 | -0.072 | 0.123 | 0.123 | 0 | 0.25 | 0.06 | 0.16 | 0 | 0.33 | 0.17 |
| 5 | 90090399 | rs4916821 | 1.872 | -0.604 | -1.147 | -0.407 | -0.042 | -0.073 | -0.073 | 0 | 0.11 | 0.01 | 0.36 | 0.37 | 0.13 | 0.28 |
| 5 | 90090437 | rs2438361 | -0.06 | -1.559 | -0.485 | -0.702 | -0.034 | 0.014 | 0.014 | 2.192 | 0.19 | 0.66 | 0.11 | 0.27 | 0.07 | 0.18 |
| 5 | 90110224 | rs2443079 | 0.186 | -1.377 | -0.397 | -0.441 | -0.123 | -0.062 | -0.062 | 0 | 0.45 | 0.08 | 0.66 | 0.57 | 0.51 | 0.54 |
| 5 | 90127959 | rs10045071 | 0.762 | -0.762 | -0.203 | -0.111 | -0.002 | -0.016 | -0.016 | 0.999 | 0.1 | 0.07 | 0.08 | 0.16 | 0.1 | 0.14 |
| 5 | 90147936 | rs10514339 | -0.595 | -0.781 | 0.094 | -0.122 | -0.007 | -0.002 | -0.002 | 0.096 | 0.21 | 0.25 | 0.14 | 0.17 | 0.16 | 0.16 |
| 5 | 90161605 | rs2438343 | -0.288 | -0.367 | 0.174 | -0.219 | 0.001 | -0.011 | -0.011 | 0 | 0.63 | 0.62 | 0.77 | 0.83 | 0.6 | 0.69 |
| 5 | 90172331 | rs6861214 | -0.661 | -0.851 | 0.526 | 0.449 | -0.078 | 0.12 | 0.12 | 0.376 | 0.18 | 0.49 | 0.1 | 0.34 | 0.07 | 0.29 |
| 5 | 90180120 | rs10942615 | -0.297 | -0.824 | 0.52 | 0.475 | -0.048 | 0.075 | 0.075 | 0.184 | 0.16 | 0.31 | 0.2 | 0.35 | 0.08 | 0.29 |
| 5 | 90188886 | rs4244205 | -1.419 | -0.546 | 0.132 | 0.071 | 0.007 | -0.011 | -0.011 | 0.235 | 0.43 | 0.63 | 0.42 | 0.51 | 0.42 | 0.52 |
| 5 | 90208481 | rs12520956 | -0.288 | -0.944 | 0.219 | -0.084 | -0.019 | 0.018 | 0.018 | 0 | 0.27 | 0.2 | 0.23 | 0.17 | 0.37 | 0.21 |
| 5 | 90209824 | rs10074525 | -2.158 | -1.076 | 0.064 | -0.523 | 0.007 | -0.051 | -0.051 | 0 | 0.79 | 0.77 | 0.85 | 1 | 0.73 | 0.86 |
| 5 | 90213250 | rs10060641 | -0.113 | -1.005 | -0.096 | -0.432 | -0.028 | 0.023 | 0.023 | 0 | 0.27 | 0.16 | 0.23 | 0.18 | 0.37 | 0.21 |
| 5 | 90226126 | rs3105792 | 0.886 | -1.225 | 0.277 | -0.211 | 0.028 | -0.009 | -0.009 | 0.055 | 0.41 | 0.31 | 0.46 | 0.59 | 0.33 | 0.49 |
| 5 | 90229426 | rs10942618 | -0.762 | -1.037 | 0.333 | -0.458 | 0.014 | -0.088 | -0.088 | 0 | 0.69 | 0.66 | 0.71 | 1 | 0.48 | 0.79 |
| 5 | 90229807 | rs11744148 | 0.86 | -0.931 | 0.222 | -0.446 | -0.019 | 0.012 | 0.012 | 0 | 0.11 | 0.09 | 0.13 | 0 | 0.26 | 0.07 |
| 5 | 90231238 | rs2367182 | 0.563 | -1.22 | 0.283 | -0.387 | 0.022 | -0.115 | -0.115 | 0.703 | 0.32 | 0.37 | 0.3 | 0.01 | 0.53 | 0.21 |
| 5 | 90233933 | rs1818350 | -0.522 | -1.395 | 0.224 | 0.367 | -0.042 | 0.092 | 0.092 | 0 | 0.18 | 0.26 | 0.2 | 0.51 | 0.07 | 0.33 |
| 5 | 90243899 | rs6891672 | -1.786 | -1.306 | -0.288 | 0.353 | -0.002 | -0.02 | -0.02 | 0.237 | 0.66 | 0.93 | 0.59 | 0.71 | 0.69 | 0.69 |
| 5 | 90248442 | rs16869247 | 0.624 | -1.334 | -0.176 | 0.3 | 0.021 | -0.007 | -0.007 | 2.083 | 0.18 | 0.09 | 0.09 | 0.18 | 0.11 | 0.16 |
| 5 | 90249900 | rs10213801 | -0.211 | -1.191 | 0.207 | 0.333 | -0.001 | 0.015 | 0.015 | 0 | 0.41 | 0.68 | 0.36 | 0.28 | 0.56 | 0.36 |
| 5 | 90252232 | rs6452916 | 0.325 | -1.27 | 0.533 | 0.631 | -0.039 | 0.078 | 0.078 | 0 | 0.11 | 0.28 | 0.07 | 0.28 | 0.05 | 0.17 |
| 5 | 90256819 | rs16869285 | 0.961 | -1.204 | 0.439 | 0.758 | -0.019 | 0.151 | 0.151 | 0 | 0.08 | 0.25 | 0.16 | 0.24 | 0.02 | 0.15 |
| 5 | 90262475 | rs12523056 | -1.304 | -0.965 | -0.211 | 0.349 | 0.012 | 0.05 | 0.05 | 0.658 | 0.46 | 0.77 | 0.43 | 0.71 | 0.42 | 0.52 |
| 5 | 90263863 | rs6879563 | -1.291 | -1.006 | -0.211 | 0.355 | 0.022 | 0.047 | 0.047 | 0.598 | 0.46 | 0.77 | 0.43 | 0.71 | 0.4 | 0.53 |
| 5 | 90263916 | rs6883495 | -1.377 | -1.44 | -0.4 | 0.321 | -0.015 | -0.016 | -0.016 | 0 | 0.66 | 0.93 | 0.58 | 0.71 | 0.68 | 0.67 |

| | | | | | | | | | | | | | | | | |
|---|---|---|---|---|---|---|---|---|---|---|---|---|---|---|---|---|
| 5 | 90264117 | rs17570760 | 0.468 | -1.119 | -0.056 | 0.472 | -0.002 | 0 | 0 | 1.859 | 0.08 | 0.08 | 0.1 | 0 | 0.18 | 0.05 |
| 5 | 90275825 | rs1995776 | -0.349 | -1.446 | 0.144 | 0.725 | -0.052 | 0.139 | 0.139 | 0.002 | 0.19 | 0.4 | 0.22 | 0.52 | 0.11 | 0.33 |
| 5 | 90286831 | rs17571029 | 1.628 | -0.912 | -0.089 | 0.059 | -0.005 | -0.007 | -0.007 | 0 | 0.24 | 0.13 | 0.16 | 0 | 0.27 | 0.21 |
| 5 | 90298632 | rs10514341 | -0.284 | -1.023 | 0.002 | 0.684 | 0.02 | -0.021 | -0.021 | 0 | 0.26 | 0.19 | 0.15 | 0.39 | 0.18 | 0.28 |
| 5 | 90301599 | rs16869330 | -0.658 | -0.459 | 0.248 | 0.51 | -0.012 | 0.125 | 0.125 | 0 | 0.1 | 0.27 | 0.22 | 0.23 | 0.09 | 0.18 |
| 5 | 90306939 | rs16869341 | -1.449 | -0.486 | 0.221 | 0.468 | -0.016 | 0.109 | 0.109 | 0 | 0.15 | 0.28 | 0.25 | 0.34 | 0.15 | 0.32 |
| 5 | 90316643 | rs10038122 | -0.926 | -0.176 | -0.324 | 0.368 | -0.042 | 0.01 | 0.01 | 0 | 0.67 | 0.86 | 0.72 | 0.67 | 0.63 | 0.67 |
| 5 | 90317558 | rs1976565 | -0.949 | -0.178 | -0.33 | 0.367 | -0.051 | 0.005 | 0.005 | 0 | 0.68 | 0.88 | 0.72 | 0.67 | 0.63 | 0.67 |
| 5 | 90348787 | rs2697542 | -0.325 | -0.364 | -0.199 | 0.426 | 0 | -0.007 | -0.007 | 0 | 0.74 | 0.8 | 0.82 | 0.71 | 0.78 | 0.85 |
| 5 | 90349562 | rs2010355 | -0.009 | -0.268 | -0.075 | 0.232 | 0.002 | 0.031 | 0.031 | 0 | 0.49 | 0.65 | 0.58 | 0.38 | 0.46 | 0.53 |
| 5 | 90350959 | rs2697543 | 1.451 | -0.621 | 1.364 | 1.3 | -0.006 | 0.005 | 0.005 | 0 | 0.62 | 0.61 | 0.5 | 0.51 | 0.68 | 0.68 |
| 5 | 90361061 | rs7706034 | 1.738 | -0.601 | 1.348 | 1.286 | -0.002 | 0 | 0 | 0 | 0.61 | 0.41 | 0.49 | 0.48 | 0.67 | 0.67 |
| 5 | 90368783 | rs16869414 | -0.188 | -0.599 | 1.363 | 1.271 | 0.032 | 0.053 | 0.053 | 2.16 | 0.14 | 0.49 | 0.1 | 0.15 | 0.1 | 0.15 |
| 5 | 90371668 | rs10037172 | 0.279 | -0.614 | 1.128 | 1.282 | 0 | -0.009 | -0.009 | 0.06 | 0.75 | 0.8 | 0.82 | 0.71 | 0.78 | 0.85 |
| 5 | 90374713 | rs2007538 | -1.043 | -0.582 | 0.812 | 1.218 | 0.002 | -0.004 | -0.004 | 1.05 | 0.12 | 0.1 | 0.32 | 0.22 | 0.1 | 0.16 |
| 5 | 90382130 | rs10454910 | 2.215 | -0.547 | 0.542 | 1.16 | -0.026 | 0.116 | 0.116 | 1.021 | 0.47 | 0.1 | 0.39 | 0.33 | 0.57 | 0.53 |
| 5 | 90395206 | rs2367279 | 0.088 | -0.518 | 0.758 | 1.297 | -0.005 | 0.003 | 0.003 | 0 | 0.79 | 0.78 | 0.89 | 0.72 | 0.9 | 0.88 |
| 5 | 90409031 | rs2973447 | -0.015 | -0.611 | 0.887 | 1.347 | 0.002 | 0.005 | 0.005 | 0.069 | 0.12 | 0.22 | 0.09 | 0.14 | 0.09 | 0.11 |
| 5 | 90410424 | rs7707023 | 0.997 | -0.692 | 0.453 | 1.132 | -0.002 | -0.003 | -0.003 | 0 | 0.54 | 0.49 | 0.61 | 0.43 | 0.64 | 0.6 |
| 5 | 90413351 | rs4916845 | 1.107 | -0.589 | -0.837 | 1.178 | 0.014 | 0.034 | 0.034 | 0.245 | 0.52 | 0.21 | 0.59 | 0.43 | 0.64 | 0.6 |
| 5 | 90415505 | rs10070771 | -0.469 | -0.743 | -1.107 | 1.212 | 0.01 | -0.007 | -0.007 | 0 | 0.08 | 0.05 | 0.13 | 0.15 | 0.04 | 0.15 |
| 5 | 90415978 | rs2367283 | 0.824 | -0.19 | -0.839 | 1.086 | -0.003 | -0.032 | -0.032 | 0 | 0.6 | 0.23 | 0.71 | 0.58 | 0.68 | 0.75 |
| 5 | 90422216 | rs2973442 | 0.574 | -0.718 | -0.654 | 0.998 | 0.002 | -0.053 | -0.053 | 0 | 0.78 | 0.93 | 0.72 | 0.78 | 0.84 | 0.9 |
| 5 | 90424279 | rs10514345 | -0.874 | -1.231 | 0.477 | 0.899 | -0.039 | 0.013 | 0.013 | 0 | 0.16 | 0.39 | 0.1 | 0.19 | 0.12 | 0.25 |
| 5 | 90426633 | rs2950858 | 1.165 | -1.586 | 1.245 | 1.041 | -0.053 | 0.023 | 0.023 | 0 | 0.79 | 0.36 | 0.87 | 0.68 | 0.88 | 0.7 |
| 5 | 90431779 | rs13158026 | -0.784 | -1.629 | 1.476 | 1.086 | 0.01 | 0.003 | 0.003 | 0 | 0.14 | 0.1 | 0.07 | 0.13 | 0.11 | 0.19 |
| 5 | 90432314 | rs2367286 | -0.692 | -1.91 | 1.437 | 0.797 | -0.002 | -0.002 | -0.002 | 1.102 | 0.17 | 0.21 | 0.11 | 0.09 | 0.14 | 0.09 |
| 5 | 90441146 | rs4458614 | 1.986 | -2 | 1.423 | 0.752 | 0.062 | -0.015 | -0.015 | 0 | 0.6 | 0.21 | 0.77 | 0.59 | 0.72 | 0.58 |
| 5 | 90447216 | rs13176728 | -0.685 | -1.798 | 1.581 | 1.018 | 0.022 | -0.002 | -0.002 | 1.44 | 0.16 | 0.02 | 0.09 | 0.12 | 0.13 | 0.21 |
| 5 | 90449031 | rs7726023 | 0.25 | -1.829 | 1.621 | 1.001 | 0.004 | 0.024 | 0.024 | 0 | 0.92 | 0.85 | 0.94 | 0.91 | 0.91 | 0.92 |
| 5 | 90449154 | rs13158963 | -1.004 | -1.851 | 1.846 | 0.987 | -0.005 | -0.001 | -0.001 | 0 | 0.18 | 0.15 | 0.09 | 0.12 | 0.12 | 0.2 |
| 5 | 90451039 | rs13169196 | -0.989 | -1.839 | 1.847 | 1.004 | 0 | -0.002 | -0.002 | 0 | 0.18 | 0.15 | 0.1 | 0.26 | 0.13 | 0.21 |
| 5 | 90453242 | rs4916704 | 1.748 | -1.812 | 2.162 | 0.964 | -0.018 | 0.106 | 0.106 | 0.545 | 0.85 | 0.36 | 0.88 | 0.72 | 0.89 | 0.84 |
| 5 | 90458126 | rs4916848 | 2.232 | -1.962 | 2.076 | 0.851 | -0.003 | -0.001 | -0.001 | 0.84 | 0.65 | 0.35 | 0.79 | 0.59 | 0.75 | 0.62 |
| 5 | 90461036 | rs10474340 | 1.245 | -1.446 | 2.528 | 0.757 | -0.003 | 0.007 | 0.007 | 0 | 0.88 | 0.82 | 0.91 | 0.89 | 0.89 | 0.92 |
| 5 | 90464235 | rs4916849 | 1.837 | -1.389 | 2.26 | 0.799 | -0.007 | 0.059 | 0.059 | 0.022 | 0.86 | 0.41 | 0.88 | 0.85 | 0.89 | 0.84 |
| 5 | 90466296 | rs4537089 | 1.878 | -1.389 | 1.722 | 0.709 | -0.033 | -0.019 | -0.019 | 0 | 0.74 | 0.47 | 0.7 | 0.79 | 0.81 | 0.8 |
| 5 | 90471610 | rs2941901 | 1.683 | -1.139 | 1.666 | 1.064 | 0.01 | 0.056 | 0.056 | 0 | 0.87 | 0.59 | 0.89 | 0.89 | 0.89 | 0.92 |
| 5 | 90478289 | rs2941876 | 1.883 | -0.97 | 0.996 | 0.996 | -0.016 | 0.027 | 0.027 | 5.314 | 0.9 | 0.29 | 0.93 | 0.88 | 0.94 | 0.9 |
| 5 | 90482854 | rs6452944 | -1.193 | -0.981 | 0.629 | 0.927 | -0.004 | -0.002 | -0.002 | 0.152 | 0.23 | 0.23 | 0.13 | 0.21 | 0.18 | 0.2 |
| 5 | 90485685 | rs1877216 | -1.757 | -1.141 | 0.993 | 1.071 | -0.01 | -0.002 | -0.002 | 0 | 0.27 | 0.71 | 0.22 | 0.34 | 0.2 | 0.27 |
| 5 | 90488667 | rs10514346 | -0.724 | -1.092 | 0.95 | 1.098 | 0.014 | -0.011 | -0.011 | 0 | 0.11 | 0.05 | 0.1 | 0.17 | 0.07 | 0.05 |
| 5 | 90493653 | rs2941886 | -1.342 | -1.048 | 1.4 | 1.129 | 0 | -0.001 | -0.001 | 0 | 0.11 | 0.14 | 0.09 | 0.1 | 0.14 | 0.14 |
| 5 | 90494075 | rs13154867 | 2.212 | -1.428 | 2.127 | 1.376 | -0.041 | 0.023 | 0.023 | 0 | 0.74 | 0.48 | 0.74 | 0.66 | 0.79 | 0.55 |
| 5 | 90494114 | rs2137009 | 0.812 | -1.495 | 2.009 | 1.332 | -0.011 | 0.002 | 0.002 | 0 | 0.9 | 0.91 | 0.86 | 0.89 | 0.93 | 0.76 |
| 5 | 90505517 | rs2941885 | 1.984 | -1.634 | 1.829 | 1.628 | -0.001 | -0.001 | -0.001 | 0 | 0.72 | 0.31 | 0.71 | 0.7 | 0.83 | 0.68 |
| 5 | 90506467 | rs2973476 | 1.946 | -1.875 | 2.504 | 1.631 | -0.074 | 0.005 | 0.005 | 0 | 0.74 | 0.49 | 0.73 | 0.73 | 0.83 | 0.67 |
| 5 | 90507623 | rs2973475 | 1.991 | -1.751 | 2.614 | 1.976 | -0.011 | 0.021 | 0.021 | 0 | 0.82 | 0.79 | 0.8 | 0.75 | 0.84 | 0.7 |
| 5 | 90510809 | rs7341115 | -2.43 | -1.669 | 3.047 | 1.881 | -0.011 | 0.03 | 0.03 | 0 | 0.15 | 0.26 | 0.25 | 0.26 | 0.13 | 0.13 |
| 5 | 90521117 | rs2941894 | 3.18 | -1.846 | 3.095 | 2.4 | -0.002 | 0.001 | 0.001 | 0 | 0.71 | 0.32 | 0.58 | 0.4 | 0.74 | 0.63 |
| 5 | 90524105 | rs7731341 | -3.181 | -1.852 | 3.441 | 2.496 | -0.013 | 0.031 | 0.031 | 0 | 0.26 | 0.36 | 0.39 | 0.58 | 0.25 | 0.38 |
| 5 | 90531378 | rs10473941 | -1.781 | -1.84 | 3.369 | 2.265 | -0.02 | -0.036 | -0.036 | 0 | 0.07 | 0.16 | 0.03 | 0 | 0.05 | 0.06 |
| 5 | 90531796 | rs2941892 | 1.35 | -1.854 | 3.441 | 2.356 | -0.007 | 0.064 | 0.064 | 0.004 | 0.93 | 0.88 | 0.87 | 0.79 | 0.95 | 0.82 |
| 5 | 90532670 | rs2973460 | -2.694 | -2.158 | 3.575 | 2.252 | 0.033 | 0.174 | 0.174 | 1.038 | 0.09 | 0.33 | 0.2 | 0.26 | 0.09 | 0.13 |
| 5 | 90533013 | rs12520306 | -0.773 | -2.122 | 3.277 | 2.257 | -0.01 | -0.026 | -0.026 | 0.055 | 0.07 | 0 | 0.06 | 0.09 | 0.06 | 0.06 |
| 5 | 90533753 | rs2941891 | -3.508 | -2.123 | 3.322 | 2.32 | -0.004 | 0.002 | 0.002 | 0 | 0.24 | 0.7 | 0.32 | 0.35 | 0.19 | 0.25 |
| 5 | 90536321 | rs2973464 | 1.386 | -1.818 | 3.303 | 2.505 | -0.007 | 0.064 | 0.064 | 0.004 | 0.93 | 0.88 | 0.87 | 0.79 | 0.95 | 0.82 |
| 5 | 90536755 | rs2941888 | 2.471 | -1.826 | 3.293 | 2.498 | -0.005 | 0.034 | 0.034 | 0 | 0.87 | 0.87 | 0.82 | 0.7 | 0.9 | 0.76 |
| 5 | 90556558 | rs1858309 | 2.903 | -1.712 | 3.386 | 2.454 | -0.001 | 0.187 | 0.187 | 1.041 | 0.86 | 0.67 | 0.7 | 0.56 | 0.87 | 0.73 |
| 5 | 90599185 | rs9293563 | -1.835 | -1.517 | 3.135 | 2.278 | 0.009 | 0.088 | 0.088 | 0 | 0.08 | 0.2 | 0.13 | 0.18 | 0.07 | 0.16 |
| 5 | 90600210 | rs7704529 | 2.043 | -1.345 | 3.141 | 2.111 | 0.069 | 0.181 | 0.181 | 0.21 | 0.94 | 0.73 | 0.83 | 0.76 | 0.93 | 0.89 |
| 5 | 90609009 | rs7727632 | 2.879 | -1.435 | 3.032 | 2.08 | -0.004 | 0.125 | 0.125 | 0.037 | 0.85 | 0.42 | 0.74 | 0.67 | 0.88 | 0.82 |
| 5 | 90609868 | rs7732832 | 2.703 | -1.383 | 2.741 | 1.968 | -0.001 | 0.074 | 0.074 | 0.976 | 0.9 | 0.73 | 0.81 | 0.76 | 0.92 | 0.85 |
| 5 | 90628300 | rs10037293 | 0.435 | -1.395 | 1.734 | 1.321 | -0.004 | -0.022 | -0.022 | 0 | 0.66 | 0.58 | 0.56 | 0.78 | 0.7 | 0.61 |
| 5 | 90642319 | rs6865416 | -0.224 | -1.253 | 1.81 | 1.386 | -0.004 | -0.017 | -0.017 | 0 | 0.29 | 0.4 | 0.49 | 0.13 | 0.24 | 0.23 |
| 5 | 90645137 | rs6452949 | -1.209 | -1.547 | 0.987 | 1.578 | 0.033 | 0.031 | 0.031 | 0.067 | 0.47 | 0.78 | 0.6 | 0.54 | 0.39 | 0.51 |
| 5 | 90645601 | rs16869685 | -1.367 | -1.185 | 0.734 | 1.86 | -0.027 | -0.099 | -0.099 | 0 | 0.13 | 0.01 | 0.07 | 0.37 | 0.13 | 0.24 |
| 5 | 90654579 | rs16869698 | -1.465 | -1.777 | 2.079 | 1.651 | -0.019 | 0.05 | 0.05 | 0 | 0.16 | 0.26 | 0.11 | 0.36 | 0.08 | 0.28 |
| 5 | 90660447 | rs1541674 | -2.489 | -1.928 | 2.4 | 1.785 | -0.012 | 0.031 | 0.031 | 0.152 | 0.24 | 0.68 | 0.23 | 0.51 | 0.14 | 0.45 |

| | | | | | | | | | | | | | | | | |
|---|---|---|---|---|---|---|---|---|---|---|---|---|---|---|---|---|
| 5 | 90676483 | rs2287875 | -0.583 | -1.584 | 2.3 | 1.679 | 0.037 | 0.054 | 0.054 | 0 | 0.06 | 0.18 | 0.09 | 0.14 | 0.06 | 0.16 |
| 5 | 90682151 | rs889213 | -1.91 | -1.582 | 2.26 | 1.595 | -0.017 | 0.085 | 0.085 | 0 | 0.22 | 0.37 | 0.2 | 0.49 | 0.14 | 0.45 |
| 5 | 90695282 | rs1363837 | -1.649 | -1.64 | 1.664 | 1.279 | -0.096 | 0.081 | 0.081 | 1.325 | 0.56 | 0.92 | 0.7 | 0.73 | 0.49 | 0.74 |
| 5 | 90713109 | rs2042339 | 0.425 | -1.637 | 1.882 | 1.264 | -0.02 | 0.004 | 0.004 | 1.098 | 0.3 | 0.12 | 0.46 | 0.23 | 0.34 | 0.29 |
| 5 | 90723034 | rs6452953 | -2.094 | -1.458 | 2.294 | 1.216 | -0.006 | 0.013 | 0.013 | 0.144 | 0.24 | 0.72 | 0.23 | 0.51 | 0.14 | 0.44 |
| 5 | 90724598 | rs6862125 | -1.703 | -1.412 | 2.189 | 1.24 | -0.021 | 0.126 | 0.126 | 0 | 0.22 | 0.45 | 0.22 | 0.51 | 0.14 | 0.44 |
| 5 | 90732225 | rs10474352 | -1.584 | -1.385 | 1.983 | 1.21 | -0.016 | 0.094 | 0.094 | 0 | 0.21 | 0.39 | 0.19 | 0.49 | 0.14 | 0.44 |
| 5 | 90750056 | rs17577253 | 0.454 | -1.093 | 1.048 | 0.419 | -0.003 | 0.004 | 0.004 | 0 | 0.08 | 0.04 | 0.06 | 0 | 0.08 | 0.02 |
| 5 | 90795928 | rs1423595 | -1.029 | -0.928 | 0.087 | 0.444 | -0.015 | -0.001 | -0.001 | 1.113 | 0.46 | 0.31 | 0.43 | 0.51 | 0.44 | 0.43 |
| 5 | 90796613 | rs16876877 | -1.382 | -0.953 | 0.047 | -0.239 | 0.005 | 0.017 | 0.017 | 0 | 0.17 | 0.16 | 0.1 | 0.08 | 0.11 | 0.14 |
| 5 | 90798611 | rs332539 | -1.403 | -1.109 | -0.805 | -0.323 | 0.016 | -0.09 | -0.09 | 0 | 0.86 | 0.99 | 0.86 | 0.73 | 0.89 | 0.82 |
| 5 | 90799333 | rs6865029 | -0.983 | -0.858 | -0.086 | -0.418 | -0.003 | -0.015 | -0.015 | 0 | 0.17 | 0.21 | 0.09 | 0.07 | 0.1 | 0.1 |
| 5 | 90805092 | rs10073149 | 0.586 | 0.115 | -0.083 | -0.63 | -0.006 | 0.007 | 0.007 | 0.004 | 0.09 | 0.07 | 0.12 | 0.04 | 0.2 | 0.25 |
| 5 | 90808954 | rs7724031 | -1.708 | 0.125 | 0.69 | -0.699 | 0.035 | -0.079 | -0.079 | 0 | 0.28 | 0.42 | 0.22 | 0.11 | 0.31 | 0.35 |
| 5 | 90810714 | rs9293569 | 0.126 | 0.116 | 0.627 | -0.688 | 0.074 | -0.035 | -0.035 | 0 | 0.11 | 0.21 | 0.13 | 0.04 | 0.2 | 0.25 |
| 5 | 90826101 | rs6870792 | 0.381 | -0.035 | 0.458 | -0.698 | -0.028 | 0.026 | 0.026 | 0.151 | 0.13 | 0.07 | 0.14 | 0.04 | 0.24 | 0.26 |
| 5 | 90830449 | rs332523 | -1.684 | -0.192 | 0.902 | -0.792 | -0.004 | 0 | 0 | 0.055 | 0.31 | 0.64 | 0.25 | 0.12 | 0.31 | 0.43 |
| 5 | 90838345 | rs1661037 | -1.684 | -0.192 | 0.902 | -0.792 | -0.004 | 0 | 0 | 0.055 | 0.31 | 0.63 | 0.25 | 0.12 | 0.31 | 0.43 |
| 5 | 90858258 | rs10942636 | -0.045 | -0.21 | 0.524 | -0.782 | -0.001 | -0.002 | -0.002 | 0 | 0.11 | 0.1 | 0.05 | 0.07 | 0.06 | 0.09 |
| 5 | 90873884 | rs6891503 | 0.387 | -0.353 | 0.548 | -0.742 | 0.018 | -0.02 | -0.02 | 0 | 0.14 | 0.13 | 0.14 | 0.04 | 0.24 | 0.26 |
| 5 | 90883485 | rs1596056 | -1.595 | -0.415 | 0.555 | -0.76 | -0.017 | 0.073 | 0.073 | 0.173 | 0.31 | 0.76 | 0.25 | 0.13 | 0.3 | 0.43 |
| 5 | 90901337 | rs17641623 | 0.534 | -0.811 | 0.207 | -0.899 | -0.103 | -0.077 | -0.077 | 0.174 | 0.36 | 0.12 | 0.55 | 0.53 | 0.48 | 0.49 |
| 5 | 90912789 | rs34113537 | -1.132 | -0.736 | 0.816 | -0.738 | -0.025 | -0.034 | -0.034 | 0 | 0.14 | 0.46 | 0.09 | 0.08 | 0.06 | 0.17 |
| 5 | 90922528 | rs6887197 | -0.081 | -0.518 | -0.018 | -0.65 | -0.002 | -0.004 | -0.004 | 0.305 | 0.1 | 0.11 | 0.04 | 0.07 | 0.05 | 0.08 |
| 5 | 90929235 | rs12188554 | -0.096 | -0.585 | -0.12 | -0.546 | 0.01 | 0.005 | 0.005 | 0 | 0.07 | 0.04 | 0.02 | 0.05 | 0.03 | 0.08 |
| 5 | 90941201 | rs2962380 | -1.035 | -1.083 | -0.482 | -0.586 | 0.047 | -0.072 | -0.072 | 0 | 0.81 | 0.96 | 0.88 | 0.7 | 0.84 | 0.82 |
| 5 | 90981532 | rs2973826 | -0.627 | -0.947 | -0.438 | -0.923 | -0.046 | -0.024 | -0.024 | 0 | 0.62 | 0.82 | 0.39 | 0.44 | 0.52 | 0.52 |
| 5 | 91006464 | rs1809814 | 0.764 | -0.758 | 0.382 | -1.092 | -0.021 | 0.101 | 0.101 | 0.512 | 0.33 | 0.18 | 0.19 | 0.08 | 0.33 | 0.18 |
| 5 | 91014718 | rs10447159 | 0.829 | -0.702 | -0.22 | -1.065 | -0.042 | 0.138 | 0.138 | 0.65 | 0.32 | 0.12 | 0.18 | 0.08 | 0.33 | 0.18 |
| 5 | 91018052 | rs2202510 | 0.04 | -0.6 | 0.077 | -0.724 | -0.007 | -0.001 | -0.001 | 0 | 0.06 | 0.11 | 0.02 | 0.05 | 0.04 | 0.1 |
| 5 | 91027493 | rs1701359 | 0.446 | -0.59 | 0.296 | -0.732 | 0.085 | 0.08 | 0.08 | 0.197 | 0.05 | 0.43 | 0.06 | 0.02 | 0.01 | 0.07 |
| 5 | 91029327 | rs1692435 | -0.476 | -0.732 | -0.234 | -0.888 | -0.043 | -0.016 | -0.016 | 0 | 0.61 | 0.8 | 0.4 | 0.45 | 0.55 | 0.53 |
| 5 | 91035641 | rs1964059 | 0.389 | -0.41 | -0.274 | -0.995 | -0.008 | 0.039 | 0.039 | 0 | 0.54 | 0.26 | 0.49 | 0.38 | 0.56 | 0.4 |
| 5 | 91036162 | rs12653151 | 1.175 | -0.602 | -0.439 | -1.085 | -0.006 | 0.065 | 0.065 | 0 | 0.33 | 0.18 | 0.18 | 0.1 | 0.32 | 0.18 |
| 5 | 91037528 | rs331403 | -0.759 | -0.579 | -0.735 | -1.025 | -0.025 | -0.029 | -0.029 | 0 | 0.65 | 0.82 | 0.58 | 0.44 | 0.63 | 0.57 |
| 5 | 91039040 | rs12516181 | 1.195 | -0.38 | -0.872 | -0.914 | -0.025 | -0.06 | -0.06 | 0.724 | 0.2 | 0.05 | 0.3 | 0.28 | 0.23 | 0.21 |
| 5 | 91053616 | rs6869800 | 0.441 | -0.381 | -0.841 | -0.916 | 0.001 | 0.002 | 0.002 | 0.039 | 0.06 | 0.04 | 0.03 | 0.05 | 0.06 | 0.1 |
| 5 | 91055657 | rs6881283 | 1.185 | -0.251 | -0.816 | -0.843 | -0.026 | -0.06 | -0.06 | 0.477 | 0.2 | 0.05 | 0.3 | 0.28 | 0.23 | 0.22 |
| 5 | 91075910 | rs7737612 | -0.052 | -0.58 | -0.566 | -1.09 | 0.004 | 0.079 | 0.079 | 0 | 0.44 | 0.31 | 0.3 | 0.16 | 0.41 | 0.3 |
| 5 | 91076695 | rs2973839 | -0.867 | -0.906 | 0.01 | -0.604 | -0.056 | -0.078 | -0.078 | 0 | 0.19 | 0.55 | 0.13 | 0.06 | 0.11 | 0.22 |
| 5 | 91080804 | rs4547954 | 0.007 | -1.032 | -0.934 | -0.249 | 0.023 | -0.045 | -0.045 | 0 | 0.3 | 0.15 | 0.34 | 0.5 | 0.27 | 0.4 |
| 5 | 91088078 | rs4916875 | -0.769 | -1.085 | -0.567 | -0.494 | 0.077 | 0.054 | 0.054 | 0 | 0.49 | 0.71 | 0.45 | 0.54 | 0.36 | 0.55 |
| 5 | 91090469 | rs4509055 | -0.691 | -1.076 | 0.312 | -0.452 | 0.017 | 0.011 | 0.011 | 0 | 0.45 | 0.35 | 0.43 | 0.53 | 0.35 | 0.57 |
| 5 | 91090726 | rs4443452 | -0.628 | -1.063 | 0.402 | -0.406 | 0.045 | 0.021 | 0.021 | 0 | 0.46 | 0.63 | 0.4 | 0.51 | 0.35 | 0.57 |
| 5 | 91093880 | rs4398667 | -0.536 | -1.101 | 0.713 | -0.613 | -0.009 | -0.002 | -0.002 | 0 | 0.38 | 0.51 | 0.37 | 0.55 | 0.33 | 0.55 |
| 5 | 91093966 | rs4270726 | -0.71 | -0.933 | 0.54 | -0.814 | 0 | 0 | 0 | 0.192 | 0.42 | 0.52 | 0.38 | 0.55 | 0.35 | 0.55 |
| 5 | 91111913 | rs2973848 | -0.526 | -1.133 | 0.75 | -0.843 | 0.008 | 0.002 | 0.002 | 0 | 0.38 | 0.58 | 0.38 | 0.56 | 0.33 | 0.55 |
| 5 | 91113824 | rs3097162 | 0.108 | -1.314 | 0.284 | -0.8 | -0.021 | -0.087 | -0.087 | 0 | 0.28 | 0.08 | 0.31 | 0.5 | 0.31 | 0.48 |
| 5 | 91119149 | rs10473954 | 0.564 | -1.365 | -0.222 | -1.278 | 0.008 | 0.006 | 0.006 | 0 | 0.32 | 0.21 | 0.37 | 0.3 | 0.27 | 0.28 |
| 5 | 91126261 | rs10074805 | 1.372 | -1.332 | -0.62 | -1.179 | -0.037 | 0.049 | 0.049 | 0 | 0.16 | 0.04 | 0.14 | 0.08 | 0.26 | 0.32 |
| 5 | 91130959 | rs10051975 | -0.232 | -1.606 | -0.223 | -1.101 | 0.073 | 0.018 | 0.018 | 0.071 | 0.42 | 0.87 | 0.44 | 0.36 | 0.29 | 0.35 |
| 5 | 91143066 | rs9293587 | 0.914 | -1.768 | -0.363 | -1.122 | 0.008 | 0.017 | 0.017 | 0 | 0.24 | 0.16 | 0.17 | 0.19 | 0.2 | 0.22 |
| 5 | 91144564 | rs16870132 | 0.704 | -1.454 | 0.699 | -0.784 | -0.016 | 0.071 | 0.071 | 0.748 | 0.09 | 0.48 | 0.05 | 0.06 | 0.02 | 0.06 |
| 5 | 91144658 | rs6887393 | 1.645 | -1.226 | 0.14 | -0.729 | 0 | -0.016 | -0.016 | 0 | 0.09 | 0.05 | 0.24 | 0.13 | 0.07 | 0.07 |
| 5 | 91155218 | rs1344690 | 0.654 | -1.274 | -0.111 | -0.701 | -0.102 | 0.095 | 0.095 | 1.912 | 0.39 | 0.08 | 0.44 | 0.24 | 0.45 | 0.27 |
| 5 | 91156283 | rs2367824 | 0.1 | -1.245 | 1.456 | 0.112 | -0.051 | 0.166 | 0.166 | 0 | 0.79 | 0.51 | 0.75 | 0.53 | 0.87 | 0.77 |
| 5 | 91166756 | rs10942647 | 0.398 | -1.27 | 1.178 | 0.1 | 0.004 | -0.006 | -0.006 | 0.078 | 0.77 | 0.23 | 0.72 | 0.53 | 0.87 | 0.77 |
| 5 | 91184723 | rs9293591 | 0.331 | -1.278 | 1.328 | 0.141 | -0.019 | 0.039 | 0.039 | 0.991 | 0.77 | 0.27 | 0.73 | 0.54 | 0.87 | 0.77 |
| 5 | 91193417 | rs17577288 | 1.414 | -0.706 | 0.862 | -0.136 | -0.055 | 0.058 | 0.058 | 0 | 0.11 | 0.01 | 0.11 | 0 | 0.19 | 0.06 |
| 5 | 91199436 | rs1863951 | 0.534 | -1.601 | 1.687 | 0.416 | -0.008 | 0.015 | 0.015 | 0.387 | 0.77 | 0.3 | 0.75 | 0.54 | 0.87 | 0.78 |
| 5 | 91204374 | rs953550 | 0.182 | -1.574 | 1.792 | 0.428 | -0.04 | 0.169 | 0.169 | 0 | 0.8 | 0.62 | 0.77 | 0.54 | 0.87 | 0.78 |
| 5 | 91227934 | rs2114980 | -0.379 | -1.568 | 1.084 | 0.509 | 0.001 | 0.015 | 0.015 | 0 | 0.84 | 0.85 | 0.78 | 0.63 | 0.87 | 0.81 |
| 5 | 91228599 | rs7705780 | -1.764 | -1.669 | 1.068 | 0.591 | -0.049 | -0.043 | -0.043 | 0 | 0.17 | 0.71 | 0.14 | 0.07 | 0.08 | 0.27 |
| 5 | 91235955 | rs825399 | 0.319 | -1.642 | 0.57 | 0.611 | -0.001 | 0.021 | 0.021 | 0 | 0.16 | 0.16 | 0.22 | 0.37 | 0.13 | 0.19 |
| 5 | 91245140 | rs825391 | -0.551 | -1.655 | 0.602 | 0.601 | -0.031 | 0.065 | 0.065 | 0 | 0.22 | 0.27 | 0.24 | 0.38 | 0.15 | 0.24 |
| 5 | 91245518 | rs13182761 | 1.492 | -1.572 | -0.147 | 0.547 | -0.011 | -0.004 | -0.004 | 0 | 0.05 | 0.04 | 0.02 | 0.02 | 0.02 | 0.05 |
| 5 | 91249160 | rs825395 | -1.2 | -1.501 | -0.463 | 0.496 | 0.08 | -0.087 | -0.087 | 0.921 | 0.33 | 0.87 | 0.35 | 0.44 | 0.21 | 0.46 |
| 5 | 91268897 | rs435195 | 0.872 | -1.202 | -0.176 | 0.593 | 0.053 | -0.083 | -0.083 | 0.035 | 0.68 | 0.14 | 0.67 | 0.56 | 0.79 | 0.59 |
| 5 | 91287437 | rs10942656 | 0.308 | -0.447 | -0.234 | 1.055 | 0.001 | -0.001 | -0.001 | 0 | 0.15 | 0.11 | 0.22 | 0.39 | 0.14 | 0.16 |

| | | | | | | | | | | | | | | | | |
|---|---|---|---|---|---|---|---|---|---|---|---|---|---|---|---|---|
| 5 | 91303719 | rs7356540 | -0.121 | -0.437 | -0.243 | 0.936 | 0.003 | -0.023 | -0.023 | 0 | 0.19 | 0.12 | 0.25 | 0.48 | 0.19 | 0.21 |
| 5 | 91321142 | rs1946485 | -0.719 | -0.584 | -0.028 | 0.843 | -0.013 | 0.039 | 0.039 | 0.654 | 0.24 | 0.65 | 0.29 | 0.5 | 0.19 | 0.21 |
| 5 | 91377829 | rs7721589 | -0.17 | -0.608 | -0.32 | -0.431 | -0.02 | -0.086 | -0.086 | 1.489 | 0.28 | 0.32 | 0.16 | 0.02 | 0.18 | 0.14 |
| 5 | 91383848 | rs7725438 | 0.11 | -0.479 | -0.392 | -0.389 | 0.001 | 0.001 | 0.001 | 0.097 | 0.14 | 0.12 | 0.06 | 0.07 | 0.11 | 0.11 |
| 5 | 91383985 | rs6868855 | 0.929 | 0.079 | -0.092 | -0.427 | 0.029 | 0.048 | 0.048 | 0.065 | 0.26 | 0.35 | 0.47 | 0.56 | 0.35 | 0.27 |
| 5 | 91390107 | rs4147091 | -1.786 | -0.336 | -0.427 | -1.204 | -0.007 | 0.016 | 0.016 | 1.185 | 0.77 | 0.83 | 0.8 | 0.94 | 0.72 | 0.81 |
| 5 | 91390316 | rs4454076 | -0.152 | -0.61 | -0.317 | -1.285 | 0.008 | 0.003 | 0.003 | 0.443 | 0.33 | 0.26 | 0.19 | 0.08 | 0.23 | 0.25 |
| 5 | 91393727 | rs6872200 | 1.248 | -0.542 | -0.349 | -1.234 | 0.006 | -0.011 | -0.011 | 0 | 0.19 | 0.14 | 0.12 | 0.01 | 0.12 | 0.14 |
| 5 | 91395086 | rs11738289 | 1.346 | -0.57 | -0.322 | -1.247 | 0.022 | 0.043 | 0.043 | 0 | 0.18 | 0.11 | 0.13 | 0.01 | 0.12 | 0.14 |
| 5 | 91396661 | rs4273628 | -0.376 | -0.54 | -0.072 | -1.28 | -0.003 | 0.009 | 0.009 | 0.013 | 0.6 | 0.68 | 0.63 | 0.63 | 0.57 | 0.52 |
| 5 | 91421138 | rs9293600 | -1.706 | -0.356 | -0.013 | -1.391 | -0.011 | 0.032 | 0.032 | 1.266 | 0.78 | 0.85 | 0.79 | 0.94 | 0.73 | 0.81 |
| 5 | 91431873 | rs7446719 | 1.794 | -0.341 | -0.098 | -1.37 | -0.024 | 0.067 | 0.067 | 0.502 | 0.21 | 0.11 | 0.21 | 0.06 | 0.27 | 0.19 |
| 5 | 91443389 | rs10942660 | -0.062 | -0.148 | 0.235 | -0.736 | 0.009 | -0.016 | -0.016 | 1.746 | 0.41 | 0.43 | 0.54 | 0.68 | 0.48 | 0.37 |
| 5 | 91452232 | rs4502864 | 1.241 | 0.236 | 0.21 | -0.276 | 0.02 | 0.075 | 0.075 | 0 | 0.16 | 0.19 | 0.29 | 0.49 | 0.26 | 0.12 |
| 5 | 91477046 | rs10040401 | -0.162 | -0.026 | -0.127 | -1.367 | 0 | -0.004 | -0.004 | 0 | 0.44 | 0.46 | 0.55 | 0.72 | 0.49 | 0.51 |
| 5 | 91489217 | rs13361289 | -0.092 | 0.035 | -0.586 | -1.343 | -0.01 | 0.039 | 0.039 | 0 | 0.45 | 0.57 | 0.55 | 0.73 | 0.49 | 0.51 |
| 5 | 91507665 | rs11744322 | 0.271 | -0.366 | -0.555 | -1.001 | -0.025 | -0.035 | -0.035 | 0 | 0.11 | 0.13 | 0.03 | 0.01 | 0.03 | 0.1 |
| 5 | 91525968 | rs7716797 | -0.527 | -0.288 | -1.451 | -1.312 | -0.08 | -0.03 | -0.03 | 0.141 | 0.72 | 0.97 | 0.68 | 0.72 | 0.68 | 0.73 |
| 5 | 91540989 | rs12651966 | 0.758 | 0.143 | -0.187 | -0.575 | 0.052 | 0.212 | 0.212 | 0.002 | 0.1 | 0.18 | 0.27 | 0.47 | 0.22 | 0.12 |
| 5 | 91557158 | rs10074207 | 0.485 | -0.034 | 0.349 | -0.582 | 0.125 | 0.423 | 0.423 | 0.108 | 0.11 | 0.4 | 0.29 | 0.47 | 0.23 | 0.12 |
| 5 | 91567488 | rs4448024 | -1.633 | -0.226 | -0.48 | -0.792 | -0.037 | -0.125 | -0.125 | 0 | 0.35 | 0.93 | 0.46 | 0.53 | 0.36 | 0.32 |
| 5 | 91575924 | rs11742941 | 0.875 | -0.585 | -0.312 | -0.673 | 0.018 | 0.018 | 0.018 | 0 | 0.09 | 0.05 | 0.03 | 0.01 | 0.03 | 0.1 |
| 5 | 91583477 | rs10942672 | 1.111 | -0.169 | -0.154 | -0.458 | 0.035 | 0.163 | 0.163 | 0 | 0.09 | 0.12 | 0.27 | 0.47 | 0.22 | 0.12 |
| 5 | 91601777 | rs13354246 | -0.067 | -0.303 | -0.2 | -0.569 | -0.003 | 0.014 | 0.014 | 0 | 0.08 | 0.05 | 0.06 | 0.01 | 0.09 | 0.07 |
| 5 | 91695086 | rs4395616 | -0.681 | -0.482 | 0.037 | -0.427 | 0.067 | 0.227 | 0.227 | 0.038 | 0.88 | 0.74 | 0.71 | 0.52 | 0.76 | 0.85 |
| 5 | 91752084 | rs10053665 | 0.103 | -0.508 | -0.836 | -0.433 | -0.005 | 0.013 | 0.013 | 0 | 0.08 | 0.04 | 0.06 | 0.01 | 0.09 | 0.07 |
| 5 | 91753965 | rs4572981 | -0.394 | -1.051 | -0.985 | -0.678 | -0.085 | -0.051 | -0.051 | 0 | 0.75 | 0.62 | 0.9 | 0.84 | 0.92 | 0.8 |
| 5 | 91754932 | rs10036746 | -0.394 | -1.051 | -0.985 | -0.678 | -0.085 | -0.051 | -0.051 | 0 | 0.75 | 0.62 | 0.9 | 0.84 | 0.92 | 0.8 |
| 5 | 91755616 | rs4260649 | 1.106 | -0.897 | -1.312 | -0.588 | -0.001 | 0.048 | 0.048 | 0.507 | 0.52 | 0.07 | 0.5 | 0.34 | 0.56 | 0.53 |
| 5 | 91782212 | rs13361467 | -0.867 | -0.682 | -1.005 | -0.378 | 0.029 | 0.118 | 0.118 | 0 | 0.89 | 0.84 | 0.75 | 0.52 | 0.74 | 0.83 |
| 5 | 91791151 | rs13168674 | 1.408 | -0.66 | -1.372 | -0.568 | -0.15 | -0.051 | -0.051 | 0 | 0.36 | 0.06 | 0.39 | 0.32 | 0.4 | 0.39 |
| 5 | 91794720 | rs10052539 | -0.572 | -1.166 | -1.098 | -0.711 | -0.005 | -0.049 | -0.049 | 0 | 0.26 | 0.65 | 0.23 | 0.07 | 0.22 | 0.27 |
| 5 | 91798227 | rs6860296 | -0.795 | -0.931 | -1.376 | -0.611 | 0.005 | -0.001 | -0.001 | 0 | 0.41 | 0.58 | 0.47 | 0.53 | 0.44 | 0.5 |
| 5 | 91804748 | rs4242231 | 0.612 | 0.384 | -1.305 | -0.11 | -0.036 | 0.078 | 0.078 | 0 | 0.14 | 0.01 | 0.13 | 0.03 | 0.15 | 0.18 |
| 5 | 91807756 | rs13155975 | -0.157 | 0.138 | -0.417 | 0.684 | 0.031 | 0.169 | 0.169 | 0.137 | 0.15 | 0.25 | 0.28 | 0.46 | 0.27 | 0.18 |
| 5 | 91822322 | rs10038166 | -0.503 | -0.276 | -1.128 | 0.041 | -0.003 | 0.003 | 0.003 | 0 | 0.06 | 0.05 | 0.04 | 0.01 | 0.07 | 0.05 |
| 5 | 91831697 | rs7710769 | -0.713 | -0.173 | -1.087 | 0.09 | 0.012 | 0.015 | 0.015 | 0.04 | 0.27 | 0.24 | 0.14 | 0.16 | 0.18 | 0.2 |
| 5 | 91856650 | rs10075367 | -1.272 | 0.094 | 0.961 | 0.703 | 0.049 | 0.068 | 0.068 | 0.37 | 0.08 | 0.48 | 0.05 | 0.08 | 0.04 | 0.12 |
| 5 | 91884247 | rs10057985 | 0.035 | 0.557 | -0.745 | 1.127 | -0.107 | 0.071 | 0.071 | 0 | 0.38 | 0.09 | 0.45 | 0.76 | 0.56 | 0.46 |
| 5 | 91893095 | rs6861029 | -0.052 | -0.286 | -1.404 | 0.3 | 0.128 | 0.149 | 0.149 | 0 | 0.43 | 0.84 | 0.19 | 0.21 | 0.18 | 0.38 |
| 5 | 91893792 | rs1027643 | -0.116 | -0.763 | -1.392 | 0.014 | -0.027 | -0.033 | -0.033 | 0.214 | 0.15 | 0.57 | 0.08 | 0.09 | 0.06 | 0.16 |
| 5 | 91902326 | rs10039860 | 1.278 | 0.103 | -2.26 | -0.313 | -0.125 | -0.171 | -0.171 | 0.034 | 0.21 | 0.02 | 0.17 | 0.49 | 0.37 | 0.22 |
| 5 | 91903948 | rs6893563 | 1.062 | -0.207 | -1.915 | -0.649 | 0.057 | 0.081 | 0.081 | 0.572 | 0.26 | 0.12 | 0.1 | 0.12 | 0.12 | 0.21 |
| 5 | 91915694 | rs6556848 | -1.136 | 0.273 | -1.818 | -0.594 | -0.031 | 0.055 | 0.055 | 0 | 0.83 | 0.94 | 0.68 | 0.98 | 0.76 | 0.85 |
| 5 | 91930882 | rs10059349 | 0.456 | 0.086 | -0.875 | -0.474 | 0 | -0.014 | -0.014 | 0 | 0.08 | 0.32 | 0.04 | 0.01 | 0.02 | 0.03 |
| 5 | 91938390 | rs2221271 | 0.263 | 0.488 | -1.978 | 0.032 | -0.138 | -0.08 | -0.08 | 1.798 | 0.36 | 0.07 | 0.3 | 0.59 | 0.49 | 0.34 |
| 5 | 91942698 | rs7709237 | -0.092 | 0.294 | -1.904 | -0.135 | 0.09 | 0.081 | 0.081 | 0 | 0.47 | 0.82 | 0.4 | 0.32 | 0.29 | 0.51 |
| 5 | 91953122 | rs9314124 | -0.324 | 0.205 | -2.263 | -0.88 | -0.02 | 0.098 | 0.098 | 0 | 0.2 | 0.06 | 0.24 | 0.04 | 0.2 | 0.17 |
| 5 | 91965961 | rs905217 | -0.919 | 0.684 | -2.853 | -0.845 | -0.06 | 0.028 | 0.028 | 0 | 0.69 | 0.88 | 0.69 | 0.76 | 0.61 | 0.7 |
| 5 | 91970189 | rs905213 | 0.001 | 0.568 | -1.822 | -0.558 | 0.005 | 0.007 | 0.007 | 1.733 | 0.43 | 0.53 | 0.41 | 0.66 | 0.37 | 0.44 |
| 5 | 91976800 | rs1501921 | -0.228 | 0.529 | -1.726 | -0.146 | -0.033 | 0.003 | 0.003 | 0 | 0.34 | 0.25 | 0.33 | 0.32 | 0.41 | 0.54 |
| 5 | 91982004 | rs1532694 | -0.434 | 0.048 | -1.081 | -0.577 | -0.002 | 0.005 | 0.005 | 0 | 0.47 | 0.47 | 0.56 | 0.76 | 0.47 | 0.66 |
| 5 | 91991628 | rs10476539 | -0.04 | 0.635 | -1.016 | -0.853 | 0.027 | -0.12 | -0.12 | 0 | 0.17 | 0.34 | 0.14 | 0.01 | 0.19 | 0.06 |
| 5 | 91996051 | rs13160867 | 0.109 | 0.522 | -1.086 | -0.793 | -0.001 | -0.023 | -0.023 | 3.541 | 0.09 | 0.09 | 0.06 | 0 | 0.09 | 0.03 |
| 5 | 92003403 | rs12655716 | 0.575 | 1.011 | -1.475 | -0.69 | -0.038 | -0.075 | -0.075 | 0.283 | 0.25 | 0.1 | 0.36 | 0.55 | 0.31 | 0.38 |
| 5 | 92005629 | rs11953547 | -0.296 | 1.171 | -0.81 | -0.766 | 0.015 | -0.097 | -0.097 | 0.147 | 0.14 | 0.33 | 0.13 | 0.01 | 0.15 | 0.05 |
| 5 | 92020006 | rs17082472 | 1.297 | 1.026 | -1.103 | -0.767 | -0.018 | -0.045 | -0.045 | 0 | 0.1 | 0 | 0.1 | 0.14 | 0.09 | 0.08 |
| 5 | 92022022 | rs7720378 | -0.406 | 1.283 | -0.863 | -0.81 | 0.072 | -0.103 | -0.103 | 0 | 0.15 | 0.33 | 0.17 | 0.01 | 0.2 | 0.06 |
| 5 | 92028637 | rs10037632 | -0.391 | 1.285 | -0.862 | -0.809 | 0.073 | -0.102 | -0.102 | 0 | 0.15 | 0.33 | 0.17 | 0.01 | 0.2 | 0.06 |
| 5 | 92029232 | rs10056094 | 0.535 | 0.943 | -0.989 | -0.818 | -0.011 | -0.021 | -0.021 | 0 | 0.11 | 0.17 | 0.04 | 0.04 | 0.05 | 0.06 |
| 5 | 92032360 | rs17082487 | 0.497 | 1.07 | -1.302 | -0.597 | -0.006 | -0.043 | -0.043 | 0 | 0.28 | 0.21 | 0.39 | 0.57 | 0.3 | 0.49 |
| 5 | 92040221 | rs10041674 | -0.377 | 1.177 | -0.863 | -0.724 | 0.035 | -0.112 | -0.112 | 2.864 | 0.19 | 0.32 | 0.19 | 0.01 | 0.24 | 0.1 |
| 5 | 92044676 | rs17082510 | 0.295 | 0.9 | -0.977 | -0.664 | -0.029 | -0.077 | -0.077 | 0 | 0.14 | 0.23 | 0.08 | 0.01 | 0.12 | 0.09 |
| 5 | 92047795 | rs10075131 | -0.419 | 0.909 | -1.351 | -0.622 | 0.004 | 0.015 | 0.015 | 0 | 0.51 | 0.56 | 0.57 | 0.62 | 0.55 | 0.65 |
| 5 | 92048984 | rs1532695 | -1.16 | 1.347 | -2.238 | -0.377 | -0.071 | 0.004 | 0.004 | 0 | 0.7 | 0.97 | 0.73 | 0.75 | 0.68 | 0.75 |
| 5 | 92055195 | rs258952 | -0.201 | 1.684 | -1.443 | 0.131 | -0.005 | -0.044 | -0.044 | 0 | 0.32 | 0.2 | 0.38 | 0.57 | 0.32 | 0.53 |
| 5 | 92063559 | rs26717 | -0.474 | 1.515 | -1.42 | 0.091 | -0.009 | -0.029 | -0.029 | 0 | 0.88 | 0.99 | 0.94 | 0.89 | 0.89 | 0.83 |
| 5 | 92064034 | rs11135445 | 1.117 | 1.697 | -1.578 | 0.835 | -0.02 | 0.022 | 0.022 | 0 | 0.07 | 0.01 | 0.21 | 0.53 | 0.07 | 0.11 |
| 5 | 92064532 | rs1501923 | -0.265 | 1.824 | -1.097 | 0.143 | 0.041 | -0.018 | -0.018 | 0 | 0.21 | 0.31 | 0.32 | 0.15 | 0.25 | 0.18 |

| | | | | | | | | | | | | | | | | |
|---|---|---|---|---|---|---|---|---|---|---|---|---|---|---|---|---|
| 5 | 92069408 | rs1160820 | 0.079 | 1.701 | -1.934 | -0.599 | -0.052 | 0.108 | 0.108 | 1.685 | 0.4 | 0.11 | 0.29 | 0.17 | 0.44 | 0.48 |
| 5 | 92083665 | rs34963 | -2.139 | 1.288 | -2.392 | -1.013 | -0.001 | 0.034 | 0.034 | 0 | 0.8 | 1 | 0.92 | 0.89 | 0.86 | 0.85 |
| 5 | 92084770 | rs6556916 | -0.869 | 1.672 | -1.506 | -0.956 | 0.021 | 0.013 | 0.013 | 0 | 0.29 | 0.6 | 0.54 | 0.67 | 0.3 | 0.21 |
| 5 | 92085529 | rs1422168 | 1.707 | 1.575 | -1.239 | -1.277 | -0.001 | -0.01 | -0.01 | 0 | 0.08 | 0.09 | 0.05 | 0 | 0.07 | 0.04 |
| 5 | 92089111 | rs6556921 | 0.318 | 1.158 | -0.926 | -0.866 | 0.08 | 0.047 | 0.047 | 0 | 0.12 | 0.32 | 0.26 | 0.15 | 0.16 | 0.08 |
| 5 | 92090548 | rs34968 | 0.258 | 1.163 | -1.08 | -0.922 | -0.003 | 0.005 | 0.005 | 2.194 | 0.13 | 0.14 | 0.1 | 0.05 | 0.17 | 0.1 |
| 5 | 92104978 | rs7734479 | -0.261 | 1.009 | -0.691 | -0.893 | 0.069 | -0.077 | -0.077 | 0 | 0.22 | 0.41 | 0.35 | 0.13 | 0.34 | 0.13 |
| 5 | 92107082 | rs2059076 | -1.055 | 0.026 | -1.268 | -0.851 | 0.019 | 0.023 | 0.023 | 0 | 0.78 | 0.82 | 0.91 | 0.87 | 0.86 | 0.82 |
| 5 | 92110386 | rs258959 | -1.32 | 0.021 | -1.386 | -0.885 | 0.004 | 0.04 | 0.04 | 0.185 | 0.78 | 0.99 | 0.92 | 0.88 | 0.84 | 0.82 |
| 5 | 92114934 | rs154809 | -0.496 | 0.162 | -1.314 | -1.176 | 0.004 | 0.017 | 0.017 | 0 | 0.32 | 0.64 | 0.38 | 0.17 | 0.33 | 0.22 |
| 5 | 92115076 | rs792466 | -0.265 | 0.134 | -0.988 | -1.201 | -0.007 | -0.054 | -0.054 | 0 | 0.27 | 0.5 | 0.34 | 0.15 | 0.31 | 0.16 |
| 5 | 92134273 | rs27540 | 0.28 | 0.193 | -1.301 | -0.978 | -0.016 | 0.094 | 0.094 | 0 | 0.55 | 0.63 | 0.52 | 0.25 | 0.56 | 0.52 |
| 5 | 92135678 | rs26720 | 0.884 | 0.159 | -1.115 | -1.001 | 0.003 | 0.048 | 0.048 | 0 | 0.5 | 0.52 | 0.49 | 0.24 | 0.53 | 0.46 |
| 5 | 92139632 | rs13165943 | 1.056 | 0.218 | -1.276 | -0.561 | -0.042 | -0.064 | -0.064 | 0 | 0.18 | 0.13 | 0.28 | 0.5 | 0.25 | 0.15 |
| 5 | 92141172 | rs17082593 | 1.705 | 0.132 | -1.278 | -0.539 | 0.006 | -0.015 | -0.015 | 0 | 0.05 | 0.03 | 0.11 | 0.12 | 0.04 | 0.15 |
| 5 | 92141841 | rs2042188 | -0.996 | 0.082 | -1.291 | -0.573 | 0.022 | 0 | 0 | 0.05 | 0.73 | 0.79 | 0.8 | 0.75 | 0.8 | 0.67 |
| 5 | 92148132 | rs258953 | -0.576 | -0.037 | -0.708 | -0.755 | 0.009 | 0 | 0 | 0 | 0.32 | 0.41 | 0.42 | 0.63 | 0.3 | 0.34 |
| 5 | 92149000 | rs1429056 | -2.003 | -0.052 | -0.846 | -0.76 | 0.02 | 0.041 | 0.041 | 0 | 0.79 | 0.97 | 0.92 | 0.87 | 0.84 | 0.82 |
| 5 | 92150839 | rs10463170 | 1.496 | 0.232 | -0.609 | -0.641 | -0.022 | -0.006 | -0.006 | 0 | 0.11 | 0 | 0.19 | 0.11 | 0.11 | 0.05 |
| 5 | 92151389 | rs1422164 | 1.235 | 0.253 | -0.148 | -0.351 | -0.016 | -0.031 | -0.031 | 0 | 0.08 | 0.24 | 0.03 | 0.02 | 0.02 | 0.04 |
| 5 | 92154579 | rs17668565 | 1.083 | 0.518 | -0.55 | -0.225 | -0.014 | 0.058 | 0.058 | 0.131 | 0.42 | 0.28 | 0.45 | 0.22 | 0.52 | 0.42 |
| 5 | 92177723 | rs17082624 | 0.627 | 0.89 | -0.733 | 0.657 | 0.002 | 0.007 | 0.007 | 0 | 0.06 | 0.05 | 0.11 | 0.16 | 0.04 | 0.13 |
| 5 | 92181633 | rs1363292 | 0.91 | 1.34 | -0.881 | 1.366 | -0.038 | -0.003 | -0.003 | 0 | 0.08 | 0.03 | 0.22 | 0.46 | 0.18 | 0.11 |
| 5 | 92184101 | rs6556969 | 0.459 | 1.114 | -0.915 | 1.304 | -0.002 | -0.033 | -0.033 | 0 | 0.1 | 0.05 | 0.11 | 0.26 | 0.09 | 0.2 |
| 5 | 92186059 | rs888812 | -1.305 | 0.757 | -0.807 | 0.875 | -0.01 | 0.08 | 0.08 | 0 | 0.54 | 0.72 | 0.57 | 0.82 | 0.49 | 0.52 |
| 5 | 92186365 | rs888813 | 0.012 | 0.586 | -0.44 | 0.818 | -0.018 | 0.128 | 0.128 | 1.474 | 0.15 | 0.51 | 0.14 | 0.28 | 0.09 | 0.21 |
| 5 | 92186429 | rs888814 | -1.372 | 0.568 | -0.456 | 0.802 | -0.006 | 0.059 | 0.059 | 0 | 0.53 | 0.66 | 0.57 | 0.82 | 0.49 | 0.52 |
| 5 | 92187932 | rs17669337 | -0.153 | 0.628 | -0.755 | 0.834 | -0.073 | -0.121 | -0.121 | 0 | 0.38 | 0.1 | 0.43 | 0.54 | 0.4 | 0.31 |
| 5 | 92201702 | rs967444 | -0.096 | 0.194 | -0.174 | 0.762 | -0.023 | 0.056 | 0.056 | 0.209 | 0.17 | 0.3 | 0.14 | 0.27 | 0.1 | 0.22 |
| 5 | 92205481 | rs1117022 | -1.271 | 0.232 | -0.622 | 0.605 | 0.01 | 0.047 | 0.047 | 0 | 0.75 | 0.9 | 0.88 | 0.84 | 0.78 | 0.79 |
| 5 | 92213757 | rs10078347 | 1.262 | 0.604 | -0.222 | 0.54 | -0.012 | 0.082 | 0.082 | 0 | 0.46 | 0.28 | 0.44 | 0.17 | 0.54 | 0.47 |
| 5 | 92215789 | rs12188632 | 1.638 | 0.718 | -0.135 | 0.543 | -0.005 | 0.014 | 0.014 | 0 | 0.36 | 0.27 | 0.36 | 0.14 | 0.4 | 0.4 |
| 5 | 92226132 | rs1010126 | 1.954 | 0.84 | -0.307 | 0.515 | 0 | 0 | 0 | 6.429 | 0.07 | 0.05 | 0.14 | 0.06 | 0.1 | 0.04 |
| 5 | 92226214 | rs1010127 | 1.429 | 0.718 | -0.04 | 0.54 | 0.007 | -0.032 | -0.032 | 0 | 0.37 | 0.45 | 0.37 | 0.17 | 0.4 | 0.39 |
| 5 | 92226608 | rs350806 | 0.199 | 0.576 | -0.18 | 0.75 | -0.074 | -0.047 | -0.047 | 0 | 0.54 | 0.87 | 0.52 | 0.44 | 0.5 | 0.61 |
| 5 | 92233582 | rs10057574 | 0.602 | 0.698 | -0.048 | 0.823 | -0.003 | 0.002 | 0.002 | 0 | 0.48 | 0.6 | 0.48 | 0.43 | 0.49 | 0.59 |
| 5 | 92236275 | rs7724557 | 1.571 | 0.709 | -0.237 | 0.865 | 0 | -0.002 | -0.002 | 0 | 0.37 | 0.33 | 0.36 | 0.17 | 0.4 | 0.39 |
| 5 | 92236350 | rs1489531 | 0.593 | 0.83 | -0.089 | 1.411 | 0.003 | 0.012 | 0.012 | 0 | 0.08 | 0.09 | 0.09 | 0.24 | 0.06 | 0.15 |
| 5 | 92242635 | rs2578495 | -0.516 | 1.107 | 0.12 | 1.396 | 0.026 | 0.117 | 0.117 | 0 | 0.15 | 0.26 | 0.34 | 0.41 | 0.17 | 0.14 |
| 5 | 92246357 | rs350802 | -0.343 | 1.112 | -0.458 | 0.154 | 0 | -0.017 | -0.017 | 0 | 0.76 | 0.78 | 0.74 | 0.94 | 0.75 | 0.89 |
| 5 | 92246775 | rs10071718 | 0.893 | 1.045 | -0.54 | 0.367 | -0.029 | 0.136 | 0.136 | 0 | 0.53 | 0.2 | 0.31 | 0.31 | 0.53 | 0.6 |
| 5 | 92282024 | rs6876638 | -1.584 | 2.396 | 0.579 | 1.071 | 0.12 | 0.044 | 0.044 | 1.667 | 0.08 | 0.44 | 0.11 | 0.04 | 0.12 | 0.03 |
| 5 | 92290203 | rs17082764 | 0.357 | 2.36 | 0.18 | 1.043 | -0.012 | -0.052 | -0.052 | 0.015 | 0.13 | 0 | 0.06 | 0.25 | 0.11 | 0.09 |
| 5 | 92313841 | rs34509 | 1.183 | 3.673 | 0.733 | 1.854 | -0.025 | -0.031 | -0.031 | 0.023 | 0.78 | 0.86 | 0.54 | 0.63 | 0.67 | 0.73 |
| 5 | 92320089 | rs10463223 | -0.873 | 3.378 | 0.671 | 1.579 | 0.007 | 0.028 | 0.028 | 0 | 0.06 | 0.06 | 0.09 | 0.31 | 0.04 | 0.2 |
| 5 | 92321477 | rs34519 | 1.509 | 3.536 | 0.843 | 1.553 | -0.009 | -0.014 | -0.014 | 0 | 0.73 | 0.77 | 0.53 | 0.64 | 0.66 | 0.7 |
| 5 | 92330332 | rs12187522 | -0.499 | 3.745 | 0.855 | 1.487 | 0.001 | -0.046 | -0.046 | 3.427 | 0.13 | 0.17 | 0.09 | 0.01 | 0.16 | 0.12 |
| 5 | 92344846 | rs2453759 | 0.128 | 3.295 | 0.357 | 1.641 | -0.045 | -0.051 | -0.051 | 0 | 0.89 | 0.97 | 0.64 | 0.65 | 0.82 | 0.84 |
| 5 | 92403727 | rs2017806 | -0.876 | 3.196 | 0.572 | 1.391 | 0.074 | 0.168 | 0.168 | 0.35 | 0.15 | 0.36 | 0.35 | 0.36 | 0.26 | 0.21 |
| 5 | 92407083 | rs17082892 | -1.121 | 3.135 | 0.418 | 1.331 | 0.023 | -0.066 | -0.066 | 0 | 0.12 | 0.02 | 0.11 | 0.29 | 0.07 | 0.26 |
| 5 | 92414916 | rs995626 | -1.177 | 3.131 | 0.496 | 1.321 | 0.089 | 0.192 | 0.192 | 0.691 | 0.17 | 0.47 | 0.37 | 0.35 | 0.26 | 0.22 |
| 5 | 92420689 | rs10078721 | -2.777 | 3.386 | 0.376 | 1.24 | 0.012 | 0.146 | 0.146 | 0.847 | 0.51 | 0.84 | 0.64 | 0.69 | 0.57 | 0.63 |
| 5 | 92428532 | rs11958890 | -0.975 | 3.327 | 0.573 | 1.475 | 0.06 | 0.113 | 0.113 | 0.479 | 0.16 | 0.31 | 0.36 | 0.35 | 0.28 | 0.21 |
| 5 | 92443100 | rs13188341 | -1.563 | 3.082 | 0.643 | 1.452 | -0.001 | 0.013 | 0.013 | 0 | 0.13 | 0.15 | 0.12 | 0.29 | 0.09 | 0.26 |
| 5 | 92477890 | rs4342312 | -1.415 | 3.233 | 0.674 | 1.375 | 0.039 | 0.061 | 0.061 | 0.07 | 0.27 | 0.57 | 0.5 | 0.39 | 0.37 | 0.28 |
| 5 | 92482188 | rs10054423 | -0.145 | 3.111 | 0.478 | 1.281 | -0.003 | 0.003 | 0.003 | 0 | 0.14 | 0.11 | 0.09 | 0.03 | 0.16 | 0.14 |
| 5 | 92487543 | rs13173682 | -2.504 | 2.928 | 0.474 | 1.37 | 0.04 | 0.122 | 0.122 | 1.408 | 0.55 | 0.84 | 0.71 | 0.73 | 0.62 | 0.68 |
| 5 | 92490451 | rs11960764 | -1.386 | 2.933 | 0.403 | 1.397 | 0.083 | -0.026 | -0.026 | 0 | 0.08 | 0.38 | 0.12 | 0.01 | 0.15 | 0.05 |
| 5 | 92491951 | rs12513458 | -0.459 | 2.918 | 0.232 | 1.496 | -0.056 | -0.106 | -0.106 | 4.521 | 0.18 | 0.03 | 0.37 | 0.39 | 0.23 | 0.27 |
| 5 | 92496410 | rs12055145 | -0.525 | 2.91 | 0.242 | 1.504 | 0.005 | -0.011 | -0.011 | 0.18 | 0.07 | 0.04 | 0.1 | 0.29 | 0.05 | 0.21 |
| 5 | 92498577 | rs12054781 | -1.513 | 2.925 | 0.291 | 1.13 | -0.007 | -0.021 | -0.021 | 0.029 | 0.28 | 0.26 | 0.49 | 0.59 | 0.29 | 0.45 |
| 5 | 92510324 | rs4554194 | -1.537 | 2.251 | 0.128 | 1.262 | -0.01 | -0.078 | -0.078 | 0.102 | 0.28 | 0.11 | 0.46 | 0.6 | 0.26 | 0.47 |
| 5 | 92515880 | rs12513551 | -0.294 | 2.306 | 0.149 | 1.293 | -0.061 | -0.142 | -0.142 | 0.019 | 0.17 | 0.02 | 0.35 | 0.43 | 0.2 | 0.24 |
| 5 | 92520141 | rs13161855 | -0.297 | 2.286 | 0.158 | 1.273 | -0.005 | 0.004 | 0.004 | 0 | 0.19 | 0.13 | 0.1 | 0.12 | 0.18 | 0.15 |
| 5 | 92528309 | rs4632783 | -2.44 | 2.698 | 0.692 | 1.766 | 0.014 | 0.021 | 0.021 | 0.993 | 0.33 | 0.55 | 0.56 | 0.6 | 0.4 | 0.48 |
| 5 | 92555377 | rs7442897 | -1.847 | 2.411 | 1.09 | 1.618 | 0.001 | -0.002 | -0.002 | 5.319 | 0.1 | 0.13 | 0.11 | 0.16 | 0.06 | 0.23 |
| 5 | 92567539 | rs4869419 | -1.861 | 2.427 | 1.037 | 1.625 | 0.015 | -0.026 | -0.026 | 0 | 0.09 | 0.03 | 0.09 | 0.16 | 0.07 | 0.16 |
| 5 | 92567901 | rs4380635 | -2.11 | 2.427 | 1.039 | 1.623 | 0.026 | -0.016 | -0.016 | 1.009 | 0.12 | 0.07 | 0.12 | 0.17 | 0.08 | 0.22 |
| 5 | 92571663 | rs12657207 | -2.197 | 2.991 | 1.314 | 1.756 | 0.084 | 0.069 | 0.069 | 0.465 | 0.11 | 0.36 | 0.18 | 0.16 | 0.17 | 0.16 |

| | | | | | | | | | | | | | | | | |
|---|---|---|---|---|---|---|---|---|---|---|---|---|---|---|---|---|
| 5 | 92586991 | rs6881733 | -1.271 | 2.995 | 0.858 | 1.664 | 0.051 | 0.129 | 0.129 | 0.035 | 0.21 | 0.53 | 0.45 | 0.41 | 0.29 | 0.24 |
| 5 | 92588936 | rs4541642 | -1.852 | 2.623 | -0.266 | 0.756 | -0.016 | -0.002 | -0.002 | 0 | 0.31 | 0.75 | 0.52 | 0.34 | 0.41 | 0.45 |
| 5 | 92594666 | rs4434363 | -0.924 | 2.746 | -0.559 | 0.821 | 0 | 0 | 0 | 0 | 0.09 | 0.15 | 0.13 | 0.14 | 0.13 | 0.2 |
| 5 | 92595749 | rs7701128 | -1.072 | 2.798 | -0.559 | 0.844 | 0.074 | 0.022 | 0.022 | 0 | 0.21 | 0.44 | 0.38 | 0.23 | 0.28 | 0.26 |
| 5 | 92595939 | rs4256329 | -0.332 | 2.527 | -1.282 | 1.002 | -0.034 | -0.066 | -0.066 | 0.044 | 0.14 | 0.01 | 0.25 | 0.22 | 0.11 | 0.18 |
| 5 | 92608801 | rs11135568 | -1.099 | 2.105 | -0.954 | 0.965 | -0.002 | -0.001 | -0.001 | 0.056 | 0.09 | 0.12 | 0.13 | 0.14 | 0.13 | 0.2 |
| 5 | 92611658 | rs6557074 | 0.546 | 1.736 | -1.088 | 0.879 | -0.031 | -0.023 | -0.023 | 0.181 | 0.89 | 0.99 | 0.96 | 0.92 | 0.92 | 0.93 |
| 5 | 92632106 | rs1545708 | -0.609 | 1.938 | -0.58 | 0.839 | 0.013 | 0.002 | 0.002 | 0 | 0.31 | 0.6 | 0.53 | 0.37 | 0.41 | 0.45 |
| 5 | 92636386 | rs13166639 | 0.728 | 2.099 | -0.322 | 0.554 | 0.035 | -0.023 | -0.023 | 0 | 0.72 | 0.61 | 0.77 | 0.81 | 0.64 | 0.62 |
| 5 | 92643805 | rs7448571 | 0.656 | 2.112 | -0.799 | 0.604 | -0.068 | -0.022 | -0.022 | 0 | 0.78 | 0.97 | 0.83 | 0.82 | 0.72 | 0.7 |
| 5 | 92665721 | rs719492 | 1.184 | 1.65 | 0.51 | 0.295 | 0.003 | -0.015 | -0.015 | 0 | 0.72 | 0.41 | 0.75 | 0.86 | 0.71 | 0.66 |
| 5 | 92677537 | rs2035053 | -1.041 | 1.731 | 0.234 | 0.754 | -0.004 | -0.026 | -0.026 | 0 | 0.36 | 0.7 | 0.49 | 0.46 | 0.33 | 0.47 |
| 5 | 92680028 | rs2343707 | -1.53 | 2.054 | 1.123 | 0.534 | 0.054 | -0.035 | -0.035 | 0.233 | 0.2 | 0.46 | 0.18 | 0.14 | 0.22 | 0.22 |
| 5 | 92705021 | rs2262637 | 1.828 | 1.554 | 1.058 | 0.719 | 0.004 | 0.02 | 0.02 | 2.208 | 0.73 | 0.21 | 0.79 | 0.8 | 0.74 | 0.81 |
| 5 | 92718948 | rs898597 | 1.457 | 1.543 | 1.273 | 0.723 | -0.006 | 0.003 | 0.003 | 0 | 0.79 | 0.86 | 0.83 | 0.81 | 0.76 | 0.82 |
| 5 | 92731616 | rs269705 | 1.284 | 1.926 | 1.195 | 0.93 | -0.001 | -0.004 | -0.004 | 0.433 | 0.83 | 0.9 | 0.87 | 0.81 | 0.82 | 0.86 |
| 5 | 92754902 | rs7715034 | -0.048 | 0.694 | 0.757 | -0.195 | -0.036 | -0.079 | -0.079 | 0.114 | 0.17 | 0.43 | 0.09 | 0.04 | 0.12 | 0.06 |
| 5 | 92776972 | rs893547 | 0.559 | 0.749 | 0.093 | -0.213 | -0.042 | -0.037 | -0.037 | 0 | 0.18 | 0.04 | 0.29 | 0.22 | 0.17 | 0.17 |
| 5 | 92815294 | rs2083227 | 0.572 | 0.728 | 0.044 | -0.188 | -0.036 | -0.075 | -0.075 | 0.498 | 0.17 | 0.43 | 0.09 | 0.04 | 0.12 | 0.06 |
| 5 | 92848652 | rs2344386 | 0.536 | 0.782 | -0.05 | -0.153 | -0.042 | -0.036 | -0.036 | 0 | 0.18 | 0.04 | 0.29 | 0.22 | 0.17 | 0.17 |
| 5 | 92849721 | rs31465 | -0.117 | 0.418 | -0.773 | -0.052 | -0.149 | -0.09 | -0.09 | 0.042 | 0.38 | 0.08 | 0.56 | 0.61 | 0.48 | 0.37 |
| 5 | 92892591 | rs6556826 | 0.796 | -0.376 | -0.272 | -0.501 | -0.085 | -0.093 | -0.093 | 0 | 0.22 | 0.46 | 0.11 | 0.05 | 0.12 | 0.13 |
| 5 | 92903582 | rs10042892 | -1.197 | 0.004 | -0.751 | -0.422 | 0.008 | 0.086 | 0.086 | 2.951 | 0.62 | 0.82 | 0.77 | 0.85 | 0.62 | 0.51 |
| 5 | 92932305 | rs2279473 | 0.977 | 0.021 | -1.038 | -0.31 | 0.034 | 0.062 | 0.062 | 0 | 0.21 | 0.12 | 0.09 | 0.04 | 0.16 | 0.08 |
| 5 | 92933747 | rs318108 | -1.222 | 0.057 | -1.446 | -0.444 | -0.043 | -0.023 | -0.023 | 0 | 0.84 | 0.99 | 0.82 | 0.85 | 0.82 | 0.85 |
| 5 | 92945156 | rs318105 | -1.108 | 0.109 | -1.448 | -0.315 | -0.047 | -0.038 | -0.038 | 1.035 | 0.84 | 0.99 | 0.82 | 0.82 | 0.81 | 0.8 |
| 5 | 92950673 | rs10064431 | 0.343 | 0.063 | -1.455 | -0.241 | -0.119 | -0.003 | -0.003 | 0 | 0.35 | 0.14 | 0.54 | 0.7 | 0.45 | 0.36 |
| 5 | 92969334 | rs7721168 | 1.017 | -0.304 | 0.095 | -0.242 | -0.032 | -0.062 | -0.062 | 0 | 0.23 | 0.33 | 0.1 | 0.04 | 0.15 | 0.08 |
| 5 | 92980916 | rs12186878 | 0.147 | -0.304 | 0.232 | 0.455 | -0.028 | 0.011 | 0.011 | 0 | 0.27 | 0.61 | 0.2 | 0.26 | 0.17 | 0.16 |
| 5 | 92987801 | rs6556829 | 0.433 | -0.298 | 0.124 | -0.15 | -0.063 | -0.105 | -0.105 | 0 | 0.24 | 0.46 | 0.11 | 0.04 | 0.14 | 0.08 |
| 5 | 92995477 | rs17083297 | 1.529 | 0.034 | -0.104 | -0.144 | -0.003 | -0.019 | -0.019 | 0 | 0.2 | 0.2 | 0.11 | 0.05 | 0.17 | 0.08 |
| 5 | 92997895 | rs700559 | 3.616 | -0.092 | -0.733 | -0.434 | -0.001 | 0.09 | 0.09 | 3.172 | 0.17 | 0.01 | 0.07 | 0.03 | 0.12 | 0.33 |
| 5 | 93004790 | rs700560 | 2.394 | 0.441 | -0.282 | 0.078 | 0.012 | 0.044 | 0.044 | 0.01 | 0.33 | 0.16 | 0.24 | 0.27 | 0.29 | 0.51 |
| 5 | 93021996 | rs9314086 | 1.124 | 0.893 | -0.263 | -0.026 | -0.004 | 0.045 | 0.045 | 0.091 | 0.22 | 0.13 | 0.12 | 0.05 | 0.19 | 0.08 |
| 5 | 93032427 | rs10041792 | 0.944 | 0.882 | -0.046 | -0.031 | -0.001 | -0.026 | -0.026 | 0 | 0.23 | 0.25 | 0.12 | 0.05 | 0.19 | 0.08 |
| 5 | 93043907 | rs17312456 | 2.187 | 0.853 | -0.277 | -0.051 | -0.023 | 0.029 | 0.029 | 0 | 0.15 | 0.01 | 0.08 | 0.05 | 0.13 | 0.07 |
| 5 | 93050453 | rs9285013 | -1.893 | 1.059 | -0.197 | 0.329 | -0.001 | -0.006 | -0.006 | 0 | 0.82 | 0.85 | 0.63 | 0.75 | 0.81 | 0.84 |
| 5 | 93054861 | rs17371926 | 1.881 | 1.559 | -0.337 | 0.338 | -0.062 | -0.049 | -0.049 | 2.724 | 0.11 | 0.02 | 0.17 | 0.17 | 0.22 | 0.13 |
| 5 | 93060781 | rs6895356 | 0.531 | 1.142 | 0.155 | 0.107 | -0.007 | -0.097 | -0.097 | 0 | 0.25 | 0.45 | 0.13 | 0.05 | 0.19 | 0.09 |
| 5 | 93071242 | rs13161885 | 2.214 | 1.203 | -0.42 | 0.128 | -0.029 | 0.046 | 0.046 | 0 | 0.14 | 0.01 | 0.07 | 0.04 | 0.13 | 0.07 |
| 5 | 93115673 | rs2119449 | 0.857 | 1.249 | -0.1 | 0.231 | 0.013 | -0.044 | -0.044 | 0 | 0.08 | 0.22 | 0.04 | 0 | 0.06 | 0.02 |
| 5 | 93133722 | rs11135392 | -0.45 | 1.301 | -0.426 | 0.342 | -0.052 | -0.048 | -0.048 | 0 | 0.34 | 0.18 | 0.55 | 0.57 | 0.44 | 0.32 |
| 5 | 93148166 | rs4242233 | -0.853 | 1.239 | -0.044 | 0.247 | 0.022 | -0.043 | -0.043 | 0.055 | 0.92 | 0.76 | 0.96 | 1 | 0.94 | 0.98 |
| 5 | 93151611 | rs1453003 | -0.896 | 1.191 | -0.107 | 0.181 | -0.002 | -0.045 | -0.045 | 0 | 0.77 | 0.72 | 0.88 | 0.95 | 0.81 | 0.91 |
| 5 | 93167165 | rs17314172 | 1.616 | 1.195 | 0.004 | 0.216 | 0.013 | -0.019 | -0.019 | 0 | 0.05 | 0.16 | 0.05 | 0 | 0.06 | 0.03 |
| 5 | 93176417 | rs4869203 | -1.577 | 1.193 | 0.136 | 0.194 | 0.01 | -0.038 | -0.038 | 0 | 0.92 | 0.86 | 0.96 | 1 | 0.94 | 0.98 |
| 5 | 93180544 | rs17594338 | 1.77 | 1.325 | 0.087 | 0.201 | 0.007 | 0.059 | 0.059 | 0 | 0.39 | 0.21 | 0.3 | 0.26 | 0.37 | 0.54 |
| 5 | 93187135 | rs11135394 | -0.076 | 1.331 | 0.312 | 0.335 | -0.008 | -0.02 | -0.02 | 0 | 0.72 | 0.43 | 0.84 | 0.83 | 0.77 | 0.86 |
| 5 | 93212833 | rs12653528 | -0.69 | 1.353 | 0.287 | 0.273 | -0.002 | -0.049 | -0.049 | 0 | 0.77 | 0.72 | 0.88 | 0.95 | 0.81 | 0.91 |
| 5 | 93214975 | rs11135397 | -0.628 | 1.355 | 0.242 | 0.288 | -0.004 | -0.067 | -0.067 | 0 | 0.76 | 0.66 | 0.87 | 0.95 | 0.81 | 0.91 |
| 5 | 93228046 | rs17314727 | 1.807 | 1.887 | -0.183 | 0.701 | -0.062 | -0.056 | -0.056 | 0.193 | 0.11 | 0.01 | 0.16 | 0.18 | 0.19 | 0.13 |
| 5 | 93257691 | rs12515593 | -0.15 | 1.822 | -0.178 | 0.676 | -0.054 | -0.06 | -0.06 | 0 | 0.36 | 0.17 | 0.56 | 0.56 | 0.43 | 0.35 |
| 5 | 93260865 | rs7735009 | -0.203 | 1.717 | 0.102 | 0.667 | -0.003 | -0.017 | -0.017 | 0 | 0.75 | 0.44 | 0.84 | 0.83 | 0.77 | 0.86 |
| 5 | 93276883 | rs1031423 | -0.218 | 1.661 | -0.014 | 0.668 | -0.007 | -0.014 | -0.014 | 0.255 | 0.75 | 0.42 | 0.84 | 0.83 | 0.79 | 0.86 |
| 5 | 93305352 | rs11135400 | 1.755 | 1.723 | -1.177 | 0.778 | 0.014 | 0.007 | 0.007 | 0 | 0.39 | 0.25 | 0.3 | 0.39 | 0.34 | 0.55 |
| 5 | 93311456 | rs7701011 | 1.92 | 1.637 | -1.396 | 0.322 | -0.012 | 0.03 | 0.03 | 0 | 0.14 | 0.05 | 0.08 | 0.05 | 0.13 | 0.07 |
| 5 | 93407803 | rs10044139 | 1.92 | 1.656 | -1.625 | 0.331 | -0.012 | 0.03 | 0.03 | 0 | 0.14 | 0.04 | 0.08 | 0.04 | 0.13 | 0.07 |
| 5 | 93427260 | rs17375167 | 1.574 | 1.913 | -1.7 | 0.391 | -0.067 | -0.054 | -0.054 | 0 | 0.11 | 0.01 | 0.17 | 0.18 | 0.22 | 0.13 |
| 5 | 93432805 | rs367971 | 1.823 | 1.597 | -1.53 | 0.233 | 0.018 | 0.048 | 0.048 | 0 | 0.37 | 0.21 | 0.26 | 0.27 | 0.3 | 0.5 |
| 5 | 93461916 | rs10476526 | 1.057 | 1.6 | -1.558 | 0.328 | 0.002 | 0.016 | 0.016 | 0 | 0.14 | 0.1 | 0.09 | 0.04 | 0.14 | 0.07 |
| 5 | 93463902 | rs6556840 | 0.211 | 1.702 | -1.803 | 0.391 | -0.079 | -0.104 | -0.104 | 1.625 | 0.2 | 0.04 | 0.21 | 0.32 | 0.31 | 0.21 |
| 5 | 93545524 | rs17376207 | 0.584 | 1.244 | -0.878 | 0.299 | 0.005 | 0.022 | 0.022 | 0 | 0.14 | 0.09 | 0.08 | 0.04 | 0.14 | 0.07 |
| 5 | 93548877 | rs13163610 | 0.584 | 1.244 | -0.878 | 0.299 | 0.005 | 0.022 | 0.022 | 0 | 0.14 | 0.09 | 0.08 | 0.04 | 0.14 | 0.06 |
| 5 | 93576698 | rs2015009 | 0.573 | 1.328 | -0.765 | 0.486 | -0.007 | -0.005 | -0.005 | 0.701 | 0.14 | 0.12 | 0.13 | 0.38 | 0.21 | 0.22 |
| 5 | 93578247 | rs5004465 | -0.143 | 1.179 | -0.618 | 0.133 | -0.002 | -0.01 | -0.01 | 0.537 | 0.88 | 0.83 | 0.86 | 0.91 | 0.87 | 0.89 |
| 5 | 93579380 | rs7444635 | -0.055 | 1.169 | -0.616 | 0.154 | -0.003 | -0.003 | -0.003 | 0 | 0.87 | 0.83 | 0.84 | 0.88 | 0.87 | 0.88 |
| 5 | 93582993 | rs1966938 | 0.667 | 1.187 | -0.972 | -0.253 | -0.002 | 0.014 | 0.014 | 0 | 0.35 | 0.34 | 0.46 | 0.16 | 0.37 | 0.24 |
| 5 | 93589609 | rs1841298 | 0.75 | 1.424 | -0.801 | -0.195 | -0.037 | -0.032 | -0.032 | 0 | 0.12 | 0.04 | 0.1 | 0.23 | 0.18 | 0.12 |
| 5 | 93615466 | rs981862 | 0.729 | 0.875 | -0.449 | -0.493 | -0.001 | -0.002 | -0.002 | 0.006 | 0.54 | 0.48 | 0.66 | 0.61 | 0.52 | 0.59 |

| | | | | | | | | | | | | | | | | |
|---|---|---|---|---|---|---|---|---|---|---|---|---|---|---|---|---|
| 5 | 93615511 | rs13154274 | 0.874 | 1.029 | -0.662 | -0.6 | -0.003 | 0.008 | 0.008 | 0 | 0.13 | 0.06 | 0.08 | 0.09 | 0.15 | 0.13 |
| 5 | 93616849 | rs2081863 | -0.358 | 1.07 | -0.372 | -0.498 | -0.004 | 0 | 0 | 0 | 0.92 | 0.87 | 0.92 | 0.91 | 0.93 | 0.91 |
| 5 | 93664162 | rs13180222 | 0.105 | 1.077 | -0.391 | -0.671 | 0.005 | -0.008 | -0.008 | 0.041 | 0.13 | 0.21 | 0.07 | 0.1 | 0.14 | 0.1 |
| 5 | 93678849 | rs160270 | 0.143 | 1.104 | -0.778 | -0.998 | -0.039 | 0.009 | 0.009 | 0 | 0.2 | 0.05 | 0.16 | 0.21 | 0.25 | 0.18 |
| 5 | 93699232 | rs13154972 | -0.648 | 0.427 | 0.234 | -0.982 | -0.009 | -0.048 | -0.048 | 0 | 0.17 | 0.49 | 0.1 | 0.09 | 0.15 | 0.1 |
| 5 | 93699673 | rs13189435 | -0.447 | 0.633 | -0.685 | -1.293 | -0.036 | 0.089 | 0.089 | 1.309 | 0.28 | 0.11 | 0.22 | 0.07 | 0.37 | 0.19 |
| 5 | 93713546 | rs29935 | -0.786 | 0.25 | -0.116 | -0.465 | -0.005 | 0 | 0 | 0 | 0.14 | 0.21 | 0.07 | 0.17 | 0.14 | 0.14 |
| 5 | 93722050 | rs29913 | -0.271 | 0.055 | -0.426 | -0.856 | 0.007 | 0.023 | 0.023 | 0.052 | 0.19 | 0.12 | 0.11 | 0 | 0.15 | 0.21 |
| 5 | 93730489 | rs29924 | -0.884 | -0.327 | 0.093 | -0.654 | -0.036 | -0.059 | -0.059 | 0 | 0.31 | 0.45 | 0.17 | 0.11 | 0.19 | 0.36 |
| 5 | 93732334 | rs308206 | -1.195 | -0.111 | -0.159 | -0.793 | 0 | 0.013 | 0.013 | 0 | 0.52 | 0.41 | 0.3 | 0.35 | 0.48 | 0.54 |
| 5 | 93733862 | rs29926 | -0.018 | -0.345 | -0.258 | -0.484 | 0.002 | -0.008 | -0.008 | 0 | 0.12 | 0.09 | 0.13 | 0.29 | 0.11 | 0.12 |
| 5 | 93742412 | rs308203 | -1.572 | -0.436 | 0.129 | -0.681 | 0.002 | 0.001 | 0.001 | 0 | 0.54 | 0.58 | 0.36 | 0.48 | 0.44 | 0.54 |
| 5 | 93751222 | rs29912 | 0.298 | -0.431 | 0.041 | -0.669 | 0.003 | -0.02 | -0.02 | 0 | 0.11 | 0.07 | 0.12 | 0.29 | 0.1 | 0.12 |
| 5 | 93751943 | rs7718517 | -0.017 | -0.506 | 0.071 | -0.72 | -0.016 | 0.078 | 0.078 | 0 | 0.09 | 0.26 | 0.04 | 0.11 | 0.04 | 0.13 |
| 5 | 93753017 | rs29910 | -0.311 | -0.358 | -0.759 | -0.695 | -0.004 | 0.129 | 0.129 | 0 | 0.31 | 0.13 | 0.17 | 0.08 | 0.29 | 0.27 |
| 5 | 93774266 | rs1861806 | -1.402 | -0.387 | -0.671 | -0.645 | 0.03 | 0.022 | 0.022 | 0 | 0.51 | 0.4 | 0.34 | 0.48 | 0.44 | 0.53 |
| 5 | 93774632 | rs1861807 | 0.393 | -0.509 | -0.605 | -0.606 | 0 | -0.018 | -0.018 | 0.006 | 0.11 | 0.07 | 0.12 | 0.29 | 0.11 | 0.11 |
| 5 | 93776023 | rs1030101 | -1.224 | -0.467 | -0.557 | -0.585 | 0.017 | 0.036 | 0.036 | 0 | 0.45 | 0.39 | 0.32 | 0.36 | 0.39 | 0.4 |
| 5 | 93783739 | rs17326006 | -1.103 | -0.103 | -0.652 | -0.45 | -0.001 | -0.008 | -0.008 | 0.075 | 0.42 | 0.48 | 0.35 | 0.34 | 0.4 | 0.38 |
| 5 | 93784383 | rs1420978 | -1.379 | -0.271 | -0.724 | -0.435 | 0.003 | 0.002 | 0.002 | 0 | 0.5 | 0.51 | 0.37 | 0.46 | 0.44 | 0.52 |
| 5 | 93788056 | rs17326131 | 0.604 | -0.303 | -0.759 | -0.562 | 0 | -0.006 | -0.006 | 0 | 0.11 | 0.09 | 0.07 | 0 | 0.11 | 0.05 |
| 5 | 93788638 | rs17326194 | 0.51 | -0.316 | -0.65 | -0.568 | -0.002 | -0.048 | -0.048 | 0.126 | 0.11 | 0.17 | 0.08 | 0 | 0.11 | 0.05 |
| 5 | 93789309 | rs10069076 | -1.105 | -0.456 | -0.777 | -0.604 | -0.003 | -0.008 | -0.008 | 0.001 | 0.37 | 0.42 | 0.24 | 0.23 | 0.33 | 0.37 |
| 5 | 93795658 | rs9314112 | -1.861 | -0.152 | -1.232 | -0.665 | -0.002 | 0.001 | 0.001 | 0 | 0.56 | 0.44 | 0.44 | 0.46 | 0.53 | 0.58 |
| 5 | 93796132 | rs9285014 | 0.322 | 0.72 | -1.55 | -0.723 | -0.062 | 0.035 | 0.035 | 0.066 | 0.17 | 0.01 | 0.13 | 0.1 | 0.19 | 0.2 |
| 5 | 93798480 | rs952664 | -0.745 | 0.479 | -0.791 | -0.229 | -0.039 | -0.035 | -0.035 | 0 | 0.14 | 0.04 | 0.16 | 0.24 | 0.18 | 0.12 |
| 5 | 93798912 | rs17379236 | -0.648 | -0.058 | -0.303 | -0.356 | 0 | 0.017 | 0.017 | 0 | 0.12 | 0.08 | 0.08 | 0.01 | 0.12 | 0.08 |
| 5 | 93810208 | rs6869388 | -1.24 | -0.33 | 0.051 | -0.42 | -0.04 | -0.107 | -0.107 | 0 | 0.25 | 0.47 | 0.2 | 0.02 | 0.2 | 0.24 |
| 5 | 93812885 | rs10057902 | -1.757 | -0.778 | 0.703 | -0.145 | -0.054 | 0.008 | 0.008 | 0 | 0.11 | 0.33 | 0.08 | 0.08 | 0.02 | 0.09 |
| 5 | 93823951 | rs13361876 | -1.605 | -0.091 | 0.295 | 0.769 | 0.003 | 0.058 | 0.058 | 0 | 0.19 | 0.28 | 0.34 | 0.33 | 0.21 | 0.21 |
| 5 | 93853266 | rs17083775 | -1.169 | -0.293 | 0.649 | 0.841 | -0.005 | -0.015 | -0.015 | 0.979 | 0.16 | 0.13 | 0.32 | 0.33 | 0.2 | 0.2 |
| 5 | 93862964 | rs1839852 | -2.179 | -0.732 | 1.574 | 0.547 | -0.01 | 0.005 | 0.005 | 0 | 0.32 | 0.66 | 0.42 | 0.32 | 0.3 | 0.26 |
| 5 | 93863813 | rs17083791 | -0.096 | -0.644 | 1.558 | 0.654 | -0.02 | -0.054 | -0.054 | 0 | 0.1 | 0.02 | 0.26 | 0.25 | 0.14 | 0.11 |
| 5 | 93864514 | rs7702344 | -2.164 | -0.623 | 1.552 | 0.649 | -0.007 | 0.003 | 0.003 | 0 | 0.32 | 0.67 | 0.42 | 0.33 | 0.3 | 0.26 |
| 5 | 93891091 | rs9314117 | 1.051 | -0.199 | 0.633 | 0.155 | -0.013 | 0.006 | 0.006 | 0 | 0.29 | 0.22 | 0.22 | 0.21 | 0.37 | 0.26 |
| 5 | 93896128 | rs6898585 | -0.461 | -0.5 | 0.622 | -0.059 | 0 | -0.005 | -0.005 | 0 | 0.49 | 0.53 | 0.35 | 0.24 | 0.52 | 0.42 |
| 5 | 93899181 | rs17083807 | 0.077 | -0.384 | 0.738 | -0.059 | 0 | -0.007 | -0.007 | 0 | 0.08 | 0.1 | 0.05 | 0.03 | 0.06 | 0.1 |
| 5 | 93918392 | rs1031389 | -0.68 | -0.164 | 0.478 | 0.111 | -0.01 | 0.008 | 0.008 | 0 | 0.44 | 0.71 | 0.48 | 0.34 | 0.47 | 0.38 |
| 5 | 93926285 | rs17327757 | 0.477 | -0.151 | 0.432 | -0.026 | 0 | 0.013 | 0.013 | 0 | 0.08 | 0.06 | 0.05 | 0.03 | 0.06 | 0.1 |
| 5 | 93936485 | rs10035650 | 0.306 | -0.151 | 0.405 | -0.014 | 0.018 | -0.122 | -0.122 | 0 | 0.32 | 0.52 | 0.23 | 0.05 | 0.34 | 0.27 |
| 5 | 93943818 | rs10043779 | 0.316 | -0.117 | 0.405 | -0.014 | 0.017 | -0.122 | -0.122 | 0.016 | 0.32 | 0.52 | 0.23 | 0.05 | 0.34 | 0.28 |
| 5 | 93947146 | rs17083818 | 0.641 | -0.234 | 0.3 | -0.07 | 0.007 | -0.054 | -0.054 | 0 | 0.3 | 0.38 | 0.22 | 0.05 | 0.34 | 0.26 |
| 5 | 93949604 | rs2047318 | -0.791 | -0.323 | 0.145 | 0.106 | 0.001 | -0.013 | -0.013 | 0.197 | 0.19 | 0.21 | 0.3 | 0.31 | 0.19 | 0.21 |
| 5 | 93956159 | rs17387486 | 0.456 | -0.329 | 0.322 | -0.014 | 0 | 0.016 | 0.016 | 0 | 0.08 | 0.07 | 0.05 | 0.03 | 0.06 | 0.11 |
| 5 | 93956937 | rs17328275 | 1.559 | -0.715 | -0.222 | -0.105 | -0.003 | -0.047 | -0.047 | 0 | 0.1 | 0 | 0.06 | 0.13 | 0.09 | 0.19 |
| 5 | 93980014 | rs10070258 | 1.712 | -0.615 | 0.03 | 0.051 | 0.035 | -0.022 | -0.022 | 0 | 0.24 | 0.38 | 0.2 | 0.25 | 0.32 | 0.25 |
| 5 | 93987532 | rs6891545 | -0.336 | -0.677 | 0.015 | 0.094 | 0.001 | -0.017 | -0.017 | 0 | 0.19 | 0.2 | 0.3 | 0.31 | 0.19 | 0.21 |
| 5 | 93991238 | rs12655521 | -0.336 | -0.677 | 0.015 | 0.073 | 0.001 | -0.017 | -0.017 | 0.562 | 0.19 | 0.2 | 0.3 | 0.31 | 0.19 | 0.21 |
| 5 | 94001174 | rs12516225 | 1.578 | -0.653 | 0.075 | 0.037 | 0.031 | -0.029 | -0.029 | 0 | 0.25 | 0.4 | 0.2 | 0.25 | 0.32 | 0.25 |
| 5 | 94001476 | rs10051931 | 2.282 | -0.648 | 0.117 | 0 | 0.025 | -0.053 | -0.053 | 0 | 0.21 | 0.26 | 0.17 | 0.02 | 0.29 | 0.16 |
| 5 | 94009546 | rs17083889 | 0.928 | -0.764 | -0.061 | 0.111 | -0.02 | -0.014 | -0.014 | 0 | 0.1 | 0 | 0.06 | 0.13 | 0.09 | 0.1 |
| 5 | 94012979 | rs1124667 | -0.205 | -0.671 | 0.209 | 0.195 | -0.012 | -0.002 | -0.002 | 0 | 0.45 | 0.66 | 0.5 | 0.56 | 0.51 | 0.46 |
| 5 | 94013241 | rs1124668 | -0.328 | -0.518 | 0.206 | 0.178 | 0.001 | -0.017 | -0.017 | 0 | 0.19 | 0.2 | 0.3 | 0.31 | 0.19 | 0.21 |
| 5 | 94019780 | rs2112449 | -0.214 | -0.514 | 0.209 | 0.154 | -0.004 | -0.001 | -0.001 | 0 | 0.45 | 0.61 | 0.5 | 0.57 | 0.51 | 0.46 |
| 5 | 94027011 | rs10078304 | 2.315 | -0.496 | 0.165 | 0.106 | 0.025 | -0.054 | -0.054 | 0 | 0.2 | 0.26 | 0.17 | 0.02 | 0.29 | 0.16 |
| 5 | 94035949 | rs11955373 | 2.356 | -0.483 | 0.187 | 0.112 | 0.024 | -0.05 | -0.05 | 0.225 | 0.2 | 0.25 | 0.17 | 0.02 | 0.29 | 0.16 |
| 5 | 94038410 | rs2112448 | -0.36 | -0.461 | -0.021 | 0.188 | -0.011 | 0.001 | 0.001 | 0 | 0.45 | 0.66 | 0.5 | 0.56 | 0.51 | 0.46 |
| 5 | 94043608 | rs7718956 | -0.601 | -0.41 | -0.191 | 0.311 | -0.001 | 0.011 | 0.011 | 0 | 0.22 | 0.27 | 0.33 | 0.65 | 0.22 | 0.32 |
| 5 | 94050447 | rs2638 | 0.217 | -0.74 | -0.268 | 0.197 | 0.011 | 0.019 | 0.019 | 0 | 0.51 | 0.19 | 0.48 | 0.43 | 0.47 | 0.51 |
| 5 | 94052733 | rs306577 | -1.969 | -0.816 | -0.744 | -0.102 | 0 | -0.013 | -0.013 | 0 | 0.83 | 0.93 | 0.88 | 0.7 | 0.86 | 0.86 |
| 5 | 94055916 | rs11135410 | 1.064 | -0.7 | -0.647 | 0.059 | 0 | 0.05 | 0.05 | 0 | 0.11 | 0.15 | 0.25 | 0.29 | 0.13 | 0.11 |
| 5 | 94056825 | rs432449 | -0.26 | -0.462 | -0.697 | 0.084 | 0.012 | -0.013 | -0.013 | 0.091 | 0.36 | 0.18 | 0.38 | 0.36 | 0.35 | 0.47 |
| 5 | 94060445 | rs306580 | -0.58 | -0.543 | -0.654 | 0.091 | 0.011 | -0.003 | -0.003 | 0.039 | 0.4 | 0.28 | 0.4 | 0.36 | 0.38 | 0.49 |
| 5 | 94082218 | rs6877393 | -0.151 | -0.569 | -0.409 | 0.181 | -0.002 | -0.019 | -0.019 | 2.567 | 0.22 | 0.35 | 0.2 | 0.17 | 0.21 | 0.29 |
| 5 | 94083237 | rs6556852 | -0.708 | -0.611 | -0.51 | 0.321 | -0.004 | 0.045 | 0.045 | 0 | 0.33 | 0.51 | 0.45 | 0.46 | 0.33 | 0.39 |
| 5 | 94083321 | rs4869220 | 0.341 | -0.689 | -1.131 | 0.347 | 0.011 | -0.036 | -0.036 | 0 | 0.34 | 0.16 | 0.33 | 0.49 | 0.31 | 0.35 |
| 5 | 94083989 | rs17084010 | 1.309 | -0.469 | -1.081 | 0.299 | -0.008 | -0.013 | -0.013 | 0 | 0.21 | 0.08 | 0.22 | 0.18 | 0.18 | 0.16 |
| 5 | 94084404 | rs6889062 | -0.342 | -0.057 | -0.174 | 0.544 | -0.002 | -0.001 | -0.001 | 0.091 | 0.19 | 0.16 | 0.19 | 0.17 | 0.18 | 0.26 |
| 5 | 94088858 | rs4869118 | -0.218 | 0.124 | -0.335 | 0.341 | 0 | -0.001 | -0.001 | 0 | 0.58 | 0.62 | 0.51 | 0.69 | 0.46 | 0.59 |

| | | | | | | | | | | | | | | | | |
|---|---|---|---|---|---|---|---|---|---|---|---|---|---|---|---|---|
| 5 | 94094647 | rs255334 | 0.285 | 0.025 | -0.428 | 0.287 | 0.014 | 0.019 | 0.019 | 0 | 0.51 | 0.44 | 0.48 | 0.69 | 0.4 | 0.57 |
| 5 | 94096010 | rs166241 | 0.997 | -0.095 | -1.14 | 0.498 | -0.009 | -0.034 | -0.034 | 5.922 | 0.36 | 0.07 | 0.31 | 0.39 | 0.29 | 0.37 |
| 5 | 94131308 | rs255344 | -0.297 | 0.348 | -0.599 | 0.819 | -0.013 | -0.031 | -0.031 | 0.067 | 0.91 | 0.95 | 0.77 | 0.72 | 0.89 | 0.88 |
| 5 | 94137963 | rs6888297 | -0.367 | 0.46 | 0.204 | 0.641 | 0.08 | 0.01 | 0.01 | 0.229 | 0.07 | 0.35 | 0.05 | 0.02 | 0.06 | 0.11 |
| 5 | 94138997 | rs255341 | 0.682 | 0.467 | 0.156 | 0.651 | 0.057 | 0.181 | 0.181 | 0 | 0.84 | 0.55 | 0.71 | 0.7 | 0.8 | 0.79 |
| 5 | 94141447 | rs11953346 | 0.735 | 0.606 | -0.556 | -0.004 | -0.06 | 0.12 | 0.12 | 0 | 0.28 | 0.1 | 0.21 | 0.02 | 0.38 | 0.22 |
| 5 | 94145255 | rs10515218 | 0.387 | 0.494 | -0.061 | 0.723 | 0.008 | 0.168 | 0.168 | 0 | 0.1 | 0.2 | 0.17 | 0.35 | 0.08 | 0.11 |
| 5 | 94149017 | rs12152974 | 0.873 | 0.485 | 0.369 | 0.4 | 0.014 | 0.097 | 0.097 | 0 | 0.08 | 0.16 | 0.12 | 0.24 | 0.07 | 0.08 |
| 5 | 94150212 | rs255332 | -1.696 | -0.194 | 0.271 | 0 | 0.008 | 0.009 | 0.009 | 0 | 0.38 | 0.64 | 0.45 | 0.88 | 0.25 | 0.47 |
| 5 | 94153513 | rs17084109 | 0.945 | -0.437 | 0.345 | 0.011 | -0.022 | -0.039 | -0.039 | 3.399 | 0.07 | 0.18 | 0.02 | 0 | 0.03 | 0.01 |
| 5 | 94153862 | rs153897 | -0.563 | -0.295 | 0.241 | 0.189 | -0.008 | 0.038 | 0.038 | 0 | 0.18 | 0.21 | 0.29 | 0.64 | 0.13 | 0.35 |
| 5 | 94154465 | rs255209 | -0.99 | -0.166 | 0.194 | 0.06 | 0.039 | -0.013 | -0.013 | 0.044 | 0.95 | 0.85 | 0.88 | 0.97 | 0.93 | 0.9 |
| 5 | 94154588 | rs17418283 | 1.818 | -0.276 | -0.284 | -0.21 | -0.043 | 0.102 | 0.102 | 0 | 0.3 | 0.02 | 0.21 | 0.07 | 0.29 | 0.2 |
| 5 | 94168290 | rs36733 | 1.317 | 0.245 | 0.615 | 0.608 | -0.001 | -0.002 | -0.002 | 0 | 0.74 | 0.76 | 0.58 | 0.33 | 0.78 | 0.49 |
| 5 | 94174183 | rs255367 | 0.705 | 0.284 | 0.675 | 0.625 | 0.035 | 0.145 | 0.145 | 0 | 0.86 | 0.56 | 0.75 | 0.73 | 0.87 | 0.83 |
| 5 | 94175381 | rs255211 | 0.465 | 0.259 | 0.441 | 0.593 | -0.037 | 0.023 | 0.023 | 0 | 0.91 | 0.99 | 0.79 | 0.93 | 0.9 | 0.8 |
| 5 | 94175811 | rs718184 | 0.097 | 0.22 | 0.549 | 0.647 | 0.011 | 0.201 | 0.201 | 0.007 | 0.1 | 0.36 | 0.13 | 0.24 | 0.06 | 0.06 |
| 5 | 94178962 | rs39715 | -0.423 | 0.354 | -0.099 | 0.688 | -0.011 | 0.081 | 0.081 | 0 | 0.14 | 0.19 | 0.19 | 0.6 | 0.1 | 0.27 |
| 5 | 94180572 | rs36724 | -1.036 | 0.452 | -0.036 | -0.119 | -0.022 | -0.043 | -0.043 | 1.242 | 0.28 | 0.34 | 0.42 | 0.88 | 0.2 | 0.46 |
| 5 | 94186517 | rs159598 | 0.167 | 1.218 | -0.034 | 0.103 | 0.049 | -0.011 | -0.011 | 0.137 | 0.94 | 0.82 | 0.87 | 0.97 | 0.93 | 0.89 |
| 5 | 94190819 | rs293061 | 0.402 | 1.147 | -0.897 | 0.152 | -0.067 | 0.106 | 0.106 | 0 | 0.23 | 0.05 | 0.15 | 0.02 | 0.31 | 0.12 |
| 5 | 94191436 | rs13184495 | -0.711 | 0.635 | -0.052 | 1.252 | 0.001 | 0.122 | 0.122 | 0 | 0.1 | 0.2 | 0.15 | 0.33 | 0.08 | 0.18 |
| 5 | 94197654 | rs27005 | -1.159 | 0.962 | -0.428 | 0.745 | -0.003 | -0.005 | -0.005 | 0 | 0.09 | 0.1 | 0.07 | 0.11 | 0.12 | 0.1 |
| 5 | 94198611 | rs149352 | 0.001 | 1.109 | -0.255 | -0.069 | 0.003 | -0.012 | -0.012 | 0 | 0.32 | 0.3 | 0.31 | 0.23 | 0.35 | 0.29 |
| 5 | 94202167 | rs27003 | -0.077 | 1.064 | -0.449 | -0.819 | -0.001 | -0.013 | -0.013 | 0 | 0.66 | 0.6 | 0.56 | 0.29 | 0.7 | 0.55 |
| 5 | 94202517 | rs6861217 | 0.783 | 1.044 | -0.606 | -1.037 | -0.008 | -0.015 | -0.015 | 0 | 0.14 | 0.22 | 0.08 | 0.07 | 0.1 | 0.11 |
| 5 | 94205837 | rs27002 | -0.952 | 0.934 | -0.654 | -1.142 | 0.013 | -0.028 | -0.028 | 0.204 | 0.76 | 0.89 | 0.65 | 0.3 | 0.81 | 0.61 |
| 5 | 94221397 | rs293056 | 1.253 | 0.96 | -0.764 | -1.262 | -0.065 | 0.007 | 0.007 | 0 | 0.19 | 0.02 | 0.21 | 0.12 | 0.25 | 0.19 |
| 5 | 94222598 | rs39988 | 0.963 | 0.789 | -0.502 | -1.456 | 0.008 | 0.083 | 0.083 | 0 | 0.3 | 0.2 | 0.22 | 0.04 | 0.3 | 0.24 |
| 5 | 94230358 | rs9885412 | 0.921 | 0.838 | -0.359 | -1.326 | 0.002 | 0.005 | 0.005 | 0 | 0.14 | 0.1 | 0.08 | 0.07 | 0.11 | 0.12 |
| 5 | 94235506 | rs26999 | -0.616 | 0.733 | -0.698 | -1.428 | -0.079 | 0.169 | 0.169 | 0 | 0.58 | 0.94 | 0.43 | 0.22 | 0.55 | 0.43 |
| 5 | 94237398 | rs26998 | 0.678 | 0.417 | -0.133 | -1.275 | 0.002 | -0.02 | -0.02 | 0.078 | 0.29 | 0.3 | 0.19 | 0.07 | 0.27 | 0.23 |
| 5 | 94239189 | rs26997 | 0.68 | 0.386 | -0.127 | -1.001 | 0.003 | 0.003 | 0.003 | 0 | 0.3 | 0.3 | 0.2 | 0.14 | 0.28 | 0.27 |
| 5 | 94247253 | rs469942 | 0.168 | 0.301 | 0.01 | -0.93 | 0.017 | 0.028 | 0.028 | 0.685 | 0.59 | 0.36 | 0.62 | 0.18 | 0.67 | 0.57 |
| 5 | 94251510 | rs159350 | -0.36 | 0.584 | 0.014 | -0.573 | -0.002 | -0.006 | -0.006 | 0.183 | 0.22 | 0.22 | 0.24 | 0.12 | 0.22 | 0.25 |
| 5 | 94252979 | rs469968 | -0.664 | 0.63 | 0.061 | -0.57 | -0.007 | -0.029 | -0.029 | 0.225 | 0.26 | 0.34 | 0.26 | 0.13 | 0.25 | 0.28 |
| 5 | 94256618 | rs17421253 | -0.051 | 1.584 | -0.679 | -0.491 | -0.058 | -0.012 | -0.012 | 0.091 | 0.17 | 0.06 | 0.29 | 0.14 | 0.23 | 0.32 |
| 5 | 94258730 | rs293049 | 0.92 | 1.08 | -1.126 | -0.138 | 0.008 | -0.07 | -0.07 | 2.87 | 0.31 | 0.17 | 0.12 | 0.49 | 0.24 | 0.25 |
| 5 | 94267120 | rs13170906 | 0.218 | 1.118 | -0.308 | -0.46 | 0.002 | -0.034 | -0.034 | 0 | 0.12 | 0.18 | 0.16 | 0.06 | 0.12 | 0.1 |
| 5 | 94268466 | rs32909 | 0.174 | 1.237 | -0.387 | -0.37 | -0.041 | -0.021 | -0.021 | 2.605 | 0.21 | 0.12 | 0.41 | 0.29 | 0.32 | 0.45 |
| 5 | 94268604 | rs17337117 | 0.57 | 1.284 | -0.475 | -0.332 | -0.085 | -0.026 | -0.026 | 0.1 | 0.18 | 0.03 | 0.37 | 0.17 | 0.28 | 0.41 |
| 5 | 94273438 | rs457891 | -1.026 | 0.646 | -0.112 | -0.571 | -0.014 | -0.026 | -0.026 | 0 | 0.72 | 0.58 | 0.75 | 0.84 | 0.73 | 0.81 |
| 5 | 94277497 | rs159022 | 0.104 | 0.617 | 0.056 | -0.424 | 0.002 | 0 | 0 | 0 | 0.48 | 0.47 | 0.33 | 0.56 | 0.37 | 0.35 |
| 5 | 94278129 | rs293035 | 1.011 | 0.639 | -0.272 | -0.17 | 0.013 | -0.069 | -0.069 | 0 | 0.32 | 0.18 | 0.13 | 0.5 | 0.24 | 0.25 |
| 5 | 94281141 | rs32913 | -0.603 | 0.643 | -0.132 | -0.44 | 0.009 | 0.032 | 0.032 | 0 | 0.6 | 0.67 | 0.61 | 0.78 | 0.61 | 0.64 |
| 5 | 94281668 | rs293033 | 0.519 | 0.62 | -0.108 | -0.202 | 0.014 | -0.016 | -0.016 | 0 | 0.37 | 0.3 | 0.23 | 0.49 | 0.29 | 0.26 |
| 5 | 94285327 | rs10866761 | -1.038 | 0.786 | 0.769 | -0.123 | 0.023 | -0.049 | -0.049 | 0 | 0.28 | 0.42 | 0.39 | 0.17 | 0.33 | 0.38 |
| 5 | 94285653 | rs11740360 | -0.466 | 0.952 | 0.726 | -0.228 | 0.003 | -0.056 | -0.056 | 1.145 | 0.22 | 0.22 | 0.25 | 0.01 | 0.24 | 0.15 |
| 5 | 94297348 | rs10043438 | 1.694 | 0.849 | 0.594 | -0.51 | 0.01 | 0.031 | 0.031 | 0.541 | 0.46 | 0.42 | 0.39 | 0.16 | 0.4 | 0.29 |
| 5 | 94301042 | rs1437118 | 1.124 | 0.814 | 0.752 | -0.31 | 0 | -0.001 | -0.001 | 0.278 | 0.56 | 0.59 | 0.58 | 0.4 | 0.52 | 0.56 |
| 5 | 94301377 | rs10051466 | -0.68 | 0.879 | 0.876 | -0.32 | -0.01 | -0.039 | -0.039 | 0.598 | 0.79 | 0.9 | 0.82 | 0.4 | 0.77 | 0.71 |
| 5 | 94302728 | rs17084340 | 0.211 | 0.866 | 1.001 | -0.321 | 0.005 | 0.071 | 0.071 | 0 | 0.11 | 0.26 | 0.19 | 0.24 | 0.13 | 0.26 |
| 5 | 94304059 | rs32922 | -1.191 | 0.879 | 0.732 | -0.179 | -0.043 | -0.126 | -0.126 | 0 | 0.82 | 0.98 | 0.85 | 0.51 | 0.81 | 0.74 |
| 5 | 94304323 | rs32921 | 0.778 | 0.896 | 0.865 | -0.182 | -0.004 | -0.027 | -0.027 | 0 | 0.62 | 0.51 | 0.6 | 0.28 | 0.62 | 0.46 |
| 5 | 94306756 | rs10079443 | 0.653 | 0.901 | 0.768 | -0.167 | 0 | 0.007 | 0.007 | 0 | 0.1 | 0.11 | 0.18 | 0.24 | 0.13 | 0.27 |
| 5 | 94308966 | rs13180001 | 1.914 | 0.903 | 0.736 | -0.252 | 0.022 | 0.108 | 0.108 | 0.698 | 0.44 | 0.26 | 0.38 | 0.16 | 0.39 | 0.29 |
| 5 | 94309166 | rs456156 | -0.198 | 0.939 | 0.907 | 0.292 | 0.013 | -0.014 | -0.014 | 0 | 0.17 | 0.21 | 0.22 | 0.12 | 0.23 | 0.17 |
| 5 | 94313365 | rs7732129 | -0.897 | 0.689 | 0.743 | 0.416 | 0 | 0 | 0 | 0.03 | 0.26 | 0.26 | 0.37 | 0.35 | 0.33 | 0.42 |
| 5 | 94313688 | rs7731439 | -0.549 | 0.697 | 0.731 | 0.429 | -0.001 | 0 | 0 | 0.07 | 0.23 | 0.22 | 0.35 | 0.24 | 0.29 | 0.38 |
| 5 | 94316007 | rs10070401 | 0.945 | 0.59 | 0.702 | -0.021 | -0.019 | 0.034 | 0.034 | 3.414 | 0.58 | 0.72 | 0.5 | 0.27 | 0.52 | 0.36 |
| 5 | 94317666 | rs13160334 | 1.502 | 0.539 | 0.865 | -0.07 | 0 | -0.014 | -0.014 | 1.044 | 0.51 | 0.55 | 0.44 | 0.16 | 0.46 | 0.31 |
| 5 | 94318833 | rs309801 | -1.68 | 0.623 | 0.818 | 0.57 | -0.008 | -0.032 | -0.032 | 0.543 | 0.36 | 0.78 | 0.51 | 0.48 | 0.38 | 0.49 |
| 5 | 94321178 | rs169822 | -0.941 | -0.017 | 1.331 | 1.197 | -0.011 | 0.032 | 0.032 | 0.031 | 0.22 | 0.29 | 0.31 | 0.36 | 0.18 | 0.32 |
| 5 | 94329800 | rs722722 | -1.589 | 0.198 | 1.336 | 1.181 | 0.039 | 0.054 | 0.054 | 0 | 0.07 | 0.3 | 0.05 | 0.11 | 0.07 | 0.05 |
| 5 | 94331398 | rs7715499 | -0.87 | 0.275 | 1.143 | 1.186 | 0.006 | 0.076 | 0.076 | 0.072 | 0.25 | 0.46 | 0.15 | 0.45 | 0.26 | 0.32 |
| 5 | 94351572 | rs7708977 | -0.023 | 0.138 | 0.928 | 0.721 | 0 | -0.022 | -0.022 | 0 | 0.12 | 0.16 | 0.06 | 0 | 0.08 | 0.06 |
| 5 | 94353905 | rs2114194 | 2.048 | -0.024 | 1.019 | 0.878 | 0.003 | -0.004 | -0.004 | 5.625 | 0.67 | 0.31 | 0.63 | 0.52 | 0.75 | 0.61 |
| 5 | 94363349 | rs7730096 | 1.746 | 0.014 | 1.045 | 0.836 | -0.012 | 0.009 | 0.009 | 0 | 0.71 | 0.33 | 0.72 | 0.61 | 0.81 | 0.65 |
| 5 | 94365306 | rs1426100 | 1.754 | 0.025 | 1.132 | 0.846 | -0.03 | 0.11 | 0.11 | 0 | 0.8 | 0.38 | 0.72 | 0.55 | 0.83 | 0.67 |

| | | | | | | | | | | | | | | | | |
|---|---|---|---|---|---|---|---|---|---|---|---|---|---|---|---|---|
| 5 | 94371947 | rs17084368 | -0.99 | 0.06 | 1.15 | 0.899 | -0.012 | 0.032 | 0.032 | 0 | 0.2 | 0.28 | 0.19 | 0.36 | 0.13 | 0.32 |
| 5 | 94377805 | rs4988879 | -1.415 | 0.018 | 1.168 | 0.912 | -0.009 | 0.04 | 0.04 | 0 | 0.23 | 0.34 | 0.23 | 0.42 | 0.19 | 0.36 |
| 5 | 94380392 | rs11135419 | -1.173 | 0.096 | 1.26 | 0.996 | -0.022 | 0.116 | 0.116 | 0.118 | 0.13 | 0.3 | 0.18 | 0.36 | 0.09 | 0.28 |
| 5 | 94380775 | rs6862790 | 1.852 | 0.24 | 1.038 | 0.867 | 0.005 | -0.007 | -0.007 | 0 | 0.61 | 0.29 | 0.65 | 0.55 | 0.67 | 0.53 |
| 5 | 94384953 | rs2881646 | 1.078 | 0.067 | 1.26 | 1.589 | 0.001 | 0.249 | 0.249 | 0 | 0.89 | 0.61 | 0.85 | 0.63 | 0.92 | 0.72 |
| 5 | 94391765 | rs13359900 | -0.58 | 0.051 | 0.63 | 1.585 | -0.002 | 0.045 | 0.045 | 0 | 0.09 | 0.13 | 0.13 | 0.36 | 0.06 | 0.28 |
| 5 | 94396805 | rs10070518 | 1.677 | -0.029 | 1.018 | 1.512 | -0.017 | 0.218 | 0.218 | 0 | 0.86 | 0.52 | 0.83 | 0.64 | 0.91 | 0.71 |
| 5 | 94399644 | rs6862865 | 2.012 | -0.238 | 1.122 | 1.5 | -0.047 | 0.126 | 0.126 | 0.264 | 0.8 | 0.4 | 0.8 | 0.59 | 0.87 | 0.69 |
| 5 | 94400017 | rs10067664 | 1.963 | -0.242 | 1.155 | 1.492 | -0.054 | 0.158 | 0.158 | 0 | 0.83 | 0.48 | 0.82 | 0.64 | 0.9 | 0.71 |
| 5 | 94407021 | rs1974960 | 0.207 | -0.161 | 1.453 | 1.633 | -0.011 | 0.164 | 0.164 | 0 | 0.93 | 0.8 | 0.92 | 0.68 | 0.96 | 0.75 |
| 5 | 94411987 | rs6865589 | 1.363 | -0.227 | 1.602 | 1.601 | -0.025 | 0.263 | 0.263 | 0 | 0.89 | 0.51 | 0.9 | 0.67 | 0.95 | 0.74 |
| 5 | 94415012 | rs12716470 | -1.142 | -0.353 | 0.969 | 1.602 | -0.034 | -0.121 | -0.121 | 0.479 | 0.24 | 0.06 | 0.14 | 0.42 | 0.26 | 0.37 |
| 5 | 94417958 | rs958492 | 0.97 | -0.347 | 0.757 | 1.606 | -0.031 | 0.228 | 0.228 | 0 | 0.88 | 0.47 | 0.89 | 0.67 | 0.95 | 0.74 |
| 5 | 94418818 | rs11744157 | -0.631 | -0.354 | 0.949 | 1.574 | 0 | -0.022 | -0.022 | 0 | 0.2 | 0.12 | 0.11 | 0.36 | 0.21 | 0.34 |
| 5 | 94424048 | rs6556858 | -1.185 | -0.244 | 0.483 | 1.178 | -0.011 | -0.034 | -0.034 | 0 | 0.34 | 0.22 | 0.19 | 0.42 | 0.33 | 0.42 |
| 5 | 94431710 | rs1895865 | 1.416 | -0.026 | 0.124 | 0.761 | 0.031 | 0.09 | 0.09 | 0 | 0.54 | 0.21 | 0.7 | 0.26 | 0.63 | 0.34 |
| 5 | 94439860 | rs1426093 | 0.893 | 0.197 | 0.534 | 0.823 | -0.004 | -0.047 | -0.047 | 0.374 | 0.81 | 0.74 | 0.91 | 0.95 | 0.87 | 0.84 |
| 5 | 94440219 | rs6894476 | 1.524 | 0.474 | 0.275 | 1.188 | -0.004 | -0.002 | -0.002 | 0 | 0.67 | 0.43 | 0.7 | 0.3 | 0.71 | 0.59 |
| 5 | 94449704 | rs1593463 | 1.164 | 0.379 | 0.187 | 1.4 | -0.009 | 0.068 | 0.068 | 0 | 0.78 | 0.62 | 0.88 | 0.61 | 0.83 | 0.66 |
| 5 | 94459053 | rs372190 | 1.354 | 0.297 | 0.423 | 1.633 | -0.016 | 0.194 | 0.194 | 0 | 0.87 | 0.63 | 0.91 | 0.62 | 0.91 | 0.69 |
| 5 | 94461607 | rs7732431 | 0.614 | 0.808 | -0.062 | 1.244 | 0.018 | 0.053 | 0.053 | 0 | 0.78 | 0.65 | 0.74 | 0.58 | 0.7 | 0.63 |
| 5 | 94466137 | rs4869238 | 0.124 | -0.24 | -0.923 | 0.372 | 0.031 | 0.067 | 0.067 | 1.596 | 0.3 | 0.15 | 0.1 | 0.12 | 0.18 | 0.17 |
| 5 | 94486474 | rs331568 | -0.323 | -0.246 | -0.756 | 1.125 | 0.022 | -0.013 | -0.013 | 0 | 0.41 | 0.21 | 0.27 | 0.6 | 0.31 | 0.47 |
| 5 | 94487600 | rs331569 | -0.33 | -0.249 | -0.758 | 1.123 | 0.022 | -0.012 | -0.012 | 0.125 | 0.41 | 0.21 | 0.27 | 0.6 | 0.31 | 0.47 |
| 5 | 94495370 | rs331573 | -0.036 | -0.328 | -0.577 | 0.574 | -0.029 | 0.041 | 0.041 | 0 | 0.78 | 0.65 | 0.72 | 0.71 | 0.83 | 0.77 |
| 5 | 94501681 | rs331576 | 0.361 | -0.113 | -0.709 | 0.641 | 0.017 | -0.014 | -0.014 | 0 | 0.41 | 0.2 | 0.28 | 0.59 | 0.32 | 0.47 |
| 5 | 94508215 | rs6885525 | -0.26 | -0.163 | -0.451 | 0.119 | -0.01 | -0.076 | -0.076 | 0.477 | 0.14 | 0.32 | 0.15 | 0.02 | 0.12 | 0.06 |
| 5 | 94509008 | rs251076 | 1.189 | -0.244 | -0.881 | -0.07 | -0.124 | 0.082 | 0.082 | 0 | 0.29 | 0.08 | 0.42 | 0.12 | 0.53 | 0.29 |
| 5 | 94511026 | rs17426610 | 1.072 | 0.951 | -1.113 | -0.284 | -0.068 | 0.037 | 0.037 | 0 | 0.2 | 0.06 | 0.31 | 0.11 | 0.35 | 0.2 |
| 5 | 94519501 | rs10064312 | 1.261 | 0.014 | -1.165 | -0.537 | 0 | 0.012 | 0.012 | 0 | 0.24 | 0.18 | 0.15 | 0.12 | 0.19 | 0.19 |
| 5 | 94519583 | rs10071202 | 0.125 | 0.236 | -1.076 | -0.836 | 0 | 0.006 | 0.006 | 0 | 0.43 | 0.38 | 0.32 | 0.21 | 0.39 | 0.4 |
| 5 | 94523865 | rs251083 | 1.149 | 0.519 | -0.87 | -0.878 | 0.027 | -0.052 | -0.052 | 1.358 | 0.1 | 0.25 | 0.09 | 0 | 0.12 | 0.04 |
| 5 | 94538011 | rs6896478 | 1.682 | 0.497 | -1.306 | -0.839 | -0.003 | 0.022 | 0.022 | 0 | 0.22 | 0.12 | 0.12 | 0.12 | 0.18 | 0.16 |
| 5 | 94550976 | rs10058958 | 1.308 | 0.361 | -1.326 | -0.79 | 0 | 0.022 | 0.022 | 0 | 0.2 | 0.12 | 0.16 | 0.09 | 0.19 | 0.2 |
| 5 | 94558809 | rs10079049 | 1.496 | 0.107 | -1.211 | -0.785 | 0.001 | 0.01 | 0.01 | 0 | 0.17 | 0.11 | 0.1 | 0.09 | 0.16 | 0.17 |
| 5 | 94572738 | rs11949033 | 0.28 | -0.139 | -1.413 | -0.967 | 0.003 | 0.008 | 0.008 | 0 | 0.46 | 0.42 | 0.28 | 0.23 | 0.38 | 0.39 |
| 5 | 94572930 | rs17084533 | 2.247 | 0.1 | -1.328 | -0.026 | -0.047 | -0.053 | -0.053 | 0 | 0.09 | 0.01 | 0.29 | 0.41 | 0.14 | 0.3 |
| 5 | 94575327 | rs385225 | 0.317 | -0.127 | -1.136 | -0.039 | 0.04 | 0.092 | 0.092 | 0 | 0.22 | 0.41 | 0.43 | 0.47 | 0.3 | 0.35 |
| 5 | 94576589 | rs10059467 | 0.331 | -0.57 | -1.338 | 0.038 | 0.004 | 0.007 | 0.007 | 0 | 0.63 | 0.44 | 0.42 | 0.36 | 0.58 | 0.49 |
| 5 | 94576946 | rs17349336 | 2.214 | -0.885 | -1.266 | -0.218 | -0.044 | -0.008 | -0.008 | 0 | 0.12 | 0.01 | 0.08 | 0.09 | 0.15 | 0.13 |
| 5 | 94578715 | rs394273 | -1.328 | -0.887 | -1.232 | -0.211 | 0.012 | -0.002 | -0.002 | 0 | 0.84 | 0.88 | 0.84 | 0.82 | 0.86 | 0.85 |
| 5 | 94586795 | rs154064 | 1.079 | -0.404 | -1.113 | -0.572 | 0.001 | 0.022 | 0.022 | 0 | 0.17 | 0.07 | 0.09 | 0.09 | 0.16 | 0.17 |
| 5 | 94606247 | rs448433 | -1.594 | -0.274 | -1.168 | -0.499 | -0.006 | 0.01 | 0.01 | 0 | 0.85 | 0.93 | 0.91 | 0.88 | 0.85 | 0.86 |
| 5 | 94619232 | rs33809 | 0.29 | -0.584 | -1.118 | -0.656 | 0.008 | 0.008 | 0.008 | 0.378 | 0.55 | 0.45 | 0.64 | 0.82 | 0.66 | 0.67 |
| 5 | 94625604 | rs26395 | -0.416 | -0.288 | -1.323 | -0.502 | -0.002 | -0.011 | -0.011 | 0.237 | 0.74 | 0.73 | 0.87 | 0.87 | 0.81 | 0.79 |
| 5 | 94637098 | rs154055 | -0.69 | -0.255 | -1.367 | -0.512 | 0.008 | 0.02 | 0.02 | 0 | 0.76 | 0.83 | 0.89 | 0.87 | 0.81 | 0.8 |
| 5 | 94642199 | rs10059732 | 1.248 | 0.074 | -1.681 | -0.337 | -0.064 | -0.044 | -0.044 | 3.936 | 0.17 | 0.02 | 0.23 | 0.17 | 0.23 | 0.11 |
| 5 | 94643766 | rs17084574 | 1.458 | 0.033 | -0.963 | 0.018 | 0.001 | 0.052 | 0.052 | 0 | 0.09 | 0.12 | 0.2 | 0.35 | 0.12 | 0.34 |
| 5 | 94654187 | rs255983 | 0.12 | -0.094 | -1 | -0.998 | -0.003 | -0.022 | -0.022 | 0.241 | 0.72 | 0.71 | 0.87 | 0.87 | 0.78 | 0.8 |
| 5 | 94666593 | rs255967 | 0.157 | 0.172 | -1.259 | -0.779 | -0.004 | 0.021 | 0.021 | 0 | 0.72 | 0.78 | 0.81 | 0.84 | 0.71 | 0.79 |
| 5 | 94668415 | rs403916 | 0.139 | 0.169 | -1.26 | -0.78 | -0.004 | 0.022 | 0.022 | 0 | 0.73 | 0.78 | 0.81 | 0.84 | 0.71 | 0.79 |
| 5 | 94681918 | rs411788 | -0.248 | 0.157 | -1.315 | -0.813 | -0.02 | 0.026 | 0.026 | 0 | 0.77 | 0.88 | 0.78 | 0.83 | 0.72 | 0.81 |
| 5 | 94688036 | rs2434250 | 0.032 | -0.151 | -1.133 | -0.927 | -0.002 | 0.019 | 0.019 | 0 | 0.75 | 0.83 | 0.77 | 0.83 | 0.73 | 0.81 |
| 5 | 94690178 | rs253710 | -0.655 | -0.102 | -1.229 | -0.919 | -0.047 | 0.019 | 0.019 | 0 | 0.8 | 0.96 | 0.8 | 0.84 | 0.75 | 0.84 |
| 5 | 94708550 | rs1363428 | -0.318 | -0.202 | -1.068 | -0.972 | -0.013 | 0.032 | 0.032 | 0 | 0.78 | 0.94 | 0.8 | 0.83 | 0.75 | 0.83 |
| 5 | 94715056 | rs2432175 | 0.699 | -0.334 | -1.537 | -1.264 | 0.003 | 0.011 | 0.011 | 0.027 | 0.39 | 0.72 | 0.41 | 0.64 | 0.36 | 0.58 |
| 5 | 94716149 | rs10077532 | 0.7 | -0.21 | -1.748 | -1.302 | -0.059 | 0.007 | 0.007 | 0 | 0.23 | 0.04 | 0.26 | 0.16 | 0.28 | 0.13 |
| 5 | 94725223 | rs181880 | -1.243 | -0.167 | -1.678 | -1.18 | -0.062 | -0.013 | -0.013 | 0 | 0.81 | 0.99 | 0.82 | 0.83 | 0.78 | 0.83 |
| 5 | 94726879 | rs2560266 | 0.005 | 0.01 | -1.457 | -1.07 | -0.011 | 0.049 | 0.049 | 0 | 0.36 | 0.2 | 0.4 | 0.2 | 0.4 | 0.24 |
| 5 | 94731347 | rs33894 | 0.963 | -0.079 | -1.316 | -1.127 | -0.001 | -0.006 | -0.006 | 0 | 0.39 | 0.52 | 0.38 | 0.64 | 0.36 | 0.56 |
| 5 | 94733704 | rs2548655 | -0.506 | 0.013 | -1.139 | -0.954 | -0.005 | 0.03 | 0.03 | 0 | 0.36 | 0.26 | 0.41 | 0.2 | 0.4 | 0.24 |
| 5 | 94734559 | rs1469393 | -0.342 | -0.393 | -0.724 | -0.892 | -0.027 | -0.075 | -0.075 | 0 | 0.24 | 0.31 | 0.2 | 0.04 | 0.21 | 0.15 |
| 5 | 94738448 | rs253720 | 0.074 | 0.051 | -0.586 | -0.481 | 0.001 | 0.041 | 0.041 | 0 | 0.52 | 0.4 | 0.58 | 0.69 | 0.49 | 0.7 |
| 5 | 94741837 | rs253715 | -0.416 | 0.319 | -0.381 | -0.63 | 0.007 | -0.013 | -0.013 | 0 | 0.8 | 0.73 | 0.82 | 0.82 | 0.81 | 0.85 |
| 5 | 94743198 | rs12522074 | 0.747 | 0.475 | -0.745 | -0.125 | -0.011 | -0.022 | -0.022 | 1.651 | 0.1 | 0.04 | 0.15 | 0.35 | 0.13 | 0.31 |
| 5 | 94744042 | rs10515225 | 0.928 | 0.659 | -0.502 | -1.058 | -0.005 | 0.01 | 0.01 | 0 | 0.26 | 0.24 | 0.23 | 0.13 | 0.32 | 0.14 |
| 5 | 94746513 | rs6870030 | 1.042 | 0.574 | -0.485 | -1.07 | -0.03 | 0.053 | 0.053 | 0 | 0.25 | 0.15 | 0.22 | 0.11 | 0.32 | 0.15 |
| 5 | 94746938 | rs6872033 | 0.903 | 0.763 | -0.578 | -0.943 | -0.008 | 0.021 | 0.021 | 0 | 0.28 | 0.24 | 0.24 | 0.11 | 0.33 | 0.16 |
| 5 | 94749288 | rs2546098 | 1.124 | 0.614 | -0.763 | -1.03 | -0.045 | 0.035 | 0.035 | 0 | 0.2 | 0.04 | 0.29 | 0.08 | 0.23 | 0.23 |

| | | | | | | | | | | | | | | | | |
|---|---|---|---|---|---|---|---|---|---|---|---|---|---|---|---|---|
| 5 | 94749723 | rs7726891 | 0.52 | 0.901 | -0.104 | -0.791 | -0.008 | 0.021 | 0.021 | 0 | 0.28 | 0.24 | 0.24 | 0.11 | 0.33 | 0.16 |
| 5 | 94751393 | rs1363430 | 0.758 | 0.956 | 0 | -0.724 | 0.006 | 0.028 | 0.028 | 0 | 0.5 | 0.43 | 0.57 | 0.73 | 0.46 | 0.69 |
| 5 | 94753303 | rs10077851 | -1.031 | 0.732 | 0.561 | 0.117 | 0.003 | -0.031 | -0.031 | 0 | 0.21 | 0.33 | 0.17 | 0.16 | 0.19 | 0.14 |
| 5 | 94756496 | rs17084686 | -1.273 | 0.762 | 0.447 | 0.076 | -0.01 | -0.05 | -0.05 | 0 | 0.23 | 0.5 | 0.19 | 0.16 | 0.18 | 0.15 |
| 5 | 94756610 | rs17084688 | -1.479 | 0.775 | 0.44 | 0.111 | 0.001 | -0.06 | -0.06 | 0 | 0.27 | 0.52 | 0.22 | 0.16 | 0.26 | 0.17 |
| 5 | 94757684 | rs9314136 | -1.487 | 0.852 | 0.289 | 0.276 | 0.017 | -0.03 | -0.03 | 0 | 0.2 | 0.42 | 0.19 | 0.14 | 0.22 | 0.16 |
| 5 | 94758774 | rs6894872 | -1.273 | 0.883 | -0.219 | 0.29 | 0.005 | -0.018 | -0.018 | 0 | 0.17 | 0.26 | 0.17 | 0.12 | 0.18 | 0.14 |
| 5 | 94759717 | rs17084706 | -0.805 | 0.644 | 0.22 | 0.421 | 0.033 | -0.015 | -0.015 | 0 | 0.06 | 0.24 | 0.03 | 0 | 0.05 | 0.04 |
| 5 | 94762306 | rs2162716 | 0.243 | 0.601 | -0.725 | 0.702 | 0.001 | -0.017 | -0.017 | 0 | 0.68 | 0.78 | 0.61 | 0.33 | 0.7 | 0.49 |
| 5 | 94778775 | rs6556871 | -0.449 | -0.812 | -1.215 | 0.564 | 0.162 | 0.013 | 0.013 | 0.054 | 0.48 | 0.86 | 0.31 | 0.41 | 0.26 | 0.4 |
| 5 | 94787954 | rs3909479 | 0.192 | -1.623 | -0.377 | 0.045 | -0.034 | -0.096 | -0.096 | 0 | 0.19 | 0.32 | 0.1 | 0.01 | 0.1 | 0.08 |
| 5 | 94789503 | rs7725219 | -0.734 | -1.885 | -0.202 | 0.185 | -0.06 | -0.081 | -0.081 | 2.174 | 0.38 | 0.47 | 0.18 | 0.12 | 0.17 | 0.24 |
| 5 | 94796041 | rs2731818 | -1.381 | -1.138 | -0.308 | 0.259 | 0.107 | 0.007 | 0.007 | 0.165 | 0.48 | 0.84 | 0.34 | 0.46 | 0.32 | 0.41 |
| 5 | 94798893 | rs9314145 | -0.663 | -1.145 | -0.218 | 0.241 | -0.055 | -0.081 | -0.081 | 0 | 0.38 | 0.47 | 0.19 | 0.12 | 0.18 | 0.24 |
| 5 | 94807358 | rs10866765 | -0.773 | -1.133 | 0.133 | 0.238 | -0.029 | -0.055 | -0.055 | 0.209 | 0.39 | 0.52 | 0.2 | 0.12 | 0.2 | 0.24 |
| 5 | 94811454 | rs2731816 | -1.171 | -0.639 | -0.787 | 0.047 | -0.011 | 0.004 | 0.004 | 1.431 | 0.8 | 0.88 | 0.78 | 0.89 | 0.78 | 0.81 |
| 5 | 94822053 | rs12516085 | 1.432 | -0.666 | -0.81 | 0.061 | -0.178 | -0.128 | -0.128 | 1.527 | 0.31 | 0.02 | 0.44 | 0.43 | 0.46 | 0.4 |
| 5 | 94826655 | rs2303650 | 0.022 | -0.935 | -0.59 | -0.105 | -0.026 | -0.043 | -0.043 | 0.008 | 0.35 | 0.4 | 0.18 | 0.12 | 0.17 | 0.2 |
| 5 | 94858574 | rs17084869 | 0.975 | -0.947 | -0.528 | -0.135 | -0.024 | -0.093 | -0.093 | 0 | 0.19 | 0.31 | 0.11 | 0.01 | 0.11 | 0.08 |
| 5 | 94865164 | rs9314148 | 0.245 | -0.883 | -0.445 | -0.08 | 0.07 | 0.22 | 0.22 | 0 | 0.09 | 0.28 | 0.14 | 0.34 | 0.14 | 0.17 |
| 5 | 94885383 | rs2881773 | -0.296 | -0.364 | -1.026 | -0.094 | 0 | -0.003 | -0.003 | 0 | 0.57 | 0.59 | 0.45 | 0.35 | 0.47 | 0.44 |
| 5 | 94890124 | rs2731826 | 0.339 | -0.955 | -0.174 | 0.049 | 0.067 | 0.184 | 0.184 | 0 | 0.08 | 0.2 | 0.13 | 0.33 | 0.14 | 0.17 |
| 5 | 94900752 | rs7737266 | 0.998 | -1.76 | -0.415 | -0.456 | 0.021 | 0.009 | 0.009 | 0 | 0.14 | 0.08 | 0.06 | 0.11 | 0.06 | 0.11 |
| 5 | 94919966 | rs6866194 | -0.491 | -1.719 | -0.488 | -0.443 | -0.041 | -0.064 | -0.064 | 0.169 | 0.38 | 0.49 | 0.18 | 0.12 | 0.18 | 0.23 |
| 5 | 94939193 | rs17084927 | 1.013 | -1.639 | -0.707 | -0.416 | 0.043 | 0.015 | 0.015 | 0 | 0.14 | 0.04 | 0.06 | 0.11 | 0.06 | 0.11 |
| 5 | 94939777 | rs10491246 | -0.571 | -1.615 | -0.693 | -0.425 | -0.03 | -0.054 | -0.054 | 6.486 | 0.39 | 0.57 | 0.2 | 0.12 | 0.18 | 0.23 |
| 5 | 94942780 | rs17084933 | 0.301 | -0.915 | -0.497 | 0.305 | -0.018 | -0.009 | -0.009 | 0 | 0.16 | 0.19 | 0.07 | 0.12 | 0.06 | 0.1 |
| 5 | 94949253 | rs10075788 | -0.244 | -0.714 | -0.544 | 0.554 | 0.008 | -0.002 | -0.002 | 0 | 0.45 | 0.59 | 0.42 | 0.57 | 0.37 | 0.55 |
| 5 | 94952136 | rs6556877 | -0.335 | -0.699 | -1.016 | 0.307 | 0.088 | 0.124 | 0.124 | 0 | 0.5 | 0.81 | 0.51 | 0.72 | 0.38 | 0.52 |
| 5 | 94955602 | rs2194051 | -0.698 | -0.306 | -0.502 | 0.605 | -0.001 | 0.043 | 0.043 | 0 | 0.19 | 0.31 | 0.26 | 0.48 | 0.19 | 0.29 |
| 5 | 94959876 | rs7731790 | -0.374 | 0.379 | -1.145 | -0.265 | -0.044 | 0.045 | 0.045 | 0 | 0.7 | 0.89 | 0.64 | 0.89 | 0.6 | 0.67 |
| 5 | 94975592 | rs6890832 | 0.15 | -0.317 | -0.437 | -0.391 | 0.001 | 0.001 | 0.001 | 0 | 0.53 | 0.57 | 0.38 | 0.34 | 0.39 | 0.37 |
| 5 | 94976651 | rs11747129 | 1.556 | -0.214 | -0.721 | -0.809 | -0.013 | 0.027 | 0.027 | 0 | 0.06 | 0 | 0.03 | 0 | 0.07 | 0.06 |
| 5 | 94987809 | rs17084946 | 1 | -0.837 | 0.15 | -1.171 | -0.02 | -0.025 | -0.025 | 0 | 0.05 | 0.09 | 0.03 | 0 | 0.03 | 0.01 |
| 5 | 94996186 | rs412026 | 0.399 | -1.149 | 0.217 | -1.361 | 0.017 | 0.025 | 0.025 | 0 | 0.43 | 0.32 | 0.34 | 0.35 | 0.36 | 0.35 |
| 5 | 94998972 | rs150548 | -0.439 | -1.187 | 0.064 | -1.377 | 0.006 | 0.025 | 0.025 | 0 | 0.57 | 0.68 | 0.43 | 0.35 | 0.43 | 0.42 |
| 5 | 95002121 | rs213512 | 0.734 | -1.282 | -0.081 | -1.409 | -0.001 | 0.024 | 0.024 | 0 | 0.39 | 0.17 | 0.31 | 0.35 | 0.35 | 0.34 |
| 5 | 95006039 | rs34902 | 1.224 | -0.216 | -0.355 | -1.459 | -0.031 | -0.009 | -0.009 | 0 | 0.12 | 0.06 | 0.11 | 0.11 | 0.18 | 0.24 |
| 5 | 95008959 | rs461873 | -0.97 | -0.59 | -0.294 | -1.576 | -0.032 | -0.034 | -0.034 | 0 | 0.68 | 0.85 | 0.55 | 0.42 | 0.61 | 0.61 |
| 5 | 95011426 | rs2548125 | -0.001 | -0.631 | -0.462 | -1.815 | -0.012 | -0.001 | -0.001 | 0.225 | 0.3 | 0.3 | 0.29 | 0.22 | 0.41 | 0.37 |
| 5 | 95018382 | rs34895 | 0.303 | -0.577 | -0.325 | -1.765 | -0.075 | -0.022 | -0.022 | 3.308 | 0.27 | 0.12 | 0.27 | 0.23 | 0.41 | 0.36 |
| 5 | 95018813 | rs2158095 | 1.486 | -0.641 | -0.342 | -1.803 | 0.013 | 0.013 | 0.013 | 0.098 | 0.25 | 0.22 | 0.16 | 0.2 | 0.12 | 0.17 |
| 5 | 95018939 | rs460697 | -0.144 | -0.596 | -0.304 | -1.778 | 0 | -0.002 | -0.002 | 0 | 0.56 | 0.53 | 0.46 | 0.42 | 0.53 | 0.55 |
| 5 | 95023545 | rs10053879 | 0.588 | -0.439 | -0.815 | -1.637 | -0.071 | -0.074 | -0.074 | 0 | 0.33 | 0.44 | 0.21 | 0.2 | 0.16 | 0.19 |
| 5 | 95026190 | rs4292466 | 0.792 | -0.369 | -1.305 | -1.602 | -0.022 | -0.024 | -0.024 | 0.835 | 0.32 | 0.37 | 0.2 | 0.2 | 0.15 | 0.19 |
| 5 | 95027958 | rs12658215 | 1.18 | 0.287 | -0.945 | -1.189 | 0.003 | 0.173 | 0.173 | 0 | 0.07 | 0.11 | 0.15 | 0.54 | 0.09 | 0.18 |
| 5 | 95033841 | rs169883 | -0.646 | 0.312 | -1.355 | -1.634 | -0.002 | 0.016 | 0.016 | 0.038 | 0.42 | 0.61 | 0.49 | 0.76 | 0.55 | 0.61 |
| 5 | 95037003 | rs150543 | 0.316 | 0.516 | -1.703 | -1.058 | -0.071 | -0.04 | -0.04 | 0.179 | 0.15 | 0.03 | 0.16 | 0.13 | 0.23 | 0.14 |
| 5 | 95041668 | rs2548130 | -0.299 | 0.584 | -1.368 | -1.365 | -0.066 | 0.152 | 0.152 | 0 | 0.68 | 0.93 | 0.62 | 0.96 | 0.63 | 0.8 |
| 5 | 95060244 | rs2545638 | -0.213 | 0.202 | -0.499 | -1.251 | 0.003 | -0.011 | -0.011 | 0 | 0.51 | 0.51 | 0.47 | 0.91 | 0.46 | 0.66 |
| 5 | 95070815 | rs185312 | -0.013 | -0.036 | -0.145 | -0.839 | 0.006 | 0.031 | 0.031 | 0 | 0.43 | 0.29 | 0.34 | 0.69 | 0.43 | 0.56 |
| 5 | 95084131 | rs34896 | 0.074 | 0.144 | -0.458 | -0.769 | -0.005 | 0.034 | 0.034 | 0 | 0.41 | 0.28 | 0.33 | 0.69 | 0.43 | 0.55 |
| 5 | 95088559 | rs1984965 | 0.413 | 0.119 | -0.438 | -0.733 | 0 | -0.005 | -0.005 | 0 | 0.05 | 0.05 | 0.02 | 0.12 | 0.04 | 0.16 |
| 5 | 95089373 | rs11745143 | 1.511 | 0.093 | -0.553 | -0.834 | -0.015 | 0.028 | 0.028 | 0 | 0.06 | 0 | 0.04 | 0 | 0.06 | 0.06 |
| 5 | 95093084 | rs34900 | 0.097 | 0.077 | -0.204 | -0.712 | 0 | 0.024 | 0.024 | 0 | 0.41 | 0.35 | 0.34 | 0.69 | 0.43 | 0.55 |
| 5 | 95094812 | rs11959859 | 0.181 | 0.258 | -0.16 | -0.65 | 0 | -0.008 | -0.008 | 0.039 | 0.49 | 0.44 | 0.61 | 0.19 | 0.53 | 0.28 |
| 5 | 95096562 | rs7729314 | 0.251 | 0.444 | -0.493 | -0.705 | 0.006 | 0.023 | 0.023 | 0 | 0.5 | 0.5 | 0.61 | 0.21 | 0.54 | 0.33 |
| 5 | 95108891 | rs13188464 | 0.197 | 0.451 | -0.542 | -0.708 | 0.001 | 0.022 | 0.022 | 0 | 0.51 | 0.51 | 0.62 | 0.19 | 0.52 | 0.28 |
| 5 | 95112866 | rs3777218 | 0.446 | 0.558 | -0.539 | -0.58 | 0.004 | -0.021 | -0.021 | 0 | 0.56 | 0.5 | 0.45 | 0.31 | 0.53 | 0.57 |
| 5 | 95116543 | rs3777217 | 0.886 | 0.062 | -0.504 | -1.153 | -0.003 | -0.028 | -0.028 | 0 | 0.41 | 0.47 | 0.36 | 0.26 | 0.37 | 0.33 |
| 5 | 95119408 | rs888808 | 0.668 | 0.115 | -0.359 | -1.15 | 0.008 | 0.038 | 0.038 | 0 | 0.44 | 0.61 | 0.38 | 0.26 | 0.38 | 0.34 |
| 5 | 95119842 | rs11738579 | -0.284 | 0.29 | -0.664 | -0.903 | 0.004 | 0.022 | 0.022 | 0 | 0.23 | 0.19 | 0.23 | 0.18 | 0.21 | 0.26 |
| 5 | 95122000 | rs3777215 | -0.236 | 0.283 | -0.722 | -0.909 | 0.002 | 0.033 | 0.033 | 0 | 0.23 | 0.16 | 0.22 | 0.19 | 0.21 | 0.27 |
| 5 | 95124328 | rs3822753 | 0.056 | 0.277 | -0.678 | -0.856 | -0.017 | -0.021 | -0.021 | 0 | 0.69 | 0.78 | 0.6 | 0.44 | 0.59 | 0.6 |
| 5 | 95133586 | rs6871090 | 0.053 | 0.281 | -0.677 | -0.892 | -0.017 | -0.02 | -0.02 | 0 | 0.69 | 0.78 | 0.6 | 0.44 | 0.59 | 0.6 |
| 5 | 95138353 | rs6556882 | -1.22 | -0.029 | -0.404 | -0.627 | 0.029 | 0.03 | 0.03 | 0.015 | 0.45 | 0.41 | 0.32 | 0.3 | 0.37 | 0.44 |
| 5 | 95140046 | rs6894498 | 0.165 | 0.156 | -0.214 | -1.334 | -0.017 | -0.06 | -0.06 | 0 | 0.17 | 0.19 | 0.07 | 0 | 0.13 | 0.04 |
| 5 | 95142318 | rs9314158 | -0.287 | 0.138 | -0.398 | -1.384 | -0.018 | 0.056 | 0.056 | 0 | 0.19 | 0.09 | 0.2 | 0.05 | 0.19 | 0.2 |
| 5 | 95143598 | rs11135434 | 1.3 | 0.464 | 0.2 | -1.318 | 0.038 | -0.015 | -0.015 | 0 | 0.65 | 0.52 | 0.44 | 0.35 | 0.56 | 0.46 |

| | | | | | | | | | | | | | | | | |
|---|---|---|---|---|---|---|---|---|---|---|---|---|---|---|---|---|
| 5 | 95143657 | rs12518592 | -0.14 | 0.295 | 0.442 | -1.308 | -0.044 | -0.085 | -0.085 | 0.387 | 0.2 | 0.27 | 0.07 | 0 | 0.12 | 0.03 |
| 5 | 95143681 | rs7736948 | -0.75 | 0.751 | -0.049 | -1.06 | 0.008 | 0.019 | 0.019 | 0.13 | 0.89 | 0.93 | 0.67 | 0.42 | 0.76 | 0.77 |
| 5 | 95144677 | rs885303 | -0.905 | -0.006 | -0.238 | -1.81 | 0.005 | 0.171 | 0.171 | 1.152 | 0.35 | 0.11 | 0.26 | 0.05 | 0.33 | 0.23 |
| 5 | 95148129 | rs7700814 | -0.91 | -0.385 | 0.398 | -1.899 | -0.057 | -0.157 | -0.157 | 0 | 0.31 | 0.57 | 0.25 | 0.01 | 0.19 | 0.11 |
| 5 | 95150245 | rs1047420 | 1.757 | 0.018 | -0.181 | -1.751 | -0.032 | 0.12 | 0.12 | 0.114 | 0.38 | 0.1 | 0.23 | 0.18 | 0.37 | 0.33 |
| 5 | 95152313 | rs4561 | 1.44 | 0.059 | 0.563 | -1.783 | 0.002 | 0.027 | 0.027 | 1.02 | 0.44 | 0.32 | 0.28 | 0.17 | 0.41 | 0.32 |
| 5 | 95156457 | rs9314160 | -1.562 | 0.55 | 0.637 | -1.717 | 0.026 | 0.058 | 0.058 | 0 | 0.52 | 0.61 | 0.55 | 0.8 | 0.59 | 0.54 |
| 5 | 95158768 | rs871775 | -0.1 | -0.537 | 0.849 | -1.494 | 0.033 | 0.04 | 0.04 | 6.21 | 0.19 | 0.07 | 0.07 | 0 | 0.13 | 0.05 |
| 5 | 95159822 | rs3756705 | -1.678 | -0.336 | 1.079 | -1.38 | -0.019 | -0.079 | -0.079 | 0 | 0.39 | 0.48 | 0.2 | 0.13 | 0.31 | 0.33 |
| 5 | 95161340 | rs6556884 | -1.123 | -0.367 | 1.052 | -1.409 | -0.025 | -0.148 | -0.148 | 0.193 | 0.31 | 0.46 | 0.17 | 0.04 | 0.25 | 0.13 |
| 5 | 95161745 | rs12513728 | -0.454 | -0.105 | 1.303 | -0.853 | -0.003 | -0.001 | -0.001 | 0 | 0.09 | 0.1 | 0.04 | 0.09 | 0.06 | 0.2 |
| 5 | 95163957 | rs6876015 | -2.006 | -0.149 | 1.448 | -0.924 | -0.016 | -0.047 | -0.047 | 0 | 0.31 | 0.37 | 0.21 | 0.11 | 0.24 | 0.35 |
| 5 | 95169909 | rs3797207 | -1.516 | 0.102 | 1.71 | -0.728 | -0.011 | -0.044 | -0.044 | 1.664 | 0.21 | 0.4 | 0.13 | 0.14 | 0.15 | 0.28 |
| 5 | 95170719 | rs10059138 | -0.091 | 0.146 | 1.628 | -0.73 | 0.011 | -0.031 | -0.031 | 0 | 0.09 | 0.18 | 0.07 | 0.04 | 0.09 | 0.08 |
| 5 | 95172281 | rs2546197 | -0.404 | 0.11 | 1.513 | -0.777 | -0.002 | -0.026 | -0.026 | 0.122 | 0.11 | 0.12 | 0.04 | 0 | 0.09 | 0.04 |
| 5 | 95173887 | rs10040697 | -1.041 | 0.217 | 1.644 | -0.649 | 0.012 | 0.065 | 0.065 | 0 | 0.15 | 0.33 | 0.31 | 0.69 | 0.16 | 0.25 |
| 5 | 95174319 | rs2547996 | -2.941 | 0.14 | 1.579 | -0.731 | 0.024 | 0.09 | 0.09 | 0 | 0.44 | 0.81 | 0.53 | 0.78 | 0.39 | 0.58 |
| 5 | 95177641 | rs6420125 | -1.883 | 0.07 | 1.53 | -0.964 | -0.015 | -0.071 | -0.071 | 0.159 | 0.23 | 0.6 | 0.14 | 0.13 | 0.17 | 0.29 |
| 5 | 95185472 | rs11135439 | -1.038 | 0.164 | 0.805 | -0.907 | 0.005 | 0.001 | 0.001 | 0.006 | 0.16 | 0.12 | 0.15 | 0 | 0.17 | 0.14 |
| 5 | 95189115 | rs10515230 | -0.356 | 0.142 | 0.892 | -0.724 | 0.013 | -0.034 | -0.034 | 0 | 0.1 | 0.23 | 0.07 | 0.04 | 0.1 | 0.08 |
| 5 | 95190176 | rs6883552 | -1.732 | 0.161 | 0.757 | -0.704 | -0.021 | -0.076 | -0.076 | 0.004 | 0.32 | 0.5 | 0.16 | 0.13 | 0.24 | 0.33 |
| 5 | 95192217 | rs889300 | -1.194 | -0.063 | 0.338 | -0.412 | 0.001 | -0.002 | -0.002 | 0 | 0.21 | 0.2 | 0.08 | 0.09 | 0.14 | 0.25 |
| 5 | 95193740 | rs2546193 | -1.635 | 0.066 | 0.779 | 0.018 | -0.029 | 0.049 | 0.049 | 0.141 | 0.23 | 0.49 | 0.33 | 0.69 | 0.19 | 0.4 |
| 5 | 95195558 | rs6873611 | -0.814 | 1.085 | 0.417 | 0.457 | -0.052 | 0.065 | 0.065 | 0 | 0.17 | 0.09 | 0.21 | 0.04 | 0.26 | 0.09 |
| 5 | 95196856 | rs12659828 | -1.712 | 1.311 | 1.117 | 0.893 | 0.028 | 0.062 | 0.062 | 0 | 0.24 | 0.36 | 0.42 | 0.6 | 0.33 | 0.24 |
| 5 | 95198252 | rs154447 | 0.867 | 1.593 | 1.161 | 0.304 | 0.022 | -0.109 | -0.109 | 1.389 | 0.78 | 0.58 | 0.76 | 0.98 | 0.71 | 0.68 |
| 5 | 95198353 | rs154448 | 0.208 | 1.65 | 1.327 | 0.284 | 0.006 | -0.044 | -0.044 | 0 | 0.85 | 0.77 | 0.81 | 0.98 | 0.8 | 0.77 |
| 5 | 95200112 | rs17464446 | 0.183 | 1.666 | 1.063 | 0.31 | 0.01 | -0.025 | -0.025 | 0 | 0.06 | 0.1 | 0.05 | 0 | 0.08 | 0.09 |
| 5 | 95201703 | rs248341 | 1.03 | 1.485 | 0.726 | 0.311 | 0.016 | 0.097 | 0.097 | 0.107 | 0.72 | 0.49 | 0.56 | 0.43 | 0.66 | 0.52 |
| 5 | 95202569 | rs248340 | 2.025 | 1.528 | 0.296 | 0.279 | 0.033 | 0.001 | 0.001 | 0 | 0.52 | 0.27 | 0.32 | 0.39 | 0.37 | 0.43 |
| 5 | 95203134 | rs1004654 | -0.839 | 1.685 | -0.215 | 0.183 | -0.066 | 0.07 | 0.07 | 0 | 0.18 | 0.1 | 0.23 | 0.04 | 0.29 | 0.09 |
| 5 | 95209382 | rs154451 | 0.709 | 0.616 | -0.321 | 0.309 | -0.001 | 0.006 | 0.006 | 1.769 | 0.46 | 0.37 | 0.41 | 0.39 | 0.5 | 0.53 |
| 5 | 95209557 | rs154452 | 0.195 | 0.651 | -0.364 | 0.444 | -0.015 | 0.018 | 0.018 | 0 | 0.84 | 0.88 | 0.79 | 0.92 | 0.77 | 0.85 |
| 5 | 95219145 | rs17085224 | 0.331 | 0.031 | -0.298 | 0.434 | -0.007 | 0.004 | 0.004 | 2.906 | 0.09 | 0.07 | 0.08 | 0.03 | 0.01 | 0.09 |
| 5 | 95219261 | rs147295 | 1.885 | -0.326 | 0.202 | 0.485 | 0.008 | 0.008 | 0.008 | 0 | 0.61 | 0.35 | 0.65 | 0.62 | 0.68 | 0.54 |
| 5 | 95221337 | rs1043381 | -1.947 | -0.285 | 0.532 | 0.544 | -0.015 | -0.005 | -0.005 | 0.165 | 0.33 | 0.56 | 0.33 | 0.38 | 0.28 | 0.44 |
| 5 | 95222156 | rs17085231 | -1.321 | -0.288 | 0.725 | 0.538 | 0.02 | 0.02 | 0.02 | 0 | 0.24 | 0.45 | 0.25 | 0.35 | 0.28 | 0.36 |
| 5 | 95222511 | rs11135442 | -1.416 | -0.368 | 0.689 | 0.5 | 0.009 | 0.009 | 0.009 | 0 | 0.26 | 0.51 | 0.25 | 0.35 | 0.28 | 0.36 |
| 5 | 95231115 | rs3777207 | -0.136 | -0.337 | 0.671 | 0.393 | -0.017 | 0.01 | 0.01 | 0 | 0.13 | 0.1 | 0.19 | 0.11 | 0.21 | 0.14 |
| 5 | 95236415 | rs17085249 | -1.444 | -0.321 | 1.092 | 0.522 | 0.012 | 0.064 | 0.064 | 0 | 0.24 | 0.37 | 0.24 | 0.43 | 0.28 | 0.35 |
| 5 | 95241127 | rs11741590 | -2.249 | -0.43 | 1.027 | 0.626 | -0.004 | 0.005 | 0.005 | 0 | 0.34 | 0.62 | 0.33 | 0.46 | 0.29 | 0.43 |
| 5 | 95245384 | rs3777194 | -2.249 | -0.43 | 1.086 | 0.626 | -0.002 | 0.001 | 0.001 | 0 | 0.34 | 0.63 | 0.33 | 0.46 | 0.29 | 0.43 |
| 5 | 95253295 | rs1458018 | -2.256 | -0.436 | 1.083 | 0.623 | -0.001 | 0.001 | 0.001 | 0 | 0.34 | 0.63 | 0.33 | 0.46 | 0.29 | 0.43 |
| 5 | 95255724 | rs1423269 | -1.59 | -0.331 | 1.021 | 0.662 | 0.016 | 0.086 | 0.086 | 0 | 0.24 | 0.43 | 0.24 | 0.43 | 0.28 | 0.35 |
| 5 | 95259862 | rs3777185 | -1.653 | -0.328 | 0.996 | 0.664 | 0.017 | 0.099 | 0.099 | 0 | 0.25 | 0.52 | 0.25 | 0.43 | 0.28 | 0.35 |
| 5 | 95268768 | rs3777179 | -0.958 | 0.063 | 1.283 | 0.801 | 0.053 | 0.136 | 0.136 | 0 | 0.17 | 0.37 | 0.22 | 0.4 | 0.24 | 0.23 |
| 5 | 95277555 | rs3777175 | -1.071 | -0.156 | 1.174 | 0.755 | 0.057 | 0.167 | 0.167 | 0 | 0.18 | 0.44 | 0.26 | 0.39 | 0.22 | 0.24 |
| 5 | 95324375 | rs7705279 | -1.957 | -0.28 | 1.213 | 0.692 | -0.002 | 0.003 | 0.003 | 0 | 0.28 | 0.66 | 0.31 | 0.42 | 0.23 | 0.33 |
| 5 | 95351767 | rs12523306 | -0.115 | -0.605 | 1.248 | 0.56 | -0.056 | -0.027 | -0.027 | 0 | 0.11 | 0.19 | 0.09 | 0.03 | 0.01 | 0.09 |
| 5 | 95357625 | rs6891871 | -1.61 | -0.754 | 1.414 | 1.133 | -0.086 | 0.169 | 0.169 | 4.502 | 0.18 | 0.54 | 0.23 | 0.37 | 0.06 | 0.23 |
| 5 | 95373029 | rs17085426 | 0.485 | -0.757 | 1.55 | 1.043 | -0.034 | -0.008 | -0.008 | 0.012 | 0.09 | 0.11 | 0.09 | 0.04 | 0.01 | 0.09 |
| 5 | 95378364 | rs9314163 | -0.19 | -0.986 | 0.852 | 0.246 | 0.006 | -0.007 | -0.007 | 0 | 0.58 | 0.62 | 0.55 | 0.53 | 0.63 | 0.65 |
| 5 | 95383587 | rs11951463 | 0.451 | -1.01 | 0.944 | 0.221 | 0.005 | 0.001 | 0.001 | 0.013 | 0.49 | 0.48 | 0.46 | 0.49 | 0.63 | 0.55 |
| 5 | 95388015 | rs17085428 | 0.778 | -0.572 | 0.542 | -0.07 | -0.034 | -0.034 | -0.034 | 0 | 0.14 | 0.1 | 0.2 | 0.37 | 0.28 | 0.2 |
| 5 | 95390839 | rs10428702 | 1.228 | -0.601 | 0.464 | -0.241 | -0.049 | -0.015 | -0.015 | 0 | 0.09 | 0.17 | 0.1 | 0.04 | 0.01 | 0.09 |
| 5 | 95400064 | rs903380 | -2.03 | -0.888 | 0.086 | -0.276 | -0.002 | 0 | 0 | 0.176 | 0.75 | 0.76 | 0.67 | 0.77 | 0.74 | 0.88 |
| 5 | 95401970 | rs4869256 | -0.884 | -0.522 | 0.336 | -0.042 | 0.012 | 0.001 | 0.001 | 0 | 0.59 | 0.66 | 0.56 | 0.6 | 0.64 | 0.66 |
| 5 | 95407719 | rs13177695 | 1.135 | -0.455 | 0.295 | -0.076 | -0.087 | 0.14 | 0.14 | 0 | 0.3 | 0.02 | 0.23 | 0.08 | 0.35 | 0.28 |
| 5 | 95407894 | rs7717348 | -0.056 | -0.405 | 0.43 | -0.043 | 0.092 | 0.184 | 0.184 | 0 | 0.19 | 0.36 | 0.23 | 0.48 | 0.29 | 0.28 |
| 5 | 95410393 | rs4128178 | -0.09 | -0.571 | 0.211 | 0.145 | 0.047 | 0.013 | 0.013 | 0 | 0.51 | 0.66 | 0.48 | 0.56 | 0.64 | 0.57 |
| 5 | 95423823 | rs13155375 | -0.687 | -0.585 | 0.427 | 0.05 | -0.003 | 0.002 | 0.002 | 0 | 0.59 | 0.6 | 0.5 | 0.52 | 0.65 | 0.67 |
| 5 | 95423944 | rs13172783 | -0.716 | -0.587 | 0.436 | 0.049 | -0.001 | 0 | 0 | 1.627 | 0.59 | 0.61 | 0.5 | 0.52 | 0.65 | 0.67 |
| 5 | 95431343 | rs6873504 | -0.708 | -0.635 | 0.732 | 0.038 | -0.013 | 0.015 | 0.015 | 0 | 0.59 | 0.54 | 0.48 | 0.5 | 0.64 | 0.68 |
| 5 | 95444969 | rs4869260 | -0.803 | -0.673 | 0.542 | 0.021 | 0.014 | -0.035 | -0.035 | 0 | 0.61 | 0.73 | 0.5 | 0.5 | 0.64 | 0.68 |
| 5 | 95447994 | rs6865159 | -1.624 | -0.669 | 0.52 | 0.064 | -0.009 | 0.012 | 0.012 | 1.447 | 0.71 | 0.9 | 0.66 | 0.75 | 0.72 | 0.84 |
| 5 | 95464398 | rs13164421 | 0.872 | -0.261 | 0.352 | -0.174 | -0.094 | 0.122 | 0.122 | 0 | 0.25 | 0.04 | 0.22 | 0.08 | 0.35 | 0.22 |
| 5 | 95470989 | rs6875999 | -0.483 | -0.415 | 0.598 | -0.187 | 0.024 | 0.062 | 0.062 | 0.014 | 0.26 | 0.43 | 0.24 | 0.4 | 0.28 | 0.37 |
| 5 | 95472410 | rs12716476 | 0.276 | -0.268 | 0.577 | -0.366 | 0.012 | -0.083 | -0.083 | 0.187 | 0.28 | 0.37 | 0.26 | 0.08 | 0.36 | 0.23 |
| 5 | 95492519 | rs4869267 | 0.009 | -0.215 | 0.353 | -0.444 | 0.009 | -0.058 | -0.058 | 0 | 0.28 | 0.31 | 0.25 | 0.08 | 0.35 | 0.22 |

| | | | | | | | | | | | | | | | | |
|---|---|---|---|---|---|---|---|---|---|---|---|---|---|---|---|---|
| 5 | 95502132 | rs4128935 | -0.277 | -0.237 | 0.347 | -0.474 | 0.002 | -0.044 | -0.044 | 0 | 0.3 | 0.3 | 0.25 | 0.08 | 0.35 | 0.25 |
| 5 | 95508726 | rs4357059 | 0.652 | -0.589 | 0.381 | -0.145 | 0.036 | 0.016 | 0.016 | 0.036 | 0.52 | 0.72 | 0.55 | 0.47 | 0.39 | 0.49 |
| 5 | 95516072 | rs13358858 | -0.904 | -0.737 | 1.108 | 0.166 | 0.003 | 0.101 | 0.101 | 0.354 | 0.11 | 0.45 | 0.15 | 0.19 | 0.07 | 0.17 |
| 5 | 95530758 | rs7706272 | 0.126 | -0.645 | 1.035 | 0.217 | 0.002 | 0.042 | 0.042 | 0 | 0.08 | 0.16 | 0.13 | 0.19 | 0.06 | 0.2 |
| 5 | 95535973 | rs6895664 | -0.246 | -0.648 | 1.031 | 0.213 | 0.033 | 0.118 | 0.118 | 0 | 0.08 | 0.31 | 0.14 | 0.19 | 0.06 | 0.17 |
| 5 | 95539448 | rs4869272 | -0.052 | -0.486 | 0.755 | 0.266 | -0.004 | -0.007 | -0.007 | 0.277 | 0.73 | 0.78 | 0.76 | 0.62 | 0.69 | 0.69 |
| 5 | 95540189 | rs10476663 | 0.783 | -0.507 | 0.611 | 0.211 | 0.006 | 0.007 | 0.007 | 0.094 | 0.54 | 0.73 | 0.58 | 0.47 | 0.42 | 0.48 |
| 5 | 95548515 | rs10866766 | 0.344 | -0.513 | 0.836 | 0.197 | 0.029 | 0.105 | 0.105 | 0 | 0.07 | 0.23 | 0.13 | 0.19 | 0.06 | 0.17 |
| 5 | 95554891 | rs4637573 | 0.123 | -0.495 | 0.423 | 0.087 | 0 | -0.013 | -0.013 | 0.253 | 0.76 | 0.74 | 0.8 | 0.85 | 0.73 | 0.75 |
| 5 | 95572369 | rs13178392 | 0.613 | -0.355 | 0.12 | 0.159 | -0.066 | -0.03 | -0.03 | 0 | 0.18 | 0.03 | 0.17 | 0.15 | 0.26 | 0.22 |
| 5 | 95610520 | rs10065122 | 0.739 | -0.516 | 0.202 | 0.142 | -0.002 | 0.015 | 0.015 | 0.446 | 0.54 | 0.72 | 0.58 | 0.46 | 0.43 | 0.48 |
| 5 | 95614860 | rs4336380 | 1.015 | -0.533 | 0.163 | 0.117 | 0.035 | 0.038 | 0.038 | 0.464 | 0.5 | 0.72 | 0.56 | 0.42 | 0.39 | 0.39 |
| 5 | 95616506 | rs6895503 | 0.027 | -0.593 | 0.621 | 0.138 | 0.033 | 0.106 | 0.106 | 0 | 0.09 | 0.33 | 0.16 | 0.19 | 0.07 | 0.2 |
| 5 | 95617095 | rs6895341 | 1.821 | -0.67 | 0.552 | 0.075 | 0.017 | 0.049 | 0.049 | 0.345 | 0.39 | 0.28 | 0.39 | 0.23 | 0.32 | 0.19 |
| 5 | 95619141 | rs4502854 | 1.689 | -0.744 | 0.739 | 0.074 | -0.019 | -0.031 | -0.031 | 0 | 0.41 | 0.5 | 0.4 | 0.23 | 0.31 | 0.19 |
| 5 | 95621789 | rs6864629 | 1.142 | -0.751 | 0.759 | 0.081 | -0.001 | -0.001 | -0.001 | 5.313 | 0.47 | 0.55 | 0.54 | 0.42 | 0.38 | 0.39 |
| 5 | 95626870 | rs13158163 | -0.366 | -0.533 | 0.774 | 0.106 | -0.002 | 0.009 | 0.009 | 0 | 0.27 | 0.21 | 0.21 | 0.19 | 0.31 | 0.32 |
| 5 | 95639072 | rs12517140 | 1.125 | -0.569 | 0.699 | 0.084 | 0.004 | 0.006 | 0.006 | 0 | 0.62 | 0.36 | 0.62 | 0.62 | 0.62 | 0.47 |
| 5 | 95658155 | rs17384294 | 2.319 | -0.695 | 0.11 | -0.056 | -0.03 | 0.058 | 0.058 | 0.815 | 0.31 | 0.09 | 0.37 | 0.19 | 0.33 | 0.13 |
| 5 | 95661486 | rs6869899 | 0.887 | 0.125 | 0.702 | 0.268 | 0.002 | 0 | 0 | 4.783 | 0.12 | 0.12 | 0.12 | 0.13 | 0.21 | 0.15 |
| 5 | 95667674 | rs10045787 | -0.526 | -0.046 | 0.576 | 0.326 | -0.002 | 0.002 | 0.002 | 0 | 0.26 | 0.21 | 0.2 | 0.22 | 0.31 | 0.29 |
| 5 | 95678301 | rs1995186 | -0.395 | -0.099 | 0.668 | 0.437 | -0.001 | 0.002 | 0.002 | 0 | 0.25 | 0.19 | 0.19 | 0.2 | 0.28 | 0.23 |
| 5 | 95684541 | rs1874929 | 0.353 | -0.141 | 0.66 | 0.48 | -0.032 | -0.052 | -0.052 | 0 | 0.73 | 0.89 | 0.68 | 0.61 | 0.65 | 0.6 |
| 5 | 95685235 | rs169240 | 1.797 | -0.204 | 0.921 | 0.386 | 0.006 | 0.015 | 0.015 | 0.082 | 0.44 | 0.42 | 0.41 | 0.24 | 0.37 | 0.28 |
| 5 | 95709132 | rs10476552 | -0.196 | -0.26 | 0.936 | 0.398 | -0.002 | -0.002 | -0.002 | 0 | 0.26 | 0.22 | 0.24 | 0.38 | 0.28 | 0.29 |
| 5 | 95715456 | rs2936428 | 0.384 | -0.236 | 0.902 | 0.405 | -0.038 | -0.062 | -0.062 | 0.427 | 0.71 | 0.92 | 0.68 | 0.61 | 0.65 | 0.57 |
| 5 | 95715558 | rs12332412 | 0.263 | -0.293 | 0.978 | 0.357 | 0.019 | 0.016 | 0.016 | 0 | 0.06 | 0.24 | 0.02 | 0.01 | 0.01 | 0.01 |
| 5 | 95719090 | rs271943 | 1.927 | -0.249 | 0.612 | 0.358 | 0.013 | 0.102 | 0.102 | 0.569 | 0.41 | 0.13 | 0.39 | 0.24 | 0.37 | 0.29 |
| 5 | 95724985 | rs1364034 | 1.608 | -0.225 | 0.739 | 0.369 | -0.004 | -0.015 | -0.015 | 0.035 | 0.43 | 0.44 | 0.41 | 0.24 | 0.37 | 0.29 |
| 5 | 95733112 | rs6233 | 1.645 | -0.221 | 0.616 | 0.371 | 0.004 | 0.007 | 0.007 | 0.067 | 0.43 | 0.41 | 0.41 | 0.24 | 0.37 | 0.29 |
| 5 | 95737362 | rs271923 | 0.296 | 0.003 | -0.329 | 0.507 | -0.068 | -0.091 | -0.091 | 0 | 0.72 | 0.93 | 0.67 | 0.61 | 0.64 | 0.57 |
| 5 | 95742313 | rs3792744 | 0.651 | 0.078 | -0.369 | 0.682 | -0.071 | -0.019 | -0.019 | 0 | 0.17 | 0.02 | 0.18 | 0.17 | 0.27 | 0.16 |
| 5 | 95744744 | rs11743265 | 0.501 | 0.095 | -0.36 | 0.587 | -0.07 | -0.061 | -0.061 | 1.532 | 0.18 | 0.02 | 0.18 | 0.23 | 0.27 | 0.19 |
| 5 | 95746355 | rs156020 | 1.816 | 0.033 | -0.271 | 0.269 | -0.015 | 0.104 | 0.104 | 1.104 | 0.4 | 0.07 | 0.41 | 0.23 | 0.37 | 0.29 |
| 5 | 95751549 | rs10515237 | -0.031 | 0.206 | -0.067 | 0.378 | 0.012 | 0.026 | 0.026 | 0 | 0.23 | 0.28 | 0.23 | 0.31 | 0.27 | 0.28 |
| 5 | 95751785 | rs6232 | 0.667 | 0.375 | -0.385 | 0.623 | 0.003 | 0.027 | 0.027 | 0 | 0.05 | 0 | 0.03 | 0 | 0.03 | 0.05 |
| 5 | 95754483 | rs436321 | -1.326 | 0.59 | -0.009 | 0.884 | 0 | 0.01 | 0.01 | 0 | 0.34 | 0.4 | 0.35 | 0.46 | 0.36 | 0.43 |
| 5 | 95757592 | rs6231 | -1.077 | 0.505 | 0.15 | 0.503 | -0.015 | -0.021 | -0.021 | 0 | 0.06 | 0.17 | 0.02 | 0.01 | 0.01 | 0.01 |
| 5 | 95767920 | rs3792747 | -0.935 | 1.228 | -0.74 | 0.645 | -0.085 | -0.047 | -0.047 | 1.178 | 0.17 | 0.02 | 0.21 | 0.15 | 0.24 | 0.17 |
| 5 | 95768847 | rs6230 | -0.86 | 1.672 | -0.036 | 1.06 | 0.041 | 0.088 | 0.088 | 0.017 | 0.12 | 0.28 | 0.16 | 0.28 | 0.16 | 0.35 |
| 5 | 95770862 | rs3762986 | -1.552 | 0.896 | 0.208 | 0.714 | -0.002 | -0.001 | -0.001 | 1.979 | 0.43 | 0.58 | 0.42 | 0.47 | 0.45 | 0.57 |
| 5 | 95779230 | rs1498924 | -1.183 | 0.415 | 0.616 | 0.359 | 0 | -0.009 | -0.009 | 0 | 0.29 | 0.26 | 0.25 | 0.36 | 0.28 | 0.49 |
| 5 | 95781172 | rs10059061 | -0.88 | 0.559 | 0.416 | 0.228 | 0.003 | -0.082 | -0.082 | 0.505 | 0.13 | 0.43 | 0.11 | 0 | 0.1 | 0.07 |
| 5 | 95782290 | rs156003 | -1.594 | 0.626 | -0.385 | 0.195 | 0.015 | 0.036 | 0.036 | 0.413 | 0.45 | 0.77 | 0.38 | 0.35 | 0.39 | 0.55 |
| 5 | 95782351 | rs6556923 | -0.684 | 1.258 | -0.322 | 0.745 | -0.006 | 0.006 | 0.006 | 0.325 | 0.08 | 0.06 | 0.11 | 0.26 | 0.17 | 0.28 |
| 5 | 95784455 | rs156000 | -0.479 | 0.257 | -0.85 | 0.204 | 0.035 | -0.056 | -0.056 | 0.198 | 0.64 | 0.86 | 0.66 | 0.52 | 0.7 | 0.46 |
| 5 | 95788810 | rs11738940 | 0.657 | 0.275 | -0.834 | -0.156 | -0.017 | -0.04 | -0.04 | 0 | 0.42 | 0.21 | 0.48 | 0.42 | 0.55 | 0.33 |
| 5 | 95790308 | rs6556924 | -0.469 | 0.211 | -0.388 | -0.153 | 0.003 | 0.025 | 0.025 | 0 | 0.18 | 0.17 | 0.14 | 0.09 | 0.16 | 0.11 |
| 5 | 95806979 | rs269853 | -0.829 | 0.45 | -0.216 | -0.549 | 0 | -0.006 | -0.006 | 0.421 | 0.43 | 0.69 | 0.48 | 0.51 | 0.43 | 0.33 |
| 5 | 95808483 | rs7700874 | -0.53 | 0.456 | -0.286 | -0.547 | -0.002 | -0.006 | -0.006 | 0.044 | 0.4 | 0.4 | 0.45 | 0.51 | 0.41 | 0.35 |
| 5 | 95809328 | rs4326166 | 0.884 | 0.497 | -0.332 | -0.557 | -0.001 | 0.008 | 0.008 | 0.081 | 0.22 | 0.22 | 0.31 | 0.42 | 0.25 | 0.23 |
| 5 | 95813277 | rs7703705 | 1.3 | 0.416 | -0.38 | -0.614 | -0.009 | 0.024 | 0.024 | 0.106 | 0.18 | 0.09 | 0.17 | 0.01 | 0.2 | 0.07 |
| 5 | 95814666 | rs155437 | -0.852 | 0.353 | -0.711 | -0.746 | -0.016 | -0.006 | -0.006 | 0.549 | 0.44 | 0.65 | 0.46 | 0.47 | 0.49 | 0.34 |
| 5 | 95815943 | rs12654207 | -0.383 | 0.618 | -1.165 | -0.647 | 0.005 | 0.1 | 0.1 | 0 | 0.19 | 0.32 | 0.27 | 0.47 | 0.2 | 0.23 |
| 5 | 95817504 | rs12153530 | 0.831 | 0.678 | -1.918 | -1.651 | 0.002 | 0.024 | 0.024 | 0.02 | 0.22 | 0.17 | 0.21 | 0.05 | 0.21 | 0.15 |
| 5 | 95818902 | rs10074809 | 0.661 | 1.081 | -2.082 | -1.267 | -0.111 | 0.134 | 0.134 | 0.194 | 0.22 | 0.01 | 0.16 | 0 | 0.29 | 0.1 |
| 5 | 95821532 | rs2350291 | -0.587 | 0.718 | -1.082 | -0.379 | -0.002 | 0.008 | 0.008 | 0 | 0.14 | 0.14 | 0.1 | 0.05 | 0.15 | 0.06 |
| 5 | 95822590 | rs7709763 | -0.201 | 0.432 | -1.046 | -0.3 | -0.003 | 0.039 | 0.039 | 0.02 | 0.22 | 0.08 | 0.19 | 0.12 | 0.21 | 0.2 |
| 5 | 95827499 | rs10067076 | 1.095 | 0.231 | 0.007 | 0.422 | 0.02 | 0.221 | 0.221 | 2.909 | 0.92 | 0.65 | 0.76 | 0.69 | 0.92 | 0.89 |
| 5 | 95836300 | rs156007 | 0.954 | -0.442 | 0.074 | -0.269 | -0.038 | 0.019 | 0.019 | 0.017 | 0.75 | 0.42 | 0.67 | 0.71 | 0.81 | 0.87 |
| 5 | 95839270 | rs156005 | 0.742 | -0.209 | -0.844 | -0.788 | -0.078 | 0.108 | 0.108 | 0 | 0.34 | 0.11 | 0.25 | 0.18 | 0.42 | 0.32 |
| 5 | 95839515 | rs156004 | 0.48 | -0.175 | -0.149 | -0.585 | 0.074 | 0.321 | 0.321 | 0.327 | 0.92 | 0.56 | 0.78 | 0.71 | 0.94 | 0.93 |
| 5 | 95842987 | rs261985 | 0.697 | -0.282 | -0.852 | -1.059 | -0.039 | 0.112 | 0.112 | 0 | 0.34 | 0.1 | 0.22 | 0.15 | 0.35 | 0.3 |
| 5 | 95844921 | rs17473246 | 0.477 | -0.38 | -0.91 | -1.209 | 0.005 | 0.111 | 0.111 | 0 | 0.2 | 0.08 | 0.13 | 0 | 0.21 | 0.08 |
| 5 | 95849587 | rs261966 | 0.678 | -0.413 | -0.155 | -1.084 | -0.002 | -0.002 | -0.002 | 0 | 0.36 | 0.38 | 0.45 | 0.47 | 0.41 | 0.37 |
| 5 | 95851693 | rs17395416 | 1.428 | -0.3 | -0.35 | -1.287 | -0.033 | 0.077 | 0.077 | 0.736 | 0.15 | 0.04 | 0.12 | 0.01 | 0.17 | 0.05 |
| 5 | 95862197 | rs261969 | 0.703 | -0.257 | 0.066 | -1.1 | 0.078 | 0.314 | 0.314 | 0 | 0.94 | 0.65 | 0.78 | 0.69 | 0.94 | 0.93 |
| 5 | 95873768 | rs11742038 | -1.035 | -0.27 | -0.118 | -1.278 | 0.003 | -0.045 | -0.045 | 0 | 0.65 | 0.57 | 0.75 | 0.85 | 0.64 | 0.7 |
| 5 | 95876525 | rs11135467 | -0.114 | -0.369 | -0.122 | -1.389 | 0.003 | -0.038 | -0.038 | 0 | 0.44 | 0.46 | 0.62 | 0.84 | 0.45 | 0.63 |

| | | | | | | | | | | | | | | | | |
|---|---|---|---|---|---|---|---|---|---|---|---|---|---|---|---|---|
| 5 | 95881444 | rs13173434 | -0.802 | -0.322 | -0.03 | -0.891 | 0.003 | -0.044 | -0.044 | 1.318 | 0.65 | 0.56 | 0.74 | 0.85 | 0.64 | 0.7 |
| 5 | 95882017 | rs6877870 | -0.746 | -0.273 | -0.276 | -0.869 | 0 | -0.05 | -0.05 | 1.689 | 0.63 | 0.44 | 0.73 | 0.85 | 0.64 | 0.7 |
| 5 | 95882162 | rs13178437 | 0.739 | -0.126 | -0.848 | -0.504 | -0.008 | -0.152 | -0.152 | 0 | 0.19 | 0.01 | 0.17 | 0.48 | 0.14 | 0.42 |
| 5 | 95884451 | rs1459839 | 0.203 | -0.162 | -0.294 | -0.834 | 0.031 | -0.005 | -0.005 | 0.109 | 0.4 | 0.31 | 0.26 | 0.58 | 0.26 | 0.51 |
| 5 | 95884775 | rs263377 | 0.442 | 0.523 | -0.274 | -0.876 | -0.013 | 0.087 | 0.087 | 0.095 | 0.37 | 0.19 | 0.26 | 0.1 | 0.39 | 0.28 |
| 5 | 95899427 | rs261999 | 0.5 | 0.528 | -0.353 | -0.94 | -0.023 | 0.14 | 0.14 | 0 | 0.35 | 0.1 | 0.19 | 0.09 | 0.36 | 0.31 |
| 5 | 95900634 | rs9918149 | -0.968 | 0.777 | 0.417 | -0.779 | 0.108 | 0.244 | 0.244 | 0 | 0.11 | 0.39 | 0.32 | 0.32 | 0.14 | 0.07 |
| 5 | 95902989 | rs261227 | 0.558 | 0.713 | 0.175 | -0.95 | -0.01 | 0.074 | 0.074 | 0.177 | 0.36 | 0.2 | 0.21 | 0.09 | 0.38 | 0.28 |
| 5 | 95906139 | rs3853202 | 0.54 | 0.675 | 0.274 | -1.005 | 0.036 | -0.005 | -0.005 | 0.434 | 0.39 | 0.28 | 0.33 | 0.58 | 0.29 | 0.52 |
| 5 | 95906373 | rs261229 | 0.382 | 0.737 | 0.646 | -1.035 | 0.007 | -0.006 | -0.006 | 1.845 | 0.42 | 0.43 | 0.49 | 0.39 | 0.47 | 0.36 |
| 5 | 95910465 | rs1459841 | 0.458 | 0.698 | 0.497 | -1.103 | 0.012 | 0.044 | 0.044 | 0 | 0.4 | 0.37 | 0.59 | 0.9 | 0.34 | 0.56 |
| 5 | 95928251 | rs7725567 | 0.121 | 1.094 | 0.124 | -0.835 | 0.015 | 0.019 | 0.019 | 0 | 0.44 | 0.65 | 0.38 | 0.63 | 0.35 | 0.54 |
| 5 | 95930564 | rs261218 | 0.012 | 1.111 | 0.282 | -0.867 | -0.001 | 0.002 | 0.002 | 0.202 | 0.44 | 0.41 | 0.5 | 0.37 | 0.46 | 0.37 |
| 5 | 95930928 | rs261217 | 0.259 | 1.141 | 0.196 | -0.829 | -0.003 | 0.012 | 0.012 | 0.004 | 0.42 | 0.34 | 0.47 | 0.37 | 0.44 | 0.37 |
| 5 | 95936428 | rs17396628 | -0.13 | 1.224 | 0.043 | -0.866 | 0.03 | -0.027 | -0.027 | 3.083 | 0.06 | 0.2 | 0.04 | 0 | 0.06 | 0.02 |
| 5 | 95941365 | rs261212 | -0.054 | 1.065 | -0.433 | -0.76 | -0.007 | 0.016 | 0.016 | 0 | 0.47 | 0.58 | 0.54 | 0.42 | 0.51 | 0.41 |
| 5 | 95948868 | rs183784 | -0.486 | 1.058 | -0.283 | -0.47 | -0.055 | 0.077 | 0.077 | 0 | 0.87 | 0.98 | 0.87 | 1 | 0.83 | 0.92 |
| 5 | 95949002 | rs6885297 | -1.552 | 1.006 | 0.137 | -0.262 | 0.075 | 0.171 | 0.171 | 0 | 0.13 | 0.52 | 0.33 | 0.35 | 0.17 | 0.13 |
| 5 | 95956828 | rs4582302 | 0.884 | 0.323 | -0.535 | -0.334 | -0.006 | -0.003 | -0.003 | 0 | 0.56 | 0.48 | 0.51 | 0.55 | 0.6 | 0.56 |
| 5 | 95957248 | rs3853206 | 0.869 | 0.945 | -0.588 | -0.327 | -0.005 | -0.011 | -0.011 | 0 | 0.56 | 0.51 | 0.48 | 0.58 | 0.54 | 0.61 |
| 5 | 95958386 | rs17086379 | 0.654 | 0.465 | -1.15 | -0.379 | 0.04 | 0.017 | 0.017 | 0.684 | 0.15 | 0.04 | 0.1 | 0.07 | 0.1 | 0.16 |
| 5 | 95958950 | rs2611713 | 0.276 | 0.673 | -0.955 | -0.238 | -0.003 | -0.004 | -0.004 | 0 | 0.75 | 0.83 | 0.63 | 0.64 | 0.7 | 0.79 |
| 5 | 95961829 | rs10515240 | 0.069 | 0.656 | -1.357 | -0.163 | -0.03 | -0.033 | -0.033 | 0.321 | 0.07 | 0.02 | 0.24 | 0.31 | 0.14 | 0.08 |
| 5 | 95962052 | rs3853203 | 0.97 | 0.441 | -1.342 | -0.493 | -0.005 | 0.015 | 0.015 | 4.279 | 0.45 | 0.39 | 0.43 | 0.56 | 0.47 | 0.4 |
| 5 | 95962344 | rs261236 | -0.547 | 0.374 | -1.897 | -0.201 | 0.018 | 0.033 | 0.033 | 0 | 0.85 | 0.93 | 0.9 | 0.95 | 0.87 | 0.88 |
| 5 | 95962417 | rs261235 | -0.552 | 0.378 | -1.896 | -0.247 | 0.018 | 0.044 | 0.044 | 0 | 0.85 | 0.93 | 0.9 | 0.96 | 0.87 | 0.88 |
| 5 | 95962591 | rs10515241 | 0.322 | 0.743 | -2.388 | 0.396 | -0.076 | -0.184 | -0.184 | 0.196 | 0.32 | 0.03 | 0.25 | 0.44 | 0.3 | 0.32 |
| 5 | 95964823 | rs155046 | 0.107 | 1.124 | -1.925 | 1.063 | -0.026 | -0.05 | -0.05 | 0 | 0.22 | 0.11 | 0.26 | 0.46 | 0.27 | 0.29 |
| 5 | 95965235 | rs17086408 | 0.353 | 1.189 | -2.542 | 0.961 | -0.063 | -0.092 | -0.092 | 0.47 | 0.22 | 0.07 | 0.16 | 0.41 | 0.25 | 0.25 |
| 5 | 95965857 | rs155044 | -0.003 | 0.968 | -1.796 | 0.74 | 0.016 | 0 | 0 | 0 | 0.28 | 0.32 | 0.29 | 0.2 | 0.28 | 0.3 |
| 5 | 95969322 | rs155041 | -0.541 | 0.815 | -1.625 | 0.718 | -0.042 | -0.042 | -0.042 | 0.759 | 0.37 | 0.72 | 0.4 | 0.38 | 0.39 | 0.37 |
| 5 | 95970970 | rs17086432 | 0.566 | 0.28 | -1.429 | 0.478 | 0.093 | 0.174 | 0.174 | 1.879 | 0.08 | 0.39 | 0.11 | 0.18 | 0.1 | 0.06 |
| 5 | 95971788 | rs155059 | -0.98 | 0.117 | -1.875 | 0.383 | 0.048 | 0.129 | 0.129 | 0.061 | 0.6 | 0.9 | 0.57 | 0.79 | 0.65 | 0.6 |
| 5 | 95975302 | rs155054 | -0.537 | 0.376 | -1.354 | 0.543 | 0.02 | 0.034 | 0.034 | 0 | 0.48 | 0.33 | 0.45 | 0.63 | 0.52 | 0.49 |
| 5 | 95976611 | rs155053 | -0.202 | 0.892 | -1.363 | 0.926 | -0.064 | -0.075 | -0.075 | 0 | 0.34 | 0.15 | 0.39 | 0.53 | 0.44 | 0.5 |
| 5 | 95977079 | rs155052 | 0.369 | 1.338 | -1.127 | 0.574 | 0.017 | 0.004 | 0.004 | 0 | 0.2 | 0.25 | 0.29 | 0.2 | 0.26 | 0.3 |
| 5 | 95977219 | rs155051 | -0.88 | 0.492 | -1.308 | 0.323 | 0.091 | 0.145 | 0.145 | 0.502 | 0.53 | 0.79 | 0.67 | 0.8 | 0.61 | 0.59 |
| 5 | 95982514 | rs6862521 | -0.57 | 1.277 | -0.758 | 0.543 | 0.028 | -0.017 | -0.017 | 0 | 0.3 | 0.38 | 0.33 | 0.2 | 0.34 | 0.33 |
| 5 | 95988787 | rs4400148 | -0.81 | 0.61 | -1.625 | 0.143 | -0.055 | 0.1 | 0.1 | 0.099 | 0.58 | 0.91 | 0.68 | 0.8 | 0.51 | 0.66 |
| 5 | 95995733 | rs2351245 | 0.015 | 0.171 | -0.826 | 0.438 | -0.053 | 0.001 | 0.001 | 0 | 0.32 | 0.54 | 0.14 | 0.31 | 0.15 | 0.29 |
| 5 | 95996712 | rs12659730 | 0.614 | 0.509 | -0.911 | 0.674 | -0.045 | -0.036 | -0.036 | 0.781 | 0.19 | 0.1 | 0.25 | 0.26 | 0.25 | 0.28 |
| 5 | 95997285 | rs4590196 | -0.617 | 0.426 | -0.431 | 0.745 | 0.018 | 0.005 | 0.005 | 0 | 0.38 | 0.68 | 0.24 | 0.33 | 0.21 | 0.33 |
| 5 | 96000947 | rs1421911 | 0.418 | 0.475 | 0.133 | 0.778 | -0.027 | 0.07 | 0.07 | 2.271 | 0.2 | 0.4 | 0.11 | 0.32 | 0.12 | 0.27 |
| 5 | 96017956 | rs26482 | -0.493 | 0.542 | 0.194 | 0.766 | -0.013 | 0.003 | 0.003 | 0.063 | 0.31 | 0.57 | 0.23 | 0.32 | 0.21 | 0.33 |
| 5 | 96025847 | rs152005 | 0.037 | 0.366 | 0.17 | 0.868 | 0.003 | -0.004 | -0.004 | 4.161 | 0.24 | 0.28 | 0.13 | 0.31 | 0.15 | 0.29 |
| 5 | 96031569 | rs9667 | 0.343 | 0.887 | 0.003 | 0.953 | -0.001 | 0.003 | 0.003 | 0 | 0.12 | 0.12 | 0.23 | 0.25 | 0.19 | 0.25 |
| 5 | 96043420 | rs10053056 | 0.504 | 0.555 | -0.044 | 0.644 | -0.001 | -0.003 | -0.003 | 0 | 0.54 | 0.42 | 0.59 | 0.42 | 0.58 | 0.38 |
| 5 | 96046039 | rs27852 | -0.078 | 0.658 | -0.179 | 0.527 | -0.004 | 0.013 | 0.013 | 3.781 | 0.65 | 0.55 | 0.64 | 0.48 | 0.72 | 0.57 |
| 5 | 96057313 | rs27991 | 0.31 | 0.77 | -0.249 | 0.799 | -0.043 | -0.067 | -0.067 | 0 | 0.12 | 0.04 | 0.22 | 0.35 | 0.19 | 0.27 |
| 5 | 96070205 | rs149313 | -0.113 | 0.618 | -0.083 | 0.278 | 0 | 0.003 | 0.003 | 0 | 0.46 | 0.48 | 0.53 | 0.47 | 0.54 | 0.51 |
| 5 | 96070982 | rs151835 | -0.656 | 0.591 | -0.435 | 0.159 | 0.059 | 0.035 | 0.035 | 0 | 0.56 | 0.68 | 0.82 | 0.67 | 0.74 | 0.63 |
| 5 | 96071194 | rs13362120 | 0.863 | 0.65 | -0.67 | -0.143 | -0.021 | 0.059 | 0.059 | 0 | 0.29 | 0.13 | 0.23 | 0.09 | 0.34 | 0.2 |
| 5 | 96079387 | rs26505 | 0.451 | 0.626 | -0.301 | 0.469 | 0.003 | 0.047 | 0.047 | 0 | 0.1 | 0.13 | 0.17 | 0.32 | 0.15 | 0.28 |
| 5 | 96079402 | rs26506 | -0.013 | 0.13 | 0.105 | 0.189 | -0.003 | -0.001 | -0.001 | 0 | 0.51 | 0.46 | 0.63 | 0.43 | 0.68 | 0.56 |
| 5 | 96080468 | rs26507 | -0.323 | -0.93 | -0.163 | -0.429 | -0.003 | 0 | 0 | 0 | 0.57 | 0.54 | 0.84 | 0.63 | 0.76 | 0.66 |
| 5 | 96080883 | rs3822683 | 0.437 | -0.648 | -0.063 | -0.372 | -0.034 | 0.024 | 0.024 | 1.982 | 0.32 | 0.44 | 0.15 | 0.38 | 0.21 | 0.29 |
| 5 | 96084431 | rs2290674 | 0.075 | -0.928 | -0.157 | -0.48 | -0.018 | -0.028 | -0.028 | 0.106 | 0.13 | 0.18 | 0.06 | 0.02 | 0.07 | 0.14 |
| 5 | 96084776 | rs10037212 | 1.456 | -0.377 | -0.479 | -0.797 | -0.059 | 0.12 | 0.12 | 1.736 | 0.31 | 0.07 | 0.23 | 0.09 | 0.34 | 0.2 |
| 5 | 96089046 | rs17401719 | 1.426 | -0.325 | -0.233 | -0.811 | -0.036 | 0.102 | 0.102 | 1.773 | 0.32 | 0.09 | 0.23 | 0.09 | 0.34 | 0.2 |
| 5 | 96089511 | rs2290675 | 0.335 | -0.386 | -0.172 | -0.801 | -0.004 | -0.011 | -0.011 | 0.016 | 0.13 | 0.16 | 0.06 | 0.02 | 0.07 | 0.14 |
| 5 | 96092990 | rs28081 | -1.951 | -0.286 | -0.025 | -0.666 | -0.012 | 0.045 | 0.045 | 0 | 0.81 | 0.74 | 0.9 | 0.66 | 0.85 | 0.82 |
| 5 | 96093222 | rs27654 | 0.045 | -0.141 | 0.058 | -0.77 | 0.012 | 0.06 | 0.06 | 0 | 0.26 | 0.4 | 0.44 | 0.53 | 0.3 | 0.38 |
| 5 | 96094694 | rs27037 | -0.888 | -0.146 | 0.026 | -0.79 | 0 | -0.002 | -0.002 | 0 | 0.71 | 0.73 | 0.78 | 0.66 | 0.7 | 0.74 |
| 5 | 96094973 | rs27429 | 1.273 | -0.124 | -0.054 | -0.823 | -0.01 | 0.053 | 0.053 | 0 | 0.09 | 0.01 | 0.04 | 0 | 0.06 | 0.09 |
| 5 | 96097895 | rs27980 | -0.265 | -0.125 | 0.002 | -0.793 | 0.013 | 0.074 | 0.074 | 0 | 0.26 | 0.42 | 0.44 | 0.53 | 0.3 | 0.38 |
| 5 | 96098214 | rs27582 | -0.239 | -0.129 | 0.003 | -0.825 | 0.014 | 0.073 | 0.073 | 0 | 0.26 | 0.42 | 0.44 | 0.53 | 0.3 | 0.38 |
| 5 | 96104835 | rs25862 | -0.047 | -0.438 | -0.016 | -0.936 | 0.003 | -0.038 | -0.038 | 0 | 0.55 | 0.43 | 0.38 | 0.13 | 0.47 | 0.45 |
| 5 | 96107774 | rs27581 | -0.044 | -0.428 | -0.171 | -0.934 | -0.001 | -0.016 | -0.016 | 0 | 0.55 | 0.42 | 0.38 | 0.13 | 0.47 | 0.45 |
| 5 | 96109244 | rs28096 | 0.741 | -0.271 | -0.512 | -0.957 | -0.024 | 0.117 | 0.117 | 1.512 | 0.41 | 0.21 | 0.29 | 0.08 | 0.4 | 0.29 |

| | | | | | | | | | | | | | | | | |
|---|---|---|---|---|---|---|---|---|---|---|---|---|---|---|---|---|
| 5 | 96109610 | rs1057569 | 1.526 | -0.232 | -0.727 | -0.976 | -0.045 | 0.109 | 0.109 | 3.047 | 0.3 | 0.08 | 0.19 | 0.06 | 0.33 | 0.14 |
| 5 | 96109935 | rs27433 | -1.776 | -0.159 | -0.479 | -0.562 | -0.01 | 0.048 | 0.048 | 0 | 0.8 | 0.74 | 0.86 | 0.63 | 0.84 | 0.79 |
| 5 | 96112083 | rs1065407 | 1.206 | -0.216 | -0.948 | -0.969 | -0.063 | 0.154 | 0.154 | 3.11 | 0.34 | 0.08 | 0.22 | 0.07 | 0.37 | 0.18 |
| 5 | 96112954 | rs27038 | -1.894 | -0.166 | -0.583 | -0.688 | -0.011 | 0.044 | 0.044 | 0 | 0.8 | 0.74 | 0.89 | 0.65 | 0.85 | 0.81 |
| 5 | 96113042 | rs27639 | 0.963 | -0.117 | -0.601 | -0.408 | -0.011 | -0.036 | -0.036 | 0.248 | 0.11 | 0.04 | 0.29 | 0.26 | 0.12 | 0.21 |
| 5 | 96114346 | rs149481 | -0.091 | -0.111 | -0.355 | -0.425 | -0.015 | -0.015 | -0.015 | 0 | 0.28 | 0.48 | 0.28 | 0.3 | 0.26 | 0.36 |
| 5 | 96114550 | rs27039 | 0.224 | -0.115 | -0.391 | -0.435 | 0.042 | 0.05 | 0.05 | 1.035 | 0.52 | 0.23 | 0.61 | 0.35 | 0.59 | 0.45 |
| 5 | 96115193 | rs149078 | -0.167 | -0.075 | -0.306 | -0.555 | -0.012 | -0.012 | -0.012 | 0 | 0.28 | 0.46 | 0.28 | 0.3 | 0.26 | 0.36 |
| 5 | 96118065 | rs27619 | -0.799 | -0.213 | -0.255 | -0.635 | -0.004 | 0.011 | 0.011 | 0.493 | 0.4 | 0.57 | 0.47 | 0.51 | 0.4 | 0.62 |
| 5 | 96120455 | rs42398 | -1.814 | -0.125 | -0.283 | -0.598 | -0.016 | 0.085 | 0.085 | 0 | 0.8 | 0.72 | 0.77 | 0.57 | 0.85 | 0.81 |
| 5 | 96122210 | rs10050860 | 1.979 | 0.639 | -0.764 | -0.707 | -0.001 | 0.005 | 0.005 | 0 | 0.1 | 0.07 | 0.13 | 0.06 | 0.23 | 0.07 |
| 5 | 96122281 | rs30380 | -0.328 | 0.091 | -0.739 | -0.919 | 0.004 | -0.003 | -0.003 | 0.01 | 0.56 | 0.6 | 0.6 | 0.55 | 0.65 | 0.59 |
| 5 | 96124330 | rs30187 | -0.328 | 0.091 | -0.726 | -0.919 | 0.005 | -0.003 | -0.003 | 0 | 0.56 | 0.6 | 0.6 | 0.55 | 0.65 | 0.59 |
| 5 | 96126308 | rs27529 | -0.328 | 0.091 | -0.726 | -0.919 | 0.005 | -0.003 | -0.003 | 1.762 | 0.56 | 0.6 | 0.6 | 0.55 | 0.65 | 0.59 |
| 5 | 96127644 | rs27690 | 1.539 | 0.09 | -0.704 | -0.984 | -0.001 | -0.022 | -0.022 | 0 | 0.09 | 0.09 | 0.04 | 0 | 0.06 | 0.11 |
| 5 | 96129512 | rs27434 | -0.958 | 0.044 | -0.729 | -1.022 | -0.035 | 0.034 | 0.034 | 0.57 | 0.66 | 0.58 | 0.71 | 0.55 | 0.79 | 0.67 |
| 5 | 96129535 | rs2287987 | 2.213 | 0.128 | -0.726 | -1.029 | 0.001 | 0.003 | 0.003 | 0 | 0.1 | 0.07 | 0.13 | 0.06 | 0.22 | 0.07 |
| 5 | 96129543 | rs27895 | 1.169 | 0.067 | -0.603 | -1.04 | -0.009 | -0.056 | -0.056 | 0 | 0.1 | 0.24 | 0.05 | 0 | 0.06 | 0.11 |
| 5 | 96130836 | rs26618 | 1.683 | 0.095 | -0.612 | -0.997 | 0.011 | 0.01 | 0.01 | 0 | 0.15 | 0.2 | 0.14 | 0.29 | 0.22 | 0.27 |
| 5 | 96132343 | rs26499 | -0.531 | 0.072 | -0.616 | -1.009 | 0.002 | 0 | 0 | 0.755 | 0.57 | 0.55 | 0.59 | 0.49 | 0.56 | 0.6 |
| 5 | 96132795 | rs998509 | 2.226 | 0.085 | -0.645 | -1.003 | 0.031 | -0.002 | -0.002 | 0 | 0.2 | 0.07 | 0.14 | 0.18 | 0.13 | 0.1 |
| 5 | 96134175 | rs2013717 | 2.168 | 0.21 | -0.649 | -0.903 | 0 | 0.004 | 0.004 | 0 | 0.1 | 0.07 | 0.13 | 0.06 | 0.22 | 0.07 |
| 5 | 96134235 | rs30333 | 0.842 | -0.057 | -0.586 | -0.827 | -0.001 | 0.041 | 0.041 | 0 | 0.11 | 0.26 | 0.13 | 0.17 | 0.1 | 0.16 |
| 5 | 96135723 | rs27036 | -0.003 | -0.051 | -0.594 | -0.823 | 0 | -0.001 | -0.001 | 0 | 0.22 | 0.28 | 0.38 | 0.2 | 0.25 | 0.28 |
| 5 | 96138861 | rs27045 | -0.734 | -0.06 | -0.481 | -0.754 | -0.027 | 0.028 | 0.028 | 0.631 | 0.66 | 0.6 | 0.72 | 0.54 | 0.79 | 0.67 |
| 5 | 96140367 | rs26654 | -0.176 | 0.034 | -0.558 | -0.693 | -0.009 | 0.001 | 0.001 | 0.916 | 0.57 | 0.48 | 0.59 | 0.52 | 0.72 | 0.53 |
| 5 | 96153624 | rs34761 | -0.306 | 0.094 | -0.752 | -0.613 | 0.009 | -0.006 | -0.006 | 2.369 | 0.58 | 0.58 | 0.74 | 0.56 | 0.72 | 0.55 |
| 5 | 96155608 | rs18036 | 1.307 | 0.265 | -0.754 | -0.871 | 0.013 | 0.014 | 0.014 | 0 | 0.16 | 0.2 | 0.16 | 0.31 | 0.23 | 0.3 |
| 5 | 96156494 | rs151908 | -0.239 | -0.267 | -0.814 | -0.76 | 0.009 | -0.003 | -0.003 | 0 | 0.58 | 0.58 | 0.61 | 0.52 | 0.72 | 0.55 |
| 5 | 96166026 | rs41135 | 0.319 | 0.555 | -0.883 | -0.504 | 0.02 | 0.004 | 0.004 | 0 | 0.27 | 0.27 | 0.37 | 0.38 | 0.47 | 0.39 |
| 5 | 96167536 | rs152468 | -1.117 | 0.474 | -0.361 | -0.273 | -0.029 | 0.142 | 0.142 | 0.06 | 0.78 | 0.57 | 0.76 | 0.59 | 0.86 | 0.81 |
| 5 | 96179346 | rs2911138 | -0.149 | 0.385 | -0.62 | -0.568 | -0.025 | 0.042 | 0.042 | 0 | 0.69 | 0.42 | 0.72 | 0.59 | 0.8 | 0.71 |
| 5 | 96180913 | rs7708451 | 0.586 | 0.49 | -0.449 | -0.435 | -0.004 | -0.03 | -0.03 | 0 | 0.09 | 0.15 | 0.04 | 0 | 0.06 | 0.1 |
| 5 | 96188666 | rs2042381 | -0.244 | 0.676 | -0.613 | -0.211 | -0.019 | 0.057 | 0.057 | 0.87 | 0.68 | 0.48 | 0.61 | 0.58 | 0.72 | 0.75 |
| 5 | 96198202 | rs2927615 | 0.541 | 1.803 | -0.956 | -0.269 | -0.015 | 0.01 | 0.01 | 0 | 0.11 | 0.08 | 0.14 | 0.03 | 0.27 | 0.12 |
| 5 | 96198692 | rs2927613 | -0.63 | 1.251 | -0.878 | -0.758 | 0 | 0.022 | 0.022 | 0.33 | 0.4 | 0.62 | 0.41 | 0.16 | 0.4 | 0.26 |
| 5 | 96199493 | rs2927612 | 1.427 | 1.109 | -0.977 | -0.825 | -0.013 | 0.006 | 0.006 | 0 | 0.09 | 0.02 | 0.04 | 0.04 | 0.12 | 0.09 |
| 5 | 96203354 | rs2911134 | -0.868 | 0.886 | -1.211 | -0.69 | 0.047 | 0.073 | 0.073 | 0.055 | 0.47 | 0.72 | 0.53 | 0.58 | 0.38 | 0.42 |
| 5 | 96204254 | rs2549778 | -1.3 | 0.598 | -1.814 | -1.2 | 0.015 | 0.001 | 0.001 | 0 | 0.81 | 0.79 | 0.69 | 0.63 | 0.59 | 0.7 |
| 5 | 96204295 | rs4092590 | 0.94 | 0.244 | -1.237 | -1.37 | -0.026 | 0.117 | 0.117 | 0 | 0.22 | 0.46 | 0.38 | 0.54 | 0.17 | 0.25 |
| 5 | 96205801 | rs6861666 | 0.762 | 0.374 | -1.372 | -1.394 | -0.001 | -0.023 | -0.023 | 0 | 0.15 | 0.13 | 0.11 | 0 | 0.09 | 0.07 |
| 5 | 96213171 | rs1230361 | 0.005 | 0.289 | -1.32 | -1.276 | 0.017 | 0.004 | 0.004 | 1.196 | 0.65 | 0.59 | 0.53 | 0.53 | 0.48 | 0.59 |
| 5 | 96216173 | rs3733904 | 1.471 | 0.071 | -1.459 | -1.363 | 0.028 | 0.17 | 0.17 | 1.203 | 0.33 | 0.02 | 0.15 | 0.06 | 0.21 | 0.28 |
| 5 | 96216387 | rs2549779 | 0.524 | 0.582 | -1.022 | -1.039 | 0.017 | 0.004 | 0.004 | 1.196 | 0.65 | 0.59 | 0.53 | 0.53 | 0.48 | 0.59 |
| 5 | 96229272 | rs4869315 | 0.987 | -0.085 | -0.979 | -1.099 | 0 | 0.001 | 0.001 | 0.087 | 0.57 | 0.57 | 0.52 | 0.42 | 0.42 | 0.46 |
| 5 | 96231000 | rs2549782 | 0.844 | 0.263 | -0.88 | -0.995 | 0.01 | 0.001 | 0.001 | 1.623 | 0.64 | 0.6 | 0.58 | 0.53 | 0.52 | 0.53 |
| 5 | 96237326 | rs2287988 | 0.844 | 0.263 | -0.878 | -0.995 | 0.009 | 0.001 | 0.001 | 1.623 | 0.64 | 0.6 | 0.58 | 0.53 | 0.52 | 0.53 |
| 5 | 96238939 | rs2303211 | 0.853 | 0.197 | -0.802 | -0.978 | -0.014 | 0.006 | 0.006 | 0 | 0.07 | 0.07 | 0.01 | 0.13 | 0.02 | 0.14 |
| 5 | 96245343 | rs2549796 | 1.082 | 0.185 | -0.615 | -1.023 | 0.012 | 0 | 0 | 2.071 | 0.63 | 0.44 | 0.57 | 0.53 | 0.52 | 0.51 |
| 5 | 96245439 | rs1056893 | 0.976 | 0.204 | -0.772 | -1.011 | 0.009 | 0.001 | 0.001 | 1.623 | 0.64 | 0.6 | 0.58 | 0.53 | 0.52 | 0.53 |
| 5 | 96249115 | rs2255546 | 0.976 | 0.204 | -0.772 | -0.987 | 0.009 | 0.001 | 0.001 | 1.623 | 0.64 | 0.6 | 0.58 | 0.53 | 0.52 | 0.53 |
| 5 | 96255017 | rs7714122 | -0.216 | 0.162 | -0.838 | -1.013 | 0.038 | 0.039 | 0.039 | 0.701 | 0.17 | 0.07 | 0.06 | 0.05 | 0.07 | 0.12 |
| 5 | 96284803 | rs10069361 | -1.024 | 0.215 | -0.768 | -1.012 | 0.005 | -0.002 | -0.002 | 0.01 | 0.36 | 0.39 | 0.37 | 0.39 | 0.47 | 0.43 |
| 5 | 96293153 | rs4360063 | -0.163 | 0.253 | -0.747 | -0.991 | 0.06 | 0.017 | 0.017 | 0.324 | 0.25 | 0.38 | 0.32 | 0.39 | 0.42 | 0.41 |
| 5 | 96320495 | rs10044354 | -0.173 | 0.256 | -0.836 | -0.99 | 0.077 | 0.022 | 0.022 | 0.394 | 0.25 | 0.4 | 0.32 | 0.39 | 0.42 | 0.41 |
| 5 | 96350710 | rs2303138 | 1.501 | 0.564 | -0.953 | -0.723 | 0.006 | 0.02 | 0.02 | 0.128 | 0.09 | 0.05 | 0.16 | 0.4 | 0.06 | 0.17 |
| 5 | 96363459 | rs11746232 | 0.175 | 1.111 | -1.48 | -1.819 | -0.012 | 0.083 | 0.083 | 0 | 0.14 | 0.03 | 0.07 | 0 | 0.13 | 0.14 |
| 5 | 96372165 | rs9127 | 0.651 | 1.89 | -0.323 | -1.073 | 0.087 | 0.013 | 0.013 | 1.287 | 0.7 | 0.52 | 0.63 | 0.58 | 0.54 | 0.67 |
| 5 | 96373981 | rs27439 | -0.183 | 1.961 | -0.851 | -0.998 | -0.115 | 0.002 | 0.002 | 0 | 0.18 | 0.02 | 0.2 | 0.11 | 0.36 | 0.1 |
| 5 | 96375671 | rs13161540 | -0.287 | 1.526 | 0.135 | -0.833 | 0.019 | -0.006 | -0.006 | 0 | 0.23 | 0.39 | 0.28 | 0.27 | 0.27 | 0.3 |
| 5 | 96380733 | rs316206 | -0.769 | 1.452 | 0.394 | -0.966 | 0.017 | -0.02 | -0.02 | 0 | 0.28 | 0.4 | 0.33 | 0.27 | 0.33 | 0.32 |
| 5 | 96383752 | rs6861399 | -1.1 | 1.38 | 0.428 | -0.934 | -0.004 | 0.011 | 0.011 | 0 | 0.33 | 0.71 | 0.41 | 0.36 | 0.34 | 0.38 |
| 5 | 96387033 | rs13175726 | -0.196 | 1.472 | 0.659 | -0.924 | 0.019 | -0.014 | -0.014 | 0.051 | 0.25 | 0.43 | 0.29 | 0.28 | 0.29 | 0.3 |
| 5 | 96391276 | rs445037 | -0.922 | 1.473 | 0.725 | -0.924 | 0.006 | -0.029 | -0.029 | 1.804 | 0.32 | 0.57 | 0.35 | 0.28 | 0.35 | 0.32 |
| 5 | 96393085 | rs10463198 | -1.133 | 1.461 | 0.68 | -0.926 | -0.006 | 0.014 | 0.014 | 0 | 0.34 | 0.72 | 0.42 | 0.35 | 0.36 | 0.37 |
| 5 | 96403620 | rs17087294 | -1.943 | 1.45 | 0.812 | -0.642 | 0.026 | 0.021 | 0.021 | 0 | 0.1 | 0.38 | 0.13 | 0.09 | 0.07 | 0.07 |
| 5 | 96413667 | rs316223 | -0.386 | 1.064 | 0.84 | -0.765 | -0.007 | -0.013 | -0.013 | 1.085 | 0.3 | 0.52 | 0.25 | 0.36 | 0.24 | 0.36 |
| 5 | 96414272 | rs316224 | -0.701 | 1.054 | 0.526 | -0.77 | 0.01 | 0.014 | 0.014 | 0.529 | 0.33 | 0.78 | 0.32 | 0.36 | 0.24 | 0.36 |
| 5 | 96417662 | rs6865929 | -0.726 | 1.006 | 0.574 | -0.767 | -0.008 | -0.017 | -0.017 | 0 | 0.1 | 0.26 | 0.03 | 0.07 | 0.06 | 0.09 |

| | | | | | | | | | | | | | | | | |
|---|---|---|---|---|---|---|---|---|---|---|---|---|---|---|---|---|
| 5 | 96427624 | rs316233 | -0.325 | 1.063 | 0.188 | -0.705 | -0.006 | -0.013 | -0.013 | 0 | 0.31 | 0.57 | 0.31 | 0.35 | 0.25 | 0.36 |
| 5 | 96427698 | rs316234 | -0.325 | 1.063 | 0.188 | -0.705 | -0.006 | -0.013 | -0.013 | 1.342 | 0.31 | 0.57 | 0.31 | 0.35 | 0.25 | 0.36 |
| 5 | 96427776 | rs17630580 | 0.99 | 1.08 | -0.768 | -0.665 | 0 | -0.018 | -0.018 | 0.19 | 0.06 | 0 | 0.01 | 0.08 | 0.05 | 0.08 |
| 5 | 96430832 | rs316187 | 0.432 | 1.126 | -0.588 | -0.703 | -0.003 | 0.008 | 0.008 | 3.331 | 0.23 | 0.23 | 0.22 | 0.27 | 0.21 | 0.27 |
| 5 | 96441391 | rs31016 | -0.574 | 1.48 | -1.478 | -0.498 | -0.031 | 0.005 | 0.005 | 0.417 | 0.48 | 0.88 | 0.44 | 0.49 | 0.47 | 0.49 |
| 5 | 96456157 | rs316218 | 0.407 | 1.454 | -0.776 | 0.185 | 0.002 | 0.011 | 0.011 | 0.302 | 0.52 | 0.58 | 0.4 | 0.35 | 0.43 | 0.5 |
| 5 | 96462244 | rs31012 | -0.49 | 1.683 | -1.614 | 0.265 | -0.006 | -0.046 | -0.046 | 4.738 | 0.66 | 0.87 | 0.57 | 0.49 | 0.67 | 0.61 |
| 5 | 96466616 | rs13167790 | -0.167 | 1.054 | -1.344 | 0.073 | 0.006 | 0.055 | 0.055 | 0 | 0.52 | 0.64 | 0.37 | 0.35 | 0.43 | 0.5 |
| 5 | 96481475 | rs17087371 | -0.483 | 1.238 | -1.792 | 0.845 | -0.036 | -0.095 | -0.095 | 0 | 0.27 | 0.05 | 0.21 | 0.47 | 0.27 | 0.36 |
| 5 | 96481613 | rs316198 | -0.393 | 2.127 | -1.298 | 1.283 | 0.042 | 0.071 | 0.071 | 0 | 0.11 | 0.14 | 0.17 | 0.32 | 0.19 | 0.19 |
| 5 | 96483480 | rs6863597 | 0.135 | 0.967 | -1.551 | 0.175 | -0.003 | 0.03 | 0.03 | 1.041 | 0.28 | 0.22 | 0.23 | 0.07 | 0.33 | 0.2 |
| 5 | 96483509 | rs316199 | -0.565 | -0.231 | -1.119 | 0.176 | -0.044 | -0.008 | -0.008 | 0.804 | 0.78 | 0.66 | 0.66 | 0.81 | 0.86 | 0.75 |
| 5 | 96484131 | rs39597 | 0.293 | -0.825 | -2.225 | 0.16 | 0.106 | -0.001 | -0.001 | 0 | 0.58 | 0.16 | 0.64 | 0.28 | 0.71 | 0.56 |
| 5 | 96497323 | rs2216709 | 0.03 | -0.066 | -2.32 | 0.195 | -0.076 | -0.1 | -0.1 | 0 | 0.28 | 0.08 | 0.42 | 0.43 | 0.41 | 0.43 |
| 5 | 96503378 | rs12188395 | 0 | -0.273 | -2.381 | -0.152 | 0.008 | 0.029 | 0.029 | 0.092 | 0.11 | 0.07 | 0.05 | 0 | 0.09 | 0.05 |
| 5 | 96503523 | rs160632 | -1.492 | -0.37 | -1.06 | -0.151 | 0.002 | 0.002 | 0.002 | 0.672 | 0.51 | 0.52 | 0.57 | 0.52 | 0.58 | 0.61 |
| 5 | 96504938 | rs27310 | -0.827 | -0.135 | -0.989 | -0.228 | -0.001 | -0.005 | -0.005 | 0 | 0.45 | 0.54 | 0.58 | 0.64 | 0.59 | 0.63 |
| 5 | 96505108 | rs27311 | -0.215 | -0.114 | -0.98 | -0.179 | 0.023 | 0.034 | 0.034 | 0 | 0.06 | 0.1 | 0.08 | 0.16 | 0.13 | 0.15 |
| 5 | 96508147 | rs316192 | 0.708 | -0.014 | -0.934 | -0.097 | -0.072 | -0.107 | -0.107 | 0.398 | 0.26 | 0.08 | 0.43 | 0.48 | 0.41 | 0.43 |
| 5 | 96512683 | rs2544771 | -1.294 | 0.101 | -0.514 | -0.126 | 0.009 | 0.012 | 0.012 | 0.949 | 0.57 | 0.68 | 0.63 | 0.64 | 0.67 | 0.68 |
| 5 | 96518792 | rs3734010 | -0.345 | 0.497 | -0.591 | 0.237 | -0.005 | 0.001 | 0.001 | 0 | 0.4 | 0.42 | 0.53 | 0.62 | 0.5 | 0.57 |
| 5 | 96521575 | rs10515255 | -0.264 | 0.229 | -0.688 | -0.34 | 0.003 | 0.004 | 0.004 | 0 | 0.17 | 0.17 | 0.09 | 0 | 0.14 | 0.08 |
| 5 | 96521669 | rs9314189 | 0.505 | -0.091 | -0.268 | -0.277 | 0.002 | 0.003 | 0.003 | 0 | 0.47 | 0.53 | 0.43 | 0.43 | 0.42 | 0.5 |
| 5 | 96522591 | rs11950429 | 0.081 | -0.719 | 0.184 | -0.559 | -0.031 | -0.078 | -0.078 | 0 | 0.16 | 0.27 | 0.07 | 0 | 0.06 | 0.08 |
| 5 | 96522675 | rs12653352 | 0.22 | -0.762 | -0.021 | -0.476 | -0.04 | -0.057 | -0.057 | 0 | 0.36 | 0.15 | 0.45 | 0.44 | 0.47 | 0.39 |
| 5 | 96523921 | rs4869151 | 1.375 | -0.881 | 0.547 | -0.351 | -0.025 | 0.004 | 0.004 | 0 | 0.4 | 0.24 | 0.42 | 0.4 | 0.43 | 0.44 |
| 5 | 96529990 | rs7703556 | 1.092 | -1.906 | 0.782 | -0.248 | -0.044 | 0.027 | 0.027 | 0.382 | 0.1 | 0.24 | 0.05 | 0.11 | 0.02 | 0.06 |
| 5 | 96531684 | rs9918187 | 0.31 | -1.617 | 0.477 | -0.259 | -0.007 | -0.051 | -0.051 | 0.735 | 0.35 | 0.14 | 0.42 | 0.42 | 0.37 | 0.39 |
| 5 | 96533469 | rs1000083 | -0.145 | -1.593 | 0.547 | -0.261 | 0 | -0.001 | -0.001 | 0 | 0.9 | 0.86 | 0.94 | 0.95 | 0.92 | 0.89 |
| 5 | 96546305 | rs10045981 | 0.267 | -1.571 | 0.402 | -0.236 | -0.006 | -0.051 | -0.051 | 0.036 | 0.35 | 0.15 | 0.42 | 0.42 | 0.37 | 0.39 |
| 5 | 96546318 | rs17630601 | 1.307 | -1.626 | 0.611 | -0.264 | -0.006 | 0.006 | 0.006 | 0 | 0.42 | 0.32 | 0.45 | 0.41 | 0.53 | 0.44 |
| 5 | 96567950 | rs10463202 | 0.727 | -1.524 | 0.493 | -0.177 | -0.071 | -0.025 | -0.025 | 0 | 0.77 | 0.52 | 0.88 | 0.84 | 0.91 | 0.83 |
| 5 | 96582060 | rs9314194 | 1.101 | -1.368 | -0.251 | -0.074 | -0.076 | -0.137 | -0.137 | 5.45 | 0.22 | 0.02 | 0.36 | 0.35 | 0.29 | 0.24 |
| 5 | 96599002 | rs931572 | 1.017 | -1.193 | 0.511 | -0.313 | -0.013 | 0.014 | 0.014 | 0 | 0.42 | 0.26 | 0.45 | 0.41 | 0.53 | 0.43 |
| 5 | 96611319 | rs13176575 | 0.494 | -1.168 | 0.564 | -0.464 | 0.004 | 0.004 | 0.004 | 0 | 0.08 | 0.07 | 0.04 | 0 | 0.05 | 0.08 |
| 5 | 96619614 | rs2113014 | 0.466 | -1.361 | 0.822 | -0.395 | -0.003 | -0.033 | -0.033 | 0.404 | 0.36 | 0.17 | 0.44 | 0.39 | 0.39 | 0.44 |
| 5 | 96621344 | rs3104168 | 0.57 | -1.4 | 1.166 | -0.415 | 0.007 | 0 | 0 | 0.207 | 0.5 | 0.54 | 0.51 | 0.55 | 0.55 | 0.45 |
| 5 | 96624676 | rs2643523 | 0.228 | -1.342 | 1.208 | -0.441 | -0.012 | -0.023 | -0.023 | 4.568 | 0.88 | 0.78 | 0.95 | 0.95 | 0.94 | 0.89 |
| 5 | 96633720 | rs1830367 | 0.176 | -1.428 | 1.355 | -0.486 | 0.02 | -0.008 | -0.008 | 0.184 | 0.41 | 0.3 | 0.46 | 0.51 | 0.36 | 0.49 |
| 5 | 96657330 | rs2643468 | 1.157 | -1.413 | 1.362 | -0.469 | -0.02 | -0.069 | -0.069 | 0 | 0.82 | 0.61 | 0.91 | 0.95 | 0.89 | 0.89 |
| 5 | 96659728 | rs1528592 | 0.57 | -1.416 | 1.3 | -0.478 | -0.016 | -0.028 | -0.028 | 0.009 | 0.87 | 0.74 | 0.95 | 0.95 | 0.94 | 0.88 |
| 5 | 96663347 | rs3108536 | 1.281 | -1.405 | 1.249 | -0.469 | -0.009 | 0.001 | 0.001 | 0 | 0.41 | 0.31 | 0.46 | 0.44 | 0.53 | 0.39 |
| 5 | 96670319 | rs2643504 | 1.317 | -0.906 | 1.649 | -0.261 | -0.019 | -0.063 | -0.063 | 0 | 0.82 | 0.62 | 0.92 | 0.95 | 0.89 | 0.89 |
| 5 | 96676099 | rs7701806 | -1.07 | -0.919 | 1.623 | -0.269 | -0.011 | -0.083 | -0.083 | 0 | 0.15 | 0.26 | 0.07 | 0 | 0.1 | 0.08 |
| 5 | 96708424 | rs10072960 | -1.361 | -0.818 | 1.259 | -0.266 | -0.016 | -0.002 | -0.002 | 0.229 | 0.58 | 0.66 | 0.54 | 0.52 | 0.53 | 0.61 |
| 5 | 96723674 | rs10063725 | -0.909 | -0.675 | 0.859 | -0.201 | -0.007 | 0.014 | 0.014 | 0.137 | 0.52 | 0.31 | 0.5 | 0.52 | 0.52 | 0.61 |
| 5 | 96723895 | rs13186882 | -0.462 | -0.673 | 0.821 | -0.19 | -0.009 | -0.027 | -0.027 | 0 | 0.44 | 0.15 | 0.46 | 0.51 | 0.45 | 0.58 |
| 5 | 96728948 | rs2220200 | 0.292 | -0.682 | 0.767 | -0.201 | -0.036 | -0.093 | -0.093 | 0 | 0.35 | 0.07 | 0.42 | 0.51 | 0.4 | 0.5 |
| 5 | 96741146 | rs291805 | 0.184 | -0.86 | 0.971 | -0.263 | -0.01 | -0.015 | -0.015 | 0 | 0.89 | 0.78 | 0.95 | 0.93 | 0.94 | 0.9 |
| 5 | 96742602 | rs10035513 | 0.155 | -0.811 | 0.679 | -0.36 | 0.025 | -0.009 | -0.009 | 0 | 0.05 | 0.15 | 0.04 | 0 | 0.04 | 0.01 |
| 5 | 96746562 | rs17087528 | 0.15 | -0.684 | 0.002 | -0.214 | -0.073 | -0.145 | -0.145 | 0.115 | 0.35 | 0.03 | 0.39 | 0.4 | 0.38 | 0.44 |
| 5 | 96747893 | rs13436264 | 0.15 | -0.646 | 0.129 | -0.193 | -0.024 | -0.082 | -0.082 | 3.149 | 0.36 | 0.1 | 0.41 | 0.4 | 0.38 | 0.45 |
| 5 | 96748665 | rs1485470 | -0.385 | -0.298 | 0.415 | -0.082 | 0 | -0.001 | -0.001 | 0.597 | 0.92 | 0.89 | 0.95 | 0.88 | 0.93 | 0.88 |
| 5 | 96770565 | rs4869166 | 0.558 | -0.343 | 0.48 | -0.197 | -0.008 | 0.013 | 0.013 | 0.15 | 0.47 | 0.73 | 0.5 | 0.48 | 0.52 | 0.4 |
| 5 | 96773996 | rs7725674 | -0.139 | -0.285 | 0.593 | -0.238 | 0.081 | 0.01 | 0.01 | 2.875 | 0.07 | 0.34 | 0.05 | 0 | 0.05 | 0.03 |
| 5 | 96782323 | rs6865802 | -0.11 | -0.278 | 0.596 | -0.235 | 0.083 | 0.012 | 0.012 | 2.896 | 0.07 | 0.34 | 0.05 | 0 | 0.05 | 0.03 |
| 5 | 96795662 | rs13181391 | 1.341 | -0.407 | 0.336 | -0.285 | -0.105 | -0.09 | -0.09 | 0.242 | 0.35 | 0.1 | 0.44 | 0.48 | 0.46 | 0.38 |
| 5 | 96797322 | rs10074597 | 0.075 | -0.439 | 0.479 | -0.287 | 0.047 | -0.012 | -0.012 | 0.84 | 0.06 | 0.26 | 0.04 | 0 | 0.05 | 0.03 |
| 5 | 96800447 | rs2351885 | -0.064 | -0.502 | 0.473 | -0.297 | -0.002 | -0.001 | -0.001 | 1.093 | 0.55 | 0.56 | 0.55 | 0.6 | 0.61 | 0.55 |
| 5 | 96817528 | rs1485469 | 1.031 | -0.032 | 0.249 | 0.023 | -0.084 | -0.076 | -0.076 | 0 | 0.35 | 0.13 | 0.44 | 0.49 | 0.46 | 0.38 |
| 5 | 96835304 | rs10056294 | -0.285 | -1.295 | 0.75 | -0.991 | 0.035 | -0.021 | -0.021 | 0.041 | 0.06 | 0.25 | 0.04 | 0 | 0.05 | 0.03 |
| 5 | 96835489 | rs2035550 | -1.224 | -1.295 | 0.087 | -1.142 | -0.022 | 0.12 | 0.12 | 0 | 0.62 | 0.88 | 0.31 | 0.16 | 0.36 | 0.48 |
| 5 | 96837053 | rs1021224 | -0.185 | -1.321 | 0.113 | -1.158 | 0.006 | -0.001 | -0.001 | 0 | 0.47 | 0.43 | 0.23 | 0.15 | 0.27 | 0.43 |
| 5 | 96840017 | rs6556983 | -0.631 | -1.122 | -0.158 | -1.048 | 0.011 | -0.056 | -0.056 | 0 | 0.1 | 0.39 | 0.06 | 0 | 0.06 | 0.03 |
| 5 | 96856112 | rs999753 | -0.764 | -0.591 | -1.265 | -0.678 | -0.044 | 0.118 | 0.118 | 0 | 0.62 | 0.91 | 0.31 | 0.16 | 0.36 | 0.49 |
| 5 | 96873781 | rs2972984 | -0.907 | -0.623 | -0.09 | -0.36 | -0.002 | 0.011 | 0.011 | 0 | 0.11 | 0.21 | 0.12 | 0.18 | 0.1 | 0.23 |
| 5 | 96882369 | rs10041715 | -1.652 | -1.064 | -0.075 | -0.616 | -0.01 | -0.009 | -0.009 | 0.858 | 0.22 | 0.29 | 0.13 | 0.2 | 0.14 | 0.23 |
| 5 | 96901049 | rs11958897 | -0.568 | -1.013 | 0.15 | -0.57 | 0 | -0.002 | -0.002 | 0 | 0.1 | 0.08 | 0.09 | 0.2 | 0.1 | 0.21 |
| 5 | 96902807 | rs10045335 | -1.597 | -1.072 | 0.542 | -0.613 | 0.002 | 0.017 | 0.017 | 0 | 0.15 | 0.26 | 0.11 | 0.21 | 0.12 | 0.22 |

| | | | | | | | | | | | | | | | | |
|---|---|---|---|---|---|---|---|---|---|---|---|---|---|---|---|---|
| 5 | 96914732 | rs6556988 | -1.891 | -1.685 | 1.021 | -0.892 | 0.026 | 0.024 | 0.024 | 0.001 | 0.38 | 0.7 | 0.28 | 0.3 | 0.23 | 0.3 |
| 5 | 96916431 | rs6894105 | -1.711 | -2.036 | 0.637 | -1.099 | 0.029 | 0.049 | 0.049 | 0.054 | 0.55 | 0.87 | 0.45 | 0.36 | 0.33 | 0.39 |
| 5 | 96940843 | rs1583087 | -1.183 | -1.541 | 1.532 | -0.571 | -0.017 | -0.005 | -0.005 | 0.089 | 0.19 | 0.24 | 0.13 | 0.2 | 0.11 | 0.21 |
| 5 | 96943871 | rs990698 | -1.107 | -1.546 | 1.533 | -0.573 | -0.008 | -0.002 | -0.002 | 0 | 0.19 | 0.21 | 0.13 | 0.2 | 0.11 | 0.21 |
| 5 | 96954005 | rs7729239 | -1.486 | -1.543 | 1.572 | -0.575 | -0.02 | -0.015 | -0.015 | 0.191 | 0.22 | 0.35 | 0.14 | 0.2 | 0.14 | 0.21 |
| 5 | 96982839 | rs17087673 | -0.575 | -1.541 | 1.506 | -0.572 | 0.011 | -0.005 | -0.005 | 5.712 | 0.16 | 0.07 | 0.11 | 0.2 | 0.11 | 0.2 |
| 5 | 96991565 | rs7733786 | -1.38 | -1.573 | 1.827 | -0.564 | -0.037 | 0.017 | 0.017 | 0 | 0.25 | 0.5 | 0.17 | 0.3 | 0.14 | 0.22 |
| 5 | 97028011 | rs1560323 | -1.138 | -1.532 | 1.373 | -0.572 | -0.022 | 0.004 | 0.004 | 0 | 0.24 | 0.36 | 0.16 | 0.29 | 0.14 | 0.24 |
| 5 | 97030575 | rs10476724 | -1.14 | -1.52 | 1.294 | -0.556 | -0.039 | 0.003 | 0.003 | 0 | 0.25 | 0.45 | 0.17 | 0.29 | 0.14 | 0.24 |
| 5 | 97047653 | rs7736841 | -1.255 | -0.553 | 0.421 | -0.82 | 0.025 | 0.084 | 0.084 | 0 | 0.45 | 0.67 | 0.28 | 0.25 | 0.35 | 0.35 |
| 5 | 97048466 | rs2914928 | -1.406 | -0.56 | 0.426 | -0.853 | 0.024 | 0.083 | 0.083 | 0 | 0.44 | 0.69 | 0.27 | 0.25 | 0.33 | 0.34 |
| 5 | 97049480 | rs2973820 | -1.283 | -0.587 | 0.54 | -1.047 | 0.014 | 0.033 | 0.033 | 0 | 0.39 | 0.66 | 0.22 | 0.1 | 0.27 | 0.21 |
| 5 | 97050542 | rs2163156 | -0.905 | -0.466 | 1.205 | -0.839 | 0.003 | 0.003 | 0.003 | 0 | 0.14 | 0.12 | 0.07 | 0.1 | 0.06 | 0.12 |
| 5 | 97050612 | rs1346699 | -2.087 | -0.473 | 1.151 | -0.927 | -0.006 | -0.032 | -0.032 | 0 | 0.28 | 0.65 | 0.16 | 0.1 | 0.18 | 0.19 |
| 5 | 97051163 | rs2115766 | -1.583 | -0.619 | 1.95 | -0.895 | 0.038 | -0.033 | -0.033 | 0 | 0.08 | 0.29 | 0.05 | 0 | 0.07 | 0.04 |
| 5 | 97064784 | rs7723898 | -2.311 | -0.884 | 1.917 | -0.817 | 0.006 | 0.006 | 0.006 | 0.195 | 0.3 | 0.68 | 0.19 | 0.21 | 0.18 | 0.3 |
| 5 | 97065386 | rs7705141 | -1.922 | -0.86 | 2.381 | -0.809 | 0.027 | -0.042 | -0.042 | 3.039 | 0.1 | 0.35 | 0.06 | 0 | 0.08 | 0.05 |
| 5 | 97070347 | rs6556998 | -1.952 | -0.864 | 2.541 | -0.811 | 0.023 | -0.044 | -0.044 | 1.107 | 0.1 | 0.36 | 0.06 | 0 | 0.08 | 0.05 |
| 5 | 97071402 | rs1429863 | -2.374 | -0.857 | 2.39 | -0.802 | -0.012 | -0.015 | -0.015 | 0 | 0.29 | 0.61 | 0.19 | 0.21 | 0.18 | 0.3 |
| 5 | 97081520 | rs10515261 | -2.222 | -0.621 | 2.387 | -0.544 | -0.006 | 0.022 | 0.022 | 0 | 0.15 | 0.27 | 0.12 | 0.2 | 0.11 | 0.22 |
| 5 | 97087390 | rs3888857 | -2.908 | -0.628 | 2.427 | -0.567 | -0.02 | -0.001 | -0.001 | 0 | 0.2 | 0.44 | 0.14 | 0.2 | 0.14 | 0.23 |
| 5 | 97096042 | rs4380692 | -2.494 | -0.64 | 2.339 | -0.604 | -0.025 | 0.012 | 0.012 | 0 | 0.16 | 0.39 | 0.13 | 0.19 | 0.1 | 0.19 |
| 5 | 97108559 | rs12659421 | -1.5 | -0.631 | 2.113 | -0.599 | 0.004 | -0.016 | -0.016 | 0 | 0.1 | 0.06 | 0.09 | 0.19 | 0.06 | 0.17 |
| 5 | 97111523 | rs12659043 | -2.735 | -0.587 | 2.133 | -0.618 | -0.02 | -0.028 | -0.028 | 1.443 | 0.22 | 0.46 | 0.16 | 0.19 | 0.16 | 0.22 |
| 5 | 97132216 | rs6884205 | -2.772 | -0.555 | 1.972 | -0.577 | -0.021 | -0.022 | -0.022 | 0 | 0.23 | 0.51 | 0.16 | 0.2 | 0.16 | 0.22 |
| 5 | 97134848 | rs6882847 | -1.906 | -0.5 | 1.919 | -0.555 | 0.037 | -0.015 | -0.015 | 0 | 0.08 | 0.32 | 0.05 | 0 | 0.06 | 0.02 |
| 5 | 97137056 | rs7701288 | -1.906 | -0.5 | 1.919 | -0.555 | 0.037 | -0.015 | -0.015 | 2.213 | 0.08 | 0.32 | 0.05 | 0 | 0.06 | 0.02 |
| 5 | 97137570 | rs11738997 | -1.906 | -0.5 | 1.919 | -0.555 | 0.037 | -0.015 | -0.015 | 0 | 0.08 | 0.32 | 0.05 | 0 | 0.06 | 0.02 |
| 5 | 97140436 | rs6881911 | -1.906 | -0.5 | 1.919 | -0.555 | 0.037 | -0.015 | -0.015 | 2.295 | 0.08 | 0.32 | 0.05 | 0 | 0.05 | 0.02 |
| 5 | 97152331 | rs4132948 | -2.085 | -0.475 | 1.911 | -0.556 | 0.039 | -0.027 | -0.027 | 0 | 0.08 | 0.31 | 0.05 | 0 | 0.06 | 0.02 |
| 5 | 97160324 | rs4597996 | -2.608 | -0.538 | 1.838 | -0.55 | -0.014 | -0.015 | -0.015 | 0.749 | 0.22 | 0.4 | 0.13 | 0.17 | 0.16 | 0.26 |
| 5 | 97165802 | rs4401597 | -2.615 | -0.544 | 1.835 | -0.369 | -0.014 | 0.01 | 0.01 | 0 | 0.22 | 0.4 | 0.15 | 0.21 | 0.16 | 0.26 |
| 5 | 97182715 | rs7719818 | -2.043 | -0.345 | 1.389 | -0.334 | -0.001 | -0.003 | -0.003 | 0 | 0.15 | 0.14 | 0.1 | 0.17 | 0.12 | 0.24 |
| 5 | 97206196 | rs6875261 | -1.774 | -0.484 | 1.838 | -0.505 | 0.014 | 0.006 | 0.006 | 0.971 | 0.1 | 0.22 | 0.07 | 0.09 | 0.07 | 0.15 |
| 5 | 97211705 | rs6557015 | -0.749 | -0.57 | 2.111 | -0.552 | -0.004 | 0.005 | 0.005 | 0 | 0.06 | 0.08 | 0.02 | 0.04 | 0.04 | 0.07 |
| 5 | 97212004 | rs11742473 | -1.592 | -0.663 | 2.449 | -0.755 | -0.005 | -0.053 | -0.053 | 0.028 | 0.09 | 0.3 | 0.05 | 0 | 0.06 | 0.01 |
| 5 | 97219737 | rs10059632 | -3.075 | -0.239 | 2.498 | -0.774 | 0 | 0.001 | 0.001 | 0.436 | 0.27 | 0.71 | 0.15 | 0.13 | 0.21 | 0.23 |
| 5 | 97242130 | rs4596407 | -1.707 | -0.399 | 2.333 | -0.726 | 0 | -0.052 | -0.052 | 0.355 | 0.1 | 0.33 | 0.05 | 0 | 0.06 | 0.01 |
| 5 | 97247756 | rs1423315 | -2.382 | -0.41 | 2.218 | -0.73 | -0.026 | -0.063 | -0.063 | 0 | 0.17 | 0.43 | 0.08 | 0.05 | 0.11 | 0.08 |
| 5 | 97258619 | rs6860141 | -2.115 | -0.498 | 2.339 | -0.748 | -0.018 | -0.073 | -0.073 | 0 | 0.12 | 0.51 | 0.06 | 0 | 0.05 | 0.01 |
| 5 | 97260047 | rs1423319 | -1.169 | -0.809 | 1.061 | -1.767 | 0.032 | 0.043 | 0.043 | 1.319 | 0.39 | 0.73 | 0.23 | 0 | 0.3 | 0.11 |
| 5 | 97260358 | rs1423320 | -2.164 | -0.889 | 0.754 | -1.831 | -0.062 | 0.114 | 0.114 | 0.072 | 0.59 | 0.87 | 0.42 | 0.17 | 0.45 | 0.41 |
| 5 | 97272105 | rs17087923 | -0.219 | -0.905 | 1.187 | -1.815 | -0.017 | -0.035 | -0.035 | 0 | 0.07 | 0.14 | 0.03 | 0 | 0.03 | 0.01 |
| 5 | 97278902 | rs7734961 | -0.743 | -0.889 | 0.836 | -1.809 | -0.007 | 0.02 | 0.02 | 0 | 0.09 | 0.17 | 0.12 | 0.08 | 0.06 | 0.1 |
| 5 | 97281641 | rs929821 | -0.048 | -0.848 | 0.722 | -1.794 | -0.017 | 0.011 | 0.011 | 0.156 | 0.12 | 0.01 | 0.06 | 0.09 | 0.09 | 0.21 |
| 5 | 97293421 | rs12658302 | -1.471 | -0.858 | 0.919 | -1.819 | 0.013 | 0.06 | 0.06 | 0 | 0.52 | 0.67 | 0.38 | 0.17 | 0.41 | 0.4 |
| 5 | 97298814 | rs6858929 | -1.537 | -0.967 | 1.068 | -1.671 | -0.012 | -0.027 | -0.027 | 0 | 0.36 | 0.41 | 0.27 | 0.17 | 0.25 | 0.31 |
| 5 | 97306269 | rs2352229 | -1.194 | -0.541 | 1.287 | -1.266 | 0.004 | -0.001 | -0.001 | 0 | 0.2 | 0.1 | 0.19 | 0.17 | 0.15 | 0.29 |
| 5 | 97308232 | rs11959302 | -1.088 | -0.229 | 1.576 | -1.062 | -0.007 | -0.037 | -0.037 | 0 | 0.08 | 0.25 | 0.05 | 0 | 0.03 | 0.01 |
| 5 | 97315701 | rs4362966 | 0.408 | 0.862 | 0.185 | -0.213 | 0.013 | 0 | 0 | 0.64 | 0.63 | 0.57 | 0.67 | 0.59 | 0.52 | 0.64 |
| 5 | 97315918 | rs1363886 | -0.98 | 0.52 | 0.005 | -0.458 | -0.009 | -0.03 | -0.03 | 0 | 0.23 | 0.4 | 0.16 | 0.17 | 0.19 | 0.09 |
| 5 | 97317922 | rs1423321 | -0.136 | 0.263 | -1.118 | -0.959 | 0.013 | 0.096 | 0.096 | 0 | 0.27 | 0.08 | 0.16 | 0.08 | 0.25 | 0.26 |
| 5 | 97318805 | rs11750435 | -0.147 | 0.162 | -0.78 | -0.99 | 0.011 | 0.039 | 0.039 | 0 | 0.27 | 0.14 | 0.16 | 0.08 | 0.24 | 0.26 |
| 5 | 97333442 | rs257239 | 0.738 | -0.175 | -0.507 | -0.391 | -0.001 | 0 | 0 | 0 | 0.43 | 0.37 | 0.6 | 0.39 | 0.55 | 0.52 |
| 5 | 97337147 | rs185385 | 1.575 | 0.577 | -0.572 | -0.594 | 0 | -0.011 | -0.011 | 0 | 0.13 | 0.13 | 0.14 | 0.01 | 0.18 | 0.11 |
| 5 | 97345454 | rs410642 | -1.1 | 0.131 | -0.168 | -0.652 | -0.014 | 0.057 | 0.057 | 1.676 | 0.8 | 0.77 | 0.85 | 0.47 | 0.89 | 0.83 |
| 5 | 97354672 | rs257246 | 0.303 | 0.064 | 0.002 | -0.796 | -0.001 | -0.001 | -0.001 | 0 | 0.53 | 0.47 | 0.66 | 0.35 | 0.62 | 0.55 |
| 5 | 97358303 | rs11957901 | 0.934 | 0.066 | -0.244 | -0.798 | 0 | -0.012 | -0.012 | 0.426 | 0.25 | 0.23 | 0.17 | 0.08 | 0.25 | 0.26 |
| 5 | 97370373 | rs257258 | -1.099 | 0.034 | -0.174 | -0.763 | 0.002 | -0.001 | -0.001 | 5.257 | 0.8 | 0.79 | 0.84 | 0.43 | 0.88 | 0.81 |
| 5 | 97372894 | rs6557027 | 0.476 | 0.344 | -0.121 | -0.852 | -0.001 | -0.012 | -0.012 | 0.003 | 0.27 | 0.27 | 0.18 | 0.08 | 0.26 | 0.28 |
| 5 | 97379184 | rs257218 | 0.672 | 0.547 | -0.242 | -0.609 | -0.004 | -0.012 | -0.012 | 0.087 | 0.43 | 0.42 | 0.32 | 0.24 | 0.37 | 0.37 |
| 5 | 97381223 | rs17706067 | 1.474 | 0.726 | -0.405 | -0.834 | -0.036 | 0.053 | 0.053 | 0 | 0.18 | 0.06 | 0.14 | 0.07 | 0.23 | 0.22 |
| 5 | 97394232 | rs257212 | 0.596 | 0.702 | -0.297 | -0.602 | -0.009 | -0.012 | -0.012 | 5.752 | 0.45 | 0.46 | 0.34 | 0.29 | 0.39 | 0.39 |
| 5 | 97402967 | rs13180770 | 1.241 | 1.28 | -0.699 | -0.239 | -0.016 | -0.004 | -0.004 | 0 | 0.14 | 0.1 | 0.28 | 0.11 | 0.16 | 0.17 |
| 5 | 97406001 | rs2432026 | -1.64 | 1.26 | -0.643 | -0.069 | -0.007 | 0.144 | 0.144 | 0 | 0.94 | 0.94 | 0.88 | 0.55 | 0.96 | 0.83 |
| 5 | 97413587 | rs13187140 | 1.001 | 1.575 | -0.797 | -0.042 | -0.016 | -0.002 | -0.002 | 0 | 0.15 | 0.11 | 0.28 | 0.13 | 0.17 | 0.17 |
| 5 | 97421946 | rs257223 | 0.571 | 1.405 | -0.65 | -0.189 | 0 | 0.003 | 0.003 | 0 | 0.37 | 0.36 | 0.32 | 0.25 | 0.38 | 0.36 |
| 5 | 97424745 | rs11956694 | -0.279 | 1.358 | -0.62 | -0.302 | -0.001 | -0.002 | -0.002 | 0 | 0.57 | 0.39 | 0.59 | 0.54 | 0.51 | 0.51 |
| 5 | 97425603 | rs17088025 | 0.562 | 0.884 | -0.653 | -0.221 | 0.008 | 0.004 | 0.004 | 0 | 0.41 | 0.32 | 0.31 | 0.41 | 0.34 | 0.32 |

| | | | | | | | | | | | | | | | | |
|---|---|---|---|---|---|---|---|---|---|---|---|---|---|---|---|---|
| 5 | 97427543 | rs7713332 | 1.165 | 1.037 | -1.45 | -0.157 | -0.034 | 0.024 | 0.024 | 0 | 0.31 | 0.43 | 0.24 | 0.41 | 0.21 | 0.29 |
| 5 | 97429223 | rs10866786 | 0.187 | 1.151 | -1.453 | -0.084 | 0.005 | -0.002 | -0.002 | 0.69 | 0.48 | 0.53 | 0.54 | 0.53 | 0.42 | 0.5 |
| 5 | 97429386 | rs7727752 | 1.188 | 1.218 | -1.735 | -0.067 | -0.015 | -0.001 | -0.001 | 0.013 | 0.15 | 0.1 | 0.28 | 0.13 | 0.19 | 0.19 |
| 5 | 97432181 | rs17088027 | 0.097 | 1.146 | -1.58 | -0.007 | -0.004 | -0.001 | -0.001 | 1.831 | 0.3 | 0.3 | 0.41 | 0.29 | 0.34 | 0.31 |
| 5 | 97435531 | rs1421804 | -0.323 | 1.397 | -1.56 | 0.075 | -0.033 | 0.013 | 0.013 | 0 | 0.37 | 0.21 | 0.33 | 0.3 | 0.43 | 0.38 |
| 5 | 97436504 | rs1421791 | 0.069 | 1.284 | -0.979 | -0.144 | -0.009 | 0.013 | 0.013 | 0 | 0.16 | 0.14 | 0.18 | 0.01 | 0.22 | 0.18 |
| 5 | 97438982 | rs7725036 | -0.486 | 0.817 | -0.546 | 0.208 | -0.005 | 0.009 | 0.009 | 0 | 0.16 | 0.27 | 0.18 | 0.2 | 0.16 | 0.14 |
| 5 | 97440934 | rs950195 | 0.318 | 0.764 | -1.123 | -0.03 | 0.076 | 0.086 | 0.086 | 0 | 0.49 | 0.69 | 0.52 | 0.7 | 0.35 | 0.56 |
| 5 | 97441007 | rs958375 | 0.312 | 1.426 | -1.428 | -0.356 | -0.031 | 0.035 | 0.035 | 0 | 0.24 | 0.19 | 0.23 | 0.01 | 0.33 | 0.26 |
| 5 | 97446209 | rs1421795 | 0.284 | 1.407 | -1.282 | -0.365 | -0.021 | 0.015 | 0.015 | 0 | 0.24 | 0.21 | 0.24 | 0.01 | 0.33 | 0.27 |
| 5 | 97449146 | rs4134469 | 0.972 | 0.496 | -0.875 | -1.004 | 0.011 | 0 | 0 | 0.444 | 0.33 | 0.27 | 0.45 | 0.69 | 0.26 | 0.47 |
| 5 | 97468211 | rs6868704 | -0.186 | 0.412 | -0.315 | -0.87 | -0.003 | 0.064 | 0.064 | 0 | 0.72 | 0.63 | 0.7 | 0.5 | 0.72 | 0.68 |
| 5 | 97469921 | rs6890789 | 0.64 | 0.46 | -0.333 | -0.852 | 0.002 | -0.002 | -0.002 | 0.007 | 0.52 | 0.45 | 0.59 | 0.49 | 0.54 | 0.6 |
| 5 | 97477227 | rs6889775 | -0.641 | 0.584 | -0.233 | -0.68 | 0.003 | 0.028 | 0.028 | 0 | 0.78 | 0.69 | 0.8 | 0.71 | 0.77 | 0.84 |
| 5 | 97478311 | rs3935679 | -0.069 | 0.437 | -0.181 | -0.532 | -0.019 | 0.067 | 0.067 | 0.476 | 0.4 | 0.24 | 0.59 | 0.2 | 0.54 | 0.43 |
| 5 | 97478555 | rs4307110 | 1.179 | 0.299 | -0.305 | -0.247 | -0.071 | -0.143 | -0.143 | 0.271 | 0.26 | 0.03 | 0.18 | 0.42 | 0.29 | 0.25 |
| 5 | 97478870 | rs10062992 | -0.761 | 0.994 | 0.479 | -0.585 | -0.003 | 0.032 | 0.032 | 0 | 0.43 | 0.42 | 0.66 | 0.23 | 0.57 | 0.44 |
| 5 | 97478920 | rs10063003 | -0.826 | 0.985 | 0.679 | -0.612 | -0.001 | 0.005 | 0.005 | 0.04 | 0.44 | 0.5 | 0.66 | 0.24 | 0.57 | 0.44 |
| 5 | 97483701 | rs4324711 | 0.02 | 1.151 | 0.565 | -0.738 | -0.028 | 0.04 | 0.04 | 0 | 0.31 | 0.24 | 0.33 | 0.01 | 0.41 | 0.25 |
| 5 | 97487315 | rs10044333 | -0.284 | 1.165 | 0.471 | -0.676 | 0.012 | 0.04 | 0.04 | 0 | 0.13 | 0.23 | 0.34 | 0.24 | 0.17 | 0.19 |
| 5 | 97488141 | rs6889798 | -0.072 | 1.145 | 0.326 | -0.68 | -0.002 | 0 | 0 | 0 | 0.05 | 0.05 | 0.08 | 0.04 | 0.1 | 0.08 |
| 5 | 97491234 | rs7725828 | -1.11 | 1.097 | 0.418 | -0.715 | 0.001 | 0.057 | 0.057 | 0.609 | 0.46 | 0.73 | 0.67 | 0.24 | 0.56 | 0.44 |
| 5 | 97497429 | rs10476736 | 1.161 | 1.195 | -0.014 | -0.587 | 0.001 | 0.034 | 0.034 | 0 | 0.07 | 0.11 | 0.24 | 0.19 | 0.07 | 0.1 |
| 5 | 97500022 | rs7702542 | 0.011 | 1.275 | -0.056 | -0.881 | -0.024 | 0.057 | 0.057 | 0 | 0.28 | 0.19 | 0.28 | 0.01 | 0.38 | 0.21 |
| 5 | 97512070 | rs4131051 | -0.068 | 1.477 | 0.399 | -0.684 | 0.03 | -0.06 | -0.06 | 0.014 | 0.2 | 0.29 | 0.26 | 0.01 | 0.38 | 0.21 |
| 5 | 97512483 | rs4330484 | -0.128 | 1.329 | 0.727 | -0.473 | -0.004 | 0.056 | 0.056 | 0 | 0.12 | 0.2 | 0.28 | 0.22 | 0.11 | 0.18 |
| 5 | 97512512 | rs4869188 | 0.611 | 0.881 | 0.897 | -0.461 | -0.028 | 0.008 | 0.008 | 1.03 | 0.67 | 0.24 | 0.46 | 0.76 | 0.53 | 0.64 |
| 5 | 97522622 | rs6872879 | 0.15 | 0.669 | 1.029 | -0.384 | -0.036 | 0.013 | 0.013 | 0 | 0.31 | 0.16 | 0.53 | 0.2 | 0.49 | 0.29 |
| 5 | 97537837 | rs4362967 | -1.175 | 0.536 | 1.173 | 0.095 | 0.001 | 0 | 0 | 0.024 | 0.69 | 0.69 | 0.76 | 0.66 | 0.71 | 0.71 |
| 5 | 97551023 | rs10066757 | 0.718 | 0.486 | 1.193 | 0.167 | -0.001 | -0.01 | -0.01 | 0.742 | 0.36 | 0.49 | 0.25 | 0.36 | 0.32 | 0.33 |
| 5 | 97555290 | rs9314227 | 0.475 | 0.641 | 0.307 | 0.239 | 0.046 | -0.074 | -0.074 | 0 | 0.31 | 0.14 | 0.2 | 0.45 | 0.19 | 0.36 |
| 5 | 97591259 | rs11749431 | -0.031 | 1.029 | 1.056 | 0.228 | -0.002 | -0.011 | -0.011 | 0 | 0.36 | 0.45 | 0.25 | 0.36 | 0.32 | 0.34 |
| 5 | 97616698 | rs10478904 | -0.28 | 0.747 | 0.118 | 0.098 | 0.006 | 0.017 | 0.017 | 0.02 | 0.39 | 0.7 | 0.3 | 0.33 | 0.34 | 0.37 |
| 5 | 97632233 | rs17655764 | -0.04 | 1.1 | -0.64 | 0.326 | -0.005 | 0.031 | 0.031 | 0.484 | 0.38 | 0.2 | 0.29 | 0.33 | 0.36 | 0.39 |
| 5 | 97655321 | rs11957758 | -0.397 | 1.083 | -0.53 | 0.471 | -0.009 | -0.013 | -0.013 | 0 | 0.42 | 0.73 | 0.38 | 0.43 | 0.4 | 0.37 |
| 5 | 97666860 | rs6882939 | -1.229 | 1.004 | 0.163 | 0.321 | 0.022 | 0.004 | 0.004 | 1.875 | 0.07 | 0.57 | 0.06 | 0.02 | 0.03 | 0.03 |
| 5 | 97674522 | rs6596729 | -0.937 | 1.01 | 1.674 | 0.323 | 0.111 | 0.094 | 0.094 | 0.12 | 0.05 | 0.39 | 0.05 | 0.02 | 0.03 | 0.02 |
| 5 | 97687754 | rs11950953 | -0.912 | 1.012 | 1.763 | 0.324 | 0.11 | 0.087 | 0.087 | 0.004 | 0.05 | 0.35 | 0.05 | 0.02 | 0.03 | 0.02 |
| 5 | 97703974 | rs1378438 | 0.077 | 0.872 | 1.2 | 0.016 | 0.005 | 0.017 | 0.017 | 1.091 | 0.11 | 0.1 | 0.07 | 0.04 | 0.13 | 0.11 |
| 5 | 97729860 | rs6594645 | -1.313 | 0.835 | 1.493 | 0.08 | 0.091 | 0.068 | 0.068 | 0.161 | 0.06 | 0.36 | 0.06 | 0.03 | 0.03 | 0.02 |
| 5 | 97740803 | rs13161187 | -0.231 | 0.467 | 1.07 | 0.131 | -0.003 | -0.011 | -0.011 | 0 | 0.44 | 0.64 | 0.43 | 0.54 | 0.44 | 0.37 |
| 5 | 97744026 | rs1378441 | 0.347 | 0.481 | 0.881 | 0.145 | -0.022 | -0.056 | -0.056 | 0.012 | 0.37 | 0.17 | 0.36 | 0.51 | 0.4 | 0.35 |
| 5 | 97757677 | rs10067255 | 0.11 | 0.473 | 0.656 | 0.16 | 0.002 | 0.012 | 0.012 | 0.041 | 0.4 | 0.43 | 0.38 | 0.51 | 0.42 | 0.35 |
| 5 | 97761687 | rs10478264 | -0.211 | 0.456 | 0.375 | 0.181 | -0.006 | -0.008 | -0.008 | 0 | 0.45 | 0.68 | 0.44 | 0.55 | 0.45 | 0.39 |
| 5 | 97763960 | rs1563320 | 0.291 | 0.741 | -0.602 | 0.193 | -0.01 | -0.034 | -0.034 | 0.399 | 0.45 | 0.12 | 0.39 | 0.44 | 0.41 | 0.5 |
| 5 | 97764522 | rs7715564 | -0.365 | 0.517 | -0.976 | 0.026 | -0.114 | -0.101 | -0.101 | 1.504 | 0.59 | 0.94 | 0.47 | 0.5 | 0.48 | 0.66 |
| 5 | 97768486 | rs1563317 | -0.221 | 0.362 | -0.698 | -0.14 | -0.025 | -0.031 | -0.031 | 0 | 0.57 | 0.83 | 0.48 | 0.43 | 0.52 | 0.61 |
| 5 | 97783491 | rs1037599 | 0.177 | 0.426 | 0.147 | -0.17 | 0.003 | 0.004 | 0.004 | 0 | 0.5 | 0.53 | 0.4 | 0.38 | 0.45 | 0.55 |
| 5 | 97784849 | rs6873376 | -0.873 | 0.545 | 0.158 | -0.17 | 0.063 | 0.019 | 0.019 | 0.022 | 0.05 | 0.21 | 0.07 | 0.04 | 0.08 | 0.06 |
| 5 | 97790692 | rs7717913 | -0.187 | 0.47 | 0.093 | -0.155 | -0.007 | -0.01 | -0.01 | 0 | 0.55 | 0.72 | 0.47 | 0.43 | 0.52 | 0.61 |
| 5 | 97793862 | rs10515274 | 0.065 | 0.431 | -0.318 | -0.172 | -0.02 | 0 | 0 | 0 | 0.19 | 0.13 | 0.28 | 0.15 | 0.26 | 0.21 |
| 5 | 97794304 | rs11241494 | -0.813 | 0.329 | 0.413 | -0.202 | -0.011 | -0.01 | -0.01 | 0.102 | 0.56 | 0.69 | 0.46 | 0.43 | 0.51 | 0.6 |
| 5 | 97794710 | rs2368554 | 0.307 | 0.31 | 0.414 | -0.211 | -0.031 | -0.014 | -0.014 | 0 | 0.36 | 0.5 | 0.18 | 0.28 | 0.25 | 0.4 |
| 5 | 97795382 | rs12655510 | -0.873 | 0.319 | 0.41 | -0.201 | -0.04 | -0.035 | -0.035 | 0.051 | 0.56 | 0.82 | 0.47 | 0.43 | 0.51 | 0.63 |
| 5 | 97809408 | rs6595256 | 0.256 | 0.211 | 0.332 | -0.27 | -0.03 | -0.013 | -0.013 | 0 | 0.36 | 0.5 | 0.18 | 0.28 | 0.25 | 0.39 |
| 5 | 97817930 | rs6861004 | -0.092 | 0.624 | 0.09 | -0.37 | -0.009 | 0.011 | 0.011 | 0 | 0.18 | 0.14 | 0.25 | 0.09 | 0.24 | 0.21 |
| 5 | 97821452 | rs11241621 | 0.673 | -0.109 | -0.029 | -0.616 | 0.003 | -0.004 | -0.004 | 0.406 | 0.27 | 0.28 | 0.12 | 0.3 | 0.17 | 0.33 |
| 5 | 97827290 | rs9327251 | 0.66 | -0.122 | -0.326 | -0.798 | 0.015 | -0.003 | -0.003 | 0 | 0.31 | 0.28 | 0.15 | 0.3 | 0.22 | 0.35 |
| 5 | 97838765 | rs891057 | -0.608 | -0.103 | -0.375 | -1.355 | -0.002 | 0 | 0 | 0 | 0.27 | 0.68 | 0.35 | 0.14 | 0.32 | 0.3 |
| 5 | 97841989 | rs6860932 | -0.643 | 0.11 | -0.217 | -1.372 | 0.002 | -0.011 | -0.011 | 0 | 0.25 | 0.66 | 0.32 | 0.14 | 0.28 | 0.27 |
| 5 | 97844408 | rs4702980 | 0.9 | -0.111 | -0.806 | -1.509 | -0.102 | -0.098 | -0.098 | 6.025 | 0.38 | 0.02 | 0.48 | 0.58 | 0.43 | 0.3 |
| 5 | 97844976 | rs2042966 | -0.833 | -0.322 | -0.377 | -1.577 | -0.033 | -0.016 | -0.016 | 0.054 | 0.64 | 0.55 | 0.83 | 0.74 | 0.78 | 0.64 |
| 5 | 97896472 | rs168654 | -0.725 | 0.278 | 0.05 | -1.538 | 0.081 | -0.053 | -0.053 | 0.098 | 0.24 | 0.46 | 0.33 | 0.15 | 0.34 | 0.27 |
| 5 | 97903158 | rs187384 | -0.198 | -0.813 | 0.384 | -1.446 | -0.002 | -0.005 | -0.005 | 0 | 0.49 | 0.51 | 0.73 | 0.68 | 0.61 | 0.51 |
| 5 | 97907147 | rs152958 | -0.161 | -0.796 | 0.458 | -1.439 | 0.008 | 0.011 | 0.011 | 0.026 | 0.48 | 0.47 | 0.71 | 0.68 | 0.61 | 0.52 |
| 5 | 97915465 | rs17716827 | 0.971 | -0.761 | 0.711 | -1.554 | 0.002 | 0 | 0 | 0 | 0.34 | 0.29 | 0.55 | 0.6 | 0.46 | 0.35 |
| 5 | 97916570 | rs11738094 | 1.057 | -0.741 | 0.69 | -1.554 | -0.043 | -0.014 | -0.014 | 0.026 | 0.33 | 0.16 | 0.54 | 0.6 | 0.46 | 0.35 |
| 5 | 97918829 | rs29661 | -1.185 | -1.054 | 0.585 | -1.738 | 0.018 | 0.018 | 0.018 | 0 | 0.68 | 0.74 | 0.85 | 0.76 | 0.77 | 0.62 |
| 5 | 97929245 | rs17132963 | 1.119 | -1.083 | 0.467 | -1.717 | 0.003 | 0.008 | 0.008 | 0.431 | 0.18 | 0.14 | 0.12 | 0.08 | 0.16 | 0.11 |

| | | | | | | | | | | | | | | | | |
|---|---|---|---|---|---|---|---|---|---|---|---|---|---|---|---|---|
| 5 | 97929333 | rs191730 | -0.946 | -1.142 | 0.599 | -1.726 | -0.003 | -0.003 | -0.003 | 0 | 0.65 | 0.64 | 0.85 | 0.76 | 0.77 | 0.62 |
| 5 | 97929966 | rs461409 | -0.653 | -1.322 | 0.795 | -1.577 | -0.005 | -0.069 | -0.069 | 0 | 0.33 | 0.51 | 0.31 | 0.16 | 0.31 | 0.28 |
| 5 | 97933888 | rs467627 | 0.094 | -1.23 | 0.476 | -1.517 | 0.006 | -0.013 | -0.013 | 0 | 0.14 | 0.19 | 0.17 | 0.09 | 0.15 | 0.15 |
| 5 | 97936210 | rs466366 | -1.348 | -1.338 | 0.328 | -1.602 | -0.067 | -0.018 | -0.018 | 1.09 | 0.65 | 0.85 | 0.46 | 0.41 | 0.52 | 0.63 |
| 5 | 97939616 | rs154201 | 0.05 | -1.109 | 0.293 | -1.481 | 0 | -0.002 | -0.002 | 0 | 0.14 | 0.16 | 0.16 | 0.09 | 0.15 | 0.15 |
| 5 | 97942014 | rs7711174 | 0.799 | -1.13 | 0.165 | -1.437 | -0.001 | -0.009 | -0.009 | 1.129 | 0.19 | 0.18 | 0.13 | 0.08 | 0.16 | 0.13 |
| 5 | 97957754 | rs25743 | 0.733 | -1.42 | 0.172 | -1.502 | 0.017 | 0.008 | 0.008 | 0 | 0.28 | 0.23 | 0.13 | 0.25 | 0.19 | 0.35 |
| 5 | 97961403 | rs29774 | -0.652 | -1.486 | 0.418 | -1.627 | 0.034 | 0.04 | 0.04 | 0 | 0.44 | 0.72 | 0.32 | 0.33 | 0.33 | 0.48 |
| 5 | 97961761 | rs6895508 | -0.505 | -1.208 | 0.577 | -1.767 | 0.025 | -0.046 | -0.046 | 0 | 0.16 | 0.48 | 0.19 | 0.08 | 0.14 | 0.14 |
| 5 | 97969137 | rs2112145 | -0.368 | -1.283 | 0.685 | -1.762 | -0.009 | -0.018 | -0.018 | 0 | 0.14 | 0.4 | 0.15 | 0.08 | 0.09 | 0.11 |
| 5 | 97969453 | rs467650 | -0.766 | -1.362 | 0.501 | -2.011 | 0.013 | 0.043 | 0.043 | 0.958 | 0.37 | 0.66 | 0.32 | 0.1 | 0.32 | 0.25 |
| 5 | 97982179 | rs27706 | 0.438 | -1.192 | 0.342 | -1.873 | -0.003 | -0.054 | -0.054 | 0.015 | 0.24 | 0.25 | 0.17 | 0.02 | 0.22 | 0.12 |
| 5 | 97985312 | rs152980 | 0.769 | -1.4 | 0.227 | -1.85 | 0.02 | 0.02 | 0.02 | 0 | 0.27 | 0.25 | 0.13 | 0.21 | 0.19 | 0.34 |
| 5 | 97991128 | rs162674 | -0.039 | -1.612 | 0.088 | -1.926 | 0.071 | 0.142 | 0.142 | 0.372 | 0.44 | 0.19 | 0.24 | 0.24 | 0.34 | 0.41 |
| 5 | 97993953 | rs2547958 | 1.535 | -1.68 | 0.377 | -1.962 | -0.044 | 0.021 | 0.021 | 0.071 | 0.36 | 0.2 | 0.58 | 0.62 | 0.52 | 0.36 |
| 5 | 97999766 | rs150822 | -1.066 | -1.804 | 0.289 | -1.985 | 0.047 | 0.039 | 0.039 | 0 | 0.75 | 0.9 | 0.88 | 0.79 | 0.83 | 0.68 |
| 5 | 97999858 | rs7729850 | -0.233 | -1.782 | 0.596 | -1.961 | 0.001 | 0.019 | 0.019 | 0.002 | 0.12 | 0.35 | 0.13 | 0.14 | 0.08 | 0.12 |
| 5 | 98011096 | rs316507 | 1.578 | -1.721 | 0.324 | -1.962 | -0.006 | 0.093 | 0.093 | 0.421 | 0.18 | 0.02 | 0.11 | 0.02 | 0.17 | 0.09 |
| 5 | 98017301 | rs6867815 | -0.056 | -1.532 | 0.733 | -1.74 | 0.041 | -0.029 | -0.029 | 0 | 0.06 | 0.19 | 0.05 | 0 | 0.06 | 0.08 |
| 5 | 98022320 | rs408005 | -0.246 | -1.638 | 0.576 | -1.613 | -0.031 | -0.019 | -0.019 | 0 | 0.7 | 0.6 | 0.86 | 0.72 | 0.83 | 0.64 |
| 5 | 98046166 | rs10479243 | 0.274 | -1.865 | 0.397 | -1.695 | -0.039 | 0.016 | 0.016 | 0 | 0.16 | 0.45 | 0.09 | 0.23 | 0.08 | 0.31 |
| 5 | 98046788 | rs2662245 | -1.469 | -1.783 | -0.501 | -1.773 | 0.01 | 0.002 | 0.002 | 0 | 0.86 | 0.92 | 0.95 | 0.98 | 0.91 | 0.93 |
| 5 | 98057031 | rs17730306 | 0.207 | -1.444 | -0.757 | -1.886 | -0.002 | 0.006 | 0.006 | 1.769 | 0.3 | 0.19 | 0.26 | 0.1 | 0.31 | 0.21 |
| 5 | 98058939 | rs2547961 | 0.995 | -0.952 | -1.142 | -1.594 | -0.173 | -0.023 | -0.023 | 0 | 0.35 | 0.02 | 0.55 | 0.67 | 0.51 | 0.35 |
| 5 | 98067832 | rs992592 | 0.81 | -0.879 | -0.781 | -1.887 | -0.043 | 0.026 | 0.026 | 0.354 | 0.35 | 0.21 | 0.58 | 0.69 | 0.55 | 0.37 |
| 5 | 98076517 | rs2662269 | 0.179 | -1.514 | -0.299 | -2.267 | 0.009 | 0.033 | 0.033 | 0.845 | 0.55 | 0.63 | 0.7 | 0.88 | 0.63 | 0.65 |
| 5 | 98078980 | rs12652843 | 0.233 | -1.638 | -0.253 | -2.396 | 0.009 | 0.068 | 0.068 | 1.918 | 0.33 | 0.2 | 0.24 | 0.1 | 0.28 | 0.26 |
| 5 | 98085065 | rs12513899 | 1.12 | -1.776 | -0.168 | -2.444 | 0.051 | -0.012 | -0.012 | 0 | 0.16 | 0.05 | 0.08 | 0.18 | 0.06 | 0.23 |
| 5 | 98093671 | rs10515283 | 1.221 | -1.864 | -0.318 | -2.623 | 0.017 | 0.137 | 0.137 | 1.654 | 0.23 | 0.03 | 0.13 | 0.01 | 0.18 | 0.08 |
| 5 | 98096735 | rs2080892 | 0.709 | -1.6 | 0.063 | -2.441 | -0.013 | -0.004 | -0.004 | 0 | 0.17 | 0.2 | 0.1 | 0.18 | 0.08 | 0.27 |
| 5 | 98099369 | rs7727348 | 0.74 | -1.644 | -0.062 | -2.439 | -0.001 | 0 | 0 | 0.116 | 0.17 | 0.15 | 0.09 | 0.18 | 0.08 | 0.27 |
| 5 | 98104977 | rs2547971 | -1.224 | -1.17 | -0.034 | -2.441 | -0.002 | -0.026 | -0.026 | 0 | 0.9 | 0.88 | 0.94 | 0.98 | 0.92 | 0.91 |
| 5 | 98107067 | rs1979980 | 0.793 | -1.078 | 0.141 | -2.327 | -0.012 | 0.008 | 0.008 | 0.967 | 0.37 | 0.29 | 0.6 | 0.67 | 0.57 | 0.41 |
| 5 | 98107876 | rs1508793 | 0.516 | -1 | -0.453 | -2.296 | -0.008 | 0.023 | 0.023 | 1.242 | 0.41 | 0.65 | 0.62 | 0.68 | 0.57 | 0.42 |
| 5 | 98115932 | rs2250807 | 0.059 | -0.862 | -1.092 | -2.338 | 0.033 | 0.076 | 0.076 | 1.861 | 0.43 | 0.14 | 0.31 | 0.31 | 0.32 | 0.46 |
| 5 | 98122187 | rs2662260 | 0.347 | -0.862 | -0.754 | -2.408 | 0.002 | -0.012 | -0.012 | 0 | 0.21 | 0.18 | 0.12 | 0.01 | 0.17 | 0.1 |
| 5 | 98123347 | rs2617515 | 0.917 | -0.85 | -0.918 | -2.401 | -0.011 | 0.092 | 0.092 | 0.369 | 0.18 | 0.05 | 0.11 | 0.01 | 0.16 | 0.06 |
| 5 | 98125530 | rs2662258 | -1.317 | -0.94 | -1.217 | -2.512 | -0.179 | -0.092 | -0.092 | 0.014 | 0.67 | 0.98 | 0.46 | 0.36 | 0.48 | 0.63 |
| 5 | 98131350 | rs2662256 | 0.981 | -0.761 | -0.873 | -2.366 | -0.011 | 0.084 | 0.084 | 0.282 | 0.17 | 0.06 | 0.11 | 0.01 | 0.16 | 0.06 |
| 5 | 98131505 | rs13541 | -1.697 | 0.41 | -1.323 | -2.689 | -0.096 | -0.103 | -0.103 | 1.112 | 0.85 | 0.98 | 0.55 | 0.35 | 0.65 | 0.75 |
| 5 | 98131883 | rs2545680 | 0.942 | -0.621 | -0.638 | -2.352 | -0.009 | 0.08 | 0.08 | 0.282 | 0.17 | 0.06 | 0.11 | 0.01 | 0.16 | 0.06 |
| 5 | 98134186 | rs2662255 | -0.22 | -0.662 | 0.096 | -2.351 | -0.02 | -0.102 | -0.102 | 0.11 | 0.25 | 0.5 | 0.13 | 0.03 | 0.17 | 0.11 |
| 5 | 98139720 | rs331934 | 0.145 | -0.428 | 0.358 | -2.274 | -0.004 | -0.083 | -0.083 | 0 | 0.68 | 0.44 | 0.77 | 0.96 | 0.72 | 0.86 |
| 5 | 98195818 | rs325203 | 0.85 | -0.433 | 0.136 | -2.238 | -0.026 | 0.111 | 0.111 | 0.289 | 0.22 | 0.04 | 0.16 | 0.03 | 0.21 | 0.08 |
| 5 | 98211890 | rs17166423 | 0.741 | -0.458 | 0.322 | -2.16 | 0.013 | -0.029 | -0.029 | 1.422 | 0.06 | 0.12 | 0.05 | 0 | 0.06 | 0.05 |
| 5 | 98212033 | rs10044279 | 0.459 | -0.615 | 0.186 | -2.266 | -0.012 | -0.024 | -0.024 | 0 | 0.05 | 0.11 | 0.02 | 0.01 | 0.02 | 0.07 |
| 5 | 98213991 | rs727637 | -0.212 | -0.578 | 0.541 | -2.234 | 0.042 | -0.012 | -0.012 | 3.08 | 0.08 | 0.3 | 0.07 | 0.01 | 0.07 | 0.06 |
| 5 | 98236831 | rs7702352 | 0.999 | -0.856 | -0.065 | -2.227 | 0.066 | 0.034 | 0.034 | 0 | 0.28 | 0.13 | 0.18 | 0.2 | 0.18 | 0.23 |
| 5 | 98250805 | rs10057320 | -0.251 | -0.516 | 0.04 | -2.276 | 0.004 | 0 | 0 | 0 | 0.12 | 0.19 | 0.09 | 0.14 | 0.09 | 0.25 |
| 5 | 98257369 | rs161952 | 0.027 | -0.584 | 0.003 | -2.432 | -0.001 | -0.033 | -0.033 | 0 | 0.32 | 0.6 | 0.23 | 0.04 | 0.28 | 0.14 |
| 5 | 98267115 | rs6860504 | 0.081 | -0.934 | 0.032 | -2.38 | 0.033 | 0.021 | 0.021 | 0 | 0.4 | 0.34 | 0.28 | 0.33 | 0.24 | 0.48 |
| 5 | 98270725 | rs9327919 | -0.178 | -0.788 | -0.062 | -2.296 | -0.005 | 0.002 | 0.002 | 0 | 0.12 | 0.18 | 0.09 | 0.13 | 0.07 | 0.24 |
| 5 | 98273862 | rs161937 | -1.682 | -0.509 | -0.404 | -2.317 | -0.141 | -0.115 | -0.115 | 0.867 | 0.75 | 0.98 | 0.52 | 0.37 | 0.56 | 0.66 |
| 5 | 98275197 | rs150992 | -0.356 | -0.218 | 0.485 | -2.053 | 0.004 | -0.067 | -0.067 | 2.336 | 0.31 | 0.37 | 0.23 | 0.06 | 0.3 | 0.15 |
| 5 | 98286819 | rs326482 | 0.268 | -0.099 | 0.16 | -2.061 | -0.053 | 0.082 | 0.082 | 0.418 | 0.24 | 0.05 | 0.18 | 0.1 | 0.25 | 0.16 |
| 5 | 98289567 | rs467728 | -0.465 | -0.134 | 0.552 | -2.078 | 0.002 | -0.029 | -0.029 | 0 | 0.32 | 0.38 | 0.24 | 0.1 | 0.31 | 0.2 |
| 5 | 98304564 | rs325220 | -0.283 | -0.132 | 0.516 | -2.1 | 0.002 | -0.031 | -0.031 | 1.085 | 0.3 | 0.33 | 0.23 | 0.08 | 0.31 | 0.19 |
| 5 | 98307825 | rs17166485 | -0.294 | -0.265 | 0.509 | -2.142 | -0.012 | -0.005 | -0.005 | 0 | 0.12 | 0.23 | 0.1 | 0.1 | 0.06 | 0.22 |
| 5 | 98318657 | rs161936 | -0.652 | -0.21 | 0.722 | -2.054 | 0.016 | -0.097 | -0.097 | 0 | 0.33 | 0.55 | 0.24 | 0.08 | 0.33 | 0.24 |
| 5 | 98330050 | rs709386 | 0.248 | -0.133 | 0.41 | -2.018 | -0.065 | 0.105 | 0.105 | 0.108 | 0.24 | 0.07 | 0.17 | 0.07 | 0.27 | 0.2 |
| 5 | 98332239 | rs377731 | -1.422 | -0.284 | 0.478 | -2.058 | -0.009 | 0.113 | 0.113 | 0 | 0.44 | 0.78 | 0.33 | 0.17 | 0.36 | 0.46 |
| 5 | 98334750 | rs416750 | -0.607 | 0.168 | 0.673 | -2.002 | -0.008 | 0.001 | 0.001 | 0 | 0.32 | 0.32 | 0.23 | 0.07 | 0.33 | 0.24 |
| 5 | 98358191 | rs6878136 | -0.122 | 0.124 | -0.193 | -1.817 | -0.018 | 0.073 | 0.073 | 0 | 0.75 | 0.9 | 0.83 | 0.93 | 0.75 | 0.85 |
| 5 | 98365881 | rs12519797 | 0.981 | -0.651 | -0.531 | -2.072 | 0.036 | 0.01 | 0.01 | 0.023 | 0.23 | 0.15 | 0.13 | 0.2 | 0.15 | 0.21 |
| 5 | 98382640 | rs10040675 | 0.174 | -0.046 | 0.177 | -2.128 | 0.036 | -0.027 | -0.027 | 1.476 | 0.05 | 0.19 | 0.05 | 0 | 0.06 | 0.04 |
| 5 | 98383404 | rs11745405 | 0.018 | -0.317 | 0.216 | -2.154 | -0.015 | 0.07 | 0.07 | 0.009 | 0.26 | 0.15 | 0.18 | 0.07 | 0.26 | 0.15 |
| 5 | 98386485 | rs12659212 | -0.63 | -0.472 | 0.634 | -2.097 | 0.001 | -0.033 | -0.033 | 1.056 | 0.34 | 0.43 | 0.3 | 0.16 | 0.32 | 0.34 |
| 5 | 98395559 | rs11242579 | -1.19 | -0.498 | 0.28 | -2.184 | 0.009 | 0.147 | 0.147 | 0.054 | 0.43 | 0.8 | 0.33 | 0.16 | 0.36 | 0.46 |
| 5 | 98411852 | rs11738286 | -0.144 | -0.738 | 0.524 | -2.198 | -0.006 | 0.02 | 0.02 | 0.448 | 0.23 | 0.18 | 0.2 | 0.1 | 0.26 | 0.29 |

| | | | | | | | | | | | | | | | | |
|---|---|---|---|---|---|---|---|---|---|---|---|---|---|---|---|---|
| 5 | 98412995 | rs6896400 | -0.188 | -0.739 | 0.89 | -2.22 | 0.001 | -0.03 | -0.03 | 1.523 | 0.23 | 0.23 | 0.19 | 0.09 | 0.26 | 0.29 |
| 5 | 98413809 | rs6596673 | -1.422 | -0.796 | 0.728 | -2.264 | 0.033 | 0.142 | 0.142 | 0.294 | 0.41 | 0.81 | 0.31 | 0.18 | 0.32 | 0.47 |
| 5 | 98418461 | rs6596679 | -1.37 | -0.833 | 1.156 | -2.268 | 0.011 | 0.032 | 0.032 | 0.014 | 0.39 | 0.65 | 0.3 | 0.18 | 0.32 | 0.47 |
| 5 | 98423627 | rs10056306 | -1.37 | -0.819 | 1.123 | -2.261 | 0.011 | 0.032 | 0.032 | 0.014 | 0.39 | 0.65 | 0.3 | 0.18 | 0.32 | 0.47 |
| 5 | 98424099 | rs17166641 | 0.034 | -0.77 | 1.343 | -2.213 | -0.019 | -0.018 | -0.018 | 0 | 0.12 | 0.18 | 0.09 | 0.08 | 0.07 | 0.16 |
| 5 | 98425232 | rs10059861 | 0.125 | -0.729 | 1.334 | -2.228 | -0.002 | -0.003 | -0.003 | 0 | 0.22 | 0.21 | 0.18 | 0.09 | 0.24 | 0.24 |
| 5 | 98427828 | rs4035624 | -0.641 | -0.772 | 1.537 | -2.239 | 0.002 | -0.1 | -0.1 | 2.884 | 0.28 | 0.44 | 0.19 | 0.1 | 0.26 | 0.31 |
| 5 | 98431058 | rs7724878 | -0.569 | -0.783 | 1.566 | -2.242 | 0.001 | -0.085 | -0.085 | 0 | 0.27 | 0.39 | 0.19 | 0.09 | 0.26 | 0.31 |
| 5 | 98435558 | rs17166666 | 0.06 | -0.852 | 1.363 | -2.184 | -0.028 | -0.014 | -0.014 | 0 | 0.12 | 0.39 | 0.11 | 0.08 | 0.05 | 0.1 |
| 5 | 98442050 | rs11748163 | 0.146 | -0.606 | 1.151 | -2.189 | -0.011 | 0.021 | 0.021 | 0.336 | 0.22 | 0.15 | 0.26 | 0.09 | 0.25 | 0.24 |
| 5 | 98442871 | rs1447379 | -0.595 | -0.626 | 1.285 | -2.203 | 0.008 | -0.088 | -0.088 | 0.057 | 0.28 | 0.42 | 0.28 | 0.09 | 0.27 | 0.32 |
| 5 | 98447107 | rs6883882 | -0.028 | -0.936 | 0.584 | -2.53 | 0.005 | 0 | 0 | 4.766 | 0.45 | 0.55 | 0.26 | 0.19 | 0.3 | 0.37 |
| 5 | 98447682 | rs12054706 | 0.846 | -0.981 | 0.612 | -2.548 | 0.026 | 0.061 | 0.061 | 3.478 | 0.33 | 0.18 | 0.15 | 0.11 | 0.23 | 0.27 |
| 5 | 98451600 | rs10464032 | 0.132 | -0.865 | 0.42 | -2.541 | 0.003 | -0.004 | -0.004 | 0.279 | 0.46 | 0.53 | 0.27 | 0.19 | 0.34 | 0.38 |
| 5 | 98452374 | rs726172 | -1.266 | -0.888 | 0.376 | -2.632 | -0.055 | 0.03 | 0.03 | 0.514 | 0.69 | 0.82 | 0.53 | 0.27 | 0.55 | 0.64 |
| 5 | 98452446 | rs1447381 | -1.014 | -0.847 | 0.346 | -2.613 | -0.017 | 0.018 | 0.018 | 1.055 | 0.67 | 0.69 | 0.51 | 0.27 | 0.55 | 0.63 |
| 5 | 98456213 | rs1447383 | 0.676 | -0.787 | 0.007 | -2.553 | 0 | -0.001 | -0.001 | 0.225 | 0.21 | 0.17 | 0.25 | 0.09 | 0.2 | 0.24 |
| 5 | 98458109 | rs7708845 | 0.78 | -0.768 | -0.002 | -2.544 | 0 | -0.001 | -0.001 | 0 | 0.21 | 0.16 | 0.25 | 0.09 | 0.2 | 0.24 |
| 5 | 98461860 | rs10039446 | 0.8 | -0.861 | -0.081 | -2.35 | 0.015 | 0.035 | 0.035 | 1.596 | 0.32 | 0.19 | 0.15 | 0.11 | 0.24 | 0.26 |
| 5 | 98476620 | rs12660041 | 0.518 | -0.322 | 0.025 | -1.871 | 0 | -0.005 | -0.005 | 0 | 0.11 | 0.12 | 0.11 | 0.08 | 0.11 | 0.12 |
| 5 | 98488261 | rs10055586 | 0.369 | -0.687 | 0.021 | -1.9 | -0.024 | -0.037 | -0.037 | 0 | 0.35 | 0.52 | 0.16 | 0.18 | 0.24 | 0.31 |
| 5 | 98496724 | rs997391 | -1.912 | 0.337 | -0.764 | -2.144 | -0.089 | -0.08 | -0.08 | 0 | 0.86 | 0.98 | 0.59 | 0.28 | 0.7 | 0.77 |
| 5 | 98497977 | rs4277937 | 0.36 | -0.278 | -0.513 | -2.125 | 0.001 | -0.011 | -0.011 | 0 | 0.33 | 0.31 | 0.31 | 0.09 | 0.3 | 0.33 |
| 5 | 98503405 | rs10071543 | 0.125 | -0.327 | -0.431 | -2.101 | 0.014 | 0.021 | 0.021 | 0 | 0.39 | 0.62 | 0.19 | 0.18 | 0.26 | 0.33 |
| 5 | 98513824 | rs10056524 | 1.081 | -0.352 | -0.4 | -2.024 | 0.007 | 0.027 | 0.027 | 0.009 | 0.28 | 0.19 | 0.13 | 0.15 | 0.22 | 0.28 |
| 5 | 98520952 | rs10479407 | 0.707 | -0.34 | -0.367 | -2.018 | 0.008 | 0.013 | 0.013 | 0 | 0.32 | 0.28 | 0.14 | 0.16 | 0.22 | 0.31 |
| 5 | 98523802 | rs12109262 | 0.585 | -0.327 | -0.419 | -1.766 | -0.001 | -0.012 | -0.012 | 1.301 | 0.29 | 0.32 | 0.29 | 0.11 | 0.28 | 0.3 |
| 5 | 98523839 | rs12108959 | 0.33 | -0.136 | -0.516 | -1.544 | 0.006 | -0.014 | -0.014 | 3.027 | 0.57 | 0.49 | 0.41 | 0.27 | 0.49 | 0.58 |
| 5 | 98524787 | rs7715226 | -1.17 | -0.092 | -0.309 | -1.57 | -0.034 | -0.071 | -0.071 | 0 | 0.16 | 0.49 | 0.09 | 0.03 | 0.06 | 0.12 |
| 5 | 98537121 | rs2121665 | -0.876 | 0.273 | -0.893 | -0.743 | -0.121 | -0.127 | -0.127 | 4.573 | 0.73 | 0.94 | 0.51 | 0.45 | 0.56 | 0.69 |
| 5 | 98552184 | rs1121306 | 0.67 | -0.04 | -0.819 | -0.848 | 0.019 | 0.006 | 0.006 | 0 | 0.3 | 0.19 | 0.3 | 0.29 | 0.27 | 0.32 |
| 5 | 98554365 | rs10057544 | 0.756 | 0.039 | -0.98 | -0.867 | 0.001 | 0.015 | 0.015 | 0 | 0.32 | 0.23 | 0.16 | 0.14 | 0.27 | 0.31 |
| 5 | 98566596 | rs1073601 | 0.176 | 0.196 | -0.882 | -0.572 | -0.018 | -0.026 | -0.026 | 0 | 0.35 | 0.46 | 0.35 | 0.29 | 0.32 | 0.34 |
| 5 | 98569709 | rs1574968 | -0.576 | 0.835 | -1.428 | -0.181 | -0.148 | -0.114 | -0.114 | 0 | 0.73 | 0.98 | 0.54 | 0.44 | 0.59 | 0.7 |
| 5 | 98576246 | rs7716477 | 0.485 | 0.162 | -0.851 | -0.432 | -0.009 | -0.01 | -0.01 | 0 | 0.22 | 0.29 | 0.21 | 0.15 | 0.2 | 0.32 |
| 5 | 98579784 | rs17166786 | -0.663 | 0.112 | -1.006 | -0.48 | 0.002 | 0.005 | 0.005 | 3.807 | 0.49 | 0.53 | 0.37 | 0.27 | 0.46 | 0.45 |
| 5 | 98583720 | rs10039243 | 1 | 0.033 | -1.025 | -0.205 | -0.001 | -0.016 | -0.016 | 0 | 0.25 | 0.33 | 0.17 | 0.16 | 0.22 | 0.28 |
| 5 | 98591861 | rs1116078 | -1.772 | 0.085 | -1.323 | -0.205 | -0.132 | -0.19 | -0.19 | 2.383 | 0.74 | 0.98 | 0.52 | 0.44 | 0.61 | 0.7 |
| 5 | 98593453 | rs1500251 | 0.717 | -0.468 | -0.597 | -0.097 | -0.004 | -0.005 | -0.005 | 0 | 0.22 | 0.23 | 0.19 | 0.15 | 0.2 | 0.31 |
| 5 | 98599968 | rs10057969 | 0.104 | -0.09 | -0.424 | 0.194 | 0.003 | -0.016 | -0.016 | 0 | 0.28 | 0.39 | 0.19 | 0.2 | 0.25 | 0.3 |
| 5 | 98644175 | rs4702966 | 0.276 | 0.354 | -1.17 | 0.604 | -0.097 | -0.103 | -0.103 | 0.181 | 0.35 | 0.12 | 0.6 | 0.53 | 0.54 | 0.37 |
| 5 | 98650208 | rs6594343 | -0.203 | -0.327 | -0.507 | 0.206 | -0.024 | -0.025 | -0.025 | 0.228 | 0.68 | 0.59 | 0.84 | 0.73 | 0.79 | 0.68 |
| 5 | 98656513 | rs6894643 | 0.771 | -0.869 | -0.117 | 0.544 | -0.013 | 0.014 | 0.014 | 0 | 0.3 | 0.37 | 0.23 | 0.3 | 0.24 | 0.32 |
| 5 | 98694849 | rs6892901 | -0.902 | -0.659 | -0.505 | 0.485 | -0.172 | -0.102 | -0.102 | 0.475 | 0.67 | 0.95 | 0.4 | 0.58 | 0.46 | 0.65 |
| 5 | 98700034 | rs1523383 | 1.361 | -0.913 | -0.428 | 0.511 | 0 | -0.016 | -0.016 | 0 | 0.24 | 0.16 | 0.18 | 0.23 | 0.22 | 0.24 |
| 5 | 98702938 | rs2461978 | -0.478 | -0.962 | -0.143 | 0.467 | 0.045 | 0.026 | 0.026 | 0.863 | 0.42 | 0.66 | 0.21 | 0.34 | 0.23 | 0.4 |
| 5 | 98713577 | rs2030605 | -0.925 | -0.384 | -0.369 | 0.26 | 0.019 | 0.061 | 0.061 | 0.664 | 0.51 | 0.73 | 0.28 | 0.37 | 0.38 | 0.49 |
| 5 | 98721872 | rs2511925 | -0.893 | -0.459 | -0.261 | 0.383 | -0.013 | -0.034 | -0.034 | 0 | 0.68 | 0.8 | 0.46 | 0.61 | 0.59 | 0.66 |
| 5 | 98725366 | rs6871534 | -0.08 | -0.756 | 0.045 | 0.056 | 0.006 | -0.012 | -0.012 | 0 | 0.34 | 0.38 | 0.21 | 0.3 | 0.33 | 0.38 |
| 5 | 98734713 | rs2963595 | -1.528 | -0.586 | -0.029 | 0.138 | -0.041 | -0.037 | -0.037 | 0.039 | 0.74 | 0.88 | 0.49 | 0.67 | 0.6 | 0.73 |
| 5 | 98779530 | rs1403660 | 1.65 | -0.717 | -0.059 | 0.088 | 0.003 | -0.025 | -0.025 | 0 | 0.24 | 0.18 | 0.22 | 0.32 | 0.22 | 0.24 |
| 5 | 98793790 | rs6863383 | -0.149 | -0.521 | -0.004 | 0.033 | 0.012 | -0.015 | -0.015 | 0.062 | 0.34 | 0.39 | 0.34 | 0.3 | 0.36 | 0.37 |
| 5 | 98802762 | rs6885370 | 0.928 | -0.404 | -0.085 | 0.048 | -0.032 | 0.023 | 0.023 | 0 | 0.07 | 0.02 | 0.06 | 0.02 | 0.14 | 0.06 |
| 5 | 98809033 | rs11954019 | 0.046 | -0.493 | -0.022 | 0.077 | -0.017 | 0.001 | 0.001 | 0 | 0.07 | 0.1 | 0.04 | 0.05 | 0.01 | 0.07 |
| 5 | 98878767 | rs1939397 | -0.328 | -0.45 | -0.027 | 0.114 | -0.004 | 0.001 | 0.001 | 0 | 0.58 | 0.57 | 0.56 | 0.58 | 0.59 | 0.6 |
| 5 | 98884847 | rs1939402 | 1.559 | -0.434 | -0.117 | 0.131 | -0.027 | -0.048 | -0.048 | 0 | 0.25 | 0.12 | 0.36 | 0.33 | 0.36 | 0.28 |
| 5 | 98888354 | rs17167031 | -0.275 | -0.457 | -0.183 | 0.042 | 0.013 | 0.008 | 0.008 | 0 | 0.1 | 0.05 | 0.04 | 0.05 | 0.05 | 0.1 |
| 5 | 98888857 | rs2155214 | -0.644 | -0.479 | -0.181 | 0.02 | -0.002 | 0.001 | 0.001 | 0.06 | 0.42 | 0.39 | 0.36 | 0.3 | 0.4 | 0.4 |
| 5 | 98899755 | rs2548103 | 1.562 | -0.366 | -0.202 | 0.125 | -0.027 | -0.048 | -0.048 | 0 | 0.25 | 0.12 | 0.36 | 0.33 | 0.36 | 0.28 |
| 5 | 98922899 | rs2682158 | -0.643 | -0.307 | -0.152 | 0.071 | -0.001 | 0.003 | 0.003 | 1.633 | 0.64 | 0.67 | 0.57 | 0.57 | 0.59 | 0.6 |
| 5 | 98923389 | rs6861784 | 0.695 | -0.072 | 0.488 | 0.133 | 0 | 0 | 0 | 0 | 0.22 | 0.25 | 0.25 | 0.27 | 0.18 | 0.27 |
| 5 | 98931151 | rs681634 | -1.617 | 0.073 | -0.055 | -0.015 | -0.095 | -0.119 | -0.119 | 0.021 | 0.78 | 0.97 | 0.65 | 0.67 | 0.64 | 0.72 |
| 5 | 98932522 | rs664978 | -0.341 | -0.076 | 0.421 | -0.125 | 0.023 | 0.007 | 0.007 | 0.516 | 0.65 | 0.48 | 0.57 | 0.65 | 0.51 | 0.64 |
| 5 | 98936840 | rs3850908 | 0.056 | -0.213 | 0.372 | -0.116 | -0.034 | -0.079 | -0.079 | 0.005 | 0.3 | 0.08 | 0.29 | 0.36 | 0.25 | 0.38 |
| 5 | 98943589 | rs4400164 | 0.103 | -0.172 | 0.402 | -0.186 | 0 | -0.001 | -0.001 | 0.506 | 0.56 | 0.47 | 0.54 | 0.65 | 0.46 | 0.62 |
| 5 | 98951156 | rs3850905 | -1.649 | -0.14 | 0.515 | -0.18 | -0.046 | -0.067 | -0.067 | 0 | 0.77 | 0.92 | 0.65 | 0.67 | 0.64 | 0.71 |
| 5 | 98954234 | rs6892152 | -1.772 | -0.135 | 0.435 | -0.178 | -0.095 | -0.119 | -0.119 | 0.091 | 0.78 | 0.97 | 0.65 | 0.67 | 0.64 | 0.71 |
| 5 | 98967037 | rs3850902 | 0.126 | -0.139 | 0.448 | -0.178 | 0 | -0.001 | -0.001 | 0 | 0.56 | 0.47 | 0.53 | 0.61 | 0.46 | 0.6 |
| 5 | 98972664 | rs6890382 | -1.59 | -0.036 | 0.158 | -0.039 | -0.06 | -0.093 | -0.093 | 0 | 0.77 | 0.96 | 0.65 | 0.63 | 0.64 | 0.69 |

| | | | | | | | | | | | | | | | | |
|---|---|---|---|---|---|---|---|---|---|---|---|---|---|---|---|---|
| 5 | 98993932 | rs10053765 | -0.197 | -0.169 | 0.121 | -0.384 | 0.021 | 0.137 | 0.137 | 1.838 | 0.49 | 0.8 | 0.37 | 0.3 | 0.4 | 0.33 |
| 5 | 98994964 | rs10036544 | 0.342 | -0.217 | 0.518 | -0.406 | -0.001 | 0.046 | 0.046 | 2.735 | 0.42 | 0.28 | 0.33 | 0.3 | 0.4 | 0.31 |
| 5 | 99001200 | rs3844093 | 1.421 | -0.143 | 0.69 | -0.225 | 0.001 | 0.002 | 0.002 | 0 | 0.29 | 0.25 | 0.26 | 0.28 | 0.26 | 0.25 |
| 5 | 99005777 | rs1462427 | 0.128 | -0.177 | 0.763 | -0.262 | 0.061 | 0.1 | 0.1 | 0.617 | 0.44 | 0.78 | 0.32 | 0.28 | 0.32 | 0.3 |
| 5 | 99012291 | rs7707171 | 1.413 | -0.108 | 0.737 | -0.197 | 0.002 | 0.003 | 0.003 | 0 | 0.29 | 0.25 | 0.26 | 0.28 | 0.26 | 0.25 |
| 5 | 99021162 | rs10078910 | 1.28 | -0.16 | 0.318 | -0.221 | -0.016 | 0.018 | 0.018 | 0 | 0.13 | 0.01 | 0.04 | 0.11 | 0.09 | 0.08 |
| 5 | 99029453 | rs12652513 | 0.632 | 0.034 | 0.756 | -0.048 | 0.014 | -0.014 | -0.014 | 1.292 | 0.08 | 0.14 | 0.06 | 0.02 | 0.12 | 0.07 |
| 5 | 99030629 | rs12187654 | 1.239 | 0.041 | 0.924 | -0.049 | -0.003 | -0.009 | -0.009 | 0 | 0.3 | 0.34 | 0.27 | 0.28 | 0.28 | 0.25 |
| 5 | 99035683 | rs9784617 | 0.158 | -0.037 | 0.33 | -0.031 | -0.055 | 0.088 | 0.088 | 0 | 0.43 | 0.91 | 0.37 | 0.29 | 0.41 | 0.32 |
| 5 | 99037694 | rs12654674 | 0.165 | -0.041 | 0.355 | -0.032 | -0.046 | 0.085 | 0.085 | 0 | 0.43 | 0.89 | 0.36 | 0.29 | 0.41 | 0.32 |
| 5 | 99049676 | rs10073413 | 0.519 | 0.18 | 0.391 | 0.141 | 0.014 | -0.07 | -0.07 | 11.365 | 0.25 | 0.05 | 0.27 | 0.32 | 0.16 | 0.35 |
| 5 | 99052298 | rs1462420 | 1.125 | 0.161 | 0.695 | 0.208 | -0.013 | -0.006 | -0.006 | 0.117 | 0.33 | 0.49 | 0.3 | 0.35 | 0.28 | 0.22 |
| 5 | 99055048 | rs1462417 | -0.422 | 0.149 | 0.452 | 0.333 | -0.049 | -0.026 | -0.026 | 0 | 0.68 | 0.89 | 0.63 | 0.69 | 0.56 | 0.64 |
| 5 | 99057627 | rs1462411 | 0.259 | 0.155 | 0.534 | 0.352 | -0.002 | 0.001 | 0.001 | 0 | 0.27 | 0.23 | 0.28 | 0.32 | 0.16 | 0.34 |
| 5 | 99063246 | rs17167155 | 0.417 | 0.347 | 0.046 | -0.21 | -0.009 | -0.048 | -0.048 | 0.073 | 0.09 | 0.17 | 0.05 | 0 | 0.06 | 0.03 |
| 5 | 99082357 | rs2118735 | -0.565 | 0.48 | -0.569 | -0.036 | -0.042 | -0.048 | -0.048 | 0 | 0.72 | 0.93 | 0.65 | 0.69 | 0.61 | 0.64 |
| 5 | 99090148 | rs10463647 | -0.573 | 0.488 | -0.566 | -0.033 | -0.042 | -0.048 | -0.048 | 0 | 0.72 | 0.93 | 0.65 | 0.69 | 0.61 | 0.64 |
| 5 | 99107961 | rs1161240 | 0.065 | 0.814 | -0.283 | 0.168 | 0.009 | 0 | 0 | 0 | 0.5 | 0.59 | 0.47 | 0.44 | 0.49 | 0.35 |
| 5 | 99110348 | rs1622879 | -0.061 | 0.871 | -0.274 | 0.103 | 0.008 | 0.001 | 0.001 | 0 | 0.5 | 0.59 | 0.44 | 0.47 | 0.47 | 0.35 |
| 5 | 99121725 | rs2028276 | 0.853 | 0.447 | -1.157 | -0.094 | -0.036 | -0.043 | -0.043 | 0.052 | 0.31 | 0.18 | 0.45 | 0.36 | 0.32 | 0.23 |
| 5 | 99140203 | rs6594760 | 0.015 | 0.355 | -1.178 | 0.102 | 0.016 | -0.02 | -0.02 | 0.078 | 0.57 | 0.78 | 0.64 | 0.4 | 0.57 | 0.42 |
| 5 | 99146425 | rs2123811 | -1.087 | 0.359 | -1.312 | 0.115 | -0.066 | -0.116 | -0.116 | 0 | 0.82 | 0.99 | 0.74 | 0.67 | 0.72 | 0.73 |
| 5 | 99146588 | rs2168002 | 0.057 | 0.418 | -1.199 | 0.097 | 0.016 | -0.02 | -0.02 | 3.97 | 0.57 | 0.78 | 0.63 | 0.4 | 0.57 | 0.42 |
| 5 | 99156362 | rs2369754 | 0.057 | 0.422 | -1.199 | 0.097 | 0.013 | -0.02 | -0.02 | 1.388 | 0.57 | 0.78 | 0.63 | 0.4 | 0.57 | 0.42 |
| 5 | 99158909 | rs4320303 | 0.057 | 0.407 | -1.188 | 0.084 | 0.016 | -0.02 | -0.02 | 0 | 0.57 | 0.78 | 0.63 | 0.39 | 0.57 | 0.42 |
| 5 | 99163402 | rs6594781 | -0.586 | 0.539 | -1.075 | 0.272 | -0.009 | -0.045 | -0.045 | 0.032 | 0.79 | 0.86 | 0.73 | 0.65 | 0.71 | 0.69 |
| 5 | 99179262 | rs7448872 | -0.878 | 0.513 | -1.126 | 0.292 | -0.066 | -0.116 | -0.116 | 0.162 | 0.82 | 0.99 | 0.74 | 0.67 | 0.72 | 0.73 |
| 5 | 99213789 | rs7709951 | 0.734 | 0.19 | -1.03 | 0.075 | 0.024 | 0.014 | 0.014 | 0 | 0.23 | 0.09 | 0.1 | 0.25 | 0.14 | 0.28 |
| 5 | 99221687 | rs3844092 | 1.169 | 0.343 | -1.2 | -0.291 | -0.051 | -0.056 | -0.056 | 0 | 0.31 | 0.15 | 0.44 | 0.35 | 0.32 | 0.23 |
| 5 | 99224816 | rs2616289 | -0.309 | 0.17 | -0.927 | -0.404 | -0.003 | -0.021 | -0.021 | 0 | 0.73 | 0.74 | 0.69 | 0.64 | 0.66 | 0.67 |
| 5 | 99226381 | rs2616293 | 0.409 | 0.186 | -0.425 | -0.43 | 0.012 | -0.093 | -0.093 | 0 | 0.2 | 0.43 | 0.14 | 0.03 | 0.19 | 0.16 |
| 5 | 99226983 | rs2602958 | -0.173 | 0.381 | -0.506 | -0.425 | 0.011 | 0.003 | 0.003 | 0.184 | 0.48 | 0.28 | 0.42 | 0.6 | 0.51 | 0.61 |
| 5 | 99229298 | rs2967107 | -1.355 | 0.408 | -0.557 | -0.409 | -0.024 | -0.04 | -0.04 | 0 | 0.69 | 0.78 | 0.56 | 0.63 | 0.7 | 0.77 |
| 5 | 99234994 | rs2616305 | 0.422 | 0.4 | -0.519 | -0.465 | 0 | -0.082 | -0.082 | 0 | 0.21 | 0.49 | 0.14 | 0.04 | 0.19 | 0.16 |
| 5 | 99235642 | rs13356795 | 0.353 | 0.27 | -0.576 | -0.396 | 0.017 | 0.019 | 0.019 | 0 | 0.3 | 0.24 | 0.14 | 0.27 | 0.2 | 0.34 |
| 5 | 99239733 | rs2924621 | -1.251 | 0.51 | -0.645 | -0.307 | -0.015 | -0.041 | -0.041 | 4.887 | 0.72 | 0.81 | 0.6 | 0.65 | 0.75 | 0.81 |
| 5 | 99248759 | rs1161319 | 0.704 | 0.629 | -0.212 | -0.238 | -0.062 | -0.115 | -0.115 | 0.171 | 0.19 | 0.01 | 0.24 | 0.35 | 0.27 | 0.26 |
| 5 | 99253302 | rs2927429 | -0.65 | 0.303 | 0.185 | -0.056 | -0.013 | 0.032 | 0.032 | 0 | 0.48 | 0.67 | 0.48 | 0.38 | 0.58 | 0.48 |
| 5 | 99258312 | rs2602970 | -0.392 | 0.742 | -0.267 | 0.297 | 0.01 | 0.006 | 0.006 | 0 | 0.39 | 0.42 | 0.44 | 0.41 | 0.54 | 0.46 |
| 5 | 99260185 | rs17167440 | -1.038 | 0.264 | -0.158 | 0.032 | -0.017 | -0.026 | -0.026 | 0 | 0.16 | 0.25 | 0.1 | 0.06 | 0.1 | 0.08 |
| 5 | 99260268 | rs17414873 | -1.342 | 0.179 | -0.398 | 0.028 | 0.01 | 0.011 | 0.011 | 0 | 0.11 | 0.09 | 0.06 | 0.04 | 0.07 | 0.03 |
| 5 | 99260992 | rs10478218 | -1.536 | 0.71 | 0.347 | -0.065 | 0.052 | -0.032 | -0.032 | 0 | 0.07 | 0.21 | 0.07 | 0 | 0.09 | 0.01 |
| 5 | 99263712 | rs13182682 | -0.472 | 1.677 | -0.203 | -0.014 | -0.053 | -0.016 | -0.016 | 1.358 | 0.14 | 0.03 | 0.3 | 0.11 | 0.24 | 0.28 |
| 5 | 99265096 | rs1184440 | -0.154 | 1.376 | -0.473 | 0.157 | -0.01 | -0.015 | -0.015 | 0 | 0.37 | 0.26 | 0.44 | 0.45 | 0.41 | 0.23 |
| 5 | 99265408 | rs2927423 | -1.291 | 0.946 | 0.195 | 0.215 | 0.048 | 0.042 | 0.042 | 0.238 | 0.58 | 0.73 | 0.78 | 0.68 | 0.68 | 0.54 |
| 5 | 99265461 | rs2924606 | -1.294 | 0.954 | 0.198 | 0.218 | 0.048 | 0.042 | 0.042 | 0.232 | 0.58 | 0.73 | 0.78 | 0.68 | 0.68 | 0.54 |
| 5 | 99268130 | rs2602972 | 0.223 | 0.245 | -0.065 | 0.033 | 0.009 | -0.063 | -0.063 | 0 | 0.42 | 0.14 | 0.38 | 0.5 | 0.36 | 0.25 |
| 5 | 99270055 | rs2259060 | 0.015 | 0.261 | -0.129 | 0.039 | -0.007 | -0.031 | -0.031 | 0 | 0.44 | 0.21 | 0.42 | 0.51 | 0.43 | 0.28 |
| 5 | 99270398 | rs2967099 | -0.324 | 0.251 | 0.335 | 0.032 | 0 | -0.001 | -0.001 | 0.051 | 0.51 | 0.45 | 0.47 | 0.51 | 0.47 | 0.28 |
| 5 | 99272740 | rs1161315 | -1.184 | 0.907 | 0.23 | 0.032 | 0.003 | -0.001 | -0.001 | 0 | 0.36 | 0.43 | 0.26 | 0.29 | 0.35 | 0.34 |
| 5 | 99279767 | rs1381988 | -1.16 | 0.907 | 0.251 | 0.032 | 0.001 | -0.001 | -0.001 | 0.116 | 0.35 | 0.41 | 0.26 | 0.29 | 0.35 | 0.34 |
| 5 | 99282835 | rs966881 | 0.537 | 0.94 | 0.32 | 0.04 | -0.001 | -0.003 | -0.003 | 0 | 0.12 | 0.15 | 0.14 | 0.11 | 0.24 | 0.23 |
| 5 | 99284570 | rs1870970 | 1.141 | 0.912 | 0.035 | -0.016 | 0.003 | 0.028 | 0.028 | 0 | 0.21 | 0.12 | 0.18 | 0.05 | 0.23 | 0.08 |
| 5 | 99285749 | rs899056 | -1.069 | 0.907 | 0.056 | 0.166 | 0.004 | 0.012 | 0.012 | 2.164 | 0.35 | 0.43 | 0.26 | 0.38 | 0.35 | 0.34 |
| 5 | 99287946 | rs10515289 | 0.373 | 0.8 | 0.028 | -0.166 | 0.007 | 0.028 | 0.028 | 0 | 0.26 | 0.18 | 0.22 | 0.05 | 0.27 | 0.08 |
| 5 | 99288390 | rs1477656 | 1.051 | 0.807 | 0.03 | -0.151 | 0.003 | 0.03 | 0.03 | 0 | 0.22 | 0.12 | 0.18 | 0.05 | 0.23 | 0.08 |
| 5 | 99290311 | rs9326958 | 0.186 | 0.69 | 0.076 | -0.098 | -0.032 | 0.008 | 0.008 | 4.622 | 0.18 | 0.28 | 0.07 | 0.18 | 0.07 | 0.11 |
| 5 | 99292308 | rs10079213 | 0.202 | 0.692 | 0.078 | -0.097 | -0.028 | 0.006 | 0.006 | 4.767 | 0.18 | 0.26 | 0.07 | 0.18 | 0.07 | 0.11 |
| 5 | 99293221 | rs4702973 | -0.879 | 0.772 | 0.058 | -0.098 | -0.002 | -0.005 | -0.005 | 0 | 0.45 | 0.44 | 0.3 | 0.23 | 0.35 | 0.19 |
| 5 | 99302219 | rs10045935 | -0.195 | 0.827 | 0.037 | -0.167 | -0.005 | -0.032 | -0.032 | 0.002 | 0.37 | 0.37 | 0.24 | 0.1 | 0.3 | 0.15 |
| 5 | 99310930 | rs1381989 | 1.163 | 0.891 | -0.079 | -0.076 | 0 | 0 | 0 | 0 | 0.42 | 0.39 | 0.55 | 0.64 | 0.4 | 0.59 |
| 5 | 99314749 | rs9326966 | 0.458 | 0.61 | -0.331 | -0.271 | -0.001 | 0.031 | 0.031 | 0 | 0.22 | 0.12 | 0.18 | 0.04 | 0.24 | 0.07 |
| 5 | 99332281 | rs2370257 | -0.037 | 0.395 | 0.252 | -0.103 | -0.029 | -0.031 | -0.031 | 0 | 0.15 | 0.34 | 0.05 | 0.06 | 0.06 | 0.06 |
| 5 | 99332696 | rs13190093 | 1.413 | 0.467 | -0.789 | -0.447 | -0.067 | 0.034 | 0.034 | 1.179 | 0.3 | 0.02 | 0.18 | 0.24 | 0.32 | 0.37 |
| 5 | 99342047 | rs10067427 | -0.66 | 0.73 | 0.045 | -0.284 | 0.011 | 0.042 | 0.042 | 2.14 | 0.42 | 0.71 | 0.33 | 0.26 | 0.38 | 0.19 |
| 5 | 99346521 | rs10057187 | -0.204 | 0.731 | 0.252 | -0.347 | 0 | -0.018 | -0.018 | 0.148 | 0.36 | 0.54 | 0.3 | 0.11 | 0.35 | 0.14 |
| 5 | 99349315 | rs9326976 | 0.567 | 0.541 | 0.379 | -0.366 | 0.024 | -0.116 | -0.116 | 0.525 | 0.25 | 0.37 | 0.25 | 0.04 | 0.3 | 0.08 |
| 5 | 99352463 | rs11241340 | 0.753 | 0.539 | 0.576 | -0.367 | -0.004 | -0.001 | -0.001 | 0 | 0.23 | 0.16 | 0.25 | 0.04 | 0.3 | 0.08 |
| 5 | 99377324 | rs13359189 | 0.912 | 0.476 | 0.514 | -0.285 | -0.017 | -0.007 | -0.007 | 0 | 0.12 | 0.18 | 0.03 | 0.06 | 0.03 | 0.06 |

| | | | | | | | | | | | | | | | | |
|---|---|---|---|---|---|---|---|---|---|---|---|---|---|---|---|---|
| 5 | 99387066 | rs10515294 | 0.584 | 0.544 | 0.839 | -0.421 | 0.003 | -0.033 | -0.033 | 0 | 0.24 | 0.22 | 0.25 | 0.05 | 0.3 | 0.08 |
| 5 | 99387360 | rs10515295 | -0.92 | 0.525 | 0.91 | -0.309 | 0.033 | 0.026 | 0.026 | 2.958 | 0.44 | 0.76 | 0.31 | 0.3 | 0.36 | 0.19 |
| 5 | 99388523 | rs17764255 | -0.967 | 0.743 | 0.907 | -0.319 | 0.014 | 0.029 | 0.029 | 2.803 | 0.44 | 0.77 | 0.33 | 0.3 | 0.41 | 0.2 |
| 5 | 99428065 | rs12332547 | -0.84 | 0.577 | 0.959 | -0.398 | -0.004 | -0.007 | -0.007 | 1.671 | 0.43 | 0.49 | 0.31 | 0.3 | 0.41 | 0.2 |
| 5 | 99451233 | rs7702667 | 0.624 | 0.614 | 0.844 | -0.374 | 0.001 | -0.024 | -0.024 | 0 | 0.24 | 0.21 | 0.25 | 0.05 | 0.3 | 0.08 |
| 5 | 99455090 | rs7737429 | -1.692 | 0.573 | 0.692 | -0.546 | -0.021 | 0.125 | 0.125 | 2.233 | 0.66 | 0.89 | 0.78 | 0.8 | 0.64 | 0.61 |
| 5 | 99458270 | rs4429827 | -1.129 | 0.519 | 1.17 | -0.347 | 0.017 | 0.029 | 0.029 | 2.66 | 0.45 | 0.81 | 0.33 | 0.3 | 0.41 | 0.2 |
| 5 | 99478428 | rs7705114 | -0.555 | 0.474 | 1.437 | -0.336 | -0.041 | 0.086 | 0.086 | 2.318 | 0.21 | 0.51 | 0.07 | 0.25 | 0.12 | 0.13 |
| 5 | 99479714 | rs4532340 | 0.716 | 0.527 | 1.066 | -0.37 | -0.014 | 0.042 | 0.042 | 0 | 0.23 | 0.12 | 0.24 | 0.05 | 0.29 | 0.09 |
| 5 | 99484502 | rs10035392 | -1.046 | 0.486 | 1.05 | -0.383 | 0.019 | 0.034 | 0.034 | 2.759 | 0.45 | 0.82 | 0.32 | 0.3 | 0.42 | 0.22 |
| 5 | 99492606 | rs4563598 | 0.664 | 0.51 | 1.016 | -0.375 | -0.014 | 0.042 | 0.042 | 0 | 0.23 | 0.12 | 0.24 | 0.05 | 0.29 | 0.09 |
| 5 | 99503303 | rs4389662 | 0.693 | 0.513 | 1.018 | -0.373 | -0.014 | 0.041 | 0.041 | 0 | 0.22 | 0.12 | 0.24 | 0.05 | 0.29 | 0.09 |
| 5 | 99507641 | rs10078007 | 0.71 | 0.537 | 1.014 | -0.377 | -0.016 | 0.04 | 0.04 | 0 | 0.22 | 0.12 | 0.24 | 0.05 | 0.29 | 0.09 |
| 5 | 99518198 | rs1580920 | -1.079 | 0.48 | 0.929 | -0.335 | 0.021 | 0.03 | 0.03 | 2.579 | 0.44 | 0.81 | 0.31 | 0.3 | 0.4 | 0.21 |
| 5 | 99522748 | rs2279855 | 0.166 | 0.632 | 1.18 | -0.184 | 0.004 | -0.008 | -0.008 | 0 | 0.15 | 0.13 | 0.05 | 0.25 | 0.1 | 0.1 |
| 5 | 99529358 | rs6895260 | 0.179 | 0.632 | 1.184 | -0.198 | 0.005 | -0.01 | -0.01 | 0 | 0.15 | 0.12 | 0.05 | 0.21 | 0.1 | 0.1 |
| 5 | 99537356 | rs2089160 | -0.61 | 0.586 | 1.491 | -0.297 | -0.003 | -0.005 | -0.005 | 0 | 0.3 | 0.61 | 0.43 | 0.24 | 0.31 | 0.14 |
| 5 | 99566704 | rs2655999 | 0.34 | 0.514 | 1.678 | -0.334 | -0.008 | 0.003 | 0.003 | 0 | 0.22 | 0.17 | 0.39 | 0.02 | 0.29 | 0.11 |
| 5 | 99582020 | rs326087 | 1.358 | 0.646 | 1.185 | -0.059 | -0.031 | 0.023 | 0.023 | 2.424 | 0.49 | 0.16 | 0.44 | 0.63 | 0.51 | 0.67 |
| 5 | 99583461 | rs167823 | 0.071 | 0.694 | 1.185 | -0.071 | 0.001 | 0.002 | 0.002 | 0.673 | 0.8 | 0.82 | 0.89 | 0.87 | 0.86 | 0.81 |
| 5 | 99598182 | rs709360 | 0.466 | 0.874 | 1.574 | -0.248 | -0.007 | -0.01 | -0.01 | 0 | 0.22 | 0.17 | 0.4 | 0.02 | 0.33 | 0.1 |
| 5 | 99601767 | rs795673 | 0.078 | 0.877 | 1.362 | -0.245 | 0.029 | -0.119 | -0.119 | 0.019 | 0.24 | 0.53 | 0.41 | 0.02 | 0.33 | 0.1 |
| 5 | 99602374 | rs1504534 | 1.465 | 1.126 | 1.024 | -0.379 | -0.026 | 0.013 | 0.013 | 0 | 0.07 | 0.01 | 0.24 | 0.02 | 0.09 | 0.06 |
| 5 | 99605709 | rs794992 | -1.106 | 1.028 | 1.094 | -0.313 | -0.001 | 0.002 | 0.002 | 0.32 | 0.31 | 0.73 | 0.44 | 0.25 | 0.35 | 0.13 |
| 5 | 99606693 | rs10515301 | -0.182 | 1.403 | 0.886 | -0.49 | -0.005 | 0.047 | 0.047 | 0 | 0.1 | 0.14 | 0.07 | 0.23 | 0.09 | 0.03 |
| 5 | 99608856 | rs610857 | 1.976 | 1.251 | 0.87 | -0.297 | -0.061 | 0.029 | 0.029 | 3.049 | 0.51 | 0.08 | 0.46 | 0.66 | 0.53 | 0.69 |
| 5 | 99609446 | rs167788 | -1.211 | 1.228 | 0.903 | -0.341 | -0.008 | -0.035 | -0.035 | 0.886 | 0.9 | 0.86 | 0.98 | 1 | 0.96 | 0.97 |
| 5 | 99612805 | rs324881 | 2.074 | 1.248 | 0.771 | -0.303 | -0.059 | 0.035 | 0.035 | 3.083 | 0.51 | 0.08 | 0.46 | 0.66 | 0.53 | 0.68 |
| 5 | 99614601 | rs1824777 | -0.475 | 1.275 | 1.124 | -0.257 | 0.021 | -0.106 | -0.106 | 0 | 0.25 | 0.54 | 0.42 | 0.02 | 0.33 | 0.1 |
| 5 | 99620958 | rs1833841 | -1.112 | 1.262 | 1.114 | -0.244 | -0.001 | 0.002 | 0.002 | 0.537 | 0.31 | 0.71 | 0.44 | 0.25 | 0.35 | 0.13 |
| 5 | 99630569 | rs794977 | -0.707 | 1.488 | 1.066 | -0.542 | 0.018 | -0.098 | -0.098 | 0.006 | 0.25 | 0.57 | 0.42 | 0.02 | 0.33 | 0.1 |
| 5 | 99636587 | rs17146237 | 0.518 | 1.431 | 0.858 | -0.5 | -0.009 | -0.004 | -0.004 | 0 | 0.08 | 0.02 | 0.08 | 0.09 | 0.08 | 0.16 |
| 5 | 99637888 | rs1973172 | -0.282 | 1.505 | 0.941 | -0.457 | 0.003 | -0.05 | -0.05 | 0 | 0.22 | 0.22 | 0.39 | 0.02 | 0.28 | 0.1 |
| 5 | 99639943 | rs1013602 | -0.076 | 1.536 | 0.604 | -0.443 | -0.043 | 0.09 | 0.09 | 0 | 0.2 | 0.04 | 0.37 | 0.02 | 0.28 | 0.1 |
| 5 | 99644635 | rs794524 | 0.204 | 1.62 | 0.542 | -0.509 | -0.021 | 0.085 | 0.085 | 0 | 0.19 | 0.03 | 0.3 | 0 | 0.21 | 0.06 |
| 5 | 99656681 | rs173467 | 0.826 | 2.311 | 1.058 | -0.531 | 0.119 | -0.059 | -0.059 | 0 | 0.83 | 0.52 | 0.85 | 0.92 | 0.77 | 0.82 |
| 5 | 99660149 | rs161321 | -0.157 | 2.373 | 1.306 | -0.528 | 0.072 | -0.01 | -0.01 | 0 | 0.11 | 0.23 | 0.13 | 0.08 | 0.19 | 0.16 |
| 5 | 99661931 | rs161320 | -0.213 | 2.373 | 1.348 | -0.528 | 0.085 | -0.009 | -0.009 | 3.954 | 0.11 | 0.26 | 0.13 | 0.08 | 0.19 | 0.16 |
| 5 | 99664527 | rs161318 | 0.513 | 2.391 | 1.23 | -0.527 | -0.023 | 0 | 0 | 0 | 0.08 | 0.03 | 0.09 | 0.08 | 0.14 | 0.15 |
| 5 | 99696831 | rs11241501 | 2.567 | 2.189 | 0.651 | -0.042 | -0.023 | -0.12 | -0.12 | 0.019 | 0.33 | 0.01 | 0.22 | 0.37 | 0.32 | 0.47 |
| 5 | 99742364 | rs898765 | -0.922 | 2.151 | 0.855 | 0.05 | 0.002 | -0.024 | -0.024 | 0 | 0.93 | 0.9 | 0.98 | 1 | 0.95 | 0.97 |
| 5 | 99772694 | rs10079469 | 0.4 | 2.092 | 0.859 | -0.066 | -0.029 | 0.129 | 0.129 | 0 | 0.24 | 0.04 | 0.36 | 0.02 | 0.27 | 0.09 |
| 5 | 99776465 | rs10068989 | 1.22 | 2.153 | 1.01 | -0.086 | 0.093 | -0.078 | -0.078 | 0.124 | 0.81 | 0.5 | 0.84 | 0.92 | 0.76 | 0.81 |
| 5 | 99782205 | rs1381315 | 2.71 | 2.166 | 0.901 | -0.024 | -0.024 | -0.092 | -0.092 | 0 | 0.35 | 0.03 | 0.24 | 0.37 | 0.34 | 0.48 |
| 5 | 99792556 | rs10900740 | 1.063 | 2.081 | 1.056 | -0.354 | 0.096 | -0.071 | -0.071 | 0.269 | 0.81 | 0.52 | 0.84 | 0.92 | 0.75 | 0.81 |
| 5 | 99819180 | rs10052212 | -0.073 | 2.059 | 1.099 | -0.373 | 0 | -0.033 | -0.033 | 0 | 0.28 | 0.23 | 0.38 | 0.02 | 0.27 | 0.08 |
| 5 | 99841999 | rs3907112 | -2.01 | 2.057 | 1.104 | -0.367 | -0.012 | 0.271 | 0.271 | 3.137 | 0.48 | 0.86 | 0.54 | 0.1 | 0.52 | 0.27 |
| 5 | 99868839 | rs32260 | -2.29 | 2.025 | 1.044 | -0.178 | -0.046 | 0.088 | 0.088 | 3.173 | 0.52 | 0.91 | 0.55 | 0.3 | 0.53 | 0.3 |
| 5 | 99869685 | rs10072409 | 0.462 | 2.229 | 1.144 | -0.698 | 0.003 | -0.029 | -0.029 | 0.02 | 0.07 | 0.13 | 0.02 | 0 | 0.05 | 0.03 |
| 5 | 99883371 | rs17447382 | 0.251 | 2.192 | 1.108 | -0.69 | -0.012 | 0.089 | 0.089 | 0 | 0.25 | 0.1 | 0.36 | 0.02 | 0.28 | 0.09 |
| 5 | 99883577 | rs26026 | 0.227 | 2.269 | 1.142 | -0.646 | 0.064 | -0.023 | -0.023 | 0 | 0.11 | 0.25 | 0.12 | 0.06 | 0.19 | 0.16 |
| 5 | 99897866 | rs6874840 | 0.105 | 2.251 | 0.68 | -0.617 | 0 | -0.04 | -0.04 | 0 | 0.26 | 0.23 | 0.37 | 0.02 | 0.27 | 0.08 |
| 5 | 99913203 | rs32236 | -0.645 | 1.969 | 0.483 | -0.551 | 0.005 | -0.041 | -0.041 | 0 | 0.36 | 0.42 | 0.49 | 0.1 | 0.46 | 0.25 |
| 5 | 99922081 | rs13876 | -1.804 | 1.978 | -0.026 | -0.569 | -0.015 | 0.274 | 0.274 | 2.342 | 0.48 | 0.87 | 0.54 | 0.1 | 0.51 | 0.27 |
| 5 | 99934829 | rs247959 | -0.899 | 2.074 | -0.176 | -0.398 | 0.014 | -0.085 | -0.085 | 0 | 0.38 | 0.51 | 0.49 | 0.09 | 0.46 | 0.25 |
| 5 | 99943991 | rs13156973 | 1.88 | 2.485 | -1.155 | 0.281 | -0.007 | 0.11 | 0.11 | 4.645 | 0.46 | 0.03 | 0.29 | 0.27 | 0.42 | 0.46 |
| 5 | 99949162 | rs9885471 | -0.3 | 2.833 | -0.336 | 0.392 | -0.049 | 0.058 | 0.058 | 0.682 | 0.14 | 0.03 | 0.32 | 0.02 | 0.21 | 0.07 |
| 5 | 99952114 | rs492938 | 1.049 | 2.446 | 0.375 | 0.865 | 0.036 | -0.013 | -0.013 | 0 | 0.77 | 0.43 | 0.84 | 0.84 | 0.76 | 0.79 |
| 5 | 99959205 | rs247932 | -0.083 | 2.015 | -0.062 | 1.427 | -0.009 | -0.011 | -0.011 | 0 | 0.12 | 0.08 | 0.25 | 0.36 | 0.17 | 0.31 |
| 5 | 99968045 | rs32282 | 0.291 | 2.633 | 0.142 | 1.196 | 0.009 | -0.002 | -0.002 | 0 | 0.82 | 0.82 | 0.84 | 0.85 | 0.71 | 0.79 |
| 5 | 99969742 | rs157436 | 1.059 | 2.482 | 0.168 | 1.2 | 0.003 | 0.003 | 0.003 | 0 | 0.68 | 0.65 | 0.57 | 0.46 | 0.54 | 0.48 |
| 5 | 99986153 | rs6880868 | 0.969 | 1.7 | 0.072 | 0.933 | -0.025 | -0.006 | -0.006 | 0 | 0.65 | 0.28 | 0.51 | 0.64 | 0.55 | 0.71 |
| 5 | 99986537 | rs6866306 | 1.432 | 1.523 | -0.417 | 0.609 | 0.004 | 0.017 | 0.017 | 0.032 | 0.55 | 0.1 | 0.46 | 0.64 | 0.46 | 0.66 |
| 5 | 99992111 | rs905829 | -0.079 | 1.855 | -0.417 | 0.537 | -0.129 | -0.008 | -0.008 | 0 | 0.8 | 0.98 | 0.8 | 0.85 | 0.68 | 0.81 |
| 5 | 100020460 | rs2590411 | -1.19 | 1.757 | 0.268 | 0.3 | -0.006 | 0.003 | 0.003 | 0 | 0.34 | 0.71 | 0.5 | 0.27 | 0.45 | 0.35 |
| 5 | 100033522 | rs6867428 | 1.427 | 1.647 | 1.042 | 0.23 | -0.001 | -0.017 | -0.017 | 0.103 | 0.6 | 0.33 | 0.49 | 0.89 | 0.5 | 0.63 |
| 5 | 100054026 | rs6595421 | -0.109 | 1.66 | 1.904 | 0.405 | 0.017 | -0.034 | -0.034 | 0 | 0.09 | 0.26 | 0.12 | 0.02 | 0.1 | 0.04 |
| 5 | 100060150 | rs13163793 | 1.047 | 1.84 | 2.085 | 0.596 | 0.007 | 0.001 | 0.001 | 3.173 | 0.7 | 0.29 | 0.54 | 0.68 | 0.64 | 0.73 |
| 5 | 100065714 | rs10062661 | -0.047 | 1.846 | 2.311 | 0.312 | 0.017 | -0.033 | -0.033 | 0 | 0.09 | 0.24 | 0.26 | 0.02 | 0.1 | 0.04 |

| | | | | | | | | | | | | | | | | |
|---|---|---|---|---|---|---|---|---|---|---|---|---|---|---|---|---|
| 5 | 100071459 | rs279091 | 0.206 | 1.854 | 2.36 | 0.305 | -0.036 | 0.009 | 0.009 | 0 | 0.17 | 0.12 | 0.17 | 0.14 | 0.26 | 0.22 |
| 5 | 100071794 | rs157178 | 1.071 | 1.8 | 2.268 | 0.28 | 0.01 | -0.016 | -0.016 | 0 | 0.69 | 0.31 | 0.55 | 0.84 | 0.64 | 0.74 |
| 5 | 100082233 | rs279109 | 0.013 | 1.82 | 2.277 | 0.265 | 0.05 | -0.027 | -0.027 | 0 | 0.81 | 0.67 | 0.81 | 0.87 | 0.74 | 0.79 |
| 5 | 100082355 | rs279108 | 1.17 | 1.817 | 1.829 | 0.264 | -0.011 | 0.009 | 0.009 | 2.414 | 0.67 | 0.23 | 0.52 | 0.85 | 0.62 | 0.73 |
| 5 | 100084878 | rs279105 | 0.012 | 1.517 | 1.472 | 0.462 | -0.005 | 0.002 | 0.002 | 0 | 0.78 | 0.78 | 0.78 | 0.84 | 0.73 | 0.79 |
| 5 | 100094210 | rs277868 | 0.017 | 1.377 | 1.217 | 0.442 | -0.041 | 0.002 | 0.002 | 0 | 0.22 | 0.18 | 0.21 | 0.19 | 0.27 | 0.22 |
| 5 | 100100157 | rs278496 | 0.566 | 1.315 | 1.804 | 0.347 | 0.012 | -0.014 | -0.014 | 0 | 0.7 | 0.28 | 0.54 | 0.8 | 0.69 | 0.75 |
| 5 | 100128913 | rs4703116 | -0.507 | 1.304 | 2.075 | 0.315 | 0.037 | -0.021 | -0.021 | 0 | 0.29 | 0.48 | 0.44 | 0.2 | 0.32 | 0.26 |
| 5 | 100145388 | rs11584 | 0.43 | 1.304 | 2.268 | 0.328 | 0.031 | -0.013 | -0.013 | 0.057 | 0.69 | 0.4 | 0.53 | 0.7 | 0.65 | 0.71 |
| 5 | 100165117 | rs2548278 | -0.173 | -0.028 | 1.258 | -0.656 | 0.014 | -0.01 | -0.01 | 0 | 0.69 | 0.39 | 0.53 | 0.7 | 0.66 | 0.72 |
| 5 | 100172595 | rs32479 | 1.029 | -1.002 | 0.026 | -0.644 | -0.028 | 0.039 | 0.039 | 0.077 | 0.36 | 0.01 | 0.22 | 0.21 | 0.32 | 0.24 |
| 5 | 100210995 | rs3776170 | 0.19 | -0.597 | 0.695 | -0.494 | -0.004 | 0.003 | 0.003 | 0.203 | 0.33 | 0.73 | 0.48 | 0.3 | 0.34 | 0.29 |
| 5 | 100214617 | rs13355889 | 1.015 | -0.574 | 0.894 | -0.489 | 0.014 | 0.001 | 0.001 | 0 | 0.25 | 0.27 | 0.24 | 0.29 | 0.3 | 0.26 |
| 5 | 100220234 | rs11241752 | 1.362 | -0.54 | 0.711 | -0.488 | -0.073 | -0.004 | -0.004 | 0 | 0.23 | 0.09 | 0.22 | 0.19 | 0.29 | 0.25 |
| 5 | 100221189 | rs1423380 | 0.205 | -0.559 | 0.864 | -0.501 | 0.018 | -0.013 | -0.013 | 0.168 | 0.32 | 0.6 | 0.47 | 0.3 | 0.34 | 0.29 |
| 5 | 100221465 | rs11241755 | 1.381 | -0.556 | 0.667 | -0.497 | -0.103 | -0.022 | -0.022 | 0.309 | 0.23 | 0.04 | 0.22 | 0.19 | 0.29 | 0.25 |
| 5 | 100238801 | rs1428439 | 0.562 | -0.587 | 0.75 | -0.523 | 0.019 | -0.009 | -0.009 | 0 | 0.3 | 0.38 | 0.28 | 0.29 | 0.33 | 0.28 |
| 5 | 100248867 | rs1152127 | -1.734 | -0.838 | 0.18 | -0.554 | -0.017 | -0.023 | -0.023 | 0 | 0.86 | 0.93 | 0.79 | 0.58 | 0.82 | 0.79 |
| 5 | 100263707 | rs11742323 | 0.84 | -0.957 | -0.17 | -0.803 | -0.135 | 0.063 | 0.063 | 0 | 0.31 | 0.04 | 0.25 | 0.16 | 0.37 | 0.35 |
| 5 | 100277256 | rs17726244 | 0.399 | -0.795 | -0.049 | -0.598 | 0.011 | -0.016 | -0.016 | 0.73 | 0.33 | 0.62 | 0.46 | 0.25 | 0.35 | 0.29 |
| 5 | 100302925 | rs12187979 | 0.052 | -0.826 | 0.048 | -0.443 | 0.009 | 0.061 | 0.061 | 0.328 | 0.23 | 0.28 | 0.28 | 0.46 | 0.25 | 0.36 |
| 5 | 100315441 | rs403409 | -0.909 | -1.202 | -0.1 | -0.885 | -0.017 | -0.092 | -0.092 | 0 | 0.75 | 0.7 | 0.67 | 0.99 | 0.79 | 0.91 |
| 5 | 100400668 | rs557953 | 1.328 | -0.998 | -0.027 | -0.662 | 0.045 | -0.004 | -0.004 | 0 | 0.4 | 0.07 | 0.28 | 0.43 | 0.37 | 0.34 |
| 5 | 100404802 | rs563361 | -0.22 | -0.927 | 0.038 | -0.644 | -0.002 | -0.002 | -0.002 | 0 | 0.66 | 0.65 | 0.55 | 0.8 | 0.59 | 0.66 |
| 5 | 100412665 | rs10041296 | -0.335 | -0.924 | 0.245 | -0.568 | 0.032 | 0.039 | 0.039 | 0.034 | 0.06 | 0.28 | 0.21 | 0.04 | 0.05 | 0.08 |
| 5 | 100420093 | rs6895082 | -0.915 | -0.941 | 0.222 | -0.588 | 0.003 | -0.016 | -0.016 | 1.32 | 0.1 | 0.29 | 0.25 | 0.04 | 0.08 | 0.09 |
| 5 | 100435429 | rs9327421 | -0.262 | -0.996 | 0.456 | -0.567 | 0.009 | 0.097 | 0.097 | 1.102 | 0.25 | 0.5 | 0.27 | 0.36 | 0.22 | 0.31 |
| 5 | 100449604 | rs10042591 | -0.039 | -1.013 | 0.676 | -0.584 | 0.008 | 0.071 | 0.071 | 0.442 | 0.23 | 0.35 | 0.26 | 0.36 | 0.22 | 0.31 |
| 5 | 100465754 | rs2400304 | -0.953 | -1.075 | 0.312 | -0.665 | -0.129 | 0.067 | 0.067 | 0 | 0.72 | 0.96 | 0.78 | 0.84 | 0.62 | 0.71 |
| 5 | 100505177 | rs10447287 | 1.208 | -1.105 | 0.612 | -0.673 | 0.001 | -0.01 | -0.01 | 0 | 0.07 | 0.05 | 0.03 | 0 | 0.03 | 0.09 |
| 5 | 100522994 | rs10062726 | -1.364 | -1.13 | 0.64 | -0.598 | 0.051 | 0.013 | 0.013 | 0 | 0.59 | 0.89 | 0.72 | 0.56 | 0.63 | 0.67 |
| 5 | 100535029 | rs10440779 | 0.032 | -1.108 | 0.699 | -0.623 | 0.012 | -0.029 | -0.029 | 0 | 0.35 | 0.41 | 0.46 | 0.2 | 0.41 | 0.36 |
| 5 | 100542852 | rs938136 | 0.279 | -1.093 | 0.518 | -0.663 | -0.026 | 0.04 | 0.04 | 0 | 0.33 | 0.25 | 0.26 | 0.16 | 0.41 | 0.3 |
| 5 | 100548406 | rs1512246 | -1.208 | -1.035 | 0.166 | -0.541 | 0.047 | 0.016 | 0.016 | 0 | 0.59 | 0.86 | 0.72 | 0.57 | 0.63 | 0.67 |
| 5 | 100557718 | rs17796189 | 0.379 | -0.993 | 0.143 | -0.65 | -0.016 | 0.019 | 0.019 | 0 | 0.31 | 0.26 | 0.45 | 0.19 | 0.4 | 0.3 |
| 5 | 100559282 | rs1588630 | -1.437 | -1.236 | 0.078 | -0.649 | 0.037 | 0.018 | 0.018 | 0 | 0.62 | 0.79 | 0.73 | 0.6 | 0.66 | 0.69 |
| 5 | 100567453 | rs11505129 | 0.188 | -1.223 | 0.468 | -0.597 | 0.005 | 0.069 | 0.069 | 0.17 | 0.24 | 0.33 | 0.25 | 0.4 | 0.21 | 0.31 |
| 5 | 100573157 | rs6869103 | -0.798 | -1.304 | 0.3 | -0.655 | 0.049 | 0.033 | 0.033 | 0.011 | 0.54 | 0.75 | 0.67 | 0.57 | 0.59 | 0.63 |
| 5 | 100578856 | rs4587054 | -0.576 | -1.367 | 0.366 | -0.701 | -0.004 | -0.003 | -0.003 | 1.636 | 0.52 | 0.49 | 0.49 | 0.59 | 0.61 | 0.64 |
| 5 | 100580114 | rs2137298 | -1.207 | -1.364 | 0.347 | -0.699 | 0.054 | 0.04 | 0.04 | 0.076 | 0.58 | 0.83 | 0.69 | 0.6 | 0.64 | 0.68 |
| 5 | 100582655 | rs10060874 | -0.584 | -1.347 | 0.257 | -0.69 | -0.004 | -0.003 | -0.003 | 0 | 0.52 | 0.49 | 0.49 | 0.59 | 0.61 | 0.64 |
| 5 | 100585407 | rs4131004 | -0.783 | -1.192 | 0.181 | -0.768 | -0.016 | -0.01 | -0.01 | 0 | 0.54 | 0.49 | 0.5 | 0.59 | 0.64 | 0.65 |
| 5 | 100586946 | rs10062319 | -1.504 | -1.223 | 0.239 | -0.729 | 0.043 | 0.013 | 0.013 | 0 | 0.62 | 0.9 | 0.74 | 0.6 | 0.66 | 0.69 |
| 5 | 100587086 | rs7733876 | -0.388 | -1.063 | 0.082 | -0.679 | 0.006 | 0.078 | 0.078 | 0.417 | 0.29 | 0.46 | 0.27 | 0.43 | 0.26 | 0.36 |
| 5 | 100587626 | rs7714143 | -0.594 | -1.053 | 0.203 | -0.639 | -0.003 | -0.003 | -0.003 | 0 | 0.52 | 0.49 | 0.49 | 0.59 | 0.61 | 0.64 |
| 5 | 100592006 | rs6862162 | 0.075 | -0.796 | 0.25 | -0.365 | 0.002 | 0.029 | 0.029 | 0.034 | 0.23 | 0.25 | 0.24 | 0.39 | 0.21 | 0.31 |
| 5 | 100592459 | rs4467676 | -0.802 | -0.972 | 0.136 | -0.459 | 0.041 | 0.01 | 0.01 | 0 | 0.58 | 0.84 | 0.72 | 0.57 | 0.63 | 0.64 |
| 5 | 100596731 | rs1599134 | -0.8 | -0.975 | 0.892 | 0.293 | -0.012 | 0.001 | 0.001 | 0 | 0.31 | 0.32 | 0.18 | 0.34 | 0.21 | 0.26 |
| 5 | 100599227 | rs2895703 | -1.013 | -0.983 | 1.126 | 0.281 | -0.044 | 0.003 | 0.003 | 0 | 0.33 | 0.46 | 0.18 | 0.34 | 0.22 | 0.29 |
| 5 | 100604495 | rs1903186 | -1.253 | -0.899 | 1.078 | 0.225 | -0.021 | -0.005 | -0.005 | 0 | 0.36 | 0.41 | 0.2 | 0.34 | 0.25 | 0.32 |
| 5 | 100605505 | rs1459574 | 0.075 | -0.699 | 0.877 | 0.393 | -0.066 | -0.127 | -0.127 | 0 | 0.16 | 0.02 | 0.27 | 0.42 | 0.22 | 0.35 |
| 5 | 100607919 | rs1107621 | -2.406 | -0.643 | 1.125 | 0.365 | -0.041 | 0.125 | 0.125 | 0 | 0.55 | 0.85 | 0.68 | 0.76 | 0.47 | 0.69 |
| 5 | 100608611 | rs4343813 | -0.084 | -0.691 | 1.213 | 0.375 | 0.003 | -0.006 | -0.006 | 3.119 | 0.25 | 0.09 | 0.16 | 0.28 | 0.22 | 0.23 |
| 5 | 100609923 | rs11949216 | -1.044 | -0.663 | 1.166 | 0.376 | -0.016 | 0 | 0 | 0.004 | 0.33 | 0.37 | 0.2 | 0.34 | 0.26 | 0.32 |
| 5 | 100618922 | rs1380391 | -0.843 | -0.469 | 1.138 | 0.505 | 0.019 | 0.125 | 0.125 | 0 | 0.2 | 0.43 | 0.49 | 0.42 | 0.22 | 0.37 |
| 5 | 100619084 | rs7724880 | -0.051 | -0.526 | 1.153 | 0.468 | -0.003 | 0 | 0 | 1.25 | 0.24 | 0.24 | 0.14 | 0.28 | 0.2 | 0.22 |
| 5 | 100620693 | rs10055969 | 0.137 | -0.502 | 1.048 | 0.49 | -0.062 | -0.124 | -0.124 | 0.097 | 0.16 | 0.02 | 0.27 | 0.42 | 0.2 | 0.34 |
| 5 | 100623751 | rs6595846 | -0.039 | -0.499 | 1.135 | 0.485 | -0.019 | 0.022 | 0.022 | 0 | 0.06 | 0.13 | 0.04 | 0.06 | 0.02 | 0.05 |
| 5 | 100632028 | rs10463847 | -2.071 | -0.504 | 0.732 | 0.451 | 0.022 | 0.131 | 0.131 | 0 | 0.51 | 0.78 | 0.67 | 0.76 | 0.44 | 0.64 |
| 5 | 100633738 | rs1459583 | -1.62 | -0.648 | 0.76 | 0.479 | -0.016 | 0.076 | 0.076 | 0.656 | 0.28 | 0.56 | 0.52 | 0.48 | 0.24 | 0.42 |
| 5 | 100637267 | rs10069443 | -1.62 | -0.648 | 0.76 | 0.479 | -0.016 | 0.076 | 0.076 | 0.656 | 0.28 | 0.56 | 0.52 | 0.48 | 0.24 | 0.42 |
| 5 | 100646372 | rs2061476 | -2.211 | -0.606 | 0.3 | 0.409 | -0.006 | 0.115 | 0.115 | 0 | 0.53 | 0.8 | 0.68 | 0.75 | 0.46 | 0.66 |
| 5 | 100660062 | rs12523560 | -0.889 | -0.394 | 0.485 | 0.003 | -0.001 | 0.012 | 0.012 | 0 | 0.2 | 0.23 | 0.29 | 0.54 | 0.21 | 0.37 |
| 5 | 100669673 | rs1599132 | 0.041 | -0.362 | 0.224 | 0.002 | 0.002 | -0.012 | -0.012 | 0 | 0.22 | 0.08 | 0.14 | 0.28 | 0.2 | 0.2 |
| 5 | 100697430 | rs346856 | -0.116 | -0.673 | 0.2 | 0.054 | 0.007 | -0.011 | -0.011 | 0 | 0.78 | 0.92 | 0.88 | 0.72 | 0.81 | 0.77 |
| 5 | 100701600 | rs12153327 | 2.087 | -0.682 | 0.088 | -0.003 | -0.017 | 0.239 | 0.239 | 0 | 0.48 | 0.14 | 0.34 | 0.14 | 0.56 | 0.35 |
| 5 | 100718992 | rs10035976 | -0.095 | -0.445 | 0.178 | 0.3 | -0.058 | -0.124 | -0.124 | 0 | 0.15 | 0.03 | 0.28 | 0.53 | 0.19 | 0.31 |
| 5 | 100721491 | rs370605 | 0.293 | -0.619 | 0.783 | 0.14 | 0.005 | 0.002 | 0.002 | 0 | 0.76 | 0.85 | 0.86 | 0.73 | 0.79 | 0.76 |
| 5 | 100759281 | rs17135633 | -0.132 | -0.599 | 0.95 | 0.097 | -0.015 | -0.028 | -0.028 | 0.097 | 0.05 | 0.12 | 0.02 | 0 | 0.02 | 0.03 |

| | | | | | | | | | | | | | | | | |
|---|---|---|---|---|---|---|---|---|---|---|---|---|---|---|---|---|
| 5 | 100760453 | rs348409 | 1.636 | -0.667 | 0.614 | 0.045 | 0.056 | 0.033 | 0.033 | 0 | 0.61 | 0.15 | 0.6 | 0.68 | 0.74 | 0.66 |
| 5 | 100765147 | rs412683 | 0.631 | -0.303 | 1.558 | 0.123 | 0.004 | -0.012 | -0.012 | 0.044 | 0.73 | 0.64 | 0.64 | 0.73 | 0.75 | 0.69 |
| 5 | 100775607 | rs6860058 | -1.177 | -0.264 | 1.772 | 0.205 | 0.002 | -0.011 | -0.011 | 0 | 0.07 | 0.28 | 0.03 | 0 | 0.01 | 0 |
| 5 | 100798650 | rs465395 | 0.446 | -0.231 | 1.828 | 0.222 | -0.002 | -0.003 | -0.003 | 0 | 0.74 | 0.72 | 0.86 | 0.73 | 0.78 | 0.72 |
| 5 | 100832707 | rs439897 | 1.801 | -0.185 | 1.525 | 0.324 | 0.002 | 0.018 | 0.018 | 0 | 0.58 | 0.34 | 0.4 | 0.25 | 0.63 | 0.48 |
| 5 | 100837326 | rs164439 | 0.428 | -0.209 | 1.453 | 0.509 | -0.004 | 0.001 | 0.001 | 0 | 0.76 | 0.74 | 0.87 | 0.72 | 0.81 | 0.79 |
| 5 | 100844041 | rs348429 | -0.122 | -0.2 | 1.146 | 0.533 | -0.005 | -0.036 | -0.036 | 1.438 | 0.8 | 0.92 | 0.89 | 0.71 | 0.82 | 0.79 |
| 5 | 100851524 | rs181161 | -0.122 | -0.2 | 1.146 | 0.527 | -0.005 | -0.034 | -0.034 | 0 | 0.8 | 0.92 | 0.89 | 0.72 | 0.82 | 0.79 |
| 5 | 100871199 | rs164619 | -1.889 | -0.22 | 1.511 | 0.487 | 0.002 | -0.031 | -0.031 | 0 | 0.39 | 0.55 | 0.59 | 0.76 | 0.37 | 0.51 |
| 5 | 100898414 | rs17137209 | -0.044 | -0.091 | 0.948 | 0.592 | -0.053 | -0.094 | -0.094 | 0 | 0.12 | 0.02 | 0.25 | 0.45 | 0.17 | 0.27 |
| 5 | 100918526 | rs17137390 | -0.13 | -0.101 | 1.027 | 0.642 | -0.007 | -0.026 | -0.026 | 1.754 | 0.19 | 0.15 | 0.29 | 0.3 | 0.18 | 0.2 |
| 5 | 100925684 | rs12109235 | -0.39 | -0.064 | 1.059 | 0.705 | 0.005 | 0.071 | 0.071 | 0 | 0.14 | 0.21 | 0.27 | 0.46 | 0.18 | 0.32 |
| 5 | 100930938 | rs11242092 | -0.407 | -0.187 | 0.589 | 0.513 | -0.028 | -0.048 | -0.048 | 0 | 0.22 | 0.05 | 0.32 | 0.3 | 0.21 | 0.22 |
| 5 | 100932981 | rs3907457 | -1.917 | -0.1 | 0.93 | 0.045 | -0.011 | 0.103 | 0.103 | 0 | 0.53 | 0.72 | 0.67 | 0.82 | 0.46 | 0.67 |
| 5 | 100948178 | rs17137638 | -1.923 | -0.144 | 1.901 | 0.22 | 0.078 | 0.057 | 0.057 | 0.426 | 0.06 | 0.39 | 0.04 | 0.02 | 0.02 | 0.11 |
| 5 | 100948545 | rs13188771 | -0.161 | -0.134 | 1.255 | 0.31 | -0.052 | -0.022 | -0.022 | 1.678 | 0.16 | 0.01 | 0.29 | 0.19 | 0.16 | 0.15 |
| 5 | 100962732 | rs17137871 | 0.095 | -0.117 | 1.575 | 0.877 | -0.037 | 0.007 | 0.007 | 0 | 0.07 | 0.01 | 0.16 | 0.49 | 0.11 | 0.22 |
| 5 | 100964192 | rs10051419 | -2.084 | -0.295 | 2.232 | 0.576 | 0.101 | 0.078 | 0.078 | 2.525 | 0.05 | 0.51 | 0.04 | 0.02 | 0.01 | 0.1 |
| 5 | 100966719 | rs6861816 | -2.901 | -0.483 | 1.548 | 0.428 | 0.001 | -0.003 | -0.003 | 0 | 0.17 | 0.77 | 0.08 | 0.06 | 0.07 | 0.15 |
| 5 | 100971002 | rs6596059 | -2.415 | -0.413 | 1.622 | 0.413 | -0.033 | -0.048 | -0.048 | 0.397 | 0.11 | 0.71 | 0.05 | 0.03 | 0.03 | 0.12 |
| 5 | 100980342 | rs1038287 | -2.507 | -0.466 | 1.604 | 0.489 | -0.029 | -0.024 | -0.024 | 0 | 0.1 | 0.82 | 0.06 | 0.03 | 0.02 | 0.11 |
| 5 | 100985751 | rs2127208 | -2.13 | -0.462 | 1.211 | 0.412 | -0.003 | 0.107 | 0.107 | 0.371 | 0.48 | 0.96 | 0.57 | 0.73 | 0.39 | 0.56 |
| 5 | 100992062 | rs17138228 | -1.651 | -0.367 | 2.092 | 0.421 | 0.031 | 0.033 | 0.033 | 0.225 | 0.07 | 0.47 | 0.04 | 0.03 | 0.02 | 0.1 |
| 5 | 100999473 | rs17138319 | -2.59 | -0.368 | 1.741 | 0.36 | -0.006 | -0.023 | -0.023 | 0 | 0.18 | 0.8 | 0.1 | 0.05 | 0.12 | 0.16 |
| 5 | 101003812 | rs1481771 | -1.471 | -0.324 | 2.037 | 0.413 | 0.042 | 0.043 | 0.043 | 0.216 | 0.07 | 0.49 | 0.04 | 0.03 | 0.02 | 0.1 |
| 5 | 101019270 | rs12654056 | -0.038 | -0.419 | 1.102 | 0.529 | 0.001 | -0.056 | -0.056 | 1.071 | 0.11 | 0.01 | 0.07 | 0.26 | 0.12 | 0.15 |
| 5 | 101019725 | rs1871332 | -2.447 | -0.153 | 1.605 | 0.567 | 0.012 | 0.02 | 0.02 | 4.09 | 0.19 | 0.84 | 0.11 | 0.04 | 0.13 | 0.18 |
| 5 | 101035851 | rs11956691 | -2.471 | -0.16 | 1.762 | 0.562 | 0.002 | -0.013 | -0.013 | 0 | 0.19 | 0.77 | 0.11 | 0.04 | 0.13 | 0.18 |
| 5 | 101039904 | rs727178 | -0.571 | -0.093 | 1.766 | 0.602 | -0.005 | -0.007 | -0.007 | 0.893 | 0.06 | 0.09 | 0.02 | 0.03 | 0.04 | 0.03 |
| 5 | 101060923 | rs12521068 | -0.336 | -0.245 | 1.386 | 0.637 | 0.029 | 0.002 | 0.002 | 0 | 0.21 | 0.07 | 0.14 | 0.2 | 0.17 | 0.19 |
| 5 | 101063450 | rs6887734 | -1.844 | -0.082 | 1.887 | 0.661 | 0.005 | -0.055 | -0.055 | 0 | 0.15 | 0.57 | 0.1 | 0.06 | 0.13 | 0.19 |
| 5 | 101067024 | rs10050964 | -0.36 | -0.032 | 1.205 | 1.041 | -0.014 | 0.016 | 0.016 | 0 | 0.09 | 0.05 | 0.34 | 0.42 | 0.13 | 0.21 |
| 5 | 101070144 | rs6895942 | -1.484 | -0.097 | 1.176 | 1.031 | -0.004 | -0.087 | -0.087 | 0 | 0.3 | 0.12 | 0.43 | 0.56 | 0.29 | 0.4 |
| 5 | 101075601 | rs11242141 | 2.247 | 0.071 | 1.604 | 0.78 | 0.028 | 0.048 | 0.048 | 0 | 0.53 | 0.26 | 0.47 | 0.32 | 0.57 | 0.42 |
| 5 | 101082433 | rs10057269 | -0.43 | 0.192 | 1.534 | 0.741 | -0.021 | -0.009 | -0.009 | 0.115 | 0.1 | 0.04 | 0.32 | 0.45 | 0.14 | 0.21 |
| 5 | 101086200 | rs12188863 | 1.772 | 0.141 | 2.042 | 0.595 | -0.003 | -0.006 | -0.006 | 0.188 | 0.58 | 0.52 | 0.48 | 0.29 | 0.59 | 0.54 |
| 5 | 101087195 | rs6864165 | -0.989 | 0.319 | 1.445 | 0.636 | -0.013 | -0.014 | -0.014 | 0 | 0.05 | 0.23 | 0.01 | 0.02 | 0.02 | 0.01 |
| 5 | 101095318 | rs1600277 | 1.306 | 0.356 | 1.309 | 0.679 | -0.001 | -0.005 | -0.005 | 0 | 0.63 | 0.56 | 0.49 | 0.34 | 0.61 | 0.45 |
| 5 | 101096015 | rs7725574 | 1.393 | 0.33 | 0.561 | 0.529 | -0.009 | 0.032 | 0.032 | 0 | 0.6 | 0.79 | 0.49 | 0.3 | 0.54 | 0.37 |
| 5 | 101097172 | rs12188461 | 1.873 | 0.519 | 0.523 | 0.638 | 0.001 | 0.01 | 0.01 | 0 | 0.55 | 0.42 | 0.49 | 0.25 | 0.52 | 0.56 |
| 5 | 101097727 | rs10070937 | -1.564 | 0.442 | 0.533 | 0.772 | 0.008 | 0.196 | 0.196 | 0 | 0.2 | 0.46 | 0.2 | 0.49 | 0.18 | 0.15 |
| 5 | 101097943 | rs7718044 | 2.008 | 0.708 | 0.396 | 0.742 | -0.001 | 0.01 | 0.01 | 0 | 0.55 | 0.42 | 0.49 | 0.25 | 0.51 | 0.56 |
| 5 | 101098660 | rs11242149 | 1.693 | 0.901 | 0.109 | 0.866 | 0.004 | -0.018 | -0.018 | 0 | 0.59 | 0.53 | 0.49 | 0.27 | 0.52 | 0.56 |
| 5 | 101100605 | rs986950 | 0.726 | 0.873 | -0.589 | 0.698 | -0.072 | -0.022 | -0.022 | 0 | 0.75 | 0.97 | 0.7 | 0.76 | 0.69 | 0.73 |
| 5 | 101105998 | rs1479073 | 0.679 | 0.912 | -0.286 | 0.661 | -0.007 | 0.026 | 0.026 | 0 | 0.75 | 0.82 | 0.73 | 0.79 | 0.69 | 0.7 |
| 5 | 101109146 | rs6870264 | 1.511 | 0.994 | 0.14 | 0.865 | 0.005 | -0.028 | -0.028 | 0 | 0.59 | 0.48 | 0.5 | 0.25 | 0.52 | 0.56 |
| 5 | 101109871 | rs4492090 | 1.089 | 0.806 | 0.232 | 0.735 | 0.001 | -0.031 | -0.031 | 3.492 | 0.67 | 0.53 | 0.64 | 0.72 | 0.63 | 0.65 |
| 5 | 101114271 | rs6863058 | -0.588 | 0.864 | -0.042 | 0.584 | 0.002 | -0.025 | -0.025 | 0 | 0.1 | 0.15 | 0.17 | 0.03 | 0.13 | 0.11 |
| 5 | 101118483 | rs4490561 | 1.813 | 0.893 | -0.843 | 0.405 | -0.064 | 0.159 | 0.159 | 0 | 0.49 | 0.08 | 0.43 | 0.24 | 0.45 | 0.47 |
| 5 | 101135059 | rs1871334 | -1.013 | 0.941 | -0.203 | 1.008 | 0 | 0.042 | 0.042 | 0 | 0.12 | 0.12 | 0.25 | 0.51 | 0.17 | 0.17 |
| 5 | 101141811 | rs1318570 | -1.514 | 0.854 | -0.19 | 0.943 | -0.017 | -0.03 | -0.03 | 0 | 0.24 | 0.18 | 0.43 | 0.54 | 0.32 | 0.29 |
| 5 | 101195531 | rs6596178 | 1.391 | 0.622 | 0.096 | 0.794 | -0.005 | -0.006 | -0.006 | 0.413 | 0.73 | 0.73 | 0.55 | 0.39 | 0.63 | 0.65 |
| 5 | 101203699 | rs1453103 | -1.709 | 0.462 | 0.637 | 0.849 | -0.003 | 0.013 | 0.013 | 0.169 | 0.35 | 0.48 | 0.42 | 0.48 | 0.37 | 0.33 |
| 5 | 101204974 | rs10036886 | -1.053 | 0.447 | 0.591 | 0.97 | -0.009 | -0.003 | -0.003 | 0 | 0.08 | 0.21 | 0.03 | 0.05 | 0.05 | 0.05 |
| 5 | 101206204 | rs10045191 | -0.974 | 0.571 | 0.676 | 1.05 | 0.006 | 0.021 | 0.021 | 0 | 0.06 | 0.21 | 0.03 | 0.05 | 0.04 | 0.05 |
| 5 | 101206369 | rs7711786 | -1.176 | 0.633 | 0.301 | 1.122 | -0.012 | -0.068 | -0.068 | 2.578 | 0.26 | 0.17 | 0.38 | 0.42 | 0.32 | 0.28 |
| 5 | 101210066 | rs10079251 | -1.275 | 0.519 | 1.108 | 0.988 | 0.004 | 0.036 | 0.036 | 0 | 0.05 | 0.21 | 0.02 | 0.06 | 0.03 | 0.05 |
| 5 | 101210703 | rs6596181 | -0.862 | 0.254 | 1.05 | -0.064 | 0 | -0.019 | -0.019 | 0 | 0.77 | 0.83 | 0.88 | 0.97 | 0.84 | 0.94 |
| 5 | 101215201 | rs10055551 | -0.579 | 0.236 | 1.104 | -0.052 | 0.009 | 0.029 | 0.029 | 0 | 0.06 | 0.22 | 0.02 | 0.05 | 0.03 | 0.05 |
| 5 | 101215710 | rs13166990 | 1.358 | 0.435 | 0.145 | -0.763 | -0.064 | 0.094 | 0.094 | 0 | 0.27 | 0.04 | 0.33 | 0.11 | 0.32 | 0.25 |
| 5 | 101216027 | rs2400660 | 0.369 | 0.283 | 0.565 | -0.142 | -0.014 | 0 | 0 | 0.24 | 0.44 | 0.27 | 0.49 | 0.57 | 0.44 | 0.36 |
| 5 | 101228173 | rs13358179 | 0.225 | 0.327 | 0.563 | -0.199 | 0.004 | 0.002 | 0.002 | 0 | 0.05 | 0.2 | 0.02 | 0.03 | 0.03 | 0.05 |
| 5 | 101234121 | rs9285932 | 1.175 | 0.307 | 0.527 | -0.282 | -0.017 | -0.087 | -0.087 | 0.184 | 0.24 | 0.38 | 0.1 | 0.03 | 0.14 | 0.07 |
| 5 | 101234570 | rs10479085 | 0.084 | 0.402 | 0.508 | -0.138 | 0.025 | 0.018 | 0.018 | 0 | 0.06 | 0.25 | 0.02 | 0.03 | 0.04 | 0.05 |
| 5 | 101237809 | rs13153577 | 0.877 | 0.385 | 0.306 | -0.089 | -0.006 | -0.011 | -0.011 | 0 | 0.22 | 0.16 | 0.38 | 0.57 | 0.27 | 0.3 |
| 5 | 101242395 | rs4642340 | -0.501 | 0.378 | 0.42 | 0.04 | 0.001 | 0.005 | 0.005 | 0 | 0.74 | 0.79 | 0.59 | 0.43 | 0.67 | 0.67 |
| 5 | 101263051 | rs9327700 | -0.19 | 0.139 | 0.795 | -0.043 | -0.007 | -0.012 | -0.012 | 0 | 0.45 | 0.51 | 0.27 | 0.29 | 0.37 | 0.4 |
| 5 | 101285347 | rs7711691 | -0.282 | 0.148 | 0.779 | -0.071 | 0.002 | 0.003 | 0.003 | 0.16 | 0.46 | 0.55 | 0.27 | 0.29 | 0.37 | 0.4 |
| 5 | 101291960 | rs6870397 | -0.012 | 0.016 | 0.728 | -0.114 | 0.002 | -0.002 | -0.002 | 0 | 0.23 | 0.21 | 0.14 | 0.26 | 0.19 | 0.34 |

| | | | | | | | | | | | | | | | | |
|---|---|---|---|---|---|---|---|---|---|---|---|---|---|---|---|---|
| 5 | 101308396 | rs2061716 | -0.067 | -0.002 | 0.784 | -0.155 | 0 | -0.001 | -0.001 | 0 | 0.41 | 0.42 | 0.22 | 0.26 | 0.31 | 0.36 |
| 5 | 101310707 | rs11242254 | -0.132 | -0.011 | 0.807 | -0.155 | -0.005 | 0 | 0 | 1.134 | 0.24 | 0.25 | 0.14 | 0.26 | 0.19 | 0.34 |
| 5 | 101318849 | rs10039287 | 2.08 | 0.071 | 0.69 | -0.157 | 0.001 | -0.028 | -0.028 | 1.718 | 0.17 | 0.16 | 0.08 | 0 | 0.11 | 0.02 |
| 5 | 101320305 | rs1011624 | 1.906 | 0.07 | 0.627 | -0.153 | -0.003 | -0.052 | -0.052 | 0.026 | 0.18 | 0.22 | 0.09 | 0 | 0.11 | 0.02 |
| 5 | 101324088 | rs7725478 | 0.144 | 0 | 0.366 | 0.026 | 0.016 | -0.01 | -0.01 | 0 | 0.22 | 0.16 | 0.14 | 0.25 | 0.2 | 0.34 |
| 5 | 101328984 | rs7356754 | -0.24 | 0.212 | 0.633 | -0.026 | 0.012 | 0.01 | 0.01 | 0 | 0.06 | 0.21 | 0.02 | 0.03 | 0.03 | 0.05 |
| 5 | 101331313 | rs1381976 | 0.231 | 0.247 | 0.57 | -0.185 | -0.004 | -0.058 | -0.058 | 0 | 0.29 | 0.6 | 0.16 | 0.09 | 0.22 | 0.14 |
| 5 | 101341123 | rs5020202 | 0.02 | 0.35 | 0.519 | -0.102 | 0.002 | 0.003 | 0.003 | 0 | 0.52 | 0.45 | 0.52 | 0.4 | 0.58 | 0.54 |
| 5 | 101343235 | rs2061714 | 0.519 | 0.39 | 0.345 | -0.229 | 0.008 | -0.026 | -0.026 | 0 | 0.2 | 0.11 | 0.15 | 0.23 | 0.2 | 0.26 |
| 5 | 101344092 | rs4569847 | -0.226 | 0.341 | 0.469 | -0.239 | -0.001 | 0.002 | 0.002 | 0 | 0.25 | 0.3 | 0.2 | 0.26 | 0.25 | 0.3 |
| 5 | 101344133 | rs4419568 | -0.385 | 0.315 | 0.511 | -0.252 | 0 | 0 | 0 | 0 | 0.53 | 0.51 | 0.52 | 0.4 | 0.58 | 0.54 |
| 5 | 101344757 | rs3846608 | 1.636 | 0.357 | 0.387 | -0.244 | -0.017 | 0.035 | 0.035 | 0 | 0.27 | 0.16 | 0.32 | 0.15 | 0.33 | 0.24 |
| 5 | 101353504 | rs9327736 | 0.5 | 0.39 | 0.35 | -0.204 | 0.008 | -0.026 | -0.026 | 0 | 0.19 | 0.11 | 0.15 | 0.22 | 0.2 | 0.26 |
| 5 | 101354219 | rs6860838 | -0.947 | 0.389 | 0.598 | -0.165 | 0 | 0.015 | 0.015 | 0 | 0.76 | 0.79 | 0.62 | 0.44 | 0.76 | 0.63 |
| 5 | 101364700 | rs10069803 | 1.434 | 0.39 | 0.679 | -0.278 | -0.003 | -0.034 | -0.034 | 0 | 0.23 | 0.25 | 0.1 | 0.04 | 0.17 | 0.1 |
| 5 | 101392473 | rs11739895 | 1.377 | 0.399 | 0.826 | -0.192 | 0.034 | 0.015 | 0.015 | 0 | 0.88 | 0.65 | 0.93 | 0.92 | 0.89 | 0.92 |
| 5 | 101394874 | rs11748160 | 0.321 | 0.424 | 0.745 | -0.041 | 0 | 0.002 | 0.002 | 0 | 0.65 | 0.39 | 0.54 | 0.4 | 0.65 | 0.55 |
| 5 | 101395391 | rs1477616 | 0.351 | 0.412 | 0.739 | -0.047 | 0 | 0.002 | 0.002 | 0 | 0.64 | 0.39 | 0.54 | 0.4 | 0.65 | 0.55 |
| 5 | 101395497 | rs1477615 | 0.32 | 0.415 | 0.741 | -0.045 | 0 | 0.002 | 0.002 | 0 | 0.65 | 0.39 | 0.54 | 0.4 | 0.65 | 0.55 |
| 5 | 101397849 | rs11739747 | 1.493 | 0.411 | 0.684 | -0.136 | -0.005 | 0.109 | 0.109 | 0.496 | 0.45 | 0.23 | 0.38 | 0.18 | 0.45 | 0.3 |
| 5 | 101403053 | rs12519005 | -0.121 | 0.287 | 0.608 | -0.05 | 0.004 | 0.008 | 0.008 | 0 | 0.56 | 0.59 | 0.75 | 0.74 | 0.63 | 0.66 |
| 5 | 101403586 | rs6861497 | 0.331 | 0.258 | 0.644 | -0.016 | 0.002 | 0.009 | 0.009 | 0 | 0.51 | 0.54 | 0.7 | 0.71 | 0.56 | 0.62 |
| 5 | 101405062 | rs17145881 | 1.59 | 0.023 | 0.491 | 0.638 | 0.006 | 0 | 0 | 0 | 0.06 | 0.04 | 0.1 | 0.2 | 0.02 | 0.14 |
| 5 | 101428403 | rs2100890 | 1.143 | 0.127 | 0.244 | 0.288 | -0.007 | -0.013 | -0.013 | 0 | 0.11 | 0.08 | 0.13 | 0.18 | 0.14 | 0.18 |
| 5 | 101432938 | rs6596344 | -2.378 | -0.003 | 0.338 | 0.105 | 0 | -0.037 | -0.037 | 0 | 0.81 | 0.84 | 0.92 | 1 | 0.86 | 0.95 |
| 5 | 101433549 | rs977664 | 0.462 | 0.045 | 0.043 | 0.141 | -0.002 | -0.005 | -0.005 | 0 | 0.21 | 0.15 | 0.17 | 0.19 | 0.22 | 0.26 |
| 5 | 101454563 | rs11745729 | 2.008 | 0.192 | -0.142 | 0.106 | -0.038 | -0.007 | -0.007 | 0 | 0.24 | 0.06 | 0.24 | 0.2 | 0.28 | 0.19 |
| 5 | 101457609 | rs382521 | -0.5 | 0.235 | 0.264 | 0.251 | 0.007 | 0.036 | 0.036 | 0 | 0.75 | 0.69 | 0.55 | 0.47 | 0.69 | 0.57 |
| 5 | 101465106 | rs428751 | 0.915 | 0.14 | 0.323 | 0.079 | -0.001 | 0.015 | 0.015 | 1.081 | 0.49 | 0.48 | 0.36 | 0.27 | 0.47 | 0.28 |
| 5 | 101482983 | rs6889035 | -0.122 | 0.223 | 0.166 | -0.332 | -0.001 | 0 | 0 | 0.326 | 0.26 | 0.27 | 0.24 | 0.24 | 0.25 | 0.29 |
| 5 | 101490022 | rs415200 | -0.903 | 0.228 | 0.485 | -0.508 | -0.001 | -0.011 | -0.011 | 0 | 0.49 | 0.51 | 0.71 | 0.8 | 0.53 | 0.72 |
| 5 | 101507854 | rs7444646 | 0.051 | 0.225 | 0.337 | -0.514 | -0.001 | 0 | 0 | 0.5 | 0.23 | 0.24 | 0.24 | 0.24 | 0.24 | 0.28 |
| 5 | 101508279 | rs453689 | -2.249 | 0.229 | 0.101 | -0.678 | -0.068 | 0.071 | 0.071 | 0 | 0.76 | 0.98 | 0.83 | 0.87 | 0.72 | 0.8 |
| 5 | 101522676 | rs2081166 | -0.393 | 0.187 | 0.357 | -0.585 | 0 | 0.023 | 0.023 | 0 | 0.27 | 0.44 | 0.28 | 0.25 | 0.26 | 0.3 |
| 5 | 101546639 | rs4703214 | 0.67 | 0.193 | -0.175 | -0.561 | -0.002 | -0.021 | -0.021 | 0 | 0.2 | 0.15 | 0.22 | 0.23 | 0.19 | 0.25 |
| 5 | 101548700 | rs1549273 | 1.803 | 0.252 | -0.199 | -0.673 | 0.002 | 0.038 | 0.038 | 0.586 | 0.2 | 0.12 | 0.09 | 0.07 | 0.14 | 0.07 |
| 5 | 101550199 | rs261116 | -1.376 | 0.502 | -0.356 | -0.659 | -0.03 | 0.088 | 0.088 | 0 | 0.74 | 0.92 | 0.84 | 0.87 | 0.69 | 0.8 |
| 5 | 101551270 | rs261115 | -0.806 | 0.618 | -0.649 | -1.097 | -0.006 | 0.001 | 0.001 | 0 | 0.66 | 0.69 | 0.75 | 0.85 | 0.56 | 0.79 |
| 5 | 101552955 | rs261114 | -1.098 | 0.534 | -0.356 | -0.947 | -0.002 | 0.076 | 0.076 | 3.518 | 0.59 | 0.75 | 0.76 | 0.87 | 0.61 | 0.78 |
| 5 | 101558083 | rs4323206 | 0.423 | 0.461 | -0.093 | -0.804 | -0.006 | 0.01 | 0.01 | 0.083 | 0.26 | 0.37 | 0.22 | 0.29 | 0.21 | 0.35 |
| 5 | 101559063 | rs12657697 | 1.15 | 0.548 | -0.157 | -0.812 | 0.018 | -0.023 | -0.023 | 0 | 0.2 | 0.14 | 0.18 | 0.23 | 0.12 | 0.25 |
| 5 | 101559119 | rs589938 | -1.427 | 0.582 | -0.143 | -0.827 | -0.001 | 0.065 | 0.065 | 3.848 | 0.59 | 0.74 | 0.76 | 0.84 | 0.6 | 0.78 |
| 5 | 101564593 | rs12658856 | 0.262 | 0.639 | 0.07 | -0.666 | -0.003 | 0.004 | 0.004 | 0.226 | 0.26 | 0.37 | 0.22 | 0.29 | 0.21 | 0.34 |
| 5 | 101566107 | rs7725428 | 0.324 | 0.658 | -0.052 | -0.724 | -0.036 | 0.01 | 0.01 | 0.558 | 0.26 | 0.48 | 0.22 | 0.23 | 0.17 | 0.3 |
| 5 | 101566775 | rs397655 | 0.375 | 0.743 | -0.205 | -0.58 | -0.005 | -0.013 | -0.013 | 0.004 | 0.3 | 0.22 | 0.5 | 0.51 | 0.37 | 0.42 |
| 5 | 101570236 | rs3811976 | -0.098 | 0.679 | -0.087 | -0.116 | 0.02 | 0.001 | 0.001 | 0 | 0.05 | 0.13 | 0.04 | 0.05 | 0.09 | 0.09 |
| 5 | 101570491 | rs4703217 | 0.531 | 0.477 | -0.181 | -0.136 | 0.005 | -0.006 | -0.006 | 0 | 0.2 | 0.2 | 0.19 | 0.23 | 0.13 | 0.27 |
| 5 | 101584499 | rs404598 | 0.394 | 0.591 | -0.167 | -0.063 | -0.003 | -0.014 | -0.014 | 0 | 0.36 | 0.31 | 0.46 | 0.52 | 0.33 | 0.43 |
| 5 | 101585209 | rs10067502 | 0.009 | 0.662 | -0.073 | -0.257 | 0.001 | -0.031 | -0.031 | 0.368 | 0.12 | 0.18 | 0.15 | 0.01 | 0.13 | 0.06 |
| 5 | 101587931 | rs10072122 | 1.636 | 0.68 | -0.235 | -0.341 | -0.024 | 0.045 | 0.045 | 2.438 | 0.26 | 0.03 | 0.14 | 0.16 | 0.28 | 0.16 |
| 5 | 101588195 | rs10067049 | 0.148 | 0.687 | -0.176 | -0.34 | 0.001 | -0.032 | -0.032 | 0 | 0.12 | 0.17 | 0.15 | 0.01 | 0.13 | 0.06 |
| 5 | 101591422 | rs431778 | -1.573 | 0.631 | 0.062 | -0.349 | 0.002 | 0.01 | 0.01 | 0 | 0.8 | 0.79 | 0.79 | 0.72 | 0.77 | 0.71 |
| 5 | 101592742 | rs174414 | -0.168 | 0.63 | 0.053 | -0.35 | 0.005 | 0 | 0 | 0 | 0.69 | 0.61 | 0.64 | 0.7 | 0.64 | 0.65 |
| 5 | 101596412 | rs10074392 | 1.755 | 0.622 | -0.038 | -0.346 | -0.043 | 0.057 | 0.057 | 2.957 | 0.26 | 0.01 | 0.13 | 0.11 | 0.27 | 0.15 |
| 5 | 101596633 | rs452857 | -1.693 | 0.628 | 0.099 | -0.351 | -0.024 | -0.045 | -0.045 | 0 | 0.81 | 0.92 | 0.79 | 0.72 | 0.77 | 0.71 |
| 5 | 101600837 | rs2113092 | 0.389 | 0.609 | 0.217 | -0.362 | 0.003 | -0.005 | -0.005 | 0 | 0.65 | 0.43 | 0.62 | 0.7 | 0.64 | 0.63 |
| 5 | 101604681 | rs2113090 | 0.558 | 0.602 | 0.227 | -0.36 | -0.001 | 0 | 0 | 0 | 0.63 | 0.37 | 0.6 | 0.68 | 0.61 | 0.58 |
| 5 | 101605641 | rs2600834 | -1.798 | 0.668 | 0.227 | -0.353 | -0.012 | -0.043 | -0.043 | 1.532 | 0.82 | 0.92 | 0.8 | 0.72 | 0.79 | 0.71 |
| 5 | 101605942 | rs2600833 | 0.393 | 0.665 | 0.228 | -0.372 | 0.003 | -0.004 | -0.004 | 0 | 0.65 | 0.43 | 0.62 | 0.7 | 0.64 | 0.63 |
| 5 | 101606002 | rs2600832 | -1.347 | 0.856 | 0.24 | -0.195 | 0.002 | 0.008 | 0.008 | 0 | 0.8 | 0.79 | 0.79 | 0.72 | 0.77 | 0.71 |
| 5 | 101608346 | rs2600827 | -1.384 | 0.866 | 0.245 | -0.191 | 0.002 | 0.01 | 0.01 | 0 | 0.8 | 0.79 | 0.79 | 0.72 | 0.78 | 0.71 |
| 5 | 101615705 | rs6884301 | 0.772 | 0.834 | -0.113 | -0.222 | 0.009 | 0.038 | 0.038 | 1.568 | 0.32 | 0.21 | 0.18 | 0.18 | 0.34 | 0.23 |
| 5 | 101620174 | rs2548724 | -1.562 | 1.046 | -0.148 | -0.043 | -0.043 | -0.091 | -0.091 | 0 | 0.82 | 0.97 | 0.81 | 0.72 | 0.79 | 0.71 |
| 5 | 101624336 | rs709369 | -0.143 | 0.942 | -0.056 | -0.14 | -0.005 | -0.021 | -0.021 | 0 | 0.45 | 0.72 | 0.58 | 0.55 | 0.41 | 0.48 |
| 5 | 101625154 | rs709372 | 0.717 | 0.942 | 0.045 | -0.155 | 0.002 | 0.007 | 0.007 | 0 | 0.36 | 0.37 | 0.47 | 0.54 | 0.34 | 0.44 |
| 5 | 101631592 | rs3114659 | -0.659 | 0.973 | -0.311 | -0.11 | -0.006 | -0.012 | -0.012 | 0 | 0.76 | 0.82 | 0.75 | 0.73 | 0.73 | 0.69 |
| 5 | 101631790 | rs10479190 | 1.355 | 0.888 | -0.441 | -0.161 | -0.033 | 0.039 | 0.039 | 1.626 | 0.28 | 0.01 | 0.14 | 0.16 | 0.28 | 0.16 |
| 5 | 101633557 | rs896544 | 0.019 | 0.947 | -0.24 | -0.081 | -0.004 | -0.022 | -0.022 | 0 | 0.55 | 0.27 | 0.42 | 0.45 | 0.59 | 0.52 |
| 5 | 101633610 | rs896545 | 0.019 | 0.947 | -0.24 | -0.081 | -0.004 | -0.022 | -0.022 | 0.511 | 0.55 | 0.27 | 0.42 | 0.45 | 0.59 | 0.52 |

| | | | | | | | | | | | | | | | | |
|---|---|---|---|---|---|---|---|---|---|---|---|---|---|---|---|---|
| 5 | 101635625 | rs10073052 | 0.019 | 0.959 | -0.24 | -0.081 | -0.002 | -0.022 | -0.022 | 0 | 0.55 | 0.27 | 0.42 | 0.45 | 0.59 | 0.52 |
| 5 | 101651873 | rs6899078 | 0.243 | 0.772 | -0.191 | -0.139 | 0.009 | 0.03 | 0.03 | 0.46 | 0.4 | 0.34 | 0.28 | 0.19 | 0.39 | 0.29 |
| 5 | 101659105 | rs17332107 | 0.668 | 0.77 | -0.223 | -0.243 | 0.003 | 0.062 | 0.062 | 0.431 | 0.34 | 0.1 | 0.19 | 0.17 | 0.34 | 0.22 |
| 5 | 101662158 | rs9327837 | 0.984 | 0.78 | -0.135 | -0.243 | 0 | -0.026 | -0.026 | 1.165 | 0.07 | 0.12 | 0.11 | 0.02 | 0.07 | 0.05 |
| 5 | 101669015 | rs12655961 | 0.735 | 0.897 | -0.031 | -0.17 | 0.001 | 0.003 | 0.003 | 0.028 | 0.23 | 0.24 | 0.25 | 0.27 | 0.26 | 0.29 |
| 5 | 101701924 | rs10075318 | -0.93 | 0.877 | -0.01 | -0.178 | -0.001 | -0.003 | -0.003 | 0.982 | 0.64 | 0.67 | 0.53 | 0.46 | 0.64 | 0.57 |
| 5 | 101707342 | rs10055753 | -0.93 | 0.899 | -0.01 | -0.178 | -0.001 | -0.003 | -0.003 | 0 | 0.64 | 0.67 | 0.53 | 0.46 | 0.64 | 0.57 |
| 5 | 101711249 | rs1901515 | -0.054 | 0.821 | -0.003 | -0.201 | 0.016 | 0.046 | 0.046 | 0.051 | 0.44 | 0.36 | 0.3 | 0.19 | 0.42 | 0.29 |
| 5 | 101711280 | rs1901514 | 0.809 | 0.842 | 0.012 | -0.199 | 0.004 | 0.063 | 0.063 | 0.431 | 0.34 | 0.11 | 0.19 | 0.17 | 0.34 | 0.22 |
| 5 | 101711558 | rs1901513 | 0.336 | 0.889 | -0.057 | -0.175 | 0.013 | 0.027 | 0.027 | 0.431 | 0.4 | 0.35 | 0.28 | 0.19 | 0.38 | 0.28 |
| 5 | 101711999 | rs7733501 | 1.15 | 0.885 | -0.052 | -0.156 | 0.008 | 0.031 | 0.031 | 0 | 0.21 | 0.31 | 0.23 | 0.27 | 0.22 | 0.28 |
| 5 | 101716620 | rs10069193 | -0.915 | 0.903 | -0.179 | -0.154 | -0.001 | -0.003 | -0.003 | 0 | 0.64 | 0.67 | 0.53 | 0.46 | 0.65 | 0.57 |
| 5 | 101719783 | rs6890130 | -0.829 | 0.898 | -0.172 | -0.156 | 0.002 | 0.014 | 0.014 | 0 | 0.64 | 0.6 | 0.53 | 0.46 | 0.65 | 0.57 |
| 5 | 101725479 | rs7728630 | 0.859 | 0.903 | -0.254 | -0.164 | -0.01 | -0.015 | -0.015 | 0 | 0.22 | 0.15 | 0.25 | 0.27 | 0.26 | 0.29 |
| 5 | 101726770 | rs10073892 | 1.112 | 0.886 | -0.523 | -0.135 | -0.046 | 0.055 | 0.055 | 0.942 | 0.31 | 0.02 | 0.17 | 0.15 | 0.31 | 0.17 |
| 5 | 101729504 | rs1562960 | -0.688 | 0.964 | -0.232 | 0.041 | -0.001 | -0.003 | -0.003 | 0 | 0.64 | 0.67 | 0.53 | 0.46 | 0.65 | 0.57 |
| 5 | 101736641 | rs6596482 | -0.698 | 0.965 | -0.232 | 0.034 | -0.001 | -0.003 | -0.003 | 0 | 0.64 | 0.66 | 0.53 | 0.46 | 0.64 | 0.57 |
| 5 | 101737777 | rs7731256 | -0.704 | 0.959 | -0.235 | 0.031 | -0.001 | -0.003 | -0.003 | 1.07 | 0.64 | 0.67 | 0.53 | 0.46 | 0.64 | 0.57 |
| 5 | 101743783 | rs1376898 | -0.697 | 0.966 | -0.232 | 0.037 | -0.001 | -0.003 | -0.003 | 0 | 0.64 | 0.67 | 0.53 | 0.46 | 0.65 | 0.57 |
| 5 | 101746708 | rs1901521 | -0.687 | 0.96 | -0.232 | 0.037 | -0.001 | -0.003 | -0.003 | 0 | 0.64 | 0.66 | 0.53 | 0.46 | 0.64 | 0.57 |
| 5 | 101746847 | rs1376896 | 0.692 | 1.288 | -0.321 | 0.353 | -0.018 | -0.013 | -0.013 | 0 | 0.09 | 0.05 | 0.16 | 0.14 | 0.14 | 0.1 |
| 5 | 101755430 | rs1542296 | -0.309 | 1.293 | -0.318 | 0.348 | -0.001 | -0.003 | -0.003 | 0.681 | 0.64 | 0.66 | 0.53 | 0.46 | 0.65 | 0.57 |
| 5 | 101762909 | rs10055023 | -0.301 | 1.287 | -0.321 | 0.32 | -0.001 | -0.003 | -0.003 | 0.018 | 0.64 | 0.67 | 0.53 | 0.46 | 0.65 | 0.57 |
| 5 | 101775394 | rs2400796 | 0.655 | 1.126 | -0.017 | 0.139 | -0.022 | -0.029 | -0.029 | 0 | 0.53 | 0.18 | 0.4 | 0.45 | 0.56 | 0.5 |
| 5 | 101776835 | rs17273466 | 0.024 | 1.118 | 0.126 | 0.135 | -0.003 | -0.017 | -0.017 | 0 | 0.6 | 0.34 | 0.51 | 0.46 | 0.64 | 0.55 |
| 5 | 101780479 | rs7734060 | 1.864 | 1.203 | -0.114 | 0.109 | -0.043 | -0.082 | -0.082 | 0 | 0.18 | 0.03 | 0.2 | 0.25 | 0.2 | 0.26 |
| 5 | 101780913 | rs4386713 | 0.836 | 1.168 | -0.154 | 0.057 | -0.015 | -0.021 | -0.021 | 0 | 0.25 | 0.18 | 0.27 | 0.3 | 0.3 | 0.34 |
| 5 | 101782930 | rs1901509 | 0.909 | 1.185 | -0.204 | 0.076 | -0.018 | -0.013 | -0.013 | 0 | 0.09 | 0.05 | 0.16 | 0.14 | 0.14 | 0.11 |
| 5 | 101788155 | rs2400798 | -0.616 | 1.203 | -0.37 | 0.084 | -0.001 | -0.003 | -0.003 | 0 | 0.64 | 0.66 | 0.53 | 0.46 | 0.64 | 0.57 |
| 5 | 101789305 | rs1452057 | -0.616 | 1.203 | -0.37 | 0.084 | -0.001 | -0.003 | -0.003 | 0.264 | 0.64 | 0.66 | 0.53 | 0.46 | 0.64 | 0.57 |
| 5 | 101795490 | rs11746217 | -0.616 | 1.173 | -0.37 | 0.084 | -0.001 | -0.003 | -0.003 | 0 | 0.64 | 0.67 | 0.53 | 0.46 | 0.66 | 0.57 |
| 5 | 101803412 | rs1597771 | -0.597 | 1.173 | -0.37 | 0.084 | -0.001 | -0.003 | -0.003 | 0 | 0.64 | 0.67 | 0.53 | 0.46 | 0.65 | 0.57 |
| 5 | 101806886 | rs7356787 | 1.646 | 1.163 | -0.442 | 0.081 | -0.028 | 0.031 | 0.031 | 1.942 | 0.28 | 0.02 | 0.14 | 0.15 | 0.27 | 0.16 |
| 5 | 101816228 | rs10041652 | 1.329 | 1.17 | -0.438 | 0.063 | -0.042 | 0.063 | 0.063 | 0.942 | 0.31 | 0.02 | 0.17 | 0.15 | 0.31 | 0.17 |
| 5 | 101817358 | rs1875634 | -0.625 | 1.128 | -0.291 | 0.116 | -0.002 | -0.008 | -0.008 | 4.745 | 0.64 | 0.69 | 0.54 | 0.46 | 0.65 | 0.57 |
| 5 | 101819981 | rs981988 | 1.851 | 0.932 | -0.477 | 0.132 | -0.042 | -0.095 | -0.095 | 0.864 | 0.22 | 0.01 | 0.24 | 0.34 | 0.18 | 0.29 |
| 5 | 101831523 | rs6860730 | 0.241 | 0.827 | -0.062 | -0.069 | -0.016 | -0.059 | -0.059 | 0 | 0.4 | 0.51 | 0.27 | 0.16 | 0.36 | 0.24 |
| 5 | 101836140 | rs6883528 | -0.525 | 0.832 | -0.004 | -0.056 | 0.035 | -0.069 | -0.069 | 0.223 | 0.12 | 0.49 | 0.13 | 0.02 | 0.09 | 0.08 |
| 5 | 101836665 | rs6898458 | -0.525 | 0.832 | -0.004 | -0.056 | 0.035 | -0.069 | -0.069 | 0 | 0.12 | 0.49 | 0.13 | 0.02 | 0.09 | 0.08 |
| 5 | 101843759 | rs953280 | 0.691 | 0.893 | -0.057 | -0.028 | -0.012 | -0.021 | -0.021 | 0 | 0.25 | 0.17 | 0.27 | 0.29 | 0.29 | 0.33 |
| 5 | 101844778 | rs10070289 | 0.911 | 0.892 | 0.03 | 0.052 | 0.011 | -0.031 | -0.031 | 0 | 0.07 | 0.15 | 0.11 | 0.02 | 0.08 | 0.05 |
| 5 | 101859136 | rs7728204 | 0.04 | 0.895 | 0.013 | 0.055 | 0.015 | -0.052 | -0.052 | 0 | 0.1 | 0.25 | 0.12 | 0.02 | 0.09 | 0.07 |
| 5 | 101860882 | rs10463983 | -0.581 | 0.877 | -0.214 | 0.149 | -0.002 | -0.009 | -0.009 | 0 | 0.64 | 0.69 | 0.54 | 0.47 | 0.65 | 0.56 |
| 5 | 101874055 | rs12657079 | 0.925 | 0.022 | -0.371 | 0.121 | 0.002 | -0.004 | -0.004 | 0 | 0.15 | 0.13 | 0.07 | 0.18 | 0.14 | 0.15 |
| 5 | 101876929 | rs1393625 | 0.778 | 0.387 | -0.235 | 0.306 | -0.032 | 0.038 | 0.038 | 0.389 | 0.33 | 0.02 | 0.17 | 0.17 | 0.32 | 0.22 |
| 5 | 101891174 | rs3105231 | -0.817 | 0.469 | 0.179 | 0.422 | -0.004 | 0.033 | 0.033 | 0 | 0.69 | 0.57 | 0.58 | 0.57 | 0.73 | 0.66 |
| 5 | 101914033 | rs17152198 | -0.43 | 0.43 | 0.342 | 0.28 | 0.005 | -0.058 | -0.058 | 0 | 0.1 | 0.33 | 0.1 | 0.02 | 0.07 | 0.07 |
| 5 | 101918306 | rs10479236 | 0.129 | 0.45 | 0.359 | 0.234 | 0.029 | 0.064 | 0.064 | 0.098 | 0.4 | 0.24 | 0.23 | 0.19 | 0.35 | 0.27 |
| 5 | 101918448 | rs10477819 | -0.433 | 0.48 | 0.414 | 0.252 | 0.006 | -0.057 | -0.057 | 6.839 | 0.1 | 0.32 | 0.1 | 0.02 | 0.07 | 0.07 |
| 5 | 101927431 | rs10479237 | -0.108 | 0.493 | 0.365 | 0.258 | -0.001 | -0.043 | -0.043 | 2.219 | 0.09 | 0.18 | 0.08 | 0.02 | 0.07 | 0.07 |
| 5 | 101929429 | rs10440756 | 1.067 | 0.443 | 0.164 | 0.264 | 0.029 | 0.044 | 0.044 | 0.413 | 0.31 | 0.1 | 0.15 | 0.18 | 0.29 | 0.2 |
| 5 | 101946798 | rs7705924 | 0.921 | 0.538 | 0.207 | 0.254 | -0.001 | -0.011 | -0.011 | 3.709 | 0.06 | 0.08 | 0.08 | 0.02 | 0.05 | 0.05 |
| 5 | 101953047 | rs3105477 | -0.652 | 0.63 | 0.248 | 0.347 | -0.035 | -0.03 | -0.03 | 0.225 | 0.72 | 0.85 | 0.66 | 0.62 | 0.61 | 0.61 |
| 5 | 101954763 | rs10070269 | 0.774 | 0.612 | 0.177 | 0.341 | -0.005 | -0.03 | -0.03 | 0 | 0.07 | 0.15 | 0.08 | 0.02 | 0.04 | 0.05 |
| 5 | 101956225 | rs10043147 | 1.11 | 0.525 | 0.16 | 0.344 | 0.027 | 0.04 | 0.04 | 0.511 | 0.31 | 0.17 | 0.15 | 0.17 | 0.28 | 0.21 |
| 5 | 101959192 | rs982094 | 0.687 | 0.69 | 0.137 | 0.529 | -0.002 | 0.011 | 0.011 | 0 | 0.31 | 0.34 | 0.42 | 0.44 | 0.27 | 0.33 |
| 5 | 101999586 | rs2657433 | -0.991 | 1.101 | 0.331 | 0.754 | -0.085 | -0.077 | -0.077 | 0.976 | 0.78 | 0.98 | 0.75 | 0.74 | 0.72 | 0.71 |
| 5 | 102004743 | rs2657445 | 0.148 | 0.858 | 0.623 | 0.643 | 0.021 | 0.016 | 0.016 | 0.288 | 0.68 | 0.46 | 0.63 | 0.62 | 0.61 | 0.6 |
| 5 | 102012989 | rs2657449 | 0.326 | 0.876 | 0.703 | 0.65 | -0.007 | -0.004 | -0.004 | 0 | 0.66 | 0.31 | 0.62 | 0.62 | 0.6 | 0.61 |
| 5 | 102044030 | rs17355759 | -0.125 | 0.726 | -0.213 | 0.672 | -0.009 | -0.01 | -0.01 | 0 | 0.34 | 0.25 | 0.38 | 0.36 | 0.37 | 0.39 |
| 5 | 102044072 | rs1895213 | -0.574 | 0.6 | 0.496 | 0.532 | 0.001 | -0.003 | -0.003 | 0 | 0.4 | 0.61 | 0.41 | 0.39 | 0.4 | 0.46 |
| 5 | 102061818 | rs258255 | -1.473 | 0.546 | -0.098 | 0.133 | -0.024 | 0.056 | 0.056 | 0 | 0.9 | 0.99 | 0.93 | 0.99 | 0.9 | 0.96 |
| 5 | 102069125 | rs1913504 | 1.452 | 0.377 | -0.446 | 0.394 | -0.083 | 0.033 | 0.033 | 0.419 | 0.26 | 0.01 | 0.18 | 0.14 | 0.29 | 0.18 |
| 5 | 102078587 | rs451819 | -0.961 | 0.49 | 0.901 | 0.915 | -0.002 | -0.002 | -0.002 | 0 | 0.4 | 0.54 | 0.41 | 0.4 | 0.39 | 0.46 |
| 5 | 102087116 | rs397605 | -0.961 | 0.49 | 0.901 | 0.915 | -0.002 | -0.002 | -0.002 | 0 | 0.4 | 0.54 | 0.41 | 0.39 | 0.39 | 0.46 |
| 5 | 102095375 | rs385827 | -0.338 | 0.455 | 0.891 | 0.887 | -0.021 | -0.032 | -0.032 | 0 | 0.33 | 0.17 | 0.38 | 0.38 | 0.37 | 0.39 |
| 5 | 102096425 | rs462957 | -1.131 | 0.485 | 0.804 | 1.373 | 0.004 | -0.001 | -0.001 | 0 | 0.43 | 0.64 | 0.45 | 0.56 | 0.41 | 0.49 |
| 5 | 102096919 | rs467186 | -0.962 | 0.591 | 0.701 | 1.196 | -0.002 | -0.001 | -0.001 | 0.809 | 0.4 | 0.61 | 0.42 | 0.41 | 0.39 | 0.46 |
| 5 | 102107466 | rs459202 | -0.666 | 0.76 | 0.678 | 1.257 | 0.011 | -0.001 | -0.001 | 0 | 0.35 | 0.54 | 0.39 | 0.38 | 0.36 | 0.4 |

| | | | | | | | | | | | | | | | | |
|---|---|---|---|---|---|---|---|---|---|---|---|---|---|---|---|---|
| 5 | 102108099 | rs2431531 | -0.665 | 0.485 | 0.886 | 1.296 | 0.001 | 0.001 | 0.001 | 0 | 0.36 | 0.48 | 0.42 | 0.39 | 0.37 | 0.45 |
| 5 | 102120269 | rs461605 | -0.813 | 0.659 | 1.125 | 1.381 | 0.003 | 0.023 | 0.023 | 0 | 0.33 | 0.57 | 0.3 | 0.42 | 0.36 | 0.32 |
| 5 | 102149795 | rs28092 | -0.086 | 0.671 | 1.18 | 1.413 | -0.011 | -0.004 | -0.004 | 0 | 0.75 | 0.8 | 0.74 | 0.73 | 0.67 | 0.82 |
| 5 | 102160785 | rs258247 | -0.062 | 0.647 | 1.229 | 1.475 | 0.006 | 0.001 | 0.001 | 0 | 0.74 | 0.7 | 0.74 | 0.73 | 0.67 | 0.82 |
| 5 | 102161751 | rs258244 | 0.51 | 0.638 | 1.293 | 1.466 | 0.012 | 0.043 | 0.043 | 0 | 0.69 | 0.58 | 0.69 | 0.58 | 0.64 | 0.8 |
| 5 | 102163325 | rs686043 | -0.065 | 0.708 | 1.124 | 1.441 | -0.001 | -0.001 | -0.001 | 0 | 0.75 | 0.75 | 0.73 | 0.74 | 0.67 | 0.82 |
| 5 | 102167154 | rs467693 | 0.738 | 0.659 | 0.958 | 1.36 | 0.004 | 0.009 | 0.009 | 0.181 | 0.65 | 0.39 | 0.67 | 0.58 | 0.62 | 0.66 |
| 5 | 102172496 | rs6886411 | -1.881 | 0.269 | 1.174 | 1.617 | -0.002 | 0.208 | 0.208 | 2.059 | 0.11 | 0.36 | 0.11 | 0.26 | 0.06 | 0.22 |
| 5 | 102196156 | rs7704560 | -1.672 | 0.209 | 1.376 | 1.593 | -0.013 | 0.171 | 0.171 | 0 | 0.07 | 0.29 | 0.05 | 0.19 | 0.01 | 0.07 |
| 5 | 102200149 | rs7708778 | -1.825 | 0.282 | 1.508 | 1.591 | 0.023 | 0.21 | 0.21 | 0 | 0.07 | 0.36 | 0.06 | 0.19 | 0.02 | 0.07 |
| 5 | 102201590 | rs249496 | 0.532 | 0.032 | 1.015 | 1.223 | 0.013 | 0.063 | 0.063 | 0 | 0.48 | 0.75 | 0.61 | 0.62 | 0.41 | 0.66 |
| 5 | 102209823 | rs6868471 | -0.938 | -0.041 | 1.251 | 1.236 | -0.009 | 0.175 | 0.175 | 0 | 0.07 | 0.3 | 0.06 | 0.18 | 0.01 | 0.07 |
| 5 | 102212963 | rs6885642 | -1.33 | -0.047 | 1.235 | 1.233 | 0.036 | 0.238 | 0.238 | 0 | 0.07 | 0.39 | 0.07 | 0.18 | 0.02 | 0.07 |
| 5 | 102225196 | rs1363786 | 0.964 | -0.257 | 1.149 | 0.774 | -0.015 | 0 | 0 | 0 | 0.89 | 0.72 | 0.84 | 0.9 | 0.92 | 0.8 |
| 5 | 102238502 | rs10038600 | 1.145 | 0.172 | 0.781 | 0.692 | 0.001 | 0.031 | 0.031 | 0 | 0.31 | 0.18 | 0.21 | 0.14 | 0.32 | 0.21 |
| 5 | 102280302 | rs6860588 | 0.313 | 0.231 | 0.876 | 1.074 | -0.021 | -0.048 | -0.048 | 0.709 | 0.2 | 0.13 | 0.23 | 0.44 | 0.29 | 0.21 |
| 5 | 102282317 | rs3776874 | 0.175 | 0.253 | 0.943 | 1.072 | -0.004 | -0.004 | -0.004 | 0.161 | 0.21 | 0.2 | 0.23 | 0.44 | 0.3 | 0.21 |
| 5 | 102287920 | rs11952361 | -0.313 | 0.222 | 1.111 | 1.055 | 0.037 | 0.1 | 0.1 | 0 | 0.24 | 0.46 | 0.26 | 0.44 | 0.3 | 0.21 |
| 5 | 102306322 | rs12652973 | 0.432 | 0.228 | 0.71 | 1.049 | -0.037 | -0.065 | -0.065 | 0.363 | 0.21 | 0.11 | 0.24 | 0.43 | 0.29 | 0.16 |
| 5 | 102310166 | rs3733939 | -0.487 | 0.114 | 1.197 | 1.067 | 0.024 | 0.075 | 0.075 | 0 | 0.28 | 0.56 | 0.36 | 0.49 | 0.34 | 0.21 |
| 5 | 102317042 | rs6596529 | -0.134 | 0.218 | 1.282 | 1.057 | 0.034 | 0.065 | 0.065 | 0 | 0.25 | 0.42 | 0.26 | 0.43 | 0.3 | 0.16 |
| 5 | 102352034 | rs3776859 | -0.133 | 0.284 | 1.356 | 1.059 | 0.036 | 0.091 | 0.091 | 0 | 0.24 | 0.49 | 0.25 | 0.43 | 0.28 | 0.16 |
| 5 | 102365186 | rs5855 | -0.195 | 0.343 | 1.261 | 0.989 | 0.017 | 0.041 | 0.041 | 0 | 0.77 | 0.69 | 0.76 | 0.55 | 0.71 | 0.86 |
| 5 | 102381641 | rs9327877 | 1.272 | 0.278 | 1.231 | 0.726 | 0.008 | 0.023 | 0.023 | 0.021 | 0.26 | 0.13 | 0.17 | 0.15 | 0.22 | 0.18 |
| 5 | 102420019 | rs185387 | 0.365 | 0.511 | 1.029 | 0.912 | 0.018 | 0.049 | 0.049 | 0 | 0.22 | 0.31 | 0.23 | 0.45 | 0.28 | 0.13 |
| 5 | 102450417 | rs34771 | 0.364 | 0.554 | 1.019 | 0.928 | 0.028 | 0.066 | 0.066 | 0 | 0.22 | 0.35 | 0.24 | 0.45 | 0.28 | 0.13 |
| 5 | 102450820 | rs34774 | 0.427 | 0.559 | 0.939 | 0.93 | 0.011 | 0.029 | 0.029 | 0 | 0.22 | 0.29 | 0.23 | 0.45 | 0.28 | 0.13 |
| 5 | 102484399 | rs26522 | -0.115 | 0.51 | 1.045 | 0.944 | 0.037 | 0.122 | 0.122 | 0 | 0.26 | 0.54 | 0.34 | 0.49 | 0.32 | 0.18 |
| 5 | 102488571 | rs27896 | 0.779 | 0.648 | 0.635 | 0.89 | -0.046 | -0.089 | -0.089 | 0 | 0.2 | 0.07 | 0.23 | 0.45 | 0.27 | 0.13 |
| 5 | 102520400 | rs26821 | 0.29 | 0.643 | 0.652 | 0.871 | 0.03 | 0.073 | 0.073 | 0 | 0.22 | 0.39 | 0.24 | 0.45 | 0.28 | 0.13 |
| 5 | 102552013 | rs26256 | -1.904 | 0.458 | -0.028 | 0.593 | -0.028 | -0.073 | -0.073 | 0 | 0.88 | 0.99 | 0.74 | 0.74 | 0.89 | 0.74 |
| 5 | 102572418 | rs17155171 | 1.235 | 0.304 | 0.281 | 0.43 | 0.003 | 0 | 0 | 0.359 | 0.27 | 0.14 | 0.18 | 0.33 | 0.23 | 0.27 |
| 5 | 102589454 | rs246903 | -0.296 | 0.276 | 0.585 | 0.636 | 0.018 | 0.013 | 0.013 | 0 | 0.26 | 0.47 | 0.24 | 0.3 | 0.29 | 0.15 |
| 5 | 102600754 | rs2288786 | -0.092 | 0.193 | 0.112 | 0.582 | -0.034 | -0.002 | -0.002 | 1.514 | 0.27 | 0.12 | 0.19 | 0.26 | 0.3 | 0.16 |
| 5 | 102614327 | rs35797 | -0.852 | 0.129 | -0.342 | 0.563 | 0.004 | 0.01 | 0.01 | 0.328 | 0.36 | 0.75 | 0.36 | 0.28 | 0.35 | 0.2 |
| 5 | 102615707 | rs386352 | -0.782 | 0.209 | -0.38 | 0.611 | 0.001 | -0.01 | -0.01 | 0.155 | 0.34 | 0.63 | 0.33 | 0.28 | 0.35 | 0.2 |
| 5 | 102634029 | rs6886092 | -0.974 | -0.049 | -1.02 | 0.543 | 0.007 | 0.016 | 0.016 | 1.235 | 0.39 | 0.77 | 0.36 | 0.32 | 0.37 | 0.21 |
| 5 | 102655261 | rs402232 | -0.935 | 0.285 | -2.134 | 0.817 | -0.047 | -0.113 | -0.113 | 0 | 0.84 | 0.99 | 0.71 | 0.69 | 0.84 | 0.71 |
| 5 | 102660400 | rs12187443 | -0.802 | 0.303 | -1.44 | 0.4 | -0.049 | -0.02 | -0.02 | 0 | 0.3 | 0.09 | 0.2 | 0.29 | 0.32 | 0.16 |
| 5 | 102677364 | rs674726 | -0.315 | 0.438 | -1.393 | 0.42 | -0.04 | -0.01 | -0.01 | 0 | 0.25 | 0.13 | 0.2 | 0.28 | 0.31 | 0.16 |
| 5 | 102681261 | rs524571 | -0.414 | 0.527 | -1.569 | 0.468 | -0.063 | -0.021 | -0.021 | 4.427 | 0.28 | 0.07 | 0.2 | 0.28 | 0.31 | 0.16 |
| 5 | 102694112 | rs2561481 | 0.332 | 0.672 | -0.684 | 0.666 | 0.002 | 0.004 | 0.004 | 0.043 | 0.66 | 0.46 | 0.56 | 0.68 | 0.62 | 0.82 |
| 5 | 102695626 | rs387569 | 0.419 | 0.632 | -0.683 | 0.478 | 0.001 | -0.017 | -0.017 | 0 | 0.59 | 0.54 | 0.56 | 0.71 | 0.54 | 0.8 |
| 5 | 102704259 | rs10068434 | 0.094 | 0.514 | -1.025 | 0.551 | -0.009 | 0.037 | 0.037 | 0.486 | 0.27 | 0.05 | 0.19 | 0.2 | 0.27 | 0.12 |
| 5 | 102709812 | rs11242490 | -0.346 | 0.796 | 0.015 | 0.707 | -0.006 | 0.004 | 0.004 | 0 | 0.33 | 0.44 | 0.35 | 0.31 | 0.36 | 0.17 |
| 5 | 102710525 | rs12654624 | -0.791 | 0.926 | -0.588 | 1.05 | -0.054 | -0.053 | -0.053 | 0 | 0.73 | 0.89 | 0.59 | 0.6 | 0.67 | 0.63 |
| 5 | 102714855 | rs11242492 | -0.547 | 0.398 | 0.403 | 1.083 | 0.012 | 0.035 | 0.035 | 0 | 0.09 | 0.16 | 0.05 | 0.09 | 0.09 | 0.04 |
| 5 | 102714963 | rs11242493 | -1.212 | 0.207 | 0.691 | 0.925 | 0.005 | 0.007 | 0.007 | 0 | 0.16 | 0.32 | 0.09 | 0.09 | 0.13 | 0.06 |
| 5 | 102721744 | rs10463994 | -0.572 | 0.265 | 0.635 | 0.969 | 0.036 | 0.078 | 0.078 | 0 | 0.09 | 0.26 | 0.06 | 0.09 | 0.09 | 0.04 |
| 5 | 102725095 | rs9327881 | -0.517 | 0.268 | 0.56 | 0.97 | 0.036 | 0.077 | 0.077 | 0 | 0.09 | 0.25 | 0.06 | 0.09 | 0.09 | 0.04 |
| 5 | 102726572 | rs17155292 | 0.551 | 0.232 | -0.179 | 1.112 | -0.011 | -0.027 | -0.027 | 0 | 0.13 | 0.07 | 0.11 | 0.22 | 0.17 | 0.13 |
| 5 | 102739759 | rs6596533 | -1.193 | -0.107 | 0.221 | 0.676 | -0.014 | -0.013 | -0.013 | 0 | 0.22 | 0.69 | 0.14 | 0.09 | 0.14 | 0.12 |
| 5 | 102753215 | rs3846615 | -1.041 | -0.043 | 0.385 | 1.223 | -0.014 | 0.017 | 0.017 | 0 | 0.3 | 0.66 | 0.34 | 0.53 | 0.25 | 0.27 |
| 5 | 102760214 | rs2068326 | -0.185 | -0.071 | 0.473 | 1.492 | 0.015 | -0.01 | -0.01 | 0 | 0.11 | 0.09 | 0.19 | 0.14 | 0.06 | 0.14 |
| 5 | 102770988 | rs2115251 | 0.125 | -0.071 | 0.727 | 1.703 | 0 | 0.009 | 0.009 | 0 | 0.14 | 0.17 | 0.12 | 0.27 | 0.17 | 0.12 |
| 5 | 102777020 | rs10479253 | -1.216 | -0.139 | 1.057 | 1.706 | -0.024 | 0.081 | 0.081 | 0 | 0.26 | 0.53 | 0.31 | 0.51 | 0.2 | 0.24 |
| 5 | 102781533 | rs1428491 | -1.292 | -0.195 | 1.069 | 1.658 | -0.024 | 0.069 | 0.069 | 0 | 0.27 | 0.57 | 0.29 | 0.41 | 0.2 | 0.24 |
| 5 | 102793568 | rs291835 | -0.137 | -0.184 | 1.124 | 1.676 | 0.003 | 0.025 | 0.025 | 0 | 0.15 | 0.22 | 0.12 | 0.27 | 0.18 | 0.12 |
| 5 | 102799858 | rs1155605 | -0.234 | -0.264 | 1.111 | 1.667 | 0.018 | -0.02 | -0.02 | 0.545 | 0.12 | 0.1 | 0.2 | 0.19 | 0.06 | 0.15 |
| 5 | 102810287 | rs294071 | 0.104 | -0.25 | 1.125 | 1.663 | -0.002 | -0.004 | -0.004 | 0 | 0.14 | 0.13 | 0.12 | 0.25 | 0.18 | 0.11 |
| 5 | 102811490 | rs150392 | -0.195 | -0.251 | 1.16 | 1.662 | 0.006 | 0.044 | 0.044 | 0 | 0.15 | 0.27 | 0.13 | 0.25 | 0.18 | 0.12 |
| 5 | 102811961 | rs291858 | -1.368 | -0.259 | 1.173 | 1.579 | -0.017 | 0.056 | 0.056 | 0 | 0.28 | 0.42 | 0.33 | 0.43 | 0.24 | 0.28 |
| 5 | 102821963 | rs158703 | -1.146 | -0.122 | 1.174 | 1.63 | -0.019 | 0.082 | 0.082 | 0 | 0.26 | 0.52 | 0.3 | 0.54 | 0.2 | 0.26 |
| 5 | 102826903 | rs432889 | 0.13 | 0.114 | 1.279 | 1.508 | 0.045 | 0.099 | 0.099 | 0 | 0.12 | 0.32 | 0.1 | 0.2 | 0.14 | 0.08 |
| 5 | 102831132 | rs460545 | -1.036 | 0.435 | 0.763 | 1.369 | -0.003 | -0.022 | -0.022 | 1.508 | 0.39 | 0.48 | 0.54 | 0.69 | 0.37 | 0.6 |
| 5 | 102834360 | rs158704 | 0.767 | 0.55 | 0.936 | 1.451 | 0.013 | 0.033 | 0.033 | 0 | 0.11 | 0.19 | 0.09 | 0.2 | 0.14 | 0.08 |
| 5 | 102840482 | rs291873 | 0.403 | 0.559 | 0.697 | 1.385 | 0.043 | 0.093 | 0.093 | 0.016 | 0.12 | 0.32 | 0.1 | 0.21 | 0.14 | 0.08 |
| 5 | 102847822 | rs378918 | 0.43 | 0.565 | 0.57 | 1.383 | 0.041 | 0.19 | 0.19 | 0 | 0.12 | 0.33 | 0.1 | 0.33 | 0.15 | 0.08 |
| 5 | 102852240 | rs12516214 | -0.193 | 0.305 | 0.408 | 1.373 | 0.002 | -0.065 | -0.065 | 0 | 0.26 | 0.11 | 0.44 | 0.36 | 0.22 | 0.48 |

| | | | | | | | | | | | | | | | | |
|---|---|---|---|---|---|---|---|---|---|---|---|---|---|---|---|---|
| 5 | 102852837 | rs291838 | -0.811 | 0.921 | 0.419 | 0.368 | 0 | -0.016 | -0.016 | 0.174 | 0.45 | 0.51 | 0.61 | 0.69 | 0.48 | 0.58 |
| 5 | 102853748 | rs1345614 | 0.563 | 1.028 | 0.291 | 0.289 | 0.003 | -0.022 | -0.022 | 0.007 | 0.35 | 0.42 | 0.28 | 0.33 | 0.38 | 0.33 |
| 5 | 102855162 | rs34229 | 0.366 | 0.912 | 0.209 | 0.549 | -0.001 | 0.017 | 0.017 | 0 | 0.21 | 0.25 | 0.41 | 0.5 | 0.2 | 0.28 |
| 5 | 102855756 | rs291840 | 0.515 | 1.104 | 0.206 | 0.214 | -0.001 | -0.077 | -0.077 | 0 | 0.26 | 0.38 | 0.21 | 0.01 | 0.27 | 0.06 |
| 5 | 102863527 | rs34225 | -1.324 | 1.149 | -0.088 | 0.131 | -0.003 | 0.016 | 0.016 | 2.499 | 0.7 | 0.82 | 0.85 | 0.76 | 0.75 | 0.71 |
| 5 | 102863759 | rs34224 | 0.676 | 1.203 | -0.395 | 0.379 | 0.002 | -0.038 | -0.038 | 0 | 0.2 | 0.16 | 0.39 | 0.4 | 0.2 | 0.24 |
| 5 | 102872552 | rs460651 | -0.197 | 1.284 | -0.597 | -0.125 | -0.001 | 0 | 0 | 0.031 | 0.46 | 0.59 | 0.62 | 0.51 | 0.47 | 0.35 |
| 5 | 102885995 | rs7724372 | -0.673 | 1.337 | -0.774 | -0.158 | 0.024 | 0.07 | 0.07 | 0 | 0.75 | 0.56 | 0.53 | 0.5 | 0.69 | 0.69 |
| 5 | 102886015 | rs10900867 | 1.021 | 1.34 | -0.734 | -0.206 | 0.001 | 0.01 | 0.01 | 2.806 | 0.33 | 0.23 | 0.17 | 0.25 | 0.27 | 0.3 |
| 5 | 102887795 | rs2271170 | 0.505 | 1.094 | -0.559 | -0.379 | 0.005 | 0.007 | 0.007 | 0.074 | 0.2 | 0.19 | 0.21 | 0.24 | 0.25 | 0.35 |
| 5 | 102895124 | rs7723689 | 0.234 | 0.594 | -0.659 | 0.018 | 0.034 | 0.048 | 0.048 | 0.222 | 0.21 | 0.14 | 0.15 | 0.01 | 0.16 | 0.04 |
| 5 | 102903747 | rs9327886 | 0.146 | 1.021 | -0.484 | 0.556 | -0.004 | -0.001 | -0.001 | 3.639 | 0.55 | 0.42 | 0.46 | 0.43 | 0.56 | 0.68 |
| 5 | 102913746 | rs371315 | -0.462 | 1.273 | -0.159 | 0.031 | -0.002 | -0.024 | -0.024 | 0 | 0.36 | 0.54 | 0.28 | 0.76 | 0.31 | 0.47 |
| 5 | 102921968 | rs6874368 | -0.373 | 1.002 | -0.509 | -0.797 | -0.003 | 0.035 | 0.035 | 1.092 | 0.38 | 0.28 | 0.35 | 0.15 | 0.35 | 0.35 |
| 5 | 102925899 | rs17155527 | -0.007 | 0.814 | -0.585 | -0.929 | 0.008 | 0.007 | 0.007 | 0.285 | 0.14 | 0.07 | 0.07 | 0.09 | 0.06 | 0.21 |
| 5 | 102926108 | rs11242497 | 0.469 | 0.881 | -0.462 | -0.944 | -0.056 | 0.071 | 0.071 | 0.021 | 0.22 | 0.1 | 0.28 | 0.01 | 0.3 | 0.15 |
| 5 | 102930270 | rs1422046 | 0.272 | 0.261 | -0.661 | -1.138 | -0.01 | -0.006 | -0.006 | 0.026 | 0.31 | 0.28 | 0.41 | 0.65 | 0.4 | 0.41 |
| 5 | 102933674 | rs4269268 | -0.69 | -0.106 | -0.658 | -1.507 | 0.006 | 0.116 | 0.116 | 0 | 0.13 | 0.56 | 0.1 | 0.26 | 0.14 | 0.15 |
| 5 | 102937084 | rs6881783 | -0.389 | -0.074 | -0.102 | -1.473 | 0.036 | 0.13 | 0.13 | 0 | 0.11 | 0.38 | 0.09 | 0.21 | 0.14 | 0.15 |
| 5 | 102945524 | rs17155558 | 0.578 | -0.329 | -0.007 | -1.607 | 0.026 | 0.019 | 0.019 | 0.022 | 0.14 | 0.07 | 0.06 | 0.09 | 0.06 | 0.2 |
| 5 | 102946979 | rs17155559 | -0.454 | -0.389 | 0.034 | -1.632 | -0.007 | 0.125 | 0.125 | 0 | 0.11 | 0.38 | 0.09 | 0.21 | 0.11 | 0.15 |
| 5 | 102949668 | rs7707391 | -0.411 | -0.487 | -0.32 | -1.898 | 0 | 0.019 | 0.019 | 0 | 0.41 | 0.35 | 0.44 | 0.73 | 0.36 | 0.57 |
| 5 | 102958282 | rs11948851 | 0.72 | -0.547 | -0.429 | -1.576 | 0.025 | 0.006 | 0.006 | 0.013 | 0.14 | 0.07 | 0.06 | 0.14 | 0.06 | 0.2 |
| 5 | 102961829 | rs10463996 | 0.36 | -0.274 | -0.484 | -1.671 | -0.032 | -0.019 | -0.019 | 0 | 0.28 | 0.16 | 0.37 | 0.66 | 0.3 | 0.4 |
| 5 | 102962282 | rs10039615 | 0.459 | 0.021 | -0.205 | -1.504 | 0.002 | 0.2 | 0.2 | 0 | 0.06 | 0.27 | 0.03 | 0.19 | 0.01 | 0.1 |
| 5 | 102970690 | rs4122274 | 1.255 | 0.04 | -0.995 | -1.751 | -0.123 | 0.163 | 0.163 | 0 | 0.25 | 0.03 | 0.35 | 0.01 | 0.32 | 0.18 |
| 5 | 102976249 | rs11952329 | -0.219 | 0.162 | -0.498 | -1.311 | 0.006 | 0.026 | 0.026 | 0 | 0.4 | 0.34 | 0.41 | 0.75 | 0.34 | 0.58 |
| 5 | 102979948 | rs12656448 | -0.465 | 0.369 | -1.079 | -1.528 | 0.032 | 0.242 | 0.242 | 0.003 | 0.5 | 0.8 | 0.37 | 0.98 | 0.43 | 0.68 |
| 5 | 102985009 | rs11242504 | -0.804 | 0.292 | -0.626 | 0.402 | 0.003 | -0.031 | -0.031 | 1.008 | 0.12 | 0.05 | 0.16 | 0.44 | 0.12 | 0.23 |
| 5 | 102985590 | rs17155627 | -1.581 | 0.633 | -0.654 | 0.435 | 0.007 | 0.09 | 0.09 | 0 | 0.19 | 0.28 | 0.36 | 0.53 | 0.26 | 0.28 |
| 5 | 102985840 | rs4145649 | 0.458 | 0.808 | -1.126 | 0.874 | -0.008 | -0.012 | -0.012 | 0 | 0.13 | 0.08 | 0.1 | 0.23 | 0.14 | 0.17 |
| 5 | 102989429 | rs3857360 | -0.896 | 0.679 | -1.159 | 0.922 | -0.018 | -0.061 | -0.061 | 1.412 | 0.15 | 0.09 | 0.2 | 0.57 | 0.21 | 0.26 |
| 5 | 102990964 | rs17155647 | -0.181 | 0.606 | -1.531 | 0.565 | -0.014 | -0.066 | -0.066 | 0 | 0.1 | 0.01 | 0.05 | 0.2 | 0.06 | 0.31 |
| 5 | 102992086 | rs2545664 | 1.361 | 0.37 | -2.196 | -0.49 | -0.091 | 0.225 | 0.225 | 0 | 0.33 | 0.04 | 0.36 | 0.01 | 0.35 | 0.16 |
| 5 | 102996464 | rs2665182 | -0.307 | -0.11 | -1.111 | -0.212 | 0.007 | 0.022 | 0.022 | 0.061 | 0.58 | 0.31 | 0.6 | 0.75 | 0.63 | 0.71 |
| 5 | 102998150 | rs10074159 | 1.204 | 0.029 | -1.817 | -0.508 | -0.032 | 0.044 | 0.044 | 1.873 | 0.28 | 0.13 | 0.39 | 0.16 | 0.29 | 0.28 |
| 5 | 102998283 | rs1288872 | -1.426 | 0.094 | -1.825 | -0.608 | -0.024 | 0.102 | 0.102 | 0.006 | 0.77 | 0.93 | 0.86 | 0.96 | 0.72 | 0.88 |
| 5 | 103001097 | rs11952040 | 1.143 | 0.063 | -1.843 | -0.299 | -0.001 | 0.063 | 0.063 | 0 | 0.07 | 0.03 | 0.12 | 0.48 | 0.08 | 0.17 |
| 5 | 103003822 | rs1288881 | -1.984 | 0.074 | -1.856 | -0.356 | 0.003 | 0.074 | 0.074 | 0.263 | 0.8 | 0.94 | 0.89 | 0.98 | 0.82 | 0.9 |
| 5 | 103003909 | rs1288882 | -0.08 | 0.02 | -1.604 | -0.293 | 0.009 | 0.014 | 0.014 | 0 | 0.42 | 0.65 | 0.35 | 0.32 | 0.35 | 0.42 |
| 5 | 103009768 | rs11950651 | 1.47 | 0.261 | -1.479 | 0.063 | -0.005 | 0.014 | 0.014 | 0 | 0.05 | 0.02 | 0.12 | 0.35 | 0.07 | 0.15 |
| 5 | 103011612 | rs1152161 | 0.261 | 0.179 | -1.184 | 0.045 | -0.011 | -0.01 | -0.01 | 0.193 | 0.36 | 0.46 | 0.32 | 0.31 | 0.31 | 0.38 |
| 5 | 103011697 | rs10060923 | 1.156 | 0.279 | -2 | 0.065 | -0.025 | -0.018 | -0.018 | 0 | 0.32 | 0.13 | 0.39 | 0.27 | 0.31 | 0.33 |
| 5 | 103013291 | rs984288 | -0.922 | 0.314 | -1.904 | 0.102 | -0.021 | -0.022 | -0.022 | 0 | 0.71 | 0.89 | 0.75 | 0.62 | 0.65 | 0.71 |
| 5 | 103014764 | rs1152168 | -0.77 | 0.297 | -1.891 | -0.513 | -0.015 | -0.09 | -0.09 | 0.033 | 0.7 | 0.87 | 0.74 | 0.47 | 0.64 | 0.7 |
| 5 | 103016305 | rs1152173 | -0.898 | 0.406 | -2.006 | -0.542 | -0.007 | -0.037 | -0.037 | 0.135 | 0.71 | 0.8 | 0.74 | 0.49 | 0.64 | 0.71 |
| 5 | 103019653 | rs320681 | 0.587 | 0.59 | -2.391 | -1.297 | -0.036 | 0.128 | 0.128 | 0 | 0.23 | 0.05 | 0.23 | 0.01 | 0.28 | 0.1 |
| 5 | 103022981 | rs3857361 | 0.195 | 0.648 | -2.136 | -1.51 | -0.063 | 0.073 | 0.073 | 1.034 | 0.37 | 0.18 | 0.44 | 0.2 | 0.41 | 0.33 |
| 5 | 103028736 | rs2545657 | -0.097 | 0.737 | -1.438 | -1.701 | -0.031 | 0.095 | 0.095 | 0 | 0.2 | 0.06 | 0.23 | 0.01 | 0.25 | 0.07 |
| 5 | 103032867 | rs634248 | 0.068 | 0.565 | -0.606 | -1.665 | 0.002 | -0.058 | -0.058 | 2.994 | 0.64 | 0.45 | 0.69 | 0.85 | 0.62 | 0.7 |
| 5 | 103038495 | rs7712812 | 0.016 | 0 | -0.586 | -2.075 | 0.029 | 0.081 | 0.081 | 0 | 0.39 | 0.28 | 0.52 | 0.12 | 0.29 | 0.27 |
| 5 | 103042597 | rs6891011 | 0.136 | -0.109 | 0.128 | -1.974 | 0.009 | 0.036 | 0.036 | 0 | 0.13 | 0.37 | 0.15 | 0.19 | 0.14 | 0.19 |
| 5 | 103042676 | rs6869108 | -0.338 | -0.167 | 0.077 | -2.029 | 0.041 | 0.108 | 0.108 | 0 | 0.51 | 0.75 | 0.64 | 0.33 | 0.43 | 0.46 |
| 5 | 103044062 | rs10041164 | -0.437 | -0.19 | 0.113 | -2.056 | 0.01 | 0.128 | 0.128 | 0 | 0.54 | 0.81 | 0.65 | 0.32 | 0.46 | 0.44 |
| 5 | 103061473 | rs11242506 | 0.654 | -0.057 | 0.866 | -1.897 | 0.004 | 0.005 | 0.005 | 0 | 0.1 | 0.16 | 0.13 | 0.19 | 0.15 | 0.17 |
| 5 | 103066438 | rs10040307 | -0.405 | -0.193 | 0.937 | -1.962 | 0.003 | 0.007 | 0.007 | 0.303 | 0.15 | 0.44 | 0.15 | 0.19 | 0.16 | 0.18 |
| 5 | 103067334 | rs4703261 | -0.483 | -0.278 | 0.02 | -2.183 | -0.016 | 0.118 | 0.118 | 0 | 0.57 | 0.82 | 0.47 | 0.3 | 0.49 | 0.49 |
| 5 | 103068468 | rs6867791 | 0.686 | -0.439 | -0.203 | -2.027 | 0.012 | 0.002 | 0.002 | 0 | 0.23 | 0.1 | 0.37 | 0.7 | 0.2 | 0.4 |
| 5 | 103069610 | rs7714087 | 0.445 | -0.195 | -0.009 | -2.038 | -0.002 | -0.065 | -0.065 | 0 | 0.33 | 0.3 | 0.27 | 0.02 | 0.3 | 0.16 |
| 5 | 103074969 | rs7720623 | -0.026 | 0.398 | -0.69 | -1.413 | 0.024 | -0.046 | -0.046 | 0.715 | 0.64 | 0.4 | 0.67 | 0.78 | 0.54 | 0.67 |
| 5 | 103078256 | rs12655517 | -0.105 | 0.701 | -0.745 | -1.285 | 0.05 | -0.06 | -0.06 | 0.766 | 0.66 | 0.49 | 0.68 | 0.77 | 0.54 | 0.68 |
| 5 | 103083642 | rs10038456 | -0.054 | 0.472 | -1.58 | -1.363 | -0.013 | 0.153 | 0.153 | 0.427 | 0.29 | 0.07 | 0.23 | 0.02 | 0.28 | 0.15 |
| 5 | 103084134 | rs10479267 | -0.057 | 0.751 | -1.103 | -2.135 | -0.001 | -0.053 | -0.053 | 0.199 | 0.7 | 0.73 | 0.81 | 1 | 0.67 | 0.83 |
| 5 | 103086147 | rs2161495 | -0.602 | 0.349 | -1.458 | -1.262 | 0.006 | -0.036 | -0.036 | 0.038 | 0.26 | 0.15 | 0.24 | 0.59 | 0.29 | 0.33 |
| 5 | 103091216 | rs10070070 | -0.611 | 0.625 | -1.458 | -0.891 | -0.002 | -0.045 | -0.045 | 0 | 0.26 | 0.16 | 0.24 | 0.56 | 0.3 | 0.33 |
| 5 | 103094133 | rs2018064 | -0.534 | 0.611 | -1.505 | -0.927 | 0.003 | -0.003 | -0.003 | 0.395 | 0.14 | 0.11 | 0.17 | 0.6 | 0.17 | 0.3 |
| 5 | 103097260 | rs320661 | 1.143 | 0.261 | -0.917 | -1.163 | 0.001 | -0.011 | -0.011 | 0.248 | 0.69 | 0.27 | 0.72 | 0.4 | 0.67 | 0.66 |
| 5 | 103104452 | rs1820121 | 0.836 | 0.249 | -1.173 | -1.54 | 0.005 | 0.012 | 0.012 | 0 | 0.32 | 0.29 | 0.44 | 0.21 | 0.27 | 0.31 |
| 5 | 103109780 | rs320654 | 0.74 | 0.286 | -0.575 | -1.536 | -0.022 | 0.036 | 0.036 | 0 | 0.4 | 0.25 | 0.3 | 0.23 | 0.41 | 0.38 |

| | | | | | | | | | | | | | | | | |
|---|---|---|---|---|---|---|---|---|---|---|---|---|---|---|---|---|
| 5 | 103116239 | rs2545618 | 0.699 | 0.169 | 0.031 | -1.578 | -0.006 | 0.031 | 0.031 | 0 | 0.46 | 0.36 | 0.35 | 0.23 | 0.47 | 0.39 |
| 5 | 103119367 | rs623961 | 0.686 | 0.202 | 0.114 | -1.572 | -0.008 | 0.041 | 0.041 | 0 | 0.46 | 0.34 | 0.35 | 0.22 | 0.48 | 0.39 |
| 5 | 103122923 | rs491507 | -0.154 | 0.165 | -0.205 | -1.489 | -0.04 | -0.095 | -0.095 | 1.219 | 0.78 | 0.93 | 0.79 | 0.43 | 0.71 | 0.71 |
| 5 | 103128418 | rs320667 | 0.203 | 0.142 | -0.087 | -1.511 | -0.014 | -0.059 | -0.059 | 0 | 0.76 | 0.86 | 0.77 | 0.43 | 0.71 | 0.68 |
| 5 | 103133258 | rs1288996 | 1.062 | 0.529 | -0.559 | -1.843 | -0.021 | 0.059 | 0.059 | 0 | 0.27 | 0.03 | 0.23 | 0.14 | 0.26 | 0.18 |
| 5 | 103136481 | rs620032 | -0.434 | 0.291 | 0.069 | -1.372 | -0.004 | 0.014 | 0.014 | 0.007 | 0.89 | 0.92 | 0.86 | 0.4 | 0.86 | 0.74 |
| 5 | 103141780 | rs1990902 | -0.971 | 0.291 | 0.206 | -1.493 | -0.035 | 0.074 | 0.074 | 0 | 0.56 | 0.73 | 0.65 | 0.76 | 0.52 | 0.64 |
| 5 | 103141923 | rs17155846 | 0.614 | 0.201 | 0.598 | -1.554 | -0.01 | -0.027 | -0.027 | 0 | 0.29 | 0.31 | 0.41 | 0.19 | 0.23 | 0.29 |
| 5 | 103142399 | rs17155849 | -0.693 | 0.172 | 0.595 | -1.517 | 0 | 0.061 | 0.061 | 0 | 0.27 | 0.43 | 0.24 | 0.56 | 0.29 | 0.33 |
| 5 | 103143532 | rs1422095 | -0.861 | 0.213 | 0.246 | -1.529 | -0.027 | 0.073 | 0.073 | 0 | 0.55 | 0.72 | 0.65 | 0.78 | 0.52 | 0.66 |
| 5 | 103158899 | rs1422099 | 0.192 | 0.29 | 0.414 | -1.521 | -0.002 | -0.139 | -0.139 | 3.627 | 0.17 | 0.48 | 0.11 | 0 | 0.15 | 0.08 |
| 5 | 103160079 | rs1039860 | -0.932 | 0.259 | 0.259 | -1.564 | -0.027 | 0.085 | 0.085 | 0.054 | 0.54 | 0.77 | 0.62 | 0.76 | 0.52 | 0.62 |
| 5 | 103160351 | rs11242508 | 0.871 | 0.492 | 0.158 | -1.395 | -0.009 | 0.04 | 0.04 | 0 | 0.07 | 0.03 | 0.11 | 0.56 | 0.11 | 0.21 |
| 5 | 103164240 | rs10056109 | 0.481 | 0.483 | 0.164 | -1.398 | -0.005 | 0.032 | 0.032 | 0 | 0.08 | 0.06 | 0.11 | 0.58 | 0.12 | 0.23 |
| 5 | 103166870 | rs2059070 | -0.946 | 0.32 | 0.285 | -0.948 | -0.007 | 0.058 | 0.058 | 0 | 0.28 | 0.51 | 0.25 | 0.58 | 0.29 | 0.39 |
| 5 | 103170089 | rs10041231 | -0.599 | 0.404 | -0.356 | -0.987 | 0.036 | 0.173 | 0.173 | 0 | 0.49 | 0.77 | 0.55 | 0.21 | 0.41 | 0.46 |
| 5 | 103172273 | rs17155896 | -0.094 | 0.803 | 0.096 | -0.139 | -0.011 | -0.024 | -0.024 | 0.236 | 0.09 | 0.03 | 0.12 | 0.58 | 0.13 | 0.23 |
| 5 | 103174685 | rs13155509 | -0.584 | 0.825 | 0.738 | -1.004 | 0.009 | -0.116 | -0.116 | 0 | 0.18 | 0.41 | 0.13 | 0 | 0.18 | 0.15 |
| 5 | 103182563 | rs7715548 | -0.922 | 1.224 | 0.476 | -0.89 | 0.038 | -0.112 | -0.112 | 0.044 | 0.21 | 0.52 | 0.16 | 0.02 | 0.23 | 0.19 |
| 5 | 103183059 | rs1363251 | -1.501 | 1.022 | -0.156 | -0.794 | -0.027 | 0.067 | 0.067 | 0 | 0.47 | 0.83 | 0.29 | 0.28 | 0.45 | 0.5 |
| 5 | 103183424 | rs6885169 | -1.116 | 1.225 | 0.529 | -1.073 | 0.016 | -0.116 | -0.116 | 0.001 | 0.28 | 0.58 | 0.18 | 0.03 | 0.3 | 0.25 |
| 5 | 103183907 | rs17155913 | 0.561 | 1.281 | 0.019 | -1.536 | 0.001 | -0.03 | -0.03 | 3.152 | 0.15 | 0.19 | 0.12 | 0.01 | 0.19 | 0.07 |
| 5 | 103196133 | rs11959903 | -0.837 | 1.325 | 0.436 | -1.59 | 0.014 | -0.08 | -0.08 | 0 | 0.3 | 0.54 | 0.27 | 0.05 | 0.4 | 0.18 |
| 5 | 103196610 | rs6871128 | -0.808 | 1.183 | 0.763 | -1.503 | 0.013 | -0.035 | -0.035 | 0 | 0.09 | 0.19 | 0.05 | 0.02 | 0.06 | 0.1 |
| 5 | 103205290 | rs17507392 | -0.792 | 1.073 | 0.435 | -1.372 | -0.001 | -0.002 | -0.002 | 0 | 0.23 | 0.18 | 0.11 | 0.23 | 0.21 | 0.31 |
| 5 | 103206340 | rs10039846 | -1.643 | 0.873 | 0.051 | -1.549 | -0.018 | 0.083 | 0.083 | 1.134 | 0.41 | 0.83 | 0.26 | 0.22 | 0.39 | 0.4 |
| 5 | 103223902 | rs7720125 | -0.921 | 0.936 | 0.316 | -1.398 | -0.004 | 0 | 0 | 0 | 0.26 | 0.31 | 0.11 | 0.23 | 0.2 | 0.3 |
| 5 | 103227422 | rs34250 | -1.197 | 1.331 | 0.056 | -1.475 | 0.022 | -0.012 | -0.012 | 0 | 0.29 | 0.37 | 0.21 | 0.22 | 0.35 | 0.31 |
| 5 | 103231731 | rs34244 | -0.521 | 1.455 | 0.276 | -1.501 | 0.108 | -0.068 | -0.068 | 0 | 0.12 | 0.4 | 0.15 | 0 | 0.21 | 0.09 |
| 5 | 103237745 | rs40054 | -0.121 | 0.928 | -1.126 | -2.121 | -0.013 | 0.101 | 0.101 | 0 | 0.65 | 0.8 | 0.72 | 0.98 | 0.72 | 0.82 |
| 5 | 103238413 | rs39653 | -0.82 | 0.856 | -1.354 | -1.98 | -0.046 | 0.115 | 0.115 | 0.036 | 0.8 | 0.98 | 0.76 | 0.98 | 0.8 | 0.87 |
| 5 | 103252374 | rs808042 | -0.113 | 0.684 | -1.824 | -1.541 | 0.012 | 0.007 | 0.007 | 0.887 | 0.49 | 0.81 | 0.32 | 0.46 | 0.49 | 0.48 |
| 5 | 103252652 | rs808041 | 0.387 | 0.346 | -1.134 | -1.568 | 0.001 | -0.064 | -0.064 | 5.287 | 0.31 | 0.47 | 0.22 | 0.17 | 0.34 | 0.35 |
| 5 | 103253764 | rs7736954 | 0.338 | 0.493 | -0.989 | -1.534 | 0.109 | -0.03 | -0.03 | 0.05 | 0.11 | 0.44 | 0.13 | 0.02 | 0.17 | 0.13 |
| 5 | 103253842 | rs1469456 | 1.011 | 0.462 | -1.684 | -1.837 | -0.059 | -0.137 | -0.137 | 0.256 | 0.22 | 0.02 | 0.42 | 0.54 | 0.26 | 0.38 |
| 5 | 103258936 | rs246373 | -0.02 | 0.38 | -0.406 | -1.501 | 0.141 | 0.196 | 0.196 | 0 | 0.11 | 0.39 | 0.14 | 0.25 | 0.2 | 0.17 |
| 5 | 103261943 | rs718377 | -0.625 | 0.252 | -0.645 | -1.596 | -0.002 | 0 | 0 | 2.356 | 0.27 | 0.7 | 0.21 | 0.28 | 0.34 | 0.27 |
| 5 | 103264987 | rs808054 | 1.581 | 0.296 | -0.463 | -1.515 | 0.001 | -0.001 | -0.001 | 0 | 0.13 | 0.09 | 0.19 | 0.3 | 0.13 | 0.11 |
| 5 | 103265452 | rs1363259 | 1.029 | 0.396 | -0.748 | -1.685 | -0.016 | -0.067 | -0.067 | 0.042 | 0.3 | 0.15 | 0.41 | 0.53 | 0.29 | 0.38 |
| 5 | 103267433 | rs971962 | -0.071 | 0.424 | -0.183 | -1.762 | -0.006 | 0.003 | 0.003 | 0 | 0.45 | 0.34 | 0.54 | 0.55 | 0.49 | 0.54 |
| 5 | 103269694 | rs1363258 | 0.997 | 0.386 | -0.207 | -1.852 | -0.018 | 0.038 | 0.038 | 0 | 0.33 | 0.19 | 0.25 | 0.15 | 0.44 | 0.34 |
| 5 | 103270358 | rs4703272 | 0.578 | 0.808 | 0.717 | -1.232 | -0.008 | -0.005 | -0.005 | 0 | 0.7 | 0.35 | 0.66 | 0.69 | 0.75 | 0.78 |
| 5 | 103272650 | rs246383 | 0.25 | 0.815 | 0.741 | -1.264 | -0.015 | -0.017 | -0.017 | 0 | 0.73 | 0.43 | 0.7 | 0.74 | 0.79 | 0.79 |
| 5 | 103275221 | rs979028 | 1.401 | 0.799 | 0.595 | -1.285 | -0.027 | 0.065 | 0.065 | 2.243 | 0.6 | 0.17 | 0.57 | 0.71 | 0.59 | 0.63 |
| 5 | 103277290 | rs173486 | -0.059 | 0.023 | 0.703 | -1.007 | 0.004 | -0.07 | -0.07 | 0.221 | 0.86 | 0.7 | 0.87 | 1 | 0.88 | 0.94 |
| 5 | 103277997 | rs34235 | 0.763 | 0.031 | 0.559 | -0.961 | -0.005 | 0.025 | 0.025 | 0 | 0.14 | 0.21 | 0.2 | 0.28 | 0.13 | 0.12 |
| 5 | 103287479 | rs2431337 | -0.247 | 0.044 | 0.435 | -1.082 | 0 | -0.024 | -0.024 | 0.162 | 0.86 | 0.81 | 0.89 | 1 | 0.88 | 0.94 |
| 5 | 103292556 | rs2195441 | -1.142 | 0.078 | 0.395 | -1.048 | -0.003 | 0.02 | 0.02 | 0 | 0.86 | 0.84 | 0.8 | 0.72 | 0.87 | 0.88 |
| 5 | 103293719 | rs7349833 | 1.127 | 0.078 | 0.248 | -1.124 | -0.006 | -0.009 | -0.009 | 0 | 0.64 | 0.38 | 0.64 | 0.7 | 0.71 | 0.75 |
| 5 | 103305737 | rs1560201 | -0.65 | 0.647 | -0.276 | -0.964 | 0 | -0.016 | -0.016 | 0.132 | 0.18 | 0.21 | 0.11 | 0.03 | 0.14 | 0.11 |
| 5 | 103309671 | rs17338777 | 0.169 | 0.39 | -0.495 | -0.717 | 0.004 | 0.004 | 0.004 | 3.589 | 0.08 | 0.04 | 0.02 | 0.02 | 0.04 | 0.07 |
| 5 | 103314475 | rs319127 | 0.565 | 0.825 | -0.343 | -0.468 | 0 | 0.001 | 0.001 | 0 | 0.13 | 0.09 | 0.18 | 0.26 | 0.12 | 0.12 |
| 5 | 103317446 | rs7727450 | 0.326 | 0.974 | -0.472 | -1.184 | 0 | -0.017 | -0.017 | 0.585 | 0.57 | 0.44 | 0.3 | 0.17 | 0.55 | 0.5 |
| 5 | 103322064 | rs319134 | -0.559 | 0.833 | -0.526 | -0.421 | 0.012 | -0.049 | -0.049 | 0 | 0.86 | 0.96 | 0.61 | 0.4 | 0.86 | 0.68 |
| 5 | 103325974 | rs319131 | 0.701 | 1.19 | -0.024 | -0.857 | 0.002 | 0.003 | 0.003 | 0 | 0.63 | 0.46 | 0.43 | 0.39 | 0.6 | 0.46 |
| 5 | 103353936 | rs950664 | 0.427 | 1.229 | 0.167 | -0.851 | -0.001 | -0.002 | -0.002 | 0 | 0.65 | 0.67 | 0.45 | 0.4 | 0.6 | 0.47 |
| 5 | 103379245 | rs6596557 | -0.568 | 1.669 | 0.194 | -1.474 | 0.028 | -0.138 | -0.138 | 0 | 0.34 | 0.38 | 0.26 | 0 | 0.42 | 0.27 |
| 5 | 103381922 | rs1990745 | -0.824 | 1.387 | -0.053 | -1.273 | 0.002 | -0.046 | -0.046 | 0 | 0.85 | 0.96 | 0.62 | 0.41 | 0.84 | 0.68 |
| 5 | 103382538 | rs10515363 | 1.356 | 0.925 | -0.436 | -1.856 | -0.019 | 0.03 | 0.03 | 0 | 0.15 | 0.08 | 0.12 | 0 | 0.19 | 0.11 |
| 5 | 103386041 | rs2403366 | -0.428 | 0.84 | -0.297 | -1.755 | -0.017 | -0.043 | -0.043 | 0 | 0.7 | 0.85 | 0.49 | 0.41 | 0.62 | 0.54 |
| 5 | 103387880 | rs6887182 | 0.905 | 1.104 | 0.164 | -2.013 | -0.009 | 0.002 | 0.002 | 0 | 0.39 | 0.45 | 0.3 | 0.41 | 0.34 | 0.35 |
| 5 | 103390933 | rs6864188 | 0.114 | 1.079 | 0.094 | -2.029 | -0.001 | -0.057 | -0.057 | 0 | 0.28 | 0.25 | 0.17 | 0 | 0.28 | 0.15 |
| 5 | 103406680 | rs1289188 | 0.379 | 1.025 | -0.048 | -1.947 | -0.018 | -0.006 | -0.006 | 0.823 | 0.6 | 0.8 | 0.47 | 0.41 | 0.58 | 0.52 |
| 5 | 103425505 | rs13190616 | -0.047 | 0.666 | 0.253 | -2.121 | -0.014 | 0.003 | 0.003 | 0 | 0.62 | 0.44 | 0.45 | 0.39 | 0.64 | 0.48 |
| 5 | 103447183 | rs7725285 | 0.898 | 0.683 | 0.16 | -2.155 | 0.003 | -0.089 | -0.089 | 2.917 | 0.22 | 0.4 | 0.18 | 0.01 | 0.25 | 0.17 |
| 5 | 103450242 | rs1997511 | 0.233 | 0.763 | 0.045 | -2.328 | -0.021 | 0.043 | 0.043 | 0.887 | 0.27 | 0.14 | 0.16 | 0 | 0.29 | 0.14 |
| 5 | 103450790 | rs1362697 | 0.974 | 0.687 | 0.026 | -2.055 | 0 | -0.003 | -0.003 | 0 | 0.35 | 0.28 | 0.29 | 0.38 | 0.35 | 0.34 |
| 5 | 103464869 | rs17156151 | 0.643 | 0.65 | -0.124 | -2.088 | 0 | 0 | 0 | 0 | 0.39 | 0.35 | 0.36 | 0.38 | 0.41 | 0.36 |
| 5 | 103465420 | rs12153622 | -0.966 | 0.333 | -0.675 | -1.671 | 0.013 | -0.021 | -0.021 | 0 | 0.86 | 0.92 | 0.65 | 0.38 | 0.89 | 0.67 |

| | | | | | | | | | | | | | | | | |
|---|---|---|---|---|---|---|---|---|---|---|---|---|---|---|---|---|
| 5 | 103472789 | rs11955569 | 0.314 | 0.571 | -1.034 | -2.037 | -0.019 | 0.075 | 0.075 | 0.084 | 0.17 | 0.04 | 0.09 | 0 | 0.14 | 0.09 |
| 5 | 103473556 | rs12189165 | 1.322 | 0.99 | -1.06 | -2.267 | -0.01 | -0.002 | -0.002 | 0 | 0.1 | 0.07 | 0.1 | 0 | 0.16 | 0.03 |
| 5 | 103475024 | rs2952549 | 1.183 | 0.691 | -0.607 | -2.674 | 0.009 | -0.164 | -0.164 | 2.255 | 0.22 | 0.48 | 0.13 | 0 | 0.21 | 0.11 |
| 5 | 103478518 | rs1922495 | 0.684 | 0.558 | -0.612 | -2.568 | -0.005 | 0.006 | 0.006 | 0 | 0.57 | 0.65 | 0.48 | 0.39 | 0.57 | 0.55 |
| 5 | 103481012 | rs3857362 | 1.221 | 0.53 | -0.654 | -2.569 | -0.002 | -0.027 | -0.027 | 0 | 0.35 | 0.15 | 0.34 | 0.39 | 0.36 | 0.44 |
| 5 | 103484564 | rs4519903 | -1.426 | 0.525 | -0.636 | -2.45 | 0 | 0.015 | 0.015 | 0 | 0.86 | 0.87 | 0.68 | 0.39 | 0.87 | 0.68 |
| 5 | 103498939 | rs1922496 | 0.739 | 0.547 | -0.628 | -2.446 | -0.006 | 0.008 | 0.008 | 0 | 0.57 | 0.67 | 0.47 | 0.38 | 0.57 | 0.56 |
| 5 | 103500168 | rs17474182 | 0.632 | 0.657 | -0.683 | -2.511 | -0.017 | 0.071 | 0.071 | 0.102 | 0.17 | 0.04 | 0.09 | 0 | 0.14 | 0.09 |
| 5 | 103500335 | rs17474342 | 1.559 | 0.694 | -0.652 | -2.529 | -0.016 | 0.007 | 0.007 | 0 | 0.1 | 0.05 | 0.1 | 0 | 0.16 | 0.03 |
| 5 | 103500778 | rs17396015 | -0.288 | 0.607 | -0.703 | -2.565 | -0.02 | 0.012 | 0.012 | 0.015 | 0.27 | 0.21 | 0.2 | 0 | 0.29 | 0.12 |
| 5 | 103508344 | rs4235637 | -0.276 | -0.145 | -0.326 | -2.433 | 0.023 | 0.088 | 0.088 | 0 | 0.51 | 0.74 | 0.34 | 0 | 0.5 | 0.24 |
| 5 | 103513066 | rs17341221 | 0.29 | -0.15 | -0.359 | -2.469 | -0.001 | 0.016 | 0.016 | 0.133 | 0.17 | 0.07 | 0.09 | 0 | 0.13 | 0.09 |
| 5 | 103515006 | rs6887495 | -0.323 | -0.134 | -0.354 | -2.477 | 0.022 | 0.098 | 0.098 | 0 | 0.51 | 0.74 | 0.34 | 0 | 0.5 | 0.24 |
| 5 | 103519842 | rs6892281 | -0.323 | -0.134 | -0.348 | -2.477 | 0.022 | 0.103 | 0.103 | 0 | 0.51 | 0.74 | 0.34 | 0 | 0.5 | 0.25 |
| 5 | 103520960 | rs1922502 | 0.531 | -0.048 | -0.104 | -2.372 | 0.01 | -0.113 | -0.113 | 0 | 0.34 | 0.64 | 0.24 | 0 | 0.37 | 0.15 |
| 5 | 103523803 | rs6596566 | -1.135 | -0.211 | -0.191 | -2.279 | 0.003 | 0.003 | 0.003 | 0 | 0.85 | 0.89 | 0.68 | 0.38 | 0.87 | 0.68 |
| 5 | 103532095 | rs11953759 | 0.622 | -0.103 | -0.169 | -2.488 | -0.001 | -0.028 | -0.028 | 0 | 0.11 | 0.11 | 0.04 | 0 | 0.06 | 0.05 |
| 5 | 103536963 | rs7707433 | -0.756 | -0.038 | -0.088 | -2.333 | 0.03 | -0.172 | -0.172 | 0 | 0.68 | 0.47 | 0.77 | 1 | 0.63 | 0.85 |
| 5 | 103540975 | rs10063272 | 0.793 | -1.071 | -0.322 | -1.588 | 0.008 | -0.039 | -0.039 | 0 | 0.15 | 0.02 | 0.2 | 0.19 | 0.1 | 0.11 |
| 5 | 103544907 | rs1445764 | -1.355 | -1.279 | 0.084 | -1.597 | 0.02 | 0.014 | 0.014 | 0 | 0.42 | 0.72 | 0.45 | 0.61 | 0.34 | 0.62 |
| 5 | 103545085 | rs1445766 | 0.382 | -1.31 | 0.116 | -1.593 | 0.004 | -0.017 | -0.017 | 0 | 0.16 | 0.11 | 0.25 | 0.6 | 0.14 | 0.39 |
| 5 | 103545551 | rs1348687 | -1.426 | -1.347 | 0.412 | -1.641 | 0.001 | 0.001 | 0.001 | 0 | 0.4 | 0.7 | 0.45 | 0.63 | 0.33 | 0.61 |
| 5 | 103562868 | rs6596568 | -1.55 | -1.409 | 0.662 | -1.386 | -0.008 | -0.004 | -0.004 | 0.115 | 0.39 | 0.62 | 0.29 | 0.63 | 0.32 | 0.6 |
| 5 | 103565928 | rs4067552 | 0.855 | -1.321 | -0.141 | -1.291 | -0.001 | -0.087 | -0.087 | 0 | 0.13 | 0.02 | 0.09 | 0.62 | 0.11 | 0.31 |
| 5 | 103575599 | rs17156302 | -0.648 | -1.043 | 0.154 | -1.455 | 0.003 | -0.012 | -0.012 | 1.131 | 0.17 | 0.16 | 0.12 | 0 | 0.14 | 0.19 |
| 5 | 103583315 | rs12523239 | -0.44 | -1.078 | -0.067 | -1.326 | 0.011 | 0.038 | 0.038 | 0 | 0.14 | 0.09 | 0.09 | 0.01 | 0.08 | 0.14 |
| 5 | 103589615 | rs2121110 | 1.036 | -1.23 | 0.131 | -1.43 | 0.004 | -0.011 | -0.011 | 0.088 | 0.76 | 0.79 | 0.84 | 0.98 | 0.82 | 0.79 |
| 5 | 103593756 | rs2715841 | 1.251 | -1.236 | 0.209 | -1.433 | -0.011 | -0.079 | -0.079 | 0.558 | 0.74 | 0.67 | 0.83 | 0.98 | 0.82 | 0.79 |
| 5 | 103598185 | rs4473744 | 0.744 | -1.616 | -0.128 | -1.421 | 0.024 | -0.047 | -0.047 | 0 | 0.14 | 0.02 | 0.2 | 0.18 | 0.11 | 0.09 |
| 5 | 103600994 | rs2591431 | 1.055 | -0.886 | 0.517 | -1.281 | 0.006 | 0.047 | 0.047 | 0 | 0.77 | 0.85 | 0.85 | 0.99 | 0.82 | 0.85 |
| 5 | 103602375 | rs1031193 | 1.412 | -0.812 | 0.526 | -1.223 | -0.01 | -0.079 | -0.079 | 0 | 0.75 | 0.67 | 0.84 | 0.98 | 0.82 | 0.81 |
| 5 | 103610468 | rs17342020 | 0.543 | -0.663 | 0.139 | -1.175 | -0.005 | 0.025 | 0.025 | 0.205 | 0.07 | 0.02 | 0.18 | 0 | 0.06 | 0.03 |
| 5 | 103612677 | rs1823031 | 0.977 | -0.651 | 0.235 | -1.191 | -0.001 | -0.032 | -0.032 | 0 | 0.8 | 0.8 | 0.86 | 0.98 | 0.85 | 0.82 |
| 5 | 103614820 | rs1838695 | 0.829 | -0.669 | 0.539 | -1.2 | -0.005 | -0.079 | -0.079 | 0 | 0.8 | 0.72 | 0.85 | 0.98 | 0.85 | 0.82 |
| 5 | 103630140 | rs2583903 | 1.49 | -0.69 | 0.555 | -1.231 | -0.012 | -0.146 | -0.146 | 0 | 0.71 | 0.61 | 0.67 | 0.98 | 0.78 | 0.78 |
| 5 | 103641154 | rs1812338 | 1.454 | -0.673 | 0.492 | -1.238 | -0.011 | -0.163 | -0.163 | 0 | 0.71 | 0.57 | 0.66 | 0.99 | 0.77 | 0.87 |
| 5 | 103643778 | rs13164145 | -0.032 | -0.498 | 0.198 | -0.997 | -0.001 | -0.021 | -0.021 | 2.041 | 0.12 | 0.13 | 0.09 | 0.01 | 0.08 | 0.08 |
| 5 | 103659607 | rs12655672 | -0.565 | -0.565 | -0.263 | -0.618 | -0.001 | 0 | 0 | 0 | 0.28 | 0.25 | 0.35 | 0.68 | 0.25 | 0.43 |
| 5 | 103669502 | rs1470209 | -0.968 | -0.415 | -0.279 | -0.198 | 0.002 | 0.005 | 0.005 | 0 | 0.39 | 0.68 | 0.44 | 0.66 | 0.34 | 0.59 |
| 5 | 103673480 | rs349587 | 0.436 | -0.644 | 0.154 | 0.407 | 0.015 | 0.093 | 0.093 | 0 | 0.74 | 0.55 | 0.52 | 0.57 | 0.72 | 0.77 |
| 5 | 103674069 | rs17156441 | 0.493 | -0.327 | 0.197 | 0.043 | 0.018 | -0.03 | -0.03 | 0 | 0.06 | 0.11 | 0.06 | 0 | 0.11 | 0.08 |
| 5 | 103676341 | rs13184907 | 0.239 | -1.071 | -0.499 | -0.625 | -0.026 | -0.091 | -0.091 | 4.029 | 0.25 | 0.31 | 0.17 | 0.03 | 0.25 | 0.2 |
| 5 | 103679255 | rs168388 | 1.173 | -1.111 | -0.946 | -0.519 | -0.093 | -0.131 | -0.131 | 0.005 | 0.38 | 0.03 | 0.28 | 0.52 | 0.35 | 0.46 |
| 5 | 103680467 | rs2963226 | 0.501 | -1.082 | 0.129 | 0.122 | -0.011 | 0.154 | 0.154 | 0 | 0.87 | 0.59 | 0.71 | 0.67 | 0.92 | 0.94 |
| 5 | 103685244 | rs17344104 | 1.4 | -1.493 | -0.084 | -0.407 | 0.004 | 0.047 | 0.047 | 0 | 0.17 | 0.07 | 0.12 | 0 | 0.21 | 0.07 |
| 5 | 103685345 | rs11749520 | 0.795 | -1.606 | 0.131 | -0.434 | -0.019 | -0.024 | -0.024 | 0.866 | 0.05 | 0.14 | 0.03 | 0.01 | 0.03 | 0.09 |
| 5 | 103692385 | rs7721449 | 0.766 | -1.533 | 0.529 | -0.519 | 0.027 | -0.043 | -0.043 | 3.269 | 0.08 | 0.13 | 0.06 | 0 | 0.12 | 0.09 |
| 5 | 103693025 | rs10036615 | 0.668 | -1.661 | 0.118 | -0.392 | -0.056 | -0.079 | -0.079 | 0.098 | 0.18 | 0.07 | 0.24 | 0.28 | 0.25 | 0.12 |
| 5 | 103701679 | rs1372501 | -0.601 | -1.69 | 0.473 | -0.236 | 0.001 | 0.124 | 0.124 | 0 | 0.92 | 0.82 | 0.76 | 0.68 | 0.94 | 0.95 |
| 5 | 103704857 | rs1899386 | 1.879 | -2.013 | 0.266 | -0.307 | -0.095 | -0.016 | -0.016 | 0 | 0.37 | 0.03 | 0.27 | 0.32 | 0.34 | 0.5 |
| 5 | 103710618 | rs2963228 | 1.296 | -1.974 | 0.755 | -0.251 | -0.023 | 0.025 | 0.025 | 0 | 0.74 | 0.43 | 0.64 | 0.66 | 0.78 | 0.75 |
| 5 | 103726321 | rs2963229 | 0.747 | -1.626 | 0.482 | -0.094 | -0.077 | -0.106 | -0.106 | 0.521 | 0.18 | 0.01 | 0.22 | 0.29 | 0.23 | 0.11 |
| 5 | 103728978 | rs10464004 | 0.04 | -1.621 | 0.555 | -0.036 | -0.012 | -0.018 | -0.018 | 0 | 0.75 | 0.82 | 0.54 | 0.39 | 0.71 | 0.82 |
| 5 | 103741435 | rs6596573 | 0.537 | -1.723 | 0.578 | -0.105 | -0.005 | -0.005 | -0.005 | 0.206 | 0.7 | 0.77 | 0.51 | 0.36 | 0.67 | 0.81 |
| 5 | 103744044 | rs2963210 | -0.672 | -1.562 | 0.678 | 0.005 | 0.009 | 0.151 | 0.151 | 0 | 0.93 | 0.84 | 0.76 | 0.68 | 0.94 | 0.95 |
| 5 | 103761407 | rs6888456 | 0.505 | -1.518 | 0.746 | -0.525 | -0.042 | -0.035 | -0.035 | 0 | 0.08 | 0.17 | 0.02 | 0.01 | 0.01 | 0.03 |
| 5 | 103766994 | rs17345277 | 0.393 | -1.397 | 0.254 | -0.585 | -0.001 | -0.053 | -0.053 | 0 | 0.25 | 0.23 | 0.17 | 0 | 0.3 | 0.14 |
| 5 | 103767504 | rs11742919 | 0.473 | -1.364 | -0.131 | -0.569 | -0.001 | 0.012 | 0.012 | 0.517 | 0.24 | 0.16 | 0.17 | 0 | 0.3 | 0.14 |
| 5 | 103771206 | rs1442111 | 1.337 | -1.176 | -0.283 | -0.261 | -0.068 | -0.007 | -0.007 | 4.788 | 0.34 | 0.08 | 0.28 | 0.33 | 0.34 | 0.48 |
| 5 | 103788551 | rs1442112 | 0.22 | -0.704 | 0.439 | 0.077 | 0.009 | 0.012 | 0.012 | 0 | 0.74 | 0.73 | 0.72 | 0.61 | 0.65 | 0.77 |
| 5 | 103828340 | rs17411887 | 1.397 | -0.986 | 0.397 | -0.174 | 0.021 | 0.028 | 0.028 | 0 | 0.14 | 0.07 | 0.1 | 0 | 0.16 | 0.04 |
| 5 | 103865167 | rs4345293 | 0.537 | -1.147 | 0.373 | -0.22 | -0.031 | -0.041 | -0.041 | 0 | 0.71 | 0.83 | 0.72 | 0.59 | 0.65 | 0.73 |
| 5 | 103867339 | rs1363098 | 1.179 | -1.216 | 0.781 | -0.241 | -0.001 | 0 | 0 | 0 | 0.51 | 0.47 | 0.59 | 0.58 | 0.48 | 0.6 |
| 5 | 103871523 | rs2403283 | 0.113 | -0.819 | 0.384 | 0.061 | -0.031 | -0.035 | -0.035 | 0 | 0.75 | 0.84 | 0.72 | 0.6 | 0.65 | 0.77 |
| 5 | 103906144 | rs11744767 | 0.354 | -0.753 | 0.272 | -0.029 | -0.013 | 0.014 | 0.014 | 0 | 0.08 | 0.04 | 0.06 | 0 | 0.13 | 0.1 |
| 5 | 103912623 | rs2447836 | 0.685 | -0.782 | 0.34 | 0.382 | -0.012 | 0.03 | 0.03 | 0 | 0.9 | 0.88 | 0.87 | 0.75 | 0.94 | 0.84 |
| 5 | 103914523 | rs13164188 | 0.08 | -0.885 | 0.532 | 0.161 | -0.003 | -0.01 | -0.01 | 3.613 | 0.36 | 0.25 | 0.31 | 0.4 | 0.4 | 0.25 |
| 5 | 103919615 | rs17156700 | 0.458 | -0.855 | 0.88 | 0.013 | 0.025 | -0.048 | -0.048 | 0 | 0.09 | 0.2 | 0.07 | 0 | 0.12 | 0.1 |
| 5 | 103920770 | rs254011 | 0.884 | -0.912 | 0.963 | 0.045 | 0.002 | 0.039 | 0.039 | 0 | 0.49 | 0.38 | 0.48 | 0.36 | 0.44 | 0.5 |

| | | | | | | | | | | | | | | | | |
|---|---|---|---|---|---|---|---|---|---|---|---|---|---|---|---|---|
| 5 | 103925012 | rs12152832 | -0.07 | -0.844 | 0.98 | -0.086 | -0.017 | -0.162 | -0.162 | 0 | 0.29 | 0.42 | 0.18 | 0 | 0.3 | 0.23 |
| 5 | 103925210 | rs2052450 | -0.109 | -0.845 | 0.979 | -0.086 | -0.018 | -0.162 | -0.162 | 0.018 | 0.29 | 0.42 | 0.18 | 0 | 0.3 | 0.23 |
| 5 | 103927774 | rs12513556 | -0.04 | -0.686 | 0.797 | 0.05 | 0.036 | -0.056 | -0.056 | 0.018 | 0.1 | 0.25 | 0.08 | 0 | 0.12 | 0.19 |
| 5 | 103936520 | rs254034 | -0.07 | -0.668 | 0.476 | 0.229 | 0.001 | 0.022 | 0.022 | 0 | 0.95 | 0.93 | 0.87 | 0.77 | 0.96 | 0.86 |
| 5 | 103940225 | rs254028 | -0.094 | -0.749 | 0.534 | -0.074 | 0.006 | 0.02 | 0.02 | 0 | 0.79 | 0.74 | 0.65 | 0.37 | 0.74 | 0.66 |
| 5 | 103944020 | rs254024 | 1.071 | -0.808 | 0.689 | -0.279 | 0 | 0.016 | 0.016 | 0 | 0.48 | 0.37 | 0.47 | 0.37 | 0.42 | 0.5 |
| 5 | 103945149 | rs2974106 | 0.603 | -0.94 | 0.672 | -0.383 | 0 | 0.004 | 0.004 | 0 | 0.13 | 0.14 | 0.28 | 0.26 | 0.1 | 0.11 |
| 5 | 103947537 | rs10515355 | 0.294 | -0.508 | 0.687 | -0.207 | -0.004 | -0.1 | -0.1 | 0 | 0.26 | 0.33 | 0.18 | 0 | 0.3 | 0.14 |
| 5 | 103950251 | rs254020 | 0.987 | -0.456 | 0.35 | -0.062 | 0.002 | 0.018 | 0.018 | 0 | 0.48 | 0.36 | 0.47 | 0.37 | 0.42 | 0.5 |
| 5 | 103973787 | rs12513686 | 0.977 | -0.44 | 0.085 | -0.286 | 0.014 | 0.012 | 0.012 | 0 | 0.18 | 0.14 | 0.11 | 0 | 0.18 | 0.04 |
| 5 | 103980993 | rs1363099 | 0.302 | -0.329 | 0.271 | 0.137 | -0.001 | 0.004 | 0.004 | 0 | 0.19 | 0.16 | 0.22 | 0.37 | 0.23 | 0.13 |
| 5 | 103989098 | rs6859797 | 0.927 | -0.436 | 0.384 | 0.033 | 0.02 | 0.021 | 0.021 | 0 | 0.19 | 0.16 | 0.12 | 0 | 0.18 | 0.04 |
| 5 | 103995368 | rs325485 | -1.445 | -0.44 | 0.469 | 0.022 | 0.01 | 0.02 | 0.02 | 0 | 0.57 | 0.74 | 0.75 | 0.65 | 0.63 | 0.56 |
| 5 | 104006833 | rs7735362 | 0.627 | -0.521 | 0.827 | -0.241 | -0.012 | -0.06 | -0.06 | 0 | 0.22 | 0.32 | 0.33 | 0.05 | 0.2 | 0.19 |
| 5 | 104019292 | rs542345 | 0.104 | -0.419 | 1.042 | -0.22 | 0.065 | -0.044 | -0.044 | 0 | 0.09 | 0.29 | 0.07 | 0 | 0.12 | 0.1 |
| 5 | 104025707 | rs428538 | -0.429 | -0.483 | 0.976 | -0.284 | -0.014 | -0.133 | -0.133 | 1.342 | 0.31 | 0.49 | 0.41 | 0.05 | 0.33 | 0.28 |
| 5 | 104028100 | rs447909 | -0.105 | -0.326 | 1.027 | -0.2 | 0.048 | -0.053 | -0.053 | 0 | 0.09 | 0.25 | 0.07 | 0 | 0.12 | 0.18 |
| 5 | 104036672 | rs17416034 | 1.142 | -0.432 | 0.796 | -0.231 | 0.022 | 0.046 | 0.046 | 0 | 0.18 | 0.07 | 0.1 | 0 | 0.18 | 0.04 |
| 5 | 104045386 | rs325523 | -1.194 | -0.335 | 0.914 | -0.023 | 0.005 | 0.016 | 0.016 | 0 | 0.52 | 0.74 | 0.73 | 0.61 | 0.58 | 0.51 |
| 5 | 104048730 | rs1875505 | -0.1 | -0.358 | 1.229 | 0.063 | 0.017 | 0.017 | 0.017 | 0 | 0.11 | 0.18 | 0.1 | 0.15 | 0.15 | 0.19 |
| 5 | 104048870 | rs106451 | 1.005 | -0.403 | 1.324 | -0.021 | -0.029 | -0.108 | -0.108 | 0 | 0.2 | 0.34 | 0.13 | 0 | 0.18 | 0.04 |
| 5 | 104052400 | rs10038581 | 0.251 | -0.368 | 1.138 | 0.08 | -0.018 | -0.038 | -0.038 | 1.204 | 0.21 | 0.15 | 0.5 | 0.47 | 0.27 | 0.21 |
| 5 | 104069562 | rs1370192 | -0.273 | -0.309 | 1.195 | 0.144 | 0.024 | 0.044 | 0.044 | 0 | 0.12 | 0.19 | 0.11 | 0.19 | 0.17 | 0.21 |
| 5 | 104069917 | rs161645 | -2.061 | -1.032 | 0.204 | -0.304 | -0.118 | 0.17 | 0.17 | 0.005 | 0.68 | 0.98 | 0.83 | 0.92 | 0.68 | 0.64 |
| 5 | 104073260 | rs161791 | 1.204 | -1.244 | 0.858 | -0.05 | -0.008 | -0.054 | -0.054 | 0 | 0.19 | 0.23 | 0.11 | 0 | 0.18 | 0.04 |
| 5 | 104073794 | rs184435 | 0.539 | -1.114 | 0.596 | 0.129 | -0.049 | 0.072 | 0.072 | 0 | 0.16 | 0.26 | 0.09 | 0.31 | 0.07 | 0.12 |
| 5 | 104079908 | rs442469 | 0.631 | -1.051 | 0.326 | 0.132 | -0.008 | 0.015 | 0.015 | 0 | 0.16 | 0.19 | 0.09 | 0.3 | 0.09 | 0.13 |
| 5 | 104086307 | rs161773 | -0.293 | -0.615 | 0.478 | -0.04 | -0.006 | -0.03 | -0.03 | 1.159 | 0.32 | 0.52 | 0.24 | 0.19 | 0.34 | 0.33 |
| 5 | 104095693 | rs1370189 | -0.001 | -0.303 | 0.212 | -0.192 | 0.025 | -0.042 | -0.042 | 0 | 0.08 | 0.18 | 0.07 | 0 | 0.12 | 0.09 |
| 5 | 104096630 | rs161641 | -0.006 | -0.672 | -0.214 | -0.632 | 0.001 | 0 | 0 | 0 | 0.36 | 0.4 | 0.22 | 0.27 | 0.29 | 0.36 |
| 5 | 104105121 | rs12657012 | 1.188 | 0.355 | -0.662 | -0.596 | -0.064 | -0.066 | -0.066 | 0 | 0.1 | 0.01 | 0.24 | 0.42 | 0.17 | 0.15 |
| 5 | 104106441 | rs13155692 | 1.799 | 0.444 | -1.46 | -1.172 | -0.092 | 0.064 | 0.064 | 0 | 0.15 | 0.02 | 0.32 | 0.05 | 0.23 | 0.07 |
| 5 | 104107038 | rs1443069 | -0.814 | 0.224 | -0.943 | -0.89 | 0 | 0.013 | 0.013 | 0 | 0.54 | 0.63 | 0.53 | 0.69 | 0.58 | 0.56 |
| 5 | 104109586 | rs750612 | 1.253 | 0.284 | -0.907 | -0.897 | 0.019 | -0.039 | -0.039 | 0.105 | 0.07 | 0.14 | 0.06 | 0 | 0.12 | 0.09 |
| 5 | 104109619 | rs2304914 | 1.195 | 0.076 | -1.086 | -0.641 | -0.04 | -0.038 | -0.038 | 0 | 0.11 | 0.03 | 0.25 | 0.42 | 0.17 | 0.15 |
| 5 | 104110119 | rs11242536 | -0.365 | 0.056 | -1.067 | -0.713 | 0.001 | 0.014 | 0.014 | 0 | 0.46 | 0.49 | 0.47 | 0.69 | 0.46 | 0.35 |
| 5 | 104115055 | rs11242537 | 1.462 | -0.825 | -1.074 | -1.237 | -0.011 | 0.095 | 0.095 | 0.4 | 0.16 | 0.01 | 0.09 | 0 | 0.16 | 0.11 |
| 5 | 104118033 | rs10050461 | -0.107 | -0.836 | -0.917 | -1.231 | 0 | 0 | 0 | 0 | 0.35 | 0.34 | 0.21 | 0.27 | 0.29 | 0.21 |
| 5 | 104123360 | rs6865855 | -0.426 | -0.48 | -0.578 | -1.268 | -0.01 | -0.012 | -0.012 | 2.276 | 0.64 | 0.49 | 0.73 | 0.74 | 0.61 | 0.82 |
| 5 | 104125475 | rs7722847 | 0.886 | -0.278 | -0.998 | -1.772 | -0.038 | -0.008 | -0.008 | 0 | 0.32 | 0.2 | 0.54 | 0.68 | 0.38 | 0.57 |
| 5 | 104125893 | rs10054555 | 0.995 | -0.137 | -0.963 | -1.772 | -0.004 | 0.025 | 0.025 | 0 | 0.07 | 0.07 | 0.06 | 0.23 | 0.12 | 0.04 |
| 5 | 104127642 | rs1020049 | 1.046 | -0.243 | -0.98 | -2.002 | 0.004 | 0.01 | 0.01 | 0 | 0.15 | 0.08 | 0.08 | 0.01 | 0.12 | 0.08 |
| 5 | 104133164 | rs9687490 | 0.436 | -0.279 | -0.962 | -2.011 | 0.009 | 0.05 | 0.05 | 0 | 0.26 | 0.16 | 0.19 | 0.08 | 0.22 | 0.24 |
| 5 | 104139803 | rs1606554 | 0.064 | -0.379 | -0.272 | -2.095 | -0.023 | -0.133 | -0.133 | 0.43 | 0.28 | 0.45 | 0.21 | 0.06 | 0.23 | 0.23 |
| 5 | 104147248 | rs12054736 | 1.272 | -0.229 | -0.474 | -2.142 | -0.053 | -0.014 | -0.014 | 0 | 0.31 | 0.17 | 0.52 | 0.69 | 0.34 | 0.57 |
| 5 | 104153592 | rs715375 | 0.329 | -0.127 | -0.349 | -1.898 | 0.003 | 0.059 | 0.059 | 0 | 0.24 | 0.15 | 0.18 | 0.06 | 0.22 | 0.22 |
| 5 | 104157752 | rs10079325 | 0.193 | -0.111 | -0.327 | -1.89 | -0.011 | -0.108 | -0.108 | 1.204 | 0.26 | 0.39 | 0.2 | 0.06 | 0.22 | 0.22 |
| 5 | 104174305 | rs9327914 | 0.819 | 0.187 | -0.62 | -1.447 | -0.005 | 0.021 | 0.021 | 0 | 0.06 | 0.07 | 0.06 | 0.22 | 0.12 | 0.04 |
| 5 | 104178282 | rs322872 | -1.42 | -0.256 | -0.797 | -1.854 | 0.034 | 0.099 | 0.099 | 0 | 0.73 | 0.9 | 0.84 | 0.97 | 0.75 | 0.85 |
| 5 | 104178598 | rs17353958 | 2.005 | -0.297 | -0.953 | -1.959 | -0.01 | 0.041 | 0.041 | 0 | 0.11 | 0.02 | 0.11 | 0.05 | 0.1 | 0.07 |
| 5 | 104179652 | rs12187200 | 1.048 | -0.219 | -0.896 | -1.889 | -0.005 | 0.02 | 0.02 | 0 | 0.06 | 0.08 | 0.06 | 0.23 | 0.11 | 0.04 |
| 5 | 104183210 | rs12517106 | 2.278 | -0.508 | -1.083 | -2.116 | 0.016 | 0.025 | 0.025 | 0 | 0.13 | 0.05 | 0.07 | 0 | 0.13 | 0.04 |
| 5 | 104198795 | rs1657365 | -0.869 | -0.534 | -0.4 | -2.13 | -0.029 | -0.116 | -0.116 | 0 | 0.65 | 0.52 | 0.8 | 0.97 | 0.72 | 0.84 |
| 5 | 104199060 | rs1632744 | 0.671 | -0.594 | -0.483 | -2.149 | 0.008 | 0.056 | 0.056 | 0 | 0.45 | 0.31 | 0.59 | 0.69 | 0.48 | 0.7 |
| 5 | 104206280 | rs2020072 | 2.117 | -0.557 | -0.478 | -2.129 | 0.003 | 0.023 | 0.023 | 0 | 0.11 | 0.08 | 0.11 | 0.05 | 0.1 | 0.07 |
| 5 | 104206398 | rs2020070 | -1.518 | -0.564 | -0.347 | -2.128 | 0.015 | 0.008 | 0.008 | 0 | 0.73 | 0.83 | 0.78 | 0.97 | 0.75 | 0.85 |
| 5 | 104208013 | rs10074959 | -0.241 | -0.38 | -0.182 | -1.895 | -0.007 | 0.019 | 0.019 | 0 | 0.26 | 0.51 | 0.21 | 0.27 | 0.25 | 0.12 |
| 5 | 104212940 | rs10069629 | -0.421 | -0.345 | -0.193 | -1.854 | -0.007 | 0.023 | 0.023 | 0 | 0.27 | 0.47 | 0.23 | 0.29 | 0.26 | 0.15 |
| 5 | 104215354 | rs6899072 | 0.204 | -0.32 | -0.362 | -1.729 | 0.001 | 0.02 | 0.02 | 0 | 0.14 | 0.14 | 0.07 | 0.24 | 0.14 | 0.07 |
| 5 | 104220584 | rs1612927 | 0.703 | -0.109 | -0.782 | -1.787 | -0.031 | 0 | 0 | 0 | 0.35 | 0.23 | 0.55 | 0.66 | 0.38 | 0.64 |
| 5 | 104224506 | rs322853 | -1.287 | 0.651 | -0.928 | -2.269 | -0.013 | 0.211 | 0.211 | 0.02 | 0.45 | 0.81 | 0.32 | 0.07 | 0.48 | 0.38 |
| 5 | 104225000 | rs1708611 | -0.729 | 0.678 | -0.526 | -2.325 | 0.001 | -0.077 | -0.077 | 0 | 0.3 | 0.59 | 0.24 | 0.06 | 0.35 | 0.31 |
| 5 | 104227288 | rs17496653 | 1.654 | 0.721 | -0.464 | -2.361 | 0.005 | 0.014 | 0.014 | 0 | 0.11 | 0.06 | 0.06 | 0.06 | 0.1 | 0.09 |
| 5 | 104227705 | rs1153996 | -0.984 | 0.724 | -0.414 | -2.381 | 0 | -0.065 | -0.065 | 0 | 0.3 | 0.62 | 0.24 | 0.06 | 0.35 | 0.28 |
| 5 | 104231294 | rs60825 | -0.28 | 0.815 | -0.301 | -2.388 | -0.004 | 0.008 | 0.008 | 0 | 0.23 | 0.18 | 0.13 | 0.01 | 0.26 | 0.14 |
| 5 | 104233022 | rs1153991 | 0.815 | 0.846 | -0.105 | -2.364 | 0.077 | -0.031 | -0.031 | 0 | 0.09 | 0.36 | 0.11 | 0 | 0.12 | 0.13 |
| 5 | 104235042 | rs319907 | -0.25 | 0.849 | -0.309 | -2.326 | -0.007 | 0.024 | 0.024 | 0 | 0.23 | 0.16 | 0.13 | 0.02 | 0.26 | 0.11 |
| 5 | 104239861 | rs1153987 | 0.007 | 0.776 | -0.168 | -2.082 | 0.008 | -0.095 | -0.095 | 0 | 0.2 | 0.42 | 0.18 | 0.06 | 0.22 | 0.21 |
| 5 | 104242877 | rs410181 | -1.415 | 0.937 | -0.864 | -1.506 | 0.006 | -0.001 | -0.001 | 0 | 0.69 | 0.69 | 0.82 | 0.65 | 0.77 | 0.58 |

| | | | | | | | | | | | | | | | | |
|---|---|---|---|---|---|---|---|---|---|---|---|---|---|---|---|---|
| 5 | 104251851 | rs319897 | -0.955 | 1.111 | -0.899 | -1.582 | -0.031 | 0.06 | 0.06 | 0 | 0.44 | 0.27 | 0.61 | 0.68 | 0.43 | 0.42 |
| 5 | 104253119 | rs319899 | 0.684 | 1.242 | -1.062 | -1.621 | -0.004 | 0.032 | 0.032 | 0 | 0.15 | 0.07 | 0.07 | 0 | 0.13 | 0.06 |
| 5 | 104269333 | rs10464009 | 0.539 | 1.409 | -0.956 | -1.43 | -0.011 | -0.007 | -0.007 | 0 | 0.14 | 0.11 | 0.46 | 0.6 | 0.17 | 0.26 |
| 5 | 104271603 | rs319914 | -1.546 | 1.232 | -0.544 | -1.673 | 0.008 | 0.075 | 0.075 | 0 | 0.49 | 0.72 | 0.66 | 0.69 | 0.46 | 0.42 |
| 5 | 104274431 | rs1821901 | -0.535 | 1.16 | -0.638 | -1.622 | -0.025 | -0.026 | -0.026 | 0 | 0.3 | 0.27 | 0.53 | 0.6 | 0.31 | 0.32 |
| 5 | 104274840 | rs1438654 | 0.637 | 1.115 | -0.666 | -1.374 | -0.032 | -0.042 | -0.042 | 0 | 0.14 | 0.1 | 0.46 | 0.6 | 0.17 | 0.26 |
| 5 | 104275994 | rs9687975 | -1.415 | 0.983 | -1.039 | -1.744 | -0.023 | 0.005 | 0.005 | 0 | 0.66 | 0.88 | 0.8 | 0.69 | 0.68 | 0.54 |
| 5 | 104278663 | rs1438656 | -0.608 | 1.058 | 0.147 | -0.85 | 0.004 | 0.004 | 0.004 | 0 | 0.27 | 0.26 | 0.54 | 0.67 | 0.28 | 0.35 |
| 5 | 104288558 | rs10479304 | -1.183 | 0.742 | 0.043 | -0.902 | 0.004 | 0.002 | 0.002 | 0 | 0.36 | 0.64 | 0.56 | 0.66 | 0.29 | 0.41 |
| 5 | 104292444 | rs17499078 | 0.72 | 0.709 | -0.169 | -0.96 | -0.028 | 0.054 | 0.054 | 0 | 0.13 | 0.02 | 0.09 | 0.04 | 0.15 | 0.09 |
| 5 | 104296661 | rs7704764 | 1.318 | 0.604 | 0.379 | -0.971 | -0.019 | -0.029 | -0.029 | 0 | 0.06 | 0.16 | 0.02 | 0 | 0.01 | 0.02 |
| 5 | 104298208 | rs17564984 | 1.215 | 0.678 | -0.489 | -1.563 | -0.064 | 0.126 | 0.126 | 0 | 0.24 | 0.05 | 0.18 | 0.06 | 0.28 | 0.16 |
| 5 | 104300273 | rs17157372 | -0.523 | 0.462 | -0.047 | -1.619 | -0.028 | -0.096 | -0.096 | 0.455 | 0.25 | 0.44 | 0.13 | 0.06 | 0.18 | 0.18 |
| 5 | 104300802 | rs1370955 | -1.534 | 0.474 | -0.634 | -1.548 | -0.024 | 0.063 | 0.063 | 0.048 | 0.6 | 0.79 | 0.71 | 0.77 | 0.53 | 0.57 |
| 5 | 104312745 | rs6866409 | 0.188 | 1.148 | -0.618 | -1.193 | -0.039 | 0.124 | 0.124 | 0 | 0.33 | 0.1 | 0.23 | 0.12 | 0.38 | 0.25 |
| 5 | 104324622 | rs13182790 | 1.151 | 1.235 | -0.358 | -0.995 | -0.024 | 0.064 | 0.064 | 0 | 0.2 | 0.09 | 0.14 | 0.06 | 0.23 | 0.15 |
| 5 | 104325297 | rs17565931 | 1.449 | 1.268 | -0.373 | -1.003 | -0.01 | 0.059 | 0.059 | 0 | 0.18 | 0.08 | 0.13 | 0.04 | 0.19 | 0.12 |
| 5 | 104334555 | rs1991012 | -0.243 | 1.107 | -0.297 | -0.884 | 0 | 0.022 | 0.022 | 0 | 0.14 | 0.18 | 0.43 | 0.6 | 0.12 | 0.27 |
| 5 | 104345468 | rs7444606 | 0.006 | 1.274 | -0.396 | -0.933 | -0.003 | 0.009 | 0.009 | 0 | 0.13 | 0.12 | 0.42 | 0.6 | 0.12 | 0.27 |
| 5 | 104348000 | rs9688255 | -0.133 | 1.253 | -0.354 | -0.94 | 0.003 | 0.047 | 0.047 | 0 | 0.14 | 0.16 | 0.42 | 0.6 | 0.12 | 0.27 |
| 5 | 104353343 | rs4235646 | -0.285 | 1.194 | -0.288 | -1.035 | -0.005 | -0.001 | -0.001 | 0 | 0.35 | 0.3 | 0.57 | 0.65 | 0.35 | 0.44 |
| 5 | 104354696 | rs6868445 | 1.021 | 1.197 | -0.339 | -0.978 | -0.007 | 0.022 | 0.022 | 0 | 0.21 | 0.14 | 0.15 | 0.06 | 0.23 | 0.16 |
| 5 | 104355383 | rs10479307 | 0.916 | 1.137 | -0.382 | -0.989 | -0.005 | 0.008 | 0.008 | 0 | 0.07 | 0.03 | 0.06 | 0.05 | 0.1 | 0.07 |
| 5 | 104362976 | rs13159678 | -0.587 | 1.031 | -0.067 | -0.731 | -0.007 | 0 | 0 | 0 | 0.35 | 0.3 | 0.57 | 0.66 | 0.35 | 0.43 |
| 5 | 104363014 | rs13176693 | 0.762 | 0.884 | -0.152 | -0.78 | -0.01 | 0.028 | 0.028 | 0 | 0.21 | 0.14 | 0.15 | 0.07 | 0.23 | 0.16 |
| 5 | 104370141 | rs13168898 | 1.453 | 0.789 | 0.404 | -0.179 | 0.001 | 0.001 | 0.001 | 0.021 | 0.65 | 0.28 | 0.58 | 0.34 | 0.67 | 0.61 |
| 5 | 104371350 | rs4259147 | 1.378 | 0.222 | 0.671 | -0.208 | -0.008 | 0.006 | 0.006 | 0.03 | 0.71 | 0.32 | 0.64 | 0.35 | 0.81 | 0.65 |
| 5 | 104380211 | rs4616867 | -0.534 | 0.32 | 0.352 | -0.08 | -0.063 | -0.129 | -0.129 | 0.103 | 0.16 | 0.01 | 0.29 | 0.59 | 0.17 | 0.28 |
| 5 | 104382896 | rs6872400 | -0.733 | 0.309 | 0.909 | -0.282 | -0.002 | 0.002 | 0.002 | 0 | 0.18 | 0.17 | 0.28 | 0.59 | 0.19 | 0.28 |
| 5 | 104394274 | rs7721060 | 0.513 | 0.303 | 1.024 | -0.351 | 0.004 | -0.031 | -0.031 | 0 | 0.06 | 0.12 | 0.03 | 0 | 0.06 | 0.01 |
| 5 | 104423734 | rs4612035 | 1.296 | 0.161 | 1.635 | -0.32 | -0.013 | 0.028 | 0.028 | 0 | 0.76 | 0.6 | 0.69 | 0.29 | 0.81 | 0.68 |
| 5 | 104426955 | rs4360015 | 0.005 | 0.183 | 1.376 | -0.294 | -0.03 | -0.081 | -0.081 | 6.152 | 0.12 | 0.01 | 0.23 | 0.6 | 0.13 | 0.24 |
| 5 | 104435148 | rs4476696 | 0.976 | 0.042 | 1.857 | -0.322 | -0.019 | 0.015 | 0.015 | 0.413 | 0.88 | 0.77 | 0.95 | 0.89 | 0.93 | 0.95 |
| 5 | 104436002 | rs4481329 | 2.028 | 0.007 | 1.929 | -0.376 | 0.029 | 0.019 | 0.019 | 0.161 | 0.65 | 0.22 | 0.63 | 0.28 | 0.73 | 0.64 |
| 5 | 104456200 | rs4388185 | 1.373 | 0.063 | 1.938 | -0.386 | -0.032 | 0 | 0 | 0 | 0.84 | 0.73 | 0.95 | 0.89 | 0.91 | 0.91 |
| 5 | 104458953 | rs294152 | 1.697 | 0.38 | 1.791 | -0.161 | -0.021 | 0 | 0 | 0 | 0.86 | 0.24 | 0.9 | 0.9 | 0.92 | 0.94 |
| 5 | 104465995 | rs472343 | -1.535 | 0.639 | 1.86 | 0.104 | -0.011 | 0.086 | 0.086 | 0.125 | 0.24 | 0.55 | 0.33 | 0.61 | 0.21 | 0.33 |
| 5 | 104483421 | rs7356635 | -1.52 | 0.965 | 1.636 | 0.043 | 0.018 | 0.104 | 0.104 | 0.312 | 0.24 | 0.49 | 0.35 | 0.61 | 0.24 | 0.36 |
| 5 | 104485484 | rs9327920 | -1.441 | 0.881 | 1.646 | 0.068 | 0.014 | 0.113 | 0.113 | 0.548 | 0.23 | 0.48 | 0.35 | 0.61 | 0.23 | 0.34 |
| 5 | 104486508 | rs10079041 | -1.167 | 0.857 | 1.705 | 0.087 | 0.01 | 0.092 | 0.092 | 0.297 | 0.21 | 0.39 | 0.34 | 0.61 | 0.22 | 0.3 |
| 5 | 104490130 | rs517484 | 1.223 | 1.038 | 0.839 | -0.465 | 0.016 | 0.136 | 0.136 | 0 | 0.39 | 0.13 | 0.19 | 0.11 | 0.35 | 0.22 |
| 5 | 104490887 | rs10043572 | -0.763 | 1.326 | 1.09 | 0.557 | -0.07 | -0.142 | -0.142 | 2.577 | 0.16 | 0.01 | 0.3 | 0.57 | 0.2 | 0.28 |
| 5 | 104494294 | rs10051385 | -0.958 | 1.321 | 1.199 | 0.554 | -0.005 | -0.003 | -0.003 | 0 | 0.17 | 0.17 | 0.32 | 0.57 | 0.2 | 0.26 |
| 5 | 104494583 | rs10051540 | -1.612 | 1.31 | 1.236 | 0.517 | 0.012 | 0.13 | 0.13 | 0.748 | 0.22 | 0.49 | 0.34 | 0.61 | 0.21 | 0.32 |
| 5 | 104500256 | rs9327922 | -1.282 | 1.334 | 1.062 | 0.528 | 0.009 | 0.071 | 0.071 | 0.177 | 0.2 | 0.33 | 0.34 | 0.61 | 0.21 | 0.28 |
| 5 | 104509754 | rs1428878 | -1.22 | 1.3 | 0.472 | 0.361 | 0.003 | 0.044 | 0.044 | 0 | 0.19 | 0.24 | 0.32 | 0.58 | 0.2 | 0.27 |
| 5 | 104509854 | rs1428877 | -1.706 | 1.303 | -0.125 | 0.332 | 0.001 | 0.057 | 0.057 | 0.274 | 0.27 | 0.51 | 0.36 | 0.62 | 0.26 | 0.32 |
| 5 | 104515164 | rs4703333 | -1.64 | 1.326 | -0.17 | 0.33 | -0.008 | 0.064 | 0.064 | 0.213 | 0.26 | 0.52 | 0.35 | 0.62 | 0.24 | 0.31 |
| 5 | 104524071 | rs9327930 | -1.629 | 1.333 | -0.166 | 0.33 | -0.007 | 0.062 | 0.062 | 0.173 | 0.26 | 0.52 | 0.35 | 0.62 | 0.24 | 0.31 |
| 5 | 104540026 | rs10070429 | -1.311 | 1.019 | -1.056 | 0.928 | -0.069 | -0.146 | -0.146 | 0 | 0.18 | 0.01 | 0.3 | 0.58 | 0.21 | 0.31 |
| 5 | 104543131 | rs11242546 | -1.11 | 1.121 | -0.938 | 0.952 | -0.072 | -0.13 | -0.13 | 0 | 0.17 | 0.03 | 0.38 | 0.61 | 0.22 | 0.3 |
| 5 | 104563176 | rs10073553 | -1.217 | 0.435 | 0.602 | 0.535 | -0.013 | -0.001 | -0.001 | 0 | 0.14 | 0.32 | 0.11 | 0.09 | 0.1 | 0.04 |
| 5 | 104582841 | rs2436365 | -2.114 | 0.183 | 1.051 | 0.8 | -0.025 | 0.033 | 0.033 | 0.356 | 0.23 | 0.66 | 0.35 | 0.65 | 0.16 | 0.21 |
| 5 | 104587852 | rs1393089 | -1.131 | 0.174 | 1.258 | 0.795 | -0.011 | 0.007 | 0.007 | 0.489 | 0.14 | 0.39 | 0.12 | 0.09 | 0.1 | 0.04 |
| 5 | 104588432 | rs2436367 | -2.598 | 0.117 | 1.071 | 0.623 | -0.007 | 0.003 | 0.003 | 0 | 0.29 | 0.69 | 0.4 | 0.67 | 0.21 | 0.26 |
| 5 | 104596490 | rs12656536 | -1.065 | 0.218 | 0.893 | 0.66 | -0.02 | -0.058 | -0.058 | 0.089 | 0.1 | 0.01 | 0.24 | 0.58 | 0.1 | 0.2 |
| 5 | 104604047 | rs249379 | -1.255 | 0.146 | 1.04 | 0.534 | -0.019 | -0.001 | -0.001 | 0 | 0.15 | 0.38 | 0.12 | 0.09 | 0.1 | 0.05 |
| 5 | 104618244 | rs173616 | -1.11 | 0.123 | 1.031 | 0.538 | 0.017 | 0.074 | 0.074 | 0.956 | 0.09 | 0.38 | 0.09 | 0.09 | 0.07 | 0.05 |
| 5 | 104620426 | rs171612 | -1.018 | 0.127 | 1.06 | 0.54 | 0.023 | 0.081 | 0.081 | 1.054 | 0.09 | 0.38 | 0.09 | 0.09 | 0.07 | 0.03 |
| 5 | 104626717 | rs512975 | -1.018 | 0.127 | 1.058 | 0.54 | 0.021 | 0.079 | 0.079 | 1.054 | 0.09 | 0.38 | 0.09 | 0.09 | 0.07 | 0.03 |
| 5 | 104688954 | rs249564 | 0.864 | 1.074 | 0.23 | -0.326 | 0.036 | -0.042 | -0.042 | 0 | 0.73 | 0.6 | 0.65 | 0.9 | 0.62 | 0.79 |
| 5 | 104690464 | rs185122 | 2.372 | 0.996 | 0.368 | -0.3 | 0.004 | 0.054 | 0.054 | 0 | 0.53 | 0.29 | 0.35 | 0.32 | 0.47 | 0.59 |
| 5 | 104692032 | rs1501668 | 1.282 | 0.986 | 0.458 | -0.31 | -0.03 | -0.012 | -0.012 | 0.322 | 0.84 | 0.43 | 0.87 | 0.91 | 0.9 | 0.95 |
| 5 | 104692551 | rs2102137 | 0.588 | 1.373 | 0.319 | -0.38 | 0.019 | -0.019 | -0.019 | 4.222 | 0.83 | 0.62 | 0.87 | 0.91 | 0.83 | 0.9 |
| 5 | 104697070 | rs2431959 | 1.258 | 1.379 | 0.311 | -0.38 | 0.029 | -0.04 | -0.04 | 4.454 | 0.72 | 0.6 | 0.64 | 0.9 | 0.62 | 0.79 |
| 5 | 104699261 | rs2455177 | 2.409 | 1.451 | 0.239 | -0.377 | 0.003 | 0.005 | 0.005 | 0 | 0.63 | 0.3 | 0.56 | 0.33 | 0.68 | 0.7 |
| 5 | 104709892 | rs7341006 | -0.245 | 1.421 | 0.729 | -0.248 | 0.016 | -0.03 | -0.03 | 0.125 | 0.12 | 0.19 | 0.25 | 0.01 | 0.21 | 0.11 |
| 5 | 104713288 | rs4440328 | 0.527 | 1.425 | 0.477 | -0.206 | -0.013 | 0.046 | 0.046 | 0 | 0.06 | 0 | 0.21 | 0.58 | 0.06 | 0.16 |
| 5 | 104713851 | rs3958113 | 1.38 | 1.435 | 0.538 | -0.232 | 0.039 | -0.052 | -0.052 | 1.267 | 0.72 | 0.54 | 0.63 | 0.89 | 0.62 | 0.79 |

| | | | | | | | | | | | | | | | | |
|---|---|---|---|---|---|---|---|---|---|---|---|---|---|---|---|---|
| 5 | 104721409 | rs1423209 | 0.831 | 1.346 | 0.645 | -0.199 | 0.02 | -0.018 | -0.018 | 0 | 0.82 | 0.52 | 0.85 | 0.91 | 0.83 | 0.9 |
| 5 | 104731689 | rs12652389 | -1.251 | 1.311 | 0.644 | -0.16 | -0.012 | 0.121 | 0.121 | 0 | 0.13 | 0.2 | 0.25 | 0.57 | 0.09 | 0.16 |
| 5 | 104734927 | rs10052529 | -1.079 | 1.337 | 0.775 | -0.473 | 0.016 | -0.022 | -0.022 | 0 | 0.18 | 0.46 | 0.14 | 0.1 | 0.17 | 0.08 |
| 5 | 104742710 | rs7702850 | -1.897 | 1.351 | 0.648 | -0.497 | 0 | -0.009 | -0.009 | 0 | 0.35 | 0.66 | 0.4 | 0.11 | 0.41 | 0.21 |
| 5 | 104747267 | rs7713672 | -1.722 | 0.947 | 0.132 | -1.056 | 0.002 | 0.067 | 0.067 | 0 | 0.66 | 0.84 | 0.81 | 0.88 | 0.67 | 0.83 |
| 5 | 104760001 | rs13177189 | 0.457 | 1.04 | 0.692 | -0.714 | 0 | -0.008 | -0.008 | 0.052 | 0.05 | 0.05 | 0.03 | 0 | 0.03 | 0 |
| 5 | 104762292 | rs10042392 | -0.668 | 1.022 | 0.972 | -0.676 | 0.034 | 0.104 | 0.104 | 0 | 0.09 | 0.42 | 0.1 | 0.09 | 0.07 | 0.04 |
| 5 | 104767470 | rs12522619 | -1.503 | 1.199 | 0.842 | -0.768 | 0.04 | -0.044 | -0.044 | 0 | 0.23 | 0.46 | 0.33 | 0.1 | 0.31 | 0.15 |
| 5 | 104794603 | rs4479819 | 1.524 | 1.089 | 1.035 | -0.944 | 0.001 | -0.012 | -0.012 | 0.218 | 0.73 | 0.34 | 0.66 | 0.9 | 0.69 | 0.83 |
| 5 | 104812752 | rs4703342 | 0.603 | 0.638 | 1.5 | -0.809 | 0.043 | 0.11 | 0.11 | 0.017 | 0.91 | 0.55 | 0.92 | 0.92 | 0.93 | 0.96 |
| 5 | 104831188 | rs6882221 | -1.832 | 0.631 | 1.481 | -0.835 | -0.017 | -0.002 | -0.002 | 0 | 0.14 | 0.55 | 0.12 | 0.08 | 0.1 | 0.04 |
| 5 | 104837273 | rs6596615 | -0.983 | 0.613 | 1.619 | -0.812 | 0.01 | 0.06 | 0.06 | 0.308 | 0.09 | 0.38 | 0.08 | 0.08 | 0.07 | 0.05 |
| 5 | 104841905 | rs7710481 | -1.377 | 0.653 | 1.317 | -0.807 | -0.018 | -0.002 | -0.002 | 0 | 0.15 | 0.54 | 0.11 | 0.08 | 0.1 | 0.06 |
| 5 | 104845257 | rs4340892 | -1.382 | 0.745 | 0.758 | -0.953 | -0.005 | -0.021 | -0.021 | 0 | 0.25 | 0.66 | 0.16 | 0.08 | 0.2 | 0.11 |
| 5 | 104863539 | rs11242553 | 0.222 | 0.501 | 0.013 | -1.386 | -0.002 | 0.022 | 0.022 | 0 | 0.37 | 0.28 | 0.26 | 0.24 | 0.37 | 0.56 |
| 5 | 104864551 | rs4128686 | -0.159 | 0.07 | 0.446 | -1.73 | -0.02 | 0.096 | 0.096 | 0 | 0.11 | 0.22 | 0.18 | 0.69 | 0.06 | 0.1 |
| 5 | 104869290 | rs7735306 | 0.264 | -0.069 | 0.671 | -1.766 | -0.005 | -0.03 | -0.03 | 0 | 0.39 | 0.47 | 0.28 | 0.25 | 0.36 | 0.56 |
| 5 | 104880515 | rs4588563 | 0.637 | -0.024 | 0.372 | -1.76 | -0.008 | 0.035 | 0.035 | 0 | 0.36 | 0.2 | 0.25 | 0.24 | 0.37 | 0.56 |
| 5 | 104892945 | rs6885620 | -0.57 | -0.138 | -0.279 | -2.211 | -0.049 | 0.192 | 0.192 | 0 | 0.58 | 0.8 | 0.55 | 0.95 | 0.51 | 0.74 |
| 5 | 104956259 | rs7716817 | 0.864 | -0.168 | -0.52 | -2.19 | 0.023 | 0.061 | 0.061 | 0 | 0.48 | 0.26 | 0.39 | 0.33 | 0.53 | 0.68 |
| 5 | 104957493 | rs13154551 | 0.296 | 0.219 | -0.552 | -2.239 | -0.014 | 0.072 | 0.072 | 0 | 0.3 | 0.16 | 0.24 | 0.03 | 0.34 | 0.14 |
| 5 | 104962218 | rs7447231 | 0.815 | 0.322 | -0.588 | -2.051 | -0.013 | 0.072 | 0.072 | 0 | 0.2 | 0.09 | 0.19 | 0.03 | 0.21 | 0.1 |
| 5 | 104965729 | rs10071907 | 0.815 | 0.322 | -0.588 | -2.051 | -0.013 | 0.072 | 0.072 | 0 | 0.2 | 0.09 | 0.19 | 0.03 | 0.21 | 0.1 |
| 5 | 104968250 | rs10079505 | 1.082 | 0.404 | -0.668 | -2.007 | -0.023 | 0.033 | 0.033 | 0 | 0.1 | 0.04 | 0.14 | 0.03 | 0.16 | 0.08 |
| 5 | 104980355 | rs6872786 | -1.44 | 0.276 | -0.852 | -2.181 | 0 | 0 | 0 | 0 | 0.69 | 0.73 | 0.7 | 0.32 | 0.67 | 0.66 |
| 5 | 104988881 | rs11960843 | -1.416 | 0.306 | -0.846 | -2.183 | -0.001 | 0.001 | 0.001 | 0 | 0.69 | 0.73 | 0.69 | 0.32 | 0.66 | 0.69 |
| 5 | 104993059 | rs2312164 | -0.326 | 0.154 | -0.974 | -2.35 | -0.011 | -0.107 | -0.107 | 0 | 0.33 | 0.39 | 0.36 | 0.05 | 0.3 | 0.21 |
| 5 | 104995351 | rs13179945 | 0.427 | 0.269 | -1.655 | -1.858 | -0.071 | -0.006 | -0.006 | 0.074 | 0.33 | 0.1 | 0.31 | 0.28 | 0.34 | 0.47 |
| 5 | 105007215 | rs4607330 | 0.386 | 0.285 | -1.575 | -1.956 | -0.017 | 0.101 | 0.101 | 0 | 0.21 | 0.06 | 0.17 | 0.03 | 0.21 | 0.13 |
| 5 | 105017111 | rs2871954 | -0.897 | 0.184 | -1.233 | -1.872 | -0.002 | 0.04 | 0.04 | 0 | 0.6 | 0.72 | 0.51 | 0.27 | 0.55 | 0.6 |
| 5 | 105022790 | rs1480583 | -0.438 | 0.367 | -1.487 | -1.914 | -0.021 | 0.132 | 0.132 | 0 | 0.36 | 0.22 | 0.27 | 0.07 | 0.35 | 0.29 |
| 5 | 105027949 | rs13157681 | -0.405 | 0.437 | -1.461 | -1.897 | -0.021 | 0.118 | 0.118 | 0 | 0.36 | 0.2 | 0.27 | 0.06 | 0.35 | 0.3 |
| 5 | 105031008 | rs1904833 | 0.948 | 0.487 | -1.524 | -1.939 | -0.005 | 0.011 | 0.011 | 0 | 0.16 | 0.14 | 0.09 | 0.03 | 0.15 | 0.18 |
| 5 | 105042740 | rs1585909 | 0.239 | 0.555 | -1.326 | -1.824 | -0.002 | 0.025 | 0.025 | 0 | 0.2 | 0.1 | 0.17 | 0.04 | 0.2 | 0.12 |
| 5 | 105055467 | rs1585911 | 0.228 | 0.572 | -1.441 | -1.819 | -0.026 | 0.09 | 0.09 | 0 | 0.19 | 0.04 | 0.17 | 0.04 | 0.21 | 0.12 |
| 5 | 105057242 | rs1600603 | -0.219 | 0.48 | -1.701 | -1.807 | -0.102 | 0.083 | 0.083 | 0 | 0.4 | 0.07 | 0.43 | 0.25 | 0.44 | 0.53 |
| 5 | 105064972 | rs4554183 | -0.79 | 0.711 | -0.897 | -1.617 | 0.004 | -0.038 | -0.038 | 0 | 0.84 | 0.81 | 0.75 | 0.96 | 0.85 | 0.8 |
| 5 | 105065283 | rs1383369 | 0.611 | 0.545 | -0.403 | -1.806 | 0.006 | -0.075 | -0.075 | 0 | 0.63 | 0.44 | 0.64 | 0.91 | 0.64 | 0.66 |
| 5 | 105070480 | rs1351403 | -0.423 | 0.578 | -0.13 | -1.17 | -0.007 | 0.005 | 0.005 | 1.729 | 0.31 | 0.28 | 0.48 | 0.24 | 0.36 | 0.46 |
| 5 | 105072874 | rs4608899 | -0.09 | 0.699 | -0.071 | -1.2 | 0.005 | -0.007 | -0.007 | 0 | 0.93 | 0.89 | 0.93 | 0.96 | 0.91 | 0.87 |
| 5 | 105078678 | rs7701292 | 0.684 | 0.68 | -0.086 | -1.182 | -0.001 | 0 | 0 | 0 | 0.08 | 0.08 | 0.09 | 0.04 | 0.13 | 0.06 |
| 5 | 105095250 | rs13182207 | -0.419 | 0.628 | -0.029 | -0.889 | -0.04 | -0.024 | -0.024 | 0 | 0.31 | 0.17 | 0.48 | 0.32 | 0.39 | 0.49 |
| 5 | 105100680 | rs10464022 | 1.011 | 0.587 | 0.311 | -0.936 | 0.04 | -0.027 | -0.027 | 0.77 | 0.91 | 0.71 | 0.93 | 0.96 | 0.93 | 0.88 |
| 5 | 105101395 | rs964986 | -0.395 | 0.612 | 0.169 | -0.924 | -0.054 | 0.007 | 0.007 | 1.383 | 0.3 | 0.12 | 0.48 | 0.24 | 0.37 | 0.46 |
| 5 | 105109427 | rs448843 | 0.976 | 0.579 | 0.38 | -0.983 | -0.001 | 0 | 0 | 0.366 | 0.62 | 0.61 | 0.45 | 0.7 | 0.53 | 0.39 |
| 5 | 105131642 | rs286717 | -1.158 | 0.614 | 0.726 | -0.986 | 0.071 | 0.016 | 0.016 | 0.195 | 0.17 | 0.42 | 0.18 | 0.15 | 0.23 | 0.21 |
| 5 | 105131707 | rs11744593 | 0.734 | 0.557 | 0.397 | -0.961 | -0.074 | 0.022 | 0.022 | 0.122 | 0.21 | 0.02 | 0.39 | 0.15 | 0.24 | 0.4 |
| 5 | 105139420 | rs286727 | -0.942 | 0.485 | 0.516 | -1.017 | 0.032 | 0.022 | 0.022 | 0.417 | 0.15 | 0.29 | 0.15 | 0.13 | 0.2 | 0.14 |
| 5 | 105144834 | rs1482900 | 1.085 | -0.445 | -0.025 | -1.547 | -0.003 | -0.005 | -0.005 | 1.84 | 0.23 | 0.19 | 0.14 | 0.67 | 0.1 | 0.13 |
| 5 | 105146129 | rs286731 | 0.672 | -0.552 | -0.18 | -1.625 | -0.009 | 0.015 | 0.015 | 0 | 0.11 | 0.04 | 0.1 | 0.05 | 0.16 | 0.08 |
| 5 | 105149353 | rs286734 | -0.808 | -0.424 | -0.011 | -1.679 | 0 | -0.004 | -0.004 | 0.006 | 0.46 | 0.52 | 0.33 | 0.76 | 0.37 | 0.37 |
| 5 | 105163415 | rs1384560 | 1.225 | -0.351 | -0.235 | -1.744 | -0.046 | -0.014 | -0.014 | 0 | 0.21 | 0.09 | 0.41 | 0.23 | 0.26 | 0.45 |
| 5 | 105163858 | rs286713 | 0.059 | -0.487 | -0.063 | -1.672 | -0.006 | -0.002 | -0.002 | 0 | 0.35 | 0.29 | 0.24 | 0.72 | 0.26 | 0.21 |
| 5 | 105174036 | rs3887065 | 0.185 | -0.127 | -0.282 | -1.635 | 0.002 | -0.012 | -0.012 | 0 | 0.08 | 0.09 | 0.06 | 0.02 | 0.08 | 0.11 |
| 5 | 105177173 | rs2086676 | 0.199 | -0.124 | -0.184 | -1.644 | 0.022 | -0.021 | -0.021 | 0 | 0.07 | 0.15 | 0.05 | 0.02 | 0.07 | 0.07 |
| 5 | 105182600 | rs10072101 | 1.488 | -0.11 | -0.174 | -1.902 | 0.021 | -0.131 | -0.131 | 0 | 0.31 | 0.53 | 0.28 | 0.03 | 0.37 | 0.21 |
| 5 | 105184914 | rs1994127 | 0.499 | -0.172 | -0.259 | -1.94 | -0.023 | 0.118 | 0.118 | 0 | 0.42 | 0.76 | 0.38 | 0.06 | 0.5 | 0.35 |
| 5 | 105185178 | rs1580395 | 1.285 | -0.271 | -0.314 | -1.486 | 0.074 | -0.093 | -0.093 | 0 | 0.22 | 0.07 | 0.12 | 0.58 | 0.1 | 0.14 |
| 5 | 105200594 | rs1384557 | 0.199 | -0.174 | -0.329 | -1.658 | 0.023 | -0.003 | -0.003 | 0 | 0.62 | 0.72 | 0.77 | 0.38 | 0.72 | 0.74 |
| 5 | 105205274 | rs10042824 | 0.015 | 0.027 | -0.23 | -1.866 | -0.01 | 0.01 | 0.01 | 0 | 0.1 | 0.08 | 0.08 | 0.01 | 0.11 | 0.11 |
| 5 | 105205829 | rs6596653 | -0.086 | -0.226 | -0.006 | -1.745 | 0.001 | -0.039 | -0.039 | 0 | 0.86 | 0.86 | 0.89 | 0.99 | 0.81 | 0.88 |
| 5 | 105206440 | rs2035437 | 1.271 | -0.227 | -0.023 | -1.862 | 0.005 | -0.093 | -0.093 | 0.87 | 0.38 | 0.53 | 0.32 | 0.03 | 0.43 | 0.25 |
| 5 | 105209225 | rs9327952 | -0.169 | -0.225 | -0.058 | -1.85 | 0 | -0.039 | -0.039 | 0 | 0.86 | 0.86 | 0.89 | 0.99 | 0.82 | 0.88 |
| 5 | 105209621 | rs10055156 | 1.24 | -0.237 | -0.015 | -1.809 | 0.011 | -0.129 | -0.129 | 5.388 | 0.37 | 0.53 | 0.32 | 0.03 | 0.43 | 0.25 |
| 5 | 105224242 | rs975290 | -0.918 | -0.648 | 0.078 | -1.534 | 0.013 | -0.054 | -0.054 | 0 | 0.19 | 0.24 | 0.16 | 0.04 | 0.25 | 0.22 |
| 5 | 105230758 | rs10070622 | 1.04 | -0.592 | -0.016 | -1.629 | 0.006 | -0.123 | -0.123 | 1.103 | 0.42 | 0.52 | 0.34 | 0.04 | 0.5 | 0.29 |
| 5 | 105246242 | rs13156240 | -1.833 | -0.583 | -0.417 | -1.175 | -0.027 | -0.097 | -0.097 | 0.208 | 0.86 | 0.97 | 0.64 | 0.64 | 0.85 | 0.65 |
| 5 | 105254305 | rs1014813 | 1.259 | -0.812 | -0.108 | -1.456 | -0.078 | 0.192 | 0.192 | 0.057 | 0.38 | 0.14 | 0.31 | 0.04 | 0.49 | 0.29 |
| 5 | 105262661 | rs10051507 | 1.336 | -0.69 | -0.061 | -1.456 | -0.064 | 0.13 | 0.13 | 0 | 0.37 | 0.2 | 0.3 | 0.04 | 0.48 | 0.29 |

| | | | | | | | | | | | | | | | | |
|---|---|---|---|---|---|---|---|---|---|---|---|---|---|---|---|---|
| 5 | 105263703 | rs6885710 | -0.173 | -0.618 | -0.287 | -1.413 | 0.063 | 0.198 | 0.198 | 0 | 0.42 | 0.17 | 0.54 | 0.94 | 0.3 | 0.53 |
| 5 | 105271209 | rs2311735 | -0.61 | -0.584 | 0.013 | -1.424 | 0.017 | 0.007 | 0.007 | 0 | 0.48 | 0.39 | 0.59 | 0.96 | 0.36 | 0.59 |
| 5 | 105282751 | rs1524355 | 0.37 | -0.467 | -0.169 | -1.185 | 0.009 | 0.047 | 0.047 | 0.158 | 0.23 | 0.32 | 0.44 | 0.39 | 0.22 | 0.44 |
| 5 | 105287914 | rs1524353 | 0.708 | -0.415 | -0.544 | -1.22 | -0.001 | 0.139 | 0.139 | 0 | 0.45 | 0.75 | 0.36 | 0.05 | 0.51 | 0.34 |
| 5 | 105306900 | rs1404163 | 1.028 | -0.695 | -0.581 | -1.052 | 0.042 | -0.055 | -0.055 | 0 | 0.22 | 0.13 | 0.14 | 0.56 | 0.1 | 0.11 |
| 5 | 105308346 | rs7380123 | -0.265 | -0.326 | -0.631 | -0.881 | -0.018 | -0.065 | -0.065 | 0 | 0.29 | 0.1 | 0.49 | 0.38 | 0.34 | 0.51 |
| 5 | 105324438 | rs13177168 | -0.829 | -0.323 | -0.106 | -0.707 | -0.001 | 0 | 0 | 0 | 0.38 | 0.47 | 0.54 | 0.37 | 0.39 | 0.56 |
| 5 | 105350426 | rs4957840 | 0.27 | -0.276 | -0.27 | -0.762 | -0.003 | 0.01 | 0.01 | 0 | 0.12 | 0.07 | 0.1 | 0.02 | 0.17 | 0.14 |
| 5 | 105355890 | rs1357718 | 1.24 | -0.66 | -0.47 | -0.88 | 0.055 | -0.087 | -0.087 | 0 | 0.19 | 0.08 | 0.14 | 0.55 | 0.09 | 0.14 |
| 5 | 105365249 | rs7725843 | 0.616 | 0.086 | 0.414 | -0.539 | 0.009 | -0.014 | -0.014 | 0 | 0.05 | 0.09 | 0.05 | 0.01 | 0.06 | 0.06 |
| 5 | 105377638 | rs13164411 | -1.37 | 0.023 | 0.625 | -0.585 | 0.025 | -0.102 | -0.102 | 0 | 0.2 | 0.44 | 0.17 | 0.03 | 0.23 | 0.2 |
| 5 | 105385709 | rs960931 | 1.463 | -0.027 | 0.4 | -0.638 | 0.005 | -0.003 | -0.003 | 0.311 | 0.55 | 0.36 | 0.43 | 0.59 | 0.56 | 0.41 |
| 5 | 105404097 | rs13162459 | 1.262 | 0.04 | 0.419 | -0.697 | 0.005 | -0.102 | -0.102 | 0 | 0.74 | 0.44 | 0.81 | 0.97 | 0.72 | 0.78 |
| 5 | 105405016 | rs12656675 | 1.301 | 0.024 | 0.423 | -0.696 | 0.009 | -0.1 | -0.1 | 0.01 | 0.74 | 0.44 | 0.81 | 0.97 | 0.72 | 0.78 |
| 5 | 105405874 | rs13165324 | 0.301 | 0.031 | -0.201 | -0.557 | -0.004 | 0.034 | 0.034 | 0 | 0.11 | 0.04 | 0.08 | 0.01 | 0.15 | 0.1 |
| 5 | 105419534 | rs1865507 | 0.64 | 0.12 | -0.049 | -0.222 | -0.013 | -0.124 | -0.124 | 0.208 | 0.18 | 0.03 | 0.38 | 0.41 | 0.16 | 0.37 |
| 5 | 105425352 | rs10059955 | 0.656 | 0.501 | 0.372 | -0.575 | 0.001 | -0.001 | -0.001 | 0.173 | 0.61 | 0.39 | 0.54 | 0.59 | 0.59 | 0.47 |
| 5 | 105453357 | rs1435194 | 0.555 | 0.601 | 0.409 | -0.54 | -0.038 | -0.083 | -0.083 | 0 | 0.22 | 0.06 | 0.32 | 0.39 | 0.24 | 0.37 |
| 5 | 105507543 | rs4457046 | 0.485 | 0.639 | 0.853 | -0.561 | 0.012 | 0.003 | 0.003 | 0 | 0.65 | 0.46 | 0.55 | 0.58 | 0.61 | 0.48 |
| 5 | 105515328 | rs10079254 | 1.879 | -0.014 | 0.386 | -1.257 | -0.019 | 0.031 | 0.031 | 0 | 0.37 | 0.25 | 0.31 | 0.01 | 0.42 | 0.28 |
| 5 | 105526457 | rs6881723 | 1.386 | -0.03 | 0.345 | -1.282 | -0.021 | -0.089 | -0.089 | 0 | 0.2 | 0.06 | 0.32 | 0.39 | 0.2 | 0.37 |
| 5 | 105527995 | rs10479360 | 1.978 | 0.418 | 0.328 | -1.322 | -0.002 | 0.025 | 0.025 | 0 | 0.36 | 0.24 | 0.26 | 0.01 | 0.34 | 0.27 |
| 5 | 105551761 | rs12719552 | -0.109 | 0.963 | 0.285 | -0.87 | -0.06 | -0.025 | -0.025 | 0 | 0.31 | 0.16 | 0.46 | 0.41 | 0.43 | 0.5 |
| 5 | 105554384 | rs6859823 | -0.132 | 0.924 | 0.24 | -0.825 | -0.064 | 0.009 | 0.009 | 0 | 0.32 | 0.14 | 0.45 | 0.37 | 0.43 | 0.53 |
| 5 | 105575013 | rs6868876 | -0.777 | 0.76 | 0.238 | -0.682 | -0.008 | -0.019 | -0.019 | 0 | 0.77 | 0.84 | 0.64 | 0.6 | 0.72 | 0.62 |
| 5 | 105613672 | rs4296777 | -1.034 | 0.924 | 0.311 | -0.58 | -0.006 | -0.007 | -0.007 | 0.378 | 0.81 | 0.85 | 0.65 | 0.65 | 0.73 | 0.64 |
| 5 | 105635825 | rs11741503 | 1.098 | 0.81 | -0.224 | -0.596 | -0.069 | -0.082 | -0.082 | 0.059 | 0.26 | 0.06 | 0.38 | 0.38 | 0.32 | 0.43 |
| 5 | 105642031 | rs12516146 | 0.177 | 0.966 | -0.137 | -0.839 | 0 | 0 | 0 | 0.269 | 0.53 | 0.5 | 0.45 | 0.6 | 0.42 | 0.42 |
| 5 | 105658285 | rs6888141 | -0.075 | 0.696 | -0.41 | -0.841 | -0.005 | -0.014 | -0.014 | 0 | 0.14 | 0.12 | 0.13 | 0.01 | 0.2 | 0.07 |
| 5 | 105668175 | rs4467679 | -0.079 | 0.701 | -0.312 | -0.718 | 0.006 | 0.016 | 0.016 | 0 | 0.45 | 0.39 | 0.53 | 0.4 | 0.57 | 0.56 |
| 5 | 105692401 | rs13158436 | -0.911 | 0.501 | -1.087 | -1.331 | -0.004 | 0.11 | 0.11 | 0 | 0.69 | 0.9 | 0.77 | 0.97 | 0.73 | 0.75 |
| 5 | 105702335 | rs12659384 | 0.247 | 0.964 | -1.49 | -0.981 | 0.009 | 0.014 | 0.014 | 0 | 0.44 | 0.31 | 0.31 | 0.43 | 0.41 | 0.52 |
| 5 | 105728438 | rs4370259 | 0.088 | 1.065 | -1.422 | -0.925 | -0.002 | -0.003 | -0.003 | 2.063 | 0.48 | 0.45 | 0.58 | 0.56 | 0.43 | 0.39 |
| 5 | 105736353 | rs7727384 | -0.965 | 1.071 | -1.534 | -0.923 | -0.011 | -0.034 | -0.034 | 0 | 0.9 | 0.87 | 0.95 | 1 | 0.95 | 0.96 |
| 5 | 105772632 | rs11951214 | 0.992 | 1.407 | -1.548 | -0.676 | 0.003 | -0.002 | -0.002 | 0 | 0.05 | 0.02 | 0.07 | 0 | 0.16 | 0.04 |
| 5 | 105780577 | rs7732321 | -0.321 | 0.71 | -1.444 | -0.524 | 0.003 | 0.004 | 0.004 | 0 | 0.49 | 0.48 | 0.58 | 0.56 | 0.42 | 0.38 |
| 5 | 105804683 | rs7706267 | -1.054 | 0.746 | -1.721 | -0.095 | -0.001 | 0.007 | 0.007 | 0 | 0.47 | 0.6 | 0.48 | 0.68 | 0.47 | 0.62 |
| 5 | 105805717 | rs9784596 | 0.593 | 1.163 | -1.619 | 0.321 | -0.033 | -0.08 | -0.08 | 0.242 | 0.11 | 0.01 | 0.13 | 0.4 | 0.17 | 0.35 |
| 5 | 105817579 | rs2135048 | 0.801 | 0.906 | -1.424 | 0.514 | -0.034 | -0.076 | -0.076 | 0 | 0.3 | 0.05 | 0.21 | 0.4 | 0.34 | 0.41 |
| 5 | 105825670 | rs10479370 | 0.237 | 1.074 | -1.102 | 0.822 | -0.007 | 0.121 | 0.121 | 0 | 0.06 | 0.11 | 0.14 | 0.26 | 0.04 | 0.1 |
| 5 | 105830691 | rs10073251 | 0.097 | 1.096 | -1.047 | 0.715 | 0.01 | 0.012 | 0.012 | 0 | 0.43 | 0.35 | 0.38 | 0.66 | 0.4 | 0.52 |
| 5 | 105846299 | rs4642355 | -0.439 | 0.954 | -0.801 | 1.322 | 0 | 0.161 | 0.161 | 0 | 0.06 | 0.15 | 0.14 | 0.25 | 0.04 | 0.1 |
| 5 | 105849895 | rs4513678 | 0.193 | 1.052 | -1.192 | 1.185 | 0.001 | -0.003 | -0.003 | 0 | 0.56 | 0.58 | 0.55 | 0.67 | 0.52 | 0.68 |
| 5 | 105854131 | rs6891659 | 0.131 | 1.104 | -1.232 | 1.21 | -0.001 | 0.002 | 0.002 | 0 | 0.56 | 0.62 | 0.55 | 0.67 | 0.52 | 0.68 |
| 5 | 105866862 | rs10038387 | -0.38 | 0.905 | -0.946 | 1.628 | 0.006 | -0.002 | -0.002 | 0 | 0.07 | 0.04 | 0.14 | 0.24 | 0.05 | 0.15 |
| 5 | 105876806 | rs11739877 | 0.321 | 1.254 | -0.783 | 1.056 | 0.003 | 0.01 | 0.01 | 0.022 | 0.63 | 0.62 | 0.61 | 0.56 | 0.61 | 0.71 |
| 5 | 105882293 | rs4551054 | 0.183 | 0.673 | -0.91 | 1.162 | -0.007 | 0.035 | 0.035 | 0.02 | 0.67 | 0.56 | 0.67 | 0.62 | 0.72 | 0.78 |
| 5 | 105910000 | rs4958081 | -0.84 | 0.741 | -1.417 | 1.391 | -0.01 | -0.056 | -0.056 | 0 | 0.16 | 0.07 | 0.07 | 0.24 | 0.16 | 0.2 |
| 5 | 105913532 | rs9327966 | -0.638 | 0.82 | -2.619 | 1.227 | -0.068 | 0.086 | 0.086 | 0 | 0.5 | 0.92 | 0.57 | 0.65 | 0.54 | 0.6 |
| 5 | 105918671 | rs6897093 | -0.695 | 0.689 | -2.676 | 1.189 | -0.075 | 0.071 | 0.071 | 0 | 0.51 | 0.93 | 0.58 | 0.65 | 0.54 | 0.6 |
| 5 | 105921679 | rs11242593 | -0.788 | 0.888 | -1.26 | 1.357 | 0.001 | 0.002 | 0.002 | 0 | 0.45 | 0.58 | 0.3 | 0.37 | 0.47 | 0.46 |
| 5 | 105923131 | rs7710861 | -0.674 | 0.296 | -0.86 | 0.969 | 0.031 | 0.064 | 0.064 | 0.111 | 0.47 | 0.35 | 0.26 | 0.25 | 0.45 | 0.44 |
| 5 | 105924416 | rs10080115 | 0.517 | 0.184 | -1.055 | 0.699 | 0.006 | 0.065 | 0.065 | 0.553 | 0.23 | 0.1 | 0.15 | 0 | 0.24 | 0.05 |
| 5 | 105936330 | rs4301221 | 0.632 | 0.483 | 0.191 | 1.214 | -0.026 | 0.006 | 0.006 | 0 | 0.73 | 0.46 | 0.88 | 0.75 | 0.8 | 0.8 |
| 5 | 105944600 | rs4467693 | 0.659 | -0.561 | -0.934 | 0.96 | 0.014 | -0.038 | -0.038 | 0.151 | 0.25 | 0.04 | 0.23 | 0.29 | 0.21 | 0.24 |
| 5 | 105944933 | rs6596684 | 0.471 | -0.146 | -0.18 | 0.97 | 0.004 | -0.002 | -0.002 | 0 | 0.58 | 0.39 | 0.66 | 0.55 | 0.66 | 0.32 |
| 5 | 105948410 | rs10040790 | 0.812 | 0.346 | -0.364 | 0.805 | -0.034 | -0.01 | -0.01 | 0 | 0.27 | 0.16 | 0.32 | 0.31 | 0.36 | 0.23 |
| 5 | 105949415 | rs1978837 | 0.425 | 0.342 | -0.243 | 0.851 | -0.03 | 0.091 | 0.091 | 0.571 | 0.87 | 0.57 | 0.95 | 0.84 | 0.93 | 0.61 |
| 5 | 105972148 | rs1437642 | 1.136 | 0.463 | -0.719 | 0.822 | -0.008 | -0.094 | -0.094 | 0.258 | 0.27 | 0.03 | 0.25 | 0.36 | 0.21 | 0.22 |
| 5 | 105980613 | rs7709573 | 1.515 | 0.593 | -0.746 | 0.706 | -0.007 | 0.152 | 0.152 | 0.956 | 0.27 | 0.03 | 0.28 | 0.05 | 0.24 | 0.06 |
| 5 | 105989479 | rs9327971 | -0.069 | 0.971 | -0.035 | 0.843 | 0.075 | 0.047 | 0.047 | 0 | 0.89 | 0.7 | 0.9 | 0.89 | 0.86 | 0.9 |
| 5 | 105989630 | rs1370473 | 0.21 | 0.91 | 0.079 | 1.193 | -0.001 | -0.001 | -0.001 | 0 | 0.65 | 0.37 | 0.57 | 0.51 | 0.55 | 0.41 |
| 5 | 105990288 | rs1370474 | 0.725 | 0.892 | 0.18 | 1.179 | 0.035 | 0.062 | 0.062 | 0 | 0.25 | 0.37 | 0.33 | 0.38 | 0.32 | 0.5 |
| 5 | 105992483 | rs11242599 | 1.732 | 0.993 | 0.074 | 1.312 | -0.096 | -0.05 | -0.05 | 0 | 0.18 | 0.02 | 0.28 | 0.22 | 0.29 | 0.24 |
| 5 | 105993785 | rs10041720 | -0.443 | 0.873 | 0.14 | 1.314 | 0.006 | 0.003 | 0.003 | 0 | 0.7 | 0.68 | 0.65 | 0.65 | 0.61 | 0.66 |
| 5 | 105995653 | rs1816641 | 0.237 | 0.742 | 0.301 | 1.226 | 0.004 | 0.04 | 0.04 | 0.076 | 0.05 | 0.14 | 0.02 | 0.1 | 0.05 | 0.11 |
| 5 | 105999196 | rs17158894 | -0.543 | 0.736 | 0.255 | 1.296 | 0.067 | 0.248 | 0.248 | 2.802 | 0.06 | 0.36 | 0.06 | 0.13 | 0.02 | 0.28 |
| 5 | 106003181 | rs11740285 | 0.628 | 0.675 | -0.035 | 1.073 | 0.017 | 0.029 | 0.029 | 0.073 | 0.36 | 0.29 | 0.31 | 0.15 | 0.29 | 0.18 |
| 5 | 106005425 | rs13359925 | -1.16 | 0.705 | 0.037 | 1.171 | -0.009 | 0.045 | 0.045 | 2.107 | 0.1 | 0.53 | 0.05 | 0.09 | 0.05 | 0.11 |

| | | | | | | | | | | | | | | | | |
|---|---|---|---|---|---|---|---|---|---|---|---|---|---|---|---|---|
| 5 | 106007557 | rs13178335 | 1.56 | 0.912 | -0.308 | 1.177 | -0.079 | -0.045 | -0.045 | 0 | 0.18 | 0.02 | 0.26 | 0.23 | 0.26 | 0.23 |
| 5 | 106009475 | rs7700912 | 0.907 | 0.97 | -0.194 | 1.122 | -0.046 | -0.003 | -0.003 | 0 | 0.23 | 0.13 | 0.32 | 0.26 | 0.32 | 0.27 |
| 5 | 106011329 | rs10068445 | -0.778 | 0.709 | 0.083 | 1.025 | 0.004 | 0.017 | 0.017 | 0.13 | 0.64 | 0.74 | 0.67 | 0.83 | 0.68 | 0.89 |
| 5 | 106014252 | rs2410869 | 0.148 | 0.669 | 0.179 | 1.125 | 0.002 | 0.02 | 0.02 | 0.709 | 0.55 | 0.36 | 0.57 | 0.69 | 0.6 | 0.56 |
| 5 | 106024211 | rs1120322 | -0.141 | 0.486 | 0.146 | 1.108 | 0.024 | 0.074 | 0.074 | 0 | 0.91 | 0.8 | 0.95 | 0.88 | 0.92 | 0.95 |
| 5 | 106033716 | rs6596693 | -0.031 | 0.74 | 0.252 | 1.015 | -0.003 | -0.006 | -0.006 | 0 | 0.57 | 0.57 | 0.6 | 0.71 | 0.61 | 0.58 |
| 5 | 106036130 | rs17158978 | 0.296 | 1.066 | 0.018 | 0.689 | -0.008 | -0.004 | -0.004 | 0 | 0.42 | 0.52 | 0.37 | 0.44 | 0.38 | 0.36 |
| 5 | 106039170 | rs10214014 | 1.849 | 1.113 | -0.243 | 0.75 | -0.071 | -0.079 | -0.079 | 0 | 0.15 | 0.02 | 0.23 | 0.27 | 0.23 | 0.23 |
| 5 | 106039296 | rs17158990 | -0.579 | 1.037 | -0.014 | 0.719 | 0.012 | 0.046 | 0.046 | 0 | 0.14 | 0.19 | 0.1 | 0.24 | 0.16 | 0.33 |
| 5 | 106039310 | rs6895448 | 1.447 | 0.766 | -0.145 | 0.324 | -0.008 | 0.145 | 0.145 | 1.449 | 0.27 | 0.03 | 0.28 | 0.05 | 0.24 | 0.08 |
| 5 | 106040216 | rs6596698 | -0.032 | 0.922 | 0.071 | 0.731 | 0.001 | 0.035 | 0.035 | 0 | 0.95 | 0.89 | 0.97 | 0.9 | 0.95 | 0.92 |
| 5 | 106047930 | rs9327973 | 0.597 | 0.676 | -0.327 | 0.55 | -0.034 | -0.044 | -0.044 | 0.292 | 0.2 | 0.12 | 0.25 | 0.3 | 0.28 | 0.25 |
| 5 | 106049034 | rs6596699 | 0.255 | 0.53 | -0.101 | 0.626 | -0.002 | 0.019 | 0.019 | 0 | 0.09 | 0.16 | 0.07 | 0.09 | 0.06 | 0.06 |
| 5 | 106056347 | rs9327976 | -0.435 | 0.189 | 0.001 | 0.348 | -0.002 | -0.002 | -0.002 | 0 | 0.59 | 0.69 | 0.5 | 0.52 | 0.62 | 0.58 |
| 5 | 106058868 | rs12109609 | -0.704 | 0.096 | 0.168 | 0.284 | -0.01 | 0.049 | 0.049 | 0 | 0.13 | 0.4 | 0.08 | 0.09 | 0.06 | 0.06 |
| 5 | 106063288 | rs293630 | -0.597 | 0.22 | 0.046 | 0.331 | -0.015 | -0.019 | -0.019 | 0 | 0.62 | 0.8 | 0.52 | 0.52 | 0.63 | 0.59 |
| 5 | 106077542 | rs6888932 | 1.121 | 0.339 | -0.29 | 0.394 | -0.033 | -0.04 | -0.04 | 0 | 0.19 | 0.11 | 0.25 | 0.3 | 0.28 | 0.25 |
| 5 | 106081893 | rs425464 | 0.267 | 0.235 | -0.09 | 0.245 | -0.002 | 0.017 | 0.017 | 0 | 0.49 | 0.37 | 0.42 | 0.43 | 0.56 | 0.52 |
| 5 | 106084494 | rs10070246 | 0.194 | 0.265 | -0.223 | 0.283 | -0.015 | -0.026 | -0.026 | 0 | 0.26 | 0.24 | 0.32 | 0.39 | 0.34 | 0.3 |
| 5 | 106089895 | rs1561354 | -1.048 | 0.234 | 0.019 | -0.319 | -0.003 | -0.015 | -0.015 | 0 | 0.68 | 0.67 | 0.82 | 0.87 | 0.72 | 0.74 |
| 5 | 106090828 | rs11952719 | -0.11 | 0.297 | -0.223 | -0.275 | 0 | -0.001 | -0.001 | 0 | 0.48 | 0.48 | 0.57 | 0.57 | 0.44 | 0.47 |
| 5 | 106094581 | rs1438243 | 0.625 | 0.482 | -0.59 | 0.156 | -0.002 | -0.001 | -0.001 | 0.054 | 0.19 | 0.16 | 0.25 | 0.3 | 0.28 | 0.24 |
| 5 | 106103975 | rs12374447 | 0.199 | 0.249 | -1.296 | -0.459 | -0.004 | -0.05 | -0.05 | 0 | 0.29 | 0.11 | 0.25 | 0.37 | 0.25 | 0.31 |
| 5 | 106104465 | rs6596703 | -0.46 | 0.021 | -1.256 | -0.754 | 0.001 | -0.008 | -0.008 | 0 | 0.71 | 0.4 | 0.62 | 0.77 | 0.69 | 0.59 |
| 5 | 106116159 | rs6596705 | -2.427 | 0.112 | -1.404 | -0.823 | -0.011 | 0.08 | 0.08 | 1.004 | 0.85 | 0.97 | 0.8 | 0.98 | 0.85 | 0.91 |
| 5 | 106116276 | rs13156926 | 1.052 | -0.039 | -1.496 | -0.917 | -0.04 | -0.044 | -0.044 | 1.047 | 0.41 | 0.14 | 0.36 | 0.38 | 0.44 | 0.29 |
| 5 | 106117058 | rs6868316 | 0.392 | 0.445 | -1.233 | -0.754 | 0.001 | -0.005 | -0.005 | 0 | 0.32 | 0.32 | 0.29 | 0.39 | 0.29 | 0.3 |
| 5 | 106123252 | rs1438255 | -1.157 | 0.679 | -0.981 | -0.648 | 0.016 | -0.098 | -0.098 | 0.09 | 0.83 | 0.68 | 0.69 | 0.97 | 0.81 | 0.87 |
| 5 | 106127543 | rs766463 | 1.536 | 0.639 | -1.197 | -0.619 | -0.054 | -0.067 | -0.067 | 0.522 | 0.27 | 0.08 | 0.21 | 0.31 | 0.33 | 0.24 |
| 5 | 106129495 | rs296275 | -0.624 | 0.758 | -0.748 | -0.42 | 0.025 | 0.016 | 0.016 | 1.209 | 0.8 | 0.64 | 0.68 | 0.77 | 0.77 | 0.62 |
| 5 | 106130200 | rs296277 | -2.27 | 0.752 | -1.259 | -0.503 | -0.004 | 0.012 | 0.012 | 0 | 0.89 | 0.94 | 0.81 | 0.98 | 0.86 | 0.9 |
| 5 | 106138146 | rs296285 | -0.11 | 0.827 | -1.334 | -0.418 | 0.029 | 0.011 | 0.011 | 0 | 0.69 | 0.51 | 0.59 | 0.66 | 0.64 | 0.57 |
| 5 | 106139212 | rs2164895 | 1.394 | 0.61 | -1.714 | -0.564 | -0.033 | -0.101 | -0.101 | 0.033 | 0.22 | 0.02 | 0.24 | 0.38 | 0.19 | 0.26 |
| 5 | 106140245 | rs6878348 | 1.176 | 0.561 | -1.591 | -0.511 | -0.021 | -0.061 | -0.061 | 0.062 | 0.25 | 0.08 | 0.27 | 0.38 | 0.23 | 0.29 |
| 5 | 106144139 | rs1898626 | 0.587 | 0.516 | -1.542 | -0.595 | -0.003 | 0.007 | 0.007 | 0.343 | 0.29 | 0.33 | 0.29 | 0.38 | 0.25 | 0.31 |
| 5 | 106144459 | rs296200 | -0.769 | 0.62 | -1.432 | -0.598 | -0.015 | -0.002 | -0.002 | 0 | 0.76 | 0.79 | 0.64 | 0.72 | 0.7 | 0.6 |
| 5 | 106146084 | rs296202 | 0.566 | 0.616 | -1.475 | -0.604 | -0.026 | -0.01 | -0.01 | 3.623 | 0.65 | 0.22 | 0.52 | 0.65 | 0.64 | 0.58 |
| 5 | 106149174 | rs296204 | -0.696 | 0.524 | -1.575 | -0.722 | 0 | 0.001 | 0.001 | 0.627 | 0.43 | 0.56 | 0.47 | 0.35 | 0.41 | 0.44 |
| 5 | 106149229 | rs296205 | 1.958 | 0.823 | -1.531 | -0.554 | -0.04 | 0.043 | 0.043 | 0 | 0.15 | 0.01 | 0.2 | 0.08 | 0.18 | 0.1 |
| 5 | 106155538 | rs296211 | -1.705 | 0.761 | -1.451 | -0.447 | -0.031 | -0.049 | -0.049 | 0 | 0.75 | 0.89 | 0.79 | 0.73 | 0.69 | 0.77 |
| 5 | 106163296 | rs6868584 | 0.557 | 0.55 | -1.222 | -0.335 | 0.026 | 0.075 | 0.075 | 0.417 | 0.06 | 0.18 | 0.06 | 0.19 | 0.08 | 0.28 |
| 5 | 106171851 | rs17159140 | 1.206 | 0.461 | -1.137 | -0.449 | 0.067 | 0.117 | 0.117 | 0 | 0.07 | 0.3 | 0.07 | 0.06 | 0.04 | 0.02 |
| 5 | 106183220 | rs467410 | 1.289 | 0.707 | -1.404 | -0.368 | -0.015 | 0.033 | 0.033 | 0 | 0.18 | 0.1 | 0.25 | 0.08 | 0.22 | 0.12 |
| 5 | 106185031 | rs159404 | 0.221 | 0.562 | -1.348 | -0.35 | 0 | -0.001 | -0.001 | 0 | 0.39 | 0.37 | 0.58 | 0.46 | 0.41 | 0.37 |
| 5 | 106191523 | rs295050 | 1.029 | 0.978 | -1.102 | -0.265 | -0.011 | 0.03 | 0.03 | 0 | 0.18 | 0.1 | 0.24 | 0.08 | 0.21 | 0.12 |
| 5 | 106198367 | rs295043 | -1.078 | 1.053 | -1.438 | -0.26 | -0.056 | -0.063 | -0.063 | 0 | 0.74 | 0.92 | 0.79 | 0.71 | 0.68 | 0.77 |
| 5 | 106199712 | rs296266 | 0.514 | 0.928 | -1.073 | -0.089 | 0.007 | -0.055 | -0.055 | 0 | 0.2 | 0.26 | 0.32 | 0.09 | 0.21 | 0.12 |
| 5 | 106208976 | rs7706073 | 0.144 | 1.003 | -1.495 | 0.562 | -0.001 | 0.038 | 0.038 | 0 | 0.06 | 0.14 | 0.05 | 0.17 | 0.07 | 0.27 |
| 5 | 106211190 | rs295018 | 0.071 | 1.048 | -1.624 | 0.313 | -0.024 | -0.038 | -0.038 | 0 | 0.63 | 0.68 | 0.72 | 0.55 | 0.57 | 0.47 |
| 5 | 106223536 | rs2454887 | 0.321 | 0.79 | -2.228 | 0.046 | -0.092 | -0.11 | -0.11 | 0 | 0.43 | 0.05 | 0.49 | 0.54 | 0.41 | 0.31 |
| 5 | 106224878 | rs11960102 | 0.228 | 0.39 | -1.473 | -0.386 | -0.052 | -0.122 | -0.122 | 0.11 | 0.2 | 0.41 | 0.09 | 0.03 | 0.11 | 0.12 |
| 5 | 106229524 | rs17159295 | 0.621 | 0.616 | -1.769 | -0.225 | -0.099 | -0.101 | -0.101 | 0 | 0.44 | 0.05 | 0.5 | 0.54 | 0.42 | 0.31 |
| 5 | 106230898 | rs1580831 | 0.629 | 0.653 | -1.788 | -0.251 | -0.1 | -0.102 | -0.102 | 0.305 | 0.44 | 0.05 | 0.5 | 0.54 | 0.42 | 0.31 |
| 5 | 106249782 | rs10058372 | 0.563 | 0.494 | -0.736 | -0.593 | -0.011 | -0.079 | -0.079 | 0 | 0.15 | 0.19 | 0.06 | 0 | 0.1 | 0.08 |
| 5 | 106256408 | rs7706863 | -0.504 | 0.428 | 0.079 | -0.488 | 0.016 | 0.073 | 0.073 | 0.112 | 0.1 | 0.35 | 0.08 | 0.15 | 0.11 | 0.23 |
| 5 | 106258427 | rs11748835 | 0.566 | 1.006 | -0.188 | -0.783 | -0.029 | 0.034 | 0.034 | 0.009 | 0.09 | 0.03 | 0.07 | 0 | 0.17 | 0.04 |
| 5 | 106260621 | rs9885083 | -0.307 | 0.878 | -0.114 | -0.735 | 0.014 | 0.082 | 0.082 | 0.037 | 0.09 | 0.36 | 0.08 | 0.15 | 0.09 | 0.22 |
| 5 | 106260687 | rs2463026 | 1.271 | 1.041 | -0.643 | -0.565 | -0.041 | -0.105 | -0.105 | 0 | 0.45 | 0.06 | 0.4 | 0.53 | 0.39 | 0.32 |
| 5 | 106263197 | rs2463031 | -0.313 | 0.845 | -0.363 | -0.712 | -0.027 | -0.031 | -0.031 | 0 | 0.77 | 0.89 | 0.69 | 0.72 | 0.7 | 0.67 |
| 5 | 106265598 | rs4421088 | 0.965 | 0.887 | -0.173 | -0.867 | -0.012 | -0.068 | -0.068 | 0.679 | 0.15 | 0.17 | 0.06 | 0 | 0.11 | 0.08 |
| 5 | 106271223 | rs7703472 | 0.286 | 1.091 | -0.396 | -1.021 | 0.007 | -0.07 | -0.07 | 0 | 0.14 | 0.17 | 0.06 | 0 | 0.13 | 0.11 |
| 5 | 106276532 | rs6880898 | 0.208 | -0.125 | -1.091 | -1.011 | -0.008 | -0.004 | -0.004 | 0 | 0.39 | 0.3 | 0.51 | 0.57 | 0.49 | 0.51 |
| 5 | 106287409 | rs4449519 | -0.467 | -0.726 | -0.949 | -1.2 | -0.071 | -0.084 | -0.084 | 0 | 0.67 | 0.53 | 0.8 | 0.86 | 0.79 | 0.87 |
| 5 | 106293263 | rs4957524 | -1.266 | -0.663 | -1.237 | -1.142 | 0.07 | 0.074 | 0.074 | 4.436 | 0.75 | 0.94 | 0.84 | 0.87 | 0.83 | 0.87 |
| 5 | 106308008 | rs4382150 | -0.455 | -0.656 | -1.015 | -1.241 | -0.052 | -0.091 | -0.091 | 0.799 | 0.65 | 0.62 | 0.8 | 0.88 | 0.78 | 0.8 |
| 5 | 106311248 | rs4245987 | 0.705 | -0.401 | -1.486 | -1.076 | -0.073 | -0.075 | -0.075 | 0.119 | 0.25 | 0.05 | 0.28 | 0.27 | 0.3 | 0.33 |
| 5 | 106328326 | rs4395620 | 0.618 | -0.12 | -0.921 | -0.966 | -0.006 | -0.005 | -0.005 | 0.099 | 0.32 | 0.31 | 0.49 | 0.6 | 0.44 | 0.46 |
| 5 | 106352714 | rs7444810 | -0.135 | 0.089 | -0.854 | -1.085 | -0.011 | 0.001 | 0.001 | 0.13 | 0.44 | 0.52 | 0.55 | 0.58 | 0.6 | 0.49 |
| 5 | 106358541 | rs12521220 | 0.421 | 0.062 | -0.935 | -1.052 | 0.018 | 0.018 | 0.018 | 0 | 0.35 | 0.39 | 0.5 | 0.56 | 0.47 | 0.42 |

| | | | | | | | | | | | | | | | | |
|---|---|---|---|---|---|---|---|---|---|---|---|---|---|---|---|---|
| 5 | 106359882 | rs11749538 | 1.017 | 0.299 | -0.592 | -0.994 | -0.04 | 0.038 | 0.038 | 0.019 | 0.09 | 0.03 | 0.07 | 0 | 0.17 | 0.04 |
| 5 | 106361340 | rs4473758 | 0.229 | 0.042 | -0.339 | -0.925 | 0.024 | -0.001 | -0.001 | 0 | 0.05 | 0.31 | 0.03 | 0 | 0.03 | 0.01 |
| 5 | 106361555 | rs4305636 | -1.263 | 0.115 | -0.54 | -0.979 | 0.013 | -0.058 | -0.058 | 0.092 | 0.12 | 0.67 | 0.1 | 0.02 | 0.09 | 0.07 |
| 5 | 106371045 | rs9327987 | -0.417 | -0.172 | -0.78 | -1.061 | 0.02 | 0.033 | 0.033 | 0 | 0.5 | 0.62 | 0.39 | 0.3 | 0.43 | 0.46 |
| 5 | 106373992 | rs11242622 | 1.354 | 0.049 | -0.95 | -0.777 | 0.003 | -0.107 | -0.107 | 0 | 0.18 | 0.01 | 0.29 | 0.29 | 0.14 | 0.32 |
| 5 | 106375215 | rs10793831 | -1.118 | 0.244 | -1.047 | -0.888 | 0.008 | 0.07 | 0.07 | 2.71 | 0.71 | 0.9 | 0.74 | 0.86 | 0.68 | 0.64 |
| 5 | 106379106 | rs4382152 | -0.338 | 0.81 | -0.324 | -0.2 | -0.02 | 0.06 | 0.06 | 1.801 | 0.39 | 0.25 | 0.54 | 0.81 | 0.4 | 0.46 |
| 5 | 106379836 | rs12656373 | -0.945 | 0.794 | -0.502 | -0.057 | 0.01 | 0.029 | 0.029 | 0.354 | 0.51 | 0.51 | 0.61 | 0.83 | 0.52 | 0.48 |
| 5 | 106384965 | rs12658882 | 1.32 | 1.436 | -1.398 | 0.364 | -0.093 | -0.116 | -0.116 | 0.505 | 0.16 | 0.03 | 0.22 | 0.43 | 0.22 | 0.14 |
| 5 | 106389374 | rs13355762 | -0.767 | 0.353 | -0.417 | 0.301 | 0.031 | -0.069 | -0.069 | 0 | 0.13 | 0.35 | 0.08 | 0.02 | 0.13 | 0.03 |
| 5 | 106390482 | rs4957684 | -0.278 | 0.525 | -0.968 | 0.543 | 0.005 | -0.009 | -0.009 | 3.367 | 0.28 | 0.15 | 0.34 | 0.29 | 0.24 | 0.31 |
| 5 | 106394177 | rs4640780 | 0.208 | 0.81 | -0.65 | 0.014 | -0.003 | -0.032 | -0.032 | 0 | 0.67 | 0.39 | 0.73 | 0.76 | 0.65 | 0.8 |
| 5 | 106394251 | rs4309958 | -0.397 | 1.416 | -0.387 | 0.564 | 0.024 | 0.084 | 0.084 | 0 | 0.25 | 0.53 | 0.26 | 0.48 | 0.22 | 0.22 |
| 5 | 106400536 | rs4413519 | 0.5 | 1.264 | -0.777 | 0.173 | -0.019 | -0.136 | -0.136 | 0.58 | 0.19 | 0.03 | 0.2 | 0.48 | 0.18 | 0.26 |
| 5 | 106415368 | rs4957527 | -0.602 | 1.253 | -1.367 | -0.746 | -0.007 | 0.034 | 0.034 | 0 | 0.58 | 0.86 | 0.57 | 0.75 | 0.62 | 0.61 |
| 5 | 106420589 | rs4957528 | -1.043 | 1.236 | -1.79 | -1.206 | 0.017 | 0.106 | 0.106 | 0 | 0.73 | 0.93 | 0.87 | 1 | 0.77 | 0.91 |
| 5 | 106428845 | rs12658379 | 0.486 | 1.286 | -0.686 | -1.364 | -0.005 | 0.038 | 0.038 | 0 | 0.46 | 0.25 | 0.42 | 0.71 | 0.48 | 0.45 |
| 5 | 106446935 | rs4512123 | 0.44 | 1.348 | -0.724 | -1.32 | 0.003 | -0.013 | -0.013 | 3.798 | 0.07 | 0.21 | 0.08 | 0.01 | 0.07 | 0.18 |
| 5 | 106449990 | rs7736860 | -0.641 | 1.434 | -0.94 | -1.392 | -0.05 | 0.032 | 0.032 | 0 | 0.59 | 0.9 | 0.57 | 0.75 | 0.6 | 0.65 |
| 5 | 106452479 | rs17450801 | 1.691 | 1.405 | -1.115 | -1.435 | -0.057 | -0.02 | -0.02 | 0 | 0.14 | 0.01 | 0.15 | 0.2 | 0.2 | 0.11 |
| 5 | 106478316 | rs6888172 | -0.294 | 1.138 | -0.064 | -1.241 | -0.013 | 0.029 | 0.029 | 0 | 0.58 | 0.78 | 0.55 | 0.72 | 0.6 | 0.63 |
| 5 | 106489108 | rs4957690 | -0.387 | 1.147 | 0.057 | -1.259 | -0.019 | 0.039 | 0.039 | 0 | 0.59 | 0.83 | 0.57 | 0.75 | 0.6 | 0.64 |
| 5 | 106493350 | rs10043242 | 0.525 | 1.092 | 0.485 | -1.322 | 0.004 | 0.01 | 0.01 | 0 | 0.48 | 0.56 | 0.44 | 0.71 | 0.48 | 0.45 |
| 5 | 106500969 | rs4957691 | -0.466 | 1.109 | 0.481 | -1.315 | -0.022 | 0.03 | 0.03 | 0 | 0.59 | 0.84 | 0.56 | 0.72 | 0.6 | 0.63 |
| 5 | 106501202 | rs4957534 | 0.446 | 1.099 | 0.512 | -1.322 | 0.005 | 0.011 | 0.011 | 0 | 0.49 | 0.56 | 0.43 | 0.71 | 0.47 | 0.45 |
| 5 | 106502006 | rs2410954 | -0.456 | 1.102 | 0.479 | -1.318 | -0.029 | 0.04 | 0.04 | 0 | 0.59 | 0.86 | 0.56 | 0.75 | 0.6 | 0.64 |
| 5 | 106504850 | rs10515368 | 0.442 | 1.037 | 0.467 | -1.302 | 0.005 | 0.012 | 0.012 | 0 | 0.49 | 0.56 | 0.43 | 0.71 | 0.47 | 0.45 |
| 5 | 106513321 | rs17159528 | -0.782 | 1.054 | 0.647 | -1.239 | 0.028 | -0.024 | -0.024 | 0 | 0.1 | 0.38 | 0.12 | 0.01 | 0.1 | 0.18 |
| 5 | 106531735 | rs7729133 | 0.629 | 1.068 | 0.222 | -1.3 | 0 | 0.002 | 0.002 | 0 | 0.47 | 0.5 | 0.42 | 0.71 | 0.47 | 0.43 |
| 5 | 106532793 | rs1549175 | 1.44 | 1.152 | 0.071 | -1.404 | -0.053 | -0.083 | -0.083 | 0 | 0.15 | 0.01 | 0.3 | 0.24 | 0.17 | 0.28 |
| 5 | 106538016 | rs2112433 | -0.749 | 0.844 | 0.528 | -1.09 | -0.042 | 0.042 | 0.042 | 0 | 0.59 | 0.9 | 0.56 | 0.75 | 0.6 | 0.63 |
| 5 | 106541842 | rs2019735 | -0.577 | 0.84 | 0.574 | -1.092 | -0.005 | 0.03 | 0.03 | 0 | 0.58 | 0.75 | 0.55 | 0.72 | 0.6 | 0.62 |
| 5 | 106542253 | rs9327992 | 0.087 | 1.029 | 0.561 | -1.003 | 0.009 | 0.023 | 0.023 | 0 | 0.49 | 0.59 | 0.44 | 0.71 | 0.48 | 0.43 |
| 5 | 106553396 | rs1159078 | 0.137 | 0.958 | 0.543 | -1.004 | -0.001 | 0.004 | 0.004 | 0 | 0.48 | 0.5 | 0.43 | 0.71 | 0.5 | 0.45 |
| 5 | 106557132 | rs6596730 | -0.569 | 0.935 | 0.484 | -1.009 | -0.011 | 0.037 | 0.037 | 0 | 0.57 | 0.8 | 0.56 | 0.73 | 0.6 | 0.63 |
| 5 | 106558682 | rs4957696 | 1.284 | 0.156 | 0.448 | -0.987 | -0.002 | 0.016 | 0.016 | 3.174 | 0.19 | 0.12 | 0.22 | 0.49 | 0.12 | 0.27 |
| 5 | 106562419 | rs7735022 | -0.185 | 0.224 | 0.39 | -0.995 | 0.013 | -0.039 | -0.039 | 1.106 | 0.07 | 0.2 | 0.12 | 0 | 0.08 | 0.13 |
| 5 | 106562841 | rs10464035 | 1.18 | 0.232 | 0.05 | -0.744 | -0.002 | 0.016 | 0.016 | 3.308 | 0.18 | 0.12 | 0.17 | 0.47 | 0.12 | 0.27 |
| 5 | 106564497 | rs4957697 | 0.056 | 0.606 | -0.307 | -0.779 | -0.001 | -0.001 | -0.001 | 0 | 0.47 | 0.47 | 0.4 | 0.69 | 0.42 | 0.47 |
| 5 | 106569691 | rs10455070 | 0.765 | 0.401 | -0.481 | -0.188 | 0.007 | -0.022 | -0.022 | 0 | 0.19 | 0.16 | 0.18 | 0.43 | 0.14 | 0.27 |
| 5 | 106569809 | rs10455071 | -0.306 | 0.349 | -0.298 | -0.613 | -0.002 | 0 | 0 | 3.161 | 0.49 | 0.58 | 0.38 | 0.46 | 0.35 | 0.51 |
| 5 | 106573961 | rs4957698 | 0.75 | 0.576 | -0.831 | -0.473 | -0.052 | -0.151 | -0.151 | 0 | 0.24 | 0.04 | 0.35 | 0.49 | 0.27 | 0.44 |
| 5 | 106574132 | rs13154130 | 0.941 | 0.171 | -0.156 | -0.781 | -0.03 | -0.076 | -0.076 | 0 | 0.2 | 0.27 | 0.09 | 0 | 0.1 | 0.06 |
| 5 | 106574196 | rs1422210 | -1.078 | 0.576 | -0.296 | -1.113 | -0.026 | -0.031 | -0.031 | 0.408 | 0.76 | 0.86 | 0.77 | 0.73 | 0.7 | 0.74 |
| 5 | 106574876 | rs11242634 | 1.625 | 0.459 | -0.18 | -1.236 | -0.021 | 0.033 | 0.033 | 0 | 0.22 | 0.08 | 0.27 | 0.21 | 0.28 | 0.19 |
| 5 | 106586798 | rs7732995 | -1.027 | 0.444 | 0.329 | -1.097 | -0.024 | -0.026 | -0.026 | 0 | 0.77 | 0.88 | 0.78 | 0.73 | 0.7 | 0.74 |
| 5 | 106587616 | rs1423336 | -0.99 | 0.446 | 0.338 | -1.096 | -0.056 | -0.058 | -0.058 | 0 | 0.77 | 0.93 | 0.78 | 0.73 | 0.7 | 0.74 |
| 5 | 106589749 | rs17159621 | 0.614 | 0.254 | 0.448 | -1.108 | -0.065 | -0.146 | -0.146 | 0 | 0.25 | 0.53 | 0.12 | 0.03 | 0.13 | 0.08 |
| 5 | 106592310 | rs2548754 | 0.373 | 0.401 | 0.404 | -0.942 | 0.003 | 0.028 | 0.028 | 0 | 0.51 | 0.35 | 0.65 | 0.7 | 0.55 | 0.65 |
| 5 | 106599519 | rs34564 | 0.212 | 0.383 | 0.617 | -0.961 | 0 | 0.004 | 0.004 | 0 | 0.52 | 0.43 | 0.65 | 0.71 | 0.56 | 0.64 |
| 5 | 106600215 | rs34566 | 0.39 | 0.433 | 0.519 | -0.931 | 0.004 | 0.034 | 0.034 | 0 | 0.5 | 0.34 | 0.63 | 0.7 | 0.56 | 0.64 |
| 5 | 106610781 | rs40083 | 0.041 | 0.418 | 0.433 | -0.959 | -0.005 | 0.027 | 0.027 | 0 | 0.55 | 0.59 | 0.65 | 0.73 | 0.56 | 0.65 |
| 5 | 106619509 | rs451833 | -0.119 | 0.392 | 0.531 | -0.94 | -0.009 | -0.009 | -0.009 | 0 | 0.59 | 0.44 | 0.69 | 0.69 | 0.67 | 0.69 |
| 5 | 106625682 | rs1363896 | -0.037 | 0.481 | 0.144 | -1.06 | 0.008 | 0.029 | 0.029 | 0 | 0.5 | 0.45 | 0.45 | 0.25 | 0.46 | 0.29 |
| 5 | 106634782 | rs34590 | -1.535 | 0.631 | -0.153 | -0.844 | -0.006 | -0.025 | -0.025 | 1.614 | 0.84 | 0.9 | 0.83 | 0.73 | 0.81 | 0.77 |
| 5 | 106636791 | rs1981851 | 0.946 | 0.527 | -0.417 | -1.084 | -0.082 | 0.046 | 0.046 | 0 | 0.28 | 0.03 | 0.29 | 0.19 | 0.35 | 0.21 |
| 5 | 106639224 | rs17343335 | 1.8 | 0.253 | -0.472 | -1.212 | 0.015 | 0.047 | 0.047 | 0 | 0.14 | 0.07 | 0.06 | 0 | 0.09 | 0.06 |
| 5 | 106641010 | rs17343546 | 0.324 | 0.14 | -0.079 | -1.282 | 0.003 | 0.066 | 0.066 | 0 | 0.38 | 0.22 | 0.33 | 0.21 | 0.38 | 0.23 |
| 5 | 106641408 | rs17159751 | 1.601 | 0.133 | 0.052 | -1.129 | 0 | -0.021 | -0.021 | 1.079 | 0.14 | 0.16 | 0.06 | 0 | 0.09 | 0.06 |
| 5 | 106645685 | rs253125 | -0.574 | 0.122 | 0.205 | -1.06 | -0.003 | -0.001 | -0.001 | 1.055 | 0.69 | 0.62 | 0.76 | 0.73 | 0.71 | 0.71 |
| 5 | 106649799 | rs252696 | 0.384 | 0.106 | 0.282 | -1.057 | -0.006 | -0.001 | -0.001 | 0 | 0.31 | 0.39 | 0.25 | 0.3 | 0.28 | 0.31 |
| 5 | 106649834 | rs252695 | 1.535 | 0.126 | -0.002 | -1.045 | -0.013 | -0.042 | -0.042 | 0 | 0.15 | 0.07 | 0.17 | 0.28 | 0.19 | 0.23 |
| 5 | 106652124 | rs17403350 | 0.966 | -0.005 | -0.039 | -1.125 | 0.043 | 0.086 | 0.086 | 0 | 0.19 | 0.07 | 0.07 | 0 | 0.11 | 0.07 |
| 5 | 106652926 | rs173757 | -0.517 | 0.11 | 0.162 | -1.148 | 0 | 0 | 0 | 0.067 | 0.54 | 0.46 | 0.51 | 0.49 | 0.56 | 0.46 |
| 5 | 106653325 | rs252691 | -0.99 | 0.089 | 0.173 | -0.983 | -0.001 | -0.003 | -0.003 | 0 | 0.7 | 0.73 | 0.62 | 0.5 | 0.69 | 0.53 |
| 5 | 106660089 | rs441650 | 1.463 | 0.069 | 0.051 | -0.983 | -0.006 | -0.024 | -0.024 | 0.046 | 0.15 | 0.09 | 0.17 | 0.28 | 0.18 | 0.22 |
| 5 | 106660124 | rs433084 | -0.966 | 0.29 | 0.244 | -1.094 | 0 | 0.02 | 0.02 | 0 | 0.68 | 0.65 | 0.66 | 0.49 | 0.69 | 0.52 |
| 5 | 106660530 | rs422261 | 0.883 | 0.212 | -0.02 | -0.978 | -0.006 | -0.032 | -0.032 | 0 | 0.19 | 0.11 | 0.19 | 0.38 | 0.21 | 0.27 |
| 5 | 106662978 | rs443277 | -0.08 | 0.227 | 0.135 | -1.219 | -0.008 | 0.006 | 0.006 | 0 | 0.4 | 0.75 | 0.34 | 0.37 | 0.41 | 0.34 |

| | | | | | | | | | | | | | | | | |
|---|---|---|---|---|---|---|---|---|---|---|---|---|---|---|---|---|
| 5 | 106664983 | rs185211 | 1.219 | 0.247 | 0.131 | -1.215 | -0.013 | -0.058 | -0.058 | 0 | 0.19 | 0.08 | 0.2 | 0.35 | 0.22 | 0.28 |
| 5 | 106665664 | rs252677 | -0.028 | 0.238 | 0.082 | -1.185 | -0.005 | 0.016 | 0.016 | 0 | 0.4 | 0.74 | 0.34 | 0.37 | 0.41 | 0.34 |
| 5 | 106665710 | rs1423341 | 0.466 | 0.164 | -0.21 | -1.245 | -0.049 | 0.103 | 0.103 | 0 | 0.33 | 0.05 | 0.36 | 0.15 | 0.34 | 0.22 |
| 5 | 106672726 | rs1423338 | 1.653 | 0.105 | 0.183 | -1.199 | -0.009 | -0.027 | -0.027 | 0 | 0.06 | 0.1 | 0.03 | 0 | 0.04 | 0.02 |
| 5 | 106673261 | rs412541 | 1.761 | 0.099 | -0.195 | -1.195 | -0.018 | -0.098 | -0.098 | 0 | 0.14 | 0 | 0.15 | 0.36 | 0.14 | 0.23 |
| 5 | 106674243 | rs7711379 | 1.511 | 0.084 | -0.182 | -1.237 | -0.033 | 0.109 | 0.109 | 0.021 | 0.18 | 0.02 | 0.09 | 0 | 0.19 | 0.07 |
| 5 | 106674516 | rs252682 | 0.805 | 0.437 | 0.327 | -1.097 | 0.001 | 0.01 | 0.01 | 0 | 0.22 | 0.28 | 0.23 | 0.36 | 0.24 | 0.27 |
| 5 | 106677407 | rs17344856 | 1.75 | 0.319 | 0.555 | -1.019 | -0.003 | -0.016 | -0.016 | 1.984 | 0.15 | 0.13 | 0.08 | 0 | 0.17 | 0.06 |
| 5 | 106678794 | rs1195947 | -1.309 | 0.188 | 0.14 | -1.032 | -0.013 | -0.097 | -0.097 | 1.486 | 0.75 | 0.91 | 0.66 | 0.51 | 0.76 | 0.55 |
| 5 | 106683271 | rs252680 | 0.126 | 0.4 | -0.522 | -1.007 | -0.016 | -0.019 | -0.019 | 0 | 0.3 | 0.25 | 0.31 | 0.37 | 0.36 | 0.29 |
| 5 | 106690481 | rs1195948 | -1.442 | 0.322 | -0.865 | -0.962 | -0.011 | -0.069 | -0.069 | 0 | 0.8 | 0.92 | 0.72 | 0.53 | 0.78 | 0.55 |
| 5 | 106697023 | rs6860893 | 0.005 | 0.227 | -0.537 | -1.587 | 0.018 | 0.118 | 0.118 | 0.562 | 0.5 | 0.64 | 0.44 | 0.16 | 0.44 | 0.25 |
| 5 | 106697485 | rs12514805 | 0.843 | 0.279 | -1.314 | -1.38 | -0.029 | 0.027 | 0.027 | 0 | 0.23 | 0.03 | 0.28 | 0.14 | 0.23 | 0.18 |
| 5 | 106701295 | rs6884177 | -0.354 | 0.317 | -1.005 | -1.148 | -0.016 | 0.101 | 0.101 | 2.57 | 0.64 | 0.83 | 0.54 | 0.17 | 0.6 | 0.38 |
| 5 | 106701494 | rs17345521 | 1.391 | 0.497 | -1.176 | -1.239 | -0.021 | 0.093 | 0.093 | 2.175 | 0.22 | 0.06 | 0.09 | 0 | 0.2 | 0.09 |
| 5 | 106702508 | rs17159829 | 0.292 | 0.107 | -0.03 | -0.561 | -0.007 | -0.053 | -0.053 | 0 | 0.59 | 0.42 | 0.47 | 0.12 | 0.59 | 0.39 |
| 5 | 106713618 | rs365807 | -0.238 | 0.237 | 0.189 | 0.45 | -0.004 | 0.115 | 0.115 | 0 | 0.25 | 0.42 | 0.38 | 0.49 | 0.26 | 0.28 |
| 5 | 106721683 | rs252815 | -0.914 | 0.276 | 0.28 | 0.242 | -0.001 | 0.028 | 0.028 | 0 | 0.85 | 0.86 | 0.83 | 0.52 | 0.83 | 0.67 |
| 5 | 106725691 | rs252818 | -0.874 | 0.283 | 0.707 | 0.226 | 0.004 | 0.068 | 0.068 | 0 | 0.85 | 0.84 | 0.77 | 0.5 | 0.81 | 0.72 |
| 5 | 106729992 | rs3797513 | 0.7 | 0.163 | 0.549 | 0.042 | -0.008 | -0.055 | -0.055 | 0 | 0.14 | 0.06 | 0.25 | 0.32 | 0.16 | 0.28 |
| 5 | 106736357 | rs252811 | -1.047 | 0.353 | 0.663 | 0.142 | 0.004 | 0.064 | 0.064 | 0 | 0.86 | 0.84 | 0.78 | 0.49 | 0.81 | 0.72 |
| 5 | 106739110 | rs252807 | 0.188 | -0.068 | 0.831 | -0.36 | 0.001 | 0.01 | 0.01 | 0 | 0.66 | 0.71 | 0.56 | 0.2 | 0.63 | 0.41 |
| 5 | 106742153 | rs682609 | 1.185 | 0.13 | 0.088 | -0.999 | -0.039 | 0.093 | 0.093 | 2.057 | 0.28 | 0.09 | 0.14 | 0 | 0.31 | 0.09 |
| 5 | 106757399 | rs252801 | 0.347 | -0.168 | 0.61 | -0.795 | 0.023 | 0.078 | 0.078 | 0 | 0.54 | 0.39 | 0.45 | 0.17 | 0.59 | 0.37 |
| 5 | 106758684 | rs13182976 | 0.882 | -0.27 | 0.585 | -0.642 | -0.003 | 0.002 | 0.002 | 0 | 0.21 | 0.24 | 0.28 | 0.29 | 0.19 | 0.33 |
| 5 | 106761136 | rs2731840 | -0.472 | -0.097 | 0.234 | -0.455 | -0.006 | 0.083 | 0.083 | 0 | 0.78 | 0.7 | 0.75 | 0.47 | 0.79 | 0.71 |
| 5 | 106762584 | rs252802 | 0.306 | 0.201 | -0.804 | -1.034 | -0.067 | 0.18 | 0.18 | 0.036 | 0.44 | 0.14 | 0.4 | 0.15 | 0.5 | 0.33 |
| 5 | 106767346 | rs252991 | -0.057 | 0.044 | -0.087 | -0.39 | -0.001 | 0.001 | 0.001 | 0 | 0.61 | 0.62 | 0.66 | 0.36 | 0.63 | 0.67 |
| 5 | 106771209 | rs152608 | 0.885 | -0.001 | -0.047 | -0.392 | -0.002 | -0.009 | -0.009 | 4.129 | 0.25 | 0.22 | 0.23 | 0.54 | 0.24 | 0.3 |
| 5 | 106773623 | rs2283 | -0.205 | 0.017 | 0.095 | -0.417 | -0.002 | 0.007 | 0.007 | 0 | 0.38 | 0.45 | 0.38 | 0.54 | 0.33 | 0.33 |
| 5 | 106778844 | rs526521 | 0.321 | -0.058 | -0.053 | -0.674 | -0.005 | -0.013 | -0.013 | 0.128 | 0.27 | 0.26 | 0.36 | 0.15 | 0.24 | 0.4 |
| 5 | 106781376 | rs152595 | 0.931 | 0.021 | -0.721 | -0.609 | -0.004 | 0.06 | 0.06 | 0 | 0.28 | 0.36 | 0.3 | 0.55 | 0.24 | 0.3 |
| 5 | 106789764 | rs1835111 | 0.834 | 0.039 | -1.7 | -0.486 | -0.021 | 0.004 | 0.004 | 0 | 0.2 | 0.01 | 0.33 | 0.15 | 0.21 | 0.38 |
| 5 | 106793773 | rs3909464 | -0.118 | 0.716 | -1.359 | -0.231 | 0 | -0.012 | -0.012 | 0 | 0.14 | 0.15 | 0.1 | 0 | 0.12 | 0.03 |
| 5 | 106795862 | rs152580 | 0.471 | 0.451 | -1.346 | -0.184 | -0.008 | 0.052 | 0.052 | 0 | 0.53 | 0.65 | 0.66 | 0.7 | 0.49 | 0.71 |
| 5 | 106800148 | rs10491390 | -0.694 | 0.321 | -0.701 | -0.117 | 0.002 | -0.015 | -0.015 | 0.067 | 0.12 | 0.13 | 0.09 | 0 | 0.1 | 0.02 |
| 5 | 106804363 | rs26247 | -0.748 | 0.155 | -0.429 | 0.332 | -0.023 | 0.011 | 0.011 | 0 | 0.29 | 0.37 | 0.46 | 0.66 | 0.2 | 0.56 |
| 5 | 106804832 | rs25998 | -1.017 | 0.614 | -1.726 | -0.753 | -0.147 | 0.06 | 0.06 | 0 | 0.67 | 0.97 | 0.75 | 0.82 | 0.54 | 0.74 |
| 5 | 106804928 | rs3890134 | 0.656 | 0.495 | -1.359 | -1.248 | -0.003 | -0.051 | -0.051 | 0 | 0.17 | 0.26 | 0.11 | 0 | 0.13 | 0.09 |
| 5 | 106806224 | rs636137 | -0.338 | 0.688 | -1.115 | -1.382 | -0.023 | -0.054 | -0.054 | 0 | 0.35 | 0.6 | 0.25 | 0.16 | 0.25 | 0.14 |
| 5 | 106806438 | rs29543 | 0.259 | 0.575 | -1.088 | -1.371 | 0.006 | 0.004 | 0.004 | 0 | 0.31 | 0.35 | 0.49 | 0.66 | 0.29 | 0.6 |
| 5 | 106810412 | rs25740 | 0.263 | 0.215 | -1.017 | -0.129 | -0.015 | 0.217 | 0.217 | 0 | 0.1 | 0.36 | 0.34 | 0.27 | 0.04 | 0.12 |
| 5 | 106811645 | rs25994 | -0.679 | 1.357 | -1.24 | -0.455 | 0.015 | 0.013 | 0.013 | 0 | 0.48 | 0.35 | 0.38 | 0.4 | 0.54 | 0.61 |
| 5 | 106818343 | rs26244 | 0.532 | 0.633 | -0.368 | -0.095 | -0.02 | 0.15 | 0.15 | 0 | 0.09 | 0.22 | 0.1 | 0.25 | 0.04 | 0.11 |
| 5 | 106820247 | rs2898622 | -0.339 | 0.547 | -0.018 | -0.122 | -0.035 | -0.017 | -0.017 | 0 | 0.09 | 0.31 | 0.04 | 0.09 | 0.03 | 0.06 |
| 5 | 106820480 | rs25966 | -1.588 | 0.785 | -0.33 | -0.287 | -0.021 | 0.077 | 0.077 | 0 | 0.57 | 0.67 | 0.53 | 0.82 | 0.48 | 0.7 |
| 5 | 106826413 | rs25970 | -0.783 | 0.95 | -0.269 | -0.257 | -0.004 | -0.002 | -0.002 | 0.765 | 0.43 | 0.5 | 0.3 | 0.4 | 0.37 | 0.54 |
| 5 | 106831574 | rs27681 | 0.154 | 0.779 | -0.241 | -0.296 | -0.003 | 0.006 | 0.006 | 0 | 0.27 | 0.31 | 0.27 | 0.4 | 0.21 | 0.45 |
| 5 | 106833151 | rs12520677 | -0.529 | 1.035 | -0.252 | -0.204 | 0.005 | -0.001 | -0.001 | 0 | 0.2 | 0.2 | 0.17 | 0.15 | 0.22 | 0.15 |
| 5 | 106834567 | rs1196516 | -0.752 | 0.795 | 0.211 | -0.477 | -0.067 | -0.027 | -0.027 | 1.475 | 0.23 | 0.41 | 0.16 | 0.18 | 0.11 | 0.19 |
| 5 | 106835010 | rs25976 | -1.681 | 0.78 | 0.524 | -0.543 | -0.023 | 0.084 | 0.084 | 0 | 0.55 | 0.76 | 0.53 | 0.73 | 0.48 | 0.7 |
| 5 | 106836319 | rs32013 | -1.984 | 0.911 | 0.53 | -0.562 | -0.026 | 0.093 | 0.093 | 0 | 0.58 | 0.81 | 0.55 | 0.75 | 0.5 | 0.7 |
| 5 | 106836774 | rs10491393 | -0.487 | 0.923 | 0.754 | -0.395 | 0.042 | 0.05 | 0.05 | 0.003 | 0.07 | 0.24 | 0.09 | 0.11 | 0.08 | 0.06 |
| 5 | 106837552 | rs26250 | 0.067 | 0.752 | 0.683 | -0.477 | 0.001 | -0.013 | -0.013 | 0 | 0.27 | 0.21 | 0.19 | 0.45 | 0.24 | 0.45 |
| 5 | 106837754 | rs27362 | 0.384 | 0.593 | 0.522 | -0.491 | -0.014 | 0.149 | 0.149 | 0.485 | 0.07 | 0.22 | 0.15 | 0.15 | 0.01 | 0.04 |
| 5 | 106838407 | rs26249 | -0.744 | 0.597 | 0.483 | -0.502 | -0.012 | 0.097 | 0.097 | 0.063 | 0.11 | 0.28 | 0.18 | 0.2 | 0.06 | 0.11 |
| 5 | 106840431 | rs17534165 | 0.874 | 0.588 | 0.386 | -0.574 | -0.006 | 0.023 | 0.023 | 0 | 0.14 | 0.08 | 0.09 | 0 | 0.13 | 0.09 |
| 5 | 106840779 | rs32006 | -0.173 | 0.567 | 0.461 | -0.566 | -0.02 | -0.009 | -0.009 | 0 | 0.21 | 0.35 | 0.18 | 0.15 | 0.14 | 0.13 |
| 5 | 106841783 | rs152562 | -1.532 | 0.825 | -0.119 | -0.809 | -0.006 | 0.025 | 0.025 | 0 | 0.55 | 0.59 | 0.52 | 0.76 | 0.48 | 0.7 |
| 5 | 106842228 | rs151943 | -0.445 | 0.929 | -0.293 | -0.811 | -0.004 | -0.034 | -0.034 | 0 | 0.37 | 0.45 | 0.4 | 0.71 | 0.33 | 0.56 |
| 5 | 106842650 | rs152560 | -1.682 | 0.942 | -1.309 | -0.67 | -0.079 | 0.097 | 0.097 | 0 | 0.59 | 0.89 | 0.55 | 0.75 | 0.5 | 0.7 |
| 5 | 106846342 | rs17534228 | 0.294 | 0.85 | -1.201 | -0.593 | -0.014 | 0.049 | 0.049 | 0 | 0.17 | 0.08 | 0.12 | 0.06 | 0.18 | 0.15 |
| 5 | 106847662 | rs152558 | -0.857 | 1.305 | -1.208 | -0.382 | -0.061 | 0.133 | 0.133 | 0 | 0.57 | 0.88 | 0.66 | 0.75 | 0.47 | 0.68 |
| 5 | 106849306 | rs153101 | -0.531 | 0.907 | -0.964 | -0.834 | 0.063 | 0.08 | 0.08 | 0 | 0.47 | 0.73 | 0.43 | 0.32 | 0.36 | 0.25 |
| 5 | 106849674 | rs152555 | 1.286 | 1.81 | -0.535 | -0.76 | -0.03 | -0.026 | -0.026 | 0 | 0.92 | 0.97 | 0.75 | 0.78 | 0.87 | 0.92 |
| 5 | 106855830 | rs250252 | 0.643 | 1.376 | -0.715 | -1.292 | -0.043 | 0.022 | 0.022 | 0 | 0.93 | 0.99 | 0.95 | 0.96 | 0.86 | 0.88 |
| 5 | 106861975 | rs395561 | -1.073 | -0.02 | -1.018 | -2.058 | 0.025 | 0.237 | 0.237 | 0.02 | 0.54 | 0.87 | 0.31 | 0.06 | 0.38 | 0.36 |
| 5 | 106863715 | rs464250 | -0.089 | -0.718 | -0.06 | -1.909 | -0.004 | -0.057 | -0.057 | 0 | 0.3 | 0.32 | 0.15 | 0.01 | 0.17 | 0.15 |
| 5 | 106875465 | rs163614 | -0.683 | -0.816 | 0.133 | -1.931 | 0.02 | -0.021 | -0.021 | 0.003 | 0.38 | 0.71 | 0.19 | 0.01 | 0.2 | 0.2 |

| | | | | | | | | | | | | | | | | |
|---|---|---|---|---|---|---|---|---|---|---|---|---|---|---|---|---|
| 5 | 106879502 | rs163895 | -1.568 | -0.4 | 0.31 | -1.656 | -0.005 | -0.037 | -0.037 | 0 | 0.2 | 0.75 | 0.13 | 0.09 | 0.14 | 0.16 |
| 5 | 106881333 | rs662627 | -0.139 | -0.013 | 0.364 | -1.28 | 0.003 | 0.024 | 0.024 | 0.154 | 0.78 | 0.69 | 0.7 | 0.71 | 0.76 | 0.61 |
| 5 | 106884480 | rs464950 | 1.587 | -0.174 | 0.307 | -1.464 | -0.001 | 0 | 0 | 0.021 | 0.4 | 0.4 | 0.47 | 0.63 | 0.45 | 0.34 |
| 5 | 106885914 | rs467089 | 1.783 | -0.128 | -0.172 | -1.461 | -0.013 | -0.001 | -0.001 | 0 | 0.36 | 0.23 | 0.46 | 0.63 | 0.43 | 0.29 |
| 5 | 106887068 | rs341272 | 0.281 | -0.014 | -0.17 | -1.468 | -0.042 | -0.022 | -0.022 | 0 | 0.71 | 0.83 | 0.64 | 0.65 | 0.63 | 0.49 |
| 5 | 106888783 | rs352610 | 0.532 | 0.532 | -0.1 | -1.176 | 0 | -0.006 | -0.006 | 0 | 0.19 | 0.12 | 0.13 | 0 | 0.18 | 0.12 |
| 5 | 106893142 | rs352605 | -0.734 | 0.743 | -0.382 | -1.024 | -0.001 | -0.025 | -0.025 | 0.022 | 0.34 | 0.68 | 0.23 | 0.08 | 0.3 | 0.28 |
| 5 | 106896484 | rs677797 | -1.098 | 0.541 | -0.334 | -0.148 | -0.003 | -0.006 | -0.006 | 0 | 0.42 | 0.47 | 0.38 | 0.32 | 0.37 | 0.57 |
| 5 | 106896617 | rs2484103 | -0.581 | 0.488 | -1.216 | 0.431 | -0.023 | -0.03 | -0.03 | 0.213 | 0.29 | 0.1 | 0.29 | 0.28 | 0.29 | 0.44 |
| 5 | 106908733 | rs164700 | 0.779 | 0.362 | -0.041 | 1.115 | -0.007 | 0.185 | 0.185 | 0 | 0.82 | 0.59 | 0.73 | 0.55 | 0.86 | 0.8 |
| 5 | 106909060 | rs6881619 | -0.987 | -0.078 | 1.114 | 0.591 | -0.027 | -0.096 | -0.096 | 0 | 0.14 | 0.38 | 0.08 | 0.01 | 0.07 | 0.03 |
| 5 | 106913484 | rs607976 | -0.306 | 0.136 | 0.515 | -0.303 | 0 | 0.064 | 0.064 | 0 | 0.5 | 0.65 | 0.33 | 0.11 | 0.49 | 0.18 |
| 5 | 106914027 | rs641406 | -0.372 | -0.111 | 0.552 | -0.452 | -0.003 | 0.024 | 0.024 | 0 | 0.51 | 0.6 | 0.33 | 0.04 | 0.49 | 0.19 |
| 5 | 106916625 | rs10061771 | 0.365 | 0.233 | 1.062 | -0.604 | 0.002 | 0.028 | 0.028 | 0 | 0.37 | 0.27 | 0.23 | 0.03 | 0.35 | 0.15 |
| 5 | 106918212 | rs872240 | -0.49 | 0.363 | 1.134 | -0.613 | 0.001 | 0.016 | 0.016 | 0 | 0.51 | 0.6 | 0.33 | 0.04 | 0.5 | 0.19 |
| 5 | 106922261 | rs9328006 | -0.601 | 0.378 | 1.137 | -0.58 | 0.017 | -0.082 | -0.082 | 0 | 0.16 | 0.49 | 0.12 | 0.04 | 0.14 | 0.06 |
| 5 | 106923569 | rs180791 | -0.189 | 0.973 | 1.038 | -0.165 | 0.09 | 0.284 | 0.284 | 0 | 0.93 | 0.79 | 0.77 | 0.56 | 0.87 | 0.8 |
| 5 | 106924829 | rs9328010 | -0.378 | 0.806 | 0.894 | -0.645 | 0 | 0.012 | 0.012 | 0 | 0.48 | 0.58 | 0.33 | 0.07 | 0.49 | 0.19 |
| 5 | 106925966 | rs3844311 | 0.44 | 0.868 | 0.594 | -0.532 | -0.016 | 0.09 | 0.09 | 0 | 0.35 | 0.14 | 0.22 | 0.03 | 0.36 | 0.15 |
| 5 | 106944011 | rs12519591 | -0.945 | 0.992 | 1.035 | -0.335 | 0.016 | -0.071 | -0.071 | 0 | 0.13 | 0.29 | 0.09 | 0.01 | 0.15 | 0.03 |
| 5 | 106944273 | rs26724 | -1.262 | 0.732 | 1.086 | -0.433 | 0.024 | -0.074 | -0.074 | 0.237 | 0.31 | 0.58 | 0.25 | 0.17 | 0.35 | 0.16 |
| 5 | 106951357 | rs17160255 | 1.418 | 0.054 | 0.269 | -0.496 | 0.019 | -0.132 | -0.132 | 0 | 0.24 | 0.01 | 0.21 | 0.38 | 0.19 | 0.47 |
| 5 | 106951925 | rs1017223 | -0.347 | 0.065 | 0.722 | -0.482 | 0.001 | -0.025 | -0.025 | 3.168 | 0.06 | 0.27 | 0.03 | 0 | 0.03 | 0.01 |
| 5 | 106955783 | rs35533 | -0.172 | -0.179 | 0.589 | -0.638 | -0.028 | 0.036 | 0.036 | 0 | 0.23 | 0.16 | 0.25 | 0.14 | 0.32 | 0.15 |
| 5 | 106956443 | rs4090594 | 1.493 | -0.02 | 0.367 | -1.125 | 0.014 | -0.001 | -0.001 | 0.338 | 0.46 | 0.12 | 0.41 | 0.4 | 0.43 | 0.59 |
| 5 | 106957027 | rs154191 | 0.736 | 0.226 | 0.395 | -1.005 | -0.008 | 0.018 | 0.018 | 0 | 0.18 | 0.12 | 0.19 | 0.13 | 0.2 | 0.13 |
| 5 | 106958958 | rs164691 | 0.377 | 0.223 | 0.244 | -1.034 | -0.048 | 0.017 | 0.017 | 0.072 | 0.19 | 0.03 | 0.18 | 0.13 | 0.2 | 0.14 |
| 5 | 106962349 | rs164840 | -0.058 | -0.173 | 0.327 | -1.068 | -0.007 | 0.055 | 0.055 | 0 | 0.71 | 0.61 | 0.61 | 0.55 | 0.76 | 0.68 |
| 5 | 106971568 | rs27550 | 0.825 | 0.075 | 0.205 | -0.955 | -0.009 | 0.015 | 0.015 | 0 | 0.16 | 0.07 | 0.17 | 0.13 | 0.18 | 0.13 |
| 5 | 106979252 | rs35419 | -0.097 | 0.126 | 0.311 | -0.99 | -0.005 | 0.003 | 0.003 | 0 | 0.23 | 0.19 | 0.25 | 0.13 | 0.33 | 0.15 |
| 5 | 106982560 | rs10074258 | 1.151 | 0.016 | 0.216 | -0.936 | 0.025 | -0.009 | -0.009 | 0 | 0.45 | 0.2 | 0.39 | 0.42 | 0.41 | 0.55 |
| 5 | 106984808 | rs17160324 | 1.657 | -0.118 | 0.045 | -0.905 | 0.028 | -0.118 | -0.118 | 0 | 0.24 | 0.01 | 0.2 | 0.33 | 0.16 | 0.45 |
| 5 | 106989475 | rs165263 | -0.375 | 0.549 | 0.753 | -0.785 | 0.084 | 0.268 | 0.268 | 0 | 0.12 | 0.31 | 0.32 | 0.55 | 0.16 | 0.22 |
| 5 | 106991031 | rs7732162 | 0.579 | 0.714 | 0.237 | -0.894 | 0.022 | -0.036 | -0.036 | 0 | 0.08 | 0.12 | 0.09 | 0.01 | 0.16 | 0.03 |
| 5 | 107000504 | rs35552 | -0.156 | 0.341 | 0.358 | -0.914 | 0.009 | 0.05 | 0.05 | 0 | 0.24 | 0.28 | 0.32 | 0.55 | 0.25 | 0.31 |
| 5 | 107004049 | rs13170272 | 1.069 | -0.119 | 0.221 | -0.915 | 0.028 | 0.013 | 0.013 | 0 | 0.43 | 0.1 | 0.41 | 0.32 | 0.39 | 0.53 |
| 5 | 107010201 | rs1490827 | 0.234 | 0.081 | 0.377 | -0.949 | 0.034 | 0.042 | 0.042 | 0 | 0.78 | 0.76 | 0.67 | 0.47 | 0.62 | 0.78 |
| 5 | 107011559 | rs6887589 | 0.036 | -0.003 | 0.607 | -1.044 | -0.036 | -0.079 | -0.079 | 0.204 | 0.34 | 0.5 | 0.23 | 0.13 | 0.22 | 0.22 |
| 5 | 107011958 | rs6596744 | 0.059 | -0.047 | 0.463 | -1.143 | 0.005 | 0.002 | 0.002 | 0 | 0.5 | 0.56 | 0.36 | 0.14 | 0.45 | 0.31 |
| 5 | 107017339 | rs17160352 | 1.159 | -0.105 | 0.268 | -1.219 | 0.028 | 0.015 | 0.015 | 0 | 0.23 | 0.14 | 0.1 | 0.01 | 0.14 | 0.1 |
| 5 | 107023656 | rs1304902 | 0.341 | 0.086 | 0.581 | -0.912 | -0.002 | 0.002 | 0.002 | 0 | 0.59 | 0.43 | 0.51 | 0.48 | 0.4 | 0.7 |
| 5 | 107024658 | rs10055714 | 2.067 | 0.249 | 0.085 | -0.737 | -0.01 | 0.015 | 0.015 | 0 | 0.16 | 0.11 | 0.14 | 0.03 | 0.22 | 0.09 |
| 5 | 107032339 | rs13160949 | 0.016 | 0.162 | 0.088 | -0.407 | 0.027 | 0.037 | 0.037 | 0 | 0.8 | 0.77 | 0.66 | 0.5 | 0.64 | 0.79 |
| 5 | 107032708 | rs7724655 | 0.871 | 0.129 | 0.037 | -0.423 | 0.073 | 0.047 | 0.047 | 0 | 0.73 | 0.63 | 0.55 | 0.38 | 0.54 | 0.66 |
| 5 | 107033576 | rs2194207 | 1.096 | 0.282 | -0.263 | -0.451 | 0.015 | -0.064 | -0.064 | 0.259 | 0.21 | 0.31 | 0.16 | 0.03 | 0.24 | 0.1 |
| 5 | 107038060 | rs1352726 | 0.785 | 0.235 | -0.316 | -0.387 | 0.002 | 0.008 | 0.008 | 0 | 0.06 | 0.09 | 0.11 | 0.12 | 0.09 | 0.13 |
| 5 | 107039550 | rs1490838 | 1.876 | 0.057 | -0.763 | -0.473 | 0.06 | -0.095 | -0.095 | 0.456 | 0.25 | 0.04 | 0.28 | 0.34 | 0.16 | 0.45 |
| 5 | 107046317 | rs6864800 | 0.65 | 0.637 | -0.396 | -0.431 | -0.001 | 0.001 | 0.001 | 0 | 0.53 | 0.46 | 0.55 | 0.51 | 0.5 | 0.69 |
| 5 | 107046588 | rs12656808 | 0.799 | 2.208 | -0.052 | 0.641 | 0.079 | 0.104 | 0.104 | 0 | 0.79 | 0.64 | 0.65 | 0.52 | 0.64 | 0.79 |
| 5 | 107050754 | rs12657057 | 0.915 | 2.432 | -0.768 | 0.236 | -0.089 | 0.093 | 0.093 | 0 | 0.17 | 0.01 | 0.13 | 0.01 | 0.25 | 0.1 |
| 5 | 107051942 | rs154167 | 0.514 | 2.771 | -0.644 | 0.552 | 0.045 | 0.002 | 0.002 | 0 | 0.86 | 0.81 | 0.82 | 0.86 | 0.68 | 0.91 |
| 5 | 107052000 | rs933066 | 0.427 | 1.851 | -0.824 | 0.697 | -0.009 | 0.006 | 0.006 | 0 | 0.18 | 0.09 | 0.14 | 0.1 | 0.24 | 0.11 |
| 5 | 107065898 | rs6894244 | -1.104 | 1.928 | -0.793 | 1.36 | 0 | 0.036 | 0.036 | 0.217 | 0.08 | 0.11 | 0.24 | 0.31 | 0.14 | 0.18 |
| 5 | 107066305 | rs154189 | 0.47 | 3.135 | -0.466 | 1.523 | 0.034 | 0.021 | 0.021 | 0 | 0.83 | 0.78 | 0.76 | 0.75 | 0.69 | 0.8 |
| 5 | 107066531 | rs154188 | -0.145 | 3.568 | -0.547 | 1.666 | -0.01 | -0.004 | -0.004 | 0.008 | 0.93 | 0.94 | 0.92 | 0.91 | 0.84 | 0.92 |
| 5 | 107072556 | rs1981953 | -0.011 | 3.634 | -0.619 | 1.598 | -0.021 | -0.021 | -0.021 | 0 | 0.09 | 0.05 | 0.16 | 0.16 | 0.15 | 0.12 |
| 5 | 107073493 | rs2411044 | 0.881 | 3.422 | -0.673 | 1.256 | -0.015 | 0.013 | 0.013 | 0 | 0.14 | 0.07 | 0.12 | 0.07 | 0.19 | 0.11 |
| 5 | 107073806 | rs35550 | 1.713 | 3.125 | -0.286 | 0.963 | -0.035 | 0.004 | 0.004 | 0.007 | 0.7 | 0.22 | 0.63 | 0.74 | 0.5 | 0.77 |
| 5 | 107081424 | rs6892045 | -0.53 | 2.735 | -0.186 | 1.058 | 0.022 | 0.127 | 0.127 | 0.107 | 0.07 | 0.13 | 0.24 | 0.32 | 0.13 | 0.2 |
| 5 | 107081478 | rs35521 | 0.785 | 1.809 | -1.027 | 0.487 | -0.002 | -0.041 | -0.041 | 0.529 | 0.27 | 0.16 | 0.27 | 0.41 | 0.23 | 0.19 |
| 5 | 107083050 | rs17160499 | 1.018 | 0.944 | -1.418 | -0.548 | 0.006 | -0.021 | -0.021 | 0 | 0.21 | 0.21 | 0.08 | 0.01 | 0.13 | 0.16 |
| 5 | 107083487 | rs1352724 | 1.272 | 1.317 | -1.616 | -0.114 | -0.061 | 0.08 | 0.08 | 4.038 | 0.24 | 0.01 | 0.13 | 0.07 | 0.22 | 0.09 |
| 5 | 107083644 | rs154212 | -0.489 | 1.69 | -0.741 | 0.476 | 0.016 | 0.009 | 0.009 | 0 | 0.17 | 0.2 | 0.26 | 0.25 | 0.32 | 0.21 |
| 5 | 107085864 | rs9328015 | 0.036 | 1.211 | -0.707 | 0.342 | 0.006 | 0.019 | 0.019 | 0.196 | 0.56 | 0.35 | 0.52 | 0.36 | 0.58 | 0.52 |
| 5 | 107086262 | rs1990918 | 0.183 | 1.063 | -0.75 | 0.377 | 0.003 | 0.016 | 0.016 | 0 | 0.57 | 0.24 | 0.51 | 0.43 | 0.57 | 0.52 |
| 5 | 107093148 | rs10052705 | -0.8 | 1.885 | -0.568 | 0.397 | 0.043 | 0.055 | 0.055 | 0 | 0.14 | 0.22 | 0.3 | 0.26 | 0.24 | 0.31 |
| 5 | 107102159 | rs35544 | -0.493 | 2.087 | -0.653 | 0.661 | -0.012 | -0.011 | -0.011 | 0 | 0.9 | 0.92 | 0.84 | 0.78 | 0.83 | 0.86 |
| 5 | 107110109 | rs249981 | -0.322 | 1.728 | -0.452 | 0.679 | -0.014 | -0.017 | -0.017 | 0.054 | 0.63 | 0.77 | 0.65 | 0.65 | 0.58 | 0.6 |
| 5 | 107110446 | rs13358604 | -0.689 | 1.597 | -0.331 | 0.652 | 0.05 | 0.078 | 0.078 | 0 | 0.13 | 0.23 | 0.29 | 0.26 | 0.21 | 0.27 |

| | | | | | | | | | | | | | | | | |
|---|---|---|---|---|---|---|---|---|---|---|---|---|---|---|---|---|
| 5 | 107110672 | rs249982 | -0.238 | 1.509 | -1.009 | 0.227 | -0.1 | 0.117 | 0.117 | 1.619 | 0.66 | 0.9 | 0.72 | 0.86 | 0.53 | 0.67 |
| 5 | 107112116 | rs17367725 | 0.769 | 1.084 | -1.38 | 0.074 | -0.05 | 0.087 | 0.087 | 0.182 | 0.33 | 0.07 | 0.28 | 0.18 | 0.36 | 0.19 |
| 5 | 107117501 | rs7708590 | 0.609 | 1.003 | -1.012 | 0.124 | -0.015 | 0.054 | 0.054 | 0.152 | 0.35 | 0.17 | 0.29 | 0.19 | 0.37 | 0.18 |
| 5 | 107118121 | rs9328019 | -0.085 | 0.925 | -0.538 | 0.283 | 0.009 | 0.032 | 0.032 | 0 | 0.17 | 0.24 | 0.23 | 0.29 | 0.2 | 0.2 |
| 5 | 107118140 | rs770182 | -1.183 | 0.954 | -0.408 | 0.264 | 0.006 | -0.039 | -0.039 | 0 | 0.72 | 0.59 | 0.72 | 0.81 | 0.69 | 0.62 |
| 5 | 107118722 | rs770180 | 0.556 | 0.637 | -0.519 | 0.402 | 0.01 | -0.074 | -0.074 | 0.028 | 0.19 | 0.06 | 0.2 | 0.33 | 0.12 | 0.23 |
| 5 | 107121419 | rs770172 | -0.678 | 0.798 | -0.394 | 0.285 | 0.002 | -0.002 | -0.002 | 0 | 0.37 | 0.39 | 0.44 | 0.62 | 0.32 | 0.44 |
| 5 | 107123947 | rs11242649 | -0.542 | 1.24 | -0.589 | 0.534 | -0.034 | -0.103 | -0.103 | 1.246 | 0.65 | 0.92 | 0.63 | 0.44 | 0.66 | 0.44 |
| 5 | 107125801 | rs1862310 | -0.435 | 1.155 | 0.043 | 0.413 | -0.011 | 0.049 | 0.049 | 1.056 | 0.22 | 0.61 | 0.27 | 0.28 | 0.2 | 0.16 |
| 5 | 107125858 | rs13159307 | 0.756 | 1.427 | 0.041 | 0.457 | -0.006 | 0.033 | 0.033 | 0 | 0.43 | 0.29 | 0.35 | 0.15 | 0.46 | 0.29 |
| 5 | 107126918 | rs770167 | -0.233 | 1.888 | 0.265 | 1.163 | -0.021 | -0.035 | -0.035 | 0 | 0.91 | 0.97 | 0.87 | 0.76 | 0.88 | 0.85 |
| 5 | 107129158 | rs10515378 | -0.366 | 1.49 | 1.034 | 0.684 | 0.064 | -0.021 | -0.021 | 4.073 | 0.1 | 0.38 | 0.08 | 0.02 | 0.1 | 0.11 |
| 5 | 107131544 | rs12518562 | -0.49 | 1.487 | 0.23 | 0.713 | 0.002 | 0.016 | 0.016 | 0 | 0.16 | 0.18 | 0.25 | 0.28 | 0.2 | 0.18 |
| 5 | 107137416 | rs1974789 | -1.134 | 1.402 | -0.32 | 0.717 | -0.001 | -0.001 | -0.001 | 0 | 0.29 | 0.7 | 0.34 | 0.31 | 0.33 | 0.3 |
| 5 | 107139388 | rs1363458 | 0.188 | 1.229 | -0.573 | -0.261 | -0.019 | 0.054 | 0.054 | 0 | 0.58 | 0.69 | 0.37 | 0.14 | 0.53 | 0.35 |
| 5 | 107139503 | rs10464043 | -0.58 | 1.22 | -0.744 | -0.551 | 0.004 | -0.021 | -0.021 | 0 | 0.41 | 0.51 | 0.3 | 0.34 | 0.42 | 0.36 |
| 5 | 107139663 | rs10515379 | 0.322 | 0.242 | -1.003 | -0.867 | -0.068 | 0.04 | 0.04 | 0.438 | 0.21 | 0.04 | 0.16 | 0.09 | 0.28 | 0.12 |
| 5 | 107140556 | rs3843492 | 1.014 | -0.267 | -0.855 | -1.37 | -0.035 | -0.133 | -0.133 | 2.297 | 0.36 | 0.47 | 0.2 | 0.03 | 0.26 | 0.17 |
| 5 | 107141733 | rs11742545 | -0.253 | -0.257 | -0.814 | -1.258 | -0.004 | 0.013 | 0.013 | 0.005 | 0.28 | 0.24 | 0.23 | 0.1 | 0.33 | 0.2 |
| 5 | 107146022 | rs4099188 | -0.379 | -0.347 | -0.774 | -1.085 | -0.015 | 0.028 | 0.028 | 0 | 0.62 | 0.68 | 0.41 | 0.15 | 0.58 | 0.36 |
| 5 | 107146674 | rs6888461 | -0.495 | -0.345 | -0.776 | -0.999 | -0.042 | 0.062 | 0.062 | 0 | 0.63 | 0.76 | 0.41 | 0.17 | 0.58 | 0.36 |
| 5 | 107156330 | rs4957544 | -0.976 | -0.412 | -0.879 | -1.172 | 0.004 | 0.067 | 0.067 | 0.002 | 0.37 | 0.76 | 0.32 | 0.14 | 0.39 | 0.2 |
| 5 | 107158158 | rs7700475 | 0.265 | -0.593 | -0.186 | -1.083 | 0.014 | -0.04 | -0.04 | 0 | 0.16 | 0.41 | 0.15 | 0.1 | 0.17 | 0.07 |
| 5 | 107164541 | rs10515381 | 0.361 | -0.267 | -0.38 | -0.256 | -0.002 | 0.016 | 0.016 | 0 | 0.32 | 0.36 | 0.25 | 0.56 | 0.25 | 0.43 |
| 5 | 107164703 | rs11955015 | 1.076 | -0.29 | -0.412 | 0.056 | -0.003 | 0.005 | 0.005 | 0 | 0.14 | 0.12 | 0.06 | 0.15 | 0.09 | 0.22 |
| 5 | 107165073 | rs6895254 | -1.014 | -0.211 | -0.554 | -0.133 | -0.05 | -0.053 | -0.053 | 0.055 | 0.85 | 0.96 | 0.57 | 0.68 | 0.72 | 0.73 |
| 5 | 107168049 | rs7708273 | 0.283 | -0.443 | -0.449 | -0.074 | 0.001 | 0.004 | 0.004 | 0 | 0.54 | 0.55 | 0.32 | 0.24 | 0.46 | 0.31 |
| 5 | 107168621 | rs11242652 | 1.535 | -0.315 | -0.43 | 0.062 | 0.004 | 0.037 | 0.037 | 0 | 0.16 | 0.19 | 0.17 | 0.4 | 0.15 | 0.21 |
| 5 | 107170088 | rs868548 | 1.603 | -0.303 | -0.446 | 0.082 | 0.004 | 0.033 | 0.033 | 0 | 0.16 | 0.18 | 0.17 | 0.39 | 0.15 | 0.21 |
| 5 | 107177381 | rs4957724 | -0.562 | -0.318 | -0.441 | 0.09 | 0.002 | 0.001 | 0.001 | 0 | 0.4 | 0.43 | 0.69 | 0.53 | 0.55 | 0.51 |
| 5 | 107190518 | rs6884480 | -1.094 | -0.376 | -0.16 | -0.01 | 0.02 | 0.003 | 0.003 | 0 | 0.56 | 0.76 | 0.77 | 0.57 | 0.64 | 0.7 |
| 5 | 107191287 | rs17160677 | 1.513 | -0.44 | -0.231 | -0.03 | 0.053 | 0.052 | 0.052 | 0 | 0.3 | 0.09 | 0.11 | 0.15 | 0.21 | 0.21 |
| 5 | 107192926 | rs7735555 | -0.282 | -0.487 | -0.157 | -0.074 | -0.001 | -0.001 | -0.001 | 0.231 | 0.37 | 0.35 | 0.62 | 0.46 | 0.46 | 0.4 |
| 5 | 107193946 | rs6882819 | 0.345 | -0.493 | -0.147 | -0.081 | 0.026 | -0.057 | -0.057 | 0 | 0.19 | 0.41 | 0.15 | 0.11 | 0.18 | 0.29 |
| 5 | 107195155 | rs12519254 | 1.579 | -0.522 | -0.702 | -0.076 | 0.052 | 0.047 | 0.047 | 0.001 | 0.3 | 0.09 | 0.11 | 0.15 | 0.21 | 0.21 |
| 5 | 107195824 | rs12522548 | 1.536 | -0.518 | -0.698 | -0.07 | 0.053 | 0.039 | 0.039 | 0 | 0.3 | 0.12 | 0.11 | 0.16 | 0.21 | 0.21 |
| 5 | 107196976 | rs1865015 | 0.285 | -0.532 | -0.37 | -0.103 | 0.02 | -0.06 | -0.06 | 0 | 0.2 | 0.54 | 0.17 | 0.11 | 0.18 | 0.29 |
| 5 | 107196998 | rs12189428 | 1.555 | -0.534 | -0.438 | -0.1 | 0.053 | 0.042 | 0.042 | 0 | 0.3 | 0.11 | 0.11 | 0.15 | 0.21 | 0.21 |
| 5 | 107202929 | rs6596759 | 0.142 | -0.412 | -0.317 | -0.044 | 0.015 | -0.065 | -0.065 | 0 | 0.2 | 0.57 | 0.17 | 0.08 | 0.18 | 0.29 |
| 5 | 107203628 | rs12152863 | 0.245 | -0.465 | -0.504 | -0.087 | -0.027 | -0.041 | -0.041 | 0.487 | 0.26 | 0.16 | 0.57 | 0.49 | 0.4 | 0.42 |
| 5 | 107205182 | rs7708343 | 1.209 | -0.293 | -0.367 | 0.138 | 0.003 | 0.103 | 0.103 | 0 | 0.1 | 0.14 | 0.14 | 0.32 | 0.11 | 0.17 |
| 5 | 107205967 | rs2900119 | -0.398 | -0.293 | -0.552 | 0.092 | -0.003 | -0.027 | -0.027 | 0.398 | 0.31 | 0.75 | 0.32 | 0.42 | 0.29 | 0.47 |
| 5 | 107209814 | rs8180457 | -2.161 | -0.08 | -0.521 | 0.158 | -0.007 | -0.008 | -0.008 | 0 | 0.87 | 0.89 | 0.88 | 0.73 | 0.85 | 0.92 |
| 5 | 107213361 | rs11748134 | 0.599 | -0.347 | -0.393 | -0.067 | -0.005 | -0.011 | -0.011 | 0 | 0.38 | 0.55 | 0.23 | 0.21 | 0.35 | 0.26 |
| 5 | 107216209 | rs1549924 | 0.843 | -0.539 | -0.412 | -0.049 | -0.007 | -0.031 | -0.031 | 0 | 0.14 | 0.17 | 0.07 | 0.04 | 0.08 | 0.21 |
| 5 | 107220280 | rs9784679 | 0.792 | -0.442 | -0.677 | 0.118 | -0.001 | -0.002 | -0.002 | 0 | 0.31 | 0.35 | 0.2 | 0.21 | 0.3 | 0.21 |
| 5 | 107222076 | rs10035487 | -1.678 | -0.461 | -0.84 | 0.34 | -0.011 | -0.013 | -0.013 | 0 | 0.87 | 0.86 | 0.87 | 0.73 | 0.86 | 0.9 |
| 5 | 107223376 | rs4957727 | -1.665 | -0.455 | -0.84 | 0.343 | -0.011 | -0.013 | -0.013 | 0 | 0.87 | 0.86 | 0.88 | 0.73 | 0.86 | 0.9 |
| 5 | 107231002 | rs13189604 | 0.309 | -0.293 | -0.835 | 0.001 | -0.052 | 0.009 | 0.009 | 2.709 | 0.44 | 0.11 | 0.5 | 0.33 | 0.49 | 0.42 |
| 5 | 107237203 | rs17160709 | 0.823 | -0.382 | -0.486 | -0.192 | -0.004 | -0.039 | -0.039 | 0 | 0.22 | 0.27 | 0.16 | 0.07 | 0.19 | 0.26 |
| 5 | 107241363 | rs998906 | 0.108 | 0.185 | -0.529 | 0.104 | 0.017 | 0.019 | 0.019 | 0 | 0.53 | 0.37 | 0.27 | 0.3 | 0.4 | 0.41 |
| 5 | 107243126 | rs12518124 | 0.798 | -0.199 | -1.772 | 0.821 | 0.019 | -0.021 | -0.021 | 1.269 | 0.22 | 0.09 | 0.1 | 0.22 | 0.18 | 0.14 |
| 5 | 107248608 | rs1433050 | -0.21 | 0.435 | -1.393 | 1.182 | 0.004 | -0.052 | -0.052 | 0 | 0.78 | 0.84 | 0.76 | 0.48 | 0.81 | 0.75 |
| 5 | 107263580 | rs11743737 | 0.304 | 0.626 | -1.252 | 0.702 | -0.012 | 0.02 | 0.02 | 0.841 | 0.57 | 0.33 | 0.39 | 0.36 | 0.51 | 0.49 |
| 5 | 107263596 | rs1821271 | -0.315 | 1.152 | -1.369 | 0.504 | -0.011 | 0.005 | 0.005 | 0 | 0.19 | 0.16 | 0.4 | 0.15 | 0.3 | 0.27 |
| 5 | 107263834 | rs1835979 | -0.331 | 0.494 | -1.494 | 0.349 | 0.009 | 0.071 | 0.071 | 0 | 0.33 | 0.15 | 0.19 | 0.1 | 0.31 | 0.32 |
| 5 | 107265051 | rs1835978 | 0.167 | 0.643 | -1.225 | 1.406 | -0.016 | -0.015 | -0.015 | 0 | 0.94 | 0.97 | 0.74 | 0.71 | 0.91 | 0.8 |
| 5 | 107265792 | rs919769 | 1.682 | 0.487 | -1.219 | 1.107 | 0.004 | -0.035 | -0.035 | 0 | 0.86 | 0.91 | 0.71 | 0.69 | 0.85 | 0.75 |
| 5 | 107265976 | rs10077567 | -0.741 | 0.555 | -0.63 | 0.692 | 0.002 | -0.049 | -0.049 | 0 | 0.13 | 0.26 | 0.1 | 0.04 | 0.12 | 0.16 |
| 5 | 107266840 | rs9328025 | -0.692 | 0.598 | -0.84 | 0.048 | -0.005 | -0.003 | -0.003 | 0 | 0.57 | 0.52 | 0.54 | 0.63 | 0.57 | 0.63 |
| 5 | 107270193 | rs13153403 | 0.608 | 0.483 | -0.972 | -0.019 | -0.027 | -0.024 | -0.024 | 0.717 | 0.4 | 0.18 | 0.41 | 0.59 | 0.4 | 0.44 |
| 5 | 107271418 | rs17437183 | -0.13 | 0.412 | -0.956 | -0.148 | -0.025 | 0.047 | 0.047 | 0 | 0.11 | 0.03 | 0.09 | 0.04 | 0.12 | 0.16 |
| 5 | 107271524 | rs13163255 | 0.958 | 0.391 | -0.773 | -0.164 | 0 | -0.023 | -0.023 | 0 | 0.37 | 0.56 | 0.21 | 0.06 | 0.34 | 0.17 |
| 5 | 107272715 | rs12657654 | 0.194 | 0.42 | -0.513 | 0.03 | -0.019 | 0.036 | 0.036 | 0.222 | 0.45 | 0.19 | 0.44 | 0.61 | 0.46 | 0.49 |
| 5 | 107276891 | rs6882731 | -0.434 | 0.401 | -0.462 | 0.144 | -0.001 | -0.003 | -0.003 | 0 | 0.92 | 0.94 | 0.73 | 0.7 | 0.9 | 0.82 |
| 5 | 107277826 | rs4957731 | -1.656 | 0.31 | -0.398 | -0.004 | -0.002 | -0.051 | -0.051 | 0 | 0.19 | 0.27 | 0.11 | 0.04 | 0.17 | 0.19 |
| 5 | 107282788 | rs7716454 | 1.02 | 0.242 | -0.091 | 0.097 | -0.003 | 0.012 | 0.012 | 0 | 0.78 | 0.7 | 0.66 | 0.66 | 0.78 | 0.67 |
| 5 | 107284514 | rs3925679 | 0.815 | 0.119 | 0.044 | 0.888 | -0.018 | 0.108 | 0.108 | 0.197 | 0.84 | 0.69 | 0.7 | 0.37 | 0.88 | 0.64 |
| 5 | 107285076 | rs1433057 | 0.377 | 0.087 | 0.129 | 0.728 | -0.044 | 0.032 | 0.032 | 0.003 | 0.43 | 0.15 | 0.41 | 0.26 | 0.45 | 0.31 |

| | | | | | | | | | | | | | | | | |
|---|---|---|---|---|---|---|---|---|---|---|---|---|---|---|---|---|
| 5 | 107288619 | rs10063386 | -0.547 | 0.068 | 0.19 | 0.667 | 0 | -0.002 | -0.002 | 3.748 | 0.56 | 0.45 | 0.5 | 0.62 | 0.55 | 0.6 |
| 5 | 107293784 | rs7701221 | -0.211 | 0.163 | 0.225 | 0.635 | -0.001 | -0.006 | -0.006 | 0 | 0.1 | 0.09 | 0.07 | 0.04 | 0.1 | 0.13 |
| 5 | 107296793 | rs17371964 | 1.269 | 0.318 | -0.131 | 0.606 | -0.061 | 0.074 | 0.074 | 0.002 | 0.27 | 0.02 | 0.18 | 0.06 | 0.23 | 0.19 |
| 5 | 107300655 | rs885624 | 0.35 | 0.286 | 0.305 | 0.852 | 0.009 | -0.004 | -0.004 | 0.23 | 0.37 | 0.35 | 0.24 | 0.4 | 0.29 | 0.39 |
| 5 | 107303576 | rs17165766 | 0.528 | 0.292 | 0.331 | 0.82 | 0.011 | -0.01 | -0.01 | 0 | 0.36 | 0.25 | 0.23 | 0.39 | 0.29 | 0.37 |
| 5 | 107304101 | rs17437979 | 0.292 | 0.353 | 0.67 | 0.83 | 0.004 | 0.018 | 0.018 | 0 | 0.1 | 0.11 | 0.2 | 0.2 | 0.15 | 0.08 |
| 5 | 107304523 | rs17165768 | 1.484 | 0.493 | 1.102 | 0.65 | -0.006 | -0.1 | -0.1 | 0.636 | 0.33 | 0.42 | 0.21 | 0.06 | 0.3 | 0.17 |
| 5 | 107305829 | rs6893313 | 1.508 | 0.493 | 1.07 | 0.65 | -0.003 | -0.071 | -0.071 | 0 | 0.33 | 0.37 | 0.2 | 0.06 | 0.3 | 0.17 |
| 5 | 107307404 | rs11242656 | -0.044 | 0.595 | 1.082 | 0.755 | -0.003 | 0.006 | 0.006 | 0 | 0.44 | 0.38 | 0.31 | 0.42 | 0.42 | 0.5 |
| 5 | 107312369 | rs1897803 | 0.663 | 0.483 | 1.394 | 0.605 | 0.001 | 0 | 0 | 0 | 0.34 | 0.31 | 0.22 | 0.06 | 0.28 | 0.22 |
| 5 | 107313403 | rs10900900 | 0.772 | 0.509 | 0.944 | 0.617 | 0.002 | 0.079 | 0.079 | 0.001 | 0.33 | 0.21 | 0.21 | 0.06 | 0.28 | 0.22 |
| 5 | 107316915 | rs172305 | -0.922 | 0.577 | 0.751 | 0.778 | 0 | 0.02 | 0.02 | 0 | 0.26 | 0.32 | 0.49 | 0.51 | 0.29 | 0.33 |
| 5 | 107318454 | rs288194 | -1.43 | 0.577 | 0.776 | 0.764 | 0.005 | 0.007 | 0.007 | 0 | 0.34 | 0.4 | 0.57 | 0.55 | 0.42 | 0.46 |
| 5 | 107318611 | rs288193 | 0.737 | 0.588 | 0.442 | 0.778 | -0.006 | -0.037 | -0.037 | 0.09 | 0.12 | 0.06 | 0.29 | 0.3 | 0.14 | 0.24 |
| 5 | 107320849 | rs10044914 | -1.058 | 0.575 | 0.679 | 1.104 | 0.006 | 0.015 | 0.015 | 0 | 0.23 | 0.3 | 0.29 | 0.37 | 0.27 | 0.23 |
| 5 | 107324595 | rs17438698 | 0.796 | 0.5 | 0.534 | 0.978 | -0.013 | 0.122 | 0.122 | 0.01 | 0.32 | 0.11 | 0.2 | 0.06 | 0.28 | 0.23 |
| 5 | 107325598 | rs10044064 | -1.113 | 0.501 | 0.623 | 0.989 | 0.009 | 0.024 | 0.024 | 0 | 0.24 | 0.31 | 0.29 | 0.37 | 0.27 | 0.23 |
| 5 | 107325759 | rs967489 | -0.895 | 0.349 | 0.191 | 1.093 | -0.023 | 0.008 | 0.008 | 0.006 | 0.6 | 0.83 | 0.53 | 0.63 | 0.57 | 0.6 |
| 5 | 107332887 | rs6880003 | -1.948 | 0 | 0.795 | 1.209 | -0.052 | 0.114 | 0.114 | 10.236 | 0.1 | 0.61 | 0.05 | 0.21 | 0.01 | 0.17 |
| 5 | 107346202 | rs6858950 | -0.058 | 0.442 | 0.334 | 1.493 | 0.012 | 0.102 | 0.102 | 0 | 0.08 | 0.09 | 0.27 | 0.34 | 0.09 | 0.13 |
| 5 | 107347834 | rs723396 | -0.225 | 0.388 | -0.148 | 0.977 | -0.043 | 0.043 | 0.043 | 0 | 0.43 | 0.18 | 0.61 | 0.58 | 0.47 | 0.39 |
| 5 | 107348180 | rs288183 | 0.495 | 0.499 | -0.045 | 0.974 | -0.008 | -0.026 | -0.026 | 0 | 0.12 | 0.09 | 0.12 | 0.35 | 0.17 | 0.38 |
| 5 | 107351517 | rs288178 | -0.681 | 0.421 | -0.115 | 0.857 | -0.006 | -0.023 | -0.023 | 0 | 0.23 | 0.2 | 0.3 | 0.56 | 0.32 | 0.47 |
| 5 | 107359523 | rs4957737 | 0.768 | 0.405 | 0.6 | 0.867 | -0.005 | 0.014 | 0.014 | 0 | 0.08 | 0.03 | 0.27 | 0.29 | 0.09 | 0.14 |
| 5 | 107364363 | rs288159 | -1.216 | 0.217 | 0.809 | 0.662 | -0.002 | 0.045 | 0.045 | 0 | 0.28 | 0.36 | 0.31 | 0.57 | 0.32 | 0.51 |
| 5 | 107374684 | rs12651724 | 0.609 | 0.28 | 0.745 | 0.782 | 0.015 | 0.038 | 0.038 | 0 | 0.07 | 0.08 | 0.18 | 0.17 | 0.14 | 0.06 |
| 5 | 107390303 | rs288147 | -0.429 | 0.203 | 0.496 | 0.636 | -0.003 | -0.057 | -0.057 | 0.167 | 0.2 | 0.08 | 0.13 | 0.39 | 0.23 | 0.47 |
| 5 | 107398866 | rs288140 | -1.585 | 0.11 | 1.074 | 0.432 | 0.007 | -0.139 | -0.139 | 4.482 | 0.7 | 0.53 | 0.77 | 0.94 | 0.71 | 0.91 |
| 5 | 107402938 | rs286751 | -1.161 | 0.097 | 1.282 | 0.457 | -0.026 | 0.032 | 0.032 | 0 | 0.64 | 0.73 | 0.62 | 0.77 | 0.57 | 0.85 |
| 5 | 107403454 | rs286749 | -0.082 | 0.058 | 1.296 | 0.454 | -0.016 | 0.097 | 0.097 | 0 | 0.18 | 0.36 | 0.13 | 0.4 | 0.18 | 0.45 |
| 5 | 107407291 | rs289232 | -0.963 | 0.078 | 0.768 | 0.454 | 0.006 | -0.012 | -0.012 | 0.208 | 0.63 | 0.44 | 0.59 | 0.77 | 0.57 | 0.85 |
| 5 | 107410205 | rs286794 | -0.305 | 0.197 | 0.754 | 0.581 | -0.005 | 0.052 | 0.052 | 0 | 0.49 | 0.64 | 0.66 | 0.54 | 0.48 | 0.42 |
| 5 | 107420876 | rs10067069 | 0.002 | 0.093 | 0.284 | 0.625 | 0.013 | 0.004 | 0.004 | 0.011 | 0.45 | 0.27 | 0.47 | 0.37 | 0.39 | 0.37 |
| 5 | 107429510 | rs286821 | -0.321 | 0.19 | 0.436 | 0.515 | -0.009 | 0.086 | 0.086 | 0.474 | 0.53 | 0.26 | 0.56 | 0.77 | 0.54 | 0.77 |
| 5 | 107437025 | rs10079990 | 0.611 | 0.075 | 0.272 | 0.627 | -0.041 | -0.072 | -0.072 | 0 | 0.37 | 0.06 | 0.42 | 0.37 | 0.35 | 0.35 |
| 5 | 107441332 | rs158185 | -1.241 | 0.108 | 0.903 | 0.404 | 0 | -0.098 | -0.098 | 0 | 0.65 | 0.58 | 0.75 | 0.93 | 0.68 | 0.88 |
| 5 | 107445351 | rs158177 | -0.804 | 0.118 | 0.906 | 0.409 | 0.002 | -0.028 | -0.028 | 0 | 0.61 | 0.35 | 0.74 | 0.93 | 0.68 | 0.87 |
| 5 | 107451983 | rs286811 | -1.459 | 0.056 | 0.97 | 0.377 | -0.005 | 0.021 | 0.021 | 0.014 | 0.68 | 0.7 | 0.77 | 0.93 | 0.71 | 0.91 |
| 5 | 107469542 | rs12655475 | 0.866 | 0.243 | 0.561 | 0.637 | -0.011 | -0.007 | -0.007 | 0 | 0.07 | 0.03 | 0.17 | 0.16 | 0.14 | 0.08 |
| 5 | 107475094 | rs10065182 | 0.419 | 0.118 | 0.51 | 0.617 | -0.013 | -0.041 | -0.041 | 0 | 0.4 | 0.12 | 0.44 | 0.37 | 0.36 | 0.35 |
| 5 | 107476501 | rs10058163 | 1.066 | -0.06 | -0.031 | 0.384 | 0.01 | 0.015 | 0.015 | 0 | 0.2 | 0.02 | 0.11 | 0.08 | 0.16 | 0.1 |
| 5 | 107485161 | rs12515105 | -0.372 | 0.015 | 0.155 | 0.464 | -0.059 | 0.013 | 0.013 | 0.041 | 0.46 | 0.12 | 0.61 | 0.52 | 0.5 | 0.44 |
| 5 | 107494277 | rs13174635 | 0.128 | -0.019 | 0.077 | 0.488 | -0.059 | -0.085 | -0.085 | 0 | 0.38 | 0.06 | 0.44 | 0.37 | 0.36 | 0.34 |
| 5 | 107495173 | rs34427 | 0.299 | 0.328 | 0.513 | 0.666 | 0.004 | 0.073 | 0.073 | 0 | 0.16 | 0.23 | 0.13 | 0.42 | 0.19 | 0.44 |
| 5 | 107500144 | rs34424 | 0.331 | 0.323 | 0.568 | 0.451 | 0.003 | 0.087 | 0.087 | 0 | 0.16 | 0.24 | 0.33 | 0.55 | 0.19 | 0.43 |
| 5 | 107504326 | rs10063866 | -0.557 | 0.383 | 0.875 | 0.237 | 0.003 | 0.059 | 0.059 | 0 | 0.51 | 0.39 | 0.64 | 0.76 | 0.48 | 0.56 |
| 5 | 107505130 | rs10036675 | -0.237 | 0.453 | 0.469 | 0.343 | 0.004 | 0.063 | 0.063 | 0 | 0.12 | 0.15 | 0.41 | 0.55 | 0.15 | 0.2 |
| 5 | 107519490 | rs158609 | 0.496 | 0.426 | 0.486 | 0.297 | -0.004 | 0.002 | 0.002 | 0 | 0.14 | 0.11 | 0.1 | 0.14 | 0.18 | 0.33 |
| 5 | 107525505 | rs286765 | 0.275 | 0.218 | 0.415 | 0.23 | 0.004 | 0.054 | 0.054 | 0 | 0.36 | 0.1 | 0.23 | 0.23 | 0.33 | 0.35 |
| 5 | 107541939 | rs2922410 | -0.123 | 0.061 | 0.862 | 0.149 | 0.018 | 0.033 | 0.033 | 0 | 0.39 | 0.26 | 0.25 | 0.23 | 0.33 | 0.35 |
| 5 | 107542118 | rs13162979 | 0.597 | 0.127 | 0.785 | 0.197 | -0.038 | -0.029 | -0.029 | 0 | 0.14 | 0.03 | 0.1 | 0.17 | 0.2 | 0.33 |
| 5 | 107557861 | rs13187816 | -0.734 | 0.209 | 1.43 | 0.38 | 0.003 | 0.001 | 0.001 | 0 | 0.25 | 0.27 | 0.28 | 0.29 | 0.34 | 0.41 |
| 5 | 107560101 | rs2044710 | -0.382 | 0.169 | 1.442 | 0.355 | -0.004 | 0.014 | 0.014 | 0 | 0.45 | 0.32 | 0.43 | 0.36 | 0.47 | 0.45 |
| 5 | 107578315 | rs10076086 | -1.092 | 0.046 | 0.998 | 0.084 | -0.002 | 0.139 | 0.139 | 0 | 0.47 | 0.71 | 0.63 | 0.79 | 0.5 | 0.65 |
| 5 | 107592304 | rs10477419 | -0.242 | 0.082 | 0.802 | 0.171 | 0.01 | 0.019 | 0.019 | 0 | 0.34 | 0.16 | 0.2 | 0.26 | 0.34 | 0.45 |
| 5 | 107593829 | rs11240960 | 1.289 | 0.079 | 0.776 | 0.144 | -0.027 | 0.015 | 0.015 | 0 | 0.16 | 0.03 | 0.13 | 0.14 | 0.18 | 0.24 |
| 5 | 107603298 | rs11742777 | -0.269 | 0.153 | 0.79 | 0.158 | 0.009 | 0.02 | 0.02 | 0 | 0.34 | 0.16 | 0.2 | 0.26 | 0.35 | 0.45 |
| 5 | 107607138 | rs13153782 | 0.522 | 0.188 | 0.894 | 0.162 | 0.001 | 0 | 0 | 0 | 0.15 | 0.18 | 0.11 | 0.17 | 0.2 | 0.34 |
| 5 | 107616085 | rs4541643 | -0.581 | 0.203 | 0.726 | 0.266 | -0.014 | 0.02 | 0.02 | 0 | 0.44 | 0.22 | 0.42 | 0.36 | 0.47 | 0.44 |
| 5 | 107649113 | rs17386875 | 0.204 | 0.513 | 0.389 | 0.309 | -0.056 | 0.018 | 0.018 | 0 | 0.21 | 0.04 | 0.26 | 0.12 | 0.26 | 0.16 |
| 5 | 107698701 | rs10515388 | 0.514 | 0.964 | 0.674 | 0.884 | -0.022 | -0.019 | -0.019 | 5.464 | 0.07 | 0.03 | 0.17 | 0.12 | 0.14 | 0.08 |
| 5 | 107721366 | rs11240962 | 0.412 | 0.753 | 0.404 | 0.42 | 0 | -0.023 | -0.023 | 0 | 0.19 | 0.25 | 0.16 | 0.06 | 0.21 | 0.1 |
| 5 | 107724017 | rs10035966 | 0.56 | 0.746 | 0.408 | 0.417 | 0 | -0.02 | -0.02 | 0.173 | 0.18 | 0.23 | 0.16 | 0.06 | 0.21 | 0.1 |
| 5 | 107731684 | rs4634342 | 0.465 | 0.726 | 0.342 | 0.408 | -0.016 | 0.02 | 0.02 | 0 | 0.7 | 0.7 | 0.76 | 0.81 | 0.64 | 0.71 |
| 5 | 107732317 | rs1122826 | 0.613 | 0.689 | 0.258 | 0.297 | 0.004 | -0.026 | -0.026 | 0 | 0.18 | 0.22 | 0.16 | 0.06 | 0.21 | 0.1 |
| 5 | 107742569 | rs10455007 | 0.926 | 0.607 | -0.073 | 0.488 | -0.02 | -0.029 | -0.029 | 0 | 0.63 | 0.16 | 0.51 | 0.41 | 0.64 | 0.6 |
| 5 | 107768782 | rs12187433 | 1.008 | 0.896 | -0.409 | -0.019 | -0.046 | 0.056 | 0.056 | 0 | 0.15 | 0.07 | 0.15 | 0.06 | 0.21 | 0.1 |
| 5 | 107781571 | rs918399 | -0.94 | 0.944 | -0.461 | 0.179 | -0.028 | -0.017 | -0.017 | 1.701 | 0.9 | 0.97 | 0.92 | 0.89 | 0.86 | 0.8 |
| 5 | 107794540 | rs748975 | 0.803 | 1.009 | -0.584 | 0.14 | -0.012 | 0.009 | 0.009 | 0 | 0.63 | 0.28 | 0.57 | 0.7 | 0.54 | 0.65 |

| | | | | | | | | | | | | | | | | |
|---|---|---|---|---|---|---|---|---|---|---|---|---|---|---|---|---|
| 5 | 107804983 | rs2080854 | -0.36 | 1.094 | -1.067 | 0.607 | -0.009 | -0.003 | -0.003 | 0 | 0.89 | 0.9 | 0.92 | 0.89 | 0.86 | 0.8 |
| 5 | 107808711 | rs2112305 | 0.467 | 0.914 | -1.203 | 0.65 | -0.057 | -0.052 | -0.052 | 0.016 | 0.78 | 0.91 | 0.72 | 0.7 | 0.68 | 0.59 |
| 5 | 107820674 | rs10066221 | 0.572 | 0.475 | -1.724 | 0.441 | 0.035 | 0.021 | 0.021 | 0 | 0.45 | 0.12 | 0.34 | 0.37 | 0.38 | 0.52 |
| 5 | 107820761 | rs6884608 | 0.36 | 0.434 | -0.814 | 0.396 | 0.02 | -0.023 | -0.023 | 0 | 0.73 | 0.63 | 0.65 | 0.8 | 0.65 | 0.74 |
| 5 | 107821228 | rs6889461 | 0.58 | 0.706 | -0.823 | 0.44 | 0.069 | 0.056 | 0.056 | 0 | 0.09 | 0.16 | 0.2 | 0.17 | 0.17 | 0.07 |
| 5 | 107842404 | rs10068143 | 0.883 | 0.368 | -1.037 | 0.292 | 0.04 | 0.022 | 0.022 | 0 | 0.45 | 0.15 | 0.34 | 0.37 | 0.38 | 0.51 |
| 5 | 107847972 | rs9326729 | 1.07 | 0.172 | -1.479 | 0.252 | 0.032 | -0.019 | -0.019 | 0.198 | 0.27 | 0.08 | 0.24 | 0.21 | 0.23 | 0.29 |
| 5 | 107853411 | rs4128352 | 0.837 | 0.732 | -1.038 | 0.37 | 0.025 | 0.018 | 0.018 | 0 | 0.08 | 0.11 | 0.18 | 0.16 | 0.16 | 0.07 |
| 5 | 107853869 | rs6861718 | 1.358 | 0.504 | -0.878 | 0.026 | -0.024 | 0.046 | 0.046 | 0.25 | 0.24 | 0.18 | 0.18 | 0.05 | 0.25 | 0.12 |
| 5 | 107866284 | rs6594316 | 0.581 | 0.222 | -1.273 | 0.143 | 0.061 | 0.061 | 0.061 | 0 | 0.48 | 0.1 | 0.35 | 0.37 | 0.39 | 0.51 |
| 5 | 107867558 | rs7446237 | -1.12 | 0.052 | -0.785 | 0.143 | -0.002 | 0.024 | 0.024 | 0 | 0.88 | 0.87 | 0.76 | 0.59 | 0.81 | 0.7 |
| 5 | 107874127 | rs11743896 | 0.745 | 0.08 | -0.416 | -0.226 | 0.005 | -0.121 | -0.121 | 0 | 0.3 | 0.56 | 0.22 | 0.05 | 0.26 | 0.12 |
| 5 | 107877657 | rs7707794 | 1.087 | 0.08 | -0.192 | -0.226 | 0.027 | -0.145 | -0.145 | 0 | 0.27 | 0.46 | 0.2 | 0.05 | 0.26 | 0.12 |
| 5 | 107881274 | rs7728461 | -0.884 | 0.271 | -0.549 | 0.524 | -0.004 | -0.005 | -0.005 | 0 | 0.88 | 0.86 | 0.85 | 0.74 | 0.8 | 0.71 |
| 5 | 107883191 | rs6594319 | -0.043 | 0.203 | -0.619 | 0.449 | -0.01 | 0.029 | 0.029 | 0.467 | 0.59 | 0.29 | 0.64 | 0.68 | 0.57 | 0.6 |
| 5 | 107896807 | rs6887376 | 0.059 | -0.167 | -1.101 | 0.717 | 0 | -0.019 | -0.019 | 0.008 | 0.64 | 0.85 | 0.53 | 0.33 | 0.64 | 0.48 |
| 5 | 107899940 | rs7447161 | 0.07 | -0.216 | -1.173 | 0.41 | -0.022 | -0.079 | -0.079 | 0 | 0.66 | 0.93 | 0.58 | 0.58 | 0.67 | 0.57 |
| 5 | 107900097 | rs7444217 | 0.119 | -0.227 | -1.036 | 0.424 | 0.007 | -0.036 | -0.036 | 0 | 0.65 | 0.87 | 0.57 | 0.58 | 0.67 | 0.57 |
| 5 | 107901554 | rs11749751 | -0.904 | -0.541 | -1.386 | 0.205 | -0.007 | -0.125 | -0.125 | 0 | 0.73 | 0.99 | 0.68 | 0.6 | 0.81 | 0.67 |
| 5 | 107915736 | rs10447248 | -0.332 | -0.459 | -1.227 | -0.041 | 0.006 | 0.002 | 0.002 | 1.122 | 0.64 | 0.91 | 0.56 | 0.31 | 0.68 | 0.38 |
| 5 | 107916909 | rs10065061 | 0.416 | -0.304 | -1.11 | -0.162 | -0.019 | 0.118 | 0.118 | 1.835 | 0.55 | 0.76 | 0.37 | 0.15 | 0.49 | 0.3 |
| 5 | 107921156 | rs12519202 | 1.459 | -0.755 | -1.503 | -0.567 | 0.002 | 0.018 | 0.018 | 0 | 0.14 | 0.03 | 0.09 | 0.07 | 0.14 | 0.16 |
| 5 | 107921232 | rs6594321 | -0.753 | -0.397 | -1.439 | -0.491 | -0.022 | 0.005 | 0.005 | 1.104 | 0.64 | 0.9 | 0.54 | 0.31 | 0.62 | 0.37 |
| 5 | 107923624 | rs7720580 | 0.302 | -0.841 | -0.606 | -0.654 | -0.035 | -0.084 | -0.084 | 3.562 | 0.3 | 0.45 | 0.18 | 0.07 | 0.25 | 0.17 |
| 5 | 107924442 | rs10477424 | 0.854 | -0.822 | -1.027 | -0.645 | 0.017 | 0.045 | 0.045 | 0 | 0.26 | 0.2 | 0.16 | 0.07 | 0.24 | 0.17 |
| 5 | 107928859 | rs6898473 | 2.066 | -0.881 | -1.338 | -0.713 | -0.072 | 0.078 | 0.078 | 1.246 | 0.21 | 0.04 | 0.16 | 0.07 | 0.24 | 0.12 |
| 5 | 107930254 | rs10044238 | -0.195 | -0.825 | -1.046 | -0.434 | -0.059 | 0.03 | 0.03 | 1.691 | 0.6 | 0.91 | 0.53 | 0.31 | 0.58 | 0.35 |
| 5 | 107932100 | rs7719547 | -1.015 | -0.845 | -0.581 | -0.53 | -0.04 | -0.066 | -0.066 | 0 | 0.11 | 0.58 | 0.08 | 0.01 | 0.04 | 0 |
| 5 | 107936020 | rs12517224 | -0.128 | -0.663 | -1.27 | -0.382 | -0.064 | 0.063 | 0.063 | 1.555 | 0.6 | 0.91 | 0.53 | 0.26 | 0.58 | 0.35 |
| 5 | 107937785 | rs7718948 | -0.043 | -1.27 | -0.781 | -0.433 | -0.004 | -0.001 | -0.001 | 0 | 0.58 | 0.61 | 0.52 | 0.57 | 0.53 | 0.44 |
| 5 | 107939906 | rs10053474 | 0.254 | -1.147 | -0.642 | -0.354 | 0.008 | 0.009 | 0.009 | 0 | 0.19 | 0.2 | 0.13 | 0.09 | 0.15 | 0.15 |
| 5 | 107951067 | rs12652136 | -0.704 | -0.79 | -0.296 | -0.144 | 0.01 | 0.048 | 0.048 | 0 | 0.8 | 0.72 | 0.64 | 0.58 | 0.75 | 0.6 |
| 5 | 107951342 | rs10214102 | -0.201 | -0.562 | -0.506 | -0.221 | -0.002 | 0.003 | 0.003 | 0 | 0.08 | 0.17 | 0.07 | 0.09 | 0.06 | 0.13 |
| 5 | 107954485 | rs6594323 | 0.384 | -0.442 | -0.508 | -0.49 | 0.002 | -0.022 | -0.022 | 0 | 0.28 | 0.22 | 0.17 | 0.02 | 0.26 | 0.17 |
| 5 | 107963472 | rs12719094 | 0.577 | -0.256 | -0.667 | 0.023 | -0.001 | -0.01 | -0.01 | 0 | 0.45 | 0.31 | 0.44 | 0.46 | 0.46 | 0.31 |
| 5 | 107979579 | rs4957775 | 0.567 | 0.062 | -0.354 | 0.212 | 0 | -0.001 | -0.001 | 0 | 0.1 | 0.09 | 0.2 | 0.11 | 0.14 | 0.05 |
| 5 | 107981747 | rs12656502 | 0.151 | -0.241 | -0.136 | -0.053 | 0.003 | -0.027 | -0.027 | 0.656 | 0.28 | 0.46 | 0.16 | 0.24 | 0.24 | 0.2 |
| 5 | 107985830 | rs10069339 | 0.83 | -0.259 | -0.195 | 0.382 | 0.011 | 0.187 | 0.187 | 0 | 0.11 | 0.17 | 0.23 | 0.41 | 0.15 | 0.13 |
| 5 | 107986411 | rs6898488 | -0.011 | -0.212 | -0.209 | 0.354 | 0.001 | -0.02 | -0.02 | 0 | 0.31 | 0.58 | 0.21 | 0.25 | 0.27 | 0.21 |
| 5 | 107987151 | rs7703870 | -1.215 | -0.368 | -0.872 | -0.118 | 0.016 | 0.111 | 0.111 | 0 | 0.49 | 0.87 | 0.46 | 0.66 | 0.42 | 0.34 |
| 5 | 107995317 | rs13187102 | 0.243 | -0.223 | -0.334 | -0.711 | -0.003 | 0.012 | 0.012 | 0 | 0.27 | 0.21 | 0.17 | 0.17 | 0.27 | 0.2 |
| 5 | 107999505 | rs1355079 | -1.556 | -0.275 | -1.014 | -1.33 | -0.027 | 0.155 | 0.155 | 0 | 0.71 | 0.91 | 0.87 | 0.98 | 0.75 | 0.84 |
| 5 | 108004312 | rs2963051 | 0.903 | -0.119 | -0.539 | -0.893 | 0 | -0.029 | -0.029 | 2.597 | 0.25 | 0.21 | 0.46 | 0.49 | 0.31 | 0.49 |
| 5 | 108008923 | rs2940516 | 0.755 | -0.081 | -0.501 | -0.882 | -0.02 | 0.023 | 0.023 | 0.119 | 0.26 | 0.17 | 0.48 | 0.76 | 0.35 | 0.55 |
| 5 | 108013210 | rs1858068 | 0.775 | -0.262 | 0.003 | -0.843 | 0.003 | -0.026 | -0.026 | 0 | 0.11 | 0.17 | 0.19 | 0.06 | 0.13 | 0.1 |
| 5 | 108013371 | rs13361304 | -1.116 | -0.287 | 0.086 | -0.837 | 0 | 0.068 | 0.068 | 0.173 | 0.2 | 0.6 | 0.26 | 0.32 | 0.19 | 0.16 |
| 5 | 108015427 | rs9968588 | 0.447 | -0.437 | -0.063 | -1.041 | 0.012 | 0.018 | 0.018 | 2.47 | 0.5 | 0.62 | 0.27 | 0.43 | 0.42 | 0.24 |
| 5 | 108016736 | rs12657495 | 1.595 | -0.731 | -0.221 | -1.184 | 0.004 | 0.005 | 0.005 | 0 | 0.19 | 0.16 | 0.07 | 0.15 | 0.11 | 0.05 |
| 5 | 108016782 | rs7737443 | -0.21 | -0.425 | -0.157 | -1.07 | -0.017 | -0.02 | -0.02 | 0 | 0.6 | 0.72 | 0.43 | 0.48 | 0.52 | 0.31 |
| 5 | 108018395 | rs11952230 | 0.069 | -0.251 | -0.277 | -0.997 | -0.001 | 0 | 0 | 0 | 0.56 | 0.47 | 0.42 | 0.5 | 0.52 | 0.31 |
| 5 | 108024159 | rs1876625 | 0.593 | 0.236 | -0.176 | -2.011 | 0.027 | -0.041 | -0.041 | 0 | 0.1 | 0.18 | 0.09 | 0.04 | 0.11 | 0.07 |
| 5 | 108029900 | rs10050689 | -0.289 | 0.041 | -0.444 | -2.385 | 0.003 | -0.035 | -0.035 | 0.169 | 0.52 | 0.52 | 0.37 | 0.12 | 0.55 | 0.41 |
| 5 | 108031617 | rs17391678 | 0.935 | 0.127 | -0.503 | -2.447 | -0.023 | 0.043 | 0.043 | 0.019 | 0.15 | 0.08 | 0.13 | 0.06 | 0.18 | 0.21 |
| 5 | 108031851 | rs7714022 | 0.026 | -0.32 | -0.195 | -2.085 | 0.004 | -0.035 | -0.035 | 0 | 0.51 | 0.51 | 0.37 | 0.12 | 0.54 | 0.41 |
| 5 | 108034255 | rs2048212 | -0.213 | -0.881 | -0.178 | -1.44 | 0.026 | -0.006 | -0.006 | 0 | 0.7 | 0.8 | 0.85 | 0.74 | 0.79 | 0.78 |
| 5 | 108034947 | rs9326745 | 0.348 | -0.686 | -0.295 | -1.285 | -0.012 | -0.011 | -0.011 | 0 | 0.58 | 0.51 | 0.76 | 0.69 | 0.68 | 0.71 |
| 5 | 108041607 | rs6859750 | -0.459 | -1.061 | -0.056 | -2.099 | -0.032 | -0.09 | -0.09 | 0 | 0.25 | 0.55 | 0.13 | 0.06 | 0.15 | 0.1 |
| 5 | 108045913 | rs7732428 | -0.316 | -1 | -0.421 | -2.263 | -0.001 | -0.004 | -0.004 | 0 | 0.14 | 0.18 | 0.09 | 0.09 | 0.11 | 0.22 |
| 5 | 108046359 | rs7718626 | -0.316 | -1 | -0.421 | -2.263 | -0.001 | -0.004 | -0.004 | 0 | 0.14 | 0.18 | 0.09 | 0.09 | 0.11 | 0.22 |
| 5 | 108048835 | rs1472916 | -1.338 | -0.806 | -1.199 | -3.008 | -0.097 | 0.156 | 0.156 | 0 | 0.64 | 0.92 | 0.36 | 0.16 | 0.54 | 0.45 |
| 5 | 108056848 | rs17161466 | 0.495 | -0.76 | -0.33 | -2.942 | 0.027 | -0.021 | -0.021 | 0 | 0.09 | 0.21 | 0.09 | 0.06 | 0.09 | 0.06 |
| 5 | 108060442 | rs11741855 | -0.562 | -1.387 | -0.247 | -3.348 | 0.007 | 0.009 | 0.009 | 3.183 | 0.53 | 0.56 | 0.27 | 0.07 | 0.42 | 0.22 |
| 5 | 108061551 | rs1423456 | -0.176 | -1.615 | -0.173 | -3.525 | -0.056 | -0.14 | -0.14 | 0 | 0.31 | 0.45 | 0.16 | 0.07 | 0.19 | 0.12 |
| 5 | 108061789 | rs4957789 | -1.642 | -1.55 | -0.045 | -3.487 | -0.03 | 0.055 | 0.055 | 0.992 | 0.66 | 0.73 | 0.35 | 0.18 | 0.54 | 0.44 |
| 5 | 108066680 | rs6594334 | -1.013 | -1.56 | 0.048 | -3.453 | 0.087 | 0.224 | 0.224 | 0.608 | 0.46 | 0.78 | 0.25 | 0.18 | 0.31 | 0.34 |
| 5 | 108066842 | rs7711892 | -0.279 | -1.39 | 0.243 | -3.355 | 0.032 | -0.057 | -0.057 | 0 | 0.14 | 0.42 | 0.12 | 0.07 | 0.13 | 0.07 |
| 5 | 108071916 | rs13169624 | 1.124 | -1.201 | -0.201 | -3.339 | -0.07 | 0.123 | 0.123 | 0 | 0.2 | 0.02 | 0.1 | 0 | 0.23 | 0.1 |
| 5 | 108085155 | rs6871143 | -1.41 | -1.235 | 0.039 | -3.35 | -0.044 | 0.079 | 0.079 | 0.261 | 0.65 | 0.76 | 0.35 | 0.19 | 0.52 | 0.44 |
| 5 | 108087810 | rs3805600 | 0.119 | -1.896 | 0.328 | -3.326 | -0.064 | -0.118 | -0.118 | 1.355 | 0.28 | 0.43 | 0.15 | 0.08 | 0.15 | 0.11 |

| | | | | | | | | | | | | | | | | |
|---|---|---|---|---|---|---|---|---|---|---|---|---|---|---|---|---|
| 5 | 108096014 | rs6882107 | -1.433 | -1.496 | 0.137 | -3.206 | -0.048 | 0.085 | 0.085 | 0.407 | 0.64 | 0.76 | 0.34 | 0.2 | 0.48 | 0.43 |
| 5 | 108099101 | rs7700630 | -1.7 | -1.553 | 0.085 | -3.212 | -0.123 | 0.117 | 0.117 | 2.574 | 0.67 | 0.89 | 0.35 | 0.18 | 0.49 | 0.44 |
| 5 | 108102347 | rs13184924 | 0.884 | -1.585 | 0.015 | -3.258 | -0.038 | 0.137 | 0.137 | 2.169 | 0.21 | 0.02 | 0.09 | 0 | 0.2 | 0.09 |
| 5 | 108107752 | rs10477933 | -0.474 | -1.902 | 0.823 | -2.743 | -0.045 | -0.112 | -0.112 | 0 | 0.35 | 0.43 | 0.2 | 0.09 | 0.23 | 0.2 |
| 5 | 108133967 | rs2229085 | 0.032 | -1.998 | 0.891 | -2.916 | -0.012 | -0.046 | -0.046 | 0 | 0.29 | 0.3 | 0.17 | 0.06 | 0.17 | 0.16 |
| 5 | 108139943 | rs10038669 | -1.087 | -2.098 | 0.87 | -3.032 | 0.064 | 0.102 | 0.102 | 2.439 | 0.46 | 0.65 | 0.27 | 0.16 | 0.29 | 0.35 |
| 5 | 108166815 | rs7710347 | -0.176 | -1.881 | 0.962 | -2.857 | 0.046 | 0.057 | 0.057 | 0.104 | 0.07 | 0.28 | 0.05 | 0.11 | 0.04 | 0.14 |
| 5 | 108168024 | rs4616948 | -0.638 | -1.887 | 0.65 | -2.863 | 0 | -0.058 | -0.058 | 0 | 0.17 | 0.36 | 0.1 | 0.11 | 0.12 | 0.19 |
| 5 | 108170607 | rs13181901 | 1.03 | -1.857 | 0.323 | -2.844 | -0.009 | 0.05 | 0.05 | 5.83 | 0.1 | 0.01 | 0.05 | 0 | 0.08 | 0.06 |
| 5 | 108185250 | rs10052531 | -0.092 | -1.89 | 0.332 | -2.873 | 0.012 | -0.013 | -0.013 | 0 | 0.34 | 0.3 | 0.19 | 0.06 | 0.22 | 0.17 |
| 5 | 108218263 | rs4560594 | -0.005 | -1.818 | 0.142 | -2.826 | 0.002 | -0.033 | -0.033 | 0 | 0.33 | 0.3 | 0.18 | 0.06 | 0.22 | 0.17 |
| 5 | 108221725 | rs12109934 | -0.274 | -1.241 | -0.485 | -2.462 | 0.007 | 0.005 | 0.005 | 0.079 | 0.08 | 0.19 | 0.05 | 0.11 | 0.05 | 0.14 |
| 5 | 108229760 | rs10079165 | -0.744 | -0.944 | -1.231 | -2.237 | 0.032 | 0.213 | 0.213 | 3.749 | 0.56 | 0.88 | 0.31 | 0.18 | 0.35 | 0.37 |
| 5 | 108243615 | rs4604227 | -0.475 | -0.795 | -1.009 | -2.144 | 0.089 | 0.199 | 0.199 | 1.603 | 0.5 | 0.73 | 0.29 | 0.16 | 0.34 | 0.36 |
| 5 | 108262478 | rs11959681 | -0.093 | -0.795 | -0.865 | -2.073 | 0.012 | 0.011 | 0.011 | 0.006 | 0.2 | 0.16 | 0.09 | 0 | 0.16 | 0.06 |
| 5 | 108262592 | rs11953198 | -0.558 | -1.044 | 0.065 | -1.89 | -0.024 | -0.055 | -0.055 | 0.111 | 0.23 | 0.39 | 0.12 | 0.04 | 0.18 | 0.06 |
| 5 | 108295480 | rs7734092 | -1.586 | -0.878 | 0.111 | -1.877 | 0.032 | 0.113 | 0.113 | 1.102 | 0.07 | 0.29 | 0.06 | 0.12 | 0.09 | 0.15 |
| 5 | 108299744 | rs7735749 | -1.17 | -0.792 | -0.328 | -2.762 | -0.004 | -0.036 | -0.036 | 5.034 | 0.45 | 0.5 | 0.29 | 0.12 | 0.42 | 0.28 |
| 5 | 108304246 | rs11948270 | -0.358 | -0.747 | -0.309 | -2.689 | -0.02 | -0.033 | -0.033 | 0 | 0.27 | 0.41 | 0.15 | 0.12 | 0.24 | 0.2 |
| 5 | 108310017 | rs13358819 | 1.049 | -0.455 | -0.699 | -2.818 | 0.002 | -0.014 | -0.014 | 0 | 0.05 | 0.04 | 0.02 | 0.01 | 0.05 | 0.1 |
| 5 | 108361566 | rs11958509 | -1.958 | -0.664 | -0.903 | -3.241 | -0.002 | -0.002 | -0.002 | 0 | 0.65 | 0.66 | 0.39 | 0.2 | 0.58 | 0.5 |
| 5 | 108371919 | rs4957795 | -1.239 | -1.386 | -1.433 | -3.543 | 0.015 | 0.216 | 0.216 | 0.101 | 0.52 | 0.84 | 0.33 | 0.15 | 0.39 | 0.33 |
| 5 | 108380419 | rs919771 | 1.503 | -1.021 | -1.193 | -3.345 | -0.005 | 0.077 | 0.077 | 0 | 0.16 | 0.06 | 0.09 | 0 | 0.16 | 0.06 |
| 5 | 108380950 | rs892009 | -1.599 | -1.114 | -1.159 | -3.404 | -0.004 | 0.017 | 0.017 | 0.654 | 0.59 | 0.64 | 0.37 | 0.21 | 0.54 | 0.47 |
| 5 | 108386660 | rs17534396 | 0.616 | -0.729 | -1.329 | -3.106 | -0.002 | 0.043 | 0.043 | 0 | 0.19 | 0.11 | 0.12 | 0.09 | 0.22 | 0.2 |
| 5 | 108389128 | rs11951976 | -1.238 | -1.05 | -1.288 | -3.321 | -0.005 | 0.071 | 0.071 | 0 | 0.43 | 0.29 | 0.24 | 0.08 | 0.43 | 0.3 |
| 5 | 108390471 | rs10062069 | -0.488 | -1.047 | -1.144 | -3.086 | -0.001 | -0.002 | -0.002 | 2.752 | 0.13 | 0.1 | 0.08 | 0.09 | 0.13 | 0.22 |
| 5 | 108398342 | rs1896582 | -0.726 | -1.054 | -1.169 | -3.109 | 0.004 | 0.018 | 0.018 | 0 | 0.18 | 0.11 | 0.1 | 0.09 | 0.19 | 0.23 |
| 5 | 108408287 | rs17450131 | 0.968 | -0.996 | -1.222 | -3.098 | 0.016 | 0.04 | 0.04 | 0 | 0.18 | 0.09 | 0.08 | 0 | 0.12 | 0.07 |
| 5 | 108409512 | rs7720808 | -0.005 | -0.799 | -1.076 | -3.061 | 0.011 | -0.016 | -0.016 | 1.264 | 0.1 | 0.15 | 0.08 | 0.07 | 0.16 | 0.08 |
| 5 | 108410880 | rs17161593 | 0.494 | -0.678 | -0.995 | -2.989 | 0.041 | -0.012 | -0.012 | 0.159 | 0.07 | 0.15 | 0.06 | 0.07 | 0.12 | 0.05 |
| 5 | 108421424 | rs10477436 | -1.872 | -1.006 | -0.719 | -2.78 | 0.013 | -0.093 | -0.093 | 0 | 0.32 | 0.46 | 0.2 | 0.16 | 0.32 | 0.26 |
| 5 | 108432970 | rs10076453 | -1.437 | -1.045 | -0.841 | -2.629 | 0.002 | -0.03 | -0.03 | 3.12 | 0.41 | 0.44 | 0.24 | 0.21 | 0.4 | 0.32 |
| 5 | 108442859 | rs17450338 | -1.365 | -1.02 | -0.808 | -2.612 | -0.009 | 0.048 | 0.048 | 0 | 0.39 | 0.28 | 0.21 | 0.12 | 0.37 | 0.22 |
| 5 | 108443391 | rs10043497 | 1.294 | -0.986 | -0.848 | -2.609 | -0.009 | 0.021 | 0.021 | 0 | 0.07 | 0.03 | 0.03 | 0 | 0.07 | 0.08 |
| 5 | 108445324 | rs17450387 | 0.038 | -1.023 | -0.802 | -2.598 | 0.002 | -0.014 | -0.014 | 0 | 0.17 | 0.16 | 0.11 | 0.07 | 0.21 | 0.13 |
| 5 | 108450242 | rs9326758 | -0.808 | -1.009 | -0.767 | -2.639 | -0.001 | -0.007 | -0.007 | 0.049 | 0.33 | 0.3 | 0.2 | 0.12 | 0.34 | 0.21 |
| 5 | 108450538 | rs7713591 | 0.664 | -1.013 | -0.81 | -2.658 | 0.001 | -0.007 | -0.007 | 0 | 0.06 | 0.05 | 0.03 | 0 | 0.05 | 0.01 |
| 5 | 108451261 | rs10477945 | -0.22 | -0.96 | -0.758 | -2.589 | -0.007 | -0.008 | -0.008 | 0 | 0.18 | 0.15 | 0.11 | 0.06 | 0.2 | 0.13 |
| 5 | 108463901 | rs17161612 | -1.616 | -1.168 | -0.639 | -2.529 | 0.003 | -0.027 | -0.027 | 0 | 0.41 | 0.42 | 0.24 | 0.22 | 0.41 | 0.31 |
| 5 | 108467319 | rs17534957 | 1.204 | -1.127 | -0.722 | -2.469 | 0.012 | 0.01 | 0.01 | 0.041 | 0.16 | 0.1 | 0.07 | 0.05 | 0.12 | 0.07 |
| 5 | 108489097 | rs17473391 | -0.325 | -0.839 | -0.802 | -2.534 | -0.01 | -0.005 | -0.005 | 0 | 0.2 | 0.17 | 0.13 | 0.07 | 0.24 | 0.15 |
| 5 | 108489944 | rs3814066 | 0.937 | -1.271 | -0.606 | -2.166 | 0.011 | 0.008 | 0.008 | 0 | 0.15 | 0.1 | 0.07 | 0.05 | 0.12 | 0.07 |
| 5 | 108502493 | rs7710527 | -0.301 | -1.248 | -0.575 | -2.156 | -0.004 | -0.002 | -0.002 | 0 | 0.23 | 0.23 | 0.11 | 0.15 | 0.19 | 0.17 |
| 5 | 108512613 | rs7735422 | 0.211 | -0.749 | -0.481 | -1.177 | -0.003 | 0 | 0 | 0.007 | 0.73 | 0.74 | 0.66 | 0.66 | 0.71 | 0.75 |
| 5 | 108517282 | rs7715602 | 0.633 | -1.001 | -0.329 | -1.326 | -0.012 | -0.009 | -0.009 | 0 | 0.19 | 0.2 | 0.06 | 0.06 | 0.12 | 0.11 |
| 5 | 108534661 | rs6594352 | 0.891 | -0.733 | -0.398 | -1.329 | 0 | 0 | 0 | 0 | 0.6 | 0.62 | 0.59 | 0.63 | 0.6 | 0.68 |
| 5 | 108542526 | rs10900654 | 0.755 | -0.696 | -0.424 | -1.308 | 0.003 | 0.019 | 0.019 | 0 | 0.53 | 0.63 | 0.6 | 0.57 | 0.59 | 0.62 |
| 5 | 108542865 | rs12187772 | 1.972 | -0.783 | -0.358 | -1.301 | -0.007 | -0.012 | -0.012 | 0 | 0.37 | 0.34 | 0.5 | 0.47 | 0.44 | 0.53 |
| 5 | 108550889 | rs13181049 | 0.861 | -1.133 | -1.136 | -1.229 | 0.009 | 0.021 | 0.021 | 0 | 0.52 | 0.6 | 0.61 | 0.57 | 0.59 | 0.6 |
| 5 | 108554780 | rs4957801 | 0.176 | -0.816 | -1.378 | -1.129 | -0.004 | -0.007 | -0.007 | 0.333 | 0.15 | 0.13 | 0.1 | 0.1 | 0.15 | 0.09 |
| 5 | 108555792 | rs6870224 | 0.182 | -0.419 | -1.147 | -0.934 | 0.038 | 0.038 | 0.038 | 1.012 | 0.32 | 0.16 | 0.13 | 0.15 | 0.26 | 0.18 |
| 5 | 108556802 | rs6875865 | 0.204 | 0.392 | -0.892 | -0.464 | -0.012 | -0.013 | -0.013 | 0.065 | 0.8 | 0.82 | 0.69 | 0.67 | 0.78 | 0.8 |
| 5 | 108556878 | rs7715933 | 0.697 | 0.379 | -0.901 | -0.408 | -0.005 | 0 | 0 | 0 | 0.72 | 0.74 | 0.66 | 0.66 | 0.71 | 0.72 |
| 5 | 108560090 | rs4593315 | 0.237 | 0.333 | -1.028 | -0.026 | 0.001 | -0.014 | -0.014 | 0 | 0.86 | 0.9 | 0.77 | 0.76 | 0.88 | 0.81 |
| 5 | 108561637 | rs6864212 | -0.452 | -0.018 | -1.585 | -0.646 | -0.046 | 0.153 | 0.153 | 0.279 | 0.55 | 0.85 | 0.52 | 0.24 | 0.49 | 0.31 |
| 5 | 108563868 | rs12515227 | -0.375 | -1.261 | -1.791 | -0.81 | 0.104 | 0.075 | 0.075 | 1.817 | 0.43 | 0.83 | 0.44 | 0.28 | 0.3 | 0.3 |
| 5 | 108566199 | rs6861689 | -0.234 | -1.564 | -1.983 | -0.989 | 0.027 | 0.055 | 0.055 | 0.728 | 0.29 | 0.83 | 0.33 | 0.2 | 0.23 | 0.2 |
| 5 | 108576515 | rs17473588 | 0.516 | -1.684 | -1.197 | -1 | -0.001 | -0.003 | -0.003 | 0 | 0.22 | 0.25 | 0.26 | 0.2 | 0.21 | 0.19 |
| 5 | 108577839 | rs10039717 | -1.015 | -1.695 | -1.392 | -1.024 | 0.059 | 0.117 | 0.117 | 0.336 | 0.52 | 0.88 | 0.46 | 0.29 | 0.39 | 0.34 |
| 5 | 108579889 | rs1968228 | -0.432 | -1.68 | -0.606 | -0.977 | -0.026 | -0.021 | -0.021 | 0 | 0.41 | 0.49 | 0.42 | 0.29 | 0.32 | 0.33 |
| 5 | 108592057 | rs9326766 | 0.352 | -1.197 | 0.003 | -0.732 | 0.005 | -0.038 | -0.038 | 0.246 | 0.07 | 0.12 | 0.03 | 0 | 0.07 | 0.02 |
| 5 | 108606564 | rs7720237 | -0.378 | -1.498 | -0.164 | -0.998 | 0.02 | 0.03 | 0.03 | 0 | 0.35 | 0.32 | 0.29 | 0.2 | 0.27 | 0.27 |
| 5 | 108609854 | rs11743452 | 0.456 | -1.559 | -0.176 | -1.026 | -0.016 | -0.017 | -0.017 | 0 | 0.2 | 0.24 | 0.18 | 0.1 | 0.16 | 0.11 |
| 5 | 108612057 | rs4645374 | 0.203 | -1.425 | -0.452 | -0.881 | 0.028 | 0.026 | 0.026 | 0 | 0.29 | 0.24 | 0.26 | 0.2 | 0.19 | 0.25 |
| 5 | 108614251 | rs2161163 | -1.529 | -1.113 | -0.76 | -0.899 | -0.103 | 0.008 | 0.008 | 0.586 | 0.65 | 0.86 | 0.51 | 0.3 | 0.52 | 0.43 |
| 5 | 108619855 | rs10058030 | -0.577 | -1.107 | -0.367 | -0.342 | 0.038 | -0.054 | -0.054 | 0 | 0.1 | 0.38 | 0.06 | 0 | 0.09 | 0.03 |
| 5 | 108620859 | rs6890213 | -1.293 | -0.82 | -1.059 | 0.351 | -0.024 | -0.073 | -0.073 | 0 | 0.89 | 0.99 | 0.85 | 0.72 | 0.91 | 0.8 |
| 5 | 108623619 | rs2416205 | -0.308 | -0.891 | -0.458 | -0.139 | 0.031 | 0.018 | 0.018 | 0 | 0.27 | 0.22 | 0.25 | 0.22 | 0.19 | 0.25 |

| | | | | | | | | | | | | | | | | |
|---|---|---|---|---|---|---|---|---|---|---|---|---|---|---|---|---|
| 5 | 108631302 | rs10056957 | -1.111 | 0.199 | -0.104 | 0.186 | 0.011 | -0.05 | -0.05 | 0 | 0.09 | 0.23 | 0.04 | 0 | 0.07 | 0.01 |
| 5 | 108633347 | rs13359409 | -0.729 | 0.244 | -0.835 | 0.455 | 0.004 | 0.002 | 0.002 | 0 | 0.27 | 0.18 | 0.23 | 0.23 | 0.27 | 0.21 |
| 5 | 108634765 | rs13168640 | -0.462 | 0.603 | -0.721 | 0.637 | -0.006 | -0.009 | -0.009 | 0 | 0.42 | 0.35 | 0.39 | 0.5 | 0.45 | 0.31 |
| 5 | 108638291 | rs1990892 | -0.506 | -0.193 | -0.381 | 0.145 | -0.003 | -0.007 | -0.007 | 0.97 | 0.36 | 0.64 | 0.2 | 0.17 | 0.25 | 0.23 |
| 5 | 108639183 | rs13166029 | -0.332 | 0.212 | -0.499 | 0.296 | 0.019 | 0.004 | 0.004 | 0 | 0.48 | 0.6 | 0.61 | 0.57 | 0.43 | 0.64 |
| 5 | 108644218 | rs11241004 | -1.07 | 0.659 | 0.667 | 0.705 | -0.004 | 0.085 | 0.085 | 2.16 | 0.18 | 0.52 | 0.21 | 0.3 | 0.11 | 0.08 |
| 5 | 108645525 | rs12657816 | -2.106 | 1.236 | 0.599 | -0.123 | -0.015 | 0.046 | 0.046 | 5.733 | 0.45 | 0.75 | 0.52 | 0.94 | 0.41 | 0.44 |
| 5 | 108648231 | rs1985666 | -0.931 | 1.074 | 0.608 | 0.292 | -0.019 | 0.119 | 0.119 | 3.467 | 0.2 | 0.54 | 0.21 | 0.38 | 0.11 | 0.08 |
| 5 | 108670421 | rs414724 | -0.377 | 1.571 | 0.394 | 0.344 | -0.037 | -0.067 | -0.067 | 0 | 0.2 | 0.06 | 0.28 | 0.56 | 0.25 | 0.35 |
| 5 | 108672946 | rs246105 | -0.493 | 1.558 | 0.522 | 0.35 | -0.016 | -0.031 | -0.031 | 0.63 | 0.21 | 0.12 | 0.28 | 0.56 | 0.25 | 0.35 |
| 5 | 108713970 | rs10042721 | 0.321 | 1.436 | 0.586 | 0.362 | -0.007 | -0.01 | -0.01 | 2.29 | 0.79 | 0.87 | 0.64 | 0.44 | 0.74 | 0.65 |
| 5 | 108721702 | rs6872547 | -0.127 | 1.551 | 0.267 | 0.483 | -0.066 | -0.089 | -0.089 | 0.021 | 0.88 | 0.99 | 0.76 | 0.52 | 0.81 | 0.71 |
| 5 | 108744375 | rs1366618 | 0.172 | 1.411 | 0.622 | 0.344 | 0.002 | -0.016 | -0.016 | 0 | 0.05 | 0.08 | 0.03 | 0 | 0.04 | 0.02 |
| 5 | 108803319 | rs10074851 | -0.071 | 1.31 | 1.067 | 0.708 | -0.021 | 0.227 | 0.227 | 5.064 | 0.11 | 0.41 | 0.06 | 0.31 | 0.04 | 0.1 |
| 5 | 108807760 | rs11741087 | -0.601 | 1.541 | 0.825 | 0.624 | -0.064 | -0.105 | -0.105 | 0.247 | 0.2 | 0.03 | 0.4 | 0.58 | 0.26 | 0.32 |
| 5 | 108826387 | rs4957817 | -0.243 | 1.535 | 1.076 | 0.724 | -0.007 | 0.257 | 0.257 | 5.578 | 0.11 | 0.45 | 0.08 | 0.37 | 0.06 | 0.1 |
| 5 | 108828810 | rs10515401 | 0.07 | 1.496 | 0.968 | 0.712 | -0.001 | 0 | 0 | 0 | 0.08 | 0.09 | 0.04 | 0.05 | 0.05 | 0.02 |
| 5 | 108833523 | rs9326773 | -0.581 | 1.83 | 0.881 | 0.618 | -0.071 | -0.105 | -0.105 | 0.037 | 0.19 | 0.04 | 0.4 | 0.53 | 0.27 | 0.33 |
| 5 | 108843934 | rs7448843 | -1.554 | 2.505 | 0.87 | 0.326 | -0.093 | -0.083 | -0.083 | 0 | 0.27 | 0.09 | 0.49 | 0.52 | 0.43 | 0.37 |
| 5 | 108844343 | rs4314456 | -0.101 | 2.441 | 0.944 | 0.543 | -0.05 | -0.063 | -0.063 | 0.093 | 0.11 | 0.01 | 0.29 | 0.42 | 0.17 | 0.28 |
| 5 | 108849336 | rs10071406 | -1.135 | 2.133 | 1.24 | 0.26 | -0.006 | 0.16 | 0.16 | 2.467 | 0.19 | 0.53 | 0.11 | 0.41 | 0.11 | 0.13 |
| 5 | 108855035 | rs17161887 | -2.186 | 1.962 | 1.205 | 0.233 | 0.002 | 0.01 | 0.01 | 2.113 | 0.26 | 0.7 | 0.22 | 0.38 | 0.26 | 0.18 |
| 5 | 108855067 | rs11747777 | -0.923 | 1.863 | 1.223 | 0.328 | -0.012 | 0.148 | 0.148 | 3.305 | 0.16 | 0.48 | 0.12 | 0.34 | 0.12 | 0.15 |
| 5 | 108911162 | rs1080895 | 0.03 | 2.066 | 1.059 | 0.147 | 0.01 | -0.02 | -0.02 | 6.527 | 0.92 | 0.85 | 0.96 | 0.96 | 0.95 | 0.97 |
| 5 | 108921039 | rs6594386 | -0.382 | 2.071 | 0.96 | 0.153 | -0.003 | 0.002 | 0.002 | 0 | 0.93 | 0.96 | 0.97 | 0.96 | 0.95 | 0.97 |
| 5 | 108922023 | rs902505 | 2.49 | 2.128 | 1.157 | 0.266 | 0.008 | 0.012 | 0.012 | 0.265 | 0.67 | 0.32 | 0.76 | 0.59 | 0.69 | 0.78 |
| 5 | 108929597 | rs9326778 | 0.199 | 1.673 | 1.764 | 0.538 | -0.003 | 0.24 | 0.24 | 5.592 | 0.09 | 0.31 | 0.06 | 0.37 | 0.05 | 0.05 |
| 5 | 108930590 | rs17161951 | -0.533 | 2.072 | 1.838 | 0.495 | 0.005 | 0.039 | 0.039 | 0.015 | 0.16 | 0.13 | 0.36 | 0.51 | 0.22 | 0.27 |
| 5 | 108932563 | rs6594388 | 0.613 | 1.998 | 1.773 | 0.444 | 0.019 | -0.027 | -0.027 | 0 | 0.91 | 0.73 | 0.95 | 0.96 | 0.94 | 0.97 |
| 5 | 108941292 | rs12719109 | 2.403 | 2.042 | 1.672 | 0.497 | 0.003 | 0.002 | 0.002 | 0 | 0.67 | 0.4 | 0.54 | 0.44 | 0.65 | 0.67 |
| 5 | 108948513 | rs1826236 | -1.511 | 1.959 | 1.738 | 0.37 | 0.049 | -0.053 | -0.053 | 2.211 | 0.13 | 0.36 | 0.09 | 0.04 | 0.11 | 0.04 |
| 5 | 108951190 | rs447849 | 1.754 | 1.981 | 1.492 | 0.61 | 0.034 | 0.072 | 0.072 | 0 | 0.73 | 0.53 | 0.59 | 0.48 | 0.71 | 0.7 |
| 5 | 108952367 | rs439182 | 0.535 | 1.888 | 1.373 | 0.772 | 0.024 | 0.125 | 0.125 | 0.002 | 0.84 | 0.78 | 0.64 | 0.52 | 0.8 | 0.74 |
| 5 | 108964907 | rs2256556 | -0.771 | 1.899 | 1.378 | 0.794 | 0.032 | 0.151 | 0.151 | 0 | 0.16 | 0.24 | 0.36 | 0.48 | 0.2 | 0.26 |
| 5 | 108980912 | rs6863703 | 0.636 | 1.949 | 1.042 | 0.847 | -0.017 | -0.002 | -0.002 | 0 | 0.08 | 0.02 | 0.13 | 0.07 | 0.12 | 0.04 |
| 5 | 108984093 | rs166181 | -1.023 | 1.887 | 1.336 | 0.688 | 0.052 | 0.228 | 0.228 | 0 | 0.17 | 0.44 | 0.38 | 0.48 | 0.2 | 0.26 |
| 5 | 108986371 | rs31593 | -2.284 | 1.903 | 1.093 | 0.441 | 0.001 | 0 | 0 | 0 | 0.36 | 0.63 | 0.5 | 0.52 | 0.37 | 0.41 |
| 5 | 108988377 | rs185088 | -1.269 | 2.143 | 0.984 | 0.153 | 0.032 | 0.078 | 0.078 | 0 | 0.19 | 0.26 | 0.4 | 0.49 | 0.27 | 0.27 |
| 5 | 108989023 | rs1490778 | 0.051 | 1.593 | 0.516 | -0.858 | 0.006 | 0.012 | 0.012 | 0.402 | 0.68 | 0.79 | 0.6 | 0.27 | 0.73 | 0.64 |
| 5 | 108990146 | rs7701590 | -0.886 | 2.023 | 0.651 | -0.727 | 0.038 | -0.044 | -0.044 | 0.328 | 0.08 | 0.18 | 0.05 | 0.03 | 0.08 | 0.03 |
| 5 | 108992465 | rs25916 | -0.86 | 1.823 | -0.057 | -0.794 | -0.001 | 0.037 | 0.037 | 0 | 0.2 | 0.29 | 0.28 | 0.62 | 0.19 | 0.45 |
| 5 | 108995505 | rs31614 | -1.719 | 2.042 | -0.397 | -0.857 | 0.004 | 0.009 | 0.009 | 0 | 0.27 | 0.33 | 0.32 | 0.63 | 0.26 | 0.48 |
| 5 | 108999666 | rs31607 | 0.347 | 2.05 | -0.433 | -0.837 | -0.008 | -0.013 | -0.013 | 1.593 | 0.11 | 0.05 | 0.24 | 0.6 | 0.12 | 0.36 |
| 5 | 109000325 | rs17162034 | -0.126 | 2.053 | -0.416 | -0.897 | -0.001 | -0.009 | -0.009 | 0 | 0.08 | 0.08 | 0.05 | 0.01 | 0.08 | 0.11 |
| 5 | 109001791 | rs845741 | -1.413 | 1.997 | -0.463 | -0.92 | 0.01 | 0.034 | 0.034 | 0 | 0.21 | 0.29 | 0.3 | 0.63 | 0.2 | 0.38 |
| 5 | 109006376 | rs31600 | -0.72 | 2.043 | -0.633 | -0.888 | -0.027 | -0.047 | -0.047 | 0 | 0.17 | 0.07 | 0.28 | 0.6 | 0.19 | 0.39 |
| 5 | 109019280 | rs10491355 | -1.679 | 2.021 | -0.65 | -0.966 | -0.004 | -0.007 | -0.007 | 0 | 0.2 | 0.18 | 0.11 | 0.04 | 0.16 | 0.13 |
| 5 | 109023410 | rs845734 | -1.824 | 2.003 | -0.55 | -0.85 | 0.004 | 0.001 | 0.001 | 0 | 0.34 | 0.37 | 0.37 | 0.65 | 0.31 | 0.5 |
| 5 | 109043183 | rs12656763 | -0.486 | 1.848 | -0.889 | -0.889 | -0.01 | -0.094 | -0.094 | 0 | 0.2 | 0.04 | 0.24 | 0.62 | 0.15 | 0.38 |
| 5 | 109044067 | rs6877440 | -0.704 | 1.792 | -0.774 | -0.952 | 0.028 | -0.071 | -0.071 | 3.396 | 0.13 | 0.3 | 0.09 | 0.04 | 0.13 | 0.12 |
| 5 | 109044558 | rs6897334 | 1.068 | 2.238 | -1.365 | -1.497 | 0.01 | 0.004 | 0.004 | 0.048 | 0.29 | 0.55 | 0.34 | 0.26 | 0.36 | 0.1 |
| 5 | 109058683 | rs1438661 | -1.413 | 2.201 | -1.551 | -1.575 | -0.005 | 0.2 | 0.2 | 0 | 0.64 | 0.92 | 0.67 | 0.92 | 0.66 | 0.62 |
| 5 | 109065708 | rs17163950 | 0.5 | 2.203 | -1.509 | -1.592 | -0.009 | -0.049 | -0.049 | 0 | 0.09 | 0.27 | 0.04 | 0.01 | 0.06 | 0.1 |
| 5 | 109066741 | rs6594396 | -1.451 | 2.216 | -1.542 | -1.603 | -0.021 | 0.199 | 0.199 | 0.248 | 0.66 | 0.95 | 0.68 | 0.92 | 0.67 | 0.62 |
| 5 | 109087406 | rs10491357 | 0.055 | 2.273 | -1.481 | -1.573 | -0.004 | -0.062 | -0.062 | 0 | 0.11 | 0.25 | 0.05 | 0.01 | 0.09 | 0.12 |
| 5 | 109088553 | rs2010653 | -1.376 | 2.651 | -1.556 | -1.645 | -0.054 | 0.223 | 0.223 | 0.688 | 0.63 | 0.94 | 0.65 | 0.94 | 0.61 | 0.63 |
| 5 | 109089550 | rs10491358 | 0.884 | 2.609 | -1.507 | -1.613 | -0.01 | -0.044 | -0.044 | 0 | 0.08 | 0.18 | 0.04 | 0.01 | 0.06 | 0.1 |
| 5 | 109095545 | rs13187428 | 1.053 | 2.652 | -1.402 | -1.579 | 0.012 | 0.009 | 0.009 | 0.018 | 0.29 | 0.48 | 0.34 | 0.28 | 0.35 | 0.12 |
| 5 | 109098354 | rs10055229 | -0.198 | 2.647 | -1.809 | -1.556 | -0.006 | -0.07 | -0.07 | 0 | 0.11 | 0.3 | 0.06 | 0.01 | 0.1 | 0.12 |
| 5 | 109108982 | rs2301004 | 1.357 | 2.665 | -1.753 | -1.56 | -0.005 | 0.014 | 0.014 | 0.382 | 0.27 | 0.54 | 0.29 | 0.26 | 0.29 | 0.1 |
| 5 | 109112145 | rs4569915 | -1.469 | 2.174 | -1.682 | -1.484 | 0.001 | 0 | 0 | 0 | 0.36 | 0.39 | 0.34 | 0.66 | 0.32 | 0.51 |
| 5 | 109113609 | rs12514321 | -0.494 | 2.158 | -1.677 | -1.467 | 0.019 | 0.067 | 0.067 | 0 | 0.54 | 0.67 | 0.63 | 0.91 | 0.58 | 0.5 |
| 5 | 109117171 | rs13189822 | -0.188 | 2.18 | -1.669 | -1.369 | -0.006 | -0.091 | -0.091 | 0 | 0.21 | 0.06 | 0.24 | 0.62 | 0.15 | 0.38 |
| 5 | 109126701 | rs2013881 | -0.699 | 2.412 | -1.821 | -1.362 | 0.04 | 0.135 | 0.135 | 0 | 0.55 | 0.76 | 0.64 | 0.92 | 0.6 | 0.51 |
| 5 | 109132461 | rs2301010 | 0.49 | 2.435 | -1.888 | -1.353 | -0.004 | -0.076 | -0.076 | 0 | 0.15 | 0.03 | 0.22 | 0.62 | 0.12 | 0.36 |
| 5 | 109136218 | rs7728733 | -1.45 | 2.642 | -1.908 | -1.395 | 0.012 | 0.23 | 0.23 | 0.253 | 0.64 | 0.98 | 0.7 | 0.92 | 0.7 | 0.63 |
| 5 | 109139335 | rs10452541 | -0.721 | 2.311 | -1.874 | -1.389 | 0.041 | 0.152 | 0.152 | 0 | 0.55 | 0.77 | 0.64 | 0.92 | 0.6 | 0.52 |
| 5 | 109143386 | rs2301015 | 1.156 | 2.36 | -1.842 | -1.359 | 0.006 | 0.009 | 0.009 | 0.013 | 0.29 | 0.54 | 0.33 | 0.27 | 0.35 | 0.1 |
| 5 | 109143602 | rs2301016 | -1.137 | 2.366 | -1.902 | -1.364 | -0.003 | 0.003 | 0.003 | 0 | 0.31 | 0.35 | 0.3 | 0.62 | 0.24 | 0.5 |

| | | | | | | | | | | | | | | | | |
|---|---|---|---|---|---|---|---|---|---|---|---|---|---|---|---|---|
| 5 | 109145646 | rs2015698 | 0.66 | 2.376 | -1.892 | -1.353 | 0.002 | -0.001 | -0.001 | 0 | 0.33 | 0.63 | 0.39 | 0.33 | 0.42 | 0.14 |
| 5 | 109159717 | rs958235 | -0.958 | 2.202 | -1.948 | -1.365 | -0.07 | 0.247 | 0.247 | 0 | 0.6 | 0.91 | 0.64 | 0.9 | 0.59 | 0.64 |
| 5 | 109172557 | rs1368358 | -1.021 | 2.192 | -1.948 | -1.358 | -0.022 | 0.245 | 0.245 | 0.017 | 0.61 | 0.93 | 0.66 | 0.92 | 0.63 | 0.6 |
| 5 | 109180817 | rs7700520 | -0.89 | 1.597 | -1.83 | -1.275 | -0.022 | 0.178 | 0.178 | 0 | 0.58 | 0.81 | 0.63 | 0.9 | 0.59 | 0.62 |
| 5 | 109187498 | rs1432805 | 0.003 | 1.496 | -1.9 | -1.263 | -0.005 | 0.042 | 0.042 | 0 | 0.47 | 0.6 | 0.59 | 0.9 | 0.52 | 0.51 |
| 5 | 109187941 | rs3797674 | 0.424 | 1.805 | -2.243 | -0.995 | -0.015 | -0.079 | -0.079 | 0 | 0.15 | 0.03 | 0.23 | 0.61 | 0.13 | 0.37 |
| 5 | 109202725 | rs1047004 | 1.691 | 1.801 | -2.032 | -1.066 | -0.014 | 0.02 | 0.02 | 0 | 0.06 | 0.01 | 0.04 | 0 | 0.08 | 0.02 |
| 5 | 109204024 | rs2241692 | -0.896 | 1.47 | -1.789 | -0.792 | -0.052 | 0.223 | 0.223 | 0 | 0.6 | 0.9 | 0.64 | 0.87 | 0.6 | 0.61 |
| 5 | 109213034 | rs9326785 | 0.593 | 1.469 | -1.822 | -0.69 | -0.001 | -0.001 | -0.001 | 2.678 | 0.29 | 0.6 | 0.33 | 0.26 | 0.31 | 0.12 |
| 5 | 109213385 | rs9326786 | -0.08 | 1.6 | -1.836 | -0.734 | -0.01 | 0.013 | 0.013 | 1.6 | 0.35 | 0.78 | 0.37 | 0.33 | 0.37 | 0.22 |
| 5 | 109213779 | rs2033463 | 0.475 | 1.471 | -1.985 | -0.818 | -0.001 | 0.011 | 0.011 | 0 | 0.64 | 0.67 | 0.54 | 0.3 | 0.66 | 0.47 |
| 5 | 109215047 | rs12519496 | 0.943 | 1.418 | -2.052 | -0.801 | -0.069 | 0.225 | 0.225 | 0 | 0.38 | 0.11 | 0.24 | 0.09 | 0.38 | 0.36 |
| 5 | 109216204 | rs6874288 | -0.08 | 1.072 | -1.942 | -0.653 | -0.017 | -0.06 | -0.06 | 0 | 0.71 | 0.94 | 0.6 | 0.37 | 0.74 | 0.56 |
| 5 | 109216361 | rs17491050 | 0.825 | 1.085 | -1.941 | -1.058 | -0.089 | 0.243 | 0.243 | 0 | 0.4 | 0.11 | 0.25 | 0.09 | 0.42 | 0.38 |
| 5 | 109221844 | rs10073312 | 0.225 | 1.346 | -1.793 | -0.86 | -0.001 | 0.008 | 0.008 | 0.195 | 0.32 | 0.68 | 0.34 | 0.2 | 0.33 | 0.11 |
| 5 | 109222729 | rs6594405 | -0.327 | 1.216 | -1.721 | -0.646 | -0.005 | -0.024 | -0.024 | 0.052 | 0.75 | 0.95 | 0.61 | 0.3 | 0.77 | 0.57 |
| 5 | 109224099 | rs2416223 | 0.689 | 1.167 | -1.557 | -0.654 | -0.019 | 0.116 | 0.116 | 0.103 | 0.43 | 0.24 | 0.25 | 0.04 | 0.42 | 0.44 |
| 5 | 109230397 | rs6884296 | -0.392 | 1.444 | -0.518 | 0.548 | 0.006 | -0.017 | -0.017 | 0 | 0.28 | 0.57 | 0.34 | 0.22 | 0.32 | 0.13 |
| 5 | 109237529 | rs1432811 | -1.17 | 1.42 | -0.65 | 1.561 | -0.012 | 0.004 | 0.004 | 0.333 | 0.07 | 0.04 | 0.2 | 0.19 | 0.1 | 0.13 |
| 5 | 109238721 | rs36835 | -1.85 | 0.803 | 0.04 | 1.609 | -0.015 | 0.105 | 0.105 | 0.272 | 0.14 | 0.33 | 0.14 | 0.25 | 0.09 | 0.17 |
| 5 | 109250630 | rs28825 | -2.015 | 0.74 | 0.607 | 1.84 | -0.004 | 0.14 | 0.14 | 0 | 0.14 | 0.37 | 0.15 | 0.25 | 0.1 | 0.17 |
| 5 | 109268575 | rs36811 | -0.978 | 0.679 | 0.47 | 0.73 | 0.009 | 0.066 | 0.066 | 0.167 | 0.49 | 0.6 | 0.34 | 0.28 | 0.43 | 0.33 |
| 5 | 109277223 | rs36804 | -1.687 | 0.075 | 0.699 | 1.466 | -0.01 | 0.177 | 0.177 | 0 | 0.1 | 0.29 | 0.09 | 0.26 | 0.07 | 0.19 |
| 5 | 109280010 | rs17596440 | -1.114 | 0.129 | 0.325 | 1.409 | -0.001 | 0.006 | 0.006 | 0 | 0.07 | 0.06 | 0.07 | 0.12 | 0.07 | 0.14 |
| 5 | 109293821 | rs36779 | -1.684 | 0.088 | 0.244 | 1.572 | 0.004 | 0.019 | 0.019 | 0 | 0.09 | 0.1 | 0.07 | 0.26 | 0.07 | 0.17 |
| 5 | 109296292 | rs10071850 | -1.118 | -0.65 | 0.244 | 0.975 | -0.038 | -0.014 | -0.014 | 0 | 0.38 | 0.51 | 0.2 | 0.3 | 0.22 | 0.29 |
| 5 | 109319962 | rs2112966 | -0.713 | -0.729 | 0.185 | 0.857 | 0.076 | 0.127 | 0.127 | 0 | 0.33 | 0.15 | 0.16 | 0.06 | 0.2 | 0.28 |
| 5 | 109322373 | rs36803 | 1.275 | -0.787 | 0.668 | 0.964 | -0.017 | -0.006 | -0.006 | 0 | 0.6 | 0.43 | 0.79 | 0.66 | 0.75 | 0.69 |
| 5 | 109323840 | rs283350 | -0.952 | -0.782 | 0.732 | 0.96 | 0.047 | 0.06 | 0.06 | 0.876 | 0.07 | 0.31 | 0.04 | 0.08 | 0.04 | 0.02 |
| 5 | 109324100 | rs13186368 | -0.579 | -0.835 | 0.763 | 0.886 | 0.066 | 0.101 | 0.101 | 0 | 0.32 | 0.17 | 0.15 | 0.05 | 0.19 | 0.27 |
| 5 | 109327087 | rs40627 | 1.341 | -0.759 | 0.602 | 0.904 | -0.011 | -0.004 | -0.004 | 0 | 0.58 | 0.43 | 0.77 | 0.66 | 0.73 | 0.68 |
| 5 | 109330195 | rs36791 | 1.328 | -0.773 | 0.416 | 0.853 | -0.01 | -0.003 | -0.003 | 0.018 | 0.58 | 0.42 | 0.79 | 0.67 | 0.74 | 0.77 |
| 5 | 109335278 | rs283358 | 1.332 | -0.776 | -0.89 | 0.322 | -0.052 | -0.162 | -0.162 | 0.018 | 0.23 | 0.02 | 0.4 | 0.53 | 0.22 | 0.35 |
| 5 | 109336913 | rs13157324 | 0.06 | -1.185 | -0.452 | 0.253 | 0.001 | 0.018 | 0.018 | 0 | 0.12 | 0.06 | 0.08 | 0.05 | 0.11 | 0.14 |
| 5 | 109343856 | rs2416227 | 0.557 | -1.137 | -0.127 | 0.416 | -0.075 | -0.023 | -0.023 | 0 | 0.71 | 0.52 | 0.86 | 0.73 | 0.85 | 0.81 |
| 5 | 109348449 | rs17162540 | 0.19 | -0.899 | -0.956 | -0.064 | 0.075 | 0.1 | 0.1 | 0.396 | 0.21 | 0.1 | 0.08 | 0 | 0.09 | 0.15 |
| 5 | 109352713 | rs1004522 | -0.458 | -0.16 | -0.687 | 0.49 | 0.003 | 0.001 | 0.001 | 0 | 0.54 | 0.64 | 0.5 | 0.38 | 0.54 | 0.52 |
| 5 | 109353601 | rs5023585 | 0.002 | -0.491 | -0.442 | -0.272 | -0.02 | -0.088 | -0.088 | 0 | 0.22 | 0.32 | 0.1 | 0 | 0.15 | 0.17 |
| 5 | 109358981 | rs2925117 | -0.561 | -1.159 | -0.757 | 0.216 | -0.035 | 0.02 | 0.02 | 0.025 | 0.42 | 0.74 | 0.61 | 0.61 | 0.44 | 0.5 |
| 5 | 109363175 | rs1455554 | -0.453 | -1.201 | -0.573 | 0.009 | 0.019 | 0.01 | 0.01 | 0 | 0.17 | 0.11 | 0.22 | 0.16 | 0.1 | 0.2 |
| 5 | 109363572 | rs1378500 | -0.453 | -1.201 | -0.573 | 0.009 | 0.019 | 0.01 | 0.01 | 0 | 0.17 | 0.11 | 0.22 | 0.16 | 0.1 | 0.2 |
| 5 | 109365711 | rs2168308 | 0.166 | -1.089 | -0.468 | 0.245 | -0.031 | -0.042 | -0.042 | 0 | 0.23 | 0.16 | 0.34 | 0.39 | 0.34 | 0.21 |
| 5 | 109367379 | rs6892037 | 0.455 | -0.914 | -0.49 | 0.271 | -0.011 | -0.024 | -0.024 | 0.015 | 0.19 | 0.15 | 0.24 | 0.38 | 0.26 | 0.2 |
| 5 | 109368613 | rs7736552 | 0.059 | -1.383 | -0.54 | 0.031 | -0.002 | -0.02 | -0.02 | 0 | 0.36 | 0.24 | 0.31 | 0.4 | 0.35 | 0.34 |
| 5 | 109369796 | rs1378507 | 1.642 | -1.571 | -0.929 | -0.625 | -0.085 | -0.085 | -0.085 | 0 | 0.32 | 0.14 | 0.49 | 0.51 | 0.47 | 0.38 |
| 5 | 109370988 | rs2900041 | -0.914 | -1.72 | -0.588 | -0.952 | 0.007 | 0.01 | 0.01 | 0.341 | 0.32 | 0.7 | 0.32 | 0.25 | 0.22 | 0.28 |
| 5 | 109381583 | rs2966765 | -0.944 | -1.789 | -0.237 | -0.967 | 0.016 | -0.002 | -0.002 | 0 | 0.78 | 0.86 | 0.84 | 0.79 | 0.79 | 0.8 |
| 5 | 109383219 | rs2966762 | 0.701 | -1.836 | -0.135 | -1.029 | 0.009 | 0.024 | 0.024 | 0 | 0.55 | 0.4 | 0.69 | 0.78 | 0.6 | 0.72 |
| 5 | 109384131 | rs17162589 | 0.284 | -1.894 | -0.023 | -1.008 | -0.008 | -0.008 | -0.008 | 0.202 | 0.23 | 0.2 | 0.2 | 0.16 | 0.11 | 0.26 |
| 5 | 109384306 | rs17162591 | 0.299 | -1.892 | -0.027 | -1.008 | -0.007 | -0.007 | -0.007 | 0 | 0.23 | 0.2 | 0.2 | 0.17 | 0.11 | 0.26 |
| 5 | 109391230 | rs1378506 | 0.627 | -1.742 | -0.223 | -1.089 | 0 | -0.005 | -0.005 | 0 | 0.54 | 0.47 | 0.6 | 0.76 | 0.56 | 0.64 |
| 5 | 109391689 | rs7701086 | -0.644 | -1.982 | -0.105 | -0.804 | -0.045 | -0.02 | -0.02 | 0 | 0.32 | 0.46 | 0.29 | 0.28 | 0.17 | 0.37 |
| 5 | 109393001 | rs17162633 | -0.933 | -1.429 | -0.38 | -0.721 | 0.001 | -0.014 | -0.014 | 0 | 0.22 | 0.28 | 0.17 | 0.16 | 0.18 | 0.17 |
| 5 | 109394587 | rs1455546 | 2.164 | -1.42 | -0.593 | -0.768 | -0.056 | -0.069 | -0.069 | 0 | 0.33 | 0.17 | 0.44 | 0.47 | 0.5 | 0.39 |
| 5 | 109398784 | rs7730450 | -0.291 | -1.333 | -0.495 | -0.815 | 0.012 | -0.021 | -0.021 | 0.317 | 0.16 | 0.25 | 0.13 | 0.08 | 0.14 | 0.11 |
| 5 | 109400991 | rs2962986 | -0.387 | -1.577 | -0.295 | -0.765 | -0.001 | 0 | 0 | 0.463 | 0.76 | 0.77 | 0.82 | 0.83 | 0.76 | 0.82 |
| 5 | 109401411 | rs17162650 | 0.522 | -1.575 | -0.27 | -0.756 | -0.021 | 0.005 | 0.005 | 0.106 | 0.11 | 0.25 | 0.07 | 0.08 | 0.06 | 0.1 |
| 5 | 109401940 | rs2966752 | 1.49 | -1.638 | -0.32 | -0.803 | -0.029 | -0.03 | -0.03 | 0 | 0.39 | 0.29 | 0.49 | 0.51 | 0.58 | 0.45 |
| 5 | 109405571 | rs10056970 | 0.452 | -1.636 | -0.248 | -0.739 | 0 | 0.003 | 0.003 | 0 | 0.63 | 0.56 | 0.58 | 0.59 | 0.67 | 0.6 |
| 5 | 109405715 | rs10079247 | 1.325 | -1.622 | -0.244 | -0.731 | 0.021 | 0 | 0 | 0 | 0.5 | 0.32 | 0.51 | 0.52 | 0.6 | 0.52 |
| 5 | 109410129 | rs1598443 | 0.377 | -1.701 | -0.307 | -0.768 | -0.004 | 0.002 | 0.002 | 0 | 0.64 | 0.58 | 0.61 | 0.59 | 0.73 | 0.66 |
| 5 | 109411047 | rs2061129 | 1.294 | -1.692 | -1.055 | -0.775 | -0.008 | -0.047 | -0.047 | 0 | 0.15 | 0.17 | 0.04 | 0 | 0.09 | 0.12 |
| 5 | 109411961 | rs1378504 | 0.034 | -1.396 | -1.303 | -0.596 | 0 | -0.032 | -0.032 | 0 | 0.13 | 0.06 | 0.23 | 0.24 | 0.09 | 0.21 |
| 5 | 109412185 | rs7718366 | 0.597 | -1.26 | -1.045 | -0.427 | 0 | -0.027 | -0.027 | 0.424 | 0.09 | 0.06 | 0.21 | 0.24 | 0.07 | 0.17 |
| 5 | 109413243 | rs6883748 | 0.876 | -1.258 | -1.039 | -0.431 | 0.03 | -0.007 | -0.007 | 0 | 0.53 | 0.34 | 0.54 | 0.51 | 0.67 | 0.56 |
| 5 | 109414387 | rs2925079 | -0.203 | -0.737 | -0.68 | -0.33 | 0 | -0.003 | -0.003 | 0 | 0.72 | 0.73 | 0.79 | 0.83 | 0.73 | 0.76 |
| 5 | 109415961 | rs2925076 | 0.185 | -0.307 | -0.474 | -0.156 | 0 | -0.002 | -0.002 | 0 | 0.67 | 0.67 | 0.61 | 0.59 | 0.73 | 0.66 |
| 5 | 109419718 | rs10067702 | 0.093 | 0.214 | -0.683 | 0.168 | 0.019 | 0.063 | 0.063 | 0 | 0.52 | 0.36 | 0.74 | 0.76 | 0.63 | 0.61 |
| 5 | 109424551 | rs10515411 | -1.133 | 0.351 | -0.051 | 0.787 | -0.001 | 0.032 | 0.032 | 0.486 | 0.25 | 0.33 | 0.28 | 0.3 | 0.28 | 0.15 |

| | | | | | | | | | | | | | | | | |
|---|---|---|---|---|---|---|---|---|---|---|---|---|---|---|---|---|
| 5 | 109428276 | rs2925068 | -1.272 | 0.118 | 0.789 | 0.893 | -0.001 | -0.078 | -0.078 | 0.102 | 0.25 | 0.36 | 0.18 | 0.06 | 0.27 | 0.15 |
| 5 | 109446858 | rs6871500 | -1.057 | 0.117 | 0.857 | 0.839 | -0.013 | 0.053 | 0.053 | 0 | 0.12 | 0.18 | 0.17 | 0.25 | 0.07 | 0.05 |
| 5 | 109447734 | rs7717231 | 2.193 | -0.18 | 1.619 | 0.705 | -0.006 | 0.012 | 0.012 | 1.029 | 0.72 | 0.4 | 0.69 | 0.7 | 0.7 | 0.85 |
| 5 | 109451963 | rs2198687 | -1.052 | -0.113 | 1.654 | 0.659 | 0.019 | -0.039 | -0.039 | 1.51 | 0.15 | 0.32 | 0.14 | 0.03 | 0.23 | 0.05 |
| 5 | 109452238 | rs13155924 | -1.274 | -0.323 | 1.773 | 0.856 | 0.023 | 0.004 | 0.004 | 0.553 | 0.09 | 0.31 | 0.09 | 0.03 | 0.12 | 0.03 |
| 5 | 109475723 | rs8180401 | 2.522 | -0.811 | 1.54 | 0.667 | -0.009 | 0.005 | 0.005 | 0.388 | 0.71 | 0.38 | 0.71 | 0.67 | 0.8 | 0.83 |
| 5 | 109477933 | rs11750108 | 1.876 | -0.796 | 1.555 | 0.239 | -0.039 | -0.031 | -0.031 | 0.547 | 0.87 | 0.39 | 0.83 | 0.94 | 0.95 | 0.91 |
| 5 | 109482778 | rs11241055 | 1.745 | -0.738 | 1.114 | 0.176 | -0.025 | -0.057 | -0.057 | 1.748 | 0.76 | 0.69 | 0.85 | 0.93 | 0.82 | 0.85 |
| 5 | 109491008 | rs7736052 | -1.175 | -0.61 | 0.96 | 0.122 | -0.003 | -0.009 | -0.009 | 10.028 | 0.15 | 0.22 | 0.1 | 0.03 | 0.14 | 0.06 |
| 5 | 109492649 | rs4615334 | 0.956 | -0.355 | 0.428 | 0.016 | 0.003 | -0.005 | -0.005 | 6.495 | 0.7 | 0.76 | 0.85 | 0.96 | 0.75 | 0.77 |
| 5 | 109497404 | rs9326794 | 0.392 | -0.346 | 0.212 | 0.53 | -0.008 | 0.021 | 0.021 | 0 | 0.69 | 0.62 | 0.56 | 0.4 | 0.73 | 0.52 |
| 5 | 109499203 | rs4429894 | 0.932 | -0.445 | 0.354 | 0.968 | 0.018 | -0.002 | -0.002 | 0.249 | 0.63 | 0.38 | 0.56 | 0.4 | 0.79 | 0.35 |
| 5 | 109502028 | rs4957876 | -0.057 | -0.463 | 0.13 | 1.613 | 0.007 | -0.006 | -0.006 | 0.025 | 0.07 | 0.04 | 0.13 | 0.24 | 0.04 | 0.11 |
| 5 | 109505808 | rs4957878 | 0.712 | -0.341 | 0.19 | 1.365 | -0.038 | 0.022 | 0.022 | 0.518 | 0.65 | 0.48 | 0.61 | 0.44 | 0.8 | 0.4 |
| 5 | 109515646 | rs4246005 | 0.476 | -0.345 | 0.149 | 1.528 | -0.064 | 0.019 | 0.019 | 0.157 | 0.68 | 0.49 | 0.55 | 0.41 | 0.87 | 0.45 |
| 5 | 109517541 | rs10062617 | 0.74 | -0.36 | -0.069 | 1.449 | 0.082 | -0.048 | -0.048 | 0.003 | 0.63 | 0.23 | 0.59 | 0.44 | 0.8 | 0.39 |
| 5 | 109520259 | rs13161437 | 0.208 | -0.012 | -0.617 | 1.297 | -0.048 | 0.012 | 0.012 | 0 | 0.1 | 0 | 0.07 | 0.05 | 0.15 | 0.05 |
| 5 | 109529109 | rs6872409 | 1.041 | -0.053 | -0.163 | 1.581 | 0.049 | -0.005 | -0.005 | 0 | 0.59 | 0.23 | 0.5 | 0.41 | 0.71 | 0.33 |
| 5 | 109536583 | rs10515412 | -2.115 | 0.845 | 0.527 | 2.406 | 0.021 | 0.071 | 0.071 | 0 | 0.2 | 0.32 | 0.27 | 0.5 | 0.27 | 0.45 |
| 5 | 109537213 | rs4529253 | -1.267 | 0.718 | 0.475 | 2.56 | -0.014 | -0.027 | -0.027 | 0 | 0.1 | 0.04 | 0.11 | 0.3 | 0.13 | 0.42 |
| 5 | 109546414 | rs6869190 | -1.96 | -0.156 | 0.694 | 2.596 | 0.002 | 0.091 | 0.091 | 0 | 0.16 | 0.26 | 0.19 | 0.48 | 0.17 | 0.35 |
| 5 | 109556518 | rs4315966 | -0.657 | -0.342 | 0.685 | 2.737 | -0.053 | 0.217 | 0.217 | 0.249 | 0.13 | 0.43 | 0.16 | 0.25 | 0.05 | 0.12 |
| 5 | 109559466 | rs11959425 | -2.136 | 0.481 | 0.527 | 2.579 | 0.041 | 0.159 | 0.159 | 0 | 0.16 | 0.36 | 0.22 | 0.46 | 0.22 | 0.36 |
| 5 | 109594268 | rs4254949 | -2.251 | 0.033 | 0.369 | 2.455 | -0.003 | 0.163 | 0.163 | 0 | 0.2 | 0.42 | 0.22 | 0.5 | 0.19 | 0.35 |
| 5 | 109601377 | rs6881487 | -1.304 | -0.075 | -0.125 | 2.007 | -0.007 | 0.053 | 0.053 | 0 | 0.09 | 0.12 | 0.08 | 0.28 | 0.07 | 0.28 |
| 5 | 109608640 | rs4532403 | -0.717 | -0.463 | -0.185 | 1.522 | -0.034 | 0.017 | 0.017 | 0 | 0.36 | 0.45 | 0.28 | 0.59 | 0.24 | 0.53 |
| 5 | 109621282 | rs7733338 | -1.101 | 0.2 | -0.179 | 1.58 | -0.017 | -0.008 | -0.008 | 2.036 | 0.06 | 0.03 | 0.05 | 0.07 | 0.11 | 0.05 |
| 5 | 109630328 | rs978830 | -1.448 | -0.641 | 1.916 | 2.313 | 0.018 | -0.025 | -0.025 | 0.102 | 0.21 | 0.81 | 0.31 | 0.4 | 0.11 | 0.24 |
| 5 | 109630971 | rs17131910 | -1.258 | -0.684 | 2.577 | 2.516 | -0.011 | 0.14 | 0.14 | 0.021 | 0.05 | 0.13 | 0.14 | 0.25 | 0.02 | 0.02 |
| 5 | 109631614 | rs12655761 | -2.2 | -0.701 | 2.81 | 2.534 | -0.019 | 0.115 | 0.115 | 0 | 0.15 | 0.32 | 0.16 | 0.34 | 0.09 | 0.2 |
| 5 | 109636606 | rs10478008 | -2.993 | -0.878 | 3.135 | 2.508 | 0.052 | -0.074 | -0.074 | 0 | 0.23 | 0.87 | 0.31 | 0.41 | 0.12 | 0.25 |
| 5 | 109643438 | rs6860004 | -2.334 | -0.822 | 3.391 | 2.546 | -0.029 | 0.106 | 0.106 | 0 | 0.16 | 0.56 | 0.18 | 0.35 | 0.09 | 0.19 |
| 5 | 109659029 | rs10038763 | -2.848 | -0.836 | 3.381 | 2.56 | 0.007 | -0.01 | -0.01 | 0 | 0.21 | 0.78 | 0.29 | 0.4 | 0.11 | 0.23 |
| 5 | 109669086 | rs13164939 | -2.008 | -0.864 | 3.897 | 2.486 | -0.019 | -0.01 | -0.01 | 0 | 0.13 | 0.28 | 0.04 | 0.12 | 0.07 | 0.18 |
| 5 | 109677736 | rs12522313 | -3.206 | -0.884 | 3.926 | 2.601 | -0.008 | 0.002 | 0.002 | 1.047 | 0.2 | 0.7 | 0.17 | 0.27 | 0.11 | 0.22 |
| 5 | 109678926 | rs6858968 | -3.211 | -0.883 | 4.021 | 2.581 | -0.037 | 0.077 | 0.077 | 0.5 | 0.2 | 0.57 | 0.21 | 0.35 | 0.11 | 0.22 |
| 5 | 109680785 | rs6889219 | -2.726 | -0.882 | 4.05 | 2.598 | -0.022 | 0.132 | 0.132 | 0 | 0.17 | 0.46 | 0.2 | 0.35 | 0.11 | 0.22 |
| 5 | 109689445 | rs13190009 | -1.635 | -0.873 | 4.041 | 2.548 | -0.008 | -0.007 | -0.007 | 0.684 | 0.11 | 0.18 | 0.03 | 0.11 | 0.07 | 0.19 |
| 5 | 109692143 | rs1004647 | -2.621 | -0.886 | 4.153 | 2.543 | -0.022 | -0.022 | -0.022 | 0.964 | 0.16 | 0.45 | 0.07 | 0.12 | 0.09 | 0.23 |
| 5 | 109703751 | rs6891701 | -3.123 | -0.86 | 4.116 | 2.592 | -0.019 | 0.032 | 0.032 | 0 | 0.19 | 0.67 | 0.22 | 0.35 | 0.11 | 0.24 |
| 5 | 109708961 | rs11955625 | -1.725 | -0.961 | 4.148 | 2.551 | -0.008 | -0.009 | -0.009 | 0 | 0.11 | 0.16 | 0.03 | 0.11 | 0.07 | 0.2 |
| 5 | 109709089 | rs12234029 | -2.692 | -0.936 | 4.273 | 2.563 | -0.015 | 0.157 | 0.157 | 0.18 | 0.07 | 0.23 | 0.15 | 0.24 | 0.02 | 0.01 |
| 5 | 109730678 | rs7714521 | -1.693 | -1.069 | 4.014 | 2.473 | -0.008 | -0.006 | -0.006 | 0.239 | 0.11 | 0.17 | 0.03 | 0.11 | 0.07 | 0.19 |
| 5 | 109732649 | rs17132048 | -3.832 | -1.092 | 4.078 | 2.494 | -0.045 | 0.116 | 0.116 | 1.392 | 0.12 | 0.63 | 0.27 | 0.29 | 0.05 | 0.07 |
| 5 | 109736454 | rs6877527 | -3.153 | -1.097 | 4.009 | 2.49 | 0.009 | -0.012 | -0.012 | 0 | 0.2 | 0.76 | 0.24 | 0.35 | 0.11 | 0.23 |
| 5 | 109743487 | rs12523032 | -3.089 | -1.076 | 4.001 | 2.499 | -0.018 | 0.202 | 0.202 | 0 | 0.1 | 0.36 | 0.26 | 0.29 | 0.05 | 0.07 |
| 5 | 109745600 | rs311721 | -3.113 | -1.06 | 3.536 | 2.466 | -0.026 | 0.038 | 0.038 | 0.425 | 0.22 | 0.67 | 0.29 | 0.39 | 0.12 | 0.27 |
| 5 | 109751913 | rs311700 | -2.948 | -1.059 | 3.61 | 2.413 | -0.032 | -0.012 | -0.012 | 0.328 | 0.18 | 0.62 | 0.15 | 0.16 | 0.1 | 0.25 |
| 5 | 109758964 | rs311689 | -3.347 | -1.075 | 3.622 | 2.52 | -0.022 | 0.029 | 0.029 | 0.558 | 0.22 | 0.69 | 0.3 | 0.41 | 0.12 | 0.26 |
| 5 | 109764417 | rs7735155 | -1.508 | -1.053 | 3.76 | 2.483 | -0.005 | -0.004 | -0.004 | 0.123 | 0.1 | 0.17 | 0.03 | 0.11 | 0.07 | 0.19 |
| 5 | 109775047 | rs1025733 | -2.922 | -1.071 | 3.859 | 2.501 | -0.025 | 0.139 | 0.139 | 0 | 0.17 | 0.48 | 0.28 | 0.33 | 0.11 | 0.23 |
| 5 | 109798056 | rs313611 | -3.373 | -1.245 | 3.701 | 2.849 | 0.032 | -0.046 | -0.046 | 0 | 0.23 | 0.8 | 0.24 | 0.41 | 0.14 | 0.24 |
| 5 | 109802199 | rs720402 | -2.82 | -1.153 | 4.099 | 2.883 | -0.025 | 0.117 | 0.117 | 0 | 0.18 | 0.32 | 0.2 | 0.41 | 0.12 | 0.22 |
| 5 | 109817549 | rs313617 | -3.724 | -1.21 | 4.019 | 2.886 | 0.058 | -0.117 | -0.117 | 0.048 | 0.24 | 0.87 | 0.25 | 0.46 | 0.14 | 0.28 |
| 5 | 109828383 | rs6860301 | -5.11 | -1.234 | 4.129 | 2.862 | 0.021 | 0.055 | 0.055 | 2.613 | 0.07 | 0.54 | 0.05 | 0.05 | 0.03 | 0.05 |
| 5 | 109833727 | rs10035983 | -3.729 | -1.247 | 4.016 | 2.909 | 0.055 | -0.087 | -0.087 | 0.497 | 0.24 | 0.85 | 0.25 | 0.44 | 0.14 | 0.27 |
| 5 | 109855077 | rs7722640 | -3.76 | -1.248 | 3.992 | 2.92 | 0.056 | -0.09 | -0.09 | 0 | 0.24 | 0.85 | 0.25 | 0.43 | 0.14 | 0.27 |
| 5 | 109858821 | rs7701890 | -2.607 | -1.235 | 4.179 | 2.859 | -0.016 | 0.006 | 0.006 | 0 | 0.15 | 0.27 | 0.07 | 0.16 | 0.12 | 0.23 |
| 5 | 109872207 | rs12189010 | -1.753 | -1.234 | 4.039 | 2.971 | -0.002 | -0.045 | -0.045 | 0 | 0.1 | 0 | 0.03 | 0.17 | 0.08 | 0.21 |
| 5 | 109880440 | rs12332108 | -3.556 | -1.242 | 4.413 | 2.974 | -0.039 | 0.012 | 0.012 | 0 | 0.18 | 0.61 | 0.09 | 0.21 | 0.11 | 0.26 |
| 5 | 109883334 | rs6861829 | -5.071 | -1.235 | 4.747 | 2.959 | 0.054 | 0.094 | 0.094 | 2.933 | 0.07 | 0.48 | 0.05 | 0.05 | 0.03 | 0.05 |
| 5 | 109885397 | rs10477457 | -4.49 | -1.233 | 4.731 | 2.96 | 0.044 | 0.073 | 0.073 | 0.188 | 0.05 | 0.31 | 0.04 | 0.05 | 0.03 | 0.05 |
| 5 | 109910067 | rs4317370 | -4.573 | -1.179 | 4.395 | 2.99 | 0.034 | -0.111 | -0.111 | 1.167 | 0.3 | 0.85 | 0.36 | 0.49 | 0.23 | 0.3 |
| 5 | 109915100 | rs4128660 | -1.485 | -1.169 | 4.386 | 2.98 | -0.001 | -0.001 | -0.001 | 0 | 0.07 | 0.07 | 0.12 | 0.06 | 0.09 | 0.07 |
| 5 | 109928666 | rs9326801 | -1.982 | -1.173 | 4.437 | 2.987 | -0.011 | 0.01 | 0.01 | 0 | 0.11 | 0.14 | 0.04 | 0.17 | 0.08 | 0.21 |
| 5 | 109946411 | rs4277958 | -2.759 | -1.181 | 4.507 | 3.013 | 0 | 0.085 | 0.085 | 1.27 | 0.12 | 0.26 | 0.21 | 0.27 | 0.12 | 0.08 |
| 5 | 109952983 | rs12186893 | -1.488 | -1.188 | 4.477 | 3.046 | -0.001 | -0.001 | -0.001 | 0.446 | 0.07 | 0.07 | 0.12 | 0.08 | 0.09 | 0.07 |
| 5 | 109955062 | rs10064687 | -4.457 | -1.117 | 4.477 | 3.05 | 0.03 | -0.115 | -0.115 | 1.769 | 0.3 | 0.84 | 0.37 | 0.49 | 0.23 | 0.3 |
| 5 | 109962107 | rs7720729 | -3.253 | -1.119 | 4.507 | 2.989 | 0.028 | 0 | 0 | 3.131 | 0.13 | 0.52 | 0.17 | 0.07 | 0.12 | 0.08 |

| | | | | | | | | | | | | | | | | |
|---|---|---|---|---|---|---|---|---|---|---|---|---|---|---|---|---|
| 5 | 109963106 | rs6886299 | -3.565 | -1.117 | 4.564 | 2.994 | -0.017 | 0.004 | 0.004 | 0 | 0.23 | 0.57 | 0.2 | 0.24 | 0.21 | 0.27 |
| 5 | 109964662 | rs12152845 | -1.901 | -1.104 | 4.373 | 2.993 | 0.001 | -0.002 | -0.002 | 0 | 0.11 | 0.07 | 0.03 | 0.17 | 0.09 | 0.21 |
| 5 | 109984076 | rs1422488 | 1.369 | -1.108 | 4.293 | 2.972 | -0.001 | -0.001 | -0.001 | 0 | 0.93 | 0.92 | 0.88 | 0.94 | 0.91 | 0.93 |
| 5 | 110021869 | rs244408 | 3.697 | -1.194 | 4.533 | 2.963 | -0.017 | 0.006 | 0.006 | 0.067 | 0.77 | 0.43 | 0.81 | 0.75 | 0.8 | 0.73 |
| 5 | 110035756 | rs13360927 | -2.037 | -1.193 | 4.511 | 2.96 | -0.011 | 0.011 | 0.011 | 0 | 0.11 | 0.14 | 0.04 | 0.17 | 0.08 | 0.19 |
| 5 | 110052401 | rs10036528 | -2.908 | -1.2 | 4.514 | 2.99 | -0.032 | 0.013 | 0.013 | 0 | 0.16 | 0.39 | 0.07 | 0.17 | 0.11 | 0.22 |
| 5 | 110070759 | rs244929 | 4.142 | -1.378 | 4.311 | 2.971 | 0.042 | -0.078 | -0.078 | 1.115 | 0.69 | 0.14 | 0.64 | 0.61 | 0.77 | 0.69 |
| 5 | 110075414 | rs665063 | 2.117 | -1.366 | 4.409 | 2.912 | 0.034 | 0.03 | 0.03 | 4.2 | 0.91 | 0.71 | 0.87 | 0.92 | 0.9 | 0.92 |
| 5 | 110079450 | rs17446534 | -0.916 | -1.364 | 4.245 | 2.907 | -0.009 | 0.012 | 0.012 | 7.062 | 0.06 | 0.02 | 0.05 | 0.02 | 0.09 | 0.03 |
| 5 | 110082585 | rs10069246 | -2.063 | -1.36 | 4.174 | 2.924 | -0.011 | 0.009 | 0.009 | 0 | 0.12 | 0.14 | 0.04 | 0.17 | 0.08 | 0.22 |
| 5 | 110085098 | rs381661 | 1.783 | -1.132 | 3.861 | 2.818 | -0.033 | 0.007 | 0.007 | 4.579 | 0.92 | 0.98 | 0.88 | 0.96 | 0.88 | 0.92 |
| 5 | 110087158 | rs456236 | 1.413 | -1.174 | 2.382 | 1.884 | -0.166 | 0.163 | 0.163 | 0 | 0.36 | 0.02 | 0.39 | 0.16 | 0.4 | 0.3 |
| 5 | 110089350 | rs12522587 | -1.985 | -1.07 | 3.182 | 2.324 | 0.01 | 0.063 | 0.063 | 1.132 | 0.12 | 0.18 | 0.21 | 0.22 | 0.12 | 0.03 |
| 5 | 110095559 | rs7718427 | -3.099 | -1.054 | 3.326 | 2.347 | 0.016 | 0.14 | 0.14 | 0 | 0.11 | 0.4 | 0.21 | 0.19 | 0.09 | 0.02 |
| 5 | 110095938 | rs27591 | 3.568 | -1.042 | 2.937 | 2.422 | 0.026 | -0.066 | -0.066 | 0 | 0.7 | 0.24 | 0.64 | 0.55 | 0.76 | 0.69 |
| 5 | 110096939 | rs7724788 | -2.183 | -1.085 | 2.891 | 2.21 | -0.028 | 0.031 | 0.031 | 0 | 0.15 | 0.46 | 0.06 | 0.17 | 0.09 | 0.23 |
| 5 | 110117283 | rs467765 | 2.53 | -0.963 | 2.198 | 2.396 | -0.001 | 0.021 | 0.021 | 0.883 | 0.86 | 0.87 | 0.78 | 0.77 | 0.87 | 0.97 |
| 5 | 110120915 | rs151809 | 2.291 | -0.978 | 2.196 | 2.331 | -0.001 | 0.037 | 0.037 | 0.546 | 0.87 | 0.86 | 0.79 | 0.75 | 0.88 | 0.97 |
| 5 | 110143168 | rs10060413 | -1.621 | -0.907 | 2.05 | 1.989 | 0.004 | -0.008 | -0.008 | 0 | 0.12 | 0.08 | 0.11 | 0.22 | 0.1 | 0.25 |
| 5 | 110170026 | rs10069077 | -1.985 | -0.943 | 2.512 | 1.987 | -0.022 | 0.037 | 0.037 | 0 | 0.14 | 0.49 | 0.12 | 0.18 | 0.08 | 0.24 |
| 5 | 110170834 | rs6594475 | -3.566 | -1.04 | 2.407 | 1.926 | 0.028 | 0.035 | 0.035 | 0.417 | 0.3 | 0.8 | 0.32 | 0.26 | 0.23 | 0.28 |
| 5 | 110175187 | rs7730010 | -2.667 | -1.038 | 2.469 | 1.935 | 0 | -0.028 | -0.028 | 0.051 | 0.16 | 0.26 | 0.2 | 0.1 | 0.15 | 0.03 |
| 5 | 110176128 | rs9885413 | -1.789 | -1.029 | 2.539 | 1.929 | -0.016 | 0.043 | 0.043 | 0 | 0.13 | 0.48 | 0.12 | 0.17 | 0.08 | 0.25 |
| 5 | 110182071 | rs1870072 | -2.16 | -1.005 | 2.189 | 1.904 | 0.001 | -0.018 | -0.018 | 0 | 0.11 | 0.13 | 0.08 | 0.01 | 0.11 | 0.02 |
| 5 | 110183740 | rs7718121 | -2.309 | -1.007 | 2.196 | 1.904 | 0.001 | -0.023 | -0.023 | 0 | 0.11 | 0.16 | 0.09 | 0.01 | 0.11 | 0.02 |
| 5 | 110193731 | rs4432941 | -3.581 | -1.041 | 2.086 | 1.919 | 0.024 | 0.03 | 0.03 | 0.306 | 0.29 | 0.77 | 0.32 | 0.26 | 0.23 | 0.28 |
| 5 | 110198463 | rs1345767 | -2.591 | -1.037 | 2.155 | 1.938 | -0.017 | 0.003 | 0.003 | 0 | 0.18 | 0.66 | 0.24 | 0.25 | 0.11 | 0.25 |
| 5 | 110207981 | rs1363797 | -3.142 | -1.117 | 1.936 | 1.908 | 0.03 | 0.027 | 0.027 | 0.477 | 0.3 | 0.79 | 0.32 | 0.27 | 0.22 | 0.28 |
| 5 | 110215781 | rs2900053 | 2.308 | -1.767 | 1.708 | 2.395 | 0.039 | -0.021 | -0.021 | 0 | 0.77 | 0.13 | 0.79 | 0.76 | 0.89 | 0.87 |
| 5 | 110237388 | rs17132484 | -2.19 | -1.558 | 2.694 | 1.974 | -0.011 | 0.03 | 0.03 | 0 | 0.12 | 0.34 | 0.11 | 0.16 | 0.08 | 0.21 |
| 5 | 110246355 | rs6876294 | -3.573 | -1.427 | 3.044 | 2.032 | 0.01 | 0 | 0 | 0.423 | 0.21 | 0.84 | 0.24 | 0.25 | 0.13 | 0.27 |
| 5 | 110253751 | rs6895612 | -2.686 | -1.43 | 3.372 | 2.026 | -0.001 | 0.002 | 0.002 | 0 | 0.14 | 0.5 | 0.14 | 0.15 | 0.1 | 0.21 |
| 5 | 110254559 | rs13340348 | -3.995 | -1.311 | 3.257 | 2.029 | 0.01 | -0.001 | -0.001 | 0 | 0.23 | 0.84 | 0.25 | 0.29 | 0.15 | 0.29 |
| 5 | 110256843 | rs12332258 | -2.403 | -1.257 | 3.667 | 1.992 | -0.002 | 0.022 | 0.022 | 0 | 0.13 | 0.25 | 0.12 | 0.18 | 0.1 | 0.21 |
| 5 | 110259077 | rs10056243 | -1.291 | -1.257 | 3.515 | 2.102 | 0.004 | -0.01 | -0.01 | 0 | 0.09 | 0.07 | 0.1 | 0.17 | 0.08 | 0.21 |
| 5 | 110287637 | rs13360174 | -1.561 | -1.241 | 3.496 | 2.088 | -0.003 | -0.02 | -0.02 | 0 | 0.1 | 0.05 | 0.11 | 0.17 | 0.1 | 0.13 |
| 5 | 110303365 | rs10043173 | -2.93 | -1.324 | 3.74 | 2.187 | -0.032 | 0.071 | 0.071 | 0 | 0.14 | 0.62 | 0.2 | 0.27 | 0.08 | 0.19 |
| 5 | 110304640 | rs12656259 | -2.454 | -1.32 | 3.79 | 2.19 | -0.02 | 0.108 | 0.108 | 0 | 0.12 | 0.37 | 0.19 | 0.27 | 0.08 | 0.19 |
| 5 | 110319359 | rs17132582 | -0.969 | -1.171 | 3.877 | 2.226 | 0.002 | 0.021 | 0.021 | 0 | 0.07 | 0.08 | 0.1 | 0.16 | 0.08 | 0.13 |
| 5 | 110331899 | rs17132590 | -3.503 | -0.995 | 4.033 | 2.193 | -0.009 | 0.044 | 0.044 | 2.101 | 0.14 | 0.69 | 0.22 | 0.26 | 0.11 | 0.2 |
| 5 | 110333886 | rs12109885 | -1.973 | -1.105 | 3.664 | 2.19 | 0.009 | 0.073 | 0.073 | 0 | 0.09 | 0.27 | 0.11 | 0.16 | 0.08 | 0.12 |
| 5 | 110345020 | rs7726472 | -3.271 | -1.17 | 3.338 | 1.809 | -0.029 | 0.043 | 0.043 | 1.781 | 0.16 | 0.72 | 0.21 | 0.27 | 0.09 | 0.2 |
| 5 | 110386641 | rs10069215 | -2.013 | -1.211 | 1.414 | 1.528 | -0.004 | 0.002 | 0.002 | 0 | 0.11 | 0.13 | 0.12 | 0.16 | 0.08 | 0.17 |
| 5 | 110389943 | rs10060511 | 0.191 | -0.573 | 0.496 | 0.735 | -0.002 | -0.073 | -0.073 | 0.859 | 0.32 | 0.56 | 0.34 | 0.12 | 0.3 | 0.11 |
| 5 | 110399033 | rs991035 | -0.625 | 0.084 | -0.612 | 1.174 | -0.163 | 0.001 | 0.001 | 0.732 | 0.65 | 0.95 | 0.62 | 0.72 | 0.55 | 0.59 |
| 5 | 110399376 | rs1545169 | -0.392 | 0.145 | 0.661 | 2.146 | -0.018 | 0.221 | 0.221 | 0 | 0.08 | 0.19 | 0.12 | 0.38 | 0.05 | 0.21 |
| 5 | 110401872 | rs1837253 | 0.872 | 1.128 | 0.032 | 1.258 | 0.007 | 0.018 | 0.018 | 0 | 0.75 | 0.69 | 0.73 | 0.39 | 0.75 | 0.54 |
| 5 | 110404185 | rs17551370 | -1.149 | 1.288 | 0.201 | 1.017 | -0.001 | 0 | 0 | 0 | 0.16 | 0.11 | 0.07 | 0.07 | 0.14 | 0.29 |
| 5 | 110404999 | rs10455025 | 0.962 | 1.418 | -0.894 | 0.1 | -0.091 | 0.15 | 0.15 | 6.929 | 0.27 | 0.02 | 0.23 | 0.04 | 0.37 | 0.06 |
| 5 | 110405675 | rs3806932 | 0.106 | 1.809 | 0.195 | 0.281 | -0.005 | 0.003 | 0.003 | 3.802 | 0.6 | 0.64 | 0.51 | 0.35 | 0.46 | 0.67 |
| 5 | 110406742 | rs3806933 | 0.588 | 1.496 | 0.476 | 0.102 | 0.004 | 0.029 | 0.029 | 1.859 | 0.55 | 0.26 | 0.48 | 0.35 | 0.45 | 0.67 |
| 5 | 110407507 | rs2289276 | 1.535 | 1.483 | 0.455 | 0.098 | 0.058 | 0.049 | 0.049 | 0.718 | 0.4 | 0.15 | 0.41 | 0.28 | 0.31 | 0.37 |
| 5 | 110412585 | rs11466749 | -0.44 | 1.67 | 0.43 | 0.08 | -0.002 | 0.001 | 0.001 | 0 | 0.16 | 0.11 | 0.08 | 0.08 | 0.17 | 0.3 |
| 5 | 110412894 | rs11466750 | -0.637 | 1.472 | 0.894 | 0.079 | -0.003 | -0.028 | -0.028 | 0 | 0.17 | 0.22 | 0.09 | 0.08 | 0.14 | 0.3 |
| 5 | 110427347 | rs7723819 | -0.023 | 1.632 | 0.943 | 0.876 | -0.009 | -0.01 | -0.01 | 0 | 0.61 | 0.69 | 0.57 | 0.42 | 0.46 | 0.69 |
| 5 | 110435490 | rs2416257 | -0.411 | 1.685 | 1.016 | 0.892 | 0 | 0.004 | 0.004 | 0 | 0.16 | 0.12 | 0.08 | 0.08 | 0.14 | 0.29 |
| 5 | 110436450 | rs10038177 | -0.214 | 1.687 | 1.122 | 0.895 | -0.02 | -0.02 | -0.02 | 0 | 0.63 | 0.72 | 0.57 | 0.42 | 0.46 | 0.69 |
| 5 | 110438357 | rs10045255 | -0.269 | 1.678 | 1.22 | 0.89 | -0.03 | -0.03 | -0.03 | 0 | 0.63 | 0.75 | 0.57 | 0.43 | 0.46 | 0.69 |
| 5 | 110439509 | rs11241095 | 1.755 | 1.375 | 1.099 | 0.93 | 0.029 | 0.019 | 0.019 | 2.172 | 0.38 | 0.06 | 0.35 | 0.27 | 0.31 | 0.39 |
| 5 | 110441439 | rs1379300 | -0.212 | 1.391 | 1.325 | 0.937 | -0.008 | -0.008 | -0.008 | 0 | 0.62 | 0.66 | 0.57 | 0.43 | 0.46 | 0.69 |
| 5 | 110447853 | rs12515367 | 1.739 | 1.396 | 1.237 | 0.943 | 0.029 | 0.019 | 0.019 | 2.02 | 0.38 | 0.06 | 0.35 | 0.27 | 0.31 | 0.39 |
| 5 | 110450584 | rs10055177 | 0.112 | 1.343 | 1.305 | 0.911 | -0.01 | -0.012 | -0.012 | 0 | 0.59 | 0.71 | 0.57 | 0.42 | 0.45 | 0.69 |
| 5 | 110452845 | rs10051830 | 0.198 | 1.338 | 1.286 | 0.911 | -0.004 | -0.003 | -0.003 | 0 | 0.58 | 0.68 | 0.57 | 0.42 | 0.45 | 0.68 |
| 5 | 110455266 | rs12522383 | 1.669 | 1.303 | 1.314 | 0.9 | 0.053 | 0.046 | 0.046 | 1.462 | 0.39 | 0.17 | 0.36 | 0.27 | 0.31 | 0.39 |
| 5 | 110462230 | rs4530809 | 1.047 | 1.345 | 1.433 | 0.903 | 0.066 | 0.049 | 0.049 | 0.643 | 0.46 | 0.28 | 0.51 | 0.35 | 0.35 | 0.44 |
| 5 | 110464008 | rs1043828 | 1 | 1.165 | 1.214 | 0.608 | -0.075 | 0.151 | 0.151 | 6.992 | 0.27 | 0.08 | 0.25 | 0.04 | 0.37 | 0.08 |
| 5 | 110467237 | rs6870356 | -1.837 | 0.45 | 1.581 | 0.812 | -0.033 | -0.045 | -0.045 | 0.506 | 0.24 | 0.5 | 0.21 | 0.14 | 0.15 | 0.3 |
| 5 | 110474161 | rs1037684 | -1.509 | 0.689 | 1.168 | 0.805 | 0.003 | 0.005 | 0.005 | 0 | 0.26 | 0.39 | 0.24 | 0.26 | 0.24 | 0.43 |
| 5 | 110478926 | rs7704730 | -1.029 | 0.667 | 1.21 | 0.817 | 0.008 | 0.066 | 0.066 | 0 | 0.13 | 0.36 | 0.1 | 0.17 | 0.09 | 0.17 |

| | | | | | | | | | | | | | | | | |
|---|---|---|---|---|---|---|---|---|---|---|---|---|---|---|---|---|
| 5 | 110481203 | rs252712 | 0.898 | 1.215 | 0.38 | 1.25 | -0.035 | -0.037 | -0.037 | 0 | 0.11 | 0.04 | 0.16 | 0.33 | 0.21 | 0.08 |
| 5 | 110482490 | rs2028258 | 0.781 | 0.798 | 0.453 | 0.761 | -0.017 | -0.036 | -0.036 | 0.139 | 0.16 | 0.22 | 0.12 | 0.02 | 0.11 | 0.05 |
| 5 | 110485574 | rs91068 | 1.634 | 0.667 | -0.277 | 0.655 | 0.004 | 0.107 | 0.107 | 3.947 | 0.39 | 0.14 | 0.38 | 0.23 | 0.36 | 0.4 |
| 5 | 110490041 | rs9632452 | -0.755 | 0.245 | 0.322 | 0.697 | -0.021 | -0.008 | -0.008 | 0 | 0.35 | 0.58 | 0.26 | 0.28 | 0.22 | 0.23 |
| 5 | 110492586 | rs10478039 | -1.106 | 0.68 | 0.204 | 0.735 | 0 | 0 | 0 | 0 | 0.27 | 0.31 | 0.28 | 0.37 | 0.25 | 0.45 |
| 5 | 110493614 | rs17459022 | 0.288 | 0.452 | 0.444 | 0.687 | -0.001 | -0.004 | -0.004 | 0 | 0.11 | 0.09 | 0.06 | 0.05 | 0.04 | 0.07 |
| 5 | 110494282 | rs151800 | -0.406 | 0.768 | 0.591 | 0.951 | -0.016 | -0.012 | -0.012 | 0 | 0.82 | 0.89 | 0.83 | 0.76 | 0.75 | 0.91 |
| 5 | 110494617 | rs17459203 | -0.278 | 1.035 | 0.701 | 0.856 | 0.002 | 0 | 0 | 0 | 0.13 | 0.17 | 0.16 | 0.14 | 0.2 | 0.21 |
| 5 | 110496071 | rs1821908 | -1.288 | 0.978 | 0.988 | 0.801 | 0.017 | -0.02 | -0.02 | 0 | 0.27 | 0.43 | 0.25 | 0.19 | 0.26 | 0.31 |
| 5 | 110498972 | rs17132862 | -0.416 | 0.992 | 1.265 | 0.89 | 0.006 | 0.094 | 0.094 | 0 | 0.06 | 0.18 | 0.09 | 0.18 | 0.04 | 0.15 |
| 5 | 110501681 | rs244515 | 2.566 | 0.626 | 1.332 | 0.807 | 0.015 | 0.042 | 0.042 | 6.34 | 0.34 | 0.16 | 0.39 | 0.27 | 0.32 | 0.4 |
| 5 | 110502044 | rs10060762 | -1.226 | 0.719 | 1.584 | 0.888 | 0.015 | -0.013 | -0.013 | 0 | 0.28 | 0.51 | 0.26 | 0.21 | 0.26 | 0.34 |
| 5 | 110502142 | rs1428644 | -1.937 | 0.725 | 1.449 | 0.896 | -0.001 | -0.004 | -0.004 | 0.058 | 0.34 | 0.69 | 0.35 | 0.39 | 0.3 | 0.47 |
| 5 | 110503378 | rs10063014 | -1.031 | 0.72 | 1.562 | 0.842 | 0.015 | -0.005 | -0.005 | 0 | 0.26 | 0.38 | 0.25 | 0.21 | 0.26 | 0.35 |
| 5 | 110504157 | rs1806353 | 1.771 | 0.755 | 1.331 | 0.873 | 0.045 | 0.061 | 0.061 | 1.369 | 0.49 | 0.28 | 0.49 | 0.37 | 0.44 | 0.45 |
| 5 | 110504412 | rs10040556 | 0.224 | 0.791 | 1.318 | 0.905 | 0.018 | 0.031 | 0.031 | 0.71 | 0.09 | 0.17 | 0.14 | 0.14 | 0.13 | 0.26 |
| 5 | 110521535 | rs17132904 | -1.471 | 0.686 | 1.386 | 0.843 | -0.002 | -0.003 | -0.003 | 0 | 0.24 | 0.61 | 0.23 | 0.24 | 0.19 | 0.24 |
| 5 | 110527505 | rs10515424 | 0.555 | 0.647 | 1.457 | 0.747 | -0.003 | -0.017 | -0.017 | 0.623 | 0.1 | 0.1 | 0.08 | 0.05 | 0.07 | 0.1 |
| 5 | 110536512 | rs10477464 | -0.592 | 0.709 | 1.537 | 0.808 | 0.004 | 0.007 | 0.007 | 0.204 | 0.18 | 0.25 | 0.2 | 0.19 | 0.18 | 0.29 |
| 5 | 110537352 | rs1370965 | 1.1 | 0.734 | 1.505 | 0.802 | 0.01 | -0.008 | -0.008 | 0 | 0.05 | 0.07 | 0.05 | 0.02 | 0.08 | 0.02 |
| 5 | 110539892 | rs184941 | -0.66 | 0.467 | 0.807 | 0.371 | -0.003 | 0 | 0 | 1.014 | 0.23 | 0.22 | 0.26 | 0.19 | 0.23 | 0.41 |
| 5 | 110542451 | rs17132949 | 1.67 | 0.354 | 0.413 | 0.368 | 0.025 | 0.007 | 0.007 | 0.511 | 0.15 | 0.03 | 0.09 | 0.1 | 0.12 | 0.1 |
| 5 | 110542854 | rs252858 | 0.64 | 0.488 | 0.319 | 0.298 | 0.003 | 0.013 | 0.013 | 0 | 0.1 | 0.14 | 0.16 | 0.14 | 0.1 | 0.28 |
| 5 | 110544623 | rs7709420 | 2.259 | 0.443 | 0.369 | 0.367 | 0.019 | 0.015 | 0.015 | 6.054 | 0.3 | 0.18 | 0.32 | 0.27 | 0.26 | 0.28 |
| 5 | 110545139 | rs1469441 | 0.565 | 0.548 | 0.203 | 0.208 | 0.01 | 0.029 | 0.029 | 0.419 | 0.08 | 0.14 | 0.12 | 0.14 | 0.11 | 0.19 |
| 5 | 110553402 | rs919334 | -0.754 | 0.198 | 0.23 | 0.236 | 0.015 | -0.031 | -0.031 | 0 | 0.34 | 0.48 | 0.34 | 0.21 | 0.34 | 0.48 |
| 5 | 110558882 | rs1457114 | 0.339 | 0.269 | -0.178 | 0.425 | -0.001 | 0.011 | 0.011 | 5.004 | 0.65 | 0.78 | 0.73 | 0.68 | 0.62 | 0.84 |
| 5 | 110561639 | rs1551565 | -0.047 | 0.824 | 0.039 | 0.311 | 0.003 | -0.009 | -0.009 | 0.056 | 0.21 | 0.22 | 0.2 | 0.07 | 0.26 | 0.1 |
| 5 | 110562604 | rs727923 | 0.153 | 0.86 | -0.181 | 0.328 | -0.006 | 0.008 | 0.008 | 0.54 | 0.2 | 0.15 | 0.19 | 0.07 | 0.26 | 0.1 |
| 5 | 110566175 | rs10057913 | 0.467 | 0.402 | 0.2 | 0.697 | -0.028 | 0.137 | 0.137 | 0.02 | 0.82 | 0.49 | 0.67 | 0.62 | 0.9 | 0.82 |
| 5 | 110566901 | rs7704970 | 0.693 | -0.01 | -0.129 | 0.442 | 0.001 | -0.001 | -0.001 | 0.342 | 0.19 | 0.22 | 0.13 | 0.14 | 0.17 | 0.32 |
| 5 | 110567598 | rs1457115 | -0.505 | -0.057 | 0.016 | 0.231 | -0.001 | -0.004 | -0.004 | 0 | 0.44 | 0.5 | 0.34 | 0.24 | 0.44 | 0.42 |
| 5 | 110568458 | rs9285875 | -2.226 | 0.149 | -0.23 | 0.032 | -0.017 | -0.043 | -0.043 | 0 | 0.72 | 0.93 | 0.76 | 0.59 | 0.7 | 0.74 |
| 5 | 110568929 | rs6875225 | -0.89 | -0.069 | -0.072 | 0.099 | 0 | 0.011 | 0.011 | 0.205 | 0.5 | 0.5 | 0.4 | 0.24 | 0.51 | 0.54 |
| 5 | 110583710 | rs2434723 | 0.682 | -0.126 | -0.216 | 0.071 | -0.02 | 0.028 | 0.028 | 2.403 | 0.22 | 0.1 | 0.2 | 0.09 | 0.28 | 0.1 |
| 5 | 110584659 | rs952733 | 1.844 | -0.035 | -0.146 | 0.16 | -0.022 | 0.007 | 0.007 | 0.1 | 0.33 | 0.16 | 0.29 | 0.39 | 0.39 | 0.28 |
| 5 | 110586438 | rs7726469 | 0.488 | -0.077 | 0.006 | 0.201 | -0.025 | -0.002 | -0.002 | 0 | 0.22 | 0.38 | 0.14 | 0.15 | 0.17 | 0.32 |
| 5 | 110595375 | rs1651404 | -0.308 | -0.225 | 0.148 | -0.038 | -0.012 | -0.015 | -0.015 | 0 | 0.3 | 0.46 | 0.24 | 0.18 | 0.26 | 0.45 |
| 5 | 110598011 | rs1560074 | -0.011 | -0.267 | -0.1 | -0.091 | 0.002 | 0.004 | 0.004 | 0 | 0.28 | 0.28 | 0.22 | 0.15 | 0.26 | 0.44 |
| 5 | 110602871 | rs11748647 | 1.794 | -0.226 | -0.371 | -0.104 | -0.042 | 0.008 | 0.008 | 0.088 | 0.34 | 0.13 | 0.3 | 0.39 | 0.4 | 0.29 |
| 5 | 110603489 | rs1650790 | 1.325 | -0.269 | -0.323 | -0.062 | -0.067 | 0.037 | 0.037 | 1.194 | 0.4 | 0.13 | 0.37 | 0.39 | 0.48 | 0.4 |
| 5 | 110606141 | rs2288397 | 0.329 | -0.273 | -0.001 | 0.239 | -0.027 | 0.004 | 0.004 | 0 | 0.24 | 0.45 | 0.16 | 0.18 | 0.19 | 0.33 |
| 5 | 110620861 | rs216535 | 0.962 | -0.174 | -0.736 | -0.123 | 0.048 | 0.035 | 0.035 | 0.348 | 0.48 | 0.29 | 0.51 | 0.43 | 0.66 | 0.44 |
| 5 | 110621221 | rs306107 | 1.312 | -0.003 | -1.058 | -0.015 | -0.051 | -0.006 | -0.006 | 0 | 0.37 | 0.19 | 0.37 | 0.43 | 0.47 | 0.4 |
| 5 | 110631265 | rs306104 | 0.991 | -0.354 | -0.547 | -0.086 | 0.025 | 0.033 | 0.033 | 0 | 0.58 | 0.28 | 0.73 | 0.74 | 0.69 | 0.53 |
| 5 | 110632243 | rs1644510 | 0.049 | -0.032 | -1.184 | -0.022 | 0 | -0.001 | -0.001 | 0 | 0.26 | 0.27 | 0.2 | 0.19 | 0.24 | 0.39 |
| 5 | 110644689 | rs306083 | -0.028 | 0.141 | -1.124 | 0.351 | 0.005 | 0.038 | 0.038 | 2.075 | 0.88 | 0.99 | 0.96 | 0.96 | 0.93 | 0.93 |
| 5 | 110650965 | rs435021 | -0.787 | 0.719 | -1.032 | 0.671 | -0.025 | -0.038 | -0.038 | 0.241 | 0.3 | 0.26 | 0.52 | 0.4 | 0.39 | 0.39 |
| 5 | 110655407 | rs306079 | -1.03 | 0.497 | 0.267 | 0.668 | 0.013 | 0.027 | 0.027 | 0.11 | 0.33 | 0.47 | 0.57 | 0.52 | 0.41 | 0.42 |
| 5 | 110656694 | rs306076 | 0.276 | 0.194 | 0.533 | 0.202 | -0.002 | -0.051 | -0.051 | 0 | 0.78 | 0.79 | 0.93 | 0.99 | 0.85 | 0.92 |
| 5 | 110658834 | rs1593864 | 1.174 | 0.435 | 1.204 | 0.133 | 0.004 | 0.014 | 0.014 | 0 | 0.61 | 0.5 | 0.47 | 0.49 | 0.6 | 0.67 |
| 5 | 110677604 | rs415322 | -0.331 | 0.431 | 1.21 | 0.19 | 0.01 | -0.07 | -0.07 | 1.848 | 0.87 | 0.79 | 0.94 | 1 | 0.9 | 0.99 |
| 5 | 110682544 | rs1650798 | 1.861 | 0.422 | 1.243 | 0.194 | 0.004 | -0.013 | -0.013 | 1.966 | 0.46 | 0.24 | 0.35 | 0.48 | 0.45 | 0.47 |
| 5 | 110708854 | rs3797755 | 0.717 | 0.626 | 1.069 | 0.67 | -0.013 | -0.043 | -0.043 | 0 | 0.07 | 0.01 | 0.17 | 0.23 | 0.07 | 0.09 |
| 5 | 110718072 | rs306119 | -0.805 | 0.701 | 1.247 | 0.454 | -0.015 | 0.048 | 0.048 | 0 | 0.36 | 0.25 | 0.45 | 0.24 | 0.4 | 0.38 |
| 5 | 110719500 | rs306117 | -0.416 | 0.762 | 1.397 | 0.355 | 0.011 | -0.066 | -0.066 | 0.277 | 0.88 | 0.8 | 0.94 | 1 | 0.9 | 0.99 |
| 5 | 110733803 | rs455531 | -0.799 | 0.727 | 1.069 | 0.379 | -0.009 | 0.013 | 0.013 | 0 | 0.36 | 0.31 | 0.48 | 0.28 | 0.41 | 0.38 |
| 5 | 110735488 | rs307548 | -0.512 | 0.827 | 0.808 | 0.104 | -0.001 | -0.02 | -0.02 | 0 | 0.89 | 0.92 | 0.95 | 0.99 | 0.9 | 0.99 |
| 5 | 110756932 | rs307532 | 0.055 | 1.014 | 0.952 | 0.544 | -0.001 | 0.016 | 0.016 | 0 | 0.73 | 0.74 | 0.67 | 0.51 | 0.77 | 0.68 |
| 5 | 110759537 | rs10052530 | 1.566 | 1.217 | 0.429 | 0.571 | 0.027 | 0.021 | 0.021 | 0 | 0.47 | 0.12 | 0.33 | 0.45 | 0.43 | 0.47 |
| 5 | 110771056 | rs1820666 | -0.319 | 1.479 | 0.838 | 0.058 | 0.011 | -0.007 | -0.007 | 0 | 0.18 | 0.23 | 0.3 | 0.07 | 0.31 | 0.17 |
| 5 | 110772942 | rs759948 | 0.392 | 1.307 | 0.881 | 0.252 | -0.001 | 0 | 0 | 0.586 | 0.63 | 0.63 | 0.59 | 0.54 | 0.64 | 0.55 |
| 5 | 110773100 | rs759947 | 1.362 | 1.145 | -0.047 | 0.243 | 0.014 | -0.042 | -0.042 | 0.821 | 0.45 | 0.12 | 0.36 | 0.49 | 0.41 | 0.46 |
| 5 | 110774225 | rs9326835 | 1.414 | 0.982 | -0.218 | -0.094 | 0.009 | -0.007 | -0.007 | 0 | 0.34 | 0.3 | 0.3 | 0.37 | 0.29 | 0.39 |
| 5 | 110779829 | rs6883539 | 0.158 | 0.869 | -0.403 | -0.229 | -0.004 | -0.025 | -0.025 | 0 | 0.06 | 0.13 | 0.03 | 0 | 0.05 | 0.05 |
| 5 | 110781537 | rs17458552 | -0.971 | 1.221 | -0.492 | -0.249 | -0.019 | 0.012 | 0.012 | 0.346 | 0.2 | 0.21 | 0.29 | 0.09 | 0.36 | 0.22 |
| 5 | 110787629 | rs971193 | 1.365 | 1.183 | -0.486 | -0.288 | 0.025 | -0.025 | -0.025 | 0.004 | 0.48 | 0.18 | 0.39 | 0.44 | 0.43 | 0.39 |
| 5 | 110788012 | rs17536705 | -0.345 | 1.357 | -0.412 | 0.05 | -0.054 | 0.003 | 0.003 | 0 | 0.14 | 0.05 | 0.27 | 0.09 | 0.32 | 0.18 |
| 5 | 110791363 | rs429219 | 0.164 | 1.345 | -0.516 | 0.057 | 0.002 | -0.002 | -0.002 | 0 | 0.68 | 0.38 | 0.67 | 0.53 | 0.79 | 0.61 |

| | | | | | | | | | | | | | | | | |
|---|---|---|---|---|---|---|---|---|---|---|---|---|---|---|---|---|
| 5 | 110791533 | rs185162 | -0.4 | 1.983 | -1.258 | 0.423 | -0.087 | -0.018 | -0.018 | 0 | 0.13 | 0.02 | 0.26 | 0.09 | 0.31 | 0.18 |
| 5 | 110799106 | rs173681 | -0.006 | 1.665 | -1.284 | 0.349 | 0.01 | -0.001 | -0.001 | 0 | 0.65 | 0.75 | 0.71 | 0.51 | 0.78 | 0.59 |
| 5 | 110815311 | rs31624 | -0.095 | 2.354 | -0.004 | 0.207 | -0.031 | 0.02 | 0.02 | 0.18 | 0.06 | 0.01 | 0.12 | 0.04 | 0.14 | 0.13 |
| 5 | 110818421 | rs3733995 | -0.836 | 1.908 | 0.466 | 0.469 | 0.118 | 0.047 | 0.047 | 0 | 0.09 | 0.24 | 0.16 | 0.05 | 0.18 | 0.06 |
| 5 | 110825128 | rs31619 | -0.748 | 1.003 | 0.123 | 0.027 | -0.009 | -0.041 | -0.041 | 0 | 0.36 | 0.53 | 0.34 | 0.21 | 0.32 | 0.31 |
| 5 | 110825198 | rs42670 | -0.965 | 1.168 | 0.185 | 0.033 | 0 | 0.006 | 0.006 | 0 | 0.19 | 0.15 | 0.16 | 0.1 | 0.2 | 0.18 |
| 5 | 110840881 | rs152883 | -0.717 | 1.162 | 0.869 | 0.261 | 0.022 | 0.032 | 0.032 | 0 | 0.11 | 0.27 | 0.22 | 0.14 | 0.14 | 0.19 |
| 5 | 110845258 | rs152880 | -0.67 | 1.248 | 0.865 | 0.26 | 0.02 | 0.022 | 0.022 | 0 | 0.1 | 0.21 | 0.22 | 0.14 | 0.15 | 0.13 |
| 5 | 110847732 | rs17458866 | -1.229 | 1.618 | 0.54 | 0.305 | 0.016 | 0.005 | 0.005 | 0 | 0.1 | 0.15 | 0.16 | 0.12 | 0.14 | 0.12 |
| 5 | 110847967 | rs40992 | -2.265 | 1.218 | -0.076 | 0.503 | -0.158 | -0.139 | -0.139 | 0.668 | 0.45 | 0.98 | 0.54 | 0.57 | 0.48 | 0.53 |
| 5 | 110855952 | rs10058699 | -1.086 | 1.042 | 0.614 | 0.298 | 0.005 | 0.007 | 0.007 | 0 | 0.09 | 0.14 | 0.13 | 0.09 | 0.07 | 0.11 |
| 5 | 110859246 | rs251133 | -1.778 | 1.069 | 1.223 | 0.392 | 0.023 | 0.01 | 0.01 | 1.057 | 0.15 | 0.7 | 0.28 | 0.16 | 0.2 | 0.14 |
| 5 | 110874123 | rs247518 | -1.825 | 0.598 | 1.777 | 0.138 | -0.016 | -0.001 | -0.001 | 1.606 | 0.25 | 0.43 | 0.33 | 0.21 | 0.21 | 0.26 |
| 5 | 110878552 | rs247527 | -2.23 | 0.553 | 1.323 | -0.011 | 0.005 | 0.022 | 0.022 | 0 | 0.32 | 0.8 | 0.41 | 0.24 | 0.29 | 0.3 |
| 5 | 110882116 | rs428995 | -1.707 | 0.749 | 1.172 | 0.067 | -0.001 | -0.001 | -0.001 | 0 | 0.24 | 0.23 | 0.33 | 0.21 | 0.23 | 0.29 |
| 5 | 110884998 | rs403237 | -1.369 | 0.688 | 1.069 | 0.059 | -0.002 | -0.008 | -0.008 | 0 | 0.31 | 0.37 | 0.38 | 0.24 | 0.31 | 0.29 |
| 5 | 110895757 | rs13356092 | -1.079 | 0.343 | 0.54 | 0.213 | 0.005 | 0.002 | 0.002 | 0 | 0.1 | 0.1 | 0.15 | 0.09 | 0.1 | 0.13 |
| 5 | 110896260 | rs247545 | -1.35 | -0.061 | 0.524 | 0.195 | 0.007 | 0.005 | 0.005 | 0 | 0.4 | 0.6 | 0.37 | 0.28 | 0.26 | 0.31 |
| 5 | 110897076 | rs6873388 | -1.704 | 0.912 | 0.989 | 0.732 | 0.042 | 0.023 | 0.023 | 0 | 0.17 | 0.25 | 0.23 | 0.23 | 0.26 | 0.15 |
| 5 | 110897159 | rs6873138 | -1.51 | 0.742 | 0.975 | 0.746 | 0.004 | 0.003 | 0.003 | 0 | 0.15 | 0.17 | 0.19 | 0.18 | 0.19 | 0.15 |
| 5 | 110897510 | rs247548 | -2.082 | -0.18 | 1.486 | 0.879 | -0.017 | 0.054 | 0.054 | 0 | 0.28 | 0.51 | 0.4 | 0.43 | 0.23 | 0.29 |
| 5 | 110906071 | rs384970 | -2.159 | -0.278 | 1.748 | 0.881 | -0.021 | 0.053 | 0.053 | 0 | 0.28 | 0.47 | 0.4 | 0.43 | 0.24 | 0.31 |
| 5 | 110908503 | rs17133427 | -1.672 | -0.309 | 1.978 | 0.878 | 0.019 | 0.057 | 0.057 | 0 | 0.17 | 0.31 | 0.34 | 0.29 | 0.19 | 0.15 |
| 5 | 110910785 | rs421880 | -0.792 | -0.351 | 1.725 | 0.699 | -0.019 | 0.001 | 0.001 | 0 | 0.11 | 0.16 | 0.07 | 0.13 | 0.04 | 0.17 |
| 5 | 110920327 | rs399503 | -0.704 | -0.344 | 1.746 | 0.702 | -0.029 | 0.006 | 0.006 | 0 | 0.11 | 0.18 | 0.07 | 0.14 | 0.04 | 0.15 |
| 5 | 110924280 | rs6886266 | 2.43 | -0.419 | 2.119 | 0.671 | 0.011 | -0.021 | -0.021 | 0 | 0.69 | 0.28 | 0.57 | 0.56 | 0.75 | 0.64 |
| 5 | 110925502 | rs4957963 | 1.577 | -0.498 | 2.477 | 0.614 | -0.015 | -0.044 | -0.044 | 0 | 0.86 | 0.54 | 0.92 | 0.94 | 0.92 | 0.8 |
| 5 | 110925663 | rs10478055 | 0.515 | -0.502 | 2.786 | 0.609 | -0.006 | -0.048 | -0.048 | 0 | 0.9 | 0.83 | 0.95 | 1 | 0.92 | 0.88 |
| 5 | 110930883 | rs1478402 | 3.164 | -0.603 | 2.673 | 1.146 | 0.02 | -0.04 | -0.04 | 0 | 0.65 | 0.27 | 0.55 | 0.51 | 0.73 | 0.6 |
| 5 | 110939390 | rs7731999 | -1.341 | -0.556 | 2.712 | 0.935 | 0.006 | 0.017 | 0.017 | 0 | 0.16 | 0.18 | 0.32 | 0.24 | 0.19 | 0.13 |
| 5 | 110939806 | rs10044756 | -2.005 | -0.554 | 2.721 | 0.927 | -0.002 | 0.027 | 0.027 | 0 | 0.2 | 0.27 | 0.37 | 0.38 | 0.19 | 0.19 |
| 5 | 110949080 | rs13187021 | 2.568 | -0.512 | 2.726 | 0.916 | -0.003 | 0.006 | 0.006 | 0 | 0.7 | 0.63 | 0.57 | 0.61 | 0.73 | 0.68 |
| 5 | 110949387 | rs2016764 | 0.455 | -0.497 | 2.671 | 0.885 | -0.001 | -0.017 | -0.017 | 0.362 | 0.9 | 0.89 | 0.95 | 1 | 0.92 | 0.87 |
| 5 | 110955100 | rs9326839 | -0.887 | -0.485 | 2.613 | 0.917 | 0.011 | 0.007 | 0.007 | 0 | 0.1 | 0.11 | 0.17 | 0.11 | 0.14 | 0.09 |
| 5 | 110958782 | rs11241130 | 2.378 | -0.528 | 2.652 | 0.909 | -0.049 | 0.007 | 0.007 | 0.306 | 0.81 | 0.49 | 0.87 | 0.8 | 0.91 | 0.74 |
| 5 | 110966946 | rs10478061 | -1.551 | -0.524 | 2.751 | 0.916 | 0.014 | 0.053 | 0.053 | 0 | 0.16 | 0.29 | 0.33 | 0.29 | 0.19 | 0.13 |
| 5 | 110968579 | rs1370969 | 2.988 | -0.523 | 2.578 | 0.892 | 0.013 | -0.019 | -0.019 | 0 | 0.66 | 0.27 | 0.55 | 0.56 | 0.72 | 0.62 |
| 5 | 110974312 | rs4957619 | 1.736 | -0.417 | 1.865 | -0.119 | -0.017 | 0.005 | 0.005 | 0 | 0.44 | 0.19 | 0.28 | 0.56 | 0.42 | 0.51 |
| 5 | 110977318 | rs7726351 | 0.89 | -0.38 | 1.691 | -0.179 | -0.003 | -0.017 | -0.017 | 0 | 0.24 | 0.22 | 0.28 | 0 | 0.3 | 0.17 |
| 5 | 110977602 | rs1457718 | -1.005 | -0.366 | 1.638 | -0.121 | 0 | -0.007 | -0.007 | 0.065 | 0.45 | 0.5 | 0.67 | 0.4 | 0.48 | 0.36 |
| 5 | 110978403 | rs1457717 | -0.099 | -0.235 | 1.261 | -0.122 | -0.002 | -0.03 | -0.03 | 0 | 0.9 | 0.86 | 0.94 | 1 | 0.92 | 0.87 |
| 5 | 110979586 | rs1457716 | 0.185 | -0.494 | 1.398 | -0.249 | -0.007 | -0.063 | -0.063 | 1.079 | 0.3 | 0.46 | 0.35 | 0.15 | 0.31 | 0.25 |
| 5 | 110983350 | rs10515433 | 0.115 | -0.577 | 1.544 | -0.279 | 0.002 | -0.049 | -0.049 | 0 | 0.29 | 0.34 | 0.31 | 0.05 | 0.34 | 0.15 |
| 5 | 110987667 | rs6885593 | 0.899 | -0.452 | 1.318 | -0.349 | 0.001 | 0.156 | 0.156 | 0.432 | 0.55 | 0.11 | 0.42 | 0.71 | 0.59 | 0.53 |
| 5 | 110991602 | rs6594525 | 1.981 | -0.427 | 1.224 | -0.304 | -0.075 | -0.052 | -0.052 | 0.016 | 0.41 | 0.06 | 0.32 | 0.56 | 0.41 | 0.45 |
| 5 | 110992525 | rs13182708 | 1.039 | -0.429 | 1.528 | -0.503 | 0.018 | -0.096 | -0.096 | 0.042 | 0.23 | 0.38 | 0.27 | 0.05 | 0.29 | 0.15 |
| 5 | 110994777 | rs6866631 | 0.814 | -0.692 | 1.245 | -0.463 | -0.051 | 0.166 | 0.166 | 0.293 | 0.54 | 0.1 | 0.4 | 0.71 | 0.53 | 0.51 |
| 5 | 111002821 | rs6860625 | -1.844 | -0.377 | 1.959 | -0.02 | -0.019 | 0.096 | 0.096 | 0 | 0.18 | 0.41 | 0.16 | 0.32 | 0.13 | 0.11 |
| 5 | 111015405 | rs1673758 | 2.115 | -0.094 | 1.597 | -0.03 | -0.017 | 0.009 | 0.009 | 0 | 0.71 | 0.51 | 0.65 | 0.62 | 0.79 | 0.67 |
| 5 | 111017793 | rs657073 | 1.152 | -0.134 | 1.609 | -0.082 | -0.015 | -0.072 | -0.072 | 1.914 | 0.82 | 0.69 | 0.77 | 0.92 | 0.89 | 0.76 |
| 5 | 111021979 | rs1154308 | 2.559 | -0.066 | 1.624 | -0.01 | -0.013 | 0.005 | 0.005 | 0 | 0.67 | 0.53 | 0.61 | 0.59 | 0.73 | 0.66 |
| 5 | 111025672 | rs1145122 | 0.788 | -0.218 | 1.578 | -0.041 | -0.005 | -0.059 | -0.059 | 0 | 0.84 | 0.72 | 0.78 | 0.92 | 0.89 | 0.76 |
| 5 | 111028566 | rs154081 | 1.835 | -0.412 | 1.402 | -0.054 | -0.06 | -0.038 | -0.038 | 0.687 | 0.72 | 0.52 | 0.66 | 0.75 | 0.85 | 0.7 |
| 5 | 111030893 | rs26050 | 1.561 | -0.424 | 1.403 | -0.088 | 0.021 | 0.02 | 0.02 | 0 | 0.79 | 0.86 | 0.76 | 0.83 | 0.86 | 0.78 |
| 5 | 111033109 | rs17133647 | -1.754 | -0.431 | 1.629 | -0.122 | -0.018 | -0.083 | -0.083 | 0 | 0.16 | 0.47 | 0.25 | 0.07 | 0.1 | 0.11 |
| 5 | 111033627 | rs17133651 | -0.328 | -0.446 | 1.549 | -0.106 | 0.003 | -0.009 | -0.009 | 0 | 0.09 | 0.04 | 0.05 | 0.13 | 0.07 | 0.06 |
| 5 | 111034217 | rs26052 | 2.953 | -0.497 | 1.682 | -0.097 | 0.048 | 0.011 | 0.011 | 1.052 | 0.64 | 0.21 | 0.58 | 0.62 | 0.79 | 0.65 |
| 5 | 111036745 | rs10515434 | -0.128 | -0.433 | 1.667 | 0.433 | -0.001 | 0.031 | 0.031 | 0.036 | 0.05 | 0.06 | 0.05 | 0.21 | 0.06 | 0.06 |
| 5 | 111036995 | rs17133655 | -0.676 | -0.444 | 1.826 | 0.395 | 0.023 | -0.057 | -0.057 | 0.974 | 0.11 | 0.35 | 0.08 | 0 | 0.07 | 0.09 |
| 5 | 111037979 | rs26054 | 1.017 | -0.423 | 1.622 | 0.41 | -0.008 | 0.011 | 0.011 | 0 | 0.87 | 0.81 | 0.87 | 0.8 | 0.89 | 0.83 |
| 5 | 111039583 | rs27461 | 0.532 | -0.857 | 0.887 | -0.176 | -0.001 | 0.015 | 0.015 | 0.319 | 0.56 | 0.71 | 0.6 | 0.61 | 0.57 | 0.59 |
| 5 | 111040064 | rs10062378 | 1.191 | -0.859 | 0.896 | -0.205 | -0.022 | 0.042 | 0.042 | 0 | 0.26 | 0.15 | 0.25 | 0.01 | 0.32 | 0.17 |
| 5 | 111040322 | rs42283 | -0.031 | -0.866 | 0.895 | -0.19 | 0.004 | 0.059 | 0.059 | 0 | 0.61 | 0.76 | 0.65 | 0.78 | 0.62 | 0.65 |
| 5 | 111040812 | rs26055 | -0.237 | -0.868 | 0.949 | -0.11 | 0.02 | -0.011 | -0.011 | 0.052 | 0.88 | 0.94 | 0.9 | 0.83 | 0.94 | 0.83 |
| 5 | 111040973 | rs17133665 | -0.24 | -0.778 | 0.96 | -0.05 | 0 | 0.046 | 0.046 | 0.003 | 0.06 | 0.08 | 0.05 | 0.25 | 0.06 | 0.06 |
| 5 | 111045446 | rs7713915 | -2.788 | -0.83 | 0.932 | -0.093 | 0.006 | -0.002 | -0.002 | 0 | 0.3 | 0.71 | 0.35 | 0.35 | 0.23 | 0.18 |
| 5 | 111054270 | rs9764455 | 0.117 | -0.841 | 0.882 | -0.076 | 0 | -0.054 | -0.054 | 1.267 | 0.09 | 0.01 | 0.04 | 0.2 | 0.04 | 0.06 |
| 5 | 111054904 | rs17132976 | -1.199 | -0.704 | 1.2 | -0.078 | 0.034 | -0.074 | -0.074 | 0 | 0.15 | 0.39 | 0.12 | 0.03 | 0.14 | 0.09 |
| 5 | 111055936 | rs1438677 | -0.477 | -0.715 | 1.028 | -0.062 | -0.009 | 0.009 | 0.009 | 0 | 0.05 | 0.22 | 0.17 | 0.07 | 0.02 | 0.02 |

| | | | | | | | | | | | | | | | | |
|---|---|---|---|---|---|---|---|---|---|---|---|---|---|---|---|---|
| 5 | 111056781 | rs17842608 | -1.876 | -0.697 | 0.964 | -0.038 | 0.016 | 0.074 | 0.074 | 0.575 | 0.2 | 0.49 | 0.16 | 0.27 | 0.18 | 0.15 |
| 5 | 111057475 | rs1145125 | 1.952 | -0.664 | 0.713 | -0.101 | -0.078 | -0.141 | -0.141 | 0.064 | 0.35 | 0.07 | 0.3 | 0.48 | 0.39 | 0.48 |
| 5 | 111061637 | rs1045903 | -2.516 | -0.826 | 1.207 | 0.165 | -0.011 | 0.021 | 0.021 | 0.799 | 0.25 | 0.58 | 0.32 | 0.32 | 0.2 | 0.17 |
| 5 | 111071297 | rs3797723 | -1.296 | -0.631 | 0.935 | 0.469 | -0.003 | 0 | 0 | 0 | 0.1 | 0.09 | 0.08 | 0.26 | 0.11 | 0.07 |
| 5 | 111074163 | rs17133684 | -0.589 | -0.614 | 1.156 | 0.391 | 0.022 | 0.009 | 0.009 | 0.287 | 0.06 | 0.22 | 0.05 | 0.04 | 0.06 | 0.02 |
| 5 | 111075728 | rs3797725 | -2.492 | -1.075 | 0.823 | 0.411 | 0.057 | -0.022 | -0.022 | 0 | 0.33 | 0.79 | 0.3 | 0.37 | 0.18 | 0.18 |
| 5 | 111081708 | rs3797730 | -1.164 | -0.855 | 1.166 | 0.487 | 0.018 | 0.091 | 0.091 | 0 | 0.07 | 0.09 | 0.07 | 0.22 | 0.07 | 0.09 |
| 5 | 111084118 | rs17133711 | -2.207 | -0.979 | 1.434 | 0.33 | -0.023 | 0.025 | 0.025 | 0.79 | 0.27 | 0.39 | 0.31 | 0.37 | 0.19 | 0.16 |
| 5 | 111084169 | rs495647 | -0.461 | -0.777 | 1.178 | 0.335 | -0.002 | -0.013 | -0.013 | 0.271 | 0.1 | 0.06 | 0.08 | 0.17 | 0.09 | 0.21 |
| 5 | 111084245 | rs17133713 | -2.46 | -0.774 | 1.454 | 0.283 | 0.015 | -0.009 | -0.009 | 0 | 0.32 | 0.73 | 0.36 | 0.37 | 0.23 | 0.19 |
| 5 | 111086273 | rs3822667 | -1.76 | -0.76 | 1.235 | 0.36 | -0.033 | 0.047 | 0.047 | 1.464 | 0.22 | 0.37 | 0.24 | 0.34 | 0.11 | 0.14 |
| 5 | 111087807 | rs11948283 | -1.291 | -0.851 | 0.708 | 0.383 | -0.009 | 0.007 | 0.007 | 1.105 | 0.2 | 0.21 | 0.2 | 0.32 | 0.09 | 0.14 |
| 5 | 111091087 | rs3822668 | -0.4 | -0.503 | 0.363 | 0.14 | -0.003 | 0.003 | 0.003 | 0.438 | 0.44 | 0.39 | 0.51 | 0.38 | 0.47 | 0.3 |
| 5 | 111092157 | rs43180 | 1.493 | -0.232 | -0.453 | 0.405 | -0.135 | -0.102 | -0.102 | 0 | 0.35 | 0.02 | 0.35 | 0.4 | 0.41 | 0.42 |
| 5 | 111094463 | rs32333 | -1.226 | -0.265 | 0.44 | 0.765 | -0.036 | 0.1 | 0.1 | 0.119 | 0.25 | 0.43 | 0.31 | 0.51 | 0.16 | 0.27 |
| 5 | 111098877 | rs529279 | -0.064 | 0.336 | -0.456 | 0.329 | -0.009 | 0.089 | 0.089 | 0 | 0.44 | 0.27 | 0.46 | 0.27 | 0.43 | 0.47 |
| 5 | 111099156 | rs872622 | 0.097 | 0.666 | 0.009 | 0.583 | -0.003 | 0.005 | 0.005 | 0 | 0.42 | 0.4 | 0.48 | 0.26 | 0.45 | 0.33 |
| 5 | 111101361 | rs4957975 | 0.359 | 0.603 | -0.002 | 0.626 | -0.003 | 0.061 | 0.061 | 0 | 0.38 | 0.25 | 0.41 | 0.2 | 0.37 | 0.29 |
| 5 | 111102229 | rs10079074 | -1.478 | 0.727 | 0.377 | 0.982 | -0.014 | 0.011 | 0.011 | 0 | 0.71 | 0.64 | 0.71 | 0.64 | 0.74 | 0.59 |
| 5 | 111103258 | rs255888 | -0.431 | 1.017 | 0.271 | 1.003 | 0.001 | -0.001 | -0.001 | 0 | 0.53 | 0.46 | 0.56 | 0.48 | 0.55 | 0.48 |
| 5 | 111109796 | rs192276 | 0.057 | 1.42 | 0.414 | 1.448 | 0.001 | -0.006 | -0.006 | 0 | 0.08 | 0.04 | 0.09 | 0.26 | 0.06 | 0.17 |
| 5 | 111116271 | rs1499272 | 0.959 | 0.827 | 0.443 | 0.778 | 0.044 | 0.046 | 0.046 | 0 | 0.43 | 0.27 | 0.34 | 0.3 | 0.33 | 0.38 |
| 5 | 111121409 | rs1948671 | -0.259 | 1.495 | -0.039 | 0.697 | -0.01 | -0.013 | -0.013 | 0.027 | 0.4 | 0.3 | 0.37 | 0.48 | 0.48 | 0.38 |
| 5 | 111124268 | rs12655884 | 1.008 | 1.61 | -0.13 | 0.763 | 0.007 | 0.024 | 0.024 | 0 | 0.1 | 0.16 | 0.24 | 0.2 | 0.14 | 0.18 |
| 5 | 111125127 | rs931684 | 0.956 | 1.352 | -0.207 | 0.762 | 0.014 | 0.015 | 0.015 | 0 | 0.5 | 0.47 | 0.39 | 0.32 | 0.39 | 0.46 |
| 5 | 111125467 | rs168871 | 0.121 | 1.728 | -1.051 | 0.889 | -0.001 | 0.026 | 0.026 | 0 | 0.05 | 0.04 | 0.1 | 0.25 | 0.13 | 0.09 |
| 5 | 111126320 | rs181918 | 0.88 | 1.567 | -1.177 | 0.653 | -0.073 | -0.035 | -0.035 | 0 | 0.23 | 0.01 | 0.2 | 0.22 | 0.25 | 0.18 |
| 5 | 111130274 | rs554514 | 0.105 | 1.216 | -0.615 | 0.63 | 0 | -0.006 | -0.006 | 0 | 0.54 | 0.71 | 0.47 | 0.54 | 0.46 | 0.53 |
| 5 | 111130474 | rs1566385 | 0.324 | 1.77 | -0.429 | 0.836 | 0.001 | 0.035 | 0.035 | 0.595 | 0.09 | 0.02 | 0.05 | 0 | 0.06 | 0.01 |
| 5 | 111135151 | rs1677316 | -1.318 | 2.251 | -0.025 | 1.353 | 0.032 | 0.066 | 0.066 | 0 | 0.06 | 0.11 | 0.1 | 0.23 | 0.13 | 0.11 |
| 5 | 111136199 | rs6890703 | -1.019 | 2.195 | 0.521 | 0.89 | 0.071 | 0.054 | 0.054 | 0 | 0.21 | 0.55 | 0.25 | 0.28 | 0.28 | 0.27 |
| 5 | 111139696 | rs10074999 | -0.337 | 1.878 | 0.782 | 0.379 | 0.026 | -0.094 | -0.094 | 0 | 0.8 | 0.63 | 0.81 | 0.96 | 0.76 | 0.9 |
| 5 | 111141556 | rs4957981 | 0.58 | 1.871 | 0.783 | 0.353 | -0.033 | 0.063 | 0.063 | 0 | 0.18 | 0.06 | 0.17 | 0.04 | 0.24 | 0.09 |
| 5 | 111143961 | rs10039269 | -0.514 | 1.864 | 1.121 | 0.35 | 0.001 | -0.019 | -0.019 | 0 | 0.81 | 0.77 | 0.82 | 0.96 | 0.76 | 0.9 |
| 5 | 111144734 | rs12520347 | -0.342 | 1.422 | 0.708 | 0.161 | -0.055 | -0.036 | -0.036 | 0 | 0.3 | 0.15 | 0.27 | 0.32 | 0.37 | 0.28 |
| 5 | 111148786 | rs10067960 | -0.287 | 1.426 | 0.751 | 0.336 | -0.055 | 0.069 | 0.069 | 0 | 0.29 | 0.14 | 0.26 | 0.09 | 0.36 | 0.2 |
| 5 | 111149480 | rs6894855 | 0.421 | 1.508 | 0.987 | 0.371 | 0.029 | 0.064 | 0.064 | 0 | 0.21 | 0.36 | 0.2 | 0.29 | 0.27 | 0.19 |
| 5 | 111161200 | rs10041953 | 1.096 | 1.311 | 1.112 | 0.371 | 0.035 | 0.01 | 0.01 | 0 | 0.49 | 0.4 | 0.54 | 0.61 | 0.37 | 0.56 |
| 5 | 111182198 | rs11952607 | -0.282 | 1.09 | 1.126 | 0.43 | -0.003 | 0.004 | 0.004 | 0 | 0.1 | 0.16 | 0.14 | 0.12 | 0.07 | 0.13 |
| 5 | 111184593 | rs6864217 | -1.778 | 1.128 | 1.25 | 0.468 | 0.037 | -0.007 | -0.007 | 0 | 0.59 | 0.7 | 0.62 | 0.52 | 0.7 | 0.52 |
| 5 | 111197930 | rs4957623 | 1.23 | 1.364 | 0.788 | 0.255 | -0.045 | 0.069 | 0.069 | 0 | 0.17 | 0.02 | 0.15 | 0.04 | 0.2 | 0.1 |
| 5 | 111204946 | rs10477473 | 0.146 | 1.046 | 1.256 | 0.482 | -0.001 | -0.013 | -0.013 | 0.032 | 0.08 | 0.08 | 0.07 | 0.01 | 0.06 | 0.03 |
| 5 | 111205268 | rs10073581 | -0.345 | 1.04 | 1.043 | 0.439 | 0.003 | 0.204 | 0.204 | 0 | 0.47 | 0.14 | 0.38 | 0.12 | 0.57 | 0.28 |
| 5 | 111215352 | rs6594555 | -0.156 | 1.033 | 1.025 | 0.436 | -0.018 | 0.19 | 0.19 | 0 | 0.45 | 0.12 | 0.38 | 0.12 | 0.57 | 0.27 |
| 5 | 111241351 | rs10056269 | -0.281 | 0.912 | 1.832 | 0.621 | 0.025 | -0.041 | -0.041 | 0 | 0.08 | 0.26 | 0.08 | 0.01 | 0.07 | 0.03 |
| 5 | 111241454 | rs1845486 | -0.97 | 1.154 | 1.782 | 0.519 | 0.015 | 0.001 | 0.001 | 0 | 0.54 | 0.55 | 0.48 | 0.38 | 0.65 | 0.39 |
| 5 | 111241954 | rs1466735 | -0.168 | 1.165 | 1.724 | 0.49 | -0.087 | 0.185 | 0.185 | 0 | 0.43 | 0.08 | 0.36 | 0.12 | 0.53 | 0.25 |
| 5 | 111246727 | rs10045343 | -0.328 | 1.197 | 1.739 | 0.504 | 0.041 | -0.044 | -0.044 | 0.758 | 0.09 | 0.26 | 0.1 | 0.01 | 0.1 | 0.04 |
| 5 | 111246794 | rs7716521 | -0.203 | 1.185 | 1.704 | 0.547 | -0.004 | 0.002 | 0.002 | 0 | 0.45 | 0.3 | 0.39 | 0.36 | 0.54 | 0.33 |
| 5 | 111250615 | rs6898801 | -0.729 | 1.162 | 1.726 | 0.495 | 0.027 | 0.059 | 0.059 | 0 | 0.52 | 0.39 | 0.46 | 0.13 | 0.63 | 0.29 |
| 5 | 111263161 | rs508089 | 0.16 | 1.073 | 1.744 | 0.73 | 0.003 | -0.006 | -0.006 | 0 | 0.54 | 0.7 | 0.59 | 0.58 | 0.45 | 0.66 |
| 5 | 111266661 | rs474807 | 0.104 | 1.018 | 1.832 | 0.719 | 0.001 | -0.01 | -0.01 | 0 | 0.55 | 0.72 | 0.61 | 0.58 | 0.45 | 0.68 |
| 5 | 111269667 | rs1493455 | -0.031 | 0.935 | 1.829 | 0.71 | 0.024 | -0.039 | -0.039 | 0 | 0.08 | 0.23 | 0.07 | 0 | 0.06 | 0.04 |
| 5 | 111275778 | rs7730997 | 0.219 | 0.94 | 1.89 | 0.721 | 0.002 | -0.002 | -0.002 | 4.488 | 0.54 | 0.63 | 0.59 | 0.57 | 0.45 | 0.66 |
| 5 | 111287944 | rs664517 | 0.476 | 0.916 | 1.929 | 0.747 | 0.048 | 0.053 | 0.053 | 0.277 | 0.91 | 0.64 | 0.9 | 0.92 | 0.93 | 0.93 |
| 5 | 111296973 | rs1994871 | -0.178 | 0.916 | 1.33 | 0.712 | -0.051 | 0.183 | 0.183 | 0 | 0.44 | 0.09 | 0.36 | 0.12 | 0.55 | 0.26 |
| 5 | 111307907 | rs490821 | -0.651 | 0.769 | 1.278 | 0.659 | 0 | 0 | 0 | 0 | 0.53 | 0.51 | 0.46 | 0.35 | 0.61 | 0.36 |
| 5 | 111332331 | rs17133979 | 0.308 | 0.636 | 1.015 | 0.558 | -0.015 | 0.021 | 0.021 | 0 | 0.05 | 0.1 | 0.11 | 0.12 | 0.02 | 0.06 |
| 5 | 111341445 | rs26563 | 0.808 | 0.6 | 1.005 | 0.324 | 0.039 | -0.039 | -0.039 | 0 | 0.43 | 0.19 | 0.49 | 0.57 | 0.37 | 0.59 |
| 5 | 111349320 | rs26540 | 0.712 | 0.477 | 1.171 | 0.361 | 0.018 | -0.008 | -0.008 | 0 | 0.42 | 0.38 | 0.47 | 0.57 | 0.32 | 0.62 |
| 5 | 111351359 | rs26539 | 0.968 | 0.498 | 0.733 | 0.363 | 0.051 | -0.049 | -0.049 | 0 | 0.4 | 0.19 | 0.45 | 0.57 | 0.32 | 0.61 |
| 5 | 111369892 | rs255910 | 0.797 | 0.235 | 0.663 | 0.233 | 0.024 | -0.061 | -0.061 | 0 | 0.33 | 0.17 | 0.48 | 0.58 | 0.29 | 0.62 |
| 5 | 111390356 | rs194359 | 0.246 | 0.221 | 1.715 | 0.063 | 0.013 | -0.01 | -0.01 | 0.071 | 0.42 | 0.33 | 0.56 | 0.57 | 0.35 | 0.62 |
| 5 | 111399885 | rs1389312 | 0.074 | 0.221 | 1.447 | 0.153 | -0.001 | -0.031 | -0.031 | 0 | 0.08 | 0.13 | 0.07 | 0 | 0.06 | 0 |
| 5 | 111400612 | rs30294 | -1.16 | 0.188 | 1.437 | 0.174 | 0.012 | -0.115 | -0.115 | 0.061 | 0.67 | 0.89 | 0.51 | 0.39 | 0.71 | 0.41 |
| 5 | 111401031 | rs154082 | -0.035 | 0.277 | 1.607 | 0.084 | 0.007 | -0.025 | -0.025 | 0 | 0.07 | 0.13 | 0.06 | 0 | 0.09 | 0.04 |
| 5 | 111402835 | rs154084 | -1.155 | 0.132 | 1.562 | 0.005 | 0.001 | -0.073 | -0.073 | 0 | 0.14 | 0.26 | 0.12 | 0 | 0.12 | 0.02 |
| 5 | 111402982 | rs39834 | -1.162 | 0.152 | 1.671 | 0.027 | 0.026 | -0.062 | -0.062 | 1.752 | 0.66 | 0.83 | 0.5 | 0.38 | 0.71 | 0.38 |
| 5 | 111408595 | rs30336 | -0.725 | 0.173 | 1.615 | 0.026 | 0 | -0.001 | -0.001 | 0.107 | 0.62 | 0.61 | 0.45 | 0.39 | 0.65 | 0.39 |

| | | | | | | | | | | | | | | | | |
|---|---|---|---|---|---|---|---|---|---|---|---|---|---|---|---|---|
| 5 | 111411516 | rs30342 | -0.583 | 0.194 | 1.449 | 0.066 | 0.015 | -0.015 | -0.015 | 1.604 | 0.59 | 0.69 | 0.43 | 0.38 | 0.64 | 0.38 |
| 5 | 111442711 | rs27536 | -0.614 | 0.359 | 0.871 | -0.619 | 0 | -0.018 | -0.018 | 1.911 | 0.28 | 0.29 | 0.17 | 0.04 | 0.23 | 0.04 |
| 5 | 111444522 | rs25796 | 0.422 | 0.644 | 0.697 | -0.815 | -0.035 | 0.067 | 0.067 | 0 | 0.36 | 0.17 | 0.26 | 0.02 | 0.43 | 0.23 |
| 5 | 111445356 | rs922431 | -0.034 | 0.667 | 0.544 | -0.506 | 0.001 | 0.026 | 0.026 | 0.852 | 0.23 | 0.17 | 0.17 | 0.09 | 0.22 | 0.06 |
| 5 | 111452329 | rs6865881 | 0.009 | 1.045 | 0.668 | -0.534 | -0.006 | 0.026 | 0.026 | 0 | 0.4 | 0.39 | 0.32 | 0.18 | 0.44 | 0.2 |
| 5 | 111452377 | rs40394 | 0.447 | 1.086 | 0.413 | -0.277 | -0.025 | -0.021 | -0.021 | 0 | 0.21 | 0.13 | 0.16 | 0.23 | 0.26 | 0.22 |
| 5 | 111454041 | rs6876779 | 1.256 | 0.481 | 0.576 | -0.287 | -0.028 | 0.036 | 0.036 | 0 | 0.28 | 0.56 | 0.35 | 0.45 | 0.21 | 0.39 |
| 5 | 111458013 | rs17252737 | 0.413 | 0.567 | 0.431 | -0.566 | -0.017 | 0.066 | 0.066 | 0 | 0.34 | 0.12 | 0.22 | 0.15 | 0.36 | 0.18 |
| 5 | 111463837 | rs27982 | -0.943 | 0.97 | 1.469 | -0.289 | 0.017 | -0.039 | -0.039 | 0 | 0.13 | 0.15 | 0.08 | 0.02 | 0.15 | 0.18 |
| 5 | 111479955 | rs17134152 | -1.462 | 0.859 | 1.744 | -0.281 | 0.032 | -0.062 | -0.062 | 0 | 0.15 | 0.32 | 0.08 | 0.06 | 0.14 | 0.17 |
| 5 | 111480534 | rs17134154 | -0.974 | 0.865 | 1.772 | -0.281 | 0.057 | -0.024 | -0.024 | 0 | 0.1 | 0.29 | 0.06 | 0.06 | 0.13 | 0.16 |
| 5 | 111481696 | rs17134155 | -1.978 | 0.835 | 1.162 | -0.384 | 0.059 | -0.088 | -0.088 | 0 | 0.17 | 0.55 | 0.13 | 0.06 | 0.18 | 0.16 |
| 5 | 111488600 | rs17651424 | -0.983 | 1.029 | 0.268 | -0.309 | -0.002 | -0.004 | -0.004 | 0 | 0.12 | 0.06 | 0.06 | 0.01 | 0.13 | 0.11 |
| 5 | 111493689 | rs7711555 | -1.881 | 1.058 | 0.286 | -0.318 | 0.004 | -0.031 | -0.031 | 0.132 | 0.18 | 0.15 | 0.12 | 0.02 | 0.2 | 0.17 |
| 5 | 111495150 | rs7730334 | -0.701 | 1.102 | 0.138 | -0.183 | 0.006 | -0.017 | -0.017 | 0 | 0.1 | 0.08 | 0.06 | 0.02 | 0.13 | 0.16 |
| 5 | 111499041 | rs12517400 | -1.854 | 0.848 | 0.428 | 0.182 | 0.012 | 0.057 | 0.057 | 0.38 | 0.23 | 0.62 | 0.3 | 0.38 | 0.23 | 0.32 |
| 5 | 111499732 | rs1054724 | -0.611 | 1.029 | -0.497 | -0.67 | -0.09 | 0.132 | 0.132 | 0.017 | 0.76 | 0.94 | 0.79 | 1 | 0.67 | 0.88 |
| 5 | 111504479 | rs7703522 | -0.535 | 0.984 | -0.469 | -0.707 | -0.081 | 0.122 | 0.122 | 0.002 | 0.75 | 0.92 | 0.78 | 1 | 0.66 | 0.88 |
| 5 | 111505578 | rs10045569 | 1.188 | 0.765 | -0.022 | -0.641 | -0.008 | 0.037 | 0.037 | 0 | 0.55 | 0.36 | 0.65 | 0.74 | 0.53 | 0.63 |
| 5 | 111506712 | rs17266727 | -1.181 | 0.758 | 0.023 | -0.57 | 0 | 0 | 0 | 0 | 0.14 | 0.07 | 0.07 | 0.01 | 0.11 | 0.09 |
| 5 | 111508960 | rs7717321 | -0.102 | 0.447 | 0.297 | -0.665 | -0.008 | -0.082 | -0.082 | 0.64 | 0.14 | 0.3 | 0.08 | 0 | 0.09 | 0.01 |
| 5 | 111515382 | rs10051634 | 1.711 | 0.553 | -0.519 | -0.365 | 0.02 | -0.031 | -0.031 | 9.117 | 0.36 | 0.1 | 0.39 | 0.66 | 0.3 | 0.57 |
| 5 | 111519018 | rs1560054 | -0.825 | 0.768 | -0.368 | -0.524 | -0.017 | 0.024 | 0.024 | 0.078 | 0.75 | 0.84 | 0.81 | 1 | 0.71 | 0.9 |
| 5 | 111519740 | rs1560058 | -0.316 | 0.704 | -0.036 | -0.548 | 0.019 | -0.176 | -0.176 | 0.071 | 0.71 | 0.5 | 0.8 | 0.99 | 0.69 | 0.84 |
| 5 | 111520646 | rs12517535 | 1.069 | 0.934 | -0.374 | -0.246 | -0.017 | -0.028 | -0.028 | 0 | 0.05 | 0.01 | 0.16 | 0.18 | 0.05 | 0.06 |
| 5 | 111530284 | rs7719346 | -1.091 | 0.886 | -0.445 | -0.52 | -0.049 | 0.077 | 0.077 | 0.049 | 0.79 | 0.92 | 0.83 | 1 | 0.72 | 0.9 |
| 5 | 111531058 | rs4958011 | 1.133 | 0.735 | -0.521 | -0.686 | 0.016 | 0.031 | 0.031 | 0 | 0.11 | 0.03 | 0.06 | 0 | 0.08 | 0.02 |
| 5 | 111531997 | rs1078129 | 0.107 | 0.778 | -0.143 | -0.174 | 0.011 | -0.005 | -0.005 | 0 | 0.69 | 0.39 | 0.75 | 0.74 | 0.64 | 0.63 |
| 5 | 111535501 | rs10515441 | 0.763 | 0.831 | -0.182 | -0.168 | -0.022 | 0.015 | 0.015 | 0 | 0.08 | 0.06 | 0.08 | 0.02 | 0.11 | 0.11 |
| 5 | 111538333 | rs13164349 | 1.751 | 0.481 | -0.333 | -0.071 | -0.004 | -0.04 | -0.04 | 0 | 0.44 | 0.04 | 0.43 | 0.44 | 0.37 | 0.46 |
| 5 | 111538602 | rs13165201 | 1.345 | 0.501 | -0.219 | -0.054 | 0.007 | 0.008 | 0.008 | 0 | 0.48 | 0.55 | 0.47 | 0.66 | 0.38 | 0.62 |
| 5 | 111545048 | rs1109590 | 0.329 | 0.744 | 0.325 | 0.127 | 0.002 | 0.022 | 0.022 | 0 | 0.15 | 0.29 | 0.1 | 0.15 | 0.16 | 0.11 |
| 5 | 111548834 | rs7722065 | -0.908 | 0.573 | 0.09 | 0.163 | -0.018 | 0.083 | 0.083 | 0 | 0.24 | 0.51 | 0.3 | 0.55 | 0.21 | 0.27 |
| 5 | 111552141 | rs7704501 | -0.129 | 1.571 | 0.404 | -0.118 | -0.014 | 0.002 | 0.002 | 0.368 | 0.71 | 0.72 | 0.78 | 0.97 | 0.64 | 0.75 |
| 5 | 111555760 | rs6873051 | 0.406 | 1.402 | 0.233 | -0.186 | 0.002 | -0.015 | -0.015 | 0 | 0.12 | 0.09 | 0.06 | 0 | 0.08 | 0.03 |
| 5 | 111556352 | rs1366145 | 0.315 | 1.537 | 0.201 | -0.188 | -0.091 | 0.137 | 0.137 | 0.213 | 0.67 | 0.83 | 0.73 | 0.97 | 0.58 | 0.74 |
| 5 | 111571207 | rs4957639 | 0.478 | 1.242 | 0.328 | 0.145 | 0 | 0.004 | 0.004 | 0 | 0.07 | 0.06 | 0.16 | 0.15 | 0.06 | 0.08 |
| 5 | 111573595 | rs7723465 | -1.498 | 1.389 | 0.227 | 0.398 | 0 | 0.024 | 0.024 | 0 | 0.46 | 0.68 | 0.48 | 0.59 | 0.53 | 0.3 |
| 5 | 111576056 | rs12186685 | 1.06 | 1.437 | 0.084 | 0.286 | -0.061 | 0.077 | 0.077 | 1.149 | 0.21 | 0.08 | 0.18 | 0 | 0.29 | 0.1 |
| 5 | 111576837 | rs4602685 | -0.237 | 1.382 | 0.171 | 0.27 | -0.03 | 0.054 | 0.054 | 0.063 | 0.32 | 0.17 | 0.24 | 0 | 0.38 | 0.13 |
| 5 | 111583782 | rs4958026 | 0.575 | 1.022 | 0.19 | 0.611 | 0.002 | 0.007 | 0.007 | 0 | 0.11 | 0.09 | 0.06 | 0.27 | 0.08 | 0.12 |
| 5 | 111586429 | rs6888754 | -1.711 | 1.095 | 0.302 | 0.579 | 0.028 | 0.029 | 0.029 | 0 | 0.52 | 0.59 | 0.53 | 0.6 | 0.59 | 0.31 |
| 5 | 111596035 | rs7732679 | -2.381 | 1.136 | 0.331 | 0.574 | 0.042 | 0.02 | 0.02 | 0.076 | 0.57 | 0.72 | 0.56 | 0.6 | 0.65 | 0.39 |
| 5 | 111597275 | rs11740678 | -0.855 | 1.709 | 0.535 | 0.541 | 0.088 | 0.055 | 0.055 | 0 | 0.23 | 0.35 | 0.33 | 0.34 | 0.34 | 0.13 |
| 5 | 111601600 | rs4957640 | 0.943 | 1.497 | 0.35 | 0.49 | 0.017 | 0.03 | 0.03 | 0 | 0.11 | 0.04 | 0.05 | 0 | 0.08 | 0.05 |
| 5 | 111602713 | rs13358904 | 0.175 | 1.475 | 0.423 | 0.546 | 0.002 | -0.003 | -0.003 | 1.358 | 0.26 | 0.23 | 0.19 | 0.27 | 0.25 | 0.2 |
| 5 | 111603606 | rs7733521 | -1.224 | 2.283 | 1.207 | 1.18 | 0.083 | 0.09 | 0.09 | 0 | 0.2 | 0.35 | 0.28 | 0.33 | 0.27 | 0.11 |
| 5 | 111607304 | rs10059647 | 2.669 | 1.99 | 0.571 | 1.002 | 0.03 | 0.04 | 0.04 | 0 | 0.52 | 0.08 | 0.5 | 0.39 | 0.44 | 0.6 |
| 5 | 111607509 | rs4311468 | 2.598 | 2.011 | 0.625 | 1.012 | 0.045 | 0.052 | 0.052 | 0 | 0.53 | 0.15 | 0.51 | 0.39 | 0.44 | 0.6 |
| 5 | 111613342 | rs1464766 | 1.944 | 2.268 | 1.505 | 1.555 | 0.045 | 0.041 | 0.041 | 0.15 | 0.77 | 0.35 | 0.66 | 0.63 | 0.68 | 0.78 |
| 5 | 111615346 | rs10478077 | 1.728 | 2.304 | 1.846 | 1.762 | 0.107 | 0.157 | 0.157 | 0.567 | 0.81 | 0.4 | 0.7 | 0.61 | 0.73 | 0.84 |
| 5 | 111620405 | rs12519569 | -2.435 | 2.158 | 2.023 | 1.756 | 0 | 0.006 | 0.006 | 0 | 0.31 | 0.63 | 0.32 | 0.47 | 0.3 | 0.31 |
| 5 | 111622713 | rs12657885 | -0.917 | 2.108 | 1.871 | 1.658 | -0.002 | 0.025 | 0.025 | 0 | 0.16 | 0.14 | 0.21 | 0.37 | 0.17 | 0.2 |
| 5 | 111626532 | rs4455607 | -2.932 | 2.191 | 2.03 | 1.701 | -0.014 | 0.014 | 0.014 | 0 | 0.37 | 0.75 | 0.39 | 0.66 | 0.36 | 0.41 |
| 5 | 111630303 | rs17134327 | -0.929 | 1.768 | 2.37 | 1.671 | 0.117 | -0.041 | -0.041 | 0 | 0.11 | 0.43 | 0.09 | 0.03 | 0.12 | 0.09 |
| 5 | 111636632 | rs6594585 | -2.366 | 1.776 | 2.067 | 1.735 | 0.062 | 0.051 | 0.051 | 0.909 | 0.19 | 0.64 | 0.17 | 0.28 | 0.24 | 0.18 |
| 5 | 111641797 | rs17134331 | 0.685 | 1.668 | 1.966 | 1.59 | -0.001 | -0.023 | -0.023 | 0 | 0.12 | 0.07 | 0.05 | 0 | 0.08 | 0.04 |
| 5 | 111642706 | rs11743178 | -1.388 | 1.821 | 2.044 | 1.649 | -0.023 | -0.022 | -0.022 | 2.539 | 0.1 | 0.07 | 0.12 | 0.24 | 0.19 | 0.09 |
| 5 | 111655010 | rs6594586 | -2.754 | 1.785 | 2.128 | 1.653 | -0.004 | -0.031 | -0.031 | 0.331 | 0.34 | 0.75 | 0.35 | 0.41 | 0.31 | 0.34 |
| 5 | 111655382 | rs1554624 | -1.48 | 1.882 | 2.264 | 1.666 | 0.063 | -0.047 | -0.047 | 2.763 | 0.12 | 0.62 | 0.11 | 0.04 | 0.12 | 0.1 |
| 5 | 111655969 | rs821749 | -0.729 | 1.892 | 2.439 | 1.663 | 0.139 | -0.041 | -0.041 | 0.469 | 0.1 | 0.44 | 0.09 | 0 | 0.12 | 0.05 |
| 5 | 111662406 | rs12514057 | -0.83 | 1.753 | 1.794 | 1.797 | -0.01 | 0.016 | 0.016 | 0 | 0.08 | 0.05 | 0.18 | 0.39 | 0.1 | 0.18 |
| 5 | 111664925 | rs885137 | 2.579 | 1.655 | 1.714 | 1.644 | -0.046 | 0.011 | 0.011 | 0 | 0.62 | 0.2 | 0.53 | 0.37 | 0.59 | 0.64 |
| 5 | 111678637 | rs1004385 | 1.184 | 1.818 | 2.457 | 1.73 | 0.07 | 0.159 | 0.159 | 0 | 0.95 | 0.86 | 0.93 | 0.8 | 0.9 | 0.97 |
| 5 | 111681843 | rs1019745 | 2.823 | 1.878 | 2.153 | 1.751 | -0.029 | -0.002 | -0.002 | 0 | 0.57 | 0.16 | 0.47 | 0.41 | 0.46 | 0.61 |
| 5 | 111682654 | rs1019743 | 1.415 | 1.8 | 2.581 | 1.687 | 0.03 | -0.004 | -0.004 | 0 | 0.75 | 0.6 | 0.79 | 0.79 | 0.68 | 0.84 |
| 5 | 111682843 | rs1560061 | -0.646 | 1.8 | 2.646 | 1.658 | -0.001 | -0.038 | -0.038 | 0.388 | 0.21 | 0.18 | 0.14 | 0 | 0.22 | 0.13 |
| 5 | 111685332 | rs2140626 | 1.568 | 1.794 | 2.675 | 1.681 | 0.009 | -0.003 | -0.003 | 0.023 | 0.71 | 0.35 | 0.7 | 0.75 | 0.68 | 0.8 |
| 5 | 111691924 | rs1962305 | 1.444 | 1.67 | 2.502 | 1.724 | 0.024 | 0 | 0 | 0 | 0.72 | 0.41 | 0.72 | 0.76 | 0.68 | 0.8 |

| | | | | | | | | | | | | | | | | |
|---|---|---|---|---|---|---|---|---|---|---|---|---|---|---|---|---|
| 5 | 111699317 | rs366180 | 2.007 | 1.657 | 2.018 | 1.354 | 0.008 | 0.002 | 0.002 | 0 | 0.63 | 0.45 | 0.52 | 0.43 | 0.54 | 0.66 |
| 5 | 111701910 | rs415229 | -2.33 | 1.728 | 2.334 | 1.159 | 0.056 | 0.019 | 0.019 | 0.013 | 0.16 | 0.48 | 0.23 | 0.16 | 0.21 | 0.15 |
| 5 | 111703577 | rs448714 | -2.462 | 1.862 | 2.538 | 0.931 | 0.043 | -0.072 | -0.072 | 0.125 | 0.16 | 0.64 | 0.17 | 0.04 | 0.22 | 0.14 |
| 5 | 111704003 | rs712682 | -1.015 | 1.533 | 2.372 | 0.596 | -0.003 | -0.037 | -0.037 | 0 | 0.23 | 0.26 | 0.2 | 0.08 | 0.21 | 0.19 |
| 5 | 111706722 | rs1094445 | -3.025 | 1.645 | 2.981 | 1.177 | 0.001 | -0.014 | -0.014 | 0.584 | 0.18 | 0.73 | 0.2 | 0.12 | 0.17 | 0.21 |
| 5 | 111737338 | rs1880332 | 2.004 | 1.574 | 2.588 | 1.46 | -0.064 | -0.025 | -0.025 | 2.062 | 0.58 | 0.14 | 0.46 | 0.42 | 0.53 | 0.63 |
| 5 | 111765962 | rs6895299 | -3.105 | 0.922 | 3.14 | 1.252 | 0.004 | 0.035 | 0.035 | 0.416 | 0.08 | 0.51 | 0.05 | 0.04 | 0.01 | 0.1 |
| 5 | 111778778 | rs9326854 | -2.574 | 1.132 | 3.081 | 1.25 | 0.044 | -0.043 | -0.043 | 0 | 0.16 | 0.6 | 0.19 | 0.1 | 0.17 | 0.21 |
| 5 | 111805307 | rs6871040 | -1.55 | 1.196 | 3.137 | 1.169 | 0.099 | -0.065 | -0.065 | 0 | 0.11 | 0.41 | 0.11 | 0 | 0.15 | 0.09 |
| 5 | 111807579 | rs1345686 | -1.834 | 1.196 | 3.07 | 1.174 | 0.12 | -0.051 | -0.051 | 0 | 0.12 | 0.52 | 0.12 | 0.04 | 0.15 | 0.16 |
| 5 | 111807744 | rs10050630 | -1.398 | 1.203 | 2.882 | 1.177 | 0.082 | -0.041 | -0.041 | 0.514 | 0.11 | 0.34 | 0.1 | 0.04 | 0.15 | 0.16 |
| 5 | 111814318 | rs6882162 | -2.005 | 1.197 | 2.177 | 1.363 | 0.078 | -0.012 | -0.012 | 1.205 | 0.13 | 0.58 | 0.12 | 0.08 | 0.15 | 0.16 |
| 5 | 111815721 | rs163618 | 1.649 | 1.235 | 0.994 | 1.433 | -0.063 | -0.016 | -0.016 | 0 | 0.54 | 0.1 | 0.44 | 0.4 | 0.51 | 0.55 |
| 5 | 111820871 | rs2067096 | -1.391 | 1.261 | 1.664 | 1.05 | 0.121 | -0.058 | -0.058 | 0 | 0.1 | 0.32 | 0.11 | 0 | 0.17 | 0.09 |
| 5 | 111822058 | rs466898 | 0.754 | 0.182 | 1.189 | 1.417 | -0.029 | 0.086 | 0.086 | 0 | 0.88 | 0.76 | 0.82 | 0.74 | 0.9 | 0.84 |
| 5 | 111822628 | rs341338 | -1.066 | 0.163 | 1.612 | 1.093 | -0.021 | -0.018 | -0.018 | 0.014 | 0.12 | 0.24 | 0.09 | 0.07 | 0.12 | 0.19 |
| 5 | 111829149 | rs341314 | 0.356 | -0.135 | 1.568 | 1.183 | 0.006 | -0.001 | -0.001 | 0 | 0.23 | 0.24 | 0.23 | 0.25 | 0.18 | 0.25 |
| 5 | 111830825 | rs341309 | -0.628 | 0.173 | 1.935 | 1.122 | 0.001 | -0.008 | -0.008 | 1.478 | 0.33 | 0.57 | 0.33 | 0.3 | 0.33 | 0.43 |
| 5 | 111836278 | rs462412 | 0.438 | -0.732 | 1.782 | 1.199 | -0.013 | 0.009 | 0.009 | 0 | 0.69 | 0.55 | 0.66 | 0.69 | 0.69 | 0.75 |
| 5 | 111837901 | rs255708 | -0.069 | -0.703 | 1.581 | 1.241 | 0.018 | -0.001 | -0.001 | 0 | 0.69 | 0.71 | 0.68 | 0.7 | 0.74 | 0.58 |
| 5 | 111848890 | rs341295 | 1.566 | -0.526 | 1.039 | 1.212 | -0.112 | -0.041 | -0.041 | 0 | 0.4 | 0.06 | 0.38 | 0.4 | 0.47 | 0.36 |
| 5 | 111869430 | rs338857 | 0.384 | -0.545 | 1.7 | 1.162 | -0.005 | 0.01 | 0.01 | 0.031 | 0.74 | 0.4 | 0.71 | 0.73 | 0.75 | 0.81 |
| 5 | 111870042 | rs163452 | 0.285 | -0.54 | 1.72 | 1.164 | -0.002 | 0.016 | 0.016 | 0 | 0.75 | 0.43 | 0.72 | 0.73 | 0.75 | 0.81 |
| 5 | 111870990 | rs11241168 | 0.801 | -0.627 | 1.221 | 0.947 | 0.027 | -0.078 | -0.078 | 1.046 | 0.21 | 0.04 | 0.17 | 0.42 | 0.16 | 0.24 |
| 5 | 111872699 | rs11241170 | 0.186 | -0.588 | 1.748 | 0.936 | -0.028 | 0.055 | 0.055 | 0 | 0.25 | 0.43 | 0.24 | 0.46 | 0.21 | 0.24 |
| 5 | 111883311 | rs6862736 | 0.408 | -0.612 | 1.666 | 0.815 | -0.018 | 0.021 | 0.021 | 0 | 0.08 | 0.11 | 0.12 | 0.11 | 0.08 | 0.2 |
| 5 | 111883877 | rs9885457 | 0.541 | -0.613 | 1.657 | 0.845 | 0.005 | -0.002 | -0.002 | 0 | 0.23 | 0.23 | 0.22 | 0.26 | 0.2 | 0.23 |
| 5 | 111886106 | rs7442992 | 0.937 | -0.623 | 1.581 | 0.89 | -0.007 | 0.007 | 0.007 | 0 | 0.65 | 0.42 | 0.61 | 0.49 | 0.69 | 0.58 |
| 5 | 111888948 | rs4574588 | 1.464 | -0.471 | 0.815 | 0.898 | -0.101 | 0.018 | 0.018 | 0.026 | 0.41 | 0.04 | 0.38 | 0.26 | 0.49 | 0.36 |
| 5 | 111890072 | rs13154524 | 0.856 | -0.602 | 0.871 | 0.868 | 0.091 | 0.035 | 0.035 | 0.064 | 0.57 | 0.12 | 0.51 | 0.29 | 0.66 | 0.44 |
| 5 | 111891417 | rs3934641 | 0.609 | -0.521 | 1.309 | 0.961 | 0.015 | 0.32 | 0.32 | 0 | 0.94 | 0.76 | 0.92 | 0.65 | 0.99 | 0.93 |
| 5 | 111893273 | rs12520993 | 0.628 | -0.459 | 0.905 | 0.834 | -0.009 | 0.045 | 0.045 | 0.343 | 0.16 | 0.09 | 0.13 | 0.04 | 0.17 | 0.09 |
| 5 | 111893837 | rs7703562 | 0.999 | -0.484 | 1.082 | 0.856 | 0.043 | 0.025 | 0.025 | 0 | 0.51 | 0.22 | 0.52 | 0.38 | 0.58 | 0.58 |
| 5 | 111894491 | rs6867731 | 0.142 | -0.221 | 0.918 | 0.636 | -0.006 | 0.039 | 0.039 | 0.333 | 0.17 | 0.1 | 0.13 | 0.03 | 0.17 | 0.09 |
| 5 | 111908927 | rs10900679 | 0.954 | -0.2 | 1.413 | 0.975 | -0.03 | 0.26 | 0.26 | 0 | 0.85 | 0.56 | 0.78 | 0.54 | 0.88 | 0.71 |
| 5 | 111911276 | rs4958063 | -0.169 | -0.235 | 0.956 | -0.062 | -0.007 | 0.001 | 0.001 | 0 | 0.22 | 0.21 | 0.22 | 0.23 | 0.13 | 0.16 |
| 5 | 111913764 | rs4958066 | -0.746 | -0.246 | 1.218 | -0.081 | -0.011 | 0.042 | 0.042 | 0 | 0.25 | 0.41 | 0.31 | 0.36 | 0.27 | 0.26 |
| 5 | 111914106 | rs13188321 | -0.381 | -0.207 | 1.241 | -0.098 | 0.002 | -0.002 | -0.002 | 0.259 | 0.14 | 0.3 | 0.14 | 0.13 | 0.19 | 0.2 |
| 5 | 111915524 | rs6594619 | 0.984 | -0.227 | 1.101 | -0.055 | -0.002 | 0.01 | 0.01 | 0.43 | 0.12 | 0.1 | 0.18 | 0.23 | 0.08 | 0.07 |
| 5 | 111915902 | rs7445902 | 1.46 | -0.314 | 1.292 | -0.212 | 0.036 | 0.025 | 0.025 | 0 | 0.58 | 0.17 | 0.59 | 0.64 | 0.64 | 0.64 |
| 5 | 111916117 | rs13170674 | -1.353 | -0.249 | 1.391 | -0.243 | -0.012 | -0.048 | -0.048 | 0 | 0.3 | 0.66 | 0.23 | 0.13 | 0.28 | 0.3 |
| 5 | 111921515 | rs6890727 | 0.322 | -0.497 | 1.188 | -0.099 | -0.022 | -0.083 | -0.083 | 0.085 | 0.13 | 0.36 | 0.06 | 0 | 0.05 | 0.08 |
| 5 | 111933873 | rs10067709 | -0.088 | -0.155 | 0.472 | 0.199 | -0.001 | 0.01 | 0.01 | 0.213 | 0.85 | 0.81 | 0.78 | 0.77 | 0.88 | 0.92 |
| 5 | 111934314 | rs4073304 | -0.087 | -0.148 | 0.475 | 0.202 | -0.001 | 0.01 | 0.01 | 0.213 | 0.85 | 0.81 | 0.78 | 0.77 | 0.88 | 0.92 |
| 5 | 111935306 | rs6594638 | 0.771 | -0.193 | 0.527 | 0.396 | -0.021 | 0.098 | 0.098 | 2.106 | 0.78 | 0.62 | 0.67 | 0.56 | 0.85 | 0.79 |
| 5 | 111937859 | rs6881630 | 0.646 | -0.306 | 0.517 | 0.441 | -0.038 | 0.145 | 0.145 | 2.02 | 0.8 | 0.6 | 0.71 | 0.56 | 0.88 | 0.79 |
| 5 | 111942769 | rs12516949 | 0.6 | -0.179 | -0.059 | 0.491 | -0.001 | 0.006 | 0.006 | 0.667 | 0.12 | 0.11 | 0.19 | 0.24 | 0.09 | 0.07 |
| 5 | 111943114 | rs11746594 | -0.122 | 0.774 | -0.772 | 0.513 | -0.014 | -0.003 | -0.003 | 0 | 0.37 | 0.18 | 0.44 | 0.64 | 0.39 | 0.43 |
| 5 | 111943552 | rs11743216 | 0.456 | 0.949 | -0.793 | 0.501 | -0.033 | -0.068 | -0.068 | 0.182 | 0.29 | 0.1 | 0.37 | 0.63 | 0.33 | 0.35 |
| 5 | 111944746 | rs7446319 | -0.06 | 0.776 | -0.792 | 0.457 | -0.009 | -0.005 | -0.005 | 4.844 | 0.36 | 0.18 | 0.43 | 0.64 | 0.38 | 0.42 |
| 5 | 111948117 | rs7732983 | -0.133 | 1.025 | -1.127 | 0.113 | 0.017 | -0.024 | -0.024 | 0.669 | 0.27 | 0.31 | 0.27 | 0.13 | 0.38 | 0.33 |
| 5 | 111949440 | rs4365835 | 0.029 | 1.111 | -1.457 | 0.796 | -0.072 | -0.137 | -0.137 | 0 | 0.26 | 0.05 | 0.28 | 0.44 | 0.33 | 0.27 |
| 5 | 111968281 | rs4355511 | -1.207 | 0.972 | -0.605 | 0.219 | 0.009 | -0.044 | -0.044 | 0 | 0.09 | 0.16 | 0.04 | 0 | 0.08 | 0.03 |
| 5 | 111977000 | rs17134832 | -0.19 | 0.59 | -0.841 | 0.39 | -0.052 | -0.168 | -0.168 | 0 | 0.26 | 0.03 | 0.22 | 0.43 | 0.29 | 0.29 |
| 5 | 111977065 | rs7707152 | 0.69 | 0.592 | -0.22 | 0.322 | 0.031 | -0.018 | -0.018 | 1.014 | 0.91 | 0.43 | 0.93 | 0.98 | 0.96 | 0.93 |
| 5 | 111978946 | rs13182407 | 0.98 | 0.22 | -1.337 | -0.171 | -0.096 | -0.082 | -0.082 | 0.063 | 0.45 | 0.08 | 0.57 | 0.52 | 0.49 | 0.42 |
| 5 | 111980106 | rs748628 | 0.29 | -0.188 | -0.48 | -0.068 | -0.029 | -0.003 | -0.003 | 0 | 0.7 | 0.56 | 0.76 | 0.72 | 0.74 | 0.7 |
| 5 | 111983052 | rs6860333 | -0.245 | 0.14 | -0.483 | -1.038 | 0.003 | -0.015 | -0.015 | 0 | 0.6 | 0.57 | 0.43 | 0.25 | 0.61 | 0.39 |
| 5 | 111983180 | rs6882143 | 0.483 | -0.106 | -1.19 | -1.546 | -0.001 | 0.003 | 0.003 | 0 | 0.24 | 0.09 | 0.19 | 0.22 | 0.24 | 0.21 |
| 5 | 111984920 | rs4705739 | 0.336 | 0.229 | -1.235 | -1.635 | -0.093 | 0.125 | 0.125 | 0.02 | 0.39 | 0.1 | 0.58 | 0.76 | 0.48 | 0.59 |
| 5 | 112006948 | rs7714497 | 0.043 | -0.008 | -0.574 | -1.73 | -0.001 | -0.017 | -0.017 | 0 | 0.16 | 0.16 | 0.11 | 0.05 | 0.19 | 0.15 |
| 5 | 112007180 | rs10428561 | 0.939 | 0.286 | -0.854 | -1.58 | -0.038 | -0.031 | -0.031 | 0.003 | 0.3 | 0.1 | 0.5 | 0.62 | 0.29 | 0.41 |
| 5 | 112008440 | rs7724805 | 0.95 | 0.283 | -0.93 | -1.684 | -0.068 | -0.041 | -0.041 | 0 | 0.3 | 0.08 | 0.5 | 0.63 | 0.29 | 0.41 |
| 5 | 112024075 | rs4705815 | -0.059 | 0.121 | 0.108 | -1.764 | 0.014 | 0.084 | 0.084 | 0 | 0.5 | 0.36 | 0.65 | 0.8 | 0.52 | 0.67 |
| 5 | 112026148 | rs1866924 | 0.465 | 0.084 | -0.038 | -1.794 | -0.123 | 0.047 | 0.047 | 0 | 0.42 | 0.08 | 0.59 | 0.68 | 0.47 | 0.56 |
| 5 | 112026480 | rs12719151 | -0.232 | 0.252 | 0.348 | -1.757 | -0.001 | -0.022 | -0.022 | 5.57 | 0.52 | 0.55 | 0.66 | 0.8 | 0.53 | 0.67 |
| 5 | 112045132 | rs6894351 | -0.298 | 0.088 | 0.707 | -1.759 | 0 | -0.011 | -0.011 | 0 | 0.54 | 0.49 | 0.66 | 0.8 | 0.53 | 0.65 |
| 5 | 112046069 | rs1734246 | 1.086 | -0.061 | 0.648 | -1.917 | 0.005 | -0.007 | -0.007 | 0 | 0.12 | 0.07 | 0.06 | 0 | 0.09 | 0.09 |
| 5 | 112057679 | rs1644242 | 1.033 | -0.072 | 0.646 | -1.922 | 0.01 | 0.011 | 0.011 | 0.008 | 0.12 | 0.05 | 0.06 | 0 | 0.1 | 0.09 |

| | | | | | | | | | | | | | | | | |
|---|---|---|---|---|---|---|---|---|---|---|---|---|---|---|---|---|
| 5 | 112060242 | rs10075281 | 0.677 | 0.08 | 0.652 | -1.935 | -0.094 | 0.061 | 0.061 | 0 | 0.44 | 0.09 | 0.6 | 0.66 | 0.47 | 0.55 |
| 5 | 112072793 | rs2464810 | 0.877 | -0.154 | 0.712 | -1.937 | -0.002 | -0.027 | -0.027 | 0.633 | 0.13 | 0.12 | 0.07 | 0 | 0.1 | 0.09 |
| 5 | 112072889 | rs2439593 | 1.027 | -0.15 | 0.682 | -1.935 | 0.005 | -0.006 | -0.006 | 0 | 0.12 | 0.07 | 0.07 | 0 | 0.1 | 0.09 |
| 5 | 112101793 | rs2464805 | -1.076 | -0.188 | 0.984 | -1.968 | -0.002 | 0.001 | 0.001 | 0 | 0.67 | 0.72 | 0.75 | 0.86 | 0.62 | 0.74 |
| 5 | 112101970 | rs2304793 | -1.03 | -0.201 | 1.188 | -1.93 | 0.015 | 0.14 | 0.14 | 0 | 0.05 | 0.22 | 0.04 | 0.13 | 0.01 | 0.09 |
| 5 | 112125419 | rs1734242 | -0.459 | -0.222 | 1.121 | -1.956 | -0.046 | -0.028 | -0.028 | 0 | 0.2 | 0.52 | 0.13 | 0.13 | 0.13 | 0.2 |
| 5 | 112146117 | rs454886 | 1.573 | -0.196 | 0.76 | -1.949 | -0.095 | -0.079 | -0.079 | 1.762 | 0.3 | 0.03 | 0.5 | 0.63 | 0.3 | 0.42 |
| 5 | 112149757 | rs2545169 | -1.218 | -0.238 | 1.292 | -1.881 | -0.017 | 0.091 | 0.091 | 0 | 0.68 | 0.86 | 0.76 | 0.91 | 0.63 | 0.75 |
| 5 | 112152206 | rs17134960 | 0.479 | -0.175 | 1.168 | -1.854 | -0.008 | -0.008 | -0.008 | 3.936 | 0.14 | 0.2 | 0.07 | 0.08 | 0.11 | 0.1 |
| 5 | 112152920 | rs17164132 | -1.566 | -0.095 | 1.29 | -1.782 | 0.033 | 0.154 | 0.154 | 0 | 0.06 | 0.27 | 0.06 | 0.16 | 0.04 | 0.11 |
| 5 | 112154303 | rs464708 | -0.987 | -0.088 | 1.207 | -1.799 | 0 | -0.02 | -0.02 | 0.032 | 0.66 | 0.64 | 0.74 | 0.82 | 0.62 | 0.75 |
| 5 | 112158431 | rs2545162 | -0.971 | -0.136 | 1.066 | -1.822 | 0.003 | -0.05 | -0.05 | 0 | 0.67 | 0.53 | 0.74 | 0.82 | 0.63 | 0.75 |
| 5 | 112164561 | rs351771 | -0.736 | 0.02 | 1.112 | -1.739 | -0.005 | -0.054 | -0.054 | 0 | 0.63 | 0.52 | 0.72 | 0.82 | 0.6 | 0.75 |
| 5 | 112175770 | rs41115 | -0.733 | 0.064 | 1.117 | -1.72 | -0.005 | -0.05 | -0.05 | 0 | 0.62 | 0.52 | 0.72 | 0.82 | 0.6 | 0.74 |
| 5 | 112176325 | rs42427 | -0.746 | 0.065 | 1.118 | -1.719 | -0.005 | -0.05 | -0.05 | 0 | 0.63 | 0.52 | 0.72 | 0.82 | 0.6 | 0.74 |
| 5 | 112176559 | rs866006 | -0.746 | 0.065 | 1.121 | -1.719 | -0.005 | -0.048 | -0.048 | 0 | 0.63 | 0.52 | 0.72 | 0.82 | 0.6 | 0.75 |
| 5 | 112177171 | rs465899 | -0.757 | 0.065 | 1.092 | -1.719 | -0.005 | -0.049 | -0.049 | 0 | 0.63 | 0.52 | 0.72 | 0.82 | 0.6 | 0.74 |
| 5 | 112179909 | rs1804197 | -1.5 | 0.054 | 1.054 | -1.643 | 0.001 | 0.124 | 0.124 | 0.018 | 0.06 | 0.23 | 0.03 | 0.16 | 0.01 | 0.1 |
| 5 | 112180257 | rs12189 | -1.5 | 0.054 | 1.054 | -1.643 | 0.001 | 0.124 | 0.124 | 0.018 | 0.06 | 0.22 | 0.03 | 0.15 | 0.01 | 0.1 |
| 5 | 112181283 | rs3733961 | -1.5 | 0.054 | 1.054 | -1.643 | 0.001 | 0.124 | 0.124 | 0.018 | 0.06 | 0.22 | 0.03 | 0.15 | 0.01 | 0.1 |
| 5 | 112185491 | rs497844 | -0.921 | 0.169 | 0.904 | -1.727 | -0.002 | -0.018 | -0.018 | 0.894 | 0.65 | 0.63 | 0.73 | 0.82 | 0.63 | 0.76 |
| 5 | 112196949 | rs460137 | 0.377 | 0.307 | 0.317 | -1.76 | -0.009 | 0.093 | 0.093 | 0 | 0.47 | 0.12 | 0.62 | 0.68 | 0.53 | 0.56 |
| 5 | 112212151 | rs3317 | 0.392 | 0.217 | 0.399 | -1.75 | -0.056 | 0.049 | 0.049 | 0 | 0.47 | 0.07 | 0.57 | 0.65 | 0.53 | 0.56 |
| 5 | 112214093 | rs2546115 | 1.549 | 0.332 | 0.485 | -1.887 | -0.086 | -0.096 | -0.096 | 0 | 0.28 | 0.03 | 0.44 | 0.61 | 0.3 | 0.39 |
| 5 | 112218700 | rs1093677 | 1.007 | 0.257 | 0.649 | -1.849 | -0.004 | -0.05 | -0.05 | 1.116 | 0.33 | 0.48 | 0.26 | 0.18 | 0.34 | 0.24 |
| 5 | 112224331 | rs458451 | 0.637 | 0.247 | 0.297 | -1.839 | 0.009 | 0.069 | 0.069 | 0.034 | 0.37 | 0.76 | 0.3 | 0.21 | 0.36 | 0.25 |
| 5 | 112228667 | rs430665 | 0.739 | 0.251 | 0.279 | -1.848 | 0.004 | 0.03 | 0.03 | 0.028 | 0.37 | 0.68 | 0.29 | 0.18 | 0.36 | 0.24 |
| 5 | 112229169 | rs466443 | -0.339 | 0.292 | 0.214 | -1.893 | 0.01 | 0.107 | 0.107 | 0 | 0.53 | 0.71 | 0.41 | 0.22 | 0.59 | 0.4 |
| 5 | 112236297 | rs153549 | -0.704 | -0.208 | 0.151 | -1.731 | -0.12 | 0.016 | 0.016 | 0 | 0.55 | 0.94 | 0.44 | 0.36 | 0.53 | 0.46 |
| 5 | 112238039 | rs153546 | 0.776 | -0.314 | 0.288 | -1.716 | 0.005 | 0.001 | 0.001 | 0 | 0.35 | 0.64 | 0.31 | 0.35 | 0.35 | 0.27 |
| 5 | 112240159 | rs10515449 | -0.336 | -0.328 | 0.495 | -1.715 | -0.036 | -0.055 | -0.055 | 0.505 | 0.19 | 0.47 | 0.16 | 0.12 | 0.13 | 0.18 |
| 5 | 112252066 | rs149192 | 0.086 | 0.026 | -0.216 | -1.515 | 0.013 | 0.099 | 0.099 | 0 | 0.48 | 0.66 | 0.36 | 0.16 | 0.46 | 0.42 |
| 5 | 112254377 | rs153560 | -1.059 | 0.312 | -0.766 | -1.186 | -0.13 | -0.067 | -0.067 | 0.044 | 0.7 | 0.96 | 0.5 | 0.36 | 0.63 | 0.61 |
| 5 | 112291861 | rs728347 | -0.692 | -0.123 | 0.602 | -0.557 | -0.02 | -0.009 | -0.009 | 0 | 0.16 | 0.39 | 0.15 | 0.15 | 0.11 | 0.22 |
| 5 | 112292127 | rs17135215 | -0.075 | -0.081 | -0.367 | -0.676 | -0.001 | -0.023 | -0.023 | 0.142 | 0.13 | 0.14 | 0.07 | 0.01 | 0.08 | 0.2 |
| 5 | 112302521 | rs9885540 | -0.864 | -0.106 | -0.754 | -0.573 | 0.027 | 0.029 | 0.029 | 0 | 0.18 | 0.08 | 0.11 | 0.15 | 0.13 | 0.12 |
| 5 | 112312528 | rs33556 | -1.285 | -0.169 | -0.707 | -0.73 | 0.018 | 0.022 | 0.022 | 0 | 0.34 | 0.22 | 0.24 | 0.3 | 0.31 | 0.29 |
| 5 | 112314281 | rs2080898 | -0.245 | 0.284 | 0.005 | -0.975 | 0.044 | -0.02 | -0.02 | 0 | 0.17 | 0.41 | 0.12 | 0.12 | 0.18 | 0.13 |
| 5 | 112317417 | rs10478101 | -0.589 | 0.191 | 0.347 | -0.766 | 0.016 | 0.006 | 0.006 | 4.516 | 0.19 | 0.6 | 0.13 | 0.17 | 0.18 | 0.14 |
| 5 | 112318928 | rs33551 | -1.84 | 0.184 | 0.426 | -0.81 | 0.002 | 0.069 | 0.069 | 0 | 0.49 | 0.68 | 0.35 | 0.34 | 0.47 | 0.41 |
| 5 | 112323738 | rs2112454 | -2.002 | 0.679 | 0.347 | -0.762 | 0.004 | 0.047 | 0.047 | 0 | 0.53 | 0.83 | 0.41 | 0.4 | 0.53 | 0.45 |
| 5 | 112324789 | rs1978430 | -0.469 | -0.106 | 0.699 | -0.719 | 0.051 | -0.106 | -0.106 | 0 | 0.19 | 0.49 | 0.13 | 0.05 | 0.19 | 0.15 |
| 5 | 112325869 | rs9784647 | 1.933 | -0.108 | 0.626 | -0.712 | -0.017 | 0.06 | 0.06 | 0 | 0.49 | 0.18 | 0.63 | 0.59 | 0.52 | 0.59 |
| 5 | 112328377 | rs3733969 | 0.589 | -0.115 | 0.591 | -0.722 | 0.031 | 0.043 | 0.043 | 0 | 0.7 | 0.8 | 0.78 | 0.76 | 0.75 | 0.75 |
| 5 | 112343544 | rs2052484 | 0.607 | -0.122 | 0.546 | -0.769 | 0.031 | 0.034 | 0.034 | 0.037 | 0.72 | 0.8 | 0.79 | 0.78 | 0.77 | 0.78 |
| 5 | 112349070 | rs9326869 | 0.81 | -0.127 | 0.609 | -0.761 | 0.032 | 0.056 | 0.056 | 0 | 0.7 | 0.84 | 0.78 | 0.78 | 0.74 | 0.75 |
| 5 | 112349297 | rs6892784 | 0.041 | -0.257 | 0.82 | -0.821 | -0.025 | 0 | 0 | 0.097 | 0.14 | 0.07 | 0.1 | 0.12 | 0.2 | 0.15 |
| 5 | 112349379 | rs4705749 | 2.29 | -0.282 | 0.834 | -0.832 | -0.066 | 0.056 | 0.056 | 0 | 0.49 | 0.08 | 0.61 | 0.61 | 0.51 | 0.59 |
| 5 | 112350707 | rs13186334 | 2.342 | -0.276 | 0.835 | -0.801 | -0.081 | 0.039 | 0.039 | 0 | 0.49 | 0.09 | 0.6 | 0.58 | 0.49 | 0.58 |
| 5 | 112356649 | rs7670 | 2.234 | -0.235 | 0.96 | -0.753 | -0.008 | 0.056 | 0.056 | 0 | 0.49 | 0.25 | 0.62 | 0.59 | 0.49 | 0.59 |
| 5 | 112358131 | rs9122 | 2.323 | -0.222 | 0.886 | -0.783 | -0.067 | 0.05 | 0.05 | 0 | 0.49 | 0.09 | 0.61 | 0.59 | 0.51 | 0.59 |
| 5 | 112358493 | rs2112452 | 0.783 | -0.205 | 0.88 | -0.776 | 0.027 | 0.046 | 0.046 | 0 | 0.69 | 0.8 | 0.77 | 0.76 | 0.73 | 0.75 |
| 5 | 112358890 | rs3209874 | 0.09 | -0.35 | 0.805 | -0.766 | -0.029 | -0.001 | -0.001 | 0.433 | 0.13 | 0.06 | 0.1 | 0.12 | 0.19 | 0.15 |
| 5 | 112362840 | rs1549181 | 2.123 | -0.382 | 0.789 | -0.751 | 0.011 | 0.065 | 0.065 | 0 | 0.5 | 0.23 | 0.64 | 0.59 | 0.54 | 0.6 |
| 5 | 112366678 | rs4705752 | 2.054 | -0.45 | 1.455 | -0.759 | -0.001 | 0.03 | 0.03 | 0 | 0.52 | 0.36 | 0.64 | 0.59 | 0.51 | 0.62 |
| 5 | 112368522 | rs2112453 | 1.19 | -0.728 | 1.4 | -0.734 | -0.013 | -0.026 | -0.026 | 2.952 | 0.67 | 0.55 | 0.74 | 0.76 | 0.71 | 0.75 |
| 5 | 112369995 | rs7447818 | -0.848 | -0.789 | 1.142 | -0.798 | 0.02 | 0.007 | 0.007 | 0 | 0.2 | 0.14 | 0.18 | 0.23 | 0.18 | 0.15 |
| 5 | 112370590 | rs878076 | 0.69 | -0.784 | 1.196 | -0.801 | -0.02 | -0.012 | -0.012 | 0 | 0.8 | 0.65 | 0.79 | 0.77 | 0.81 | 0.85 |
| 5 | 112374562 | rs6594659 | 1.653 | -0.786 | 1.482 | -0.796 | -0.024 | -0.022 | -0.022 | 0.806 | 0.67 | 0.42 | 0.74 | 0.7 | 0.72 | 0.75 |
| 5 | 112378836 | rs13360862 | 0.065 | -0.501 | 1.21 | -0.42 | -0.02 | 0.003 | 0.003 | 0.015 | 0.12 | 0.07 | 0.1 | 0.1 | 0.19 | 0.13 |
| 5 | 112379659 | rs7732198 | -2.403 | -0.774 | 1.295 | -0.558 | -0.001 | 0.056 | 0.056 | 0 | 0.43 | 0.28 | 0.33 | 0.34 | 0.45 | 0.4 |
| 5 | 112384802 | rs6594664 | 1.573 | -0.711 | 1.726 | -0.506 | 0.121 | 0.143 | 0.143 | 4.815 | 0.95 | 0.59 | 0.95 | 0.94 | 0.96 | 0.98 |
| 5 | 112386217 | rs7731890 | -2.686 | -0.784 | 1.716 | -0.44 | 0.007 | 0.032 | 0.032 | 0 | 0.48 | 0.71 | 0.37 | 0.43 | 0.45 | 0.41 |
| 5 | 112388225 | rs6594665 | -2.572 | -0.718 | 1.829 | -0.523 | -0.005 | -0.005 | -0.005 | 0.088 | 0.39 | 0.56 | 0.29 | 0.4 | 0.36 | 0.38 |
| 5 | 112390951 | rs4457065 | -0.641 | -0.851 | 1.777 | -0.593 | -0.005 | -0.048 | -0.048 | 0 | 0.18 | 0.24 | 0.09 | 0.01 | 0.15 | 0.18 |
| 5 | 112391033 | rs4705758 | 3.18 | -1.051 | 1.665 | -0.598 | 0.015 | 0.05 | 0.05 | 0 | 0.51 | 0.27 | 0.64 | 0.61 | 0.55 | 0.61 |
| 5 | 112393300 | rs6892724 | -2.102 | -1.056 | 1.662 | -0.572 | -0.021 | 0.084 | 0.084 | 0 | 0.15 | 0.43 | 0.17 | 0.28 | 0.1 | 0.06 |
| 5 | 112393483 | rs6893389 | -1.939 | -1.001 | 1.348 | -0.589 | 0.002 | 0.003 | 0.003 | 0 | 0.3 | 0.22 | 0.26 | 0.33 | 0.3 | 0.21 |
| 5 | 112394380 | rs6879589 | -1.889 | -1.1 | 1.468 | -0.426 | 0.003 | 0.014 | 0.014 | 0.065 | 0.36 | 0.3 | 0.3 | 0.34 | 0.35 | 0.3 |

| | | | | | | | | | | | | | | | | |
|---|---|---|---|---|---|---|---|---|---|---|---|---|---|---|---|---|
| 5 | 112396174 | rs3922410 | -0.049 | -1.581 | 1.565 | -0.069 | 0.014 | -0.038 | -0.038 | 0.018 | 0.93 | 0.81 | 0.95 | 1 | 0.91 | 0.9 |
| 5 | 112398161 | rs6890872 | 2.42 | -1.806 | 1.355 | -0.057 | 0.019 | 0.006 | 0.006 | 0.59 | 0.68 | 0.21 | 0.8 | 0.69 | 0.76 | 0.74 |
| 5 | 112398203 | rs9688048 | -2.281 | -2.159 | 2.103 | 0.233 | -0.034 | -0.02 | -0.02 | 0 | 0.24 | 0.34 | 0.15 | 0.2 | 0.14 | 0.15 |
| 5 | 112399791 | rs2227947 | 3.095 | -2.215 | 1.449 | -0.069 | 0.052 | 0.048 | 0.048 | 0 | 0.56 | 0.14 | 0.67 | 0.63 | 0.66 | 0.63 |
| 5 | 112417363 | rs4235789 | 0.502 | -1.906 | 1.875 | -0.165 | -0.02 | 0.012 | 0.012 | 0 | 0.93 | 0.85 | 0.91 | 0.89 | 0.94 | 0.97 |
| 5 | 112417907 | rs6859485 | -0.943 | -1.934 | 2.206 | -0.199 | 0.005 | 0.03 | 0.03 | 0 | 0.11 | 0.23 | 0.1 | 0.19 | 0.09 | 0.09 |
| 5 | 112420569 | rs11743316 | 0.028 | -1.943 | 2.179 | -0.198 | 0.003 | 0.001 | 0.001 | 0 | 0.08 | 0.07 | 0.06 | 0.07 | 0.03 | 0.06 |
| 5 | 112422167 | rs7708215 | -0.748 | -1.987 | 2.268 | -0.239 | -0.007 | 0.042 | 0.042 | 0.009 | 0.14 | 0.09 | 0.09 | 0.04 | 0.16 | 0.21 |
| 5 | 112424337 | rs7736823 | -2.92 | -1.95 | 2.488 | -0.231 | -0.001 | -0.002 | -0.002 | 0.033 | 0.33 | 0.4 | 0.29 | 0.34 | 0.32 | 0.35 |
| 5 | 112427271 | rs11744828 | -3.649 | -1.968 | 2.593 | -0.243 | 0.017 | 0.038 | 0.038 | 0 | 0.4 | 0.75 | 0.32 | 0.34 | 0.34 | 0.36 |
| 5 | 112428869 | rs7731125 | -2.046 | -1.952 | 2.616 | -0.236 | 0.076 | 0.037 | 0.037 | 1.915 | 0.06 | 0.36 | 0.03 | 0 | 0.03 | 0.02 |
| 5 | 112432454 | rs6594680 | -3.19 | -1.922 | 2.407 | -0.214 | 0.018 | 0.036 | 0.036 | 0 | 0.39 | 0.75 | 0.32 | 0.35 | 0.34 | 0.36 |
| 5 | 112432680 | rs10077067 | -2.004 | -1.93 | 2.447 | -0.217 | -0.001 | 0.028 | 0.028 | 0 | 0.15 | 0.22 | 0.17 | 0.26 | 0.15 | 0.12 |
| 5 | 112434312 | rs7724258 | -2.602 | -1.887 | 2.209 | -0.19 | 0.007 | 0.008 | 0.008 | 0 | 0.36 | 0.65 | 0.29 | 0.27 | 0.28 | 0.33 |
| 5 | 112435864 | rs6861336 | -1.372 | -1.852 | 1.717 | -0.161 | 0.002 | 0.025 | 0.025 | 0 | 0.12 | 0.26 | 0.11 | 0.19 | 0.09 | 0.1 |
| 5 | 112440142 | rs6869399 | -2.074 | -1.966 | 1.338 | 0.156 | 0.021 | 0.013 | 0.013 | 0.85 | 0.36 | 0.76 | 0.4 | 0.33 | 0.31 | 0.34 |
| 5 | 112452529 | rs13156718 | 1.505 | -2.278 | 0.797 | -0.265 | 0.056 | 0.106 | 0.106 | 0.241 | 0.52 | 0.15 | 0.56 | 0.69 | 0.61 | 0.54 |
| 5 | 112453010 | rs4705542 | 1.519 | -2.286 | 1.266 | -0.263 | 0.059 | 0.099 | 0.099 | 0.348 | 0.51 | 0.21 | 0.56 | 0.68 | 0.6 | 0.53 |
| 5 | 112456258 | rs7701321 | -0.988 | -2.389 | 1.622 | -0.266 | -0.024 | 0.061 | 0.061 | 0 | 0.08 | 0.28 | 0.12 | 0.16 | 0.05 | 0.06 |
| 5 | 112460633 | rs6883605 | 2.076 | -2.59 | 1.176 | -0.34 | 0.003 | 0.039 | 0.039 | 0 | 0.45 | 0.08 | 0.52 | 0.62 | 0.6 | 0.51 |
| 5 | 112461641 | rs6594683 | 1.463 | -2.554 | 1.469 | -0.373 | 0.051 | 0.073 | 0.073 | 0.148 | 0.53 | 0.26 | 0.61 | 0.69 | 0.64 | 0.55 |
| 5 | 112464953 | rs1443693 | 0.638 | -2.442 | 1.353 | -0.435 | -0.001 | -0.006 | -0.006 | 0.157 | 0.64 | 0.65 | 0.74 | 0.87 | 0.71 | 0.67 |
| 5 | 112478397 | rs17135341 | 0.706 | -2.203 | 0.866 | -0.314 | 0.006 | 0.013 | 0.013 | 0 | 0.06 | 0.01 | 0.07 | 0.05 | 0.04 | 0.12 |
| 5 | 112482033 | rs2416303 | 1.823 | -2.483 | 0.924 | -0.386 | 0.078 | 0.093 | 0.093 | 0 | 0.5 | 0.21 | 0.52 | 0.69 | 0.65 | 0.52 |
| 5 | 112483604 | rs2416305 | 1.369 | -2.298 | 1.493 | -0.308 | -0.018 | -0.014 | -0.014 | 1.833 | 0.57 | 0.51 | 0.6 | 0.72 | 0.68 | 0.58 |
| 5 | 112486175 | rs10068491 | -0.922 | -2.044 | 1.723 | -0.234 | 0.015 | 0.007 | 0.007 | 0 | 0.34 | 0.25 | 0.31 | 0.25 | 0.29 | 0.37 |
| 5 | 112490295 | rs10477488 | -2.314 | -1.899 | 2.145 | 0.036 | 0.062 | 0.041 | 0.041 | 0.024 | 0.08 | 0.26 | 0.1 | 0.11 | 0.07 | 0.1 |
| 5 | 112492839 | rs13156236 | 1.997 | -2.134 | 1.973 | -0.085 | 0.02 | 0.009 | 0.009 | 0.979 | 0.6 | 0.26 | 0.63 | 0.67 | 0.68 | 0.59 |
| 5 | 112494411 | rs4705798 | 1.718 | -2.023 | 2.302 | 0.052 | 0.016 | 0.059 | 0.059 | 0 | 0.92 | 0.75 | 0.91 | 0.87 | 0.96 | 0.92 |
| 5 | 112494583 | rs1867322 | 2.845 | -2.022 | 2.522 | 0.037 | 0.128 | 0.123 | 0.123 | 1.609 | 0.94 | 0.59 | 0.96 | 0.92 | 0.97 | 0.95 |
| 5 | 112495421 | rs7732695 | -1.015 | -2.026 | 2.264 | 0.046 | -0.017 | 0.061 | 0.061 | 0 | 0.06 | 0.1 | 0.04 | 0.17 | 0.03 | 0.05 |
| 5 | 112496354 | rs935162 | -1.949 | -2.345 | 2.041 | -0.144 | 0.003 | 0.001 | 0.001 | 0 | 0.37 | 0.58 | 0.33 | 0.34 | 0.3 | 0.33 |
| 5 | 112496622 | rs1443687 | 2.17 | -2.345 | 1.813 | -0.144 | 0.031 | 0.013 | 0.013 | 0.109 | 0.6 | 0.25 | 0.65 | 0.66 | 0.7 | 0.67 |
| 5 | 112500410 | rs11241195 | -0.709 | -2.199 | 2.059 | -0.183 | -0.003 | -0.022 | -0.022 | 0 | 0.24 | 0.25 | 0.21 | 0.12 | 0.22 | 0.22 |
| 5 | 112501783 | rs12654269 | -0.671 | -2.256 | 2.021 | -0.21 | -0.002 | -0.012 | -0.012 | 0.99 | 0.24 | 0.24 | 0.21 | 0.12 | 0.22 | 0.22 |
| 5 | 112501801 | rs12652257 | -1.974 | -2.336 | 2.132 | -0.219 | 0.003 | 0.003 | 0.003 | 0 | 0.37 | 0.56 | 0.28 | 0.27 | 0.3 | 0.32 |
| 5 | 112502941 | rs11952815 | -0.615 | -1.805 | 2.181 | -0.295 | -0.008 | -0.028 | -0.028 | 0.649 | 0.22 | 0.28 | 0.1 | 0.11 | 0.2 | 0.22 |
| 5 | 112504775 | rs17135372 | -1.386 | -1.665 | 1.662 | -0.157 | -0.009 | 0.023 | 0.023 | 0 | 0.07 | 0.1 | 0.04 | 0.12 | 0.05 | 0.08 |
| 5 | 112512499 | rs7719681 | 2.047 | -1.76 | 1.654 | -0.217 | 0.025 | 0.043 | 0.043 | 0 | 0.61 | 0.27 | 0.76 | 0.72 | 0.7 | 0.66 |
| 5 | 112516709 | rs6594693 | 1.02 | -1.565 | 1.33 | -0.108 | 0.005 | -0.004 | -0.004 | 0 | 0.93 | 0.93 | 0.93 | 0.9 | 0.95 | 0.94 |
| 5 | 112517571 | rs4398626 | -0.962 | -1.488 | 1.287 | -0.09 | -0.002 | -0.002 | -0.002 | 0 | 0.05 | 0.04 | 0.02 | 0.08 | 0.06 | 0.07 |
| 5 | 112519471 | rs1005746 | -1.547 | -1.777 | 1.244 | -0.268 | 0.003 | 0.014 | 0.014 | 0 | 0.11 | 0.07 | 0.05 | 0.04 | 0.1 | 0.07 |
| 5 | 112521248 | rs7718971 | -0.303 | -1.937 | 1.56 | -0.329 | -0.001 | -0.007 | -0.007 | 0 | 0.19 | 0.23 | 0.09 | 0.11 | 0.18 | 0.18 |
| 5 | 112521347 | rs1348431 | 0.673 | -1.901 | 1.174 | -0.308 | -0.017 | -0.075 | -0.075 | 0.111 | 0.74 | 0.53 | 0.85 | 0.89 | 0.78 | 0.79 |
| 5 | 112523526 | rs2044241 | 0.906 | -1.834 | 0.805 | -0.295 | -0.007 | -0.014 | -0.014 | 0 | 0.72 | 0.42 | 0.81 | 0.81 | 0.76 | 0.76 |
| 5 | 112530417 | rs2416315 | 1.814 | -1.881 | 0.181 | -0.301 | 0.029 | 0.035 | 0.035 | 0 | 0.59 | 0.25 | 0.72 | 0.69 | 0.67 | 0.67 |
| 5 | 112530947 | rs2416313 | 1.025 | -1.541 | 0.321 | -0.225 | 0.003 | 0.002 | 0.002 | 0 | 0.7 | 0.34 | 0.76 | 0.73 | 0.77 | 0.74 |
| 5 | 112532935 | rs2120349 | 1.568 | -1.167 | -0.462 | -0.498 | -0.002 | 0.089 | 0.089 | 0.426 | 0.56 | 0.04 | 0.51 | 0.75 | 0.67 | 0.69 |
| 5 | 112535345 | rs12659754 | 1.128 | -1.065 | 0.465 | -0.588 | -0.004 | -0.015 | -0.015 | 0.009 | 0.64 | 0.43 | 0.53 | 0.72 | 0.67 | 0.69 |
| 5 | 112542527 | rs1822489 | 1.203 | -1.108 | 0.71 | -0.563 | -0.01 | -0.02 | -0.02 | 3.528 | 0.64 | 0.43 | 0.53 | 0.72 | 0.68 | 0.68 |
| 5 | 112550247 | rs509237 | 1.5 | -1.378 | -0.161 | -0.734 | 0.048 | 0.052 | 0.052 | 0.871 | 0.64 | 0.08 | 0.53 | 0.75 | 0.79 | 0.77 |
| 5 | 112563686 | rs7717906 | -1.791 | -1.377 | 0.065 | -0.761 | 0.026 | 0.032 | 0.032 | 2.16 | 0.21 | 0.79 | 0.42 | 0.14 | 0.11 | 0.09 |
| 5 | 112563944 | rs557438 | -1.613 | -1.653 | 0.275 | -0.983 | 0.008 | 0.009 | 0.009 | 0 | 0.32 | 0.33 | 0.12 | 0.15 | 0.23 | 0.24 |
| 5 | 112565996 | rs496977 | -1.05 | -1.66 | -0.269 | -0.995 | -0.004 | 0.061 | 0.061 | 0 | 0.27 | 0.05 | 0.09 | 0.15 | 0.23 | 0.24 |
| 5 | 112566758 | rs12521391 | 0.592 | -1.623 | 0.118 | -0.97 | -0.01 | -0.006 | -0.006 | 0.122 | 0.07 | 0.15 | 0.31 | 0.07 | 0.05 | 0.04 |
| 5 | 112567834 | rs569540 | -0.227 | -1.835 | -0.287 | -0.601 | -0.011 | -0.064 | -0.064 | 0 | 0.18 | 0.01 | 0.06 | 0.2 | 0.15 | 0.17 |
| 5 | 112568212 | rs680503 | -2.224 | -1.606 | 0.017 | -0.85 | 0.065 | 0.149 | 0.149 | 0.101 | 0.48 | 0.88 | 0.53 | 0.29 | 0.33 | 0.33 |
| 5 | 112577696 | rs11740353 | 0.257 | -1.415 | 0.454 | -0.719 | -0.011 | -0.006 | -0.006 | 0.159 | 0.07 | 0.18 | 0.32 | 0.07 | 0.05 | 0.04 |
| 5 | 112578543 | rs2091803 | -1.812 | -1.691 | 1.116 | -0.926 | -0.035 | -0.075 | -0.075 | 0 | 0.21 | 0.62 | 0.4 | 0.08 | 0.15 | 0.12 |
| 5 | 112579347 | rs637634 | -2.787 | -1.166 | 0.852 | -0.917 | -0.037 | 0.095 | 0.095 | 0.068 | 0.48 | 0.96 | 0.53 | 0.31 | 0.37 | 0.3 |
| 5 | 112584120 | rs619146 | -1.2 | -1.084 | 1.022 | -0.799 | 0.002 | -0.003 | -0.003 | 0 | 0.26 | 0.32 | 0.13 | 0.24 | 0.22 | 0.19 |
| 5 | 112587455 | rs665034 | 0.449 | -1.092 | 0.854 | -0.817 | -0.007 | -0.016 | -0.016 | 0 | 0.12 | 0.01 | 0.03 | 0.12 | 0.08 | 0.11 |
| 5 | 112592606 | rs10519343 | 0.484 | -1.095 | 0.955 | -0.824 | -0.01 | -0.005 | -0.005 | 0 | 0.07 | 0.19 | 0.32 | 0.07 | 0.05 | 0.04 |
| 5 | 112593347 | rs10519347 | -1.994 | -1.121 | 1.327 | -0.86 | -0.001 | -0.012 | -0.012 | 0 | 0.33 | 0.34 | 0.16 | 0.23 | 0.29 | 0.24 |
| 5 | 112601878 | rs10519348 | -0.715 | -0.883 | 1.251 | -0.912 | -0.05 | 0.059 | 0.059 | 0.009 | 0.14 | 0.02 | 0.06 | 0.03 | 0.18 | 0.12 |
| 5 | 112608343 | rs17135534 | -0.58 | -0.908 | 1.528 | -0.861 | -0.015 | -0.021 | -0.021 | 0 | 0.11 | 0.28 | 0.34 | 0.09 | 0.08 | 0.07 |
| 5 | 112612123 | rs11749864 | 0.02 | -0.926 | 1.59 | -0.865 | -0.013 | -0.008 | -0.008 | 0 | 0.08 | 0.26 | 0.32 | 0.07 | 0.05 | 0.04 |
| 5 | 112615044 | rs13190054 | -2.724 | 1.186 | 0.607 | -0.895 | -0.126 | 0.076 | 0.076 | 1.5 | 0.43 | 0.97 | 0.58 | 0.28 | 0.49 | 0.32 |
| 5 | 112624552 | rs11956139 | -1.966 | 1.19 | 0.749 | -0.81 | -0.048 | 0.041 | 0.041 | 1.149 | 0.25 | 0.83 | 0.49 | 0.18 | 0.27 | 0.19 |

| | | | | | | | | | | | | | | | | |
|---|---|---|---|---|---|---|---|---|---|---|---|---|---|---|---|---|
| 5 | 112635794 | rs10434754 | 0.886 | 1.26 | 0.785 | -0.848 | 0.03 | 0.001 | 0.001 | 0 | 0.08 | 0.07 | 0.11 | 0.08 | 0.18 | 0.11 |
| 5 | 112641650 | rs11241198 | -1.865 | 1.209 | 0.762 | -0.849 | -0.017 | 0.023 | 0.023 | 0.708 | 0.25 | 0.78 | 0.49 | 0.19 | 0.27 | 0.21 |
| 5 | 112655413 | rs2068923 | 0.176 | 1.233 | 0.844 | -0.732 | 0 | 0.009 | 0.009 | 0 | 0.09 | 0.27 | 0.34 | 0.08 | 0.07 | 0.04 |
| 5 | 112674497 | rs4705812 | -2.081 | 1.33 | 0.772 | -0.794 | -0.091 | 0.119 | 0.119 | 3.037 | 0.39 | 0.8 | 0.55 | 0.2 | 0.46 | 0.31 |
| 5 | 112679373 | rs1965429 | -1.802 | 1.464 | 0.949 | -0.73 | 0.017 | -0.02 | -0.02 | 0 | 0.26 | 0.65 | 0.21 | 0.14 | 0.35 | 0.24 |
| 5 | 112681830 | rs6876848 | -0.846 | 1.559 | 1.011 | -0.686 | 0.121 | 0.025 | 0.025 | 0.078 | 0.15 | 0.38 | 0.18 | 0.18 | 0.26 | 0.19 |
| 5 | 112696768 | rs1850412 | -2.004 | 1.509 | 0.677 | -0.682 | -0.007 | 0.003 | 0.003 | 0 | 0.3 | 0.71 | 0.25 | 0.22 | 0.41 | 0.26 |
| 5 | 112709282 | rs17135581 | -1.428 | 1.528 | 0.83 | -0.639 | 0.029 | -0.022 | -0.022 | 0 | 0.18 | 0.59 | 0.11 | 0.13 | 0.17 | 0.11 |
| 5 | 112718730 | rs7737155 | -1.412 | 1.531 | 0.832 | -0.637 | 0.029 | -0.022 | -0.022 | 0.469 | 0.17 | 0.59 | 0.11 | 0.13 | 0.17 | 0.11 |
| 5 | 112719722 | rs17135597 | -0.11 | 1.29 | 0.661 | -0.79 | -0.028 | -0.038 | -0.038 | 0 | 0.16 | 0.26 | 0.36 | 0.08 | 0.13 | 0.1 |
| 5 | 112723567 | rs1318772 | -0.151 | 1.318 | 0.409 | -1.008 | 0.061 | -0.034 | -0.034 | 0.339 | 0.11 | 0.33 | 0.05 | 0.03 | 0.13 | 0.07 |
| 5 | 112735231 | rs7714760 | 0.351 | 1.475 | -0.034 | -0.983 | 0.018 | -0.001 | -0.001 | 0 | 0.1 | 0.12 | 0.13 | 0.09 | 0.22 | 0.13 |
| 5 | 112739017 | rs26965 | 0.47 | 1.396 | 0.078 | -0.957 | 0.031 | -0.023 | -0.023 | 0.333 | 0.08 | 0.17 | 0.04 | 0.03 | 0.12 | 0.07 |
| 5 | 112739130 | rs27566 | -1.996 | 1.324 | -0.479 | -0.951 | -0.119 | 0.032 | 0.032 | 2.15 | 0.42 | 0.97 | 0.6 | 0.3 | 0.51 | 0.33 |
| 5 | 112755990 | rs36611 | 0.603 | 1.376 | -0.232 | -0.819 | -0.03 | 0.017 | 0.017 | 0 | 0.08 | 0.02 | 0.03 | 0.03 | 0.12 | 0.07 |
| 5 | 112759591 | rs10053157 | -0.773 | 1.318 | 0.142 | -0.819 | 0.036 | 0.05 | 0.05 | 0 | 0.08 | 0.33 | 0.06 | 0.1 | 0.05 | 0.04 |
| 5 | 112768385 | rs17135618 | -0.738 | 1.334 | 0.149 | -0.818 | 0.04 | 0.049 | 0.049 | 0 | 0.07 | 0.32 | 0.06 | 0.1 | 0.05 | 0.04 |
| 5 | 112770027 | rs26980 | 0.626 | 1.375 | 0.064 | -0.843 | -0.03 | 0.02 | 0.02 | 0 | 0.08 | 0.02 | 0.03 | 0.03 | 0.12 | 0.06 |
| 5 | 112771824 | rs7727449 | -1.612 | 1.546 | 0.225 | -0.741 | -0.021 | 0.011 | 0.011 | 0.134 | 0.33 | 0.8 | 0.54 | 0.29 | 0.36 | 0.27 |
| 5 | 112780914 | rs7709818 | -0.765 | 1.294 | 0.206 | -0.675 | 0.039 | 0.053 | 0.053 | 0 | 0.07 | 0.33 | 0.06 | 0.1 | 0.05 | 0.04 |
| 5 | 112783791 | rs17388817 | 0.713 | 1.149 | -0.339 | -0.568 | -0.025 | -0.021 | -0.021 | 0.28 | 0.11 | 0 | 0.04 | 0.1 | 0.09 | 0.1 |
| 5 | 112790404 | rs6594718 | -1.072 | 1.153 | 0.063 | -0.637 | -0.009 | -0.03 | -0.03 | 0 | 0.24 | 0.6 | 0.44 | 0.19 | 0.21 | 0.17 |
| 5 | 112796598 | rs17389165 | 0.961 | 1.635 | -0.056 | -0.644 | -0.038 | -0.016 | -0.016 | 0 | 0.07 | 0.03 | 0.09 | 0.08 | 0.17 | 0.12 |
| 5 | 112825032 | rs348938 | 1.051 | 1.566 | -0.198 | -0.802 | 0.023 | -0.021 | -0.021 | 0.92 | 0.7 | 0.63 | 0.83 | 0.77 | 0.66 | 0.74 |
| 5 | 112825677 | rs348937 | -2.107 | 1.52 | 0.058 | -0.829 | 0.011 | 0.065 | 0.065 | 0.085 | 0.48 | 0.67 | 0.55 | 0.35 | 0.5 | 0.41 |
| 5 | 112829305 | rs6594721 | -1.085 | 1.477 | 0.006 | -0.719 | -0.009 | -0.008 | -0.008 | 0.958 | 0.25 | 0.59 | 0.44 | 0.21 | 0.21 | 0.17 |
| 5 | 112842992 | rs3906371 | -1.737 | 1.496 | 0.094 | -0.727 | -0.042 | 0.03 | 0.03 | 0.663 | 0.35 | 0.83 | 0.54 | 0.28 | 0.38 | 0.29 |
| 5 | 112848767 | rs2416282 | 2.545 | 1.378 | -0.097 | -0.81 | -0.046 | -0.096 | -0.096 | 0.011 | 0.43 | 0.02 | 0.37 | 0.57 | 0.43 | 0.56 |
| 5 | 112853189 | rs1833678 | -0.052 | 1.536 | 0.392 | -0.631 | -0.008 | -0.024 | -0.024 | 0 | 0.17 | 0.27 | 0.38 | 0.1 | 0.16 | 0.13 |
| 5 | 112868693 | rs6594732 | -1.043 | 1.642 | 0.337 | -0.602 | -0.002 | -0.004 | -0.004 | 0.452 | 0.24 | 0.58 | 0.45 | 0.21 | 0.21 | 0.17 |
| 5 | 112885560 | rs6594735 | 1.231 | 1.688 | 0.241 | -0.638 | -0.005 | 0.018 | 0.018 | 3.184 | 0.69 | 0.22 | 0.54 | 0.76 | 0.7 | 0.77 |
| 5 | 112899573 | rs6867579 | 1.021 | 1.782 | 0.275 | -0.591 | -0.043 | -0.019 | -0.019 | 0.028 | 0.08 | 0.03 | 0.09 | 0.08 | 0.18 | 0.12 |
| 5 | 112899702 | rs6886628 | -0.704 | 1.758 | 0.421 | -0.587 | -0.006 | 0.001 | 0.001 | 0.752 | 0.22 | 0.62 | 0.43 | 0.2 | 0.19 | 0.16 |
| 5 | 112911165 | rs10065790 | 0.274 | 1.752 | 0.264 | -0.611 | 0.024 | -0.001 | -0.001 | 0 | 0.09 | 0.12 | 0.11 | 0.09 | 0.18 | 0.13 |
| 5 | 112916780 | rs6864369 | -0.171 | 1.504 | 0.373 | -0.586 | -0.008 | -0.024 | -0.024 | 0.443 | 0.17 | 0.27 | 0.38 | 0.1 | 0.17 | 0.13 |
| 5 | 112917278 | rs11748794 | -0.204 | 1.503 | 0.37 | -0.587 | -0.009 | -0.025 | -0.025 | 0 | 0.18 | 0.28 | 0.38 | 0.1 | 0.17 | 0.13 |
| 5 | 112926755 | rs2303718 | -0.771 | 1.535 | 0.523 | -0.547 | 0.049 | 0.072 | 0.072 | 0 | 0.07 | 0.32 | 0.06 | 0.1 | 0.05 | 0.04 |
| 5 | 112926982 | rs2303717 | -2.223 | 1.633 | 0.059 | -0.673 | -0.097 | 0.005 | 0.005 | 2.31 | 0.44 | 0.97 | 0.59 | 0.32 | 0.5 | 0.36 |
| 5 | 112929694 | rs9552 | -2.023 | 1.619 | 0.036 | -0.691 | -0.124 | -0.019 | -0.019 | 0 | 0.41 | 0.96 | 0.59 | 0.32 | 0.49 | 0.35 |
| 5 | 112930203 | rs7730949 | -1.123 | 1.544 | 0.25 | -0.673 | -0.006 | -0.008 | -0.008 | 0 | 0.25 | 0.65 | 0.45 | 0.2 | 0.21 | 0.17 |
| 5 | 112930565 | rs7713451 | -1.123 | 1.544 | 0.25 | -0.673 | -0.006 | -0.008 | -0.008 | 0 | 0.25 | 0.65 | 0.45 | 0.2 | 0.21 | 0.17 |
| 5 | 112942820 | rs10076131 | -1.042 | 1.566 | 0.251 | -0.621 | -0.002 | -0.006 | -0.006 | 0 | 0.24 | 0.62 | 0.45 | 0.21 | 0.22 | 0.18 |
| 5 | 112945992 | rs2914200 | 2.414 | 1.448 | -0.174 | -0.668 | -0.043 | -0.09 | -0.09 | 4.766 | 0.45 | 0.02 | 0.39 | 0.59 | 0.45 | 0.57 |
| 5 | 112947050 | rs17333442 | 0.28 | 1.725 | 0.313 | -0.549 | 0.018 | -0.023 | -0.023 | 0.066 | 0.08 | 0.15 | 0.04 | 0.03 | 0.11 | 0.07 |
| 5 | 112963488 | rs4538600 | -1.696 | 1.685 | 0.445 | -0.599 | 0.063 | -0.014 | -0.014 | 0 | 0.24 | 0.57 | 0.22 | 0.19 | 0.32 | 0.22 |
| 5 | 112965145 | rs6889348 | -1.279 | 1.766 | 0.58 | -0.606 | 0.068 | -0.004 | -0.004 | 0 | 0.16 | 0.35 | 0.16 | 0.11 | 0.2 | 0.14 |
| 5 | 112965930 | rs13160538 | 0.545 | 1.748 | 0.549 | -0.611 | 0.005 | -0.006 | -0.006 | 0.29 | 0.07 | 0.08 | 0.04 | 0.02 | 0.11 | 0.07 |
| 5 | 112969775 | rs10061922 | -1.821 | 1.721 | 0.498 | -0.608 | 0.059 | -0.032 | -0.032 | 2.429 | 0.24 | 0.57 | 0.21 | 0.13 | 0.31 | 0.21 |
| 5 | 112970054 | rs6877548 | -1.81 | 1.726 | 0.5 | -0.605 | 0.06 | -0.032 | -0.032 | 2.607 | 0.24 | 0.57 | 0.21 | 0.13 | 0.31 | 0.21 |
| 5 | 112973977 | rs2914177 | 0.594 | 1.378 | 0.58 | -0.67 | -0.012 | -0.012 | -0.012 | 2.095 | 0.73 | 0.68 | 0.61 | 0.83 | 0.81 | 0.79 |
| 5 | 112979910 | rs2914173 | 0.05 | 1.523 | 0.751 | -0.563 | -0.002 | 0.009 | 0.009 | 0 | 0.86 | 0.82 | 0.9 | 0.78 | 0.9 | 0.84 |
| 5 | 112981528 | rs12189067 | 0.114 | 1.538 | 0.769 | -0.583 | 0.006 | -0.001 | -0.001 | 1.78 | 0.08 | 0.1 | 0.11 | 0.02 | 0.16 | 0.11 |
| 5 | 112989901 | rs2972664 | 2.815 | 1.215 | 0.352 | -0.888 | -0.102 | -0.011 | -0.011 | 0 | 0.55 | 0.09 | 0.67 | 0.63 | 0.58 | 0.65 |
| 5 | 112990207 | rs6594744 | -2.02 | 1.289 | 0.593 | -0.942 | -0.008 | 0.009 | 0.009 | 0.142 | 0.3 | 0.7 | 0.26 | 0.2 | 0.35 | 0.22 |
| 5 | 112995542 | rs11747724 | -0.758 | 1.411 | 0.957 | -0.944 | 0 | -0.01 | -0.01 | 0.201 | 0.14 | 0.16 | 0.06 | 0.09 | 0.14 | 0.09 |
| 5 | 112996654 | rs17135859 | 0 | 1.539 | 0.856 | -0.94 | 0.056 | -0.016 | -0.016 | 0.144 | 0.08 | 0.23 | 0.12 | 0.02 | 0.16 | 0.11 |
| 5 | 112998285 | rs2115207 | -1.662 | 1.361 | 0.467 | -0.905 | -0.001 | -0.008 | -0.008 | 0 | 0.28 | 0.68 | 0.21 | 0.11 | 0.31 | 0.21 |
| 5 | 113007302 | rs12515653 | 2.029 | 1.351 | -0.343 | -0.347 | -0.056 | -0.07 | -0.07 | 0 | 0.15 | 0.01 | 0.18 | 0.26 | 0.22 | 0.25 |
| 5 | 113007423 | rs6859521 | -1.25 | 1.88 | 0.744 | -0.483 | 0.055 | -0.082 | -0.082 | 0 | 0.22 | 0.48 | 0.19 | 0.05 | 0.28 | 0.2 |
| 5 | 113007517 | rs17419034 | 0.677 | 2.223 | 0.497 | -0.101 | -0.032 | -0.045 | -0.045 | 0.02 | 0.11 | 0.03 | 0.3 | 0.15 | 0.09 | 0.14 |
| 5 | 113009026 | rs6891711 | -1.566 | 1.869 | 0.978 | -0.323 | 0.025 | -0.038 | -0.038 | 1.406 | 0.25 | 0.64 | 0.2 | 0.1 | 0.29 | 0.17 |
| 5 | 113013752 | rs2972693 | 1.993 | 1.761 | 0.134 | -0.265 | -0.039 | 0.067 | 0.067 | 0.252 | 0.61 | 0.2 | 0.72 | 0.79 | 0.56 | 0.7 |
| 5 | 113018177 | rs890924 | 2.133 | 1.762 | -0.052 | 0.027 | -0.082 | -0.037 | -0.037 | 0 | 0.58 | 0.09 | 0.7 | 0.63 | 0.58 | 0.72 |
| 5 | 113018908 | rs2416325 | 0.407 | 2.059 | 0.632 | 0.076 | 0.086 | 0.004 | 0.004 | 0 | 0.06 | 0.14 | 0.09 | 0.02 | 0.15 | 0.07 |
| 5 | 113019365 | rs2416326 | -0.186 | 2.11 | 0.719 | 0.074 | -0.004 | -0.01 | -0.01 | 0 | 0.08 | 0.14 | 0.04 | 0.06 | 0.04 | 0.09 |
| 5 | 113019569 | rs2162846 | -1.757 | 2.055 | 0.982 | 0.041 | 0.005 | -0.022 | -0.022 | 0 | 0.21 | 0.71 | 0.17 | 0.08 | 0.2 | 0.17 |
| 5 | 113022960 | rs2416328 | -2.166 | 2.089 | 0.914 | -0.142 | -0.011 | -0.022 | -0.022 | 0 | 0.23 | 0.81 | 0.19 | 0.26 | 0.22 | 0.19 |
| 5 | 113023069 | rs10519361 | -0.937 | 2.126 | 1.237 | -0.129 | 0.009 | 0.15 | 0.15 | 0 | 0.11 | 0.33 | 0.06 | 0.24 | 0.06 | 0.11 |
| 5 | 113023119 | rs2416329 | -0.043 | 2.144 | 1.391 | -0.19 | 0.106 | -0.018 | -0.018 | 0.791 | 0.11 | 0.36 | 0.12 | 0.02 | 0.16 | 0.08 |

| | | | | | | | | | | | | | | | | |
|---|---|---|---|---|---|---|---|---|---|---|---|---|---|---|---|---|
| 5 | 113030164 | rs4621553 | 0.192 | 2.005 | 0.75 | -0.468 | -0.001 | -0.017 | -0.017 | 0 | 0.77 | 0.8 | 0.59 | 0.93 | 0.76 | 0.83 |
| 5 | 113042748 | rs4274969 | -0.561 | 2.013 | 0.416 | -0.432 | 0 | -0.046 | -0.046 | 0 | 0.92 | 0.81 | 0.96 | 1 | 0.94 | 0.97 |
| 5 | 113053253 | rs1482347 | -0.053 | 2.093 | 0.367 | -0.334 | 0.011 | 0.067 | 0.067 | 0.051 | 0.79 | 0.91 | 0.63 | 0.93 | 0.8 | 0.86 |
| 5 | 113053318 | rs1482346 | 0.964 | 2.094 | 0.511 | -0.413 | -0.016 | -0.103 | -0.103 | 1.389 | 0.69 | 0.61 | 0.58 | 0.93 | 0.76 | 0.77 |
| 5 | 113053462 | rs13188669 | -0.347 | 2.321 | 0.322 | -0.488 | -0.014 | 0.02 | 0.02 | 0 | 0.12 | 0.07 | 0.07 | 0.06 | 0.14 | 0.06 |
| 5 | 113054380 | rs6885608 | 0.166 | 2.268 | 0.394 | -0.423 | 0.035 | -0.039 | -0.039 | 0 | 0.08 | 0.17 | 0.06 | 0.01 | 0.08 | 0.12 |
| 5 | 113055010 | rs1871909 | 1.03 | 2.269 | 0.411 | -0.412 | 0 | -0.002 | -0.002 | 0 | 0.69 | 0.64 | 0.54 | 0.69 | 0.71 | 0.72 |
| 5 | 113058627 | rs1871908 | 0.336 | 2.158 | -0.195 | -0.348 | 0.009 | -0.012 | -0.012 | 0 | 0.76 | 0.8 | 0.6 | 0.7 | 0.78 | 0.84 |
| 5 | 113067189 | rs7737958 | 0.512 | 2.155 | -0.159 | -0.349 | -0.01 | 0.007 | 0.007 | 0.02 | 0.75 | 0.66 | 0.59 | 0.7 | 0.78 | 0.84 |
| 5 | 113072475 | rs11241230 | 2.175 | 2.315 | -0.684 | -0.474 | -0.031 | 0.004 | 0.004 | 1.116 | 0.65 | 0.13 | 0.74 | 0.74 | 0.68 | 0.77 |
| 5 | 113072836 | rs11241231 | -0.167 | 2.18 | -0.853 | -0.262 | -0.001 | 0.033 | 0.033 | 0 | 0.91 | 0.87 | 0.94 | 0.75 | 0.92 | 0.95 |
| 5 | 113076095 | rs10064445 | 1.452 | 2.393 | -0.731 | -0.543 | -0.005 | -0.089 | -0.089 | 0 | 0.64 | 0.61 | 0.55 | 0.93 | 0.7 | 0.74 |
| 5 | 113076176 | rs6879991 | -0.41 | 2.375 | -0.719 | -0.557 | 0 | 0.041 | 0.041 | 0 | 0.23 | 0.14 | 0.4 | 0.07 | 0.21 | 0.23 |
| 5 | 113076866 | rs4705582 | 1.49 | 2.361 | -0.689 | -0.545 | -0.015 | -0.126 | -0.126 | 0 | 0.8 | 0.64 | 0.93 | 0.99 | 0.87 | 0.87 |
| 5 | 113078335 | rs6594752 | -0.153 | 2.532 | -1.208 | -0.526 | 0.01 | -0.007 | -0.007 | 0 | 0.11 | 0.14 | 0.07 | 0.07 | 0.16 | 0.15 |
| 5 | 113078457 | rs6594753 | 0.092 | 2.399 | -1.047 | -0.404 | -0.001 | -0.025 | -0.025 | 0 | 0.88 | 0.89 | 0.95 | 1 | 0.92 | 0.96 |
| 5 | 113079354 | rs11740330 | -0.308 | 2.417 | -1.081 | -0.425 | -0.033 | 0.065 | 0.065 | 0 | 0.28 | 0.16 | 0.45 | 0.13 | 0.32 | 0.29 |
| 5 | 113079424 | rs1384286 | 1.274 | 2.411 | -1.093 | -0.407 | -0.019 | 0.068 | 0.068 | 0 | 0.6 | 0.73 | 0.51 | 0.87 | 0.59 | 0.68 |
| 5 | 113083726 | rs1871916 | 1.257 | 2.317 | -0.902 | -0.305 | -0.015 | 0.008 | 0.008 | 0.162 | 0.6 | 0.68 | 0.49 | 0.63 | 0.58 | 0.66 |
| 5 | 113094772 | rs10059162 | 1.926 | 2.377 | -0.904 | -0.284 | 0.026 | 0.255 | 0.255 | 3.584 | 0.47 | 0.11 | 0.35 | 0.85 | 0.42 | 0.6 |
| 5 | 113096508 | rs1482371 | 2.006 | 2.379 | -0.894 | -0.243 | 0.049 | -0.009 | -0.009 | 0.089 | 0.46 | 0.1 | 0.34 | 0.61 | 0.39 | 0.58 |
| 5 | 113102185 | rs7720273 | -0.245 | 1.911 | -0.83 | -0.197 | 0.073 | -0.049 | -0.049 | 0 | 0.13 | 0.54 | 0.14 | 0.02 | 0.17 | 0.08 |
| 5 | 113106735 | rs7732090 | 0.002 | 1.904 | -1.481 | -0.106 | -0.004 | 0.053 | 0.053 | 0 | 0.88 | 0.78 | 0.93 | 0.76 | 0.89 | 0.94 |
| 5 | 113115477 | rs1905135 | -0.35 | 2.039 | -1.765 | -0.169 | -0.048 | 0.103 | 0.103 | 0 | 0.27 | 0.08 | 0.44 | 0.13 | 0.3 | 0.28 |
| 5 | 113118026 | rs2170528 | 1.318 | 1.671 | -1.724 | -0.072 | -0.094 | -0.038 | -0.038 | 1.027 | 0.48 | 0.02 | 0.36 | 0.61 | 0.45 | 0.63 |
| 5 | 113118069 | rs2128313 | 0.762 | 1.44 | -0.962 | 0.257 | -0.002 | -0.003 | -0.003 | 1.115 | 0.65 | 0.69 | 0.51 | 0.63 | 0.63 | 0.7 |
| 5 | 113128872 | rs1871915 | 0.704 | 1.226 | -1.259 | 0.22 | -0.081 | 0.018 | 0.018 | 0.019 | 0.26 | 0.07 | 0.23 | 0.19 | 0.35 | 0.24 |
| 5 | 113129950 | rs900303 | 0.86 | 0.345 | -0.839 | -0.334 | -0.018 | -0.059 | -0.059 | 0.009 | 0.76 | 0.41 | 0.83 | 0.95 | 0.82 | 0.8 |
| 5 | 113133844 | rs17336742 | 0.653 | 0.301 | -0.949 | -0.353 | 0.009 | 0.018 | 0.018 | 0 | 0.08 | 0.02 | 0.05 | 0.03 | 0.07 | 0.12 |
| 5 | 113134734 | rs17135999 | 0.267 | 0.271 | -0.321 | -0.386 | 0.005 | -0.038 | -0.038 | 0 | 0.1 | 0.17 | 0.09 | 0.02 | 0.1 | 0.07 |
| 5 | 113135019 | rs6885793 | 1.31 | 0.139 | -0.341 | -0.259 | -0.009 | -0.03 | -0.03 | 0 | 0.35 | 0.42 | 0.36 | 0.18 | 0.3 | 0.16 |
| 5 | 113137884 | rs10477505 | 1.293 | 0.074 | -0.451 | 0.209 | -0.033 | -0.092 | -0.092 | 0 | 0.13 | 0.02 | 0.13 | 0.36 | 0.11 | 0.25 |
| 5 | 113141831 | rs6881055 | 0.786 | 0.086 | -0.289 | 0.199 | -0.002 | -0.015 | -0.015 | 0 | 0.43 | 0.55 | 0.41 | 0.18 | 0.39 | 0.21 |
| 5 | 113143478 | rs17337236 | 0.858 | 0.223 | -0.695 | 0.391 | -0.098 | -0.087 | -0.087 | 2.696 | 0.3 | 0.08 | 0.31 | 0.41 | 0.42 | 0.4 |
| 5 | 113145310 | rs17136012 | 0.665 | 0.316 | -0.631 | 0.45 | 0.009 | 0.013 | 0.013 | 0 | 0.08 | 0.04 | 0.05 | 0.03 | 0.07 | 0.1 |
| 5 | 113149124 | rs1384282 | -0.14 | 0.573 | -0.266 | 0.647 | -0.004 | -0.001 | -0.001 | 0.165 | 0.55 | 0.43 | 0.56 | 0.53 | 0.5 | 0.47 |
| 5 | 113150200 | rs17136028 | -0.41 | 0.736 | -0.249 | 0.686 | 0 | 0 | 0 | 0.122 | 0.58 | 0.46 | 0.56 | 0.54 | 0.51 | 0.49 |
| 5 | 113150598 | rs12719168 | -0.024 | 0.785 | -0.508 | 0.355 | -0.015 | -0.052 | -0.052 | 0 | 0.64 | 0.54 | 0.64 | 0.82 | 0.7 | 0.83 |
| 5 | 113160490 | rs1482358 | -0.194 | 0.722 | -0.689 | 0.666 | -0.003 | -0.001 | -0.001 | 0.394 | 0.19 | 0.2 | 0.33 | 0.39 | 0.24 | 0.3 |
| 5 | 113160543 | rs17338434 | 0.107 | 0.14 | -0.812 | 0.144 | 0.009 | -0.002 | -0.002 | 0 | 0.29 | 0.23 | 0.18 | 0.29 | 0.26 | 0.25 |
| 5 | 113161020 | rs13182734 | 0.07 | 0.102 | -0.692 | 0.126 | 0.005 | -0.001 | -0.001 | 0 | 0.29 | 0.26 | 0.18 | 0.29 | 0.26 | 0.25 |
| 5 | 113161178 | rs17136043 | 0.049 | -0.285 | -0.967 | -0.525 | 0.001 | -0.015 | -0.015 | 1.734 | 0.44 | 0.31 | 0.45 | 0.43 | 0.43 | 0.48 |
| 5 | 113163228 | rs7725392 | -0.719 | -0.243 | -0.877 | -0.538 | 0.028 | -0.02 | -0.02 | 0.087 | 0.71 | 0.53 | 0.62 | 0.71 | 0.69 | 0.72 |
| 5 | 113164028 | rs4537053 | -0.961 | -0.269 | -0.716 | -0.55 | 0.036 | -0.012 | -0.012 | 0 | 0.73 | 0.6 | 0.63 | 0.71 | 0.69 | 0.73 |
| 5 | 113170445 | rs1482349 | 0.087 | -0.253 | -0.75 | -0.544 | 0.001 | 0.043 | 0.043 | 0 | 0.46 | 0.38 | 0.49 | 0.67 | 0.5 | 0.53 |
| 5 | 113171928 | rs13189273 | 1.034 | 0.244 | -0.373 | -0.586 | 0.104 | 0.046 | 0.046 | 0 | 0.05 | 0.22 | 0.15 | 0.09 | 0.06 | 0.07 |
| 5 | 113179090 | rs7734201 | 0.939 | 0.147 | -0.609 | -0.587 | 0.02 | 0.057 | 0.057 | 0 | 0.25 | 0.13 | 0.12 | 0.04 | 0.19 | 0.19 |
| 5 | 113181289 | rs4299731 | 0.499 | 0.463 | -0.591 | -0.415 | -0.04 | -0.055 | -0.055 | 0.098 | 0.26 | 0.16 | 0.4 | 0.39 | 0.36 | 0.32 |
| 5 | 113185123 | rs7711822 | 0.907 | 0.712 | -0.352 | -0.16 | 0 | 0.007 | 0.007 | 0.623 | 0.11 | 0.08 | 0.05 | 0.25 | 0.08 | 0.16 |
| 5 | 113191311 | rs12516340 | 0.763 | 0.726 | -0.32 | -0.522 | -0.053 | -0.01 | -0.01 | 0 | 0.22 | 0.05 | 0.22 | 0.21 | 0.26 | 0.23 |
| 5 | 113207613 | rs13186060 | 1.095 | 0.864 | 0.055 | -0.481 | 0.036 | 0.095 | 0.095 | 0 | 0.37 | 0.2 | 0.17 | 0.04 | 0.26 | 0.21 |
| 5 | 113210609 | rs7713629 | -0.681 | 0.968 | 0.408 | -0.462 | -0.001 | -0.046 | -0.046 | 0 | 0.57 | 0.48 | 0.68 | 0.91 | 0.69 | 0.71 |
| 5 | 113211212 | rs1482483 | 0.945 | 0.973 | 0.297 | -0.39 | -0.002 | 0.007 | 0.007 | 0 | 0.16 | 0.22 | 0.2 | 0.26 | 0.19 | 0.21 |
| 5 | 113213560 | rs2416342 | -0.809 | 1.014 | 0.296 | -0.441 | -0.002 | -0.046 | -0.046 | 0.138 | 0.57 | 0.48 | 0.68 | 0.91 | 0.7 | 0.71 |
| 5 | 113225701 | rs1482377 | 0.698 | 0.759 | -0.897 | 0.073 | 0.001 | -0.034 | -0.034 | 0.252 | 0.2 | 0.14 | 0.15 | 0.32 | 0.22 | 0.23 |
| 5 | 113227379 | rs1384338 | -1.022 | 0.327 | -1.096 | -0.018 | 0.005 | -0.037 | -0.037 | 7.347 | 0.6 | 0.43 | 0.8 | 0.97 | 0.73 | 0.78 |
| 5 | 113230902 | rs1482484 | -0.161 | 0.485 | -0.544 | 0.893 | 0.003 | 0.011 | 0.011 | 0 | 0.24 | 0.29 | 0.31 | 0.38 | 0.28 | 0.29 |
| 5 | 113237638 | rs11950195 | -0.201 | 0.558 | -1.115 | 0.538 | -0.004 | 0.007 | 0.007 | 0.331 | 0.42 | 0.67 | 0.5 | 0.66 | 0.46 | 0.5 |
| 5 | 113246018 | rs7379658 | -0.35 | 0.03 | -1.402 | 1.056 | 0.028 | -0.063 | -0.063 | 0 | 0.17 | 0.08 | 0.22 | 0.41 | 0.12 | 0.28 |
| 5 | 113249232 | rs6893845 | -0.026 | -0.393 | -0.761 | 0.273 | -0.017 | -0.098 | -0.098 | 0 | 0.84 | 0.72 | 0.93 | 1 | 0.89 | 0.94 |
| 5 | 113249296 | rs6893860 | 0.512 | 0.285 | -1.166 | 0.09 | 0.004 | -0.034 | -0.034 | 0 | 0.69 | 0.65 | 0.75 | 0.85 | 0.71 | 0.76 |
| 5 | 113250798 | rs1384334 | 0.147 | 0.791 | -1.381 | 0.258 | -0.043 | 0.009 | 0.009 | 0 | 0.18 | 0.09 | 0.2 | 0.14 | 0.23 | 0.19 |
| 5 | 113254757 | rs1384337 | -0.772 | 0.619 | -1.314 | 0.23 | -0.03 | -0.082 | -0.082 | 0 | 0.87 | 0.73 | 0.95 | 1 | 0.94 | 0.95 |
| 5 | 113258058 | rs6594771 | -0.196 | 0.429 | -1.614 | -0.372 | 0.023 | 0.008 | 0.008 | 0 | 0.64 | 0.32 | 0.69 | 0.69 | 0.69 | 0.69 |
| 5 | 113258218 | rs6874345 | 0.54 | 0.484 | -1.674 | -0.374 | -0.067 | -0.023 | -0.023 | 0.28 | 0.21 | 0.1 | 0.29 | 0.25 | 0.28 | 0.26 |
| 5 | 113275375 | rs4705607 | 1.319 | 0.364 | -1.883 | -0.505 | 0.009 | -0.12 | -0.12 | 5.099 | 0.18 | 0.02 | 0.22 | 0.4 | 0.12 | 0.25 |
| 5 | 113279043 | rs7732200 | -0.175 | 0.408 | -1.739 | -0.555 | -0.001 | -0.001 | -0.001 | 0.175 | 0.46 | 0.57 | 0.56 | 0.56 | 0.53 | 0.55 |
| 5 | 113279595 | rs6894635 | -0.851 | 0.493 | -1.71 | -0.551 | -0.008 | -0.089 | -0.089 | 0 | 0.63 | 0.6 | 0.78 | 0.95 | 0.66 | 0.8 |
| 5 | 113280827 | rs2416344 | -1.096 | 0.587 | -1.659 | -0.447 | -0.037 | -0.094 | -0.094 | 0 | 0.85 | 0.7 | 0.95 | 0.99 | 0.93 | 0.91 |

| | | | | | | | | | | | | | | | | |
|---|---|---|---|---|---|---|---|---|---|---|---|---|---|---|---|---|
| 5 | 113282554 | rs10053880 | 0.916 | 0.666 | -1.842 | -0.315 | -0.008 | -0.054 | -0.054 | 0 | 0.2 | 0.12 | 0.24 | 0.34 | 0.24 | 0.27 |
| 5 | 113288654 | rs17136126 | 0.129 | 1.074 | -1.441 | -0.272 | 0 | 0 | 0 | 0.016 | 0.22 | 0.24 | 0.3 | 0.26 | 0.27 | 0.25 |
| 5 | 113288869 | rs1487215 | 0.129 | 1.074 | -1.441 | -0.272 | 0 | 0 | 0 | 0 | 0.22 | 0.24 | 0.3 | 0.26 | 0.27 | 0.25 |
| 5 | 113290107 | rs543136 | 0.88 | 0.957 | -1.635 | -0.234 | -0.033 | 0.074 | 0.074 | 0 | 0.22 | 0.07 | 0.17 | 0.04 | 0.28 | 0.18 |
| 5 | 113296006 | rs2931490 | 0.314 | 1.283 | -2.05 | -0.095 | -0.116 | -0.121 | -0.121 | 1.098 | 0.33 | 0.04 | 0.31 | 0.36 | 0.4 | 0.31 |
| 5 | 113296159 | rs17136136 | 0.202 | 0.879 | -1.32 | -0.275 | -0.018 | 0.002 | 0.002 | 0.034 | 0.33 | 0.41 | 0.34 | 0.37 | 0.28 | 0.35 |
| 5 | 113298993 | rs10519366 | 0.298 | 1.149 | -1.086 | -0.224 | -0.024 | 0.002 | 0.002 | 0 | 0.16 | 0.12 | 0.23 | 0.17 | 0.22 | 0.22 |
| 5 | 113306851 | rs1482487 | 0.309 | 0.976 | -0.802 | 0.021 | -0.005 | -0.018 | -0.018 | 0 | 0.17 | 0.09 | 0.19 | 0.31 | 0.19 | 0.17 |
| 5 | 113307417 | rs1482486 | 0.072 | 0.404 | 0.147 | -0.437 | 0.001 | -0.039 | -0.039 | 0 | 0.1 | 0.48 | 0.08 | 0 | 0.02 | 0.02 |
| 5 | 113321924 | rs12658199 | 0.732 | 0.579 | -0.903 | -0.369 | 0.005 | 0.027 | 0.027 | 0.478 | 0.23 | 0.1 | 0.17 | 0.17 | 0.23 | 0.35 |
| 5 | 113327484 | rs2974558 | 0.822 | 0.768 | -0.812 | -0.693 | 0.002 | -0.056 | -0.056 | 0.667 | 0.43 | 0.61 | 0.26 | 0.03 | 0.31 | 0.24 |
| 5 | 113327561 | rs4342322 | 0.822 | 0.768 | -0.815 | -0.693 | 0.003 | -0.052 | -0.052 | 0 | 0.43 | 0.61 | 0.26 | 0.03 | 0.32 | 0.24 |
| 5 | 113330738 | rs1487220 | 0.835 | 0.838 | -0.777 | -0.694 | 0.001 | -0.06 | -0.06 | 0 | 0.42 | 0.61 | 0.27 | 0.03 | 0.32 | 0.24 |
| 5 | 113340036 | rs12187494 | 0.863 | 0.806 | -0.389 | -0.702 | -0.014 | -0.118 | -0.118 | 0.161 | 0.42 | 0.51 | 0.22 | 0.01 | 0.32 | 0.22 |
| 5 | 113342732 | rs4422524 | 0.816 | 0.919 | -0.181 | -0.669 | -0.009 | -0.023 | -0.023 | 0.056 | 0.81 | 0.29 | 0.83 | 0.89 | 0.88 | 0.88 |
| 5 | 113347652 | rs4518384 | 0.008 | 1.157 | 0.187 | -0.071 | 0.028 | 0.075 | 0.075 | 0 | 0.75 | 0.58 | 0.72 | 0.58 | 0.73 | 0.73 |
| 5 | 113355447 | rs7735190 | 1.415 | 1.108 | -0.304 | 0.006 | -0.01 | -0.025 | -0.025 | 0.075 | 0.07 | 0.11 | 0.03 | 0 | 0.02 | 0.02 |
| 5 | 113356277 | rs10043271 | 0.862 | 1.369 | -0.643 | 0.361 | -0.04 | 0 | 0 | 0 | 0.07 | 0.01 | 0.32 | 0.36 | 0.15 | 0.12 |
| 5 | 113367177 | rs13355051 | 0.304 | 1.266 | -0.678 | 0.48 | -0.006 | -0.04 | -0.04 | 0.014 | 0.27 | 0.18 | 0.44 | 0.48 | 0.33 | 0.45 |
| 5 | 113368278 | rs11241257 | 0.135 | 1.378 | -0.565 | 0.522 | -0.003 | -0.001 | -0.001 | 0.072 | 0.88 | 0.88 | 0.86 | 0.84 | 0.83 | 0.86 |
| 5 | 113369605 | rs11241258 | -0.597 | 1.211 | -0.473 | 0.462 | 0.011 | 0.002 | 0.002 | 0 | 0.93 | 0.89 | 0.93 | 0.91 | 0.91 | 0.91 |
| 5 | 113385553 | rs2721314 | 0.488 | 1.072 | -0.522 | 0.66 | -0.013 | -0.014 | -0.014 | 0 | 0.67 | 0.71 | 0.68 | 0.61 | 0.64 | 0.71 |
| 5 | 113389336 | rs17136225 | 0.024 | 0.478 | -0.815 | 0.771 | 0.008 | -0.001 | -0.001 | 0.059 | 0.32 | 0.25 | 0.21 | 0.37 | 0.28 | 0.47 |
| 5 | 113389560 | rs1487227 | 1.947 | 0.725 | -1.099 | 0.49 | -0.041 | 0.216 | 0.216 | 0.692 | 0.34 | 0.04 | 0.19 | 0.03 | 0.31 | 0.22 |
| 5 | 113390625 | rs873512 | 0.587 | 1.482 | -0.399 | 1.056 | -0.002 | -0.002 | -0.002 | 0 | 0.17 | 0.22 | 0.14 | 0.16 | 0.16 | 0.16 |
| 5 | 113393781 | rs2632151 | -1.222 | 1.347 | -0.567 | 1.217 | -0.027 | -0.015 | -0.015 | 5.613 | 0.27 | 0.21 | 0.54 | 0.6 | 0.41 | 0.33 |
| 5 | 113394815 | rs12523313 | 0.083 | 0.993 | -0.618 | 1.44 | -0.045 | -0.011 | -0.011 | 0.495 | 0.07 | 0.01 | 0.36 | 0.4 | 0.17 | 0.16 |
| 5 | 113396378 | rs12520827 | 0.529 | 0.104 | -0.433 | 0.828 | -0.002 | 0.005 | 0.005 | 0 | 0.15 | 0.15 | 0.26 | 0.28 | 0.11 | 0.14 |
| 5 | 113402762 | rs17348806 | 1.135 | -0.039 | -0.802 | -0.257 | 0.014 | 0.042 | 0.042 | 0 | 0.2 | 0.12 | 0.1 | 0.01 | 0.17 | 0.17 |
| 5 | 113403389 | rs901583 | -0.443 | -0.682 | -0.555 | -0.948 | -0.018 | -0.063 | -0.063 | 0 | 0.15 | 0.17 | 0.13 | 0 | 0.15 | 0.13 |
| 5 | 113404933 | rs2974491 | -0.641 | -0.538 | -0.702 | -0.94 | -0.011 | -0.034 | -0.034 | 0 | 0.75 | 0.83 | 0.65 | 0.64 | 0.74 | 0.83 |
| 5 | 113407106 | rs11952942 | 2.274 | -0.62 | -0.7 | -0.996 | 0.033 | 0.068 | 0.068 | 0 | 0.18 | 0.05 | 0.05 | 0.01 | 0.12 | 0.16 |
| 5 | 113409641 | rs35703899 | 1.802 | -0.665 | -0.626 | -1.019 | 0.013 | 0.032 | 0.032 | 0 | 0.21 | 0.14 | 0.11 | 0.01 | 0.17 | 0.14 |
| 5 | 113414351 | rs2632138 | 0.099 | -0.636 | -0.624 | -0.987 | -0.017 | -0.059 | -0.059 | 0 | 0.15 | 0.17 | 0.13 | 0 | 0.15 | 0.13 |
| 5 | 113418512 | rs2931491 | -1.279 | -0.769 | -0.954 | -0.965 | 0.011 | -0.014 | -0.014 | 0 | 0.82 | 0.83 | 0.75 | 0.66 | 0.9 | 0.83 |
| 5 | 113429984 | rs1158117 | 0.785 | -0.714 | -0.428 | -1.04 | 0.046 | 0.036 | 0.036 | 0.153 | 0.44 | 0.3 | 0.51 | 0.66 | 0.59 | 0.56 |
| 5 | 113432832 | rs4705623 | 1.435 | -0.539 | -0.271 | -0.949 | -0.05 | -0.016 | -0.016 | 0 | 0.37 | 0.23 | 0.39 | 0.61 | 0.42 | 0.52 |
| 5 | 113433547 | rs2220318 | -0.528 | -1.045 | 0.178 | -0.984 | -0.063 | -0.016 | -0.016 | 0 | 0.32 | 0.45 | 0.37 | 0.3 | 0.24 | 0.27 |
| 5 | 113434432 | rs2721304 | 1.106 | -0.3 | 0.172 | -1.153 | -0.001 | -0.036 | -0.036 | 0 | 0.23 | 0.29 | 0.13 | 0.02 | 0.21 | 0.14 |
| 5 | 113434724 | rs2632112 | 1.25 | -0.536 | 0.213 | -1.143 | -0.001 | -0.027 | -0.027 | 0.469 | 0.22 | 0.25 | 0.11 | 0.02 | 0.18 | 0.13 |
| 5 | 113435957 | rs13190004 | 1.279 | -0.564 | 0.346 | -1.165 | -0.005 | -0.038 | -0.038 | 0 | 0.21 | 0.26 | 0.11 | 0.02 | 0.18 | 0.12 |
| 5 | 113436206 | rs1487216 | 0.194 | -0.389 | 0.34 | -1.295 | -0.002 | -0.065 | -0.065 | 0.434 | 0.29 | 0.39 | 0.24 | 0.02 | 0.32 | 0.19 |
| 5 | 113440892 | rs1487222 | -0.892 | -0.693 | 0.514 | -1.087 | 0.091 | 0.093 | 0.093 | 0 | 0.5 | 0.77 | 0.5 | 0.31 | 0.43 | 0.39 |
| 5 | 113451025 | rs6860602 | 1.182 | -0.637 | 0.61 | -1.063 | -0.026 | -0.12 | -0.12 | 0.046 | 0.23 | 0.45 | 0.13 | 0.02 | 0.18 | 0.12 |
| 5 | 113453870 | rs12522520 | 1.531 | -0.767 | 0.124 | -1.01 | 0.019 | -0.081 | -0.081 | 0 | 0.14 | 0.01 | 0.24 | 0.26 | 0.1 | 0.18 |
| 5 | 113459893 | rs2954363 | -0.667 | -0.599 | 0.383 | -1.07 | 0.094 | 0.093 | 0.093 | 0.009 | 0.5 | 0.79 | 0.51 | 0.31 | 0.43 | 0.39 |
| 5 | 113463029 | rs2954359 | 1.6 | -0.534 | 0.496 | -1.17 | -0.021 | -0.052 | -0.052 | 0 | 0.2 | 0.26 | 0.07 | 0.02 | 0.13 | 0.12 |
| 5 | 113466023 | rs2954358 | -0.855 | 0.075 | -0.093 | -1.105 | -0.023 | 0.014 | 0.014 | 0 | 0.57 | 0.73 | 0.58 | 0.36 | 0.55 | 0.44 |
| 5 | 113466377 | rs7726250 | 0.366 | 0.21 | 0.087 | -0.753 | 0.007 | -0.028 | -0.028 | 0 | 0.08 | 0.08 | 0.13 | 0 | 0.14 | 0.07 |
| 5 | 113470953 | rs10463656 | 0.718 | 0.025 | -0.07 | -0.845 | -0.019 | -0.004 | -0.004 | 0 | 0.32 | 0.41 | 0.32 | 0.32 | 0.23 | 0.32 |
| 5 | 113476277 | rs2974470 | 0.068 | 0.732 | -0.263 | -1.186 | -0.002 | 0.003 | 0.003 | 0.274 | 0.25 | 0.22 | 0.2 | 0.08 | 0.27 | 0.11 |
| 5 | 113486177 | rs10044576 | 0.584 | 0.799 | -0.306 | -1.246 | -0.013 | 0.012 | 0.012 | 0 | 0.05 | 0.03 | 0.11 | 0 | 0.11 | 0.07 |
| 5 | 113486329 | rs13180608 | 1.524 | 0.83 | -0.784 | -1.311 | -0.017 | 0.004 | 0.004 | 3.118 | 0.19 | 0.02 | 0.13 | 0.11 | 0.16 | 0.13 |
| 5 | 113488823 | rs17350454 | 1.554 | 0.923 | -0.726 | -1.435 | 0.005 | 0.009 | 0.009 | 1.481 | 0.14 | 0.09 | 0.1 | 0.05 | 0.11 | 0.06 |
| 5 | 113489245 | rs4429843 | 0.739 | 1.092 | -0.283 | -1.724 | 0 | -0.003 | -0.003 | 0 | 0.31 | 0.33 | 0.43 | 0.73 | 0.33 | 0.5 |
| 5 | 113489353 | rs13158827 | -0.005 | 1.09 | -0.245 | -1.736 | 0.018 | 0.165 | 0.165 | 0.529 | 0.52 | 0.21 | 0.44 | 0.22 | 0.55 | 0.44 |
| 5 | 113503422 | rs11747037 | 0.5 | 1.211 | -0.209 | -1.644 | -0.056 | 0.107 | 0.107 | 0.296 | 0.32 | 0.16 | 0.28 | 0.1 | 0.39 | 0.27 |
| 5 | 113505686 | rs7719809 | 1.036 | 1.555 | 0.321 | -0.905 | 0.005 | 0.01 | 0.01 | 0 | 0.23 | 0.07 | 0.31 | 0.21 | 0.22 | 0.3 |
| 5 | 113506795 | rs6594788 | 0.026 | 1.566 | 0.67 | -0.915 | -0.003 | -0.01 | -0.01 | 0.213 | 0.33 | 0.48 | 0.45 | 0.73 | 0.33 | 0.5 |
| 5 | 113524524 | rs4705634 | 0.256 | 1.516 | 0.579 | -1.03 | 0.009 | 0.019 | 0.019 | 0 | 0.57 | 0.45 | 0.55 | 0.21 | 0.65 | 0.5 |
| 5 | 113525477 | rs6594793 | -1.486 | 1.507 | 0.453 | -1.026 | -0.01 | -0.021 | -0.021 | 0 | 0.85 | 0.75 | 0.89 | 0.95 | 0.88 | 0.94 |
| 5 | 113526724 | rs7738011 | 0.478 | 1.48 | 0.651 | -0.889 | 0.004 | -0.003 | -0.003 | 0 | 0.27 | 0.22 | 0.33 | 0.74 | 0.24 | 0.44 |
| 5 | 113531516 | rs7726566 | 1.21 | 1.344 | 0.517 | -0.854 | 0.017 | 0.005 | 0.005 | 0 | 0.21 | 0.16 | 0.22 | 0.21 | 0.13 | 0.25 |
| 5 | 113550042 | rs6893764 | -0.218 | 1.685 | -0.181 | -1.104 | -0.023 | 0.043 | 0.043 | 0 | 0.72 | 0.84 | 0.77 | 0.88 | 0.7 | 0.8 |
| 5 | 113552328 | rs4512129 | 0.045 | 1.271 | 0.157 | -0.583 | 0.001 | -0.004 | -0.004 | 0 | 0.3 | 0.34 | 0.44 | 0.74 | 0.34 | 0.5 |
| 5 | 113561408 | rs9637896 | 1.758 | 1.073 | -0.381 | -0.522 | 0.032 | -0.025 | -0.025 | 0.277 | 0.17 | 0.03 | 0.2 | 0.22 | 0.11 | 0.23 |
| 5 | 113563230 | rs13186908 | 0.855 | 1.255 | -0.284 | -1.269 | -0.012 | 0.149 | 0.149 | 0 | 0.47 | 0.24 | 0.38 | 0.16 | 0.51 | 0.37 |
| 5 | 113570190 | rs6594804 | -0.243 | 1.345 | -0.131 | -0.996 | -0.001 | -0.005 | -0.005 | 0 | 0.79 | 0.73 | 0.83 | 0.9 | 0.85 | 0.87 |
| 5 | 113570293 | rs7708130 | -0.261 | 1.355 | -0.126 | -0.991 | -0.001 | -0.005 | -0.005 | 2.022 | 0.79 | 0.73 | 0.83 | 0.9 | 0.85 | 0.87 |

| | | | | | | | | | | | | | | | | |
|---|---|---|---|---|---|---|---|---|---|---|---|---|---|---|---|---|
| 5 | 113591317 | rs4610451 | 0.174 | 1.879 | -0.312 | -1.054 | 0.001 | -0.006 | -0.006 | 0 | 0.75 | 0.77 | 0.78 | 0.88 | 0.73 | 0.86 |
| 5 | 113591334 | rs4466147 | 0.818 | 1.422 | 0.105 | -1.241 | -0.02 | 0.001 | 0.001 | 0.152 | 0.25 | 0.33 | 0.3 | 0.75 | 0.2 | 0.43 |
| 5 | 113591476 | rs4293905 | 0.436 | 1.35 | 0.099 | -1.338 | -0.006 | -0.011 | -0.011 | 0 | 0.31 | 0.36 | 0.41 | 0.76 | 0.3 | 0.5 |
| 5 | 113596392 | rs13184901 | 0.648 | 1.492 | -0.29 | -1.299 | -0.035 | 0.178 | 0.178 | 0.491 | 0.45 | 0.19 | 0.37 | 0.15 | 0.49 | 0.37 |
| 5 | 113601371 | rs10065785 | -0.068 | 1.485 | -0.277 | -1.273 | -0.008 | -0.037 | -0.037 | 0.156 | 0.42 | 0.53 | 0.5 | 0.8 | 0.37 | 0.56 |
| 5 | 113602162 | rs6594807 | -0.58 | 1.428 | -0.043 | -0.674 | 0.014 | -0.02 | -0.02 | 1.119 | 0.88 | 0.78 | 0.87 | 0.95 | 0.86 | 0.92 |
| 5 | 113604682 | rs4705653 | -0.333 | 1.575 | -0.593 | -0.571 | -0.03 | -0.004 | -0.004 | 0 | 0.1 | 0.03 | 0.14 | 0.09 | 0.13 | 0.1 |
| 5 | 113604806 | rs4582264 | -0.258 | 1.191 | -0.471 | -0.666 | -0.002 | 0.002 | 0.002 | 0 | 0.69 | 0.78 | 0.78 | 0.68 | 0.65 | 0.69 |
| 5 | 113604818 | rs4412104 | -0.076 | 0.657 | 0.383 | -0.017 | 0.002 | 0.348 | 0.348 | 0 | 0.08 | 0.23 | 0.25 | 0.51 | 0.04 | 0.19 |
| 5 | 113605976 | rs7737235 | 0.562 | 0.532 | 0.575 | -1.285 | -0.011 | -0.015 | -0.015 | 0 | 0.88 | 0.6 | 0.94 | 0.94 | 0.95 | 0.93 |
| 5 | 113616993 | rs1457759 | 0.225 | 0.533 | 0.785 | -1.271 | -0.01 | -0.008 | -0.008 | 0.117 | 0.89 | 0.74 | 0.95 | 0.93 | 0.95 | 0.93 |
| 5 | 113622824 | rs13172051 | 0.449 | 0.9 | 0.413 | -2.281 | 0.008 | -0.047 | -0.047 | 0 | 0.38 | 0.42 | 0.37 | 0.1 | 0.42 | 0.34 |
| 5 | 113623738 | rs1037719 | 0.841 | 0.766 | 0.255 | -2.532 | -0.005 | -0.046 | -0.046 | 0.006 | 0.1 | 0.15 | 0.06 | 0.01 | 0.11 | 0.05 |
| 5 | 113633321 | rs6894645 | 0.141 | 0.551 | 0.449 | -2.603 | 0 | -0.014 | -0.014 | 0 | 0.1 | 0.3 | 0.05 | 0.04 | 0.05 | 0.06 |
| 5 | 113633599 | rs12522842 | 0.191 | 0.557 | 0.392 | -2.64 | 0 | 0.013 | 0.013 | 0 | 0.46 | 0.43 | 0.4 | 0.15 | 0.47 | 0.4 |
| 5 | 113640906 | rs7706180 | 1.627 | 0.156 | -0.03 | -2.815 | 0.03 | -0.002 | -0.002 | 0 | 0.22 | 0.12 | 0.3 | 0.81 | 0.13 | 0.39 |
| 5 | 113641145 | rs11958855 | 1.245 | 0.452 | -0.316 | -3.257 | 0.01 | -0.016 | -0.016 | 0 | 0.2 | 0.24 | 0.27 | 0.1 | 0.22 | 0.27 |
| 5 | 113643828 | rs2125143 | 0.941 | 0.403 | -0.422 | -3.347 | -0.039 | 0.119 | 0.119 | 0 | 0.25 | 0.08 | 0.14 | 0.01 | 0.31 | 0.12 |
| 5 | 113644030 | rs2416366 | 0.229 | 0.834 | -0.236 | -3.476 | 0.012 | -0.094 | -0.094 | 0 | 0.58 | 0.47 | 0.66 | 0.93 | 0.47 | 0.74 |
| 5 | 113645426 | rs1379560 | 0.657 | 0.765 | 0.042 | -3.208 | 0.072 | -0.033 | -0.033 | 0.102 | 0.06 | 0.21 | 0.15 | 0 | 0.17 | 0.08 |
| 5 | 113645899 | rs1124188 | 1.305 | 0.692 | -0.041 | -3.244 | 0.002 | 0.003 | 0.003 | 0 | 0.27 | 0.23 | 0.42 | 0.81 | 0.29 | 0.47 |
| 5 | 113650445 | rs656942 | -0.877 | 0.555 | -0.018 | -3.31 | -0.003 | -0.005 | -0.005 | 0 | 0.4 | 0.38 | 0.35 | 0.09 | 0.47 | 0.27 |
| 5 | 113651694 | rs672894 | 0.659 | 0.702 | -0.024 | -3.071 | -0.017 | -0.044 | -0.044 | 0 | 0.38 | 0.49 | 0.49 | 0.84 | 0.39 | 0.52 |
| 5 | 113653803 | rs3112767 | 0.418 | 0.703 | 0.034 | -3.052 | -0.008 | -0.075 | -0.075 | 0.314 | 0.18 | 0.3 | 0.12 | 0.02 | 0.14 | 0.07 |
| 5 | 113658141 | rs36963 | 0.259 | 0.689 | -0.145 | -2.854 | -0.002 | -0.026 | -0.026 | 0.582 | 0.46 | 0.45 | 0.53 | 0.87 | 0.42 | 0.57 |
| 5 | 113659909 | rs12520028 | 0.255 | 0.529 | -0.291 | -2.496 | -0.005 | 0.013 | 0.013 | 0 | 0.23 | 0.19 | 0.28 | 0.1 | 0.23 | 0.27 |
| 5 | 113673427 | rs36952 | -0.059 | 0.364 | -0.104 | -2.951 | -0.008 | -0.083 | -0.083 | 0 | 0.66 | 0.64 | 0.85 | 0.96 | 0.73 | 0.84 |
| 5 | 113674308 | rs1457765 | 0.035 | 0.52 | 0.081 | -3.082 | -0.001 | -0.006 | -0.006 | 0 | 0.29 | 0.27 | 0.33 | 0.11 | 0.33 | 0.32 |
| 5 | 113679153 | rs36935 | 0.754 | 0.32 | 0.268 | -3.223 | 0 | -0.001 | -0.001 | 0 | 0.46 | 0.44 | 0.54 | 0.85 | 0.46 | 0.58 |
| 5 | 113679204 | rs36936 | -0.866 | 0.301 | 0.231 | -3.202 | -0.012 | -0.07 | -0.07 | 1.514 | 0.76 | 0.71 | 0.88 | 0.96 | 0.79 | 0.9 |
| 5 | 113680739 | rs13181125 | 1.75 | 0.619 | 0.023 | -3.302 | -0.022 | 0.017 | 0.017 | 0 | 0.15 | 0.02 | 0.2 | 0.08 | 0.14 | 0.23 |
| 5 | 113682972 | rs257468 | 1.735 | 0.491 | 0.217 | -3.304 | -0.038 | 0.092 | 0.092 | 0 | 0.35 | 0.13 | 0.5 | 0.83 | 0.41 | 0.51 |
| 5 | 113688284 | rs12189350 | 0.136 | 0.5 | 0.266 | -3.312 | -0.015 | 0.024 | 0.024 | 0 | 0.28 | 0.21 | 0.33 | 0.11 | 0.32 | 0.32 |
| 5 | 113690176 | rs163303 | 1.501 | 0.53 | 0.282 | -3.255 | -0.013 | 0.055 | 0.055 | 0 | 0.36 | 0.29 | 0.51 | 0.85 | 0.41 | 0.52 |
| 5 | 113697587 | rs36945 | 0.716 | 0.553 | 0.282 | -3.271 | -0.007 | 0.084 | 0.084 | 0.972 | 0.45 | 0.32 | 0.53 | 0.85 | 0.46 | 0.57 |
| 5 | 113700382 | rs36946 | 1.481 | 0.526 | 0.253 | -3.265 | -0.005 | 0.02 | 0.02 | 0 | 0.37 | 0.28 | 0.5 | 0.83 | 0.4 | 0.51 |
| 5 | 113703273 | rs36949 | 0.539 | 0.548 | 0.216 | -3.216 | 0.001 | 0.023 | 0.023 | 0.088 | 0.48 | 0.58 | 0.54 | 0.84 | 0.46 | 0.58 |
| 5 | 113716532 | rs388084 | -1.771 | 0.533 | 0.446 | -2.739 | -0.013 | 0.231 | 0.231 | 0 | 0.49 | 0.84 | 0.43 | 0.19 | 0.45 | 0.42 |
| 5 | 113726539 | rs715730 | 0.38 | 0.158 | 0.948 | -2.419 | -0.008 | -0.058 | -0.058 | 3.352 | 0.1 | 0.19 | 0.03 | 0 | 0.06 | 0.06 |
| 5 | 113731423 | rs337701 | 0.624 | 0.107 | 1.006 | -2.435 | -0.016 | -0.036 | -0.036 | 0 | 0.1 | 0.43 | 0.06 | 0.04 | 0.06 | 0.03 |
| 5 | 113737062 | rs337705 | -1.621 | 0.401 | 0.345 | -2.456 | 0.014 | 0.169 | 0.169 | 0 | 0.43 | 0.83 | 0.38 | 0.17 | 0.39 | 0.38 |
| 5 | 113742869 | rs13183393 | 0.298 | 0.384 | 0.359 | -2.456 | -0.03 | 0.045 | 0.045 | 0 | 0.21 | 0.08 | 0.27 | 0.1 | 0.27 | 0.26 |
| 5 | 113743653 | rs4705665 | 0.726 | 0.445 | 0.384 | -2.452 | -0.002 | 0.008 | 0.008 | 0 | 0.18 | 0.16 | 0.23 | 0.1 | 0.17 | 0.24 |
| 5 | 113745738 | rs163210 | -1.441 | 0.417 | 0.041 | -2.433 | 0.015 | 0.104 | 0.104 | 0 | 0.42 | 0.7 | 0.37 | 0.15 | 0.39 | 0.37 |
| 5 | 113755797 | rs337715 | -0.681 | 0.342 | 0.118 | -2.351 | -0.046 | -0.101 | -0.101 | 0 | 0.2 | 0.56 | 0.09 | 0.04 | 0.12 | 0.09 |
| 5 | 113762810 | rs338625 | 1.115 | 0.69 | -0.512 | -2.297 | -0.015 | 0.138 | 0.138 | 0 | 0.58 | 0.19 | 0.59 | 0.81 | 0.58 | 0.61 |
| 5 | 113771811 | rs338660 | -0.181 | 0.528 | -0.383 | -2.22 | -0.002 | -0.037 | -0.037 | 1.383 | 0.79 | 0.81 | 0.89 | 0.95 | 0.76 | 0.86 |
| 5 | 113775870 | rs2416371 | 0.152 | 0.464 | -0.579 | -1.748 | -0.007 | 0.025 | 0.025 | 0 | 0.19 | 0.1 | 0.25 | 0.11 | 0.18 | 0.22 |
| 5 | 113779421 | rs10050725 | 0.618 | 0.239 | -1.121 | -1.939 | -0.06 | 0.078 | 0.078 | 0 | 0.36 | 0.15 | 0.57 | 0.77 | 0.43 | 0.53 |
| 5 | 113783989 | rs2169095 | 0.444 | -0.164 | -1.588 | -2.503 | -0.002 | 0.093 | 0.093 | 0 | 0.26 | 0.12 | 0.16 | 0 | 0.23 | 0.17 |
| 5 | 113785673 | rs13169372 | 0.621 | -0.337 | -1.559 | -2.331 | -0.026 | 0.02 | 0.02 | 0.012 | 0.22 | 0.06 | 0.17 | 0.17 | 0.25 | 0.16 |
| 5 | 113788755 | rs12153735 | 1.121 | -0.447 | -1.912 | -1.681 | -0.086 | 0.024 | 0.024 | 0 | 0.33 | 0.1 | 0.34 | 0.26 | 0.45 | 0.3 |
| 5 | 113794394 | rs1459782 | -0.684 | -0.468 | -0.954 | -1.301 | -0.015 | -0.02 | -0.02 | 0.526 | 0.8 | 0.69 | 0.87 | 0.91 | 0.83 | 0.89 |
| 5 | 113798016 | rs898354 | -0.291 | -0.573 | -0.601 | -1.333 | -0.037 | -0.05 | -0.05 | 2.257 | 0.77 | 0.47 | 0.85 | 0.91 | 0.83 | 0.88 |
| 5 | 113799403 | rs12516818 | 0.518 | -0.485 | -0.694 | -1.296 | 0.013 | 0.11 | 0.11 | 0 | 0.16 | 0.28 | 0.38 | 0.6 | 0.18 | 0.38 |
| 5 | 113804486 | rs17458304 | 1.618 | -0.873 | -1.128 | -1.488 | 0 | 0.068 | 0.068 | 0 | 0.16 | 0.06 | 0.07 | 0 | 0.1 | 0.09 |
| 5 | 113808031 | rs6870038 | -0.153 | -1 | -0.918 | -1.539 | -0.022 | -0.079 | -0.079 | 0.88 | 0.74 | 0.4 | 0.83 | 0.95 | 0.8 | 0.89 |
| 5 | 113810634 | rs1993657 | -0.408 | -0.847 | -0.898 | -1.256 | -0.002 | -0.009 | -0.009 | 0.192 | 0.32 | 0.3 | 0.46 | 0.62 | 0.3 | 0.49 |
| 5 | 113834663 | rs3891371 | 0.504 | -0.944 | -1.052 | -1.362 | -0.003 | -0.02 | -0.02 | 0 | 0.61 | 0.58 | 0.82 | 0.92 | 0.73 | 0.77 |
| 5 | 113834780 | rs4256337 | -0.083 | -1.016 | -1.045 | -1.367 | -0.045 | -0.129 | -0.129 | 3.638 | 0.76 | 0.58 | 0.9 | 1 | 0.89 | 0.89 |
| 5 | 113837198 | rs1469249 | 1.537 | -0.563 | -1.741 | -1.123 | 0.016 | 0.051 | 0.051 | 0 | 0.23 | 0.07 | 0.21 | 0.13 | 0.23 | 0.14 |
| 5 | 113846922 | rs404025 | 0.467 | -0.566 | -1.819 | -0.944 | 0.014 | 0.007 | 0.007 | 0 | 0.38 | 0.11 | 0.28 | 0.31 | 0.39 | 0.22 |
| 5 | 113847912 | rs1366088 | 1.5 | -0.248 | -1.723 | -0.95 | -0.004 | 0.009 | 0.009 | 0 | 0.14 | 0.09 | 0.08 | 0.07 | 0.16 | 0.08 |
| 5 | 113849294 | rs400028 | -1.63 | -0.176 | -1.591 | -0.919 | 0.012 | -0.008 | -0.008 | 2.893 | 0.75 | 0.74 | 0.85 | 0.82 | 0.72 | 0.73 |
| 5 | 113854743 | rs17459453 | -0.357 | -0.253 | -2.067 | -0.837 | 0.016 | 0.103 | 0.103 | 0 | 0.31 | 0.15 | 0.16 | 0.08 | 0.28 | 0.21 |
| 5 | 113874136 | rs17362782 | 1.317 | -0.091 | -1.988 | -0.603 | -0.005 | -0.005 | -0.005 | 0 | 0.14 | 0.03 | 0.06 | 0.08 | 0.14 | 0.08 |
| 5 | 113898201 | rs1427867 | 1.934 | 0.168 | -2.143 | -0.163 | -0.01 | 0.037 | 0.037 | 0 | 0.08 | 0.07 | 0.36 | 0.39 | 0.14 | 0.29 |
| 5 | 113910679 | rs3096548 | -1.835 | -0.025 | -2.727 | -0.513 | -0.094 | -0.028 | -0.028 | 0 | 0.76 | 0.95 | 0.86 | 0.77 | 0.73 | 0.72 |
| 5 | 113914493 | rs403133 | 0.805 | -0.34 | -2.967 | -0.839 | -0.029 | 0.021 | 0.021 | 0.452 | 0.34 | 0.04 | 0.26 | 0.23 | 0.35 | 0.24 |

| | | | | | | | | | | | | | | | | |
|---|---|---|---|---|---|---|---|---|---|---|---|---|---|---|---|---|
| 5 | 113918545 | rs17461363 | 1.255 | -0.454 | -2.528 | -0.849 | 0.006 | 0.012 | 0.012 | 0.195 | 0.18 | 0.1 | 0.1 | 0.08 | 0.18 | 0.09 |
| 5 | 113919964 | rs13189108 | 1.018 | -0.459 | -2.392 | -0.883 | 0.002 | 0.003 | 0.003 | 0.022 | 0.19 | 0.18 | 0.11 | 0.08 | 0.18 | 0.1 |
| 5 | 113928284 | rs674588 | 3.098 | -0.431 | -2.406 | -0.785 | -0.023 | -0.042 | -0.042 | 0.273 | 0.08 | 0.01 | 0.09 | 0.14 | 0.15 | 0.05 |
| 5 | 113929984 | rs11745459 | 2.969 | -0.423 | -2.078 | -0.824 | -0.002 | 0.001 | 0.001 | 0 | 0.07 | 0.04 | 0.07 | 0 | 0.07 | 0.02 |
| 5 | 113930285 | rs402641 | 0.521 | -0.46 | -1.127 | -0.633 | -0.019 | -0.033 | -0.033 | 0 | 0.38 | 0.52 | 0.3 | 0.23 | 0.34 | 0.25 |
| 5 | 113940187 | rs10071490 | 1.757 | -0.007 | -1.509 | -0.246 | -0.013 | 0.025 | 0.025 | 0 | 0.08 | 0.06 | 0.35 | 0.39 | 0.14 | 0.3 |
| 5 | 113957120 | rs618550 | 0.855 | -0.096 | -1.773 | -0.479 | -0.002 | 0.035 | 0.035 | 0 | 0.35 | 0.13 | 0.27 | 0.24 | 0.35 | 0.24 |
| 5 | 113961148 | rs10478173 | 1.413 | 0.083 | -1.463 | -0.269 | 0.016 | 0.123 | 0.123 | 0 | 0.09 | 0.13 | 0.36 | 0.41 | 0.14 | 0.27 |
| 5 | 113980360 | rs2553476 | -0.996 | -0.102 | -1.671 | -0.714 | -0.02 | 0.027 | 0.027 | 0 | 0.66 | 0.75 | 0.77 | 0.71 | 0.66 | 0.63 |
| 5 | 113984242 | rs283777 | 2.027 | -0.156 | -2.109 | -0.646 | -0.041 | -0.071 | -0.071 | 0 | 0.22 | 0.01 | 0.2 | 0.23 | 0.25 | 0.14 |
| 5 | 113986223 | rs1364931 | 0.791 | -0.127 | -1.921 | -0.606 | 0.006 | 0.046 | 0.046 | 0 | 0.36 | 0.09 | 0.27 | 0.24 | 0.36 | 0.25 |
| 5 | 113993771 | rs285890 | 0.566 | 0.675 | -0.566 | -0.193 | 0.003 | 0.012 | 0.012 | 0 | 0.6 | 0.57 | 0.45 | 0.5 | 0.56 | 0.51 |
| 5 | 113997778 | rs287652 | -0.051 | 1.306 | -0.405 | 0.122 | -0.025 | 0.009 | 0.009 | 0 | 0.89 | 0.98 | 0.94 | 0.9 | 0.84 | 0.88 |
| 5 | 114001677 | rs6876724 | -0.701 | 1.106 | 0.075 | -0.242 | -0.009 | -0.077 | -0.077 | 0.695 | 0.21 | 0.68 | 0.18 | 0.06 | 0.15 | 0.12 |
| 5 | 114004560 | rs1520248 | 0.682 | 1.078 | 0.692 | -0.243 | 0.035 | -0.055 | -0.055 | 0 | 0.11 | 0.39 | 0.12 | 0.01 | 0.09 | 0.04 |
| 5 | 114015223 | rs11241287 | 2.264 | 1.007 | 0.162 | -0.289 | 0.045 | 0.01 | 0.01 | 0.321 | 0.42 | 0.02 | 0.25 | 0.3 | 0.34 | 0.3 |
| 5 | 114015591 | rs10491498 | 2.198 | 1.084 | 0.288 | 0.126 | -0.026 | -0.031 | -0.031 | 1.595 | 0.06 | 0 | 0.1 | 0.1 | 0.05 | 0.06 |
| 5 | 114018075 | rs7732393 | 0.078 | 1.197 | 0.904 | 0.289 | 0.029 | -0.07 | -0.07 | 0 | 0.12 | 0.47 | 0.13 | 0.01 | 0.09 | 0.05 |
| 5 | 114023787 | rs10491499 | 0.963 | 1.184 | 1.159 | 0.293 | -0.003 | -0.053 | -0.053 | 0 | 0.09 | 0.24 | 0.07 | 0 | 0.07 | 0.02 |
| 5 | 114032769 | rs7721917 | 0.838 | 1.202 | 1.144 | 0.27 | 0.005 | -0.039 | -0.039 | 0 | 0.07 | 0.2 | 0.04 | 0 | 0.05 | 0.01 |
| 5 | 114036919 | rs958240 | -0.702 | 1.205 | 1.121 | 0.25 | 0.003 | -0.135 | -0.135 | 0.118 | 0.18 | 0.51 | 0.15 | 0.01 | 0.14 | 0.06 |
| 5 | 114046141 | rs2161182 | -0.285 | 1.577 | 1.194 | 0.404 | 0.006 | -0.003 | -0.003 | 0 | 0.24 | 0.25 | 0.21 | 0.22 | 0.27 | 0.29 |
| 5 | 114047808 | rs157980 | 0.191 | 1.75 | 1.292 | 0.514 | -0.003 | 0.012 | 0.012 | 0 | 0.84 | 0.86 | 0.62 | 0.55 | 0.75 | 0.65 |
| 5 | 114047868 | rs285879 | 1.719 | 1.733 | 1.17 | 0.5 | -0.012 | -0.022 | -0.022 | 0 | 0.65 | 0.26 | 0.46 | 0.54 | 0.61 | 0.59 |
| 5 | 114048498 | rs285878 | 1.221 | 1.727 | 1.264 | 0.556 | 0.01 | 0.03 | 0.03 | 0 | 0.71 | 0.37 | 0.49 | 0.54 | 0.66 | 0.6 |
| 5 | 114063419 | rs2394 | 0.092 | 1.647 | 1.447 | 0.476 | -0.017 | -0.05 | -0.05 | 0 | 0.12 | 0.31 | 0.09 | 0.08 | 0.08 | 0.09 |
| 5 | 114067361 | rs12109407 | 0.293 | 1.752 | 1.521 | 0.524 | -0.009 | -0.046 | -0.046 | 0.021 | 0.11 | 0.32 | 0.09 | 0.07 | 0.08 | 0.09 |
| 5 | 114075021 | rs34311 | 0.181 | 1.848 | 1.464 | 0.777 | -0.017 | -0.02 | -0.02 | 0 | 0.84 | 0.9 | 0.62 | 0.56 | 0.75 | 0.65 |
| 5 | 114075239 | rs287656 | 1.34 | 1.905 | 1.603 | 0.819 | 0.005 | -0.007 | -0.007 | 0.618 | 0.6 | 0.57 | 0.41 | 0.33 | 0.48 | 0.35 |
| 5 | 114075585 | rs34312 | 0.049 | 1.874 | 1.418 | 0.77 | -0.015 | 0.01 | 0.01 | 0 | 0.21 | 0.14 | 0.18 | 0.16 | 0.26 | 0.23 |
| 5 | 114078632 | rs158240 | 0.494 | 1.76 | 1.038 | 0.924 | -0.003 | 0.012 | 0.012 | 0 | 0.83 | 0.84 | 0.62 | 0.51 | 0.74 | 0.63 |
| 5 | 114079298 | rs6878057 | 0.806 | 1.36 | 1.473 | 0.654 | -0.005 | -0.051 | -0.051 | 2.12 | 0.09 | 0.25 | 0.08 | 0 | 0.07 | 0.02 |
| 5 | 114082720 | rs158359 | -2.434 | 1.25 | 0.947 | 0.665 | 0.051 | 0.04 | 0.04 | 0.456 | 0.58 | 0.94 | 0.75 | 0.72 | 0.65 | 0.66 |
| 5 | 114104210 | rs463926 | -1.445 | 1.323 | 1.638 | 0.671 | -0.003 | -0.02 | -0.02 | 0 | 0.43 | 0.49 | 0.63 | 0.74 | 0.54 | 0.59 |
| 5 | 114108659 | rs289026 | -0.637 | 1.578 | 1.871 | 1.026 | -0.017 | -0.014 | -0.014 | 0 | 0.28 | 0.18 | 0.23 | 0.36 | 0.35 | 0.3 |
| 5 | 114109101 | rs289027 | -2.677 | 1.525 | 1.997 | 0.983 | 0.073 | 0.096 | 0.096 | 0.655 | 0.56 | 0.85 | 0.74 | 0.75 | 0.65 | 0.65 |
| 5 | 114116285 | rs10085128 | 0.172 | 1.635 | 2.183 | 1.087 | 0.058 | 0.125 | 0.125 | 4.15 | 0.13 | 0.25 | 0.39 | 0.33 | 0.18 | 0.27 |
| 5 | 114117381 | rs4705511 | 0.397 | 1.674 | 2.147 | 1.105 | -0.008 | 0.002 | 0.002 | 0 | 0.12 | 0.1 | 0.35 | 0.32 | 0.16 | 0.25 |
| 5 | 114120596 | rs289031 | -1.72 | 1.252 | 2.068 | 0.948 | 0.012 | 0.016 | 0.016 | 0 | 0.49 | 0.43 | 0.68 | 0.69 | 0.6 | 0.59 |
| 5 | 114127793 | rs10478183 | -0.378 | 1.368 | 2.163 | 0.952 | 0.026 | 0.03 | 0.03 | 0 | 0.2 | 0.3 | 0.46 | 0.28 | 0.25 | 0.28 |
| 5 | 114128978 | rs542263 | -0.72 | 1.446 | 2.079 | 0.92 | 0.002 | 0.004 | 0.004 | 0 | 0.26 | 0.27 | 0.22 | 0.34 | 0.3 | 0.28 |
| 5 | 114130420 | rs2247496 | 0.292 | 1.483 | 1.761 | 0.925 | -0.038 | 0.004 | 0.004 | 0 | 0.17 | 0.05 | 0.13 | 0.13 | 0.18 | 0.24 |
| 5 | 114133676 | rs11955050 | -1.809 | 1.433 | 1.939 | 0.637 | 0.028 | 0.039 | 0.039 | 0 | 0.48 | 0.69 | 0.68 | 0.62 | 0.55 | 0.57 |
| 5 | 114138605 | rs755759 | -0.304 | 1.416 | 2.137 | 0.449 | 0.058 | 0.073 | 0.073 | 1.423 | 0.16 | 0.31 | 0.43 | 0.29 | 0.2 | 0.27 |
| 5 | 114139169 | rs6870296 | -2.179 | 1.365 | 1.995 | 0.404 | 0.066 | 0.018 | 0.018 | 0.245 | 0.53 | 0.86 | 0.72 | 0.64 | 0.61 | 0.62 |
| 5 | 114139525 | rs11745694 | -0.462 | 1.367 | 2.107 | 0.376 | -0.032 | -0.027 | -0.027 | 0.471 | 0.25 | 0.11 | 0.2 | 0.29 | 0.31 | 0.29 |
| 5 | 114140181 | rs1020992 | -2.222 | 1.375 | 2.061 | 0.363 | 0.066 | 0.018 | 0.018 | 0.245 | 0.53 | 0.86 | 0.72 | 0.64 | 0.61 | 0.63 |
| 5 | 114140694 | rs2416380 | 0.191 | 1.382 | 2.096 | 0.37 | 0.012 | 0.015 | 0.015 | 0 | 0.13 | 0.15 | 0.39 | 0.25 | 0.18 | 0.26 |
| 5 | 114141842 | rs10519398 | 0.387 | 1.303 | 2.019 | 0.325 | 0.005 | -0.053 | -0.053 | 1.265 | 0.1 | 0.17 | 0.09 | 0 | 0.11 | 0.06 |
| 5 | 114141956 | rs1479347 | -2.184 | 1.295 | 2.022 | 0.323 | 0.038 | -0.02 | -0.02 | 0 | 0.54 | 0.95 | 0.72 | 0.64 | 0.61 | 0.62 |
| 5 | 114145524 | rs6894095 | -0.318 | 1.226 | 2.091 | 0.539 | 0.005 | -0.076 | -0.076 | 0 | 0.12 | 0.31 | 0.1 | 0 | 0.11 | 0.06 |
| 5 | 114162202 | rs6879582 | -0.084 | 1.232 | 2.06 | 0.542 | 0.007 | -0.069 | -0.069 | 0 | 0.11 | 0.29 | 0.09 | 0 | 0.11 | 0.06 |
| 5 | 114166068 | rs10069405 | -0.203 | 1.258 | 2.04 | 0.543 | 0.009 | -0.072 | -0.072 | 0.987 | 0.12 | 0.29 | 0.09 | 0 | 0.11 | 0.06 |
| 5 | 114202263 | rs7718032 | 0.28 | 1.185 | 2.034 | 0.554 | 0.001 | -0.052 | -0.052 | 0 | 0.09 | 0.28 | 0.08 | 0 | 0.07 | 0.02 |
| 5 | 114208426 | rs13360560 | -1.479 | 1.254 | 1.66 | 0.614 | -0.012 | 0.002 | 0.002 | 0.087 | 0.28 | 0.74 | 0.51 | 0.3 | 0.3 | 0.34 |
| 5 | 114218797 | rs11748926 | -0.467 | 1.353 | 1.729 | 0.682 | 0 | 0.008 | 0.008 | 0 | 0.22 | 0.23 | 0.19 | 0.33 | 0.23 | 0.25 |
| 5 | 114241746 | rs2416384 | -1.887 | 1.148 | 1.82 | 0.627 | 0.021 | 0.019 | 0.019 | 0.15 | 0.46 | 0.75 | 0.68 | 0.63 | 0.53 | 0.55 |
| 5 | 114279914 | rs4292441 | -1.108 | 1.145 | 1.827 | 0.609 | 0.019 | 0.015 | 0.015 | 0 | 0.32 | 0.35 | 0.57 | 0.43 | 0.43 | 0.5 |
| 5 | 114283042 | rs11241298 | -0.979 | 1.147 | 1.762 | 0.627 | -0.006 | -0.005 | -0.005 | 0 | 0.31 | 0.29 | 0.56 | 0.43 | 0.42 | 0.5 |
| 5 | 114284436 | rs988700 | -0.013 | 0.676 | 1.856 | 0.705 | 0.021 | 0.021 | 0.021 | 0 | 0.13 | 0.16 | 0.38 | 0.23 | 0.18 | 0.26 |
| 5 | 114284539 | rs1453013 | 0.955 | 0.622 | 1.897 | 0.685 | -0.004 | -0.032 | -0.032 | 0.02 | 0.08 | 0.13 | 0.07 | 0 | 0.07 | 0.02 |
| 5 | 114287552 | rs10477524 | -0.49 | 0.744 | 1.779 | 0.766 | 0.026 | 0.018 | 0.018 | 0 | 0.15 | 0.18 | 0.39 | 0.23 | 0.23 | 0.3 |
| 5 | 114289334 | rs11749427 | -0.577 | 0.717 | 1.696 | 0.737 | 0.097 | 0.071 | 0.071 | 0 | 0.16 | 0.35 | 0.4 | 0.24 | 0.23 | 0.32 |
| 5 | 114294204 | rs9326939 | -0.391 | 0.725 | 1.439 | 0.749 | -0.027 | -0.022 | -0.022 | 0 | 0.14 | 0.12 | 0.38 | 0.24 | 0.23 | 0.32 |
| 5 | 114323981 | rs7701456 | 1.881 | 0.561 | 1.233 | 0.617 | -0.029 | 0.002 | 0.002 | 0.022 | 0.5 | 0.12 | 0.36 | 0.38 | 0.5 | 0.39 |
| 5 | 114324954 | rs1453027 | -1.722 | 0.557 | 1.502 | 0.618 | -0.004 | 0.006 | 0.006 | 0 | 0.46 | 0.62 | 0.61 | 0.57 | 0.49 | 0.59 |
| 5 | 114337809 | rs1377333 | -0.201 | 0.281 | 1.55 | 0.759 | 0.011 | -0.024 | -0.024 | 0.111 | 0.22 | 0.11 | 0.14 | 0.33 | 0.2 | 0.26 |
| 5 | 114338758 | rs11954151 | -1.091 | 0.274 | 1.358 | 0.758 | 0.021 | 0.001 | 0.001 | 0.801 | 0.34 | 0.68 | 0.25 | 0.38 | 0.28 | 0.29 |
| 5 | 114364916 | rs6869551 | -1.772 | 0.405 | 1.023 | 0.745 | -0.016 | 0.008 | 0.008 | 0.016 | 0.46 | 0.67 | 0.62 | 0.63 | 0.46 | 0.57 |

| | | | | | | | | | | | | | | | | |
|---|---|---|---|---|---|---|---|---|---|---|---|---|---|---|---|---|
| 5 | 114381312 | rs4487481 | -1.836 | 0.322 | 0.855 | 0.85 | -0.006 | 0.023 | 0.023 | 0.042 | 0.49 | 0.7 | 0.63 | 0.62 | 0.48 | 0.59 |
| 5 | 114383501 | rs10055057 | -0.521 | 0.592 | 0.726 | 1.012 | 0.085 | 0.06 | 0.06 | 0 | 0.14 | 0.37 | 0.38 | 0.22 | 0.18 | 0.27 |
| 5 | 114386109 | rs11956229 | 0.17 | 0.514 | -0.286 | 0.628 | -0.001 | -0.021 | -0.021 | 0 | 0.19 | 0.08 | 0.16 | 0.21 | 0.22 | 0.28 |
| 5 | 114404806 | rs4705516 | 1.186 | 0.272 | -0.446 | 0.762 | 0.018 | -0.103 | -0.103 | 0 | 0.2 | 0.01 | 0.11 | 0.25 | 0.2 | 0.1 |
| 5 | 114405806 | rs11951180 | -0.775 | 1.28 | 0.322 | 1.185 | 0.011 | 0.029 | 0.029 | 0.001 | 0.34 | 0.21 | 0.22 | 0.25 | 0.32 | 0.3 |
| 5 | 114407438 | rs12517339 | -0.075 | 1.951 | 0.51 | 1.461 | 0.021 | 0.071 | 0.071 | 0 | 0.07 | 0.09 | 0.37 | 0.36 | 0.14 | 0.28 |
| 5 | 114423353 | rs1026006 | -0.47 | 1.913 | 0.198 | 1.411 | 0.001 | 0.012 | 0.012 | 0 | 0.3 | 0.21 | 0.22 | 0.26 | 0.32 | 0.3 |
| 5 | 114434126 | rs966791 | 0.664 | 1.316 | -0.541 | 0.883 | -0.02 | -0.008 | -0.008 | 0 | 0.58 | 0.28 | 0.38 | 0.39 | 0.53 | 0.43 |
| 5 | 114451513 | rs17137452 | -0.447 | 1.105 | -0.52 | 0.858 | 0 | -0.011 | -0.011 | 0.007 | 0.06 | 0.07 | 0.02 | 0 | 0.05 | 0.01 |
| 5 | 114451587 | rs2034246 | -0.69 | 1.449 | -0.85 | 1.041 | -0.076 | -0.067 | -0.067 | 0 | 0.34 | 0.16 | 0.58 | 0.58 | 0.43 | 0.48 |
| 5 | 114452634 | rs2974520 | 1.68 | 1.185 | 0.529 | 2.074 | 0.018 | 0.436 | 0.436 | 4.276 | 0.94 | 0.7 | 0.76 | 0.5 | 0.97 | 0.8 |
| 5 | 114466145 | rs6866580 | -0.906 | 1.57 | 1.151 | 2.2 | -0.002 | 0.017 | 0.017 | 0 | 0.33 | 0.35 | 0.48 | 0.59 | 0.34 | 0.36 |
| 5 | 114497708 | rs1422068 | -1.03 | 1.489 | 0.948 | 2.112 | -0.001 | -0.003 | -0.003 | 0 | 0.36 | 0.66 | 0.53 | 0.69 | 0.38 | 0.4 |
| 5 | 114497972 | rs11738500 | -0.689 | 1.683 | 0.844 | 1.493 | 0.01 | -0.033 | -0.033 | 0 | 0.31 | 0.54 | 0.3 | 0.28 | 0.34 | 0.2 |
| 5 | 114506461 | rs973589 | -0.831 | 1.594 | -0.239 | 1.538 | 0.012 | 0.004 | 0.004 | 5.596 | 0.33 | 0.7 | 0.27 | 0.34 | 0.3 | 0.19 |
| 5 | 114508952 | rs3805590 | -0.862 | 1.34 | -0.451 | 1.278 | 0.005 | 0.031 | 0.031 | 6.48 | 0.34 | 0.76 | 0.32 | 0.3 | 0.32 | 0.19 |
| 5 | 114527392 | rs10051090 | -0.181 | 1.303 | -0.757 | 1.543 | 0.001 | -0.008 | -0.008 | 0 | 0.28 | 0.22 | 0.35 | 0.51 | 0.28 | 0.29 |
| 5 | 114531072 | rs10059320 | 0.544 | 1.46 | -1.112 | 1.478 | -0.039 | 0.125 | 0.125 | 0.091 | 0.2 | 0.03 | 0.15 | 0.03 | 0.22 | 0.09 |
| 5 | 114533832 | rs10519400 | -0.688 | 1.33 | -0.827 | 1.7 | 0.002 | 0.16 | 0.16 | 0 | 0.06 | 0.07 | 0.19 | 0.49 | 0.06 | 0.22 |
| 5 | 114546906 | rs265421 | -0.226 | 0.914 | -1.723 | 1.075 | -0.089 | 0.072 | 0.072 | 0 | 0.66 | 0.96 | 0.61 | 0.84 | 0.63 | 0.7 |
| 5 | 114552350 | rs2053697 | 0.273 | 0.02 | -1.904 | -0.056 | -0.012 | 0.033 | 0.033 | 0 | 0.4 | 0.35 | 0.53 | 0.83 | 0.51 | 0.53 |
| 5 | 114554974 | rs265425 | -1.009 | 0.067 | -1.986 | -0.02 | -0.089 | 0.076 | 0.076 | 0 | 0.66 | 0.96 | 0.61 | 0.84 | 0.63 | 0.7 |
| 5 | 114564400 | rs4705731 | 0.485 | 0.308 | -1.633 | 0.247 | 0.009 | 0.243 | 0.243 | 0 | 0.05 | 0.09 | 0.2 | 0.47 | 0.06 | 0.24 |
| 5 | 114567027 | rs3805586 | 0.789 | 0.101 | -2.103 | -0.148 | -0.122 | -0.058 | -0.058 | 1.449 | 0.23 | 0.07 | 0.27 | 0.36 | 0.39 | 0.25 |
| 5 | 114576598 | rs10076833 | -0.119 | -0.038 | -2.034 | 0.721 | -0.051 | -0.052 | -0.052 | 0.03 | 0.34 | 0.2 | 0.37 | 0.52 | 0.47 | 0.29 |
| 5 | 114580652 | rs10077453 | 2.073 | -0.311 | -2.35 | 0.014 | -0.038 | 0.17 | 0.17 | 0 | 0.25 | 0.02 | 0.16 | 0.03 | 0.26 | 0.1 |
| 5 | 114580742 | rs1005056 | 1.008 | -0.021 | -2.054 | 0.357 | 0.002 | 0.165 | 0.165 | 0 | 0.05 | 0.06 | 0.2 | 0.46 | 0.05 | 0.24 |
| 5 | 114591853 | rs7700478 | 1.34 | -0.506 | -1.39 | -0.077 | -0.01 | -0.043 | -0.043 | 0 | 0.16 | 0.2 | 0.09 | 0.01 | 0.09 | 0.12 |
| 5 | 114593335 | rs17137584 | 1.083 | -0.556 | -0.969 | -0.088 | -0.046 | -0.096 | -0.096 | 0.448 | 0.17 | 0.32 | 0.1 | 0.01 | 0.07 | 0.12 |
| 5 | 114617706 | rs10055125 | 2.017 | -0.299 | -1.114 | -0.04 | 0.003 | -0.031 | -0.031 | 0.018 | 0.06 | 0.08 | 0.03 | 0 | 0.05 | 0.04 |
| 5 | 114618289 | rs6594883 | 0.27 | -0.45 | -1.295 | 0.092 | -0.066 | -0.025 | -0.025 | 0 | 0.29 | 0.16 | 0.32 | 0.4 | 0.47 | 0.29 |
| 5 | 114622705 | rs267300 | -2.298 | -0.354 | -1.239 | 0.271 | -0.037 | -0.044 | -0.044 | 0 | 0.91 | 0.98 | 0.84 | 0.87 | 0.9 | 0.8 |
| 5 | 114635340 | rs267291 | 0.751 | -0.465 | -1.134 | 0.415 | 0.003 | 0.129 | 0.129 | 0 | 0.07 | 0.06 | 0.21 | 0.52 | 0.08 | 0.25 |
| 5 | 114638625 | rs2100521 | 0.32 | -0.677 | -1.102 | 0.11 | -0.004 | 0.035 | 0.035 | 0 | 0.22 | 0.3 | 0.29 | 0.47 | 0.17 | 0.33 |
| 5 | 114640464 | rs2161196 | 1.126 | -0.736 | -1.098 | -0.248 | -0.017 | -0.061 | -0.061 | 0 | 0.15 | 0.24 | 0.08 | 0.01 | 0.1 | 0.12 |
| 5 | 114645507 | rs267286 | -2.623 | -0.367 | -1.234 | -0.132 | -0.038 | -0.041 | -0.041 | 0.054 | 0.92 | 0.98 | 0.84 | 0.87 | 0.9 | 0.84 |
| 5 | 114647578 | rs2925160 | 1.544 | -0.738 | -1.429 | -0.586 | 0.053 | 0.177 | 0.177 | 0 | 0.34 | 0.11 | 0.17 | 0.05 | 0.28 | 0.12 |
| 5 | 114653622 | rs2925178 | 0.381 | -0.246 | -0.906 | -0.428 | -0.007 | 0.039 | 0.039 | 0.203 | 0.55 | 0.23 | 0.54 | 0.63 | 0.48 | 0.47 |
| 5 | 114654042 | rs6894010 | 1.09 | -0.213 | -1.205 | -0.321 | 0.005 | 0.002 | 0.002 | 0.01 | 0.25 | 0.25 | 0.25 | 0.21 | 0.2 | 0.32 |
| 5 | 114673479 | rs1350106 | 1.494 | -0.201 | -1.202 | -0.343 | 0.004 | 0.007 | 0.007 | 0 | 0.22 | 0.22 | 0.25 | 0.15 | 0.19 | 0.31 |
| 5 | 114676772 | rs2925180 | 0.864 | 0.217 | -1.251 | -0.419 | 0.05 | 0.147 | 0.147 | 0 | 0.45 | 0.17 | 0.31 | 0.19 | 0.4 | 0.24 |
| 5 | 114677729 | rs269513 | -0.856 | 0.54 | -1.197 | -0.572 | -0.042 | -0.001 | -0.001 | 0 | 0.85 | 0.9 | 0.84 | 0.79 | 0.7 | 0.84 |
| 5 | 114678852 | rs2925179 | 1.278 | -0.117 | -1.092 | -0.478 | 0.035 | 0.061 | 0.061 | 0 | 0.32 | 0.14 | 0.16 | 0.04 | 0.28 | 0.14 |
| 5 | 114679296 | rs269515 | 0.121 | 0.232 | -0.755 | -0.616 | -0.008 | 0.009 | 0.009 | 0 | 0.6 | 0.55 | 0.59 | 0.65 | 0.53 | 0.53 |
| 5 | 114683867 | rs17137641 | -0.003 | 0.493 | -0.473 | -0.247 | -0.005 | -0.002 | -0.002 | 0 | 0.09 | 0.1 | 0.14 | 0.12 | 0.11 | 0.06 |
| 5 | 114685541 | rs269505 | -0.606 | 0.673 | -0.693 | -0.577 | -0.091 | -0.102 | -0.102 | 2.221 | 0.17 | 0.06 | 0.29 | 0.59 | 0.33 | 0.41 |
| 5 | 114687463 | rs2925175 | -0.284 | 0.607 | -0.355 | -0.751 | -0.012 | -0.015 | -0.015 | 0 | 0.13 | 0.2 | 0.13 | 0.08 | 0.12 | 0.06 |
| 5 | 114687642 | rs1455845 | 0.424 | 0.674 | -0.328 | -0.682 | -0.007 | 0 | 0 | 0.01 | 0.1 | 0.08 | 0.11 | 0.11 | 0.1 | 0.09 |
| 5 | 114688671 | rs269511 | -2.039 | 0.548 | -0.038 | -0.723 | 0.026 | -0.013 | -0.013 | 0 | 0.37 | 0.52 | 0.44 | 0.73 | 0.47 | 0.53 |
| 5 | 114691287 | rs10519406 | -0.634 | 0.603 | -0.078 | -0.694 | -0.088 | -0.077 | -0.077 | 0 | 0.19 | 0.06 | 0.31 | 0.66 | 0.35 | 0.44 |
| 5 | 114693074 | rs2963765 | 1.672 | 0.462 | 0.12 | -0.781 | -0.001 | -0.014 | -0.014 | 0 | 0.58 | 0.57 | 0.46 | 0.12 | 0.46 | 0.4 |
| 5 | 114700699 | rs12657417 | -0.123 | 0.739 | 0.17 | -0.647 | -0.073 | -0.075 | -0.075 | 0 | 0.15 | 0.06 | 0.3 | 0.63 | 0.34 | 0.42 |
| 5 | 114702503 | rs11241322 | 1.882 | 0.417 | 0.211 | -0.728 | 0.003 | 0.054 | 0.054 | 0 | 0.55 | 0.36 | 0.44 | 0.11 | 0.46 | 0.42 |
| 5 | 114709028 | rs17137669 | 0.465 | 0.498 | 0.217 | -0.674 | 0.003 | -0.024 | -0.024 | 0.28 | 0.11 | 0.15 | 0.11 | 0.07 | 0.1 | 0.08 |
| 5 | 114712055 | rs7715232 | -0.081 | 0.58 | 0.036 | -0.62 | -0.073 | -0.075 | -0.075 | 0 | 0.14 | 0.05 | 0.29 | 0.65 | 0.32 | 0.42 |
| 5 | 114713898 | rs10478230 | 1.207 | 0.364 | 0.053 | -0.617 | -0.004 | -0.035 | -0.035 | 0 | 0.09 | 0.01 | 0.1 | 0.11 | 0.08 | 0.05 |
| 5 | 114718506 | rs13357967 | -0.402 | 0.31 | 0.129 | -0.713 | -0.005 | -0.062 | -0.062 | 0.596 | 0.14 | 0.3 | 0.14 | 0.06 | 0.1 | 0.09 |
| 5 | 114719148 | rs751485 | 1.822 | 0.437 | -0.007 | -0.684 | -0.001 | -0.025 | -0.025 | 0.588 | 0.56 | 0.42 | 0.45 | 0.11 | 0.46 | 0.42 |
| 5 | 114721062 | rs269542 | 0.899 | 0.541 | 0.122 | -0.521 | -0.021 | -0.053 | -0.053 | 0 | 0.81 | 0.54 | 0.83 | 0.87 | 0.86 | 0.88 |
| 5 | 114722599 | rs2166339 | -0.232 | 0.871 | -0.337 | -0.466 | -0.007 | -0.017 | -0.017 | 0 | 0.23 | 0.27 | 0.2 | 0.07 | 0.11 | 0.16 |
| 5 | 114723502 | rs269539 | -1.604 | 1.141 | -0.819 | -0.45 | 0.004 | 0.003 | 0.003 | 0.025 | 0.52 | 0.57 | 0.62 | 0.89 | 0.52 | 0.65 |
| 5 | 114724345 | rs1809211 | 1.251 | 0.594 | -0.925 | -0.316 | 0.004 | -0.047 | -0.047 | 0.19 | 0.64 | 0.5 | 0.6 | 0.18 | 0.62 | 0.55 |
| 5 | 114728073 | rs269537 | 0.263 | 0.663 | -1.246 | -0.315 | -0.004 | -0.015 | -0.015 | 0.617 | 0.76 | 0.76 | 0.76 | 0.82 | 0.77 | 0.86 |
| 5 | 114733752 | rs2605190 | 0.94 | 0.736 | -1.676 | -0.212 | -0.004 | 0.022 | 0.022 | 0 | 0.07 | 0.01 | 0.08 | 0.47 | 0.06 | 0.28 |
| 5 | 114743006 | rs17137761 | 0.969 | 0.29 | -1.603 | -1.238 | 0.003 | -0.003 | -0.003 | 0 | 0.1 | 0.02 | 0.1 | 0.08 | 0.08 | 0.05 |
| 5 | 114743023 | rs17470150 | 0.891 | 0.677 | -1.544 | -1.102 | -0.018 | -0.011 | -0.011 | 0 | 0.14 | 0.02 | 0.18 | 0.1 | 0.08 | 0.15 |
| 5 | 114743497 | rs2964546 | -1.102 | 0.677 | -1.499 | -1.121 | 0.03 | 0.048 | 0.048 | 0.158 | 0.49 | 0.42 | 0.47 | 0.31 | 0.34 | 0.33 |
| 5 | 114744304 | rs2963750 | 1.154 | 0.616 | -2.185 | -1.242 | 0.002 | 0.019 | 0.019 | 0.207 | 0.4 | 0.32 | 0.3 | 0.04 | 0.43 | 0.32 |
| 5 | 114745089 | rs13189365 | -0.254 | 2.057 | -2.512 | -0.09 | -0.023 | 0.054 | 0.054 | 0 | 0.13 | 0.02 | 0.13 | 0.07 | 0.13 | 0.1 |

| | | | | | | | | | | | | | | | | |
|---|---|---|---|---|---|---|---|---|---|---|---|---|---|---|---|---|
| 5 | 114745394 | rs269500 | -0.852 | 2.161 | -1.826 | 0.649 | -0.101 | -0.029 | -0.029 | 0 | 0.25 | 0.07 | 0.42 | 0.74 | 0.33 | 0.5 |
| 5 | 114751410 | rs2605200 | -0.345 | 0.586 | -1.504 | 0.141 | 0.038 | 0.015 | 0.015 | 0 | 0.39 | 0.32 | 0.33 | 0.6 | 0.25 | 0.46 |
| 5 | 114754424 | rs269526 | -0.229 | 0.998 | -1.09 | -0.492 | -0.015 | 0.004 | 0.004 | 0 | 0.68 | 0.71 | 0.6 | 0.7 | 0.55 | 0.71 |
| 5 | 114759167 | rs269528 | -0.154 | 1.052 | -0.866 | -0.307 | 0.003 | -0.004 | -0.004 | 0.48 | 0.68 | 0.68 | 0.67 | 0.75 | 0.58 | 0.68 |
| 5 | 114759195 | rs3852203 | -0.418 | 0.47 | -0.373 | -0.814 | -0.005 | -0.001 | -0.001 | 0.026 | 0.17 | 0.19 | 0.26 | 0.17 | 0.15 | 0.11 |
| 5 | 114771127 | rs2546477 | -0.713 | 0.136 | -0.227 | -1.011 | 0.013 | -0.009 | -0.009 | 0 | 0.76 | 0.75 | 0.77 | 0.82 | 0.6 | 0.73 |
| 5 | 114776341 | rs174908 | -0.641 | 0.146 | 0.124 | -1.041 | 0.089 | -0.055 | -0.055 | 0 | 0.75 | 0.58 | 0.74 | 0.81 | 0.6 | 0.7 |
| 5 | 114785465 | rs790669 | 0.742 | -0.15 | 0.096 | -1.129 | -0.006 | 0.056 | 0.056 | 0 | 0.55 | 0.27 | 0.45 | 0.63 | 0.45 | 0.59 |
| 5 | 114792198 | rs2900095 | 0.517 | 0.297 | 0.464 | -1.427 | 0.07 | -0.057 | -0.057 | 0.483 | 0.14 | 0.21 | 0.16 | 0.03 | 0.29 | 0.19 |
| 5 | 114797458 | rs17137802 | -0.015 | 0.21 | 0.319 | -1.237 | -0.022 | -0.042 | -0.042 | 0 | 0.14 | 0.04 | 0.22 | 0.22 | 0.15 | 0.08 |
| 5 | 114799354 | rs6888963 | 0.662 | 0.208 | 0.952 | -1.258 | 0.052 | -0.034 | -0.034 | 0 | 0.71 | 0.5 | 0.74 | 0.77 | 0.61 | 0.63 |
| 5 | 114802697 | rs10478234 | 0.885 | 0.239 | 0.96 | -1.251 | 0.068 | -0.044 | -0.044 | 0 | 0.12 | 0.18 | 0.13 | 0.02 | 0.26 | 0.21 |
| 5 | 114811667 | rs2112613 | 0.903 | 0.01 | 1.312 | -1.346 | -0.006 | 0.007 | 0.007 | 0 | 0.69 | 0.63 | 0.64 | 0.58 | 0.72 | 0.75 |
| 5 | 114812584 | rs2112614 | 0.199 | 0.007 | 1.333 | -1.344 | -0.025 | 0.009 | 0.009 | 0 | 0.82 | 0.65 | 0.86 | 0.81 | 0.87 | 0.84 |
| 5 | 114813155 | rs2546480 | 1.504 | 0.025 | 1.361 | -1.36 | 0.004 | 0.016 | 0.016 | 0 | 0.56 | 0.29 | 0.5 | 0.57 | 0.46 | 0.55 |
| 5 | 114813639 | rs10037359 | -0.574 | 0.09 | 1.366 | -1.315 | 0.042 | -0.014 | -0.014 | 1.626 | 0.27 | 0.37 | 0.36 | 0.24 | 0.41 | 0.29 |
| 5 | 114819995 | rs1833716 | 0.673 | 0.209 | 1.271 | -1.316 | 0.206 | -0.042 | -0.042 | 0 | 0.1 | 0.3 | 0.13 | 0.01 | 0.25 | 0.07 |
| 5 | 114821140 | rs12517357 | 1.52 | -0.621 | 0.486 | -1.595 | 0.026 | 0.008 | 0.008 | 0 | 0.43 | 0.25 | 0.47 | 0.55 | 0.38 | 0.48 |
| 5 | 114822737 | rs10040274 | -1.287 | -1.313 | 0.612 | -1.239 | -0.029 | 0.091 | 0.091 | 0.005 | 0.23 | 0.48 | 0.31 | 0.46 | 0.19 | 0.34 |
| 5 | 114824202 | rs9687406 | -0.459 | -1.357 | 0.815 | -1.237 | -0.002 | 0.01 | 0.01 | 0 | 0.15 | 0.18 | 0.22 | 0.21 | 0.14 | 0.18 |
| 5 | 114826454 | rs13180979 | 1.383 | -1.709 | 0.219 | -1.492 | -0.014 | 0 | 0 | 0 | 0.59 | 0.5 | 0.63 | 0.54 | 0.7 | 0.55 |
| 5 | 114827270 | rs1422640 | 0.137 | -1.456 | -0.482 | -1.38 | 0.034 | 0.11 | 0.11 | 0 | 0.81 | 0.98 | 0.94 | 0.99 | 0.89 | 0.88 |
| 5 | 114828491 | rs9687184 | 0.236 | -1.312 | -0.366 | -1.282 | 0.024 | 0.022 | 0.022 | 0 | 0.78 | 0.9 | 0.88 | 0.86 | 0.84 | 0.76 |
| 5 | 114835146 | rs1363550 | 0.212 | -1.719 | -0.693 | -1.484 | -0.001 | -0.003 | -0.003 | 0 | 0.85 | 0.87 | 0.92 | 0.88 | 0.89 | 0.9 |
| 5 | 114844882 | rs2546484 | -0.22 | -0.742 | -1.004 | -0.632 | 0.004 | 0.077 | 0.077 | 0 | 0.51 | 0.74 | 0.6 | 0.69 | 0.55 | 0.61 |
| 5 | 114856307 | rs7712026 | 1.447 | -0.963 | -0.415 | -0.993 | -0.02 | -0.058 | -0.058 | 0 | 0.16 | 0.22 | 0.07 | 0.03 | 0.07 | 0.07 |
| 5 | 114858437 | rs12187973 | 0.139 | -1.249 | -0.295 | -1.095 | -0.108 | -0.074 | -0.074 | 1.679 | 0.33 | 0.12 | 0.49 | 0.54 | 0.47 | 0.51 |
| 5 | 114859653 | rs3776801 | 1.475 | -1.205 | -0.093 | -1.066 | -0.017 | -0.05 | -0.05 | 0 | 0.15 | 0.21 | 0.06 | 0.03 | 0.07 | 0.07 |
| 5 | 114868337 | rs256938 | 1.139 | -1.199 | -0.123 | -1.092 | 0.008 | 0.068 | 0.068 | 0 | 0.49 | 0.28 | 0.4 | 0.31 | 0.45 | 0.39 |
| 5 | 114878065 | rs256958 | 0.502 | -1.233 | 0.122 | -1.034 | -0.018 | -0.023 | -0.023 | 0 | 0.16 | 0.18 | 0.09 | 0.12 | 0.11 | 0.09 |
| 5 | 114885896 | rs256978 | -1.137 | -0.904 | -0.024 | -0.978 | -0.049 | -0.055 | -0.055 | 0 | 0.82 | 0.9 | 0.55 | 0.45 | 0.61 | 0.56 |
| 5 | 114890507 | rs256976 | -0.283 | -0.962 | 0.042 | -0.998 | -0.072 | -0.044 | -0.044 | 0 | 0.68 | 0.84 | 0.49 | 0.42 | 0.55 | 0.49 |
| 5 | 114893223 | rs256975 | 0.326 | -0.902 | 0.549 | -0.999 | -0.027 | -0.04 | -0.04 | 0 | 0.14 | 0.29 | 0.09 | 0.12 | 0.11 | 0.1 |
| 5 | 114893518 | rs256974 | 0.326 | -0.902 | 0.549 | -0.999 | -0.027 | -0.04 | -0.04 | 0 | 0.14 | 0.29 | 0.09 | 0.12 | 0.11 | 0.1 |
| 5 | 114915460 | rs17473484 | 1.628 | -0.86 | 0.095 | -1.064 | 0.015 | 0.053 | 0.053 | 1.373 | 0.13 | 0.04 | 0.05 | 0.03 | 0.06 | 0.07 |
| 5 | 114916028 | rs2288384 | 1.256 | -0.873 | 0.196 | -1.067 | -0.005 | -0.003 | -0.003 | 0 | 0.1 | 0.11 | 0.05 | 0.12 | 0.08 | 0.13 |
| 5 | 114928381 | rs256943 | 0.147 | -1.192 | 0.005 | -1.28 | -0.046 | -0.002 | -0.002 | 0 | 0.58 | 0.77 | 0.42 | 0.43 | 0.5 | 0.46 |
| 5 | 114942711 | rs256962 | 0.616 | -1.276 | 0.36 | -1.18 | -0.003 | 0.001 | 0.001 | 0.762 | 0.52 | 0.39 | 0.4 | 0.48 | 0.49 | 0.46 |
| 5 | 114949680 | rs17474216 | 1.544 | -1.291 | 0.131 | -1.215 | 0.019 | 0.023 | 0.023 | 1.762 | 0.15 | 0.08 | 0.03 | 0 | 0.06 | 0.07 |
| 5 | 114951080 | rs698365 | -0.498 | -1.196 | 0.354 | -1.16 | -0.012 | -0.017 | -0.017 | 0 | 0.71 | 0.76 | 0.45 | 0.48 | 0.55 | 0.53 |
| 5 | 114951140 | rs698366 | -0.486 | -1.196 | 0.354 | -1.16 | -0.013 | -0.018 | -0.018 | 0 | 0.71 | 0.75 | 0.45 | 0.48 | 0.55 | 0.54 |
| 5 | 114951553 | rs400570 | -0.498 | -1.195 | 0.354 | -1.16 | -0.012 | -0.017 | -0.017 | 0 | 0.71 | 0.76 | 0.45 | 0.48 | 0.55 | 0.53 |
| 5 | 114967205 | rs256942 | -0.537 | -1.191 | 0.163 | -1.129 | -0.051 | -0.072 | -0.072 | 0 | 0.72 | 0.87 | 0.45 | 0.49 | 0.55 | 0.53 |
| 5 | 114971118 | rs10051170 | 0.527 | -1.349 | 0.116 | -1.272 | -0.011 | 0.035 | 0.035 | 0 | 0.52 | 0.28 | 0.34 | 0.35 | 0.44 | 0.4 |
| 5 | 114973120 | rs10055339 | 1.496 | -1.174 | 0.189 | -1.185 | -0.014 | -0.005 | -0.005 | 0 | 0.38 | 0.24 | 0.31 | 0.35 | 0.38 | 0.34 |
| 5 | 114973671 | rs10055677 | 0.376 | -0.551 | 0.725 | -0.657 | 0.021 | 0.039 | 0.039 | 0 | 0.67 | 0.49 | 0.43 | 0.48 | 0.55 | 0.52 |
| 5 | 114983982 | rs7727624 | -0.02 | -0.362 | -0.102 | -0.706 | -0.055 | -0.053 | -0.053 | 0.008 | 0.67 | 0.88 | 0.43 | 0.42 | 0.53 | 0.55 |
| 5 | 114989990 | rs1820631 | 0.181 | -0.43 | 0.004 | -0.674 | -0.029 | -0.023 | -0.023 | 0 | 0.65 | 0.74 | 0.41 | 0.42 | 0.5 | 0.48 |
| 5 | 114994084 | rs1698099 | 0.957 | 0.212 | -0.179 | -0.571 | -0.057 | -0.076 | -0.076 | 0 | 0.16 | 0.05 | 0.47 | 0.46 | 0.32 | 0.21 |
| 5 | 114995781 | rs6863960 | 0.446 | 0.142 | 0.053 | -0.623 | -0.006 | -0.004 | -0.004 | 1.955 | 0.64 | 0.66 | 0.4 | 0.41 | 0.5 | 0.47 |
| 5 | 114999751 | rs11745035 | 0.882 | -0.083 | 0.16 | -0.72 | -0.001 | -0.001 | -0.001 | 0 | 0.55 | 0.56 | 0.38 | 0.42 | 0.48 | 0.43 |
| 5 | 115001810 | rs249712 | 1.018 | 0.065 | -0.131 | -0.663 | -0.055 | -0.074 | -0.074 | 0 | 0.16 | 0.05 | 0.47 | 0.44 | 0.32 | 0.22 |
| 5 | 115017389 | rs365818 | 0.036 | -0.185 | -0.058 | -0.802 | 0.011 | 0.007 | 0.007 | 0 | 0.74 | 0.75 | 0.85 | 0.87 | 0.8 | 0.67 |
| 5 | 115030677 | rs166082 | 0.925 | 0.01 | -0.302 | -0.614 | -0.056 | -0.082 | -0.082 | 1.68 | 0.17 | 0.06 | 0.46 | 0.45 | 0.32 | 0.23 |
| 5 | 115038523 | rs465437 | 0.001 | -0.248 | -0.472 | -0.963 | 0.013 | -0.002 | -0.002 | 0.087 | 0.37 | 0.39 | 0.55 | 0.59 | 0.49 | 0.47 |
| 5 | 115041671 | rs4921010 | 1.322 | -0.104 | -1 | -0.952 | -0.019 | -0.033 | -0.033 | 0 | 0.31 | 0.07 | 0.28 | 0.28 | 0.29 | 0.19 |
| 5 | 115044230 | rs403039 | -1.591 | 0.007 | -0.8 | -0.881 | 0.004 | 0.016 | 0.016 | 0 | 0.85 | 0.89 | 0.9 | 0.87 | 0.86 | 0.72 |
| 5 | 115050362 | rs251891 | 1.186 | 0.912 | -0.678 | -0.734 | -0.01 | -0.002 | -0.002 | 0 | 0.15 | 0.13 | 0.42 | 0.44 | 0.26 | 0.2 |
| 5 | 115051490 | rs251890 | 0.297 | 0.765 | -0.414 | -0.728 | -0.001 | -0.014 | -0.014 | 0 | 0.52 | 0.44 | 0.74 | 0.72 | 0.57 | 0.4 |
| 5 | 115059411 | rs173635 | -0.987 | 0.82 | -0.282 | -0.443 | -0.013 | 0.011 | 0.011 | 0 | 0.83 | 0.87 | 0.84 | 0.83 | 0.73 | 0.68 |
| 5 | 115059538 | rs13360899 | 0.974 | 0.534 | -0.299 | -0.324 | 0.006 | 0.007 | 0.007 | 0 | 0.36 | 0.25 | 0.31 | 0.26 | 0.33 | 0.27 |
| 5 | 115060230 | rs1422289 | -0.211 | 0.568 | -0.285 | -0.262 | 0 | 0.003 | 0.003 | 0 | 0.71 | 0.7 | 0.47 | 0.39 | 0.6 | 0.58 |
| 5 | 115068172 | rs251867 | 1.204 | 0.296 | -0.64 | 0.329 | 0 | -0.015 | -0.015 | 0 | 0.13 | 0.07 | 0.39 | 0.45 | 0.17 | 0.14 |
| 5 | 115068238 | rs10478249 | -0.335 | 0.514 | -0.465 | 0.135 | 0.041 | 0.024 | 0.024 | 0 | 0.39 | 0.16 | 0.27 | 0.25 | 0.28 | 0.23 |
| 5 | 115068867 | rs13356510 | 0.183 | 0.554 | -0.558 | 0.146 | 0.011 | -0.014 | -0.014 | 0.666 | 0.32 | 0.12 | 0.26 | 0.25 | 0.27 | 0.23 |
| 5 | 115070403 | rs10477538 | -0.401 | 0.722 | -0.464 | 0.39 | 0.023 | 0.027 | 0.027 | 0 | 0.43 | 0.32 | 0.32 | 0.26 | 0.33 | 0.27 |
| 5 | 115091586 | rs10060156 | -0.062 | 0.863 | -0.308 | 0.029 | -0.002 | -0.021 | -0.021 | 0 | 0.25 | 0.23 | 0.33 | 0.04 | 0.29 | 0.14 |
| 5 | 115091714 | rs4921023 | -0.088 | 0.854 | -0.312 | 0.024 | -0.001 | -0.024 | -0.024 | 2.911 | 0.25 | 0.23 | 0.34 | 0.04 | 0.29 | 0.14 |
| 5 | 115092509 | rs1529453 | 0.139 | 0.795 | -0.579 | 0.109 | -0.024 | -0.033 | -0.033 | 0 | 0.06 | 0.15 | 0.01 | 0 | 0.01 | 0.02 |

| | | | | | | | | | | | | | | | | |
|---|---|---|---|---|---|---|---|---|---|---|---|---|---|---|---|---|
| 5 | 115096570 | rs154769 | -0.106 | 1.475 | -0.74 | 0.929 | -0.002 | 0.012 | 0.012 | 0 | 0.15 | 0.14 | 0.21 | 0.45 | 0.17 | 0.16 |
| 5 | 115110090 | rs154763 | -0.823 | 1.402 | -0.999 | 0.826 | -0.001 | 0.008 | 0.008 | 0 | 0.48 | 0.65 | 0.57 | 0.49 | 0.48 | 0.42 |
| 5 | 115112928 | rs6869363 | 0.042 | 1.48 | -0.856 | 0.751 | -0.009 | -0.138 | -0.138 | 0 | 0.34 | 0.5 | 0.39 | 0.04 | 0.32 | 0.27 |
| 5 | 115114235 | rs71186 | 0.569 | 1.737 | -1.634 | 0.841 | 0.001 | 0.021 | 0.021 | 0 | 0.53 | 0.34 | 0.46 | 0.57 | 0.58 | 0.59 |
| 5 | 115114515 | rs1864122 | -0.338 | 2.091 | -1.982 | 1.601 | -0.045 | -0.044 | -0.044 | 0.105 | 0.08 | 0.01 | 0.14 | 0.2 | 0.15 | 0.18 |
| 5 | 115117353 | rs249487 | -0.765 | 1.273 | -1.582 | 0.933 | -0.008 | -0.011 | -0.011 | 0 | 0.15 | 0.09 | 0.15 | 0.28 | 0.16 | 0.14 |
| 5 | 115120408 | rs34849 | -0.552 | 0.323 | -2.058 | 0.152 | 0.019 | 0.095 | 0.095 | 0 | 0.22 | 0.03 | 0.14 | 0.05 | 0.16 | 0.07 |
| 5 | 115122951 | rs12189241 | 0.045 | 0.229 | -3.007 | 0.409 | -0.033 | -0.12 | -0.12 | 0 | 0.3 | 0.03 | 0.36 | 0.31 | 0.26 | 0.35 |
| 5 | 115128540 | rs411148 | 0.049 | 1.034 | -2.869 | 0.661 | -0.002 | 0.053 | 0.053 | 0 | 0.91 | 0.92 | 0.86 | 0.51 | 0.84 | 0.8 |
| 5 | 115130926 | rs258737 | 0.41 | -0.073 | -2.681 | -0.068 | 0.022 | 0.105 | 0.105 | 0 | 0.56 | 0.24 | 0.39 | 0.24 | 0.61 | 0.51 |
| 5 | 115137144 | rs154759 | 0.192 | -0.859 | -2.496 | -0.925 | 0.04 | 0.03 | 0.03 | 0.057 | 0.45 | 0.18 | 0.48 | 0.32 | 0.4 | 0.44 |
| 5 | 115138748 | rs13174348 | -0.414 | -1.179 | -1.909 | -0.9 | 0.036 | -0.001 | -0.001 | 0 | 0.37 | 0.69 | 0.45 | 0.32 | 0.23 | 0.31 |
| 5 | 115157476 | rs6872406 | 0.511 | -1.234 | -1.049 | -0.919 | 0.005 | -0.007 | -0.007 | 0 | 0.27 | 0.2 | 0.38 | 0.32 | 0.19 | 0.29 |
| 5 | 115166713 | rs1058600 | -0.39 | -1.201 | -1.088 | -0.908 | 0.048 | -0.001 | -0.001 | 0 | 0.37 | 0.7 | 0.45 | 0.32 | 0.23 | 0.31 |
| 5 | 115175503 | rs6594908 | -0.094 | -0.685 | -0.563 | -0.637 | 0.05 | 0.033 | 0.033 | 4.352 | 0.07 | 0.48 | 0.05 | 0 | 0.01 | 0 |
| 5 | 115186899 | rs171809 | -0.16 | -0.769 | -0.917 | -0.997 | -0.01 | -0.007 | -0.007 | 0 | 0.45 | 0.5 | 0.48 | 0.33 | 0.35 | 0.48 |
| 5 | 115189500 | rs152367 | -0.694 | -0.133 | -1.222 | -0.889 | -0.016 | -0.005 | -0.005 | 0 | 0.67 | 0.78 | 0.71 | 0.63 | 0.64 | 0.77 |
| 5 | 115196792 | rs174046 | 1.549 | 0.14 | -1.698 | -0.772 | -0.054 | 0.115 | 0.115 | 0 | 0.19 | 0.02 | 0.11 | 0 | 0.24 | 0.16 |
| 5 | 115206937 | rs11241337 | -0.54 | 0.049 | -1.04 | -0.311 | -0.019 | 0.005 | 0.005 | 0 | 0.35 | 0.47 | 0.47 | 0.32 | 0.28 | 0.35 |
| 5 | 115229725 | rs6882704 | 0.143 | 0.108 | -0.909 | -0.428 | 0.001 | -0.112 | -0.112 | 0 | 0.19 | 0.35 | 0.16 | 0 | 0.2 | 0.07 |
| 5 | 115233116 | rs252738 | 1.036 | 0.138 | -1.242 | -0.285 | -0.082 | -0.017 | -0.017 | 0 | 0.42 | 0.08 | 0.27 | 0.38 | 0.42 | 0.36 |
| 5 | 115236690 | rs9326977 | 0.366 | -0.344 | -0.21 | -0.04 | -0.004 | -0.053 | -0.053 | 0.001 | 0.14 | 0.19 | 0.12 | 0 | 0.16 | 0.06 |
| 5 | 115242142 | rs3733902 | -0.322 | -0.051 | -0.394 | 0.264 | -0.024 | 0.01 | 0.01 | 0 | 0.36 | 0.5 | 0.46 | 0.35 | 0.31 | 0.37 |
| 5 | 115260315 | rs156661 | 0.314 | -0.366 | 0.114 | 0.684 | -0.024 | 0.027 | 0.027 | 0 | 0.67 | 0.57 | 0.66 | 0.47 | 0.8 | 0.65 |
| 5 | 115263853 | rs156666 | 0.536 | -0.127 | 0.366 | 0.582 | -0.007 | 0.007 | 0.007 | 0 | 0.64 | 0.43 | 0.64 | 0.47 | 0.77 | 0.61 |
| 5 | 115270867 | rs890785 | 0.835 | 0.269 | 0.029 | 0.275 | -0.011 | 0.014 | 0.014 | 0.048 | 0.43 | 0.24 | 0.43 | 0.41 | 0.43 | 0.5 |
| 5 | 115272911 | rs2416419 | -0.215 | 0.298 | 0.504 | 0.529 | 0.023 | 0.004 | 0.004 | 0 | 0.36 | 0.46 | 0.4 | 0.38 | 0.42 | 0.24 |
| 5 | 115285514 | rs9326979 | 0.01 | -0.018 | 0.791 | 0.576 | 0.027 | -0.005 | -0.005 | 0 | 0.3 | 0.34 | 0.28 | 0.26 | 0.4 | 0.17 |
| 5 | 115285650 | rs12189092 | 1.318 | -0.146 | 0.528 | 0.614 | -0.026 | 0.013 | 0.013 | 0 | 0.42 | 0.18 | 0.42 | 0.38 | 0.43 | 0.49 |
| 5 | 115291291 | rs7705750 | -1.097 | -0.311 | 0.448 | 0.462 | 0.016 | -0.049 | -0.049 | 0 | 0.83 | 0.96 | 0.84 | 0.78 | 0.87 | 0.76 |
| 5 | 115293612 | rs10071062 | 0.073 | 0.019 | 0.782 | 0.555 | 0.001 | 0 | 0 | 0.092 | 0.29 | 0.25 | 0.26 | 0.22 | 0.38 | 0.17 |
| 5 | 115298475 | rs12520255 | -0.167 | -0.282 | 0.344 | 0.339 | 0.043 | 0.006 | 0.006 | 0 | 0.7 | 0.85 | 0.8 | 0.75 | 0.78 | 0.66 |
| 5 | 115302066 | rs6896619 | 1.159 | -0.023 | 0.468 | 0.251 | -0.009 | 0.011 | 0.011 | 0 | 0.2 | 0.12 | 0.18 | 0.04 | 0.31 | 0.06 |
| 5 | 115307266 | rs6594921 | 1.234 | -0.114 | 0.712 | 0.277 | -0.007 | -0.019 | -0.019 | 1.12 | 0.34 | 0.3 | 0.49 | 0.53 | 0.37 | 0.49 |
| 5 | 115307292 | rs6895571 | -0.226 | -0.169 | 0.877 | 0.271 | 0.05 | 0.107 | 0.107 | 0 | 0.13 | 0.49 | 0.12 | 0.2 | 0.08 | 0.12 |
| 5 | 115309222 | rs6866413 | 0.339 | -0.18 | 0.774 | 0.215 | -0.005 | 0.061 | 0.061 | 0.02 | 0.47 | 0.75 | 0.61 | 0.73 | 0.45 | 0.61 |
| 5 | 115310760 | rs7341066 | 0.042 | -0.267 | 0.614 | 0.206 | -0.001 | -0.008 | -0.008 | 0.054 | 0.08 | 0.08 | 0.04 | 0.02 | 0.05 | 0.06 |
| 5 | 115312671 | rs6594925 | -0.892 | -0.208 | 0.43 | 0.217 | 0.023 | -0.038 | -0.038 | 0 | 0.76 | 0.96 | 0.83 | 0.79 | 0.81 | 0.72 |
| 5 | 115314458 | rs17482037 | 1.549 | -0.102 | 0.401 | -0.127 | -0.015 | 0.044 | 0.044 | 0 | 0.26 | 0.21 | 0.17 | 0.03 | 0.28 | 0.25 |
| 5 | 115315701 | rs10519434 | 0.718 | -0.078 | 0.663 | 0.152 | 0.007 | 0.038 | 0.038 | 0 | 0.09 | 0.13 | 0.07 | 0.2 | 0.04 | 0.09 |
| 5 | 115318303 | rs6594927 | -1.143 | -0.016 | 0.577 | -0.252 | -0.01 | 0.007 | 0.007 | 0 | 0.78 | 0.83 | 0.8 | 0.97 | 0.64 | 0.87 |
| 5 | 115318418 | rs6594928 | -0.534 | -0.086 | 0.796 | -0.321 | -0.01 | 0.007 | 0.007 | 0 | 0.73 | 0.79 | 0.78 | 0.96 | 0.6 | 0.84 |
| 5 | 115320107 | rs12515495 | -1.252 | -0.072 | 0.646 | -0.311 | -0.025 | 0.039 | 0.039 | 0 | 0.78 | 0.88 | 0.81 | 0.97 | 0.64 | 0.87 |
| 5 | 115321124 | rs6864491 | 0.48 | -0.329 | 0.76 | -0.002 | -0.024 | 0.035 | 0.035 | 0 | 0.56 | 0.35 | 0.47 | 0.28 | 0.48 | 0.52 |
| 5 | 115321748 | rs1366199 | -0.257 | -0.316 | 0.917 | 0.045 | -0.003 | 0.108 | 0.108 | 0 | 0.22 | 0.4 | 0.33 | 0.69 | 0.17 | 0.35 |
| 5 | 115322822 | rs10519435 | 1.651 | -0.472 | 0.639 | -0.049 | -0.015 | 0.034 | 0.034 | 0 | 0.27 | 0.23 | 0.26 | 0.05 | 0.31 | 0.25 |
| 5 | 115323234 | rs17138562 | 1.353 | -0.514 | 0.603 | -0.04 | 0.041 | -0.056 | -0.056 | 0.34 | 0.21 | 0.02 | 0.16 | 0.22 | 0.12 | 0.23 |
| 5 | 115323724 | rs10519437 | 0.824 | -0.413 | 0.833 | 0.263 | -0.004 | 0.045 | 0.045 | 0.005 | 0.12 | 0.16 | 0.17 | 0.42 | 0.12 | 0.26 |
| 5 | 115325279 | rs2195299 | 0.165 | -0.382 | 0.854 | 0.265 | 0.005 | -0.007 | -0.007 | 0 | 0.61 | 0.52 | 0.6 | 0.69 | 0.53 | 0.73 |
| 5 | 115328436 | rs10042431 | 0.534 | -0.351 | 0.855 | 0.342 | -0.011 | 0.114 | 0.114 | 0 | 0.13 | 0.24 | 0.26 | 0.5 | 0.12 | 0.25 |
| 5 | 115330927 | rs10044122 | 0.645 | -0.394 | 0.768 | 0.282 | 0.009 | 0.015 | 0.015 | 0 | 0.49 | 0.46 | 0.51 | 0.35 | 0.42 | 0.49 |
| 5 | 115332461 | rs4921045 | 0.522 | -0.462 | 0.608 | 0.395 | -0.009 | -0.005 | -0.005 | 0 | 0.43 | 0.4 | 0.45 | 0.46 | 0.45 | 0.52 |
| 5 | 115334143 | rs4921049 | 1.421 | -0.402 | 0.495 | 0.327 | -0.026 | 0.064 | 0.064 | 0 | 0.3 | 0.2 | 0.28 | 0.04 | 0.33 | 0.27 |
| 5 | 115334893 | rs7710003 | 0.803 | -0.103 | 0.623 | 0.391 | 0.006 | -0.044 | -0.044 | 0 | 0.24 | 0.37 | 0.29 | 0.12 | 0.32 | 0.08 |
| 5 | 115339086 | rs6594929 | -0.553 | 0.007 | 0.193 | 0.625 | 0.011 | -0.03 | -0.03 | 0 | 0.76 | 0.94 | 0.8 | 0.77 | 0.81 | 0.68 |
| 5 | 115341028 | rs7717144 | -0.265 | 0.243 | 0.396 | 0.668 | -0.001 | 0 | 0 | 0.038 | 0.32 | 0.63 | 0.37 | 0.32 | 0.37 | 0.15 |
| 5 | 115344224 | rs1593872 | 0.586 | 0.098 | 0.189 | 0.828 | 0.004 | 0.001 | 0.001 | 0 | 0.5 | 0.38 | 0.48 | 0.49 | 0.49 | 0.56 |
| 5 | 115345115 | rs2081862 | -0.067 | 0.166 | -0.34 | 0.833 | -0.009 | -0.002 | -0.002 | 0 | 0.73 | 0.68 | 0.78 | 0.78 | 0.8 | 0.71 |
| 5 | 115348816 | rs17138656 | 0.285 | 0.04 | -1.035 | 0.945 | 0.024 | -0.007 | -0.007 | 0.02 | 0.23 | 0.13 | 0.17 | 0.22 | 0.13 | 0.22 |
| 5 | 115349375 | rs1529442 | -0.454 | 1.152 | -0.967 | 1.321 | -0.018 | 0.019 | 0.019 | 0 | 0.81 | 0.87 | 0.8 | 0.96 | 0.67 | 0.87 |
| 5 | 115349682 | rs7732824 | -0.365 | 0.602 | -0.813 | 2.1 | -0.002 | 0.012 | 0.012 | 0 | 0.06 | 0.05 | 0.14 | 0.25 | 0.04 | 0.09 |
| 5 | 115350534 | rs11954031 | -0.684 | 0.458 | -0.791 | 1.744 | -0.001 | 0.028 | 0.028 | 0 | 0.09 | 0.11 | 0.07 | 0.28 | 0.08 | 0.16 |
| 5 | 115356672 | rs17138679 | -0.595 | 0.826 | -1.205 | 1.85 | -0.041 | -0.08 | -0.08 | 0 | 0.11 | 0.01 | 0.11 | 0.31 | 0.13 | 0.15 |
| 5 | 115356834 | rs11957856 | -0.787 | 1.172 | -1.445 | 1.021 | -0.028 | -0.032 | -0.032 | 0 | 0.38 | 0.2 | 0.3 | 0.4 | 0.46 | 0.41 |
| 5 | 115358663 | rs2662482 | 0.315 | 0.879 | -0.833 | 0.539 | 0.001 | 0.036 | 0.036 | 0.074 | 0.27 | 0.19 | 0.19 | 0.09 | 0.26 | 0.16 |
| 5 | 115361188 | rs1428089 | -0.771 | 0.455 | -0.764 | 0.824 | -0.029 | 0.113 | 0.113 | 0 | 0.54 | 0.77 | 0.6 | 0.68 | 0.47 | 0.54 |
| 5 | 115363934 | rs2560684 | -0.765 | 0.453 | -0.761 | 0.859 | -0.028 | 0.129 | 0.129 | 0.003 | 0.54 | 0.77 | 0.6 | 0.74 | 0.47 | 0.54 |
| 5 | 115364344 | rs17138687 | 0.404 | 0.418 | 0.02 | 1.345 | 0.005 | 0.061 | 0.061 | 0.337 | 0.08 | 0.17 | 0.1 | 0.17 | 0.07 | 0.15 |
| 5 | 115364622 | rs11241349 | 1.104 | 0.598 | -0.102 | 1.055 | 0 | 0 | 0 | 1.677 | 0.09 | 0.1 | 0.11 | 0.03 | 0.21 | 0.07 |

| | | | | | | | | | | | | | | | | |
|---|---|---|---|---|---|---|---|---|---|---|---|---|---|---|---|---|
| 5 | 115366925 | rs2662447 | -0.916 | -0.053 | 0.151 | 1.368 | 0.018 | 0.122 | 0.122 | 0 | 0.49 | 0.77 | 0.51 | 0.64 | 0.4 | 0.44 |
| 5 | 115369915 | rs1428090 | -0.365 | -0.107 | 0.561 | 1.48 | -0.002 | 0.007 | 0.007 | 0.007 | 0.42 | 0.48 | 0.3 | 0.42 | 0.38 | 0.29 |
| 5 | 115370311 | rs11241350 | 0.269 | -0.068 | 0.55 | 1.551 | -0.007 | 0.138 | 0.138 | 0 | 0.06 | 0.2 | 0.19 | 0.21 | 0.02 | 0.1 |
| 5 | 115371978 | rs969558 | -0.864 | -0.266 | 0.124 | 1.385 | 0.018 | 0.093 | 0.093 | 0 | 0.48 | 0.76 | 0.49 | 0.57 | 0.4 | 0.43 |
| 5 | 115373181 | rs1030465 | -0.324 | 0.15 | -0.156 | 1.103 | -0.001 | -0.002 | -0.002 | 0 | 0.39 | 0.34 | 0.28 | 0.39 | 0.37 | 0.29 |
| 5 | 115377150 | rs1396489 | 0.449 | -0.525 | -0.4 | 0.954 | 0.022 | 0.017 | 0.017 | 1.062 | 0.77 | 0.91 | 0.86 | 0.77 | 0.85 | 0.73 |
| 5 | 115378103 | rs2063254 | -0.882 | -0.853 | 0.578 | 1.352 | 0 | 0.14 | 0.14 | 0 | 0.1 | 0.4 | 0.09 | 0.21 | 0.06 | 0.09 |
| 5 | 115378630 | rs1355100 | -0.502 | -0.985 | 0.238 | 1.197 | 0.026 | -0.014 | -0.014 | 0.345 | 0.88 | 1 | 0.96 | 0.88 | 0.96 | 0.83 |
| 5 | 115382097 | rs2662468 | 1.137 | -1.224 | 0.961 | 1.6 | -0.005 | 0.004 | 0.004 | 1.889 | 0.69 | 0.34 | 0.62 | 0.55 | 0.82 | 0.64 |
| 5 | 115384346 | rs2136210 | 0.574 | -1.231 | 1.295 | 1.617 | 0.006 | -0.004 | -0.004 | 0 | 0.75 | 0.78 | 0.67 | 0.69 | 0.85 | 0.66 |
| 5 | 115385416 | rs10060276 | -2.217 | -1.111 | 1.656 | 1.381 | 0.075 | 0.206 | 0.206 | 9.786 | 0.06 | 0.46 | 0.06 | 0.14 | 0.02 | 0.06 |
| 5 | 115387119 | rs1541802 | -0.46 | -1.046 | 1.257 | 0.963 | 0.007 | -0.042 | -0.042 | 0.099 | 0.12 | 0.17 | 0.06 | 0.04 | 0.1 | 0.17 |
| 5 | 115388544 | rs2560696 | 1.407 | -1.659 | 1.076 | 1.308 | 0.02 | 0.012 | 0.012 | 0 | 0.56 | 0.34 | 0.43 | 0.32 | 0.69 | 0.57 |
| 5 | 115390957 | rs2560685 | 0.667 | -1.693 | 1.271 | 1.444 | 0.052 | -0.069 | -0.069 | 0.026 | 0.68 | 0.85 | 0.58 | 0.5 | 0.79 | 0.67 |
| 5 | 115393280 | rs2457142 | 0.694 | -1.699 | 1.454 | 1.436 | 0.024 | -0.029 | -0.029 | 0.737 | 0.67 | 0.78 | 0.57 | 0.5 | 0.79 | 0.67 |
| 5 | 115399346 | rs6878141 | 1.688 | -1.712 | 1.51 | 1.411 | 0.006 | 0.003 | 0.003 | 0 | 0.58 | 0.38 | 0.43 | 0.32 | 0.69 | 0.58 |
| 5 | 115402108 | rs1508879 | 0.344 | -1.834 | 1.651 | 1.661 | 0.023 | -0.017 | -0.017 | 0 | 0.73 | 0.81 | 0.6 | 0.66 | 0.82 | 0.68 |
| 5 | 115402958 | rs17138784 | 0.723 | -1.847 | 1.687 | 1.652 | -0.018 | 0.032 | 0.032 | 0 | 0.07 | 0.09 | 0.03 | 0.16 | 0.03 | 0.03 |
| 5 | 115403924 | rs6862008 | -2.476 | -1.917 | 1.542 | 1.574 | 0.031 | 0.165 | 0.165 | 0.503 | 0.05 | 0.33 | 0.04 | 0.14 | 0.01 | 0.02 |
| 5 | 115406154 | rs1105724 | 0.593 | -1.995 | 1.234 | 1.362 | -0.003 | 0.003 | 0.003 | 0.478 | 0.71 | 0.72 | 0.57 | 0.54 | 0.81 | 0.69 |
| 5 | 115408118 | rs10037346 | 1.424 | -1.907 | 0.859 | 1.2 | 0.009 | 0 | 0 | 0.116 | 0.6 | 0.37 | 0.52 | 0.37 | 0.77 | 0.63 |
| 5 | 115412283 | rs11241352 | 1.775 | -1.749 | -0.054 | 0.859 | 0.079 | -0.037 | -0.037 | 0.614 | 0.55 | 0.19 | 0.54 | 0.38 | 0.67 | 0.57 |
| 5 | 115412737 | rs6863734 | 0.158 | -1.825 | 0.153 | 0.862 | 0.019 | -0.022 | -0.022 | 0 | 0.78 | 0.82 | 0.8 | 0.61 | 0.85 | 0.74 |
| 5 | 115413956 | rs2292266 | -0.577 | -1.81 | 0.589 | 0.819 | 0.034 | -0.025 | -0.025 | 0 | 0.08 | 0.25 | 0.18 | 0.04 | 0.11 | 0.1 |
| 5 | 115414612 | rs1508884 | -0.743 | -1.95 | 0.443 | 0.919 | 0.024 | -0.035 | -0.035 | 5.923 | 0.86 | 0.94 | 0.85 | 0.75 | 0.9 | 0.83 |
| 5 | 115419968 | rs10054121 | 1.409 | -1.899 | 0.485 | 0.898 | -0.016 | 0.011 | 0.011 | 0.195 | 0.68 | 0.57 | 0.62 | 0.57 | 0.74 | 0.64 |
| 5 | 115420415 | rs1133186 | 1.393 | -1.888 | 0.491 | 0.9 | -0.017 | 0.013 | 0.013 | 0.211 | 0.69 | 0.58 | 0.62 | 0.56 | 0.74 | 0.64 |
| 5 | 115423531 | rs12517505 | -0.811 | -1.934 | 0.747 | 0.843 | 0.02 | -0.016 | -0.016 | 0 | 0.12 | 0.45 | 0.2 | 0.06 | 0.11 | 0.11 |
| 5 | 115425203 | rs9326985 | -0.811 | -1.953 | 0.703 | 0.87 | 0.017 | -0.037 | -0.037 | 0.025 | 0.86 | 0.94 | 0.84 | 0.75 | 0.9 | 0.83 |
| 5 | 115425396 | rs6896147 | 0.868 | -1.961 | 0.769 | 0.776 | -0.013 | 0.017 | 0.017 | 0 | 0.74 | 0.64 | 0.78 | 0.6 | 0.81 | 0.71 |
| 5 | 115425628 | rs17138884 | 0.572 | -1.956 | 0.777 | 0.772 | 0.006 | -0.012 | -0.012 | 1.544 | 0.06 | 0.15 | 0.17 | 0.04 | 0.07 | 0.08 |
| 5 | 115428334 | rs1129494 | 2.583 | -2.025 | 0.735 | 0.798 | 0.063 | -0.037 | -0.037 | 0 | 0.55 | 0.17 | 0.53 | 0.38 | 0.66 | 0.57 |
| 5 | 115428444 | rs10069656 | 0.312 | -1.967 | 0.681 | 0.811 | 0.006 | -0.011 | -0.011 | 0 | 0.06 | 0.15 | 0.17 | 0.04 | 0.07 | 0.08 |
| 5 | 115429249 | rs13186125 | 1.19 | -1.761 | 0.267 | 0.455 | -0.034 | 0.058 | 0.058 | 0 | 0.1 | 0.01 | 0.08 | 0.01 | 0.11 | 0.06 |
| 5 | 115429757 | rs10477550 | -1.122 | -1.65 | 0.557 | 0.69 | -0.03 | 0.022 | 0.022 | 0 | 0.19 | 0.43 | 0.24 | 0.21 | 0.14 | 0.13 |
| 5 | 115450657 | rs1876672 | -1.356 | -1.575 | 0.41 | 0.704 | -0.004 | -0.003 | -0.003 | 1.211 | 0.22 | 0.66 | 0.26 | 0.2 | 0.18 | 0.15 |
| 5 | 115453671 | rs4463206 | -0.94 | -1.557 | 0.386 | 0.748 | 0.016 | -0.041 | -0.041 | 0.785 | 0.86 | 0.94 | 0.84 | 0.73 | 0.9 | 0.82 |
| 5 | 115465899 | rs7704903 | 0.285 | -1.502 | 0.382 | 0.749 | 0.02 | -0.042 | -0.042 | 2.76 | 0.78 | 0.85 | 0.8 | 0.58 | 0.84 | 0.73 |
| 5 | 115471015 | rs17138924 | 1.147 | -1.455 | 0.394 | 0.734 | -0.008 | -0.035 | -0.035 | 0 | 0.08 | 0 | 0.03 | 0.15 | 0.05 | 0.09 |
| 5 | 115472493 | rs10519443 | 0.913 | -1.438 | 0.584 | 0.713 | 0.021 | 0.014 | 0.014 | 0.15 | 0.07 | 0.25 | 0.03 | 0.04 | 0.04 | 0.03 |
| 5 | 115473423 | rs10519444 | -0.553 | -1.435 | 0.362 | 0.723 | -0.007 | 0.017 | 0.017 | 0.169 | 0.16 | 0.52 | 0.1 | 0.17 | 0.12 | 0.07 |
| 5 | 115494919 | rs2136208 | 0.607 | -1.403 | 0.262 | 0.707 | 0.012 | -0.014 | -0.014 | 1.879 | 0.06 | 0.15 | 0.17 | 0.02 | 0.07 | 0.08 |
| 5 | 115496470 | rs1508875 | 2.279 | -1.446 | -0.033 | 0.74 | 0.039 | -0.064 | -0.064 | 1.104 | 0.55 | 0.11 | 0.54 | 0.38 | 0.66 | 0.57 |
| 5 | 115501792 | rs1567335 | 1.433 | -1.539 | -0.019 | 0.756 | 0.024 | -0.036 | -0.036 | 0 | 0.66 | 0.24 | 0.58 | 0.53 | 0.72 | 0.67 |
| 5 | 115514834 | rs1567337 | -2.07 | -1.612 | 0.063 | 0.665 | 0.039 | 0.005 | 0.005 | 1.795 | 0.33 | 0.86 | 0.31 | 0.35 | 0.24 | 0.25 |
| 5 | 115517399 | rs6878871 | -1.248 | -1.454 | 0.17 | 0.691 | 0 | 0 | 0 | 0.163 | 0.22 | 0.7 | 0.26 | 0.2 | 0.19 | 0.15 |
| 5 | 115519292 | rs9686231 | -1.248 | -1.454 | 0.17 | 0.691 | 0 | 0 | 0 | 0 | 0.22 | 0.7 | 0.26 | 0.2 | 0.19 | 0.15 |
| 5 | 115525539 | rs7733187 | -0.054 | -1.467 | 0.487 | 0.711 | -0.021 | 0.031 | 0.031 | 0 | 0.13 | 0.27 | 0.07 | 0.18 | 0.07 | 0.06 |
| 5 | 115540041 | rs11241377 | -2.028 | -1.546 | 0.2 | 0.695 | 0.014 | -0.001 | -0.001 | 0 | 0.3 | 0.73 | 0.3 | 0.35 | 0.24 | 0.25 |
| 5 | 115542005 | rs11241378 | -1.402 | -1.541 | 0.324 | 0.695 | -0.016 | 0.013 | 0.013 | 0.114 | 0.24 | 0.59 | 0.13 | 0.33 | 0.17 | 0.17 |
| 5 | 115542982 | rs13362436 | -1.054 | -1.548 | 0.012 | 0.684 | -0.009 | -0.004 | -0.004 | 0 | 0.18 | 0.71 | 0.11 | 0.18 | 0.12 | 0.11 |
| 5 | 115547937 | rs17139182 | 1.534 | -1.528 | -0.017 | 0.705 | 0.009 | -0.014 | -0.014 | 0.288 | 0.06 | 0.02 | 0.04 | 0.14 | 0.03 | 0.03 |
| 5 | 115548435 | rs7723648 | -1.33 | -1.534 | -0.017 | 0.697 | 0.001 | 0.001 | 0.001 | 0.828 | 0.22 | 0.73 | 0.26 | 0.2 | 0.19 | 0.15 |
| 5 | 115552945 | rs712561 | 2.025 | -1.554 | -0.249 | 0.717 | 0.043 | -0.079 | -0.079 | 1.609 | 0.55 | 0.08 | 0.53 | 0.38 | 0.66 | 0.58 |
| 5 | 115562150 | rs1422498 | 0.337 | -1.508 | -0.034 | 0.727 | 0.001 | 0.003 | 0.003 | 0 | 0.1 | 0.12 | 0.05 | 0.15 | 0.09 | 0.11 |
| 5 | 115568093 | rs4283832 | 0.951 | -1.484 | -0.287 | 0.719 | 0.002 | 0.003 | 0.003 | 1.11 | 0.77 | 0.25 | 0.74 | 0.79 | 0.79 | 0.78 |
| 5 | 115568712 | rs12515329 | 0.985 | -1.482 | -0.42 | 0.72 | 0.007 | 0.011 | 0.011 | 1.157 | 0.76 | 0.22 | 0.73 | 0.79 | 0.79 | 0.79 |
| 5 | 115571703 | rs250331 | 2.254 | -1.586 | -0.901 | 0.672 | -0.139 | -0.058 | -0.058 | 0.03 | 0.41 | 0.03 | 0.43 | 0.26 | 0.55 | 0.45 |
| 5 | 115591333 | rs6594961 | -1.482 | -1.574 | -0.401 | 0.709 | 0.014 | 0 | 0 | 0 | 0.3 | 0.72 | 0.3 | 0.35 | 0.23 | 0.25 |
| 5 | 115592141 | rs4921085 | -0.019 | -1.627 | 0.043 | 0.713 | -0.021 | 0.03 | 0.03 | 0 | 0.13 | 0.28 | 0.07 | 0.17 | 0.07 | 0.06 |
| 5 | 115594144 | rs37183 | -0.115 | -1.777 | 0.075 | 0.758 | -0.001 | 0.001 | 0.001 | 0.045 | 0.19 | 0.2 | 0.32 | 0.29 | 0.17 | 0.26 |
| 5 | 115601291 | rs10066072 | 1.005 | -1.753 | 0.113 | 0.732 | 0 | -0.001 | -0.001 | 0 | 0.75 | 0.28 | 0.86 | 0.68 | 0.83 | 0.83 |
| 5 | 115607289 | rs6888823 | 0.018 | -1.65 | 0.023 | 0.756 | -0.02 | 0.023 | 0.023 | 0 | 0.13 | 0.28 | 0.07 | 0.17 | 0.07 | 0.06 |
| 5 | 115616288 | rs784482 | 0.796 | -1.63 | -0.06 | 0.763 | 0.013 | -0.034 | -0.034 | 3.335 | 0.13 | 0.06 | 0.17 | 0.26 | 0.1 | 0.18 |
| 5 | 115616807 | rs804199 | 2.331 | -1.426 | -1.055 | -0.081 | -0.127 | 0.168 | 0.168 | 0 | 0.25 | 0.03 | 0.23 | 0 | 0.38 | 0.13 |
| 5 | 115617297 | rs784480 | -0.954 | -1.498 | 0.027 | 0.987 | -0.027 | 0.033 | 0.033 | 0 | 0.27 | 0.37 | 0.24 | 0.44 | 0.17 | 0.24 |
| 5 | 115627570 | rs2068456 | 0.141 | -1.458 | -0.244 | 0.954 | 0.003 | -0.01 | -0.01 | 4.703 | 0.1 | 0.05 | 0.19 | 0.16 | 0.1 | 0.07 |
| 5 | 115633474 | rs153613 | -1.351 | -1.483 | -0.069 | 0.99 | -0.006 | 0.006 | 0.006 | 0 | 0.31 | 0.63 | 0.26 | 0.46 | 0.23 | 0.32 |
| 5 | 115641393 | rs254153 | -0.638 | -1.409 | -0.431 | 1.044 | 0.032 | -0.066 | -0.066 | 0 | 0.24 | 0.08 | 0.24 | 0.41 | 0.18 | 0.23 |

| | | | | | | | | | | | | | | | | |
|---|---|---|---|---|---|---|---|---|---|---|---|---|---|---|---|---|
| 5 | 115642365 | rs11956927 | 0.602 | -1.336 | 0.219 | 1.18 | 0.015 | 0.008 | 0.008 | 0.186 | 0.07 | 0.26 | 0.03 | 0.04 | 0.04 | 0.03 |
| 5 | 115642513 | rs185263 | -2.292 | -1.547 | 0.406 | 1.105 | 0.074 | -0.02 | -0.02 | 0 | 0.44 | 0.74 | 0.46 | 0.61 | 0.3 | 0.42 |
| 5 | 115643769 | rs11948029 | -0.684 | -1.539 | 0.47 | 1.133 | -0.004 | 0.031 | 0.031 | 0.043 | 0.14 | 0.37 | 0.08 | 0.19 | 0.09 | 0.12 |
| 5 | 115645582 | rs712574 | -1.707 | -1.517 | 0.47 | 1.107 | -0.039 | 0.033 | 0.033 | 0 | 0.37 | 0.45 | 0.29 | 0.58 | 0.23 | 0.34 |
| 5 | 115649941 | rs712577 | -1.214 | -1.5 | 0.187 | 1.299 | 0.038 | -0.054 | -0.054 | 0 | 0.29 | 0.16 | 0.25 | 0.55 | 0.19 | 0.31 |
| 5 | 115655198 | rs785372 | -0.37 | -1.459 | 0.184 | 1.353 | 0.017 | -0.03 | -0.03 | 0 | 0.21 | 0.15 | 0.21 | 0.41 | 0.16 | 0.28 |
| 5 | 115664057 | rs7721593 | -0.65 | -1.387 | 0.631 | 1.193 | -0.015 | -0.006 | -0.006 | 0 | 0.17 | 0.4 | 0.22 | 0.16 | 0.14 | 0.13 |
| 5 | 115670932 | rs804152 | -1.135 | -1.458 | 0.498 | 1.31 | -0.008 | 0.007 | 0.007 | 0 | 0.27 | 0.29 | 0.36 | 0.43 | 0.19 | 0.3 |
| 5 | 115678956 | rs804163 | 0.353 | -1.173 | 0.339 | 1.832 | 0.004 | -0.069 | -0.069 | 0.458 | 0.15 | 0.07 | 0.27 | 0.4 | 0.13 | 0.2 |
| 5 | 115693166 | rs7727761 | -0.272 | -1.357 | 0.718 | 1.239 | 0.042 | -0.001 | -0.001 | 0.015 | 0.1 | 0.39 | 0.06 | 0.04 | 0.08 | 0.05 |
| 5 | 115695077 | rs27634 | -1.087 | -1.482 | 0.339 | 1.266 | 0.017 | -0.012 | -0.012 | 0 | 0.3 | 0.27 | 0.35 | 0.43 | 0.19 | 0.27 |
| 5 | 115698959 | rs12521635 | 0.245 | -1.436 | 0.042 | 1.261 | 0.01 | 0.001 | 0.001 | 0 | 0.15 | 0.12 | 0.19 | 0.17 | 0.1 | 0.1 |
| 5 | 115700756 | rs5027681 | -1.889 | -1.558 | 0.256 | 1.27 | -0.017 | -0.01 | -0.01 | 0 | 0.35 | 0.56 | 0.3 | 0.31 | 0.24 | 0.24 |
| 5 | 115701098 | rs1421847 | -2.08 | -1.351 | 0.126 | 1.242 | 0.091 | -0.02 | -0.02 | 0 | 0.48 | 0.79 | 0.47 | 0.58 | 0.31 | 0.43 |
| 5 | 115707457 | rs4921096 | -1.047 | -1.438 | 0.097 | 1.225 | -0.019 | -0.02 | -0.02 | 0.152 | 0.19 | 0.49 | 0.1 | 0.17 | 0.13 | 0.14 |
| 5 | 115710803 | rs6594967 | -1.881 | -1.336 | -0.783 | 1.278 | 0.11 | -0.041 | -0.041 | 0 | 0.49 | 0.89 | 0.47 | 0.6 | 0.32 | 0.43 |
| 5 | 115713000 | rs26333 | 2.227 | -0.837 | -1.177 | 1.073 | -0.175 | -0.098 | -0.098 | 1.462 | 0.37 | 0.03 | 0.39 | 0.32 | 0.5 | 0.42 |
| 5 | 115718006 | rs17139652 | 0.514 | -0.546 | -0.566 | 1.398 | -0.001 | 0.006 | 0.006 | 0.303 | 0.13 | 0.15 | 0.19 | 0.26 | 0.11 | 0.17 |
| 5 | 115718845 | rs26331 | -0.434 | -0.512 | -0.855 | 1.205 | -0.029 | 0.015 | 0.015 | 0 | 0.87 | 0.93 | 0.89 | 0.92 | 0.84 | 0.86 |
| 5 | 115720094 | rs10053028 | -1.559 | -1.686 | -0.247 | 1.53 | 0.011 | -0.007 | -0.007 | 0 | 0.4 | 0.62 | 0.44 | 0.46 | 0.29 | 0.37 |
| 5 | 115721584 | rs17139655 | 0.339 | -1.529 | -0.645 | 1.231 | 0.022 | -0.016 | -0.016 | 0 | 0.09 | 0.02 | 0.04 | 0.15 | 0.03 | 0.04 |
| 5 | 115725791 | rs10477554 | -1.483 | -1.631 | -0.591 | 1.399 | 0.032 | -0.048 | -0.048 | 0 | 0.38 | 0.8 | 0.42 | 0.59 | 0.29 | 0.41 |
| 5 | 115727838 | rs7702057 | -1.078 | -1.758 | 0.682 | 1.371 | -0.001 | 0.076 | 0.076 | 1.42 | 0.11 | 0.43 | 0.04 | 0.14 | 0.05 | 0.13 |
| 5 | 115731448 | rs1990877 | -1.326 | -1.821 | 0.561 | 1.389 | -0.022 | 0.053 | 0.053 | 0.209 | 0.34 | 0.49 | 0.3 | 0.56 | 0.23 | 0.26 |
| 5 | 115733327 | rs28019 | 2.595 | -1.992 | -0.03 | 1.207 | -0.038 | 0.047 | 0.047 | 1.775 | 0.42 | 0.06 | 0.45 | 0.2 | 0.62 | 0.44 |
| 5 | 115735483 | rs6896985 | -1.058 | -2.2 | 0.622 | 0.96 | -0.083 | -0.083 | -0.083 | 0 | 0.21 | 0.48 | 0.09 | 0.05 | 0.08 | 0.19 |
| 5 | 115737646 | rs26325 | 1.085 | -1.972 | 0.548 | 1.129 | -0.008 | -0.002 | -0.002 | 0 | 0.72 | 0.69 | 0.75 | 0.75 | 0.81 | 0.77 |
| 5 | 115739315 | rs17139708 | 0.022 | -1.998 | 0.388 | 1.205 | 0.009 | 0.023 | 0.023 | 0 | 0.06 | 0.1 | 0.03 | 0.09 | 0.04 | 0.05 |
| 5 | 115740185 | rs26322 | -0.787 | -1.99 | 0.177 | 1.178 | 0.014 | -0.012 | -0.012 | 5.043 | 0.91 | 0.98 | 0.96 | 0.87 | 0.94 | 0.95 |
| 5 | 115744515 | rs17139712 | 0.049 | -1.85 | 0.229 | 1.332 | 0.001 | -0.016 | -0.016 | 1.945 | 0.16 | 0.13 | 0.21 | 0.42 | 0.14 | 0.14 |
| 5 | 115746451 | rs27953 | 2.667 | -1.98 | -0.061 | 1.158 | -0.119 | 0.028 | 0.028 | 1.058 | 0.41 | 0.05 | 0.4 | 0.2 | 0.55 | 0.41 |
| 5 | 115746567 | rs12521753 | -0.887 | -2.092 | 0.478 | 1.146 | -0.003 | 0.001 | 0.001 | 0 | 0.28 | 0.3 | 0.4 | 0.46 | 0.21 | 0.28 |
| 5 | 115746776 | rs26581 | -0.779 | -2.072 | 0.886 | 1.197 | -0.011 | 0.014 | 0.014 | 0 | 0.91 | 0.9 | 0.96 | 0.87 | 0.94 | 0.96 |
| 5 | 115751889 | rs254082 | -0.728 | -2.117 | 0.772 | 1.176 | 0.028 | -0.033 | -0.033 | 0.241 | 0.26 | 0.17 | 0.38 | 0.46 | 0.2 | 0.27 |
| 5 | 115752385 | rs6861280 | -0.52 | -2.111 | 1.153 | 1.166 | -0.002 | -0.004 | -0.004 | 0 | 0.18 | 0.21 | 0.1 | 0.1 | 0.12 | 0.15 |
| 5 | 115754261 | rs33071 | 1.471 | -2.018 | 0.955 | 1.195 | 0.002 | -0.014 | -0.014 | 0 | 0.58 | 0.36 | 0.61 | 0.39 | 0.64 | 0.68 |
| 5 | 115756397 | rs11241390 | 0.498 | -1.864 | 0.616 | 1.226 | -0.001 | -0.036 | -0.036 | 0 | 0.14 | 0.08 | 0.21 | 0.41 | 0.12 | 0.13 |
| 5 | 115761175 | rs26600 | 2.509 | -1.932 | 0.465 | 0.947 | -0.134 | -0.025 | -0.025 | 0.258 | 0.41 | 0.04 | 0.4 | 0.27 | 0.56 | 0.47 |
| 5 | 115764462 | rs154610 | 0.153 | -1.882 | 0.883 | 0.731 | -0.017 | -0.022 | -0.022 | 0 | 0.1 | 0.17 | 0.18 | 0.03 | 0.08 | 0.07 |
| 5 | 115764968 | rs153744 | 1.592 | -1.817 | 1.031 | 0.796 | -0.022 | -0.017 | -0.017 | 0 | 0.61 | 0.5 | 0.68 | 0.75 | 0.73 | 0.74 |
| 5 | 115769681 | rs40303 | -1.532 | -1.822 | 1.193 | 0.514 | -0.046 | -0.053 | -0.053 | 0 | 0.26 | 0.51 | 0.24 | 0.1 | 0.18 | 0.18 |
| 5 | 115771509 | rs33064 | 0.696 | -1.492 | 0.85 | 0.663 | -0.014 | -0.026 | -0.026 | 2.997 | 0.65 | 0.8 | 0.65 | 0.51 | 0.7 | 0.63 |
| 5 | 115778852 | rs17431689 | -0.659 | -1.436 | 0.847 | 0.44 | -0.005 | -0.002 | -0.002 | 0 | 0.13 | 0.16 | 0.07 | 0.08 | 0.08 | 0.12 |
| 5 | 115783553 | rs17139805 | -0.872 | -1.483 | 1.015 | 0.396 | -0.031 | 0.005 | 0.005 | 0 | 0.21 | 0.33 | 0.1 | 0.16 | 0.12 | 0.13 |
| 5 | 115785512 | rs2303752 | -0.591 | -1.406 | 1.128 | 0.376 | 0.028 | 0.076 | 0.076 | 0 | 0.14 | 0.34 | 0.09 | 0.16 | 0.12 | 0.06 |
| 5 | 115785522 | rs2303753 | 0.165 | -1.416 | 0.98 | 0.362 | 0.007 | 0.004 | 0.004 | 0 | 0.09 | 0.07 | 0.05 | 0.01 | 0.04 | 0.07 |
| 5 | 115787223 | rs10447236 | 0.863 | -1.487 | 0.868 | 0.418 | -0.014 | -0.033 | -0.033 | 0 | 0.2 | 0.1 | 0.18 | 0.39 | 0.2 | 0.23 |
| 5 | 115788104 | rs26602 | 1.586 | -1.545 | 1.045 | 0.462 | -0.001 | 0 | 0 | 1.254 | 0.72 | 0.34 | 0.71 | 0.65 | 0.8 | 0.73 |
| 5 | 115788412 | rs17139820 | -1.315 | -1.573 | 1.065 | 0.442 | -0.011 | 0.01 | 0.01 | 0 | 0.23 | 0.57 | 0.23 | 0.25 | 0.17 | 0.17 |
| 5 | 115789599 | rs1963858 | -1.972 | -1.723 | 0.949 | 0.346 | -0.031 | -0.016 | -0.016 | 0.013 | 0.56 | 0.79 | 0.54 | 0.65 | 0.45 | 0.45 |
| 5 | 115791050 | rs728529 | -1.18 | -1.704 | 0.986 | 0.354 | -0.015 | -0.004 | -0.004 | 0 | 0.23 | 0.53 | 0.15 | 0.17 | 0.16 | 0.13 |
| 5 | 115791669 | rs3733725 | -0.726 | -1.807 | 0.848 | 0.389 | -0.01 | -0.001 | -0.001 | 0 | 0.2 | 0.21 | 0.16 | 0.17 | 0.12 | 0.12 |
| 5 | 115793437 | rs2288476 | 0.88 | -1.722 | 0.747 | 0.405 | -0.011 | -0.032 | -0.032 | 0.042 | 0.2 | 0.13 | 0.26 | 0.47 | 0.2 | 0.26 |
| 5 | 115795101 | rs40104 | -0.027 | -1.716 | 0.863 | 0.411 | 0.042 | 0.083 | 0.083 | 0 | 0.94 | 0.73 | 0.93 | 0.9 | 0.97 | 0.9 |
| 5 | 115795188 | rs34971 | 0.805 | -1.618 | 0.857 | 0.056 | -0.012 | 0.009 | 0.009 | 0 | 0.61 | 0.47 | 0.54 | 0.42 | 0.68 | 0.57 |
| 5 | 115796103 | rs187600 | 0.079 | -1.62 | 0.84 | 0.054 | 0 | 0.051 | 0.051 | 0 | 0.7 | 0.54 | 0.65 | 0.43 | 0.77 | 0.64 |
| 5 | 115798501 | rs153742 | 1.906 | -1.479 | 0.595 | 0.144 | -0.025 | 0.031 | 0.031 | 0 | 0.41 | 0.24 | 0.44 | 0.24 | 0.55 | 0.44 |
| 5 | 115800189 | rs10077506 | 0.147 | -1.307 | 0.418 | 0.543 | -0.008 | 0.027 | 0.027 | 0 | 0.11 | 0.13 | 0.04 | 0.16 | 0.08 | 0.06 |
| 5 | 115807688 | rs258019 | -0.515 | -1.099 | 0.219 | 0.573 | 0 | -0.001 | -0.001 | 0 | 0.31 | 0.33 | 0.27 | 0.51 | 0.24 | 0.38 |
| 5 | 115811220 | rs423834 | 0.167 | -0.773 | 0.332 | 0.573 | -0.005 | -0.008 | -0.008 | 0.007 | 0.37 | 0.67 | 0.4 | 0.51 | 0.34 | 0.4 |
| 5 | 115814106 | rs34967 | -1.487 | -0.751 | 0.282 | 0.606 | -0.053 | 0.013 | 0.013 | 0 | 0.65 | 0.8 | 0.62 | 0.78 | 0.55 | 0.62 |
| 5 | 115815607 | rs469035 | -1.295 | -0.769 | 0.32 | 0.557 | 0.005 | -0.004 | -0.004 | 0 | 0.63 | 0.42 | 0.57 | 0.77 | 0.5 | 0.62 |
| 5 | 115816823 | rs2278264 | -0.043 | -0.829 | 0.297 | 0.596 | 0.007 | -0.002 | -0.002 | 0 | 0.39 | 0.26 | 0.31 | 0.5 | 0.33 | 0.42 |
| 5 | 115821215 | rs17139903 | -1.076 | -0.742 | 0.387 | 0.65 | -0.024 | 0.03 | 0.03 | 0 | 0.22 | 0.38 | 0.2 | 0.26 | 0.18 | 0.14 |
| 5 | 115822392 | rs2059874 | -0.747 | -0.69 | -0.646 | 0.751 | 0.036 | -0.04 | -0.04 | 0 | 0.19 | 0.08 | 0.16 | 0.26 | 0.14 | 0.14 |
| 5 | 115822858 | rs17433317 | -0.811 | -0.477 | -0.678 | 0.625 | 0.015 | 0.031 | 0.031 | 0 | 0.09 | 0.03 | 0.05 | 0.01 | 0.04 | 0.08 |
| 5 | 115826299 | rs11956022 | -1.099 | -0.332 | -0.297 | 0.907 | -0.021 | 0.027 | 0.027 | 0.115 | 0.26 | 0.44 | 0.23 | 0.26 | 0.23 | 0.17 |
| 5 | 115827356 | rs154599 | 0.897 | -0.655 | -1.404 | 0.819 | 0.037 | -0.009 | -0.009 | 0 | 0.74 | 0.87 | 0.8 | 0.8 | 0.82 | 0.81 |
| 5 | 115827945 | rs154601 | 0.931 | 0.081 | -1.272 | 0.904 | -0.003 | -0.025 | -0.025 | 1.283 | 0.92 | 0.98 | 0.98 | 0.92 | 0.94 | 0.93 |

| | | | | | | | | | | | | | | | | |
|---|---|---|---|---|---|---|---|---|---|---|---|---|---|---|---|---|
| 5 | 115830469 | rs1014643 | -0.674 | -0.249 | -1.723 | 0.724 | 0.024 | -0.028 | -0.028 | 0 | 0.24 | 0.11 | 0.24 | 0.4 | 0.21 | 0.28 |
| 5 | 115831092 | rs32972 | -0.277 | 0.335 | -1.681 | -0.248 | 0.001 | 0.03 | 0.03 | 0 | 0.46 | 0.35 | 0.4 | 0.32 | 0.48 | 0.39 |
| 5 | 115831305 | rs32973 | 0.022 | 0.16 | -1.93 | -0.821 | 0.003 | 0.004 | 0.004 | 0 | 0.64 | 0.71 | 0.59 | 0.37 | 0.64 | 0.53 |
| 5 | 115831330 | rs2287686 | 0.005 | 0.702 | -2.064 | -0.827 | -0.022 | -0.008 | -0.008 | 0 | 0.25 | 0.06 | 0.3 | 0.21 | 0.2 | 0.3 |
| 5 | 115831574 | rs1991504 | -0.354 | 0.973 | -1.617 | -0.811 | -0.014 | 0.023 | 0.023 | 0.008 | 0.3 | 0.16 | 0.37 | 0.21 | 0.3 | 0.29 |
| 5 | 115831873 | rs32974 | 0.21 | 0.934 | -1.4 | -0.815 | 0.001 | 0.007 | 0.007 | 0 | 0.13 | 0.11 | 0.1 | 0.07 | 0.17 | 0.06 |
| 5 | 115832056 | rs41099 | -0.763 | 0.619 | -1.05 | -0.994 | -0.02 | -0.051 | -0.051 | 0.037 | 0.23 | 0.31 | 0.14 | 0.07 | 0.2 | 0.09 |
| 5 | 115833644 | rs258014 | 1.391 | 0.618 | -1.165 | -1.011 | 0 | 0 | 0 | 0.06 | 0.36 | 0.35 | 0.44 | 0.64 | 0.38 | 0.51 |
| 5 | 115834599 | rs258016 | 0.486 | 0.629 | -1.434 | -0.723 | -0.002 | -0.026 | -0.026 | 0 | 0.78 | 0.71 | 0.86 | 0.92 | 0.76 | 0.91 |
| 5 | 115837772 | rs154466 | -0.184 | 1.412 | -1.964 | -0.285 | -0.042 | 0.051 | 0.051 | 0 | 0.81 | 0.97 | 0.86 | 0.92 | 0.76 | 0.91 |
| 5 | 115837789 | rs6885854 | -0.622 | 1.548 | -1.203 | 0.57 | -0.016 | -0.026 | -0.026 | 0 | 0.22 | 0.15 | 0.31 | 0.25 | 0.22 | 0.28 |
| 5 | 115840528 | rs153644 | -0.638 | 1.239 | -1.43 | 0.535 | -0.049 | -0.047 | -0.047 | 1.643 | 0.22 | 0.03 | 0.28 | 0.3 | 0.22 | 0.28 |
| 5 | 115842768 | rs153599 | -1.13 | 0.893 | -1.081 | 0.365 | 0.001 | 0.009 | 0.009 | 0.095 | 0.37 | 0.73 | 0.4 | 0.38 | 0.31 | 0.33 |
| 5 | 115845251 | rs7701234 | -1.441 | 0.213 | -0.472 | -0.005 | -0.037 | -0.056 | -0.056 | 0 | 0.2 | 0.63 | 0.12 | 0.06 | 0.12 | 0.22 |
| 5 | 115845585 | rs7721479 | -1.79 | 0.765 | -1.023 | 0.358 | -0.002 | -0.021 | -0.021 | 0 | 0.38 | 0.82 | 0.46 | 0.4 | 0.36 | 0.53 |
| 5 | 115845811 | rs918425 | -1.496 | 0.694 | -0.633 | 0.355 | 0 | 0.002 | 0.002 | 0 | 0.28 | 0.7 | 0.37 | 0.38 | 0.28 | 0.39 |
| 5 | 115847241 | rs4920916 | -1.777 | 0.171 | -0.428 | 0.04 | -0.021 | -0.039 | -0.039 | 3.442 | 0.24 | 0.69 | 0.21 | 0.08 | 0.18 | 0.23 |
| 5 | 115848046 | rs6897947 | -2.151 | 0.467 | -0.577 | 0.138 | 0.032 | 0.021 | 0.021 | 0.044 | 0.39 | 0.74 | 0.45 | 0.33 | 0.35 | 0.49 |
| 5 | 115849242 | rs6882841 | -1.86 | 1.045 | 0.575 | 0.16 | -0.009 | 0.001 | 0.001 | 0.412 | 0.27 | 0.51 | 0.38 | 0.27 | 0.27 | 0.34 |
| 5 | 115851237 | rs11956300 | 0.102 | 1.029 | 0.555 | 0.241 | 0.011 | -0.032 | -0.032 | 0 | 0.11 | 0.04 | 0.21 | 0.25 | 0.11 | 0.23 |
| 5 | 115852553 | rs9327005 | -0.836 | 0.929 | 1.017 | 0.244 | -0.021 | 0.012 | 0.012 | 0 | 0.09 | 0.24 | 0.04 | 0.08 | 0.06 | 0.14 |
| 5 | 115852960 | rs6860340 | -1.692 | 1.315 | 0.901 | 0.478 | 0.001 | 0.011 | 0.011 | 0 | 0.28 | 0.66 | 0.31 | 0.32 | 0.28 | 0.3 |
| 5 | 115857491 | rs154577 | 1.365 | 0.872 | -0.201 | -0.705 | -0.047 | -0.093 | -0.093 | 0 | 0.39 | 0.07 | 0.39 | 0.59 | 0.38 | 0.33 |
| 5 | 115861057 | rs7702266 | 0.434 | 0.066 | 0.581 | -1.141 | 0.006 | 0 | 0 | 0.03 | 0.12 | 0.1 | 0.07 | 0 | 0.09 | 0.09 |
| 5 | 115861632 | rs154576 | 1.267 | 0.11 | 0.729 | -1.205 | 0.001 | -0.048 | -0.048 | 0 | 0.67 | 0.34 | 0.77 | 0.99 | 0.73 | 0.84 |
| 5 | 115868611 | rs17140038 | -1.756 | 0.087 | 0.749 | -0.515 | 0.032 | -0.039 | -0.039 | 0 | 0.1 | 0.3 | 0.11 | 0.02 | 0.09 | 0.03 |
| 5 | 115877786 | rs11746151 | -1.624 | 0.193 | 0.675 | -0.567 | 0.046 | -0.039 | -0.039 | 0 | 0.08 | 0.26 | 0.05 | 0 | 0.08 | 0.03 |
| 5 | 115879239 | rs17140078 | -0.431 | -0.809 | 0.068 | -1.196 | 0.011 | -0.029 | -0.029 | 0 | 0.31 | 0.27 | 0.14 | 0 | 0.17 | 0.16 |
| 5 | 115880797 | rs439891 | -0.462 | -0.496 | 0.34 | -0.756 | 0.03 | 0.08 | 0.08 | 0.075 | 0.18 | 0.29 | 0.36 | 0.31 | 0.23 | 0.35 |
| 5 | 115881429 | rs10077439 | -0.315 | -0.7 | 0.09 | -1.069 | 0.02 | -0.001 | -0.001 | 0 | 0.34 | 0.27 | 0.18 | 0 | 0.23 | 0.19 |
| 5 | 115885282 | rs254226 | 0.453 | -0.665 | 0.73 | -0.314 | -0.007 | -0.019 | -0.019 | 0 | 0.77 | 0.59 | 0.79 | 0.86 | 0.76 | 0.68 |
| 5 | 115885574 | rs254227 | 0.142 | -0.689 | 0.72 | -0.32 | -0.008 | -0.01 | -0.01 | 0 | 0.79 | 0.62 | 0.8 | 0.85 | 0.79 | 0.68 |
| 5 | 115885858 | rs254230 | 1.261 | -0.696 | 0.854 | -0.16 | 0.001 | -0.003 | -0.003 | 2.007 | 0.64 | 0.34 | 0.55 | 0.62 | 0.6 | 0.46 |
| 5 | 115887109 | rs254232 | 2.019 | -1.204 | 0.547 | -0.327 | 0.018 | 0.016 | 0.016 | 0 | 0.47 | 0.08 | 0.51 | 0.63 | 0.56 | 0.4 |
| 5 | 115888522 | rs7716209 | -0.489 | -1.083 | 0.724 | -0.324 | -0.007 | -0.05 | -0.05 | 0 | 0.09 | 0.21 | 0.05 | 0 | 0.07 | 0.03 |
| 5 | 115891083 | rs254079 | 1.003 | -1.079 | 0.376 | -0.215 | -0.02 | -0.076 | -0.076 | 0 | 0.64 | 0.16 | 0.54 | 0.56 | 0.61 | 0.46 |
| 5 | 115892410 | rs153640 | 0.83 | -0.866 | 0.488 | -0.152 | -0.01 | -0.03 | -0.03 | 0.09 | 0.7 | 0.34 | 0.74 | 0.86 | 0.74 | 0.64 |
| 5 | 115894579 | rs254209 | 0.717 | -0.531 | 0.649 | 0.04 | -0.014 | -0.021 | -0.021 | 0.372 | 0.62 | 0.37 | 0.75 | 0.85 | 0.75 | 0.62 |
| 5 | 115896636 | rs5016403 | -1.251 | 0.256 | 1.437 | 0.64 | 0.024 | 0.072 | 0.072 | 0.828 | 0.14 | 0.22 | 0.26 | 0.3 | 0.22 | 0.22 |
| 5 | 115899132 | rs9327007 | -0.577 | 0.272 | 1.411 | 0.65 | 0.026 | 0.089 | 0.089 | 1.687 | 0.12 | 0.19 | 0.24 | 0.3 | 0.19 | 0.21 |
| 5 | 115910233 | rs1862372 | -0.586 | -0.18 | 1.32 | 0.193 | -0.016 | -0.031 | -0.031 | 0 | 0.12 | 0.04 | 0.23 | 0.23 | 0.18 | 0.2 |
| 5 | 115922669 | rs9327012 | -1.709 | -0.217 | 1.533 | 0.135 | 0.002 | -0.009 | -0.009 | 0 | 0.24 | 0.5 | 0.3 | 0.22 | 0.24 | 0.24 |
| 5 | 115939341 | rs682684 | -1.458 | -0.007 | 0.646 | 0.665 | 0.015 | 0.1 | 0.1 | 0.046 | 0.2 | 0.42 | 0.28 | 0.4 | 0.22 | 0.22 |
| 5 | 115939896 | rs17140201 | -0.574 | -0.034 | 0.09 | 0.851 | 0 | 0.03 | 0.03 | 0.77 | 0.11 | 0.06 | 0.22 | 0.4 | 0.16 | 0.19 |
| 5 | 115940710 | rs684730 | -1.012 | -0.709 | 0.542 | -0.179 | -0.051 | -0.131 | -0.131 | 1.164 | 0.18 | 0.56 | 0.09 | 0 | 0.11 | 0.06 |
| 5 | 115942140 | rs17140205 | 0.199 | -0.897 | -0.613 | -0.089 | 0.008 | -0.032 | -0.032 | 0.495 | 0.35 | 0.22 | 0.43 | 0.54 | 0.34 | 0.6 |
| 5 | 115943194 | rs607930 | -0.901 | -1.484 | -0.911 | -0.442 | -0.011 | -0.052 | -0.052 | 0 | 0.6 | 0.87 | 0.54 | 0.53 | 0.47 | 0.72 |
| 5 | 115944749 | rs9327014 | 0.071 | -1.337 | -0.751 | -0.21 | 0 | -0.002 | -0.002 | 0.058 | 0.34 | 0.31 | 0.42 | 0.52 | 0.3 | 0.53 |
| 5 | 115947220 | rs589869 | 0.162 | -1.176 | -1.836 | -0.398 | -0.025 | -0.077 | -0.077 | 0 | 0.41 | 0.09 | 0.43 | 0.47 | 0.54 | 0.4 |
| 5 | 115947887 | rs592965 | 0.194 | -1.51 | -1.991 | -0.909 | 0.026 | 0.079 | 0.079 | 0 | 0.68 | 0.91 | 0.82 | 0.84 | 0.75 | 0.68 |
| 5 | 115948109 | rs632993 | 0.456 | -0.223 | -1.514 | -0.576 | 0.006 | -0.002 | -0.002 | 1.118 | 0.46 | 0.58 | 0.49 | 0.57 | 0.46 | 0.4 |
| 5 | 115949578 | rs1873853 | -0.459 | 0.273 | -0.868 | -0.835 | 0.015 | -0.02 | -0.02 | 0 | 0.25 | 0.28 | 0.2 | 0.08 | 0.35 | 0.22 |
| 5 | 115952274 | rs680878 | 1.416 | 0.128 | -1.493 | -0.924 | 0.01 | -0.044 | -0.044 | 0 | 0.4 | 0.1 | 0.5 | 0.6 | 0.39 | 0.52 |
| 5 | 115954404 | rs2546511 | 0.408 | 0.145 | -0.908 | -0.927 | 0.003 | -0.005 | -0.005 | 0.023 | 0.68 | 0.3 | 0.76 | 0.84 | 0.61 | 0.76 |
| 5 | 115957300 | rs6896713 | 0.892 | 0.848 | -0.911 | -1.076 | -0.045 | -0.003 | -0.003 | 0 | 0.14 | 0.08 | 0.13 | 0.09 | 0.25 | 0.22 |
| 5 | 115958978 | rs680404 | 0.517 | 0.483 | -0.593 | -1.066 | 0.007 | 0.011 | 0.011 | 0.27 | 0.63 | 0.32 | 0.7 | 0.79 | 0.72 | 0.76 |
| 5 | 115961967 | rs635958 | 0.901 | 0.337 | -0.68 | -0.791 | 0.005 | 0.055 | 0.055 | 0.427 | 0.51 | 0.24 | 0.47 | 0.69 | 0.52 | 0.56 |
| 5 | 115966256 | rs10463685 | -0.099 | 0.43 | -0.483 | -0.735 | -0.001 | 0 | 0 | 0 | 0.18 | 0.21 | 0.23 | 0.19 | 0.16 | 0.18 |
| 5 | 115970828 | rs595117 | -0.221 | 0.513 | -0.523 | -0.735 | 0.019 | -0.059 | -0.059 | 4.763 | 0.7 | 0.47 | 0.71 | 0.89 | 0.69 | 0.73 |
| 5 | 115973077 | rs881020 | -1.034 | 0.194 | -0.227 | -0.283 | -0.005 | -0.006 | -0.006 | 0.068 | 0.23 | 0.26 | 0.25 | 0.19 | 0.2 | 0.18 |
| 5 | 115975656 | rs6898653 | -0.812 | 0.394 | 0.236 | -0.321 | 0 | -0.031 | -0.031 | 0 | 0.25 | 0.43 | 0.18 | 0.19 | 0.21 | 0.22 |
| 5 | 115975963 | rs11957113 | -0.712 | 0.248 | 0.22 | -0.525 | -0.005 | -0.015 | -0.015 | 0 | 0.57 | 0.56 | 0.49 | 0.32 | 0.58 | 0.48 |
| 5 | 115976839 | rs622996 | 1.194 | 0.468 | 0.722 | -0.602 | 0.02 | -0.066 | -0.066 | 0.216 | 0.89 | 0.71 | 0.95 | 1 | 0.92 | 0.91 |
| 5 | 115979013 | rs643995 | 0.528 | 0.652 | 1.134 | -0.735 | 0.018 | -0.034 | -0.034 | 0 | 0.67 | 0.46 | 0.68 | 0.74 | 0.65 | 0.74 |
| 5 | 115979152 | rs10455011 | 0.616 | 0.64 | 0.673 | -0.73 | -0.08 | 0.014 | 0.014 | 0 | 0.17 | 0.02 | 0.15 | 0.12 | 0.23 | 0.18 |
| 5 | 115987168 | rs6594974 | -0.404 | 0.648 | 1.256 | -0.831 | 0.03 | -0.054 | -0.054 | 0 | 0.24 | 0.4 | 0.19 | 0.11 | 0.28 | 0.18 |
| 5 | 115988836 | rs12189229 | -0.207 | 0.67 | 1.107 | -0.854 | -0.027 | 0.027 | 0.027 | 0 | 0.23 | 0.19 | 0.18 | 0.13 | 0.29 | 0.22 |
| 5 | 115990628 | rs7707446 | 0.021 | 0.408 | 1.176 | -0.619 | 0.003 | -0.023 | -0.023 | 0 | 0.71 | 0.64 | 0.73 | 0.75 | 0.72 | 0.74 |
| 5 | 115994775 | rs1948429 | 0.215 | 0.377 | 1.13 | -0.254 | 0.002 | -0.002 | -0.002 | 0 | 0.67 | 0.66 | 0.7 | 0.64 | 0.67 | 0.64 |

| | | | | | | | | | | | | | | | | |
|---|---|---|---|---|---|---|---|---|---|---|---|---|---|---|---|---|
| 5 | 115998514 | rs1353046 | 0.108 | 0.448 | 1.197 | -0.284 | 0.008 | -0.008 | -0.008 | 0 | 0.69 | 0.63 | 0.7 | 0.68 | 0.65 | 0.67 |
| 5 | 115999520 | rs1389164 | 0.048 | 0.441 | 1.164 | -0.294 | 0.004 | -0.003 | -0.003 | 0 | 0.69 | 0.66 | 0.71 | 0.67 | 0.67 | 0.67 |
| 5 | 116011292 | rs12521259 | 0.555 | 0.263 | 0.998 | -0.39 | -0.001 | -0.003 | -0.003 | 0 | 0.62 | 0.41 | 0.68 | 0.67 | 0.63 | 0.66 |
| 5 | 116019372 | rs7719129 | 0.172 | 0.1 | 1.212 | -0.298 | 0.02 | 0.003 | 0.003 | 0 | 0.68 | 0.43 | 0.69 | 0.6 | 0.65 | 0.68 |
| 5 | 116019417 | rs10072088 | -0.018 | -0.041 | 1.138 | -0.547 | -0.001 | -0.001 | -0.001 | 0 | 0.24 | 0.25 | 0.21 | 0.1 | 0.26 | 0.19 |
| 5 | 116019604 | rs10040214 | 1.59 | -0.074 | 0.898 | -0.458 | 0.009 | -0.058 | -0.058 | 1.142 | 0.38 | 0.08 | 0.47 | 0.52 | 0.37 | 0.45 |
| 5 | 116019920 | rs17140312 | 1.358 | 0.29 | 1.207 | -0.256 | 0.047 | 0.004 | 0.004 | 0.416 | 0.44 | 0.18 | 0.48 | 0.52 | 0.39 | 0.48 |
| 5 | 116020849 | rs2964663 | 1.053 | 0.387 | 1.487 | -0.327 | 0.053 | -0.078 | -0.078 | 0 | 0.88 | 0.5 | 0.92 | 1 | 0.92 | 0.92 |
| 5 | 116030080 | rs2933608 | -0.741 | 0.259 | 1.171 | -0.324 | 0.002 | 0.016 | 0.016 | 1.462 | 0.48 | 0.31 | 0.45 | 0.48 | 0.53 | 0.48 |
| 5 | 116049340 | rs2933630 | -0.866 | 0.179 | 1.569 | -0.381 | 0.009 | 0.011 | 0.011 | 0.891 | 0.51 | 0.57 | 0.49 | 0.48 | 0.58 | 0.52 |
| 5 | 116055123 | rs2933624 | 0.485 | 0.295 | 1.295 | -0.395 | 0.002 | 0.014 | 0.014 | 2.204 | 0.24 | 0.3 | 0.26 | 0.44 | 0.28 | 0.33 |
| 5 | 116055533 | rs2933622 | 0.213 | 0.276 | 1.278 | -0.404 | 0.001 | 0.022 | 0.022 | 1.016 | 0.27 | 0.36 | 0.26 | 0.45 | 0.28 | 0.33 |
| 5 | 116064934 | rs2909823 | 1.766 | 0.216 | 1.194 | -0.47 | 0.029 | -0.009 | -0.009 | 0 | 0.4 | 0.17 | 0.46 | 0.52 | 0.35 | 0.43 |
| 5 | 116066910 | rs1076985 | -0.325 | 0.48 | 0.692 | -0.389 | -0.039 | 0.104 | 0.104 | 0 | 0.27 | 0.11 | 0.21 | 0.04 | 0.3 | 0.17 |
| 5 | 116066998 | rs2909821 | 0.163 | 0.74 | 0.827 | 0.008 | 0.003 | 0.018 | 0.018 | 1.922 | 0.25 | 0.35 | 0.26 | 0.48 | 0.27 | 0.34 |
| 5 | 116069685 | rs2964690 | 0.678 | 0.873 | 0.618 | 0.125 | 0.005 | 0.017 | 0.017 | 0.989 | 0.21 | 0.26 | 0.24 | 0.4 | 0.24 | 0.24 |
| 5 | 116070510 | rs10056291 | 0.189 | 1.088 | -0.071 | 0.12 | -0.093 | 0.054 | 0.054 | 0 | 0.18 | 0.02 | 0.2 | 0.04 | 0.29 | 0.15 |
| 5 | 116080029 | rs26674 | 0.547 | 0.727 | 0.285 | 0.17 | -0.002 | -0.007 | -0.007 | 0 | 0.63 | 0.32 | 0.69 | 0.52 | 0.67 | 0.63 |
| 5 | 116080067 | rs26673 | 0.559 | 0.728 | 0.285 | 0.17 | -0.001 | -0.007 | -0.007 | 0 | 0.63 | 0.32 | 0.69 | 0.52 | 0.67 | 0.63 |
| 5 | 116088403 | rs26670 | -0.177 | 0.964 | 0.034 | 0.231 | 0.005 | 0.018 | 0.018 | 0.476 | 0.27 | 0.39 | 0.25 | 0.39 | 0.27 | 0.31 |
| 5 | 116088853 | rs2068933 | -0.682 | 1.222 | -0.713 | -0.498 | -0.055 | 0.05 | 0.05 | 0 | 0.22 | 0.08 | 0.24 | 0.05 | 0.35 | 0.18 |
| 5 | 116090613 | rs6883782 | -1.132 | 1.117 | -0.871 | -0.427 | -0.039 | 0.013 | 0.013 | 0 | 0.19 | 0.12 | 0.21 | 0.09 | 0.31 | 0.22 |
| 5 | 116098004 | rs2933639 | -1.026 | 0.835 | -0.541 | -0.452 | 0.025 | 0.008 | 0.008 | 0.241 | 0.52 | 0.7 | 0.48 | 0.44 | 0.61 | 0.55 |
| 5 | 116099168 | rs2937054 | 0.915 | 0.888 | -0.441 | -0.541 | 0.004 | -0.016 | -0.016 | 0 | 0.19 | 0.11 | 0.15 | 0.3 | 0.17 | 0.23 |
| 5 | 116101059 | rs2933643 | -1.257 | 0.807 | -0.259 | -0.562 | 0.022 | 0.002 | 0.002 | 0.299 | 0.52 | 0.71 | 0.47 | 0.45 | 0.61 | 0.56 |
| 5 | 116106226 | rs4920935 | -0.256 | 1.036 | -0.368 | -0.522 | -0.037 | -0.008 | -0.008 | 0.004 | 0.19 | 0.07 | 0.18 | 0.12 | 0.28 | 0.27 |
| 5 | 116107372 | rs2242410 | 0.945 | 0.557 | -0.577 | -0.544 | 0.013 | -0.024 | -0.024 | 0 | 0.23 | 0.09 | 0.16 | 0.31 | 0.18 | 0.25 |
| 5 | 116109222 | rs2933646 | -0.835 | 0.382 | -0.507 | -0.54 | -0.008 | -0.005 | -0.005 | 0 | 0.56 | 0.79 | 0.47 | 0.42 | 0.57 | 0.56 |
| 5 | 116124578 | rs924453 | 0.167 | -0.309 | -0.246 | -0.545 | -0.013 | -0.003 | -0.003 | 0.005 | 0.61 | 0.54 | 0.79 | 0.62 | 0.73 | 0.6 |
| 5 | 116125078 | rs3097167 | -0.831 | -0.31 | -0.692 | -0.527 | 0.045 | -0.038 | -0.038 | 0 | 0.76 | 0.94 | 0.85 | 0.68 | 0.84 | 0.68 |
| 5 | 116131757 | rs2964671 | -1.07 | 0.286 | -1.071 | -0.983 | -0.059 | 0.166 | 0.166 | 0.447 | 0.48 | 0.92 | 0.49 | 0.17 | 0.57 | 0.35 |
| 5 | 116133051 | rs2133759 | -0.65 | 0.253 | -0.096 | -1.213 | 0 | -0.014 | -0.014 | 0.036 | 0.33 | 0.35 | 0.26 | 0.06 | 0.45 | 0.3 |
| 5 | 116138341 | rs2913600 | -0.922 | 0.163 | -0.091 | -1.281 | 0.002 | -0.027 | -0.027 | 0 | 0.37 | 0.41 | 0.29 | 0.06 | 0.49 | 0.33 |
| 5 | 116140038 | rs2937067 | -1.641 | 0.115 | -0.18 | -1.307 | -0.061 | 0.291 | 0.291 | 0.031 | 0.48 | 0.92 | 0.37 | 0.06 | 0.56 | 0.35 |
| 5 | 116143444 | rs13181946 | -1.264 | 0.225 | 0.477 | -1.475 | 0.018 | -0.094 | -0.094 | 0 | 0.35 | 0.51 | 0.3 | 0.05 | 0.44 | 0.3 |
| 5 | 116146765 | rs10043156 | -1.454 | 0.371 | 0.317 | -1.512 | -0.009 | 0.094 | 0.094 | 1.457 | 0.36 | 0.78 | 0.3 | 0.05 | 0.42 | 0.29 |
| 5 | 116149999 | rs2034849 | 2.011 | 0.29 | -0.043 | -1.545 | -0.081 | -0.147 | -0.147 | 0 | 0.27 | 0.03 | 0.37 | 0.52 | 0.28 | 0.31 |
| 5 | 116150016 | rs9885302 | -0.998 | 0.868 | 1.138 | -1.561 | 0.006 | -0.02 | -0.02 | 0 | 0.24 | 0.25 | 0.21 | 0.05 | 0.33 | 0.25 |
| 5 | 116153277 | rs698377 | -0.169 | 0.772 | 0.817 | -1.451 | -0.001 | -0.002 | -0.002 | 0.926 | 0.89 | 0.89 | 0.83 | 0.88 | 0.88 | 0.95 |
| 5 | 116156020 | rs725151 | -0.369 | 0.421 | 0.847 | -1.397 | -0.089 | -0.001 | -0.001 | 0.138 | 0.15 | 0.01 | 0.17 | 0.05 | 0.3 | 0.16 |
| 5 | 116160862 | rs153577 | 0.061 | 0.374 | 1.061 | -1.313 | -0.001 | -0.008 | -0.008 | 0 | 0.1 | 0.11 | 0.06 | 0.01 | 0.12 | 0.06 |
| 5 | 116160951 | rs6870730 | -1.96 | 0.308 | 1.26 | -1.377 | -0.003 | 0.029 | 0.029 | 0.594 | 0.33 | 0.73 | 0.38 | 0.16 | 0.36 | 0.25 |
| 5 | 116165532 | rs4920937 | -1.684 | 0.343 | 1.406 | -1.368 | 0.011 | -0.032 | -0.032 | 0 | 0.31 | 0.51 | 0.36 | 0.16 | 0.36 | 0.25 |
| 5 | 116176431 | rs6860399 | -0.997 | 0.154 | 1.488 | -1.327 | 0.013 | -0.018 | -0.018 | 0 | 0.08 | 0.36 | 0.04 | 0 | 0.03 | 0.01 |
| 5 | 116178814 | rs10069564 | -1.927 | 0.172 | 1.511 | -1.33 | -0.007 | 0.048 | 0.048 | 0.99 | 0.33 | 0.8 | 0.37 | 0.16 | 0.34 | 0.25 |
| 5 | 116180180 | rs6882638 | -1.388 | 0.083 | 1.607 | -1.293 | 0.027 | -0.017 | -0.017 | 0 | 0.25 | 0.45 | 0.32 | 0.16 | 0.31 | 0.24 |
| 5 | 116183712 | rs253897 | -2.156 | 0.044 | 1.475 | -1.394 | -0.002 | 0.062 | 0.062 | 0 | 0.41 | 0.7 | 0.42 | 0.15 | 0.45 | 0.32 |
| 5 | 116184752 | rs12521549 | -0.836 | 0.296 | 1.031 | -1.262 | 0.015 | -0.005 | -0.005 | 0 | 0.21 | 0.26 | 0.28 | 0.14 | 0.27 | 0.17 |
| 5 | 116185523 | rs253896 | -0.746 | 0.243 | 0.335 | -1.075 | 0.021 | -0.036 | -0.036 | 0.002 | 0.3 | 0.58 | 0.36 | 0.15 | 0.39 | 0.29 |
| 5 | 116188453 | rs13179097 | -1.101 | -0.234 | 0.18 | -0.754 | -0.029 | -0.132 | -0.132 | 0.001 | 0.18 | 0.46 | 0.12 | 0 | 0.14 | 0.05 |
| 5 | 116197865 | rs2080867 | -0.448 | -0.149 | -0.576 | -0.176 | -0.013 | -0.014 | -0.014 | 0.126 | 0.32 | 0.48 | 0.35 | 0.24 | 0.28 | 0.4 |
| 5 | 116199838 | rs10058021 | -0.139 | 0.111 | -1.226 | -0.095 | -0.003 | -0.003 | -0.003 | 0 | 0.24 | 0.23 | 0.3 | 0.25 | 0.26 | 0.31 |
| 5 | 116200854 | rs698384 | 0.058 | 0.634 | -1.996 | 0.379 | -0.132 | -0.126 | -0.126 | 0.311 | 0.36 | 0.08 | 0.38 | 0.58 | 0.47 | 0.43 |
| 5 | 116208147 | rs4607382 | -0.099 | 0.297 | -1.617 | 0.16 | -0.001 | 0.027 | 0.027 | 0 | 0.42 | 0.31 | 0.45 | 0.75 | 0.58 | 0.55 |
| 5 | 116211296 | rs6859942 | 0.056 | 0.462 | -1.74 | -0.068 | -0.034 | 0.022 | 0.022 | 0 | 0.34 | 0.25 | 0.4 | 0.77 | 0.51 | 0.53 |
| 5 | 116218493 | rs13161202 | -0.255 | 0.135 | -1.765 | 0.74 | -0.033 | -0.048 | -0.048 | 0 | 0.38 | 0.2 | 0.4 | 0.59 | 0.56 | 0.55 |
| 5 | 116218547 | rs13161243 | -0.256 | 0.118 | -1.543 | 0.732 | -0.025 | -0.037 | -0.037 | 0.46 | 0.38 | 0.23 | 0.4 | 0.59 | 0.56 | 0.56 |
| 5 | 116219459 | rs6897267 | -0.34 | -1.129 | -0.906 | 0.013 | 0.012 | 0.007 | 0.007 | 0.02 | 0.56 | 0.65 | 0.63 | 0.67 | 0.66 | 0.7 |
| 5 | 116219745 | rs6876340 | 0.366 | -1.072 | -0.741 | 0.066 | -0.021 | -0.035 | -0.035 | 0.537 | 0.39 | 0.26 | 0.41 | 0.6 | 0.56 | 0.56 |
| 5 | 116222219 | rs1972623 | -0.188 | -1.744 | -0.289 | -0.383 | 0.01 | 0.011 | 0.011 | 0 | 0.51 | 0.5 | 0.52 | 0.21 | 0.36 | 0.38 |
| 5 | 116223322 | rs2034850 | 0.089 | -2.303 | -0.456 | -0.681 | -0.004 | -0.011 | -0.011 | 0.05 | 0.26 | 0.26 | 0.27 | 0.07 | 0.15 | 0.19 |
| 5 | 116225123 | rs1643889 | -0.555 | -1.81 | -0.306 | -0.425 | -0.003 | 0.068 | 0.068 | 0 | 0.88 | 0.76 | 0.93 | 0.8 | 0.92 | 0.93 |
| 5 | 116228312 | rs6594987 | -0.682 | -1.868 | -0.324 | -0.471 | -0.007 | -0.016 | -0.016 | 0 | 0.79 | 0.77 | 0.77 | 0.84 | 0.82 | 0.84 |
| 5 | 116231413 | rs1643897 | 0.552 | -1.435 | -0.442 | -0.67 | 0 | -0.005 | -0.005 | 0 | 0.61 | 0.47 | 0.63 | 0.8 | 0.68 | 0.8 |
| 5 | 116232263 | rs7718938 | -0.587 | -1.692 | -0.362 | -0.529 | -0.042 | 0.074 | 0.074 | 0 | 0.14 | 0.28 | 0.04 | 0.19 | 0.05 | 0.06 |
| 5 | 116233236 | rs2112342 | -0.084 | -0.506 | -0.317 | -0.64 | 0.008 | -0.012 | -0.012 | 0 | 0.71 | 0.72 | 0.64 | 0.8 | 0.63 | 0.8 |
| 5 | 116234449 | rs1366156 | 0.667 | -1.184 | -0.884 | -0.497 | -0.002 | 0.029 | 0.029 | 0 | 0.07 | 0.05 | 0.03 | 0.19 | 0.04 | 0.05 |
| 5 | 116234868 | rs10053736 | 1.139 | -0.953 | -1.056 | -0.495 | -0.001 | -0.002 | -0.002 | 0 | 0.14 | 0.17 | 0.28 | 0.09 | 0.14 | 0.21 |
| 5 | 116245826 | rs712623 | 1.437 | -0.733 | -1.382 | -0.323 | -0.14 | -0.124 | -0.124 | 0 | 0.37 | 0.08 | 0.3 | 0.52 | 0.49 | 0.52 |

| | | | | | | | | | | | | | | | | |
|---|---|---|---|---|---|---|---|---|---|---|---|---|---|---|---|---|
| 5 | 116248504 | rs712624 | 1.616 | -0.525 | -1.284 | -0.128 | -0.108 | -0.114 | -0.114 | 0 | 0.33 | 0.08 | 0.26 | 0.5 | 0.4 | 0.48 |
| 5 | 116249713 | rs712626 | -0.582 | 0.067 | -0.408 | 0.253 | -0.009 | -0.008 | -0.008 | 0 | 0.13 | 0.07 | 0.18 | 0.24 | 0.18 | 0.08 |
| 5 | 116249992 | rs17140799 | 0.03 | -0.249 | -0.192 | 0.324 | 0.013 | 0.086 | 0.086 | 0 | 0.05 | 0.06 | 0.03 | 0.19 | 0.04 | 0.05 |
| 5 | 116252792 | rs1284011 | -0.115 | -0.523 | -0.072 | -0.332 | -0.035 | -0.045 | -0.045 | 0 | 0.78 | 0.55 | 0.81 | 0.84 | 0.84 | 0.85 |
| 5 | 116255188 | rs993764 | -0.367 | -0.399 | -0.196 | -0.662 | -0.003 | -0.043 | -0.043 | 0.031 | 0.2 | 0.19 | 0.18 | 0.02 | 0.19 | 0.05 |
| 5 | 116261947 | rs2416451 | 0.024 | -0.497 | -0.088 | -1.162 | -0.003 | -0.052 | -0.052 | 0.265 | 0.27 | 0.36 | 0.32 | 0.06 | 0.23 | 0.32 |
| 5 | 116264097 | rs1459089 | 0.619 | -0.748 | -0.284 | -0.636 | 0.004 | -0.03 | -0.03 | 0 | 0.21 | 0.18 | 0.19 | 0.46 | 0.16 | 0.14 |
| 5 | 116265087 | rs7706388 | -0.739 | -0.775 | -0.264 | -0.705 | 0.001 | -0.002 | -0.002 | 0 | 0.43 | 0.53 | 0.39 | 0.48 | 0.35 | 0.18 |
| 5 | 116275461 | rs749977 | -0.884 | -1.062 | -0.137 | -0.622 | 0.017 | 0.082 | 0.082 | 0 | 0.48 | 0.58 | 0.44 | 0.12 | 0.4 | 0.34 |
| 5 | 116276837 | rs4920862 | 0.3 | -1.286 | 0.031 | -0.799 | -0.025 | 0.064 | 0.064 | 0.409 | 0.31 | 0.14 | 0.36 | 0.11 | 0.34 | 0.32 |
| 5 | 116279182 | rs17140860 | -0.615 | -1.482 | 0.047 | -0.867 | -0.093 | -0.157 | -0.157 | 0 | 0.2 | 0.44 | 0.08 | 0 | 0.06 | 0.03 |
| 5 | 116279948 | rs10077835 | 0.175 | -0.887 | -0.014 | -0.288 | -0.02 | 0.055 | 0.055 | 0.127 | 0.23 | 0.4 | 0.21 | 0.44 | 0.14 | 0.12 |
| 5 | 116280537 | rs4920942 | -0.657 | 0.128 | -0.203 | 0.143 | -0.037 | -0.038 | -0.038 | 0 | 0.65 | 0.78 | 0.62 | 0.55 | 0.53 | 0.48 |
| 5 | 116291904 | rs7447358 | -0.45 | -0.217 | -0.446 | 0.019 | -0.055 | -0.032 | -0.032 | 0.433 | 0.63 | 0.82 | 0.49 | 0.41 | 0.45 | 0.37 |
| 5 | 116300347 | rs6866207 | 0.006 | -0.424 | 0.195 | 0.337 | 0.004 | 0 | 0 | 0.556 | 0.6 | 0.43 | 0.48 | 0.43 | 0.48 | 0.38 |
| 5 | 116319808 | rs4920944 | -0.298 | -0.516 | 0.318 | 0.433 | 0.001 | 0.001 | 0.001 | 0.023 | 0.63 | 0.59 | 0.52 | 0.47 | 0.49 | 0.4 |
| 5 | 116329875 | rs1459085 | -0.011 | -0.306 | -0.643 | 0.322 | -0.019 | -0.011 | -0.011 | 0 | 0.61 | 0.71 | 0.49 | 0.44 | 0.47 | 0.37 |
| 5 | 116355602 | rs1350574 | -0.258 | -0.022 | -0.851 | 0.353 | -0.081 | -0.035 | -0.035 | 0.854 | 0.65 | 0.84 | 0.51 | 0.4 | 0.48 | 0.4 |
| 5 | 116369531 | rs17140968 | -0.765 | -0.778 | 0.238 | 0.034 | -0.033 | -0.063 | -0.063 | 3.031 | 0.27 | 0.39 | 0.3 | 0.11 | 0.16 | 0.24 |
| 5 | 116375473 | rs930922 | -0.397 | -0.531 | 0.936 | 0.056 | -0.001 | 0.025 | 0.025 | 0.267 | 0.26 | 0.12 | 0.2 | 0.01 | 0.24 | 0.22 |
| 5 | 116376690 | rs2061448 | 0.615 | -0.586 | 1.18 | 0.068 | -0.006 | -0.06 | -0.06 | 0.195 | 0.7 | 0.58 | 0.72 | 0.84 | 0.73 | 0.73 |
| 5 | 116396722 | rs6595005 | -0.545 | -0.396 | 0.874 | 0.446 | -0.015 | -0.047 | -0.047 | 0 | 0.4 | 0.48 | 0.32 | 0.16 | 0.27 | 0.24 |
| 5 | 116401118 | rs6869550 | 0.154 | -0.326 | 0.892 | 0.461 | 0.009 | -0.02 | -0.02 | 0 | 0.3 | 0.26 | 0.19 | 0.05 | 0.16 | 0.2 |
| 5 | 116403002 | rs11241410 | 0.36 | -0.286 | 0.847 | 0.655 | 0.001 | 0.144 | 0.144 | 0 | 0.29 | 0.03 | 0.18 | 0.06 | 0.2 | 0.21 |
| 5 | 116403902 | rs6882410 | -1.815 | 0.417 | 2.21 | 1.63 | 0.028 | 0.001 | 0.001 | 0 | 0.08 | 0.23 | 0.07 | 0.06 | 0.09 | 0.09 |
| 5 | 116406053 | rs4266447 | -2.706 | 0.391 | 2.271 | 1.601 | 0.017 | 0.058 | 0.058 | 2.355 | 0.14 | 0.42 | 0.14 | 0.17 | 0.12 | 0.12 |
| 5 | 116413397 | rs12521805 | -1.594 | 0.652 | 2.264 | 1.709 | -0.006 | 0.08 | 0.08 | 5.712 | 0.07 | 0.24 | 0.08 | 0.11 | 0.03 | 0.03 |
| 5 | 116414003 | rs11241411 | 0.182 | 0.581 | 2.105 | 2.051 | 0.012 | -0.017 | -0.017 | 0 | 0.95 | 0.97 | 0.98 | 0.86 | 0.97 | 0.95 |
| 5 | 116416022 | rs7733755 | -2.55 | 0.623 | 2.179 | 1.911 | 0.025 | 0.066 | 0.066 | 4.109 | 0.13 | 0.56 | 0.16 | 0.17 | 0.12 | 0.1 |
| 5 | 116418850 | rs6888475 | 0.71 | 0.972 | 1.388 | 1.989 | 0 | 0.001 | 0.001 | 0 | 0.83 | 0.82 | 0.88 | 0.77 | 0.84 | 0.85 |
| 5 | 116419511 | rs6872138 | -1.315 | 0.861 | 1.189 | 1.947 | 0.036 | 0.076 | 0.076 | 0 | 0.05 | 0.18 | 0.06 | 0.15 | 0.09 | 0.07 |
| 5 | 116421862 | rs6595008 | -2.531 | 1.02 | 0.854 | 1.952 | 0.062 | 0.142 | 0.142 | 3.051 | 0.13 | 0.4 | 0.18 | 0.26 | 0.18 | 0.1 |
| 5 | 116423771 | rs7716590 | -1.525 | 0.581 | -0.307 | 1.507 | 0 | 0.037 | 0.037 | 4.163 | 0.06 | 0.07 | 0.15 | 0.22 | 0.07 | 0.04 |
| 5 | 116436198 | rs11241414 | 1.329 | 0.289 | 0.152 | 0.835 | -0.007 | -0.003 | -0.003 | 0 | 0.9 | 0.73 | 0.94 | 0.92 | 0.91 | 0.97 |
| 5 | 116438019 | rs12332394 | -0.817 | -0.326 | -0.454 | 0.709 | 0.021 | -0.009 | -0.009 | 0 | 0.14 | 0.04 | 0.04 | 0.17 | 0.11 | 0.14 |
| 5 | 116439083 | rs13356416 | -0.93 | -0.3 | -0.425 | 0.523 | -0.01 | -0.014 | -0.014 | 0.576 | 0.07 | 0.02 | 0.04 | 0.08 | 0.09 | 0.03 |
| 5 | 116446390 | rs4537091 | 1.97 | -0.463 | 0.48 | 0.477 | -0.01 | 0.033 | 0.033 | 8.133 | 0.9 | 0.57 | 0.94 | 0.92 | 0.95 | 0.98 |
| 5 | 116465497 | rs6862540 | -2.217 | -0.838 | 0.057 | -0.178 | -0.018 | -0.05 | -0.05 | 0 | 0.32 | 0.42 | 0.13 | 0.15 | 0.23 | 0.17 |
| 5 | 116468054 | rs10478330 | -1.047 | -0.786 | -0.073 | -0.135 | 0.012 | -0.007 | -0.007 | 0.037 | 0.08 | 0.04 | 0.04 | 0.08 | 0.06 | 0.02 |
| 5 | 116482190 | rs7710231 | -2.355 | -1.661 | -0.216 | -0.049 | 0.027 | 0.014 | 0.014 | 0 | 0.49 | 0.61 | 0.33 | 0.45 | 0.34 | 0.37 |
| 5 | 116485063 | rs6595012 | -0.842 | -0.694 | -0.626 | -1.426 | 0.001 | -0.052 | -0.052 | 2.364 | 0.29 | 0.38 | 0.27 | 0.07 | 0.31 | 0.12 |
| 5 | 116486485 | rs17141187 | -0.543 | -0.924 | -0.696 | -1.299 | 0.004 | -0.007 | -0.007 | 0.592 | 0.1 | 0.07 | 0.1 | 0.17 | 0.11 | 0.06 |
| 5 | 116486522 | rs2036023 | -1.31 | -0.805 | -0.47 | -1.996 | -0.005 | 0.019 | 0.019 | 0 | 0.46 | 0.65 | 0.34 | 0.08 | 0.43 | 0.25 |
| 5 | 116490357 | rs6897500 | -1.493 | -0.388 | -0.697 | -2.249 | -0.001 | 0.062 | 0.062 | 0 | 0.49 | 0.65 | 0.34 | 0.08 | 0.44 | 0.27 |
| 5 | 116497698 | rs6886341 | -1.321 | 0.152 | -0.573 | -2.279 | -0.001 | -0.084 | -0.084 | 0 | 0.47 | 0.57 | 0.33 | 0.08 | 0.42 | 0.27 |
| 5 | 116500367 | rs13154048 | 0.074 | -0.207 | -0.407 | -2.386 | 0.007 | -0.096 | -0.096 | 0.421 | 0.24 | 0.35 | 0.23 | 0.03 | 0.25 | 0.12 |
| 5 | 116500651 | rs7704021 | -1.33 | -0.23 | -0.414 | -2.415 | -0.001 | -0.079 | -0.079 | 0 | 0.42 | 0.48 | 0.3 | 0.08 | 0.37 | 0.24 |
| 5 | 116505127 | rs11241417 | -0.051 | -0.226 | -0.644 | -2.392 | -0.001 | 0.038 | 0.038 | 0 | 0.16 | 0.1 | 0.06 | 0.05 | 0.11 | 0.11 |
| 5 | 116513164 | rs7706447 | -0.005 | -0.292 | -0.654 | -2.421 | 0 | -0.04 | -0.04 | 0 | 0.07 | 0.18 | 0.04 | 0 | 0.06 | 0.03 |
| 5 | 116515829 | rs10067550 | -1.416 | -0.33 | -0.816 | -2.474 | 0 | -0.077 | -0.077 | 0 | 0.42 | 0.48 | 0.3 | 0.08 | 0.38 | 0.25 |
| 5 | 116517515 | rs13168914 | -0.131 | -0.058 | -0.694 | -2.268 | 0.011 | -0.078 | -0.078 | 0 | 0.23 | 0.31 | 0.23 | 0.04 | 0.26 | 0.14 |
| 5 | 116519696 | rs7711251 | -0.595 | 0.146 | -0.926 | -2.147 | 0.017 | -0.089 | -0.089 | 0 | 0.25 | 0.32 | 0.25 | 0.04 | 0.31 | 0.14 |
| 5 | 116525783 | rs1494826 | -0.505 | 0.056 | -1.09 | -1.93 | 0.005 | -0.009 | -0.009 | 0.167 | 0.14 | 0.1 | 0.15 | 0.17 | 0.15 | 0.13 |
| 5 | 116528042 | rs10068492 | 0.71 | 0.326 | -0.96 | -2.011 | -0.078 | 0.076 | 0.076 | 0 | 0.15 | 0.02 | 0.18 | 0.03 | 0.23 | 0.11 |
| 5 | 116529909 | rs7718941 | -0.957 | -0.029 | -0.472 | -1.606 | 0.001 | -0.126 | -0.126 | 0.924 | 0.19 | 0.32 | 0.09 | 0 | 0.15 | 0.13 |
| 5 | 116544172 | rs1021516 | -1.036 | 0.244 | -1.646 | -2.04 | -0.049 | 0.268 | 0.268 | 0 | 0.56 | 0.88 | 0.38 | 0.09 | 0.44 | 0.34 |
| 5 | 116549669 | rs7710998 | -0.075 | -0.264 | -1.072 | -2.472 | -0.007 | -0.129 | -0.129 | 1.149 | 0.29 | 0.38 | 0.15 | 0.01 | 0.25 | 0.19 |
| 5 | 116549781 | rs6595018 | -0.52 | -0.526 | -0.826 | -2.613 | -0.016 | -0.095 | -0.095 | 0 | 0.32 | 0.54 | 0.16 | 0.02 | 0.22 | 0.21 |
| 5 | 116553253 | rs11739552 | -0.275 | -0.45 | -1.05 | -2.564 | -0.001 | -0.014 | -0.014 | 1.194 | 0.12 | 0.12 | 0.07 | 0.05 | 0.09 | 0.1 |
| 5 | 116553503 | rs11746186 | -0.225 | -0.444 | -1.071 | -2.562 | -0.001 | -0.012 | -0.012 | 0 | 0.11 | 0.12 | 0.07 | 0.05 | 0.09 | 0.1 |
| 5 | 116563077 | rs6595023 | -0.539 | -0.576 | -1.027 | -2.636 | 0.021 | 0.081 | 0.081 | 1.46 | 0.38 | 0.72 | 0.2 | 0.01 | 0.3 | 0.23 |
| 5 | 116565409 | rs6860917 | -1.419 | -0.569 | -1.164 | -2.606 | 0.047 | 0.338 | 0.338 | 0.123 | 0.51 | 0.88 | 0.26 | 0.07 | 0.37 | 0.32 |
| 5 | 116574744 | rs1021517 | -1.687 | -0.391 | -1.207 | -2.393 | 0.006 | 0.041 | 0.041 | 0 | 0.2 | 0.16 | 0.26 | 0.08 | 0.18 | 0.13 |
| 5 | 116581841 | rs10073120 | -0.32 | -0.656 | -0.781 | -2.435 | -0.015 | -0.137 | -0.137 | 0.048 | 0.35 | 0.47 | 0.17 | 0.01 | 0.27 | 0.21 |
| 5 | 116588233 | rs1494820 | -1.122 | -0.705 | -0.829 | -2.432 | 0.049 | 0.263 | 0.263 | 3.316 | 0.47 | 0.82 | 0.25 | 0.06 | 0.36 | 0.31 |
| 5 | 116600516 | rs12719228 | -0.139 | -0.584 | -0.81 | -2.424 | -0.009 | -0.103 | -0.103 | 0 | 0.36 | 0.53 | 0.18 | 0.01 | 0.28 | 0.22 |
| 5 | 116607979 | rs6595026 | -1.295 | -0.652 | -0.934 | -2.437 | 0.046 | 0.324 | 0.324 | 0.246 | 0.52 | 0.86 | 0.27 | 0.07 | 0.37 | 0.31 |
| 5 | 116619728 | rs17141409 | -0.042 | -0.698 | -0.747 | -2.46 | -0.015 | -0.142 | -0.142 | 0 | 0.34 | 0.39 | 0.17 | 0.01 | 0.27 | 0.21 |
| 5 | 116624982 | rs7710852 | -1.269 | -0.662 | -0.778 | -2.348 | 0.056 | 0.139 | 0.139 | 0 | 0.49 | 0.66 | 0.29 | 0.1 | 0.36 | 0.31 |

| | | | | | | | | | | | | | | | | |
|---|---|---|---|---|---|---|---|---|---|---|---|---|---|---|---|---|
| 5 | 116625850 | rs7712165 | -1.22 | -0.636 | -0.816 | -2.336 | 0.043 | 0.326 | 0.326 | 0.226 | 0.51 | 0.88 | 0.3 | 0.1 | 0.37 | 0.31 |
| 5 | 116626388 | rs10463690 | -1.205 | -0.64 | -0.818 | -2.337 | 0.045 | 0.328 | 0.328 | 0.204 | 0.51 | 0.88 | 0.3 | 0.1 | 0.37 | 0.31 |
| 5 | 116630974 | rs7701246 | -0.031 | -0.558 | -0.765 | -2.322 | -0.012 | -0.142 | -0.142 | 0.203 | 0.35 | 0.48 | 0.18 | 0 | 0.27 | 0.22 |
| 5 | 116647725 | rs7735145 | -0.335 | -0.378 | -0.943 | -2.26 | -0.022 | -0.021 | -0.021 | 1.741 | 0.14 | 0.24 | 0.12 | 0.09 | 0.09 | 0.08 |
| 5 | 116653901 | rs10068349 | -1.244 | -0.445 | -1.403 | -2.3 | 0.004 | 0.307 | 0.307 | 0.058 | 0.53 | 0.89 | 0.34 | 0.09 | 0.39 | 0.31 |
| 5 | 116657759 | rs10058376 | -0.021 | -0.497 | -1.219 | -2.271 | -0.015 | -0.154 | -0.154 | 1.23 | 0.35 | 0.41 | 0.18 | 0 | 0.27 | 0.22 |
| 5 | 116661710 | rs1565893 | 0.051 | -0.435 | -1.964 | -1.634 | 0 | -0.002 | -0.002 | 0 | 0.06 | 0.05 | 0.06 | 0.01 | 0.03 | 0.02 |
| 5 | 116663400 | rs1845560 | -0.994 | 0.28 | -2.218 | -0.6 | -0.044 | -0.105 | -0.105 | 0 | 0.65 | 0.95 | 0.69 | 0.4 | 0.68 | 0.58 |
| 5 | 116666584 | rs10040906 | -0.521 | 0.416 | -1.831 | -0.261 | -0.057 | -0.062 | -0.062 | 0 | 0.68 | 0.91 | 0.66 | 0.59 | 0.64 | 0.54 |
| 5 | 116669707 | rs6880664 | -0.836 | -0.21 | -0.806 | -0.725 | -0.033 | -0.044 | -0.044 | 0 | 0.16 | 0.29 | 0.07 | 0.05 | 0.08 | 0.03 |
| 5 | 116671103 | rs10076399 | -0.409 | 0.188 | -1.025 | -1.155 | -0.018 | 0.009 | 0.009 | 0 | 0.13 | 0.12 | 0.14 | 0 | 0.21 | 0.12 |
| 5 | 116671392 | rs1032491 | -0.204 | 0.371 | -1.03 | -1.16 | -0.007 | 0.003 | 0.003 | 0 | 0.12 | 0.11 | 0.09 | 0 | 0.14 | 0.1 |
| 5 | 116672520 | rs4513729 | -0.543 | 0.373 | -1.074 | -1.176 | -0.003 | 0.044 | 0.044 | 0 | 0.14 | 0.07 | 0.08 | 0.05 | 0.12 | 0.03 |
| 5 | 116680760 | rs4457117 | -2.106 | 0.045 | -0.471 | -1.3 | 0.008 | 0.076 | 0.076 | 0.93 | 0.42 | 0.7 | 0.3 | 0.13 | 0.39 | 0.18 |
| 5 | 116691333 | rs7716581 | -2.111 | 0.016 | -0.358 | -1.307 | 0.009 | 0.086 | 0.086 | 1.824 | 0.42 | 0.72 | 0.3 | 0.13 | 0.39 | 0.19 |
| 5 | 116703668 | rs6896597 | -0.045 | -0.542 | -0.225 | -1.436 | 0 | -0.018 | -0.018 | 0 | 0.11 | 0.15 | 0.07 | 0.01 | 0.09 | 0.08 |
| 5 | 116710914 | rs17335992 | -0.202 | -0.546 | -0.265 | -1.445 | -0.005 | 0.039 | 0.039 | 0.303 | 0.13 | 0.06 | 0.07 | 0.04 | 0.11 | 0.03 |
| 5 | 116711571 | rs17336048 | 0.079 | -0.593 | 0.211 | -1.464 | -0.006 | -0.033 | -0.033 | 0.06 | 0.12 | 0.2 | 0.06 | 0.05 | 0.1 | 0.04 |
| 5 | 116713132 | rs10519507 | 0.079 | -0.593 | 0.211 | -1.477 | -0.006 | -0.036 | -0.036 | 0.225 | 0.12 | 0.2 | 0.06 | 0.05 | 0.1 | 0.04 |
| 5 | 116713295 | rs2217261 | -1.982 | -0.578 | 0.448 | -1.468 | -0.014 | -0.085 | -0.085 | 0 | 0.39 | 0.57 | 0.22 | 0.11 | 0.32 | 0.17 |
| 5 | 116713465 | rs2195294 | -1.395 | -0.532 | 0.139 | -1.135 | -0.015 | -0.074 | -0.074 | 0.227 | 0.29 | 0.55 | 0.16 | 0.1 | 0.23 | 0.09 |
| 5 | 116719741 | rs2560611 | -1.839 | -0.817 | 0.176 | -1.1 | -0.036 | -0.078 | -0.078 | 0.013 | 0.17 | 0.64 | 0.1 | 0.03 | 0.09 | 0.11 |
| 5 | 116723208 | rs2560606 | 0.436 | -0.641 | -0.497 | -0.873 | -0.008 | 0.013 | 0.013 | 0 | 0.4 | 0.34 | 0.5 | 0.35 | 0.51 | 0.56 |
| 5 | 116727172 | rs2624495 | 0.49 | -0.775 | -0.554 | -0.703 | -0.008 | 0.004 | 0.004 | 0 | 0.41 | 0.35 | 0.51 | 0.4 | 0.52 | 0.56 |
| 5 | 116727621 | rs10900718 | -0.889 | -1.035 | -0.621 | -0.706 | 0.027 | -0.081 | -0.081 | 0 | 0.61 | 0.87 | 0.62 | 0.55 | 0.65 | 0.68 |
| 5 | 116728965 | rs6887206 | -0.292 | -0.857 | -0.193 | -0.603 | 0.004 | -0.001 | -0.001 | 0.123 | 0.52 | 0.54 | 0.59 | 0.55 | 0.62 | 0.62 |
| 5 | 116730016 | rs2624489 | -0.169 | -0.822 | -0.495 | -0.602 | 0 | 0.001 | 0.001 | 0 | 0.1 | 0.1 | 0.07 | 0.16 | 0.1 | 0.06 |
| 5 | 116732988 | rs2560600 | 1.243 | -0.475 | -0.591 | -0.787 | -0.018 | -0.006 | -0.006 | 0 | 0.31 | 0.26 | 0.45 | 0.38 | 0.42 | 0.53 |
| 5 | 116738589 | rs10061995 | 1.411 | -1.189 | -0.913 | -1.059 | 0.027 | 0.064 | 0.064 | 0 | 0.17 | 0.05 | 0.07 | 0.02 | 0.13 | 0.1 |
| 5 | 116750291 | rs1030461 | -0.653 | -0.603 | -0.586 | -0.582 | -0.014 | 0.011 | 0.011 | 0 | 0.6 | 0.7 | 0.5 | 0.62 | 0.48 | 0.44 |
| 5 | 116751170 | rs13184765 | 0.531 | -0.576 | -0.638 | -0.592 | 0.006 | 0.022 | 0.022 | 0.017 | 0.09 | 0.06 | 0.04 | 0 | 0.07 | 0.02 |
| 5 | 116752778 | rs10065401 | 0.115 | -0.593 | -0.546 | -0.538 | 0 | 0 | 0 | 0 | 0.49 | 0.48 | 0.45 | 0.6 | 0.41 | 0.39 |
| 5 | 116757581 | rs2624542 | -1.089 | -0.344 | -0.762 | -0.426 | -0.026 | -0.007 | -0.007 | 0 | 0.68 | 0.73 | 0.54 | 0.63 | 0.55 | 0.46 |
| 5 | 116759655 | rs17141747 | -0.195 | -0.46 | -1.062 | -0.543 | 0.01 | 0.038 | 0.038 | 1.153 | 0.12 | 0.07 | 0.04 | 0 | 0.08 | 0.03 |
| 5 | 116762741 | rs2560532 | 0.175 | -0.505 | -0.961 | -0.532 | 0.004 | -0.001 | -0.001 | 0 | 0.06 | 0.07 | 0.04 | 0.05 | 0.06 | 0.04 |
| 5 | 116772093 | rs13164296 | -0.538 | -0.623 | -0.783 | -0.6 | -0.004 | -0.032 | -0.032 | 0 | 0.14 | 0.16 | 0.05 | 0.01 | 0.08 | 0.05 |
| 5 | 116777116 | rs17141780 | 0.704 | -1.02 | -1.109 | -0.626 | 0.035 | -0.091 | -0.091 | 0 | 0.42 | 0.11 | 0.39 | 0.44 | 0.36 | 0.35 |
| 5 | 116783660 | rs17141799 | -0.058 | -0.928 | -0.824 | -0.746 | 0.006 | 0.006 | 0.006 | 0.32 | 0.11 | 0.07 | 0.04 | 0 | 0.08 | 0.03 |
| 5 | 116786213 | rs2929729 | -1.455 | -0.94 | -0.793 | -0.753 | 0.007 | 0.024 | 0.024 | 0.125 | 0.21 | 0.17 | 0.11 | 0.15 | 0.18 | 0.08 |
| 5 | 116790756 | rs2560519 | -1.037 | -0.912 | -0.121 | -0.823 | -0.002 | -0.071 | -0.071 | 0 | 0.15 | 0.29 | 0.08 | 0.02 | 0.14 | 0.06 |
| 5 | 116799810 | rs17141817 | 0.289 | -0.552 | -0.698 | -0.784 | -0.002 | 0.008 | 0.008 | 0 | 0.08 | 0.05 | 0.05 | 0.01 | 0.08 | 0.04 |
| 5 | 116804254 | rs7718223 | -0.35 | -0.904 | -0.412 | -0.852 | -0.008 | 0.037 | 0.037 | 0 | 0.11 | 0.41 | 0.09 | 0.13 | 0.07 | 0.07 |
| 5 | 116809323 | rs2162748 | -0.162 | -0.815 | -0.814 | -0.952 | 0.002 | -0.036 | -0.036 | 0 | 0.1 | 0.15 | 0.06 | 0 | 0.1 | 0.03 |
| 5 | 116809749 | rs1025275 | -1.537 | -1.277 | -1.54 | -0.715 | -0.169 | -0.153 | -0.153 | 0.103 | 0.68 | 0.95 | 0.53 | 0.59 | 0.51 | 0.44 |
| 5 | 116816776 | rs12652139 | 1.395 | -1.099 | -1.624 | -0.511 | -0.027 | -0.021 | -0.021 | 0 | 0.07 | 0.01 | 0.12 | 0.23 | 0.08 | 0.18 |
| 5 | 116818239 | rs2560635 | 0.792 | -1.884 | -1.664 | -1.586 | -0.003 | -0.021 | -0.021 | 0 | 0.22 | 0.2 | 0.08 | 0.04 | 0.15 | 0.11 |
| 5 | 116818810 | rs2560634 | -1.302 | -1.671 | -2.218 | -1.474 | 0.087 | 0.305 | 0.305 | 0.363 | 0.49 | 0.9 | 0.25 | 0.17 | 0.36 | 0.26 |
| 5 | 116821562 | rs2624569 | -1.015 | -1.302 | -1.591 | -1.039 | -0.005 | -0.007 | -0.007 | 0 | 0.21 | 0.69 | 0.12 | 0.16 | 0.13 | 0.14 |
| 5 | 116824450 | rs890779 | -1.667 | -0.238 | -1.962 | -0.827 | -0.104 | -0.129 | -0.129 | 0.509 | 0.83 | 0.99 | 0.65 | 0.61 | 0.69 | 0.64 |
| 5 | 116825352 | rs1863977 | -0.642 | -1.473 | -1.97 | -2.04 | 0.085 | 0.327 | 0.327 | 0.695 | 0.46 | 0.88 | 0.22 | 0.06 | 0.32 | 0.31 |
| 5 | 116828604 | rs2624564 | -0.529 | -1.658 | -1.718 | -2.189 | 0.004 | -0.057 | -0.057 | 0.124 | 0.14 | 0.64 | 0.1 | 0.04 | 0.13 | 0.09 |
| 5 | 116828752 | rs12332110 | 0.085 | -1.815 | -1.611 | -2.045 | 0.03 | 0.01 | 0.01 | 0 | 0.32 | 0.25 | 0.12 | 0.02 | 0.19 | 0.22 |
| 5 | 116840907 | rs10519513 | -0.493 | -1.126 | -1.455 | -1.65 | -0.022 | 0.026 | 0.026 | 0.5 | 0.16 | 0.07 | 0.12 | 0.16 | 0.19 | 0.1 |
| 5 | 116843143 | rs10071606 | 0.791 | -1.673 | -0.879 | -1.976 | -0.024 | -0.057 | -0.057 | 0.062 | 0.18 | 0.18 | 0.07 | 0.02 | 0.11 | 0.05 |
| 5 | 116843574 | rs265871 | -0.569 | -2.027 | -0.804 | -2.052 | -0.044 | -0.098 | -0.098 | 0.057 | 0.16 | 0.62 | 0.1 | 0.02 | 0.1 | 0.14 |
| 5 | 116849573 | rs17141968 | 0.343 | -1.814 | -1.03 | -1.986 | -0.025 | 0.021 | 0.021 | 1.653 | 0.11 | 0.02 | 0.09 | 0.11 | 0.14 | 0.06 |
| 5 | 116855837 | rs388865 | -1.512 | -1.932 | -0.959 | -2.102 | 0 | -0.011 | -0.011 | 0.323 | 0.3 | 0.69 | 0.22 | 0.15 | 0.29 | 0.25 |
| 5 | 116857589 | rs390708 | -0.24 | -2.012 | -0.424 | -2.064 | -0.002 | -0.055 | -0.055 | 0.068 | 0.15 | 0.24 | 0.1 | 0.03 | 0.14 | 0.18 |
| 5 | 116869940 | rs4920974 | 0.867 | -2.144 | -0.966 | -2.133 | -0.031 | -0.056 | -0.056 | 0 | 0.18 | 0.19 | 0.07 | 0.02 | 0.09 | 0.07 |
| 5 | 116885018 | rs265893 | -1.713 | -2.064 | -1.207 | -2.126 | 0.046 | 0.289 | 0.289 | 0.758 | 0.53 | 0.9 | 0.36 | 0.19 | 0.39 | 0.34 |
| 5 | 116888640 | rs12659561 | -0.194 | -1.894 | -0.963 | -2.072 | 0.026 | 0.053 | 0.053 | 0.372 | 0.32 | 0.18 | 0.18 | 0.15 | 0.25 | 0.15 |
| 5 | 116895323 | rs187859 | -1.191 | -1.834 | -0.891 | -2.018 | 0.102 | 0.211 | 0.211 | 0 | 0.47 | 0.8 | 0.28 | 0.17 | 0.36 | 0.3 |
| 5 | 116910855 | rs265918 | -0.135 | -1.698 | -0.252 | -1.906 | -0.034 | -0.06 | -0.06 | 0 | 0.11 | 0.25 | 0.03 | 0.02 | 0.06 | 0.11 |
| 5 | 116914436 | rs10478363 | 0.733 | -1.901 | -0.448 | -2.092 | 0.007 | -0.009 | -0.009 | 0 | 0.21 | 0.16 | 0.11 | 0.02 | 0.15 | 0.1 |
| 5 | 116915749 | rs449716 | -1.861 | -2.015 | -0.693 | -2.295 | -0.056 | 0.224 | 0.224 | 0.113 | 0.58 | 0.93 | 0.4 | 0.19 | 0.44 | 0.35 |
| 5 | 116915880 | rs17142077 | 0.786 | -1.894 | -0.629 | -2.141 | -0.026 | -0.015 | -0.015 | 0.778 | 0.1 | 0.02 | 0.13 | 0.15 | 0.11 | 0.08 |
| 5 | 116916674 | rs1553557 | 0.035 | -2.054 | -0.25 | -1.973 | 0.01 | 0.015 | 0.015 | 0.02 | 0.27 | 0.17 | 0.17 | 0.02 | 0.22 | 0.13 |
| 5 | 116933316 | rs398793 | -0.916 | -1.956 | 0.107 | -1.762 | -0.044 | -0.104 | -0.104 | 0.34 | 0.16 | 0.61 | 0.09 | 0.02 | 0.11 | 0.14 |
| 5 | 116948591 | rs267070 | 0.1 | -1.739 | -0.319 | -1.67 | -0.006 | -0.033 | -0.033 | 0 | 0.09 | 0.13 | 0.05 | 0.02 | 0.09 | 0.13 |

| | | | | | | | | | | | | | | | | |
|---|---|---|---|---|---|---|---|---|---|---|---|---|---|---|---|---|
| 5 | 116950104 | rs267069 | -0.192 | -1.785 | -0.267 | -1.691 | -0.013 | -0.052 | -0.052 | 0 | 0.1 | 0.24 | 0.06 | 0.02 | 0.08 | 0.13 |
| 5 | 116962366 | rs434097 | -0.948 | -1.982 | -0.552 | -1.912 | 0.057 | 0.127 | 0.127 | 0 | 0.43 | 0.74 | 0.27 | 0.03 | 0.34 | 0.28 |
| 5 | 116968077 | rs389286 | -1.067 | -1.894 | -1.011 | -1.871 | 0.076 | 0.37 | 0.37 | 0.117 | 0.46 | 0.93 | 0.29 | 0.03 | 0.34 | 0.28 |
| 5 | 116979988 | rs1686413 | 0.716 | -1.541 | -0.511 | -1.372 | 0.001 | 0.011 | 0.011 | 0 | 0.07 | 0.03 | 0.04 | 0 | 0.08 | 0.12 |
| 5 | 116991351 | rs842009 | -0.836 | -1.273 | -1.214 | -0.785 | -0.035 | -0.145 | -0.145 | 1.286 | 0.77 | 0.97 | 0.79 | 0.52 | 0.77 | 0.67 |
| 5 | 116993837 | rs841999 | -0.124 | -0.905 | -0.829 | -0.992 | -0.003 | -0.023 | -0.023 | 0 | 0.09 | 0.11 | 0.05 | 0.02 | 0.08 | 0.13 |
| 5 | 117005392 | rs17346050 | -0.834 | -1.076 | -1.116 | -1.106 | 0.019 | 0.034 | 0.034 | 1.336 | 0.14 | 0.07 | 0.11 | 0 | 0.11 | 0.07 |
| 5 | 117005977 | rs17411921 | -0.834 | -1.432 | -1.008 | -1.687 | 0.008 | -0.012 | -0.012 | 1.757 | 0.36 | 0.29 | 0.24 | 0.03 | 0.33 | 0.28 |
| 5 | 117006071 | rs951482 | -0.577 | -1.67 | -0.63 | -1.712 | -0.015 | -0.078 | -0.078 | 0.043 | 0.31 | 0.6 | 0.16 | 0.03 | 0.22 | 0.23 |
| 5 | 117007686 | rs10043348 | -0.059 | -1.689 | -1.322 | -1.595 | 0.009 | 0.018 | 0.018 | 0 | 0.14 | 0.08 | 0.08 | 0 | 0.12 | 0.13 |
| 5 | 117011725 | rs10051273 | 1.056 | -1.897 | -2.093 | -1.081 | -0.064 | -0.087 | -0.087 | 0 | 0.44 | 0.06 | 0.51 | 0.55 | 0.54 | 0.45 |
| 5 | 117012266 | rs17412135 | 1.144 | -1.963 | -2.327 | -1.372 | -0.063 | 0.001 | 0.001 | 0 | 0.15 | 0.02 | 0.08 | 0.08 | 0.18 | 0.2 |
| 5 | 117013294 | rs1526881 | -1.257 | -2.024 | -2.202 | -1.435 | 0.041 | 0.22 | 0.22 | 0.405 | 0.41 | 0.87 | 0.26 | 0.06 | 0.33 | 0.32 |
| 5 | 117016265 | rs1526880 | -1.512 | -2.045 | -2.143 | -1.442 | 0.052 | 0.19 | 0.19 | 0.036 | 0.43 | 0.84 | 0.27 | 0.07 | 0.36 | 0.33 |
| 5 | 117017440 | rs10085098 | 0.132 | -1.45 | -1.406 | -0.984 | 0.002 | -0.019 | -0.019 | 0.212 | 0.16 | 0.18 | 0.07 | 0.04 | 0.15 | 0.13 |
| 5 | 117026872 | rs1526882 | -0.324 | -1.465 | -1.38 | -0.884 | -0.011 | -0.043 | -0.043 | 0 | 0.09 | 0.16 | 0.07 | 0 | 0.07 | 0.06 |
| 5 | 117037025 | rs6882497 | -1.808 | -1.75 | -1.882 | -1.057 | 0.014 | -0.005 | -0.005 | 0 | 0.22 | 0.79 | 0.16 | 0.01 | 0.16 | 0.12 |
| 5 | 117048409 | rs9327050 | 0.194 | -1.291 | -2.05 | -0.817 | 0.006 | 0.001 | 0.001 | 0 | 0.9 | 0.91 | 0.93 | 1 | 0.9 | 0.93 |
| 5 | 117048725 | rs1830169 | -0.19 | -1.398 | -2.535 | -0.414 | -0.037 | -0.111 | -0.111 | 0 | 0.22 | 0.05 | 0.19 | 0.44 | 0.24 | 0.3 |
| 5 | 117055145 | rs1358226 | 0.26 | -1.006 | -2.717 | -0.15 | -0.034 | -0.105 | -0.105 | 0 | 0.6 | 0.91 | 0.59 | 0.44 | 0.61 | 0.69 |
| 5 | 117063252 | rs4532391 | 0.154 | -0.8 | -2.631 | -0.328 | 0.002 | 0.046 | 0.046 | 0 | 0.78 | 0.89 | 0.85 | 0.96 | 0.78 | 0.75 |
| 5 | 117069229 | rs4076522 | 0.451 | -0.58 | -2.406 | 0.305 | -0.028 | -0.118 | -0.118 | 0 | 0.61 | 0.91 | 0.6 | 0.48 | 0.63 | 0.73 |
| 5 | 117085745 | rs12520471 | -0.688 | -0.551 | -1.683 | 0.673 | -0.003 | -0.032 | -0.032 | 0 | 0.1 | 0.06 | 0.2 | 0.28 | 0.11 | 0.23 |
| 5 | 117087858 | rs4482937 | -0.497 | -0.563 | -0.964 | 0.337 | -0.007 | 0.024 | 0.024 | 0 | 0.67 | 0.58 | 0.65 | 0.53 | 0.71 | 0.61 |
| 5 | 117088421 | rs12514675 | -1.054 | -0.604 | -0.785 | 0.261 | -0.012 | 0.052 | 0.052 | 0.622 | 0.1 | 0.12 | 0.18 | 0.28 | 0.07 | 0.17 |
| 5 | 117089085 | rs7736745 | -1.07 | -0.067 | -0.898 | 0 | -0.004 | -0.005 | -0.005 | 0 | 0.2 | 0.16 | 0.29 | 0.31 | 0.28 | 0.3 |
| 5 | 117096806 | rs12187403 | 0.336 | 0.086 | -0.948 | -0.127 | -0.066 | 0.01 | 0.01 | 0.002 | 0.11 | 0.01 | 0.12 | 0.04 | 0.2 | 0.19 |
| 5 | 117105363 | rs4327633 | 0.315 | -0.696 | -0.971 | -0.512 | 0.036 | 0.196 | 0.196 | 0 | 0.46 | 0.17 | 0.36 | 0.14 | 0.59 | 0.41 |
| 5 | 117109134 | rs10477564 | -1.201 | -0.928 | -0.399 | -0.447 | 0.004 | -0.011 | -0.011 | 0 | 0.66 | 0.73 | 0.74 | 0.52 | 0.68 | 0.51 |
| 5 | 117114675 | rs4437413 | -1.305 | -0.979 | -0.197 | -0.486 | -0.054 | 0.148 | 0.148 | 0 | 0.63 | 0.84 | 0.74 | 0.9 | 0.58 | 0.79 |
| 5 | 117118798 | rs7732707 | -0.985 | -1.088 | 0.092 | -0.463 | -0.048 | 0.081 | 0.081 | 0 | 0.16 | 0.34 | 0.23 | 0.24 | 0.08 | 0.2 |
| 5 | 117120444 | rs6863742 | -1.489 | -0.825 | -0.194 | -0.527 | -0.098 | 0.165 | 0.165 | 0.008 | 0.65 | 0.93 | 0.76 | 0.9 | 0.57 | 0.79 |
| 5 | 117132151 | rs4244396 | 0.429 | -1.172 | 0.045 | -0.405 | -0.02 | 0.013 | 0.013 | 0.814 | 0.39 | 0.47 | 0.28 | 0.49 | 0.32 | 0.45 |
| 5 | 117139928 | rs10061470 | -0.032 | -1.148 | 0.113 | -0.478 | 0.002 | -0.001 | -0.001 | 0 | 0.46 | 0.5 | 0.47 | 0.72 | 0.37 | 0.54 |
| 5 | 117143718 | rs9327056 | -0.443 | -0.9 | -0.138 | -0.402 | 0.005 | 0.019 | 0.019 | 0 | 0.5 | 0.4 | 0.68 | 0.9 | 0.55 | 0.72 |
| 5 | 117143804 | rs10054762 | -0.443 | -0.9 | -0.138 | -0.402 | 0.005 | 0.019 | 0.019 | 0 | 0.5 | 0.4 | 0.68 | 0.9 | 0.55 | 0.72 |
| 5 | 117146271 | rs12655706 | -0.096 | -0.896 | -0.31 | -0.379 | -0.002 | 0.073 | 0.073 | 0 | 0.47 | 0.32 | 0.55 | 0.76 | 0.54 | 0.7 |
| 5 | 117146651 | rs6595061 | -0.358 | -0.885 | -0.304 | -0.418 | 0.007 | 0.071 | 0.071 | 0 | 0.49 | 0.34 | 0.68 | 0.9 | 0.55 | 0.72 |
| 5 | 117146864 | rs4304104 | 1.025 | -0.599 | -0.316 | -0.191 | 0.001 | -0.007 | -0.007 | 0 | 0.31 | 0.27 | 0.25 | 0.48 | 0.3 | 0.47 |
| 5 | 117168998 | rs7730998 | -0.218 | -0.349 | -0.534 | -0.259 | -0.079 | 0.023 | 0.023 | 0 | 0.14 | 0.02 | 0.15 | 0.05 | 0.24 | 0.14 |
| 5 | 117186295 | rs6879319 | 1.072 | -0.57 | -0.316 | -0.224 | -0.002 | 0.005 | 0.005 | 0 | 0.32 | 0.33 | 0.25 | 0.49 | 0.29 | 0.51 |
| 5 | 117212923 | rs6887429 | 0.218 | -0.813 | -0.472 | -0.301 | -0.031 | 0.018 | 0.018 | 0 | 0.46 | 0.78 | 0.52 | 0.66 | 0.49 | 0.64 |
| 5 | 117224933 | rs1904134 | -0.855 | -0.64 | -0.617 | -0.309 | -0.007 | -0.001 | -0.001 | 0 | 0.95 | 1 | 0.81 | 0.78 | 0.94 | 0.9 |
| 5 | 117225247 | rs1904135 | -0.855 | -0.64 | -0.617 | -0.309 | -0.007 | -0.001 | -0.001 | 0.118 | 0.95 | 1 | 0.81 | 0.78 | 0.94 | 0.9 |
| 5 | 117227611 | rs10076425 | 0.292 | -0.773 | -0.511 | -0.521 | 0.038 | 0.016 | 0.016 | 0.222 | 0.47 | 0.7 | 0.48 | 0.65 | 0.4 | 0.56 |
| 5 | 117231418 | rs11956306 | 1.113 | -0.683 | -0.657 | -0.599 | -0.12 | 0.171 | 0.171 | 0.299 | 0.32 | 0.04 | 0.21 | 0.07 | 0.39 | 0.21 |
| 5 | 117235680 | rs12186546 | -0.788 | -0.75 | -0.414 | -0.625 | -0.003 | 0.011 | 0.011 | 0 | 0.22 | 0.2 | 0.12 | 0.06 | 0.21 | 0.2 |
| 5 | 117239454 | rs4546421 | 1.018 | -0.791 | -0.132 | -0.498 | -0.007 | 0.02 | 0.02 | 0 | 0.35 | 0.51 | 0.42 | 0.62 | 0.34 | 0.49 |
| 5 | 117241490 | rs7720835 | 0.637 | -0.745 | -0.598 | -0.632 | 0.018 | -0.009 | -0.009 | 0 | 0.39 | 0.66 | 0.47 | 0.64 | 0.34 | 0.51 |
| 5 | 117255401 | rs4389718 | 1.124 | -0.602 | -0.885 | -0.671 | -0.005 | -0.009 | -0.009 | 0.107 | 0.67 | 0.81 | 0.67 | 0.74 | 0.72 | 0.69 |
| 5 | 117260126 | rs4301259 | 0.704 | -0.025 | -0.448 | -0.261 | 0.003 | 0.049 | 0.049 | 0 | 0.78 | 0.61 | 0.69 | 0.73 | 0.8 | 0.74 |
| 5 | 117263731 | rs11951689 | 0.626 | 0.597 | -1.143 | -0.815 | -0.087 | 0.142 | 0.142 | 0.408 | 0.28 | 0.04 | 0.19 | 0.08 | 0.32 | 0.2 |
| 5 | 117265658 | rs7726540 | 0.433 | 0.64 | -1.078 | -0.817 | -0.034 | 0.1 | 0.1 | 0.034 | 0.31 | 0.14 | 0.23 | 0.08 | 0.39 | 0.21 |
| 5 | 117272473 | rs4302625 | -0.029 | 0.406 | -1.184 | -0.65 | -0.078 | 0.124 | 0.124 | 1.146 | 0.38 | 0.16 | 0.27 | 0.13 | 0.46 | 0.33 |
| 5 | 117278010 | rs718228 | -1.079 | 0.321 | -0.27 | -0.281 | 0.016 | -0.02 | -0.02 | 0 | 0.22 | 0.36 | 0.26 | 0.2 | 0.21 | 0.15 |
| 5 | 117282234 | rs17143339 | -0.778 | 0.334 | 0.366 | -0.543 | 0.036 | -0.022 | -0.022 | 0 | 0.07 | 0.33 | 0.07 | 0 | 0.05 | 0.02 |
| 5 | 117283938 | rs17143349 | -0.411 | -0.18 | -0.168 | -0.04 | -0.009 | 0.038 | 0.038 | 0.098 | 0.11 | 0.15 | 0.22 | 0.26 | 0.08 | 0.11 |
| 5 | 117286831 | rs10071777 | -1.157 | -0.244 | 0.084 | -0.472 | 0.006 | -0.07 | -0.07 | 0 | 0.27 | 0.61 | 0.19 | 0.08 | 0.23 | 0.2 |
| 5 | 117292776 | rs10061408 | -1.516 | -0.015 | -0.176 | -0.546 | 0 | -0.026 | -0.026 | 0.862 | 0.31 | 0.69 | 0.21 | 0.08 | 0.28 | 0.21 |
| 5 | 117297258 | rs17414326 | 0.744 | -0.104 | -0.336 | -0.467 | -0.047 | -0.017 | -0.017 | 0 | 0.08 | 0.01 | 0.07 | 0.06 | 0.14 | 0.09 |
| 5 | 117298452 | rs17143450 | -1.158 | -0.418 | -0.141 | -0.634 | -0.042 | -0.113 | -0.113 | 0 | 0.21 | 0.65 | 0.14 | 0.03 | 0.11 | 0.09 |
| 5 | 117299721 | rs10519530 | -0.507 | -0.211 | -0.715 | -0.181 | -0.017 | -0.031 | -0.031 | 0 | 0.15 | 0.08 | 0.3 | 0.26 | 0.19 | 0.21 |
| 5 | 117300672 | rs472296 | 1.21 | -0.39 | -1.712 | -0.728 | -0.013 | -0.174 | -0.174 | 0 | 0.26 | 0.02 | 0.21 | 0.52 | 0.24 | 0.44 |
| 5 | 117301623 | rs553374 | 1.258 | -0.284 | -1.398 | -0.575 | -0.007 | -0.022 | -0.022 | 0.531 | 0.63 | 0.23 | 0.44 | 0.56 | 0.66 | 0.67 |
| 5 | 117306795 | rs684884 | -0.209 | -0.458 | -1.415 | -0.661 | -0.003 | 0.006 | 0.006 | 0 | 0.84 | 0.91 | 0.78 | 0.81 | 0.87 | 0.89 |
| 5 | 117315725 | rs6595079 | 1.443 | -0.316 | -1.67 | -0.56 | 0.007 | -0.043 | -0.043 | 0.179 | 0.58 | 0.17 | 0.43 | 0.54 | 0.66 | 0.64 |
| 5 | 117319372 | rs4921000 | 0.676 | -0.491 | -1.15 | -1.204 | -0.01 | -0.017 | -0.017 | 0 | 0.77 | 0.28 | 0.76 | 0.85 | 0.88 | 0.89 |
| 5 | 117330989 | rs1996778 | 0.774 | -0.289 | -1.823 | -1.776 | -0.029 | -0.065 | -0.065 | 0 | 0.25 | 0.03 | 0.4 | 0.79 | 0.24 | 0.45 |
| 5 | 117331544 | rs13162213 | 0.156 | -0.083 | -1.101 | -1.896 | 0.026 | 0.041 | 0.041 | 0 | 0.47 | 0.32 | 0.45 | 0.23 | 0.64 | 0.47 |

| | | | | | | | | | | | | | | | | |
|---|---|---|---|---|---|---|---|---|---|---|---|---|---|---|---|---|
| 5 | 117336746 | rs2900110 | 0.513 | 0.694 | -1.31 | -2.329 | 0.018 | 0.005 | 0.005 | 0.053 | 0.17 | 0.19 | 0.31 | 0.17 | 0.29 | 0.21 |
| 5 | 117343784 | rs2900111 | 0.184 | -0.195 | -1.148 | -2.346 | 0.029 | 0.046 | 0.046 | 0 | 0.48 | 0.34 | 0.5 | 0.24 | 0.65 | 0.47 |
| 5 | 117345425 | rs9327065 | -0.045 | -0.148 | -1.558 | -2.482 | 0.045 | 0.116 | 0.116 | 0 | 0.56 | 0.85 | 0.55 | 0.19 | 0.66 | 0.48 |
| 5 | 117347530 | rs2416494 | -0.553 | 0.072 | -0.712 | -4.243 | -0.001 | -0.059 | -0.059 | 0 | 0.36 | 0.61 | 0.23 | 0.02 | 0.27 | 0.19 |
| 5 | 117353851 | rs4340948 | -0.572 | -0.388 | -0.542 | -4.179 | -0.031 | -0.065 | -0.065 | 1.259 | 0.86 | 0.84 | 0.96 | 1 | 0.92 | 0.91 |
| 5 | 117354366 | rs10037996 | 1.595 | 0.166 | -1.044 | -4.715 | -0.072 | 0.176 | 0.176 | 0 | 0.36 | 0.13 | 0.64 | 0.98 | 0.47 | 0.58 |
| 5 | 117360819 | rs2416498 | 1.119 | 0.097 | -0.439 | -4.85 | -0.004 | -0.066 | -0.066 | 0.001 | 0.68 | 0.37 | 0.85 | 0.99 | 0.77 | 0.75 |
| 5 | 117364600 | rs7710446 | 0.154 | 0.211 | -0.203 | -4.754 | 0.079 | -0.012 | -0.012 | 0 | 0.92 | 0.55 | 0.94 | 1 | 0.96 | 0.99 |
| 5 | 117371148 | rs4435901 | -0.052 | 0.252 | 0.157 | -4.74 | 0.089 | 0.001 | 0.001 | 1.373 | 0.93 | 0.64 | 0.95 | 1 | 0.95 | 0.99 |
| 5 | 117375545 | rs6595087 | 1.63 | 0.228 | -0.269 | -4.927 | -0.074 | 0.194 | 0.194 | 0 | 0.39 | 0.15 | 0.68 | 0.98 | 0.53 | 0.59 |
| 5 | 117376303 | rs2416500 | -0.498 | 0.204 | -0.061 | -4.847 | 0.003 | -0.022 | -0.022 | 0 | 0.7 | 0.76 | 0.83 | 0.99 | 0.75 | 0.81 |
| 5 | 117421203 | rs17143539 | 0.386 | 0.413 | 0 | -4.758 | 0.015 | -0.088 | -0.088 | 4.705 | 0.16 | 0.28 | 0.11 | 0 | 0.17 | 0.12 |
| 5 | 117434840 | rs10074918 | -0.283 | 0.215 | -0.008 | -4.744 | 0 | -0.041 | -0.041 | 0.01 | 0.26 | 0.22 | 0.14 | 0.01 | 0.24 | 0.21 |
| 5 | 117441642 | rs4632836 | -0.242 | 0.194 | 0.259 | -4.707 | 0.082 | -0.005 | -0.005 | 0 | 0.07 | 0.4 | 0.05 | 0 | 0.05 | 0.01 |
| 5 | 117445263 | rs11955772 | 0.779 | -0.024 | -0.058 | -4.869 | 0.002 | -0.02 | -0.02 | 1.037 | 0.24 | 0.17 | 0.13 | 0.01 | 0.17 | 0.14 |
| 5 | 117451628 | rs4559034 | -0.642 | 0.127 | 0.053 | -4.816 | 0 | -0.133 | -0.133 | 0 | 0.3 | 0.43 | 0.16 | 0.01 | 0.29 | 0.25 |
| 5 | 117455495 | rs4389719 | 0.221 | 0.125 | 0.179 | -4.831 | -0.009 | -0.078 | -0.078 | 0 | 0.3 | 0.37 | 0.19 | 0.01 | 0.23 | 0.16 |
| 5 | 117492799 | rs1479207 | -2.056 | 0.373 | -0.107 | -4.91 | -0.101 | 0.212 | 0.212 | 0 | 0.66 | 0.94 | 0.38 | 0.02 | 0.55 | 0.41 |
| 5 | 117492859 | rs1382723 | 0.059 | 0.389 | -0.027 | -4.936 | 0.031 | -0.026 | -0.026 | 2.453 | 0.05 | 0.18 | 0.02 | 0 | 0.05 | 0.03 |
| 5 | 117494640 | rs10079352 | -2.053 | 0.397 | -0.125 | -4.912 | -0.11 | 0.218 | 0.218 | 0 | 0.66 | 0.95 | 0.38 | 0.02 | 0.55 | 0.41 |
| 5 | 117499364 | rs1479202 | 0.074 | 0.316 | 0.36 | -4.589 | -0.019 | -0.106 | -0.106 | 0 | 0.26 | 0.38 | 0.17 | 0.01 | 0.19 | 0.12 |
| 5 | 117499480 | rs1479201 | -0.522 | 0.366 | 0.225 | -4.592 | -0.007 | -0.095 | -0.095 | 7.496 | 0.32 | 0.6 | 0.2 | 0.01 | 0.24 | 0.18 |
| 5 | 117506665 | rs1479196 | -1.903 | 0.396 | -0.057 | -4.573 | -0.091 | 0.236 | 0.236 | 0 | 0.61 | 0.9 | 0.36 | 0.02 | 0.5 | 0.39 |
| 5 | 117508036 | rs13187513 | -0.346 | 0.3 | 0.101 | -4.695 | 0 | -0.064 | -0.064 | 0.128 | 0.24 | 0.24 | 0.14 | 0.01 | 0.22 | 0.17 |
| 5 | 117509503 | rs4526142 | -1.566 | 0.284 | 0.126 | -4.722 | -0.01 | 0.107 | 0.107 | 0 | 0.55 | 0.7 | 0.33 | 0.02 | 0.45 | 0.35 |
| 5 | 117510153 | rs10060461 | -0.355 | 0.31 | 0.261 | -4.655 | -0.001 | -0.027 | -0.027 | 0 | 0.24 | 0.21 | 0.14 | 0.01 | 0.22 | 0.17 |
| 5 | 117512698 | rs10519551 | -0.507 | 0.375 | 0.159 | -4.648 | -0.001 | -0.168 | -0.168 | 2.233 | 0.26 | 0.47 | 0.15 | 0.01 | 0.22 | 0.17 |
| 5 | 117515575 | rs7709033 | -0.509 | 0.366 | 0.154 | -4.648 | -0.004 | -0.153 | -0.153 | 0 | 0.26 | 0.5 | 0.16 | 0.01 | 0.22 | 0.17 |
| 5 | 117518502 | rs899467 | -0.066 | 0.347 | -0.342 | -4.684 | 0.012 | 0.011 | 0.011 | 0 | 0.29 | 0.22 | 0.18 | 0.01 | 0.22 | 0.18 |
| 5 | 117520174 | rs10078247 | -2.04 | 0.324 | -0.392 | -4.695 | -0.1 | 0.258 | 0.258 | 0 | 0.6 | 0.89 | 0.36 | 0.02 | 0.5 | 0.39 |
| 5 | 117521059 | rs6861306 | -1.085 | 0.565 | 0.017 | -4.353 | -0.001 | -0.045 | -0.045 | 0 | 0.31 | 0.66 | 0.18 | 0.01 | 0.28 | 0.21 |
| 5 | 117524559 | rs6879786 | -0.927 | 0.383 | 0.053 | -4.308 | 0 | -0.047 | -0.047 | 0.16 | 0.3 | 0.65 | 0.17 | 0.01 | 0.25 | 0.2 |
| 5 | 117528371 | rs1382713 | -1.607 | 0.103 | 0.408 | -4.305 | -0.018 | -0.109 | -0.109 | 0.293 | 0.33 | 0.59 | 0.16 | 0.01 | 0.22 | 0.13 |
| 5 | 117538971 | rs13188829 | -1.61 | 0.089 | 0.42 | -4.313 | -0.021 | -0.115 | -0.115 | 1.459 | 0.33 | 0.59 | 0.16 | 0.01 | 0.22 | 0.13 |
| 5 | 117551523 | rs1479230 | -1.202 | 0.098 | 0.317 | -4.307 | 0.004 | -0.022 | -0.022 | 0.043 | 0.26 | 0.22 | 0.13 | 0.01 | 0.19 | 0.11 |
| 5 | 117560263 | rs6883098 | -1.927 | 0.229 | -0.076 | -4.539 | -0.015 | 0.137 | 0.137 | 0 | 0.55 | 0.73 | 0.32 | 0.02 | 0.41 | 0.28 |
| 5 | 117561189 | rs10519544 | -0.375 | 0.086 | -0.148 | -4.556 | 0.027 | 0.073 | 0.073 | 0 | 0.14 | 0.03 | 0.04 | 0.01 | 0.08 | 0.07 |
| 5 | 117566290 | rs1479211 | -1.66 | 0.2 | -0.104 | -4.61 | 0.039 | 0.16 | 0.16 | 0.127 | 0.51 | 0.7 | 0.29 | 0.02 | 0.37 | 0.22 |
| 5 | 117568527 | rs17143900 | -1.171 | 0.149 | 0.322 | -4.501 | -0.018 | -0.12 | -0.12 | 0 | 0.36 | 0.43 | 0.22 | 0.03 | 0.27 | 0.18 |
| 5 | 117572000 | rs1479227 | -1.83 | 1.213 | 0.56 | -4.27 | -0.008 | 0.002 | 0.002 | 0 | 0.21 | 0.19 | 0.11 | 0.01 | 0.23 | 0.14 |
| 5 | 117577409 | rs1479198 | -1.211 | 1.22 | 0.597 | -4.252 | 0 | -0.021 | -0.021 | 0.361 | 0.15 | 0.18 | 0.05 | 0.01 | 0.13 | 0.11 |
| 5 | 117590121 | rs17143910 | -0.42 | 1.185 | 0.586 | -4.232 | -0.006 | -0.023 | -0.023 | 0 | 0.11 | 0.15 | 0.03 | 0.01 | 0.07 | 0.08 |
| 5 | 117591619 | rs17368151 | 0.504 | 1.194 | 0.693 | -4.337 | 0 | -0.092 | -0.092 | 1.527 | 0.18 | 0.28 | 0.16 | 0.01 | 0.16 | 0.07 |
| 5 | 117592341 | rs7341174 | -1.513 | 1.346 | 0.727 | -4.277 | -0.015 | 0.116 | 0.116 | 1.002 | 0.41 | 0.73 | 0.28 | 0.03 | 0.39 | 0.21 |
| 5 | 117592453 | rs17143923 | -0.647 | 0.911 | 0.752 | -4.105 | -0.016 | -0.139 | -0.139 | 2.702 | 0.29 | 0.45 | 0.18 | 0.03 | 0.23 | 0.15 |
| 5 | 117603886 | rs6872263 | -0.481 | 0.822 | 0.774 | -4.092 | -0.025 | -0.14 | -0.14 | 2.353 | 0.3 | 0.49 | 0.19 | 0.02 | 0.23 | 0.15 |
| 5 | 117610910 | rs6860479 | -0.487 | 0.914 | 0.708 | -4.191 | -0.014 | -0.096 | -0.096 | 0 | 0.29 | 0.36 | 0.19 | 0.02 | 0.23 | 0.15 |
| 5 | 117634630 | rs1479224 | -0.114 | 0.659 | 0.71 | -4.075 | -0.008 | -0.027 | -0.027 | 0.492 | 0.05 | 0.13 | 0.02 | 0 | 0.02 | 0.02 |
| 5 | 117649329 | rs10057536 | 0.175 | 0.623 | 0.647 | -4.139 | -0.02 | -0.094 | -0.094 | 0 | 0.24 | 0.3 | 0.16 | 0.01 | 0.18 | 0.08 |
| 5 | 117649475 | rs6880706 | -1.973 | 0.567 | 0.295 | -4.193 | -0.081 | 0.368 | 0.368 | 0.643 | 0.52 | 0.94 | 0.32 | 0.03 | 0.41 | 0.26 |
| 5 | 117662282 | rs7446931 | 0.162 | 0.554 | 0.651 | -4.02 | -0.019 | -0.096 | -0.096 | 0.736 | 0.23 | 0.3 | 0.17 | 0.01 | 0.18 | 0.08 |
| 5 | 117669302 | rs7448606 | -0.649 | 0.579 | 0.516 | -4.05 | -0.006 | -0.082 | -0.082 | 0.004 | 0.35 | 0.56 | 0.23 | 0.01 | 0.28 | 0.14 |
| 5 | 117675855 | rs11241454 | 1.189 | 0.679 | 0.532 | -3.856 | 0.015 | 0.045 | 0.045 | 0 | 0.54 | 0.41 | 0.68 | 0.9 | 0.64 | 0.78 |
| 5 | 117676469 | rs6595113 | 1.139 | 0.672 | 0.526 | -3.853 | 0.013 | 0.035 | 0.035 | 0 | 0.55 | 0.42 | 0.68 | 0.9 | 0.64 | 0.78 |
| 5 | 117678736 | rs4920888 | 1.136 | 0.558 | 0.514 | -3.814 | 0.007 | 0.001 | 0.001 | 3.007 | 0.57 | 0.44 | 0.67 | 0.89 | 0.65 | 0.81 |
| 5 | 117683871 | rs7443402 | -0.199 | 0.425 | 0.669 | -3.647 | -0.04 | -0.155 | -0.155 | 0 | 0.27 | 0.41 | 0.17 | 0.01 | 0.19 | 0.1 |
| 5 | 117690833 | rs7707505 | 0.027 | 0.402 | 0.771 | -3.415 | 0.016 | -0.151 | -0.151 | 0.923 | 0.3 | 0.4 | 0.26 | 0.02 | 0.31 | 0.16 |
| 5 | 117695003 | rs6882527 | -0.978 | 0.491 | 0.081 | -3.499 | 0.007 | 0.11 | 0.11 | 0.844 | 0.51 | 0.69 | 0.4 | 0.11 | 0.5 | 0.3 |
| 5 | 117698479 | rs7701305 | -1.345 | 0.513 | -0.022 | -3.436 | 0.007 | 0.074 | 0.074 | 1.31 | 0.55 | 0.68 | 0.44 | 0.12 | 0.55 | 0.3 |
| 5 | 117701230 | rs7736991 | -0.626 | 0.587 | -0.01 | -3.613 | 0 | -0.055 | -0.055 | 0.019 | 0.41 | 0.54 | 0.33 | 0.1 | 0.39 | 0.25 |
| 5 | 117701468 | rs7445762 | -0.339 | 0.615 | 0.266 | -3.39 | 0.002 | -0.076 | -0.076 | 0.315 | 0.36 | 0.53 | 0.29 | 0.09 | 0.33 | 0.23 |
| 5 | 117706786 | rs6876064 | -0.135 | 0.56 | -0.144 | -3.302 | -0.001 | 0.003 | 0.003 | 0.004 | 0.16 | 0.12 | 0.09 | 0 | 0.16 | 0.06 |
| 5 | 117711064 | rs7704260 | -1.011 | 0.762 | -0.611 | -3.135 | -0.014 | 0.262 | 0.262 | 0.437 | 0.5 | 0.77 | 0.31 | 0.01 | 0.43 | 0.28 |
| 5 | 117714620 | rs7721152 | -0.865 | 0.486 | 0.332 | -2.34 | -0.017 | -0.155 | -0.155 | 0 | 0.21 | 0.48 | 0.12 | 0 | 0.16 | 0.1 |
| 5 | 117715695 | rs6878247 | -1.219 | 1.067 | 0.506 | -2.169 | 0.073 | 0.012 | 0.012 | 0 | 0.05 | 0.19 | 0.06 | 0.02 | 0.09 | 0.02 |
| 5 | 117730127 | rs7711344 | -1.878 | 1.599 | 0.906 | -2.53 | 0.018 | -0.023 | -0.023 | 0.03 | 0.23 | 0.69 | 0.23 | 0.11 | 0.28 | 0.11 |
| 5 | 117740545 | rs7446054 | -1.462 | 1.59 | 1.482 | -2.577 | 0.076 | 0.112 | 0.112 | 1.621 | 0.06 | 0.4 | 0.1 | 0.08 | 0.05 | 0.04 |
| 5 | 117747508 | rs7708440 | -1.401 | 1.36 | -0.107 | -3.393 | -0.129 | 0.046 | 0.046 | 3.429 | 0.69 | 0.97 | 0.5 | 0.18 | 0.57 | 0.4 |
| 5 | 117754341 | rs11951938 | -0.677 | 1.521 | 0.112 | -3.418 | 0.037 | -0.026 | -0.026 | 0 | 0.12 | 0.18 | 0.11 | 0.03 | 0.21 | 0.08 |

| | | | | | | | | | | | | | | | | |
|---|---|---|---|---|---|---|---|---|---|---|---|---|---|---|---|---|
| 5 | 117754697 | rs9327072 | 0.498 | 0.736 | 0.205 | -3.821 | 0.023 | 0.151 | 0.151 | 1.603 | 0.38 | 0.17 | 0.28 | 0.06 | 0.31 | 0.18 |
| 5 | 117763178 | rs6887550 | -0.794 | 0.276 | 0.854 | -3.927 | -0.007 | 0.028 | 0.028 | 0 | 0.09 | 0.25 | 0.09 | 0.08 | 0.06 | 0.03 |
| 5 | 117763622 | rs6595129 | -1.42 | 0.298 | 0.978 | -3.917 | 0.079 | 0.129 | 0.129 | 1.059 | 0.06 | 0.38 | 0.09 | 0.08 | 0.04 | 0.04 |
| 5 | 117770863 | rs10056223 | -0.115 | 0.531 | 0.816 | -3.932 | 0.046 | -0.016 | -0.016 | 3.933 | 0.91 | 0.82 | 0.91 | 0.97 | 0.84 | 0.92 |
| 5 | 117772695 | rs7448017 | 0.962 | 0.345 | 0.906 | -4.019 | 0.047 | -0.072 | -0.072 | 0 | 0.84 | 0.61 | 0.89 | 0.98 | 0.83 | 0.9 |
| 5 | 117793509 | rs4329039 | -0.788 | 0.616 | 0.025 | -4.234 | 0.04 | 0.184 | 0.184 | 1.063 | 0.53 | 0.8 | 0.42 | 0.16 | 0.4 | 0.31 |
| 5 | 117795324 | rs12719243 | -2.252 | 0.634 | -0.32 | -4.158 | -0.103 | -0.004 | -0.004 | 0 | 0.72 | 0.98 | 0.56 | 0.19 | 0.65 | 0.41 |
| 5 | 117795654 | rs10046044 | 0.378 | 0.332 | 0.115 | -4.007 | 0.041 | 0.092 | 0.092 | 0.966 | 0.33 | 0.17 | 0.23 | 0.05 | 0.24 | 0.16 |
| 5 | 117799807 | rs6885949 | 0.212 | 0.64 | 0.071 | -3.773 | -0.017 | -0.064 | -0.064 | 0 | 0.36 | 0.36 | 0.26 | 0.06 | 0.27 | 0.17 |
| 5 | 117805637 | rs4073021 | -1.225 | 1.163 | 0.216 | -3.437 | 0.02 | -0.052 | -0.052 | 0 | 0.23 | 0.41 | 0.18 | 0.1 | 0.22 | 0.14 |
| 5 | 117810106 | rs9327074 | -0.302 | 1.165 | 0.178 | -3.593 | 0.014 | 0.071 | 0.071 | 0.365 | 0.46 | 0.67 | 0.36 | 0.07 | 0.4 | 0.32 |
| 5 | 117814294 | rs13187374 | -1.203 | 1.305 | 0.512 | -3.302 | 0.015 | -0.047 | -0.047 | 0 | 0.1 | 0.14 | 0.18 | 0 | 0.14 | 0.09 |
| 5 | 117815175 | rs10077519 | 0.143 | 1.323 | 0.075 | -3.905 | 0.026 | 0.14 | 0.14 | 1.224 | 0.34 | 0.14 | 0.18 | 0.05 | 0.25 | 0.17 |
| 5 | 117823265 | rs6859213 | -1.31 | 1.61 | 0.412 | -3.973 | -0.029 | 0.165 | 0.165 | 1.712 | 0.58 | 0.84 | 0.48 | 0.13 | 0.53 | 0.31 |
| 5 | 117823346 | rs6859532 | -1.174 | 1.628 | 1.273 | -4.014 | 0.081 | -0.148 | -0.148 | 0 | 0.19 | 0.41 | 0.24 | 0 | 0.24 | 0.11 |
| 5 | 117824287 | rs12654169 | -0.101 | 1.588 | 1.214 | -4.009 | -0.005 | -0.001 | -0.001 | 0 | 0.08 | 0.06 | 0.17 | 0 | 0.11 | 0.07 |
| 5 | 117825449 | rs12109441 | -0.554 | 1.551 | 1.598 | -4.036 | -0.002 | -0.039 | -0.039 | 1.628 | 0.47 | 0.49 | 0.4 | 0.13 | 0.39 | 0.28 |
| 5 | 117825588 | rs12110298 | 0.329 | 1.356 | 1.456 | -4.031 | 0.028 | 0.154 | 0.154 | 1.869 | 0.35 | 0.13 | 0.15 | 0.04 | 0.25 | 0.18 |
| 5 | 117843869 | rs3924079 | -0.159 | 1.278 | 1.514 | -3.975 | 0.082 | -0.066 | -0.066 | 0 | 0.11 | 0.31 | 0.07 | 0 | 0.16 | 0.05 |
| 5 | 117845579 | rs3923756 | -1.245 | 1.222 | 0.862 | -4.003 | -0.042 | 0.17 | 0.17 | 1.369 | 0.58 | 0.82 | 0.47 | 0.13 | 0.51 | 0.31 |
| 5 | 117847097 | rs10058844 | -1.23 | 1.228 | 0.869 | -3.999 | -0.037 | 0.163 | 0.163 | 1.381 | 0.58 | 0.81 | 0.47 | 0.13 | 0.51 | 0.31 |
| 5 | 117861849 | rs5000240 | -1.824 | 1.037 | 0.293 | -4.091 | -0.029 | 0.117 | 0.117 | 1.015 | 0.67 | 0.9 | 0.57 | 0.11 | 0.65 | 0.43 |
| 5 | 117862376 | rs6880637 | 0.59 | 0.584 | 0.303 | -4.004 | 0.044 | 0.167 | 0.167 | 2.625 | 0.31 | 0.08 | 0.13 | 0.01 | 0.18 | 0.17 |
| 5 | 117863779 | rs6868178 | 0.568 | 0.585 | 0.304 | -4.003 | 0.045 | 0.127 | 0.127 | 2.425 | 0.31 | 0.08 | 0.18 | 0.1 | 0.19 | 0.18 |
| 5 | 117869104 | rs6870931 | -0.21 | 0.608 | 0.473 | -3.954 | -0.001 | -0.035 | -0.035 | 0 | 0.08 | 0.11 | 0.14 | 0 | 0.08 | 0.1 |
| 5 | 117869174 | rs6595143 | -1.094 | 0.628 | 0.759 | -4.034 | -0.002 | -0.005 | -0.005 | 0 | 0.55 | 0.58 | 0.46 | 0.1 | 0.47 | 0.39 |
| 5 | 117884604 | rs10058989 | 0.424 | 0.2 | 0.141 | -3.837 | 0.04 | 0.115 | 0.115 | 0.696 | 0.31 | 0.14 | 0.14 | 0.02 | 0.2 | 0.18 |
| 5 | 117888912 | rs11739499 | -0.761 | 0.059 | 0.16 | -4.031 | 0.003 | -0.042 | -0.042 | 0.045 | 0.17 | 0.18 | 0.11 | 0.02 | 0.19 | 0.12 |
| 5 | 117891968 | rs1012695 | -0.739 | -0.497 | -0.073 | -3.795 | 0.027 | 0.039 | 0.039 | 0.176 | 0.42 | 0.68 | 0.25 | 0.12 | 0.27 | 0.24 |
| 5 | 117893138 | rs7737007 | 0.016 | -0.207 | -0.67 | -3.501 | 0 | -0.047 | -0.047 | 0 | 0.3 | 0.29 | 0.13 | 0.01 | 0.2 | 0.19 |
| 5 | 117894902 | rs7705821 | 0.311 | -0.106 | -1.586 | -3.413 | 0.022 | 0.069 | 0.069 | 0 | 0.27 | 0.14 | 0.12 | 0.01 | 0.18 | 0.18 |
| 5 | 117899908 | rs6595144 | -0.979 | 0.407 | -1.693 | -2.985 | -0.05 | 0.139 | 0.139 | 0.752 | 0.63 | 0.92 | 0.43 | 0.16 | 0.58 | 0.43 |
| 5 | 117901357 | rs6894915 | -0.618 | -0.676 | -1.573 | -2.806 | 0.055 | 0.28 | 0.28 | 1.313 | 0.51 | 0.93 | 0.29 | 0.14 | 0.32 | 0.33 |
| 5 | 117906849 | rs12654918 | -0.557 | -0.749 | -1.385 | -2.7 | 0.097 | 0.278 | 0.278 | 1.011 | 0.48 | 0.83 | 0.24 | 0.05 | 0.3 | 0.29 |
| 5 | 117907292 | rs10213999 | -0.574 | -0.748 | -1.415 | -2.712 | 0.099 | 0.29 | 0.29 | 1.058 | 0.49 | 0.83 | 0.23 | 0.05 | 0.29 | 0.29 |
| 5 | 117910620 | rs13164702 | -0.567 | -0.654 | -1.415 | -2.532 | 0.092 | 0.281 | 0.281 | 0.631 | 0.52 | 0.85 | 0.26 | 0.08 | 0.3 | 0.3 |
| 5 | 117912524 | rs13157174 | -0.774 | -0.623 | -1.462 | -2.509 | 0.012 | 0.199 | 0.199 | 0.778 | 0.55 | 0.95 | 0.32 | 0.18 | 0.32 | 0.34 |
| 5 | 117914813 | rs750569 | -0.081 | -0.675 | -1.302 | -2.599 | 0.042 | 0.11 | 0.11 | 0.105 | 0.4 | 0.21 | 0.25 | 0.17 | 0.29 | 0.28 |
| 5 | 117916235 | rs1429723 | -0.465 | -0.282 | -0.761 | -2.349 | 0.012 | 0.013 | 0.013 | 0.166 | 0.51 | 0.54 | 0.29 | 0.18 | 0.32 | 0.34 |
| 5 | 117929951 | rs6862770 | -0.386 | -0.096 | -1.67 | -1.794 | 0.033 | 0.122 | 0.122 | 0 | 0.33 | 0.16 | 0.21 | 0.1 | 0.26 | 0.22 |
| 5 | 117935041 | rs1002916 | -0.75 | -0.712 | -1.993 | -2.205 | 0.032 | 0.029 | 0.029 | 2.863 | 0.13 | 0.08 | 0.05 | 0 | 0.04 | 0.03 |
| 5 | 117937429 | rs6595148 | 0.815 | -0.385 | -2.146 | -2.884 | 0.001 | -0.047 | -0.047 | 0 | 0.17 | 0.18 | 0.14 | 0 | 0.21 | 0.15 |
| 5 | 117939969 | rs17371061 | 0.101 | -0.444 | -2.153 | -2.902 | 0.029 | 0.133 | 0.133 | 0 | 0.32 | 0.13 | 0.21 | 0.1 | 0.26 | 0.21 |
| 5 | 117943909 | rs10519559 | 1.088 | 0.9 | -2.241 | -2.801 | -0.059 | 0.029 | 0.029 | 0 | 0.09 | 0.01 | 0.11 | 0.01 | 0.2 | 0.1 |
| 5 | 117946207 | rs2043048 | 0.102 | 0.61 | -2.2 | -2.833 | -0.047 | 0.045 | 0.045 | 0 | 0.26 | 0.14 | 0.24 | 0.02 | 0.41 | 0.25 |
| 5 | 117950885 | rs6881807 | 0.296 | -0.151 | -1.408 | -2.499 | -0.011 | -0.023 | -0.023 | 0 | 0.65 | 0.6 | 0.48 | 0.19 | 0.73 | 0.56 |
| 5 | 117951540 | rs1429739 | -0.151 | -0.229 | -1.539 | -0.888 | 0.022 | -0.044 | -0.044 | 0 | 0.76 | 0.86 | 0.71 | 0.59 | 0.8 | 0.62 |
| 5 | 117955915 | rs4519946 | -0.172 | -0.156 | -1.908 | -0.936 | -0.051 | -0.114 | -0.114 | 0.757 | 0.7 | 0.92 | 0.66 | 0.55 | 0.66 | 0.58 |
| 5 | 117967472 | rs4895236 | -0.15 | -0.301 | -0.708 | -1.069 | -0.023 | -0.1 | -0.1 | 0 | 0.14 | 0.33 | 0.11 | 0 | 0.11 | 0.08 |
| 5 | 117967649 | rs1429734 | -0.564 | -0.406 | -0.604 | -1.007 | -0.015 | -0.022 | -0.022 | 0 | 0.16 | 0.37 | 0.22 | 0.1 | 0.12 | 0.13 |
| 5 | 117968633 | rs13153473 | -1.047 | -0.02 | -0.723 | -0.795 | 0.037 | 0.031 | 0.031 | 0 | 0.08 | 0.2 | 0.11 | 0.11 | 0.13 | 0.07 |
| 5 | 117976258 | rs2099154 | -0.29 | 0.625 | -0.869 | -1.042 | -0.003 | 0.043 | 0.043 | 0 | 0.41 | 0.63 | 0.4 | 0.11 | 0.42 | 0.28 |
| 5 | 117981154 | rs6885983 | -0.048 | 0.207 | -0.649 | -1.142 | -0.008 | -0.071 | -0.071 | 0 | 0.2 | 0.25 | 0.11 | 0 | 0.14 | 0.08 |
| 5 | 117982046 | rs2914908 | 0.186 | 0.511 | -0.985 | -0.6 | -0.015 | -0.01 | -0.01 | 0 | 0.44 | 0.68 | 0.49 | 0.51 | 0.48 | 0.45 |
| 5 | 117983612 | rs2972219 | -1.299 | 0.633 | -1.171 | -0.638 | -0.071 | 0.097 | 0.097 | 0 | 0.79 | 0.98 | 0.78 | 0.97 | 0.74 | 0.8 |
| 5 | 117989334 | rs734514 | -0.655 | 0.955 | -0.938 | -0.625 | -0.021 | 0.269 | 0.269 | 0 | 0.5 | 0.83 | 0.56 | 0.88 | 0.45 | 0.59 |
| 5 | 117990821 | rs7442981 | -0.23 | 0.958 | -0.792 | -0.544 | 0.038 | 0.175 | 0.175 | 0 | 0.17 | 0.57 | 0.28 | 0.43 | 0.19 | 0.18 |
| 5 | 118009472 | rs2376682 | 0.358 | 0.842 | -0.84 | -0.569 | 0.005 | -0.003 | -0.003 | 0 | 0.34 | 0.33 | 0.29 | 0.43 | 0.26 | 0.4 |
| 5 | 118018091 | rs7728143 | -0.124 | 0.977 | -0.279 | -0.37 | 0.042 | 0.17 | 0.17 | 0 | 0.15 | 0.29 | 0.26 | 0.48 | 0.19 | 0.18 |
| 5 | 118032245 | rs7737088 | 0.091 | 1.012 | -0.255 | -0.412 | 0.021 | -0.001 | -0.001 | 0 | 0.13 | 0.21 | 0.1 | 0.08 | 0.16 | 0.14 |
| 5 | 118045916 | rs10069027 | 0.286 | 0.702 | -0.452 | -0.524 | 0.014 | -0.006 | -0.006 | 0 | 0.34 | 0.27 | 0.28 | 0.41 | 0.26 | 0.39 |
| 5 | 118049776 | rs10077150 | 0.639 | 1.003 | -0.641 | -0.92 | -0.004 | -0.044 | -0.044 | 0 | 0.42 | 0.43 | 0.32 | 0.1 | 0.41 | 0.43 |
| 5 | 118058065 | rs735620 | 0.416 | 1.024 | -0.683 | -0.884 | -0.001 | -0.062 | -0.062 | 0 | 0.45 | 0.48 | 0.35 | 0.1 | 0.47 | 0.44 |
| 5 | 118058631 | rs730907 | -0.313 | 1.137 | -0.918 | -0.5 | -0.008 | -0.012 | -0.012 | 0.101 | 0.79 | 0.81 | 0.64 | 0.57 | 0.73 | 0.84 |
| 5 | 118058726 | rs730906 | 0.526 | 0.774 | -1.061 | -1.036 | -0.011 | 0.101 | 0.101 | 0 | 0.3 | 0.1 | 0.22 | 0.08 | 0.3 | 0.27 |
| 5 | 118059534 | rs728676 | -0.013 | 0.824 | -1.379 | -1.125 | -0.033 | 0.191 | 0.191 | 0 | 0.45 | 0.22 | 0.34 | 0.1 | 0.51 | 0.45 |
| 5 | 118060890 | rs17144476 | -0.865 | 0.585 | -1.023 | -1.025 | 0 | -0.011 | -0.011 | 0.289 | 0.18 | 0.17 | 0.12 | 0.03 | 0.18 | 0.15 |
| 5 | 118062597 | rs2193965 | -1.239 | 0.138 | -0.419 | -1.221 | -0.025 | -0.086 | -0.086 | 0 | 0.21 | 0.61 | 0.12 | 0.03 | 0.15 | 0.15 |
| 5 | 118064125 | rs7727675 | -1.196 | 0.118 | -0.373 | -1.23 | -0.021 | -0.075 | -0.075 | 0 | 0.21 | 0.65 | 0.12 | 0.03 | 0.15 | 0.15 |

| | | | | | | | | | | | | | | | | |
|---|---|---|---|---|---|---|---|---|---|---|---|---|---|---|---|---|
| 5 | 118070512 | rs11949509 | 0.155 | 0.315 | -0.712 | -0.488 | 0 | 0.026 | 0.026 | 0 | 0.86 | 0.88 | 0.62 | 0.43 | 0.81 | 0.81 |
| 5 | 118071881 | rs1544740 | 1.285 | 0.28 | -0.609 | -0.495 | 0.002 | 0.009 | 0.009 | 0 | 0.71 | 0.66 | 0.53 | 0.39 | 0.67 | 0.66 |
| 5 | 118073157 | rs981763 | 0.397 | 0.283 | -0.454 | -0.521 | 0.065 | 0.054 | 0.054 | 3.619 | 0.06 | 0.43 | 0.03 | 0 | 0.01 | 0.01 |
| 5 | 118073885 | rs6870403 | -0.05 | 0.383 | -0.57 | -0.525 | 0.004 | -0.023 | -0.023 | 0.933 | 0.11 | 0.14 | 0.08 | 0.03 | 0.13 | 0.14 |
| 5 | 118080021 | rs17144507 | 1.351 | 0.128 | -0.906 | -0.824 | -0.001 | -0.037 | -0.037 | 0 | 0.07 | 0.15 | 0.04 | 0 | 0.05 | 0.03 |
| 5 | 118082030 | rs7728873 | -0.017 | 0.115 | -0.853 | -0.33 | -0.018 | -0.038 | -0.038 | 0 | 0.86 | 0.91 | 0.68 | 0.54 | 0.8 | 0.82 |
| 5 | 118085428 | rs17144520 | -0.954 | -0.024 | -0.817 | -0.521 | -0.04 | -0.112 | -0.112 | 0 | 0.19 | 0.49 | 0.1 | 0.03 | 0.11 | 0.14 |
| 5 | 118087436 | rs584789 | 2.122 | -0.048 | -0.843 | -0.421 | 0.02 | -0.021 | -0.021 | 0 | 0.65 | 0.22 | 0.52 | 0.38 | 0.69 | 0.65 |
| 5 | 118089798 | rs1510973 | -0.718 | -0.038 | -0.714 | -0.38 | -0.06 | -0.068 | -0.068 | 0 | 0.18 | 0.49 | 0.1 | 0.06 | 0.07 | 0.15 |
| 5 | 118094406 | rs17144556 | 0.499 | 0.273 | -0.864 | -0.215 | 0.002 | 0.002 | 0.002 | 0 | 0.08 | 0.05 | 0.06 | 0.06 | 0.06 | 0.14 |
| 5 | 118094523 | rs586056 | 1.051 | 0.515 | -1.507 | -0.749 | -0.011 | 0.021 | 0.021 | 0.502 | 0.42 | 0.2 | 0.35 | 0.35 | 0.41 | 0.45 |
| 5 | 118122504 | rs7728676 | -0.576 | 0.806 | -0.793 | -0.027 | 0.003 | 0.023 | 0.023 | 0 | 0.13 | 0.21 | 0.11 | 0.14 | 0.11 | 0.18 |
| 5 | 118129948 | rs365096 | -0.015 | 0.953 | -0.926 | 0.044 | -0.054 | -0.077 | -0.077 | 0 | 0.88 | 0.97 | 0.63 | 0.47 | 0.78 | 0.84 |
| 5 | 118134414 | rs17144660 | -0.521 | 0.584 | -0.43 | -0.452 | 0.01 | 0.039 | 0.039 | 0 | 0.1 | 0.26 | 0.08 | 0.16 | 0.08 | 0.16 |
| 5 | 118159871 | rs1567520 | 0.249 | 0.133 | -0.654 | -1.092 | 0.03 | 0.05 | 0.05 | 0 | 0.16 | 0.05 | 0.06 | 0.03 | 0.06 | 0.16 |
| 5 | 118161392 | rs1510976 | 0.823 | 0.72 | -1.039 | -1.812 | -0.093 | 0.146 | 0.146 | 0.088 | 0.26 | 0.06 | 0.21 | 0.01 | 0.36 | 0.16 |
| 5 | 118167525 | rs2081957 | -0.797 | 0.726 | -0.131 | -1.548 | 0.001 | -0.042 | -0.042 | 3.638 | 0.25 | 0.26 | 0.22 | 0.06 | 0.25 | 0.25 |
| 5 | 118168648 | rs4895347 | -1.514 | 0.402 | 0.157 | -1.71 | 0.008 | 0.035 | 0.035 | 2.257 | 0.33 | 0.75 | 0.29 | 0.09 | 0.29 | 0.27 |
| 5 | 118175020 | rs17144716 | -1.505 | 0.403 | 0.173 | -1.709 | 0.004 | 0.005 | 0.005 | 2.236 | 0.33 | 0.69 | 0.29 | 0.08 | 0.28 | 0.27 |
| 5 | 118175674 | rs17132677 | -0.673 | 0.376 | 0.08 | -1.695 | -0.076 | -0.117 | -0.117 | 0 | 0.25 | 0.52 | 0.12 | 0.05 | 0.1 | 0.19 |
| 5 | 118179104 | rs6892437 | -0.321 | 0.34 | 0.107 | -1.683 | -0.049 | -0.064 | -0.064 | 0 | 0.23 | 0.33 | 0.1 | 0.05 | 0.07 | 0.19 |
| 5 | 118180935 | rs7700373 | -0.166 | 0.633 | -0.058 | -1.551 | 0.033 | -0.016 | -0.016 | 0 | 0.08 | 0.09 | 0.16 | 0.03 | 0.18 | 0.08 |
| 5 | 118191798 | rs7722194 | 1.339 | 0.167 | -0.599 | -1.314 | -0.026 | -0.062 | -0.062 | 0 | 0.17 | 0.04 | 0.19 | 0.25 | 0.17 | 0.35 |
| 5 | 118210835 | rs6870923 | 0.139 | 0.052 | -0.174 | -1.706 | -0.025 | -0.06 | -0.06 | 0.306 | 0.2 | 0.22 | 0.12 | 0.04 | 0.11 | 0.18 |
| 5 | 118216272 | rs10434765 | -0.767 | 0.222 | -0.214 | -1.746 | 0.051 | -0.025 | -0.025 | 0 | 0.09 | 0.29 | 0.16 | 0.03 | 0.14 | 0.08 |
| 5 | 118216318 | rs4895183 | -1.166 | -0.026 | -0.323 | -1.871 | 0.012 | 0.01 | 0.01 | 3.352 | 0.32 | 0.73 | 0.25 | 0.08 | 0.2 | 0.26 |
| 5 | 118218002 | rs6863802 | -0.826 | -0.082 | -0.286 | -1.832 | -0.007 | -0.026 | -0.026 | 0.604 | 0.28 | 0.71 | 0.17 | 0.06 | 0.14 | 0.2 |
| 5 | 118225616 | rs6874399 | -0.419 | -0.077 | -0.191 | -1.759 | -0.049 | -0.076 | -0.076 | 0.023 | 0.24 | 0.58 | 0.12 | 0.05 | 0.1 | 0.18 |
| 5 | 118241854 | rs4895184 | -1.412 | 0.285 | -0.273 | -1.591 | 0.054 | -0.02 | -0.02 | 4.216 | 0.13 | 0.29 | 0.17 | 0.06 | 0.23 | 0.18 |
| 5 | 118245193 | rs6868097 | 0.17 | -0.004 | -0.727 | -1.145 | -0.089 | -0.126 | -0.126 | 0.026 | 0.28 | 0.04 | 0.44 | 0.38 | 0.31 | 0.39 |
| 5 | 118264866 | rs17144824 | -0.406 | -0.415 | 0.235 | -1.361 | -0.036 | -0.056 | -0.056 | 0 | 0.24 | 0.66 | 0.12 | 0.06 | 0.09 | 0.17 |
| 5 | 118267633 | rs12109252 | 0.616 | -0.425 | 0.243 | -1.365 | -0.024 | -0.042 | -0.042 | 0 | 0.17 | 0.19 | 0.06 | 0.04 | 0.06 | 0.16 |
| 5 | 118275079 | rs17144836 | -0.651 | -0.444 | 0.152 | -1.396 | -0.012 | -0.027 | -0.027 | 0 | 0.27 | 0.7 | 0.13 | 0.05 | 0.11 | 0.17 |
| 5 | 118275869 | rs2029036 | 0.736 | -0.184 | -0.003 | -1.244 | -0.001 | -0.009 | -0.009 | 0.486 | 0.18 | 0.14 | 0.06 | 0.04 | 0.08 | 0.16 |
| 5 | 118285396 | rs13156156 | 0.496 | 0.131 | -0.26 | -1.195 | 0.003 | 0.004 | 0.004 | 0.01 | 0.61 | 0.35 | 0.74 | 0.8 | 0.67 | 0.75 |
| 5 | 118293501 | rs2442123 | -0.159 | 0.19 | -0.49 | -2.003 | -0.013 | 0.003 | 0.003 | 0 | 0.7 | 0.75 | 0.76 | 0.99 | 0.66 | 0.81 |
| 5 | 118295726 | rs6894429 | 0.771 | 0.382 | -0.272 | -2.074 | -0.001 | 0.093 | 0.093 | 0.076 | 0.48 | 0.31 | 0.64 | 0.94 | 0.52 | 0.65 |
| 5 | 118299216 | rs6894997 | 0.475 | 0.433 | -0.269 | -2.081 | 0.001 | -0.007 | -0.007 | 0.56 | 0.52 | 0.59 | 0.66 | 0.96 | 0.52 | 0.67 |
| 5 | 118303481 | rs299201 | -0.721 | 0.513 | -0.576 | -2.026 | -0.079 | 0.127 | 0.127 | 0 | 0.75 | 0.92 | 0.78 | 0.99 | 0.67 | 0.82 |
| 5 | 118320318 | rs11959589 | 0.004 | 0.492 | 0.139 | -1.559 | -0.032 | -0.138 | -0.138 | 0 | 0.24 | 0.47 | 0.13 | 0.03 | 0.16 | 0.16 |
| 5 | 118325429 | rs299208 | -1.296 | 0.822 | -0.195 | -1.5 | -0.052 | 0.061 | 0.061 | 0 | 0.83 | 0.95 | 0.82 | 0.99 | 0.73 | 0.86 |
| 5 | 118326072 | rs976586 | -1.004 | 0.344 | 0.573 | -0.962 | -0.024 | -0.006 | -0.006 | 0.011 | 0.1 | 0.5 | 0.05 | 0.02 | 0.01 | 0.12 |
| 5 | 118349058 | rs17144869 | 1.033 | 0.838 | -0.132 | -0.397 | -0.032 | -0.029 | -0.029 | 0 | 0.1 | 0.03 | 0.13 | 0.38 | 0.19 | 0.26 |
| 5 | 118349195 | rs17144871 | -0.367 | 0.519 | 0.22 | -0.755 | -0.016 | -0.031 | -0.031 | 0 | 0.11 | 0.33 | 0.07 | 0.03 | 0.05 | 0.13 |
| 5 | 118357780 | rs7706037 | -0.701 | 0.263 | -0.729 | -1.005 | -0.037 | -0.112 | -0.112 | 0.976 | 0.28 | 0.57 | 0.13 | 0.03 | 0.16 | 0.2 |
| 5 | 118368516 | rs17144883 | 0.706 | 0.764 | -1.588 | -0.673 | 0.018 | 0.053 | 0.053 | 0.157 | 0.17 | 0.07 | 0.07 | 0.02 | 0.11 | 0.1 |
| 5 | 118394717 | rs11740134 | 0.48 | 1.21 | -1.573 | -0.193 | -0.06 | -0.073 | -0.073 | 0.573 | 0.12 | 0.03 | 0.17 | 0.34 | 0.22 | 0.36 |
| 5 | 118396228 | rs299216 | 0.486 | 0.75 | -1.5 | -0.984 | 0.007 | 0.035 | 0.035 | 0 | 0.21 | 0.12 | 0.1 | 0.01 | 0.17 | 0.1 |
| 5 | 118404296 | rs13189818 | 0.451 | 0.429 | -2.052 | -1.734 | -0.008 | 0.053 | 0.053 | 0 | 0.33 | 0.23 | 0.26 | 0.05 | 0.35 | 0.23 |
| 5 | 118405195 | rs12188483 | 0.363 | 0.093 | -1.796 | -2.073 | -0.031 | 0.121 | 0.121 | 0 | 0.37 | 0.19 | 0.57 | 0.94 | 0.43 | 0.58 |
| 5 | 118431709 | rs12516300 | 0.51 | 0.021 | -2.414 | -2.374 | -0.105 | 0.156 | 0.156 | 0 | 0.31 | 0.09 | 0.56 | 0.94 | 0.44 | 0.54 |
| 5 | 118453771 | rs4576176 | -1.869 | -0.035 | -2.384 | -2.353 | -0.04 | 0.114 | 0.114 | 0.053 | 0.82 | 0.99 | 0.9 | 0.99 | 0.84 | 0.89 |
| 5 | 118474109 | rs4895185 | 0.973 | -0.368 | -2.619 | -2.618 | -0.034 | 0.169 | 0.169 | 0 | 0.33 | 0.1 | 0.22 | 0.03 | 0.33 | 0.21 |
| 5 | 118474205 | rs4895361 | -1.95 | -0.324 | -2.637 | -2.581 | -0.039 | 0.116 | 0.116 | 0.261 | 0.82 | 0.99 | 0.9 | 0.99 | 0.84 | 0.9 |
| 5 | 118481205 | rs7717384 | 0.308 | -0.214 | -2.754 | -1.841 | 0.001 | -0.112 | -0.112 | 0 | 0.25 | 0.13 | 0.27 | 0.55 | 0.24 | 0.2 |
| 5 | 118495671 | rs11742202 | 1.308 | 0.048 | -2.495 | -1.565 | -0.052 | -0.077 | -0.077 | 1.537 | 0.13 | 0.03 | 0.32 | 0.38 | 0.22 | 0.38 |
| 5 | 118507250 | rs6866260 | 0.323 | 0.156 | -2.469 | -1.565 | -0.037 | -0.088 | -0.088 | 1.227 | 0.28 | 0.11 | 0.32 | 0.59 | 0.33 | 0.3 |
| 5 | 118516913 | rs7727906 | 0.128 | -0.553 | -1.057 | -1.85 | -0.039 | 0.103 | 0.103 | 1.312 | 0.39 | 0.22 | 0.62 | 0.94 | 0.52 | 0.62 |
| 5 | 118530872 | rs3992620 | 1.32 | -0.547 | -0.523 | -1.826 | 0.003 | -0.034 | -0.034 | 0.382 | 0.07 | 0.16 | 0.05 | 0 | 0.05 | 0.09 |
| 5 | 118532034 | rs6595178 | -0.431 | -0.567 | -0.655 | -1.934 | 0.004 | -0.052 | -0.052 | 0 | 0.51 | 0.58 | 0.71 | 0.96 | 0.61 | 0.74 |
| 5 | 118534517 | rs10064311 | 0.838 | -0.295 | -0.662 | -1.589 | -0.025 | 0.126 | 0.126 | 0 | 0.27 | 0.09 | 0.18 | 0.03 | 0.27 | 0.17 |
| 5 | 118553470 | rs3797337 | -0.07 | -0.514 | -0.146 | -1.291 | 0.003 | 0.102 | 0.102 | 0 | 0.91 | 0.65 | 0.93 | 0.87 | 0.94 | 0.87 |
| 5 | 118570172 | rs9327091 | 0.943 | -0.48 | -1.017 | -1.47 | 0.019 | 0.089 | 0.089 | 0 | 0.26 | 0.09 | 0.17 | 0.02 | 0.22 | 0.14 |
| 5 | 118583487 | rs2896840 | -0.985 | -0.161 | -0.697 | -1.145 | -0.005 | -0.039 | -0.039 | 0.246 | 0.93 | 0.86 | 0.95 | 1 | 0.95 | 0.91 |
| 5 | 118589485 | rs4895187 | 0.733 | 0.007 | -1.43 | -0.944 | -0.002 | 0.029 | 0.029 | 0 | 0.12 | 0.13 | 0.2 | 0.46 | 0.15 | 0.11 |
| 5 | 118597487 | rs1553544 | 0.789 | -0.387 | -1.645 | -2.102 | -0.113 | 0.051 | 0.051 | 0 | 0.33 | 0.1 | 0.56 | 0.8 | 0.45 | 0.56 |
| 5 | 118601775 | rs9327093 | -0.123 | -0.695 | -0.617 | -1.614 | -0.008 | -0.077 | -0.077 | 2.08 | 0.31 | 0.34 | 0.2 | 0.05 | 0.27 | 0.18 |
| 5 | 118605023 | rs1004531 | 0.527 | -0.54 | -1.102 | -1.488 | 0.045 | 0.13 | 0.13 | 0 | 0.59 | 0.22 | 0.71 | 0.86 | 0.67 | 0.73 |
| 5 | 118607342 | rs7704111 | 0.038 | -0.282 | -0.892 | -1.402 | 0.015 | 0.028 | 0.028 | 0.141 | 0.69 | 0.28 | 0.79 | 0.87 | 0.75 | 0.84 |

| | | | | | | | | | | | | | | | | |
|---|---|---|---|---|---|---|---|---|---|---|---|---|---|---|---|---|
| 5 | 118608427 | rs7709638 | -0.428 | 0.008 | -0.875 | -1.634 | 0.003 | -0.001 | -0.001 | 0 | 0.4 | 0.32 | 0.27 | 0.06 | 0.37 | 0.31 |
| 5 | 118610519 | rs1504977 | -0.528 | 0.125 | -0.534 | -1.422 | 0.002 | -0.031 | -0.031 | 0.719 | 0.29 | 0.35 | 0.32 | 0.06 | 0.34 | 0.21 |
| 5 | 118611118 | rs17145066 | 0.07 | -0.019 | -0.249 | -0.991 | 0.004 | 0.002 | 0.002 | 0 | 0.1 | 0.17 | 0.07 | 0.06 | 0.08 | 0.1 |
| 5 | 118619289 | rs4895188 | 0.04 | -0.72 | -0.173 | -1.171 | -0.042 | -0.133 | -0.133 | 1.031 | 0.8 | 0.49 | 0.88 | 0.98 | 0.87 | 0.85 |
| 5 | 118621164 | rs888772 | -1.057 | -0.764 | -0.235 | -1.191 | 0.003 | 0.062 | 0.062 | 0.066 | 0.87 | 0.96 | 0.94 | 0.99 | 0.9 | 0.85 |
| 5 | 118621672 | rs4895364 | 1.329 | -0.762 | -0.801 | -0.858 | -0.033 | -0.078 | -0.078 | 0 | 0.22 | 0.08 | 0.28 | 0.41 | 0.27 | 0.14 |
| 5 | 118624190 | rs905831 | 0.831 | -0.572 | -0.573 | -1.187 | -0.003 | -0.059 | -0.059 | 0.478 | 0.62 | 0.5 | 0.7 | 0.92 | 0.65 | 0.66 |
| 5 | 118624941 | rs6890983 | 1.11 | -0.637 | -0.609 | -1.462 | 0.041 | 0.098 | 0.098 | 0 | 0.54 | 0.25 | 0.65 | 0.84 | 0.65 | 0.64 |
| 5 | 118625333 | rs888769 | 0.438 | -0.375 | -0.508 | -1.467 | -0.018 | -0.072 | -0.072 | 0 | 0.67 | 0.51 | 0.75 | 0.93 | 0.74 | 0.68 |
| 5 | 118636227 | rs6897511 | -0.48 | -0.402 | -0.556 | -1.36 | -0.018 | -0.005 | -0.005 | 0 | 0.2 | 0.71 | 0.13 | 0.12 | 0.09 | 0.08 |
| 5 | 118639499 | rs10519581 | 0.867 | -0.346 | -0.596 | -1.345 | -0.014 | -0.058 | -0.058 | 0.321 | 0.11 | 0.22 | 0.06 | 0.02 | 0.06 | 0.02 |
| 5 | 118641096 | rs4895365 | 0.502 | -0.372 | -0.7 | -1.439 | -0.052 | 0.061 | 0.061 | 0 | 0.17 | 0.04 | 0.19 | 0.03 | 0.22 | 0.14 |
| 5 | 118642703 | rs10054006 | 1.131 | -0.28 | -0.669 | -1.325 | 0.025 | 0.161 | 0.161 | 0 | 0.5 | 0.22 | 0.62 | 0.84 | 0.58 | 0.63 |
| 5 | 118645320 | rs17145135 | 0.578 | -0.544 | -0.114 | -1.072 | -0.012 | -0.057 | -0.057 | 0 | 0.11 | 0.22 | 0.06 | 0.02 | 0.06 | 0.02 |
| 5 | 118649282 | rs6595183 | -0.67 | -0.375 | -0.694 | -0.929 | 0.017 | 0.049 | 0.049 | 0.247 | 0.33 | 0.74 | 0.27 | 0.08 | 0.24 | 0.2 |
| 5 | 118653933 | rs10066380 | 0.149 | -0.233 | -0.334 | -0.244 | 0.01 | 0 | 0 | 4.163 | 0.46 | 0.4 | 0.38 | 0.46 | 0.4 | 0.48 |
| 5 | 118658527 | rs7721976 | -0.41 | -0.116 | 0.475 | 0.069 | 0.06 | 0.105 | 0.105 | 1.063 | 0.06 | 0.44 | 0.06 | 0.03 | 0.01 | 0.01 |
| 5 | 118659579 | rs6888116 | -0.32 | -0.551 | -0.367 | -0.194 | 0.034 | -0.006 | -0.006 | 0 | 0.56 | 0.71 | 0.72 | 0.5 | 0.62 | 0.49 |
| 5 | 118660289 | rs11956215 | 0.319 | -0.519 | -0.622 | -0.352 | -0.009 | -0.017 | -0.017 | 0 | 0.08 | 0.17 | 0.08 | 0.05 | 0.06 | 0.04 |
| 5 | 118665393 | rs1876736 | 0.216 | 0.308 | -1.076 | -0.687 | -0.011 | -0.026 | -0.026 | 0 | 0.31 | 0.2 | 0.37 | 0.45 | 0.34 | 0.3 |
| 5 | 118668050 | rs986649 | 0.384 | 0.531 | -0.97 | -0.581 | -0.084 | 0.077 | 0.077 | 0.217 | 0.2 | 0.04 | 0.32 | 0.05 | 0.31 | 0.17 |
| 5 | 118669094 | rs250306 | 1.039 | -0.329 | -1.216 | -0.495 | -0.04 | -0.142 | -0.142 | 1.278 | 0.18 | 0.01 | 0.15 | 0.41 | 0.19 | 0.15 |
| 5 | 118671062 | rs11951771 | -0.071 | -0.172 | -0.334 | -0.695 | -0.007 | -0.012 | -0.012 | 0 | 0.08 | 0.21 | 0.08 | 0.03 | 0.06 | 0.03 |
| 5 | 118676529 | rs10519585 | -0.584 | 0.248 | -0.331 | -1.069 | -0.001 | -0.096 | -0.096 | 0 | 0.28 | 0.55 | 0.31 | 0.06 | 0.3 | 0.21 |
| 5 | 118677093 | rs250303 | 1.189 | 0.355 | -1.165 | -0.841 | -0.124 | -0.129 | -0.129 | 0 | 0.22 | 0.02 | 0.36 | 0.42 | 0.31 | 0.27 |
| 5 | 118677323 | rs10060971 | -0.578 | 0.019 | -0.418 | -0.946 | -0.001 | -0.011 | -0.011 | 0 | 0.1 | 0.43 | 0.12 | 0.03 | 0.06 | 0.08 |
| 5 | 118681650 | rs408095 | -0.799 | -0.072 | -0.664 | -1.049 | 0.033 | 0.004 | 0.004 | 0 | 0.54 | 0.69 | 0.73 | 0.47 | 0.62 | 0.48 |
| 5 | 118682789 | rs6878879 | 0.723 | -0.005 | -0.443 | -1.208 | -0.002 | 0.001 | 0.001 | 0.109 | 0.18 | 0.17 | 0.23 | 0.05 | 0.2 | 0.16 |
| 5 | 118685885 | rs1396609 | -0.907 | 0.1 | -0.608 | -1.208 | 0 | -0.008 | -0.008 | 0.02 | 0.31 | 0.67 | 0.37 | 0.08 | 0.31 | 0.24 |
| 5 | 118691888 | rs3797343 | -0.796 | -0.083 | -0.592 | -1.227 | 0 | -0.016 | -0.016 | 0 | 0.3 | 0.66 | 0.29 | 0.08 | 0.27 | 0.23 |
| 5 | 118696911 | rs6870253 | -0.846 | -0.148 | -0.621 | -1.249 | 0 | -0.015 | -0.015 | 0 | 0.31 | 0.66 | 0.36 | 0.08 | 0.29 | 0.24 |
| 5 | 118698583 | rs10064783 | -1.009 | -0.258 | -0.283 | -1.22 | -0.014 | -0.07 | -0.07 | 0 | 0.11 | 0.48 | 0.13 | 0 | 0.06 | 0.03 |
| 5 | 118699108 | rs13155880 | 0.398 | -0.187 | -0.645 | -1.158 | 0.034 | -0.007 | -0.007 | 0 | 0.48 | 0.32 | 0.28 | 0.49 | 0.41 | 0.52 |
| 5 | 118729327 | rs1045242 | -0.218 | -0.097 | -0.101 | -1.192 | 0.005 | -0.042 | -0.042 | 0.97 | 0.25 | 0.4 | 0.28 | 0.12 | 0.26 | 0.25 |
| 5 | 118733464 | rs17386422 | 1.394 | -0.487 | -0.419 | -1.537 | -0.007 | 0.056 | 0.056 | 0 | 0.14 | 0.06 | 0.09 | 0 | 0.15 | 0.06 |
| 5 | 118734731 | rs3813311 | -0.632 | -0.441 | -0.265 | -1.468 | 0.055 | 0.03 | 0.03 | 0 | 0.06 | 0.32 | 0.1 | 0.02 | 0.05 | 0.08 |
| 5 | 118740848 | rs17386541 | 0.489 | -0.54 | -0.661 | -1.598 | 0.008 | 0.025 | 0.025 | 0.033 | 0.3 | 0.24 | 0.18 | 0.06 | 0.27 | 0.36 |
| 5 | 118752111 | rs461108 | -0.572 | -0.482 | -0.471 | -1.299 | 0.001 | 0.002 | 0.002 | 0.053 | 0.57 | 0.48 | 0.4 | 0.49 | 0.51 | 0.6 |
| 5 | 118753515 | rs467076 | 0.922 | -0.443 | -0.96 | -1.274 | -0.062 | 0.055 | 0.055 | 0 | 0.2 | 0.04 | 0.21 | 0.08 | 0.25 | 0.36 |
| 5 | 118766143 | rs258047 | -2.155 | 0.031 | -0.615 | -0.933 | -0.065 | -0.087 | -0.087 | 0 | 0.79 | 0.94 | 0.64 | 0.57 | 0.7 | 0.79 |
| 5 | 118789338 | rs6897978 | -0.193 | -0.462 | -0.097 | -0.909 | -0.038 | 0.097 | 0.097 | 0 | 0.26 | 0.44 | 0.3 | 0.5 | 0.17 | 0.28 |
| 5 | 118789413 | rs6898518 | 1.338 | -0.453 | -0.048 | -0.904 | -0.001 | -0.002 | -0.002 | 0 | 0.15 | 0.15 | 0.13 | 0.1 | 0.12 | 0.15 |
| 5 | 118802221 | rs257977 | 1.79 | -0.57 | 0.045 | -1.017 | -0.001 | -0.022 | -0.022 | 0 | 0.11 | 0.1 | 0.06 | 0 | 0.08 | 0.03 |
| 5 | 118810964 | rs2287712 | -0.763 | -0.392 | 0.31 | -0.777 | -0.026 | 0.277 | 0.277 | 0.021 | 0.15 | 0.45 | 0.12 | 0.4 | 0.07 | 0.16 |
| 5 | 118811533 | rs25640 | 1.068 | -0.378 | 0.133 | -0.944 | -0.102 | -0.111 | -0.111 | 0.046 | 0.36 | 0.11 | 0.51 | 0.51 | 0.45 | 0.36 |
| 5 | 118812180 | rs2678070 | -0.03 | -0.715 | 0.26 | -0.94 | -0.001 | -0.035 | -0.035 | 0 | 0.49 | 0.46 | 0.56 | 0.91 | 0.49 | 0.52 |
| 5 | 118812383 | rs3756512 | 0.459 | -0.685 | 0.262 | -0.916 | -0.042 | 0.254 | 0.254 | 0.116 | 0.08 | 0.21 | 0.03 | 0.4 | 0.01 | 0.11 |
| 5 | 118812648 | rs3850200 | -0.83 | -0.683 | 0.232 | -0.919 | -0.026 | 0.266 | 0.266 | 0.067 | 0.15 | 0.43 | 0.12 | 0.4 | 0.07 | 0.15 |
| 5 | 118813240 | rs457106 | 1.09 | -0.618 | -0.077 | -0.806 | -0.194 | -0.17 | -0.17 | 0 | 0.36 | 0.03 | 0.51 | 0.55 | 0.47 | 0.37 |
| 5 | 118827601 | rs2546210 | 0.174 | -0.572 | 0.098 | -0.882 | -0.003 | 0 | 0 | 0.061 | 0.47 | 0.49 | 0.63 | 0.51 | 0.55 | 0.46 |
| 5 | 118836954 | rs3797363 | 0.986 | -0.603 | -0.006 | -0.741 | -0.024 | 0.128 | 0.128 | 0 | 0.06 | 0.08 | 0.02 | 0.4 | 0.01 | 0.11 |
| 5 | 118837658 | rs257969 | 1.79 | -0.496 | -0.228 | -0.837 | -0.148 | -0.135 | -0.135 | 0.983 | 0.24 | 0.02 | 0.38 | 0.41 | 0.37 | 0.22 |
| 5 | 118844486 | rs32662 | 0.545 | -0.57 | -0.03 | -0.883 | -0.029 | -0.032 | -0.032 | 0 | 0.4 | 0.26 | 0.54 | 0.51 | 0.48 | 0.37 |
| 5 | 118857497 | rs10069875 | -0.154 | -0.563 | 0.266 | -0.72 | -0.053 | 0.249 | 0.249 | 0.037 | 0.09 | 0.23 | 0.03 | 0.41 | 0.01 | 0.16 |
| 5 | 118861227 | rs1155777 | 1.804 | -0.33 | -0.025 | -0.775 | -0.103 | -0.101 | -0.101 | 0 | 0.23 | 0.05 | 0.37 | 0.41 | 0.35 | 0.21 |
| 5 | 118869440 | rs17145475 | -0.101 | -0.377 | 0.406 | -0.777 | 0.046 | -0.025 | -0.025 | 0 | 0.09 | 0.36 | 0.1 | 0 | 0.06 | 0.05 |
| 5 | 118890915 | rs246965 | -0.159 | -0.577 | -0.132 | -0.827 | 0.02 | -0.005 | -0.005 | 0 | 0.52 | 0.36 | 0.36 | 0.49 | 0.46 | 0.51 |
| 5 | 118891682 | rs246966 | -0.404 | -0.505 | 0.049 | -0.733 | 0.008 | -0.035 | -0.035 | 0 | 0.59 | 0.54 | 0.5 | 0.18 | 0.57 | 0.59 |
| 5 | 118900602 | rs92631 | -0.461 | -0.589 | -0.586 | -0.752 | 0.033 | 0.036 | 0.036 | 0 | 0.49 | 0.31 | 0.42 | 0.59 | 0.43 | 0.65 |
| 5 | 118906769 | rs17454921 | 1.7 | -0.529 | -0.885 | -0.796 | -0.005 | -0.009 | -0.009 | 0 | 0.12 | 0.01 | 0.12 | 0.07 | 0.1 | 0.19 |
| 5 | 118907117 | rs455355 | -0.107 | -0.709 | -0.492 | -0.887 | -0.02 | 0.025 | 0.025 | 0 | 0.36 | 0.51 | 0.28 | 0.49 | 0.32 | 0.49 |
| 5 | 118908168 | rs412069 | -1.365 | -0.271 | -1.048 | -0.745 | -0.008 | -0.012 | -0.012 | 0 | 0.71 | 0.76 | 0.63 | 0.59 | 0.68 | 0.78 |
| 5 | 118908524 | rs6864002 | 1.142 | 0.268 | -1.583 | -1.043 | -0.006 | 0.035 | 0.035 | 0.387 | 0.15 | 0.06 | 0.16 | 0.07 | 0.15 | 0.18 |
| 5 | 118908855 | rs405031 | 0.024 | -0.152 | -1.645 | -1.157 | -0.06 | 0.125 | 0.125 | 0 | 0.38 | 0.19 | 0.39 | 0.1 | 0.48 | 0.42 |
| 5 | 118909006 | rs1388107 | 0.639 | -0.105 | -1.322 | -0.566 | 0.015 | 0.032 | 0.032 | 0.124 | 0.15 | 0.06 | 0.15 | 0.07 | 0.12 | 0.18 |
| 5 | 118921139 | rs2896965 | 0.812 | 0.1 | -1.344 | -0.669 | -0.009 | 0.06 | 0.06 | 0 | 0.15 | 0.05 | 0.09 | 0 | 0.15 | 0.05 |
| 5 | 118922563 | rs40150 | -1.22 | 0.076 | -1.835 | -0.661 | -0.094 | -0.028 | -0.028 | 3.744 | 0.64 | 0.92 | 0.56 | 0.61 | 0.61 | 0.75 |
| 5 | 118923303 | rs10463719 | 0.234 | 0.145 | -1.79 | -0.686 | -0.035 | 0.126 | 0.126 | 0.438 | 0.31 | 0.1 | 0.28 | 0.08 | 0.32 | 0.25 |
| 5 | 118925050 | rs162198 | 0.431 | -0.303 | -0.972 | -0.777 | -0.011 | -0.075 | -0.075 | 0 | 0.31 | 0.52 | 0.29 | 0.1 | 0.33 | 0.42 |

| | | | | | | | | | | | | | | | | |
|---|---|---|---|---|---|---|---|---|---|---|---|---|---|---|---|---|
| 5 | 118926618 | rs328680 | -1.631 | -0.115 | -1.026 | -0.105 | -0.135 | -0.161 | -0.161 | 0.079 | 0.73 | 0.97 | 0.61 | 0.52 | 0.67 | 0.79 |
| 5 | 118931912 | rs1428432 | -0.659 | -0.162 | -0.302 | -0.52 | 0.006 | -0.049 | -0.049 | 0 | 0.63 | 0.53 | 0.57 | 0.18 | 0.65 | 0.68 |
| 5 | 118942612 | rs10059784 | 0.444 | -0.585 | 0.31 | -0.199 | -0.02 | 0.386 | 0.386 | 0 | 0.07 | 0.21 | 0.03 | 0.43 | 0.01 | 0.11 |
| 5 | 118943108 | rs7712974 | -0.608 | -0.617 | 0.164 | -0.205 | -0.037 | 0.485 | 0.485 | 1.439 | 0.1 | 0.43 | 0.03 | 0.43 | 0.01 | 0.11 |
| 5 | 118947467 | rs721676 | -0.115 | 0.19 | -0.806 | -0.841 | -0.084 | 0.146 | 0.146 | 0.488 | 0.35 | 0.09 | 0.29 | 0.09 | 0.41 | 0.28 |
| 5 | 118953519 | rs184497 | -0.532 | 0.142 | -0.101 | -0.673 | -0.002 | -0.003 | -0.003 | 0 | 0.54 | 0.45 | 0.5 | 0.54 | 0.56 | 0.61 |
| 5 | 118962192 | rs1318067 | -1.156 | -0.195 | -0.04 | -0.692 | -0.004 | 0.002 | 0.002 | 0 | 0.67 | 0.61 | 0.6 | 0.62 | 0.71 | 0.78 |
| 5 | 118962245 | rs1574641 | 0.435 | -0.131 | 0.097 | -0.784 | -0.015 | 0.011 | 0.011 | 0 | 0.21 | 0.15 | 0.21 | 0.12 | 0.27 | 0.22 |
| 5 | 118963199 | rs2115237 | -0.075 | -0.089 | 0.102 | -0.695 | -0.002 | 0 | 0 | 0 | 0.46 | 0.45 | 0.39 | 0.5 | 0.44 | 0.56 |
| 5 | 118968423 | rs328698 | -0.035 | -0.045 | -0.179 | -0.692 | 0.016 | -0.009 | -0.009 | 0 | 0.45 | 0.33 | 0.38 | 0.5 | 0.45 | 0.56 |
| 5 | 118969960 | rs328694 | -0.091 | -0.054 | -0.194 | -0.699 | 0.014 | -0.007 | -0.007 | 1.168 | 0.46 | 0.35 | 0.39 | 0.5 | 0.45 | 0.56 |
| 5 | 118972089 | rs2431378 | 0.138 | -0.134 | -0.338 | -0.935 | -0.009 | 0.107 | 0.107 | 0 | 0.38 | 0.21 | 0.36 | 0.09 | 0.43 | 0.4 |
| 5 | 118972113 | rs17145587 | -0.331 | 0.392 | 0.032 | -0.291 | 0.041 | -0.046 | -0.046 | 0.095 | 0.24 | 0.38 | 0.23 | 0.12 | 0.26 | 0.22 |
| 5 | 118972185 | rs2443743 | 1.088 | 0.704 | -0.763 | -0.005 | -0.016 | -0.015 | -0.015 | 0 | 0.06 | 0.02 | 0.15 | 0.08 | 0.13 | 0.19 |
| 5 | 118977870 | rs715715 | -0.993 | 0.186 | -1.467 | -0.31 | -0.027 | -0.025 | -0.025 | 0 | 0.68 | 0.86 | 0.6 | 0.64 | 0.63 | 0.71 |
| 5 | 118982484 | rs328647 | -1.139 | 0.281 | -1.718 | -0.283 | -0.07 | -0.059 | -0.059 | 0 | 0.68 | 0.93 | 0.59 | 0.63 | 0.59 | 0.69 |
| 5 | 118982698 | rs17145616 | 0.687 | -0.156 | -2.233 | -0.351 | -0.021 | -0.159 | -0.159 | 2.312 | 0.22 | 0.02 | 0.14 | 0.44 | 0.14 | 0.14 |
| 5 | 118986107 | rs10519604 | 0.242 | 0.05 | -2.34 | -0.457 | -0.04 | -0.165 | -0.165 | 1.78 | 0.26 | 0.03 | 0.19 | 0.44 | 0.2 | 0.16 |
| 5 | 118987108 | rs179274 | -1.453 | -0.062 | -1.993 | -0.401 | -0.01 | 0.017 | 0.017 | 0 | 0.6 | 0.75 | 0.55 | 0.63 | 0.54 | 0.62 |
| 5 | 118988269 | rs1820725 | 2.164 | -0.07 | -2.18 | -0.501 | -0.054 | -0.004 | -0.004 | 0 | 0.11 | 0 | 0.1 | 0.08 | 0.16 | 0.22 |
| 5 | 118989772 | rs329144 | -1.356 | -0.111 | -2.003 | -0.423 | -0.019 | 0.027 | 0.027 | 0 | 0.57 | 0.73 | 0.5 | 0.63 | 0.48 | 0.6 |
| 5 | 118990871 | rs780433 | 0.254 | 0.213 | -1.531 | -0.222 | -0.014 | 0.001 | 0.001 | 0 | 0.21 | 0.15 | 0.27 | 0.18 | 0.28 | 0.4 |
| 5 | 118995809 | rs330192 | -0.332 | 0.003 | -1.551 | 0.003 | -0.014 | -0.03 | -0.03 | 0 | 0.27 | 0.14 | 0.31 | 0.59 | 0.34 | 0.48 |
| 5 | 118996115 | rs780429 | -1.719 | -0.363 | -0.873 | -0.332 | 0.003 | 0.008 | 0.008 | 0.02 | 0.44 | 0.64 | 0.46 | 0.6 | 0.43 | 0.56 |
| 5 | 119002347 | rs1597713 | 0.354 | -0.415 | -0.886 | -0.07 | -0.004 | -0.037 | -0.037 | 0 | 0.19 | 0.11 | 0.13 | 0.5 | 0.2 | 0.28 |
| 5 | 119011525 | rs330206 | -0.986 | -0.558 | -0.854 | 0.032 | -0.074 | 0.16 | 0.16 | 0.038 | 0.17 | 0.42 | 0.11 | 0.43 | 0.06 | 0.11 |
| 5 | 119017300 | rs330201 | -0.856 | -0.067 | -1.665 | -0.444 | -0.052 | 0.004 | 0.004 | 0.413 | 0.77 | 0.96 | 0.7 | 0.86 | 0.76 | 0.73 |
| 5 | 119019195 | rs330200 | -0.841 | -0.053 | -1.659 | -0.422 | -0.051 | 0.005 | 0.005 | 0.272 | 0.77 | 0.96 | 0.7 | 0.86 | 0.76 | 0.73 |
| 5 | 119029069 | rs1451809 | -0.456 | -0.04 | -0.9 | -0.503 | 0 | -0.026 | -0.026 | 0.139 | 0.26 | 0.7 | 0.23 | 0.08 | 0.25 | 0.28 |
| 5 | 119046220 | rs17145711 | 1.264 | -0.073 | -1.424 | 0.208 | -0.042 | -0.033 | -0.033 | 0.087 | 0.29 | 0.05 | 0.2 | 0.34 | 0.35 | 0.29 |
| 5 | 119046825 | rs330173 | -0.19 | -0.657 | -0.759 | -0.183 | -0.056 | -0.062 | -0.062 | 0 | 0.18 | 0.27 | 0.16 | 0.05 | 0.06 | 0.08 |
| 5 | 119064355 | rs6890105 | 0.332 | -0.153 | -0.725 | 0.317 | -0.01 | 0.007 | 0.007 | 0.205 | 0.39 | 0.16 | 0.26 | 0.32 | 0.38 | 0.42 |
| 5 | 119065358 | rs162608 | 0.993 | -0.108 | -0.537 | 0.244 | 0.006 | 0.013 | 0.013 | 0 | 0.32 | 0.27 | 0.23 | 0.35 | 0.34 | 0.32 |
| 5 | 119067213 | rs259188 | 1.318 | -0.007 | -0.742 | 0.288 | -0.015 | -0.003 | -0.003 | 0 | 0.29 | 0.12 | 0.24 | 0.34 | 0.33 | 0.29 |
| 5 | 119082631 | rs162602 | 1.157 | 0.032 | -0.493 | 0.298 | 0.001 | 0.022 | 0.022 | 0 | 0.31 | 0.12 | 0.23 | 0.33 | 0.34 | 0.29 |
| 5 | 119092956 | rs259175 | 1.355 | 0.043 | -0.566 | 0.303 | -0.026 | 0 | 0 | 0 | 0.3 | 0.07 | 0.22 | 0.32 | 0.34 | 0.29 |
| 5 | 119122508 | rs2115138 | -0.137 | 0.64 | -0.3 | 0.533 | 0 | -0.039 | -0.039 | 0 | 0.42 | 0.46 | 0.3 | 0.14 | 0.4 | 0.44 |
| 5 | 119124054 | rs2973940 | -1.513 | 0.48 | -1.488 | 0.445 | -0.004 | -0.021 | -0.021 | 0 | 0.37 | 0.8 | 0.54 | 0.56 | 0.34 | 0.33 |
| 5 | 119126638 | rs1125859 | -1.51 | 0.401 | -1.616 | 0.454 | 0.001 | 0.03 | 0.03 | 0 | 0.62 | 0.95 | 0.76 | 0.65 | 0.65 | 0.68 |
| 5 | 119133696 | rs11747748 | 0.174 | 0.46 | -1.441 | 0.397 | 0.006 | -0.003 | -0.003 | 0 | 0.14 | 0.11 | 0.05 | 0.02 | 0.07 | 0.09 |
| 5 | 119145275 | rs7709933 | 0.529 | 0.838 | -1.793 | 0.898 | -0.09 | -0.08 | -0.08 | 1.717 | 0.31 | 0.06 | 0.25 | 0.36 | 0.39 | 0.4 |
| 5 | 119159236 | rs10793801 | -1.222 | 1.179 | -1.37 | 0.662 | -0.007 | -0.012 | -0.012 | 0 | 0.36 | 0.77 | 0.54 | 0.54 | 0.34 | 0.42 |
| 5 | 119165688 | rs2914045 | 1.111 | 0.919 | -1.764 | 0.683 | 0.019 | -0.033 | -0.033 | 0.032 | 0.64 | 0.15 | 0.47 | 0.45 | 0.7 | 0.57 |
| 5 | 119177725 | rs12518342 | 0.031 | 0.901 | -1.969 | 0.56 | -0.082 | 0.024 | 0.024 | 0.244 | 0.38 | 0.07 | 0.66 | 0.27 | 0.56 | 0.5 |
| 5 | 119182542 | rs2973996 | 0.933 | 0.983 | -1.972 | 0.805 | -0.026 | -0.132 | -0.132 | 0 | 0.38 | 0.05 | 0.24 | 0.5 | 0.36 | 0.35 |
| 5 | 119183940 | rs12719264 | 1.005 | 0.952 | -1.939 | 0.807 | -0.019 | -0.1 | -0.1 | 1.416 | 0.37 | 0.06 | 0.24 | 0.45 | 0.35 | 0.34 |
| 5 | 119195837 | rs4131101 | -0.034 | 1.137 | -1.298 | 0.67 | -0.007 | -0.014 | -0.014 | 0 | 0.58 | 0.33 | 0.58 | 0.5 | 0.56 | 0.53 |
| 5 | 119196820 | rs3935993 | 0.089 | 1.126 | -0.862 | 0.647 | -0.001 | -0.005 | -0.005 | 0.034 | 0.55 | 0.34 | 0.37 | 0.49 | 0.46 | 0.51 |
| 5 | 119199265 | rs2974025 | -0.272 | 0.549 | -0.231 | 0.649 | 0.013 | 0.325 | 0.325 | 1.534 | 0.91 | 0.59 | 0.97 | 0.71 | 0.97 | 0.97 |
| 5 | 119203911 | rs7701785 | 1.057 | 0.873 | -0.708 | 0.263 | 0.003 | -0.001 | -0.001 | 0 | 0.34 | 0.13 | 0.27 | 0.34 | 0.31 | 0.32 |
| 5 | 119210995 | rs2973958 | 0.45 | 0.584 | -0.024 | 0.074 | -0.002 | -0.03 | -0.03 | 0 | 0.8 | 0.8 | 0.75 | 0.98 | 0.83 | 0.83 |
| 5 | 119228851 | rs2973963 | -0.983 | 0.872 | -0.244 | 0.149 | -0.042 | 0.097 | 0.097 | 0.073 | 0.72 | 0.92 | 0.74 | 0.93 | 0.68 | 0.81 |
| 5 | 119228879 | rs11241518 | -0.188 | 0.655 | 0.431 | -0.057 | -0.003 | -0.066 | -0.066 | 0.076 | 0.13 | 0.26 | 0.22 | 0.01 | 0.13 | 0.17 |
| 5 | 119231326 | rs2974041 | -0.583 | 0.344 | 0.669 | 0.209 | -0.043 | 0.13 | 0.13 | 0.197 | 0.13 | 0.45 | 0.04 | 0.23 | 0.04 | 0.06 |
| 5 | 119232050 | rs2974039 | -0.946 | 0.671 | 0.234 | 0.422 | -0.09 | -0.039 | -0.039 | 0 | 0.68 | 0.91 | 0.47 | 0.67 | 0.59 | 0.69 |
| 5 | 119233362 | rs4272156 | 0.563 | 0.672 | 0.342 | 0.416 | -0.001 | -0.022 | -0.022 | 0 | 0.11 | 0.11 | 0.06 | 0.01 | 0.11 | 0.17 |
| 5 | 119234983 | rs4352622 | 1.012 | 0.627 | 0.386 | 0.42 | -0.005 | -0.03 | -0.03 | 1.545 | 0.42 | 0.16 | 0.36 | 0.45 | 0.45 | 0.45 |
| 5 | 119235327 | rs12188729 | -0.949 | 0.593 | 1.438 | 0.871 | -0.046 | 0.1 | 0.1 | 0.077 | 0.13 | 0.43 | 0.04 | 0.2 | 0.04 | 0.06 |
| 5 | 119243084 | rs7725436 | -1.616 | 0.612 | 1.331 | 0.879 | -0.028 | -0.011 | -0.011 | 2.429 | 0.25 | 0.61 | 0.25 | 0.22 | 0.17 | 0.24 |
| 5 | 119253621 | rs7713766 | -2.161 | 0.683 | 1.222 | 0.877 | 0.021 | -0.05 | -0.05 | 0 | 0.3 | 0.78 | 0.37 | 0.47 | 0.2 | 0.36 |
| 5 | 119270840 | rs4295414 | -0.191 | 0.6 | 1.02 | 0.69 | -0.002 | -0.036 | -0.036 | 0 | 0.13 | 0.19 | 0.21 | 0.02 | 0.13 | 0.17 |
| 5 | 119276292 | rs7713770 | -1.284 | 0.569 | 1.159 | 0.841 | -0.053 | 0.049 | 0.049 | 0.705 | 0.14 | 0.59 | 0.05 | 0.2 | 0.04 | 0.06 |
| 5 | 119290814 | rs4496729 | 0.681 | 0.736 | 0.585 | 0.791 | 0.023 | -0.031 | -0.031 | 0 | 0.35 | 0.15 | 0.32 | 0.42 | 0.32 | 0.47 |
| 5 | 119298330 | rs10068616 | -0.602 | 0.951 | 0.615 | 0.519 | -0.002 | 0.085 | 0.085 | 0.476 | 0.56 | 0.8 | 0.69 | 0.75 | 0.57 | 0.78 |
| 5 | 119316941 | rs12186828 | -0.568 | -0.005 | 1.432 | 0.669 | 0.005 | 0.051 | 0.051 | 0.662 | 0.16 | 0.26 | 0.29 | 0.29 | 0.16 | 0.29 |
| 5 | 119317372 | rs931452 | -0.426 | -0.006 | 1.572 | 0.598 | 0 | 0.001 | 0.001 | 0.175 | 0.52 | 0.53 | 0.61 | 0.74 | 0.5 | 0.76 |
| 5 | 119319410 | rs11241528 | -0.344 | 0.011 | 1.245 | 0.623 | 0.003 | 0.027 | 0.027 | 0.596 | 0.15 | 0.19 | 0.28 | 0.28 | 0.16 | 0.29 |
| 5 | 119328679 | rs10447240 | -0.616 | -0.034 | 1.136 | 0.396 | -0.034 | 0.107 | 0.107 | 0.749 | 0.55 | 0.86 | 0.63 | 0.74 | 0.49 | 0.77 |
| 5 | 119328777 | rs17145952 | 0.658 | 0.05 | 1.032 | 0.495 | -0.006 | 0.005 | 0.005 | 0 | 0.37 | 0.36 | 0.33 | 0.47 | 0.33 | 0.47 |

| | | | | | | | | | | | | | | | | |
|---|---|---|---|---|---|---|---|---|---|---|---|---|---|---|---|---|
| 5 | 119329324 | rs6875179 | -0.342 | 0.056 | 0.981 | 0.365 | 0 | 0.001 | 0.001 | 0.078 | 0.52 | 0.47 | 0.61 | 0.76 | 0.49 | 0.76 |
| 5 | 119332670 | rs9327121 | 0.478 | 0.05 | 0.198 | 0.444 | -0.014 | 0.016 | 0.016 | 0.109 | 0.37 | 0.52 | 0.33 | 0.49 | 0.33 | 0.47 |
| 5 | 119342045 | rs7718303 | -0.073 | 0.03 | -0.046 | 0.519 | 0.038 | 0.017 | 0.017 | 0.773 | 0.43 | 0.73 | 0.51 | 0.61 | 0.37 | 0.59 |
| 5 | 119368801 | rs7735851 | 0.354 | -0.134 | -0.485 | 0.557 | -0.007 | 0.004 | 0.004 | 0 | 0.41 | 0.53 | 0.34 | 0.49 | 0.33 | 0.49 |
| 5 | 119372015 | rs13186435 | 0.238 | 0.018 | -1.738 | 0.755 | 0.014 | -0.057 | -0.057 | 0 | 0.43 | 0.12 | 0.34 | 0.49 | 0.4 | 0.48 |
| 5 | 119372212 | rs173219 | -0.247 | 0.269 | -1.238 | 0.839 | 0.008 | -0.006 | -0.006 | 0 | 0.94 | 0.95 | 0.94 | 0.83 | 0.98 | 0.97 |
| 5 | 119392433 | rs1209532 | 0.179 | 0.519 | -1.799 | 0.068 | -0.118 | 0.133 | 0.133 | 0.122 | 0.38 | 0.06 | 0.39 | 0.14 | 0.52 | 0.21 |
| 5 | 119399177 | rs11959439 | -0.44 | -0.752 | -0.557 | 0.385 | -0.063 | 0.093 | 0.093 | 1.117 | 0.19 | 0.46 | 0.16 | 0.3 | 0.1 | 0.23 |
| 5 | 119410495 | rs10052192 | -0.546 | -0.981 | -0.159 | 0.2 | -0.021 | 0.007 | 0.007 | 3.081 | 0.26 | 0.68 | 0.21 | 0.34 | 0.15 | 0.24 |
| 5 | 119412658 | rs1191746 | 0.963 | -1.003 | -0.594 | 0.082 | -0.173 | 0.165 | 0.165 | 1.175 | 0.37 | 0.03 | 0.38 | 0.1 | 0.5 | 0.18 |
| 5 | 119413039 | rs1191745 | -0.212 | -1.159 | -0.236 | 0.215 | -0.041 | 0.077 | 0.077 | 0 | 0.8 | 0.7 | 0.82 | 0.6 | 0.87 | 0.76 |
| 5 | 119413303 | rs1191743 | 0.303 | -1.156 | -0.326 | 0.216 | 0.072 | -0.072 | -0.072 | 0 | 0.75 | 0.13 | 0.79 | 0.6 | 0.87 | 0.75 |
| 5 | 119434193 | rs1348478 | -0.48 | -1.146 | -0.347 | 0.182 | 0.043 | -0.028 | -0.028 | 0 | 0.25 | 0.77 | 0.2 | 0.34 | 0.13 | 0.24 |
| 5 | 119437585 | rs4895232 | 1.154 | -1.212 | -0.79 | -0.185 | -0.02 | -0.117 | -0.117 | 2.07 | 0.23 | 0.06 | 0.32 | 0.49 | 0.21 | 0.45 |
| 5 | 119439819 | rs7721883 | -1.402 | -0.946 | -0.68 | -0.115 | -0.173 | 0.092 | 0.092 | 0.8 | 0.63 | 0.97 | 0.61 | 0.84 | 0.49 | 0.76 |
| 5 | 119488924 | rs1981819 | -0.34 | -0.786 | -0.668 | -0.259 | 0.096 | 0.014 | 0.014 | 1.484 | 0.39 | 0.86 | 0.27 | 0.34 | 0.25 | 0.34 |
| 5 | 119502826 | rs982476 | -0.298 | -0.781 | -0.233 | -0.257 | 0.095 | 0.005 | 0.005 | 1.364 | 0.39 | 0.88 | 0.27 | 0.33 | 0.25 | 0.34 |
| 5 | 119503363 | rs4354076 | -0.354 | -0.589 | -0.268 | -0.269 | -0.005 | 0.025 | 0.025 | 0 | 0.62 | 0.51 | 0.58 | 0.49 | 0.66 | 0.57 |
| 5 | 119507243 | rs1348473 | 0.87 | -0.086 | -0.451 | -0.061 | -0.158 | -0.007 | -0.007 | 0.015 | 0.27 | 0.02 | 0.35 | 0.19 | 0.42 | 0.23 |
| 5 | 119526332 | rs2112498 | 0.496 | 0.449 | 0.319 | 0.97 | -0.044 | -0.118 | -0.118 | 0 | 0.21 | 0.06 | 0.41 | 0.47 | 0.23 | 0.4 |
| 5 | 119526884 | rs1422377 | 0.101 | 0.184 | 0.57 | 0.783 | -0.006 | 0.131 | 0.131 | 2.168 | 0.37 | 0.12 | 0.25 | 0.18 | 0.33 | 0.29 |
| 5 | 119528887 | rs11951942 | -0.533 | 0.167 | 0.622 | 0.825 | 0.008 | 0.007 | 0.007 | 0.578 | 0.44 | 0.61 | 0.34 | 0.38 | 0.36 | 0.35 |
| 5 | 119532996 | rs17146259 | -0.471 | 0.19 | 0.524 | 0.798 | -0.003 | -0.002 | -0.002 | 0 | 0.44 | 0.54 | 0.34 | 0.38 | 0.36 | 0.35 |
| 5 | 119544266 | rs13156712 | -0.175 | 0.257 | 0.265 | 0.622 | -0.008 | -0.042 | -0.042 | 0 | 0.41 | 0.47 | 0.28 | 0.18 | 0.35 | 0.31 |
| 5 | 119564388 | rs2161264 | -0.052 | 0.237 | 0.315 | 0.641 | 0.003 | 0.006 | 0.006 | 1.321 | 0.4 | 0.38 | 0.38 | 0.21 | 0.35 | 0.33 |
| 5 | 119577574 | rs1422384 | -0.471 | 0.19 | 0.469 | 0.838 | 0.006 | 0.003 | 0.003 | 0.179 | 0.46 | 0.58 | 0.43 | 0.41 | 0.38 | 0.37 |
| 5 | 119580326 | rs1363438 | -0.093 | 0.28 | 0.164 | 0.354 | 0.004 | 0.009 | 0.009 | 0.671 | 0.41 | 0.38 | 0.46 | 0.32 | 0.34 | 0.33 |
| 5 | 119587219 | rs7711327 | 0.326 | 0.371 | -0.097 | -0.418 | -0.006 | 0.059 | 0.059 | 4.864 | 0.24 | 0.15 | 0.23 | 0.14 | 0.25 | 0.14 |
| 5 | 119588486 | rs17146327 | 0.266 | 0.323 | -0.037 | -0.464 | -0.001 | 0.044 | 0.044 | 5.401 | 0.25 | 0.16 | 0.24 | 0.17 | 0.25 | 0.14 |
| 5 | 119593371 | rs1422392 | -0.537 | 0.324 | 0.362 | -0.529 | 0.007 | 0.014 | 0.014 | 0.794 | 0.51 | 0.41 | 0.53 | 0.56 | 0.46 | 0.42 |
| 5 | 119596306 | rs2112495 | -1.505 | 0.531 | 0.15 | -0.767 | -0.039 | 0.044 | 0.044 | 0.006 | 0.73 | 0.91 | 0.8 | 0.81 | 0.65 | 0.8 |
| 5 | 119600027 | rs17146331 | 0.595 | 0.559 | 0.146 | -0.675 | -0.013 | 0.033 | 0.033 | 0 | 0.26 | 0.3 | 0.29 | 0.38 | 0.19 | 0.26 |
| 5 | 119600902 | rs870009 | -0.3 | 0.61 | 0.172 | -0.847 | 0.002 | 0.004 | 0.004 | 0.682 | 0.51 | 0.48 | 0.54 | 0.56 | 0.46 | 0.42 |
| 5 | 119603547 | rs1001012 | 0.055 | 0.602 | -0.079 | -0.849 | -0.008 | -0.01 | -0.01 | 3.213 | 0.44 | 0.17 | 0.49 | 0.56 | 0.42 | 0.38 |
| 5 | 119614401 | rs4498277 | -0.147 | 0.446 | 0.174 | -0.839 | 0.003 | -0.024 | -0.024 | 0.404 | 0.94 | 0.82 | 0.97 | 1 | 0.96 | 0.97 |
| 5 | 119615110 | rs12108988 | 1.16 | 0.416 | -0.002 | -0.807 | 0.017 | -0.061 | -0.061 | 0.384 | 0.21 | 0.1 | 0.26 | 0.42 | 0.16 | 0.23 |
| 5 | 119627087 | rs4379229 | -0.142 | 0.71 | 0.211 | -0.811 | 0 | 0 | 0 | 0 | 0.43 | 0.41 | 0.48 | 0.39 | 0.42 | 0.49 |
| 5 | 119635847 | rs10478459 | 0.066 | 0.706 | 0.193 | -1.084 | -0.008 | -0.085 | -0.085 | 0.938 | 0.68 | 0.53 | 0.71 | 0.85 | 0.7 | 0.8 |
| 5 | 119636391 | rs9327136 | 0.257 | 0.59 | 0.184 | -1.208 | -0.018 | -0.031 | -0.031 | 0 | 0.39 | 0.49 | 0.51 | 0.66 | 0.3 | 0.52 |
| 5 | 119636402 | rs9327137 | -0.345 | 0.328 | 0.394 | -1.334 | -0.001 | -0.01 | -0.01 | 0 | 0.6 | 0.47 | 0.46 | 0.32 | 0.65 | 0.49 |
| 5 | 119640500 | rs12653659 | 0.031 | 0.621 | 0.015 | -1.248 | 0.003 | 0.012 | 0.012 | 0 | 0.45 | 0.62 | 0.52 | 0.57 | 0.41 | 0.45 |
| 5 | 119641791 | rs1368188 | -0.069 | 0.627 | -0.063 | -1.239 | 0.008 | 0.025 | 0.025 | 0 | 0.46 | 0.69 | 0.53 | 0.57 | 0.41 | 0.45 |
| 5 | 119641970 | rs1368187 | -1.109 | 0.569 | -0.204 | -1.345 | -0.004 | 0.005 | 0.005 | 0 | 0.74 | 0.8 | 0.71 | 0.76 | 0.75 | 0.67 |
| 5 | 119657733 | rs10519630 | 0.214 | 0.778 | -0.345 | -1.356 | 0.004 | -0.007 | -0.007 | 0 | 0.11 | 0.14 | 0.17 | 0.13 | 0.14 | 0.2 |
| 5 | 119660331 | rs17146446 | 0.547 | 0.937 | -0.707 | -1.275 | -0.013 | 0.023 | 0.023 | 0.011 | 0.36 | 0.5 | 0.35 | 0.46 | 0.28 | 0.26 |
| 5 | 119661088 | rs2405806 | 0.062 | 0.862 | -0.685 | -1.224 | -0.037 | 0.05 | 0.05 | 0 | 0.26 | 0.05 | 0.18 | 0.17 | 0.33 | 0.2 |
| 5 | 119666172 | rs17480640 | -0.615 | 0.844 | -0.284 | -1.19 | -0.02 | 0.039 | 0.039 | 0 | 0.4 | 0.31 | 0.44 | 0.23 | 0.52 | 0.42 |
| 5 | 119668237 | rs7728770 | -0.532 | 0.476 | -0.326 | -1.171 | 0.007 | -0.036 | -0.036 | 0 | 0.37 | 0.59 | 0.41 | 0.25 | 0.41 | 0.43 |
| 5 | 119669457 | rs1368183 | -1.019 | 0.138 | -0.738 | -1.558 | 0.001 | 0.017 | 0.017 | 0 | 0.18 | 0.12 | 0.13 | 0 | 0.16 | 0.09 |
| 5 | 119672998 | rs7717041 | -0.968 | -0.983 | -0.508 | -2.238 | -0.023 | -0.075 | -0.075 | 0 | 0.36 | 0.43 | 0.26 | 0.08 | 0.25 | 0.28 |
| 5 | 119675960 | rs6860067 | -0.071 | -1.192 | -0.728 | -2.361 | -0.041 | 0.013 | 0.013 | 0 | 0.12 | 0.03 | 0.16 | 0.08 | 0.15 | 0.19 |
| 5 | 119682607 | rs17164648 | -0.422 | -1.309 | -0.668 | -2.383 | 0.007 | 0.005 | 0.005 | 0 | 0.08 | 0.06 | 0.05 | 0 | 0.07 | 0.06 |
| 5 | 119682668 | rs6899092 | -2.073 | -1.709 | -0.131 | -2.659 | -0.009 | -0.056 | -0.056 | 0 | 0.42 | 0.53 | 0.29 | 0.09 | 0.29 | 0.35 |
| 5 | 119686188 | rs6863052 | -2.531 | -2.187 | -0.921 | -3.075 | -0.074 | 0.242 | 0.242 | 0.98 | 0.59 | 0.97 | 0.39 | 0.1 | 0.42 | 0.41 |
| 5 | 119690431 | rs17146503 | 1.119 | -2.223 | -0.662 | -2.976 | -0.018 | -0.023 | -0.023 | 2.697 | 0.11 | 0.11 | 0.02 | 0 | 0.01 | 0.01 |
| 5 | 119696973 | rs4895244 | -0.079 | -2.155 | -0.725 | -2.957 | -0.017 | 0.029 | 0.029 | 0 | 0.15 | 0.08 | 0.2 | 0.08 | 0.19 | 0.2 |
| 5 | 119698794 | rs4895155 | -0.202 | -2.119 | -0.727 | -3.085 | -0.05 | 0.086 | 0.086 | 0 | 0.15 | 0.04 | 0.19 | 0.01 | 0.18 | 0.19 |
| 5 | 119709256 | rs17146550 | 0.282 | -2.169 | -0.439 | -3.074 | -0.025 | -0.026 | -0.026 | 0 | 0.05 | 0.1 | 0.01 | 0 | 0.01 | 0.02 |
| 5 | 119720613 | rs4895248 | -1.373 | -2.242 | -0.433 | -3.136 | 0.036 | 0.092 | 0.092 | 0.316 | 0.33 | 0.18 | 0.25 | 0 | 0.26 | 0.26 |
| 5 | 119741331 | rs6864618 | -0.009 | -2.138 | -0.235 | -3.004 | -0.009 | -0.033 | -0.033 | 0.836 | 0.08 | 0.11 | 0.05 | 0 | 0.07 | 0.04 |
| 5 | 119771119 | rs17146596 | -0.571 | -2.225 | -0.278 | -3.061 | -0.01 | -0.022 | -0.022 | 0.217 | 0.09 | 0.08 | 0.02 | 0 | 0.02 | 0.07 |
| 5 | 119771549 | rs17146599 | -0.216 | -2.141 | -0.315 | -3.01 | -0.005 | -0.03 | -0.03 | 0 | 0.18 | 0.18 | 0.06 | 0 | 0.08 | 0.04 |
| 5 | 119782404 | rs10478465 | -1.113 | -2.235 | 0.076 | -3.046 | -0.056 | -0.104 | -0.104 | 0 | 0.24 | 0.34 | 0.09 | 0.02 | 0.12 | 0.08 |
| 5 | 119783715 | rs10043459 | 0.77 | -2.196 | 0.036 | -3.109 | -0.007 | 0.029 | 0.029 | 0 | 0.09 | 0.04 | 0.07 | 0.02 | 0.1 | 0.05 |
| 5 | 119786093 | rs12109029 | -2.089 | -2.37 | 0.187 | -3.348 | -0.013 | -0.115 | -0.115 | 0 | 0.46 | 0.49 | 0.31 | 0.03 | 0.31 | 0.35 |
| 5 | 119786253 | rs2973133 | -2.758 | -2.171 | -0.083 | -3.353 | -0.13 | 0.297 | 0.297 | 1.536 | 0.6 | 0.97 | 0.42 | 0.06 | 0.44 | 0.42 |
| 5 | 119789615 | rs17424128 | -1.19 | -1.961 | 0.158 | -3.211 | 0 | -0.037 | -0.037 | 0 | 0.23 | 0.22 | 0.22 | 0.01 | 0.2 | 0.27 |
| 5 | 119803829 | rs4895254 | -2.276 | -1.618 | 0.285 | -3.106 | 0.004 | -0.016 | -0.016 | 0 | 0.53 | 0.56 | 0.4 | 0.09 | 0.45 | 0.4 |
| 5 | 119806872 | rs7715549 | -1.531 | -1.563 | 0.265 | -3.188 | -0.005 | -0.078 | -0.078 | 0.904 | 0.24 | 0.31 | 0.25 | 0.04 | 0.22 | 0.28 |

| | | | | | | | | | | | | | | | | |
|---|---|---|---|---|---|---|---|---|---|---|---|---|---|---|---|---|
| 5 | 119809193 | rs2601208 | -1.589 | -1.567 | 0.254 | -3.187 | -0.027 | -0.135 | -0.135 | 0 | 0.38 | 0.49 | 0.2 | 0.03 | 0.27 | 0.17 |
| 5 | 119811722 | rs17146665 | -1.232 | -1.477 | 0.114 | -2.987 | -0.003 | -0.045 | -0.045 | 0 | 0.21 | 0.23 | 0.21 | 0.02 | 0.16 | 0.21 |
| 5 | 119814488 | rs6595236 | -0.808 | -1.467 | 0.192 | -2.85 | -0.012 | -0.087 | -0.087 | 0 | 0.27 | 0.27 | 0.15 | 0.01 | 0.2 | 0.13 |
| 5 | 119816120 | rs2655067 | 1.461 | -1.472 | 0.186 | -2.819 | -0.024 | -0.118 | -0.118 | 0 | 0.64 | 0.5 | 0.8 | 0.96 | 0.75 | 0.81 |
| 5 | 119838578 | rs10478474 | -1.547 | -1.708 | 0.206 | -2.818 | -0.049 | -0.155 | -0.155 | 0.38 | 0.34 | 0.5 | 0.17 | 0.01 | 0.2 | 0.15 |
| 5 | 119842665 | rs13356782 | -0.511 | -1.804 | 0.235 | -2.865 | 0 | -0.025 | -0.025 | 2.279 | 0.17 | 0.15 | 0.12 | 0.01 | 0.16 | 0.11 |
| 5 | 119843757 | rs9327146 | -1.437 | -1.663 | 0.074 | -2.791 | -0.047 | -0.155 | -0.155 | 0 | 0.34 | 0.5 | 0.17 | 0.01 | 0.2 | 0.15 |
| 5 | 119846247 | rs16766 | -1.451 | -1.62 | 0.048 | -2.786 | -0.044 | -0.149 | -0.149 | 1.38 | 0.36 | 0.46 | 0.21 | 0.04 | 0.25 | 0.19 |
| 5 | 119866504 | rs4895161 | -1.59 | -1.389 | -0.112 | -2.659 | 0.028 | 0.094 | 0.094 | 0.404 | 0.42 | 0.71 | 0.32 | 0.06 | 0.35 | 0.3 |
| 5 | 119868284 | rs10066512 | -1.086 | -1.232 | 0.252 | -2.315 | -0.008 | -0.052 | -0.052 | 0 | 0.2 | 0.24 | 0.13 | 0.03 | 0.18 | 0.1 |
| 5 | 119869631 | rs13161840 | -0.166 | -0.62 | -0.754 | -1.856 | -0.04 | -0.031 | -0.031 | 0.936 | 0.26 | 0.11 | 0.32 | 0.67 | 0.32 | 0.32 |
| 5 | 119869763 | rs300974 | 0.411 | -0.934 | -0.955 | -2.456 | 0.007 | 0.004 | 0.004 | 0 | 0.65 | 0.38 | 0.8 | 0.86 | 0.8 | 0.68 |
| 5 | 119876209 | rs300970 | 1.217 | -0.241 | -0.531 | -2.139 | 0.005 | 0.025 | 0.025 | 0 | 0.5 | 0.44 | 0.45 | 0.08 | 0.52 | 0.43 |
| 5 | 119877678 | rs10074464 | 0.283 | 0.08 | -0.467 | -1.692 | -0.006 | 0.006 | 0.006 | 0 | 0.18 | 0.12 | 0.13 | 0.15 | 0.21 | 0.19 |
| 5 | 119879082 | rs10063554 | 0.146 | 0.073 | -0.333 | -1.693 | 0 | -0.002 | -0.002 | 1.477 | 0.19 | 0.19 | 0.13 | 0.15 | 0.21 | 0.18 |
| 5 | 119884420 | rs2174236 | 1.388 | 0.078 | -0.373 | -0.712 | 0.002 | 0.013 | 0.013 | 0 | 0.51 | 0.44 | 0.45 | 0.12 | 0.52 | 0.44 |
| 5 | 119885348 | rs1157177 | 0.417 | 0.094 | -0.254 | -0.476 | 0.021 | -0.05 | -0.05 | 0 | 0.19 | 0.12 | 0.27 | 0.6 | 0.16 | 0.3 |
| 5 | 119887839 | rs12109575 | -0.331 | 0.688 | 0.167 | -0.175 | 0.001 | -0.004 | -0.004 | 1.001 | 0.19 | 0.22 | 0.14 | 0.15 | 0.21 | 0.18 |
| 5 | 119892906 | rs10077688 | -0.134 | 0.705 | 0.167 | -0.162 | -0.005 | 0.004 | 0.004 | 0 | 0.19 | 0.14 | 0.13 | 0.15 | 0.21 | 0.18 |
| 5 | 119901028 | rs10478478 | -1.396 | 0.721 | 0.107 | -0.195 | 0.019 | 0.001 | 0.001 | 0 | 0.27 | 0.37 | 0.21 | 0.25 | 0.31 | 0.27 |
| 5 | 119903794 | rs17146783 | -0.811 | 0.749 | 0.11 | -0.21 | 0.015 | 0.019 | 0.019 | 0 | 0.08 | 0.08 | 0.07 | 0.11 | 0.1 | 0.09 |
| 5 | 119903803 | rs10478480 | -0.343 | 0.566 | 0.061 | -0.433 | 0.001 | 0.001 | 0.001 | 0 | 0.19 | 0.21 | 0.13 | 0.19 | 0.22 | 0.18 |
| 5 | 119911812 | rs13171462 | 0.564 | 0.533 | -0.281 | -0.561 | 0.026 | -0.025 | -0.025 | 11.902 | 0.77 | 0.58 | 0.83 | 0.84 | 0.76 | 0.79 |
| 5 | 119918341 | rs17427447 | 0.262 | 0.115 | -1.279 | -1.281 | 0.048 | 0.137 | 0.137 | 0 | 0.34 | 0.14 | 0.16 | 0.03 | 0.26 | 0.3 |
| 5 | 119919047 | rs9765600 | -0.6 | 0.308 | -0.743 | -1.331 | -0.003 | 0.013 | 0.013 | 0.548 | 0.58 | 0.61 | 0.47 | 0.22 | 0.44 | 0.49 |
| 5 | 119920720 | rs9285891 | -1.221 | 1.036 | -0.928 | -1.321 | -0.023 | 0.002 | 0.002 | 0 | 0.88 | 0.94 | 0.91 | 0.91 | 0.82 | 0.87 |
| 5 | 119924532 | rs6865072 | 0.61 | 0.694 | -0.943 | -1.46 | -0.005 | 0.003 | 0.003 | 0 | 0.34 | 0.32 | 0.47 | 0.73 | 0.4 | 0.3 |
| 5 | 119925692 | rs6881097 | 0.885 | 0.548 | -0.818 | -1.754 | -0.001 | -0.03 | -0.03 | 0.655 | 0.73 | 0.39 | 0.83 | 0.84 | 0.75 | 0.78 |
| 5 | 119926305 | rs1037569 | 0.135 | 0.75 | -0.047 | -1.762 | 0.013 | -0.038 | -0.038 | 0 | 0.82 | 0.67 | 0.91 | 0.91 | 0.81 | 0.85 |
| 5 | 119928674 | rs13167682 | 2.102 | 1.334 | -0.439 | -1.963 | -0.084 | 0.115 | 0.115 | 0.119 | 0.19 | 0.01 | 0.16 | 0.01 | 0.29 | 0.05 |
| 5 | 119932276 | rs2405955 | 0.284 | 0.94 | -0.332 | -2.024 | -0.012 | 0.025 | 0.025 | 0.969 | 0.38 | 0.32 | 0.63 | 0.82 | 0.45 | 0.4 |
| 5 | 119935830 | rs9327155 | -1.555 | 1.011 | -0.02 | -1.798 | 0.007 | -0.003 | -0.003 | 0 | 0.89 | 0.89 | 0.93 | 0.91 | 0.83 | 0.88 |
| 5 | 119937691 | rs1584468 | 0.467 | 0.884 | 0.262 | -1.687 | 0.032 | -0.008 | -0.008 | 0 | 0.06 | 0.29 | 0.04 | 0 | 0.02 | 0.03 |
| 5 | 119940108 | rs1597285 | -0.566 | 0.841 | 0.207 | -1.742 | 0.005 | -0.028 | -0.028 | 0.567 | 0.12 | 0.29 | 0.1 | 0.06 | 0.09 | 0.1 |
| 5 | 119951494 | rs12514071 | 1.173 | 0.91 | -0.089 | -1.647 | -0.036 | -0.022 | -0.022 | 0 | 0.1 | 0.01 | 0.26 | 0.11 | 0.09 | 0.1 |
| 5 | 119953529 | rs10042938 | 0.838 | 0.771 | 0.071 | -1.713 | -0.02 | -0.006 | -0.006 | 0.356 | 0.07 | 0 | 0.06 | 0.06 | 0.07 | 0.07 |
| 5 | 119956764 | rs2218717 | 0.364 | 0.653 | 0.182 | -1.934 | -0.011 | 0.016 | 0.016 | 0.139 | 0.37 | 0.32 | 0.63 | 0.82 | 0.45 | 0.49 |
| 5 | 119957977 | rs1524559 | 0.113 | 0.563 | 0.046 | -1.958 | -0.014 | 0.078 | 0.078 | 0.009 | 0.55 | 0.33 | 0.33 | 0.18 | 0.52 | 0.48 |
| 5 | 119962441 | rs10463729 | -0.633 | 0.21 | 0.342 | -2.017 | -0.021 | -0.099 | -0.099 | 0.021 | 0.2 | 0.54 | 0.11 | 0.05 | 0.15 | 0.15 |
| 5 | 119966585 | rs1524561 | 0.242 | 0.229 | 0.507 | -2.251 | 0.017 | -0.076 | -0.076 | 1.522 | 0.38 | 0.49 | 0.63 | 0.86 | 0.44 | 0.38 |
| 5 | 119978197 | rs7711106 | -0.559 | 0.275 | 0.358 | -2.228 | 0.007 | -0.035 | -0.035 | 0 | 0.63 | 0.62 | 0.38 | 0.14 | 0.57 | 0.53 |
| 5 | 119979333 | rs2122193 | 1.389 | 0.158 | 0.09 | -2.332 | 0.002 | 0.21 | 0.21 | 0 | 0.38 | 0.09 | 0.21 | 0.03 | 0.35 | 0.4 |
| 5 | 119987933 | rs2405963 | 0.781 | 0.158 | 0.164 | -2.323 | -0.014 | -0.063 | -0.063 | 0 | 0.73 | 0.41 | 0.83 | 0.89 | 0.78 | 0.78 |
| 5 | 119989370 | rs17429498 | 1.666 | -0.123 | -0.151 | -2.493 | -0.023 | 0.184 | 0.184 | 0 | 0.33 | 0.07 | 0.19 | 0.03 | 0.34 | 0.3 |
| 5 | 119997409 | rs716815 | 1.188 | 0.095 | -0.01 | -2.315 | -0.012 | 0.012 | 0.012 | 0 | 0.27 | 0.23 | 0.52 | 0.8 | 0.36 | 0.37 |
| 5 | 120009805 | rs4314428 | 0.662 | 0.072 | 0.135 | -2.25 | -0.009 | -0.047 | -0.047 | 0.491 | 0.74 | 0.59 | 0.84 | 0.89 | 0.78 | 0.79 |
| 5 | 120016536 | rs6898545 | 1.436 | -0.003 | -0.083 | -1.941 | -0.01 | -0.006 | -0.006 | 2.012 | 0.26 | 0.23 | 0.5 | 0.65 | 0.36 | 0.27 |
| 5 | 120016963 | rs12187844 | 1.164 | -0.024 | -0.386 | -2.016 | 0.01 | 0.015 | 0.015 | 0 | 0.36 | 0.24 | 0.21 | 0.03 | 0.34 | 0.41 |
| 5 | 120020074 | rs17146934 | 0.895 | 0.229 | -0.58 | -1.494 | -0.013 | -0.025 | -0.025 | 0 | 0.09 | 0.04 | 0.12 | 0.19 | 0.08 | 0.11 |
| 5 | 120022759 | rs10069230 | 0.538 | 0.171 | -0.6 | -1.784 | -0.019 | -0.008 | -0.008 | 0.612 | 0.07 | 0 | 0.06 | 0.06 | 0.07 | 0.06 |
| 5 | 120023393 | rs4895272 | 0.396 | 0.199 | -0.455 | -1.608 | -0.019 | -0.004 | -0.004 | 0 | 0.08 | 0.06 | 0.08 | 0.06 | 0.15 | 0.09 |
| 5 | 120026332 | rs17502163 | 1.107 | 0.185 | -0.506 | -1.632 | -0.024 | 0.014 | 0.014 | 0 | 0.09 | 0.04 | 0.1 | 0.04 | 0.09 | 0.11 |
| 5 | 120028001 | rs1449145 | 0.279 | -0.036 | -0.126 | -1.577 | 0.035 | -0.008 | -0.008 | 0.211 | 0.89 | 0.71 | 0.92 | 0.94 | 0.85 | 0.9 |
| 5 | 120043003 | rs10075522 | -0.128 | -0.483 | -0.237 | -1.849 | 0.003 | -0.036 | -0.036 | 0 | 0.56 | 0.56 | 0.35 | 0.16 | 0.53 | 0.51 |
| 5 | 120047147 | rs1449157 | -0.617 | -1.63 | 0.077 | -0.887 | -0.001 | 0.018 | 0.018 | 0.102 | 0.19 | 0.28 | 0.46 | 0.64 | 0.19 | 0.31 |
| 5 | 120049065 | rs9285894 | -0.519 | -1.665 | 0.097 | -0.854 | -0.001 | 0.022 | 0.022 | 0.049 | 0.18 | 0.28 | 0.46 | 0.64 | 0.18 | 0.3 |
| 5 | 120052870 | rs1449154 | -0.368 | -1.656 | 0.091 | -0.766 | -0.001 | 0.01 | 0.01 | 0.28 | 0.17 | 0.21 | 0.45 | 0.6 | 0.18 | 0.29 |
| 5 | 120065986 | rs7710640 | -0.024 | -1.568 | -0.129 | -1.294 | -0.002 | 0.001 | 0.001 | 0 | 0.79 | 0.83 | 0.82 | 0.9 | 0.72 | 0.78 |
| 5 | 120066820 | rs1449148 | 1.886 | -1.937 | -0.611 | -1.336 | -0.148 | 0.113 | 0.113 | 0 | 0.29 | 0.02 | 0.22 | 0.1 | 0.41 | 0.32 |
| 5 | 120084554 | rs10073004 | -0.129 | -1.707 | 0.46 | -1.083 | -0.007 | -0.015 | -0.015 | 0.011 | 0.1 | 0.19 | 0.07 | 0.06 | 0.06 | 0.06 |
| 5 | 120100225 | rs7705890 | -1.051 | -1.854 | 0.595 | -1.161 | -0.04 | -0.039 | -0.039 | 0 | 0.14 | 0.33 | 0.09 | 0.06 | 0.07 | 0.07 |
| 5 | 120103173 | rs11241565 | 1.47 | -1.976 | 0.74 | -1.228 | -0.003 | 0.013 | 0.013 | 0 | 0.6 | 0.39 | 0.63 | 0.79 | 0.63 | 0.65 |
| 5 | 120109243 | rs13358641 | -1.633 | -1.959 | 0.723 | -1.222 | 0.006 | -0.037 | -0.037 | 0.277 | 0.4 | 0.52 | 0.38 | 0.17 | 0.38 | 0.35 |
| 5 | 120114265 | rs4605820 | -1.892 | -2.235 | 0.424 | -1.29 | 0 | 0.059 | 0.059 | 0 | 0.54 | 0.65 | 0.42 | 0.21 | 0.43 | 0.43 |
| 5 | 120114673 | rs13357432 | -0.422 | -1.921 | 0.067 | -1.125 | -0.033 | -0.021 | -0.021 | 0 | 0.11 | 0.18 | 0.07 | 0.06 | 0.05 | 0.07 |
| 5 | 120115857 | rs6891234 | -0.248 | -1.91 | 0.027 | -1.274 | -0.028 | 0.019 | 0.019 | 0 | 0.24 | 0.18 | 0.26 | 0.14 | 0.29 | 0.28 |
| 5 | 120120064 | rs6877942 | -1.528 | -1.931 | -0.047 | -1.241 | 0.001 | -0.016 | -0.016 | 0 | 0.38 | 0.41 | 0.37 | 0.18 | 0.37 | 0.35 |
| 5 | 120123994 | rs12523559 | -0.647 | -1.42 | -0.856 | -0.887 | -0.048 | 0.041 | 0.041 | 0 | 0.27 | 0.17 | 0.29 | 0.14 | 0.31 | 0.29 |
| 5 | 120140600 | rs9327167 | 0.122 | -1.847 | -0.795 | -0.91 | 0.047 | 0.035 | 0.035 | 0 | 0.22 | 0.1 | 0.1 | 0 | 0.09 | 0.1 |

| | | | | | | | | | | | | | | | | |
|---|---|---|---|---|---|---|---|---|---|---|---|---|---|---|---|---|
| 5 | 120140803 | rs4340941 | -0.448 | -1.604 | -0.812 | -0.913 | -0.064 | 0.039 | 0.039 | 0 | 0.24 | 0.11 | 0.31 | 0.14 | 0.31 | 0.27 |
| 5 | 120141792 | rs12520572 | -0.726 | -1.641 | -0.663 | -0.881 | -0.021 | 0.018 | 0.018 | 0 | 0.27 | 0.22 | 0.32 | 0.18 | 0.33 | 0.3 |
| 5 | 120154484 | rs6595265 | -1.832 | -1.881 | -0.378 | -0.177 | 0.005 | 0.002 | 0.002 | 0 | 0.6 | 0.58 | 0.58 | 0.57 | 0.46 | 0.61 |
| 5 | 120159013 | rs12523326 | 0.053 | -2.026 | -0.7 | -0.08 | 0.052 | -0.037 | -0.037 | 2.513 | 0.26 | 0.1 | 0.21 | 0.36 | 0.12 | 0.3 |
| 5 | 120175344 | rs4895169 | -0.259 | -2.026 | -0.532 | -0.083 | 0.022 | -0.017 | -0.017 | 1.93 | 0.29 | 0.21 | 0.23 | 0.41 | 0.14 | 0.33 |
| 5 | 120177018 | rs13170286 | 0.474 | -1.945 | -0.692 | -0.019 | -0.027 | -0.005 | -0.005 | 0 | 0.1 | 0.01 | 0.2 | 0.06 | 0.13 | 0.02 |
| 5 | 120178687 | rs11958476 | 0.156 | -1.853 | -0.46 | 0.02 | -0.011 | 0.007 | 0.007 | 0.149 | 0.11 | 0.08 | 0.17 | 0.06 | 0.16 | 0.16 |
| 5 | 120183320 | rs10054920 | -0.427 | -2.054 | -0.434 | 0.106 | 0.066 | -0.039 | -0.039 | 0 | 0.3 | 0.13 | 0.23 | 0.41 | 0.14 | 0.32 |
| 5 | 120198311 | rs10041997 | -1.575 | -1.758 | -0.303 | 0.127 | 0.004 | 0.001 | 0.001 | 0 | 0.56 | 0.51 | 0.55 | 0.58 | 0.43 | 0.58 |
| 5 | 120207152 | rs12522426 | 0.201 | -1.856 | -0.601 | 0.263 | 0.063 | -0.062 | -0.062 | 0 | 0.23 | 0.06 | 0.18 | 0.35 | 0.09 | 0.24 |
| 5 | 120209700 | rs4383741 | 0.621 | -1.825 | -0.397 | 0.245 | 0.003 | -0.001 | -0.001 | 0 | 0.11 | 0.09 | 0.12 | 0.13 | 0.11 | 0.09 |
| 5 | 120219340 | rs10041968 | -1.505 | -1.713 | 0.367 | 0.296 | -0.009 | 0.009 | 0.009 | 0 | 0.55 | 0.65 | 0.53 | 0.6 | 0.43 | 0.56 |
| 5 | 120244182 | rs7732798 | -0.918 | -2.145 | 0.553 | 0.267 | -0.026 | 0.19 | 0.19 | 0.376 | 0.13 | 0.37 | 0.15 | 0.38 | 0.05 | 0.21 |
| 5 | 120244495 | rs4895287 | 1.315 | -2.156 | 0.635 | 0.192 | 0.004 | 0.01 | 0.01 | 0 | 0.48 | 0.44 | 0.49 | 0.41 | 0.61 | 0.45 |
| 5 | 120245440 | rs6595276 | 0.454 | -2.112 | 0.569 | 0.128 | 0.004 | 0.027 | 0.027 | 0 | 0.59 | 0.72 | 0.64 | 0.79 | 0.66 | 0.67 |
| 5 | 120246666 | rs7733696 | 0.427 | -2.432 | 0.554 | 0.108 | 0.02 | 0.044 | 0.044 | 0 | 0.26 | 0.1 | 0.17 | 0.12 | 0.18 | 0.16 |
| 5 | 120247295 | rs4895290 | -1.25 | -2.634 | 0.737 | 0.006 | -0.041 | -0.049 | -0.049 | 1.083 | 0.12 | 0.42 | 0.07 | 0.04 | 0.05 | 0.05 |
| 5 | 120253083 | rs1469049 | 1.995 | -2.548 | 0.398 | 0.076 | -0.061 | -0.055 | -0.055 | 0 | 0.39 | 0.1 | 0.44 | 0.4 | 0.6 | 0.44 |
| 5 | 120255182 | rs4621580 | -0.401 | -2.249 | 0.733 | 0.337 | -0.035 | 0.001 | 0.001 | 0 | 0.88 | 0.98 | 0.82 | 0.91 | 0.87 | 0.84 |
| 5 | 120256243 | rs6861434 | -0.454 | -2.248 | 0.752 | 0.337 | -0.029 | 0.003 | 0.003 | 0 | 0.88 | 0.98 | 0.82 | 0.91 | 0.87 | 0.84 |
| 5 | 120258577 | rs6892749 | -1.017 | -2.294 | 0.941 | 0.309 | -0.035 | 0.085 | 0.085 | 1.005 | 0.15 | 0.3 | 0.18 | 0.35 | 0.05 | 0.2 |
| 5 | 120273930 | rs17147116 | -0.967 | -2.273 | 1.009 | 0.319 | -0.036 | 0.085 | 0.085 | 0.872 | 0.15 | 0.31 | 0.18 | 0.35 | 0.05 | 0.19 |
| 5 | 120275942 | rs6889233 | -0.119 | -2.223 | 0.862 | 0.247 | 0.005 | -0.012 | -0.012 | 0 | 0.87 | 0.87 | 0.82 | 0.92 | 0.87 | 0.84 |
| 5 | 120280194 | rs2909938 | -0.895 | -2.23 | 0.817 | 0.216 | 0.002 | -0.031 | -0.031 | 0.294 | 0.18 | 0.34 | 0.2 | 0.12 | 0.16 | 0.2 |
| 5 | 120310093 | rs4895293 | -1.998 | -2.325 | 0.687 | 0.213 | 0.012 | -0.017 | -0.017 | 0 | 0.34 | 0.77 | 0.38 | 0.49 | 0.2 | 0.39 |
| 5 | 120317035 | rs2909919 | -1.367 | -1.869 | 0.171 | -0.39 | 0.002 | -0.017 | -0.017 | 0 | 0.24 | 0.28 | 0.37 | 0.13 | 0.27 | 0.24 |
| 5 | 120318488 | rs2909918 | 0.001 | -1.788 | -0.556 | -0.329 | -0.012 | 0.016 | 0.016 | 0 | 0.12 | 0.03 | 0.16 | 0.08 | 0.11 | 0.16 |
| 5 | 120328270 | rs10053084 | -0.969 | -1.711 | -0.383 | -0.333 | 0.005 | 0.009 | 0.009 | 0 | 0.15 | 0.17 | 0.22 | 0.05 | 0.16 | 0.1 |
| 5 | 120331625 | rs2973163 | -1.265 | -2.003 | -0.799 | -0.543 | -0.019 | 0.161 | 0.161 | 0.014 | 0.58 | 0.85 | 0.67 | 0.85 | 0.39 | 0.61 |
| 5 | 120331891 | rs2973162 | -2.02 | -2.508 | -0.844 | 0.089 | -0.05 | -0.017 | -0.017 | 1.124 | 0.64 | 0.87 | 0.63 | 0.59 | 0.46 | 0.76 |
| 5 | 120334802 | rs1422601 | -1.416 | -3.177 | -0.218 | -0.346 | -0.018 | -0.01 | -0.01 | 0 | 0.37 | 0.5 | 0.29 | 0.63 | 0.29 | 0.58 |
| 5 | 120339865 | rs1833646 | -2.169 | -3.23 | -0.035 | -0.378 | -0.001 | -0.003 | -0.003 | 0 | 0.62 | 0.62 | 0.42 | 0.63 | 0.44 | 0.75 |
| 5 | 120345122 | rs17147214 | 0.284 | -3.33 | -0.146 | -0.101 | 0.001 | 0.016 | 0.016 | 0.157 | 0.07 | 0.03 | 0.1 | 0.42 | 0.05 | 0.22 |
| 5 | 120359319 | rs1862280 | 0.291 | -3.351 | -0.136 | -0.437 | 0.051 | 0.044 | 0.044 | 0 | 0.25 | 0.12 | 0.12 | 0 | 0.15 | 0.17 |
| 5 | 120363673 | rs13169397 | -0.055 | -3.112 | -0.122 | -0.297 | -0.003 | -0.02 | -0.02 | 0 | 0.13 | 0.01 | 0.08 | 0.12 | 0.13 | 0.23 |
| 5 | 120367551 | rs1422606 | -1.772 | -3.014 | 0.177 | -0.163 | 0.006 | 0.006 | 0.006 | 0 | 0.59 | 0.4 | 0.41 | 0.62 | 0.42 | 0.73 |
| 5 | 120369453 | rs17147260 | 0.23 | -3.204 | 0.125 | -0.083 | -0.008 | 0.001 | 0.001 | 0 | 0.11 | 0.14 | 0.06 | 0.13 | 0.11 | 0.19 |
| 5 | 120369695 | rs7704984 | -2.054 | -3.007 | -0.003 | -0.049 | -0.019 | -0.014 | -0.014 | 0.108 | 0.64 | 0.87 | 0.41 | 0.63 | 0.43 | 0.72 |
| 5 | 120371760 | rs1422607 | -1.073 | -2.513 | 0.247 | 0.245 | 0.019 | 0.048 | 0.048 | 0 | 0.3 | 0.2 | 0.19 | 0.11 | 0.23 | 0.28 |
| 5 | 120373318 | rs10038211 | -2.02 | -2.588 | 0.296 | 0.222 | 0.006 | -0.002 | -0.002 | 0 | 0.44 | 0.59 | 0.32 | 0.49 | 0.29 | 0.52 |
| 5 | 120374562 | rs17446625 | 0.061 | -2.454 | -0.069 | 0.157 | 0.026 | 0.083 | 0.083 | 0 | 0.17 | 0.03 | 0.09 | 0 | 0.13 | 0.15 |
| 5 | 120381682 | rs17446950 | -0.081 | -2.237 | 0.097 | 0.325 | -0.001 | 0 | 0 | 0.015 | 0.1 | 0.11 | 0.07 | 0.1 | 0.09 | 0.1 |
| 5 | 120388440 | rs1820068 | -1.91 | -2.471 | 0.2 | 0.294 | -0.015 | 0.015 | 0.015 | 0.252 | 0.34 | 0.4 | 0.26 | 0.49 | 0.21 | 0.4 |
| 5 | 120388986 | rs7727775 | -0.381 | -2.469 | 0.079 | 0.315 | 0.006 | -0.004 | -0.004 | 0 | 0.16 | 0.16 | 0.13 | 0.39 | 0.08 | 0.25 |
| 5 | 120392422 | rs10463734 | -0.766 | -2.442 | 0.072 | 0.297 | 0 | 0.004 | 0.004 | 0 | 0.14 | 0.08 | 0.1 | 0.1 | 0.12 | 0.12 |
| 5 | 120396412 | rs10519660 | -1.326 | -2.365 | 0.08 | 0.358 | 0.02 | -0.036 | -0.036 | 0 | 0.3 | 0.23 | 0.23 | 0.49 | 0.2 | 0.37 |
| 5 | 120412220 | rs17447768 | 0.37 | -2.473 | -0.105 | 0.258 | 0.01 | 0.026 | 0.026 | 0 | 0.11 | 0.07 | 0.07 | 0 | 0.1 | 0.13 |
| 5 | 120424662 | rs11960097 | 0.388 | -2.387 | -0.077 | 0.344 | -0.031 | -0.037 | -0.037 | 2.016 | 0.08 | 0.01 | 0.06 | 0.1 | 0.09 | 0.1 |
| 5 | 120425411 | rs966452 | -1.343 | -2.38 | 0.164 | 0.3 | -0.037 | 0.062 | 0.062 | 0 | 0.31 | 0.48 | 0.25 | 0.49 | 0.2 | 0.37 |
| 5 | 120440686 | rs6886868 | -1.49 | -2.415 | 0.011 | 0.284 | -0.001 | -0.001 | -0.001 | 0 | 0.37 | 0.66 | 0.26 | 0.39 | 0.21 | 0.42 |
| 5 | 120445673 | rs7724896 | -2.014 | -2.362 | -0.152 | 0.244 | 0.061 | 0.003 | 0.003 | 1.143 | 0.48 | 0.71 | 0.33 | 0.54 | 0.31 | 0.69 |
| 5 | 120449584 | rs4440396 | -1.836 | -2.096 | 0.102 | 0.597 | 0 | -0.002 | -0.002 | 0.342 | 0.37 | 0.67 | 0.25 | 0.54 | 0.2 | 0.56 |
| 5 | 120457296 | rs11954476 | -1.167 | -1.981 | 0.316 | 0.56 | 0.043 | 0.012 | 0.012 | 0 | 0.14 | 0.42 | 0.11 | 0.09 | 0.12 | 0.14 |
| 5 | 120468093 | rs11959708 | -0.978 | -2.005 | 0.258 | 0.563 | 0.013 | 0.012 | 0.012 | 0 | 0.13 | 0.36 | 0.1 | 0.1 | 0.1 | 0.13 |
| 5 | 120471719 | rs7731784 | -0.891 | -2.043 | -0.182 | 0.674 | -0.018 | 0.01 | 0.01 | 0.098 | 0.23 | 0.29 | 0.14 | 0.45 | 0.08 | 0.41 |
| 5 | 120474531 | rs13436259 | -1.192 | -1.062 | -0.279 | 0.598 | 0.017 | 0 | 0 | 3.15 | 0.14 | 0.29 | 0.11 | 0.1 | 0.12 | 0.14 |
| 5 | 120477945 | rs10055647 | 1.001 | -0.858 | -1.539 | 0.503 | 0.001 | 0.011 | 0.011 | 0.023 | 0.6 | 0.84 | 0.56 | 0.35 | 0.65 | 0.45 |
| 5 | 120482741 | rs9327182 | 0.664 | -0.634 | -1.525 | 0.81 | -0.011 | -0.094 | -0.094 | 0 | 0.81 | 0.94 | 0.76 | 0.55 | 0.81 | 0.62 |
| 5 | 120484783 | rs4305661 | -0.798 | -0.419 | -1.282 | 0.455 | -0.014 | -0.009 | -0.009 | 0 | 0.08 | 0.04 | 0.16 | 0.1 | 0.09 | 0.06 |
| 5 | 120490664 | rs4585482 | -0.997 | -0.763 | -1.089 | 0.258 | -0.002 | -0.003 | -0.003 | 1.259 | 0.11 | 0.15 | 0.18 | 0.08 | 0.1 | 0.07 |
| 5 | 120501442 | rs7724812 | -0.801 | -0.267 | -1.027 | -0.445 | 0.003 | -0.035 | -0.035 | 0.151 | 0.23 | 0.25 | 0.12 | 0.02 | 0.15 | 0.22 |
| 5 | 120511120 | rs12109237 | -1.077 | -0.691 | -1.015 | -0.479 | 0.062 | 0.03 | 0.03 | 0 | 0.5 | 0.73 | 0.51 | 0.48 | 0.38 | 0.63 |
| 5 | 120511792 | rs10053662 | 1.458 | -0.741 | -1.25 | -0.625 | -0.032 | -0.036 | -0.036 | 0 | 0.37 | 0.26 | 0.3 | 0.45 | 0.46 | 0.28 |
| 5 | 120515118 | rs2914084 | -0.624 | -0.172 | -1.074 | -0.543 | 0.003 | 0.091 | 0.091 | 0 | 0.94 | 0.96 | 0.71 | 0.59 | 0.89 | 0.87 |
| 5 | 120517296 | rs2115143 | 0.437 | -0.3 | -0.958 | -0.879 | 0.036 | 0.087 | 0.087 | 0 | 0.89 | 0.79 | 0.68 | 0.67 | 0.83 | 0.82 |
| 5 | 120524720 | rs1896656 | 1.63 | -0.441 | -1.061 | -1.147 | -0.024 | -0.005 | -0.005 | 0 | 0.36 | 0.25 | 0.3 | 0.58 | 0.43 | 0.31 |
| 5 | 120529121 | rs17147484 | -0.23 | -0.671 | -0.362 | -1.344 | 0.019 | -0.018 | -0.018 | 0.312 | 0.42 | 0.4 | 0.21 | 0.01 | 0.28 | 0.47 |
| 5 | 120529790 | rs10478509 | 1.313 | -0.478 | -0.256 | -1.346 | -0.014 | 0.018 | 0.018 | 0 | 0.42 | 0.31 | 0.34 | 0.59 | 0.5 | 0.34 |
| 5 | 120536562 | rs2973419 | 0.727 | -0.612 | -0.232 | -1.417 | -0.013 | -0.081 | -0.081 | 0 | 0.22 | 0.32 | 0.13 | 0 | 0.15 | 0.27 |

| | | | | | | | | | | | | | | | | |
|---|---|---|---|---|---|---|---|---|---|---|---|---|---|---|---|---|
| 5 | 120540854 | rs12153288 | -0.363 | -0.611 | -0.359 | -1.455 | 0.049 | 0.183 | 0.183 | 0 | 0.41 | 0.16 | 0.19 | 0.02 | 0.28 | 0.45 |
| 5 | 120542415 | rs17515002 | 1.121 | -0.291 | -0.237 | -1.136 | -0.022 | 0.032 | 0.032 | 0 | 0.08 | 0.05 | 0.07 | 0 | 0.1 | 0.13 |
| 5 | 120565606 | rs919288 | 0.182 | -0.371 | 0.066 | -1.017 | 0.046 | 0.18 | 0.18 | 0 | 0.87 | 0.52 | 0.71 | 0.66 | 0.86 | 0.83 |
| 5 | 120569108 | rs4895298 | 0.117 | -0.48 | 0.084 | -1.016 | 0.038 | 0.187 | 0.187 | 0 | 0.87 | 0.52 | 0.71 | 0.66 | 0.88 | 0.84 |
| 5 | 120570124 | rs11738220 | 0.127 | -0.496 | 0.14 | -1.022 | 0.043 | 0.202 | 0.202 | 0 | 0.87 | 0.53 | 0.71 | 0.65 | 0.88 | 0.84 |
| 5 | 120579984 | rs4895300 | 2.194 | 0.012 | -0.347 | -1.622 | -0.006 | -0.013 | -0.013 | 0 | 0.28 | 0.24 | 0.25 | 0.57 | 0.36 | 0.28 |
| 5 | 120580857 | rs17449200 | 1.426 | 0.026 | -0.337 | -1.608 | -0.01 | 0.012 | 0.012 | 0 | 0.09 | 0.07 | 0.07 | 0 | 0.1 | 0.13 |
| 5 | 120581128 | rs2406422 | 0.148 | -0.039 | -0.436 | -1.493 | -0.011 | -0.043 | -0.043 | 0 | 0.69 | 0.46 | 0.63 | 0.85 | 0.75 | 0.71 |
| 5 | 120589282 | rs747244 | 0.853 | 0.053 | -0.672 | -1.491 | 0.013 | 0.018 | 0.018 | 0 | 0.56 | 0.34 | 0.42 | 0.6 | 0.6 | 0.58 |
| 5 | 120599222 | rs11948980 | 1.058 | 0.042 | -0.714 | -1.43 | -0.001 | 0.024 | 0.024 | 0 | 0.51 | 0.35 | 0.37 | 0.57 | 0.51 | 0.54 |
| 5 | 120608476 | rs12188286 | 0.432 | -0.05 | -0.98 | -1.754 | 0.027 | 0.058 | 0.058 | 0 | 0.23 | 0.08 | 0.12 | 0 | 0.15 | 0.29 |
| 5 | 120615810 | rs11744147 | -0.276 | 0.434 | -0.641 | -1.261 | -0.002 | -0.005 | -0.005 | 0 | 0.13 | 0.13 | 0.1 | 0.04 | 0.18 | 0.09 |
| 5 | 120621269 | rs2052555 | -0.498 | -0.2 | -0.957 | -1.919 | 0.029 | 0.1 | 0.1 | 0 | 0.65 | 0.85 | 0.82 | 1 | 0.73 | 0.65 |
| 5 | 120622200 | rs730464 | -0.394 | -0.664 | -0.268 | -2.25 | -0.011 | -0.12 | -0.12 | 0.041 | 0.63 | 0.57 | 0.79 | 1 | 0.74 | 0.68 |
| 5 | 120626646 | rs4409129 | 1.008 | -0.397 | -0.112 | -1.866 | 0.015 | 0.07 | 0.07 | 0 | 0.07 | 0.13 | 0.28 | 0.2 | 0.11 | 0.11 |
| 5 | 120626777 | rs4895176 | 0.383 | -0.526 | 0.014 | -1.945 | 0.027 | 0.074 | 0.074 | 0 | 0.5 | 0.39 | 0.49 | 0.79 | 0.59 | 0.5 |
| 5 | 120634862 | rs17456449 | 1.009 | -0.549 | -0.069 | -2.024 | -0.002 | 0.128 | 0.128 | 0 | 0.31 | 0.09 | 0.17 | 0 | 0.26 | 0.36 |
| 5 | 120635617 | rs2216616 | -0.641 | -0.557 | 0.295 | -1.95 | -0.013 | 0.013 | 0.013 | 0 | 0.19 | 0.37 | 0.33 | 0.21 | 0.18 | 0.17 |
| 5 | 120636776 | rs6859471 | -1.173 | -0.426 | 0.085 | -1.91 | -0.004 | -0.003 | -0.003 | 0 | 0.41 | 0.49 | 0.57 | 0.4 | 0.42 | 0.43 |
| 5 | 120638202 | rs4146402 | 0.573 | -0.403 | 0.107 | -1.885 | -0.005 | 0.007 | 0.007 | 0 | 0.22 | 0.14 | 0.24 | 0.21 | 0.24 | 0.26 |
| 5 | 120638576 | rs1812546 | -1.473 | -0.526 | 0.076 | -1.982 | 0.023 | 0.024 | 0.024 | 0 | 0.47 | 0.71 | 0.61 | 0.41 | 0.44 | 0.43 |
| 5 | 120645496 | rs6861011 | -1.074 | -0.54 | 0.224 | -1.903 | 0.002 | -0.001 | -0.001 | 0 | 0.38 | 0.66 | 0.57 | 0.41 | 0.39 | 0.4 |
| 5 | 120646062 | rs10051149 | 1.432 | -0.45 | 0.203 | -1.917 | 0.001 | 0.055 | 0.055 | 0 | 0.3 | 0.16 | 0.17 | 0 | 0.26 | 0.34 |
| 5 | 120653398 | rs6885327 | 0.298 | -0.293 | 0.365 | -1.719 | 0.017 | 0.094 | 0.094 | 0.618 | 0.12 | 0.36 | 0.3 | 0.21 | 0.13 | 0.15 |
| 5 | 120657454 | rs17456960 | -0.352 | -0.174 | 0.242 | -1.803 | -0.003 | 0.014 | 0.014 | 0.029 | 0.34 | 0.54 | 0.53 | 0.4 | 0.36 | 0.42 |
| 5 | 120659585 | rs1592893 | -0.268 | -0.094 | 0.003 | -1.766 | -0.001 | 0.015 | 0.015 | 0 | 0.33 | 0.56 | 0.53 | 0.4 | 0.36 | 0.4 |
| 5 | 120661074 | rs9885119 | -1.162 | -0.091 | -0.476 | -1.693 | -0.018 | -0.021 | -0.021 | 0 | 0.69 | 0.79 | 0.73 | 0.41 | 0.64 | 0.76 |
| 5 | 120662660 | rs9327190 | 0.18 | 0.01 | -0.173 | -1.596 | -0.003 | -0.041 | -0.041 | 0 | 0.4 | 0.44 | 0.41 | 0.18 | 0.36 | 0.46 |
| 5 | 120683069 | rs3900188 | -0.912 | 0.181 | 0.005 | -1.469 | 0.03 | -0.054 | -0.054 | 0 | 0.19 | 0.44 | 0.22 | 0.12 | 0.21 | 0.2 |
| 5 | 120689867 | rs6861838 | -1.318 | 0.846 | -0.775 | -1.186 | -0.077 | -0.105 | -0.105 | 0 | 0.86 | 0.97 | 0.82 | 0.41 | 0.77 | 0.85 |
| 5 | 120690354 | rs17457815 | 1.634 | 0.333 | -1.427 | -1.939 | -0.051 | 0.274 | 0.274 | 0.052 | 0.38 | 0.07 | 0.2 | 0.01 | 0.34 | 0.4 |
| 5 | 120692891 | rs11241595 | -1.755 | 0.326 | -1.389 | -1.826 | -0.077 | -0.087 | -0.087 | 0 | 0.86 | 0.97 | 0.69 | 0.34 | 0.77 | 0.84 |
| 5 | 120694906 | rs13159763 | 1.058 | 0.145 | -1.073 | -2.033 | 0.018 | -0.008 | -0.008 | 3.173 | 0.24 | 0.16 | 0.35 | 0.27 | 0.19 | 0.29 |
| 5 | 120708748 | rs246210 | 0.959 | 0.359 | -0.459 | -1.727 | -0.021 | 0.023 | 0.023 | 0.161 | 0.63 | 0.27 | 0.57 | 0.28 | 0.55 | 0.68 |
| 5 | 120712417 | rs711423 | 0.791 | 0.355 | -0.178 | -1.728 | 0.012 | -0.012 | -0.012 | 0.019 | 0.65 | 0.43 | 0.59 | 0.28 | 0.55 | 0.69 |
| 5 | 120713370 | rs700604 | 0.869 | 0.284 | -0.692 | -2.027 | 0.007 | 0.04 | 0.04 | 0 | 0.37 | 0.22 | 0.45 | 0.27 | 0.34 | 0.45 |
| 5 | 120715592 | rs1592892 | -1.118 | 0.672 | -0.066 | -1.482 | 0.012 | 0.039 | 0.039 | 0 | 0.84 | 0.77 | 0.81 | 0.41 | 0.77 | 0.85 |
| 5 | 120716442 | rs11948148 | 0.621 | 0.537 | -0.117 | -1.622 | -0.001 | -0.018 | -0.018 | 0.978 | 0.06 | 0.06 | 0.05 | 0.01 | 0.07 | 0.08 |
| 5 | 120721018 | rs17147676 | 0.988 | 0.701 | -0.356 | -1.562 | 0.071 | 0.038 | 0.038 | 0.091 | 0.07 | 0.18 | 0.15 | 0.08 | 0.13 | 0.07 |
| 5 | 120722413 | rs160122 | -1.442 | 0.945 | -0.46 | -1.236 | -0.012 | 0.037 | 0.037 | 0 | 0.91 | 0.9 | 0.85 | 0.46 | 0.82 | 0.91 |
| 5 | 120722596 | rs160121 | 1.469 | 0.207 | -0.648 | -1.538 | 0.006 | -0.007 | -0.007 | 0.347 | 0.58 | 0.52 | 0.57 | 0.37 | 0.5 | 0.67 |
| 5 | 120723763 | rs17147694 | 0.829 | 0.165 | -0.653 | -1.552 | 0.021 | 0.002 | 0.002 | 1.198 | 0.19 | 0.13 | 0.08 | 0.01 | 0.11 | 0.07 |
| 5 | 120727051 | rs33584 | 0.851 | 1.225 | -0.142 | -0.936 | 0.059 | 0.05 | 0.05 | 0.122 | 0.77 | 0.66 | 0.65 | 0.38 | 0.61 | 0.75 |
| 5 | 120732139 | rs711418 | -0.037 | 1.186 | -0.787 | -0.834 | -0.01 | -0.002 | -0.002 | 0 | 0.84 | 0.84 | 0.8 | 0.45 | 0.74 | 0.82 |
| 5 | 120744968 | rs4493702 | -1.287 | 0.713 | -1.23 | -0.897 | -0.008 | -0.024 | -0.024 | 0.832 | 0.61 | 0.93 | 0.54 | 0.65 | 0.69 | 0.56 |
| 5 | 120752908 | rs4481374 | -0.2 | 1.058 | -0.364 | -0.585 | -0.007 | -0.06 | -0.06 | 0 | 0.15 | 0.21 | 0.09 | 0.02 | 0.16 | 0.09 |
| 5 | 120757262 | rs1428702 | -0.723 | 1.719 | -0.684 | -0.197 | -0.076 | -0.074 | -0.074 | 0.005 | 0.22 | 0.09 | 0.33 | 0.57 | 0.38 | 0.26 |
| 5 | 120760608 | rs10054289 | -1.702 | 1.424 | 0.066 | -0.402 | 0.021 | 0.003 | 0.003 | 0 | 0.36 | 0.44 | 0.42 | 0.59 | 0.5 | 0.33 |
| 5 | 120767312 | rs17147802 | 0.413 | 1.797 | -0.104 | -0.16 | 0.021 | 0.007 | 0.007 | 0.035 | 0.07 | 0.1 | 0.15 | 0.09 | 0.14 | 0.08 |
| 5 | 120776375 | rs2217318 | 0.531 | 1.809 | -0.125 | -0.138 | 0.008 | 0.003 | 0.003 | 0 | 0.07 | 0.08 | 0.16 | 0.09 | 0.14 | 0.08 |
| 5 | 120776767 | rs10065963 | -0.17 | 1.681 | -0.329 | -0.046 | -0.076 | -0.126 | -0.126 | 0.034 | 0.16 | 0.02 | 0.19 | 0.51 | 0.24 | 0.18 |
| 5 | 120792844 | rs9885550 | 0.8 | 1.841 | -0.317 | -0.016 | -0.048 | -0.027 | -0.027 | 4.376 | 0.09 | 0.02 | 0.16 | 0.53 | 0.18 | 0.11 |
| 5 | 120800320 | rs6891688 | 1.139 | 1.699 | 0.556 | -0.036 | 0.039 | 0.06 | 0.06 | 0 | 0.79 | 0.6 | 0.76 | 0.33 | 0.71 | 0.74 |
| 5 | 120804862 | rs6874603 | 1.047 | 1.693 | 0.556 | -0.037 | 0.043 | 0.067 | 0.067 | 0 | 0.79 | 0.6 | 0.76 | 0.33 | 0.71 | 0.74 |
| 5 | 120804959 | rs6896409 | 0.745 | 1.698 | 0.55 | -0.07 | -0.004 | 0.002 | 0.002 | 0 | 0.9 | 0.89 | 0.94 | 0.83 | 0.92 | 0.84 |
| 5 | 120804992 | rs35560164 | 0.794 | 1.694 | 0.579 | 0.002 | -0.053 | -0.035 | -0.035 | 0.843 | 0.09 | 0.03 | 0.17 | 0.53 | 0.21 | 0.11 |
| 5 | 120817358 | rs1428696 | 0.91 | 1.279 | 0.681 | -0.065 | -0.009 | 0.063 | 0.063 | 0 | 0.95 | 0.84 | 0.97 | 0.84 | 0.97 | 0.92 |
| 5 | 120821261 | rs11748934 | 0.435 | 1.413 | 0.584 | -0.22 | 0.011 | -0.046 | -0.046 | 0.17 | 0.1 | 0.22 | 0.07 | 0.02 | 0.11 | 0.07 |
| 5 | 120826561 | rs7709711 | 1.373 | 1.126 | -0.39 | -0.718 | 0.019 | 0.036 | 0.036 | 0 | 0.37 | 0.06 | 0.43 | 0.22 | 0.27 | 0.35 |
| 5 | 120827799 | rs11241599 | -0.573 | 1.605 | -0.116 | -0.542 | -0.003 | -0.003 | -0.003 | 0 | 0.88 | 0.89 | 0.83 | 0.73 | 0.82 | 0.83 |
| 5 | 120828366 | rs12523300 | -0.701 | 1.752 | -0.096 | -0.62 | -0.041 | -0.033 | -0.033 | 0 | 0.24 | 0.16 | 0.36 | 0.59 | 0.4 | 0.28 |
| 5 | 120831491 | rs2115332 | -0.916 | 1.211 | 0.259 | -0.761 | -0.017 | -0.008 | -0.008 | 0.345 | 0.56 | 0.49 | 0.53 | 0.62 | 0.69 | 0.54 |
| 5 | 120833832 | rs4394153 | 0.995 | 1.339 | -0.244 | -0.737 | -0.003 | 0 | 0 | 0.312 | 0.1 | 0.09 | 0.16 | 0.1 | 0.16 | 0.1 |
| 5 | 120839597 | rs4895310 | 1.053 | 1.003 | -0.499 | -0.834 | 0.03 | 0.134 | 0.134 | 0 | 0.26 | 0.06 | 0.13 | 0.02 | 0.22 | 0.27 |
| 5 | 120843516 | rs1594006 | -1.609 | 1.152 | -0.26 | -0.877 | -0.015 | -0.002 | -0.002 | 0 | 0.38 | 0.75 | 0.47 | 0.62 | 0.55 | 0.36 |
| 5 | 120844396 | rs11750797 | -0.167 | 1.226 | 0.186 | -0.901 | 0.008 | -0.091 | -0.091 | 3.753 | 0.15 | 0.34 | 0.12 | 0.03 | 0.17 | 0.14 |
| 5 | 120846718 | rs11954404 | -0.207 | 1.232 | 0.287 | -0.899 | 0.004 | -0.076 | -0.076 | 0 | 0.14 | 0.27 | 0.11 | 0.03 | 0.16 | 0.13 |
| 5 | 120847814 | rs10519672 | 1.737 | 1.137 | -0.143 | -0.945 | 0.013 | 0.139 | 0.139 | 0 | 0.22 | 0.01 | 0.09 | 0.01 | 0.17 | 0.18 |
| 5 | 120850224 | rs1428710 | -0.079 | 1.155 | 0.066 | -0.931 | -0.024 | -0.129 | -0.129 | 0 | 0.37 | 0.42 | 0.21 | 0.04 | 0.33 | 0.32 |

| | | | | | | | | | | | | | | | | |
|---|---|---|---|---|---|---|---|---|---|---|---|---|---|---|---|---|
| 5 | 120850360 | rs17462291 | 2.053 | 0.805 | -0.521 | -1.254 | 0.002 | -0.062 | -0.062 | 0 | 0.21 | 0.01 | 0.36 | 0.21 | 0.17 | 0.13 |
| 5 | 120853669 | rs17147959 | 0.206 | 1.836 | 0.371 | -0.594 | 0.02 | -0.008 | -0.008 | 0 | 0.14 | 0.25 | 0.18 | 0.1 | 0.17 | 0.11 |
| 5 | 120854632 | rs4895178 | -1.21 | 1.834 | 0.382 | -0.615 | 0.083 | 0.129 | 0.129 | 0.341 | 0.24 | 0.48 | 0.29 | 0.51 | 0.37 | 0.24 |
| 5 | 120856183 | rs17147988 | -1.373 | 1.721 | 0.425 | -0.689 | 0.001 | -0.012 | -0.012 | 0 | 0.28 | 0.66 | 0.3 | 0.13 | 0.34 | 0.24 |
| 5 | 120859113 | rs4473796 | -0.833 | 1.599 | 0.422 | -0.631 | 0.018 | -0.096 | -0.096 | 0.648 | 0.15 | 0.41 | 0.12 | 0.03 | 0.16 | 0.13 |
| 5 | 120873947 | rs11956612 | -0.698 | 1.669 | 0.49 | -0.635 | 0.025 | -0.087 | -0.087 | 0.352 | 0.14 | 0.31 | 0.12 | 0.02 | 0.17 | 0.12 |
| 5 | 120881440 | rs9327199 | -0.729 | 1.71 | 0.343 | -0.773 | -0.021 | -0.012 | -0.012 | 0 | 0.44 | 0.69 | 0.38 | 0.52 | 0.49 | 0.34 |
| 5 | 120883097 | rs17148019 | -0.268 | 1.762 | 0.426 | -0.749 | -0.016 | -0.124 | -0.124 | 0.868 | 0.38 | 0.46 | 0.22 | 0.04 | 0.35 | 0.32 |
| 5 | 120883308 | rs17148024 | 1.195 | 1.785 | 0.413 | -0.725 | -0.002 | 0.048 | 0.048 | 0 | 0.09 | 0.08 | 0.17 | 0.49 | 0.21 | 0.11 |
| 5 | 120883361 | rs13189608 | 0.04 | 1.71 | 0.452 | -0.735 | 0.043 | -0.052 | -0.052 | 0 | 0.1 | 0.29 | 0.08 | 0.01 | 0.12 | 0.06 |
| 5 | 120886161 | rs10072279 | -0.931 | 1.683 | 0.347 | -0.713 | 0.016 | 0.005 | 0.005 | 0 | 0.49 | 0.72 | 0.41 | 0.52 | 0.56 | 0.42 |
| 5 | 120886375 | rs12518214 | 1.651 | 1.627 | 0.276 | -0.752 | 0.045 | 0.029 | 0.029 | 0 | 0.36 | 0.04 | 0.41 | 0.21 | 0.25 | 0.34 |
| 5 | 120887552 | rs11738537 | -1.219 | 2.201 | 0.936 | -0.189 | 0.01 | -0.101 | -0.101 | 0.099 | 0.16 | 0.55 | 0.14 | 0.02 | 0.17 | 0.12 |
| 5 | 120894559 | rs7725716 | 1.274 | 2.143 | 0.736 | -0.179 | 0.026 | 0.015 | 0.015 | 0.234 | 0.71 | 0.43 | 0.67 | 0.33 | 0.59 | 0.61 |
| 5 | 120894696 | rs4895318 | -0.235 | 1.79 | 0.724 | -0.693 | -0.007 | -0.011 | -0.011 | 0 | 0.72 | 0.66 | 0.87 | 0.83 | 0.81 | 0.7 |
| 5 | 120906150 | rs12108891 | 0.619 | 2.084 | 0.38 | -0.592 | 0.059 | 0.105 | 0.105 | 0 | 0.95 | 0.74 | 0.82 | 0.85 | 0.93 | 0.85 |
| 5 | 120908778 | rs4559029 | 0.902 | 2.213 | 0.398 | -0.705 | 0.022 | 0.096 | 0.096 | 0 | 0.56 | 0.3 | 0.58 | 0.81 | 0.59 | 0.54 |
| 5 | 120911071 | rs4128972 | -0.136 | 1.946 | 0.211 | -0.431 | 0.003 | 0.001 | 0.001 | 0 | 0.72 | 0.74 | 0.72 | 0.84 | 0.77 | 0.66 |
| 5 | 120917249 | rs6885430 | -1.849 | 2.938 | 0.61 | 0.725 | -0.017 | -0.013 | -0.013 | 0 | 0.32 | 0.74 | 0.45 | 0.53 | 0.4 | 0.37 |
| 5 | 120919205 | rs13166683 | -0.87 | 3.113 | 1.044 | 1.097 | 0.004 | 0.073 | 0.073 | 0 | 0.1 | 0.34 | 0.15 | 0.15 | 0.08 | 0.08 |
| 5 | 120919796 | rs4348243 | -0.228 | 3.177 | 0.851 | 1.126 | -0.046 | -0.046 | -0.046 | 0 | 0.13 | 0.06 | 0.24 | 0.32 | 0.24 | 0.2 |
| 5 | 120922897 | rs10055397 | -1.662 | 3.127 | 0.994 | 0.929 | 0.082 | 0.067 | 0.067 | 0 | 0.25 | 0.47 | 0.38 | 0.55 | 0.35 | 0.28 |
| 5 | 120930705 | rs7729331 | 0.07 | 3.225 | 1.025 | 0.94 | 0.01 | 0.04 | 0.04 | 0.152 | 0.11 | 0.12 | 0.23 | 0.35 | 0.22 | 0.17 |
| 5 | 120931849 | rs7734808 | -1.357 | 3.205 | 1.058 | 0.938 | 0.03 | 0.078 | 0.078 | 0 | 0.14 | 0.36 | 0.15 | 0.2 | 0.13 | 0.1 |
| 5 | 120934256 | rs7712083 | 1.631 | 2.93 | 1.359 | 1.08 | 0.11 | 0.218 | 0.218 | 0.311 | 0.95 | 0.56 | 0.81 | 0.88 | 0.95 | 0.91 |
| 5 | 120942971 | rs11948809 | -0.78 | 2.768 | 1.015 | 0.948 | 0.067 | 0.139 | 0.139 | 0.039 | 0.13 | 0.31 | 0.26 | 0.35 | 0.22 | 0.19 |
| 5 | 120945421 | rs4343869 | -0.658 | 1.808 | 0.713 | 0.215 | 0.067 | 0.001 | 0.001 | 0 | 0.07 | 0.3 | 0.04 | 0 | 0.05 | 0.01 |
| 5 | 120947041 | rs4895324 | -0.827 | 1.913 | -0.143 | 0.569 | -0.028 | -0.09 | -0.09 | 0 | 0.16 | 0.06 | 0.24 | 0.5 | 0.2 | 0.27 |
| 5 | 120948472 | rs11241609 | -0.1 | 1.834 | 0.097 | 0.363 | -0.012 | -0.025 | -0.025 | 0 | 0.23 | 0.37 | 0.35 | 0.16 | 0.19 | 0.24 |
| 5 | 120948875 | rs11241611 | -0.711 | 2.098 | -0.071 | 1.191 | -0.029 | -0.056 | -0.056 | 0.193 | 0.12 | 0.06 | 0.16 | 0.5 | 0.17 | 0.18 |
| 5 | 120951703 | rs10793805 | -1.513 | 2.333 | 0.5 | 1.331 | 0.015 | 0.061 | 0.061 | 0 | 0.16 | 0.3 | 0.38 | 0.62 | 0.19 | 0.3 |
| 5 | 120952441 | rs10044992 | -0.868 | 2.864 | 0.502 | 0.842 | 0.034 | -0.076 | -0.076 | 0 | 0.19 | 0.32 | 0.11 | 0.02 | 0.23 | 0.07 |
| 5 | 120952522 | rs11959611 | 0.208 | 2.806 | 0.22 | 0.853 | -0.011 | 0.017 | 0.017 | 0 | 0.06 | 0.03 | 0.02 | 0 | 0.06 | 0.01 |
| 5 | 120954676 | rs13155638 | 0.048 | 2.617 | 0.228 | 0.974 | -0.032 | 0.067 | 0.067 | 0 | 0.05 | 0.01 | 0.11 | 0.44 | 0.12 | 0.12 |
| 5 | 120955493 | rs9686533 | -1.333 | 2.632 | 0.47 | 0.879 | 0.04 | -0.044 | -0.044 | 2.878 | 0.3 | 0.49 | 0.41 | 0.21 | 0.33 | 0.26 |
| 5 | 120958689 | rs7700795 | -0.977 | 1.935 | -0.023 | 0.445 | 0.066 | 0.018 | 0.018 | 0 | 0.27 | 0.46 | 0.46 | 0.63 | 0.36 | 0.28 |
| 5 | 120965472 | rs11742173 | -0.551 | 1.152 | -0.764 | -0.255 | -0.009 | 0.098 | 0.098 | 0 | 0.48 | 0.67 | 0.54 | 0.72 | 0.48 | 0.45 |
| 5 | 120965891 | rs6888393 | -0.882 | 0.407 | -0.596 | -0.713 | -0.01 | -0.027 | -0.027 | 0 | 0.26 | 0.62 | 0.13 | 0.05 | 0.16 | 0.09 |
| 5 | 120966235 | rs1911920 | 0.381 | 0.351 | -0.379 | -1.06 | 0.009 | -0.085 | -0.085 | 0 | 0.77 | 0.65 | 0.83 | 0.87 | 0.81 | 0.84 |
| 5 | 120968757 | rs4437402 | 0.927 | -0.054 | -0.625 | -0.721 | 0.032 | 0.036 | 0.036 | 0.464 | 0.61 | 0.23 | 0.74 | 0.64 | 0.68 | 0.67 |
| 5 | 120969316 | rs9327207 | -0.409 | 0.244 | -0.875 | -0.661 | -0.006 | 0 | 0 | 0 | 0.86 | 0.88 | 0.87 | 0.79 | 0.85 | 0.86 |
| 5 | 120970448 | rs1508805 | -0.042 | 0.214 | -0.635 | -0.408 | -0.018 | -0.057 | -0.057 | 0.561 | 0.68 | 0.48 | 0.78 | 0.77 | 0.72 | 0.76 |
| 5 | 120974416 | rs4612103 | 0.158 | 0.697 | -1.134 | 0.496 | -0.101 | -0.187 | -0.187 | 0.084 | 0.28 | 0.02 | 0.5 | 0.49 | 0.33 | 0.39 |
| 5 | 120988738 | rs4895332 | -0.065 | 0.565 | -0.419 | 0.231 | -0.02 | 0.028 | 0.028 | 0 | 0.05 | 0 | 0.04 | 0 | 0.08 | 0.02 |
| 5 | 121007018 | rs10519677 | -1.064 | 0.028 | 0.159 | 0.525 | -0.019 | 0.021 | 0.021 | 0 | 0.11 | 0.16 | 0.12 | 0.16 | 0.06 | 0.22 |
| 5 | 121007574 | rs9327213 | -0.715 | -0.317 | -0.128 | 0.029 | 0.013 | 0.083 | 0.083 | 0 | 0.54 | 0.8 | 0.46 | 0.34 | 0.57 | 0.58 |
| 5 | 121018550 | rs10053041 | 0.476 | -0.619 | 0.614 | -0.274 | -0.055 | 0.062 | 0.062 | 0 | 0.32 | 0.2 | 0.28 | 0.17 | 0.46 | 0.27 |
| 5 | 121027720 | rs1876648 | -0.904 | -0.865 | 1.1 | 0.4 | 0.002 | 0 | 0 | 0 | 0.52 | 0.55 | 0.42 | 0.44 | 0.55 | 0.55 |
| 5 | 121027934 | rs7712025 | 0.23 | -0.614 | 1.143 | 0.227 | -0.047 | 0.007 | 0.007 | 0 | 0.33 | 0.2 | 0.25 | 0.26 | 0.42 | 0.3 |
| 5 | 121030389 | rs7727002 | 0.472 | -0.567 | 1.149 | 0.255 | -0.035 | -0.003 | -0.003 | 0 | 0.3 | 0.17 | 0.23 | 0.26 | 0.39 | 0.3 |
| 5 | 121031130 | rs7732370 | 0.466 | -0.613 | 1.16 | 0.263 | -0.045 | 0.001 | 0.001 | 0 | 0.31 | 0.18 | 0.25 | 0.26 | 0.42 | 0.3 |
| 5 | 121032073 | rs17148198 | 0.013 | -0.574 | 1.318 | 0.317 | 0.005 | -0.034 | -0.034 | 0.018 | 0.08 | 0.15 | 0.05 | 0 | 0.07 | 0.02 |
| 5 | 121038138 | rs2662302 | -0.584 | -0.661 | 1.289 | 0.291 | -0.003 | -0.003 | -0.003 | 0 | 0.46 | 0.41 | 0.37 | 0.45 | 0.49 | 0.53 |
| 5 | 121043540 | rs173154 | -1.097 | -0.628 | 1.257 | 0.353 | 0.007 | -0.006 | -0.006 | 0.073 | 0.54 | 0.65 | 0.44 | 0.45 | 0.56 | 0.55 |
| 5 | 121062939 | rs10056717 | -1.92 | -0.852 | 2.377 | 0.973 | 0 | 0.021 | 0.021 | 0 | 0.15 | 0.41 | 0.17 | 0.16 | 0.11 | 0.16 |
| 5 | 121063011 | rs10058293 | -1.525 | -0.878 | 2.403 | 0.976 | -0.016 | 0.031 | 0.031 | 0 | 0.08 | 0.13 | 0.1 | 0.16 | 0.04 | 0.14 |
| 5 | 121064424 | rs7710680 | -0.549 | -0.784 | 2.42 | 0.965 | 0.068 | -0.033 | -0.033 | 0.018 | 0.07 | 0.28 | 0.07 | 0 | 0.08 | 0.02 |
| 5 | 121064897 | rs2567819 | 2.077 | -0.895 | 2.385 | 1.059 | -0.044 | 0.049 | 0.049 | 0 | 0.84 | 0.69 | 0.83 | 0.77 | 0.92 | 0.76 |
| 5 | 121066267 | rs2662288 | -2.727 | -0.832 | 2.48 | 0.978 | -0.018 | 0.09 | 0.09 | 0 | 0.14 | 0.41 | 0.18 | 0.2 | 0.1 | 0.26 |
| 5 | 121067170 | rs7735996 | 2.009 | -0.81 | 2.32 | 0.99 | -0.002 | 0.11 | 0.11 | 0 | 0.89 | 0.67 | 0.83 | 0.79 | 0.92 | 0.79 |
| 5 | 121079701 | rs2249574 | -2.915 | -1.243 | 1.816 | 0.822 | 0.032 | 0.025 | 0.025 | 0.026 | 0.3 | 0.9 | 0.27 | 0.21 | 0.19 | 0.33 |
| 5 | 121082692 | rs2662295 | -1.77 | -1.16 | 2.233 | 0.883 | 0.03 | 0.067 | 0.067 | 0.196 | 0.16 | 0.53 | 0.23 | 0.21 | 0.14 | 0.21 |
| 5 | 121087457 | rs1560553 | -2.69 | -1.216 | 1.529 | 0.998 | 0.042 | 0.007 | 0.007 | 0.017 | 0.31 | 0.89 | 0.29 | 0.25 | 0.2 | 0.32 |
| 5 | 121100211 | rs1431944 | -1.996 | -1.218 | 1.904 | 0.718 | -0.068 | 0.02 | 0.02 | 0.222 | 0.15 | 0.48 | 0.12 | 0.1 | 0.05 | 0.13 |
| 5 | 121100484 | rs6595320 | -3.039 | -1.286 | 1.315 | 1.145 | 0.022 | -0.167 | -0.167 | 0.745 | 0.34 | 0.96 | 0.31 | 0.36 | 0.19 | 0.36 |
| 5 | 121103216 | rs6862426 | -1.932 | -1.461 | 1.888 | 0.704 | -0.008 | 0.004 | 0.004 | 0 | 0.09 | 0.12 | 0.03 | 0.1 | 0.04 | 0.12 |
| 5 | 121104400 | rs1431945 | -2.448 | -1.545 | 2.357 | 0.663 | -0.064 | 0.012 | 0.012 | 0.058 | 0.15 | 0.39 | 0.05 | 0.1 | 0.06 | 0.21 |
| 5 | 121106601 | rs10053356 | -0.948 | -1.556 | 2.345 | 0.643 | -0.014 | -0.021 | -0.021 | 5.01 | 0.06 | 0.22 | 0.02 | 0 | 0.02 | 0.07 |
| 5 | 121110144 | rs17148400 | -3.001 | -1.54 | 2.448 | 0.708 | -0.076 | 0.069 | 0.069 | 0.456 | 0.19 | 0.55 | 0.14 | 0.2 | 0.08 | 0.22 |

| | | | | | | | | | | | | | | | | |
|---|---|---|---|---|---|---|---|---|---|---|---|---|---|---|---|---|
| 5 | 121112267 | rs1431948 | -3.135 | -1.614 | 2.176 | 0.708 | 0.022 | -0.012 | -0.012 | 0.561 | 0.22 | 0.82 | 0.16 | 0.21 | 0.08 | 0.21 |
| 5 | 121129445 | rs17148460 | -1.785 | -1.216 | 2.038 | 0.188 | -0.002 | 0.084 | 0.084 | 0.009 | 0.08 | 0.33 | 0.1 | 0.1 | 0.03 | 0.04 |
| 5 | 121136126 | rs6595322 | 2.369 | -1.309 | 1.777 | 0.087 | -0.063 | -0.011 | -0.011 | 0 | 0.84 | 0.44 | 0.87 | 0.9 | 0.93 | 0.91 |
| 5 | 121153924 | rs10037268 | -3.043 | -1.379 | 1.86 | 0.105 | -0.077 | -0.02 | -0.02 | 0 | 0.19 | 0.62 | 0.12 | 0.1 | 0.05 | 0.1 |
| 5 | 121155521 | rs7736854 | -2.433 | -1.374 | 1.893 | 0.109 | -0.068 | 0.05 | 0.05 | 1.572 | 0.12 | 0.51 | 0.11 | 0.1 | 0.02 | 0.06 |
| 5 | 121167471 | rs10478541 | -2.549 | -1.168 | 1.797 | 0.207 | -0.067 | 0.004 | 0.004 | 0.829 | 0.15 | 0.6 | 0.12 | 0.1 | 0.05 | 0.1 |
| 5 | 121173834 | rs17148538 | -0.971 | -0.733 | 0.879 | 0.314 | -0.012 | -0.028 | -0.028 | 0 | 0.06 | 0.13 | 0.01 | 0 | 0.03 | 0.03 |
| 5 | 121174252 | rs11959845 | -0.971 | -0.733 | 0.877 | 0.314 | -0.012 | -0.029 | -0.029 | 0 | 0.06 | 0.12 | 0.01 | 0 | 0.03 | 0.03 |
| 5 | 121178515 | rs6887649 | -1.918 | -0.645 | 1.138 | 0.4 | 0.007 | 0.069 | 0.069 | 0.019 | 0.09 | 0.3 | 0.12 | 0.1 | 0.05 | 0.04 |
| 5 | 121182996 | rs2116427 | -1.205 | -0.752 | -0.257 | -0.562 | -0.006 | 0.091 | 0.091 | 0.472 | 0.34 | 0.92 | 0.28 | 0.18 | 0.26 | 0.3 |
| 5 | 121189496 | rs1560550 | -1.019 | -0.361 | -0.899 | -1.257 | -0.142 | 0.044 | 0.044 | 0.07 | 0.61 | 0.98 | 0.45 | 0.26 | 0.51 | 0.53 |
| 5 | 121197292 | rs17148577 | -1.584 | -0.396 | 0.087 | -1.843 | -0.003 | -0.024 | -0.024 | 0 | 0.15 | 0.2 | 0.17 | 0.1 | 0.11 | 0.13 |
| 5 | 121197663 | rs1982052 | -0.971 | -0.404 | 0.07 | -1.854 | 0.015 | 0.032 | 0.032 | 0 | 0.1 | 0.05 | 0.05 | 0 | 0.06 | 0.04 |
| 5 | 121207152 | rs4481375 | 1.785 | -0.609 | 0.416 | -2.503 | 0.031 | 0.103 | 0.103 | 0 | 0.55 | 0.29 | 0.65 | 0.83 | 0.63 | 0.73 |
| 5 | 121211925 | rs4632823 | 0.026 | -0.72 | 1.06 | -2.564 | -0.004 | -0.049 | -0.049 | 0.61 | 0.76 | 0.72 | 0.84 | 0.96 | 0.78 | 0.88 |
| 5 | 121212154 | rs4301249 | 1.59 | -0.726 | 1.189 | -2.569 | -0.021 | -0.115 | -0.115 | 0 | 0.6 | 0.45 | 0.76 | 0.94 | 0.68 | 0.82 |
| 5 | 121213868 | rs7728244 | -0.299 | -0.792 | 1.003 | -2.567 | 0.004 | 0.047 | 0.047 | 0 | 0.79 | 0.89 | 0.85 | 0.96 | 0.78 | 0.88 |
| 5 | 121214164 | rs13182775 | -0.243 | -0.739 | 0.996 | -2.504 | 0.004 | -0.003 | -0.003 | 0.669 | 0.08 | 0.08 | 0.05 | 0 | 0.06 | 0.04 |
| 5 | 121226853 | rs4637581 | -0.286 | -0.764 | 0.778 | -2.545 | 0.003 | 0.063 | 0.063 | 0 | 0.79 | 0.9 | 0.85 | 0.96 | 0.78 | 0.88 |
| 5 | 121229399 | rs6595333 | 1.803 | -0.706 | 0.84 | -2.577 | 0.024 | 0.067 | 0.067 | 0 | 0.55 | 0.31 | 0.66 | 0.84 | 0.63 | 0.74 |
| 5 | 121234474 | rs4343870 | 1.617 | -0.732 | 0.769 | -2.574 | 0.006 | -0.028 | -0.028 | 0.373 | 0.58 | 0.36 | 0.75 | 0.89 | 0.69 | 0.78 |
| 5 | 121249306 | rs4621583 | 1.691 | -0.594 | 1.021 | -2.296 | 0.002 | -0.011 | -0.011 | 0 | 0.61 | 0.35 | 0.68 | 0.85 | 0.67 | 0.8 |
| 5 | 121252616 | rs9637910 | -1.644 | -0.457 | 1.123 | -2.156 | 0.002 | 0.004 | 0.004 | 0 | 0.06 | 0.11 | 0.1 | 0.13 | 0.05 | 0.08 |
| 5 | 121260543 | rs12518706 | 0.733 | -0.477 | 1.205 | -2.176 | -0.029 | -0.081 | -0.081 | 0 | 0.72 | 0.47 | 0.81 | 0.9 | 0.78 | 0.86 |
| 5 | 121260607 | rs12516371 | 1.464 | -0.469 | 1.136 | -2.177 | 0.004 | 0.008 | 0.008 | 0 | 0.64 | 0.32 | 0.69 | 0.81 | 0.69 | 0.81 |
| 5 | 121265925 | rs2972720 | 0.929 | -0.375 | 1.766 | -1.641 | -0.03 | -0.077 | -0.077 | 0 | 0.79 | 0.42 | 0.88 | 0.94 | 0.86 | 0.91 |
| 5 | 121269498 | rs2914598 | 1.844 | -0.313 | 1.428 | -1.922 | -0.087 | 0.192 | 0.192 | 0 | 0.46 | 0.11 | 0.52 | 0.74 | 0.55 | 0.56 |
| 5 | 121270012 | rs6869790 | -1.335 | -0.328 | 1.604 | -1.914 | -0.007 | 0.042 | 0.042 | 0.195 | 0.08 | 0.45 | 0.03 | 0.06 | 0.01 | 0.03 |
| 5 | 121271817 | rs11750012 | 1.046 | -0.262 | 1.65 | -1.89 | -0.025 | -0.047 | -0.047 | 0 | 0.78 | 0.35 | 0.87 | 0.9 | 0.86 | 0.88 |
| 5 | 121273302 | rs11241627 | 0.391 | -0.069 | 0.798 | -1.811 | -0.001 | -0.024 | -0.024 | 0 | 0.83 | 0.75 | 0.87 | 0.96 | 0.84 | 0.9 |
| 5 | 121279253 | rs12153375 | 1.754 | -0.036 | 0.782 | -1.812 | -0.088 | 0.183 | 0.183 | 0 | 0.45 | 0.1 | 0.51 | 0.74 | 0.55 | 0.54 |
| 5 | 121297115 | rs2972350 | 0.353 | 0.506 | 1.301 | -1.108 | 0.001 | 0.019 | 0.019 | 0 | 0.86 | 0.91 | 0.91 | 0.96 | 0.88 | 0.9 |
| 5 | 121308199 | rs12657394 | 1.34 | 0.558 | 1.377 | -1.108 | -0.006 | -0.035 | -0.035 | 0.009 | 0.81 | 0.68 | 0.85 | 0.92 | 0.84 | 0.86 |
| 5 | 121309324 | rs13172533 | 0.515 | 0.556 | 1.367 | -1.091 | -0.002 | 0.022 | 0.022 | 0 | 0.86 | 0.91 | 0.9 | 0.95 | 0.87 | 0.9 |
| 5 | 121314168 | rs2914610 | 1.62 | 0.553 | 1.462 | -1.1 | -0.015 | -0.07 | -0.07 | 0 | 0.8 | 0.55 | 0.84 | 0.92 | 0.84 | 0.86 |
| 5 | 121354221 | rs10078419 | -0.635 | 0.48 | 1.073 | -1.058 | 0.002 | 0.02 | 0.02 | 0 | 0.14 | 0.1 | 0.09 | 0.04 | 0.13 | 0.1 |
| 5 | 121402597 | rs10053942 | -1.42 | 0.451 | 0.628 | -1.043 | -0.018 | -0.082 | -0.082 | 0 | 0.2 | 0.49 | 0.14 | 0.08 | 0.16 | 0.15 |
| 5 | 121403659 | rs3792802 | -1.42 | 0.451 | 0.628 | -1.043 | -0.018 | -0.082 | -0.082 | 0 | 0.2 | 0.49 | 0.14 | 0.08 | 0.16 | 0.15 |
| 5 | 121413208 | rs1800449 | -0.611 | 0.689 | -0.59 | -0.62 | 0.003 | -0.005 | -0.005 | 0.002 | 0.17 | 0.14 | 0.16 | 0.19 | 0.16 | 0.14 |
| 5 | 121418427 | rs17431081 | -0.32 | 0.73 | -0.73 | -0.772 | -0.019 | 0.029 | 0.029 | 0 | 0.19 | 0.15 | 0.17 | 0.09 | 0.24 | 0.21 |
| 5 | 121418545 | rs17352686 | -0.306 | 0.725 | -0.755 | -0.765 | -0.038 | 0.05 | 0.05 | 0 | 0.19 | 0.11 | 0.17 | 0.09 | 0.24 | 0.21 |
| 5 | 121419259 | rs840461 | 1.249 | 1.04 | -0.891 | -0.875 | -0.05 | 0.074 | 0.074 | 0.957 | 0.59 | 0.74 | 0.67 | 0.74 | 0.56 | 0.66 |
| 5 | 121422057 | rs995689 | 0.465 | 1.069 | -0.998 | -0.691 | -0.042 | -0.032 | -0.032 | 1.636 | 0.81 | 0.98 | 0.84 | 0.84 | 0.82 | 0.88 |
| 5 | 121426675 | rs3900446 | -0.126 | 1.254 | -0.382 | -0.523 | 0.001 | 0 | 0 | 0.167 | 0.17 | 0.15 | 0.16 | 0.07 | 0.23 | 0.16 |
| 5 | 121426712 | rs711413 | 1.37 | 0.039 | 0.443 | -0.412 | 0.006 | 0.013 | 0.013 | 0.009 | 0.68 | 0.24 | 0.63 | 0.76 | 0.74 | 0.75 |
| 5 | 121427230 | rs763497 | -1.525 | -0.052 | 1.401 | -0.358 | 0.069 | 0.031 | 0.031 | 0.389 | 0.13 | 0.48 | 0.23 | 0.16 | 0.11 | 0.16 |
| 5 | 121428330 | rs2560303 | 0.512 | -0.557 | 1.295 | -0.399 | 0.023 | 0.017 | 0.017 | 0.323 | 0.74 | 0.82 | 0.83 | 0.84 | 0.82 | 0.86 |
| 5 | 121428395 | rs2560302 | 1.832 | -0.589 | 1.088 | -0.457 | 0.032 | 0.049 | 0.049 | 1.255 | 0.57 | 0.13 | 0.57 | 0.68 | 0.68 | 0.67 |
| 5 | 121429808 | rs2560301 | 1.545 | -0.326 | 1.46 | -0.354 | -0.017 | -0.016 | -0.016 | 0 | 0.62 | 0.46 | 0.61 | 0.68 | 0.71 | 0.8 |
| 5 | 121452158 | rs984279 | 0.568 | -0.277 | 1.028 | -0.055 | 0.022 | 0.032 | 0.032 | 0.455 | 0.75 | 0.92 | 0.82 | 0.85 | 0.8 | 0.85 |
| 5 | 121480802 | rs6595344 | -1.258 | 0.047 | 1.295 | 0.273 | 0.047 | 0.056 | 0.056 | 0 | 0.08 | 0.28 | 0.18 | 0.15 | 0.07 | 0.05 |
| 5 | 121488203 | rs2560306 | -1.545 | -0.205 | 1.334 | 0.363 | 0.026 | 0.097 | 0.097 | 0 | 0.07 | 0.32 | 0.03 | 0.1 | 0.05 | 0.08 |
| 5 | 121492603 | rs11951324 | -1.332 | -1.208 | 1.446 | -0.28 | -0.079 | -0.098 | -0.098 | 0 | 0.23 | 0.46 | 0.09 | 0.09 | 0.09 | 0.16 |
| 5 | 121497094 | rs10519697 | -0.876 | -1.348 | 1.337 | -0.352 | 0.026 | 0.103 | 0.103 | 0.014 | 0.17 | 0.01 | 0.05 | 0 | 0.09 | 0.15 |
| 5 | 121497706 | rs17148903 | -1.778 | -1.29 | 1.41 | -0.369 | -0.07 | -0.098 | -0.098 | 0 | 0.25 | 0.49 | 0.1 | 0.09 | 0.1 | 0.16 |
| 5 | 121505407 | rs10900755 | -1.874 | -2.007 | 1.329 | -0.306 | -0.003 | -0.002 | -0.002 | 0 | 0.26 | 0.72 | 0.26 | 0.13 | 0.17 | 0.39 |
| 5 | 121506678 | rs1427936 | -1.965 | -1.998 | 1.337 | -0.301 | -0.009 | -0.007 | -0.007 | 0 | 0.24 | 0.72 | 0.26 | 0.12 | 0.17 | 0.38 |
| 5 | 121510586 | rs17148926 | -1.872 | -2.539 | 1.577 | -0.726 | -0.007 | -0.025 | -0.025 | 0.276 | 0.26 | 0.26 | 0.2 | 0.07 | 0.2 | 0.21 |
| 5 | 121522316 | rs11948266 | -1.802 | -2.524 | 1.326 | -0.669 | 0.036 | 0.081 | 0.081 | 0 | 0.24 | 0.07 | 0.19 | 0.07 | 0.19 | 0.22 |
| 5 | 121525621 | rs9968686 | -2.128 | -2.505 | 1.651 | -0.652 | -0.029 | -0.017 | -0.017 | 0 | 0.13 | 0.28 | 0.14 | 0.07 | 0.08 | 0.11 |
| 5 | 121532424 | rs992788 | -2.132 | -2.881 | 1.27 | -0.497 | 0.05 | 0.02 | 0.02 | 0 | 0.4 | 0.78 | 0.41 | 0.3 | 0.28 | 0.47 |
| 5 | 121550921 | rs6893318 | -1.824 | -2.487 | 2.098 | -0.771 | -0.009 | -0.066 | -0.066 | 0 | 0.28 | 0.64 | 0.22 | 0.09 | 0.23 | 0.24 |
| 5 | 121595053 | rs11241634 | -1.272 | -2.739 | 1.646 | -0.892 | -0.01 | 0.086 | 0.086 | 0.002 | 0.25 | 0.08 | 0.19 | 0.09 | 0.23 | 0.31 |
| 5 | 121601488 | rs10085169 | 1.341 | -2.43 | 0.53 | -0.656 | -0.065 | -0.009 | -0.009 | 0 | 0.42 | 0.13 | 0.41 | 0.44 | 0.49 | 0.29 |
| 5 | 121622291 | rs12655730 | -1.332 | -2.268 | 0.77 | -0.784 | -0.003 | -0.017 | -0.017 | 0 | 0.32 | 0.7 | 0.22 | 0.11 | 0.23 | 0.33 |
| 5 | 121633067 | rs995140 | -1.037 | -2.332 | 0.639 | -0.753 | -0.025 | -0.061 | -0.061 | 0 | 0.27 | 0.59 | 0.24 | 0.1 | 0.2 | 0.31 |
| 5 | 121635375 | rs17434434 | -0.839 | -1.52 | -0.087 | -0.315 | 0.013 | 0.018 | 0.018 | 0 | 0.12 | 0.02 | 0.11 | 0.06 | 0.12 | 0.12 |
| 5 | 121644052 | rs4461665 | -1.312 | -1.266 | 0.432 | -0.097 | 0.03 | -0.011 | -0.011 | 0 | 0.18 | 0.42 | 0.23 | 0.13 | 0.18 | 0.28 |
| 5 | 121651149 | rs4895350 | -1.461 | -1.23 | 0.519 | -0.141 | 0.022 | -0.013 | -0.013 | 0.154 | 0.21 | 0.48 | 0.18 | 0.18 | 0.19 | 0.29 |

| | | | | | | | | | | | | | | | | |
|---|---|---|---|---|---|---|---|---|---|---|---|---|---|---|---|---|
| 5 | 121704974 | rs11241642 | -1.368 | -0.748 | 0.553 | -0.128 | -0.005 | 0.019 | 0.019 | 0.506 | 0.16 | 0.41 | 0.15 | 0.19 | 0.13 | 0.25 |
| 5 | 121707541 | rs10065900 | -0.167 | -0.113 | -0.365 | 0.36 | -0.049 | -0.022 | -0.022 | 0 | 0.8 | 0.96 | 0.78 | 0.79 | 0.79 | 0.68 |
| 5 | 121726123 | rs17149115 | -0.825 | -0.085 | 0.138 | 0.591 | -0.001 | 0.002 | 0.002 | 0.354 | 0.13 | 0.12 | 0.07 | 0.17 | 0.12 | 0.25 |
| 5 | 121740909 | rs17149141 | -0.773 | -0.098 | -0.152 | 0.604 | -0.001 | -0.004 | -0.004 | 0 | 0.13 | 0.11 | 0.08 | 0.18 | 0.12 | 0.25 |
| 5 | 121750329 | rs10066937 | -0.013 | 0.155 | -0.375 | 0.427 | -0.044 | -0.005 | -0.005 | 0 | 0.78 | 0.96 | 0.72 | 0.8 | 0.76 | 0.68 |
| 5 | 121753782 | rs304391 | 0.766 | 0.172 | 0.272 | 0.56 | -0.001 | -0.001 | -0.001 | 0 | 0.47 | 0.39 | 0.47 | 0.61 | 0.39 | 0.44 |
| 5 | 121755954 | rs304388 | 0.666 | 0.207 | 0.462 | 0.536 | -0.013 | 0.024 | 0.024 | 0 | 0.29 | 0.25 | 0.25 | 0.1 | 0.37 | 0.22 |
| 5 | 121761461 | rs2242223 | 0.29 | 0.205 | 0.487 | 0.78 | -0.026 | -0.048 | -0.048 | 0 | 0.19 | 0.08 | 0.31 | 0.3 | 0.24 | 0.34 |
| 5 | 121765801 | rs3811891 | 0.29 | 0.1 | 0.44 | 0.802 | -0.044 | -0.068 | -0.068 | 0.051 | 0.19 | 0.04 | 0.31 | 0.28 | 0.23 | 0.34 |
| 5 | 121768671 | rs304379 | -0.895 | 0.064 | 0.638 | 0.762 | 0.005 | 0.006 | 0.006 | 0 | 0.52 | 0.63 | 0.53 | 0.37 | 0.6 | 0.55 |
| 5 | 121772722 | rs3828578 | -0.3 | 0.573 | 0.646 | 0.99 | 0.013 | 0.023 | 0.023 | 0 | 0.22 | 0.38 | 0.29 | 0.26 | 0.24 | 0.34 |
| 5 | 121776301 | rs2290987 | -0.01 | 0.69 | 0.493 | 1.054 | -0.001 | -0.001 | -0.001 | 0 | 0.19 | 0.16 | 0.27 | 0.26 | 0.23 | 0.34 |
| 5 | 121777426 | rs1369044 | -0.198 | 0.683 | 0.477 | 1.011 | 0.001 | 0.018 | 0.018 | 0 | 0.21 | 0.29 | 0.28 | 0.27 | 0.22 | 0.32 |
| 5 | 121779262 | rs2737094 | 0.087 | 0.804 | 0.179 | 0.952 | -0.01 | -0.01 | -0.01 | 0 | 0.09 | 0.04 | 0.13 | 0.12 | 0.13 | 0.09 |
| 5 | 121788166 | rs3828576 | -0.061 | 0.864 | 0.134 | 0.964 | -0.035 | -0.044 | -0.044 | 0 | 0.19 | 0.05 | 0.22 | 0.25 | 0.23 | 0.33 |
| 5 | 121793477 | rs3811873 | -0.154 | 0.86 | 0.156 | 0.962 | -0.025 | -0.035 | -0.035 | 0 | 0.21 | 0.09 | 0.22 | 0.25 | 0.23 | 0.32 |
| 5 | 121793874 | rs6862890 | 1.209 | 0.9 | 0.323 | 0.789 | -0.004 | -0.002 | -0.002 | 0 | 0.2 | 0.2 | 0.18 | 0.08 | 0.27 | 0.12 |
| 5 | 121796012 | rs1478370 | -0.523 | 0.302 | 0.617 | 1.384 | 0.002 | 0.007 | 0.007 | 0 | 0.19 | 0.21 | 0.22 | 0.24 | 0.21 | 0.3 |
| 5 | 121808701 | rs10067389 | -1.382 | 0.373 | 0.636 | 1.104 | 0.01 | 0.043 | 0.043 | 0 | 0.28 | 0.32 | 0.41 | 0.45 | 0.34 | 0.4 |
| 5 | 121826721 | rs1000590 | -1.404 | 0.444 | 0.338 | 1.066 | 0.006 | 0.024 | 0.024 | 0 | 0.31 | 0.34 | 0.43 | 0.44 | 0.38 | 0.39 |
| 5 | 121826941 | rs2407209 | -1.355 | 0.499 | 0.136 | 1.087 | 0.002 | 0.01 | 0.01 | 0 | 0.3 | 0.31 | 0.43 | 0.44 | 0.38 | 0.39 |
| 5 | 121829635 | rs6866750 | -1.501 | 0.635 | -0.243 | 0.846 | -0.002 | -0.004 | -0.004 | 0 | 0.3 | 0.3 | 0.44 | 0.4 | 0.38 | 0.38 |
| 5 | 121832058 | rs747013 | -0.484 | 0.734 | -0.699 | 0.962 | -0.031 | -0.057 | -0.057 | 0 | 0.17 | 0.06 | 0.26 | 0.3 | 0.22 | 0.29 |
| 5 | 121838912 | rs10519702 | -0.806 | 0.355 | -0.103 | 0.237 | -0.003 | -0.032 | -0.032 | 0.135 | 0.13 | 0.21 | 0.13 | 0.03 | 0.15 | 0.14 |
| 5 | 121844320 | rs17149268 | -1.39 | 0.115 | 0.146 | -0.228 | -0.001 | -0.066 | -0.066 | 0 | 0.24 | 0.38 | 0.26 | 0.04 | 0.24 | 0.26 |
| 5 | 121852531 | rs4835803 | 0.698 | -0.204 | -0.54 | -0.111 | 0.008 | -0.032 | -0.032 | 0.256 | 0.17 | 0.07 | 0.19 | 0.27 | 0.14 | 0.1 |
| 5 | 121854525 | rs6867820 | -1.176 | -0.124 | -0.243 | -0.369 | 0 | -0.02 | -0.02 | 0 | 0.26 | 0.27 | 0.25 | 0.04 | 0.26 | 0.29 |
| 5 | 121856506 | rs10519706 | -0.081 | -0.008 | -0.186 | -0.106 | -0.002 | -0.017 | -0.017 | 0.292 | 0.11 | 0.14 | 0.09 | 0.03 | 0.1 | 0.14 |
| 5 | 121860590 | rs6893509 | -0.315 | -0.279 | -0.041 | -0.237 | -0.001 | -0.008 | -0.008 | 0 | 0.12 | 0.13 | 0.18 | 0.04 | 0.12 | 0.13 |
| 5 | 121861856 | rs1478374 | -1.151 | -0.76 | 0.68 | -0.488 | -0.011 | -0.078 | -0.078 | 0.451 | 0.24 | 0.37 | 0.28 | 0.07 | 0.2 | 0.25 |
| 5 | 121862860 | rs12518676 | -1.158 | -0.759 | 0.704 | -0.193 | -0.011 | -0.046 | -0.046 | 0 | 0.24 | 0.36 | 0.28 | 0.09 | 0.2 | 0.26 |
| 5 | 121866583 | rs17149301 | 0.885 | -0.952 | 0.505 | -0.12 | -0.003 | 0.005 | 0.005 | 0 | 0.17 | 0.15 | 0.2 | 0.24 | 0.14 | 0.08 |
| 5 | 121871938 | rs10519707 | -1.4 | -1.213 | 0.133 | -0.246 | 0.057 | 0.029 | 0.029 | 0.896 | 0.48 | 0.89 | 0.52 | 0.32 | 0.35 | 0.35 |
| 5 | 121874834 | rs1904139 | 0.026 | -1.002 | -0.221 | -0.269 | -0.002 | 0.031 | 0.031 | 0 | 0.68 | 0.82 | 0.73 | 0.73 | 0.71 | 0.74 |
| 5 | 121879366 | rs6595377 | 0.165 | -1.02 | -0.168 | -0.449 | 0.003 | 0.058 | 0.058 | 0 | 0.67 | 0.78 | 0.73 | 0.82 | 0.71 | 0.73 |
| 5 | 121883682 | rs2407210 | 0.404 | -1.051 | 0.141 | -0.406 | 0.004 | 0.017 | 0.017 | 0 | 0.65 | 0.7 | 0.73 | 0.78 | 0.71 | 0.73 |
| 5 | 121884439 | rs2407211 | 0.116 | -1.054 | 0.142 | -0.41 | 0.002 | 0.054 | 0.054 | 0 | 0.67 | 0.81 | 0.77 | 0.81 | 0.72 | 0.74 |
| 5 | 121884501 | rs2407212 | 0.372 | -1.051 | 0.137 | -0.415 | 0.003 | 0.013 | 0.013 | 0 | 0.65 | 0.7 | 0.77 | 0.84 | 0.71 | 0.72 |
| 5 | 121887537 | rs4836045 | 0.716 | -1.158 | 0.286 | -0.456 | -0.039 | 0.011 | 0.011 | 1.319 | 0.22 | 0.55 | 0.23 | 0.22 | 0.14 | 0.09 |
| 5 | 121889627 | rs4836049 | 0.116 | -1.058 | 0.153 | -0.467 | 0.002 | 0.055 | 0.055 | 0 | 0.67 | 0.8 | 0.77 | 0.81 | 0.72 | 0.74 |
| 5 | 121898571 | rs7700476 | 0.341 | -1.055 | 0.144 | -0.469 | 0.004 | 0.022 | 0.022 | 0 | 0.65 | 0.73 | 0.73 | 0.82 | 0.71 | 0.74 |
| 5 | 121899110 | rs17149348 | 1.75 | -1.092 | 0.293 | -0.425 | 0.01 | -0.021 | -0.021 | 0 | 0.16 | 0.07 | 0.18 | 0.22 | 0.12 | 0.08 |
| 5 | 121905828 | rs4629597 | 0.895 | -0.967 | 0.241 | -0.379 | -0.002 | -0.001 | -0.001 | 0 | 0.39 | 0.63 | 0.43 | 0.57 | 0.49 | 0.51 |
| 5 | 121907730 | rs1478369 | 0.784 | -1.012 | 0.113 | -0.479 | 0.011 | 0.02 | 0.02 | 0 | 0.12 | 0.04 | 0.14 | 0.06 | 0.1 | 0.13 |
| 5 | 121907799 | rs17149379 | 0.342 | -1.062 | 0.251 | -0.308 | 0.004 | 0.005 | 0.005 | 0 | 0.28 | 0.24 | 0.31 | 0.22 | 0.22 | 0.22 |
| 5 | 121908577 | rs1478367 | 0.008 | -0.8 | -0.404 | -0.342 | 0.003 | 0.067 | 0.067 | 0 | 0.67 | 0.83 | 0.73 | 0.82 | 0.71 | 0.74 |
| 5 | 121910555 | rs17380695 | 0.625 | -0.186 | -0.56 | 0.133 | -0.031 | -0.024 | -0.024 | 1.472 | 0.31 | 0.15 | 0.35 | 0.58 | 0.46 | 0.49 |
| 5 | 121914456 | rs10477635 | -1.508 | 0.227 | -0.8 | 0.333 | -0.051 | -0.011 | -0.011 | 0 | 0.82 | 0.91 | 0.73 | 0.79 | 0.68 | 0.75 |
| 5 | 121916555 | rs4836062 | 0.261 | -0.653 | -0.884 | -0.439 | 0.023 | 0.108 | 0.108 | 0 | 0.41 | 0.16 | 0.31 | 0.21 | 0.32 | 0.28 |
| 5 | 121916936 | rs7702503 | -0.213 | -0.801 | -0.24 | -0.61 | 0.006 | 0.01 | 0.01 | 0.157 | 0.5 | 0.57 | 0.4 | 0.21 | 0.39 | 0.32 |
| 5 | 121917603 | rs899181 | -0.253 | -0.81 | -0.159 | -0.495 | 0.003 | -0.003 | -0.003 | 0 | 0.48 | 0.57 | 0.38 | 0.19 | 0.37 | 0.3 |
| 5 | 121921114 | rs7727904 | -0.237 | -1.108 | -0.267 | -0.841 | -0.009 | -0.085 | -0.085 | 0.04 | 0.29 | 0.4 | 0.29 | 0.01 | 0.23 | 0.21 |
| 5 | 121924581 | rs12656417 | -0.778 | -1.286 | 0.01 | -0.092 | 0.001 | -0.002 | -0.002 | 0 | 0.37 | 0.69 | 0.31 | 0.43 | 0.27 | 0.25 |
| 5 | 121925160 | rs13362479 | -0.265 | -0.656 | -0.125 | 0.078 | 0.004 | 0.016 | 0.016 | 0 | 0.15 | 0.23 | 0.18 | 0.23 | 0.15 | 0.18 |
| 5 | 121934906 | rs11739754 | -0.856 | -1.342 | -0.193 | -0.498 | 0.001 | -0.035 | -0.035 | 0 | 0.31 | 0.67 | 0.33 | 0.04 | 0.22 | 0.33 |
| 5 | 121943904 | rs6874307 | -1.391 | -1.529 | -0.383 | -0.114 | 0.045 | 0.028 | 0.028 | 0 | 0.42 | 0.18 | 0.44 | 0.34 | 0.35 | 0.47 |
| 5 | 121965806 | rs7349770 | -0.632 | -1.461 | -0.786 | -0.02 | 0 | -0.097 | -0.097 | 0 | 0.17 | 0.01 | 0.12 | 0.27 | 0.14 | 0.18 |
| 5 | 121987792 | rs13156690 | 0.039 | -1.307 | -0.501 | -0.128 | -0.003 | 0.096 | 0.096 | 0.273 | 0.2 | 0.07 | 0.21 | 0.01 | 0.18 | 0.24 |
| 5 | 121990722 | rs9327261 | -1.422 | -1.441 | -0.075 | 0.169 | 0.04 | 0 | 0 | 3.126 | 0.42 | 0.66 | 0.49 | 0.37 | 0.33 | 0.46 |
| 5 | 121991807 | rs17149450 | -0.996 | -0.907 | -0.19 | 0.346 | -0.001 | 0 | 0 | 0 | 0.35 | 0.36 | 0.33 | 0.33 | 0.31 | 0.41 |
| 5 | 121992016 | rs17149451 | -0.954 | -0.87 | -0.449 | 0.364 | 0.007 | -0.006 | -0.006 | 1.213 | 0.34 | 0.29 | 0.32 | 0.33 | 0.31 | 0.41 |
| 5 | 122004636 | rs4240383 | 0.21 | -0.786 | -0.756 | -0.19 | 0.009 | 0.028 | 0.028 | 0 | 0.82 | 0.89 | 0.91 | 0.99 | 0.84 | 0.86 |
| 5 | 122005153 | rs7706930 | 0.209 | -1.24 | -0.534 | 0.203 | -0.005 | 0.004 | 0.004 | 0 | 0.16 | 0.18 | 0.2 | 0.28 | 0.1 | 0.22 |
| 5 | 122007862 | rs13186938 | -0.577 | -1.738 | -0.527 | 0.761 | 0.001 | -0.002 | -0.002 | 0 | 0.27 | 0.25 | 0.33 | 0.53 | 0.18 | 0.38 |
| 5 | 122010221 | rs4835860 | -0.894 | -1.674 | -0.277 | 0.38 | -0.01 | -0.028 | -0.028 | 0 | 0.42 | 0.43 | 0.44 | 0.8 | 0.31 | 0.63 |
| 5 | 122025478 | rs10079345 | 1.435 | -1.742 | -0.193 | 0.137 | -0.047 | 0.068 | 0.068 | 0.108 | 0.35 | 0.19 | 0.37 | 0.12 | 0.47 | 0.19 |
| 5 | 122029790 | rs6863273 | 0.215 | -1.86 | 0.196 | 0.616 | -0.002 | 0.098 | 0.098 | 0 | 0.16 | 0.28 | 0.14 | 0.33 | 0.16 | 0.26 |
| 5 | 122030711 | rs7709626 | 1.436 | -1.455 | 0.117 | 0.213 | 0.011 | -0.017 | -0.017 | 0.403 | 0.27 | 0.3 | 0.3 | 0.11 | 0.37 | 0.17 |
| 5 | 122036899 | rs10042243 | 0.103 | -1.605 | -0.224 | 0.146 | 0.007 | 0.007 | 0.007 | 0 | 0.48 | 0.66 | 0.51 | 0.43 | 0.55 | 0.48 |

| | | | | | | | | | | | | | | | | |
|---|---|---|---|---|---|---|---|---|---|---|---|---|---|---|---|---|
| 5 | 122042452 | rs6866004 | 0.343 | -1.884 | -0.398 | 0.11 | 0.009 | 0.002 | 0.002 | 0 | 0.42 | 0.54 | 0.5 | 0.41 | 0.48 | 0.33 |
| 5 | 122050927 | rs10050385 | 1.573 | -1.743 | -0.22 | -0.294 | 0.042 | -0.042 | -0.042 | 0.53 | 0.25 | 0.4 | 0.35 | 0.1 | 0.35 | 0.12 |
| 5 | 122051019 | rs4836191 | -0.172 | -2.056 | -0.643 | -0.626 | 0.002 | -0.002 | -0.002 | 0 | 0.39 | 0.32 | 0.37 | 0.57 | 0.28 | 0.58 |
| 5 | 122059215 | rs7722933 | -1.63 | -2.097 | -0.992 | -0.686 | -0.099 | 0.129 | 0.129 | 0 | 0.8 | 0.99 | 0.86 | 0.99 | 0.74 | 0.87 |
| 5 | 122060284 | rs17149587 | 0.906 | -2.407 | -0.797 | -0.596 | 0 | 0 | 0 | 0 | 0.2 | 0.2 | 0.24 | 0.25 | 0.11 | 0.29 |
| 5 | 122072632 | rs17149596 | 0.503 | -2.327 | -0.85 | -0.583 | 0.019 | -0.038 | -0.038 | 0 | 0.18 | 0.12 | 0.13 | 0.32 | 0.14 | 0.26 |
| 5 | 122076870 | rs9327274 | 0.62 | -2.224 | -0.772 | -0.645 | 0.007 | 0.007 | 0.007 | 0 | 0.39 | 0.48 | 0.49 | 0.42 | 0.45 | 0.28 |
| 5 | 122096103 | rs4836231 | 1.158 | -1.677 | -0.511 | -1.078 | 0.05 | -0.034 | -0.034 | 0.313 | 0.28 | 0.49 | 0.38 | 0.16 | 0.39 | 0.15 |
| 5 | 122098688 | rs28603112 | 1.158 | -1.677 | -0.511 | -1.078 | 0.05 | -0.034 | -0.034 | 0 | 0.28 | 0.49 | 0.38 | 0.16 | 0.39 | 0.15 |
| 5 | 122102442 | rs1545696 | 0.322 | -1.881 | -0.534 | -1.115 | -0.001 | 0.003 | 0.003 | 0 | 0.56 | 0.67 | 0.7 | 0.64 | 0.56 | 0.56 |
| 5 | 122105232 | rs1460040 | -1.334 | -1.758 | -0.527 | -1.107 | -0.044 | 0.064 | 0.064 | 0 | 0.78 | 0.91 | 0.85 | 0.99 | 0.73 | 0.85 |
| 5 | 122109207 | rs6887101 | -0.12 | -1.651 | -0.429 | -0.982 | -0.019 | -0.012 | -0.012 | 0 | 0.6 | 0.79 | 0.72 | 0.64 | 0.58 | 0.58 |
| 5 | 122109923 | rs4835894 | 1.395 | -1.463 | -0.548 | -0.925 | 0.067 | -0.052 | -0.052 | 0 | 0.26 | 0.52 | 0.36 | 0.11 | 0.36 | 0.14 |
| 5 | 122132725 | rs10071705 | -0.078 | -1.512 | -0.625 | -0.899 | -0.007 | -0.003 | -0.003 | 0 | 0.59 | 0.73 | 0.72 | 0.64 | 0.58 | 0.58 |
| 5 | 122162857 | rs3797071 | 0.549 | -1.63 | -0.636 | -0.935 | 0.03 | -0.03 | -0.03 | 0 | 0.29 | 0.21 | 0.32 | 0.49 | 0.17 | 0.41 |
| 5 | 122169545 | rs7726915 | 1.57 | -1.673 | -0.62 | -1.094 | 0.079 | -0.078 | -0.078 | 0.227 | 0.24 | 0.49 | 0.36 | 0.06 | 0.35 | 0.12 |
| 5 | 122206447 | rs26372 | -0.705 | -1.742 | -1.132 | -1.257 | -0.108 | 0.187 | 0.187 | 0 | 0.72 | 0.97 | 0.82 | 0.98 | 0.68 | 0.82 |
| 5 | 122213846 | rs165976 | 0.621 | -1.71 | -0.99 | -1.074 | 0.047 | -0.047 | -0.047 | 0.705 | 0.28 | 0.18 | 0.31 | 0.49 | 0.15 | 0.4 |
| 5 | 122253304 | rs193541 | 0.017 | -1.727 | -1.014 | -1.201 | -0.01 | -0.004 | -0.004 | 0 | 0.61 | 0.73 | 0.72 | 0.63 | 0.61 | 0.6 |
| 5 | 122260654 | rs3897978 | 1.977 | -1.833 | -0.773 | -1.282 | -0.012 | 0.008 | 0.008 | 1.297 | 0.23 | 0.17 | 0.33 | 0.04 | 0.36 | 0.14 |
| 5 | 122267367 | rs17149748 | 0.886 | -1.892 | -0.869 | -1.308 | 0.007 | -0.079 | -0.079 | 0 | 0.14 | 0.04 | 0.11 | 0.35 | 0.12 | 0.24 |
| 5 | 122314954 | rs4836353 | 1.371 | -1.608 | -0.556 | -1.383 | 0.06 | -0.092 | -0.092 | 0.461 | 0.26 | 0.51 | 0.35 | 0.05 | 0.36 | 0.14 |
| 5 | 122328421 | rs1551937 | -0.026 | -1.763 | -0.646 | -1.479 | 0.032 | 0.041 | 0.041 | 0.116 | 0.58 | 0.66 | 0.55 | 0.06 | 0.69 | 0.32 |
| 5 | 122343870 | rs1060353 | 0.382 | -1.715 | -0.7 | -1.467 | -0.108 | 0.191 | 0.191 | 0 | 0.3 | 0.05 | 0.2 | 0.01 | 0.35 | 0.19 |
| 5 | 122348548 | rs1871180 | 1.288 | -1.538 | -0.673 | -1.477 | -0.016 | 0.016 | 0.016 | 0.881 | 0.27 | 0.18 | 0.34 | 0.15 | 0.38 | 0.18 |
| 5 | 122351493 | rs10080107 | 0.329 | -1.268 | -0.82 | -1.501 | 0.036 | 0.044 | 0.044 | 0 | 0.52 | 0.59 | 0.52 | 0.16 | 0.68 | 0.34 |
| 5 | 122353194 | rs7731806 | 0.875 | -1.379 | -0.754 | -1.468 | 0.006 | -0.035 | -0.035 | 0 | 0.3 | 0.62 | 0.37 | 0.05 | 0.38 | 0.16 |
| 5 | 122354298 | rs3813313 | -0.429 | -1.43 | -0.773 | -1.461 | 0.014 | 0.015 | 0.015 | 0 | 0.63 | 0.67 | 0.59 | 0.17 | 0.77 | 0.38 |
| 5 | 122357713 | rs448442 | 0.222 | -1.384 | -0.715 | -1.498 | 0.015 | -0.003 | -0.003 | 2.677 | 0.55 | 0.6 | 0.56 | 0.06 | 0.71 | 0.33 |
| 5 | 122358015 | rs6891052 | 1.248 | -1.304 | -0.785 | -1.261 | -0.005 | 0.045 | 0.045 | 0 | 0.12 | 0.12 | 0.11 | 0.34 | 0.1 | 0.23 |
| 5 | 122359190 | rs15577 | 0.407 | -1.174 | -0.911 | -1.373 | -0.122 | 0.192 | 0.192 | 0 | 0.29 | 0.04 | 0.19 | 0.01 | 0.34 | 0.18 |
| 5 | 122359640 | rs451195 | 0.417 | -1.179 | -0.929 | -1.376 | -0.127 | 0.196 | 0.196 | 0 | 0.29 | 0.03 | 0.19 | 0.01 | 0.34 | 0.18 |
| 5 | 122367347 | rs10076327 | -1.399 | -0.999 | -0.846 | -1.267 | -0.127 | 0.14 | 0.14 | 0 | 0.78 | 0.99 | 0.85 | 0.99 | 0.71 | 0.85 |
| 5 | 122368076 | rs10455038 | -0.229 | -0.744 | -0.669 | -1.555 | 0.012 | -0.018 | -0.018 | 0 | 0.5 | 0.47 | 0.59 | 0.95 | 0.43 | 0.72 |
| 5 | 122374233 | rs10061195 | 0.744 | -0.672 | -0.57 | -1.074 | 0.002 | 0.206 | 0.206 | 0 | 0.13 | 0.32 | 0.14 | 0.35 | 0.13 | 0.23 |
| 5 | 122381363 | rs10079941 | -0.69 | -0.559 | -0.877 | -1.208 | -0.063 | 0.092 | 0.092 | 0 | 0.73 | 0.92 | 0.82 | 0.93 | 0.67 | 0.8 |
| 5 | 122391626 | rs4349713 | -0.706 | -0.547 | -0.872 | -1.202 | -0.063 | 0.091 | 0.091 | 0 | 0.73 | 0.92 | 0.82 | 0.93 | 0.67 | 0.8 |
| 5 | 122398510 | rs4333309 | 0.582 | -1.184 | -0.544 | -0.758 | 0.047 | -0.063 | -0.063 | 0.601 | 0.26 | 0.16 | 0.32 | 0.49 | 0.14 | 0.39 |
| 5 | 122405090 | rs7700388 | 0.266 | -0.496 | -0.612 | -1.717 | 0.004 | -0.001 | -0.001 | 0 | 0.34 | 0.35 | 0.45 | 0.87 | 0.26 | 0.58 |
| 5 | 122406325 | rs3924912 | 0.069 | 0.31 | -0.561 | -1.789 | -0.002 | -0.044 | -0.044 | 0 | 0.33 | 0.61 | 0.19 | 0.06 | 0.24 | 0.21 |
| 5 | 122409434 | rs11747907 | -0.262 | 1.203 | -0.386 | -0.876 | 0.009 | -0.033 | -0.033 | 0 | 0.25 | 0.29 | 0.26 | 0.08 | 0.32 | 0.16 |
| 5 | 122418806 | rs4288112 | 0.861 | 0.831 | -0.274 | -0.376 | -0.002 | 0.006 | 0.006 | 0 | 0.93 | 0.89 | 0.96 | 0.92 | 0.91 | 0.9 |
| 5 | 122420169 | rs929772 | 1.199 | 1.157 | 0.376 | -0.611 | 0.078 | -0.013 | -0.013 | 0 | 0.91 | 0.7 | 0.9 | 0.96 | 0.86 | 0.95 |
| 5 | 122421661 | rs2302991 | -0.056 | 1.656 | 0.061 | -1.157 | 0 | -0.026 | -0.026 | 0 | 0.24 | 0.25 | 0.22 | 0.03 | 0.34 | 0.13 |
| 5 | 122422693 | rs918441 | 0.346 | 0.248 | -0.965 | -1.125 | 0.017 | -0.054 | -0.054 | 0 | 0.25 | 0.04 | 0.33 | 0.69 | 0.16 | 0.44 |
| 5 | 122423281 | rs888857 | 0.577 | 0.038 | -1.043 | -0.323 | 0.01 | -0.071 | -0.071 | 1.483 | 0.12 | 0.01 | 0.12 | 0.27 | 0.07 | 0.23 |
| 5 | 122424162 | rs6896708 | 0.282 | 0.175 | -0.718 | -0.927 | 0.019 | -0.091 | -0.091 | 0 | 0.3 | 0.36 | 0.26 | 0.03 | 0.39 | 0.14 |
| 5 | 122434666 | rs2546343 | 0.692 | -0.368 | -0.429 | -0.587 | -0.004 | -0.009 | -0.009 | 0.004 | 0.63 | 0.68 | 0.56 | 0.2 | 0.7 | 0.39 |
| 5 | 122435627 | rs1008058 | 0.442 | -0.265 | -0.879 | 0.254 | 0.01 | -0.045 | -0.045 | 2.742 | 0.12 | 0.06 | 0.23 | 0.39 | 0.11 | 0.19 |
| 5 | 122442322 | rs335143 | -0.722 | -0.781 | -0.869 | -0.589 | 0.038 | -0.009 | -0.009 | 0 | 0.28 | 0.19 | 0.21 | 0.37 | 0.18 | 0.4 |
| 5 | 122442627 | rs163189 | -0.094 | -1.352 | -0.919 | -0.848 | 0.019 | -0.048 | -0.048 | 0 | 0.77 | 0.85 | 0.74 | 0.52 | 0.82 | 0.73 |
| 5 | 122446619 | rs151283 | -0.478 | -1.031 | -1.135 | -1.02 | 0.009 | -0.1 | -0.1 | 0.007 | 0.28 | 0.14 | 0.34 | 0.5 | 0.25 | 0.49 |
| 5 | 122447774 | rs632678 | -0.833 | -0.355 | -0.975 | -1.459 | -0.004 | -0.001 | -0.001 | 0 | 0.48 | 0.4 | 0.51 | 0.52 | 0.5 | 0.6 |
| 5 | 122448244 | rs17149910 | 1.606 | -0.413 | -0.841 | -1.547 | 0.012 | -0.153 | -0.153 | 0 | 0.32 | 0.49 | 0.23 | 0.01 | 0.32 | 0.16 |
| 5 | 122448701 | rs17389042 | 0.474 | -0.428 | -1.294 | -1.439 | -0.02 | -0.05 | -0.05 | 0 | 0.12 | 0.01 | 0.19 | 0.14 | 0.11 | 0.17 |
| 5 | 122448927 | rs335183 | 0.247 | -0.594 | -1.135 | -1.425 | -0.019 | 0.008 | 0.008 | 0 | 0.59 | 0.67 | 0.57 | 0.42 | 0.53 | 0.58 |
| 5 | 122449650 | rs570682 | -1.342 | -0.47 | -1.05 | -1.247 | 0 | 0.018 | 0.018 | 0 | 0.84 | 0.85 | 0.75 | 0.53 | 0.83 | 0.78 |
| 5 | 122450771 | rs10077410 | 1.319 | -0.475 | -1.107 | -1.252 | -0.002 | 0.068 | 0.068 | 0 | 0.14 | 0.14 | 0.24 | 0.43 | 0.14 | 0.21 |
| 5 | 122450824 | rs335182 | 1.082 | -0.403 | -1.289 | -1.298 | -0.068 | 0.094 | 0.094 | 0 | 0.17 | 0.04 | 0.14 | 0.01 | 0.24 | 0.09 |
| 5 | 122453652 | rs466712 | -1.249 | -0.268 | -0.793 | -1.178 | 0 | -0.028 | -0.028 | 2.523 | 0.91 | 0.88 | 0.96 | 0.99 | 0.92 | 0.97 |
| 5 | 122453933 | rs335180 | 0.318 | -0.472 | -0.641 | -1.232 | 0.001 | -0.002 | -0.002 | 0 | 0.7 | 0.71 | 0.63 | 0.63 | 0.74 | 0.59 |
| 5 | 122454305 | rs186749 | 1.331 | -0.673 | -0.226 | -1.336 | 0.007 | -0.004 | -0.004 | 0 | 0.58 | 0.37 | 0.58 | 0.53 | 0.67 | 0.47 |
| 5 | 122454550 | rs398028 | 0.609 | -0.435 | -0.49 | -1.377 | 0.001 | -0.079 | -0.079 | 0 | 0.18 | 0.05 | 0.31 | 0.35 | 0.17 | 0.37 |
| 5 | 122455147 | rs335162 | 0.173 | -0.331 | -0.44 | -1.36 | 0.016 | -0.098 | -0.098 | 0 | 0.23 | 0.06 | 0.32 | 0.43 | 0.19 | 0.42 |
| 5 | 122457047 | rs431518 | 0.175 | -0.147 | -0.152 | -1.603 | -0.011 | -0.066 | -0.066 | 0.048 | 0.86 | 0.73 | 0.94 | 0.99 | 0.91 | 0.95 |
| 5 | 122459869 | rs577774 | -0.214 | -0.039 | -0.164 | -1.55 | -0.005 | -0.06 | -0.06 | 3.52 | 0.88 | 0.76 | 0.95 | 0.99 | 0.91 | 0.95 |
| 5 | 122461778 | rs465305 | -0.214 | -0.039 | -0.175 | -1.55 | -0.006 | -0.059 | -0.059 | 0 | 0.88 | 0.76 | 0.95 | 0.99 | 0.92 | 0.97 |
| 5 | 122471860 | rs375185 | 1.044 | 0.099 | -0.418 | -1.751 | -0.011 | 0.081 | 0.081 | 0 | 0.37 | 0.25 | 0.23 | 0.02 | 0.35 | 0.2 |
| 5 | 122472792 | rs393761 | -0.427 | -0.004 | -0.119 | -1.7 | -0.006 | 0.014 | 0.014 | 0.164 | 0.76 | 0.73 | 0.67 | 0.62 | 0.78 | 0.57 |

| | | | | | | | | | | | | | | | | |
|---|---|---|---|---|---|---|---|---|---|---|---|---|---|---|---|---|
| 5 | 122473679 | rs419201 | 1.171 | -0.009 | -0.033 | -1.693 | -0.002 | 0.021 | 0.021 | 0 | 0.39 | 0.34 | 0.24 | 0.02 | 0.36 | 0.2 |
| 5 | 122474938 | rs4836465 | 0.861 | -0.104 | -0.068 | -1.391 | 0 | 0.007 | 0.007 | 0 | 0.18 | 0.14 | 0.27 | 0.59 | 0.17 | 0.29 |
| 5 | 122483513 | rs335169 | -0.033 | 0.042 | 0.143 | -1.056 | -0.012 | 0.029 | 0.029 | 0.117 | 0.75 | 0.67 | 0.66 | 0.62 | 0.78 | 0.58 |
| 5 | 122496931 | rs459743 | -0.182 | 0.333 | 0.631 | -1.47 | 0.009 | -0.108 | -0.108 | 0.048 | 0.18 | 0.36 | 0.09 | 0.02 | 0.18 | 0.11 |
| 5 | 122501476 | rs4836483 | 0.463 | 0.321 | 0.297 | -1.637 | -0.004 | -0.024 | -0.024 | 2.151 | 0.32 | 0.2 | 0.37 | 0.58 | 0.32 | 0.34 |
| 5 | 122503511 | rs10077908 | 0.345 | 0.31 | 0.535 | -1.642 | -0.001 | 0.003 | 0.003 | 0 | 0.33 | 0.34 | 0.38 | 0.58 | 0.32 | 0.34 |
| 5 | 122504566 | rs335206 | -0.714 | 0.238 | 0.547 | -1.758 | -0.004 | 0.029 | 0.029 | 0 | 0.53 | 0.64 | 0.48 | 0.6 | 0.51 | 0.46 |
| 5 | 122506237 | rs2112487 | 1.957 | 0.259 | 0.242 | -1.954 | -0.028 | 0.083 | 0.083 | 0 | 0.26 | 0.11 | 0.2 | 0.02 | 0.29 | 0.14 |
| 5 | 122514120 | rs335178 | -0.723 | 0.096 | -0.033 | -1.72 | 0.014 | -0.048 | -0.048 | 0.014 | 0.82 | 0.92 | 0.7 | 0.66 | 0.85 | 0.64 |
| 5 | 122514588 | rs163186 | 0.276 | -0.008 | -0.023 | -1.971 | 0.006 | 0.009 | 0.009 | 0.005 | 0.69 | 0.73 | 0.79 | 0.77 | 0.73 | 0.69 |
| 5 | 122524977 | rs10519720 | -0.602 | 0.024 | 0.219 | -2.094 | -0.033 | -0.008 | -0.008 | 0 | 0.1 | 0.27 | 0.03 | 0.06 | 0.04 | 0.05 |
| 5 | 122526666 | rs629818 | -0.158 | 0.286 | 0.695 | -2.174 | 0.051 | -0.01 | -0.01 | 0 | 0.09 | 0.27 | 0.07 | 0.05 | 0.12 | 0.06 |
| 5 | 122526819 | rs335168 | -0.825 | 0.371 | 0.054 | -2.786 | -0.062 | 0.227 | 0.227 | 0.028 | 0.59 | 0.89 | 0.34 | 0.13 | 0.55 | 0.36 |
| 5 | 122528994 | rs337113 | -0.27 | 0.345 | 0.331 | -2.806 | 0.004 | 0.058 | 0.058 | 0.579 | 0.53 | 0.62 | 0.32 | 0.07 | 0.52 | 0.32 |
| 5 | 122532390 | rs17470253 | 2.407 | 0.457 | -0.097 | -2.898 | -0.022 | 0.03 | 0.03 | 0 | 0.06 | 0 | 0.03 | 0 | 0.08 | 0.03 |
| 5 | 122532869 | rs1644317 | 0.87 | 0.38 | 0.333 | -2.823 | 0 | 0.067 | 0.067 | 1.292 | 0.32 | 0.15 | 0.21 | 0.01 | 0.34 | 0.18 |
| 5 | 122546510 | rs337100 | -0.44 | 0.423 | 0.442 | -2.798 | 0.008 | 0.043 | 0.043 | 0 | 0.49 | 0.59 | 0.28 | 0.08 | 0.44 | 0.31 |
| 5 | 122548721 | rs7702622 | -0.349 | 0.71 | 0.536 | -2.818 | 0.015 | -0.028 | -0.028 | 0 | 0.19 | 0.28 | 0.13 | 0.07 | 0.23 | 0.12 |
| 5 | 122551276 | rs10519721 | 0.068 | 0.697 | 0.552 | -2.798 | 0.121 | -0.02 | -0.02 | 0.155 | 0.12 | 0.41 | 0.13 | 0.04 | 0.17 | 0.06 |
| 5 | 122552158 | rs417825 | -0.963 | 0.189 | 0.122 | -2.583 | 0.012 | 0.136 | 0.136 | 0 | 0.6 | 0.81 | 0.38 | 0.13 | 0.61 | 0.37 |
| 5 | 122552265 | rs448166 | -0.231 | 0.246 | 0.148 | -2.558 | 0.022 | 0.046 | 0.046 | 0.183 | 0.52 | 0.62 | 0.36 | 0.07 | 0.58 | 0.31 |
| 5 | 122554856 | rs6895930 | -0.423 | 0.768 | -0.459 | -1.574 | -0.013 | 0.012 | 0.012 | 0 | 0.06 | 0.16 | 0.02 | 0.05 | 0.03 | 0.04 |
| 5 | 122555432 | rs1565059 | 0.646 | 1.123 | -0.873 | -1.537 | -0.018 | 0.028 | 0.028 | 0 | 0.08 | 0.04 | 0.02 | 0.03 | 0.06 | 0.06 |
| 5 | 122559120 | rs2450508 | -0.744 | 0.803 | -1.054 | -1.635 | 0.021 | 0.065 | 0.065 | 0 | 0.6 | 0.78 | 0.56 | 0.22 | 0.67 | 0.51 |
| 5 | 122559169 | rs6874105 | -0.388 | 1.088 | -1.216 | -2.031 | -0.001 | 0.011 | 0.011 | 1.328 | 0.35 | 0.69 | 0.35 | 0.2 | 0.32 | 0.3 |
| 5 | 122560170 | rs17405383 | 1.674 | 1.196 | -1.204 | -1.963 | 0 | -0.012 | -0.012 | 0 | 0.08 | 0.06 | 0.05 | 0 | 0.1 | 0.03 |
| 5 | 122562800 | rs338796 | 0.714 | 1.266 | -0.868 | -1.972 | -0.009 | 0.001 | 0.001 | 0 | 0.18 | 0.13 | 0.17 | 0.02 | 0.26 | 0.17 |
| 5 | 122569039 | rs337096 | -0.386 | 1.089 | -0.641 | -2.07 | 0.001 | -0.05 | -0.05 | 0 | 0.3 | 0.32 | 0.22 | 0.03 | 0.36 | 0.27 |
| 5 | 122571493 | rs255664 | 0.438 | 1.005 | -0.638 | -1.977 | 0.012 | 0.005 | 0.005 | 0 | 0.17 | 0.19 | 0.28 | 0.16 | 0.25 | 0.16 |
| 5 | 122572116 | rs17150263 | 0.724 | 0.892 | -0.357 | -2.136 | 0.006 | -0.077 | -0.077 | 1.021 | 0.14 | 0.27 | 0.04 | 0.02 | 0.08 | 0.1 |
| 5 | 122578507 | rs378377 | 0.021 | 0.703 | -0.508 | -1.974 | 0.02 | -0.137 | -0.137 | 0.766 | 0.34 | 0.53 | 0.21 | 0.05 | 0.34 | 0.27 |
| 5 | 122580617 | rs453473 | -0.007 | 0.644 | -0.522 | -1.757 | 0.002 | -0.079 | -0.079 | 0 | 0.41 | 0.59 | 0.26 | 0.05 | 0.43 | 0.31 |
| 5 | 122596593 | rs380164 | -1.02 | 0.782 | -0.71 | -1.527 | 0.031 | 0.047 | 0.047 | 0 | 0.61 | 0.73 | 0.53 | 0.22 | 0.69 | 0.47 |
| 5 | 122598142 | rs35824 | 0.183 | 1.017 | -0.489 | -0.572 | -0.011 | 0.004 | 0.004 | 0 | 0.1 | 0.03 | 0.14 | 0.07 | 0.13 | 0.13 |
| 5 | 122603725 | rs35830 | -0.933 | 0.914 | -0.366 | -0.186 | 0.022 | 0.067 | 0.067 | 0 | 0.24 | 0.49 | 0.31 | 0.29 | 0.31 | 0.29 |
| 5 | 122607023 | rs255628 | -0.173 | 0.843 | -1.045 | -0.526 | -0.019 | 0.04 | 0.04 | 0.25 | 0.72 | 0.77 | 0.73 | 0.86 | 0.67 | 0.76 |
| 5 | 122607156 | rs35832 | 0.761 | 0.459 | -1.231 | -0.384 | 0.004 | 0 | 0 | 0 | 0.15 | 0.12 | 0.08 | 0.14 | 0.13 | 0.19 |
| 5 | 122614010 | rs28045 | 0.769 | 0.278 | -1.006 | -0.136 | 0.004 | 0.018 | 0.018 | 0 | 0.31 | 0.35 | 0.43 | 0.48 | 0.33 | 0.3 |
| 5 | 122617574 | rs7714132 | 0.319 | 0.208 | -0.927 | -0.344 | 0.003 | -0.034 | -0.034 | 4.025 | 0.33 | 0.42 | 0.3 | 0.13 | 0.35 | 0.24 |
| 5 | 122622141 | rs255670 | -1.226 | 0.264 | -1.33 | -0.123 | -0.016 | -0.051 | -0.051 | 1.442 | 0.77 | 0.87 | 0.62 | 0.53 | 0.76 | 0.76 |
| 5 | 122626463 | rs10455039 | 1.748 | 0.196 | -1.666 | -0.435 | -0.077 | 0.1 | 0.1 | 0.029 | 0.18 | 0.04 | 0.15 | 0 | 0.26 | 0.09 |
| 5 | 122627059 | rs890930 | 0.169 | -0.287 | -1.207 | -0.314 | -0.007 | 0.027 | 0.027 | 0 | 0.25 | 0.39 | 0.22 | 0.23 | 0.22 | 0.27 |
| 5 | 122627814 | rs13188307 | 0.373 | -0.26 | -1.501 | -0.276 | 0.004 | -0.011 | -0.011 | 0 | 0.14 | 0.06 | 0.17 | 0.15 | 0.16 | 0.2 |
| 5 | 122629266 | rs1366337 | 0.053 | -0.637 | -1.431 | -0.202 | 0.018 | 0.015 | 0.015 | 0 | 0.31 | 0.71 | 0.13 | 0.18 | 0.17 | 0.31 |
| 5 | 122630230 | rs10055144 | -0.434 | -0.023 | -1.295 | 0.193 | -0.053 | -0.018 | -0.018 | 0 | 0.67 | 0.86 | 0.45 | 0.34 | 0.61 | 0.6 |
| 5 | 122632423 | rs2897737 | -0.484 | 0.058 | -0.842 | 0.295 | 0.009 | 0.044 | 0.044 | 0 | 0.18 | 0.29 | 0.2 | 0.17 | 0.17 | 0.17 |
| 5 | 122640214 | rs890933 | -0.956 | 0.065 | -1.264 | 0.278 | -0.01 | -0.048 | -0.048 | 1.505 | 0.77 | 0.84 | 0.62 | 0.5 | 0.76 | 0.75 |
| 5 | 122642381 | rs4836527 | 0.275 | -0.253 | -0.994 | -0.247 | -0.004 | 0.017 | 0.017 | 1.311 | 0.43 | 0.55 | 0.24 | 0.17 | 0.42 | 0.28 |
| 5 | 122644698 | rs675443 | -0.37 | -0.413 | -1.452 | 0.35 | 0.011 | -0.094 | -0.094 | 0 | 0.25 | 0.09 | 0.36 | 0.38 | 0.26 | 0.37 |
| 5 | 122646088 | rs6886384 | 0.836 | -0.273 | -1.085 | -0.129 | 0.005 | 0.005 | 0.005 | 0 | 0.18 | 0.15 | 0.06 | 0.16 | 0.14 | 0.17 |
| 5 | 122659109 | rs1428393 | -0.499 | -0.2 | -0.836 | 0.214 | -0.004 | -0.001 | -0.001 | 0 | 0.48 | 0.58 | 0.51 | 0.45 | 0.51 | 0.53 |
| 5 | 122676525 | rs7708474 | -0.358 | 0.093 | -0.236 | 0.166 | 0 | 0.067 | 0.067 | 0 | 0.22 | 0.3 | 0.27 | 0.36 | 0.22 | 0.3 |
| 5 | 122680189 | rs17472832 | 0.534 | 0.141 | -0.599 | -0.029 | -0.012 | 0.147 | 0.147 | 12.282 | 0.46 | 0.22 | 0.32 | 0.15 | 0.51 | 0.25 |
| 5 | 122680667 | rs1047484 | 1.584 | 0.182 | -0.592 | -0.125 | -0.05 | 0.069 | 0.069 | 5.787 | 0.19 | 0.07 | 0.15 | 0 | 0.26 | 0.09 |
| 5 | 122681834 | rs1047440 | -0.06 | 0.108 | -0.479 | -0.033 | 0.001 | -0.006 | -0.006 | 0.008 | 0.43 | 0.31 | 0.42 | 0.5 | 0.45 | 0.45 |
| 5 | 122682154 | rs2303719 | -1.235 | 0.119 | -0.394 | -0.012 | -0.005 | 0.005 | 0.005 | 1.193 | 0.7 | 0.61 | 0.67 | 0.59 | 0.74 | 0.67 |
| 5 | 122682348 | rs1047438 | -1.199 | 0.107 | -0.222 | -0.035 | 0.003 | -0.008 | -0.008 | 0.173 | 0.71 | 0.72 | 0.67 | 0.59 | 0.74 | 0.67 |
| 5 | 122697009 | rs17473342 | 1.45 | 0.161 | -0.219 | -0.08 | -0.005 | -0.023 | -0.023 | 0 | 0.2 | 0.18 | 0.16 | 0 | 0.26 | 0.09 |
| 5 | 122714843 | rs11951466 | 0.645 | 0.039 | 0.267 | 0.342 | -0.014 | -0.005 | -0.005 | 0.014 | 0.07 | 0.11 | 0.04 | 0.09 | 0.05 | 0.06 |
| 5 | 122714972 | rs3990341 | -0.493 | 0.129 | 0.129 | 0.408 | 0.005 | -0.013 | -0.013 | 0 | 0.66 | 0.65 | 0.59 | 0.51 | 0.71 | 0.53 |
| 5 | 122744178 | rs13154356 | -0.128 | 0.262 | 0.429 | 0.842 | 0.003 | 0.037 | 0.037 | 0 | 0.13 | 0.11 | 0.22 | 0.26 | 0.16 | 0.22 |
| 5 | 122759862 | rs11241696 | -1.104 | 0.156 | 0.466 | 0.776 | -0.005 | 0.071 | 0.071 | 0 | 0.2 | 0.26 | 0.26 | 0.35 | 0.21 | 0.28 |
| 5 | 122769296 | rs13165542 | 0.038 | 0.332 | 0.316 | 0.65 | 0.003 | 0.033 | 0.033 | 0 | 0.11 | 0.09 | 0.23 | 0.24 | 0.16 | 0.22 |
| 5 | 122776642 | rs13155399 | -1.718 | 0.325 | 0.683 | 0.573 | 0.007 | 0.089 | 0.089 | 0.069 | 0.25 | 0.4 | 0.35 | 0.39 | 0.28 | 0.41 |
| 5 | 122778705 | rs7735409 | -1.786 | 0.321 | 0.681 | 0.571 | 0.008 | 0.092 | 0.092 | 0.048 | 0.25 | 0.4 | 0.35 | 0.39 | 0.28 | 0.41 |
| 5 | 122788602 | rs6876223 | -0.087 | 0.34 | 0.664 | 0.574 | 0.009 | 0.015 | 0.015 | 0 | 0.06 | 0.1 | 0.07 | 0.08 | 0.06 | 0.13 |
| 5 | 122789901 | rs12109037 | 1.224 | -0.032 | 0.246 | 0.164 | 0.023 | 0.057 | 0.057 | 0.973 | 0.28 | 0.1 | 0.16 | 0.12 | 0.21 | 0.13 |
| 5 | 122791137 | rs4572998 | -0.815 | 0.002 | 0.378 | 0.169 | -0.001 | 0.012 | 0.012 | 0 | 0.53 | 0.6 | 0.52 | 0.52 | 0.49 | 0.55 |
| 5 | 122798261 | rs7703080 | -1.148 | 0.286 | 0.181 | 0.204 | -0.004 | 0.003 | 0.003 | 0.319 | 0.14 | 0.19 | 0.11 | 0.16 | 0.12 | 0.2 |

| | | | | | | | | | | | | | | | | |
|---|---|---|---|---|---|---|---|---|---|---|---|---|---|---|---|---|
| 5 | 122814733 | rs13171367 | -0.17 | 0.709 | 0.388 | 0.387 | 0.007 | 0.032 | 0.032 | 0 | 0.13 | 0.13 | 0.18 | 0.23 | 0.18 | 0.24 |
| 5 | 122836911 | rs10477646 | 0.003 | 1.006 | 0.6 | -0.059 | 0.032 | -0.022 | -0.022 | 0 | 0.27 | 0.35 | 0.31 | 0.13 | 0.35 | 0.23 |
| 5 | 122837204 | rs13170938 | 1.701 | 1.091 | 0.631 | -0.018 | 0.007 | 0.033 | 0.033 | 0 | 0.05 | 0.07 | 0.09 | 0.15 | 0.07 | 0.17 |
| 5 | 122847622 | rs11745493 | 0.732 | 1.058 | 0.73 | -0.074 | 0.023 | -0.033 | -0.033 | 0 | 0.21 | 0.28 | 0.17 | 0.05 | 0.27 | 0.1 |
| 5 | 122860462 | rs7731839 | -0.326 | 0.735 | 0.514 | -0.223 | -0.028 | -0.016 | -0.016 | 0 | 0.61 | 0.81 | 0.5 | 0.34 | 0.58 | 0.42 |
| 5 | 122863961 | rs17164879 | 1.321 | 0.597 | 0.798 | -0.28 | 0.002 | 0.002 | 0.002 | 0.147 | 0.33 | 0.27 | 0.17 | 0.19 | 0.22 | 0.19 |
| 5 | 122864567 | rs6595456 | 0.297 | 0.609 | 0.85 | -0.27 | -0.005 | 0.031 | 0.031 | 2.524 | 0.54 | 0.61 | 0.34 | 0.25 | 0.49 | 0.29 |
| 5 | 122884414 | rs6887967 | 1.058 | 0.594 | 0.925 | -0.225 | 0.018 | -0.021 | -0.021 | 0.116 | 0.2 | 0.22 | 0.17 | 0.06 | 0.27 | 0.1 |
| 5 | 122890729 | rs1972627 | 0.674 | 0.716 | 0.911 | -0.192 | 0.009 | 0.015 | 0.015 | 0 | 0.43 | 0.33 | 0.32 | 0.29 | 0.35 | 0.33 |
| 5 | 122893560 | rs10519732 | 0.172 | 0.734 | 0.899 | -0.169 | 0.006 | 0.011 | 0.011 | 2.431 | 0.06 | 0.09 | 0.14 | 0.1 | 0.09 | 0.13 |
| 5 | 122899301 | rs6884946 | 1.315 | 0.706 | 0.89 | -0.2 | 0.003 | 0.002 | 0.002 | 0.863 | 0.37 | 0.34 | 0.18 | 0.2 | 0.26 | 0.19 |
| 5 | 122905683 | rs7728845 | 0.912 | 0.749 | 0.815 | -0.21 | 0.033 | -0.028 | -0.028 | 0.297 | 0.17 | 0.29 | 0.16 | 0.05 | 0.23 | 0.1 |
| 5 | 122905812 | rs10052983 | 0.968 | 0.68 | 0.673 | -0.191 | -0.003 | -0.11 | -0.11 | 2.228 | 0.28 | 0.08 | 0.34 | 0.49 | 0.25 | 0.33 |
| 5 | 122921103 | rs1579036 | 0.668 | 0.682 | 0.794 | -0.257 | -0.006 | -0.01 | -0.01 | 0 | 0.43 | 0.47 | 0.33 | 0.29 | 0.35 | 0.33 |
| 5 | 122943249 | rs2407856 | -0.256 | 0.9 | 0.791 | -0.183 | 0.052 | -0.048 | -0.048 | 0 | 0.8 | 0.58 | 0.8 | 0.94 | 0.73 | 0.88 |
| 5 | 122950589 | rs17150513 | 1.995 | 0.637 | 0.237 | -0.324 | 0.007 | -0.04 | -0.04 | 0 | 0.1 | 0 | 0.06 | 0.13 | 0.06 | 0.08 |
| 5 | 122950911 | rs1059259 | -0.541 | 0.854 | 0.473 | -0.332 | 0.038 | -0.032 | -0.032 | 0 | 0.82 | 0.69 | 0.83 | 0.95 | 0.74 | 0.88 |
| 5 | 122957239 | rs12659062 | 0.386 | 0.851 | 0.39 | -0.294 | -0.004 | -0.006 | -0.006 | 0.608 | 0.07 | 0.05 | 0.15 | 0.09 | 0.09 | 0.13 |
| 5 | 122959130 | rs10463762 | 1.11 | 0.81 | 0.471 | -0.252 | -0.001 | -0.001 | -0.001 | 0 | 0.09 | 0.07 | 0.15 | 0.15 | 0.13 | 0.22 |
| 5 | 122976045 | rs7720106 | 0.515 | 0.708 | 0.62 | -0.465 | 0.022 | -0.023 | -0.023 | 0 | 0.17 | 0.28 | 0.15 | 0.05 | 0.23 | 0.09 |
| 5 | 122993059 | rs1973174 | -0.096 | 0.743 | 0.728 | -0.473 | 0.009 | -0.005 | -0.005 | 0 | 0.21 | 0.25 | 0.28 | 0.13 | 0.26 | 0.2 |
| 5 | 122994255 | rs1834887 | 0.392 | 0.862 | 0.838 | -0.48 | -0.019 | -0.018 | -0.018 | 0.23 | 0.38 | 0.19 | 0.49 | 0.67 | 0.39 | 0.58 |
| 5 | 122997026 | rs10478586 | 0.921 | 0.939 | 1.025 | -0.202 | 0.024 | -0.016 | -0.016 | 0 | 0.14 | 0.22 | 0.14 | 0.05 | 0.2 | 0.09 |
| 5 | 122998955 | rs958883 | 0.116 | 0.848 | 1.174 | -0.276 | 0.006 | -0.005 | -0.005 | 0 | 0.73 | 0.67 | 0.78 | 0.82 | 0.69 | 0.79 |
| 5 | 123002130 | rs1862265 | -1.262 | 0.729 | 0.946 | -0.212 | -0.002 | 0.001 | 0.001 | 0 | 0.87 | 0.86 | 0.92 | 0.87 | 0.9 | 0.88 |
| 5 | 123003913 | rs13175487 | -1.211 | 0.73 | 0.946 | -0.212 | -0.002 | 0.001 | 0.001 | 0 | 0.87 | 0.86 | 0.92 | 0.87 | 0.9 | 0.88 |
| 5 | 123008100 | rs12658122 | 0.354 | 0.66 | 0.609 | -0.358 | -0.001 | 0.01 | 0.01 | 0 | 0.22 | 0.22 | 0.36 | 0.63 | 0.2 | 0.47 |
| 5 | 123009413 | rs4569864 | 1.478 | 0.735 | 0.565 | -0.623 | -0.008 | 0.085 | 0.085 | 0.163 | 0.31 | 0.13 | 0.22 | 0.06 | 0.36 | 0.12 |
| 5 | 123018374 | rs6890830 | 1.458 | 0.751 | 0.875 | -0.517 | -0.003 | 0.001 | 0.001 | 0 | 0.13 | 0.13 | 0.13 | 0.08 | 0.19 | 0.09 |
| 5 | 123024195 | rs2194046 | -0.731 | 0.665 | 1.051 | -0.546 | 0 | 0.014 | 0.014 | 0 | 0.4 | 0.68 | 0.51 | 0.72 | 0.39 | 0.57 |
| 5 | 123026041 | rs1820031 | 0.013 | 0.568 | 1.135 | -0.428 | -0.012 | 0.037 | 0.037 | 0 | 0.25 | 0.32 | 0.36 | 0.64 | 0.2 | 0.48 |
| 5 | 123026867 | rs12521632 | 1.696 | 0.594 | 1.048 | -0.439 | 0.006 | -0.005 | -0.005 | 0 | 0.11 | 0.09 | 0.07 | 0.13 | 0.08 | 0.08 |
| 5 | 123030802 | rs6889596 | -0.419 | 0.575 | 1.125 | -0.615 | -0.017 | -0.017 | -0.017 | 0 | 0.36 | 0.43 | 0.44 | 0.76 | 0.28 | 0.56 |
| 5 | 123042675 | rs4835808 | 1.682 | 0.612 | 0.82 | -0.543 | 0.006 | -0.001 | -0.001 | 0 | 0.11 | 0.09 | 0.07 | 0.11 | 0.08 | 0.08 |
| 5 | 123044859 | rs1422262 | 0.385 | 0.593 | 0.922 | -0.561 | -0.001 | 0.005 | 0.005 | 0 | 0.22 | 0.22 | 0.36 | 0.65 | 0.2 | 0.48 |
| 5 | 123046778 | rs4836033 | 1.628 | 0.596 | 0.638 | -0.567 | 0.002 | -0.001 | -0.001 | 0 | 0.11 | 0.1 | 0.07 | 0.11 | 0.08 | 0.07 |
| 5 | 123050342 | rs12515947 | -0.707 | 0.748 | 0.643 | -0.607 | 0.006 | 0.05 | 0.05 | 0 | 0.54 | 0.62 | 0.68 | 0.83 | 0.53 | 0.76 |
| 5 | 123054717 | rs17150658 | 0.868 | 0.562 | 0.733 | -0.425 | -0.001 | -0.033 | -0.033 | 0.056 | 0.19 | 0.34 | 0.2 | 0.1 | 0.14 | 0.19 |
| 5 | 123055659 | rs187683 | -1.132 | 0.519 | 1.01 | -0.301 | 0.007 | -0.013 | -0.013 | 1.178 | 0.81 | 0.75 | 0.79 | 0.89 | 0.82 | 0.77 |
| 5 | 123057964 | rs331580 | -1.49 | 0.508 | 0.976 | -0.299 | -0.017 | 0.008 | 0.008 | 0.655 | 0.83 | 0.92 | 0.8 | 0.89 | 0.83 | 0.77 |
| 5 | 123058338 | rs10478591 | 0.32 | 0.628 | 0.859 | -0.351 | -0.021 | -0.071 | -0.071 | 1.065 | 0.23 | 0.11 | 0.41 | 0.65 | 0.29 | 0.48 |
| 5 | 123072033 | rs51777 | 0.604 | 0.57 | 1.081 | -0.492 | -0.001 | -0.001 | -0.001 | 0 | 0.56 | 0.42 | 0.36 | 0.22 | 0.54 | 0.28 |
| 5 | 123082557 | rs17150713 | 1.291 | 0.608 | 0.837 | -0.28 | 0.013 | -0.005 | -0.005 | 0 | 0.12 | 0.08 | 0.21 | 0.57 | 0.08 | 0.25 |
| 5 | 123083753 | rs9327305 | 1.112 | 0.6 | 0.912 | -0.284 | -0.021 | 0.123 | 0.123 | 0 | 0.13 | 0.21 | 0.22 | 0.57 | 0.08 | 0.25 |
| 5 | 123085026 | rs1017699 | 0.671 | 0.611 | 0.934 | -0.49 | 0 | -0.005 | -0.005 | 0.046 | 0.25 | 0.27 | 0.2 | 0.16 | 0.26 | 0.17 |
| 5 | 123086916 | rs29559 | 1.565 | 0.602 | 0.903 | -0.505 | 0.012 | 0.045 | 0.045 | 1.109 | 0.32 | 0.18 | 0.17 | 0.05 | 0.27 | 0.11 |
| 5 | 123090139 | rs3901873 | 1.032 | 0.695 | 0.561 | -0.456 | -0.021 | 0.004 | 0.004 | 0 | 0.11 | 0.06 | 0.2 | 0.11 | 0.21 | 0.24 |
| 5 | 123092028 | rs11746044 | 0.437 | 0.55 | 0.643 | -0.448 | -0.001 | -0.012 | -0.012 | 0 | 0.26 | 0.33 | 0.2 | 0.16 | 0.27 | 0.17 |
| 5 | 123115485 | rs28132 | 0.221 | 0.594 | 0.528 | -0.4 | -0.016 | 0.01 | 0.01 | 0.334 | 0.29 | 0.27 | 0.41 | 0.2 | 0.38 | 0.46 |
| 5 | 123117953 | rs3909335 | 1.291 | 0.423 | 0.473 | -0.336 | -0.027 | -0.012 | -0.012 | 0 | 0.13 | 0.25 | 0.08 | 0.1 | 0.07 | 0.08 |
| 5 | 123119565 | rs11241713 | 1.404 | 0.403 | 0.155 | -0.221 | 0.012 | 0 | 0 | 0 | 0.12 | 0.08 | 0.21 | 0.56 | 0.08 | 0.25 |
| 5 | 123120882 | rs12518486 | 1.341 | 0.397 | 0.226 | -0.279 | -0.011 | -0.005 | -0.005 | 0 | 0.13 | 0.19 | 0.08 | 0.1 | 0.07 | 0.08 |
| 5 | 123121022 | rs25771 | -1.488 | 0.499 | 0.305 | -0.254 | 0 | -0.025 | -0.025 | 0 | 0.68 | 0.73 | 0.83 | 0.95 | 0.73 | 0.89 |
| 5 | 123124233 | rs29614 | -0.679 | 0.548 | 0.072 | -0.171 | -0.003 | -0.001 | -0.001 | 0 | 0.55 | 0.56 | 0.61 | 0.39 | 0.66 | 0.64 |
| 5 | 123129656 | rs10043442 | -0.23 | 0.681 | 0.054 | -0.226 | 0.005 | -0.008 | -0.008 | 0.022 | 0.37 | 0.45 | 0.41 | 0.28 | 0.48 | 0.41 |
| 5 | 123131730 | rs255277 | 1.406 | 0.8 | -0.221 | -0.312 | -0.017 | 0.006 | 0.006 | 0.523 | 0.17 | 0.1 | 0.2 | 0.12 | 0.18 | 0.23 |
| 5 | 123131879 | rs11746266 | -0.765 | 0.908 | -0.089 | -0.011 | 0.001 | 0.023 | 0.023 | 0 | 0.5 | 0.58 | 0.62 | 0.83 | 0.55 | 0.66 |
| 5 | 123132945 | rs4835811 | -0.107 | 0.992 | -0.702 | -0.03 | -0.005 | 0.009 | 0.009 | 0 | 0.23 | 0.17 | 0.27 | 0.19 | 0.28 | 0.31 |
| 5 | 123136377 | rs257140 | 0.811 | 0.972 | -0.927 | 0.196 | -0.028 | 0.003 | 0.003 | 0 | 0.21 | 0.14 | 0.19 | 0.22 | 0.2 | 0.23 |
| 5 | 123136656 | rs6891344 | 0.906 | 1.038 | -0.369 | 0.065 | 0.007 | -0.017 | -0.017 | 0.008 | 0.16 | 0.3 | 0.12 | 0.11 | 0.19 | 0.12 |
| 5 | 123138998 | rs257139 | -0.112 | 0.972 | -0.385 | 0.043 | -0.001 | -0.001 | -0.001 | 0.943 | 0.38 | 0.43 | 0.32 | 0.34 | 0.39 | 0.35 |
| 5 | 123140147 | rs27735 | -1.675 | 0.863 | -0.926 | -0.528 | 0.007 | 0.119 | 0.119 | 0.048 | 0.75 | 0.94 | 0.87 | 0.98 | 0.79 | 0.9 |
| 5 | 123144113 | rs6895877 | 0.901 | 0.652 | -0.717 | 0.014 | -0.026 | -0.016 | -0.016 | 0.296 | 0.23 | 0.15 | 0.33 | 0.25 | 0.33 | 0.37 |
| 5 | 123147064 | rs6882579 | 1.302 | 0.511 | -0.348 | 0.016 | -0.021 | 0.096 | 0.096 | 0.002 | 0.11 | 0.12 | 0.19 | 0.4 | 0.05 | 0.17 |
| 5 | 123147945 | rs29644 | -1.328 | 0.714 | -0.03 | -0.274 | 0.011 | 0.009 | 0.009 | 0 | 0.72 | 0.8 | 0.85 | 0.98 | 0.76 | 0.89 |
| 5 | 123148436 | rs29645 | -0.754 | 0.738 | 0.105 | -0.106 | -0.004 | -0.032 | -0.032 | 0 | 0.57 | 0.48 | 0.73 | 0.87 | 0.61 | 0.78 |
| 5 | 123149884 | rs25767 | 0.773 | 0.691 | -0.26 | -0.04 | -0.001 | -0.034 | -0.034 | 0 | 0.31 | 0.15 | 0.38 | 0.61 | 0.25 | 0.4 |
| 5 | 123152620 | rs13162753 | 0.38 | 0.992 | -0.328 | -0.137 | 0.057 | 0.077 | 0.077 | 0 | 0.14 | 0.31 | 0.3 | 0.27 | 0.26 | 0.32 |
| 5 | 123152672 | rs153884 | -0.737 | 0.343 | -0.624 | -0.426 | -0.007 | 0.185 | 0.185 | 0.846 | 0.64 | 0.9 | 0.77 | 0.98 | 0.62 | 0.83 |

| | | | | | | | | | | | | | | | | |
|---|---|---|---|---|---|---|---|---|---|---|---|---|---|---|---|---|
| 5 | 123168242 | rs3909329 | -0.737 | 0.239 | -0.276 | -0.297 | -0.004 | 0.047 | 0.047 | 1.88 | 0.55 | 0.63 | 0.33 | 0.22 | 0.49 | 0.34 |
| 5 | 123169574 | rs7708937 | 0.944 | -0.009 | -0.21 | 0.318 | -0.027 | 0.166 | 0.166 | 0.053 | 0.07 | 0.18 | 0.18 | 0.33 | 0.02 | 0.12 |
| 5 | 123170153 | rs153880 | -1.338 | 0.022 | -0.365 | 0.229 | 0.05 | 0.016 | 0.016 | 0.85 | 0.35 | 0.83 | 0.33 | 0.63 | 0.23 | 0.34 |
| 5 | 123170490 | rs10477650 | 1.828 | 0.043 | -0.239 | 0.193 | 0.006 | 0.006 | 0.006 | 0 | 0.12 | 0.08 | 0.06 | 0.09 | 0.09 | 0.1 |
| 5 | 123171342 | rs3909326 | 1.288 | 0.059 | -0.135 | 0.215 | 0.001 | 0.051 | 0.051 | 0 | 0.07 | 0.22 | 0.04 | 0.11 | 0.04 | 0.08 |
| 5 | 123176408 | rs11241716 | -0.019 | 0.011 | -0.088 | 0.187 | -0.041 | -0.001 | -0.001 | 0 | 0.22 | 0.51 | 0.12 | 0.2 | 0.13 | 0.18 |
| 5 | 123179118 | rs10477652 | 0.552 | 0.053 | -0.251 | 0.239 | -0.007 | 0.03 | 0.03 | 0 | 0.17 | 0.32 | 0.1 | 0.19 | 0.13 | 0.14 |
| 5 | 123179249 | rs6875311 | 0.089 | 0.069 | -0.389 | 0.129 | 0.006 | 0.002 | 0.002 | 0 | 0.42 | 0.37 | 0.39 | 0.35 | 0.39 | 0.29 |
| 5 | 123179734 | rs6876166 | -1.718 | 0.106 | -0.508 | 0.111 | -0.142 | -0.002 | -0.002 | 0 | 0.67 | 0.95 | 0.53 | 0.66 | 0.56 | 0.5 |
| 5 | 123179990 | rs11747272 | 0.834 | 0.061 | -0.447 | 0.049 | -0.008 | 0.125 | 0.125 | 2.977 | 0.32 | 0.11 | 0.2 | 0.02 | 0.33 | 0.16 |
| 5 | 123182944 | rs17150847 | -0.051 | 0.031 | -0.324 | 0.231 | 0 | 0.098 | 0.098 | 0 | 0.11 | 0.31 | 0.06 | 0.21 | 0.08 | 0.13 |
| 5 | 123187446 | rs1592906 | -0.757 | 0.005 | -0.983 | 0.205 | 0.051 | -0.039 | -0.039 | 0 | 0.32 | 0.8 | 0.32 | 0.53 | 0.19 | 0.32 |
| 5 | 123188566 | rs10052257 | 0.562 | 0.585 | -1.348 | 0.305 | 0 | -0.004 | -0.004 | 2.983 | 0.34 | 0.32 | 0.21 | 0.14 | 0.28 | 0.17 |
| 5 | 123188660 | rs3909324 | -0.661 | 0.801 | -1.35 | 0.759 | -0.099 | -0.002 | -0.002 | 0 | 0.7 | 0.95 | 0.56 | 0.68 | 0.62 | 0.51 |
| 5 | 123192434 | rs1021769 | -0.266 | -0.012 | -1.161 | 0.716 | -0.071 | -0.046 | -0.046 | 0.02 | 0.64 | 0.83 | 0.53 | 0.54 | 0.53 | 0.47 |
| 5 | 123192945 | rs1030179 | 0.321 | -0.078 | -1.142 | 0.203 | -0.023 | 0.078 | 0.078 | 1.027 | 0.57 | 0.7 | 0.35 | 0.22 | 0.49 | 0.35 |
| 5 | 123193433 | rs1021770 | -0.323 | 0.045 | -1.543 | -0.066 | -0.104 | -0.042 | -0.042 | 0 | 0.69 | 0.95 | 0.55 | 0.65 | 0.6 | 0.51 |
| 5 | 123204222 | rs7708319 | 0.257 | -0.559 | -0.135 | -0.571 | 0.004 | -0.082 | -0.082 | 3.332 | 0.25 | 0.31 | 0.21 | 0.03 | 0.24 | 0.2 |
| 5 | 123204325 | rs11748635 | 0.321 | -0.81 | -0.431 | -0.661 | 0.009 | 0.014 | 0.014 | 0.48 | 0.08 | 0.07 | 0.04 | 0 | 0.07 | 0.01 |
| 5 | 123206264 | rs330681 | 0.891 | -1.026 | -0.342 | -0.546 | 0 | -0.002 | -0.002 | 0 | 0.39 | 0.38 | 0.54 | 0.37 | 0.41 | 0.49 |
| 5 | 123208563 | rs330683 | -0.511 | -0.883 | -0.571 | -0.462 | 0 | -0.008 | -0.008 | 0 | 0.73 | 0.8 | 0.8 | 0.4 | 0.72 | 0.69 |
| 5 | 123210364 | rs1422415 | 0.252 | -0.683 | -0.9 | -0.368 | -0.003 | -0.001 | -0.001 | 0 | 0.6 | 0.43 | 0.74 | 0.41 | 0.65 | 0.67 |
| 5 | 123212430 | rs11948130 | 0.773 | -0.483 | -1.389 | -0.877 | -0.004 | 0.032 | 0.032 | 0 | 0.25 | 0.16 | 0.2 | 0.03 | 0.25 | 0.2 |
| 5 | 123218166 | rs4836051 | -0.82 | -0.052 | -1.215 | -0.402 | -0.04 | -0.051 | -0.051 | 2.59 | 0.65 | 0.81 | 0.44 | 0.47 | 0.57 | 0.58 |
| 5 | 123223532 | rs9687603 | 0.086 | 0.722 | -1.375 | -0.038 | 0.003 | 0.062 | 0.062 | 0 | 0.3 | 0.17 | 0.22 | 0.03 | 0.28 | 0.2 |
| 5 | 123238611 | rs2546850 | 0.418 | 1.02 | -0.745 | 1.337 | 0 | 0.011 | 0.011 | 0 | 0.79 | 0.82 | 0.82 | 0.58 | 0.75 | 0.69 |
| 5 | 123242975 | rs1501762 | -0.695 | 0.693 | -0.917 | 1.203 | 0.01 | -0.024 | -0.024 | 0.149 | 0.08 | 0.01 | 0.07 | 0.07 | 0.06 | 0.1 |
| 5 | 123246694 | rs1990962 | 0.726 | 1.732 | -0.04 | 1.755 | 0.116 | 0.14 | 0.14 | 0.202 | 0.81 | 0.53 | 0.64 | 0.64 | 0.71 | 0.71 |
| 5 | 123247587 | rs2255981 | -1.138 | 1.333 | -0.227 | 1.423 | 0.01 | -0.028 | -0.028 | 0 | 0.26 | 0.16 | 0.33 | 0.55 | 0.25 | 0.4 |
| 5 | 123248573 | rs2859487 | 0.07 | 1.849 | 0.631 | 1.952 | 0.116 | 0.197 | 0.197 | 0 | 0.93 | 0.7 | 0.76 | 0.75 | 0.84 | 0.93 |
| 5 | 123261616 | rs2652267 | -1.354 | 1.371 | 0.674 | 1.907 | -0.013 | 0.04 | 0.04 | 0 | 0.25 | 0.32 | 0.31 | 0.54 | 0.22 | 0.37 |
| 5 | 123272896 | rs6884110 | -0.35 | 1.438 | 0.465 | 1.969 | -0.002 | -0.04 | -0.04 | 0 | 0.16 | 0.08 | 0.24 | 0.44 | 0.18 | 0.26 |
| 5 | 123279262 | rs10054053 | -0.129 | 1.411 | 0.503 | 1.942 | 0.001 | 0.054 | 0.054 | 0.069 | 0.12 | 0.13 | 0.24 | 0.45 | 0.17 | 0.26 |
| 5 | 123287402 | rs330454 | -1.418 | 1.403 | 0.651 | 2.119 | -0.003 | 0.024 | 0.024 | 0 | 0.24 | 0.25 | 0.32 | 0.5 | 0.24 | 0.36 |
| 5 | 123292142 | rs11952019 | -0.092 | 1.45 | 0.514 | 1.847 | 0.004 | -0.006 | -0.006 | 0 | 0.08 | 0.04 | 0.06 | 0.11 | 0.07 | 0.12 |
| 5 | 123293132 | rs2652257 | -0.439 | 1.245 | 0.4 | 1.841 | -0.005 | -0.029 | -0.029 | 0.472 | 0.17 | 0.11 | 0.26 | 0.53 | 0.19 | 0.29 |
| 5 | 123297332 | rs2407999 | -0.509 | 1.245 | 0.455 | 1.697 | -0.008 | -0.005 | -0.005 | 0 | 0.06 | 0.05 | 0.22 | 0.2 | 0.12 | 0.04 |
| 5 | 123298624 | rs4835814 | -1.145 | 1.705 | 1.112 | 1.511 | 0.079 | 0.057 | 0.057 | 0 | 0.21 | 0.45 | 0.36 | 0.29 | 0.31 | 0.31 |
| 5 | 123302367 | rs1363666 | -0.607 | 1.564 | 0.81 | 1.683 | 0.003 | -0.056 | -0.056 | 0 | 0.21 | 0.12 | 0.27 | 0.53 | 0.2 | 0.3 |
| 5 | 123332534 | rs10054840 | -1.849 | 1.95 | 1.575 | 1.477 | 0.098 | 0.121 | 0.121 | 1.287 | 0.1 | 0.37 | 0.28 | 0.26 | 0.15 | 0.06 |
| 5 | 123333374 | rs179354 | 1.24 | 1.734 | 1.493 | 1.668 | 0.012 | 0.085 | 0.085 | 0 | 0.86 | 0.71 | 0.63 | 0.61 | 0.82 | 0.8 |
| 5 | 123335648 | rs330439 | -0.65 | 2.013 | 1.275 | 1.491 | -0.003 | -0.047 | -0.047 | 0 | 0.26 | 0.15 | 0.28 | 0.53 | 0.26 | 0.29 |
| 5 | 123340716 | rs950392 | 0.552 | 2.194 | 1.39 | 1.886 | -0.012 | -0.032 | -0.032 | 0 | 0.06 | 0 | 0.08 | 0.11 | 0.07 | 0.09 |
| 5 | 123343862 | rs4836057 | 0.543 | 2.216 | 1.525 | 1.907 | 0.015 | 0.078 | 0.078 | 0 | 0.93 | 0.86 | 0.72 | 0.72 | 0.89 | 0.89 |
| 5 | 123347900 | rs4836058 | 1.587 | 2.424 | 1.367 | 1.709 | 0.011 | 0.018 | 0.018 | 0 | 0.76 | 0.77 | 0.54 | 0.52 | 0.62 | 0.6 |
| 5 | 123350243 | rs1862404 | -1.378 | 2.463 | 1.422 | 1.668 | 0.042 | 0.05 | 0.05 | 0 | 0.18 | 0.23 | 0.39 | 0.38 | 0.31 | 0.3 |
| 5 | 123360908 | rs10463770 | -1.2 | 2.487 | 1.3 | 1.666 | 0.03 | 0.036 | 0.036 | 0 | 0.17 | 0.18 | 0.38 | 0.38 | 0.31 | 0.31 |
| 5 | 123374257 | rs6595498 | -0.66 | 2.093 | 1.275 | 1.509 | 0 | 0.015 | 0.015 | 0 | 0.94 | 0.91 | 0.92 | 0.9 | 0.93 | 0.91 |
| 5 | 123375432 | rs6595499 | -0.696 | 2.078 | 1.259 | 1.513 | -0.001 | 0.011 | 0.011 | 0 | 0.94 | 0.93 | 0.92 | 0.9 | 0.93 | 0.89 |
| 5 | 123383613 | rs17151170 | -0.403 | 2.304 | 1.315 | 1.544 | 0.018 | 0 | 0 | 0 | 0.1 | 0.1 | 0.1 | 0.11 | 0.2 | 0.15 |
| 5 | 123384579 | rs17151176 | -0.579 | 2.065 | 1.421 | 1.511 | -0.004 | -0.022 | -0.022 | 0 | 0.21 | 0.14 | 0.23 | 0.49 | 0.24 | 0.34 |
| 5 | 123388957 | rs6885003 | 0.365 | 2.005 | 1.486 | 1.847 | -0.015 | -0.015 | -0.015 | 0 | 0.84 | 0.9 | 0.82 | 0.7 | 0.73 | 0.74 |
| 5 | 123399421 | rs10045249 | 1.765 | 1.912 | 1.454 | 1.727 | -0.047 | 0.022 | 0.022 | 0 | 0.61 | 0.23 | 0.59 | 0.37 | 0.53 | 0.43 |
| 5 | 123401074 | rs7447381 | 1.737 | 2.218 | 1.583 | 1.775 | 0.017 | -0.016 | -0.016 | 0 | 0.61 | 0.51 | 0.62 | 0.37 | 0.54 | 0.4 |
| 5 | 123402920 | rs7733161 | 0.88 | 2.344 | 1.46 | 2.179 | 0.026 | 0.01 | 0.01 | 0 | 0.81 | 0.66 | 0.81 | 0.8 | 0.76 | 0.72 |
| 5 | 123406035 | rs10066505 | -0.26 | 2.432 | 1.156 | 1.974 | -0.024 | -0.018 | -0.018 | 0 | 0.08 | 0.04 | 0.36 | 0.35 | 0.15 | 0.12 |
| 5 | 123408472 | rs7718242 | 0.671 | 1.99 | 1.364 | 1.812 | 0.032 | 0.014 | 0.014 | 0 | 0.82 | 0.71 | 0.82 | 0.8 | 0.75 | 0.73 |
| 5 | 123414747 | rs17151197 | -0.508 | 1.257 | 0.742 | 1.588 | -0.003 | 0.07 | 0.07 | 0 | 0.23 | 0.31 | 0.29 | 0.46 | 0.23 | 0.35 |
| 5 | 123416065 | rs6871469 | 1.074 | 1.457 | 0.978 | 1.329 | 0.024 | 0.002 | 0.002 | 0 | 0.76 | 0.41 | 0.82 | 0.8 | 0.76 | 0.78 |
| 5 | 123418270 | rs6883321 | -0.478 | 1.609 | 0.973 | 1.471 | 0.069 | 0.051 | 0.051 | 0 | 0.09 | 0.24 | 0.11 | 0.11 | 0.12 | 0.13 |
| 5 | 123425663 | rs7732928 | -0.161 | 2.212 | -0.219 | 1.227 | -0.028 | -0.027 | -0.027 | 0.23 | 0.77 | 0.84 | 0.54 | 0.58 | 0.59 | 0.8 |
| 5 | 123449266 | rs4240372 | -0.518 | 0.561 | -0.666 | 0.571 | -0.003 | -0.001 | -0.001 | 0 | 0.42 | 0.28 | 0.35 | 0.54 | 0.37 | 0.53 |
| 5 | 123452759 | rs10038458 | -0.857 | 0.377 | -0.013 | 0.888 | 0.037 | 0.062 | 0.062 | 0 | 0.23 | 0.5 | 0.18 | 0.32 | 0.23 | 0.36 |
| 5 | 123462031 | rs17151233 | 0.469 | 0.392 | 0.014 | 1.273 | 0.014 | 0.021 | 0.021 | 0 | 0.15 | 0.24 | 0.13 | 0.23 | 0.17 | 0.28 |
| 5 | 123462858 | rs1004891 | -1.332 | -0.093 | -0.35 | 0.755 | 0.008 | 0.02 | 0.02 | 0 | 0.31 | 0.53 | 0.24 | 0.34 | 0.34 | 0.44 |
| 5 | 123463805 | rs11950574 | -1.417 | -0.438 | -0.622 | 0.615 | -0.001 | -0.001 | -0.001 | 0 | 0.34 | 0.32 | 0.25 | 0.34 | 0.37 | 0.47 |
| 5 | 123466714 | rs4836066 | -0.031 | -0.516 | -0.55 | 0.596 | 0.011 | 0.056 | 0.056 | 0 | 0.19 | 0.27 | 0.2 | 0.39 | 0.21 | 0.33 |
| 5 | 123472576 | rs7719325 | 0.464 | -0.907 | -0.964 | 0.366 | 0.053 | 0.027 | 0.027 | 0.883 | 0.33 | 0.2 | 0.22 | 0.23 | 0.2 | 0.23 |
| 5 | 123474979 | rs2125127 | -1.087 | -0.513 | -0.805 | 0.258 | -0.064 | 0.021 | 0.021 | 3.585 | 0.65 | 0.83 | 0.49 | 0.64 | 0.52 | 0.64 |

| | | | | | | | | | | | | | | | | |
|---|---|---|---|---|---|---|---|---|---|---|---|---|---|---|---|---|
| 5 | 123493686 | rs2408118 | -1.145 | -0.465 | -0.926 | 0.218 | -0.081 | 0.015 | 0.015 | 2.997 | 0.66 | 0.88 | 0.49 | 0.64 | 0.51 | 0.63 |
| 5 | 123502228 | rs17151277 | 0.251 | -1.012 | -0.071 | 0.07 | -0.002 | 0.004 | 0.004 | 0 | 0.2 | 0.2 | 0.15 | 0.37 | 0.1 | 0.28 |
| 5 | 123510199 | rs17151283 | 1.216 | -1.03 | 0.106 | -0.11 | 0.009 | -0.046 | -0.046 | 0.017 | 0.09 | 0.23 | 0.06 | 0 | 0.09 | 0.04 |
| 5 | 123515763 | rs12186497 | -1.428 | -1.162 | -0.503 | -0.306 | 0.056 | 0.011 | 0.011 | 0.072 | 0.44 | 0.85 | 0.31 | 0.38 | 0.34 | 0.42 |
| 5 | 123517721 | rs1428417 | -1.524 | -1.233 | -0.758 | -0.313 | 0.055 | 0.001 | 0.001 | 0.386 | 0.45 | 0.9 | 0.32 | 0.38 | 0.35 | 0.42 |
| 5 | 123521221 | rs1007191 | 0.695 | -1.388 | -0.265 | 0.034 | -0.001 | -0.006 | -0.006 | 0 | 0.11 | 0.07 | 0.06 | 0.01 | 0.14 | 0.11 |
| 5 | 123524842 | rs10519752 | -1.021 | -1.409 | -0.165 | 0.02 | -0.014 | 0.005 | 0.005 | 0 | 0.34 | 0.41 | 0.23 | 0.37 | 0.26 | 0.37 |
| 5 | 123525965 | rs37391 | -0.621 | -0.913 | -1.474 | 0.186 | -0.083 | -0.004 | -0.004 | 0.291 | 0.69 | 0.98 | 0.74 | 0.8 | 0.68 | 0.74 |
| 5 | 123534021 | rs7702700 | -1.76 | -1.557 | -0.313 | 0.957 | 0.075 | -0.033 | -0.033 | 0 | 0.38 | 0.9 | 0.27 | 0.37 | 0.24 | 0.37 |
| 5 | 123545499 | rs12188854 | -0.526 | -1.169 | -0.052 | 0.43 | -0.001 | -0.045 | -0.045 | 0 | 0.13 | 0.15 | 0.07 | 0.01 | 0.15 | 0.12 |
| 5 | 123561794 | rs9687273 | -1.886 | -1.237 | -0.253 | 0.497 | 0.1 | 0.021 | 0.021 | 0 | 0.5 | 0.87 | 0.4 | 0.55 | 0.36 | 0.56 |
| 5 | 123564714 | rs1428431 | -1.417 | -1.326 | 0.299 | 0.993 | 0.017 | -0.007 | -0.007 | 0 | 0.37 | 0.69 | 0.33 | 0.54 | 0.2 | 0.47 |
| 5 | 123572141 | rs316389 | -0.927 | -0.948 | 0.325 | 0.659 | -0.029 | -0.02 | -0.02 | 2.297 | 0.18 | 0.25 | 0.09 | 0.1 | 0.07 | 0.15 |
| 5 | 123575862 | rs149533 | -1.965 | -0.731 | 0.078 | 0.869 | 0.023 | -0.047 | -0.047 | 0 | 0.46 | 0.91 | 0.4 | 0.53 | 0.4 | 0.6 |
| 5 | 123576004 | rs160705 | -0.837 | -0.748 | 0.568 | 1.031 | -0.04 | -0.013 | -0.013 | 0 | 0.16 | 0.31 | 0.09 | 0.12 | 0.06 | 0.17 |
| 5 | 123587152 | rs160851 | -0.958 | -0.735 | 0.544 | 1.039 | -0.042 | -0.018 | -0.018 | 0 | 0.17 | 0.33 | 0.11 | 0.12 | 0.06 | 0.16 |
| 5 | 123587452 | rs7713532 | -0.33 | -0.278 | 0.415 | 1.42 | -0.005 | 0.061 | 0.061 | 0 | 0.1 | 0.16 | 0.12 | 0.27 | 0.08 | 0.23 |
| 5 | 123589859 | rs17151495 | -0.713 | -0.164 | 0.188 | 1.074 | -0.005 | 0.004 | 0.004 | 0 | 0.11 | 0.37 | 0.07 | 0.08 | 0.05 | 0.11 |
| 5 | 123594293 | rs170316 | -1.048 | -0.073 | 0.046 | 1.112 | -0.005 | 0.035 | 0.035 | 0 | 0.19 | 0.43 | 0.16 | 0.24 | 0.17 | 0.23 |
| 5 | 123596603 | rs318008 | -1.208 | -0.216 | -1.13 | 0.948 | 0.017 | -0.071 | -0.071 | 1.746 | 0.42 | 0.91 | 0.39 | 0.51 | 0.37 | 0.57 |
| 5 | 123599714 | rs7711913 | -1.129 | 0.05 | -0.458 | 0.839 | -0.006 | 0.011 | 0.011 | 0 | 0.3 | 0.57 | 0.21 | 0.37 | 0.28 | 0.31 |
| 5 | 123603595 | rs31333 | -1.284 | -0.23 | -0.177 | 1.036 | 0.005 | 0.145 | 0.145 | 0.535 | 0.2 | 0.6 | 0.17 | 0.49 | 0.2 | 0.28 |
| 5 | 123603631 | rs31332 | -1.284 | -0.23 | -0.177 | 1.061 | 0.005 | 0.147 | 0.147 | 0.535 | 0.2 | 0.6 | 0.17 | 0.49 | 0.2 | 0.28 |
| 5 | 123603674 | rs31331 | -1.478 | -0.044 | -0.199 | 1.098 | 0.015 | 0.135 | 0.135 | 0.138 | 0.22 | 0.6 | 0.21 | 0.5 | 0.23 | 0.3 |
| 5 | 123605840 | rs161014 | -1.215 | -0.075 | -0.106 | 1.141 | 0.028 | 0.071 | 0.071 | 0.357 | 0.15 | 0.49 | 0.1 | 0.16 | 0.14 | 0.19 |
| 5 | 123614481 | rs31318 | -1.447 | -0.054 | 0.015 | 1.111 | -0.017 | -0.022 | -0.022 | 0 | 0.25 | 0.58 | 0.15 | 0.16 | 0.19 | 0.23 |
| 5 | 123617110 | rs149556 | -0.651 | 0.145 | -0.924 | 0.137 | -0.01 | -0.042 | -0.042 | 0.004 | 0.7 | 0.82 | 0.49 | 0.49 | 0.66 | 0.6 |
| 5 | 123618908 | rs11953978 | 1.444 | 0.145 | -1.245 | 0.081 | -0.04 | 0.179 | 0.179 | 0 | 0.37 | 0.1 | 0.26 | 0.01 | 0.4 | 0.21 |
| 5 | 123618933 | rs319634 | -0.305 | -0.126 | -0.405 | 0.86 | 0.038 | 0.083 | 0.083 | 0.297 | 0.1 | 0.41 | 0.06 | 0.11 | 0.09 | 0.12 |
| 5 | 123620112 | rs809039 | -0.942 | 0.027 | -1.073 | 0.767 | 0.038 | -0.067 | -0.067 | 4.146 | 0.33 | 0.85 | 0.2 | 0.49 | 0.24 | 0.38 |
| 5 | 123621707 | rs187678 | -0.967 | -0.038 | -1.056 | 0.72 | 0.022 | -0.076 | -0.076 | 0.236 | 0.33 | 0.85 | 0.26 | 0.5 | 0.26 | 0.4 |
| 5 | 123623533 | rs149509 | -0.535 | 0.093 | -1.068 | 0.623 | 0.02 | -0.002 | -0.002 | 0 | 0.51 | 0.55 | 0.28 | 0.44 | 0.41 | 0.45 |
| 5 | 123633249 | rs2032806 | -0.068 | -0.016 | -1.249 | 0.098 | 0.017 | 0.047 | 0.047 | 0 | 0.58 | 0.76 | 0.66 | 0.75 | 0.63 | 0.7 |
| 5 | 123633753 | rs922558 | -0.117 | -0.07 | -1.203 | 0.072 | 0.014 | 0.064 | 0.064 | 0 | 0.59 | 0.83 | 0.66 | 0.74 | 0.63 | 0.7 |
| 5 | 123634176 | rs2619943 | -0.119 | -0.156 | -1.254 | 0.085 | -0.009 | 0.111 | 0.111 | 0 | 0.6 | 0.91 | 0.7 | 0.81 | 0.65 | 0.7 |
| 5 | 123648021 | rs920997 | 0.188 | -0.535 | -0.295 | -0.315 | -0.001 | -0.011 | -0.011 | 0 | 0.52 | 0.53 | 0.66 | 0.81 | 0.6 | 0.67 |
| 5 | 123659824 | rs256857 | 0.325 | -0.608 | 0.265 | -0.053 | 0.016 | 0.022 | 0.022 | 0 | 0.51 | 0.36 | 0.72 | 0.7 | 0.61 | 0.69 |
| 5 | 123664565 | rs698185 | 0.324 | -0.617 | 0.264 | -0.057 | 0.015 | 0.021 | 0.021 | 0 | 0.51 | 0.36 | 0.71 | 0.7 | 0.6 | 0.69 |
| 5 | 123690883 | rs1904463 | -0.809 | -0.7 | -0.107 | -0.252 | -0.045 | 0.172 | 0.172 | 0 | 0.67 | 0.93 | 0.79 | 0.94 | 0.68 | 0.75 |
| 5 | 123704134 | rs10063748 | 0.393 | -0.714 | -0.198 | -0.302 | -0.013 | 0.178 | 0.178 | 0 | 0.35 | 0.1 | 0.21 | 0.06 | 0.31 | 0.24 |
| 5 | 123712903 | rs17151710 | -1.456 | -0.973 | 0.536 | -0.126 | -0.052 | 0.08 | 0.08 | 1.513 | 0.16 | 0.68 | 0.14 | 0.28 | 0.07 | 0.09 |
| 5 | 123717687 | rs11956092 | -0.271 | -0.948 | 0.746 | -0.102 | -0.032 | 0.022 | 0.022 | 0 | 0.09 | 0.19 | 0.09 | 0.14 | 0.03 | 0.07 |
| 5 | 123717754 | rs7712066 | -0.209 | -1.039 | 0.545 | -0.215 | -0.007 | -0.006 | -0.006 | 0 | 0.44 | 0.53 | 0.31 | 0.33 | 0.35 | 0.33 |
| 5 | 123730483 | rs988409 | -0.491 | -0.975 | 0.608 | -0.185 | 0.009 | 0.008 | 0.008 | 0 | 0.49 | 0.6 | 0.31 | 0.33 | 0.35 | 0.33 |
| 5 | 123731118 | rs185401 | -1.193 | -0.497 | 0.085 | -0.444 | -0.041 | 0.009 | 0.009 | 0.274 | 0.87 | 0.99 | 0.88 | 0.93 | 0.83 | 0.83 |
| 5 | 123734441 | rs895298 | -0.819 | -0.536 | -0.092 | -0.322 | -0.051 | 0.023 | 0.023 | 0.9 | 0.16 | 0.68 | 0.07 | 0.19 | 0.07 | 0.09 |
| 5 | 123735241 | rs469053 | 0.468 | 0.09 | -0.58 | -0.141 | -0.037 | -0.052 | -0.052 | 0 | 0.24 | 0.16 | 0.48 | 0.53 | 0.38 | 0.43 |
| 5 | 123736406 | rs6878727 | 0.506 | -0.278 | -0.985 | -0.343 | -0.004 | -0.003 | -0.003 | 0.032 | 0.44 | 0.59 | 0.24 | 0.3 | 0.34 | 0.31 |
| 5 | 123737922 | rs257701 | 0.368 | 0.413 | -1.485 | -0.264 | -0.131 | -0.087 | -0.087 | 4.354 | 0.23 | 0.02 | 0.47 | 0.7 | 0.37 | 0.46 |
| 5 | 123741065 | rs467485 | 0.68 | -0.428 | -1.159 | -0.995 | 0 | -0.026 | -0.026 | 0 | 0.78 | 0.77 | 0.86 | 0.98 | 0.81 | 0.86 |
| 5 | 123741332 | rs12656478 | -0.139 | -0.445 | -1.053 | -0.994 | 0.012 | 0.024 | 0.024 | 0 | 0.14 | 0.09 | 0.07 | 0.02 | 0.09 | 0.12 |
| 5 | 123741914 | rs419936 | 0.929 | 0.262 | -1.186 | -1.775 | 0.002 | -0.015 | -0.015 | 0 | 0.21 | 0.17 | 0.19 | 0.03 | 0.32 | 0.14 |
| 5 | 123744036 | rs12518621 | 0.985 | -0.213 | -2.05 | -1.794 | -0.051 | -0.103 | -0.103 | 0.268 | 0.27 | 0.03 | 0.43 | 0.69 | 0.27 | 0.44 |
| 5 | 123744093 | rs187702 | 0.244 | -0.105 | -0.987 | -1.72 | -0.003 | -0.005 | -0.005 | 0 | 0.78 | 0.75 | 0.86 | 0.9 | 0.81 | 0.85 |
| 5 | 123746171 | rs7705687 | 0.162 | 0.002 | -0.695 | -1.609 | 0.015 | -0.021 | -0.021 | 0 | 0.7 | 0.62 | 0.73 | 0.9 | 0.57 | 0.83 |
| 5 | 123746753 | rs27399 | 0.311 | 0.183 | -0.656 | -1.534 | 0.004 | -0.022 | -0.022 | 0 | 0.22 | 0.26 | 0.2 | 0.03 | 0.32 | 0.14 |
| 5 | 123750873 | rs17151784 | 0.294 | -0.022 | -0.627 | -1.245 | -0.033 | -0.027 | -0.027 | 0 | 0.12 | 0.22 | 0.03 | 0.03 | 0.04 | 0.11 |
| 5 | 123750902 | rs33890 | 0.718 | -0.162 | -0.773 | -1.417 | -0.004 | -0.018 | -0.018 | 0.092 | 0.27 | 0.17 | 0.5 | 0.71 | 0.3 | 0.38 |
| 5 | 123751831 | rs33887 | -1.258 | -0.204 | -0.498 | -1.671 | 0.002 | 0.004 | 0.004 | 0 | 0.66 | 0.69 | 0.76 | 0.83 | 0.71 | 0.67 |
| 5 | 123754638 | rs154463 | 0.443 | 0.086 | -0.235 | -1.524 | 0.005 | -0.04 | -0.04 | 0 | 0.23 | 0.22 | 0.23 | 0.02 | 0.36 | 0.14 |
| 5 | 123757079 | rs154444 | -0.44 | 0.083 | -0.304 | -1.447 | 0.005 | -0.054 | -0.054 | 0 | 0.34 | 0.46 | 0.22 | 0.12 | 0.35 | 0.24 |
| 5 | 123764757 | rs459380 | 0.757 | 0.287 | -0.656 | -1.418 | 0 | -0.026 | -0.026 | 0 | 0.2 | 0.22 | 0.2 | 0.02 | 0.32 | 0.11 |
| 5 | 123772792 | rs35137 | 1.057 | 0.313 | -1.052 | -1.373 | -0.099 | 0.101 | 0.101 | 0 | 0.17 | 0.02 | 0.18 | 0.02 | 0.3 | 0.11 |
| 5 | 123776023 | rs253417 | -0.11 | -0.206 | -0.817 | -1.069 | 0.015 | 0.047 | 0.047 | 0 | 0.36 | 0.14 | 0.51 | 0.75 | 0.32 | 0.47 |
| 5 | 123778418 | rs594434 | 0.762 | -0.189 | -0.942 | -1.039 | -0.075 | -0.092 | -0.092 | 0 | 0.25 | 0.03 | 0.49 | 0.72 | 0.29 | 0.35 |
| 5 | 123779300 | rs393547 | -0.727 | -0.477 | -0.391 | -1.225 | -0.015 | -0.047 | -0.047 | 0.353 | 0.42 | 0.49 | 0.54 | 0.81 | 0.34 | 0.49 |
| 5 | 123782602 | rs1199746 | 0.051 | -0.027 | -0.464 | -0.925 | -0.006 | 0.03 | 0.03 | 0 | 0.29 | 0.13 | 0.48 | 0.78 | 0.29 | 0.39 |
| 5 | 123784733 | rs17151866 | 0.887 | 0.041 | -0.877 | -0.182 | -0.067 | -0.174 | -0.174 | 3.455 | 0.22 | 0.03 | 0.42 | 0.53 | 0.27 | 0.29 |
| 5 | 123786852 | rs4568357 | 0.177 | 0.553 | -0.713 | -0.662 | 0.027 | -0.049 | -0.049 | 0 | 0.75 | 0.51 | 0.8 | 0.82 | 0.74 | 0.85 |

| | | | | | | | | | | | | | | | | |
|---|---|---|---|---|---|---|---|---|---|---|---|---|---|---|---|---|
| 5 | 123787030 | rs10900771 | 0.134 | 0.228 | -0.98 | -0.209 | -0.126 | -0.104 | -0.104 | 0 | 0.31 | 0.03 | 0.51 | 0.63 | 0.41 | 0.52 |
| 5 | 123793306 | rs4836083 | -0.547 | 0.046 | 0.296 | 0.601 | 0.015 | 0.09 | 0.09 | 0 | 0.17 | 0.32 | 0.26 | 0.31 | 0.18 | 0.26 |
| 5 | 123796107 | rs6890381 | -0.126 | 0.097 | 0.165 | 0.58 | 0.019 | 0.059 | 0.059 | 0 | 0.15 | 0.24 | 0.23 | 0.29 | 0.2 | 0.26 |
| 5 | 123796615 | rs6891277 | -0.205 | 0.365 | -0.219 | 0.602 | -0.013 | 0.026 | 0.026 | 0.394 | 0.9 | 0.78 | 0.91 | 0.84 | 0.93 | 0.92 |
| 5 | 123796639 | rs6890828 | 0.386 | 0.385 | -1.073 | 0.427 | 0.042 | -0.028 | -0.028 | 0 | 0.58 | 0.3 | 0.64 | 0.52 | 0.7 | 0.58 |
| 5 | 123798436 | rs4836084 | 0.475 | 1.719 | -0.655 | 0.064 | 0.027 | -0.005 | -0.005 | 0 | 0.84 | 0.76 | 0.8 | 0.83 | 0.76 | 0.76 |
| 5 | 123798978 | rs4574539 | 0.619 | 1.339 | -1.307 | -0.533 | -0.001 | 0.082 | 0.082 | 0 | 0.51 | 0.2 | 0.55 | 0.62 | 0.47 | 0.69 |
| 5 | 123801827 | rs11241745 | -0.398 | 1.468 | -0.804 | -1.511 | 0.005 | -0.002 | -0.002 | 0 | 0.9 | 0.86 | 0.88 | 0.88 | 0.85 | 0.97 |
| 5 | 123811602 | rs10067065 | 1.003 | 1.416 | -1.354 | -1.885 | -0.031 | -0.056 | -0.056 | 0.188 | 0.22 | 0.08 | 0.36 | 0.58 | 0.28 | 0.51 |
| 5 | 123814026 | rs6862843 | 0.786 | 0.796 | -1.936 | -2.502 | -0.027 | 0.112 | 0.112 | 0.066 | 0.45 | 0.19 | 0.42 | 0.28 | 0.45 | 0.36 |
| 5 | 123815748 | rs13188115 | 0.938 | 0.641 | -2.123 | -2.13 | -0.07 | -0.004 | -0.004 | 0.064 | 0.13 | 0.01 | 0.13 | 0.13 | 0.2 | 0.14 |
| 5 | 123816089 | rs4457071 | 0.777 | 0.347 | -1.519 | -1.754 | -0.018 | -0.039 | -0.039 | 0 | 0.22 | 0.13 | 0.39 | 0.62 | 0.28 | 0.5 |
| 5 | 123819489 | rs4397127 | 0.241 | 0.334 | -0.75 | -1.815 | 0.005 | -0.008 | -0.008 | 0 | 0.91 | 0.83 | 0.89 | 0.93 | 0.86 | 0.98 |
| 5 | 123828524 | rs6883157 | 0.119 | 0.317 | -0.723 | -1.841 | -0.001 | -0.006 | -0.006 | 0 | 0.84 | 0.8 | 0.86 | 0.93 | 0.83 | 0.98 |
| 5 | 123850956 | rs6887119 | -0.393 | -0.398 | -0.482 | -1.932 | -0.018 | -0.068 | -0.068 | 0 | 0.21 | 0.32 | 0.06 | 0 | 0.11 | 0.08 |
| 5 | 123851734 | rs10076865 | -0.931 | -0.24 | -0.624 | -2.145 | -0.005 | 0.049 | 0.049 | 0 | 0.45 | 0.67 | 0.26 | 0.27 | 0.44 | 0.25 |
| 5 | 123853534 | rs949973 | 0.193 | -0.198 | -0.861 | -2.108 | -0.063 | -0.018 | -0.018 | 0 | 0.23 | 0.09 | 0.18 | 0.27 | 0.33 | 0.17 |
| 5 | 123853591 | rs949974 | -1.321 | -0.337 | -0.997 | -2.16 | -0.067 | 0.072 | 0.072 | 0 | 0.58 | 0.9 | 0.37 | 0.29 | 0.55 | 0.31 |
| 5 | 123863647 | rs10478625 | -0.591 | -0.241 | -0.269 | -1.948 | 0.023 | -0.04 | -0.04 | 0 | 0.34 | 0.51 | 0.26 | 0.19 | 0.41 | 0.24 |
| 5 | 123867087 | rs10046065 | -0.13 | -0.614 | -0.283 | -1.954 | 0.039 | -0.059 | -0.059 | 0 | 0.22 | 0.41 | 0.15 | 0.12 | 0.24 | 0.2 |
| 5 | 123870271 | rs6880412 | 1.441 | -0.878 | -1.154 | -1.937 | -0.116 | -0.08 | -0.08 | 0 | 0.31 | 0.04 | 0.5 | 0.63 | 0.41 | 0.55 |
| 5 | 123873685 | rs1494672 | -1.194 | -0.418 | 0.025 | -1.584 | -0.001 | -0.019 | -0.019 | 0 | 0.37 | 0.44 | 0.27 | 0.29 | 0.39 | 0.27 |
| 5 | 123873998 | rs1494673 | 0.716 | -0.17 | 0.036 | -1.503 | -0.039 | -0.009 | -0.009 | 0 | 0.12 | 0.01 | 0.09 | 0.12 | 0.17 | 0.11 |
| 5 | 123875127 | rs10478626 | -0.329 | -0.137 | 0.081 | -1.481 | 0.003 | -0.009 | -0.009 | 0.661 | 0.19 | 0.29 | 0.16 | 0.17 | 0.22 | 0.14 |
| 5 | 123880851 | rs1494671 | -1.392 | -0.206 | -0.231 | -1.451 | -0.004 | 0.027 | 0.027 | 0 | 0.43 | 0.66 | 0.3 | 0.28 | 0.44 | 0.32 |
| 5 | 123886769 | rs9327331 | -1.071 | -0.143 | -0.188 | -1.686 | -0.001 | -0.003 | -0.003 | 1.107 | 0.41 | 0.57 | 0.3 | 0.26 | 0.44 | 0.3 |
| 5 | 123888331 | rs10061885 | 0.41 | -0.103 | -0.148 | -1.665 | 0 | -0.008 | -0.008 | 0.043 | 0.15 | 0.14 | 0.11 | 0.04 | 0.16 | 0.09 |
| 5 | 123888562 | rs3924161 | -0.231 | 0.078 | 0.319 | -0.899 | 0.006 | 0 | 0 | 0 | 0.2 | 0.25 | 0.16 | 0.22 | 0.24 | 0.17 |
| 5 | 123896355 | rs1018047 | -0.408 | 0.121 | -0.464 | -0.697 | 0.005 | -0.031 | -0.031 | 0 | 0.29 | 0.43 | 0.19 | 0.22 | 0.29 | 0.24 |
| 5 | 123899147 | rs11958144 | -0.435 | 0.312 | -0.283 | -0.206 | -0.015 | -0.056 | -0.056 | 0 | 0.1 | 0.36 | 0.05 | 0 | 0.04 | 0 |
| 5 | 123909500 | rs6873640 | 0.257 | 0.964 | -1.546 | 0.142 | -0.031 | 0.03 | 0.03 | 0 | 0.63 | 0.87 | 0.38 | 0.34 | 0.59 | 0.52 |
| 5 | 123914126 | rs7719941 | 0.432 | 1.43 | -1.102 | 0.047 | -0.004 | 0.007 | 0.007 | 0.43 | 0.63 | 0.69 | 0.37 | 0.24 | 0.52 | 0.43 |
| 5 | 123915337 | rs4323228 | -0.716 | 1.007 | -0.721 | 0.513 | -0.012 | 0.026 | 0.026 | 0 | 0.35 | 0.52 | 0.19 | 0.44 | 0.28 | 0.36 |
| 5 | 123918548 | rs1389881 | 0.127 | 1.332 | -0.65 | 0.81 | 0.014 | 0.043 | 0.043 | 0 | 0.95 | 0.88 | 0.78 | 0.87 | 0.92 | 0.94 |
| 5 | 123919181 | rs10071778 | -0.025 | 1.101 | -0.224 | 1.288 | -0.001 | 0.002 | 0.002 | 0 | 0.16 | 0.18 | 0.09 | 0.26 | 0.14 | 0.2 |
| 5 | 123919931 | rs11743981 | 1.057 | 1.092 | -1.098 | 0.387 | -0.052 | -0.044 | -0.044 | 0.856 | 0.3 | 0.07 | 0.37 | 0.39 | 0.38 | 0.46 |
| 5 | 123922314 | rs10070372 | 0.166 | 1.056 | -0.497 | 0.39 | -0.005 | 0.028 | 0.028 | 0 | 0.18 | 0.23 | 0.1 | 0.3 | 0.15 | 0.24 |
| 5 | 123924246 | rs7707575 | -0.53 | 0.699 | -0.61 | 0.234 | -0.002 | 0.007 | 0.007 | 0 | 0.49 | 0.42 | 0.51 | 0.44 | 0.5 | 0.44 |
| 5 | 123930829 | rs10463782 | 0.348 | 0.428 | -1.26 | 0.183 | -0.058 | -0.015 | -0.015 | 0 | 0.3 | 0.01 | 0.21 | 0.25 | 0.28 | 0.33 |
| 5 | 123933446 | rs10038823 | -0.67 | 0.612 | -0.745 | 0.716 | 0.01 | -0.029 | -0.029 | 0 | 0.59 | 0.73 | 0.62 | 0.52 | 0.64 | 0.63 |
| 5 | 123939264 | rs4835839 | 0.75 | 0.91 | -0.863 | 0.619 | -0.018 | -0.02 | -0.02 | 0 | 0.11 | 0.06 | 0.31 | 0.26 | 0.24 | 0.21 |
| 5 | 123941250 | rs4132580 | 0.657 | 0.837 | -0.012 | 0.45 | 0.031 | -0.054 | -0.054 | 0 | 0.16 | 0.46 | 0.12 | 0.07 | 0.18 | 0.09 |
| 5 | 123941736 | rs12109090 | -0.251 | 0.85 | -0.229 | 0.451 | 0.021 | 0.08 | 0.08 | 0 | 0.47 | 0.67 | 0.31 | 0.13 | 0.38 | 0.38 |
| 5 | 123948093 | rs4240375 | -1.036 | 0.832 | -0.168 | 0.528 | 0.006 | 0.005 | 0.005 | 0 | 0.58 | 0.72 | 0.62 | 0.39 | 0.62 | 0.59 |
| 5 | 123948730 | rs11956421 | 0.08 | 0.835 | -0.386 | 0.53 | -0.026 | 0.027 | 0.027 | 0.531 | 0.41 | 0.18 | 0.49 | 0.33 | 0.44 | 0.5 |
| 5 | 123950146 | rs4626333 | 0.316 | 0.799 | 0.197 | 0.305 | 0.042 | -0.067 | -0.067 | 0.089 | 0.17 | 0.53 | 0.13 | 0.08 | 0.19 | 0.11 |
| 5 | 123950609 | rs4568359 | 1.075 | 0.257 | -0.438 | 0.254 | -0.022 | 0.089 | 0.089 | 0.427 | 0.26 | 0.01 | 0.16 | 0.05 | 0.18 | 0.2 |
| 5 | 123950880 | rs10066524 | 1.056 | 0.012 | -0.206 | 0.65 | 0.003 | -0.017 | -0.017 | 0 | 0.25 | 0.23 | 0.3 | 0.43 | 0.22 | 0.32 |
| 5 | 123954634 | rs4304076 | 0.508 | -0.05 | -0.081 | 0.411 | 0.067 | 0.061 | 0.061 | 0.817 | 0.49 | 0.3 | 0.4 | 0.36 | 0.34 | 0.44 |
| 5 | 123958110 | rs6859251 | 1.387 | -0.101 | -0.091 | 0.398 | 0.005 | -0.001 | -0.001 | 0.783 | 0.25 | 0.26 | 0.3 | 0.3 | 0.21 | 0.29 |
| 5 | 123958159 | rs11957228 | 1.124 | -0.005 | -0.174 | 0.328 | -0.005 | 0.095 | 0.095 | 0.102 | 0.27 | 0.07 | 0.18 | 0.08 | 0.22 | 0.23 |
| 5 | 123959513 | rs10074057 | -1.362 | -0.016 | 0.41 | 0.094 | 0.003 | -0.083 | -0.083 | 1.187 | 0.8 | 0.78 | 0.91 | 1 | 0.83 | 0.84 |
| 5 | 123960447 | rs6871217 | 1.667 | 0.06 | 0.228 | 0.168 | 0.004 | -0.022 | -0.022 | 0 | 0.23 | 0.11 | 0.29 | 0.3 | 0.2 | 0.28 |
| 5 | 123962891 | rs10035209 | -0.225 | -0.015 | 0.556 | 0.159 | 0.018 | 0.044 | 0.044 | 0 | 0.52 | 0.43 | 0.59 | 0.67 | 0.63 | 0.53 |
| 5 | 123964252 | rs9687477 | -0.449 | 0.001 | 0.706 | 0.132 | -0.005 | -0.018 | -0.018 | 0 | 0.54 | 0.54 | 0.61 | 0.7 | 0.64 | 0.55 |
| 5 | 123966399 | rs11953210 | 1.417 | -0.136 | 0.386 | 0.363 | 0.032 | -0.024 | -0.024 | 0 | 0.26 | 0.21 | 0.14 | 0.27 | 0.17 | 0.22 |
| 5 | 123966937 | rs12522836 | 1.771 | -0.086 | 0.194 | 0.436 | 0.009 | -0.043 | -0.043 | 0 | 0.23 | 0.11 | 0.43 | 0.32 | 0.19 | 0.28 |
| 5 | 123967000 | rs10045786 | 0.122 | 0.658 | 0.639 | 0.268 | 0.045 | -0.029 | -0.029 | 0 | 0.06 | 0.22 | 0.07 | 0.01 | 0.07 | 0.06 |
| 5 | 123968495 | rs4836101 | -1.129 | 0.722 | 0.337 | 0.303 | -0.002 | -0.081 | -0.081 | 0 | 0.79 | 0.78 | 0.91 | 1 | 0.83 | 0.83 |
| 5 | 123971932 | rs4541656 | 0.431 | 0.676 | -0.242 | 0.805 | -0.064 | -0.041 | -0.041 | 0 | 0.24 | 0.08 | 0.25 | 0.35 | 0.4 | 0.25 |
| 5 | 123980872 | rs6866116 | 0.329 | 0.692 | 0.083 | 0.478 | -0.008 | -0.011 | -0.011 | 4.469 | 0.26 | 0.22 | 0.25 | 0.46 | 0.38 | 0.26 |
| 5 | 123983915 | rs6862252 | -0.684 | 0.079 | -0.157 | 0.076 | -0.007 | 0.12 | 0.12 | 0 | 0.51 | 0.71 | 0.45 | 0.75 | 0.55 | 0.61 |
| 5 | 123984763 | rs7708070 | -1.463 | 0.096 | -0.471 | -0.072 | -0.022 | -0.007 | -0.007 | 0 | 0.77 | 0.89 | 0.6 | 0.76 | 0.75 | 0.72 |
| 5 | 123986316 | rs7721111 | 1.216 | 0.044 | -0.012 | -0.139 | -0.037 | -0.081 | -0.081 | 0 | 0.14 | 0.22 | 0.05 | 0.01 | 0.06 | 0.12 |
| 5 | 123988063 | rs6871146 | 0.264 | 0.067 | -0.223 | 0.091 | -0.032 | 0.055 | 0.055 | 0 | 0.21 | 0.39 | 0.19 | 0.27 | 0.16 | 0.22 |
| 5 | 123989154 | rs10042766 | -0.76 | 0.808 | -0.029 | 0.5 | 0.005 | 0.002 | 0.002 | 0 | 0.73 | 0.66 | 0.55 | 0.73 | 0.72 | 0.6 |
| 5 | 123990972 | rs10040793 | 0.024 | 1.301 | -0.322 | 1.017 | -0.018 | -0.039 | -0.039 | 0.198 | 0.26 | 0.19 | 0.23 | 0.45 | 0.36 | 0.26 |
| 5 | 124001206 | rs4836104 | -1.35 | 1.603 | 0.044 | 0.983 | -0.001 | 0.035 | 0.035 | 0 | 0.09 | 0.12 | 0.26 | 0.32 | 0.19 | 0.22 |
| 5 | 124002342 | rs6595536 | -1.967 | 1.669 | 0.66 | 0.88 | 0.097 | 0.198 | 0.198 | 0.251 | 0.15 | 0.46 | 0.32 | 0.33 | 0.24 | 0.26 |

| | | | | | | | | | | | | | | | | |
|---|---|---|---|---|---|---|---|---|---|---|---|---|---|---|---|---|
| 5 | 124004404 | rs13162151 | -0.5 | 1.684 | 0.4 | 0.965 | -0.001 | 0.023 | 0.023 | 0 | 0.07 | 0.09 | 0.17 | 0.22 | 0.17 | 0.1 |
| 5 | 124006906 | rs6895878 | -1.656 | 1.647 | 0.508 | 0.839 | 0.112 | 0.195 | 0.195 | 0.073 | 0.16 | 0.55 | 0.34 | 0.31 | 0.25 | 0.26 |
| 5 | 124015270 | rs4835842 | 0.737 | 1.413 | -0.423 | 0.262 | -0.03 | 0.098 | 0.098 | 0 | 0.16 | 0.01 | 0.08 | 0.01 | 0.18 | 0.06 |
| 5 | 124018279 | rs6871443 | -1.332 | 1.333 | -0.133 | 0.43 | -0.012 | -0.033 | -0.033 | 0 | 0.34 | 0.8 | 0.38 | 0.35 | 0.4 | 0.28 |
| 5 | 124022189 | rs4616890 | -0.465 | 1.372 | 0.251 | 0.493 | 0 | 0 | 0 | 0.474 | 0.24 | 0.25 | 0.23 | 0.2 | 0.33 | 0.13 |
| 5 | 124031011 | rs12515071 | 0.911 | 1.16 | -0.305 | -0.138 | 0.006 | -0.005 | -0.005 | 2.005 | 0.43 | 0.28 | 0.53 | 0.66 | 0.42 | 0.63 |
| 5 | 124033113 | rs4621556 | 0.471 | 1.057 | -0.447 | -0.821 | 0.001 | -0.008 | -0.008 | 0 | 0.52 | 0.46 | 0.69 | 0.84 | 0.62 | 0.7 |
| 5 | 124036555 | rs6873550 | -0.889 | 1.509 | -0.941 | -0.285 | 0 | 0.139 | 0.139 | 0.016 | 0.79 | 0.99 | 0.92 | 1 | 0.88 | 0.93 |
| 5 | 124038251 | rs12513661 | -0.773 | 1.811 | -0.071 | 0.434 | 0.016 | 0.039 | 0.039 | 0 | 0.05 | 0.1 | 0.12 | 0.13 | 0.06 | 0.15 |
| 5 | 124043721 | rs4836112 | 0.81 | 1.679 | -0.875 | 0.314 | -0.053 | 0.031 | 0.031 | 0 | 0.69 | 0.81 | 0.62 | 0.23 | 0.56 | 0.42 |
| 5 | 124045025 | rs12514819 | 0.679 | 1.015 | -1.157 | -0.079 | 0.008 | 0.025 | 0.025 | 0 | 0.23 | 0.1 | 0.33 | 0.14 | 0.16 | 0.27 |
| 5 | 124050437 | rs4389685 | 0.327 | 1.216 | -1.267 | 0.282 | -0.046 | -0.053 | -0.053 | 0 | 0.73 | 0.87 | 0.72 | 0.38 | 0.6 | 0.56 |
| 5 | 124055918 | rs4623140 | 0.493 | 0.795 | -1.203 | 0.264 | -0.005 | 0.037 | 0.037 | 0 | 0.23 | 0.07 | 0.32 | 0.12 | 0.17 | 0.26 |
| 5 | 124073861 | rs4629598 | 0.492 | 1.027 | -0.175 | 1.048 | -0.001 | -0.001 | -0.001 | 0 | 0.83 | 0.79 | 0.88 | 0.86 | 0.81 | 0.83 |
| 5 | 124084849 | rs4421098 | 0.878 | 0.748 | 0.253 | 1.248 | -0.032 | 0.19 | 0.19 | 0 | 0.82 | 0.58 | 0.79 | 0.5 | 0.82 | 0.67 |
| 5 | 124097421 | rs11739274 | -1.269 | 1.504 | -0.63 | 0.674 | -0.03 | -0.004 | -0.004 | 0 | 0.05 | 0.01 | 0.04 | 0.05 | 0.12 | 0.15 |
| 5 | 124110646 | rs10069971 | -0.649 | 1.199 | -0.532 | 0.745 | 0 | 0 | 0 | 0.158 | 0.4 | 0.52 | 0.37 | 0.45 | 0.42 | 0.58 |
| 5 | 124112498 | rs4836120 | 0.332 | 1.346 | 0.056 | 0.711 | 0.028 | 0.03 | 0.03 | 0.001 | 0.75 | 0.62 | 0.69 | 0.6 | 0.64 | 0.6 |
| 5 | 124125793 | rs6595545 | -1.502 | 0.91 | 0.629 | 0.646 | 0.003 | -0.043 | -0.043 | 0 | 0.09 | 0.34 | 0.05 | 0.01 | 0.04 | 0.05 |
| 5 | 124129139 | rs6882715 | -0.107 | 0.706 | 0.364 | -0.151 | -0.001 | 0 | 0 | 0 | 0.6 | 0.63 | 0.55 | 0.59 | 0.45 | 0.54 |
| 5 | 124132647 | rs10045691 | -0.307 | 0.715 | 0.236 | -0.13 | 0.009 | -0.031 | -0.031 | 0 | 0.07 | 0.27 | 0.04 | 0 | 0.04 | 0.04 |
| 5 | 124132673 | rs4835845 | 0.952 | 0.5 | -0.72 | 0.283 | -0.028 | -0.052 | -0.052 | 0.078 | 0.34 | 0.02 | 0.3 | 0.26 | 0.27 | 0.25 |
| 5 | 124139895 | rs6595551 | -1.076 | 0.576 | 0.16 | 0.827 | 0.029 | 0.191 | 0.191 | 0 | 0.17 | 0.44 | 0.11 | 0.36 | 0.18 | 0.13 |
| 5 | 124146728 | rs13168430 | 0.119 | -0.118 | -0.133 | 0.038 | -0.001 | -0.001 | -0.001 | 0 | 0.55 | 0.43 | 0.43 | 0.49 | 0.5 | 0.57 |
| 5 | 124146768 | rs13168447 | -0.003 | -0.119 | -0.134 | 0.037 | -0.001 | -0.001 | -0.001 | 0 | 0.55 | 0.43 | 0.43 | 0.49 | 0.5 | 0.57 |
| 5 | 124147560 | rs4357032 | -0.968 | -0.194 | -0.304 | -0.102 | -0.045 | 0.114 | 0.114 | 0.335 | 0.71 | 0.89 | 0.67 | 0.92 | 0.68 | 0.79 |
| 5 | 124154976 | rs715393 | 0.003 | -0.15 | 0.057 | 0.029 | 0.005 | 0.003 | 0.003 | 0 | 0.07 | 0.17 | 0.03 | 0.08 | 0.04 | 0.1 |
| 5 | 124157377 | rs17152077 | 0.147 | -0.117 | -0.121 | 0.103 | 0.005 | 0.087 | 0.087 | 0 | 0.08 | 0.11 | 0.19 | 0.33 | 0.14 | 0.13 |
| 5 | 124177701 | rs10041012 | -0.338 | -0.135 | -0.097 | -0.06 | 0.008 | -0.08 | -0.08 | 0.841 | 0.62 | 0.51 | 0.63 | 0.84 | 0.64 | 0.69 |
| 5 | 124179292 | rs11241771 | 0.909 | -0.502 | -0.514 | -0.002 | -0.02 | -0.009 | -0.009 | 0.221 | 0.28 | 0.22 | 0.34 | 0.39 | 0.4 | 0.39 |
| 5 | 124188887 | rs930606 | -0.825 | -1.433 | 0.089 | -0.355 | -0.043 | -0.03 | -0.03 | 0 | 0.35 | 0.53 | 0.15 | 0.17 | 0.14 | 0.3 |
| 5 | 124193973 | rs13166738 | -0.326 | -1.141 | -0.14 | -0.129 | -0.006 | 0 | 0 | 0 | 0.25 | 0.19 | 0.09 | 0.17 | 0.1 | 0.25 |
| 5 | 124217262 | rs1078675 | -0.115 | -0.604 | -0.079 | -0.246 | -0.004 | 0 | 0 | 0.008 | 0.56 | 0.67 | 0.39 | 0.56 | 0.53 | 0.48 |
| 5 | 124222305 | rs2408183 | 0.979 | -0.769 | -0.265 | -0.56 | -0.015 | 0.128 | 0.128 | 0 | 0.33 | 0.15 | 0.24 | 0.07 | 0.32 | 0.25 |
| 5 | 124223035 | rs17448433 | 0.629 | -0.744 | -0.162 | -0.61 | -0.03 | 0.131 | 0.131 | 0 | 0.38 | 0.19 | 0.29 | 0.07 | 0.42 | 0.32 |
| 5 | 124224444 | rs11241775 | 0.647 | -0.8 | 0.106 | -0.594 | 0.001 | 0.016 | 0.016 | 0 | 0.13 | 0.13 | 0.07 | 0.48 | 0.11 | 0.15 |
| 5 | 124224634 | rs745085 | 0.132 | -0.823 | 0.378 | -0.624 | 0 | -0.026 | -0.026 | 0 | 0.43 | 0.45 | 0.32 | 0.07 | 0.42 | 0.34 |
| 5 | 124238375 | rs6880908 | 0.199 | -1.002 | 0.415 | -0.658 | -0.004 | -0.007 | -0.007 | 0 | 0.42 | 0.35 | 0.3 | 0.55 | 0.42 | 0.39 |
| 5 | 124245657 | rs2897883 | -0.285 | -1.069 | 0.316 | -0.597 | -0.015 | 0.004 | 0.004 | 0 | 0.85 | 0.77 | 0.91 | 0.84 | 0.93 | 0.89 |
| 5 | 124250018 | rs13362134 | -0.102 | -1.083 | 0.667 | -0.641 | -0.001 | 0.001 | 0.001 | 0 | 0.48 | 0.52 | 0.36 | 0.55 | 0.51 | 0.46 |
| 5 | 124253737 | rs10078386 | 0.228 | -1.104 | 0.552 | -0.552 | -0.014 | 0.116 | 0.116 | 0 | 0.16 | 0.27 | 0.09 | 0.48 | 0.11 | 0.16 |
| 5 | 124254518 | rs1562258 | -0.322 | -1.301 | 0.608 | -0.68 | -0.001 | -0.004 | -0.004 | 0 | 0.54 | 0.44 | 0.7 | 0.76 | 0.64 | 0.65 |
| 5 | 124257374 | rs1026009 | 1.134 | -1.222 | 0.052 | -0.667 | -0.048 | 0.013 | 0.013 | 0 | 0.36 | 0.15 | 0.57 | 0.3 | 0.44 | 0.46 |
| 5 | 124273899 | rs6869863 | -0.235 | -0.735 | 0.349 | -0.682 | 0.026 | 0.196 | 0.196 | 0 | 0.19 | 0.39 | 0.13 | 0.46 | 0.2 | 0.16 |
| 5 | 124277064 | rs4458563 | -0.282 | -0.759 | -0.32 | -0.79 | 0.024 | 0.033 | 0.033 | 0.014 | 0.87 | 0.92 | 0.96 | 0.98 | 0.95 | 0.96 |
| 5 | 124278397 | rs2408226 | 1.229 | -0.69 | -0.526 | -0.823 | -0.043 | 0.125 | 0.125 | 0.059 | 0.29 | 0.1 | 0.22 | 0.08 | 0.31 | 0.28 |
| 5 | 124279194 | rs13361521 | 1.261 | -0.683 | -0.543 | -0.82 | -0.052 | 0.127 | 0.127 | 0.078 | 0.29 | 0.08 | 0.22 | 0.08 | 0.31 | 0.28 |
| 5 | 124287674 | rs4836131 | 1.144 | -0.451 | -0.062 | -0.414 | 0 | -0.004 | -0.004 | 0.123 | 0.68 | 0.38 | 0.81 | 0.52 | 0.74 | 0.78 |
| 5 | 124288960 | rs7707774 | -0.256 | -0.223 | -0.436 | -0.41 | 0 | 0.015 | 0.015 | 0 | 0.88 | 0.92 | 0.95 | 0.98 | 0.9 | 0.97 |
| 5 | 124290967 | rs723187 | 0.703 | -0.565 | -0.807 | -0.739 | 0.06 | 0.11 | 0.11 | 0 | 0.38 | 0.18 | 0.2 | 0.09 | 0.28 | 0.34 |
| 5 | 124294448 | rs17152193 | 0.581 | -0.16 | -0.997 | -0.596 | 0.01 | 0.033 | 0.033 | 0 | 0.69 | 0.48 | 0.76 | 0.56 | 0.65 | 0.8 |
| 5 | 124298997 | rs6595565 | 0.785 | -0.293 | -1.966 | -0.328 | 0.032 | 0.161 | 0.161 | 0.627 | 0.35 | 0.05 | 0.19 | 0.09 | 0.28 | 0.34 |
| 5 | 124302354 | rs6875187 | 0.261 | -0.092 | -1.373 | 0.041 | 0.001 | -0.034 | -0.034 | 0 | 0.77 | 0.81 | 0.87 | 0.56 | 0.78 | 0.83 |
| 5 | 124318270 | rs10073969 | 0.374 | -0.326 | -1.79 | -0.79 | -0.005 | 0.222 | 0.222 | 0 | 0.37 | 0.07 | 0.23 | 0.07 | 0.32 | 0.34 |
| 5 | 124322471 | rs1374131 | -0.188 | -0.558 | -0.703 | 0.235 | -0.013 | 0.159 | 0.159 | 0.172 | 0.09 | 0.3 | 0.06 | 0.19 | 0.04 | 0.11 |
| 5 | 124327862 | rs4518389 | -0.72 | -0.542 | -1.866 | 0.082 | -0.011 | -0.015 | -0.015 | 0 | 0.48 | 0.86 | 0.67 | 0.47 | 0.46 | 0.56 |
| 5 | 124329306 | rs10440778 | -0.823 | -0.656 | -1.915 | 0.037 | -0.007 | -0.015 | -0.015 | 0 | 0.5 | 0.87 | 0.68 | 0.48 | 0.48 | 0.59 |
| 5 | 124330522 | rs6864049 | -1.105 | -0.411 | -1.833 | -0.201 | -0.066 | 0.086 | 0.086 | 0 | 0.59 | 0.87 | 0.71 | 0.75 | 0.53 | 0.64 |
| 5 | 124332738 | rs4240386 | -0.456 | -0.517 | -0.976 | -0.457 | -0.009 | -0.03 | -0.03 | 0.956 | 0.34 | 0.43 | 0.56 | 0.73 | 0.31 | 0.47 |
| 5 | 124340703 | rs6891054 | -0.669 | -1.203 | -0.491 | -0.465 | -0.028 | -0.02 | -0.02 | 0 | 0.34 | 0.53 | 0.53 | 0.72 | 0.22 | 0.43 |
| 5 | 124341945 | rs4240388 | -1.652 | -1.25 | -0.516 | -0.749 | 0.009 | 0.093 | 0.093 | 0 | 0.57 | 0.8 | 0.65 | 0.74 | 0.43 | 0.58 |
| 5 | 124342918 | rs4472259 | 0.551 | -1.479 | -0.324 | -0.578 | -0.042 | 0.034 | 0.034 | 0 | 0.14 | 0.23 | 0.06 | 0.2 | 0.06 | 0.13 |
| 5 | 124343052 | rs7707465 | 0.384 | -1.222 | -0.264 | -1.06 | 0.008 | 0.038 | 0.038 | 0.091 | 0.22 | 0.15 | 0.12 | 0.02 | 0.21 | 0.17 |
| 5 | 124343550 | rs7708412 | -0.083 | -1.201 | -0.268 | -1.051 | -0.006 | -0.065 | -0.065 | 0.841 | 0.25 | 0.35 | 0.14 | 0.02 | 0.21 | 0.17 |
| 5 | 124351539 | rs6883400 | -0.486 | -1.504 | -0.597 | -0.907 | -0.012 | -0.007 | -0.007 | 1.9 | 0.32 | 0.3 | 0.51 | 0.72 | 0.22 | 0.45 |
| 5 | 124353599 | rs6595569 | -1.359 | -1.74 | -0.573 | -1.025 | 0.055 | 0.046 | 0.046 | 0 | 0.42 | 0.66 | 0.54 | 0.72 | 0.25 | 0.48 |
| 5 | 124354175 | rs6859102 | -0.521 | -1.591 | -1.01 | -0.937 | -0.01 | -0.007 | -0.007 | 0.346 | 0.33 | 0.3 | 0.52 | 0.72 | 0.25 | 0.46 |
| 5 | 124358513 | rs11241783 | -1.482 | -1.576 | -1.894 | -0.986 | -0.043 | 0.094 | 0.094 | 0 | 0.58 | 0.91 | 0.66 | 0.74 | 0.43 | 0.6 |
| 5 | 124359018 | rs4836134 | 1.019 | -0.79 | -1.258 | -1.182 | -0.006 | 0.007 | 0.007 | 0 | 0.08 | 0.05 | 0.42 | 0.22 | 0.13 | 0.24 |

| | | | | | | | | | | | | | | | | |
|---|---|---|---|---|---|---|---|---|---|---|---|---|---|---|---|---|
| 5 | 124361588 | rs6869617 | -0.89 | -1.2 | -1.316 | -2.661 | 0.043 | 0.218 | 0.218 | 0.783 | 0.35 | 0.93 | 0.19 | 0.03 | 0.23 | 0.19 |
| 5 | 124361999 | rs6874416 | 0.191 | -1.293 | -0.477 | -2.557 | -0.018 | -0.02 | -0.02 | 0 | 0.07 | 0.26 | 0.02 | 0 | 0.01 | 0.02 |
| 5 | 124374180 | rs6872503 | -0.465 | -1.267 | -0.894 | -2.611 | 0.005 | -0.022 | -0.022 | 0 | 0.28 | 0.73 | 0.15 | 0.02 | 0.21 | 0.17 |
| 5 | 124376698 | rs6876152 | -0.785 | -1.331 | -1.065 | -2.676 | 0.037 | 0.053 | 0.053 | 1.283 | 0.33 | 0.78 | 0.16 | 0.02 | 0.21 | 0.19 |
| 5 | 124378785 | rs6595571 | -0.924 | -0.347 | -1.19 | -2.457 | -0.072 | -0.044 | -0.044 | 0 | 0.87 | 0.99 | 0.9 | 0.86 | 0.82 | 0.89 |
| 5 | 124386991 | rs11950485 | -1.147 | -1.252 | -0.202 | -2.888 | -0.044 | -0.09 | -0.09 | 0 | 0.18 | 0.6 | 0.08 | 0.01 | 0.06 | 0.07 |
| 5 | 124392917 | rs7444307 | -0.589 | -1.486 | -0.603 | -3.269 | 0.024 | 0.201 | 0.201 | 0 | 0.45 | 0.81 | 0.37 | 0.09 | 0.44 | 0.39 |
| 5 | 124397677 | rs6866873 | -0.582 | -1.483 | -0.603 | -3.267 | 0.023 | 0.209 | 0.209 | 0 | 0.45 | 0.82 | 0.37 | 0.09 | 0.43 | 0.38 |
| 5 | 124407926 | rs6872841 | 0.833 | -1.202 | -0.38 | -3.116 | -0.021 | 0.04 | 0.04 | 0.021 | 0.24 | 0.19 | 0.29 | 0.08 | 0.36 | 0.29 |
| 5 | 124411766 | rs6595573 | -0.599 | -1.189 | -0.448 | -3.075 | 0.008 | 0.137 | 0.137 | 0 | 0.42 | 0.8 | 0.37 | 0.09 | 0.42 | 0.37 |
| 5 | 124432527 | rs12152877 | 0.281 | -1.473 | -0.151 | -2.636 | 0.011 | 0.003 | 0.003 | 0.657 | 0.1 | 0.09 | 0.04 | 0 | 0.05 | 0.05 |
| 5 | 124436615 | rs4643951 | -0.771 | -1.288 | 0.018 | -2.749 | -0.007 | 0.012 | 0.012 | 0.076 | 0.78 | 0.8 | 0.74 | 0.92 | 0.67 | 0.77 |
| 5 | 124451809 | rs875331 | 0.654 | -1.501 | 0.183 | -2.808 | 0.033 | 0.071 | 0.071 | 0 | 0.57 | 0.28 | 0.67 | 0.92 | 0.62 | 0.68 |
| 5 | 124452279 | rs875332 | 0.604 | -1.463 | 0.293 | -2.793 | 0.019 | 0.036 | 0.036 | 0 | 0.58 | 0.27 | 0.67 | 0.91 | 0.62 | 0.67 |
| 5 | 124459547 | rs7735615 | 0.709 | -1.337 | 0.495 | -2.734 | -0.001 | -0.04 | -0.04 | 0 | 0.47 | 0.41 | 0.52 | 0.95 | 0.48 | 0.65 |
| 5 | 124471691 | rs7733868 | 0.278 | -1.199 | 0.501 | -2.616 | -0.02 | 0.044 | 0.044 | 0 | 0.33 | 0.24 | 0.36 | 0.05 | 0.48 | 0.27 |
| 5 | 124477401 | rs1550357 | 0.004 | -1.192 | 0.224 | -2.611 | -0.001 | -0.083 | -0.083 | 0 | 0.6 | 0.42 | 0.62 | 0.95 | 0.52 | 0.7 |
| 5 | 124480529 | rs1550363 | -0.043 | -1.287 | 0.284 | -2.672 | -0.019 | -0.048 | -0.048 | 1.806 | 0.84 | 0.76 | 0.94 | 1 | 0.93 | 0.92 |
| 5 | 124483173 | rs5016082 | 0.285 | -1.171 | 0.317 | -2.606 | 0.012 | -0.135 | -0.135 | 0 | 0.35 | 0.47 | 0.38 | 0.05 | 0.47 | 0.27 |
| 5 | 124485073 | rs7724689 | 0.147 | -1.122 | -0.014 | -2.589 | 0.009 | -0.109 | -0.109 | 1.892 | 0.35 | 0.51 | 0.38 | 0.05 | 0.47 | 0.27 |
| 5 | 124488884 | rs1030954 | -0.02 | -0.608 | -0.196 | -2.109 | -0.018 | -0.025 | -0.025 | 0 | 0.07 | 0.12 | 0.02 | 0 | 0.01 | 0.03 |
| 5 | 124488902 | rs11741331 | 0.419 | -0.677 | -0.318 | -2.147 | -0.024 | 0.167 | 0.167 | 0.211 | 0.45 | 0.19 | 0.52 | 0.92 | 0.45 | 0.62 |
| 5 | 124489336 | rs6868078 | -1.117 | -0.66 | 0.262 | -0.975 | -0.032 | -0.041 | -0.041 | 3.116 | 0.08 | 0.19 | 0.03 | 0 | 0.01 | 0.03 |
| 5 | 124490816 | rs1866543 | 0.506 | -0.679 | 0.222 | -1.2 | -0.052 | -0.129 | -0.129 | 0.015 | 0.79 | 0.58 | 0.91 | 0.97 | 0.88 | 0.94 |
| 5 | 124491364 | rs1016476 | -0.459 | 0.182 | 0.099 | -0.404 | 0.01 | 0.004 | 0.004 | 0 | 0.26 | 0.35 | 0.3 | 0.22 | 0.3 | 0.37 |
| 5 | 124491386 | rs1016475 | -0.569 | 0.19 | 0.157 | 0.011 | 0.009 | 0.033 | 0.033 | 0 | 0.19 | 0.27 | 0.27 | 0.31 | 0.21 | 0.29 |
| 5 | 124491958 | rs11241791 | 0.141 | 0.088 | -0.113 | 0.133 | -0.009 | 0.006 | 0.006 | 0 | 0.75 | 0.62 | 0.87 | 0.7 | 0.78 | 0.67 |
| 5 | 124500651 | rs4370270 | -0.644 | 0.151 | -0.002 | 0.373 | 0.003 | 0.007 | 0.007 | 0 | 0.46 | 0.42 | 0.44 | 0.34 | 0.55 | 0.62 |
| 5 | 124507426 | rs17152354 | 0.544 | -0.191 | -0.445 | 0.147 | -0.004 | -0.008 | -0.008 | 0 | 0.12 | 0.15 | 0.07 | 0.04 | 0.09 | 0.09 |
| 5 | 124510131 | rs4836157 | 0.57 | 0.051 | -0.638 | 0.041 | 0.006 | -0.01 | -0.01 | 0 | 0.37 | 0.25 | 0.47 | 0.43 | 0.35 | 0.21 |
| 5 | 124510520 | rs17457509 | 0.997 | -0.28 | -0.769 | -0.427 | -0.002 | 0.039 | 0.039 | 5.134 | 0.13 | 0.06 | 0.08 | 0 | 0.11 | 0.03 |
| 5 | 124512345 | rs11743924 | 1.514 | -0.302 | -0.686 | -0.24 | -0.025 | 0.02 | 0.02 | 0 | 0.12 | 0.04 | 0.07 | 0.09 | 0.17 | 0.18 |
| 5 | 124518252 | rs6595591 | 0.458 | -0.359 | -1.058 | -0.391 | -0.018 | 0.09 | 0.09 | 0.754 | 0.42 | 0.24 | 0.42 | 0.21 | 0.43 | 0.43 |
| 5 | 124520411 | rs6873604 | -0.402 | 0.365 | -0.851 | -0.128 | -0.004 | -0.01 | -0.01 | 0 | 0.64 | 0.46 | 0.52 | 0.36 | 0.65 | 0.73 |
| 5 | 124525027 | rs6892878 | 0.694 | 1.05 | -0.953 | 0.229 | -0.009 | 0.008 | 0.008 | 0 | 0.13 | 0.07 | 0.07 | 0.09 | 0.17 | 0.18 |
| 5 | 124526594 | rs9637914 | 0.299 | 0.533 | -1.24 | 1.284 | -0.015 | -0.043 | -0.043 | 1.461 | 0.1 | 0.04 | 0.27 | 0.29 | 0.12 | 0.05 |
| 5 | 124526673 | rs9327354 | -0.142 | 0.31 | -1.08 | 0.643 | 0 | 0.001 | 0.001 | 0 | 0.49 | 0.45 | 0.65 | 0.61 | 0.45 | 0.58 |
| 5 | 124526865 | rs17517907 | -0.128 | 0.13 | -0.944 | 0.182 | 0.003 | 0.004 | 0.004 | 0 | 0.26 | 0.24 | 0.16 | 0.09 | 0.25 | 0.15 |
| 5 | 124528073 | rs1550370 | 0.363 | -0.277 | -0.498 | 0.432 | -0.035 | 0.091 | 0.091 | 0 | 0.77 | 0.55 | 0.67 | 0.57 | 0.86 | 0.7 |
| 5 | 124529484 | rs6895024 | 0.219 | -0.253 | -0.89 | -0.471 | -0.007 | 0.092 | 0.092 | 0 | 0.45 | 0.25 | 0.39 | 0.21 | 0.47 | 0.34 |
| 5 | 124530606 | rs2119001 | -0.597 | -0.249 | -0.121 | -0.404 | -0.008 | -0.109 | -0.109 | 0.452 | 0.16 | 0.34 | 0.1 | 0.01 | 0.12 | 0.16 |
| 5 | 124530708 | rs17152394 | 0.822 | 0.115 | -0.486 | -0.293 | 0.021 | -0.008 | -0.008 | 0 | 0.16 | 0.22 | 0.17 | 0.12 | 0.23 | 0.14 |
| 5 | 124532423 | rs10519804 | 0.664 | 0.11 | -0.705 | -0.515 | -0.004 | 0.063 | 0.063 | 0 | 0.22 | 0.08 | 0.18 | 0.07 | 0.22 | 0.22 |
| 5 | 124534055 | rs920386 | -0.467 | -0.09 | -0.572 | -0.323 | 0.004 | 0.014 | 0.014 | 0 | 0.46 | 0.32 | 0.37 | 0.39 | 0.46 | 0.66 |
| 5 | 124534114 | rs920387 | -0.068 | -0.037 | -0.504 | -0.159 | -0.003 | 0.019 | 0.019 | 0 | 0.39 | 0.25 | 0.32 | 0.28 | 0.42 | 0.54 |
| 5 | 124534697 | rs10058654 | -0.182 | 0.129 | -0.345 | 0.007 | -0.008 | 0.007 | 0.007 | 1.086 | 0.07 | 0.15 | 0.05 | 0.11 | 0.05 | 0.07 |
| 5 | 124536079 | rs6863765 | 0.585 | 0.267 | -0.296 | 0.02 | 0.046 | 0.036 | 0.036 | 0 | 0.16 | 0.31 | 0.18 | 0.23 | 0.2 | 0.16 |
| 5 | 124538332 | rs1439576 | 1.13 | 0.023 | -0.191 | 0.167 | 0.055 | 0.097 | 0.097 | 0.076 | 0.06 | 0.22 | 0.03 | 0.12 | 0.05 | 0.01 |
| 5 | 124540711 | rs11241795 | 0.572 | -0.137 | -0.441 | 0.142 | 0.002 | -0.07 | -0.07 | 0.006 | 0.23 | 0.11 | 0.18 | 0.32 | 0.23 | 0.42 |
| 5 | 124541162 | rs11745996 | -0.244 | -0.168 | -0.082 | 0.173 | 0.002 | 0.023 | 0.023 | 0 | 0.73 | 0.59 | 0.58 | 0.63 | 0.72 | 0.7 |
| 5 | 124543797 | rs1439573 | 0.508 | -0.099 | -0.187 | 0.181 | 0.001 | -0.001 | -0.001 | 0 | 0.54 | 0.42 | 0.41 | 0.44 | 0.55 | 0.53 |
| 5 | 124544884 | rs10075641 | 0.45 | -0.163 | -0.176 | 0.14 | 0 | -0.001 | -0.001 | 0 | 0.56 | 0.45 | 0.43 | 0.54 | 0.55 | 0.58 |
| 5 | 124545732 | rs1439568 | -0.273 | 0.37 | -0.054 | -0.706 | 0.004 | -0.056 | -0.056 | 0 | 0.37 | 0.4 | 0.35 | 0.09 | 0.41 | 0.32 |
| 5 | 124553329 | rs6595594 | -0.322 | 0.72 | 0.118 | -0.302 | 0.003 | -0.006 | -0.006 | 0 | 0.26 | 0.38 | 0.23 | 0.2 | 0.26 | 0.35 |
| 5 | 124555847 | rs1898910 | 1.019 | 0.536 | 0.067 | -0.131 | 0 | -0.001 | -0.001 | 0 | 0.12 | 0.09 | 0.14 | 0.08 | 0.16 | 0.12 |
| 5 | 124556118 | rs13185379 | 0.679 | 0.461 | -0.02 | -0.293 | 0.004 | 0.178 | 0.178 | 0 | 0.48 | 0.22 | 0.39 | 0.19 | 0.5 | 0.38 |
| 5 | 124557411 | rs4836164 | -0.632 | 0.509 | 0.068 | -0.196 | -0.006 | 0.017 | 0.017 | 0.05 | 0.7 | 0.62 | 0.61 | 0.42 | 0.71 | 0.61 |
| 5 | 124559651 | rs1439564 | 0.524 | 0.682 | 0.043 | -0.205 | -0.003 | 0.16 | 0.16 | 0 | 0.46 | 0.22 | 0.4 | 0.13 | 0.51 | 0.28 |
| 5 | 124561232 | rs10214269 | -0.601 | 0.574 | 0.323 | 0.565 | -0.001 | 0.077 | 0.077 | 0 | 0.07 | 0.21 | 0.06 | 0.15 | 0.03 | 0.11 |
| 5 | 124563598 | rs1439612 | -1.169 | 0.506 | -0.376 | 0.577 | -0.021 | 0.028 | 0.028 | 0 | 0.11 | 0.21 | 0.06 | 0.19 | 0.03 | 0.21 |
| 5 | 124568300 | rs7715337 | 0.359 | 1.027 | -0.83 | 0.407 | -0.006 | 0.003 | 0.003 | 2.629 | 0.7 | 0.29 | 0.72 | 0.73 | 0.68 | 0.66 |
| 5 | 124569338 | rs1439609 | -0.094 | 0.469 | -1.067 | 0.741 | 0.013 | -0.053 | -0.053 | 0 | 0.73 | 0.83 | 0.7 | 0.52 | 0.73 | 0.61 |
| 5 | 124569471 | rs1439607 | 0.155 | 0.697 | -0.622 | 0.504 | 0.001 | 0.007 | 0.007 | 0.2 | 0.47 | 0.45 | 0.4 | 0.36 | 0.39 | 0.21 |
| 5 | 124573609 | rs1439605 | 0.653 | 0.352 | -0.566 | 1.089 | -0.001 | 0.006 | 0.006 | 0 | 0.72 | 0.39 | 0.68 | 0.56 | 0.73 | 0.76 |
| 5 | 124580839 | rs1118619 | 0.2 | -0.071 | -1.427 | 0.867 | 0.018 | -0.053 | -0.053 | 0 | 0.24 | 0.12 | 0.19 | 0.35 | 0.23 | 0.45 |
| 5 | 124581782 | rs719829 | -0.761 | -0.629 | -1.828 | 0.316 | 0.054 | -0.057 | -0.057 | 0.051 | 0.4 | 0.2 | 0.33 | 0.54 | 0.31 | 0.58 |
| 5 | 124585022 | rs12055008 | -0.347 | -0.632 | -1.26 | 0.423 | 0 | 0.116 | 0.116 | 0.03 | 0.77 | 0.5 | 0.7 | 0.53 | 0.76 | 0.81 |
| 5 | 124593173 | rs1347924 | 1.143 | -0.764 | -0.412 | 0.581 | 0.002 | 0.022 | 0.022 | 0 | 0.46 | 0.57 | 0.41 | 0.12 | 0.5 | 0.29 |
| 5 | 124593936 | rs11738947 | 0.856 | -0.989 | -0.466 | 0.889 | 0.012 | -0.041 | -0.041 | 0 | 0.21 | 0.15 | 0.33 | 0.56 | 0.21 | 0.49 |

| | | | | | | | | | | | | | | | | |
|---|---|---|---|---|---|---|---|---|---|---|---|---|---|---|---|---|
| 5 | 124594637 | rs2033841 | 0.468 | -0.986 | -0.322 | 0.897 | 0 | 0.011 | 0.011 | 0 | 0.76 | 0.56 | 0.84 | 0.77 | 0.74 | 0.86 |
| 5 | 124597571 | rs6880511 | 1.118 | -1.099 | -0.43 | 1.159 | 0.02 | -0.026 | -0.026 | 0 | 0.11 | 0.05 | 0.13 | 0.19 | 0.05 | 0.07 |
| 5 | 124598619 | rs17152498 | 0.514 | -1.061 | -0.383 | 1.155 | 0.013 | 0.075 | 0.075 | 0.075 | 0.09 | 0.27 | 0.04 | 0.1 | 0.05 | 0.06 |
| 5 | 124600022 | rs17152509 | -0.945 | -0.983 | -0.855 | 1.199 | 0.054 | 0.057 | 0.057 | 0 | 0.4 | 0.24 | 0.49 | 0.76 | 0.31 | 0.62 |
| 5 | 124608570 | rs17152551 | 0.267 | -0.872 | -0.85 | 1.231 | 0.018 | -0.015 | -0.015 | 0 | 0.14 | 0.06 | 0.14 | 0.18 | 0.08 | 0.07 |
| 5 | 124611833 | rs17152569 | 0.251 | -0.683 | -0.428 | 1.075 | -0.006 | 0.023 | 0.023 | 0.112 | 0.07 | 0.12 | 0.03 | 0.11 | 0.05 | 0.06 |
| 5 | 124615537 | rs1562125 | -0.207 | -0.833 | -0.516 | 1.097 | 0.046 | -0.068 | -0.068 | 2.858 | 0.32 | 0.17 | 0.28 | 0.51 | 0.24 | 0.51 |
| 5 | 124621156 | rs767177 | 0.236 | -0.463 | -0.339 | 0.911 | 0.003 | -0.008 | -0.008 | 0 | 0.27 | 0.2 | 0.27 | 0.43 | 0.23 | 0.44 |
| 5 | 124622296 | rs11744285 | 1.461 | -0.115 | -0.29 | 0.326 | -0.025 | 0.068 | 0.068 | 0 | 0.38 | 0.27 | 0.29 | 0.02 | 0.43 | 0.2 |
| 5 | 124622536 | rs7722013 | 0.354 | 0.157 | 0.146 | 0.799 | 0.004 | 0.082 | 0.082 | 0 | 0.77 | 0.55 | 0.75 | 0.63 | 0.79 | 0.74 |
| 5 | 124624902 | rs1530121 | -0.299 | 0.201 | 0.109 | 0.825 | 0.005 | -0.014 | -0.014 | 0 | 0.83 | 0.64 | 0.92 | 0.88 | 0.84 | 0.81 |
| 5 | 124625976 | rs7710220 | 0.552 | 0.458 | -0.19 | 0.763 | 0.007 | 0.023 | 0.023 | 0 | 0.06 | 0.09 | 0.17 | 0.18 | 0.1 | 0.05 |
| 5 | 124626392 | rs1371370 | 0.555 | 0.149 | -0.339 | 0.372 | -0.005 | 0.034 | 0.034 | 0 | 0.47 | 0.37 | 0.46 | 0.27 | 0.48 | 0.26 |
| 5 | 124627321 | rs10519810 | 0.515 | -0.237 | -0.544 | 0.4 | -0.006 | 0.013 | 0.013 | 0.375 | 0.35 | 0.38 | 0.27 | 0.51 | 0.29 | 0.59 |
| 5 | 124629015 | rs17521821 | 0.808 | 0.477 | -1.231 | -0.436 | -0.027 | 0.07 | 0.07 | 0 | 0.19 | 0.06 | 0.16 | 0.03 | 0.23 | 0.07 |
| 5 | 124631460 | rs1439598 | -0.318 | 1.171 | -1.365 | -0.812 | -0.041 | 0.058 | 0.058 | 0 | 0.74 | 0.52 | 0.71 | 0.66 | 0.83 | 0.78 |
| 5 | 124635329 | rs11740072 | -0.151 | 0.955 | -1.477 | -0.91 | -0.066 | 0.137 | 0.137 | 0 | 0.36 | 0.13 | 0.45 | 0.13 | 0.56 | 0.19 |
| 5 | 124636443 | rs17152620 | 1.31 | 0.826 | -1.38 | -0.941 | -0.012 | 0.01 | 0.01 | 0.314 | 0.36 | 0.34 | 0.24 | 0.52 | 0.26 | 0.59 |
| 5 | 124639468 | rs1444649 | -0.568 | 0.578 | -1.195 | -0.905 | -0.025 | 0.066 | 0.066 | 0 | 0.76 | 0.58 | 0.71 | 0.65 | 0.83 | 0.81 |
| 5 | 124643194 | rs13189771 | -0.495 | 0.815 | -1.345 | -0.94 | -0.042 | 0.139 | 0.139 | 0 | 0.39 | 0.2 | 0.47 | 0.13 | 0.55 | 0.19 |
| 5 | 124644516 | rs4640788 | 0.94 | 0.992 | -1.205 | -0.725 | -0.032 | -0.013 | -0.013 | 0 | 0.1 | 0.06 | 0.2 | 0.13 | 0.22 | 0.05 |
| 5 | 124655412 | rs1371383 | 0.967 | 0.757 | -1.271 | -0.719 | -0.056 | -0.006 | -0.006 | 4.419 | 0.14 | 0.07 | 0.22 | 0.13 | 0.26 | 0.06 |
| 5 | 124655934 | rs2408382 | -0.561 | 0.597 | -1.04 | -0.701 | -0.018 | 0.11 | 0.11 | 0 | 0.81 | 0.54 | 0.72 | 0.66 | 0.86 | 0.82 |
| 5 | 124660883 | rs10519815 | 0.512 | 0.415 | -0.554 | -0.633 | 0.001 | 0 | 0 | 0.534 | 0.55 | 0.53 | 0.43 | 0.53 | 0.51 | 0.69 |
| 5 | 124663527 | rs17523117 | 1.241 | 0.488 | -0.762 | -0.87 | -0.045 | 0.078 | 0.078 | 0 | 0.17 | 0.04 | 0.17 | 0 | 0.23 | 0.08 |
| 5 | 124666245 | rs7701571 | -0.696 | 0.421 | -0.415 | -0.592 | 0.009 | 0.108 | 0.108 | 0 | 0.83 | 0.7 | 0.73 | 0.54 | 0.78 | 0.84 |
| 5 | 124672949 | rs1822758 | 1.457 | 0.355 | -0.223 | -0.643 | 0.01 | -0.005 | -0.005 | 0 | 0.36 | 0.33 | 0.24 | 0.4 | 0.26 | 0.59 |
| 5 | 124676033 | rs13354270 | 0.065 | 0.51 | -0.497 | -0.658 | -0.054 | 0.085 | 0.085 | 0 | 0.34 | 0.15 | 0.39 | 0.13 | 0.42 | 0.15 |
| 5 | 124676142 | rs17463846 | 1.378 | 0.475 | -0.54 | -0.673 | -0.058 | -0.004 | -0.004 | 0.016 | 0.16 | 0.03 | 0.22 | 0.12 | 0.19 | 0.07 |
| 5 | 124676698 | rs2678180 | 0.711 | 0.673 | -0.085 | -0.363 | 0.003 | 0.001 | 0.001 | 0 | 0.57 | 0.57 | 0.48 | 0.54 | 0.46 | 0.69 |
| 5 | 124676830 | rs2678181 | 0.112 | 0.945 | -0.068 | -0.119 | 0.004 | 0.039 | 0.039 | 0 | 0.74 | 0.67 | 0.65 | 0.55 | 0.69 | 0.76 |
| 5 | 124679857 | rs1444653 | 0.025 | 0.862 | -0.607 | -0.272 | 0.013 | 0.024 | 0.024 | 0 | 0.71 | 0.7 | 0.65 | 0.56 | 0.6 | 0.81 |
| 5 | 124681086 | rs895042 | -1.062 | 0.878 | -0.616 | -0.221 | 0.025 | 0.151 | 0.151 | 1.236 | 0.87 | 0.72 | 0.8 | 0.68 | 0.83 | 0.88 |
| 5 | 124683175 | rs2678201 | 1.619 | 0.108 | -0.78 | -0.294 | -0.007 | 0.031 | 0.031 | 0 | 0.22 | 0.33 | 0.29 | 0.38 | 0.2 | 0.42 |
| 5 | 124685311 | rs17523724 | 1.521 | -0.003 | -0.977 | -0.514 | -0.022 | 0.116 | 0.116 | 0.937 | 0.21 | 0.04 | 0.08 | 0.01 | 0.18 | 0.15 |
| 5 | 124685783 | rs2678200 | 1.002 | 0.005 | -0.984 | -0.548 | 0.003 | -0.007 | -0.007 | 0 | 0.31 | 0.32 | 0.36 | 0.39 | 0.28 | 0.49 |
| 5 | 124686839 | rs6887011 | 0.831 | 0.098 | -0.928 | -0.537 | 0.008 | 0.009 | 0.009 | 0 | 0.3 | 0.29 | 0.42 | 0.4 | 0.37 | 0.2 |
| 5 | 124690313 | rs2678196 | -0.09 | -0.235 | -0.927 | -0.584 | -0.001 | -0.002 | -0.002 | 0.323 | 0.59 | 0.45 | 0.6 | 0.52 | 0.53 | 0.7 |
| 5 | 124692073 | rs7701782 | 1.473 | 0.105 | -0.92 | -0.515 | 0.006 | 0.003 | 0.003 | 1.101 | 0.24 | 0.24 | 0.27 | 0.27 | 0.32 | 0.14 |
| 5 | 124704493 | rs10063763 | 0.556 | -0.125 | -1.629 | -0.458 | -0.05 | 0.075 | 0.075 | 0.874 | 0.28 | 0.07 | 0.25 | 0.14 | 0.26 | 0.23 |
| 5 | 124706208 | rs1517808 | 1.299 | 0.441 | -0.798 | 0.493 | 0.016 | 0.026 | 0.026 | 0 | 0.44 | 0.32 | 0.36 | 0.38 | 0.39 | 0.58 |
| 5 | 124714338 | rs6864886 | -1.03 | 0.684 | -0.383 | -0.058 | -0.003 | 0.009 | 0.009 | 0.632 | 0.42 | 0.56 | 0.39 | 0.31 | 0.45 | 0.3 |
| 5 | 124723432 | rs925894 | -0.148 | 1.065 | 0.139 | 0.687 | 0.033 | -0.009 | -0.009 | 0.296 | 0.28 | 0.33 | 0.29 | 0.24 | 0.36 | 0.16 |
| 5 | 124730924 | rs12653164 | 0.49 | 1.173 | -0.223 | 0.77 | -0.002 | -0.004 | -0.004 | 0 | 0.05 | 0.16 | 0.04 | 0.04 | 0.04 | 0.05 |
| 5 | 124737623 | rs4836190 | -0.274 | 1.289 | -0.934 | 0.885 | 0.014 | -0.016 | -0.016 | 0.433 | 0.35 | 0.51 | 0.33 | 0.28 | 0.41 | 0.25 |
| 5 | 124751860 | rs6875251 | 0.406 | 0.981 | -1.561 | 0.253 | -0.034 | 0.1 | 0.1 | 0 | 0.16 | 0.02 | 0.08 | 0 | 0.15 | 0.06 |
| 5 | 124754302 | rs13180857 | 0.464 | 0.97 | -2.576 | 0.069 | -0.044 | 0.011 | 0.011 | 0 | 0.41 | 0.13 | 0.38 | 0.6 | 0.42 | 0.52 |
| 5 | 124769855 | rs17152760 | 0.488 | 0.959 | -3.037 | 0.271 | -0.019 | -0.101 | -0.101 | 0.223 | 0.2 | 0.07 | 0.2 | 0.51 | 0.21 | 0.36 |
| 5 | 124807122 | rs1517805 | 0.714 | 1.134 | -2.167 | -0.238 | -0.028 | -0.012 | -0.012 | 0.612 | 0.05 | 0.01 | 0.14 | 0.05 | 0.11 | 0.11 |
| 5 | 124807348 | rs1400870 | 0.348 | 0.822 | -1.82 | -0.841 | -0.013 | 0.005 | 0.005 | 0 | 0.37 | 0.31 | 0.52 | 0.38 | 0.44 | 0.35 |
| 5 | 124811582 | rs1826262 | 0.403 | 0.318 | -1.573 | -0.638 | 0.028 | 0.03 | 0.03 | 0 | 0.54 | 0.31 | 0.6 | 0.42 | 0.6 | 0.4 |
| 5 | 124815907 | rs13187817 | 1.431 | 0.282 | -1.72 | -0.578 | -0.004 | 0.063 | 0.063 | 0 | 0.15 | 0.02 | 0.06 | 0.05 | 0.11 | 0.13 |
| 5 | 124817800 | rs2046908 | -0.196 | 0.152 | -1.264 | -0.65 | 0 | -0.009 | -0.009 | 0.165 | 0.67 | 0.77 | 0.68 | 0.39 | 0.69 | 0.51 |
| 5 | 124820042 | rs4240391 | -1.551 | 0.187 | -1.439 | 0.268 | -0.043 | -0.037 | -0.037 | 0.928 | 0.92 | 0.99 | 0.79 | 0.7 | 0.87 | 0.72 |
| 5 | 124820663 | rs1491070 | 1.258 | 0.201 | -1.415 | 0.09 | -0.007 | -0.033 | -0.033 | 0.491 | 0.17 | 0.11 | 0.28 | 0.36 | 0.17 | 0.11 |
| 5 | 124824336 | rs17152833 | 1.74 | 0.148 | -1.404 | 0.234 | 0.017 | -0.019 | -0.019 | 0.043 | 0.11 | 0.07 | 0.03 | 0.19 | 0.08 | 0.1 |
| 5 | 124825014 | rs13340392 | 1.362 | 0.395 | -1.15 | 0.191 | 0.014 | -0.014 | -0.014 | 0 | 0.25 | 0.19 | 0.11 | 0.31 | 0.18 | 0.22 |
| 5 | 124826243 | rs331702 | 1.32 | 0.399 | -1.098 | 0.074 | -0.028 | 0.105 | 0.105 | 0 | 0.37 | 0.21 | 0.32 | 0.06 | 0.44 | 0.29 |
| 5 | 124828696 | rs331696 | 0.987 | 0.497 | -0.993 | 0.194 | -0.002 | -0.011 | -0.011 | 0 | 0.18 | 0.16 | 0.28 | 0.35 | 0.16 | 0.11 |
| 5 | 124832592 | rs230736 | 0.668 | 0.467 | -0.699 | 0.18 | -0.001 | 0.017 | 0.017 | 0 | 0.19 | 0.23 | 0.28 | 0.34 | 0.16 | 0.12 |
| 5 | 124843659 | rs4428402 | -0.843 | 0.261 | -0.602 | 0.172 | 0.048 | 0.093 | 0.093 | 0.786 | 0.86 | 0.73 | 0.74 | 0.7 | 0.79 | 0.62 |
| 5 | 124844266 | rs331757 | 0.053 | -0.099 | -0.699 | 0.477 | -0.013 | 0.021 | 0.021 | 0 | 0.23 | 0.31 | 0.32 | 0.35 | 0.16 | 0.12 |
| 5 | 124847603 | rs331755 | -0.489 | 0.068 | -0.936 | 0.357 | -0.005 | -0.003 | -0.003 | 0 | 0.47 | 0.49 | 0.42 | 0.63 | 0.34 | 0.33 |
| 5 | 124848906 | rs17468244 | 0.72 | 0.939 | -1.495 | 0.644 | -0.027 | -0.01 | -0.01 | 0 | 0.05 | 0.01 | 0.13 | 0.05 | 0.12 | 0.11 |
| 5 | 124849125 | rs17152861 | 0.581 | 0.54 | -1.301 | 0.845 | -0.046 | -0.084 | -0.084 | 0.266 | 0.15 | 0.01 | 0.16 | 0.29 | 0.15 | 0.1 |
| 5 | 124853564 | rs331744 | -0.122 | 0.489 | -0.857 | 0.576 | 0.012 | 0.014 | 0.014 | 0 | 0.49 | 0.45 | 0.43 | 0.64 | 0.4 | 0.48 |
| 5 | 124854425 | rs10041548 | -0.254 | 1.134 | -0.321 | 0.786 | 0 | -0.001 | -0.001 | 0 | 0.39 | 0.38 | 0.37 | 0.36 | 0.29 | 0.5 |
| 5 | 124856792 | rs7712191 | 0.138 | 1.385 | -0.1 | 1.017 | -0.007 | -0.005 | -0.005 | 0.009 | 0.77 | 0.8 | 0.56 | 0.56 | 0.65 | 0.82 |
| 5 | 124858464 | rs331785 | -0.642 | 1.634 | -0.129 | 0.969 | -0.017 | -0.055 | -0.055 | 0.307 | 0.84 | 0.92 | 0.65 | 0.57 | 0.81 | 0.84 |

| | | | | | | | | | | | | | | | | |
|---|---|---|---|---|---|---|---|---|---|---|---|---|---|---|---|---|
| 5 | 124863567 | rs2125139 | -0.437 | 1.422 | 0.328 | 0.577 | -0.002 | 0.008 | 0.008 | 0.164 | 0.47 | 0.56 | 0.42 | 0.33 | 0.47 | 0.51 |
| 5 | 124866783 | rs331743 | -0.77 | 1.342 | 0.359 | 0.425 | 0.032 | 0.011 | 0.011 | 0 | 0.31 | 0.5 | 0.35 | 0.32 | 0.35 | 0.47 |
| 5 | 124871683 | rs331732 | -1.027 | 1.246 | 0.309 | 0.417 | -0.005 | -0.009 | -0.009 | 0.003 | 0.83 | 0.88 | 0.65 | 0.57 | 0.81 | 0.84 |
| 5 | 124875058 | rs2408428 | 0.155 | 1.292 | 0.72 | 0.428 | -0.007 | 0.011 | 0.011 | 0 | 0.22 | 0.21 | 0.23 | 0.3 | 0.13 | 0.45 |
| 5 | 124877035 | rs331728 | -0.655 | 1.479 | 0.761 | 0.39 | 0.124 | -0.023 | -0.023 | 0 | 0.09 | 0.27 | 0.11 | 0.03 | 0.2 | 0.03 |
| 5 | 124877051 | rs1117076 | 0.326 | 1.38 | 0.621 | 0.384 | 0.001 | -0.003 | -0.003 | 0 | 0.19 | 0.19 | 0.22 | 0.3 | 0.13 | 0.43 |
| 5 | 124880974 | rs331726 | 0.847 | 1.369 | 0.596 | -0.09 | 0.038 | 0.027 | 0.027 | 0 | 0.9 | 0.82 | 0.81 | 0.85 | 0.82 | 0.92 |
| 5 | 124881304 | rs331723 | 0.342 | 1.383 | 0.498 | -0.139 | 0.01 | -0.01 | -0.01 | 0 | 0.06 | 0.07 | 0.09 | 0 | 0.17 | 0.02 |
| 5 | 124885971 | rs331773 | 0.03 | 1.228 | 0.493 | -0.139 | -0.01 | -0.003 | -0.003 | 0 | 0.67 | 0.81 | 0.53 | 0.38 | 0.69 | 0.47 |
| 5 | 124890054 | rs331706 | -0.855 | 1.497 | 0.575 | -0.084 | 0.068 | 0.06 | 0.06 | 0 | 0.08 | 0.22 | 0.17 | 0.12 | 0.15 | 0.07 |
| 5 | 124896039 | rs331717 | 0.695 | 1.217 | 0.468 | -0.137 | 0.004 | -0.029 | -0.029 | 0.121 | 0.23 | 0.26 | 0.21 | 0.04 | 0.29 | 0.08 |
| 5 | 124900197 | rs331721 | -0.304 | 1.173 | 0.13 | 0.015 | -0.013 | -0.015 | -0.015 | 0 | 0.77 | 0.83 | 0.55 | 0.37 | 0.72 | 0.51 |
| 5 | 124910047 | rs330338 | 0.091 | 1.202 | 0.186 | 0.016 | -0.013 | -0.017 | -0.017 | 2.274 | 0.23 | 0.18 | 0.46 | 0.61 | 0.28 | 0.48 |
| 5 | 124910196 | rs330339 | -1.284 | 0.967 | 0.275 | -0.078 | 0.014 | 0.079 | 0.079 | 0 | 0.48 | 0.63 | 0.61 | 0.79 | 0.4 | 0.68 |
| 5 | 124914520 | rs330343 | 0.379 | 1.037 | 0.207 | 0.145 | -0.008 | -0.013 | -0.013 | 0 | 0.21 | 0.17 | 0.44 | 0.57 | 0.25 | 0.45 |
| 5 | 124916977 | rs12651765 | -0.25 | 1.088 | 0.379 | 0.146 | -0.058 | -0.004 | -0.004 | 0 | 0.26 | 0.48 | 0.16 | 0.2 | 0.13 | 0.21 |
| 5 | 124927547 | rs330349 | -0.89 | 1.112 | 0.328 | 0.062 | 0 | -0.005 | -0.005 | 0 | 0.46 | 0.5 | 0.57 | 0.69 | 0.36 | 0.65 |
| 5 | 124933851 | rs10044325 | -0.624 | 1.071 | 0.378 | 0.08 | -0.03 | 0.049 | 0.049 | 0 | 0.07 | 0.16 | 0.07 | 0.1 | 0.01 | 0.08 |
| 5 | 124937427 | rs330354 | -0.533 | 1.162 | 0.225 | 0.111 | 0 | 0.004 | 0.004 | 0 | 0.39 | 0.35 | 0.5 | 0.59 | 0.35 | 0.57 |
| 5 | 124946532 | rs1490996 | -0.709 | 1.36 | -0.265 | 0.016 | -0.006 | 0.013 | 0.013 | 0.211 | 0.73 | 0.77 | 0.84 | 0.94 | 0.71 | 0.89 |
| 5 | 124958205 | rs10037649 | 0.958 | 0.641 | -0.631 | 0.032 | -0.024 | -0.026 | -0.026 | 0.058 | 0.18 | 0.22 | 0.12 | 0.06 | 0.09 | 0.14 |
| 5 | 124963890 | rs10213918 | 1.071 | 1.222 | -1.477 | 0.268 | -0.002 | 0.015 | 0.015 | 0.099 | 0.3 | 0.21 | 0.27 | 0.33 | 0.34 | 0.29 |
| 5 | 124964532 | rs10214334 | 1.257 | 1.223 | -1.829 | -0.137 | 0.002 | 0.047 | 0.047 | 0.067 | 0.28 | 0.14 | 0.23 | 0.23 | 0.31 | 0.28 |
| 5 | 124969756 | rs1490974 | 1.181 | 2.28 | -1.411 | 0.722 | 0.023 | 0.032 | 0.032 | 0 | 0.54 | 0.62 | 0.42 | 0.32 | 0.42 | 0.52 |
| 5 | 124974976 | rs7702671 | 0.118 | 2.357 | -1.762 | 0.288 | -0.088 | -0.055 | -0.055 | 0.64 | 0.14 | 0.01 | 0.28 | 0.18 | 0.22 | 0.09 |
| 5 | 124977016 | rs1602402 | -0.114 | 1.168 | -1.246 | -0.131 | 0 | 0.026 | 0.026 | 1.193 | 0.8 | 0.87 | 0.87 | 1 | 0.77 | 0.94 |
| 5 | 124989000 | rs7736175 | 0.341 | 1.134 | -1.156 | 0.339 | 0.017 | 0.029 | 0.029 | 0 | 0.55 | 0.64 | 0.39 | 0.34 | 0.44 | 0.54 |
| 5 | 124989579 | rs6595621 | 1.031 | 2.304 | -0.498 | 0.149 | 0.081 | 0.006 | 0.006 | 0.758 | 0.9 | 0.77 | 0.82 | 0.93 | 0.76 | 0.9 |
| 5 | 124993133 | rs10057318 | 0.192 | 2.22 | -1.325 | -0.515 | -0.07 | 0.05 | 0.05 | 0 | 0.16 | 0.02 | 0.15 | 0.04 | 0.17 | 0.14 |
| 5 | 124996496 | rs2172102 | 0.591 | 2.27 | -1.58 | 0.096 | -0.015 | -0.019 | -0.019 | 0 | 0.19 | 0.12 | 0.26 | 0.26 | 0.21 | 0.14 |
| 5 | 125001052 | rs10085075 | 0.067 | 1.651 | -0.902 | -0.426 | 0 | -0.002 | -0.002 | 0 | 0.71 | 0.44 | 0.68 | 0.73 | 0.71 | 0.78 |
| 5 | 125003197 | rs10059503 | 0.47 | 1.043 | -1.164 | -0.765 | 0 | 0.14 | 0.14 | 4.85 | 0.33 | 0.14 | 0.22 | 0.04 | 0.26 | 0.16 |
| 5 | 125007308 | rs4836202 | 0.33 | 0.537 | -0.58 | -0.096 | -0.027 | 0.012 | 0.012 | 0.225 | 0.3 | 0.39 | 0.29 | 0.33 | 0.2 | 0.33 |
| 5 | 125007623 | rs902596 | 0.005 | 0.224 | -0.116 | -0.18 | -0.002 | 0.002 | 0.002 | 0 | 0.84 | 0.77 | 0.89 | 0.88 | 0.81 | 0.87 |
| 5 | 125008619 | rs13358753 | 1.36 | 0.301 | -0.103 | -0.172 | -0.023 | 0.012 | 0.012 | 0.13 | 0.33 | 0.2 | 0.35 | 0.32 | 0.41 | 0.4 |
| 5 | 125011212 | rs1994665 | 1.588 | 0.524 | -0.331 | -0.057 | -0.022 | -0.027 | -0.027 | 0 | 0.16 | 0.01 | 0.21 | 0.16 | 0.13 | 0.09 |
| 5 | 125014143 | rs9327364 | 0.317 | 0.502 | -0.22 | -0.037 | 0 | -0.001 | -0.001 | 0 | 0.63 | 0.64 | 0.64 | 0.64 | 0.61 | 0.72 |
| 5 | 125014575 | rs7700496 | 0.236 | 0.503 | -0.267 | -0.026 | -0.003 | -0.006 | -0.006 | 2.254 | 0.64 | 0.72 | 0.64 | 0.64 | 0.6 | 0.72 |
| 5 | 125017426 | rs10519829 | 0.006 | 0.384 | -0.319 | 0.03 | 0.042 | 0.091 | 0.091 | 0.214 | 0.49 | 0.3 | 0.43 | 0.31 | 0.45 | 0.3 |
| 5 | 125017930 | rs2408482 | 0.215 | 0.414 | -0.192 | 0.148 | -0.034 | 0.192 | 0.192 | 0 | 0.82 | 0.5 | 0.85 | 0.65 | 0.88 | 0.69 |
| 5 | 125018570 | rs2172101 | 1.196 | 0.412 | -0.238 | 0.166 | -0.025 | 0.006 | 0.006 | 0.026 | 0.33 | 0.21 | 0.42 | 0.31 | 0.41 | 0.39 |
| 5 | 125018595 | rs13161952 | 1.058 | 0.449 | -0.398 | 0.232 | -0.028 | -0.006 | -0.006 | 1.89 | 0.21 | 0.05 | 0.25 | 0.23 | 0.22 | 0.14 |
| 5 | 125019413 | rs2408483 | 0.529 | 0.447 | -0.378 | 0.226 | -0.008 | 0.007 | 0.007 | 0 | 0.62 | 0.53 | 0.6 | 0.42 | 0.67 | 0.55 |
| 5 | 125019633 | rs721194 | 2.617 | 0.119 | -0.795 | -0.1 | -0.07 | -0.073 | -0.073 | 0.036 | 0.18 | 0.01 | 0.25 | 0.27 | 0.22 | 0.23 |
| 5 | 125022679 | rs13183847 | 0.913 | 0.084 | -0.516 | 0.02 | -0.015 | 0.008 | 0.008 | 0 | 0.21 | 0.07 | 0.25 | 0.21 | 0.21 | 0.14 |
| 5 | 125023190 | rs11960504 | -0.574 | -0.028 | -0.156 | 0.324 | -0.022 | 0.184 | 0.184 | 0 | 0.17 | 0.44 | 0.15 | 0.34 | 0.12 | 0.31 |
| 5 | 125023618 | rs7702793 | 0.518 | -0.032 | -0.161 | 0.322 | -0.028 | 0.166 | 0.166 | 0 | 0.83 | 0.55 | 0.85 | 0.65 | 0.88 | 0.69 |
| 5 | 125031051 | rs6595623 | -0.759 | 0.508 | -0.479 | 0.561 | -0.001 | 0.002 | 0.002 | 0.063 | 0.44 | 0.53 | 0.52 | 0.62 | 0.51 | 0.7 |
| 5 | 125032464 | rs902594 | 0.73 | 0.481 | -0.436 | 0.543 | -0.003 | 0.017 | 0.017 | 0.023 | 0.55 | 0.36 | 0.47 | 0.39 | 0.49 | 0.3 |
| 5 | 125039645 | rs17153120 | 0.962 | 0.377 | -0.631 | 0.26 | 0.015 | -0.001 | -0.001 | 0.914 | 0.3 | 0.22 | 0.25 | 0.32 | 0.2 | 0.18 |
| 5 | 125039981 | rs1490989 | 0.211 | 0.648 | -0.366 | 0.398 | 0.01 | 0.015 | 0.015 | 0 | 0.33 | 0.38 | 0.38 | 0.37 | 0.37 | 0.31 |
| 5 | 125043721 | rs2029547 | -0.785 | 0.673 | -0.316 | 0.336 | 0.004 | -0.028 | -0.028 | 0 | 0.95 | 0.85 | 0.95 | 1 | 0.94 | 0.91 |
| 5 | 125044634 | rs2029543 | 1.756 | 0.662 | -0.367 | 0.315 | -0.001 | 0.072 | 0.072 | 0 | 0.25 | 0.14 | 0.22 | 0.03 | 0.3 | 0.11 |
| 5 | 125047409 | rs1490986 | 1.497 | 0.674 | -0.304 | 0.415 | 0.012 | 0.057 | 0.057 | 0.056 | 0.07 | 0.11 | 0.09 | 0.24 | 0.07 | 0.3 |
| 5 | 125051385 | rs7729374 | 0.219 | 0.65 | -0.341 | 0.278 | 0.01 | 0.018 | 0.018 | 0 | 0.32 | 0.38 | 0.38 | 0.39 | 0.37 | 0.31 |
| 5 | 125054298 | rs6885482 | 1.547 | 0.74 | -0.702 | 0.246 | 0.012 | 0.072 | 0.072 | 0 | 0.26 | 0.14 | 0.21 | 0.02 | 0.27 | 0.1 |
| 5 | 125055261 | rs11950923 | 1.567 | 0.677 | -0.685 | 0.265 | 0.006 | -0.004 | -0.004 | 0 | 0.31 | 0.23 | 0.27 | 0.35 | 0.24 | 0.2 |
| 5 | 125056207 | rs17485515 | 1.818 | 0.562 | -0.707 | 0.141 | 0.007 | 0.09 | 0.09 | 0.14 | 0.24 | 0.1 | 0.2 | 0.02 | 0.26 | 0.1 |
| 5 | 125057918 | rs17485634 | 1.852 | 0.788 | -0.97 | 0.191 | -0.048 | -0.033 | -0.033 | 0 | 0.08 | 0.01 | 0.13 | 0.09 | 0.15 | 0.06 |
| 5 | 125059228 | rs13154003 | 0.732 | 0.212 | -0.628 | 0.078 | 0.002 | 0.014 | 0.014 | 0 | 0.32 | 0.24 | 0.31 | 0.24 | 0.34 | 0.39 |
| 5 | 125059432 | rs13154204 | 1.194 | -0.302 | -0.932 | -0.369 | 0.002 | 0.121 | 0.121 | 0.003 | 0.26 | 0.09 | 0.21 | 0.03 | 0.27 | 0.1 |
| 5 | 125060278 | rs17153195 | 0.555 | 0.18 | -0.232 | 0.295 | 0.007 | -0.003 | -0.003 | 1.889 | 0.31 | 0.23 | 0.27 | 0.35 | 0.24 | 0.2 |
| 5 | 125080061 | rs162309 | 0.066 | 0.061 | 0.573 | 0.156 | 0.002 | 0.027 | 0.027 | 0 | 0.12 | 0.17 | 0.13 | 0.24 | 0.16 | 0.14 |
| 5 | 125080313 | rs329185 | 1.109 | -0.207 | 0.697 | -0.27 | -0.004 | 0.079 | 0.079 | 0 | 0.51 | 0.19 | 0.5 | 0.72 | 0.5 | 0.62 |
| 5 | 125082217 | rs330238 | -0.577 | -0.447 | 1.157 | -0.305 | -0.023 | -0.159 | -0.159 | 0 | 0.73 | 0.56 | 0.81 | 1 | 0.79 | 0.92 |
| 5 | 125082295 | rs329187 | 0.315 | -0.403 | 1.107 | -0.297 | 0.007 | 0.033 | 0.033 | 0 | 0.12 | 0.18 | 0.14 | 0.23 | 0.19 | 0.14 |
| 5 | 125084717 | rs13176171 | 0.407 | -0.463 | 1.073 | -0.34 | -0.059 | 0.112 | 0.112 | 0.435 | 0.27 | 0.04 | 0.29 | 0.05 | 0.26 | 0.14 |
| 5 | 125092500 | rs329195 | 1.241 | -0.399 | 1.052 | -0.304 | -0.004 | 0.073 | 0.073 | 0 | 0.52 | 0.19 | 0.51 | 0.73 | 0.51 | 0.68 |
| 5 | 125099344 | rs2910090 | 0.859 | -0.219 | 1.285 | -0.162 | 0.065 | 0.065 | 0.065 | 0.174 | 0.91 | 0.51 | 0.84 | 0.95 | 0.95 | 0.94 |

| | | | | | | | | | | | | | | | | |
|---|---|---|---|---|---|---|---|---|---|---|---|---|---|---|---|---|
| 5 | 125099527 | rs407595 | 0.994 | -0.217 | 1.045 | -0.162 | -0.001 | 0.005 | 0.005 | 0 | 0.78 | 0.25 | 0.71 | 0.73 | 0.79 | 0.81 |
| 5 | 125103347 | rs329162 | -1.493 | -0.216 | 1.043 | -0.156 | -0.001 | 0 | 0 | 0.012 | 0.23 | 0.76 | 0.32 | 0.29 | 0.26 | 0.2 |
| 5 | 125111363 | rs17153274 | 0.526 | -0.179 | 1.127 | -0.168 | 0.028 | 0.029 | 0.029 | 0 | 0.06 | 0.19 | 0.14 | 0.05 | 0.05 | 0.06 |
| 5 | 125119750 | rs228869 | -0.737 | -0.526 | 1.028 | -0.365 | -0.005 | -0.023 | -0.023 | 2.223 | 0.39 | 0.41 | 0.35 | 0.23 | 0.41 | 0.32 |
| 5 | 125121113 | rs2431597 | 0.168 | -0.292 | 0.257 | -0.153 | 0 | 0.007 | 0.007 | 0 | 0.13 | 0.14 | 0.15 | 0.24 | 0.19 | 0.14 |
| 5 | 125124027 | rs17153316 | 0.447 | -0.245 | -0.304 | -0.19 | 0.012 | 0.025 | 0.025 | 0 | 0.26 | 0.19 | 0.19 | 0 | 0.24 | 0.14 |
| 5 | 125124150 | rs329154 | -0.865 | -0.128 | 0.44 | 1.002 | 0.017 | 0.095 | 0.095 | 0 | 0.17 | 0.51 | 0.18 | 0.24 | 0.2 | 0.15 |
| 5 | 125127240 | rs2434572 | 0.422 | -0.895 | -0.172 | 1.139 | -0.026 | 0.019 | 0.019 | 0 | 0.68 | 0.41 | 0.67 | 0.57 | 0.83 | 0.55 |
| 5 | 125127876 | rs2431598 | 0.829 | -1.269 | 0.044 | 1.525 | -0.04 | 0.036 | 0.036 | 0 | 0.8 | 0.73 | 0.88 | 0.7 | 0.91 | 0.75 |
| 5 | 125130065 | rs2161342 | -0.389 | -0.427 | -0.033 | 1.384 | 0.014 | 0.012 | 0.012 | 0 | 0.12 | 0.16 | 0.25 | 0.2 | 0.18 | 0.11 |
| 5 | 125131156 | rs329175 | -0.206 | -0.319 | -0.074 | 0.931 | 0 | 0 | 0 | 0 | 0.55 | 0.53 | 0.58 | 0.42 | 0.61 | 0.42 |
| 5 | 125132831 | rs329173 | 0.776 | -0.231 | -0.109 | 0.585 | 0.003 | 0.005 | 0.005 | 0.062 | 0.27 | 0.17 | 0.22 | 0.22 | 0.3 | 0.24 |
| 5 | 125133971 | rs329168 | -0.18 | -0.306 | 0.005 | 0.525 | -0.001 | 0.013 | 0.013 | 0 | 0.41 | 0.37 | 0.34 | 0.23 | 0.43 | 0.33 |
| 5 | 125134363 | rs11958977 | 0.652 | -0.334 | 0.038 | 0.551 | 0.001 | -0.054 | -0.054 | 0.016 | 0.14 | 0.2 | 0.12 | 0 | 0.13 | 0.08 |
| 5 | 125134665 | rs328992 | 0.194 | -0.599 | -0.274 | 0.797 | 0.014 | 0.021 | 0.021 | 0 | 0.13 | 0.02 | 0.06 | 0.05 | 0.08 | 0.17 |
| 5 | 125137593 | rs388910 | 0.398 | -0.297 | -0.009 | 1.238 | 0.004 | -0.05 | -0.05 | 0 | 0.81 | 0.88 | 0.76 | 0.61 | 0.84 | 0.63 |
| 5 | 125139660 | rs1432338 | 0.841 | -0.461 | 0.163 | 1.042 | -0.022 | 0.052 | 0.052 | 0.451 | 0.73 | 0.4 | 0.71 | 0.43 | 0.8 | 0.44 |
| 5 | 125145154 | rs17153375 | 0.743 | -0.371 | -0.201 | 0.684 | -0.006 | 0.049 | 0.049 | 0 | 0.33 | 0.2 | 0.39 | 0.2 | 0.36 | 0.17 |
| 5 | 125147841 | rs327826 | -0.389 | -0.368 | 0.115 | 0.678 | 0.017 | 0.028 | 0.028 | 0 | 0.51 | 0.72 | 0.5 | 0.4 | 0.46 | 0.43 |
| 5 | 125154277 | rs179241 | -0.162 | -0.196 | 0.177 | 0.353 | -0.026 | -0.065 | -0.065 | 4.052 | 0.15 | 0.44 | 0.08 | 0.02 | 0.08 | 0.11 |
| 5 | 125159259 | rs327838 | 0.154 | -0.031 | -0.308 | 0.42 | 0.001 | 0.002 | 0.002 | 0 | 0.51 | 0.49 | 0.4 | 0.27 | 0.54 | 0.52 |
| 5 | 125160316 | rs985130 | 0.101 | 0.239 | -0.502 | 0.952 | -0.023 | -0.025 | -0.025 | 0 | 0.05 | 0 | 0.04 | 0.17 | 0.08 | 0.17 |
| 5 | 125161198 | rs1991488 | -0.039 | 0.697 | -0.441 | 0.558 | 0.002 | 0.017 | 0.017 | 0 | 0.49 | 0.49 | 0.58 | 0.33 | 0.55 | 0.36 |
| 5 | 125162446 | rs10519853 | 1.007 | 0.922 | -0.568 | 0.557 | 0.046 | 0.061 | 0.061 | 0.469 | 0.31 | 0.16 | 0.34 | 0.19 | 0.26 | 0.13 |
| 5 | 125166979 | rs12519875 | -0.167 | 0.861 | -0.542 | 0.635 | 0.008 | 0.076 | 0.076 | 0 | 0.5 | 0.65 | 0.58 | 0.34 | 0.55 | 0.36 |
| 5 | 125172583 | rs327832 | 0.533 | 0.78 | -0.843 | 0.486 | 0.007 | -0.003 | -0.003 | 0 | 0.42 | 0.21 | 0.3 | 0.34 | 0.36 | 0.4 |
| 5 | 125188931 | rs401425 | -0.552 | 0.556 | -0.902 | 0.229 | 0.002 | -0.031 | -0.031 | 3.919 | 0.84 | 0.91 | 0.85 | 0.68 | 0.87 | 0.76 |
| 5 | 125189406 | rs411361 | -0.294 | 0.241 | -1.016 | 0.004 | 0.001 | 0.012 | 0.012 | 0 | 0.57 | 0.63 | 0.59 | 0.29 | 0.56 | 0.41 |
| 5 | 125189968 | rs6895419 | 1.001 | 0.11 | -1.088 | -0.049 | 0 | -0.006 | -0.006 | 1.226 | 0.33 | 0.28 | 0.29 | 0.37 | 0.35 | 0.35 |
| 5 | 125197317 | rs13354717 | 0.756 | 0.109 | -0.566 | -0.124 | -0.005 | 0.003 | 0.003 | 1.125 | 0.27 | 0.21 | 0.27 | 0.14 | 0.35 | 0.26 |
| 5 | 125198949 | rs432111 | -0.083 | 0.08 | -0.404 | -0.191 | 0.001 | -0.005 | -0.005 | 0 | 0.39 | 0.32 | 0.31 | 0.17 | 0.42 | 0.35 |
| 5 | 125199582 | rs7703325 | -1.043 | 0.085 | -0.486 | -0.141 | -0.003 | 0.003 | 0.003 | 0 | 0.73 | 0.69 | 0.6 | 0.56 | 0.75 | 0.7 |
| 5 | 125199923 | rs1035954 | -1.133 | 0.095 | -0.483 | -0.136 | -0.001 | 0.018 | 0.018 | 0 | 0.92 | 0.91 | 0.88 | 0.68 | 0.92 | 0.76 |
| 5 | 125203280 | rs389333 | -1.133 | 0.095 | -0.483 | -0.136 | -0.001 | 0.018 | 0.018 | 0 | 0.92 | 0.91 | 0.88 | 0.68 | 0.92 | 0.76 |
| 5 | 125203610 | rs13172859 | 2.091 | 0.053 | -0.487 | -0.157 | -0.008 | -0.008 | -0.008 | 0 | 0.19 | 0.23 | 0.29 | 0.12 | 0.17 | 0.06 |
| 5 | 125204801 | rs1946598 | 1.42 | 0.034 | -0.159 | -0.137 | 0 | 0.003 | 0.003 | 0 | 0.29 | 0.34 | 0.26 | 0.3 | 0.28 | 0.3 |
| 5 | 125209744 | rs1368131 | 1.138 | 0.224 | -0.505 | -0.011 | -0.002 | -0.023 | -0.023 | 0 | 0.07 | 0.02 | 0.04 | 0.32 | 0.05 | 0.22 |
| 5 | 125211527 | rs1036161 | 0.973 | 0.221 | -0.477 | -0.013 | 0 | 0.027 | 0.027 | 0 | 0.07 | 0.08 | 0.04 | 0.32 | 0.05 | 0.22 |
| 5 | 125220878 | rs2116565 | 0.887 | 0.785 | -0.499 | -0.11 | 0.001 | 0.013 | 0.013 | 0 | 0.27 | 0.33 | 0.36 | 0.44 | 0.3 | 0.29 |
| 5 | 125225989 | rs17153597 | 1.043 | 0.761 | -0.492 | -0.1 | -0.016 | 0.018 | 0.018 | 0 | 0.25 | 0.32 | 0.33 | 0.44 | 0.22 | 0.29 |
| 5 | 125226570 | rs327864 | 1.563 | 0.871 | -0.797 | -0.083 | -0.007 | 0.053 | 0.053 | 0 | 0.11 | 0 | 0.04 | 0.01 | 0.07 | 0.09 |
| 5 | 125226999 | rs6868678 | 0.611 | 1.007 | -0.581 | -0.005 | 0.001 | -0.001 | -0.001 | 0 | 0.26 | 0.21 | 0.24 | 0.16 | 0.28 | 0.25 |
| 5 | 125229157 | rs2562262 | -0.528 | 1.233 | -0.708 | 0.182 | -0.042 | -0.038 | -0.038 | 0 | 0.92 | 0.97 | 0.86 | 0.68 | 0.84 | 0.75 |
| 5 | 125231983 | rs1432345 | 0.814 | 0.345 | -0.719 | 0.1 | 0.002 | -0.025 | -0.025 | 0 | 0.11 | 0.05 | 0.21 | 0.44 | 0.11 | 0.25 |
| 5 | 125232864 | rs34035 | -1.274 | 0.097 | -0.528 | -0.156 | -0.005 | -0.012 | -0.012 | 0.752 | 0.56 | 0.45 | 0.56 | 0.69 | 0.58 | 0.65 |
| 5 | 125246304 | rs6595650 | 0.954 | 0.29 | -0.543 | -0.19 | -0.01 | -0.01 | -0.01 | 0.06 | 0.45 | 0.54 | 0.44 | 0.32 | 0.37 | 0.35 |
| 5 | 125274491 | rs465730 | 0.931 | 0.222 | -0.479 | -0.124 | -0.002 | 0.016 | 0.016 | 0 | 0.12 | 0.1 | 0.3 | 0.45 | 0.15 | 0.26 |
| 5 | 125276922 | rs4460136 | 1.731 | 0.04 | -0.48 | -0.351 | 0.005 | -0.02 | -0.02 | 0 | 0.27 | 0.22 | 0.24 | 0.32 | 0.24 | 0.29 |
| 5 | 125279529 | rs327819 | 1.114 | 0.063 | -0.512 | -0.335 | -0.011 | -0.026 | -0.026 | 0 | 0.11 | 0.06 | 0.3 | 0.43 | 0.15 | 0.26 |
| 5 | 125284770 | rs17153713 | 2.472 | -0.006 | -0.374 | -0.386 | -0.018 | -0.056 | -0.056 | 0.019 | 0.14 | 0.21 | 0.11 | 0 | 0.11 | 0.04 |
| 5 | 125287589 | rs399295 | -0.986 | -0.082 | -0.463 | -0.491 | -0.089 | 0.007 | 0.007 | 0.474 | 0.73 | 0.95 | 0.74 | 0.84 | 0.65 | 0.73 |
| 5 | 125287868 | rs10045999 | 0.623 | -0.211 | -0.533 | -0.53 | 0.024 | 0.017 | 0.017 | 0.02 | 0.51 | 0.61 | 0.39 | 0.4 | 0.44 | 0.4 |
| 5 | 125293366 | rs327816 | 0.201 | 0.154 | -0.638 | -0.439 | 0.027 | 0.011 | 0.011 | 0 | 0.67 | 0.62 | 0.46 | 0.4 | 0.53 | 0.43 |
| 5 | 125295228 | rs17153717 | 2.407 | -0.063 | -0.532 | -0.242 | 0.008 | -0.008 | -0.008 | 0 | 0.14 | 0.12 | 0.11 | 0 | 0.11 | 0.04 |
| 5 | 125296857 | rs188986 | 1.072 | -0.011 | -0.689 | -0.185 | 0.018 | -0.008 | -0.008 | 0 | 0.35 | 0.21 | 0.26 | 0.31 | 0.27 | 0.32 |
| 5 | 125297247 | rs326020 | 0.039 | 0.172 | -0.572 | -0.124 | 0.001 | 0.105 | 0.105 | 0 | 0.51 | 0.31 | 0.6 | 0.8 | 0.51 | 0.63 |
| 5 | 125301304 | rs34029 | 0.331 | 0.096 | -0.28 | -0.082 | -0.016 | 0.035 | 0.035 | 0 | 0.25 | 0.44 | 0.42 | 0.41 | 0.24 | 0.29 |
| 5 | 125304547 | rs1910003 | -0.105 | 0.048 | -0.228 | -0.137 | -0.004 | -0.006 | -0.006 | 0 | 0.12 | 0.12 | 0.05 | 0.12 | 0.13 | 0.12 |
| 5 | 125305451 | rs2568399 | -0.813 | 0.085 | -0.358 | -0.103 | -0.014 | 0.008 | 0.008 | 0.021 | 0.38 | 0.57 | 0.47 | 0.53 | 0.37 | 0.4 |
| 5 | 125308509 | rs42783 | 0.185 | 0.376 | -0.711 | 0.34 | -0.001 | 0.02 | 0.02 | 0 | 0.12 | 0.17 | 0.3 | 0.41 | 0.14 | 0.24 |
| 5 | 125310154 | rs10434776 | 0.372 | 0.388 | -0.93 | 0.283 | -0.01 | -0.001 | -0.001 | 0 | 0.06 | 0.07 | 0.01 | 0.09 | 0.03 | 0.06 |
| 5 | 125312031 | rs994228 | 0.546 | 0.281 | -0.81 | 0.23 | -0.006 | 0.016 | 0.016 | 0 | 0.29 | 0.59 | 0.44 | 0.36 | 0.3 | 0.29 |
| 5 | 125312789 | rs326016 | 0.649 | -0.137 | -1.352 | 0.104 | 0.022 | -0.014 | -0.014 | 0 | 0.26 | 0.1 | 0.18 | 0.26 | 0.12 | 0.31 |
| 5 | 125314685 | rs4836239 | 0.924 | -0.085 | -1.497 | 0.055 | 0 | 0 | 0 | 0 | 0.14 | 0.12 | 0.16 | 0.14 | 0.15 | 0.13 |
| 5 | 125315488 | rs426136 | 0.688 | -0.094 | -1.425 | 0.056 | 0.009 | -0.004 | -0.004 | 0 | 0.26 | 0.17 | 0.18 | 0.26 | 0.12 | 0.31 |
| 5 | 125315960 | rs434265 | -0.806 | -0.088 | -1.863 | 0.018 | -0.076 | -0.023 | -0.023 | 2.223 | 0.66 | 0.82 | 0.7 | 0.65 | 0.55 | 0.6 |
| 5 | 125324472 | rs326038 | -1.614 | 0.765 | -1.973 | -0.405 | -0.103 | 0.094 | 0.094 | 0 | 0.83 | 0.98 | 0.81 | 0.93 | 0.74 | 0.87 |
| 5 | 125325978 | rs463757 | 0.626 | 0.145 | -1.035 | 0.297 | -0.013 | -0.024 | -0.024 | 0.697 | 0.25 | 0.57 | 0.36 | 0.19 | 0.27 | 0.12 |
| 5 | 125328001 | rs12659015 | -0.441 | 0.454 | -1.046 | 0.612 | -0.001 | -0.001 | -0.001 | 0 | 0.14 | 0.14 | 0.16 | 0.14 | 0.15 | 0.12 |

| | | | | | | | | | | | | | | | | |
|---|---|---|---|---|---|---|---|---|---|---|---|---|---|---|---|---|
| 5 | 125338324 | rs6859472 | 0.392 | 0.464 | -1.618 | 0.547 | -0.02 | 0.062 | 0.062 | 0 | 0.48 | 0.16 | 0.44 | 0.32 | 0.48 | 0.43 |
| 5 | 125339876 | rs1393144 | 0.469 | 0.271 | -0.625 | 0.959 | -0.005 | 0.054 | 0.054 | 0 | 0.18 | 0.34 | 0.12 | 0.29 | 0.19 | 0.28 |
| 5 | 125340731 | rs17566147 | 0.706 | 0.154 | -1.053 | 0.278 | -0.014 | -0.001 | -0.001 | 0 | 0.36 | 0.19 | 0.34 | 0.32 | 0.35 | 0.4 |
| 5 | 125343531 | rs9327373 | 1.182 | 0.18 | -0.57 | 0.62 | 0.004 | -0.004 | -0.004 | 0 | 0.13 | 0.11 | 0.09 | 0.2 | 0.13 | 0.16 |
| 5 | 125345974 | rs10075656 | -0.108 | -0.375 | -0.611 | 0.343 | -0.017 | -0.014 | -0.014 | 1.111 | 0.69 | 0.58 | 0.59 | 0.7 | 0.73 | 0.69 |
| 5 | 125348776 | rs1393141 | -0.051 | -0.417 | -0.633 | 0.323 | -0.006 | -0.028 | -0.028 | 0.545 | 0.71 | 0.49 | 0.65 | 0.7 | 0.72 | 0.76 |
| 5 | 125350072 | rs3849056 | -0.671 | -0.58 | -0.384 | 0.358 | -0.009 | -0.012 | -0.012 | 0.318 | 0.77 | 0.67 | 0.69 | 0.8 | 0.79 | 0.86 |
| 5 | 125350909 | rs9327375 | -0.143 | -0.586 | 0.005 | 0.493 | 0.002 | -0.007 | -0.007 | 0 | 0.25 | 0.24 | 0.2 | 0.34 | 0.23 | 0.38 |
| 5 | 125354082 | rs17153843 | 0.817 | -0.442 | 0.005 | 0.533 | 0.003 | -0.009 | -0.009 | 0.278 | 0.08 | 0.01 | 0.1 | 0.06 | 0.06 | 0.11 |
| 5 | 125356046 | rs3913480 | 1.289 | -0.69 | -0.091 | 0.835 | -0.011 | -0.004 | -0.004 | 0 | 0.31 | 0.25 | 0.29 | 0.38 | 0.38 | 0.27 |
| 5 | 125356973 | rs440187 | -1.46 | -0.88 | 0.095 | 0.567 | 0.009 | -0.001 | -0.001 | 0.071 | 0.83 | 0.91 | 0.72 | 0.79 | 0.83 | 0.87 |
| 5 | 125359766 | rs12653591 | 2.064 | -0.983 | 0.175 | 0.599 | 0.002 | 0.002 | 0.002 | 0 | 0.23 | 0.2 | 0.24 | 0.09 | 0.18 | 0.2 |
| 5 | 125360438 | rs10463812 | 0.104 | -0.94 | 0.261 | 0.625 | 0.001 | -0.007 | -0.007 | 0 | 0.71 | 0.6 | 0.65 | 0.69 | 0.68 | 0.77 |
| 5 | 125368345 | rs17507502 | 1.253 | -0.899 | 0.203 | 0.612 | -0.037 | -0.005 | -0.005 | 0 | 0.06 | 0.01 | 0.05 | 0.05 | 0.12 | 0.09 |
| 5 | 125368536 | rs6859953 | 0.322 | -0.973 | 0.265 | 0.604 | -0.007 | 0.011 | 0.011 | 0 | 0.44 | 0.27 | 0.39 | 0.55 | 0.48 | 0.5 |
| 5 | 125369113 | rs12719343 | 0.072 | -0.844 | 0.322 | 0.647 | 0 | -0.004 | -0.004 | 0 | 0.71 | 0.64 | 0.65 | 0.69 | 0.68 | 0.77 |
| 5 | 125370720 | rs7712273 | -0.632 | -0.898 | 0.406 | 0.655 | -0.013 | -0.012 | -0.012 | 0.904 | 0.77 | 0.62 | 0.69 | 0.75 | 0.8 | 0.86 |
| 5 | 125370834 | rs11746120 | -0.232 | -0.973 | 0.538 | 0.686 | -0.027 | 0.031 | 0.031 | 0 | 0.74 | 0.53 | 0.59 | 0.65 | 0.79 | 0.7 |
| 5 | 125371267 | rs3909300 | -1.126 | -0.974 | 0.204 | 0.63 | -0.004 | -0.015 | -0.015 | 0 | 0.8 | 0.93 | 0.7 | 0.73 | 0.8 | 0.85 |
| 5 | 125373297 | rs12521931 | 1.488 | -0.743 | 0.153 | 0.572 | -0.004 | 0.003 | 0.003 | 0 | 0.29 | 0.25 | 0.27 | 0.21 | 0.32 | 0.19 |
| 5 | 125380430 | rs3849064 | 1.999 | -0.677 | 0.403 | 0.707 | -0.017 | 0.134 | 0.134 | 0.064 | 0.47 | 0.22 | 0.37 | 0.2 | 0.47 | 0.22 |
| 5 | 125382324 | rs10519881 | 0.299 | -0.465 | 0.422 | 0.959 | 0.007 | 0.074 | 0.074 | 0 | 0.71 | 0.51 | 0.64 | 0.45 | 0.68 | 0.65 |
| 5 | 125382449 | rs6868912 | -0.907 | -0.529 | 0.153 | 0.923 | -0.001 | -0.013 | -0.013 | 0.862 | 0.8 | 0.92 | 0.7 | 0.73 | 0.8 | 0.85 |
| 5 | 125385126 | rs955626 | -0.339 | -0.49 | 0.094 | 0.896 | -0.022 | -0.003 | -0.003 | 0 | 0.14 | 0.04 | 0.14 | 0.11 | 0.15 | 0.2 |
| 5 | 125385248 | rs7708364 | 1.115 | -0.502 | 0.172 | 0.817 | 0 | -0.001 | -0.001 | 0 | 0.59 | 0.38 | 0.46 | 0.53 | 0.63 | 0.5 |
| 5 | 125387794 | rs10064418 | 0.917 | -0.303 | 0.238 | 0.869 | -0.001 | 0.003 | 0.003 | 0 | 0.63 | 0.39 | 0.47 | 0.53 | 0.65 | 0.53 |
| 5 | 125388544 | rs3849065 | 1.121 | -0.308 | 0.237 | 0.863 | 0 | -0.001 | -0.001 | 0 | 0.61 | 0.39 | 0.46 | 0.48 | 0.64 | 0.53 |
| 5 | 125390294 | rs3849066 | 1.103 | -0.081 | 0.166 | 0.795 | 0.002 | -0.007 | -0.007 | 0 | 0.58 | 0.36 | 0.42 | 0.48 | 0.6 | 0.53 |
| 5 | 125402989 | rs1175281 | 1.421 | -0.179 | 0.219 | 0.671 | -0.006 | 0.019 | 0.019 | 0 | 0.08 | 0.03 | 0.07 | 0 | 0.09 | 0.03 |
| 5 | 125429205 | rs3860121 | 0.561 | -0.126 | 0.539 | 0.87 | -0.018 | 0.01 | 0.01 | 0 | 0.64 | 0.45 | 0.52 | 0.57 | 0.72 | 0.59 |
| 5 | 125430870 | rs3843817 | -0.173 | -0.199 | 0.489 | 0.927 | -0.001 | -0.015 | -0.015 | 0 | 0.72 | 0.92 | 0.64 | 0.67 | 0.73 | 0.76 |
| 5 | 125432585 | rs10042227 | -0.855 | -0.196 | 0.639 | 0.977 | -0.012 | -0.017 | -0.017 | 0.116 | 0.79 | 0.52 | 0.7 | 0.8 | 0.81 | 0.88 |
| 5 | 125436331 | rs3909382 | 1.224 | -0.195 | 0.578 | 0.932 | 0.01 | -0.012 | -0.012 | 0 | 0.58 | 0.35 | 0.46 | 0.48 | 0.62 | 0.57 |
| 5 | 125441453 | rs10070545 | -0.517 | 0.023 | 0.508 | 1.008 | 0.006 | 0.039 | 0.039 | 0 | 0.08 | 0.1 | 0.14 | 0.15 | 0.11 | 0.18 |
| 5 | 125450818 | rs1600681 | 0.963 | 0.277 | 0.237 | 0.636 | -0.082 | 0.047 | 0.047 | 0 | 0.29 | 0.04 | 0.28 | 0.18 | 0.33 | 0.31 |
| 5 | 125451265 | rs11747637 | 1.182 | 0.284 | 0.639 | 0.977 | 0.01 | 0.04 | 0.04 | 0 | 0.55 | 0.23 | 0.43 | 0.3 | 0.56 | 0.45 |
| 5 | 125455992 | rs11960577 | -0.484 | 0.312 | 1.011 | 0.961 | 0.088 | 0.074 | 0.074 | 0 | 0.05 | 0.23 | 0.05 | 0.08 | 0.1 | 0.05 |
| 5 | 125470423 | rs17154090 | 0.816 | 0.233 | 0.552 | 0.983 | -0.011 | -0.02 | -0.02 | 0 | 0.1 | 0.01 | 0.09 | 0.12 | 0.08 | 0.11 |
| 5 | 125471402 | rs10463817 | 0.899 | 0.284 | 0.757 | 1.073 | 0.004 | -0.005 | -0.005 | 0 | 0.58 | 0.37 | 0.46 | 0.5 | 0.59 | 0.59 |
| 5 | 125474036 | rs3849076 | 0.196 | 0.522 | 0.84 | 1.171 | 0 | 0.011 | 0.011 | 0 | 0.71 | 0.69 | 0.57 | 0.61 | 0.67 | 0.7 |
| 5 | 125475599 | rs2408573 | -0.619 | 0.552 | 0.601 | 1.201 | 0.007 | -0.032 | -0.032 | 1.048 | 0.77 | 0.91 | 0.62 | 0.69 | 0.77 | 0.75 |
| 5 | 125507307 | rs6890677 | -0.618 | 0.555 | 0.604 | 1.197 | 0.009 | -0.031 | -0.031 | 0 | 0.77 | 0.91 | 0.62 | 0.65 | 0.78 | 0.74 |
| 5 | 125520876 | rs6895621 | 0.232 | 1.103 | 0.724 | 1.111 | 0.01 | 0.009 | 0.009 | 0 | 0.71 | 0.69 | 0.5 | 0.65 | 0.62 | 0.69 |
| 5 | 125528830 | rs7730830 | -0.124 | 0.975 | 0.653 | 0.983 | -0.001 | -0.001 | -0.001 | 0 | 0.41 | 0.37 | 0.36 | 0.49 | 0.41 | 0.5 |
| 5 | 125531647 | rs7356716 | -0.536 | 0.965 | 0.824 | 1.005 | 0.003 | -0.002 | -0.002 | 0 | 0.46 | 0.59 | 0.42 | 0.56 | 0.51 | 0.56 |
| 5 | 125533130 | rs17584191 | 1.659 | 0.934 | 0.768 | 1.034 | 0.017 | 0.05 | 0.05 | 0 | 0.3 | 0.23 | 0.24 | 0.12 | 0.27 | 0.17 |
| 5 | 125534820 | rs7704934 | -0.183 | 0.984 | 0.972 | 1.057 | 0.005 | 0.001 | 0.001 | 0.014 | 0.41 | 0.42 | 0.37 | 0.49 | 0.41 | 0.5 |
| 5 | 125537459 | rs1427885 | -0.088 | 0.958 | 0.881 | 1.064 | -0.001 | 0 | 0 | 0 | 0.4 | 0.4 | 0.3 | 0.53 | 0.35 | 0.5 |
| 5 | 125539560 | rs10065670 | -0.142 | 0.967 | 1.005 | 1.06 | -0.003 | -0.005 | -0.005 | 0.067 | 0.39 | 0.34 | 0.3 | 0.48 | 0.35 | 0.49 |
| 5 | 125547517 | rs2036675 | -0.77 | 1.199 | 0.933 | 1.112 | 0.061 | 0.066 | 0.066 | 0 | 0.06 | 0.21 | 0.05 | 0.07 | 0.1 | 0.08 |
| 5 | 125567968 | rs7712923 | -0.718 | 1.11 | 0.576 | 1.123 | 0.007 | -0.025 | -0.025 | 0 | 0.77 | 0.88 | 0.66 | 0.64 | 0.78 | 0.74 |
| 5 | 125576242 | rs17584877 | 1.494 | 1.487 | 0.113 | 0.865 | -0.086 | 0.069 | 0.069 | 0 | 0.15 | 0.01 | 0.12 | 0.07 | 0.2 | 0.23 |
| 5 | 125580205 | rs11241878 | -0.257 | 1.388 | 0.755 | 1.32 | 0.005 | 0.03 | 0.03 | 0 | 0.73 | 0.71 | 0.57 | 0.51 | 0.67 | 0.7 |
| 5 | 125592690 | rs7704232 | -0.81 | 1.406 | 0.814 | 1.344 | -0.002 | -0.008 | -0.008 | 0 | 0.77 | 0.85 | 0.69 | 0.68 | 0.72 | 0.77 |
| 5 | 125603060 | rs1366106 | 0.33 | 1.397 | 0.962 | 1.292 | 0.012 | 0.021 | 0.021 | 0 | 0.68 | 0.47 | 0.51 | 0.44 | 0.57 | 0.62 |
| 5 | 125609245 | rs1303821 | -0.696 | 1.189 | 0.42 | 1.235 | -0.005 | -0.024 | -0.024 | 0 | 0.76 | 0.89 | 0.7 | 0.68 | 0.72 | 0.76 |
| 5 | 125615229 | rs1156854 | -0.786 | 1.265 | 0.398 | 1.192 | -0.006 | -0.034 | -0.034 | 0.356 | 0.77 | 0.9 | 0.69 | 0.69 | 0.72 | 0.76 |
| 5 | 125616401 | rs10478719 | 0.253 | 1.297 | 0.666 | 1.132 | 0.007 | 0.016 | 0.016 | 0 | 0.68 | 0.42 | 0.52 | 0.39 | 0.59 | 0.64 |
| 5 | 125619900 | rs10477668 | -0.78 | 1.312 | 0.534 | 1.139 | 0.011 | -0.028 | -0.028 | 0.157 | 0.77 | 0.89 | 0.75 | 0.63 | 0.78 | 0.76 |
| 5 | 125621479 | rs7718098 | 1.083 | 1.163 | 0.437 | 0.827 | -0.01 | -0.021 | -0.021 | 0 | 0.44 | 0.51 | 0.37 | 0.21 | 0.35 | 0.36 |
| 5 | 125633592 | rs7448802 | 1.619 | 1.21 | 0.251 | 0.616 | -0.085 | 0.049 | 0.049 | 0 | 0.14 | 0.01 | 0.13 | 0.07 | 0.21 | 0.18 |
| 5 | 125641993 | rs12517548 | 1.329 | 1.227 | 0.315 | 0.472 | 0.018 | 0.013 | 0.013 | 0 | 0.38 | 0.36 | 0.26 | 0.12 | 0.26 | 0.2 |
| 5 | 125648541 | rs882216 | 0.42 | 1.161 | 0.369 | 0.573 | -0.002 | -0.002 | -0.002 | 1.054 | 0.26 | 0.22 | 0.33 | 0.36 | 0.32 | 0.33 |
| 5 | 125649162 | rs17527858 | 2.958 | 0.166 | 0.24 | 0.254 | 0.031 | 0.117 | 0.117 | 1.631 | 0.22 | 0.04 | 0.12 | 0 | 0.16 | 0.03 |
| 5 | 125655221 | rs1567249 | 1.253 | 0.344 | 0.364 | 0.372 | -0.011 | -0.045 | -0.045 | 0 | 0.38 | 0.43 | 0.26 | 0.12 | 0.26 | 0.19 |
| 5 | 125661113 | rs1896588 | 0.911 | 0.364 | 0.403 | 0.424 | -0.029 | -0.047 | -0.047 | 0 | 0.41 | 0.54 | 0.27 | 0.19 | 0.27 | 0.26 |
| 5 | 125663367 | rs7733611 | -0.238 | 0.474 | 0.258 | 0.525 | -0.002 | -0.011 | -0.011 | 0 | 0.67 | 0.73 | 0.65 | 0.5 | 0.65 | 0.5 |
| 5 | 125663900 | rs10055040 | -0.032 | 0.492 | 0.205 | 0.426 | -0.005 | -0.002 | -0.002 | 0 | 0.31 | 0.28 | 0.41 | 0.37 | 0.42 | 0.37 |
| 5 | 125664939 | rs10900787 | -0.211 | 0.827 | 0.353 | 1.077 | -0.016 | -0.061 | -0.061 | 0 | 0.71 | 0.87 | 0.67 | 0.53 | 0.67 | 0.57 |

| | | | | | | | | | | | | | | | | |
|---|---|---|---|---|---|---|---|---|---|---|---|---|---|---|---|---|
| 5 | 125665285 | rs1427895 | 1.401 | 0.817 | 0.353 | 0.381 | -0.003 | 0.126 | 0.126 | 0.114 | 0.5 | 0.35 | 0.32 | 0.18 | 0.46 | 0.26 |
| 5 | 125670036 | rs17154410 | 1.525 | 1.178 | 0.329 | 0.911 | -0.033 | -0.032 | -0.032 | 0 | 0.05 | 0 | 0.07 | 0.08 | 0.11 | 0.16 |
| 5 | 125672801 | rs10045325 | -1.292 | 1.006 | 0.653 | 0.57 | 0.071 | 0.071 | 0.071 | 0 | 0.08 | 0.31 | 0.22 | 0.1 | 0.11 | 0.12 |
| 5 | 125674421 | rs918440 | 0.86 | 1.096 | 0.385 | 0.465 | 0.005 | -0.021 | -0.021 | 0 | 0.59 | 0.47 | 0.42 | 0.3 | 0.53 | 0.39 |
| 5 | 125676464 | rs9327402 | -0.111 | 1.129 | -0.626 | 0.632 | -0.03 | -0.02 | -0.02 | 0 | 0.69 | 0.87 | 0.65 | 0.39 | 0.65 | 0.51 |
| 5 | 125678354 | rs6595693 | -1.075 | 1.271 | 0.7 | 0.794 | 0.043 | -0.033 | -0.033 | 0 | 0.12 | 0.39 | 0.23 | 0.05 | 0.14 | 0.15 |
| 5 | 125679739 | rs11738463 | 0.154 | 1.785 | -0.008 | 1.274 | -0.063 | -0.066 | -0.066 | 0 | 0.13 | 0.02 | 0.16 | 0.17 | 0.2 | 0.23 |
| 5 | 125680888 | rs7730928 | -0.931 | 1.536 | 0.564 | 1.209 | 0.005 | -0.012 | -0.012 | 0 | 0.33 | 0.6 | 0.43 | 0.33 | 0.36 | 0.23 |
| 5 | 125683958 | rs10068159 | -0.911 | 1.514 | 1.461 | 1.406 | 0.029 | 0.007 | 0.007 | 0.553 | 0.37 | 0.56 | 0.38 | 0.43 | 0.45 | 0.39 |
| 5 | 125685374 | rs10057227 | -0.034 | 1.566 | 1.732 | 1.384 | 0.01 | -0.014 | -0.014 | 0 | 0.23 | 0.35 | 0.18 | 0.26 | 0.26 | 0.18 |
| 5 | 125685898 | rs12513492 | -0.817 | 1.557 | 1.684 | 1.336 | 0.041 | 0.063 | 0.063 | 0 | 0.06 | 0.22 | 0.22 | 0.09 | 0.09 | 0.03 |
| 5 | 125685909 | rs10478724 | 0.276 | 1.549 | 1.297 | 1.334 | -0.038 | 0.023 | 0.023 | 0 | 0.22 | 0.12 | 0.17 | 0.24 | 0.26 | 0.18 |
| 5 | 125686381 | rs7727523 | -1.896 | 1.558 | 1.262 | 1.348 | 0.005 | -0.032 | -0.032 | 0.204 | 0.45 | 0.83 | 0.63 | 0.51 | 0.55 | 0.44 |
| 5 | 125688296 | rs1549189 | -0.16 | 1.683 | 0.943 | 1.234 | 0.001 | -0.005 | -0.005 | 0 | 0.24 | 0.29 | 0.18 | 0.25 | 0.26 | 0.18 |
| 5 | 125693382 | rs10519907 | 0.779 | 1.647 | 0.715 | 1.304 | -0.059 | -0.061 | -0.061 | 0.044 | 0.11 | 0.02 | 0.19 | 0.16 | 0.19 | 0.2 |
| 5 | 125694196 | rs10519908 | -0.152 | 1.38 | 0.584 | 1.271 | -0.017 | -0.036 | -0.036 | 0.322 | 0.06 | 0 | 0.11 | 0.11 | 0.06 | 0.27 |
| 5 | 125696690 | rs7733479 | -0.969 | 1.244 | 1.029 | 0.825 | 0.031 | -0.022 | -0.022 | 0 | 0.32 | 0.51 | 0.48 | 0.28 | 0.39 | 0.22 |
| 5 | 125697656 | rs7700582 | -0.576 | 1.206 | 0.992 | 0.838 | 0.011 | -0.006 | -0.006 | 0 | 0.27 | 0.33 | 0.27 | 0.26 | 0.33 | 0.18 |
| 5 | 125704211 | rs17595984 | 0.205 | 0.776 | 1.074 | 0.594 | 0.021 | -0.006 | -0.006 | 1.261 | 0.2 | 0.22 | 0.24 | 0.2 | 0.29 | 0.17 |
| 5 | 125705163 | rs10519910 | 1.459 | 0.315 | 0.905 | 0.729 | 0.054 | 0.005 | 0.005 | 0 | 0.46 | 0.27 | 0.26 | 0.48 | 0.34 | 0.6 |
| 5 | 125705876 | rs10478726 | -0.579 | 0.62 | 1.024 | 0.51 | 0.042 | -0.015 | -0.015 | 0 | 0.25 | 0.39 | 0.44 | 0.23 | 0.34 | 0.22 |
| 5 | 125708877 | rs10069151 | 1.165 | -0.78 | 1.297 | 0.793 | -0.022 | -0.023 | -0.023 | 1.211 | 0.73 | 0.61 | 0.84 | 0.81 | 0.82 | 0.54 |
| 5 | 125709909 | rs10078989 | -0.595 | -0.769 | 1.384 | 0.805 | -0.006 | -0.034 | -0.034 | 0.007 | 0.93 | 0.79 | 0.96 | 1 | 0.97 | 0.98 |
| 5 | 125712172 | rs1023613 | 1.469 | -0.806 | 1.085 | 0.8 | -0.028 | -0.025 | -0.025 | 0.27 | 0.82 | 0.43 | 0.86 | 0.88 | 0.89 | 0.67 |
| 5 | 125726117 | rs7732081 | -0.164 | -0.604 | 0.755 | 0.912 | 0.004 | -0.004 | -0.004 | 0.178 | 0.18 | 0.14 | 0.19 | 0.23 | 0.09 | 0.19 |
| 5 | 125727454 | rs10037420 | -0.685 | -0.416 | 0.431 | 0.78 | 0.002 | 0.01 | 0.01 | 0 | 0.07 | 0.16 | 0.06 | 0.12 | 0.07 | 0.24 |
| 5 | 125731213 | rs1422271 | 0.656 | -0.169 | 0.636 | 0.426 | -0.037 | 0.006 | 0.006 | 0 | 0.74 | 0.66 | 0.77 | 0.73 | 0.87 | 0.78 |
| 5 | 125738273 | rs1422270 | 0.742 | -0.163 | 0.525 | 0.413 | -0.039 | 0.002 | 0.002 | 0 | 0.73 | 0.65 | 0.75 | 0.74 | 0.85 | 0.76 |
| 5 | 125740942 | rs1422269 | 0.962 | -0.145 | 0.373 | 0.403 | -0.029 | -0.015 | -0.015 | 0 | 0.69 | 0.45 | 0.71 | 0.75 | 0.81 | 0.73 |
| 5 | 125754678 | rs17154542 | -0.351 | -0.348 | -1.28 | 0.698 | 0.024 | -0.03 | -0.03 | 0 | 0.19 | 0.07 | 0.13 | 0.25 | 0.1 | 0.22 |
| 5 | 125758373 | rs4388209 | -0.483 | -0.288 | -1.33 | 0.667 | 0.077 | -0.033 | -0.033 | 0.538 | 0.59 | 0.81 | 0.76 | 0.5 | 0.69 | 0.53 |
| 5 | 125768020 | rs11241891 | -0.137 | 0.375 | -1.04 | 0.12 | -0.003 | 0.007 | 0.007 | 0.017 | 0.3 | 0.24 | 0.35 | 0.19 | 0.26 | 0.32 |
| 5 | 125773331 | rs11241892 | -1.02 | 0.5 | -2.078 | 0.039 | 0.001 | -0.094 | -0.094 | 0 | 0.63 | 0.88 | 0.68 | 0.44 | 0.62 | 0.45 |
| 5 | 125773570 | rs4413529 | -0.018 | 0.444 | -1.441 | -0.246 | 0.044 | 0.017 | 0.017 | 0.61 | 0.08 | 0.46 | 0.12 | 0.01 | 0.04 | 0.04 |
| 5 | 125774155 | rs12186425 | 0.416 | 0.48 | -1.865 | -0.289 | -0.058 | -0.015 | -0.015 | 0.34 | 0.3 | 0.02 | 0.33 | 0.2 | 0.23 | 0.31 |
| 5 | 125778935 | rs4426911 | -0.706 | 0.792 | -0.839 | -0.37 | -0.02 | -0.063 | -0.063 | 0 | 0.13 | 0.55 | 0.15 | 0.01 | 0.07 | 0.05 |
| 5 | 125779180 | rs11744511 | -0.387 | 1.271 | -1.479 | 0.08 | -0.036 | 0.005 | 0.005 | 0 | 0.19 | 0.11 | 0.19 | 0.18 | 0.31 | 0.09 |
| 5 | 125791140 | rs13162396 | -0.235 | 0.765 | -1.737 | 0.017 | -0.011 | -0.02 | -0.02 | 0 | 0.34 | 0.2 | 0.36 | 0.58 | 0.36 | 0.4 |
| 5 | 125795515 | rs6886050 | -0.013 | 0.194 | -0.895 | -0.499 | -0.027 | -0.084 | -0.084 | 2.38 | 0.87 | 0.53 | 0.91 | 1 | 0.94 | 0.94 |
| 5 | 125795563 | rs12516622 | 0.329 | -0.129 | -2.118 | -1.272 | -0.054 | 0.094 | 0.094 | 0 | 0.33 | 0.01 | 0.38 | 0.17 | 0.33 | 0.39 |
| 5 | 125795898 | rs4240399 | -0.321 | -0.115 | -0.649 | -0.842 | -0.01 | -0.075 | -0.075 | 0 | 0.88 | 0.6 | 0.92 | 1 | 0.94 | 0.94 |
| 5 | 125797951 | rs9327407 | 0.015 | -0.118 | -1.384 | -0.834 | -0.04 | -0.081 | -0.081 | 0.225 | 0.86 | 0.42 | 0.9 | 0.99 | 0.94 | 0.94 |
| 5 | 125800646 | rs9687567 | -0.019 | 0.252 | -2.1 | -0.754 | 0 | 0.03 | 0.03 | 0.213 | 0.42 | 0.21 | 0.37 | 0.61 | 0.38 | 0.44 |
| 5 | 125800836 | rs6595711 | 0.114 | 0.618 | -2.07 | -0.297 | -0.018 | -0.028 | -0.028 | 0 | 0.24 | 0.18 | 0.33 | 0.5 | 0.32 | 0.33 |
| 5 | 125801394 | rs6860957 | -0.131 | -0.087 | -2.192 | -1.04 | 0.052 | 0.119 | 0.119 | 0 | 0.58 | 0.76 | 0.51 | 0.81 | 0.65 | 0.55 |
| 5 | 125815602 | rs11740694 | -0.259 | 0.537 | -2.087 | -1.006 | -0.098 | 0.034 | 0.034 | 0.403 | 0.32 | 0.1 | 0.46 | 0.77 | 0.4 | 0.6 |
| 5 | 125818698 | rs4526101 | 0.279 | 0.28 | -1.524 | -0.929 | -0.025 | 0.029 | 0.029 | 0.004 | 0.64 | 0.69 | 0.52 | 0.23 | 0.59 | 0.39 |
| 5 | 125822005 | rs10037307 | 1.136 | -0.157 | -1.388 | -1.465 | -0.007 | 0.013 | 0.013 | 0 | 0.28 | 0.21 | 0.18 | 0.21 | 0.29 | 0.21 |
| 5 | 125824819 | rs4836266 | 0.095 | -0.087 | -0.797 | -1.582 | -0.004 | -0.056 | -0.056 | 0 | 0.71 | 0.68 | 0.71 | 0.98 | 0.71 | 0.84 |
| 5 | 125825824 | rs10214090 | 0.129 | -0.105 | -0.906 | -1.591 | 0.002 | -0.028 | -0.028 | 1.738 | 0.71 | 0.7 | 0.7 | 0.98 | 0.71 | 0.83 |
| 5 | 125826191 | rs10213948 | -0.063 | 0.125 | -1.356 | -1.006 | -0.043 | 0.002 | 0.002 | 0 | 0.8 | 0.87 | 0.76 | 0.81 | 0.75 | 0.88 |
| 5 | 125826789 | rs9327410 | -0.028 | -0.276 | -0.971 | -1.307 | 0.039 | 0.083 | 0.083 | 0 | 0.76 | 0.88 | 0.7 | 0.96 | 0.8 | 0.71 |
| 5 | 125828803 | rs4358517 | -0.541 | 0.223 | -0.641 | -0.965 | -0.004 | 0.028 | 0.028 | 0 | 0.83 | 0.88 | 0.69 | 0.93 | 0.79 | 0.73 |
| 5 | 125838022 | rs6888676 | 0.116 | 0.003 | -0.303 | -1.09 | -0.018 | 0.083 | 0.083 | 0 | 0.47 | 0.28 | 0.42 | 0.73 | 0.46 | 0.5 |
| 5 | 125845095 | rs1552243 | 0.522 | -0.014 | -0.426 | -1.11 | -0.013 | 0.029 | 0.029 | 0 | 0.41 | 0.24 | 0.41 | 0.72 | 0.43 | 0.48 |
| 5 | 125859181 | rs6595718 | -0.538 | 0.145 | -1.253 | -0.83 | -0.003 | 0.016 | 0.016 | 0 | 0.81 | 0.87 | 0.71 | 0.89 | 0.77 | 0.7 |
| 5 | 125861056 | rs17597145 | 0.788 | 0.301 | -1.057 | -0.754 | -0.013 | 0.038 | 0.038 | 0 | 0.31 | 0.21 | 0.23 | 0.22 | 0.32 | 0.21 |
| 5 | 125862026 | rs1552240 | 0.018 | 0.34 | -1.257 | -0.48 | 0.008 | 0.036 | 0.036 | 0 | 0.75 | 0.81 | 0.69 | 0.88 | 0.75 | 0.7 |
| 5 | 125866090 | rs11953659 | -0.001 | 0.543 | -0.962 | -0.292 | -0.01 | 0.025 | 0.025 | 0 | 0.57 | 0.7 | 0.53 | 0.33 | 0.52 | 0.52 |
| 5 | 125872017 | rs4836271 | -0.25 | 0.649 | -1.696 | -0.143 | -0.021 | 0.003 | 0.003 | 0 | 0.39 | 0.18 | 0.54 | 0.6 | 0.41 | 0.39 |
| 5 | 125874334 | rs1384880 | 0.153 | 0.559 | -1.267 | -0.067 | -0.001 | 0 | 0 | 0 | 0.59 | 0.42 | 0.44 | 0.42 | 0.6 | 0.59 |
| 5 | 125874525 | rs2408700 | 0.847 | 0.483 | -1.88 | -0.512 | -0.036 | -0.049 | -0.049 | 0 | 0.21 | 0.04 | 0.17 | 0.31 | 0.18 | 0.17 |
| 5 | 125874993 | rs730870 | 0.172 | 0.58 | -1.23 | -0.085 | -0.004 | 0 | 0 | 1.123 | 0.59 | 0.37 | 0.44 | 0.45 | 0.6 | 0.61 |
| 5 | 125879549 | rs1483780 | 0.182 | 0.726 | -1.318 | 0.01 | -0.002 | -0.001 | -0.001 | 0 | 0.59 | 0.39 | 0.42 | 0.46 | 0.58 | 0.63 |
| 5 | 125880109 | rs1060856 | -0.197 | 0.955 | -1.612 | 0.317 | -0.048 | -0.031 | -0.031 | 0 | 0.12 | 0.01 | 0.08 | 0.12 | 0.15 | 0.15 |
| 5 | 125883228 | rs11241901 | 0.15 | 0.797 | -1.398 | 0.046 | -0.007 | 0.018 | 0.018 | 0 | 0.57 | 0.64 | 0.37 | 0.46 | 0.5 | 0.61 |
| 5 | 125883341 | rs4836272 | 0.091 | 0.826 | -0.912 | 0.183 | 0 | 0 | 0 | 0 | 0.61 | 0.57 | 0.62 | 0.63 | 0.62 | 0.66 |
| 5 | 125884934 | rs13179664 | -0.029 | 0.835 | -1.046 | -0.485 | -0.021 | 0.019 | 0.019 | 0 | 0.43 | 0.17 | 0.46 | 0.58 | 0.41 | 0.49 |
| 5 | 125885214 | rs1038378 | 0.512 | 0.763 | -1.32 | -0.58 | -0.018 | -0.085 | -0.085 | 0.343 | 0.2 | 0.06 | 0.29 | 0.48 | 0.17 | 0.25 |

| | | | | | | | | | | | | | | | | |
|---|---|---|---|---|---|---|---|---|---|---|---|---|---|---|---|---|
| 5 | 125885803 | rs1038380 | 0.699 | 0.832 | -1.288 | -0.548 | -0.021 | -0.081 | -0.081 | 0.006 | 0.19 | 0.06 | 0.29 | 0.48 | 0.17 | 0.24 |
| 5 | 125886071 | rs1038381 | 1.18 | 0.82 | -1.304 | -0.564 | -0.032 | 0.1 | 0.1 | 0 | 0.24 | 0.01 | 0.17 | 0.09 | 0.24 | 0.22 |
| 5 | 125887496 | rs4379190 | -0.793 | 0.72 | -1.26 | -0.86 | 0.009 | 0.026 | 0.026 | 0.101 | 0.7 | 0.82 | 0.81 | 0.76 | 0.72 | 0.62 |
| 5 | 125887685 | rs2306617 | -0.645 | 0.778 | -0.913 | -0.833 | -0.007 | -0.015 | -0.015 | 0 | 0.68 | 0.66 | 0.79 | 0.76 | 0.72 | 0.62 |
| 5 | 125887856 | rs2306619 | 0.464 | 0.629 | -0.942 | -0.187 | -0.021 | -0.082 | -0.082 | 0 | 0.19 | 0.06 | 0.29 | 0.47 | 0.17 | 0.24 |
| 5 | 125890559 | rs12513847 | -0.51 | 0.484 | -0.566 | -0.101 | -0.003 | 0.05 | 0.05 | 0 | 0.48 | 0.38 | 0.62 | 0.72 | 0.51 | 0.55 |
| 5 | 125896918 | rs4836276 | -0.028 | 0.512 | -1.751 | 0.116 | -0.099 | -0.133 | -0.133 | 0 | 0.39 | 0.02 | 0.42 | 0.52 | 0.38 | 0.43 |
| 5 | 125897227 | rs4626334 | 0.671 | 0.503 | -0.287 | 0.645 | -0.014 | -0.023 | -0.023 | 0.424 | 0.74 | 0.5 | 0.71 | 0.78 | 0.75 | 0.73 |
| 5 | 125897411 | rs4626335 | 0.522 | 0.348 | -0.324 | 0.548 | -0.001 | -0.002 | -0.002 | 0.072 | 0.59 | 0.5 | 0.58 | 0.46 | 0.62 | 0.51 |
| 5 | 125905569 | rs6870785 | -1.742 | -0.223 | -0.284 | 0.733 | 0.015 | 0.005 | 0.005 | 0.914 | 0.23 | 0.86 | 0.17 | 0.21 | 0.19 | 0.21 |
| 5 | 125908090 | rs7709115 | -1.382 | -0.274 | 0.554 | 0.708 | 0.001 | 0.067 | 0.067 | 1.035 | 0.12 | 0.6 | 0.1 | 0.15 | 0.09 | 0.07 |
| 5 | 125908203 | rs17154671 | -1.208 | -0.238 | 0.497 | 0.726 | 0 | -0.037 | -0.037 | 0 | 0.12 | 0.26 | 0.07 | 0.06 | 0.1 | 0.14 |
| 5 | 125918148 | rs13182402 | -1.219 | -0.284 | 0.527 | 0.702 | 0.003 | -0.037 | -0.037 | 0 | 0.12 | 0.26 | 0.07 | 0.06 | 0.1 | 0.15 |
| 5 | 125921768 | rs6595719 | -1.325 | -0.301 | 0.719 | 0.75 | 0.006 | 0.111 | 0.111 | 0.139 | 0.11 | 0.58 | 0.1 | 0.16 | 0.08 | 0.06 |
| 5 | 125928737 | rs6861395 | -1.013 | -0.294 | 0.559 | 0.517 | 0 | -0.034 | -0.034 | 0 | 0.09 | 0.11 | 0.06 | 0 | 0.09 | 0.1 |
| 5 | 125931022 | rs900641 | -2.074 | -0.408 | 1.272 | 0.498 | -0.021 | -0.06 | -0.06 | 0 | 0.15 | 0.64 | 0.11 | 0.06 | 0.1 | 0.15 |
| 5 | 125931041 | rs900640 | -2.017 | -0.427 | 1.792 | 0.511 | 0.04 | 0.015 | 0.015 | 1.244 | 0.06 | 0.35 | 0.04 | 0 | 0.02 | 0 |
| 5 | 125936399 | rs2306620 | -1.037 | -1.041 | 1.226 | 0.909 | -0.037 | 0.049 | 0.049 | 0 | 0.19 | 0.3 | 0.22 | 0.35 | 0.09 | 0.21 |
| 5 | 125937190 | rs6882015 | -1.705 | -1.067 | 1.38 | 0.876 | -0.05 | 0.018 | 0.018 | 0 | 0.26 | 0.45 | 0.27 | 0.3 | 0.15 | 0.32 |
| 5 | 125938092 | rs7726892 | -1.937 | -1.093 | 1.376 | 0.872 | -0.056 | 0.025 | 0.025 | 0 | 0.28 | 0.52 | 0.27 | 0.36 | 0.15 | 0.34 |
| 5 | 125943743 | rs10063061 | -1.852 | -1.095 | 1.383 | 0.871 | -0.054 | 0.028 | 0.028 | 0 | 0.27 | 0.52 | 0.28 | 0.36 | 0.15 | 0.34 |
| 5 | 125945742 | rs11960347 | -1.001 | -1.095 | 1.388 | 0.871 | -0.044 | 0.039 | 0.039 | 0 | 0.19 | 0.3 | 0.21 | 0.3 | 0.08 | 0.2 |
| 5 | 125946011 | rs900646 | -1.725 | -1.048 | 1.191 | 0.86 | -0.056 | 0.025 | 0.025 | 0 | 0.28 | 0.52 | 0.28 | 0.36 | 0.15 | 0.34 |
| 5 | 125953144 | rs6883308 | -1.725 | -1.048 | 1.191 | 0.86 | -0.056 | 0.025 | 0.025 | 0 | 0.28 | 0.52 | 0.28 | 0.36 | 0.15 | 0.34 |
| 5 | 125967653 | rs12655861 | -1.425 | -0.876 | 1.187 | 0.511 | -0.042 | 0.069 | 0.069 | 0.735 | 0.13 | 0.43 | 0.07 | 0.15 | 0.06 | 0.02 |
| 5 | 125968843 | rs34599807 | -0.706 | -0.803 | 0.856 | 0.545 | 0 | -0.001 | -0.001 | 0.036 | 0.1 | 0.09 | 0.05 | 0.15 | 0.06 | 0.02 |
| 5 | 125969749 | rs35726580 | -0.137 | -0.826 | 0.867 | 0.508 | -0.002 | -0.001 | -0.001 | 0 | 0.06 | 0.05 | 0.04 | 0 | 0.07 | 0.1 |
| 5 | 125972624 | rs12652642 | -0.775 | -0.779 | 0.864 | 0.577 | -0.003 | 0.001 | 0.001 | 0.507 | 0.1 | 0.1 | 0.05 | 0.15 | 0.06 | 0.02 |
| 5 | 125981300 | rs4836280 | -1.268 | -0.956 | 0.277 | -0.176 | 0.048 | 0.096 | 0.096 | 0.063 | 0.5 | 0.67 | 0.46 | 0.77 | 0.41 | 0.71 |
| 5 | 125984043 | rs7707878 | 1.685 | -0.353 | -0.444 | -0.659 | -0.117 | 0.222 | 0.222 | 0.738 | 0.34 | 0.04 | 0.25 | 0.07 | 0.39 | 0.16 |
| 5 | 125986096 | rs7701818 | 0.357 | -0.489 | -0.04 | -0.667 | 0.013 | -0.043 | -0.043 | 0 | 0.87 | 0.75 | 0.88 | 0.94 | 0.84 | 0.92 |
| 5 | 125987855 | rs13176922 | 1.144 | -1.015 | -0.418 | 0.042 | -0.035 | -0.08 | -0.08 | 0.09 | 0.13 | 0.01 | 0.1 | 0.2 | 0.15 | 0.25 |
| 5 | 125989661 | rs7718574 | 0.064 | -0.868 | -0.013 | -0.61 | 0.01 | 0.006 | 0.006 | 0.023 | 0.29 | 0.22 | 0.33 | 0.73 | 0.25 | 0.52 |
| 5 | 126003817 | rs7727701 | -1.349 | -0.622 | -0.013 | -0.641 | 0.001 | 0.002 | 0.002 | 0 | 0.36 | 0.67 | 0.41 | 0.22 | 0.32 | 0.35 |
| 5 | 126011133 | rs9637915 | 0.428 | -0.711 | -0.385 | -0.582 | 0.019 | -0.013 | -0.013 | 0 | 0.28 | 0.15 | 0.32 | 0.65 | 0.24 | 0.46 |
| 5 | 126014766 | rs7720178 | -0.458 | -0.355 | -0.232 | -0.476 | 0.011 | 0.022 | 0.022 | 0 | 0.52 | 0.57 | 0.48 | 0.72 | 0.42 | 0.69 |
| 5 | 126016244 | rs6595726 | -1.168 | -0.298 | -0.239 | -0.846 | -0.045 | 0.059 | 0.059 | 0 | 0.64 | 0.75 | 0.72 | 0.95 | 0.54 | 0.84 |
| 5 | 126017826 | rs6888913 | -0.045 | -0.092 | -0.596 | -0.596 | 0.006 | -0.002 | -0.002 | 3.701 | 0.11 | 0.07 | 0.07 | 0.01 | 0.08 | 0.04 |
| 5 | 126041342 | rs12521148 | -0.334 | 0.172 | -0.535 | -0.213 | 0.004 | 0 | 0 | 0 | 0.46 | 0.44 | 0.46 | 0.83 | 0.4 | 0.69 |
| 5 | 126041754 | rs6884533 | -0.334 | 0.172 | -0.535 | -0.213 | 0.004 | 0 | 0 | 0.218 | 0.46 | 0.44 | 0.46 | 0.83 | 0.4 | 0.69 |
| 5 | 126055067 | rs907195 | -0.721 | 1.345 | -0.82 | 0.472 | -0.004 | -0.002 | -0.002 | 0.02 | 0.24 | 0.21 | 0.35 | 0.27 | 0.31 | 0.17 |
| 5 | 126085712 | rs2036846 | -0.371 | 0.943 | -1.153 | 0.091 | 0.018 | 0.072 | 0.072 | 0 | 0.5 | 0.66 | 0.55 | 0.66 | 0.46 | 0.53 |
| 5 | 126086748 | rs2175359 | -0.25 | 1.211 | -1.122 | -0.654 | 0 | -0.006 | -0.006 | 0.207 | 0.64 | 0.4 | 0.64 | 0.71 | 0.65 | 0.57 |
| 5 | 126087378 | rs10478752 | 0.016 | 1.224 | -0.246 | -0.628 | 0.101 | -0.014 | -0.014 | 2.236 | 0.07 | 0.29 | 0.06 | 0 | 0.09 | 0.04 |
| 5 | 126089190 | rs7716198 | 0.353 | 0.672 | -0.744 | -0.388 | 0.024 | 0.009 | 0.009 | 0.022 | 0.33 | 0.2 | 0.19 | 0.34 | 0.24 | 0.29 |
| 5 | 126091664 | rs2271353 | 0.317 | 0.942 | -0.649 | -0.186 | -0.049 | -0.014 | -0.014 | 0 | 0.43 | 0.21 | 0.52 | 0.57 | 0.48 | 0.58 |
| 5 | 126095011 | rs12523362 | -0.404 | 0.254 | -0.379 | -0.686 | -0.015 | -0.046 | -0.046 | 0 | 0.28 | 0.34 | 0.18 | 0.08 | 0.21 | 0.15 |
| 5 | 126112283 | rs3812062 | 0.893 | 0.349 | -0.534 | -0.61 | -0.006 | 0.008 | 0.008 | 0 | 0.05 | 0.02 | 0.05 | 0.02 | 0.07 | 0.07 |
| 5 | 126125209 | rs2973588 | 1.367 | 0.558 | -0.469 | -0.54 | -0.008 | 0.006 | 0.006 | 0.227 | 0.31 | 0.21 | 0.29 | 0.29 | 0.33 | 0.4 |
| 5 | 126129243 | rs2036847 | 0.46 | 0.541 | -0.374 | -0.424 | 0.003 | 0.027 | 0.027 | 0 | 0.16 | 0.2 | 0.28 | 0.3 | 0.2 | 0.19 |
| 5 | 126130101 | rs2973586 | -0.671 | 0.407 | -0.356 | -0.543 | 0.002 | -0.008 | -0.008 | 0 | 0.64 | 0.53 | 0.66 | 0.62 | 0.63 | 0.65 |
| 5 | 126135382 | rs17598783 | -0.304 | 0.39 | -0.398 | -0.584 | -0.005 | -0.013 | -0.013 | 0 | 0.25 | 0.25 | 0.36 | 0.35 | 0.31 | 0.25 |
| 5 | 126140350 | rs968355 | 0.105 | 0.424 | -0.519 | -0.686 | 0.017 | 0.045 | 0.045 | 0 | 0.5 | 0.36 | 0.4 | 0.34 | 0.43 | 0.47 |
| 5 | 126140522 | rs3749830 | 1.095 | 0.637 | -0.605 | -0.546 | -0.006 | 0.009 | 0.009 | 0 | 0.05 | 0.02 | 0.05 | 0.02 | 0.06 | 0.07 |
| 5 | 126159441 | rs2941673 | 1.468 | 0.464 | -0.52 | -0.507 | -0.018 | 0.003 | 0.003 | 0.041 | 0.3 | 0.16 | 0.28 | 0.29 | 0.33 | 0.39 |
| 5 | 126161629 | rs3763156 | 0.925 | -0.07 | -0.031 | -0.623 | -0.008 | 0.008 | 0.008 | 0 | 0.08 | 0.05 | 0.08 | 0.04 | 0.1 | 0.06 |
| 5 | 126166191 | rs3805739 | 1.133 | 0.007 | -0.223 | -0.588 | -0.027 | -0.007 | -0.007 | 0 | 0.08 | 0.01 | 0.07 | 0.05 | 0.1 | 0.06 |
| 5 | 126171999 | rs1051643 | -0.647 | -0.139 | 0.013 | -0.581 | 0.001 | -0.005 | -0.005 | 0.265 | 0.67 | 0.67 | 0.67 | 0.66 | 0.67 | 0.69 |
| 5 | 126172195 | rs1051644 | 0.648 | -0.111 | -0.452 | -0.541 | -0.026 | -0.006 | -0.006 | 1.93 | 0.4 | 0.26 | 0.47 | 0.56 | 0.46 | 0.55 |
| 5 | 126181862 | rs959573 | -0.744 | -0.372 | -0.198 | -0.418 | 0.001 | -0.001 | -0.001 | 1.797 | 0.7 | 0.77 | 0.69 | 0.63 | 0.71 | 0.6 |
| 5 | 126183486 | rs17533927 | 1.261 | -0.238 | -0.54 | -0.307 | -0.027 | 0.019 | 0.019 | 0 | 0.08 | 0.01 | 0.07 | 0.03 | 0.1 | 0.06 |
| 5 | 126183502 | rs17165293 | 1.434 | -0.519 | -0.342 | -0.484 | -0.026 | -0.049 | -0.049 | 0.024 | 0.12 | 0.16 | 0.05 | 0 | 0.06 | 0.01 |
| 5 | 126186957 | rs884241 | 0.289 | -0.447 | -0.055 | -0.432 | -0.006 | -0.008 | -0.008 | 0 | 0.39 | 0.43 | 0.39 | 0.42 | 0.36 | 0.5 |
| 5 | 126188634 | rs6869763 | -0.28 | 0.225 | -0.234 | 0.124 | -0.014 | 0.009 | 0.009 | 0 | 0.58 | 0.61 | 0.5 | 0.43 | 0.53 | 0.53 |
| 5 | 126188989 | rs11740069 | -0.47 | 0.24 | -0.597 | 0.131 | -0.046 | 0.006 | 0.006 | 0 | 0.61 | 0.74 | 0.51 | 0.43 | 0.54 | 0.55 |
| 5 | 126191551 | rs12652098 | -0.335 | 0.401 | -0.602 | 0.219 | -0.023 | 0.009 | 0.009 | 0.019 | 0.59 | 0.68 | 0.5 | 0.43 | 0.53 | 0.53 |
| 5 | 126200982 | rs2055873 | 0.247 | -0.367 | -0.928 | 0.124 | 0.073 | 0.078 | 0.078 | 1.027 | 0.39 | 0.18 | 0.28 | 0.21 | 0.32 | 0.3 |
| 5 | 126201232 | rs11749282 | 0.316 | -0.429 | -0.557 | 0.277 | 0.054 | 0.044 | 0.044 | 0.357 | 0.4 | 0.26 | 0.29 | 0.25 | 0.33 | 0.3 |
| 5 | 126203506 | rs3533 | -0.619 | 0.105 | -1.41 | 0.66 | -0.142 | -0.175 | -0.175 | 0 | 0.64 | 0.97 | 0.68 | 0.54 | 0.54 | 0.54 |

| | | | | | | | | | | | | | | | | |
|---|---|---|---|---|---|---|---|---|---|---|---|---|---|---|---|---|
| 5 | 126211262 | rs9637916 | 0.377 | -0.119 | -0.234 | 0.877 | 0.007 | 0.018 | 0.018 | 0.012 | 0.13 | 0.23 | 0.31 | 0.18 | 0.16 | 0.09 |
| 5 | 126214855 | rs4836300 | 0.665 | 0.724 | -0.018 | 0.321 | 0.016 | 0.143 | 0.143 | 0.056 | 0.51 | 0.2 | 0.36 | 0.25 | 0.49 | 0.33 |
| 5 | 126220091 | rs7701976 | -1.026 | 0.654 | 1.068 | 0.271 | 0.076 | 0.176 | 0.176 | 1.038 | 0.06 | 0.45 | 0.05 | 0.1 | 0.02 | 0.14 |
| 5 | 126224390 | rs10040840 | -1.384 | 0.644 | 1.068 | 0.266 | 0.055 | 0.15 | 0.15 | 0.715 | 0.07 | 0.46 | 0.06 | 0.1 | 0.02 | 0.15 |
| 5 | 126267351 | rs12655258 | -0.668 | 0.678 | 0.935 | 0.278 | 0.053 | 0.141 | 0.141 | 0.377 | 0.06 | 0.32 | 0.04 | 0.1 | 0.02 | 0.15 |
| 5 | 126282246 | rs11949302 | -0.168 | 0.725 | 0.609 | 0.263 | -0.008 | 0.01 | 0.01 | 0.018 | 0.42 | 0.38 | 0.58 | 0.68 | 0.47 | 0.52 |
| 5 | 126307610 | rs4836307 | -0.664 | 0.7 | 0.368 | 0.214 | -0.006 | 0.105 | 0.105 | 0.015 | 0.48 | 0.69 | 0.62 | 0.78 | 0.49 | 0.67 |
| 5 | 126325510 | rs12659476 | -0.661 | 0.714 | 0.384 | 0.386 | -0.001 | 0.047 | 0.047 | 0.042 | 0.05 | 0.16 | 0.02 | 0.1 | 0.02 | 0.15 |
| 5 | 126340086 | rs10491285 | 1.062 | 0.362 | -0.197 | 0.015 | -0.041 | 0.048 | 0.048 | 0 | 0.3 | 0.06 | 0.26 | 0.18 | 0.31 | 0.28 |
| 5 | 126365228 | rs3828668 | 0.776 | 0.523 | 0.507 | 1.311 | -0.035 | -0.076 | -0.076 | 0.631 | 0.11 | 0.01 | 0.29 | 0.32 | 0.16 | 0.09 |
| 5 | 126376954 | rs6877692 | -1.041 | 0.493 | 1.076 | 0.865 | 0.021 | 0.094 | 0.094 | 0 | 0.05 | 0.22 | 0.03 | 0.1 | 0.01 | 0.15 |
| 5 | 126377240 | rs6860581 | -1.618 | 0.509 | 1.13 | 0.859 | 0.047 | 0.106 | 0.106 | 0.211 | 0.06 | 0.28 | 0.05 | 0.1 | 0.04 | 0.16 |
| 5 | 126394484 | rs17165251 | -1.32 | 0.517 | 1.085 | 0.863 | 0.015 | 0.059 | 0.059 | 0.013 | 0.06 | 0.18 | 0.04 | 0.1 | 0.04 | 0.16 |
| 5 | 126419094 | rs6870711 | -2.232 | 0.484 | 1.085 | 0.837 | 0.035 | 0.077 | 0.077 | 1.121 | 0.07 | 0.36 | 0.05 | 0.08 | 0.04 | 0.16 |
| 5 | 126429749 | rs6898415 | -0.715 | 0.526 | 0.67 | 0.706 | -0.004 | 0.159 | 0.159 | 0 | 0.51 | 0.83 | 0.64 | 0.73 | 0.51 | 0.7 |
| 5 | 126433164 | rs3936305 | -0.413 | 0.547 | 0.946 | 0.705 | -0.005 | 0.009 | 0.009 | 0 | 0.44 | 0.59 | 0.6 | 0.65 | 0.47 | 0.54 |
| 5 | 126436308 | rs758628 | -1.117 | 0.375 | 0.808 | 0.611 | -0.05 | 0.157 | 0.157 | 0 | 0.56 | 0.89 | 0.66 | 0.78 | 0.55 | 0.73 |
| 5 | 126436942 | rs727038 | -0.494 | 0.515 | 1.316 | 0.659 | -0.003 | 0.005 | 0.005 | 0 | 0.44 | 0.57 | 0.6 | 0.65 | 0.46 | 0.58 |
| 5 | 126449925 | rs887372 | -0.25 | 0.496 | 1.537 | 0.67 | -0.011 | 0.013 | 0.013 | 0.007 | 0.42 | 0.36 | 0.58 | 0.65 | 0.46 | 0.58 |
| 5 | 126469930 | rs6884988 | -0.266 | 0.504 | 1.454 | 0.668 | 0.002 | -0.004 | -0.004 | 0 | 0.42 | 0.45 | 0.59 | 0.64 | 0.46 | 0.58 |
| 5 | 126475572 | rs13178353 | -0.61 | 0.508 | 1.349 | 0.671 | -0.034 | 0.06 | 0.06 | 0.011 | 0.46 | 0.8 | 0.63 | 0.64 | 0.5 | 0.59 |
| 5 | 126528653 | rs173555 | 0.059 | 0.508 | 1.222 | 0.706 | -0.002 | 0 | 0 | 0 | 0.59 | 0.6 | 0.41 | 0.36 | 0.53 | 0.42 |
| 5 | 126541479 | rs6894083 | -0.838 | 0.505 | 0.771 | 0.962 | 0 | 0.001 | 0.001 | 0.429 | 0.22 | 0.71 | 0.37 | 0.35 | 0.2 | 0.22 |
| 5 | 126542592 | rs726990 | 0.895 | 0.497 | 0.796 | 0.875 | -0.005 | 0.103 | 0.103 | 0 | 0.51 | 0.25 | 0.36 | 0.3 | 0.49 | 0.33 |
| 5 | 126547326 | rs6897890 | 0.167 | 0.742 | 1.091 | 0.973 | -0.002 | 0.132 | 0.132 | 0.032 | 0.15 | 0.4 | 0.34 | 0.3 | 0.15 | 0.12 |
| 5 | 126551732 | rs7720606 | 0.945 | 0.68 | 0.297 | 0.62 | -0.036 | 0.107 | 0.107 | 0.335 | 0.52 | 0.17 | 0.52 | 0.26 | 0.49 | 0.36 |
| 5 | 126568929 | rs10078274 | 0.864 | 0.815 | 0.704 | 0.564 | 0.001 | 0.027 | 0.027 | 0 | 0.52 | 0.56 | 0.52 | 0.3 | 0.46 | 0.36 |
| 5 | 126569039 | rs10065046 | 0.932 | 0.8 | 0.859 | 0.557 | 0.005 | 0.116 | 0.116 | 0 | 0.5 | 0.24 | 0.5 | 0.3 | 0.45 | 0.36 |
| 5 | 126587180 | rs12186854 | 0.264 | 0.812 | 0.796 | 0.575 | -0.016 | 0.001 | 0.001 | 0 | 0.62 | 0.67 | 0.56 | 0.38 | 0.53 | 0.57 |
| 5 | 126591501 | rs6899068 | -0.664 | 0.819 | 0.56 | 0.636 | -0.129 | -0.118 | -0.118 | 1.869 | 0.75 | 0.98 | 0.71 | 0.63 | 0.67 | 0.69 |
| 5 | 126617387 | rs17165119 | -0.374 | 0.488 | 1.113 | -0.056 | -0.005 | -0.031 | -0.031 | 0 | 0.06 | 0.1 | 0.03 | 0 | 0.06 | 0.1 |
| 5 | 126625548 | rs2408868 | 0.024 | 0.639 | 1.102 | -0.067 | 0.012 | 0.001 | 0.001 | 0 | 0.6 | 0.51 | 0.42 | 0.45 | 0.53 | 0.57 |
| 5 | 126625758 | rs10519929 | -0.395 | 0.617 | 0.955 | -0.082 | -0.008 | -0.022 | -0.022 | 0 | 0.06 | 0.13 | 0.03 | 0.03 | 0.05 | 0.08 |
| 5 | 126626044 | rs1363387 | 0.696 | 0.579 | 0.985 | -0.086 | -0.002 | 0.086 | 0.086 | 0.174 | 0.51 | 0.23 | 0.37 | 0.34 | 0.46 | 0.38 |
| 5 | 126626645 | rs2032834 | 0.821 | 0.56 | 0.975 | -0.095 | -0.004 | 0.092 | 0.092 | 0 | 0.49 | 0.21 | 0.37 | 0.34 | 0.46 | 0.38 |
| 5 | 126628098 | rs2194079 | -0.051 | 0.653 | 0.959 | -0.039 | -0.005 | -0.001 | -0.001 | 4.527 | 0.6 | 0.4 | 0.41 | 0.45 | 0.53 | 0.57 |
| 5 | 126629170 | rs12519784 | 0.275 | 0.748 | 0.88 | -0.054 | -0.016 | 0.031 | 0.031 | 0 | 0.55 | 0.34 | 0.4 | 0.37 | 0.51 | 0.46 |
| 5 | 126631841 | rs2585200 | 0.486 | 0.391 | 0.7 | -0.182 | 0.006 | 0.023 | 0.023 | 0 | 0.57 | 0.39 | 0.48 | 0.31 | 0.59 | 0.4 |
| 5 | 126639591 | rs2431496 | 0.369 | 0.352 | 1.019 | -0.178 | 0.001 | 0.005 | 0.005 | 0 | 0.59 | 0.42 | 0.44 | 0.3 | 0.61 | 0.4 |
| 5 | 126641654 | rs2546082 | -0.137 | 0.518 | 1.067 | -0.034 | 0.003 | 0.001 | 0.001 | 0 | 0.69 | 0.65 | 0.49 | 0.33 | 0.68 | 0.51 |
| 5 | 126641770 | rs2585193 | -0.156 | 0.53 | 1.064 | 0.043 | 0.002 | 0.013 | 0.013 | 0 | 0.71 | 0.65 | 0.5 | 0.43 | 0.69 | 0.63 |
| 5 | 126646864 | rs2161283 | 1.306 | 0.29 | 0.464 | -0.428 | -0.109 | 0.094 | 0.094 | 0.031 | 0.45 | 0.07 | 0.35 | 0.23 | 0.48 | 0.36 |
| 5 | 126650182 | rs1345663 | -0.062 | -0.302 | 0.878 | -0.025 | 0.007 | 0 | 0 | 0 | 0.68 | 0.55 | 0.48 | 0.33 | 0.67 | 0.51 |
| 5 | 126655749 | rs246887 | 0.548 | -0.295 | 0.531 | 0.042 | -0.002 | 0.03 | 0.03 | 0.218 | 0.59 | 0.31 | 0.41 | 0.28 | 0.58 | 0.4 |
| 5 | 126668683 | rs2546078 | 1.015 | -0.416 | 0.196 | -0.035 | 0.009 | 0.09 | 0.09 | 0.191 | 0.51 | 0.15 | 0.36 | 0.27 | 0.51 | 0.36 |
| 5 | 126679697 | rs246896 | 1.207 | -0.369 | 0.166 | -0.006 | -0.041 | 0.037 | 0.037 | 0 | 0.45 | 0.16 | 0.35 | 0.29 | 0.48 | 0.36 |
| 5 | 126683809 | rs27388 | 0.078 | -0.323 | 0.173 | -0.183 | 0.011 | 0.014 | 0.014 | 0.352 | 0.65 | 0.52 | 0.43 | 0.42 | 0.57 | 0.52 |
| 5 | 126683883 | rs27652 | 1.289 | -0.291 | 0.118 | -0.15 | -0.008 | 0.034 | 0.034 | 0 | 0.46 | 0.33 | 0.36 | 0.31 | 0.48 | 0.36 |
| 5 | 126687687 | rs43148 | 0.09 | 0.689 | -0.08 | 0.247 | -0.003 | -0.004 | -0.004 | 0.07 | 0.7 | 0.71 | 0.46 | 0.45 | 0.63 | 0.61 |
| 5 | 126699347 | rs35525 | 0.76 | 0.597 | -0.088 | 0.19 | 0.002 | 0 | 0 | 0 | 0.58 | 0.51 | 0.38 | 0.4 | 0.5 | 0.48 |
| 5 | 126701190 | rs17165035 | -0.925 | 0.439 | -0.056 | 0.076 | -0.002 | -0.016 | -0.016 | 0 | 0.12 | 0.16 | 0.08 | 0.06 | 0.12 | 0.14 |
| 5 | 126701622 | rs35524 | -0.713 | -0.083 | -0.062 | -0.09 | 0.039 | 0.02 | 0.02 | 2.063 | 0.48 | 0.42 | 0.55 | 0.63 | 0.39 | 0.62 |
| 5 | 126713606 | rs1363388 | -1.957 | -0.094 | 1.285 | 0.026 | 0.031 | 0.002 | 0.002 | 0.005 | 0.15 | 0.41 | 0.19 | 0.12 | 0.17 | 0.23 |
| 5 | 126722859 | rs6881894 | -1.947 | -0.138 | 1.971 | 0.015 | 0.027 | -0.004 | -0.004 | 0 | 0.16 | 0.52 | 0.2 | 0.12 | 0.17 | 0.23 |
| 5 | 126726669 | rs6874521 | -1.955 | -0.127 | 1.969 | -0.029 | 0.059 | 0.004 | 0.004 | 0.296 | 0.13 | 0.43 | 0.13 | 0.1 | 0.16 | 0.22 |
| 5 | 126727418 | rs1153432 | 0.174 | -0.601 | 1.632 | -0.249 | -0.003 | 0.008 | 0.008 | 1.473 | 0.75 | 0.73 | 0.61 | 0.44 | 0.8 | 0.66 |
| 5 | 126732399 | rs3812055 | -1.357 | -0.459 | 1.822 | -0.263 | 0.044 | 0.01 | 0.01 | 0.202 | 0.1 | 0.25 | 0.09 | 0.1 | 0.14 | 0.21 |
| 5 | 126732427 | rs3812054 | -0.715 | -0.457 | 1.571 | -0.265 | -0.011 | 0.012 | 0.012 | 0.131 | 0.07 | 0.06 | 0.07 | 0.05 | 0.12 | 0.13 |
| 5 | 126740316 | rs17165090 | -1.995 | -0.487 | 1.714 | -0.12 | 0.032 | 0.017 | 0.017 | 0 | 0.14 | 0.4 | 0.13 | 0.15 | 0.17 | 0.23 |
| 5 | 126744988 | rs17673474 | -0.8 | -0.485 | 1.607 | -0.124 | -0.028 | 0.029 | 0.029 | 0 | 0.09 | 0.04 | 0.06 | 0.05 | 0.12 | 0.14 |
| 5 | 126760822 | rs6897163 | -1.536 | -0.472 | 1.78 | -0.098 | 0.088 | 0.161 | 0.161 | 0 | 0.05 | 0.33 | 0.06 | 0.11 | 0.05 | 0.1 |
| 5 | 126767961 | rs6866678 | 0.189 | -0.649 | 1.676 | -0.05 | -0.014 | 0.045 | 0.045 | 2.42 | 0.75 | 0.67 | 0.61 | 0.45 | 0.81 | 0.65 |
| 5 | 126780212 | rs12516897 | 1.725 | -0.677 | 1.743 | -0.086 | 0.022 | 0.015 | 0.015 | 3.857 | 0.59 | 0.22 | 0.48 | 0.3 | 0.64 | 0.42 |
| 5 | 126791282 | rs17164935 | -1.11 | -0.666 | 1.793 | -0.084 | -0.006 | 0.008 | 0.008 | 0 | 0.11 | 0.1 | 0.1 | 0.1 | 0.15 | 0.15 |
| 5 | 126795159 | rs3756721 | -1.929 | -0.659 | 1.811 | -0.102 | 0.033 | 0.015 | 0.015 | 0 | 0.15 | 0.44 | 0.13 | 0.15 | 0.17 | 0.23 |
| 5 | 126797292 | rs4836323 | 1.005 | -0.674 | 1.648 | -0.091 | 0.015 | -0.011 | -0.011 | 0 | 0.67 | 0.3 | 0.55 | 0.37 | 0.75 | 0.55 |
| 5 | 126798031 | rs17164923 | -1.421 | -0.597 | 1.591 | -0.07 | 0.071 | 0.155 | 0.155 | 1.635 | 0.07 | 0.33 | 0.06 | 0.11 | 0.05 | 0.1 |
| 5 | 126805804 | rs6894441 | 1.765 | -0.604 | 0.867 | -0.48 | 0.005 | 0.063 | 0.063 | 2.76 | 0.53 | 0.15 | 0.44 | 0.29 | 0.58 | 0.38 |
| 5 | 126811978 | rs2898044 | -1.987 | -0.701 | 1.43 | -0.253 | 0.012 | -0.023 | -0.023 | 0.382 | 0.16 | 0.49 | 0.13 | 0.12 | 0.16 | 0.17 |

| | | | | | | | | | | | | | | | | |
|---|---|---|---|---|---|---|---|---|---|---|---|---|---|---|---|---|
| 5 | 126814254 | rs35522 | -1.846 | -0.873 | 1.252 | -0.259 | 0.018 | 0.012 | 0.012 | 4.613 | 0.4 | 0.75 | 0.52 | 0.67 | 0.36 | 0.58 |
| 5 | 126817110 | rs2910219 | -0.272 | -0.772 | 0.957 | -0.201 | 0.006 | -0.05 | -0.05 | 1.286 | 0.22 | 0.16 | 0.39 | 0.54 | 0.2 | 0.35 |
| 5 | 126822042 | rs7718640 | -0.915 | -0.695 | 0.979 | -0.4 | -0.002 | -0.002 | -0.002 | 0.199 | 0.09 | 0.07 | 0.07 | 0.04 | 0.12 | 0.13 |
| 5 | 126835863 | rs151794 | -0.47 | -0.985 | 1.145 | -0.214 | 0.004 | -0.008 | -0.008 | 0.164 | 0.23 | 0.24 | 0.39 | 0.57 | 0.18 | 0.34 |
| 5 | 126838945 | rs6892265 | -1.716 | -1.494 | 1.61 | -0.119 | 0.025 | 0.001 | 0.001 | 0 | 0.1 | 0.28 | 0.05 | 0.08 | 0.06 | 0.1 |
| 5 | 126841120 | rs151796 | -1.718 | -1.525 | 1.58 | -0.139 | -0.008 | 0.001 | 0.001 | 0.093 | 0.34 | 0.63 | 0.46 | 0.64 | 0.25 | 0.43 |
| 5 | 126847729 | rs182844 | -1.223 | -1.354 | 1.391 | -0.028 | -0.01 | 0.01 | 0.01 | 0.895 | 0.3 | 0.64 | 0.44 | 0.63 | 0.23 | 0.42 |
| 5 | 126865295 | rs233031 | -0.51 | -1.575 | 1.237 | -0.173 | 0.022 | -0.014 | -0.014 | 1.67 | 0.39 | 0.31 | 0.45 | 0.55 | 0.29 | 0.37 |
| 5 | 126868653 | rs233038 | -0.463 | -1.576 | 1.251 | -0.173 | 0.051 | -0.039 | -0.039 | 0 | 0.38 | 0.24 | 0.45 | 0.55 | 0.29 | 0.37 |
| 5 | 126887569 | rs7522 | -0.513 | -1.571 | 1.257 | -0.171 | 0.019 | -0.013 | -0.013 | 2.13 | 0.39 | 0.32 | 0.45 | 0.55 | 0.29 | 0.37 |
| 5 | 126887706 | rs7729354 | 0.453 | -1.576 | 1.263 | -0.179 | -0.003 | 0.004 | 0.004 | 0 | 0.05 | 0.06 | 0.03 | 0.02 | 0.04 | 0.04 |
| 5 | 126889640 | rs418657 | -0.687 | -1.403 | 1.384 | -0.024 | 0.019 | -0.013 | -0.013 | 0.033 | 0.39 | 0.32 | 0.45 | 0.55 | 0.29 | 0.37 |
| 5 | 126889986 | rs432672 | -0.687 | -1.403 | 1.39 | -0.024 | 0.019 | -0.013 | -0.013 | 0 | 0.39 | 0.32 | 0.45 | 0.55 | 0.29 | 0.37 |
| 5 | 126890541 | rs384556 | -0.427 | -1.418 | 1.201 | -0.03 | 0.033 | -0.026 | -0.026 | 0.04 | 0.37 | 0.27 | 0.44 | 0.55 | 0.27 | 0.34 |
| 5 | 126897060 | rs374499 | -1.081 | -1.42 | 1.168 | -0.035 | 0.006 | 0.001 | 0.001 | 1.13 | 0.43 | 0.6 | 0.49 | 0.63 | 0.32 | 0.42 |
| 5 | 126899205 | rs4835927 | -0.483 | -1.829 | 1.208 | 0.026 | 0.031 | -0.022 | -0.022 | 3.47 | 0.36 | 0.32 | 0.41 | 0.55 | 0.23 | 0.33 |
| 5 | 126901904 | rs2408952 | -0.815 | -1.793 | 0.903 | 0.038 | -0.016 | 0 | 0 | 1.239 | 0.4 | 0.6 | 0.45 | 0.59 | 0.25 | 0.4 |
| 5 | 126918504 | rs3909885 | -0.996 | -1.485 | 0.539 | 0.279 | 0.008 | -0.001 | -0.001 | 3.152 | 0.4 | 0.65 | 0.44 | 0.61 | 0.25 | 0.42 |
| 5 | 126921292 | rs13180404 | -0.239 | -1.35 | -0.407 | 0.356 | 0.047 | -0.067 | -0.067 | 0.876 | 0.31 | 0.17 | 0.4 | 0.54 | 0.22 | 0.35 |
| 5 | 126930433 | rs6595779 | -0.893 | -1.128 | -0.686 | 0.308 | 0.019 | 0.008 | 0.008 | 0 | 0.43 | 0.69 | 0.48 | 0.65 | 0.3 | 0.47 |
| 5 | 126936096 | rs2447822 | 0.288 | -0.273 | -0.851 | 0.833 | -0.01 | -0.035 | -0.035 | 0 | 0.11 | 0.02 | 0.1 | 0.12 | 0.09 | 0.06 |
| 5 | 126938115 | rs791333 | 0.859 | -0.336 | -0.343 | 0.64 | -0.001 | 0.012 | 0.012 | 0 | 0.88 | 0.57 | 0.9 | 0.89 | 0.93 | 0.82 |
| 5 | 126940058 | rs17685437 | -0.707 | -0.26 | -1.518 | 1.035 | 0.021 | -0.057 | -0.057 | 0 | 0.17 | 0.04 | 0.33 | 0.29 | 0.13 | 0.1 |
| 5 | 126943278 | rs17685455 | -0.91 | -0.091 | -1.433 | 1.061 | 0.004 | -0.019 | -0.019 | 0 | 0.24 | 0.18 | 0.34 | 0.63 | 0.22 | 0.33 |
| 5 | 126944070 | rs3906536 | 0.282 | 0.186 | -1.94 | 0.052 | 0.002 | 0.003 | 0.003 | 0.316 | 0.35 | 0.65 | 0.23 | 0.15 | 0.28 | 0.22 |
| 5 | 126946236 | rs10035170 | 0.19 | 0.099 | -1.593 | 0.121 | -0.006 | -0.029 | -0.029 | 0 | 0.34 | 0.58 | 0.23 | 0.16 | 0.28 | 0.22 |
| 5 | 126953752 | rs7723560 | 0.584 | 0.068 | -1.507 | -0.024 | -0.019 | 0.058 | 0.058 | 0 | 0.21 | 0.1 | 0.13 | 0 | 0.19 | 0.12 |
| 5 | 126958230 | rs791326 | 0.804 | 0.261 | -2.262 | -0.68 | -0.065 | 0.058 | 0.058 | 0.407 | 0.33 | 0.02 | 0.23 | 0.16 | 0.33 | 0.32 |
| 5 | 126963224 | rs17764057 | 0.657 | 0.425 | -2.059 | -0.243 | 0.011 | 0.054 | 0.054 | 0 | 0.05 | 0 | 0.16 | 0.6 | 0.03 | 0.27 |
| 5 | 126965630 | rs17764063 | -0.155 | 0.455 | -1.985 | 0.071 | -0.018 | -0.045 | -0.045 | 0.107 | 0.15 | 0.06 | 0.11 | 0.45 | 0.18 | 0.31 |
| 5 | 126967831 | rs1031847 | 0.257 | 0.668 | -1.353 | -0.816 | -0.004 | -0.052 | -0.052 | 0 | 0.26 | 0.61 | 0.17 | 0.05 | 0.19 | 0.13 |
| 5 | 126972005 | rs2408996 | -0.411 | 1.075 | -1.253 | -0.901 | 0.005 | -0.073 | -0.073 | 0.019 | 0.31 | 0.6 | 0.23 | 0.02 | 0.32 | 0.17 |
| 5 | 126975509 | rs2898063 | 0.143 | 1.001 | -1.03 | -0.389 | 0.015 | -0.12 | -0.12 | 0 | 0.22 | 0.55 | 0.15 | 0.02 | 0.18 | 0.12 |
| 5 | 126982737 | rs6595787 | 0.487 | 0.924 | -1.54 | -0.39 | -0.006 | 0.01 | 0.01 | 7.862 | 0.22 | 0.16 | 0.35 | 0.17 | 0.26 | 0.19 |
| 5 | 126984115 | rs10041400 | -0.391 | 0.961 | -0.786 | -0.209 | 0 | -0.075 | -0.075 | 0.009 | 0.22 | 0.63 | 0.15 | 0.02 | 0.17 | 0.12 |
| 5 | 126986357 | rs791077 | 1.014 | 1.468 | -0.313 | 0.637 | 0.038 | 0.058 | 0.058 | 0 | 0.81 | 0.64 | 0.62 | 0.77 | 0.75 | 0.81 |
| 5 | 126987061 | rs6595788 | -0.609 | 1.43 | 0.019 | 0.17 | 0.001 | -0.03 | -0.03 | 6.652 | 0.25 | 0.25 | 0.18 | 0.11 | 0.26 | 0.29 |
| 5 | 126988749 | rs17764136 | 0.526 | 1.284 | -1.172 | 0.208 | -0.036 | -0.016 | -0.016 | 0 | 0.31 | 0.09 | 0.21 | 0.24 | 0.3 | 0.18 |
| 5 | 126990095 | rs791076 | -0.041 | 0.785 | -1.168 | -0.367 | -0.001 | 0.09 | 0.09 | 0 | 0.32 | 0.16 | 0.21 | 0.09 | 0.31 | 0.36 |
| 5 | 126990589 | rs791075 | -1.151 | 0.493 | -1.324 | 0.001 | -0.018 | -0.014 | -0.014 | 0 | 0.79 | 0.88 | 0.86 | 0.8 | 0.79 | 0.81 |
| 5 | 126992734 | rs248710 | -0.637 | 0.228 | -0.576 | -0.3 | -0.007 | -0.006 | -0.006 | 0.319 | 0.35 | 0.39 | 0.39 | 0.38 | 0.35 | 0.38 |
| 5 | 126999937 | rs248694 | 0.682 | -0.182 | -0.348 | -0.095 | -0.004 | 0.012 | 0.012 | 1.913 | 0.42 | 0.35 | 0.28 | 0.24 | 0.4 | 0.23 |
| 5 | 127003242 | rs2673765 | 0.08 | -0.353 | -0.287 | 0.03 | 0.003 | -0.024 | -0.024 | 0.094 | 0.18 | 0.21 | 0.16 | 0.12 | 0.24 | 0.25 |
| 5 | 127006007 | rs11241937 | 0.612 | -0.327 | -0.269 | 0.076 | -0.025 | 0.025 | 0.025 | 0.565 | 0.08 | 0.15 | 0.16 | 0.26 | 0.01 | 0.09 |
| 5 | 127062190 | rs12515892 | 1.035 | -0.308 | -0.34 | 0.035 | -0.034 | -0.081 | -0.081 | 0.238 | 0.31 | 0.17 | 0.39 | 0.39 | 0.34 | 0.44 |
| 5 | 127069127 | rs2673748 | 0.294 | -0.381 | -0.275 | -0.012 | 0.005 | 0.007 | 0.007 | 3.255 | 0.51 | 0.62 | 0.46 | 0.51 | 0.41 | 0.31 |
| 5 | 127069197 | rs2569152 | -0.24 | -0.352 | -0.399 | -0.017 | 0.002 | -0.027 | -0.027 | 0.014 | 0.81 | 0.79 | 0.84 | 0.9 | 0.76 | 0.76 |
| 5 | 127078541 | rs10054323 | 0.764 | -0.219 | -1.08 | -0.008 | -0.061 | 0.059 | 0.059 | 3.097 | 0.4 | 0.11 | 0.27 | 0.25 | 0.39 | 0.23 |
| 5 | 127090699 | rs2673772 | 0.82 | -0.407 | -0.912 | 0.192 | -0.037 | -0.097 | -0.097 | 1.105 | 0.31 | 0.09 | 0.35 | 0.39 | 0.33 | 0.45 |
| 5 | 127110486 | rs10463832 | 0.678 | -0.367 | -0.739 | 0.288 | -0.016 | 0.024 | 0.024 | 0.06 | 0.06 | 0.09 | 0.15 | 0.27 | 0.01 | 0.07 |
| 5 | 127122492 | rs11241942 | 0.688 | -0.445 | -0.887 | 0.175 | -0.025 | -0.09 | -0.09 | 0.005 | 0.34 | 0.09 | 0.36 | 0.39 | 0.34 | 0.45 |
| 5 | 127139616 | rs1515641 | 0.487 | -0.336 | -0.438 | 0.222 | -0.014 | 0.021 | 0.021 | 0.06 | 0.06 | 0.09 | 0.14 | 0.27 | 0.01 | 0.07 |
| 5 | 127147382 | rs10051423 | 0.588 | 0.21 | -0.849 | -0.054 | -0.038 | 0.027 | 0.027 | 0 | 0.31 | 0.09 | 0.23 | 0.22 | 0.32 | 0.14 |
| 5 | 127147547 | rs790480 | 0.737 | 0.082 | -0.273 | -0.1 | -0.009 | 0 | 0 | 0.153 | 0.75 | 0.42 | 0.63 | 0.68 | 0.75 | 0.65 |
| 5 | 127148845 | rs790477 | 0.137 | 0.157 | -0.162 | -0.177 | 0.001 | -0.025 | -0.025 | 0.107 | 0.82 | 0.82 | 0.8 | 0.94 | 0.77 | 0.72 |
| 5 | 127149861 | rs790475 | 1.14 | 0.126 | -0.372 | -0.221 | -0.046 | -0.091 | -0.091 | 0.142 | 0.42 | 0.11 | 0.39 | 0.46 | 0.41 | 0.51 |
| 5 | 127151003 | rs790474 | -0.672 | 0.073 | -0.245 | -0.293 | -0.004 | 0.019 | 0.019 | 0 | 0.89 | 0.93 | 0.85 | 0.95 | 0.84 | 0.82 |
| 5 | 127151976 | rs1421746 | 0.143 | 0.098 | -0.218 | -0.247 | -0.003 | -0.014 | -0.014 | 0 | 0.37 | 0.71 | 0.41 | 0.48 | 0.36 | 0.22 |
| 5 | 127155001 | rs711359 | 0.178 | -0.136 | -0.109 | -0.406 | 0 | -0.014 | -0.014 | 0.605 | 0.82 | 0.84 | 0.8 | 0.95 | 0.77 | 0.72 |
| 5 | 127156760 | rs698171 | 0.312 | -0.757 | -0.032 | -0.345 | 0.013 | 0.022 | 0.022 | 0 | 0.43 | 0.36 | 0.46 | 0.77 | 0.3 | 0.53 |
| 5 | 127169212 | rs245201 | 0.52 | -0.98 | -0.373 | -0.194 | 0.052 | -0.026 | -0.026 | 0 | 0.45 | 0.22 | 0.35 | 0.42 | 0.31 | 0.49 |
| 5 | 127170643 | rs151849 | 0.52 | -0.968 | -0.373 | -0.194 | 0.056 | -0.026 | -0.026 | 0 | 0.45 | 0.22 | 0.35 | 0.42 | 0.31 | 0.49 |
| 5 | 127196066 | rs6864687 | 0.811 | -0.98 | -0.321 | -0.689 | -0.02 | -0.025 | -0.025 | 0 | 0.33 | 0.31 | 0.4 | 0.53 | 0.42 | 0.24 |
| 5 | 127203192 | rs245178 | -0.859 | -1.244 | -0.52 | -0.588 | -0.002 | -0.114 | -0.114 | 0 | 0.6 | 0.9 | 0.68 | 0.58 | 0.69 | 0.51 |
| 5 | 127212228 | rs1421750 | -0.364 | -1.245 | 0.389 | -0.69 | 0.012 | -0.003 | -0.003 | 0 | 0.53 | 0.6 | 0.61 | 0.54 | 0.67 | 0.59 |
| 5 | 127218995 | rs6894681 | 0.159 | -1.183 | 0.583 | -0.646 | -0.018 | -0.028 | -0.028 | 0.743 | 0.06 | 0.2 | 0.04 | 0 | 0.01 | 0 |
| 5 | 127219305 | rs245237 | 0.573 | -1.269 | 0.524 | -0.7 | -0.002 | 0 | 0 | 0 | 0.4 | 0.38 | 0.36 | 0.46 | 0.29 | 0.39 |
| 5 | 127225652 | rs151848 | -0.513 | -0.393 | 0.629 | -0.173 | -0.005 | 0.024 | 0.024 | 0 | 0.22 | 0.39 | 0.3 | 0.31 | 0.21 | 0.31 |
| 5 | 127234944 | rs193495 | -1.086 | -0.444 | 0.138 | -0.209 | 0.028 | -0.029 | -0.029 | 0 | 0.58 | 0.79 | 0.64 | 0.54 | 0.68 | 0.6 |

| | | | | | | | | | | | | | | | | |
|---|---|---|---|---|---|---|---|---|---|---|---|---|---|---|---|---|
| 5 | 127236014 | rs2147866 | 0.48 | -0.447 | 0.354 | -0.258 | 0.02 | -0.035 | -0.035 | 0 | 0.37 | 0.5 | 0.35 | 0.23 | 0.47 | 0.28 |
| 5 | 127236220 | rs245161 | -0.514 | -0.494 | 0.502 | -0.085 | -0.002 | 0.007 | 0.007 | 0 | 0.22 | 0.29 | 0.3 | 0.31 | 0.21 | 0.31 |
| 5 | 127243524 | rs6891675 | 0.245 | -1.21 | 0.459 | -0.06 | 0.004 | -0.033 | -0.033 | 0 | 0.38 | 0.5 | 0.42 | 0.23 | 0.56 | 0.35 |
| 5 | 127248485 | rs734642 | -0.966 | -1.107 | 0.889 | 0.427 | -0.013 | 0.016 | 0.016 | 0 | 0.18 | 0.25 | 0.24 | 0.31 | 0.16 | 0.25 |
| 5 | 127257406 | rs7711172 | -2.12 | -0.931 | 1.312 | 0.733 | 0.039 | 0.424 | 0.424 | 0.948 | 0.07 | 0.52 | 0.18 | 0.31 | 0.01 | 0.06 |
| 5 | 127260499 | rs10477678 | -2.003 | -0.928 | 1.188 | 0.735 | 0.046 | 0.432 | 0.432 | 1.004 | 0.06 | 0.46 | 0.17 | 0.31 | 0.01 | 0.06 |
| 5 | 127266725 | rs1520246 | -1.244 | -0.959 | 0.942 | 0.607 | -0.013 | 0.168 | 0.168 | 0 | 0.06 | 0.22 | 0.19 | 0.3 | 0.02 | 0.09 |
| 5 | 127267482 | rs9327458 | 0.695 | -1.176 | 0.134 | -0.029 | -0.027 | 0.039 | 0.039 | 0.017 | 0.4 | 0.16 | 0.52 | 0.63 | 0.57 | 0.54 |
| 5 | 127272483 | rs11738398 | -0.557 | -1.606 | 0.36 | -0.307 | -0.011 | -0.048 | -0.048 | 1.683 | 0.67 | 0.39 | 0.83 | 0.95 | 0.84 | 0.8 |
| 5 | 127274447 | rs7717741 | -0.814 | -1.633 | 0.558 | -0.32 | -0.042 | -0.102 | -0.102 | 0 | 0.7 | 0.61 | 0.84 | 0.95 | 0.84 | 0.8 |
| 5 | 127277217 | rs1013375 | 0.531 | -1.463 | 0.52 | -0.368 | 0.001 | -0.034 | -0.034 | 0.065 | 0.48 | 0.54 | 0.7 | 0.92 | 0.6 | 0.67 |
| 5 | 127285152 | rs12517005 | -0.342 | -1.236 | 0.54 | -0.076 | -0.058 | -0.013 | -0.013 | 0 | 0.66 | 0.51 | 0.78 | 0.66 | 0.82 | 0.71 |
| 5 | 127294777 | rs12523625 | 0.85 | -1.154 | 0.404 | -0.056 | -0.005 | 0.029 | 0.029 | 0 | 0.44 | 0.24 | 0.64 | 0.61 | 0.58 | 0.57 |
| 5 | 127304700 | rs17606751 | 1.01 | -0.932 | -0.098 | 0.435 | -0.032 | 0.093 | 0.093 | 0 | 0.43 | 0.2 | 0.41 | 0.25 | 0.56 | 0.48 |
| 5 | 127314400 | rs9327462 | 1.048 | -0.725 | -0.246 | 0.477 | -0.036 | 0.08 | 0.08 | 0 | 0.42 | 0.23 | 0.46 | 0.25 | 0.54 | 0.48 |
| 5 | 127317581 | rs17839662 | -0.496 | -0.375 | -0.665 | 0.133 | -0.021 | 0.044 | 0.044 | 0 | 0.2 | 0.12 | 0.14 | 0.04 | 0.24 | 0.15 |
| 5 | 127337086 | rs4835937 | 0.165 | -0.46 | -0.76 | -0.594 | 0.011 | -0.023 | -0.023 | 0.256 | 0.67 | 0.72 | 0.82 | 0.98 | 0.77 | 0.7 |
| 5 | 127347815 | rs1003136 | -0.33 | -0.134 | -0.975 | 0.14 | 0.02 | 0.004 | 0.004 | 0 | 0.4 | 0.26 | 0.26 | 0.43 | 0.33 | 0.44 |
| 5 | 127350549 | rs3749748 | 0.357 | -0.412 | -1.327 | 0.073 | -0.027 | -0.094 | -0.094 | 0 | 0.25 | 0.02 | 0.16 | 0.35 | 0.21 | 0.37 |
| 5 | 127357526 | rs17764730 | 0.357 | -0.412 | -1.327 | 0.073 | -0.027 | -0.094 | -0.094 | 0 | 0.25 | 0.02 | 0.16 | 0.35 | 0.21 | 0.37 |
| 5 | 127362543 | rs12188497 | 0.874 | -0.146 | -1.636 | -0.236 | -0.022 | 0.032 | 0.032 | 0.07 | 0.06 | 0 | 0.04 | 0 | 0.07 | 0.02 |
| 5 | 127371588 | rs10063647 | -0.606 | -0.122 | -1.446 | 0.074 | 0.027 | 0.132 | 0.132 | 0.05 | 0.51 | 0.81 | 0.57 | 0.73 | 0.42 | 0.58 |
| 5 | 127378828 | rs722888 | -1.399 | 0.515 | -2.128 | -0.456 | -0.053 | -0.032 | -0.032 | 0 | 0.82 | 0.95 | 0.73 | 0.78 | 0.73 | 0.83 |
| 5 | 127420700 | rs1864922 | -0.171 | 0.306 | -1.068 | -0.139 | 0 | 0.001 | 0.001 | 0.988 | 0.24 | 0.74 | 0.4 | 0.33 | 0.19 | 0.2 |
| 5 | 127468177 | rs806100 | 0.067 | 0.298 | -1.062 | -0.199 | 0.037 | 0.091 | 0.091 | 0.037 | 0.51 | 0.25 | 0.45 | 0.29 | 0.6 | 0.43 |
| 5 | 127480998 | rs3805608 | 0.934 | 0.262 | -0.65 | -0.206 | 0 | -0.016 | -0.016 | 0 | 0.12 | 0.25 | 0.08 | 0.07 | 0.1 | 0.06 |
| 5 | 127493206 | rs2054444 | 0.144 | 0.216 | -0.216 | -0.085 | -0.002 | 0.07 | 0.07 | 0.079 | 0.22 | 0.54 | 0.38 | 0.31 | 0.19 | 0.21 |
| 5 | 127498380 | rs790155 | -1.534 | 0.231 | -0.5 | -0.117 | -0.021 | -0.134 | -0.134 | 0 | 0.74 | 0.98 | 0.85 | 0.59 | 0.8 | 0.64 |
| 5 | 127519049 | rs3805615 | 0.238 | 0.302 | -0.262 | -0.099 | 0.008 | 0.086 | 0.086 | 0 | 0.21 | 0.41 | 0.37 | 0.32 | 0.19 | 0.22 |
| 5 | 127522543 | rs10089 | 0.467 | 0.306 | -0.574 | -0.091 | 0.001 | 0.016 | 0.016 | 0 | 0.19 | 0.19 | 0.35 | 0.32 | 0.19 | 0.21 |
| 5 | 127528880 | rs10519976 | 0.801 | 0.45 | -1.063 | -0.805 | -0.054 | 0.109 | 0.109 | 0 | 0.22 | 0.01 | 0.13 | 0.04 | 0.25 | 0.15 |
| 5 | 127543622 | rs251215 | -0.229 | 0.236 | -0.625 | -0.757 | 0.031 | 0.153 | 0.153 | 0.233 | 0.5 | 0.21 | 0.45 | 0.25 | 0.58 | 0.42 |
| 5 | 127548700 | rs36693 | -1.401 | 0.257 | -0.768 | -0.482 | 0.016 | -0.097 | -0.097 | 1.149 | 0.73 | 0.93 | 0.71 | 0.43 | 0.78 | 0.65 |
| 5 | 127550055 | rs1351141 | 0.439 | 0.271 | -0.097 | -0.502 | 0.025 | -0.011 | -0.011 | 0 | 0.2 | 0.45 | 0.26 | 0.2 | 0.21 | 0.19 |
| 5 | 127552827 | rs36703 | 1.267 | 0.492 | -0.829 | -0.721 | -0.059 | -0.016 | -0.016 | 0 | 0.18 | 0.05 | 0.27 | 0.19 | 0.26 | 0.15 |
| 5 | 127557814 | rs11955148 | 0.462 | 0.273 | -0.065 | -0.404 | 0.021 | 0.005 | 0.005 | 0 | 0.19 | 0.34 | 0.24 | 0.23 | 0.2 | 0.19 |
| 5 | 127562880 | rs36715 | -0.952 | 0.061 | -0.158 | -0.414 | -0.041 | 0.056 | 0.056 | 0 | 0.69 | 0.53 | 0.67 | 0.47 | 0.78 | 0.65 |
| 5 | 127563563 | rs36716 | 0.471 | 0.158 | -1.12 | -0.525 | -0.016 | 0.122 | 0.122 | 0.236 | 0.46 | 0.21 | 0.4 | 0.24 | 0.52 | 0.37 |
| 5 | 127574115 | rs6891025 | -1.317 | 0.764 | -0.696 | -0.706 | -0.001 | -0.024 | -0.024 | 2.208 | 0.13 | 0.74 | 0.13 | 0.05 | 0.13 | 0.08 |
| 5 | 127576726 | rs17608067 | 0.245 | 1.154 | -1.264 | -0.541 | -0.078 | -0.012 | -0.012 | 0 | 0.22 | 0.03 | 0.25 | 0.19 | 0.26 | 0.26 |
| 5 | 127592925 | rs17676260 | 0.502 | 0.885 | -0.569 | -0.368 | -0.035 | -0.01 | -0.01 | 0 | 0.06 | 0 | 0.06 | 0.05 | 0.11 | 0.07 |
| 5 | 127593870 | rs7288 | -0.48 | 0.877 | -0.354 | -0.37 | 0.024 | -0.011 | -0.011 | 0.039 | 0.09 | 0.2 | 0.09 | 0.05 | 0.13 | 0.08 |
| 5 | 127609633 | rs2291628 | 0.702 | 0.258 | -0.999 | -0.696 | 0.012 | 0.015 | 0.015 | 0.058 | 0.12 | 0.08 | 0.06 | 0.07 | 0.07 | 0.1 |
| 5 | 127610002 | rs2307108 | -0.624 | 0.391 | -1.326 | -1.231 | 0.001 | 0.081 | 0.081 | 0.59 | 0.55 | 0.71 | 0.62 | 0.3 | 0.55 | 0.55 |
| 5 | 127614472 | rs190450 | -0.531 | 0.098 | -0.824 | -1.466 | 0.007 | -0.056 | -0.056 | 0 | 0.71 | 0.61 | 0.71 | 0.85 | 0.7 | 0.79 |
| 5 | 127617336 | rs32206 | -1.17 | 0.12 | -0.904 | -1.638 | 0.003 | -0.027 | -0.027 | 0 | 0.84 | 0.84 | 0.83 | 0.96 | 0.78 | 0.86 |
| 5 | 127622747 | rs11746935 | 1.534 | -0.184 | -1.373 | -2.542 | -0.022 | 0.244 | 0.244 | 0 | 0.36 | 0.04 | 0.24 | 0.02 | 0.33 | 0.23 |
| 5 | 127638759 | rs11955288 | 1.331 | 0.264 | -0.964 | -2.654 | 0.015 | 0.02 | 0.02 | 0 | 0.14 | 0.1 | 0.07 | 0.07 | 0.1 | 0.12 |
| 5 | 127640076 | rs32216 | -0.226 | 0.351 | -0.923 | -2.766 | 0 | -0.011 | -0.011 | 0.733 | 0.72 | 0.72 | 0.71 | 0.85 | 0.7 | 0.76 |
| 5 | 127640861 | rs27855 | 0.607 | 0.408 | -1.186 | -2.713 | 0.02 | 0.196 | 0.196 | 0.003 | 0.53 | 0.12 | 0.61 | 0.79 | 0.61 | 0.64 |
| 5 | 127641916 | rs32220 | 0.978 | 0.896 | -0.491 | -2.454 | -0.026 | 0.037 | 0.037 | 1.772 | 0.17 | 0.01 | 0.13 | 0.07 | 0.12 | 0.13 |
| 5 | 127661976 | rs32215 | 1.409 | 0.957 | -0.187 | -2.598 | 0.021 | 0.042 | 0.042 | 0 | 0.42 | 0.36 | 0.48 | 0.79 | 0.37 | 0.52 |
| 5 | 127666565 | rs469722 | -0.047 | 1.078 | -0.013 | -2.665 | -0.008 | 0.016 | 0.016 | 0 | 0.33 | 0.24 | 0.29 | 0.13 | 0.32 | 0.25 |
| 5 | 127667613 | rs467610 | 1.754 | 1.075 | -0.12 | -2.533 | 0.027 | 0.012 | 0.012 | 0 | 0.35 | 0.28 | 0.25 | 0.67 | 0.3 | 0.36 |
| 5 | 127680818 | rs468026 | -0.934 | 1.153 | -0.373 | -2.728 | 0.014 | -0.039 | -0.039 | 0.2 | 0.84 | 0.67 | 0.83 | 0.92 | 0.8 | 0.86 |
| 5 | 127682184 | rs11746746 | 2.378 | 1.11 | -0.573 | -2.816 | 0.088 | 0.196 | 0.196 | 0 | 0.3 | 0.02 | 0.15 | 0.03 | 0.2 | 0.17 |
| 5 | 127685135 | rs154001 | 0.367 | 1.193 | -0.44 | -2.746 | 0.013 | -0.077 | -0.077 | 0 | 0.73 | 0.6 | 0.76 | 0.92 | 0.67 | 0.76 |
| 5 | 127685624 | rs255690 | 0.007 | 1.516 | -0.631 | -2.423 | -0.074 | 0.121 | 0.121 | 0.116 | 0.31 | 0.06 | 0.27 | 0.12 | 0.32 | 0.25 |
| 5 | 127691272 | rs3792871 | 0.512 | 1.388 | -0.469 | -2.204 | -0.018 | 0.021 | 0.021 | 0 | 0.09 | 0.06 | 0.07 | 0.01 | 0.12 | 0.09 |
| 5 | 127694805 | rs27754 | -0.599 | 1.371 | -0.429 | -2.365 | -0.011 | 0.087 | 0.087 | 0.008 | 0.31 | 0.16 | 0.26 | 0.13 | 0.29 | 0.3 |
| 5 | 127705251 | rs331098 | -0.839 | 1.505 | -0.777 | -2.481 | -0.071 | 0.118 | 0.118 | 0 | 0.4 | 0.11 | 0.47 | 0.19 | 0.43 | 0.3 |
| 5 | 127706046 | rs331097 | -2.263 | 1.486 | -0.235 | -2.6 | 0.018 | -0.002 | -0.002 | 0 | 0.62 | 0.7 | 0.73 | 0.98 | 0.69 | 0.72 |
| 5 | 127707075 | rs331096 | 0.283 | 1.374 | -0.442 | -2.481 | 0.003 | 0.058 | 0.058 | 0.534 | 0.24 | 0.08 | 0.32 | 0.12 | 0.19 | 0.23 |
| 5 | 127712939 | rs331094 | -1.023 | 1.374 | -0.033 | -2.486 | 0.005 | -0.06 | -0.06 | 0.224 | 0.44 | 0.48 | 0.49 | 0.13 | 0.45 | 0.34 |
| 5 | 127714784 | rs17676891 | 1.131 | 1.223 | -0.016 | -2.427 | -0.008 | -0.023 | -0.023 | 0 | 0.06 | 0.02 | 0.05 | 0.07 | 0.09 | 0.04 |
| 5 | 127716424 | rs331089 | -1.299 | 1.232 | 0.197 | -2.483 | 0.013 | 0.072 | 0.072 | 0 | 0.5 | 0.35 | 0.43 | 0.19 | 0.54 | 0.38 |
| 5 | 127722735 | rs331074 | -1.125 | 1.261 | 0.246 | -2.504 | -0.008 | -0.044 | -0.044 | 0 | 0.42 | 0.52 | 0.34 | 0.19 | 0.39 | 0.34 |
| 5 | 127725467 | rs331069 | -0.913 | 1.3 | -0.055 | -2.508 | -0.017 | 0.06 | 0.06 | 0 | 0.44 | 0.27 | 0.5 | 0.13 | 0.46 | 0.34 |
| 5 | 127739700 | rs10041869 | -0.234 | 1.098 | 0.757 | -1.542 | 0.023 | -0.05 | -0.05 | 0.204 | 0.09 | 0.25 | 0.07 | 0.01 | 0.09 | 0.09 |

| | | | | | | | | | | | | | | | | |
|---|---|---|---|---|---|---|---|---|---|---|---|---|---|---|---|---|
| 5 | 127740576 | rs10463844 | 0.78 | 1.233 | 0.78 | -1.392 | 0.008 | 0.064 | 0.064 | 0 | 0.06 | 0.07 | 0.1 | 0.69 | 0.11 | 0.22 |
| 5 | 127742721 | rs6868664 | -0.225 | 1.176 | 0.839 | -1.535 | 0.052 | -0.004 | -0.004 | 0 | 0.14 | 0.52 | 0.2 | 0.11 | 0.13 | 0.19 |
| 5 | 127755378 | rs331087 | -2.584 | 0.517 | 0.493 | -1.405 | -0.097 | 0.078 | 0.078 | 0 | 0.56 | 0.93 | 0.66 | 0.29 | 0.52 | 0.55 |
| 5 | 127756829 | rs3805651 | -0.909 | 0.569 | 0.49 | -1.389 | -0.005 | 0.013 | 0.013 | 0 | 0.15 | 0.11 | 0.11 | 0.11 | 0.19 | 0.16 |
| 5 | 127768601 | rs331082 | 0.153 | 0.546 | 0.73 | -1.357 | -0.002 | -0.007 | -0.007 | 0 | 0.18 | 0.14 | 0.28 | 0.06 | 0.11 | 0.11 |
| 5 | 127771987 | rs1848501 | -0.238 | 0.852 | 1.176 | -0.898 | 0.014 | -0.046 | -0.046 | 0.619 | 0.08 | 0.19 | 0.07 | 0.01 | 0.09 | 0.09 |
| 5 | 127780460 | rs6883276 | -0.34 | 0.852 | 1.175 | -0.898 | 0.016 | -0.048 | -0.048 | 0 | 0.09 | 0.22 | 0.07 | 0.01 | 0.09 | 0.09 |
| 5 | 127787964 | rs11241959 | -2.642 | 0.852 | 0.76 | -0.904 | -0.019 | 0.188 | 0.188 | 0 | 0.48 | 0.9 | 0.41 | 0.23 | 0.46 | 0.48 |
| 5 | 127801323 | rs17677244 | -0.081 | 0.085 | 1.283 | -0.439 | -0.003 | -0.009 | -0.009 | 1.767 | 0.08 | 0.05 | 0.06 | 0.01 | 0.1 | 0.09 |
| 5 | 127803816 | rs6595827 | -2.166 | 0.056 | 1.439 | -0.457 | -0.003 | -0.003 | -0.003 | 0 | 0.32 | 0.68 | 0.26 | 0.22 | 0.27 | 0.34 |
| 5 | 127819132 | rs10520002 | 0.146 | 0.017 | 1.419 | -0.488 | -0.021 | -0.021 | -0.021 | 0 | 0.06 | 0.01 | 0.05 | 0.07 | 0.09 | 0.04 |
| 5 | 127822876 | rs4836375 | -2.002 | -0.078 | 1.606 | -0.511 | 0.008 | -0.036 | -0.036 | 0 | 0.22 | 0.3 | 0.18 | 0.12 | 0.28 | 0.25 |
| 5 | 127823986 | rs6864057 | -1.295 | -0.372 | 1.68 | -0.448 | -0.047 | -0.031 | -0.031 | 0 | 0.19 | 0.58 | 0.14 | 0.11 | 0.09 | 0.18 |
| 5 | 127825485 | rs968008 | -0.248 | -0.37 | 1.656 | -0.448 | 0.018 | -0.044 | -0.044 | 0.014 | 0.08 | 0.2 | 0.05 | 0.01 | 0.08 | 0.09 |
| 5 | 127832856 | rs11960575 | -2.123 | -0.358 | 1.431 | -0.44 | 0.034 | 0.054 | 0.054 | 0 | 0.28 | 0.82 | 0.2 | 0.12 | 0.18 | 0.27 |
| 5 | 127850855 | rs17164094 | -0.738 | -0.37 | 1.772 | -0.779 | 0.036 | 0.087 | 0.087 | 0 | 0.1 | 0.46 | 0.12 | 0.11 | 0.06 | 0.15 |
| 5 | 127856474 | rs17164083 | -0.227 | -0.323 | 1.404 | -0.795 | 0.003 | -0.038 | -0.038 | 0.652 | 0.08 | 0.11 | 0.05 | 0.01 | 0.08 | 0.09 |
| 5 | 127856494 | rs13155917 | -0.965 | -0.22 | 1.165 | -0.739 | 0.002 | 0.01 | 0.01 | 0 | 0.19 | 0.14 | 0.14 | 0.11 | 0.22 | 0.17 |
| 5 | 127860887 | rs6595836 | -0.492 | 0.001 | 1.463 | -0.45 | 0.031 | 0.08 | 0.08 | 0 | 0.08 | 0.32 | 0.11 | 0.12 | 0.05 | 0.15 |
| 5 | 127871750 | rs6860901 | -1.81 | -0.13 | 0.755 | -0.63 | 0.026 | 0.074 | 0.074 | 0 | 0.35 | 0.86 | 0.27 | 0.19 | 0.31 | 0.33 |
| 5 | 127895438 | rs10040056 | -0.35 | 0.034 | 1.333 | -0.626 | 0.041 | 0.087 | 0.087 | 0 | 0.08 | 0.27 | 0.1 | 0.11 | 0.06 | 0.15 |
| 5 | 127915543 | rs4345355 | -0.914 | -0.009 | 1.618 | -0.715 | 0.008 | -0.059 | -0.059 | 0 | 0.11 | 0.3 | 0.09 | 0.01 | 0.13 | 0.09 |
| 5 | 127923763 | rs11956881 | -0.914 | -0.009 | 1.618 | -0.715 | 0.008 | -0.059 | -0.059 | 1.23 | 0.11 | 0.3 | 0.09 | 0.01 | 0.12 | 0.09 |
| 5 | 127932658 | rs6859278 | -1.958 | -0.115 | 1.38 | -0.706 | 0.026 | 0.049 | 0.049 | 0 | 0.31 | 0.86 | 0.24 | 0.18 | 0.26 | 0.24 |
| 5 | 127982590 | rs10044764 | -1.254 | -0.056 | 1.833 | -0.589 | 0.084 | 0.2 | 0.2 | 0 | 0.05 | 0.33 | 0.07 | 0.11 | 0.02 | 0.07 |
| 5 | 127985881 | rs7730549 | -1.287 | -0.244 | 1.061 | -0.703 | -0.041 | -0.019 | -0.019 | 0.095 | 0.23 | 0.53 | 0.22 | 0.16 | 0.13 | 0.15 |
| 5 | 127999225 | rs10520006 | -1.396 | -0.222 | 0.482 | -0.693 | 0.004 | 0.003 | 0.003 | 1.304 | 0.24 | 0.69 | 0.23 | 0.16 | 0.14 | 0.15 |
| 5 | 128010071 | rs10053477 | -1.169 | 0.354 | 0.898 | -0.341 | 0.04 | 0.075 | 0.075 | 0 | 0.12 | 0.45 | 0.11 | 0.13 | 0.1 | 0.15 |
| 5 | 128011688 | rs4836390 | -0.285 | 0.316 | 0.115 | -0.473 | -0.029 | 0.037 | 0.037 | 0 | 0.13 | 0.03 | 0.17 | 0.05 | 0.16 | 0.1 |
| 5 | 128015559 | rs4835955 | -0.081 | 0.35 | 0.012 | -0.749 | -0.003 | 0.01 | 0.01 | 0 | 0.77 | 0.77 | 0.77 | 0.27 | 0.79 | 0.56 |
| 5 | 128017544 | rs10072361 | -0.941 | -0.292 | 0.363 | -1.239 | -0.017 | -0.002 | -0.002 | 0 | 0.25 | 0.33 | 0.24 | 0.21 | 0.18 | 0.23 |
| 5 | 128030391 | rs17211718 | -0.239 | 1.084 | 0.631 | -1.194 | 0.043 | 0.015 | 0.015 | 0 | 0.12 | 0.18 | 0.16 | 0.14 | 0.24 | 0.16 |
| 5 | 128033561 | rs4440414 | -0.25 | 1.054 | 0.569 | -1.209 | -0.013 | -0.06 | -0.06 | 0 | 0.79 | 0.74 | 0.82 | 0.93 | 0.84 | 0.73 |
| 5 | 128040308 | rs1422450 | 0.492 | 0.915 | 0.622 | -1.159 | 0.009 | -0.044 | -0.044 | 0 | 0.69 | 0.66 | 0.7 | 0.91 | 0.62 | 0.64 |
| 5 | 128042790 | rs10463845 | 0.813 | 0.834 | 0.235 | -1.168 | -0.021 | 0.026 | 0.026 | 0 | 0.66 | 0.36 | 0.64 | 0.79 | 0.6 | 0.56 |
| 5 | 128060951 | rs1422453 | 0.6 | 0.265 | 0.231 | -1.592 | -0.01 | -0.068 | -0.068 | 0 | 0.72 | 0.69 | 0.81 | 0.95 | 0.78 | 0.7 |
| 5 | 128072198 | rs247179 | -0.516 | -0.252 | 0.018 | -2.11 | -0.009 | -0.02 | -0.02 | 0.098 | 0.66 | 0.58 | 0.73 | 0.79 | 0.74 | 0.72 |
| 5 | 128080556 | rs247197 | -0.545 | 0.22 | 0.036 | -2.253 | 0.035 | -0.063 | -0.063 | 0 | 0.3 | 0.37 | 0.33 | 0.13 | 0.4 | 0.38 |
| 5 | 128085409 | rs247206 | -0.629 | 0.11 | 0.051 | -2.412 | -0.005 | -0.031 | -0.031 | 0 | 0.62 | 0.45 | 0.67 | 0.79 | 0.69 | 0.72 |
| 5 | 128093877 | rs247096 | 0.416 | 0.169 | 0.249 | -2.282 | 0.049 | -0.06 | -0.06 | 0.264 | 0.17 | 0.24 | 0.23 | 0.05 | 0.29 | 0.21 |
| 5 | 128100444 | rs4836402 | -1.504 | 0.031 | 0.189 | -2.321 | -0.03 | 0.089 | 0.089 | 0.062 | 0.74 | 0.99 | 0.8 | 0.91 | 0.79 | 0.81 |
| 5 | 128100622 | rs4836403 | 0.731 | 0.157 | 0.537 | -2.213 | -0.001 | -0.011 | -0.011 | 0 | 0.41 | 0.46 | 0.48 | 0.78 | 0.39 | 0.4 |
| 5 | 128106619 | rs10059650 | -0.03 | -0.005 | 0.217 | -2.212 | 0.02 | 0.031 | 0.031 | 0.805 | 0.57 | 0.64 | 0.69 | 0.83 | 0.66 | 0.59 |
| 5 | 128110085 | rs1422446 | -0.479 | 0.11 | 0.634 | -1.568 | 0.006 | -0.024 | -0.024 | 0 | 0.66 | 0.58 | 0.65 | 0.76 | 0.69 | 0.67 |
| 5 | 128111864 | rs7449021 | -0.265 | 0.092 | 0.731 | -1.735 | 0.019 | -0.04 | -0.04 | 0.041 | 0.22 | 0.32 | 0.21 | 0.07 | 0.32 | 0.27 |
| 5 | 128114425 | rs11742783 | 1.203 | 0.053 | 0.902 | -1.711 | 0.033 | 0.022 | 0.022 | 0.078 | 0.4 | 0.19 | 0.42 | 0.66 | 0.34 | 0.35 |
| 5 | 128118088 | rs10491268 | -0.099 | 0.287 | 1.235 | -1.515 | -0.017 | 0.017 | 0.017 | 0 | 0.19 | 0.13 | 0.19 | 0.07 | 0.31 | 0.27 |
| 5 | 128123652 | rs28870456 | -1.144 | 0.23 | 1.117 | -1.552 | 0.012 | -0.01 | -0.01 | 0.505 | 0.32 | 0.6 | 0.37 | 0.21 | 0.44 | 0.42 |
| 5 | 128129794 | rs1019137 | -0.87 | 0.24 | 1.107 | -1.555 | 0.041 | -0.08 | -0.08 | 0.144 | 0.28 | 0.39 | 0.29 | 0.07 | 0.41 | 0.35 |
| 5 | 128138057 | rs6595861 | -1.694 | 0.256 | 0.499 | -1.522 | -0.061 | 0.063 | 0.063 | 0 | 0.38 | 0.83 | 0.39 | 0.22 | 0.48 | 0.47 |
| 5 | 128149665 | rs10039077 | 0.057 | 0.228 | 0.755 | -1.379 | 0.054 | 0.053 | 0.053 | 0 | 0.11 | 0.3 | 0.14 | 0.15 | 0.11 | 0.14 |
| 5 | 128150215 | rs1390615 | -1.518 | 0.25 | 0.294 | -1.415 | -0.025 | 0.039 | 0.039 | 0 | 0.37 | 0.76 | 0.34 | 0.22 | 0.44 | 0.45 |
| 5 | 128172850 | rs2158174 | -0.669 | 0.196 | 0.514 | -1.259 | -0.009 | -0.102 | -0.102 | 0 | 0.15 | 0.52 | 0.11 | 0.02 | 0.09 | 0.13 |
| 5 | 128173339 | rs17616087 | -1.278 | 0.229 | 0.446 | -1.335 | 0.007 | -0.047 | -0.047 | 0 | 0.31 | 0.64 | 0.27 | 0.07 | 0.37 | 0.34 |
| 5 | 128179151 | rs6891265 | 0.076 | 0.018 | 0.063 | -1.648 | 0.035 | 0.096 | 0.096 | 0 | 0.5 | 0.72 | 0.51 | 0.71 | 0.42 | 0.5 |
| 5 | 128193116 | rs1422458 | -0.599 | 0.22 | 0.522 | -1.393 | -0.011 | -0.013 | -0.013 | 0.227 | 0.71 | 0.55 | 0.7 | 0.76 | 0.77 | 0.74 |
| 5 | 128196492 | rs1476085 | 0.612 | 0.217 | 0.262 | -1.394 | 0.002 | -0.046 | -0.046 | 6.601 | 0.1 | 0.17 | 0.07 | 0.02 | 0.08 | 0.12 |
| 5 | 128198631 | rs2106980 | 0.608 | 0.225 | 0.258 | -1.415 | 0.014 | 0.021 | 0.021 | 0.214 | 0.5 | 0.54 | 0.5 | 0.69 | 0.41 | 0.41 |
| 5 | 128198862 | rs11745830 | 1.065 | 0.254 | 0.156 | -1.399 | 0.027 | 0.057 | 0.057 | 0.114 | 0.45 | 0.17 | 0.46 | 0.69 | 0.4 | 0.4 |
| 5 | 128200502 | rs10050439 | 0.26 | 0.493 | -0.159 | -1.205 | -0.042 | 0.034 | 0.034 | 0 | 0.16 | 0.09 | 0.16 | 0.05 | 0.29 | 0.21 |
| 5 | 128201893 | rs9327493 | -0.59 | 0.476 | -0.151 | -1.186 | 0.004 | -0.017 | -0.017 | 0 | 0.25 | 0.27 | 0.23 | 0.07 | 0.36 | 0.32 |
| 5 | 128220643 | rs1421892 | 0.052 | 0.514 | -0.486 | -0.933 | -0.04 | 0.018 | 0.018 | 0 | 0.62 | 0.71 | 0.57 | 0.71 | 0.51 | 0.54 |
| 5 | 128232901 | rs6873372 | -0.129 | 0.576 | -0.279 | -0.943 | -0.03 | 0.04 | 0.04 | 0 | 0.2 | 0.13 | 0.21 | 0.04 | 0.32 | 0.2 |
| 5 | 128238649 | rs3851461 | -0.863 | 0.352 | -0.185 | -0.882 | -0.016 | 0.014 | 0.014 | 0 | 0.35 | 0.74 | 0.4 | 0.22 | 0.43 | 0.39 |
| 5 | 128244740 | rs2577470 | 1.043 | 0.145 | -0.31 | -1.022 | 0.03 | 0.038 | 0.038 | 0 | 0.42 | 0.14 | 0.38 | 0.69 | 0.36 | 0.4 |
| 5 | 128249256 | rs2522378 | 0.87 | 0.102 | 0.169 | -1.046 | 0.011 | 0.014 | 0.014 | 0 | 0.44 | 0.36 | 0.4 | 0.68 | 0.37 | 0.4 |
| 5 | 128250479 | rs2577472 | 0.955 | 0.098 | 0.166 | -1.049 | 0.025 | 0.033 | 0.033 | 0 | 0.43 | 0.22 | 0.39 | 0.69 | 0.37 | 0.4 |
| 5 | 128255056 | rs2526256 | 0.658 | 0.113 | 0.53 | -1.073 | 0.007 | 0.025 | 0.025 | 0 | 0.46 | 0.38 | 0.47 | 0.82 | 0.4 | 0.46 |
| 5 | 128261415 | rs2577487 | 1.143 | 0.11 | 0.485 | -1.069 | 0.021 | 0.034 | 0.034 | 0 | 0.42 | 0.12 | 0.38 | 0.68 | 0.36 | 0.4 |

| | | | | | | | | | | | | | | | | |
|---|---|---|---|---|---|---|---|---|---|---|---|---|---|---|---|---|
| 5 | 128262334 | rs10520015 | 1.264 | 0.129 | 0.381 | -1.231 | 0.001 | 0.028 | 0.028 | 0 | 0.21 | 0.18 | 0.21 | 0.1 | 0.2 | 0.21 |
| 5 | 128264677 | rs2526185 | 0.345 | 0.176 | 0.416 | -1.222 | 0.015 | -0.067 | -0.067 | 0.519 | 0.68 | 0.62 | 0.69 | 0.92 | 0.61 | 0.67 |
| 5 | 128291354 | rs2577448 | 1.201 | 0.08 | 0.239 | -0.816 | 0.015 | 0.092 | 0.092 | 0 | 0.36 | 0.17 | 0.19 | 0.05 | 0.3 | 0.17 |
| 5 | 128291364 | rs2577447 | 1.201 | 0.08 | 0.239 | -0.816 | 0.015 | 0.092 | 0.092 | 0 | 0.36 | 0.17 | 0.19 | 0.05 | 0.3 | 0.17 |
| 5 | 128313024 | rs185411 | -0.595 | 0.257 | 0.18 | -0.39 | 0.005 | 0.028 | 0.028 | 0.022 | 0.68 | 0.75 | 0.7 | 0.88 | 0.74 | 0.65 |
| 5 | 128323346 | rs258007 | -0.443 | 0.303 | 0.037 | -0.391 | -0.003 | -0.003 | -0.003 | 0 | 0.25 | 0.22 | 0.35 | 0.6 | 0.39 | 0.4 |
| 5 | 128323644 | rs3886286 | 0.099 | 0.358 | -0.006 | -0.531 | -0.027 | 0.029 | 0.029 | 0 | 0.19 | 0.12 | 0.21 | 0.03 | 0.34 | 0.2 |
| 5 | 128325025 | rs7730969 | 0.061 | 0.363 | -0.062 | -0.529 | 0.001 | -0.015 | -0.015 | 0.661 | 0.19 | 0.19 | 0.22 | 0.03 | 0.34 | 0.2 |
| 5 | 128330093 | rs1181983 | 0.871 | 0.101 | 0.113 | -0.34 | 0.007 | 0.033 | 0.033 | 0.136 | 0.41 | 0.28 | 0.34 | 0.28 | 0.34 | 0.24 |
| 5 | 128330723 | rs6859755 | -0.609 | -0.01 | -0.246 | -0.357 | 0.024 | -0.011 | -0.011 | 0.301 | 0.61 | 0.73 | 0.69 | 0.4 | 0.69 | 0.64 |
| 5 | 128336024 | rs3851465 | 0.383 | -0.421 | 0 | -0.51 | -0.003 | -0.002 | -0.002 | 0 | 0.27 | 0.26 | 0.22 | 0.24 | 0.18 | 0.25 |
| 5 | 128339246 | rs4836430 | -0.622 | -0.704 | 0.15 | -0.801 | -0.007 | 0.009 | 0.009 | 0 | 0.19 | 0.28 | 0.36 | 0.69 | 0.17 | 0.39 |
| 5 | 128340794 | rs1181975 | -0.452 | 0.019 | 0.165 | -1.005 | 0.019 | -0.03 | -0.03 | 0.323 | 0.86 | 0.84 | 0.86 | 0.99 | 0.72 | 0.84 |
| 5 | 128354053 | rs1181968 | 1.713 | -0.236 | 0.3 | -1.054 | -0.021 | 0.15 | 0.15 | 0 | 0.36 | 0.09 | 0.23 | 0.04 | 0.36 | 0.18 |
| 5 | 128367988 | rs1181966 | 1.665 | -0.255 | 0.091 | -1.061 | -0.075 | 0.225 | 0.225 | 0 | 0.35 | 0.03 | 0.23 | 0.04 | 0.36 | 0.17 |
| 5 | 128373968 | rs2538017 | 1.084 | -0.348 | 0.456 | -1.057 | 0.017 | 0.016 | 0.016 | 0 | 0.26 | 0.16 | 0.21 | 0.24 | 0.18 | 0.25 |
| 5 | 128386801 | rs1181962 | 1.717 | -0.278 | 0.325 | -1.07 | -0.032 | 0.181 | 0.181 | 0 | 0.36 | 0.08 | 0.23 | 0.04 | 0.36 | 0.17 |
| 5 | 128395295 | rs257921 | -0.455 | -0.252 | 0.486 | -0.64 | 0.012 | 0.018 | 0.018 | 0.774 | 0.2 | 0.22 | 0.4 | 0.5 | 0.35 | 0.32 |
| 5 | 128397565 | rs10064916 | -0.433 | -0.312 | 0.622 | -0.654 | -0.03 | 0.033 | 0.033 | 0.147 | 0.17 | 0.43 | 0.14 | 0.19 | 0.11 | 0.26 |
| 5 | 128400356 | rs42563 | -0.347 | -0.242 | 0.448 | -0.386 | -0.008 | -0.004 | -0.004 | 1.568 | 0.19 | 0.17 | 0.38 | 0.47 | 0.34 | 0.33 |
| 5 | 128404226 | rs42265 | -1.423 | -0.264 | 0.403 | -0.504 | 0.016 | -0.02 | -0.02 | 0.465 | 0.37 | 0.57 | 0.53 | 0.67 | 0.45 | 0.57 |
| 5 | 128404326 | rs42562 | -0.912 | -0.293 | 0.396 | -0.503 | -0.016 | -0.013 | -0.013 | 0.419 | 0.73 | 0.67 | 0.77 | 0.74 | 0.82 | 0.75 |
| 5 | 128408502 | rs13181777 | 1.473 | -0.217 | 0.036 | -0.571 | -0.032 | 0.172 | 0.172 | 0 | 0.37 | 0.1 | 0.25 | 0.07 | 0.38 | 0.18 |
| 5 | 128412653 | rs7735174 | -0.564 | -0.325 | 0.575 | -0.267 | -0.008 | 0.007 | 0.007 | 0.788 | 0.41 | 0.59 | 0.59 | 0.68 | 0.48 | 0.51 |
| 5 | 128413093 | rs192299 | 0.076 | -0.02 | 0.608 | -0.15 | -0.027 | -0.006 | -0.006 | 0.109 | 0.11 | 0.06 | 0.32 | 0.42 | 0.25 | 0.24 |
| 5 | 128414577 | rs6867156 | 0.932 | -0.325 | 0.996 | -0.203 | -0.015 | -0.013 | -0.013 | 0.006 | 0.22 | 0.28 | 0.22 | 0.26 | 0.16 | 0.21 |
| 5 | 128417940 | rs17839625 | -0.243 | -0.56 | 0.841 | -0.347 | -0.012 | 0.028 | 0.028 | 0 | 0.13 | 0.27 | 0.12 | 0.18 | 0.1 | 0.23 |
| 5 | 128422167 | rs4836437 | 0.007 | -0.539 | 0.982 | -0.298 | -0.003 | 0.057 | 0.057 | 0.016 | 0.12 | 0.34 | 0.11 | 0.17 | 0.09 | 0.18 |
| 5 | 128428851 | rs3756746 | 0.058 | -0.538 | 0.987 | -0.298 | -0.006 | 0.046 | 0.046 | 0 | 0.12 | 0.29 | 0.11 | 0.17 | 0.09 | 0.18 |
| 5 | 128434448 | rs17163708 | 1.074 | -0.634 | 1.17 | -0.387 | 0 | -0.002 | -0.002 | 0 | 0.22 | 0.21 | 0.23 | 0.18 | 0.17 | 0.21 |
| 5 | 128438147 | rs3798111 | 0.108 | -0.607 | 1.182 | -0.37 | -0.018 | 0.039 | 0.039 | 0 | 0.12 | 0.3 | 0.11 | 0.17 | 0.08 | 0.19 |
| 5 | 128441227 | rs976607 | 0.779 | -0.418 | 1.324 | -0.405 | 0.024 | -0.007 | -0.007 | 0 | 0.08 | 0.2 | 0.05 | 0.02 | 0.08 | 0.06 |
| 5 | 128442038 | rs17679811 | 0.513 | -0.47 | 1.483 | -0.407 | 0.003 | -0.006 | -0.006 | 0.071 | 0.11 | 0.11 | 0.11 | 0.17 | 0.09 | 0.18 |
| 5 | 128442644 | rs3734161 | -0.04 | -0.591 | 1.336 | -0.567 | -0.031 | -0.029 | -0.029 | 0 | 0.09 | 0.19 | 0.03 | 0.04 | 0.02 | 0.02 |
| 5 | 128442948 | rs17165184 | 1.117 | -0.601 | 1.282 | -0.323 | -0.004 | -0.011 | -0.011 | 0.478 | 0.22 | 0.22 | 0.23 | 0.18 | 0.18 | 0.21 |
| 5 | 128448016 | rs2059124 | -0.092 | -0.516 | 0.507 | -0.235 | -0.031 | -0.029 | -0.029 | 0.762 | 0.09 | 0.18 | 0.03 | 0.04 | 0.02 | 0.02 |
| 5 | 128454132 | rs7714427 | -1.964 | -0.336 | -0.104 | -0.246 | -0.078 | 0.202 | 0.202 | 0 | 0.62 | 0.97 | 0.74 | 0.89 | 0.61 | 0.8 |
| 5 | 128458834 | rs10520025 | 0.811 | -0.501 | 0.191 | -0.194 | -0.004 | -0.005 | -0.005 | 1.758 | 0.2 | 0.22 | 0.26 | 0.2 | 0.13 | 0.15 |
| 5 | 128460713 | rs10071265 | -2.322 | -0.376 | -0.304 | -0.181 | -0.088 | 0.166 | 0.166 | 0 | 0.63 | 0.97 | 0.74 | 0.88 | 0.61 | 0.81 |
| 5 | 128463532 | rs4835970 | -1.329 | -0.444 | 0.108 | -0.018 | 0.053 | 0.053 | 0.053 | 0 | 0.48 | 0.77 | 0.47 | 0.43 | 0.38 | 0.49 |
| 5 | 128465327 | rs2312243 | -0.587 | -0.483 | 0.336 | -0.033 | -0.03 | -0.044 | -0.044 | 0.355 | 0.11 | 0.2 | 0.05 | 0.03 | 0.06 | 0.08 |
| 5 | 128466832 | rs17163689 | -0.657 | -0.678 | 0.175 | -0.064 | -0.01 | 0.001 | 0.001 | 0 | 0.18 | 0.23 | 0.15 | 0.17 | 0.12 | 0.29 |
| 5 | 128469491 | rs963677 | -2.394 | -0.591 | -0.271 | -0.101 | -0.047 | 0.101 | 0.101 | 0 | 0.63 | 0.92 | 0.74 | 0.83 | 0.62 | 0.81 |
| 5 | 128473773 | rs1345722 | -0.35 | -0.354 | 0.299 | 0.024 | -0.025 | -0.014 | -0.014 | 0 | 0.33 | 0.24 | 0.54 | 0.61 | 0.46 | 0.52 |
| 5 | 128474742 | rs7707761 | -0.402 | -0.458 | 0.57 | 0.116 | 0.004 | -0.003 | -0.003 | 0 | 0.17 | 0.18 | 0.13 | 0.19 | 0.11 | 0.22 |
| 5 | 128477058 | rs7732209 | -1.231 | -0.474 | 0.707 | 0.105 | -0.023 | -0.005 | -0.005 | 0 | 0.23 | 0.34 | 0.17 | 0.2 | 0.14 | 0.25 |
| 5 | 128477439 | rs17839618 | 0.771 | -0.042 | 0.744 | 0.331 | -0.008 | -0.025 | -0.025 | 0 | 0.06 | 0.15 | 0.04 | 0 | 0.03 | 0.03 |
| 5 | 128489453 | rs1422802 | 0.258 | 0.609 | 0.302 | 0.777 | 0.032 | 0.164 | 0.164 | 0 | 0.05 | 0.06 | 0.21 | 0.42 | 0.15 | 0.22 |
| 5 | 128504564 | rs17618074 | -0.317 | 0.496 | 0.259 | 0.437 | -0.002 | -0.001 | -0.001 | 0 | 0.28 | 0.27 | 0.49 | 0.59 | 0.35 | 0.47 |
| 5 | 128506589 | rs7356681 | -0.521 | 0.544 | 0.505 | -0.302 | 0.009 | -0.033 | -0.033 | 0 | 0.12 | 0.24 | 0.07 | 0.02 | 0.1 | 0.09 |
| 5 | 128509412 | rs10063241 | -1.436 | 0.956 | 0.33 | -0.137 | 0.04 | -0.018 | -0.018 | 0 | 0.14 | 0.28 | 0.16 | 0.06 | 0.24 | 0.21 |
| 5 | 128529794 | rs4509075 | 0.574 | 0.706 | -0.167 | 0.133 | -0.016 | -0.037 | -0.037 | 1.806 | 0.13 | 0.07 | 0.2 | 0.6 | 0.18 | 0.29 |
| 5 | 128531752 | rs4314458 | -1.718 | 0.306 | -0.579 | -0.799 | 0 | -0.01 | -0.01 | 0 | 0.58 | 0.62 | 0.62 | 0.85 | 0.58 | 0.72 |
| 5 | 128532085 | rs13157663 | -0.744 | 0.611 | -0.372 | -1.179 | 0.006 | -0.038 | -0.038 | 0.263 | 0.21 | 0.23 | 0.15 | 0.04 | 0.25 | 0.22 |
| 5 | 128532350 | rs10074559 | -1.56 | 0.451 | -0.433 | -1.222 | -0.004 | 0.014 | 0.014 | 0.323 | 0.47 | 0.62 | 0.53 | 0.64 | 0.48 | 0.66 |
| 5 | 128535398 | rs10054567 | -1.317 | 0.294 | -0.563 | -1.104 | 0.008 | -0.032 | -0.032 | 0.743 | 0.24 | 0.26 | 0.19 | 0.07 | 0.28 | 0.27 |
| 5 | 128541597 | rs35100037 | -0.115 | -0.352 | -0.31 | -0.748 | -0.002 | -0.026 | -0.026 | 0.587 | 0.11 | 0.24 | 0.06 | 0.04 | 0.09 | 0.19 |
| 5 | 128552706 | rs1181841 | 1.43 | -0.434 | -1.136 | -1.038 | -0.077 | -0.008 | -0.008 | 0.473 | 0.4 | 0.05 | 0.41 | 0.38 | 0.43 | 0.3 |
| 5 | 128553667 | rs1184334 | 1.26 | -0.352 | -0.651 | -0.958 | -0.004 | 0.02 | 0.02 | 0 | 0.44 | 0.25 | 0.53 | 0.39 | 0.46 | 0.38 |
| 5 | 128554088 | rs1181842 | -0.407 | -0.062 | -0.771 | -0.685 | -0.029 | -0.05 | -0.05 | 0.826 | 0.74 | 0.86 | 0.69 | 0.45 | 0.67 | 0.66 |
| 5 | 128554789 | rs11746085 | 0.602 | -0.016 | -0.778 | -0.587 | 0 | -0.004 | -0.004 | 0 | 0.07 | 0.08 | 0.03 | 0.02 | 0.05 | 0.03 |
| 5 | 128556817 | rs1183063 | 0.595 | -0.09 | -0.711 | -0.633 | -0.001 | 0.003 | 0.003 | 0.114 | 0.53 | 0.55 | 0.59 | 0.39 | 0.49 | 0.41 |
| 5 | 128565893 | rs2312255 | -0.641 | 0.167 | -0.236 | -0.833 | 0.052 | -0.003 | -0.003 | 0 | 0.11 | 0.55 | 0.17 | 0.06 | 0.12 | 0.14 |
| 5 | 128573779 | rs13189185 | -0.74 | 0.122 | -0.358 | -0.768 | 0.035 | 0.031 | 0.031 | 1.034 | 0.12 | 0.58 | 0.17 | 0.12 | 0.12 | 0.14 |
| 5 | 128582254 | rs10072264 | 1.377 | -0.267 | -0.744 | -1.072 | 0.024 | 0.054 | 0.054 | 1.335 | 0.14 | 0.04 | 0.11 | 0.02 | 0.09 | 0.06 |
| 5 | 128590209 | rs2189752 | -1.872 | -0.523 | -1.355 | -1.533 | -0.086 | 0.132 | 0.132 | 0 | 0.68 | 0.99 | 0.71 | 0.92 | 0.69 | 0.81 |
| 5 | 128592181 | rs10477700 | -1.292 | -0.488 | -1.211 | -0.699 | 0.006 | -0.001 | -0.001 | 0 | 0.46 | 0.68 | 0.45 | 0.53 | 0.42 | 0.43 |
| 5 | 128595494 | rs7705332 | -0.696 | -0.528 | -1.238 | -0.753 | 0.014 | 0.047 | 0.047 | 0.254 | 0.28 | 0.59 | 0.33 | 0.51 | 0.31 | 0.32 |
| 5 | 128596232 | rs13160758 | 1.185 | -0.539 | -1.381 | -0.755 | 0.033 | 0.072 | 0.072 | 0.459 | 0.17 | 0.06 | 0.12 | 0.02 | 0.11 | 0.08 |

| | | | | | | | | | | | | | | | | |
|---|---|---|---|---|---|---|---|---|---|---|---|---|---|---|---|---|
| 5 | 128607170 | rs10057716 | 1.779 | -0.477 | -1.354 | -0.711 | 0.026 | 0.056 | 0.056 | 1.113 | 0.14 | 0.04 | 0.11 | 0.02 | 0.09 | 0.06 |
| 5 | 128615051 | rs252671 | -1.568 | -0.478 | -1.308 | -0.717 | 0.017 | 0.002 | 0.002 | 0.442 | 0.48 | 0.78 | 0.46 | 0.53 | 0.43 | 0.43 |
| 5 | 128615902 | rs2068046 | 1.132 | -0.396 | -1.714 | -0.677 | -0.018 | -0.035 | -0.035 | 0 | 0.08 | 0 | 0.06 | 0.37 | 0.06 | 0.06 |
| 5 | 128616356 | rs7722731 | -2.106 | -0.597 | -1.699 | -0.92 | -0.053 | 0.097 | 0.097 | 0 | 0.73 | 0.97 | 0.73 | 0.92 | 0.73 | 0.82 |
| 5 | 128618834 | rs10077978 | 1.7 | -0.567 | -1.607 | -0.795 | 0.024 | 0.054 | 0.054 | 1 | 0.14 | 0.04 | 0.11 | 0.02 | 0.09 | 0.07 |
| 5 | 128627794 | rs252668 | 0.645 | -0.559 | -1.409 | -0.767 | -0.014 | -0.033 | -0.033 | 0 | 0.18 | 0.1 | 0.24 | 0.46 | 0.21 | 0.21 |
| 5 | 128630825 | rs6893462 | 1.279 | -0.625 | -1.393 | -0.811 | 0.005 | -0.002 | -0.002 | 1.438 | 0.16 | 0.17 | 0.12 | 0.02 | 0.1 | 0.07 |
| 5 | 128636024 | rs245786 | -0.179 | -0.629 | -1.311 | -0.812 | 0.005 | -0.006 | -0.006 | 0 | 0.35 | 0.3 | 0.36 | 0.48 | 0.3 | 0.28 |
| 5 | 128638883 | rs6869011 | 0.896 | -0.647 | -0.739 | -0.487 | 0.003 | -0.005 | -0.005 | 0 | 0.16 | 0.17 | 0.12 | 0.02 | 0.1 | 0.07 |
| 5 | 128643236 | rs7717089 | -1.24 | -0.525 | -0.792 | -0.492 | -0.046 | -0.035 | -0.035 | 0 | 0.57 | 0.85 | 0.49 | 0.46 | 0.51 | 0.65 |
| 5 | 128644746 | rs4836453 | 0.917 | -0.026 | -0.978 | -0.093 | -0.022 | -0.045 | -0.045 | 0 | 0.21 | 0.09 | 0.27 | 0.38 | 0.29 | 0.36 |
| 5 | 128645304 | rs13168004 | -0.051 | -0.375 | -0.521 | -0.384 | 0.014 | 0.008 | 0.008 | 0.872 | 0.19 | 0.17 | 0.12 | 0.02 | 0.1 | 0.1 |
| 5 | 128650612 | rs13174910 | -0.036 | -0.363 | -0.542 | -0.378 | 0.022 | 0.02 | 0.02 | 0.894 | 0.19 | 0.15 | 0.12 | 0.02 | 0.1 | 0.1 |
| 5 | 128651624 | rs17163509 | -0.036 | -0.363 | -0.547 | -0.378 | 0.016 | 0.011 | 0.011 | 0.894 | 0.19 | 0.17 | 0.12 | 0.02 | 0.1 | 0.1 |
| 5 | 128658431 | rs10079852 | 0.288 | -0.096 | -0.827 | -0.235 | 0.029 | 0.036 | 0.036 | 0.036 | 0.17 | 0.1 | 0.12 | 0.02 | 0.1 | 0.1 |
| 5 | 128660264 | rs13161848 | 0.288 | -0.096 | -0.827 | -0.235 | 0.029 | 0.036 | 0.036 | 0.036 | 0.17 | 0.1 | 0.12 | 0.02 | 0.1 | 0.1 |
| 5 | 128661339 | rs10056165 | -0.972 | 0.923 | -0.241 | 0.431 | 0.015 | 0 | 0 | 0 | 0.09 | 0.24 | 0.06 | 0.06 | 0.1 | 0.15 |
| 5 | 128674953 | rs6595902 | 0.568 | 0.766 | -0.157 | 0.369 | 0.041 | 0.016 | 0.016 | 0 | 0.92 | 0.74 | 0.95 | 0.94 | 0.92 | 0.86 |
| 5 | 128677982 | rs17680011 | 0.346 | 0.91 | -1.281 | 0.418 | -0.117 | -0.042 | -0.042 | 0 | 0.26 | 0.02 | 0.39 | 0.25 | 0.37 | 0.25 |
| 5 | 128692695 | rs17680057 | 0.222 | 0.857 | -1.334 | 0.459 | -0.12 | -0.043 | -0.043 | 6.266 | 0.27 | 0.02 | 0.39 | 0.25 | 0.38 | 0.27 |
| 5 | 128692877 | rs11959939 | 0.494 | 0.923 | -0.473 | 0.974 | 0.042 | 0.092 | 0.092 | 0.053 | 0.15 | 0.24 | 0.24 | 0.35 | 0.27 | 0.35 |
| 5 | 128694141 | rs12109162 | -0.315 | 0.178 | -0.566 | 0.529 | 0.003 | 0.001 | 0.001 | 0 | 0.31 | 0.27 | 0.37 | 0.22 | 0.31 | 0.26 |
| 5 | 128694216 | rs12108748 | 0.149 | -0.016 | -0.89 | 0.507 | -0.018 | -0.018 | -0.018 | 0 | 0.68 | 0.78 | 0.56 | 0.68 | 0.61 | 0.69 |
| 5 | 128696341 | rs10038633 | -0.329 | -1.333 | -0.822 | 0.091 | -0.084 | -0.04 | -0.04 | 0 | 0.26 | 0.51 | 0.16 | 0.16 | 0.1 | 0.19 |
| 5 | 128699833 | rs7725892 | 0.008 | -1.464 | -0.875 | 0.062 | 0.01 | 0.055 | 0.055 | 0.203 | 0.1 | 0.42 | 0.09 | 0.12 | 0.05 | 0.02 |
| 5 | 128702357 | rs11241996 | 1.307 | -1.498 | -1.034 | 0.137 | 0.012 | 0.005 | 0.005 | 2.025 | 0.36 | 0.43 | 0.36 | 0.43 | 0.4 | 0.54 |
| 5 | 128704748 | rs13182736 | 0.124 | -1.52 | -1.06 | 0.125 | -0.07 | -0.051 | -0.051 | 0 | 0.38 | 0.11 | 0.49 | 0.39 | 0.5 | 0.3 |
| 5 | 128707863 | rs6861188 | 1.46 | -1.367 | -0.967 | 0.251 | 0.007 | 0 | 0 | 0 | 0.33 | 0.35 | 0.34 | 0.35 | 0.39 | 0.5 |
| 5 | 128709025 | rs2012795 | -0.702 | -0.76 | -0.635 | 1.042 | -0.08 | -0.064 | -0.064 | 0 | 0.23 | 0.44 | 0.13 | 0.12 | 0.09 | 0.16 |
| 5 | 128709872 | rs10478880 | 0.189 | -1.011 | -0.737 | 0.998 | -0.046 | -0.093 | -0.093 | 0.359 | 0.24 | 0.11 | 0.4 | 0.45 | 0.38 | 0.26 |
| 5 | 128718147 | rs6873564 | 0.65 | -0.987 | -0.683 | 0.586 | 0.001 | 0.022 | 0.022 | 0 | 0.65 | 0.46 | 0.49 | 0.47 | 0.58 | 0.72 |
| 5 | 128718653 | rs6893553 | -0.542 | -0.696 | -0.629 | 0.764 | -0.084 | -0.037 | -0.037 | 0 | 0.23 | 0.42 | 0.14 | 0.14 | 0.08 | 0.16 |
| 5 | 128718832 | rs17165195 | -0.288 | -0.589 | -0.91 | 0.779 | -0.006 | -0.001 | -0.001 | 0 | 0.21 | 0.19 | 0.12 | 0.12 | 0.07 | 0.15 |
| 5 | 128722696 | rs10077875 | -0.547 | 0.029 | -0.679 | 0.888 | 0.005 | 0.001 | 0.001 | 0 | 0.47 | 0.55 | 0.54 | 0.55 | 0.45 | 0.41 |
| 5 | 128730528 | rs10053585 | -0.679 | 0.199 | -0.569 | 0.908 | -0.014 | 0.046 | 0.046 | 0.386 | 0.07 | 0.2 | 0.09 | 0.11 | 0.04 | 0.02 |
| 5 | 128732672 | rs13171514 | -0.418 | 0.565 | -1.956 | 0.624 | -0.056 | -0.031 | -0.031 | 1.496 | 0.32 | 0.05 | 0.45 | 0.21 | 0.43 | 0.31 |
| 5 | 128735170 | rs17618372 | -0.305 | -0.129 | -2.284 | -0.039 | 0 | 0.071 | 0.071 | 0 | 0.27 | 0.07 | 0.17 | 0.16 | 0.21 | 0.14 |
| 5 | 128737229 | rs12515721 | -0.323 | -0.135 | -2.287 | -0.042 | 0 | 0.072 | 0.072 | 0 | 0.27 | 0.07 | 0.17 | 0.16 | 0.21 | 0.14 |
| 5 | 128738407 | rs13174854 | 0.242 | -0.042 | -1.462 | -0.236 | 0.027 | -0.029 | -0.029 | 0 | 0.33 | 0.46 | 0.46 | 0.17 | 0.41 | 0.37 |
| 5 | 128742671 | rs10067879 | -0.17 | -0.548 | -1.391 | -0.211 | 0.009 | 0.064 | 0.064 | 0 | 0.2 | 0.08 | 0.09 | 0.02 | 0.15 | 0.1 |
| 5 | 128743894 | rs17163440 | -1.448 | -0.926 | -0.707 | 0.251 | -0.003 | -0.002 | -0.002 | 0 | 0.24 | 0.7 | 0.16 | 0.21 | 0.17 | 0.16 |
| 5 | 128744872 | rs12520926 | -1.238 | -1.25 | -0.357 | 0.3 | 0.001 | -0.001 | -0.001 | 0 | 0.16 | 0.13 | 0.1 | 0.19 | 0.14 | 0.24 |
| 5 | 128747841 | rs17163425 | -1.168 | -1.222 | -0.342 | 0.317 | 0 | -0.001 | -0.001 | 0 | 0.16 | 0.13 | 0.1 | 0.2 | 0.14 | 0.2 |
| 5 | 128760076 | rs17163418 | -1.272 | -1.257 | -0.199 | 0.3 | -0.008 | 0.004 | 0.004 | 0 | 0.16 | 0.2 | 0.11 | 0.2 | 0.14 | 0.2 |
| 5 | 128768047 | rs6886494 | -1.306 | -1.237 | -0.077 | 0.283 | -0.01 | 0.003 | 0.003 | 0 | 0.17 | 0.22 | 0.11 | 0.19 | 0.13 | 0.2 |
| 5 | 128779359 | rs11241998 | -0.607 | -1.768 | 0.021 | 0.858 | -0.026 | 0.088 | 0.088 | 0 | 0.16 | 0.22 | 0.13 | 0.35 | 0.11 | 0.16 |
| 5 | 128787411 | rs9686134 | -1.773 | -2.041 | 0.952 | 0.785 | -0.013 | 0.009 | 0.009 | 0.059 | 0.27 | 0.72 | 0.19 | 0.37 | 0.16 | 0.23 |
| 5 | 128789554 | rs11241999 | -1.287 | -1.651 | 1.547 | 0.721 | -0.02 | 0.076 | 0.076 | 0 | 0.2 | 0.28 | 0.16 | 0.37 | 0.15 | 0.18 |
| 5 | 128794080 | rs3866223 | -0.795 | -1.685 | 1.718 | 0.719 | -0.029 | 0.068 | 0.068 | 0 | 0.17 | 0.22 | 0.13 | 0.34 | 0.11 | 0.19 |
| 5 | 128802515 | rs11242001 | -0.309 | -1.631 | 1.298 | 0.76 | 0.019 | -0.08 | -0.08 | 0 | 0.13 | 0.03 | 0.09 | 0.32 | 0.1 | 0.17 |
| 5 | 128803345 | rs6866231 | 0.101 | -2.687 | 1.324 | 0.433 | -0.08 | -0.056 | -0.056 | 0 | 0.17 | 0.4 | 0.19 | 0.09 | 0.08 | 0.12 |
| 5 | 128806861 | rs7733098 | -1.699 | -2.76 | 1.327 | 0.442 | 0.084 | -0.031 | -0.031 | 0 | 0.41 | 0.79 | 0.36 | 0.44 | 0.27 | 0.37 |
| 5 | 128813527 | rs1862164 | -1.85 | -2.974 | 1.295 | 0.341 | 0.129 | -0.041 | -0.041 | 1.279 | 0.43 | 0.87 | 0.36 | 0.44 | 0.28 | 0.38 |
| 5 | 128831307 | rs10214155 | 0.179 | -2.41 | 1.805 | 0.239 | -0.048 | -0.026 | -0.026 | 0.751 | 0.15 | 0.24 | 0.18 | 0.09 | 0.08 | 0.13 |
| 5 | 128847172 | rs7724326 | -0.584 | -1.743 | 1.026 | 0.333 | -0.034 | -0.04 | -0.04 | 0 | 0.81 | 0.94 | 0.81 | 0.71 | 0.77 | 0.71 |
| 5 | 128856669 | rs7707594 | -0.797 | -1.909 | 1.639 | 0.033 | 0.021 | -0.026 | -0.026 | 0.326 | 0.12 | 0.33 | 0.07 | 0.04 | 0.1 | 0.08 |
| 5 | 128865864 | rs10478885 | -0.599 | -1.798 | 0.777 | 0.152 | -0.083 | -0.087 | -0.087 | 0.197 | 0.22 | 0.45 | 0.22 | 0.1 | 0.14 | 0.19 |
| 5 | 128876639 | rs10072248 | -0.29 | -1.813 | 0.48 | 0.804 | -0.037 | 0.026 | 0.026 | 0 | 0.19 | 0.22 | 0.21 | 0.31 | 0.12 | 0.15 |
| 5 | 128877617 | rs6866597 | -1.313 | -1.722 | -0.38 | 0.176 | -0.092 | 0.114 | 0.114 | 0.853 | 0.64 | 0.94 | 0.58 | 0.78 | 0.51 | 0.61 |
| 5 | 128888039 | rs13158524 | 0.686 | -2.074 | 0.778 | 0.348 | 0.056 | -0.014 | -0.014 | 0 | 0.54 | 0.29 | 0.6 | 0.52 | 0.64 | 0.55 |
| 5 | 128891688 | rs7707487 | 0.046 | -2.148 | 0.89 | 0.344 | -0.004 | -0.027 | -0.027 | 0.448 | 0.26 | 0.34 | 0.18 | 0.12 | 0.23 | 0.3 |
| 5 | 128900524 | rs7380441 | -0.546 | -2.12 | 1.001 | 0.348 | -0.073 | 0.093 | 0.093 | 0 | 0.79 | 0.45 | 0.77 | 0.64 | 0.88 | 0.84 |
| 5 | 128926595 | rs10043738 | -0.755 | -2.131 | 1.082 | 0.347 | -0.032 | 0.033 | 0.033 | 0 | 0.81 | 0.78 | 0.79 | 0.67 | 0.87 | 0.85 |
| 5 | 128940618 | rs1017201 | 0.795 | -2.138 | 0.887 | 0.324 | 0.092 | -0.051 | -0.051 | 0 | 0.54 | 0.11 | 0.59 | 0.53 | 0.64 | 0.55 |
| 5 | 128951136 | rs1946289 | -0.002 | -2.166 | 0.941 | 0.274 | -0.017 | 0.019 | 0.019 | 0 | 0.75 | 0.34 | 0.69 | 0.61 | 0.79 | 0.74 |
| 5 | 128951428 | rs1946290 | 0.724 | -2.177 | 0.899 | 0.261 | 0.045 | -0.01 | -0.01 | 0 | 0.59 | 0.31 | 0.62 | 0.58 | 0.67 | 0.61 |
| 5 | 128960082 | rs30708 | -0.551 | -2.182 | 1.042 | 0.254 | -0.041 | 0.015 | 0.015 | 1.744 | 0.37 | 0.44 | 0.29 | 0.44 | 0.25 | 0.33 |
| 5 | 128969615 | rs3909548 | 0.958 | -2.083 | 0.892 | 0.212 | 0 | 0.048 | 0.048 | 5.101 | 0.16 | 0.02 | 0.07 | 0.05 | 0.12 | 0.16 |
| 5 | 128977732 | rs17163231 | 0.075 | -2.079 | 0.978 | 0.208 | -0.003 | -0.055 | -0.055 | 3.202 | 0.21 | 0.33 | 0.16 | 0.08 | 0.2 | 0.26 |

| | | | | | | | | | | | | | | | | |
|---|---|---|---|---|---|---|---|---|---|---|---|---|---|---|---|---|
| 5 | 128993273 | rs25807 | -0.939 | -2.199 | 0.837 | 0.136 | 0.064 | 0.087 | 0.087 | 0.717 | 0.46 | 0.75 | 0.48 | 0.66 | 0.38 | 0.44 |
| 5 | 129010769 | rs30656 | -0.45 | -1.652 | 0.932 | -0.039 | -0.047 | 0.087 | 0.087 | 0 | 0.78 | 0.54 | 0.73 | 0.68 | 0.8 | 0.77 |
| 5 | 129015452 | rs30645 | 1.526 | -1.607 | 0.589 | -0.084 | -0.037 | 0.119 | 0.119 | 0.955 | 0.38 | 0.08 | 0.46 | 0.24 | 0.53 | 0.41 |
| 5 | 129018092 | rs30642 | 0.67 | -1.176 | 1.225 | 0.228 | -0.014 | 0.014 | 0.014 | 0 | 0.17 | 0.17 | 0.19 | 0.32 | 0.12 | 0.14 |
| 5 | 129018855 | rs30640 | -0.027 | -1.189 | 1.248 | 0.223 | -0.035 | 0.033 | 0.033 | 0.019 | 0.22 | 0.32 | 0.2 | 0.35 | 0.15 | 0.19 |
| 5 | 129043518 | rs30302 | 0.761 | -0.504 | 0.899 | 0.019 | -0.006 | -0.002 | -0.002 | 0 | 0.58 | 0.45 | 0.63 | 0.63 | 0.62 | 0.55 |
| 5 | 129047454 | rs32816 | 0.573 | -0.404 | 0.522 | -0.047 | -0.017 | 0.052 | 0.052 | 0.708 | 0.17 | 0.05 | 0.1 | 0.05 | 0.17 | 0.2 |
| 5 | 129062364 | rs7735351 | 1.112 | -0.413 | 0.743 | 0.125 | 0.051 | -0.002 | -0.002 | 0 | 0.52 | 0.25 | 0.59 | 0.55 | 0.59 | 0.54 |
| 5 | 129064393 | rs1363475 | 1.097 | 0.018 | 0.83 | 0.088 | 0.033 | 0.007 | 0.007 | 0 | 0.5 | 0.25 | 0.5 | 0.56 | 0.54 | 0.52 |
| 5 | 129070945 | rs1422466 | 0.121 | -0.156 | 1.317 | 0.62 | -0.006 | 0.004 | 0.004 | 0 | 0.2 | 0.2 | 0.23 | 0.29 | 0.17 | 0.14 |
| 5 | 129081099 | rs17163010 | 0.148 | 0.892 | 1.673 | 0.419 | 0.086 | -0.012 | -0.012 | 0.005 | 0.06 | 0.24 | 0.05 | 0 | 0.06 | 0.03 |
| 5 | 129097638 | rs6866940 | 1.113 | 0.614 | 0.914 | 0.61 | 0.038 | 0.03 | 0.03 | 1.155 | 0.9 | 0.67 | 0.91 | 0.91 | 0.9 | 0.87 |
| 5 | 129107137 | rs888899 | -0.645 | 0.335 | 0.439 | 0.858 | -0.002 | 0.009 | 0.009 | 0 | 0.21 | 0.25 | 0.25 | 0.37 | 0.21 | 0.23 |
| 5 | 129126598 | rs4836474 | -1.218 | -0.086 | 0.461 | 0.185 | 0.014 | -0.006 | -0.006 | 0 | 0.43 | 0.35 | 0.42 | 0.41 | 0.4 | 0.48 |
| 5 | 129147396 | rs6595926 | 1.229 | 0.364 | 1.134 | 0.07 | -0.002 | -0.002 | -0.002 | 0 | 0.56 | 0.58 | 0.53 | 0.59 | 0.52 | 0.49 |
| 5 | 129159544 | rs2112564 | 1.324 | 0.419 | 1.429 | 0.135 | 0.002 | -0.003 | -0.003 | 0.439 | 0.54 | 0.38 | 0.52 | 0.58 | 0.52 | 0.49 |
| 5 | 129168399 | rs1006611 | -0.951 | 0.968 | 1.288 | -0.012 | 0.021 | -0.041 | -0.041 | 0 | 0.39 | 0.16 | 0.39 | 0.41 | 0.38 | 0.47 |
| 5 | 129170412 | rs458490 | 1.057 | 0.979 | 1.317 | -0.043 | 0.164 | 0.042 | 0.042 | 2.677 | 0.93 | 0.55 | 0.91 | 0.98 | 0.9 | 0.97 |
| 5 | 129180529 | rs445027 | 0.37 | 0.988 | 0.794 | -0.139 | 0.017 | 0.04 | 0.04 | 0.284 | 0.18 | 0.07 | 0.12 | 0.03 | 0.14 | 0.21 |
| 5 | 129181051 | rs6882935 | 1.337 | 1.024 | 1.225 | 0.092 | 0.008 | 0.002 | 0.002 | 0 | 0.52 | 0.23 | 0.49 | 0.59 | 0.52 | 0.52 |
| 5 | 129182833 | rs1422479 | 1.824 | 0.579 | 0.69 | 0.004 | -0.097 | 0.057 | 0.057 | 0.447 | 0.31 | 0.07 | 0.31 | 0.25 | 0.38 | 0.32 |
| 5 | 129199151 | rs1432784 | 0.742 | 0.647 | 0.939 | 0.048 | 0.003 | -0.001 | -0.001 | 0 | 0.54 | 0.32 | 0.52 | 0.59 | 0.52 | 0.51 |
| 5 | 129218180 | rs33910 | 0.791 | 0.555 | 0.887 | 0.058 | 0.007 | -0.002 | -0.002 | 0.24 | 0.53 | 0.29 | 0.53 | 0.59 | 0.54 | 0.52 |
| 5 | 129242068 | rs26417 | 0.887 | 0.601 | 0.844 | 0.071 | 0.008 | 0.002 | 0.002 | 0 | 0.52 | 0.23 | 0.51 | 0.59 | 0.54 | 0.52 |
| 5 | 129277660 | rs13168872 | -0.479 | 0.541 | 0.907 | 0.275 | 0.018 | -0.025 | -0.025 | 0 | 0.39 | 0.24 | 0.39 | 0.43 | 0.37 | 0.47 |
| 5 | 129279813 | rs4530793 | 1.93 | 0.53 | 0.599 | 0.235 | -0.114 | 0.048 | 0.048 | 0.188 | 0.3 | 0.03 | 0.23 | 0.22 | 0.37 | 0.3 |
| 5 | 129280920 | rs10036904 | 0.741 | 0.69 | 1.279 | 0.398 | -0.003 | -0.002 | -0.002 | 0 | 0.55 | 0.29 | 0.51 | 0.57 | 0.53 | 0.49 |
| 5 | 129315115 | rs6864481 | 0.747 | 0.344 | 1.701 | 0.433 | 0.001 | -0.001 | -0.001 | 0 | 0.56 | 0.59 | 0.54 | 0.57 | 0.55 | 0.52 |
| 5 | 129327874 | rs12513704 | 0.823 | 0.158 | 1.337 | 0.414 | 0.017 | -0.07 | -0.07 | 0 | 0.17 | 0.06 | 0.24 | 0.32 | 0.13 | 0.16 |
| 5 | 129351499 | rs4836487 | -0.579 | -0.038 | 1.098 | 0.385 | 0.03 | 0.001 | 0.001 | 0 | 0.41 | 0.24 | 0.39 | 0.39 | 0.37 | 0.46 |
| 5 | 129364691 | rs4323260 | 0.573 | -0.115 | 0.838 | 0.348 | -0.043 | -0.075 | -0.075 | 0.736 | 0.15 | 0.01 | 0.22 | 0.23 | 0.18 | 0.2 |
| 5 | 129368358 | rs4836488 | 0.92 | 0.007 | 1.208 | 0.369 | 0.004 | 0.031 | 0.031 | 0 | 0.55 | 0.34 | 0.52 | 0.69 | 0.56 | 0.53 |
| 5 | 129372263 | rs4146658 | 1.04 | 0.042 | 1.224 | 0.384 | 0.001 | 0.007 | 0.007 | 0 | 0.55 | 0.41 | 0.52 | 0.69 | 0.56 | 0.52 |
| 5 | 129377047 | rs1000140 | 1.04 | 0.042 | 1.224 | 0.384 | 0.001 | 0.007 | 0.007 | 0 | 0.55 | 0.41 | 0.52 | 0.69 | 0.56 | 0.52 |
| 5 | 129385674 | rs1541450 | 0.863 | 0.059 | 1.096 | 0.392 | -0.002 | 0.052 | 0.052 | 0 | 0.57 | 0.72 | 0.54 | 0.69 | 0.57 | 0.53 |
| 5 | 129395456 | rs10491272 | 0.709 | -0.067 | 0.863 | 0.338 | -0.02 | 0.083 | 0.083 | 0.048 | 0.2 | 0.01 | 0.14 | 0.06 | 0.16 | 0.21 |
| 5 | 129412126 | rs4836491 | -0.283 | 0.365 | 0.877 | 0.659 | 0.029 | 0.042 | 0.042 | 0 | 0.41 | 0.23 | 0.4 | 0.32 | 0.37 | 0.44 |
| 5 | 129439574 | rs6858978 | -1.099 | 0.249 | 0.63 | 0.77 | -0.008 | 0.049 | 0.049 | 0 | 0.58 | 0.72 | 0.65 | 0.68 | 0.48 | 0.58 |
| 5 | 129442887 | rs2052007 | 0.072 | 0.135 | 0.733 | 1.122 | -0.008 | 0.032 | 0.032 | 0 | 0.18 | 0.26 | 0.26 | 0.38 | 0.11 | 0.17 |
| 5 | 129452658 | rs6862930 | -0.586 | 0.477 | 0.977 | 1.058 | 0 | 0.029 | 0.029 | 0 | 0.47 | 0.57 | 0.52 | 0.27 | 0.46 | 0.43 |
| 5 | 129453234 | rs2190815 | -1.186 | 0.367 | 0.632 | 1.449 | -0.043 | 0.043 | 0.043 | 0 | 0.55 | 0.85 | 0.72 | 0.59 | 0.49 | 0.65 |
| 5 | 129469043 | rs13167635 | -0.289 | 0.522 | 0.637 | 1.26 | 0.018 | 0.06 | 0.06 | 0 | 0.43 | 0.2 | 0.45 | 0.27 | 0.38 | 0.37 |
| 5 | 129486905 | rs30264 | 0.986 | 0.48 | 1.188 | 1.83 | 0.023 | 0.097 | 0.097 | 0 | 0.91 | 0.82 | 0.85 | 0.73 | 0.9 | 0.73 |
| 5 | 129517194 | rs2051802 | 1.628 | 0.504 | 1.226 | 1.815 | 0.052 | 0.175 | 0.175 | 0 | 0.9 | 0.64 | 0.84 | 0.75 | 0.89 | 0.73 |
| 5 | 129520679 | rs10068403 | -1.52 | 0.006 | 1.388 | 1.961 | 0.005 | 0.307 | 0.307 | 0.953 | 0.06 | 0.3 | 0.16 | 0.26 | 0.01 | 0.08 |
| 5 | 129522136 | rs10057029 | -1.52 | 0.006 | 1.388 | 2.022 | 0.005 | 0.327 | 0.327 | 0.953 | 0.06 | 0.29 | 0.17 | 0.26 | 0.01 | 0.08 |
| 5 | 129524245 | rs40564 | 1.16 | 0.227 | 1.077 | 1.734 | 0.014 | 0.095 | 0.095 | 0 | 0.86 | 0.69 | 0.77 | 0.73 | 0.86 | 0.87 |
| 5 | 129525635 | rs30268 | 1.119 | 0.229 | 1.078 | 1.718 | 0.016 | 0.095 | 0.095 | 0 | 0.86 | 0.7 | 0.77 | 0.74 | 0.86 | 0.87 |
| 5 | 129544292 | rs4326178 | -0.201 | -0.167 | 0.463 | 1.681 | 0.009 | -0.019 | -0.019 | 0.37 | 0.17 | 0.08 | 0.2 | 0.17 | 0.14 | 0.21 |
| 5 | 129552816 | rs6861345 | -1.898 | -0.43 | 0.488 | 1.452 | 0.007 | -0.037 | -0.037 | 1.441 | 0.31 | 0.83 | 0.34 | 0.37 | 0.25 | 0.36 |
| 5 | 129555759 | rs10075474 | 0.053 | -0.381 | 0.569 | 1.224 | 0.017 | -0.056 | -0.056 | 0 | 0.11 | 0.18 | 0.08 | 0 | 0.13 | 0.06 |
| 5 | 129566101 | rs6860728 | -0.455 | -0.49 | 0.471 | 1.285 | -0.001 | -0.006 | -0.006 | 0 | 0.14 | 0.14 | 0.05 | 0.08 | 0.1 | 0.2 |
| 5 | 129579206 | rs6883532 | -1.694 | -0.367 | 0.156 | 1.516 | 0.003 | -0.01 | -0.01 | 0 | 0.3 | 0.78 | 0.34 | 0.31 | 0.25 | 0.34 |
| 5 | 129587463 | rs10520068 | -0.17 | -0.438 | 0.109 | 1.34 | 0.006 | -0.024 | -0.024 | 0 | 0.16 | 0.08 | 0.2 | 0.21 | 0.14 | 0.21 |
| 5 | 129591296 | rs2418536 | -1.139 | -0.735 | 0.073 | 0.974 | 0.005 | 0.004 | 0.004 | 0 | 0.88 | 0.87 | 0.94 | 0.85 | 0.92 | 0.95 |
| 5 | 129614109 | rs3919478 | 0.516 | -0.855 | 0.558 | 1.105 | -0.003 | 0 | 0 | 0 | 0.78 | 0.28 | 0.87 | 0.75 | 0.84 | 0.8 |
| 5 | 129615610 | rs2418539 | 0.118 | -0.853 | 1.123 | 1.123 | 0.004 | 0.01 | 0.01 | 0 | 0.08 | 0.15 | 0.05 | 0.07 | 0.08 | 0.15 |
| 5 | 129616513 | rs13173386 | -1.271 | -0.816 | 1.141 | 1.1 | 0.004 | 0.003 | 0.003 | 0 | 0.88 | 0.87 | 0.94 | 0.85 | 0.92 | 0.95 |
| 5 | 129632890 | rs10478920 | 0.662 | -1.207 | 0.394 | 1.355 | 0.002 | -0.087 | -0.087 | 1.35 | 0.13 | 0 | 0.13 | 0.28 | 0.1 | 0.06 |
| 5 | 129633301 | rs7711634 | -0.961 | -1.102 | 1.082 | 1.202 | -0.031 | 0.112 | 0.112 | 0 | 0.25 | 0.46 | 0.37 | 0.41 | 0.18 | 0.35 |
| 5 | 129647781 | rs264122 | -1.88 | -1.102 | 1.009 | 1.107 | 0.062 | 0.065 | 0.065 | 0 | 0.46 | 0.87 | 0.5 | 0.66 | 0.35 | 0.55 |
| 5 | 129651668 | rs17713413 | -0.475 | -1.137 | 0.81 | 1.185 | 0.01 | -0.009 | -0.009 | 0.088 | 0.22 | 0.12 | 0.21 | 0.22 | 0.17 | 0.27 |
| 5 | 129659729 | rs2108425 | -1.843 | -1.069 | 0.735 | 1.211 | 0.054 | 0.069 | 0.069 | 0 | 0.46 | 0.87 | 0.5 | 0.67 | 0.35 | 0.55 |
| 5 | 129669944 | rs1473134 | -0.016 | -1.116 | 0.839 | 1.187 | -0.003 | -0.006 | -0.006 | 0 | 0.16 | 0.13 | 0.21 | 0.26 | 0.17 | 0.23 |
| 5 | 129673206 | rs6876137 | -2.29 | -0.908 | 0.606 | 1.021 | -0.027 | -0.036 | -0.036 | 0 | 0.53 | 0.88 | 0.43 | 0.51 | 0.45 | 0.52 |
| 5 | 129673919 | rs6898857 | -1.69 | -0.961 | 0.919 | 1.023 | -0.002 | 0 | 0 | 0 | 0.33 | 0.65 | 0.31 | 0.36 | 0.25 | 0.42 |
| 5 | 129684723 | rs12519614 | -0.164 | -0.92 | 0.679 | 0.63 | -0.003 | -0.003 | -0.003 | 0 | 0.17 | 0.15 | 0.34 | 0.46 | 0.18 | 0.3 |
| 5 | 129684777 | rs6869219 | -2.44 | -0.876 | 0.445 | 0.614 | -0.061 | 0.086 | 0.086 | 0 | 0.54 | 0.9 | 0.57 | 0.71 | 0.47 | 0.6 |
| 5 | 129684909 | rs12516966 | -1.121 | -0.586 | 0.786 | 0.773 | -0.009 | 0.146 | 0.146 | 0 | 0.23 | 0.45 | 0.37 | 0.49 | 0.19 | 0.35 |

| | | | | | | | | | | | | | | | | |
|---|---|---|---|---|---|---|---|---|---|---|---|---|---|---|---|---|
| 5 | 129693875 | rs11741204 | 0.623 | -0.155 | 0.255 | 0.511 | -0.038 | 0.047 | 0.047 | 0 | 0.08 | 0.01 | 0.06 | 0 | 0.13 | 0.05 |
| 5 | 129703802 | rs6870318 | -0.275 | -0.455 | 1.349 | 0.78 | -0.045 | -0.026 | -0.026 | 0 | 0.19 | 0.44 | 0.14 | 0.19 | 0.1 | 0.09 |
| 5 | 129708350 | rs7726515 | -0.585 | 0.07 | 1.241 | 0.91 | 0.033 | 0.001 | 0.001 | 0 | 0.14 | 0.16 | 0.15 | 0.19 | 0.21 | 0.2 |
| 5 | 129710445 | rs17165472 | -1.307 | -0.061 | 1.44 | 1.219 | -0.042 | 0.166 | 0.166 | 0 | 0.17 | 0.38 | 0.27 | 0.34 | 0.11 | 0.28 |
| 5 | 129723516 | rs7356549 | -0.773 | 0.195 | 1.253 | 0.807 | 0.017 | -0.007 | -0.007 | 0.097 | 0.19 | 0.19 | 0.16 | 0.19 | 0.21 | 0.2 |
| 5 | 129732849 | rs17165462 | -0.981 | -0.036 | 1.721 | 1.089 | -0.03 | 0.26 | 0.26 | 1.416 | 0.07 | 0.2 | 0.22 | 0.31 | 0.02 | 0.1 |
| 5 | 129740524 | rs12188544 | -0.685 | -0.152 | 1.959 | 0.922 | -0.01 | -0.018 | -0.018 | 0.063 | 0.12 | 0.18 | 0.06 | 0.04 | 0.08 | 0.16 |
| 5 | 129741850 | rs12516185 | -0.283 | -0.119 | 1.993 | 0.905 | 0.03 | 0.016 | 0.016 | 0 | 0.09 | 0.11 | 0.13 | 0.19 | 0.17 | 0.19 |
| 5 | 129751475 | rs10068281 | 0.629 | -0.116 | 1.879 | 0.711 | -0.003 | -0.024 | -0.024 | 1.109 | 0.1 | 0.1 | 0.03 | 0 | 0.04 | 0 |
| 5 | 129759493 | rs9285919 | -2.569 | -0.251 | 1.984 | 0.656 | 0.013 | 0.008 | 0.008 | 0 | 0.41 | 0.82 | 0.49 | 0.54 | 0.32 | 0.47 |
| 5 | 129781272 | rs10478926 | -1.299 | -0.261 | 1.977 | 0.531 | -0.043 | -0.117 | -0.117 | 2.687 | 0.23 | 0.48 | 0.14 | 0.04 | 0.13 | 0.18 |
| 5 | 129797129 | rs10463875 | -2.493 | -0.242 | 1.816 | 0.568 | 0.01 | 0.006 | 0.006 | 0 | 0.4 | 0.77 | 0.49 | 0.54 | 0.32 | 0.47 |
| 5 | 129818555 | rs6893341 | -0.496 | -0.12 | 1.766 | 0.606 | 0.004 | 0.006 | 0.006 | 0 | 0.12 | 0.08 | 0.06 | 0.04 | 0.08 | 0.17 |
| 5 | 129834311 | rs10520081 | -0.681 | -0.241 | 1.959 | 0.831 | 0.011 | 0.032 | 0.032 | 0 | 0.12 | 0.06 | 0.06 | 0.04 | 0.08 | 0.16 |
| 5 | 129846173 | rs6595951 | -1.422 | -0.089 | 2.067 | 0.901 | 0.047 | 0.181 | 0.181 | 0.232 | 0.16 | 0.33 | 0.35 | 0.5 | 0.19 | 0.29 |
| 5 | 129863694 | rs1009686 | -2.365 | -0.121 | 1.91 | 0.87 | 0.003 | 0.021 | 0.021 | 0 | 0.29 | 0.56 | 0.45 | 0.54 | 0.28 | 0.44 |
| 5 | 129873659 | rs11242026 | -1.255 | 0.068 | 1.991 | 0.971 | 0.001 | 0.028 | 0.028 | 0 | 0.27 | 0.32 | 0.4 | 0.6 | 0.27 | 0.33 |
| 5 | 129877045 | rs2215258 | -2.516 | 0.068 | 1.855 | 0.609 | -0.02 | 0.038 | 0.038 | 0 | 0.5 | 0.78 | 0.51 | 0.63 | 0.45 | 0.52 |
| 5 | 129885843 | rs13359516 | 0.776 | 0.076 | 1.913 | 0.559 | 0.004 | 0.009 | 0.009 | 1.837 | 0.09 | 0.04 | 0.02 | 0 | 0.04 | 0 |
| 5 | 129888054 | rs7729880 | -1.594 | 0.05 | 2.097 | 0.567 | 0.009 | 0.039 | 0.039 | 0.038 | 0.31 | 0.39 | 0.41 | 0.59 | 0.32 | 0.37 |
| 5 | 129891761 | rs10035791 | -0.394 | 0.065 | 2.099 | 0.67 | 0.003 | -0.002 | -0.002 | 0 | 0.09 | 0.07 | 0.13 | 0.17 | 0.18 | 0.19 |
| 5 | 129907414 | rs9327564 | 0.668 | 0.052 | 2.241 | 0.655 | -0.004 | -0.052 | -0.052 | 1.71 | 0.1 | 0.18 | 0.03 | 0 | 0.04 | 0 |
| 5 | 129909600 | rs11242029 | -2.739 | 0.017 | 2.248 | 0.646 | -0.03 | -0.014 | -0.014 | 0.03 | 0.41 | 0.78 | 0.49 | 0.62 | 0.39 | 0.49 |
| 5 | 129911049 | rs17839718 | -0.239 | 0.019 | 2.267 | 0.662 | 0 | -0.005 | -0.005 | 0 | 0.08 | 0.09 | 0.06 | 0.04 | 0.07 | 0.15 |
| 5 | 129912666 | rs1468550 | -1.762 | 0.003 | 2.241 | 0.681 | -0.013 | 0.188 | 0.188 | 3.078 | 0.12 | 0.29 | 0.25 | 0.41 | 0.09 | 0.16 |
| 5 | 129925322 | rs2215257 | -2.167 | 0.05 | 2.195 | 0.646 | -0.002 | 0 | 0 | 0.036 | 0.42 | 0.6 | 0.48 | 0.62 | 0.4 | 0.52 |
| 5 | 129984085 | rs1073833 | -2.271 | 0.275 | 2.337 | 0.773 | 0.005 | 0.023 | 0.023 | 0 | 0.34 | 0.57 | 0.46 | 0.44 | 0.32 | 0.41 |
| 5 | 129997196 | rs17714209 | -1.808 | -0.079 | 2.197 | 0.791 | 0.007 | 0.037 | 0.037 | 0.142 | 0.22 | 0.24 | 0.4 | 0.45 | 0.27 | 0.4 |
| 5 | 130005745 | rs6595954 | -2.445 | -0.131 | 2.252 | 0.75 | 0.002 | -0.027 | -0.027 | 0.829 | 0.36 | 0.72 | 0.48 | 0.45 | 0.31 | 0.41 |
| 5 | 130017287 | rs4836519 | -1.049 | -0.017 | 1.903 | 0.846 | -0.031 | -0.016 | -0.016 | 0.239 | 0.16 | 0.08 | 0.16 | 0.21 | 0.22 | 0.33 |
| 5 | 130051646 | rs11954837 | -1.154 | -0.005 | 1.985 | 0.734 | -0.016 | 0.116 | 0.116 | 0 | 0.05 | 0.11 | 0.24 | 0.22 | 0.04 | 0.08 |
| 5 | 130091947 | rs10478939 | -1.04 | -0.03 | 1.944 | 0.74 | -0.029 | -0.018 | -0.018 | 0 | 0.16 | 0.09 | 0.17 | 0.22 | 0.22 | 0.33 |
| 5 | 130096260 | rs12518350 | -2.348 | -0.049 | 2.044 | 0.724 | 0.007 | 0.051 | 0.051 | 0.059 | 0.24 | 0.32 | 0.43 | 0.44 | 0.27 | 0.4 |
| 5 | 130102023 | rs1862647 | -2.184 | -0.224 | 2.038 | 0.668 | -0.001 | 0.007 | 0.007 | 0 | 0.36 | 0.36 | 0.45 | 0.44 | 0.32 | 0.41 |
| 5 | 130113901 | rs10044304 | -2.232 | -0.096 | 2.1 | 0.732 | 0.002 | 0.037 | 0.037 | 0 | 0.35 | 0.47 | 0.44 | 0.44 | 0.31 | 0.38 |
| 5 | 130131711 | rs12517046 | -0.807 | -0.109 | 2.018 | 0.746 | -0.029 | 0.068 | 0.068 | 1.63 | 0.15 | 0.3 | 0.26 | 0.22 | 0.08 | 0.05 |
| 5 | 130140680 | rs12652579 | -1.311 | -0.006 | 1.581 | 0.747 | -0.035 | -0.013 | -0.013 | 0 | 0.19 | 0.08 | 0.17 | 0.22 | 0.24 | 0.33 |
| 5 | 130152348 | rs10052692 | -2.201 | -0.587 | 1.757 | 0.726 | 0.01 | -0.024 | -0.024 | 0 | 0.37 | 0.69 | 0.47 | 0.45 | 0.28 | 0.38 |
| 5 | 130164018 | rs6867080 | -2.271 | -0.604 | 1.81 | 0.734 | 0.015 | -0.032 | -0.032 | 0 | 0.37 | 0.73 | 0.47 | 0.45 | 0.29 | 0.38 |
| 5 | 130164900 | rs3981405 | -1.331 | -0.567 | 1.68 | 0.748 | -0.025 | -0.015 | -0.015 | 0 | 0.19 | 0.07 | 0.17 | 0.23 | 0.21 | 0.33 |
| 5 | 130167659 | rs4492106 | -1.354 | -0.555 | 1.659 | 0.811 | -0.026 | -0.036 | -0.036 | 0 | 0.2 | 0.07 | 0.17 | 0.37 | 0.22 | 0.33 |
| 5 | 130176202 | rs4705896 | -1.631 | -0.479 | 1.881 | 0.765 | 0.007 | 0.007 | 0.007 | 0 | 0.19 | 0.26 | 0.16 | 0.24 | 0.2 | 0.28 |
| 5 | 130185380 | rs10478953 | -1.086 | -0.524 | 1.714 | 0.707 | -0.041 | -0.025 | -0.025 | 0 | 0.17 | 0.07 | 0.17 | 0.23 | 0.23 | 0.28 |
| 5 | 130195731 | rs12522693 | 0.255 | -0.515 | 1.687 | 0.723 | -0.029 | -0.02 | -0.02 | 0 | 0.08 | 0.03 | 0.1 | 0.14 | 0.15 | 0.15 |
| 5 | 130195812 | rs10900798 | -1.518 | -0.676 | 1.944 | 0.669 | 0.003 | 0.023 | 0.023 | 0 | 0.29 | 0.34 | 0.42 | 0.45 | 0.28 | 0.34 |
| 5 | 130202197 | rs10520092 | -0.48 | -0.585 | 1.509 | 0.472 | -0.046 | -0.028 | -0.028 | 0 | 0.17 | 0.06 | 0.17 | 0.23 | 0.23 | 0.28 |
| 5 | 130215911 | rs6888236 | -1.585 | -0.695 | 1.446 | 0.409 | -0.001 | -0.003 | -0.003 | 1.652 | 0.34 | 0.64 | 0.26 | 0.23 | 0.27 | 0.3 |
| 5 | 130217998 | rs6863693 | -1.585 | -0.695 | 1.446 | 0.409 | -0.001 | -0.003 | -0.003 | 0.127 | 0.34 | 0.64 | 0.26 | 0.23 | 0.27 | 0.3 |
| 5 | 130219200 | rs4705925 | -0.588 | -0.687 | 1.508 | 0.41 | 0.003 | 0 | 0 | 0 | 0.21 | 0.22 | 0.2 | 0.23 | 0.23 | 0.3 |
| 5 | 130225121 | rs6873442 | 0.544 | -0.671 | 1.201 | 0.417 | 0.004 | -0.008 | -0.008 | 0 | 0.09 | 0.07 | 0.01 | 0 | 0.03 | 0 |
| 5 | 130242429 | rs2522031 | -1.489 | -0.631 | 1.066 | 0.435 | -0.003 | -0.014 | -0.014 | 0 | 0.33 | 0.59 | 0.26 | 0.22 | 0.26 | 0.29 |
| 5 | 130247078 | rs6898224 | 1.341 | -0.85 | 1.174 | 0.557 | -0.001 | -0.003 | -0.003 | 0 | 0.67 | 0.36 | 0.74 | 0.78 | 0.74 | 0.71 |
| 5 | 130249954 | rs6870590 | 0.567 | -0.847 | 1.05 | 0.559 | 0.029 | 0.001 | 0.001 | 0 | 0.78 | 0.59 | 0.77 | 0.78 | 0.77 | 0.71 |
| 5 | 130272586 | rs12187094 | 2.156 | -0.967 | -0.007 | 0.507 | -0.101 | -0.095 | -0.095 | 0 | 0.45 | 0.03 | 0.51 | 0.4 | 0.58 | 0.48 |
| 5 | 130300594 | rs10040560 | -1.404 | -0.39 | 0.346 | -0.029 | -0.003 | -0.015 | -0.015 | 1.62 | 0.33 | 0.59 | 0.26 | 0.22 | 0.26 | 0.29 |
| 5 | 130330467 | rs17692694 | 0.039 | 0.017 | -0.441 | 0.275 | -0.023 | -0.042 | -0.042 | 0 | 0.14 | 0.01 | 0.04 | 0.21 | 0.13 | 0.09 |
| 5 | 130359215 | rs6595989 | 0.769 | 0.297 | 0.399 | 0.288 | 0.02 | 0.004 | 0.004 | 0 | 0.8 | 0.73 | 0.8 | 0.77 | 0.77 | 0.71 |
| 5 | 130378070 | rs1363696 | 1.22 | 0.195 | -0.848 | 0.254 | 0.004 | 0.049 | 0.049 | 1.021 | 0.64 | 0.12 | 0.72 | 0.76 | 0.74 | 0.67 |
| 5 | 130407827 | rs7709727 | -1.124 | -0.176 | 0.161 | -0.021 | -0.019 | -0.048 | -0.048 | 0 | 0.22 | 0.69 | 0.13 | 0.01 | 0.09 | 0.07 |
| 5 | 130418134 | rs12153520 | -0.812 | 0.006 | 0.039 | 0.081 | -0.024 | -0.036 | -0.036 | 0 | 0.14 | 0.04 | 0.14 | 0.2 | 0.17 | 0.22 |
| 5 | 130433358 | rs7715572 | 0.107 | -0.099 | 0.568 | 0.01 | -0.002 | -0.021 | -0.021 | 0 | 0.11 | 0.09 | 0.02 | 0 | 0.03 | 0.01 |
| 5 | 130439091 | rs3891636 | -1.751 | -0.098 | 1.199 | 0.021 | 0.005 | 0.008 | 0.008 | 1.318 | 0.23 | 0.53 | 0.23 | 0.21 | 0.23 | 0.27 |
| 5 | 130463135 | rs6874948 | -2.168 | -0.155 | 1.283 | -0.039 | 0.024 | 0.04 | 0.04 | 0.972 | 0.36 | 0.8 | 0.26 | 0.21 | 0.27 | 0.28 |
| 5 | 130487957 | rs10070034 | 0.015 | -0.181 | 1.311 | -0.034 | 0.003 | -0.013 | -0.013 | 0 | 0.11 | 0.09 | 0.03 | 0 | 0.03 | 0.01 |
| 5 | 130492613 | rs9327595 | -0.52 | -0.189 | 1.5 | -0.038 | -0.035 | -0.068 | -0.068 | 0.235 | 0.13 | 0.25 | 0.05 | 0 | 0.04 | 0.01 |
| 5 | 130494365 | rs10062233 | -0.615 | -0.187 | 2.162 | -0.037 | -0.048 | -0.092 | -0.092 | 0 | 0.14 | 0.38 | 0.06 | 0 | 0.04 | 0.01 |
| 5 | 130498228 | rs17165688 | -0.615 | -0.187 | 2.195 | -0.037 | -0.048 | -0.092 | -0.092 | 0 | 0.14 | 0.38 | 0.06 | 0 | 0.04 | 0.01 |
| 5 | 130505761 | rs7735116 | -1.416 | -0.238 | 2.644 | -0.064 | -0.076 | -0.123 | -0.123 | 0.869 | 0.19 | 0.61 | 0.1 | 0 | 0.05 | 0.06 |
| 5 | 130506144 | rs9327597 | -1.416 | -0.238 | 2.644 | -0.064 | -0.076 | -0.123 | -0.123 | 0 | 0.19 | 0.61 | 0.1 | 0 | 0.05 | 0.06 |

| | | | | | | | | | | | | | | | | |
|---|---|---|---|---|---|---|---|---|---|---|---|---|---|---|---|---|
| 5 | 130515072 | rs7721068 | -1.734 | -0.228 | 2.681 | -0.045 | 0 | -0.009 | -0.009 | 1.514 | 0.16 | 0.55 | 0.14 | 0.11 | 0.14 | 0.19 |
| 5 | 130516368 | rs10052199 | -2.148 | -0.227 | 2.619 | -0.045 | 0.02 | 0.026 | 0.026 | 3.095 | 0.29 | 0.8 | 0.18 | 0.11 | 0.18 | 0.2 |
| 5 | 130529122 | rs17165964 | -0.671 | -0.199 | 2.559 | -0.005 | -0.022 | -0.031 | -0.031 | 0 | 0.06 | 0 | 0.08 | 0.1 | 0.07 | 0.08 |
| 5 | 130533828 | rs2108392 | -2.159 | -0.259 | 2.619 | -0.044 | 0.003 | 0 | 0 | 1.296 | 0.3 | 0.75 | 0.19 | 0.11 | 0.18 | 0.21 |
| 5 | 130536044 | rs4706009 | -1.508 | -0.336 | 2.619 | -0.081 | 0.002 | -0.007 | -0.007 | 0 | 0.15 | 0.3 | 0.14 | 0.11 | 0.15 | 0.2 |
| 5 | 130541807 | rs7729291 | 0.211 | -0.327 | 2.671 | -0.077 | -0.013 | -0.038 | -0.038 | 0 | 0.11 | 0.16 | 0.03 | 0 | 0.03 | 0.01 |
| 5 | 130543648 | rs6882231 | -2.033 | -0.308 | 2.594 | -0.065 | 0.006 | 0.014 | 0.014 | 0.255 | 0.21 | 0.42 | 0.22 | 0.21 | 0.21 | 0.27 |
| 5 | 130547165 | rs6596004 | -2.047 | -0.295 | 2.238 | -0.06 | -0.019 | -0.028 | -0.028 | 0.001 | 0.32 | 0.52 | 0.25 | 0.21 | 0.24 | 0.28 |
| 5 | 130548380 | rs4706011 | -1.817 | -0.348 | 2.271 | -0.075 | -0.03 | -0.055 | -0.055 | 0 | 0.28 | 0.58 | 0.18 | 0.11 | 0.18 | 0.21 |
| 5 | 130550343 | rs7710461 | -1.979 | -0.355 | 2.262 | -0.078 | 0.022 | 0.028 | 0.028 | 1.453 | 0.3 | 0.79 | 0.19 | 0.11 | 0.18 | 0.21 |
| 5 | 130551178 | rs763812 | -1.968 | -0.355 | 2.262 | -0.078 | 0.021 | 0.028 | 0.028 | 1.523 | 0.3 | 0.79 | 0.19 | 0.11 | 0.18 | 0.21 |
| 5 | 130553378 | rs7726567 | -1.654 | -0.345 | 2.235 | -0.074 | -0.034 | -0.065 | -0.065 | 3.602 | 0.27 | 0.52 | 0.17 | 0.11 | 0.18 | 0.21 |
| 5 | 130560128 | rs2079511 | -0.06 | -0.299 | 1.73 | -0.047 | -0.018 | -0.015 | -0.015 | 0 | 0.08 | 0.03 | 0.08 | 0.11 | 0.13 | 0.15 |
| 5 | 130561257 | rs2420147 | 0.282 | -0.328 | 1.126 | 0.076 | -0.049 | -0.071 | -0.071 | 0 | 0.83 | 0.96 | 0.86 | 0.69 | 0.79 | 0.83 |
| 5 | 130567793 | rs6867367 | -1.535 | -0.371 | 1.318 | -0.031 | 0.019 | 0.024 | 0.024 | 1.39 | 0.31 | 0.78 | 0.19 | 0.11 | 0.18 | 0.21 |
| 5 | 130575413 | rs6596006 | -1.601 | -0.444 | 1.198 | -0.063 | 0.056 | 0.09 | 0.09 | 2.963 | 0.39 | 0.82 | 0.2 | 0.11 | 0.22 | 0.23 |
| 5 | 130588008 | rs6895586 | 0.857 | -0.554 | 1.25 | 0.706 | 0.018 | 0.12 | 0.12 | 0.115 | 0.85 | 0.82 | 0.64 | 0.48 | 0.77 | 0.74 |
| 5 | 130598318 | rs11242069 | -1.955 | -0.727 | 1.183 | 0.608 | 0.017 | -0.149 | -0.149 | 0 | 0.34 | 0.92 | 0.46 | 0.51 | 0.27 | 0.33 |
| 5 | 130616792 | rs12513768 | 1.62 | -0.714 | 1.394 | 0.62 | 0.024 | -0.122 | -0.122 | 0.027 | 0.67 | 0.12 | 0.55 | 0.49 | 0.73 | 0.67 |
| 5 | 130635566 | rs10042016 | -0.454 | -0.69 | 1.639 | 0.524 | -0.064 | -0.11 | -0.11 | 0 | 0.17 | 0.65 | 0.09 | 0 | 0.04 | 0.06 |
| 5 | 130736800 | rs7735957 | 0.242 | -0.178 | 2.216 | 0.112 | -0.045 | -0.087 | -0.087 | 0 | 0.15 | 0.34 | 0.06 | 0 | 0.04 | 0.05 |
| 5 | 130739895 | rs6596018 | 0.229 | -0.178 | 2.203 | 0.112 | -0.06 | -0.112 | -0.112 | 0 | 0.16 | 0.44 | 0.07 | 0 | 0.04 | 0.05 |
| 5 | 130761470 | rs4996522 | -0.802 | -0.05 | 2.209 | 0.222 | 0.013 | 0.101 | 0.101 | 0.1 | 0.15 | 0.2 | 0.32 | 0.51 | 0.23 | 0.26 |
| 5 | 130766662 | rs1291602 | -0.542 | -0.181 | 1.479 | 0.11 | -0.016 | 0.038 | 0.038 | 2.233 | 0.81 | 0.98 | 0.9 | 0.9 | 0.83 | 0.85 |
| 5 | 130788698 | rs10054168 | -1.204 | -0.361 | 1.867 | 0.169 | -0.011 | 0.051 | 0.051 | 0 | 0.29 | 0.53 | 0.39 | 0.52 | 0.26 | 0.28 |
| 5 | 130813377 | rs31239 | 1.072 | -0.3 | 1.87 | 0.203 | 0.003 | -0.008 | -0.008 | 0 | 0.69 | 0.36 | 0.61 | 0.48 | 0.74 | 0.72 |
| 5 | 130833946 | rs31251 | 2.116 | -0.476 | 1.533 | 0.036 | -0.118 | -0.062 | -0.062 | 0.005 | 0.44 | 0.03 | 0.46 | 0.38 | 0.53 | 0.48 |
| 5 | 130840385 | rs10063129 | 0.725 | -0.388 | 1.853 | 0.075 | -0.041 | -0.077 | -0.077 | 3.087 | 0.14 | 0.25 | 0.04 | 0 | 0.04 | 0.01 |
| 5 | 130850938 | rs3776019 | 0.713 | -0.406 | 1.704 | 0.242 | -0.045 | -0.02 | -0.02 | 0.646 | 0.18 | 0.01 | 0.09 | 0.15 | 0.19 | 0.18 |
| 5 | 130857727 | rs11746464 | 1.816 | -0.381 | 1.717 | 0.212 | -0.069 | -0.009 | -0.009 | 0 | 0.48 | 0.05 | 0.48 | 0.37 | 0.55 | 0.52 |
| 5 | 130861874 | rs2189383 | -0.684 | -0.363 | 1.902 | 0.342 | 0.001 | 0.027 | 0.027 | 0 | 0.94 | 0.94 | 0.82 | 0.8 | 0.93 | 0.87 |
| 5 | 130879181 | rs6878488 | 0.799 | -0.39 | 2.011 | 0.32 | -0.042 | -0.078 | -0.078 | 0 | 0.14 | 0.27 | 0.04 | 0 | 0.04 | 0.01 |
| 5 | 130909732 | rs10477734 | -0.365 | -0.379 | 1.775 | 0.348 | 0 | 0.025 | 0.025 | 0 | 0.94 | 0.93 | 0.82 | 0.79 | 0.93 | 0.86 |
| 5 | 130936652 | rs10054049 | 0.532 | -0.379 | 1.926 | 0.325 | -0.086 | -0.119 | -0.119 | 0 | 0.15 | 0.47 | 0.06 | 0 | 0.03 | 0.02 |
| 5 | 130942624 | rs1422870 | 1.165 | -0.349 | 1.847 | 0.381 | 0.007 | -0.018 | -0.018 | 5.414 | 0.69 | 0.34 | 0.6 | 0.48 | 0.74 | 0.72 |
| 5 | 130962033 | rs1295873 | -0.612 | -0.296 | 1.789 | 0.417 | 0 | 0.034 | 0.034 | 0 | 0.94 | 0.93 | 0.82 | 0.79 | 0.93 | 0.86 |
| 5 | 130980027 | rs2042252 | 1.048 | -0.308 | 1.761 | 0.405 | 0.003 | -0.009 | -0.009 | 0 | 0.69 | 0.35 | 0.61 | 0.49 | 0.75 | 0.72 |
| 5 | 131008194 | rs26008 | -0.562 | -0.302 | 1.736 | 0.427 | -0.003 | 0.067 | 0.067 | 0 | 0.94 | 0.9 | 0.81 | 0.79 | 0.94 | 0.87 |
| 5 | 131040552 | rs32108 | 2.004 | -0.478 | 1.361 | 0.143 | -0.117 | -0.06 | -0.06 | 1.456 | 0.44 | 0.03 | 0.47 | 0.39 | 0.54 | 0.49 |
| 5 | 131050780 | rs6596034 | 0.564 | -0.315 | 1.729 | 0.206 | -0.063 | -0.11 | -0.11 | 2.029 | 0.16 | 0.36 | 0.05 | 0 | 0.04 | 0.02 |
| 5 | 131107692 | rs6895631 | 0.688 | -0.312 | 1.773 | 0.002 | -0.059 | -0.105 | -0.105 | 0 | 0.15 | 0.34 | 0.05 | 0 | 0.04 | 0.02 |
| 5 | 131125988 | rs6861579 | 0.45 | -0.313 | 1.706 | 0.002 | -0.072 | -0.124 | -0.124 | 0 | 0.16 | 0.44 | 0.06 | 0 | 0.04 | 0.03 |
| 5 | 131195002 | rs10054618 | 0.575 | -0.233 | 1.571 | 0.048 | 0.014 | 0.094 | 0.094 | 0.189 | 0.85 | 0.82 | 0.68 | 0.49 | 0.78 | 0.74 |
| 5 | 131258378 | rs630044 | -1.459 | -0.251 | 1.089 | 0.044 | 0.015 | -0.143 | -0.143 | 0.441 | 0.34 | 0.9 | 0.43 | 0.5 | 0.26 | 0.33 |
| 5 | 131271022 | rs2108450 | 1.843 | -0.267 | 0.966 | -0.029 | -0.106 | -0.037 | -0.037 | 4.477 | 0.46 | 0.03 | 0.47 | 0.4 | 0.55 | 0.51 |
| 5 | 131288468 | rs676944 | -1.231 | 0.004 | 1.272 | 0.185 | 0.008 | -0.02 | -0.02 | 0 | 0.3 | 0.67 | 0.36 | 0.52 | 0.24 | 0.28 |
| 5 | 131294966 | rs10078535 | 0.466 | -0.02 | 1.153 | 0.166 | -0.059 | -0.109 | -0.109 | 2.397 | 0.16 | 0.34 | 0.04 | 0 | 0.04 | 0.01 |
| 5 | 131312327 | rs370652 | -0.879 | 0.237 | 0.305 | 0.357 | -0.046 | 0.021 | 0.021 | 0.295 | 0.83 | 0.99 | 0.91 | 0.92 | 0.81 | 0.82 |
| 5 | 131316164 | rs559971 | -1.136 | 0.358 | 0.909 | 0.734 | 0.012 | 0.104 | 0.104 | 0.124 | 0.15 | 0.18 | 0.32 | 0.52 | 0.21 | 0.25 |
| 5 | 131319036 | rs380182 | -1.259 | 0.359 | 0.868 | 0.734 | 0.027 | 0.189 | 0.189 | 0.244 | 0.16 | 0.27 | 0.32 | 0.53 | 0.21 | 0.25 |
| 5 | 131322494 | rs77938 | 1.121 | 0.171 | 0.166 | 0.657 | 0.028 | -0.084 | -0.084 | 0 | 0.7 | 0.17 | 0.59 | 0.43 | 0.77 | 0.76 |
| 5 | 131322976 | rs253949 | 1.071 | 0.191 | 0.778 | 0.975 | 0.01 | 0.096 | 0.096 | 0 | 0.85 | 0.75 | 0.68 | 0.39 | 0.81 | 0.78 |
| 5 | 131329591 | rs2074808 | -0.49 | 0.215 | 0.658 | 1.181 | 0.002 | 0.015 | 0.015 | 0 | 0.09 | 0.07 | 0.12 | 0.32 | 0.15 | 0.12 |
| 5 | 131336287 | rs440970 | 1.24 | 0.636 | 0.255 | 0.622 | -0.007 | -0.044 | -0.044 | 0 | 0.56 | 0.2 | 0.37 | 0.43 | 0.42 | 0.68 |
| 5 | 131344358 | rs247004 | 0.885 | 0.577 | 0.43 | 0.799 | 0.011 | 0.114 | 0.114 | 0 | 0.85 | 0.74 | 0.65 | 0.43 | 0.82 | 0.78 |
| 5 | 131354094 | rs6596048 | -1.069 | 0.846 | 0.677 | 0.876 | 0.006 | 0.005 | 0.005 | 0.024 | 0.39 | 0.63 | 0.6 | 0.57 | 0.54 | 0.3 |
| 5 | 131359614 | rs247011 | 1.704 | 0.314 | 0.097 | 0.545 | 0.076 | -0.047 | -0.047 | 0 | 0.34 | 0.11 | 0.35 | 0.35 | 0.2 | 0.39 |
| 5 | 131369241 | rs3916441 | -1.074 | 0.755 | 0.761 | 0.872 | -0.004 | -0.003 | -0.003 | 0.002 | 0.39 | 0.66 | 0.49 | 0.57 | 0.54 | 0.31 |
| 5 | 131370827 | rs6868554 | -0.98 | 0.753 | 0.715 | 0.874 | 0.014 | 0.011 | 0.011 | 0 | 0.38 | 0.53 | 0.49 | 0.57 | 0.54 | 0.3 |
| 5 | 131395115 | rs181781 | 0.25 | 1.026 | 0.162 | 1.47 | -0.031 | -0.014 | -0.014 | 0 | 0.06 | 0.02 | 0.08 | 0.33 | 0.13 | 0.11 |
| 5 | 131401334 | rs246844 | -1.247 | 0.956 | 0.509 | 1.356 | 0.013 | 0.072 | 0.072 | 0.041 | 0.13 | 0.23 | 0.24 | 0.59 | 0.21 | 0.23 |
| 5 | 131407044 | rs25879 | -1.625 | 0.96 | 0.731 | 1.342 | 0.029 | 0.107 | 0.107 | 0.218 | 0.16 | 0.31 | 0.28 | 0.61 | 0.25 | 0.25 |
| 5 | 131408077 | rs2069616 | -0.39 | 0.797 | 0.704 | 0.506 | 0.042 | -0.184 | -0.184 | 3.773 | 0.32 | 0.48 | 0.3 | 0 | 0.4 | 0.11 |
| 5 | 131408842 | rs1469149 | -0.225 | 0.967 | 0.711 | 0.533 | 0.125 | -0.159 | -0.159 | 0 | 0.21 | 0.49 | 0.29 | 0 | 0.37 | 0.1 |
| 5 | 131411138 | rs25881 | -0.854 | 0.881 | 0.619 | 1.084 | 0.026 | 0.084 | 0.084 | 0 | 0.13 | 0.21 | 0.25 | 0.62 | 0.21 | 0.24 |
| 5 | 131413255 | rs27438 | -1.213 | 0.578 | 1.238 | 0.874 | -0.001 | 0 | 0 | 0.107 | 0.28 | 0.62 | 0.29 | 0.63 | 0.25 | 0.28 |
| 5 | 131413722 | rs27349 | -0.972 | 0.657 | 1.132 | 0.918 | 0.023 | 0.078 | 0.078 | 0.01 | 0.13 | 0.22 | 0.25 | 0.62 | 0.21 | 0.25 |
| 5 | 131416061 | rs25887 | -1.038 | 0.702 | 1.146 | 0.807 | 0.027 | 0.007 | 0.007 | 1.798 | 0.31 | 0.43 | 0.51 | 0.62 | 0.58 | 0.32 |
| 5 | 131436486 | rs10065787 | -0.19 | 0.69 | 0.956 | 1.574 | -0.086 | -0.02 | -0.02 | 0 | 0.27 | 0.12 | 0.27 | 0.19 | 0.44 | 0.13 |

| | | | | | | | | | | | | | | | | |
|---|---|---|---|---|---|---|---|---|---|---|---|---|---|---|---|---|
| 5 | 131436838 | rs31467 | -1.165 | 1.267 | 1.489 | 1.72 | 0.013 | 0.009 | 0.009 | 0 | 0.34 | 0.56 | 0.36 | 0.37 | 0.52 | 0.23 |
| 5 | 131436896 | rs39897 | -0.201 | 1.067 | 1.631 | 1.881 | -0.02 | -0.019 | -0.019 | 0 | 0.22 | 0.2 | 0.32 | 0.34 | 0.54 | 0.22 |
| 5 | 131447104 | rs247008 | 0.442 | -0.009 | 1.227 | 1.535 | -0.001 | -0.006 | -0.006 | 0 | 0.65 | 0.4 | 0.65 | 0.45 | 0.66 | 0.61 |
| 5 | 131472553 | rs154734 | -1.598 | 0.008 | 1.607 | 1.59 | 0.062 | 0.123 | 0.123 | 1.078 | 0.14 | 0.55 | 0.27 | 0.34 | 0.19 | 0.2 |
| 5 | 131526190 | rs156031 | -1.844 | -0.34 | 0.989 | 1.375 | 0.053 | -0.01 | -0.01 | 0.336 | 0.69 | 0.99 | 0.65 | 0.76 | 0.8 | 0.64 |
| 5 | 131527321 | rs156033 | -1.365 | -0.208 | 1.447 | 1.47 | 0.083 | 0.155 | 0.155 | 0.361 | 0.13 | 0.49 | 0.25 | 0.34 | 0.19 | 0.2 |
| 5 | 131534499 | rs3749756 | 0.329 | -0.21 | 1.095 | 1.402 | -0.091 | -0.006 | -0.006 | 0 | 0.3 | 0.13 | 0.28 | 0.19 | 0.45 | 0.13 |
| 5 | 131540875 | rs128738 | -0.346 | -0.193 | 1.249 | 1.433 | 0.034 | 0.069 | 0.069 | 0 | 0.1 | 0.18 | 0.22 | 0.33 | 0.18 | 0.17 |
| 5 | 131540884 | rs156025 | 1.433 | -0.437 | 0.941 | 1.364 | 0.062 | 0.017 | 0.017 | 0.059 | 0.33 | 0.06 | 0.36 | 0.24 | 0.2 | 0.43 |
| 5 | 131541833 | rs75189 | -0.894 | -0.141 | 1.298 | 1.456 | 0.005 | -0.002 | -0.002 | 0 | 0.43 | 0.6 | 0.53 | 0.53 | 0.64 | 0.33 |
| 5 | 131546280 | rs154483 | -0.91 | -0.165 | 1.286 | 1.461 | 0.002 | -0.001 | -0.001 | 0.351 | 0.43 | 0.59 | 0.53 | 0.53 | 0.65 | 0.31 |
| 5 | 131556309 | rs7733814 | 0.552 | -0.516 | 1.024 | 1.365 | 0.001 | -0.001 | -0.001 | 0.128 | 0.16 | 0.15 | 0.06 | 0.2 | 0.08 | 0.05 |
| 5 | 131562015 | rs154487 | -0.556 | -0.818 | 0.442 | 0.958 | -0.001 | -0.058 | -0.058 | 0 | 0.75 | 0.93 | 0.84 | 0.73 | 0.8 | 0.67 |
| 5 | 131567924 | rs11955347 | 0.242 | -0.609 | 0.669 | 0.507 | -0.032 | 0.066 | 0.066 | 0 | 0.36 | 0.22 | 0.31 | 0 | 0.42 | 0.11 |
| 5 | 131568617 | rs2077380 | -0.634 | -0.671 | 0.729 | 0.582 | 0.025 | 0.114 | 0.114 | 0.241 | 0.12 | 0.14 | 0.23 | 0.54 | 0.17 | 0.23 |
| 5 | 131573529 | rs2089855 | 0.272 | -0.708 | 0.698 | 0.393 | -0.05 | 0.129 | 0.129 | 0 | 0.37 | 0.17 | 0.3 | 0 | 0.42 | 0.11 |
| 5 | 131575338 | rs12521097 | -0.091 | -0.732 | 1.015 | 0.38 | 0.005 | -0.111 | -0.111 | 0 | 0.39 | 0.45 | 0.32 | 0 | 0.43 | 0.11 |
| 5 | 131577894 | rs2136187 | -0.35 | -0.762 | 1.049 | 0.594 | 0 | -0.007 | -0.007 | 0 | 0.45 | 0.62 | 0.39 | 0.27 | 0.53 | 0.17 |
| 5 | 131580152 | rs162907 | -0.994 | -0.761 | 0.99 | 0.586 | 0.075 | -0.022 | -0.022 | 0 | 0.53 | 0.82 | 0.57 | 0.55 | 0.61 | 0.34 |
| 5 | 131590534 | rs7727544 | -0.261 | -0.634 | 0.987 | 0.463 | -0.017 | 0.095 | 0.095 | 0 | 0.43 | 0.31 | 0.44 | 0.1 | 0.49 | 0.24 |
| 5 | 131592405 | rs7720230 | 0.287 | -0.645 | 0.968 | 0.45 | -0.041 | 0.107 | 0.107 | 0 | 0.37 | 0.19 | 0.31 | 0 | 0.42 | 0.11 |
| 5 | 131592870 | rs3900945 | -0.678 | -0.695 | 1.089 | 0.591 | 0.019 | 0.017 | 0.017 | 0 | 0.5 | 0.61 | 0.51 | 0.38 | 0.6 | 0.3 |
| 5 | 131602166 | rs1007602 | -0.69 | -0.747 | 1.07 | 0.595 | 0.012 | 0.011 | 0.011 | 0 | 0.5 | 0.59 | 0.51 | 0.38 | 0.61 | 0.3 |
| 5 | 131606936 | rs2292260 | -0.12 | -0.892 | 1.371 | 0.953 | 0.086 | 0.184 | 0.184 | 0 | 0.08 | 0.29 | 0.07 | 0.28 | 0.1 | 0.06 |
| 5 | 131607721 | rs10479001 | 0.665 | -0.994 | 1.199 | 0.832 | 0.019 | -0.004 | -0.004 | 0.981 | 0.22 | 0.17 | 0.08 | 0 | 0.05 | 0.02 |
| 5 | 131608794 | rs162881 | -0.997 | -0.9 | 1.333 | 1.027 | 0.091 | 0.396 | 0.396 | 0 | 0.1 | 0.53 | 0.1 | 0.47 | 0.11 | 0.11 |
| 5 | 131615593 | rs162899 | -0.936 | -0.922 | 0.868 | 0.859 | 0.024 | -0.023 | -0.023 | 0.219 | 0.38 | 0.75 | 0.32 | 0.56 | 0.24 | 0.26 |
| 5 | 131615966 | rs7721296 | -0.428 | -0.207 | 0.957 | 1.289 | 0.115 | 0.23 | 0.23 | 0 | 0.08 | 0.35 | 0.07 | 0.28 | 0.1 | 0.06 |
| 5 | 131620366 | rs162901 | -0.922 | -0.373 | 0.2 | 1.042 | 0.021 | 0.138 | 0.138 | 0 | 0.08 | 0.23 | 0.16 | 0.28 | 0.08 | 0.17 |
| 5 | 131625169 | rs2662314 | -0.645 | -0.747 | -0.312 | 0.206 | 0.001 | -0.007 | -0.007 | 0.828 | 0.8 | 0.83 | 0.85 | 0.92 | 0.84 | 0.79 |
| 5 | 131626611 | rs156109 | 0.463 | -1.253 | -0.424 | -0.122 | 0.021 | 0.027 | 0.027 | 3.195 | 0.52 | 0.58 | 0.62 | 0.81 | 0.69 | 0.63 |
| 5 | 131633355 | rs419291 | -0.425 | -0.581 | -0.436 | 0.602 | 0.038 | 0.01 | 0.01 | 0 | 0.52 | 0.8 | 0.69 | 0.35 | 0.59 | 0.33 |
| 5 | 131637338 | rs270612 | -0.895 | -0.539 | -0.456 | 0.731 | 0.078 | 0.039 | 0.039 | 0 | 0.58 | 0.88 | 0.75 | 0.66 | 0.68 | 0.41 |
| 5 | 131640583 | rs270613 | -0.298 | -0.488 | -0.437 | 0.657 | 0.027 | 0.021 | 0.021 | 0 | 0.51 | 0.68 | 0.68 | 0.33 | 0.58 | 0.33 |
| 5 | 131644965 | rs3792880 | 0.935 | -0.114 | -0.18 | 0.896 | 0.07 | 0.153 | 0.153 | 0 | 0.05 | 0.14 | 0.06 | 0.3 | 0.09 | 0.07 |
| 5 | 131650867 | rs270606 | -0.8 | -0.485 | -0.179 | 0.751 | 0.084 | 0.048 | 0.048 | 0 | 0.57 | 0.85 | 0.75 | 0.66 | 0.68 | 0.41 |
| 5 | 131651261 | rs3792884 | 0.776 | -0.401 | -0.354 | 0.74 | -0.032 | -0.034 | -0.034 | 0 | 0.06 | 0 | 0.12 | 0.1 | 0.09 | 0.15 |
| 5 | 131651811 | rs270605 | -0.333 | -0.399 | -0.286 | 0.764 | 0.029 | 0.021 | 0.021 | 0 | 0.51 | 0.68 | 0.67 | 0.33 | 0.58 | 0.33 |
| 5 | 131653925 | rs156322 | -0.896 | -0.401 | -0.289 | 0.763 | 0.084 | 0.049 | 0.049 | 0 | 0.57 | 0.83 | 0.75 | 0.65 | 0.68 | 0.41 |
| 5 | 131656517 | rs272842 | -0.333 | -0.4 | -0.285 | 0.764 | 0.029 | 0.021 | 0.021 | 0 | 0.51 | 0.68 | 0.67 | 0.33 | 0.58 | 0.33 |
| 5 | 131656700 | rs270602 | -0.481 | -0.41 | -0.301 | 0.758 | 0.041 | 0.008 | 0.008 | 0 | 0.52 | 0.77 | 0.68 | 0.35 | 0.59 | 0.34 |
| 5 | 131656997 | rs270601 | -0.893 | -0.479 | -0.354 | 0.72 | 0.084 | 0.049 | 0.049 | 0 | 0.57 | 0.83 | 0.75 | 0.66 | 0.68 | 0.41 |
| 5 | 131660119 | rs273915 | -0.893 | -0.494 | -0.36 | 0.696 | 0.084 | 0.049 | 0.049 | 0 | 0.57 | 0.83 | 0.75 | 0.66 | 0.68 | 0.41 |
| 5 | 131661156 | rs273913 | -0.451 | -0.494 | -0.361 | 0.697 | 0.036 | 0.023 | 0.023 | 0 | 0.52 | 0.79 | 0.68 | 0.33 | 0.58 | 0.34 |
| 5 | 131661349 | rs273912 | -0.958 | -0.502 | -0.365 | 0.694 | 0.082 | 0.044 | 0.044 | 0 | 0.57 | 0.87 | 0.75 | 0.66 | 0.68 | 0.41 |
| 5 | 131662460 | rs272894 | -0.335 | -0.502 | -0.348 | 0.693 | 0.029 | 0.021 | 0.021 | 0 | 0.51 | 0.68 | 0.67 | 0.33 | 0.58 | 0.33 |
| 5 | 131663062 | rs272893 | -0.475 | -0.5 | -0.371 | 0.693 | 0.041 | 0.008 | 0.008 | 0 | 0.52 | 0.77 | 0.68 | 0.35 | 0.59 | 0.34 |
| 5 | 131664349 | rs272892 | -0.89 | -0.501 | -0.372 | 0.693 | 0.084 | 0.049 | 0.049 | 0 | 0.57 | 0.83 | 0.75 | 0.65 | 0.68 | 0.41 |
| 5 | 131665378 | rs272889 | -0.475 | -0.5 | -0.371 | 0.694 | 0.041 | 0.008 | 0.008 | 0 | 0.52 | 0.78 | 0.68 | 0.35 | 0.59 | 0.34 |
| 5 | 131665423 | rs272888 | -0.885 | -0.515 | -0.383 | 0.676 | 0.084 | 0.049 | 0.049 | 0 | 0.57 | 0.83 | 0.75 | 0.66 | 0.68 | 0.41 |
| 5 | 131665713 | rs272887 | -0.935 | -0.534 | -0.387 | 0.667 | 0.082 | 0.045 | 0.045 | 0 | 0.57 | 0.84 | 0.75 | 0.66 | 0.68 | 0.41 |
| 5 | 131665840 | rs272886 | -0.324 | -0.52 | -0.357 | 0.668 | 0.029 | 0.021 | 0.021 | 0 | 0.51 | 0.68 | 0.67 | 0.33 | 0.58 | 0.33 |
| 5 | 131667736 | rs272885 | -0.88 | -0.556 | 0.166 | 0.604 | 0.084 | 0.049 | 0.049 | 0 | 0.57 | 0.83 | 0.75 | 0.66 | 0.68 | 0.41 |
| 5 | 131668698 | rs272883 | -0.88 | -0.554 | 0.17 | 0.605 | 0.084 | 0.049 | 0.049 | 0 | 0.57 | 0.83 | 0.75 | 0.65 | 0.68 | 0.41 |
| 5 | 131669264 | rs272881 | -0.308 | -0.565 | 0.19 | 0.601 | 0.03 | 0.022 | 0.022 | 0 | 0.51 | 0.68 | 0.67 | 0.33 | 0.58 | 0.33 |
| 5 | 131670178 | rs272880 | -1.012 | -0.566 | 0.201 | 0.6 | 0.078 | 0.039 | 0.039 | 0 | 0.58 | 0.87 | 0.75 | 0.65 | 0.68 | 0.41 |
| 5 | 131674888 | rs272875 | -0.242 | -0.56 | 0.218 | 0.603 | 0.019 | 0.015 | 0.015 | 0 | 0.5 | 0.62 | 0.67 | 0.33 | 0.58 | 0.33 |
| 5 | 131675046 | rs272874 | -0.532 | -0.558 | 0.187 | 0.604 | 0.041 | 0.008 | 0.008 | 0 | 0.52 | 0.77 | 0.68 | 0.35 | 0.59 | 0.34 |
| 5 | 131677997 | rs272869 | -0.789 | -0.838 | -0.141 | 0.339 | 0.041 | 0.008 | 0.008 | 0 | 0.52 | 0.77 | 0.68 | 0.35 | 0.59 | 0.34 |
| 5 | 131681057 | rs272867 | -0.614 | -0.85 | -0.162 | 0.329 | 0.028 | 0.021 | 0.021 | 0 | 0.51 | 0.68 | 0.67 | 0.33 | 0.58 | 0.33 |
| 5 | 131686146 | rs10058074 | 0.722 | -1.283 | -0.466 | 0.026 | -0.169 | 0.208 | 0.208 | 0 | 0.29 | 0.04 | 0.28 | 0 | 0.43 | 0.1 |
| 5 | 131699867 | rs274546 | -0.955 | -1.152 | -0.328 | 0.157 | 0.043 | 0.012 | 0.012 | 0 | 0.52 | 0.76 | 0.68 | 0.35 | 0.59 | 0.33 |
| 5 | 131704519 | rs2569265 | 1.252 | -1.155 | -0.294 | 0.18 | 0.087 | 0.053 | 0.053 | 0 | 0.44 | 0.18 | 0.26 | 0.35 | 0.32 | 0.6 |
| 5 | 131707095 | rs2631363 | 0.884 | -1.146 | -0.256 | 0.187 | 0.041 | 0.012 | 0.012 | 0 | 0.48 | 0.26 | 0.32 | 0.65 | 0.41 | 0.67 |
| 5 | 131707293 | rs2631362 | 1.508 | -1.16 | -0.258 | 0.181 | 0.075 | 0.039 | 0.039 | 0 | 0.42 | 0.15 | 0.25 | 0.34 | 0.32 | 0.59 |
| 5 | 131707380 | rs2631361 | 0.871 | -1.16 | -0.258 | 0.181 | 0.041 | 0.012 | 0.012 | 0 | 0.48 | 0.26 | 0.32 | 0.65 | 0.41 | 0.67 |
| 5 | 131707892 | rs2631359 | 1.507 | -1.16 | -0.26 | 0.18 | 0.075 | 0.039 | 0.039 | 0 | 0.42 | 0.15 | 0.25 | 0.34 | 0.32 | 0.59 |
| 5 | 131711100 | rs4646301 | 1.438 | -1.142 | -0.258 | 0.183 | 0.015 | 0.055 | 0.055 | 0 | 0.05 | 0.06 | 0.07 | 0.31 | 0.09 | 0.07 |
| 5 | 131712125 | rs274571 | 1.507 | -1.152 | -0.26 | 0.18 | 0.075 | 0.039 | 0.039 | 0 | 0.42 | 0.15 | 0.25 | 0.34 | 0.32 | 0.59 |

| | | | | | | | | | | | | | | | | |
|---|---|---|---|---|---|---|---|---|---|---|---|---|---|---|---|---|
| 5 | 131713226 | rs274570 | 1.489 | -1.156 | -0.262 | 0.178 | 0.076 | 0.04 | 0.04 | 0 | 0.42 | 0.15 | 0.25 | 0.34 | 0.32 | 0.59 |
| 5 | 131713774 | rs2073642 | 1.384 | -1.148 | -0.258 | 0.183 | 0.012 | 0.053 | 0.053 | 0 | 0.05 | 0.08 | 0.07 | 0.31 | 0.09 | 0.07 |
| 5 | 131714409 | rs274567 | 0.872 | -1.16 | -0.06 | 0.175 | 0.041 | 0.012 | 0.012 | 0 | 0.48 | 0.26 | 0.32 | 0.65 | 0.41 | 0.67 |
| 5 | 131716902 | rs183898 | 1.507 | -1.162 | -0.069 | 0.173 | 0.075 | 0.039 | 0.039 | 0 | 0.42 | 0.15 | 0.25 | 0.34 | 0.32 | 0.59 |
| 5 | 131717050 | rs17622208 | 0.2 | -1.245 | 0.182 | 0.107 | -0.081 | 0.16 | 0.16 | 0 | 0.35 | 0.14 | 0.31 | 0 | 0.44 | 0.1 |
| 5 | 131718539 | rs4646305 | 1.393 | -1.189 | 0.171 | 0.174 | 0.012 | 0.053 | 0.053 | 0 | 0.05 | 0.08 | 0.07 | 0.31 | 0.09 | 0.07 |
| 5 | 131719228 | rs274561 | 1.023 | -1.202 | 0.17 | 0.168 | 0.036 | 0.003 | 0.003 | 0 | 0.47 | 0.22 | 0.32 | 0.65 | 0.41 | 0.67 |
| 5 | 131719450 | rs274560 | 0.893 | -1.204 | 0.172 | 0.167 | 0.042 | 0.012 | 0.012 | 0 | 0.48 | 0.26 | 0.32 | 0.65 | 0.41 | 0.67 |
| 5 | 131720265 | rs635619 | 1.537 | -1.208 | 0.155 | 0.163 | 0.075 | 0.039 | 0.039 | 0 | 0.42 | 0.15 | 0.25 | 0.34 | 0.32 | 0.59 |
| 5 | 131721174 | rs274558 | 0.746 | -1.199 | 0.216 | 0.154 | 0.027 | 0.02 | 0.02 | 0 | 0.49 | 0.34 | 0.33 | 0.67 | 0.41 | 0.67 |
| 5 | 131721204 | rs274557 | 0.746 | -1.199 | 0.216 | 0.154 | 0.027 | 0.02 | 0.02 | 0 | 0.49 | 0.34 | 0.33 | 0.67 | 0.41 | 0.67 |
| 5 | 131722951 | rs274555 | 0.663 | -1.292 | 0.199 | -0.078 | 0.017 | 0.109 | 0.109 | 0.083 | 0.52 | 0.31 | 0.52 | 0.89 | 0.45 | 0.75 |
| 5 | 131723065 | rs17689550 | 1.48 | -1.657 | -0.219 | -0.31 | -0.033 | 0.054 | 0.054 | 0.686 | 0.12 | 0 | 0.11 | 0.04 | 0.14 | 0.21 |
| 5 | 131723288 | rs2073643 | 0.03 | -1.653 | -0.095 | -0.305 | 0 | 0.011 | 0.011 | 0 | 0.42 | 0.36 | 0.39 | 0.33 | 0.53 | 0.17 |
| 5 | 131724814 | rs2073645 | 1.969 | -1.212 | -0.453 | -0.038 | 0.007 | 0.04 | 0.04 | 0 | 0.05 | 0.04 | 0.07 | 0.3 | 0.09 | 0.07 |
| 5 | 131724950 | rs274554 | -0.156 | -1.313 | -0.449 | -0.254 | -0.038 | -0.054 | -0.054 | 1.296 | 0.66 | 0.46 | 0.78 | 0.9 | 0.85 | 0.84 |
| 5 | 131727033 | rs11746555 | 2.147 | -0.922 | -0.49 | -0.275 | -0.102 | 0.087 | 0.087 | 1.814 | 0.14 | 0.04 | 0.25 | 0 | 0.37 | 0.08 |
| 5 | 131727680 | rs274551 | -0.143 | -1.029 | -0.414 | -0.219 | -0.038 | -0.054 | -0.054 | 0 | 0.66 | 0.47 | 0.78 | 0.9 | 0.85 | 0.84 |
| 5 | 131728712 | rs274550 | 0.015 | -1.009 | -0.408 | -0.214 | 0.014 | 0.014 | 0.014 | 2.032 | 0.65 | 0.36 | 0.78 | 0.88 | 0.85 | 0.83 |
| 5 | 131729118 | rs274549 | -0.144 | -1.032 | -0.397 | -0.216 | -0.038 | -0.054 | -0.054 | 0.013 | 0.66 | 0.47 | 0.78 | 0.9 | 0.85 | 0.84 |
| 5 | 131730807 | rs274548 | -0.139 | -1.119 | -0.385 | -0.207 | -0.038 | -0.054 | -0.054 | 0 | 0.66 | 0.46 | 0.78 | 0.9 | 0.85 | 0.83 |
| 5 | 131731304 | rs274547 | -0.139 | -1.119 | -0.385 | -0.207 | -0.038 | -0.054 | -0.054 | 1.44 | 0.66 | 0.46 | 0.78 | 0.9 | 0.85 | 0.83 |
| 5 | 131732456 | rs7705826 | -0.014 | -1.123 | -0.662 | -0.198 | 0.014 | 0.014 | 0.014 | 2.032 | 0.65 | 0.36 | 0.78 | 0.88 | 0.85 | 0.83 |
| 5 | 131732774 | rs17689595 | 2.038 | -1.273 | -0.916 | -0.297 | 0.024 | 0.164 | 0.164 | 0 | 0.26 | 0.01 | 0.08 | 0.03 | 0.18 | 0.35 |
| 5 | 131743465 | rs6897597 | 0.304 | -1.007 | -0.676 | -0.042 | 0.037 | 0.006 | 0.006 | 0.407 | 0.6 | 0.31 | 0.71 | 0.58 | 0.76 | 0.75 |
| 5 | 131744202 | rs17771891 | 1.6 | -0.974 | -0.575 | 0.148 | 0.05 | 0.002 | 0.002 | 0 | 0.29 | 0.15 | 0.1 | 0.27 | 0.2 | 0.42 |
| 5 | 131744574 | rs1016988 | 1.529 | -0.95 | -0.563 | 0.055 | 0.052 | -0.014 | -0.014 | 0 | 0.3 | 0.15 | 0.14 | 0.32 | 0.2 | 0.42 |
| 5 | 131752620 | rs10071051 | 0.5 | -0.852 | -0.758 | -0.011 | -0.078 | 0.013 | 0.013 | 0 | 0.19 | 0.03 | 0.24 | 0.12 | 0.24 | 0.32 |
| 5 | 131755004 | rs4705943 | 0.247 | -1.165 | -0.503 | 0.083 | 0.039 | 0.204 | 0.204 | 0 | 0.09 | 0.19 | 0.28 | 0.52 | 0.13 | 0.15 |
| 5 | 131755651 | rs1003533 | 1.449 | -1.243 | -0.322 | 0.13 | 0.025 | -0.004 | -0.004 | 0 | 0.3 | 0.23 | 0.15 | 0.33 | 0.2 | 0.43 |
| 5 | 131758314 | rs2269821 | -0.698 | -1.164 | -0.331 | 0.213 | -0.048 | -0.022 | -0.022 | 0 | 0.69 | 0.55 | 0.85 | 0.69 | 0.86 | 0.92 |
| 5 | 131770805 | rs2188962 | 1.59 | -0.722 | -0.677 | -0.08 | -0.106 | 0.09 | 0.09 | 2.06 | 0.14 | 0.04 | 0.25 | 0 | 0.39 | 0.09 |
| 5 | 131778413 | rs10062371 | -0.42 | -0.877 | -0.508 | -0.04 | -0.016 | 0.028 | 0.028 | 0 | 0.79 | 0.85 | 0.77 | 0.88 | 0.77 | 0.68 |
| 5 | 131781288 | rs7730247 | 1.267 | -1.046 | -0.41 | 0.083 | -0.024 | 0.008 | 0.008 | 0 | 0.31 | 0.32 | 0.16 | 0.36 | 0.2 | 0.44 |
| 5 | 131784393 | rs12521868 | 1.448 | -0.242 | -0.894 | 0.166 | -0.103 | 0.087 | 0.087 | 1.343 | 0.14 | 0.04 | 0.24 | 0 | 0.39 | 0.08 |
| 5 | 131786608 | rs7703230 | 0.475 | -0.714 | -0.859 | 0.284 | 0.015 | -0.044 | -0.044 | 3.913 | 0.61 | 0.79 | 0.32 | 0.65 | 0.34 | 0.5 |
| 5 | 131792058 | rs7734115 | 0.875 | -0.971 | -0.502 | -0.007 | 0.004 | 0.003 | 0.003 | 1.719 | 0.24 | 0.27 | 0.13 | 0.27 | 0.13 | 0.08 |
| 5 | 131796922 | rs11745587 | -0.651 | -0.882 | -0.23 | -0.068 | -0.004 | -0.023 | -0.023 | 0 | 0.45 | 0.49 | 0.39 | 0.38 | 0.36 | 0.38 |
| 5 | 131797578 | rs2522051 | -0.21 | -0.918 | -0.346 | -0.11 | 0.005 | -0.022 | -0.022 | 0 | 0.6 | 0.42 | 0.6 | 0.71 | 0.48 | 0.81 |
| 5 | 131801158 | rs10077785 | 0.12 | -0.098 | -0.657 | 0.138 | -0.067 | 0.056 | 0.056 | 0 | 0.2 | 0.02 | 0.24 | 0.11 | 0.23 | 0.31 |
| 5 | 131804347 | rs2248116 | -1.479 | -0.047 | -0.605 | 0.142 | -0.047 | 0.034 | 0.034 | 1.851 | 0.85 | 0.88 | 0.75 | 0.98 | 0.62 | 0.89 |
| 5 | 131808869 | rs2548993 | 0.527 | -0.289 | -0.338 | 0.495 | 0.007 | 0.021 | 0.021 | 0 | 0.75 | 0.73 | 0.54 | 0.66 | 0.71 | 0.59 |
| 5 | 131809316 | rs2057657 | -0.022 | -0.222 | -1.185 | -0.237 | -0.069 | 0.045 | 0.045 | 0 | 0.2 | 0.02 | 0.23 | 0.12 | 0.23 | 0.31 |
| 5 | 131813350 | rs13165038 | 0.123 | -0.876 | -0.865 | -0.428 | -0.003 | -0.011 | -0.011 | 0 | 0.44 | 0.47 | 0.35 | 0.38 | 0.34 | 0.43 |
| 5 | 131820997 | rs17622656 | 1.147 | -0.012 | -1.241 | -0.438 | -0.074 | 0.062 | 0.062 | 0.942 | 0.13 | 0.04 | 0.24 | 0 | 0.35 | 0.07 |
| 5 | 131839618 | rs4705952 | 0.124 | -1.739 | -0.972 | -0.333 | 0.101 | 0.044 | 0.044 | 0 | 0.58 | 0.31 | 0.78 | 0.45 | 0.73 | 0.55 |
| 5 | 131847892 | rs10074713 | 0.728 | -1.195 | -1.327 | -0.645 | -0.122 | 0.154 | 0.154 | 0 | 0.34 | 0.1 | 0.48 | 0.17 | 0.48 | 0.47 |
| 5 | 131862977 | rs4143832 | -1.014 | -1.397 | -0.354 | -0.305 | -0.023 | -0.043 | -0.043 | 0 | 0.78 | 0.6 | 0.86 | 0.88 | 0.83 | 0.78 |
| 5 | 131864152 | rs11739623 | 1.013 | -1.539 | -0.934 | -0.068 | -0.035 | -0.122 | -0.122 | 0 | 0.19 | 0.03 | 0.17 | 0.34 | 0.26 | 0.19 |
| 5 | 131864506 | rs2079103 | -0.184 | -1.624 | -0.665 | -0.069 | 0.006 | 0.001 | 0.001 | 0 | 0.64 | 0.36 | 0.72 | 0.61 | 0.76 | 0.75 |
| 5 | 131867835 | rs17690122 | 1.06 | -1.36 | -0.926 | -0.001 | -0.003 | -0.005 | -0.005 | 3.116 | 0.19 | 0.2 | 0.13 | 0.13 | 0.17 | 0.22 |
| 5 | 131872383 | rs743562 | 0.198 | -1.239 | -1 | -0.009 | -0.009 | -0.021 | -0.021 | 5.012 | 0.37 | 0.14 | 0.29 | 0.46 | 0.43 | 0.41 |
| 5 | 131872865 | rs739719 | 0.029 | -1.565 | -0.262 | 0.121 | -0.042 | -0.002 | -0.002 | 0 | 0.8 | 0.35 | 0.88 | 0.75 | 0.92 | 0.92 |
| 5 | 131879916 | rs2069812 | 0.692 | -1.929 | -1.028 | -0.004 | 0.105 | 0.076 | 0.076 | 0 | 0.55 | 0.12 | 0.61 | 0.32 | 0.69 | 0.69 |
| 5 | 131901225 | rs2244012 | -0.064 | -1.852 | -0.521 | -0.403 | 0.001 | 0 | 0 | 1.23 | 0.27 | 0.66 | 0.19 | 0.19 | 0.2 | 0.23 |
| 5 | 131951996 | rs2301713 | 0.393 | -1.905 | 0.438 | -0.428 | -0.007 | -0.013 | -0.013 | 0 | 0.24 | 0.29 | 0.14 | 0.19 | 0.19 | 0.22 |
| 5 | 131952405 | rs17772565 | 1.355 | -2.225 | 0.379 | -0.114 | -0.005 | 0.01 | 0.01 | 0 | 0.12 | 0.13 | 0.14 | 0.34 | 0.06 | 0.08 |
| 5 | 131953510 | rs17772583 | 1.333 | -2.288 | 0.041 | -0.176 | -0.041 | -0.137 | -0.137 | 0 | 0.19 | 0.03 | 0.16 | 0.34 | 0.24 | 0.21 |
| 5 | 131965179 | rs3798134 | -0.062 | -1.916 | 0.505 | -0.483 | -0.025 | -0.038 | -0.038 | 0.906 | 0.27 | 0.43 | 0.16 | 0.19 | 0.19 | 0.23 |
| 5 | 131970885 | rs2237060 | 1.514 | -1.53 | 0.079 | -0.69 | -0.163 | 0.138 | 0.138 | 0.132 | 0.27 | 0.02 | 0.28 | 0.05 | 0.42 | 0.24 |
| 5 | 131992409 | rs1881457 | -0.231 | -1.975 | 1.057 | 0.519 | 0.009 | -0.001 | -0.001 | 0.92 | 0.25 | 0.2 | 0.2 | 0.24 | 0.18 | 0.21 |
| 5 | 131995843 | rs1295686 | 1.892 | -1.58 | 1.636 | 0.78 | 0.005 | 0 | 0 | 0 | 0.68 | 0.29 | 0.56 | 0.64 | 0.79 | 0.71 |
| 5 | 131995964 | rs20541 | 1.251 | -1.234 | 1.606 | 0.956 | -0.002 | -0.019 | -0.019 | 0.169 | 0.78 | 0.81 | 0.62 | 0.64 | 0.79 | 0.73 |
| 5 | 131996445 | rs1295685 | 1.128 | -1.227 | 1.492 | 1.039 | -0.039 | -0.085 | -0.085 | 1.531 | 0.78 | 0.97 | 0.62 | 0.64 | 0.79 | 0.73 |
| 5 | 131996500 | rs848 | 1.692 | -1.231 | 1.825 | 1.037 | -0.013 | 0.023 | 0.023 | 0.347 | 0.73 | 0.44 | 0.59 | 0.64 | 0.79 | 0.73 |
| 5 | 131999494 | rs2243204 | -1.62 | -1.203 | 1.792 | 0.634 | -0.022 | -0.025 | -0.025 | 3.426 | 0.16 | 0.67 | 0.21 | 0.15 | 0.08 | 0.14 |
| 5 | 132001151 | rs2243208 | -0.897 | -0.95 | 0.913 | 0.741 | 0 | 0 | 0 | 0.609 | 0.13 | 0.19 | 0.18 | 0.15 | 0.09 | 0.13 |
| 5 | 132001386 | rs2243210 | -0.604 | -0.916 | 0.864 | 0.442 | -0.006 | -0.03 | -0.03 | 0 | 0.12 | 0.25 | 0.14 | 0.04 | 0.07 | 0.13 |
| 5 | 132008644 | rs2243248 | -0.804 | -0.819 | 1.137 | 0.553 | -0.002 | -0.004 | -0.004 | 0 | 0.13 | 0.15 | 0.13 | 0.07 | 0.07 | 0.12 |

| | | | | | | | | | | | | | | | | |
|---|---|---|---|---|---|---|---|---|---|---|---|---|---|---|---|---|
| 5 | 132013179 | rs2227282 | -1.676 | -1.157 | 0.727 | 0.515 | 0.021 | 0.089 | 0.089 | 0 | 0.38 | 0.05 | 0.48 | 0.15 | 0.72 | 0.66 |
| 5 | 132024166 | rs11747814 | 1.231 | -1.134 | 1.856 | 0.617 | -0.008 | -0.017 | -0.017 | 0 | 0.68 | 0.38 | 0.51 | 0.16 | 0.73 | 0.67 |
| 5 | 132032806 | rs2023823 | 1.41 | -1.136 | 1.907 | 0.555 | 0.017 | 0.044 | 0.044 | 0 | 0.65 | 0.3 | 0.51 | 0.17 | 0.71 | 0.67 |
| 5 | 132034346 | rs2023822 | 0.13 | -1.017 | 1.756 | 0.628 | 0.019 | 0.056 | 0.056 | 0 | 0.79 | 0.9 | 0.85 | 0.92 | 0.89 | 0.84 |
| 5 | 132036252 | rs1468216 | 0.199 | -0.994 | 1.71 | 0.639 | -0.004 | -0.015 | -0.015 | 0 | 0.79 | 0.76 | 0.85 | 0.92 | 0.89 | 0.84 |
| 5 | 132043032 | rs2299007 | -0.784 | -0.422 | 2.004 | 1.957 | -0.01 | 0.043 | 0.043 | 0 | 0.08 | 0.05 | 0.27 | 0.5 | 0.16 | 0.15 |
| 5 | 132135372 | rs256869 | -0.475 | -0.584 | 1.819 | 1.396 | -0.008 | 0.076 | 0.076 | 3.184 | 0.11 | 0.24 | 0.2 | 0.18 | 0.1 | 0.17 |
| 5 | 132138129 | rs4705869 | -1.024 | -0.306 | 1.44 | 1.851 | 0.012 | 0.169 | 0.169 | 0 | 0.11 | 0.24 | 0.25 | 0.58 | 0.12 | 0.15 |
| 5 | 132138324 | rs803136 | 0.212 | -0.738 | 0.921 | 1.051 | -0.011 | 0.01 | 0.01 | 0 | 0.19 | 0.22 | 0.33 | 0.2 | 0.13 | 0.18 |
| 5 | 132151071 | rs757537 | 0.21 | -0.645 | 0.407 | 0.61 | 0.006 | 0.049 | 0.049 | 0.477 | 0.2 | 0.07 | 0.08 | 0.12 | 0.12 | 0.18 |
| 5 | 132154270 | rs12523593 | -0.599 | -1.208 | 1.367 | 0.318 | -0.039 | -0.074 | -0.074 | 0 | 0.28 | 0.56 | 0.12 | 0.12 | 0.13 | 0.19 |
| 5 | 132159134 | rs4705870 | -0.008 | -1.116 | 1.158 | 0.312 | 0.022 | 0.075 | 0.075 | 0.468 | 0.23 | 0.05 | 0.09 | 0.12 | 0.13 | 0.19 |
| 5 | 132167542 | rs254289 | 0.407 | -1.054 | 1.474 | 0.983 | 0.029 | 0.247 | 0.247 | 0 | 0.91 | 0.75 | 0.72 | 0.55 | 0.88 | 0.84 |
| 5 | 132174492 | rs803217 | 1.412 | -0.86 | 0.815 | 0.697 | 0.006 | 0.082 | 0.082 | 0 | 0.51 | 0.09 | 0.47 | 0.29 | 0.59 | 0.62 |
| 5 | 132180990 | rs17166268 | 0.524 | -0.889 | 1.282 | 0.71 | -0.017 | -0.044 | -0.044 | 0 | 0.22 | 0.3 | 0.11 | 0.12 | 0.12 | 0.19 |
| 5 | 132181800 | rs1557852 | -0.729 | -0.918 | 1.496 | 0.711 | -0.019 | -0.031 | -0.031 | 0.017 | 0.37 | 0.51 | 0.23 | 0.27 | 0.28 | 0.22 |
| 5 | 132196383 | rs10491280 | 0.539 | -0.961 | 1.486 | 0.711 | -0.017 | -0.044 | -0.044 | 0.063 | 0.22 | 0.3 | 0.11 | 0.12 | 0.12 | 0.19 |
| 5 | 132198199 | rs254286 | 1.192 | -0.973 | 1.494 | 0.73 | 0.016 | 0.033 | 0.033 | 0 | 0.54 | 0.33 | 0.49 | 0.29 | 0.6 | 0.62 |
| 5 | 132200943 | rs11748063 | -0.722 | -0.971 | 1.465 | 0.719 | -0.02 | -0.031 | -0.031 | 0 | 0.38 | 0.51 | 0.23 | 0.26 | 0.28 | 0.22 |
| 5 | 132208168 | rs708459 | 1.208 | -0.739 | 1.466 | 0.601 | 0.003 | 0.009 | 0.009 | 0 | 0.51 | 0.46 | 0.48 | 0.29 | 0.54 | 0.59 |
| 5 | 132208363 | rs803223 | 0.047 | -0.741 | 1.588 | 0.594 | -0.006 | -0.065 | -0.065 | 0.199 | 0.71 | 0.82 | 0.63 | 0.44 | 0.72 | 0.65 |
| 5 | 132208861 | rs765760 | 0.589 | -0.753 | 1.608 | 0.588 | -0.007 | -0.001 | -0.001 | 0 | 0.16 | 0.28 | 0.14 | 0.14 | 0.18 | 0.06 |
| 5 | 132210436 | rs803221 | 0.07 | -0.9 | 1.543 | 0.542 | 0 | -0.064 | -0.064 | 0.014 | 0.69 | 0.83 | 0.63 | 0.43 | 0.73 | 0.65 |
| 5 | 132214330 | rs42395 | 1.369 | -0.603 | 1.434 | 0.67 | 0.015 | 0.079 | 0.079 | 0 | 0.48 | 0.22 | 0.45 | 0.29 | 0.54 | 0.59 |
| 5 | 132214931 | rs803220 | 1.199 | -0.605 | 1.434 | 0.666 | 0.003 | 0.01 | 0.01 | 0 | 0.51 | 0.46 | 0.48 | 0.28 | 0.54 | 0.59 |
| 5 | 132215807 | rs803219 | 1.37 | -0.603 | 1.414 | 0.667 | 0.015 | 0.084 | 0.084 | 0 | 0.48 | 0.22 | 0.45 | 0.28 | 0.54 | 0.59 |
| 5 | 132216264 | rs803218 | 0.683 | -0.531 | 0.997 | 0.725 | -0.001 | 0.056 | 0.056 | 0 | 0.74 | 0.55 | 0.58 | 0.39 | 0.69 | 0.78 |
| 5 | 132245943 | rs4380643 | -0.896 | -0.507 | 1.003 | 0.707 | -0.002 | 0.058 | 0.058 | 0 | 0.26 | 0.44 | 0.42 | 0.6 | 0.31 | 0.24 |
| 5 | 132279016 | rs7710795 | 0.526 | -0.344 | 0.532 | 0.735 | -0.007 | -0.001 | -0.001 | 0 | 0.16 | 0.29 | 0.14 | 0.14 | 0.18 | 0.06 |
| 5 | 132284251 | rs17691584 | 0.241 | -0.261 | -0.382 | 0.942 | 0.016 | 0.062 | 0.062 | 0.725 | 0.24 | 0.09 | 0.1 | 0.12 | 0.15 | 0.19 |
| 5 | 132304222 | rs7723827 | -0.816 | 0.262 | 0.107 | 1.564 | 0.004 | 0.103 | 0.103 | 0 | 0.09 | 0.17 | 0.28 | 0.45 | 0.13 | 0.17 |
| 5 | 132334252 | rs3087646 | -0.893 | -0.406 | 0.23 | 0.988 | -0.02 | 0.03 | 0.03 | 0.72 | 0.31 | 0.61 | 0.24 | 0.4 | 0.26 | 0.16 |
| 5 | 132335165 | rs6859346 | -0.594 | -0.457 | 0.472 | 0.955 | -0.039 | 0.109 | 0.109 | 0.108 | 0.27 | 0.52 | 0.21 | 0.4 | 0.23 | 0.16 |
| 5 | 132355147 | rs12517292 | 1.405 | -0.491 | -0.306 | 0.487 | 0.016 | 0.093 | 0.093 | 0.912 | 0.3 | 0.03 | 0.22 | 0.18 | 0.17 | 0.25 |
| 5 | 132358914 | rs10053483 | -0.789 | -0.764 | 0.128 | 0.588 | -0.02 | 0.028 | 0.028 | 0.672 | 0.31 | 0.61 | 0.23 | 0.39 | 0.26 | 0.16 |
| 5 | 132361894 | rs3885730 | -0.058 | -0.794 | 0.447 | 0.682 | -0.006 | -0.014 | -0.014 | 0 | 0.23 | 0.24 | 0.17 | 0.1 | 0.23 | 0.15 |
| 5 | 132384689 | rs4574536 | 0.693 | -0.328 | 0.681 | 0.214 | -0.007 | -0.056 | -0.056 | 0 | 0.28 | 0.36 | 0.12 | 0.09 | 0.23 | 0.22 |
| 5 | 132386105 | rs9327640 | -0.898 | -0.41 | 0.679 | 0.231 | -0.008 | -0.019 | -0.019 | 0 | 0.55 | 0.81 | 0.34 | 0.39 | 0.49 | 0.38 |
| 5 | 132387260 | rs3088225 | -0.328 | -0.306 | 1.126 | 0.969 | 0.005 | -0.008 | -0.008 | 0 | 0.36 | 0.67 | 0.19 | 0.39 | 0.26 | 0.23 |
| 5 | 132407688 | rs13161158 | 0.04 | 0.369 | 0.008 | 0.456 | -0.02 | -0.035 | -0.035 | 0 | 0.06 | 0 | 0.24 | 0.23 | 0.07 | 0.1 |
| 5 | 132441302 | rs13718 | 0.057 | 0.301 | -0.163 | -0.033 | 0.001 | 0 | 0 | 1.327 | 0.27 | 0.25 | 0.11 | 0.09 | 0.23 | 0.22 |
| 5 | 132463441 | rs4515334 | -0.082 | 0.001 | 0.184 | 0.457 | 0.008 | 0.019 | 0.019 | 0.75 | 0.44 | 0.42 | 0.37 | 0.16 | 0.38 | 0.45 |
| 5 | 132464549 | rs12513436 | -0.105 | -0.014 | 0.13 | 0.435 | -0.011 | 0 | 0 | 0 | 0.44 | 0.5 | 0.37 | 0.38 | 0.35 | 0.46 |
| 5 | 132465165 | rs6897881 | 0.496 | -0.142 | -0.921 | 0.211 | -0.001 | -0.001 | -0.001 | 0 | 0.26 | 0.29 | 0.26 | 0.16 | 0.23 | 0.4 |
| 5 | 132465841 | rs6899003 | -0.314 | 0.436 | -0.951 | 0.557 | -0.093 | -0.064 | -0.064 | 0 | 0.6 | 0.91 | 0.52 | 0.38 | 0.49 | 0.54 |
| 5 | 132466886 | rs10477746 | 0.132 | 0.371 | -0.687 | 0.031 | 0.007 | 0.014 | 0.014 | 1.952 | 0.43 | 0.38 | 0.34 | 0.16 | 0.35 | 0.45 |
| 5 | 132475314 | rs10051096 | 0.244 | 0.694 | -0.929 | 0.721 | -0.003 | -0.006 | -0.006 | 0 | 0.66 | 0.68 | 0.71 | 0.39 | 0.61 | 0.64 |
| 5 | 132482518 | rs2059805 | -0.666 | 0.611 | -1.334 | 0.118 | -0.006 | 0.003 | 0.003 | 0.554 | 0.24 | 0.21 | 0.28 | 0.11 | 0.38 | 0.36 |
| 5 | 132484208 | rs4958125 | 1.258 | -0.184 | -2.092 | 0.327 | -0.142 | 0.036 | 0.036 | 0.272 | 0.42 | 0.04 | 0.26 | 0.35 | 0.44 | 0.26 |
| 5 | 132487330 | rs12186846 | 0.395 | 0.178 | -1.948 | 0.278 | -0.019 | 0.018 | 0.018 | 0 | 0.16 | 0.13 | 0.19 | 0.08 | 0.3 | 0.2 |
| 5 | 132490991 | rs4358577 | -0.068 | 0.391 | -2.216 | 0.59 | -0.003 | -0.049 | -0.049 | 0 | 0.18 | 0.08 | 0.42 | 0.3 | 0.19 | 0.41 |
| 5 | 132491562 | rs7717377 | 1.22 | 0.316 | -2.078 | 0.618 | -0.062 | 0.043 | 0.043 | 0.728 | 0.41 | 0.09 | 0.25 | 0.35 | 0.39 | 0.25 |
| 5 | 132495185 | rs13178251 | -0.583 | -0.143 | -1.654 | 0.671 | 0.025 | -0.047 | -0.047 | 2.653 | 0.85 | 0.94 | 0.93 | 0.73 | 0.91 | 0.86 |
| 5 | 132500078 | rs7718731 | -0.089 | 0.104 | -0.89 | 0.476 | -0.004 | 0 | 0 | 0 | 0.22 | 0.28 | 0.17 | 0.2 | 0.18 | 0.17 |
| 5 | 132500250 | rs1428142 | -0.338 | 0.23 | -0.181 | 0.447 | -0.008 | 0.065 | 0.065 | 0 | 0.24 | 0.34 | 0.44 | 0.4 | 0.25 | 0.36 |
| 5 | 132506685 | rs2864676 | 0.078 | 0.217 | -0.12 | 0.434 | 0.001 | -0.061 | -0.061 | 0.206 | 0.21 | 0.15 | 0.41 | 0.4 | 0.23 | 0.35 |
| 5 | 132511553 | rs1896634 | -0.142 | 0.214 | -0.013 | 0.433 | 0.006 | -0.045 | -0.045 | 0 | 0.22 | 0.16 | 0.41 | 0.4 | 0.23 | 0.35 |
| 5 | 132526551 | rs7715671 | 0.214 | -0.002 | -0.663 | -0.327 | 0.01 | 0.028 | 0.028 | 0 | 0.38 | 0.35 | 0.3 | 0.2 | 0.34 | 0.24 |
| 5 | 132533135 | rs3749819 | -1.053 | -0.391 | -0.982 | -0.473 | 0.006 | 0 | 0 | 0 | 0.69 | 0.7 | 0.75 | 0.61 | 0.59 | 0.62 |
| 5 | 132533517 | rs13183672 | -0.003 | -0.495 | -0.927 | -0.517 | 0.02 | -0.032 | -0.032 | 0.004 | 0.3 | 0.25 | 0.43 | 0.4 | 0.25 | 0.38 |
| 5 | 132534021 | rs3798104 | -0.988 | -0.291 | -1.221 | -0.422 | 0.005 | -0.001 | -0.001 | 5.332 | 0.69 | 0.71 | 0.75 | 0.61 | 0.59 | 0.62 |
| 5 | 132534059 | rs3798103 | 0.376 | -0.378 | -1.427 | -0.41 | 0.007 | -0.127 | -0.127 | 0 | 0.25 | 0.06 | 0.41 | 0.4 | 0.23 | 0.35 |
| 5 | 132535046 | rs3749817 | 0.427 | -0.373 | -1.445 | -0.408 | 0.003 | -0.13 | -0.13 | 0 | 0.24 | 0.06 | 0.4 | 0.39 | 0.23 | 0.34 |
| 5 | 132536038 | rs720622 | -0.974 | 0.638 | -1.304 | -0.588 | -0.024 | -0.033 | -0.033 | 0 | 0.77 | 0.86 | 0.82 | 0.6 | 0.7 | 0.64 |
| 5 | 132540109 | rs2162769 | -0.434 | 0.483 | -1.296 | -0.562 | 0.028 | 0.004 | 0.004 | 0 | 0.69 | 0.64 | 0.74 | 0.59 | 0.58 | 0.62 |
| 5 | 132543001 | rs4958168 | 0.085 | 0.159 | -1.845 | -0.201 | 0.014 | -0.111 | -0.111 | 0 | 0.27 | 0.09 | 0.41 | 0.43 | 0.25 | 0.34 |
| 5 | 132546059 | rs3828692 | 0.107 | 0.371 | -1.547 | -0.26 | 0.011 | 0.002 | 0.002 | 0 | 0.46 | 0.44 | 0.47 | 0.38 | 0.38 | 0.39 |
| 5 | 132550830 | rs6871976 | -0.001 | 0.152 | -2.526 | -0.094 | 0.138 | 0.004 | 0.004 | 0 | 0.48 | 0.83 | 0.46 | 0.39 | 0.3 | 0.41 |
| 5 | 132558754 | rs736324 | -0.095 | 0.985 | -2.153 | 0.452 | -0.043 | -0.04 | -0.04 | 0 | 0.89 | 0.97 | 0.82 | 0.79 | 0.81 | 0.88 |

| | | | | | | | | | | | | | | | | |
|---|---|---|---|---|---|---|---|---|---|---|---|---|---|---|---|---|
| 5 | 132559950 | rs2303671 | 0.102 | 0.061 | -2.365 | 0.397 | 0.048 | -0.005 | -0.005 | 0.173 | 0.38 | 0.79 | 0.4 | 0.35 | 0.26 | 0.29 |
| 5 | 132562731 | rs10038863 | -0.181 | 0.627 | -2.443 | 0.241 | -0.073 | 0.03 | 0.03 | 0 | 0.61 | 0.86 | 0.6 | 0.64 | 0.49 | 0.6 |
| 5 | 132566211 | rs959727 | -0.046 | 0.094 | -1.445 | -0.098 | -0.001 | -0.007 | -0.007 | 0 | 0.45 | 0.5 | 0.55 | 0.7 | 0.43 | 0.48 |
| 5 | 132572752 | rs11242145 | -0.428 | 0.059 | -1.068 | -0.429 | -0.001 | 0.004 | 0.004 | 0 | 0.47 | 0.51 | 0.68 | 0.85 | 0.51 | 0.62 |
| 5 | 132573216 | rs4958109 | 0.065 | -0.107 | -1.096 | -0.27 | -0.001 | -0.003 | -0.003 | 0 | 0.41 | 0.49 | 0.51 | 0.69 | 0.36 | 0.46 |
| 5 | 132579659 | rs11741756 | -0.138 | -0.271 | -0.71 | 0.094 | 0 | 0.008 | 0.008 | 0 | 0.07 | 0.1 | 0.14 | 0.16 | 0.11 | 0.15 |
| 5 | 132580990 | rs7736339 | 0.584 | -0.74 | -0.605 | 0.502 | -0.025 | 0.072 | 0.072 | 0.159 | 0.21 | 0.36 | 0.31 | 0.48 | 0.13 | 0.26 |
| 5 | 132585528 | rs17681747 | 0.738 | -0.486 | -1.397 | 0.018 | -0.031 | -0.038 | -0.038 | 0.645 | 0.25 | 0.01 | 0.14 | 0.2 | 0.22 | 0.28 |
| 5 | 132586717 | rs890815 | 0.68 | -1.053 | -0.889 | 0.058 | 0.044 | -0.064 | -0.064 | 0 | 0.22 | 0.11 | 0.34 | 0.38 | 0.12 | 0.22 |
| 5 | 132587049 | rs6890910 | 0.272 | 0.017 | 0.14 | -0.778 | 0.013 | -0.069 | -0.069 | 2.283 | 0.64 | 0.53 | 0.74 | 0.84 | 0.6 | 0.8 |
| 5 | 132588939 | rs17166462 | -0.942 | -0.114 | -0.037 | -0.222 | -0.009 | -0.031 | -0.031 | 0.018 | 0.07 | 0.12 | 0.05 | 0 | 0.04 | 0.02 |
| 5 | 132605601 | rs3798102 | -0.967 | -0.075 | -1.323 | -0.145 | -0.032 | -0.024 | -0.024 | 0 | 0.12 | 0.7 | 0.1 | 0.07 | 0.03 | 0.04 |
| 5 | 132609517 | rs4958112 | 0.137 | 0.463 | -2.574 | -0.398 | -0.086 | 0.209 | 0.209 | 0 | 0.54 | 0.09 | 0.61 | 0.8 | 0.49 | 0.74 |
| 5 | 132612432 | rs10069497 | 0.325 | 0.973 | -2.917 | 0.168 | -0.128 | -0.107 | -0.107 | 0 | 0.62 | 0.96 | 0.54 | 0.39 | 0.59 | 0.55 |
| 5 | 132612643 | rs4958087 | -0.657 | 1.12 | -2.843 | 0.341 | -0.114 | -0.138 | -0.138 | 0 | 0.3 | 0.06 | 0.41 | 0.57 | 0.36 | 0.42 |
| 5 | 132619333 | rs4365881 | 0.456 | 1.621 | -1.675 | 0.863 | 0.001 | 0 | 0 | 0.317 | 0.76 | 0.79 | 0.81 | 0.75 | 0.73 | 0.71 |
| 5 | 132619938 | rs1428146 | 0.371 | 1.619 | -1.583 | 1.056 | -0.03 | -0.029 | -0.029 | 0.307 | 0.76 | 0.9 | 0.8 | 0.71 | 0.72 | 0.71 |
| 5 | 132621489 | rs6874084 | 0.155 | 2.613 | -0.546 | 0.701 | 0.172 | -0.034 | -0.034 | 0 | 0.08 | 0.33 | 0.16 | 0 | 0.19 | 0.04 |
| 5 | 132622724 | rs6880078 | 1.637 | 2.454 | -0.452 | 1.004 | 0.058 | 0.131 | 0.131 | 3.339 | 0.51 | 0.18 | 0.37 | 0.27 | 0.44 | 0.39 |
| 5 | 132623165 | rs17166490 | -0.168 | 2.779 | -0.394 | 1.587 | -0.012 | -0.042 | -0.042 | 0 | 0.09 | 0.03 | 0.16 | 0.26 | 0.09 | 0.16 |
| 5 | 132623423 | rs13155634 | 2.114 | 2.277 | -0.476 | 1.356 | 0.075 | 0.105 | 0.105 | 0.084 | 0.45 | 0.09 | 0.34 | 0.27 | 0.38 | 0.42 |
| 5 | 132626033 | rs4958114 | 0.178 | 2.203 | -0.216 | 1.618 | 0.014 | -0.029 | -0.029 | 0 | 0.08 | 0.03 | 0.15 | 0.27 | 0.06 | 0.15 |
| 5 | 132626997 | rs6596118 | 0.513 | 2.187 | -0.053 | 1.486 | 0.044 | -0.012 | -0.012 | 0 | 0.69 | 0.46 | 0.66 | 0.73 | 0.62 | 0.68 |
| 5 | 132630270 | rs12187702 | 0.857 | 1.988 | 0.268 | 1.596 | -0.016 | 0.026 | 0.026 | 0.216 | 0.66 | 0.57 | 0.65 | 0.44 | 0.72 | 0.54 |
| 5 | 132631659 | rs2162768 | 1.221 | 2.089 | 0.298 | 1.522 | -0.004 | -0.025 | -0.025 | 0 | 0.6 | 0.57 | 0.53 | 0.27 | 0.63 | 0.44 |
| 5 | 132632476 | rs11242148 | 0.02 | 2.159 | 0.397 | 1.56 | -0.026 | -0.016 | -0.016 | 0 | 0.76 | 0.89 | 0.8 | 0.73 | 0.74 | 0.68 |
| 5 | 132633880 | rs10061620 | -0.154 | 2.092 | 0.4 | 1.565 | -0.01 | -0.022 | -0.022 | 0.018 | 0.77 | 0.89 | 0.82 | 0.72 | 0.78 | 0.69 |
| 5 | 132635935 | rs7733176 | -0.579 | 2.064 | 0.506 | 1.633 | 0.024 | 0.102 | 0.102 | 0 | 0.08 | 0.17 | 0.13 | 0.19 | 0.08 | 0.1 |
| 5 | 132636493 | rs6859098 | -0.151 | 1.924 | 0.458 | 1.662 | 0.008 | 0.153 | 0.153 | 0 | 0.06 | 0.16 | 0.1 | 0.18 | 0.04 | 0.09 |
| 5 | 132637766 | rs2864680 | -0.642 | 1.927 | 0.234 | 1.658 | 0.032 | 0.204 | 0.204 | 0 | 0.08 | 0.32 | 0.11 | 0.17 | 0.04 | 0.09 |
| 5 | 132638993 | rs10900816 | 1.548 | 2.037 | -0.43 | 1.555 | -0.023 | 0.14 | 0.14 | 1.505 | 0.52 | 0.13 | 0.43 | 0.27 | 0.5 | 0.41 |
| 5 | 132641670 | rs6868719 | -0.144 | 2.291 | -0.471 | 1.482 | -0.021 | -0.043 | -0.043 | 0 | 0.82 | 0.94 | 0.87 | 0.73 | 0.81 | 0.71 |
| 5 | 132642611 | rs7714610 | -0.335 | 2.172 | -0.46 | 1.474 | -0.002 | -0.044 | -0.044 | 0.093 | 0.83 | 0.94 | 0.88 | 0.72 | 0.85 | 0.72 |
| 5 | 132645814 | rs1366229 | 0.749 | 2.2 | 0.101 | 1.028 | -0.009 | -0.004 | -0.004 | 0.527 | 0.61 | 0.6 | 0.52 | 0.41 | 0.53 | 0.5 |
| 5 | 132645858 | rs6880509 | 0.04 | 2.367 | 0.527 | 0.724 | 0.109 | -0.03 | -0.03 | 0 | 0.06 | 0.25 | 0.14 | 0 | 0.16 | 0.03 |
| 5 | 132650871 | rs10515457 | -0.11 | 0.965 | 0.168 | 0.68 | 0 | -0.001 | -0.001 | 0 | 0.19 | 0.17 | 0.16 | 0.2 | 0.16 | 0.12 |
| 5 | 132654848 | rs7730752 | 1.076 | 0.66 | -0.557 | 0.303 | -0.036 | 0.089 | 0.089 | 0.707 | 0.42 | 0.16 | 0.41 | 0.24 | 0.48 | 0.53 |
| 5 | 132655046 | rs1079229 | 0.204 | 0.867 | 0.192 | 0.977 | 0.006 | -0.051 | -0.051 | 0.02 | 0.7 | 0.81 | 0.63 | 0.51 | 0.76 | 0.72 |
| 5 | 132655625 | rs733023 | 0.651 | 0.812 | 0.529 | 0.951 | 0.001 | -0.005 | -0.005 | 0 | 0.64 | 0.63 | 0.59 | 0.48 | 0.71 | 0.67 |
| 5 | 132655919 | rs10455054 | 0.148 | 0.836 | 0.305 | 0.964 | -0.028 | -0.137 | -0.137 | 0 | 0.71 | 0.94 | 0.63 | 0.51 | 0.74 | 0.71 |
| 5 | 132657126 | rs9686988 | 0.902 | 0.74 | 0.366 | 0.903 | 0.011 | -0.013 | -0.013 | 0 | 0.61 | 0.67 | 0.59 | 0.48 | 0.71 | 0.7 |
| 5 | 132659915 | rs7729554 | 1.056 | 1.165 | 0.151 | 0.995 | -0.004 | -0.022 | -0.022 | 0 | 0.6 | 0.58 | 0.49 | 0.24 | 0.65 | 0.6 |
| 5 | 132663664 | rs2864246 | 0.35 | 0.903 | -1.409 | 1.317 | -0.074 | -0.184 | -0.184 | 0 | 0.74 | 0.98 | 0.58 | 0.47 | 0.73 | 0.7 |
| 5 | 132669374 | rs17166584 | -0.911 | 0.946 | -0.651 | 1.493 | 0.001 | 0.037 | 0.037 | 0 | 0.12 | 0.14 | 0.1 | 0.3 | 0.13 | 0.18 |
| 5 | 132669425 | rs7716072 | -0.572 | 0.544 | 0.041 | 1.457 | -0.008 | 0.028 | 0.028 | 0 | 0.12 | 0.28 | 0.05 | 0.15 | 0.08 | 0.12 |
| 5 | 132673660 | rs10515459 | -0.91 | 1.321 | 0.839 | 0.937 | 0.096 | -0.029 | -0.029 | 1.694 | 0.08 | 0.28 | 0.15 | 0 | 0.13 | 0.09 |
| 5 | 132677170 | rs4958118 | -1.239 | 1.067 | -0.128 | 0.118 | -0.024 | -0.05 | -0.05 | 0.713 | 0.68 | 0.83 | 0.5 | 0.52 | 0.64 | 0.55 |
| 5 | 132682130 | rs6596123 | -1.696 | 0.587 | 0.01 | 0.291 | 0.007 | -0.02 | -0.02 | 0 | 0.4 | 0.8 | 0.33 | 0.53 | 0.35 | 0.43 |
| 5 | 132699846 | rs17166642 | 0.062 | 0.738 | 0.308 | 0.213 | 0.054 | -0.023 | -0.023 | 0 | 0.06 | 0.18 | 0.12 | 0 | 0.11 | 0.09 |
| 5 | 132703700 | rs4958121 | -1.877 | 0.692 | 0.192 | 0.258 | 0.006 | -0.03 | -0.03 | 2.115 | 0.43 | 0.81 | 0.36 | 0.51 | 0.38 | 0.48 |
| 5 | 132706629 | rs29550 | 0.632 | 0.572 | 0.649 | 0.41 | -0.008 | 0.062 | 0.062 | 0 | 0.74 | 0.6 | 0.82 | 0.5 | 0.79 | 0.66 |
| 5 | 132710063 | rs40255 | 0.445 | 0.325 | 1.019 | 0.442 | -0.03 | 0.106 | 0.106 | 0 | 0.79 | 0.66 | 0.84 | 0.49 | 0.85 | 0.7 |
| 5 | 132713811 | rs700711 | 0.659 | 0.429 | 0.808 | 0.442 | -0.024 | 0.078 | 0.078 | 0 | 0.77 | 0.6 | 0.83 | 0.46 | 0.82 | 0.69 |
| 5 | 132716935 | rs250858 | 1.304 | 0.442 | 0.595 | 0.354 | -0.008 | 0.012 | 0.012 | 0 | 0.65 | 0.41 | 0.68 | 0.48 | 0.69 | 0.57 |
| 5 | 132722340 | rs1428912 | 0.765 | 0.078 | 0.124 | -0.446 | 0.004 | 0.063 | 0.063 | 2.069 | 0.36 | 0.19 | 0.18 | 0.01 | 0.31 | 0.16 |
| 5 | 132723075 | rs700712 | 0.456 | -0.034 | 0.5 | -0.331 | -0.003 | 0.019 | 0.019 | 1.394 | 0.42 | 0.43 | 0.67 | 0.81 | 0.54 | 0.6 |
| 5 | 132723619 | rs700713 | 0.155 | -0.374 | 0.626 | -0.161 | -0.004 | -0.001 | -0.001 | 0 | 0.83 | 0.85 | 0.89 | 0.86 | 0.88 | 0.78 |
| 5 | 132726731 | rs700715 | -0.246 | -0.494 | 0.713 | -0.342 | 0.02 | 0.018 | 0.018 | 1.079 | 0.5 | 0.62 | 0.74 | 0.86 | 0.65 | 0.61 |
| 5 | 132727291 | rs153196 | 0.825 | -0.326 | 0.59 | 0.149 | -0.066 | -0.073 | -0.073 | 0 | 0.34 | 0.19 | 0.57 | 0.49 | 0.53 | 0.41 |
| 5 | 132727692 | rs153197 | 0.005 | -0.321 | 1.287 | 0.887 | -0.04 | 0.046 | 0.046 | 0 | 0.92 | 0.81 | 0.97 | 0.88 | 0.98 | 0.85 |
| 5 | 132728928 | rs250856 | 1.267 | -0.311 | 0.88 | 0.832 | -0.045 | 0.156 | 0.156 | 0.661 | 0.8 | 0.49 | 0.82 | 0.48 | 0.89 | 0.68 |
| 5 | 132732713 | rs17166677 | 0.168 | 0.226 | 0.093 | 0.868 | -0.034 | -0.027 | -0.027 | 0.784 | 0.07 | 0.01 | 0.11 | 0.27 | 0.08 | 0.17 |
| 5 | 132734362 | rs6875606 | -0.646 | -0.214 | 0.631 | 0.191 | -0.052 | -0.115 | -0.115 | 0.016 | 0.17 | 0.39 | 0.12 | 0.01 | 0.1 | 0.11 |
| 5 | 132735155 | rs25735 | 1.007 | 0.975 | 0.123 | 1.029 | 0.014 | 0.054 | 0.054 | 0 | 0.82 | 0.66 | 0.71 | 0.69 | 0.78 | 0.77 |
| 5 | 132739326 | rs17685465 | -1.128 | 1.329 | -0.38 | 0.582 | -0.002 | 0 | 0 | 0 | 0.19 | 0.21 | 0.23 | 0.12 | 0.25 | 0.11 |
| 5 | 132742450 | rs11242152 | 0.504 | 1.201 | -0.879 | 0.635 | -0.047 | -0.009 | -0.009 | 0.15 | 0.37 | 0.2 | 0.31 | 0.4 | 0.41 | 0.25 |
| 5 | 132742865 | rs10793814 | -0.759 | 0.471 | -1.027 | 0.476 | -0.008 | -0.027 | -0.027 | 0.081 | 0.71 | 0.89 | 0.66 | 0.68 | 0.68 | 0.58 |
| 5 | 132746820 | rs17629149 | 0.231 | 0.482 | -1.121 | 0.506 | -0.041 | -0.029 | -0.029 | 0.528 | 0.13 | 0.06 | 0.1 | 0.2 | 0.17 | 0.13 |
| 5 | 132747063 | rs6888605 | 0.478 | 0.089 | -0.841 | 0.444 | 0.007 | 0.002 | 0.002 | 0 | 0.12 | 0.07 | 0.19 | 0.09 | 0.08 | 0.1 |

| | | | | | | | | | | | | | | | | |
|---|---|---|---|---|---|---|---|---|---|---|---|---|---|---|---|---|
| 5 | 132747765 | rs7729507 | 0.252 | 0.326 | 0.017 | 0.499 | -0.004 | -0.004 | -0.004 | 0 | 0.53 | 0.56 | 0.52 | 0.46 | 0.48 | 0.35 |
| 5 | 132748007 | rs10077572 | 1.117 | 0.362 | 0.026 | 0.497 | -0.013 | -0.004 | -0.004 | 0 | 0.39 | 0.27 | 0.31 | 0.37 | 0.39 | 0.26 |
| 5 | 132751172 | rs17696621 | -0.645 | 0.371 | 0.034 | 0.489 | -0.009 | -0.001 | -0.001 | 0 | 0.28 | 0.19 | 0.32 | 0.29 | 0.29 | 0.22 |
| 5 | 132752066 | rs166112 | -0.901 | 0.323 | 0.24 | 0.295 | -0.021 | -0.001 | -0.001 | 0 | 0.69 | 0.49 | 0.64 | 0.69 | 0.71 | 0.57 |
| 5 | 132758353 | rs250850 | 1.208 | 0.669 | -0.131 | 0.121 | -0.026 | -0.048 | -0.048 | 0 | 0.36 | 0.2 | 0.28 | 0.44 | 0.36 | 0.24 |
| 5 | 132765214 | rs7730256 | -0.165 | 0.054 | 0.005 | -0.1 | 0.029 | 0 | 0 | 0 | 0.55 | 0.32 | 0.56 | 0.53 | 0.61 | 0.44 |
| 5 | 132783187 | rs25869 | -0.088 | 0.333 | -0.064 | -0.543 | -0.04 | 0.069 | 0.069 | 0 | 0.22 | 0.09 | 0.29 | 0.08 | 0.23 | 0.17 |
| 5 | 132791882 | rs27435 | -1.043 | 0.06 | 0.223 | -0.878 | -0.004 | 0.033 | 0.033 | 0 | 0.39 | 0.3 | 0.41 | 0.12 | 0.4 | 0.45 |
| 5 | 132801509 | rs31341 | 0.531 | 0.366 | 0.026 | -0.835 | -0.037 | 0.03 | 0.03 | 0 | 0.1 | 0.01 | 0.07 | 0.03 | 0.1 | 0.09 |
| 5 | 132805024 | rs12186611 | 1.545 | 0.499 | 0.304 | -0.738 | -0.002 | -0.005 | -0.005 | 0 | 0.31 | 0.29 | 0.25 | 0.47 | 0.36 | 0.22 |
| 5 | 132805442 | rs1494666 | -0.309 | 0.378 | 0.48 | -0.723 | -0.018 | -0.102 | -0.102 | 0 | 0.29 | 0.5 | 0.35 | 0.08 | 0.28 | 0.23 |
| 5 | 132807320 | rs31345 | -1.086 | 0.673 | 0.489 | -0.291 | 0.112 | 0.028 | 0.028 | 1.842 | 0.07 | 0.33 | 0.07 | 0.03 | 0.05 | 0.08 |
| 5 | 132807672 | rs718410 | 0.214 | 0.647 | 0.184 | -0.266 | -0.002 | -0.017 | -0.017 | 0 | 0.06 | 0.12 | 0.05 | 0.02 | 0.04 | 0.13 |
| 5 | 132809967 | rs190317 | -0.674 | 0.607 | -0.475 | -0.214 | -0.01 | -0.088 | -0.088 | 0 | 0.76 | 0.93 | 0.71 | 0.59 | 0.78 | 0.68 |
| 5 | 132811282 | rs246960 | -1.399 | 0.844 | 0.221 | -0.582 | 0.07 | -0.029 | -0.029 | 0.148 | 0.13 | 0.46 | 0.12 | 0.05 | 0.1 | 0.22 |
| 5 | 132811978 | rs246961 | -1.01 | 0.414 | -0.995 | -0.794 | 0.014 | 0.029 | 0.029 | 0.319 | 0.24 | 0.77 | 0.15 | 0.04 | 0.15 | 0.15 |
| 5 | 132815696 | rs12522909 | -0.321 | 0.645 | -1.142 | -0.626 | -0.021 | 0.079 | 0.079 | 1.011 | 0.26 | 0.08 | 0.14 | 0.05 | 0.22 | 0.33 |
| 5 | 132819346 | rs17630080 | 0.854 | 0.698 | -1.72 | -0.795 | 0.008 | 0.052 | 0.052 | 0.422 | 0.27 | 0.01 | 0.19 | 0.22 | 0.21 | 0.23 |
| 5 | 132820319 | rs27986 | -1.06 | 1.81 | -0.391 | -0.316 | 0 | -0.049 | -0.049 | 0 | 0.17 | 0.66 | 0.15 | 0.05 | 0.19 | 0.13 |
| 5 | 132821596 | rs31325 | -1.098 | 1.829 | -0.405 | -0.296 | 0.001 | -0.043 | -0.043 | 1.384 | 0.18 | 0.66 | 0.15 | 0.06 | 0.2 | 0.13 |
| 5 | 132831435 | rs31323 | -0.066 | 1.947 | -0.36 | 0.535 | 0 | 0.01 | 0.01 | 0 | 0.15 | 0.16 | 0.29 | 0.39 | 0.28 | 0.23 |
| 5 | 132837218 | rs10040489 | -0.544 | 1.688 | -0.145 | 0.152 | 0.024 | -0.056 | -0.056 | 0.114 | 0.14 | 0.25 | 0.12 | 0.02 | 0.21 | 0.1 |
| 5 | 132839137 | rs1366604 | -0.941 | 1.347 | -0.307 | 0.383 | -0.01 | 0 | 0 | 0.09 | 0.37 | 0.63 | 0.43 | 0.37 | 0.51 | 0.28 |
| 5 | 132844488 | rs457886 | -0.377 | 1.307 | -0.104 | 0.335 | 0.012 | -0.007 | -0.007 | 0 | 0.1 | 0.12 | 0.1 | 0.03 | 0.15 | 0.07 |
| 5 | 132852187 | rs397649 | -1.26 | 0.788 | 0.1 | 0.322 | 0.009 | -0.029 | -0.029 | 0.858 | 0.23 | 0.53 | 0.18 | 0.16 | 0.21 | 0.14 |
| 5 | 132859669 | rs155581 | 0.014 | 0.276 | -0.586 | 0.705 | 0.029 | 0.001 | 0.001 | 0 | 0.46 | 0.38 | 0.54 | 0.56 | 0.64 | 0.56 |
| 5 | 132863198 | rs1318103 | 0.458 | 0.45 | -1.049 | 0.935 | 0.024 | -0.055 | -0.055 | 2.611 | 0.3 | 0.12 | 0.37 | 0.42 | 0.28 | 0.31 |
| 5 | 132863381 | rs700703 | -0.424 | 0.595 | -0.71 | 0.788 | -0.002 | -0.018 | -0.018 | 0 | 0.44 | 0.33 | 0.53 | 0.52 | 0.46 | 0.39 |
| 5 | 132864295 | rs700706 | 0.077 | 0.379 | -0.51 | 0.864 | 0.001 | -0.002 | -0.002 | 0.225 | 0.37 | 0.33 | 0.43 | 0.42 | 0.37 | 0.39 |
| 5 | 132866748 | rs42567 | 0.777 | 1.222 | -0.661 | 1.24 | -0.017 | -0.016 | -0.016 | 0.031 | 0.13 | 0.08 | 0.23 | 0.33 | 0.24 | 0.23 |
| 5 | 132885741 | rs187411 | -0.909 | 1.003 | -0.427 | 1.021 | 0.023 | 0 | 0 | 0 | 0.35 | 0.57 | 0.45 | 0.33 | 0.48 | 0.48 |
| 5 | 132889644 | rs26367 | 0.641 | 1.085 | -0.457 | 0.995 | -0.032 | -0.023 | -0.023 | 0 | 0.05 | 0 | 0.05 | 0.08 | 0.09 | 0.06 |
| 5 | 132892497 | rs30487 | -1.504 | 0.743 | -0.614 | 0.922 | 0.019 | -0.001 | -0.001 | 0.253 | 0.69 | 0.82 | 0.82 | 0.69 | 0.77 | 0.76 |
| 5 | 132894431 | rs9327655 | -0.335 | 0.564 | -0.462 | 0.935 | 0.043 | -0.002 | -0.002 | 0 | 0.28 | 0.43 | 0.38 | 0.25 | 0.38 | 0.42 |
| 5 | 132895893 | rs10479044 | -0.62 | 0.577 | -0.556 | 0.924 | 0 | -0.002 | -0.002 | 0 | 0.62 | 0.7 | 0.76 | 0.62 | 0.67 | 0.7 |
| 5 | 132916408 | rs256257 | -0.777 | 0.762 | -0.592 | 0.995 | -0.002 | -0.002 | -0.002 | 0 | 0.65 | 0.66 | 0.52 | 0.58 | 0.6 | 0.68 |
| 5 | 132917106 | rs256259 | 0.824 | 0.513 | -1.165 | 0.79 | -0.003 | -0.011 | -0.011 | 0 | 0.26 | 0.14 | 0.17 | 0.28 | 0.23 | 0.23 |
| 5 | 132927454 | rs171742 | -0.196 | 0.522 | -1.199 | 0.721 | -0.021 | 0.066 | 0.066 | 0.943 | 0.49 | 0.23 | 0.4 | 0.6 | 0.5 | 0.46 |
| 5 | 132929310 | rs369853 | -0.577 | 0.376 | 0.258 | 0.589 | 0 | -0.006 | -0.006 | 4.556 | 0.57 | 0.6 | 0.47 | 0.66 | 0.57 | 0.51 |
| 5 | 132929702 | rs377434 | -0.457 | 0.368 | 0.007 | 0.755 | 0 | 0.002 | 0.002 | 0.612 | 0.3 | 0.36 | 0.29 | 0.35 | 0.34 | 0.28 |
| 5 | 132930325 | rs11948984 | 0.998 | 0.229 | -0.135 | 0.721 | 0.003 | -0.025 | -0.025 | 0 | 0.27 | 0.15 | 0.17 | 0.32 | 0.23 | 0.23 |
| 5 | 132933128 | rs381879 | -0.299 | 0.339 | 0.146 | 0.429 | -0.012 | -0.017 | -0.017 | 0.01 | 0.11 | 0.2 | 0.07 | 0.06 | 0.08 | 0.05 |
| 5 | 132935420 | rs407758 | 0.736 | 0.363 | 0.134 | 0.193 | 0.003 | -0.002 | -0.002 | 0 | 0.31 | 0.28 | 0.33 | 0.37 | 0.31 | 0.29 |
| 5 | 132936076 | rs256249 | 0.505 | 0.427 | 0.142 | 0.156 | -0.007 | 0.009 | 0.009 | 0 | 0.44 | 0.42 | 0.39 | 0.6 | 0.49 | 0.45 |
| 5 | 132941617 | rs7722071 | 1.186 | 0.596 | 0.217 | 0.313 | 0.044 | 0.065 | 0.065 | 0 | 0.12 | 0.22 | 0.15 | 0.25 | 0.25 | 0.22 |
| 5 | 132941794 | rs17642974 | 1.348 | 0.571 | 0.064 | 0.273 | 0.01 | -0.018 | -0.018 | 0 | 0.26 | 0.12 | 0.35 | 0.34 | 0.27 | 0.25 |
| 5 | 132942656 | rs405954 | -1.173 | 0.85 | 0.709 | 0.057 | 0.001 | -0.002 | -0.002 | 0 | 0.37 | 0.55 | 0.28 | 0.32 | 0.4 | 0.5 |
| 5 | 132944069 | rs17166869 | 0.726 | 0.844 | 0.572 | -0.104 | 0 | -0.015 | -0.015 | 0 | 0.1 | 0.07 | 0.05 | 0 | 0.07 | 0.2 |
| 5 | 132949899 | rs256241 | -1.293 | 0.891 | 0.509 | -0.012 | 0.04 | -0.013 | -0.013 | 0 | 0.93 | 0.87 | 0.79 | 0.97 | 0.89 | 0.97 |
| 5 | 132959138 | rs453376 | -0.925 | -0.041 | 0.392 | -0.282 | -0.024 | -0.059 | -0.059 | 0 | 0.25 | 0.32 | 0.12 | 0.07 | 0.15 | 0.28 |
| 5 | 132971131 | rs256209 | 0.024 | -0.103 | -0.104 | -0.348 | 0.004 | 0.003 | 0.003 | 0 | 0.34 | 0.35 | 0.5 | 0.48 | 0.49 | 0.4 |
| 5 | 132971697 | rs256214 | 1.132 | 0.017 | -0.244 | -0.328 | 0.009 | 0.002 | 0.002 | 0.059 | 0.18 | 0.18 | 0.27 | 0.25 | 0.34 | 0.25 |
| 5 | 132975474 | rs256229 | -1.45 | -0.156 | -0.547 | -0.49 | 0.019 | -0.05 | -0.05 | 0.144 | 0.64 | 0.83 | 0.78 | 0.62 | 0.7 | 0.74 |
| 5 | 132975698 | rs256231 | 1.066 | 0.125 | -0.45 | -0.343 | 0.013 | 0.005 | 0.005 | 0.408 | 0.2 | 0.2 | 0.28 | 0.33 | 0.37 | 0.3 |
| 5 | 132978751 | rs4958130 | 1.202 | -0.663 | -0.398 | -0.177 | 0.005 | -0.012 | -0.012 | 0 | 0.13 | 0.03 | 0.27 | 0.19 | 0.13 | 0.11 |
| 5 | 132990323 | rs10491294 | -0.146 | -0.77 | -0.406 | -0.36 | -0.015 | 0.047 | 0.047 | 0 | 0.25 | 0.09 | 0.34 | 0.19 | 0.24 | 0.3 |
| 5 | 132991517 | rs4958133 | -1.581 | -0.832 | -0.625 | -0.591 | -0.085 | -0.011 | -0.011 | 0.607 | 0.75 | 0.89 | 0.68 | 0.76 | 0.56 | 0.71 |
| 5 | 132992201 | rs10054346 | -0.493 | -0.979 | -0.13 | -0.651 | -0.017 | 0.003 | 0.003 | 0.95 | 0.12 | 0.32 | 0.08 | 0.08 | 0.09 | 0.09 |
| 5 | 132992973 | rs17694216 | 1.754 | -0.968 | -0.828 | -0.624 | 0.006 | -0.003 | -0.003 | 0 | 0.07 | 0.06 | 0.1 | 0.1 | 0.05 | 0.12 |
| 5 | 132993189 | rs10479054 | 0.815 | -0.89 | -0.924 | -0.62 | -0.018 | 0.032 | 0.032 | 0 | 0.11 | 0.05 | 0.07 | 0.01 | 0.11 | 0.18 |
| 5 | 132996659 | rs7706203 | -1.371 | -0.959 | -0.847 | -0.63 | 0.035 | 0.05 | 0.05 | 0 | 0.44 | 0.69 | 0.53 | 0.38 | 0.37 | 0.5 |
| 5 | 132998708 | rs4958136 | -0.793 | -0.96 | -0.775 | -0.58 | 0.003 | 0.001 | 0.001 | 0.053 | 0.34 | 0.64 | 0.47 | 0.37 | 0.27 | 0.31 |
| 5 | 133001245 | rs2867327 | -0.779 | -0.886 | -0.919 | -0.522 | -0.041 | -0.006 | -0.006 | 0.686 | 0.62 | 0.79 | 0.43 | 0.58 | 0.44 | 0.6 |
| 5 | 133002123 | rs9327661 | 0.405 | -0.708 | -0.825 | -0.474 | -0.001 | 0 | 0 | 0 | 0.15 | 0.12 | 0.1 | 0.08 | 0.16 | 0.22 |
| 5 | 133003070 | rs11242167 | 0.024 | -0.649 | -0.755 | -0.394 | 0.033 | 0.015 | 0.015 | 0 | 0.52 | 0.6 | 0.48 | 0.59 | 0.35 | 0.36 |
| 5 | 133005912 | rs17694351 | 0.861 | -0.494 | -1.624 | -0.264 | 0.019 | -0.061 | -0.061 | 0.298 | 0.32 | 0.17 | 0.16 | 0.39 | 0.21 | 0.28 |
| 5 | 133006274 | rs10068769 | -0.891 | 0.337 | -1.058 | -0.119 | -0.091 | -0.023 | -0.023 | 0 | 0.76 | 0.9 | 0.69 | 0.76 | 0.56 | 0.71 |
| 5 | 133009547 | rs2270088 | -0.275 | -0.92 | -0.387 | -0.577 | -0.018 | -0.023 | -0.023 | 0 | 0.34 | 0.39 | 0.32 | 0.2 | 0.19 | 0.19 |
| 5 | 133011262 | rs10479055 | -0.576 | 0.065 | -0.036 | 0.11 | -0.005 | -0.007 | -0.007 | 0 | 0.33 | 0.61 | 0.42 | 0.25 | 0.32 | 0.26 |

| | | | | | | | | | | | | | | | | |
|---|---|---|---|---|---|---|---|---|---|---|---|---|---|---|---|---|
| 5 | 133011503 | rs17699116 | -0.294 | 0.165 | -0.365 | 0.046 | -0.015 | -0.001 | -0.001 | 0 | 0.28 | 0.11 | 0.38 | 0.28 | 0.3 | 0.27 |
| 5 | 133012034 | rs17642519 | -0.181 | 0.138 | -0.416 | 0.054 | -0.03 | -0.015 | -0.015 | 0 | 0.27 | 0.06 | 0.37 | 0.29 | 0.29 | 0.24 |
| 5 | 133013103 | rs13359225 | -0.83 | 0.27 | 0.358 | -0.058 | 0.12 | 0.058 | 0.058 | 3.861 | 0.06 | 0.51 | 0.07 | 0 | 0.03 | 0.01 |
| 5 | 133014915 | rs13359919 | -0.83 | 0.27 | 0.358 | -0.058 | 0.12 | 0.058 | 0.058 | 3.861 | 0.06 | 0.51 | 0.07 | 0 | 0.03 | 0.01 |
| 5 | 133016054 | rs4958095 | 0.883 | 0.055 | -0.42 | -0.269 | 0.01 | 0.06 | 0.06 | 0 | 0.58 | 0.26 | 0.53 | 0.71 | 0.64 | 0.58 |
| 5 | 133017882 | rs758309 | -0.627 | 0.092 | -0.299 | -0.054 | 0.003 | 0.013 | 0.013 | 0 | 0.34 | 0.73 | 0.45 | 0.29 | 0.32 | 0.26 |
| 5 | 133018603 | rs4141220 | 0.028 | -0.139 | -0.397 | 0.143 | 0.009 | -0.006 | -0.006 | 0 | 0.56 | 0.65 | 0.51 | 0.52 | 0.64 | 0.75 |
| 5 | 133019116 | rs6596137 | -1.251 | -0.333 | 0.634 | -0.011 | 0.04 | -0.001 | -0.001 | 0 | 0.08 | 0.5 | 0.06 | 0 | 0.02 | 0.13 |
| 5 | 133019542 | rs13185268 | 0.075 | -0.514 | 0.408 | 0.583 | -0.011 | 0.055 | 0.055 | 0 | 0.07 | 0.13 | 0.07 | 0.16 | 0.06 | 0.04 |
| 5 | 133021143 | rs10793820 | 0.393 | -0.455 | 0.081 | 0.523 | -0.002 | -0.003 | -0.003 | 0 | 0.41 | 0.52 | 0.41 | 0.36 | 0.42 | 0.33 |
| 5 | 133021590 | rs4958144 | 0.046 | -0.49 | 0.168 | 0.58 | -0.002 | -0.001 | -0.001 | 0 | 0.46 | 0.54 | 0.45 | 0.51 | 0.46 | 0.36 |
| 5 | 133021940 | rs6596142 | 0.249 | -0.455 | -0.221 | 0.359 | -0.002 | -0.003 | -0.003 | 0 | 0.35 | 0.37 | 0.43 | 0.38 | 0.33 | 0.41 |
| 5 | 133022935 | rs7713865 | 0.184 | -0.494 | 0.186 | 0.379 | -0.003 | 0.21 | 0.21 | 0 | 0.08 | 0.39 | 0.07 | 0.16 | 0.04 | 0.03 |
| 5 | 133023122 | rs10751462 | -0.505 | -0.134 | -0.587 | 0.218 | -0.032 | 0.052 | 0.052 | 0 | 0.78 | 0.92 | 0.86 | 0.9 | 0.78 | 0.76 |
| 5 | 133028493 | rs6596146 | 0.054 | -0.709 | -0.302 | 0.229 | -0.009 | 0 | 0 | 0.835 | 0.33 | 0.56 | 0.35 | 0.35 | 0.26 | 0.28 |
| 5 | 133029003 | rs7703486 | 0.063 | -0.711 | 0.051 | 0.3 | -0.025 | 0.005 | 0.005 | 0.833 | 0.33 | 0.52 | 0.34 | 0.38 | 0.26 | 0.28 |
| 5 | 133029196 | rs17166927 | -0.191 | -1.069 | -0.444 | -0.134 | 0.068 | 0.032 | 0.032 | 1.847 | 0.5 | 0.64 | 0.42 | 0.41 | 0.35 | 0.33 |
| 5 | 133039590 | rs876932 | 0.716 | -1.303 | -0.095 | -0.573 | -0.05 | -0.018 | -0.018 | 1.399 | 0.28 | 0.42 | 0.14 | 0.21 | 0.18 | 0.14 |
| 5 | 133045139 | rs11749241 | 1.118 | -1.344 | -0.237 | -0.618 | 0.028 | 0.048 | 0.048 | 0 | 0.18 | 0.06 | 0.07 | 0.04 | 0.09 | 0.05 |
| 5 | 133047775 | rs6596147 | -1.642 | -1.255 | -0.726 | -0.748 | -0.068 | 0.03 | 0.03 | 0 | 0.79 | 0.98 | 0.87 | 0.9 | 0.78 | 0.78 |
| 5 | 133048578 | rs10068258 | 0.659 | -1.377 | -0.264 | -0.644 | -0.034 | -0.014 | -0.014 | 0 | 0.28 | 0.51 | 0.14 | 0.22 | 0.18 | 0.14 |
| 5 | 133049626 | rs10900821 | 1.039 | -1.713 | 0.145 | -0.794 | 0.057 | -0.036 | -0.036 | 0.582 | 0.1 | 0.27 | 0.11 | 0.03 | 0.14 | 0.09 |
| 5 | 133051079 | rs10052064 | -0.464 | -1.688 | -0.236 | -0.815 | 0.05 | 0.046 | 0.046 | 3.387 | 0.46 | 0.6 | 0.23 | 0.25 | 0.27 | 0.23 |
| 5 | 133056044 | rs17643897 | 1.075 | -1.525 | -0.642 | -0.759 | 0.032 | 0.063 | 0.063 | 0 | 0.18 | 0.08 | 0.08 | 0.04 | 0.09 | 0.05 |
| 5 | 133059581 | rs740048 | -0.04 | -0.875 | -0.288 | -0.41 | 0.104 | 0.091 | 0.091 | 4.681 | 0.47 | 0.78 | 0.25 | 0.26 | 0.27 | 0.18 |
| 5 | 133064960 | rs6870951 | 0.096 | 0.243 | 0.423 | -0.266 | 0.048 | -0.043 | -0.043 | 0 | 0.09 | 0.36 | 0.08 | 0.01 | 0.08 | 0.05 |
| 5 | 133072525 | rs4958148 | 0.055 | 1.189 | -1.363 | 0.32 | -0.084 | -0.068 | -0.068 | 0 | 0.28 | 0.14 | 0.52 | 0.49 | 0.53 | 0.5 |
| 5 | 133073611 | rs1981608 | 0.297 | 0.675 | -1.285 | -0.552 | 0.098 | 0.079 | 0.079 | 1.288 | 0.49 | 0.26 | 0.43 | 0.65 | 0.35 | 0.48 |
| 5 | 133074478 | rs17700949 | -0.893 | 2.553 | -0.784 | -0.12 | -0.045 | 0.028 | 0.028 | 0 | 0.15 | 0.09 | 0.23 | 0.11 | 0.33 | 0.17 |
| 5 | 133075934 | rs17166985 | -0.149 | 1.806 | -0.36 | -0.03 | -0.006 | -0.001 | -0.001 | 0 | 0.34 | 0.35 | 0.29 | 0.67 | 0.45 | 0.51 |
| 5 | 133076556 | rs17166987 | -0.892 | 1.119 | 0.509 | 0.274 | 0.008 | -0.012 | -0.012 | 0 | 0.1 | 0.29 | 0.3 | 0.07 | 0.12 | 0.1 |
| 5 | 133078832 | rs6890528 | -0.175 | 1.326 | 0.234 | 0.156 | 0.013 | 0.013 | 0.013 | 0.021 | 0.42 | 0.66 | 0.36 | 0.69 | 0.38 | 0.47 |
| 5 | 133079344 | rs1109309 | -0.794 | 1.384 | 0.111 | 0.071 | -0.005 | 0.114 | 0.114 | 0 | 0.52 | 0.73 | 0.72 | 0.92 | 0.53 | 0.66 |
| 5 | 133082666 | rs10515462 | -0.221 | 1.332 | 0.115 | 0.674 | -0.02 | -0.004 | -0.004 | 0 | 0.07 | 0.02 | 0.21 | 0.05 | 0.11 | 0.11 |
| 5 | 133082746 | rs7712990 | -1.069 | 1.305 | 0.388 | 0.65 | 0.022 | 0.093 | 0.093 | 0.433 | 0.12 | 0.3 | 0.38 | 0.23 | 0.15 | 0.18 |
| 5 | 133089667 | rs11739843 | 0.569 | 1.33 | -0.115 | -0.08 | -0.018 | 0.162 | 0.162 | 0.193 | 0.47 | 0.25 | 0.38 | 0.1 | 0.44 | 0.29 |
| 5 | 133089768 | rs12188394 | 0.693 | 1.194 | 0.191 | 0.709 | 0.022 | -0.027 | -0.027 | 0 | 0.27 | 0.16 | 0.21 | 0.39 | 0.21 | 0.35 |
| 5 | 133098198 | rs258074 | 0.066 | 1.313 | 0.905 | 0.851 | -0.015 | 0.013 | 0.013 | 0 | 0.63 | 0.61 | 0.55 | 0.51 | 0.66 | 0.52 |
| 5 | 133100553 | rs258071 | 0.541 | 1.418 | 1.259 | 0.821 | -0.002 | 0.001 | 0.001 | 0 | 0.58 | 0.46 | 0.51 | 0.51 | 0.6 | 0.5 |
| 5 | 133101137 | rs258068 | -0.017 | 1.419 | 1.36 | 0.824 | 0.016 | -0.018 | -0.018 | 0 | 0.63 | 0.7 | 0.55 | 0.51 | 0.66 | 0.52 |
| 5 | 133101257 | rs6891832 | 0.772 | 1.436 | 1.146 | 0.857 | 0.032 | -0.044 | -0.044 | 0.053 | 0.27 | 0.14 | 0.21 | 0.38 | 0.21 | 0.35 |
| 5 | 133101506 | rs10068902 | -0.859 | 2.042 | 1.512 | 1.295 | 0.064 | -0.004 | -0.004 | 1.131 | 0.05 | 0.22 | 0.04 | 0 | 0.06 | 0.02 |
| 5 | 133112458 | rs6596148 | -0.419 | 1.93 | 0.731 | 1.454 | 0.013 | 0.058 | 0.058 | 0 | 0.11 | 0.15 | 0.12 | 0.4 | 0.16 | 0.22 |
| 5 | 133114228 | rs1859856 | 0.269 | 1.582 | 0.414 | 1.058 | -0.022 | -0.029 | -0.029 | 0 | 0.91 | 0.98 | 0.77 | 0.89 | 0.87 | 0.88 |
| 5 | 133117988 | rs17167038 | -0.441 | 1.743 | 0.771 | 1.288 | 0.004 | 0.03 | 0.03 | 0 | 0.11 | 0.15 | 0.11 | 0.35 | 0.16 | 0.19 |
| 5 | 133120623 | rs3853752 | 0.475 | 2.197 | 1.34 | 0.667 | 0.025 | -0.024 | -0.024 | 0.179 | 0.82 | 0.71 | 0.68 | 0.92 | 0.75 | 0.79 |
| 5 | 133131036 | rs731432 | 1.354 | 2.269 | 1.478 | 0.865 | -0.002 | 0.001 | 0.001 | 0.005 | 0.73 | 0.31 | 0.56 | 0.75 | 0.63 | 0.7 |
| 5 | 133131129 | rs17646132 | -0.844 | 2.185 | 1.406 | 0.969 | 0.005 | 0.042 | 0.042 | 0 | 0.07 | 0.11 | 0.04 | 0.16 | 0.07 | 0.14 |
| 5 | 133134016 | rs3909368 | 1.378 | 2.155 | 1.793 | 0.927 | 0.037 | 0.113 | 0.113 | 0 | 0.84 | 0.49 | 0.85 | 0.68 | 0.82 | 0.78 |
| 5 | 133134183 | rs1644299 | 2.005 | 2.129 | 1.28 | 0.868 | -0.107 | -0.054 | -0.054 | 0 | 0.65 | 0.12 | 0.53 | 0.6 | 0.57 | 0.57 |
| 5 | 133138196 | rs12653634 | -0.942 | 2.408 | 1.185 | 0.811 | -0.03 | -0.039 | -0.039 | 0 | 0.1 | 0.05 | 0.19 | 0.18 | 0.16 | 0.2 |
| 5 | 133138979 | rs2052496 | -1.942 | 2.476 | 1.874 | 0.87 | 0.049 | 0.101 | 0.101 | 0 | 0.19 | 0.37 | 0.29 | 0.39 | 0.27 | 0.29 |
| 5 | 133139439 | rs2873324 | -0.403 | 2.204 | 1.935 | 0.573 | 0.016 | -0.058 | -0.058 | 0 | 0.14 | 0.25 | 0.17 | 0.01 | 0.18 | 0.11 |
| 5 | 133140038 | rs7712156 | 0.472 | 2.188 | 1.867 | 0.702 | 0.01 | 0.023 | 0.023 | 0 | 0.07 | 0.14 | 0.07 | 0.11 | 0.08 | 0.02 |
| 5 | 133144207 | rs6895072 | -0.461 | 2.149 | 1.959 | 0.724 | 0.016 | 0.072 | 0.072 | 0 | 0.09 | 0.25 | 0.09 | 0.22 | 0.1 | 0.11 |
| 5 | 133146042 | rs184977 | -0.973 | 1.45 | 1.372 | 0.385 | -0.015 | 0.047 | 0.047 | 0 | 0.62 | 0.78 | 0.63 | 0.81 | 0.58 | 0.77 |
| 5 | 133155469 | rs4958231 | 0.068 | 1.381 | 0.821 | 0.22 | 0.04 | 0.045 | 0.045 | 0 | 0.51 | 0.4 | 0.53 | 0.3 | 0.65 | 0.31 |
| 5 | 133158325 | rs245582 | 0.831 | 1.439 | 0.261 | 0.137 | -0.071 | 0.106 | 0.106 | 0 | 0.37 | 0.08 | 0.36 | 0.18 | 0.42 | 0.22 |
| 5 | 133160484 | rs11242190 | 0.087 | 1.442 | 0.491 | 0.471 | -0.028 | -0.027 | -0.027 | 0 | 0.06 | 0.01 | 0.15 | 0.07 | 0.09 | 0.12 |
| 5 | 133161452 | rs17167074 | -0.104 | 1.432 | 0.9 | 0.286 | 0.03 | 0.087 | 0.087 | 0.222 | 0.08 | 0.21 | 0.17 | 0.18 | 0.1 | 0.13 |
| 5 | 133161907 | rs12514637 | 0.067 | 1.425 | 0.916 | 0.282 | 0.021 | 0.072 | 0.072 | 0.243 | 0.08 | 0.18 | 0.16 | 0.18 | 0.09 | 0.13 |
| 5 | 133163039 | rs1644311 | 1.209 | 1.318 | 0.402 | -0.345 | -0.023 | 0.15 | 0.15 | 0.176 | 0.3 | 0.1 | 0.18 | 0.01 | 0.31 | 0.13 |
| 5 | 133164490 | rs7729087 | -0.257 | 0.912 | 0.953 | -0.115 | -0.018 | -0.013 | -0.013 | 0 | 0.58 | 0.51 | 0.56 | 0.4 | 0.65 | 0.53 |
| 5 | 133168558 | rs11242192 | -0.185 | 1.177 | 0.591 | 0.066 | -0.003 | -0.003 | -0.003 | 1.203 | 0.07 | 0.07 | 0.17 | 0.1 | 0.12 | 0.12 |
| 5 | 133168839 | rs11242193 | -0.082 | 1.189 | 0.596 | 0.072 | -0.002 | -0.002 | -0.002 | 0 | 0.06 | 0.07 | 0.17 | 0.1 | 0.12 | 0.12 |
| 5 | 133177588 | rs2457175 | -0.986 | 1.26 | 0.632 | -0.16 | -0.017 | -0.011 | -0.011 | 0.012 | 0.22 | 0.18 | 0.38 | 0.3 | 0.32 | 0.33 |
| 5 | 133178199 | rs2431561 | 0.06 | 1.292 | 0.708 | -0.07 | -0.012 | -0.006 | -0.006 | 0.041 | 0.14 | 0.08 | 0.2 | 0.2 | 0.18 | 0.18 |
| 5 | 133179080 | rs2457174 | 0.993 | 1.024 | 0.648 | -0.34 | -0.031 | 0.084 | 0.084 | 0 | 0.35 | 0.19 | 0.28 | 0.05 | 0.41 | 0.18 |
| 5 | 133179726 | rs1681138 | 1.033 | 1.046 | 0.981 | -0.274 | 0.054 | 0.087 | 0.087 | 0.169 | 0.77 | 0.46 | 0.66 | 0.59 | 0.68 | 0.72 |

| | | | | | | | | | | | | | | | | |
|---|---|---|---|---|---|---|---|---|---|---|---|---|---|---|---|---|
| 5 | 133180735 | rs1703425 | 0.328 | 1.124 | 1.014 | -0.24 | 0.061 | 0.078 | 0.078 | 0.115 | 0.82 | 0.57 | 0.76 | 0.69 | 0.79 | 0.8 |
| 5 | 133181613 | rs2436447 | 0.13 | 0.992 | 0.932 | -0.349 | -0.001 | -0.008 | -0.008 | 0 | 0.47 | 0.5 | 0.44 | 0.26 | 0.56 | 0.45 |
| 5 | 133183441 | rs1681127 | -0.649 | 1.142 | 0.675 | -0.184 | -0.052 | -0.043 | -0.043 | 0.01 | 0.19 | 0.08 | 0.28 | 0.31 | 0.3 | 0.28 |
| 5 | 133183870 | rs10078111 | -0.592 | 1.028 | 0.836 | -0.168 | -0.003 | -0.021 | -0.021 | 0.035 | 0.06 | 0.17 | 0.05 | 0.05 | 0.03 | 0.12 |
| 5 | 133184986 | rs4958262 | 0.912 | 1.056 | 1.085 | -0.098 | 0.09 | 0.203 | 0.203 | 1.077 | 0.91 | 0.48 | 0.86 | 0.8 | 0.86 | 0.91 |
| 5 | 133186592 | rs10062785 | 1.894 | 1.105 | 1.023 | -0.181 | 0.051 | 0.104 | 0.104 | 0.214 | 0.86 | 0.47 | 0.82 | 0.75 | 0.83 | 0.79 |
| 5 | 133195356 | rs1644312 | 0.181 | 1.142 | 0.414 | -0.165 | -0.044 | -0.014 | -0.014 | 0.238 | 0.14 | 0.04 | 0.13 | 0.19 | 0.18 | 0.2 |
| 5 | 133199252 | rs1644309 | -0.029 | 0.939 | 0.212 | -0.328 | 0.023 | 0.069 | 0.069 | 0 | 0.51 | 0.11 | 0.53 | 0.65 | 0.41 | 0.6 |
| 5 | 133199549 | rs1644308 | 0.875 | 1.087 | 0.426 | -0.401 | -0.002 | 0.006 | 0.006 | 0 | 0.37 | 0.3 | 0.29 | 0.14 | 0.41 | 0.28 |
| 5 | 133201441 | rs12519506 | -0.052 | 0.939 | -0.002 | -0.373 | 0.091 | 0.071 | 0.071 | 0 | 0.51 | 0.29 | 0.53 | 0.65 | 0.36 | 0.59 |
| 5 | 133202612 | rs12515157 | 0.796 | 0.775 | -0.196 | -0.396 | 0.073 | -0.117 | -0.117 | 0.075 | 0.34 | 0.08 | 0.32 | 0.42 | 0.22 | 0.35 |
| 5 | 133211490 | rs2640684 | 0.252 | 0.728 | 0.238 | -1.039 | -0.002 | 0.039 | 0.039 | 0 | 0.47 | 0.36 | 0.37 | 0.06 | 0.5 | 0.23 |
| 5 | 133215213 | rs4958200 | 0.978 | 0.573 | 0.165 | -0.994 | 0.021 | -0.037 | -0.037 | 0 | 0.3 | 0.15 | 0.38 | 0.58 | 0.24 | 0.48 |
| 5 | 133217659 | rs4077649 | 0.915 | 0.727 | 0.336 | -1.535 | -0.01 | 0.084 | 0.084 | 0.016 | 0.32 | 0.17 | 0.25 | 0.04 | 0.37 | 0.14 |
| 5 | 133219341 | rs11745938 | 0.463 | 0.721 | 0.408 | -1.519 | -0.007 | -0.008 | -0.008 | 0.186 | 0.38 | 0.32 | 0.31 | 0.04 | 0.42 | 0.17 |
| 5 | 133220528 | rs10751463 | -0.088 | 1.114 | 0.507 | -1.467 | 0.006 | -0.152 | -0.152 | 0 | 0.63 | 0.51 | 0.68 | 0.96 | 0.59 | 0.8 |
| 5 | 133221460 | rs7378772 | 1.184 | 1.607 | 1.041 | -0.504 | 0.041 | -0.05 | -0.05 | 0 | 0.9 | 0.7 | 0.89 | 0.97 | 0.9 | 0.9 |
| 5 | 133222833 | rs6890778 | 1.146 | 1.691 | 1.045 | -0.37 | -0.008 | 0.005 | 0.005 | 0.801 | 0.73 | 0.42 | 0.71 | 0.71 | 0.71 | 0.64 |
| 5 | 133223085 | rs11242196 | 1.09 | 1.6 | 1.131 | -0.367 | 0.067 | -0.046 | -0.046 | 0 | 0.89 | 0.57 | 0.88 | 0.97 | 0.9 | 0.9 |
| 5 | 133223945 | rs10055533 | -1.4 | 1.911 | 0.851 | -0.318 | 0.019 | -0.003 | -0.003 | 0 | 0.33 | 0.64 | 0.45 | 0.31 | 0.43 | 0.41 |
| 5 | 133224404 | rs903077 | 1.264 | 1.002 | -0.356 | -0.261 | 0.016 | 0.013 | 0.013 | 0.286 | 0.58 | 0.26 | 0.55 | 0.62 | 0.6 | 0.54 |
| 5 | 133226649 | rs10479058 | -1.501 | 1.424 | -0.031 | -0.156 | -0.001 | -0.006 | -0.006 | 0 | 0.24 | 0.69 | 0.25 | 0.17 | 0.22 | 0.25 |
| 5 | 133227257 | rs2452857 | -1.728 | 1.569 | -0.065 | -0.2 | 0.001 | -0.003 | -0.003 | 0 | 0.27 | 0.69 | 0.29 | 0.16 | 0.27 | 0.28 |
| 5 | 133240927 | rs1618181 | -1.576 | 1.027 | 0.136 | 0.065 | 0 | -0.009 | -0.009 | 0.129 | 0.24 | 0.65 | 0.24 | 0.18 | 0.21 | 0.24 |
| 5 | 133243168 | rs11242198 | -1.064 | 1.371 | -0.189 | 0.3 | 0.025 | -0.002 | -0.002 | 0 | 0.13 | 0.24 | 0.15 | 0.13 | 0.15 | 0.2 |
| 5 | 133246201 | rs6874189 | -0.425 | 1.057 | -0.711 | 0.191 | -0.016 | 0.022 | 0.022 | 0 | 0.42 | 0.32 | 0.45 | 0.3 | 0.49 | 0.34 |
| 5 | 133246615 | rs3936585 | -0.746 | 0.505 | -0.029 | 0.369 | -0.017 | -0.009 | -0.009 | 0 | 0.19 | 0.6 | 0.22 | 0.14 | 0.13 | 0.22 |
| 5 | 133249555 | rs3892476 | -0.728 | 0.274 | -0.195 | 0.537 | -0.006 | 0.003 | 0.003 | 0 | 0.43 | 0.46 | 0.43 | 0.43 | 0.36 | 0.53 |
| 5 | 133251538 | rs6863158 | -0.577 | 0.083 | 0.337 | 0.459 | -0.002 | 0.007 | 0.007 | 0 | 0.37 | 0.34 | 0.43 | 0.43 | 0.34 | 0.5 |
| 5 | 133252333 | rs1961427 | -0.237 | -0.018 | -0.179 | 0.778 | 0.003 | -0.007 | -0.007 | 0 | 0.25 | 0.17 | 0.22 | 0.3 | 0.23 | 0.33 |
| 5 | 133253849 | rs17683751 | -1.242 | 0.537 | -0.977 | 0.287 | -0.033 | 0.013 | 0.013 | 0 | 0.54 | 0.81 | 0.58 | 0.55 | 0.51 | 0.69 |
| 5 | 133255854 | rs13185944 | -0.253 | 0.508 | -1.298 | 0.031 | 0.006 | -0.072 | -0.072 | 0.786 | 0.37 | 0.11 | 0.43 | 0.44 | 0.39 | 0.5 |
| 5 | 133256102 | rs6596163 | -1.003 | 0.377 | -1.411 | 0.324 | -0.025 | -0.002 | -0.002 | 0 | 0.58 | 0.83 | 0.61 | 0.55 | 0.58 | 0.71 |
| 5 | 133260485 | rs6865260 | -0.877 | 0.378 | -2.016 | 0.438 | -0.015 | -0.059 | -0.059 | 0 | 0.45 | 0.88 | 0.59 | 0.55 | 0.55 | 0.48 |
| 5 | 133266198 | rs6887853 | -0.63 | 0.971 | -1.515 | -0.159 | 0.001 | -0.014 | -0.014 | 0.038 | 0.2 | 0.23 | 0.15 | 0.08 | 0.21 | 0.2 |
| 5 | 133268881 | rs2869178 | -0.35 | 1.565 | -1.745 | 0.092 | -0.042 | -0.027 | -0.027 | 0.127 | 0.24 | 0.14 | 0.32 | 0.32 | 0.29 | 0.4 |
| 5 | 133273053 | rs4544869 | -0.367 | 1.552 | -1.619 | -0.321 | -0.035 | 0.074 | 0.074 | 0 | 0.17 | 0.07 | 0.12 | 0.04 | 0.17 | 0.06 |
| 5 | 133287464 | rs10064104 | -1.237 | 1.714 | -0.591 | -0.397 | 0.058 | -0.083 | -0.083 | 1.988 | 0.27 | 0.41 | 0.33 | 0.09 | 0.37 | 0.24 |
| 5 | 133290851 | rs7449072 | -0.363 | 1.838 | -0.953 | -0.358 | -0.072 | 0.075 | 0.075 | 0 | 0.17 | 0.08 | 0.28 | 0.04 | 0.27 | 0.11 |
| 5 | 133295892 | rs3733946 | -0.489 | 1.911 | -0.949 | -0.358 | -0.076 | 0.078 | 0.078 | 0 | 0.18 | 0.07 | 0.29 | 0.04 | 0.29 | 0.12 |
| 5 | 133303063 | rs2112585 | -1.968 | 0.927 | -1.081 | -0.061 | 0.013 | -0.012 | -0.012 | 0 | 0.7 | 0.87 | 0.8 | 0.72 | 0.73 | 0.82 |
| 5 | 133303819 | rs13169435 | -0.493 | 1.438 | -0.455 | -0.46 | 0.028 | -0.061 | -0.061 | 0.031 | 0.28 | 0.41 | 0.38 | 0.1 | 0.4 | 0.29 |
| 5 | 133307106 | rs4279383 | -0.92 | 0.748 | -0.104 | -0.161 | -0.005 | 0.015 | 0.015 | 0 | 0.45 | 0.56 | 0.47 | 0.22 | 0.48 | 0.38 |
| 5 | 133307337 | rs13354887 | 1.263 | 0.793 | -0.094 | -0.055 | -0.007 | 0.01 | 0.01 | 0.03 | 0.06 | 0.06 | 0.06 | 0.13 | 0.04 | 0.03 |
| 5 | 133310029 | rs11952145 | 0.826 | 0.701 | -0.041 | 0.354 | 0.002 | 0.032 | 0.032 | 0.122 | 0.12 | 0.14 | 0.14 | 0.34 | 0.14 | 0.2 |
| 5 | 133310248 | rs10068191 | -1.293 | 0.676 | -0.079 | 0.311 | 0.011 | -0.005 | -0.005 | 0 | 0.54 | 0.58 | 0.61 | 0.55 | 0.61 | 0.57 |
| 5 | 133322024 | rs10074308 | 1.211 | 0.67 | -0.12 | 0.323 | -0.009 | 0.008 | 0.008 | 0.371 | 0.06 | 0.07 | 0.06 | 0.13 | 0.03 | 0.03 |
| 5 | 133324558 | rs6878448 | -1.473 | 0.613 | -0.019 | 0.261 | 0.013 | -0.017 | -0.017 | 0 | 0.57 | 0.66 | 0.64 | 0.55 | 0.63 | 0.58 |
| 5 | 133325781 | rs11242200 | 0.844 | 0.715 | -0.264 | 0.328 | 0.005 | 0.05 | 0.05 | 0.146 | 0.12 | 0.16 | 0.14 | 0.34 | 0.14 | 0.2 |
| 5 | 133328825 | rs6893145 | 1.473 | 0.823 | -0.261 | 0.205 | -0.007 | -0.001 | -0.001 | 0 | 0.1 | 0.1 | 0.08 | 0.03 | 0.13 | 0.21 |
| 5 | 133329931 | rs10078324 | -0.037 | 0.82 | -0.213 | 0.195 | -0.009 | -0.003 | -0.003 | 0.157 | 0.35 | 0.33 | 0.4 | 0.08 | 0.43 | 0.35 |
| 5 | 133330926 | rs10050682 | 1.446 | 0.684 | -0.266 | 0.232 | -0.006 | 0.006 | 0.006 | 0.453 | 0.06 | 0.05 | 0.06 | 0.13 | 0.03 | 0.03 |
| 5 | 133331466 | rs17517610 | 0.974 | 0.827 | -0.17 | 0.132 | -0.029 | 0.051 | 0.051 | 0 | 0.23 | 0.13 | 0.31 | 0.06 | 0.29 | 0.14 |
| 5 | 133346511 | rs10061925 | -1.125 | 0.858 | -0.128 | 0.226 | 0.011 | -0.004 | -0.004 | 0 | 0.54 | 0.58 | 0.63 | 0.55 | 0.65 | 0.59 |
| 5 | 133356534 | rs10079663 | -1.096 | 0.873 | -0.431 | 0.223 | -0.001 | -0.001 | -0.001 | 0 | 0.54 | 0.53 | 0.62 | 0.55 | 0.65 | 0.57 |
| 5 | 133364258 | rs13173424 | -0.083 | 0.164 | -0.515 | 0.416 | -0.017 | -0.007 | -0.007 | 0 | 0.35 | 0.26 | 0.44 | 0.4 | 0.42 | 0.32 |
| 5 | 133368048 | rs11956750 | -0.675 | -0.737 | 0.189 | 0.996 | 0 | 0.096 | 0.096 | 0 | 0.16 | 0.24 | 0.19 | 0.45 | 0.15 | 0.24 |
| 5 | 133370196 | rs247461 | 1.542 | -0.702 | -0.577 | 0.558 | 0.061 | 0.012 | 0.012 | 0 | 0.5 | 0.26 | 0.47 | 0.38 | 0.59 | 0.31 |
| 5 | 133374511 | rs247456 | 0.255 | -0.578 | -0.436 | 0.324 | 0.006 | 0.019 | 0.019 | 0 | 0.81 | 0.89 | 0.89 | 0.87 | 0.84 | 0.74 |
| 5 | 133382456 | rs247459 | 0.82 | -0.933 | -0.209 | 0.57 | -0.018 | -0.002 | -0.002 | 0 | 0.68 | 0.47 | 0.79 | 0.65 | 0.76 | 0.56 |
| 5 | 133384491 | rs13357750 | -0.379 | -1.16 | -0.209 | 0.779 | -0.005 | 0.008 | 0.008 | 0.042 | 0.16 | 0.15 | 0.16 | 0.31 | 0.14 | 0.27 |
| 5 | 133384656 | rs11740710 | 0.026 | -1.128 | -0.416 | 0.798 | 0.002 | -0.051 | -0.051 | 0 | 0.13 | 0.04 | 0.12 | 0.31 | 0.11 | 0.27 |
| 5 | 133391516 | rs247472 | 0.302 | -1.083 | -0.399 | 0.367 | 0.001 | 0.031 | 0.031 | 0 | 0.72 | 0.8 | 0.7 | 0.88 | 0.7 | 0.76 |
| 5 | 133392291 | rs247475 | 0.179 | -1.073 | -0.491 | 0.371 | -0.011 | 0.072 | 0.072 | 0.001 | 0.73 | 0.9 | 0.71 | 0.88 | 0.7 | 0.76 |
| 5 | 133394797 | rs2429080 | 0.778 | -1.271 | 0.173 | 0.351 | -0.013 | -0.05 | -0.05 | 0 | 0.67 | 0.56 | 0.64 | 0.79 | 0.69 | 0.71 |
| 5 | 133404705 | rs11242202 | -0.157 | -1.182 | -0.058 | 0.406 | -0.034 | 0.058 | 0.058 | 0 | 0.77 | 0.95 | 0.73 | 0.88 | 0.72 | 0.76 |
| 5 | 133408988 | rs10463908 | -0.668 | -1.053 | 0.378 | 0.587 | -0.005 | 0.042 | 0.042 | 0.168 | 0.11 | 0.38 | 0.11 | 0.12 | 0.06 | 0.19 |
| 5 | 133410192 | rs10059571 | 0.407 | -1.071 | -0.019 | 0.582 | 0.002 | -0.001 | -0.001 | 0.013 | 0.51 | 0.45 | 0.5 | 0.45 | 0.54 | 0.29 |
| 5 | 133415061 | rs1541907 | 0.775 | -0.803 | -0.352 | 0.676 | -0.013 | -0.002 | -0.002 | 0 | 0.44 | 0.22 | 0.45 | 0.35 | 0.52 | 0.25 |

| | | | | | | | | | | | | | | | | |
|---|---|---|---|---|---|---|---|---|---|---|---|---|---|---|---|---|
| 5 | 133416258 | rs244669 | 0.482 | -0.617 | -0.369 | 0.707 | 0 | 0 | 0 | 0 | 0.47 | 0.47 | 0.49 | 0.44 | 0.54 | 0.3 |
| 5 | 133417805 | rs10900826 | -0.758 | -0.679 | -0.667 | 1.413 | -0.007 | -0.047 | -0.047 | 0 | 0.12 | 0.04 | 0.28 | 0.32 | 0.12 | 0.15 |
| 5 | 133422816 | rs244689 | 0.964 | -1.611 | -0.202 | 1.343 | -0.01 | 0.001 | 0.001 | 0 | 0.76 | 0.72 | 0.73 | 0.63 | 0.84 | 0.7 |
| 5 | 133427110 | rs244684 | 0.527 | -1.787 | 0.511 | 1.071 | -0.025 | -0.015 | -0.015 | 0 | 0.86 | 0.69 | 0.92 | 0.9 | 0.92 | 0.86 |
| 5 | 133428082 | rs244680 | 1.772 | -1.97 | 0.816 | 1.025 | -0.054 | -0.028 | -0.028 | 0 | 0.73 | 0.34 | 0.72 | 0.82 | 0.85 | 0.72 |
| 5 | 133431834 | rs7726414 | -1.24 | -1.941 | 0.984 | 1.033 | -0.058 | -0.015 | -0.015 | 0 | 0.13 | 0.36 | 0.2 | 0.08 | 0.07 | 0.16 |
| 5 | 133434914 | rs244660 | -0.486 | -1.777 | 0.601 | 1.068 | -0.007 | -0.014 | -0.014 | 0 | 0.9 | 1 | 0.94 | 0.9 | 0.92 | 0.85 |
| 5 | 133439876 | rs30500 | 1.12 | -2.162 | 0.504 | 1.316 | -0.035 | 0.081 | 0.081 | 0 | 0.77 | 0.63 | 0.78 | 0.52 | 0.83 | 0.72 |
| 5 | 133441726 | rs30499 | 1.811 | -2.097 | 0.523 | 1.311 | -0.057 | 0.052 | 0.052 | 0 | 0.67 | 0.48 | 0.61 | 0.45 | 0.76 | 0.59 |
| 5 | 133445061 | rs244948 | 0.838 | -1.911 | 0.465 | 0.786 | -0.031 | -0.018 | -0.018 | 1.715 | 0.8 | 0.66 | 0.76 | 0.85 | 0.85 | 0.75 |
| 5 | 133473697 | rs249610 | -1.356 | -2.002 | 0.405 | 1.205 | -0.063 | 0.146 | 0.146 | 0 | 0.21 | 0.46 | 0.41 | 0.46 | 0.17 | 0.21 |
| 5 | 133481467 | rs30489 | 0.643 | -2.041 | 0.297 | 1.212 | -0.011 | -0.003 | -0.003 | 0.432 | 0.07 | 0 | 0.22 | 0.42 | 0.09 | 0.11 |
| 5 | 133486263 | rs4958216 | 0.149 | -1.74 | -0.357 | 0.501 | -0.041 | 0.042 | 0.042 | 0 | 0.15 | 0.04 | 0.1 | 0.06 | 0.17 | 0.24 |
| 5 | 133509294 | rs17167353 | -0.052 | -1.76 | -0.551 | 0.699 | -0.006 | -0.004 | -0.004 | 0.016 | 0.1 | 0.12 | 0.16 | 0.05 | 0.08 | 0.09 |
| 5 | 133514972 | rs3776841 | 0.594 | -1.578 | -0.685 | 0.692 | -0.02 | -0.017 | -0.017 | 0 | 0.1 | 0.01 | 0.06 | 0.1 | 0.08 | 0.15 |
| 5 | 133531449 | rs10051026 | 0.934 | -1.568 | -0.333 | 0.833 | 0.022 | -0.029 | -0.029 | 0 | 0.74 | 0.77 | 0.6 | 0.54 | 0.79 | 0.75 |
| 5 | 133551445 | rs13187105 | 0.893 | -1.573 | -0.318 | 0.838 | 0.021 | -0.029 | -0.029 | 0 | 0.74 | 0.77 | 0.6 | 0.54 | 0.79 | 0.75 |
| 5 | 133552454 | rs7704116 | -0.144 | -1.585 | -0.32 | 0.835 | -0.009 | -0.007 | -0.007 | 0 | 0.1 | 0.16 | 0.17 | 0.06 | 0.09 | 0.1 |
| 5 | 133557876 | rs4431386 | 0.982 | -1.596 | -0.317 | 0.838 | 0.021 | -0.028 | -0.028 | 2.237 | 0.74 | 0.75 | 0.6 | 0.54 | 0.79 | 0.75 |
| 5 | 133563318 | rs2292283 | -0.234 | -1.437 | -0.41 | 0.914 | 0.019 | -0.049 | -0.049 | 0 | 0.85 | 0.94 | 0.77 | 0.59 | 0.88 | 0.84 |
| 5 | 133576837 | rs17167399 | -1.207 | -1.621 | -0.078 | 0.55 | -0.053 | -0.052 | -0.052 | 1.249 | 0.18 | 0.34 | 0.19 | 0.06 | 0.1 | 0.11 |
| 5 | 133591127 | rs734620 | -0.788 | -1.643 | 0.071 | 0.577 | -0.043 | -0.036 | -0.036 | 0 | 0.15 | 0.29 | 0.19 | 0.06 | 0.11 | 0.11 |
| 5 | 133638034 | rs17165388 | 0.671 | -0.816 | -0.446 | 0.27 | -0.044 | 0.004 | 0.004 | 0 | 0.12 | 0.02 | 0.12 | 0.07 | 0.16 | 0.1 |
| 5 | 133665121 | rs10061825 | -0.915 | -1.318 | 0.249 | 0.136 | -0.073 | -0.064 | -0.064 | 0.588 | 0.17 | 0.49 | 0.21 | 0.06 | 0.11 | 0.11 |
| 5 | 133706612 | rs1379247 | -0.365 | -1.345 | 0.425 | 0.123 | -0.037 | -0.03 | -0.03 | 0 | 0.14 | 0.26 | 0.19 | 0.06 | 0.11 | 0.11 |
| 5 | 133706994 | rs17167484 | -0.885 | -1.362 | 0.418 | 0.115 | -0.05 | -0.044 | -0.044 | 2.317 | 0.17 | 0.34 | 0.19 | 0.06 | 0.11 | 0.11 |
| 5 | 133719752 | rs6863127 | -0.957 | -1.422 | 0.422 | 0.086 | -0.053 | -0.046 | -0.046 | 0.654 | 0.17 | 0.36 | 0.2 | 0.06 | 0.11 | 0.11 |
| 5 | 133727303 | rs3777373 | -0.381 | -1.377 | 0.374 | 0.107 | -0.037 | -0.03 | -0.03 | 0.037 | 0.14 | 0.26 | 0.19 | 0.06 | 0.11 | 0.11 |
| 5 | 133730455 | rs4637596 | 0.825 | -1.34 | 0.004 | 0.125 | -0.068 | -0.061 | -0.061 | 0.013 | 0.81 | 0.39 | 0.77 | 0.94 | 0.89 | 0.89 |
| 5 | 133741881 | rs729800 | 1.609 | -1.339 | -0.149 | 0.147 | 0.029 | -0.028 | -0.028 | 0 | 0.6 | 0.27 | 0.47 | 0.49 | 0.68 | 0.55 |
| 5 | 133745398 | rs328043 | 0.768 | -1.385 | -0.277 | 0.336 | -0.005 | -0.092 | -0.092 | 0 | 0.13 | 0 | 0.24 | 0.4 | 0.11 | 0.16 |
| 5 | 133772648 | rs6872005 | 0.712 | -2.087 | -0.343 | -0.283 | -0.123 | -0.108 | -0.108 | 0 | 0.74 | 0.48 | 0.76 | 0.95 | 0.87 | 0.87 |
| 5 | 133774168 | rs328040 | 0.672 | -1.82 | -0.369 | 0.035 | -0.061 | 0.003 | 0.003 | 0 | 0.64 | 0.46 | 0.54 | 0.61 | 0.78 | 0.66 |
| 5 | 133777231 | rs12514418 | 0.524 | -1.135 | -0.65 | -0.467 | 0.019 | -0.012 | -0.012 | 0 | 0.39 | 0.46 | 0.51 | 0.6 | 0.5 | 0.54 |
| 5 | 133781587 | rs10491288 | 0.216 | -1.603 | -0.578 | -0.455 | -0.001 | -0.001 | -0.001 | 0 | 0.25 | 0.28 | 0.4 | 0.38 | 0.19 | 0.36 |
| 5 | 133781618 | rs328834 | 0.362 | -1.882 | -0.67 | -0.637 | -0.001 | 0.021 | 0.021 | 0 | 0.2 | 0.16 | 0.11 | 0.11 | 0.17 | 0.18 |
| 5 | 133782153 | rs10056003 | 0.635 | -1.774 | -0.403 | -0.207 | -0.04 | -0.002 | -0.002 | 0 | 0.12 | 0.27 | 0.19 | 0.08 | 0.08 | 0.11 |
| 5 | 133782963 | rs328833 | -0.871 | -1.478 | 0.112 | -0.38 | 0.036 | -0.005 | -0.005 | 0 | 0.44 | 0.74 | 0.52 | 0.47 | 0.33 | 0.49 |
| 5 | 133784798 | rs12516971 | 0.546 | -1.439 | -0.002 | -0.361 | 0.005 | -0.008 | -0.008 | 0 | 0.28 | 0.28 | 0.43 | 0.42 | 0.23 | 0.37 |
| 5 | 133784852 | rs328831 | 0.386 | -1.327 | -0.038 | -0.798 | 0.006 | -0.014 | -0.014 | 0 | 0.09 | 0.15 | 0.07 | 0.05 | 0.09 | 0.12 |
| 5 | 133788713 | rs328828 | 0.406 | -1.312 | -0.03 | -0.825 | 0.007 | -0.018 | -0.018 | 0 | 0.09 | 0.15 | 0.06 | 0.05 | 0.09 | 0.11 |
| 5 | 133796095 | rs739953 | -0.074 | -1.258 | -0.297 | -0.508 | -0.002 | 0.002 | 0.002 | 0 | 0.37 | 0.44 | 0.5 | 0.52 | 0.33 | 0.47 |
| 5 | 133799374 | rs7706779 | -0.536 | -1.15 | -0.386 | -0.473 | -0.012 | 0.003 | 0.003 | 0 | 0.43 | 0.55 | 0.53 | 0.54 | 0.34 | 0.48 |
| 5 | 133809801 | rs258514 | -0.573 | -0.476 | -0.683 | -0.373 | 0.05 | -0.016 | -0.016 | 0.001 | 0.74 | 0.59 | 0.76 | 0.78 | 0.61 | 0.67 |
| 5 | 133812710 | rs1476095 | 0.24 | -0.47 | -0.911 | -0.548 | 0.002 | 0.009 | 0.009 | 0 | 0.55 | 0.56 | 0.59 | 0.74 | 0.47 | 0.6 |
| 5 | 133817247 | rs258522 | 0.775 | -0.149 | -0.69 | -0.398 | 0.003 | -0.002 | -0.002 | 0.062 | 0.21 | 0.17 | 0.18 | 0.04 | 0.14 | 0.08 |
| 5 | 133819314 | rs7719606 | 0.666 | -0.461 | -0.647 | 0.011 | -0.012 | 0.025 | 0.025 | 0 | 0.29 | 0.33 | 0.46 | 0.5 | 0.25 | 0.35 |
| 5 | 133823078 | rs7702415 | 0.328 | -0.176 | -0.799 | 0.21 | -0.002 | 0 | 0 | 0 | 0.22 | 0.23 | 0.13 | 0.24 | 0.22 | 0.24 |
| 5 | 133826527 | rs6889807 | 0.299 | -0.287 | -0.864 | 0.061 | 0.017 | 0.033 | 0.033 | 0 | 0.16 | 0.07 | 0.14 | 0.05 | 0.1 | 0.06 |
| 5 | 133839534 | rs12519914 | 0.257 | -0.476 | -0.177 | 0.816 | -0.028 | 0.13 | 0.13 | 0 | 0.21 | 0.41 | 0.2 | 0.51 | 0.15 | 0.36 |
| 5 | 133841675 | rs17644077 | -0.075 | -0.098 | -0.557 | 0.354 | -0.003 | -0.02 | -0.02 | 0.025 | 0.24 | 0.19 | 0.35 | 0.35 | 0.25 | 0.27 |
| 5 | 133845380 | rs4958244 | -1.538 | -0.068 | -0.391 | -0.235 | -0.013 | 0.02 | 0.02 | 0 | 0.69 | 0.76 | 0.74 | 0.92 | 0.59 | 0.74 |
| 5 | 133848760 | rs11242219 | -1.533 | -0.087 | -0.4 | -0.233 | -0.013 | 0.019 | 0.019 | 0 | 0.69 | 0.76 | 0.74 | 0.92 | 0.59 | 0.74 |
| 5 | 133851599 | rs6891492 | 0.887 | -0.181 | -0.169 | -0.293 | 0.001 | -0.019 | -0.019 | 2.026 | 0.23 | 0.16 | 0.17 | 0.04 | 0.18 | 0.13 |
| 5 | 133853332 | rs10040679 | 1.055 | -0.16 | -0.273 | -0.15 | 0.005 | 0.013 | 0.013 | 0.015 | 0.1 | 0.11 | 0.12 | 0.15 | 0.14 | 0.13 |
| 5 | 133858719 | rs329115 | -0.166 | -0.249 | -0.189 | -0.178 | 0.011 | 0.022 | 0.022 | 0 | 0.36 | 0.27 | 0.41 | 0.23 | 0.29 | 0.25 |
| 5 | 133860101 | rs329117 | -1.732 | -0.12 | -0.13 | -0.196 | -0.011 | 0.016 | 0.016 | 0 | 0.69 | 0.75 | 0.74 | 0.92 | 0.59 | 0.74 |
| 5 | 133861716 | rs329119 | 0.717 | -0.02 | -0.364 | 0.034 | -0.004 | -0.044 | -0.044 | 0 | 0.23 | 0.23 | 0.17 | 0.04 | 0.18 | 0.13 |
| 5 | 133861756 | rs329120 | -0.666 | 0.048 | -0.435 | 0.229 | 0.001 | 0.005 | 0.005 | 0 | 0.46 | 0.42 | 0.53 | 0.37 | 0.43 | 0.39 |
| 5 | 133869935 | rs11949657 | 0.216 | -0.074 | -0.741 | 0.8 | 0.003 | -0.028 | -0.028 | 0.271 | 0.11 | 0.04 | 0.32 | 0.26 | 0.12 | 0.12 |
| 5 | 133871101 | rs329125 | 0.595 | 0.004 | -0.449 | 0.762 | 0 | 0.004 | 0.004 | 0 | 0.22 | 0.2 | 0.22 | 0.56 | 0.18 | 0.35 |
| 5 | 133881485 | rs2569340 | -0.648 | -0.178 | -0.422 | 0.037 | -0.001 | -0.023 | -0.023 | 0 | 0.78 | 0.8 | 0.82 | 0.96 | 0.84 | 0.89 |
| 5 | 133886166 | rs11242224 | 0.36 | 0.158 | -0.563 | 0.431 | -0.011 | -0.019 | -0.019 | 0 | 0.1 | 0.05 | 0.2 | 0.21 | 0.14 | 0.21 |
| 5 | 133887780 | rs12163993 | 0.324 | 0.233 | -0.598 | 0.444 | -0.015 | -0.024 | -0.024 | 0 | 0.1 | 0.05 | 0.21 | 0.21 | 0.16 | 0.22 |
| 5 | 133891282 | rs2084007 | 0.525 | 0.391 | -0.23 | 0.237 | -0.004 | 0.002 | 0.002 | 0 | 0.38 | 0.35 | 0.45 | 0.33 | 0.51 | 0.37 |
| 5 | 133894781 | rs12054866 | 0.71 | 0.438 | -0.538 | 0.286 | -0.012 | -0.02 | -0.02 | 0 | 0.09 | 0.04 | 0.2 | 0.25 | 0.14 | 0.19 |
| 5 | 133897829 | rs329304 | 0.734 | 0.069 | -0.407 | 0.091 | -0.049 | 0.011 | 0.011 | 0 | 0.37 | 0.2 | 0.44 | 0.33 | 0.5 | 0.37 |
| 5 | 133898136 | rs10056247 | 0.393 | -0.098 | -0.065 | 0.12 | -0.001 | -0.045 | -0.045 | 0.081 | 0.31 | 0.36 | 0.3 | 0.1 | 0.28 | 0.24 |
| 5 | 133900111 | rs329311 | -0.914 | -0.119 | -0.284 | 0.827 | -0.016 | 0.015 | 0.015 | 0 | 0.89 | 0.88 | 0.98 | 0.78 | 0.98 | 0.74 |

| | | | | | | | | | | | | | | | | |
|---|---|---|---|---|---|---|---|---|---|---|---|---|---|---|---|---|
| 5 | 133900498 | rs329312 | -0.011 | -0.162 | -0.309 | 0.814 | -0.001 | 0 | 0 | 0 | 0.69 | 0.69 | 0.75 | 0.44 | 0.79 | 0.61 |
| 5 | 133900576 | rs4958191 | 0.491 | -0.061 | -0.308 | 0.576 | -0.003 | -0.064 | -0.064 | 0 | 0.32 | 0.43 | 0.31 | 0.12 | 0.29 | 0.24 |
| 5 | 133900974 | rs17167621 | 0.84 | -0.268 | -0.417 | 1.091 | 0.001 | -0.032 | -0.032 | 0 | 0.19 | 0.11 | 0.21 | 0.34 | 0.19 | 0.13 |
| 5 | 133903814 | rs329317 | 0.418 | -0.043 | -0.363 | 0.428 | 0.003 | 0.016 | 0.016 | 0 | 0.6 | 0.68 | 0.54 | 0.19 | 0.63 | 0.41 |
| 5 | 133906527 | rs11953143 | 0.78 | 0.38 | -0.502 | 0.644 | -0.012 | -0.004 | -0.004 | 0.043 | 0.06 | 0.03 | 0.1 | 0.05 | 0.09 | 0.07 |
| 5 | 133906609 | rs329319 | 0.159 | 0.281 | -0.393 | 0.38 | 0.004 | 0.021 | 0.021 | 0 | 0.47 | 0.47 | 0.57 | 0.7 | 0.57 | 0.5 |
| 5 | 133908361 | rs329320 | 0.583 | 0.121 | 0.032 | 0.631 | 0.005 | 0.066 | 0.066 | 0 | 0.81 | 0.6 | 0.87 | 0.71 | 0.86 | 0.57 |
| 5 | 133913553 | rs2589400 | 1.481 | -0.355 | -0.218 | 0.306 | 0.015 | -0.017 | -0.017 | 0 | 0.62 | 0.24 | 0.65 | 0.42 | 0.69 | 0.39 |
| 5 | 133930528 | rs17167632 | 0.021 | -0.331 | 0.021 | 0.615 | 0 | 0.044 | 0.044 | 0 | 0.09 | 0.09 | 0.19 | 0.28 | 0.14 | 0.18 |
| 5 | 133942492 | rs7728741 | -0.689 | -0.39 | 0.376 | 0.547 | -0.013 | 0.058 | 0.058 | 0.653 | 0.16 | 0.19 | 0.22 | 0.34 | 0.15 | 0.19 |
| 5 | 133943937 | rs2305049 | -0.689 | -0.477 | 0.315 | 0.483 | -0.013 | 0.058 | 0.058 | 0.653 | 0.16 | 0.19 | 0.21 | 0.34 | 0.15 | 0.19 |
| 5 | 133973889 | rs4958192 | 1.004 | -0.344 | 0.63 | 0.288 | -0.016 | 0.049 | 0.049 | 2.676 | 0.74 | 0.4 | 0.84 | 0.61 | 0.85 | 0.6 |
| 5 | 133976554 | rs11747528 | 1.746 | -0.265 | -0.07 | -0.204 | -0.109 | 0.113 | 0.113 | 0 | 0.26 | 0.02 | 0.25 | 0.08 | 0.31 | 0.14 |
| 5 | 134007166 | rs4958263 | 0.606 | -0.23 | 0.162 | -0.136 | -0.009 | 0.077 | 0.077 | 1.008 | 0.76 | 0.5 | 0.85 | 0.61 | 0.85 | 0.61 |
| 5 | 134043468 | rs246344 | -2.165 | -0.231 | -0.277 | -0.47 | -0.037 | 0.107 | 0.107 | 0.008 | 0.82 | 0.99 | 0.86 | 0.99 | 0.81 | 0.87 |
| 5 | 134051404 | rs1820677 | 0.739 | -0.207 | -0.048 | -0.325 | 0 | 0.016 | 0.016 | 0.03 | 0.13 | 0.13 | 0.09 | 0.32 | 0.13 | 0.37 |
| 5 | 134076812 | rs12657663 | 1.029 | -0.149 | -0.239 | -0.274 | -0.008 | -0.031 | -0.031 | 0 | 0.09 | 0.01 | 0.18 | 0.28 | 0.13 | 0.18 |
| 5 | 134090360 | rs6880257 | -1.367 | -0.225 | -0.032 | -0.334 | -0.005 | 0.003 | 0.003 | 1.108 | 0.32 | 0.51 | 0.33 | 0.68 | 0.28 | 0.57 |
| 5 | 134093932 | rs7718958 | 0.563 | 0.132 | -0.018 | -0.361 | 0.016 | 0.04 | 0.04 | 0.042 | 0.13 | 0.19 | 0.11 | 0.33 | 0.16 | 0.37 |
| 5 | 134096294 | rs246347 | -2.268 | -0.116 | -0.089 | -0.446 | -0.019 | 0.093 | 0.093 | 0 | 0.82 | 0.98 | 0.86 | 0.99 | 0.82 | 0.88 |
| 5 | 134097290 | rs10067976 | -1.937 | 0.377 | 0.825 | 0.254 | 0.013 | 0.006 | 0.006 | 1.529 | 0.36 | 0.75 | 0.35 | 0.68 | 0.29 | 0.57 |
| 5 | 134111769 | rs254709 | 0.391 | 0.33 | 0.741 | 0.183 | -0.037 | -0.012 | -0.012 | 0 | 0.1 | 0.23 | 0.03 | 0.07 | 0.01 | 0.03 |
| 5 | 134130064 | rs435397 | 1.899 | 0.387 | 0.746 | 0.205 | -0.017 | -0.007 | -0.007 | 0 | 0.05 | 0.09 | 0.02 | 0.05 | 0.01 | 0.02 |
| 5 | 134132269 | rs403005 | -1.216 | 0.386 | 0.693 | 0.21 | 0.002 | 0.028 | 0.028 | 0.817 | 0.6 | 0.7 | 0.57 | 0.79 | 0.58 | 0.71 |
| 5 | 134136333 | rs2108665 | 0.325 | 0.508 | 0.656 | 0.64 | -0.007 | -0.02 | -0.02 | 0.019 | 0.09 | 0.04 | 0.18 | 0.31 | 0.14 | 0.18 |
| 5 | 134144277 | rs254712 | 1.321 | 0.479 | 0.717 | 0.635 | -0.019 | -0.008 | -0.008 | 0 | 0.06 | 0.1 | 0.02 | 0.05 | 0.01 | 0.02 |
| 5 | 134161929 | rs10463924 | 0.808 | 0.728 | 0.174 | 0.756 | 0.009 | 0.003 | 0.003 | 0 | 0.07 | 0.01 | 0.08 | 0.29 | 0.06 | 0.14 |
| 5 | 134175339 | rs11242228 | 0.239 | 0.808 | 0.065 | 0.745 | 0.014 | 0.054 | 0.054 | 0.016 | 0.12 | 0.19 | 0.1 | 0.33 | 0.13 | 0.36 |
| 5 | 134190225 | rs7717860 | 0.239 | 0.808 | -0.023 | 0.745 | 0.008 | 0.038 | 0.038 | 0.016 | 0.12 | 0.16 | 0.1 | 0.33 | 0.13 | 0.36 |
| 5 | 134199105 | rs11950533 | 0.586 | 0.752 | -0.108 | 0.694 | -0.007 | -0.019 | -0.019 | 0 | 0.09 | 0.04 | 0.18 | 0.31 | 0.14 | 0.18 |
| 5 | 134215127 | rs2548966 | -0.131 | 0.871 | -0.096 | 0.371 | -0.021 | -0.022 | -0.022 | 0.799 | 0.64 | 0.77 | 0.61 | 0.54 | 0.61 | 0.53 |
| 5 | 134222164 | rs319602 | -0.202 | 0.848 | -0.106 | 0.36 | -0.021 | -0.022 | -0.022 | 0.096 | 0.64 | 0.77 | 0.61 | 0.54 | 0.61 | 0.56 |
| 5 | 134223593 | rs3733897 | 0.959 | 0.758 | -0.233 | 0.344 | -0.021 | -0.066 | -0.066 | 0.034 | 0.13 | 0.04 | 0.14 | 0.27 | 0.15 | 0.21 |
| 5 | 134235613 | rs319600 | 1.312 | 0.669 | 0.11 | 0.278 | 0.002 | 0.064 | 0.064 | 0.076 | 0.41 | 0.23 | 0.42 | 0.21 | 0.45 | 0.34 |
| 5 | 134239350 | rs319599 | -1.747 | 0.916 | -0.281 | 0.398 | -0.069 | 0.041 | 0.041 | 0 | 0.8 | 0.98 | 0.78 | 0.9 | 0.75 | 0.93 |
| 5 | 134240235 | rs319598 | -0.444 | 1.117 | -0.153 | 0.518 | -0.02 | -0.017 | -0.017 | 0.46 | 0.64 | 0.77 | 0.6 | 0.55 | 0.61 | 0.56 |
| 5 | 134244534 | rs6874344 | 1.191 | 1.173 | 0.031 | 0.507 | 0.002 | 0 | 0 | 0 | 0.06 | 0.23 | 0.03 | 0.02 | 0.01 | 0.01 |
| 5 | 134244946 | rs319596 | 0.913 | 1.163 | 0.036 | 0.504 | -0.003 | -0.015 | -0.015 | 0 | 0.44 | 0.47 | 0.43 | 0.2 | 0.45 | 0.32 |
| 5 | 134250146 | rs319594 | -1.269 | 1.27 | -0.43 | 0.57 | -0.01 | 0.049 | 0.049 | 0.851 | 0.74 | 0.91 | 0.76 | 0.85 | 0.74 | 0.89 |
| 5 | 134253500 | rs4345357 | 0.875 | 1.138 | -0.57 | 0.486 | 0.002 | 0 | 0 | 0 | 0.06 | 0.23 | 0.03 | 0.02 | 0.01 | 0.01 |
| 5 | 134257519 | rs9285933 | 0.733 | 1.174 | -0.953 | 0.504 | -0.03 | -0.074 | -0.074 | 0 | 0.13 | 0.03 | 0.14 | 0.27 | 0.15 | 0.2 |
| 5 | 134258183 | rs17167693 | 0.764 | 1.03 | -0.273 | -0.044 | -0.008 | 0.017 | 0.017 | 0 | 0.06 | 0.19 | 0.03 | 0.05 | 0.01 | 0.01 |
| 5 | 134272525 | rs6862828 | -0.243 | 0.568 | -0.861 | -0.205 | 0.003 | 0.034 | 0.034 | 0 | 0.59 | 0.7 | 0.58 | 0.79 | 0.55 | 0.67 |
| 5 | 134275189 | rs6889478 | 0.36 | 1.086 | -0.795 | 1.083 | -0.009 | -0.01 | -0.01 | 0 | 0.05 | 0.11 | 0.02 | 0.02 | 0.01 | 0.01 |
| 5 | 134305197 | rs12515798 | -0.504 | 1.531 | -0.981 | 1.254 | -0.026 | -0.035 | -0.035 | 0 | 0.1 | 0.03 | 0.21 | 0.19 | 0.18 | 0.17 |
| 5 | 134331921 | rs299364 | -0.43 | 1.215 | -0.906 | 1.042 | -0.008 | -0.043 | -0.043 | 0 | 0.32 | 0.15 | 0.39 | 0.64 | 0.32 | 0.59 |
| 5 | 134343799 | rs177252 | -0.803 | 0.684 | 0.297 | 1.05 | -0.001 | 0.005 | 0.005 | 2.589 | 0.34 | 0.64 | 0.41 | 0.56 | 0.32 | 0.54 |
| 5 | 134348503 | rs12516279 | -0.585 | 1.685 | 1.232 | 1.056 | 0.057 | 0.043 | 0.043 | 0 | 0.05 | 0.11 | 0.2 | 0.12 | 0.1 | 0.11 |
| 5 | 134348564 | rs660273 | -0.053 | 1.665 | 1.242 | 1.05 | -0.01 | -0.031 | -0.031 | 0 | 0.28 | 0.17 | 0.36 | 0.52 | 0.31 | 0.54 |
| 5 | 134352534 | rs28330 | -0.565 | 1.615 | 1.218 | 0.997 | -0.004 | -0.003 | -0.003 | 0 | 0.33 | 0.26 | 0.56 | 0.64 | 0.41 | 0.62 |
| 5 | 134361020 | rs31210 | 0.221 | 1.735 | 0.952 | 1.037 | -0.016 | -0.022 | -0.022 | 0.19 | 0.23 | 0.11 | 0.3 | 0.33 | 0.27 | 0.42 |
| 5 | 134364996 | rs1131611 | -0.978 | 1.75 | 1.375 | 0.698 | 0.162 | -0.015 | -0.015 | 0 | 0.08 | 0.46 | 0.21 | 0 | 0.11 | 0.05 |
| 5 | 134368169 | rs6596189 | -0.978 | 1.75 | 1.375 | 0.698 | 0.162 | -0.015 | -0.015 | 0.038 | 0.08 | 0.46 | 0.21 | 0 | 0.11 | 0.05 |
| 5 | 134372685 | rs31198 | 0.003 | 1.563 | 0.282 | 0.963 | -0.004 | -0.043 | -0.043 | 0.518 | 0.26 | 0.15 | 0.33 | 0.48 | 0.27 | 0.46 |
| 5 | 134379311 | rs39881 | -0.743 | 0.704 | -0.553 | 0.053 | 0.027 | 0.079 | 0.079 | 0 | 0.69 | 0.86 | 0.79 | 0.99 | 0.74 | 0.91 |
| 5 | 134380531 | rs254551 | -0.304 | 1.449 | -0.339 | 0.82 | -0.027 | -0.029 | -0.029 | 0.963 | 0.34 | 0.28 | 0.54 | 0.51 | 0.45 | 0.55 |
| 5 | 134385234 | rs713563 | -0.964 | 0.94 | -0.105 | 0.79 | 0.001 | -0.004 | -0.004 | 0 | 0.21 | 0.24 | 0.29 | 0.77 | 0.22 | 0.45 |
| 5 | 134390581 | rs4976242 | -0.604 | 1.021 | -0.327 | 1.587 | -0.043 | -0.072 | -0.072 | 0 | 0.12 | 0.02 | 0.22 | 0.5 | 0.17 | 0.23 |
| 5 | 134404390 | rs10065996 | -1.189 | 0.674 | 0.021 | 1.376 | -0.008 | 0.223 | 0.223 | 0 | 0.11 | 0.23 | 0.15 | 0.49 | 0.11 | 0.31 |
| 5 | 134406317 | rs11242232 | 0.249 | 1.262 | -0.715 | -0.174 | -0.029 | 0.055 | 0.055 | 0 | 0.26 | 0.14 | 0.24 | 0.19 | 0.32 | 0.38 |
| 5 | 134431163 | rs6596197 | -0.421 | 0.621 | -0.439 | -0.428 | -0.007 | -0.038 | -0.038 | 0 | 0.3 | 0.43 | 0.37 | 0.75 | 0.27 | 0.37 |
| 5 | 134441457 | rs254562 | 0.579 | 0.936 | -1.324 | -0.827 | -0.066 | 0.056 | 0.056 | 0 | 0.31 | 0.07 | 0.36 | 0.24 | 0.42 | 0.44 |
| 5 | 134444120 | rs495406 | 0.203 | 0.479 | -1.179 | -0.944 | 0.006 | 0.008 | 0.008 | 0 | 0.44 | 0.55 | 0.37 | 0.63 | 0.39 | 0.35 |
| 5 | 134445988 | rs6891074 | 0.701 | 0.647 | -2.025 | -0.386 | -0.032 | -0.106 | -0.106 | 0.006 | 0.19 | 0.04 | 0.16 | 0.52 | 0.19 | 0.19 |
| 5 | 134452308 | rs779848 | -0.544 | 0.849 | -1.605 | -1.043 | -0.012 | 0.009 | 0.009 | 0 | 0.82 | 0.88 | 0.77 | 0.88 | 0.82 | 0.78 |
| 5 | 134456005 | rs540917 | -0.23 | 0.526 | -1.376 | -1.006 | -0.015 | 0.061 | 0.061 | 0 | 0.55 | 0.75 | 0.44 | 0.64 | 0.48 | 0.42 |
| 5 | 134467700 | rs477687 | 0.247 | 0.582 | -0.525 | -1.274 | 0 | -0.06 | -0.06 | 0 | 0.19 | 0.25 | 0.12 | 0 | 0.16 | 0.1 |
| 5 | 134467751 | rs639933 | -0.931 | 0.641 | -1.326 | -1.554 | -0.068 | 0.003 | 0.003 | 0 | 0.68 | 0.94 | 0.59 | 0.65 | 0.63 | 0.5 |
| 5 | 134470030 | rs1559050 | 0.365 | 0.426 | -0.773 | -1.627 | -0.007 | -0.005 | -0.005 | 0 | 0.23 | 0.18 | 0.31 | 0.64 | 0.29 | 0.3 |

| | | | | | | | | | | | | | | | | |
|---|---|---|---|---|---|---|---|---|---|---|---|---|---|---|---|---|
| 5 | 134472374 | rs641455 | -1.059 | 0.353 | -0.08 | -1.872 | 0.004 | 0.04 | 0.04 | 0 | 0.52 | 0.66 | 0.49 | 0.7 | 0.53 | 0.47 |
| 5 | 134475494 | rs11745068 | 0.925 | 0.704 | -0.1 | -1.194 | -0.054 | -0.068 | -0.068 | 0.841 | 0.11 | 0 | 0.14 | 0.56 | 0.18 | 0.23 |
| 5 | 134485285 | rs566750 | -1.004 | 0.472 | 0.6 | -1.579 | 0.005 | -0.016 | -0.016 | 0 | 0.33 | 0.58 | 0.36 | 0.69 | 0.34 | 0.36 |
| 5 | 134493323 | rs17660170 | 0.002 | 0.438 | 0.48 | -1.395 | -0.004 | -0.003 | -0.003 | 0 | 0.06 | 0.04 | 0.07 | 0 | 0.08 | 0.07 |
| 5 | 134509987 | rs1122171 | -0.827 | 0.211 | 0.798 | -1.648 | -0.002 | -0.014 | -0.014 | 0.75 | 0.55 | 0.49 | 0.48 | 0.69 | 0.57 | 0.53 |
| 5 | 134510303 | rs9327705 | 0.533 | 0.257 | 0.916 | -1.927 | 0 | -0.022 | -0.022 | 0 | 0.24 | 0.18 | 0.13 | 0 | 0.22 | 0.11 |
| 5 | 134512608 | rs9327706 | -1.069 | 0.239 | 0.965 | -1.932 | -0.002 | -0.008 | -0.008 | 0 | 0.54 | 0.5 | 0.48 | 0.7 | 0.58 | 0.52 |
| 5 | 134514658 | rs6869259 | -0.807 | -0.818 | 1.797 | -1.782 | 0 | -0.003 | -0.003 | 0.847 | 0.35 | 0.37 | 0.38 | 0.66 | 0.36 | 0.43 |
| 5 | 134526551 | rs2162709 | -0.453 | -1.083 | 1.754 | -1.796 | 0.01 | -0.017 | -0.017 | 0 | 0.3 | 0.48 | 0.35 | 0.67 | 0.32 | 0.37 |
| 5 | 134530116 | rs1346441 | -1.261 | -1.174 | 1.307 | -1.823 | 0.021 | 0.079 | 0.079 | 0 | 0.41 | 0.82 | 0.42 | 0.68 | 0.38 | 0.44 |
| 5 | 134543927 | rs10515467 | -0.31 | -0.919 | 0.97 | -1.731 | 0.004 | -0.004 | -0.004 | 0 | 0.28 | 0.33 | 0.34 | 0.68 | 0.31 | 0.37 |
| 5 | 134550777 | rs683756 | -1.316 | -1.194 | 1.294 | -1.126 | 0.001 | -0.021 | -0.021 | 0.338 | 0.39 | 0.58 | 0.45 | 0.69 | 0.43 | 0.46 |
| 5 | 134555620 | rs2059779 | -1.131 | -1.627 | 1.225 | -1.032 | -0.033 | -0.021 | -0.021 | 0 | 0.26 | 0.39 | 0.26 | 0.12 | 0.16 | 0.13 |
| 5 | 134556555 | rs11950409 | -0.841 | -1.349 | 0.486 | -0.919 | 0.041 | -0.038 | -0.038 | 0 | 0.09 | 0.38 | 0.09 | 0 | 0.1 | 0.06 |
| 5 | 134557037 | rs11951162 | -0.308 | -1.551 | 0.491 | -0.839 | 0.031 | -0.025 | -0.025 | 0 | 0.08 | 0.37 | 0.08 | 0 | 0.07 | 0.02 |
| 5 | 134558028 | rs12514014 | -0.758 | -1.318 | -0.566 | -0.39 | -0.005 | 0 | 0 | 0 | 0.18 | 0.21 | 0.2 | 0.12 | 0.09 | 0.11 |
| 5 | 134563157 | rs634308 | 0.346 | -1.187 | -1.002 | -0.531 | 0.001 | 0.001 | 0.001 | 0.156 | 0.62 | 0.63 | 0.66 | 0.68 | 0.68 | 0.48 |
| 5 | 134563752 | rs17717975 | 0.081 | -0.85 | -0.867 | -0.555 | 0.008 | 0.027 | 0.027 | 0 | 0.11 | 0.06 | 0.06 | 0.01 | 0.07 | 0.06 |
| 5 | 134563817 | rs631607 | 0.45 | -1.336 | -0.569 | -0.471 | 0.038 | 0.05 | 0.05 | 0 | 0.55 | 0.42 | 0.61 | 0.81 | 0.71 | 0.56 |
| 5 | 134564405 | rs688622 | 0.119 | -1.098 | -1.19 | 0.017 | 0.044 | 0.037 | 0.037 | 0 | 0.68 | 0.8 | 0.68 | 0.79 | 0.8 | 0.61 |
| 5 | 134569607 | rs6880486 | -1.103 | -0.197 | -0.702 | 0.471 | -0.001 | 0.017 | 0.017 | 0 | 0.05 | 0.04 | 0.12 | 0.09 | 0.07 | 0.15 |
| 5 | 134571150 | rs10044699 | -0.771 | -0.365 | -0.178 | 0.027 | 0.004 | -0.048 | -0.048 | 0 | 0.08 | 0.2 | 0.04 | 0 | 0.07 | 0.01 |
| 5 | 134572460 | rs1025262 | 0.194 | 0.246 | -0.159 | 0.274 | 0.006 | 0.006 | 0.006 | 0 | 0.68 | 0.62 | 0.64 | 0.66 | 0.61 | 0.6 |
| 5 | 134572661 | rs606995 | -0.055 | 0.53 | -0.166 | 0.271 | -0.059 | -0.017 | -0.017 | 0 | 0.8 | 0.96 | 0.82 | 0.84 | 0.74 | 0.77 |
| 5 | 134572865 | rs1016344 | 0.942 | -0.576 | -0.115 | -0.496 | 0.006 | 0.013 | 0.013 | 0 | 0.57 | 0.39 | 0.63 | 0.76 | 0.61 | 0.69 |
| 5 | 134573007 | rs1346440 | 1.166 | -0.614 | -0.736 | -0.813 | -0.069 | -0.037 | -0.037 | 3.887 | 0.36 | 0.14 | 0.38 | 0.68 | 0.44 | 0.51 |
| 5 | 134573992 | rs2019973 | 0.96 | -0.376 | -0.357 | -0.946 | -0.014 | 0.035 | 0.035 | 0 | 0.42 | 0.29 | 0.5 | 0.76 | 0.53 | 0.55 |
| 5 | 134574232 | rs2114961 | -0.427 | -0.474 | -0.372 | -1.028 | 0.042 | 0.049 | 0.049 | 0 | 0.4 | 0.69 | 0.42 | 0.15 | 0.28 | 0.25 |
| 5 | 134574406 | rs890731 | 1.275 | -0.437 | -0.264 | -1.04 | -0.058 | -0.022 | -0.022 | 0.075 | 0.38 | 0.19 | 0.39 | 0.69 | 0.45 | 0.52 |
| 5 | 134575601 | rs919233 | -0.803 | -0.69 | -0.085 | -0.949 | -0.002 | -0.001 | -0.001 | 0 | 0.81 | 0.81 | 0.84 | 0.84 | 0.81 | 0.8 |
| 5 | 134577757 | rs6896456 | 0.261 | -0.516 | 0.389 | -0.725 | -0.012 | 0.034 | 0.034 | 0 | 0.18 | 0.46 | 0.21 | 0.15 | 0.13 | 0.15 |
| 5 | 134579744 | rs1019736 | 0.008 | -0.713 | -0.123 | -0.877 | 0.028 | 0.06 | 0.06 | 0 | 0.58 | 0.71 | 0.64 | 0.85 | 0.65 | 0.69 |
| 5 | 134586211 | rs17168022 | -0.495 | -0.365 | -0.087 | -0.652 | -0.006 | 0.01 | 0.01 | 0 | 0.09 | 0.11 | 0.06 | 0.05 | 0.06 | 0.1 |
| 5 | 134588881 | rs1834902 | -0.656 | -0.289 | -0.15 | -0.43 | -0.018 | 0.011 | 0.011 | 0 | 0.63 | 0.87 | 0.5 | 0.67 | 0.63 | 0.61 |
| 5 | 134589155 | rs1834903 | -0.485 | -0.294 | -0.185 | -0.453 | -0.028 | 0.025 | 0.025 | 0 | 0.59 | 0.86 | 0.47 | 0.67 | 0.6 | 0.59 |
| 5 | 134589923 | rs10038188 | 0.02 | 0.177 | 0.41 | -0.059 | 0.005 | 0.003 | 0.003 | 3.147 | 0.05 | 0.34 | 0.03 | 0 | 0 | 0 |
| 5 | 134590912 | rs11242237 | 0.352 | 0.108 | -0.241 | -0.106 | -0.03 | -0.025 | -0.025 | 0.873 | 0.4 | 0.23 | 0.35 | 0.61 | 0.51 | 0.51 |
| 5 | 134597267 | rs609385 | -0.076 | 0.419 | -0.81 | -0.171 | -0.01 | 0.03 | 0.03 | 0 | 0.45 | 0.72 | 0.36 | 0.66 | 0.46 | 0.53 |
| 5 | 134599783 | rs6891343 | -0.571 | 0.494 | -1.267 | -0.294 | 0.002 | 0.048 | 0.048 | 0 | 0.34 | 0.25 | 0.21 | 0.2 | 0.3 | 0.36 |
| 5 | 134599977 | rs652371 | -0.725 | 0.495 | -1.617 | -0.577 | 0.023 | 0.073 | 0.073 | 1.655 | 0.38 | 0.78 | 0.29 | 0.16 | 0.29 | 0.27 |
| 5 | 134600902 | rs666601 | -1.022 | 0.56 | -0.646 | -0.773 | 0.004 | 0.037 | 0.037 | 0 | 0.48 | 0.65 | 0.44 | 0.26 | 0.44 | 0.47 |
| 5 | 134601755 | rs7703593 | 1.308 | 0.597 | -1.463 | -0.653 | -0.081 | -0.053 | -0.053 | 0.411 | 0.37 | 0.11 | 0.47 | 0.51 | 0.44 | 0.44 |
| 5 | 134602537 | rs595269 | -0.472 | 0.467 | -0.892 | -0.647 | 0.017 | 0.004 | 0.004 | 0 | 0.05 | 0.12 | 0.04 | 0.03 | 0.06 | 0.1 |
| 5 | 134603531 | rs2162708 | 1.186 | 0.417 | -0.763 | -0.651 | -0.054 | -0.008 | -0.008 | 0 | 0.4 | 0.19 | 0.5 | 0.49 | 0.49 | 0.46 |
| 5 | 134608010 | rs11242240 | 1.104 | 0.525 | -0.862 | -0.68 | -0.111 | -0.04 | -0.04 | 0 | 0.39 | 0.11 | 0.5 | 0.49 | 0.49 | 0.47 |
| 5 | 134609822 | rs4976285 | -0.147 | 0.084 | 0.057 | -0.826 | -0.002 | 0.001 | 0.001 | 0.094 | 0.25 | 0.27 | 0.29 | 0.38 | 0.28 | 0.24 |
| 5 | 134611916 | rs630231 | -1.578 | -0.038 | 0.306 | -1.305 | 0 | -0.026 | -0.026 | 0 | 0.86 | 0.87 | 0.94 | 1 | 0.91 | 0.92 |
| 5 | 134612072 | rs10515469 | 1.04 | -0.148 | 0.355 | -1.355 | 0 | -0.006 | -0.006 | 0 | 0.14 | 0.09 | 0.11 | 0.06 | 0.16 | 0.06 |
| 5 | 134612693 | rs2652080 | 1.052 | -0.114 | 0 | -1.88 | -0.016 | 0.026 | 0.026 | 0 | 0.4 | 0.32 | 0.45 | 0.63 | 0.42 | 0.49 |
| 5 | 134615015 | rs2603021 | -1.016 | -0.275 | 0.303 | -1.259 | 0.012 | 0.01 | 0.01 | 0.503 | 0.49 | 0.56 | 0.49 | 0.47 | 0.47 | 0.44 |
| 5 | 134615858 | rs2652093 | -0.873 | -0.902 | 0.787 | -1.539 | -0.021 | -0.017 | -0.017 | 1.662 | 0.29 | 0.57 | 0.24 | 0.19 | 0.22 | 0.24 |
| 5 | 134623699 | rs10067160 | -1.107 | -1.018 | 0.768 | -1.721 | 0.018 | 0.025 | 0.025 | 0 | 0.34 | 0.69 | 0.22 | 0.12 | 0.21 | 0.19 |
| 5 | 134629551 | rs2262634 | -0.711 | -0.996 | 0.874 | -1.687 | -0.006 | -0.019 | -0.019 | 0 | 0.29 | 0.65 | 0.21 | 0.13 | 0.21 | 0.14 |
| 5 | 134633657 | rs10515470 | 0.69 | -1.297 | 0.989 | -1.769 | -0.001 | -0.036 | -0.036 | 0 | 0.17 | 0.18 | 0.1 | 0.05 | 0.16 | 0.1 |
| 5 | 134638707 | rs2652085 | -1.576 | -1.471 | 0.825 | -1.942 | 0.035 | 0.095 | 0.095 | 0.517 | 0.51 | 0.72 | 0.43 | 0.23 | 0.41 | 0.34 |
| 5 | 134639915 | rs2603019 | -0.77 | -1.555 | 1.245 | -2.073 | -0.028 | -0.042 | -0.042 | 0 | 0.27 | 0.57 | 0.2 | 0.12 | 0.2 | 0.16 |
| 5 | 134642115 | rs2603017 | -0.146 | -1.513 | 1.265 | -2.103 | -0.044 | -0.048 | -0.048 | 0 | 0.21 | 0.43 | 0.22 | 0.08 | 0.14 | 0.11 |
| 5 | 134643116 | rs2603014 | -1.504 | -1.434 | 0.758 | -2.089 | 0.042 | 0.094 | 0.094 | 1.342 | 0.48 | 0.68 | 0.4 | 0.13 | 0.38 | 0.28 |
| 5 | 134644855 | rs2603013 | -1.096 | -1.753 | 0.868 | -1.851 | 0.03 | 0.05 | 0.05 | 3.847 | 0.4 | 0.67 | 0.34 | 0.13 | 0.3 | 0.22 |
| 5 | 134652489 | rs2652092 | -1.155 | -1.514 | 1.696 | -1.064 | -0.059 | -0.011 | -0.011 | 0 | 0.19 | 0.3 | 0.12 | 0.12 | 0.08 | 0.13 |
| 5 | 134652996 | rs11738420 | -1.746 | -1.356 | 1.59 | -1.135 | 0.004 | 0.006 | 0.006 | 0 | 0.39 | 0.57 | 0.26 | 0.17 | 0.28 | 0.25 |
| 5 | 134655895 | rs10068105 | -1.743 | -1.268 | 1.224 | -1.137 | 0.02 | 0.052 | 0.052 | 0 | 0.41 | 0.65 | 0.26 | 0.16 | 0.3 | 0.27 |
| 5 | 134658634 | rs7706804 | -0.745 | -0.542 | 1.202 | -0.605 | 0.004 | -0.032 | -0.032 | 0.032 | 0.2 | 0.19 | 0.12 | 0.06 | 0.21 | 0.14 |
| 5 | 134663097 | rs7712997 | -0.685 | -0.968 | 0.996 | -0.472 | -0.053 | -0.057 | -0.057 | 0 | 0.24 | 0.49 | 0.16 | 0.09 | 0.13 | 0.16 |
| 5 | 134665082 | rs13163460 | -1.092 | -0.993 | 0.368 | -0.469 | -0.001 | -0.004 | -0.004 | 0 | 0.31 | 0.73 | 0.21 | 0.14 | 0.19 | 0.2 |
| 5 | 134670741 | rs3733728 | -1.407 | -0.623 | 0.087 | -0.098 | -0.008 | -0.001 | -0.001 | 0 | 0.06 | 0.01 | 0.02 | 0.05 | 0.05 | 0.06 |
| 5 | 134677927 | rs1393081 | -2.227 | -0.636 | 1.362 | 0.018 | 0.095 | 0.145 | 0.145 | 1.1 | 0.06 | 0.41 | 0.12 | 0.1 | 0.05 | 0.04 |
| 5 | 134679247 | rs2292011 | -2.193 | -0.485 | 1.808 | -0.014 | 0.032 | 0.022 | 0.022 | 0.797 | 0.17 | 0.58 | 0.26 | 0.18 | 0.17 | 0.22 |
| 5 | 134692112 | rs3776209 | -2.232 | -0.487 | 2.737 | -0.016 | 0.08 | 0.134 | 0.134 | 0 | 0.06 | 0.37 | 0.11 | 0.09 | 0.04 | 0.06 |

| | | | | | | | | | | | | | | | | |
|---|---|---|---|---|---|---|---|---|---|---|---|---|---|---|---|---|
| 5 | 134707073 | rs1393082 | -1.703 | -1.477 | 1.743 | 0.013 | 0.021 | -0.024 | -0.024 | 0 | 0.51 | 0.94 | 0.39 | 0.38 | 0.33 | 0.49 |
| 5 | 134708765 | rs3776203 | -0.509 | -0.705 | 2.5 | 0.594 | -0.022 | -0.01 | -0.01 | 0 | 0.1 | 0.03 | 0.13 | 0.11 | 0.12 | 0.17 |
| 5 | 134729522 | rs779562 | -2.563 | -0.79 | 2.636 | 0.596 | 0.055 | 0.105 | 0.105 | 1.94 | 0.07 | 0.57 | 0.13 | 0.09 | 0.04 | 0.04 |
| 5 | 134756056 | rs6870570 | 1.621 | -0.854 | 2.636 | 0.731 | 0.033 | 0.115 | 0.115 | 0 | 0.94 | 0.76 | 0.9 | 0.88 | 0.95 | 0.95 |
| 5 | 134759783 | rs4246779 | 2.529 | -0.873 | 2.302 | 0.722 | 0.01 | 0.095 | 0.095 | 1.176 | 0.9 | 0.39 | 0.87 | 0.88 | 0.95 | 0.95 |
| 5 | 134768329 | rs7710100 | 1.789 | -0.845 | 2.007 | 1.249 | 0.015 | 0.191 | 0.191 | 0.661 | 0.9 | 0.65 | 0.87 | 0.76 | 0.93 | 0.92 |
| 5 | 134768688 | rs4976296 | -0.39 | -1.236 | 1.644 | 1.076 | -0.016 | 0.003 | 0.003 | 0.037 | 0.15 | 0.19 | 0.13 | 0.13 | 0.09 | 0.06 |
| 5 | 134769827 | rs4976297 | 1.787 | -1.164 | 0.68 | 1.123 | 0.023 | -0.05 | -0.05 | 0.018 | 0.66 | 0.26 | 0.69 | 0.56 | 0.75 | 0.69 |
| 5 | 134773951 | rs11242246 | 1.642 | -0.909 | 0.247 | 0.688 | -0.023 | -0.011 | -0.011 | 0.012 | 0.45 | 0.18 | 0.56 | 0.61 | 0.59 | 0.42 |
| 5 | 134775617 | rs874021 | 1.751 | -0.818 | 0.739 | 0.907 | -0.026 | 0.064 | 0.064 | 0.832 | 0.82 | 0.55 | 0.83 | 0.79 | 0.87 | 0.81 |
| 5 | 134775697 | rs874020 | 0.919 | -0.549 | -0.313 | 0.877 | -0.011 | 0.008 | 0.008 | 0.12 | 0.89 | 0.86 | 0.94 | 0.85 | 0.92 | 0.91 |
| 5 | 134778470 | rs6896596 | 0.931 | -0.425 | -0.796 | 1.129 | 0.008 | -0.044 | -0.044 | 0 | 0.66 | 0.85 | 0.64 | 0.58 | 0.72 | 0.63 |
| 5 | 134779242 | rs4976254 | -1.145 | -0.158 | -0.288 | 1.554 | -0.008 | -0.024 | -0.024 | 0 | 0.35 | 0.23 | 0.39 | 0.38 | 0.32 | 0.4 |
| 5 | 134779417 | rs874158 | -1.351 | -0.534 | 0.479 | 1.632 | -0.014 | 0.02 | 0.02 | 0.529 | 0.12 | 0.24 | 0.08 | 0.16 | 0.1 | 0.19 |
| 5 | 134780909 | rs17168353 | -1.519 | -0.524 | 0.849 | 1.492 | -0.014 | 0.008 | 0.008 | 0 | 0.27 | 0.34 | 0.17 | 0.32 | 0.28 | 0.31 |
| 5 | 134781400 | rs744247 | -0.291 | -0.422 | 0.744 | 1.5 | -0.008 | -0.014 | -0.014 | 2.57 | 0.25 | 0.15 | 0.32 | 0.25 | 0.21 | 0.22 |
| 5 | 134781992 | rs17168355 | -0.31 | -0.474 | 0.805 | 1.471 | -0.002 | -0.002 | -0.002 | 0 | 0.15 | 0.18 | 0.11 | 0.16 | 0.18 | 0.12 |
| 5 | 134782414 | rs12518053 | -2.572 | -0.613 | 0.647 | 1.307 | -0.136 | -0.057 | -0.057 | 0.078 | 0.57 | 0.96 | 0.55 | 0.56 | 0.51 | 0.54 |
| 5 | 134782450 | rs12520799 | -2.536 | -0.615 | 0.745 | 1.307 | -0.12 | -0.044 | -0.044 | 0.03 | 0.56 | 0.93 | 0.55 | 0.56 | 0.51 | 0.54 |
| 5 | 134782510 | rs12520809 | -2.585 | -0.615 | 0.745 | 1.306 | -0.122 | -0.047 | -0.047 | 0.368 | 0.57 | 0.93 | 0.55 | 0.56 | 0.51 | 0.54 |
| 5 | 134782576 | rs1031844 | -2.536 | -0.612 | 0.747 | 1.308 | -0.12 | -0.044 | -0.044 | 0.111 | 0.56 | 0.93 | 0.55 | 0.56 | 0.52 | 0.54 |
| 5 | 134784086 | rs12519481 | -0.246 | -0.562 | 0.916 | 1.296 | -0.005 | -0.012 | -0.012 | 0 | 0.25 | 0.17 | 0.33 | 0.25 | 0.21 | 0.22 |
| 5 | 134786259 | rs13173226 | -0.521 | -0.696 | 1.503 | 1.315 | -0.024 | 0 | 0 | 0.018 | 0.3 | 0.17 | 0.39 | 0.26 | 0.33 | 0.24 |
| 5 | 134794909 | rs899162 | -0.584 | -0.597 | 1.467 | 1.32 | 0.007 | -0.015 | -0.015 | 1.135 | 0.11 | 0.07 | 0.06 | 0.16 | 0.1 | 0.19 |
| 5 | 134798723 | rs251562 | -1.486 | -0.528 | 1.6 | 1.277 | -0.006 | 0.009 | 0.009 | 0.566 | 0.15 | 0.27 | 0.09 | 0.16 | 0.14 | 0.2 |
| 5 | 134799411 | rs251565 | -1.833 | -0.599 | 1.657 | 1.328 | -0.039 | 0.03 | 0.03 | 0.232 | 0.25 | 0.47 | 0.14 | 0.31 | 0.2 | 0.31 |
| 5 | 134801438 | rs251567 | -1.07 | -0.572 | 1.425 | 1.353 | 0 | -0.011 | -0.011 | 0.126 | 0.13 | 0.08 | 0.07 | 0.16 | 0.14 | 0.2 |
| 5 | 134807383 | rs6862436 | -0.292 | -0.612 | 1.509 | 1.352 | -0.033 | 0.04 | 0.04 | 0.048 | 0.1 | 0.27 | 0.04 | 0.14 | 0.05 | 0.11 |
| 5 | 134808026 | rs11738925 | 2.055 | -0.605 | 1.522 | 1.456 | 0.01 | 0.002 | 0.002 | 0 | 0.51 | 0.27 | 0.55 | 0.48 | 0.54 | 0.46 |
| 5 | 134812100 | rs171654 | 0.375 | -0.513 | 1.22 | 1.522 | -0.028 | -0.006 | -0.006 | 2.616 | 0.72 | 0.84 | 0.6 | 0.74 | 0.68 | 0.71 |
| 5 | 134824264 | rs17168440 | -1.502 | -0.443 | 1.627 | 1.612 | -0.009 | 0.141 | 0.141 | 0 | 0.14 | 0.46 | 0.15 | 0.23 | 0.1 | 0.15 |
| 5 | 134825122 | rs17597227 | -0.232 | -0.548 | 1.298 | 1.737 | -0.044 | -0.021 | -0.021 | 0 | 0.24 | 0.1 | 0.32 | 0.23 | 0.28 | 0.31 |
| 5 | 134826809 | rs1076379 | 1.913 | -0.564 | 1.296 | 1.786 | 0 | 0.016 | 0.016 | 0.039 | 0.54 | 0.35 | 0.54 | 0.45 | 0.57 | 0.43 |
| 5 | 134828617 | rs740657 | -1.03 | -0.313 | 1.491 | 1.684 | -0.016 | 0.071 | 0.071 | 0.177 | 0.13 | 0.39 | 0.12 | 0.19 | 0.1 | 0.12 |
| 5 | 134829420 | rs17644674 | 0.046 | -0.061 | 1.052 | 1.605 | 0.001 | -0.03 | -0.03 | 2.33 | 0.07 | 0.03 | 0.05 | 0.14 | 0.08 | 0.15 |
| 5 | 134830781 | rs7714565 | -1.434 | -0.16 | 0.884 | 1.115 | -0.001 | 0 | 0 | 0 | 0.41 | 0.61 | 0.39 | 0.37 | 0.38 | 0.47 |
| 5 | 134832884 | rs2058169 | 1.464 | -0.828 | 0.995 | 0.2 | -0.017 | -0.006 | -0.006 | 0 | 0.8 | 0.34 | 0.85 | 0.81 | 0.87 | 0.72 |
| 5 | 134834401 | rs13165759 | 1.951 | -0.741 | 0.524 | 0.099 | 0.037 | 0.039 | 0.039 | 0.697 | 0.57 | 0.2 | 0.64 | 0.61 | 0.62 | 0.43 |
| 5 | 134846824 | rs12374511 | -0.455 | -0.692 | 0.701 | 0.102 | -0.006 | 0.015 | 0.015 | 0 | 0.22 | 0.12 | 0.21 | 0.2 | 0.25 | 0.29 |
| 5 | 134861178 | rs6878746 | -1.905 | -0.797 | 1.734 | 0.038 | -0.004 | -0.005 | -0.005 | 0 | 0.41 | 0.49 | 0.35 | 0.39 | 0.39 | 0.57 |
| 5 | 134864640 | rs6875282 | -1.483 | -0.936 | 0.727 | -0.048 | 0.006 | 0.021 | 0.021 | 0 | 0.34 | 0.2 | 0.28 | 0.35 | 0.35 | 0.51 |
| 5 | 134868339 | rs3860110 | -0.582 | -0.686 | 0.543 | 0.099 | -0.007 | -0.002 | -0.002 | 0 | 0.12 | 0.14 | 0.05 | 0.14 | 0.08 | 0.22 |
| 5 | 134876123 | rs13356152 | -1.325 | -0.577 | -0.161 | 0.061 | 0.021 | 0.059 | 0.059 | 1.642 | 0.42 | 0.83 | 0.33 | 0.36 | 0.36 | 0.58 |
| 5 | 134893439 | rs3860112 | 0.648 | -0.197 | -0.441 | -0.269 | -0.022 | 0.017 | 0.017 | 0.367 | 0.41 | 0.2 | 0.47 | 0.35 | 0.46 | 0.36 |
| 5 | 134896455 | rs6868474 | -0.473 | -1.059 | 0.612 | -0.861 | 0.023 | 0.011 | 0.011 | 0 | 0.48 | 0.58 | 0.26 | 0.43 | 0.27 | 0.38 |
| 5 | 134896833 | rs10038760 | 0.717 | -1.291 | 0.44 | -0.917 | 0.079 | -0.009 | -0.009 | 0.669 | 0.3 | 0.11 | 0.14 | 0.32 | 0.15 | 0.22 |
| 5 | 134901156 | rs10045413 | -0.063 | -1.06 | 0.34 | -1.012 | 0.024 | 0.019 | 0.019 | 1.331 | 0.39 | 0.11 | 0.23 | 0.34 | 0.33 | 0.33 |
| 5 | 134904639 | rs13357245 | 0.541 | -0.962 | 0.588 | -0.796 | 0.079 | -0.026 | -0.026 | 0 | 0.31 | 0.1 | 0.15 | 0.33 | 0.16 | 0.24 |
| 5 | 134904975 | rs11749604 | 0.556 | -0.883 | 0.688 | -0.754 | 0.004 | -0.01 | -0.01 | 0 | 0.09 | 0.06 | 0.05 | 0.11 | 0.06 | 0.15 |
| 5 | 134907031 | rs1046092 | 0.315 | -0.836 | 0.736 | -0.732 | -0.001 | 0 | 0 | 1.115 | 0.1 | 0.08 | 0.04 | 0.11 | 0.07 | 0.17 |
| 5 | 134908402 | rs1016666 | -0.438 | -0.89 | 0.936 | -0.772 | -0.015 | -0.003 | -0.003 | 0 | 0.45 | 0.52 | 0.23 | 0.45 | 0.26 | 0.41 |
| 5 | 134912330 | rs2237061 | 0.243 | -0.907 | 0.899 | -0.736 | 0.003 | -0.001 | -0.001 | 0 | 0.1 | 0.08 | 0.04 | 0.11 | 0.07 | 0.16 |
| 5 | 134914233 | rs11950696 | -0.413 | -0.906 | 0.979 | -0.751 | 0.022 | -0.028 | -0.028 | 0 | 0.08 | 0.28 | 0.07 | 0 | 0.05 | 0.01 |
| 5 | 134914350 | rs2072347 | 0.694 | -0.899 | 1.02 | -0.735 | 0.073 | -0.011 | -0.011 | 0.033 | 0.31 | 0.14 | 0.15 | 0.33 | 0.16 | 0.21 |
| 5 | 134925099 | rs17168697 | -0.85 | -0.684 | 0.52 | -0.657 | 0.004 | 0.002 | 0.002 | 0 | 0.53 | 0.56 | 0.31 | 0.45 | 0.4 | 0.46 |
| 5 | 134926532 | rs17168700 | -0.514 | -0.979 | 0.596 | -0.717 | -0.024 | -0.005 | -0.005 | 0 | 0.45 | 0.5 | 0.24 | 0.44 | 0.25 | 0.4 |
| 5 | 134934087 | rs10515473 | -0.794 | -0.497 | 0.617 | -0.545 | 0.002 | -0.051 | -0.051 | 0 | 0.25 | 0.38 | 0.15 | 0.11 | 0.26 | 0.23 |
| 5 | 134938615 | rs1848474 | -1.083 | -0.479 | 0.352 | -0.532 | 0 | 0.003 | 0.003 | 0 | 0.57 | 0.65 | 0.34 | 0.46 | 0.45 | 0.47 |
| 5 | 134939703 | rs7732567 | -1.165 | -0.486 | 0.03 | -0.535 | -0.003 | 0.005 | 0.005 | 0 | 0.57 | 0.71 | 0.34 | 0.46 | 0.45 | 0.47 |
| 5 | 134939990 | rs2344483 | 0.458 | -0.207 | -1.084 | -0.36 | 0.055 | -0.004 | -0.004 | 0.063 | 0.31 | 0.17 | 0.18 | 0.34 | 0.19 | 0.24 |
| 5 | 134942818 | rs1876623 | 0.518 | -0.161 | -1.194 | -0.339 | 0.062 | -0.015 | -0.015 | 0.336 | 0.3 | 0.12 | 0.17 | 0.34 | 0.19 | 0.24 |
| 5 | 134957413 | rs6873508 | 0.29 | 0.438 | -0.807 | 0.271 | 0.019 | 0.029 | 0.029 | 0.338 | 0.46 | 0.38 | 0.28 | 0.34 | 0.39 | 0.31 |
| 5 | 134961400 | rs1396439 | -1.029 | 0.732 | -0.752 | 0.607 | 0.006 | -0.01 | -0.01 | 0 | 0.09 | 0.02 | 0.03 | 0.12 | 0.06 | 0.16 |
| 5 | 134966642 | rs31265 | -1.239 | 0.909 | -0.667 | 0.436 | -0.002 | 0.005 | 0.005 | 0 | 0.15 | 0.12 | 0.09 | 0.1 | 0.14 | 0.19 |
| 5 | 134967868 | rs7701856 | -1.362 | 0.951 | -0.396 | 1.124 | 0.001 | 0.019 | 0.019 | 0 | 0.07 | 0.09 | 0.07 | 0.25 | 0.08 | 0.09 |
| 5 | 134971621 | rs249592 | -1.065 | 1.121 | -0.539 | 1.612 | -0.007 | -0.034 | -0.034 | 0 | 0.22 | 0.17 | 0.38 | 0.61 | 0.27 | 0.38 |
| 5 | 134973802 | rs2521968 | 1.308 | 1.14 | -0.226 | 0.499 | -0.01 | 0.003 | 0.003 | 0 | 0.55 | 0.39 | 0.48 | 0.38 | 0.5 | 0.5 |
| 5 | 134976095 | rs2521964 | 1.256 | 1.088 | 0.063 | 0.462 | -0.005 | 0 | 0 | 0 | 0.56 | 0.45 | 0.48 | 0.38 | 0.49 | 0.52 |
| 5 | 134977833 | rs11738809 | 1.265 | 1.082 | 0.05 | 0.429 | -0.007 | 0 | 0 | 0 | 0.56 | 0.44 | 0.48 | 0.38 | 0.48 | 0.5 |

| | | | | | | | | | | | | | | | | |
|---|---|---|---|---|---|---|---|---|---|---|---|---|---|---|---|---|
| 5 | 134979444 | rs801569 | 0.125 | 1.088 | 0.514 | 0.401 | 0.007 | -0.12 | -0.12 | 0 | 0.76 | 0.67 | 0.87 | 1 | 0.75 | 0.87 |
| 5 | 134987770 | rs11738518 | 1.297 | 0.93 | 0.784 | 0.352 | -0.02 | 0.01 | 0.01 | 0 | 0.54 | 0.27 | 0.47 | 0.38 | 0.49 | 0.49 |
| 5 | 134993246 | rs801553 | -0.71 | 1.085 | 0.696 | 0.372 | 0.007 | 0.038 | 0.038 | 0 | 0.23 | 0.34 | 0.41 | 0.62 | 0.3 | 0.37 |
| 5 | 134999199 | rs17736119 | -0.423 | 0.315 | 1.024 | -0.06 | 0 | 0.015 | 0.015 | 0 | 0.25 | 0.16 | 0.11 | 0 | 0.23 | 0.12 |
| 5 | 135007644 | rs2188919 | 0.393 | 0.358 | 1.694 | 0.05 | 0.002 | 0.088 | 0.088 | 0 | 0.79 | 0.65 | 0.62 | 0.39 | 0.75 | 0.64 |
| 5 | 135013095 | rs6873251 | 0.483 | 0.266 | 1.716 | 0.65 | -0.002 | 0.112 | 0.112 | 0 | 0.84 | 0.66 | 0.67 | 0.6 | 0.83 | 0.73 |
| 5 | 135015599 | rs10045021 | 0.426 | 0.246 | 1.791 | 0.646 | -0.008 | 0.049 | 0.049 | 0 | 0.86 | 0.79 | 0.68 | 0.6 | 0.86 | 0.74 |
| 5 | 135028674 | rs11242261 | 0.328 | 0.174 | 1.921 | 0.627 | -0.005 | 0.122 | 0.122 | 0 | 0.86 | 0.71 | 0.68 | 0.6 | 0.85 | 0.74 |
| 5 | 135029394 | rs4246783 | 1.349 | 0.038 | 2.102 | 0.503 | -0.002 | -0.002 | -0.002 | 0.286 | 0.59 | 0.44 | 0.55 | 0.6 | 0.62 | 0.62 |
| 5 | 135035085 | rs6596257 | 0.25 | 0.026 | 1.892 | 0.558 | -0.003 | 0.03 | 0.03 | 0 | 0.86 | 0.84 | 0.69 | 0.61 | 0.85 | 0.74 |
| 5 | 135035711 | rs6887694 | 1.665 | 0.265 | 1.586 | 0.566 | 0.014 | 0.03 | 0.03 | 0 | 0.55 | 0.31 | 0.53 | 0.6 | 0.59 | 0.62 |
| 5 | 135039790 | rs4613698 | -0.495 | 0.345 | 1.227 | 0.374 | 0.004 | -0.013 | -0.013 | 0 | 0.24 | 0.19 | 0.1 | 0.01 | 0.19 | 0.11 |
| 5 | 135047254 | rs4976321 | 1.805 | 0.083 | 0.79 | 0.835 | 0.009 | 0.068 | 0.068 | 0 | 0.48 | 0.15 | 0.39 | 0.34 | 0.52 | 0.45 |
| 5 | 135050175 | rs6861738 | 0.919 | -0.181 | 1.663 | 0.931 | -0.037 | 0.046 | 0.046 | 0 | 0.76 | 0.57 | 0.59 | 0.63 | 0.78 | 0.65 |
| 5 | 135050965 | rs17168967 | -0.525 | -0.199 | 1.828 | 0.924 | -0.001 | -0.007 | -0.007 | 1.027 | 0.21 | 0.21 | 0.16 | 0.28 | 0.23 | 0.19 |
| 5 | 135052055 | rs4976323 | 2.137 | -0.077 | 1.861 | 0.938 | 0.052 | 0.072 | 0.072 | 0 | 0.51 | 0.2 | 0.41 | 0.35 | 0.54 | 0.46 |
| 5 | 135053005 | rs13186945 | 0.589 | -0.081 | 1.75 | 1.069 | -0.007 | 0.011 | 0.011 | 0 | 0.05 | 0.01 | 0.07 | 0.29 | 0.07 | 0.09 |
| 5 | 135055456 | rs4976324 | 0.136 | 0.013 | 1.688 | 1.067 | -0.032 | -0.048 | -0.048 | 0 | 0.92 | 0.99 | 0.68 | 0.82 | 0.88 | 0.89 |
| 5 | 135061332 | rs10515477 | 2.097 | -0.29 | 1.603 | 1.017 | 0.041 | 0.08 | 0.08 | 0 | 0.48 | 0.11 | 0.4 | 0.31 | 0.55 | 0.46 |
| 5 | 135071512 | rs4285212 | -1.214 | -0.411 | 1.605 | 0.916 | 0.022 | -0.019 | -0.019 | 0 | 0.34 | 0.27 | 0.23 | 0.46 | 0.3 | 0.33 |
| 5 | 135085098 | rs7732275 | -1.264 | -0.451 | 1.846 | 0.852 | -0.01 | 0.01 | 0.01 | 0 | 0.31 | 0.34 | 0.23 | 0.47 | 0.28 | 0.28 |
| 5 | 135123055 | rs7445602 | -0.938 | -0.438 | 1.549 | 0.853 | 0.016 | -0.066 | -0.066 | 0 | 0.28 | 0.12 | 0.21 | 0.44 | 0.29 | 0.27 |
| 5 | 135144329 | rs17169129 | -0.171 | -0.402 | 1.592 | 0.835 | -0.011 | -0.032 | -0.032 | 1.253 | 0.07 | 0.16 | 0.02 | 0 | 0.02 | 0.05 |
| 5 | 135148372 | rs1465291 | -1.159 | -0.533 | 1.864 | 0.814 | -0.061 | 0.047 | 0.047 | 0 | 0.27 | 0.43 | 0.16 | 0.44 | 0.15 | 0.27 |
| 5 | 135148655 | rs3930698 | -1.882 | -0.48 | 1.801 | 0.752 | -0.007 | 0.001 | 0.001 | 0 | 0.38 | 0.44 | 0.25 | 0.44 | 0.34 | 0.32 |
| 5 | 135148662 | rs2881156 | -1.882 | -0.48 | 1.801 | 0.769 | -0.007 | 0.001 | 0.001 | 0 | 0.38 | 0.44 | 0.25 | 0.45 | 0.34 | 0.32 |
| 5 | 135149952 | rs4521465 | -1.873 | -0.479 | 1.801 | 0.765 | -0.007 | 0.001 | 0.001 | 0 | 0.38 | 0.44 | 0.25 | 0.44 | 0.34 | 0.33 |
| 5 | 135150915 | rs4976465 | 0.935 | -0.289 | 1.913 | 0.944 | 0.041 | 0.202 | 0.202 | 0.247 | 0.86 | 0.56 | 0.64 | 0.71 | 0.84 | 0.76 |
| 5 | 135153111 | rs6877036 | 0.76 | -0.286 | 2.092 | 0.947 | 0.03 | 0.154 | 0.154 | 0.181 | 0.87 | 0.63 | 0.66 | 0.72 | 0.84 | 0.76 |
| 5 | 135160208 | rs721834 | -0.11 | -0.306 | 1.804 | 0.918 | 0.023 | -0.031 | -0.031 | 0 | 0.18 | 0.1 | 0.13 | 0.44 | 0.13 | 0.22 |
| 5 | 135161055 | rs17169180 | -0.895 | -0.224 | 1.705 | 0.909 | 0.019 | -0.024 | -0.024 | 0 | 0.24 | 0.22 | 0.16 | 0.44 | 0.17 | 0.27 |
| 5 | 135161418 | rs877356 | -0.895 | -0.224 | 1.705 | 0.923 | 0.019 | -0.022 | -0.022 | 0.113 | 0.24 | 0.22 | 0.16 | 0.43 | 0.17 | 0.27 |
| 5 | 135166905 | rs714091 | -1.183 | -0.132 | 1.681 | 0.864 | -0.002 | -0.024 | -0.024 | 0 | 0.19 | 0.21 | 0.12 | 0.01 | 0.21 | 0.11 |
| 5 | 135168344 | rs17738592 | 3.066 | -0.293 | 1.547 | 0.882 | -0.101 | 0.067 | 0.067 | 0 | 0.46 | 0.04 | 0.35 | 0.26 | 0.47 | 0.44 |
| 5 | 135173188 | rs6890039 | -1.244 | -0.257 | 1.627 | 0.753 | -0.002 | -0.022 | -0.022 | 0 | 0.19 | 0.21 | 0.12 | 0.01 | 0.21 | 0.11 |
| 5 | 135178124 | rs2304075 | 0.662 | -0.303 | 2.165 | 0.778 | 0.021 | 0.121 | 0.121 | 0.21 | 0.88 | 0.73 | 0.66 | 0.71 | 0.83 | 0.76 |
| 5 | 135178142 | rs6596270 | -1.217 | -0.302 | 2.138 | 0.771 | -0.002 | -0.022 | -0.022 | 0.986 | 0.19 | 0.21 | 0.12 | 0.01 | 0.21 | 0.11 |
| 5 | 135182123 | rs4976493 | 1.307 | -0.26 | 1.953 | 0.708 | 0.016 | 0.014 | 0.014 | 0 | 0.75 | 0.69 | 0.57 | 0.71 | 0.63 | 0.69 |
| 5 | 135191461 | rs4976375 | 2.203 | -0.052 | 1.894 | 0.704 | -0.024 | 0.007 | 0.007 | 1.517 | 0.64 | 0.16 | 0.5 | 0.7 | 0.59 | 0.64 |
| 5 | 135197638 | rs17169232 | 2.724 | 0.158 | 0.051 | 0.245 | -0.05 | 0.039 | 0.039 | 0 | 0.29 | 0.01 | 0.21 | 0.15 | 0.26 | 0.21 |
| 5 | 135199709 | rs4246802 | 0.479 | 0.27 | 0.872 | 0.436 | 0.006 | 0.12 | 0.12 | 0.889 | 0.89 | 0.73 | 0.78 | 0.71 | 0.84 | 0.78 |
| 5 | 135203513 | rs4272127 | 1.562 | 0.298 | 0.285 | 0.249 | 0.004 | 0.059 | 0.059 | 2.316 | 0.67 | 0.18 | 0.68 | 0.83 | 0.66 | 0.67 |
| 5 | 135204686 | rs4246806 | -0.244 | -0.067 | -0.357 | 0.033 | 0.02 | -0.116 | -0.116 | 0 | 0.3 | 0.08 | 0.28 | 0.51 | 0.3 | 0.23 |
| 5 | 135206025 | rs4582270 | 0.037 | 1.205 | 0.531 | -0.339 | -0.014 | -0.007 | -0.007 | 0 | 0.85 | 0.89 | 0.8 | 0.81 | 0.76 | 0.79 |
| 5 | 135207870 | rs11960025 | -2.021 | 0.973 | 1.066 | -0.513 | -0.031 | -0.131 | -0.131 | 0 | 0.19 | 0.54 | 0.1 | 0.01 | 0.13 | 0.17 |
| 5 | 135211466 | rs7717673 | -1.635 | 0.925 | 1.253 | -0.524 | -0.02 | -0.101 | -0.101 | 0 | 0.15 | 0.52 | 0.09 | 0.01 | 0.09 | 0.05 |
| 5 | 135212048 | rs10477775 | 2.303 | 1.001 | 0.909 | -0.506 | -0.013 | 0.043 | 0.043 | 0.917 | 0.65 | 0.27 | 0.58 | 0.8 | 0.62 | 0.62 |
| 5 | 135216112 | rs31565 | 0.202 | 1.058 | 0.406 | 0.002 | 0.002 | 0 | 0 | 0 | 0.85 | 0.86 | 0.72 | 0.8 | 0.8 | 0.8 |
| 5 | 135216503 | rs10077832 | -1.071 | 0.631 | 0.589 | -0.243 | 0 | -0.014 | -0.014 | 0 | 0.08 | 0.31 | 0.05 | 0.01 | 0.01 | 0.02 |
| 5 | 135217384 | rs1557918 | 1.21 | 1.094 | -0.135 | -0.262 | 0.014 | -0.051 | -0.051 | 0 | 0.66 | 0.53 | 0.6 | 0.84 | 0.65 | 0.74 |
| 5 | 135218585 | rs17169306 | -1.084 | 1.124 | -0.232 | -0.218 | -0.001 | -0.016 | -0.016 | 0 | 0.09 | 0.1 | 0.05 | 0 | 0.11 | 0.03 |
| 5 | 135220829 | rs6596271 | -0.732 | 0.972 | 0.088 | -0.179 | -0.004 | -0.029 | -0.029 | 0 | 0.07 | 0.14 | 0.04 | 0.04 | 0.06 | 0.03 |
| 5 | 135226137 | rs11741137 | 0.109 | 0.71 | 0.014 | -0.646 | -0.001 | -0.03 | -0.03 | 0.009 | 0.13 | 0.13 | 0.09 | 0 | 0.13 | 0.12 |
| 5 | 135228165 | rs2069885 | 0.082 | 0.717 | 0.017 | -0.629 | -0.001 | -0.025 | -0.025 | 0 | 0.13 | 0.13 | 0.09 | 0 | 0.13 | 0.11 |
| 5 | 135230253 | rs31564 | -1.47 | 0.709 | -0.48 | -0.542 | 0.009 | -0.011 | -0.011 | 0 | 0.6 | 0.79 | 0.64 | 0.45 | 0.64 | 0.53 |
| 5 | 135230513 | rs1859430 | -0.529 | 0.509 | 0.157 | -0.44 | -0.007 | -0.072 | -0.072 | 1.488 | 0.2 | 0.28 | 0.13 | 0.04 | 0.19 | 0.14 |
| 5 | 135235606 | rs31563 | 0.407 | 0.978 | 0.272 | -0.299 | 0.011 | 0.014 | 0.014 | 0 | 0.08 | 0.13 | 0.23 | 0.16 | 0.14 | 0.08 |
| 5 | 135236015 | rs17169344 | -0.228 | 0.455 | 0.226 | -0.542 | 0.001 | -0.001 | -0.001 | 0 | 0.11 | 0.05 | 0.08 | 0 | 0.12 | 0.03 |
| 5 | 135239560 | rs10477776 | -0.382 | -0.093 | 0.022 | -0.191 | 0.028 | 0.032 | 0.032 | 0.731 | 0.46 | 0.59 | 0.26 | 0.23 | 0.29 | 0.31 |
| 5 | 135240813 | rs11739800 | -0.342 | 0.045 | -0.431 | -0.159 | 0.013 | 0.016 | 0.016 | 0.519 | 0.45 | 0.41 | 0.27 | 0.2 | 0.31 | 0.35 |
| 5 | 135248015 | rs10059744 | 0.868 | 0.085 | -0.243 | 0.705 | -0.011 | 0.039 | 0.039 | 0 | 0.11 | 0.22 | 0.08 | 0.13 | 0.07 | 0.13 |
| 5 | 135251107 | rs1859428 | -0.451 | 0.097 | -0.055 | 0.336 | 0 | -0.001 | -0.001 | 0 | 0.44 | 0.57 | 0.28 | 0.38 | 0.29 | 0.32 |
| 5 | 135259846 | rs6861170 | -0.468 | -0.045 | -0.04 | 0.308 | 0.002 | 0 | 0 | 0.325 | 0.44 | 0.6 | 0.28 | 0.37 | 0.28 | 0.31 |
| 5 | 135261332 | rs6890950 | 0.114 | -0.113 | 0.357 | 0.285 | -0.012 | -0.016 | -0.016 | 0 | 0.34 | 0.43 | 0.21 | 0.24 | 0.21 | 0.19 |
| 5 | 135266859 | rs185069 | 0.362 | 0.143 | 0.327 | 0.299 | 0.003 | 0 | 0 | 0 | 0.56 | 0.39 | 0.67 | 0.63 | 0.69 | 0.69 |
| 5 | 135267337 | rs17701996 | 1.688 | 0.074 | 0.163 | 0.387 | 0.011 | -0.025 | -0.025 | 0.983 | 0.1 | 0.06 | 0.07 | 0.14 | 0.07 | 0.12 |
| 5 | 135274148 | rs31553 | -0.968 | 0.388 | -0.364 | -0.094 | -0.132 | -0.009 | -0.009 | 0.034 | 0.73 | 0.98 | 0.77 | 0.8 | 0.68 | 0.75 |
| 5 | 135275330 | rs31551 | -0.593 | -0.119 | -0.251 | 0.168 | -0.001 | -0.002 | -0.002 | 0 | 0.33 | 0.35 | 0.47 | 0.39 | 0.38 | 0.38 |
| 5 | 135276049 | rs31549 | -0.862 | -0.174 | -0.19 | 0.161 | 0.005 | 0.001 | 0.001 | 0.277 | 0.44 | 0.44 | 0.54 | 0.52 | 0.45 | 0.5 |

| | | | | | | | | | | | | | | | | |
|---|---|---|---|---|---|---|---|---|---|---|---|---|---|---|---|---|
| 5 | 135276314 | rs40986 | -1.7 | -0.054 | 0.065 | 0.15 | -0.048 | 0.006 | 0.006 | 0.049 | 0.76 | 0.92 | 0.8 | 0.78 | 0.72 | 0.82 |
| 5 | 135276847 | rs31547 | -1.15 | -0.099 | 0.164 | 0.128 | 0.015 | -0.022 | -0.022 | 0 | 0.71 | 0.64 | 0.76 | 0.78 | 0.71 | 0.81 |
| 5 | 135279666 | rs31537 | -1.634 | -0.086 | 0.216 | 0.134 | -0.005 | 0.002 | 0.002 | 0 | 0.75 | 0.8 | 0.79 | 0.78 | 0.72 | 0.82 |
| 5 | 135280552 | rs31535 | -0.141 | 0.06 | 0.022 | 0.171 | 0 | 0 | 0 | 0 | 0.55 | 0.5 | 0.62 | 0.51 | 0.52 | 0.56 |
| 5 | 135282514 | rs31531 | -1.94 | 0.07 | 0.009 | 0.17 | -0.095 | -0.021 | -0.021 | 1.29 | 0.77 | 0.96 | 0.8 | 0.78 | 0.72 | 0.82 |
| 5 | 135286437 | rs31519 | 0.228 | 0.063 | 0.025 | 0.17 | -0.001 | 0.021 | 0.021 | 0 | 0.45 | 0.41 | 0.55 | 0.37 | 0.45 | 0.45 |
| 5 | 135287029 | rs31517 | -0.637 | 0.07 | 0.038 | 0.17 | 0.002 | 0.008 | 0.008 | 0.019 | 0.62 | 0.55 | 0.7 | 0.65 | 0.64 | 0.7 |
| 5 | 135287201 | rs2073615 | 2.199 | 0.066 | 0.039 | 0.175 | 0.007 | -0.008 | -0.008 | 0 | 0.1 | 0.09 | 0.07 | 0.13 | 0.07 | 0.12 |
| 5 | 135296362 | rs2526150 | -0.557 | 0.052 | -0.058 | 0.176 | 0 | 0.01 | 0.01 | 0.015 | 0.61 | 0.55 | 0.69 | 0.63 | 0.63 | 0.7 |
| 5 | 135300039 | rs2158440 | -1.755 | 0.075 | -0.043 | 0.187 | -0.004 | 0.001 | 0.001 | 0 | 0.75 | 0.79 | 0.79 | 0.78 | 0.72 | 0.82 |
| 5 | 135302002 | rs12652910 | 1.132 | 0.071 | -0.04 | 0.188 | -0.032 | 0.044 | 0.044 | 0 | 0.14 | 0.24 | 0.1 | 0.15 | 0.09 | 0.12 |
| 5 | 135303001 | rs2214080 | -1.743 | 0.098 | -0.065 | 0.169 | -0.004 | 0.001 | 0.001 | 0 | 0.75 | 0.79 | 0.79 | 0.78 | 0.72 | 0.82 |
| 5 | 135307373 | rs2526166 | -1.743 | 0.098 | -0.065 | 0.169 | -0.004 | 0.001 | 0.001 | 0 | 0.75 | 0.79 | 0.79 | 0.78 | 0.72 | 0.82 |
| 5 | 135307505 | rs2526165 | -0.674 | 0.023 | -0.099 | 0.147 | -0.011 | -0.011 | -0.011 | 0.628 | 0.29 | 0.26 | 0.44 | 0.35 | 0.38 | 0.37 |
| 5 | 135310491 | rs7710932 | 0.567 | -0.012 | -0.103 | 0.16 | -0.038 | 0.013 | 0.013 | 0 | 0.17 | 0.26 | 0.12 | 0.13 | 0.08 | 0.12 |
| 5 | 135311863 | rs2526156 | 0.031 | 0.025 | -0.455 | 0.165 | -0.014 | 0.014 | 0.014 | 0 | 0.13 | 0.12 | 0.29 | 0.07 | 0.19 | 0.12 |
| 5 | 135312001 | rs11750675 | 2.246 | 0.132 | -0.537 | 0.22 | 0.015 | -0.027 | -0.027 | 0 | 0.1 | 0.05 | 0.07 | 0.14 | 0.07 | 0.12 |
| 5 | 135312174 | rs2526154 | -1.739 | 0.185 | -0.585 | 0.21 | -0.004 | 0.001 | 0.001 | 0 | 0.75 | 0.79 | 0.79 | 0.78 | 0.72 | 0.82 |
| 5 | 135313852 | rs2428158 | 1.291 | 0.015 | -1.328 | 0.212 | 0.003 | 0.011 | 0.011 | 0 | 0.29 | 0.22 | 0.23 | 0.26 | 0.26 | 0.31 |
| 5 | 135323434 | rs30737 | -0.634 | 0.42 | -1.768 | 0.341 | -0.015 | 0.008 | 0.008 | 0 | 0.9 | 0.94 | 0.72 | 0.93 | 0.82 | 0.88 |
| 5 | 135326590 | rs30740 | -0.002 | 0.26 | -1.711 | 0.359 | -0.007 | -0.01 | -0.01 | 0 | 0.75 | 0.77 | 0.69 | 0.67 | 0.69 | 0.66 |
| 5 | 135327491 | rs1859295 | -0.11 | -0.293 | -1.324 | 0.355 | 0.066 | -0.015 | -0.015 | 0.047 | 0.46 | 0.74 | 0.44 | 0.44 | 0.35 | 0.45 |
| 5 | 135329749 | rs728479 | -0.199 | 0.526 | -0.85 | 0.343 | 0 | 0 | 0 | 0 | 0.38 | 0.6 | 0.28 | 0.29 | 0.27 | 0.32 |
| 5 | 135331900 | rs40277 | 0.698 | 1.408 | -1.298 | 0.726 | -0.012 | -0.004 | -0.004 | 0 | 0.86 | 0.88 | 0.78 | 0.84 | 0.77 | 0.82 |
| 5 | 135332571 | rs30743 | 0.045 | -0.049 | -1.327 | 0.621 | 0.008 | 0.037 | 0.037 | 0.795 | 0.72 | 0.86 | 0.68 | 0.84 | 0.76 | 0.68 |
| 5 | 135333749 | rs30745 | 0.032 | -0.203 | -1.304 | 0.666 | 0.014 | 0.035 | 0.035 | 0.717 | 0.73 | 0.87 | 0.68 | 0.83 | 0.78 | 0.71 |
| 5 | 135335229 | rs10079806 | -0.59 | -0.35 | -1.268 | 0.646 | 0.006 | 0.003 | 0.003 | 0 | 0.18 | 0.15 | 0.12 | 0.14 | 0.16 | 0.16 |
| 5 | 135338599 | rs916950 | -0.382 | -0.095 | -1.274 | 1.189 | 0.02 | -0.058 | -0.058 | 1.149 | 0.3 | 0.12 | 0.29 | 0.42 | 0.23 | 0.29 |
| 5 | 135343547 | rs9327738 | 0.633 | -0.343 | -0.167 | 0.951 | -0.033 | 0.027 | 0.027 | 0.138 | 0.87 | 0.62 | 0.91 | 0.88 | 0.94 | 0.93 |
| 5 | 135348376 | rs2106491 | 0.269 | 0.063 | -1.2 | 0.926 | -0.073 | -0.056 | -0.056 | 0.923 | 0.2 | 0.01 | 0.17 | 0.18 | 0.22 | 0.36 |
| 5 | 135355038 | rs6894906 | -0.431 | -0.171 | -0.278 | 1.028 | -0.002 | -0.004 | -0.004 | 0 | 0.3 | 0.3 | 0.21 | 0.33 | 0.31 | 0.42 |
| 5 | 135363875 | rs2282790 | -0.548 | -0.242 | -0.905 | 0.866 | -0.038 | 0.009 | 0.009 | 0 | 0.54 | 0.87 | 0.6 | 0.61 | 0.49 | 0.52 |
| 5 | 135367757 | rs756463 | -0.109 | -0.721 | -0.76 | 0.621 | -0.01 | 0.009 | 0.009 | 0 | 0.3 | 0.44 | 0.41 | 0.36 | 0.26 | 0.19 |
| 5 | 135375042 | rs11738979 | -0.51 | -0.133 | -0.335 | 0.584 | -0.003 | -0.001 | -0.001 | 0.049 | 0.57 | 0.76 | 0.53 | 0.58 | 0.57 | 0.54 |
| 5 | 135377349 | rs2107331 | -0.237 | -0.472 | 0.038 | 0.742 | 0.004 | 0.017 | 0.017 | 0 | 0.51 | 0.66 | 0.47 | 0.58 | 0.48 | 0.49 |
| 5 | 135378781 | rs3805700 | 0.354 | -0.505 | 0.138 | 0.75 | -0.006 | -0.001 | -0.001 | 0 | 0.33 | 0.6 | 0.37 | 0.34 | 0.28 | 0.37 |
| 5 | 135383357 | rs2073508 | 0.676 | -0.514 | 0.228 | 0.78 | 0.002 | 0.01 | 0.01 | 0.021 | 0.2 | 0.28 | 0.12 | 0.24 | 0.2 | 0.12 |
| 5 | 135391326 | rs6860369 | 0.305 | -0.544 | 0.379 | 0.724 | 0.006 | 0.001 | 0.001 | 0 | 0.33 | 0.67 | 0.25 | 0.34 | 0.26 | 0.37 |
| 5 | 135392426 | rs4669 | 0.292 | -0.553 | 0.272 | 0.709 | 0.006 | 0.001 | 0.001 | 0 | 0.33 | 0.67 | 0.37 | 0.34 | 0.26 | 0.37 |
| 5 | 135393138 | rs17689879 | 0.589 | -0.563 | 0.013 | 0.607 | -0.134 | -0.081 | -0.081 | 0.47 | 0.44 | 0.05 | 0.47 | 0.42 | 0.51 | 0.49 |
| 5 | 135395626 | rs10036667 | 0.466 | -0.553 | 0.406 | 0.633 | 0.018 | -0.024 | -0.024 | 3.359 | 0.2 | 0.34 | 0.28 | 0.11 | 0.22 | 0.06 |
| 5 | 135396669 | rs6880837 | -0.396 | -0.606 | 0.373 | 0.88 | -0.007 | 0.037 | 0.037 | 0 | 0.52 | 0.76 | 0.51 | 0.59 | 0.5 | 0.52 |
| 5 | 135397702 | rs6886556 | 0.273 | -0.593 | 0.471 | 0.896 | -0.005 | 0.011 | 0.011 | 0 | 0.35 | 0.61 | 0.27 | 0.47 | 0.29 | 0.46 |
| 5 | 135398279 | rs3749782 | 0.822 | -0.551 | 0.48 | 0.902 | 0.006 | -0.007 | -0.007 | 0 | 0.18 | 0.22 | 0.13 | 0.12 | 0.22 | 0.06 |
| 5 | 135403529 | rs6899012 | 0.011 | -0.526 | 0.507 | 0.936 | 0.015 | -0.022 | -0.022 | 0 | 0.61 | 0.31 | 0.7 | 0.53 | 0.69 | 0.54 |
| 5 | 135411281 | rs11749522 | 0.256 | -0.187 | 0.734 | 0.86 | -0.006 | 0.014 | 0.014 | 0 | 0.36 | 0.49 | 0.39 | 0.37 | 0.31 | 0.46 |
| 5 | 135414280 | rs2190622 | -0.026 | -0.196 | 0.567 | 0.862 | 0.01 | -0.002 | -0.002 | 0 | 0.59 | 0.31 | 0.55 | 0.63 | 0.66 | 0.54 |
| 5 | 135419046 | rs7728385 | 0.264 | 0.022 | 0.552 | 0.641 | 0.052 | -0.026 | -0.026 | 0 | 0.06 | 0.19 | 0.06 | 0.01 | 0.11 | 0.02 |
| 5 | 135419341 | rs13159052 | 0.259 | -0.325 | 0.409 | 0.493 | -0.002 | 0.067 | 0.067 | 0 | 0.5 | 0.19 | 0.47 | 0.68 | 0.54 | 0.48 |
| 5 | 135424352 | rs6596283 | -0.01 | -0.294 | 0.494 | 0.491 | 0 | 0.008 | 0.008 | 0 | 0.05 | 0.09 | 0.06 | 0.09 | 0.06 | 0.08 |
| 5 | 135426282 | rs10056462 | 0.436 | -0.27 | 0.6 | 0.496 | 0.047 | -0.027 | -0.027 | 1.309 | 0.06 | 0.18 | 0.06 | 0.01 | 0.11 | 0.02 |
| 5 | 135429020 | rs7715300 | -0.408 | -0.598 | 0.642 | 0.434 | -0.006 | 0.023 | 0.023 | 0 | 0.12 | 0.34 | 0.09 | 0.09 | 0.08 | 0.08 |
| 5 | 135434609 | rs12654032 | 1.252 | -0.703 | 0.066 | 0.459 | -0.012 | -0.059 | -0.059 | 0.55 | 0.13 | 0.01 | 0.06 | 0.23 | 0.09 | 0.07 |
| 5 | 135439740 | rs17691375 | 0.069 | -0.28 | 0.207 | 0.156 | -0.003 | -0.003 | -0.003 | 0.189 | 0.4 | 0.45 | 0.28 | 0.26 | 0.31 | 0.47 |
| 5 | 135440176 | rs10052731 | 0.126 | -0.154 | -0.097 | 0.109 | 0.035 | -0.037 | -0.037 | 0 | 0.09 | 0.17 | 0.07 | 0.03 | 0.16 | 0.02 |
| 5 | 135441313 | rs17748071 | 1.487 | -0.299 | -0.479 | 0.138 | -0.03 | -0.006 | -0.006 | 0.196 | 0.25 | 0.01 | 0.17 | 0.18 | 0.2 | 0.39 |
| 5 | 135442965 | rs2006225 | 0.541 | -0.181 | -0.136 | 0.244 | -0.031 | 0.066 | 0.066 | 0 | 0.47 | 0.18 | 0.61 | 0.69 | 0.52 | 0.5 |
| 5 | 135448505 | rs2346361 | -0.092 | -0.488 | 0.017 | 0.257 | 0.001 | 0.001 | 0.001 | 0 | 0.51 | 0.58 | 0.32 | 0.32 | 0.43 | 0.49 |
| 5 | 135456085 | rs2906829 | -1.575 | -0.317 | 0.241 | 0.6 | 0.015 | 0.038 | 0.038 | 2.521 | 0.3 | 0.55 | 0.24 | 0.39 | 0.33 | 0.19 |
| 5 | 135469500 | rs3764942 | 1.414 | 0.035 | 0.321 | 0.543 | 0 | 0.021 | 0.021 | 1.107 | 0.12 | 0.12 | 0.07 | 0.28 | 0.1 | 0.09 |
| 5 | 135473762 | rs9327744 | -0.481 | 0.053 | 0.265 | 0.544 | 0.008 | -0.001 | -0.001 | 0 | 0.13 | 0.17 | 0.11 | 0.09 | 0.21 | 0.09 |
| 5 | 135494074 | rs13166063 | -1.458 | 0.023 | 0.259 | 0.512 | 0.009 | 0.025 | 0.025 | 3.257 | 0.31 | 0.59 | 0.24 | 0.38 | 0.33 | 0.19 |
| 5 | 135514055 | rs12719481 | -1.458 | 0.023 | 0.264 | 0.512 | 0.009 | 0.026 | 0.026 | 3.257 | 0.31 | 0.58 | 0.24 | 0.38 | 0.33 | 0.19 |
| 5 | 135514216 | rs12719482 | -0.474 | 0.024 | 0.32 | 0.518 | 0.006 | -0.001 | -0.001 | 0 | 0.13 | 0.17 | 0.11 | 0.09 | 0.21 | 0.09 |
| 5 | 135515708 | rs6865297 | -1.263 | -0.011 | 0.145 | 0.485 | 0.009 | 0.026 | 0.026 | 3.257 | 0.31 | 0.58 | 0.24 | 0.38 | 0.33 | 0.19 |
| 5 | 135516203 | rs3206634 | -1.263 | -0.011 | 0.145 | 0.485 | 0.009 | 0.026 | 0.026 | 3.257 | 0.31 | 0.58 | 0.24 | 0.38 | 0.33 | 0.19 |
| 5 | 135516282 | rs3206635 | -1.263 | -0.011 | 0.145 | 0.485 | 0.009 | 0.026 | 0.026 | 3.257 | 0.31 | 0.58 | 0.24 | 0.38 | 0.33 | 0.19 |
| 5 | 135516707 | rs6596293 | -1.263 | -0.011 | 0.145 | 0.485 | 0.009 | 0.026 | 0.026 | 3.257 | 0.31 | 0.58 | 0.24 | 0.38 | 0.33 | 0.19 |
| 5 | 135516836 | rs6596294 | -1.252 | -0.009 | 0.143 | 0.487 | 0.01 | 0.027 | 0.027 | 3.098 | 0.31 | 0.58 | 0.24 | 0.38 | 0.33 | 0.19 |

| | | | | | | | | | | | | | | | | |
|---|---|---|---|---|---|---|---|---|---|---|---|---|---|---|---|---|
| 5 | 135544695 | rs7728953 | 0.31 | -0.225 | 0.064 | 0.414 | -0.002 | -0.002 | -0.002 | 0 | 0.45 | 0.43 | 0.34 | 0.48 | 0.45 | 0.47 |
| 5 | 135545093 | rs7705351 | 0.591 | -0.468 | -0.066 | 0.138 | 0.02 | 0.013 | 0.013 | 0 | 0.16 | 0.12 | 0.07 | 0.07 | 0.1 | 0.08 |
| 5 | 135570567 | rs6596299 | -1.059 | -0.487 | 0.065 | 0.392 | -0.039 | -0.011 | -0.011 | 0 | 0.68 | 0.87 | 0.47 | 0.63 | 0.57 | 0.58 |
| 5 | 135574111 | rs10463951 | -0.776 | -0.48 | 0.306 | 0.388 | -0.011 | 0.002 | 0.002 | 0.831 | 0.65 | 0.77 | 0.46 | 0.63 | 0.56 | 0.58 |
| 5 | 135579422 | rs13157486 | 0.69 | -0.589 | 0.262 | 0.348 | -0.002 | 0 | 0 | 0 | 0.47 | 0.42 | 0.35 | 0.56 | 0.46 | 0.5 |
| 5 | 135581659 | rs11740657 | 0.196 | -0.468 | 0.309 | 0.412 | 0.003 | 0.003 | 0.003 | 0.539 | 0.18 | 0.18 | 0.09 | 0.06 | 0.11 | 0.08 |
| 5 | 135583912 | rs2042243 | 0.43 | -0.192 | 0.071 | 0.492 | 0.013 | -0.021 | -0.021 | 0.362 | 0.22 | 0.06 | 0.2 | 0.3 | 0.15 | 0.09 |
| 5 | 135589692 | rs7724982 | -0.073 | 0.048 | 0.527 | 0.406 | -0.008 | -0.005 | -0.005 | 0 | 0.05 | 0.19 | 0.03 | 0.01 | 0.02 | 0 |
| 5 | 135605350 | rs2546651 | -0.918 | 0.282 | -0.406 | 0.425 | -0.028 | -0.016 | -0.016 | 0 | 0.7 | 0.85 | 0.52 | 0.68 | 0.63 | 0.65 |
| 5 | 135611442 | rs6868895 | 0.473 | 0.244 | -0.294 | 0.398 | 0.029 | -0.004 | -0.004 | 0.683 | 0.48 | 0.69 | 0.35 | 0.54 | 0.36 | 0.52 |
| 5 | 135631449 | rs1392171 | 0.281 | 0.63 | -0.05 | 0.307 | 0.004 | 0.01 | 0.01 | 0 | 0.09 | 0.13 | 0.06 | 0.07 | 0.13 | 0.1 |
| 5 | 135631588 | rs1392172 | -0.443 | 0.573 | 0.007 | 0.192 | 0.014 | 0.05 | 0.05 | 0 | 0.58 | 0.82 | 0.48 | 0.68 | 0.6 | 0.67 |
| 5 | 135633741 | rs6894680 | 0.665 | 0.685 | 0.294 | 0.197 | 0.036 | -0.01 | -0.01 | 1.05 | 0.08 | 0.15 | 0.08 | 0.08 | 0.13 | 0.06 |
| 5 | 135650298 | rs4976485 | -1.334 | 0.502 | 0.06 | 0.325 | -0.003 | 0 | 0 | 0 | 0.67 | 0.79 | 0.78 | 0.73 | 0.66 | 0.48 |
| 5 | 135651243 | rs2277052 | 0.517 | 0.444 | 0.164 | 0.331 | -0.017 | 0.123 | 0.123 | 0.329 | 0.17 | 0.43 | 0.17 | 0.36 | 0.12 | 0.1 |
| 5 | 135663592 | rs2881486 | 0.714 | 0.435 | 0.146 | 0.294 | -0.001 | 0.05 | 0.05 | 0 | 0.39 | 0.11 | 0.5 | 0.29 | 0.39 | 0.32 |
| 5 | 135668524 | rs10053931 | 0.791 | 0.414 | 0.378 | 0.318 | -0.01 | 0.107 | 0.107 | 0.149 | 0.16 | 0.35 | 0.16 | 0.36 | 0.12 | 0.1 |
| 5 | 135678957 | rs4976489 | -1.414 | 0.46 | 0.125 | 0.32 | -0.002 | -0.004 | -0.004 | 0 | 0.67 | 0.78 | 0.77 | 0.71 | 0.66 | 0.48 |
| 5 | 135680239 | rs950715 | 0.494 | 0.507 | -0.139 | 0.325 | 0.009 | 0.039 | 0.039 | 0 | 0.42 | 0.28 | 0.53 | 0.27 | 0.41 | 0.32 |
| 5 | 135691678 | rs2673930 | -0.939 | 0.51 | 0.009 | 0.419 | -0.001 | 0.018 | 0.018 | 3.955 | 0.34 | 0.51 | 0.29 | 0.45 | 0.29 | 0.16 |
| 5 | 135692575 | rs2546661 | -0.005 | 0.468 | 0.025 | 0.435 | -0.03 | 0.028 | 0.028 | 3.867 | 0.26 | 0.43 | 0.21 | 0.36 | 0.16 | 0.1 |
| 5 | 135699853 | rs2649693 | 0.046 | 0.462 | -0.061 | 0.423 | -0.026 | 0.025 | 0.025 | 3.62 | 0.26 | 0.41 | 0.21 | 0.37 | 0.16 | 0.1 |
| 5 | 135705674 | rs2673925 | -1.261 | 0.585 | -0.11 | 0.371 | -0.003 | 0.005 | 0.005 | 1.597 | 0.37 | 0.6 | 0.31 | 0.46 | 0.31 | 0.2 |
| 5 | 135709277 | rs1587436 | -1.703 | 0.81 | -0.265 | 0.247 | -0.001 | -0.001 | -0.001 | 0 | 0.7 | 0.73 | 0.82 | 0.72 | 0.68 | 0.55 |
| 5 | 135722467 | rs184704 | 0.906 | 0.857 | -0.333 | 0.241 | -0.034 | 0.023 | 0.023 | 0 | 0.33 | 0.12 | 0.51 | 0.26 | 0.38 | 0.35 |
| 5 | 135724921 | rs346650 | 0.536 | 0.845 | -0.377 | 0.246 | 0.008 | -0.003 | -0.003 | 0 | 0.41 | 0.47 | 0.29 | 0.37 | 0.46 | 0.55 |
| 5 | 135727158 | rs346652 | -0.13 | 1.133 | -1.092 | 0.309 | -0.03 | 0.048 | 0.048 | 0 | 0.43 | 0.15 | 0.61 | 0.35 | 0.52 | 0.45 |
| 5 | 135753008 | rs7705405 | -0.229 | 1.562 | -0.818 | 0.654 | -0.059 | 0.007 | 0.007 | 0 | 0.1 | 0.01 | 0.09 | 0.07 | 0.15 | 0.1 |
| 5 | 135755386 | rs11748540 | -0.331 | 1.303 | -1.114 | 1.092 | -0.087 | -0.141 | -0.141 | 0 | 0.18 | 0.03 | 0.34 | 0.49 | 0.24 | 0.16 |
| 5 | 135756606 | rs346656 | 0.722 | 0.757 | -0.923 | 1.052 | -0.001 | -0.027 | -0.027 | 0 | 0.16 | 0.06 | 0.33 | 0.22 | 0.18 | 0.17 |
| 5 | 135758057 | rs346658 | -1.087 | 0.601 | -0.694 | 0.991 | 0.017 | 0.008 | 0.008 | 0.309 | 0.42 | 0.72 | 0.59 | 0.65 | 0.37 | 0.34 |
| 5 | 135764923 | rs7736962 | -0.352 | 0.905 | 0.066 | 0.896 | -0.002 | 0 | 0 | 0 | 0.32 | 0.66 | 0.27 | 0.44 | 0.21 | 0.18 |
| 5 | 135772456 | rs6872555 | -0.01 | 1.113 | -0.066 | 0.994 | 0.002 | 0.022 | 0.022 | 0 | 0.28 | 0.32 | 0.27 | 0.44 | 0.26 | 0.19 |
| 5 | 135795159 | rs346653 | -0.527 | 1.355 | -0.121 | 0.637 | -0.032 | 0.048 | 0.048 | 0 | 0.14 | 0.02 | 0.09 | 0.05 | 0.15 | 0.09 |
| 5 | 135842661 | rs10042003 | -0.906 | 1.339 | 0.568 | 1.143 | 0.02 | 0.031 | 0.031 | 0.254 | 0.24 | 0.48 | 0.44 | 0.27 | 0.25 | 0.22 |
| 5 | 135851739 | rs4270686 | -0.207 | 1.704 | -0.306 | 0.907 | -0.029 | 0 | 0 | 0.185 | 0.18 | 0.08 | 0.38 | 0.15 | 0.24 | 0.2 |
| 5 | 135864565 | rs4423993 | -1.106 | 1.793 | 0.735 | 1.169 | 0.075 | 0.002 | 0.002 | 0 | 0.19 | 0.5 | 0.22 | 0.14 | 0.26 | 0.13 |
| 5 | 135865077 | rs11952460 | -0.71 | 1.741 | 0.887 | 1.165 | -0.008 | 0.005 | 0.005 | 0 | 0.15 | 0.15 | 0.17 | 0.06 | 0.26 | 0.07 |
| 5 | 135895603 | rs7705446 | 1.384 | 1.383 | 1.48 | 1.838 | 0.015 | 0.039 | 0.039 | 0.098 | 0.78 | 0.35 | 0.76 | 0.63 | 0.73 | 0.85 |
| 5 | 135896392 | rs28378599 | -0.222 | 1.494 | 0.84 | 1.867 | -0.004 | -0.005 | -0.005 | 0.064 | 0.2 | 0.21 | 0.26 | 0.39 | 0.29 | 0.24 |
| 5 | 135916094 | rs13355974 | 1.257 | 1.427 | 1.235 | 1.905 | 0.023 | 0.059 | 0.059 | 0.098 | 0.78 | 0.38 | 0.76 | 0.63 | 0.73 | 0.85 |
| 5 | 135922353 | rs7735135 | 0.588 | 1.208 | 1.408 | 1.902 | 0.039 | 0.295 | 0.295 | 1.391 | 0.95 | 0.68 | 0.97 | 0.77 | 1 | 0.98 |
| 5 | 135929255 | rs4569863 | 1.049 | 1.444 | 1.274 | 1.863 | 0.024 | 0.062 | 0.062 | 0 | 0.78 | 0.41 | 0.76 | 0.63 | 0.73 | 0.85 |
| 5 | 135952213 | rs4141140 | -0.265 | 1.747 | 1.176 | 2.022 | 0.012 | 0.046 | 0.046 | 2.281 | 0.15 | 0.22 | 0.23 | 0.39 | 0.23 | 0.22 |
| 5 | 135956384 | rs11242336 | 0.1 | 1.703 | 1.743 | 1.944 | 0.109 | -0.033 | -0.033 | 0 | 0.88 | 0.71 | 0.83 | 0.94 | 0.76 | 0.94 |
| 5 | 135962502 | rs11744460 | 1.45 | 1.669 | 1.646 | 1.926 | 0.016 | 0.044 | 0.044 | 0.346 | 0.8 | 0.32 | 0.76 | 0.63 | 0.75 | 0.85 |
| 5 | 135973083 | rs2347092 | -1.556 | 1.72 | 1.678 | 1.93 | 0.055 | 0.062 | 0.062 | 1.684 | 0.3 | 0.54 | 0.44 | 0.47 | 0.49 | 0.36 |
| 5 | 135975093 | rs4976516 | -1.391 | 1.744 | 1.728 | 1.986 | 0.06 | 0.031 | 0.031 | 1.708 | 0.27 | 0.4 | 0.4 | 0.39 | 0.48 | 0.29 |
| 5 | 135978129 | rs4976518 | -1.641 | 1.748 | 1.688 | 1.967 | 0.068 | 0.075 | 0.075 | 1.464 | 0.29 | 0.46 | 0.44 | 0.48 | 0.48 | 0.36 |
| 5 | 135980919 | rs1331980 | -0.272 | 1.639 | 1.203 | 1.806 | 0.017 | 0.047 | 0.047 | 2.834 | 0.15 | 0.24 | 0.23 | 0.34 | 0.24 | 0.24 |
| 5 | 135990317 | rs10050402 | -0.708 | 1.655 | 1.151 | 1.835 | 0.018 | 0.006 | 0.006 | 0.605 | 0.23 | 0.25 | 0.32 | 0.28 | 0.37 | 0.23 |
| 5 | 135993207 | rs11950654 | 0.495 | 0.807 | 0.294 | 1.133 | -0.024 | 0.062 | 0.062 | 0 | 0.31 | 0.12 | 0.21 | 0.23 | 0.3 | 0.39 |
| 5 | 135993372 | rs11242338 | -0.03 | 0.82 | 0.681 | 1.161 | 0.052 | 0.017 | 0.017 | 0.425 | 0.76 | 0.66 | 0.68 | 0.72 | 0.63 | 0.77 |
| 5 | 136000937 | rs10075735 | -0.072 | 0.659 | 0.955 | 1.546 | 0.035 | 0.012 | 0.012 | 0.464 | 0.23 | 0.32 | 0.32 | 0.28 | 0.37 | 0.23 |
| 5 | 136003152 | rs13155995 | 0.718 | 0.174 | 0.92 | 1.309 | 0.05 | 0.006 | 0.006 | 0.26 | 0.49 | 0.28 | 0.52 | 0.51 | 0.4 | 0.44 |
| 5 | 136004552 | rs2188467 | -0.863 | 0.175 | 0.824 | 1.383 | 0.011 | -0.002 | -0.002 | 0 | 0.55 | 0.43 | 0.53 | 0.52 | 0.68 | 0.63 |
| 5 | 136010114 | rs981123 | 1.38 | -0.094 | 0.52 | 1.398 | 0.01 | 0.12 | 0.12 | 0 | 0.36 | 0.03 | 0.38 | 0.16 | 0.3 | 0.21 |
| 5 | 136014505 | rs7718740 | -0.581 | 0.112 | 0.545 | 1.389 | -0.003 | 0.058 | 0.058 | 0 | 0.68 | 0.87 | 0.8 | 0.76 | 0.69 | 0.59 |
| 5 | 136017011 | rs10073753 | 0.407 | -0.123 | 0.642 | 1.454 | -0.002 | -0.001 | -0.001 | 0 | 0.45 | 0.56 | 0.48 | 0.47 | 0.32 | 0.39 |
| 5 | 136019657 | rs6596333 | 0.407 | -0.123 | 0.642 | 1.454 | -0.002 | -0.001 | -0.001 | 0 | 0.45 | 0.56 | 0.48 | 0.46 | 0.32 | 0.39 |
| 5 | 136021586 | rs4476715 | -1.188 | -0.115 | 0.851 | 1.625 | 0.001 | 0.475 | 0.475 | 3.878 | 0.08 | 0.52 | 0.09 | 0.31 | 0.02 | 0.18 |
| 5 | 136027793 | rs13163027 | -0.901 | -0.07 | 0.281 | 1.634 | -0.004 | 0.406 | 0.406 | 2.974 | 0.07 | 0.35 | 0.08 | 0.31 | 0.02 | 0.18 |
| 5 | 136027890 | rs2188464 | 1.247 | -0.063 | -0.916 | 1.097 | -0.001 | 0.116 | 0.116 | 0 | 0.33 | 0.04 | 0.38 | 0.16 | 0.31 | 0.2 |
| 5 | 136029849 | rs4380648 | -0.497 | 0.073 | -0.907 | 0.852 | -0.036 | -0.048 | -0.048 | 3.918 | 0.37 | 0.76 | 0.41 | 0.5 | 0.43 | 0.34 |
| 5 | 136034863 | rs2188472 | -0.456 | 0.307 | -0.904 | 0.654 | -0.018 | 0.026 | 0.026 | 0 | 0.7 | 0.87 | 0.81 | 0.74 | 0.69 | 0.6 |
| 5 | 136035580 | rs4976524 | -0.076 | 0.649 | -0.212 | 0.686 | 0.031 | 0.012 | 0.012 | 0 | 0.23 | 0.3 | 0.33 | 0.28 | 0.35 | 0.18 |
| 5 | 136036257 | rs193493 | -0.08 | -0.15 | -0.492 | -0.126 | 0.004 | 0.061 | 0.061 | 0.047 | 0.47 | 0.68 | 0.42 | 0.27 | 0.54 | 0.37 |
| 5 | 136038199 | rs193489 | 0.79 | -0.121 | -0.136 | -0.424 | -0.001 | 0.03 | 0.03 | 0 | 0.44 | 0.37 | 0.44 | 0.81 | 0.45 | 0.43 |
| 5 | 136039745 | rs2188469 | -0.712 | -0.3 | -0.144 | -0.589 | -0.031 | 0.025 | 0.025 | 0 | 0.2 | 0.07 | 0.14 | 0.12 | 0.25 | 0.34 |

| | | | | | | | | | | | | | | | | |
|---|---|---|---|---|---|---|---|---|---|---|---|---|---|---|---|---|
| 5 | 136046520 | rs7701533 | 1.668 | -0.469 | 0.002 | -0.751 | -0.023 | -0.068 | -0.068 | 0.988 | 0.67 | 0.5 | 0.81 | 0.85 | 0.72 | 0.59 |
| 5 | 136055430 | rs7446328 | -0.272 | -0.883 | 0.166 | -0.612 | -0.068 | -0.072 | -0.072 | 0 | 0.87 | 0.55 | 0.94 | 0.97 | 0.98 | 0.93 |
| 5 | 136061610 | rs4976526 | 1.558 | -0.853 | 0.281 | -0.628 | -0.027 | -0.052 | -0.052 | 0 | 0.69 | 0.54 | 0.83 | 0.85 | 0.75 | 0.6 |
| 5 | 136068650 | rs12518838 | -0.325 | -0.815 | 0.355 | -0.599 | -0.063 | -0.066 | -0.066 | 0 | 0.87 | 0.61 | 0.95 | 0.97 | 0.98 | 0.93 |
| 5 | 136072105 | rs726847 | 0.08 | -0.807 | 0.231 | -0.451 | -0.009 | -0.053 | -0.053 | 0 | 0.63 | 0.47 | 0.58 | 0.82 | 0.69 | 0.75 |
| 5 | 136079775 | rs11242352 | 0.802 | -0.665 | -0.191 | -0.376 | -0.007 | 0.007 | 0.007 | 0 | 0.24 | 0.21 | 0.38 | 0.15 | 0.29 | 0.19 |
| 5 | 136083077 | rs12188559 | 0.512 | -0.737 | -0.437 | -0.445 | -0.02 | 0.041 | 0.041 | 0 | 0.27 | 0.15 | 0.37 | 0.15 | 0.29 | 0.18 |
| 5 | 136100390 | rs13176455 | -0.393 | -0.367 | -0.413 | -0.193 | -0.037 | 0.006 | 0.006 | 0 | 0.17 | 0.03 | 0.13 | 0.12 | 0.22 | 0.31 |
| 5 | 136109982 | rs13159721 | 0.286 | -0.581 | -0.435 | -0.143 | -0.029 | -0.018 | -0.018 | 1.052 | 0.08 | 0.01 | 0.06 | 0.09 | 0.13 | 0.07 |
| 5 | 136114752 | rs2157916 | 0.016 | -0.68 | -0.108 | -0.278 | -0.003 | 0.006 | 0.006 | 1.953 | 0.33 | 0.36 | 0.4 | 0.16 | 0.32 | 0.2 |
| 5 | 136115243 | rs929197 | 0.16 | -0.675 | -0.213 | -0.275 | -0.013 | 0.038 | 0.038 | 2.896 | 0.32 | 0.25 | 0.39 | 0.16 | 0.32 | 0.2 |
| 5 | 136133321 | rs10035385 | -0.618 | -0.56 | -0.024 | -0.164 | 0.002 | 0.016 | 0.016 | 0 | 0.17 | 0.06 | 0.12 | 0.12 | 0.19 | 0.3 |
| 5 | 136136002 | rs12517066 | 0.9 | -0.591 | -0.081 | -0.26 | -0.042 | 0.025 | 0.025 | 0.287 | 0.23 | 0.11 | 0.38 | 0.16 | 0.31 | 0.23 |
| 5 | 136141652 | rs12518384 | -0.78 | -0.59 | 0.051 | -0.206 | -0.004 | 0.013 | 0.013 | 0 | 0.45 | 0.43 | 0.53 | 0.3 | 0.52 | 0.56 |
| 5 | 136172926 | rs17712407 | -0.224 | -0.451 | -0.172 | 0.28 | -0.012 | -0.001 | -0.001 | 0 | 0.1 | 0.03 | 0.06 | 0.1 | 0.13 | 0.07 |
| 5 | 136179240 | rs4289554 | -1.565 | -0.62 | -0.471 | 0.888 | -0.12 | -0.039 | -0.039 | 0 | 0.56 | 0.93 | 0.56 | 0.63 | 0.52 | 0.66 |
| 5 | 136181754 | rs2190580 | -2.254 | -0.747 | 0.371 | 1.004 | 0.042 | -0.029 | -0.029 | 0 | 0.33 | 0.76 | 0.2 | 0.47 | 0.23 | 0.44 |
| 5 | 136189846 | rs2041064 | -1.969 | -0.632 | 0.563 | 1.059 | -0.022 | 0.019 | 0.019 | 0 | 0.3 | 0.6 | 0.18 | 0.47 | 0.22 | 0.41 |
| 5 | 136201487 | rs10515486 | 0.287 | -0.849 | 0.795 | 1.046 | -0.02 | -0.021 | -0.021 | 0 | 0.1 | 0.16 | 0.04 | 0.02 | 0.02 | 0.04 |
| 5 | 136208388 | rs2214993 | -1.195 | -0.781 | -0.115 | 1.082 | -0.039 | 0.037 | 0.037 | 0 | 0.5 | 0.81 | 0.54 | 0.62 | 0.48 | 0.55 |
| 5 | 136220872 | rs13188072 | -0.604 | -0.186 | 0.271 | 1.213 | -0.001 | -0.008 | -0.008 | 0 | 0.13 | 0.14 | 0.08 | 0.09 | 0.14 | 0.09 |
| 5 | 136221763 | rs6421910 | -0.754 | -0.011 | 0.414 | 0.903 | -0.011 | -0.04 | -0.04 | 0 | 0.33 | 0.75 | 0.44 | 0.54 | 0.31 | 0.29 |
| 5 | 136224357 | rs7702626 | 0.945 | 0.029 | 0.342 | 0.903 | 0.003 | -0.003 | -0.003 | 2.473 | 0.08 | 0.07 | 0.01 | 0 | 0.01 | 0 |
| 5 | 136228845 | rs7448833 | 0.083 | 0.247 | 0.445 | 1.029 | 0.028 | 0.065 | 0.065 | 0 | 0.24 | 0.58 | 0.4 | 0.51 | 0.29 | 0.25 |
| 5 | 136261046 | rs13179633 | 0.189 | -0.125 | 0.306 | 0.207 | 0.001 | -0.006 | -0.006 | 0.015 | 0.12 | 0.12 | 0.08 | 0.09 | 0.13 | 0.06 |
| 5 | 136261052 | rs11749072 | 0.23 | -0.199 | 0.387 | 0.299 | 0.011 | -0.001 | -0.001 | 0 | 0.68 | 0.43 | 0.5 | 0.29 | 0.67 | 0.67 |
| 5 | 136273789 | rs7710457 | 1.105 | -0.184 | 0.083 | 0.252 | -0.047 | 0.128 | 0.128 | 0.224 | 0.55 | 0.12 | 0.4 | 0.2 | 0.54 | 0.6 |
| 5 | 136274796 | rs740366 | 1.284 | -0.212 | -0.057 | 0.136 | 0.024 | 0.153 | 0.153 | 0.271 | 0.49 | 0.12 | 0.36 | 0.2 | 0.47 | 0.58 |
| 5 | 136311770 | rs6596361 | -0.723 | -0.017 | 0.609 | 0.098 | 0 | -0.033 | -0.033 | 0 | 0.43 | 0.45 | 0.34 | 0.1 | 0.42 | 0.19 |
| 5 | 136312039 | rs10043215 | 1.498 | 0.007 | 0.593 | 0.179 | 0.007 | 0.119 | 0.119 | 0.934 | 0.45 | 0.11 | 0.35 | 0.2 | 0.47 | 0.56 |
| 5 | 136313051 | rs3192243 | -1.058 | 0.039 | 0.65 | -0.16 | 0.005 | -0.061 | -0.061 | 0 | 0.44 | 0.51 | 0.35 | 0.13 | 0.43 | 0.25 |
| 5 | 136313801 | rs1051853 | -0.308 | 0.147 | 0.68 | -0.122 | 0.023 | -0.028 | -0.028 | 0 | 0.14 | 0.37 | 0.1 | 0.09 | 0.13 | 0.08 |
| 5 | 136319856 | rs6872714 | -0.911 | 0 | 0.451 | -0.142 | 0.003 | -0.056 | -0.056 | 0 | 0.42 | 0.48 | 0.33 | 0.14 | 0.41 | 0.23 |
| 5 | 136327205 | rs3756709 | 0.497 | 0.064 | 0.147 | 0.107 | -0.045 | 0 | 0 | 0.142 | 0.1 | 0.01 | 0.06 | 0.08 | 0.13 | 0.09 |
| 5 | 136334314 | rs4308478 | 1.027 | -0.472 | 0.405 | 0.176 | -0.029 | 0.001 | 0.001 | 0 | 0.82 | 0.73 | 0.9 | 0.87 | 0.9 | 0.8 |
| 5 | 136335318 | rs7703118 | 0.554 | -0.492 | 0.85 | 0.141 | -0.019 | 0.074 | 0.074 | 0 | 0.88 | 0.61 | 0.92 | 0.87 | 0.92 | 0.82 |
| 5 | 136335948 | rs6863244 | -1.062 | -0.462 | 0.662 | 0.031 | 0.004 | 0.018 | 0.018 | 0.817 | 0.52 | 0.54 | 0.63 | 0.79 | 0.53 | 0.45 |
| 5 | 136345401 | rs17521284 | 0.284 | -0.223 | 0.444 | 0.372 | -0.01 | 0.018 | 0.018 | 0.112 | 0.26 | 0.18 | 0.23 | 0.24 | 0.28 | 0.28 |
| 5 | 136345669 | rs10067912 | 0.062 | -0.213 | 0.69 | 0.627 | 0.017 | 0.318 | 0.318 | 0 | 0.87 | 0.46 | 0.67 | 0.6 | 0.88 | 0.96 |
| 5 | 136351987 | rs13158068 | 0.078 | 0.028 | 0.172 | 0.386 | -0.067 | 0.093 | 0.093 | 0.047 | 0.29 | 0.12 | 0.26 | 0.11 | 0.34 | 0.22 |
| 5 | 136354227 | rs10491300 | -1.11 | 0.068 | 0.264 | 0.623 | -0.024 | 0.077 | 0.077 | 0.065 | 0.14 | 0.22 | 0.09 | 0.32 | 0.1 | 0.3 |
| 5 | 136358263 | rs17703026 | -0.726 | 0.349 | 0.056 | 0.145 | 0.023 | 0.126 | 0.126 | 0 | 0.07 | 0.21 | 0.04 | 0.23 | 0.06 | 0.19 |
| 5 | 136368258 | rs3849044 | 0.879 | 0.364 | -0.741 | 0.009 | -0.07 | 0.16 | 0.16 | 0 | 0.37 | 0.05 | 0.3 | 0.14 | 0.41 | 0.36 |
| 5 | 136368354 | rs6596363 | 0.24 | 0.117 | -0.359 | -0.054 | 0 | -0.005 | -0.005 | 0 | 0.27 | 0.23 | 0.18 | 0.2 | 0.23 | 0.25 |
| 5 | 136371511 | rs17521509 | 0.079 | 0.072 | -0.553 | 0.108 | 0.003 | 0.143 | 0.143 | 0 | 0.38 | 0.13 | 0.26 | 0.13 | 0.38 | 0.37 |
| 5 | 136380195 | rs6898165 | -0.674 | 0.868 | -0.98 | 0.814 | -0.038 | 0.038 | 0.038 | 0 | 0.2 | 0.06 | 0.23 | 0.11 | 0.23 | 0.15 |
| 5 | 136394804 | rs6880363 | -0.177 | 0.217 | -0.433 | 0.257 | -0.019 | -0.052 | -0.052 | 0 | 0.84 | 0.72 | 0.94 | 0.93 | 0.89 | 0.94 |
| 5 | 136402958 | rs718704 | -0.051 | 0.243 | 0.036 | 0.958 | -0.08 | 0.004 | 0.004 | 0 | 0.18 | 0.01 | 0.22 | 0.11 | 0.22 | 0.16 |
| 5 | 136408030 | rs10434747 | 1.451 | -1.353 | -0.89 | 1.324 | -0.075 | -0.04 | -0.04 | 0 | 0.43 | 0.05 | 0.43 | 0.37 | 0.57 | 0.57 |
| 5 | 136409585 | rs2189599 | 0.353 | -1.657 | 0.049 | 1.506 | -0.044 | 0.006 | 0.006 | 0.278 | 0.74 | 0.48 | 0.75 | 0.69 | 0.83 | 0.78 |
| 5 | 136415192 | rs7708798 | -0.307 | -1.592 | 0.299 | 1.605 | -0.026 | 0.23 | 0.23 | 0 | 0.1 | 0.25 | 0.18 | 0.37 | 0.06 | 0.16 |
| 5 | 136416957 | rs713584 | 1.409 | -1.654 | 0.019 | 1.524 | -0.061 | -0.016 | -0.016 | 0 | 0.45 | 0.06 | 0.44 | 0.36 | 0.57 | 0.58 |
| 5 | 136417084 | rs713583 | 0.105 | -1.354 | 0.459 | 1.564 | -0.018 | 0.121 | 0.121 | 0 | 0.08 | 0.12 | 0.17 | 0.26 | 0.06 | 0.16 |
| 5 | 136422213 | rs2107443 | -0.98 | -1.661 | 0.947 | 1.394 | -0.005 | -0.005 | -0.005 | 0 | 0.31 | 0.44 | 0.49 | 0.38 | 0.31 | 0.37 |
| 5 | 136422358 | rs2158409 | -0.571 | -1.279 | 0.431 | 0.328 | -0.009 | -0.097 | -0.097 | 0 | 0.75 | 0.66 | 0.86 | 1 | 0.77 | 0.88 |
| 5 | 136424167 | rs2348188 | 1.24 | -1.077 | 0.028 | 0.964 | -0.034 | -0.016 | -0.016 | 0.191 | 0.41 | 0.23 | 0.29 | 0.39 | 0.43 | 0.43 |
| 5 | 136426367 | rs11743969 | 0.303 | -1.194 | -0.043 | 0.824 | 0 | 0.04 | 0.04 | 0 | 0.2 | 0.07 | 0.22 | 0.13 | 0.18 | 0.14 |
| 5 | 136427154 | rs10040966 | -0.764 | -1.259 | 0.254 | 0.92 | 0.004 | -0.006 | -0.006 | 0 | 0.44 | 0.35 | 0.49 | 0.45 | 0.43 | 0.39 |
| 5 | 136427927 | rs4976407 | -0.506 | -0.801 | 0.102 | 0.628 | 0.015 | 0.002 | 0.002 | 0 | 0.35 | 0.43 | 0.48 | 0.43 | 0.4 | 0.37 |
| 5 | 136430647 | rs1116919 | 0.778 | -0.773 | -0.075 | -0.061 | -0.032 | -0.007 | -0.007 | 0 | 0.28 | 0.17 | 0.28 | 0.23 | 0.34 | 0.22 |
| 5 | 136431150 | rs6888621 | -0.068 | -0.994 | -0.329 | -0.028 | -0.005 | 0.017 | 0.017 | 0 | 0.66 | 0.64 | 0.7 | 0.51 | 0.69 | 0.65 |
| 5 | 136435483 | rs7701979 | 0.991 | -0.909 | -0.791 | -0.051 | -0.009 | 0.077 | 0.077 | 0 | 0.3 | 0.11 | 0.18 | 0.16 | 0.29 | 0.29 |
| 5 | 136443115 | rs11741632 | 1.137 | -0.536 | -0.872 | 0.215 | -0.049 | -0.07 | -0.07 | 0 | 0.14 | 0.02 | 0.17 | 0.24 | 0.17 | 0.13 |
| 5 | 136444759 | rs11742757 | 1.092 | -0.641 | -0.547 | 0.255 | -0.013 | -0.026 | -0.026 | 1.953 | 0.17 | 0.12 | 0.21 | 0.36 | 0.19 | 0.18 |
| 5 | 136448486 | rs757422 | -0.23 | -0.825 | -0.21 | 0.187 | 0.005 | 0.002 | 0.002 | 0 | 0.87 | 0.87 | 0.81 | 0.85 | 0.89 | 0.83 |
| 5 | 136449299 | rs757423 | 1.326 | 0.034 | 0.068 | 0.05 | 0.024 | 0.05 | 0.05 | 1.561 | 0.26 | 0.13 | 0.16 | 0 | 0.24 | 0.13 |
| 5 | 136449339 | rs6869576 | 1.055 | 0.161 | -0.154 | 0.241 | -0.039 | -0.058 | -0.058 | 0.018 | 0.15 | 0.04 | 0.18 | 0.22 | 0.16 | 0.2 |
| 5 | 136452619 | rs715187 | 1.238 | 0.153 | 0.048 | 0.034 | 0.002 | 0.054 | 0.054 | 0 | 0.28 | 0.13 | 0.19 | 0.13 | 0.26 | 0.28 |
| 5 | 136452856 | rs2905964 | -0.318 | 0.181 | 0.231 | 0.066 | -0.014 | -0.04 | -0.04 | 0 | 0.78 | 0.83 | 0.69 | 0.63 | 0.75 | 0.63 |

| | | | | | | | | | | | | | | | | |
|---|---|---|---|---|---|---|---|---|---|---|---|---|---|---|---|---|
| 5 | 136453078 | rs2966736 | -1.187 | 0.229 | 0.226 | 0.074 | 0 | -0.002 | -0.002 | 0 | 0.94 | 0.93 | 0.83 | 0.85 | 0.9 | 0.83 |
| 5 | 136460641 | rs916980 | 0.442 | 0.184 | 0.715 | 0.038 | -0.001 | -0.001 | -0.001 | 0 | 0.47 | 0.51 | 0.48 | 0.5 | 0.49 | 0.35 |
| 5 | 136460855 | rs10072929 | 0.925 | 0.278 | 0.596 | 0.096 | 0 | -0.007 | -0.007 | 0 | 0.3 | 0.3 | 0.21 | 0.12 | 0.26 | 0.28 |
| 5 | 136475061 | rs7725922 | 0.951 | 0.434 | 0.602 | 0.43 | 0.016 | 0.048 | 0.048 | 0 | 0.21 | 0.28 | 0.3 | 0.37 | 0.22 | 0.17 |
| 5 | 136478798 | rs7705445 | -0.222 | 0.424 | 0.341 | 0.316 | 0.011 | 0.087 | 0.087 | 0 | 0.92 | 0.79 | 0.79 | 0.72 | 0.88 | 0.78 |
| 5 | 136484839 | rs1434660 | 0.164 | 0.314 | 0.229 | -0.026 | -0.038 | 0.068 | 0.068 | 0.521 | 0.6 | 0.75 | 0.41 | 0.26 | 0.52 | 0.46 |
| 5 | 136485859 | rs10515487 | 0.573 | 0.346 | 0.504 | 0.626 | -0.004 | 0.058 | 0.058 | 0 | 0.28 | 0.11 | 0.19 | 0.12 | 0.27 | 0.28 |
| 5 | 136513869 | rs17720840 | 0.863 | 0.311 | 0.135 | 0.976 | -0.042 | -0.061 | -0.061 | 4.615 | 0.15 | 0.03 | 0.13 | 0.23 | 0.15 | 0.2 |
| 5 | 136518169 | rs17720952 | 0.966 | 0.105 | 0.339 | 0.855 | -0.017 | -0.014 | -0.014 | 0 | 0.47 | 0.24 | 0.52 | 0.51 | 0.54 | 0.37 |
| 5 | 136521281 | rs10072007 | 0.109 | 0.305 | 0.332 | 0.925 | -0.008 | -0.014 | -0.014 | 0 | 0.9 | 0.92 | 0.81 | 0.86 | 0.88 | 0.84 |
| 5 | 136521684 | rs1434666 | 0.291 | 0.16 | 1.181 | 1.025 | 0.016 | 0.116 | 0.116 | 0.189 | 0.18 | 0.4 | 0.18 | 0.36 | 0.17 | 0.22 |
| 5 | 136522085 | rs930275 | 0.467 | 0.161 | 1.405 | 0.871 | -0.005 | -0.011 | -0.011 | 0.652 | 0.61 | 0.47 | 0.65 | 0.74 | 0.67 | 0.53 |
| 5 | 136527370 | rs6883260 | 1.199 | 0.092 | 1.292 | 0.854 | 0.001 | 0.017 | 0.017 | 0.466 | 0.68 | 0.33 | 0.6 | 0.49 | 0.68 | 0.59 |
| 5 | 136536194 | rs11957994 | 0.889 | 0.099 | 1.079 | 0.817 | 0.007 | 0.044 | 0.044 | 0 | 0.23 | 0.08 | 0.12 | 0.11 | 0.18 | 0.29 |
| 5 | 136536309 | rs1560932 | 1.725 | 0.101 | 1.096 | 0.822 | -0.037 | 0.026 | 0.026 | 0 | 0.44 | 0.16 | 0.48 | 0.38 | 0.5 | 0.3 |
| 5 | 136550402 | rs10479149 | 1.69 | 0.106 | 1.336 | 0.835 | -0.01 | 0.025 | 0.025 | 0 | 0.44 | 0.24 | 0.49 | 0.38 | 0.49 | 0.3 |
| 5 | 136550583 | rs11743006 | 1.226 | 0.141 | 1.121 | 0.882 | -0.044 | -0.064 | -0.064 | 1.207 | 0.14 | 0.01 | 0.12 | 0.22 | 0.15 | 0.18 |
| 5 | 136564577 | rs10065544 | 0.831 | 0.227 | 0.62 | 0.943 | 0.006 | 0.031 | 0.031 | 0 | 0.23 | 0.09 | 0.13 | 0.13 | 0.19 | 0.29 |
| 5 | 136571412 | rs10900848 | 1.705 | 0.202 | 0.658 | 0.947 | -0.033 | 0.035 | 0.035 | 0 | 0.44 | 0.17 | 0.48 | 0.38 | 0.49 | 0.3 |
| 5 | 136572793 | rs17779700 | 0.585 | 0.383 | 0.429 | 0.963 | 0.009 | 0.031 | 0.031 | 0 | 0.23 | 0.09 | 0.12 | 0.13 | 0.19 | 0.29 |
| 5 | 136573355 | rs1434637 | 0.243 | 0.527 | -0.363 | 0.789 | -0.015 | -0.017 | -0.017 | 0 | 0.89 | 0.91 | 0.8 | 0.85 | 0.85 | 0.81 |
| 5 | 136576623 | rs4976425 | 0.795 | 0.2 | -0.476 | 0.492 | 0.019 | 0.055 | 0.055 | 0.827 | 0.17 | 0.03 | 0.16 | 0.05 | 0.12 | 0.1 |
| 5 | 136593823 | rs4976428 | 1.493 | 0.028 | -0.174 | 0.412 | -0.018 | -0.017 | -0.017 | 0.086 | 0.45 | 0.28 | 0.53 | 0.51 | 0.52 | 0.34 |
| 5 | 136596050 | rs10055051 | 1.25 | 0.02 | -0.437 | 0.486 | -0.05 | -0.072 | -0.072 | 0 | 0.13 | 0.01 | 0.13 | 0.25 | 0.16 | 0.18 |
| 5 | 136597822 | rs2961624 | -0.219 | -0.127 | -0.255 | 0.346 | -0.027 | -0.033 | -0.033 | 0 | 0.81 | 0.92 | 0.86 | 0.79 | 0.8 | 0.63 |
| 5 | 136603181 | rs6596370 | 0.886 | -0.224 | 0.114 | 0.288 | 0.005 | 0.008 | 0.008 | 0.385 | 0.59 | 0.4 | 0.67 | 0.76 | 0.67 | 0.52 |
| 5 | 136605421 | rs2060426 | 0.533 | 0.013 | -0.024 | 0.555 | -0.018 | -0.041 | -0.041 | 0.508 | 0.68 | 0.78 | 0.73 | 0.46 | 0.65 | 0.42 |
| 5 | 136613156 | rs2961639 | -0.257 | 0.321 | -0.269 | 0.363 | -0.058 | -0.03 | -0.03 | 0 | 0.81 | 0.97 | 0.85 | 0.82 | 0.74 | 0.63 |
| 5 | 136614606 | rs1865405 | 0.599 | 0.063 | -0.056 | 0.537 | -0.001 | 0.004 | 0.004 | 0 | 0.17 | 0.18 | 0.17 | 0.37 | 0.18 | 0.22 |
| 5 | 136615395 | rs4976348 | 0.981 | 0.002 | 0.157 | 0.512 | 0.011 | 0.049 | 0.049 | 0.509 | 0.19 | 0.06 | 0.2 | 0.08 | 0.15 | 0.12 |
| 5 | 136619518 | rs988856 | 1.342 | 0.017 | 0.567 | 0.556 | -0.057 | -0.074 | -0.074 | 0 | 0.13 | 0.01 | 0.13 | 0.25 | 0.16 | 0.18 |
| 5 | 136620224 | rs2905560 | 1.352 | 0.038 | 0.931 | 0.615 | 0 | -0.005 | -0.005 | 0.193 | 0.46 | 0.48 | 0.5 | 0.38 | 0.5 | 0.31 |
| 5 | 136620998 | rs2905559 | 0.109 | 0.068 | 0.407 | 0.614 | -0.001 | 0.002 | 0.002 | 0 | 0.81 | 0.79 | 0.85 | 0.71 | 0.81 | 0.61 |
| 5 | 136624058 | rs2961638 | -0.393 | 0.105 | 0.333 | 0.506 | -0.002 | -0.008 | -0.008 | 0 | 0.83 | 0.88 | 0.89 | 0.81 | 0.83 | 0.65 |
| 5 | 136630810 | rs9327782 | -0.41 | 0.11 | 0.281 | 0.587 | 0.002 | -0.016 | -0.016 | 0.099 | 0.37 | 0.27 | 0.37 | 0.45 | 0.33 | 0.34 |
| 5 | 136637321 | rs2961637 | -0.309 | 0.38 | 0.29 | 0.663 | -0.003 | -0.008 | -0.008 | 0 | 0.84 | 0.89 | 0.89 | 0.83 | 0.83 | 0.64 |
| 5 | 136638780 | rs2961634 | -0.197 | 0.33 | 0.283 | 0.64 | -0.009 | -0.019 | -0.019 | 0 | 0.84 | 0.91 | 0.89 | 0.79 | 0.83 | 0.69 |
| 5 | 136639892 | rs2916647 | 1.25 | 0.175 | 0.849 | 0.647 | 0 | -0.001 | -0.001 | 0 | 0.6 | 0.4 | 0.68 | 0.4 | 0.61 | 0.46 |
| 5 | 136642189 | rs7733981 | -0.326 | 0.244 | 1.214 | 0.769 | 0.013 | 0.07 | 0.07 | 0.153 | 0.21 | 0.36 | 0.19 | 0.38 | 0.22 | 0.23 |
| 5 | 136643868 | rs9327783 | -0.936 | 0.197 | 1.202 | 0.677 | 0 | 0.069 | 0.069 | 0 | 0.27 | 0.5 | 0.21 | 0.39 | 0.23 | 0.25 |
| 5 | 136646649 | rs11745363 | 0.093 | 0.193 | 1.288 | 0.701 | -0.003 | -0.004 | -0.004 | 0 | 0.21 | 0.16 | 0.16 | 0.27 | 0.22 | 0.21 |
| 5 | 136648381 | rs739697 | -0.14 | 0.187 | 1.513 | 0.698 | 0.004 | 0.011 | 0.011 | 0 | 0.22 | 0.3 | 0.17 | 0.27 | 0.22 | 0.21 |
| 5 | 136650942 | rs17171354 | -2.321 | 0.193 | 1.394 | 0.72 | 0.054 | 0.211 | 0.211 | 3.848 | 0.06 | 0.41 | 0.06 | 0.13 | 0.01 | 0.05 |
| 5 | 136652348 | rs11741229 | 0.162 | 0.272 | 0.919 | 0.628 | -0.03 | -0.038 | -0.038 | 0 | 0.2 | 0.06 | 0.15 | 0.27 | 0.21 | 0.21 |
| 5 | 136665962 | rs11740211 | -0.344 | 0.305 | 1.076 | 0.62 | -0.001 | 0.005 | 0.005 | 0.27 | 0.23 | 0.29 | 0.16 | 0.26 | 0.21 | 0.2 |
| 5 | 136681134 | rs1229734 | -0.158 | 0.264 | 1.479 | 0.469 | 0.002 | 0.001 | 0.001 | 0 | 0.21 | 0.25 | 0.17 | 0.26 | 0.22 | 0.31 |
| 5 | 136683860 | rs2698233 | -1.411 | 0.035 | 1.447 | 0.381 | 0.001 | 0.001 | 0.001 | 0 | 0.42 | 0.46 | 0.32 | 0.6 | 0.42 | 0.57 |
| 5 | 136684506 | rs12517481 | 0.191 | 0.041 | 1.415 | 0.378 | -0.017 | -0.012 | -0.012 | 0 | 0.19 | 0.11 | 0.16 | 0.26 | 0.23 | 0.31 |
| 5 | 136684768 | rs1229732 | -1.408 | 0.108 | 1.371 | 0.373 | 0.003 | 0.001 | 0.001 | 0 | 0.42 | 0.5 | 0.32 | 0.6 | 0.42 | 0.57 |
| 5 | 136686608 | rs1229729 | -1.381 | 0.081 | 1.265 | 0.366 | 0.003 | 0.002 | 0.002 | 0 | 0.42 | 0.47 | 0.33 | 0.6 | 0.42 | 0.57 |
| 5 | 136698758 | rs1229708 | -1.164 | 0.142 | 0.676 | -0.001 | 0.004 | 0.011 | 0.011 | 0 | 0.53 | 0.38 | 0.56 | 0.66 | 0.52 | 0.66 |
| 5 | 136700372 | rs1229707 | 2.069 | 0.112 | -0.435 | -0.719 | -0.007 | 0.128 | 0.128 | 0 | 0.26 | 0.06 | 0.15 | 0 | 0.27 | 0.17 |
| 5 | 136706160 | rs11740681 | 1.449 | 0.411 | -0.37 | -0.47 | -0.046 | -0.077 | -0.077 | 0 | 0.18 | 0.01 | 0.13 | 0.28 | 0.18 | 0.2 |
| 5 | 136708619 | rs17171376 | 1.785 | 0.728 | 0.2 | -0.128 | 0.007 | 0.063 | 0.063 | 2.266 | 0.05 | 0.17 | 0.03 | 0.07 | 0.02 | 0.06 |
| 5 | 136710361 | rs6596374 | -1.526 | 0.968 | -0.343 | -0.067 | 0.003 | 0.024 | 0.024 | 0 | 0.85 | 0.84 | 0.75 | 0.67 | 0.8 | 0.83 |
| 5 | 136715749 | rs1229742 | -0.306 | 0.961 | -0.629 | -0.135 | -0.011 | 0.022 | 0.022 | 0 | 0.32 | 0.19 | 0.44 | 0.31 | 0.32 | 0.43 |
| 5 | 136732394 | rs1229726 | 0.121 | 1.125 | -0.675 | -0.134 | -0.003 | -0.003 | -0.003 | 0 | 0.22 | 0.16 | 0.19 | 0.27 | 0.22 | 0.32 |
| 5 | 136733948 | rs6873075 | 0.466 | 0.932 | -0.453 | -0.213 | -0.002 | -0.082 | -0.082 | 0.828 | 0.16 | 0.35 | 0.27 | 0.04 | 0.12 | 0.13 |
| 5 | 136736272 | rs17703757 | 0.929 | 1.119 | -1.151 | 0.241 | -0.075 | -0.103 | -0.103 | 0 | 0.14 | 0.01 | 0.24 | 0.33 | 0.24 | 0.18 |
| 5 | 136738113 | rs1229724 | -0.919 | 0.609 | -1.115 | -0.27 | -0.025 | 0.015 | 0.015 | 0.491 | 0.46 | 0.87 | 0.5 | 0.58 | 0.46 | 0.62 |
| 5 | 136739650 | rs739701 | 0.457 | 0.671 | -0.143 | -0.397 | 0 | -0.002 | -0.002 | 0 | 0.61 | 0.39 | 0.55 | 0.38 | 0.58 | 0.43 |
| 5 | 136739724 | rs739700 | -0.135 | 0.648 | 0.318 | -0.467 | -0.009 | -0.005 | -0.005 | 0 | 0.33 | 0.25 | 0.4 | 0.63 | 0.39 | 0.56 |
| 5 | 136742178 | rs739699 | -0.505 | 0.871 | 0.326 | -0.662 | 0.005 | -0.025 | -0.025 | 0 | 0.82 | 0.71 | 0.71 | 0.9 | 0.84 | 0.84 |
| 5 | 136772683 | rs2974501 | -0.128 | 0.703 | 0.347 | -0.284 | -0.016 | -0.009 | -0.009 | 0.085 | 0.32 | 0.23 | 0.4 | 0.63 | 0.39 | 0.56 |
| 5 | 136783523 | rs11948441 | 1.104 | 0.045 | 0.111 | 0.027 | -0.011 | -0.009 | -0.009 | 0 | 0.35 | 0.35 | 0.21 | 0.25 | 0.29 | 0.16 |
| 5 | 136787898 | rs17727364 | 1.241 | 0.518 | -0.298 | -0.115 | -0.044 | 0.052 | 0.052 | 0 | 0.14 | 0.03 | 0.3 | 0.05 | 0.2 | 0.07 |
| 5 | 136793148 | rs1528961 | -0.121 | 0.234 | -0.12 | -0.12 | -0.045 | 0.016 | 0.016 | 0 | 0.38 | 0.2 | 0.43 | 0.65 | 0.43 | 0.64 |
| 5 | 136797494 | rs4976447 | 0.099 | 0.415 | 0.093 | 0.006 | 0.008 | -0.049 | -0.049 | 0 | 0.76 | 0.65 | 0.65 | 0.89 | 0.74 | 0.81 |
| 5 | 136817159 | rs1108547 | 0.023 | 0.654 | -0.425 | 0.763 | -0.06 | -0.059 | -0.059 | 0 | 0.31 | 0.15 | 0.35 | 0.38 | 0.35 | 0.48 |

| | | | | | | | | | | | | | | | | |
|---|---|---|---|---|---|---|---|---|---|---|---|---|---|---|---|---|
| 5 | 136832279 | rs1104978 | 0.392 | 0.738 | -0.096 | 0.935 | 0.018 | 0.066 | 0.066 | 0.749 | 0.11 | 0.18 | 0.16 | 0.21 | 0.16 | 0.17 |
| 5 | 136839964 | rs1544028 | -0.221 | 0.527 | -0.303 | 0.89 | -0.051 | -0.05 | -0.05 | 0.008 | 0.69 | 0.84 | 0.65 | 0.62 | 0.65 | 0.52 |
| 5 | 136858359 | rs4976442 | 0.387 | 0.172 | 0.178 | 1.036 | 0.004 | 0.028 | 0.028 | 0.519 | 0.1 | 0.09 | 0.15 | 0.2 | 0.15 | 0.17 |
| 5 | 136882278 | rs11740609 | 0.035 | 0.002 | 0.016 | 0.813 | 0.039 | -0.015 | -0.015 | 0 | 0.45 | 0.21 | 0.62 | 0.47 | 0.59 | 0.55 |
| 5 | 136882334 | rs10056463 | -0.203 | -0.083 | 0.448 | 0.831 | -0.028 | -0.013 | -0.013 | 0.146 | 0.07 | 0.17 | 0.02 | 0.04 | 0.02 | 0.11 |
| 5 | 136893576 | rs10058264 | -0.532 | -0.097 | 0.581 | 0.824 | -0.03 | -0.011 | -0.011 | 0.055 | 0.09 | 0.3 | 0.03 | 0.04 | 0.02 | 0.11 |
| 5 | 136893719 | rs1989916 | 0.488 | 0.024 | 0.389 | 0.876 | 0.007 | -0.004 | -0.004 | 0 | 0.43 | 0.34 | 0.33 | 0.49 | 0.38 | 0.34 |
| 5 | 136897592 | rs1989919 | 1.229 | -0.041 | 0.09 | 0.872 | 0.037 | 0.013 | 0.013 | 0.305 | 0.32 | 0.2 | 0.18 | 0.29 | 0.24 | 0.17 |
| 5 | 136900203 | rs10463961 | 0.049 | 0.405 | 0.582 | 1.059 | 0.008 | 0.05 | 0.05 | 0.59 | 0.1 | 0.18 | 0.15 | 0.21 | 0.14 | 0.17 |
| 5 | 136901419 | rs10491336 | -0.461 | 0.766 | 0.307 | 1.173 | 0.018 | 0 | 0 | 0.038 | 0.25 | 0.39 | 0.25 | 0.24 | 0.32 | 0.41 |
| 5 | 136902696 | rs1541364 | 0.993 | 0.83 | -0.273 | 1.129 | -0.015 | -0.031 | -0.031 | 0 | 0.62 | 0.31 | 0.63 | 0.55 | 0.54 | 0.44 |
| 5 | 136909613 | rs4240364 | 0.922 | 0.748 | -0.326 | 1.05 | -0.015 | -0.019 | -0.019 | 0 | 0.6 | 0.32 | 0.63 | 0.53 | 0.53 | 0.43 |
| 5 | 136914326 | rs2189082 | -1.199 | 0.756 | -0.558 | 1.003 | -0.014 | -0.014 | -0.014 | 0 | 0.51 | 0.72 | 0.62 | 0.45 | 0.54 | 0.62 |
| 5 | 136930564 | rs6892472 | -0.51 | 1.011 | 0.287 | 1.011 | 0.051 | 0.105 | 0.105 | 1.119 | 0.1 | 0.29 | 0.11 | 0.12 | 0.13 | 0.09 |
| 5 | 136934484 | rs2189087 | 0.225 | 1.026 | -0.008 | 0.522 | 0.006 | -0.045 | -0.045 | 0 | 0.75 | 0.62 | 0.58 | 0.82 | 0.77 | 0.8 |
| 5 | 136940216 | rs6878439 | -1.057 | 0.948 | 0.144 | 1.053 | 0.031 | 0.082 | 0.082 | 0.232 | 0.13 | 0.35 | 0.29 | 0.16 | 0.13 | 0.11 |
| 5 | 136954719 | rs3813314 | 0.397 | 1.087 | -0.512 | 0.936 | 0.017 | 0.061 | 0.061 | 0 | 0.71 | 0.57 | 0.68 | 0.54 | 0.65 | 0.74 |
| 5 | 136955707 | rs3813318 | 0.206 | 1.721 | -1.533 | -0.085 | -0.103 | 0.125 | 0.125 | 0 | 0.19 | 0.01 | 0.24 | 0.02 | 0.26 | 0.16 |
| 5 | 136956683 | rs7444370 | 0.823 | 0.868 | -1.142 | -0.324 | -0.004 | 0.012 | 0.012 | 0.86 | 0.08 | 0.3 | 0.03 | 0.04 | 0.03 | 0.07 |
| 5 | 136961400 | rs2074344 | 0.414 | 1.032 | -1.441 | -0.581 | 0.006 | 0.148 | 0.148 | 0.016 | 0.06 | 0.08 | 0.18 | 0.46 | 0.08 | 0.24 |
| 5 | 136965249 | rs3777376 | 0.53 | 0.524 | -2.267 | -1.564 | -0.011 | 0.103 | 0.103 | 0 | 0.21 | 0.08 | 0.18 | 0.03 | 0.21 | 0.2 |
| 5 | 136969793 | rs2301708 | 0.215 | 0.428 | -2.473 | -1.693 | -0.002 | 0.052 | 0.052 | 0.473 | 0.1 | 0.09 | 0.21 | 0.46 | 0.18 | 0.24 |
| 5 | 136969953 | rs2301709 | 0.9 | -0.259 | -1.988 | -2.143 | -0.009 | 0.058 | 0.058 | 0.032 | 0.32 | 0.51 | 0.29 | 0.49 | 0.31 | 0.31 |
| 5 | 136972187 | rs17459939 | 0.974 | -0.249 | -1.967 | -2.156 | 0.002 | 0.072 | 0.072 | 0 | 0.21 | 0.13 | 0.19 | 0.03 | 0.21 | 0.2 |
| 5 | 136973913 | rs3777379 | 0.991 | -0.555 | -2.13 | -2.306 | -0.083 | 0.229 | 0.229 | 0 | 0.32 | 0.03 | 0.29 | 0.02 | 0.29 | 0.25 |
| 5 | 136996852 | rs11242395 | 1.088 | -0.563 | -2.071 | -2.252 | -0.011 | 0.102 | 0.102 | 0 | 0.21 | 0.08 | 0.18 | 0.03 | 0.21 | 0.2 |
| 5 | 136998182 | rs10037465 | -0.063 | -0.521 | -2.024 | -1.791 | -0.002 | 0.002 | 0.002 | 0 | 0.56 | 0.62 | 0.48 | 0.63 | 0.53 | 0.51 |
| 5 | 137000965 | rs11242396 | 1.266 | -0.623 | -1.967 | -1.663 | -0.004 | -0.018 | -0.018 | 2.138 | 0.25 | 0.25 | 0.28 | 0.52 | 0.3 | 0.26 |
| 5 | 137005673 | rs2040694 | 0.655 | -1.274 | -1.903 | -2.246 | 0.001 | -0.065 | -0.065 | 0.1 | 0.39 | 0.41 | 0.47 | 0.9 | 0.36 | 0.52 |
| 5 | 137016586 | rs4371746 | -0.118 | -1.222 | -1.997 | -2.25 | -0.015 | 0.237 | 0.237 | 0 | 0.54 | 0.77 | 0.58 | 0.93 | 0.52 | 0.6 |
| 5 | 137017681 | rs12188115 | 0.768 | -0.522 | -2.083 | -2.322 | -0.02 | 0.058 | 0.058 | 0.37 | 0.2 | 0.13 | 0.18 | 0.05 | 0.23 | 0.21 |
| 5 | 137037269 | rs2967806 | -1.018 | -0.486 | -1.977 | -2.202 | 0.008 | 0.043 | 0.043 | 0 | 0.74 | 0.82 | 0.75 | 0.98 | 0.76 | 0.81 |
| 5 | 137044526 | rs10515496 | 0.662 | -0.28 | -1.221 | -2.042 | 0.015 | -0.06 | -0.06 | 0.862 | 0.14 | 0.26 | 0.11 | 0.03 | 0.16 | 0.08 |
| 5 | 137135984 | rs17171590 | 1.381 | 0.138 | -1.55 | -2.108 | -0.01 | -0.055 | -0.055 | 0 | 0.15 | 0.06 | 0.13 | 0.52 | 0.16 | 0.17 |
| 5 | 137142853 | rs12652106 | 1.439 | 0.126 | -1.586 | -2.121 | -0.029 | -0.045 | -0.045 | 0 | 0.11 | 0.02 | 0.22 | 0.36 | 0.12 | 0.29 |
| 5 | 137164863 | rs10039139 | 0.578 | -0.175 | -1.345 | -2.445 | -0.017 | -0.057 | -0.057 | 0 | 0.23 | 0.37 | 0.18 | 0.05 | 0.19 | 0.17 |
| 5 | 137179924 | rs847821 | 0.805 | -0.219 | -1.379 | -2.468 | -0.023 | 0 | 0 | 0 | 0.53 | 0.19 | 0.49 | 0.56 | 0.53 | 0.45 |
| 5 | 137194762 | rs7722600 | 1.217 | -0.279 | -1.127 | -2.458 | 0.009 | -0.044 | -0.044 | 0 | 0.14 | 0.21 | 0.11 | 0.04 | 0.16 | 0.08 |
| 5 | 137201693 | rs3756687 | 1.239 | -0.275 | -1.145 | -2.456 | 0.007 | -0.04 | -0.04 | 0 | 0.13 | 0.2 | 0.11 | 0.04 | 0.16 | 0.08 |
| 5 | 137210651 | rs7713839 | -0.359 | -0.095 | -0.517 | -2.494 | 0.001 | -0.015 | -0.015 | 2.34 | 0.35 | 0.68 | 0.29 | 0.05 | 0.35 | 0.24 |
| 5 | 137241856 | rs700605 | -0.535 | -0.232 | -0.184 | -2.333 | 0.004 | -0.067 | -0.067 | 0 | 0.82 | 0.57 | 0.84 | 0.95 | 0.82 | 0.93 |
| 5 | 137244311 | rs2003491 | 0.811 | 0.517 | -0.392 | -1.291 | -0.005 | 0.019 | 0.019 | 0 | 0.53 | 0.24 | 0.43 | 0.58 | 0.5 | 0.56 |
| 5 | 137248721 | rs2349556 | 0.912 | 0.582 | -0.262 | -1.308 | 0.046 | 0.077 | 0.077 | 0 | 0.22 | 0.35 | 0.34 | 0.38 | 0.27 | 0.35 |
| 5 | 137255967 | rs6899035 | -0.197 | 0.34 | -0.13 | -1.43 | -0.019 | -0.125 | -0.125 | 0.592 | 0.75 | 0.52 | 0.77 | 0.97 | 0.77 | 0.9 |
| 5 | 137257848 | rs6891280 | 0.212 | 0.344 | 0.107 | -1.317 | -0.002 | -0.023 | -0.023 | 0 | 0.23 | 0.28 | 0.22 | 0.04 | 0.23 | 0.1 |
| 5 | 137271340 | rs11743778 | 3.333 | -0.126 | -0.446 | -1.835 | 0.009 | 0.004 | 0.004 | 0 | 0.1 | 0.08 | 0.11 | 0 | 0.08 | 0.03 |
| 5 | 137271884 | rs13357883 | 2.297 | 0.049 | -0.313 | -1.82 | -0.001 | -0.007 | -0.007 | 0 | 0.13 | 0.15 | 0.17 | 0.03 | 0.18 | 0.07 |
| 5 | 137275115 | rs3777118 | -0.419 | -0.024 | -0.277 | -1.819 | -0.014 | -0.097 | -0.097 | 0.474 | 0.75 | 0.59 | 0.78 | 0.97 | 0.76 | 0.9 |
| 5 | 137275199 | rs1141538 | -0.419 | -0.024 | -0.277 | -1.819 | -0.014 | -0.097 | -0.097 | 0.995 | 0.75 | 0.59 | 0.78 | 0.97 | 0.76 | 0.9 |
| 5 | 137281708 | rs1125979 | -0.419 | -0.024 | -0.277 | -1.819 | -0.014 | -0.097 | -0.097 | 0 | 0.75 | 0.59 | 0.78 | 0.97 | 0.76 | 0.9 |
| 5 | 137283812 | rs4557401 | 2.312 | -0.013 | -0.287 | -1.813 | -0.001 | -0.007 | -0.007 | 0 | 0.13 | 0.15 | 0.17 | 0.03 | 0.18 | 0.07 |
| 5 | 137298890 | rs4835662 | 2.102 | 0.031 | -0.916 | -2.4 | -0.048 | 0.068 | 0.068 | 0 | 0.13 | 0.01 | 0.17 | 0.02 | 0.18 | 0.19 |
| 5 | 137334791 | rs12516717 | 3.175 | 0.651 | -0.493 | -2.133 | 0.004 | 0.049 | 0.049 | 0.043 | 0.11 | 0.12 | 0.24 | 0.39 | 0.11 | 0.28 |
| 5 | 137359757 | rs13169864 | -1.04 | 0.565 | -0.668 | -2.091 | -0.014 | -0.097 | -0.097 | 2.065 | 0.75 | 0.59 | 0.78 | 0.97 | 0.76 | 0.9 |
| 5 | 137373502 | rs17171718 | 3.272 | 0.282 | -1.137 | -2.252 | -0.025 | -0.085 | -0.085 | 0 | 0.13 | 0.02 | 0.08 | 0.54 | 0.15 | 0.2 |
| 5 | 137412786 | rs11242415 | 0.89 | 0.542 | -0.655 | -2.23 | -0.004 | 0.012 | 0.012 | 0 | 0.53 | 0.32 | 0.43 | 0.58 | 0.5 | 0.56 |
| 5 | 137412963 | rs217259 | 1.443 | 0.593 | -0.583 | -2.272 | 0.027 | 0.047 | 0.047 | 0 | 0.22 | 0.28 | 0.34 | 0.39 | 0.27 | 0.35 |
| 5 | 137419494 | rs10041787 | 2.731 | 0.756 | -1.025 | -2.248 | -0.013 | 0.021 | 0.021 | 0.033 | 0.09 | 0.08 | 0.11 | 0.02 | 0.13 | 0.07 |
| 5 | 137419725 | rs10036244 | 2.299 | 0.759 | -0.981 | -2.193 | -0.004 | 0.001 | 0.001 | 0 | 0.14 | 0.14 | 0.17 | 0.03 | 0.18 | 0.08 |
| 5 | 137419989 | rs2040862 | 2.808 | 0.775 | -0.952 | -2.137 | -0.015 | 0.011 | 0.011 | 0 | 0.1 | 0.05 | 0.09 | 0 | 0.15 | 0.07 |
| 5 | 137435730 | rs11741240 | 0.462 | 0.788 | -0.894 | -2.005 | -0.002 | 0.011 | 0.011 | 0 | 0.53 | 0.32 | 0.43 | 0.58 | 0.49 | 0.56 |
| 5 | 137442353 | rs7720650 | 1.72 | 1.348 | -0.748 | -1.78 | 0 | -0.013 | -0.013 | 0 | 0.14 | 0.18 | 0.17 | 0.03 | 0.18 | 0.07 |
| 5 | 137445000 | rs12516561 | -0.234 | 1.716 | -0.935 | -1.371 | -0.035 | -0.045 | -0.045 | 1.229 | 0.13 | 0.06 | 0.28 | 0.32 | 0.17 | 0.38 |
| 5 | 137457418 | rs3849045 | 0.694 | 1.456 | -0.573 | -2.246 | -0.001 | 0.001 | 0.001 | 0 | 0.46 | 0.53 | 0.38 | 0.59 | 0.48 | 0.51 |
| 5 | 137475787 | rs423258 | 0.807 | 1.612 | -0.882 | -2.33 | -0.046 | -0.087 | -0.087 | 0 | 0.29 | 0.09 | 0.4 | 0.38 | 0.31 | 0.4 |
| 5 | 137486426 | rs217272 | 1.439 | 1.754 | -0.945 | -2.379 | -0.044 | -0.15 | -0.15 | 0.062 | 0.23 | 0.01 | 0.34 | 0.38 | 0.19 | 0.35 |
| 5 | 137493528 | rs17171767 | 0.303 | 1.666 | -0.566 | -2.426 | -0.001 | -0.007 | -0.007 | 1.755 | 0.33 | 0.27 | 0.42 | 0.38 | 0.32 | 0.4 |
| 5 | 137500665 | rs11750814 | 1.287 | 1.931 | -0.584 | -2.506 | -0.029 | 0.028 | 0.028 | 0 | 0.11 | 0.06 | 0.1 | 0.02 | 0.17 | 0.25 |
| 5 | 137506434 | rs13155519 | 1.243 | 1.649 | -0.37 | -2.625 | 0.002 | 0.001 | 0.001 | 0 | 0.39 | 0.38 | 0.35 | 0.59 | 0.43 | 0.47 |

| | | | | | | | | | | | | | | | | |
|---|---|---|---|---|---|---|---|---|---|---|---|---|---|---|---|---|
| 5 | 137523430 | rs2864 | 1.074 | 1.582 | -0.41 | -2.533 | -0.049 | -0.088 | -0.088 | 2.697 | 0.28 | 0.09 | 0.4 | 0.38 | 0.31 | 0.4 |
| 5 | 137527917 | rs2231478 | 0.593 | 1.304 | -0.358 | -2.491 | -0.003 | -0.056 | -0.056 | 0 | 0.28 | 0.33 | 0.23 | 0.02 | 0.25 | 0.13 |
| 5 | 137543135 | rs2107682 | 1.024 | 1.355 | -0.559 | -2.333 | -0.053 | -0.094 | -0.094 | 0.004 | 0.29 | 0.06 | 0.4 | 0.38 | 0.31 | 0.4 |
| 5 | 137552970 | rs1468333 | 0.531 | 1.382 | -0.524 | -2.332 | -0.002 | -0.011 | -0.011 | 0 | 0.33 | 0.26 | 0.42 | 0.38 | 0.32 | 0.41 |
| 5 | 137568867 | rs4835672 | 2.121 | 1.319 | -0.396 | -2.227 | -0.005 | 0.023 | 0.023 | 0 | 0.27 | 0.27 | 0.26 | 0.58 | 0.27 | 0.21 |
| 5 | 137573725 | rs10040989 | 1.997 | 1.05 | -1.009 | -2.234 | 0.006 | -0.017 | -0.017 | 0.034 | 0.06 | 0.07 | 0.06 | 0 | 0.12 | 0.05 |
| 5 | 137586330 | rs10515498 | 1.277 | 1.026 | -1.156 | -2.238 | -0.034 | -0.073 | -0.073 | 0 | 0.25 | 0.01 | 0.36 | 0.29 | 0.2 | 0.35 |
| 5 | 137587806 | rs9327809 | 0.796 | 0.822 | -0.997 | -2.139 | -0.037 | -0.004 | -0.004 | 0 | 0.3 | 0.08 | 0.42 | 0.29 | 0.32 | 0.4 |
| 5 | 137590786 | rs28704868 | 0.753 | 0.819 | -0.999 | -2.14 | -0.036 | -0.002 | -0.002 | 0 | 0.31 | 0.08 | 0.42 | 0.29 | 0.32 | 0.4 |
| 5 | 137599334 | rs11242417 | 1.923 | 0.966 | -0.917 | -2.076 | -0.046 | 0.094 | 0.094 | 0.02 | 0.16 | 0.02 | 0.15 | 0.02 | 0.16 | 0.09 |
| 5 | 137618772 | rs10054835 | 0.526 | 0.929 | -0.793 | -2.041 | -0.006 | 0.039 | 0.039 | 0 | 0.37 | 0.14 | 0.45 | 0.29 | 0.38 | 0.45 |
| 5 | 137619591 | rs4457069 | 1.358 | 0.951 | -0.515 | -2.012 | -0.029 | 0.022 | 0.022 | 0 | 0.33 | 0.48 | 0.27 | 0.61 | 0.27 | 0.19 |
| 5 | 137631073 | rs7716275 | -0.111 | 1.016 | -0.629 | -1.946 | -0.008 | -0.001 | -0.001 | 1.284 | 0.14 | 0.1 | 0.11 | 0.02 | 0.18 | 0.26 |
| 5 | 137646221 | rs10072318 | 0.55 | 1.017 | -0.701 | -1.969 | -0.008 | 0.025 | 0.025 | 0 | 0.35 | 0.11 | 0.42 | 0.29 | 0.34 | 0.45 |
| 5 | 137648679 | rs13361619 | 0.55 | 1.017 | -0.701 | -1.969 | -0.008 | 0.025 | 0.025 | 0 | 0.35 | 0.11 | 0.42 | 0.29 | 0.34 | 0.45 |
| 5 | 137665323 | rs3734166 | 1.403 | 0.976 | -0.396 | -1.954 | -0.025 | 0.026 | 0.026 | 0 | 0.33 | 0.48 | 0.3 | 0.61 | 0.31 | 0.2 |
| 5 | 137667200 | rs11567957 | 0.239 | 1.079 | -0.229 | -1.879 | -0.012 | 0.006 | 0.006 | 0 | 0.12 | 0.08 | 0.11 | 0.02 | 0.18 | 0.29 |
| 5 | 137667345 | rs1042124 | 0.946 | 1.098 | -0.26 | -1.87 | -0.063 | 0.046 | 0.046 | 0 | 0.1 | 0.01 | 0.1 | 0.02 | 0.17 | 0.29 |
| 5 | 137671058 | rs3756761 | 1.968 | 0.851 | -0.018 | -1.896 | -0.019 | 0.065 | 0.065 | 0 | 0.25 | 0.37 | 0.23 | 0.55 | 0.22 | 0.17 |
| 5 | 137673167 | rs3756766 | -0.769 | 0.863 | -0.046 | -1.929 | 0 | -0.037 | -0.037 | 2.15 | 0.73 | 0.58 | 0.76 | 0.85 | 0.73 | 0.89 |
| 5 | 137681986 | rs10900851 | 0.884 | 0.863 | -0.01 | -1.968 | 0 | -0.003 | -0.003 | 0 | 0.26 | 0.27 | 0.23 | 0.14 | 0.26 | 0.11 |
| 5 | 137682714 | rs2269953 | 0.599 | 0.886 | -0.059 | -2.001 | 0.006 | 0.008 | 0.008 | 0 | 0.38 | 0.46 | 0.34 | 0.58 | 0.4 | 0.41 |
| 5 | 137683305 | rs2269952 | 0.573 | 0.9 | -0.053 | -1.994 | 0.005 | 0.007 | 0.007 | 0 | 0.38 | 0.46 | 0.34 | 0.58 | 0.4 | 0.47 |
| 5 | 137684330 | rs740074 | 2.019 | 0.898 | -0.036 | -2.007 | -0.019 | 0.061 | 0.061 | 0 | 0.25 | 0.38 | 0.24 | 0.56 | 0.22 | 0.18 |
| 5 | 137707315 | rs757647 | 1.43 | 0.955 | -0.052 | -2.012 | -0.024 | 0.047 | 0.047 | 0 | 0.3 | 0.45 | 0.26 | 0.58 | 0.26 | 0.31 |
| 5 | 137708434 | rs4835678 | -0.065 | 0.957 | -0.054 | -2.033 | 0.014 | -0.039 | -0.039 | 0 | 0.65 | 0.56 | 0.67 | 0.85 | 0.59 | 0.74 |
| 5 | 137717213 | rs10073922 | 0.602 | 0.952 | -0.064 | -2.031 | -0.008 | 0.028 | 0.028 | 0 | 0.35 | 0.11 | 0.42 | 0.27 | 0.34 | 0.42 |
| 5 | 137718687 | rs3822837 | 2.153 | 0.95 | -0.097 | -1.993 | -0.011 | 0.043 | 0.043 | 0 | 0.24 | 0.32 | 0.23 | 0.56 | 0.22 | 0.18 |
| 5 | 137728774 | rs2269951 | 0.587 | 1.032 | -0.279 | -1.969 | 0.005 | 0.008 | 0.008 | 0 | 0.38 | 0.46 | 0.34 | 0.58 | 0.4 | 0.47 |
| 5 | 137733891 | rs17294563 | 0.586 | 1.086 | -0.327 | -1.932 | -0.066 | 0.047 | 0.047 | 0 | 0.11 | 0.01 | 0.1 | 0.02 | 0.17 | 0.29 |
| 5 | 137759010 | rs10040792 | -0.421 | 0.243 | -0.037 | -1.693 | 0.011 | -0.032 | -0.032 | 0 | 0.65 | 0.57 | 0.67 | 0.85 | 0.59 | 0.74 |
| 5 | 137763798 | rs17599026 | 2.175 | 0.101 | -0.315 | -1.713 | 0.001 | 0.008 | 0.008 | 0.231 | 0.15 | 0.01 | 0.12 | 0.1 | 0.08 | 0.13 |
| 5 | 137773525 | rs12659034 | 0.886 | 0.471 | 0.177 | -1.546 | -0.025 | 0.04 | 0.04 | 0 | 0.31 | 0.48 | 0.25 | 0.58 | 0.26 | 0.31 |
| 5 | 137776931 | rs2269947 | 0.083 | 0.41 | -0.024 | -1.582 | 0.002 | 0.072 | 0.072 | 0 | 0.39 | 0.17 | 0.25 | 0.24 | 0.37 | 0.56 |
| 5 | 137777269 | rs12516269 | 0.035 | 0.425 | -0.017 | -1.576 | 0.004 | 0.076 | 0.076 | 0 | 0.39 | 0.17 | 0.25 | 0.25 | 0.37 | 0.56 |
| 5 | 137782115 | rs4835796 | 0.772 | 1.348 | 0.911 | -1.664 | 0.01 | -0.028 | -0.028 | 0 | 0.61 | 0.53 | 0.48 | 0.78 | 0.57 | 0.6 |
| 5 | 137782204 | rs3798153 | 0.813 | 1.35 | 0.805 | -1.663 | 0.01 | -0.031 | -0.031 | 0 | 0.61 | 0.49 | 0.47 | 0.79 | 0.57 | 0.59 |
| 5 | 137788607 | rs740076 | -0.643 | 1.73 | 0.412 | -1.701 | -0.002 | -0.002 | -0.002 | 0 | 0.36 | 0.35 | 0.51 | 0.21 | 0.42 | 0.38 |
| 5 | 137797380 | rs7729723 | -0.729 | 1.638 | 0.153 | -1.913 | -0.018 | 0.088 | 0.088 | 0.603 | 0.41 | 0.29 | 0.51 | 0.18 | 0.45 | 0.29 |
| 5 | 137800024 | rs3813321 | -0.792 | 1.696 | 0.143 | -1.919 | -0.026 | 0.099 | 0.099 | 0.71 | 0.41 | 0.23 | 0.51 | 0.18 | 0.46 | 0.29 |
| 5 | 137802404 | rs11743810 | -0.969 | 1.654 | 0.167 | -2.025 | -0.003 | 0.071 | 0.071 | 0 | 0.43 | 0.28 | 0.57 | 0.18 | 0.54 | 0.32 |
| 5 | 137805652 | rs11741807 | -1.443 | 1.726 | 0.276 | -2.021 | 0.012 | 0.026 | 0.026 | 0 | 0.5 | 0.45 | 0.59 | 0.19 | 0.58 | 0.35 |
| 5 | 137818916 | rs6870458 | -1.094 | 1.607 | 0.372 | -1.998 | -0.001 | -0.003 | -0.003 | 0 | 0.44 | 0.45 | 0.57 | 0.19 | 0.53 | 0.33 |
| 5 | 137822866 | rs11242430 | -1.275 | 1.644 | 0.322 | -1.978 | 0.001 | -0.002 | -0.002 | 1.028 | 0.48 | 0.49 | 0.57 | 0.19 | 0.55 | 0.35 |
| 5 | 137834139 | rs11745596 | -0.333 | 1.729 | -0.092 | -1.878 | -0.064 | 0.085 | 0.085 | 0.016 | 0.35 | 0.16 | 0.5 | 0.19 | 0.44 | 0.27 |
| 5 | 137844274 | rs2242599 | 0.9 | 1.556 | 0.404 | -1.719 | -0.001 | 0.01 | 0.01 | 0 | 0.47 | 0.43 | 0.38 | 0.78 | 0.45 | 0.58 |
| 5 | 137849526 | rs10077407 | 0.9 | 1.554 | 0.491 | -1.72 | -0.001 | 0.01 | 0.01 | 0 | 0.47 | 0.43 | 0.38 | 0.78 | 0.45 | 0.58 |
| 5 | 137872056 | rs4835687 | 0.8 | 1.552 | 0.663 | -1.667 | 0.001 | 0.07 | 0.07 | 0 | 0.44 | 0.67 | 0.35 | 0.79 | 0.4 | 0.57 |
| 5 | 137876920 | rs154069 | 0.141 | 1.516 | 0.567 | -1.705 | -0.03 | 0.151 | 0.151 | 0 | 0.54 | 0.82 | 0.37 | 0.8 | 0.44 | 0.67 |
| 5 | 137881104 | rs188731 | 0.132 | 1.515 | 0.566 | -1.705 | -0.031 | 0.151 | 0.151 | 0 | 0.54 | 0.82 | 0.38 | 0.79 | 0.44 | 0.67 |
| 5 | 137883734 | rs11740501 | 0.616 | 1.761 | 0.457 | -1.817 | -0.052 | -0.01 | -0.01 | 0 | 0.15 | 0.04 | 0.2 | 0.13 | 0.2 | 0.19 |
| 5 | 137892170 | rs10117 | -0.343 | 1.193 | 0.713 | -1.224 | -0.03 | 0.152 | 0.152 | 0 | 0.54 | 0.82 | 0.38 | 0.79 | 0.44 | 0.67 |
| 5 | 137902339 | rs1042665 | 0.998 | 1.235 | 0.774 | -1.18 | 0.002 | 0.001 | 0.001 | 0 | 0.36 | 0.35 | 0.25 | 0.76 | 0.25 | 0.47 |
| 5 | 137924729 | rs7443140 | 0.026 | 0.974 | -0.801 | -1.063 | -0.008 | -0.018 | -0.018 | 0 | 0.91 | 0.9 | 0.98 | 0.99 | 0.96 | 0.94 |
| 5 | 137945037 | rs13159624 | -0.026 | 1.134 | -0.905 | -1.034 | -0.037 | 0.173 | 0.173 | 0 | 0.46 | 0.21 | 0.52 | 0.17 | 0.55 | 0.37 |
| 5 | 137947980 | rs6421919 | 0.897 | 1.072 | -0.047 | -0.098 | -0.016 | -0.031 | -0.031 | 0.243 | 0.91 | 0.89 | 0.97 | 0.99 | 0.96 | 0.92 |
| 5 | 137958314 | rs2060214 | -1.67 | 1.635 | -0.049 | 0.143 | 0.001 | -0.002 | -0.002 | 0 | 0.06 | 0.1 | 0.17 | 0.12 | 0.07 | 0.15 |
| 5 | 137965309 | rs6596445 | -1.791 | 2.04 | 0.134 | -0.031 | -0.002 | 0.001 | 0.001 | 0.582 | 0.22 | 0.73 | 0.38 | 0.26 | 0.25 | 0.23 |
| 5 | 137967734 | rs11743323 | -0.531 | 2.647 | -0.138 | -0.037 | -0.012 | 0.007 | 0.007 | 0 | 0.09 | 0.05 | 0.12 | 0.04 | 0.14 | 0.1 |
| 5 | 137967937 | rs7716228 | -0.79 | 2.522 | 0.878 | -0.049 | 0.044 | 0.011 | 0.011 | 0 | 0.12 | 0.27 | 0.18 | 0.14 | 0.18 | 0.07 |
| 5 | 137972584 | rs4835696 | -1.904 | 2.772 | 1.05 | -0.005 | 0.043 | -0.018 | -0.018 | 0 | 0.29 | 0.54 | 0.37 | 0.23 | 0.39 | 0.21 |
| 5 | 137976852 | rs7379760 | -1.617 | 2.147 | 1.228 | 0.715 | 0.02 | 0.052 | 0.052 | 0.659 | 0.3 | 0.51 | 0.39 | 0.39 | 0.37 | 0.42 |
| 5 | 137977109 | rs17657378 | -1.412 | 2.053 | 1.063 | 0.409 | -0.013 | 0.009 | 0.009 | 0 | 0.16 | 0.13 | 0.18 | 0.1 | 0.22 | 0.16 |
| 5 | 137977864 | rs17171904 | -0.853 | 2.058 | 0.846 | 0.306 | -0.005 | 0.01 | 0.01 | 0 | 0.14 | 0.09 | 0.1 | 0.11 | 0.14 | 0.12 |
| 5 | 137979158 | rs4835646 | -1.506 | 2.087 | 1.241 | 0.251 | 0.007 | -0.054 | -0.054 | 0 | 0.2 | 0.46 | 0.16 | 0.12 | 0.2 | 0.13 |
| 5 | 137980476 | rs3849047 | -1.56 | 2.303 | 0.561 | 0.57 | -0.02 | 0.006 | 0.006 | 0 | 0.29 | 0.2 | 0.28 | 0.28 | 0.35 | 0.35 |
| 5 | 137987439 | rs1651031 | -1.237 | 2.212 | 0.771 | 1.109 | 0.036 | 0.051 | 0.051 | 0 | 0.27 | 0.42 | 0.5 | 0.5 | 0.38 | 0.24 |
| 5 | 137995762 | rs10062943 | -1.283 | 2.338 | 0.557 | 0.224 | -0.015 | 0.024 | 0.024 | 0 | 0.16 | 0.12 | 0.21 | 0.02 | 0.21 | 0.17 |
| 5 | 138012666 | rs11957671 | -1.39 | 2.21 | 1.101 | 0.177 | 0.004 | -0.041 | -0.041 | 0 | 0.17 | 0.2 | 0.2 | 0.01 | 0.21 | 0.19 |

| | | | | | | | | | | | | | | | |
|---|---|---|---|---|---|---|---|---|---|---|---|---|---|---|---|
| 5 | 138020898 | rs10078499 | -2.115 | 1.931 | 1.966 | 0.133 | 0.01 | -0.12 | -0.12 | 0 | 0.23 | 0.53 | 0.24 | 0.03 | 0.23 | 0.2 |
| 5 | 138031907 | rs11742397 | -1.237 | 1.963 | 1.693 | 0.113 | -0.011 | 0.012 | 0.012 | 0 | 0.16 | 0.12 | 0.2 | 0.01 | 0.21 | 0.17 |
| 5 | 138053059 | rs311602 | -1.157 | 1.31 | 1.583 | -0.31 | -0.004 | -0.084 | -0.084 | 0 | 0.32 | 0.55 | 0.29 | 0.11 | 0.27 | 0.26 |
| 5 | 138055308 | rs17286171 | 0.038 | 2.87 | 1.22 | -0.605 | -0.025 | 0.028 | 0.028 | 0 | 0.05 | 0.01 | 0.09 | 0.04 | 0.18 | 0.1 |
| 5 | 138061341 | rs160401 | -0.851 | 2.663 | 1.5 | -0.216 | -0.001 | 0.001 | 0.001 | 0 | 0.29 | 0.36 | 0.26 | 0.16 | 0.27 | 0.26 |
| 5 | 138084269 | rs160404 | 0.709 | 2.657 | 1.554 | -0.566 | -0.001 | -0.054 | -0.054 | 0 | 0.69 | 0.57 | 0.73 | 0.92 | 0.73 | 0.74 |
| 5 | 138087873 | rs700616 | 0.649 | 2.655 | 1.574 | -0.557 | 0 | -0.04 | -0.04 | 0 | 0.7 | 0.61 | 0.73 | 0.92 | 0.73 | 0.74 |
| 5 | 138091697 | rs12653845 | 0.851 | 1.924 | 0.505 | -0.735 | 0.015 | -0.037 | -0.037 | 1.098 | 0.14 | 0.01 | 0.1 | 0.13 | 0.08 | 0.1 |
| 5 | 138115219 | rs825770 | 0.234 | 1.991 | 0.615 | -0.969 | 0 | -0.035 | -0.035 | 0 | 0.7 | 0.61 | 0.73 | 0.92 | 0.73 | 0.74 |
| 5 | 138167497 | rs6596456 | 0.499 | 1.959 | 0.731 | -0.999 | -0.004 | -0.057 | -0.057 | 0.029 | 0.68 | 0.4 | 0.71 | 0.88 | 0.72 | 0.74 |
| 5 | 138206200 | rs288039 | -0.416 | 2.14 | 0.693 | -1.015 | -0.001 | -0.054 | -0.054 | 0 | 0.31 | 0.43 | 0.28 | 0.07 | 0.27 | 0.26 |
| 5 | 138210895 | rs288036 | -0.08 | 2.129 | 0.769 | -1.339 | -0.015 | 0.049 | 0.049 | 0 | 0.27 | 0.19 | 0.25 | 0.01 | 0.27 | 0.23 |
| 5 | 138222330 | rs288027 | -0.903 | 1.702 | 0.142 | -2.119 | -0.038 | 0.077 | 0.077 | 0 | 0.87 | 1 | 0.91 | 1 | 0.83 | 0.97 |
| 5 | 138256139 | rs10078079 | 0.184 | 1.476 | 0.532 | -1.808 | -0.028 | 0.079 | 0.079 | 0 | 0.26 | 0.13 | 0.24 | 0.01 | 0.27 | 0.23 |
| 5 | 138266546 | rs1059110 | -0.313 | 1.429 | 0.515 | -1.819 | 0.001 | -0.103 | -0.103 | 0 | 0.29 | 0.43 | 0.27 | 0.01 | 0.27 | 0.22 |
| 5 | 138272536 | rs9942422 | -0.429 | 1.43 | 0.47 | -1.811 | 0.002 | -0.129 | -0.129 | 0 | 0.3 | 0.54 | 0.28 | 0.05 | 0.27 | 0.22 |
| 5 | 138277847 | rs10037071 | -0.509 | 1.283 | 0.48 | -1.821 | -0.008 | -0.135 | -0.135 | 0 | 0.32 | 0.54 | 0.28 | 0.05 | 0.27 | 0.24 |
| 5 | 138278251 | rs7722413 | -0.538 | 1.29 | 0.47 | -1.818 | -0.008 | -0.126 | -0.126 | 0 | 0.32 | 0.55 | 0.28 | 0.05 | 0.27 | 0.24 |
| 5 | 138281261 | rs6878910 | 0.338 | 1.366 | 0.715 | -1.908 | -0.011 | -0.08 | -0.08 | 0 | 0.21 | 0.31 | 0.16 | 0.02 | 0.15 | 0.14 |
| 5 | 138299630 | rs12108892 | -0.249 | 1.571 | 1.329 | -1.502 | -0.001 | -0.014 | -0.014 | 2.516 | 0.42 | 0.61 | 0.36 | 0.14 | 0.33 | 0.28 |
| 5 | 138307306 | rs12055237 | -0.137 | 1.586 | 0.963 | -1.508 | 0.014 | 0.066 | 0.066 | 0 | 0.4 | 0.3 | 0.34 | 0.12 | 0.33 | 0.29 |
| 5 | 138308466 | rs2060222 | 0.099 | 1.683 | 0.899 | -1.336 | 0.009 | 0.091 | 0.091 | 0 | 0.39 | 0.21 | 0.34 | 0.16 | 0.33 | 0.31 |
| 5 | 138314106 | rs2882667 | 0.202 | 1.687 | 0.844 | -1.234 | 0.004 | 0.078 | 0.078 | 0 | 0.38 | 0.16 | 0.34 | 0.17 | 0.33 | 0.32 |
| 5 | 138318158 | rs11747454 | 1.511 | 1.908 | 0.386 | -1.604 | -0.03 | 0.094 | 0.094 | 0 | 0.22 | 0.01 | 0.11 | 0.06 | 0.2 | 0.18 |
| 5 | 138324706 | rs10070021 | -0.195 | 1.695 | 1.558 | -1.269 | 0.024 | 0.061 | 0.061 | 0.16 | 0.48 | 0.69 | 0.39 | 0.16 | 0.35 | 0.35 |
| 5 | 138357211 | rs3749665 | -0.21 | 1.849 | 1.64 | -1.205 | 0.051 | 0.159 | 0.159 | 0.484 | 0.48 | 0.81 | 0.4 | 0.19 | 0.35 | 0.32 |
| 5 | 138361135 | rs6863347 | -0.009 | 1.858 | 1.728 | -1.185 | -0.002 | -0.018 | -0.018 | 0 | 0.44 | 0.57 | 0.38 | 0.16 | 0.34 | 0.32 |
| 5 | 138401773 | rs10057932 | -0.92 | 1.26 | 1.87 | -1.082 | 0.066 | 0.031 | 0.031 | 1.804 | 0.05 | 0.34 | 0.05 | 0.01 | 0.03 | 0 |
| 5 | 138406078 | rs7701116 | 0.057 | 1.479 | 1.581 | -1.141 | -0.014 | 0.08 | 0.08 | 0 | 0.44 | 0.27 | 0.41 | 0.19 | 0.41 | 0.37 |
| 5 | 138436392 | rs6596462 | -0.95 | 1.444 | 1.662 | -1.067 | 0.077 | 0.068 | 0.068 | 1.574 | 0.05 | 0.34 | 0.05 | 0.03 | 0.03 | 0 |
| 5 | 138444170 | rs13174479 | 0.768 | 1.59 | 0.732 | -1.238 | -0.016 | 0.142 | 0.142 | 0.045 | 0.38 | 0.09 | 0.32 | 0.15 | 0.32 | 0.32 |
| 5 | 138454192 | rs7731574 | -0.252 | 1.707 | 1.014 | -1.233 | -0.005 | 0.161 | 0.161 | 0.071 | 0.5 | 0.75 | 0.46 | 0.19 | 0.44 | 0.37 |
| 5 | 138461547 | rs11951084 | -0.257 | 1.548 | 0.957 | -1.235 | 0.007 | 0.124 | 0.124 | 0.136 | 0.49 | 0.71 | 0.44 | 0.18 | 0.42 | 0.34 |
| 5 | 138468344 | rs1549203 | 0.301 | 1.482 | 0.986 | -1.235 | -0.003 | 0.032 | 0.032 | 0 | 0.42 | 0.36 | 0.36 | 0.17 | 0.37 | 0.36 |
| 5 | 138471330 | rs2351905 | -0.296 | 1.334 | 0.716 | -1.193 | 0.005 | 0.152 | 0.152 | 0.068 | 0.49 | 0.74 | 0.44 | 0.18 | 0.42 | 0.34 |
| 5 | 138483565 | rs13354264 | -1.361 | 1.555 | 1.132 | -0.774 | 0.13 | 0.031 | 0.031 | 0.936 | 0.08 | 0.39 | 0.11 | 0.02 | 0.09 | 0.02 |
| 5 | 138486543 | rs11958425 | 0.596 | 0.978 | -0.023 | -0.921 | -0.013 | 0.253 | 0.253 | 0 | 0.5 | 0.19 | 0.41 | 0.13 | 0.45 | 0.5 |
| 5 | 138487253 | rs10900858 | 0.492 | 0.826 | 0.206 | -0.897 | 0.01 | 0.092 | 0.092 | 0 | 0.53 | 0.34 | 0.44 | 0.16 | 0.5 | 0.53 |
| 5 | 138647766 | rs13178336 | 0.994 | 0.331 | -0.399 | -0.345 | -0.002 | 0.045 | 0.045 | 1.614 | 0.08 | 0 | 0.03 | 0 | 0.05 | 0.03 |
| 5 | 138714474 | rs11950646 | 1.408 | -0.006 | -0.893 | 0.067 | -0.022 | 0.049 | 0.049 | 1.12 | 0.57 | 0.05 | 0.6 | 0.29 | 0.62 | 0.5 |
| 5 | 138719526 | rs10063949 | -1.331 | 0.012 | -0.53 | 0.182 | 0.008 | 0.036 | 0.036 | 0.788 | 0.41 | 0.83 | 0.39 | 0.67 | 0.38 | 0.48 |
| 5 | 138719658 | rs6596475 | 1.163 | -0.061 | -0.535 | 0.224 | 0.045 | 0.023 | 0.023 | 0.392 | 0.61 | 0.21 | 0.63 | 0.33 | 0.65 | 0.52 |
| 5 | 138721164 | rs12516732 | 1.396 | -0.073 | -0.582 | 0.204 | -0.014 | 0.035 | 0.035 | 0.847 | 0.57 | 0.07 | 0.6 | 0.31 | 0.62 | 0.5 |
| 5 | 138732579 | rs10900862 | 1.389 | -0.069 | -0.581 | 0.205 | -0.015 | 0.035 | 0.035 | 0.705 | 0.57 | 0.07 | 0.6 | 0.31 | 0.62 | 0.5 |
| 5 | 138733811 | rs4835726 | -1.296 | -0.055 | -0.555 | 0.237 | 0.006 | 0.027 | 0.027 | 0.631 | 0.4 | 0.83 | 0.39 | 0.66 | 0.38 | 0.49 |
| 5 | 138747460 | rs11477 | 0.868 | -0.055 | -0.44 | 0.243 | 0.023 | 0.009 | 0.009 | 0.107 | 0.63 | 0.26 | 0.64 | 0.35 | 0.67 | 0.52 |
| 5 | 138762305 | rs11242465 | 0.416 | -0.215 | -0.102 | 0.464 | 0.009 | -0.043 | -0.043 | 0.672 | 0.71 | 0.84 | 0.72 | 0.39 | 0.7 | 0.53 |
| 5 | 138762998 | rs11748320 | 1.055 | -0.058 | -0.373 | 0.477 | 0.034 | 0.02 | 0.02 | 0.26 | 0.62 | 0.21 | 0.57 | 0.35 | 0.65 | 0.52 |
| 5 | 138767565 | rs11242467 | 1.263 | -0.054 | -0.385 | 0.579 | -0.004 | 0.031 | 0.031 | 0.918 | 0.59 | 0.1 | 0.6 | 0.32 | 0.63 | 0.5 |
| 5 | 138772180 | rs10900863 | 1.091 | -0.264 | -0.349 | 0.616 | 0.041 | -0.005 | -0.005 | 0 | 0.61 | 0.21 | 0.63 | 0.37 | 0.66 | 0.52 |
| 5 | 138773090 | rs10900864 | 1.152 | -0.289 | -0.53 | 0.522 | 0.032 | 0.038 | 0.038 | 0.878 | 0.6 | 0.12 | 0.61 | 0.31 | 0.65 | 0.51 |
| 5 | 138775552 | rs7445264 | 1.134 | -0.267 | -0.527 | 0.485 | 0.026 | 0.032 | 0.032 | 0.771 | 0.6 | 0.11 | 0.61 | 0.31 | 0.65 | 0.51 |
| 5 | 138844599 | rs28419191 | 0.042 | -0.467 | 0.198 | 1.014 | 0.002 | 0.008 | 0.008 | 0 | 0.9 | 0.9 | 0.86 | 0.78 | 0.86 | 0.9 |
| 5 | 138847901 | rs7380062 | 0.208 | -0.568 | 0.468 | 1.34 | -0.005 | -0.094 | -0.094 | 0.008 | 0.14 | 0.01 | 0.19 | 0.34 | 0.16 | 0.17 |
| 5 | 138850905 | rs13181561 | 1.29 | -0.651 | 1.147 | 1.243 | -0.007 | 0.023 | 0.023 | 0 | 0.71 | 0.38 | 0.61 | 0.37 | 0.69 | 0.63 |
| 5 | 138856982 | rs7380824 | -0.342 | -0.725 | 1.248 | 0.884 | -0.024 | 0.047 | 0.047 | 0 | 0.18 | 0.27 | 0.29 | 0.4 | 0.15 | 0.26 |
| 5 | 138857919 | rs1131769 | -0.077 | -0.691 | 1.285 | 0.836 | 0.005 | -0.001 | -0.001 | 0.338 | 0.9 | 0.89 | 0.85 | 0.86 | 0.85 | 0.9 |
| 5 | 138860604 | rs7380272 | -0.294 | -1.138 | 1.21 | 0.742 | -0.016 | 0.029 | 0.029 | 0 | 0.18 | 0.24 | 0.29 | 0.4 | 0.15 | 0.26 |
| 5 | 138919078 | rs2336901 | -1.492 | -1.288 | 0.841 | -0.408 | 0.071 | 0.162 | 0.162 | 0.508 | 0.4 | 0.79 | 0.26 | 0.16 | 0.31 | 0.45 |
| 5 | 138933151 | rs261537 | -0.561 | -1.337 | 1.158 | -0.441 | -0.013 | -0.04 | -0.04 | 1.521 | 0.29 | 0.56 | 0.2 | 0.1 | 0.23 | 0.34 |
| 5 | 138949362 | rs261532 | -1.596 | -1.298 | 0.872 | -0.444 | 0.049 | 0.212 | 0.212 | 1.746 | 0.41 | 0.95 | 0.27 | 0.16 | 0.31 | 0.45 |
| 5 | 138985694 | rs165179 | -1.685 | -1.069 | 0.543 | -0.307 | -0.002 | -0.013 | -0.013 | 0 | 0.1 | 0.15 | 0.05 | 0.06 | 0.08 | 0.12 |
| 5 | 138992703 | rs356428 | -0.109 | -1.094 | 0.659 | -0.315 | 0.015 | 0.024 | 0.024 | 0 | 0.26 | 0.17 | 0.17 | 0.1 | 0.22 | 0.33 |
| 5 | 139029000 | rs187167 | -2.015 | -0.679 | 0.368 | -0.005 | -0.003 | -0.026 | -0.026 | 0 | 0.1 | 0.16 | 0.06 | 0.05 | 0.09 | 0.14 |
| 5 | 139033279 | rs356447 | -1.746 | -0.678 | 0.347 | 0.06 | -0.006 | -0.016 | -0.016 | 0 | 0.08 | 0.21 | 0.06 | 0.06 | 0.06 | 0.14 |
| 5 | 139045003 | rs6580190 | -1.252 | -1.25 | -0.882 | -0.081 | -0.01 | -0.177 | -0.177 | 0.509 | 0.78 | 1 | 0.88 | 0.56 | 0.87 | 0.77 |
| 5 | 139053563 | rs356453 | -0.598 | -1.083 | 0.129 | -1.129 | -0.012 | -0.06 | -0.06 | 0.55 | 0.33 | 0.44 | 0.24 | 0.09 | 0.32 | 0.35 |
| 5 | 139058454 | rs585101 | -0.907 | -1.158 | 0.008 | -1.175 | -0.002 | -0.03 | -0.03 | 0.153 | 0.4 | 0.44 | 0.26 | 0.1 | 0.34 | 0.35 |
| 5 | 139063728 | rs2336880 | -0.961 | -0.926 | -0.744 | -1.126 | 0.078 | 0.137 | 0.137 | 1.173 | 0.43 | 0.75 | 0.29 | 0.13 | 0.33 | 0.45 |

| | | | | | | | | | | | | | | | | |
|---|---|---|---|---|---|---|---|---|---|---|---|---|---|---|---|---|
| 5 | 139088077 | rs165193 | -0.926 | -0.057 | -0.237 | -0.67 | 0.008 | -0.029 | -0.029 | 0.413 | 0.06 | 0.11 | 0.06 | 0.01 | 0.08 | 0.07 |
| 5 | 139103881 | rs260735 | 0.377 | -0.385 | -0.205 | -0.564 | -0.002 | 0.001 | 0.001 | 0 | 0.9 | 0.87 | 0.93 | 0.91 | 0.91 | 0.93 |
| 5 | 139106190 | rs260736 | 0.067 | -0.396 | -0.395 | -0.552 | -0.004 | -0.014 | -0.014 | 0 | 0.91 | 0.95 | 0.94 | 0.9 | 0.9 | 0.93 |
| 5 | 139110879 | rs262787 | -0.667 | -0.363 | -0.019 | -0.724 | 0.018 | -0.005 | -0.005 | 4.703 | 0.08 | 0.2 | 0.08 | 0.08 | 0.12 | 0.15 |
| 5 | 139120826 | rs356472 | -0.07 | -0.814 | -0.199 | -1.262 | 0.001 | -0.013 | -0.013 | 0 | 0.23 | 0.24 | 0.13 | 0.04 | 0.16 | 0.3 |
| 5 | 139132796 | rs260718 | 1.361 | -0.467 | 0.286 | -0.876 | -0.04 | -0.021 | -0.021 | 0.764 | 0.82 | 0.55 | 0.89 | 0.88 | 0.87 | 0.86 |
| 5 | 139140709 | rs260727 | 0.06 | -0.2 | -0.406 | -0.716 | 0.001 | -0.011 | -0.011 | 0.015 | 0.91 | 0.95 | 0.94 | 0.9 | 0.9 | 0.93 |
| 5 | 139144766 | rs165185 | 0.948 | 0.241 | -0.858 | -1.315 | -0.018 | 0.128 | 0.128 | 0 | 0.53 | 0.24 | 0.61 | 0.78 | 0.45 | 0.48 |
| 5 | 139145049 | rs165187 | 1.589 | -0.484 | -0.105 | -0.559 | -0.01 | 0.002 | 0.002 | 1.509 | 0.7 | 0.39 | 0.8 | 0.74 | 0.73 | 0.71 |
| 5 | 139145084 | rs262792 | -1.229 | -0.525 | 0.317 | -0.576 | -0.016 | -0.015 | -0.015 | 0.24 | 0.1 | 0.23 | 0.05 | 0.08 | 0.06 | 0.19 |
| 5 | 139147918 | rs356478 | -1.88 | -0.426 | 0.57 | -0.483 | 0.014 | -0.015 | -0.015 | 0 | 0.12 | 0.32 | 0.1 | 0.11 | 0.11 | 0.2 |
| 5 | 139150252 | rs2336977 | -0.735 | -1.036 | 0.08 | -0.921 | -0.009 | -0.014 | -0.014 | 0 | 0.53 | 0.56 | 0.77 | 0.78 | 0.7 | 0.59 |
| 5 | 139166034 | rs2878853 | -0.748 | -0.587 | -0.159 | -0.546 | 0.006 | 0.054 | 0.054 | 0 | 0.16 | 0.34 | 0.11 | 0.26 | 0.17 | 0.27 |
| 5 | 139166808 | rs749125 | 0.266 | -0.563 | -0.831 | -0.462 | -0.132 | -0.125 | -0.125 | 0 | 0.32 | 0.1 | 0.58 | 0.55 | 0.45 | 0.27 |
| 5 | 139173441 | rs2194141 | -1.287 | -0.983 | 0.106 | -0.487 | 0 | 0.002 | 0.002 | 0 | 0.14 | 0.16 | 0.12 | 0.21 | 0.16 | 0.17 |
| 5 | 139185509 | rs1422780 | 1.235 | -2.078 | -0.009 | -0.965 | 0.054 | 0.062 | 0.062 | 0 | 0.6 | 0.73 | 0.75 | 0.74 | 0.69 | 0.46 |
| 5 | 139188856 | rs3797901 | -1.647 | -1.595 | 1.055 | -0.492 | -0.025 | 0.025 | 0.025 | 0.533 | 0.22 | 0.5 | 0.16 | 0.25 | 0.19 | 0.31 |
| 5 | 139195160 | rs17118443 | -1.772 | -1.592 | 1.089 | -0.53 | -0.038 | -0.003 | -0.003 | 0 | 0.26 | 0.51 | 0.19 | 0.25 | 0.2 | 0.31 |
| 5 | 139201064 | rs7723670 | -1.722 | -1.588 | 1.15 | -0.573 | -0.032 | -0.007 | -0.007 | 0 | 0.26 | 0.45 | 0.19 | 0.24 | 0.2 | 0.31 |
| 5 | 139211723 | rs1006967 | -2.086 | -1.295 | 1.038 | -0.515 | 0.023 | 0.065 | 0.065 | 0.027 | 0.07 | 0.26 | 0.06 | 0.06 | 0.06 | 0.04 |
| 5 | 139222208 | rs3797904 | -2.055 | -1.276 | 0.929 | -0.521 | 0.014 | 0.047 | 0.047 | 0.08 | 0.07 | 0.26 | 0.06 | 0.07 | 0.07 | 0.12 |
| 5 | 139222591 | rs10078529 | -1.544 | -1.267 | 0.872 | -0.517 | 0.004 | 0.023 | 0.023 | 0 | 0.06 | 0.15 | 0.05 | 0.06 | 0.05 | 0.04 |
| 5 | 139223085 | rs17286536 | -0.117 | -1.236 | 0.927 | -0.581 | -0.001 | 0.002 | 0.002 | 0 | 0.12 | 0.08 | 0.1 | 0.09 | 0.13 | 0.14 |
| 5 | 139223145 | rs10061228 | -2.193 | -1.291 | 1.015 | -0.471 | 0.018 | 0.055 | 0.055 | 0.131 | 0.07 | 0.28 | 0.06 | 0.07 | 0.07 | 0.12 |
| 5 | 139224735 | rs17118468 | -0.239 | -2.093 | 0.642 | -0.722 | -0.021 | -0.014 | -0.014 | 1.466 | 0.2 | 0.25 | 0.06 | 0.07 | 0.07 | 0.06 |
| 5 | 139238893 | rs3756675 | -0.496 | -2.18 | 0.895 | -0.79 | -0.07 | -0.065 | -0.065 | 0.852 | 0.26 | 0.53 | 0.11 | 0.09 | 0.1 | 0.17 |
| 5 | 139240190 | rs12517577 | 0.148 | -2.099 | 0.865 | -0.752 | 0.035 | 0.031 | 0.031 | 1.904 | 0.21 | 0.12 | 0.06 | 0.07 | 0.09 | 0.07 |
| 5 | 139245086 | rs1800954 | -0.574 | -2.124 | 0.928 | -1.031 | -0.061 | -0.116 | -0.116 | 0 | 0.27 | 0.53 | 0.12 | 0.09 | 0.1 | 0.19 |
| 5 | 139247971 | rs3777106 | -0.263 | -2.067 | 0.637 | -1.001 | 0.015 | 0.014 | 0.014 | 4.058 | 0.31 | 0.2 | 0.12 | 0.13 | 0.13 | 0.18 |
| 5 | 139257603 | rs3777102 | -0.463 | -1.794 | -0.021 | -0.854 | -0.046 | -0.077 | -0.077 | 0 | 0.36 | 0.45 | 0.15 | 0.13 | 0.16 | 0.29 |
| 5 | 139262117 | rs6860347 | -0.304 | -0.149 | -0.886 | 0.149 | -0.04 | -0.041 | -0.041 | 0 | 0.72 | 0.77 | 0.39 | 0.49 | 0.5 | 0.69 |
| 5 | 139276664 | rs1422188 | -0.729 | -0.907 | -0.825 | -0.355 | 0.026 | 0.035 | 0.035 | 0.102 | 0.24 | 0.14 | 0.1 | 0.17 | 0.16 | 0.27 |
| 5 | 139276748 | rs1422187 | -1.223 | -1.056 | -0.461 | -0.59 | 0.006 | 0.004 | 0.004 | 0.215 | 0.37 | 0.66 | 0.21 | 0.21 | 0.24 | 0.37 |
| 5 | 139290054 | rs10515509 | -0.577 | -0.83 | -0.746 | -0.37 | 0.018 | 0.016 | 0.016 | 0.416 | 0.2 | 0.15 | 0.11 | 0.2 | 0.13 | 0.25 |
| 5 | 139293166 | rs4912936 | -0.647 | -0.348 | -0.889 | -0.498 | -0.009 | 0.034 | 0.034 | 0.718 | 0.21 | 0.08 | 0.14 | 0.17 | 0.23 | 0.27 |
| 5 | 139295821 | rs6895139 | -0.174 | -0.636 | -1.049 | -0.549 | 0.011 | 0.042 | 0.042 | 0 | 0.12 | 0.02 | 0.06 | 0.03 | 0.07 | 0.07 |
| 5 | 139303341 | rs17118539 | -0.933 | -0.579 | -0.185 | -0.575 | -0.028 | -0.046 | -0.046 | 0 | 0.23 | 0.34 | 0.13 | 0.18 | 0.15 | 0.28 |
| 5 | 139323257 | rs2060296 | 0.739 | -0.118 | -1.427 | -0.242 | -0.174 | -0.181 | -0.181 | 0.794 | 0.27 | 0.02 | 0.55 | 0.51 | 0.44 | 0.26 |
| 5 | 139329066 | rs9686629 | -0.533 | -0.742 | -0.453 | -0.426 | -0.01 | -0.013 | -0.013 | 0 | 0.19 | 0.23 | 0.11 | 0.18 | 0.12 | 0.27 |
| 5 | 139335948 | rs7736276 | -0.378 | -0.698 | -0.546 | -0.405 | -0.005 | -0.007 | -0.007 | 0.831 | 0.19 | 0.18 | 0.11 | 0.18 | 0.12 | 0.26 |
| 5 | 139348915 | rs197197 | -0.257 | -0.532 | -0.228 | -0.071 | -0.001 | -0.003 | -0.003 | 0 | 0.56 | 0.54 | 0.49 | 0.7 | 0.38 | 0.57 |
| 5 | 139353857 | rs11746363 | 0.425 | -1.144 | -0.665 | 0.047 | 0.073 | -0.149 | -0.149 | 0 | 0.33 | 0.04 | 0.25 | 0.51 | 0.2 | 0.25 |
| 5 | 139360590 | rs264347 | -1.078 | -0.871 | -0.698 | -0.18 | -0.067 | 0.042 | 0.042 | 0 | 0.61 | 0.87 | 0.57 | 0.74 | 0.48 | 0.47 |
| 5 | 139362321 | rs264350 | -0.921 | -0.058 | 0.449 | 0.321 | 0.01 | -0.02 | -0.02 | 0.045 | 0.25 | 0.54 | 0.3 | 0.23 | 0.29 | 0.21 |
| 5 | 139372149 | rs171951 | -1.04 | 0.08 | 0.693 | 0.544 | 0.026 | 0.08 | 0.08 | 0 | 0.27 | 0.43 | 0.35 | 0.61 | 0.32 | 0.23 |
| 5 | 139372521 | rs6891114 | -0.258 | 0.08 | 1.005 | 0.575 | 0 | 0.131 | 0.131 | 0 | 0.09 | 0.11 | 0.11 | 0.49 | 0.09 | 0.15 |
| 5 | 139375692 | rs17118604 | 0.025 | 0.08 | 0.987 | 0.531 | -0.002 | 0.057 | 0.057 | 0 | 0.08 | 0.04 | 0.1 | 0.45 | 0.09 | 0.15 |
| 5 | 139375932 | rs6896087 | 1.257 | 0.278 | 0.908 | 0.395 | 0.013 | 0.035 | 0.035 | 0 | 0.7 | 0.49 | 0.59 | 0.42 | 0.59 | 0.75 |
| 5 | 139378986 | rs265151 | 0.68 | 0.4 | 0.948 | 0.448 | 0.004 | 0.077 | 0.077 | 0 | 0.87 | 0.87 | 0.83 | 0.54 | 0.83 | 0.82 |
| 5 | 139384602 | rs11167875 | 1.578 | 0.093 | 1.006 | 0.329 | -0.003 | 0.008 | 0.008 | 0 | 0.58 | 0.43 | 0.38 | 0.27 | 0.5 | 0.6 |
| 5 | 139390467 | rs265158 | -0.086 | 0.048 | 1.11 | -0.356 | -0.009 | -0.011 | -0.011 | 0 | 0.84 | 0.94 | 0.79 | 0.83 | 0.81 | 0.84 |
| 5 | 139390914 | rs265160 | -0.5 | -0.063 | 1.151 | -0.902 | 0.031 | -0.061 | -0.061 | 0.581 | 0.18 | 0.43 | 0.25 | 0.12 | 0.22 | 0.07 |
| 5 | 139394941 | rs2560711 | -0.203 | -0.047 | 1.16 | -0.894 | 0.019 | -0.041 | -0.041 | 0 | 0.17 | 0.3 | 0.24 | 0.12 | 0.22 | 0.07 |
| 5 | 139400779 | rs265148 | -0.315 | 0.025 | 0.726 | -0.885 | 0.026 | -0.046 | -0.046 | 0 | 0.17 | 0.36 | 0.23 | 0.11 | 0.22 | 0.06 |
| 5 | 139405072 | rs12520398 | 0.719 | 0.025 | 0.398 | -0.777 | 0.009 | -0.081 | -0.081 | 0.301 | 0.67 | 0.55 | 0.64 | 0.81 | 0.61 | 0.79 |
| 5 | 139405701 | rs7734418 | 0.828 | -0.227 | 0.116 | -0.655 | -0.021 | -0.061 | -0.061 | 1.758 | 0.13 | 0.23 | 0.07 | 0 | 0.07 | 0.01 |
| 5 | 139407044 | rs4404729 | -0.724 | 0.312 | -0.445 | -0.249 | 0.005 | -0.039 | -0.039 | 0 | 0.28 | 0.33 | 0.3 | 0.19 | 0.31 | 0.19 |
| 5 | 139410510 | rs13173983 | -0.632 | -0.009 | -1.084 | -0.128 | 0.017 | 0.064 | 0.064 | 0.09 | 0.33 | 0.19 | 0.2 | 0.21 | 0.31 | 0.22 |
| 5 | 139410687 | rs6580353 | -0.807 | 0.114 | -0.47 | 0.065 | 0.013 | 0.024 | 0.024 | 0 | 0.25 | 0.51 | 0.31 | 0.3 | 0.26 | 0.17 |
| 5 | 139411703 | rs7723119 | -0.322 | 0.127 | -1.727 | -0.375 | 0.005 | 0.088 | 0.088 | 0.096 | 0.57 | 0.89 | 0.57 | 0.33 | 0.63 | 0.5 |
| 5 | 139414752 | rs2916092 | -0.416 | -0.236 | -1.961 | -0.05 | 0.012 | 0.073 | 0.073 | 0 | 0.67 | 0.9 | 0.73 | 0.85 | 0.76 | 0.63 |
| 5 | 139419985 | rs2436387 | 0.134 | -0.456 | -0.842 | -0.088 | 0.002 | 0.002 | 0.002 | 0 | 0.36 | 0.37 | 0.3 | 0.15 | 0.25 | 0.36 |
| 5 | 139437110 | rs155939 | 0.103 | -0.502 | -0.788 | -0.068 | 0.012 | 0.024 | 0.024 | 0 | 0.35 | 0.3 | 0.29 | 0.16 | 0.25 | 0.35 |
| 5 | 139438106 | rs6877423 | 1.052 | -0.849 | -1.09 | -0.292 | 0.026 | 0.039 | 0.039 | 0 | 0.15 | 0.06 | 0.06 | 0.03 | 0.1 | 0.19 |
| 5 | 139445100 | rs1347142 | 1.247 | -1.28 | -1.588 | -0.833 | 0.024 | 0.042 | 0.042 | 0 | 0.23 | 0.05 | 0.12 | 0.19 | 0.15 | 0.31 |
| 5 | 139460236 | rs1623177 | -0.392 | -1.099 | -0.86 | -1.115 | 0.046 | 0.078 | 0.078 | 0 | 0.45 | 0.67 | 0.27 | 0.23 | 0.31 | 0.51 |
| 5 | 139470240 | rs2974421 | 1.269 | -1.089 | -0.78 | -1.108 | -0.002 | 0.026 | 0.026 | 0.1 | 0.18 | 0.12 | 0.21 | 0.03 | 0.16 | 0.24 |
| 5 | 139489185 | rs13187159 | 1.356 | -0.94 | -0.895 | -1.108 | 0.018 | 0.043 | 0.043 | 0 | 0.23 | 0.05 | 0.12 | 0.18 | 0.15 | 0.32 |
| 5 | 139501147 | rs10060173 | 0.67 | -0.721 | -0.45 | -1.016 | -0.006 | -0.077 | -0.077 | 0 | 0.21 | 0.38 | 0.23 | 0.03 | 0.16 | 0.24 |

| | | | | | | | | | | | | | | | | |
|---|---|---|---|---|---|---|---|---|---|---|---|---|---|---|---|---|
| 5 | 139508705 | rs2429310 | 0.73 | -0.704 | -0.482 | -1.008 | -0.005 | -0.07 | -0.07 | 0 | 0.2 | 0.37 | 0.23 | 0.03 | 0.16 | 0.24 |
| 5 | 139514964 | rs13156549 | 0.659 | -0.97 | -0.715 | -0.514 | 0.044 | -0.044 | -0.044 | 0 | 0.33 | 0.14 | 0.27 | 0.55 | 0.23 | 0.35 |
| 5 | 139515394 | rs269783 | 0.879 | 0.237 | -0.334 | -0.659 | 0.001 | 0 | 0 | 0 | 0.19 | 0.27 | 0.2 | 0.23 | 0.19 | 0.2 |
| 5 | 139517197 | rs185123 | -0.606 | 0.12 | -0.195 | -0.807 | -0.001 | -0.004 | -0.004 | 0 | 0.64 | 0.66 | 0.67 | 0.83 | 0.51 | 0.82 |
| 5 | 139517561 | rs156094 | 0.208 | 0.074 | -0.39 | -0.657 | 0.017 | 0.01 | 0.01 | 0 | 0.43 | 0.31 | 0.46 | 0.61 | 0.33 | 0.57 |
| 5 | 139522328 | rs156089 | 0.351 | 0.088 | -0.483 | -0.651 | 0.022 | 0.011 | 0.011 | 0.09 | 0.42 | 0.25 | 0.45 | 0.61 | 0.33 | 0.57 |
| 5 | 139533563 | rs2436396 | -1.86 | 0.265 | -0.563 | -0.716 | -0.015 | 0.013 | 0.013 | 0.268 | 0.85 | 0.93 | 0.75 | 0.93 | 0.72 | 0.86 |
| 5 | 139557385 | rs2436391 | -0.119 | -0.23 | -0.22 | -1.081 | 0.01 | -0.005 | -0.005 | 0 | 0.39 | 0.47 | 0.52 | 0.56 | 0.55 | 0.24 |
| 5 | 139564473 | rs10080146 | 1.208 | -0.487 | -0.241 | -1.289 | 0.006 | -0.001 | -0.001 | 0.162 | 0.23 | 0.33 | 0.22 | 0.24 | 0.22 | 0.25 |
| 5 | 139565028 | rs10038443 | 1.247 | -0.495 | -0.122 | -1.292 | 0.002 | -0.001 | -0.001 | 0.013 | 0.23 | 0.29 | 0.22 | 0.24 | 0.22 | 0.25 |
| 5 | 139570731 | rs37355 | -1.117 | -0.506 | -0.134 | -1.307 | 0.061 | 0.083 | 0.083 | 0 | 0.62 | 0.8 | 0.74 | 0.8 | 0.77 | 0.49 |
| 5 | 139577219 | rs2436399 | -1.183 | -0.519 | -0.124 | -1.312 | 0.077 | 0.115 | 0.115 | 0.099 | 0.63 | 0.85 | 0.75 | 0.8 | 0.77 | 0.49 |
| 5 | 139581205 | rs17286613 | 1.428 | -0.562 | -0.123 | -1.294 | -0.016 | -0.008 | -0.008 | 0.022 | 0.21 | 0.13 | 0.21 | 0.24 | 0.22 | 0.26 |
| 5 | 139617885 | rs10477333 | 1.254 | -0.543 | -0.204 | -1.284 | 0.013 | 0.004 | 0.004 | 0 | 0.21 | 0.33 | 0.22 | 0.24 | 0.22 | 0.25 |
| 5 | 139625224 | rs11558407 | 2.179 | -0.969 | -0.711 | -1.506 | 0.044 | 0.054 | 0.054 | 0.329 | 0.14 | 0.01 | 0.12 | 0.04 | 0.07 | 0.18 |
| 5 | 139658794 | rs1030667 | 1.334 | -0.799 | -0.651 | -1.464 | 0.02 | -0.071 | -0.071 | 0 | 0.23 | 0.47 | 0.19 | 0.12 | 0.24 | 0.09 |
| 5 | 139664820 | rs12187444 | 1.57 | -0.683 | -0.727 | -1.387 | -0.045 | -0.027 | -0.027 | 0 | 0.2 | 0.06 | 0.2 | 0.24 | 0.21 | 0.25 |
| 5 | 139669386 | rs1347143 | -0.126 | -0.621 | -0.632 | -1.383 | 0.001 | -0.002 | -0.002 | 0 | 0.58 | 0.44 | 0.46 | 0.44 | 0.44 | 0.76 |
| 5 | 139684364 | rs13190290 | -2.073 | -0.564 | -0.955 | -1.489 | 0.002 | 0.091 | 0.091 | 0 | 0.8 | 0.98 | 0.9 | 0.88 | 0.86 | 0.75 |
| 5 | 139695230 | rs11738439 | 1.562 | -0.033 | -0.405 | -0.976 | -0.021 | -0.024 | -0.024 | 0.209 | 0.16 | 0.08 | 0.35 | 0.43 | 0.31 | 0.19 |
| 5 | 139697220 | rs2189659 | 0.131 | -0.979 | -0.363 | -1.358 | 0.008 | -0.005 | -0.005 | 0 | 0.4 | 0.48 | 0.53 | 0.56 | 0.55 | 0.28 |
| 5 | 139708382 | rs6884244 | 0.444 | -0.308 | -0.637 | -0.935 | -0.002 | 0.013 | 0.013 | 0 | 0.56 | 0.75 | 0.57 | 0.55 | 0.57 | 0.44 |
| 5 | 139711132 | rs10875633 | 0.405 | -0.332 | -0.648 | -0.946 | 0.002 | 0.018 | 0.018 | 0 | 0.55 | 0.75 | 0.57 | 0.55 | 0.57 | 0.44 |
| 5 | 139712550 | rs7268 | 0.544 | -0.216 | -0.509 | -0.883 | 0.004 | 0.008 | 0.008 | 0 | 0.53 | 0.55 | 0.56 | 0.54 | 0.57 | 0.41 |
| 5 | 139712878 | rs13385 | -0.077 | 0.307 | -0.671 | -0.387 | -0.05 | -0.062 | -0.062 | 0.185 | 0.19 | 0.05 | 0.2 | 0.31 | 0.23 | 0.25 |
| 5 | 139713761 | rs4150237 | 0.683 | 0.261 | -0.38 | -0.527 | 0.01 | 0 | 0 | 1.044 | 0.1 | 0.08 | 0.04 | 0 | 0.05 | 0.01 |
| 5 | 139720400 | rs2237077 | -0.791 | 0.365 | -0.102 | -0.571 | -0.025 | -0.061 | -0.061 | 1.352 | 0.31 | 0.43 | 0.24 | 0.15 | 0.25 | 0.29 |
| 5 | 139730444 | rs1862176 | 0.511 | 0.987 | 0.027 | -0.57 | -0.014 | -0.058 | -0.058 | 0 | 0.7 | 0.57 | 0.75 | 0.83 | 0.71 | 0.73 |
| 5 | 139745154 | rs6860077 | 0.501 | 0.954 | 0.02 | -0.592 | -0.018 | -0.06 | -0.06 | 0 | 0.7 | 0.57 | 0.75 | 0.84 | 0.72 | 0.74 |
| 5 | 139747914 | rs11168051 | 0.49 | 0.956 | 0.021 | -0.591 | -0.018 | -0.06 | -0.06 | 0.323 | 0.7 | 0.57 | 0.75 | 0.83 | 0.72 | 0.73 |
| 5 | 139749028 | rs17286676 | 0.423 | 1.051 | -0.408 | -0.485 | -0.032 | -0.027 | -0.027 | 0 | 0.19 | 0.07 | 0.21 | 0.27 | 0.23 | 0.26 |
| 5 | 139775839 | rs6872579 | -0.269 | 0.77 | -0.081 | -0.531 | 0.01 | 0.022 | 0.022 | 1.286 | 0.78 | 0.86 | 0.83 | 0.94 | 0.83 | 0.89 |
| 5 | 139790364 | rs975243 | 0.544 | 0.766 | -0.084 | -0.528 | -0.019 | -0.06 | -0.06 | 0 | 0.7 | 0.57 | 0.75 | 0.84 | 0.72 | 0.73 |
| 5 | 139805611 | rs10042299 | 1.388 | 0.666 | 0.116 | -0.661 | -0.025 | 0.036 | 0.036 | 0 | 0.46 | 0.22 | 0.51 | 0.56 | 0.48 | 0.47 |
| 5 | 139820931 | rs10476907 | 0.365 | 0.722 | 0.318 | -0.636 | -0.021 | -0.064 | -0.064 | 0 | 0.69 | 0.55 | 0.75 | 0.84 | 0.72 | 0.73 |
| 5 | 139862774 | rs10041762 | 0.405 | 0.705 | 0.31 | -0.663 | -0.022 | -0.066 | -0.066 | 0.479 | 0.69 | 0.55 | 0.75 | 0.83 | 0.72 | 0.73 |
| 5 | 139920311 | rs1835959 | -1.074 | 0.8 | 0.582 | -0.643 | -0.01 | 0 | 0 | 0 | 0.94 | 0.95 | 0.87 | 0.9 | 0.88 | 0.89 |
| 5 | 139928128 | rs250425 | 0.63 | 0.738 | 0.548 | -0.675 | -0.036 | -0.031 | -0.031 | 0 | 0.18 | 0.07 | 0.21 | 0.26 | 0.23 | 0.26 |
| 5 | 139941228 | rs250430 | -0.978 | 0.57 | 0.313 | -0.612 | -0.01 | 0 | 0 | 0 | 0.94 | 0.95 | 0.88 | 0.9 | 0.88 | 0.89 |
| 5 | 139944826 | rs250429 | 0.649 | 0.586 | 0.284 | -0.611 | -0.019 | -0.068 | -0.068 | 0 | 0.7 | 0.55 | 0.75 | 0.83 | 0.72 | 0.74 |
| 5 | 139950138 | rs917302 | 0.248 | 0.364 | -0.079 | -0.628 | 0.007 | 0.01 | 0.01 | 1.975 | 0.21 | 0.16 | 0.1 | 0.06 | 0.15 | 0.08 |
| 5 | 139974473 | rs1864255 | -1.225 | 0.502 | 0.257 | -0.656 | 0.002 | 0.016 | 0.016 | 0 | 0.49 | 0.44 | 0.45 | 0.45 | 0.51 | 0.5 |
| 5 | 139987897 | rs11167532 | 1.374 | -0.024 | -0.571 | -0.798 | -0.158 | -0.135 | -0.135 | 0 | 0.39 | 0.02 | 0.45 | 0.53 | 0.42 | 0.49 |
| 5 | 140005110 | rs778583 | -0.205 | 0.143 | 0.089 | -0.772 | -0.003 | -0.027 | -0.027 | 0 | 0.26 | 0.32 | 0.22 | 0.11 | 0.27 | 0.22 |
| 5 | 140005786 | rs7721577 | 0.9 | 0.22 | 0.037 | -0.734 | -0.021 | -0.034 | -0.034 | 0 | 0.17 | 0.06 | 0.2 | 0.3 | 0.24 | 0.25 |
| 5 | 140006430 | rs778587 | -0.533 | 0.28 | 0.884 | -0.692 | -0.018 | -0.076 | -0.076 | 0 | 0.3 | 0.5 | 0.26 | 0.13 | 0.27 | 0.22 |
| 5 | 140011315 | rs2563298 | 0.024 | 0.261 | 0.96 | -0.701 | 0.004 | 0.028 | 0.028 | 0 | 0.25 | 0.21 | 0.23 | 0.13 | 0.26 | 0.22 |
| 5 | 140012916 | rs2569190 | -1.47 | 0.219 | 1.176 | -0.713 | 0.008 | 0.04 | 0.04 | 0 | 0.5 | 0.7 | 0.47 | 0.43 | 0.51 | 0.47 |
| 5 | 140013307 | rs5744455 | 0.91 | 0.294 | 1.092 | -0.67 | -0.021 | -0.033 | -0.033 | 0.52 | 0.18 | 0.06 | 0.2 | 0.3 | 0.24 | 0.25 |
| 5 | 140019921 | rs12517200 | -0.188 | 0.27 | 1.271 | -0.689 | 0.005 | 0.018 | 0.018 | 0 | 0.26 | 0.25 | 0.25 | 0.15 | 0.27 | 0.24 |
| 5 | 140023818 | rs753279 | 1.156 | 0.623 | 1.376 | -0.475 | -0.004 | 0.002 | 0.002 | 0 | 0.48 | 0.33 | 0.48 | 0.51 | 0.45 | 0.46 |
| 5 | 140025484 | rs778591 | -0.666 | 0.714 | 1.595 | -0.424 | 0.004 | 0.01 | 0.01 | 0 | 0.34 | 0.42 | 0.35 | 0.45 | 0.41 | 0.45 |
| 5 | 140026527 | rs778592 | 0.341 | 0.826 | 1.594 | -0.375 | 0.002 | 0.001 | 0.001 | 0 | 0.13 | 0.19 | 0.14 | 0.15 | 0.18 | 0.21 |
| 5 | 140047600 | rs2302104 | 0.568 | 0.74 | 1.47 | -0.373 | -0.017 | -0.031 | -0.031 | 0 | 0.22 | 0.05 | 0.22 | 0.3 | 0.24 | 0.24 |
| 5 | 140048544 | rs2286394 | 0.568 | 0.74 | 1.47 | -0.382 | -0.017 | -0.033 | -0.033 | 0 | 0.22 | 0.05 | 0.22 | 0.3 | 0.24 | 0.24 |
| 5 | 140050265 | rs702397 | -1.093 | 0.774 | 1.576 | -0.409 | -0.002 | -0.002 | -0.002 | 0 | 0.41 | 0.7 | 0.41 | 0.45 | 0.44 | 0.45 |
| 5 | 140058893 | rs2073512 | 0.643 | 0.474 | 1.596 | -0.344 | -0.067 | -0.085 | -0.085 | 0 | 0.22 | 0.01 | 0.21 | 0.3 | 0.24 | 0.24 |
| 5 | 140079051 | rs2563302 | -0.678 | 0.478 | 1.546 | -0.342 | 0.001 | 0.003 | 0.003 | 0 | 0.37 | 0.38 | 0.36 | 0.45 | 0.43 | 0.45 |
| 5 | 140119534 | rs2531352 | -1.205 | 0.322 | 1.284 | -0.335 | 0.006 | 0.012 | 0.012 | 0 | 0.45 | 0.72 | 0.44 | 0.47 | 0.46 | 0.47 |
| 5 | 140123428 | rs2262572 | -0.189 | 0.485 | 1.147 | -0.261 | 0.011 | -0.008 | -0.008 | 0 | 0.17 | 0.36 | 0.19 | 0.15 | 0.21 | 0.21 |
| 5 | 140137644 | rs2563263 | -0.181 | 0.57 | 1.156 | -0.081 | -0.004 | -0.003 | -0.003 | 0 | 0.41 | 0.31 | 0.4 | 0.47 | 0.46 | 0.48 |
| 5 | 140165568 | rs12055222 | 0.042 | 0.59 | 1.222 | -0.089 | -0.006 | -0.004 | -0.004 | 0 | 0.59 | 0.69 | 0.6 | 0.53 | 0.54 | 0.52 |
| 5 | 140180561 | rs3756338 | 0.014 | 0.54 | 1.488 | -0.117 | -0.01 | -0.007 | -0.007 | 0.001 | 0.6 | 0.72 | 0.61 | 0.53 | 0.54 | 0.52 |
| 5 | 140181892 | rs3733707 | -0.004 | 0.538 | 1.527 | -0.119 | -0.018 | -0.013 | -0.013 | 0 | 0.6 | 0.76 | 0.61 | 0.53 | 0.54 | 0.52 |
| 5 | 140185132 | rs1030163 | 0.015 | 0.541 | 1.474 | -0.117 | -0.01 | -0.007 | -0.007 | 0 | 0.6 | 0.72 | 0.61 | 0.53 | 0.54 | 0.52 |
| 5 | 140185473 | rs1030166 | 0.007 | 0.538 | 1.473 | -0.118 | -0.01 | -0.007 | -0.007 | 0 | 0.6 | 0.72 | 0.61 | 0.53 | 0.54 | 0.52 |
| 5 | 140187322 | rs3822346 | 0.017 | 0.538 | 1.473 | -0.118 | -0.01 | -0.007 | -0.007 | 0.085 | 0.6 | 0.72 | 0.61 | 0.53 | 0.54 | 0.52 |
| 5 | 140192643 | rs10223116 | 0.497 | 0.49 | 1.537 | -0.146 | 0 | 0.005 | 0.005 | 0.176 | 0.55 | 0.33 | 0.57 | 0.53 | 0.53 | 0.52 |
| 5 | 140197222 | rs10037757 | 0.361 | 0.496 | 1.54 | -0.144 | 0 | 0.005 | 0.005 | 0.169 | 0.55 | 0.33 | 0.57 | 0.53 | 0.53 | 0.52 |

| | | | | | | | | | | | | | | | | |
|---|---|---|---|---|---|---|---|---|---|---|---|---|---|---|---|---|
| 5 | 140204650 | rs10050455 | 0.199 | 0.576 | 1.06 | -0.076 | 0.002 | 0.001 | 0.001 | 0 | 0.58 | 0.51 | 0.59 | 0.53 | 0.54 | 0.52 |
| 5 | 140210616 | rs10038174 | 0.016 | 0.655 | 0.311 | -0.039 | -0.017 | -0.012 | -0.012 | 0 | 0.6 | 0.75 | 0.61 | 0.53 | 0.54 | 0.52 |
| 5 | 140246287 | rs251370 | 0.193 | 0.311 | 0.677 | 0.078 | 0.027 | 0.022 | 0.022 | 0 | 0.5 | 0.31 | 0.41 | 0.42 | 0.48 | 0.62 |
| 5 | 140246511 | rs17208327 | 0.406 | 0.629 | 0.688 | -0.074 | -0.008 | 0.028 | 0.028 | 0 | 0.15 | 0.09 | 0.13 | 0.02 | 0.14 | 0.05 |
| 5 | 140255792 | rs251377 | 0.442 | 0.608 | 0.771 | -0.038 | 0.031 | 0.023 | 0.023 | 0 | 0.5 | 0.29 | 0.42 | 0.43 | 0.48 | 0.62 |
| 5 | 140258922 | rs251379 | -0.854 | 0.487 | 1.517 | 0.01 | -0.01 | -0.018 | -0.018 | 0.57 | 0.72 | 0.72 | 0.6 | 0.44 | 0.66 | 0.67 |
| 5 | 140282431 | rs1119032 | 0.194 | 0.474 | 1.839 | -0.005 | 0 | -0.024 | -0.024 | 0 | 0.16 | 0.19 | 0.13 | 0.02 | 0.14 | 0.05 |
| 5 | 140298815 | rs56002 | 0.833 | 0.48 | 1.677 | 0.003 | -0.015 | -0.028 | -0.028 | 0 | 0.28 | 0.23 | 0.39 | 0.55 | 0.33 | 0.33 |
| 5 | 140307284 | rs155361 | -0.705 | 0.252 | 1.92 | 0.004 | 0.024 | 0.032 | 0.032 | 0 | 0.51 | 0.72 | 0.59 | 0.61 | 0.52 | 0.38 |
| 5 | 140309016 | rs155360 | -0.785 | 0.246 | 1.892 | 0.002 | 0.021 | 0.041 | 0.041 | 0.01 | 0.52 | 0.79 | 0.6 | 0.61 | 0.52 | 0.38 |
| 5 | 140309488 | rs11747154 | 0.508 | 0.249 | 1.852 | 0 | -0.02 | 0.051 | 0.051 | 0 | 0.15 | 0.05 | 0.12 | 0.02 | 0.14 | 0.05 |
| 5 | 140321593 | rs155814 | 0.953 | 0.24 | 1.985 | 0.004 | -0.033 | -0.063 | -0.063 | 0 | 0.27 | 0.15 | 0.38 | 0.55 | 0.33 | 0.32 |
| 5 | 140324431 | rs269554 | -0.675 | 0.201 | 1.987 | -0.019 | -0.015 | -0.069 | -0.069 | 0 | 0.21 | 0.45 | 0.16 | 0.06 | 0.15 | 0.06 |
| 5 | 140327590 | rs246049 | -0.771 | 0.201 | 1.937 | -0.011 | 0.019 | 0.027 | 0.027 | 0 | 0.52 | 0.71 | 0.6 | 0.61 | 0.52 | 0.39 |
| 5 | 140331867 | rs12187982 | 0.438 | 0.375 | 1.446 | 0.058 | -0.023 | 0.057 | 0.057 | 0 | 0.15 | 0.05 | 0.12 | 0.02 | 0.14 | 0.06 |
| 5 | 140336168 | rs155799 | 0.957 | 0.391 | 1.517 | 0.082 | -0.072 | -0.123 | -0.123 | 0 | 0.27 | 0.09 | 0.38 | 0.55 | 0.33 | 0.32 |
| 5 | 140343598 | rs155811 | 0.548 | 0.42 | 1.458 | 0.09 | 0.006 | 0.011 | 0.011 | 2.303 | 0.31 | 0.33 | 0.44 | 0.55 | 0.37 | 0.33 |
| 5 | 140344242 | rs2067599 | -0.343 | 0.014 | 0.934 | 0.149 | -0.002 | -0.023 | -0.023 | 0 | 0.18 | 0.29 | 0.14 | 0.06 | 0.15 | 0.07 |
| 5 | 140346712 | rs155808 | 0.601 | 0.133 | 0.061 | 0.153 | -0.019 | -0.038 | -0.038 | 0.966 | 0.28 | 0.22 | 0.4 | 0.55 | 0.34 | 0.34 |
| 5 | 140347053 | rs155807 | 0.69 | 0.151 | 0.074 | 0.166 | -0.018 | -0.036 | -0.036 | 0.057 | 0.28 | 0.22 | 0.39 | 0.55 | 0.33 | 0.33 |
| 5 | 140347398 | rs12153295 | -0.055 | 0.272 | -0.118 | -0.085 | -0.026 | 0.058 | 0.058 | 0 | 0.15 | 0.05 | 0.13 | 0.02 | 0.15 | 0.06 |
| 5 | 140349840 | rs155806 | 0.079 | 0.249 | -0.43 | -0.113 | -0.023 | 0.047 | 0.047 | 0 | 0.13 | 0.01 | 0.13 | 0.02 | 0.1 | 0.05 |
| 5 | 140353907 | rs11743888 | 0.439 | 0.472 | -0.207 | -0.076 | -0.025 | 0.028 | 0.028 | 0 | 0.06 | 0.01 | 0.03 | 0 | 0.09 | 0.01 |
| 5 | 140366901 | rs13153056 | 0.986 | -0.13 | -0.136 | -0.135 | 0.018 | 0.005 | 0.005 | 0 | 0.13 | 0.05 | 0.04 | 0.11 | 0.06 | 0.11 |
| 5 | 140368859 | rs3756325 | 1.106 | -0.141 | -0.169 | 0.121 | -0.02 | -0.027 | -0.027 | 0.904 | 0.26 | 0.23 | 0.39 | 0.45 | 0.31 | 0.34 |
| 5 | 140370339 | rs3822343 | 2.221 | -0.127 | -0.216 | 0.108 | 0.009 | 0.004 | 0.004 | 0 | 0.1 | 0 | 0.03 | 0.07 | 0.06 | 0.11 |
| 5 | 140372222 | rs31872 | -1.547 | -0.736 | -0.271 | -0.02 | -0.037 | -0.021 | -0.021 | 0 | 0.7 | 0.9 | 0.76 | 0.68 | 0.6 | 0.55 |
| 5 | 140373888 | rs13163241 | 1.476 | -1.118 | -0.135 | 0.024 | 0.016 | 0.005 | 0.005 | 0 | 0.13 | 0.05 | 0.04 | 0.11 | 0.06 | 0.11 |
| 5 | 140419991 | rs31746 | 0.165 | -0.581 | 0.527 | -0.108 | 0.002 | -0.006 | -0.006 | 0 | 0.39 | 0.59 | 0.42 | 0.53 | 0.32 | 0.44 |
| 5 | 140436063 | rs246584 | 0.691 | -0.652 | 0.645 | -0.176 | -0.037 | -0.052 | -0.052 | 0.063 | 0.17 | 0.39 | 0.1 | 0.08 | 0.09 | 0.11 |
| 5 | 140442209 | rs702390 | 1.392 | -0.66 | 0.175 | -0.229 | -0.097 | -0.044 | -0.044 | 1.037 | 0.34 | 0.06 | 0.34 | 0.34 | 0.45 | 0.45 |
| 5 | 140464559 | rs12521934 | 0.307 | -0.682 | 0.483 | -0.097 | 0.003 | 0.004 | 0.004 | 0 | 0.2 | 0.15 | 0.18 | 0.12 | 0.14 | 0.11 |
| 5 | 140474756 | rs31853 | 2.094 | -0.609 | 0.511 | -0.032 | -0.007 | -0.007 | -0.007 | 0 | 0.1 | 0.15 | 0.05 | 0.08 | 0.07 | 0.11 |
| 5 | 140476894 | rs3776103 | 2.126 | -0.577 | 0.387 | -0.037 | -0.008 | -0.001 | -0.001 | 5.887 | 0.1 | 0 | 0.04 | 0.08 | 0.08 | 0.11 |
| 5 | 140480852 | rs12515688 | 0.352 | -0.655 | 0.476 | -0.039 | 0.002 | 0.003 | 0.003 | 0 | 0.21 | 0.17 | 0.2 | 0.12 | 0.16 | 0.11 |
| 5 | 140482871 | rs7722330 | 1.602 | -0.649 | 0.528 | -0.058 | -0.009 | -0.029 | -0.029 | 0.245 | 0.22 | 0.11 | 0.32 | 0.45 | 0.23 | 0.33 |
| 5 | 140498598 | rs13171965 | 2.203 | -0.63 | 0.211 | -0.05 | -0.002 | 0.001 | 0.001 | 0 | 0.1 | 0.01 | 0.03 | 0.08 | 0.07 | 0.11 |
| 5 | 140498780 | rs13171205 | 2.203 | -0.63 | 0.211 | -0.05 | -0.002 | 0.001 | 0.001 | 0 | 0.1 | 0 | 0.03 | 0.08 | 0.07 | 0.1 |
| 5 | 140501183 | rs4151691 | 2.203 | -0.63 | 0.211 | -0.05 | -0.002 | 0.001 | 0.001 | 0 | 0.1 | 0.01 | 0.03 | 0.08 | 0.07 | 0.1 |
| 5 | 140502177 | rs3776100 | 1.743 | -0.699 | 0.204 | -0.05 | -0.081 | -0.151 | -0.151 | 0.205 | 0.21 | 0.01 | 0.32 | 0.45 | 0.23 | 0.33 |
| 5 | 140502344 | rs3733697 | 0.365 | -0.736 | 0.238 | -0.038 | 0.002 | 0.003 | 0.003 | 0 | 0.21 | 0.17 | 0.2 | 0.12 | 0.16 | 0.11 |
| 5 | 140502841 | rs3776099 | 2.09 | -0.598 | -0.198 | 0.057 | 0.002 | 0.001 | 0.001 | 0 | 0.1 | 0.01 | 0.03 | 0.08 | 0.07 | 0.1 |
| 5 | 140504310 | rs3088331 | 1.621 | -0.569 | -0.218 | 0.064 | -0.084 | -0.151 | -0.151 | 0 | 0.21 | 0.01 | 0.32 | 0.45 | 0.23 | 0.33 |
| 5 | 140504432 | rs3822339 | 1.798 | -0.429 | 0.13 | -0.046 | -0.011 | 0.003 | 0.003 | 0 | 0.1 | 0 | 0.03 | 0.08 | 0.08 | 0.11 |
| 5 | 140517431 | rs246725 | 0.708 | -0.552 | 0.226 | -0.03 | 0.006 | 0.006 | 0.006 | 0 | 0.15 | 0.15 | 0.07 | 0.08 | 0.08 | 0.11 |
| 5 | 140519095 | rs6864827 | 0.688 | -0.514 | 0.057 | -0.029 | -0.034 | 0.016 | 0.016 | 0 | 0.4 | 0.26 | 0.38 | 0.35 | 0.52 | 0.45 |
| 5 | 140519166 | rs246723 | -0.07 | -0.109 | 0.245 | 0.256 | -0.003 | 0.001 | 0.001 | 0 | 0.58 | 0.73 | 0.48 | 0.42 | 0.61 | 0.56 |
| 5 | 140522425 | rs246718 | -0.305 | -0.279 | 0.426 | 0.046 | 0.003 | 0.002 | 0.002 | 0 | 0.45 | 0.59 | 0.56 | 0.57 | 0.41 | 0.44 |
| 5 | 140527753 | rs13355454 | -0.049 | -0.435 | 0.446 | 0.336 | -0.01 | 0.006 | 0.006 | 0.028 | 0.19 | 0.21 | 0.2 | 0.19 | 0.15 | 0.1 |
| 5 | 140528906 | rs12653888 | 1.3 | -0.462 | 0.235 | -0.358 | -0.089 | -0.167 | -0.167 | 2.341 | 0.21 | 0.01 | 0.31 | 0.53 | 0.24 | 0.34 |
| 5 | 140529711 | rs4151693 | 1.313 | -0.454 | 0.238 | -0.354 | -0.089 | -0.167 | -0.167 | 0 | 0.21 | 0.01 | 0.31 | 0.53 | 0.24 | 0.34 |
| 5 | 140530529 | rs3776096 | 1.313 | -0.454 | 0.238 | -0.354 | -0.089 | -0.167 | -0.167 | 0.008 | 0.21 | 0.01 | 0.31 | 0.53 | 0.24 | 0.34 |
| 5 | 140532409 | rs17629216 | 1.189 | -0.332 | 0.331 | -0.416 | -0.003 | 0 | 0 | 0 | 0.06 | 0.02 | 0.03 | 0 | 0.08 | 0.01 |
| 5 | 140544992 | rs246690 | -1.245 | -0.868 | 0.424 | -0.688 | -0.018 | 0.001 | 0.001 | 0 | 0.64 | 0.69 | 0.64 | 0.66 | 0.55 | 0.58 |
| 5 | 140552774 | rs17286891 | 0.829 | -0.951 | 0.34 | -0.735 | -0.019 | 0.021 | 0.021 | 0 | 0.08 | 0 | 0.12 | 0.02 | 0.1 | 0.04 |
| 5 | 140553664 | rs3733695 | 0.25 | -0.806 | 0.69 | -0.634 | -0.018 | -0.048 | -0.048 | 0.102 | 0.19 | 0.26 | 0.19 | 0.04 | 0.15 | 0.09 |
| 5 | 140561768 | rs2740584 | 1.038 | -0.771 | 0.804 | -0.579 | -0.005 | -0.016 | -0.016 | 0 | 0.23 | 0.25 | 0.35 | 0.53 | 0.25 | 0.32 |
| 5 | 140561983 | rs2204704 | 1.222 | -0.856 | 0.956 | -0.67 | -0.056 | -0.009 | -0.009 | 0 | 0.36 | 0.2 | 0.35 | 0.34 | 0.45 | 0.42 |
| 5 | 140562073 | rs4151695 | 0.328 | -0.888 | 1.001 | -0.721 | -0.016 | -0.048 | -0.048 | 0 | 0.19 | 0.26 | 0.19 | 0.04 | 0.16 | 0.09 |
| 5 | 140566244 | rs6580182 | 1.154 | -0.89 | 1.057 | -0.713 | -0.019 | 0.002 | 0.002 | 0 | 0.36 | 0.31 | 0.36 | 0.34 | 0.45 | 0.42 |
| 5 | 140570973 | rs12654953 | -0.154 | -0.842 | 0.887 | -0.674 | -0.003 | -0.005 | -0.005 | 0 | 0.39 | 0.31 | 0.5 | 0.57 | 0.39 | 0.41 |
| 5 | 140571526 | rs4912742 | -0.447 | -0.836 | 0.886 | -0.67 | -0.002 | -0.002 | -0.002 | 0 | 0.43 | 0.51 | 0.54 | 0.57 | 0.41 | 0.42 |
| 5 | 140579358 | rs3756323 | -0.379 | -1.164 | 1.223 | -1.152 | -0.016 | -0.048 | -0.048 | 0.442 | 0.19 | 0.26 | 0.19 | 0.04 | 0.16 | 0.09 |
| 5 | 140588373 | rs2910326 | -1.396 | -1.154 | 0.784 | -1.168 | 0.079 | 0.144 | 0.144 | 0.146 | 0.44 | 0.73 | 0.34 | 0.13 | 0.32 | 0.27 |
| 5 | 140590766 | rs2910006 | -1.463 | -1.119 | 0.571 | -1.152 | 0.098 | 0.192 | 0.192 | 0.264 | 0.45 | 0.75 | 0.34 | 0.13 | 0.32 | 0.27 |
| 5 | 140596704 | rs2910333 | -1.266 | -1.112 | 0.574 | -1.166 | 0.101 | 0.197 | 0.197 | 0.253 | 0.45 | 0.75 | 0.34 | 0.13 | 0.32 | 0.27 |
| 5 | 140602119 | rs2910004 | -1.266 | -1.112 | 0.564 | -1.166 | 0.1 | 0.193 | 0.193 | 0.253 | 0.45 | 0.75 | 0.34 | 0.13 | 0.32 | 0.27 |
| 5 | 140606802 | rs2907320 | -0.5 | -0.523 | 1.211 | -0.545 | -0.057 | -0.106 | -0.106 | 0.524 | 0.25 | 0.44 | 0.15 | 0.09 | 0.16 | 0.17 |
| 5 | 140614038 | rs2907309 | -0.487 | -0.523 | 1.21 | -0.545 | -0.057 | -0.106 | -0.106 | 0 | 0.25 | 0.44 | 0.15 | 0.09 | 0.16 | 0.17 |

| | | | | | | | | | | | | | | | | |
|---|---|---|---|---|---|---|---|---|---|---|---|---|---|---|---|---|
| 5 | 140614830 | rs2907308 | -0.43 | -0.509 | 1.025 | -0.539 | -0.05 | -0.091 | -0.091 | 1.97 | 0.25 | 0.48 | 0.15 | 0.09 | 0.16 | 0.17 |
| 5 | 140615269 | rs2907307 | -0.43 | -0.509 | 1.025 | -0.539 | -0.05 | -0.091 | -0.091 | 0 | 0.25 | 0.48 | 0.15 | 0.09 | 0.16 | 0.17 |
| 5 | 140631925 | rs2910337 | -0.493 | -0.44 | 0.312 | -0.506 | -0.058 | -0.108 | -0.108 | 0 | 0.25 | 0.42 | 0.15 | 0.09 | 0.16 | 0.17 |
| 5 | 140653258 | rs10044936 | 1.029 | -0.011 | -0.217 | -0.329 | 0.014 | 0.012 | 0.012 | 0 | 0.07 | 0.03 | 0.03 | 0.04 | 0.04 | 0.07 |
| 5 | 140659538 | rs618578 | -1.103 | 0.582 | -0.05 | -0.614 | -0.003 | 0.005 | 0.005 | 0 | 0.36 | 0.67 | 0.36 | 0.32 | 0.38 | 0.34 |
| 5 | 140663663 | rs619268 | 0.064 | 0.721 | 0.545 | -0.688 | -0.003 | -0.006 | -0.006 | 0 | 0.14 | 0.2 | 0.15 | 0.1 | 0.09 | 0.1 |
| 5 | 140668375 | rs2879202 | -0.071 | 0.82 | 0.665 | -0.668 | 0.137 | 0.021 | 0.021 | 0 | 0.12 | 0.37 | 0.15 | 0.09 | 0.21 | 0.11 |
| 5 | 140672089 | rs1422411 | 0.025 | 0.814 | 0.631 | -0.667 | 0 | 0 | 0 | 0 | 0.84 | 0.77 | 0.83 | 0.8 | 0.87 | 0.84 |
| 5 | 140674609 | rs1076361 | 0.167 | 0.791 | 1.097 | -0.67 | 0.01 | -0.002 | -0.002 | 0 | 0.1 | 0.14 | 0.14 | 0.07 | 0.2 | 0.11 |
| 5 | 140675316 | rs11740372 | 1.885 | 1.228 | 0.886 | -0.876 | -0.014 | 0.023 | 0.023 | 0 | 0.59 | 0.32 | 0.6 | 0.68 | 0.55 | 0.63 |
| 5 | 140676273 | rs6580184 | 1.931 | 1.214 | 0.924 | -0.443 | -0.018 | 0 | 0 | 0.081 | 0.58 | 0.28 | 0.49 | 0.59 | 0.54 | 0.6 |
| 5 | 140677114 | rs13187825 | 0.57 | 1.102 | 1.133 | -0.457 | 0.002 | 0.001 | 0.001 | 0 | 0.76 | 0.73 | 0.79 | 0.77 | 0.82 | 0.77 |
| 5 | 140678474 | rs10491308 | 1.087 | 1.106 | 1.177 | -0.482 | -0.001 | 0.01 | 0.01 | 0 | 0.09 | 0.29 | 0.04 | 0.06 | 0.04 | 0.06 |
| 5 | 140679023 | rs10491309 | 0.905 | 0.841 | 1.141 | -0.775 | -0.007 | -0.019 | -0.019 | 0 | 0.08 | 0.07 | 0.05 | 0.01 | 0.07 | 0.02 |
| 5 | 140679359 | rs7735993 | -0.675 | 0.868 | 1.073 | -0.724 | 0.048 | -0.034 | -0.034 | 0 | 0.19 | 0.44 | 0.18 | 0.13 | 0.25 | 0.19 |
| 5 | 140682757 | rs3749780 | 0.336 | 0.925 | 0.912 | -0.71 | 0.011 | -0.003 | -0.003 | 0 | 0.1 | 0.13 | 0.13 | 0.06 | 0.2 | 0.11 |
| 5 | 140682958 | rs10075302 | 2.274 | 0.875 | 0.8 | -0.703 | 0.01 | 0.011 | 0.011 | 0 | 0.07 | 0.03 | 0.04 | 0.04 | 0.05 | 0.06 |
| 5 | 140683265 | rs11952797 | 1.443 | 0.852 | 0.765 | -0.712 | -0.005 | -0.005 | -0.005 | 0 | 0.63 | 0.32 | 0.53 | 0.59 | 0.61 | 0.59 |
| 5 | 140695905 | rs17097145 | 0.222 | 0.612 | 1.442 | -0.627 | -0.039 | 0.005 | 0.005 | 0 | 0.15 | 0.2 | 0.18 | 0.15 | 0.11 | 0.14 |
| 5 | 140700489 | rs10875595 | 0.395 | 0.807 | 1.507 | -0.686 | 0.015 | 0 | 0 | 0 | 0.08 | 0.09 | 0.12 | 0.06 | 0.18 | 0.12 |
| 5 | 140701730 | rs13359820 | -0.275 | 0.731 | 1.518 | -0.681 | -0.035 | 0.035 | 0.035 | 0 | 0.18 | 0.34 | 0.2 | 0.24 | 0.14 | 0.18 |
| 5 | 140711853 | rs17097187 | 0.445 | 0.705 | 1.169 | -1.031 | 0.001 | 0.002 | 0.002 | 0 | 0.07 | 0.04 | 0.05 | 0 | 0.07 | 0.01 |
| 5 | 140717739 | rs10491311 | -0.006 | 0.729 | 1.245 | -0.788 | -0.026 | 0.034 | 0.034 | 0 | 0.16 | 0.25 | 0.2 | 0.24 | 0.14 | 0.18 |
| 5 | 140719090 | rs6861047 | 0.321 | 1.036 | 0.714 | -0.936 | 0.008 | 0 | 0 | 0 | 0.08 | 0.08 | 0.11 | 0.06 | 0.17 | 0.11 |
| 5 | 140723174 | rs3806837 | -0.963 | 0.383 | 1.013 | -1.168 | -0.043 | -0.078 | -0.078 | 0.297 | 0.2 | 0.31 | 0.1 | 0.04 | 0.11 | 0.08 |
| 5 | 140728534 | rs17097224 | 0.293 | 0.395 | 1.246 | -1.145 | -0.035 | -0.044 | -0.044 | 0 | 0.09 | 0.12 | 0.03 | 0 | 0.03 | 0.02 |
| 5 | 140733878 | rs10214288 | -0.688 | 0.39 | 1.321 | -1.156 | -0.027 | -0.054 | -0.054 | 0.143 | 0.13 | 0.31 | 0.07 | 0.04 | 0.09 | 0.06 |
| 5 | 140742483 | rs17097238 | -0.661 | 0.735 | 1.435 | -1.433 | -0.028 | -0.055 | -0.055 | 0 | 0.13 | 0.31 | 0.07 | 0.04 | 0.09 | 0.06 |
| 5 | 140753245 | rs4151698 | -0.446 | 0.72 | 1.617 | -1.446 | -0.021 | -0.044 | -0.044 | 0.076 | 0.13 | 0.21 | 0.07 | 0.04 | 0.09 | 0.07 |
| 5 | 140763029 | rs2072315 | -1.954 | 0.846 | 1.416 | -1.635 | 0.036 | 0.076 | 0.076 | 0.336 | 0.47 | 0.85 | 0.63 | 0.33 | 0.42 | 0.42 |
| 5 | 140765000 | rs2024141 | 0.389 | 0.874 | 1.65 | -1.604 | -0.004 | -0.002 | -0.002 | 0.092 | 0.11 | 0.12 | 0.15 | 0.12 | 0.06 | 0.13 |
| 5 | 140771392 | rs3806831 | 0.011 | 0.856 | 1.807 | -1.622 | -0.022 | -0.045 | -0.045 | 0 | 0.13 | 0.21 | 0.07 | 0.04 | 0.09 | 0.07 |
| 5 | 140772427 | rs726684 | 0.701 | 0.886 | 1.876 | -1.626 | 0.009 | 0.006 | 0.006 | 0 | 0.09 | 0.1 | 0.33 | 0.13 | 0.19 | 0.13 |
| 5 | 140777577 | rs10045443 | -1.605 | 0.783 | 1.912 | -1.654 | -0.033 | -0.034 | -0.034 | 0 | 0.34 | 0.53 | 0.26 | 0.2 | 0.19 | 0.28 |
| 5 | 140778256 | rs6867460 | -0.21 | 0.784 | 1.9 | -1.649 | -0.033 | -0.017 | -0.017 | 0 | 0.2 | 0.32 | 0.19 | 0.15 | 0.1 | 0.21 |
| 5 | 140782160 | rs4151700 | 2.176 | 0.735 | 1.85 | -1.665 | 0.015 | 0.005 | 0.005 | 0 | 0.08 | 0.05 | 0.03 | 0.04 | 0.04 | 0.08 |
| 5 | 140784055 | rs3749770 | -1.643 | 0.79 | 1.901 | -1.666 | -0.031 | -0.035 | -0.035 | 0 | 0.34 | 0.53 | 0.26 | 0.2 | 0.2 | 0.28 |
| 5 | 140784427 | rs3749769 | 0.187 | 0.771 | 1.917 | -1.659 | -0.016 | -0.006 | -0.006 | 0 | 0.12 | 0.18 | 0.15 | 0.12 | 0.06 | 0.13 |
| 5 | 140789933 | rs3749767 | -1.597 | 0.778 | 1.922 | -1.666 | -0.033 | -0.034 | -0.034 | 2.267 | 0.33 | 0.53 | 0.26 | 0.19 | 0.19 | 0.28 |
| 5 | 140796099 | rs3806830 | 2.54 | 0.756 | 1.682 | -1.851 | 0.012 | 0.005 | 0.005 | 0 | 0.07 | 0.05 | 0.03 | 0.04 | 0.04 | 0.08 |
| 5 | 140800708 | rs7732160 | 2.18 | 0.752 | 1.719 | -1.853 | -0.006 | -0.003 | -0.003 | 0 | 0.08 | 0.13 | 0.04 | 0.04 | 0.04 | 0.08 |
| 5 | 140801104 | rs11167744 | 0.604 | 0.752 | 1.739 | -1.854 | -0.004 | -0.002 | -0.002 | 0 | 0.11 | 0.12 | 0.15 | 0.12 | 0.06 | 0.13 |
| 5 | 140805771 | rs10046053 | 0.218 | 0.756 | 1.625 | -1.853 | -0.017 | -0.007 | -0.007 | 2.25 | 0.12 | 0.19 | 0.15 | 0.12 | 0.06 | 0.13 |
| 5 | 140807256 | rs3749765 | -0.047 | 0.772 | 1.591 | -1.846 | -0.037 | -0.019 | -0.019 | 0 | 0.2 | 0.34 | 0.19 | 0.15 | 0.1 | 0.21 |
| 5 | 140815348 | rs3828680 | -2.16 | 0.831 | 1.594 | -1.859 | 0.021 | 0.068 | 0.068 | 0.424 | 0.46 | 0.85 | 0.62 | 0.33 | 0.42 | 0.42 |
| 5 | 140817231 | rs10035418 | -1.432 | 0.722 | 1.518 | -1.835 | -0.033 | -0.035 | -0.035 | 0 | 0.33 | 0.53 | 0.26 | 0.19 | 0.19 | 0.28 |
| 5 | 140822723 | rs1059029 | -1.414 | 0.664 | 1.482 | -1.832 | -0.048 | -0.05 | -0.05 | 0.238 | 0.33 | 0.48 | 0.26 | 0.19 | 0.19 | 0.28 |
| 5 | 140833077 | rs11958830 | -0.067 | 0.682 | 1.466 | -1.827 | -0.014 | -0.007 | -0.007 | 0.03 | 0.2 | 0.31 | 0.2 | 0.19 | 0.13 | 0.23 |
| 5 | 140839695 | rs10051366 | -1.123 | 0.722 | 1.392 | -1.807 | -0.05 | -0.052 | -0.052 | 0 | 0.32 | 0.45 | 0.25 | 0.19 | 0.19 | 0.27 |
| 5 | 140853856 | rs11956411 | 0.352 | 0.9 | 1.329 | -1.748 | 0.046 | 0.045 | 0.045 | 0 | 0.1 | 0.2 | 0.35 | 0.15 | 0.21 | 0.16 |
| 5 | 140857798 | rs1423148 | -1.939 | 0.783 | 1.255 | -1.846 | 0.003 | 0.06 | 0.06 | 0.665 | 0.46 | 0.89 | 0.62 | 0.33 | 0.42 | 0.43 |
| 5 | 140858245 | rs3805695 | -0.733 | 0.864 | 1.293 | -2.087 | 0.003 | -0.036 | -0.036 | 0 | 0.18 | 0.4 | 0.21 | 0.11 | 0.14 | 0.16 |
| 5 | 140861801 | rs11748256 | -0.961 | 0.857 | 1.272 | -2.099 | -0.006 | -0.046 | -0.046 | 0 | 0.26 | 0.43 | 0.26 | 0.17 | 0.22 | 0.27 |
| 5 | 140863674 | rs13361997 | -0.814 | 0.78 | 1.097 | -2.044 | -0.058 | -0.11 | -0.11 | 1.734 | 0.25 | 0.55 | 0.12 | 0.06 | 0.15 | 0.16 |
| 5 | 140868499 | rs2233598 | 0.258 | 1.028 | 0.693 | -1.915 | 0.045 | 0.025 | 0.025 | 0 | 0.11 | 0.19 | 0.36 | 0.13 | 0.22 | 0.16 |
| 5 | 140869003 | rs1002519 | -0.285 | 0.984 | 0.56 | -1.955 | -0.002 | 0.005 | 0.005 | 0 | 0.19 | 0.12 | 0.37 | 0.13 | 0.23 | 0.19 |
| 5 | 140870516 | rs2074912 | 0.426 | 1.17 | 0.618 | -1.855 | 0.028 | 0.014 | 0.014 | 0 | 0.1 | 0.14 | 0.35 | 0.13 | 0.21 | 0.16 |
| 5 | 140870949 | rs918558 | 0.426 | 1.17 | 0.618 | -1.855 | 0.028 | 0.014 | 0.014 | 0 | 0.1 | 0.14 | 0.35 | 0.13 | 0.21 | 0.16 |
| 5 | 140871249 | rs11952292 | 2.066 | 0.994 | 0.333 | -1.86 | -0.007 | 0.011 | 0.011 | 0 | 0.08 | 0.02 | 0.04 | 0.06 | 0.07 | 0.11 |
| 5 | 140877905 | rs2237079 | -1.281 | 1.092 | 0.645 | -1.886 | 0.004 | -0.026 | -0.026 | 0 | 0.39 | 0.49 | 0.57 | 0.25 | 0.42 | 0.43 |
| 5 | 140881474 | rs4912761 | -1.734 | 1.049 | 0.382 | -1.915 | -0.017 | 0.11 | 0.11 | 0.279 | 0.5 | 0.84 | 0.64 | 0.3 | 0.5 | 0.47 |
| 5 | 140881691 | rs4912762 | -1.585 | 1.065 | 0.391 | -1.906 | -0.004 | 0.045 | 0.045 | 0 | 0.48 | 0.7 | 0.62 | 0.3 | 0.49 | 0.47 |
| 5 | 140882770 | rs10477147 | 2.502 | 1.104 | -0.114 | -1.753 | 0.004 | 0.034 | 0.034 | 0 | 0.07 | 0.01 | 0.02 | 0.01 | 0.04 | 0.08 |
| 5 | 140887429 | rs17286954 | 0.134 | 1.233 | 0.009 | -1.752 | -0.007 | 0.002 | 0.002 | 0 | 0.14 | 0.12 | 0.18 | 0.11 | 0.12 | 0.15 |
| 5 | 140888248 | rs970069 | 0.408 | 1.329 | 0.125 | -1.787 | 0.004 | 0 | 0 | 0.093 | 0.1 | 0.09 | 0.34 | 0.12 | 0.2 | 0.16 |
| 5 | 140889376 | rs13163163 | -0.269 | 1.335 | 0.191 | -1.791 | 0.013 | 0.005 | 0.005 | 0 | 0.13 | 0.15 | 0.37 | 0.19 | 0.26 | 0.23 |
| 5 | 140892643 | rs251022 | -1.465 | 0.308 | -0.398 | -2.213 | -0.038 | 0.054 | 0.054 | 0.062 | 0.56 | 0.79 | 0.65 | 0.3 | 0.5 | 0.54 |
| 5 | 140895377 | rs6892185 | 0.451 | 0.093 | -0.121 | -2.392 | -0.047 | -0.136 | -0.136 | 0 | 0.17 | 0.47 | 0.07 | 0 | 0.09 | 0.04 |
| 5 | 140899268 | rs251021 | 1.536 | 0.073 | -0.244 | -2.442 | 0.005 | 0.025 | 0.025 | 0 | 0.13 | 0.01 | 0.05 | 0.05 | 0.07 | 0.15 |

| | | | | | | | | | | | | | | | | |
|---|---|---|---|---|---|---|---|---|---|---|---|---|---|---|---|---|
| 5 | 140902116 | rs251018 | -1.822 | -0.085 | -0.578 | -2.725 | -0.056 | 0.117 | 0.117 | 0 | 0.6 | 0.9 | 0.42 | 0.21 | 0.43 | 0.47 |
| 5 | 140914459 | rs6859379 | 0.646 | -0.003 | -0.392 | -2.631 | -0.045 | -0.134 | -0.134 | 0 | 0.17 | 0.47 | 0.07 | 0 | 0.1 | 0.04 |
| 5 | 140921411 | rs11954658 | 1.176 | 0.142 | -0.456 | -2.569 | 0.013 | 0.007 | 0.007 | 0 | 0.08 | 0.09 | 0.33 | 0.14 | 0.2 | 0.18 |
| 5 | 140930162 | rs11748064 | 0.66 | 0.052 | -0.514 | -2.604 | -0.004 | 0.006 | 0.006 | 0 | 0.14 | 0.13 | 0.18 | 0.1 | 0.11 | 0.16 |
| 5 | 140933573 | rs250791 | 3.459 | 0.486 | -0.883 | -2.811 | -0.026 | 0.026 | 0.026 | 0.161 | 0.05 | 0.01 | 0.04 | 0 | 0.09 | 0.03 |
| 5 | 140937935 | rs10044763 | 3.108 | -0.239 | -0.493 | -2.507 | 0.006 | 0.013 | 0.013 | 0 | 0.07 | 0.02 | 0.03 | 0.02 | 0.04 | 0.07 |
| 5 | 140938633 | rs7703688 | 1.749 | -0.163 | -0.47 | -2.424 | -0.001 | 0 | 0 | 0 | 0.06 | 0.06 | 0.07 | 0.11 | 0.07 | 0.11 |
| 5 | 140965528 | rs6580191 | 0.049 | -0.295 | -0.373 | -2.447 | -0.041 | -0.122 | -0.122 | 0.007 | 0.26 | 0.51 | 0.1 | 0.06 | 0.16 | 0.15 |
| 5 | 140983565 | rs1035468 | 0.608 | -0.287 | -0.356 | -2.425 | -0.057 | -0.145 | -0.145 | 0 | 0.18 | 0.46 | 0.07 | 0 | 0.1 | 0.04 |
| 5 | 140989410 | rs11167753 | 0.629 | -0.177 | -0.425 | -2.392 | -0.001 | 0 | 0 | 0 | 0.18 | 0.14 | 0.36 | 0.15 | 0.27 | 0.28 |
| 5 | 140992235 | rs3844598 | -0.997 | -0.168 | -0.423 | -2.346 | -0.008 | 0.016 | 0.016 | 0 | 0.45 | 0.39 | 0.61 | 0.35 | 0.48 | 0.6 |
| 5 | 141005687 | rs2547547 | 1.494 | -0.082 | -0.856 | -2.22 | 0.003 | 0.006 | 0.006 | 0 | 0.13 | 0.12 | 0.07 | 0.1 | 0.1 | 0.17 |
| 5 | 141011346 | rs2545025 | 1.55 | -0.081 | -0.867 | -2.218 | 0.002 | 0.006 | 0.006 | 0 | 0.13 | 0.09 | 0.07 | 0.1 | 0.11 | 0.17 |
| 5 | 141014494 | rs2530223 | -1.703 | -0.108 | -0.932 | -2.195 | -0.057 | -0.012 | -0.012 | 0.837 | 0.64 | 0.89 | 0.69 | 0.35 | 0.57 | 0.64 |
| 5 | 141015242 | rs7702834 | 0.815 | -0.352 | -0.323 | -2.176 | -0.025 | -0.073 | -0.073 | 0.95 | 0.15 | 0.25 | 0.05 | 0 | 0.09 | 0.04 |
| 5 | 141019110 | rs14251 | -1.165 | -0.173 | -0.535 | -2.27 | 0 | -0.013 | -0.013 | 1.821 | 0.57 | 0.58 | 0.38 | 0.16 | 0.38 | 0.43 |
| 5 | 141020100 | rs41098 | -0.312 | -0.218 | -0.209 | -2.107 | -0.043 | -0.166 | -0.166 | 0.086 | 0.3 | 0.46 | 0.16 | 0.01 | 0.22 | 0.12 |
| 5 | 141022848 | rs702379 | -0.558 | -0.373 | -0.227 | -2.13 | -0.009 | -0.04 | -0.04 | 0.037 | 0.34 | 0.39 | 0.15 | 0.06 | 0.2 | 0.17 |
| 5 | 141022882 | rs468650 | 0.086 | -0.323 | -0.183 | -2.085 | -0.04 | -0.113 | -0.113 | 0.016 | 0.23 | 0.38 | 0.12 | 0 | 0.15 | 0.04 |
| 5 | 141025581 | rs2052455 | 0.424 | 0.322 | -0.271 | -1.463 | 0.054 | 0.066 | 0.066 | 0 | 0.09 | 0.19 | 0.31 | 0.17 | 0.17 | 0.2 |
| 5 | 141025929 | rs7711960 | -0.038 | 0.21 | -0.495 | -1.517 | -0.01 | 0.006 | 0.006 | 0 | 0.21 | 0.33 | 0.36 | 0.23 | 0.19 | 0.32 |
| 5 | 141026884 | rs251177 | 0.136 | 0.3 | -0.531 | -1.648 | -0.019 | -0.113 | -0.113 | 0 | 0.3 | 0.41 | 0.15 | 0.01 | 0.23 | 0.13 |
| 5 | 141027399 | rs468968 | -1.029 | 0.415 | -1.494 | -1.422 | -0.007 | 0.12 | 0.12 | 0 | 0.53 | 0.8 | 0.56 | 0.24 | 0.53 | 0.47 |
| 5 | 141035955 | rs2306339 | -0.255 | 0.569 | -1.176 | -1.023 | -0.002 | 0.007 | 0.007 | 0 | 0.24 | 0.18 | 0.26 | 0.19 | 0.19 | 0.37 |
| 5 | 141041148 | rs6891143 | 0.775 | 0.193 | -1.988 | -0.21 | 0.024 | -0.037 | -0.037 | 0.059 | 0.2 | 0.09 | 0.09 | 0.25 | 0.14 | 0.1 |
| 5 | 141041435 | rs3749759 | -1.401 | 0.521 | -2.175 | -0.133 | -0.025 | -0.043 | -0.043 | 0 | 0.65 | 0.85 | 0.75 | 0.59 | 0.63 | 0.68 |
| 5 | 141046936 | rs17097438 | 0.499 | 0.461 | -2.234 | 0.013 | 0 | -0.001 | -0.001 | 2.401 | 0.15 | 0.13 | 0.29 | 0.14 | 0.17 | 0.21 |
| 5 | 141049407 | rs3763120 | -0.363 | 0.479 | -2.228 | -0.214 | 0.022 | 0.002 | 0.002 | 0 | 0.47 | 0.33 | 0.5 | 0.44 | 0.45 | 0.45 |
| 5 | 141054240 | rs4912610 | 0.521 | 0.58 | -2.997 | 0.209 | -0.012 | -0.033 | -0.033 | 2.968 | 0.11 | 0.07 | 0.31 | 0.26 | 0.17 | 0.21 |
| 5 | 141058487 | rs11957761 | 0.362 | 0.216 | -3.009 | 0.015 | 0.041 | 0.088 | 0.088 | 0.103 | 0.29 | 0.04 | 0.1 | 0.14 | 0.19 | 0.14 |
| 5 | 141059649 | rs417503 | -0.989 | 1.01 | -2.658 | 0.538 | -0.103 | -0.141 | -0.141 | 0 | 0.75 | 0.99 | 0.81 | 0.61 | 0.71 | 0.75 |
| 5 | 141064627 | rs349132 | -0.373 | 0.856 | -2.494 | -0.007 | -0.011 | 0.125 | 0.125 | 0 | 0.52 | 0.82 | 0.45 | 0.34 | 0.49 | 0.44 |
| 5 | 141071879 | rs1651423 | 0.311 | 0.744 | -2.245 | 0.185 | -0.017 | -0.035 | -0.035 | 0 | 0.11 | 0.04 | 0.1 | 0.24 | 0.14 | 0.11 |
| 5 | 141073166 | rs9324855 | -0.004 | 1.067 | -2.205 | -0.243 | -0.022 | 0.087 | 0.087 | 0 | 0.42 | 0.71 | 0.41 | 0.15 | 0.47 | 0.27 |
| 5 | 141073377 | rs396682 | -0.469 | 1.014 | -1.928 | -0.395 | -0.001 | -0.001 | -0.001 | 0.572 | 0.19 | 0.22 | 0.16 | 0.24 | 0.22 | 0.11 |
| 5 | 141075430 | rs248590 | -1.105 | 0.941 | -1.879 | -0.394 | 0.021 | 0.026 | 0.026 | 0.092 | 0.57 | 0.78 | 0.54 | 0.38 | 0.64 | 0.39 |
| 5 | 141079168 | rs11750900 | 0.91 | 1.029 | -1.446 | -0.463 | -0.028 | 0.033 | 0.033 | 0 | 0.27 | 0.14 | 0.19 | 0.15 | 0.32 | 0.19 |
| 5 | 141080897 | rs349128 | -0.556 | 0.967 | -1.55 | -0.282 | 0.014 | 0.046 | 0.046 | 0.221 | 0.49 | 0.61 | 0.48 | 0.37 | 0.56 | 0.38 |
| 5 | 141082821 | rs466157 | -1.216 | 0.903 | -1.937 | 0.381 | -0.021 | -0.06 | -0.06 | 1.124 | 0.74 | 0.93 | 0.83 | 0.71 | 0.78 | 0.78 |
| 5 | 141084320 | rs248589 | 0.664 | 1.178 | -1.528 | 0.531 | -0.013 | -0.021 | -0.021 | 0.441 | 0.08 | 0.06 | 0.07 | 0.23 | 0.12 | 0.08 |
| 5 | 141086579 | rs4912768 | -0.781 | 1.089 | -1.376 | -0.056 | -0.002 | 0.081 | 0.081 | 0 | 0.49 | 0.71 | 0.51 | 0.73 | 0.5 | 0.66 |
| 5 | 141087689 | rs17097481 | -0.267 | 1.05 | -0.627 | 0.1 | 0.08 | 0.126 | 0.126 | 0 | 0.1 | 0.36 | 0.16 | 0.16 | 0.12 | 0.12 |
| 5 | 141091311 | rs248616 | -1.339 | 0.674 | -1.38 | 0.063 | -0.029 | -0.064 | -0.064 | 0.021 | 0.71 | 0.96 | 0.64 | 0.72 | 0.75 | 0.76 |
| 5 | 141092705 | rs248618 | -0.64 | 0.648 | -0.781 | 0.222 | 0.013 | -0.018 | -0.018 | 0.018 | 0.38 | 0.32 | 0.33 | 0.57 | 0.35 | 0.52 |
| 5 | 141098864 | rs248628 | 0.528 | 0.323 | -0.689 | 0.417 | 0.008 | 0.001 | 0.001 | 0 | 0.12 | 0.09 | 0.05 | 0.15 | 0.06 | 0.07 |
| 5 | 141101086 | rs755772 | 0.134 | 0.569 | -0.455 | 0.42 | 0.006 | 0.023 | 0.023 | 0 | 0.2 | 0.24 | 0.32 | 0.33 | 0.26 | 0.45 |
| 5 | 141103117 | rs248638 | -0.498 | 0.766 | -0.575 | 0.15 | 0 | -0.007 | -0.007 | 0.097 | 0.4 | 0.55 | 0.42 | 0.54 | 0.42 | 0.57 |
| 5 | 141107382 | rs32929 | -0.758 | 0.455 | -1.007 | 0.077 | 0.023 | -0.011 | -0.011 | 0 | 0.72 | 0.89 | 0.67 | 0.72 | 0.79 | 0.83 |
| 5 | 141109570 | rs10515511 | -0.446 | 0.907 | 0.041 | 0.046 | 0.069 | -0.023 | -0.023 | 0 | 0.15 | 0.49 | 0.19 | 0.13 | 0.2 | 0.24 |
| 5 | 141110907 | rs9686896 | -0.639 | 0.731 | 0.118 | -0.304 | 0.018 | -0.096 | -0.096 | 0.03 | 0.21 | 0.43 | 0.22 | 0.03 | 0.24 | 0.24 |
| 5 | 141112713 | rs11954808 | -0.757 | 0.799 | 0.301 | -0.314 | 0.003 | -0.13 | -0.13 | 0 | 0.21 | 0.55 | 0.22 | 0.03 | 0.21 | 0.23 |
| 5 | 141112927 | rs32930 | 0.619 | 0.226 | -0.434 | -0.692 | -0.006 | 0.067 | 0.067 | 0 | 0.46 | 0.17 | 0.39 | 0.68 | 0.5 | 0.58 |
| 5 | 141114617 | rs887269 | 1.917 | -0.075 | -0.638 | -0.69 | 0.01 | -0.013 | -0.013 | 0 | 0.2 | 0.15 | 0.33 | 0.29 | 0.17 | 0.1 |
| 5 | 141116062 | rs32923 | -1.069 | 0.144 | -0.562 | -0.524 | 0.011 | 0.002 | 0.002 | 0 | 0.69 | 0.74 | 0.63 | 0.69 | 0.74 | 0.85 |
| 5 | 141119040 | rs32926 | -0.294 | 0.199 | -0.515 | -0.541 | 0.001 | 0.001 | 0.001 | 0 | 0.58 | 0.54 | 0.55 | 0.71 | 0.64 | 0.77 |
| 5 | 141122067 | rs32927 | -0.567 | 0.976 | -0.929 | 0.294 | 0.022 | 0.009 | 0.009 | 0 | 0.18 | 0.25 | 0.2 | 0.26 | 0.26 | 0.26 |
| 5 | 141127231 | rs12518743 | 0.942 | 1.099 | -1.28 | 0.936 | -0.022 | -0.069 | -0.069 | 0 | 0.14 | 0.01 | 0.13 | 0.2 | 0.12 | 0.25 |
| 5 | 141127665 | rs6580202 | 0.802 | 1.291 | -0.325 | 0.832 | 0.021 | -0.029 | -0.029 | 0 | 0.09 | 0.23 | 0.13 | 0.03 | 0.11 | 0.16 |
| 5 | 141128076 | rs13158321 | 0.452 | 1.11 | -1.158 | 1.201 | 0.036 | -0.07 | -0.07 | 0.383 | 0.13 | 0.01 | 0.04 | 0.2 | 0.06 | 0.07 |
| 5 | 141129204 | rs32946 | -1.103 | 1.167 | -1.095 | 1.203 | -0.005 | -0.014 | -0.014 | 0 | 0.24 | 0.18 | 0.19 | 0.48 | 0.27 | 0.31 |
| 5 | 141133945 | rs32935 | -0.993 | 1.763 | -1.051 | 0.993 | -0.022 | -0.021 | -0.021 | 0 | 0.09 | 0.03 | 0.12 | 0.27 | 0.19 | 0.2 |
| 5 | 141147245 | rs32928 | -0.315 | 1.461 | -1.401 | 1.117 | -0.049 | -0.021 | -0.021 | 0.469 | 0.08 | 0.01 | 0.2 | 0.35 | 0.18 | 0.17 |
| 5 | 141148482 | rs740666 | 0.633 | -0.71 | -1.395 | -0.07 | 0.061 | 0.019 | 0.019 | 1.925 | 0.3 | 0.19 | 0.15 | 0.2 | 0.17 | 0.16 |
| 5 | 141153269 | rs173568 | 0.311 | -0.229 | -0.444 | -0.012 | 0.008 | 0.007 | 0.007 | 0 | 0.48 | 0.39 | 0.69 | 0.78 | 0.64 | 0.63 |
| 5 | 141155858 | rs152273 | 0.361 | -0.166 | -0.305 | -0.007 | 0.009 | 0.009 | 0.009 | 0 | 0.47 | 0.38 | 0.69 | 0.72 | 0.62 | 0.58 |
| 5 | 141156299 | rs152355 | 0.327 | -0.154 | -0.453 | -0.001 | -0.003 | -0.003 | -0.003 | 0 | 0.47 | 0.45 | 0.69 | 0.71 | 0.62 | 0.57 |
| 5 | 141165875 | rs248492 | 0.714 | 0.477 | -0.132 | -0.048 | 0.008 | -0.047 | -0.047 | 0 | 0.78 | 0.65 | 0.83 | 0.91 | 0.78 | 0.76 |
| 5 | 141167461 | rs152276 | 0.488 | 0.457 | -0.085 | -0.062 | 0 | -0.007 | -0.007 | 0.792 | 0.8 | 0.83 | 0.84 | 0.91 | 0.79 | 0.79 |
| 5 | 141171742 | rs457639 | -0.493 | 0.415 | -0.178 | -0.111 | 0.012 | 0.075 | 0.075 | 0.224 | 0.88 | 0.98 | 0.95 | 0.99 | 0.93 | 0.86 |

| | | | | | | | | | | | | | | | | |
|---|---|---|---|---|---|---|---|---|---|---|---|---|---|---|---|---|
| 5 | 141188315 | rs17097654 | -1.776 | 0.598 | 0.048 | -0.043 | 0.097 | 0.019 | 0.019 | 0 | 0.09 | 0.32 | 0.13 | 0.08 | 0.14 | 0.09 |
| 5 | 141189168 | rs248471 | 0.591 | 0.347 | -0.772 | -0.486 | 0.009 | -0.005 | -0.005 | 0.219 | 0.35 | 0.58 | 0.58 | 0.68 | 0.48 | 0.39 |
| 5 | 141189817 | rs11746161 | 1.049 | -0.475 | -1.121 | -0.606 | 0.074 | -0.007 | -0.007 | 0.01 | 0.15 | 0.01 | 0.03 | 0.16 | 0.03 | 0.12 |
| 5 | 141191724 | rs187454 | -0.366 | -0.042 | -0.661 | -0.652 | -0.008 | -0.022 | -0.022 | 0.118 | 0.84 | 0.82 | 0.9 | 0.92 | 0.88 | 0.8 |
| 5 | 141198969 | rs4912778 | 0.662 | 0.302 | -0.281 | -0.354 | -0.007 | 0.011 | 0.011 | 0 | 0.31 | 0.2 | 0.47 | 0.25 | 0.31 | 0.38 |
| 5 | 141203031 | rs248516 | 0.393 | 0.261 | -0.025 | -0.447 | 0 | 0 | 0 | 0 | 0.48 | 0.44 | 0.39 | 0.73 | 0.51 | 0.45 |
| 5 | 141204945 | rs248514 | -0.617 | 0.253 | -0.007 | -0.378 | -0.006 | -0.083 | -0.083 | 3.104 | 0.85 | 0.69 | 0.92 | 0.99 | 0.89 | 0.86 |
| 5 | 141206585 | rs13162537 | 0.806 | 0.34 | -0.242 | -0.029 | -0.005 | 0.026 | 0.026 | 0 | 0.29 | 0.21 | 0.25 | 0.16 | 0.27 | 0.31 |
| 5 | 141206685 | rs11746146 | 0.353 | 0.081 | -0.061 | -0.248 | 0.04 | 0.081 | 0.081 | 0.05 | 0.42 | 0.28 | 0.27 | 0.18 | 0.3 | 0.32 |
| 5 | 141207740 | rs248512 | 0.438 | -0.097 | 0.46 | 0.2 | -0.046 | 0.002 | 0.002 | 0 | 0.68 | 0.5 | 0.65 | 0.65 | 0.81 | 0.7 |
| 5 | 141209467 | rs248504 | -0.051 | 0.251 | 0.26 | 0.279 | 0 | 0.006 | 0.006 | 0.323 | 0.82 | 0.63 | 0.7 | 0.82 | 0.86 | 0.79 |
| 5 | 141210753 | rs166040 | -0.314 | 0.383 | 0.129 | 0.279 | -0.005 | 0.001 | 0.001 | 0.142 | 0.84 | 0.7 | 0.73 | 0.83 | 0.88 | 0.8 |
| 5 | 141216310 | rs10491313 | 0.666 | 0.156 | -0.717 | 0.144 | 0.05 | -0.02 | -0.02 | 0 | 0.17 | 0.02 | 0.07 | 0.16 | 0.07 | 0.11 |
| 5 | 141216732 | rs415388 | 0.528 | 0.953 | -0.496 | 0.225 | 0 | 0 | 0 | 0 | 0.54 | 0.53 | 0.53 | 0.53 | 0.6 | 0.57 |
| 5 | 141219628 | rs17649125 | -0.087 | 1.686 | -0.571 | 0.77 | 0.067 | 0.052 | 0.052 | 0 | 0.13 | 0.29 | 0.34 | 0.28 | 0.23 | 0.16 |
| 5 | 141220006 | rs3815355 | 0.58 | 1.699 | -0.968 | 0.947 | -0.001 | 0.005 | 0.005 | 0 | 0.19 | 0.16 | 0.25 | 0.35 | 0.27 | 0.22 |
| 5 | 141222253 | rs17097801 | -0.089 | 1.503 | -1.191 | 0.984 | 0.038 | 0.044 | 0.044 | 0 | 0.12 | 0.23 | 0.32 | 0.28 | 0.2 | 0.14 |
| 5 | 141226970 | rs17208551 | 0.701 | 0.739 | -2.054 | 0.986 | -0.007 | -0.027 | -0.027 | 0 | 0.26 | 0.09 | 0.23 | 0.24 | 0.25 | 0.2 |
| 5 | 141227042 | rs2974704 | 0.102 | 0.358 | -1.605 | 0.781 | 0.042 | 0.029 | 0.029 | 0 | 0.76 | 0.88 | 0.94 | 1 | 0.91 | 0.86 |
| 5 | 141228889 | rs17097812 | -1.441 | 1.866 | -1.129 | 2.117 | 0.001 | 0.054 | 0.054 | 0 | 0.08 | 0.07 | 0.18 | 0.34 | 0.08 | 0.17 |
| 5 | 141241883 | rs12653948 | -0.835 | 2.382 | -1.449 | 1.361 | -0.058 | -0.041 | -0.041 | 0 | 0.1 | 0.02 | 0.22 | 0.2 | 0.2 | 0.11 |
| 5 | 141244356 | rs3822357 | -0.959 | 1.42 | -1.256 | 1.458 | -0.003 | -0.03 | -0.03 | 2.281 | 0.07 | 0.03 | 0.15 | 0.14 | 0.07 | 0.1 |
| 5 | 141246646 | rs10063472 | 0.467 | 1.571 | -0.904 | 0.725 | 0.005 | -0.006 | -0.006 | 0 | 0.79 | 0.76 | 0.85 | 0.8 | 0.78 | 0.87 |
| 5 | 141248963 | rs12515587 | -0.763 | 0.934 | -0.567 | -0.022 | -0.011 | -0.05 | -0.05 | 3.544 | 0.17 | 0.74 | 0.15 | 0.04 | 0.13 | 0.1 |
| 5 | 141254063 | rs6888135 | 0.092 | 0.941 | -1.763 | -0.808 | -0.024 | 0.208 | 0.208 | 0 | 0.5 | 0.21 | 0.61 | 0.81 | 0.5 | 0.66 |
| 5 | 141279350 | rs9324859 | -0.666 | 0.843 | -0.483 | -0.911 | -0.039 | 0.086 | 0.086 | 0 | 0.65 | 0.83 | 0.73 | 0.85 | 0.59 | 0.73 |
| 5 | 141293587 | rs17097850 | 0.558 | 0.197 | -0.878 | -1.517 | 0.022 | 0.019 | 0.019 | 0 | 0.17 | 0.12 | 0.06 | 0.01 | 0.1 | 0.12 |
| 5 | 141296502 | rs4141206 | -0.594 | -0.006 | -0.299 | -1.918 | 0 | -0.034 | -0.034 | 0.111 | 0.54 | 0.54 | 0.62 | 0.82 | 0.49 | 0.66 |
| 5 | 141298977 | rs3855086 | -1.173 | 0.151 | 0.053 | -1.961 | -0.062 | 0.128 | 0.128 | 0 | 0.61 | 0.88 | 0.69 | 0.86 | 0.54 | 0.73 |
| 5 | 141299444 | rs3861853 | 0.16 | 0.181 | 0.132 | -1.884 | -0.009 | 0.035 | 0.035 | 0 | 0.34 | 0.3 | 0.55 | 0.81 | 0.38 | 0.54 |
| 5 | 141300369 | rs3861854 | 0.583 | 0.321 | 0.406 | -1.713 | -0.003 | -0.017 | -0.017 | 0.476 | 0.07 | 0.13 | 0.05 | 0.04 | 0.06 | 0.07 |
| 5 | 141300906 | rs445310 | 0.198 | 0.705 | 0.245 | -2.148 | 0.018 | -0.056 | -0.056 | 0 | 0.83 | 0.77 | 0.88 | 0.98 | 0.79 | 0.88 |
| 5 | 141301096 | rs378361 | 0.715 | 1.034 | -0.631 | -1.505 | -0.098 | -0.064 | -0.064 | 0 | 0.35 | 0.05 | 0.39 | 0.38 | 0.4 | 0.31 |
| 5 | 141303633 | rs13183870 | 0.622 | 0.546 | 0.289 | -1.854 | -0.039 | -0.115 | -0.115 | 0.409 | 0.16 | 0.42 | 0.08 | 0.01 | 0.09 | 0.14 |
| 5 | 141307833 | rs10036567 | 0.73 | 0.597 | 0.007 | -1.832 | -0.035 | -0.106 | -0.106 | 0 | 0.16 | 0.38 | 0.07 | 0.01 | 0.09 | 0.14 |
| 5 | 141309824 | rs351260 | 0.658 | 1.12 | -0.767 | -1.42 | -0.053 | 0.075 | 0.075 | 1.09 | 0.3 | 0.02 | 0.22 | 0.13 | 0.31 | 0.2 |
| 5 | 141324623 | rs153148 | -0.079 | 1.148 | -0.694 | -1.485 | -0.079 | 0.141 | 0.141 | 0.373 | 0.61 | 0.87 | 0.69 | 0.86 | 0.54 | 0.73 |
| 5 | 141330544 | rs164514 | 0.704 | 1.145 | -1.086 | -1.321 | -0.056 | 0.052 | 0.052 | 1.115 | 0.3 | 0.02 | 0.22 | 0.17 | 0.31 | 0.2 |
| 5 | 141334921 | rs2434322 | 1.156 | 0.753 | -1.165 | -0.768 | -0.014 | -0.024 | -0.024 | 0 | 0.08 | 0.02 | 0.07 | 0.27 | 0.12 | 0.16 |
| 5 | 141335241 | rs164073 | -0.748 | 0.903 | -0.901 | -1.485 | -0.121 | 0.114 | 0.114 | 0 | 0.67 | 0.94 | 0.72 | 0.86 | 0.58 | 0.74 |
| 5 | 141335284 | rs164074 | -0.32 | 1.124 | -0.623 | -1.359 | -0.015 | 0.054 | 0.054 | 0 | 0.59 | 0.32 | 0.66 | 0.86 | 0.55 | 0.7 |
| 5 | 141337126 | rs164076 | -0.071 | 1.24 | -0.49 | -1.202 | -0.01 | 0.081 | 0.081 | 0.007 | 0.54 | 0.35 | 0.64 | 0.85 | 0.51 | 0.71 |
| 5 | 141338771 | rs252108 | 1.303 | 2.065 | -0.077 | -0.907 | 0.026 | -0.057 | -0.057 | 0 | 0.85 | 0.78 | 0.87 | 1 | 0.79 | 0.89 |
| 5 | 141338980 | rs164078 | -0.287 | 1.758 | 0.025 | -0.644 | 0 | -0.009 | -0.009 | 0 | 0.39 | 0.36 | 0.51 | 0.62 | 0.39 | 0.42 |
| 5 | 141339522 | rs252110 | 0.351 | 2.156 | 0.526 | -1.016 | -0.028 | 0.026 | 0.026 | 0 | 0.92 | 0.98 | 0.92 | 0.99 | 0.83 | 0.89 |
| 5 | 141343853 | rs252143 | 1.193 | 2.045 | 1.055 | -1.066 | 0.035 | -0.05 | -0.05 | 0 | 0.88 | 0.76 | 0.9 | 0.98 | 0.83 | 0.88 |
| 5 | 141344123 | rs6580209 | -1.477 | 2.209 | 1.689 | -1.293 | 0.042 | -0.09 | -0.09 | 0.489 | 0.14 | 0.31 | 0.14 | 0 | 0.19 | 0.13 |
| 5 | 141345062 | rs252141 | 2.324 | 2.125 | 1.597 | -1.337 | 0.006 | -0.067 | -0.067 | 0.12 | 0.7 | 0.35 | 0.75 | 0.98 | 0.63 | 0.73 |
| 5 | 141348708 | rs252133 | 0.299 | 2.174 | 1.193 | -1.254 | -0.032 | 0.03 | 0.03 | 0 | 0.91 | 0.96 | 0.92 | 0.98 | 0.83 | 0.9 |
| 5 | 141354040 | rs164495 | 1.726 | 2.156 | 1.445 | -1.295 | 0.028 | -0.067 | -0.067 | 0.183 | 0.81 | 0.76 | 0.86 | 0.98 | 0.74 | 0.84 |
| 5 | 141355948 | rs740662 | -0.776 | 2.13 | 1.314 | -1.283 | 0.001 | -0.037 | -0.037 | 0 | 0.1 | 0.11 | 0.06 | 0 | 0.09 | 0.06 |
| 5 | 141367202 | rs374840 | 1.573 | 2.158 | 1.28 | -1.293 | 0.024 | -0.074 | -0.074 | 0.015 | 0.8 | 0.71 | 0.84 | 0.98 | 0.74 | 0.84 |
| 5 | 141367898 | rs433623 | 2.713 | 2.313 | 1.375 | -1.456 | -0.014 | -0.016 | -0.016 | 0.573 | 0.65 | 0.28 | 0.69 | 0.98 | 0.55 | 0.71 |
| 5 | 141369635 | rs17097934 | -0.61 | 2.34 | 1.142 | -1.433 | 0.002 | -0.053 | -0.053 | 0 | 0.1 | 0.19 | 0.06 | 0 | 0.09 | 0.05 |
| 5 | 141380348 | rs7408 | 2.036 | 2.28 | 0.496 | -1.384 | 0.013 | -0.095 | -0.095 | 0.007 | 0.66 | 0.39 | 0.7 | 0.98 | 0.56 | 0.71 |
| 5 | 141382584 | rs252114 | 0.426 | 2.206 | 0.068 | -1.114 | -0.041 | 0.033 | 0.033 | 0 | 0.91 | 0.97 | 0.92 | 0.98 | 0.83 | 0.9 |
| 5 | 141386638 | rs108593 | 1.053 | 1.842 | 0.151 | -1.039 | 0.03 | -0.127 | -0.127 | 0 | 0.7 | 0.4 | 0.77 | 0.98 | 0.63 | 0.75 |
| 5 | 141391532 | rs164080 | 0.191 | 1.406 | -0.497 | -0.342 | -0.008 | 0.031 | 0.031 | 0 | 0.56 | 0.67 | 0.44 | 0.63 | 0.46 | 0.46 |
| 5 | 141395998 | rs6580215 | -0.274 | 1.48 | -0.153 | 0.327 | 0.005 | 0.054 | 0.054 | 0 | 0.25 | 0.32 | 0.25 | 0.43 | 0.23 | 0.37 |
| 5 | 141398031 | rs10078983 | -0.294 | 1.501 | -0.232 | 0.355 | -0.006 | -0.027 | -0.027 | 0 | 0.24 | 0.19 | 0.24 | 0.43 | 0.22 | 0.35 |
| 5 | 141400028 | rs252200 | -0.066 | 1.378 | -0.433 | 0.216 | -0.008 | 0.033 | 0.033 | 0 | 0.3 | 0.15 | 0.37 | 0.2 | 0.27 | 0.28 |
| 5 | 141405507 | rs163918 | 0.897 | 1.256 | 0.135 | 0.711 | 0.068 | 0.12 | 0.12 | 0 | 0.94 | 0.75 | 0.93 | 0.86 | 0.92 | 0.87 |
| 5 | 141414460 | rs492620 | -0.57 | 1.039 | -0.982 | 0.543 | -0.01 | -0.016 | -0.016 | 0 | 0.23 | 0.11 | 0.21 | 0.27 | 0.21 | 0.31 |
| 5 | 141417287 | rs617384 | -0.995 | 1.247 | -0.436 | 0.74 | -0.001 | 0 | 0 | 0 | 0.46 | 0.56 | 0.38 | 0.52 | 0.49 | 0.69 |
| 5 | 141425905 | rs4912795 | -1.03 | 1.273 | -0.002 | 0.899 | 0.008 | 0.113 | 0.113 | 0.015 | 0.2 | 0.24 | 0.37 | 0.49 | 0.21 | 0.28 |
| 5 | 141428100 | rs252106 | 0.043 | 1.532 | -0.439 | 0.815 | -0.033 | -0.035 | -0.035 | 0 | 0.13 | 0.03 | 0.16 | 0.21 | 0.18 | 0.19 |
| 5 | 141428161 | rs7729341 | -0.26 | 1.243 | 0.112 | 0.593 | 0.014 | -0.029 | -0.029 | 0 | 0.16 | 0.35 | 0.1 | 0.09 | 0.14 | 0.17 |
| 5 | 141432269 | rs153145 | -0.037 | 1.238 | -0.183 | 0.733 | -0.03 | -0.036 | -0.036 | 0 | 0.13 | 0.03 | 0.17 | 0.21 | 0.18 | 0.19 |
| 5 | 141439358 | rs153149 | -0.048 | 1.22 | -0.223 | 0.749 | -0.052 | -0.06 | -0.06 | 0 | 0.13 | 0.01 | 0.17 | 0.22 | 0.18 | 0.21 |

| | | | | | | | | | | | | | | | | |
|---|---|---|---|---|---|---|---|---|---|---|---|---|---|---|---|---|
| 5 | 141439441 | rs252144 | -0.648 | 1.236 | 0.151 | 0.675 | -0.002 | -0.002 | -0.002 | 0.019 | 0.17 | 0.16 | 0.21 | 0.22 | 0.22 | 0.23 |
| 5 | 141441170 | rs252149 | -0.733 | 1.223 | 0.356 | 0.651 | 0.01 | 0.002 | 0.002 | 0.554 | 0.17 | 0.22 | 0.21 | 0.22 | 0.23 | 0.23 |
| 5 | 141442133 | rs1438699 | 0.417 | 0.858 | -0.018 | 0.193 | -0.007 | 0.013 | 0.013 | 0 | 0.77 | 0.88 | 0.78 | 0.79 | 0.75 | 0.77 |
| 5 | 141445774 | rs252152 | 1.071 | 0.617 | 0.379 | 0.16 | 0.001 | 0.037 | 0.037 | 0 | 0.7 | 0.43 | 0.51 | 0.49 | 0.67 | 0.65 |
| 5 | 141445980 | rs6867913 | 0.024 | 0.58 | 0.37 | 0.182 | -0.021 | -0.041 | -0.041 | 0 | 0.14 | 0.06 | 0.36 | 0.5 | 0.2 | 0.32 |
| 5 | 141446303 | rs12654522 | 1.704 | 0.516 | 0.654 | 0.09 | -0.008 | 0.073 | 0.073 | 0 | 0.54 | 0.32 | 0.36 | 0.25 | 0.49 | 0.41 |
| 5 | 141453291 | rs4912798 | 0.689 | 0.456 | 0.703 | 0.017 | -0.005 | 0.013 | 0.013 | 0 | 0.64 | 0.69 | 0.72 | 0.71 | 0.63 | 0.64 |
| 5 | 141467856 | rs12655443 | 1.172 | 0.276 | 0.693 | -0.268 | -0.001 | -0.003 | -0.003 | 0 | 0.57 | 0.58 | 0.39 | 0.27 | 0.51 | 0.38 |
| 5 | 141468254 | rs1821263 | -1.409 | 0.311 | 1.008 | -0.465 | -0.001 | -0.011 | -0.011 | 0 | 0.43 | 0.34 | 0.57 | 0.81 | 0.47 | 0.66 |
| 5 | 141469000 | rs9324864 | 0.367 | 0.389 | 1.004 | -0.042 | 0.008 | 0.083 | 0.083 | 0 | 0.76 | 0.61 | 0.56 | 0.48 | 0.73 | 0.68 |
| 5 | 141476168 | rs2879271 | -1.09 | -0.697 | 0.438 | -0.325 | 0.004 | 0.007 | 0.007 | 0 | 0.46 | 0.45 | 0.51 | 0.64 | 0.37 | 0.51 |
| 5 | 141482700 | rs1835967 | 2.74 | -0.67 | -0.065 | -0.486 | -0.12 | 0.061 | 0.061 | 0 | 0.3 | 0.02 | 0.26 | 0.18 | 0.37 | 0.25 |
| 5 | 141484349 | rs6887599 | -0.598 | -0.712 | 0.3 | -0.47 | 0.002 | -0.007 | -0.007 | 0 | 0.81 | 0.81 | 0.83 | 0.83 | 0.79 | 0.8 |
| 5 | 141485168 | rs735683 | -0.38 | -0.939 | 0.28 | -0.571 | 0.019 | -0.035 | -0.035 | 0 | 0.29 | 0.14 | 0.41 | 0.55 | 0.24 | 0.4 |
| 5 | 141487203 | rs36061508 | 0.777 | -0.922 | 0.575 | -0.582 | 0.004 | -0.044 | -0.044 | 0 | 0.15 | 0.31 | 0.07 | 0.07 | 0.1 | 0.1 |
| 5 | 141487530 | rs12517169 | 1.136 | -0.961 | 0.366 | -0.264 | -0.006 | -0.014 | -0.014 | 0 | 0.16 | 0.07 | 0.15 | 0.19 | 0.17 | 0.26 |
| 5 | 141490587 | rs4912622 | 0.273 | -0.967 | 0.383 | -0.271 | -0.01 | -0.007 | -0.007 | 0 | 0.66 | 0.8 | 0.69 | 0.63 | 0.62 | 0.54 |
| 5 | 141500436 | rs11749731 | 0.273 | -0.967 | 0.383 | -0.276 | -0.01 | -0.005 | -0.005 | 0.729 | 0.66 | 0.8 | 0.69 | 0.64 | 0.62 | 0.54 |
| 5 | 141513204 | rs6863411 | 0.273 | -0.967 | 0.396 | -0.276 | -0.012 | -0.006 | -0.006 | 0 | 0.66 | 0.81 | 0.69 | 0.64 | 0.62 | 0.54 |
| 5 | 141516727 | rs7725157 | 0.823 | -1.041 | 0.385 | -0.287 | 0.014 | -0.012 | -0.012 | 0 | 0.13 | 0.07 | 0.29 | 0.35 | 0.08 | 0.13 |
| 5 | 141518104 | rs12655675 | 2.999 | -1.035 | 0.369 | -0.316 | -0.045 | 0.078 | 0.078 | 0 | 0.32 | 0.11 | 0.26 | 0.19 | 0.37 | 0.27 |
| 5 | 141523000 | rs1062158 | 0.263 | -0.977 | 0.387 | -0.307 | -0.012 | -0.006 | -0.006 | 0 | 0.66 | 0.81 | 0.69 | 0.64 | 0.62 | 0.54 |
| 5 | 141525620 | rs13185299 | 0.731 | -0.965 | 0.459 | -0.305 | 0.005 | -0.046 | -0.046 | 1.27 | 0.15 | 0.32 | 0.07 | 0.07 | 0.1 | 0.1 |
| 5 | 141531735 | rs249637 | -2.316 | -0.985 | 0.124 | -0.306 | 0.002 | 0.005 | 0.005 | 0.36 | 0.92 | 0.96 | 0.94 | 1 | 0.92 | 0.98 |
| 5 | 141540017 | rs2338819 | 0.004 | -0.968 | -0.312 | -0.375 | -0.005 | -0.013 | -0.013 | 0 | 0.26 | 0.16 | 0.24 | 0.36 | 0.28 | 0.44 |
| 5 | 141540911 | rs17705913 | 1.174 | -1.084 | -0.343 | -0.335 | -0.027 | -0.052 | -0.052 | 0.014 | 0.15 | 0.02 | 0.13 | 0.19 | 0.15 | 0.25 |
| 5 | 141543351 | rs2961694 | -0.945 | -1.212 | -0.025 | -0.352 | 0.017 | -0.035 | -0.035 | 0.305 | 0.84 | 0.71 | 0.88 | 0.91 | 0.84 | 0.86 |
| 5 | 141544204 | rs2915866 | 1.273 | -1.238 | -0.424 | -0.315 | 0.016 | -0.023 | -0.023 | 0.196 | 0.65 | 0.27 | 0.57 | 0.55 | 0.75 | 0.72 |
| 5 | 141548024 | rs2288820 | -0.124 | -1.135 | -0.132 | 0.14 | -0.002 | 0.016 | 0.016 | 0 | 0.15 | 0.15 | 0.38 | 0.35 | 0.13 | 0.18 |
| 5 | 141548194 | rs2288819 | -1.901 | -1.161 | -0.104 | 0.068 | -0.004 | -0.012 | -0.012 | 0 | 0.44 | 0.5 | 0.65 | 0.72 | 0.41 | 0.61 |
| 5 | 141549536 | rs17649451 | -0.118 | -1.144 | -0.342 | 0.022 | -0.044 | -0.103 | -0.103 | 0 | 0.26 | 0.04 | 0.23 | 0.36 | 0.28 | 0.43 |
| 5 | 141552745 | rs11745706 | -0.113 | -1.107 | -0.278 | 0.043 | -0.007 | -0.042 | -0.042 | 0 | 0.26 | 0.11 | 0.24 | 0.36 | 0.28 | 0.43 |
| 5 | 141555464 | rs10491314 | 0.906 | -0.997 | -0.295 | 0.03 | -0.047 | -0.056 | -0.056 | 0 | 0.14 | 0.02 | 0.16 | 0.18 | 0.18 | 0.27 |
| 5 | 141557236 | rs3910205 | -1.407 | -1.002 | -0.153 | -0.028 | -0.004 | -0.011 | -0.011 | 0.033 | 0.41 | 0.41 | 0.6 | 0.71 | 0.36 | 0.57 |
| 5 | 141564645 | rs2961693 | -0.027 | -0.912 | -0.116 | 0.539 | -0.032 | 0.151 | 0.151 | 0.074 | 0.12 | 0.29 | 0.34 | 0.34 | 0.07 | 0.09 |
| 5 | 141564948 | rs11948854 | 0.438 | -0.598 | -0.62 | 0.425 | -0.025 | -0.058 | -0.058 | 0 | 0.1 | 0.01 | 0.07 | 0.2 | 0.1 | 0.15 |
| 5 | 141566785 | rs11738456 | 0.433 | -0.88 | -0.456 | -0.262 | -0.016 | -0.082 | -0.082 | 2.472 | 0.63 | 0.47 | 0.8 | 0.92 | 0.74 | 0.79 |
| 5 | 141567415 | rs17098224 | -0.216 | -0.494 | -0.677 | 0.009 | 0.005 | 0.071 | 0.071 | 0.69 | 0.25 | 0.48 | 0.27 | 0.38 | 0.26 | 0.42 |
| 5 | 141573265 | rs3910203 | 0.785 | -0.546 | -1.297 | 0.063 | 0 | -0.003 | -0.003 | 0 | 0.57 | 0.42 | 0.43 | 0.56 | 0.6 | 0.62 |
| 5 | 141574172 | rs249663 | -1.124 | -0.732 | -1.269 | 0.183 | -0.019 | 0.07 | 0.07 | 0 | 0.25 | 0.52 | 0.48 | 0.57 | 0.25 | 0.31 |
| 5 | 141576973 | rs13354387 | -0.51 | -0.692 | -1.121 | 0.135 | 0.007 | 0.042 | 0.042 | 0 | 0.18 | 0.43 | 0.18 | 0.21 | 0.18 | 0.2 |
| 5 | 141577017 | rs11952091 | 0.87 | -0.693 | -1.179 | 0.336 | -0.035 | 0.003 | 0.003 | 0 | 0.2 | 0.4 | 0.18 | 0.18 | 0.15 | 0.34 |
| 5 | 141577451 | rs249660 | 1.675 | -0.467 | -1.918 | 0.612 | -0.001 | 0.114 | 0.114 | 0 | 0.05 | 0.05 | 0.31 | 0.35 | 0.07 | 0.11 |
| 5 | 141580122 | rs2906067 | -1.17 | -0.297 | -2.192 | -0.276 | -0.058 | 0.038 | 0.038 | 0 | 0.86 | 1 | 0.94 | 0.94 | 0.89 | 0.9 |
| 5 | 141581995 | rs17539696 | 0.499 | -0.241 | -1.894 | 0.019 | -0.005 | -0.008 | -0.008 | 0 | 0.08 | 0.05 | 0.1 | 0.17 | 0.13 | 0.11 |
| 5 | 141584972 | rs249646 | 0.084 | -0.264 | -1.859 | -0.199 | 0 | -0.004 | -0.004 | 0 | 0.19 | 0.19 | 0.18 | 0.17 | 0.21 | 0.13 |
| 5 | 141586530 | rs2906061 | 1.309 | 0.102 | -1.948 | -0.032 | -0.102 | 0.103 | 0.103 | 0 | 0.36 | 0.07 | 0.26 | 0.22 | 0.47 | 0.23 |
| 5 | 141590358 | rs2906057 | 1.145 | 0 | -1.761 | 0.035 | -0.067 | 0.107 | 0.107 | 0 | 0.38 | 0.16 | 0.28 | 0.22 | 0.48 | 0.25 |
| 5 | 141593628 | rs4912823 | 0.83 | 0.404 | -1.405 | 0.23 | -0.033 | 0.063 | 0.063 | 0 | 0.66 | 0.85 | 0.79 | 0.75 | 0.7 | 0.68 |
| 5 | 141594907 | rs10075594 | 0.099 | 0.623 | -1.202 | 0.31 | -0.018 | 0.008 | 0.008 | 0 | 0.77 | 0.83 | 0.83 | 0.76 | 0.76 | 0.78 |
| 5 | 141605252 | rs2339024 | -0.482 | 0.677 | -0.873 | 0.438 | 0.007 | -0.086 | -0.086 | 0 | 0.14 | 0.4 | 0.26 | 0.01 | 0.09 | 0.12 |
| 5 | 141611957 | rs17649631 | 0.407 | 0.884 | -0.855 | 1.466 | -0.017 | -0.026 | -0.026 | 0 | 0.05 | 0.02 | 0.09 | 0.13 | 0.08 | 0.12 |
| 5 | 141613468 | rs1860991 | 0.582 | 0.853 | -0.764 | 1.395 | 0.001 | 0.004 | 0.004 | 0 | 0.63 | 0.41 | 0.64 | 0.47 | 0.56 | 0.67 |
| 5 | 141624346 | rs17649655 | 0.495 | 1.228 | -0.889 | 1.567 | -0.025 | -0.026 | -0.026 | 0.078 | 0.15 | 0.1 | 0.2 | 0.29 | 0.3 | 0.17 |
| 5 | 141628405 | rs10072521 | -0.195 | 1.069 | -0.959 | 1.515 | -0.049 | 0.011 | 0.011 | 0.003 | 0.76 | 0.45 | 0.79 | 0.74 | 0.85 | 0.82 |
| 5 | 141633461 | rs2908528 | 0.317 | 1.319 | -1.804 | 1.472 | -0.088 | -0.084 | -0.084 | 0 | 0.14 | 0.02 | 0.19 | 0.28 | 0.31 | 0.17 |
| 5 | 141637058 | rs4912636 | 0.221 | 1.181 | -1.482 | 1.089 | 0 | -0.023 | -0.023 | 0 | 0.56 | 0.19 | 0.58 | 0.45 | 0.52 | 0.64 |
| 5 | 141642966 | rs4244027 | -0.801 | 2.012 | -1.029 | 1.149 | -0.003 | 0.006 | 0.006 | 0 | 0.19 | 0.14 | 0.48 | 0.49 | 0.35 | 0.26 |
| 5 | 141646188 | rs7710406 | 0.723 | 1.246 | -1.384 | -0.443 | 0.014 | 0.169 | 0.169 | 0.083 | 0.52 | 0.18 | 0.53 | 0.21 | 0.51 | 0.5 |
| 5 | 141668625 | rs11167773 | -0.298 | 1.164 | -1.485 | -0.576 | -0.079 | -0.049 | -0.049 | 0 | 0.21 | 0.05 | 0.3 | 0.64 | 0.28 | 0.31 |
| 5 | 141669599 | rs7734788 | -0.248 | 1.043 | -0.253 | -0.683 | 0.111 | 0.074 | 0.074 | 0.854 | 0.06 | 0.47 | 0.06 | 0.02 | 0.07 | 0.04 |
| 5 | 141676257 | rs11744608 | 1.337 | 0.973 | -0.072 | -0.826 | 0.03 | 0.092 | 0.092 | 0 | 0.51 | 0.31 | 0.51 | 0.29 | 0.47 | 0.47 |
| 5 | 141681788 | rs4624820 | 0.98 | 1.059 | -0.083 | -0.798 | 0.002 | 0.036 | 0.036 | 0.022 | 0.56 | 0.63 | 0.58 | 0.3 | 0.54 | 0.51 |
| 5 | 141685730 | rs11745862 | 0.018 | 1.132 | -0.182 | -0.745 | -0.058 | -0.054 | -0.054 | 0.022 | 0.18 | 0.07 | 0.28 | 0.6 | 0.26 | 0.29 |
| 5 | 141687732 | rs6893780 | -0.544 | 1.073 | -0.171 | -0.786 | 0 | -0.03 | -0.03 | 0 | 0.77 | 0.76 | 0.87 | 0.93 | 0.81 | 0.82 |
| 5 | 141689777 | rs7723984 | 0.695 | 1.081 | -0.181 | -0.783 | -0.013 | 0.039 | 0.039 | 0 | 0.6 | 0.72 | 0.59 | 0.33 | 0.55 | 0.54 |
| 5 | 141690524 | rs6874376 | 0.122 | 0.861 | -0.068 | -0.91 | 0 | -0.037 | -0.037 | 4.068 | 0.74 | 0.69 | 0.85 | 0.91 | 0.8 | 0.8 |
| 5 | 141691471 | rs6880007 | -1.421 | 0.821 | 0.411 | -0.912 | 0.027 | -0.018 | -0.018 | 0.092 | 0.08 | 0.35 | 0.07 | 0.01 | 0.08 | 0.06 |
| 5 | 141693593 | rs10040443 | 0.492 | 0.79 | 0.079 | -0.917 | 0.002 | -0.09 | -0.09 | 0 | 0.71 | 0.45 | 0.8 | 0.9 | 0.74 | 0.78 |

| | | | | | | | | | | | | | | | | |
|---|---|---|---|---|---|---|---|---|---|---|---|---|---|---|---|---|
| 5 | 141700952 | rs6897690 | -0.636 | 0.493 | -0.736 | -1.063 | 0.008 | 0.092 | 0.092 | 0.624 | 0.68 | 0.85 | 0.84 | 0.98 | 0.78 | 0.84 |
| 5 | 141703671 | rs12516550 | -0.14 | 0.863 | -1.142 | -0.865 | -0.078 | -0.059 | -0.059 | 0 | 0.2 | 0.05 | 0.42 | 0.63 | 0.3 | 0.38 |
| 5 | 141706638 | rs9324876 | 0.212 | 0.041 | -1.179 | -0.967 | 0.065 | 0.152 | 0.152 | 0.293 | 0.51 | 0.91 | 0.57 | 0.34 | 0.48 | 0.48 |
| 5 | 141707606 | rs959662 | 0.247 | -0.007 | -0.919 | -0.956 | 0.099 | 0.142 | 0.142 | 0 | 0.51 | 0.82 | 0.53 | 0.3 | 0.4 | 0.45 |
| 5 | 141710499 | rs10463352 | -0.611 | 0.453 | -0.672 | -0.754 | -0.047 | -0.001 | -0.001 | 12.5 | 0.23 | 0.16 | 0.35 | 0.73 | 0.43 | 0.42 |
| 5 | 141722865 | rs1198963 | -0.512 | 0.025 | -0.258 | -1.039 | 0.002 | -0.027 | -0.027 | 3.264 | 0.65 | 0.66 | 0.82 | 0.95 | 0.74 | 0.8 |
| 5 | 141737196 | rs1198965 | 0.056 | 0.189 | -0.726 | -0.815 | -0.073 | -0.048 | -0.048 | 0.05 | 0.19 | 0.07 | 0.32 | 0.63 | 0.34 | 0.36 |
| 5 | 141744512 | rs10061008 | 0.86 | -0.498 | -0.575 | -1.024 | 0.036 | 0.095 | 0.095 | 0.354 | 0.37 | 0.13 | 0.44 | 0.24 | 0.34 | 0.37 |
| 5 | 141755921 | rs11954937 | 0.345 | -0.78 | 0.284 | -0.518 | -0.001 | -0.004 | -0.004 | 0 | 0.38 | 0.6 | 0.2 | 0.22 | 0.29 | 0.25 |
| 5 | 141758788 | rs450988 | -0.207 | -0.942 | -0.483 | -0.576 | 0.097 | 0.013 | 0.013 | 0 | 0.42 | 0.81 | 0.51 | 0.42 | 0.28 | 0.42 |
| 5 | 141764656 | rs254441 | 0.066 | -0.086 | 0.04 | 0.1 | 0.034 | 0.043 | 0.043 | 0 | 0.72 | 0.66 | 0.69 | 0.54 | 0.53 | 0.71 |
| 5 | 141765054 | rs254439 | -0.169 | -0.245 | -0.802 | 0.337 | -0.168 | -0.136 | -0.136 | 0 | 0.73 | 0.96 | 0.71 | 0.63 | 0.52 | 0.7 |
| 5 | 141765951 | rs254437 | 0.26 | -0.686 | -0.451 | 0.046 | 0.009 | -0.013 | -0.013 | 0 | 0.29 | 0.25 | 0.46 | 0.41 | 0.26 | 0.3 |
| 5 | 141775868 | rs6883937 | 0.355 | -0.939 | 0.191 | -0.367 | 0.022 | 0.082 | 0.082 | 1.671 | 0.39 | 0.23 | 0.21 | 0.07 | 0.26 | 0.22 |
| 5 | 141779528 | rs7716114 | 0.197 | -1.014 | 0.681 | -0.313 | -0.002 | -0.016 | -0.016 | 0 | 0.42 | 0.41 | 0.26 | 0.13 | 0.28 | 0.33 |
| 5 | 141783536 | rs1864003 | -0.915 | -0.813 | 0.013 | -0.585 | -0.088 | 0.118 | 0.118 | 0.314 | 0.63 | 0.87 | 0.44 | 0.74 | 0.46 | 0.64 |
| 5 | 141805088 | rs10062749 | 0.332 | -0.804 | 0.514 | -0.581 | -0.001 | -0.027 | -0.027 | 0 | 0.4 | 0.39 | 0.22 | 0.08 | 0.26 | 0.32 |
| 5 | 141807365 | rs42243 | 0.546 | -0.777 | 0.552 | -0.351 | -0.012 | 0.102 | 0.102 | 0 | 0.86 | 0.72 | 0.85 | 0.43 | 0.85 | 0.65 |
| 5 | 141810877 | rs254424 | 0.126 | -0.752 | 0.459 | -0.402 | 0.015 | 0.039 | 0.039 | 0.436 | 0.5 | 0.58 | 0.68 | 0.37 | 0.6 | 0.42 |
| 5 | 141813945 | rs153423 | -1.011 | -0.967 | 0.518 | -0.434 | -0.027 | 0.034 | 0.034 | 0.929 | 0.21 | 0.4 | 0.18 | 0.68 | 0.18 | 0.37 |
| 5 | 141815211 | rs4912850 | -0.636 | -0.581 | 0.203 | -0.406 | -0.028 | 0.115 | 0.115 | 0.499 | 0.59 | 0.78 | 0.41 | 0.75 | 0.44 | 0.65 |
| 5 | 141821010 | rs6873267 | 0.093 | -0.249 | -0.139 | -0.152 | 0.032 | 0.05 | 0.05 | 1.821 | 0.48 | 0.3 | 0.28 | 0.58 | 0.36 | 0.45 |
| 5 | 141823848 | rs6865028 | -1.27 | 0.172 | 0.471 | 0.438 | -0.003 | 0.038 | 0.038 | 0.199 | 0.1 | 0.22 | 0.1 | 0.14 | 0.09 | 0.09 |
| 5 | 141826254 | rs2560868 | -0.816 | 0.411 | -0.022 | -0.174 | -0.001 | -0.015 | -0.015 | 0 | 0.48 | 0.53 | 0.48 | 0.7 | 0.44 | 0.63 |
| 5 | 141829010 | rs6862405 | -0.987 | 0.221 | 0.286 | -0.263 | 0.014 | 0.072 | 0.072 | 0 | 0.48 | 0.67 | 0.51 | 0.74 | 0.42 | 0.67 |
| 5 | 141835595 | rs2042896 | -0.47 | -0.01 | 0.339 | 0.277 | -0.005 | -0.021 | -0.021 | 0 | 0.32 | 0.22 | 0.37 | 0.51 | 0.31 | 0.45 |
| 5 | 141839507 | rs10064191 | -0.265 | -0.113 | -0.12 | 0.275 | -0.027 | -0.075 | -0.075 | 0 | 0.3 | 0.1 | 0.36 | 0.51 | 0.3 | 0.45 |
| 5 | 141841925 | rs4912639 | -1.264 | -0.126 | 0.052 | 0.246 | -0.004 | -0.019 | -0.019 | 1.81 | 0.43 | 0.55 | 0.46 | 0.72 | 0.34 | 0.66 |
| 5 | 141842845 | rs10477181 | -0.508 | 0.496 | -0.691 | 0.486 | -0.016 | -0.052 | -0.052 | 0 | 0.31 | 0.15 | 0.36 | 0.51 | 0.3 | 0.48 |
| 5 | 141851826 | rs4244028 | -0.81 | 0.207 | -0.544 | 0.01 | 0.002 | 0.002 | 0.002 | 0.042 | 0.39 | 0.62 | 0.38 | 0.73 | 0.27 | 0.65 |
| 5 | 141854131 | rs17208817 | 0.537 | 0.135 | -1.049 | -0.704 | 0.004 | 0.022 | 0.022 | 0 | 0.18 | 0.11 | 0.09 | 0.04 | 0.16 | 0.12 |
| 5 | 141854163 | rs11167777 | -0.46 | 0.447 | -0.772 | -0.781 | 0.015 | 0.061 | 0.061 | 0 | 0.5 | 0.67 | 0.41 | 0.7 | 0.46 | 0.5 |
| 5 | 141855300 | rs17098699 | -0.289 | 0.406 | -0.19 | -0.491 | 0.05 | 0.097 | 0.097 | 0.03 | 0.07 | 0.33 | 0.07 | 0.07 | 0.06 | 0.07 |
| 5 | 141856176 | rs7716375 | -0.021 | 0.45 | -0.173 | -0.558 | 0.076 | 0.01 | 0.01 | 0.02 | 0.06 | 0.31 | 0.06 | 0 | 0.05 | 0.01 |
| 5 | 141857865 | rs10463262 | 1.153 | 0.666 | -1.093 | -1.058 | -0.13 | 0.042 | 0.042 | 0 | 0.26 | 0.02 | 0.31 | 0.1 | 0.37 | 0.15 |
| 5 | 141858677 | rs975385 | 0.238 | 0.594 | -0.574 | -0.676 | 0.009 | -0.009 | -0.009 | 0 | 0.4 | 0.29 | 0.31 | 0.63 | 0.37 | 0.43 |
| 5 | 141859640 | rs4912856 | -0.088 | 0.6 | -1.074 | -0.523 | 0.004 | 0.003 | 0.003 | 0 | 0.46 | 0.59 | 0.37 | 0.63 | 0.42 | 0.44 |
| 5 | 141862148 | rs17638714 | 0.273 | 0.589 | -1.479 | -0.32 | -0.023 | 0.039 | 0.039 | 0 | 0.37 | 0.13 | 0.36 | 0.27 | 0.4 | 0.37 |
| 5 | 141865255 | rs10155516 | -0.415 | 0.428 | -0.328 | -0.512 | -0.014 | 0.06 | 0.06 | 0 | 0.63 | 0.77 | 0.67 | 0.93 | 0.65 | 0.8 |
| 5 | 141865513 | rs4632826 | 0.232 | 0.238 | 0.41 | 0.221 | 0 | 0.003 | 0.003 | 0 | 0.49 | 0.45 | 0.46 | 0.34 | 0.54 | 0.48 |
| 5 | 141868481 | rs10062025 | -1.428 | -0.06 | 0.549 | 0.156 | -0.048 | 0.031 | 0.031 | 0 | 0.2 | 0.51 | 0.07 | 0.19 | 0.1 | 0.2 |
| 5 | 141869664 | rs10491399 | 1.289 | 0.09 | -0.025 | -0.054 | -0.114 | 0.06 | 0.06 | 0.01 | 0.35 | 0.03 | 0.43 | 0.15 | 0.44 | 0.35 |
| 5 | 141869933 | rs10491398 | 0.352 | 0.007 | 0.117 | 0.16 | -0.001 | -0.004 | -0.004 | 0 | 0.08 | 0.05 | 0.02 | 0.07 | 0.07 | 0.09 |
| 5 | 141870200 | rs4912859 | -0.944 | -0.05 | 0.01 | -0.011 | -0.076 | 0.178 | 0.178 | 0 | 0.66 | 0.92 | 0.68 | 0.93 | 0.67 | 0.8 |
| 5 | 141870609 | rs17639134 | 0.986 | 0.152 | 0.128 | 0.188 | -0.047 | 0.081 | 0.081 | 0 | 0.37 | 0.13 | 0.43 | 0.15 | 0.45 | 0.34 |
| 5 | 141875313 | rs249725 | -1.119 | 0.23 | 1.452 | 1.106 | -0.003 | -0.017 | -0.017 | 0.262 | 0.12 | 0.23 | 0.06 | 0.06 | 0.09 | 0.1 |
| 5 | 141876649 | rs10491296 | -0.343 | 0.263 | 1.085 | 1.378 | -0.01 | -0.022 | -0.022 | 0 | 0.08 | 0 | 0.14 | 0.54 | 0.07 | 0.19 |
| 5 | 141887551 | rs249732 | -2.18 | 0.034 | 2.488 | 1.282 | -0.027 | 0.111 | 0.111 | 0 | 0.26 | 0.44 | 0.27 | 0.6 | 0.23 | 0.4 |
| 5 | 141888038 | rs10491297 | 0.024 | 0.086 | 2.351 | 1.119 | 0 | -0.019 | -0.019 | 0 | 0.08 | 0.08 | 0.1 | 0 | 0.1 | 0.07 |
| 5 | 141891293 | rs249736 | -1.877 | -0.041 | 2.645 | 1.182 | -0.044 | 0.178 | 0.178 | 0 | 0.21 | 0.38 | 0.2 | 0.58 | 0.12 | 0.34 |
| 5 | 141891695 | rs10476838 | -0.782 | -0.075 | 2.366 | 1.316 | 0.002 | 0.031 | 0.031 | 0 | 0.1 | 0.1 | 0.13 | 0.66 | 0.06 | 0.23 |
| 5 | 141892009 | rs249738 | -0.592 | -0.056 | 2.441 | 1.226 | -0.004 | -0.01 | -0.01 | 0 | 0.09 | 0.13 | 0.05 | 0.06 | 0.07 | 0.08 |
| 5 | 141893763 | rs17649915 | 0.044 | -0.048 | 2.212 | 1.28 | 0.001 | -0.003 | -0.003 | 0 | 0.07 | 0 | 0.12 | 0.52 | 0.05 | 0.19 |
| 5 | 141894261 | rs249740 | -2.178 | -0.05 | 2.333 | 1.025 | -0.006 | -0.031 | -0.031 | 1.375 | 0.25 | 0.61 | 0.19 | 0.09 | 0.19 | 0.21 |
| 5 | 141910036 | rs249749 | -2.588 | -0.086 | 2.098 | 1.26 | -0.015 | 0.047 | 0.047 | 0 | 0.29 | 0.55 | 0.3 | 0.57 | 0.2 | 0.37 |
| 5 | 141910393 | rs249750 | -2.151 | 0.12 | 2.194 | 0.795 | -0.011 | -0.048 | -0.048 | 1.682 | 0.22 | 0.54 | 0.18 | 0.09 | 0.15 | 0.19 |
| 5 | 141912050 | rs17098939 | -0.884 | 0.127 | 1.999 | 0.784 | 0.003 | -0.044 | -0.044 | 0 | 0.13 | 0.16 | 0.12 | 0.01 | 0.12 | 0.13 |
| 5 | 141912239 | rs249752 | -2.587 | 0.116 | 1.739 | 0.817 | -0.017 | 0.043 | 0.043 | 0 | 0.29 | 0.55 | 0.3 | 0.56 | 0.2 | 0.37 |
| 5 | 141912334 | rs249754 | -0.934 | -0.12 | 1.689 | 1.198 | -0.003 | 0.015 | 0.015 | 0 | 0.07 | 0.14 | 0.04 | 0.07 | 0.03 | 0.06 |
| 5 | 141912841 | rs4244029 | -1.618 | 0.014 | 0.999 | 1.266 | -0.017 | 0.109 | 0.109 | 0.13 | 0.16 | 0.3 | 0.2 | 0.5 | 0.15 | 0.27 |
| 5 | 141920095 | rs152442 | -1.82 | 0.015 | 0.478 | 1.044 | -0.027 | 0.257 | 0.257 | 0.104 | 0.15 | 0.5 | 0.11 | 0.52 | 0.08 | 0.25 |
| 5 | 141924395 | rs152439 | -1.94 | -0.073 | 0.377 | 0.893 | -0.023 | -0.03 | -0.03 | 0 | 0.15 | 0.71 | 0.1 | 0.06 | 0.07 | 0.1 |
| 5 | 141928868 | rs33961 | -1.505 | 0.199 | 0.441 | 0.748 | 0.006 | -0.028 | -0.028 | 0 | 0.14 | 0.23 | 0.11 | 0.06 | 0.16 | 0.15 |
| 5 | 141929551 | rs33962 | -2.528 | 0.176 | 0.321 | 0.818 | 0.025 | -0.063 | -0.063 | 0 | 0.29 | 0.81 | 0.22 | 0.54 | 0.22 | 0.41 |
| 5 | 141936472 | rs11750296 | -1.594 | 0.526 | 0.127 | 1.014 | -0.007 | 0.204 | 0.204 | 0.433 | 0.18 | 0.5 | 0.16 | 0.47 | 0.17 | 0.35 |
| 5 | 141945329 | rs152422 | -0.569 | 0.973 | -0.443 | 0.96 | -0.072 | 0.062 | 0.062 | 0 | 0.81 | 0.98 | 0.74 | 0.95 | 0.73 | 0.86 |
| 5 | 141945931 | rs17706715 | -1.149 | 0.413 | 0.481 | 1.626 | 0.03 | 0.33 | 0.33 | 0.033 | 0.09 | 0.32 | 0.1 | 0.46 | 0.08 | 0.25 |
| 5 | 141952510 | rs33988 | -1.175 | 0.47 | 0.425 | 1.431 | -0.02 | 0.263 | 0.263 | 0 | 0.16 | 0.4 | 0.12 | 0.54 | 0.09 | 0.28 |
| 5 | 141958831 | rs41123 | -1.047 | 0.458 | 0.027 | 1.436 | 0.009 | 0.259 | 0.259 | 0 | 0.16 | 0.42 | 0.14 | 0.54 | 0.12 | 0.29 |

| | | | | | | | | | | | | | | | | |
|---|---|---|---|---|---|---|---|---|---|---|---|---|---|---|---|---|
| 5 | 141971281 | rs13179022 | 0.271 | 0.348 | -1.835 | 0.584 | 0.013 | 0.021 | 0.021 | 0.013 | 0.55 | 0.18 | 0.58 | 0.34 | 0.56 | 0.48 |
| 5 | 141972713 | rs4912868 | 0.557 | 0.711 | -2.383 | 0.257 | -0.127 | 0.002 | 0.002 | 0 | 0.22 | 0.02 | 0.35 | 0.14 | 0.32 | 0.17 |
| 5 | 141973501 | rs34000 | -0.169 | -0.209 | -1.632 | 0.028 | -0.002 | -0.005 | -0.005 | 2.84 | 0.43 | 0.52 | 0.34 | 0.61 | 0.42 | 0.49 |
| 5 | 141974533 | rs17099029 | 0.385 | -0.616 | -1.188 | -0.107 | -0.018 | -0.024 | -0.024 | 0.006 | 0.13 | 0.23 | 0.06 | 0.08 | 0.08 | 0.08 |
| 5 | 141975067 | rs34003 | -0.849 | -0.979 | -0.993 | 0.182 | 0.021 | 0.046 | 0.046 | 0 | 0.52 | 0.77 | 0.38 | 0.61 | 0.46 | 0.5 |
| 5 | 141979016 | rs2299020 | -0.389 | -0.855 | 0.097 | 0.858 | -0.025 | 0.036 | 0.036 | 0 | 0.08 | 0.26 | 0.11 | 0.13 | 0.03 | 0.07 |
| 5 | 141979200 | rs2299022 | -0.156 | -0.975 | -0.003 | 0.829 | 0.001 | -0.025 | -0.025 | 0 | 0.08 | 0.23 | 0.06 | 0.04 | 0.07 | 0.06 |
| 5 | 141981709 | rs2282794 | -0.278 | -0.885 | -0.312 | 1.216 | -0.025 | 0.026 | 0.026 | 0 | 0.07 | 0.16 | 0.1 | 0.15 | 0.03 | 0.07 |
| 5 | 141982923 | rs152523 | 0.597 | -0.753 | -0.516 | 1.368 | -0.074 | -0.09 | -0.09 | 0.007 | 0.15 | 0.03 | 0.12 | 0.3 | 0.23 | 0.29 |
| 5 | 141988810 | rs921001 | -0.59 | -0.808 | -0.247 | 1.374 | -0.023 | 0.078 | 0.078 | 0 | 0.08 | 0.21 | 0.11 | 0.18 | 0.03 | 0.08 |
| 5 | 141992229 | rs17099100 | 0.255 | -0.792 | -0.26 | 1.33 | -0.021 | 0.029 | 0.029 | 0.048 | 0.09 | 0.23 | 0.05 | 0.13 | 0.05 | 0.04 |
| 5 | 141994473 | rs250108 | -0.363 | -0.137 | -1.011 | 1.005 | -0.024 | -0.042 | -0.042 | 0.055 | 0.23 | 0.14 | 0.22 | 0.36 | 0.27 | 0.38 |
| 5 | 141998966 | rs34016 | -0.658 | -0.949 | -0.559 | 0.906 | -0.026 | 0.019 | 0.019 | 0 | 0.28 | 0.57 | 0.22 | 0.42 | 0.15 | 0.28 |
| 5 | 142004422 | rs6891913 | -0.124 | 0.243 | -0.475 | 0.012 | 0.006 | -0.063 | -0.063 | 0 | 0.33 | 0.38 | 0.32 | 0.06 | 0.4 | 0.33 |
| 5 | 142005577 | rs10069453 | -0.504 | -0.422 | -0.242 | 0.967 | -0.024 | 0.1 | 0.1 | 0 | 0.06 | 0.11 | 0.03 | 0.26 | 0.01 | 0.11 |
| 5 | 142008525 | rs249917 | -0.019 | -0.223 | -0.48 | 0.687 | -0.002 | -0.005 | -0.005 | 0 | 0.56 | 0.4 | 0.45 | 0.5 | 0.57 | 0.54 |
| 5 | 142008657 | rs6884797 | -0.048 | -0.532 | -0.722 | 0.55 | -0.024 | -0.002 | -0.002 | 0.032 | 0.25 | 0.29 | 0.12 | 0.18 | 0.12 | 0.24 |
| 5 | 142015389 | rs249923 | 1.127 | -0.379 | -0.939 | 0.545 | -0.126 | -0.006 | -0.006 | 0 | 0.29 | 0.04 | 0.3 | 0.24 | 0.42 | 0.31 |
| 5 | 142015715 | rs4912870 | -0.702 | -0.482 | -0.483 | 0.245 | -0.004 | -0.072 | -0.072 | 0 | 0.22 | 0.36 | 0.26 | 0.06 | 0.23 | 0.16 |
| 5 | 142017860 | rs152528 | 1.559 | -0.407 | -1.025 | 0.276 | -0.117 | -0.027 | -0.027 | 3.351 | 0.27 | 0.04 | 0.26 | 0.26 | 0.38 | 0.29 |
| 5 | 142018424 | rs1609763 | -1.317 | -0.648 | -0.828 | 0.42 | -0.008 | -0.044 | -0.044 | 0 | 0.68 | 0.84 | 0.66 | 0.52 | 0.7 | 0.62 |
| 5 | 142019596 | rs17099156 | -0.373 | -0.292 | -0.349 | 0.489 | 0.072 | -0.025 | -0.025 | 0 | 0.11 | 0.51 | 0.25 | 0.05 | 0.16 | 0.07 |
| 5 | 142020380 | rs10070885 | -0.91 | -0.377 | -0.481 | 0.519 | -0.012 | -0.013 | -0.013 | 0 | 0.26 | 0.68 | 0.29 | 0.18 | 0.25 | 0.21 |
| 5 | 142020438 | rs12523052 | 1.189 | -0.278 | -1.493 | 0.579 | -0.131 | 0.001 | 0.001 | 0 | 0.32 | 0.05 | 0.3 | 0.26 | 0.41 | 0.32 |
| 5 | 142020508 | rs12523624 | 0.661 | 0.279 | -0.954 | 0.784 | -0.024 | 0.034 | 0.034 | 0 | 0.48 | 0.23 | 0.34 | 0.39 | 0.49 | 0.45 |
| 5 | 142021377 | rs17223381 | -0.155 | 0.084 | -0.933 | 0.351 | -0.024 | 0.027 | 0.027 | 0 | 0.07 | 0.02 | 0.14 | 0 | 0.14 | 0.05 |
| 5 | 142021586 | rs17216995 | -0.344 | -0.473 | -0.875 | 0.401 | -0.025 | -0.003 | -0.003 | 2.211 | 0.11 | 0.19 | 0.14 | 0.09 | 0.05 | 0.11 |
| 5 | 142022777 | rs6893408 | -0.974 | -0.009 | -0.834 | 0.499 | 0.001 | -0.017 | -0.017 | 0 | 0.1 | 0.16 | 0.17 | 0.04 | 0.14 | 0.11 |
| 5 | 142024256 | rs2070715 | 0.359 | -0.063 | -1.48 | 0.888 | 0.044 | -0.043 | -0.043 | 0 | 0.53 | 0.14 | 0.47 | 0.5 | 0.57 | 0.51 |
| 5 | 142030342 | rs17099157 | -1.118 | -0.3 | -0.511 | 1.162 | -0.009 | 0.003 | 0.003 | 0 | 0.09 | 0.08 | 0.08 | 0.11 | 0.04 | 0.04 |
| 5 | 142030968 | rs4912872 | -0.55 | -0.822 | -0.556 | 0.895 | 0.012 | 0.014 | 0.014 | 0.233 | 0.17 | 0.12 | 0.09 | 0.1 | 0.11 | 0.08 |
| 5 | 142031751 | rs17099158 | -1.435 | -0.868 | 0.296 | 0.978 | -0.051 | 0.053 | 0.053 | 0 | 0.11 | 0.39 | 0.12 | 0.15 | 0.05 | 0.04 |
| 5 | 142043344 | rs11747068 | -1.078 | -0.038 | 0.388 | 1.149 | 0.052 | -0.053 | -0.053 | 0 | 0.11 | 0.25 | 0.15 | 0.02 | 0.19 | 0.06 |
| 5 | 142047191 | rs11742776 | -1.964 | -0.025 | -0.307 | 0.931 | -0.084 | -0.045 | -0.045 | 0.017 | 0.53 | 0.94 | 0.44 | 0.55 | 0.53 | 0.38 |
| 5 | 142047909 | rs9324888 | -0.917 | 0 | 0.363 | 0.826 | 0.059 | -0.056 | -0.056 | 0 | 0.1 | 0.25 | 0.15 | 0 | 0.19 | 0.06 |
| 5 | 142059978 | rs17208908 | -1.401 | 0.074 | -0.016 | 0.797 | -0.038 | 0.035 | 0.035 | 0 | 0.15 | 0.1 | 0.18 | 0 | 0.25 | 0.15 |
| 5 | 142064203 | rs11167786 | -2.152 | -0.153 | 0.268 | 0.861 | 0.01 | -0.018 | -0.018 | 0 | 0.24 | 0.28 | 0.28 | 0.12 | 0.29 | 0.16 |
| 5 | 142066991 | rs11955552 | -0.151 | 0.308 | -0.593 | 0.698 | -0.015 | 0.034 | 0.034 | 0 | 0.06 | 0.02 | 0.03 | 0 | 0.06 | 0.09 |
| 5 | 142068366 | rs11167790 | -1.257 | 0.28 | -0.575 | 0.714 | -0.044 | 0.043 | 0.043 | 0 | 0.14 | 0.08 | 0.17 | 0 | 0.25 | 0.15 |
| 5 | 142073939 | rs4912876 | -0.883 | -0.952 | -0.431 | 1.202 | 0.045 | -0.009 | -0.009 | 0 | 0.46 | 0.77 | 0.32 | 0.5 | 0.36 | 0.29 |
| 5 | 142076661 | rs6580256 | -2.103 | -1.76 | -0.298 | 1.919 | 0.108 | -0.123 | -0.123 | 0 | 0.3 | 0.93 | 0.18 | 0.47 | 0.14 | 0.29 |
| 5 | 142077863 | rs1860230 | -2.41 | -1.919 | -0.261 | 1.896 | 0.123 | -0.101 | -0.101 | 0 | 0.3 | 0.93 | 0.23 | 0.43 | 0.12 | 0.29 |
| 5 | 142080602 | rs6875059 | -2.477 | -1.978 | -0.149 | 1.966 | 0.13 | -0.098 | -0.098 | 2.145 | 0.29 | 0.93 | 0.23 | 0.42 | 0.1 | 0.29 |
| 5 | 142080756 | rs10069883 | -1.917 | -2.063 | 0.237 | 2.054 | -0.016 | 0.003 | 0.003 | 0 | 0.07 | 0.11 | 0.02 | 0.09 | 0.01 | 0.11 |
| 5 | 142081528 | rs1860228 | -2.116 | -2.109 | 0.606 | 2.074 | -0.028 | 0.002 | 0.002 | 1.039 | 0.18 | 0.76 | 0.18 | 0.21 | 0.06 | 0.15 |
| 5 | 142081559 | rs7724522 | -2.456 | -2.11 | 0.653 | 2.079 | -0.033 | 0.055 | 0.055 | 0 | 0.09 | 0.17 | 0.04 | 0.22 | 0.02 | 0.14 |
| 5 | 142085997 | rs7723040 | -2.861 | -2.136 | 0.547 | 2.175 | 0.112 | -0.103 | -0.103 | 0.175 | 0.28 | 0.91 | 0.24 | 0.51 | 0.12 | 0.3 |
| 5 | 142090022 | rs9324895 | -2.76 | -2.153 | 0.867 | 2.24 | 0.059 | -0.046 | -0.046 | 0.686 | 0.27 | 0.85 | 0.19 | 0.45 | 0.1 | 0.25 |
| 5 | 142095250 | rs17099388 | -2.891 | -2.202 | 1.6 | 2.234 | -0.062 | 0.158 | 0.158 | 0 | 0.14 | 0.28 | 0.14 | 0.49 | 0.05 | 0.22 |
| 5 | 142095411 | rs12516796 | -3.341 | -2.224 | 1.409 | 2.216 | 0.123 | -0.105 | -0.105 | 0.166 | 0.3 | 0.91 | 0.24 | 0.51 | 0.13 | 0.31 |
| 5 | 142096793 | rs12517068 | -2.643 | -2.17 | 1.448 | 2.157 | -0.049 | 0.098 | 0.098 | 0 | 0.13 | 0.23 | 0.1 | 0.43 | 0.03 | 0.16 |
| 5 | 142099658 | rs17099399 | -3.324 | -2.074 | 1.551 | 2.122 | 0.07 | -0.051 | -0.051 | 0.073 | 0.3 | 0.82 | 0.25 | 0.49 | 0.14 | 0.3 |
| 5 | 142100354 | rs17099403 | -2.36 | -1.771 | 1.525 | 2.139 | 0.023 | -0.026 | -0.026 | 0.121 | 0.09 | 0.05 | 0.03 | 0.3 | 0.03 | 0.15 |
| 5 | 142121774 | rs17099428 | -2.551 | -1.891 | 2.071 | 1.574 | -0.031 | 0.002 | 0.002 | 0 | 0.16 | 0.26 | 0.17 | 0.2 | 0.07 | 0.16 |
| 5 | 142125177 | rs7732591 | -3.203 | -2.105 | 1.824 | 1.731 | 0.123 | -0.069 | -0.069 | 0.048 | 0.28 | 0.95 | 0.25 | 0.37 | 0.11 | 0.26 |
| 5 | 142129500 | rs6580258 | -2.282 | -2.127 | 2.575 | 1.71 | -0.028 | -0.044 | -0.044 | 0 | 0.1 | 0.22 | 0.06 | 0.01 | 0.06 | 0.09 |
| 5 | 142131369 | rs930206 | -3.318 | -2.119 | 2.902 | 1.728 | -0.051 | 0.009 | 0.009 | 1.182 | 0.24 | 0.67 | 0.23 | 0.33 | 0.09 | 0.26 |
| 5 | 142142015 | rs7736417 | -3.369 | -2.104 | 2.049 | 1.65 | 0.123 | -0.06 | -0.06 | 0 | 0.28 | 0.95 | 0.25 | 0.34 | 0.12 | 0.26 |
| 5 | 142151659 | rs6898675 | -2.286 | -1.793 | 2.486 | 1.672 | -0.041 | 0.073 | 0.073 | 1.617 | 0.12 | 0.48 | 0.15 | 0.18 | 0.04 | 0.13 |
| 5 | 142159867 | rs11749540 | -2.055 | -1.642 | 2.206 | 1.582 | -0.001 | 0.092 | 0.092 | 0 | 0.06 | 0.21 | 0.03 | 0.12 | 0.03 | 0.07 |
| 5 | 142162633 | rs7728894 | -2.563 | -1.559 | 1.307 | 1.992 | 0.041 | -0.089 | -0.089 | 0 | 0.31 | 0.95 | 0.24 | 0.33 | 0.24 | 0.28 |
| 5 | 142165841 | rs724301 | -2.246 | -1.526 | 2.33 | 2.081 | -0.056 | -0.033 | -0.033 | 0 | 0.21 | 0.59 | 0.1 | 0.12 | 0.11 | 0.15 |
| 5 | 142170174 | rs1155226 | -2.698 | -1.94 | 2.527 | 1.93 | -0.061 | -0.01 | -0.01 | 1.036 | 0.18 | 0.68 | 0.1 | 0.15 | 0.07 | 0.14 |
| 5 | 142174079 | rs185205 | -1.958 | -2.087 | 2.321 | 2.333 | -0.03 | 0.023 | 0.023 | 0 | 0.14 | 0.21 | 0.13 | 0.25 | 0.07 | 0.2 |
| 5 | 142176206 | rs388940 | -2.395 | -2.656 | 2.631 | 2.436 | -0.009 | 0 | 0 | 0 | 0.2 | 0.21 | 0.16 | 0.25 | 0.08 | 0.22 |
| 5 | 142176721 | rs439140 | -3.05 | -2.672 | 2.896 | 2.674 | -0.059 | 0.072 | 0.072 | 0 | 0.25 | 0.57 | 0.26 | 0.48 | 0.1 | 0.25 |
| 5 | 142177490 | rs11167792 | 0.989 | -2.666 | 1.7 | 2.006 | -0.167 | -0.111 | -0.111 | 0.132 | 0.31 | 0.02 | 0.3 | 0.35 | 0.43 | 0.34 |
| 5 | 142183930 | rs429465 | -2.593 | -3.66 | 2.241 | 2.346 | -0.096 | 0.028 | 0.028 | 0 | 0.24 | 0.46 | 0.18 | 0.3 | 0.09 | 0.21 |
| 5 | 142188049 | rs371846 | -1.949 | -3.24 | 1.978 | 2.289 | -0.042 | 0.071 | 0.071 | 0 | 0.19 | 0.28 | 0.23 | 0.41 | 0.07 | 0.18 |

| | | | | | | | | | | | | | | | | |
|---|---|---|---|---|---|---|---|---|---|---|---|---|---|---|---|---|
| 5 | 142194451 | rs2436379 | -2.002 | -3.288 | 1.958 | 2.131 | -0.081 | 0.03 | 0.03 | 0 | 0.3 | 0.42 | 0.23 | 0.45 | 0.12 | 0.24 |
| 5 | 142202317 | rs417020 | -2.144 | -3.216 | 2 | 2.163 | -0.026 | 0.01 | 0.01 | 0.066 | 0.33 | 0.6 | 0.27 | 0.49 | 0.15 | 0.29 |
| 5 | 142202417 | rs398067 | -1.499 | -3.169 | 2.002 | 2.172 | -0.032 | 0.035 | 0.035 | 0 | 0.17 | 0.26 | 0.19 | 0.3 | 0.07 | 0.15 |
| 5 | 142205359 | rs17099585 | 0.47 | -2.775 | 0.944 | 1.927 | -0.005 | -0.011 | -0.011 | 0 | 0.27 | 0.26 | 0.31 | 0.42 | 0.35 | 0.37 |
| 5 | 142217070 | rs35297 | -1.906 | -3.311 | 1.617 | 2.395 | -0.038 | 0.116 | 0.116 | 0 | 0.15 | 0.25 | 0.21 | 0.45 | 0.07 | 0.17 |
| 5 | 142221460 | rs6883314 | -1.495 | -3.234 | 1.529 | 2.177 | -0.063 | -0.05 | -0.05 | 0 | 0.19 | 0.37 | 0.09 | 0.12 | 0.07 | 0.13 |
| 5 | 142228026 | rs185207 | -2.168 | -2.977 | 0.567 | 2.394 | -0.029 | 0.009 | 0.009 | 0.365 | 0.32 | 0.59 | 0.3 | 0.63 | 0.14 | 0.31 |
| 5 | 142228057 | rs17099608 | -1.375 | -1.989 | -0.054 | 2.697 | -0.019 | 0.103 | 0.103 | 0 | 0.06 | 0.12 | 0.14 | 0.34 | 0.03 | 0.08 |
| 5 | 142230251 | rs40127 | -1.333 | -1.943 | -0.07 | 2.261 | -0.002 | 0.023 | 0.023 | 0 | 0.13 | 0.13 | 0.19 | 0.49 | 0.07 | 0.17 |
| 5 | 142238119 | rs10044036 | -0.824 | -2.28 | 0.298 | 1.763 | -0.036 | 0.033 | 0.033 | 0 | 0.23 | 0.33 | 0.19 | 0.33 | 0.15 | 0.36 |
| 5 | 142242139 | rs13159551 | -1.024 | -2.16 | 0.117 | 2.17 | 0.004 | -0.005 | -0.005 | 0 | 0.08 | 0.08 | 0.06 | 0.15 | 0.04 | 0.09 |
| 5 | 142243575 | rs13185255 | -1.784 | -2.675 | 0.351 | 2.233 | -0.062 | 0.124 | 0.124 | 0 | 0.2 | 0.32 | 0.2 | 0.51 | 0.09 | 0.19 |
| 5 | 142244277 | rs12520828 | -1.799 | -2.679 | 0.327 | 2.201 | -0.061 | 0.116 | 0.116 | 0 | 0.2 | 0.32 | 0.2 | 0.5 | 0.09 | 0.19 |
| 5 | 142246574 | rs3733733 | -1.881 | -2.545 | -0.11 | 1.565 | -0.003 | -0.001 | -0.001 | 0 | 0.2 | 0.22 | 0.21 | 0.22 | 0.11 | 0.17 |
| 5 | 142248702 | rs12519983 | -0.749 | -2.178 | -0.301 | 1.288 | 0.012 | 0.006 | 0.006 | 0 | 0.06 | 0.01 | 0.11 | 0.07 | 0.03 | 0.07 |
| 5 | 142250890 | rs463717 | -1.35 | -3.041 | -1.171 | 0.469 | 0.146 | 0.302 | 0.302 | 0 | 0.53 | 0.9 | 0.54 | 0.9 | 0.3 | 0.67 |
| 5 | 142251807 | rs6896092 | -1.29 | -3.544 | -0.276 | 1.904 | -0.035 | 0.023 | 0.023 | 0.007 | 0.2 | 0.19 | 0.15 | 0.68 | 0.07 | 0.19 |
| 5 | 142251892 | rs707177 | -1.77 | -3.328 | -0.582 | 1.57 | 0.071 | 0.317 | 0.317 | 0.119 | 0.52 | 0.91 | 0.48 | 0.91 | 0.33 | 0.71 |
| 5 | 142252549 | rs10039856 | -1.896 | -3.654 | 0.709 | 1.805 | -0.103 | 0.039 | 0.039 | 0.149 | 0.24 | 0.44 | 0.15 | 0.68 | 0.09 | 0.21 |
| 5 | 142262294 | rs6887217 | -0.998 | -3.547 | 0.826 | 2.048 | 0.03 | 0.463 | 0.463 | 3.529 | 0.07 | 0.42 | 0.09 | 0.39 | 0.02 | 0.05 |
| 5 | 142264406 | rs6580260 | -1.519 | -3.582 | 1.002 | 2.053 | -0.031 | 0.381 | 0.381 | 2.587 | 0.11 | 0.46 | 0.12 | 0.44 | 0.04 | 0.08 |
| 5 | 142277132 | rs153169 | -1.161 | -3.698 | 1.252 | 1.949 | -0.001 | -0.002 | -0.002 | 0 | 0.22 | 0.24 | 0.33 | 0.47 | 0.2 | 0.54 |
| 5 | 142278636 | rs37214 | -0.634 | -3.7 | 1.414 | 1.95 | -0.015 | -0.002 | -0.002 | 0 | 0.18 | 0.12 | 0.31 | 0.19 | 0.19 | 0.43 |
| 5 | 142288545 | rs13187932 | -1.029 | -3.677 | 1.553 | 1.809 | -0.02 | 0.06 | 0.06 | 0.406 | 0.09 | 0.31 | 0.06 | 0.13 | 0.03 | 0.04 |
| 5 | 142298353 | rs712166 | -3.063 | -3.797 | 1.824 | 1.629 | 0.073 | 0.306 | 0.306 | 0.262 | 0.51 | 0.89 | 0.53 | 0.9 | 0.33 | 0.73 |
| 5 | 142300493 | rs37191 | -0.448 | -3.881 | 2.038 | 1.756 | -0.013 | 0.13 | 0.13 | 0 | 0.06 | 0.16 | 0.05 | 0.26 | 0.03 | 0.1 |
| 5 | 142307908 | rs17099709 | -1.848 | -3.895 | 2.173 | 1.722 | -0.077 | 0.021 | 0.021 | 1.864 | 0.13 | 0.48 | 0.06 | 0.14 | 0.03 | 0.03 |
| 5 | 142311970 | rs37184 | -0.883 | -3.882 | 2.07 | 1.768 | -0.02 | -0.068 | -0.068 | 0 | 0.19 | 0.09 | 0.31 | 0.47 | 0.19 | 0.48 |
| 5 | 142319301 | rs712178 | -2.201 | -3.853 | 2.152 | 1.735 | -0.006 | -0.02 | -0.02 | 0 | 0.35 | 0.44 | 0.4 | 0.76 | 0.26 | 0.65 |
| 5 | 142329145 | rs252234 | -1.55 | -3.832 | 2.187 | 1.745 | -0.001 | -0.005 | -0.005 | 0 | 0.26 | 0.29 | 0.36 | 0.73 | 0.22 | 0.58 |
| 5 | 142337083 | rs7723229 | -0.81 | -3.77 | 3.181 | 1.775 | 0.001 | 0.042 | 0.042 | 0.532 | 0.06 | 0.3 | 0.04 | 0.05 | 0.02 | 0.02 |
| 5 | 142353097 | rs37199 | -1.823 | -3.74 | 2.967 | 1.803 | -0.04 | 0.062 | 0.062 | 0 | 0.17 | 0.28 | 0.08 | 0.27 | 0.07 | 0.16 |
| 5 | 142357116 | rs35295 | -2.212 | -4.132 | 2.968 | 1.729 | -0.013 | -0.026 | -0.026 | 1.794 | 0.29 | 0.42 | 0.36 | 0.77 | 0.21 | 0.54 |
| 5 | 142357758 | rs37195 | -1.126 | -4.151 | 2.805 | 1.867 | -0.022 | 0.084 | 0.084 | 0 | 0.08 | 0.12 | 0.03 | 0.26 | 0.02 | 0.13 |
| 5 | 142359037 | rs35294 | -3.639 | -4.211 | 2.799 | 1.275 | 0.127 | 0.253 | 0.253 | 0.476 | 0.46 | 0.85 | 0.43 | 0.91 | 0.27 | 0.64 |
| 5 | 142371467 | rs11167795 | 3.444 | -4.193 | 2.771 | 1.284 | 0.111 | 0.2 | 0.2 | 0.387 | 0.54 | 0.19 | 0.58 | 0.09 | 0.73 | 0.36 |
| 5 | 142372435 | rs6898510 | -1.494 | -3.959 | 2.83 | 1.437 | -0.033 | 0.032 | 0.032 | 0 | 0.09 | 0.27 | 0.05 | 0.14 | 0.04 | 0.11 |
| 5 | 142378861 | rs13161500 | -1.599 | -3.954 | 2.832 | 1.44 | -0.037 | 0.026 | 0.026 | 0 | 0.1 | 0.27 | 0.05 | 0.14 | 0.04 | 0.11 |
| 5 | 142379626 | rs7721905 | 1.432 | -3.772 | 2.238 | 1.531 | 0.001 | 0.003 | 0.003 | 0.299 | 0.79 | 0.73 | 0.67 | 0.28 | 0.81 | 0.55 |
| 5 | 142384160 | rs728465 | 1.349 | -3.766 | 2.143 | 1.534 | 0.001 | 0.002 | 0.002 | 0.321 | 0.8 | 0.75 | 0.67 | 0.28 | 0.81 | 0.56 |
| 5 | 142393562 | rs3822397 | 1.384 | -3.75 | 2.07 | 1.542 | 0.002 | 0.004 | 0.004 | 0.205 | 0.79 | 0.73 | 0.68 | 0.28 | 0.81 | 0.54 |
| 5 | 142398857 | rs3776362 | -0.435 | -3.129 | 1.764 | 1.677 | -0.052 | -0.038 | -0.038 | 0.887 | 0.08 | 0.01 | 0.2 | 0.14 | 0.14 | 0.24 |
| 5 | 142400119 | rs7721749 | -2.049 | -3.295 | 2.168 | 1.392 | -0.098 | -0.016 | -0.016 | 0.65 | 0.16 | 0.49 | 0.08 | 0.14 | 0.05 | 0.12 |
| 5 | 142403036 | rs3756387 | -1.871 | -3.228 | 2.002 | 1.72 | -0.033 | 0.012 | 0.012 | 0.407 | 0.25 | 0.67 | 0.15 | 0.28 | 0.1 | 0.29 |
| 5 | 142406143 | rs171538 | -0.911 | -3.109 | 1.379 | 1.411 | -0.012 | 0.027 | 0.027 | 1.423 | 0.21 | 0.09 | 0.31 | 0.18 | 0.21 | 0.36 |
| 5 | 142410183 | rs246637 | -1.619 | -3.236 | 2.082 | 1.516 | -0.071 | 0.025 | 0.025 | 0 | 0.22 | 0.59 | 0.12 | 0.25 | 0.09 | 0.3 |
| 5 | 142410922 | rs187419 | -0.982 | -3.251 | 2.421 | 1.508 | -0.013 | 0.012 | 0.012 | 0 | 0.16 | 0.24 | 0.08 | 0.25 | 0.07 | 0.29 |
| 5 | 142411430 | rs246636 | -2.51 | -3.261 | 2.492 | 1.603 | -0.004 | -0.001 | -0.001 | 0 | 0.27 | 0.75 | 0.16 | 0.26 | 0.11 | 0.3 |
| 5 | 142414146 | rs245828 | -3.206 | -3.282 | 2.473 | 1.588 | 0.077 | 0.07 | 0.07 | 0.63 | 0.48 | 0.84 | 0.47 | 0.39 | 0.32 | 0.66 |
| 5 | 142414548 | rs1036596 | -2.525 | -3.204 | 2.483 | 1.631 | -0.004 | 0 | 0 | 0 | 0.27 | 0.75 | 0.16 | 0.22 | 0.11 | 0.3 |
| 5 | 142415898 | rs13162227 | -1.452 | -3.192 | 2.495 | 1.497 | -0.028 | -0.061 | -0.061 | 0 | 0.11 | 0.51 | 0.08 | 0 | 0.04 | 0.01 |
| 5 | 142416115 | rs11960805 | 0.03 | -3.173 | 2.131 | 1.506 | -0.001 | -0.023 | -0.023 | 0 | 0.06 | 0.17 | 0.04 | 0 | 0.02 | 0 |
| 5 | 142417494 | rs17707048 | -0.555 | -3.124 | 1.848 | 1.586 | 0.036 | -0.01 | -0.01 | 0.016 | 0.13 | 0.07 | 0.05 | 0.11 | 0.05 | 0.17 |
| 5 | 142420072 | rs190313 | -1.664 | -3.113 | 1.94 | 1.595 | -0.058 | 0.041 | 0.041 | 0 | 0.21 | 0.62 | 0.12 | 0.25 | 0.09 | 0.31 |
| 5 | 142429811 | rs3822394 | -1.925 | -2.657 | 1.703 | 1.75 | 0.035 | 0.051 | 0.051 | 0.581 | 0.37 | 0.21 | 0.37 | 0.25 | 0.27 | 0.51 |
| 5 | 142431193 | rs3776346 | 2.045 | -2.619 | 1.892 | 1.773 | -0.021 | -0.048 | -0.048 | 1.633 | 0.59 | 0.53 | 0.53 | 0.19 | 0.72 | 0.46 |
| 5 | 142433727 | rs6580265 | -1.866 | -2.264 | 1.021 | 1.973 | 0.025 | 0.042 | 0.042 | 0.642 | 0.34 | 0.17 | 0.36 | 0.24 | 0.26 | 0.5 |
| 5 | 142438222 | rs10057800 | -1.857 | -2.336 | 1.056 | 1.96 | 0.024 | 0.04 | 0.04 | 0.575 | 0.34 | 0.17 | 0.36 | 0.24 | 0.26 | 0.5 |
| 5 | 142454472 | rs17541441 | -1.044 | -1.548 | 0.516 | 2.108 | -0.033 | 0.017 | 0.017 | 1.401 | 0.15 | 0.03 | 0.28 | 0.13 | 0.18 | 0.32 |
| 5 | 142463668 | rs17707291 | -1.413 | -1.418 | 1.388 | 2.324 | 0.009 | 0.248 | 0.248 | 0 | 0.06 | 0.21 | 0.05 | 0.29 | 0.03 | 0.19 |
| 5 | 142465816 | rs7727006 | -1.87 | -1.504 | 1.576 | 2.338 | 0.006 | 0.274 | 0.274 | 0 | 0.07 | 0.28 | 0.06 | 0.37 | 0.03 | 0.2 |
| 5 | 142466623 | rs3776317 | 1.766 | -2.222 | 2.21 | 2.161 | -0.059 | 0.092 | 0.092 | 1.093 | 0.79 | 0.38 | 0.88 | 0.56 | 0.93 | 0.66 |
| 5 | 142477096 | rs10060327 | 0.146 | -2.207 | 2.309 | 1.943 | 0.006 | -0.002 | -0.002 | 2.983 | 0.9 | 0.9 | 0.98 | 1 | 0.96 | 0.92 |
| 5 | 142488144 | rs3776309 | 1.878 | -2.155 | 2.558 | 2.012 | -0.046 | 0.008 | 0.008 | 0 | 0.81 | 0.64 | 0.93 | 0.8 | 0.92 | 0.72 |
| 5 | 142490003 | rs17099984 | -1.191 | -2.113 | 2.52 | 2.036 | 0.008 | 0.101 | 0.101 | 0 | 0.06 | 0.13 | 0.05 | 0.2 | 0.04 | 0.19 |
| 5 | 142501187 | rs7705013 | -2.141 | -2.182 | 1.86 | 1.54 | -0.019 | 0.012 | 0.012 | 1.268 | 0.3 | 0.67 | 0.17 | 0.42 | 0.15 | 0.43 |
| 5 | 142503573 | rs187418 | -1.214 | -2.184 | 2.087 | 1.487 | -0.041 | -0.059 | -0.059 | 1.523 | 0.16 | 0.3 | 0.04 | 0.09 | 0.05 | 0.11 |
| 5 | 142507171 | rs3797074 | -0.688 | -2.21 | 2.226 | 1.468 | -0.005 | 0.026 | 0.026 | 0 | 0.12 | 0.17 | 0.07 | 0.2 | 0.09 | 0.24 |
| 5 | 142507651 | rs3776299 | 1.405 | -1.183 | 1.381 | 0.539 | -0.045 | 0.101 | 0.101 | 0 | 0.35 | 0.15 | 0.33 | 0.13 | 0.48 | 0.22 |

| | | | | | | | | | | | | | | | | |
|---|---|---|---|---|---|---|---|---|---|---|---|---|---|---|---|---|
| 5 | 142508981 | rs6876462 | -1.409 | -1.109 | 0.85 | 0.376 | -0.062 | 0.031 | 0.031 | 0.54 | 0.27 | 0.48 | 0.11 | 0.36 | 0.13 | 0.34 |
| 5 | 142514131 | rs10075062 | -0.437 | -0.574 | 0.738 | 0.804 | 0 | 0.003 | 0.003 | 0 | 0.63 | 0.65 | 0.43 | 0.47 | 0.63 | 0.64 |
| 5 | 142516670 | rs246602 | 0.007 | -0.49 | 0.349 | 0.75 | 0.002 | 0 | 0 | 0 | 0.58 | 0.49 | 0.39 | 0.38 | 0.58 | 0.57 |
| 5 | 142517601 | rs3756374 | -0.986 | -0.246 | 0.479 | 1.146 | -0.019 | 0.137 | 0.137 | 0.01 | 0.16 | 0.39 | 0.12 | 0.39 | 0.12 | 0.32 |
| 5 | 142521025 | rs246599 | 0.202 | 0.248 | -0.226 | 0.592 | -0.001 | -0.01 | -0.01 | 0.046 | 0.58 | 0.51 | 0.38 | 0.36 | 0.61 | 0.51 |
| 5 | 142529960 | rs258799 | 0.559 | 1.14 | -0.115 | 0.275 | 0.029 | -0.037 | -0.037 | 0.952 | 0.7 | 0.54 | 0.54 | 0.79 | 0.62 | 0.61 |
| 5 | 142533985 | rs174049 | 0.809 | 2.578 | -0.426 | 0.439 | 0.109 | 0.052 | 0.052 | 0 | 0.76 | 0.52 | 0.55 | 0.67 | 0.6 | 0.71 |
| 5 | 142535913 | rs6887887 | 0.56 | 1.322 | -1.103 | -0.702 | 0.013 | -0.01 | -0.01 | 1.08 | 0.19 | 0.17 | 0.11 | 0 | 0.12 | 0.06 |
| 5 | 142537474 | rs853170 | 1.513 | 1.012 | -1.078 | -0.268 | 0.036 | -0.069 | -0.069 | 0 | 0.39 | 0.08 | 0.26 | 0.57 | 0.28 | 0.52 |
| 5 | 142538959 | rs6888245 | -0.575 | 0.021 | -0.774 | -0.074 | 0.02 | -0.005 | -0.005 | 0 | 0.4 | 0.69 | 0.29 | 0.42 | 0.34 | 0.42 |
| 5 | 142540017 | rs258795 | 1.624 | -0.096 | -1.031 | -0.264 | -0.015 | -0.104 | -0.104 | 0.528 | 0.31 | 0.09 | 0.24 | 0.4 | 0.27 | 0.2 |
| 5 | 142541417 | rs258791 | -0.335 | -0.33 | -0.939 | -0.404 | -0.008 | -0.045 | -0.045 | 2.688 | 0.81 | 0.93 | 0.64 | 0.62 | 0.81 | 0.55 |
| 5 | 142541977 | rs258790 | 1.11 | 0.108 | -0.616 | -0.389 | -0.052 | -0.043 | -0.043 | 0 | 0.38 | 0.17 | 0.29 | 0.48 | 0.39 | 0.22 |
| 5 | 142544155 | rs17287641 | 1.27 | -0.041 | -0.565 | -0.447 | -0.016 | 0.076 | 0.076 | 0 | 0.19 | 0.07 | 0.13 | 0.03 | 0.2 | 0.16 |
| 5 | 142546222 | rs3776249 | 1.574 | 0.019 | -0.218 | -0.069 | -0.006 | -0.007 | -0.007 | 0 | 0.07 | 0.04 | 0.05 | 0.08 | 0.13 | 0.03 |
| 5 | 142550251 | rs853165 | 0.381 | -0.293 | 0.299 | -0.505 | 0.005 | 0.006 | 0.006 | 0 | 0.46 | 0.6 | 0.33 | 0.51 | 0.38 | 0.26 |
| 5 | 142551397 | rs2107622 | 0.358 | -0.535 | 0.29 | -0.325 | 0.01 | -0.018 | -0.018 | 0 | 0.15 | 0.1 | 0.24 | 0.36 | 0.12 | 0.42 |
| 5 | 142552526 | rs17209139 | 1.045 | -0.553 | 0.455 | -0.382 | -0.011 | -0.035 | -0.035 | 0 | 0.09 | 0.1 | 0.07 | 0 | 0.06 | 0.03 |
| 5 | 142553082 | rs853163 | 0.474 | -0.683 | 0.831 | -0.368 | -0.015 | 0.008 | 0.008 | 0 | 0.39 | 0.39 | 0.27 | 0.45 | 0.24 | 0.24 |
| 5 | 142556593 | rs258759 | 1.33 | -0.628 | 0.298 | -0.271 | 0.03 | -0.095 | -0.095 | 0 | 0.28 | 0.05 | 0.19 | 0.4 | 0.18 | 0.19 |
| 5 | 142559092 | rs853160 | -0.864 | -0.471 | 0.274 | -0.484 | -0.09 | 0.128 | 0.128 | 0 | 0.66 | 0.88 | 0.64 | 0.93 | 0.54 | 0.81 |
| 5 | 142561332 | rs258769 | 0.336 | -0.692 | 0.426 | -0.45 | 0.031 | -0.005 | -0.005 | 0 | 0.41 | 0.75 | 0.35 | 0.55 | 0.33 | 0.35 |
| 5 | 142562196 | rs258772 | 0.928 | -0.742 | 0.553 | -0.441 | 0.017 | -0.02 | -0.02 | 0 | 0.32 | 0.7 | 0.24 | 0.45 | 0.16 | 0.2 |
| 5 | 142564672 | rs258774 | 1.361 | -0.71 | 0.624 | -0.411 | 0.027 | -0.061 | -0.061 | 0.413 | 0.28 | 0.12 | 0.2 | 0.41 | 0.17 | 0.2 |
| 5 | 142571720 | rs13169445 | 0.193 | -0.476 | 1.266 | -0.333 | 0.077 | 0.045 | 0.045 | 0.901 | 0.11 | 0.45 | 0.13 | 0.1 | 0.16 | 0.14 |
| 5 | 142576453 | rs258764 | -0.906 | -0.344 | 0.908 | -0.483 | -0.074 | 0.128 | 0.128 | 0 | 0.65 | 0.87 | 0.66 | 0.93 | 0.56 | 0.8 |
| 5 | 142579766 | rs6864988 | 0.04 | -0.023 | 1.38 | 0.017 | 0.066 | 0.079 | 0.079 | 0.62 | 0.11 | 0.41 | 0.12 | 0.13 | 0.16 | 0.17 |
| 5 | 142579907 | rs11167807 | 0.182 | -0.084 | 0.836 | 0.007 | -0.003 | -0.038 | -0.038 | 0 | 0.16 | 0.07 | 0.26 | 0.35 | 0.17 | 0.4 |
| 5 | 142581364 | rs12234019 | -0.453 | -0.074 | 0.913 | 0.014 | 0.036 | 0.08 | 0.08 | 2.117 | 0.13 | 0.59 | 0.15 | 0.17 | 0.16 | 0.17 |
| 5 | 142586382 | rs2074637 | -0.154 | 0.014 | 0.3 | 0.144 | 0 | 0.005 | 0.005 | 0.02 | 0.17 | 0.13 | 0.26 | 0.35 | 0.17 | 0.4 |
| 5 | 142589474 | rs258815 | -0.184 | -0.206 | -0.621 | 0.34 | 0.017 | 0.007 | 0.007 | 0.029 | 0.39 | 0.19 | 0.43 | 0.7 | 0.32 | 0.58 |
| 5 | 142590233 | rs3776221 | -0.891 | 0.103 | -0.022 | 1.59 | 0.061 | 0.195 | 0.195 | 0 | 0.17 | 0.39 | 0.34 | 0.44 | 0.27 | 0.22 |
| 5 | 142590414 | rs258816 | -0.201 | 0.612 | -0.59 | 1.439 | -0.002 | -0.004 | -0.004 | 0 | 0.36 | 0.35 | 0.29 | 0.53 | 0.4 | 0.33 |
| 5 | 142591027 | rs258818 | 0.291 | 0.782 | -0.721 | 1.185 | 0.002 | 0.006 | 0.006 | 0.01 | 0.3 | 0.33 | 0.18 | 0.42 | 0.27 | 0.21 |
| 5 | 142592978 | rs7705435 | -0.525 | -0.117 | -1.229 | 0.914 | 0.012 | 0.008 | 0.008 | 0.502 | 0.31 | 0.71 | 0.31 | 0.29 | 0.28 | 0.45 |
| 5 | 142594521 | rs258820 | -0.382 | 0.087 | -1.626 | 0.911 | -0.03 | -0.021 | -0.021 | 0 | 0.56 | 0.85 | 0.38 | 0.66 | 0.47 | 0.66 |
| 5 | 142595995 | rs258821 | 0.507 | 0.066 | -1.788 | 0.989 | -0.064 | -0.12 | -0.12 | 0 | 0.27 | 0.05 | 0.2 | 0.5 | 0.32 | 0.29 |
| 5 | 142596013 | rs258822 | 0.004 | 0.441 | -1.009 | 0.774 | 0.006 | -0.008 | -0.008 | 0 | 0.36 | 0.29 | 0.44 | 0.75 | 0.44 | 0.39 |
| 5 | 142596624 | rs1438734 | -0.458 | -0.073 | -0.581 | 0.843 | -0.035 | 0.013 | 0.013 | 0 | 0.29 | 0.56 | 0.33 | 0.32 | 0.25 | 0.42 |
| 5 | 142597290 | rs258824 | 0.555 | 0.089 | -1.518 | 0.834 | -0.065 | -0.142 | -0.142 | 0 | 0.28 | 0.02 | 0.18 | 0.43 | 0.24 | 0.2 |
| 5 | 142599442 | rs9324909 | -0.539 | -0.104 | -1.225 | 0.828 | 0.032 | 0 | 0 | 0 | 0.32 | 0.74 | 0.36 | 0.34 | 0.25 | 0.43 |
| 5 | 142605172 | rs853158 | 0.737 | -0.595 | -0.591 | 0.427 | -0.015 | 0.003 | 0.003 | 0 | 0.4 | 0.48 | 0.25 | 0.43 | 0.29 | 0.24 |
| 5 | 142606971 | rs154786 | -0.576 | -0.539 | -1.032 | 0.442 | -0.146 | 0.056 | 0.056 | 0.226 | 0.67 | 0.94 | 0.58 | 0.79 | 0.52 | 0.63 |
| 5 | 142609642 | rs33385 | 0.733 | -0.623 | -0.664 | 0.403 | -0.001 | 0.011 | 0.011 | 0 | 0.12 | 0.14 | 0.24 | 0.34 | 0.15 | 0.34 |
| 5 | 142611567 | rs40596 | 0.101 | -0.722 | -0.614 | 0.475 | -0.026 | 0.017 | 0.017 | 1.848 | 0.24 | 0.38 | 0.29 | 0.31 | 0.2 | 0.36 |
| 5 | 142612488 | rs258796 | 0.702 | -0.467 | -1.662 | 0.655 | 0.014 | 0.059 | 0.059 | 0 | 0.39 | 0.17 | 0.22 | 0.29 | 0.29 | 0.23 |
| 5 | 142612938 | rs13359372 | -0.204 | 0.589 | -1.569 | 0.366 | -0.166 | -0.026 | -0.026 | 0 | 0.26 | 0.05 | 0.42 | 0.19 | 0.49 | 0.38 |
| 5 | 142623288 | rs174047 | 0.139 | 0.82 | -1.073 | 0.169 | 0.002 | 0.01 | 0.01 | 0 | 0.5 | 0.46 | 0.48 | 0.34 | 0.53 | 0.41 |
| 5 | 142623471 | rs258753 | 1.25 | 1.048 | -1.089 | -0.22 | -0.002 | 0.011 | 0.011 | 0 | 0.3 | 0.27 | 0.2 | 0.11 | 0.31 | 0.18 |
| 5 | 142639062 | rs17209216 | 0.545 | 1.592 | -0.985 | -0.126 | -0.048 | -0.028 | -0.028 | 0 | 0.13 | 0.01 | 0.16 | 0.14 | 0.18 | 0.17 |
| 5 | 142645903 | rs244465 | -1.401 | 0.719 | -0.476 | 0.197 | -0.029 | 0.04 | 0.04 | 0.547 | 0.86 | 0.78 | 0.78 | 0.74 | 0.95 | 0.83 |
| 5 | 142655015 | rs17287745 | 0.818 | 0.829 | -0.446 | -0.001 | -0.055 | 0.157 | 0.157 | 0 | 0.44 | 0.09 | 0.29 | 0.19 | 0.42 | 0.35 |
| 5 | 142656813 | rs258747 | 0.504 | 0.818 | -0.308 | -0.006 | 0.007 | 0.028 | 0.028 | 0 | 0.48 | 0.41 | 0.37 | 0.3 | 0.56 | 0.36 |
| 5 | 142657021 | rs17287758 | 2.148 | 0.85 | -0.38 | -0.043 | 0.022 | 0.094 | 0.094 | 0 | 0.28 | 0.07 | 0.1 | 0 | 0.17 | 0.15 |
| 5 | 142657212 | rs17209237 | 0.822 | 0.81 | -0.397 | -0.04 | -0.079 | -0.05 | -0.05 | 0.208 | 0.16 | 0.01 | 0.19 | 0.18 | 0.25 | 0.2 |
| 5 | 142657621 | rs6198 | 2.183 | 0.706 | -0.323 | -0.062 | 0.007 | 0.138 | 0.138 | 0 | 0.27 | 0.02 | 0.1 | 0 | 0.17 | 0.15 |
| 5 | 142661889 | rs258750 | 1.549 | 0.7 | -0.293 | -0.072 | -0.001 | -0.007 | -0.007 | 0 | 0.32 | 0.3 | 0.21 | 0.12 | 0.32 | 0.18 |
| 5 | 142669223 | rs17209251 | 0.872 | 0.69 | -0.295 | 0.035 | -0.071 | -0.053 | -0.053 | 5.711 | 0.15 | 0.01 | 0.17 | 0.19 | 0.23 | 0.18 |
| 5 | 142673397 | rs17209258 | 0.871 | 0.689 | -0.287 | 0.012 | -0.071 | -0.048 | -0.048 | 0 | 0.15 | 0.01 | 0.17 | 0.18 | 0.23 | 0.18 |
| 5 | 142674690 | rs258813 | 1.597 | 0.659 | -0.265 | -0.073 | -0.001 | -0.007 | -0.007 | 5.616 | 0.32 | 0.3 | 0.2 | 0.12 | 0.32 | 0.18 |
| 5 | 142680344 | rs6188 | 1.651 | 0.433 | -0.42 | -0.235 | -0.001 | -0.011 | -0.011 | 0 | 0.32 | 0.3 | 0.2 | 0.12 | 0.32 | 0.18 |
| 5 | 142687494 | rs852977 | 1.589 | 0.429 | -0.409 | -0.238 | -0.001 | -0.009 | -0.009 | 0.345 | 0.32 | 0.3 | 0.2 | 0.12 | 0.32 | 0.18 |
| 5 | 142688323 | rs860457 | 1.692 | 0.473 | -0.408 | -0.238 | -0.003 | 0 | 0 | 0 | 0.32 | 0.27 | 0.2 | 0.09 | 0.32 | 0.18 |
| 5 | 142692533 | rs10482672 | 0.785 | 0.327 | -0.568 | -0.067 | 0.015 | 0.001 | 0.001 | 0 | 0.21 | 0.18 | 0.11 | 0.16 | 0.13 | 0.28 |
| 5 | 142693846 | rs4986593 | 1.002 | 0.311 | -0.615 | -0.088 | -0.075 | -0.049 | -0.049 | 0 | 0.16 | 0.01 | 0.17 | 0.18 | 0.24 | 0.19 |
| 5 | 142697295 | rs33388 | -0.976 | 0.307 | -0.607 | -0.091 | 0.007 | 0.037 | 0.037 | 0 | 0.52 | 0.6 | 0.63 | 0.73 | 0.45 | 0.64 |
| 5 | 142705421 | rs852982 | 2.168 | 0.344 | -0.581 | -0.091 | -0.003 | 0.001 | 0.001 | 0 | 0.31 | 0.26 | 0.2 | 0.09 | 0.32 | 0.18 |
| 5 | 142705895 | rs10482655 | 1.021 | 0.343 | -0.586 | -0.091 | -0.071 | -0.031 | -0.031 | 0 | 0.15 | 0.01 | 0.16 | 0.16 | 0.23 | 0.18 |
| 5 | 142709986 | rs33383 | -1.026 | 0.382 | -0.543 | -0.076 | 0.003 | 0.012 | 0.012 | 0 | 0.53 | 0.57 | 0.63 | 0.73 | 0.45 | 0.64 |

| | | | | | | | | | | | | | | | | |
|---|---|---|---|---|---|---|---|---|---|---|---|---|---|---|---|---|
| 5 | 142716357 | rs11745958 | 1.062 | 0.389 | -0.579 | -0.056 | -0.073 | -0.028 | -0.028 | 0.128 | 0.15 | 0.01 | 0.16 | 0.16 | 0.24 | 0.18 |
| 5 | 142720726 | rs17100236 | 0.587 | 0.366 | -0.472 | -0.054 | -0.017 | -0.002 | -0.002 | 0 | 0.22 | 0.31 | 0.28 | 0.16 | 0.14 | 0.27 |
| 5 | 142723937 | rs4912650 | 1.361 | 0.622 | -0.729 | 0.096 | -0.057 | -0.064 | -0.064 | 1.835 | 0.11 | 0.04 | 0.28 | 0.31 | 0.24 | 0.19 |
| 5 | 142726170 | rs2918417 | 1.862 | 0.631 | -0.623 | -0.101 | -0.001 | 0.002 | 0.002 | 0 | 0.32 | 0.28 | 0.2 | 0.12 | 0.32 | 0.18 |
| 5 | 142727193 | rs6877893 | -0.719 | 0.605 | -0.624 | -0.109 | 0.008 | 0.024 | 0.024 | 0 | 0.51 | 0.57 | 0.62 | 0.72 | 0.44 | 0.64 |
| 5 | 142728031 | rs10482642 | 2.396 | 0.613 | -0.539 | -0.076 | 0.02 | 0.09 | 0.09 | 0 | 0.28 | 0.08 | 0.1 | 0 | 0.17 | 0.15 |
| 5 | 142740166 | rs10515521 | 2.452 | 0.612 | -0.584 | -0.282 | 0.018 | 0.087 | 0.087 | 0 | 0.27 | 0.08 | 0.1 | 0 | 0.17 | 0.15 |
| 5 | 142741027 | rs2918416 | 1.818 | 0.649 | -0.556 | -0.265 | -0.001 | -0.008 | -0.008 | 0 | 0.32 | 0.3 | 0.2 | 0.12 | 0.32 | 0.18 |
| 5 | 142750449 | rs10482634 | 0.947 | 0.651 | -0.627 | -0.243 | -0.071 | -0.027 | -0.027 | 2.356 | 0.16 | 0.02 | 0.16 | 0.16 | 0.23 | 0.18 |
| 5 | 142750533 | rs10482633 | 2.443 | 0.649 | -0.623 | -0.263 | 0.018 | 0.087 | 0.087 | 0 | 0.27 | 0.08 | 0.1 | 0 | 0.17 | 0.15 |
| 5 | 142756004 | rs2963155 | 0.754 | 0.564 | -0.613 | -0.251 | 0.004 | 0.005 | 0.005 | 0.691 | 0.17 | 0.26 | 0.17 | 0.2 | 0.22 | 0.24 |
| 5 | 142758496 | rs2963156 | -0.909 | 0.795 | -0.66 | -0.141 | -0.024 | 0.001 | 0.001 | 0 | 0.84 | 0.9 | 0.83 | 0.85 | 0.77 | 0.82 |
| 5 | 142767740 | rs9324921 | 1.417 | 0.367 | -0.632 | 0.118 | -0.036 | 0.043 | 0.043 | 0.512 | 0.14 | 0.22 | 0.05 | 0.26 | 0.05 | 0.16 |
| 5 | 142776684 | rs10041520 | -0.838 | 0.718 | -1.056 | -0.495 | 0.002 | 0.013 | 0.013 | 0 | 0.53 | 0.58 | 0.63 | 0.73 | 0.45 | 0.64 |
| 5 | 142781567 | rs10482616 | 0.247 | 0.33 | -1.069 | -0.52 | 0.015 | 0.007 | 0.007 | 0 | 0.21 | 0.18 | 0.27 | 0.14 | 0.13 | 0.28 |
| 5 | 142786701 | rs10052957 | 0.636 | 1.143 | -1.015 | -0.461 | -0.002 | 0 | 0 | 0 | 0.31 | 0.27 | 0.2 | 0.12 | 0.32 | 0.18 |
| 5 | 142791677 | rs11749561 | 0.108 | 1.102 | -1.044 | -0.554 | -0.002 | 0.064 | 0.064 | 0 | 0.44 | 0.34 | 0.38 | 0.13 | 0.54 | 0.3 |
| 5 | 142792484 | rs9324924 | -0.407 | 0.671 | -1.104 | 0.199 | 0.035 | -0.011 | -0.011 | 0 | 0.38 | 0.77 | 0.43 | 0.41 | 0.33 | 0.36 |
| 5 | 142792650 | rs7701443 | -0.41 | 0.826 | -0.944 | 0.429 | -0.004 | -0.002 | -0.002 | 0 | 0.46 | 0.48 | 0.42 | 0.57 | 0.39 | 0.42 |
| 5 | 142792990 | rs6865292 | 0.488 | 0.453 | -1.458 | 0.246 | -0.027 | -0.024 | -0.024 | 0 | 0.21 | 0.06 | 0.18 | 0.2 | 0.26 | 0.11 |
| 5 | 142796532 | rs4607376 | -0.017 | 0.607 | -0.915 | 0.389 | -0.02 | -0.007 | -0.007 | 0 | 0.45 | 0.21 | 0.44 | 0.54 | 0.51 | 0.37 |
| 5 | 142798757 | rs12653301 | -0.334 | 0.867 | -0.726 | 0.307 | 0.007 | 0.033 | 0.033 | 0 | 0.24 | 0.35 | 0.4 | 0.44 | 0.27 | 0.45 |
| 5 | 142801480 | rs13182800 | 0.779 | 0.33 | -0.938 | -0.51 | 0.004 | -0.006 | -0.006 | 0 | 0.27 | 0.19 | 0.14 | 0.03 | 0.22 | 0.17 |
| 5 | 142801934 | rs13184611 | 0.801 | 0.338 | -0.935 | -0.493 | 0.003 | -0.005 | -0.005 | 1.518 | 0.27 | 0.19 | 0.14 | 0.03 | 0.22 | 0.17 |
| 5 | 142805874 | rs10477211 | 0.524 | 0.175 | -0.975 | -0.187 | -0.008 | 0.003 | 0.003 | 0 | 0.83 | 0.84 | 0.7 | 0.84 | 0.74 | 0.74 |
| 5 | 142806890 | rs4912910 | 0.569 | -1.118 | -0.947 | -0.134 | 0.05 | -0.003 | -0.003 | 0 | 0.56 | 0.29 | 0.58 | 0.54 | 0.67 | 0.57 |
| 5 | 142807150 | rs4912912 | 0.673 | -0.214 | -1.056 | 0.108 | -0.084 | -0.034 | -0.034 | 0 | 0.37 | 0.11 | 0.45 | 0.42 | 0.44 | 0.4 |
| 5 | 142817607 | rs12521436 | -0.107 | -0.008 | -0.657 | 0.14 | 0.024 | 0.129 | 0.129 | 0 | 0.15 | 0.28 | 0.29 | 0.42 | 0.17 | 0.22 |
| 5 | 142827239 | rs4128755 | 0.165 | -0.503 | -0.955 | -0.008 | -0.038 | 0.063 | 0.063 | 0.672 | 0.45 | 0.2 | 0.35 | 0.27 | 0.53 | 0.32 |
| 5 | 142827853 | rs6863190 | 0.016 | -0.512 | -0.722 | 0.001 | -0.004 | 0.013 | 0.013 | 0.236 | 0.47 | 0.41 | 0.36 | 0.3 | 0.53 | 0.32 |
| 5 | 142835748 | rs4912916 | -0.101 | -0.722 | -0.476 | -0.052 | -0.07 | 0.052 | 0.052 | 1.283 | 0.17 | 0.35 | 0.21 | 0.28 | 0.07 | 0.3 |
| 5 | 142839717 | rs6871464 | -1.241 | -0.625 | -0.62 | -0.026 | -0.04 | -0.021 | -0.021 | 0.009 | 0.64 | 0.81 | 0.58 | 0.58 | 0.59 | 0.61 |
| 5 | 142849104 | rs11167813 | 1.148 | -0.681 | -0.635 | -0.049 | -0.072 | -0.042 | -0.042 | 0 | 0.34 | 0.11 | 0.41 | 0.42 | 0.41 | 0.39 |
| 5 | 142849317 | rs7713008 | -0.412 | -0.608 | -0.684 | -0.18 | 0.036 | 0.007 | 0.007 | 3.848 | 0.54 | 0.67 | 0.42 | 0.53 | 0.37 | 0.52 |
| 5 | 142850814 | rs7700825 | -1.251 | -0.595 | -0.696 | -0.189 | -0.056 | 0.042 | 0.042 | 0 | 0.9 | 0.99 | 0.85 | 0.97 | 0.78 | 0.91 |
| 5 | 142852051 | rs2121152 | 0.919 | -0.622 | -0.631 | -0.201 | -0.016 | -0.002 | -0.002 | 0 | 0.72 | 0.58 | 0.63 | 0.7 | 0.71 | 0.61 |
| 5 | 142852148 | rs7714402 | 0.861 | -0.657 | -0.529 | -0.161 | 0.007 | 0.008 | 0.008 | 1.953 | 0.36 | 0.24 | 0.2 | 0.27 | 0.3 | 0.22 |
| 5 | 142857495 | rs12055197 | -0.126 | -0.683 | -0.49 | -0.107 | -0.072 | 0.053 | 0.053 | 1.423 | 0.17 | 0.35 | 0.21 | 0.28 | 0.06 | 0.3 |
| 5 | 142859231 | rs12054921 | -0.425 | -0.896 | -0.958 | -0.773 | 0.039 | -0.006 | -0.006 | 5.431 | 0.39 | 0.73 | 0.28 | 0.34 | 0.17 | 0.23 |
| 5 | 142859606 | rs7724289 | 1.492 | -0.905 | -0.947 | -0.747 | 0.02 | -0.02 | -0.02 | 0 | 0.21 | 0.15 | 0.07 | 0.26 | 0.11 | 0.07 |
| 5 | 142866564 | rs17100330 | 0.204 | -0.858 | -0.982 | -0.72 | -0.074 | -0.022 | -0.022 | 0.844 | 0.17 | 0.39 | 0.2 | 0.08 | 0.05 | 0.16 |
| 5 | 142868277 | rs6879079 | -1.699 | -0.832 | -2.003 | -0.863 | -0.034 | -0.077 | -0.077 | 0.034 | 0.87 | 0.98 | 0.71 | 0.47 | 0.84 | 0.63 |
| 5 | 142870815 | rs921017 | 0.09 | -1.044 | -1.59 | -0.435 | -0.066 | 0.005 | 0.005 | 1.816 | 0.15 | 0.57 | 0.21 | 0.09 | 0.05 | 0.16 |
| 5 | 142873977 | rs1374001 | 1.288 | -0.763 | -1.489 | -0.31 | -0.016 | 0.012 | 0.012 | 0 | 0.15 | 0.06 | 0.13 | 0.12 | 0.21 | 0.14 |
| 5 | 142877286 | rs4428446 | 0.366 | -0.907 | -1.367 | -0.155 | -0.031 | 0.032 | 0.032 | 0 | 0.26 | 0.15 | 0.28 | 0.13 | 0.44 | 0.22 |
| 5 | 142881689 | rs4912922 | -0.6 | -1.418 | -1.268 | -0.001 | 0.013 | -0.003 | -0.003 | 0 | 0.39 | 0.67 | 0.28 | 0.34 | 0.17 | 0.23 |
| 5 | 142883695 | rs11167816 | 0.28 | -0.275 | -1.936 | -0.039 | 0.043 | -0.025 | -0.025 | 0 | 0.26 | 0.1 | 0.21 | 0.24 | 0.16 | 0.09 |
| 5 | 142892428 | rs6876238 | -0.385 | 0.742 | -1.664 | -0.237 | -0.065 | 0.081 | 0.081 | 0 | 0.26 | 0.1 | 0.42 | 0.13 | 0.42 | 0.22 |
| 5 | 142895811 | rs1445873 | -0.353 | 0.259 | -0.527 | 0.375 | -0.025 | 0.071 | 0.071 | 0 | 0.23 | 0.42 | 0.17 | 0.32 | 0.16 | 0.21 |
| 5 | 142900260 | rs17342419 | 1.291 | 0.399 | -1.006 | 0.143 | 0.013 | 0.04 | 0.04 | 0 | 0.49 | 0.35 | 0.24 | 0.2 | 0.38 | 0.29 |
| 5 | 142904949 | rs7700977 | 1.071 | 0.45 | -0.884 | 0.144 | 0 | -0.002 | -0.002 | 0.057 | 0.53 | 0.43 | 0.35 | 0.22 | 0.55 | 0.32 |
| 5 | 142905423 | rs7718505 | 1.307 | -0.225 | -1.36 | -0.462 | 0.05 | -0.031 | -0.031 | 0.545 | 0.22 | 0.07 | 0.06 | 0.21 | 0.11 | 0.1 |
| 5 | 142905928 | rs6879513 | 1.572 | -0.211 | -1.334 | -0.472 | 0.02 | -0.082 | -0.082 | 0 | 0.3 | 0.31 | 0.29 | 0.02 | 0.44 | 0.22 |
| 5 | 142915171 | rs7730230 | 0.668 | -0.228 | -1.123 | -0.447 | -0.002 | -0.005 | -0.005 | 0 | 0.27 | 0.16 | 0.18 | 0.22 | 0.29 | 0.13 |
| 5 | 142916732 | rs17402447 | 1.968 | -0.257 | -0.994 | -0.501 | 0.004 | -0.088 | -0.088 | 0 | 0.27 | 0.29 | 0.18 | 0 | 0.27 | 0.19 |
| 5 | 142917007 | rs6893954 | -0.293 | -0.383 | -0.98 | -0.326 | -0.014 | -0.025 | -0.025 | 0 | 0.75 | 0.88 | 0.53 | 0.59 | 0.71 | 0.53 |
| 5 | 142922087 | rs4635964 | 0.181 | -0.356 | -1.026 | -0.435 | -0.002 | -0.008 | -0.008 | 0.255 | 0.63 | 0.44 | 0.53 | 0.27 | 0.65 | 0.37 |
| 5 | 142925752 | rs4912924 | 0.531 | -0.126 | -1.311 | -0.359 | 0.012 | 0.03 | 0.03 | 0 | 0.12 | 0.05 | 0.18 | 0.02 | 0.11 | 0.05 |
| 5 | 142930730 | rs7712869 | -0.649 | -0.003 | -1.439 | -0.227 | -0.034 | -0.036 | -0.036 | 0.246 | 0.82 | 0.89 | 0.62 | 0.5 | 0.7 | 0.53 |
| 5 | 142934649 | rs682490 | -1.959 | -0.049 | -1.404 | -0.201 | 0.013 | 0.003 | 0.003 | 0 | 0.94 | 0.93 | 0.9 | 0.88 | 0.87 | 0.93 |
| 5 | 142938663 | rs1037100 | -0.165 | -0.058 | -1.458 | -0.255 | -0.017 | -0.014 | -0.014 | 0 | 0.77 | 0.82 | 0.49 | 0.48 | 0.51 | 0.49 |
| 5 | 142938964 | rs4392659 | 2.108 | -0.271 | -1.202 | -0.43 | -0.004 | -0.024 | -0.024 | 0.072 | 0.26 | 0.19 | 0.17 | 0 | 0.27 | 0.19 |
| 5 | 142941194 | rs17100400 | 1.225 | -0.814 | -1.428 | -0.301 | 0.027 | -0.038 | -0.038 | 0 | 0.14 | 0.04 | 0.19 | 0.18 | 0.1 | 0.09 |
| 5 | 142942400 | rs888993 | -0.052 | -0.897 | -1.336 | -0.359 | -0.092 | 0.03 | 0.03 | 0.063 | 0.28 | 0.5 | 0.11 | 0.25 | 0.12 | 0.19 |
| 5 | 142951094 | rs4912927 | -0.655 | -0.51 | -1.441 | -0.33 | -0.045 | -0.023 | -0.023 | 0 | 0.67 | 0.86 | 0.6 | 0.5 | 0.67 | 0.52 |
| 5 | 142951947 | rs7722813 | 0.219 | -0.722 | -1.415 | -0.481 | 0.003 | 0.005 | 0.005 | 0 | 0.18 | 0.22 | 0.32 | 0.23 | 0.27 | 0.12 |
| 5 | 142954968 | rs149485 | 1.233 | -0.631 | -1.624 | -0.411 | -0.042 | -0.034 | -0.034 | 0 | 0.26 | 0.03 | 0.31 | 0.36 | 0.2 | 0.39 |
| 5 | 142956828 | rs153521 | -0.111 | -0.838 | -1.571 | -0.38 | -0.024 | 0.004 | 0.004 | 0.745 | 0.56 | 0.86 | 0.57 | 0.47 | 0.62 | 0.49 |
| 5 | 142962119 | rs185057 | 2.133 | -0.459 | -1.535 | -0.505 | -0.03 | 0.049 | 0.049 | 0.184 | 0.26 | 0.12 | 0.19 | 0 | 0.32 | 0.21 |

| | | | | | | | | | | | | | | | | |
|---|---|---|---|---|---|---|---|---|---|---|---|---|---|---|---|---|
| 5 | 142962809 | rs17100410 | 0.233 | -0.793 | -0.787 | -0.102 | -0.054 | 0.091 | 0.091 | 0 | 0.19 | 0.39 | 0.09 | 0.26 | 0.09 | 0.19 |
| 5 | 142969131 | rs153512 | 0.281 | -0.274 | -1.291 | -0.389 | 0.007 | 0.087 | 0.087 | 0 | 0.47 | 0.27 | 0.5 | 0.25 | 0.6 | 0.36 |
| 5 | 142972345 | rs152227 | 1.192 | 0.267 | -1.401 | -0.761 | 0.005 | 0.005 | 0.005 | 0.737 | 0.19 | 0.17 | 0.14 | 0 | 0.18 | 0.17 |
| 5 | 142972893 | rs10515523 | 0.55 | 0.672 | -1.158 | -0.062 | -0.016 | -0.056 | -0.056 | 0 | 0.1 | 0.02 | 0.21 | 0.18 | 0.11 | 0.1 |
| 5 | 142974644 | rs11167821 | -0.161 | 1.04 | -0.337 | -0.414 | 0.124 | -0.074 | -0.074 | 0.787 | 0.13 | 0.33 | 0.15 | 0.02 | 0.26 | 0.06 |
| 5 | 142979183 | rs153516 | -0.229 | 0.374 | -1.222 | 0.146 | 0.034 | -0.061 | -0.061 | 1.868 | 0.42 | 0.14 | 0.48 | 0.55 | 0.33 | 0.55 |
| 5 | 142982181 | rs246430 | -0.939 | 0.477 | -0.452 | -0.095 | -0.005 | -0.043 | -0.043 | 0.527 | 0.87 | 0.8 | 0.94 | 0.96 | 0.92 | 0.91 |
| 5 | 142984844 | rs17100459 | 0.204 | 0.508 | -0.757 | 0.689 | -0.005 | 0.096 | 0.096 | 0 | 0.08 | 0.13 | 0.04 | 0.26 | 0.05 | 0.13 |
| 5 | 142986271 | rs246537 | 1.325 | 0.756 | -1.298 | 0.058 | 0.013 | 0.198 | 0.198 | 0 | 0.41 | 0.1 | 0.44 | 0.08 | 0.37 | 0.32 |
| 5 | 142991165 | rs152225 | 0.972 | 1.392 | -0.93 | 0.477 | -0.028 | 0.041 | 0.041 | 0 | 0.62 | 0.22 | 0.65 | 0.33 | 0.57 | 0.62 |
| 5 | 142991422 | rs247638 | 0.547 | 2.118 | -1.033 | 1.099 | -0.012 | -0.01 | -0.01 | 0 | 0.05 | 0.03 | 0.06 | 0.11 | 0.1 | 0.09 |
| 5 | 142994641 | rs444218 | 0.882 | 1.924 | -1.206 | 0.951 | -0.021 | 0.132 | 0.132 | 0 | 0.31 | 0.05 | 0.38 | 0.08 | 0.3 | 0.22 |
| 5 | 143003729 | rs17100500 | -1.365 | 2.25 | -0.597 | 2.149 | 0.027 | 0.075 | 0.075 | 0 | 0.14 | 0.22 | 0.19 | 0.53 | 0.24 | 0.18 |
| 5 | 143004996 | rs13181739 | -1.406 | 2.086 | -0.594 | 2.152 | 0.02 | 0.04 | 0.04 | 0 | 0.17 | 0.22 | 0.2 | 0.53 | 0.3 | 0.23 |
| 5 | 143006537 | rs11948121 | -0.656 | 2.288 | -1.04 | 1.562 | -0.029 | -0.029 | -0.029 | 0 | 0.11 | 0.06 | 0.16 | 0.29 | 0.25 | 0.23 |
| 5 | 143015669 | rs11741557 | -0.808 | 1.369 | -0.651 | 1.065 | 0.027 | 0.033 | 0.033 | 0.049 | 0.06 | 0.16 | 0.07 | 0.12 | 0.1 | 0.1 |
| 5 | 143019997 | rs6898504 | 0 | 0.838 | -1.625 | 0.68 | -0.035 | -0.004 | -0.004 | 0 | 0.75 | 0.98 | 0.8 | 0.75 | 0.78 | 0.79 |
| 5 | 143024627 | rs12657648 | 0.275 | 0.344 | -1.874 | 0.812 | 0.015 | -0.05 | -0.05 | 0 | 0.3 | 0.15 | 0.32 | 0.34 | 0.27 | 0.29 |
| 5 | 143030387 | rs1213707 | 0.726 | 0.091 | -0.493 | 0.482 | 0.013 | 0.002 | 0.002 | 0 | 0.36 | 0.36 | 0.42 | 0.41 | 0.52 | 0.48 |
| 5 | 143031025 | rs1213705 | 0.015 | 0.029 | -0.38 | 0.43 | -0.015 | -0.023 | -0.023 | 0 | 0.66 | 0.56 | 0.75 | 0.75 | 0.78 | 0.77 |
| 5 | 143034513 | rs2910256 | 0.121 | 0.329 | -0.23 | 0.794 | 0.008 | 0.014 | 0.014 | 0 | 0.35 | 0.63 | 0.44 | 0.49 | 0.46 | 0.52 |
| 5 | 143036860 | rs1427864 | -0.399 | -0.823 | -0.429 | 0.649 | 0.038 | 0.001 | 0.001 | 0 | 0.47 | 0.61 | 0.41 | 0.33 | 0.29 | 0.34 |
| 5 | 143039972 | rs10070905 | -0.293 | -0.313 | -0.279 | 0.905 | -0.002 | 0.001 | 0.001 | 0 | 0.3 | 0.3 | 0.29 | 0.3 | 0.23 | 0.24 |
| 5 | 143045017 | rs1366084 | -0.512 | 0.389 | 0.218 | 0.799 | 0.051 | 0.031 | 0.031 | 0.028 | 0.22 | 0.47 | 0.38 | 0.28 | 0.26 | 0.21 |
| 5 | 143046538 | rs890710 | -0.326 | 0.388 | 0.436 | 0.798 | 0.012 | 0.007 | 0.007 | 0.03 | 0.21 | 0.27 | 0.37 | 0.28 | 0.26 | 0.19 |
| 5 | 143053517 | rs11738753 | -0.719 | 0.185 | -0.298 | 0.517 | -0.004 | 0.004 | 0.004 | 0 | 0.53 | 0.71 | 0.49 | 0.54 | 0.44 | 0.49 |
| 5 | 143058402 | rs17346995 | 0.669 | 0.378 | -0.334 | 0.52 | -0.034 | 0.004 | 0.004 | 0 | 0.14 | 0.08 | 0.19 | 0.14 | 0.19 | 0.11 |
| 5 | 143069305 | rs10477215 | -0.225 | 0.321 | -0.176 | 0.54 | -0.003 | 0 | 0 | 0.798 | 0.39 | 0.54 | 0.31 | 0.42 | 0.26 | 0.37 |
| 5 | 143074342 | rs12717904 | -0.029 | 0.015 | 0.189 | 0.476 | -0.007 | 0.006 | 0.006 | 0.838 | 0.32 | 0.34 | 0.26 | 0.39 | 0.23 | 0.28 |
| 5 | 143076516 | rs7707734 | -1.144 | 0.024 | 0.091 | 0.446 | -0.026 | 0.003 | 0.003 | 0 | 0.53 | 0.82 | 0.5 | 0.56 | 0.44 | 0.49 |
| 5 | 143083967 | rs1895344 | 0.481 | 0.147 | 0.018 | 0.461 | 0.007 | 0.005 | 0.005 | 0.012 | 0.16 | 0.04 | 0.21 | 0.11 | 0.19 | 0.15 |
| 5 | 143085460 | rs6861643 | -1.764 | -0.116 | -0.217 | 0.264 | -0.045 | 0.021 | 0.021 | 0.012 | 0.6 | 0.92 | 0.69 | 0.67 | 0.59 | 0.62 |
| 5 | 143086037 | rs4912940 | 0.364 | -0.1 | 0.169 | 0.352 | 0.005 | 0.005 | 0.005 | 0 | 0.16 | 0.11 | 0.22 | 0.11 | 0.19 | 0.15 |
| 5 | 143087656 | rs867924 | 0.272 | -0.031 | 0.13 | 0.362 | -0.011 | 0.078 | 0.078 | 0.001 | 0.24 | 0.43 | 0.24 | 0.38 | 0.19 | 0.25 |
| 5 | 143093377 | rs17100697 | 0.384 | 0.033 | -0.467 | 0.451 | -0.008 | -0.036 | -0.036 | 0 | 0.15 | 0.03 | 0.21 | 0.17 | 0.2 | 0.16 |
| 5 | 143100276 | rs2161416 | 0.393 | -0.058 | 0.08 | 0.386 | 0.011 | -0.029 | -0.029 | 0 | 0.24 | 0.36 | 0.24 | 0.15 | 0.24 | 0.25 |
| 5 | 143103853 | rs2636096 | -1.483 | 0.054 | -0.067 | 0.358 | -0.006 | 0.046 | 0.046 | 0 | 0.6 | 0.86 | 0.68 | 0.7 | 0.6 | 0.64 |
| 5 | 143110862 | rs325260 | 0.478 | 0.034 | 0.26 | 0.344 | 0.02 | -0.046 | -0.046 | 0.678 | 0.24 | 0.4 | 0.24 | 0.15 | 0.24 | 0.25 |
| 5 | 143112659 | rs17414396 | 0.374 | -1.49 | 0.286 | 0.402 | 0.005 | 0.101 | 0.101 | 0.334 | 0.21 | 0.32 | 0.21 | 0.38 | 0.17 | 0.27 |
| 5 | 143114091 | rs17100742 | 0.497 | -1.37 | -0.03 | 0.394 | -0.023 | -0.029 | -0.029 | 0 | 0.14 | 0.03 | 0.22 | 0.14 | 0.2 | 0.12 |
| 5 | 143115442 | rs7723052 | 1.584 | -0.952 | -0.106 | 0.304 | 0.094 | 0.025 | 0.025 | 0.26 | 0.38 | 0.08 | 0.24 | 0.29 | 0.24 | 0.35 |
| 5 | 143115696 | rs325253 | -0.975 | -0.736 | 0.129 | 0.424 | -0.002 | 0.007 | 0.007 | 0 | 0.37 | 0.51 | 0.44 | 0.54 | 0.37 | 0.39 |
| 5 | 143126066 | rs17348968 | 0.986 | -1.002 | 0.07 | 0.556 | -0.022 | -0.044 | -0.044 | 0 | 0.12 | 0.04 | 0.17 | 0.3 | 0.12 | 0.1 |
| 5 | 143129033 | rs325231 | -0.044 | -0.852 | 0.178 | 0.593 | 0.006 | 0.108 | 0.108 | 0.395 | 0.22 | 0.36 | 0.21 | 0.4 | 0.17 | 0.28 |
| 5 | 143134575 | rs2546879 | -0.136 | -0.85 | -0.153 | 0.501 | -0.035 | 0.013 | 0.013 | 0.648 | 0.65 | 0.78 | 0.55 | 0.69 | 0.43 | 0.65 |
| 5 | 143138340 | rs1862407 | -0.057 | -0.948 | 0.258 | 0.625 | -0.025 | 0.037 | 0.037 | 0.462 | 0.25 | 0.53 | 0.23 | 0.32 | 0.16 | 0.24 |
| 5 | 143145881 | rs844301 | -0.375 | -1.705 | 0.067 | 0.535 | -0.033 | -0.037 | -0.037 | 0.02 | 0.29 | 0.56 | 0.16 | 0.18 | 0.16 | 0.24 |
| 5 | 143146139 | rs17100840 | 0.267 | -1.645 | -0.242 | 0.661 | -0.01 | -0.036 | -0.036 | 0 | 0.16 | 0.03 | 0.21 | 0.18 | 0.2 | 0.15 |
| 5 | 143160187 | rs2636112 | -0.022 | -1.791 | 0.117 | 0.614 | -0.047 | -0.048 | -0.048 | 0 | 0.28 | 0.45 | 0.14 | 0.18 | 0.16 | 0.27 |
| 5 | 143160225 | rs11167828 | 1.599 | -1.899 | -0.389 | 0.679 | 0.049 | -0.047 | -0.047 | 0 | 0.29 | 0.07 | 0.28 | 0.3 | 0.22 | 0.28 |
| 5 | 143161132 | rs7704419 | 1.247 | -1.865 | -0.236 | 0.638 | 0.012 | -0.014 | -0.014 | 0.059 | 0.33 | 0.27 | 0.41 | 0.49 | 0.25 | 0.38 |
| 5 | 143166433 | rs4912948 | 0.7 | -1.926 | -0.246 | 0.6 | 0.025 | -0.013 | -0.013 | 0 | 0.4 | 0.32 | 0.43 | 0.5 | 0.28 | 0.4 |
| 5 | 143168435 | rs7715118 | 1.162 | -1.848 | -0.191 | 0.653 | 0.002 | -0.003 | -0.003 | 0 | 0.34 | 0.32 | 0.41 | 0.5 | 0.25 | 0.38 |
| 5 | 143173023 | rs833748 | -0.529 | -1.866 | -0.134 | 0.618 | -0.008 | -0.001 | -0.001 | 0 | 0.39 | 0.55 | 0.34 | 0.38 | 0.34 | 0.39 |
| 5 | 143174218 | rs4912956 | 0.592 | -1.749 | -0.581 | 0.696 | -0.013 | -0.043 | -0.043 | 0 | 0.15 | 0.04 | 0.2 | 0.18 | 0.2 | 0.13 |
| 5 | 143176424 | rs17101011 | 1.485 | -1.866 | -0.643 | 0.622 | 0.026 | -0.086 | -0.086 | 0.512 | 0.29 | 0.04 | 0.26 | 0.34 | 0.22 | 0.28 |
| 5 | 143183845 | rs1024961 | 0.934 | -1.946 | -0.714 | 0.65 | 0.063 | -0.02 | -0.02 | 0 | 0.37 | 0.04 | 0.23 | 0.33 | 0.25 | 0.3 |
| 5 | 143192211 | rs3792815 | 0.58 | -1.788 | -0.169 | 0.75 | -0.039 | -0.027 | -0.027 | 0 | 0.23 | 0.39 | 0.13 | 0.18 | 0.14 | 0.25 |
| 5 | 143193206 | rs6876125 | 1.695 | -1.838 | -0.727 | 0.456 | 0.053 | -0.09 | -0.09 | 0 | 0.25 | 0.04 | 0.17 | 0.37 | 0.16 | 0.21 |
| 5 | 143197848 | rs918377 | -1.27 | -1.247 | -0.388 | 0.539 | -0.076 | 0.057 | 0.057 | 1.064 | 0.79 | 0.89 | 0.77 | 0.93 | 0.61 | 0.78 |
| 5 | 143200053 | rs161557 | -0.034 | -1.401 | 0.178 | 0.733 | -0.006 | 0.08 | 0.08 | 0 | 0.19 | 0.3 | 0.33 | 0.39 | 0.22 | 0.23 |
| 5 | 143201705 | rs918378 | -0.032 | -1.503 | 0.126 | 0.748 | -0.012 | 0.098 | 0.098 | 0 | 0.08 | 0.18 | 0.13 | 0.22 | 0.07 | 0.16 |
| 5 | 143212029 | rs7718152 | 0.267 | -1.65 | 0.089 | 0.249 | -0.014 | -0.016 | -0.016 | 0 | 0.26 | 0.28 | 0.14 | 0.13 | 0.15 | 0.26 |
| 5 | 143216173 | rs10059157 | -0.001 | -1.659 | 0.036 | 0.26 | -0.055 | -0.039 | -0.039 | 0.169 | 0.28 | 0.46 | 0.21 | 0.17 | 0.16 | 0.26 |
| 5 | 143222173 | rs323982 | 1.079 | -1.241 | -0.193 | 0.538 | 0.06 | 0.022 | 0.022 | 0 | 0.35 | 0.13 | 0.22 | 0.34 | 0.25 | 0.3 |
| 5 | 143223005 | rs323985 | 1.203 | -1.23 | -0.614 | 0.544 | 0.04 | -0.023 | -0.023 | 0 | 0.33 | 0.03 | 0.21 | 0.34 | 0.25 | 0.3 |
| 5 | 143224078 | rs13165784 | 0.252 | -1.327 | -0.479 | 0.437 | 0.015 | 0.017 | 0.017 | 0 | 0.28 | 0.2 | 0.13 | 0.13 | 0.18 | 0.32 |
| 5 | 143226759 | rs323990 | -0.62 | -0.751 | -0.836 | 0.463 | 0.011 | -0.004 | -0.004 | 2.355 | 0.76 | 0.75 | 0.71 | 0.85 | 0.61 | 0.78 |
| 5 | 143230325 | rs2636111 | -0.62 | -0.751 | -0.836 | 0.451 | 0.011 | -0.004 | -0.004 | 0 | 0.76 | 0.75 | 0.71 | 0.85 | 0.61 | 0.78 |

| | | | | | | | | | | | | | | | | |
|---|---|---|---|---|---|---|---|---|---|---|---|---|---|---|---|---|
| 5 | 143230480 | rs2644832 | 0.939 | -0.672 | -1.682 | 0.37 | 0.038 | -0.021 | -0.021 | 0 | 0.33 | 0.03 | 0.21 | 0.33 | 0.25 | 0.3 |
| 5 | 143240844 | rs321202 | -0.26 | 0.736 | -0.532 | 0.413 | -0.131 | 0.061 | 0.061 | 1.489 | 0.78 | 0.93 | 0.78 | 0.88 | 0.62 | 0.78 |
| 5 | 143249879 | rs1421776 | -1.306 | 0.225 | 0.579 | 0.958 | 0.005 | 0.06 | 0.06 | 0 | 0.14 | 0.28 | 0.16 | 0.19 | 0.13 | 0.22 |
| 5 | 143258738 | rs10035652 | -1.711 | -0.261 | 0.977 | 1.211 | -0.008 | 0.025 | 0.025 | 0 | 0.27 | 0.64 | 0.3 | 0.39 | 0.22 | 0.46 |
| 5 | 143266118 | rs17101166 | -0.434 | 0.116 | 0.967 | 1.062 | -0.001 | -0.011 | -0.011 | 0 | 0.08 | 0.06 | 0.04 | 0 | 0.1 | 0.11 |
| 5 | 143287833 | rs2042157 | -0.994 | -0.007 | 0.894 | 1.132 | -0.001 | -0.001 | -0.001 | 1.749 | 0.13 | 0.12 | 0.07 | 0 | 0.13 | 0.11 |
| 5 | 143287847 | rs315176 | -0.355 | -0.062 | 1.27 | 1.238 | 0.006 | 0.04 | 0.04 | 0 | 0.06 | 0.15 | 0.05 | 0.13 | 0.07 | 0.18 |
| 5 | 143288412 | rs315177 | -1.89 | -0.063 | 1.273 | 1.237 | -0.003 | 0.084 | 0.084 | 0.036 | 0.21 | 0.38 | 0.22 | 0.36 | 0.2 | 0.38 |
| 5 | 143298046 | rs456304 | -0.851 | -0.061 | 1.424 | 1.315 | 0.004 | 0.167 | 0.167 | 0 | 0.06 | 0.15 | 0.14 | 0.32 | 0.07 | 0.25 |
| 5 | 143312556 | rs315212 | -1.817 | -0.217 | 1.558 | 1.263 | -0.031 | 0.14 | 0.14 | 0.3 | 0.2 | 0.56 | 0.21 | 0.37 | 0.15 | 0.38 |
| 5 | 143319871 | rs318373 | -0.814 | -0.516 | 0.536 | 0.59 | -0.157 | 0.148 | 0.148 | 0.344 | 0.66 | 0.97 | 0.62 | 0.86 | 0.53 | 0.8 |
| 5 | 143324439 | rs1603418 | -1.279 | -0.221 | 1.766 | 1.288 | -0.003 | 0.039 | 0.039 | 0 | 0.14 | 0.27 | 0.16 | 0.19 | 0.12 | 0.18 |
| 5 | 143332894 | rs312557 | -1.892 | -0.227 | 1.75 | 1.291 | -0.031 | 0.138 | 0.138 | 0.34 | 0.2 | 0.56 | 0.21 | 0.37 | 0.14 | 0.38 |
| 5 | 143345026 | rs312570 | -0.853 | -0.192 | 1.377 | 1.298 | 0.003 | 0.168 | 0.168 | 0 | 0.06 | 0.31 | 0.04 | 0.18 | 0.02 | 0.2 |
| 5 | 143348795 | rs17101282 | -1.283 | 0.187 | 0.648 | 1.012 | -0.002 | 0.019 | 0.019 | 0 | 0.14 | 0.2 | 0.15 | 0.19 | 0.12 | 0.18 |
| 5 | 143364448 | rs12717907 | -0.987 | 0.681 | 1.001 | 0.527 | 0.035 | -0.065 | -0.065 | 0 | 0.11 | 0.34 | 0.08 | 0 | 0.1 | 0.06 |
| 5 | 143372963 | rs312612 | -0.907 | 0.98 | 1.06 | 0.482 | 0.011 | -0.01 | -0.01 | 0 | 0.37 | 0.62 | 0.49 | 0.28 | 0.44 | 0.48 |
| 5 | 143386014 | rs312588 | 0.038 | 0.978 | 0.917 | 0.515 | 0.009 | 0 | 0 | 0.082 | 0.26 | 0.29 | 0.42 | 0.28 | 0.34 | 0.41 |
| 5 | 143395577 | rs3913197 | 0.139 | 0.949 | 0.928 | 0.484 | -0.002 | -0.001 | -0.001 | 0 | 0.25 | 0.24 | 0.41 | 0.28 | 0.34 | 0.42 |
| 5 | 143396282 | rs160617 | -0.852 | 0.767 | 0.771 | 0.72 | -0.016 | -0.007 | -0.007 | 6.456 | 0.39 | 0.7 | 0.6 | 0.43 | 0.47 | 0.55 |
| 5 | 143406683 | rs6861682 | -0.289 | 0.225 | 0.818 | 0.645 | -0.017 | 0.006 | 0.006 | 0 | 0.14 | 0.31 | 0.16 | 0.13 | 0.1 | 0.22 |
| 5 | 143412790 | rs588129 | -0.673 | 0.189 | 1.295 | 0.554 | -0.019 | 0.003 | 0.003 | 0 | 0.17 | 0.57 | 0.19 | 0.13 | 0.12 | 0.23 |
| 5 | 143414743 | rs316056 | -1.007 | -0.107 | 0.546 | 0.409 | -0.044 | -0.045 | -0.045 | 0.481 | 0.65 | 0.9 | 0.57 | 0.52 | 0.58 | 0.77 |
| 5 | 143416814 | rs372853 | -0.486 | -0.107 | 0.614 | 0.046 | -0.024 | 0.035 | 0.035 | 0 | 0.17 | 0.66 | 0.19 | 0.22 | 0.12 | 0.22 |
| 5 | 143417210 | rs391535 | -0.315 | -0.106 | 0.613 | 0.044 | -0.017 | 0.006 | 0.006 | 0 | 0.17 | 0.57 | 0.19 | 0.13 | 0.12 | 0.22 |
| 5 | 143426260 | rs6889431 | 0.819 | -0.041 | -0.169 | -0.246 | 0 | 0.096 | 0.096 | 1.042 | 0.2 | 0.07 | 0.1 | 0 | 0.17 | 0.13 |
| 5 | 143464943 | rs10061858 | 0.189 | -0.214 | 0.015 | -0.214 | -0.024 | -0.071 | -0.071 | 0 | 0.24 | 0.44 | 0.13 | 0.09 | 0.18 | 0.14 |
| 5 | 143467659 | rs1318584 | -0.816 | -0.269 | 0.032 | -0.219 | 0.013 | 0.018 | 0.018 | 0.034 | 0.39 | 0.74 | 0.29 | 0.22 | 0.28 | 0.35 |
| 5 | 143467718 | rs11951400 | 1.023 | -0.098 | -0.826 | -0.029 | -0.109 | -0.042 | -0.042 | 0.734 | 0.21 | 0.01 | 0.24 | 0.17 | 0.3 | 0.19 |
| 5 | 143474131 | rs9285658 | -0.923 | -0.977 | 0.071 | -0.215 | -0.011 | -0.011 | -0.011 | 1.192 | 0.22 | 0.3 | 0.19 | 0.12 | 0.15 | 0.31 |
| 5 | 143479040 | rs7717773 | -0.659 | -0.624 | 0.312 | 0.299 | -0.023 | 0.078 | 0.078 | 0 | 0.16 | 0.48 | 0.16 | 0.21 | 0.1 | 0.22 |
| 5 | 143482130 | rs10069436 | -0.528 | -0.551 | -0.301 | 0.033 | -0.022 | 0.006 | 0.006 | 0 | 0.15 | 0.37 | 0.16 | 0.12 | 0.09 | 0.21 |
| 5 | 143483204 | rs2398717 | -0.228 | 0.021 | -1.318 | -0.005 | 0.009 | 0.087 | 0.087 | 0.024 | 0.19 | 0.07 | 0.1 | 0 | 0.16 | 0.13 |
| 5 | 143484405 | rs1895151 | 0.767 | 0.197 | -0.793 | 0.786 | 0.005 | -0.008 | -0.008 | 0.163 | 0.86 | 0.9 | 0.85 | 0.84 | 0.9 | 0.85 |
| 5 | 143496926 | rs2544605 | -1.207 | 0.314 | 0.159 | 0.767 | 0.005 | 0.032 | 0.032 | 0 | 0.16 | 0.39 | 0.18 | 0.17 | 0.13 | 0.18 |
| 5 | 143504102 | rs4513726 | -0.158 | 0.308 | -0.325 | 0.941 | 0.003 | 0.004 | 0.004 | 0 | 0.74 | 0.65 | 0.73 | 0.69 | 0.7 | 0.61 |
| 5 | 143510587 | rs3924119 | 1.16 | 0.199 | -0.457 | 0.899 | -0.025 | -0.067 | -0.067 | 0 | 0.56 | 0.11 | 0.54 | 0.51 | 0.57 | 0.43 |
| 5 | 143524766 | rs7716084 | -0.68 | -0.189 | -0.98 | 0.806 | 0.05 | 0.004 | 0.004 | 0 | 0.5 | 0.64 | 0.52 | 0.5 | 0.39 | 0.4 |
| 5 | 143527974 | rs7704225 | 1.013 | -0.502 | -1.048 | 0.397 | 0.022 | 0.023 | 0.023 | 0 | 0.23 | 0.12 | 0.27 | 0.03 | 0.16 | 0.17 |
| 5 | 143536329 | rs13354411 | -0.78 | -0.506 | -1.005 | 0.666 | -0.014 | 0.045 | 0.045 | 0 | 0.28 | 0.52 | 0.25 | 0.42 | 0.23 | 0.25 |
| 5 | 143536704 | rs17101605 | 1.012 | -0.466 | -1.018 | 0.644 | 0.021 | 0.02 | 0.02 | 0 | 0.23 | 0.12 | 0.27 | 0.01 | 0.16 | 0.14 |
| 5 | 143539785 | rs12109284 | 1.042 | -0.495 | -0.495 | 0.659 | 0.033 | 0.052 | 0.052 | 0 | 0.22 | 0.1 | 0.27 | 0.02 | 0.16 | 0.14 |
| 5 | 143540718 | rs6864640 | -1.784 | -0.504 | -0.753 | 1.002 | -0.055 | -0.049 | -0.049 | 0.699 | 0.84 | 0.97 | 0.8 | 0.85 | 0.77 | 0.87 |
| 5 | 143549711 | rs1862140 | -0.951 | -0.751 | -0.463 | 0.943 | 0.047 | 0.034 | 0.034 | 0 | 0.5 | 0.64 | 0.52 | 0.43 | 0.39 | 0.4 |
| 5 | 143555740 | rs6650972 | 0.338 | -0.827 | 0.002 | 0.902 | -0.011 | -0.017 | -0.017 | 0.285 | 0.06 | 0.24 | 0.02 | 0 | 0.01 | 0 |
| 5 | 143561329 | rs13357041 | -0.969 | -0.808 | -0.174 | 0.91 | -0.015 | 0.046 | 0.046 | 0 | 0.27 | 0.52 | 0.25 | 0.43 | 0.23 | 0.25 |
| 5 | 143563483 | rs1609849 | 0.825 | -0.852 | -0.216 | 0.891 | 0.023 | 0.039 | 0.039 | 0.001 | 0.25 | 0.12 | 0.28 | 0.02 | 0.18 | 0.14 |
| 5 | 143570841 | rs9324943 | -0.97 | -0.816 | -0.411 | 0.925 | 0.048 | 0.035 | 0.035 | 0 | 0.51 | 0.66 | 0.52 | 0.44 | 0.39 | 0.39 |
| 5 | 143579812 | rs6860529 | 0.139 | -0.403 | -0.516 | 1.23 | -0.028 | -0.086 | -0.086 | 0 | 0.14 | 0.02 | 0.17 | 0.38 | 0.15 | 0.2 |
| 5 | 143587990 | rs879717 | -0.732 | -0.304 | -0.21 | 0.523 | -0.004 | -0.033 | -0.033 | 0 | 0.42 | 0.5 | 0.4 | 0.18 | 0.37 | 0.3 |
| 5 | 143589949 | rs6868068 | -0.543 | -0.529 | -0.204 | 0.912 | -0.031 | 0.028 | 0.028 | 0 | 0.17 | 0.31 | 0.12 | 0.22 | 0.12 | 0.16 |
| 5 | 143594031 | rs17438740 | 1.274 | -0.469 | -0.616 | 0.573 | -0.017 | 0.032 | 0.032 | 0 | 0.06 | 0 | 0.04 | 0 | 0.06 | 0.04 |
| 5 | 143595812 | rs6580315 | 0.627 | -0.881 | -0.588 | 0.289 | 0.02 | 0.078 | 0.078 | 0 | 0.21 | 0.08 | 0.06 | 0.01 | 0.14 | 0.14 |
| 5 | 143605039 | rs17438802 | 0.216 | -0.741 | -0.313 | 0.52 | 0.016 | -0.01 | -0.01 | 0 | 0.13 | 0.03 | 0.08 | 0.12 | 0.11 | 0.1 |
| 5 | 143620425 | rs2112157 | 0.961 | -0.237 | -0.551 | 0.333 | -0.068 | -0.008 | -0.008 | 1.099 | 0.19 | 0.03 | 0.24 | 0.18 | 0.3 | 0.12 |
| 5 | 143621838 | rs1422711 | -1.729 | -0.686 | -0.954 | 0.724 | -0.13 | -0.043 | -0.043 | 0 | 0.71 | 0.97 | 0.6 | 0.73 | 0.62 | 0.63 |
| 5 | 143644780 | rs13182980 | 0.665 | -0.431 | -0.38 | 1.021 | -0.017 | 0.009 | 0.009 | 0 | 0.2 | 0.1 | 0.25 | 0.18 | 0.28 | 0.13 |
| 5 | 143656916 | rs13357117 | 0.587 | -0.459 | -0.135 | 1.008 | -0.007 | 0.007 | 0.007 | 0 | 0.21 | 0.16 | 0.25 | 0.18 | 0.28 | 0.12 |
| 5 | 143669068 | rs7711546 | -0.855 | -0.718 | 0.211 | 1.05 | 0.003 | 0.001 | 0.001 | 0 | 0.19 | 0.15 | 0.2 | 0.53 | 0.16 | 0.3 |
| 5 | 143683902 | rs4912969 | 0.653 | -0.43 | 0.269 | 0.666 | -0.019 | 0.007 | 0.007 | 0 | 0.2 | 0.09 | 0.24 | 0.18 | 0.28 | 0.12 |
| 5 | 143694706 | rs10515538 | 0.653 | -0.43 | 0.315 | 0.666 | -0.012 | 0.005 | 0.005 | 0 | 0.2 | 0.11 | 0.24 | 0.18 | 0.28 | 0.12 |
| 5 | 143708458 | rs29893 | -0.072 | -0.563 | 1.139 | 0.595 | 0.001 | -0.01 | -0.01 | 0 | 0.26 | 0.29 | 0.29 | 0.19 | 0.31 | 0.14 |
| 5 | 143718288 | rs831419 | 0.672 | -0.525 | 1.058 | 0.616 | -0.005 | 0.003 | 0.003 | 0 | 0.2 | 0.14 | 0.25 | 0.18 | 0.28 | 0.12 |
| 5 | 143727167 | rs244526 | 0.65 | -0.54 | 1.107 | 0.609 | -0.002 | 0.001 | 0.001 | 0 | 0.2 | 0.15 | 0.25 | 0.18 | 0.28 | 0.12 |
| 5 | 143728206 | rs244524 | 0.288 | -0.547 | 1.105 | 0.607 | 0.002 | -0.005 | -0.005 | 1.437 | 0.22 | 0.25 | 0.25 | 0.19 | 0.28 | 0.12 |
| 5 | 143736618 | rs17101828 | 0.675 | -0.459 | 0.922 | 0.929 | -0.009 | -0.04 | -0.04 | 1.121 | 0.07 | 0 | 0.11 | 0.18 | 0.06 | 0.15 |
| 5 | 143746369 | rs3095960 | -0.412 | -0.341 | 1.532 | 0.491 | -0.001 | -0.035 | -0.035 | 0 | 0.27 | 0.54 | 0.31 | 0.18 | 0.31 | 0.13 |
| 5 | 143758377 | rs10515542 | -0.714 | -0.191 | 1.436 | 0.571 | -0.012 | 0.005 | 0.005 | 0 | 0.81 | 0.89 | 0.75 | 0.82 | 0.72 | 0.89 |
| 5 | 143775494 | rs4345344 | -0.057 | -0.225 | 1.377 | 0.594 | 0.025 | -0.016 | -0.016 | 0 | 0.76 | 0.52 | 0.72 | 0.8 | 0.72 | 0.81 |

| | | | | | | | | | | | | | | | | |
|---|---|---|---|---|---|---|---|---|---|---|---|---|---|---|---|---|
| 5 | 143819327 | rs2017862 | 0.412 | -0.119 | 0.986 | 0.582 | -0.044 | -0.007 | -0.007 | 0.027 | 0.2 | 0.08 | 0.23 | 0.2 | 0.28 | 0.2 |
| 5 | 143837325 | rs9324959 | -0.813 | -0.494 | 0.971 | 0.684 | 0.005 | -0.01 | -0.01 | 0 | 0.37 | 0.41 | 0.59 | 0.68 | 0.42 | 0.42 |
| 5 | 143849358 | rs349682 | -1.154 | -0.16 | 0.633 | -0.422 | 0.024 | 0.061 | 0.061 | 0.222 | 0.6 | 0.68 | 0.84 | 0.94 | 0.71 | 0.6 |
| 5 | 143854604 | rs187034 | -0.224 | -0.211 | 0.651 | -0.466 | 0.008 | -0.011 | -0.011 | 0.229 | 0.36 | 0.43 | 0.52 | 0.67 | 0.41 | 0.42 |
| 5 | 143855031 | rs349704 | 0.001 | -0.272 | 0.872 | -0.49 | -0.029 | 0.003 | 0.003 | 0 | 0.33 | 0.2 | 0.5 | 0.67 | 0.41 | 0.42 |
| 5 | 143855082 | rs349703 | -0.203 | -0.218 | 0.612 | -0.487 | 0.009 | -0.011 | -0.011 | 0 | 0.36 | 0.43 | 0.52 | 0.67 | 0.41 | 0.42 |
| 5 | 143855606 | rs358651 | -2 | -0.314 | 0.629 | -0.552 | -0.048 | -0.038 | -0.038 | 1.703 | 0.82 | 0.97 | 0.73 | 0.73 | 0.73 | 0.84 |
| 5 | 143855915 | rs349700 | -0.169 | -0.294 | 0.592 | -0.542 | 0.009 | -0.011 | -0.011 | 0.033 | 0.36 | 0.43 | 0.52 | 0.67 | 0.41 | 0.42 |
| 5 | 143856528 | rs349698 | -0.198 | -0.297 | 0.571 | -0.556 | 0.005 | -0.007 | -0.007 | 0 | 0.36 | 0.41 | 0.53 | 0.67 | 0.42 | 0.42 |
| 5 | 143860437 | rs358650 | 0.846 | -0.236 | 0.317 | -0.562 | 0.035 | 0.072 | 0.072 | 0 | 0.44 | 0.34 | 0.2 | 0.06 | 0.32 | 0.42 |
| 5 | 143869004 | rs6882650 | 1.245 | -0.163 | -0.092 | -0.523 | 0.023 | 0.063 | 0.063 | 0.189 | 0.4 | 0.31 | 0.21 | 0.07 | 0.29 | 0.4 |
| 5 | 143880746 | rs467019 | 1.788 | 0.402 | -0.764 | -0.187 | -0.041 | -0.007 | -0.007 | 0 | 0.15 | 0.05 | 0.2 | 0.16 | 0.24 | 0.12 |
| 5 | 143881073 | rs358673 | -1.501 | 0.362 | -0.519 | -0.217 | -0.029 | -0.023 | -0.023 | 0.013 | 0.82 | 0.93 | 0.73 | 0.74 | 0.73 | 0.84 |
| 5 | 143898188 | rs463245 | -0.574 | 0.306 | -0.395 | -0.402 | -0.008 | -0.005 | -0.005 | 0 | 0.29 | 0.25 | 0.52 | 0.61 | 0.41 | 0.4 |
| 5 | 143901489 | rs358661 | -0.301 | -0.707 | -0.485 | -1.123 | -0.002 | 0.002 | 0.002 | 0 | 0.36 | 0.37 | 0.52 | 0.59 | 0.41 | 0.56 |
| 5 | 143909677 | rs17691826 | 1.794 | -1.634 | -0.511 | -1.59 | -0.01 | 0.121 | 0.121 | 0 | 0.29 | 0.14 | 0.11 | 0.01 | 0.24 | 0.2 |
| 5 | 143948732 | rs358094 | 0.454 | -1.593 | -0.549 | -1.318 | -0.016 | 0.06 | 0.06 | 0 | 0.2 | 0.32 | 0.26 | 0.5 | 0.14 | 0.38 |
| 5 | 143956168 | rs1559057 | 1.764 | -1.624 | -0.213 | -1.385 | -0.004 | -0.014 | -0.014 | 0 | 0.07 | 0.06 | 0.04 | 0.06 | 0.04 | 0.06 |
| 5 | 143957988 | rs358093 | 1.883 | -1.638 | -0.225 | -1.377 | -0.001 | -0.003 | -0.003 | 0.016 | 0.12 | 0.16 | 0.04 | 0.05 | 0.06 | 0.25 |
| 5 | 143959674 | rs11953106 | 1.714 | -1.442 | -0.202 | -1.389 | 0 | 0.083 | 0.083 | 0 | 0.29 | 0.19 | 0.12 | 0.01 | 0.25 | 0.2 |
| 5 | 143965621 | rs1432672 | 1.652 | -1.483 | 0.205 | -1.408 | 0.004 | 0.012 | 0.012 | 0 | 0.3 | 0.26 | 0.12 | 0.01 | 0.24 | 0.21 |
| 5 | 143966681 | rs165445 | -0.565 | -1.332 | 0.123 | -1.528 | -0.001 | 0.009 | 0.009 | 0 | 0.48 | 0.44 | 0.6 | 0.65 | 0.51 | 0.67 |
| 5 | 143999885 | rs258928 | 1.614 | -1.097 | 0.173 | -1.346 | 0.015 | 0.036 | 0.036 | 0 | 0.13 | 0.12 | 0.23 | 0.34 | 0.21 | 0.1 |
| 5 | 144013825 | rs165463 | 1.122 | -1.129 | 0.285 | -1.361 | 0.05 | 0.109 | 0.109 | 0 | 0.16 | 0.39 | 0.25 | 0.34 | 0.21 | 0.11 |
| 5 | 144017979 | rs165400 | -1.809 | -1.108 | -0.042 | -1.394 | -0.073 | 0.024 | 0.024 | 0 | 0.79 | 0.96 | 0.71 | 0.91 | 0.68 | 0.78 |
| 5 | 144018068 | rs165401 | -2.63 | -0.969 | 0.006 | -1.348 | -0.06 | 0.001 | 0.001 | 0 | 0.84 | 0.96 | 0.75 | 0.91 | 0.72 | 0.89 |
| 5 | 144022041 | rs165451 | 1.452 | -0.686 | 0.314 | -1.14 | 0.017 | 0.035 | 0.035 | 0 | 0.13 | 0.12 | 0.23 | 0.32 | 0.21 | 0.1 |
| 5 | 144025843 | rs171818 | 0.097 | -0.843 | 0.168 | -1.21 | 0.001 | 0.035 | 0.035 | 0 | 0.51 | 0.58 | 0.4 | 0.41 | 0.5 | 0.37 |
| 5 | 144026983 | rs17385747 | 1.077 | -1.147 | -0.219 | -1.303 | 0.003 | 0.005 | 0.005 | 0 | 0.07 | 0.03 | 0.04 | 0.06 | 0.04 | 0.06 |
| 5 | 144030993 | rs17451739 | 1.326 | -1.13 | -0.299 | -1.294 | -0.001 | 0.024 | 0.024 | 0 | 0.06 | 0 | 0.04 | 0.06 | 0.04 | 0.06 |
| 5 | 144035490 | rs984827 | 0.684 | -1.418 | -0.173 | -1.547 | -0.001 | -0.063 | -0.063 | 0.061 | 0.34 | 0.4 | 0.17 | 0.07 | 0.28 | 0.26 |
| 5 | 144045754 | rs3910188 | -0.349 | -2.108 | -0.568 | -1.528 | 0.018 | 0.08 | 0.08 | 0 | 0.54 | 0.66 | 0.49 | 0.83 | 0.44 | 0.65 |
| 5 | 144048517 | rs10042924 | 0.512 | -2.123 | -0.477 | -1.564 | -0.003 | 0.024 | 0.024 | 0 | 0.41 | 0.4 | 0.47 | 0.17 | 0.52 | 0.25 |
| 5 | 144050938 | rs7720947 | 0.891 | -2.665 | -0.856 | -1.146 | 0.003 | 0.002 | 0.002 | 0 | 0.19 | 0.18 | 0.18 | 0.49 | 0.11 | 0.37 |
| 5 | 144052047 | rs17387218 | 0.825 | -2.458 | -0.775 | -1.256 | 0.017 | 0.098 | 0.098 | 0.196 | 0.24 | 0.07 | 0.13 | 0.01 | 0.22 | 0.33 |
| 5 | 144055534 | rs2108647 | -0.809 | -2.251 | -0.17 | -0.983 | 0.006 | 0.126 | 0.126 | 0.039 | 0.83 | 0.46 | 0.76 | 0.66 | 0.85 | 0.95 |
| 5 | 144064654 | rs7704625 | -0.23 | -2.155 | -0.646 | -1.204 | 0 | 0 | 0 | 0 | 0.69 | 0.32 | 0.66 | 0.59 | 0.67 | 0.82 |
| 5 | 144065162 | rs724382 | -0.202 | -2.164 | -0.651 | -1.208 | 0 | 0 | 0 | 0 | 0.69 | 0.33 | 0.67 | 0.59 | 0.67 | 0.82 |
| 5 | 144092494 | rs34970666 | 1.913 | -2.198 | -0.769 | -1.341 | 0.006 | 0.078 | 0.078 | 0.039 | 0.16 | 0.04 | 0.05 | 0 | 0.12 | 0.18 |
| 5 | 144098872 | rs13357155 | 1.852 | -2.227 | -0.655 | -1.034 | 0.006 | 0.241 | 0.241 | 0 | 0.07 | 0.16 | 0.15 | 0.43 | 0.06 | 0.16 |
| 5 | 144099791 | rs6863555 | 2.231 | -2.251 | -0.843 | -1.156 | -0.05 | 0.057 | 0.057 | 0.121 | 0.13 | 0.01 | 0.08 | 0.07 | 0.18 | 0.13 |
| 5 | 144103375 | rs2041070 | 2.259 | -2.501 | -0.68 | -1.058 | -0.073 | 0.019 | 0.019 | 0 | 0.19 | 0.04 | 0.33 | 0.1 | 0.31 | 0.11 |
| 5 | 144103408 | rs17102282 | 0.339 | -2.722 | -0.145 | -1.155 | -0.071 | 0.082 | 0.082 | 0 | 0.27 | 0.56 | 0.23 | 0.48 | 0.12 | 0.38 |
| 5 | 144116622 | rs13154970 | -0.15 | -2.644 | -0.131 | -1.123 | 0 | 0.016 | 0.016 | 0 | 0.46 | 0.61 | 0.57 | 0.58 | 0.43 | 0.49 |
| 5 | 144132644 | rs6880076 | 1.804 | -2.498 | -0.147 | -0.962 | 0 | 0.149 | 0.149 | 0 | 0.07 | 0.11 | 0.14 | 0.43 | 0.06 | 0.16 |
| 5 | 144132727 | rs10491258 | 1.077 | -2.535 | 0.028 | -0.961 | -0.039 | -0.036 | -0.036 | 0 | 0.2 | 0.28 | 0.09 | 0.06 | 0.06 | 0.22 |
| 5 | 144136931 | rs461599 | 0.571 | -2.601 | 0.182 | -1.097 | 0 | 0.001 | 0.001 | 0 | 0.45 | 0.45 | 0.36 | 0.51 | 0.48 | 0.62 |
| 5 | 144138146 | rs459807 | -0.596 | -2.413 | 0.176 | -1.012 | -0.024 | -0.011 | -0.011 | 0 | 0.66 | 0.74 | 0.51 | 0.6 | 0.55 | 0.84 |
| 5 | 144147675 | rs247571 | 0.213 | -2.486 | 0.002 | -0.968 | -0.001 | 0 | 0 | 0 | 0.54 | 0.45 | 0.46 | 0.55 | 0.52 | 0.65 |
| 5 | 144148044 | rs247572 | -0.563 | -2.355 | -0.129 | -0.94 | -0.006 | -0.002 | -0.002 | 0.075 | 0.67 | 0.72 | 0.51 | 0.61 | 0.54 | 0.83 |
| 5 | 144151495 | rs171585 | -0.59 | -2.368 | -0.143 | -0.938 | -0.009 | -0.003 | -0.003 | 4.77 | 0.67 | 0.72 | 0.51 | 0.61 | 0.55 | 0.83 |
| 5 | 144167718 | rs4912999 | 1.043 | -2.529 | -0.355 | -1.094 | 0.016 | 0.086 | 0.086 | 2.583 | 0.2 | 0.03 | 0.14 | 0.01 | 0.19 | 0.21 |
| 5 | 144168303 | rs11952422 | 2.035 | -2.395 | -0.079 | -0.957 | -0.043 | -0.029 | -0.029 | 0 | 0.12 | 0.25 | 0.04 | 0.06 | 0.03 | 0.18 |
| 5 | 144181430 | rs693443 | -0.466 | -1.877 | -0.192 | -0.654 | 0.007 | 0.002 | 0.002 | 0 | 0.66 | 0.66 | 0.51 | 0.6 | 0.55 | 0.83 |
| 5 | 144193857 | rs2400161 | -1.181 | -1.84 | -0.334 | -0.701 | -0.001 | 0.035 | 0.035 | 1.306 | 0.86 | 0.83 | 0.84 | 0.69 | 0.86 | 0.94 |
| 5 | 144215915 | rs1347121 | 2.001 | -1.756 | -0.55 | -0.678 | -0.069 | 0.031 | 0.031 | 0 | 0.18 | 0.05 | 0.32 | 0.09 | 0.31 | 0.11 |
| 5 | 144219439 | rs2163748 | 0.22 | -1.864 | -0.281 | -0.682 | 0.043 | 0.012 | 0.012 | 0 | 0.34 | 0.42 | 0.49 | 0.41 | 0.46 | 0.18 |
| 5 | 144220904 | rs12515080 | 0.718 | -1.677 | -0.217 | -0.383 | -0.003 | 0.037 | 0.037 | 0.139 | 0.13 | 0.2 | 0.14 | 0.23 | 0.12 | 0.06 |
| 5 | 144223076 | rs1347120 | -0.878 | -1.768 | -0.321 | -0.418 | -0.003 | -0.017 | -0.017 | 0 | 0.67 | 0.66 | 0.67 | 0.48 | 0.67 | 0.58 |
| 5 | 144242950 | rs13181482 | -1.048 | -1.672 | -0.377 | -0.426 | 0.022 | 0.046 | 0.046 | 1.599 | 0.7 | 0.49 | 0.68 | 0.45 | 0.67 | 0.61 |
| 5 | 144243287 | rs1368338 | -1.183 | -1.673 | -0.332 | -0.426 | 0.001 | 0.004 | 0.004 | 1.483 | 0.7 | 0.64 | 0.68 | 0.45 | 0.67 | 0.61 |
| 5 | 144255145 | rs13357903 | 2.037 | -1.713 | -0.547 | -0.582 | -0.067 | 0.022 | 0.022 | 0.32 | 0.18 | 0.04 | 0.33 | 0.1 | 0.32 | 0.14 |
| 5 | 144255465 | rs12518187 | 0.898 | -2.02 | -0.377 | -0.703 | -0.007 | 0.049 | 0.049 | 2.819 | 0.15 | 0.26 | 0.17 | 0.29 | 0.14 | 0.06 |
| 5 | 144259773 | rs17454394 | 1.082 | -2.195 | -0.558 | -0.914 | 0.005 | 0.049 | 0.049 | 0 | 0.16 | 0.04 | 0.1 | 0.01 | 0.16 | 0.2 |
| 5 | 144262743 | rs1549896 | 1.706 | -2.37 | -0.514 | -0.941 | -0.047 | -0.035 | -0.035 | 0 | 0.18 | 0.24 | 0.07 | 0.06 | 0.04 | 0.2 |
| 5 | 144262851 | rs1549897 | 2.16 | -2.387 | -0.567 | -0.993 | -0.005 | 0.002 | 0.002 | 0 | 0.05 | 0.03 | 0.06 | 0.01 | 0.12 | 0.08 |
| 5 | 144267106 | rs7712842 | 0.678 | -2.169 | -0.561 | -0.89 | -0.001 | 0.026 | 0.026 | 0 | 0.36 | 0.27 | 0.39 | 0.18 | 0.34 | 0.35 |
| 5 | 144275245 | rs1368335 | 1.833 | -2.136 | -0.6 | -0.869 | -0.01 | -0.005 | -0.005 | 0 | 0.17 | 0.16 | 0.06 | 0.06 | 0.04 | 0.2 |
| 5 | 144277803 | rs13158888 | -1.159 | -2.231 | -0.587 | -0.806 | -0.002 | -0.003 | -0.003 | 2.805 | 0.64 | 0.61 | 0.66 | 0.49 | 0.64 | 0.6 |

| | | | | | | | | | | | | | | | | |
|---|---|---|---|---|---|---|---|---|---|---|---|---|---|---|---|---|
| 5 | 144292791 | rs4912684 | -0.949 | -2.286 | -0.687 | -0.827 | -0.002 | -0.005 | -0.005 | 0 | 0.62 | 0.62 | 0.66 | 0.49 | 0.64 | 0.6 |
| 5 | 144296502 | rs11167874 | -1.06 | -2.162 | -0.63 | -0.767 | -0.003 | -0.004 | -0.004 | 0 | 0.64 | 0.62 | 0.66 | 0.49 | 0.64 | 0.6 |
| 5 | 144296874 | rs10044798 | 0.993 | -2.19 | -0.655 | -0.811 | -0.004 | 0.053 | 0.053 | 0 | 0.36 | 0.2 | 0.39 | 0.18 | 0.35 | 0.35 |
| 5 | 144298836 | rs4583918 | 1.539 | -2.216 | -0.715 | -0.127 | 0 | -0.008 | -0.008 | 1.526 | 0.1 | 0.09 | 0.2 | 0.24 | 0.09 | 0.13 |
| 5 | 144299821 | rs10070261 | -1.213 | -1.697 | -0.488 | -0.487 | 0 | 0.001 | 0.001 | 0.229 | 0.64 | 0.56 | 0.66 | 0.49 | 0.64 | 0.6 |
| 5 | 144327414 | rs767665 | -1.01 | -0.369 | -0.588 | 0.111 | -0.003 | -0.004 | -0.004 | 0 | 0.47 | 0.46 | 0.56 | 0.44 | 0.61 | 0.41 |
| 5 | 144329947 | rs17613806 | 1.486 | -0.348 | -0.608 | 0.095 | -0.061 | 0.012 | 0.012 | 0 | 0.18 | 0.05 | 0.28 | 0.13 | 0.31 | 0.15 |
| 5 | 144333462 | rs1476736 | -0.099 | -0.355 | -0.568 | 0.029 | 0.025 | 0.01 | 0.01 | 0 | 0.32 | 0.38 | 0.46 | 0.42 | 0.45 | 0.22 |
| 5 | 144337292 | rs4913013 | -0.677 | -0.61 | -0.321 | 0.324 | -0.003 | -0.005 | -0.005 | 0 | 0.47 | 0.47 | 0.56 | 0.43 | 0.61 | 0.41 |
| 5 | 144338833 | rs10463267 | -0.598 | -0.172 | 0.086 | 0.909 | 0.012 | 0.08 | 0.08 | 1.15 | 0.13 | 0.34 | 0.17 | 0.29 | 0.13 | 0.06 |
| 5 | 144339411 | rs917378 | -0.489 | -0.155 | -0.315 | 0.836 | 0.004 | 0.061 | 0.061 | 1.615 | 0.13 | 0.23 | 0.17 | 0.3 | 0.13 | 0.06 |
| 5 | 144341743 | rs248014 | -0.519 | -0.144 | -0.393 | 0.818 | 0.005 | 0.067 | 0.067 | 1.442 | 0.13 | 0.26 | 0.17 | 0.31 | 0.13 | 0.06 |
| 5 | 144358716 | rs11741672 | -0.603 | -0.507 | -1.206 | 0.747 | 0.024 | -0.028 | -0.028 | 0 | 0.68 | 0.82 | 0.82 | 0.52 | 0.79 | 0.68 |
| 5 | 144369950 | rs370906 | -0.739 | -0.495 | -1.211 | 0.263 | 0.022 | 0.066 | 0.066 | 0.208 | 0.69 | 0.83 | 0.89 | 0.84 | 0.8 | 0.71 |
| 5 | 144373766 | rs13188843 | 0.343 | -0.58 | -1.499 | 0.137 | -0.056 | 0.115 | 0.115 | 0 | 0.4 | 0.14 | 0.44 | 0.17 | 0.56 | 0.25 |
| 5 | 144397224 | rs740477 | 0.345 | -0.564 | -1.299 | 0.127 | -0.052 | 0.104 | 0.104 | 0.104 | 0.4 | 0.14 | 0.44 | 0.17 | 0.55 | 0.24 |
| 5 | 144397607 | rs740475 | 0.857 | -0.611 | -1.705 | -0.245 | -0.068 | 0.015 | 0.015 | 0 | 0.24 | 0.09 | 0.34 | 0.16 | 0.4 | 0.16 |
| 5 | 144405640 | rs17102624 | 1.624 | -0.968 | -0.976 | -0.505 | -0.008 | -0.029 | -0.029 | 0.049 | 0.09 | 0.11 | 0.03 | 0 | 0.06 | 0.1 |
| 5 | 144412335 | rs12655091 | -0.065 | -0.993 | -0.805 | -0.448 | -0.03 | 0.069 | 0.069 | 0.078 | 0.41 | 0.21 | 0.44 | 0.25 | 0.55 | 0.25 |
| 5 | 144416863 | rs248017 | -1.876 | -1.226 | -0.792 | -0.563 | 0.025 | 0.077 | 0.077 | 0.192 | 0.7 | 0.83 | 0.89 | 0.9 | 0.8 | 0.72 |
| 5 | 144433331 | rs6860134 | -0.033 | -0.895 | -0.695 | -0.413 | -0.021 | 0.072 | 0.072 | 0.159 | 0.41 | 0.19 | 0.44 | 0.24 | 0.56 | 0.25 |
| 5 | 144437410 | rs13154577 | 1.074 | -0.991 | -0.742 | -0.366 | 0.015 | 0.005 | 0.005 | 1.616 | 0.17 | 0.1 | 0.1 | 0.14 | 0.15 | 0.09 |
| 5 | 144441278 | rs6892136 | 0.24 | -0.875 | -0.699 | -0.294 | -0.045 | 0.085 | 0.085 | 0 | 0.41 | 0.15 | 0.43 | 0.24 | 0.55 | 0.25 |
| 5 | 144441794 | rs952583 | 1.237 | -1.045 | -0.35 | -0.204 | 0.031 | 0.068 | 0.068 | 0 | 0.38 | 0.29 | 0.16 | 0.15 | 0.25 | 0.39 |
| 5 | 144443517 | rs966180 | 0.276 | -0.948 | -0.506 | -0.174 | -0.062 | 0.084 | 0.084 | 0 | 0.4 | 0.11 | 0.43 | 0.23 | 0.55 | 0.25 |
| 5 | 144451833 | rs1422838 | -0.468 | -0.25 | -0.534 | -0.27 | -0.064 | 0.113 | 0.113 | 0.3 | 0.65 | 0.88 | 0.59 | 0.89 | 0.51 | 0.81 |
| 5 | 144456420 | rs6580340 | 1.456 | -0.249 | 0.147 | 0.554 | 0.014 | 0.202 | 0.202 | 0 | 0.05 | 0.14 | 0.11 | 0.37 | 0.04 | 0.06 |
| 5 | 144461473 | rs10875610 | -1.208 | -0.38 | 0.321 | 0.454 | 0.013 | -0.009 | -0.009 | 0.251 | 0.57 | 0.64 | 0.59 | 0.52 | 0.67 | 0.41 |
| 5 | 144461614 | rs11167894 | -0.328 | -0.253 | 0.309 | 0.343 | -0.008 | 0.001 | 0.001 | 0 | 0.24 | 0.48 | 0.22 | 0.28 | 0.21 | 0.25 |
| 5 | 144462121 | rs1363645 | -0.858 | -0.149 | 0.156 | 0.197 | 0.004 | 0.037 | 0.037 | 0 | 0.48 | 0.62 | 0.52 | 0.38 | 0.54 | 0.35 |
| 5 | 144464260 | rs9324976 | 1.283 | -0.149 | 0.246 | 0.179 | 0.004 | 0.001 | 0.001 | 0.017 | 0.14 | 0.09 | 0.06 | 0 | 0.12 | 0.19 |
| 5 | 144465810 | rs1895336 | -0.046 | -0.067 | 0.31 | 0.246 | 0.024 | 0.041 | 0.041 | 0.329 | 0.34 | 0.46 | 0.46 | 0.5 | 0.42 | 0.2 |
| 5 | 144466643 | rs4286710 | 0.938 | -0.063 | 0.427 | 0.247 | 0.023 | 0.106 | 0.106 | 1.734 | 0.08 | 0.21 | 0.15 | 0.28 | 0.08 | 0.06 |
| 5 | 144467735 | rs11740769 | 0.27 | -0.052 | -0.005 | 0.213 | -0.054 | 0.095 | 0.095 | 0.239 | 0.32 | 0.1 | 0.37 | 0.11 | 0.45 | 0.14 |
| 5 | 144470886 | rs166609 | 1.056 | -0.05 | 0.007 | 0.239 | -0.03 | 0.047 | 0.047 | 1.086 | 0.24 | 0.09 | 0.3 | 0.1 | 0.34 | 0.11 |
| 5 | 144474779 | rs10875612 | -0.753 | -0.029 | 0.062 | 0.245 | 0.001 | 0.001 | 0.001 | 0 | 0.43 | 0.53 | 0.53 | 0.52 | 0.54 | 0.21 |
| 5 | 144475467 | rs271586 | 0.924 | 0.055 | -0.149 | 0.01 | -0.022 | 0.041 | 0.041 | 2.222 | 0.25 | 0.12 | 0.31 | 0.11 | 0.34 | 0.11 |
| 5 | 144480425 | rs10043443 | 1.122 | 0.098 | 0.031 | 0.217 | 0.002 | -0.004 | -0.004 | 0 | 0.13 | 0.1 | 0.06 | 0 | 0.12 | 0.16 |
| 5 | 144483226 | rs6885281 | -0.572 | -0.005 | 0.415 | 0.682 | 0.036 | 0.067 | 0.067 | 0 | 0.17 | 0.41 | 0.23 | 0.3 | 0.2 | 0.09 |
| 5 | 144490550 | rs11949054 | 1.478 | 0.006 | 0.033 | 0.364 | -0.008 | 0.016 | 0.016 | 0 | 0.06 | 0.02 | 0.06 | 0 | 0.1 | 0.03 |
| 5 | 144494659 | rs13359856 | -0.5 | -0.392 | 0.421 | 0.582 | -0.028 | 0.066 | 0.066 | 0 | 0.27 | 0.47 | 0.31 | 0.4 | 0.24 | 0.2 |
| 5 | 144528455 | rs1981626 | -1.29 | -0.506 | -0.134 | 0.578 | 0.006 | 0.013 | 0.013 | 0 | 0.89 | 1 | 0.95 | 0.95 | 0.93 | 0.82 |
| 5 | 144528984 | rs1002807 | -0.853 | -0.361 | 0.016 | 0.394 | -0.01 | 0.012 | 0.012 | 0 | 0.43 | 0.63 | 0.53 | 0.38 | 0.53 | 0.26 |
| 5 | 144532826 | rs271072 | -1.305 | -0.486 | 0.009 | 0.291 | 0.031 | 0.091 | 0.091 | 0 | 0.51 | 0.8 | 0.79 | 0.63 | 0.63 | 0.42 |
| 5 | 144541531 | rs6890604 | 0.775 | -0.212 | 0.019 | -0.395 | 0.022 | -0.016 | -0.016 | 0.543 | 0.18 | 0.24 | 0.24 | 0.13 | 0.32 | 0.1 |
| 5 | 144545930 | rs2400170 | -0.917 | -0.197 | -0.02 | -0.362 | -0.015 | 0.024 | 0.024 | 0 | 0.44 | 0.72 | 0.55 | 0.39 | 0.53 | 0.25 |
| 5 | 144549093 | rs11949752 | 0.87 | -0.211 | -0.316 | -0.408 | -0.004 | -0.028 | -0.028 | 0.005 | 0.09 | 0.13 | 0.07 | 0.02 | 0.1 | 0.11 |
| 5 | 144551333 | rs2400171 | 1.636 | -0.714 | -0.562 | -0.579 | -0.118 | 0.063 | 0.063 | 0.701 | 0.3 | 0.01 | 0.14 | 0.18 | 0.29 | 0.34 |
| 5 | 144572513 | rs2400173 | 0.308 | -0.011 | 0.433 | -0.212 | -0.017 | 0.046 | 0.046 | 2.264 | 0.17 | 0.32 | 0.23 | 0.23 | 0.12 | 0.05 |
| 5 | 144578788 | rs13179777 | 0.28 | -0.253 | -0.282 | -0.048 | -0.018 | 0.056 | 0.056 | 2.65 | 0.17 | 0.38 | 0.23 | 0.23 | 0.12 | 0.06 |
| 5 | 144582213 | rs7736228 | 0.053 | 0.515 | -1.084 | -0.556 | -0.017 | 0.013 | 0.013 | 0 | 0.17 | 0.12 | 0.23 | 0.12 | 0.31 | 0.09 |
| 5 | 144584265 | rs17470789 | 0.613 | 0.296 | -1.134 | -0.436 | -0.011 | 0.047 | 0.047 | 0.131 | 0.09 | 0.01 | 0.07 | 0.01 | 0.12 | 0.04 |
| 5 | 144585367 | rs6580355 | -0.369 | 0.516 | -0.585 | -0.836 | 0.005 | -0.027 | -0.027 | 0 | 0.34 | 0.37 | 0.38 | 0.11 | 0.57 | 0.17 |
| 5 | 144587447 | rs2400176 | 0.265 | -0.005 | -0.365 | -0.451 | -0.003 | -0.009 | -0.009 | 0 | 0.63 | 0.42 | 0.46 | 0.28 | 0.7 | 0.45 |
| 5 | 144593285 | rs2895634 | -0.035 | -0.035 | 0.234 | -0.001 | -0.004 | 0.011 | 0.011 | 0 | 0.71 | 0.69 | 0.63 | 0.53 | 0.8 | 0.52 |
| 5 | 144597373 | rs3851498 | 0.428 | 0.251 | 0.026 | -0.395 | -0.01 | 0.008 | 0.008 | 0 | 0.15 | 0.14 | 0.23 | 0.08 | 0.32 | 0.07 |
| 5 | 144598309 | rs13170885 | 1.314 | 0.315 | 0.005 | -0.416 | 0.046 | 0.104 | 0.104 | 0 | 0.47 | 0.27 | 0.22 | 0.21 | 0.36 | 0.37 |
| 5 | 144599586 | rs2895635 | 0.47 | 0.102 | 0.208 | -0.184 | -0.003 | -0.009 | -0.009 | 0 | 0.64 | 0.42 | 0.46 | 0.28 | 0.71 | 0.49 |
| 5 | 144603982 | rs13362029 | 0.933 | 0.038 | 0.403 | -0.171 | 0.001 | 0.005 | 0.005 | 0 | 0.54 | 0.4 | 0.27 | 0.22 | 0.43 | 0.42 |
| 5 | 144605231 | rs2547002 | 1.132 | -0.108 | 0.358 | 0.044 | -0.002 | 0 | 0 | 0 | 0.57 | 0.36 | 0.48 | 0.47 | 0.5 | 0.44 |
| 5 | 144605745 | rs12717935 | 0.236 | -0.177 | -0.143 | 0.409 | -0.006 | -0.046 | -0.046 | 0 | 0.84 | 0.89 | 0.78 | 0.55 | 0.85 | 0.54 |
| 5 | 144615100 | rs11745248 | -0.247 | 0.114 | -0.067 | -0.006 | -0.059 | 0.054 | 0.054 | 0 | 0.18 | 0.04 | 0.3 | 0.07 | 0.34 | 0.1 |
| 5 | 144619630 | rs12655715 | 0.572 | -0.467 | -0.316 | 0.198 | 0.001 | -0.101 | -0.101 | 0 | 0.14 | 0.01 | 0.18 | 0.34 | 0.08 | 0.08 |
| 5 | 144620356 | rs13180918 | 0.774 | -0.099 | 0.674 | -0.54 | -0.027 | -0.042 | -0.042 | 0 | 0.09 | 0.13 | 0.04 | 0 | 0.02 | 0 |
| 5 | 144625079 | rs13169495 | -0.245 | -0.167 | 1.108 | -0.521 | -0.052 | -0.019 | -0.019 | 0.152 | 0.12 | 0.46 | 0.07 | 0.03 | 0.03 | 0.02 |
| 5 | 144655047 | rs17103008 | -0.922 | -0.046 | 1.059 | -0.525 | 0.05 | -0.043 | -0.043 | 1.377 | 0.22 | 0.57 | 0.29 | 0.12 | 0.34 | 0.12 |
| 5 | 144660484 | rs6871262 | -0.883 | -0.004 | 1.068 | -0.492 | 0.051 | -0.036 | -0.036 | 0.386 | 0.22 | 0.57 | 0.29 | 0.14 | 0.34 | 0.12 |
| 5 | 144665996 | rs2400181 | -1.39 | -0.031 | 0.687 | -0.491 | 0.045 | -0.066 | -0.066 | 1 | 0.27 | 0.47 | 0.31 | 0.13 | 0.39 | 0.19 |
| 5 | 144666680 | rs6890617 | -0.38 | 0.134 | 0.633 | -0.432 | 0.015 | 0.025 | 0.025 | 0.087 | 0.07 | 0.26 | 0.11 | 0.05 | 0.05 | 0.09 |

| | | | | | | | | | | | | | | | | |
|---|---|---|---|---|---|---|---|---|---|---|---|---|---|---|---|---|
| 5 | 144672163 | rs7708225 | -1.312 | 0.155 | -0.135 | -0.424 | 0.024 | -0.034 | -0.034 | 0 | 0.26 | 0.62 | 0.31 | 0.14 | 0.38 | 0.19 |
| 5 | 144684867 | rs13172414 | -0.206 | 0.23 | -0.279 | -0.342 | -0.063 | -0.071 | -0.071 | 0 | 0.17 | 0.47 | 0.17 | 0.05 | 0.07 | 0.1 |
| 5 | 144688529 | rs12522659 | -0.829 | 1.369 | -1.079 | 0.235 | -0.02 | 0.015 | 0.015 | 0 | 0.18 | 0.14 | 0.19 | 0.1 | 0.34 | 0.09 |
| 5 | 144697159 | rs26174 | -0.319 | 0.079 | -1.254 | 0.701 | 0.033 | -0.018 | -0.018 | 0 | 0.36 | 0.24 | 0.27 | 0.53 | 0.25 | 0.31 |
| 5 | 144697489 | rs10515549 | 0.44 | 0.541 | -2.026 | -0.096 | -0.093 | 0.108 | 0.108 | 0 | 0.35 | 0.12 | 0.45 | 0.15 | 0.48 | 0.39 |
| 5 | 144726896 | rs2442760 | -0.558 | 0.676 | -1.187 | 0.173 | 0.038 | -0.044 | -0.044 | 0 | 0.76 | 0.66 | 0.62 | 0.88 | 0.62 | 0.87 |
| 5 | 144735126 | rs1450332 | 0.907 | 0.274 | -1.45 | 0.022 | -0.012 | 0.064 | 0.064 | 0 | 0.27 | 0.08 | 0.23 | 0.13 | 0.26 | 0.37 |
| 5 | 144769376 | rs32190 | -0.647 | 0.118 | -1.041 | -0.081 | 0.039 | -0.055 | -0.055 | 0 | 0.74 | 0.6 | 0.61 | 0.88 | 0.59 | 0.87 |
| 5 | 144773424 | rs282457 | 0.252 | -0.051 | -0.299 | 0.251 | 0.042 | 0.047 | 0.047 | 0 | 0.45 | 0.27 | 0.36 | 0.72 | 0.34 | 0.49 |
| 5 | 144778846 | rs711168 | 0.268 | -0.984 | -0.675 | 0.466 | -0.005 | 0.038 | 0.038 | 0 | 0.24 | 0.28 | 0.21 | 0.59 | 0.22 | 0.27 |
| 5 | 144781346 | rs711170 | 0.117 | -0.997 | -0.476 | 0.46 | -0.013 | 0.086 | 0.086 | 0 | 0.25 | 0.39 | 0.22 | 0.59 | 0.22 | 0.27 |
| 5 | 144782407 | rs813188 | -0.527 | -1.207 | -0.505 | 0.218 | -0.016 | 0.021 | 0.021 | 0 | 0.65 | 0.67 | 0.5 | 0.79 | 0.55 | 0.82 |
| 5 | 144809625 | rs1946246 | -0.256 | -1.243 | -0.521 | 0.509 | 0.01 | -0.008 | -0.008 | 0 | 0.63 | 0.72 | 0.75 | 0.41 | 0.71 | 0.57 |
| 5 | 144810692 | rs7721549 | 0.333 | -1.155 | -0.628 | 0.546 | -0.017 | 0.022 | 0.022 | 0 | 0.35 | 0.33 | 0.5 | 0.21 | 0.45 | 0.19 |
| 5 | 144812460 | rs7718608 | 1.223 | -0.997 | -0.463 | 0.249 | -0.052 | 0.062 | 0.062 | 0 | 0.25 | 0.03 | 0.22 | 0.11 | 0.25 | 0.38 |
| 5 | 144818318 | rs17103175 | 0.973 | -0.931 | -0.397 | 0.234 | -0.002 | 0.033 | 0.033 | 0 | 0.28 | 0.17 | 0.24 | 0.1 | 0.25 | 0.39 |
| 5 | 144820417 | rs889064 | -1.417 | -1.055 | -0.481 | 0.309 | -0.017 | -0.035 | -0.035 | 0 | 0.89 | 0.97 | 0.91 | 0.71 | 0.84 | 0.84 |
| 5 | 144821606 | rs10477276 | 0.984 | -0.921 | -0.407 | -0.538 | -0.002 | 0.023 | 0.023 | 0 | 0.28 | 0.18 | 0.24 | 0.14 | 0.25 | 0.39 |
| 5 | 144829826 | rs10068420 | 1.121 | -0.916 | -0.455 | -0.545 | -0.013 | 0.069 | 0.069 | 0 | 0.27 | 0.13 | 0.24 | 0.1 | 0.26 | 0.39 |
| 5 | 144845050 | rs2974572 | 0.626 | -1.018 | -0.401 | -0.603 | 0.008 | -0.013 | -0.013 | 0 | 0.32 | 0.39 | 0.42 | 0.17 | 0.43 | 0.17 |
| 5 | 144849881 | rs1422922 | 1.195 | -1.048 | -0.424 | -0.619 | -0.04 | 0.094 | 0.094 | 0 | 0.27 | 0.04 | 0.23 | 0.1 | 0.26 | 0.39 |
| 5 | 144851525 | rs2910272 | 0.52 | -1.037 | -0.31 | -0.572 | 0.021 | -0.017 | -0.017 | 0 | 0.33 | 0.43 | 0.43 | 0.23 | 0.44 | 0.18 |
| 5 | 144855154 | rs450337 | -0.038 | -1.037 | -0.278 | -0.548 | -0.018 | -0.016 | -0.016 | 0 | 0.61 | 0.57 | 0.66 | 0.27 | 0.69 | 0.56 |
| 5 | 144856948 | rs2400194 | -0.379 | -1.017 | -0.251 | -0.555 | 0.019 | 0.001 | 0.001 | 0 | 0.64 | 0.73 | 0.71 | 0.32 | 0.72 | 0.57 |
| 5 | 144857230 | rs427979 | 0.473 | -0.893 | -0.292 | -1.042 | -0.013 | 0.014 | 0.014 | 0 | 0.33 | 0.35 | 0.43 | 0.17 | 0.46 | 0.2 |
| 5 | 144861632 | rs717076 | 0.082 | -0.76 | -0.203 | -0.95 | -0.005 | -0.007 | -0.007 | 0 | 0.4 | 0.4 | 0.34 | 0.72 | 0.29 | 0.42 |
| 5 | 144868408 | rs11167904 | 0.035 | -0.611 | -0.172 | -1.704 | -0.006 | -0.036 | -0.036 | 0 | 0.4 | 0.37 | 0.46 | 0.88 | 0.3 | 0.48 |
| 5 | 144874969 | rs1345698 | -0.934 | -0.395 | -0.228 | -1.826 | -0.033 | -0.003 | -0.003 | 1.041 | 0.72 | 0.74 | 0.69 | 1 | 0.51 | 0.85 |
| 5 | 144877201 | rs10463375 | 0.765 | -0.193 | -0.115 | -1.723 | 0.017 | 0.038 | 0.038 | 0 | 0.27 | 0.15 | 0.2 | 0.11 | 0.2 | 0.38 |
| 5 | 144878860 | rs13177421 | 0.788 | -0.13 | -0.124 | -1.719 | -0.013 | 0.083 | 0.083 | 0 | 0.27 | 0.07 | 0.23 | 0.11 | 0.24 | 0.39 |
| 5 | 144880889 | rs7718967 | 0.325 | -0.145 | -0.099 | -1.727 | -0.008 | -0.042 | -0.042 | 0 | 0.3 | 0.36 | 0.26 | 0.11 | 0.24 | 0.4 |
| 5 | 144883076 | rs13179436 | 0.855 | -0.088 | -0.365 | -1.697 | 0.02 | 0.079 | 0.079 | 0 | 0.26 | 0.07 | 0.19 | 0.11 | 0.2 | 0.39 |
| 5 | 144883233 | rs12657244 | 0.743 | -0.088 | -0.365 | -1.697 | 0.006 | 0.008 | 0.008 | 0 | 0.27 | 0.19 | 0.21 | 0.11 | 0.2 | 0.39 |
| 5 | 144888814 | rs17411333 | 1.134 | -0.163 | -0.823 | -1.613 | -0.054 | -0.059 | -0.059 | 0 | 0.13 | 0.01 | 0.09 | 0.19 | 0.17 | 0.13 |
| 5 | 144890837 | rs12657926 | 0.36 | 0.714 | -0.025 | -2.188 | -0.011 | 0.081 | 0.081 | 0 | 0.27 | 0.06 | 0.23 | 0.1 | 0.24 | 0.39 |
| 5 | 144906456 | rs1363556 | -0.005 | 0.568 | 0.378 | -2.609 | -0.006 | 0.001 | 0.001 | 0 | 0.6 | 0.61 | 0.6 | 0.92 | 0.44 | 0.74 |
| 5 | 144906846 | rs11167905 | 0.438 | 0.534 | 0.908 | -1.26 | -0.015 | 0.032 | 0.032 | 0 | 0.13 | 0.04 | 0.13 | 0.07 | 0.14 | 0.12 |
| 5 | 144924760 | rs7727300 | 0.001 | 0.456 | 0.693 | -1.268 | 0 | 0.026 | 0.026 | 0 | 0.16 | 0.09 | 0.14 | 0.07 | 0.16 | 0.15 |
| 5 | 144924901 | rs7714479 | -1.165 | 0.332 | 0.511 | -1.294 | 0.004 | 0.003 | 0.003 | 0.059 | 0.34 | 0.67 | 0.3 | 0.12 | 0.27 | 0.39 |
| 5 | 144939151 | rs17103387 | -0.751 | 0.016 | 1.175 | -0.692 | -0.031 | -0.046 | -0.046 | 0.939 | 0.19 | 0.66 | 0.16 | 0.08 | 0.11 | 0.12 |
| 5 | 144943205 | rs6866539 | -0.43 | -0.017 | 1.071 | -0.679 | -0.053 | -0.084 | -0.084 | 3.234 | 0.2 | 0.54 | 0.16 | 0.08 | 0.11 | 0.14 |
| 5 | 144944054 | rs964923 | -2.279 | 0.569 | 1.852 | -0.041 | 0.051 | 0.141 | 0.141 | 2.137 | 0.08 | 0.46 | 0.27 | 0.13 | 0.08 | 0.12 |
| 5 | 144944159 | rs964924 | -1.576 | 0.572 | 1.878 | -0.037 | -0.002 | 0.006 | 0.006 | 0.436 | 0.06 | 0.08 | 0.23 | 0.12 | 0.05 | 0.11 |
| 5 | 144944249 | rs867715 | -0.125 | 1.486 | 1.574 | -0.377 | -0.054 | 0.051 | 0.051 | 0.223 | 0.1 | 0.01 | 0.16 | 0 | 0.27 | 0.03 |
| 5 | 144945649 | rs17103398 | -1.504 | 0.869 | 1.848 | 0.002 | -0.001 | 0.005 | 0.005 | 0.276 | 0.05 | 0.07 | 0.23 | 0.12 | 0.05 | 0.12 |
| 5 | 144972449 | rs11953390 | -2.788 | 0.866 | 2.451 | -0.048 | 0.058 | 0.14 | 0.14 | 1.933 | 0.08 | 0.49 | 0.26 | 0.13 | 0.08 | 0.11 |
| 5 | 144989671 | rs1865010 | -2.368 | 0.868 | 2.653 | -0.046 | 0.024 | 0.091 | 0.091 | 0.659 | 0.07 | 0.32 | 0.25 | 0.13 | 0.07 | 0.11 |
| 5 | 145003254 | rs4339405 | -1.899 | 0.867 | 2.612 | -0.019 | 0.026 | 0.098 | 0.098 | 0.533 | 0.06 | 0.27 | 0.24 | 0.15 | 0.07 | 0.12 |
| 5 | 145007570 | rs12653168 | -1.974 | 0.861 | 2.487 | 0.004 | 0.005 | 0.069 | 0.069 | 0.29 | 0.07 | 0.21 | 0.23 | 0.16 | 0.07 | 0.11 |
| 5 | 145041065 | rs7733723 | -2.671 | 0.774 | 2.784 | -0.051 | -0.006 | 0.047 | 0.047 | 4.438 | 0.1 | 0.63 | 0.29 | 0.14 | 0.08 | 0.11 |
| 5 | 145052223 | rs4509079 | -2.594 | 0.797 | 2.263 | -0.039 | 0.006 | 0.082 | 0.082 | 4.095 | 0.1 | 0.6 | 0.28 | 0.15 | 0.07 | 0.11 |
| 5 | 145062940 | rs11948853 | -2.229 | 0.86 | 2.035 | -0.12 | 0.022 | 0.125 | 0.125 | 3.677 | 0.09 | 0.52 | 0.27 | 0.13 | 0.05 | 0.11 |
| 5 | 145068327 | rs1368434 | -0.775 | 0.693 | 0.301 | -0.409 | -0.096 | 0.145 | 0.145 | 0.598 | 0.6 | 0.95 | 0.73 | 0.79 | 0.63 | 0.64 |
| 5 | 145080340 | rs972351 | -2.234 | 0.6 | 1.73 | 0.349 | 0.046 | 0.15 | 0.15 | 2.679 | 0.08 | 0.47 | 0.26 | 0.15 | 0.06 | 0.12 |
| 5 | 145091210 | rs1654135 | -2.118 | 0.203 | 1.367 | 0.002 | -0.015 | -0.023 | -0.023 | 0.014 | 0.21 | 0.32 | 0.18 | 0.13 | 0.14 | 0.24 |
| 5 | 145109218 | rs2400197 | 1.915 | 0.173 | 1.389 | 0.029 | -0.017 | -0.022 | -0.022 | 0.005 | 0.78 | 0.68 | 0.82 | 0.87 | 0.86 | 0.76 |
| 5 | 145131986 | rs7734685 | 1.923 | 0.368 | 1.902 | 0.092 | -0.015 | -0.033 | -0.033 | 0 | 0.8 | 0.57 | 0.81 | 0.87 | 0.87 | 0.78 |
| 5 | 145139816 | rs6867440 | 1.624 | 0.397 | 1.825 | -0.083 | 0 | -0.018 | -0.018 | 0.153 | 0.85 | 0.85 | 0.91 | 0.98 | 0.89 | 0.87 |
| 5 | 145146015 | rs338896 | 1.838 | 0.207 | 2.622 | -0.154 | -0.016 | -0.116 | -0.116 | 0.051 | 0.8 | 0.49 | 0.86 | 0.95 | 0.87 | 0.81 |
| 5 | 145149318 | rs1821250 | 2.22 | 0.072 | 2.759 | 0 | -0.026 | -0.054 | -0.054 | 0 | 0.77 | 0.38 | 0.8 | 0.9 | 0.87 | 0.79 |
| 5 | 145149940 | rs1897545 | 2.187 | 0.073 | 2.851 | -0.004 | -0.036 | -0.089 | -0.089 | 0 | 0.77 | 0.46 | 0.81 | 0.91 | 0.87 | 0.79 |
| 5 | 145168203 | rs12173238 | 2.209 | -0.019 | 2.876 | -0.05 | -0.048 | -0.083 | -0.083 | 0 | 0.77 | 0.48 | 0.81 | 0.9 | 0.88 | 0.8 |
| 5 | 145170910 | rs13157947 | 2.058 | -0.265 | 2.77 | -0.179 | -0.076 | -0.092 | -0.092 | 0 | 0.78 | 0.47 | 0.84 | 0.89 | 0.94 | 0.84 |
| 5 | 145174947 | rs1014292 | -2.367 | 0.037 | 2.548 | 0.246 | -0.003 | 0.006 | 0.006 | 0.763 | 0.12 | 0.15 | 0.16 | 0.21 | 0.1 | 0.21 |
| 5 | 145201132 | rs12658465 | -2.643 | 0.121 | 2.937 | 0.298 | 0.001 | 0.025 | 0.025 | 0.79 | 0.13 | 0.3 | 0.18 | 0.22 | 0.11 | 0.2 |
| 5 | 145202461 | rs6898776 | 2.418 | 0.323 | 2.659 | 0.826 | 0.019 | 0.071 | 0.071 | 1.05 | 0.57 | 0.17 | 0.57 | 0.28 | 0.69 | 0.51 |
| 5 | 145204975 | rs13155845 | -1.989 | 0.327 | 2.682 | 0.826 | -0.001 | 0.01 | 0.01 | 0.492 | 0.12 | 0.2 | 0.17 | 0.22 | 0.11 | 0.19 |
| 5 | 145205480 | rs340046 | 2.325 | 0.335 | 2.666 | 0.918 | 0.022 | 0.049 | 0.049 | 0.843 | 0.58 | 0.21 | 0.58 | 0.31 | 0.69 | 0.51 |
| 5 | 145210041 | rs443033 | 1.147 | 0.508 | 2.706 | 1.062 | 0.002 | 0.093 | 0.093 | 0 | 0.75 | 0.48 | 0.76 | 0.51 | 0.81 | 0.74 |

| | | | | | | | | | | | | | | | | |
|---|---|---|---|---|---|---|---|---|---|---|---|---|---|---|---|---|
| 5 | 145212145 | rs681591 | 1.59 | 1.113 | 2.646 | 1.012 | 0.012 | 0.032 | 0.032 | 0.172 | 0.7 | 0.45 | 0.69 | 0.44 | 0.69 | 0.64 |
| 5 | 145212518 | rs1984956 | 1.626 | 0.617 | 2.648 | 1.062 | -0.001 | 0.001 | 0.001 | 0.035 | 0.7 | 0.35 | 0.71 | 0.41 | 0.77 | 0.65 |
| 5 | 145219445 | rs13359571 | -1.344 | 0.594 | 2.317 | 1.031 | -0.015 | -0.017 | -0.017 | 0.153 | 0.08 | 0.05 | 0.13 | 0.13 | 0.09 | 0.12 |
| 5 | 145220936 | rs1001432 | -1.669 | 0.657 | 2.417 | 1.012 | 0.027 | 0.019 | 0.019 | 0.056 | 0.07 | 0.35 | 0.08 | 0.07 | 0.07 | 0.14 |
| 5 | 145226615 | rs722714 | -1.299 | 0.676 | 1.864 | 1.023 | 0.002 | 0.002 | 0.002 | 0 | 0.08 | 0.1 | 0.13 | 0.13 | 0.08 | 0.12 |
| 5 | 145230253 | rs1490366 | 0.944 | 0.764 | 1.279 | 1.223 | -0.012 | -0.07 | -0.07 | 0 | 0.75 | 0.88 | 0.76 | 0.54 | 0.75 | 0.78 |
| 5 | 145231621 | rs6892053 | -2.611 | 0.852 | 1.335 | 0.793 | 0.003 | 0 | 0 | 0 | 0.21 | 0.81 | 0.24 | 0.24 | 0.17 | 0.28 |
| 5 | 145235121 | rs340051 | 0.701 | 0.89 | 0.783 | 1.188 | -0.071 | 0.022 | 0.022 | 0 | 0.81 | 0.99 | 0.89 | 0.93 | 0.78 | 0.87 |
| 5 | 145235669 | rs3995495 | -1.397 | 0.967 | 1.295 | 1.158 | 0.03 | 0.157 | 0.157 | 1.418 | 0.06 | 0.24 | 0.1 | 0.14 | 0.03 | 0.04 |
| 5 | 145235874 | rs13180312 | -1.7 | 1.454 | 0.148 | 0.807 | -0.002 | 0.008 | 0.008 | 0.02 | 0.1 | 0.1 | 0.14 | 0.07 | 0.12 | 0.14 |
| 5 | 145238464 | rs381658 | 1.3 | 1.234 | 0.016 | 1.069 | -0.007 | -0.022 | -0.022 | 0 | 0.58 | 0.32 | 0.66 | 0.47 | 0.59 | 0.5 |
| 5 | 145241164 | rs2195929 | -0.865 | 1.18 | -0.204 | 1.158 | 0 | 0.006 | 0.006 | 0 | 0.21 | 0.2 | 0.38 | 0.43 | 0.22 | 0.34 |
| 5 | 145244646 | rs2569008 | -0.49 | -0.157 | 0 | -0.302 | -0.055 | -0.095 | -0.095 | 0.406 | 0.2 | 0.53 | 0.1 | 0.05 | 0.1 | 0.1 |
| 5 | 145246085 | rs2569006 | 0.152 | 0.333 | -1.307 | -0.646 | -0.102 | -0.001 | -0.001 | 0.108 | 0.69 | 0.9 | 0.73 | 0.73 | 0.58 | 0.77 |
| 5 | 145249934 | rs1347127 | 0.55 | -0.431 | -1.033 | -1.723 | -0.011 | -0.006 | -0.006 | 0 | 0.32 | 0.43 | 0.47 | 0.69 | 0.27 | 0.55 |
| 5 | 145252574 | rs10037373 | 0.116 | -1.23 | -0.992 | -2.236 | -0.004 | 0.016 | 0.016 | 0 | 0.54 | 0.61 | 0.69 | 0.72 | 0.55 | 0.7 |
| 5 | 145255328 | rs7701544 | 0.01 | -1.614 | -0.967 | -2.322 | -0.069 | -0.106 | -0.106 | 0 | 0.17 | 0.32 | 0.05 | 0 | 0.04 | 0.08 |
| 5 | 145266174 | rs10074608 | 1.327 | -1.894 | -0.925 | -2.25 | -0.002 | -0.023 | -0.023 | 0 | 0.21 | 0.14 | 0.21 | 0.62 | 0.22 | 0.46 |
| 5 | 145270339 | rs1821251 | 0.293 | -1.767 | -0.73 | -2.146 | -0.008 | -0.014 | -0.014 | 0.283 | 0.36 | 0.42 | 0.49 | 0.71 | 0.33 | 0.58 |
| 5 | 145281565 | rs1907062 | -0.513 | -1.54 | -0.909 | -2.046 | 0.075 | 0.131 | 0.131 | 0 | 0.52 | 0.79 | 0.54 | 0.72 | 0.36 | 0.64 |
| 5 | 145282109 | rs4913050 | 1 | -1.401 | -0.138 | -1.915 | 0.003 | 0.028 | 0.028 | 0 | 0.08 | 0.28 | 0.26 | 0.08 | 0.07 | 0.09 |
| 5 | 145285134 | rs13169102 | 0.953 | -1.49 | -0.081 | -1.96 | 0.002 | -0.007 | -0.007 | 0 | 0.08 | 0.04 | 0.02 | 0 | 0.02 | 0.05 |
| 5 | 145285452 | rs2569011 | -0.532 | -1.378 | -0.198 | -1.906 | -0.062 | -0.059 | -0.059 | 0 | 0.23 | 0.6 | 0.31 | 0.08 | 0.1 | 0.16 |
| 5 | 145297971 | rs970456 | -0.434 | 0.025 | -0.534 | -1.317 | -0.006 | -0.029 | -0.029 | 0 | 0.17 | 0.33 | 0.38 | 0.12 | 0.15 | 0.11 |
| 5 | 145310336 | rs6580395 | 0.082 | 0.561 | -0.625 | -1.086 | -0.009 | 0.038 | 0.038 | 0 | 0.18 | 0.13 | 0.19 | 0.03 | 0.22 | 0.15 |
| 5 | 145313920 | rs13436709 | 0.147 | 1.205 | -0.59 | -1.128 | 0.029 | -0.049 | -0.049 | 0 | 0.2 | 0.3 | 0.18 | 0.05 | 0.28 | 0.11 |
| 5 | 145314878 | rs7734655 | 0.343 | 1.162 | -0.745 | -1.066 | -0.008 | 0.021 | 0.021 | 0 | 0.17 | 0.12 | 0.18 | 0.1 | 0.2 | 0.16 |
| 5 | 145315962 | rs7725688 | -0.376 | 1.31 | -0.353 | -0.507 | 0.038 | -0.056 | -0.056 | 0 | 0.21 | 0.34 | 0.19 | 0.05 | 0.29 | 0.11 |
| 5 | 145316958 | rs17104037 | -0.971 | 1.098 | -0.914 | 0.513 | -0.05 | -0.075 | -0.075 | 0.098 | 0.12 | 0.01 | 0.13 | 0.26 | 0.16 | 0.11 |
| 5 | 145322071 | rs6869382 | -0.195 | 0.449 | -0.065 | 0.795 | 0.004 | 0.002 | 0.002 | 0 | 0.42 | 0.61 | 0.4 | 0.44 | 0.34 | 0.45 |
| 5 | 145341353 | rs17491399 | 0.53 | -0.124 | 0.225 | 0.547 | 0.024 | 0.037 | 0.037 | 0 | 0.21 | 0.04 | 0.11 | 0.15 | 0.16 | 0.16 |
| 5 | 145342331 | rs4912701 | -0.202 | -0.166 | 0.414 | 0.619 | -0.009 | 0.107 | 0.107 | 0.027 | 0.14 | 0.45 | 0.24 | 0.29 | 0.1 | 0.23 |
| 5 | 145356042 | rs886949 | -0.949 | -0.157 | 0.542 | 0.643 | -0.017 | -0.001 | -0.001 | 0.164 | 0.35 | 0.61 | 0.36 | 0.44 | 0.25 | 0.37 |
| 5 | 145361499 | rs6883675 | -0.624 | 0.426 | 0.933 | 1.031 | -0.007 | 0.125 | 0.125 | 0 | 0.14 | 0.54 | 0.24 | 0.28 | 0.1 | 0.22 |
| 5 | 145361864 | rs4913057 | -0.223 | 0.483 | 0.834 | 1.058 | -0.001 | 0.025 | 0.025 | 0.053 | 0.09 | 0.13 | 0.21 | 0.28 | 0.09 | 0.22 |
| 5 | 145376268 | rs6876373 | 0.008 | 1.353 | 0.875 | 1.277 | 0.093 | 0.067 | 0.067 | 0 | 0.78 | 0.61 | 0.57 | 0.72 | 0.61 | 0.78 |
| 5 | 145386138 | rs1859894 | -0.026 | 1.52 | 0.935 | 1.253 | 0.079 | 0.063 | 0.063 | 0 | 0.78 | 0.63 | 0.58 | 0.72 | 0.62 | 0.79 |
| 5 | 145396585 | rs168712 | 0.251 | 1.641 | 0.656 | 1.343 | -0.051 | -0.045 | -0.045 | 0.084 | 0.18 | 0.12 | 0.39 | 0.28 | 0.35 | 0.22 |
| 5 | 145401360 | rs2215010 | 0.066 | 1.648 | 0.664 | 1.347 | 0.006 | 0.003 | 0.003 | 0 | 0.19 | 0.23 | 0.39 | 0.28 | 0.35 | 0.22 |
| 5 | 145415118 | rs6580404 | 0.011 | 1.606 | 0.482 | 1.312 | -0.029 | -0.027 | -0.027 | 0.002 | 0.19 | 0.21 | 0.39 | 0.28 | 0.35 | 0.22 |
| 5 | 145422043 | rs3733937 | -0.032 | 1.603 | 0.552 | 1.325 | 0.003 | 0.001 | 0.001 | 0 | 0.19 | 0.26 | 0.38 | 0.28 | 0.35 | 0.22 |
| 5 | 145436874 | rs248771 | -0.065 | 1.492 | 0.69 | 1.269 | 0.032 | 0.026 | 0.026 | 0 | 0.2 | 0.33 | 0.38 | 0.28 | 0.35 | 0.22 |
| 5 | 145442133 | rs11435 | -0.424 | 1.377 | 0.368 | 1.448 | -0.033 | -0.075 | -0.075 | 0 | 0.12 | 0.01 | 0.22 | 0.3 | 0.13 | 0.27 |
| 5 | 145442203 | rs1056149 | 0.208 | 1.498 | 1.045 | 1.293 | 0.115 | 0.093 | 0.093 | 0 | 0.78 | 0.46 | 0.59 | 0.72 | 0.65 | 0.78 |
| 5 | 145450586 | rs2962540 | 0.077 | 2.183 | -0.076 | 1.493 | -0.104 | -0.083 | -0.083 | 0 | 0.14 | 0.01 | 0.41 | 0.23 | 0.32 | 0.21 |
| 5 | 145453440 | rs12374457 | -0.233 | 2.432 | 0.014 | 1.433 | -0.035 | -0.023 | -0.023 | 0.102 | 0.17 | 0.11 | 0.4 | 0.24 | 0.3 | 0.2 |
| 5 | 145477855 | rs6877277 | 0.015 | 2.331 | 0.141 | 1.273 | -0.014 | -0.01 | -0.01 | 5.334 | 0.16 | 0.13 | 0.4 | 0.2 | 0.25 | 0.19 |
| 5 | 145483703 | rs2962544 | 0.996 | 1.32 | -0.877 | 1.002 | -0.012 | -0.046 | -0.046 | 0.284 | 0.26 | 0.01 | 0.2 | 0.24 | 0.2 | 0.16 |
| 5 | 145489771 | rs13174055 | 0.765 | 1.428 | -0.746 | 0.961 | -0.095 | -0.072 | -0.072 | 0 | 0.14 | 0.01 | 0.38 | 0.2 | 0.25 | 0.19 |
| 5 | 145496311 | rs7728381 | -0.928 | 1.018 | -0.061 | 0.687 | 0.065 | -0.011 | -0.011 | 0 | 0.26 | 0.54 | 0.46 | 0.2 | 0.33 | 0.23 |
| 5 | 145499996 | rs10988 | 0.868 | 1.033 | -0.14 | 0.694 | -0.08 | -0.059 | -0.059 | 0 | 0.14 | 0.02 | 0.38 | 0.2 | 0.25 | 0.19 |
| 5 | 145505337 | rs17427286 | 0.868 | 1.033 | -0.14 | 0.694 | -0.08 | -0.059 | -0.059 | 0 | 0.14 | 0.02 | 0.38 | 0.2 | 0.25 | 0.19 |
| 5 | 145509573 | rs11953517 | 0.93 | 1.033 | -0.141 | 0.694 | -0.095 | -0.072 | -0.072 | 0 | 0.14 | 0.01 | 0.38 | 0.2 | 0.25 | 0.19 |
| 5 | 145509728 | rs17104268 | 0.099 | 1.01 | -0.052 | 0.683 | 0 | -0.012 | -0.012 | 0 | 0.06 | 0.06 | 0.03 | 0 | 0.07 | 0.04 |
| 5 | 145515379 | rs2963919 | 0.597 | 1.008 | 0.048 | 0.682 | 0.004 | -0.001 | -0.001 | 0 | 0.76 | 0.72 | 0.55 | 0.8 | 0.67 | 0.77 |
| 5 | 145519821 | rs17493851 | 0.933 | 1.008 | 0.045 | 0.682 | -0.095 | -0.072 | -0.072 | 0 | 0.14 | 0.01 | 0.38 | 0.2 | 0.25 | 0.19 |
| 5 | 145532859 | rs17500483 | -0.556 | 1.008 | 0.046 | 0.682 | -0.03 | -0.001 | -0.001 | 0 | 0.23 | 0.17 | 0.42 | 0.2 | 0.33 | 0.23 |
| 5 | 145543972 | rs3763373 | 0.865 | 1.075 | -0.217 | 0.737 | -0.095 | -0.072 | -0.072 | 0.061 | 0.14 | 0.01 | 0.38 | 0.2 | 0.25 | 0.19 |
| 5 | 145551181 | rs3828935 | 0.904 | 1.078 | -0.215 | 0.738 | -0.095 | -0.072 | -0.072 | 1.094 | 0.14 | 0.01 | 0.38 | 0.2 | 0.25 | 0.19 |
| 5 | 145583623 | rs2279895 | 0.786 | 1.054 | -0.174 | 0.736 | -0.031 | -0.02 | -0.02 | 0 | 0.15 | 0.1 | 0.38 | 0.2 | 0.25 | 0.18 |
| 5 | 145588483 | rs6881180 | -0.887 | 1.026 | -0.083 | 0.705 | 0.062 | -0.008 | -0.008 | 0 | 0.25 | 0.51 | 0.43 | 0.21 | 0.3 | 0.21 |
| 5 | 145612822 | rs17104331 | 0.002 | 1.02 | -0.116 | 0.701 | 0.005 | -0.03 | -0.03 | 0 | 0.06 | 0.08 | 0.03 | 0 | 0.07 | 0.04 |
| 5 | 145617707 | rs10515565 | -0.702 | 1.028 | -0.381 | 0.705 | 0.056 | -0.009 | -0.009 | 0 | 0.25 | 0.52 | 0.43 | 0.21 | 0.3 | 0.21 |
| 5 | 145630737 | rs1567057 | -0.818 | 1.046 | -0.374 | 0.713 | 0.057 | -0.008 | -0.008 | 0 | 0.25 | 0.52 | 0.43 | 0.21 | 0.3 | 0.21 |
| 5 | 145636671 | rs11953090 | 1.002 | 1.051 | -0.389 | 0.717 | -0.088 | -0.07 | -0.07 | 0.073 | 0.14 | 0.01 | 0.37 | 0.21 | 0.23 | 0.17 |
| 5 | 145642007 | rs13155119 | 1.002 | 1.051 | -0.389 | 0.722 | -0.088 | -0.067 | -0.067 | 0 | 0.14 | 0.01 | 0.37 | 0.2 | 0.23 | 0.17 |
| 5 | 145643048 | rs7729098 | -0.798 | 1.054 | -0.379 | 0.709 | 0.059 | -0.008 | -0.008 | 0 | 0.25 | 0.52 | 0.43 | 0.21 | 0.3 | 0.21 |
| 5 | 145647151 | rs2963925 | 0.727 | 1.04 | -1.466 | 0.818 | 0.001 | 0.006 | 0.006 | 0.437 | 0.31 | 0.27 | 0.19 | 0.23 | 0.29 | 0.43 |
| 5 | 145668298 | rs11954297 | 0.792 | 1.322 | -1.714 | 1.012 | -0.062 | -0.047 | -0.047 | 0.021 | 0.14 | 0.02 | 0.37 | 0.21 | 0.23 | 0.17 |

| | | | | | | | | | | | | | | | | |
|---|---|---|---|---|---|---|---|---|---|---|---|---|---|---|---|---|
| 5 | 145701961 | rs12515446 | 0.405 | 0.569 | -1.502 | 0.901 | -0.01 | -0.048 | -0.048 | 0 | 0.28 | 0.01 | 0.21 | 0.25 | 0.21 | 0.18 |
| 5 | 145710848 | rs1201598 | 0.641 | 1.083 | -0.714 | 0.986 | -0.027 | 0.074 | 0.074 | 0 | 0.55 | 0.19 | 0.48 | 0.7 | 0.57 | 0.46 |
| 5 | 145713164 | rs6866292 | -2.323 | 1.141 | 0.759 | 1.516 | 0.009 | 0.006 | 0.006 | 0.69 | 0.23 | 0.72 | 0.21 | 0.21 | 0.27 | 0.14 |
| 5 | 145717285 | rs1368402 | -2.267 | 1.266 | 0.916 | 1.546 | 0.054 | 0.06 | 0.06 | 0.481 | 0.2 | 0.57 | 0.19 | 0.21 | 0.24 | 0.11 |
| 5 | 145720431 | rs891970 | -1.614 | 1.444 | 0.152 | 1.749 | -0.024 | -0.025 | -0.025 | 2.617 | 0.12 | 0.07 | 0.12 | 0.21 | 0.22 | 0.12 |
| 5 | 145732560 | rs10515569 | -1.893 | 1.114 | 0.455 | 1.204 | 0.019 | -0.041 | -0.041 | 0.19 | 0.07 | 0.21 | 0.05 | 0 | 0.09 | 0.03 |
| 5 | 145742435 | rs9325005 | -1.746 | 0.965 | 0.49 | 1.741 | 0.006 | 0.031 | 0.031 | 0.285 | 0.27 | 0.66 | 0.21 | 0.51 | 0.26 | 0.19 |
| 5 | 145749535 | rs7718446 | -2.107 | 0.924 | 0.629 | 1.652 | 0.002 | 0.008 | 0.008 | 0.768 | 0.29 | 0.67 | 0.29 | 0.48 | 0.31 | 0.17 |
| 5 | 145753001 | rs1991800 | -1.211 | 1.202 | 0.619 | 1.976 | 0.01 | 0.01 | 0.01 | 0.012 | 0.09 | 0.07 | 0.12 | 0.2 | 0.15 | 0.09 |
| 5 | 145766174 | rs879943 | -1.327 | 1.131 | 0.788 | 1.954 | 0.023 | 0.072 | 0.072 | 0 | 0.08 | 0.14 | 0.1 | 0.26 | 0.12 | 0.06 |
| 5 | 145771821 | rs997011 | 1.085 | 1.049 | 0.763 | 1.701 | -0.019 | -0.019 | -0.019 | 0 | 0.9 | 0.91 | 0.88 | 0.8 | 0.85 | 0.91 |
| 5 | 145772645 | rs1368404 | -2.129 | 0.887 | 1.395 | 1.741 | 0.017 | 0.07 | 0.07 | 1.378 | 0.18 | 0.47 | 0.15 | 0.26 | 0.16 | 0.09 |
| 5 | 145772967 | rs9325006 | -1.68 | 0.943 | 1.515 | 1.601 | 0.016 | -0.063 | -0.063 | 1.086 | 0.13 | 0.35 | 0.1 | 0.02 | 0.11 | 0.05 |
| 5 | 145775892 | rs12655536 | 1.156 | 0.19 | 0.45 | 1.368 | 0.039 | -0.017 | -0.017 | 0 | 0.22 | 0.01 | 0.09 | 0.2 | 0.13 | 0.32 |
| 5 | 145776261 | rs6868971 | 1.785 | 1.036 | 1.439 | 1.25 | 0.059 | 0.161 | 0.161 | 0 | 0.85 | 0.66 | 0.83 | 0.55 | 0.8 | 0.87 |
| 5 | 145776488 | rs6580418 | 1.516 | 1.055 | 1.195 | 1.256 | 0.017 | 0.061 | 0.061 | 0 | 0.86 | 0.81 | 0.84 | 0.55 | 0.8 | 0.87 |
| 5 | 145777478 | rs6886412 | 2.355 | 0.99 | 1.106 | 1.209 | -0.015 | -0.029 | -0.029 | 0.051 | 0.7 | 0.28 | 0.72 | 0.53 | 0.7 | 0.81 |
| 5 | 145784922 | rs11747475 | 2.052 | 0.863 | 0.836 | 0.585 | 0.015 | 0.01 | 0.01 | 0.237 | 0.77 | 0.4 | 0.79 | 0.71 | 0.78 | 0.85 |
| 5 | 145789106 | rs4705295 | 0.324 | 1.379 | 0.082 | 0.823 | -0.042 | -0.035 | -0.035 | 0 | 0.93 | 0.99 | 0.91 | 0.72 | 0.88 | 0.89 |
| 5 | 145795417 | rs2116800 | 0.394 | 1.383 | -0.435 | 0.579 | -0.133 | 0.154 | 0.154 | 0.777 | 0.32 | 0.01 | 0.26 | 0.09 | 0.43 | 0.26 |
| 5 | 145797020 | rs979893 | 1.634 | 1.57 | 1.023 | 1.588 | 0.062 | 0.057 | 0.057 | 0 | 0.85 | 0.69 | 0.83 | 0.71 | 0.8 | 0.85 |
| 5 | 145804118 | rs10056189 | 1.842 | 1.355 | 0.826 | 1.465 | 0.036 | 0.037 | 0.037 | 0 | 0.88 | 0.75 | 0.85 | 0.83 | 0.87 | 0.9 |
| 5 | 145804417 | rs9325009 | 1.73 | 0.964 | 1.047 | 1.615 | 0.011 | 0.057 | 0.057 | 0.035 | 0.8 | 0.63 | 0.82 | 0.66 | 0.84 | 0.94 |
| 5 | 145808408 | rs7712762 | 2.247 | 0.68 | 1.643 | 1.169 | 0.038 | -0.039 | -0.039 | 0 | 0.9 | 0.72 | 0.9 | 0.94 | 0.85 | 0.94 |
| 5 | 145822073 | rs1978708 | 2.598 | 0.568 | 2.756 | 1.237 | 0.023 | -0.028 | -0.028 | 0 | 0.94 | 0.82 | 0.95 | 1 | 0.92 | 0.99 |
| 5 | 145822515 | rs4705334 | 1.994 | 0.313 | 2.274 | 0.719 | -0.001 | -0.041 | -0.041 | 1.877 | 0.82 | 0.81 | 0.86 | 1 | 0.84 | 0.94 |
| 5 | 145834529 | rs10044956 | -2.061 | 0.379 | 2.482 | 0.494 | 0 | -0.026 | -0.026 | 0 | 0.17 | 0.26 | 0.23 | 0.84 | 0.19 | 0.29 |
| 5 | 145838440 | rs11743333 | 3.24 | 0.346 | 2.538 | 0.476 | 0.007 | 0.05 | 0.05 | 0 | 0.62 | 0.27 | 0.61 | 0.16 | 0.65 | 0.64 |
| 5 | 145857166 | rs9986288 | -1.021 | 0.379 | 2.534 | 0.5 | -0.001 | -0.031 | -0.031 | 0 | 0.11 | 0.03 | 0.12 | 0.71 | 0.09 | 0.22 |
| 5 | 145873365 | rs3797299 | -2.176 | 0.358 | 2.547 | 0.487 | 0 | -0.023 | -0.023 | 0 | 0.17 | 0.25 | 0.23 | 0.84 | 0.19 | 0.3 |
| 5 | 145890228 | rs1802027 | -0.683 | 0.33 | 2.44 | 0.47 | 0.004 | 0.006 | 0.006 | 0 | 0.12 | 0.09 | 0.09 | 0 | 0.08 | 0.05 |
| 5 | 145890455 | rs3822506 | -1.041 | 0.331 | 2.361 | 0.474 | -0.001 | -0.03 | -0.03 | 0 | 0.12 | 0.03 | 0.12 | 0.72 | 0.09 | 0.22 |
| 5 | 145893765 | rs6580423 | 2.975 | 0.399 | 2.319 | 0.458 | -0.002 | -0.053 | -0.053 | 0 | 0.63 | 0.47 | 0.67 | 0.22 | 0.65 | 0.66 |
| 5 | 145895558 | rs17104742 | -0.586 | 0.484 | 1.849 | 0.502 | -0.003 | 0.05 | 0.05 | 0 | 0.12 | 0.02 | 0.09 | 0 | 0.08 | 0.05 |
| 5 | 145897895 | rs7735001 | -2.759 | 0.51 | 1.887 | 0.493 | 0.001 | -0.04 | -0.04 | 0 | 0.22 | 0.5 | 0.22 | 0.77 | 0.22 | 0.28 |
| 5 | 145904225 | rs11956501 | 2.642 | 0.453 | 1.836 | 1.267 | 0.001 | 0.005 | 0.005 | 0 | 0.65 | 0.57 | 0.68 | 0.44 | 0.66 | 0.67 |
| 5 | 145915073 | rs4705423 | 1.176 | 0.395 | 1.666 | 1.259 | -0.018 | 0.068 | 0.068 | 0 | 0.84 | 0.95 | 0.87 | 1 | 0.85 | 0.94 |
| 5 | 145917201 | rs918533 | 2.831 | 0.355 | 1.828 | 1.387 | -0.004 | 0.009 | 0.009 | 0 | 0.66 | 0.48 | 0.69 | 0.46 | 0.7 | 0.69 |
| 5 | 145918304 | rs10035912 | -1.081 | 0.357 | 1.667 | 1.409 | -0.003 | -0.015 | -0.015 | 0 | 0.09 | 0.03 | 0.1 | 0.49 | 0.06 | 0.19 |
| 5 | 145920072 | rs10037468 | -1.995 | 0.357 | 1.751 | 1.45 | 0 | 0.038 | 0.038 | 0 | 0.13 | 0.17 | 0.14 | 0.51 | 0.15 | 0.24 |
| 5 | 145921524 | rs7717186 | 2.413 | 0.375 | 1.655 | 1.342 | -0.005 | -0.002 | -0.002 | 0 | 0.66 | 0.47 | 0.61 | 0.4 | 0.7 | 0.68 |
| 5 | 145924124 | rs4260721 | 1.902 | 0.143 | 1.386 | 1.481 | -0.009 | 0.015 | 0.015 | 0 | 0.71 | 0.65 | 0.67 | 0.46 | 0.81 | 0.74 |
| 5 | 145925989 | rs986061 | 1.391 | 0.232 | 0.807 | 1.43 | -0.032 | -0.132 | -0.132 | 0.032 | 0.81 | 0.95 | 0.84 | 0.56 | 0.86 | 0.8 |
| 5 | 145929602 | rs2216657 | -1.333 | -0.204 | 0.49 | 1.034 | 0.024 | 0.088 | 0.088 | 0.122 | 0.19 | 0.05 | 0.12 | 0.04 | 0.18 | 0.14 |
| 5 | 145930544 | rs1423029 | 2.654 | 0.719 | 0.917 | 0.843 | -0.001 | -0.002 | -0.002 | 0 | 0.53 | 0.55 | 0.62 | 0.51 | 0.54 | 0.62 |
| 5 | 145931611 | rs11743811 | -1.685 | 0.927 | 0.328 | 0.636 | -0.007 | 0.053 | 0.053 | 0.152 | 0.26 | 0.19 | 0.2 | 0.05 | 0.31 | 0.17 |
| 5 | 145949061 | rs2014944 | 1.595 | 0.994 | 0.564 | 1.397 | -0.015 | -0.096 | -0.096 | 0 | 0.76 | 0.91 | 0.78 | 0.53 | 0.79 | 0.69 |
| 5 | 145949309 | rs6892770 | -1.868 | 0.655 | 1.567 | 1.268 | 0.009 | 0.006 | 0.006 | 0 | 0.06 | 0.32 | 0.03 | 0 | 0.01 | 0.01 |
| 5 | 145953954 | rs1159045 | 2.758 | 0.689 | 2.128 | 1.208 | -0.005 | -0.012 | -0.012 | 0 | 0.61 | 0.35 | 0.64 | 0.55 | 0.64 | 0.66 |
| 5 | 145957498 | rs1363745 | 1.736 | 0.725 | 2.161 | 1.336 | 0.006 | 0.015 | 0.015 | 0 | 0.74 | 0.7 | 0.72 | 0.56 | 0.71 | 0.76 |
| 5 | 145978224 | rs3096089 | 1.924 | 0.738 | 2.31 | 1.358 | 0.021 | 0.044 | 0.044 | 0 | 0.74 | 0.66 | 0.72 | 0.56 | 0.7 | 0.76 |
| 5 | 145985870 | rs48625 | -2.471 | 0.676 | 2.279 | 1.794 | 0.004 | 0.159 | 0.159 | 0 | 0.08 | 0.3 | 0.14 | 0.23 | 0.03 | 0.11 |
| 5 | 145986083 | rs715891 | -1.903 | 0.791 | 2.049 | 1.789 | -0.001 | 0.027 | 0.027 | 0 | 0.07 | 0.08 | 0.14 | 0.22 | 0.06 | 0.15 |
| 5 | 146008385 | rs322487 | -2.627 | 0.373 | 2.359 | 1.748 | -0.06 | 0.064 | 0.064 | 0 | 0.18 | 0.49 | 0.21 | 0.22 | 0.09 | 0.2 |
| 5 | 146009812 | rs322485 | -2.562 | 0.439 | 2.224 | 1.807 | -0.004 | 0.188 | 0.188 | 0 | 0.09 | 0.35 | 0.15 | 0.22 | 0.03 | 0.14 |
| 5 | 146015615 | rs17448496 | -1.567 | 0.708 | 1.641 | 1.677 | -0.018 | -0.014 | -0.014 | 0 | 0.18 | 0.13 | 0.12 | 0.23 | 0.23 | 0.14 |
| 5 | 146025090 | rs1838336 | 1.766 | -0.249 | 2.224 | 1.581 | -0.021 | -0.02 | -0.02 | 0 | 0.79 | 0.62 | 0.86 | 0.82 | 0.85 | 0.85 |
| 5 | 146029906 | rs6580434 | 2.035 | -0.321 | 2.454 | 1.447 | -0.032 | -0.036 | -0.036 | 0.159 | 0.75 | 0.53 | 0.77 | 0.81 | 0.83 | 0.79 |
| 5 | 146033182 | rs404167 | 2.542 | -0.848 | 2.267 | 1.475 | -0.025 | 0.013 | 0.013 | 0 | 0.68 | 0.5 | 0.7 | 0.62 | 0.79 | 0.71 |
| 5 | 146033408 | rs403151 | 1.906 | -0.571 | 2.156 | 1.337 | -0.006 | -0.044 | -0.044 | 0 | 0.73 | 0.59 | 0.74 | 0.82 | 0.78 | 0.81 |
| 5 | 146034535 | rs460413 | 1.372 | -0.751 | 2.587 | 1.68 | 0.003 | 0.029 | 0.029 | 0 | 0.86 | 0.71 | 0.92 | 0.83 | 0.91 | 0.9 |
| 5 | 146034749 | rs460881 | 2.334 | -0.851 | 2.337 | 1.654 | -0.039 | 0.051 | 0.051 | 0 | 0.74 | 0.49 | 0.75 | 0.62 | 0.86 | 0.76 |
| 5 | 146048377 | rs17105086 | -2.477 | -0.947 | 2.35 | 1.681 | -0.024 | 0.045 | 0.045 | 0 | 0.21 | 0.3 | 0.16 | 0.36 | 0.12 | 0.2 |
| 5 | 146049511 | rs1833773 | -1.182 | -0.957 | 2.378 | 1.691 | 0.013 | -0.015 | -0.015 | 0 | 0.11 | 0.07 | 0.09 | 0.2 | 0.04 | 0.1 |
| 5 | 146052888 | rs458725 | 1.919 | -0.917 | 1.676 | 1.552 | -0.043 | 0.053 | 0.053 | 0 | 0.76 | 0.51 | 0.82 | 0.65 | 0.86 | 0.81 |
| 5 | 146055403 | rs661408 | 1.945 | -0.914 | 1.653 | 1.611 | -0.045 | 0.069 | 0.069 | 0 | 0.75 | 0.51 | 0.75 | 0.61 | 0.86 | 0.77 |
| 5 | 146062112 | rs386039 | -2.084 | -0.919 | 1.442 | 1.586 | -0.032 | 0.034 | 0.034 | 0.016 | 0.25 | 0.58 | 0.19 | 0.36 | 0.14 | 0.2 |
| 5 | 146068662 | rs7736604 | -1.601 | -1.188 | 0.87 | 1.468 | 0.005 | -0.001 | -0.001 | 0.016 | 0.34 | 0.28 | 0.36 | 0.63 | 0.23 | 0.33 |
| 5 | 146069598 | rs2400228 | -2.233 | -1.663 | 0.034 | 0.957 | -0.05 | 0.277 | 0.277 | 0.003 | 0.56 | 0.97 | 0.56 | 0.9 | 0.41 | 0.49 |

| | | | | | | | | | | | | | | | | |
|---|---|---|---|---|---|---|---|---|---|---|---|---|---|---|---|---|
| 5 | 146069910 | rs10038136 | -1.623 | -1.57 | 1.548 | 1.691 | -0.035 | 0.155 | 0.155 | 0.541 | 0.12 | 0.47 | 0.13 | 0.26 | 0.08 | 0.1 |
| 5 | 146077526 | rs6879694 | -1.866 | -1.226 | 1.858 | 2.335 | 0.02 | 0.079 | 0.079 | 0 | 0.13 | 0.25 | 0.18 | 0.27 | 0.14 | 0.17 |
| 5 | 146080012 | rs7716683 | -2.599 | -1.351 | 2.188 | 2.329 | -0.003 | 0.078 | 0.078 | 0 | 0.18 | 0.52 | 0.21 | 0.29 | 0.15 | 0.18 |
| 5 | 146083136 | rs10477307 | -2.148 | -1.27 | 2.165 | 2.327 | 0.041 | 0.132 | 0.132 | 0.15 | 0.15 | 0.49 | 0.21 | 0.27 | 0.15 | 0.17 |
| 5 | 146085978 | rs1194995 | -1.139 | -1.5 | 1.345 | 2.243 | 0.017 | -0.018 | -0.018 | 0 | 0.06 | 0.01 | 0.02 | 0.15 | 0.01 | 0.07 |
| 5 | 146094785 | rs3851489 | -1.76 | -1.373 | 1.604 | 2.666 | 0.004 | 0.042 | 0.042 | 0 | 0.15 | 0.2 | 0.22 | 0.37 | 0.19 | 0.17 |
| 5 | 146099634 | rs1194998 | -2.927 | -2.035 | 1.765 | 2.674 | 0.045 | 0.076 | 0.076 | 0 | 0.41 | 0.72 | 0.42 | 0.81 | 0.29 | 0.4 |
| 5 | 146104452 | rs13340393 | -2.09 | -1.826 | 1.892 | 2.71 | 0.006 | 0.102 | 0.102 | 0 | 0.18 | 0.36 | 0.23 | 0.38 | 0.19 | 0.18 |
| 5 | 146105578 | rs3844538 | -0.896 | -1.798 | 1.577 | 2.902 | 0.009 | -0.097 | -0.097 | 0.54 | 0.13 | 0.01 | 0.14 | 0.3 | 0.08 | 0.17 |
| 5 | 146110880 | rs7446531 | -1.754 | -1.746 | 1.881 | 2.919 | -0.007 | 0.092 | 0.092 | 0 | 0.09 | 0.12 | 0.15 | 0.39 | 0.06 | 0.14 |
| 5 | 146111707 | rs3910815 | -2.689 | -1.771 | 2.342 | 2.89 | -0.011 | 0.231 | 0.231 | 0.197 | 0.13 | 0.43 | 0.2 | 0.41 | 0.09 | 0.15 |
| 5 | 146117992 | rs17105326 | -0.981 | -1.826 | 2.391 | 3.109 | -0.006 | 0.004 | 0.004 | 0 | 0.16 | 0.17 | 0.15 | 0.19 | 0.1 | 0.19 |
| 5 | 146118962 | rs11167944 | 2.356 | -1.69 | 2.672 | 3.191 | -0.007 | 0.008 | 0.008 | 0 | 0.69 | 0.36 | 0.63 | 0.43 | 0.73 | 0.66 |
| 5 | 146125038 | rs17105330 | -2.471 | -1.621 | 2.797 | 3.202 | 0.031 | 0.184 | 0.184 | 0 | 0.12 | 0.33 | 0.22 | 0.4 | 0.15 | 0.16 |
| 5 | 146126203 | rs11949312 | -1.787 | -1.637 | 2.797 | 2.969 | 0.005 | -0.051 | -0.051 | 0 | 0.09 | 0.21 | 0.07 | 0 | 0.09 | 0.04 |
| 5 | 146126245 | rs7715375 | 1.156 | -1.73 | 2.757 | 3.041 | -0.012 | 0.006 | 0.006 | 0 | 0.82 | 0.76 | 0.85 | 0.8 | 0.88 | 0.81 |
| 5 | 146128342 | rs4091215 | 2.555 | -1.621 | 2.549 | 3.124 | 0.007 | -0.001 | -0.001 | 0 | 0.64 | 0.31 | 0.58 | 0.38 | 0.7 | 0.62 |
| 5 | 146150857 | rs249907 | 2.031 | -1.717 | 2.54 | 3.059 | -0.034 | -0.041 | -0.041 | 0.723 | 0.7 | 0.37 | 0.76 | 0.88 | 0.8 | 0.78 |
| 5 | 146153055 | rs249904 | 2.431 | -1.709 | 2.342 | 3.008 | 0.012 | -0.012 | -0.012 | 5.397 | 0.63 | 0.29 | 0.58 | 0.46 | 0.71 | 0.62 |
| 5 | 146165753 | rs161034 | 2.401 | -2.132 | 2.155 | 2.627 | 0.009 | -0.005 | -0.005 | 0 | 0.6 | 0.33 | 0.58 | 0.44 | 0.71 | 0.61 |
| 5 | 146166378 | rs161036 | 1.532 | -2.121 | 2.206 | 2.361 | -0.044 | -0.059 | -0.059 | 0 | 0.73 | 0.59 | 0.78 | 0.89 | 0.8 | 0.8 |
| 5 | 146169511 | rs160974 | 2.177 | -1.826 | 1.682 | 2.668 | 0.08 | 0.049 | 0.049 | 1.676 | 0.55 | 0.16 | 0.54 | 0.33 | 0.67 | 0.45 |
| 5 | 146175331 | rs11748900 | 2.454 | -1.669 | 1.117 | 2.533 | 0.043 | 0.059 | 0.059 | 1.633 | 0.52 | 0.04 | 0.5 | 0.32 | 0.63 | 0.45 |
| 5 | 146184132 | rs161040 | 0.285 | -1.631 | 1.331 | 2.555 | 0.017 | 0.01 | 0.01 | 0 | 0.87 | 0.94 | 0.86 | 0.9 | 0.9 | 0.85 |
| 5 | 146184951 | rs161041 | -2.3 | -1.461 | 1.752 | 2.689 | 0.071 | 0.079 | 0.079 | 0 | 0.06 | 0.18 | 0.13 | 0.14 | 0.11 | 0.08 |
| 5 | 146197381 | rs12513869 | -1.526 | -1.637 | 1.62 | 2.695 | 0.004 | -0.006 | -0.006 | 0 | 0.22 | 0.23 | 0.26 | 0.43 | 0.13 | 0.29 |
| 5 | 146201876 | rs10067441 | -0.805 | -1.623 | 1.532 | 2.738 | 0.034 | -0.059 | -0.059 | 0.41 | 0.17 | 0.08 | 0.23 | 0.41 | 0.1 | 0.26 |
| 5 | 146205786 | rs9325024 | -1.189 | -1.655 | 1.525 | 2.711 | 0.048 | -0.073 | -0.073 | 1 | 0.2 | 0.1 | 0.23 | 0.44 | 0.1 | 0.3 |
| 5 | 146208681 | rs1835951 | 2.629 | -1.611 | 2.02 | 2.67 | 0.031 | 0.038 | 0.038 | 0.311 | 0.53 | 0.34 | 0.46 | 0.29 | 0.62 | 0.45 |
| 5 | 146215708 | rs160971 | -1.504 | -1.226 | 2.222 | 2.791 | 0.057 | 0.103 | 0.103 | 0 | 0.06 | 0.2 | 0.11 | 0.17 | 0.1 | 0.07 |
| 5 | 146218226 | rs173638 | -2.621 | -1.24 | 2.236 | 2.782 | 0.017 | 0.029 | 0.029 | 0 | 0.11 | 0.22 | 0.13 | 0.17 | 0.13 | 0.12 |
| 5 | 146218243 | rs452399 | -2.109 | -1.226 | 2.209 | 2.803 | -0.033 | 0.03 | 0.03 | 0 | 0.31 | 0.47 | 0.37 | 0.61 | 0.22 | 0.39 |
| 5 | 146222814 | rs10515577 | -1.165 | -1.596 | 1.881 | 2.805 | 0.034 | -0.032 | -0.032 | 0 | 0.22 | 0.16 | 0.2 | 0.35 | 0.09 | 0.3 |
| 5 | 146225178 | rs609325 | -2.26 | -1.445 | 2.523 | 2.616 | 0.075 | 0.012 | 0.012 | 0 | 0.05 | 0.28 | 0.06 | 0 | 0.06 | 0.02 |
| 5 | 146225244 | rs609412 | -2.299 | -1.464 | 2.596 | 2.851 | 0.007 | -0.007 | -0.007 | 0 | 0.33 | 0.64 | 0.34 | 0.57 | 0.2 | 0.39 |
| 5 | 146229428 | rs2125443 | -1.086 | -1.421 | 2.512 | 2.714 | -0.001 | 0.009 | 0.009 | 0 | 0.17 | 0.2 | 0.19 | 0.34 | 0.18 | 0.22 |
| 5 | 146230996 | rs4515335 | -2.492 | -1.433 | 2.836 | 2.862 | 0.067 | 0.081 | 0.081 | 0.339 | 0.45 | 0.73 | 0.49 | 0.69 | 0.33 | 0.54 |
| 5 | 146233639 | rs10477313 | -0.922 | -1.496 | 3.079 | 2.888 | -0.039 | 0.036 | 0.036 | 0 | 0.21 | 0.27 | 0.21 | 0.33 | 0.1 | 0.27 |
| 5 | 146242647 | rs17105535 | -1.34 | -1.498 | 3.152 | 2.899 | -0.001 | 0.013 | 0.013 | 0 | 0.2 | 0.21 | 0.25 | 0.34 | 0.21 | 0.26 |
| 5 | 146242996 | rs7718587 | -1.661 | -1.498 | 3.189 | 2.973 | -0.001 | 0.012 | 0.012 | 0 | 0.23 | 0.29 | 0.27 | 0.34 | 0.24 | 0.27 |
| 5 | 146249101 | rs10037715 | -1.544 | -1.698 | 2.694 | 2.357 | 0.004 | -0.005 | -0.005 | 1.109 | 0.33 | 0.27 | 0.32 | 0.34 | 0.31 | 0.28 |
| 5 | 146255753 | rs319215 | -1.239 | -1.085 | 3.218 | 2.844 | -0.069 | 0.084 | 0.084 | 0 | 0.23 | 0.41 | 0.23 | 0.35 | 0.11 | 0.29 |
| 5 | 146258596 | rs2082405 | -2.506 | -1.091 | 3.153 | 2.649 | 0.003 | 0.003 | 0.003 | 0.106 | 0.44 | 0.54 | 0.48 | 0.68 | 0.34 | 0.54 |
| 5 | 146262489 | rs319204 | -2.418 | -1.084 | 2.774 | 2.639 | 0.055 | 0.065 | 0.065 | 0.468 | 0.46 | 0.69 | 0.49 | 0.69 | 0.34 | 0.55 |
| 5 | 146272403 | rs319185 | -2.536 | -0.746 | 2.536 | 2.63 | 0.04 | 0.056 | 0.056 | 0.433 | 0.44 | 0.69 | 0.55 | 0.68 | 0.34 | 0.52 |
| 5 | 146282105 | rs9325026 | -1.669 | -0.831 | 2.746 | 2.826 | 0 | 0.021 | 0.021 | 0.025 | 0.23 | 0.26 | 0.34 | 0.38 | 0.24 | 0.26 |
| 5 | 146286561 | rs2963075 | -0.836 | -0.842 | 2.829 | 2.828 | 0.004 | -0.006 | -0.006 | 1.318 | 0.19 | 0.19 | 0.19 | 0.31 | 0.09 | 0.26 |
| 5 | 146287183 | rs2436401 | -2.629 | -0.817 | 2.546 | 2.906 | 0.037 | 0.03 | 0.03 | 0.492 | 0.43 | 0.67 | 0.52 | 0.64 | 0.33 | 0.53 |
| 5 | 146291010 | rs319168 | -2.614 | -0.873 | 2.48 | 2.899 | 0.074 | 0.101 | 0.101 | 0.602 | 0.46 | 0.78 | 0.56 | 0.7 | 0.33 | 0.54 |
| 5 | 146295990 | rs178549 | -2.584 | -0.84 | 2.486 | 2.915 | 0.069 | 0.104 | 0.104 | 0.529 | 0.45 | 0.81 | 0.56 | 0.7 | 0.34 | 0.54 |
| 5 | 146302106 | rs10068414 | -1.242 | -0.829 | 2.611 | 2.97 | -0.001 | 0.033 | 0.033 | 0.012 | 0.2 | 0.21 | 0.32 | 0.37 | 0.21 | 0.26 |
| 5 | 146305665 | rs319142 | -2.764 | -0.855 | 2.552 | 2.968 | 0.045 | 0.063 | 0.063 | 0.164 | 0.44 | 0.71 | 0.55 | 0.7 | 0.33 | 0.55 |
| 5 | 146311042 | rs6878648 | -1.311 | -0.971 | 2.6 | 3.102 | -0.005 | 0.029 | 0.029 | 0.002 | 0.2 | 0.24 | 0.32 | 0.38 | 0.19 | 0.27 |
| 5 | 146315393 | rs2004709 | -1.547 | -0.976 | 2.596 | 3.091 | -0.012 | 0.066 | 0.066 | 0.115 | 0.22 | 0.31 | 0.35 | 0.44 | 0.19 | 0.28 |
| 5 | 146320823 | rs1864982 | -1.193 | -0.952 | 2.436 | 3.069 | -0.041 | 0.064 | 0.064 | 0 | 0.19 | 0.41 | 0.2 | 0.26 | 0.12 | 0.21 |
| 5 | 146334385 | rs1125900 | -0.871 | -0.833 | 1.798 | 3.013 | -0.011 | -0.089 | -0.089 | 2.711 | 0.16 | 0.03 | 0.28 | 0.38 | 0.16 | 0.24 |
| 5 | 146336387 | rs6580446 | 2.406 | -0.744 | 1.95 | 2.66 | 0.04 | 0.063 | 0.063 | 0.245 | 0.57 | 0.26 | 0.51 | 0.28 | 0.67 | 0.47 |
| 5 | 146339841 | rs4705449 | 2.176 | -0.233 | 2.292 | 2.739 | -0.011 | -0.018 | -0.018 | 0 | 0.62 | 0.5 | 0.55 | 0.33 | 0.63 | 0.5 |
| 5 | 146344847 | rs17105791 | -0.945 | -0.269 | 2.284 | 3.006 | -0.031 | -0.056 | -0.056 | 0 | 0.16 | 0.08 | 0.32 | 0.34 | 0.24 | 0.2 |
| 5 | 146347495 | rs13182712 | 2.404 | -0.268 | 2.377 | 3.035 | 0.025 | 0.062 | 0.062 | 0.313 | 0.59 | 0.26 | 0.5 | 0.24 | 0.63 | 0.49 |
| 5 | 146351658 | rs2400235 | 1.137 | 0.01 | 2.407 | 3.101 | -0.017 | 0.117 | 0.117 | 0 | 0.78 | 0.58 | 0.85 | 0.66 | 0.82 | 0.71 |
| 5 | 146370029 | rs10077240 | 0.822 | 0.036 | 2.346 | 3.18 | -0.01 | -0.021 | -0.021 | 1.295 | 0.84 | 0.89 | 0.68 | 0.71 | 0.76 | 0.81 |
| 5 | 146374689 | rs1383168 | 0.702 | 0.09 | 2.291 | 3.205 | -0.025 | -0.04 | -0.04 | 0 | 0.86 | 0.93 | 0.68 | 0.71 | 0.76 | 0.81 |
| 5 | 146375787 | rs10875618 | -1.393 | 0.087 | 2.43 | 3.398 | -0.018 | 0.078 | 0.078 | 0 | 0.22 | 0.42 | 0.15 | 0.29 | 0.18 | 0.29 |
| 5 | 146407693 | rs7713487 | 2.02 | 0.19 | 1.996 | 3.316 | 0 | -0.001 | -0.001 | 2.909 | 0.65 | 0.41 | 0.53 | 0.36 | 0.59 | 0.51 |
| 5 | 146417802 | rs13183391 | 1.59 | 0.19 | 1.821 | 3.344 | 0.029 | 0.183 | 0.183 | 0.586 | 0.8 | 0.52 | 0.59 | 0.54 | 0.74 | 0.79 |
| 5 | 146418746 | rs1480158 | -0.889 | 0.335 | 0.623 | 3.34 | 0.004 | -0.053 | -0.053 | 4.482 | 0.16 | 0.05 | 0.09 | 0.3 | 0.16 | 0.29 |
| 5 | 146429843 | rs1031915 | -1.199 | 0.623 | 0.838 | 3.216 | -0.062 | -0.082 | -0.082 | 0 | 0.14 | 0.02 | 0.31 | 0.33 | 0.22 | 0.19 |
| 5 | 146434800 | rs1383169 | -1.824 | 0.128 | 1.408 | 2.869 | -0.048 | 0.197 | 0.197 | 1.133 | 0.17 | 0.54 | 0.2 | 0.42 | 0.08 | 0.23 |

| | | | | | | | | | | | | | | | |
|---|---|---|---|---|---|---|---|---|---|---|---|---|---|---|---|
| 5 | 146435163 | rs11958187 | 1.832 | 1.03 | 1.213 | 2.502 | 0.001 | -0.015 | -0.015 | 0 | 0.66 | 0.38 | 0.54 | 0.32 | 0.62 | 0.57 |
| 5 | 146436523 | rs11167957 | 1.899 | 1.17 | 1.148 | 2.35 | 0.008 | -0.016 | -0.016 | 0.004 | 0.67 | 0.4 | 0.54 | 0.32 | 0.61 | 0.58 |
| 5 | 146438035 | rs6580448 | 1.51 | 1.129 | 1.151 | 2.5 | 0.018 | 0.067 | 0.067 | 0.006 | 0.73 | 0.61 | 0.59 | 0.48 | 0.67 | 0.59 |
| 5 | 146448551 | rs1480149 | 1.376 | 1.359 | 0.978 | 2.44 | 0.039 | 0.059 | 0.059 | 0 | 0.74 | 0.56 | 0.58 | 0.46 | 0.62 | 0.65 |
| 5 | 146457870 | rs1156700 | 1.363 | 1.375 | 1.068 | 2.501 | -0.014 | -0.025 | -0.025 | 0.395 | 0.74 | 0.76 | 0.58 | 0.5 | 0.62 | 0.61 |
| 5 | 146460474 | rs1480151 | 1.564 | 1.478 | 1.074 | 2.579 | 0.04 | 0.076 | 0.076 | 0 | 0.74 | 0.56 | 0.56 | 0.5 | 0.61 | 0.64 |
| 5 | 146472853 | rs6895471 | -1.663 | 1.043 | 0.783 | 2.512 | 0.02 | 0.114 | 0.114 | 0.586 | 0.08 | 0.26 | 0.07 | 0.17 | 0.06 | 0.08 |
| 5 | 146475879 | rs1480159 | 0.992 | 1.302 | -0.05 | 2.59 | -0.005 | -0.022 | -0.022 | 0 | 0.8 | 0.84 | 0.68 | 0.64 | 0.79 | 0.69 |
| 5 | 146475990 | rs1480160 | 0.835 | 1.281 | -0.06 | 2.46 | -0.005 | -0.025 | -0.025 | 0.518 | 0.8 | 0.82 | 0.67 | 0.56 | 0.78 | 0.68 |
| 5 | 146482041 | rs6880853 | 0.926 | 0.786 | -1.34 | 0.93 | -0.104 | 0.114 | 0.114 | 0.183 | 0.44 | 0.04 | 0.47 | 0.3 | 0.58 | 0.49 |
| 5 | 146483985 | rs6867092 | 0.973 | 0.859 | -0.47 | 0.698 | 0.017 | -0.022 | -0.022 | 0 | 0.17 | 0.28 | 0.13 | 0.17 | 0.2 | 0.11 |
| 5 | 146485682 | rs728937 | 1.253 | 0.323 | -1.328 | 0.219 | -0.078 | 0.068 | 0.068 | 1.06 | 0.42 | 0.05 | 0.48 | 0.32 | 0.55 | 0.54 |
| 5 | 146492274 | rs989817 | 0.046 | -0.05 | -0.638 | 0.136 | 0.034 | -0.06 | -0.06 | 9.887 | 0.69 | 0.84 | 0.66 | 0.5 | 0.77 | 0.69 |
| 5 | 146494880 | rs12659167 | -0.463 | 0.033 | -0.232 | 0.215 | 0.008 | 0.002 | 0.002 | 0 | 0.06 | 0.21 | 0.03 | 0.04 | 0.01 | 0.04 |
| 5 | 146496726 | rs1480163 | 1.575 | 0.071 | -0.054 | 0.239 | 0.012 | -0.018 | -0.018 | 0 | 0.17 | 0.25 | 0.12 | 0.17 | 0.2 | 0.11 |
| 5 | 146512747 | rs12374531 | 1.059 | 0.045 | 0.127 | 0.283 | 0.012 | -0.014 | -0.014 | 0 | 0.2 | 0.38 | 0.13 | 0.21 | 0.2 | 0.12 |
| 5 | 146523279 | rs13360969 | 1.625 | 0.034 | -0.085 | 0.29 | 0.015 | -0.013 | -0.013 | 0.004 | 0.17 | 0.25 | 0.12 | 0.17 | 0.2 | 0.11 |
| 5 | 146527326 | rs11957305 | 0.57 | -0.184 | -0.52 | 0.395 | -0.006 | -0.14 | -0.14 | 0 | 0.17 | 0.01 | 0.26 | 0.44 | 0.11 | 0.25 |
| 5 | 146527589 | rs10079730 | 0.432 | -0.12 | -0.054 | 0.093 | -0.009 | -0.02 | -0.02 | 0 | 0.24 | 0.65 | 0.15 | 0.21 | 0.21 | 0.14 |
| 5 | 146538989 | rs11949165 | -0.423 | 0.229 | -1.507 | 0.053 | -0.015 | 0.029 | 0.029 | 0 | 0.39 | 0.32 | 0.44 | 0.7 | 0.39 | 0.41 |
| 5 | 146539138 | rs11958839 | -0.015 | 0.311 | -1.699 | 0.319 | 0.009 | -0.017 | -0.017 | 0 | 0.2 | 0.21 | 0.3 | 0.55 | 0.14 | 0.3 |
| 5 | 146540899 | rs2913326 | 0.72 | 0.561 | -2.008 | 0.059 | -0.067 | 0.046 | 0.046 | 0.518 | 0.41 | 0.04 | 0.45 | 0.29 | 0.47 | 0.5 |
| 5 | 146553369 | rs40474 | -0.506 | 0.628 | -0.917 | 0.039 | 0.001 | 0.01 | 0.01 | 0.023 | 0.71 | 0.72 | 0.76 | 0.83 | 0.68 | 0.87 |
| 5 | 146554234 | rs10075422 | 1.282 | 0.489 | -1.342 | 0.877 | 0.006 | 0.003 | 0.003 | 0 | 0.06 | 0.02 | 0.05 | 0.29 | 0.04 | 0.09 |
| 5 | 146554363 | rs30950 | 0.204 | 0.953 | -1.509 | 0.971 | -0.013 | -0.029 | -0.029 | 0 | 0.17 | 0.11 | 0.12 | 0.3 | 0.18 | 0.22 |
| 5 | 146554431 | rs4705021 | 0.459 | 1.179 | -1.32 | 0.499 | -0.002 | 0 | 0 | 0.271 | 0.25 | 0.25 | 0.49 | 0.28 | 0.36 | 0.42 |
| 5 | 146554649 | rs30949 | -0.247 | 0.793 | -0.926 | 0.612 | -0.007 | -0.006 | -0.006 | 0 | 0.47 | 0.49 | 0.29 | 0.58 | 0.31 | 0.45 |
| 5 | 146556930 | rs1119704 | 0.478 | 1.846 | -1.259 | -0.068 | 0.01 | 0.014 | 0.014 | 0 | 0.77 | 0.71 | 0.81 | 0.66 | 0.65 | 0.78 |
| 5 | 146559069 | rs40473 | 0.023 | 1.709 | -1.535 | -0.185 | -0.033 | 0.041 | 0.041 | 0.749 | 0.92 | 0.99 | 0.94 | 1 | 0.89 | 0.95 |
| 5 | 146561416 | rs17582364 | 0.283 | 1.35 | -1.64 | -0.006 | -0.028 | -0.037 | -0.037 | 0.008 | 0.16 | 0.01 | 0.25 | 0.21 | 0.15 | 0.43 |
| 5 | 146566438 | rs10037981 | -0.006 | 1.411 | -1.268 | -0.22 | 0.009 | 0.016 | 0.016 | 0 | 0.24 | 0.32 | 0.41 | 0.36 | 0.35 | 0.19 |
| 5 | 146571012 | rs890742 | 1.076 | 1.283 | -1.271 | -0.046 | 0 | 0.012 | 0.012 | 0 | 0.34 | 0.27 | 0.21 | 0.05 | 0.39 | 0.15 |
| 5 | 146578854 | rs30983 | 0.459 | 1.377 | -1.507 | 0.398 | -0.013 | 0.001 | 0.001 | 0.023 | 0.63 | 0.74 | 0.56 | 0.36 | 0.63 | 0.65 |
| 5 | 146580239 | rs2053030 | 0.111 | 1.086 | -1.386 | -0.565 | -0.002 | -0.01 | -0.01 | 0 | 0.23 | 0.17 | 0.31 | 0.31 | 0.23 | 0.19 |
| 5 | 146587911 | rs30979 | 0.186 | 1.196 | -1.046 | -0.573 | -0.005 | -0.098 | -0.098 | 0 | 0.42 | 0.5 | 0.29 | 0.05 | 0.45 | 0.18 |
| 5 | 146591566 | rs31021 | 0.175 | 1.102 | -1.018 | -0.545 | -0.006 | -0.096 | -0.096 | 0 | 0.43 | 0.5 | 0.29 | 0.05 | 0.44 | 0.18 |
| 5 | 146610003 | rs4705024 | -0.497 | 1.377 | -1.269 | 0.567 | -0.014 | -0.019 | -0.019 | 1.249 | 0.26 | 0.22 | 0.38 | 0.36 | 0.34 | 0.25 |
| 5 | 146615260 | rs31036 | -0.009 | 1.167 | -1.624 | 0.512 | -0.012 | -0.002 | -0.002 | 0 | 0.68 | 0.77 | 0.71 | 0.64 | 0.67 | 0.56 |
| 5 | 146617405 | rs31039 | 0.293 | 0.924 | -2.365 | 0.641 | -0.021 | -0.081 | -0.081 | 0 | 0.2 | 0.03 | 0.12 | 0.33 | 0.2 | 0.23 |
| 5 | 146618078 | rs17590508 | 0.607 | 0.317 | -2.542 | 0.213 | -0.044 | 0.031 | 0.031 | 0 | 0.28 | 0.12 | 0.43 | 0.19 | 0.36 | 0.27 |
| 5 | 146620082 | rs891898 | -0.45 | 0.237 | -1.967 | -0.235 | -0.007 | 0.018 | 0.018 | 0.201 | 0.57 | 0.64 | 0.73 | 0.31 | 0.56 | 0.38 |
| 5 | 146623577 | rs4705028 | 0.433 | -0.057 | -1.108 | -0.382 | 0.001 | -0.009 | -0.009 | 0 | 0.37 | 0.41 | 0.49 | 0.26 | 0.41 | 0.3 |
| 5 | 146625346 | rs17106248 | 0.489 | -0.061 | -1.324 | -0.389 | -0.007 | -0.03 | -0.03 | 0 | 0.19 | 0.22 | 0.24 | 0.05 | 0.15 | 0.07 |
| 5 | 146628427 | rs1368284 | 0.393 | -0.066 | -1.327 | -0.392 | -0.008 | -0.031 | -0.031 | 0.091 | 0.19 | 0.23 | 0.24 | 0.05 | 0.15 | 0.07 |
| 5 | 146630049 | rs10515579 | 0.919 | -0.029 | -1.373 | -0.352 | -0.026 | -0.083 | -0.083 | 0 | 0.19 | 0.03 | 0.11 | 0.35 | 0.2 | 0.21 |
| 5 | 146630091 | rs4705140 | -0.103 | 0.152 | -1.689 | -0.574 | 0.002 | 0.006 | 0.006 | 0 | 0.3 | 0.25 | 0.3 | 0.13 | 0.26 | 0.19 |
| 5 | 146630580 | rs7701791 | 1.39 | 0.324 | -1.642 | -0.44 | -0.023 | -0.02 | -0.02 | 0 | 0.18 | 0.11 | 0.37 | 0.19 | 0.26 | 0.15 |
| 5 | 146632594 | rs11167971 | -1.723 | 0.375 | -1.566 | -0.456 | -0.027 | -0.045 | -0.045 | 0 | 0.77 | 0.9 | 0.86 | 0.66 | 0.77 | 0.58 |
| 5 | 146632842 | rs12654852 | 0.982 | 0.329 | -1.257 | -0.556 | -0.001 | -0.002 | -0.002 | 2.364 | 0.11 | 0.11 | 0.07 | 0.07 | 0.11 | 0.12 |
| 5 | 146635260 | rs11167975 | -0.393 | 0.227 | -0.88 | -0.557 | -0.002 | -0.011 | -0.011 | 0 | 0.34 | 0.36 | 0.36 | 0.15 | 0.3 | 0.21 |
| 5 | 146641238 | rs4705034 | 0.79 | 0.477 | -0.576 | -0.402 | 0.017 | 0.004 | 0.004 | 0 | 0.2 | 0.4 | 0.39 | 0.17 | 0.26 | 0.15 |
| 5 | 146641615 | rs6890102 | -1.671 | 0.406 | -0.728 | -0.209 | -0.028 | -0.048 | -0.048 | 0 | 0.78 | 0.91 | 0.86 | 0.66 | 0.77 | 0.57 |
| 5 | 146642893 | rs1432648 | 0.681 | 0.242 | -0.488 | -0.536 | -0.005 | -0.052 | -0.052 | 0 | 0.17 | 0.16 | 0.17 | 0 | 0.13 | 0.08 |
| 5 | 146643572 | rs2895679 | 0.47 | 0.516 | -0.441 | -0.063 | 0.02 | 0.007 | 0.007 | 0 | 0.2 | 0.4 | 0.38 | 0.17 | 0.26 | 0.15 |
| 5 | 146647500 | rs17106354 | 0.242 | 0.428 | -0.792 | 0.08 | 0.002 | -0.01 | -0.01 | 0.263 | 0.11 | 0.06 | 0.07 | 0.19 | 0.1 | 0.12 |
| 5 | 146649327 | rs4307135 | -0.016 | 0.708 | -0.938 | -0.469 | 0.008 | 0.054 | 0.054 | 0 | 0.49 | 0.69 | 0.67 | 0.65 | 0.56 | 0.33 |
| 5 | 146652132 | rs6877196 | -0.349 | 0.786 | -1.034 | -0.637 | -0.011 | 0.035 | 0.035 | 0 | 0.58 | 0.79 | 0.69 | 0.66 | 0.62 | 0.41 |
| 5 | 146652943 | rs987651 | 0.615 | 0.775 | -0.995 | -0.67 | 0.001 | 0.018 | 0.018 | 0 | 0.13 | 0.12 | 0.16 | 0.54 | 0.16 | 0.16 |
| 5 | 146655511 | rs17468085 | 1.197 | 0.729 | -1.134 | -0.79 | 0.001 | -0.021 | -0.021 | 0 | 0.09 | 0.1 | 0.02 | 0 | 0.06 | 0.08 |
| 5 | 146656123 | rs1368285 | -2.159 | 0.667 | -1.333 | -0.817 | -0.036 | -0.045 | -0.045 | 0.551 | 0.81 | 0.97 | 0.89 | 0.74 | 0.83 | 0.59 |
| 5 | 146657838 | rs6580459 | -0.315 | 0.238 | -0.839 | -0.568 | 0.001 | 0.01 | 0.01 | 0 | 0.43 | 0.68 | 0.6 | 0.61 | 0.5 | 0.3 |
| 5 | 146671021 | rs7724577 | -0.047 | 0.474 | -0.799 | -0.563 | -0.021 | 0.002 | 0.002 | 0 | 0.35 | 0.54 | 0.52 | 0.62 | 0.31 | 0.28 |
| 5 | 146671385 | rs2161747 | 1.141 | 0.428 | -1.033 | -0.688 | -0.007 | 0.017 | 0.017 | 0 | 0.39 | 0.27 | 0.2 | 0.31 | 0.34 | 0.63 |
| 5 | 146675710 | rs4705158 | 0.804 | 0.755 | -1.161 | -1.178 | 0.002 | -0.008 | -0.008 | 0 | 0.14 | 0.08 | 0.14 | 0 | 0.14 | 0.05 |
| 5 | 146676419 | rs11956052 | 0.17 | 0.926 | -0.844 | -0.502 | -0.013 | -0.078 | -0.078 | 0 | 0.23 | 0.36 | 0.36 | 0.05 | 0.18 | 0.13 |
| 5 | 146678112 | rs4705159 | 0.627 | 1.433 | -1.2 | 0.253 | -0.07 | 0.041 | 0.041 | 0.886 | 0.86 | 0.99 | 0.86 | 0.93 | 0.75 | 0.93 |
| 5 | 146685292 | rs4705036 | -0.858 | 0.302 | -0.564 | 0.269 | 0.008 | 0 | 0 | 0.081 | 0.21 | 0.19 | 0.21 | 0.21 | 0.16 | 0.15 |
| 5 | 146687393 | rs6880199 | -0.066 | 1.285 | -1.078 | 0.947 | -0.055 | -0.062 | -0.062 | 2.064 | 0.87 | 0.95 | 0.77 | 0.65 | 0.8 | 0.63 |
| 5 | 146687914 | rs1582611 | 1.311 | 1.081 | -0.887 | 0.932 | -0.006 | -0.04 | -0.04 | 4.375 | 0.66 | 0.74 | 0.54 | 0.46 | 0.65 | 0.49 |

| | | | | | | | | | | | | | | | | |
|---|---|---|---|---|---|---|---|---|---|---|---|---|---|---|---|---|
| 5 | 146689412 | rs12659800 | -0.133 | 0.913 | -0.403 | 0.781 | 0.008 | 0.142 | 0.142 | 0.177 | 0.07 | 0.21 | 0.03 | 0.22 | 0.03 | 0.02 |
| 5 | 146689428 | rs4705160 | 0.096 | 0.979 | -0.153 | 0.766 | 0.001 | 0.005 | 0.005 | 0 | 0.86 | 0.83 | 0.77 | 0.73 | 0.83 | 0.63 |
| 5 | 146690030 | rs6866338 | 1.202 | 0.82 | -0.282 | 0.068 | -0.018 | 0.025 | 0.025 | 0.432 | 0.28 | 0.23 | 0.31 | 0.13 | 0.4 | 0.15 |
| 5 | 146690712 | rs12652074 | 0.789 | 0.967 | 0.171 | 0.111 | 0.009 | -0.017 | -0.017 | 0 | 0.61 | 0.52 | 0.52 | 0.63 | 0.6 | 0.49 |
| 5 | 146695681 | rs2400293 | -0.178 | 0.815 | 0.424 | -0.088 | 0.002 | -0.027 | -0.027 | 0 | 0.24 | 0.28 | 0.24 | 0.1 | 0.23 | 0.15 |
| 5 | 146697763 | rs17547243 | 0.386 | 0.988 | 0.493 | 0.08 | 0.001 | -0.017 | -0.017 | 0.293 | 0.07 | 0.06 | 0.05 | 0 | 0.08 | 0.02 |
| 5 | 146701867 | rs17106430 | 0.225 | 0.725 | 0.85 | 0.075 | 0.04 | -0.014 | -0.014 | 0.065 | 0.06 | 0.26 | 0.04 | 0 | 0.04 | 0.01 |
| 5 | 146704598 | rs10064995 | -0.377 | 0.331 | 0.983 | -0.284 | -0.01 | -0.032 | -0.032 | 1.136 | 0.2 | 0.24 | 0.19 | 0.09 | 0.17 | 0.18 |
| 5 | 146707117 | rs7707452 | -1.266 | 0.3 | 1.047 | 0.196 | -0.022 | -0.007 | -0.007 | 0 | 0.28 | 0.53 | 0.31 | 0.27 | 0.21 | 0.2 |
| 5 | 146720355 | rs11955126 | -0.819 | 0.27 | 1.09 | 0.138 | -0.001 | -0.061 | -0.061 | 2.048 | 0.19 | 0.37 | 0.19 | 0.08 | 0.16 | 0.14 |
| 5 | 146722890 | rs6884181 | -1.308 | 0.233 | 1.219 | 0.168 | -0.006 | 0 | 0 | 0 | 0.25 | 0.66 | 0.28 | 0.26 | 0.18 | 0.18 |
| 5 | 146724726 | rs889829 | -0.655 | -0.016 | 0.582 | 0.143 | 0.027 | 0.053 | 0.053 | 0 | 0.52 | 0.67 | 0.6 | 0.6 | 0.38 | 0.47 |
| 5 | 146729164 | rs4705165 | -0.221 | 0.095 | 0.43 | 0.051 | -0.022 | 0.057 | 0.057 | 0 | 0.19 | 0.52 | 0.24 | 0.26 | 0.14 | 0.17 |
| 5 | 146736875 | rs12332485 | -0.315 | -0.071 | 0.348 | 0.054 | 0.004 | 0.049 | 0.049 | 0.008 | 0.21 | 0.42 | 0.23 | 0.27 | 0.16 | 0.18 |
| 5 | 146737467 | rs9325041 | -0.291 | -0.216 | 0.051 | -0.108 | 0.017 | 0.029 | 0.029 | 0.014 | 0.51 | 0.54 | 0.59 | 0.61 | 0.38 | 0.47 |
| 5 | 146753004 | rs4235731 | -0.354 | -0.149 | -0.225 | -0.201 | 0.008 | 0.014 | 0.014 | 0 | 0.49 | 0.53 | 0.59 | 0.85 | 0.38 | 0.5 |
| 5 | 146758154 | rs12657525 | 0.68 | -0.329 | -0.981 | 0.106 | 0.048 | -0.072 | -0.072 | 0 | 0.3 | 0.05 | 0.34 | 0.59 | 0.21 | 0.32 |
| 5 | 146759867 | rs37227 | -1.294 | -0.305 | -1.057 | -0.023 | -0.026 | -0.022 | -0.022 | 0 | 0.87 | 0.98 | 0.92 | 0.85 | 0.86 | 0.69 |
| 5 | 146767470 | rs729530 | 1.156 | -0.47 | -0.94 | 0.248 | 0.045 | -0.086 | -0.086 | 0.059 | 0.26 | 0.06 | 0.33 | 0.59 | 0.19 | 0.32 |
| 5 | 146774592 | rs3805535 | 0.068 | -0.146 | -0.186 | 0.117 | 0.012 | 0.006 | 0.006 | 0.07 | 0.17 | 0.3 | 0.25 | 0.18 | 0.15 | 0.14 |
| 5 | 146775475 | rs2288806 | -0.159 | -0.395 | -0.267 | 0.734 | 0.015 | -0.001 | -0.001 | 0 | 0.45 | 0.6 | 0.5 | 0.55 | 0.37 | 0.48 |
| 5 | 146779986 | rs3213858 | 0.353 | -0.547 | -0.684 | 1.242 | -0.013 | -0.063 | -0.063 | 0 | 0.17 | 0.03 | 0.11 | 0.25 | 0.16 | 0.34 |
| 5 | 146780521 | rs13168262 | 0.557 | -0.728 | 0.04 | 1.335 | 0.02 | -0.052 | -0.052 | 0.14 | 0.27 | 0.11 | 0.27 | 0.38 | 0.23 | 0.35 |
| 5 | 146810610 | rs10064194 | -0.178 | -0.992 | -0.175 | 0.993 | 0.04 | -0.015 | -0.015 | 0 | 0.41 | 0.65 | 0.36 | 0.56 | 0.22 | 0.41 |
| 5 | 146813845 | rs7725771 | -0.976 | -0.105 | -0.174 | 0.624 | 0.011 | -0.014 | -0.014 | 0 | 0.55 | 0.76 | 0.54 | 0.56 | 0.55 | 0.42 |
| 5 | 146814432 | rs10079318 | -0.443 | -0.064 | -0.143 | 0.454 | 0.017 | 0.014 | 0.014 | 0 | 0.51 | 0.38 | 0.46 | 0.45 | 0.54 | 0.39 |
| 5 | 146839364 | rs10072652 | 0.336 | -0.401 | -0.558 | -0.213 | -0.061 | -0.041 | -0.041 | 0 | 0.28 | 0.37 | 0.23 | 0.19 | 0.11 | 0.23 |
| 5 | 146846799 | rs17106769 | -0.288 | -0.324 | -0.352 | 0.002 | 0.001 | 0.022 | 0.022 | 0 | 0.46 | 0.41 | 0.44 | 0.21 | 0.51 | 0.32 |
| 5 | 146866283 | rs1368366 | -0.401 | -0.368 | -0.522 | 0.01 | 0.023 | 0.136 | 0.136 | 0.259 | 0.52 | 0.73 | 0.53 | 0.26 | 0.51 | 0.35 |
| 5 | 146881381 | rs4705040 | 0.554 | -0.267 | -0.429 | -0.237 | -0.002 | -0.002 | -0.002 | 0 | 0.17 | 0.19 | 0.19 | 0.11 | 0.14 | 0.11 |
| 5 | 146890363 | rs7706630 | -0.817 | -0.351 | -0.634 | -0.214 | 0.019 | 0.109 | 0.109 | 0.758 | 0.53 | 0.74 | 0.53 | 0.29 | 0.51 | 0.37 |
| 5 | 146896383 | rs6580467 | 0.259 | -0.334 | -0.652 | -0.393 | 0.019 | -0.033 | -0.033 | 0 | 0.31 | 0.36 | 0.33 | 0.17 | 0.37 | 0.26 |
| 5 | 146896655 | rs6860091 | 0.817 | -0.358 | -0.639 | -0.404 | 0.003 | 0.021 | 0.021 | 0 | 0.17 | 0.1 | 0.11 | 0.03 | 0.14 | 0.06 |
| 5 | 146896967 | rs2032856 | 0.14 | -0.384 | -0.851 | -0.279 | 0.019 | -0.033 | -0.033 | 0 | 0.31 | 0.36 | 0.33 | 0.17 | 0.37 | 0.26 |
| 5 | 146898773 | rs4235732 | -0.541 | -0.291 | -1.259 | 0.278 | -0.036 | -0.013 | -0.013 | 0 | 0.81 | 0.98 | 0.86 | 0.84 | 0.77 | 0.8 |
| 5 | 146906766 | rs17106850 | -0.451 | -0.685 | -0.193 | 0.782 | -0.009 | -0.008 | -0.008 | 0 | 0.23 | 0.46 | 0.22 | 0.17 | 0.17 | 0.1 |
| 5 | 146912107 | rs7716144 | -0.286 | -0.434 | -0.25 | 0.624 | 0.006 | -0.084 | -0.084 | 0.003 | 0.2 | 0.31 | 0.12 | 0.02 | 0.17 | 0.08 |
| 5 | 146934785 | rs2194156 | -0.386 | -1.135 | -1.096 | 1.006 | 0.013 | 0.009 | 0.009 | 0 | 0.41 | 0.33 | 0.48 | 0.67 | 0.3 | 0.51 |
| 5 | 146938734 | rs6881655 | -0.088 | -0.554 | -1.428 | 0.947 | 0.014 | -0.011 | -0.011 | 0.012 | 0.28 | 0.17 | 0.27 | 0.27 | 0.22 | 0.34 |
| 5 | 146944239 | rs10075595 | -1.267 | -0.145 | -1.101 | 1.404 | 0.007 | -0.036 | -0.036 | 0 | 0.23 | 0.16 | 0.23 | 0.34 | 0.18 | 0.27 |
| 5 | 146946871 | rs6867633 | -0.626 | -0.097 | -1.016 | 1.044 | 0.005 | 0.041 | 0.041 | 0 | 0.12 | 0.03 | 0.06 | 0.01 | 0.1 | 0.08 |
| 5 | 146951712 | rs7706398 | -0.635 | -0.395 | 0.407 | 2.106 | 0.003 | 0.024 | 0.024 | 0 | 0.08 | 0.11 | 0.14 | 0.16 | 0.08 | 0.15 |
| 5 | 146961115 | rs17106906 | -0.056 | -1.535 | -0.259 | 2.29 | 0.049 | -0.037 | -0.037 | 0.008 | 0.16 | 0.02 | 0.15 | 0.19 | 0.05 | 0.17 |
| 5 | 147014141 | rs17106977 | -1.364 | -1.776 | 0.593 | 2.587 | -0.06 | 0.05 | 0.05 | 0 | 0.32 | 0.45 | 0.27 | 0.52 | 0.16 | 0.27 |
| 5 | 147017322 | rs17106994 | -1.16 | -1.978 | 1.019 | 2.627 | 0.012 | -0.008 | -0.008 | 0 | 0.3 | 0.31 | 0.36 | 0.61 | 0.16 | 0.29 |
| 5 | 147018084 | rs17106995 | -1.382 | -2.004 | 1.051 | 2.605 | -0.064 | 0.031 | 0.031 | 0 | 0.32 | 0.47 | 0.36 | 0.6 | 0.16 | 0.29 |
| 5 | 147019114 | rs10515588 | -1.14 | -1.997 | 0.973 | 2.597 | 0.011 | -0.016 | -0.016 | 0 | 0.3 | 0.31 | 0.35 | 0.47 | 0.16 | 0.28 |
| 5 | 147024509 | rs2116765 | -1.403 | -1.994 | 0.888 | 2.596 | -0.058 | 0.057 | 0.057 | 0 | 0.32 | 0.51 | 0.36 | 0.47 | 0.16 | 0.28 |
| 5 | 147027930 | rs7735403 | -0.937 | -1.856 | 0.42 | 2.108 | -0.048 | 0.031 | 0.031 | 0.65 | 0.31 | 0.51 | 0.27 | 0.49 | 0.17 | 0.21 |
| 5 | 147029505 | rs2053052 | 1.169 | -1.856 | 0.377 | 2.105 | -0.025 | -0.015 | -0.015 | 0.077 | 0.11 | 0 | 0.07 | 0.13 | 0.11 | 0.04 |
| 5 | 147041185 | rs9325048 | -0.985 | -1.889 | 0.481 | 2.211 | -0.057 | 0.041 | 0.041 | 0.612 | 0.31 | 0.49 | 0.27 | 0.49 | 0.17 | 0.21 |
| 5 | 147041517 | rs17495305 | 1.206 | -1.899 | 0.473 | 2.201 | -0.029 | -0.019 | -0.019 | 0.088 | 0.11 | 0 | 0.07 | 0.12 | 0.11 | 0.04 |
| 5 | 147041720 | rs12658788 | -0.978 | -1.893 | 0.477 | 2.21 | -0.056 | 0.041 | 0.041 | 0.597 | 0.31 | 0.49 | 0.27 | 0.49 | 0.17 | 0.21 |
| 5 | 147046695 | rs6883792 | -1.749 | -1.872 | 0.998 | 2.191 | 0.011 | 0.007 | 0.007 | 0.125 | 0.46 | 0.5 | 0.46 | 0.65 | 0.31 | 0.43 |
| 5 | 147050646 | rs7737054 | 0.237 | -1.868 | 0.855 | 2.217 | 0.028 | -0.014 | -0.014 | 10.13 | 0.16 | 0.1 | 0.16 | 0.22 | 0.05 | 0.15 |
| 5 | 147053189 | rs1106894 | -1.38 | -1.81 | 0.904 | 1.983 | -0.016 | 0 | 0 | 0.023 | 0.42 | 0.46 | 0.43 | 0.59 | 0.27 | 0.32 |
| 5 | 147056194 | rs12651994 | -0.834 | -1.956 | 1.287 | 1.945 | -0.023 | 0.078 | 0.078 | 0 | 0.18 | 0.25 | 0.31 | 0.43 | 0.11 | 0.28 |
| 5 | 147059075 | rs10463279 | 0.351 | -2.162 | 1.122 | 1.801 | -0.004 | 0.032 | 0.032 | 2.897 | 0.11 | 0 | 0.03 | 0.07 | 0.05 | 0.13 |
| 5 | 147059645 | rs6580493 | -1.142 | -2.171 | 1.572 | 1.701 | -0.016 | -0.093 | -0.093 | 0 | 0.22 | 0.38 | 0.12 | 0.07 | 0.15 | 0.15 |
| 5 | 147061995 | rs6895574 | -2.183 | -2.214 | 1.626 | 1.727 | 0.051 | -0.028 | -0.028 | 0.182 | 0.41 | 0.72 | 0.44 | 0.5 | 0.26 | 0.42 |
| 5 | 147063674 | rs6881313 | -1.563 | -2.268 | 1.626 | 1.712 | -0.009 | -0.06 | -0.06 | 0 | 0.35 | 0.57 | 0.19 | 0.15 | 0.26 | 0.19 |
| 5 | 147064881 | rs17107181 | 0.885 | -2.214 | 1.491 | 1.271 | 0 | -0.001 | -0.001 | 0 | 0.13 | 0.14 | 0.07 | 0.09 | 0.11 | 0.04 |
| 5 | 147065573 | rs2288826 | -2.116 | -2.176 | 1.338 | 1.29 | 0.043 | 0.023 | 0.023 | 0 | 0.52 | 0.73 | 0.5 | 0.59 | 0.37 | 0.47 |
| 5 | 147068247 | rs6877552 | 0.542 | -2.047 | 1.276 | 1.329 | -0.029 | 0.029 | 0.029 | 0 | 0.07 | 0 | 0.06 | 0 | 0.09 | 0.02 |
| 5 | 147078815 | rs1432830 | -1.594 | -2.318 | 1.53 | 1.374 | -0.037 | 0.029 | 0.029 | 0.287 | 0.37 | 0.46 | 0.31 | 0.49 | 0.21 | 0.39 |
| 5 | 147079910 | rs1835943 | -1.686 | -2.321 | 1.562 | 1.372 | 0.004 | -0.005 | -0.005 | 0 | 0.38 | 0.6 | 0.32 | 0.5 | 0.21 | 0.39 |
| 5 | 147082525 | rs17107218 | -1.408 | -2.32 | 1.639 | 1.416 | -0.036 | -0.02 | -0.02 | 1.067 | 0.28 | 0.45 | 0.14 | 0.17 | 0.17 | 0.24 |
| 5 | 147087337 | rs17107228 | 0.496 | -2.285 | 1.604 | 1.47 | -0.019 | 0.058 | 0.058 | 0 | 0.09 | 0.12 | 0.18 | 0.33 | 0.04 | 0.15 |
| 5 | 147088555 | rs17107233 | -0.817 | -2.318 | 1.641 | 1.438 | -0.009 | -0.052 | -0.052 | 3.313 | 0.22 | 0.27 | 0.11 | 0.06 | 0.15 | 0.16 |

| | | | | | | | | | | | | | | | | |
|---|---|---|---|---|---|---|---|---|---|---|---|---|---|---|---|---|
| 5 | 147088798 | rs11167996 | 0.897 | -1.887 | 1.189 | 1.275 | -0.039 | 0.048 | 0.048 | 0 | 0.14 | 0.01 | 0.15 | 0.07 | 0.16 | 0.09 |
| 5 | 147089438 | rs1549911 | -0.26 | -1.9 | 1.261 | 1.288 | -0.002 | 0.005 | 0.005 | 0 | 0.23 | 0.23 | 0.34 | 0.4 | 0.2 | 0.25 |
| 5 | 147092355 | rs2288825 | -1.752 | -1.795 | 0.894 | 1.294 | 0.032 | 0.013 | 0.013 | 0 | 0.52 | 0.7 | 0.49 | 0.57 | 0.38 | 0.49 |
| 5 | 147101920 | rs3734122 | 0.185 | -1.804 | 1.103 | 1.346 | -0.028 | 0.125 | 0.125 | 0 | 0.11 | 0.28 | 0.19 | 0.37 | 0.06 | 0.16 |
| 5 | 147104275 | rs7713010 | 1.366 | -1.431 | 0.367 | 0.889 | -0.023 | -0.017 | -0.017 | 0 | 0.12 | 0.05 | 0.21 | 0.17 | 0.16 | 0.1 |
| 5 | 147111031 | rs17497858 | 1.102 | -1.277 | 0.482 | 0.892 | -0.009 | 0.027 | 0.027 | 0 | 0.15 | 0.08 | 0.16 | 0.06 | 0.15 | 0.1 |
| 5 | 147118153 | rs4705192 | -1.316 | -1.289 | 0.484 | 0.94 | 0.026 | 0.027 | 0.027 | 0 | 0.52 | 0.65 | 0.48 | 0.59 | 0.38 | 0.51 |
| 5 | 147122020 | rs6895278 | -1.809 | -1.146 | 0.074 | 0.661 | -0.013 | -0.002 | -0.002 | 0 | 0.74 | 0.81 | 0.8 | 0.76 | 0.66 | 0.66 |
| 5 | 147138729 | rs1016104 | 0.44 | -1.237 | 0.261 | 0.743 | 0.001 | 0.004 | 0.004 | 0 | 0.29 | 0.3 | 0.39 | 0.28 | 0.32 | 0.26 |
| 5 | 147139449 | rs12657336 | -0.084 | -1.331 | 0.814 | 1.025 | -0.028 | 0.101 | 0.101 | 0 | 0.12 | 0.25 | 0.14 | 0.4 | 0.06 | 0.17 |
| 5 | 147151179 | rs11746607 | -0.247 | -1.035 | 0.919 | 0.143 | -0.028 | 0.04 | 0.04 | 1.502 | 0.17 | 0.25 | 0.15 | 0.39 | 0.07 | 0.17 |
| 5 | 147152885 | rs994025 | -0.876 | -0.946 | 0.665 | 0.078 | -0.019 | 0.01 | 0.01 | 0.082 | 0.33 | 0.58 | 0.33 | 0.47 | 0.21 | 0.27 |
| 5 | 147165953 | rs7729260 | -0.033 | -0.546 | -0.08 | -0.402 | -0.006 | -0.061 | -0.061 | 1.363 | 0.12 | 0.19 | 0.09 | 0.01 | 0.1 | 0.08 |
| 5 | 147170949 | rs7447800 | 0.163 | -0.923 | -0.305 | -0.278 | -0.039 | 0.057 | 0.057 | 2.167 | 0.14 | 0.26 | 0.14 | 0.34 | 0.06 | 0.18 |
| 5 | 147171592 | rs4705195 | -0.154 | 0.121 | -1.176 | -0.411 | -0.082 | 0.108 | 0.108 | 0.384 | 0.79 | 0.98 | 0.79 | 0.95 | 0.78 | 0.88 |
| 5 | 147174874 | rs10477349 | 1.491 | 0.139 | -0.754 | -0.136 | 0.018 | 0.05 | 0.05 | 0 | 0.49 | 0.35 | 0.55 | 0.59 | 0.61 | 0.49 |
| 5 | 147177325 | rs1594671 | 0.005 | 1.359 | -0.907 | -0.176 | -0.061 | 0.055 | 0.055 | 0 | 0.86 | 0.98 | 0.83 | 0.95 | 0.84 | 0.9 |
| 5 | 147180762 | rs12110175 | 0.489 | 1.298 | -0.743 | -0.05 | -0.006 | 0.004 | 0.004 | 0 | 0.74 | 0.8 | 0.75 | 0.95 | 0.73 | 0.82 |
| 5 | 147183123 | rs4705198 | -1.181 | 1.574 | -0.366 | 1.12 | 0.018 | 0.063 | 0.063 | 0 | 0.17 | 0.31 | 0.35 | 0.43 | 0.19 | 0.25 |
| 5 | 147183694 | rs1368412 | -0.571 | 1.653 | -0.739 | 0.67 | -0.01 | 0.003 | 0.003 | 0 | 0.35 | 0.31 | 0.56 | 0.71 | 0.52 | 0.59 |
| 5 | 147188475 | rs1025489 | -0.873 | 1.54 | -0.514 | 0.196 | 0.013 | -0.042 | -0.042 | 0 | 0.32 | 0.52 | 0.38 | 0.18 | 0.35 | 0.41 |
| 5 | 147192121 | rs4705201 | -0.654 | 0.748 | -0.191 | -0.076 | -0.004 | -0.014 | -0.014 | 0 | 0.38 | 0.55 | 0.37 | 0.66 | 0.34 | 0.47 |
| 5 | 147203033 | rs17107298 | -0.763 | 0.521 | -0.341 | 0.003 | -0.015 | -0.006 | -0.006 | 0 | 0.26 | 0.64 | 0.32 | 0.24 | 0.2 | 0.36 |
| 5 | 147204592 | rs4705202 | -0.478 | 0.795 | -0.507 | -0.01 | -0.004 | -0.003 | -0.003 | 2.002 | 0.33 | 0.32 | 0.35 | 0.58 | 0.42 | 0.48 |
| 5 | 147208043 | rs6580502 | -1.836 | 0.741 | -0.289 | -0.072 | 0.009 | 0.049 | 0.049 | 1.398 | 0.59 | 0.7 | 0.66 | 0.81 | 0.62 | 0.84 |
| 5 | 147211393 | rs4705203 | -0.048 | 0.88 | -0.642 | 0.157 | 0.008 | 0.029 | 0.029 | 0.827 | 0.19 | 0.1 | 0.11 | 0.13 | 0.15 | 0.16 |
| 5 | 147215120 | rs4705204 | 0.246 | 0.933 | -0.594 | 0.36 | 0.006 | 0.055 | 0.055 | 0.379 | 0.18 | 0.23 | 0.24 | 0.53 | 0.19 | 0.29 |
| 5 | 147225514 | rs891992 | -0.436 | 0.784 | -0.676 | 0.568 | -0.006 | 0.004 | 0.004 | 0.864 | 0.21 | 0.22 | 0.24 | 0.31 | 0.15 | 0.18 |
| 5 | 147234371 | rs13154671 | 0.061 | 1.36 | -0.89 | 0.098 | -0.027 | 0.029 | 0.029 | 0 | 0.14 | 0.1 | 0.12 | 0.05 | 0.23 | 0.18 |
| 5 | 147236299 | rs4705209 | -0.676 | 0.626 | -1.356 | -0.395 | -0.041 | 0.09 | 0.09 | 1.411 | 0.72 | 0.93 | 0.78 | 0.88 | 0.7 | 0.82 |
| 5 | 147236546 | rs1002182 | 0.55 | 0.319 | -1.454 | -0.432 | -0.027 | 0.035 | 0.035 | 0 | 0.22 | 0.12 | 0.16 | 0.09 | 0.28 | 0.23 |
| 5 | 147241696 | rs9654488 | -0.608 | -0.36 | -1.34 | 0.065 | -0.031 | -0.02 | -0.02 | 0 | 0.59 | 0.77 | 0.56 | 0.41 | 0.58 | 0.55 |
| 5 | 147242621 | rs17703536 | 1.072 | -0.632 | -1.418 | 0.22 | 0 | -0.029 | -0.029 | 0 | 0.1 | 0 | 0.22 | 0.08 | 0.1 | 0.08 |
| 5 | 147243641 | rs6889194 | 0.637 | -0.68 | -0.638 | 0.239 | -0.003 | 0.007 | 0.007 | 0 | 0.16 | 0.16 | 0.25 | 0.46 | 0.18 | 0.31 |
| 5 | 147255920 | rs7727031 | 0.595 | -0.498 | 0.325 | -0.014 | -0.011 | 0.028 | 0.028 | 0 | 0.11 | 0.33 | 0.08 | 0.07 | 0.09 | 0.06 |
| 5 | 147258162 | rs1368408 | -0.277 | -0.51 | 0.537 | 0.109 | -0.014 | 0.074 | 0.074 | 0 | 0.15 | 0.34 | 0.34 | 0.22 | 0.14 | 0.15 |
| 5 | 147260274 | rs13355726 | -0.906 | -0.764 | 0.743 | -0.038 | -0.029 | 0.002 | 0.002 | 0 | 0.19 | 0.44 | 0.35 | 0.15 | 0.16 | 0.14 |
| 5 | 147261557 | rs6890965 | 0.515 | -0.752 | 0.345 | -0.033 | -0.009 | -0.027 | -0.027 | 0 | 0.06 | 0.21 | 0.03 | 0 | 0.02 | 0.04 |
| 5 | 147261812 | rs3843496 | 0.027 | -0.926 | -0.442 | -0.277 | 0.034 | 0.077 | 0.077 | 0 | 0.53 | 0.14 | 0.49 | 0.69 | 0.6 | 0.65 |
| 5 | 147262978 | rs1144986 | 1.397 | -0.885 | -0.501 | -0.313 | -0.061 | 0.078 | 0.078 | 0.158 | 0.23 | 0.08 | 0.17 | 0.06 | 0.33 | 0.2 |
| 5 | 147266794 | rs2400439 | 0.792 | -1.119 | 0.226 | 0.101 | -0.003 | 0 | 0 | 0 | 0.1 | 0.11 | 0.07 | 0.07 | 0.1 | 0.06 |
| 5 | 147267123 | rs17107391 | 0.193 | -1.395 | 0.251 | 0.134 | -0.001 | 0 | 0 | 0 | 0.12 | 0.12 | 0.31 | 0.15 | 0.14 | 0.11 |
| 5 | 147271465 | rs1549204 | -1.998 | -1.066 | 0.289 | 0.175 | -0.032 | 0.032 | 0.032 | 0.194 | 0.14 | 0.67 | 0.33 | 0.17 | 0.08 | 0.13 |
| 5 | 147273776 | rs6889356 | -1.025 | -1.059 | 0.301 | 0.131 | -0.019 | -0.023 | -0.023 | 0 | 0.1 | 0.52 | 0.06 | 0.01 | 0.02 | 0.04 |
| 5 | 147275849 | rs1153084 | -1.137 | -0.908 | 0.21 | 0.156 | -0.01 | 0.042 | 0.042 | 0.638 | 0.77 | 0.88 | 0.88 | 0.9 | 0.76 | 0.84 |
| 5 | 147277031 | rs6862197 | -0.472 | -1.039 | 0.158 | 0.197 | -0.065 | -0.042 | -0.042 | 0 | 0.22 | 0.54 | 0.13 | 0.13 | 0.09 | 0.14 |
| 5 | 147279371 | rs11958481 | -0.472 | -1.039 | 0.158 | 0.197 | -0.065 | -0.042 | -0.042 | 0.145 | 0.22 | 0.54 | 0.13 | 0.13 | 0.09 | 0.14 |
| 5 | 147286054 | rs2250145 | 0.745 | -0.907 | -0.43 | 0.184 | 0.053 | 0.075 | 0.075 | 0 | 0.5 | 0.18 | 0.47 | 0.62 | 0.62 | 0.61 |
| 5 | 147286458 | rs7706789 | -1.016 | -0.954 | 0.517 | 1.041 | -0.044 | 0.007 | 0.007 | 3.24 | 0.22 | 0.64 | 0.14 | 0.22 | 0.09 | 0.18 |
| 5 | 147289666 | rs7719581 | -0.995 | -0.488 | 0.086 | 1.359 | -0.049 | 0.054 | 0.054 | 0.35 | 0.25 | 0.46 | 0.37 | 0.38 | 0.14 | 0.28 |
| 5 | 147289799 | rs7715716 | -0.679 | -0.538 | -0.252 | 1.35 | -0.03 | 0.012 | 0.012 | 0.214 | 0.21 | 0.3 | 0.11 | 0.29 | 0.09 | 0.22 |
| 5 | 147290321 | rs6884703 | -0.878 | -0.352 | -0.435 | 1.424 | -0.005 | 0.005 | 0.005 | 0.295 | 0.26 | 0.3 | 0.21 | 0.48 | 0.16 | 0.33 |
| 5 | 147290867 | rs4100170 | -1.279 | 0.334 | -0.742 | 1.275 | -0.014 | -0.03 | -0.03 | 0 | 0.09 | 0.03 | 0.14 | 0.21 | 0.11 | 0.12 |
| 5 | 147291200 | rs7704889 | -0.616 | 0.667 | -0.539 | 1.188 | 0 | 0.004 | 0.004 | 0 | 0.32 | 0.34 | 0.44 | 0.39 | 0.33 | 0.37 |
| 5 | 147297470 | rs2112599 | -0.75 | 0.062 | -0.274 | 0.752 | 0.013 | 0.047 | 0.047 | 0.582 | 0.09 | 0.02 | 0.04 | 0 | 0.06 | 0.08 |
| 5 | 147297817 | rs7709676 | -1.25 | 0.321 | 0.073 | 1.05 | 0.056 | 0.188 | 0.188 | 0 | 0.15 | 0.5 | 0.24 | 0.34 | 0.18 | 0.29 |
| 5 | 147298497 | rs2400440 | -2.272 | 0.377 | -0.055 | 1.366 | 0.004 | -0.053 | -0.053 | 0 | 0.29 | 0.8 | 0.33 | 0.42 | 0.29 | 0.42 |
| 5 | 147303051 | rs721570 | -1.916 | 0.162 | 1.243 | 1.231 | -0.001 | -0.016 | -0.016 | 0 | 0.24 | 0.22 | 0.27 | 0.38 | 0.25 | 0.42 |
| 5 | 147304361 | rs6867813 | -1.707 | 0.156 | 1.287 | 1.228 | -0.009 | -0.038 | -0.038 | 11.348 | 0.22 | 0.16 | 0.26 | 0.38 | 0.25 | 0.41 |
| 5 | 147309468 | rs7731860 | -1.929 | 0.23 | 1.554 | 1.231 | -0.001 | -0.006 | -0.006 | 0 | 0.24 | 0.24 | 0.28 | 0.38 | 0.26 | 0.42 |
| 5 | 147310577 | rs6873510 | -2.191 | 0.276 | 1.456 | 1.253 | 0.001 | -0.004 | -0.004 | 0.013 | 0.28 | 0.72 | 0.31 | 0.38 | 0.26 | 0.41 |
| 5 | 147311794 | rs13187585 | 1.372 | 0.143 | 0.355 | 0.681 | -0.009 | 0.061 | 0.061 | 0.38 | 0.5 | 0.13 | 0.52 | 0.43 | 0.47 | 0.47 |
| 5 | 147326902 | rs13361066 | -0.799 | 0.283 | 0.638 | 0.475 | -0.013 | -0.051 | -0.051 | 0 | 0.14 | 0.22 | 0.09 | 0.03 | 0.1 | 0.16 |
| 5 | 147333945 | rs6580511 | 0.773 | 0.625 | 0.775 | 0.207 | -0.002 | 0.033 | 0.033 | 0 | 0.57 | 0.68 | 0.67 | 0.76 | 0.51 | 0.65 |
| 5 | 147334005 | rs1422587 | -0.504 | 0.633 | 0.707 | 0.278 | -0.065 | 0.125 | 0.125 | 1.458 | 0.72 | 0.96 | 0.82 | 0.92 | 0.66 | 0.83 |
| 5 | 147335875 | rs10515600 | 1.584 | 0.459 | 0.538 | 0.358 | 0 | 0.026 | 0.026 | 0 | 0.42 | 0.21 | 0.46 | 0.41 | 0.36 | 0.4 |
| 5 | 147336286 | rs10515601 | -0.572 | 0.421 | 0.53 | 0.4 | -0.012 | 0.026 | 0.026 | 0.099 | 0.15 | 0.29 | 0.16 | 0.17 | 0.14 | 0.18 |
| 5 | 147337164 | rs17704205 | 0.133 | 0.451 | 0.165 | 0.41 | 0.001 | 0.048 | 0.048 | 1.032 | 0.12 | 0.04 | 0.07 | 0.04 | 0.1 | 0.16 |
| 5 | 147345190 | rs1345689 | -0.94 | 0.458 | 0.133 | 0.306 | -0.052 | 0.101 | 0.101 | 0.402 | 0.75 | 0.99 | 0.86 | 0.92 | 0.72 | 0.84 |

| | | | | | | | | | | | | | | | | |
|---|---|---|---|---|---|---|---|---|---|---|---|---|---|---|---|---|
| 5 | 147347842 | rs4493662 | -1.091 | 0.288 | 0.338 | 0.526 | 0.005 | 0.054 | 0.054 | 1.61 | 0.26 | 0.39 | 0.28 | 0.38 | 0.25 | 0.42 |
| 5 | 147360904 | rs2080085 | 0.956 | 0.218 | -0.12 | 0.337 | 0.015 | 0.027 | 0.027 | 0.017 | 0.46 | 0.21 | 0.44 | 0.45 | 0.4 | 0.48 |
| 5 | 147363251 | rs7700964 | -0.595 | 0.324 | -0.342 | 0.165 | -0.104 | 0.199 | 0.199 | 0.768 | 0.7 | 0.97 | 0.78 | 0.97 | 0.64 | 0.8 |
| 5 | 147382111 | rs6882483 | 0.893 | 0.472 | -0.382 | 0.451 | 0.01 | 0.031 | 0.031 | 0.004 | 0.47 | 0.19 | 0.44 | 0.45 | 0.4 | 0.49 |
| 5 | 147388234 | rs7707803 | -0.668 | 0.587 | 0.126 | 1.269 | 0.01 | 0.089 | 0.089 | 0 | 0.1 | 0.17 | 0.18 | 0.35 | 0.14 | 0.18 |
| 5 | 147393664 | rs1422997 | 0.284 | 0.618 | 0.13 | 0.456 | 0.028 | -0.028 | -0.028 | 1.604 | 0.66 | 0.43 | 0.66 | 0.78 | 0.54 | 0.8 |
| 5 | 147394348 | rs2216655 | -0.444 | 0.493 | 0.089 | 0.401 | -0.131 | 0.122 | 0.122 | 2.602 | 0.73 | 0.93 | 0.76 | 0.9 | 0.58 | 0.86 |
| 5 | 147400155 | rs17641748 | 0.868 | -0.172 | -0.421 | 0.698 | 0 | -0.088 | -0.088 | 0 | 0.16 | 0.01 | 0.06 | 0.27 | 0.1 | 0.27 |
| 5 | 147404281 | rs10477360 | 0.825 | -0.159 | 0.089 | 0.877 | 0.022 | -0.005 | -0.005 | 0 | 0.49 | 0.29 | 0.59 | 0.51 | 0.44 | 0.52 |
| 5 | 147413500 | rs11948836 | -0.804 | -0.001 | 0.984 | 1.041 | 0.068 | 0.223 | 0.223 | 0.054 | 0.05 | 0.39 | 0.09 | 0.12 | 0.04 | 0.06 |
| 5 | 147438586 | rs11168017 | -0.049 | -0.081 | 0.174 | 0.937 | -0.036 | 0.043 | 0.043 | 0.143 | 0.59 | 0.77 | 0.74 | 0.68 | 0.53 | 0.67 |
| 5 | 147442779 | rs3756688 | -0.817 | -0.113 | -0.026 | 0.768 | -0.174 | 0.088 | 0.088 | 3.239 | 0.75 | 0.97 | 0.79 | 0.91 | 0.6 | 0.86 |
| 5 | 147442868 | rs1423007 | -0.723 | -0.113 | 0.113 | 0.768 | -0.042 | 0.036 | 0.036 | 3.156 | 0.74 | 0.8 | 0.78 | 0.91 | 0.6 | 0.86 |
| 5 | 147456843 | rs6864564 | -0.346 | -0.226 | 0.378 | 0.777 | 0.047 | -0.019 | -0.019 | 0 | 0.7 | 0.57 | 0.69 | 0.79 | 0.57 | 0.8 |
| 5 | 147471811 | rs4519913 | 0.802 | -0.422 | 0.172 | 0.841 | 0.019 | 0.001 | 0.001 | 0.391 | 0.5 | 0.28 | 0.63 | 0.51 | 0.48 | 0.51 |
| 5 | 147474639 | rs6580519 | -0.816 | -0.237 | 0.263 | 0.793 | -0.098 | 0.007 | 0.007 | 3.609 | 0.74 | 0.91 | 0.73 | 0.79 | 0.57 | 0.8 |
| 5 | 147475386 | rs6892205 | 0.823 | -0.523 | 0.831 | 0.8 | 0.003 | 0.003 | 0.003 | 0 | 0.5 | 0.47 | 0.65 | 0.48 | 0.49 | 0.51 |
| 5 | 147475798 | rs11948461 | 0.955 | -0.562 | 0.638 | 0.805 | 0.006 | 0.003 | 0.003 | 0 | 0.48 | 0.27 | 0.63 | 0.48 | 0.48 | 0.51 |
| 5 | 147477551 | rs34482796 | 0.969 | -0.562 | 0.619 | 0.805 | 0.004 | 0.002 | 0.002 | 0 | 0.48 | 0.25 | 0.63 | 0.48 | 0.48 | 0.51 |
| 5 | 147480007 | rs2303062 | 0.98 | -0.553 | 0.666 | 0.813 | -0.009 | -0.012 | -0.012 | 0.762 | 0.47 | 0.2 | 0.62 | 0.49 | 0.49 | 0.51 |
| 5 | 147480027 | rs2303063 | 0.98 | -0.553 | 0.666 | 0.813 | -0.009 | -0.012 | -0.012 | 0 | 0.47 | 0.2 | 0.61 | 0.48 | 0.49 | 0.51 |
| 5 | 147480080 | rs2303064 | -0.125 | -0.566 | 0.926 | 0.802 | -0.07 | 0.081 | 0.081 | 0 | 0.25 | 0.56 | 0.16 | 0.42 | 0.12 | 0.35 |
| 5 | 147480509 | rs1862445 | -1.07 | -0.566 | 0.736 | 0.8 | -0.159 | 0.098 | 0.098 | 3.15 | 0.75 | 0.97 | 0.8 | 0.91 | 0.61 | 0.86 |
| 5 | 147480955 | rs2303067 | 0.994 | -0.53 | 0.778 | 0.808 | -0.009 | -0.012 | -0.012 | 2.009 | 0.47 | 0.2 | 0.62 | 0.49 | 0.49 | 0.51 |
| 5 | 147488367 | rs2303071 | 0.463 | -0.537 | 0.921 | 0.805 | -0.01 | -0.013 | -0.013 | 0 | 0.54 | 0.81 | 0.67 | 0.49 | 0.49 | 0.51 |
| 5 | 147488654 | rs17704764 | 1.163 | -1.358 | 0.159 | 0.837 | 0.008 | 0.001 | 0.001 | 0 | 0.2 | 0.06 | 0.14 | 0.18 | 0.17 | 0.15 |
| 5 | 147491218 | rs2287770 | 0.342 | -1.365 | 0.751 | 1.021 | -0.042 | 0.02 | 0.02 | 0 | 0.23 | 0.29 | 0.08 | 0.31 | 0.09 | 0.29 |
| 5 | 147497871 | rs6891892 | -1.445 | -0.305 | 0.262 | 0.699 | -0.088 | 0.079 | 0.079 | 1.095 | 0.78 | 0.98 | 0.81 | 0.9 | 0.72 | 0.87 |
| 5 | 147497932 | rs3777135 | 0.553 | -0.62 | 0.764 | 0.774 | 0.038 | 0.004 | 0.004 | 0 | 0.51 | 0.32 | 0.64 | 0.48 | 0.59 | 0.52 |
| 5 | 147498019 | rs3777134 | 0.574 | -0.616 | 0.76 | 0.772 | 0.041 | 0.007 | 0.007 | 0 | 0.51 | 0.31 | 0.63 | 0.48 | 0.59 | 0.52 |
| 5 | 147499426 | rs6580523 | 0.429 | -0.618 | 1.28 | 0.784 | -0.069 | 0.102 | 0.102 | 0 | 0.25 | 0.42 | 0.16 | 0.42 | 0.13 | 0.35 |
| 5 | 147499891 | rs2303070 | 0.962 | -0.633 | 1.18 | 0.845 | 0.027 | -0.019 | -0.019 | 0 | 0.21 | 0.12 | 0.06 | 0.29 | 0.09 | 0.3 |
| 5 | 147505116 | rs3764930 | 0.577 | -0.626 | 0.963 | 0.823 | 0.05 | -0.006 | -0.006 | 0 | 0.49 | 0.19 | 0.62 | 0.48 | 0.59 | 0.52 |
| 5 | 147510866 | rs2400478 | 0.464 | -0.548 | 0.752 | 0.859 | 0.027 | 0.004 | 0.004 | 0 | 0.51 | 0.39 | 0.64 | 0.48 | 0.59 | 0.52 |
| 5 | 147516165 | rs13185274 | 1.081 | -0.295 | 0.208 | 0.871 | -0.03 | -0.006 | -0.006 | 0 | 0.24 | 0.12 | 0.26 | 0.22 | 0.32 | 0.18 |
| 5 | 147518389 | rs4362936 | -1.272 | 0.474 | 0.092 | 1.31 | -0.022 | -0.023 | -0.023 | 0 | 0.94 | 0.98 | 0.73 | 0.87 | 0.89 | 0.87 |
| 5 | 147520710 | rs12332183 | 0.227 | 0.204 | 0.242 | 1.222 | 0.082 | 0.071 | 0.071 | 0 | 0.5 | 0.79 | 0.3 | 0.39 | 0.37 | 0.44 |
| 5 | 147522547 | rs10463397 | -1.662 | -0.2 | 1.101 | 1.691 | -0.064 | 0.167 | 0.167 | 0.401 | 0.15 | 0.59 | 0.12 | 0.29 | 0.06 | 0.16 |
| 5 | 147528249 | rs1016256 | 1.337 | -0.066 | 0.675 | 1.294 | -0.071 | 0.294 | 0.294 | 0.823 | 0.47 | 0.13 | 0.34 | 0.09 | 0.53 | 0.35 |
| 5 | 147531776 | rs10038045 | -1.358 | -0.076 | 0.827 | 1.362 | -0.063 | 0.063 | 0.063 | 0.149 | 0.14 | 0.57 | 0.07 | 0.18 | 0.05 | 0.15 |
| 5 | 147536452 | rs10077953 | 1.189 | -0.121 | 1.013 | 1.39 | -0.022 | 0.034 | 0.034 | 0 | 0.06 | 0.13 | 0.02 | 0.13 | 0.03 | 0.05 |
| 5 | 147537156 | rs1117507 | 1.34 | -0.098 | 0.9 | 1.311 | -0.034 | 0.241 | 0.241 | 0.883 | 0.47 | 0.19 | 0.34 | 0.09 | 0.53 | 0.35 |
| 5 | 147538478 | rs17599675 | 0.785 | -0.175 | 0.703 | 1.595 | -0.078 | -0.081 | -0.081 | 0.144 | 0.17 | 0.02 | 0.12 | 0.3 | 0.23 | 0.14 |
| 5 | 147547917 | rs10068916 | 0.357 | -0.176 | 0.546 | 1.207 | -0.026 | -0.021 | -0.021 | 0.147 | 0.57 | 0.78 | 0.42 | 0.4 | 0.53 | 0.47 |
| 5 | 147547978 | rs10068988 | 0.386 | -0.571 | 0.892 | 1.416 | -0.009 | -0.018 | -0.018 | 0 | 0.61 | 0.75 | 0.45 | 0.4 | 0.6 | 0.5 |
| 5 | 147548328 | rs6881658 | 0.356 | -0.596 | 0.865 | 1.417 | -0.016 | -0.028 | -0.028 | 0 | 0.61 | 0.79 | 0.46 | 0.4 | 0.6 | 0.49 |
| 5 | 147550938 | rs1156195 | 0.472 | -0.607 | 1.216 | 1.409 | -0.003 | -0.007 | -0.007 | 0 | 0.6 | 0.62 | 0.44 | 0.4 | 0.6 | 0.5 |
| 5 | 147551346 | rs9325081 | -0.195 | -0.631 | 1.539 | 1.442 | -0.004 | 0.135 | 0.135 | 0 | 0.08 | 0.31 | 0.04 | 0.14 | 0.03 | 0.06 |
| 5 | 147556458 | rs767570 | 0.341 | -0.584 | 1.05 | 1.425 | -0.017 | -0.029 | -0.029 | 0 | 0.62 | 0.79 | 0.46 | 0.4 | 0.6 | 0.5 |
| 5 | 147572841 | rs7732604 | -0.945 | -0.422 | 1.088 | 1.283 | 0.007 | 0.221 | 0.221 | 0.113 | 0.48 | 0.8 | 0.63 | 0.87 | 0.42 | 0.52 |
| 5 | 147578262 | rs1432689 | -0.785 | -0.397 | 1.082 | 1.384 | 0.001 | 0.029 | 0.029 | 0.131 | 0.45 | 0.33 | 0.57 | 0.75 | 0.42 | 0.52 |
| 5 | 147586789 | rs10477364 | 0.661 | 0.129 | 2.24 | 2.272 | 0 | 0.002 | 0.002 | 0 | 0.95 | 0.9 | 0.96 | 0.95 | 0.93 | 0.87 |
| 5 | 147593231 | rs10038416 | 2.911 | 0.089 | 2.429 | 2.331 | -0.003 | 0.13 | 0.13 | 0.467 | 0.9 | 0.37 | 0.89 | 0.83 | 0.92 | 0.86 |
| 5 | 147629534 | rs13179719 | 2.856 | 0.106 | 2.445 | 2.379 | -0.004 | 0.136 | 0.136 | 0.392 | 0.9 | 0.37 | 0.89 | 0.83 | 0.93 | 0.86 |
| 5 | 147635072 | rs2400503 | 0.877 | 0.107 | 2.478 | 2.374 | -0.002 | 0.001 | 0.001 | 0 | 0.94 | 0.88 | 0.95 | 0.95 | 0.93 | 0.86 |
| 5 | 147639229 | rs4705243 | -0.442 | 0.253 | 1.939 | 2.324 | -0.01 | -0.005 | -0.005 | 0 | 0.24 | 0.16 | 0.36 | 0.32 | 0.29 | 0.19 |
| 5 | 147648510 | rs7734352 | -2.816 | -0.07 | 2.988 | 2.438 | 0.077 | 0.519 | 0.519 | 2.764 | 0.06 | 0.48 | 0.06 | 0.26 | 0.02 | 0.03 |
| 5 | 147685952 | rs7734458 | -0.457 | 0.161 | 1.381 | 1.705 | 0.006 | 0.064 | 0.064 | 0 | 0.27 | 0.54 | 0.3 | 0.42 | 0.25 | 0.25 |
| 5 | 147713645 | rs6580545 | -0.883 | -0.639 | 1.995 | 2.225 | 0.001 | -0.004 | -0.004 | 0.197 | 0.3 | 0.7 | 0.38 | 0.6 | 0.26 | 0.31 |
| 5 | 147715820 | rs6894032 | -0.55 | -0.64 | 1.997 | 2.264 | 0 | 0.005 | 0.005 | 0.233 | 0.28 | 0.69 | 0.32 | 0.42 | 0.25 | 0.25 |
| 5 | 147720723 | rs10476892 | -0.249 | -0.64 | 1.966 | 2.264 | 0.011 | 0.056 | 0.056 | 0 | 0.25 | 0.37 | 0.28 | 0.42 | 0.25 | 0.25 |
| 5 | 147724086 | rs1422977 | -1.094 | -0.821 | 1.71 | 2.08 | 0.013 | -0.017 | -0.017 | 0 | 0.35 | 0.77 | 0.4 | 0.6 | 0.29 | 0.37 |
| 5 | 147761285 | rs4705252 | -0.154 | -0.749 | 1.688 | 2.09 | -0.001 | -0.002 | -0.002 | 0 | 0.25 | 0.24 | 0.33 | 0.33 | 0.25 | 0.3 |
| 5 | 147773318 | rs10068216 | -0.428 | -0.862 | 1.916 | 1.777 | -0.002 | 0.006 | 0.006 | 0 | 0.28 | 0.23 | 0.26 | 0.25 | 0.23 | 0.29 |
| 5 | 147785009 | rs4705254 | -1.629 | -0.902 | 1.91 | 1.824 | 0.022 | 0.526 | 0.526 | 1.921 | 0.07 | 0.46 | 0.12 | 0.35 | 0.02 | 0.1 |
| 5 | 147788798 | rs10077690 | 0.398 | -0.886 | 1.892 | 1.825 | -0.017 | -0.005 | -0.005 | 0 | 0.21 | 0.13 | 0.25 | 0.25 | 0.21 | 0.24 |
| 5 | 147789721 | rs17720191 | -0.923 | -1.24 | 1.384 | 1.455 | -0.028 | -0.014 | -0.014 | 0 | 0.32 | 0.56 | 0.43 | 0.66 | 0.29 | 0.37 |
| 5 | 147794582 | rs17108251 | -0.122 | -1.21 | 1.289 | 1.585 | 0.001 | -0.009 | -0.009 | 0 | 0.28 | 0.31 | 0.27 | 0.25 | 0.23 | 0.29 |
| 5 | 147797539 | rs10055430 | -0.687 | -1.2 | 1.18 | 1.556 | -0.01 | 0.013 | 0.013 | 0.385 | 0.33 | 0.51 | 0.35 | 0.6 | 0.26 | 0.39 |

| | | | | | | | | | | | | | | | | |
|---|---|---|---|---|---|---|---|---|---|---|---|---|---|---|---|---|
| 5 | 147827921 | rs7721661 | -1.073 | -1.123 | 0.332 | 1.539 | -0.003 | -0.067 | -0.067 | 0 | 0.87 | 0.99 | 0.76 | 0.78 | 0.86 | 0.84 |
| 5 | 147832486 | rs7727161 | -0.289 | -0.825 | 0.591 | 1.519 | -0.001 | 0.009 | 0.009 | 0 | 0.29 | 0.23 | 0.26 | 0.25 | 0.23 | 0.29 |
| 5 | 147835912 | rs2037997 | -0.774 | -0.712 | 0.27 | 0.837 | 0.033 | -0.012 | -0.012 | 0.207 | 0.38 | 0.8 | 0.38 | 0.6 | 0.27 | 0.38 |
| 5 | 147836880 | rs12374521 | 0.337 | 0.122 | 0.923 | 0.387 | 0.023 | 0.15 | 0.15 | 3.021 | 0.53 | 0.23 | 0.36 | 0.18 | 0.59 | 0.51 |
| 5 | 147839537 | rs10075211 | -0.402 | 0.144 | 0.933 | 0.398 | 0.023 | 0.145 | 0.145 | 3.602 | 0.46 | 0.77 | 0.64 | 0.81 | 0.4 | 0.47 |
| 5 | 147842353 | rs11168048 | 0.363 | 0.107 | 1.144 | 0.435 | -0.01 | 0.031 | 0.031 | 0.346 | 0.37 | 0.19 | 0.55 | 0.72 | 0.39 | 0.41 |
| 5 | 147847273 | rs4597955 | -1.175 | 0.238 | 0.677 | 0.487 | -0.139 | 0.284 | 0.284 | 2.293 | 0.58 | 0.98 | 0.72 | 0.93 | 0.57 | 0.62 |
| 5 | 147849759 | rs3995091 | -0.683 | 0.05 | 1.334 | 1.03 | 0.008 | 0.038 | 0.038 | 3.174 | 0.45 | 0.58 | 0.61 | 0.81 | 0.4 | 0.47 |
| 5 | 147854970 | rs10037493 | -0.678 | 0.288 | 1.027 | 1.129 | -0.002 | -0.014 | -0.014 | 0 | 0.43 | 0.48 | 0.61 | 0.73 | 0.4 | 0.48 |
| 5 | 147856232 | rs6580550 | -0.885 | 0.296 | 0.687 | 1.145 | 0.009 | 0.035 | 0.035 | 3.734 | 0.45 | 0.68 | 0.62 | 0.74 | 0.4 | 0.46 |
| 5 | 147856333 | rs7733088 | -0.435 | 0.673 | -0.155 | 1.317 | 0.001 | -0.003 | -0.003 | 0 | 0.36 | 0.29 | 0.55 | 0.64 | 0.37 | 0.41 |
| 5 | 147867479 | rs10076060 | -0.531 | 1.046 | 0.117 | 1.575 | 0.009 | 0.016 | 0.016 | 0.144 | 0.37 | 0.58 | 0.51 | 0.52 | 0.44 | 0.33 |
| 5 | 147877724 | rs7723153 | -0.422 | 0.87 | 0.144 | 1.384 | 0.027 | 0.045 | 0.045 | 0.056 | 0.32 | 0.6 | 0.5 | 0.52 | 0.42 | 0.3 |
| 5 | 147897161 | rs1368388 | -0.974 | 0.755 | 1.25 | 1.327 | -0.001 | -0.001 | -0.001 | 0 | 0.38 | 0.67 | 0.52 | 0.52 | 0.44 | 0.33 |
| 5 | 147902703 | rs7725785 | -1.319 | 0.785 | 1.241 | 1.528 | 0.019 | 0.167 | 0.167 | 0.037 | 0.11 | 0.45 | 0.19 | 0.27 | 0.07 | 0.1 |
| 5 | 147908345 | rs12152801 | -0.082 | 0.683 | 1.032 | 1.459 | -0.089 | -0.033 | -0.033 | 0 | 0.22 | 0.06 | 0.3 | 0.22 | 0.34 | 0.19 |
| 5 | 147927899 | rs13359903 | -1.792 | 1.058 | 1.24 | 1.217 | 0.029 | 0.103 | 0.103 | 0 | 0.14 | 0.46 | 0.22 | 0.27 | 0.12 | 0.16 |
| 5 | 147932201 | rs1422636 | 0.398 | 1.362 | 0.448 | 0.617 | -0.039 | 0.068 | 0.068 | 0 | 0.8 | 0.92 | 0.86 | 0.95 | 0.75 | 0.87 |
| 5 | 147936120 | rs12332417 | -0.885 | 1.385 | 1.006 | 0.846 | 0.138 | 0.273 | 0.273 | 0 | 0.07 | 0.44 | 0.2 | 0.25 | 0.1 | 0.13 |
| 5 | 147940042 | rs1820075 | 0.159 | 1.278 | 1.274 | 1.11 | 0.02 | 0.007 | 0.007 | 0 | 0.23 | 0.35 | 0.31 | 0.25 | 0.32 | 0.18 |
| 5 | 147942238 | rs1345697 | -1.73 | 0.904 | 1.107 | 0.839 | -0.046 | -0.055 | -0.055 | 0.111 | 0.45 | 0.84 | 0.58 | 0.52 | 0.5 | 0.47 |
| 5 | 147943769 | rs17720691 | -1.063 | 1.067 | 1.393 | 0.944 | -0.042 | -0.043 | -0.043 | 0.178 | 0.36 | 0.76 | 0.55 | 0.51 | 0.47 | 0.35 |
| 5 | 147964658 | rs10040819 | -1.995 | 0.939 | 1.7 | 0.934 | 0.017 | 0.063 | 0.063 | 0.098 | 0.14 | 0.61 | 0.26 | 0.24 | 0.16 | 0.15 |
| 5 | 147967746 | rs1833710 | -1.829 | 1.02 | 1.918 | 1.042 | 0.022 | 0.062 | 0.062 | 0.021 | 0.1 | 0.27 | 0.27 | 0.17 | 0.13 | 0.24 |
| 5 | 147972619 | rs4280857 | -0.148 | 0.77 | 1.667 | 0.729 | 0.004 | 0.003 | 0.003 | 0 | 0.09 | 0.06 | 0.03 | 0 | 0.04 | 0.06 |
| 5 | 147973265 | rs17108477 | -0.297 | 0.841 | 1.671 | 0.844 | 0.029 | 0.004 | 0.004 | 0 | 0.05 | 0.14 | 0.13 | 0.06 | 0.1 | 0.09 |
| 5 | 147982855 | rs7711800 | -0.652 | 0.403 | 1.3 | 0.787 | 0.01 | 0.01 | 0.01 | 0 | 0.39 | 0.52 | 0.54 | 0.47 | 0.47 | 0.52 |
| 5 | 147995605 | rs13182913 | 0.396 | 0.439 | 0.56 | 0.862 | -0.076 | -0.077 | -0.077 | 0.218 | 0.28 | 0.05 | 0.31 | 0.34 | 0.34 | 0.3 |
| 5 | 147996087 | rs10223307 | -1.743 | -0.283 | 0.015 | 1.387 | -0.121 | 0.01 | 0.01 | 1.503 | 0.6 | 0.98 | 0.63 | 0.73 | 0.56 | 0.68 |
| 5 | 147998579 | rs7701432 | -1.743 | -0.283 | 0.015 | 1.387 | -0.121 | 0.01 | 0.01 | 1.503 | 0.6 | 0.98 | 0.63 | 0.73 | 0.56 | 0.68 |
| 5 | 148001470 | rs10078551 | 0.513 | -0.318 | 0.371 | 1.565 | -0.047 | 0.019 | 0.019 | 0.112 | 0.28 | 0.11 | 0.2 | 0.24 | 0.33 | 0.26 |
| 5 | 148007013 | rs2068190 | -1.008 | -0.379 | 0.842 | 1.345 | 0.009 | 0.04 | 0.04 | 0.098 | 0.49 | 0.62 | 0.42 | 0.67 | 0.43 | 0.57 |
| 5 | 148010913 | rs9686886 | 0.648 | 0.088 | 0.329 | 1.454 | -0.05 | -0.087 | -0.087 | 0 | 0.24 | 0.04 | 0.16 | 0.41 | 0.28 | 0.21 |
| 5 | 148016537 | rs17639735 | 0.372 | 0.182 | 0.391 | 1.174 | -0.013 | -0.029 | -0.029 | 0 | 0.1 | 0.01 | 0.06 | 0.08 | 0.09 | 0.07 |
| 5 | 148024701 | rs1833704 | -0.332 | 0.231 | 0.812 | 1.146 | 0.003 | 0.009 | 0.009 | 0 | 0.2 | 0.17 | 0.12 | 0.13 | 0.16 | 0.2 |
| 5 | 148025070 | rs7737338 | -0.764 | 0.299 | 1.022 | 1.251 | 0 | -0.005 | -0.005 | 0 | 0.43 | 0.31 | 0.36 | 0.53 | 0.4 | 0.39 |
| 5 | 148025158 | rs7713886 | -1.467 | 0.268 | 0.287 | 1.145 | -0.068 | -0.02 | -0.02 | 0 | 0.87 | 0.99 | 0.84 | 0.9 | 0.78 | 0.9 |
| 5 | 148030064 | rs1862342 | -1.493 | -0.612 | 1.45 | 1.157 | 0.008 | 0.023 | 0.023 | 2.304 | 0.53 | 0.72 | 0.46 | 0.59 | 0.43 | 0.56 |
| 5 | 148054452 | rs1820074 | -1.272 | -0.657 | 1.704 | 1.199 | 0 | 0 | 0 | 1.986 | 0.49 | 0.57 | 0.41 | 0.59 | 0.42 | 0.57 |
| 5 | 148056656 | rs10476898 | -0.906 | -0.656 | 1.752 | 1.194 | 0.005 | 0.003 | 0.003 | 0.289 | 0.45 | 0.46 | 0.37 | 0.56 | 0.4 | 0.44 |
| 5 | 148063894 | rs6882807 | -1.268 | -0.651 | 1.639 | 1.16 | 0.004 | 0.001 | 0.001 | 0.316 | 0.49 | 0.58 | 0.41 | 0.51 | 0.42 | 0.55 |
| 5 | 148065581 | rs12521854 | -0.718 | -0.627 | 1.654 | 1.182 | 0.002 | -0.027 | -0.027 | 0 | 0.42 | 0.23 | 0.35 | 0.4 | 0.4 | 0.31 |
| 5 | 148065877 | rs12521922 | -1.014 | -0.624 | 1.673 | 1.156 | 0 | 0 | 0 | 0 | 0.46 | 0.51 | 0.38 | 0.49 | 0.4 | 0.42 |
| 5 | 148067337 | rs13159091 | -0.337 | -0.713 | 1.591 | 1.116 | 0.004 | 0.006 | 0.006 | 0 | 0.19 | 0.2 | 0.12 | 0.09 | 0.14 | 0.12 |
| 5 | 148073605 | rs3923307 | -1.041 | -0.741 | 1.712 | 1.279 | 0.018 | 0.008 | 0.008 | 0.745 | 0.47 | 0.62 | 0.39 | 0.53 | 0.39 | 0.44 |
| 5 | 148074640 | rs13177640 | -0.808 | -0.717 | 1.623 | 1.29 | 0.013 | -0.015 | -0.015 | 0.835 | 0.43 | 0.23 | 0.35 | 0.5 | 0.39 | 0.4 |
| 5 | 148088972 | rs11953006 | -0.873 | -0.719 | 1.676 | 1.295 | -0.003 | 0.001 | 0.001 | 0.862 | 0.45 | 0.51 | 0.38 | 0.45 | 0.38 | 0.32 |
| 5 | 148093644 | rs764414 | -1.39 | -0.725 | 1.708 | 1.294 | 0.049 | 0.036 | 0.036 | 1.187 | 0.5 | 0.73 | 0.42 | 0.55 | 0.41 | 0.56 |
| 5 | 148099993 | rs13188076 | 0.868 | -0.745 | 1.571 | 1.24 | -0.03 | -0.09 | -0.09 | 0.016 | 0.21 | 0.04 | 0.21 | 0.35 | 0.24 | 0.19 |
| 5 | 148104167 | rs11168056 | -0.755 | -0.744 | 1.459 | 1.225 | -0.026 | -0.023 | -0.023 | 0 | 0.24 | 0.4 | 0.14 | 0.15 | 0.15 | 0.22 |
| 5 | 148105558 | rs17108677 | -0.578 | -0.71 | 0.976 | 1.212 | -0.023 | -0.043 | -0.043 | 0 | 0.22 | 0.38 | 0.14 | 0.09 | 0.15 | 0.14 |
| 5 | 148110028 | rs9325115 | -1.163 | -0.66 | 0.283 | 1.278 | 0.042 | 0.004 | 0.004 | 1.172 | 0.5 | 0.73 | 0.41 | 0.48 | 0.41 | 0.57 |
| 5 | 148111919 | rs1432749 | -0.525 | -0.17 | -0.038 | 1.188 | -0.003 | -0.006 | -0.006 | 0 | 0.2 | 0.26 | 0.13 | 0.12 | 0.15 | 0.16 |
| 5 | 148114806 | rs2400642 | 0.073 | 0.094 | -0.413 | 1.31 | -0.002 | 0.008 | 0.008 | 0.202 | 0.27 | 0.25 | 0.26 | 0.44 | 0.23 | 0.23 |
| 5 | 148116202 | rs1465405 | 0.131 | 0.144 | -0.446 | 1.169 | -0.001 | 0.003 | 0.003 | 0.233 | 0.26 | 0.22 | 0.26 | 0.37 | 0.26 | 0.22 |
| 5 | 148116653 | rs10059336 | -0.471 | 0.27 | -0.724 | 1.003 | 0.004 | -0.003 | -0.003 | 0 | 0.46 | 0.55 | 0.39 | 0.46 | 0.38 | 0.35 |
| 5 | 148118029 | rs17777882 | -0.818 | 0.248 | -1.04 | 0.794 | -0.006 | -0.045 | -0.045 | 0 | 0.11 | 0.03 | 0.18 | 0.24 | 0.09 | 0.28 |
| 5 | 148120217 | rs888956 | 0.407 | -0.16 | -1.304 | -0.375 | 0.024 | 0.068 | 0.068 | 0.926 | 0.28 | 0.13 | 0.22 | 0.03 | 0.23 | 0.2 |
| 5 | 148135026 | rs2116719 | -0.148 | 0.477 | -1.471 | 0.212 | -0.107 | -0.046 | -0.046 | 0 | 0.64 | 0.88 | 0.54 | 0.35 | 0.51 | 0.54 |
| 5 | 148138790 | rs6883606 | -0.162 | 0.463 | -1.477 | 0.205 | -0.122 | -0.057 | -0.057 | 3.26 | 0.64 | 0.91 | 0.55 | 0.35 | 0.52 | 0.54 |
| 5 | 148145138 | rs2163752 | 0.036 | 0.423 | -0.699 | 0.185 | -0.013 | -0.002 | -0.002 | 0 | 0.62 | 0.67 | 0.53 | 0.35 | 0.51 | 0.54 |
| 5 | 148156653 | rs30312 | -0.102 | 0.418 | -0.63 | 0.041 | -0.073 | 0.001 | 0.001 | 0.088 | 0.63 | 0.86 | 0.54 | 0.31 | 0.51 | 0.48 |
| 5 | 148168332 | rs30330 | 0.021 | 0.501 | -0.792 | 0.108 | 0 | -0.001 | -0.001 | 0 | 0.59 | 0.6 | 0.5 | 0.28 | 0.49 | 0.45 |
| 5 | 148172689 | rs171551 | 0.265 | 1.022 | -1.095 | 0.153 | -0.082 | 0.032 | 0.032 | 0 | 0.69 | 0.87 | 0.6 | 0.75 | 0.53 | 0.77 |
| 5 | 148172928 | rs9285673 | -1.679 | 0.178 | 0.325 | 0.321 | 0.014 | -0.02 | -0.02 | 0 | 0.14 | 0.54 | 0.09 | 0.06 | 0.08 | 0.06 |
| 5 | 148175047 | rs9325117 | -1.821 | -0.345 | 1.133 | 0.755 | 0.052 | 0.166 | 0.166 | 2.437 | 0.06 | 0.5 | 0.05 | 0.12 | 0.01 | 0.05 |
| 5 | 148178371 | rs17778143 | -0.261 | -0.908 | 0.614 | 1.19 | 0.019 | -0.049 | -0.049 | 0.782 | 0.26 | 0.16 | 0.33 | 0.5 | 0.22 | 0.25 |
| 5 | 148182123 | rs10059242 | -1.039 | -0.88 | 0.697 | 1.217 | 0.083 | 0.143 | 0.143 | 2.738 | 0.05 | 0.5 | 0.05 | 0.05 | 0.01 | 0.01 |
| 5 | 148183832 | rs11742884 | -0.266 | -0.862 | 0.487 | 1.176 | 0.017 | -0.046 | -0.046 | 0.843 | 0.25 | 0.16 | 0.33 | 0.51 | 0.22 | 0.25 |

| | | | | | | | | | | | | | | | | |
|---|---|---|---|---|---|---|---|---|---|---|---|---|---|---|---|---|
| 5 | 148191261 | rs2116713 | -1.54 | -0.926 | 1.145 | 1.46 | -0.028 | 0.303 | 0.303 | 1.204 | 0.1 | 0.46 | 0.07 | 0.24 | 0.01 | 0.19 |
| 5 | 148191398 | rs2116714 | -1.544 | -0.939 | 1.008 | 1.454 | -0.047 | 0.254 | 0.254 | 2.619 | 0.11 | 0.57 | 0.08 | 0.24 | 0.01 | 0.19 |
| 5 | 148200600 | rs2082395 | -0.005 | -0.357 | 0.306 | 0.99 | 0.047 | 0.004 | 0.004 | 0 | 0.7 | 0.54 | 0.7 | 0.74 | 0.59 | 0.74 |
| 5 | 148205741 | rs12654778 | 0.914 | -0.999 | 0.027 | 1.057 | 0.015 | 0.009 | 0.009 | 0.218 | 0.39 | 0.2 | 0.4 | 0.37 | 0.38 | 0.39 |
| 5 | 148206348 | rs1042711 | -0.771 | -1.146 | -0.106 | 0.635 | -0.067 | 0.049 | 0.049 | 0 | 0.75 | 0.86 | 0.76 | 0.92 | 0.59 | 0.81 |
| 5 | 148206440 | rs1042713 | 0.881 | -1.174 | -0.024 | 0.689 | -0.007 | 0.002 | 0.002 | 0 | 0.43 | 0.52 | 0.46 | 0.55 | 0.39 | 0.45 |
| 5 | 148206646 | rs1042717 | -0.69 | -1.394 | 0.383 | 1.217 | -0.009 | 0 | 0 | 0 | 0.33 | 0.34 | 0.3 | 0.36 | 0.2 | 0.36 |
| 5 | 148206917 | rs1042718 | -0.26 | -1.367 | 0.434 | 1.272 | -0.02 | 0.019 | 0.019 | 0 | 0.26 | 0.34 | 0.29 | 0.37 | 0.17 | 0.31 |
| 5 | 148211991 | rs4705271 | -0.365 | -1.035 | -0.073 | 0.886 | -0.053 | 0.01 | 0.01 | 0 | 0.36 | 0.45 | 0.29 | 0.6 | 0.2 | 0.47 |
| 5 | 148213082 | rs3857420 | -0.313 | -0.996 | -0.31 | 0.801 | -0.038 | 0.025 | 0.025 | 0 | 0.36 | 0.49 | 0.29 | 0.52 | 0.2 | 0.43 |
| 5 | 148214229 | rs12652757 | -0.329 | -0.149 | -0.551 | 0.404 | -0.008 | 0.012 | 0.012 | 0 | 0.07 | 0.16 | 0.03 | 0.06 | 0.03 | 0.08 |
| 5 | 148216187 | rs11957757 | -0.36 | 0.657 | -0.987 | -0.042 | -0.025 | 0.021 | 0.021 | 0 | 0.28 | 0.19 | 0.26 | 0.07 | 0.43 | 0.23 |
| 5 | 148216402 | rs6888329 | -0.166 | 0.491 | -1.071 | 0.475 | -0.001 | -0.004 | -0.004 | 0.597 | 0.66 | 0.75 | 0.63 | 0.39 | 0.65 | 0.63 |
| 5 | 148216692 | rs11957895 | -0.75 | 0.749 | -1.001 | 0.6 | 0.002 | 0 | 0 | 0.073 | 0.28 | 0.18 | 0.24 | 0.26 | 0.26 | 0.19 |
| 5 | 148221473 | rs6888011 | 0.332 | 1.056 | -0.516 | 0.771 | 0.001 | 0.038 | 0.038 | 0 | 0.7 | 0.63 | 0.57 | 0.53 | 0.69 | 0.74 |
| 5 | 148232677 | rs10070291 | -0.714 | 1.311 | -1.062 | 0.533 | -0.021 | -0.001 | -0.001 | 0 | 0.25 | 0.82 | 0.27 | 0.27 | 0.26 | 0.21 |
| 5 | 148236409 | rs17707884 | 1.222 | 1.19 | -1.679 | -0.002 | 0.009 | 0.062 | 0.062 | 0.679 | 0.43 | 0.08 | 0.29 | 0.27 | 0.37 | 0.53 |
| 5 | 148238456 | rs10214302 | -0.115 | 1.357 | -0.412 | 0.423 | -0.041 | 0.019 | 0.019 | 0.183 | 0.14 | 0.62 | 0.1 | 0.16 | 0.07 | 0.08 |
| 5 | 148240456 | rs13189358 | 0.587 | 1.397 | -0.715 | 0.405 | -0.037 | -0.112 | -0.112 | 0 | 0.19 | 0.04 | 0.37 | 0.44 | 0.25 | 0.17 |
| 5 | 148242723 | rs6580586 | -0.577 | 1.393 | 0.12 | 0.242 | -0.031 | -0.006 | -0.006 | 0 | 0.17 | 0.67 | 0.14 | 0.16 | 0.12 | 0.12 |
| 5 | 148243473 | rs6884617 | -1.483 | 1.513 | -0.151 | 0.132 | -0.053 | 0.091 | 0.091 | 0.491 | 0.45 | 0.89 | 0.64 | 0.71 | 0.51 | 0.38 |
| 5 | 148248818 | rs9325124 | -0.732 | 1.611 | 0.201 | 0.138 | -0.001 | 0.007 | 0.007 | 0 | 0.41 | 0.67 | 0.54 | 0.62 | 0.43 | 0.29 |
| 5 | 148250573 | rs6893517 | -1.136 | 1.6 | 0.114 | 0.151 | 0.027 | 0.083 | 0.083 | 0 | 0.46 | 0.71 | 0.67 | 0.75 | 0.57 | 0.4 |
| 5 | 148256109 | rs1368277 | -0.09 | 1.762 | 0.272 | 0.423 | -0.025 | -0.074 | -0.074 | 2.091 | 0.28 | 0.16 | 0.44 | 0.47 | 0.34 | 0.2 |
| 5 | 148260007 | rs2082383 | 0.383 | 1.872 | 0.088 | 0.445 | -0.025 | -0.018 | -0.018 | 0 | 0.06 | 0.01 | 0.1 | 0.1 | 0.13 | 0.11 |
| 5 | 148261996 | rs919725 | -0.077 | 1.807 | 0.805 | 0.468 | 0.007 | 0.035 | 0.035 | 0 | 0.28 | 0.38 | 0.46 | 0.47 | 0.34 | 0.18 |
| 5 | 148267406 | rs1864932 | -0.643 | 1.342 | 0.324 | 0.396 | -0.017 | -0.008 | -0.008 | 0 | 0.37 | 0.69 | 0.58 | 0.58 | 0.47 | 0.3 |
| 5 | 148278925 | rs4705280 | -0.48 | 1.334 | 0.008 | 0.446 | -0.068 | -0.031 | -0.031 | 0 | 0.38 | 0.16 | 0.56 | 0.59 | 0.47 | 0.35 |
| 5 | 148280187 | rs1181141 | -0.283 | 1.079 | 0.678 | -0.158 | 0.011 | -0.062 | -0.062 | 0.684 | 0.11 | 0.25 | 0.07 | 0 | 0.1 | 0.07 |
| 5 | 148280902 | rs1181139 | -0.918 | 1.569 | 0.781 | -0.083 | 0.001 | 0.025 | 0.025 | 0 | 0.49 | 0.44 | 0.59 | 0.65 | 0.48 | 0.35 |
| 5 | 148283322 | rs17108911 | 0.286 | 1.548 | 0.866 | -0.019 | -0.001 | 0.013 | 0.013 | 0 | 0.07 | 0.11 | 0.04 | 0.06 | 0.04 | 0.04 |
| 5 | 148283944 | rs4705283 | -0.899 | 1.416 | 0.712 | -0.158 | 0.002 | 0.163 | 0.163 | 0 | 0.51 | 0.85 | 0.68 | 0.7 | 0.52 | 0.41 |
| 5 | 148286432 | rs17600134 | 0.282 | 1.429 | 0.717 | -0.098 | 0 | -0.021 | -0.021 | 0 | 0.34 | 0.21 | 0.5 | 0.59 | 0.35 | 0.25 |
| 5 | 148287096 | rs4705284 | 0.607 | 1.325 | 0.651 | 0.034 | -0.031 | -0.002 | -0.002 | 0 | 0.05 | 0.01 | 0.09 | 0.05 | 0.13 | 0.11 |
| 5 | 148287177 | rs4705285 | -1.217 | 0.863 | 0.736 | 0.017 | 0.026 | 0.091 | 0.091 | 0 | 0.59 | 0.89 | 0.73 | 0.7 | 0.62 | 0.47 |
| 5 | 148293429 | rs10075995 | -0.372 | 0.983 | 0.714 | 0.091 | 0.023 | 0.047 | 0.047 | 0 | 0.47 | 0.31 | 0.64 | 0.64 | 0.57 | 0.42 |
| 5 | 148294678 | rs2400709 | -0.446 | 1.018 | 0.731 | 0.167 | -0.006 | 0.015 | 0.015 | 0 | 0.07 | 0.12 | 0.04 | 0.05 | 0.03 | 0.03 |
| 5 | 148297564 | rs11957970 | -1.296 | 1.315 | 1.143 | 0.524 | 0.051 | 0.095 | 0.095 | 0 | 0.13 | 0.43 | 0.15 | 0.19 | 0.16 | 0.22 |
| 5 | 148302641 | rs4705286 | -2.275 | 1.323 | 1.208 | 0.406 | 0 | 0.012 | 0.012 | 0.823 | 0.23 | 0.72 | 0.22 | 0.25 | 0.22 | 0.29 |
| 5 | 148305690 | rs11740127 | 1.728 | 0.672 | 0.043 | -0.494 | 0.068 | 0.144 | 0.144 | 0 | 0.41 | 0.16 | 0.22 | 0.01 | 0.31 | 0.37 |
| 5 | 148306141 | rs11740851 | -0.103 | 0.71 | 0.003 | -0.441 | -0.118 | 0.002 | 0.002 | 0 | 0.64 | 0.92 | 0.44 | 0.24 | 0.53 | 0.65 |
| 5 | 148307111 | rs741146 | 0.774 | 0.708 | 0.333 | -0.553 | 0.017 | 0.006 | 0.006 | 0 | 0.5 | 0.45 | 0.3 | 0.06 | 0.38 | 0.42 |
| 5 | 148310151 | rs1029942 | -0.739 | 0.519 | 0.446 | -0.379 | 0.008 | 0.023 | 0.023 | 0 | 0.09 | 0.29 | 0.07 | 0.05 | 0.07 | 0.07 |
| 5 | 148311053 | rs12652493 | 0.235 | 0.939 | 0.309 | 0.224 | 0 | 0 | 0 | 2.039 | 0.61 | 0.62 | 0.48 | 0.35 | 0.55 | 0.71 |
| 5 | 148312767 | rs7729953 | 1.27 | 1 | -0.123 | -0.045 | 0.016 | 0.18 | 0.18 | 0 | 0.45 | 0.18 | 0.26 | 0.01 | 0.41 | 0.42 |
| 5 | 148313933 | rs4705064 | -1.017 | 1.013 | 0.064 | 0.142 | 0.019 | 0.04 | 0.04 | 0 | 0.47 | 0.61 | 0.63 | 0.7 | 0.6 | 0.48 |
| 5 | 148316076 | rs917875 | -0.107 | 0.917 | 0.481 | -0.117 | 0.001 | -0.023 | -0.023 | 0.238 | 0.07 | 0.16 | 0.04 | 0.01 | 0.05 | 0.12 |
| 5 | 148318559 | rs7729246 | -0.07 | 0.919 | 0.288 | -0.116 | -0.002 | -0.016 | -0.016 | 1.941 | 0.07 | 0.13 | 0.04 | 0.01 | 0.05 | 0.12 |
| 5 | 148329991 | rs973057 | 1.276 | 0.814 | -0.427 | -0.577 | 0.024 | 0.179 | 0.179 | 0.045 | 0.4 | 0.14 | 0.26 | 0.11 | 0.34 | 0.27 |
| 5 | 148331046 | rs10491337 | 1.083 | 0.815 | -0.412 | -0.537 | 0.004 | 0.005 | 0.005 | 1.099 | 0.07 | 0.04 | 0.03 | 0.08 | 0.04 | 0.15 |
| 5 | 148331571 | rs759135 | -0.536 | 0.96 | -0.46 | -0.271 | 0.027 | 0.04 | 0.04 | 0.145 | 0.72 | 0.6 | 0.5 | 0.37 | 0.65 | 0.75 |
| 5 | 148333934 | rs2192256 | 0.86 | 0.673 | -1.157 | -0.555 | 0.023 | 0.178 | 0.178 | 0.045 | 0.4 | 0.14 | 0.26 | 0.12 | 0.34 | 0.27 |
| 5 | 148341110 | rs11168076 | 0.63 | 0.694 | -0.911 | -0.606 | 0.012 | 0.046 | 0.046 | 0 | 0.42 | 0.28 | 0.23 | 0.11 | 0.38 | 0.33 |
| 5 | 148342810 | rs7703500 | -0.67 | 1.017 | -0.445 | 0.531 | -0.002 | -0.004 | -0.004 | 0 | 0.13 | 0.13 | 0.16 | 0.18 | 0.14 | 0.31 |
| 5 | 148345447 | rs7732060 | -0.656 | 1.019 | -0.509 | 0.532 | -0.004 | -0.011 | -0.011 | 0.03 | 0.13 | 0.12 | 0.16 | 0.18 | 0.14 | 0.31 |
| 5 | 148346443 | rs7737361 | -1.32 | 0.546 | -0.323 | 0.331 | -0.026 | 0.023 | 0.023 | 0 | 0.25 | 0.63 | 0.42 | 0.27 | 0.19 | 0.41 |
| 5 | 148348395 | rs994446 | -0.959 | 0.672 | -0.533 | 0.294 | 0.002 | 0.023 | 0.023 | 0 | 0.19 | 0.25 | 0.37 | 0.32 | 0.19 | 0.29 |
| 5 | 148352990 | rs10045726 | -0.511 | 0.891 | -0.895 | -0.432 | 0.002 | -0.109 | -0.109 | 0 | 0.75 | 0.61 | 0.87 | 0.98 | 0.77 | 0.79 |
| 5 | 148357632 | rs733032 | 0.961 | 0.969 | -0.812 | -0.405 | 0 | -0.002 | -0.002 | 0.004 | 0.57 | 0.4 | 0.49 | 0.66 | 0.58 | 0.5 |
| 5 | 148365234 | rs930205 | 0.894 | 0.913 | -0.85 | -0.396 | 0 | 0.012 | 0.012 | 0 | 0.55 | 0.37 | 0.49 | 0.65 | 0.58 | 0.49 |
| 5 | 148367643 | rs4705303 | 1.173 | 1.021 | -0.858 | -0.453 | -0.01 | 0.056 | 0.056 | 0 | 0.52 | 0.22 | 0.47 | 0.65 | 0.55 | 0.49 |
| 5 | 148373306 | rs13166730 | 1.49 | 0.572 | -1.319 | -0.898 | -0.005 | 0.081 | 0.081 | 0 | 0.15 | 0.02 | 0.13 | 0.01 | 0.16 | 0.07 |
| 5 | 148381078 | rs3763022 | -0.148 | 0.789 | -0.863 | -0.398 | -0.005 | -0.034 | -0.034 | 0 | 0.3 | 0.64 | 0.35 | 0.14 | 0.25 | 0.28 |
| 5 | 148383555 | rs10050933 | -0.223 | 0.86 | -0.886 | -0.378 | -0.003 | -0.01 | -0.01 | 0 | 0.17 | 0.14 | 0.17 | 0.2 | 0.18 | 0.23 |
| 5 | 148389763 | rs1025476 | -0.913 | 0.875 | -0.834 | -0.408 | -0.001 | 0 | 0 | 0 | 0.42 | 0.47 | 0.3 | 0.22 | 0.4 | 0.44 |
| 5 | 148392565 | rs1549907 | -0.058 | 0.895 | -0.853 | -0.305 | -0.023 | -0.035 | -0.035 | 0 | 0.16 | 0.07 | 0.17 | 0.2 | 0.18 | 0.23 |
| 5 | 148406032 | rs10075404 | -0.424 | 0.904 | -0.84 | -0.276 | 0.002 | 0.033 | 0.033 | 0 | 0.18 | 0.25 | 0.35 | 0.33 | 0.18 | 0.3 |
| 5 | 148407893 | rs6875902 | 0.381 | 0.816 | -0.939 | -0.378 | -0.001 | -0.069 | -0.069 | 0 | 0.26 | 0.34 | 0.13 | 0.02 | 0.22 | 0.21 |
| 5 | 148408101 | rs1432793 | -0.994 | 1.082 | -0.997 | -0.369 | -0.002 | 0.015 | 0.015 | 0.001 | 0.46 | 0.62 | 0.35 | 0.22 | 0.47 | 0.45 |

| | | | | | | | | | | | | | | | | |
|---|---|---|---|---|---|---|---|---|---|---|---|---|---|---|---|---|
| 5 | 148410920 | rs2304034 | -0.21 | 0.254 | -1.151 | -0.076 | -0.01 | -0.018 | -0.018 | 0 | 0.14 | 0.08 | 0.33 | 0.21 | 0.12 | 0.28 |
| 5 | 148416105 | rs17109268 | 1.503 | 0.101 | -1.135 | -0.077 | -0.008 | 0.014 | 0.014 | 3.069 | 0.06 | 0.13 | 0.03 | 0.05 | 0.04 | 0.09 |
| 5 | 148416952 | rs17795259 | -0.065 | 0.154 | -1.171 | -0.081 | -0.041 | -0.034 | -0.034 | 0 | 0.14 | 0.05 | 0.14 | 0.18 | 0.14 | 0.32 |
| 5 | 148417553 | rs36067 | 0.263 | 0.086 | -1.301 | 0.042 | -0.006 | -0.039 | -0.039 | 0 | 0.27 | 0.4 | 0.16 | 0.07 | 0.22 | 0.2 |
| 5 | 148431964 | rs36077 | -1.345 | 0.048 | -1.269 | 0.062 | 0.001 | -0.001 | -0.001 | 0.99 | 0.52 | 0.74 | 0.55 | 0.43 | 0.42 | 0.67 |
| 5 | 148441128 | rs36044 | 0.311 | -0.151 | -1.07 | 0.192 | -0.014 | -0.008 | -0.008 | 0.154 | 0.25 | 0.42 | 0.33 | 0.2 | 0.19 | 0.27 |
| 5 | 148441321 | rs36047 | -0.904 | -0.157 | -1.064 | 0.192 | -0.002 | -0.002 | -0.002 | 0 | 0.41 | 0.6 | 0.49 | 0.38 | 0.34 | 0.58 |
| 5 | 148446523 | rs250671 | 0.248 | -0.245 | -1.011 | 0.223 | -0.014 | -0.008 | -0.008 | 0.053 | 0.25 | 0.42 | 0.33 | 0.2 | 0.19 | 0.27 |
| 5 | 148454394 | rs250665 | -0.288 | -0.243 | -1.016 | 0.222 | -0.025 | -0.002 | -0.002 | 0 | 0.31 | 0.58 | 0.37 | 0.25 | 0.24 | 0.35 |
| 5 | 148468467 | rs2112598 | -0.386 | -0.126 | -0.981 | 0.286 | -0.007 | -0.001 | -0.001 | 0 | 0.74 | 0.54 | 0.67 | 0.77 | 0.77 | 0.71 |
| 5 | 148472410 | rs1120540 | -0.386 | -0.126 | -0.981 | 0.289 | -0.007 | -0.003 | -0.003 | 0 | 0.74 | 0.54 | 0.67 | 0.77 | 0.77 | 0.71 |
| 5 | 148484187 | rs11168086 | 0.549 | -0.026 | -0.983 | 0.342 | 0 | 0.001 | 0.001 | 0.032 | 0.59 | 0.4 | 0.53 | 0.59 | 0.63 | 0.41 |
| 5 | 148484757 | rs1422579 | 1.183 | 0.111 | -1.101 | 0.344 | 0 | -0.002 | -0.002 | 0 | 0.51 | 0.25 | 0.48 | 0.53 | 0.54 | 0.32 |
| 5 | 148486371 | rs1422577 | 1.068 | -0.06 | -1.086 | 0.36 | 0.006 | -0.006 | -0.006 | 0 | 0.52 | 0.26 | 0.49 | 0.53 | 0.58 | 0.32 |
| 5 | 148499460 | rs11746996 | 0.944 | -0.047 | -1.106 | 0.339 | 0.013 | -0.012 | -0.012 | 0.348 | 0.53 | 0.26 | 0.5 | 0.54 | 0.62 | 0.37 |
| 5 | 148501158 | rs13153796 | 0.249 | -0.085 | -1.048 | 0.383 | -0.007 | -0.003 | -0.003 | 0 | 0.65 | 0.38 | 0.64 | 0.71 | 0.71 | 0.62 |
| 5 | 148505628 | rs11168091 | -0.483 | 0.077 | -1.018 | 0.462 | -0.003 | -0.025 | -0.025 | 0 | 0.74 | 0.64 | 0.86 | 0.92 | 0.78 | 0.78 |
| 5 | 148516012 | rs17795445 | 0.262 | 0.107 | -1.565 | 0.565 | -0.027 | -0.036 | -0.036 | 0 | 0.12 | 0.06 | 0.14 | 0.19 | 0.13 | 0.29 |
| 5 | 148520213 | rs4579247 | -0.074 | 0.082 | -1.326 | 0.61 | -0.017 | -0.023 | -0.023 | 0.249 | 0.71 | 0.48 | 0.68 | 0.73 | 0.77 | 0.71 |
| 5 | 148520669 | rs11168092 | 1.174 | -0.149 | -1.632 | 0.483 | -0.143 | -0.15 | -0.15 | 5.232 | 0.31 | 0.02 | 0.35 | 0.4 | 0.44 | 0.25 |
| 5 | 148528107 | rs4147470 | 1.129 | -0.616 | -1.087 | 0.642 | -0.026 | -0.012 | -0.012 | 0 | 0.34 | 0.24 | 0.63 | 0.41 | 0.51 | 0.25 |
| 5 | 148532533 | rs3922512 | 0.454 | -0.799 | -1.119 | 0.584 | -0.001 | -0.073 | -0.073 | 0 | 0.25 | 0.4 | 0.12 | 0.07 | 0.21 | 0.23 |
| 5 | 148539100 | rs4546368 | 0.438 | -0.888 | -1.159 | 0.555 | 0.003 | 0.009 | 0.009 | 0.092 | 0.49 | 0.41 | 0.68 | 0.64 | 0.57 | 0.42 |
| 5 | 148540904 | rs10077985 | 1.177 | -0.73 | -1.315 | 0.488 | 0.003 | 0.004 | 0.004 | 8.214 | 0.14 | 0.12 | 0.05 | 0.06 | 0.06 | 0.11 |
| 5 | 148544809 | rs4361503 | -0.495 | -0.673 | -1.413 | 0.44 | 0.004 | -0.051 | -0.051 | 0.484 | 0.74 | 0.9 | 0.81 | 0.7 | 0.78 | 0.65 |
| 5 | 148544856 | rs4314386 | 0.801 | 0.16 | -1.032 | 0.182 | -0.011 | -0.003 | -0.003 | 0.019 | 0.35 | 0.27 | 0.46 | 0.39 | 0.51 | 0.25 |
| 5 | 148545567 | rs10053271 | -0.708 | 0.605 | -0.384 | 0.057 | 0.003 | -0.082 | -0.082 | 0 | 0.24 | 0.38 | 0.11 | 0.06 | 0.21 | 0.23 |
| 5 | 148548007 | rs9325132 | -0.608 | 0.237 | -0.351 | 0.097 | -0.013 | -0.079 | -0.079 | 0 | 0.23 | 0.37 | 0.16 | 0.06 | 0.18 | 0.23 |
| 5 | 148548523 | rs9765483 | 0.467 | 0.044 | -1.01 | 0.459 | 0.027 | 0.013 | 0.013 | 0 | 0.7 | 0.84 | 0.82 | 0.72 | 0.78 | 0.63 |
| 5 | 148551105 | rs10052843 | -0.084 | 0.819 | 0.593 | 1.127 | 0.028 | 0.089 | 0.089 | 0 | 0.21 | 0.33 | 0.3 | 0.53 | 0.32 | 0.38 |
| 5 | 148554730 | rs6871775 | -0.007 | 0.852 | 0.587 | 1.175 | -0.001 | 0.01 | 0.01 | 0 | 0.19 | 0.2 | 0.28 | 0.52 | 0.32 | 0.37 |
| 5 | 148562981 | rs6887628 | -0.143 | 0.749 | 0.788 | 0.927 | 0.012 | 0.041 | 0.041 | 0 | 0.19 | 0.24 | 0.28 | 0.52 | 0.32 | 0.37 |
| 5 | 148576253 | rs7720260 | 0.853 | 0.286 | -0.051 | 0.25 | -0.022 | -0.029 | -0.029 | 0 | 0.6 | 0.73 | 0.45 | 0.39 | 0.55 | 0.49 |
| 5 | 148576697 | rs4705331 | -0.022 | 0.844 | 0.366 | 0.662 | 0.01 | 0.027 | 0.027 | 0 | 0.78 | 0.74 | 0.7 | 0.47 | 0.67 | 0.61 |
| 5 | 148581063 | rs10463419 | -0.078 | 0.895 | 0.193 | 0.704 | 0.009 | 0.024 | 0.024 | 0 | 0.21 | 0.24 | 0.29 | 0.53 | 0.33 | 0.39 |
| 5 | 148593958 | rs2963489 | -0.522 | 0.309 | 0.321 | 0.079 | 0.011 | 0.003 | 0.003 | 0 | 0.32 | 0.62 | 0.41 | 0.65 | 0.4 | 0.44 |
| 5 | 148615987 | rs4295373 | -0.192 | 0.217 | 0.671 | 0.046 | 0.002 | 0 | 0 | 0 | 0.29 | 0.32 | 0.37 | 0.59 | 0.39 | 0.42 |
| 5 | 148620535 | rs748066 | -0.403 | 0.145 | 0.716 | 0.028 | 0.017 | 0.003 | 0.003 | 0 | 0.32 | 0.4 | 0.4 | 0.64 | 0.4 | 0.49 |
| 5 | 148623098 | rs10067047 | -0.521 | 0.216 | 0.758 | 0.029 | 0.042 | 0.014 | 0.014 | 0 | 0.33 | 0.54 | 0.41 | 0.62 | 0.42 | 0.44 |
| 5 | 148624898 | rs3816159 | -0.265 | 0.222 | 0.635 | 0.041 | -0.008 | -0.007 | -0.007 | 0 | 0.3 | 0.28 | 0.37 | 0.57 | 0.4 | 0.4 |
| 5 | 148631252 | rs2289285 | 0.878 | 0.019 | 0.138 | -0.273 | -0.006 | -0.009 | -0.009 | 0.42 | 0.11 | 0.07 | 0.07 | 0.12 | 0.14 | 0.11 |
| 5 | 148631433 | rs1438689 | -0.033 | 0.14 | 0.615 | -0.166 | 0.077 | 0.047 | 0.047 | 0 | 0.28 | 0.42 | 0.36 | 0.61 | 0.39 | 0.44 |
| 5 | 148639062 | rs1438688 | 0.439 | 0.184 | -0.017 | -0.214 | -0.109 | -0.098 | -0.098 | 0 | 0.24 | 0.08 | 0.33 | 0.57 | 0.39 | 0.39 |
| 5 | 148641917 | rs6887404 | -0.82 | -0.271 | 0.369 | -0.612 | 0.038 | -0.01 | -0.01 | 0 | 0.08 | 0.42 | 0.06 | 0 | 0.03 | 0.08 |
| 5 | 148646778 | rs4100758 | -0.011 | -0.796 | -1.147 | -1.017 | 0.013 | 0.019 | 0.019 | 0.037 | 0.31 | 0.26 | 0.45 | 0.19 | 0.26 | 0.28 |
| 5 | 148651711 | rs12054749 | -0.464 | -0.816 | -1.418 | -1.317 | -0.026 | 0.098 | 0.098 | 0 | 0.45 | 0.7 | 0.67 | 0.77 | 0.47 | 0.52 |
| 5 | 148653731 | rs4705335 | -1.304 | -0.587 | -1.19 | -1.096 | 0.001 | 0.043 | 0.043 | 0 | 0.79 | 0.85 | 0.89 | 0.97 | 0.8 | 0.87 |
| 5 | 148656344 | rs17109880 | 0.274 | -0.29 | -0.991 | -0.555 | -0.007 | -0.034 | -0.034 | 0.434 | 0.08 | 0.16 | 0.05 | 0.01 | 0.05 | 0.1 |
| 5 | 148657910 | rs1438693 | -0.876 | -0.145 | -0.785 | -0.542 | 0 | 0.037 | 0.037 | 0 | 0.49 | 0.44 | 0.65 | 0.78 | 0.49 | 0.6 |
| 5 | 148659664 | rs1438692 | -0.306 | 0.807 | -0.731 | 0.213 | -0.007 | -0.008 | -0.008 | 0.361 | 0.27 | 0.25 | 0.46 | 0.59 | 0.35 | 0.43 |
| 5 | 148663788 | rs352360 | 0.35 | 0.16 | -0.764 | -0.41 | -0.034 | 0.084 | 0.084 | 0 | 0.41 | 0.22 | 0.54 | 0.83 | 0.47 | 0.66 |
| 5 | 148663992 | rs352358 | -0.558 | -0.36 | -0.358 | -0.85 | 0.009 | -0.091 | -0.091 | 0 | 0.64 | 0.47 | 0.69 | 0.95 | 0.6 | 0.77 |
| 5 | 148664244 | rs352357 | 0.591 | -0.814 | 0.977 | 0.012 | 0.018 | 0.009 | 0.009 | 0 | 0.91 | 0.62 | 0.95 | 0.96 | 0.97 | 0.96 |
| 5 | 148665430 | rs352355 | 1.353 | -0.907 | 1.006 | -0.031 | -0.054 | -0.021 | -0.021 | 0 | 0.74 | 0.43 | 0.86 | 0.83 | 0.89 | 0.86 |
| 5 | 148666133 | rs352354 | -0.158 | -0.811 | 0.433 | 0.022 | 0.003 | -0.005 | -0.005 | 0 | 0.92 | 0.93 | 0.94 | 0.87 | 0.94 | 0.93 |
| 5 | 148668174 | rs1623904 | 1.813 | -0.865 | 0.75 | 0.004 | -0.042 | -0.012 | -0.012 | 0.251 | 0.76 | 0.56 | 0.86 | 0.84 | 0.89 | 0.87 |
| 5 | 148671814 | rs1621136 | 1.034 | -0.818 | 0.453 | 0.594 | -0.005 | 0.008 | 0.008 | 0.165 | 0.83 | 0.78 | 0.83 | 0.72 | 0.91 | 0.91 |
| 5 | 148672177 | rs1617849 | 1.778 | -0.84 | 0.461 | -0.177 | -0.069 | -0.042 | -0.042 | 0.103 | 0.69 | 0.45 | 0.81 | 0.83 | 0.85 | 0.83 |
| 5 | 148675225 | rs1659091 | 1.393 | -0.363 | 0.197 | 0.126 | -0.052 | -0.007 | -0.007 | 0 | 0.77 | 0.52 | 0.84 | 0.83 | 0.87 | 0.85 |
| 5 | 148683087 | rs17642472 | -0.429 | 0.005 | -0.585 | 0.175 | 0.01 | 0.008 | 0.008 | 0.02 | 0.08 | 0.06 | 0.05 | 0.04 | 0.05 | 0.03 |
| 5 | 148685500 | rs352338 | 0.888 | 0.391 | -0.399 | 0.253 | -0.001 | 0.002 | 0.002 | 0 | 0.79 | 0.79 | 0.84 | 0.79 | 0.81 | 0.84 |
| 5 | 148690879 | rs920317 | -0.896 | 0.561 | -0.463 | 0.077 | -0.005 | -0.038 | -0.038 | 0 | 0.1 | 0.11 | 0.06 | 0 | 0.08 | 0.01 |
| 5 | 148694097 | rs2438491 | -1.503 | 1.009 | -0.702 | -0.056 | -0.002 | -0.004 | -0.004 | 0.125 | 0.15 | 0.12 | 0.15 | 0.03 | 0.22 | 0.06 |
| 5 | 148696042 | rs10072778 | -1.373 | 0.629 | -0.265 | 0.127 | -0.016 | 0.019 | 0.019 | 0.03 | 0.12 | 0.25 | 0.09 | 0.12 | 0.08 | 0.03 |
| 5 | 148696383 | rs10079672 | -1.632 | 0.84 | -0.658 | -0.259 | -0.003 | -0.025 | -0.025 | 0.086 | 0.2 | 0.27 | 0.12 | 0.12 | 0.15 | 0.08 |
| 5 | 148699951 | rs17642594 | -0.893 | 0.844 | -1.097 | -0.355 | 0.003 | 0.009 | 0.009 | 0.019 | 0.11 | 0.06 | 0.07 | 0 | 0.09 | 0.01 |
| 5 | 148700714 | rs10515626 | -0.177 | 1.12 | -1.977 | 0.084 | -0.102 | -0.179 | -0.179 | 0 | 0.28 | 0.02 | 0.45 | 0.53 | 0.32 | 0.44 |
| 5 | 148712150 | rs2270682 | -0.299 | 1.421 | -1.347 | -0.653 | 0.022 | -0.057 | -0.057 | 0 | 0.28 | 0.5 | 0.32 | 0.16 | 0.32 | 0.26 |
| 5 | 148718498 | rs11743308 | 1.375 | 0.991 | -2.168 | -1.166 | 0.028 | 0.13 | 0.13 | 0 | 0.29 | 0.07 | 0.18 | 0.07 | 0.22 | 0.2 |

| | | | | | | | | | | | | | | | | |
|---|---|---|---|---|---|---|---|---|---|---|---|---|---|---|---|---|
| 5 | 148720087 | rs813035 | -0.879 | 0.934 | -2.006 | -0.918 | 0.009 | 0.067 | 0.067 | 0 | 0.52 | 0.76 | 0.46 | 0.31 | 0.54 | 0.37 |
| 5 | 148725199 | rs352347 | -0.772 | 0.879 | -1.948 | -0.87 | -0.021 | 0.055 | 0.055 | 0 | 0.46 | 0.76 | 0.44 | 0.31 | 0.49 | 0.34 |
| 5 | 148725821 | rs352346 | -0.741 | 0.872 | -1.825 | -0.873 | -0.012 | 0.04 | 0.04 | 0 | 0.45 | 0.7 | 0.43 | 0.31 | 0.49 | 0.34 |
| 5 | 148731218 | rs783777 | 0.973 | 0.84 | -1.886 | -0.843 | -0.049 | -0.034 | -0.034 | 1.809 | 0.11 | 0 | 0.1 | 0.08 | 0.17 | 0.05 |
| 5 | 148748728 | rs353255 | -0.686 | 0.808 | -1.614 | -1.039 | -0.013 | 0.021 | 0.021 | 0 | 0.41 | 0.66 | 0.43 | 0.24 | 0.5 | 0.31 |
| 5 | 148749852 | rs173296 | 0.997 | 0.844 | -1.623 | -0.977 | -0.044 | -0.027 | -0.027 | 0.215 | 0.11 | 0.01 | 0.1 | 0.08 | 0.17 | 0.05 |
| 5 | 148756848 | rs2227456 | 0.942 | 0.742 | -1.572 | -0.97 | -0.013 | 0.003 | 0.003 | 0 | 0.05 | 0.06 | 0.03 | 0.07 | 0.03 | 0.03 |
| 5 | 148758027 | rs2227447 | 0.185 | 0.776 | -1.477 | -1.014 | 0.022 | -0.062 | -0.062 | 0 | 0.28 | 0.52 | 0.31 | 0.16 | 0.33 | 0.25 |
| 5 | 148759014 | rs353272 | 0.956 | 1 | -1.823 | -0.992 | -0.044 | -0.027 | -0.027 | 0 | 0.11 | 0.01 | 0.1 | 0.08 | 0.17 | 0.05 |
| 5 | 148765904 | rs353275 | 0.8 | 1.013 | -1.775 | -0.999 | 0.009 | 0 | 0 | 0 | 0.12 | 0.17 | 0.11 | 0.08 | 0.17 | 0.05 |
| 5 | 148768604 | rs353282 | -0.425 | 0.622 | -2.004 | -0.838 | -0.026 | 0.067 | 0.067 | 0 | 0.46 | 0.76 | 0.45 | 0.3 | 0.52 | 0.34 |
| 5 | 148775566 | rs353265 | -0.372 | 0.671 | -1.375 | -0.08 | -0.01 | -0.034 | -0.034 | 0.916 | 0.77 | 0.85 | 0.64 | 0.38 | 0.76 | 0.55 |
| 5 | 148779127 | rs10054004 | -0.577 | 0.339 | -0.507 | -0.094 | -0.034 | -0.013 | -0.013 | 0 | 0.18 | 0.49 | 0.15 | 0.16 | 0.12 | 0.21 |
| 5 | 148780576 | rs387936 | 0.221 | 0.28 | -1.286 | -0.255 | 0.052 | 0.033 | 0.033 | 0 | 0.61 | 0.75 | 0.53 | 0.28 | 0.73 | 0.49 |
| 5 | 148780713 | rs10515627 | -0.2 | 1.487 | -0.756 | 0.34 | 0.008 | 0.044 | 0.044 | 0 | 0.07 | 0.08 | 0.1 | 0.22 | 0.15 | 0.04 |
| 5 | 148783268 | rs6861564 | -0.66 | 1.368 | -0.888 | -0.294 | 0.011 | 0.127 | 0.127 | 1.316 | 0.52 | 0.7 | 0.48 | 0.84 | 0.53 | 0.5 |
| 5 | 148783767 | rs6887452 | -0.355 | 1.169 | -0.496 | 0.026 | 0.053 | 0.041 | 0.041 | 0.03 | 0.46 | 0.29 | 0.33 | 0.58 | 0.32 | 0.48 |
| 5 | 148784809 | rs726953 | 0.863 | 1.773 | 0.331 | -0.043 | 0.032 | 0.071 | 0.071 | 0.537 | 0.5 | 0.37 | 0.39 | 0.14 | 0.35 | 0.42 |
| 5 | 148785449 | rs17796636 | -0.577 | 1.806 | 0.555 | 1.104 | 0.027 | 0.112 | 0.112 | 0 | 0.07 | 0.12 | 0.1 | 0.3 | 0.14 | 0.04 |
| 5 | 148787662 | rs12652986 | -0.613 | 1.802 | 0.629 | 1.102 | 0.039 | 0.142 | 0.142 | 0 | 0.07 | 0.16 | 0.1 | 0.3 | 0.14 | 0.03 |
| 5 | 148792761 | rs353286 | 0.989 | 1.954 | -0.105 | 0.912 | -0.118 | -0.072 | -0.072 | 0.685 | 0.54 | 0.04 | 0.54 | 0.5 | 0.52 | 0.37 |
| 5 | 148792853 | rs10476909 | -1.457 | 2.1 | 0.967 | 0.952 | 0.119 | 0.24 | 0.24 | 0.048 | 0.1 | 0.37 | 0.12 | 0.28 | 0.15 | 0.04 |
| 5 | 148798732 | rs2042249 | -0.666 | 1.775 | 0.594 | 0.776 | -0.07 | -0.101 | -0.101 | 0.377 | 0.26 | 0.43 | 0.21 | 0.12 | 0.16 | 0.17 |
| 5 | 148799117 | rs353303 | -0.241 | 2.119 | 0.186 | 1.028 | 0.03 | 0.058 | 0.058 | 0 | 0.82 | 0.74 | 0.79 | 0.61 | 0.69 | 0.58 |
| 5 | 148801101 | rs353299 | 1.534 | 1.803 | -0.203 | 0.814 | 0.006 | 0.187 | 0.187 | 0 | 0.44 | 0.12 | 0.47 | 0.16 | 0.38 | 0.33 |
| 5 | 148801826 | rs353296 | -1.451 | 1.871 | 0.712 | 1.298 | 0.005 | 0.011 | 0.011 | 0 | 0.31 | 0.56 | 0.34 | 0.45 | 0.36 | 0.21 |
| 5 | 148801876 | rs353295 | -1.039 | 1.577 | 0.254 | 1.217 | 0.003 | -0.042 | -0.042 | 0 | 0.17 | 0.33 | 0.21 | 0.13 | 0.19 | 0.16 |
| 5 | 148805005 | rs12659504 | -1.726 | 1.551 | -0.182 | 1.87 | 0.011 | 0.078 | 0.078 | 0 | 0.11 | 0.21 | 0.11 | 0.33 | 0.14 | 0.08 |
| 5 | 148805051 | rs878008 | -1.451 | 1.698 | -0.672 | 1.94 | -0.005 | 0.011 | 0.011 | 0 | 0.1 | 0.07 | 0.1 | 0.33 | 0.15 | 0.08 |
| 5 | 148810746 | rs353291 | -0.743 | 1.773 | -0.958 | 1.634 | -0.045 | -0.044 | -0.044 | 0 | 0.3 | 0.19 | 0.32 | 0.41 | 0.38 | 0.56 |
| 5 | 148815204 | rs476741 | -1.05 | 1.367 | -0.187 | 1.851 | 0.015 | 0.007 | 0.007 | 0 | 0.39 | 0.53 | 0.39 | 0.49 | 0.41 | 0.54 |
| 5 | 148818629 | rs452366 | -1.105 | 1.212 | -0.364 | 1.553 | -0.007 | 0.034 | 0.034 | 0 | 0.47 | 0.83 | 0.43 | 0.48 | 0.48 | 0.61 |
| 5 | 148818868 | rs443402 | 0.432 | 0.33 | 0.143 | 1.26 | -0.035 | -0.003 | -0.003 | 1.471 | 0.83 | 0.65 | 0.9 | 0.91 | 0.91 | 0.92 |
| 5 | 148819714 | rs403084 | 1.369 | 0.673 | 0.646 | 1.467 | 0.002 | -0.005 | -0.005 | 0 | 0.72 | 0.29 | 0.77 | 0.53 | 0.77 | 0.86 |
| 5 | 148821990 | rs2454104 | -0.112 | 0.6 | 0.24 | 0.92 | -0.029 | -0.008 | -0.008 | 0 | 0.32 | 0.24 | 0.42 | 0.44 | 0.4 | 0.57 |
| 5 | 148822118 | rs4705344 | 1.087 | 0.608 | 0.709 | 0.914 | 0.019 | 0.008 | 0.008 | 0 | 0.5 | 0.32 | 0.46 | 0.43 | 0.46 | 0.36 |
| 5 | 148823007 | rs734510 | -0.207 | 0.802 | 1.007 | 1.005 | -0.027 | -0.008 | -0.008 | 0 | 0.31 | 0.23 | 0.42 | 0.44 | 0.4 | 0.55 |
| 5 | 148824756 | rs353249 | 0.354 | 0.704 | 1.409 | 1.355 | 0.059 | 0.069 | 0.069 | 0 | 0.93 | 0.68 | 0.92 | 0.9 | 0.91 | 0.97 |
| 5 | 148825669 | rs10062536 | -1.025 | 0.657 | 1.374 | 1.414 | 0.065 | 0.371 | 0.371 | 0 | 0.07 | 0.33 | 0.07 | 0.34 | 0.07 | 0.09 |
| 5 | 148828478 | rs353241 | 1.514 | 0.669 | 1.354 | 0.971 | 0.04 | -0.036 | -0.036 | 0.32 | 0.88 | 0.39 | 0.89 | 0.96 | 0.9 | 0.92 |
| 5 | 148832326 | rs4705073 | -1.175 | 0.772 | 1.064 | 1.045 | 0.006 | -0.005 | -0.005 | 0.405 | 0.38 | 0.42 | 0.54 | 0.38 | 0.45 | 0.59 |
| 5 | 148835789 | rs191618 | -0.859 | 0.393 | 1.216 | 1.423 | -0.027 | 0.087 | 0.087 | 0 | 0.93 | 0.88 | 0.98 | 0.88 | 0.99 | 0.97 |
| 5 | 148839424 | rs11167468 | 1.506 | 0.444 | 1.296 | 1.471 | -0.031 | 0.101 | 0.101 | 0.174 | 0.81 | 0.46 | 0.88 | 0.48 | 0.89 | 0.9 |
| 5 | 148846426 | rs447950 | -1.286 | 0.945 | 0.838 | 1.361 | -0.01 | -0.019 | -0.019 | 0 | 0.24 | 0.18 | 0.43 | 0.4 | 0.3 | 0.45 |
| 5 | 148846518 | rs11750184 | -0.877 | 1.057 | 0.628 | 0.345 | -0.014 | 0.032 | 0.032 | 0 | 0.25 | 0.21 | 0.18 | 0.17 | 0.28 | 0.2 |
| 5 | 148846557 | rs386763 | -2.059 | 0.972 | 0.68 | 0.35 | 0.028 | 0.059 | 0.059 | 0 | 0.5 | 0.43 | 0.6 | 0.96 | 0.58 | 0.67 |
| 5 | 148851149 | rs414582 | -1.026 | 0.637 | 1.017 | 0.626 | 0.017 | -0.032 | -0.032 | 0 | 0.13 | 0.49 | 0.11 | 0.07 | 0.09 | 0.06 |
| 5 | 148861298 | rs241280 | -2.11 | 0.31 | 0.652 | 0.522 | 0.014 | 0.054 | 0.054 | 0.009 | 0.4 | 0.75 | 0.32 | 0.73 | 0.33 | 0.3 |
| 5 | 148869387 | rs10045427 | -1.413 | 0.375 | 0.729 | 0.279 | 0.021 | -0.073 | -0.073 | 0 | 0.25 | 0.34 | 0.21 | 0.1 | 0.25 | 0.16 |
| 5 | 148873854 | rs10057083 | -0.628 | 0.455 | 0.414 | 0.279 | 0.002 | -0.044 | -0.044 | 0.143 | 0.11 | 0.14 | 0.08 | 0 | 0.09 | 0.02 |
| 5 | 148883634 | rs6886243 | -0.578 | 0.915 | 0.421 | 0.56 | -0.001 | -0.008 | -0.008 | 0 | 0.21 | 0.24 | 0.42 | 0.63 | 0.26 | 0.44 |
| 5 | 148887713 | rs13170358 | -0.437 | 0.906 | 0.438 | 0.861 | -0.007 | -0.013 | -0.013 | 0 | 0.2 | 0.18 | 0.42 | 0.31 | 0.25 | 0.42 |
| 5 | 148889018 | rs752822 | -0.511 | 0.906 | 0.401 | 0.861 | -0.019 | -0.029 | -0.029 | 0.969 | 0.2 | 0.13 | 0.42 | 0.31 | 0.25 | 0.42 |
| 5 | 148893101 | rs7712431 | 0.229 | -0.209 | 0.036 | 0.259 | -0.02 | 0.033 | 0.033 | 0 | 0.15 | 0.07 | 0.11 | 0.1 | 0.16 | 0.15 |
| 5 | 148917992 | rs7719315 | -0.723 | -0.334 | 0.584 | 0.076 | 0.006 | -0.046 | -0.046 | 0 | 0.27 | 0.33 | 0.2 | 0.1 | 0.25 | 0.16 |
| 5 | 148932568 | rs13184089 | -2.415 | -0.461 | -0.191 | -0.039 | -0.056 | 0.182 | 0.182 | 0 | 0.6 | 0.89 | 0.71 | 0.99 | 0.53 | 0.72 |
| 5 | 148956032 | rs921978 | -0.877 | -0.802 | 0.175 | 0.131 | -0.007 | -0.097 | -0.097 | 0.509 | 0.28 | 0.38 | 0.17 | 0.1 | 0.2 | 0.15 |
| 5 | 148958653 | rs6861618 | -0.998 | -0.818 | 0.102 | 0.515 | -0.023 | -0.011 | -0.011 | 0 | 0.32 | 0.43 | 0.23 | 0.28 | 0.21 | 0.21 |
| 5 | 148960686 | rs13184845 | 0.575 | -0.729 | -0.546 | 0.658 | -0.058 | -0.056 | -0.056 | 0 | 0.1 | 0.01 | 0.36 | 0.22 | 0.19 | 0.37 |
| 5 | 148964897 | rs4705353 | -1.885 | -1.269 | -0.698 | 0.276 | -0.058 | 0.191 | 0.191 | 0.283 | 0.56 | 0.87 | 0.69 | 0.89 | 0.46 | 0.65 |
| 5 | 148974261 | rs6882002 | -0.725 | -1.208 | -0.026 | 0.482 | -0.011 | 0.034 | 0.034 | 0.013 | 0.29 | 0.35 | 0.19 | 0.38 | 0.18 | 0.26 |
| 5 | 148978105 | rs2400889 | -0.245 | -1.117 | -0.205 | 0.729 | -0.002 | -0.012 | -0.012 | 0 | 0.27 | 0.23 | 0.46 | 0.34 | 0.28 | 0.44 |
| 5 | 148981046 | rs1379544 | -1.86 | -1.107 | -0.097 | 0.615 | 0.003 | -0.012 | -0.012 | 0.084 | 0.58 | 0.58 | 0.68 | 0.72 | 0.49 | 0.7 |
| 5 | 148984001 | rs4705357 | -1.813 | -1.036 | -0.103 | 0.591 | 0.002 | -0.017 | -0.017 | 0 | 0.58 | 0.54 | 0.68 | 0.72 | 0.49 | 0.7 |
| 5 | 148985791 | rs7715047 | 2.31 | -1.063 | -0.376 | 0.519 | -0.107 | 0.229 | 0.229 | 0.06 | 0.36 | 0.06 | 0.28 | 0.01 | 0.49 | 0.26 |
| 5 | 148989122 | rs2400891 | 1.357 | -1.117 | -0.284 | 0.544 | -0.007 | 0.021 | 0.021 | 0 | 0.47 | 0.28 | 0.33 | 0.36 | 0.55 | 0.37 |
| 5 | 148989358 | rs17654288 | -0.396 | -1.043 | -0.05 | 0.67 | 0.002 | 0.048 | 0.048 | 0 | 0.09 | 0.21 | 0.05 | 0.08 | 0.05 | 0.08 |
| 5 | 148998847 | rs7728256 | -0.683 | -1.135 | -0.144 | 0.589 | -0.013 | 0.029 | 0.029 | 0 | 0.12 | 0.23 | 0.05 | 0.08 | 0.06 | 0.13 |
| 5 | 149001551 | rs4629585 | -1.424 | -1.177 | -0.174 | 0.648 | 0.004 | 0.021 | 0.021 | 0 | 0.32 | 0.38 | 0.48 | 0.5 | 0.34 | 0.52 |

| | | | | | | | | | | | | | | | | |
|---|---|---|---|---|---|---|---|---|---|---|---|---|---|---|---|---|
| 5 | 149001950 | rs4409073 | 1.773 | -1.43 | -0.422 | 0.404 | -0.034 | 0.147 | 0.147 | 0 | 0.42 | 0.2 | 0.34 | 0.17 | 0.5 | 0.28 |
| 5 | 149006640 | rs9324624 | 1.458 | -1.424 | -0.26 | 0.406 | -0.001 | -0.036 | -0.036 | 0 | 0.45 | 0.49 | 0.36 | 0.17 | 0.51 | 0.31 |
| 5 | 149006842 | rs7732874 | -0.196 | -1.386 | -0.231 | 0.504 | -0.002 | 0.05 | 0.05 | 0 | 0.11 | 0.27 | 0.05 | 0.08 | 0.06 | 0.1 |
| 5 | 149007059 | rs6894391 | 0.718 | -1.407 | -0.21 | 0.486 | -0.001 | 0.005 | 0.005 | 0 | 0.54 | 0.43 | 0.39 | 0.26 | 0.57 | 0.41 |
| 5 | 149007672 | rs9885369 | -0.454 | -1.391 | -0.295 | 0.475 | -0.015 | 0.021 | 0.021 | 0 | 0.34 | 0.4 | 0.21 | 0.42 | 0.21 | 0.3 |
| 5 | 149008403 | rs1056993 | -0.712 | -1.526 | -0.211 | 0.805 | 0.041 | -0.017 | -0.017 | 0 | 0.8 | 0.6 | 0.55 | 0.84 | 0.72 | 0.64 |
| 5 | 149008467 | rs3733662 | 0.968 | -1.517 | -0.219 | 0.803 | -0.029 | -0.083 | -0.083 | 0 | 0.23 | 0.02 | 0.14 | 0.33 | 0.15 | 0.2 |
| 5 | 149008521 | rs1135093 | -0.68 | -1.414 | -0.128 | 0.784 | 0.032 | -0.014 | -0.014 | 0 | 0.8 | 0.68 | 0.55 | 0.84 | 0.72 | 0.64 |
| 5 | 149012782 | rs3733658 | 1.528 | -1.336 | -0.779 | 0.358 | -0.073 | 0.131 | 0.131 | 0 | 0.4 | 0.12 | 0.29 | 0.12 | 0.51 | 0.34 |
| 5 | 149013587 | rs2279132 | 0.686 | -1.21 | -0.611 | 0.418 | 0.002 | -0.037 | -0.037 | 0 | 0.21 | 0.08 | 0.14 | 0.3 | 0.14 | 0.19 |
| 5 | 149020020 | rs13356972 | 0.084 | -1.212 | -0.488 | 0.287 | -0.005 | -0.001 | -0.001 | 0 | 0.24 | 0.27 | 0.12 | 0.25 | 0.15 | 0.2 |
| 5 | 149020986 | rs17724303 | 0.87 | -1.229 | -1.12 | 0.406 | 0.006 | -0.003 | -0.003 | 1.428 | 0.13 | 0.14 | 0.08 | 0.17 | 0.05 | 0.07 |
| 5 | 149023879 | rs10061434 | 0.277 | -0.756 | -1.229 | 0.12 | -0.008 | -0.036 | -0.036 | 0 | 0.23 | 0.07 | 0.11 | 0.27 | 0.18 | 0.23 |
| 5 | 149024328 | rs4705360 | 0.168 | -0.403 | -0.653 | 0.192 | -0.01 | -0.02 | -0.02 | 0 | 0.61 | 0.6 | 0.45 | 0.28 | 0.68 | 0.43 |
| 5 | 149026533 | rs6866304 | 0.1 | 0.055 | -0.784 | 0.496 | 0 | -0.001 | -0.001 | 1.347 | 0.42 | 0.54 | 0.5 | 0.39 | 0.51 | 0.2 |
| 5 | 149031582 | rs10074889 | -0.168 | 0.337 | -0.885 | 0.702 | -0.003 | -0.012 | -0.012 | 0 | 0.45 | 0.36 | 0.44 | 0.49 | 0.47 | 0.62 |
| 5 | 149043308 | rs6874724 | 0.36 | 0.865 | -0.526 | 0.422 | 0.011 | -0.017 | -0.017 | 2.938 | 0.15 | 0.2 | 0.13 | 0.1 | 0.18 | 0.12 |
| 5 | 149045703 | rs1478366 | -1.364 | 0.727 | -1 | 0.269 | -0.044 | 0.008 | 0.008 | 0.373 | 0.79 | 0.93 | 0.83 | 0.8 | 0.79 | 0.92 |
| 5 | 149047786 | rs728837 | 0.48 | 0.614 | -0.056 | 0.965 | 0.002 | -0.005 | -0.005 | 0 | 0.4 | 0.35 | 0.43 | 0.46 | 0.41 | 0.51 |
| 5 | 149051489 | rs12658991 | 0.528 | 0.631 | 0.488 | 1.132 | 0.068 | 0.204 | 0.204 | 0 | 0.05 | 0.26 | 0.1 | 0.17 | 0.05 | 0.12 |
| 5 | 149055331 | rs17110245 | 0.182 | 0.486 | 0.162 | 1.279 | -0.01 | 0.036 | 0.036 | 0.017 | 0.12 | 0.27 | 0.24 | 0.15 | 0.12 | 0.25 |
| 5 | 149057371 | rs12153727 | 0.922 | 0.388 | -0.116 | 0.901 | -0.019 | -0.018 | -0.018 | 0 | 0.28 | 0.14 | 0.19 | 0.3 | 0.29 | 0.25 |
| 5 | 149058864 | rs899174 | 0.827 | 0.377 | -0.025 | 0.896 | -0.005 | -0.004 | -0.004 | 0.45 | 0.29 | 0.21 | 0.19 | 0.3 | 0.29 | 0.25 |
| 5 | 149059130 | rs12652107 | 1.08 | 0.375 | 0.049 | 0.916 | -0.003 | 0.002 | 0.002 | 0.121 | 0.15 | 0.13 | 0.11 | 0.11 | 0.18 | 0.12 |
| 5 | 149061387 | rs17110264 | -0.588 | 0.383 | 0.258 | 1.055 | 0.008 | 0.157 | 0.157 | 0.075 | 0.17 | 0.43 | 0.33 | 0.32 | 0.17 | 0.38 |
| 5 | 149061981 | rs2061765 | -0.576 | 0.375 | 0.243 | 1.057 | 0.004 | 0.153 | 0.153 | 0.06 | 0.16 | 0.42 | 0.33 | 0.32 | 0.17 | 0.38 |
| 5 | 149064858 | rs12332370 | -0.016 | 0.402 | -0.285 | 0.601 | 0 | 0.003 | 0.003 | 0.454 | 0.2 | 0.16 | 0.3 | 0.15 | 0.19 | 0.26 |
| 5 | 149066552 | rs17797412 | 0.466 | 0.629 | -0.874 | 0.563 | -0.082 | -0.062 | -0.062 | 0 | 0.28 | 0.05 | 0.17 | 0.29 | 0.35 | 0.31 |
| 5 | 149070220 | rs6878452 | -0.325 | 0.626 | -0.316 | 0.609 | 0.006 | -0.002 | -0.002 | 0 | 0.28 | 0.28 | 0.28 | 0.27 | 0.38 | 0.39 |
| 5 | 149076956 | rs7731599 | -0.863 | 0.269 | -0.299 | 0.537 | 0.005 | 0.009 | 0.009 | 6.148 | 0.6 | 0.74 | 0.74 | 0.6 | 0.65 | 0.76 |
| 5 | 149082092 | rs2340623 | -0.29 | 0.185 | 0.132 | 0.604 | -0.004 | 0.024 | 0.024 | 4.257 | 0.48 | 0.35 | 0.58 | 0.35 | 0.55 | 0.54 |
| 5 | 149082704 | rs10078478 | -2.863 | 0.055 | -0.429 | 0.504 | 0.001 | -0.065 | -0.065 | 0 | 0.88 | 1 | 0.92 | 0.77 | 0.91 | 0.94 |
| 5 | 149082878 | rs10515634 | 0.3 | 0.289 | -0.019 | 0.741 | -0.01 | -0.016 | -0.016 | 0 | 0.13 | 0.08 | 0.18 | 0.18 | 0.19 | 0.31 |
| 5 | 149083207 | rs7721676 | 0.65 | -0.133 | 0.219 | 0.718 | -0.001 | 0.006 | 0.006 | 0 | 0.26 | 0.27 | 0.41 | 0.32 | 0.24 | 0.32 |
| 5 | 149096383 | rs4705079 | 1.195 | -0.273 | -0.104 | 0.47 | 0.029 | 0.021 | 0.021 | 0 | 0.23 | 0.11 | 0.3 | 0.16 | 0.21 | 0.28 |
| 5 | 149103165 | rs11740140 | 2.174 | -0.257 | 0.132 | 0.49 | 0.003 | 0.003 | 0.003 | 0 | 0.17 | 0.13 | 0.28 | 0.09 | 0.15 | 0.16 |
| 5 | 149105093 | rs17797652 | 0.66 | -0.137 | 0.161 | 0.847 | 0.003 | 0.041 | 0.041 | 0 | 0.1 | 0.12 | 0.13 | 0.28 | 0.1 | 0.16 |
| 5 | 149113096 | rs9986116 | -0.321 | -0.194 | 0.491 | 0.626 | -0.001 | 0.078 | 0.078 | 0 | 0.25 | 0.46 | 0.22 | 0.41 | 0.22 | 0.32 |
| 5 | 149114224 | rs10060424 | -0.684 | -0.257 | 0.424 | 0.541 | 0.02 | -0.012 | -0.012 | 0.236 | 0.44 | 0.67 | 0.5 | 0.49 | 0.37 | 0.48 |
| 5 | 149117004 | rs1124995 | -2.102 | -0.129 | 0.251 | 0.467 | -0.01 | 0 | 0 | 0 | 0.75 | 0.86 | 0.81 | 0.76 | 0.77 | 0.92 |
| 5 | 149118841 | rs10080122 | -2.068 | -0.132 | 0.249 | 0.466 | -0.009 | 0.001 | 0.001 | 1.835 | 0.75 | 0.85 | 0.81 | 0.76 | 0.77 | 0.92 |
| 5 | 149119815 | rs6650970 | 2.415 | -0.113 | 0.481 | 0.513 | -0.003 | 0.001 | 0.001 | 0.815 | 0.1 | 0.25 | 0.06 | 0.08 | 0.06 | 0.04 |
| 5 | 149124228 | rs17110375 | 1.294 | -0.101 | -0.008 | 0.538 | 0.01 | 0.008 | 0.008 | 0 | 0.09 | 0.05 | 0.04 | 0.08 | 0.07 | 0.14 |
| 5 | 149125449 | rs6880142 | 0.572 | -0.097 | 0.002 | 0.528 | -0.023 | 0.008 | 0.008 | 0.491 | 0.28 | 0.16 | 0.3 | 0.27 | 0.39 | 0.37 |
| 5 | 149133734 | rs11740247 | -2.049 | -0.042 | 0.028 | 0.528 | 0.002 | 0.005 | 0.005 | 1.636 | 0.74 | 0.83 | 0.8 | 0.76 | 0.77 | 0.9 |
| 5 | 149133874 | rs11746929 | 0.075 | -0.191 | -0.064 | 0.496 | 0.033 | -0.032 | -0.032 | 0 | 0.34 | 0.14 | 0.39 | 0.32 | 0.24 | 0.32 |
| 5 | 149136662 | rs4705089 | -0.075 | -0.212 | 0.061 | 0.525 | -0.014 | -0.015 | -0.015 | 0.526 | 0.38 | 0.31 | 0.43 | 0.5 | 0.49 | 0.54 |
| 5 | 149136958 | rs4705369 | -0.846 | -0.067 | -0.065 | 0.52 | 0.012 | 0.014 | 0.014 | 0.145 | 0.64 | 0.72 | 0.75 | 0.68 | 0.7 | 0.75 |
| 5 | 149142316 | rs2340621 | -0.908 | -0.04 | -0.379 | 0.493 | 0.015 | 0.011 | 0.011 | 0.266 | 0.65 | 0.82 | 0.76 | 0.67 | 0.7 | 0.75 |
| 5 | 149143963 | rs11746690 | 2.506 | 0.1 | -0.11 | 0.497 | -0.007 | -0.004 | -0.004 | 0 | 0.09 | 0.17 | 0.04 | 0.08 | 0.06 | 0.04 |
| 5 | 149152531 | rs10515638 | 0.408 | 0.074 | -0.336 | 0.467 | 0.002 | 0.002 | 0.002 | 0 | 0.1 | 0.09 | 0.04 | 0.09 | 0.07 | 0.19 |
| 5 | 149152691 | rs6894584 | 1.69 | -0.014 | -0.748 | 0.483 | -0.008 | -0.025 | -0.025 | 0 | 0.27 | 0.09 | 0.2 | 0.27 | 0.23 | 0.11 |
| 5 | 149153570 | rs13190217 | -0.224 | 0.12 | -0.285 | 0.507 | 0.017 | -0.009 | -0.009 | 0 | 0.64 | 0.53 | 0.65 | 0.66 | 0.55 | 0.64 |
| 5 | 149154877 | rs880769 | 1.411 | -0.105 | -0.854 | 0.382 | -0.001 | -0.04 | -0.04 | 12.285 | 0.36 | 0.1 | 0.25 | 0.4 | 0.3 | 0.15 |
| 5 | 149161437 | rs6579757 | 0.307 | 0.194 | -0.226 | 0.553 | 0.001 | -0.015 | -0.015 | 0.014 | 0.56 | 0.74 | 0.59 | 0.56 | 0.46 | 0.45 |
| 5 | 149163414 | rs4705377 | -0.391 | 0.408 | -0.376 | 0.404 | -0.044 | 0.047 | 0.047 | 0 | 0.66 | 0.86 | 0.72 | 0.79 | 0.57 | 0.63 |
| 5 | 149166434 | rs1422429 | 0.067 | 0.5 | -0.048 | 0.545 | 0.001 | 0.014 | 0.014 | 0 | 0.48 | 0.47 | 0.42 | 0.72 | 0.42 | 0.46 |
| 5 | 149172257 | rs13158830 | 0.596 | 0.726 | -0.191 | 1.048 | -0.01 | -0.004 | -0.004 | 0 | 0.09 | 0.07 | 0.13 | 0.23 | 0.11 | 0.21 |
| 5 | 149173134 | rs10875551 | 0.221 | 0.979 | 0.101 | 0.957 | 0.094 | 0.145 | 0.145 | 0 | 0.82 | 0.59 | 0.78 | 0.68 | 0.75 | 0.63 |
| 5 | 149174952 | rs4705380 | 2.178 | 0.461 | -0.531 | 0.508 | 0.005 | 0.008 | 0.008 | 0 | 0.16 | 0.1 | 0.27 | 0.06 | 0.15 | 0.11 |
| 5 | 149179725 | rs17461842 | 1.523 | 0.449 | -1.282 | 0.188 | -0.024 | 0.05 | 0.05 | 0 | 0.16 | 0 | 0.12 | 0.05 | 0.1 | 0.04 |
| 5 | 149181366 | rs4705382 | -0.418 | 0.5 | -0.572 | 0.429 | -0.004 | 0.005 | 0.005 | 0 | 0.43 | 0.51 | 0.56 | 0.41 | 0.37 | 0.36 |
| 5 | 149181386 | rs759813 | 2.687 | 0.502 | -0.649 | 0.534 | -0.017 | -0.009 | -0.009 | 0 | 0.14 | 0 | 0.27 | 0.06 | 0.15 | 0.09 |
| 5 | 149182447 | rs12717846 | 1.919 | 0.528 | -0.717 | 0.386 | -0.019 | -0.004 | -0.004 | 0.09 | 0.28 | 0.16 | 0.27 | 0.32 | 0.32 | 0.12 |
| 5 | 149186077 | rs4705094 | -1.882 | 0.549 | -0.602 | 0.401 | 0.03 | 0.019 | 0.019 | 0.001 | 0.91 | 0.83 | 0.92 | 0.91 | 0.86 | 0.85 |
| 5 | 149187038 | rs1544744 | -1.974 | 0.561 | -0.694 | 0.407 | 0.021 | 0.013 | 0.013 | 0.009 | 0.92 | 0.86 | 0.92 | 0.91 | 0.86 | 0.85 |
| 5 | 149187134 | rs9324628 | 1.729 | 0.188 | -0.86 | 0.354 | -0.018 | -0.003 | -0.003 | 0 | 0.28 | 0.17 | 0.27 | 0.32 | 0.33 | 0.12 |
| 5 | 149187457 | rs17110439 | 0.718 | 0.275 | -0.663 | 0.109 | 0.015 | 0.02 | 0.02 | 0 | 0.2 | 0.14 | 0.09 | 0.16 | 0.16 | 0.37 |
| 5 | 149189489 | rs10875552 | -0.933 | 0.429 | -0.695 | -0.102 | -0.011 | 0.074 | 0.074 | 0 | 0.67 | 0.81 | 0.78 | 0.88 | 0.64 | 0.59 |

| | | | | | | | | | | | | | | | | |
|---|---|---|---|---|---|---|---|---|---|---|---|---|---|---|---|---|
| 5 | 149189997 | rs2161257 | -1.356 | 0.651 | -0.57 | 0.258 | 0.001 | 0.013 | 0.013 | 1.116 | 0.45 | 0.59 | 0.61 | 0.66 | 0.4 | 0.56 |
| 5 | 149191652 | rs2010994 | -1.98 | 0.505 | -0.543 | 0.252 | -0.083 | -0.052 | -0.052 | 0 | 0.8 | 0.99 | 0.65 | 0.85 | 0.72 | 0.74 |
| 5 | 149194485 | rs251468 | -0.458 | 0.154 | 0.508 | 0.471 | -0.02 | -0.022 | -0.022 | 0 | 0.32 | 0.42 | 0.41 | 0.22 | 0.26 | 0.32 |
| 5 | 149194923 | rs109075 | -0.982 | 0.149 | 0.618 | 0.703 | 0.016 | -0.002 | -0.002 | 0 | 0.38 | 0.7 | 0.46 | 0.38 | 0.29 | 0.39 |
| 5 | 149196329 | rs251463 | -0.366 | 0.148 | 0.774 | 0.595 | -0.023 | -0.062 | -0.062 | 0 | 0.3 | 0.43 | 0.22 | 0.14 | 0.22 | 0.28 |
| 5 | 149199889 | rs32589 | 1.073 | 0.061 | 0.769 | 0.549 | -0.024 | -0.035 | -0.035 | 0.005 | 0.19 | 0.25 | 0.08 | 0.09 | 0.14 | 0.22 |
| 5 | 149200043 | rs32588 | 0.996 | 0.067 | 0.727 | 0.551 | -0.042 | -0.062 | -0.062 | 0 | 0.19 | 0.38 | 0.09 | 0.09 | 0.14 | 0.22 |
| 5 | 149200402 | rs17462080 | 1.081 | 0.265 | 0.07 | 0.582 | -0.022 | 0.017 | 0.017 | 0 | 0.1 | 0.02 | 0.13 | 0.06 | 0.08 | 0.06 |
| 5 | 149201670 | rs741580 | -0.152 | 0.335 | 0.177 | 0.686 | 0.005 | 0.039 | 0.039 | 0 | 0.06 | 0.15 | 0.06 | 0.16 | 0.05 | 0.08 |
| 5 | 149205630 | rs32581 | -0.583 | 0.278 | 0.293 | 0.671 | -0.003 | 0.008 | 0.008 | 0 | 0.31 | 0.62 | 0.45 | 0.36 | 0.28 | 0.36 |
| 5 | 149205940 | rs2003604 | 0.424 | 0.41 | 0.303 | 0.79 | -0.001 | 0.018 | 0.018 | 0 | 0.05 | 0.07 | 0.03 | 0.15 | 0.03 | 0.07 |
| 5 | 149206104 | rs11749124 | -0.606 | 0.347 | 0.27 | 0.665 | -0.006 | 0.03 | 0.03 | 0 | 0.07 | 0.14 | 0.04 | 0.13 | 0.03 | 0.07 |
| 5 | 149208768 | rs10491361 | 0.889 | 0.454 | 0.426 | 0.669 | 0.019 | -0.02 | -0.02 | 0.449 | 0.11 | 0.25 | 0.15 | 0.06 | 0.08 | 0.07 |
| 5 | 149212510 | rs11959820 | 0.022 | 0.66 | -0.113 | 0.787 | 0.026 | 0.092 | 0.092 | 0 | 0.05 | 0.2 | 0.05 | 0.13 | 0.03 | 0.07 |
| 5 | 149213456 | rs32576 | -0.288 | 0.61 | -0.098 | 0.743 | -0.029 | 0.032 | 0.032 | 0 | 0.25 | 0.49 | 0.31 | 0.29 | 0.22 | 0.33 |
| 5 | 149216987 | rs10783180 | -1.221 | 0.512 | -0.107 | 0.746 | 0.034 | 0.218 | 0.218 | 0 | 0.09 | 0.34 | 0.26 | 0.27 | 0.08 | 0.12 |
| 5 | 149218886 | rs32574 | -0.387 | 0.505 | -0.207 | 0.737 | -0.028 | 0.016 | 0.016 | 0 | 0.27 | 0.55 | 0.33 | 0.28 | 0.23 | 0.34 |
| 5 | 149230730 | rs25846 | -0.907 | 1.068 | -1.592 | 0.241 | -0.088 | -0.07 | -0.07 | 0.138 | 0.76 | 0.99 | 0.64 | 0.78 | 0.7 | 0.75 |
| 5 | 149231519 | rs26122 | -0.092 | 0.634 | 0.067 | 0.897 | -0.017 | -0.009 | -0.009 | 0 | 0.21 | 0.27 | 0.28 | 0.16 | 0.19 | 0.27 |
| 5 | 149233555 | rs3733665 | 0.763 | 1.084 | -0.279 | 0.855 | -0.031 | -0.029 | -0.029 | 0 | 0.15 | 0.01 | 0.07 | 0.14 | 0.13 | 0.05 |
| 5 | 149237916 | rs30884 | -0.07 | 1.093 | 0.113 | 0.914 | -0.024 | 0.002 | 0.002 | 0.406 | 0.33 | 0.24 | 0.43 | 0.38 | 0.41 | 0.22 |
| 5 | 149239093 | rs888895 | 0.406 | 0.859 | 0.791 | 1.199 | 0.02 | -0.007 | -0.007 | 0 | 0.58 | 0.67 | 0.62 | 0.53 | 0.6 | 0.37 |
| 5 | 149240767 | rs30887 | -0.007 | 0.853 | 0.673 | 0.387 | -0.011 | -0.053 | -0.053 | 0.177 | 0.77 | 0.57 | 0.75 | 0.86 | 0.78 | 0.77 |
| 5 | 149241601 | rs10515637 | 0.671 | 1.166 | -0.103 | 0.313 | -0.065 | 0.097 | 0.097 | 2.079 | 0.41 | 0.1 | 0.29 | 0.19 | 0.4 | 0.16 |
| 5 | 149251139 | rs245082 | -0.938 | 0.424 | -0.128 | 0.613 | -0.016 | -0.04 | -0.04 | 1.511 | 0.43 | 0.82 | 0.42 | 0.59 | 0.41 | 0.71 |
| 5 | 149251539 | rs6886532 | -1.93 | 0.041 | 0.663 | 0.361 | 0.003 | -0.004 | -0.004 | 0 | 0.05 | 0.05 | 0.09 | 0.09 | 0.04 | 0.07 |
| 5 | 149253914 | rs10223176 | -1.252 | 0.096 | 0.831 | 0.199 | 0.036 | 0.038 | 0.038 | 0 | 0.06 | 0.35 | 0.12 | 0.01 | 0.03 | 0.05 |
| 5 | 149266279 | rs17096125 | -1.346 | 0.389 | -0.207 | 0.234 | -0.007 | -0.031 | -0.031 | 2.781 | 0.13 | 0.17 | 0.09 | 0.05 | 0.08 | 0.26 |
| 5 | 149276567 | rs30818 | 0.047 | 0.151 | -1.478 | -0.177 | 0.022 | 0.133 | 0.133 | 0 | 0.48 | 0.12 | 0.36 | 0.27 | 0.55 | 0.33 |
| 5 | 149279976 | rs245073 | 0.054 | -0.039 | -1.565 | -0.084 | 0.021 | 0.125 | 0.125 | 0.039 | 0.48 | 0.12 | 0.36 | 0.28 | 0.55 | 0.33 |
| 5 | 149285033 | rs2569200 | -0.226 | -0.758 | -2.133 | 0.663 | 0.015 | -0.133 | -0.133 | 0.351 | 0.75 | 0.94 | 0.76 | 0.57 | 0.78 | 0.73 |
| 5 | 149291275 | rs245068 | 0.916 | -1.039 | -2.617 | 0.196 | -0.026 | 0.102 | 0.102 | 0.094 | 0.44 | 0.14 | 0.28 | 0.26 | 0.48 | 0.31 |
| 5 | 149293060 | rs30825 | 0.297 | -1.243 | -1.131 | 0.188 | -0.028 | 0.029 | 0.029 | 0.011 | 0.63 | 0.51 | 0.6 | 0.49 | 0.68 | 0.53 |
| 5 | 149301335 | rs11167487 | -1.215 | -1.341 | -0.385 | 0.193 | 0.002 | 0.03 | 0.03 | 0.003 | 0.19 | 0.7 | 0.35 | 0.26 | 0.17 | 0.33 |
| 5 | 149310447 | rs3756305 | 0.06 | -1.351 | 0.564 | 0.198 | 0.071 | 0.241 | 0.241 | 0 | 0.09 | 0.37 | 0.21 | 0.22 | 0.08 | 0.2 |
| 5 | 149315287 | rs3776069 | -0.872 | -1.567 | 1.107 | 0.128 | 0.014 | 0.047 | 0.047 | 0 | 0.16 | 0.22 | 0.22 | 0.25 | 0.16 | 0.3 |
| 5 | 149323772 | rs2277926 | -1.065 | -1.565 | 1.272 | 0.126 | 0.012 | 0.055 | 0.055 | 0 | 0.17 | 0.28 | 0.23 | 0.25 | 0.17 | 0.31 |
| 5 | 149325721 | rs12109815 | -1.252 | -1.668 | 0.892 | 0.363 | -0.003 | 0.012 | 0.012 | 0 | 0.34 | 0.61 | 0.49 | 0.49 | 0.34 | 0.6 |
| 5 | 149326063 | rs13357787 | -0.97 | -1.684 | 1.243 | 0.468 | 0 | 0.024 | 0.024 | 0 | 0.29 | 0.38 | 0.47 | 0.46 | 0.33 | 0.53 |
| 5 | 149326249 | rs10040644 | -1.044 | -1.713 | 1.151 | 0.471 | -0.003 | 0.045 | 0.045 | 0 | 0.31 | 0.57 | 0.48 | 0.47 | 0.33 | 0.55 |
| 5 | 149327532 | rs245048 | 1.463 | -2.057 | 0.585 | 0.37 | 0.009 | 0.056 | 0.056 | 0 | 0.55 | 0.28 | 0.39 | 0.25 | 0.57 | 0.4 |
| 5 | 149331638 | rs446426 | -0.244 | -2.234 | 0.285 | 0.309 | 0.013 | -0.034 | -0.034 | 4.188 | 0.33 | 0.24 | 0.53 | 0.44 | 0.34 | 0.43 |
| 5 | 149335379 | rs6579764 | -0.132 | -1.753 | 0.484 | 0.503 | 0.048 | 0.01 | 0.01 | 0 | 0.08 | 0.24 | 0.06 | 0.08 | 0.11 | 0.13 |
| 5 | 149337416 | rs173438 | 1.144 | -1.883 | 0.014 | 0.264 | -0.021 | 0.039 | 0.039 | 0 | 0.38 | 0.26 | 0.27 | 0.21 | 0.42 | 0.29 |
| 5 | 149338157 | rs245057 | 0.461 | -2.468 | -0.306 | 0.038 | -0.001 | -0.001 | -0.001 | 0 | 0.47 | 0.52 | 0.39 | 0.49 | 0.51 | 0.35 |
| 5 | 149341279 | rs245055 | -0.501 | -2.852 | -1.359 | 0.585 | -0.019 | -0.119 | -0.119 | 0 | 0.29 | 0.03 | 0.24 | 0.38 | 0.28 | 0.38 |
| 5 | 149345975 | rs245051 | -1.697 | -1.514 | -1.176 | 0.259 | 0.016 | -0.016 | -0.016 | 0 | 0.55 | 0.76 | 0.55 | 0.48 | 0.44 | 0.49 |
| 5 | 149349161 | rs11739934 | 1.322 | -1.636 | -0.824 | 0.186 | 0.01 | 0.013 | 0.013 | 4.503 | 0.19 | 0.1 | 0.26 | 0.1 | 0.16 | 0.06 |
| 5 | 149350648 | rs245076 | 0.582 | -1.647 | -0.04 | 0.188 | 0.014 | -0.018 | -0.018 | 0 | 0.32 | 0.28 | 0.26 | 0.38 | 0.28 | 0.4 |
| 5 | 149356486 | rs3756307 | 1.043 | -1.54 | 0.014 | 0.096 | 0.001 | 0.003 | 0.003 | 0 | 0.16 | 0.1 | 0.26 | 0.1 | 0.15 | 0.06 |
| 5 | 149357060 | rs17110742 | 0.78 | -1.55 | 0.123 | 0.092 | -0.005 | -0.008 | -0.008 | 0 | 0.17 | 0.27 | 0.28 | 0.1 | 0.15 | 0.06 |
| 5 | 149362400 | rs8073 | 0.441 | -1.708 | 0.645 | 0.256 | 0.013 | 0.005 | 0.005 | 0 | 0.33 | 0.27 | 0.24 | 0.3 | 0.23 | 0.32 |
| 5 | 149366714 | rs17711997 | 0.933 | -1.638 | 0.302 | 0.271 | 0.001 | 0.003 | 0.003 | 0 | 0.17 | 0.14 | 0.26 | 0.1 | 0.16 | 0.06 |
| 5 | 149369776 | rs10038824 | 0.295 | -1.801 | 0.44 | 0.249 | -0.022 | -0.011 | -0.011 | 0.704 | 0.33 | 0.54 | 0.25 | 0.29 | 0.23 | 0.29 |
| 5 | 149373140 | rs973077 | 0.991 | -1.647 | 0.371 | 0.271 | 0.002 | 0.003 | 0.003 | 0.007 | 0.16 | 0.09 | 0.26 | 0.1 | 0.15 | 0.06 |
| 5 | 149379897 | rs6579765 | 0.711 | -1.641 | 0.528 | 0.337 | -0.009 | -0.005 | -0.005 | 0 | 0.16 | 0.29 | 0.27 | 0.1 | 0.13 | 0.04 |
| 5 | 149380493 | rs4705403 | 0.966 | -1.636 | 0.304 | 0.339 | 0.002 | 0 | 0 | 0.02 | 0.14 | 0.09 | 0.26 | 0.1 | 0.13 | 0.04 |
| 5 | 149381952 | rs4705404 | 1.475 | -1.779 | 0.251 | 0.367 | 0.075 | 0.02 | 0.02 | 0.009 | 0.5 | 0.19 | 0.48 | 0.59 | 0.64 | 0.62 |
| 5 | 149390148 | rs2241698 | 1.038 | -1.732 | 0.181 | 0.384 | 0.002 | 0 | 0 | 0.02 | 0.14 | 0.09 | 0.26 | 0.1 | 0.13 | 0.04 |
| 5 | 149393006 | rs1962603 | -0.887 | -1.713 | -0.082 | 0.38 | -0.025 | -0.074 | -0.074 | 0 | 0.72 | 0.99 | 0.8 | 0.69 | 0.78 | 0.69 |
| 5 | 149397862 | rs17655812 | 2 | -1.284 | -0.279 | 0.249 | -0.036 | 0.014 | 0.014 | 0 | 0.07 | 0.01 | 0.08 | 0.03 | 0.15 | 0.14 |
| 5 | 149398480 | rs7720470 | 1.595 | -2.08 | 0.066 | 0.633 | 0.014 | -0.043 | -0.043 | 0 | 0.46 | 0.07 | 0.46 | 0.49 | 0.6 | 0.55 |
| 5 | 149410153 | rs2278393 | -0.542 | -2.081 | 0.384 | 0.577 | -0.018 | -0.014 | -0.014 | 0 | 0.2 | 0.64 | 0.3 | 0.1 | 0.13 | 0.05 |
| 5 | 149414899 | rs4705409 | -0.842 | -2.095 | 0.371 | 0.584 | 0.016 | -0.021 | -0.021 | 0.754 | 0.71 | 0.92 | 0.79 | 0.7 | 0.77 | 0.71 |
| 5 | 149417138 | rs6887999 | 0.864 | -2.027 | 0.639 | 0.572 | -0.007 | -0.002 | -0.002 | 0 | 0.16 | 0.27 | 0.27 | 0.1 | 0.12 | 0.04 |
| 5 | 149420698 | rs2304070 | 0.692 | -2.021 | 0.453 | 0.575 | -0.007 | 0 | 0 | 0.021 | 0.17 | 0.4 | 0.29 | 0.1 | 0.13 | 0.04 |
| 5 | 149421502 | rs2276982 | -0.377 | -1.98 | -0.492 | 0.594 | -0.019 | -0.014 | -0.014 | 0 | 0.2 | 0.63 | 0.3 | 0.1 | 0.13 | 0.05 |
| 5 | 149424508 | rs13178760 | -0.096 | -1.951 | -0.607 | 0.542 | -0.013 | 0.011 | 0.011 | 0 | 0.05 | 0.11 | 0.03 | 0.07 | 0.05 | 0.08 |
| 5 | 149429209 | rs216128 | 0.675 | -1.818 | -0.526 | 0.715 | 0.054 | 0.024 | 0.024 | 0 | 0.51 | 0.34 | 0.49 | 0.6 | 0.65 | 0.67 |

| | | | | | | | | | | | | | | | | |
|---|---|---|---|---|---|---|---|---|---|---|---|---|---|---|---|---|
| 5 | 149431025 | rs4705411 | -1.088 | -1.322 | -0.792 | 0.798 | 0.047 | -0.003 | -0.003 | 0 | 0.25 | 0.89 | 0.34 | 0.18 | 0.17 | 0.16 |
| 5 | 149435759 | rs216136 | 0.792 | -1.079 | -0.806 | 0.792 | 0.007 | 0.006 | 0.006 | 0 | 0.48 | 0.1 | 0.44 | 0.46 | 0.58 | 0.51 |
| 5 | 149442474 | rs2282806 | 0.084 | -0.903 | -0.331 | 0.845 | -0.002 | 0.002 | 0.002 | 0 | 0.13 | 0.08 | 0.26 | 0.1 | 0.13 | 0.05 |
| 5 | 149443298 | rs216140 | 0.019 | -1.136 | 0.025 | 0.954 | 0.003 | -0.005 | -0.005 | 0 | 0.63 | 0.38 | 0.72 | 0.54 | 0.72 | 0.56 |
| 5 | 149445921 | rs216146 | 0.567 | -1.107 | 0.139 | 0.966 | 0.005 | -0.001 | -0.001 | 0 | 0.49 | 0.49 | 0.47 | 0.48 | 0.58 | 0.55 |
| 5 | 149445987 | rs7719910 | -0.384 | -1.293 | 0.203 | 1.223 | 0.003 | 0.001 | 0.001 | 1.083 | 0.18 | 0.16 | 0.1 | 0.18 | 0.13 | 0.07 |
| 5 | 149446487 | rs216148 | 0.151 | -1.75 | 0.316 | 1.634 | -0.07 | 0.194 | 0.194 | 0 | 0.21 | 0.5 | 0.29 | 0.5 | 0.09 | 0.23 |
| 5 | 149450132 | rs10079250 | 0.952 | -1.82 | -0.111 | 1.571 | 0.015 | -0.016 | -0.016 | 0 | 0.15 | 0.11 | 0.1 | 0.38 | 0.07 | 0.11 |
| 5 | 149453450 | rs1549919 | -1.127 | -1.906 | -0.17 | 1.219 | 0.005 | 0.02 | 0.02 | 0.013 | 0.73 | 0.78 | 0.74 | 0.79 | 0.75 | 0.74 |
| 5 | 149454918 | rs12651699 | -0.337 | -1.999 | 0.34 | 1.142 | -0.086 | 0.13 | 0.13 | 0.037 | 0.26 | 0.52 | 0.16 | 0.53 | 0.1 | 0.16 |
| 5 | 149455029 | rs1076869 | -0.426 | -2.006 | -0.027 | 1.132 | -0.031 | 0.031 | 0.031 | 0.081 | 0.27 | 0.67 | 0.17 | 0.53 | 0.1 | 0.16 |
| 5 | 149456771 | rs2855781 | 0.951 | -1.809 | -0.07 | 0.868 | -0.022 | 0.022 | 0.022 | 0 | 0.32 | 0.23 | 0.34 | 0.07 | 0.52 | 0.44 |
| 5 | 149456811 | rs3829987 | -0.283 | -1.813 | 0.026 | 1.057 | -0.025 | 0.145 | 0.145 | 1.881 | 0.12 | 0.4 | 0.21 | 0.22 | 0.03 | 0.05 |
| 5 | 149457678 | rs2228422 | 0.393 | -1.609 | -0.698 | 0.555 | -0.015 | 0.025 | 0.025 | 0 | 0.37 | 0.28 | 0.37 | 0.21 | 0.55 | 0.57 |
| 5 | 149458829 | rs2569075 | 0.257 | -1.029 | -0.928 | 0.293 | -0.017 | 0.101 | 0.101 | 0.451 | 0.42 | 0.24 | 0.35 | 0.07 | 0.54 | 0.45 |
| 5 | 149460176 | rs1986027 | 0.802 | -1.151 | -0.994 | 0.532 | 0.03 | -0.047 | -0.047 | 0.101 | 0.14 | 0.06 | 0.1 | 0.34 | 0.07 | 0.12 |
| 5 | 149460553 | rs216123 | -0.129 | -0.476 | -0.681 | -0.17 | -0.007 | 0.085 | 0.085 | 0.1 | 0.43 | 0.26 | 0.35 | 0.09 | 0.55 | 0.46 |
| 5 | 149461912 | rs6870146 | 0.067 | 1.176 | -0.676 | 0.041 | -0.03 | 0.028 | 0.028 | 0 | 0.14 | 0.07 | 0.22 | 0.02 | 0.25 | 0.09 |
| 5 | 149462009 | rs216124 | -0.694 | 0.275 | -0.541 | 0.29 | -0.003 | 0.009 | 0.009 | 0.045 | 0.82 | 0.92 | 0.68 | 0.83 | 0.81 | 0.86 |
| 5 | 149465491 | rs2237084 | 0.256 | 0.208 | -0.316 | 0.319 | 0.005 | 0.008 | 0.008 | 0 | 0.26 | 0.17 | 0.26 | 0.18 | 0.27 | 0.21 |
| 5 | 149467663 | rs6874087 | -0.049 | 0.165 | 0.077 | 0.3 | -0.021 | -0.029 | -0.029 | 0.403 | 0.3 | 0.52 | 0.3 | 0.18 | 0.3 | 0.28 |
| 5 | 149479647 | rs11746309 | 0.641 | 0.44 | -0.29 | 0.367 | 0.005 | -0.021 | -0.021 | 0 | 0.34 | 0.29 | 0.19 | 0.47 | 0.28 | 0.26 |
| 5 | 149479852 | rs4337842 | 0.641 | 0.399 | -0.29 | 0.367 | 0.006 | -0.021 | -0.021 | 0 | 0.34 | 0.28 | 0.19 | 0.47 | 0.28 | 0.27 |
| 5 | 149480879 | rs13362413 | 0.671 | 0.992 | -0.019 | 0.484 | 0.007 | -0.009 | -0.009 | 0 | 0.61 | 0.56 | 0.52 | 0.63 | 0.59 | 0.56 |
| 5 | 149483141 | rs11743220 | 0.248 | 1.476 | -0.649 | 0.751 | -0.009 | -0.002 | -0.002 | 0 | 0.68 | 0.74 | 0.58 | 0.63 | 0.64 | 0.56 |
| 5 | 149487768 | rs17798680 | -0.906 | 1.342 | -0.158 | 1.22 | 0.021 | 0.087 | 0.087 | 0 | 0.09 | 0.17 | 0.1 | 0.28 | 0.12 | 0.05 |
| 5 | 149487941 | rs7727195 | -1.444 | 1.668 | 0.106 | 1.119 | 0.033 | 0.048 | 0.048 | 0 | 0.19 | 0.36 | 0.13 | 0.32 | 0.21 | 0.1 |
| 5 | 149492455 | rs11740298 | -0.149 | 1.044 | -1.187 | 0.119 | -0.075 | 0.187 | 0.187 | 0.227 | 0.35 | 0.03 | 0.23 | 0.11 | 0.3 | 0.27 |
| 5 | 149494521 | rs1017375 | -0.165 | 0.821 | -0.204 | 0.003 | 0.035 | -0.003 | -0.003 | 0 | 0.07 | 0.22 | 0.11 | 0.01 | 0.05 | 0.06 |
| 5 | 149494984 | rs6887454 | 0.121 | 0.818 | -0.278 | 0.002 | -0.001 | 0 | 0 | 0 | 0.06 | 0.07 | 0.1 | 0.01 | 0.05 | 0.06 |
| 5 | 149495770 | rs165979 | -0.792 | 0.558 | 0.057 | -0.137 | -0.015 | 0.037 | 0.037 | 0.236 | 0.56 | 0.74 | 0.5 | 0.37 | 0.45 | 0.45 |
| 5 | 149497177 | rs246391 | -0.166 | 0.68 | 0.046 | -0.217 | -0.007 | 0.071 | 0.071 | 1.127 | 0.46 | 0.35 | 0.32 | 0.11 | 0.4 | 0.31 |
| 5 | 149498821 | rs3733676 | 1.314 | 0.047 | -0.664 | 0.458 | 0.02 | -0.07 | -0.07 | 0 | 0.11 | 0.01 | 0.04 | 0.22 | 0.06 | 0.1 |
| 5 | 149499672 | rs246395 | 0.537 | 0.161 | -0.428 | 0.15 | -0.018 | 0.112 | 0.112 | 0.375 | 0.37 | 0.15 | 0.3 | 0.15 | 0.29 | 0.33 |
| 5 | 149501803 | rs17656204 | 0.624 | 0.181 | -0.516 | 0.127 | -0.057 | 0.182 | 0.182 | 0.806 | 0.35 | 0.04 | 0.23 | 0.1 | 0.28 | 0.26 |
| 5 | 149504158 | rs1075846 | -0.269 | 0.249 | -0.016 | 0.114 | -0.001 | 0.004 | 0.004 | 0.826 | 0.42 | 0.6 | 0.39 | 0.38 | 0.33 | 0.35 |
| 5 | 149505489 | rs919751 | 1.511 | 0.435 | -0.832 | -0.446 | -0.068 | 0.031 | 0.031 | 0 | 0.18 | 0.08 | 0.41 | 0.13 | 0.4 | 0.31 |
| 5 | 149505624 | rs3756309 | 0.124 | 0.351 | -0.591 | -0.403 | 0.013 | 0.056 | 0.056 | 1.191 | 0.39 | 0.26 | 0.35 | 0.18 | 0.28 | 0.32 |
| 5 | 149509111 | rs1864971 | -0.701 | 0.053 | -0.655 | -0.427 | -0.01 | 0.113 | 0.113 | 2.486 | 0.63 | 0.87 | 0.74 | 0.85 | 0.66 | 0.8 |
| 5 | 149512678 | rs246396 | 1.715 | -0.048 | -0.941 | -0.683 | -0.058 | 0.059 | 0.059 | 0.472 | 0.1 | 0.01 | 0.09 | 0 | 0.15 | 0.05 |
| 5 | 149515074 | rs740750 | 0.097 | -0.411 | -0.674 | -0.378 | -0.024 | -0.001 | -0.001 | 0.094 | 0.42 | 0.26 | 0.57 | 0.43 | 0.56 | 0.57 |
| 5 | 149515888 | rs740751 | -0.751 | -0.344 | -0.962 | -0.428 | -0.006 | 0.107 | 0.107 | 2.222 | 0.64 | 0.87 | 0.75 | 0.85 | 0.67 | 0.8 |
| 5 | 149521238 | rs17708574 | 0.77 | -1.18 | -0.94 | 0.072 | -0.012 | -0.068 | -0.068 | 0 | 0.13 | 0.01 | 0.07 | 0.22 | 0.11 | 0.19 |
| 5 | 149527844 | rs9324641 | -0.578 | -1.646 | -0.3 | -0.096 | -0.002 | 0.001 | 0.001 | 0 | 0.61 | 0.57 | 0.42 | 0.55 | 0.37 | 0.4 |
| 5 | 149531832 | rs4358508 | 0.581 | -1.957 | -0.468 | -0.164 | -0.065 | 0.099 | 0.099 | 0 | 0.24 | 0.45 | 0.11 | 0.37 | 0.11 | 0.15 |
| 5 | 149532107 | rs3776081 | -0.09 | -1.953 | -1.189 | -0.299 | 0.082 | 0.025 | 0.025 | 0.071 | 0.51 | 0.33 | 0.33 | 0.55 | 0.32 | 0.34 |
| 5 | 149535255 | rs2302273 | 0.753 | -1.832 | -2.13 | -0.192 | 0.029 | 0.102 | 0.102 | 0.34 | 0.27 | 0.02 | 0.17 | 0.13 | 0.19 | 0.13 |
| 5 | 149537080 | rs3756311 | -0.631 | -1.528 | -1.833 | -0.018 | -0.001 | -0.022 | -0.022 | 0 | 0.63 | 0.76 | 0.45 | 0.56 | 0.38 | 0.42 |
| 5 | 149538885 | rs2342256 | -0.398 | -1.251 | -0.766 | 0.692 | -0.016 | 0.01 | 0.01 | 0 | 0.27 | 0.66 | 0.19 | 0.42 | 0.12 | 0.24 |
| 5 | 149544045 | rs3776082 | -0.01 | -0.279 | -0.665 | 0.502 | -0.001 | 0.001 | 0.001 | 0.133 | 0.6 | 0.58 | 0.43 | 0.6 | 0.43 | 0.47 |
| 5 | 149548823 | rs887343 | 0.466 | -0.788 | -0.381 | 0.223 | 0.088 | 0.007 | 0.007 | 0 | 0.48 | 0.22 | 0.26 | 0.47 | 0.34 | 0.36 |
| 5 | 149549022 | rs887344 | -0.668 | -0.828 | -0.035 | 0.15 | -0.015 | -0.049 | -0.049 | 0 | 0.63 | 0.78 | 0.4 | 0.59 | 0.45 | 0.51 |
| 5 | 149552699 | rs2282810 | 0.01 | -1.135 | -0.313 | 0.04 | 0.021 | -0.009 | -0.009 | 0 | 0.44 | 0.39 | 0.28 | 0.58 | 0.34 | 0.33 |
| 5 | 149559937 | rs10463230 | 2.011 | -1.258 | -0.314 | 0.146 | 0.015 | -0.004 | -0.004 | 0.116 | 0.05 | 0.01 | 0.01 | 0.1 | 0.03 | 0.03 |
| 5 | 149566309 | rs10063514 | 0.176 | -1.227 | 0.016 | 0.114 | 0.028 | -0.025 | -0.025 | 0 | 0.06 | 0.22 | 0.06 | 0 | 0.07 | 0.02 |
| 5 | 149568093 | rs3776084 | 0.426 | -0.678 | -0.301 | 0.143 | 0.007 | -0.029 | -0.029 | 0 | 0.41 | 0.24 | 0.29 | 0.5 | 0.39 | 0.32 |
| 5 | 149575702 | rs2240784 | 0.115 | -0.523 | -0.527 | 0.123 | -0.001 | 0.018 | 0.018 | 0 | 0.47 | 0.27 | 0.33 | 0.49 | 0.48 | 0.42 |
| 5 | 149575855 | rs13360134 | 0.039 | -0.532 | 0.113 | 0.214 | -0.008 | 0 | 0 | 0 | 0.11 | 0.28 | 0.07 | 0.14 | 0.08 | 0.09 |
| 5 | 149577919 | rs758593 | -0.221 | -0.176 | 0.297 | 0.286 | -0.007 | -0.005 | -0.005 | 0 | 0.58 | 0.63 | 0.4 | 0.51 | 0.58 | 0.54 |
| 5 | 149581230 | rs6579777 | -1.329 | 0.503 | -0.094 | 0.712 | -0.013 | -0.029 | -0.029 | 0.606 | 0.82 | 0.89 | 0.79 | 0.69 | 0.76 | 0.81 |
| 5 | 149581252 | rs13171447 | 0.434 | 0.139 | -0.361 | 0.557 | -0.005 | 0.087 | 0.087 | 0 | 0.47 | 0.25 | 0.32 | 0.32 | 0.48 | 0.42 |
| 5 | 149581321 | rs13153971 | 0.434 | 0.139 | -0.361 | 0.557 | -0.005 | 0.087 | 0.087 | 0 | 0.47 | 0.25 | 0.32 | 0.32 | 0.48 | 0.42 |
| 5 | 149586369 | rs12652602 | 0.345 | 0.171 | -1.79 | 0.138 | -0.034 | 0.058 | 0.058 | 0 | 0.29 | 0.06 | 0.28 | 0.13 | 0.24 | 0.34 |
| 5 | 149588609 | rs4705430 | -0.025 | 0.05 | -1.912 | -0.175 | -0.059 | 0.02 | 0.02 | 0 | 0.33 | 0.08 | 0.18 | 0.29 | 0.35 | 0.34 |
| 5 | 149589647 | rs2240795 | -1.432 | -0.071 | -1.982 | -0.114 | -0.082 | 0.048 | 0.048 | 0 | 0.45 | 0.92 | 0.47 | 0.37 | 0.46 | 0.46 |
| 5 | 149590133 | rs13153325 | -0.704 | -0.078 | -0.982 | 0.072 | -0.003 | 0.001 | 0.001 | 0 | 0.34 | 0.34 | 0.38 | 0.34 | 0.38 | 0.36 |
| 5 | 149594257 | rs17111051 | -1.021 | -0.521 | 0.451 | 0.416 | 0.142 | 0.046 | 0.046 | 0.125 | 0.06 | 0.46 | 0.06 | 0 | 0.07 | 0.06 |
| 5 | 149596208 | rs6895585 | -1.119 | -0.529 | 0.447 | 0.413 | 0.139 | 0.044 | 0.044 | 0.28 | 0.06 | 0.46 | 0.06 | 0 | 0.07 | 0.06 |
| 5 | 149596241 | rs887347 | -1.976 | -0.93 | -0.5 | 0.503 | -0.035 | -0.026 | -0.026 | 0.021 | 0.49 | 0.91 | 0.47 | 0.4 | 0.42 | 0.53 |

| | | | | | | | | | | | | | | | | |
|---|---|---|---|---|---|---|---|---|---|---|---|---|---|---|---|---|
| 5 | 149596955 | rs980272 | -0.872 | -0.47 | -0.076 | 0.673 | 0.038 | -0.004 | -0.004 | 0 | 0.14 | 0.2 | 0.26 | 0.13 | 0.24 | 0.14 |
| 5 | 149602608 | rs2241694 | -0.952 | -0.515 | -0.352 | 0.855 | -0.026 | -0.033 | -0.033 | 0 | 0.94 | 1 | 0.97 | 0.84 | 0.92 | 0.82 |
| 5 | 149602824 | rs2241695 | 2.225 | -0.685 | -0.394 | 0.864 | -0.016 | 0.022 | 0.022 | 0.052 | 0.51 | 0.17 | 0.54 | 0.63 | 0.57 | 0.48 |
| 5 | 149605116 | rs4958469 | -0.533 | -0.73 | -0.483 | 0.643 | -0.115 | 0.031 | 0.031 | 0.309 | 0.22 | 0.05 | 0.25 | 0.13 | 0.28 | 0.17 |
| 5 | 149608779 | rs6885505 | -1.921 | -0.834 | -0.278 | 0.619 | -0.013 | 0.062 | 0.062 | 1.12 | 0.38 | 0.76 | 0.43 | 0.16 | 0.34 | 0.31 |
| 5 | 149609393 | rs2053053 | -1.795 | -0.759 | -0.702 | 0.653 | -0.024 | 0.078 | 0.078 | 0.099 | 0.38 | 0.18 | 0.38 | 0.16 | 0.34 | 0.33 |
| 5 | 149611688 | rs919741 | -0.496 | -0.34 | -0.915 | 0.717 | -0.068 | 0.036 | 0.036 | 0.825 | 0.2 | 0.05 | 0.22 | 0.11 | 0.21 | 0.13 |
| 5 | 149615681 | rs10066581 | -0.72 | -0.283 | -0.78 | 0.859 | 0.025 | 0.024 | 0.024 | 0 | 0.93 | 0.9 | 0.93 | 0.8 | 0.86 | 0.78 |
| 5 | 149616620 | rs6869180 | -0.951 | -0.474 | -0.441 | 0.971 | 0.027 | 0.123 | 0.123 | 0.552 | 0.06 | 0.25 | 0.05 | 0.15 | 0.02 | 0.06 |
| 5 | 149616719 | rs6869634 | -1.03 | -0.425 | -0.487 | 0.954 | -0.001 | -0.008 | -0.008 | 0.064 | 0.25 | 0.3 | 0.26 | 0.25 | 0.19 | 0.18 |
| 5 | 149623365 | rs4958456 | -0.526 | -0.652 | -0.745 | 1.153 | 0.007 | 0.026 | 0.026 | 0 | 0.1 | 0.1 | 0.1 | 0.34 | 0.14 | 0.28 |
| 5 | 149626752 | rs4958454 | -2.143 | -1.248 | -1.128 | 0.855 | -0.064 | -0.083 | -0.083 | 1.49 | 0.56 | 0.87 | 0.53 | 0.49 | 0.43 | 0.55 |
| 5 | 149627423 | rs7711562 | -2.087 | -1.023 | -1.184 | 0.948 | -0.104 | -0.081 | -0.081 | 0.014 | 0.55 | 0.9 | 0.51 | 0.52 | 0.43 | 0.55 |
| 5 | 149628644 | rs3756577 | -0.541 | -0.608 | -0.589 | 1.272 | -0.027 | -0.022 | -0.022 | 0.019 | 0.12 | 0.04 | 0.23 | 0.21 | 0.14 | 0.26 |
| 5 | 149631413 | rs2288799 | -2.038 | -0.73 | -0.396 | 1.345 | -0.009 | 0.04 | 0.04 | 0 | 0.51 | 0.82 | 0.45 | 0.6 | 0.45 | 0.57 |
| 5 | 149632955 | rs7701427 | -0.728 | -0.46 | -0.68 | 1.138 | -0.003 | 0.002 | 0.002 | 0.016 | 0.2 | 0.22 | 0.1 | 0.24 | 0.17 | 0.09 |
| 5 | 149638360 | rs4958445 | -1.257 | -0.143 | -1.107 | 1.264 | -0.062 | -0.019 | -0.019 | 0.565 | 0.27 | 0.05 | 0.3 | 0.31 | 0.28 | 0.47 |
| 5 | 149648780 | rs4958902 | -0.544 | -0.476 | -0.636 | 1.701 | -0.037 | -0.046 | -0.046 | 0.718 | 0.14 | 0.03 | 0.21 | 0.33 | 0.15 | 0.31 |
| 5 | 149650925 | rs874083 | -1.258 | -1.489 | -0.111 | 1.577 | 0.014 | -0.017 | -0.017 | 2.097 | 0.2 | 0.11 | 0.09 | 0.23 | 0.19 | 0.1 |
| 5 | 149657367 | rs34087853 | -1.786 | -1.278 | 0.644 | 1.549 | -0.03 | 0.03 | 0.03 | 0 | 0.3 | 0.43 | 0.23 | 0.47 | 0.28 | 0.25 |
| 5 | 149658711 | rs2228706 | -1.68 | -1.237 | 0.643 | 1.489 | -0.025 | 0.025 | 0.025 | 0 | 0.28 | 0.62 | 0.35 | 0.39 | 0.28 | 0.18 |
| 5 | 149659180 | rs6894342 | -0.994 | -0.691 | -0.017 | 1.777 | 0.005 | -0.042 | -0.042 | 0 | 0.11 | 0.05 | 0.07 | 0.29 | 0.13 | 0.08 |
| 5 | 149671251 | rs3806948 | 0.842 | -0.946 | 0.56 | 1.056 | -0.032 | 0.001 | 0.001 | 0 | 0.81 | 0.65 | 0.72 | 0.82 | 0.87 | 0.86 |
| 5 | 149675169 | rs7714832 | 1.637 | -1.138 | 0.24 | 0.99 | 0.023 | 0.003 | 0.003 | 0.189 | 0.65 | 0.32 | 0.64 | 0.67 | 0.77 | 0.77 |
| 5 | 149675349 | rs10037049 | -1.679 | -0.554 | 0.137 | 0.885 | -0.015 | -0.004 | -0.004 | 0 | 0.16 | 0.2 | 0.06 | 0.19 | 0.09 | 0.11 |
| 5 | 149680259 | rs4286653 | -1.302 | -0.101 | -0.252 | 0.63 | 0.002 | 0.01 | 0.01 | 0 | 0.12 | 0.08 | 0.05 | 0.08 | 0.11 | 0.06 |
| 5 | 149687113 | rs891943 | -1.021 | 0.81 | -0.228 | 0.923 | 0.001 | 0 | 0 | 0 | 0.34 | 0.66 | 0.32 | 0.43 | 0.42 | 0.3 |
| 5 | 149689401 | rs9688182 | -1.691 | 1.322 | 0.073 | 0.492 | 0.015 | -0.025 | -0.025 | 0 | 0.3 | 0.57 | 0.28 | 0.23 | 0.36 | 0.23 |
| 5 | 149695449 | rs6888551 | -1.42 | 1.362 | 0.804 | 0.473 | 0.002 | 0.001 | 0.001 | 0 | 0.21 | 0.26 | 0.23 | 0.22 | 0.29 | 0.15 |
| 5 | 149699621 | rs4588582 | 1.822 | 1.446 | 0.495 | 0.277 | 0.005 | 0.008 | 0.008 | 0.716 | 0.46 | 0.43 | 0.33 | 0.2 | 0.38 | 0.43 |
| 5 | 149700006 | rs891942 | -0.044 | 1.399 | 0.738 | 0.464 | -0.002 | -0.008 | -0.008 | 0 | 0.74 | 0.76 | 0.59 | 0.45 | 0.73 | 0.67 |
| 5 | 149704728 | rs13161248 | 0.93 | 1.226 | 0.335 | 0.515 | 0.011 | -0.021 | -0.021 | 0 | 0.58 | 0.29 | 0.65 | 0.49 | 0.67 | 0.58 |
| 5 | 149706449 | rs12651800 | -1.151 | 1.126 | 0.623 | 1.002 | 0.005 | 0.017 | 0.017 | 0.021 | 0.11 | 0.16 | 0.08 | 0.34 | 0.16 | 0.17 |
| 5 | 149707778 | rs11750343 | -0.002 | 1.12 | 0.614 | 1 | 0.009 | 0.03 | 0.03 | 0 | 0.06 | 0.1 | 0.04 | 0.22 | 0.1 | 0.08 |
| 5 | 149708571 | rs11167499 | 1.353 | 1.307 | 0.484 | 0.702 | 0.028 | 0.079 | 0.079 | 0 | 0.45 | 0.26 | 0.32 | 0.21 | 0.41 | 0.47 |
| 5 | 149709644 | rs891946 | 0.108 | 1.112 | 0.691 | 0.637 | -0.004 | -0.012 | -0.012 | 0 | 0.59 | 0.7 | 0.43 | 0.56 | 0.57 | 0.63 |
| 5 | 149712217 | rs6579788 | -1.091 | 1.188 | 0.442 | 0.673 | -0.004 | -0.006 | -0.006 | 0 | 0.12 | 0.12 | 0.09 | 0.27 | 0.16 | 0.12 |
| 5 | 149713921 | rs1012173 | -1.83 | 1.187 | 0.679 | 0.682 | 0.012 | 0.024 | 0.024 | 0.225 | 0.18 | 0.24 | 0.13 | 0.39 | 0.22 | 0.21 |
| 5 | 149718858 | rs10039050 | -0.925 | 1.182 | 0.573 | 0.613 | 0.01 | 0.003 | 0.003 | 0.045 | 0.11 | 0.18 | 0.09 | 0.15 | 0.13 | 0.13 |
| 5 | 149742236 | rs1104959 | 1.028 | 0.988 | 0.593 | 0.41 | -0.002 | -0.022 | -0.022 | 0 | 0.07 | 0.07 | 0.04 | 0 | 0.03 | 0.13 |
| 5 | 149743976 | rs891941 | -1.549 | 0.977 | 0.555 | 0.544 | 0.003 | -0.002 | -0.002 | 0 | 0.13 | 0.2 | 0.08 | 0.15 | 0.13 | 0.14 |
| 5 | 149753035 | rs17656747 | 0.369 | 0.998 | 0.557 | 0.553 | 0 | -0.007 | -0.007 | 0 | 0.05 | 0.07 | 0.05 | 0.03 | 0.06 | 0.04 |
| 5 | 149755421 | rs2071239 | -1.766 | 0.997 | 0.601 | 0.518 | 0.007 | -0.007 | -0.007 | 0 | 0.13 | 0.25 | 0.09 | 0.14 | 0.13 | 0.14 |
| 5 | 149759096 | rs7713638 | -1.766 | 1.073 | 0.601 | 0.518 | 0.008 | -0.007 | -0.007 | 0.015 | 0.13 | 0.25 | 0.09 | 0.14 | 0.13 | 0.14 |
| 5 | 149764293 | rs2748220 | 1.205 | 1.231 | 0.57 | 0.62 | 0.003 | 0.006 | 0.006 | 0 | 0.44 | 0.36 | 0.33 | 0.43 | 0.46 | 0.5 |
| 5 | 149766790 | rs1864959 | 1.233 | 1.235 | 0.388 | 0.648 | 0.003 | 0.009 | 0.009 | 0 | 0.43 | 0.28 | 0.33 | 0.43 | 0.46 | 0.5 |
| 5 | 149772932 | rs11743855 | -0.521 | 1.373 | 0.222 | 0.965 | 0 | -0.019 | -0.019 | 0.053 | 0.06 | 0.12 | 0.05 | 0.02 | 0.06 | 0.04 |
| 5 | 149779030 | rs7702237 | -0.709 | 1.394 | 0.224 | 0.974 | -0.001 | -0.019 | -0.019 | 0 | 0.07 | 0.11 | 0.05 | 0.02 | 0.06 | 0.03 |
| 5 | 149781627 | rs1056400 | -0.211 | 1.492 | -0.007 | 1.171 | -0.032 | -0.031 | -0.031 | 0 | 0.06 | 0.01 | 0.05 | 0.23 | 0.12 | 0.08 |
| 5 | 149787068 | rs2288817 | -0.942 | 1.206 | 0.23 | 0.884 | 0.033 | 0.031 | 0.031 | 0 | 0.05 | 0.18 | 0.04 | 0.12 | 0.07 | 0.06 |
| 5 | 149792600 | rs2748249 | -0.254 | 1.324 | -0.615 | 0.381 | -0.044 | -0.008 | -0.008 | 0.885 | 0.18 | 0.03 | 0.16 | 0.12 | 0.18 | 0.15 |
| 5 | 149793457 | rs1560661 | -0.608 | 0.952 | -0.694 | 0.79 | -0.016 | -0.014 | -0.014 | 0 | 0.64 | 0.76 | 0.44 | 0.6 | 0.55 | 0.69 |
| 5 | 149795052 | rs2569094 | -0.149 | 0.751 | -0.383 | 0.779 | -0.002 | -0.004 | -0.004 | 0 | 0.55 | 0.43 | 0.39 | 0.49 | 0.48 | 0.56 |
| 5 | 149799339 | rs891956 | 0.459 | 0.005 | -0.11 | 1.449 | -0.017 | 0.043 | 0.043 | 0 | 0.88 | 0.85 | 0.95 | 0.65 | 0.96 | 0.87 |
| 5 | 149800334 | rs13175409 | -0.153 | 0.061 | -0.321 | 1.21 | -0.016 | 0.066 | 0.066 | 0.536 | 0.3 | 0.08 | 0.25 | 0.16 | 0.3 | 0.36 |
| 5 | 149802111 | rs2569103 | -1.405 | -0.039 | 0.41 | 1.566 | -0.029 | 0.14 | 0.14 | 0 | 0.14 | 0.32 | 0.13 | 0.36 | 0.07 | 0.15 |
| 5 | 149816968 | rs3901731 | -1.796 | -0.343 | 0.371 | 1.583 | 0.005 | 0 | 0 | 0 | 0.41 | 0.62 | 0.3 | 0.63 | 0.3 | 0.62 |
| 5 | 149821847 | rs2569107 | -1.844 | -0.494 | 0.398 | 1.617 | -0.049 | 0.027 | 0.027 | 0.732 | 0.19 | 0.69 | 0.11 | 0.36 | 0.05 | 0.21 |
| 5 | 149822596 | rs13181401 | -0.379 | -0.471 | 0.219 | 1.645 | -0.051 | -0.045 | -0.045 | 0.432 | 0.23 | 0.06 | 0.2 | 0.27 | 0.26 | 0.41 |
| 5 | 149823785 | rs2070844 | -0.379 | -0.454 | 0.074 | 1.678 | -0.054 | -0.047 | -0.047 | 0 | 0.23 | 0.06 | 0.2 | 0.27 | 0.26 | 0.41 |
| 5 | 149827494 | rs2748241 | -1.583 | -0.169 | 0.155 | 1.525 | 0.043 | -0.002 | -0.002 | 0.221 | 0.42 | 0.8 | 0.31 | 0.63 | 0.3 | 0.62 |
| 5 | 149843942 | rs6579790 | -0.749 | 0.059 | 0.445 | 1.549 | -0.004 | -0.008 | -0.008 | 0 | 0.33 | 0.39 | 0.26 | 0.24 | 0.28 | 0.41 |
| 5 | 149845446 | rs2983102 | -1.149 | 0.296 | 0.092 | 1.544 | 0.004 | -0.002 | -0.002 | 2.07 | 0.42 | 0.61 | 0.31 | 0.59 | 0.34 | 0.66 |
| 5 | 149857460 | rs3846703 | -0.629 | 0.648 | 0.058 | 1.854 | -0.016 | -0.031 | -0.031 | 0 | 0.06 | 0 | 0.05 | 0.25 | 0.07 | 0.16 |
| 5 | 149859235 | rs10061076 | -1.773 | 0.538 | 0.668 | 1.673 | 0.05 | 0.258 | 0.258 | 1.293 | 0.06 | 0.36 | 0.03 | 0.13 | 0.03 | 0.04 |
| 5 | 149878840 | rs2343813 | -1.305 | 0.598 | 0.468 | 1.679 | -0.001 | 0.061 | 0.061 | 0 | 0.09 | 0.12 | 0.26 | 0.27 | 0.1 | 0.16 |
| 5 | 149886437 | rs9324645 | -1.8 | 1.003 | 0.439 | 1.484 | 0.03 | 0.097 | 0.097 | 0.157 | 0.18 | 0.43 | 0.34 | 0.27 | 0.21 | 0.27 |
| 5 | 149893287 | rs2983101 | 1.32 | 0.476 | -0.213 | 0.981 | 0.013 | 0.065 | 0.065 | 0 | 0.46 | 0.32 | 0.38 | 0.25 | 0.51 | 0.37 |
| 5 | 149893372 | rs10463299 | -0.821 | 0.991 | -0.293 | 1.37 | -0.04 | 0 | 0 | 0 | 0.13 | 0.04 | 0.13 | 0.13 | 0.19 | 0.24 |

| | | | | | | | | | | | | | | | | |
|---|---|---|---|---|---|---|---|---|---|---|---|---|---|---|---|---|
| 5 | 149895492 | rs2748216 | 0.362 | 0.435 | -0.549 | 1.559 | 0.022 | -0.095 | -0.095 | 0 | 0.75 | 0.93 | 0.83 | 0.6 | 0.78 | 0.78 |
| 5 | 149897291 | rs10074650 | -1.397 | 1.917 | 0.36 | 1.601 | 0.102 | 0.084 | 0.084 | 0.595 | 0.19 | 0.45 | 0.38 | 0.26 | 0.29 | 0.27 |
| 5 | 149899836 | rs4409074 | 0.756 | 1.769 | -0.416 | 1.285 | -0.071 | -0.059 | -0.059 | 0 | 0.15 | 0.02 | 0.2 | 0.21 | 0.21 | 0.25 |
| 5 | 149901834 | rs12516924 | -0.487 | 1.985 | -0.31 | 1.183 | -0.12 | 0.016 | 0.016 | 0.121 | 0.2 | 0.04 | 0.38 | 0.16 | 0.36 | 0.3 |
| 5 | 149906520 | rs3797620 | -0.422 | 1.792 | -0.225 | 1.095 | -0.077 | 0.033 | 0.033 | 0.988 | 0.21 | 0.12 | 0.41 | 0.14 | 0.4 | 0.34 |
| 5 | 149907533 | rs2273235 | -1.127 | 1.644 | -0.045 | 1.195 | -0.024 | -0.005 | -0.005 | 0 | 0.28 | 0.24 | 0.5 | 0.28 | 0.47 | 0.42 |
| 5 | 149907602 | rs2273234 | -0.993 | 1.659 | -0.297 | 1.068 | -0.098 | 0.07 | 0.07 | 1.44 | 0.27 | 0.12 | 0.49 | 0.15 | 0.46 | 0.41 |
| 5 | 149922208 | rs3095905 | 0.384 | 2.007 | 0.288 | 1.404 | -0.014 | -0.006 | -0.006 | 0 | 0.17 | 0.09 | 0.21 | 0.21 | 0.21 | 0.25 |
| 5 | 149926998 | rs3095902 | -0.687 | 1.735 | 0.543 | 1.666 | -0.001 | 0.07 | 0.07 | 1.262 | 0.25 | 0.65 | 0.28 | 0.48 | 0.25 | 0.35 |
| 5 | 149935680 | rs11542921 | -0.257 | 1.846 | 0.364 | 1.587 | -0.133 | 0.022 | 0.022 | 0.676 | 0.2 | 0.04 | 0.4 | 0.15 | 0.4 | 0.34 |
| 5 | 149937739 | rs11994 | -0.848 | 1.725 | 0.38 | 1.577 | -0.108 | -0.048 | -0.048 | 0.907 | 0.25 | 0.1 | 0.41 | 0.29 | 0.44 | 0.43 |
| 5 | 149946821 | rs253290 | 0.29 | 1.475 | 0.503 | 1.49 | 0.023 | -0.039 | -0.039 | 0 | 0.74 | 0.43 | 0.72 | 0.8 | 0.74 | 0.72 |
| 5 | 149949806 | rs888714 | 1.647 | 1.481 | 0.459 | 1.531 | 0.097 | 0.039 | 0.039 | 0 | 0.5 | 0.27 | 0.32 | 0.39 | 0.34 | 0.31 |
| 5 | 149952749 | rs253296 | -0.346 | 1.74 | 0.09 | 1.616 | -0.103 | 0.018 | 0.018 | 1.115 | 0.19 | 0.05 | 0.37 | 0.15 | 0.35 | 0.29 |
| 5 | 149955482 | rs7734132 | 0.046 | 0.122 | 0.602 | 1.852 | -0.027 | -0.008 | -0.008 | 0 | 0.64 | 0.94 | 0.41 | 0.65 | 0.42 | 0.41 |
| 5 | 149955709 | rs7716678 | -1.233 | -0.233 | 1.377 | 1.998 | -0.051 | 0.221 | 0.221 | 3.892 | 0.11 | 0.58 | 0.07 | 0.27 | 0.05 | 0.1 |
| 5 | 149971345 | rs10463300 | 0.661 | 0.248 | 0.377 | 1.536 | 0.067 | 0 | 0 | 0 | 0.5 | 0.21 | 0.32 | 0.47 | 0.39 | 0.36 |
| 5 | 149972167 | rs11739838 | 0.483 | 0.292 | 0.379 | 1.559 | 0.018 | -0.002 | -0.002 | 0 | 0.52 | 0.34 | 0.33 | 0.47 | 0.39 | 0.35 |
| 5 | 149986473 | rs9324646 | 0.496 | -0.036 | -0.227 | 1.087 | 0.085 | -0.009 | -0.009 | 0.03 | 0.48 | 0.26 | 0.36 | 0.47 | 0.36 | 0.33 |
| 5 | 149991012 | rs1560657 | 0.569 | -0.445 | 0.147 | 0.866 | -0.002 | 0.001 | 0.001 | 0 | 0.47 | 0.5 | 0.35 | 0.35 | 0.33 | 0.26 |
| 5 | 149991853 | rs752921 | -1.281 | 1.179 | 1.182 | 0.487 | 0.107 | -0.132 | -0.132 | 0.12 | 0.18 | 0.5 | 0.16 | 0.03 | 0.24 | 0.22 |
| 5 | 149992045 | rs752922 | -1.267 | 1.112 | 1.405 | 0.59 | 0.044 | -0.087 | -0.087 | 0 | 0.19 | 0.49 | 0.35 | 0.09 | 0.22 | 0.24 |
| 5 | 150000145 | rs7719412 | -2.355 | 0.601 | 1.538 | 1.362 | -0.027 | 0.223 | 0.223 | 0.789 | 0.08 | 0.28 | 0.05 | 0.28 | 0.01 | 0.19 |
| 5 | 150001729 | rs9918250 | -1.695 | 0.652 | 1.333 | 1.386 | -0.022 | 0.155 | 0.155 | 0.036 | 0.05 | 0.13 | 0.03 | 0.28 | 0.01 | 0.21 |
| 5 | 150010099 | rs4958728 | -1.24 | 1.112 | 1.224 | 1.207 | 0.015 | -0.014 | -0.014 | 0.369 | 0.24 | 0.26 | 0.41 | 0.14 | 0.29 | 0.27 |
| 5 | 150011387 | rs10039597 | -0.338 | 1.56 | 0.7 | 0.996 | -0.004 | 0.005 | 0.005 | 0 | 0.21 | 0.2 | 0.15 | 0.1 | 0.23 | 0.18 |
| 5 | 150012917 | rs17111261 | -1.247 | 1.814 | 0.412 | 0.991 | 0.026 | -0.027 | -0.027 | 1.1 | 0.29 | 0.36 | 0.44 | 0.16 | 0.35 | 0.38 |
| 5 | 150016115 | rs253345 | -1.675 | 1.273 | -0.129 | 1.282 | 0.014 | -0.047 | -0.047 | 0.107 | 0.51 | 0.89 | 0.6 | 0.52 | 0.5 | 0.59 |
| 5 | 150022919 | rs1351119 | -1.571 | 1.173 | 0.593 | 1.459 | -0.002 | 0.004 | 0.004 | 0.468 | 0.29 | 0.36 | 0.41 | 0.38 | 0.24 | 0.28 |
| 5 | 150025994 | rs253349 | -0.385 | 1.289 | 0.36 | 0.962 | -0.05 | 0.057 | 0.057 | 0.119 | 0.17 | 0.05 | 0.15 | 0.07 | 0.22 | 0.14 |
| 5 | 150033470 | rs33412 | -1.212 | 1.085 | 0.576 | 1.099 | -0.004 | 0 | 0 | 0 | 0.34 | 0.27 | 0.45 | 0.32 | 0.35 | 0.35 |
| 5 | 150038265 | rs33409 | -1.287 | 0.538 | 0.901 | 1.024 | 0.024 | -0.009 | -0.009 | 0 | 0.37 | 0.47 | 0.49 | 0.33 | 0.42 | 0.41 |
| 5 | 150044701 | rs10038115 | -1.409 | -0.334 | 1.259 | 0.59 | -0.052 | -0.056 | -0.056 | 0 | 0.16 | 0.41 | 0.08 | 0.05 | 0.07 | 0.09 |
| 5 | 150048067 | rs10036981 | -1.376 | -0.325 | 1.297 | 0.594 | -0.049 | -0.052 | -0.052 | 0 | 0.15 | 0.42 | 0.08 | 0.05 | 0.07 | 0.09 |
| 5 | 150049655 | rs10064957 | -1.592 | -0.378 | 1.448 | 0.896 | -0.059 | 0.027 | 0.027 | 1.601 | 0.16 | 0.47 | 0.08 | 0.14 | 0.07 | 0.11 |
| 5 | 150087245 | rs252160 | -1.638 | -0.874 | 1.491 | 0.876 | -0.009 | 0.117 | 0.117 | 0.037 | 0.53 | 0.82 | 0.33 | 0.68 | 0.48 | 0.6 |
| 5 | 150090344 | rs3733925 | -0.351 | -1.062 | 1.459 | 0.906 | 0.004 | -0.045 | -0.045 | 0.264 | 0.08 | 0 | 0.03 | 0.1 | 0.06 | 0.09 |
| 5 | 150102244 | rs17111302 | -1.63 | -1.083 | 1.674 | 0.777 | -0.046 | 0.081 | 0.081 | 0.148 | 0.15 | 0.34 | 0.07 | 0.26 | 0.06 | 0.15 |
| 5 | 150107876 | rs10053702 | -1.935 | -1.097 | 1.602 | 0.77 | -0.069 | 0.103 | 0.103 | 0.26 | 0.17 | 0.4 | 0.07 | 0.26 | 0.06 | 0.16 |
| 5 | 150116980 | rs12332423 | -2.036 | -1.147 | 1.579 | 0.749 | -0.073 | 0.093 | 0.093 | 0.538 | 0.18 | 0.4 | 0.07 | 0.26 | 0.06 | 0.16 |
| 5 | 150138196 | rs2278395 | -0.885 | -1.063 | 1.52 | 0.788 | -0.001 | 0.009 | 0.009 | 0 | 0.12 | 0.12 | 0.04 | 0.26 | 0.06 | 0.15 |
| 5 | 150138634 | rs7706089 | -1.785 | -1.095 | 1.716 | 0.773 | -0.05 | 0.093 | 0.093 | 0.149 | 0.15 | 0.34 | 0.06 | 0.26 | 0.06 | 0.15 |
| 5 | 150140140 | rs10071236 | -1.805 | -1.096 | 1.734 | 0.772 | -0.052 | 0.1 | 0.1 | 0.196 | 0.15 | 0.35 | 0.07 | 0.26 | 0.06 | 0.15 |
| 5 | 150151271 | rs6868496 | -1.611 | -1.249 | 1.241 | 0.537 | -0.037 | 0.106 | 0.106 | 0.626 | 0.53 | 0.87 | 0.37 | 0.66 | 0.5 | 0.56 |
| 5 | 150158164 | rs12653537 | -1.759 | -1.489 | 1.984 | 1.025 | -0.038 | 0.068 | 0.068 | 0.068 | 0.14 | 0.2 | 0.05 | 0.26 | 0.06 | 0.16 |
| 5 | 150161038 | rs11952896 | -2.079 | -1.417 | 1.645 | 1.08 | -0.071 | 0.105 | 0.105 | 0.911 | 0.17 | 0.44 | 0.07 | 0.25 | 0.06 | 0.2 |
| 5 | 150167118 | rs1529476 | -1.868 | -1.252 | 1.509 | 1.121 | -0.053 | 0.124 | 0.124 | 0.499 | 0.15 | 0.39 | 0.06 | 0.26 | 0.06 | 0.18 |
| 5 | 150170609 | rs9324656 | -0.81 | -1.66 | 0.598 | 0.615 | 0.037 | -0.017 | -0.017 | 0 | 0.26 | 0.17 | 0.16 | 0.63 | 0.15 | 0.29 |
| 5 | 150176331 | rs9885376 | -0.669 | -1.706 | 0.68 | 0.878 | -0.006 | 0.058 | 0.058 | 2.36 | 0.08 | 0.1 | 0.04 | 0.2 | 0.05 | 0.16 |
| 5 | 150182434 | rs2112637 | -1.145 | -1.852 | 1.016 | 0.765 | -0.013 | 0 | 0 | 0.01 | 0.3 | 0.33 | 0.18 | 0.63 | 0.16 | 0.29 |
| 5 | 150187805 | rs6873063 | -1.374 | -1.986 | 1.692 | 0.708 | -0.054 | 0.001 | 0.001 | 0.299 | 0.33 | 0.52 | 0.19 | 0.63 | 0.15 | 0.29 |
| 5 | 150228833 | rs10052080 | -1.236 | -1.855 | 1.8 | 0.722 | -0.062 | 0.006 | 0.006 | 0.484 | 0.31 | 0.53 | 0.19 | 0.6 | 0.15 | 0.28 |
| 5 | 150230950 | rs11957134 | -0.149 | -1.784 | 1.608 | 0.718 | -0.034 | 0.033 | 0.033 | 0 | 0.19 | 0.24 | 0.13 | 0.4 | 0.1 | 0.12 |
| 5 | 150233004 | rs6869426 | 0.263 | -0.977 | 1.019 | 0.167 | -0.109 | 0.089 | 0.089 | 0.424 | 0.19 | 0.04 | 0.24 | 0.01 | 0.43 | 0.2 |
| 5 | 150240076 | rs1000113 | -0.044 | -1.472 | 1.52 | 0.607 | -0.043 | 0.046 | 0.046 | 0 | 0.19 | 0.25 | 0.13 | 0.41 | 0.1 | 0.12 |
| 5 | 150244041 | rs10077333 | -1.142 | -1.558 | 1.733 | 0.612 | -0.065 | 0.007 | 0.007 | 0.497 | 0.31 | 0.52 | 0.19 | 0.6 | 0.15 | 0.28 |
| 5 | 150253104 | rs2344182 | -1.168 | -1.564 | 1.807 | 0.609 | -0.061 | 0.006 | 0.006 | 0.514 | 0.31 | 0.53 | 0.19 | 0.6 | 0.15 | 0.28 |
| 5 | 150254620 | rs10463307 | -0.199 | -1.533 | 1.596 | 0.516 | -0.078 | 0.103 | 0.103 | 0 | 0.21 | 0.36 | 0.14 | 0.43 | 0.11 | 0.21 |
| 5 | 150257616 | rs1428554 | -0.432 | -1.539 | 1.596 | 0.514 | -0.091 | 0.106 | 0.106 | 0 | 0.22 | 0.51 | 0.15 | 0.43 | 0.11 | 0.22 |
| 5 | 150260382 | rs6888154 | -1.158 | -1.558 | 1.652 | 0.538 | -0.046 | 0.011 | 0.011 | 0.464 | 0.31 | 0.54 | 0.19 | 0.57 | 0.15 | 0.28 |
| 5 | 150270420 | rs7714584 | -0.409 | -1.543 | 1.653 | 0.542 | -0.091 | 0.106 | 0.106 | 0 | 0.22 | 0.51 | 0.15 | 0.43 | 0.11 | 0.22 |
| 5 | 150274079 | rs10515643 | 0.865 | -1.453 | 1.321 | 0.562 | 0.001 | -0.019 | -0.019 | 0 | 0.08 | 0.03 | 0.04 | 0.14 | 0.04 | 0.06 |
| 5 | 150274374 | rs12653571 | 0.865 | -1.453 | 1.321 | 0.562 | 0.001 | -0.019 | -0.019 | 0 | 0.08 | 0.03 | 0.05 | 0.14 | 0.04 | 0.06 |
| 5 | 150277111 | rs17800777 | 0.865 | -1.453 | 1.321 | 0.562 | 0.001 | -0.019 | -0.019 | 0 | 0.08 | 0.03 | 0.04 | 0.14 | 0.04 | 0.06 |
| 5 | 150278587 | rs4958427 | -1.401 | -1.76 | 1.381 | 0.385 | 0.101 | -0.015 | -0.015 | 0.141 | 0.35 | 0.82 | 0.22 | 0.57 | 0.15 | 0.28 |
| 5 | 150284808 | rs7724036 | -0.56 | -1.601 | 1.587 | 0.499 | -0.095 | 0.127 | 0.127 | 0 | 0.23 | 0.49 | 0.15 | 0.44 | 0.12 | 0.23 |
| 5 | 150293820 | rs1600151 | 0.16 | -1.644 | 1.006 | 0.022 | -0.02 | 0.051 | 0.051 | 0.019 | 0.63 | 0.77 | 0.68 | 0.84 | 0.48 | 0.63 |
| 5 | 150299621 | rs6868828 | 0.112 | -1.671 | 0.955 | 0.009 | -0.055 | 0.085 | 0.085 | 0.521 | 0.63 | 0.89 | 0.69 | 0.84 | 0.48 | 0.63 |
| 5 | 150328866 | rs7724558 | 0.105 | -1.685 | 1.206 | 0.155 | 0.005 | -0.011 | -0.011 | 0.549 | 0.19 | 0.15 | 0.11 | 0.41 | 0.11 | 0.11 |

| | | | | | | | | | | | | | | | | |
|---|---|---|---|---|---|---|---|---|---|---|---|---|---|---|---|---|
| 5 | 150331600 | rs6579821 | 1.455 | -1.84 | 1.524 | 0.067 | -0.006 | -0.01 | -0.01 | 0 | 0.4 | 0.52 | 0.53 | 0.4 | 0.35 | 0.42 |
| 5 | 150332481 | rs11167526 | 0.063 | -2.114 | 1.557 | 0.323 | 0.044 | -0.054 | -0.054 | 3.978 | 0.19 | 0.08 | 0.11 | 0.41 | 0.1 | 0.12 |
| 5 | 150350990 | rs10050426 | -0.267 | -2 | 1.642 | -0.297 | -0.026 | -0.011 | -0.011 | 0 | 0.22 | 0.24 | 0.12 | 0.13 | 0.13 | 0.08 |
| 5 | 150353775 | rs977696 | 1.365 | -1.838 | 1.808 | -0.475 | 0.035 | 0.049 | 0.049 | 0.279 | 0.44 | 0.28 | 0.56 | 0.7 | 0.35 | 0.52 |
| 5 | 150358459 | rs11950220 | 0.113 | -1.79 | 1.617 | -0.469 | 0.003 | 0.003 | 0.003 | 2.318 | 0.22 | 0.16 | 0.11 | 0.12 | 0.13 | 0.09 |
| 5 | 150358957 | rs1038074 | 0.315 | -1.46 | 1.634 | -0.5 | -0.003 | -0.003 | -0.003 | 0 | 0.74 | 0.73 | 0.85 | 0.85 | 0.81 | 0.81 |
| 5 | 150362371 | rs10064261 | 0.259 | -1.156 | 1.39 | -0.442 | 0.004 | -0.017 | -0.017 | 0.103 | 0.26 | 0.25 | 0.26 | 0.05 | 0.45 | 0.27 |
| 5 | 150372984 | rs13158099 | 0.359 | -1.049 | 1.167 | -0.383 | -0.048 | 0.038 | 0.038 | 0 | 0.25 | 0.15 | 0.26 | 0.06 | 0.45 | 0.27 |
| 5 | 150376966 | rs6579833 | 1.249 | -1.012 | 1.657 | -0.21 | 0.075 | 0.092 | 0.092 | 0 | 0.92 | 0.57 | 0.94 | 0.88 | 0.92 | 0.86 |
| 5 | 150381243 | rs17111637 | 1.021 | -1.158 | 1.521 | -0.249 | 0.021 | 0.028 | 0.028 | 0.766 | 0.47 | 0.48 | 0.58 | 0.78 | 0.36 | 0.53 |
| 5 | 150383403 | rs12109402 | -0.819 | -1.052 | 0.78 | -0.266 | 0 | -0.005 | -0.005 | 0 | 0.48 | 0.39 | 0.38 | 0.19 | 0.57 | 0.37 |
| 5 | 150384959 | rs2054440 | 0.387 | -0.986 | -0.071 | -0.36 | -0.017 | 0.098 | 0.098 | 0 | 0.62 | 0.8 | 0.66 | 0.91 | 0.42 | 0.62 |
| 5 | 150386198 | rs707141 | 0.072 | -0.941 | -0.153 | 0.05 | -0.034 | -0.021 | -0.021 | 0 | 0.34 | 0.21 | 0.32 | 0.36 | 0.47 | 0.36 |
| 5 | 150387141 | rs707143 | -0.964 | -0.724 | -0.753 | -0.185 | -0.015 | -0.005 | -0.005 | 0 | 0.61 | 0.83 | 0.47 | 0.57 | 0.61 | 0.48 |
| 5 | 150387736 | rs6869405 | 0.679 | -0.839 | -0.799 | -0.156 | -0.015 | -0.036 | -0.036 | 0.107 | 0.1 | 0.16 | 0.07 | 0 | 0.07 | 0.01 |
| 5 | 150390517 | rs707149 | -0.611 | -0.141 | -1.046 | 0.186 | -0.025 | 0 | 0 | 2.293 | 0.6 | 0.8 | 0.47 | 0.57 | 0.59 | 0.48 |
| 5 | 150397914 | rs2042235 | 0.424 | -0.201 | -0.298 | -0.296 | -0.011 | -0.075 | -0.075 | 0 | 0.72 | 0.48 | 0.8 | 0.9 | 0.76 | 0.81 |
| 5 | 150398115 | rs2042236 | -0.169 | -1.064 | -0.59 | -0.014 | -0.024 | 0.001 | 0.001 | 0.283 | 0.19 | 0.28 | 0.09 | 0.15 | 0.11 | 0.1 |
| 5 | 150399210 | rs1946234 | -0.131 | -1.236 | -0.643 | -0.049 | -0.016 | -0.001 | -0.001 | 0 | 0.22 | 0.25 | 0.09 | 0.17 | 0.12 | 0.15 |
| 5 | 150399225 | rs1946235 | -0.146 | -1.24 | -0.645 | -0.051 | -0.016 | -0.001 | -0.001 | 0.074 | 0.22 | 0.25 | 0.09 | 0.17 | 0.12 | 0.15 |
| 5 | 150399584 | rs8177404 | -0.102 | -1.255 | -0.571 | -0.057 | -0.012 | -0.001 | -0.001 | 0 | 0.21 | 0.24 | 0.09 | 0.15 | 0.12 | 0.15 |
| 5 | 150399634 | rs8177406 | -0.102 | -1.255 | -0.571 | -0.057 | -0.012 | -0.001 | -0.001 | 0 | 0.21 | 0.24 | 0.09 | 0.15 | 0.12 | 0.15 |
| 5 | 150400087 | rs8177412 | -0.116 | -1.244 | -0.614 | -0.052 | -0.013 | -0.001 | -0.001 | 0 | 0.22 | 0.24 | 0.09 | 0.17 | 0.12 | 0.16 |
| 5 | 150400601 | rs870406 | -0.075 | -1.17 | -0.982 | -0.053 | 0.037 | -0.027 | -0.027 | 0 | 0.2 | 0.03 | 0.07 | 0.15 | 0.12 | 0.14 |
| 5 | 150401296 | rs3805435 | 0.276 | -1.158 | -1.101 | 0.654 | 0.004 | -0.036 | -0.036 | 0 | 0.09 | 0 | 0.18 | 0.42 | 0.09 | 0.2 |
| 5 | 150401765 | rs3792798 | -0.255 | -1.461 | -0.701 | 0.383 | -0.019 | 0.111 | 0.111 | 0 | 0.12 | 0.2 | 0.19 | 0.43 | 0.09 | 0.31 |
| 5 | 150401796 | rs3828599 | 0.779 | -1.311 | -0.041 | -0.204 | 0 | 0.092 | 0.092 | 0 | 0.71 | 0.49 | 0.68 | 0.54 | 0.73 | 0.53 |
| 5 | 150401891 | rs3792797 | 0.298 | -1.488 | 0.681 | -0.284 | 0.004 | -0.089 | -0.089 | 0 | 0.78 | 0.36 | 0.83 | 0.93 | 0.81 | 0.72 |
| 5 | 150402334 | rs4958872 | 0.879 | -1.491 | 0.728 | -0.274 | -0.002 | 0.012 | 0.012 | 0 | 0.7 | 0.29 | 0.66 | 0.52 | 0.73 | 0.53 |
| 5 | 150402490 | rs3792796 | -1.405 | -1.858 | 0.75 | -0.426 | 0.066 | 0.124 | 0.124 | 0.016 | 0.5 | 0.17 | 0.59 | 0.35 | 0.64 | 0.39 |
| 5 | 150402570 | rs4958434 | -0.431 | -1.807 | 0.831 | -0.423 | 0.042 | -0.099 | -0.099 | 0 | 0.82 | 0.55 | 0.86 | 0.97 | 0.82 | 0.83 |
| 5 | 150402940 | rs8177426 | -0.011 | -1.761 | 0.803 | -0.465 | 0.015 | -0.113 | -0.113 | 0 | 0.79 | 0.52 | 0.84 | 0.95 | 0.81 | 0.72 |
| 5 | 150403802 | rs3792795 | 1.559 | -1.752 | 0.777 | -0.432 | 0.004 | 0.017 | 0.017 | 0.153 | 0.08 | 0.01 | 0.16 | 0.41 | 0.08 | 0.19 |
| 5 | 150404096 | rs4958873 | -0.534 | -1.74 | 0.841 | -0.481 | 0.041 | -0.091 | -0.091 | 0.085 | 0.83 | 0.6 | 0.86 | 0.97 | 0.82 | 0.83 |
| 5 | 150404311 | rs8177433 | -0.078 | -1.804 | 0.808 | -0.497 | 0.048 | -0.02 | -0.02 | 0 | 0.28 | 0.14 | 0.23 | 0.59 | 0.17 | 0.34 |
| 5 | 150404383 | rs8177435 | 1.329 | -1.859 | 0.919 | -0.529 | 0.065 | 0.119 | 0.119 | 0 | 0.5 | 0.21 | 0.6 | 0.35 | 0.64 | 0.39 |
| 5 | 150404657 | rs3763012 | -0.411 | -1.86 | 0.93 | -0.503 | 0.01 | 0 | 0 | 0 | 0.31 | 0.25 | 0.25 | 0.62 | 0.18 | 0.44 |
| 5 | 150406437 | rs869976 | 0.769 | -1.726 | 0.975 | -0.469 | 0.007 | 0.019 | 0.019 | 0.186 | 0.05 | 0.27 | 0.04 | 0.04 | 0.01 | 0.11 |
| 5 | 150407456 | rs8177447 | -0.52 | -1.745 | 0.93 | -0.511 | 0.045 | -0.088 | -0.088 | 0.233 | 0.83 | 0.6 | 0.86 | 0.97 | 0.82 | 0.83 |
| 5 | 150407783 | rs11548 | 1.035 | -1.749 | 0.773 | -0.415 | 0.003 | 0.055 | 0.055 | 0.192 | 0.08 | 0.09 | 0.17 | 0.41 | 0.08 | 0.19 |
| 5 | 150409477 | rs2277940 | 1.347 | -1.683 | 0.319 | -0.429 | 0.005 | 0.013 | 0.013 | 0.196 | 0.08 | 0.01 | 0.16 | 0.42 | 0.08 | 0.2 |
| 5 | 150409989 | rs8177834 | 0.699 | -1.848 | 0.224 | -0.595 | 0.05 | -0.019 | -0.019 | 0.216 | 0.2 | 0.05 | 0.06 | 0.17 | 0.09 | 0.14 |
| 5 | 150412140 | rs3763009 | 1.544 | -1.004 | -0.573 | -1.272 | -0.13 | -0.061 | -0.061 | 0.65 | 0.19 | 0.01 | 0.4 | 0.29 | 0.35 | 0.29 |
| 5 | 150413406 | rs2233305 | -0.254 | -1.046 | -0.047 | -1.331 | 0.007 | -0.126 | -0.126 | 1.822 | 0.78 | 0.51 | 0.84 | 0.95 | 0.82 | 0.72 |
| 5 | 150422765 | rs4958876 | -1.472 | -0.918 | 0.073 | -1.32 | 0.016 | -0.038 | -0.038 | 0 | 0.88 | 0.75 | 0.9 | 0.97 | 0.87 | 0.87 |
| 5 | 150425467 | rs2233299 | 1.311 | -1.312 | -0.162 | -1.642 | 0 | 0.137 | 0.137 | 0 | 0.29 | 0.07 | 0.18 | 0.06 | 0.28 | 0.1 |
| 5 | 150426868 | rs11747787 | -0.134 | -1.298 | 0.34 | -1.573 | 0 | -0.009 | -0.009 | 0 | 0.6 | 0.56 | 0.32 | 0.28 | 0.45 | 0.38 |
| 5 | 150428195 | rs4292439 | -0.442 | -1.184 | 0.314 | -1.64 | 0.063 | -0.119 | -0.119 | 0.721 | 0.81 | 0.47 | 0.84 | 0.97 | 0.81 | 0.83 |
| 5 | 150428871 | rs6862024 | 1.472 | -0.87 | -0.404 | -1.254 | -0.067 | -0.013 | -0.013 | 0 | 0.18 | 0.05 | 0.22 | 0.26 | 0.34 | 0.28 |
| 5 | 150430696 | rs7713223 | -1.2 | -1.246 | -0.51 | -1.18 | 0.015 | -0.063 | -0.063 | 3.546 | 0.8 | 0.99 | 0.93 | 0.81 | 0.89 | 0.85 |
| 5 | 150432153 | rs2112635 | -0.83 | -1.73 | -0.795 | -1.612 | 0.033 | 0.092 | 0.092 | 0 | 0.62 | 0.92 | 0.81 | 0.85 | 0.7 | 0.74 |
| 5 | 150432388 | rs871269 | 0.093 | -0.151 | -0.213 | -0.948 | 0.018 | 0.042 | 0.042 | 0 | 0.31 | 0.42 | 0.55 | 0.54 | 0.34 | 0.42 |
| 5 | 150432446 | rs17111695 | -0.134 | -0.271 | 0.24 | -0.379 | 0.013 | 0.005 | 0.005 | 0 | 0.14 | 0.19 | 0.12 | 0.23 | 0.17 | 0.18 |
| 5 | 150434722 | rs3792794 | 0.757 | -0.457 | -0.084 | -0.17 | 0.006 | -0.045 | -0.045 | 2.056 | 0.1 | 0.01 | 0.29 | 0.48 | 0.07 | 0.19 |
| 5 | 150435645 | rs12516176 | -0.727 | -0.595 | 0.386 | -0.399 | 0.005 | -0.008 | -0.008 | 0 | 0.24 | 0.34 | 0.22 | 0.27 | 0.25 | 0.32 |
| 5 | 150438284 | rs4958435 | -1.562 | -0.771 | 0.539 | -0.576 | 0.008 | 0.125 | 0.125 | 0.005 | 0.52 | 0.73 | 0.62 | 0.76 | 0.45 | 0.6 |
| 5 | 150438988 | rs1422673 | -0.379 | -1.162 | 1.139 | 0.081 | -0.042 | 0.071 | 0.071 | 0 | 0.28 | 0.39 | 0.39 | 0.5 | 0.19 | 0.29 |
| 5 | 150440097 | rs2233287 | 0.999 | -1.168 | 1.314 | 0.034 | -0.024 | -0.065 | -0.065 | 0.128 | 0.14 | 0.21 | 0.09 | 0 | 0.08 | 0.05 |
| 5 | 150443005 | rs17728260 | 0.771 | -1.213 | 1.238 | 0.114 | 0.009 | -0.032 | -0.032 | 0 | 0.1 | 0.03 | 0.31 | 0.49 | 0.09 | 0.2 |
| 5 | 150444212 | rs7732451 | -0.015 | -1.213 | 1.348 | 0.052 | -0.041 | -0.137 | -0.137 | 0.156 | 0.19 | 0.46 | 0.11 | 0.01 | 0.12 | 0.09 |
| 5 | 150445609 | rs1422674 | 0.738 | -1.012 | 1.105 | 0.253 | -0.005 | -0.046 | -0.046 | 0 | 0.1 | 0.01 | 0.3 | 0.49 | 0.1 | 0.2 |
| 5 | 150448861 | rs12109187 | 0.007 | -0.95 | 1.179 | 0.047 | 0.033 | 0.004 | 0.004 | 0 | 0.05 | 0.25 | 0.02 | 0.01 | 0.04 | 0.05 |
| 5 | 150451650 | rs3792785 | 0.723 | -1.073 | 0.946 | 0.061 | -0.014 | -0.005 | -0.005 | 0 | 0.16 | 0.21 | 0.09 | 0.18 | 0.09 | 0.14 |
| 5 | 150455732 | rs3792783 | -0.34 | -1.203 | 1.18 | 0.114 | -0.046 | -0.014 | -0.014 | 0.065 | 0.24 | 0.52 | 0.14 | 0.25 | 0.16 | 0.2 |
| 5 | 150458146 | rs10036748 | -1.205 | -1.292 | 0.604 | 0.208 | 0.081 | 0.03 | 0.03 | 0.005 | 0.37 | 0.85 | 0.46 | 0.73 | 0.26 | 0.4 |
| 5 | 150459788 | rs918498 | -0.296 | -1.202 | 0.957 | 0.299 | -0.039 | 0.234 | 0.234 | 0 | 0.16 | 0.44 | 0.35 | 0.54 | 0.12 | 0.2 |
| 5 | 150461049 | rs960709 | -1.148 | -1.206 | -0.099 | 0.216 | 0.093 | 0.043 | 0.043 | 0 | 0.38 | 0.85 | 0.47 | 0.73 | 0.26 | 0.4 |
| 5 | 150461069 | rs4958438 | 0.355 | -0.673 | 0.278 | 0.56 | -0.005 | 0.245 | 0.245 | 0 | 0.11 | 0.3 | 0.32 | 0.48 | 0.1 | 0.19 |
| 5 | 150462638 | rs13168551 | -0.762 | -0.507 | -0.869 | -0.221 | 0.043 | 0.147 | 0.147 | 0.046 | 0.5 | 0.84 | 0.58 | 0.73 | 0.43 | 0.51 |

| | | | | | | | | | | | | | | | | |
|---|---|---|---|---|---|---|---|---|---|---|---|---|---|---|---|---|
| 5 | 150469426 | rs3762999 | -0.908 | -0.392 | -0.181 | -0.447 | -0.001 | -0.03 | -0.03 | 0 | 0.54 | 0.54 | 0.59 | 0.73 | 0.45 | 0.6 |
| 5 | 150473674 | rs999556 | -0.749 | -0.61 | -0.08 | -0.556 | -0.001 | -0.008 | -0.008 | 0 | 0.38 | 0.45 | 0.5 | 0.72 | 0.32 | 0.56 |
| 5 | 150476004 | rs1024995 | -0.691 | -0.458 | -0.138 | -0.103 | -0.023 | 0.061 | 0.061 | 0 | 0.19 | 0.36 | 0.11 | 0.39 | 0.14 | 0.31 |
| 5 | 150477087 | rs4958890 | -1.428 | -0.278 | -0.586 | -0.202 | -0.051 | 0.111 | 0.111 | 0.095 | 0.62 | 0.82 | 0.66 | 0.83 | 0.51 | 0.77 |
| 5 | 150478318 | rs17728338 | 0.183 | 0.112 | -0.685 | 0.308 | 0.005 | 0.001 | 0.001 | 0 | 0.07 | 0.1 | 0.05 | 0.1 | 0.07 | 0.15 |
| 5 | 150479351 | rs999011 | 0.341 | 0.087 | -0.989 | 0.427 | 0 | 0.008 | 0.008 | 0.024 | 0.13 | 0.1 | 0.13 | 0.04 | 0.13 | 0.17 |
| 5 | 150480520 | rs11960458 | 0.198 | 0.432 | -0.904 | 0.971 | 0.004 | -0.041 | -0.041 | 0 | 0.22 | 0.11 | 0.37 | 0.38 | 0.19 | 0.19 |
| 5 | 150483095 | rs4958892 | -0.21 | 0.605 | -0.725 | 0.515 | -0.008 | -0.007 | -0.007 | 0 | 0.4 | 0.4 | 0.48 | 0.63 | 0.34 | 0.39 |
| 5 | 150483825 | rs6896621 | 0.311 | 1.322 | -1.509 | 0.874 | -0.071 | -0.073 | -0.073 | 0 | 0.78 | 0.95 | 0.78 | 0.62 | 0.7 | 0.79 |
| 5 | 150485241 | rs868641 | 0.49 | 1.346 | -1.07 | 0.909 | -0.003 | -0.007 | -0.007 | 0 | 0.78 | 0.8 | 0.77 | 0.67 | 0.75 | 0.75 |
| 5 | 150486991 | rs4958893 | 0.653 | 1.569 | -1.025 | 0.799 | -0.012 | -0.029 | -0.029 | 0 | 0.77 | 0.88 | 0.77 | 0.67 | 0.76 | 0.73 |
| 5 | 150489390 | rs1133202 | 0.596 | 1.723 | -0.806 | 0.777 | -0.011 | -0.025 | -0.025 | 0 | 0.77 | 0.84 | 0.76 | 0.67 | 0.76 | 0.74 |
| 5 | 150489817 | rs9324675 | -0.44 | 1.062 | -0.189 | 0.746 | -0.008 | -0.004 | -0.004 | 0 | 0.17 | 0.34 | 0.1 | 0.18 | 0.14 | 0.28 |
| 5 | 150490751 | rs7706525 | 0.454 | 0.848 | -0.896 | 0.288 | 0 | 0.001 | 0.001 | 0.328 | 0.52 | 0.47 | 0.58 | 0.58 | 0.49 | 0.47 |
| 5 | 150490805 | rs7724316 | 0.006 | 0.74 | -0.601 | 0.209 | 0.002 | 0.003 | 0.003 | 0 | 0.61 | 0.63 | 0.62 | 0.67 | 0.59 | 0.61 |
| 5 | 150490958 | rs7724949 | 0.633 | 0.84 | -0.731 | 0.308 | -0.014 | -0.031 | -0.031 | 0.245 | 0.09 | 0.04 | 0.06 | 0.18 | 0.12 | 0.16 |
| 5 | 150492562 | rs4958441 | 0.035 | 0.801 | -0.587 | 0.346 | -0.038 | 0.025 | 0.025 | 3.321 | 0.15 | 0.55 | 0.09 | 0.18 | 0.08 | 0.22 |
| 5 | 150493523 | rs4526094 | 0.553 | 1.127 | -1.118 | 0.385 | 0.003 | -0.031 | -0.031 | 0.027 | 0.46 | 0.17 | 0.53 | 0.49 | 0.5 | 0.39 |
| 5 | 150493725 | rs12651780 | 1.197 | 1.172 | -1.369 | 0.394 | -0.064 | -0.044 | -0.044 | 0 | 0.38 | 0.07 | 0.49 | 0.4 | 0.4 | 0.24 |
| 5 | 150495808 | rs11167536 | 0.589 | 1.25 | -0.835 | 0.648 | -0.034 | -0.025 | -0.025 | 0 | 0.08 | 0 | 0.04 | 0.09 | 0.11 | 0.14 |
| 5 | 150499733 | rs4958897 | 0.048 | 1.298 | -0.615 | 0.694 | -0.004 | 0.028 | 0.028 | 0 | 0.27 | 0.35 | 0.48 | 0.68 | 0.28 | 0.41 |
| 5 | 150499884 | rs728052 | 1.118 | 1.348 | -0.762 | 0.388 | 0.004 | 0.046 | 0.046 | 1.73 | 0.35 | 0.26 | 0.18 | 0.17 | 0.29 | 0.18 |
| 5 | 150503337 | rs3792775 | 0.692 | 1.184 | -0.872 | 0.662 | -0.005 | 0.042 | 0.042 | 0 | 0.24 | 0.28 | 0.46 | 0.5 | 0.24 | 0.21 |
| 5 | 150503675 | rs2303025 | -0.24 | 1.228 | -0.807 | 0.513 | 0 | 0 | 0 | 0 | 0.43 | 0.53 | 0.64 | 0.59 | 0.39 | 0.41 |
| 5 | 150504416 | rs17659044 | 1.564 | 1.189 | -0.772 | 0.499 | 0.001 | 0.03 | 0.03 | 0.467 | 0.17 | 0.09 | 0.15 | 0.06 | 0.15 | 0.18 |
| 5 | 150504585 | rs10476965 | -0.177 | 1.186 | -0.715 | 0.499 | -0.006 | 0.004 | 0.004 | 0 | 0.42 | 0.47 | 0.63 | 0.56 | 0.39 | 0.38 |
| 5 | 150505516 | rs10037814 | -0.513 | 1.16 | -0.78 | 0.543 | -0.003 | -0.005 | -0.005 | 0 | 0.44 | 0.52 | 0.64 | 0.76 | 0.41 | 0.58 |
| 5 | 150505673 | rs17659056 | 1.826 | 1.342 | -0.904 | 0.528 | -0.039 | -0.022 | -0.022 | 1.324 | 0.08 | 0 | 0.06 | 0.06 | 0.13 | 0.12 |
| 5 | 150505794 | rs2303028 | 1.46 | 1.263 | -0.853 | 0.72 | -0.005 | 0.046 | 0.046 | 0 | 0.19 | 0.24 | 0.35 | 0.39 | 0.2 | 0.15 |
| 5 | 150506086 | rs2303029 | 0.796 | 0.836 | -0.73 | 1.012 | 0.008 | 0.003 | 0.003 | 0.184 | 0.2 | 0.12 | 0.24 | 0.2 | 0.17 | 0.21 |
| 5 | 150506894 | rs7706092 | -0.062 | 0.861 | -0.725 | 1.019 | -0.005 | 0.001 | 0.001 | 2.467 | 0.28 | 0.19 | 0.3 | 0.26 | 0.29 | 0.33 |
| 5 | 150507435 | rs1012415 | -0.903 | 0.942 | -0.661 | 0.907 | 0.003 | 0.018 | 0.018 | 0 | 0.67 | 0.6 | 0.42 | 0.48 | 0.6 | 0.75 |
| 5 | 150508602 | rs6886358 | 0.227 | 0.928 | -0.891 | 0.726 | 0 | 0.01 | 0.01 | 0 | 0.52 | 0.46 | 0.33 | 0.22 | 0.42 | 0.35 |
| 5 | 150510233 | rs3762993 | 0.254 | 0.974 | -1.383 | 0.439 | -0.009 | 0.068 | 0.068 | 0 | 0.53 | 0.26 | 0.33 | 0.31 | 0.46 | 0.58 |
| 5 | 150513738 | rs9324677 | -0.396 | 0.349 | -0.812 | 0.584 | -0.011 | -0.001 | -0.001 | 0 | 0.63 | 0.76 | 0.47 | 0.52 | 0.56 | 0.54 |
| 5 | 150513847 | rs28705982 | -0.621 | -0.273 | -0.481 | 0.5 | 0.002 | -0.001 | -0.001 | 0 | 0.21 | 0.23 | 0.15 | 0.07 | 0.13 | 0.2 |
| 5 | 150516492 | rs11952969 | -0.849 | -0.316 | -0.603 | 0.78 | -0.002 | -0.001 | -0.001 | 0 | 0.25 | 0.3 | 0.27 | 0.25 | 0.17 | 0.26 |
| 5 | 150517899 | rs10039049 | -0.735 | -0.215 | -0.95 | 0.858 | -0.005 | -0.05 | -0.05 | 0 | 0.32 | 0.23 | 0.42 | 0.54 | 0.31 | 0.56 |
| 5 | 150517935 | rs12519495 | -0.72 | 0.097 | -1.233 | 0.763 | -0.053 | -0.116 | -0.116 | 0 | 0.29 | 0.03 | 0.33 | 0.45 | 0.29 | 0.55 |
| 5 | 150518532 | rs3815725 | -1.096 | 0.068 | -0.653 | 0.391 | 0 | -0.01 | -0.01 | 0.552 | 0.47 | 0.5 | 0.56 | 0.77 | 0.51 | 0.74 |
| 5 | 150518988 | rs2228458 | 0.574 | 0.071 | -1.008 | 0.443 | -0.043 | -0.124 | -0.124 | 0.175 | 0.21 | 0.01 | 0.31 | 0.45 | 0.22 | 0.5 |
| 5 | 150522559 | rs13185706 | 0.049 | 0.036 | -0.818 | 0.351 | 0 | 0 | 0 | 0 | 0.1 | 0.16 | 0.11 | 0.09 | 0.09 | 0.06 |
| 5 | 150523163 | rs883887 | 0.265 | 0.029 | -0.352 | 0.601 | -0.006 | 0.03 | 0.03 | 0.491 | 0.23 | 0.27 | 0.37 | 0.63 | 0.21 | 0.55 |
| 5 | 150523187 | rs2161358 | -2.042 | -0.553 | -0.296 | 0.202 | 0.004 | -0.031 | -0.031 | 0 | 0.65 | 0.68 | 0.69 | 0.88 | 0.62 | 0.9 |
| 5 | 150523987 | rs884083 | -1.398 | -0.627 | -0.343 | 0.155 | 0.013 | 0.035 | 0.035 | 2.385 | 0.52 | 0.66 | 0.55 | 0.72 | 0.4 | 0.76 |
| 5 | 150525901 | rs10515644 | -1.092 | 0.078 | -0.285 | 0.606 | -0.031 | 0.014 | 0.014 | 0 | 0.31 | 0.21 | 0.28 | 0.25 | 0.38 | 0.26 |
| 5 | 150530679 | rs1862362 | -0.759 | 0.196 | -0.179 | 0.654 | -0.002 | 0.031 | 0.031 | 0.075 | 0.39 | 0.32 | 0.52 | 0.78 | 0.43 | 0.68 |
| 5 | 150531247 | rs951224 | -0.666 | 0.182 | -0.16 | 0.678 | -0.002 | 0.018 | 0.018 | 0.168 | 0.38 | 0.35 | 0.53 | 0.78 | 0.43 | 0.68 |
| 5 | 150534671 | rs12517644 | 1.067 | 0.08 | -0.292 | 0.861 | 0.022 | -0.038 | -0.038 | 2.346 | 0.12 | 0.03 | 0.23 | 0.44 | 0.08 | 0.34 |
| 5 | 150534953 | rs990995 | -1.961 | 0.086 | -0.242 | 0.809 | -0.002 | -0.039 | -0.039 | 0 | 0.54 | 0.58 | 0.66 | 0.88 | 0.59 | 0.79 |
| 5 | 150535013 | rs2303038 | 0.824 | 0.086 | -0.265 | 0.819 | -0.058 | -0.04 | -0.04 | 0.547 | 0.14 | 0.01 | 0.14 | 0.15 | 0.22 | 0.14 |
| 5 | 150537629 | rs1030199 | -1.979 | 0.224 | -0.189 | 0.833 | -0.001 | -0.027 | -0.027 | 0 | 0.54 | 0.58 | 0.66 | 0.88 | 0.58 | 0.78 |
| 5 | 150544571 | rs7731328 | 0.565 | 0.207 | -0.068 | 1.025 | 0 | 0 | 0 | 0 | 0.08 | 0.13 | 0.11 | 0.09 | 0.09 | 0.07 |
| 5 | 150544620 | rs371586 | 0.071 | 0.166 | -0.043 | 1.062 | -0.003 | 0.023 | 0.023 | 0.704 | 0.77 | 0.71 | 0.62 | 0.36 | 0.79 | 0.46 |
| 5 | 150548231 | rs10476966 | -0.863 | 0.931 | 0.102 | 1.144 | -0.01 | 0.016 | 0.016 | 0 | 0.16 | 0.11 | 0.11 | 0.09 | 0.15 | 0.11 |
| 5 | 150548526 | rs7724685 | -1.594 | 1.094 | 0.228 | 1.158 | 0.012 | 0.011 | 0.011 | 0 | 0.28 | 0.39 | 0.28 | 0.28 | 0.31 | 0.31 |
| 5 | 150551827 | rs10515645 | -1.521 | 1.224 | -0.168 | 1.146 | 0.007 | 0.005 | 0.005 | 0 | 0.27 | 0.36 | 0.27 | 0.3 | 0.31 | 0.31 |
| 5 | 150553419 | rs891899 | -0.994 | 1.226 | -0.169 | 1.15 | 0.012 | 0.062 | 0.062 | 0 | 0.17 | 0.36 | 0.18 | 0.29 | 0.15 | 0.14 |
| 5 | 150555579 | rs13157589 | -1.655 | 1.198 | -0.211 | 1.228 | 0.003 | 0.004 | 0.004 | 0 | 0.4 | 0.48 | 0.43 | 0.46 | 0.51 | 0.43 |
| 5 | 150556327 | rs7349812 | 0.304 | 1.206 | -0.284 | 1.206 | -0.043 | 0.038 | 0.038 | 0 | 0.1 | 0.02 | 0.09 | 0.02 | 0.16 | 0.17 |
| 5 | 150557223 | rs6579839 | -0.98 | 0.609 | -0.113 | 1.351 | 0 | 0.001 | 0.001 | 0 | 0.4 | 0.46 | 0.43 | 0.46 | 0.51 | 0.43 |
| 5 | 150559991 | rs7717659 | -0.945 | 0.633 | -0.55 | 1.276 | 0.003 | 0.001 | 0.001 | 0 | 0.42 | 0.56 | 0.45 | 0.52 | 0.52 | 0.44 |
| 5 | 150560007 | rs7717803 | -1.059 | 0.435 | -0.526 | 1.556 | -0.001 | 0.13 | 0.13 | 0 | 0.1 | 0.25 | 0.15 | 0.27 | 0.06 | 0.09 |
| 5 | 150563302 | rs12518395 | -0.352 | 0.2 | -1.713 | 0.766 | -0.014 | -0.017 | -0.017 | 0 | 0.25 | 0.01 | 0.37 | 0.17 | 0.24 | 0.18 |
| 5 | 150565724 | rs248426 | 0.236 | 0.682 | -1.39 | 0.594 | -0.001 | 0.007 | 0.007 | 0.019 | 0.45 | 0.45 | 0.4 | 0.35 | 0.44 | 0.38 |
| 5 | 150566792 | rs448503 | 0.005 | 0.829 | -1.448 | 0.712 | 0.009 | 0.05 | 0.05 | 0 | 0.71 | 0.46 | 0.77 | 0.51 | 0.69 | 0.67 |
| 5 | 150572439 | rs248441 | -0.329 | 1.061 | -0.962 | 0.982 | 0.04 | 0.127 | 0.127 | 0 | 0.87 | 0.74 | 0.84 | 0.64 | 0.78 | 0.81 |
| 5 | 150574632 | rs248448 | 0.624 | 0.67 | -1.074 | 0.907 | -0.001 | 0.064 | 0.064 | 0 | 0.9 | 0.89 | 0.93 | 0.6 | 0.92 | 0.78 |
| 5 | 150576767 | rs248450 | -0.117 | 0.713 | -1.013 | 0.008 | -0.006 | -0.008 | -0.008 | 0 | 0.53 | 0.46 | 0.69 | 0.28 | 0.69 | 0.47 |

| | | | | | | | | | | | | | | | | |
|---|---|---|---|---|---|---|---|---|---|---|---|---|---|---|---|---|
| 5 | 150579350 | rs248452 | -0.031 | 1.711 | -0.657 | -0.264 | 0.092 | -0.052 | -0.052 | 0.387 | 0.22 | 0.4 | 0.29 | 0.1 | 0.4 | 0.11 |
| 5 | 150582456 | rs248457 | -0.424 | 1.111 | -0.759 | -0.181 | 0.024 | 0.001 | 0.001 | 0 | 0.26 | 0.42 | 0.34 | 0.31 | 0.33 | 0.3 |
| 5 | 150587346 | rs6865418 | -0.039 | 0.144 | -1.587 | -0.65 | 0.038 | 0.075 | 0.075 | 0 | 0.24 | 0.12 | 0.13 | 0 | 0.17 | 0.11 |
| 5 | 150598381 | rs153470 | 0.065 | 0.193 | -1.714 | 0.238 | 0.018 | -0.064 | -0.064 | 0.021 | 0.3 | 0.17 | 0.47 | 0.52 | 0.23 | 0.44 |
| 5 | 150599974 | rs152265 | -1.051 | 1.28 | -2.191 | -0.415 | -0.088 | 0.081 | 0.081 | 0 | 0.87 | 0.96 | 0.83 | 1 | 0.68 | 0.94 |
| 5 | 150605070 | rs153462 | 0.217 | 1.026 | -1.867 | 0.027 | -0.001 | -0.026 | -0.026 | 0.797 | 0.3 | 0.26 | 0.3 | 0.46 | 0.27 | 0.43 |
| 5 | 150612568 | rs6866925 | 0.93 | 0.502 | -1.316 | -0.286 | -0.008 | -0.008 | -0.008 | 0 | 0.12 | 0.19 | 0.1 | 0.09 | 0.11 | 0.08 |
| 5 | 150612849 | rs1004736 | 0.636 | 0.23 | -1.082 | -0.381 | 0.005 | 0.017 | 0.017 | 0 | 0.18 | 0.13 | 0.09 | 0.09 | 0.15 | 0.13 |
| 5 | 150617370 | rs3097793 | 0.228 | -0.105 | -0.986 | -0.149 | -0.002 | 0.019 | 0.019 | 0 | 0.26 | 0.31 | 0.26 | 0.47 | 0.2 | 0.46 |
| 5 | 150618829 | rs173567 | 0.069 | -0.121 | -0.968 | -0.146 | -0.002 | 0.01 | 0.01 | 0 | 0.27 | 0.36 | 0.26 | 0.57 | 0.21 | 0.54 |
| 5 | 150619092 | rs1991798 | 1.072 | -0.189 | -0.867 | -0.337 | 0.044 | 0.045 | 0.045 | 0.064 | 0.39 | 0.16 | 0.44 | 0.31 | 0.32 | 0.26 |
| 5 | 150626940 | rs6579844 | 0.448 | -0.083 | -0.744 | -0.144 | 0.004 | 0.006 | 0.006 | 0 | 0.52 | 0.57 | 0.61 | 0.71 | 0.41 | 0.5 |
| 5 | 150630845 | rs153483 | 0.027 | 0.211 | -0.342 | -0.387 | 0.017 | -0.072 | -0.072 | 0 | 0.67 | 0.62 | 0.73 | 0.92 | 0.53 | 0.81 |
| 5 | 150635429 | rs152266 | 0.857 | -0.063 | -0.514 | -0.32 | -0.003 | 0.04 | 0.04 | 0 | 0.56 | 0.33 | 0.6 | 0.63 | 0.48 | 0.63 |
| 5 | 150638537 | rs153474 | 0.102 | 0.511 | 0.337 | -0.222 | 0.072 | 0.011 | 0.011 | 0.001 | 0.75 | 0.56 | 0.73 | 0.66 | 0.63 | 0.75 |
| 5 | 150639409 | rs153477 | -0.002 | 0.544 | 0.404 | -0.223 | 0.067 | 0.016 | 0.016 | 0 | 0.76 | 0.57 | 0.74 | 0.66 | 0.65 | 0.76 |
| 5 | 150639778 | rs153479 | 0.511 | 0.206 | 0.243 | -0.342 | 0.044 | -0.018 | -0.018 | 0 | 0.67 | 0.5 | 0.69 | 0.65 | 0.56 | 0.67 |
| 5 | 150642265 | rs6579845 | 1.023 | 0.015 | 0.011 | -0.013 | 0.006 | 0.011 | 0.011 | 0 | 0.32 | 0.23 | 0.48 | 0.27 | 0.32 | 0.31 |
| 5 | 150647156 | rs9324686 | -0.459 | 0.017 | 0.019 | -0.026 | 0.023 | 0.129 | 0.129 | 0.274 | 0.06 | 0.24 | 0.07 | 0.25 | 0.02 | 0.09 |
| 5 | 150648878 | rs3806953 | 1.106 | -0.098 | -0.971 | -0.38 | -0.049 | 0.016 | 0.016 | 0 | 0.31 | 0.02 | 0.44 | 0.21 | 0.31 | 0.29 |
| 5 | 150651775 | rs1065018 | 0.937 | -0.12 | -0.769 | -0.34 | 0.016 | 0.009 | 0.009 | 0 | 0.33 | 0.11 | 0.45 | 0.3 | 0.3 | 0.37 |
| 5 | 150652985 | rs248469 | -1.301 | 0.126 | -0.568 | -0.698 | 0.019 | -0.048 | -0.048 | 0 | 0.83 | 0.81 | 0.78 | 1 | 0.65 | 0.92 |
| 5 | 150653407 | rs153445 | -0.486 | 0.062 | -0.646 | -0.69 | 0.053 | 0.11 | 0.11 | 0 | 0.52 | 0.82 | 0.34 | 0.7 | 0.35 | 0.55 |
| 5 | 150654481 | rs7717132 | 0.048 | -0.188 | -0.461 | -0.691 | 0.001 | 0.024 | 0.024 | 0 | 0.53 | 0.64 | 0.61 | 0.47 | 0.43 | 0.53 |
| 5 | 150656548 | rs17659864 | 1.401 | -0.237 | -0.969 | -0.386 | -0.043 | -0.009 | -0.009 | 0.02 | 0.29 | 0.03 | 0.44 | 0.23 | 0.29 | 0.28 |
| 5 | 150658260 | rs17802052 | 0.613 | -0.166 | -0.832 | -0.475 | 0.016 | 0.019 | 0.019 | 0.337 | 0.41 | 0.18 | 0.51 | 0.45 | 0.37 | 0.41 |
| 5 | 150659204 | rs1347144 | -1.296 | -0.441 | 0.065 | -0.454 | 0.037 | 0.114 | 0.114 | 4.561 | 0.06 | 0.56 | 0.06 | 0.09 | 0.01 | 0.08 |
| 5 | 150660769 | rs1368393 | -0.111 | -0.205 | -0.182 | -0.578 | 0.039 | 0.025 | 0.025 | 1.78 | 0.49 | 0.77 | 0.57 | 0.53 | 0.39 | 0.53 |
| 5 | 150660794 | rs1368394 | -0.019 | -0.209 | -0.174 | -0.579 | 0.033 | 0.021 | 0.021 | 1.634 | 0.49 | 0.73 | 0.57 | 0.53 | 0.39 | 0.53 |
| 5 | 150664002 | rs969835 | 1.621 | -0.246 | -0.083 | -0.608 | 0.024 | 0.032 | 0.032 | 1.012 | 0.31 | 0.13 | 0.42 | 0.24 | 0.26 | 0.29 |
| 5 | 150664183 | rs17111902 | -0.625 | -0.244 | 0.024 | -0.607 | 0.064 | 0.01 | 0.01 | 0.994 | 0.05 | 0.3 | 0.04 | 0 | 0.04 | 0.04 |
| 5 | 150666361 | rs10055434 | -0.625 | -0.244 | 0.024 | -0.607 | 0.064 | 0.01 | 0.01 | 0.994 | 0.05 | 0.3 | 0.04 | 0 | 0.04 | 0.04 |
| 5 | 150666962 | rs12520516 | 1.504 | -0.239 | -0.024 | -0.599 | -0.025 | -0.036 | -0.036 | 0 | 0.09 | 0.02 | 0.06 | 0.22 | 0.08 | 0.12 |
| 5 | 150667016 | rs978012 | -0.174 | -0.241 | -0.034 | -0.599 | 0.04 | 0.027 | 0.027 | 1.823 | 0.5 | 0.77 | 0.57 | 0.53 | 0.4 | 0.54 |
| 5 | 150667775 | rs6579847 | 0.395 | -0.372 | 0.011 | -0.533 | 0.015 | 0.08 | 0.08 | 0.204 | 0.13 | 0.33 | 0.11 | 0.3 | 0.09 | 0.21 |
| 5 | 150673387 | rs190309 | 0.806 | -0.242 | -0.168 | -0.621 | -0.001 | 0.007 | 0.007 | 0 | 0.22 | 0.25 | 0.15 | 0.45 | 0.18 | 0.28 |
| 5 | 150677182 | rs6876589 | 0.93 | -0.198 | -0.389 | -0.608 | -0.005 | -0.004 | -0.004 | 0 | 0.43 | 0.52 | 0.53 | 0.43 | 0.35 | 0.41 |
| 5 | 150680283 | rs246498 | -0.684 | -0.094 | -0.726 | -0.67 | -0.15 | 0.175 | 0.175 | 0.268 | 0.73 | 0.97 | 0.73 | 0.98 | 0.58 | 0.84 |
| 5 | 150683463 | rs7356742 | 0.315 | 0.033 | -0.551 | -0.418 | 0.041 | -0.001 | -0.001 | 0 | 0.48 | 0.77 | 0.57 | 0.54 | 0.39 | 0.5 |
| 5 | 150695700 | rs9324688 | -0.927 | 0.017 | -0.014 | -0.294 | 0.02 | 0.064 | 0.064 | 0.017 | 0.05 | 0.21 | 0.03 | 0.14 | 0.04 | 0.07 |
| 5 | 150695724 | rs383915 | -0.537 | -0.063 | -0.338 | -0.547 | -0.057 | 0.088 | 0.088 | 0 | 0.68 | 0.8 | 0.72 | 0.99 | 0.58 | 0.82 |
| 5 | 150698806 | rs3846710 | 0.913 | -0.195 | -0.478 | -0.13 | 0.006 | -0.125 | -0.125 | 0 | 0.2 | 0.04 | 0.1 | 0.41 | 0.17 | 0.33 |
| 5 | 150702281 | rs408316 | 0.062 | -0.055 | -0.196 | -0.278 | 0.015 | -0.059 | -0.059 | 0 | 0.62 | 0.53 | 0.68 | 0.85 | 0.54 | 0.75 |
| 5 | 150708723 | rs246491 | -0.472 | 0.107 | -0.847 | -0.488 | -0.03 | 0.03 | 0.03 | 0 | 0.67 | 0.8 | 0.71 | 0.99 | 0.58 | 0.8 |
| 5 | 150716364 | rs17111939 | -1 | 0.031 | -0.766 | -0.393 | -0.012 | -0.055 | -0.055 | 0 | 0.09 | 0.7 | 0.09 | 0 | 0.04 | 0.02 |
| 5 | 150722675 | rs2431079 | 1.075 | 0.104 | -1.581 | -0.049 | -0.05 | -0.165 | -0.165 | 0 | 0.38 | 0.02 | 0.53 | 0.46 | 0.36 | 0.42 |
| 5 | 150723806 | rs192192 | 0.506 | 0.633 | -0.666 | -0.717 | -0.023 | 0.034 | 0.034 | 0 | 0.59 | 0.31 | 0.64 | 0.98 | 0.53 | 0.77 |
| 5 | 150743405 | rs1605712 | -1.006 | 1.107 | 0.691 | 0.003 | 0.015 | -0.07 | -0.07 | 1.788 | 0.18 | 0.71 | 0.19 | 0.01 | 0.23 | 0.14 |
| 5 | 150760282 | rs11167542 | -1.323 | 0.941 | 0.243 | 0.351 | -0.024 | -0.038 | -0.038 | 0.01 | 0.35 | 0.81 | 0.3 | 0.43 | 0.4 | 0.47 |
| 5 | 150764714 | rs7702931 | 0.159 | 0.904 | 0.2 | 0.432 | -0.003 | -0.046 | -0.046 | 4.081 | 0.18 | 0.12 | 0.1 | 0.38 | 0.17 | 0.33 |
| 5 | 150781849 | rs12659858 | -1.078 | 0.627 | -0.638 | -0.094 | -0.074 | 0.036 | 0.036 | 0 | 0.74 | 0.99 | 0.64 | 0.9 | 0.73 | 0.83 |
| 5 | 150789051 | rs7708940 | -0.958 | 0.822 | 0.659 | -0.118 | 0.011 | 0.019 | 0.019 | 6.989 | 0.34 | 0.46 | 0.26 | 0.39 | 0.39 | 0.48 |
| 5 | 150795275 | rs1116037 | 0 | 0.867 | 0.89 | -0.166 | 0.072 | -0.078 | -0.078 | 2.94 | 0.12 | 0.26 | 0.16 | 0 | 0.22 | 0.12 |
| 5 | 150795336 | rs11167546 | 0.19 | 0.844 | 0.778 | -0.159 | 0.004 | -0.061 | -0.061 | 0 | 0.2 | 0.13 | 0.11 | 0.39 | 0.17 | 0.37 |
| 5 | 150801545 | rs6871318 | -1.218 | 0.815 | 0.913 | -0.072 | -0.021 | -0.045 | -0.045 | 0 | 0.37 | 0.8 | 0.31 | 0.56 | 0.4 | 0.49 |
| 5 | 150808555 | rs17111995 | -0.7 | 1.019 | 1.183 | -0.049 | 0.076 | -0.009 | -0.009 | 0 | 0.14 | 0.53 | 0.19 | 0.14 | 0.22 | 0.12 |
| 5 | 150820233 | rs1876620 | -0.7 | 1.019 | 1.228 | -0.05 | 0.076 | -0.009 | -0.009 | 0 | 0.14 | 0.53 | 0.19 | 0.14 | 0.22 | 0.12 |
| 5 | 150825329 | rs3905908 | -0.433 | 1.077 | 0.843 | -0.129 | 0.086 | -0.089 | -0.089 | 3.25 | 0.13 | 0.34 | 0.16 | 0.01 | 0.22 | 0.11 |
| 5 | 150831352 | rs17740674 | -0.204 | 0.62 | -0.068 | -0.35 | 0.059 | -0.02 | -0.02 | 0 | 0.45 | 0.24 | 0.16 | 0.43 | 0.3 | 0.48 |
| 5 | 150831989 | rs918421 | -0.953 | 1.076 | 0.149 | -0.034 | -0.015 | -0.013 | -0.013 | 0 | 0.35 | 0.7 | 0.33 | 0.54 | 0.46 | 0.51 |
| 5 | 150836145 | rs11954054 | -0.328 | 0.956 | 0.07 | 0.057 | -0.001 | -0.015 | -0.015 | 0 | 0.22 | 0.17 | 0.14 | 0.4 | 0.23 | 0.38 |
| 5 | 150836580 | rs17802828 | -0.365 | 0.991 | 0.151 | 0.138 | -0.001 | 0.013 | 0.013 | 0.778 | 0.19 | 0.17 | 0.13 | 0.4 | 0.18 | 0.37 |
| 5 | 150838085 | rs35060330 | -0.474 | 0.729 | 0.432 | 0.009 | -0.007 | -0.015 | -0.015 | 0 | 0.42 | 0.49 | 0.29 | 0.64 | 0.33 | 0.61 |
| 5 | 150840380 | rs357608 | -0.626 | 0.79 | 0.005 | -0.024 | 0.027 | 0.077 | 0.077 | 0.157 | 0.48 | 0.25 | 0.38 | 0.65 | 0.52 | 0.71 |
| 5 | 150845369 | rs357616 | 0.011 | 0.888 | 0.125 | -0.22 | -0.028 | 0.018 | 0.018 | 0 | 0.32 | 0.23 | 0.29 | 0.26 | 0.4 | 0.34 |
| 5 | 150847187 | rs11949709 | -1.138 | 1.262 | 1.271 | 0.145 | -0.009 | 0.065 | 0.065 | 0.983 | 0.23 | 0.35 | 0.16 | 0.49 | 0.19 | 0.39 |
| 5 | 150853893 | rs165350 | 0.068 | 1.361 | 1.149 | 0.03 | -0.036 | 0.03 | 0.03 | 0.182 | 0.09 | 0.04 | 0.12 | 0.03 | 0.2 | 0.1 |
| 5 | 150856107 | rs165352 | -1.674 | 1.314 | 1.157 | 0.032 | 0.008 | 0.015 | 0.015 | 2.138 | 0.32 | 0.39 | 0.29 | 0.52 | 0.39 | 0.5 |
| 5 | 150861701 | rs17112022 | -0.986 | 1.42 | 1.141 | 0.049 | 0.01 | -0.03 | -0.03 | 0 | 0.06 | 0.12 | 0.04 | 0.01 | 0.07 | 0.03 |

| | | | | | | | | | | | | | | | | |
|---|---|---|---|---|---|---|---|---|---|---|---|---|---|---|---|---|
| 5 | 150868490 | rs2278373 | 0.029 | 1.381 | 1.269 | 0.09 | -0.003 | -0.113 | -0.113 | 0.009 | 0.15 | 0.01 | 0.09 | 0.35 | 0.11 | 0.35 |
| 5 | 150868745 | rs2278372 | -0.071 | 1.463 | 1.306 | 0.129 | -0.006 | -0.109 | -0.109 | 0.125 | 0.15 | 0.02 | 0.09 | 0.35 | 0.11 | 0.34 |
| 5 | 150869352 | rs165369 | 0.012 | 1.646 | 1.533 | -0.616 | -0.042 | 0.032 | 0.032 | 0.014 | 0.09 | 0.03 | 0.12 | 0.03 | 0.2 | 0.1 |
| 5 | 150871280 | rs14160 | -2.097 | 1.543 | 1.656 | -0.663 | -0.031 | 0.025 | 0.025 | 0.962 | 0.2 | 0.84 | 0.25 | 0.18 | 0.28 | 0.15 |
| 5 | 150871795 | rs1175 | -0.289 | 1.59 | 1.751 | -0.639 | 0.021 | -0.018 | -0.018 | 0 | 0.11 | 0.13 | 0.13 | 0.03 | 0.2 | 0.09 |
| 5 | 150875468 | rs165330 | -1.409 | 1.55 | 1.308 | -0.651 | 0.041 | -0.011 | -0.011 | 0 | 0.15 | 0.69 | 0.21 | 0.17 | 0.21 | 0.11 |
| 5 | 150878177 | rs165335 | -2.041 | 1.531 | 1.236 | -0.613 | -0.016 | 0.014 | 0.014 | 1.008 | 0.21 | 0.82 | 0.25 | 0.17 | 0.28 | 0.15 |
| 5 | 150884089 | rs165344 | -0.209 | 1.87 | 0.932 | -0.485 | -0.036 | 0.03 | 0.03 | 0.114 | 0.1 | 0.04 | 0.13 | 0.03 | 0.2 | 0.1 |
| 5 | 150887925 | rs11750722 | 0.933 | 1.27 | 0.901 | -0.709 | -0.011 | 0.03 | 0.03 | 0 | 0.64 | 0.29 | 0.59 | 0.82 | 0.48 | 0.8 |
| 5 | 150891733 | rs2304028 | 0.181 | 1.39 | 1.21 | -0.678 | 0.046 | -0.055 | -0.055 | 0 | 0.81 | 0.68 | 0.8 | 0.95 | 0.72 | 0.88 |
| 5 | 150892489 | rs10476987 | 0.838 | 1.35 | 1.212 | -0.765 | -0.002 | -0.011 | -0.011 | 0 | 0.64 | 0.6 | 0.64 | 0.82 | 0.52 | 0.83 |
| 5 | 150894920 | rs12717853 | -0.197 | 1.356 | 1.317 | -0.746 | -0.02 | 0.016 | 0.016 | 0 | 0.83 | 0.9 | 0.82 | 0.95 | 0.72 | 0.88 |
| 5 | 150896898 | rs17741008 | 0.784 | 1.273 | 1.01 | -0.42 | 0.003 | -0.053 | -0.053 | 0 | 0.08 | 0 | 0.02 | 0.18 | 0.03 | 0.25 |
| 5 | 150901111 | rs3822699 | 0.495 | 1.85 | 1.602 | -1.007 | 0.056 | -0.073 | -0.073 | 0.18 | 0.8 | 0.63 | 0.8 | 0.95 | 0.72 | 0.85 |
| 5 | 150901613 | rs2053028 | 0.407 | 1.851 | 1.532 | -1.007 | 0.064 | -0.075 | -0.075 | 0.017 | 0.81 | 0.62 | 0.8 | 0.95 | 0.72 | 0.85 |
| 5 | 150901630 | rs3734049 | 0.655 | 1.87 | 0.727 | -1.004 | -0.053 | 0.024 | 0.024 | 0 | 0.16 | 0.01 | 0.16 | 0.13 | 0.19 | 0.05 |
| 5 | 150901699 | rs3734050 | -0.569 | 1.979 | 0.843 | -0.92 | 0.068 | 0.011 | 0.011 | 0.001 | 0.05 | 0.17 | 0.11 | 0.04 | 0.11 | 0.06 |
| 5 | 150903392 | rs6879158 | 1.343 | 1.714 | -0.094 | -0.905 | 0.024 | 0.132 | 0.132 | 0 | 0.57 | 0.23 | 0.51 | 0.78 | 0.41 | 0.73 |
| 5 | 150903601 | rs13360326 | -0.777 | 1.81 | 0.099 | -0.833 | 0.106 | 0.022 | 0.022 | 0 | 0.05 | 0.33 | 0.05 | 0.01 | 0.07 | 0.03 |
| 5 | 150906198 | rs1009091 | 1.055 | 1.562 | -0.682 | -0.774 | 0.041 | 0.126 | 0.126 | 0 | 0.54 | 0.26 | 0.51 | 0.78 | 0.4 | 0.72 |
| 5 | 150909672 | rs6864752 | 0.703 | 1.847 | -0.211 | -0.242 | 0.052 | -0.067 | -0.067 | 0 | 0.76 | 0.47 | 0.7 | 0.92 | 0.66 | 0.81 |
| 5 | 150920566 | rs2288778 | -0.005 | 1.495 | -1.224 | 0.423 | -0.038 | -0.032 | -0.032 | 0 | 0.22 | 0.09 | 0.19 | 0.29 | 0.32 | 0.25 |
| 5 | 150925144 | rs3797875 | -0.11 | 1.754 | -1.809 | 0.853 | -0.074 | -0.087 | -0.087 | 0 | 0.85 | 0.99 | 0.84 | 0.73 | 0.82 | 0.64 |
| 5 | 150928486 | rs6890712 | 0.323 | 1.002 | -1.639 | -0.067 | 0 | 0.056 | 0.056 | 0 | 0.47 | 0.3 | 0.46 | 0.3 | 0.59 | 0.36 |
| 5 | 150930186 | rs2278370 | 0.978 | 0.099 | -1.143 | 0.188 | 0.009 | -0.008 | -0.008 | 0 | 0.09 | 0.08 | 0.22 | 0.21 | 0.04 | 0.08 |
| 5 | 150930345 | rs2278371 | 0.945 | 0.092 | -1.093 | 0.201 | 0.006 | 0 | 0 | 0.463 | 0.09 | 0.08 | 0.22 | 0.23 | 0.04 | 0.08 |
| 5 | 150933411 | rs3097775 | -0.109 | -0.235 | -0.898 | 0.049 | -0.01 | 0.018 | 0.018 | 0.194 | 0.22 | 0.27 | 0.3 | 0.36 | 0.13 | 0.16 |
| 5 | 150935419 | rs3846714 | 0.635 | -0.308 | -0.362 | -0.221 | -0.003 | -0.045 | -0.045 | 0.231 | 0.08 | 0.23 | 0.05 | 0 | 0.08 | 0.02 |
| 5 | 150942969 | rs2304053 | 1.088 | -0.148 | -0.897 | -0.167 | -0.007 | 0.019 | 0.019 | 0 | 0.41 | 0.3 | 0.43 | 0.31 | 0.49 | 0.33 |
| 5 | 150943085 | rs2304054 | 0.871 | -0.131 | -0.856 | -0.16 | -0.001 | 0.005 | 0.005 | 0 | 0.43 | 0.41 | 0.45 | 0.36 | 0.49 | 0.36 |
| 5 | 150945483 | rs3734055 | 0.901 | -0.131 | -0.86 | -0.16 | -0.002 | 0.008 | 0.008 | 0 | 0.43 | 0.38 | 0.45 | 0.36 | 0.49 | 0.36 |
| 5 | 150945699 | rs10085060 | 0.835 | -0.123 | -0.872 | -0.267 | 0 | -0.004 | -0.004 | 0 | 0.44 | 0.44 | 0.45 | 0.36 | 0.49 | 0.36 |
| 5 | 150946436 | rs9324700 | 0.835 | -0.123 | -0.872 | -0.285 | 0 | -0.005 | -0.005 | 0 | 0.44 | 0.44 | 0.45 | 0.36 | 0.49 | 0.36 |
| 5 | 150946773 | rs1432862 | 0.845 | -0.123 | -0.866 | -0.285 | 0 | -0.002 | -0.002 | 0 | 0.44 | 0.44 | 0.45 | 0.36 | 0.49 | 0.36 |
| 5 | 150946966 | rs1465690 | -1.356 | -0.129 | -0.928 | -0.175 | -0.003 | 0.015 | 0.015 | 0.421 | 0.89 | 0.9 | 0.77 | 0.78 | 0.94 | 0.91 |
| 5 | 150948016 | rs3734062 | 1.265 | -0.042 | -0.992 | -0.184 | -0.026 | -0.01 | -0.01 | 0 | 0.1 | 0 | 0.06 | 0.08 | 0.09 | 0.05 |
| 5 | 150952884 | rs10072841 | 0.673 | -0.153 | -0.938 | -0.236 | -0.002 | 0.01 | 0.01 | 0 | 0.44 | 0.38 | 0.37 | 0.38 | 0.48 | 0.38 |
| 5 | 150955813 | rs11750443 | 1.421 | 0.019 | -1.395 | -0.41 | -0.001 | -0.018 | -0.018 | 0.742 | 0.1 | 0.1 | 0.12 | 0 | 0.13 | 0.05 |
| 5 | 150963802 | rs9688110 | -0.111 | 0.124 | -1.312 | 0.041 | 0 | 0 | 0 | 2.593 | 0.3 | 0.34 | 0.29 | 0.33 | 0.34 | 0.49 |
| 5 | 150964777 | rs7736409 | -0.234 | 0.318 | -1.693 | 0.092 | 0.001 | 0.001 | 0.001 | 0.035 | 0.35 | 0.5 | 0.31 | 0.39 | 0.35 | 0.54 |
| 5 | 150967836 | rs766206 | 0.634 | 0.711 | -1.287 | 0.312 | 0.002 | 0.009 | 0.009 | 0 | 0.53 | 0.4 | 0.51 | 0.27 | 0.61 | 0.48 |
| 5 | 150968303 | rs1368369 | 0.53 | 0.767 | -1.534 | 0.556 | 0.004 | -0.007 | -0.007 | 0 | 0.87 | 0.88 | 0.76 | 0.78 | 0.91 | 0.91 |
| 5 | 150971110 | rs13157373 | -0.51 | 1.538 | -2.068 | -0.236 | -0.095 | 0.103 | 0.103 | 0.172 | 0.26 | 0.05 | 0.3 | 0.07 | 0.36 | 0.17 |
| 5 | 150971682 | rs4958462 | 0.125 | 0.847 | -1.901 | -0.224 | 0.011 | 0.014 | 0.014 | 1.353 | 0.67 | 0.72 | 0.81 | 0.76 | 0.71 | 0.84 |
| 5 | 150972154 | rs11953148 | -0.581 | 1.117 | -1.389 | -0.81 | -0.008 | -0.086 | -0.086 | 0.653 | 0.29 | 0.39 | 0.24 | 0.07 | 0.24 | 0.25 |
| 5 | 150972177 | rs2278382 | -0.834 | 1.777 | -1.75 | -0.15 | -0.044 | 0.006 | 0.006 | 0 | 0.15 | 0.05 | 0.13 | 0.17 | 0.28 | 0.11 |
| 5 | 150972319 | rs2278383 | -1.024 | 1.436 | -1.619 | 0.394 | -0.022 | -0.021 | -0.021 | 0 | 0.16 | 0.06 | 0.11 | 0.2 | 0.21 | 0.08 |
| 5 | 150974701 | rs17717970 | -0.849 | 1.495 | -1.404 | 0.558 | 0.008 | 0.002 | 0.002 | 0 | 0.06 | 0.06 | 0.07 | 0.08 | 0.13 | 0.11 |
| 5 | 150975571 | rs7728335 | -0.919 | 0.674 | -2.1 | 0.706 | -0.029 | 0.068 | 0.068 | 0.478 | 0.53 | 0.86 | 0.55 | 0.65 | 0.48 | 0.47 |
| 5 | 150978726 | rs1529690 | -0.458 | 0.037 | -0.805 | 1.173 | 0 | 0.006 | 0.006 | 0 | 0.23 | 0.21 | 0.37 | 0.38 | 0.25 | 0.27 |
| 5 | 150981259 | rs753458 | -0.988 | -0.526 | -0.163 | 1.669 | -0.013 | 0.008 | 0.008 | 0.517 | 0.35 | 0.36 | 0.22 | 0.44 | 0.26 | 0.33 |
| 5 | 150981304 | rs753459 | -1.476 | -0.838 | 0.069 | 1.649 | 0.022 | -0.003 | -0.003 | 0 | 0.47 | 0.59 | 0.42 | 0.56 | 0.35 | 0.4 |
| 5 | 150982156 | rs12109416 | -0.554 | -0.511 | -0.013 | 1.401 | -0.007 | -0.018 | -0.018 | 0.162 | 0.12 | 0.05 | 0.16 | 0.17 | 0.12 | 0.1 |
| 5 | 150987402 | rs10071849 | -1.734 | -0.87 | 0.67 | 1.248 | -0.009 | 0.001 | 0.001 | 0.417 | 0.22 | 0.42 | 0.27 | 0.22 | 0.22 | 0.19 |
| 5 | 150987979 | rs11744028 | 0.494 | -0.411 | 0.523 | 1.554 | -0.038 | -0.026 | -0.026 | 0 | 0.18 | 0.09 | 0.13 | 0.19 | 0.24 | 0.11 |
| 5 | 150992585 | rs10052562 | -1.868 | -0.698 | 0.899 | 1.399 | -0.021 | -0.013 | -0.013 | 0 | 0.25 | 0.4 | 0.27 | 0.22 | 0.22 | 0.2 |
| 5 | 150998029 | rs7706276 | -2.096 | -1.171 | 1.595 | 1.739 | 0.003 | 0.025 | 0.025 | 0.627 | 0.49 | 0.65 | 0.55 | 0.63 | 0.49 | 0.45 |
| 5 | 151000157 | rs17741337 | 0.292 | -1.208 | 1.484 | 1.763 | -0.024 | 0.023 | 0.023 | 0.056 | 0.1 | 0.15 | 0.15 | 0.22 | 0.04 | 0.15 |
| 5 | 151004356 | rs17661732 | -0.575 | -1.184 | 1.209 | 1.755 | -0.022 | -0.028 | -0.028 | 0 | 0.15 | 0.03 | 0.2 | 0.17 | 0.18 | 0.17 |
| 5 | 151004915 | rs11167551 | -2.029 | -1.195 | 1.699 | 1.786 | -0.016 | 0.039 | 0.039 | 0.543 | 0.54 | 0.81 | 0.57 | 0.61 | 0.5 | 0.47 |
| 5 | 151006449 | rs11959894 | 0.255 | -1.164 | 1.473 | 1.674 | -0.022 | -0.031 | -0.031 | 0.061 | 0.06 | 0.12 | 0.03 | 0 | 0.02 | 0.01 |
| 5 | 151011600 | rs4958474 | -2.066 | -1.309 | 0.945 | 1.707 | -0.046 | 0.04 | 0.04 | 0.995 | 0.56 | 0.9 | 0.58 | 0.63 | 0.51 | 0.46 |
| 5 | 151012257 | rs1036196 | 0.117 | -1.188 | 0.929 | 1.543 | 0.017 | -0.018 | -0.018 | 0 | 0.11 | 0.08 | 0.16 | 0.22 | 0.06 | 0.18 |
| 5 | 151013080 | rs11744940 | -0.066 | -1.084 | 0.797 | 1.533 | -0.067 | 0.001 | 0.001 | 0 | 0.28 | 0.09 | 0.19 | 0.21 | 0.36 | 0.18 |
| 5 | 151020581 | rs4958476 | -1.043 | -0.654 | 0.927 | 1.296 | 0.002 | -0.01 | -0.01 | 0 | 0.44 | 0.66 | 0.4 | 0.46 | 0.44 | 0.37 |
| 5 | 151027147 | rs11167552 | -1.284 | -0.661 | 0.487 | 1.289 | 0.034 | -0.009 | -0.009 | 0 | 0.51 | 0.8 | 0.43 | 0.45 | 0.46 | 0.38 |
| 5 | 151034010 | rs6878184 | -1.186 | -0.785 | 0.037 | 1.054 | -0.031 | 0.002 | 0.002 | 0.342 | 0.55 | 0.93 | 0.61 | 0.38 | 0.51 | 0.47 |
| 5 | 151037329 | rs11747938 | 0.392 | -1.005 | -0.015 | 0.606 | -0.074 | 0.132 | 0.132 | 0 | 0.27 | 0.06 | 0.18 | 0.05 | 0.36 | 0.14 |
| 5 | 151039626 | rs2033467 | -1.179 | -0.836 | 0.686 | 0.573 | -0.001 | 0.003 | 0.003 | 0 | 0.5 | 0.38 | 0.57 | 0.5 | 0.55 | 0.5 |

| | | | | | | | | | | | | | | | | |
|---|---|---|---|---|---|---|---|---|---|---|---|---|---|---|---|---|
| 5 | 151041932 | rs3210714 | -0.763 | -0.878 | 0.567 | 0.612 | -0.014 | -0.011 | -0.011 | 0 | 0.43 | 0.25 | 0.49 | 0.43 | 0.47 | 0.37 |
| 5 | 151042029 | rs1059829 | -0.782 | -0.88 | 0.587 | 0.611 | -0.013 | -0.01 | -0.01 | 0 | 0.43 | 0.26 | 0.5 | 0.43 | 0.47 | 0.37 |
| 5 | 151045061 | rs2288810 | -0.4 | -0.951 | 1.134 | 0.569 | -0.035 | -0.003 | -0.003 | 0 | 0.13 | 0.51 | 0.12 | 0.07 | 0.09 | 0.13 |
| 5 | 151046928 | rs729853 | 0.021 | -0.948 | 1.06 | 0.597 | -0.001 | 0.013 | 0.013 | 0 | 0.15 | 0.15 | 0.32 | 0.38 | 0.11 | 0.23 |
| 5 | 151048688 | rs4958279 | -1.727 | -0.662 | 1.022 | 0.746 | -0.065 | -0.028 | -0.028 | 0.68 | 0.63 | 0.94 | 0.67 | 0.65 | 0.61 | 0.59 |
| 5 | 151049404 | rs7719521 | -1.16 | -0.617 | 1.15 | 0.635 | -0.034 | -0.031 | -0.031 | 0 | 0.56 | 0.81 | 0.62 | 0.52 | 0.57 | 0.54 |
| 5 | 151054227 | rs2304052 | -0.41 | -0.513 | 1.268 | 0.541 | -0.031 | 0 | 0 | 0.911 | 0.12 | 0.47 | 0.12 | 0.07 | 0.09 | 0.13 |
| 5 | 151056711 | rs725937 | -0.521 | -0.453 | 0.715 | 0.496 | -0.004 | 0.005 | 0.005 | 0.003 | 0.52 | 0.7 | 0.59 | 0.45 | 0.56 | 0.47 |
| 5 | 151057686 | rs2881558 | -1.168 | 0.006 | -0.587 | -0.085 | -0.068 | -0.007 | -0.007 | 0 | 0.62 | 0.94 | 0.64 | 0.66 | 0.62 | 0.58 |
| 5 | 151058893 | rs17718347 | -0.269 | 0.594 | -0.214 | -0.956 | -0.002 | -0.058 | -0.058 | 0.37 | 0.67 | 0.66 | 0.79 | 0.99 | 0.59 | 0.84 |
| 5 | 151059231 | rs11745387 | 0.413 | 0.218 | -0.717 | -0.883 | -0.039 | 0.16 | 0.16 | 0.873 | 0.29 | 0.06 | 0.16 | 0.01 | 0.33 | 0.13 |
| 5 | 151063674 | rs4958487 | -0.253 | -0.217 | -0.359 | -0.994 | 0.013 | 0.013 | 0.013 | 0 | 0.46 | 0.64 | 0.38 | 0.06 | 0.41 | 0.28 |
| 5 | 151067412 | rs1545030 | -0.452 | -0.21 | -0.381 | -0.741 | 0.012 | 0.034 | 0.034 | 0 | 0.52 | 0.63 | 0.42 | 0.17 | 0.46 | 0.36 |
| 5 | 151071719 | rs6579888 | -1.009 | -0.571 | -0.533 | -0.593 | 0.061 | 0.087 | 0.087 | 2.18 | 0.42 | 0.81 | 0.5 | 0.7 | 0.3 | 0.46 |
| 5 | 151078359 | rs7734196 | -1.087 | -0.512 | 0.519 | -0.513 | 0.044 | 0.082 | 0.082 | 1.028 | 0.07 | 0.47 | 0.1 | 0.02 | 0.01 | 0.02 |
| 5 | 151080003 | rs13153997 | -0.997 | -0.571 | -0.15 | -0.533 | 0.055 | 0.005 | 0.005 | 1.617 | 0.4 | 0.81 | 0.44 | 0.6 | 0.29 | 0.39 |
| 5 | 151088968 | rs12656024 | -0.377 | -0.207 | -0.078 | -0.407 | -0.006 | -0.031 | -0.031 | 0 | 0.31 | 0.23 | 0.34 | 0.56 | 0.29 | 0.34 |
| 5 | 151097281 | rs2011050 | -0.708 | -0.262 | 0.017 | -1.246 | 0.01 | 0.059 | 0.059 | 0 | 0.44 | 0.65 | 0.48 | 0.68 | 0.4 | 0.51 |
| 5 | 151110223 | rs4958493 | 0.659 | 0.116 | -0.02 | -1.547 | -0.001 | -0.014 | -0.014 | 0 | 0.26 | 0.29 | 0.25 | 0.03 | 0.29 | 0.13 |
| 5 | 151142878 | rs2964584 | 1.09 | -0.014 | 0.162 | -1.346 | -0.022 | -0.018 | -0.018 | 0 | 0.2 | 0.09 | 0.26 | 0.31 | 0.26 | 0.33 |
| 5 | 151150745 | rs2915876 | 1.138 | -0.012 | 0.11 | -1.345 | -0.069 | -0.059 | -0.059 | 0.037 | 0.19 | 0.02 | 0.24 | 0.31 | 0.26 | 0.33 |
| 5 | 151175748 | rs10477000 | 0.062 | -0.425 | 0.459 | -1.896 | -0.01 | -0.065 | -0.065 | 0 | 0.23 | 0.34 | 0.12 | 0.02 | 0.16 | 0.16 |
| 5 | 151184701 | rs1062177 | 1.503 | -0.365 | 0.465 | -1.878 | -0.071 | -0.058 | -0.058 | 0.135 | 0.19 | 0.02 | 0.25 | 0.31 | 0.27 | 0.3 |
| 5 | 151193637 | rs10044368 | -0.296 | -0.434 | 1.008 | -1.903 | -0.001 | 0.005 | 0.005 | 0.057 | 0.42 | 0.38 | 0.39 | 0.36 | 0.42 | 0.48 |
| 5 | 151198031 | rs7720579 | -0.296 | -0.47 | 1.008 | -1.903 | -0.002 | 0.005 | 0.005 | 0.057 | 0.42 | 0.38 | 0.39 | 0.35 | 0.42 | 0.48 |
| 5 | 151198772 | rs11958614 | 0.598 | -0.62 | 0.963 | -2.007 | -0.001 | 0.003 | 0.003 | 0 | 0.15 | 0.15 | 0.09 | 0.01 | 0.14 | 0.06 |
| 5 | 151201487 | rs2964611 | -0.433 | -0.646 | 1.034 | -1.966 | 0 | -0.011 | -0.011 | 0 | 0.42 | 0.43 | 0.38 | 0.34 | 0.42 | 0.48 |
| 5 | 151201689 | rs10066206 | 1.063 | -0.627 | 1.038 | -1.972 | 0.002 | -0.001 | -0.001 | 2.822 | 0.07 | 0.05 | 0.02 | 0.02 | 0.02 | 0.1 |
| 5 | 151221643 | rs7715874 | -0.312 | -0.662 | 1.09 | -1.932 | -0.007 | 0.041 | 0.041 | 0.113 | 0.42 | 0.32 | 0.35 | 0.33 | 0.42 | 0.46 |
| 5 | 151237970 | rs6879622 | -1.242 | -1.049 | 0.986 | -2.327 | -0.046 | 0.059 | 0.059 | 0.284 | 0.71 | 0.82 | 0.75 | 0.99 | 0.59 | 0.86 |
| 5 | 151259975 | rs2913894 | -0.014 | -0.782 | 0.584 | -2.046 | -0.022 | 0.062 | 0.062 | 0.151 | 0.59 | 0.74 | 0.65 | 0.68 | 0.58 | 0.54 |
| 5 | 151270122 | rs4467698 | 0.638 | -0.657 | 0.557 | -1.885 | 0.001 | 0.006 | 0.006 | 0.548 | 0.47 | 0.45 | 0.52 | 0.54 | 0.45 | 0.45 |
| 5 | 151270330 | rs890832 | -0.168 | -0.641 | 0.451 | -1.94 | -0.011 | 0.047 | 0.047 | 0 | 0.75 | 0.79 | 0.82 | 0.92 | 0.71 | 0.86 |
| 5 | 151273630 | rs2964565 | 0.574 | -0.609 | 0.293 | -1.955 | -0.101 | -0.047 | -0.047 | 1.013 | 0.31 | 0.04 | 0.33 | 0.36 | 0.4 | 0.37 |
| 5 | 151274117 | rs6877199 | 0.659 | -0.571 | 0.466 | -1.918 | 0.002 | 0.009 | 0.009 | 0.393 | 0.47 | 0.43 | 0.51 | 0.54 | 0.45 | 0.45 |
| 5 | 151279255 | rs1864001 | 1.166 | -0.31 | 0.266 | -1.83 | -0.025 | 0.066 | 0.066 | 0.099 | 0.12 | 0.01 | 0.1 | 0 | 0.14 | 0.07 |
| 5 | 151289698 | rs1465555 | -0.758 | 0.005 | 0.322 | -1.755 | -0.019 | -0.022 | -0.022 | 0 | 0.69 | 0.91 | 0.67 | 0.64 | 0.65 | 0.61 |
| 5 | 151290725 | rs7709656 | 0.053 | -0.087 | 0.433 | -1.749 | -0.004 | -0.037 | -0.037 | 0 | 0.19 | 0.31 | 0.12 | 0.1 | 0.15 | 0.17 |
| 5 | 151304306 | rs2071221 | 0.674 | -0.042 | 0.331 | -1.802 | 0.01 | 0.026 | 0.026 | 0.36 | 0.47 | 0.28 | 0.49 | 0.51 | 0.45 | 0.43 |
| 5 | 151304760 | rs2071220 | 0.037 | -0.043 | 0.28 | -1.832 | -0.014 | -0.001 | -0.001 | 0.66 | 0.57 | 0.76 | 0.57 | 0.49 | 0.43 | 0.52 |
| 5 | 151306610 | rs1428157 | 0.523 | -0.09 | 0.18 | -2.131 | 0.01 | -0.011 | -0.011 | 0 | 0.33 | 0.25 | 0.33 | 0.59 | 0.3 | 0.37 |
| 5 | 151309666 | rs11740566 | 0.986 | -0.148 | 0.237 | -2.186 | -0.004 | 0.044 | 0.044 | 0 | 0.32 | 0.2 | 0.26 | 0.02 | 0.31 | 0.18 |
| 5 | 151312478 | rs13183768 | 0.697 | -0.144 | 0.433 | -2.167 | -0.004 | -0.146 | -0.146 | 0.444 | 0.34 | 0.43 | 0.3 | 0.04 | 0.31 | 0.2 |
| 5 | 151317924 | rs919260 | -0.101 | -0.063 | -0.006 | -2.141 | -0.003 | 0.013 | 0.013 | 0.138 | 0.6 | 0.67 | 0.6 | 0.63 | 0.57 | 0.51 |
| 5 | 151321248 | rs302401 | -0.804 | -0.412 | 0.107 | -2.111 | 0.02 | 0.03 | 0.03 | 0 | 0.72 | 0.6 | 0.67 | 0.41 | 0.69 | 0.68 |
| 5 | 151322875 | rs12521103 | 0.857 | -0.324 | -0.251 | -2.073 | -0.002 | 0.017 | 0.017 | 0 | 0.33 | 0.27 | 0.26 | 0.02 | 0.31 | 0.18 |
| 5 | 151323905 | rs10463321 | 1.832 | -0.356 | -0.449 | -2.092 | -0.036 | -0.086 | -0.086 | 0 | 0.18 | 0.02 | 0.23 | 0.36 | 0.23 | 0.32 |
| 5 | 151326579 | rs186217 | 0.096 | -0.413 | 0.028 | -1.783 | 0.003 | 0.001 | 0.001 | 0 | 0.63 | 0.38 | 0.63 | 0.4 | 0.69 | 0.59 |
| 5 | 151328528 | rs182488 | 0.737 | -0.497 | -0.069 | -1.859 | 0.052 | 0.093 | 0.093 | 1.369 | 0.51 | 0.16 | 0.5 | 0.37 | 0.63 | 0.48 |
| 5 | 151330830 | rs13186058 | 1.757 | -0.283 | -0.118 | -1.827 | -0.02 | 0.011 | 0.011 | 0 | 0.2 | 0.1 | 0.16 | 0 | 0.26 | 0.09 |
| 5 | 151332812 | rs302412 | 0.571 | -0.363 | 0.243 | -1.796 | -0.002 | -0.003 | -0.003 | 0 | 0.53 | 0.42 | 0.52 | 0.37 | 0.63 | 0.48 |
| 5 | 151333011 | rs10067788 | -0.66 | -0.445 | 0.141 | -1.729 | -0.089 | -0.123 | -0.123 | 0 | 0.21 | 0.45 | 0.16 | 0.02 | 0.06 | 0.19 |
| 5 | 151333124 | rs7711643 | 0.244 | 0.229 | -0.276 | -1.389 | -0.028 | -0.044 | -0.044 | 0 | 0.13 | 0.27 | 0.11 | 0.01 | 0.05 | 0.1 |
| 5 | 151338950 | rs748463 | 0.393 | 1.05 | -0.443 | -0.95 | 0.02 | -0.101 | -0.101 | 0.51 | 0.22 | 0.25 | 0.17 | 0 | 0.26 | 0.09 |
| 5 | 151344558 | rs170020 | 0.006 | 0.92 | -1.217 | -0.519 | -0.008 | 0.074 | 0.074 | 0.901 | 0.54 | 0.77 | 0.5 | 0.36 | 0.54 | 0.56 |
| 5 | 151347364 | rs4958506 | -0.488 | 1.343 | -0.954 | -0.271 | 0.002 | 0.001 | 0.001 | 0 | 0.25 | 0.24 | 0.38 | 0.42 | 0.43 | 0.42 |
| 5 | 151349490 | rs255522 | -0.695 | 0.609 | -1.056 | -1.157 | 0.004 | -0.009 | -0.009 | 0.048 | 0.72 | 0.78 | 0.84 | 0.94 | 0.79 | 0.72 |
| 5 | 151351067 | rs2194139 | 0.751 | 1.567 | -1.42 | -0.818 | -0.055 | -0.076 | -0.076 | 0.12 | 0.13 | 0.02 | 0.24 | 0.44 | 0.23 | 0.29 |
| 5 | 151352448 | rs159922 | 0.166 | 0.833 | -1.091 | -1.129 | -0.005 | 0.157 | 0.157 | 0 | 0.5 | 0.27 | 0.63 | 0.87 | 0.53 | 0.56 |
| 5 | 151354752 | rs4958508 | 1.583 | 0.356 | -1.525 | -1.408 | 0.024 | -0.143 | -0.143 | 0.042 | 0.22 | 0.03 | 0.28 | 0.41 | 0.12 | 0.2 |
| 5 | 151357064 | rs426783 | 0.714 | 0.809 | -1.009 | -1.412 | 0.003 | -0.072 | -0.072 | 0 | 0.38 | 0.13 | 0.41 | 0.43 | 0.35 | 0.3 |
| 5 | 151377281 | rs159933 | 0.049 | 0.762 | -0.953 | -1.422 | 0.021 | 0.152 | 0.152 | 0 | 0.52 | 0.27 | 0.66 | 0.89 | 0.57 | 0.6 |
| 5 | 151377631 | rs2161387 | 0.888 | 0.7 | -0.901 | -1.531 | -0.02 | 0.054 | 0.054 | 0 | 0.16 | 0.08 | 0.14 | 0.01 | 0.22 | 0.14 |
| 5 | 151379310 | rs29664 | -1.179 | 0.699 | -0.257 | -1.481 | -0.014 | -0.097 | -0.097 | 0 | 0.71 | 0.63 | 0.84 | 0.94 | 0.79 | 0.74 |
| 5 | 151383115 | rs7735270 | 0.901 | 0.871 | -0.221 | -1.31 | 0.03 | 0.09 | 0.09 | 0.07 | 0.14 | 0.18 | 0.26 | 0.46 | 0.23 | 0.31 |
| 5 | 151383692 | rs12523233 | 0.837 | 1.094 | -0.226 | -1.385 | 0.042 | 0.08 | 0.08 | 0.124 | 0.14 | 0.18 | 0.28 | 0.45 | 0.26 | 0.32 |
| 5 | 151384425 | rs29667 | 0.025 | 0.758 | -0.409 | -1.632 | 0.005 | 0.006 | 0.006 | 0 | 0.55 | 0.4 | 0.69 | 0.93 | 0.59 | 0.6 |
| 5 | 151385578 | rs1445675 | 0.485 | 0.655 | -0.387 | -1.656 | 0.001 | -0.049 | -0.049 | 0 | 0.19 | 0.24 | 0.17 | 0.01 | 0.22 | 0.14 |
| 5 | 151390009 | rs11744717 | 1.07 | 0.659 | -0.682 | -1.492 | 0.03 | 0.098 | 0.098 | 0 | 0.14 | 0.18 | 0.26 | 0.49 | 0.23 | 0.3 |

| | | | | | | | | | | | | | | | |
|---|---|---|---|---|---|---|---|---|---|---|---|---|---|---|---|
| 5 | 151392480 | rs159926 | 0.7 | 0.633 | -0.707 | -1.51 | -0.021 | -0.128 | -0.128 | 0 | 0.4 | 0.07 | 0.41 | 0.44 | 0.36 | 0.29 |
| 5 | 151398349 | rs255493 | -0.191 | 0.63 | -0.68 | -1.514 | 0.007 | 0.03 | 0.03 | 0 | 0.55 | 0.38 | 0.68 | 0.93 | 0.59 | 0.58 |
| 5 | 151410888 | rs29658 | 0.487 | 0.374 | -0.98 | -1.338 | 0.042 | -0.048 | -0.048 | 1.703 | 0.41 | 0.22 | 0.41 | 0.44 | 0.35 | 0.28 |
| 5 | 151426516 | rs185321 | 0.47 | 0.355 | -1.472 | -1.307 | -0.021 | -0.128 | -0.128 | 0.095 | 0.4 | 0.07 | 0.41 | 0.43 | 0.36 | 0.29 |
| 5 | 151435875 | rs564489 | 0.611 | -0.312 | -1.421 | -1.372 | 0.094 | -0.092 | -0.092 | 0.081 | 0.34 | 0.15 | 0.35 | 0.45 | 0.18 | 0.25 |
| 5 | 151438217 | rs9324717 | 0.309 | -0.37 | -1.107 | -1.504 | -0.021 | -0.069 | -0.069 | 0.736 | 0.19 | 0.64 | 0.19 | 0.01 | 0.17 | 0.13 |
| 5 | 151441395 | rs154111 | 0.831 | -0.119 | -1.432 | -1.616 | -0.042 | -0.076 | -0.076 | 0 | 0.41 | 0.12 | 0.45 | 0.5 | 0.43 | 0.3 |
| 5 | 151444352 | rs169953 | -1.164 | 0.142 | -1.535 | -1.48 | 0 | 0.028 | 0.028 | 0 | 0.77 | 0.82 | 0.87 | 0.94 | 0.81 | 0.77 |
| 5 | 151449400 | rs29848 | 0.091 | 0.145 | -1.507 | -1.444 | -0.027 | -0.035 | -0.035 | 0.204 | 0.6 | 0.75 | 0.64 | 0.5 | 0.6 | 0.43 |
| 5 | 151453383 | rs10515655 | 0.599 | 0.47 | -1.246 | -1.09 | -0.006 | -0.003 | -0.003 | 0 | 0.16 | 0.14 | 0.24 | 0.42 | 0.2 | 0.34 |
| 5 | 151463491 | rs29849 | -0.589 | -0.036 | -1.297 | -1.522 | 0.015 | 0.01 | 0.01 | 0.523 | 0.51 | 0.63 | 0.55 | 0.43 | 0.63 | 0.58 |
| 5 | 151464176 | rs29851 | 0.122 | -0.028 | -1.449 | -1.613 | -0.177 | -0.149 | -0.149 | 0 | 0.29 | 0.03 | 0.36 | 0.41 | 0.46 | 0.44 |
| 5 | 151471642 | rs10477008 | 0.005 | -0.378 | -1.14 | -1.41 | -0.042 | -0.102 | -0.102 | 0 | 0.2 | 0.6 | 0.17 | 0.01 | 0.14 | 0.14 |
| 5 | 151472918 | rs891164 | 0.796 | -0.43 | -1.124 | -1.277 | 0.006 | -0.011 | -0.011 | 0 | 0.31 | 0.25 | 0.3 | 0.51 | 0.23 | 0.21 |
| 5 | 151482147 | rs10075826 | -0.369 | 0.001 | -0.835 | -0.827 | -0.017 | 0.049 | 0.049 | 0.511 | 0.23 | 0.67 | 0.29 | 0.43 | 0.19 | 0.27 |
| 5 | 151483769 | rs297449 | 0.349 | -0.131 | -1.342 | -1.007 | 0.01 | -0.031 | -0.031 | 0 | 0.44 | 0.18 | 0.39 | 0.41 | 0.41 | 0.44 |
| 5 | 151483806 | rs10515656 | 0.209 | -0.033 | -0.138 | -1.001 | 0.005 | 0.341 | 0.341 | 0 | 0.11 | 0.35 | 0.18 | 0.43 | 0.08 | 0.22 |
| 5 | 151495545 | rs154095 | -0.013 | -0.108 | -0.182 | -1.35 | -0.004 | 0.022 | 0.022 | 0 | 0.68 | 0.41 | 0.59 | 0.47 | 0.76 | 0.58 |
| 5 | 151500945 | rs40812 | -1.193 | -0.017 | -0.107 | -1.347 | 0.003 | -0.038 | -0.038 | 0 | 0.8 | 0.73 | 0.78 | 0.89 | 0.83 | 0.79 |
| 5 | 151503347 | rs10053232 | -0.246 | -0.288 | -0.591 | -1.595 | -0.012 | 0.074 | 0.074 | 0 | 0.27 | 0.39 | 0.37 | 0.44 | 0.21 | 0.36 |
| 5 | 151507977 | rs4958295 | 1.161 | -0.479 | -1.344 | -1.536 | -0.018 | -0.109 | -0.109 | 0.676 | 0.3 | 0.05 | 0.2 | 0.34 | 0.28 | 0.31 |
| 5 | 151513383 | rs40266 | 1.695 | -0.32 | -1.39 | -1.608 | -0.063 | 0.076 | 0.076 | 0 | 0.13 | 0.01 | 0.12 | 0 | 0.22 | 0.07 |
| 5 | 151514171 | rs29826 | 0.054 | -0.3 | -1.159 | -1.586 | -0.009 | -0.018 | -0.018 | 0 | 0.53 | 0.27 | 0.39 | 0.44 | 0.6 | 0.41 |
| 5 | 151522878 | rs11167571 | 1.271 | -0.291 | -1.326 | -1.436 | -0.03 | -0.132 | -0.132 | 0.605 | 0.27 | 0.03 | 0.2 | 0.37 | 0.27 | 0.28 |
| 5 | 151524210 | rs154690 | -0.862 | -0.289 | -1.175 | -1.616 | -0.001 | -0.016 | -0.016 | 0 | 0.68 | 0.63 | 0.69 | 0.82 | 0.71 | 0.72 |
| 5 | 151525147 | rs154689 | 0.712 | -0.281 | -1.081 | -1.654 | -0.008 | -0.002 | -0.002 | 0.261 | 0.42 | 0.26 | 0.32 | 0.37 | 0.47 | 0.35 |
| 5 | 151527585 | rs6874463 | 0.157 | -0.282 | -1.059 | -1.642 | 0.001 | 0.071 | 0.071 | 0 | 0.27 | 0.38 | 0.38 | 0.45 | 0.25 | 0.38 |
| 5 | 151527831 | rs6579920 | 0.259 | -0.398 | -0.525 | -1.187 | -0.004 | 0.013 | 0.013 | 0 | 0.54 | 0.52 | 0.58 | 0.82 | 0.51 | 0.66 |
| 5 | 151529317 | rs3857428 | -0.434 | -0.332 | -0.639 | -1.211 | -0.005 | -0.06 | -0.06 | 0 | 0.68 | 0.55 | 0.69 | 0.82 | 0.71 | 0.73 |
| 5 | 151532898 | rs3843498 | 1.172 | -0.215 | -0.642 | -0.998 | -0.018 | 0.02 | 0.02 | 0 | 0.14 | 0.11 | 0.11 | 0 | 0.2 | 0.06 |
| 5 | 151540970 | rs10223081 | 0.375 | -0.307 | -0.449 | -0.839 | 0.003 | 0.262 | 0.262 | 0 | 0.12 | 0.31 | 0.18 | 0.43 | 0.08 | 0.22 |
| 5 | 151551733 | rs294952 | -1.22 | -0.371 | -1.061 | -1.134 | -0.079 | -0.075 | -0.075 | 0 | 0.59 | 0.93 | 0.61 | 0.63 | 0.57 | 0.57 |
| 5 | 151556303 | rs787751 | 0.072 | -0.456 | -0.898 | -1.086 | -0.003 | 0.023 | 0.023 | 0 | 0.33 | 0.51 | 0.4 | 0.61 | 0.27 | 0.42 |
| 5 | 151558198 | rs294968 | -0.594 | -0.489 | -1.266 | -1.022 | -0.012 | 0.002 | 0.002 | 0.104 | 0.49 | 0.78 | 0.53 | 0.55 | 0.47 | 0.51 |
| 5 | 151561278 | rs10515652 | 1.307 | 0.08 | -0.941 | -0.713 | 0.003 | 0.019 | 0.019 | 0 | 0.09 | 0.05 | 0.07 | 0.08 | 0.1 | 0.04 |
| 5 | 151568099 | rs294958 | -1.06 | -0.074 | -1.259 | -0.709 | -0.028 | 0.013 | 0.013 | 0.12 | 0.52 | 0.92 | 0.58 | 0.61 | 0.54 | 0.5 |
| 5 | 151568481 | rs10072147 | 1.418 | 0.067 | -0.789 | -0.9 | 0.009 | -0.06 | -0.06 | 0 | 0.13 | 0.18 | 0.1 | 0 | 0.18 | 0.06 |
| 5 | 151575731 | rs1156008 | -1.259 | 0.104 | -0.819 | -0.778 | -0.03 | 0.106 | 0.106 | 0.12 | 0.52 | 0.92 | 0.58 | 0.73 | 0.54 | 0.5 |
| 5 | 151576474 | rs11951697 | 1.563 | 0.116 | -0.763 | -0.736 | -0.011 | 0.022 | 0.022 | 0 | 0.11 | 0.06 | 0.1 | 0.05 | 0.14 | 0.05 |
| 5 | 151578349 | rs190903 | -0.504 | 0.116 | -0.493 | -0.652 | 0.008 | 0.024 | 0.024 | 0 | 0.26 | 0.55 | 0.38 | 0.69 | 0.23 | 0.39 |
| 5 | 151582211 | rs1422929 | -0.667 | -0.031 | -0.614 | -0.884 | -0.021 | 0.019 | 0.019 | 1.503 | 0.63 | 0.66 | 0.48 | 0.69 | 0.53 | 0.59 |
| 5 | 151583410 | rs4639211 | 0.946 | -0.179 | -0.621 | -0.846 | 0.079 | -0.049 | -0.049 | 0.01 | 0.33 | 0.05 | 0.23 | 0.63 | 0.2 | 0.44 |
| 5 | 151586580 | rs12522523 | 1.587 | -0.058 | -0.485 | -0.834 | 0.01 | -0.062 | -0.062 | 0 | 0.13 | 0.2 | 0.1 | 0 | 0.18 | 0.06 |
| 5 | 151603058 | rs10065797 | 0.796 | -0.108 | -0.947 | -0.759 | 0.095 | -0.024 | -0.024 | 0 | 0.36 | 0.1 | 0.25 | 0.61 | 0.2 | 0.44 |
| 5 | 151607001 | rs17805030 | 1.938 | -0.054 | -1.086 | -0.662 | -0.015 | 0.013 | 0.013 | 0 | 0.07 | 0 | 0.12 | 0.4 | 0.06 | 0.2 |
| 5 | 151608073 | rs17112911 | 1.254 | 0.343 | -0.66 | -1.392 | 0.019 | -0.081 | -0.081 | 0 | 0.14 | 0.26 | 0.11 | 0 | 0.18 | 0.06 |
| 5 | 151612227 | rs4958296 | 0.845 | 0.11 | -0.491 | -1.116 | 0.022 | 0.005 | 0.005 | 0.043 | 0.36 | 0.25 | 0.27 | 0.63 | 0.2 | 0.44 |
| 5 | 151617817 | rs2112693 | -0.045 | 1.162 | -0.208 | -0.471 | 0.018 | 0.02 | 0.02 | 0 | 0.12 | 0.19 | 0.19 | 0.27 | 0.15 | 0.14 |
| 5 | 151623192 | rs11167574 | 0.153 | 1.164 | -0.216 | -0.506 | 0.008 | -0.061 | -0.061 | 0.026 | 0.25 | 0.32 | 0.21 | 0.05 | 0.32 | 0.11 |
| 5 | 151627889 | rs17805132 | -0.679 | 1.244 | -0.36 | 0.497 | 0.019 | 0.021 | 0.021 | 0 | 0.12 | 0.18 | 0.19 | 0.27 | 0.15 | 0.14 |
| 5 | 151640402 | rs1316849 | -0.019 | 0.686 | -0.996 | 0.208 | -0.01 | 0.027 | 0.027 | 0 | 0.05 | 0.01 | 0.03 | 0 | 0.05 | 0.03 |
| 5 | 151665028 | rs1997520 | 0.226 | 0.678 | -0.902 | 0.165 | -0.004 | 0.018 | 0.018 | 0 | 0.56 | 0.36 | 0.51 | 0.75 | 0.55 | 0.51 |
| 5 | 151673546 | rs13176984 | 0.414 | -0.167 | -1.33 | 0.605 | 0.018 | -0.102 | -0.102 | 0 | 0.22 | 0.01 | 0.12 | 0.32 | 0.16 | 0.24 |
| 5 | 151674065 | rs6579926 | -0.762 | -0.179 | -0.798 | 0.16 | 0 | -0.001 | -0.001 | 0 | 0.85 | 0.81 | 0.92 | 0.84 | 0.91 | 0.79 |
| 5 | 151682559 | rs1422358 | -0.662 | -0.299 | -0.203 | 0.144 | 0.002 | -0.017 | -0.017 | 0 | 0.44 | 0.51 | 0.48 | 0.26 | 0.45 | 0.49 |
| 5 | 151686722 | rs1422361 | -0.374 | 0.291 | -0.284 | -0.034 | 0.001 | 0.013 | 0.013 | 0 | 0.32 | 0.3 | 0.19 | 0.23 | 0.27 | 0.34 |
| 5 | 151715667 | rs1422350 | 0.438 | 0.289 | -0.3 | -0.016 | 0.007 | 0 | 0 | 0 | 0.17 | 0.18 | 0.08 | 0.16 | 0.09 | 0.21 |
| 5 | 151716974 | rs1363414 | 0.127 | 0.26 | -0.144 | -0.03 | -0.023 | -0.005 | -0.005 | 0 | 0.19 | 0.33 | 0.09 | 0.16 | 0.09 | 0.22 |
| 5 | 151719391 | rs6876461 | -0.552 | 0.27 | -0.083 | -0.07 | 0.02 | -0.03 | -0.03 | 0 | 0.3 | 0.54 | 0.38 | 0.19 | 0.28 | 0.36 |
| 5 | 151731277 | rs17113138 | -0.543 | 0.269 | -0.082 | -0.068 | 0.018 | -0.03 | -0.03 | 0 | 0.3 | 0.54 | 0.38 | 0.19 | 0.28 | 0.36 |
| 5 | 151734109 | rs1010254 | -0.567 | 0.243 | -0.105 | -0.081 | 0.014 | -0.027 | -0.027 | 0 | 0.31 | 0.56 | 0.38 | 0.19 | 0.28 | 0.37 |
| 5 | 151737056 | rs1002798 | 0.432 | 0.231 | -0.082 | -0.08 | -0.004 | -0.001 | -0.001 | 10.577 | 0.17 | 0.22 | 0.08 | 0.16 | 0.09 | 0.22 |
| 5 | 151739218 | rs2016865 | -1.25 | 0.108 | -0.297 | -0.178 | 0.003 | 0.024 | 0.024 | 0 | 0.48 | 0.62 | 0.49 | 0.27 | 0.46 | 0.5 |
| 5 | 151742571 | rs918462 | -0.661 | 0.107 | -0.211 | -0.18 | -0.01 | -0.029 | -0.029 | 0 | 0.36 | 0.5 | 0.2 | 0.25 | 0.27 | 0.36 |
| 5 | 151748808 | rs991637 | 1.268 | 0.31 | -0.55 | -0.208 | -0.026 | 0.013 | 0.013 | 0 | 0.11 | 0.05 | 0.29 | 0.03 | 0.19 | 0.14 |
| 5 | 151751433 | rs17447280 | 0.604 | 0.295 | -0.511 | -0.125 | -0.023 | 0.061 | 0.061 | 0.826 | 0.17 | 0.05 | 0.11 | 0.09 | 0.18 | 0.13 |
| 5 | 151756408 | rs6866565 | -0.819 | -0.034 | -0.514 | 0.098 | -0.001 | -0.01 | -0.01 | 0 | 0.53 | 0.52 | 0.49 | 0.26 | 0.48 | 0.53 |
| 5 | 151765489 | rs2348424 | -0.93 | -0.521 | -0.177 | 0.48 | 0.003 | 0.002 | 0.002 | 0 | 0.57 | 0.51 | 0.61 | 0.41 | 0.5 | 0.59 |
| 5 | 151768955 | rs10476783 | -0.93 | -0.47 | -0.177 | 0.48 | 0.003 | 0.002 | 0.002 | 0 | 0.57 | 0.51 | 0.61 | 0.41 | 0.49 | 0.58 |

| | | | | | | | | | | | | | | | | |
|---|---|---|---|---|---|---|---|---|---|---|---|---|---|---|---|---|
| 5 | 151771103 | rs982714 | 1.178 | -0.494 | -0.493 | 0.297 | -0.069 | 0.032 | 0.032 | 0 | 0.14 | 0.01 | 0.3 | 0.03 | 0.2 | 0.17 |
| 5 | 151771186 | rs982715 | 1.178 | -0.494 | -0.493 | 0.297 | -0.069 | 0.032 | 0.032 | 0 | 0.14 | 0.01 | 0.3 | 0.03 | 0.2 | 0.17 |
| 5 | 151771356 | rs982716 | 1.134 | -0.514 | -0.502 | 0.288 | -0.069 | 0.033 | 0.033 | 0 | 0.14 | 0.01 | 0.3 | 0.03 | 0.2 | 0.17 |
| 5 | 151771838 | rs4958531 | 1.181 | -0.503 | -0.497 | 0.293 | -0.069 | 0.032 | 0.032 | 0 | 0.14 | 0.01 | 0.3 | 0.03 | 0.2 | 0.17 |
| 5 | 151771852 | rs4958532 | 1.181 | -0.503 | -0.497 | 0.293 | -0.069 | 0.032 | 0.032 | 0 | 0.14 | 0.01 | 0.3 | 0.03 | 0.2 | 0.17 |
| 5 | 151772055 | rs1895245 | 1.181 | -0.503 | -0.497 | 0.293 | -0.069 | 0.032 | 0.032 | 0 | 0.14 | 0.01 | 0.3 | 0.03 | 0.2 | 0.17 |
| 5 | 151778214 | rs10434724 | 0.663 | -0.767 | -0.287 | 0.335 | 0 | -0.001 | -0.001 | 2.047 | 0.2 | 0.16 | 0.24 | 0.3 | 0.19 | 0.17 |
| 5 | 151791969 | rs9687964 | -0.441 | -0.755 | -0.235 | 0.299 | -0.001 | 0.063 | 0.063 | 0 | 0.37 | 0.23 | 0.37 | 0.12 | 0.31 | 0.42 |
| 5 | 151792075 | rs9688095 | -0.441 | -0.755 | -0.235 | 0.299 | -0.001 | 0.063 | 0.063 | 0 | 0.37 | 0.23 | 0.37 | 0.12 | 0.31 | 0.42 |
| 5 | 151801207 | rs1422337 | 0.585 | -0.619 | -0.378 | 0.371 | -0.002 | 0.008 | 0.008 | 0.462 | 0.21 | 0.21 | 0.24 | 0.3 | 0.19 | 0.17 |
| 5 | 151803033 | rs13188352 | 0.66 | -0.631 | -0.547 | 0.363 | 0.017 | 0.015 | 0.015 | 0 | 0.19 | 0.11 | 0.07 | 0.09 | 0.11 | 0.25 |
| 5 | 151810876 | rs4958541 | 1.18 | -0.357 | -0.441 | 0.409 | -0.074 | 0.027 | 0.027 | 0 | 0.14 | 0.01 | 0.3 | 0.03 | 0.2 | 0.17 |
| 5 | 151813000 | rs2080930 | 1.039 | -0.449 | -0.456 | 0.374 | -0.037 | 0.034 | 0.034 | 0 | 0.14 | 0.06 | 0.3 | 0.03 | 0.2 | 0.17 |
| 5 | 151834529 | rs1019475 | -0.393 | -0.501 | -0.452 | 0.411 | -0.027 | 0.005 | 0.005 | 0.019 | 0.34 | 0.18 | 0.53 | 0.32 | 0.4 | 0.33 |
| 5 | 151846236 | rs1422363 | -0.333 | -0.683 | -0.535 | 0.326 | -0.026 | 0.01 | 0.01 | 0.014 | 0.35 | 0.18 | 0.54 | 0.33 | 0.4 | 0.33 |
| 5 | 151847116 | rs11956510 | 0.235 | -0.783 | -0.555 | 0.663 | -0.03 | 0.059 | 0.059 | 0 | 0.06 | 0.12 | 0.14 | 0.22 | 0.01 | 0.06 |
| 5 | 151853233 | rs1363407 | 1.124 | -0.68 | -0.604 | 0.513 | -0.027 | 0.025 | 0.025 | 0 | 0.14 | 0.06 | 0.3 | 0.03 | 0.2 | 0.19 |
| 5 | 151854624 | rs17113267 | 0.114 | -0.784 | -0.769 | 0.433 | 0.001 | -0.015 | -0.015 | 0 | 0.2 | 0.12 | 0.23 | 0.28 | 0.19 | 0.16 |
| 5 | 151855232 | rs6897802 | -1.284 | -1.192 | -0.68 | 0.653 | 0 | -0.006 | -0.006 | 0 | 0.55 | 0.41 | 0.61 | 0.47 | 0.5 | 0.59 |
| 5 | 151855467 | rs17113268 | 0.616 | -0.96 | -0.807 | 0.268 | 0.023 | -0.014 | -0.014 | 0 | 0.19 | 0.08 | 0.07 | 0.16 | 0.11 | 0.25 |
| 5 | 151855626 | rs11750622 | 0.893 | -0.607 | -1.121 | 0.019 | -0.024 | -0.042 | -0.042 | 0.774 | 0.13 | 0.03 | 0.12 | 0.18 | 0.16 | 0.12 |
| 5 | 151866663 | rs10039253 | 1.205 | -0.614 | -0.893 | -0.391 | -0.003 | 0 | 0 | 0 | 0.14 | 0.11 | 0.3 | 0.03 | 0.2 | 0.19 |
| 5 | 151875652 | rs7731937 | 0.38 | -1.227 | -0.901 | -0.715 | 0.001 | 0.018 | 0.018 | 0.55 | 0.51 | 0.56 | 0.49 | 0.76 | 0.51 | 0.49 |
| 5 | 151876462 | rs7717092 | -0.436 | -1.063 | -0.836 | -0.948 | -0.008 | -0.057 | -0.057 | 0.013 | 0.83 | 0.72 | 0.9 | 0.95 | 0.85 | 0.92 |
| 5 | 151879912 | rs17451303 | 1.768 | -0.946 | -0.967 | -0.961 | -0.076 | 0.055 | 0.055 | 0 | 0.17 | 0.01 | 0.34 | 0.04 | 0.22 | 0.23 |
| 5 | 151880020 | rs7705865 | 0.751 | -0.618 | -1.053 | -1.06 | -0.001 | -0.004 | -0.004 | 0 | 0.43 | 0.55 | 0.43 | 0.76 | 0.43 | 0.45 |
| 5 | 151885658 | rs2961738 | -0.366 | -0.382 | -1.59 | -0.992 | -0.055 | 0.137 | 0.137 | 2.158 | 0.68 | 0.91 | 0.57 | 0.9 | 0.63 | 0.69 |
| 5 | 151886147 | rs2909950 | -0.094 | -0.539 | -1.303 | -1.027 | -0.013 | 0.052 | 0.052 | 0 | 0.42 | 0.24 | 0.47 | 0.18 | 0.42 | 0.45 |
| 5 | 151886485 | rs2112536 | 0.738 | -0.34 | -0.927 | -0.652 | 0.002 | 0.009 | 0.009 | 0 | 0.59 | 0.66 | 0.78 | 0.8 | 0.64 | 0.67 |
| 5 | 151887435 | rs2909951 | -0.87 | -0.279 | -0.485 | -0.321 | -0.01 | -0.046 | -0.046 | 2.368 | 0.92 | 0.82 | 0.96 | 1 | 0.95 | 0.96 |
| 5 | 151888663 | rs7725174 | -0.003 | -0.211 | -0.905 | -0.274 | -0.002 | 0.003 | 0.003 | 0 | 0.18 | 0.1 | 0.1 | 0.1 | 0.16 | 0.21 |
| 5 | 151889058 | rs11748577 | -0.021 | -0.262 | -0.914 | -0.302 | 0 | 0.006 | 0.006 | 0 | 0.18 | 0.09 | 0.09 | 0.09 | 0.15 | 0.21 |
| 5 | 151893373 | rs765674 | -0.154 | -0.24 | -0.707 | -0.594 | -0.003 | -0.027 | -0.027 | 0 | 0.85 | 0.82 | 0.93 | 0.95 | 0.86 | 0.92 |
| 5 | 151894477 | rs4585434 | -0.746 | -0.213 | -0.665 | -0.43 | 0.038 | 0.073 | 0.073 | 0 | 0.76 | 0.5 | 0.77 | 0.69 | 0.72 | 0.7 |
| 5 | 151900203 | rs12654017 | 1.477 | -0.487 | -1.12 | -0.549 | -0.006 | 0.053 | 0.053 | 0 | 0.15 | 0.18 | 0.23 | 0.53 | 0.13 | 0.12 |
| 5 | 151902735 | rs13436634 | 0.255 | -0.447 | -1.256 | -0.538 | -0.006 | -0.001 | -0.001 | 0 | 0.55 | 0.4 | 0.69 | 0.62 | 0.55 | 0.44 |
| 5 | 151902834 | rs6875911 | 1.251 | -0.364 | -0.46 | -0.426 | 0.013 | 0.01 | 0.01 | 0 | 0.19 | 0.26 | 0.2 | 0.2 | 0.27 | 0.25 |
| 5 | 151903277 | rs10463245 | -0.103 | -0.36 | -0.756 | -0.408 | -0.012 | 0.029 | 0.029 | 0.256 | 0.3 | 0.37 | 0.3 | 0.58 | 0.27 | 0.2 |
| 5 | 151903680 | rs13190288 | -0.103 | -0.36 | -0.764 | -0.408 | -0.011 | 0.026 | 0.026 | 0.256 | 0.3 | 0.36 | 0.3 | 0.58 | 0.27 | 0.2 |
| 5 | 151907086 | rs4958556 | 1.196 | -0.263 | -0.975 | -1.049 | 0 | 0.008 | 0.008 | 0 | 0.07 | 0.08 | 0.03 | 0.05 | 0.07 | 0.03 |
| 5 | 151910392 | rs10053527 | 0.304 | -0.386 | -1.074 | -1.215 | 0.006 | 0.045 | 0.045 | 0.004 | 0.22 | 0.1 | 0.09 | 0.07 | 0.17 | 0.26 |
| 5 | 151912745 | rs4569857 | 0.88 | -0.228 | -0.961 | -1.17 | -0.027 | 0.123 | 0.123 | 0 | 0.32 | 0.15 | 0.4 | 0.09 | 0.34 | 0.28 |
| 5 | 151914803 | rs4958559 | 1.049 | -0.177 | -1.133 | -1.145 | -0.072 | 0.152 | 0.152 | 0 | 0.31 | 0.09 | 0.39 | 0.09 | 0.34 | 0.28 |
| 5 | 151921873 | rs2964243 | 0.332 | -0.119 | -0.492 | -0.863 | -0.001 | 0.001 | 0.001 | 0 | 0.46 | 0.44 | 0.47 | 0.21 | 0.44 | 0.48 |
| 5 | 151929836 | rs4958560 | 1.553 | 0.035 | -0.631 | -1.012 | -0.018 | 0.087 | 0.087 | 0 | 0.27 | 0.12 | 0.35 | 0.03 | 0.28 | 0.27 |
| 5 | 151931892 | rs2961766 | 1.402 | -0.011 | -0.448 | -1.034 | -0.006 | 0.049 | 0.049 | 0 | 0.28 | 0.18 | 0.36 | 0.03 | 0.28 | 0.27 |
| 5 | 151939650 | rs2961759 | -0.287 | -0.342 | -0.033 | -1.184 | -0.002 | 0.005 | 0.005 | 0 | 0.54 | 0.57 | 0.49 | 0.3 | 0.48 | 0.54 |
| 5 | 151939905 | rs2961760 | 1.161 | -0.304 | 0.058 | -1.121 | 0.003 | 0.014 | 0.014 | 0 | 0.31 | 0.28 | 0.37 | 0.03 | 0.28 | 0.27 |
| 5 | 151950825 | rs6868331 | 1.125 | -0.258 | 0.013 | -1.057 | -0.014 | 0.023 | 0.023 | 0 | 0.31 | 0.24 | 0.37 | 0.2 | 0.33 | 0.4 |
| 5 | 151964513 | rs10056525 | 0.613 | -0.132 | -0.219 | -1.069 | -0.027 | -0.081 | -0.081 | 0 | 0.24 | 0.1 | 0.32 | 0.54 | 0.25 | 0.23 |
| 5 | 151965983 | rs17113468 | 0.204 | 0.005 | -0.158 | -1.263 | 0.056 | 0.095 | 0.095 | 0 | 0.05 | 0.17 | 0.04 | 0.18 | 0.07 | 0.14 |
| 5 | 151981337 | rs4958563 | 1.358 | -0.028 | 0.26 | -1.324 | 0.004 | -0.012 | -0.012 | 2.418 | 0.33 | 0.36 | 0.4 | 0.22 | 0.34 | 0.4 |
| 5 | 151981696 | rs12188451 | 1.341 | -0.068 | 0.705 | -1.377 | 0 | -0.04 | -0.04 | 0 | 0.2 | 0.19 | 0.13 | 0.01 | 0.25 | 0.18 |
| 5 | 151985487 | rs7735160 | -0.483 | -0.027 | 0.565 | -1.237 | 0.105 | 0.151 | 0.151 | 0.397 | 0.07 | 0.31 | 0.06 | 0.18 | 0.08 | 0.14 |
| 5 | 152010642 | rs10515674 | 0.879 | -0.191 | 0.04 | -1.276 | 0.011 | 0.023 | 0.023 | 0 | 0.15 | 0.01 | 0.06 | 0.11 | 0.12 | 0.11 |
| 5 | 152011505 | rs7733725 | -0.017 | -0.144 | 0.179 | -1.327 | -0.019 | -0.007 | -0.007 | 0.008 | 0.29 | 0.13 | 0.38 | 0.67 | 0.3 | 0.31 |
| 5 | 152020604 | rs4958321 | 1.32 | -0.198 | 0.666 | -1.383 | 0.009 | -0.035 | -0.035 | 1.027 | 0.33 | 0.43 | 0.41 | 0.22 | 0.34 | 0.4 |
| 5 | 152024757 | rs6579952 | 0.076 | -0.234 | 0.661 | -1.422 | -0.012 | -0.066 | -0.066 | 0 | 0.6 | 0.54 | 0.79 | 0.88 | 0.63 | 0.71 |
| 5 | 152028550 | rs10477025 | 0.308 | -0.23 | 0.616 | -1.422 | -0.038 | -0.025 | -0.025 | 0 | 0.27 | 0.1 | 0.37 | 0.66 | 0.29 | 0.31 |
| 5 | 152030714 | rs4958571 | 1.47 | -0.145 | 0.661 | -1.343 | 0.01 | -0.027 | -0.027 | 0.054 | 0.32 | 0.41 | 0.41 | 0.22 | 0.33 | 0.4 |
| 5 | 152054181 | rs17565170 | 1.244 | -0.087 | 0.545 | -1.403 | -0.006 | 0.019 | 0.019 | 0 | 0.21 | 0.14 | 0.12 | 0.01 | 0.25 | 0.18 |
| 5 | 152054744 | rs12517092 | 1.249 | -0.52 | 0.644 | -1.338 | 0.005 | -0.023 | -0.023 | 0 | 0.33 | 0.37 | 0.37 | 0.21 | 0.34 | 0.44 |
| 5 | 152057492 | rs2974554 | -0.169 | -0.631 | 0.709 | -1.113 | -0.03 | -0.036 | -0.036 | 0 | 0.8 | 0.68 | 0.91 | 0.89 | 0.88 | 0.89 |
| 5 | 152059348 | rs2964254 | 0.031 | -0.131 | 0.758 | -0.946 | 0.001 | -0.009 | -0.009 | 0 | 0.25 | 0.16 | 0.34 | 0.67 | 0.22 | 0.3 |
| 5 | 152067929 | rs2964252 | -0.492 | -0.172 | 0.799 | -1.163 | 0.001 | -0.014 | -0.014 | 4.035 | 0.29 | 0.29 | 0.38 | 0.85 | 0.29 | 0.46 |
| 5 | 152094555 | rs11950313 | 0.082 | -0.624 | 0.461 | -1.001 | -0.001 | -0.005 | -0.005 | 0 | 0.75 | 0.74 | 0.66 | 0.15 | 0.76 | 0.56 |
| 5 | 152117400 | rs11746937 | -0.032 | -0.701 | 0.033 | -0.889 | -0.005 | 0.002 | 0.002 | 0 | 0.75 | 0.85 | 0.65 | 0.3 | 0.74 | 0.72 |
| 5 | 152130453 | rs6879032 | -0.015 | -0.659 | 0.035 | -0.889 | -0.007 | 0 | 0 | 0.057 | 0.76 | 0.85 | 0.65 | 0.3 | 0.74 | 0.73 |
| 5 | 152137936 | rs7702336 | 0.081 | -0.665 | 0.137 | -0.89 | -0.001 | -0.001 | -0.001 | 1.999 | 0.75 | 0.76 | 0.64 | 0.3 | 0.74 | 0.73 |

| | | | | | | | | | | | | | | | | |
|---|---|---|---|---|---|---|---|---|---|---|---|---|---|---|---|---|
| 5 | 152159184 | rs4610449 | -0.387 | -0.25 | 0.478 | -0.212 | -0.008 | 0.043 | 0.043 | 0 | 0.93 | 0.91 | 0.87 | 0.85 | 0.96 | 0.92 |
| 5 | 152200109 | rs7445606 | -0.308 | -0.248 | 0.437 | -0.546 | 0.011 | -0.006 | -0.006 | 0.188 | 0.35 | 0.46 | 0.33 | 0.35 | 0.4 | 0.42 |
| 5 | 152206046 | rs11958854 | -0.536 | -0.374 | -0.036 | -0.18 | 0.005 | -0.024 | -0.024 | 0.002 | 0.69 | 0.78 | 0.81 | 0.46 | 0.72 | 0.73 |
| 5 | 152219802 | rs4623205 | 0.492 | -0.425 | -0.269 | -0.124 | 0.005 | -0.035 | -0.035 | 0 | 0.3 | 0.19 | 0.19 | 0.54 | 0.28 | 0.27 |
| 5 | 152236125 | rs17113719 | -0.106 | -0.323 | -0.297 | 0.115 | 0.002 | 0.015 | 0.015 | 1.448 | 0.1 | 0.08 | 0.19 | 0.37 | 0.05 | 0.25 |
| 5 | 152244389 | rs2069118 | -1.048 | -0.56 | 0.03 | -0.102 | -0.024 | 0.025 | 0.025 | 1.298 | 0.17 | 0.24 | 0.21 | 0.36 | 0.09 | 0.26 |
| 5 | 152249065 | rs4246037 | 0.269 | -0.604 | 0.194 | -0.523 | 0.048 | 0.027 | 0.027 | 0.132 | 0.53 | 0.29 | 0.44 | 0.54 | 0.63 | 0.43 |
| 5 | 152251201 | rs4958587 | 1.559 | -0.659 | 0.077 | -0.571 | -0.075 | 0.076 | 0.076 | 0 | 0.26 | 0.02 | 0.3 | 0.11 | 0.28 | 0.31 |
| 5 | 152261044 | rs4524565 | 1.103 | -0.603 | -0.015 | -0.744 | -0.053 | 0.053 | 0.053 | 1.201 | 0.19 | 0.12 | 0.21 | 0.02 | 0.34 | 0.15 |
| 5 | 152265018 | rs4958591 | 1.162 | -0.811 | 0.764 | -0.062 | -0.039 | 0.034 | 0.034 | 1.161 | 0.82 | 0.7 | 0.78 | 0.65 | 0.91 | 0.74 |
| 5 | 152279502 | rs9285647 | 0.728 | -0.862 | 0.533 | -0.038 | 0.048 | 0.024 | 0.024 | 0.151 | 0.53 | 0.28 | 0.45 | 0.56 | 0.63 | 0.43 |
| 5 | 152286477 | rs10056647 | -0.401 | -0.869 | 0.81 | -0.05 | -0.009 | -0.091 | -0.091 | 0 | 0.65 | 0.49 | 0.64 | 0.92 | 0.68 | 0.68 |
| 5 | 152288508 | rs4262151 | -0.148 | -0.851 | 0.604 | -0.041 | 0.006 | 0.003 | 0.003 | 0 | 0.63 | 0.35 | 0.63 | 0.92 | 0.68 | 0.69 |
| 5 | 152291650 | rs4478368 | 0.12 | -0.85 | 0.611 | 0.027 | 0.01 | 0.019 | 0.019 | 0 | 0.61 | 0.35 | 0.63 | 0.73 | 0.67 | 0.51 |
| 5 | 152302912 | rs3924314 | 0.743 | -0.848 | 0.577 | -0.092 | 0.048 | 0.025 | 0.025 | 0.12 | 0.52 | 0.29 | 0.44 | 0.55 | 0.63 | 0.43 |
| 5 | 152312444 | rs17489682 | 0.942 | -0.56 | 0.172 | 0.091 | -0.074 | 0.067 | 0.067 | 0 | 0.27 | 0.02 | 0.29 | 0.11 | 0.28 | 0.31 |
| 5 | 152318758 | rs17113811 | -0.336 | -0.4 | 0.477 | 0.345 | 0.013 | -0.009 | -0.009 | 0 | 0.1 | 0.06 | 0.18 | 0.33 | 0.04 | 0.26 |
| 5 | 152327668 | rs4958596 | -0.686 | -0.042 | 0.709 | 0.187 | 0.006 | -0.082 | -0.082 | 0.892 | 0.75 | 0.62 | 0.7 | 0.92 | 0.76 | 0.75 |
| 5 | 152340416 | rs11167606 | 0.226 | 0.147 | -0.075 | 0.635 | -0.003 | 0.024 | 0.024 | 5.331 | 0.07 | 0.05 | 0.19 | 0.41 | 0.06 | 0.25 |
| 5 | 152343943 | rs17113831 | 0.211 | 0.194 | -0.073 | 0.634 | -0.006 | 0.016 | 0.016 | 4.74 | 0.07 | 0.05 | 0.19 | 0.4 | 0.08 | 0.25 |
| 5 | 152347463 | rs6864972 | 0.91 | 0.148 | 0.098 | 0.382 | -0.001 | 0.004 | 0.004 | 0 | 0.21 | 0.16 | 0.26 | 0.06 | 0.2 | 0.21 |
| 5 | 152352546 | rs4484483 | -0.255 | 0.309 | -0.557 | 0.441 | -0.023 | -0.031 | -0.031 | 0 | 0.1 | 0.05 | 0.21 | 0.39 | 0.12 | 0.26 |
| 5 | 152354943 | rs1010100 | 0.697 | 0.24 | -0.521 | 0.295 | -0.002 | 0.003 | 0.003 | 0.329 | 0.61 | 0.44 | 0.5 | 0.55 | 0.64 | 0.49 |
| 5 | 152357770 | rs17568427 | 0.614 | 0.448 | -1.34 | -0.1 | -0.049 | 0.074 | 0.074 | 0 | 0.21 | 0.01 | 0.24 | 0.04 | 0.2 | 0.21 |
| 5 | 152367546 | rs1438935 | 0.784 | 0.508 | -1.203 | -0.037 | -0.009 | 0.046 | 0.046 | 0 | 0.15 | 0.07 | 0.19 | 0.03 | 0.16 | 0.15 |
| 5 | 152370448 | rs1438937 | -0.735 | 0.114 | -1.024 | 0.044 | -0.002 | 0 | 0 | 0 | 0.47 | 0.56 | 0.59 | 0.48 | 0.41 | 0.6 |
| 5 | 152373129 | rs1438940 | -0.832 | 0.367 | -1.717 | -0.45 | 0.011 | 0.073 | 0.073 | 0 | 0.75 | 0.63 | 0.8 | 0.53 | 0.72 | 0.78 |
| 5 | 152373633 | rs10515679 | 0.155 | 0.034 | -1.261 | -0.239 | 0.001 | -0.005 | -0.005 | 0 | 0.29 | 0.29 | 0.36 | 0.41 | 0.22 | 0.45 |
| 5 | 152375973 | rs17491087 | 2.205 | 0.038 | -1.308 | -0.239 | -0.009 | 0.046 | 0.046 | 0 | 0.15 | 0.07 | 0.19 | 0.03 | 0.16 | 0.15 |
| 5 | 152389065 | rs1866419 | 0.251 | -0.177 | -1.093 | -0.193 | -0.027 | -0.094 | -0.094 | 0 | 0.26 | 0.08 | 0.34 | 0.41 | 0.22 | 0.45 |
| 5 | 152402347 | rs1595586 | 0.098 | -0.17 | -1.063 | -0.129 | 0 | -0.038 | -0.038 | 0.416 | 0.27 | 0.19 | 0.35 | 0.43 | 0.22 | 0.46 |
| 5 | 152409030 | rs966088 | -0.194 | -0.19 | -1.005 | -0.11 | 0 | 0 | 0 | 0 | 0.29 | 0.32 | 0.37 | 0.43 | 0.22 | 0.46 |
| 5 | 152426881 | rs1583692 | 0.42 | 0.179 | -1.324 | 0.256 | -0.038 | -0.057 | -0.057 | 0 | 0.12 | 0.03 | 0.28 | 0.33 | 0.14 | 0.34 |
| 5 | 152460375 | rs12658477 | 0.595 | -0.105 | -1.586 | 0.398 | 0.021 | -0.043 | -0.043 | 0.285 | 0.07 | 0 | 0.02 | 0.21 | 0.02 | 0.06 |
| 5 | 152466290 | rs11744443 | -0.61 | -0.003 | -1.614 | 0.06 | 0.003 | -0.006 | -0.006 | 0.431 | 0.56 | 0.56 | 0.6 | 0.7 | 0.47 | 0.69 |
| 5 | 152474181 | rs17493227 | 0.777 | 0.16 | -1.603 | 0.277 | -0.053 | -0.091 | -0.091 | 0 | 0.19 | 0.01 | 0.15 | 0.25 | 0.24 | 0.1 |
| 5 | 152488891 | rs11746889 | -1.161 | 0.128 | -0.903 | 0.338 | 0.004 | -0.01 | -0.01 | 0 | 0.33 | 0.33 | 0.47 | 0.44 | 0.29 | 0.49 |
| 5 | 152495144 | rs987369 | 0.091 | 0.112 | -1.03 | 0.374 | -0.038 | -0.059 | -0.059 | 0.12 | 0.11 | 0.03 | 0.34 | 0.35 | 0.15 | 0.33 |
| 5 | 152498173 | rs6881504 | -0.853 | 0.08 | -0.877 | 0.372 | -0.002 | 0.005 | 0.005 | 0 | 0.33 | 0.41 | 0.47 | 0.46 | 0.3 | 0.49 |
| 5 | 152504929 | rs1438949 | -0.495 | 0.277 | -0.816 | 0.17 | -0.003 | -0.001 | -0.001 | 0 | 0.93 | 0.89 | 0.95 | 0.93 | 0.93 | 0.93 |
| 5 | 152509001 | rs1438946 | -0.025 | 0.23 | -1.028 | 0.248 | 0.007 | -0.025 | -0.025 | 0 | 0.57 | 0.74 | 0.72 | 0.43 | 0.56 | 0.63 |
| 5 | 152531583 | rs6580003 | -0.003 | 0.445 | -0.46 | -0.847 | 0.004 | -0.058 | -0.058 | 0 | 0.34 | 0.42 | 0.55 | 0.83 | 0.34 | 0.52 |
| 5 | 152535502 | rs10073138 | 0.739 | 0.196 | -0.4 | -1.27 | -0.006 | 0.104 | 0.104 | 0.275 | 0.38 | 0.21 | 0.23 | 0.1 | 0.4 | 0.29 |
| 5 | 152536787 | rs2910031 | 0.802 | 0.335 | -0.742 | -1.225 | -0.024 | -0.057 | -0.057 | 0 | 0.13 | 0.05 | 0.32 | 0.63 | 0.15 | 0.35 |
| 5 | 152540118 | rs2973145 | -0.893 | 0.357 | -0.735 | -1.222 | -0.003 | -0.071 | -0.071 | 0 | 0.72 | 0.63 | 0.79 | 0.94 | 0.74 | 0.81 |
| 5 | 152546146 | rs2973141 | 1.343 | 0.331 | -0.662 | -1.201 | -0.011 | 0.008 | 0.008 | 0 | 0.19 | 0.1 | 0.21 | 0.2 | 0.19 | 0.17 |
| 5 | 152548821 | rs2973139 | 0.525 | 0.255 | -0.686 | -1.024 | -0.009 | 0.003 | 0.003 | 0 | 0.77 | 0.57 | 0.6 | 0.3 | 0.79 | 0.6 |
| 5 | 152576916 | rs13156828 | 1.637 | 0.382 | -1.275 | -0.959 | -0.059 | 0.115 | 0.115 | 0 | 0.18 | 0.01 | 0.13 | 0 | 0.2 | 0.14 |
| 5 | 152589262 | rs2615160 | 0.327 | -0.167 | -0.689 | -0.826 | -0.019 | 0.001 | 0.001 | 0 | 0.1 | 0.31 | 0.07 | 0.1 | 0.07 | 0.06 |
| 5 | 152590591 | rs2615159 | 0.346 | -0.039 | -0.766 | -1.034 | -0.005 | 0.019 | 0.019 | 0 | 0.58 | 0.56 | 0.46 | 0.26 | 0.58 | 0.45 |
| 5 | 152592621 | rs296178 | 0.831 | -0.303 | -0.695 | -1.118 | -0.019 | -0.058 | -0.058 | 0.126 | 0.22 | 0.41 | 0.24 | 0.16 | 0.18 | 0.17 |
| 5 | 152633770 | rs2168966 | -0.973 | -0.615 | -1.445 | -1.07 | 0.067 | 0.018 | 0.018 | 0 | 0.44 | 0.91 | 0.39 | 0.4 | 0.33 | 0.34 |
| 5 | 152637000 | rs2926289 | -1.1 | -0.619 | -1.622 | -1.222 | -0.14 | -0.124 | -0.124 | 0.808 | 0.71 | 0.98 | 0.56 | 0.46 | 0.65 | 0.53 |
| 5 | 152644679 | rs1350900 | 1.075 | -0.732 | -1.23 | -0.972 | -0.004 | 0.003 | 0.003 | 0 | 0.07 | 0.07 | 0.07 | 0.02 | 0.1 | 0.06 |
| 5 | 152649498 | rs2926286 | -0.968 | -0.822 | -0.83 | -0.958 | -0.005 | 0.07 | 0.07 | 0 | 0.9 | 0.62 | 0.92 | 0.86 | 0.92 | 0.94 |
| 5 | 152651765 | rs2199123 | 0.057 | -0.896 | -0.653 | -0.993 | -0.036 | 0.059 | 0.059 | 0 | 0.86 | 0.31 | 0.87 | 0.7 | 0.92 | 0.91 |
| 5 | 152655834 | rs1824 | 0.949 | -0.872 | -0.683 | -1.053 | 0.038 | 0.07 | 0.07 | 0 | 0.22 | 0.09 | 0.18 | 0.08 | 0.14 | 0.19 |
| 5 | 152696107 | rs2615152 | 1.552 | -0.657 | -0.564 | -1.029 | 0.019 | 0.059 | 0.059 | 0 | 0.17 | 0.06 | 0.17 | 0 | 0.13 | 0.16 |
| 5 | 152725255 | rs160163 | -0.484 | -0.404 | -0.065 | -0.595 | 0.003 | 0.085 | 0.085 | 0 | 0.9 | 0.6 | 0.92 | 0.86 | 0.92 | 0.94 |
| 5 | 152725480 | rs159974 | 0.68 | -0.393 | -0.103 | -0.606 | 0.005 | 0.026 | 0.026 | 0 | 0.82 | 0.47 | 0.84 | 0.81 | 0.82 | 0.88 |
| 5 | 152729750 | rs13354091 | -0.685 | -0.082 | -0.026 | -0.462 | 0.038 | 0.088 | 0.088 | 0.065 | 0.12 | 0.41 | 0.11 | 0.19 | 0.1 | 0.09 |
| 5 | 152734724 | rs17114473 | -0.66 | -0.078 | -0.214 | -0.475 | 0.034 | 0.072 | 0.072 | 0.046 | 0.12 | 0.41 | 0.11 | 0.19 | 0.1 | 0.09 |
| 5 | 152749009 | rs7722746 | 0.918 | -0.107 | -0.928 | -0.781 | -0.07 | 0.098 | 0.098 | 0.504 | 0.28 | 0.09 | 0.19 | 0.13 | 0.39 | 0.21 |
| 5 | 152760340 | rs9324747 | -0.342 | -0.313 | -0.823 | -0.904 | -0.01 | -0.001 | -0.001 | 0 | 0.65 | 0.57 | 0.48 | 0.43 | 0.72 | 0.5 |
| 5 | 152770561 | rs304844 | 0.225 | -0.019 | -0.535 | -0.468 | 0.026 | 0.062 | 0.062 | 0 | 0.58 | 0.27 | 0.68 | 0.72 | 0.64 | 0.7 |
| 5 | 152772852 | rs160063 | 0.128 | -0.02 | -0.504 | -0.469 | 0.017 | 0.04 | 0.04 | 0 | 0.59 | 0.34 | 0.69 | 0.72 | 0.64 | 0.7 |
| 5 | 152776865 | rs922426 | 0.195 | 0.055 | -0.808 | 0.52 | -0.034 | -0.053 | -0.053 | 0 | 0.09 | 0 | 0.29 | 0.28 | 0.1 | 0.09 |
| 5 | 152777855 | rs6877008 | -0.434 | -0.088 | -0.342 | -0.511 | -0.031 | 0.029 | 0.029 | 0.064 | 0.1 | 0.31 | 0.08 | 0.12 | 0.04 | 0.06 |
| 5 | 152783598 | rs304842 | 0.165 | 0.218 | -1.36 | -0.487 | 0.026 | 0.035 | 0.035 | 0 | 0.61 | 0.24 | 0.69 | 0.69 | 0.67 | 0.71 |

| | | | | | | | | | | | | | | | | |
|---|---|---|---|---|---|---|---|---|---|---|---|---|---|---|---|---|
| 5 | 152785000 | rs11167633 | 0.414 | 0.535 | -1.541 | -0.918 | -0.036 | 0.097 | 0.097 | 0 | 0.31 | 0.13 | 0.21 | 0.14 | 0.41 | 0.23 |
| 5 | 152786520 | rs1423466 | 0.238 | 0.321 | -0.831 | -0.885 | -0.021 | -0.022 | -0.022 | 0 | 0.63 | 0.43 | 0.66 | 0.73 | 0.7 | 0.7 |
| 5 | 152787454 | rs1019738 | 1.283 | 0.968 | -1.242 | -1.246 | -0.009 | 0.061 | 0.061 | 0 | 0.24 | 0.09 | 0.15 | 0.12 | 0.25 | 0.24 |
| 5 | 152789621 | rs2927585 | -0.467 | 0.504 | -0.853 | -1.412 | -0.025 | -0.007 | -0.007 | 0 | 0.87 | 0.72 | 0.93 | 0.89 | 0.94 | 0.89 |
| 5 | 152789711 | rs4958644 | 0.819 | 0.383 | -0.995 | -1.537 | -0.015 | 0.032 | 0.032 | 0 | 0.1 | 0.25 | 0.16 | 0.16 | 0.07 | 0.06 |
| 5 | 152790163 | rs4958645 | 0.614 | 0.359 | -0.912 | -1.594 | -0.004 | -0.009 | -0.009 | 0.399 | 0.35 | 0.42 | 0.32 | 0.28 | 0.32 | 0.29 |
| 5 | 152791465 | rs1871164 | 0.53 | 0.551 | -1.413 | -1.644 | -0.045 | 0.009 | 0.009 | 0 | 0.39 | 0.16 | 0.51 | 0.61 | 0.48 | 0.46 |
| 5 | 152795131 | rs13159257 | 1.858 | 0.643 | -1.492 | -1.626 | -0.068 | 0.071 | 0.071 | 0 | 0.23 | 0.01 | 0.14 | 0.12 | 0.24 | 0.21 |
| 5 | 152797679 | rs13361457 | 0.955 | 1.372 | -1.395 | -1.656 | -0.014 | -0.018 | -0.018 | 0 | 0.19 | 0.12 | 0.39 | 0.59 | 0.26 | 0.31 |
| 5 | 152798385 | rs1478364 | 1.315 | 1.237 | -1.186 | -1.704 | 0.006 | 0.011 | 0.011 | 0.021 | 0.19 | 0.12 | 0.21 | 0 | 0.2 | 0.15 |
| 5 | 152798614 | rs1019507 | -0.11 | 1.411 | -1.099 | -1.528 | -0.044 | -0.009 | -0.009 | 0 | 0.34 | 0.21 | 0.51 | 0.6 | 0.45 | 0.45 |
| 5 | 152802553 | rs13169527 | 0.816 | 1.236 | -0.664 | -1.63 | -0.001 | -0.016 | -0.016 | 0 | 0.42 | 0.39 | 0.26 | 0.11 | 0.41 | 0.37 |
| 5 | 152806220 | rs13190209 | 1.508 | 1.248 | -0.655 | -1.657 | 0.011 | 0.065 | 0.065 | 0 | 0.31 | 0.21 | 0.17 | 0.08 | 0.28 | 0.2 |
| 5 | 152817363 | rs6883921 | -0.514 | 1.102 | -0.694 | -1.466 | 0.013 | 0.022 | 0.022 | 0.003 | 0.72 | 0.83 | 0.75 | 1 | 0.74 | 0.75 |
| 5 | 152839315 | rs530376 | 0.548 | 1.058 | -0.848 | -1.286 | 0.021 | 0.133 | 0.133 | 0 | 0.5 | 0.72 | 0.3 | 0.12 | 0.44 | 0.43 |
| 5 | 152857519 | rs472477 | 0.157 | 0.866 | -0.407 | -0.227 | -0.012 | -0.01 | -0.01 | 0 | 0.07 | 0.29 | 0.05 | 0.03 | 0.03 | 0.04 |
| 5 | 152861509 | rs6882575 | -0.224 | 1.378 | -1.293 | 0.108 | -0.019 | -0.034 | -0.034 | 0 | 0.2 | 0.11 | 0.41 | 0.6 | 0.24 | 0.31 |
| 5 | 152867348 | rs578772 | -0.035 | 0.915 | -0.908 | -1.003 | 0.001 | -0.058 | -0.058 | 3.482 | 0.78 | 0.77 | 0.84 | 1 | 0.74 | 0.84 |
| 5 | 152868437 | rs548294 | 0.248 | 0.41 | -0.976 | -0.912 | 0.004 | 0.014 | 0.014 | 0 | 0.58 | 0.64 | 0.65 | 0.73 | 0.6 | 0.71 |
| 5 | 152869624 | rs3792766 | 0.736 | 1.284 | -1.361 | -1.137 | -0.042 | 0.023 | 0.023 | 0 | 0.18 | 0.05 | 0.16 | 0.13 | 0.27 | 0.11 |
| 5 | 152874549 | rs4145160 | 0.449 | 0.476 | -1.008 | -0.782 | -0.002 | 0.052 | 0.052 | 0 | 0.09 | 0.11 | 0.36 | 0.6 | 0.08 | 0.22 |
| 5 | 152876121 | rs509555 | 0.626 | 0.02 | -0.682 | -1.389 | 0 | -0.006 | -0.006 | 0 | 0.53 | 0.45 | 0.4 | 0.24 | 0.59 | 0.35 |
| 5 | 152877008 | rs1428920 | 0.552 | 0.366 | -0.793 | -1.635 | 0.021 | 0.047 | 0.047 | 0 | 0.46 | 0.35 | 0.58 | 0.7 | 0.35 | 0.53 |
| 5 | 152878727 | rs540375 | -0.521 | 0.256 | -0.774 | -1.706 | -0.041 | 0.11 | 0.11 | 0 | 0.71 | 0.95 | 0.83 | 0.88 | 0.7 | 0.74 |
| 5 | 152879456 | rs17515563 | 1.642 | 0.392 | -0.778 | -1.66 | -0.108 | -0.017 | -0.017 | 0 | 0.18 | 0.01 | 0.19 | 0.14 | 0.34 | 0.08 |
| 5 | 152883381 | rs7707530 | 1.583 | 0.404 | -0.744 | -1.822 | -0.05 | 0.024 | 0.024 | 0.893 | 0.18 | 0.04 | 0.2 | 0.12 | 0.34 | 0.08 |
| 5 | 152887567 | rs17114681 | 0.533 | 0.404 | -0.89 | -1.786 | 0.019 | 0.059 | 0.059 | 0 | 0.46 | 0.34 | 0.58 | 0.74 | 0.35 | 0.54 |
| 5 | 152892304 | rs7727515 | 0.956 | 0.366 | -1.336 | -2.257 | 0 | -0.016 | -0.016 | 0 | 0.11 | 0.29 | 0.06 | 0.11 | 0.1 | 0.14 |
| 5 | 152892800 | rs515887 | -0.623 | 0.412 | -1.961 | -2.447 | -0.032 | 0.113 | 0.113 | 0 | 0.81 | 0.97 | 0.88 | 1 | 0.81 | 0.87 |
| 5 | 152895409 | rs512983 | 0.086 | 0.08 | -1.162 | -2.337 | 0.008 | -0.003 | -0.003 | 0 | 0.35 | 0.37 | 0.19 | 0.12 | 0.23 | 0.25 |
| 5 | 152901395 | rs17588123 | 0.093 | 0.791 | -1.413 | -2.108 | -0.06 | 0.091 | 0.091 | 0 | 0.25 | 0.13 | 0.22 | 0.05 | 0.41 | 0.14 |
| 5 | 152907442 | rs549100 | -0.539 | -0.247 | -0.944 | -2.345 | -0.006 | -0.052 | -0.052 | 0 | 0.39 | 0.49 | 0.39 | 0.17 | 0.34 | 0.43 |
| 5 | 152913504 | rs1463747 | 1.398 | -0.415 | -1.273 | -2.582 | -0.115 | 0.222 | 0.222 | 0 | 0.4 | 0.08 | 0.27 | 0.13 | 0.49 | 0.33 |
| 5 | 152913770 | rs1463746 | 0.122 | -0.336 | -1.249 | -2.571 | -0.006 | -0.052 | -0.052 | 0 | 0.61 | 0.51 | 0.6 | 0.83 | 0.66 | 0.57 |
| 5 | 152915962 | rs17114771 | 0.417 | -0.716 | -1.224 | -2.689 | 0.005 | 0.006 | 0.006 | 0 | 0.1 | 0.08 | 0.12 | 0.08 | 0.07 | 0.16 |
| 5 | 152916115 | rs10041179 | 0.412 | -0.744 | -0.696 | -2.737 | 0.022 | -0.02 | -0.02 | 0 | 0.06 | 0.18 | 0.14 | 0.01 | 0.07 | 0.07 |
| 5 | 152923433 | rs6893720 | 1.32 | -0.876 | -0.737 | -2.815 | 0.01 | 0.172 | 0.172 | 0 | 0.47 | 0.24 | 0.33 | 0.13 | 0.56 | 0.37 |
| 5 | 152927244 | rs4958344 | -0.502 | -1.074 | -0.696 | -2.931 | -0.006 | -0.035 | -0.035 | 0.174 | 0.35 | 0.59 | 0.51 | 0.71 | 0.31 | 0.4 |
| 5 | 152935392 | rs10515691 | 1.408 | -0.967 | -1.441 | -2.869 | 0.012 | 0.172 | 0.172 | 0 | 0.47 | 0.23 | 0.33 | 0.13 | 0.56 | 0.35 |
| 5 | 152940405 | rs10065813 | 1.144 | -0.752 | -2.114 | -2.755 | 0 | -0.031 | -0.031 | 1.424 | 0.15 | 0.13 | 0.1 | 0 | 0.15 | 0.13 |
| 5 | 152940728 | rs12656103 | 0.889 | -0.583 | -2.568 | -2.53 | -0.035 | -0.058 | -0.058 | 0 | 0.12 | 0.01 | 0.26 | 0.68 | 0.11 | 0.22 |
| 5 | 152942223 | rs12522802 | -0.325 | -0.618 | -2.487 | -2.512 | 0.006 | 0.038 | 0.038 | 0 | 0.18 | 0.06 | 0.25 | 0.16 | 0.16 | 0.25 |
| 5 | 152942869 | rs12520266 | 0.321 | -0.619 | -2.489 | -2.515 | 0.005 | 0.021 | 0.021 | 0 | 0.15 | 0.06 | 0.15 | 0.15 | 0.12 | 0.21 |
| 5 | 152956050 | rs12515622 | -0.216 | -0.608 | -2.481 | -2.493 | 0.013 | 0.046 | 0.046 | 0 | 0.2 | 0.06 | 0.26 | 0.16 | 0.17 | 0.31 |
| 5 | 152958550 | rs17519383 | 1.005 | -0.085 | -2.363 | -2.329 | -0.156 | 0.225 | 0.225 | 0 | 0.38 | 0.04 | 0.26 | 0.1 | 0.48 | 0.27 |
| 5 | 152968158 | rs778816 | 0.44 | -0.055 | -2.196 | -2.006 | -0.034 | -0.061 | -0.061 | 0.155 | 0.12 | 0.01 | 0.26 | 0.71 | 0.11 | 0.21 |
| 5 | 152969250 | rs778819 | 0.088 | -0.533 | -1.508 | -2.533 | -0.022 | -0.124 | -0.124 | 4.664 | 0.7 | 0.59 | 0.74 | 0.94 | 0.72 | 0.76 |
| 5 | 152972201 | rs778821 | 0.38 | -0.132 | -2.586 | -2.237 | 0.023 | 0.278 | 0.278 | 0.934 | 0.53 | 0.04 | 0.53 | 0.79 | 0.62 | 0.5 |
| 5 | 152991398 | rs1493383 | -0.532 | 0.21 | -2.113 | -2.098 | -0.014 | 0.064 | 0.064 | 0 | 0.2 | 0.06 | 0.12 | 0 | 0.2 | 0.15 |
| 5 | 152992110 | rs6866738 | -0.847 | 0.622 | -2.247 | -2.397 | -0.026 | 0.065 | 0.065 | 0 | 0.25 | 0.12 | 0.18 | 0 | 0.3 | 0.23 |
| 5 | 152993589 | rs2910268 | 0.611 | 1.493 | -2.287 | -2.653 | -0.001 | 0.002 | 0.002 | 0 | 0.08 | 0.07 | 0.07 | 0.03 | 0.14 | 0.08 |
| 5 | 153003764 | rs1559157 | 0.781 | 1.459 | -2.462 | -2.163 | -0.044 | -0.099 | -0.099 | 0 | 0.14 | 0.01 | 0.25 | 0.65 | 0.15 | 0.26 |
| 5 | 153004943 | rs11748500 | -0.177 | 1.449 | -2.467 | -2.199 | -0.026 | 0.065 | 0.065 | 0 | 0.25 | 0.13 | 0.18 | 0 | 0.3 | 0.23 |
| 5 | 153009489 | rs1422884 | -0.66 | 0.767 | -1.967 | -2.642 | -0.011 | -0.034 | -0.034 | 1.777 | 0.69 | 0.54 | 0.75 | 0.78 | 0.74 | 0.72 |
| 5 | 153014862 | rs2910266 | 1.566 | -0.074 | -1.782 | -2.743 | -0.031 | 0.05 | 0.05 | 0 | 0.18 | 0.06 | 0.25 | 0.02 | 0.27 | 0.1 |
| 5 | 153019167 | rs2910261 | -0.318 | -0.432 | -1.708 | -2.68 | 0 | -0.018 | -0.018 | 0 | 0.54 | 0.53 | 0.49 | 0.13 | 0.59 | 0.46 |
| 5 | 153019210 | rs2216649 | 1.155 | -0.594 | -1.696 | -2.815 | 0.037 | 0.032 | 0.032 | 0 | 0.2 | 0.11 | 0.08 | 0.11 | 0.12 | 0.23 |
| 5 | 153020030 | rs2910260 | 0.182 | 0.443 | -1.369 | -2.362 | -0.027 | 0.042 | 0.042 | 0 | 0.29 | 0.14 | 0.32 | 0.02 | 0.37 | 0.15 |
| 5 | 153020606 | rs1422894 | -0.325 | 0.444 | -1.304 | -2.362 | -0.006 | -0.037 | -0.037 | 0.017 | 0.34 | 0.28 | 0.39 | 0.02 | 0.47 | 0.23 |
| 5 | 153021549 | rs17114887 | 1.156 | 0.443 | -1.302 | -2.357 | 0.027 | -0.029 | -0.029 | 0.903 | 0.05 | 0.14 | 0.07 | 0 | 0.1 | 0.08 |
| 5 | 153023165 | rs1363673 | -0.293 | 0.456 | -1.358 | -2.346 | -0.006 | -0.037 | -0.037 | 0 | 0.34 | 0.28 | 0.39 | 0.02 | 0.47 | 0.23 |
| 5 | 153027222 | rs729329 | -0.919 | 0.508 | -1.568 | -2.276 | -0.06 | -0.045 | -0.045 | 1.076 | 0.77 | 0.92 | 0.67 | 0.32 | 0.71 | 0.67 |
| 5 | 153029960 | rs707176 | 0.424 | 1.007 | -0.894 | -2.269 | -0.019 | 0.013 | 0.013 | 0 | 0.21 | 0.1 | 0.28 | 0.02 | 0.32 | 0.11 |
| 5 | 153039315 | rs2963941 | 0.018 | 0.61 | -1.104 | -2.165 | 0.055 | -0.124 | -0.124 | 0 | 0.27 | 0.38 | 0.37 | 0.02 | 0.42 | 0.19 |
| 5 | 153044440 | rs17114975 | -0.627 | -0.22 | -1.338 | -1.104 | 0.013 | 0.013 | 0.013 | 2.202 | 0.11 | 0.08 | 0.06 | 0 | 0.06 | 0.08 |
| 5 | 153044744 | rs13173910 | -0.485 | -0.182 | -1.378 | -1.087 | 0.014 | 0.023 | 0.023 | 2.584 | 0.11 | 0.05 | 0.05 | 0 | 0.06 | 0.08 |
| 5 | 153054941 | rs10036436 | -0.395 | -0.771 | -1.281 | -1.517 | 0.014 | -0.008 | -0.008 | 0 | 0.23 | 0.16 | 0.12 | 0.01 | 0.13 | 0.2 |
| 5 | 153055075 | rs7714428 | -0.582 | -0.162 | -1.416 | -2.623 | -0.021 | 0.017 | 0.017 | 0 | 0.73 | 0.8 | 0.72 | 0.95 | 0.67 | 0.84 |
| 5 | 153055356 | rs13186237 | 0.779 | -0.562 | -1.241 | -1.418 | -0.003 | -0.013 | -0.013 | 0.68 | 0.23 | 0.18 | 0.4 | 0.42 | 0.3 | 0.2 |

| | | | | | | | | | | | | | | | | |
|---|---|---|---|---|---|---|---|---|---|---|---|---|---|---|---|---|
| 5 | 153056350 | rs6580028 | 1.25 | -0.538 | -1.377 | -1.604 | -0.001 | 0.018 | 0.018 | 0 | 0.16 | 0.14 | 0.34 | 0.42 | 0.19 | 0.18 |
| 5 | 153057548 | rs12189362 | 1.193 | -1.071 | -1.089 | -1.751 | 0 | 0.003 | 0.003 | 0.023 | 0.14 | 0.13 | 0.06 | 0.02 | 0.13 | 0.24 |
| 5 | 153058142 | rs11741511 | 0.733 | -1.119 | -1.007 | -1.779 | 0.031 | 0.053 | 0.053 | 0 | 0.31 | 0.17 | 0.18 | 0.01 | 0.24 | 0.23 |
| 5 | 153060357 | rs10070086 | 1.436 | -1.106 | -0.99 | -1.746 | 0.014 | -0.008 | -0.008 | 0 | 0.23 | 0.16 | 0.12 | 0.01 | 0.13 | 0.2 |
| 5 | 153061732 | rs10057486 | 0.939 | -1.447 | -1.072 | -1.946 | 0.02 | -0.036 | -0.036 | 0 | 0.07 | 0.15 | 0.06 | 0.01 | 0.06 | 0.11 |
| 5 | 153061830 | rs7719292 | 0.999 | -1.642 | -1.033 | -2.095 | -0.026 | 0.046 | 0.046 | 0 | 0.17 | 0.07 | 0.11 | 0.01 | 0.24 | 0.15 |
| 5 | 153062161 | rs10041275 | -0.123 | -1.897 | -0.549 | -1.959 | -0.043 | 0.012 | 0.012 | 0 | 0.39 | 0.49 | 0.26 | 0.52 | 0.23 | 0.43 |
| 5 | 153064231 | rs9324756 | 1.242 | -1.815 | -0.311 | -1.882 | -0.009 | 0.066 | 0.066 | 0 | 0.13 | 0.18 | 0.12 | 0.51 | 0.1 | 0.19 |
| 5 | 153070392 | rs13189947 | 1.721 | -1.639 | -0.267 | -1.835 | -0.003 | 0.005 | 0.005 | 0 | 0.08 | 0.04 | 0.06 | 0.01 | 0.11 | 0.04 |
| 5 | 153070876 | rs12717878 | 0.177 | -1.703 | -0.232 | -1.901 | 0.014 | -0.051 | -0.051 | 0.112 | 0.16 | 0.2 | 0.14 | 0.02 | 0.19 | 0.16 |
| 5 | 153072199 | rs13359392 | -0.617 | -1.724 | -0.152 | -1.943 | 0.023 | 0.006 | 0.006 | 0 | 0.46 | 0.61 | 0.33 | 0.52 | 0.34 | 0.42 |
| 5 | 153083431 | rs17523488 | 0.562 | -1.352 | -0.125 | -1.742 | 0.017 | -0.036 | -0.036 | 2.866 | 0.07 | 0.11 | 0.07 | 0 | 0.08 | 0.12 |
| 5 | 153087153 | rs12523051 | 0.469 | -1.504 | -0.079 | -1.867 | 0 | 0.001 | 0.001 | 0.271 | 0.26 | 0.28 | 0.39 | 0.41 | 0.31 | 0.36 |
| 5 | 153093323 | rs4336409 | -0.545 | -1.472 | -0.088 | -1.951 | 0.008 | -0.002 | -0.002 | 0 | 0.4 | 0.42 | 0.51 | 0.43 | 0.49 | 0.51 |
| 5 | 153093591 | rs13180912 | 1.449 | -1.347 | -0.134 | -2.232 | 0.002 | -0.016 | -0.016 | 0.292 | 0.08 | 0.09 | 0.06 | 0.01 | 0.11 | 0.04 |
| 5 | 153122863 | rs11741924 | -0.144 | -1.664 | 0.072 | -2.54 | -0.007 | -0.063 | -0.063 | 0 | 0.37 | 0.45 | 0.48 | 0.93 | 0.35 | 0.63 |
| 5 | 153123488 | rs6890057 | 0.849 | -2.108 | 0.052 | -2.866 | 0.013 | -0.011 | -0.011 | 0.019 | 0.25 | 0.18 | 0.14 | 0 | 0.18 | 0.1 |
| 5 | 153123729 | rs6890276 | -1.27 | -2.11 | 0.037 | -2.875 | 0.001 | -0.026 | -0.026 | 0 | 0.64 | 0.65 | 0.64 | 0.93 | 0.55 | 0.79 |
| 5 | 153128715 | rs4128572 | -1.318 | -2.117 | 0.157 | -2.878 | 0.002 | -0.028 | -0.028 | 0 | 0.64 | 0.66 | 0.64 | 0.93 | 0.55 | 0.79 |
| 5 | 153133722 | rs13168358 | 1.209 | -2.219 | 0.073 | -2.943 | 0.004 | 0.009 | 0.009 | 0 | 0.22 | 0.19 | 0.41 | 0.9 | 0.23 | 0.35 |
| 5 | 153142662 | rs6891142 | 2.296 | -2.331 | -0.298 | -2.947 | 0.043 | 0.082 | 0.082 | 0 | 0.14 | 0.02 | 0.04 | 0 | 0.07 | 0.07 |
| 5 | 153146891 | rs11749754 | 0.866 | -2.227 | -0.179 | -2.9 | 0.02 | 0.014 | 0.014 | 0 | 0.25 | 0.16 | 0.14 | 0 | 0.19 | 0.1 |
| 5 | 153148582 | rs6580040 | -2.12 | -2.445 | 0.001 | -3.015 | -0.043 | 0.055 | 0.055 | 0 | 0.73 | 0.87 | 0.72 | 0.94 | 0.67 | 0.82 |
| 5 | 153148731 | rs10155614 | 1.198 | -2.422 | -0.02 | -2.915 | 0.022 | 0 | 0 | 0.441 | 0.22 | 0.16 | 0.09 | 0 | 0.13 | 0.14 |
| 5 | 153158686 | rs1381119 | 0.663 | -2.388 | 0.078 | -2.87 | 0.01 | -0.024 | -0.024 | 0 | 0.27 | 0.24 | 0.13 | 0.01 | 0.19 | 0.19 |
| 5 | 153169740 | rs1461226 | -1.593 | -2.027 | 0.221 | -2.816 | -0.025 | 0.031 | 0.031 | 0 | 0.72 | 0.82 | 0.71 | 0.95 | 0.67 | 0.82 |
| 5 | 153175750 | rs764604 | 0.239 | -1.945 | 0.096 | -2.832 | 0.007 | 0.067 | 0.067 | 0 | 0.38 | 0.23 | 0.21 | 0.02 | 0.36 | 0.35 |
| 5 | 153224301 | rs34054798 | 2.119 | -1.834 | 0.125 | -2.743 | -0.003 | 0.001 | 0.001 | 0 | 0.07 | 0.06 | 0.06 | 0.01 | 0.12 | 0.03 |
| 5 | 153238093 | rs2926858 | -1.552 | -2.123 | 0.017 | -2.921 | -0.026 | 0.031 | 0.031 | 0 | 0.73 | 0.8 | 0.71 | 0.94 | 0.67 | 0.83 |
| 5 | 153238323 | rs1461237 | 0.81 | -2.164 | 0.245 | -2.478 | -0.03 | -0.013 | -0.013 | 0.61 | 0.23 | 0.32 | 0.4 | 0.84 | 0.18 | 0.34 |
| 5 | 153238466 | rs17115298 | 1.732 | -1.902 | -0.041 | -2.325 | 0.013 | -0.028 | -0.028 | 0.091 | 0.09 | 0.17 | 0.07 | 0.01 | 0.12 | 0.03 |
| 5 | 153243573 | rs17115312 | 1.46 | -1.993 | -0.584 | -2.45 | -0.024 | -0.001 | -0.001 | 0.074 | 0.08 | 0.13 | 0.03 | 0.07 | 0.02 | 0.08 |
| 5 | 153245475 | rs1461234 | 0.2 | -2.4 | -0.842 | -2.59 | 0.011 | 0.127 | 0.127 | 0 | 0.39 | 0.18 | 0.24 | 0.04 | 0.38 | 0.36 |
| 5 | 153250819 | rs2926830 | -0.762 | -2.052 | -1.085 | -2.107 | -0.039 | 0.02 | 0.02 | 0 | 0.75 | 0.86 | 0.73 | 0.85 | 0.68 | 0.78 |
| 5 | 153263479 | rs2964029 | 0.173 | -1.512 | -0.701 | -0.915 | -0.007 | 0.005 | 0.005 | 0 | 0.4 | 0.64 | 0.54 | 0.7 | 0.49 | 0.52 |
| 5 | 153266148 | rs10476798 | 0.941 | -0.486 | -0.761 | -1.054 | -0.006 | -0.038 | -0.038 | 0 | 0.09 | 0.14 | 0.09 | 0 | 0.1 | 0.02 |
| 5 | 153267753 | rs2614125 | 0.279 | -0.468 | -0.805 | -1.015 | -0.007 | 0.003 | 0.003 | 0 | 0.4 | 0.63 | 0.54 | 0.69 | 0.49 | 0.52 |
| 5 | 153268620 | rs2560036 | -1.216 | -0.435 | -0.938 | -1.164 | -0.042 | 0.018 | 0.018 | 0 | 0.77 | 0.92 | 0.76 | 0.84 | 0.75 | 0.77 |
| 5 | 153269843 | rs10061978 | 0.601 | -0.549 | -0.612 | -1.354 | 0.016 | 0.013 | 0.013 | 0 | 0.32 | 0.24 | 0.18 | 0.02 | 0.22 | 0.22 |
| 5 | 153273168 | rs6873055 | 0.806 | -0.369 | -0.314 | -1.364 | 0.002 | -0.074 | -0.074 | 0.421 | 0.14 | 0.36 | 0.13 | 0.01 | 0.15 | 0.04 |
| 5 | 153277759 | rs2434220 | -1.454 | -0.184 | -0.808 | -1.247 | -0.046 | 0.04 | 0.04 | 0 | 0.77 | 0.92 | 0.77 | 0.86 | 0.75 | 0.77 |
| 5 | 153281406 | rs2560030 | 0.731 | -0.201 | -0.588 | -1.233 | 0 | -0.003 | -0.003 | 4.283 | 0.31 | 0.31 | 0.44 | 0.83 | 0.37 | 0.46 |
| 5 | 153286964 | rs898709 | 0.863 | -0.178 | -0.564 | -0.943 | -0.004 | -0.011 | -0.011 | 0.985 | 0.28 | 0.23 | 0.4 | 0.61 | 0.34 | 0.42 |
| 5 | 153289425 | rs2614119 | -0.089 | -0.331 | -0.555 | -1.01 | 0 | -0.001 | -0.001 | 0 | 0.45 | 0.55 | 0.55 | 0.63 | 0.51 | 0.49 |
| 5 | 153305438 | rs1461241 | 1.039 | -0.14 | -0.513 | -1.336 | -0.015 | 0.021 | 0.021 | 0.051 | 0.34 | 0.25 | 0.47 | 0.81 | 0.41 | 0.44 |
| 5 | 153305796 | rs970751 | -0.245 | -0.159 | -0.48 | -1.385 | 0.017 | 0.099 | 0.099 | 0 | 0.53 | 0.7 | 0.66 | 0.91 | 0.62 | 0.58 |
| 5 | 153312122 | rs17115409 | 0.688 | -0.171 | -0.5 | -1.335 | -0.014 | 0.047 | 0.047 | 0.034 | 0.18 | 0.41 | 0.18 | 0.23 | 0.19 | 0.1 |
| 5 | 153316866 | rs508306 | 1.044 | -0.098 | -0.628 | -1.292 | -0.017 | -0.007 | -0.007 | 0 | 0.33 | 0.25 | 0.47 | 0.68 | 0.41 | 0.44 |
| 5 | 153321768 | rs286969 | -0.095 | -0.169 | -1.041 | -1.515 | -0.001 | -0.024 | -0.024 | 0.047 | 0.66 | 0.63 | 0.69 | 0.92 | 0.65 | 0.71 |
| 5 | 153339577 | rs13179753 | 1.515 | -0.513 | -1.087 | -1.121 | 0.042 | 0.07 | 0.07 | 0.005 | 0.2 | 0.07 | 0.08 | 0 | 0.11 | 0.1 |
| 5 | 153358226 | rs17115481 | 0.244 | -0.376 | -0.849 | -1.034 | -0.002 | -0.014 | -0.014 | 0.608 | 0.37 | 0.39 | 0.23 | 0.14 | 0.29 | 0.19 |
| 5 | 153371463 | rs707179 | -0.019 | 0.001 | -0.761 | -0.885 | 0.022 | 0.033 | 0.033 | 0.636 | 0.56 | 0.62 | 0.72 | 0.86 | 0.63 | 0.79 |
| 5 | 153372525 | rs1057772 | 0.043 | 0.018 | -0.559 | -0.662 | -0.002 | -0.028 | -0.028 | 0.117 | 0.54 | 0.5 | 0.69 | 0.86 | 0.62 | 0.73 |
| 5 | 153377113 | rs9324761 | -1.395 | -0.113 | -0.61 | -0.74 | 0.045 | 0.088 | 0.088 | 0 | 0.73 | 0.93 | 0.85 | 0.88 | 0.81 | 0.85 |
| 5 | 153407397 | rs7704562 | 0.761 | 0.06 | -0.563 | -0.882 | -0.012 | 0.004 | 0.004 | 0.1 | 0.38 | 0.28 | 0.51 | 0.67 | 0.48 | 0.52 |
| 5 | 153412529 | rs720985 | -0.898 | -0.03 | -0.63 | -0.791 | 0.011 | 0.029 | 0.029 | 0 | 0.65 | 0.74 | 0.78 | 0.86 | 0.69 | 0.82 |
| 5 | 153413390 | rs2578377 | -0.267 | 0.172 | -0.565 | -0.681 | 0.022 | 0.033 | 0.033 | 0.636 | 0.56 | 0.63 | 0.72 | 0.86 | 0.63 | 0.79 |
| 5 | 153417721 | rs920309 | -0.267 | 0.172 | -0.565 | -0.681 | 0.022 | 0.033 | 0.033 | 0.636 | 0.56 | 0.63 | 0.72 | 0.86 | 0.63 | 0.79 |
| 5 | 153429169 | rs478020 | 0.22 | 0.178 | -0.55 | -0.682 | -0.003 | -0.005 | -0.005 | 0 | 0.52 | 0.42 | 0.68 | 0.73 | 0.61 | 0.73 |
| 5 | 153430783 | rs1370912 | 1.735 | 0.524 | -0.841 | -0.285 | -0.051 | -0.085 | -0.085 | 0.044 | 0.12 | 0.01 | 0.23 | 0.41 | 0.14 | 0.17 |
| 5 | 153432970 | rs2255493 | -0.111 | 0.4 | -0.472 | -1.265 | 0.022 | 0.034 | 0.034 | 0.716 | 0.55 | 0.62 | 0.72 | 0.86 | 0.63 | 0.79 |
| 5 | 153434407 | rs1382278 | -0.034 | 0.438 | -0.716 | -1.237 | -0.002 | -0.027 | -0.027 | 0 | 0.53 | 0.5 | 0.69 | 0.86 | 0.62 | 0.74 |
| 5 | 153435497 | rs11958069 | 1.615 | 0.76 | -0.746 | -0.816 | 0.025 | -0.023 | -0.023 | 0 | 0.08 | 0.19 | 0.08 | 0.02 | 0.12 | 0.03 |
| 5 | 153452002 | rs816028 | 0.093 | 0.384 | -1.377 | -0.914 | -0.002 | -0.005 | -0.005 | 0 | 0.52 | 0.42 | 0.67 | 0.74 | 0.61 | 0.73 |
| 5 | 153468889 | rs13173354 | 1.185 | 0.197 | -1.485 | -1.05 | 0.032 | 0.086 | 0.086 | 0 | 0.27 | 0.07 | 0.15 | 0.13 | 0.2 | 0.16 |
| 5 | 153471737 | rs707183 | 0.103 | 0.63 | -1.39 | -0.947 | 0 | -0.002 | -0.002 | 0.139 | 0.48 | 0.57 | 0.35 | 0.27 | 0.44 | 0.28 |
| 5 | 153477465 | rs984078 | 0.693 | -0.017 | -1.152 | -0.504 | 0 | 0.011 | 0.011 | 0 | 0.19 | 0.15 | 0.26 | 0.57 | 0.19 | 0.21 |
| 5 | 153480536 | rs1093167 | -0.431 | -0.042 | -1.201 | -1.282 | -0.001 | -0.013 | -0.013 | 0 | 0.85 | 0.84 | 0.83 | 0.94 | 0.86 | 0.79 |
| 5 | 153486785 | rs815630 | -0.236 | 0.093 | -1.182 | -0.837 | 0.03 | 0.049 | 0.049 | 0 | 0.09 | 0.01 | 0.05 | 0 | 0.06 | 0.03 |

| | | | | | | | | | | | | | | | | |
|---|---|---|---|---|---|---|---|---|---|---|---|---|---|---|---|---|
| 5 | 153495869 | rs890798 | -0.066 | 0.459 | -0.887 | -1.328 | -0.001 | -0.071 | -0.071 | 0.332 | 0.37 | 0.55 | 0.58 | 0.91 | 0.34 | 0.38 |
| 5 | 153497048 | rs7721733 | -0.516 | 0.276 | -0.711 | -1.812 | 0.011 | -0.074 | -0.074 | 0 | 0.16 | 0.51 | 0.11 | 0.06 | 0.11 | 0.15 |
| 5 | 153497780 | rs1366220 | 0.439 | 0.302 | -0.959 | -2.089 | -0.055 | 0.129 | 0.129 | 0 | 0.29 | 0.06 | 0.55 | 0.9 | 0.33 | 0.36 |
| 5 | 153513403 | rs17625484 | 0.859 | 0.286 | -0.229 | -2.108 | 0.024 | -0.049 | -0.049 | 0 | 0.09 | 0.2 | 0.05 | 0 | 0.07 | 0.01 |
| 5 | 153518766 | rs815611 | 0.747 | 0.447 | -0.189 | -2.3 | -0.001 | -0.07 | -0.07 | 0 | 0.52 | 0.42 | 0.33 | 0.04 | 0.54 | 0.47 |
| 5 | 153526051 | rs17536714 | 1.362 | 0.278 | -0.427 | -3.279 | 0 | -0.04 | -0.04 | 0 | 0.26 | 0.22 | 0.15 | 0.02 | 0.3 | 0.26 |
| 5 | 153531842 | rs9324766 | 1.146 | 0.364 | -0.763 | -3.366 | 0.006 | 0.027 | 0.027 | 0 | 0.21 | 0.14 | 0.14 | 0.02 | 0.18 | 0.19 |
| 5 | 153535126 | rs7719786 | -0.789 | 0.237 | -0.83 | -3.402 | -0.072 | 0.18 | 0.18 | 0 | 0.66 | 0.92 | 0.41 | 0.04 | 0.6 | 0.53 |
| 5 | 153537026 | rs13177936 | -0.981 | 0.328 | -0.936 | -2.909 | -0.084 | 0.092 | 0.092 | 0.029 | 0.68 | 0.95 | 0.41 | 0.15 | 0.62 | 0.55 |
| 5 | 153538241 | rs7719067 | -0.308 | 0.357 | -0.768 | -3.236 | -0.012 | 0.045 | 0.045 | 0 | 0.57 | 0.68 | 0.36 | 0.04 | 0.56 | 0.52 |
| 5 | 153546023 | rs11747752 | 0.908 | 0.148 | -0.818 | -3.215 | -0.003 | 0.037 | 0.037 | 0 | 0.22 | 0.14 | 0.15 | 0.02 | 0.22 | 0.19 |
| 5 | 153547306 | rs11957885 | 2.169 | 0 | -1.013 | -3.445 | 0.014 | 0.009 | 0.009 | 0 | 0.11 | 0.07 | 0.04 | 0 | 0.06 | 0.11 |
| 5 | 153550364 | rs11747325 | 0.089 | -0.224 | -1.19 | -3.613 | -0.001 | -0.032 | -0.032 | 0 | 0.24 | 0.24 | 0.17 | 0.02 | 0.25 | 0.17 |
| 5 | 153551892 | rs12658319 | 1.622 | -0.189 | -1.18 | -3.413 | -0.023 | 0.086 | 0.086 | 0 | 0.28 | 0.12 | 0.53 | 0.87 | 0.32 | 0.36 |
| 5 | 153555164 | rs10052189 | -0.486 | -0.226 | -1.021 | -3.44 | 0.002 | -0.024 | -0.024 | 0 | 0.56 | 0.53 | 0.39 | 0.08 | 0.61 | 0.52 |
| 5 | 153557619 | rs2351227 | 0.647 | -0.048 | -1.291 | -3.481 | -0.016 | 0.082 | 0.082 | 0 | 0.15 | 0.01 | 0.11 | 0 | 0.15 | 0.14 |
| 5 | 153557907 | rs890794 | -1.178 | -0.016 | -1.198 | -3.454 | -0.008 | 0.083 | 0.083 | 0.032 | 0.65 | 0.76 | 0.43 | 0.08 | 0.63 | 0.53 |
| 5 | 153573579 | rs6868044 | 0.216 | 0.033 | -1.136 | -3.453 | -0.004 | -0.126 | -0.126 | 0 | 0.31 | 0.46 | 0.22 | 0.07 | 0.28 | 0.25 |
| 5 | 153573938 | rs890793 | 1.357 | -0.073 | -1.335 | -3.263 | -0.045 | 0.115 | 0.115 | 0 | 0.32 | 0.14 | 0.58 | 0.87 | 0.38 | 0.38 |
| 5 | 153577521 | rs6860445 | -0.718 | 0.031 | -1.251 | -3.11 | -0.072 | 0.244 | 0.244 | 0.941 | 0.58 | 0.87 | 0.37 | 0.08 | 0.53 | 0.43 |
| 5 | 153580181 | rs6876928 | 0.293 | 0.003 | -0.787 | -2.865 | -0.006 | -0.075 | -0.075 | 0.207 | 0.15 | 0.28 | 0.1 | 0.02 | 0.12 | 0.13 |
| 5 | 153584363 | rs4958711 | 0.506 | 0.147 | -1.144 | -2.388 | -0.024 | 0.104 | 0.104 | 0 | 0.4 | 0.26 | 0.62 | 0.8 | 0.45 | 0.43 |
| 5 | 153586443 | rs13162836 | 0.238 | 0.082 | -1.157 | -2.473 | 0.005 | 0.132 | 0.132 | 0 | 0.45 | 0.3 | 0.65 | 0.8 | 0.47 | 0.44 |
| 5 | 153591020 | rs2161393 | -1.08 | 0.244 | -1.021 | -2.502 | 0.007 | 0.001 | 0.001 | 0 | 0.94 | 0.93 | 0.96 | 0.93 | 0.94 | 0.85 |
| 5 | 153592852 | rs7716370 | 0.03 | 0.229 | -0.909 | -2.53 | 0.004 | 0.083 | 0.083 | 0 | 0.48 | 0.6 | 0.3 | 0.13 | 0.47 | 0.4 |
| 5 | 153593934 | rs12187228 | 0.419 | 0.272 | -0.915 | -2.257 | -0.01 | 0.086 | 0.086 | 0 | 0.39 | 0.26 | 0.62 | 0.8 | 0.42 | 0.42 |
| 5 | 153596545 | rs11744052 | -0.618 | 1.014 | -0.989 | -1.68 | 0 | 0.001 | 0.001 | 0 | 0.18 | 0.22 | 0.31 | 0.73 | 0.3 | 0.33 |
| 5 | 153597924 | rs4958715 | 0.445 | 0.158 | -0.688 | -1.509 | -0.01 | 0.044 | 0.044 | 0 | 0.36 | 0.25 | 0.63 | 0.84 | 0.42 | 0.4 |
| 5 | 153601658 | rs13169185 | 0.628 | 0.131 | -1.047 | -1.486 | -0.095 | 0.095 | 0.095 | 0 | 0.35 | 0.07 | 0.62 | 0.83 | 0.42 | 0.39 |
| 5 | 153614213 | rs6880803 | -0.199 | -0.233 | -0.947 | -1.902 | -0.008 | -0.03 | -0.03 | 0 | 0.59 | 0.44 | 0.77 | 0.84 | 0.7 | 0.61 |
| 5 | 153616916 | rs11739888 | 0.569 | -0.308 | -1.2 | -1.977 | -0.029 | -0.03 | -0.03 | 0 | 0.08 | 0.01 | 0.07 | 0.08 | 0.11 | 0.08 |
| 5 | 153629068 | rs11951742 | -0.072 | -0.365 | -0.728 | -2.056 | -0.015 | -0.059 | -0.059 | 0 | 0.6 | 0.46 | 0.78 | 0.86 | 0.7 | 0.62 |
| 5 | 153638793 | rs532270 | 0.875 | -0.176 | -0.649 | -2.022 | 0.077 | -0.052 | -0.052 | 0.379 | 0.17 | 0.46 | 0.13 | 0.09 | 0.21 | 0.16 |
| 5 | 153645178 | rs7737215 | 0.573 | -0.298 | -1.009 | -1.996 | -0.073 | 0.159 | 0.159 | 0 | 0.45 | 0.07 | 0.66 | 0.77 | 0.49 | 0.44 |
| 5 | 153650262 | rs753685 | 0.805 | -0.61 | -0.767 | -2.027 | 0.014 | 0.03 | 0.03 | 0.564 | 0.21 | 0.13 | 0.12 | 0.06 | 0.18 | 0.15 |
| 5 | 153654930 | rs192256 | 0.392 | -0.466 | -0.736 | -1.796 | -0.017 | -0.012 | -0.012 | 0 | 0.1 | 0.17 | 0.08 | 0.07 | 0.06 | 0.07 |
| 5 | 153663103 | rs576374 | 0.46 | -0.455 | -0.783 | -1.791 | -0.001 | 0 | 0 | 0 | 0.1 | 0.11 | 0.07 | 0.07 | 0.06 | 0.07 |
| 5 | 153665854 | rs17553077 | 0.409 | -0.343 | -0.908 | -1.957 | -0.002 | 0.149 | 0.149 | 0 | 0.48 | 0.25 | 0.69 | 0.84 | 0.5 | 0.49 |
| 5 | 153667124 | rs12514791 | 1.195 | -0.189 | -0.824 | -1.622 | -0.021 | 0.028 | 0.028 | 0 | 0.31 | 0.13 | 0.56 | 0.77 | 0.35 | 0.39 |
| 5 | 153667787 | rs4958723 | 0.517 | -0.126 | -0.748 | -1.103 | -0.001 | -0.007 | -0.007 | 0 | 0.41 | 0.26 | 0.64 | 0.62 | 0.43 | 0.42 |
| 5 | 153668942 | rs13169724 | 0.954 | -0.303 | -0.655 | -1.343 | 0.009 | 0.004 | 0.004 | 0.752 | 0.2 | 0.17 | 0.1 | 0.03 | 0.15 | 0.16 |
| 5 | 153668985 | rs11950904 | 1.091 | -0.298 | -0.661 | -1.351 | -0.002 | -0.035 | -0.035 | 0 | 0.09 | 0.12 | 0.08 | 0 | 0.09 | 0.02 |
| 5 | 153669038 | rs10057248 | 0.939 | -0.311 | -0.493 | -1.357 | -0.001 | -0.053 | -0.053 | 0.789 | 0.1 | 0.19 | 0.08 | 0 | 0.09 | 0.02 |
| 5 | 153671061 | rs729987 | 0.244 | -0.236 | -0.313 | -1.473 | -0.001 | -0.028 | -0.028 | 0.565 | 0.32 | 0.31 | 0.18 | 0.05 | 0.36 | 0.27 |
| 5 | 153673621 | rs6580068 | -1.134 | -0.103 | -0.34 | -1.282 | -0.001 | 0.001 | 0.001 | 0 | 0.74 | 0.69 | 0.83 | 0.74 | 0.79 | 0.7 |
| 5 | 153673897 | rs13158652 | 1.521 | -0.159 | -0.402 | -1.258 | -0.038 | -0.056 | -0.056 | 0.111 | 0.3 | 0.09 | 0.55 | 0.69 | 0.34 | 0.36 |
| 5 | 153674032 | rs7707954 | 0.176 | -0.166 | -0.353 | -1.613 | -0.012 | -0.058 | -0.058 | 0 | 0.29 | 0.31 | 0.17 | 0.03 | 0.23 | 0.18 |
| 5 | 153676440 | rs7722581 | -0.774 | -0.103 | -0.643 | -1.537 | 0.001 | -0.02 | -0.02 | 0 | 0.44 | 0.57 | 0.28 | 0.05 | 0.45 | 0.34 |
| 5 | 153693475 | rs10057369 | 1.177 | -0.497 | -0.506 | -0.994 | -0.059 | -0.013 | -0.013 | 0.241 | 0.4 | 0.16 | 0.62 | 0.62 | 0.52 | 0.48 |
| 5 | 153695382 | rs13180928 | -0.355 | -0.537 | -0.444 | -1.069 | -0.036 | -0.12 | -0.12 | 0.062 | 0.32 | 0.48 | 0.19 | 0.05 | 0.25 | 0.19 |
| 5 | 153698362 | rs889031 | 1.064 | -0.456 | -0.782 | -0.974 | -0.121 | -0.067 | -0.067 | 0 | 0.4 | 0.08 | 0.62 | 0.61 | 0.52 | 0.48 |
| 5 | 153702252 | rs889032 | -0.605 | -0.442 | -0.332 | -0.983 | -0.002 | 0.022 | 0.022 | 0 | 0.72 | 0.49 | 0.81 | 0.67 | 0.77 | 0.66 |
| 5 | 153708963 | rs1004977 | 0.432 | -0.676 | -0.637 | -1.14 | 0.041 | 0.037 | 0.037 | 0 | 0.56 | 0.13 | 0.73 | 0.65 | 0.7 | 0.58 |
| 5 | 153710692 | rs7736159 | -0.651 | -0.609 | -0.279 | -1.094 | -0.004 | 0.025 | 0.025 | 0 | 0.72 | 0.45 | 0.8 | 0.67 | 0.77 | 0.66 |
| 5 | 153712738 | rs10075400 | -0.617 | -0.717 | -0.434 | -1.165 | -0.018 | -0.07 | -0.07 | 0 | 0.32 | 0.6 | 0.21 | 0.05 | 0.25 | 0.18 |
| 5 | 153716363 | rs17115956 | 1.21 | -0.579 | -0.551 | -1.083 | -0.027 | -0.076 | -0.076 | 0 | 0.14 | 0.25 | 0.06 | 0 | 0.07 | 0.09 |
| 5 | 153718007 | rs10463257 | 0.922 | -0.246 | -1.117 | -0.861 | -0.125 | -0.062 | -0.062 | 0 | 0.4 | 0.05 | 0.61 | 0.61 | 0.53 | 0.49 |
| 5 | 153731210 | rs7733050 | -0.738 | -0.297 | 0.021 | -0.869 | -0.015 | -0.077 | -0.077 | 0.037 | 0.17 | 0.43 | 0.13 | 0.02 | 0.15 | 0.1 |
| 5 | 153734373 | rs7711600 | -0.046 | -0.101 | -0.457 | -0.942 | -0.01 | -0.037 | -0.037 | 2.957 | 0.15 | 0.22 | 0.09 | 0.03 | 0.13 | 0.1 |
| 5 | 153734582 | rs27152 | -0.116 | 0.656 | -0.867 | -0.573 | 0.006 | 0 | 0 | 0 | 0.74 | 0.73 | 0.78 | 0.71 | 0.68 | 0.68 |
| 5 | 153735484 | rs918510 | 0.542 | 0.576 | -1.489 | -0.408 | -0.021 | -0.008 | -0.008 | 0 | 0.53 | 0.17 | 0.53 | 0.55 | 0.54 | 0.52 |
| 5 | 153737288 | rs7730696 | 0.092 | 1.244 | -1.005 | -0.388 | -0.003 | 0 | 0 | 0 | 0.78 | 0.79 | 0.8 | 0.78 | 0.67 | 0.68 |
| 5 | 153741139 | rs3886611 | 0.81 | 0.423 | -1.966 | -0.837 | 0.017 | -0.062 | -0.062 | 0 | 0.37 | 0.1 | 0.45 | 0.53 | 0.34 | 0.3 |
| 5 | 153741843 | rs27154 | -0.178 | 0.699 | -1.26 | -1.037 | 0.008 | 0.024 | 0.024 | 0 | 0.66 | 0.74 | 0.76 | 0.85 | 0.7 | 0.66 |
| 5 | 153744999 | rs462515 | 0.068 | 1.04 | -0.675 | -0.882 | 0.001 | -0.057 | -0.057 | 0 | 0.66 | 0.51 | 0.73 | 0.85 | 0.67 | 0.65 |
| 5 | 153749697 | rs153415 | -0.396 | 1.746 | -0.771 | -0.182 | 0.036 | 0.031 | 0.031 | 1.539 | 0.88 | 0.79 | 0.87 | 0.85 | 0.83 | 0.7 |
| 5 | 153753481 | rs153411 | -0.283 | 1.642 | -0.981 | -0.299 | -0.015 | 0.001 | 0.001 | 1.087 | 0.18 | 0.15 | 0.23 | 0.21 | 0.28 | 0.29 |
| 5 | 153756128 | rs149128 | -0.389 | 1.519 | -1.054 | -0.373 | -0.017 | 0.001 | 0.001 | 0.034 | 0.19 | 0.11 | 0.22 | 0.23 | 0.25 | 0.29 |
| 5 | 153760329 | rs17116042 | 0.327 | 1.199 | -1.524 | -0.576 | -0.025 | -0.013 | -0.013 | 0 | 0.22 | 0.12 | 0.2 | 0.25 | 0.3 | 0.28 |

| | | | | | | | | | | | | | | | | |
|---|---|---|---|---|---|---|---|---|---|---|---|---|---|---|---|---|
| 5 | 153760704 | rs750063 | 0.754 | 0.81 | -1.276 | -1.241 | 0.015 | -0.017 | -0.017 | 0 | 0.35 | 0.23 | 0.43 | 0.51 | 0.3 | 0.31 |
| 5 | 153762653 | rs918509 | -0.829 | 1.102 | -1.176 | -1.321 | -0.012 | -0.002 | -0.002 | 0 | 0.83 | 0.89 | 0.8 | 0.76 | 0.8 | 0.69 |
| 5 | 153765431 | rs6870010 | 0.9 | 0.875 | -0.942 | -1.339 | -0.028 | -0.003 | -0.003 | 0 | 0.2 | 0.11 | 0.2 | 0.18 | 0.29 | 0.22 |
| 5 | 153765462 | rs6889241 | -0.307 | 0.862 | -0.941 | -1.362 | -0.01 | 0.004 | 0.004 | 0.338 | 0.79 | 0.84 | 0.78 | 0.76 | 0.76 | 0.63 |
| 5 | 153766356 | rs2034577 | 1.508 | 0.299 | -0.818 | -1.404 | 0.01 | 0 | 0 | 3.282 | 0.17 | 0.1 | 0.12 | 0 | 0.16 | 0.05 |
| 5 | 153767845 | rs6871105 | -0.46 | -0.162 | -0.692 | -1.297 | -0.018 | -0.019 | -0.019 | 0 | 0.34 | 0.53 | 0.25 | 0.24 | 0.32 | 0.4 |
| 5 | 153768664 | rs12520590 | 0.275 | 0.155 | -1.088 | -1.25 | -0.019 | -0.008 | -0.008 | 0.168 | 0.22 | 0.15 | 0.22 | 0.2 | 0.31 | 0.2 |
| 5 | 153769293 | rs13154706 | 0.286 | 0.153 | -1.089 | -1.25 | -0.019 | -0.008 | -0.008 | 0.003 | 0.22 | 0.15 | 0.22 | 0.2 | 0.31 | 0.21 |
| 5 | 153770059 | rs10073540 | -0.202 | 0.165 | -1.058 | -1.067 | 0.017 | 0.048 | 0.048 | 0.131 | 0.5 | 0.61 | 0.58 | 0.63 | 0.47 | 0.47 |
| 5 | 153772081 | rs10072289 | -0.738 | 0.393 | -1.243 | -0.965 | -0.044 | 0.079 | 0.079 | 0 | 0.58 | 0.79 | 0.63 | 0.67 | 0.54 | 0.57 |
| 5 | 153772289 | rs10064618 | -0.433 | 0.161 | -1.123 | -0.659 | 0.039 | 0.11 | 0.11 | 0.239 | 0.47 | 0.77 | 0.55 | 0.66 | 0.42 | 0.54 |
| 5 | 153774171 | rs17116065 | 0.17 | 0.279 | -0.719 | 0.006 | 0.002 | -0.004 | -0.004 | 0 | 0.08 | 0.05 | 0.05 | 0.14 | 0.07 | 0.12 |
| 5 | 153774496 | rs17637310 | 0.32 | 0.648 | -1.228 | -0.075 | -0.031 | 0.052 | 0.052 | 0 | 0.33 | 0.11 | 0.31 | 0.23 | 0.33 | 0.32 |
| 5 | 153775672 | rs744938 | -0.532 | 0.671 | 0.048 | 0.488 | 0.016 | 0.092 | 0.092 | 0 | 0.12 | 0.5 | 0.09 | 0.14 | 0.1 | 0.12 |
| 5 | 153775699 | rs744939 | 0.271 | 0.996 | -0.3 | 0.723 | 0.011 | 0.071 | 0.071 | 0.044 | 0.2 | 0.27 | 0.34 | 0.47 | 0.21 | 0.24 |
| 5 | 153780104 | rs2161343 | -0.248 | 1.096 | -0.37 | 0.534 | 0.025 | 0.024 | 0.024 | 0.279 | 0.36 | 0.49 | 0.41 | 0.44 | 0.42 | 0.45 |
| 5 | 153781229 | rs3756645 | -0.829 | 0.387 | -0.847 | 0.331 | -0.014 | -0.097 | -0.097 | 0 | 0.64 | 0.89 | 0.56 | 0.45 | 0.61 | 0.7 |
| 5 | 153783587 | rs4958733 | 0.463 | 0.411 | -0.59 | 0.145 | -0.022 | 0.021 | 0.021 | 0.036 | 0.33 | 0.19 | 0.3 | 0.25 | 0.35 | 0.32 |
| 5 | 153783753 | rs6580076 | -0.181 | 0.297 | -0.097 | 0.306 | -0.042 | -0.02 | -0.02 | 0 | 0.27 | 0.51 | 0.16 | 0.16 | 0.16 | 0.33 |
| 5 | 153785497 | rs7703034 | 0.607 | 0.75 | -0.314 | 0.71 | 0.023 | 0.071 | 0.071 | 0 | 0.2 | 0.25 | 0.48 | 0.54 | 0.24 | 0.23 |
| 5 | 153785632 | rs13174523 | 0.17 | 0.686 | -0.843 | 0.621 | -0.083 | -0.028 | -0.028 | 0 | 0.34 | 0.06 | 0.25 | 0.32 | 0.38 | 0.34 |
| 5 | 153787189 | rs2882672 | 0.12 | 0.723 | -0.652 | 0.687 | -0.033 | -0.024 | -0.024 | 0.07 | 0.33 | 0.19 | 0.26 | 0.37 | 0.37 | 0.34 |
| 5 | 153788339 | rs12658875 | 0.16 | 0.746 | -0.711 | 0.725 | -0.034 | -0.028 | -0.028 | 1.524 | 0.33 | 0.16 | 0.26 | 0.37 | 0.37 | 0.34 |
| 5 | 153789590 | rs4958735 | -0.983 | 0.483 | -1.032 | 0.33 | 0.019 | 0.033 | 0.033 | 0 | 0.77 | 0.91 | 0.83 | 0.87 | 0.76 | 0.8 |
| 5 | 153790531 | rs1465417 | -0.721 | 0.568 | -1.201 | 0.296 | -0.007 | 0.057 | 0.057 | 0 | 0.74 | 0.89 | 0.8 | 0.87 | 0.66 | 0.76 |
| 5 | 153790664 | rs2034578 | 0.129 | 0.837 | -1.072 | 0.8 | -0.053 | -0.057 | -0.057 | 0 | 0.31 | 0.17 | 0.29 | 0.41 | 0.38 | 0.33 |
| 5 | 153796875 | rs888979 | 0.486 | 1.003 | -1.547 | 0.963 | -0.058 | -0.032 | -0.032 | 0 | 0.23 | 0.05 | 0.45 | 0.2 | 0.28 | 0.27 |
| 5 | 153797049 | rs3776996 | 0.409 | 0.418 | -1.596 | 0.816 | 0.014 | -0.016 | -0.016 | 0 | 0.27 | 0.14 | 0.14 | 0.41 | 0.21 | 0.3 |
| 5 | 153797880 | rs3172941 | -0.069 | 0.696 | -0.711 | 0.802 | 0 | 0.022 | 0.022 | 0 | 0.32 | 0.37 | 0.2 | 0.68 | 0.32 | 0.33 |
| 5 | 153799165 | rs2277937 | 0.185 | 0.732 | -0.605 | 0.729 | -0.003 | -0.014 | -0.014 | 1.434 | 0.28 | 0.24 | 0.17 | 0.53 | 0.29 | 0.31 |
| 5 | 153800513 | rs7244 | 0.547 | 0.425 | -0.564 | 0.202 | 0.002 | 0.002 | 0.002 | 0 | 0.25 | 0.23 | 0.12 | 0.26 | 0.18 | 0.26 |
| 5 | 153803086 | rs10477100 | 0.823 | 0.481 | -0.449 | 0.079 | -0.029 | -0.125 | -0.125 | 0 | 0.19 | 0.37 | 0.15 | 0.01 | 0.18 | 0.09 |
| 5 | 153810063 | rs11167680 | 1.268 | 0.635 | -0.811 | 0.299 | 0 | -0.002 | -0.002 | 0 | 0.18 | 0.12 | 0.11 | 0.26 | 0.16 | 0.21 |
| 5 | 153811490 | rs10037670 | 0.574 | 0.519 | -0.622 | 0.229 | -0.001 | -0.005 | -0.005 | 0.623 | 0.24 | 0.27 | 0.2 | 0.1 | 0.19 | 0.26 |
| 5 | 153814835 | rs13169441 | 0.016 | 0.973 | -0.169 | 0.664 | -0.028 | -0.022 | -0.022 | 0 | 0.21 | 0.32 | 0.15 | 0.16 | 0.21 | 0.13 |
| 5 | 153822342 | rs11167682 | -0.596 | 1.497 | -0.002 | 1.19 | 0.025 | 0.152 | 0.152 | 0 | 0.18 | 0.39 | 0.15 | 0.45 | 0.27 | 0.14 |
| 5 | 153824197 | rs972596 | -0.772 | 1.444 | -0.328 | 1.086 | 0.044 | 0.11 | 0.11 | 0 | 0.11 | 0.29 | 0.19 | 0.27 | 0.18 | 0.14 |
| 5 | 153831867 | rs3340 | 0.861 | 1.099 | -1.44 | 0.531 | -0.004 | -0.055 | -0.055 | 0 | 0.21 | 0.06 | 0.42 | 0.27 | 0.21 | 0.21 |
| 5 | 153832289 | rs11747607 | 0.086 | 1.162 | -1.223 | -0.226 | 0.028 | -0.012 | -0.012 | 0 | 0.63 | 0.46 | 0.67 | 0.28 | 0.54 | 0.63 |
| 5 | 153840472 | rs6884266 | -0.335 | 1.066 | -1.806 | -0.302 | -0.044 | -0.001 | -0.001 | 0 | 0.92 | 0.99 | 0.94 | 0.96 | 0.87 | 0.89 |
| 5 | 153844993 | rs17116175 | 1.127 | 1.141 | -1.772 | -0.293 | -0.022 | -0.005 | -0.005 | 0 | 0.06 | 0.02 | 0.04 | 0.04 | 0.11 | 0.08 |
| 5 | 153846868 | rs10037652 | -0.208 | 0.874 | -1.901 | -0.425 | -0.003 | -0.021 | -0.021 | 3.206 | 0.65 | 0.79 | 0.51 | 0.65 | 0.65 | 0.63 |
| 5 | 153850649 | rs4958740 | -1.129 | 0.694 | -2.224 | -0.221 | -0.035 | -0.048 | -0.048 | 0.012 | 0.79 | 0.97 | 0.88 | 0.69 | 0.81 | 0.76 |
| 5 | 153852352 | rs10058323 | 1.529 | 0.487 | -2.229 | -0.53 | 0.041 | 0.061 | 0.061 | 0.028 | 0.24 | 0.09 | 0.1 | 0 | 0.12 | 0.22 |
| 5 | 153861047 | rs880083 | 0.309 | 0.717 | -2.073 | -0.313 | 0.004 | -0.023 | -0.023 | 0.008 | 0.32 | 0.19 | 0.48 | 0.35 | 0.32 | 0.34 |
| 5 | 153864515 | rs2351485 | 1.594 | 0.555 | -2.225 | -0.23 | 0.007 | -0.072 | -0.072 | 0 | 0.19 | 0.05 | 0.4 | 0.31 | 0.17 | 0.22 |
| 5 | 153865729 | rs6580083 | 0.079 | 0.707 | -2.105 | -0.277 | -0.024 | 0.045 | 0.045 | 0 | 0.47 | 0.75 | 0.43 | 0.37 | 0.51 | 0.47 |
| 5 | 153867221 | rs13178677 | 0.164 | 0.722 | -1.935 | -0.27 | -0.014 | 0.028 | 0.028 | 0 | 0.46 | 0.67 | 0.42 | 0.37 | 0.51 | 0.47 |
| 5 | 153869040 | rs13165478 | 0.342 | 1.05 | -1.549 | 0.064 | 0.069 | 0.071 | 0.071 | 0.205 | 0.22 | 0.57 | 0.31 | 0.37 | 0.38 | 0.21 |
| 5 | 153872961 | rs7735256 | 0.472 | 0.76 | -1.787 | 0.357 | 0.027 | -0.036 | -0.036 | 0.166 | 0.26 | 0.11 | 0.41 | 0.32 | 0.2 | 0.26 |
| 5 | 153873560 | rs283438 | -0.708 | 1.141 | -1.779 | 0.429 | -0.038 | 0.026 | 0.026 | 0 | 0.83 | 0.96 | 0.9 | 0.88 | 0.85 | 0.81 |
| 5 | 153874518 | rs2055422 | 0.865 | 0.859 | -1.677 | 0.708 | 0.005 | -0.038 | -0.038 | 0.009 | 0.2 | 0.12 | 0.42 | 0.32 | 0.2 | 0.2 |
| 5 | 153875961 | rs4958742 | 0.795 | 0.873 | -1.736 | 0.715 | 0.003 | -0.05 | -0.05 | 0 | 0.2 | 0.07 | 0.42 | 0.32 | 0.2 | 0.2 |
| 5 | 153877690 | rs10477102 | 0.035 | 0.739 | -1.195 | 0.675 | 0.003 | 0.022 | 0.022 | 0 | 0.56 | 0.72 | 0.46 | 0.37 | 0.59 | 0.5 |
| 5 | 153884609 | rs922535 | 0.142 | 1.354 | -1.266 | 0.4 | 0.001 | -0.004 | -0.004 | 0 | 0.07 | 0.08 | 0.12 | 0.11 | 0.07 | 0.11 |
| 5 | 153885015 | rs7733449 | 0.725 | 0.908 | -1.285 | -0.481 | 0.002 | -0.034 | -0.034 | 0 | 0.27 | 0.25 | 0.16 | 0.01 | 0.24 | 0.27 |
| 5 | 153887991 | rs374418 | 0.423 | 0.916 | -1.067 | -0.206 | -0.018 | 0.021 | 0.021 | 0 | 0.17 | 0.08 | 0.14 | 0.12 | 0.24 | 0.1 |
| 5 | 153891017 | rs4958377 | 0.488 | 0.448 | -0.739 | 0.324 | -0.009 | -0.04 | -0.04 | 0 | 0.26 | 0.11 | 0.42 | 0.31 | 0.24 | 0.38 |
| 5 | 153892083 | rs7708891 | -0.217 | 0.528 | -0.56 | 0.387 | 0.002 | 0.02 | 0.02 | 0 | 0.09 | 0.26 | 0.14 | 0.11 | 0.07 | 0.11 |
| 5 | 153894704 | rs6861375 | 0.592 | 0.267 | -0.522 | 0.358 | -0.025 | 0.01 | 0.01 | 0 | 0.06 | 0.23 | 0.02 | 0.07 | 0.01 | 0 |
| 5 | 153896160 | rs6871767 | -0.142 | 0.485 | -0.576 | 0.395 | -0.008 | 0.022 | 0.022 | 0 | 0.35 | 0.54 | 0.57 | 0.42 | 0.31 | 0.36 |
| 5 | 153899307 | rs10056769 | -0.268 | 0.614 | -0.704 | 0.269 | 0.008 | 0.029 | 0.029 | 0 | 0.29 | 0.19 | 0.23 | 0.23 | 0.28 | 0.25 |
| 5 | 153900209 | rs283433 | -0.55 | 1.278 | -0.088 | 0.938 | 0.018 | 0.017 | 0.017 | 0 | 0.47 | 0.27 | 0.74 | 0.53 | 0.63 | 0.54 |
| 5 | 153902050 | rs10076602 | 1.141 | 1.301 | 0.36 | 0.907 | 0.011 | 0.005 | 0.005 | 0 | 0.42 | 0.33 | 0.17 | 0.39 | 0.26 | 0.34 |
| 5 | 153904423 | rs2351749 | 0.518 | 1.597 | 0.304 | 0.897 | 0 | 0.005 | 0.005 | 0 | 0.53 | 0.4 | 0.3 | 0.59 | 0.46 | 0.42 |
| 5 | 153907246 | rs9324786 | 1.598 | 1.426 | 0.322 | 0.965 | 0.031 | 0.074 | 0.074 | 0 | 0.37 | 0.19 | 0.14 | 0.19 | 0.23 | 0.3 |
| 5 | 153912076 | rs7703330 | -0.179 | 1.449 | 0.543 | 0.964 | -0.029 | 0.027 | 0.027 | 0 | 0.67 | 0.72 | 0.44 | 0.74 | 0.59 | 0.49 |
| 5 | 153913156 | rs1032741 | 0.048 | 1.448 | 0.628 | 0.966 | 0.014 | -0.03 | -0.03 | 0 | 0.64 | 0.46 | 0.42 | 0.73 | 0.58 | 0.49 |
| 5 | 153919809 | rs11749542 | -1.038 | 1.539 | 0.455 | 0.886 | 0.016 | 0.015 | 0.015 | 0 | 0.51 | 0.65 | 0.5 | 0.61 | 0.66 | 0.62 |

| | | | | | | | | | | | | | | | | |
|---|---|---|---|---|---|---|---|---|---|---|---|---|---|---|---|---|
| 5 | 153921460 | rs17639612 | 0.037 | 1.655 | 0.109 | 1.203 | -0.01 | -0.007 | -0.007 | 0 | 0.12 | 0.09 | 0.14 | 0.17 | 0.23 | 0.07 |
| 5 | 153949908 | rs7443454 | -1.768 | 1.405 | 0.02 | 1.004 | 0.018 | 0.013 | 0.013 | 0 | 0.66 | 0.75 | 0.63 | 0.72 | 0.8 | 0.73 |
| 5 | 153955820 | rs283458 | 0.7 | 1.517 | -0.56 | 0.832 | -0.068 | 0.027 | 0.027 | 0 | 0.25 | 0.03 | 0.19 | 0.16 | 0.28 | 0.4 |
| 5 | 153960485 | rs12655409 | -1.573 | 1.361 | -0.212 | 0.696 | 0.025 | -0.01 | -0.01 | 0 | 0.6 | 0.76 | 0.57 | 0.47 | 0.75 | 0.66 |
| 5 | 153968022 | rs2351752 | 0.779 | 1.575 | -1.47 | 0.463 | -0.03 | 0.081 | 0.081 | 0 | 0.22 | 0.1 | 0.17 | 0.1 | 0.27 | 0.33 |
| 5 | 153970243 | rs13173061 | -0.174 | 1.281 | -1.479 | 0.54 | -0.12 | -0.031 | -0.031 | 0 | 0.35 | 0.11 | 0.31 | 0.33 | 0.51 | 0.45 |
| 5 | 153972821 | rs13184629 | 0.395 | 1.46 | -1.447 | 0.417 | -0.008 | 0.016 | 0.016 | 0 | 0.09 | 0.06 | 0.1 | 0 | 0.1 | 0.12 |
| 5 | 153973305 | rs10057687 | -1.206 | 1.402 | -1.286 | 0.7 | -0.018 | 0.058 | 0.058 | 0 | 0.57 | 0.86 | 0.52 | 0.68 | 0.67 | 0.6 |
| 5 | 153986044 | rs1824936 | -0.836 | 1.678 | -0.757 | 0.692 | 0.003 | -0.005 | -0.005 | 0 | 0.39 | 0.41 | 0.26 | 0.39 | 0.42 | 0.33 |
| 5 | 153986462 | rs1824933 | 0.085 | 1.892 | -1.634 | 0.175 | -0.094 | 0.12 | 0.12 | 0 | 0.2 | 0.02 | 0.11 | 0.01 | 0.28 | 0.18 |
| 5 | 153990543 | rs6876806 | -0.383 | 1.413 | -1.229 | 0.72 | 0.025 | 0.01 | 0.01 | 0.097 | 0.51 | 0.75 | 0.48 | 0.38 | 0.61 | 0.45 |
| 5 | 153991222 | rs1975985 | 0.756 | 0.606 | -1.648 | 0.784 | 0.083 | -0.03 | -0.03 | 0.019 | 0.34 | 0.1 | 0.35 | 0.36 | 0.17 | 0.22 |
| 5 | 153991379 | rs1975984 | 0.26 | 0.876 | -0.918 | 0.487 | -0.002 | -0.001 | -0.001 | 0 | 0.47 | 0.47 | 0.52 | 0.4 | 0.3 | 0.46 |
| 5 | 153992798 | rs7708052 | 0.124 | 1.694 | -1.101 | 0.209 | -0.01 | 0.008 | 0.008 | 0 | 0.2 | 0.14 | 0.24 | 0.13 | 0.32 | 0.26 |
| 5 | 153995314 | rs4958751 | 1.499 | 1.479 | -1.307 | 0.169 | 0.077 | -0.011 | -0.011 | 2.57 | 0.32 | 0.08 | 0.35 | 0.31 | 0.17 | 0.2 |
| 5 | 153997708 | rs6580092 | -0.183 | 1.614 | -0.542 | 0.219 | 0.048 | -0.018 | -0.018 | 0.727 | 0.13 | 0.39 | 0.15 | 0.08 | 0.14 | 0.23 |
| 5 | 153998704 | rs12522617 | 0.495 | 1.748 | -1.157 | 0.403 | -0.036 | 0.045 | 0.045 | 0 | 0.16 | 0.07 | 0.16 | 0.05 | 0.26 | 0.12 |
| 5 | 153999695 | rs13168119 | 0.261 | 1.841 | -1.22 | 0.852 | 0.004 | 0.044 | 0.044 | 0.137 | 0.91 | 0.88 | 0.94 | 0.84 | 0.92 | 0.91 |
| 5 | 154001174 | rs12374480 | 0.47 | 1.904 | -1.303 | 0.648 | -0.036 | 0.043 | 0.043 | 0 | 0.16 | 0.07 | 0.15 | 0.05 | 0.27 | 0.12 |
| 5 | 154005049 | rs2056429 | -0.222 | 1.334 | -1.444 | 1.016 | -0.01 | -0.032 | -0.032 | 0 | 0.16 | 0.09 | 0.15 | 0.33 | 0.19 | 0.2 |
| 5 | 154007669 | rs6892346 | 0.189 | 0.903 | -1.177 | 0.823 | -0.019 | 0.065 | 0.065 | 0 | 0.71 | 0.49 | 0.6 | 0.54 | 0.77 | 0.62 |
| 5 | 154010041 | rs4958379 | 0.306 | 0.443 | -1.34 | 0.815 | 0.009 | -0.002 | -0.002 | 0 | 0.44 | 0.27 | 0.33 | 0.37 | 0.42 | 0.34 |
| 5 | 154023119 | rs11739775 | -0.441 | 0.217 | -1.199 | 0.615 | -0.003 | -0.015 | -0.015 | 0 | 0.52 | 0.48 | 0.63 | 0.78 | 0.4 | 0.55 |
| 5 | 154045825 | rs4958756 | -0.523 | 0.063 | -1.23 | 0.675 | 0.006 | 0.012 | 0.012 | 0 | 0.49 | 0.43 | 0.54 | 0.73 | 0.36 | 0.45 |
| 5 | 154050914 | rs11167699 | -0.423 | 0.077 | -1.525 | 1.446 | -0.022 | -0.07 | -0.07 | 0 | 0.13 | 0.01 | 0.07 | 0.21 | 0.12 | 0.12 |
| 5 | 154053249 | rs7723312 | 0.344 | 0.067 | -1.411 | 1.609 | 0 | -0.039 | -0.039 | 0.427 | 0.86 | 0.93 | 0.92 | 0.57 | 0.89 | 0.87 |
| 5 | 154056002 | rs13186731 | -0.526 | 0.077 | -1.398 | 1.628 | -0.001 | -0.042 | -0.042 | 0 | 0.13 | 0.06 | 0.08 | 0.42 | 0.11 | 0.14 |
| 5 | 154060180 | rs918365 | -0.221 | 0.004 | -1.492 | 1.369 | -0.002 | -0.031 | -0.031 | 0 | 0.7 | 0.89 | 0.64 | 0.72 | 0.73 | 0.73 |
| 5 | 154063295 | rs965235 | -0.218 | -0.017 | -1.514 | 1.422 | 0.002 | -0.064 | -0.064 | 0 | 0.69 | 0.92 | 0.63 | 0.69 | 0.75 | 0.73 |
| 5 | 154063416 | rs17116366 | 0.401 | -0.048 | -1.192 | 1.39 | -0.002 | 0.023 | 0.023 | 0 | 0.07 | 0.14 | 0.16 | 0.12 | 0.03 | 0.08 |
| 5 | 154065758 | rs4958760 | 0.342 | -0.071 | -1.159 | 1.343 | 0.011 | -0.008 | -0.008 | 0.005 | 0.61 | 0.7 | 0.47 | 0.57 | 0.72 | 0.65 |
| 5 | 154080958 | rs6868098 | -0.232 | -0.119 | -1.361 | 1.336 | 0.011 | -0.044 | -0.044 | 3.634 | 0.69 | 0.9 | 0.62 | 0.67 | 0.75 | 0.73 |
| 5 | 154091904 | rs12055015 | 0.317 | 0.082 | -1.62 | 1.729 | -0.037 | -0.056 | -0.056 | 0 | 0.09 | 0.01 | 0.07 | 0.19 | 0.11 | 0.13 |
| 5 | 154092202 | rs3867429 | 1.42 | 0.155 | -1.54 | 0.801 | -0.057 | 0.148 | 0.148 | 0 | 0.38 | 0.18 | 0.27 | 0 | 0.5 | 0.28 |
| 5 | 154096447 | rs9324794 | 0.188 | 0.212 | -0.758 | 1.514 | 0.009 | 0.05 | 0.05 | 0 | 0.07 | 0.16 | 0.06 | 0.17 | 0.08 | 0.22 |
| 5 | 154101875 | rs4117784 | -0.452 | -0.084 | -0.934 | 1.173 | 0.002 | -0.028 | -0.028 | 0 | 0.8 | 0.9 | 0.65 | 0.68 | 0.81 | 0.75 |
| 5 | 154106976 | rs6875019 | -0.564 | 0.037 | -0.645 | 0.875 | 0 | 0.002 | 0.002 | 0.76 | 0.34 | 0.33 | 0.35 | 0.25 | 0.3 | 0.33 |
| 5 | 154111128 | rs1233078 | -1.078 | 0.228 | 0.077 | 0.843 | -0.013 | 0.004 | 0.004 | 2.86 | 0.27 | 0.2 | 0.41 | 0.8 | 0.33 | 0.45 |
| 5 | 154121691 | rs17116409 | -0.242 | 0.044 | 0.234 | 0.972 | -0.023 | 0.018 | 0.018 | 0 | 0.14 | 0.25 | 0.21 | 0.14 | 0.1 | 0.21 |
| 5 | 154142057 | rs17116411 | -0.348 | 0.134 | 0.659 | 0.952 | 0.006 | 0.006 | 0.006 | 0 | 0.14 | 0.41 | 0.23 | 0.11 | 0.12 | 0.11 |
| 5 | 154145474 | rs17116417 | 0.474 | 0.115 | 0.645 | 0.938 | 0.005 | 0.009 | 0.009 | 0 | 0.1 | 0.06 | 0.18 | 0.05 | 0.09 | 0.09 |
| 5 | 154183497 | rs816717 | -0.555 | 0.012 | 0.819 | 1.007 | 0 | -0.004 | -0.004 | 0 | 0.13 | 0.32 | 0.22 | 0.11 | 0.12 | 0.11 |
| 5 | 154199368 | rs39614 | -0.578 | 0.012 | 0.822 | 0.668 | 0.001 | -0.003 | -0.003 | 2.635 | 0.13 | 0.33 | 0.22 | 0.11 | 0.12 | 0.11 |
| 5 | 154203121 | rs17641488 | 0.988 | 0.053 | 0.697 | 0.473 | -0.013 | 0.033 | 0.033 | 0 | 0.09 | 0 | 0.05 | 0.01 | 0.07 | 0.05 |
| 5 | 154203264 | rs6895741 | -0.058 | 0.122 | 0.842 | 0.58 | -0.011 | -0.015 | -0.015 | 0.035 | 0.11 | 0.16 | 0.19 | 0.05 | 0.09 | 0.09 |
| 5 | 154209800 | rs12187921 | -0.058 | 0.122 | 0.842 | 0.583 | -0.011 | -0.006 | -0.006 | 0 | 0.11 | 0.17 | 0.19 | 0.09 | 0.09 | 0.09 |
| 5 | 154210129 | rs2278223 | 0.438 | 0.129 | 0.762 | 0.563 | -0.01 | -0.017 | -0.017 | 0 | 0.11 | 0.07 | 0.12 | 0.19 | 0.12 | 0.22 |
| 5 | 154224949 | rs17096232 | 0.153 | 0.131 | 0.803 | 0.478 | -0.004 | -0.006 | -0.006 | 0.028 | 0.11 | 0.13 | 0.19 | 0.05 | 0.09 | 0.09 |
| 5 | 154236694 | rs3776984 | 0.324 | 0.111 | 0.779 | 0.458 | 0.006 | 0.011 | 0.011 | 0 | 0.1 | 0.06 | 0.18 | 0.05 | 0.09 | 0.09 |
| 5 | 154254818 | rs17116534 | -0.617 | 0.154 | 0.892 | 0.477 | 0 | -0.025 | -0.025 | 0 | 0.13 | 0.33 | 0.22 | 0.08 | 0.11 | 0.11 |
| 5 | 154256175 | rs12300 | -0.617 | 0.157 | 0.867 | 0.478 | 0 | -0.025 | -0.025 | 1.122 | 0.13 | 0.33 | 0.22 | 0.08 | 0.11 | 0.11 |
| 5 | 154265119 | rs2544880 | -0.616 | 0.273 | 0.836 | 0.448 | -0.002 | -0.005 | -0.005 | 0 | 0.14 | 0.33 | 0.22 | 0.11 | 0.11 | 0.11 |
| 5 | 154275786 | rs6865950 | -0.163 | 0.272 | 0.882 | 0.448 | -0.014 | -0.02 | -0.02 | 0 | 0.12 | 0.18 | 0.2 | 0.05 | 0.09 | 0.09 |
| 5 | 154282407 | rs1421815 | 0.085 | 0.271 | 0.873 | 0.447 | -0.005 | -0.007 | -0.007 | 0.945 | 0.11 | 0.14 | 0.19 | 0.05 | 0.09 | 0.09 |
| 5 | 154291409 | rs1974777 | 0.375 | 0.277 | 0.688 | 0.45 | -0.002 | -0.006 | -0.006 | 0 | 0.86 | 0.67 | 0.78 | 0.89 | 0.89 | 0.89 |
| 5 | 154300940 | rs348739 | 0.567 | 0.269 | 0.686 | 0.446 | 0 | -0.005 | -0.005 | 2.967 | 0.86 | 0.57 | 0.77 | 0.89 | 0.89 | 0.89 |
| 5 | 154308307 | rs1035368 | 0.548 | 0.285 | 0.687 | 0.453 | 0.002 | -0.003 | -0.003 | 0.474 | 0.86 | 0.57 | 0.77 | 0.89 | 0.89 | 0.89 |
| 5 | 154336832 | rs1421819 | -0.205 | 0.211 | 0.378 | 0.334 | -0.017 | -0.024 | -0.024 | 0.007 | 0.12 | 0.22 | 0.2 | 0.05 | 0.09 | 0.09 |
| 5 | 154345987 | rs13156669 | 0.773 | 0.48 | 0.141 | 0.684 | -0.036 | 0.013 | 0.013 | 0.023 | 0.07 | 0.01 | 0.22 | 0.5 | 0.09 | 0.12 |
| 5 | 154353012 | rs13165424 | -0.818 | 0.486 | 0.184 | -0.15 | -0.031 | -0.007 | -0.007 | 0 | 0.2 | 0.1 | 0.35 | 0.81 | 0.22 | 0.47 |
| 5 | 154357463 | rs2431184 | 1.768 | 0.382 | 0.418 | -0.553 | -0.01 | 0.067 | 0.067 | 0 | 0.57 | 0.64 | 0.51 | 0.11 | 0.58 | 0.37 |
| 5 | 154359159 | rs158134 | 1.183 | 0.473 | 0.375 | -0.505 | -0.015 | 0.04 | 0.04 | 0.456 | 0.69 | 0.77 | 0.57 | 0.12 | 0.66 | 0.43 |
| 5 | 154369436 | rs286621 | 1.076 | 0.535 | 0.886 | -0.155 | 0.008 | -0.004 | -0.004 | 0 | 0.67 | 0.46 | 0.56 | 0.35 | 0.69 | 0.71 |
| 5 | 154375465 | rs286593 | 0.867 | 0.831 | 0.627 | -0.26 | 0.007 | 0 | 0 | 0.1 | 0.69 | 0.62 | 0.56 | 0.36 | 0.65 | 0.69 |
| 5 | 154389411 | rs17116707 | 0.065 | 0.475 | 0.019 | -0.132 | 0.009 | 0.032 | 0.032 | 0 | 0.13 | 0.05 | 0.2 | 0.05 | 0.12 | 0.13 |
| 5 | 154395158 | rs6580126 | 0.453 | 0.535 | -0.044 | -0.103 | -0.033 | -0.047 | -0.047 | 0 | 0.77 | 0.91 | 0.64 | 0.41 | 0.76 | 0.81 |
| 5 | 154403211 | rs9324797 | 0.048 | 0.429 | 0.104 | -0.222 | 0.002 | 0.001 | 0.001 | 0 | 0.14 | 0.17 | 0.2 | 0.05 | 0.12 | 0.18 |
| 5 | 154404176 | rs7736342 | 0.215 | 0.436 | -0.05 | -0.211 | 0.015 | 0.03 | 0.03 | 0 | 0.13 | 0.04 | 0.2 | 0.06 | 0.12 | 0.18 |
| 5 | 154406132 | rs176273 | 2.282 | 0.384 | -0.167 | -0.538 | -0.045 | 0.142 | 0.142 | 2.169 | 0.37 | 0.09 | 0.24 | 0.12 | 0.42 | 0.33 |

| | | | | | | | | | | | | | | | | |
|---|---|---|---|---|---|---|---|---|---|---|---|---|---|---|---|---|
| 5 | 154423527 | rs286632 | 1.639 | 0.355 | 0.407 | -0.499 | 0.012 | 0.121 | 0.121 | 0.023 | 0.52 | 0.18 | 0.35 | 0.32 | 0.61 | 0.55 |
| 5 | 154425547 | rs287125 | 1.662 | 0.366 | 0.413 | -0.493 | 0.016 | 0.115 | 0.115 | 0.004 | 0.52 | 0.22 | 0.36 | 0.32 | 0.61 | 0.55 |
| 5 | 154436799 | rs101271 | 0.229 | 0.375 | 0.533 | -0.316 | 0.017 | 0.035 | 0.035 | 0 | 0.78 | 0.71 | 0.54 | 0.43 | 0.76 | 0.78 |
| 5 | 154440574 | rs286612 | 0.023 | 0.383 | 0.126 | -0.175 | -0.016 | -0.029 | -0.029 | 1.097 | 0.8 | 0.84 | 0.64 | 0.43 | 0.77 | 0.79 |
| 5 | 154443069 | rs286607 | 1.542 | 0.377 | 0.157 | -0.238 | 0.006 | 0.037 | 0.037 | 0.045 | 0.57 | 0.33 | 0.38 | 0.34 | 0.63 | 0.58 |
| 5 | 154443639 | rs286605 | 1.474 | 0.436 | 0.008 | -0.442 | -0.007 | 0.099 | 0.099 | 2.358 | 0.49 | 0.1 | 0.52 | 0.62 | 0.57 | 0.44 |
| 5 | 154444008 | rs7730481 | -1.355 | 0.29 | 0.17 | -0.583 | -0.003 | -0.02 | -0.02 | 0.005 | 0.29 | 0.67 | 0.31 | 0.09 | 0.21 | 0.27 |
| 5 | 154444377 | rs11750225 | 0.461 | 1.027 | 0.277 | 0.15 | 0.011 | -0.027 | -0.027 | 0 | 0.05 | 0.1 | 0.04 | 0 | 0.05 | 0.06 |
| 5 | 154446956 | rs444984 | 1.428 | 1.065 | -0.163 | 0.161 | -0.039 | 0.179 | 0.179 | 1.459 | 0.51 | 0.14 | 0.43 | 0.21 | 0.56 | 0.52 |
| 5 | 154447535 | rs351291 | 0.19 | 0.948 | 0.031 | 0.52 | 0.005 | 0.074 | 0.074 | 0 | 0.13 | 0.15 | 0.31 | 0.48 | 0.14 | 0.1 |
| 5 | 154448263 | rs351292 | 0.067 | 0.91 | -0.274 | 0.41 | -0.003 | -0.003 | -0.003 | 0 | 0.69 | 0.62 | 0.77 | 0.68 | 0.72 | 0.68 |
| 5 | 154448739 | rs351294 | -0.358 | 0.799 | -0.132 | 0.605 | 0.056 | 0.434 | 0.434 | 0 | 0.09 | 0.42 | 0.28 | 0.42 | 0.07 | 0.06 |
| 5 | 154448827 | rs13162193 | 0.219 | 0.972 | -1.264 | -0.432 | 0.009 | -0.018 | -0.018 | 0 | 0.13 | 0.13 | 0.09 | 0.09 | 0.13 | 0.12 |
| 5 | 154448863 | rs970029 | 0.473 | 0.961 | -1.666 | -0.342 | -0.061 | 0.18 | 0.18 | 1.854 | 0.48 | 0.08 | 0.42 | 0.18 | 0.55 | 0.49 |
| 5 | 154449947 | rs2314305 | -0.001 | 0.745 | -1.097 | -0.579 | 0.011 | 0.042 | 0.042 | 0 | 0.47 | 0.68 | 0.42 | 0.19 | 0.42 | 0.37 |
| 5 | 154454291 | rs721259 | -0.338 | 0.367 | -0.931 | -0.703 | 0.007 | 0.019 | 0.019 | 0 | 0.23 | 0.16 | 0.13 | 0.05 | 0.18 | 0.28 |
| 5 | 154457905 | rs1319694 | -1.351 | 0.093 | -0.425 | 0.123 | 0.002 | -0.002 | -0.002 | 0 | 0.4 | 0.7 | 0.26 | 0.43 | 0.28 | 0.44 |
| 5 | 154459454 | rs9324802 | -1.045 | -0.206 | -0.072 | 0.236 | -0.028 | -0.081 | -0.081 | 0 | 0.34 | 0.58 | 0.2 | 0.08 | 0.23 | 0.37 |
| 5 | 154477576 | rs7722634 | -0.242 | 0.103 | 0.031 | 0.936 | 0.039 | 0.15 | 0.15 | 0 | 0.94 | 0.86 | 0.68 | 0.78 | 0.93 | 0.92 |
| 5 | 154478636 | rs1025300 | 0.46 | -0.069 | 0.342 | 0.721 | -0.02 | -0.025 | -0.025 | 0 | 0.05 | 0.18 | 0.02 | 0 | 0.02 | 0.01 |
| 5 | 154486109 | rs6893762 | -1.24 | -0.093 | 0.352 | 0.782 | -0.029 | 0.158 | 0.158 | 0 | 0.1 | 0.28 | 0.09 | 0.35 | 0.04 | 0.08 |
| 5 | 154487768 | rs10454998 | 0.396 | -0.241 | 0.385 | 0.698 | 0.021 | 0.093 | 0.093 | 0 | 0.21 | 0.09 | 0.12 | 0.02 | 0.16 | 0.21 |
| 5 | 154488947 | rs6897414 | 0.851 | -0.255 | -0.176 | 0.275 | 0 | 0.002 | 0.002 | 0 | 0.52 | 0.57 | 0.37 | 0.51 | 0.51 | 0.36 |
| 5 | 154489393 | rs1549647 | 1.963 | -0.195 | -0.164 | 0.297 | -0.061 | -0.082 | -0.082 | 0 | 0.38 | 0.15 | 0.29 | 0.45 | 0.43 | 0.22 |
| 5 | 154493253 | rs7703759 | 0.263 | 0.019 | 0.047 | 0.627 | 0.016 | 0.044 | 0.044 | 0 | 0.73 | 0.71 | 0.49 | 0.52 | 0.67 | 0.59 |
| 5 | 154493790 | rs277311 | 0.864 | 0.089 | -0.12 | 0.6 | -0.035 | -0.019 | -0.019 | 0 | 0.1 | 0 | 0.11 | 0.14 | 0.14 | 0.21 |
| 5 | 154496905 | rs7720827 | -0.83 | 0.027 | 0.228 | 0.545 | -0.01 | -0.005 | -0.005 | 1.45 | 0.11 | 0.33 | 0.08 | 0.06 | 0.07 | 0.14 |
| 5 | 154499047 | rs10066223 | 0.688 | 0.093 | 0.063 | 0.267 | 0.015 | 0.032 | 0.032 | 0 | 0.67 | 0.6 | 0.44 | 0.49 | 0.61 | 0.47 |
| 5 | 154509643 | rs7715630 | 0.359 | 0.192 | 0.174 | 0.268 | -0.028 | -0.029 | -0.029 | 0 | 0.05 | 0.12 | 0.02 | 0 | 0.02 | 0.01 |
| 5 | 154510634 | rs13190138 | 1.568 | 0.057 | -0.102 | 0.245 | -0.032 | 0.007 | 0.007 | 0.488 | 0.32 | 0.2 | 0.21 | 0.25 | 0.35 | 0.21 |
| 5 | 154511881 | rs6866348 | 0.472 | -0.34 | 0.194 | 0.033 | -0.021 | -0.122 | -0.122 | 0 | 0.25 | 0.36 | 0.13 | 0.01 | 0.17 | 0.22 |
| 5 | 154513646 | rs12518198 | 1.659 | -0.225 | -0.206 | 0.049 | -0.041 | 0.012 | 0.012 | 0.66 | 0.33 | 0.21 | 0.23 | 0.25 | 0.37 | 0.22 |
| 5 | 154515885 | rs17652057 | 0.73 | -0.662 | -0.4 | -0.09 | 0.004 | 0.123 | 0.123 | 0.016 | 0.23 | 0.07 | 0.12 | 0.01 | 0.17 | 0.22 |
| 5 | 154523910 | rs6863955 | 1.266 | -0.168 | -0.22 | 0.177 | -0.012 | -0.014 | -0.014 | 0 | 0.37 | 0.31 | 0.27 | 0.48 | 0.38 | 0.29 |
| 5 | 154529948 | rs4958400 | -0.519 | -0.253 | 0.177 | 0.164 | -0.034 | -0.009 | -0.009 | 0.407 | 0.35 | 0.54 | 0.2 | 0.26 | 0.23 | 0.32 |
| 5 | 154530330 | rs3851503 | 0.818 | -0.23 | -0.285 | 0.183 | -0.001 | 0 | 0 | 0 | 0.56 | 0.56 | 0.36 | 0.33 | 0.51 | 0.42 |
| 5 | 154546402 | rs986179 | -0.928 | -0.928 | -0.288 | 0.257 | 0.041 | 0.011 | 0.011 | 0.047 | 0.35 | 0.33 | 0.19 | 0.26 | 0.22 | 0.33 |
| 5 | 154559401 | rs2872502 | -0.936 | -0.927 | -0.324 | 0.258 | 0.03 | 0.009 | 0.009 | 0.043 | 0.35 | 0.35 | 0.19 | 0.26 | 0.22 | 0.33 |
| 5 | 154564146 | rs17574895 | 0.051 | -0.479 | -0.894 | 0.13 | 0.003 | 0.102 | 0.102 | 0.053 | 0.22 | 0.06 | 0.11 | 0.06 | 0.17 | 0.21 |
| 5 | 154569150 | rs1864125 | -0.131 | -0.155 | -0.778 | 0.417 | -0.024 | -0.025 | -0.025 | 0 | 0.13 | 0.01 | 0.14 | 0.15 | 0.17 | 0.15 |
| 5 | 154572428 | rs753943 | 0.063 | 0.173 | -0.376 | 0.811 | 0.005 | 0.055 | 0.055 | 0 | 0.06 | 0.1 | 0.05 | 0.16 | 0.06 | 0.07 |
| 5 | 154578318 | rs1422834 | -1.536 | -0.274 | -0.337 | 0.5 | -0.039 | -0.033 | -0.033 | 0 | 0.56 | 0.79 | 0.64 | 0.53 | 0.49 | 0.56 |
| 5 | 154594301 | rs2062400 | -0.787 | 0.363 | -0.042 | 1.107 | 0.033 | 0.143 | 0.143 | 0.075 | 0.09 | 0.16 | 0.35 | 0.38 | 0.14 | 0.19 |
| 5 | 154595914 | rs17653560 | -0.614 | -0.587 | -0.957 | -0.427 | 0 | 0.097 | 0.097 | 2.721 | 0.19 | 0.05 | 0.1 | 0.03 | 0.15 | 0.18 |
| 5 | 154601557 | rs1422832 | -0.694 | -0.684 | -0.26 | -0.274 | -0.012 | -0.054 | -0.054 | 0 | 0.18 | 0.22 | 0.09 | 0 | 0.08 | 0.1 |
| 5 | 154609080 | rs17653878 | -0.883 | -0.542 | -0.14 | 0.023 | -0.001 | 0.01 | 0.01 | 0 | 0.06 | 0.08 | 0.07 | 0.09 | 0.05 | 0.15 |
| 5 | 154609902 | rs961548 | -0.549 | -1.201 | 0.309 | 0.538 | -0.024 | 0.095 | 0.095 | 0 | 0.16 | 0.26 | 0.19 | 0.37 | 0.1 | 0.11 |
| 5 | 154614881 | rs17117138 | -0.166 | -1.25 | 0.364 | 0.516 | -0.007 | 0.086 | 0.086 | 0.242 | 0.1 | 0.15 | 0.17 | 0.37 | 0.08 | 0.09 |
| 5 | 154627585 | rs12655293 | -1.891 | -1.743 | 0.59 | 0.187 | 0.109 | -0.052 | -0.052 | 0.555 | 0.42 | 0.82 | 0.36 | 0.46 | 0.23 | 0.4 |
| 5 | 154631839 | rs10071781 | -0.679 | -1.693 | 1.063 | 0.18 | -0.039 | -0.071 | -0.071 | 2.705 | 0.23 | 0.38 | 0.16 | 0.09 | 0.13 | 0.25 |
| 5 | 154635358 | rs1155578 | -1.385 | -1.405 | 0.377 | 0.25 | 0.06 | -0.036 | -0.036 | 0 | 0.4 | 0.75 | 0.36 | 0.44 | 0.23 | 0.39 |
| 5 | 154643606 | rs11748833 | -0.714 | -0.721 | 0.504 | -0.166 | 0 | -0.02 | -0.02 | 0 | 0.09 | 0.08 | 0.06 | 0 | 0.12 | 0.13 |
| 5 | 154648174 | rs4958401 | -1.82 | -0.57 | 1.189 | -0.444 | 0.016 | -0.054 | -0.054 | 0 | 0.24 | 0.36 | 0.2 | 0.16 | 0.22 | 0.26 |
| 5 | 154648481 | rs12514142 | -1.153 | -0.596 | 1.401 | -0.452 | 0.002 | 0.006 | 0.006 | 0 | 0.12 | 0.16 | 0.13 | 0.16 | 0.12 | 0.19 |
| 5 | 154650537 | rs12110234 | -2.079 | -0.648 | 1.459 | -0.898 | -0.001 | -0.03 | -0.03 | 0 | 0.36 | 0.53 | 0.28 | 0.18 | 0.32 | 0.38 |
| 5 | 154650883 | rs17117188 | -0.303 | -0.52 | 1.021 | -0.766 | 0 | -0.005 | -0.005 | 0.416 | 0.11 | 0.13 | 0.07 | 0.02 | 0.09 | 0.12 |
| 5 | 154668311 | rs12187859 | -0.316 | -0.53 | 1.017 | -0.772 | -0.003 | -0.008 | -0.008 | 0.289 | 0.09 | 0.12 | 0.07 | 0.02 | 0.08 | 0.12 |
| 5 | 154678693 | rs42673 | 1.494 | -0.69 | 1.21 | -0.848 | -0.007 | -0.029 | -0.029 | 11.005 | 0.76 | 0.68 | 0.8 | 0.84 | 0.82 | 0.76 |
| 5 | 154689635 | rs822470 | 1.506 | -0.336 | 0.022 | -0.845 | -0.089 | -0.155 | -0.155 | 0.043 | 0.32 | 0.01 | 0.38 | 0.46 | 0.33 | 0.33 |
| 5 | 154690482 | rs13173883 | 0.255 | -0.119 | 0.605 | -0.432 | -0.002 | -0.009 | -0.009 | 0 | 0.09 | 0.11 | 0.07 | 0.01 | 0.09 | 0.11 |
| 5 | 154691146 | rs17117230 | 0.416 | -0.113 | 0.634 | -0.433 | 0 | 0.002 | 0.002 | 0 | 0.08 | 0.09 | 0.07 | 0.02 | 0.09 | 0.12 |
| 5 | 154694954 | rs1428887 | 1.618 | -0.161 | 0.577 | -0.742 | -0.011 | 0.064 | 0.064 | 0 | 0.16 | 0.03 | 0.12 | 0 | 0.19 | 0.11 |
| 5 | 154699695 | rs154966 | 0.753 | 0.016 | 1.635 | 0.082 | -0.01 | -0.019 | -0.019 | 0 | 0.79 | 0.82 | 0.75 | 0.65 | 0.76 | 0.84 |
| 5 | 154703277 | rs697144 | 2.026 | -0.029 | 1.842 | 0.084 | 0.001 | 0.024 | 0.024 | 0 | 0.66 | 0.56 | 0.61 | 0.47 | 0.66 | 0.63 |
| 5 | 154709107 | rs12658964 | 0.471 | 0.103 | 1.841 | 0.173 | -0.006 | 0.026 | 0.026 | 0 | 0.09 | 0.07 | 0.18 | 0.35 | 0.14 | 0.1 |
| 5 | 154716929 | rs10039324 | -0.933 | 0.067 | 1.895 | 0.154 | -0.002 | -0.036 | -0.036 | 0 | 0.1 | 0.26 | 0.07 | 0.02 | 0.09 | 0.13 |
| 5 | 154719256 | rs245568 | 2.143 | 0.084 | 1.963 | 0.163 | 0.001 | 0.012 | 0.012 | 0 | 0.64 | 0.45 | 0.61 | 0.47 | 0.65 | 0.63 |
| 5 | 154721185 | rs702744 | 1.651 | 0.521 | 1.912 | 0.379 | 0.035 | 0.075 | 0.075 | 0 | 0.74 | 0.55 | 0.69 | 0.63 | 0.7 | 0.79 |
| 5 | 154738333 | rs10477131 | -1.699 | 0.249 | 2.382 | 0.858 | 0.038 | 0.214 | 0.214 | 0.03 | 0.16 | 0.45 | 0.25 | 0.36 | 0.21 | 0.14 |

| | | | | | | | | | | | | | | | | |
|---|---|---|---|---|---|---|---|---|---|---|---|---|---|---|---|---|
| 5 | 154746704 | rs10058975 | -0.878 | 0.03 | 2.206 | 0.977 | 0.022 | 0.218 | 0.218 | 0.02 | 0.12 | 0.31 | 0.2 | 0.35 | 0.13 | 0.11 |
| 5 | 154749572 | rs3929283 | -1.692 | 0.006 | 1.957 | 1.008 | -0.012 | 0.235 | 0.235 | 0.887 | 0.17 | 0.49 | 0.27 | 0.5 | 0.15 | 0.2 |
| 5 | 154757969 | rs9324810 | -0.495 | 0.394 | 0.853 | 0.896 | 0.009 | 0.114 | 0.114 | 0 | 0.11 | 0.14 | 0.18 | 0.34 | 0.13 | 0.11 |
| 5 | 154759008 | rs17117358 | -0.495 | 0.394 | 0.835 | 0.896 | 0.008 | 0.11 | 0.11 | 0 | 0.11 | 0.14 | 0.18 | 0.34 | 0.13 | 0.12 |
| 5 | 154762555 | rs822405 | 1.135 | 0.347 | 0.738 | 0.253 | 0.009 | -0.003 | -0.003 | 0 | 0.89 | 0.74 | 0.87 | 0.85 | 0.89 | 0.89 |
| 5 | 154771346 | rs13361944 | -1.4 | 0.282 | 0.531 | 0.039 | -0.002 | -0.046 | -0.046 | 0.532 | 0.13 | 0.22 | 0.07 | 0.06 | 0.1 | 0.05 |
| 5 | 154772970 | rs10072751 | -0.93 | 0.132 | -0.482 | 0.381 | -0.005 | -0.006 | -0.006 | 0 | 0.53 | 0.9 | 0.66 | 0.55 | 0.44 | 0.57 |
| 5 | 154773775 | rs17117397 | -0.439 | -0.205 | -0.243 | 0.06 | -0.004 | -0.048 | -0.048 | 1.295 | 0.14 | 0.17 | 0.07 | 0 | 0.1 | 0.11 |
| 5 | 154774815 | rs10054652 | -1.372 | -0.244 | -0.052 | 0.106 | 0.007 | 0.016 | 0.016 | 0 | 0.53 | 0.84 | 0.66 | 0.56 | 0.45 | 0.57 |
| 5 | 154778196 | rs9324815 | -0.447 | -0.247 | 0.264 | 0.238 | -0.001 | 0 | 0 | 0 | 0.38 | 0.59 | 0.58 | 0.57 | 0.34 | 0.47 |
| 5 | 154781106 | rs17117435 | -0.778 | -0.327 | 0.84 | 0.458 | 0.027 | 0.065 | 0.065 | 0.075 | 0.14 | 0.43 | 0.15 | 0.19 | 0.09 | 0.17 |
| 5 | 154794480 | rs4958803 | 1.551 | -0.138 | 0.986 | 0.447 | 0.003 | 0.019 | 0.019 | 0 | 0.47 | 0.81 | 0.64 | 0.56 | 0.45 | 0.57 |
| 5 | 154803797 | rs10515719 | -0.075 | -0.175 | 1.622 | 0.454 | -0.007 | 0.014 | 0.014 | 0 | 0.1 | 0.18 | 0.09 | 0.16 | 0.05 | 0.13 |
| 5 | 154805541 | rs10063605 | -1.082 | -0.127 | 1.984 | 0.445 | 0.009 | -0.004 | -0.004 | 0 | 0.43 | 0.64 | 0.56 | 0.4 | 0.39 | 0.44 |
| 5 | 154806830 | rs10065961 | -0.809 | -0.132 | 2.208 | 0.471 | -0.017 | 0.001 | 0.001 | 0 | 0.39 | 0.52 | 0.51 | 0.36 | 0.35 | 0.38 |
| 5 | 154810653 | rs13357562 | -0.312 | -0.033 | 1.802 | 0.452 | 0.02 | -0.014 | -0.014 | 0.26 | 0.09 | 0.19 | 0.08 | 0.04 | 0.1 | 0.08 |
| 5 | 154818499 | rs1471744 | -1.716 | -0.078 | 1.765 | 0.43 | -0.01 | -0.043 | -0.043 | 2.255 | 0.28 | 0.5 | 0.2 | 0.2 | 0.2 | 0.29 |
| 5 | 154819446 | rs11167724 | -1.576 | -0.178 | 1.303 | 0.402 | 0.007 | 0.017 | 0.017 | 0 | 0.52 | 0.89 | 0.66 | 0.56 | 0.44 | 0.56 |
| 5 | 154820449 | rs10515717 | -0.926 | -0.083 | 1.453 | 0.582 | 0.001 | -0.022 | -0.022 | 0.327 | 0.18 | 0.27 | 0.12 | 0.08 | 0.15 | 0.16 |
| 5 | 154821619 | rs11167725 | -0.926 | -0.083 | 1.453 | 0.582 | 0.001 | -0.022 | -0.022 | 0 | 0.18 | 0.27 | 0.12 | 0.08 | 0.15 | 0.16 |
| 5 | 154831511 | rs267004 | -1.67 | 1.143 | 0.724 | 0.125 | -0.021 | -0.013 | -0.013 | 0 | 0.61 | 0.91 | 0.77 | 0.6 | 0.6 | 0.64 |
| 5 | 154840829 | rs461355 | -0.794 | 1.208 | 0.902 | 0.032 | 0.012 | -0.052 | -0.052 | 0.183 | 0.27 | 0.34 | 0.23 | 0.03 | 0.32 | 0.23 |
| 5 | 154841170 | rs12652707 | 0.921 | 0.966 | 1.09 | 0.223 | 0.003 | -0.007 | -0.007 | 2.349 | 0.23 | 0.22 | 0.43 | 0.35 | 0.24 | 0.27 |
| 5 | 154841939 | rs10515716 | 0.992 | 0.966 | 1.06 | 0.223 | 0.014 | -0.039 | -0.039 | 0 | 0.22 | 0.15 | 0.42 | 0.35 | 0.24 | 0.27 |
| 5 | 154845771 | rs448373 | -0.52 | 1.039 | 0.392 | -0.259 | -0.009 | 0.042 | 0.042 | 0.018 | 0.42 | 0.33 | 0.31 | 0.2 | 0.47 | 0.39 |
| 5 | 154853461 | rs1148124 | -0.524 | 1.045 | 0.742 | -0.476 | 0.029 | 0.002 | 0.002 | 0 | 0.47 | 0.7 | 0.61 | 0.52 | 0.4 | 0.52 |
| 5 | 154866721 | rs17117712 | 1.098 | 1.096 | 0.67 | -0.437 | 0.002 | -0.007 | -0.007 | 0 | 0.09 | 0.08 | 0.03 | 0 | 0.05 | 0.08 |
| 5 | 154888314 | rs17117822 | 1.023 | 1.072 | 0.655 | -0.449 | 0.003 | -0.006 | -0.006 | 7.985 | 0.09 | 0.08 | 0.03 | 0 | 0.05 | 0.08 |
| 5 | 154889665 | rs9324826 | -0.082 | 1.07 | 0.9 | -0.455 | -0.006 | -0.04 | -0.04 | 0.905 | 0.15 | 0.25 | 0.1 | 0.04 | 0.12 | 0.15 |
| 5 | 154891822 | rs6884002 | 1.147 | 1.116 | 0.893 | -0.419 | 0.023 | 0.017 | 0.017 | 0 | 0.06 | 0.18 | 0.08 | 0.05 | 0.07 | 0.07 |
| 5 | 154894169 | rs1295242 | 1.124 | 1.041 | 1.014 | -0.457 | -0.01 | 0.037 | 0.037 | 2.018 | 0.25 | 0.34 | 0.49 | 0.43 | 0.25 | 0.34 |
| 5 | 154895569 | rs10515714 | 1.11 | 1.548 | 0.803 | -0.582 | 0.013 | 0.003 | 0.003 | 0 | 0.39 | 0.11 | 0.21 | 0.41 | 0.37 | 0.34 |
| 5 | 154895902 | rs1295233 | -0.054 | 1.814 | 0.809 | -0.563 | 0.031 | -0.064 | -0.064 | 1.397 | 0.78 | 0.7 | 0.8 | 0.97 | 0.73 | 0.8 |
| 5 | 154896454 | rs1295232 | -0.9 | 1.867 | 0.819 | -0.5 | 0.055 | -0.044 | -0.044 | 0 | 0.86 | 0.75 | 0.83 | 0.97 | 0.78 | 0.87 |
| 5 | 154906291 | rs10040300 | 0.572 | 1.932 | 0.777 | -0.54 | 0.005 | -0.016 | -0.016 | 0.1 | 0.16 | 0.18 | 0.17 | 0.03 | 0.22 | 0.13 |
| 5 | 154907699 | rs12515228 | 1.877 | 1.624 | 0.44 | -0.528 | 0.009 | -0.038 | -0.038 | 0 | 0.17 | 0.12 | 0.38 | 0.33 | 0.16 | 0.2 |
| 5 | 154910252 | rs17117884 | 1.233 | 1.881 | 0.561 | -0.471 | 0.003 | -0.002 | -0.002 | 0 | 0.08 | 0.04 | 0.02 | 0 | 0.05 | 0.08 |
| 5 | 154911215 | rs17117886 | 0.113 | 1.881 | 0.728 | -0.509 | -0.004 | -0.064 | -0.064 | 0.877 | 0.14 | 0.2 | 0.08 | 0 | 0.12 | 0.11 |
| 5 | 154913937 | rs10068643 | -1.295 | 1.752 | 0.633 | -0.264 | -0.004 | 0.014 | 0.014 | 0 | 0.61 | 0.82 | 0.78 | 0.6 | 0.61 | 0.66 |
| 5 | 154916462 | rs12516532 | 1.285 | 1.683 | 0.694 | -0.168 | 0.001 | -0.001 | -0.001 | 0 | 0.24 | 0.26 | 0.47 | 0.45 | 0.22 | 0.34 |
| 5 | 154917681 | rs12521335 | 0.608 | 1.699 | 0.723 | -0.169 | -0.004 | 0.005 | 0.005 | 0.737 | 0.31 | 0.34 | 0.5 | 0.53 | 0.27 | 0.38 |
| 5 | 154936212 | rs12655584 | 0.607 | 1.7 | 0.722 | -0.169 | -0.003 | 0.003 | 0.003 | 0.648 | 0.31 | 0.34 | 0.5 | 0.53 | 0.27 | 0.38 |
| 5 | 154942738 | rs12517952 | 1.237 | 1.665 | 0.673 | -0.193 | -0.005 | 0.014 | 0.014 | 0.857 | 0.24 | 0.29 | 0.48 | 0.49 | 0.23 | 0.37 |
| 5 | 154944885 | rs725883 | 0.044 | 1.665 | 0.762 | -0.219 | -0.015 | 0.013 | 0.013 | 0.105 | 0.37 | 0.52 | 0.56 | 0.53 | 0.33 | 0.41 |
| 5 | 154956642 | rs17117987 | -0.356 | 1.668 | 0.698 | -0.248 | -0.002 | -0.002 | -0.002 | 0 | 0.45 | 0.53 | 0.6 | 0.57 | 0.38 | 0.52 |
| 5 | 154971987 | rs7710000 | 1.314 | 0.958 | 0.978 | -0.448 | -0.001 | -0.004 | -0.004 | 0.138 | 0.07 | 0.07 | 0.04 | 0.08 | 0.04 | 0.04 |
| 5 | 154972936 | rs10515709 | 0.494 | 0.989 | 0.92 | -0.465 | -0.029 | 0.033 | 0.033 | 0 | 0.16 | 0.08 | 0.15 | 0.03 | 0.22 | 0.15 |
| 5 | 154975163 | rs10477135 | -1.454 | 0.945 | 0.981 | -0.489 | 0.005 | -0.034 | -0.034 | 0 | 0.39 | 0.53 | 0.31 | 0.15 | 0.38 | 0.33 |
| 5 | 154975827 | rs7731279 | -0.577 | 0.92 | 1.063 | -0.448 | 0.026 | -0.052 | -0.052 | 0 | 0.24 | 0.33 | 0.21 | 0.11 | 0.27 | 0.19 |
| 5 | 154980966 | rs6580169 | -0.519 | 0.901 | 0.792 | -0.447 | 0.033 | -0.049 | -0.049 | 0 | 0.22 | 0.36 | 0.23 | 0.07 | 0.29 | 0.21 |
| 5 | 154986042 | rs10072112 | 1.36 | 0.921 | 0.721 | -0.426 | 0.001 | -0.024 | -0.024 | 0 | 0.05 | 0.09 | 0.05 | 0 | 0.06 | 0.03 |
| 5 | 154988618 | rs6865003 | -1.075 | 0.909 | 0.696 | -0.423 | 0.003 | -0.005 | -0.005 | 0 | 0.32 | 0.63 | 0.33 | 0.24 | 0.34 | 0.36 |
| 5 | 154989490 | rs17118094 | 0.257 | 0.982 | 0.715 | -0.504 | 0.035 | -0.051 | -0.051 | 1.823 | 0.17 | 0.26 | 0.17 | 0.03 | 0.22 | 0.15 |
| 5 | 154992510 | rs6868624 | -0.934 | 0.768 | 0.461 | -0.375 | -0.02 | -0.004 | -0.004 | 0 | 0.53 | 0.8 | 0.73 | 0.5 | 0.5 | 0.54 |
| 5 | 155000148 | rs2431754 | -1.43 | 0.822 | 0.489 | -0.253 | -0.041 | -0.048 | -0.048 | 0 | 0.62 | 0.85 | 0.76 | 0.5 | 0.57 | 0.62 |
| 5 | 155001507 | rs7728980 | -0.41 | 0.637 | 1.235 | 0.187 | 0 | 0.029 | 0.029 | 0 | 0.16 | 0.38 | 0.16 | 0.2 | 0.12 | 0.21 |
| 5 | 155006293 | rs11957223 | -1.093 | 0.771 | 0.797 | 0.156 | 0.011 | -0.011 | -0.011 | 0 | 0.27 | 0.54 | 0.27 | 0.24 | 0.27 | 0.32 |
| 5 | 155010605 | rs3863202 | -0.514 | 0.469 | 1.084 | 0.316 | -0.004 | 0.01 | 0.01 | 0 | 0.16 | 0.42 | 0.15 | 0.17 | 0.12 | 0.17 |
| 5 | 155018265 | rs6885497 | 0.43 | 0.397 | 0.7 | 0.258 | 0.005 | -0.005 | -0.005 | 0 | 0.25 | 0.24 | 0.44 | 0.25 | 0.24 | 0.23 |
| 5 | 155023174 | rs2544746 | -1.102 | 0.437 | 0.687 | 0.245 | 0.001 | -0.014 | -0.014 | 0.364 | 0.28 | 0.54 | 0.26 | 0.24 | 0.26 | 0.32 |
| 5 | 155027916 | rs7726153 | -1.385 | 0.486 | 0.585 | 0.309 | 0.004 | 0.007 | 0.007 | 0 | 0.53 | 0.76 | 0.68 | 0.5 | 0.48 | 0.55 |
| 5 | 155036508 | rs6877767 | 0.87 | 0.404 | 0.48 | 0.26 | 0.025 | -0.027 | -0.027 | 0 | 0.22 | 0.14 | 0.41 | 0.26 | 0.19 | 0.21 |
| 5 | 155042560 | rs2552976 | 0.279 | 0.567 | 0.518 | 0.179 | 0.015 | -0.046 | -0.046 | 0 | 0.13 | 0.26 | 0.11 | 0.03 | 0.13 | 0.12 |
| 5 | 155046264 | rs2544756 | -1.202 | 0.544 | 0.339 | 0.222 | 0.006 | -0.018 | -0.018 | 0 | 0.31 | 0.57 | 0.27 | 0.24 | 0.3 | 0.34 |
| 5 | 155049089 | rs11167733 | -1.364 | 0.579 | 0.414 | 0.286 | 0.017 | -0.018 | -0.018 | 0 | 0.3 | 0.59 | 0.26 | 0.23 | 0.32 | 0.34 |
| 5 | 155057677 | rs4958416 | -1.369 | 0.444 | 0.318 | 0.329 | -0.002 | 0.013 | 0.013 | 0 | 0.52 | 0.82 | 0.67 | 0.5 | 0.49 | 0.55 |
| 5 | 155060719 | rs4631234 | -0.239 | 0.339 | 0.45 | 0.421 | -0.056 | 0.056 | 0.056 | 0.069 | 0.32 | 0.42 | 0.49 | 0.46 | 0.23 | 0.36 |
| 5 | 155060775 | rs4610514 | 0.604 | 0.38 | -0.004 | 0.455 | -0.016 | 0.009 | 0.009 | 0.724 | 0.22 | 0.24 | 0.4 | 0.27 | 0.17 | 0.21 |
| 5 | 155060978 | rs10477140 | -0.141 | 1.358 | -0.004 | 0.298 | 0.025 | -0.03 | -0.03 | 0 | 0.11 | 0.19 | 0.09 | 0.03 | 0.14 | 0.12 |

| | | | | | | | | | | | | | | | | |
|---|---|---|---|---|---|---|---|---|---|---|---|---|---|---|---|---|
| 5 | 155066004 | rs4310017 | -0.992 | 1.154 | -0.198 | 0.375 | 0.031 | -0.026 | -0.026 | 0 | 0.29 | 0.49 | 0.25 | 0.23 | 0.31 | 0.34 |
| 5 | 155066776 | rs4585515 | 0.074 | 0.566 | -0.842 | 0.549 | 0.022 | -0.019 | -0.019 | 0 | 0.72 | 0.82 | 0.57 | 0.73 | 0.76 | 0.78 |
| 5 | 155072951 | rs1363121 | 0.727 | 0.896 | -1.14 | 0.449 | -0.038 | -0.048 | -0.048 | 0 | 0.19 | 0.14 | 0.14 | 0.41 | 0.27 | 0.22 |
| 5 | 155074865 | rs1579282 | -0.021 | -0.13 | -0.74 | -0.1 | -0.001 | -0.002 | -0.002 | 0.011 | 0.35 | 0.4 | 0.22 | 0.43 | 0.34 | 0.33 |
| 5 | 155080747 | rs1862147 | -0.823 | -0.455 | -0.478 | -0.445 | 0.004 | 0.002 | 0.002 | 0.225 | 0.51 | 0.68 | 0.31 | 0.5 | 0.5 | 0.44 |
| 5 | 155087160 | rs716651 | -0.222 | -0.465 | -0.499 | -0.46 | -0.036 | -0.019 | -0.019 | 0.399 | 0.4 | 0.25 | 0.23 | 0.4 | 0.4 | 0.35 |
| 5 | 155088339 | rs11954814 | 1.191 | -0.462 | -0.54 | -0.438 | -0.022 | 0.029 | 0.029 | 0 | 0.06 | 0.01 | 0.02 | 0 | 0.06 | 0.02 |
| 5 | 155089792 | rs4532407 | -0.848 | -0.968 | -0.68 | -1.119 | 0 | -0.033 | -0.033 | 0 | 0.77 | 0.82 | 0.89 | 1 | 0.83 | 0.78 |
| 5 | 155102440 | rs3922737 | -0.6 | -1.175 | -0.347 | -1.163 | -0.026 | -0.16 | -0.16 | 2.399 | 0.75 | 0.59 | 0.86 | 1 | 0.83 | 0.78 |
| 5 | 155110509 | rs4958418 | 1.526 | -1.062 | -0.964 | -1.151 | -0.075 | -0.19 | -0.19 | 0 | 0.26 | 0.01 | 0.56 | 0.53 | 0.32 | 0.34 |
| 5 | 155114977 | rs10077093 | 2.154 | -1.112 | -0.846 | -1.095 | -0.103 | -0.117 | -0.117 | 0 | 0.18 | 0.03 | 0.14 | 0.38 | 0.3 | 0.22 |
| 5 | 155115300 | rs7715469 | 0.138 | -1.816 | -0.364 | -1.393 | -0.049 | -0.061 | -0.061 | 0.816 | 0.27 | 0.65 | 0.16 | 0.09 | 0.15 | 0.19 |
| 5 | 155120001 | rs9324841 | -0.385 | -1.142 | -0.33 | -1.155 | -0.008 | -0.068 | -0.068 | 0 | 0.77 | 0.78 | 0.89 | 1 | 0.84 | 0.78 |
| 5 | 155122182 | rs11167741 | 0.108 | -1.145 | -0.387 | -0.775 | 0.005 | -0.023 | -0.023 | 0 | 0.58 | 0.52 | 0.63 | 0.7 | 0.5 | 0.58 |
| 5 | 155124320 | rs6580180 | -0.101 | -1.225 | -0.357 | -0.922 | -0.004 | -0.068 | -0.068 | 0 | 0.86 | 0.81 | 0.94 | 1 | 0.89 | 0.83 |
| 5 | 155128570 | rs10044666 | -0.569 | -1.203 | -0.526 | -0.877 | 0 | -0.002 | -0.002 | 0.09 | 0.24 | 0.2 | 0.21 | 0.23 | 0.25 | 0.46 |
| 5 | 155129755 | rs7448810 | -0.117 | -1.664 | -0.931 | -0.832 | 0.072 | 0.038 | 0.038 | 0.097 | 0.48 | 0.22 | 0.41 | 0.64 | 0.63 | 0.72 |
| 5 | 155197081 | rs6556604 | -0.64 | -1.537 | -1.035 | -0.882 | 0.003 | -0.012 | -0.012 | 0 | 0.82 | 0.9 | 0.92 | 1 | 0.87 | 0.82 |
| 5 | 155202050 | rs12173249 | 0.768 | -1.95 | -1.104 | -1.046 | -0.016 | -0.014 | -0.014 | 0.076 | 0.41 | 0.34 | 0.29 | 0.47 | 0.48 | 0.35 |
| 5 | 155214166 | rs17524681 | 1.005 | -2.172 | -1.414 | -0.96 | 0.005 | 0.007 | 0.007 | 0 | 0.3 | 0.15 | 0.35 | 0.25 | 0.29 | 0.18 |
| 5 | 155236287 | rs17052889 | 1.947 | -1.88 | -1.854 | -0.849 | -0.019 | 0.009 | 0.009 | 0 | 0.1 | 0.02 | 0.14 | 0.08 | 0.12 | 0.23 |
| 5 | 155238565 | rs17052899 | 0.938 | -1.943 | -1.814 | -0.849 | 0.005 | -0.001 | -0.001 | 0 | 0.14 | 0.13 | 0.17 | 0.15 | 0.14 | 0.28 |
| 5 | 155239884 | rs1545773 | 2.314 | -1.857 | -2.029 | -0.888 | -0.132 | -0.139 | -0.139 | 0 | 0.19 | 0.02 | 0.18 | 0.44 | 0.37 | 0.27 |
| 5 | 155247903 | rs2619722 | 1.823 | -1.948 | -2.078 | -0.932 | -0.049 | -0.08 | -0.08 | 0 | 0.22 | 0.02 | 0.43 | 0.26 | 0.23 | 0.12 |
| 5 | 155253747 | rs1432816 | -0.285 | -2.158 | -2.002 | -1.167 | 0.002 | 0.056 | 0.056 | 0 | 0.73 | 0.88 | 0.79 | 0.91 | 0.75 | 0.6 |
| 5 | 155253880 | rs6875111 | 0.686 | -1.876 | -1.251 | -0.698 | -0.015 | 0.114 | 0.114 | 0.07 | 0.08 | 0.33 | 0.06 | 0.19 | 0.03 | 0.1 |
| 5 | 155273228 | rs2619728 | -1.527 | -1.926 | -1.837 | -0.308 | -0.018 | -0.019 | -0.019 | 0 | 0.62 | 0.78 | 0.74 | 0.57 | 0.5 | 0.55 |
| 5 | 155274700 | rs1432812 | 0.676 | -1.225 | -2.183 | -0.256 | -0.048 | -0.09 | -0.09 | 0 | 0.23 | 0.01 | 0.43 | 0.27 | 0.23 | 0.12 |
| 5 | 155281159 | rs11738514 | 0.216 | -1.306 | -1.029 | -0.681 | -0.016 | 0.011 | 0.011 | 0 | 0.1 | 0.03 | 0.14 | 0.08 | 0.12 | 0.22 |
| 5 | 155290198 | rs17052933 | -0.243 | -1.416 | -0.403 | -0.508 | 0.007 | 0.007 | 0.007 | 0.076 | 0.13 | 0.12 | 0.05 | 0.03 | 0.05 | 0.05 |
| 5 | 155292787 | rs7726529 | -0.883 | -0.837 | 0.279 | -0.335 | 0.03 | -0.011 | -0.011 | 0 | 0.11 | 0.33 | 0.07 | 0.04 | 0.09 | 0.16 |
| 5 | 155302582 | rs1432723 | -0.215 | -1.172 | -0.597 | -0.531 | 0 | 0.007 | 0.007 | 0 | 0.23 | 0.06 | 0.42 | 0.2 | 0.2 | 0.13 |
| 5 | 155305036 | rs244990 | 0.463 | -1.445 | -0.726 | -0.562 | 0.006 | -0.001 | -0.001 | 0 | 0.16 | 0.1 | 0.06 | 0.2 | 0.1 | 0.2 |
| 5 | 155310585 | rs244982 | -1.548 | -1.895 | -0.543 | -0.745 | -0.018 | -0.013 | -0.013 | 0 | 0.56 | 0.82 | 0.62 | 0.47 | 0.41 | 0.52 |
| 5 | 155316740 | rs244977 | -0.851 | -1.969 | -0.583 | -0.746 | 0.005 | -0.001 | -0.001 | 0 | 0.85 | 0.9 | 0.94 | 0.8 | 0.9 | 0.81 |
| 5 | 155317884 | rs9313417 | -0.832 | -1.991 | -0.533 | -0.751 | 0.022 | 0.032 | 0.032 | 0 | 0.39 | 0.69 | 0.55 | 0.28 | 0.31 | 0.32 |
| 5 | 155319175 | rs244976 | 1.405 | -1.859 | -0.881 | -0.708 | -0.067 | -0.075 | -0.075 | 0 | 0.43 | 0.07 | 0.37 | 0.52 | 0.59 | 0.48 |
| 5 | 155323385 | rs244975 | -0.867 | -1.999 | -0.685 | -0.977 | 0.004 | -0.012 | -0.012 | 0 | 0.86 | 0.97 | 0.95 | 0.8 | 0.91 | 0.82 |
| 5 | 155325538 | rs244972 | 0.895 | -2.4 | 0.273 | -1.064 | 0.005 | 0.005 | 0.005 | 0.16 | 0.58 | 0.4 | 0.45 | 0.62 | 0.71 | 0.68 |
| 5 | 155326812 | rs1864956 | -1.049 | -2.36 | 0.727 | -0.999 | -0.027 | -0.037 | -0.037 | 0 | 0.34 | 0.49 | 0.5 | 0.22 | 0.25 | 0.24 |
| 5 | 155328492 | rs7724912 | -0.14 | -2.29 | 0.673 | -0.973 | -0.006 | -0.027 | -0.027 | 0 | 0.12 | 0.31 | 0.06 | 0.05 | 0.09 | 0.17 |
| 5 | 155331080 | rs1432734 | 0.064 | -1.919 | 0.052 | -0.789 | 0.01 | 0.014 | 0.014 | 0 | 0.21 | 0.14 | 0.43 | 0.16 | 0.16 | 0.08 |
| 5 | 155333254 | rs3097821 | 1.286 | -2.246 | -0.231 | -1.07 | -0.039 | -0.036 | -0.036 | 0 | 0.32 | 0.17 | 0.33 | 0.51 | 0.51 | 0.41 |
| 5 | 155333533 | rs244955 | -0.852 | -2.893 | -0.715 | -1.252 | 0.017 | 0 | 0 | 0.087 | 0.82 | 0.88 | 0.91 | 0.8 | 0.88 | 0.78 |
| 5 | 155344802 | rs244953 | -0.882 | -2.866 | -0.701 | -1.259 | 0.015 | -0.005 | -0.005 | 0 | 0.82 | 0.88 | 0.91 | 0.78 | 0.87 | 0.78 |
| 5 | 155346383 | rs1368331 | 0.576 | -3.146 | -0.667 | -1.454 | 0.001 | -0.016 | -0.016 | 0 | 0.14 | 0.16 | 0.04 | 0.03 | 0.09 | 0.17 |
| 5 | 155348825 | rs17053001 | 0.571 | -3.298 | -0.775 | -1.672 | 0.002 | -0.017 | -0.017 | 0 | 0.13 | 0.13 | 0.06 | 0 | 0.09 | 0.03 |
| 5 | 155351357 | rs244959 | 1.376 | -3.423 | -0.662 | -1.734 | 0.002 | 0 | 0 | 0 | 0.42 | 0.34 | 0.36 | 0.53 | 0.55 | 0.44 |
| 5 | 155359757 | rs13178470 | 1.832 | -3.341 | -0.994 | -1.585 | -0.129 | -0.134 | -0.134 | 0 | 0.33 | 0.02 | 0.33 | 0.52 | 0.55 | 0.44 |
| 5 | 155361116 | rs17053011 | 0.934 | -2.315 | -0.368 | -0.949 | -0.035 | -0.064 | -0.064 | 0.006 | 0.12 | 0.01 | 0.42 | 0.23 | 0.15 | 0.15 |
| 5 | 155362564 | rs6556055 | -0.818 | -2.307 | -0.305 | -0.951 | 0.012 | -0.006 | -0.006 | 0 | 0.83 | 0.88 | 0.91 | 0.78 | 0.88 | 0.79 |
| 5 | 155367923 | rs4704876 | -0.974 | -1.682 | 0.033 | -0.858 | 0.009 | 0.009 | 0.009 | 0 | 0.27 | 0.2 | 0.47 | 0.26 | 0.25 | 0.32 |
| 5 | 155369080 | rs17053035 | 0.034 | -1.432 | 0.151 | -0.985 | -0.006 | -0.028 | -0.028 | 0.134 | 0.1 | 0.13 | 0.04 | 0 | 0.06 | 0.02 |
| 5 | 155370467 | rs17053040 | -0.999 | -0.884 | 0.002 | -0.497 | 0.001 | -0.017 | -0.017 | 0 | 0.15 | 0.15 | 0.04 | 0.03 | 0.09 | 0.18 |
| 5 | 155371739 | rs6894361 | -0.933 | -0.826 | -0.055 | -0.47 | -0.026 | -0.068 | -0.068 | 0 | 0.16 | 0.28 | 0.06 | 0.03 | 0.09 | 0.17 |
| 5 | 155373213 | rs6556104 | -1.023 | -0.554 | -0.242 | -0.163 | -0.011 | -0.011 | -0.011 | 0.345 | 0.12 | 0.21 | 0.08 | 0.07 | 0.09 | 0.15 |
| 5 | 155378761 | rs6556145 | -0.451 | -0.102 | 0.04 | 0.473 | 0.009 | 0.032 | 0.032 | 0 | 0.15 | 0.18 | 0.39 | 0.27 | 0.17 | 0.05 |
| 5 | 155379999 | rs10062650 | -0.58 | 0.308 | -0.63 | 0.49 | -0.007 | 0.049 | 0.049 | 0 | 0.56 | 0.75 | 0.7 | 0.67 | 0.54 | 0.59 |
| 5 | 155384557 | rs4704749 | -0.079 | 0.204 | 0.397 | 0.49 | 0.035 | 0.106 | 0.106 | 0 | 0.17 | 0.38 | 0.41 | 0.28 | 0.18 | 0.05 |
| 5 | 155394230 | rs17053082 | 0.282 | 0.081 | 0.224 | 0.521 | -0.003 | -0.003 | -0.003 | 0 | 0.08 | 0.08 | 0.05 | 0.03 | 0.05 | 0.1 |
| 5 | 155400562 | rs2310879 | -0.251 | 0.552 | 0.141 | 0.811 | 0 | 0 | 0 | 0 | 0.4 | 0.64 | 0.32 | 0.28 | 0.37 | 0.52 |
| 5 | 155410970 | rs17053102 | 0.671 | 0.167 | -0.705 | 0.39 | 0 | -0.058 | -0.058 | 0 | 0.17 | 0.03 | 0.06 | 0.25 | 0.1 | 0.14 |
| 5 | 155413224 | rs735331 | 0.922 | 0.195 | -0.612 | 0.401 | 0.006 | 0.01 | 0.01 | 0 | 0.08 | 0.03 | 0.04 | 0.01 | 0.05 | 0.09 |
| 5 | 155415603 | rs17053110 | 0.537 | 0.273 | -0.561 | 0.61 | -0.007 | -0.057 | -0.057 | 0 | 0.15 | 0.03 | 0.06 | 0.25 | 0.1 | 0.13 |
| 5 | 155415607 | rs10866706 | 0.451 | 0.163 | -0.039 | 0.657 | -0.002 | -0.004 | -0.004 | 2.021 | 0.28 | 0.45 | 0.26 | 0.25 | 0.32 | 0.41 |
| 5 | 155422838 | rs1864943 | 0.876 | 0.059 | -0.21 | 0.618 | -0.073 | -0.073 | -0.073 | 0 | 0.24 | 0.01 | 0.2 | 0.24 | 0.29 | 0.41 |
| 5 | 155427558 | rs10041446 | 0.471 | -0.056 | -0.089 | 0.629 | -0.012 | 0.002 | 0.002 | 1.487 | 0.26 | 0.45 | 0.25 | 0.24 | 0.26 | 0.4 |
| 5 | 155443062 | rs11750885 | -0.324 | 0.112 | -1.186 | 0.821 | 0.004 | -0.026 | -0.026 | 0.303 | 0.18 | 0.08 | 0.4 | 0.28 | 0.16 | 0.05 |
| 5 | 155448823 | rs256635 | 0.052 | 0.305 | -1.436 | 0.861 | 0.004 | -0.065 | -0.065 | 0 | 0.39 | 0.09 | 0.27 | 0.44 | 0.4 | 0.49 |

| | | | | | | | | | | | | | | | | |
|---|---|---|---|---|---|---|---|---|---|---|---|---|---|---|---|---|
| 5 | 155450424 | rs2116737 | -0.029 | 0.355 | -1.129 | 0.59 | 0.04 | 0.014 | 0.014 | 0.043 | 0.56 | 0.3 | 0.55 | 0.59 | 0.67 | 0.64 |
| 5 | 155453475 | rs3097811 | -0.243 | 0.532 | -0.77 | 0.51 | 0 | 0.001 | 0.001 | 0 | 0.57 | 0.54 | 0.66 | 0.73 | 0.53 | 0.58 |
| 5 | 155453603 | rs7721978 | -0.074 | 0.512 | -0.754 | 0.822 | -0.008 | 0.017 | 0.017 | 0 | 0.2 | 0.27 | 0.37 | 0.28 | 0.12 | 0.09 |
| 5 | 155455954 | rs256625 | 0.413 | 0.59 | -0.761 | 0.732 | -0.015 | -0.024 | -0.024 | 0.707 | 0.38 | 0.25 | 0.3 | 0.45 | 0.43 | 0.5 |
| 5 | 155456437 | rs6862821 | -0.624 | 0.183 | -0.473 | 0.605 | -0.01 | 0.017 | 0.017 | 0 | 0.19 | 0.28 | 0.37 | 0.26 | 0.11 | 0.09 |
| 5 | 155458803 | rs17537018 | -0.376 | 0.486 | -0.71 | -0.098 | -0.002 | -0.023 | -0.023 | 0.026 | 0.16 | 0.18 | 0.14 | 0 | 0.21 | 0.09 |
| 5 | 155460299 | rs12188985 | -0.295 | -0.212 | -0.573 | -0.192 | -0.016 | 0 | 0 | 0 | 0.18 | 0.21 | 0.23 | 0.15 | 0.12 | 0.16 |
| 5 | 155462260 | rs1347118 | -0.314 | -0.238 | -0.601 | -0.207 | -0.022 | 0 | 0 | 0 | 0.18 | 0.21 | 0.22 | 0.15 | 0.1 | 0.16 |
| 5 | 155468133 | rs17537158 | 0.012 | -0.134 | -0.706 | -0.335 | -0.033 | 0.042 | 0.042 | 0 | 0.15 | 0.06 | 0.13 | 0 | 0.21 | 0.1 |
| 5 | 155471714 | rs6556352 | -1.361 | -1.408 | -1.439 | 0.089 | -0.008 | -0.083 | -0.083 | 0 | 0.44 | 0.97 | 0.44 | 0.56 | 0.33 | 0.42 |
| 5 | 155500631 | rs12187915 | -0.393 | -1.323 | -0.559 | -0.13 | -0.021 | -0.005 | -0.005 | 0 | 0.19 | 0.24 | 0.23 | 0.14 | 0.12 | 0.18 |
| 5 | 155519933 | rs7702370 | 0.216 | -1.729 | -0.126 | 0.02 | -0.089 | 0.218 | 0.218 | 0.92 | 0.17 | 0.44 | 0.22 | 0.42 | 0.06 | 0.26 |
| 5 | 155520303 | rs891909 | 0.293 | -1.729 | -0.119 | 0.019 | -0.082 | 0.197 | 0.197 | 0.673 | 0.16 | 0.41 | 0.23 | 0.42 | 0.06 | 0.26 |
| 5 | 155528034 | rs1368321 | 0.054 | -1.754 | -0.374 | -0.552 | -0.013 | 0.025 | 0.025 | 0 | 0.24 | 0.19 | 0.16 | 0 | 0.26 | 0.12 |
| 5 | 155530223 | rs1549887 | -1.023 | -1.105 | -0.636 | -0.774 | -0.003 | 0 | 0 | 0 | 0.39 | 0.6 | 0.4 | 0.19 | 0.41 | 0.33 |
| 5 | 155532735 | rs1030659 | 0.561 | -1.074 | -0.951 | -0.748 | -0.035 | 0.068 | 0.068 | 0 | 0.22 | 0.11 | 0.16 | 0 | 0.26 | 0.12 |
| 5 | 155540311 | rs1368319 | -0.194 | -0.691 | -0.935 | -0.444 | 0.02 | -0.012 | -0.012 | 0 | 0.43 | 0.77 | 0.41 | 0.42 | 0.36 | 0.38 |
| 5 | 155564123 | rs256773 | 0.383 | -0.515 | -1.092 | -1.122 | -0.027 | -0.002 | -0.002 | 0 | 0.07 | 0.01 | 0.04 | 0.04 | 0.08 | 0.12 |
| 5 | 155566544 | rs256771 | -0.145 | -0.486 | -1.873 | -1.088 | -0.007 | 0.021 | 0.021 | 0 | 0.12 | 0.06 | 0.07 | 0.04 | 0.1 | 0.17 |
| 5 | 155566924 | rs256770 | -0.397 | -0.426 | -1.748 | -0.631 | -0.035 | -0.005 | -0.005 | 5.346 | 0.54 | 0.84 | 0.48 | 0.56 | 0.46 | 0.56 |
| 5 | 155573559 | rs10078170 | 0.243 | -0.073 | -1.677 | -1.243 | -0.1 | 0.182 | 0.182 | 1.19 | 0.65 | 0.93 | 0.7 | 0.95 | 0.64 | 0.8 |
| 5 | 155603358 | rs4526092 | 0.222 | -0.246 | -0.806 | -1.338 | -0.077 | 0.157 | 0.157 | 0.888 | 0.66 | 0.91 | 0.7 | 0.95 | 0.64 | 0.8 |
| 5 | 155615071 | rs991611 | 0.465 | -0.305 | -0.529 | -0.911 | -0.017 | 0.007 | 0.007 | 0.603 | 0.34 | 0.37 | 0.21 | 0.6 | 0.22 | 0.35 |
| 5 | 155615982 | rs4704988 | -0.459 | -0.033 | -0.824 | -0.806 | -0.008 | -0.015 | -0.015 | 0 | 0.71 | 0.81 | 0.54 | 0.65 | 0.61 | 0.56 |
| 5 | 155620464 | rs35426260 | -0.081 | 0.447 | -1.033 | -1.153 | -0.085 | 0.14 | 0.14 | 0.648 | 0.31 | 0.06 | 0.28 | 0.06 | 0.35 | 0.2 |
| 5 | 155642709 | rs249881 | -0.287 | 0.596 | -0.792 | -1.104 | -0.023 | 0.028 | 0.028 | 0 | 0.24 | 0.16 | 0.28 | 0.08 | 0.31 | 0.18 |
| 5 | 155651950 | rs447817 | 0.317 | 1.03 | -0.945 | -1.16 | -0.066 | -0.006 | -0.006 | 1.55 | 0.13 | 0.02 | 0.24 | 0.09 | 0.23 | 0.14 |
| 5 | 155666794 | rs249888 | -0.8 | 0.933 | -0.766 | -1.198 | -0.014 | 0.015 | 0.015 | 0 | 0.25 | 0.18 | 0.28 | 0.08 | 0.31 | 0.19 |
| 5 | 155673942 | rs4704996 | 1.024 | 0.144 | -1.022 | -1.32 | 0.014 | -0.065 | -0.065 | 1.293 | 0.27 | 0.13 | 0.17 | 0.62 | 0.23 | 0.3 |
| 5 | 155678185 | rs2033208 | 0.756 | -0.265 | -0.357 | -1.547 | -0.001 | 0.01 | 0.01 | 0.364 | 0.38 | 0.32 | 0.47 | 0.3 | 0.43 | 0.46 |
| 5 | 155686871 | rs12653936 | 0.505 | 0.625 | -0.012 | -1.719 | 0 | -0.001 | -0.001 | 0 | 0.13 | 0.1 | 0.22 | 0.09 | 0.18 | 0.14 |
| 5 | 155691924 | rs6866397 | -1.051 | 0.505 | 0.653 | -1.761 | 0.035 | -0.062 | -0.062 | 0 | 0.11 | 0.38 | 0.09 | 0 | 0.11 | 0.04 |
| 5 | 155703142 | rs10068407 | 0.253 | 0.372 | 0.567 | -1.795 | -0.035 | -0.037 | -0.037 | 0 | 0.07 | 0.19 | 0.03 | 0 | 0 | 0.03 |
| 5 | 155703396 | rs10065545 | -1.626 | 0.401 | 0.576 | -1.836 | -0.02 | -0.13 | -0.13 | 0 | 0.17 | 0.52 | 0.11 | 0 | 0.12 | 0.07 |
| 5 | 155710837 | rs10064971 | -1.278 | 0.288 | 0.423 | -1.835 | 0 | -0.059 | -0.059 | 0 | 0.32 | 0.6 | 0.32 | 0.09 | 0.31 | 0.22 |
| 5 | 155714159 | rs10050429 | 0.192 | 0.053 | 0.45 | -1.762 | -0.035 | -0.037 | -0.037 | 0 | 0.07 | 0.18 | 0.03 | 0 | 0 | 0.03 |
| 5 | 155714344 | rs10052410 | -1.295 | 0.202 | 0.107 | -1.791 | 0.004 | -0.051 | -0.051 | 0.517 | 0.37 | 0.6 | 0.37 | 0.09 | 0.38 | 0.22 |
| 5 | 155717149 | rs10077764 | -0.962 | -0.112 | 0.331 | -1.434 | -0.014 | -0.062 | -0.062 | 2.628 | 0.11 | 0.45 | 0.07 | 0 | 0.03 | 0.03 |
| 5 | 155729890 | rs1580927 | 0.14 | -0.072 | -1.054 | -1.673 | 0.032 | 0.135 | 0.135 | 0 | 0.44 | 0.23 | 0.32 | 0.18 | 0.37 | 0.46 |
| 5 | 155741238 | rs7737496 | -0.536 | -0.094 | -0.393 | -1.892 | -0.014 | -0.099 | -0.099 | 0.417 | 0.23 | 0.32 | 0.18 | 0.02 | 0.18 | 0.24 |
| 5 | 155742820 | rs6898100 | -1.249 | -0.372 | -0.123 | -1.848 | 0.006 | -0.009 | -0.009 | 1.885 | 0.44 | 0.6 | 0.28 | 0.11 | 0.3 | 0.28 |
| 5 | 155743144 | rs7710388 | -0.705 | -0.29 | -0.091 | -1.841 | -0.01 | 0.003 | 0.003 | 0.259 | 0.08 | 0.2 | 0.03 | 0.07 | 0.04 | 0.02 |
| 5 | 155752007 | rs10057826 | -0.416 | -0.423 | -0.27 | -1.847 | 0.012 | 0.19 | 0.19 | 0 | 0.34 | 0.11 | 0.21 | 0.03 | 0.27 | 0.25 |
| 5 | 155753058 | rs10058119 | 1.533 | 0.162 | -0.328 | -1.66 | 0.002 | 0.005 | 0.005 | 0 | 0.45 | 0.39 | 0.48 | 0.79 | 0.44 | 0.54 |
| 5 | 155756299 | rs11744154 | 1.546 | 0.149 | -0.087 | -1.653 | 0.005 | 0.033 | 0.033 | 0 | 0.45 | 0.35 | 0.48 | 0.79 | 0.44 | 0.53 |
| 5 | 155757872 | rs13358340 | 1.241 | 0.08 | -0.267 | -1.645 | 0.004 | 0.01 | 0.01 | 0.148 | 0.09 | 0.08 | 0.1 | 0.03 | 0.06 | 0.17 |
| 5 | 155759846 | rs1394606 | 1.239 | 0.059 | -0.336 | -1.669 | 0.012 | 0.122 | 0.122 | 0.042 | 0.19 | 0.02 | 0.07 | 0 | 0.13 | 0.08 |
| 5 | 155763769 | rs10061194 | -0.357 | 0.38 | 0.382 | -1.386 | -0.038 | -0.035 | -0.035 | 2.942 | 0.16 | 0.36 | 0.13 | 0.11 | 0.09 | 0.22 |
| 5 | 155764666 | rs10039835 | -0.269 | 0.395 | 0.35 | -1.379 | -0.032 | -0.029 | -0.029 | 0 | 0.16 | 0.31 | 0.13 | 0.11 | 0.09 | 0.21 |
| 5 | 155774155 | rs4345305 | -1.145 | 0.774 | 0.3 | -1.325 | -0.037 | 0.001 | 0.001 | 0 | 0.89 | 0.95 | 0.73 | 0.91 | 0.73 | 0.84 |
| 5 | 155778744 | rs10037772 | -0.412 | 0.765 | 0.305 | -1.344 | -0.032 | 0.016 | 0.016 | 0 | 0.85 | 0.9 | 0.68 | 0.91 | 0.65 | 0.83 |
| 5 | 155779301 | rs1504946 | 1.581 | 0.424 | 0.726 | -1.556 | 0.016 | 0.109 | 0.109 | 0 | 0.46 | 0.24 | 0.46 | 0.79 | 0.45 | 0.54 |
| 5 | 155779927 | rs10059872 | 0.686 | 0.365 | 0.83 | -1.57 | -0.012 | -0.05 | -0.05 | 0 | 0.12 | 0.28 | 0.11 | 0.03 | 0.09 | 0.2 |
| 5 | 155780885 | rs17053465 | 0.155 | 0.361 | 0.783 | -1.555 | -0.04 | -0.069 | -0.069 | 0 | 0.26 | 0.37 | 0.11 | 0.08 | 0.12 | 0.09 |
| 5 | 155784924 | rs11743063 | 0.908 | 0.453 | 0.812 | -1.517 | 0.002 | 0.026 | 0.026 | 0 | 0.53 | 0.57 | 0.51 | 0.87 | 0.47 | 0.58 |
| 5 | 155784983 | rs4704795 | 0.585 | 0.423 | 0.784 | -1.511 | 0.016 | 0.124 | 0.124 | 0.854 | 0.24 | 0.05 | 0.11 | 0 | 0.17 | 0.06 |
| 5 | 155789169 | rs7714201 | 0.24 | 0.434 | 0.823 | -1.498 | 0.003 | -0.038 | -0.038 | 0 | 0.27 | 0.25 | 0.14 | 0 | 0.19 | 0.08 |
| 5 | 155790030 | rs10043401 | 0.449 | 0.533 | 0.79 | -1.335 | -0.005 | -0.056 | -0.056 | 0 | 0.12 | 0.38 | 0.12 | 0.04 | 0.09 | 0.2 |
| 5 | 155799466 | rs7732608 | 0.624 | 0.44 | 0.56 | -1.366 | -0.001 | -0.012 | -0.012 | 0.234 | 0.58 | 0.42 | 0.57 | 0.83 | 0.53 | 0.75 |
| 5 | 155800253 | rs9313900 | 0.627 | 0.435 | 0.27 | -1.38 | -0.008 | -0.03 | -0.03 | 0 | 0.12 | 0.18 | 0.11 | 0.03 | 0.08 | 0.19 |
| 5 | 155804271 | rs2055609 | 0.889 | 0.832 | 0.435 | -0.267 | -0.009 | 0.026 | 0.026 | 0 | 0.44 | 0.13 | 0.33 | 0.61 | 0.45 | 0.51 |
| 5 | 155810954 | rs140616 | 1.307 | 2.141 | 1.677 | 1.047 | 0.001 | -0.002 | -0.002 | 0 | 0.65 | 0.4 | 0.58 | 0.71 | 0.6 | 0.58 |
| 5 | 155813177 | rs7723660 | -0.049 | 2.08 | 1.542 | 1.186 | -0.005 | 0.003 | 0.003 | 0 | 0.16 | 0.21 | 0.12 | 0.18 | 0.11 | 0.22 |
| 5 | 155813531 | rs4438860 | -0.049 | 2.067 | 1.499 | 1.186 | 0.005 | -0.002 | -0.002 | 0 | 0.16 | 0.13 | 0.11 | 0.18 | 0.09 | 0.22 |
| 5 | 155822589 | rs6869537 | -0.112 | 2.218 | 1.429 | 1.181 | 0 | 0.002 | 0.002 | 0 | 0.17 | 0.21 | 0.14 | 0.19 | 0.14 | 0.25 |
| 5 | 155824774 | rs4454042 | 0.254 | 2.494 | 1.505 | 1.269 | 0.015 | 0.009 | 0.009 | 0.01 | 0.87 | 0.83 | 0.6 | 0.73 | 0.73 | 0.79 |
| 5 | 155834919 | rs7731517 | -0.229 | 2.42 | 1.391 | 1.229 | -0.003 | 0.002 | 0.002 | 0 | 0.18 | 0.26 | 0.13 | 0.19 | 0.13 | 0.24 |
| 5 | 155859368 | rs6556615 | -0.006 | 2.556 | 1.147 | 1.547 | -0.014 | -0.016 | -0.016 | 0.744 | 0.88 | 0.94 | 0.62 | 0.68 | 0.71 | 0.8 |
| 5 | 155862225 | rs905788 | -0.829 | 2.296 | 1.474 | 1.541 | -0.04 | -0.013 | -0.013 | 0.863 | 0.25 | 0.34 | 0.14 | 0.22 | 0.12 | 0.26 |

| | | | | | | | | | | | | | | | | |
|---|---|---|---|---|---|---|---|---|---|---|---|---|---|---|---|---|
| 5 | 155863846 | rs1504931 | 1.658 | 2.469 | 1.234 | 1.443 | 0.004 | 0.009 | 0.009 | 0 | 0.59 | 0.5 | 0.42 | 0.45 | 0.49 | 0.54 |
| 5 | 155868442 | rs2312182 | 0.794 | 2.501 | 1.055 | 1.476 | 0.103 | 0.068 | 0.068 | 0 | 0.82 | 0.65 | 0.54 | 0.65 | 0.65 | 0.8 |
| 5 | 155875529 | rs6883722 | -1.352 | 2.144 | 0.5 | 1.244 | 0.021 | -0.005 | -0.005 | 0.186 | 0.29 | 0.78 | 0.2 | 0.36 | 0.19 | 0.28 |
| 5 | 155898139 | rs7721082 | 0.064 | 2.678 | 0.379 | 1.765 | -0.078 | -0.07 | -0.07 | 0 | 0.89 | 0.99 | 0.76 | 0.77 | 0.77 | 0.84 |
| 5 | 155907487 | rs4704804 | 1.248 | 2.921 | 0.928 | 1.67 | 0.032 | 0.013 | 0.013 | 0.035 | 0.67 | 0.43 | 0.55 | 0.58 | 0.55 | 0.6 |
| 5 | 155908785 | rs980149 | 1.34 | 2.948 | 0.925 | 1.788 | 0.019 | 0.027 | 0.027 | 0.122 | 0.67 | 0.42 | 0.53 | 0.44 | 0.5 | 0.6 |
| 5 | 155909073 | rs17638584 | 0.027 | 3.003 | 0.708 | 1.439 | -0.076 | 0.004 | 0.004 | 0 | 0.17 | 0.01 | 0.16 | 0.15 | 0.23 | 0.23 |
| 5 | 155912152 | rs2055613 | 1.289 | 2.66 | 1.429 | 1.772 | 0.01 | 0.002 | 0.002 | 0 | 0.65 | 0.65 | 0.59 | 0.59 | 0.56 | 0.61 |
| 5 | 155914451 | rs4307078 | -0.409 | 2.632 | 1.41 | 1.621 | 0.025 | 0.019 | 0.019 | 0 | 0.2 | 0.48 | 0.18 | 0.21 | 0.21 | 0.21 |
| 5 | 155914878 | rs2312185 | 1.404 | 2.409 | 1.389 | 1.682 | -0.025 | -0.021 | -0.021 | 0.038 | 0.62 | 0.28 | 0.52 | 0.38 | 0.49 | 0.61 |
| 5 | 155922368 | rs256841 | -0.663 | 2.019 | 1.644 | 1.757 | 0.028 | 0.067 | 0.067 | 0 | 0.16 | 0.22 | 0.27 | 0.42 | 0.25 | 0.17 |
| 5 | 155926544 | rs256821 | -1.378 | 2.115 | 1.442 | 1.488 | 0.02 | 0 | 0 | 0 | 0.46 | 0.61 | 0.51 | 0.61 | 0.6 | 0.42 |
| 5 | 155927543 | rs6880345 | 0.194 | 2.226 | 0.809 | 1.476 | -0.13 | 0.049 | 0.049 | 0 | 0.25 | 0.01 | 0.2 | 0.15 | 0.32 | 0.24 |
| 5 | 155932280 | rs256822 | -0.262 | 2.302 | 0.924 | 1.592 | -0.068 | -0.083 | -0.083 | 0 | 0.14 | 0.01 | 0.22 | 0.25 | 0.2 | 0.16 |
| 5 | 155933860 | rs256824 | -0.529 | 2.256 | 1.203 | 1.605 | 0.074 | 0.12 | 0.12 | 0 | 0.16 | 0.3 | 0.29 | 0.42 | 0.27 | 0.17 |
| 5 | 155938322 | rs1368299 | -0.376 | 2.036 | 0.723 | 1.196 | -0.01 | 0 | 0 | 0 | 0.29 | 0.22 | 0.25 | 0.35 | 0.38 | 0.28 |
| 5 | 155957439 | rs7715923 | -0.903 | 1.64 | 0.767 | 1.104 | 0.005 | -0.014 | -0.014 | 0 | 0.4 | 0.5 | 0.45 | 0.38 | 0.5 | 0.42 |
| 5 | 155960089 | rs253604 | -0.874 | 1.771 | 0.817 | 1.167 | 0.007 | -0.032 | -0.032 | 0.271 | 0.4 | 0.47 | 0.44 | 0.2 | 0.48 | 0.41 |
| 5 | 155966087 | rs253602 | -1.339 | 1.442 | 0.988 | 1.331 | 0.037 | 0.02 | 0.02 | 0 | 0.49 | 0.73 | 0.56 | 0.46 | 0.57 | 0.51 |
| 5 | 155967030 | rs6885454 | 0.066 | 1.589 | 1.441 | 1.257 | 0.04 | -0.008 | -0.008 | 0.153 | 0.28 | 0.42 | 0.23 | 0.34 | 0.34 | 0.25 |
| 5 | 155968271 | rs253598 | -0.931 | 1.567 | 1.491 | 1.277 | 0.008 | -0.026 | -0.026 | 0.481 | 0.4 | 0.52 | 0.44 | 0.35 | 0.49 | 0.4 |
| 5 | 155970607 | rs253594 | -1.736 | 0.907 | 1.089 | 1.308 | 0.023 | 0.016 | 0.016 | 0.106 | 0.58 | 0.87 | 0.58 | 0.67 | 0.61 | 0.61 |
| 5 | 155975264 | rs7717533 | -0.873 | 1.239 | 1.33 | 0.939 | -0.002 | -0.001 | -0.001 | 0 | 0.42 | 0.62 | 0.46 | 0.2 | 0.5 | 0.41 |
| 5 | 155982551 | rs2312187 | -0.948 | 1.227 | 1.279 | 0.944 | -0.002 | 0.01 | 0.01 | 0 | 0.42 | 0.64 | 0.46 | 0.39 | 0.5 | 0.43 |
| 5 | 155986366 | rs447938 | 0.001 | 1.242 | 1.093 | 0.927 | -0.006 | -0.005 | -0.005 | 0.816 | 0.12 | 0.1 | 0.2 | 0.19 | 0.15 | 0.16 |
| 5 | 156000686 | rs10052818 | 0.25 | 1.262 | 1.257 | 0.923 | -0.005 | 0.004 | 0.004 | 0 | 0.27 | 0.23 | 0.21 | 0.15 | 0.32 | 0.24 |
| 5 | 156013831 | rs7710178 | -0.624 | 1.03 | 1.368 | 1.154 | 0.014 | -0.025 | -0.025 | 0 | 0.13 | 0.23 | 0.22 | 0.05 | 0.16 | 0.16 |
| 5 | 156019919 | rs10515738 | -1.83 | 0.98 | 1.395 | 1.204 | 0.032 | 0.052 | 0.052 | 0 | 0.22 | 0.53 | 0.36 | 0.29 | 0.25 | 0.27 |
| 5 | 156039751 | rs2194036 | -0.443 | 0.995 | 1.227 | 1.295 | 0.002 | -0.01 | -0.01 | 0.471 | 0.07 | 0.05 | 0.1 | 0.06 | 0.06 | 0.09 |
| 5 | 156052194 | rs1363743 | -1.786 | 0.989 | 1.25 | 1.231 | 0.002 | 0.009 | 0.009 | 0 | 0.2 | 0.3 | 0.22 | 0.29 | 0.22 | 0.24 |
| 5 | 156060370 | rs11743091 | -0.074 | 1.005 | 1.134 | 1.1 | -0.008 | 0.006 | 0.006 | 0.266 | 0.1 | 0.08 | 0.08 | 0.01 | 0.14 | 0.11 |
| 5 | 156065646 | rs6877490 | -1.743 | 0.937 | 1.184 | 1.14 | -0.003 | 0.011 | 0.011 | 0 | 0.12 | 0.24 | 0.14 | 0.1 | 0.1 | 0.13 |
| 5 | 156067682 | rs32060 | -1.456 | 0.926 | 1.494 | 1.188 | 0.054 | 0.105 | 0.105 | 0 | 0.16 | 0.44 | 0.18 | 0.25 | 0.2 | 0.2 |
| 5 | 156069442 | rs32058 | -2.5 | 0.845 | 1.44 | 1.157 | 0.01 | 0.033 | 0.033 | 0 | 0.24 | 0.57 | 0.24 | 0.28 | 0.24 | 0.25 |
| 5 | 156082997 | rs281046 | 0.738 | 0.859 | 1.521 | 1.123 | 0.021 | -0.053 | -0.053 | 0 | 0.87 | 0.77 | 0.9 | 0.99 | 0.86 | 0.88 |
| 5 | 156116412 | rs39926 | 1.567 | 0.763 | 1.418 | 1.172 | 0.024 | 0.037 | 0.037 | 0 | 0.82 | 0.56 | 0.87 | 0.77 | 0.83 | 0.83 |
| 5 | 156116642 | rs1353249 | -1.325 | 0.794 | 1.189 | 1.218 | -0.002 | 0.014 | 0.014 | 0 | 0.06 | 0.1 | 0.11 | 0.06 | 0.07 | 0.08 |
| 5 | 156121269 | rs32094 | 0.699 | 0.825 | 1.093 | 1.157 | 0.033 | 0.047 | 0.047 | 0.034 | 0.88 | 0.73 | 0.89 | 0.77 | 0.84 | 0.86 |
| 5 | 156133899 | rs157341 | -0.349 | 0.865 | 0.605 | 1.03 | -0.019 | 0.022 | 0.022 | 0.316 | 0.91 | 0.97 | 0.92 | 0.99 | 0.87 | 0.9 |
| 5 | 156141917 | rs282474 | 1.296 | 0.874 | 0.067 | 1.107 | -0.084 | -0.025 | -0.025 | 0 | 0.42 | 0.08 | 0.52 | 0.37 | 0.45 | 0.47 |
| 5 | 156144783 | rs157676 | -1.771 | 0.759 | 0.732 | 0.956 | 0.01 | 0.038 | 0.038 | 0 | 0.23 | 0.54 | 0.23 | 0.29 | 0.24 | 0.23 |
| 5 | 156151046 | rs459055 | 1.134 | 0.519 | 1.129 | 0.686 | 0.026 | -0.025 | -0.025 | 0.917 | 0.83 | 0.63 | 0.84 | 0.9 | 0.83 | 0.84 |
| 5 | 156153399 | rs282466 | 1.263 | 0.528 | 0.746 | 0.704 | 0.014 | 0.02 | 0.02 | 0 | 0.46 | 0.21 | 0.51 | 0.57 | 0.42 | 0.53 |
| 5 | 156154739 | rs462058 | -0.141 | 0.745 | 0.686 | 0.915 | -0.013 | 0.012 | 0.012 | 0 | 0.89 | 0.94 | 0.92 | 0.94 | 0.87 | 0.91 |
| 5 | 156158356 | rs284444 | 1.043 | 0.642 | 1.053 | 0.946 | 0.03 | 0.018 | 0.018 | 0 | 0.84 | 0.62 | 0.84 | 0.85 | 0.82 | 0.85 |
| 5 | 156164978 | rs284442 | 1.418 | 0.616 | 1.104 | 0.984 | 0.026 | 0.056 | 0.056 | 0 | 0.83 | 0.53 | 0.83 | 0.82 | 0.83 | 0.83 |
| 5 | 156172418 | rs284439 | 1.401 | 0.692 | 1.169 | 0.991 | 0.002 | 0.034 | 0.034 | 0.285 | 0.85 | 0.57 | 0.87 | 0.85 | 0.87 | 0.87 |
| 5 | 156174011 | rs381258 | -0.533 | 0.727 | 0.93 | 0.987 | -0.029 | -0.006 | -0.006 | 0 | 0.92 | 0.99 | 0.96 | 0.93 | 0.92 | 0.93 |
| 5 | 156187928 | rs284445 | 0.603 | 0.063 | 0.476 | 0.433 | -0.146 | 0.024 | 0.024 | 0.058 | 0.31 | 0.02 | 0.25 | 0.23 | 0.37 | 0.24 |
| 5 | 156188019 | rs1389818 | -1.905 | 0.067 | 0.621 | 0.444 | 0.06 | 0.149 | 0.149 | 0.034 | 0.06 | 0.4 | 0.09 | 0.08 | 0.04 | 0.06 |
| 5 | 156191214 | rs1845479 | -1.716 | 0.067 | 0.585 | 0.444 | 0.069 | 0.161 | 0.161 | 0 | 0.06 | 0.38 | 0.09 | 0.08 | 0.04 | 0.06 |
| 5 | 156192264 | rs11953631 | -2.315 | 0.063 | 0.518 | 0.45 | 0.035 | 0.186 | 0.186 | 0.164 | 0.08 | 0.45 | 0.1 | 0.12 | 0.04 | 0.1 |
| 5 | 156226550 | rs2436424 | 0.545 | 0.137 | 0.171 | 0.439 | -0.133 | 0.04 | 0.04 | 0.043 | 0.31 | 0.05 | 0.27 | 0.23 | 0.37 | 0.23 |
| 5 | 156273481 | rs4704810 | 0.622 | -0.295 | 0.158 | 0.703 | -0.126 | -0.121 | -0.121 | 1.302 | 0.34 | 0.02 | 0.48 | 0.38 | 0.39 | 0.47 |
| 5 | 156279593 | rs10054293 | -3.037 | 0.071 | 1.226 | 0.245 | 0.012 | 0.044 | 0.044 | 0 | 0.06 | 0.13 | 0.08 | 0.11 | 0.07 | 0.14 |
| 5 | 156280556 | rs10060710 | -3.114 | 0.067 | 1.232 | 0.22 | 0.012 | 0.041 | 0.041 | 0 | 0.06 | 0.13 | 0.08 | 0.11 | 0.07 | 0.14 |
| 5 | 156283415 | rs3909320 | -3.221 | 0.12 | 1.333 | 0.245 | 0.118 | 0.194 | 0.194 | 0.027 | 0.06 | 0.35 | 0.1 | 0.11 | 0.07 | 0.14 |
| 5 | 156295198 | rs17570186 | -0.269 | 0.142 | 0.974 | 0.106 | -0.025 | 0.019 | 0.019 | 0 | 0.1 | 0.03 | 0.08 | 0.04 | 0.09 | 0.07 |
| 5 | 156295372 | rs17053996 | -1.145 | 0.124 | 1.142 | 0.105 | 0 | -0.001 | -0.001 | 0.396 | 0.14 | 0.21 | 0.17 | 0.11 | 0.13 | 0.19 |
| 5 | 156312678 | rs12187637 | -1.251 | 0.175 | 0.446 | 0.129 | 0.008 | 0.002 | 0.002 | 0 | 0.15 | 0.3 | 0.17 | 0.11 | 0.13 | 0.2 |
| 5 | 156323717 | rs4704815 | -0.676 | 0.426 | -0.49 | 0.065 | -0.015 | 0.011 | 0.011 | 0 | 0.12 | 0.04 | 0.15 | 0.07 | 0.13 | 0.16 |
| 5 | 156336987 | rs12187482 | -0.373 | 0.223 | -0.575 | -0.061 | 0.005 | 0.025 | 0.025 | 0.178 | 0.21 | 0.11 | 0.32 | 0.14 | 0.23 | 0.15 |
| 5 | 156345735 | rs10070224 | -0.842 | -0.042 | 0.439 | -0.023 | -0.018 | -0.022 | -0.022 | 0 | 0.25 | 0.53 | 0.35 | 0.16 | 0.23 | 0.18 |
| 5 | 156351347 | rs1345618 | -0.641 | 0.053 | 1.077 | -0.126 | -0.003 | -0.04 | -0.04 | 0 | 0.26 | 0.32 | 0.33 | 0.11 | 0.27 | 0.23 |
| 5 | 156351674 | rs1345616 | -0.731 | 0.056 | 1.134 | -0.123 | -0.005 | -0.059 | -0.059 | 0 | 0.27 | 0.41 | 0.34 | 0.11 | 0.27 | 0.23 |
| 5 | 156366181 | rs7700944 | -0.883 | 0.037 | 1.126 | -0.1 | -0.005 | -0.014 | -0.014 | 0 | 0.28 | 0.32 | 0.34 | 0.16 | 0.27 | 0.28 |
| 5 | 156390297 | rs6882076 | -0.14 | -0.325 | 0.428 | -0.505 | 0.042 | 0.043 | 0.043 | 0 | 0.52 | 0.32 | 0.7 | 0.73 | 0.65 | 0.7 |
| 5 | 156398253 | rs6883317 | 1.273 | -0.891 | 0.066 | -0.947 | 0.002 | 0.003 | 0.003 | 0 | 0.2 | 0.23 | 0.15 | 0.16 | 0.15 | 0.16 |
| 5 | 156405784 | rs10059560 | 0.374 | -1.07 | 0.052 | -1.085 | -0.004 | -0.001 | -0.001 | 0 | 0.27 | 0.34 | 0.17 | 0.21 | 0.18 | 0.22 |

| | | | | | | | | | | | | | | | | |
|---|---|---|---|---|---|---|---|---|---|---|---|---|---|---|---|---|
| 5 | 156418790 | rs1501910 | -1.472 | -0.278 | 0.246 | -0.831 | -0.006 | 0.016 | 0.016 | 0 | 0.9 | 0.95 | 0.92 | 0.99 | 0.84 | 0.96 |
| 5 | 156419765 | rs17054104 | 1.468 | 0.619 | -0.011 | -0.427 | -0.038 | 0.03 | 0.03 | 0 | 0.06 | 0 | 0.24 | 0.5 | 0.12 | 0.21 |
| 5 | 156422887 | rs13360569 | 0.243 | -0.264 | 0.114 | -1.133 | 0 | -0.006 | -0.006 | 0 | 0.64 | 0.44 | 0.56 | 0.83 | 0.54 | 0.72 |
| 5 | 156432018 | rs2102151 | 0.624 | -0.57 | 0.296 | -1.198 | -0.035 | -0.02 | -0.02 | 0 | 0.37 | 0.58 | 0.19 | 0.21 | 0.17 | 0.22 |
| 5 | 156432215 | rs2088917 | -1.619 | -0.42 | 0.059 | -1.164 | -0.002 | 0.004 | 0.004 | 0 | 0.89 | 0.93 | 0.92 | 0.99 | 0.84 | 0.96 |
| 5 | 156449748 | rs7707445 | 0.911 | -0.598 | 0.239 | -1.214 | 0.024 | 0.017 | 0.017 | 0 | 0.33 | 0.31 | 0.17 | 0.2 | 0.17 | 0.22 |
| 5 | 156460095 | rs2277025 | 0.955 | -0.346 | -0.236 | -1.138 | -0.113 | -0.112 | -0.112 | 0 | 0.28 | 0.05 | 0.37 | 0.61 | 0.37 | 0.5 |
| 5 | 156464242 | rs2036402 | 0.85 | -0.269 | -0.273 | -1.324 | -0.069 | -0.001 | -0.001 | 0.164 | 0.21 | 0.04 | 0.31 | 0.12 | 0.29 | 0.2 |
| 5 | 156469783 | rs2270926 | 1.572 | -0.426 | -0.268 | -1.396 | 0.048 | 0.051 | 0.051 | 0.117 | 0.28 | 0.17 | 0.14 | 0.15 | 0.16 | 0.13 |
| 5 | 156469808 | rs13173581 | 0.477 | -0.433 | -0.081 | -1.399 | -0.001 | -0.001 | -0.001 | 0.153 | 0.24 | 0.25 | 0.35 | 0.15 | 0.3 | 0.24 |
| 5 | 156470401 | rs10038271 | 1.427 | -0.286 | 0.012 | -1.27 | 0.048 | 0.051 | 0.051 | 0.117 | 0.28 | 0.17 | 0.14 | 0.15 | 0.16 | 0.13 |
| 5 | 156475398 | rs7706174 | 0.995 | 0.224 | 0.09 | -1.143 | -0.034 | -0.026 | -0.026 | 1.428 | 0.34 | 0.39 | 0.19 | 0.2 | 0.19 | 0.19 |
| 5 | 156477540 | rs1393209 | -1.434 | 0.402 | -0.285 | -1.099 | -0.003 | 0.01 | 0.01 | 0 | 0.89 | 0.93 | 0.92 | 0.99 | 0.84 | 0.96 |
| 5 | 156479204 | rs2279804 | -0.036 | 0.237 | -0.41 | -1.243 | -0.004 | -0.002 | -0.002 | 0 | 0.65 | 0.65 | 0.55 | 0.35 | 0.5 | 0.46 |
| 5 | 156479424 | rs1553317 | -0.752 | 0.724 | -0.223 | -1.267 | -0.001 | -0.005 | -0.005 | 0 | 0.88 | 0.87 | 0.92 | 0.99 | 0.84 | 0.96 |
| 5 | 156479426 | rs12522248 | -0.55 | 0.723 | -0.287 | -1.105 | 0 | 0.046 | 0.046 | 0.005 | 0.3 | 0.14 | 0.34 | 0.15 | 0.3 | 0.27 |
| 5 | 156488903 | rs6873777 | 1.085 | 1.412 | 0.39 | -0.73 | 0.001 | -0.017 | -0.017 | 0.008 | 0.69 | 0.6 | 0.62 | 0.85 | 0.61 | 0.8 |
| 5 | 156493740 | rs6555828 | -0.375 | 1.445 | -0.103 | -0.753 | -0.012 | 0.023 | 0.023 | 0 | 0.86 | 0.91 | 0.89 | 0.99 | 0.79 | 0.96 |
| 5 | 156494601 | rs4704844 | 1.348 | 1.412 | 0.007 | -0.759 | 0 | 0.015 | 0.015 | 0 | 0.68 | 0.27 | 0.58 | 0.85 | 0.61 | 0.8 |
| 5 | 156499981 | rs2116787 | -0.09 | 1.464 | -0.529 | -0.433 | -0.023 | -0.027 | -0.027 | 0 | 0.11 | 0.05 | 0.25 | 0.12 | 0.16 | 0.15 |
| 5 | 156503767 | rs12152865 | 0.261 | 1.639 | -0.421 | 0.146 | -0.057 | -0.034 | -0.034 | 0 | 0.11 | 0.01 | 0.06 | 0.13 | 0.15 | 0.13 |
| 5 | 156505233 | rs11134551 | -1.308 | 1.307 | 0.172 | 0.066 | 0.01 | 0.014 | 0.014 | 0 | 0.13 | 0.29 | 0.26 | 0.13 | 0.14 | 0.21 |
| 5 | 156513282 | rs3087616 | 0.29 | 1.073 | 0.149 | -0.221 | 0 | -0.025 | -0.025 | 0 | 0.84 | 0.8 | 0.9 | 0.98 | 0.82 | 0.97 |
| 5 | 156531330 | rs919747 | 0.161 | 0.947 | 0.48 | -0.195 | -0.007 | 0.035 | 0.035 | 0 | 0.85 | 0.89 | 0.91 | 0.98 | 0.82 | 0.97 |
| 5 | 156531736 | rs1036199 | 0.273 | 0.939 | 0.528 | -0.198 | 0 | -0.028 | -0.028 | 0.133 | 0.84 | 0.79 | 0.89 | 0.98 | 0.82 | 0.97 |
| 5 | 156543695 | rs2033475 | 1.203 | 0.739 | 0.915 | -0.27 | -0.009 | -0.031 | -0.031 | 0 | 0.76 | 0.64 | 0.72 | 0.93 | 0.76 | 0.83 |
| 5 | 156546464 | rs422450 | 0.465 | 0.754 | 0.709 | -0.423 | 0 | 0.016 | 0.016 | 0 | 0.83 | 0.84 | 0.9 | 0.98 | 0.82 | 0.97 |
| 5 | 156554132 | rs420586 | -1.621 | 1.185 | 0.838 | -0.37 | 0.001 | 0.03 | 0.03 | 0 | 0.37 | 0.7 | 0.22 | 0.18 | 0.33 | 0.26 |
| 5 | 156563909 | rs190314 | -0.057 | 1.133 | 0.841 | -0.371 | 0.004 | 0.005 | 0.005 | 0 | 0.14 | 0.13 | 0.09 | 0.02 | 0.14 | 0.03 |
| 5 | 156564882 | rs13182119 | -1.023 | 1.192 | 1.126 | -0.318 | 0.019 | -0.009 | -0.009 | 0.838 | 0.22 | 0.54 | 0.13 | 0.17 | 0.19 | 0.23 |
| 5 | 156565372 | rs3797851 | -0.76 | 1.215 | 1.415 | -0.307 | 0.03 | 0.005 | 0.005 | 0 | 0.19 | 0.34 | 0.11 | 0.17 | 0.18 | 0.23 |
| 5 | 156565612 | rs9007 | 0.254 | 1.259 | 1.306 | -0.317 | -0.045 | -0.035 | -0.035 | 0 | 0.12 | 0.03 | 0.07 | 0.17 | 0.15 | 0.2 |
| 5 | 156573140 | rs13158986 | -0.619 | 1.215 | 1.542 | -0.34 | 0.019 | 0.003 | 0.003 | 1.646 | 0.17 | 0.22 | 0.1 | 0.17 | 0.19 | 0.22 |
| 5 | 156579780 | rs442773 | -2.021 | 1.304 | 1.419 | -0.666 | -0.003 | 0.016 | 0.016 | 0.034 | 0.41 | 0.63 | 0.25 | 0.2 | 0.37 | 0.3 |
| 5 | 156585154 | rs17054294 | -0.938 | 1.308 | 1.471 | -0.676 | 0.022 | -0.002 | -0.002 | 0 | 0.21 | 0.28 | 0.14 | 0.19 | 0.23 | 0.24 |
| 5 | 156589188 | rs31207 | -2.19 | 1.224 | 1.472 | -0.72 | -0.006 | 0.05 | 0.05 | 0.075 | 0.41 | 0.71 | 0.26 | 0.2 | 0.37 | 0.3 |
| 5 | 156589399 | rs2278385 | 0.051 | 1.232 | 1.344 | -0.708 | -0.067 | -0.054 | -0.054 | 1.885 | 0.14 | 0.02 | 0.08 | 0.18 | 0.18 | 0.21 |
| 5 | 156589585 | rs31208 | -0.398 | 1.104 | 1.279 | -0.673 | -0.005 | -0.051 | -0.051 | 0 | 0.17 | 0.24 | 0.09 | 0.01 | 0.12 | 0.05 |
| 5 | 156590355 | rs31220 | -1.061 | 1.107 | 1.148 | -0.672 | -0.018 | -0.085 | -0.085 | 0 | 0.2 | 0.36 | 0.11 | 0.01 | 0.14 | 0.05 |
| 5 | 156602286 | rs865332 | -0.204 | 0.907 | 0.354 | -0.793 | -0.03 | 0.077 | 0.077 | 0 | 0.16 | 0.02 | 0.08 | 0.01 | 0.13 | 0.05 |
| 5 | 156609000 | rs411174 | -1.519 | 1.159 | 0.68 | -0.693 | 0.006 | -0.028 | -0.028 | 0 | 0.35 | 0.46 | 0.21 | 0.22 | 0.34 | 0.31 |
| 5 | 156618003 | rs163310 | -0.363 | 1.09 | 0.748 | -0.683 | -0.003 | -0.068 | -0.068 | 0 | 0.16 | 0.27 | 0.1 | 0.01 | 0.13 | 0.05 |
| 5 | 156619180 | rs10079961 | 0.032 | 0.912 | 0.751 | -0.148 | 0.056 | 0.157 | 0.157 | 0.042 | 0.06 | 0.27 | 0.07 | 0.18 | 0.07 | 0.05 |
| 5 | 156620451 | rs2436384 | 0.149 | 0.6 | -0.397 | -0.627 | 0.053 | 0.025 | 0.025 | 2.588 | 0.51 | 0.77 | 0.55 | 0.43 | 0.47 | 0.35 |
| 5 | 156621333 | rs163425 | 0.615 | 0.806 | -0.277 | -0.456 | -0.011 | -0.076 | -0.076 | 0.072 | 0.15 | 0.36 | 0.11 | 0.01 | 0.13 | 0.03 |
| 5 | 156623904 | rs25856 | -0.867 | 1.078 | -1.196 | -0.281 | -0.041 | -0.057 | -0.057 | 2.945 | 0.8 | 0.94 | 0.73 | 0.67 | 0.81 | 0.77 |
| 5 | 156625279 | rs31223 | 0.504 | 0.64 | -0.949 | -0.475 | -0.009 | 0.001 | 0.001 | 0.072 | 0.39 | 0.41 | 0.37 | 0.43 | 0.38 | 0.39 |
| 5 | 156625763 | rs163433 | 0.252 | 0.957 | -1.721 | -0.067 | 0.014 | 0.007 | 0.007 | 0 | 0.54 | 0.69 | 0.44 | 0.44 | 0.5 | 0.41 |
| 5 | 156626573 | rs17054348 | -0.222 | 0.905 | -1.688 | 0.022 | -0.046 | -0.006 | -0.006 | 0 | 0.16 | 0.02 | 0.17 | 0.16 | 0.15 | 0.17 |
| 5 | 156628287 | rs246852 | 0.395 | 0.887 | -1.213 | 0.499 | -0.001 | -0.006 | -0.006 | 0 | 0.69 | 0.72 | 0.71 | 0.62 | 0.67 | 0.69 |
| 5 | 156637347 | rs40563 | -0.164 | 0.647 | -0.532 | -0.553 | 0.051 | -0.092 | -0.092 | 0 | 0.81 | 0.66 | 0.81 | 0.98 | 0.72 | 0.86 |
| 5 | 156644624 | rs152112 | 0.761 | 0.38 | -1.646 | -1.068 | -0.081 | 0.041 | 0.041 | 0.075 | 0.27 | 0.06 | 0.33 | 0.2 | 0.31 | 0.2 |
| 5 | 156644649 | rs2288504 | 0.525 | -0.41 | -1.467 | -0.982 | -0.009 | 0.007 | 0.007 | 0.251 | 0.37 | 0.45 | 0.39 | 0.56 | 0.35 | 0.41 |
| 5 | 156650052 | rs2288502 | 1.329 | -0.442 | -1.654 | -1.1 | 0.009 | -0.041 | -0.041 | 0 | 0.09 | 0.01 | 0.07 | 0.17 | 0.06 | 0.05 |
| 5 | 156656955 | rs4122648 | 1.338 | -0.759 | -1.755 | -1.187 | -0.015 | -0.127 | -0.127 | 0.008 | 0.22 | 0.04 | 0.31 | 0.54 | 0.24 | 0.28 |
| 5 | 156664822 | rs13355489 | 0.283 | -0.723 | -1.593 | -1.227 | -0.045 | -0.007 | -0.007 | 0.014 | 0.31 | 0.41 | 0.14 | 0.25 | 0.14 | 0.46 |
| 5 | 156664955 | rs13361568 | 1.391 | -0.844 | -1.478 | -1.992 | -0.021 | -0.042 | -0.042 | 0 | 0.16 | 0.25 | 0.07 | 0.03 | 0.11 | 0.2 |
| 5 | 156666799 | rs30118 | -0.153 | -0.7 | -1.734 | -1.954 | -0.026 | 0.114 | 0.114 | 0.147 | 0.62 | 0.19 | 0.72 | 0.95 | 0.58 | 0.66 |
| 5 | 156670047 | rs6876898 | 0.469 | 0.025 | -1.63 | -1.738 | -0.02 | -0.025 | -0.025 | 0 | 0.33 | 0.43 | 0.26 | 0.8 | 0.23 | 0.6 |
| 5 | 156671873 | rs13190673 | 0.615 | 0.447 | -1.495 | -1.502 | 0.017 | 0.059 | 0.059 | 2.731 | 0.22 | 0.08 | 0.23 | 0 | 0.19 | 0.08 |
| 5 | 156671918 | rs25775 | 0.179 | 1.128 | -0.704 | -0.707 | 0.041 | -0.112 | -0.112 | 2.775 | 0.78 | 0.6 | 0.79 | 0.98 | 0.7 | 0.86 |
| 5 | 156680629 | rs3892245 | 0.33 | 0.484 | -1.15 | -0.353 | -0.004 | -0.017 | -0.017 | 0 | 0.14 | 0.19 | 0.2 | 0.06 | 0.13 | 0.1 |
| 5 | 156680685 | rs27988 | -1.055 | 0.311 | -1.5 | 0.169 | -0.008 | -0.057 | -0.057 | 0 | 0.9 | 0.99 | 0.9 | 0.55 | 0.9 | 0.79 |
| 5 | 156680876 | rs17054397 | 0.351 | 0.57 | -1.207 | -0.217 | -0.005 | -0.021 | -0.021 | 0 | 0.14 | 0.2 | 0.21 | 0.06 | 0.13 | 0.1 |
| 5 | 156684911 | rs246750 | 0.295 | 1.368 | -1.745 | -0.362 | -0.082 | 0.089 | 0.089 | 0 | 0.2 | 0.06 | 0.28 | 0.02 | 0.3 | 0.16 |
| 5 | 156686040 | rs17054414 | -0.495 | 0.594 | -1.545 | 0.154 | -0.033 | -0.071 | -0.071 | 0.233 | 0.14 | 0.01 | 0.09 | 0.25 | 0.14 | 0.18 |
| 5 | 156699129 | rs727588 | -0.24 | 0.036 | -0.934 | -0.248 | 0.023 | 0.023 | 0.023 | 0 | 0.5 | 0.35 | 0.48 | 0.58 | 0.56 | 0.55 |
| 5 | 156699305 | rs714158 | 0.364 | 0.196 | -1.277 | -0.592 | 0.005 | 0.002 | 0.002 | 0.014 | 0.36 | 0.34 | 0.41 | 0.57 | 0.45 | 0.48 |
| 5 | 156703572 | rs1030270 | 0.178 | 0.111 | -0.521 | -0.697 | 0.016 | 0.176 | 0.176 | 0.195 | 0.07 | 0.21 | 0.05 | 0.26 | 0.06 | 0.19 |

| | | | | | | | | | | | | | | | | |
|---|---|---|---|---|---|---|---|---|---|---|---|---|---|---|---|---|
| 5 | 156705880 | rs10463005 | -0.783 | -0.11 | -0.479 | -0.741 | -0.006 | 0.062 | 0.062 | 0 | 0.17 | 0.29 | 0.14 | 0.3 | 0.17 | 0.3 |
| 5 | 156707948 | rs12654973 | 0.383 | -0.047 | -0.836 | -0.755 | -0.008 | 0.03 | 0.03 | 0 | 0.1 | 0.01 | 0.09 | 0.03 | 0.11 | 0.11 |
| 5 | 156714137 | rs2288068 | 0.394 | -0.099 | -0.671 | -0.772 | 0.001 | -0.001 | -0.001 | 0 | 0.1 | 0.08 | 0.09 | 0.03 | 0.1 | 0.11 |
| 5 | 156717629 | rs411720 | 0.137 | -0.574 | -0.776 | -1.186 | -0.033 | -0.02 | -0.02 | 0 | 0.62 | 0.47 | 0.8 | 0.73 | 0.73 | 0.71 |
| 5 | 156721986 | rs7724099 | 0.499 | -0.524 | -1.564 | -1.611 | -0.059 | 0.054 | 0.054 | 0 | 0.25 | 0.13 | 0.26 | 0.02 | 0.37 | 0.21 |
| 5 | 156722813 | rs889827 | -0.429 | -0.845 | -1.119 | -1.244 | 0.003 | 0.001 | 0.001 | 0.163 | 0.69 | 0.35 | 0.83 | 0.73 | 0.77 | 0.77 |
| 5 | 156724968 | rs1236180 | -0.566 | -0.608 | -1.685 | -1.275 | 0.048 | 0.158 | 0.158 | 3.638 | 0.71 | 0.92 | 0.87 | 0.99 | 0.79 | 0.88 |
| 5 | 156736310 | rs392512 | 0.587 | -0.443 | -1.861 | -0.914 | -0.048 | -0.143 | -0.143 | 0.564 | 0.29 | 0.01 | 0.42 | 0.57 | 0.27 | 0.42 |
| 5 | 156742186 | rs6897056 | 0.209 | -0.802 | -1.036 | -1.292 | -0.037 | -0.021 | -0.021 | 0 | 0.38 | 0.52 | 0.2 | 0.27 | 0.27 | 0.29 |
| 5 | 156747669 | rs1823035 | 0.235 | -0.803 | -0.679 | -1.265 | -0.037 | -0.02 | -0.02 | 2.167 | 0.38 | 0.53 | 0.2 | 0.27 | 0.27 | 0.29 |
| 5 | 156754251 | rs11743919 | 1.058 | -0.94 | -0.346 | -1.333 | 0.041 | 0.058 | 0.058 | 3.899 | 0.3 | 0.21 | 0.13 | 0.01 | 0.2 | 0.12 |
| 5 | 156754659 | rs13154298 | 2.037 | -1.421 | -0.687 | -1.613 | 0.018 | 0.075 | 0.075 | 0.429 | 0.16 | 0.04 | 0.06 | 0.01 | 0.12 | 0.05 |
| 5 | 156761070 | rs12656549 | -0.496 | -1.671 | -0.605 | -1.807 | 0.095 | 0.061 | 0.061 | 0.11 | 0.45 | 0.79 | 0.28 | 0.31 | 0.32 | 0.39 |
| 5 | 156765981 | rs3734026 | -0.751 | -1.486 | -0.651 | -1.847 | 0.083 | 0.076 | 0.076 | 0.186 | 0.47 | 0.8 | 0.31 | 0.31 | 0.38 | 0.39 |
| 5 | 156769008 | rs11739823 | -0.542 | -1.522 | -0.281 | -1.866 | -0.015 | 0.091 | 0.091 | 0.006 | 0.19 | 0.39 | 0.16 | 0.3 | 0.17 | 0.28 |
| 5 | 156769668 | rs7715719 | -0.771 | -1.526 | -0.304 | -1.872 | 0.083 | 0.076 | 0.076 | 0.169 | 0.47 | 0.79 | 0.31 | 0.31 | 0.38 | 0.39 |
| 5 | 156770133 | rs10037485 | 0.641 | -1.423 | -0.242 | -1.905 | -0.029 | -0.117 | -0.117 | 0.381 | 0.29 | 0.38 | 0.15 | 0.01 | 0.2 | 0.11 |
| 5 | 156772844 | rs13156380 | 0.601 | -1.423 | -0.165 | -1.905 | -0.044 | -0.153 | -0.153 | 0.06 | 0.29 | 0.43 | 0.15 | 0.01 | 0.2 | 0.11 |
| 5 | 156782843 | rs11952897 | 0.556 | -1.427 | -0.159 | -1.939 | -0.002 | -0.054 | -0.054 | 0.334 | 0.3 | 0.31 | 0.16 | 0.01 | 0.21 | 0.11 |
| 5 | 156785067 | rs11738062 | 0.658 | -1.357 | -0.202 | -1.897 | 0.004 | 0.019 | 0.019 | 0 | 0.11 | 0.07 | 0.09 | 0.03 | 0.11 | 0.12 |
| 5 | 156795204 | rs652267 | 0.971 | -1.087 | -0.259 | -1.934 | -0.012 | 0.008 | 0.008 | 0 | 0.22 | 0.2 | 0.26 | 0.12 | 0.33 | 0.18 |
| 5 | 156795719 | rs2910160 | 1.02 | -1.092 | -0.241 | -1.937 | -0.035 | 0.031 | 0.031 | 0 | 0.22 | 0.16 | 0.26 | 0.12 | 0.33 | 0.18 |
| 5 | 156796765 | rs684291 | -1.011 | -1.457 | -0.343 | -2.116 | 0 | 0.033 | 0.033 | 0 | 0.68 | 0.76 | 0.52 | 0.16 | 0.68 | 0.49 |
| 5 | 156798576 | rs2431686 | -0.808 | -1.366 | -0.215 | -2.117 | 0.003 | 0.037 | 0.037 | 0 | 0.66 | 0.75 | 0.52 | 0.16 | 0.67 | 0.49 |
| 5 | 156801180 | rs10076465 | -0.652 | -1.618 | -0.227 | -1.938 | 0.022 | 0.012 | 0.012 | 0 | 0.51 | 0.59 | 0.35 | 0.41 | 0.43 | 0.52 |
| 5 | 156802019 | rs12522498 | -0.285 | -1.578 | -0.279 | -2.235 | -0.001 | -0.024 | -0.024 | 0 | 0.45 | 0.47 | 0.31 | 0.13 | 0.36 | 0.32 |
| 5 | 156805787 | rs10515749 | -0.349 | -1.551 | -0.17 | -2.132 | -0.014 | -0.019 | -0.019 | 1.975 | 0.22 | 0.29 | 0.19 | 0.13 | 0.16 | 0.26 |
| 5 | 156809050 | rs889034 | -1.249 | 0.352 | -0.235 | -2.522 | 0.008 | -0.047 | -0.047 | 0 | 0.8 | 0.76 | 0.72 | 0.98 | 0.7 | 0.82 |
| 5 | 156809094 | rs884623 | 1.183 | 0.206 | -0.16 | -2.521 | -0.003 | -0.027 | -0.027 | 0 | 0.12 | 0.15 | 0.09 | 0 | 0.09 | 0.06 |
| 5 | 156809248 | rs3822691 | -0.197 | 0.223 | -0.158 | -2.46 | 0 | 0 | 0 | 0 | 0.09 | 0.07 | 0.1 | 0.13 | 0.07 | 0.19 |
| 5 | 156810072 | rs10515750 | 0.937 | 0.087 | -0.167 | -2.574 | 0.003 | -0.001 | -0.001 | 0 | 0.09 | 0.09 | 0.05 | 0 | 0.08 | 0.01 |
| 5 | 156811045 | rs10054755 | -0.284 | 0.079 | -0.165 | -2.576 | 0.003 | 0.004 | 0.004 | 0 | 0.15 | 0.13 | 0.11 | 0.11 | 0.12 | 0.1 |
| 5 | 156816522 | rs889035 | 1.537 | -0.234 | -0.083 | -2.487 | -0.005 | -0.009 | -0.009 | 0.002 | 0.3 | 0.18 | 0.37 | 0.67 | 0.3 | 0.39 |
| 5 | 156817266 | rs17599222 | -0.075 | -0.264 | -0.031 | -2.51 | 0.003 | -0.028 | -0.028 | 0 | 0.55 | 0.46 | 0.57 | 0.8 | 0.49 | 0.65 |
| 5 | 156817376 | rs7724666 | 2.169 | -0.239 | -0.057 | -2.499 | -0.011 | -0.045 | -0.045 | 1.538 | 0.14 | 0.18 | 0.07 | 0.01 | 0.1 | 0.05 |
| 5 | 156819669 | rs12518740 | 0.059 | -0.399 | -0.105 | -2.616 | 0.004 | -0.124 | -0.124 | 0 | 0.55 | 0.44 | 0.59 | 0.97 | 0.52 | 0.79 |
| 5 | 156820957 | rs6555992 | 0.848 | -0.335 | -0.142 | -2.768 | -0.013 | -0.111 | -0.111 | 0 | 0.24 | 0.4 | 0.13 | 0.01 | 0.17 | 0.18 |
| 5 | 156821217 | rs3734034 | 0.019 | -0.745 | -0.404 | -2.847 | -0.017 | -0.008 | -0.008 | 0 | 0.73 | 0.77 | 0.65 | 0.7 | 0.63 | 0.77 |
| 5 | 156821313 | rs6862302 | 2.433 | -0.382 | -0.391 | -2.733 | -0.001 | -0.004 | -0.004 | 0 | 0.14 | 0.11 | 0.07 | 0 | 0.12 | 0.05 |
| 5 | 156822230 | rs10073822 | 1.004 | -0.375 | -0.452 | -2.729 | -0.012 | -0.109 | -0.109 | 0 | 0.24 | 0.4 | 0.13 | 0.01 | 0.17 | 0.18 |
| 5 | 156828010 | rs1895311 | -0.962 | -0.239 | -0.765 | -2.722 | -0.031 | 0.029 | 0.029 | 0 | 0.83 | 0.91 | 0.72 | 0.92 | 0.73 | 0.87 |
| 5 | 156836438 | rs1157344 | 1.117 | -0.2 | -1.007 | -2.579 | -0.001 | 0.015 | 0.015 | 0 | 0.51 | 0.41 | 0.48 | 0.7 | 0.47 | 0.39 |
| 5 | 156836513 | rs11134736 | 2.405 | -0.311 | -1.019 | -2.638 | 0 | -0.016 | -0.016 | 0 | 0.16 | 0.16 | 0.18 | 0 | 0.13 | 0.06 |
| 5 | 156849722 | rs1833736 | 0.609 | -0.29 | -0.974 | -2.544 | 0.003 | -0.012 | -0.012 | 0 | 0.61 | 0.44 | 0.56 | 0.71 | 0.56 | 0.6 |
| 5 | 156876151 | rs3844 | 1.05 | -0.335 | -0.99 | -2.602 | 0.002 | -0.005 | -0.005 | 0.246 | 0.25 | 0.25 | 0.27 | 0.03 | 0.22 | 0.28 |
| 5 | 156876784 | rs11951889 | 1.861 | -0.411 | -1.045 | -2.632 | -0.001 | -0.008 | -0.008 | 0 | 0.29 | 0.17 | 0.25 | 0.69 | 0.27 | 0.34 |
| 5 | 156879727 | rs4704863 | 0.926 | -0.312 | -0.998 | -2.532 | -0.001 | 0.004 | 0.004 | 0 | 0.54 | 0.42 | 0.51 | 0.73 | 0.49 | 0.63 |
| 5 | 156891353 | rs4704742 | 0.985 | -0.33 | -1.107 | -2.65 | 0 | -0.003 | -0.003 | 0 | 0.52 | 0.43 | 0.47 | 0.72 | 0.47 | 0.43 |
| 5 | 156898690 | rs6860507 | 0.992 | -0.324 | -1.106 | -2.638 | -0.001 | -0.002 | -0.002 | 0 | 0.52 | 0.43 | 0.47 | 0.72 | 0.47 | 0.43 |
| 5 | 156900047 | rs9313608 | 2.089 | -0.473 | -1.089 | -2.637 | 0.001 | 0.003 | 0.003 | 0 | 0.08 | 0.06 | 0.05 | 0 | 0.08 | 0.01 |
| 5 | 156900534 | rs10476052 | 2.089 | -0.473 | -1.089 | -2.637 | 0.001 | 0.003 | 0.003 | 0 | 0.08 | 0.06 | 0.05 | 0 | 0.08 | 0.01 |
| 5 | 156901615 | rs11749762 | 1.822 | -0.453 | -1.071 | -2.642 | 0 | -0.014 | -0.014 | 0 | 0.08 | 0.09 | 0.06 | 0 | 0.08 | 0.01 |
| 5 | 156901701 | rs11739062 | 2.305 | -0.484 | -1.161 | -2.639 | -0.013 | 0.039 | 0.039 | 0 | 0.07 | 0 | 0.05 | 0 | 0.08 | 0.01 |
| 5 | 156904633 | rs10404 | 1.875 | -0.52 | -1.185 | -2.716 | 0 | -0.001 | -0.001 | 0.954 | 0.3 | 0.22 | 0.25 | 0.72 | 0.27 | 0.37 |
| 5 | 156905517 | rs11466822 | 1.008 | -0.494 | -1.195 | -2.687 | -0.001 | 0.004 | 0.004 | 0.028 | 0.52 | 0.43 | 0.36 | 0.72 | 0.46 | 0.42 |
| 5 | 156906819 | rs6895849 | 1.559 | -0.627 | -1.089 | -2.795 | -0.014 | -0.047 | -0.047 | 0.28 | 0.1 | 0.19 | 0.06 | 0 | 0.06 | 0.12 |
| 5 | 156906986 | rs3822585 | 0.914 | -0.381 | -1.51 | -2.596 | -0.001 | 0.004 | 0.004 | 0.04 | 0.51 | 0.42 | 0.36 | 0.72 | 0.46 | 0.42 |
| 5 | 156907110 | rs11466816 | 2.654 | -0.358 | -1.815 | -2.549 | -0.03 | 0.079 | 0.079 | 0.519 | 0.13 | 0.02 | 0.05 | 0 | 0.12 | 0.05 |
| 5 | 156907171 | rs11466815 | 2.166 | -0.369 | -1.792 | -2.535 | -0.005 | 0.031 | 0.031 | 0 | 0.09 | 0.03 | 0.08 | 0.03 | 0.09 | 0.2 |
| 5 | 156917340 | rs10067096 | 1.333 | -0.343 | -1.685 | -2.572 | 0.001 | -0.013 | -0.013 | 0.011 | 0.22 | 0.21 | 0.11 | 0.01 | 0.2 | 0.05 |
| 5 | 156918850 | rs2287749 | 2.632 | -0.324 | -1.722 | -2.556 | -0.015 | 0.057 | 0.057 | 0.519 | 0.13 | 0.05 | 0.05 | 0 | 0.12 | 0.05 |
| 5 | 156920756 | rs1990950 | 1.788 | -0.369 | -1.615 | -2.547 | 0 | 0.021 | 0.021 | 0 | 0.42 | 0.29 | 0.31 | 0.72 | 0.38 | 0.41 |
| 5 | 156920911 | rs1990951 | 0.894 | -0.334 | -1.609 | -2.502 | -0.001 | 0.007 | 0.007 | 0.065 | 0.52 | 0.44 | 0.37 | 0.73 | 0.46 | 0.42 |
| 5 | 156927875 | rs17054657 | -0.047 | -0.495 | -1.589 | -2.621 | -0.036 | -0.041 | -0.041 | 0 | 0.24 | 0.62 | 0.32 | 0.1 | 0.18 | 0.26 |
| 5 | 156932376 | rs2277027 | -1.07 | -0.531 | -1.769 | -2.759 | 0.008 | 0.009 | 0.009 | 0 | 0.41 | 0.64 | 0.44 | 0.13 | 0.36 | 0.45 |
| 5 | 156936364 | rs1422795 | -1.051 | -0.532 | -1.769 | -2.759 | 0.007 | 0.008 | 0.008 | 0 | 0.41 | 0.64 | 0.44 | 0.13 | 0.36 | 0.45 |
| 5 | 156936549 | rs1422794 | -1.005 | -0.468 | -1.894 | -2.807 | -0.093 | 0.119 | 0.119 | 0 | 0.71 | 0.88 | 0.71 | 0.92 | 0.62 | 0.81 |
| 5 | 156938327 | rs6860540 | 1.231 | -0.465 | -1.772 | -2.814 | 0 | -0.001 | -0.001 | 0 | 0.38 | 0.29 | 0.31 | 0.79 | 0.34 | 0.37 |
| 5 | 156940760 | rs3734031 | 1.812 | -1.366 | -1.168 | -2.341 | -0.015 | 0.022 | 0.022 | 0 | 0.09 | 0.01 | 0.06 | 0.02 | 0.09 | 0.17 |

| | | | | | | | | | | | | | | | | |
|---|---|---|---|---|---|---|---|---|---|---|---|---|---|---|---|---|
| 5 | 156943285 | rs6899205 | -0.682 | -1.312 | -1.098 | -2.078 | -0.029 | -0.062 | -0.062 | 0 | 0.34 | 0.53 | 0.39 | 0.14 | 0.28 | 0.45 |
| 5 | 156946049 | rs2042247 | -1.498 | -0.905 | -1.023 | -1.819 | -0.026 | 0.034 | 0.034 | 0 | 0.63 | 0.79 | 0.53 | 0.22 | 0.53 | 0.58 |
| 5 | 156953340 | rs11742401 | -0.378 | -1.03 | -0.858 | -2.041 | -0.031 | -0.071 | -0.071 | 0.06 | 0.32 | 0.48 | 0.36 | 0.11 | 0.27 | 0.29 |
| 5 | 156957113 | rs11750135 | -1.128 | -1.118 | -0.915 | -2.091 | -0.026 | -0.047 | -0.047 | 0 | 0.38 | 0.49 | 0.38 | 0.17 | 0.31 | 0.37 |
| 5 | 156957741 | rs3822696 | -1.61 | -0.85 | -1.008 | -1.879 | -0.012 | 0.022 | 0.022 | 0 | 0.61 | 0.73 | 0.51 | 0.2 | 0.52 | 0.58 |
| 5 | 156957784 | rs3734032 | -0.125 | -0.593 | -1.048 | -1.343 | 0.011 | 0.027 | 0.027 | 0 | 0.15 | 0.11 | 0.09 | 0.08 | 0.14 | 0.24 |
| 5 | 156962401 | rs17054692 | -0.05 | -0.414 | -0.969 | -1.089 | 0.008 | -0.01 | -0.01 | 0.051 | 0.07 | 0.03 | 0.05 | 0.09 | 0.08 | 0.19 |
| 5 | 156964594 | rs9313634 | -1.415 | -0.417 | -1.126 | -1.515 | -0.045 | 0.072 | 0.072 | 0.073 | 0.61 | 0.8 | 0.53 | 0.21 | 0.51 | 0.56 |
| 5 | 156966595 | rs6875485 | 0.126 | -0.768 | -0.534 | -1.793 | -0.011 | -0.078 | -0.078 | 1.333 | 0.21 | 0.3 | 0.14 | 0 | 0.17 | 0.14 |
| 5 | 156970878 | rs11739504 | 1.904 | -0.776 | -0.827 | -1.828 | -0.02 | 0.08 | 0.08 | 1.147 | 0.13 | 0.02 | 0.05 | 0 | 0.1 | 0.03 |
| 5 | 156972294 | rs10056240 | 0.6 | -1.005 | -0.337 | -1.194 | -0.001 | 0 | 0 | 0 | 0.21 | 0.21 | 0.14 | 0.21 | 0.17 | 0.24 |
| 5 | 156973288 | rs17054709 | 0.059 | -0.9 | -0.388 | -1.112 | 0.002 | 0.014 | 0.014 | 0 | 0.07 | 0.04 | 0.08 | 0.21 | 0.07 | 0.21 |
| 5 | 156975016 | rs6896064 | 0.734 | -0.986 | 0.078 | -1.17 | -0.003 | -0.005 | -0.005 | 0.302 | 0.17 | 0.16 | 0.28 | 0.12 | 0.18 | 0.2 |
| 5 | 156977853 | rs12189024 | -0.018 | -0.916 | -0.342 | -1.375 | 0.019 | 0.017 | 0.017 | 0 | 0.3 | 0.18 | 0.17 | 0.21 | 0.23 | 0.38 |
| 5 | 156981913 | rs4461616 | -0.009 | -0.955 | -0.349 | -1.371 | 0.021 | 0.018 | 0.018 | 0 | 0.3 | 0.17 | 0.17 | 0.21 | 0.23 | 0.38 |
| 5 | 156986576 | rs11749951 | -0.514 | -1.22 | -0.195 | -1.605 | 0.006 | -0.027 | -0.027 | 0.01 | 0.42 | 0.64 | 0.22 | 0.06 | 0.29 | 0.24 |
| 5 | 156989120 | rs17054724 | 0.815 | -1.194 | -0.253 | -1.564 | -0.002 | -0.003 | -0.003 | 0.87 | 0.07 | 0.03 | 0.1 | 0.15 | 0.09 | 0.17 |
| 5 | 156989757 | rs7725069 | -0.494 | -1.219 | -0.117 | -1.775 | -0.016 | -0.099 | -0.099 | 1.38 | 0.42 | 0.55 | 0.21 | 0.06 | 0.29 | 0.24 |
| 5 | 156991508 | rs11134819 | 0.968 | -1.24 | -0.073 | -1.817 | -0.007 | -0.106 | -0.106 | 1.671 | 0.16 | 0.4 | 0.12 | 0 | 0.13 | 0.02 |
| 5 | 156994437 | rs6876128 | 0.378 | -1.233 | -0.238 | -1.814 | -0.008 | -0.139 | -0.139 | 2.917 | 0.18 | 0.47 | 0.15 | 0 | 0.15 | 0.04 |
| 5 | 156999340 | rs7719224 | 0.663 | -1.041 | -0.63 | -1.638 | 0.029 | 0.073 | 0.073 | 0 | 0.23 | 0.15 | 0.07 | 0.06 | 0.14 | 0.21 |
| 5 | 157001302 | rs11134822 | 0.728 | -1.033 | -0.657 | -1.634 | 0.027 | 0.085 | 0.085 | 0 | 0.23 | 0.11 | 0.07 | 0.06 | 0.14 | 0.21 |
| 5 | 157030395 | rs10035961 | -0.555 | -1.073 | -0.557 | -1.695 | 0.001 | -0.007 | -0.007 | 0.419 | 0.37 | 0.62 | 0.43 | 0.11 | 0.34 | 0.24 |
| 5 | 157031235 | rs13153959 | -0.485 | -1.087 | -0.337 | -1.704 | -0.02 | -0.072 | -0.072 | 0 | 0.36 | 0.51 | 0.41 | 0.11 | 0.33 | 0.23 |
| 5 | 157034655 | rs6556121 | -2.383 | -0.687 | -0.807 | -1.34 | -0.071 | -0.042 | -0.042 | 0 | 0.68 | 0.9 | 0.6 | 0.34 | 0.59 | 0.62 |
| 5 | 157036921 | rs17054764 | 0.473 | -0.576 | -0.791 | -1.433 | 0.007 | 0.055 | 0.055 | 0 | 0.25 | 0.18 | 0.11 | 0.06 | 0.18 | 0.2 |
| 5 | 157037412 | rs10063242 | -1.368 | -0.42 | -0.623 | -1.195 | -0.019 | 0.036 | 0.036 | 0 | 0.61 | 0.72 | 0.51 | 0.17 | 0.5 | 0.44 |
| 5 | 157039737 | rs889056 | -1.164 | -0.648 | -0.602 | -0.971 | -0.015 | 0.038 | 0.038 | 0 | 0.55 | 0.78 | 0.36 | 0.29 | 0.48 | 0.41 |
| 5 | 157054080 | rs17054782 | 1.383 | -0.413 | -0.303 | -0.978 | 0.003 | -0.017 | -0.017 | 0.137 | 0.13 | 0.13 | 0.06 | 0 | 0.09 | 0.03 |
| 5 | 157055047 | rs6895071 | -0.281 | -0.611 | 0.174 | -0.889 | -0.055 | -0.055 | -0.055 | 0.334 | 0.26 | 0.55 | 0.17 | 0.1 | 0.12 | 0.2 |
| 5 | 157069796 | rs11134860 | -0.149 | -0.553 | 0.132 | -0.882 | -0.026 | -0.028 | -0.028 | 0 | 0.24 | 0.26 | 0.14 | 0.1 | 0.12 | 0.2 |
| 5 | 157070998 | rs6866972 | -0.137 | -0.539 | 0.136 | -0.875 | -0.029 | -0.031 | -0.031 | 0 | 0.24 | 0.28 | 0.15 | 0.1 | 0.12 | 0.2 |
| 5 | 157087764 | rs13165191 | 0.978 | -0.591 | -0.235 | -0.91 | -0.016 | -0.031 | -0.031 | 0 | 0.13 | 0.01 | 0.07 | 0.12 | 0.13 | 0.16 |
| 5 | 157095798 | rs6880381 | -1.392 | -0.879 | -0.351 | -1.013 | 0.028 | 0.055 | 0.055 | 0 | 0.53 | 0.91 | 0.38 | 0.29 | 0.39 | 0.46 |
| 5 | 157098233 | rs2277948 | -0.038 | -0.589 | -0.073 | -0.915 | -0.01 | -0.02 | -0.02 | 0 | 0.23 | 0.3 | 0.15 | 0.08 | 0.18 | 0.18 |
| 5 | 157098756 | rs11740603 | 0.47 | -0.597 | -0.071 | -0.9 | -0.012 | -0.022 | -0.022 | 0 | 0.2 | 0.29 | 0.13 | 0.07 | 0.16 | 0.13 |
| 5 | 157102159 | rs10051838 | -0.925 | -0.484 | 0.031 | -0.779 | 0.027 | 0.027 | 0.027 | 0 | 0.12 | 0.31 | 0.11 | 0.12 | 0.13 | 0.19 |
| 5 | 157114139 | rs6887040 | -1.608 | -0.019 | -0.28 | -0.784 | -0.035 | 0.013 | 0.013 | 0 | 0.59 | 0.77 | 0.37 | 0.32 | 0.53 | 0.53 |
| 5 | 157117920 | rs11748404 | 2.063 | -0.063 | -0.353 | -0.823 | -0.007 | 0.059 | 0.059 | 0.001 | 0.11 | 0.02 | 0.04 | 0 | 0.09 | 0.03 |
| 5 | 157118579 | rs12188530 | 0.056 | 0.208 | 0.254 | -0.393 | 0.003 | 0.094 | 0.094 | 0 | 0.07 | 0.27 | 0.1 | 0.11 | 0.03 | 0.08 |
| 5 | 157120226 | rs10045580 | -0.401 | 0.159 | 0.319 | -0.467 | -0.001 | -0.011 | -0.011 | 0 | 0.43 | 0.5 | 0.36 | 0.31 | 0.42 | 0.37 |
| 5 | 157122324 | rs2133732 | -1.723 | 0.119 | 0.307 | -0.487 | -0.011 | -0.012 | -0.012 | 0 | 0.64 | 0.72 | 0.44 | 0.43 | 0.55 | 0.62 |
| 5 | 157126005 | rs2418851 | -0.568 | 0.141 | 0.269 | -0.486 | 0.001 | -0.002 | -0.002 | 0 | 0.47 | 0.58 | 0.38 | 0.42 | 0.45 | 0.42 |
| 5 | 157130208 | rs1909701 | -1.757 | 0.138 | 0.304 | -0.489 | -0.064 | -0.068 | -0.068 | 0.009 | 0.65 | 0.87 | 0.46 | 0.43 | 0.55 | 0.62 |
| 5 | 157130903 | rs11744274 | 0.147 | 0.268 | 0.161 | -0.425 | -0.016 | -0.034 | -0.034 | 0.373 | 0.37 | 0.45 | 0.29 | 0.19 | 0.31 | 0.22 |
| 5 | 157131079 | rs13436754 | 0.362 | 0.45 | -0.528 | -0.553 | -0.013 | -0.013 | -0.013 | 0 | 0.06 | 0.05 | 0.07 | 0.13 | 0.11 | 0.15 |
| 5 | 157131941 | rs13155976 | 0.948 | 0.403 | -0.541 | -0.575 | 0.008 | 0.07 | 0.07 | 0 | 0.13 | 0.02 | 0.05 | 0 | 0.08 | 0.13 |
| 5 | 157136938 | rs4704894 | 0.235 | 0.623 | -0.456 | -0.37 | -0.013 | -0.013 | -0.013 | 0 | 0.06 | 0.05 | 0.07 | 0.13 | 0.11 | 0.15 |
| 5 | 157140461 | rs6864538 | 0.406 | 0.142 | -0.412 | -0.446 | 0.005 | 0.036 | 0.036 | 0 | 0.3 | 0.24 | 0.19 | 0.08 | 0.28 | 0.14 |
| 5 | 157154518 | rs1846664 | -0.815 | 0.129 | 0.044 | -0.146 | -0.008 | 0.089 | 0.089 | 0 | 0.06 | 0.19 | 0.03 | 0.12 | 0.03 | 0.05 |
| 5 | 157158372 | rs2270820 | -1.554 | 0.067 | -0.126 | -0.152 | -0.002 | 0 | 0 | 0 | 0.37 | 0.63 | 0.3 | 0.37 | 0.32 | 0.47 |
| 5 | 157158439 | rs2270818 | -0.393 | 0.491 | -0.427 | 0.049 | -0.04 | -0.046 | -0.046 | 0.016 | 0.11 | 0.01 | 0.08 | 0.17 | 0.14 | 0.25 |
| 5 | 157158553 | rs2270817 | -0.838 | 0.487 | -0.05 | 0.076 | -0.014 | 0.067 | 0.067 | 0 | 0.06 | 0.19 | 0.03 | 0.12 | 0.03 | 0.05 |
| 5 | 157164889 | rs2270813 | -0.744 | 0.978 | -0.409 | -0.256 | 0.027 | -0.044 | -0.044 | 0 | 0.68 | 0.56 | 0.5 | 0.82 | 0.65 | 0.72 |
| 5 | 157164962 | rs2270812 | -0.772 | 1.053 | -0.556 | -0.173 | -0.015 | 0.013 | 0.013 | 0 | 0.7 | 0.77 | 0.51 | 0.81 | 0.65 | 0.72 |
| 5 | 157166598 | rs3194515 | 0.846 | 1.044 | -0.884 | -0.219 | 0.006 | -0.004 | -0.004 | 1.03 | 0.41 | 0.37 | 0.34 | 0.57 | 0.35 | 0.32 |
| 5 | 157169968 | rs6867670 | 0.895 | 1.013 | -0.898 | -0.234 | 0.005 | -0.005 | -0.005 | 0.48 | 0.41 | 0.37 | 0.33 | 0.57 | 0.34 | 0.31 |
| 5 | 157178219 | rs13162531 | 0.148 | 1.107 | -0.888 | -0.308 | 0.007 | 0.008 | 0.008 | 0 | 0.52 | 0.42 | 0.41 | 0.59 | 0.47 | 0.42 |
| 5 | 157182897 | rs6882516 | -0.034 | 1.364 | -0.535 | 0.115 | 0.013 | 0.044 | 0.044 | 0.075 | 0.17 | 0.35 | 0.1 | 0.24 | 0.18 | 0.3 |
| 5 | 157183739 | rs2279770 | -0.17 | 1.178 | -0.916 | 0.282 | 0.006 | -0.002 | -0.002 | 0 | 0.41 | 0.37 | 0.33 | 0.59 | 0.35 | 0.32 |
| 5 | 157189618 | rs11738432 | -0.155 | 1.089 | -0.928 | 0.284 | 0.012 | -0.006 | -0.006 | 0 | 0.41 | 0.3 | 0.33 | 0.59 | 0.35 | 0.32 |
| 5 | 157191954 | rs7713308 | -0.16 | 1.128 | -1.146 | 0.101 | 0.002 | -0.003 | -0.003 | 4.392 | 0.41 | 0.4 | 0.34 | 0.52 | 0.35 | 0.31 |
| 5 | 157196835 | rs17605098 | -0.701 | 1.887 | -1.457 | -0.593 | -0.048 | 0.038 | 0.038 | 0 | 0.15 | 0.07 | 0.17 | 0 | 0.24 | 0.08 |
| 5 | 157201095 | rs11134939 | -0.189 | 1.127 | -2.042 | -0.985 | -0.021 | 0.102 | 0.102 | 0 | 0.64 | 0.88 | 0.76 | 0.83 | 0.63 | 0.6 |
| 5 | 157201700 | rs10050757 | -0.166 | 0.871 | -1.299 | -1.093 | 0 | -0.002 | -0.002 | 0.665 | 0.68 | 0.69 | 0.78 | 0.79 | 0.66 | 0.62 |
| 5 | 157207502 | rs254680 | 0.604 | 0.887 | -0.828 | -1.171 | 0.02 | 0.024 | 0.024 | 0.156 | 0.47 | 0.28 | 0.55 | 0.64 | 0.41 | 0.39 |
| 5 | 157211542 | rs17054884 | 0.71 | 0.34 | -1.001 | -1.632 | 0.005 | -0.006 | -0.006 | 0.325 | 0.15 | 0.1 | 0.14 | 0.02 | 0.12 | 0.13 |
| 5 | 157213497 | rs254682 | 0.029 | 0.402 | -1.03 | -1.702 | 0.021 | 0.031 | 0.031 | 0.44 | 0.48 | 0.27 | 0.56 | 0.63 | 0.43 | 0.39 |
| 5 | 157214434 | rs12283 | 1.166 | 0.321 | -0.587 | -1.787 | -0.017 | 0.071 | 0.071 | 0 | 0.1 | 0.29 | 0.08 | 0.16 | 0.08 | 0.15 |

| | | | | | | | | | | | | | | | | |
|---|---|---|---|---|---|---|---|---|---|---|---|---|---|---|---|---|
| 5 | 157221489 | rs17679725 | 1.6 | 0.193 | -0.806 | -2.011 | -0.04 | 0.045 | 0.045 | 0.998 | 0.08 | 0.02 | 0.08 | 0 | 0.09 | 0.05 |
| 5 | 157225195 | rs406150 | 0.04 | 0.228 | -0.71 | -1.918 | 0.02 | 0.03 | 0.03 | 1.005 | 0.48 | 0.27 | 0.56 | 0.63 | 0.43 | 0.39 |
| 5 | 157232642 | rs254670 | -0.587 | 0.266 | -0.721 | -1.904 | -0.004 | 0.006 | 0.006 | 1.636 | 0.59 | 0.63 | 0.65 | 0.79 | 0.51 | 0.54 |
| 5 | 157232764 | rs10515755 | 1.13 | 0.273 | -1.342 | -1.856 | -0.028 | 0.052 | 0.052 | 0 | 0.11 | 0.29 | 0.09 | 0.16 | 0.08 | 0.16 |
| 5 | 157233289 | rs254671 | -1.775 | 0.144 | -1.802 | -2.261 | -0.035 | -0.014 | -0.014 | 0 | 0.8 | 0.98 | 0.85 | 0.85 | 0.82 | 0.76 |
| 5 | 157233521 | rs4704753 | 1.119 | 0.476 | -1.322 | -1.791 | -0.028 | 0.052 | 0.052 | 0 | 0.11 | 0.29 | 0.09 | 0.16 | 0.08 | 0.16 |
| 5 | 157235372 | rs3097836 | 1.669 | 0.415 | -1.251 | -1.82 | 0.005 | -0.084 | -0.084 | 0 | 0.23 | 0.08 | 0.32 | 0.6 | 0.21 | 0.21 |
| 5 | 157248847 | rs10515753 | 0.831 | 0.666 | -1.166 | -1.976 | -0.026 | 0.033 | 0.033 | 0 | 0.06 | 0.01 | 0.11 | 0 | 0.16 | 0.07 |
| 5 | 157259513 | rs17607769 | 0.606 | 0.662 | -1.159 | -1.975 | -0.014 | 0.017 | 0.017 | 0 | 0.07 | 0.03 | 0.12 | 0 | 0.17 | 0.07 |
| 5 | 157260015 | rs254663 | 0.476 | -0.126 | -1.035 | -1.783 | -0.001 | -0.001 | -0.001 | 0.679 | 0.34 | 0.37 | 0.41 | 0.76 | 0.3 | 0.37 |
| 5 | 157298694 | rs7735412 | 0.936 | 0.378 | -0.973 | -1.306 | -0.007 | 0.02 | 0.02 | 0 | 0.11 | 0.03 | 0.06 | 0.06 | 0.13 | 0.14 |
| 5 | 157317932 | rs11738052 | 1.205 | 0.294 | -0.875 | -1.391 | 0 | 0.002 | 0.002 | 0.135 | 0.09 | 0.07 | 0.05 | 0 | 0.1 | 0.01 |
| 5 | 157318560 | rs10052561 | 0.288 | -0.183 | -1.053 | -2.022 | 0.021 | 0.089 | 0.089 | 3.923 | 0.39 | 0.25 | 0.33 | 0.14 | 0.31 | 0.42 |
| 5 | 157320276 | rs1186919 | -1.348 | -0.278 | -0.996 | -1.529 | 0.013 | -0.057 | -0.057 | 0 | 0.77 | 0.87 | 0.66 | 0.39 | 0.79 | 0.79 |
| 5 | 157329451 | rs2430586 | 0.645 | -0.135 | -0.413 | -1.338 | -0.015 | 0.012 | 0.012 | 0 | 0.12 | 0.21 | 0.09 | 0.12 | 0.1 | 0.16 |
| 5 | 157335926 | rs12716331 | -0.252 | -0.06 | -0.401 | -1.649 | 0.031 | 0.044 | 0.044 | 5.947 | 0.54 | 0.37 | 0.51 | 0.27 | 0.6 | 0.64 |
| 5 | 157340772 | rs17608675 | 0.023 | 0.149 | -0.728 | -1.731 | -0.06 | 0.028 | 0.028 | 1.888 | 0.37 | 0.19 | 0.33 | 0.28 | 0.46 | 0.53 |
| 5 | 157341414 | rs1211159 | -0.022 | -0.252 | -0.431 | -2.143 | 0.006 | -0.119 | -0.119 | 0 | 0.23 | 0.39 | 0.16 | 0.03 | 0.18 | 0.2 |
| 5 | 157341463 | rs17608755 | 1.159 | -0.208 | -0.785 | -2.303 | -0.003 | 0.057 | 0.057 | 3.918 | 0.23 | 0.03 | 0.16 | 0.11 | 0.2 | 0.31 |
| 5 | 157342660 | rs1209063 | -0.84 | -0.463 | -0.181 | -2.142 | -0.018 | -0.013 | -0.013 | 0.152 | 0.59 | 0.45 | 0.48 | 0.3 | 0.66 | 0.74 |
| 5 | 157342764 | rs899541 | -0.919 | -0.433 | -0.376 | -2.129 | 0.006 | 0.005 | 0.005 | 6.347 | 0.6 | 0.59 | 0.51 | 0.3 | 0.66 | 0.73 |
| 5 | 157343390 | rs1186905 | -0.384 | -0.308 | -0.025 | -1.888 | 0.006 | -0.122 | -0.122 | 0.019 | 0.23 | 0.39 | 0.17 | 0.03 | 0.18 | 0.2 |
| 5 | 157346367 | rs7728721 | -0.349 | -0.285 | 0.215 | -1.952 | -0.023 | -0.148 | -0.148 | 0 | 0.27 | 0.49 | 0.18 | 0.02 | 0.19 | 0.15 |
| 5 | 157357100 | rs3095702 | 0.824 | 0.373 | -0.569 | -1.806 | -0.096 | 0.024 | 0.024 | 0.014 | 0.29 | 0.02 | 0.36 | 0.16 | 0.29 | 0.4 |
| 5 | 157357767 | rs1109985 | -1.534 | 0.524 | -0.479 | -1.935 | 0.004 | -0.014 | -0.014 | 1.992 | 0.72 | 0.85 | 0.67 | 0.31 | 0.71 | 0.72 |
| 5 | 157369504 | rs2418849 | 0.41 | 0.54 | -0.069 | -1.949 | 0.015 | -0.071 | -0.071 | 0.299 | 0.26 | 0.36 | 0.17 | 0.05 | 0.28 | 0.16 |
| 5 | 157369533 | rs3095997 | -1.484 | 0.549 | 0.189 | -1.95 | -0.001 | 0 | 0 | 0 | 0.71 | 0.74 | 0.66 | 0.31 | 0.72 | 0.72 |
| 5 | 157370372 | rs3095996 | -0.062 | 0.536 | 0.154 | -1.937 | -0.013 | 0.062 | 0.062 | 1.445 | 0.44 | 0.33 | 0.48 | 0.26 | 0.45 | 0.55 |
| 5 | 157383522 | rs10515756 | -0.687 | 0.555 | 0.531 | -1.824 | -0.01 | -0.049 | -0.049 | 3.371 | 0.32 | 0.53 | 0.22 | 0.18 | 0.34 | 0.27 |
| 5 | 157385701 | rs4454041 | -1.124 | 0.585 | 0.245 | -1.779 | 0.001 | 0.054 | 0.054 | 0 | 0.41 | 0.7 | 0.31 | 0.18 | 0.43 | 0.33 |
| 5 | 157387156 | rs9313764 | 0.621 | 0.738 | 0.47 | -1.512 | 0.012 | -0.044 | -0.044 | 0.331 | 0.17 | 0.36 | 0.12 | 0.05 | 0.2 | 0.1 |
| 5 | 157390840 | rs949602 | -0.624 | 0.759 | 0.216 | -1.044 | -0.003 | 0.001 | 0.001 | 0 | 0.62 | 0.63 | 0.65 | 0.32 | 0.53 | 0.63 |
| 5 | 157390960 | rs1894964 | -0.031 | 0.796 | 0.149 | -1.137 | 0.005 | 0.016 | 0.016 | 0 | 0.52 | 0.57 | 0.57 | 0.32 | 0.43 | 0.55 |
| 5 | 157394806 | rs3095710 | -1.478 | 0.707 | -0.526 | -0.906 | -0.038 | -0.167 | -0.167 | 0 | 0.74 | 0.99 | 0.78 | 0.45 | 0.75 | 0.77 |
| 5 | 157402820 | rs2900797 | -0.802 | 0.66 | 0.069 | -0.979 | 0.008 | 0.005 | 0.005 | 0 | 0.64 | 0.75 | 0.68 | 0.32 | 0.61 | 0.64 |
| 5 | 157404028 | rs2080462 | 0.845 | 0.733 | -0.335 | -1.219 | -0.027 | 0.034 | 0.034 | 0.537 | 0.31 | 0.13 | 0.3 | 0.18 | 0.31 | 0.3 |
| 5 | 157412103 | rs9686690 | 0.576 | 0.738 | 0.459 | -1.225 | 0.015 | -0.074 | -0.074 | 0.081 | 0.13 | 0.26 | 0.1 | 0 | 0.1 | 0.1 |
| 5 | 157422405 | rs12332160 | 1.104 | 0.66 | 0.313 | -1.287 | 0.014 | 0.014 | 0.014 | 0 | 0.17 | 0.08 | 0.24 | 0.14 | 0.11 | 0.22 |
| 5 | 157425198 | rs11949394 | 0.594 | 0.768 | 0.596 | -1.265 | 0.002 | -0.037 | -0.037 | 0 | 0.13 | 0.14 | 0.08 | 0 | 0.1 | 0.1 |
| 5 | 157437035 | rs6884132 | -0.503 | 0.785 | 0.6 | -1.285 | 0.046 | -0.124 | -0.124 | 1.223 | 0.18 | 0.57 | 0.15 | 0 | 0.18 | 0.11 |
| 5 | 157451580 | rs10044900 | -0.754 | 0.732 | 0.513 | -1.31 | 0.027 | -0.125 | -0.125 | 0.477 | 0.19 | 0.62 | 0.16 | 0 | 0.18 | 0.11 |
| 5 | 157455284 | rs3117740 | -0.507 | 0.654 | 0.141 | -1.444 | -0.029 | 0.059 | 0.059 | 0 | 0.38 | 0.77 | 0.36 | 0.12 | 0.41 | 0.34 |
| 5 | 157456191 | rs17229224 | 0.657 | 0.399 | -0.039 | -1.328 | 0.034 | 0.012 | 0.012 | 0 | 0.27 | 0.14 | 0.34 | 0.21 | 0.19 | 0.24 |
| 5 | 157467325 | rs3117730 | -0.183 | 0.531 | 0.013 | -1.345 | -0.011 | 0.01 | 0.01 | 0 | 0.37 | 0.69 | 0.36 | 0.12 | 0.42 | 0.33 |
| 5 | 157478630 | rs1957562 | 1.155 | 0.587 | 0.275 | -1.345 | 0.01 | -0.019 | -0.019 | 0.013 | 0.21 | 0.3 | 0.21 | 0.12 | 0.24 | 0.23 |
| 5 | 157483485 | rs2962392 | 1.929 | 0.458 | 0.034 | -1.298 | -0.012 | -0.029 | -0.029 | 0 | 0.13 | 0.01 | 0.22 | 0.16 | 0.09 | 0.19 |
| 5 | 157486849 | rs3117722 | -0.913 | 0.423 | 0.098 | -1.339 | 0.009 | 0.062 | 0.062 | 0 | 0.49 | 0.68 | 0.47 | 0.17 | 0.51 | 0.36 |
| 5 | 157487568 | rs1002566 | -0.884 | 0.421 | 0.097 | -1.34 | 0.008 | 0.06 | 0.06 | 0 | 0.48 | 0.68 | 0.46 | 0.17 | 0.51 | 0.36 |
| 5 | 157488424 | rs17055248 | 1.184 | 0.37 | 0.159 | -1.305 | 0.002 | 0.004 | 0.004 | 0.249 | 0.1 | 0.1 | 0.06 | 0.15 | 0.14 | 0.18 |
| 5 | 157488655 | rs3117721 | -1.514 | 0.278 | 0.17 | -1.33 | 0.011 | 0.005 | 0.005 | 0 | 0.62 | 0.84 | 0.7 | 0.33 | 0.6 | 0.55 |
| 5 | 157489027 | rs17229238 | 2.1 | 0.388 | 0.032 | -1.291 | 0.024 | -0.004 | -0.004 | 0 | 0.12 | 0.03 | 0.12 | 0.06 | 0.1 | 0.04 |
| 5 | 157494859 | rs2962383 | 1.173 | 0.533 | 0.193 | -1.298 | 0.013 | -0.016 | -0.016 | 0 | 0.21 | 0.29 | 0.23 | 0.12 | 0.29 | 0.24 |
| 5 | 157496046 | rs2962384 | -0.546 | 0.497 | 0.224 | -1.303 | -0.012 | 0.062 | 0.062 | 0 | 0.43 | 0.35 | 0.39 | 0.17 | 0.43 | 0.33 |
| 5 | 157505115 | rs17242471 | 0.702 | 0.49 | 0.215 | -1.328 | -0.007 | 0.065 | 0.065 | 0 | 0.23 | 0.08 | 0.18 | 0.06 | 0.19 | 0.1 |
| 5 | 157507491 | rs3096010 | -0.849 | 0.488 | 0.312 | -1.193 | 0.005 | 0.038 | 0.038 | 0 | 0.48 | 0.64 | 0.47 | 0.19 | 0.51 | 0.35 |
| 5 | 157509770 | rs2025215 | 1.433 | 0.498 | 0.152 | -1.186 | -0.016 | 0.012 | 0.012 | 0 | 0.19 | 0.15 | 0.2 | 0.13 | 0.24 | 0.25 |
| 5 | 157513404 | rs2475777 | -2.187 | 0.549 | 0.061 | -1.14 | -0.012 | -0.026 | -0.026 | 0 | 0.87 | 0.99 | 0.78 | 0.85 | 0.91 | 0.82 |
| 5 | 157513737 | rs7732107 | -0.087 | 0.814 | 0.469 | -1.046 | 0.005 | -0.049 | -0.049 | 0.593 | 0.27 | 0.36 | 0.25 | 0.06 | 0.27 | 0.11 |
| 5 | 157515153 | rs882086 | -0.615 | 0.66 | 0.161 | -0.638 | -0.009 | 0.048 | 0.048 | 0.025 | 0.42 | 0.75 | 0.41 | 0.19 | 0.44 | 0.3 |
| 5 | 157519669 | rs12514994 | -0.642 | 0.183 | 0.277 | -0.349 | -0.016 | -0.027 | -0.027 | 0.261 | 0.3 | 0.57 | 0.27 | 0.19 | 0.3 | 0.2 |
| 5 | 157521976 | rs1744344 | -1.124 | 0.197 | -0.041 | 0.016 | -0.001 | -0.025 | -0.025 | 0.152 | 0.87 | 0.95 | 0.74 | 0.79 | 0.89 | 0.78 |
| 5 | 157522100 | rs1744343 | 0.249 | 0.158 | 0.109 | -0.163 | 0.001 | -0.003 | -0.003 | 0.181 | 0.12 | 0.18 | 0.11 | 0.13 | 0.18 | 0.14 |
| 5 | 157523151 | rs10045606 | 0.326 | 0.453 | 0.283 | -0.016 | 0.036 | -0.023 | -0.023 | 0.014 | 0.26 | 0.38 | 0.25 | 0.15 | 0.32 | 0.3 |
| 5 | 157523642 | rs6878263 | 0.449 | -0.148 | -0.057 | -0.061 | 0.002 | 0.001 | 0.001 | 0 | 0.6 | 0.6 | 0.57 | 0.61 | 0.65 | 0.61 |
| 5 | 157525060 | rs13159558 | 0.274 | -0.728 | 0.199 | -0.132 | -0.079 | -0.057 | -0.057 | 0.594 | 0.19 | 0.39 | 0.08 | 0.09 | 0.06 | 0.05 |
| 5 | 157525853 | rs7716011 | 1.33 | -0.279 | -0.653 | -0.239 | -0.03 | 0.015 | 0.015 | 0 | 0.15 | 0.09 | 0.22 | 0.09 | 0.25 | 0.23 |
| 5 | 157526367 | rs13182707 | -0.382 | -0.611 | -0.693 | 0.153 | -0.039 | 0.007 | 0.007 | 0.618 | 0.31 | 0.57 | 0.16 | 0.35 | 0.19 | 0.18 |
| 5 | 157526827 | rs17294992 | 1.704 | -0.385 | -1.581 | 0.063 | -0.099 | -0.126 | -0.126 | 0 | 0.28 | 0.01 | 0.4 | 0.38 | 0.3 | 0.26 |
| 5 | 157528286 | rs17229266 | 0.836 | -0.746 | -1.226 | 0.309 | 0.011 | -0.048 | -0.048 | 0 | 0.11 | 0.02 | 0.06 | 0.24 | 0.12 | 0.14 |

| | | | | | | | | | | | | | | | | |
|---|---|---|---|---|---|---|---|---|---|---|---|---|---|---|---|---|
| 5 | 157528485 | rs13159354 | 0.484 | -0.464 | -0.996 | 0.177 | 0.074 | -0.001 | -0.001 | 0.142 | 0.52 | 0.29 | 0.67 | 0.47 | 0.66 | 0.57 |
| 5 | 157530478 | rs7726988 | -0.127 | -0.165 | -0.27 | 0.312 | -0.005 | 0.001 | 0.001 | 0.946 | 0.44 | 0.46 | 0.37 | 0.43 | 0.44 | 0.44 |
| 5 | 157531969 | rs4704916 | 0.635 | -0.621 | -0.671 | 0.238 | 0.035 | -0.057 | -0.057 | 0 | 0.33 | 0.09 | 0.4 | 0.42 | 0.23 | 0.28 |
| 5 | 157535313 | rs2984629 | -0.919 | -0.397 | -0.27 | -0.504 | 0.005 | -0.006 | -0.006 | 0.269 | 0.87 | 0.89 | 0.94 | 0.79 | 0.89 | 0.89 |
| 5 | 157538994 | rs1628560 | 0.674 | -0.514 | 0.109 | -0.783 | 0.028 | 0.003 | 0.003 | 0.017 | 0.39 | 0.28 | 0.47 | 0.6 | 0.25 | 0.39 |
| 5 | 157539010 | rs1613292 | 0.831 | -0.598 | 0.164 | -0.874 | -0.034 | -0.001 | -0.001 | 0.423 | 0.37 | 0.57 | 0.46 | 0.6 | 0.21 | 0.34 |
| 5 | 157541101 | rs1650551 | 0.575 | -0.576 | 0.149 | -0.99 | -0.005 | -0.002 | -0.002 | 0.114 | 0.41 | 0.6 | 0.49 | 0.62 | 0.26 | 0.39 |
| 5 | 157545316 | rs1744359 | 1.075 | -0.339 | 0.811 | -0.836 | 0 | 0.029 | 0.029 | 0.933 | 0.54 | 0.64 | 0.71 | 0.69 | 0.55 | 0.61 |
| 5 | 157550899 | rs2065138 | 0.442 | -0.795 | 1.061 | -0.578 | 0 | -0.001 | -0.001 | 5.768 | 0.78 | 0.76 | 0.85 | 0.78 | 0.8 | 0.8 |
| 5 | 157551324 | rs1744357 | 0.362 | -0.833 | 1 | -0.609 | -0.002 | -0.006 | -0.006 | 0 | 0.19 | 0.2 | 0.11 | 0.1 | 0.18 | 0.15 |
| 5 | 157551411 | rs1744356 | 1.382 | -0.842 | 0.785 | -0.605 | 0.036 | 0.096 | 0.096 | 3.181 | 0.55 | 0.09 | 0.71 | 0.69 | 0.61 | 0.65 |
| 5 | 157559979 | rs1173481 | 1.411 | -0.552 | 0.68 | -0.54 | 0.016 | 0.039 | 0.039 | 1.43 | 0.56 | 0.29 | 0.72 | 0.67 | 0.62 | 0.64 |
| 5 | 157560130 | rs1173482 | 1.357 | -0.505 | 0.724 | -0.518 | 0.01 | 0.025 | 0.025 | 1.264 | 0.57 | 0.34 | 0.73 | 0.67 | 0.62 | 0.65 |
| 5 | 157564869 | rs4639206 | 0.959 | -0.315 | 0.032 | -0.428 | 0 | 0.007 | 0.007 | 0.012 | 0.75 | 0.8 | 0.84 | 0.78 | 0.78 | 0.79 |
| 5 | 157565338 | rs10055302 | -0.17 | 0.106 | -0.042 | -0.202 | 0.003 | -0.018 | -0.018 | 0.049 | 0.09 | 0.08 | 0.07 | 0 | 0.1 | 0.08 |
| 5 | 157570419 | rs6860826 | 0.157 | 0.091 | -0.187 | -0.076 | 0.008 | -0.008 | -0.008 | 0 | 0.77 | 0.64 | 0.83 | 0.78 | 0.79 | 0.81 |
| 5 | 157579932 | rs1173455 | -0.617 | 0.024 | -0.645 | 0.049 | -0.002 | -0.006 | -0.006 | 0 | 0.19 | 0.22 | 0.1 | 0.11 | 0.15 | 0.16 |
| 5 | 157582088 | rs1173454 | -0.555 | 0.028 | -0.844 | 0.019 | 0.01 | 0.015 | 0.015 | 0 | 0.2 | 0.19 | 0.1 | 0.11 | 0.16 | 0.17 |
| 5 | 157588271 | rs2419398 | -0.149 | 0.368 | -1.196 | 0.207 | -0.023 | 0.003 | 0.003 | 0 | 0.53 | 0.73 | 0.38 | 0.46 | 0.48 | 0.66 |
| 5 | 157588362 | rs4512122 | 0.064 | 0.177 | -1.339 | 0.053 | 0.004 | 0.01 | 0.01 | 0 | 0.19 | 0.09 | 0.1 | 0.2 | 0.15 | 0.18 |
| 5 | 157589544 | rs2181233 | -0.209 | 0.621 | -0.187 | -0.055 | 0.008 | -0.019 | -0.019 | 0 | 0.21 | 0.32 | 0.14 | 0.2 | 0.2 | 0.22 |
| 5 | 157590857 | rs6886021 | -0.801 | 0.497 | 0.559 | -0.197 | 0 | 0.001 | 0.001 | 0 | 0.44 | 0.45 | 0.32 | 0.46 | 0.42 | 0.55 |
| 5 | 157592608 | rs1027355 | 0.553 | 0.468 | 0.632 | -0.147 | -0.001 | -0.002 | -0.002 | 0 | 0.23 | 0.15 | 0.19 | 0.24 | 0.25 | 0.36 |
| 5 | 157593214 | rs12186870 | 0.017 | 0.294 | 0.673 | -0.308 | -0.017 | 0.005 | 0.005 | 0 | 0.19 | 0.05 | 0.11 | 0.2 | 0.16 | 0.18 |
| 5 | 157596056 | rs927465 | 0.436 | 0.55 | 0.639 | -0.369 | -0.006 | 0.018 | 0.018 | 1.275 | 0.08 | 0.05 | 0.05 | 0 | 0.07 | 0.07 |
| 5 | 157600753 | rs1173447 | -0.446 | -0.212 | 0.911 | -0.636 | -0.009 | -0.006 | -0.006 | 0 | 0.21 | 0.36 | 0.13 | 0.22 | 0.17 | 0.28 |
| 5 | 157614896 | rs11748189 | 0.334 | -0.076 | 1.121 | -0.584 | 0.01 | -0.049 | -0.049 | 0 | 0.09 | 0.21 | 0.07 | 0 | 0.07 | 0.07 |
| 5 | 157615710 | rs1874161 | 0.533 | -0.063 | 0.96 | -0.578 | -0.002 | 0.002 | 0.002 | 1.079 | 0.09 | 0.07 | 0.06 | 0 | 0.07 | 0.07 |
| 5 | 157625299 | rs202821 | -1.036 | -0.444 | 1.507 | -0.639 | -0.007 | -0.031 | -0.031 | 0 | 0.31 | 0.44 | 0.17 | 0.25 | 0.24 | 0.31 |
| 5 | 157637018 | rs169370 | -0.327 | -0.39 | 1.751 | -0.632 | -0.02 | 0.001 | 0.001 | 0 | 0.22 | 0.29 | 0.11 | 0.24 | 0.15 | 0.25 |
| 5 | 157637276 | rs202793 | -0.265 | -0.401 | 1.714 | -0.631 | -0.01 | 0 | 0 | 0 | 0.21 | 0.23 | 0.1 | 0.24 | 0.15 | 0.25 |
| 5 | 157645207 | rs1145634 | -1.419 | -0.408 | 1.828 | -0.777 | -0.006 | -0.028 | -0.028 | 0 | 0.34 | 0.58 | 0.22 | 0.25 | 0.29 | 0.33 |
| 5 | 157645927 | rs1896603 | -0.297 | -0.417 | 1.71 | -0.784 | -0.001 | -0.021 | -0.021 | 0.193 | 0.12 | 0.18 | 0.1 | 0 | 0.14 | 0.06 |
| 5 | 157646252 | rs1145632 | -1.418 | -0.454 | 1.653 | -0.8 | 0.006 | 0.016 | 0.016 | 0 | 0.35 | 0.71 | 0.24 | 0.25 | 0.3 | 0.33 |
| 5 | 157652081 | rs1145625 | -0.135 | -0.418 | 1.196 | -0.666 | -0.009 | 0.001 | 0.001 | 0 | 0.2 | 0.21 | 0.1 | 0.24 | 0.15 | 0.26 |
| 5 | 157671789 | rs1177762 | -0.45 | 0.048 | 0.646 | -1.006 | 0.006 | 0.031 | 0.031 | 0.046 | 0.3 | 0.21 | 0.28 | 0.21 | 0.31 | 0.3 |
| 5 | 157677264 | rs11739307 | 0.336 | 0.046 | 0.986 | -1.036 | 0.028 | -0.044 | -0.044 | 0 | 0.1 | 0.17 | 0.07 | 0.01 | 0.09 | 0.06 |
| 5 | 157683676 | rs961585 | 1.079 | 0.069 | 1.029 | -1.025 | 0.024 | -0.029 | -0.029 | 0 | 0.08 | 0.1 | 0.07 | 0.01 | 0.09 | 0.06 |
| 5 | 157689852 | rs1650609 | -1.042 | -0.022 | 0.839 | -0.816 | -0.002 | 0.066 | 0.066 | 0 | 0.41 | 0.69 | 0.36 | 0.25 | 0.4 | 0.35 |
| 5 | 157694866 | rs17612017 | 0.985 | 0.079 | -0.022 | -0.685 | -0.066 | 0.014 | 0.014 | 0 | 0.27 | 0.01 | 0.11 | 0.17 | 0.26 | 0.41 |
| 5 | 157703679 | rs1145608 | -0.854 | 0.064 | 0.346 | -0.583 | 0 | -0.012 | -0.012 | 0 | 0.4 | 0.53 | 0.35 | 0.21 | 0.4 | 0.35 |
| 5 | 157704332 | rs1145605 | 0.756 | 0.41 | -0.502 | 0.539 | -0.033 | -0.043 | -0.043 | 0.666 | 0.09 | 0.01 | 0.18 | 0.4 | 0.1 | 0.15 |
| 5 | 157710122 | rs1565867 | -0.12 | 0.048 | -0.453 | -0.237 | 0.007 | 0.018 | 0.018 | 1.058 | 0.29 | 0.21 | 0.22 | 0.21 | 0.25 | 0.31 |
| 5 | 157713315 | rs1038993 | 0.065 | 0.172 | -0.474 | -0.115 | -0.019 | 0.024 | 0.024 | 0 | 0.41 | 0.25 | 0.66 | 0.66 | 0.52 | 0.6 |
| 5 | 157717906 | rs2172745 | 0.8 | -0.352 | -0.367 | -0.487 | 0.009 | 0.024 | 0.024 | 0 | 0.3 | 0.2 | 0.17 | 0.1 | 0.25 | 0.25 |
| 5 | 157721041 | rs9765118 | 0.22 | -0.465 | 0.043 | -0.653 | -0.015 | -0.008 | -0.008 | 0 | 0.4 | 0.47 | 0.21 | 0.34 | 0.26 | 0.31 |
| 5 | 157722151 | rs12173246 | 0.652 | -0.404 | 0.424 | -0.42 | -0.023 | 0.102 | 0.102 | 0.088 | 0.08 | 0.32 | 0.04 | 0.13 | 0.01 | 0.06 |
| 5 | 157727005 | rs11953309 | 0.967 | -0.395 | 0.292 | -0.415 | -0.035 | 0.04 | 0.04 | 0 | 0.07 | 0.17 | 0.03 | 0.13 | 0.01 | 0.06 |
| 5 | 157727123 | rs11950350 | 0.967 | -0.395 | 0.292 | -0.394 | -0.035 | 0.043 | 0.043 | 0 | 0.07 | 0.17 | 0.03 | 0.13 | 0.01 | 0.06 |
| 5 | 157727648 | rs882884 | 0.318 | -0.295 | 0.268 | -0.423 | -0.014 | -0.005 | -0.005 | 0 | 0.72 | 0.57 | 0.85 | 0.76 | 0.77 | 0.85 |
| 5 | 157730917 | rs10515760 | 0.631 | -0.19 | 0.142 | -0.517 | -0.025 | -0.049 | -0.049 | 0 | 0.32 | 0.53 | 0.2 | 0.21 | 0.25 | 0.25 |
| 5 | 157735216 | rs4992528 | 0.408 | -0.058 | 0.119 | -0.558 | -0.003 | -0.013 | -0.013 | 0 | 0.33 | 0.39 | 0.21 | 0.22 | 0.28 | 0.27 |
| 5 | 157740602 | rs6870603 | 0.664 | -0.036 | -0.09 | -0.769 | -0.003 | -0.013 | -0.013 | 0.098 | 0.33 | 0.39 | 0.21 | 0.22 | 0.29 | 0.27 |
| 5 | 157743781 | rs2964330 | -0.145 | -0.257 | 0.218 | -1.016 | 0.015 | 0.045 | 0.045 | 0 | 0.55 | 0.4 | 0.72 | 0.78 | 0.59 | 0.65 |
| 5 | 157745474 | rs12655968 | 1.821 | 0.632 | -0.193 | -0.44 | -0.021 | -0.025 | -0.025 | 0 | 0.19 | 0.01 | 0.27 | 0.18 | 0.15 | 0.3 |
| 5 | 157752386 | rs10515761 | 1.069 | 1.08 | -0.197 | 0.213 | -0.059 | -0.068 | -0.068 | 0.023 | 0.12 | 0.03 | 0.32 | 0.3 | 0.25 | 0.2 |
| 5 | 157753540 | rs6896221 | 1.414 | 0.2 | 0.124 | -0.445 | 0.023 | 0.176 | 0.176 | 0.174 | 0.37 | 0.09 | 0.2 | 0.1 | 0.32 | 0.32 |
| 5 | 157758162 | rs4516837 | 1.349 | 0.224 | 0.176 | -0.435 | 0.031 | 0.157 | 0.157 | 0.11 | 0.37 | 0.12 | 0.19 | 0.1 | 0.32 | 0.32 |
| 5 | 157765934 | rs6894827 | 0.611 | 0.192 | 0.418 | -0.352 | -0.005 | -0.026 | -0.026 | 0 | 0.43 | 0.44 | 0.24 | 0.21 | 0.35 | 0.34 |
| 5 | 157766167 | rs2988321 | -1.148 | 0.233 | 0.337 | -0.349 | -0.036 | 0.087 | 0.087 | 0.579 | 0.86 | 0.99 | 0.92 | 1 | 0.84 | 0.93 |
| 5 | 157788342 | rs2773218 | 0.646 | 0.222 | 0.423 | -0.201 | -0.038 | 0.004 | 0.004 | 0 | 0.38 | 0.24 | 0.63 | 0.62 | 0.48 | 0.56 |
| 5 | 157789025 | rs13357130 | -0.827 | 0.343 | 0.524 | -0.306 | 0.096 | 0.162 | 0.162 | 1.479 | 0.07 | 0.48 | 0.06 | 0.11 | 0.04 | 0.02 |
| 5 | 157797077 | rs1015527 | 0.341 | 0.516 | -0.055 | -0.309 | 0.007 | 0.018 | 0.018 | 0 | 0.45 | 0.58 | 0.28 | 0.21 | 0.39 | 0.34 |
| 5 | 157803588 | rs13436194 | -0.029 | 0.514 | -0.126 | 0.102 | 0.001 | 0.008 | 0.008 | 0 | 0.59 | 0.64 | 0.45 | 0.36 | 0.42 | 0.47 |
| 5 | 157817634 | rs11960210 | 0.55 | 0.188 | -0.131 | -0.061 | 0.021 | 0.029 | 0.029 | 0 | 0.51 | 0.55 | 0.4 | 0.3 | 0.36 | 0.42 |
| 5 | 157822886 | rs10045385 | 1.207 | 0.197 | -0.133 | -0.181 | 0.012 | 0.003 | 0.003 | 0.673 | 0.09 | 0.05 | 0.04 | 0.06 | 0.06 | 0.06 |
| 5 | 157827241 | rs9313774 | 0.283 | 0.211 | 0.152 | -0.112 | 0.005 | 0.015 | 0.015 | 0.058 | 0.55 | 0.4 | 0.31 | 0.25 | 0.41 | 0.46 |
| 5 | 157861965 | rs10036622 | 0.241 | 0.221 | 0.008 | -0.176 | 0.011 | 0.039 | 0.039 | 0.176 | 0.54 | 0.35 | 0.31 | 0.25 | 0.43 | 0.46 |
| 5 | 157864334 | rs2901007 | 0.241 | 0.221 | 0.008 | -0.176 | 0.011 | 0.039 | 0.039 | 0.176 | 0.54 | 0.35 | 0.31 | 0.25 | 0.43 | 0.46 |

| | | | | | | | | | | | | | | | | |
|---|---|---|---|---|---|---|---|---|---|---|---|---|---|---|---|---|
| 5 | 157875231 | rs6556348 | 0.215 | 0.225 | 0.018 | -0.174 | 0.01 | 0.038 | 0.038 | 0.231 | 0.55 | 0.37 | 0.31 | 0.25 | 0.43 | 0.46 |
| 5 | 157877838 | rs10069494 | 0.226 | 0.288 | -0.077 | -0.238 | -0.001 | -0.01 | -0.01 | 0.322 | 0.56 | 0.43 | 0.3 | 0.24 | 0.41 | 0.46 |
| 5 | 157878160 | rs10045014 | -0.033 | 0.632 | -0.037 | -0.035 | 0.016 | -0.016 | -0.016 | 0 | 0.61 | 0.54 | 0.44 | 0.37 | 0.51 | 0.56 |
| 5 | 157879489 | rs6556350 | 0.453 | 0.45 | -0.155 | -0.457 | 0.005 | -0.009 | -0.009 | 0 | 0.49 | 0.42 | 0.36 | 0.12 | 0.44 | 0.44 |
| 5 | 157889391 | rs1316379 | 1.366 | 0.309 | -0.466 | -0.649 | 0.028 | 0.194 | 0.194 | 0.743 | 0.36 | 0.02 | 0.18 | 0.1 | 0.29 | 0.33 |
| 5 | 157891878 | rs2946160 | -0.786 | 1.087 | -0.385 | -0.086 | 0.002 | 0.003 | 0.003 | 0 | 0.71 | 0.67 | 0.74 | 0.65 | 0.73 | 0.66 |
| 5 | 157895459 | rs12332652 | 0.483 | 1.09 | -0.228 | -0.052 | -0.001 | -0.006 | -0.006 | 0 | 0.55 | 0.36 | 0.58 | 0.51 | 0.6 | 0.53 |
| 5 | 157899616 | rs2901003 | 0.401 | 0.915 | -0.121 | -0.016 | 0.001 | 0.002 | 0.002 | 0 | 0.12 | 0.14 | 0.14 | 0.14 | 0.12 | 0.11 |
| 5 | 157900829 | rs11135035 | 0.401 | 1.16 | -0.278 | -0.077 | 0.007 | 0.016 | 0.016 | 0.147 | 0.58 | 0.36 | 0.61 | 0.69 | 0.63 | 0.58 |
| 5 | 157901024 | rs11135036 | -0.931 | 1.171 | -0.227 | -0.055 | -0.002 | 0.003 | 0.003 | 0 | 0.73 | 0.67 | 0.77 | 0.72 | 0.76 | 0.68 |
| 5 | 157901746 | rs11135037 | 0.193 | 1.241 | -0.316 | 0.014 | 0.037 | 0.081 | 0.081 | 0 | 0.19 | 0.29 | 0.41 | 0.42 | 0.31 | 0.21 |
| 5 | 157917275 | rs4563609 | -1.34 | 0.619 | -0.975 | -0.29 | 0.005 | 0.086 | 0.086 | 0.075 | 0.61 | 0.94 | 0.79 | 0.72 | 0.67 | 0.63 |
| 5 | 157927591 | rs1317146 | -0.511 | 0.383 | -0.591 | -0.185 | 0.001 | 0.006 | 0.006 | 0.747 | 0.43 | 0.58 | 0.46 | 0.31 | 0.35 | 0.41 |
| 5 | 157947537 | rs10515765 | 2.205 | 0.518 | -0.785 | -0.273 | -0.037 | 0.009 | 0.009 | 0 | 0.06 | 0 | 0.1 | 0.02 | 0.1 | 0.09 |
| 5 | 157952205 | rs1317414 | -0.333 | 0.257 | -0.568 | -0.431 | 0.002 | -0.012 | -0.012 | 0 | 0.44 | 0.42 | 0.57 | 0.72 | 0.48 | 0.48 |
| 5 | 157953614 | rs17055794 | 1.393 | 0.168 | -0.598 | -0.421 | 0.001 | -0.006 | -0.006 | 1.187 | 0.08 | 0.08 | 0.04 | 0 | 0.03 | 0.01 |
| 5 | 157953813 | rs2963436 | 0.912 | 0.177 | -0.639 | -0.432 | -0.012 | 0.01 | 0.01 | 0 | 0.26 | 0.31 | 0.27 | 0.28 | 0.2 | 0.35 |
| 5 | 157956327 | rs10074759 | -0.239 | 0.306 | -0.781 | -0.37 | -0.082 | -0.099 | -0.099 | 0 | 0.27 | 0.13 | 0.4 | 0.45 | 0.38 | 0.25 |
| 5 | 157957642 | rs7732691 | -0.426 | 0.303 | -0.657 | -0.372 | -0.003 | -0.003 | -0.003 | 0 | 0.28 | 0.29 | 0.41 | 0.45 | 0.38 | 0.25 |
| 5 | 157969475 | rs17055818 | 1.08 | 0.345 | -0.868 | -0.308 | -0.04 | -0.069 | -0.069 | 0 | 0.16 | 0.08 | 0.27 | 0.44 | 0.26 | 0.17 |
| 5 | 157973399 | rs2963429 | -0.045 | 0.235 | -0.628 | -0.452 | 0.008 | -0.014 | -0.014 | 5.196 | 0.37 | 0.45 | 0.41 | 0.3 | 0.32 | 0.43 |
| 5 | 157978668 | rs6898621 | -0.133 | 0.387 | -0.634 | -0.399 | -0.006 | -0.055 | -0.055 | 0.581 | 0.24 | 0.19 | 0.31 | 0.44 | 0.29 | 0.17 |
| 5 | 157983114 | rs17055847 | 1.137 | 0.344 | -0.68 | -0.325 | -0.044 | -0.059 | -0.059 | 0 | 0.15 | 0.07 | 0.31 | 0.44 | 0.3 | 0.13 |
| 5 | 157996329 | rs2963475 | 0.181 | 0.169 | -0.479 | -0.365 | -0.007 | -0.024 | -0.024 | 0 | 0.31 | 0.41 | 0.3 | 0.31 | 0.21 | 0.4 |
| 5 | 158007165 | rs2914228 | -0.744 | 0.033 | -0.394 | -0.438 | 0.029 | 0.044 | 0.044 | 1.007 | 0.43 | 0.63 | 0.35 | 0.31 | 0.24 | 0.42 |
| 5 | 158016237 | rs1122081 | 0.177 | 0.163 | -0.133 | -0.344 | -0.069 | -0.078 | -0.078 | 1.685 | 0.13 | 0.37 | 0.06 | 0 | 0.03 | 0.01 |
| 5 | 158020161 | rs1629585 | -1.382 | 0.584 | -0.31 | -0.281 | 0.013 | 0.036 | 0.036 | 0 | 0.61 | 0.73 | 0.75 | 0.76 | 0.6 | 0.63 |
| 5 | 158020790 | rs975630 | -0.039 | 0.937 | -0.075 | -0.135 | 0 | -0.003 | -0.003 | 0 | 0.28 | 0.24 | 0.28 | 0.31 | 0.21 | 0.4 |
| 5 | 158029734 | rs1650505 | -0.032 | 0.939 | -0.041 | -0.135 | -0.001 | -0.001 | -0.001 | 0.043 | 0.27 | 0.24 | 0.28 | 0.31 | 0.21 | 0.4 |
| 5 | 158030482 | rs1650506 | -0.569 | 0.918 | -0.299 | -0.09 | 0.012 | -0.015 | -0.015 | 0 | 0.35 | 0.41 | 0.4 | 0.33 | 0.31 | 0.49 |
| 5 | 158033070 | rs1896695 | -0.244 | 0.19 | -0.863 | -0.161 | 0.007 | -0.008 | -0.008 | 0 | 0.27 | 0.23 | 0.35 | 0.28 | 0.24 | 0.33 |
| 5 | 158038931 | rs824872 | -0.966 | 0.118 | -0.77 | -0.371 | 0.019 | 0.017 | 0.017 | 0.488 | 0.39 | 0.7 | 0.48 | 0.29 | 0.34 | 0.44 |
| 5 | 158040346 | rs716451 | 0.324 | -0.423 | -0.372 | -0.477 | 0.026 | -0.061 | -0.061 | 0.113 | 0.11 | 0.38 | 0.12 | 0 | 0.09 | 0.07 |
| 5 | 158040399 | rs699082 | 0.262 | -0.38 | -0.432 | -0.404 | 0.002 | -0.002 | -0.002 | 0 | 0.22 | 0.22 | 0.23 | 0.25 | 0.15 | 0.28 |
| 5 | 158041829 | rs824879 | -1.372 | -0.636 | -1.04 | -0.686 | -0.002 | 0.056 | 0.056 | 0.886 | 0.7 | 0.99 | 0.88 | 0.83 | 0.76 | 0.69 |
| 5 | 158043052 | rs824881 | 0.369 | -0.416 | -0.71 | -0.444 | -0.021 | 0.005 | 0.005 | 0 | 0.22 | 0.29 | 0.24 | 0.25 | 0.15 | 0.28 |
| 5 | 158044089 | rs12654386 | 0.281 | -0.009 | -0.666 | -0.281 | 0.02 | 0.002 | 0.002 | 0 | 0.07 | 0.11 | 0.08 | 0.09 | 0.11 | 0.12 |
| 5 | 158044628 | rs972341 | 0.663 | -0.238 | -0.768 | -0.163 | -0.036 | -0.044 | -0.044 | 0 | 0.32 | 0.22 | 0.45 | 0.49 | 0.42 | 0.24 |
| 5 | 158045955 | rs824883 | 0.689 | -0.559 | -0.884 | -0.175 | 0.013 | -0.027 | -0.027 | 0 | 0.2 | 0.01 | 0.11 | 0.17 | 0.13 | 0.15 |
| 5 | 158048225 | rs4270681 | 0.452 | -0.548 | -0.618 | -0.161 | 0.015 | 0.077 | 0.077 | 0.053 | 0.05 | 0.32 | 0.14 | 0.09 | 0.04 | 0.14 |
| 5 | 158048398 | rs824893 | -0.023 | -0.523 | -1.126 | -0.152 | -0.011 | -0.001 | -0.001 | 0 | 0.26 | 0.3 | 0.25 | 0.25 | 0.17 | 0.29 |
| 5 | 158052650 | rs824890 | 0.434 | -0.092 | -1.369 | -0.322 | 0.035 | 0.012 | 0.012 | 0.67 | 0.22 | 0.07 | 0.12 | 0.17 | 0.12 | 0.16 |
| 5 | 158057822 | rs824888 | 0.031 | 0.483 | -0.886 | -0.408 | -0.001 | -0.031 | -0.031 | 0 | 0.6 | 0.51 | 0.7 | 0.74 | 0.6 | 0.53 |
| 5 | 158059982 | rs1428449 | -0.68 | 0.199 | -1.52 | -0.265 | 0.024 | 0.022 | 0.022 | 2.424 | 0.74 | 0.93 | 0.86 | 0.77 | 0.81 | 0.72 |
| 5 | 158079394 | rs184946 | -0.659 | 0.787 | -1.056 | -0.883 | -0.007 | -0.008 | -0.008 | 0 | 0.27 | 0.27 | 0.29 | 0.04 | 0.32 | 0.19 |
| 5 | 158083398 | rs824854 | 1.307 | 0.369 | -0.42 | -1.319 | -0.004 | -0.023 | -0.023 | 0 | 0.58 | 0.31 | 0.6 | 0.92 | 0.56 | 0.66 |
| 5 | 158090351 | rs13178063 | 0.086 | 0.46 | -0.509 | -1.211 | -0.016 | -0.056 | -0.056 | 0 | 0.13 | 0.32 | 0.09 | 0.03 | 0.1 | 0.15 |
| 5 | 158096651 | rs403334 | -0.391 | 0.826 | -1.173 | -1.278 | -0.012 | 0.014 | 0.014 | 0 | 0.84 | 0.86 | 0.84 | 0.96 | 0.81 | 0.87 |
| 5 | 158096975 | rs10515769 | 1.341 | 0.583 | -1.655 | -0.646 | -0.054 | 0.012 | 0.012 | 0.936 | 0.45 | 0.07 | 0.28 | 0.35 | 0.46 | 0.39 |
| 5 | 158104515 | rs1991493 | -0.785 | 1.007 | -0.617 | -0.488 | -0.006 | -0.008 | -0.008 | 0 | 0.19 | 0.18 | 0.18 | 0.03 | 0.24 | 0.2 |
| 5 | 158114288 | rs6867864 | -0.31 | 0.828 | -0.199 | -0.394 | -0.004 | -0.062 | -0.062 | 0 | 0.18 | 0.24 | 0.17 | 0.01 | 0.17 | 0.07 |
| 5 | 158115878 | rs17056092 | -0.354 | 0.72 | -0.197 | 0.193 | -0.002 | 0.055 | 0.055 | 0 | 0.08 | 0.05 | 0.31 | 0.64 | 0.1 | 0.31 |
| 5 | 158122679 | rs2161140 | 0.163 | 1.188 | 0.31 | -0.393 | 0.049 | -0.056 | -0.056 | 3.63 | 0.87 | 0.66 | 0.82 | 0.97 | 0.85 | 0.89 |
| 5 | 158131574 | rs6883772 | 1.336 | 1.075 | 0.357 | -0.475 | 0.007 | -0.145 | -0.145 | 0 | 0.67 | 0.46 | 0.67 | 0.96 | 0.69 | 0.79 |
| 5 | 158137439 | rs17056109 | -0.019 | 0.802 | -0.234 | 0.555 | -0.009 | 0.066 | 0.066 | 0 | 0.05 | 0 | 0.07 | 0.44 | 0.07 | 0.18 |
| 5 | 158162106 | rs2018466 | -0.098 | 0.777 | -0.113 | -0.167 | 0.008 | 0.04 | 0.04 | 0.175 | 0.17 | 0.09 | 0.15 | 0.01 | 0.16 | 0.07 |
| 5 | 158173509 | rs17712592 | -0.672 | 1.323 | -0.191 | 0.001 | -0.008 | 0.034 | 0.034 | 0 | 0.23 | 0.16 | 0.29 | 0.15 | 0.25 | 0.12 |
| 5 | 158174315 | rs17056159 | -0.863 | 1.267 | 0.525 | -0.006 | -0.001 | -0.032 | -0.032 | 0.573 | 0.24 | 0.39 | 0.31 | 0.15 | 0.25 | 0.12 |
| 5 | 158179152 | rs10476265 | -0.753 | 1.388 | 0.22 | 0.072 | -0.005 | -0.004 | -0.004 | 0 | 0.07 | 0.09 | 0.11 | 0.1 | 0.09 | 0.06 |
| 5 | 158182330 | rs17712788 | -0.102 | 1.883 | 0.105 | -0.497 | -0.019 | 0.012 | 0.012 | 0 | 0.18 | 0.09 | 0.13 | 0 | 0.22 | 0.15 |
| 5 | 158183996 | rs2277041 | 0.429 | 1.759 | 1.087 | -0.159 | -0.002 | -0.018 | -0.018 | 0 | 0.16 | 0.1 | 0.18 | 0.05 | 0.16 | 0.06 |
| 5 | 158203512 | rs17713494 | 0.714 | 1.872 | 1.051 | 0.32 | 0 | -0.014 | -0.014 | 0.032 | 0.09 | 0.08 | 0.06 | 0.01 | 0.1 | 0.12 |
| 5 | 158204347 | rs12657410 | -0.249 | 1.593 | 1.549 | 0.671 | -0.01 | 0.035 | 0.035 | 0.948 | 0.08 | 0.15 | 0.04 | 0.13 | 0.05 | 0.07 |
| 5 | 158204425 | rs1368298 | -2.108 | 1.544 | 1.503 | 0.644 | -0.002 | 0.021 | 0.021 | 0.911 | 0.45 | 0.85 | 0.61 | 0.62 | 0.45 | 0.35 |
| 5 | 158220193 | rs31864 | -2.341 | 1.609 | 1.451 | 0.645 | -0.003 | 0.019 | 0.019 | 0.583 | 0.45 | 0.85 | 0.61 | 0.61 | 0.45 | 0.35 |
| 5 | 158220492 | rs31863 | -0.913 | 1.571 | 1.718 | 0.617 | -0.005 | 0.021 | 0.021 | 0 | 0.31 | 0.46 | 0.47 | 0.38 | 0.32 | 0.22 |
| 5 | 158222797 | rs31862 | 0.526 | 1.682 | 1.154 | 0.402 | -0.005 | 0.022 | 0.022 | 0 | 0.17 | 0.07 | 0.21 | 0.07 | 0.21 | 0.07 |
| 5 | 158229781 | rs11750796 | 0.804 | 1.671 | 1.033 | 0.479 | -0.001 | -0.012 | -0.012 | 0.172 | 0.09 | 0.08 | 0.06 | 0.01 | 0.1 | 0.12 |
| 5 | 158246650 | rs7720275 | -0.15 | 1.744 | 1.211 | 0.546 | 0.026 | -0.009 | -0.009 | 0 | 0.12 | 0.24 | 0.15 | 0.1 | 0.18 | 0.17 |

| | | | | | | | | | | | | | | | | |
|---|---|---|---|---|---|---|---|---|---|---|---|---|---|---|---|---|
| 5 | 158256767 | rs13359166 | 0.123 | 1.616 | 1.267 | 0.619 | -0.005 | 0.018 | 0.018 | 0.003 | 0.77 | 0.77 | 0.6 | 0.63 | 0.77 | 0.89 |
| 5 | 158261163 | rs17715065 | 1.824 | 1.979 | 0.923 | 0.352 | -0.046 | 0.034 | 0.034 | 0.708 | 0.53 | 0.14 | 0.37 | 0.38 | 0.49 | 0.64 |
| 5 | 158263144 | rs6885822 | -0.026 | 1.823 | 1.075 | -0.207 | -0.002 | 0.001 | 0.001 | 0 | 0.12 | 0.14 | 0.15 | 0.09 | 0.18 | 0.17 |
| 5 | 158269081 | rs10052777 | 1.779 | 1.917 | 1.334 | -0.048 | 0.024 | 0.058 | 0.058 | 0.204 | 0.49 | 0.37 | 0.35 | 0.28 | 0.37 | 0.59 |
| 5 | 158269734 | rs17715535 | 1.889 | 1.917 | 1.265 | -0.048 | 0.035 | 0.121 | 0.121 | 0.438 | 0.48 | 0.19 | 0.33 | 0.28 | 0.37 | 0.59 |
| 5 | 158273665 | rs11747709 | 0.224 | 2.165 | 1.243 | 0.022 | -0.01 | 0.004 | 0.004 | 0 | 0.13 | 0.11 | 0.1 | 0.01 | 0.21 | 0.17 |
| 5 | 158282326 | rs7715522 | 0.154 | 2.162 | 1.346 | 0.02 | 0.024 | -0.054 | -0.054 | 0 | 0.14 | 0.21 | 0.1 | 0.01 | 0.21 | 0.17 |
| 5 | 158295127 | rs31193 | 0.132 | 2.182 | 1.334 | 0.029 | 0.023 | -0.054 | -0.054 | 0 | 0.14 | 0.21 | 0.1 | 0.01 | 0.21 | 0.17 |
| 5 | 158310631 | rs929626 | 1.452 | 2.012 | 1.252 | -0.027 | -0.001 | 0.056 | 0.056 | 2.744 | 0.49 | 0.31 | 0.41 | 0.36 | 0.44 | 0.64 |
| 5 | 158332380 | rs17543752 | 1.824 | 1.976 | 1.268 | -0.033 | 0.017 | 0.095 | 0.095 | 1.171 | 0.46 | 0.2 | 0.33 | 0.28 | 0.37 | 0.59 |
| 5 | 158338397 | rs1862385 | 0.111 | 1.743 | 1.377 | 0.023 | -0.013 | 0.019 | 0.019 | 0.042 | 0.74 | 0.73 | 0.59 | 0.47 | 0.79 | 0.9 |
| 5 | 158345923 | rs6869051 | -1.083 | 1.849 | 1.003 | 0.077 | 0.003 | 0.02 | 0.02 | 0 | 0.84 | 0.92 | 0.79 | 0.95 | 0.83 | 0.94 |
| 5 | 158359476 | rs6556373 | 1.23 | 1.847 | 0.839 | 0.063 | -0.005 | 0.015 | 0.015 | 0 | 0.56 | 0.34 | 0.4 | 0.39 | 0.51 | 0.66 |
| 5 | 158366263 | rs4921537 | 0.906 | 1.763 | 0.822 | 0.091 | -0.005 | 0.005 | 0.005 | 0.562 | 0.61 | 0.43 | 0.5 | 0.46 | 0.6 | 0.74 |
| 5 | 158386181 | rs11135050 | 1.591 | 0.962 | 0.296 | 0.141 | 0.076 | 0.05 | 0.05 | 0.341 | 0.42 | 0.18 | 0.24 | 0.33 | 0.28 | 0.54 |
| 5 | 158387921 | rs6420100 | -0.272 | 1.039 | 0.329 | 0.399 | -0.013 | 0.132 | 0.132 | 0 | 0.94 | 0.9 | 0.82 | 0.58 | 0.99 | 0.96 |
| 5 | 158393842 | rs12109219 | 0.213 | 1.505 | 0.323 | -0.06 | 0.009 | -0.014 | -0.014 | 0 | 0.17 | 0.17 | 0.19 | 0.03 | 0.3 | 0.23 |
| 5 | 158404129 | rs2161356 | 2.198 | 0.987 | -0.183 | -0.342 | 0.008 | -0.032 | -0.032 | 2.839 | 0.31 | 0.01 | 0.21 | 0.29 | 0.23 | 0.43 |
| 5 | 158418490 | rs10447210 | 0.594 | 1.329 | 0.167 | -0.152 | 0.031 | 0.012 | 0.012 | 3.049 | 0.6 | 0.3 | 0.47 | 0.39 | 0.66 | 0.8 |
| 5 | 158423745 | rs12055028 | 1.789 | 1.336 | 0.157 | -0.17 | 0.069 | 0.074 | 0.074 | 0.318 | 0.41 | 0.11 | 0.24 | 0.29 | 0.29 | 0.53 |
| 5 | 158424391 | rs10040979 | 0.622 | 1.069 | 0.543 | -0.152 | -0.021 | 0.002 | 0.002 | 3.077 | 0.62 | 0.49 | 0.52 | 0.44 | 0.67 | 0.82 |
| 5 | 158442109 | rs4244439 | -0.494 | 1.577 | 0.292 | -0.477 | 0 | -0.011 | -0.011 | 1.749 | 0.14 | 0.2 | 0.08 | 0.1 | 0.15 | 0.09 |
| 5 | 158443352 | rs10515785 | -0.297 | 1.757 | -0.124 | -0.374 | 0.008 | -0.014 | -0.014 | 0 | 0.17 | 0.17 | 0.19 | 0.03 | 0.3 | 0.22 |
| 5 | 158451492 | rs6890049 | -0.211 | 0.508 | -0.307 | -0.43 | 0 | -0.003 | -0.003 | 0.321 | 0.32 | 0.68 | 0.34 | 0.05 | 0.31 | 0.21 |
| 5 | 158452056 | rs10475669 | -0.107 | 0.296 | -0.351 | 0.576 | -0.02 | 0.22 | 0.22 | 0 | 0.05 | 0.14 | 0.19 | 0.31 | 0.02 | 0.09 |
| 5 | 158459721 | rs7720283 | -1.569 | -0.21 | -1.246 | 0.075 | 0.052 | -0.048 | -0.048 | 0 | 0.73 | 0.99 | 0.85 | 0.74 | 0.85 | 0.72 |
| 5 | 158460212 | rs7734385 | 0.217 | -0.093 | -1.165 | 0.004 | -0.07 | 0.022 | 0.022 | 0 | 0.29 | 0.14 | 0.31 | 0.23 | 0.45 | 0.35 |
| 5 | 158462476 | rs13174549 | -0.118 | -0.435 | -0.753 | -0.037 | -0.002 | 0.002 | 0.002 | 0 | 0.11 | 0.11 | 0.05 | 0.08 | 0.06 | 0.08 |
| 5 | 158467401 | rs1422672 | -0.335 | -0.311 | -0.332 | -0.144 | 0.01 | -0.013 | -0.013 | 0 | 0.34 | 0.34 | 0.43 | 0.5 | 0.28 | 0.28 |
| 5 | 158475986 | rs10515786 | -0.414 | -0.315 | 0.103 | -0.154 | -0.038 | 0.031 | 0.031 | 0.918 | 0.36 | 0.5 | 0.44 | 0.51 | 0.28 | 0.28 |
| 5 | 158476709 | rs17645325 | -0.219 | 0.597 | -0.13 | 0.267 | 0.02 | -0.019 | -0.019 | 0 | 0.06 | 0.07 | 0.06 | 0 | 0.1 | 0.08 |
| 5 | 158476891 | rs6877077 | 0.709 | 0.51 | -0.094 | 0.319 | 0.001 | -0.002 | -0.002 | 0 | 0.22 | 0.18 | 0.37 | 0.42 | 0.21 | 0.2 |
| 5 | 158486993 | rs4921121 | 0.479 | 0.232 | -0.353 | 0.344 | -0.004 | -0.031 | -0.031 | 0 | 0.26 | 0.2 | 0.42 | 0.43 | 0.26 | 0.22 |
| 5 | 158488198 | rs929779 | -0.454 | 0.22 | -0.515 | 0.254 | -0.008 | 0.023 | 0.023 | 0 | 0.37 | 0.26 | 0.36 | 0.23 | 0.55 | 0.52 |
| 5 | 158489315 | rs7711823 | -0.247 | 0.513 | -1.171 | 0.685 | 0.139 | 0.022 | 0.022 | 0 | 0.21 | 0.48 | 0.23 | 0.22 | 0.38 | 0.35 |
| 5 | 158490202 | rs17056506 | -0.012 | 0.542 | -1.384 | 0.843 | 0.008 | 0.175 | 0.175 | 0 | 0.08 | 0.16 | 0.22 | 0.39 | 0.07 | 0.16 |
| 5 | 158490797 | rs6883429 | -1.07 | 0.161 | -2.138 | -0.008 | 0.106 | 0.03 | 0.03 | 0.034 | 0.68 | 0.88 | 0.83 | 0.74 | 0.83 | 0.78 |
| 5 | 158492279 | rs13181905 | -0.298 | 1.111 | -1.706 | -0.146 | -0.02 | 0.107 | 0.107 | 0.042 | 0.39 | 0.21 | 0.39 | 0.13 | 0.42 | 0.34 |
| 5 | 158501850 | rs17056526 | -0.712 | 1.516 | -1.73 | 0.945 | -0.076 | -0.093 | -0.093 | 0.696 | 0.16 | 0.03 | 0.33 | 0.31 | 0.23 | 0.38 |
| 5 | 158502378 | rs17056534 | -0.796 | 1.535 | -1.773 | 1.236 | -0.079 | -0.099 | -0.099 | 0 | 0.15 | 0.01 | 0.3 | 0.34 | 0.2 | 0.44 |
| 5 | 158504140 | rs7726961 | 0.203 | 0.929 | -2.102 | 0.451 | 0.057 | 0.026 | 0.026 | 0 | 0.51 | 0.78 | 0.39 | 0.56 | 0.39 | 0.43 |
| 5 | 158506415 | rs10515789 | 0.215 | 0.647 | -1.274 | 0.905 | -0.009 | 0.197 | 0.197 | 0 | 0.09 | 0.16 | 0.13 | 0.34 | 0.06 | 0.1 |
| 5 | 158507676 | rs12054738 | 0.199 | 0.638 | -1.278 | 0.901 | -0.01 | 0.194 | 0.194 | 0 | 0.09 | 0.16 | 0.13 | 0.34 | 0.06 | 0.1 |
| 5 | 158510054 | rs6866195 | 0.491 | -0.134 | -0.89 | 0.484 | 0 | -0.004 | -0.004 | 0 | 0.32 | 0.29 | 0.22 | 0.16 | 0.29 | 0.25 |
| 5 | 158513125 | rs7707897 | -0.274 | -0.1 | -0.655 | 0.537 | 0.013 | -0.012 | -0.012 | 0 | 0.42 | 0.6 | 0.35 | 0.5 | 0.35 | 0.34 |
| 5 | 158513831 | rs6898651 | 0.406 | 0.117 | -0.814 | 0.005 | 0 | 0.006 | 0.006 | 0 | 0.3 | 0.25 | 0.2 | 0.15 | 0.29 | 0.25 |
| 5 | 158514771 | rs10035751 | 0.137 | 0.096 | -0.788 | 0.006 | -0.001 | -0.012 | -0.012 | 0 | 0.33 | 0.4 | 0.22 | 0.16 | 0.31 | 0.26 |
| 5 | 158515199 | rs6879995 | 0.137 | 0.096 | -0.788 | 0.006 | -0.001 | -0.012 | -0.012 | 0.989 | 0.33 | 0.4 | 0.22 | 0.16 | 0.31 | 0.26 |
| 5 | 158530015 | rs7732511 | 0.4 | 0.339 | -1.336 | -0.058 | -0.014 | 0.04 | 0.04 | 0 | 0.15 | 0.03 | 0.07 | 0.06 | 0.15 | 0.12 |
| 5 | 158537119 | rs2161357 | 0.089 | 0.586 | -1.124 | -0.141 | 0.028 | -0.023 | -0.023 | 0 | 0.07 | 0.11 | 0.19 | 0.02 | 0.18 | 0.08 |
| 5 | 158538277 | rs4921317 | -1.398 | 0.458 | -1.384 | 0.52 | -0.018 | -0.024 | -0.024 | 0.678 | 0.37 | 0.21 | 0.4 | 0.54 | 0.42 | 0.45 |
| 5 | 158541200 | rs6897374 | 0.615 | 0.492 | -1.334 | 0.711 | 0 | 0.061 | 0.061 | 0 | 0.08 | 0.07 | 0.12 | 0.39 | 0.08 | 0.11 |
| 5 | 158543969 | rs11950065 | 0.307 | 0.48 | -1.402 | 0.561 | 0.019 | -0.018 | -0.018 | 0.027 | 0.07 | 0.1 | 0.19 | 0.02 | 0.18 | 0.08 |
| 5 | 158556738 | rs270664 | 1.267 | 0.963 | -1.453 | 0.372 | -0.059 | 0 | 0 | 0 | 0.6 | 0.81 | 0.65 | 0.44 | 0.52 | 0.53 |
| 5 | 158560154 | rs270661 | -0.165 | 1.038 | -1.546 | 0.295 | 0.008 | 0.028 | 0.028 | 0 | 0.24 | 0.17 | 0.11 | 0.13 | 0.18 | 0.23 |
| 5 | 158560181 | rs270660 | 1.516 | 0.576 | -2.041 | -0.388 | 0.008 | 0.034 | 0.034 | 0 | 0.14 | 0.09 | 0.09 | 0.04 | 0.11 | 0.04 |
| 5 | 158562019 | rs6556396 | 0.179 | 0.648 | -2.131 | -0.227 | -0.036 | -0.057 | -0.057 | 0 | 0.82 | 0.88 | 0.75 | 0.59 | 0.7 | 0.77 |
| 5 | 158565109 | rs270654 | 1.939 | 0.54 | -2.277 | -0.389 | -0.025 | 0.043 | 0.043 | 0 | 0.08 | 0 | 0.05 | 0 | 0.09 | 0.06 |
| 5 | 158568110 | rs2420355 | 1.051 | 0.245 | -2.063 | -0.446 | 0.022 | 0.025 | 0.025 | 0 | 0.52 | 0.72 | 0.57 | 0.48 | 0.43 | 0.62 |
| 5 | 158579712 | rs6556401 | 0.55 | 0.923 | -2.579 | -0.699 | -0.167 | -0.137 | -0.137 | 0 | 0.69 | 0.94 | 0.66 | 0.51 | 0.53 | 0.62 |
| 5 | 158584991 | rs10035989 | -0.978 | 1.022 | -2.578 | -0.604 | -0.045 | -0.021 | -0.021 | 0 | 0.91 | 0.98 | 0.89 | 0.93 | 0.82 | 0.9 |
| 5 | 158589858 | rs10515774 | 2.202 | 0.94 | -2.632 | -0.68 | 0.008 | 0.027 | 0.027 | 0 | 0.12 | 0.06 | 0.07 | 0.04 | 0.09 | 0.03 |
| 5 | 158590454 | rs10515775 | 1.324 | 0.99 | -2.657 | -0.675 | -0.037 | -0.026 | -0.026 | 0 | 0.09 | 0.01 | 0.14 | 0.36 | 0.12 | 0.11 |
| 5 | 158597936 | rs2043269 | 2.263 | 0.792 | -2.608 | -0.771 | 0.008 | 0.031 | 0.031 | 0 | 0.12 | 0.04 | 0.07 | 0.04 | 0.09 | 0.03 |
| 5 | 158602694 | rs12655958 | 2.263 | 0.792 | -2.606 | -0.771 | 0.008 | 0.034 | 0.034 | 0 | 0.12 | 0.04 | 0.07 | 0.04 | 0.09 | 0.03 |
| 5 | 158603271 | rs13158488 | 1.032 | 0.663 | -2.499 | -0.729 | -0.008 | 0.035 | 0.035 | 0 | 0.17 | 0.08 | 0.1 | 0.07 | 0.17 | 0.11 |
| 5 | 158604963 | rs10076782 | 1.179 | 0.582 | -2.426 | -0.678 | 0.088 | 0.017 | 0.017 | 0 | 0.39 | 0.74 | 0.49 | 0.4 | 0.26 | 0.47 |
| 5 | 158624751 | rs6888950 | 1.527 | 0.571 | -2.345 | -0.683 | 0.044 | 0.008 | 0.008 | 0 | 0.38 | 0.67 | 0.48 | 0.4 | 0.26 | 0.47 |
| 5 | 158626111 | rs17663989 | 0.915 | 0.849 | -2.46 | -0.541 | -0.072 | 0.008 | 0.008 | 0 | 0.14 | 0.01 | 0.09 | 0.09 | 0.17 | 0.18 |

| | | | | | | | | | | | | | | | | |
|---|---|---|---|---|---|---|---|---|---|---|---|---|---|---|---|---|
| 5 | 158626347 | rs2163775 | 0.624 | 0.751 | -2.463 | -0.59 | 0.026 | 0.017 | 0.017 | 0 | 0.55 | 0.76 | 0.58 | 0.46 | 0.43 | 0.59 |
| 5 | 158650814 | rs7721001 | 0.579 | 0.789 | -2.384 | -0.44 | 0.016 | 0.008 | 0.008 | 0 | 0.57 | 0.81 | 0.59 | 0.45 | 0.44 | 0.59 |
| 5 | 158658012 | rs10515778 | 0.488 | 1.096 | -2.174 | -0.162 | -0.019 | 0.018 | 0.018 | 0 | 0.14 | 0.06 | 0.1 | 0.08 | 0.17 | 0.18 |
| 5 | 158659707 | rs17056627 | 0.729 | 0.013 | -2.701 | -0.198 | 0.029 | -0.071 | -0.071 | 0.475 | 0.11 | 0 | 0.26 | 0.37 | 0.07 | 0.35 |
| 5 | 158666731 | rs254850 | 0.569 | -0.085 | -2.555 | -0.464 | -0.035 | 0.062 | 0.062 | 0 | 0.18 | 0.01 | 0.1 | 0.07 | 0.18 | 0.12 |
| 5 | 158679579 | rs3846687 | 0.084 | 0.299 | -2.312 | 0.048 | -0.141 | -0.142 | -0.142 | 1.291 | 0.22 | 0.02 | 0.32 | 0.38 | 0.34 | 0.31 |
| 5 | 158687153 | rs11744690 | 0.394 | -0.069 | -1.853 | -0.126 | -0.028 | -0.022 | -0.022 | 0.319 | 0.2 | 0.15 | 0.32 | 0.31 | 0.3 | 0.32 |
| 5 | 158690951 | rs4921437 | -0.779 | -0.404 | -1.354 | -0.571 | 0.002 | -0.084 | -0.084 | 0.544 | 0.8 | 0.74 | 0.89 | 1 | 0.83 | 0.87 |
| 5 | 158698068 | rs4147384 | -0.47 | -0.715 | -1.662 | -0.642 | -0.057 | -0.059 | -0.059 | 0.103 | 0.31 | 0.2 | 0.44 | 0.45 | 0.48 | 0.44 |
| 5 | 158699263 | rs11745659 | 1.062 | -0.648 | -1.637 | -0.682 | -0.03 | -0.024 | -0.024 | 0 | 0.2 | 0.15 | 0.32 | 0.31 | 0.3 | 0.32 |
| 5 | 158705370 | rs12520035 | 1.604 | -1.326 | -1.695 | -0.877 | 0.042 | 0.049 | 0.049 | 0 | 0.2 | 0.1 | 0.11 | 0.07 | 0.12 | 0.06 |
| 5 | 158711219 | rs4921209 | 1.338 | -1.249 | -1.666 | -0.843 | -0.074 | -0.06 | -0.06 | 0 | 0.2 | 0.09 | 0.31 | 0.31 | 0.3 | 0.32 |
| 5 | 158715663 | rs4921458 | -1.491 | -1.238 | -1.435 | -0.887 | -0.006 | 0.006 | 0.006 | 0 | 0.82 | 0.92 | 0.91 | 1 | 0.83 | 0.87 |
| 5 | 158722094 | rs12652431 | 1.471 | -1.418 | -1.361 | -0.723 | 0.038 | 0.045 | 0.045 | 0 | 0.2 | 0.1 | 0.11 | 0.07 | 0.12 | 0.06 |
| 5 | 158735933 | rs4921468 | 0.288 | -1.427 | -1.388 | -0.661 | 0.021 | 0.014 | 0.014 | 0 | 0.31 | 0.19 | 0.25 | 0.22 | 0.31 | 0.18 |
| 5 | 158740623 | rs3181225 | 0.689 | -1.356 | -1.363 | -0.72 | -0.016 | -0.026 | -0.026 | 0 | 0.11 | 0.05 | 0.11 | 0.15 | 0.17 | 0.12 |
| 5 | 158740850 | rs3181224 | 1.453 | -1.393 | -1.351 | -0.735 | 0.03 | 0.041 | 0.041 | 0 | 0.2 | 0.11 | 0.14 | 0.04 | 0.13 | 0.06 |
| 5 | 158742950 | rs3212227 | 1.846 | -1.601 | -1.053 | -0.801 | -0.016 | 0.029 | 0.029 | 0 | 0.28 | 0.35 | 0.34 | 0.5 | 0.22 | 0.37 |
| 5 | 158743846 | rs11574790 | 1.538 | -1.575 | -0.926 | -0.816 | -0.005 | -0.018 | -0.018 | 0 | 0.21 | 0.22 | 0.15 | 0.04 | 0.13 | 0.06 |
| 5 | 158749088 | rs2853694 | -0.129 | -1.58 | -0.894 | -0.796 | -0.028 | 0.027 | 0.027 | 0.619 | 0.6 | 0.78 | 0.61 | 0.69 | 0.53 | 0.55 |
| 5 | 158750013 | rs2288831 | 1.743 | -1.437 | -0.941 | -0.654 | -0.016 | 0.029 | 0.029 | 0 | 0.28 | 0.36 | 0.34 | 0.51 | 0.22 | 0.37 |
| 5 | 158750769 | rs3213094 | 1.743 | -1.532 | -0.96 | -0.699 | -0.016 | 0.029 | 0.029 | 0 | 0.28 | 0.36 | 0.34 | 0.51 | 0.22 | 0.38 |
| 5 | 158751249 | rs2569254 | 0.683 | -1.523 | -0.969 | -0.736 | -0.016 | -0.025 | -0.025 | 0 | 0.11 | 0.05 | 0.11 | 0.14 | 0.17 | 0.12 |
| 5 | 158754195 | rs3212220 | 1.63 | -1.491 | -0.907 | -0.716 | -0.016 | 0.03 | 0.03 | 0 | 0.28 | 0.36 | 0.34 | 0.51 | 0.23 | 0.37 |
| 5 | 158755130 | rs3212217 | 1.641 | -1.487 | -0.937 | -0.714 | -0.016 | 0.029 | 0.029 | 0 | 0.28 | 0.34 | 0.34 | 0.51 | 0.23 | 0.37 |
| 5 | 158755566 | rs1003199 | 0.15 | -1.135 | -1.302 | -0.603 | -0.034 | -0.013 | -0.013 | 0.245 | 0.31 | 0.22 | 0.39 | 0.37 | 0.47 | 0.44 |
| 5 | 158755960 | rs2546893 | -0.062 | -0.822 | -1.246 | -0.81 | -0.043 | -0.049 | -0.049 | 0 | 0.35 | 0.21 | 0.41 | 0.45 | 0.47 | 0.45 |
| 5 | 158756227 | rs730691 | -0.843 | -1.138 | -1.04 | -0.661 | 0.008 | -0.008 | -0.008 | 0 | 0.59 | 0.35 | 0.56 | 0.5 | 0.62 | 0.53 |
| 5 | 158759900 | rs2546890 | 0.532 | -0.996 | -0.676 | -0.419 | 0.017 | 0.001 | 0.001 | 0.191 | 0.52 | 0.65 | 0.59 | 0.59 | 0.51 | 0.44 |
| 5 | 158764177 | rs7709212 | 1.501 | -0.799 | -0.404 | -0.351 | 0 | 0.001 | 0.001 | 0 | 0.29 | 0.3 | 0.46 | 0.51 | 0.35 | 0.4 |
| 5 | 158764420 | rs6868898 | 0.545 | -0.491 | -0.819 | -0.163 | -0.035 | -0.028 | -0.028 | 0 | 0.27 | 0.2 | 0.46 | 0.35 | 0.32 | 0.32 |
| 5 | 158770755 | rs17056706 | -0.057 | -0.456 | -0.727 | -0.282 | 0.004 | 0.008 | 0.008 | 0.161 | 0.37 | 0.54 | 0.5 | 0.46 | 0.35 | 0.45 |
| 5 | 158772582 | rs953861 | -1.15 | -0.565 | -0.834 | -0.42 | -0.037 | 0.098 | 0.098 | 0 | 0.84 | 0.99 | 0.92 | 1 | 0.85 | 0.88 |
| 5 | 158773779 | rs11746138 | 1.487 | -0.467 | -0.658 | -0.333 | 0.012 | 0.009 | 0.009 | 0 | 0.16 | 0.05 | 0.11 | 0.08 | 0.16 | 0.05 |
| 5 | 158781604 | rs6869411 | -0.045 | -0.353 | -0.551 | -0.236 | 0.004 | 0.007 | 0.007 | 0.175 | 0.37 | 0.54 | 0.5 | 0.45 | 0.34 | 0.45 |
| 5 | 158806244 | rs4262088 | 0.733 | -0.295 | -1.542 | -0.138 | -0.024 | 0.028 | 0.028 | 0 | 0.11 | 0.23 | 0.22 | 0.18 | 0.05 | 0.15 |
| 5 | 158814533 | rs10045431 | -0.359 | -0.289 | -1.683 | -0.185 | -0.001 | 0.083 | 0.083 | 0 | 0.71 | 0.91 | 0.83 | 0.92 | 0.72 | 0.85 |
| 5 | 158820822 | rs10463070 | 0.885 | -0.126 | -1.688 | -0.177 | -0.042 | -0.018 | -0.018 | 0.506 | 0.25 | 0.16 | 0.28 | 0.27 | 0.3 | 0.3 |
| 5 | 158821493 | rs7704367 | 0.788 | 0.221 | -1.196 | -0.1 | -0.005 | 0.034 | 0.034 | 0 | 0.32 | 0.48 | 0.31 | 0.46 | 0.34 | 0.39 |
| 5 | 158823786 | rs6556417 | 0.234 | 0.443 | -1.106 | -0.283 | -0.041 | 0.083 | 0.083 | 0 | 0.86 | 1 | 0.93 | 1 | 0.86 | 0.89 |
| 5 | 158824979 | rs17056737 | -1.05 | 0.302 | -0.503 | 0.297 | -0.006 | -0.001 | -0.001 | 0 | 0.14 | 0.17 | 0.07 | 0.12 | 0.07 | 0.17 |
| 5 | 158826357 | rs918519 | 0.383 | 1.221 | -0.123 | 0.035 | 0.063 | -0.072 | -0.072 | 0 | 0.86 | 0.73 | 0.88 | 1 | 0.82 | 0.88 |
| 5 | 158830819 | rs4921226 | 0.169 | 1.136 | -1.533 | 0.021 | -0.075 | -0.013 | -0.013 | 0.008 | 0.35 | 0.06 | 0.35 | 0.24 | 0.41 | 0.36 |
| 5 | 158836107 | rs4921493 | 0.751 | 0.432 | -1.118 | 0.334 | -0.012 | 0 | 0 | 0 | 0.34 | 0.26 | 0.29 | 0.38 | 0.34 | 0.37 |
| 5 | 158843388 | rs17056747 | -0.303 | 0.156 | -1.285 | 0.198 | 0.007 | 0.005 | 0.005 | 0.026 | 0.14 | 0.11 | 0.08 | 0.08 | 0.08 | 0.11 |
| 5 | 158848071 | rs4921496 | -0.352 | 0.205 | -0.579 | 0.452 | 0.004 | -0.001 | -0.001 | 0 | 0.82 | 0.77 | 0.87 | 0.87 | 0.8 | 0.81 |
| 5 | 158848492 | rs17454570 | 0.672 | 0.191 | -0.743 | 0.544 | -0.032 | -0.05 | -0.05 | 0 | 0.33 | 0.05 | 0.35 | 0.36 | 0.38 | 0.37 |
| 5 | 158849491 | rs10866712 | 0.429 | 0.16 | -0.715 | 0.55 | 0.001 | 0.001 | 0.001 | 0 | 0.7 | 0.66 | 0.61 | 0.74 | 0.71 | 0.72 |
| 5 | 158849837 | rs4921227 | 1.012 | 0.204 | -0.397 | 0.52 | 0.001 | -0.005 | -0.005 | 0.515 | 0.36 | 0.52 | 0.26 | 0.38 | 0.33 | 0.35 |
| 5 | 158858447 | rs12055208 | -0.013 | 0.618 | -1.25 | 0.432 | -0.074 | -0.052 | -0.052 | 0 | 0.43 | 0.14 | 0.64 | 0.47 | 0.53 | 0.55 |
| 5 | 158860636 | rs10515802 | -0.199 | 0.589 | -0.43 | 0.359 | 0.004 | -0.01 | -0.01 | 2.283 | 0.18 | 0.41 | 0.12 | 0.11 | 0.15 | 0.15 |
| 5 | 158860939 | rs10060059 | -0.176 | 0.525 | -1.306 | 0.179 | -0.045 | -0.021 | -0.021 | 0.251 | 0.65 | 0.91 | 0.62 | 0.62 | 0.57 | 0.54 |
| 5 | 158865362 | rs1422882 | 0.42 | 0.269 | -0.89 | 0.031 | -0.001 | -0.009 | -0.009 | 0 | 0.46 | 0.34 | 0.55 | 0.54 | 0.51 | 0.47 |
| 5 | 158867212 | rs7736632 | 0.533 | 0.229 | -0.869 | -0.311 | 0.003 | -0.003 | -0.003 | 0 | 0.53 | 0.35 | 0.51 | 0.5 | 0.51 | 0.5 |
| 5 | 158868175 | rs2059131 | -0.165 | 0.178 | -0.514 | -0.251 | 0.043 | 0.067 | 0.067 | 0 | 0.74 | 0.51 | 0.68 | 0.66 | 0.68 | 0.7 |
| 5 | 158879069 | rs6896881 | 0.353 | 0.194 | -0.136 | -0.262 | 0.008 | 0.016 | 0.016 | 0.373 | 0.13 | 0.38 | 0.09 | 0.11 | 0.11 | 0.1 |
| 5 | 158880032 | rs7703043 | 1.355 | 0.039 | -0.642 | -0.441 | -0.004 | 0.025 | 0.025 | 0 | 0.16 | 0.1 | 0.27 | 0.05 | 0.16 | 0.13 |
| 5 | 158881497 | rs6882695 | 0.993 | -0.023 | -0.67 | -0.313 | 0.035 | 0.002 | 0.002 | 0 | 0.38 | 0.22 | 0.25 | 0.34 | 0.34 | 0.35 |
| 5 | 158884535 | rs7720046 | 0.078 | 0.012 | -0.29 | -0.297 | -0.009 | -0.013 | -0.013 | 0.048 | 0.54 | 0.73 | 0.37 | 0.46 | 0.5 | 0.46 |
| 5 | 158888106 | rs10057242 | -0.485 | 0.121 | -0.144 | -0.258 | 0.004 | -0.01 | -0.01 | 0 | 0.36 | 0.57 | 0.44 | 0.29 | 0.37 | 0.31 |
| 5 | 158901965 | rs6889446 | -0.353 | 0.21 | -0.085 | -0.163 | 0.032 | 0.029 | 0.029 | 0.278 | 0.2 | 0.49 | 0.16 | 0.26 | 0.21 | 0.19 |
| 5 | 158904313 | rs12657996 | 1.587 | 0.019 | -0.06 | -0.271 | -0.008 | 0.021 | 0.021 | 0 | 0.27 | 0.45 | 0.2 | 0.33 | 0.25 | 0.27 |
| 5 | 158904390 | rs7706354 | 1.09 | 0.05 | -0.065 | -0.23 | 0.008 | 0.025 | 0.025 | 0 | 0.12 | 0.24 | 0.08 | 0.12 | 0.11 | 0.1 |
| 5 | 158907016 | rs2421048 | -0.328 | 0.022 | -0.419 | -0.321 | -0.029 | -0.05 | -0.05 | 0 | 0.63 | 0.91 | 0.46 | 0.61 | 0.61 | 0.58 |
| 5 | 158908227 | rs10515794 | 0.832 | -0.149 | -0.297 | -0.777 | 0.014 | 0.038 | 0.038 | 0 | 0.26 | 0.19 | 0.14 | 0 | 0.22 | 0.14 |
| 5 | 158908650 | rs6556425 | -0.262 | 0.038 | -0.451 | -0.51 | -0.038 | -0.078 | -0.078 | 0 | 0.67 | 0.89 | 0.45 | 0.46 | 0.63 | 0.52 |
| 5 | 158915287 | rs10515797 | -0.179 | 0.433 | 0.304 | -0.095 | 0.046 | 0.016 | 0.016 | 0 | 0.14 | 0.38 | 0.12 | 0.12 | 0.15 | 0.11 |
| 5 | 158932186 | rs2421049 | -0.255 | 0.459 | 0.029 | -0.103 | -0.016 | -0.007 | -0.007 | 0.228 | 0.2 | 0.23 | 0.29 | 0.16 | 0.15 | 0.19 |
| 5 | 158935479 | rs11747171 | 0.446 | 0.664 | -0.257 | -0.02 | -0.011 | -0.004 | -0.004 | 0 | 0.14 | 0.17 | 0.07 | 0.12 | 0.11 | 0.1 |

| | | | | | | | | | | | | | | | | |
|---|---|---|---|---|---|---|---|---|---|---|---|---|---|---|---|---|
| 5 | 158943044 | rs2421050 | -0.351 | 0.821 | -1.491 | 0.148 | -0.005 | -0.036 | -0.036 | 0 | 0.67 | 0.79 | 0.67 | 0.49 | 0.67 | 0.59 |
| 5 | 158944085 | rs11959014 | -0.256 | 0.847 | -1.493 | -0.089 | 0.005 | -0.013 | -0.013 | 0.025 | 0.24 | 0.22 | 0.13 | 0 | 0.21 | 0.15 |
| 5 | 158946495 | rs6881547 | 0.636 | 1.483 | -1.497 | 1.033 | -0.015 | -0.02 | -0.02 | 0 | 0.77 | 0.79 | 0.82 | 0.31 | 0.73 | 0.61 |
| 5 | 158948962 | rs10037360 | -0.034 | 1.453 | -1.198 | 1.384 | -0.001 | 0.005 | 0.005 | 0.248 | 0.79 | 0.76 | 0.68 | 0.75 | 0.8 | 0.65 |
| 5 | 158948994 | rs12659608 | 0.437 | 1.567 | -1.48 | 0.97 | 0.003 | -0.023 | -0.023 | 0 | 0.41 | 0.48 | 0.24 | 0.34 | 0.36 | 0.31 |
| 5 | 158949047 | rs10045568 | -0.056 | 1.442 | -2.316 | 0.989 | -0.063 | -0.102 | -0.102 | 0 | 0.62 | 0.93 | 0.48 | 0.48 | 0.54 | 0.46 |
| 5 | 158958254 | rs2421052 | -0.329 | 1.133 | -2.215 | 1.028 | -0.002 | -0.02 | -0.02 | 0 | 0.62 | 0.92 | 0.53 | 0.36 | 0.63 | 0.36 |
| 5 | 158965775 | rs10476298 | -0.477 | 1.15 | -2.216 | 1.086 | -0.019 | -0.03 | -0.03 | 0 | 0.65 | 0.94 | 0.59 | 0.68 | 0.67 | 0.5 |
| 5 | 158972473 | rs4921508 | -0.282 | 0.617 | -1.501 | 0.424 | -0.022 | -0.102 | -0.102 | 0.321 | 0.22 | 0.36 | 0.1 | 0.03 | 0.17 | 0.12 |
| 5 | 158975663 | rs4493656 | -0.133 | 1.308 | -1.725 | 0.139 | 0.002 | -0.004 | -0.004 | 0 | 0.43 | 0.54 | 0.45 | 0.61 | 0.49 | 0.38 |
| 5 | 158976183 | rs4304068 | 0.793 | 1.148 | -1.223 | 0.352 | 0 | -0.002 | -0.002 | 0 | 0.31 | 0.29 | 0.36 | 0.36 | 0.32 | 0.22 |
| 5 | 158984093 | rs7718001 | -0.927 | 1.283 | -1.081 | 0.218 | -0.006 | -0.037 | -0.037 | 0 | 0.27 | 0.64 | 0.16 | 0.15 | 0.21 | 0.2 |
| 5 | 158984239 | rs4576132 | -0.526 | 1.118 | -1.036 | 0.258 | 0.032 | 0.022 | 0.022 | 2.281 | 0.49 | 0.6 | 0.41 | 0.58 | 0.37 | 0.4 |
| 5 | 158984287 | rs4334868 | 0.685 | 1.389 | -0.979 | 0.462 | 0.001 | -0.01 | -0.01 | 0.048 | 0.28 | 0.25 | 0.34 | 0.5 | 0.29 | 0.24 |
| 5 | 158986637 | rs10515799 | -0.678 | 1.403 | -0.886 | 0.487 | -0.004 | 0.001 | 0.001 | 0 | 0.26 | 0.33 | 0.14 | 0.32 | 0.21 | 0.25 |
| 5 | 158987588 | rs6895787 | -0.981 | 1.218 | -0.899 | 0.492 | -0.001 | -0.001 | -0.001 | 0 | 0.32 | 0.35 | 0.19 | 0.4 | 0.33 | 0.34 |
| 5 | 159001299 | rs4315909 | -0.438 | 1.13 | -0.633 | 0.719 | 0.019 | 0 | 0 | 1.367 | 0.41 | 0.65 | 0.39 | 0.38 | 0.36 | 0.21 |
| 5 | 159004095 | rs7717259 | 0.07 | 1.458 | -0.771 | 0.51 | -0.039 | 0.04 | 0.04 | 0.002 | 0.08 | 0.01 | 0.07 | 0.02 | 0.14 | 0.08 |
| 5 | 159005439 | rs34618866 | -0.341 | 1.402 | -0.43 | 0.476 | -0.002 | -0.004 | -0.004 | 0.176 | 0.09 | 0.09 | 0.03 | 0.07 | 0.06 | 0.08 |
| 5 | 159007778 | rs10065906 | -1.856 | 1.821 | -0.937 | 0.44 | 0.015 | -0.005 | -0.005 | 0 | 0.64 | 0.93 | 0.61 | 0.68 | 0.7 | 0.55 |
| 5 | 159009273 | rs11948965 | -1.3 | 1.913 | -0.344 | 0.389 | 0.004 | -0.043 | -0.043 | 0.042 | 0.14 | 0.45 | 0.09 | 0.07 | 0.1 | 0.09 |
| 5 | 159016266 | rs11135065 | -1.849 | 1.962 | -0.359 | 0.485 | 0.024 | 0.022 | 0.022 | 0 | 0.35 | 0.52 | 0.38 | 0.57 | 0.5 | 0.44 |
| 5 | 159021900 | rs6860449 | -1.941 | 0.812 | -0.301 | 0.51 | 0.03 | 0.028 | 0.028 | 0 | 0.34 | 0.47 | 0.34 | 0.57 | 0.42 | 0.43 |
| 5 | 159022275 | rs2421059 | 0.69 | 0.928 | -0.263 | 0.593 | -0.018 | -0.021 | -0.021 | 1.42 | 0.11 | 0.07 | 0.16 | 0.19 | 0.17 | 0.13 |
| 5 | 159025716 | rs2421057 | -1.808 | 0.915 | -0.28 | 0.569 | 0.029 | 0.066 | 0.066 | 0 | 0.29 | 0.5 | 0.3 | 0.55 | 0.31 | 0.36 |
| 5 | 159036930 | rs9313812 | -0.558 | 0.808 | -0.336 | 0.515 | -0.002 | -0.001 | -0.001 | 0.462 | 0.42 | 0.45 | 0.33 | 0.41 | 0.4 | 0.35 |
| 5 | 159038109 | rs11958768 | -1.108 | 0.821 | -0.332 | 0.521 | 0.042 | 0.044 | 0.044 | 0 | 0.23 | 0.46 | 0.25 | 0.26 | 0.26 | 0.21 |
| 5 | 159038698 | rs2421056 | -2.256 | 0.66 | -0.409 | 0.428 | -0.024 | -0.008 | -0.008 | 0.422 | 0.42 | 0.73 | 0.4 | 0.57 | 0.47 | 0.5 |
| 5 | 159041812 | rs17056839 | 1.193 | 0.639 | -0.391 | 0.419 | 0.003 | 0.001 | 0.001 | 0.883 | 0.23 | 0.21 | 0.18 | 0.1 | 0.19 | 0.06 |
| 5 | 159044278 | rs2421055 | -2.267 | 0.69 | -0.518 | 0.446 | -0.061 | -0.034 | -0.034 | 0 | 0.65 | 0.94 | 0.59 | 0.67 | 0.66 | 0.56 |
| 5 | 159056499 | rs11949241 | -0.072 | 0.461 | -0.23 | 0.439 | 0.02 | 0.016 | 0.016 | 0.022 | 0.09 | 0.37 | 0.05 | 0.07 | 0.05 | 0.07 |
| 5 | 159057330 | rs1422858 | -2.311 | 0.494 | -0.258 | 0.476 | -0.001 | 0 | 0 | 1.091 | 0.31 | 0.66 | 0.24 | 0.39 | 0.3 | 0.37 |
| 5 | 159060317 | rs7725393 | -1.076 | 0.623 | -0.274 | 0.371 | -0.001 | -0.001 | -0.001 | 0 | 0.51 | 0.51 | 0.51 | 0.59 | 0.54 | 0.43 |
| 5 | 159062530 | rs13163267 | 0.864 | 0.631 | -0.409 | 0.401 | -0.02 | 0.027 | 0.027 | 0.053 | 0.06 | 0.02 | 0.06 | 0.02 | 0.1 | 0.06 |
| 5 | 159065624 | rs6873462 | -1.293 | 0.493 | -0.259 | 0.341 | 0.007 | 0 | 0 | 0.321 | 0.3 | 0.64 | 0.3 | 0.29 | 0.36 | 0.26 |
| 5 | 159071666 | rs6556439 | -0.566 | 0.497 | 0.014 | 0.359 | 0.039 | 0.038 | 0.038 | 0 | 0.23 | 0.54 | 0.23 | 0.27 | 0.27 | 0.2 |
| 5 | 159072960 | rs17056872 | 0.216 | 0.455 | -0.545 | 0.454 | 0.026 | -0.018 | -0.018 | 1.158 | 0.31 | 0.24 | 0.25 | 0.36 | 0.27 | 0.22 |
| 5 | 159076347 | rs11135070 | -2.322 | 0.456 | -0.57 | 0.44 | -0.083 | -0.044 | -0.044 | 0 | 0.66 | 0.94 | 0.58 | 0.66 | 0.65 | 0.55 |
| 5 | 159078346 | rs7709089 | 0.558 | 0.248 | -0.466 | 0.494 | 0.018 | -0.022 | -0.022 | 0 | 0.26 | 0.19 | 0.21 | 0.36 | 0.23 | 0.2 |
| 5 | 159084481 | rs11135072 | -1.033 | 0.2 | -0.509 | 0.501 | -0.026 | 0.064 | 0.064 | 1.156 | 0.51 | 0.85 | 0.45 | 0.62 | 0.5 | 0.41 |
| 5 | 159087744 | rs10065065 | -1.854 | -1.248 | -0.464 | 0.588 | -0.044 | -0.038 | -0.038 | 1.131 | 0.6 | 0.93 | 0.5 | 0.6 | 0.47 | 0.46 |
| 5 | 159087887 | rs6872351 | -0.683 | -1.4 | -0.114 | 0.625 | -0.027 | 0.057 | 0.057 | 0 | 0.23 | 0.49 | 0.23 | 0.31 | 0.17 | 0.25 |
| 5 | 159091105 | rs10042565 | -0.587 | -1.409 | -0.163 | 0.616 | -0.027 | -0.012 | -0.012 | 0 | 0.38 | 0.57 | 0.28 | 0.29 | 0.29 | 0.21 |
| 5 | 159092417 | rs4921515 | 0.037 | -1.365 | -0.182 | 0.654 | -0.013 | 0.079 | 0.079 | 0 | 0.18 | 0.36 | 0.2 | 0.3 | 0.15 | 0.17 |
| 5 | 159097894 | rs17056921 | -1.2 | -1.396 | -0.121 | 0.629 | 0.001 | 0.001 | 0.001 | 0.007 | 0.5 | 0.48 | 0.39 | 0.58 | 0.36 | 0.4 |
| 5 | 159098864 | rs12655809 | 0.288 | -1.319 | -0.607 | 0.638 | 0.07 | -0.023 | -0.023 | 0.001 | 0.28 | 0.12 | 0.17 | 0.27 | 0.19 | 0.15 |
| 5 | 159101288 | rs7720446 | -0.024 | -1.266 | -0.67 | 0.644 | 0.036 | 0.007 | 0.007 | 0 | 0.31 | 0.1 | 0.26 | 0.24 | 0.3 | 0.21 |
| 5 | 159101706 | rs7705455 | -1.507 | -1.324 | -0.583 | 0.683 | 0.012 | 0 | 0 | 0 | 0.36 | 0.69 | 0.33 | 0.38 | 0.28 | 0.31 |
| 5 | 159101936 | rs1030206 | -1.681 | -1.078 | -0.478 | 0.762 | -0.003 | 0 | 0 | 0.138 | 0.6 | 0.75 | 0.51 | 0.6 | 0.49 | 0.46 |
| 5 | 159102593 | rs974805 | -1.62 | -1.429 | -0.439 | 0.512 | 0.017 | 0.02 | 0.02 | 0.11 | 0.38 | 0.69 | 0.31 | 0.28 | 0.32 | 0.31 |
| 5 | 159102717 | rs12652868 | 0.499 | -1.213 | -0.363 | 0.596 | -0.008 | -0.001 | -0.001 | 0.001 | 0.24 | 0.24 | 0.18 | 0.31 | 0.15 | 0.15 |
| 5 | 159102814 | rs974807 | -1.314 | -1.157 | -0.182 | 0.684 | -0.003 | 0 | 0 | 0 | 0.54 | 0.55 | 0.47 | 0.44 | 0.47 | 0.44 |
| 5 | 159103367 | rs1422853 | -0.17 | -1.155 | -0.451 | 0.002 | -0.005 | -0.153 | -0.153 | 0 | 0.21 | 0.45 | 0.12 | 0.01 | 0.18 | 0.19 |
| 5 | 159104799 | rs7720413 | -0.295 | -1.189 | -0.643 | 0.119 | 0.047 | 0 | 0 | 0 | 0.33 | 0.19 | 0.25 | 0.34 | 0.28 | 0.25 |
| 5 | 159124404 | rs13355113 | 0.048 | -1.212 | -0.68 | 0.119 | 0.052 | -0.002 | -0.002 | 0 | 0.29 | 0.17 | 0.2 | 0.25 | 0.18 | 0.22 |
| 5 | 159124879 | rs10476306 | 0.108 | -1.244 | -0.663 | 0.149 | 0.027 | 0.028 | 0.028 | 0 | 0.28 | 0.2 | 0.19 | 0.08 | 0.17 | 0.15 |
| 5 | 159125765 | rs1363659 | -0.768 | -1.146 | -0.598 | 0.122 | 0.003 | -0.021 | -0.021 | 0 | 0.27 | 0.43 | 0.19 | 0.27 | 0.29 | 0.35 |
| 5 | 159129383 | rs17057016 | 0.826 | -1.169 | -1 | -0.273 | 0.055 | 0.063 | 0.063 | 0.072 | 0.23 | 0.03 | 0.16 | 0.07 | 0.15 | 0.06 |
| 5 | 159129977 | rs16750 | 0.42 | -1.123 | -1.039 | -0.28 | -0.045 | 0.085 | 0.085 | 0.043 | 0.14 | 0.01 | 0.09 | 0.01 | 0.16 | 0.14 |
| 5 | 159130117 | rs17057017 | -0.911 | -1.427 | -0.875 | -0.024 | 0.162 | 0.107 | 0.107 | 1.377 | 0.41 | 0.9 | 0.35 | 0.19 | 0.22 | 0.21 |
| 5 | 159130222 | rs17057019 | -2.067 | -0.404 | 0.187 | 0.505 | -0.029 | -0.007 | -0.007 | 2.079 | 0.15 | 0.73 | 0.17 | 0.11 | 0.05 | 0.07 |
| 5 | 159130866 | rs11951956 | -1.386 | -0.36 | 0.234 | 0.538 | -0.036 | 0.042 | 0.042 | 0.127 | 0.09 | 0.29 | 0.14 | 0.11 | 0.02 | 0.02 |
| 5 | 159132395 | rs12652367 | -1.28 | 0.12 | 0.187 | 0.512 | 0.035 | -0.034 | -0.034 | 0 | 0.12 | 0.42 | 0.09 | 0.09 | 0.13 | 0.09 |
| 5 | 159137011 | rs10063205 | -0.994 | -0.198 | -0.627 | 0.86 | 0.021 | -0.004 | -0.004 | 0.533 | 0.3 | 0.72 | 0.21 | 0.29 | 0.21 | 0.35 |
| 5 | 159139029 | rs6864613 | -0.172 | 0.358 | -0.641 | 0.665 | 0.001 | 0.003 | 0.003 | 0.288 | 0.49 | 0.56 | 0.55 | 0.48 | 0.47 | 0.65 |
| 5 | 159141973 | rs6890497 | -0.367 | 0.289 | -0.616 | 0.651 | 0.002 | 0.08 | 0.08 | 0.227 | 0.54 | 0.74 | 0.66 | 0.78 | 0.51 | 0.85 |
| 5 | 159155592 | rs10035228 | -0.383 | 0.393 | -0.549 | 0.536 | 0.003 | -0.035 | -0.035 | 0 | 0.12 | 0.19 | 0.08 | 0.01 | 0.11 | 0.06 |
| 5 | 159170752 | rs6885423 | 0.389 | 1.064 | -0.587 | 0.543 | 0 | 0.002 | 0.002 | 0.86 | 0.43 | 0.44 | 0.48 | 0.44 | 0.41 | 0.57 |
| 5 | 159170813 | rs1592968 | -0.945 | 1.08 | -1.461 | 0.75 | 0.012 | 0.03 | 0.03 | 1.542 | 0.65 | 0.84 | 0.73 | 0.79 | 0.64 | 0.9 |

| | | | | | | | | | | | | | | | | |
|---|---|---|---|---|---|---|---|---|---|---|---|---|---|---|---|---|
| 5 | 159173642 | rs11744947 | 0.118 | 1.334 | -1.498 | 0.08 | -0.078 | -0.104 | -0.104 | 0 | 0.28 | 0.03 | 0.39 | 0.43 | 0.28 | 0.38 |
| 5 | 159183922 | rs1432714 | -1.506 | 1.361 | 0.027 | -0.127 | 0.021 | -0.013 | -0.013 | 0 | 0.18 | 0.62 | 0.16 | 0.13 | 0.17 | 0.14 |
| 5 | 159192346 | rs365804 | 1.213 | 1.009 | 0.172 | -0.153 | 0.025 | -0.029 | -0.029 | 0 | 0.83 | 0.41 | 0.85 | 0.91 | 0.81 | 0.84 |
| 5 | 159209041 | rs12518680 | -1.3 | 0.888 | -0.028 | 0.128 | 0.01 | -0.031 | -0.031 | 1.619 | 0.15 | 0.09 | 0.37 | 0.63 | 0.11 | 0.49 |
| 5 | 159214593 | rs1549888 | -1.38 | 0.895 | 0.966 | -0.417 | 0.041 | -0.032 | -0.032 | 0 | 0.09 | 0.43 | 0.06 | 0 | 0.05 | 0.08 |
| 5 | 159214702 | rs1003068 | -1.752 | 0.84 | 1.11 | -0.443 | -0.002 | -0.065 | -0.065 | 0 | 0.11 | 0.56 | 0.08 | 0 | 0.05 | 0.08 |
| 5 | 159215298 | rs6891014 | -1.375 | 0.867 | 1.018 | -0.43 | 0.041 | -0.032 | -0.032 | 0 | 0.09 | 0.43 | 0.06 | 0 | 0.05 | 0.08 |
| 5 | 159220255 | rs17733689 | -0.331 | 0.47 | 0.533 | -0.632 | -0.022 | 0.048 | 0.048 | 0 | 0.15 | 0.01 | 0.08 | 0.04 | 0.15 | 0.09 |
| 5 | 159237381 | rs4558971 | -0.807 | 0.773 | 0.377 | -0.679 | -0.001 | 0.062 | 0.062 | 0.594 | 0.36 | 0.83 | 0.3 | 0.24 | 0.38 | 0.24 |
| 5 | 159261636 | rs12513480 | -0.858 | 0.695 | 1.096 | -0.324 | -0.03 | -0.039 | -0.039 | 4.211 | 0.15 | 0.02 | 0.18 | 0.17 | 0.17 | 0.17 |
| 5 | 159261646 | rs10044331 | -1.025 | 0.528 | 1.578 | -0.659 | 0.024 | -0.04 | -0.04 | 0.522 | 0.08 | 0.22 | 0.05 | 0 | 0.07 | 0.09 |
| 5 | 159265860 | rs1835928 | -1.285 | 0.477 | 1.279 | -0.687 | -0.031 | -0.037 | -0.037 | 0 | 0.2 | 0.37 | 0.13 | 0.14 | 0.19 | 0.18 |
| 5 | 159268879 | rs6881745 | -1.639 | 0.765 | 0.661 | -0.716 | -0.001 | -0.075 | -0.075 | 3.023 | 0.3 | 0.44 | 0.19 | 0.15 | 0.3 | 0.29 |
| 5 | 159270713 | rs959931 | -0.952 | 0.932 | 0.946 | -0.268 | 0.02 | -0.038 | -0.038 | 0 | 0.18 | 0.39 | 0.15 | 0.14 | 0.22 | 0.19 |
| 5 | 159283689 | rs12657061 | 0.126 | -0.154 | 0.114 | -1.082 | 0.065 | 0.106 | 0.106 | 0 | 0.4 | 0.21 | 0.38 | 0.82 | 0.26 | 0.44 |
| 5 | 159283738 | rs713580 | 0.205 | -0.717 | -0.16 | -1.293 | 0.007 | 0.003 | 0.003 | 0 | 0.16 | 0.11 | 0.13 | 0.02 | 0.11 | 0.08 |
| 5 | 159284708 | rs31687 | 0.595 | -0.406 | 0.28 | -1.692 | 0.003 | -0.042 | -0.042 | 0.848 | 0.1 | 0.18 | 0.05 | 0.01 | 0.06 | 0.09 |
| 5 | 159288177 | rs1432674 | -0.135 | -0.755 | 0.742 | -1.826 | 0.007 | 0.003 | 0.003 | 0 | 0.42 | 0.38 | 0.4 | 0.82 | 0.27 | 0.44 |
| 5 | 159288452 | rs31689 | -0.889 | 0.739 | 0.22 | -2.043 | 0.01 | 0.001 | 0.001 | 0 | 0.84 | 0.83 | 0.63 | 0.83 | 0.66 | 0.8 |
| 5 | 159289295 | rs11135085 | 0.59 | 0.524 | 0.117 | -2.24 | -0.016 | 0.027 | 0.027 | 0 | 0.27 | 0.13 | 0.15 | 0.03 | 0.28 | 0.27 |
| 5 | 159290037 | rs17052550 | 0.043 | 0.088 | -0.136 | -1.734 | -0.01 | -0.034 | -0.034 | 0 | 0.16 | 0.28 | 0.1 | 0.07 | 0.14 | 0.03 |
| 5 | 159291373 | rs13436303 | -0.017 | 0.08 | -0.291 | -1.738 | -0.008 | -0.025 | -0.025 | 0 | 0.16 | 0.19 | 0.1 | 0.07 | 0.14 | 0.03 |
| 5 | 159292703 | rs31694 | -0.349 | 0.452 | -0.797 | -1.576 | -0.036 | -0.015 | -0.015 | 0 | 0.86 | 0.94 | 0.69 | 0.84 | 0.74 | 0.79 |
| 5 | 159294343 | rs6556458 | 0.084 | -0.018 | -1.349 | -1.581 | -0.056 | -0.001 | -0.001 | 0 | 0.65 | 0.82 | 0.52 | 0.39 | 0.54 | 0.61 |
| 5 | 159297656 | rs6865665 | 0.175 | -0.15 | -0.795 | -1.888 | 0.007 | -0.024 | -0.024 | 0 | 0.07 | 0.23 | 0.08 | 0.03 | 0.07 | 0.07 |
| 5 | 159298474 | rs1432675 | 0.861 | -0.132 | -0.546 | -2.2 | 0.005 | -0.061 | -0.061 | 0 | 0.16 | 0.24 | 0.12 | 0.04 | 0.19 | 0.2 |
| 5 | 159303793 | rs10045099 | 0.729 | -0.16 | -0.494 | -2.175 | 0.002 | -0.021 | -0.021 | 0.335 | 0.07 | 0.16 | 0.07 | 0.03 | 0.07 | 0.07 |
| 5 | 159308536 | rs1432676 | 0.773 | -0.191 | -0.958 | -2.32 | 0.004 | -0.025 | -0.025 | 0 | 0.24 | 0.27 | 0.18 | 0.04 | 0.3 | 0.19 |
| 5 | 159310105 | rs10060486 | -1.516 | -0.356 | -1.693 | -2.422 | -0.05 | 0.035 | 0.035 | 0 | 0.7 | 0.91 | 0.5 | 0.23 | 0.7 | 0.53 |
| 5 | 159310695 | rs6861020 | -1.516 | -0.356 | -1.693 | -2.422 | -0.05 | 0.035 | 0.035 | 0 | 0.7 | 0.91 | 0.5 | 0.23 | 0.7 | 0.53 |
| 5 | 159314858 | rs13176344 | 1.123 | -0.299 | -1.235 | -1.729 | 0.019 | 0.036 | 0.036 | 0.241 | 0.15 | 0.02 | 0.08 | 0.11 | 0.08 | 0.06 |
| 5 | 159320525 | rs13179580 | -0.076 | -0.226 | -1.53 | -1.722 | -0.001 | -0.018 | -0.018 | 1.475 | 0.31 | 0.27 | 0.21 | 0.14 | 0.28 | 0.25 |
| 5 | 159323677 | rs12656915 | 0.675 | -0.155 | -1.613 | -1.713 | 0.002 | 0.004 | 0.004 | 0.112 | 0.07 | 0.23 | 0.07 | 0.05 | 0.07 | 0.06 |
| 5 | 159325351 | rs7712049 | -0.702 | -0.067 | -1.542 | -1.657 | -0.018 | -0.067 | -0.067 | 1.119 | 0.39 | 0.48 | 0.28 | 0.19 | 0.34 | 0.32 |
| 5 | 159327355 | rs4921240 | 0.038 | 0.678 | -1.588 | -0.985 | 0 | -0.059 | -0.059 | 1.771 | 0.22 | 0.23 | 0.16 | 0.04 | 0.24 | 0.19 |
| 5 | 159333536 | rs1025466 | 0.006 | 0.502 | -1.448 | -0.635 | 0.006 | 0.089 | 0.089 | 0 | 0.54 | 0.28 | 0.41 | 0.2 | 0.59 | 0.42 |
| 5 | 159335852 | rs185289 | -0.481 | 1.029 | -1.6 | 0.066 | -0.022 | -0.001 | -0.001 | 0 | 0.84 | 0.94 | 0.69 | 0.83 | 0.81 | 0.71 |
| 5 | 159339533 | rs1019920 | -0.767 | 0.48 | -0.902 | 0.208 | 0 | -0.006 | -0.006 | 0 | 0.4 | 0.44 | 0.36 | 0.75 | 0.36 | 0.38 |
| 5 | 159341032 | rs752266 | -0.362 | 0.455 | -0.656 | 1.042 | 0.008 | 0.108 | 0.108 | 0 | 0.08 | 0.21 | 0.11 | 0.27 | 0.06 | 0.1 |
| 5 | 159344461 | rs3729604 | 0.318 | 0.52 | -0.59 | 1.404 | -0.001 | -0.039 | -0.039 | 3.825 | 0.21 | 0.17 | 0.19 | 0.35 | 0.18 | 0.13 |
| 5 | 159347481 | rs2030373 | -0.459 | 0.532 | -0.577 | 1.267 | 0 | -0.023 | -0.023 | 0 | 0.28 | 0.23 | 0.3 | 0.61 | 0.25 | 0.23 |
| 5 | 159348443 | rs6884105 | -1.36 | 0.551 | -0.613 | 1.278 | -0.002 | -0.028 | -0.028 | 0 | 0.41 | 0.52 | 0.37 | 0.73 | 0.36 | 0.38 |
| 5 | 159348898 | rs17455628 | 0.028 | 0.71 | -0.716 | 1.299 | 0.001 | -0.008 | -0.008 | 0 | 0.08 | 0.06 | 0.03 | 0.12 | 0.06 | 0.12 |
| 5 | 159351460 | rs756275 | -0.135 | 0.855 | -0.736 | 1.288 | -0.004 | -0.01 | -0.01 | 0.326 | 0.09 | 0.04 | 0.04 | 0.1 | 0.08 | 0.13 |
| 5 | 159354775 | rs11952941 | -1.666 | 0.915 | -0.879 | 0.988 | 0.001 | 0.019 | 0.019 | 0 | 0.5 | 0.51 | 0.47 | 0.72 | 0.5 | 0.44 |
| 5 | 159360485 | rs6892282 | -1.509 | 0.672 | -0.92 | 1.275 | -0.002 | 0.018 | 0.018 | 0 | 0.45 | 0.45 | 0.41 | 0.73 | 0.45 | 0.42 |
| 5 | 159365567 | rs10515805 | -1.224 | 0.471 | -0.812 | 1.44 | -0.007 | 0.017 | 0.017 | 0 | 0.12 | 0.07 | 0.07 | 0.06 | 0.17 | 0.08 |
| 5 | 159366698 | rs4921242 | -0.833 | 0.472 | -0.813 | 1.439 | -0.013 | 0.026 | 0.026 | 0 | 0.1 | 0.03 | 0.07 | 0.04 | 0.16 | 0.06 |
| 5 | 159367114 | rs6888306 | -1.839 | 0.452 | -0.76 | 1.622 | 0.005 | 0.046 | 0.046 | 0 | 0.22 | 0.3 | 0.2 | 0.39 | 0.25 | 0.24 |
| 5 | 159368286 | rs11743425 | -1.249 | 0.459 | -0.776 | 1.34 | -0.007 | 0.017 | 0.017 | 0 | 0.12 | 0.07 | 0.07 | 0.06 | 0.17 | 0.08 |
| 5 | 159369429 | rs7737796 | -1.71 | 0.394 | -0.752 | 1.399 | 0 | 0.003 | 0.003 | 0 | 0.43 | 0.45 | 0.38 | 0.73 | 0.4 | 0.4 |
| 5 | 159377721 | rs34467921 | -0.827 | 0.29 | -1.212 | 1.145 | -0.013 | 0.024 | 0.024 | 0 | 0.1 | 0.03 | 0.07 | 0.05 | 0.16 | 0.06 |
| 5 | 159378159 | rs13171967 | -0.932 | 0.262 | -1.179 | 1.266 | -0.002 | 0.013 | 0.013 | 0 | 0.21 | 0.27 | 0.13 | 0.31 | 0.23 | 0.24 |
| 5 | 159381526 | rs3896275 | -0.034 | 0.253 | -1.055 | 1.446 | -0.032 | -0.102 | -0.102 | 0 | 0.21 | 0.01 | 0.17 | 0.3 | 0.16 | 0.16 |
| 5 | 159383271 | rs12653825 | -0.687 | 0.416 | -0.864 | 0.849 | 0.005 | 0.002 | 0.002 | 0 | 0.38 | 0.32 | 0.26 | 0.62 | 0.32 | 0.38 |
| 5 | 159390877 | rs952037 | -0.54 | 0.128 | -0.437 | 0.883 | -0.017 | 0.015 | 0.015 | 0 | 0.37 | 0.5 | 0.23 | 0.45 | 0.32 | 0.43 |
| 5 | 159400511 | rs10214211 | -0.877 | -0.216 | 0.216 | 0.454 | -0.029 | -0.043 | -0.043 | 0 | 0.16 | 0.38 | 0.1 | 0.15 | 0.07 | 0.11 |
| 5 | 159407164 | rs10053468 | -0.879 | -0.286 | 0.6 | 0.447 | -0.03 | -0.036 | -0.036 | 0.319 | 0.16 | 0.33 | 0.09 | 0.15 | 0.07 | 0.1 |
| 5 | 159412045 | rs13177334 | -1.345 | -0.474 | 0.67 | 0.95 | -0.042 | 0.035 | 0.035 | 0.059 | 0.23 | 0.43 | 0.2 | 0.36 | 0.15 | 0.17 |
| 5 | 159418490 | rs10068999 | -0.812 | -0.331 | -0.114 | 0.049 | -0.051 | 0.022 | 0.022 | 1.156 | 0.61 | 0.81 | 0.56 | 0.41 | 0.56 | 0.48 |
| 5 | 159425524 | rs6878309 | -1.185 | -0.712 | 0.056 | -0.131 | -0.035 | 0.024 | 0.024 | 0 | 0.2 | 0.43 | 0.31 | 0.31 | 0.15 | 0.21 |
| 5 | 159428322 | rs12519460 | -0.121 | -0.703 | -0.241 | -0.124 | 0 | -0.013 | -0.013 | 0 | 0.06 | 0.02 | 0.05 | 0.14 | 0.03 | 0.09 |
| 5 | 159436546 | rs10044313 | 1.924 | -0.808 | -0.23 | -0.146 | 0.017 | 0.015 | 0.015 | 0.664 | 0.07 | 0.06 | 0.03 | 0 | 0.01 | 0.01 |
| 5 | 159446499 | rs7712634 | -0.67 | -0.79 | 0.087 | 0.008 | 0.006 | -0.009 | -0.009 | 0.031 | 0.15 | 0.14 | 0.28 | 0.29 | 0.14 | 0.19 |
| 5 | 159447213 | rs10515804 | -1.228 | -0.85 | -0.408 | 0.298 | -0.075 | -0.1 | -0.1 | 0 | 0.76 | 0.93 | 0.78 | 0.6 | 0.7 | 0.71 |
| 5 | 159448136 | rs2176830 | 0.938 | -1.17 | -0.176 | 0.405 | 0.006 | -0.019 | -0.019 | 0 | 0.08 | 0.04 | 0.22 | 0.17 | 0.1 | 0.09 |
| 5 | 159466751 | rs6870694 | -0.597 | -1.166 | -0.045 | 0.39 | -0.002 | 0 | 0 | 0 | 0.07 | 0.1 | 0.06 | 0.12 | 0.05 | 0.1 |
| 5 | 159473094 | rs11749533 | 0.932 | -1.167 | -0.027 | 0.4 | 0.003 | -0.007 | -0.007 | 0 | 0.08 | 0.06 | 0.22 | 0.17 | 0.1 | 0.09 |
| 5 | 159475476 | rs4385243 | -0.456 | -1.79 | 0.025 | -0.325 | 0 | 0.009 | 0.009 | 0 | 0.68 | 0.74 | 0.66 | 0.72 | 0.66 | 0.71 |

| | | | | | | | | | | | | | | | | |
|---|---|---|---|---|---|---|---|---|---|---|---|---|---|---|---|---|
| 5 | 159478541 | rs3733870 | -1.213 | -1.852 | 0.083 | -0.355 | -0.007 | 0.023 | 0.023 | 0 | 0.77 | 0.84 | 0.88 | 0.89 | 0.76 | 0.81 |
| 5 | 159478640 | rs12516548 | -0.158 | -1.867 | 0 | -0.352 | -0.005 | 0.001 | 0.001 | 0 | 0.07 | 0.12 | 0.06 | 0.12 | 0.05 | 0.1 |
| 5 | 159487953 | rs7704770 | 0.333 | -1.824 | 0.321 | -0.369 | -0.001 | -0.003 | -0.003 | 0 | 0.61 | 0.61 | 0.59 | 0.61 | 0.61 | 0.59 |
| 5 | 159488365 | rs12514103 | -0.254 | -1.818 | 0.304 | -0.356 | -0.005 | -0.002 | -0.002 | 0.014 | 0.07 | 0.12 | 0.05 | 0.1 | 0.05 | 0.11 |
| 5 | 159488697 | rs12514671 | -0.033 | -1.821 | 0.302 | -0.358 | -0.001 | 0 | 0 | 0 | 0.07 | 0.08 | 0.05 | 0.1 | 0.05 | 0.11 |
| 5 | 159489455 | rs3733868 | 1.255 | -2.007 | 0.139 | -0.381 | -0.013 | -0.038 | -0.038 | 0 | 0.08 | 0.01 | 0.03 | 0.16 | 0.08 | 0.14 |
| 5 | 159489550 | rs3733869 | -0.421 | -2.086 | 0.289 | -0.344 | 0 | -0.006 | -0.006 | 0 | 0.69 | 0.69 | 0.64 | 0.7 | 0.66 | 0.72 |
| 5 | 159490247 | rs6861719 | -1.098 | -2.084 | 0.285 | -0.342 | 0.003 | -0.018 | -0.018 | 0 | 0.77 | 0.78 | 0.86 | 0.87 | 0.76 | 0.81 |
| 5 | 159491520 | rs1106055 | 0.273 | -2.035 | 0.301 | -0.326 | -0.003 | -0.011 | -0.011 | 0.878 | 0.61 | 0.57 | 0.59 | 0.61 | 0.61 | 0.6 |
| 5 | 159513867 | rs10037844 | 0.355 | -2.021 | 0.313 | -0.327 | -0.005 | -0.018 | -0.018 | 0 | 0.6 | 0.51 | 0.58 | 0.61 | 0.61 | 0.59 |
| 5 | 159522488 | rs2546963 | 0.521 | -1.956 | 0.584 | -0.243 | 0.033 | 0.067 | 0.067 | 0.014 | 0.52 | 0.29 | 0.53 | 0.61 | 0.6 | 0.59 |
| 5 | 159560569 | rs11952950 | 0.907 | -1.616 | 0.415 | -0.168 | 0.012 | -0.024 | -0.024 | 0.136 | 0.16 | 0.23 | 0.12 | 0.13 | 0.18 | 0.12 |
| 5 | 159569779 | rs17057459 | -0.2 | -1.816 | 0.239 | 0.181 | -0.002 | 0.005 | 0.005 | 0 | 0.26 | 0.34 | 0.18 | 0.33 | 0.24 | 0.25 |
| 5 | 159577813 | rs11954055 | -0.138 | -1.696 | 0.208 | 0.2 | 0.004 | 0.037 | 0.037 | 0 | 0.24 | 0.43 | 0.17 | 0.36 | 0.23 | 0.28 |
| 5 | 159591003 | rs7378736 | -0.615 | -1.685 | -0.445 | -0.1 | -0.014 | 0.009 | 0.009 | 0.119 | 0.86 | 0.99 | 0.91 | 0.91 | 0.86 | 0.81 |
| 5 | 159592857 | rs2546371 | -0.617 | -1.667 | -0.087 | -0.107 | 0.03 | 0.04 | 0.04 | 0 | 0.7 | 0.77 | 0.85 | 0.91 | 0.79 | 0.73 |
| 5 | 159596509 | rs7734683 | 0.478 | -1.616 | 0.228 | -0.051 | 0.029 | -0.011 | -0.011 | 0 | 0.22 | 0.46 | 0.37 | 0.21 | 0.26 | 0.2 |
| 5 | 159602367 | rs918476 | 1.017 | -1.866 | 0.114 | 0.683 | -0.034 | 0.032 | 0.032 | 0 | 0.34 | 0.25 | 0.41 | 0.14 | 0.44 | 0.23 |
| 5 | 159607979 | rs2546375 | 1.54 | -1.323 | 0.045 | 0.546 | -0.04 | 0.016 | 0.016 | 0.086 | 0.25 | 0.13 | 0.34 | 0.14 | 0.34 | 0.21 |
| 5 | 159608980 | rs4921110 | 0.056 | -1.101 | 0.261 | 1.089 | 0.039 | 0.009 | 0.009 | 0 | 0.57 | 0.72 | 0.78 | 0.54 | 0.69 | 0.54 |
| 5 | 159613503 | rs2901520 | 1.065 | -0.962 | 0.195 | 0.965 | -0.023 | 0.027 | 0.027 | 0 | 0.33 | 0.25 | 0.4 | 0.14 | 0.42 | 0.23 |
| 5 | 159618158 | rs7701539 | 0.453 | -0.711 | 0.193 | 0.392 | 0.011 | 0.002 | 0.002 | 0 | 0.49 | 0.51 | 0.53 | 0.39 | 0.61 | 0.46 |
| 5 | 159619826 | rs7707569 | 1.166 | -0.572 | -0.017 | 0.134 | 0.011 | -0.021 | -0.021 | 0.059 | 0.14 | 0.16 | 0.1 | 0.07 | 0.19 | 0.12 |
| 5 | 159622419 | rs1465403 | 0.492 | -0.748 | -0.226 | -0.093 | 0.023 | 0.041 | 0.041 | 0 | 0.3 | 0.23 | 0.15 | 0.09 | 0.22 | 0.26 |
| 5 | 159633484 | rs4568401 | -1.348 | -0.636 | -0.017 | 0.125 | -0.003 | 0.03 | 0.03 | 0 | 0.91 | 0.81 | 0.73 | 0.85 | 0.91 | 0.91 |
| 5 | 159633681 | rs10866717 | 0.213 | -0.719 | -0.462 | 0.207 | 0.01 | -0.041 | -0.041 | 0 | 0.8 | 0.83 | 0.69 | 0.49 | 0.85 | 0.72 |
| 5 | 159637815 | rs17057519 | 0.689 | -0.712 | -0.409 | 0.278 | 0.011 | -0.02 | -0.02 | 0 | 0.1 | 0.06 | 0.04 | 0.36 | 0.05 | 0.16 |
| 5 | 159638777 | rs2421730 | -1.961 | -0.686 | -0.421 | 0.285 | -0.001 | -0.005 | -0.005 | 0 | 0.92 | 0.92 | 0.74 | 0.86 | 0.91 | 0.91 |
| 5 | 159640062 | rs11135094 | 0.422 | -0.668 | -0.38 | -0.078 | 0.054 | -0.009 | -0.009 | 1.046 | 0.53 | 0.67 | 0.55 | 0.4 | 0.63 | 0.49 |
| 5 | 159644037 | rs6877438 | 0.477 | -0.489 | -0.761 | -0.061 | 0.034 | 0.093 | 0.093 | 0 | 0.3 | 0.16 | 0.14 | 0.09 | 0.23 | 0.26 |
| 5 | 159646094 | rs918480 | 0.499 | -0.221 | -0.477 | 0.425 | -0.008 | -0.007 | -0.007 | 0 | 0.37 | 0.33 | 0.42 | 0.36 | 0.45 | 0.36 |
| 5 | 159654109 | rs10041333 | 0.208 | -0.201 | -0.342 | 0.031 | -0.003 | -0.056 | -0.056 | 0 | 0.38 | 0.45 | 0.25 | 0.1 | 0.35 | 0.28 |
| 5 | 159655116 | rs11746179 | -0.26 | -0.06 | -0.604 | 0.688 | -0.008 | -0.011 | -0.011 | 4.292 | 0.61 | 0.69 | 0.63 | 0.52 | 0.57 | 0.58 |
| 5 | 159665599 | rs2277954 | 0.535 | -0.796 | -0.059 | 0.87 | -0.059 | 0.038 | 0.038 | 0 | 0.75 | 0.45 | 0.68 | 0.64 | 0.89 | 0.85 |
| 5 | 159667613 | rs3926393 | 0.429 | -0.815 | 0.143 | 0.866 | -0.053 | 0.051 | 0.051 | 0 | 0.77 | 0.57 | 0.69 | 0.64 | 0.9 | 0.86 |
| 5 | 159677743 | rs6874570 | -1.188 | -1.164 | 0.233 | 0.461 | -0.028 | -0.022 | -0.022 | 0.174 | 0.23 | 0.32 | 0.12 | 0.15 | 0.12 | 0.21 |
| 5 | 159678995 | rs3749798 | -0.911 | -1.267 | 0.13 | 0.396 | 0.011 | -0.021 | -0.021 | 0 | 0.1 | 0.03 | 0.03 | 0.15 | 0.05 | 0.2 |
| 5 | 159679636 | rs4921270 | -1.658 | -1.286 | 0.238 | 0.386 | 0.032 | 0.021 | 0.021 | 0 | 0.21 | 0.1 | 0.08 | 0.15 | 0.12 | 0.21 |
| 5 | 159691302 | rs17057575 | -1.064 | -1.269 | 0.344 | 0.465 | -0.005 | 0.004 | 0.004 | 0 | 0.11 | 0.13 | 0.04 | 0.16 | 0.05 | 0.2 |
| 5 | 159693842 | rs10064968 | -0.504 | -1.486 | -0.423 | -0.306 | -0.004 | 0.056 | 0.056 | 0 | 0.8 | 0.9 | 0.75 | 0.93 | 0.8 | 0.82 |
| 5 | 159694357 | rs17472426 | -0.195 | -1.641 | -0.321 | -0.142 | 0.02 | 0.073 | 0.073 | 0 | 0.12 | 0.01 | 0.05 | 0 | 0.07 | 0.01 |
| 5 | 159696554 | rs10071116 | -1.182 | -1.574 | -0.208 | -0.061 | -0.026 | 0.002 | 0.002 | 0 | 0.16 | 0.27 | 0.07 | 0.15 | 0.07 | 0.26 |
| 5 | 159696951 | rs10071377 | -0.885 | -1.572 | -0.284 | -0.06 | 0.017 | -0.02 | -0.02 | 0 | 0.14 | 0.03 | 0.03 | 0.15 | 0.07 | 0.26 |
| 5 | 159699112 | rs12658758 | -1.43 | -1.799 | -0.836 | 0.618 | 0.006 | -0.004 | -0.004 | 0 | 0.38 | 0.39 | 0.23 | 0.51 | 0.27 | 0.46 |
| 5 | 159699709 | rs4921273 | 0.013 | -1.784 | -0.847 | 0.626 | 0.022 | -0.091 | -0.091 | 5.089 | 0.22 | 0.07 | 0.15 | 0.36 | 0.2 | 0.2 |
| 5 | 159702834 | rs17057594 | -1.172 | -1.15 | -0.299 | 0.597 | -0.031 | 0.001 | 0.001 | 0 | 0.16 | 0.31 | 0.07 | 0.15 | 0.07 | 0.26 |
| 5 | 159714528 | rs2303059 | -1.388 | -1.153 | 0.046 | 0.754 | -0.039 | 0.032 | 0.032 | 0.241 | 0.18 | 0.54 | 0.1 | 0.19 | 0.09 | 0.27 |
| 5 | 159720780 | rs6899125 | 0.443 | 0.181 | -0.823 | 0.564 | -0.037 | -0.001 | -0.001 | 0 | 0.11 | 0.04 | 0.21 | 0.15 | 0.24 | 0.1 |
| 5 | 159721672 | rs754111 | -0.147 | 0.504 | 0.039 | 0.693 | 0.112 | 0.13 | 0.13 | 0.046 | 0.08 | 0.43 | 0.15 | 0.09 | 0.12 | 0.13 |
| 5 | 159723358 | rs2421778 | -1.38 | 1.062 | -0.044 | 1.022 | 0.035 | 0.054 | 0.054 | 1.062 | 0.19 | 0.6 | 0.24 | 0.26 | 0.23 | 0.35 |
| 5 | 159732443 | rs12658296 | -0.778 | 1.312 | -0.015 | 0.952 | -0.015 | -0.005 | -0.005 | 0.371 | 0.21 | 0.19 | 0.26 | 0.23 | 0.3 | 0.27 |
| 5 | 159734986 | rs13362118 | -1.298 | 0.913 | -0.373 | 0.868 | -0.004 | -0.001 | -0.001 | 0.009 | 0.32 | 0.7 | 0.43 | 0.3 | 0.42 | 0.43 |
| 5 | 159735012 | rs17057631 | -0.024 | 0.94 | -0.477 | 0.895 | -0.023 | -0.036 | -0.036 | 0 | 0.12 | 0.06 | 0.08 | 0.18 | 0.11 | 0.22 |
| 5 | 159736411 | rs2421780 | -1.508 | 1.264 | -1.285 | 0.953 | -0.037 | -0.043 | -0.043 | 0 | 0.82 | 0.96 | 0.71 | 0.78 | 0.85 | 0.79 |
| 5 | 159738709 | rs6556490 | 0.064 | 1.104 | -0.849 | 0.769 | 0.059 | -0.02 | -0.02 | 0 | 0.19 | 0.55 | 0.34 | 0.13 | 0.3 | 0.21 |
| 5 | 159740138 | rs10038395 | -0.159 | 0.509 | -1.172 | 0.601 | -0.005 | 0.006 | 0.006 | 0.072 | 0.39 | 0.65 | 0.45 | 0.3 | 0.44 | 0.43 |
| 5 | 159748167 | rs10063519 | -0.117 | 0.614 | -1.191 | 0.65 | -0.001 | -0.031 | -0.031 | 0 | 0.64 | 0.78 | 0.59 | 0.52 | 0.66 | 0.53 |
| 5 | 159766954 | rs4921132 | 0.16 | 0.622 | -1.103 | 0.478 | -0.002 | 0.002 | 0.002 | 0.003 | 0.56 | 0.69 | 0.52 | 0.59 | 0.59 | 0.57 |
| 5 | 159772467 | rs13157941 | 0.851 | 0.297 | -1.385 | 0.504 | -0.016 | 0.013 | 0.013 | 0.042 | 0.41 | 0.17 | 0.4 | 0.32 | 0.48 | 0.34 |
| 5 | 159774217 | rs4276447 | -0.587 | 0.469 | -0.727 | 0.618 | 0.018 | 0.014 | 0.014 | 0 | 0.74 | 0.69 | 0.65 | 0.69 | 0.75 | 0.63 |
| 5 | 159775741 | rs6556492 | -0.527 | 0.463 | -0.419 | 0.614 | 0.024 | 0.019 | 0.019 | 0 | 0.73 | 0.66 | 0.64 | 0.68 | 0.75 | 0.63 |
| 5 | 159776047 | rs9216 | 0.701 | 0.067 | -0.37 | 0.331 | -0.004 | -0.017 | -0.017 | 0.103 | 0.45 | 0.33 | 0.49 | 0.47 | 0.56 | 0.38 |
| 5 | 159786594 | rs13185269 | 0.267 | 0.277 | -0.371 | 0.397 | 0.001 | 0 | 0 | 0 | 0.57 | 0.62 | 0.51 | 0.53 | 0.54 | 0.59 |
| 5 | 159787106 | rs4921133 | -0.903 | 0.311 | 0.276 | 0.861 | 0.011 | 0.01 | 0.01 | 0 | 0.75 | 0.66 | 0.66 | 0.74 | 0.78 | 0.67 |
| 5 | 159789092 | rs12519408 | 1.058 | 0.272 | 0.319 | 0.87 | -0.008 | -0.01 | -0.01 | 0 | 0.39 | 0.23 | 0.39 | 0.34 | 0.48 | 0.38 |
| 5 | 159791206 | rs1105788 | 0.306 | 0.177 | 0.599 | 0.836 | -0.002 | 0.01 | 0.01 | 0 | 0.52 | 0.48 | 0.53 | 0.55 | 0.62 | 0.48 |
| 5 | 159791231 | rs1105789 | 0.782 | 0.21 | 0.594 | 0.838 | 0.001 | 0.002 | 0.002 | 0 | 0.44 | 0.47 | 0.44 | 0.34 | 0.51 | 0.43 |
| 5 | 159791334 | rs1574468 | 0.82 | 0.309 | 0.293 | 0.932 | 0 | 0.016 | 0.016 | 0 | 0.09 | 0.1 | 0.1 | 0.21 | 0.11 | 0.05 |
| 5 | 159794511 | rs2961923 | 0.624 | 0.243 | 0.204 | 0.626 | -0.019 | 0.026 | 0.026 | 0 | 0.47 | 0.25 | 0.49 | 0.54 | 0.59 | 0.44 |

| | | | | | | | | | | | | | | | | |
|---|---|---|---|---|---|---|---|---|---|---|---|---|---|---|---|---|
| 5 | 159796182 | rs2961924 | -1.556 | 0.57 | 0.255 | 0.898 | -0.036 | -0.045 | -0.045 | 0 | 0.84 | 0.95 | 0.72 | 0.76 | 0.83 | 0.71 |
| 5 | 159818125 | rs2961932 | -0.306 | 0.399 | 0.472 | 0.646 | -0.005 | 0.02 | 0.02 | 0.614 | 0.37 | 0.32 | 0.43 | 0.17 | 0.4 | 0.48 |
| 5 | 159820931 | rs10515808 | 0.892 | 0.26 | 0.359 | 0.6 | 0.001 | 0.042 | 0.042 | 0 | 0.12 | 0.02 | 0.07 | 0 | 0.11 | 0.06 |
| 5 | 159823099 | rs2910177 | -0.317 | 0.242 | 0.484 | 0.725 | -0.009 | 0.052 | 0.052 | 0 | 0.41 | 0.3 | 0.45 | 0.17 | 0.45 | 0.48 |
| 5 | 159824464 | rs4921136 | 1.424 | 0.22 | 0.432 | 0.953 | -0.008 | 0.006 | 0.006 | 0.02 | 0.34 | 0.53 | 0.35 | 0.34 | 0.32 | 0.31 |
| 5 | 159825798 | rs1075059 | -0.135 | 0.279 | 0.194 | 0.981 | -0.009 | 0.05 | 0.05 | 0 | 0.41 | 0.3 | 0.45 | 0.18 | 0.45 | 0.48 |
| 5 | 159826674 | rs7717016 | 1.582 | 0.146 | 0.043 | 1.04 | -0.001 | -0.029 | -0.029 | 0.657 | 0.32 | 0.21 | 0.31 | 0.34 | 0.32 | 0.31 |
| 5 | 159842673 | rs2910191 | 0.394 | 0.11 | 0.604 | 1.401 | -0.012 | 0.09 | 0.09 | 0 | 0.76 | 0.65 | 0.8 | 0.52 | 0.82 | 0.82 |
| 5 | 159843996 | rs11949572 | -0.375 | 0.165 | 0.537 | 1.669 | 0.013 | 0.11 | 0.11 | 0.109 | 0.12 | 0.33 | 0.19 | 0.24 | 0.12 | 0.13 |
| 5 | 159844275 | rs2910195 | -0.592 | -0.031 | -0.061 | 1.727 | 0.003 | -0.015 | -0.015 | 0 | 0.87 | 0.9 | 0.86 | 0.52 | 0.88 | 0.9 |
| 5 | 159846074 | rs17057781 | -0.055 | 0.082 | -0.198 | 1.4 | -0.002 | 0.011 | 0.011 | 0 | 0.2 | 0.12 | 0.1 | 0.21 | 0.17 | 0.21 |
| 5 | 159846872 | rs3811999 | 1.187 | 0.183 | -0.458 | 1.362 | -0.001 | 0.084 | 0.084 | 0 | 0.34 | 0.17 | 0.32 | 0.15 | 0.38 | 0.28 |
| 5 | 159847080 | rs2910199 | 1.055 | 0.185 | -0.341 | 1.369 | -0.019 | 0.009 | 0.009 | 0.467 | 0.21 | 0.12 | 0.33 | 0.12 | 0.25 | 0.31 |
| 5 | 159850372 | rs2910201 | 0.895 | 0.027 | 0.043 | 1.646 | 0.019 | 0.042 | 0.042 | 0 | 0.55 | 0.29 | 0.65 | 0.28 | 0.63 | 0.59 |
| 5 | 159857528 | rs6897560 | 0.331 | 0.141 | -0.078 | 1.262 | -0.018 | 0.136 | 0.136 | 0 | 0.07 | 0.28 | 0.02 | 0.25 | 0.01 | 0.17 |
| 5 | 159861575 | rs2961911 | 0.214 | 0.724 | -0.465 | 0.723 | 0.002 | -0.005 | -0.005 | 0 | 0.68 | 0.57 | 0.77 | 0.72 | 0.7 | 0.85 |
| 5 | 159865579 | rs11135102 | 0.186 | 0.524 | -0.718 | 0.501 | -0.002 | 0.01 | 0.01 | 2.235 | 0.57 | 0.43 | 0.47 | 0.25 | 0.64 | 0.34 |
| 5 | 159870611 | rs4921283 | 0.115 | 0.033 | -1.045 | 0.422 | 0.03 | 0.04 | 0.04 | 0 | 0.5 | 0.29 | 0.47 | 0.63 | 0.63 | 0.54 |
| 5 | 159876888 | rs2910211 | -0.298 | -0.159 | -0.9 | 0.4 | -0.018 | -0.009 | -0.009 | 0 | 0.7 | 0.4 | 0.8 | 0.76 | 0.81 | 0.84 |
| 5 | 159879258 | rs2431696 | 0.904 | 0.037 | -1.085 | 0.357 | -0.056 | -0.01 | -0.01 | 0 | 0.19 | 0.01 | 0.32 | 0.13 | 0.18 | 0.3 |
| 5 | 159879978 | rs2431697 | -0.344 | 0.005 | -0.87 | -0.069 | -0.001 | -0.013 | -0.013 | 0 | 0.46 | 0.46 | 0.49 | 0.11 | 0.43 | 0.39 |
| 5 | 159880356 | rs11951834 | 1.178 | -0.082 | -0.47 | -0.18 | -0.006 | -0.038 | -0.038 | 0 | 0.14 | 0.19 | 0.07 | 0 | 0.11 | 0.01 |
| 5 | 159883652 | rs2909733 | -0.623 | 0.1 | -0.501 | 0.296 | 0 | 0.019 | 0.019 | 0 | 0.93 | 0.93 | 0.94 | 0.77 | 0.95 | 0.87 |
| 5 | 159886620 | rs2431099 | 0.613 | 0.105 | -0.645 | 0.164 | -0.001 | 0.001 | 0.001 | 0 | 0.46 | 0.46 | 0.45 | 0.66 | 0.52 | 0.48 |
| 5 | 159890885 | rs2431097 | 0.512 | 0.084 | -0.993 | 0.133 | -0.028 | 0.069 | 0.069 | 0 | 0.43 | 0.27 | 0.39 | 0.25 | 0.51 | 0.42 |
| 5 | 159892111 | rs17476918 | 0.541 | 0.25 | -0.601 | 0.427 | -0.029 | 0.035 | 0.035 | 0.351 | 0.08 | 0.02 | 0.04 | 0 | 0.07 | 0.11 |
| 5 | 159896897 | rs17057833 | 0.786 | -0.126 | -0.469 | 0.339 | 0.005 | -0.008 | -0.008 | 0 | 0.13 | 0.13 | 0.06 | 0 | 0.09 | 0.01 |
| 5 | 159902313 | rs17057846 | -0.113 | 0.009 | -0.44 | 0.628 | -0.001 | -0.001 | -0.001 | 0.884 | 0.14 | 0.15 | 0.12 | 0.21 | 0.12 | 0.27 |
| 5 | 159904729 | rs883517 | -0.344 | -0.09 | -0.103 | 1.092 | 0.02 | 0.265 | 0.265 | 0.342 | 0.13 | 0.44 | 0.23 | 0.39 | 0.11 | 0.17 |
| 5 | 159911506 | rs2961920 | 0.388 | -0.267 | -0.208 | 0.931 | -0.035 | 0.052 | 0.052 | 0 | 0.69 | 0.56 | 0.68 | 0.37 | 0.77 | 0.7 |
| 5 | 159920295 | rs4921291 | -1.149 | -0.181 | -1.315 | 0.596 | 0.018 | 0.162 | 0.162 | 0 | 0.47 | 0.94 | 0.47 | 0.84 | 0.34 | 0.53 |
| 5 | 159923720 | rs1895330 | 1.068 | 0.763 | -1.508 | 0.415 | -0.064 | -0.021 | -0.021 | 0 | 0.16 | 0.02 | 0.3 | 0.1 | 0.21 | 0.2 |
| 5 | 159926960 | rs17057882 | 0.06 | 0.39 | -0.662 | 0.443 | 0.031 | 0.129 | 0.129 | 0 | 0.11 | 0.26 | 0.22 | 0.36 | 0.14 | 0.16 |
| 5 | 159928372 | rs2431103 | -1.274 | 0.257 | -1.459 | 0.21 | -0.077 | 0.216 | 0.216 | 0.092 | 0.64 | 0.96 | 0.77 | 0.96 | 0.58 | 0.83 |
| 5 | 159936883 | rs4921145 | 0.275 | -0.129 | -0.571 | 0.714 | -0.028 | 0.003 | 0.003 | 0.415 | 0.16 | 0.22 | 0.11 | 0.2 | 0.08 | 0.13 |
| 5 | 159938903 | rs11742500 | 0.329 | -0.048 | -0.641 | 0.74 | -0.013 | 0.002 | 0.002 | 0.417 | 0.16 | 0.2 | 0.1 | 0.2 | 0.09 | 0.13 |
| 5 | 159942144 | rs1592743 | -0.759 | 0.195 | -0.75 | 0.485 | -0.001 | -0.003 | -0.003 | 0 | 0.49 | 0.54 | 0.53 | 0.61 | 0.43 | 0.62 |
| 5 | 159961053 | rs11741878 | 0.252 | -0.433 | -1.242 | 0.132 | -0.007 | 0.002 | 0.002 | 0.994 | 0.13 | 0.17 | 0.09 | 0.18 | 0.07 | 0.11 |
| 5 | 159966924 | rs6897982 | -0.189 | -0.47 | -1.399 | 0.128 | 0.011 | 0.002 | 0.002 | 0.368 | 0.22 | 0.23 | 0.13 | 0.21 | 0.11 | 0.13 |
| 5 | 159967244 | rs12187077 | 0.116 | 0.396 | -1.625 | 0.146 | -0.047 | 0.023 | 0.023 | 2.335 | 0.36 | 0.25 | 0.61 | 0.68 | 0.44 | 0.62 |
| 5 | 159972064 | rs1592750 | -0.059 | 0.159 | -1.165 | 0.007 | 0.016 | 0.037 | 0.037 | 0 | 0.6 | 0.38 | 0.72 | 0.86 | 0.61 | 0.76 |
| 5 | 159975042 | rs7717590 | -0.287 | 0.203 | -1.031 | 0.561 | -0.011 | 0.046 | 0.046 | 0 | 0.53 | 0.61 | 0.59 | 0.65 | 0.48 | 0.59 |
| 5 | 159976464 | rs10515809 | -0.705 | -0.404 | -0.015 | 1.311 | -0.011 | 0.245 | 0.245 | 0 | 0.12 | 0.29 | 0.24 | 0.46 | 0.1 | 0.22 |
| 5 | 159976624 | rs17057955 | 0.098 | -0.439 | -0.178 | 1.592 | 0.002 | -0.002 | -0.002 | 0.511 | 0.07 | 0.05 | 0.04 | 0.25 | 0.08 | 0.16 |
| 5 | 159979651 | rs3926946 | -0.754 | -0.597 | 0.147 | 1.24 | 0.003 | 0.001 | 0.001 | 0 | 0.38 | 0.59 | 0.55 | 0.62 | 0.33 | 0.43 |
| 5 | 159980580 | rs10515811 | -0.187 | -0.562 | 0.134 | 1.498 | -0.001 | 0.11 | 0.11 | 0 | 0.07 | 0.11 | 0.21 | 0.37 | 0.08 | 0.19 |
| 5 | 159984054 | rs7712888 | -0.686 | -0.584 | 0.028 | 1.39 | -0.004 | 0.005 | 0.005 | 0 | 0.37 | 0.52 | 0.53 | 0.54 | 0.33 | 0.44 |
| 5 | 159985757 | rs12656515 | -0.721 | -0.24 | 0.023 | 1.754 | -0.001 | 0.107 | 0.107 | 0 | 0.07 | 0.12 | 0.18 | 0.36 | 0.08 | 0.2 |
| 5 | 159990141 | rs11959870 | 0.226 | -0.284 | -0.167 | 0.91 | 0.002 | 0 | 0 | 0 | 0.12 | 0.11 | 0.07 | 0.01 | 0.08 | 0.14 |
| 5 | 159990237 | rs1053468 | 0.226 | -0.284 | -0.167 | 0.916 | 0.002 | 0.001 | 0.001 | 0 | 0.12 | 0.11 | 0.07 | 0.01 | 0.08 | 0.14 |
| 5 | 159990279 | rs1053467 | -0.739 | -0.671 | 0.302 | 0.728 | 0.002 | 0.01 | 0.01 | 0 | 0.34 | 0.33 | 0.4 | 0.17 | 0.31 | 0.34 |
| 5 | 159991367 | rs11953731 | 0.483 | -0.656 | 0.23 | 0.73 | 0.002 | 0.001 | 0.001 | 0 | 0.11 | 0.1 | 0.07 | 0.01 | 0.07 | 0.14 |
| 5 | 159992655 | rs3812005 | -0.963 | -0.793 | 0.866 | 0.653 | -0.013 | -0.044 | -0.044 | 0.5 | 0.39 | 0.48 | 0.43 | 0.17 | 0.33 | 0.36 |
| 5 | 159992751 | rs4921150 | -0.185 | -0.792 | 0.894 | 0.64 | -0.021 | -0.086 | -0.086 | 0 | 0.16 | 0.29 | 0.1 | 0.01 | 0.1 | 0.16 |
| 5 | 159995502 | rs1895191 | 0.275 | -0.655 | 0.734 | 0.771 | 0 | 0.005 | 0.005 | 0 | 0.22 | 0.18 | 0.33 | 0.16 | 0.23 | 0.2 |
| 5 | 160003685 | rs4921305 | -0.412 | -0.655 | 0.734 | 0.771 | 0 | 0.005 | 0.005 | 0 | 0.78 | 0.82 | 0.67 | 0.84 | 0.77 | 0.8 |
| 5 | 160016007 | rs2052459 | -0.172 | -0.687 | 1.016 | 0.775 | -0.005 | -0.009 | -0.009 | 0 | 0.76 | 0.66 | 0.65 | 0.83 | 0.77 | 0.8 |
| 5 | 160026777 | rs4921307 | 0.319 | -0.938 | 1.202 | 0.655 | 0.02 | 0.024 | 0.024 | 0 | 0.8 | 0.55 | 0.66 | 0.82 | 0.79 | 0.77 |
| 5 | 160034484 | rs11135108 | 1.808 | -1.456 | 0.149 | 0.303 | -0.145 | 0.166 | 0.166 | 0 | 0.32 | 0.04 | 0.25 | 0.09 | 0.44 | 0.22 |
| 5 | 160034720 | rs1019455 | -0.136 | -1.252 | 0.71 | 0.704 | 0.008 | 0.011 | 0.011 | 0.038 | 0.82 | 0.72 | 0.68 | 0.82 | 0.79 | 0.77 |
| 5 | 160040461 | rs6556505 | -0.951 | -1.237 | 0.648 | 0.724 | -0.006 | -0.006 | -0.006 | 1.153 | 0.87 | 0.88 | 0.71 | 0.83 | 0.81 | 0.81 |
| 5 | 160043071 | rs4921308 | 1.518 | -1.474 | 1.048 | 0.575 | -0.01 | 0.167 | 0.167 | 0 | 0.44 | 0.2 | 0.32 | 0.09 | 0.51 | 0.34 |
| 5 | 160045164 | rs888747 | 0.359 | -0.676 | 1.348 | 1.039 | 0.021 | 0.025 | 0.025 | 0 | 0.8 | 0.55 | 0.66 | 0.82 | 0.79 | 0.77 |
| 5 | 160046184 | rs17058066 | 0.221 | -0.765 | 0.645 | 1.168 | -0.012 | -0.008 | -0.008 | 0 | 0.09 | 0.03 | 0.06 | 0.06 | 0.07 | 0.07 |
| 5 | 160050162 | rs10476328 | 0.168 | -0.665 | 0.886 | 1.09 | 0.002 | 0.003 | 0.003 | 0 | 0.82 | 0.79 | 0.68 | 0.82 | 0.79 | 0.78 |
| 5 | 160064605 | rs6888425 | 0.16 | -0.453 | 0.896 | 1.325 | -0.002 | -0.002 | -0.002 | 0 | 0.89 | 0.91 | 0.73 | 0.89 | 0.87 | 0.9 |
| 5 | 160069545 | rs1363197 | 0.992 | -0.489 | 1.206 | 1.378 | 0.014 | 0.068 | 0.068 | 0 | 0.85 | 0.56 | 0.7 | 0.82 | 0.85 | 0.84 |
| 5 | 160091381 | rs1549905 | 0.503 | -0.325 | 1.04 | 1.232 | 0.02 | 0.025 | 0.025 | 0 | 0.89 | 0.74 | 0.72 | 0.88 | 0.87 | 0.9 |
| 5 | 160093769 | rs2421533 | 0.379 | -0.308 | 0.937 | 1.256 | 0.016 | 0.023 | 0.023 | 0 | 0.89 | 0.79 | 0.73 | 0.88 | 0.87 | 0.9 |

| | | | | | | | | | | | | | | | | |
|---|---|---|---|---|---|---|---|---|---|---|---|---|---|---|---|---|
| 5 | 160097496 | rs958912 | 0.71 | 0.171 | 0.724 | 1.174 | -0.018 | -0.002 | -0.002 | 0.003 | 0.88 | 0.93 | 0.71 | 0.89 | 0.81 | 0.9 |
| 5 | 160097632 | rs958911 | 0.338 | 0.101 | 0.748 | 1.199 | -0.007 | -0.005 | -0.005 | 2.302 | 0.9 | 0.93 | 0.74 | 0.89 | 0.87 | 0.9 |
| 5 | 160099440 | rs11135109 | 1.376 | -0.17 | 1.286 | 1.77 | 0.011 | 0.129 | 0.129 | 0.082 | 0.74 | 0.51 | 0.59 | 0.53 | 0.74 | 0.69 |
| 5 | 160100301 | rs7704503 | 0.134 | -0.177 | 1.37 | 1.75 | 0.014 | -0.003 | -0.003 | 0 | 0.9 | 0.85 | 0.73 | 0.91 | 0.87 | 0.91 |
| 5 | 160101765 | rs2024524 | 0.142 | -0.159 | 1.293 | 1.758 | 0.001 | -0.001 | -0.001 | 0 | 0.9 | 0.9 | 0.73 | 0.91 | 0.87 | 0.91 |
| 5 | 160106482 | rs17058110 | -0.864 | -0.218 | 1.312 | 1.774 | 0.002 | 0.202 | 0.202 | 0.12 | 0.16 | 0.39 | 0.15 | 0.39 | 0.13 | 0.22 |
| 5 | 160114020 | rs12652215 | 0.064 | -0.161 | 1.355 | 1.2 | 0.001 | -0.002 | -0.002 | 0 | 0.9 | 0.87 | 0.72 | 0.91 | 0.88 | 0.9 |
| 5 | 160115779 | rs6894908 | -0.961 | -0.415 | 1.176 | 1.363 | -0.013 | 0.052 | 0.052 | 1.257 | 0.2 | 0.34 | 0.25 | 0.66 | 0.16 | 0.35 |
| 5 | 160134369 | rs2033462 | 0.813 | 0.026 | 1.187 | 1.302 | 0.038 | 0.006 | 0.006 | 0 | 0.88 | 0.72 | 0.72 | 0.9 | 0.86 | 0.91 |
| 5 | 160135873 | rs6866731 | 0.255 | 0.074 | 0.869 | 1.308 | 0.003 | -0.001 | -0.001 | 0.3 | 0.9 | 0.88 | 0.73 | 0.9 | 0.86 | 0.9 |
| 5 | 160150822 | rs10062528 | 0.944 | 0.042 | 0.556 | 1.175 | -0.012 | 0.003 | 0.003 | 0 | 0.79 | 0.85 | 0.65 | 0.82 | 0.73 | 0.85 |
| 5 | 160152931 | rs11742097 | 0.274 | 0.057 | 0.592 | 1.169 | 0.018 | 0 | 0 | 0 | 0.89 | 0.81 | 0.72 | 0.9 | 0.86 | 0.91 |
| 5 | 160157153 | rs2421535 | 0.205 | -0.001 | 0.486 | 1.134 | 0.001 | 0.005 | 0.005 | 0.468 | 0.11 | 0.07 | 0.06 | 0.09 | 0.1 | 0.08 |
| 5 | 160172536 | rs1432788 | -0.161 | -0.043 | 0.758 | 1.124 | -0.001 | -0.01 | -0.01 | 0 | 0.1 | 0.17 | 0.07 | 0.08 | 0.09 | 0.07 |
| 5 | 160173130 | rs7737399 | -0.747 | -0.11 | 1.328 | 1.092 | 0.01 | -0.007 | -0.007 | 0 | 0.12 | 0.4 | 0.09 | 0.08 | 0.09 | 0.07 |
| 5 | 160173283 | rs9313857 | -0.66 | 0.292 | 1.072 | 1.422 | 0.004 | 0.055 | 0.055 | 0 | 0.2 | 0.24 | 0.3 | 0.46 | 0.23 | 0.26 |
| 5 | 160174315 | rs1432791 | -1.364 | 0.106 | 0.816 | 0.957 | -0.004 | -0.003 | -0.003 | 0 | 0.35 | 0.34 | 0.57 | 0.58 | 0.41 | 0.41 |
| 5 | 160174947 | rs1005790 | -2.087 | 0.049 | 0.482 | 0.899 | -0.013 | 0.066 | 0.066 | 0.243 | 0.48 | 0.74 | 0.67 | 0.66 | 0.51 | 0.49 |
| 5 | 160184267 | rs4292483 | -2.045 | 0.052 | 0.263 | 0.901 | -0.005 | 0.023 | 0.023 | 0.174 | 0.47 | 0.67 | 0.66 | 0.64 | 0.51 | 0.48 |
| 5 | 160187121 | rs11135114 | -1.207 | -0.061 | 0.361 | 1.359 | -0.011 | 0.007 | 0.007 | 0 | 0.3 | 0.24 | 0.26 | 0.23 | 0.35 | 0.26 |
| 5 | 160187406 | rs7725968 | -0.891 | 0.096 | 0.291 | 1.524 | -0.002 | 0.154 | 0.154 | 0 | 0.17 | 0.43 | 0.4 | 0.41 | 0.16 | 0.23 |
| 5 | 160187458 | rs12517584 | -0.567 | 0.148 | -0.417 | 1.361 | -0.008 | -0.002 | -0.002 | 0.15 | 0.2 | 0.18 | 0.19 | 0.17 | 0.26 | 0.19 |
| 5 | 160196305 | rs4479880 | -0.727 | 0.221 | -1.109 | 0.94 | -0.016 | 0.017 | 0.017 | 0 | 0.57 | 0.35 | 0.6 | 0.63 | 0.51 | 0.53 |
| 5 | 160196465 | rs11135118 | -0.222 | -0.041 | -1.17 | 1.348 | 0 | -0.002 | -0.002 | 0 | 0.15 | 0.17 | 0.27 | 0.22 | 0.12 | 0.13 |
| 5 | 160196659 | rs11958424 | -0.894 | -0.193 | -1.236 | 1.301 | -0.003 | 0.022 | 0.022 | 0.319 | 0.21 | 0.27 | 0.33 | 0.5 | 0.15 | 0.28 |
| 5 | 160197921 | rs12517197 | -1.04 | -0.401 | -0.99 | 1.295 | -0.028 | 0.108 | 0.108 | 0 | 0.14 | 0.27 | 0.29 | 0.43 | 0.05 | 0.23 |
| 5 | 160200108 | rs4594897 | -0.675 | -0.265 | -1.159 | 1.058 | 0.033 | 0.005 | 0.005 | 0 | 0.49 | 0.34 | 0.56 | 0.54 | 0.4 | 0.46 |
| 5 | 160201741 | rs17058247 | -0.366 | -0.282 | -0.936 | 0.976 | -0.003 | -0.002 | -0.002 | 0.193 | 0.43 | 0.43 | 0.53 | 0.59 | 0.36 | 0.48 |
| 5 | 160202116 | rs13160719 | -0.302 | -0.37 | -1.339 | 0.852 | 0 | 0.001 | 0.001 | 0 | 0.25 | 0.27 | 0.38 | 0.32 | 0.26 | 0.24 |
| 5 | 160205012 | rs7724015 | 1.242 | -0.536 | -1.424 | 0.657 | 0.001 | 0.073 | 0.073 | 0 | 0.18 | 0.04 | 0.15 | 0.01 | 0.15 | 0.14 |
| 5 | 160208277 | rs4921336 | -1.019 | -0.624 | -1.336 | 0.769 | 0.005 | 0.004 | 0.004 | 0 | 0.47 | 0.44 | 0.42 | 0.59 | 0.4 | 0.54 |
| 5 | 160209733 | rs7712586 | -0.047 | -0.664 | -1.33 | 0.832 | 0.003 | 0.013 | 0.013 | 0 | 0.29 | 0.23 | 0.29 | 0.2 | 0.26 | 0.27 |
| 5 | 160211319 | rs4921338 | 1.009 | -0.542 | -1.065 | 1.245 | 0.009 | -0.018 | -0.018 | 0.213 | 0.12 | 0.05 | 0.08 | 0.13 | 0.12 | 0.1 |
| 5 | 160212103 | rs11953648 | -0.108 | -0.546 | -1.075 | 1.244 | 0.001 | 0.019 | 0.019 | 0 | 0.11 | 0.17 | 0.15 | 0.19 | 0.11 | 0.12 |
| 5 | 160212237 | rs11135121 | -0.805 | -0.537 | -1.059 | 1.223 | -0.01 | 0.028 | 0.028 | 0 | 0.25 | 0.35 | 0.24 | 0.31 | 0.23 | 0.25 |
| 5 | 160212590 | rs13190543 | -0.765 | -0.529 | -1.151 | 1.227 | -0.003 | 0.007 | 0.007 | 0 | 0.25 | 0.29 | 0.24 | 0.31 | 0.23 | 0.25 |
| 5 | 160217933 | rs6862853 | 0.593 | -0.404 | -1.191 | 0.99 | 0.005 | -0.003 | -0.003 | 0.008 | 0.13 | 0.12 | 0.09 | 0.13 | 0.12 | 0.12 |
| 5 | 160219464 | rs10036919 | -0.418 | 0.093 | -1.442 | 0.781 | 0 | -0.019 | -0.019 | 0 | 0.32 | 0.28 | 0.31 | 0.54 | 0.32 | 0.35 |
| 5 | 160219844 | rs10046095 | -0.161 | -0.075 | -1.465 | 0.555 | 0.007 | -0.033 | -0.033 | 0 | 0.27 | 0.34 | 0.26 | 0.18 | 0.31 | 0.18 |
| 5 | 160220268 | rs10037124 | -1.191 | -0.138 | -1.399 | 0.679 | -0.001 | -0.002 | -0.002 | 0 | 0.44 | 0.5 | 0.42 | 0.63 | 0.43 | 0.47 |
| 5 | 160226968 | rs13175076 | -0.933 | -0.335 | -1.312 | 0.29 | -0.001 | -0.011 | -0.011 | 0 | 0.45 | 0.5 | 0.4 | 0.38 | 0.44 | 0.3 |
| 5 | 160233213 | rs4921342 | -0.811 | 0.158 | -1.657 | -0.191 | 0 | -0.005 | -0.005 | 0 | 0.48 | 0.48 | 0.47 | 0.78 | 0.48 | 0.53 |
| 5 | 160233794 | rs10062727 | 1.076 | 0.146 | -1.56 | 0.282 | -0.001 | -0.008 | -0.008 | 0 | 0.18 | 0.15 | 0.21 | 0.34 | 0.19 | 0.17 |
| 5 | 160235125 | rs4569911 | 0.367 | -0.355 | -1.607 | -0.294 | -0.048 | -0.032 | -0.032 | 0 | 0.33 | 0.14 | 0.38 | 0.6 | 0.43 | 0.46 |
| 5 | 160264007 | rs10078103 | 0.6 | -0.963 | -0.872 | -0.694 | 0.002 | -0.013 | -0.013 | 0.184 | 0.12 | 0.17 | 0.11 | 0.11 | 0.12 | 0.11 |
| 5 | 160271059 | rs17058343 | 0.887 | -1.177 | -0.776 | -0.8 | 0.007 | -0.001 | -0.001 | 0 | 0.09 | 0.06 | 0.04 | 0 | 0.04 | 0.08 |
| 5 | 160275493 | rs17058351 | 2.463 | -1.357 | -0.919 | -0.991 | 0.003 | 0.047 | 0.047 | 0 | 0.15 | 0.05 | 0.06 | 0.01 | 0.13 | 0.15 |
| 5 | 160276677 | rs4559036 | -0.889 | -1.231 | -0.731 | -0.907 | -0.05 | -0.147 | -0.147 | 0.125 | 0.73 | 0.51 | 0.73 | 0.98 | 0.83 | 0.72 |
| 5 | 160286514 | rs4921347 | -0.885 | -1.227 | -0.802 | -0.904 | -0.045 | -0.134 | -0.134 | 1.382 | 0.73 | 0.53 | 0.73 | 0.98 | 0.83 | 0.72 |
| 5 | 160312017 | rs7706812 | -0.161 | -1.096 | -1.242 | -0.844 | 0.04 | 0.082 | 0.082 | 0 | 0.59 | 0.27 | 0.61 | 0.86 | 0.71 | 0.61 |
| 5 | 160317284 | rs1104769 | 0.385 | -0.984 | -1.595 | -0.425 | -0.043 | -0.053 | -0.053 | 0.311 | 0.39 | 0.15 | 0.42 | 0.51 | 0.56 | 0.32 |
| 5 | 160318038 | rs6859868 | 0.288 | -1.37 | -1.629 | -0.607 | 0.023 | -0.045 | -0.045 | 0 | 0.26 | 0.11 | 0.22 | 0.36 | 0.19 | 0.36 |
| 5 | 160318988 | rs12658840 | -0.392 | -1.155 | -1.433 | -0.593 | 0.023 | 0.01 | 0.01 | 0 | 0.54 | 0.39 | 0.53 | 0.62 | 0.69 | 0.47 |
| 5 | 160326374 | rs4921349 | 1.03 | -0.722 | -1.405 | -0.21 | -0.003 | -0.069 | -0.069 | 0.369 | 0.18 | 0.05 | 0.18 | 0.36 | 0.16 | 0.27 |
| 5 | 160328128 | rs1516651 | -0.275 | -0.241 | -0.963 | 0.125 | -0.015 | -0.021 | -0.021 | 0 | 0.4 | 0.27 | 0.43 | 0.51 | 0.56 | 0.32 |
| 5 | 160330420 | rs7700972 | 1.504 | -0.014 | -0.698 | 0.122 | 0.007 | -0.036 | -0.036 | 0.113 | 0.08 | 0.15 | 0.09 | 0 | 0.12 | 0.05 |
| 5 | 160331001 | rs885466 | 1.088 | 0.138 | -0.779 | 0.89 | -0.035 | -0.029 | -0.029 | 0.005 | 0.07 | 0.01 | 0.31 | 0.29 | 0.1 | 0.12 |
| 5 | 160331497 | rs867358 | 0.348 | 0.112 | -0.504 | 1.048 | 0.005 | 0 | 0 | 0.068 | 0.64 | 0.59 | 0.43 | 0.37 | 0.54 | 0.54 |
| 5 | 160333436 | rs6871718 | 1.981 | -0.25 | -0.568 | 0.87 | 0.001 | -0.128 | -0.128 | 0 | 0.34 | 0.44 | 0.23 | 0.04 | 0.32 | 0.24 |
| 5 | 160337021 | rs949719 | -0.415 | -0.189 | -0.779 | 0.903 | -0.061 | -0.031 | -0.031 | 1.769 | 0.69 | 0.85 | 0.47 | 0.63 | 0.56 | 0.68 |
| 5 | 160337126 | rs949718 | -0.057 | -0.069 | -0.945 | 0.26 | 0.009 | -0.071 | -0.071 | 2.097 | 0.24 | 0.05 | 0.17 | 0.33 | 0.2 | 0.23 |
| 5 | 160343115 | rs6898138 | 0.394 | 0.187 | -0.372 | 0.782 | 0.011 | 0.003 | 0.003 | 0 | 0.64 | 0.58 | 0.43 | 0.38 | 0.54 | 0.54 |
| 5 | 160350132 | rs6872549 | 1.002 | 0.307 | -0.176 | 0.869 | 0.003 | 0.015 | 0.015 | 0 | 0.52 | 0.35 | 0.39 | 0.39 | 0.53 | 0.48 |
| 5 | 160353356 | rs962102 | 0.393 | 0.369 | -0.256 | 1.124 | 0.001 | 0.014 | 0.014 | 0 | 0.63 | 0.59 | 0.62 | 0.42 | 0.62 | 0.58 |
| 5 | 160353471 | rs17058469 | 0.675 | 0.561 | -0.676 | 1.094 | -0.019 | -0.036 | -0.036 | 0.173 | 0.06 | 0.01 | 0.04 | 0.12 | 0.09 | 0.13 |
| 5 | 160356255 | rs10051325 | -0.993 | 0.347 | -0.412 | 1.134 | -0.003 | 0.019 | 0.019 | 0 | 0.38 | 0.49 | 0.45 | 0.55 | 0.38 | 0.49 |
| 5 | 160357338 | rs17058486 | 0.686 | 0.37 | -0.422 | 1.3 | -0.001 | 0.012 | 0.012 | 0 | 0.08 | 0.09 | 0.09 | 0.21 | 0.07 | 0.15 |
| 5 | 160359601 | rs10079953 | -0.753 | 0.346 | -0.262 | 1.249 | -0.004 | 0.02 | 0.02 | 0 | 0.3 | 0.4 | 0.37 | 0.36 | 0.31 | 0.32 |
| 5 | 160360749 | rs2090374 | 0.26 | 0.367 | -0.321 | 1.331 | -0.001 | 0.004 | 0.004 | 0.148 | 0.17 | 0.15 | 0.13 | 0.26 | 0.2 | 0.19 |

| | | | | | | | | | | | | | | | |
|---|---|---|---|---|---|---|---|---|---|---|---|---|---|---|---|
| 5 | 160367797 | rs13166904 | -0.282 | 0.382 | -0.348 | 1.133 | -0.002 | 0.001 | 0.001 | 0 | 0.2 | 0.16 | 0.18 | 0.2 | 0.25 | 0.22 |
| 5 | 160375241 | rs17501698 | 1.036 | 0.633 | -0.324 | 1.15 | 0 | -0.001 | -0.001 | 0 | 0.07 | 0.06 | 0.08 | 0.09 | 0.11 | 0.13 |
| 5 | 160375908 | rs869339 | -1.049 | 0.56 | -0.055 | 1.187 | -0.002 | 0.024 | 0.024 | 0 | 0.3 | 0.42 | 0.37 | 0.37 | 0.31 | 0.33 |
| 5 | 160375995 | rs938540 | -1.503 | 0.562 | -0.081 | 1.134 | -0.003 | 0.016 | 0.016 | 0 | 0.38 | 0.5 | 0.45 | 0.56 | 0.38 | 0.48 |
| 5 | 160380004 | rs6872185 | 1.919 | 0.236 | -0.388 | -0.038 | -0.017 | 0.086 | 0.086 | 0 | 0.42 | 0.27 | 0.29 | 0.05 | 0.43 | 0.27 |
| 5 | 160380401 | rs954929 | -1.498 | 0.175 | -0.546 | 0.456 | -0.003 | -0.055 | -0.055 | 0 | 0.37 | 0.46 | 0.48 | 0.8 | 0.37 | 0.48 |
| 5 | 160382733 | rs17058520 | 0.84 | 0.172 | -0.558 | 0.779 | -0.001 | 0.059 | 0.059 | 0 | 0.08 | 0.08 | 0.12 | 0.41 | 0.07 | 0.15 |
| 5 | 160382872 | rs999341 | 0.351 | 0.187 | -0.547 | -0.465 | 0.005 | -0.012 | -0.012 | 0 | 0.12 | 0.08 | 0.16 | 0.69 | 0.09 | 0.27 |
| 5 | 160384176 | rs1124418 | 0.512 | 0.321 | -0.365 | -0.5 | 0.001 | -0.038 | -0.038 | 0 | 0.56 | 0.45 | 0.59 | 0.82 | 0.51 | 0.58 |
| 5 | 160384249 | rs4513732 | 0.827 | 0.31 | -0.29 | -0.542 | 0.011 | -0.032 | -0.032 | 0.197 | 0.06 | 0.18 | 0.04 | 0 | 0.03 | 0.05 |
| 5 | 160387968 | rs1516655 | 1.612 | 0.315 | -0.73 | -0.546 | -0.035 | 0.047 | 0.047 | 0 | 0.41 | 0.12 | 0.34 | 0.7 | 0.41 | 0.43 |
| 5 | 160388599 | rs2176297 | 1.232 | 0.43 | -0.788 | -0.485 | -0.001 | -0.001 | -0.001 | 0.064 | 0.06 | 0.03 | 0.19 | 0.11 | 0.06 | 0.07 |
| 5 | 160388643 | rs2176298 | -0.259 | 0.449 | -0.938 | -0.489 | -0.003 | -0.034 | -0.034 | 0 | 0.68 | 0.58 | 0.69 | 0.82 | 0.69 | 0.64 |
| 5 | 160390549 | rs2222123 | 0.032 | 0.887 | 0.01 | -0.147 | -0.001 | -0.017 | -0.017 | 0 | 0.12 | 0.13 | 0.1 | 0 | 0.18 | 0.06 |
| 5 | 160390969 | rs1400334 | -0.778 | 0.522 | 0.183 | 0.433 | 0.006 | 0.002 | 0.002 | 0 | 0.14 | 0.33 | 0.24 | 0.1 | 0.1 | 0.12 |
| 5 | 160396816 | rs13154439 | -0.645 | 0.5 | -0.439 | 0.252 | -0.002 | 0.015 | 0.015 | 0 | 0.26 | 0.23 | 0.22 | 0.13 | 0.26 | 0.31 |
| 5 | 160398857 | rs7730983 | -0.759 | -0.054 | -0.079 | 0.488 | -0.024 | 0.004 | 0.004 | 0 | 0.09 | 0.15 | 0.05 | 0.12 | 0.03 | 0.14 |
| 5 | 160405064 | rs4244414 | -0.821 | -0.055 | 0.145 | 0.762 | 0.002 | 0.001 | 0.001 | 0 | 0.48 | 0.48 | 0.39 | 0.59 | 0.41 | 0.47 |
| 5 | 160412665 | rs1516660 | -0.013 | -0.121 | 0.525 | 0.688 | 0.004 | 0.01 | 0.01 | 0 | 0.37 | 0.45 | 0.34 | 0.47 | 0.35 | 0.34 |
| 5 | 160416534 | rs2421858 | 1.452 | -0.172 | 0.609 | 0.264 | -0.003 | 0.016 | 0.016 | 0 | 0.28 | 0.2 | 0.17 | 0.06 | 0.28 | 0.24 |
| 5 | 160420365 | rs2421857 | 0.385 | -0.172 | 0.577 | 0.277 | 0.003 | -0.014 | -0.014 | 0 | 0.65 | 0.65 | 0.5 | 0.54 | 0.63 | 0.59 |
| 5 | 160432975 | rs4921364 | -0.166 | -0.214 | 0.663 | 0.261 | -0.008 | -0.001 | -0.001 | 0 | 0.12 | 0.14 | 0.07 | 0.11 | 0.06 | 0.15 |
| 5 | 160436102 | rs11135140 | 0.606 | -0.159 | 0.642 | 0.306 | 0 | 0 | 0 | 0 | 0.37 | 0.38 | 0.33 | 0.47 | 0.35 | 0.34 |
| 5 | 160436777 | rs2138573 | 0.973 | -0.097 | 0.75 | 0.166 | -0.005 | -0.03 | -0.03 | 0 | 0.31 | 0.2 | 0.21 | 0.47 | 0.27 | 0.3 |
| 5 | 160442842 | rs1464144 | 1.454 | -0.072 | 0.387 | 0.467 | -0.03 | -0.039 | -0.039 | 0 | 0.15 | 0.06 | 0.37 | 0.2 | 0.25 | 0.19 |
| 5 | 160447362 | rs6894598 | 0.157 | -0.046 | 0.312 | 0.103 | -0.001 | 0.006 | 0.006 | 0 | 0.69 | 0.65 | 0.53 | 0.6 | 0.66 | 0.63 |
| 5 | 160448976 | rs17503515 | 1.445 | -0.128 | 0.32 | 0.142 | -0.039 | 0.1 | 0.1 | 0 | 0.26 | 0.07 | 0.15 | 0.06 | 0.28 | 0.24 |
| 5 | 160471514 | rs9313867 | 0.82 | 0.006 | 0.61 | 0.243 | 0.001 | 0.018 | 0.018 | 0 | 0.33 | 0.28 | 0.22 | 0.12 | 0.32 | 0.28 |
| 5 | 160488378 | rs10053266 | 0.566 | 0.209 | 0.619 | 0.687 | 0.023 | 0.062 | 0.062 | 0 | 0.2 | 0.32 | 0.5 | 0.35 | 0.33 | 0.27 |
| 5 | 160493675 | rs6861122 | 0.075 | -0.037 | 0.263 | 1.017 | -0.018 | 0.07 | 0.07 | 0 | 0.09 | 0.14 | 0.06 | 0.35 | 0.04 | 0.16 |
| 5 | 160494759 | rs10075958 | -0.537 | -0.203 | 0.37 | 0.672 | -0.004 | -0.007 | -0.007 | 0 | 0.56 | 0.54 | 0.67 | 0.66 | 0.61 | 0.68 |
| 5 | 160505992 | rs1489712 | 0.706 | -0.249 | 0.327 | 0.675 | -0.01 | -0.004 | -0.004 | 0 | 0.08 | 0.1 | 0.02 | 0.04 | 0.03 | 0.11 |
| 5 | 160506084 | rs9313868 | 0.665 | -0.111 | 0.41 | 0.722 | 0.015 | 0.05 | 0.05 | 0 | 0.22 | 0.3 | 0.5 | 0.46 | 0.33 | 0.32 |
| 5 | 160510960 | rs10866726 | 0.999 | -0.163 | -0.048 | 0.652 | -0.006 | 0.044 | 0.044 | 0 | 0.26 | 0.12 | 0.14 | 0.16 | 0.23 | 0.24 |
| 5 | 160517449 | rs7734700 | -0.051 | -0.097 | 0.055 | 0.78 | -0.013 | 0.002 | 0.002 | 0 | 0.1 | 0.14 | 0.03 | 0.14 | 0.05 | 0.18 |
| 5 | 160517753 | rs384943 | 0.59 | 0.266 | 0.392 | 0.887 | 0.061 | 0.112 | 0.112 | 0 | 0.21 | 0.43 | 0.51 | 0.37 | 0.33 | 0.26 |
| 5 | 160519563 | rs1387611 | -0.345 | -0.689 | -0.058 | 0.573 | 0.001 | -0.005 | -0.005 | 0 | 0.61 | 0.83 | 0.76 | 0.63 | 0.63 | 0.62 |
| 5 | 160521734 | rs4921374 | -0.37 | -0.694 | -0.015 | 0.475 | 0.003 | 0.019 | 0.019 | 0 | 0.62 | 0.82 | 0.76 | 0.66 | 0.63 | 0.62 |
| 5 | 160522668 | rs7733472 | -0.589 | -0.829 | 0.642 | 0.192 | -0.031 | -0.044 | -0.044 | 0 | 0.15 | 0.35 | 0.11 | 0.08 | 0.07 | 0.1 |
| 5 | 160536882 | rs7725291 | -0.567 | -1.734 | 0.584 | 0.204 | 0.013 | 0.01 | 0.01 | 0 | 0.26 | 0.26 | 0.15 | 0.16 | 0.18 | 0.18 |
| 5 | 160537025 | rs406195 | 0.788 | -1.753 | 0.304 | -0.246 | 0.002 | 0.012 | 0.012 | 0 | 0.47 | 0.56 | 0.57 | 0.65 | 0.52 | 0.5 |
| 5 | 160537180 | rs1489707 | -0.1 | -1.75 | 0.314 | -0.255 | 0.017 | 0.013 | 0.013 | 0 | 0.26 | 0.24 | 0.15 | 0.16 | 0.18 | 0.18 |
| 5 | 160537832 | rs861320 | -0.948 | -1.929 | 0.028 | -0.392 | -0.002 | -0.001 | -0.001 | 0.399 | 0.77 | 0.8 | 0.75 | 0.9 | 0.73 | 0.73 |
| 5 | 160541427 | rs17058779 | -0.056 | -1.878 | 0.01 | -0.336 | 0.014 | 0.01 | 0.01 | 0 | 0.25 | 0.24 | 0.15 | 0.17 | 0.17 | 0.19 |
| 5 | 160545841 | rs437220 | 1.538 | -2.359 | -0.296 | -0.3 | -0.083 | -0.082 | -0.082 | 0 | 0.29 | 0.13 | 0.29 | 0.44 | 0.42 | 0.37 |
| 5 | 160551087 | rs1038704 | -0.075 | -1.96 | 0.21 | 0.062 | 0.048 | 0.03 | 0.03 | 0 | 0.25 | 0.17 | 0.14 | 0.17 | 0.17 | 0.19 |
| 5 | 160556882 | rs1875526 | -0.075 | -1.96 | 0.219 | 0.062 | 0.045 | 0.028 | 0.028 | 0 | 0.25 | 0.17 | 0.14 | 0.17 | 0.18 | 0.19 |
| 5 | 160567322 | rs624722 | 0.978 | -1.76 | 0.383 | 0.075 | -0.007 | -0.019 | -0.019 | 0 | 0.42 | 0.29 | 0.37 | 0.45 | 0.51 | 0.44 |
| 5 | 160573507 | rs349753 | -0.764 | -2.107 | 0.838 | -0.719 | -0.029 | -0.097 | -0.097 | 0 | 0.28 | 0.44 | 0.28 | 0.07 | 0.22 | 0.21 |
| 5 | 160575832 | rs11956273 | -0.015 | -2.435 | 0.789 | -0.727 | 0.025 | -0.013 | -0.013 | 0 | 0.16 | 0.13 | 0.07 | 0.22 | 0.08 | 0.21 |
| 5 | 160577798 | rs375314 | -1.783 | -2.082 | 0.99 | -0.766 | 0.046 | 0.066 | 0.066 | 0 | 0.46 | 0.7 | 0.38 | 0.28 | 0.35 | 0.44 |
| 5 | 160581147 | rs349746 | 0.006 | -2.147 | 1.125 | -0.813 | 0.003 | -0.012 | -0.012 | 0 | 0.13 | 0.15 | 0.17 | 0.06 | 0.14 | 0.21 |
| 5 | 160585163 | rs411005 | -2.045 | -2.196 | 1.062 | -0.829 | 0.061 | 0.088 | 0.088 | 0 | 0.48 | 0.75 | 0.39 | 0.28 | 0.35 | 0.44 |
| 5 | 160585904 | rs10042476 | 0.079 | -2.193 | 1.035 | -0.817 | 0.024 | -0.013 | -0.013 | 0 | 0.16 | 0.13 | 0.07 | 0.22 | 0.08 | 0.21 |
| 5 | 160588561 | rs2546621 | 2.152 | -2.114 | 0.952 | -0.914 | -0.045 | 0.042 | 0.042 | 0 | 0.4 | 0.19 | 0.51 | 0.71 | 0.49 | 0.48 |
| 5 | 160591276 | rs349732 | -0.96 | -2.119 | 1.123 | -0.811 | 0.008 | -0.003 | -0.003 | 0.921 | 0.25 | 0.24 | 0.19 | 0.27 | 0.17 | 0.35 |
| 5 | 160593748 | rs164043 | 0.768 | -1.925 | 1.027 | -0.725 | -0.003 | 0.001 | 0.001 | 0.158 | 0.08 | 0.07 | 0.12 | 0.05 | 0.09 | 0.15 |
| 5 | 160604424 | rs467746 | 1.529 | -1.99 | 1.207 | -0.758 | 0.007 | 0.007 | 0.007 | 0 | 0.55 | 0.4 | 0.65 | 0.71 | 0.7 | 0.57 |
| 5 | 160613194 | rs455707 | -1.098 | -1.888 | 1.328 | -0.702 | -0.04 | -0.036 | -0.036 | 0 | 0.34 | 0.55 | 0.21 | 0.22 | 0.21 | 0.24 |
| 5 | 160622725 | rs180781 | -1.002 | -1.684 | 1.27 | -0.612 | -0.004 | 0 | 0 | 0.666 | 0.92 | 0.95 | 0.89 | 0.95 | 0.91 | 0.86 |
| 5 | 160623076 | rs164369 | 1.299 | -1.763 | 1.14 | -0.743 | -0.014 | 0.035 | 0.035 | 0 | 0.14 | 0.06 | 0.12 | 0.01 | 0.19 | 0.1 |
| 5 | 160624458 | rs164368 | 1.376 | -1.912 | 1.236 | -0.7 | 0.02 | 0.044 | 0.044 | 0 | 0.63 | 0.27 | 0.66 | 0.78 | 0.7 | 0.63 |
| 5 | 160635051 | rs248358 | -0.236 | -1.955 | 1.383 | -0.716 | 0.019 | -0.007 | -0.007 | 0 | 0.89 | 0.69 | 0.86 | 0.95 | 0.91 | 0.86 |
| 5 | 160639336 | rs463778 | 0.541 | -1.931 | 1.356 | -0.706 | -0.023 | -0.078 | -0.078 | 4.035 | 0.72 | 0.56 | 0.73 | 0.93 | 0.77 | 0.78 |
| 5 | 160641385 | rs386388 | 0.552 | -1.932 | 1.356 | -0.706 | -0.023 | -0.078 | -0.078 | 0.063 | 0.72 | 0.56 | 0.73 | 0.93 | 0.77 | 0.78 |
| 5 | 160645939 | rs449969 | 0.54 | -1.931 | 1.356 | -0.706 | -0.023 | -0.078 | -0.078 | 0 | 0.72 | 0.56 | 0.73 | 0.93 | 0.77 | 0.78 |
| 5 | 160649086 | rs17059065 | 0.296 | -1.644 | 1.235 | -0.464 | -0.015 | 0.046 | 0.046 | 0 | 0.06 | 0.12 | 0.03 | 0.15 | 0.03 | 0.15 |
| 5 | 160658223 | rs193722 | 1.244 | -1.607 | 1.547 | -0.262 | 0.01 | 0.042 | 0.042 | 0.216 | 0.82 | 0.37 | 0.81 | 0.76 | 0.86 | 0.74 |
| 5 | 160660200 | rs4921381 | 2.083 | -1.734 | 1.101 | -0.425 | -0.064 | 0.038 | 0.038 | 0 | 0.42 | 0.14 | 0.52 | 0.67 | 0.49 | 0.51 |

| | | | | | | | | | | | | | | | | |
|---|---|---|---|---|---|---|---|---|---|---|---|---|---|---|---|---|
| 5 | 160661108 | rs1818429 | 1.504 | -1.59 | 1.647 | 0.087 | -0.005 | -0.029 | -0.029 | 1.673 | 0.73 | 0.35 | 0.72 | 0.87 | 0.77 | 0.71 |
| 5 | 160665145 | rs10045194 | -0.618 | -1.329 | 1.455 | -0.215 | -0.038 | -0.045 | -0.045 | 1.12 | 0.15 | 0.34 | 0.12 | 0.08 | 0.1 | 0.13 |
| 5 | 160674476 | rs687501 | -0.342 | -0.95 | 1.016 | -0.025 | -0.001 | 0 | 0 | 0 | 0.92 | 0.94 | 0.89 | 0.95 | 0.91 | 0.85 |
| 5 | 160680142 | rs430279 | 0.173 | -0.956 | 1.052 | -0.027 | 0.055 | -0.004 | -0.004 | 0 | 0.9 | 0.81 | 0.86 | 0.95 | 0.86 | 0.84 |
| 5 | 160689448 | rs173564 | 1.392 | -1.531 | 1.12 | 0.251 | -0.001 | 0 | 0 | 0.176 | 0.62 | 0.36 | 0.51 | 0.74 | 0.7 | 0.67 |
| 5 | 160689685 | rs248367 | 0.099 | -1.386 | 1.177 | 0.346 | 0.006 | 0.05 | 0.05 | 0.308 | 0.91 | 0.84 | 0.85 | 0.83 | 0.91 | 0.86 |
| 5 | 160692454 | rs248368 | -0.549 | -1.519 | 0.833 | -0.01 | 0.008 | 0.068 | 0.068 | 0 | 0.24 | 0.11 | 0.16 | 0.01 | 0.23 | 0.09 |
| 5 | 160696308 | rs4618471 | -0.119 | -1.514 | 1.376 | 0.659 | -0.042 | 0.003 | 0.003 | 0 | 0.17 | 0.31 | 0.1 | 0.17 | 0.09 | 0.06 |
| 5 | 160696361 | rs248372 | 1.307 | -1.576 | 0.954 | 0.457 | 0.019 | -0.003 | -0.003 | 0.631 | 0.58 | 0.39 | 0.48 | 0.61 | 0.68 | 0.63 |
| 5 | 160701994 | rs687524 | -1.057 | -1.381 | 1.225 | 0.255 | -0.045 | -0.094 | -0.094 | 0 | 0.24 | 0.37 | 0.31 | 0.04 | 0.18 | 0.15 |
| 5 | 160702411 | rs664806 | -0.45 | -1.425 | 1.244 | 0.218 | -0.049 | -0.09 | -0.09 | 0 | 0.16 | 0.31 | 0.26 | 0.01 | 0.1 | 0.14 |
| 5 | 160703125 | rs10072922 | -1.221 | -1.424 | 1.257 | 0.212 | -0.065 | -0.134 | -0.134 | 1.764 | 0.26 | 0.43 | 0.31 | 0.04 | 0.18 | 0.15 |
| 5 | 160705005 | rs640476 | -1.762 | -1.453 | 0.816 | 0.41 | 0.043 | 0.022 | 0.022 | 0 | 0.37 | 0.76 | 0.53 | 0.27 | 0.28 | 0.26 |
| 5 | 160711978 | rs252955 | -1.1 | -1.342 | 1.049 | 0.339 | -0.026 | -0.07 | -0.07 | 0.216 | 0.26 | 0.37 | 0.34 | 0.06 | 0.24 | 0.21 |
| 5 | 160712407 | rs252954 | -1.025 | -1.366 | 1.24 | 0.307 | 0.005 | 0.003 | 0.003 | 0 | 0.24 | 0.21 | 0.33 | 0.05 | 0.24 | 0.15 |
| 5 | 160712947 | rs13178594 | -0.468 | -1.319 | 1.548 | 0.475 | 0.022 | 0.114 | 0.114 | 0 | 0.09 | 0.24 | 0.2 | 0.23 | 0.09 | 0.1 |
| 5 | 160716628 | rs17059231 | 0.023 | -1.534 | 1.386 | 0.974 | -0.002 | 0.094 | 0.094 | 0 | 0.07 | 0.18 | 0.17 | 0.15 | 0.05 | 0.09 |
| 5 | 160719244 | rs12523641 | -0.219 | -1.605 | 1.092 | 0.956 | -0.028 | -0.031 | -0.031 | 0 | 0.15 | 0.21 | 0.23 | 0.03 | 0.08 | 0.11 |
| 5 | 160719984 | rs592403 | -2.276 | -1.658 | 1.129 | 0.931 | 0.053 | -0.049 | -0.049 | 0.877 | 0.41 | 0.79 | 0.57 | 0.44 | 0.33 | 0.37 |
| 5 | 160721319 | rs2229944 | -0.143 | -1.472 | 1.085 | 0.8 | -0.021 | -0.025 | -0.025 | 0.803 | 0.14 | 0.16 | 0.23 | 0.03 | 0.08 | 0.11 |
| 5 | 160722863 | rs252965 | -0.348 | -1.106 | 1.124 | 0.963 | -0.008 | 0.014 | 0.014 | 0.015 | 0.1 | 0.08 | 0.09 | 0.03 | 0.14 | 0.11 |
| 5 | 160725431 | rs980161 | 0.949 | -1.132 | 1.051 | 0.871 | -0.002 | 0.02 | 0.02 | 0 | 0.07 | 0.03 | 0.05 | 0.03 | 0.08 | 0.01 |
| 5 | 160725513 | rs6556546 | -1.903 | -1.297 | 1.371 | 1.061 | -0.012 | 0.054 | 0.054 | 0 | 0.18 | 0.65 | 0.42 | 0.4 | 0.12 | 0.21 |
| 5 | 160726954 | rs9313879 | -1.506 | -1.308 | 1.831 | 1.099 | -0.045 | 0.204 | 0.204 | 0.242 | 0.14 | 0.42 | 0.2 | 0.29 | 0.05 | 0.2 |
| 5 | 160730392 | rs679633 | -1.294 | -1.222 | 1.769 | 1.04 | 0.046 | 0.001 | 0.001 | 0 | 0.15 | 0.36 | 0.32 | 0.14 | 0.21 | 0.11 |
| 5 | 160730639 | rs252977 | 0.316 | -1.213 | 1.656 | 1.04 | -0.029 | 0.012 | 0.012 | 0.252 | 0.08 | 0.03 | 0.09 | 0.03 | 0.14 | 0.04 |
| 5 | 160732593 | rs173767 | -0.086 | -1.269 | 1.63 | 1.013 | -0.015 | 0.017 | 0.017 | 0.087 | 0.09 | 0.06 | 0.11 | 0.03 | 0.14 | 0.04 |
| 5 | 160733095 | rs10050588 | -0.028 | -1.24 | 1.889 | 1.139 | 0.001 | 0.119 | 0.119 | 0 | 0.07 | 0.19 | 0.16 | 0.18 | 0.05 | 0.1 |
| 5 | 160738317 | rs3850733 | -0.524 | -1.199 | 1.798 | 1.104 | 0.085 | 0.091 | 0.091 | 0 | 0.06 | 0.26 | 0.21 | 0.11 | 0.07 | 0.08 |
| 5 | 160749825 | rs10515826 | -0.518 | -1.231 | 1.458 | 1.119 | -0.006 | 0.117 | 0.117 | 0 | 0.09 | 0.24 | 0.17 | 0.18 | 0.05 | 0.16 |
| 5 | 160754549 | rs1644522 | -2.301 | -1.36 | 1.042 | 0.978 | 0.024 | -0.04 | -0.04 | 0 | 0.36 | 0.76 | 0.55 | 0.43 | 0.31 | 0.37 |
| 5 | 160755835 | rs173766 | -1.838 | -1.203 | 1.214 | 0.848 | -0.001 | -0.002 | -0.002 | 0 | 0.33 | 0.38 | 0.37 | 0.22 | 0.32 | 0.25 |
| 5 | 160757462 | rs10052351 | 0.921 | -1.155 | 1.279 | 0.892 | -0.002 | -0.014 | -0.014 | 0 | 0.07 | 0.09 | 0.06 | 0.03 | 0.09 | 0.01 |
| 5 | 160758655 | rs194072 | -0.606 | -1.095 | 1.202 | 0.907 | 0.002 | 0.024 | 0.024 | 0 | 0.09 | 0.12 | 0.21 | 0.18 | 0.14 | 0.17 |
| 5 | 160759656 | rs1816072 | -2.316 | -1.141 | 1.396 | 0.906 | 0.014 | -0.005 | -0.005 | 0.004 | 0.4 | 0.71 | 0.43 | 0.4 | 0.36 | 0.3 |
| 5 | 160759956 | rs1816071 | -1.751 | -1.063 | 1.352 | 0.894 | 0 | 0 | 0 | 0.109 | 0.33 | 0.39 | 0.39 | 0.29 | 0.36 | 0.26 |
| 5 | 160765206 | rs2617503 | -2.561 | -1.017 | 1.012 | 0.894 | 0.04 | -0.05 | -0.05 | 0 | 0.44 | 0.85 | 0.63 | 0.45 | 0.38 | 0.36 |
| 5 | 160786065 | rs13362062 | -0.576 | -0.928 | 1.462 | 0.818 | 0.007 | 0.054 | 0.054 | 0 | 0.13 | 0.4 | 0.13 | 0.19 | 0.11 | 0.05 |
| 5 | 160795065 | rs2910295 | 1.841 | -0.972 | 1.48 | 0.746 | -0.009 | -0.025 | -0.025 | 0 | 0.68 | 0.43 | 0.71 | 0.77 | 0.73 | 0.8 |
| 5 | 160801723 | rs11949658 | 0.118 | -0.999 | 1.489 | 0.714 | -0.001 | -0.029 | -0.029 | 0 | 0.08 | 0.14 | 0.07 | 0.01 | 0.07 | 0.09 |
| 5 | 160802427 | rs4921390 | -0.497 | -1.003 | 1.471 | 0.724 | 0.004 | 0.051 | 0.051 | 0 | 0.13 | 0.41 | 0.14 | 0.19 | 0.11 | 0.05 |
| 5 | 160812482 | rs2910291 | 1.433 | -1.003 | 1.617 | 0.724 | -0.016 | -0.02 | -0.02 | 0 | 0.76 | 0.61 | 0.78 | 0.78 | 0.79 | 0.89 |
| 5 | 160831867 | rs17521304 | 1.839 | -0.265 | 0.351 | 0.382 | -0.091 | -0.022 | -0.022 | 0 | 0.2 | 0.01 | 0.13 | 0.13 | 0.23 | 0.08 |
| 5 | 160833871 | rs2962393 | -0.49 | -0.168 | 0.691 | 0.467 | -0.02 | 0.03 | 0.03 | 0.008 | 0.9 | 0.99 | 0.92 | 0.97 | 0.9 | 0.94 |
| 5 | 160836620 | rs1946247 | 0.569 | -0.178 | 0.721 | 0.458 | 0.01 | 0.009 | 0.009 | 0 | 0.82 | 0.85 | 0.87 | 0.97 | 0.88 | 0.85 |
| 5 | 160847964 | rs10043074 | -0.496 | -0.033 | 0.646 | 0.55 | 0.023 | 0.026 | 0.026 | 0 | 0.16 | 0.56 | 0.17 | 0.21 | 0.15 | 0.06 |
| 5 | 160859923 | rs7714930 | -1.046 | -0.02 | 0.61 | 0.519 | 0.023 | -0.013 | -0.013 | 0 | 0.2 | 0.53 | 0.21 | 0.21 | 0.2 | 0.07 |
| 5 | 160872062 | rs3923156 | 0.98 | 0.646 | 0.062 | 0.932 | -0.036 | -0.035 | -0.035 | 0 | 0.06 | 0.01 | 0.07 | 0.17 | 0.12 | 0.13 |
| 5 | 160882045 | rs10515829 | -0.53 | -0.037 | 0.263 | 0.818 | 0 | -0.001 | -0.001 | 0 | 0.19 | 0.21 | 0.23 | 0.22 | 0.13 | 0.15 |
| 5 | 160894631 | rs7724146 | 1.469 | -0.055 | -0.474 | 0.556 | 0.025 | 0.012 | 0.012 | 0 | 0.2 | 0.08 | 0.08 | 0.19 | 0.17 | 0.34 |
| 5 | 160898328 | rs10051667 | 0.586 | 0.573 | 0.273 | 0.825 | 0.004 | -0.034 | -0.034 | 0 | 0.1 | 0.19 | 0.09 | 0.04 | 0.11 | 0.02 |
| 5 | 160908195 | rs2962407 | 1.232 | 0.602 | -0.159 | 0.9 | 0.001 | -0.003 | -0.003 | 0 | 0.67 | 0.3 | 0.59 | 0.58 | 0.7 | 0.77 |
| 5 | 160910212 | rs2964773 | 0.367 | 1.02 | -0.304 | 0.635 | -0.004 | 0.005 | 0.005 | 0 | 0.81 | 0.83 | 0.73 | 0.89 | 0.78 | 0.86 |
| 5 | 160911968 | rs6881515 | -0.458 | 1.107 | -0.513 | 0.577 | -0.02 | 0.026 | 0.026 | 0 | 0.21 | 0.09 | 0.24 | 0.11 | 0.25 | 0.16 |
| 5 | 160917953 | rs10069900 | -1.504 | 1.132 | 0.364 | 0.801 | 0.011 | -0.002 | -0.002 | 0.041 | 0.29 | 0.64 | 0.51 | 0.29 | 0.3 | 0.23 |
| 5 | 160952734 | rs17522737 | -0.929 | 0.949 | 0.057 | 1.117 | -0.029 | -0.015 | -0.015 | 3.262 | 0.24 | 0.15 | 0.46 | 0.29 | 0.29 | 0.23 |
| 5 | 160953031 | rs4426954 | 1.706 | 0.964 | -0.2 | 1.159 | -0.025 | -0.028 | -0.028 | 0.38 | 0.58 | 0.18 | 0.34 | 0.55 | 0.55 | 0.65 |
| 5 | 160975332 | rs3816596 | -1.265 | 1.489 | 0.482 | 1.145 | 0.02 | 0.023 | 0.023 | 0 | 0.33 | 0.48 | 0.55 | 0.38 | 0.38 | 0.4 |
| 5 | 160981706 | rs9313889 | -0.916 | 1.558 | 0.296 | 1.185 | -0.001 | 0 | 0 | 0 | 0.3 | 0.32 | 0.34 | 0.23 | 0.35 | 0.38 |
| 5 | 160984243 | rs1002067 | -1.369 | 1.636 | 0.238 | 1.351 | 0 | 0 | 0 | 0 | 0.34 | 0.38 | 0.58 | 0.43 | 0.42 | 0.43 |
| 5 | 161001591 | rs2009913 | 1.169 | 0.921 | -0.386 | 1.096 | -0.001 | -0.002 | -0.002 | 0 | 0.36 | 0.32 | 0.22 | 0.24 | 0.36 | 0.27 |
| 5 | 161008001 | rs4623189 | 0.624 | 0.602 | -0.682 | 1.037 | 0.019 | -0.025 | -0.025 | 0 | 0.28 | 0.18 | 0.19 | 0.51 | 0.19 | 0.4 |
| 5 | 161012011 | rs1899928 | -0.114 | 0.654 | -0.735 | 1.024 | 0.023 | -0.015 | -0.015 | 0 | 0.35 | 0.21 | 0.45 | 0.68 | 0.3 | 0.44 |
| 5 | 161018271 | rs11956866 | -0.914 | 0.538 | -0.633 | 0.943 | 0.002 | -0.008 | -0.008 | 0 | 0.54 | 0.26 | 0.48 | 0.59 | 0.53 | 0.65 |
| 5 | 161020788 | rs9313891 | 0.123 | 0.629 | -0.645 | 0.896 | -0.07 | 0.075 | 0.075 | 0 | 0.25 | 0.11 | 0.29 | 0.08 | 0.34 | 0.24 |
| 5 | 161025848 | rs12519172 | 0.869 | 0.226 | -0.489 | 1.111 | -0.013 | -0.025 | -0.025 | 0.082 | 0.06 | 0.03 | 0.26 | 0.17 | 0.1 | 0.04 |
| 5 | 161028766 | rs17059579 | 0.539 | 0.081 | -0.505 | 1.205 | 0.031 | -0.017 | -0.017 | 2.096 | 0.28 | 0.14 | 0.18 | 0.38 | 0.18 | 0.41 |
| 5 | 161031824 | rs6888242 | 0.605 | 0.128 | -0.329 | 1.055 | -0.006 | -0.002 | -0.002 | 0 | 0.57 | 0.46 | 0.44 | 0.32 | 0.64 | 0.47 |
| 5 | 161052120 | rs7706918 | 1.738 | 0.508 | -0.454 | 0.802 | -0.002 | -0.001 | -0.001 | 1.649 | 0.32 | 0.31 | 0.15 | 0.24 | 0.26 | 0.26 |

| | | | | | | | | | | | | | | | | |
|---|---|---|---|---|---|---|---|---|---|---|---|---|---|---|---|---|
| 5 | 161055576 | rs1037786 | 0.88 | 0.033 | -0.583 | 0.993 | 0.028 | -0.027 | -0.027 | 0 | 0.27 | 0.15 | 0.18 | 0.43 | 0.18 | 0.41 |
| 5 | 161056055 | rs2422006 | 1.188 | 0.175 | -0.707 | 0.994 | -0.032 | -0.051 | -0.051 | 0 | 0.1 | 0.03 | 0.33 | 0.25 | 0.17 | 0.07 |
| 5 | 161077036 | rs1822784 | 0.938 | -0.017 | -0.661 | 0.89 | 0.036 | -0.036 | -0.036 | 0 | 0.27 | 0.08 | 0.17 | 0.38 | 0.18 | 0.4 |
| 5 | 161097702 | rs1373599 | 1.012 | 0.639 | -0.504 | 0.774 | -0.006 | -0.021 | -0.021 | 0 | 0.43 | 0.48 | 0.26 | 0.24 | 0.39 | 0.32 |
| 5 | 161103142 | rs6893149 | 0.942 | 0.257 | -0.568 | 0.723 | -0.009 | 0.006 | 0.006 | 0 | 0.26 | 0.27 | 0.18 | 0.38 | 0.17 | 0.48 |
| 5 | 161108140 | rs11948174 | 0.906 | 0.256 | -0.443 | 0.731 | 0.013 | 0.001 | 0.001 | 0 | 0.26 | 0.19 | 0.17 | 0.27 | 0.17 | 0.47 |
| 5 | 161111588 | rs4921195 | -1.086 | 0.39 | -0.396 | 0.894 | 0.016 | 0.104 | 0.104 | 0 | 0.88 | 0.76 | 0.64 | 0.64 | 0.82 | 0.93 |
| 5 | 161112693 | rs3811995 | 0.922 | 0.142 | -0.398 | 0.671 | 0.017 | 0.017 | 0.017 | 0.068 | 0.54 | 0.33 | 0.37 | 0.32 | 0.57 | 0.4 |
| 5 | 161113004 | rs13188991 | 2.846 | -0.656 | -0.972 | 0.545 | -0.096 | 0.039 | 0.039 | 0.706 | 0.22 | 0.01 | 0.11 | 0.07 | 0.27 | 0.08 |
| 5 | 161114522 | rs4454083 | 1.833 | -0.293 | -0.707 | 0.703 | 0 | 0.001 | 0.001 | 0 | 0.34 | 0.31 | 0.17 | 0.24 | 0.3 | 0.24 |
| 5 | 161115761 | rs12518088 | 1.429 | -0.271 | -0.686 | 0.748 | -0.004 | 0.009 | 0.009 | 0 | 0.1 | 0.09 | 0.34 | 0.36 | 0.18 | 0.07 |
| 5 | 161127822 | rs11956731 | 1.072 | -0.287 | -0.696 | 0.786 | 0.003 | -0.001 | -0.001 | 0.042 | 0.24 | 0.2 | 0.17 | 0.27 | 0.17 | 0.46 |
| 5 | 161128914 | rs3219151 | 0.868 | -0.258 | -0.893 | 0.676 | -0.001 | -0.001 | -0.001 | 0 | 0.55 | 0.43 | 0.39 | 0.32 | 0.58 | 0.42 |
| 5 | 161131600 | rs13178296 | 0.068 | -0.237 | -0.85 | 0.727 | -0.013 | 0.004 | 0.004 | 0.964 | 0.63 | 0.47 | 0.46 | 0.32 | 0.65 | 0.47 |
| 5 | 161140141 | rs4921405 | 0.531 | -0.499 | -0.926 | 0.663 | 0.008 | -0.014 | -0.014 | 0 | 0.21 | 0.27 | 0.23 | 0.08 | 0.28 | 0.18 |
| 5 | 161143459 | rs13156569 | -1.346 | -0.587 | -1.291 | 0.719 | -0.001 | 0.004 | 0.004 | 0.069 | 0.57 | 0.62 | 0.74 | 0.76 | 0.62 | 0.74 |
| 5 | 161154713 | rs12658134 | 1.464 | -0.5 | -1.34 | 0.838 | -0.038 | -0.073 | -0.073 | 1.42 | 0.11 | 0.01 | 0.32 | 0.34 | 0.17 | 0.06 |
| 5 | 161172541 | rs4921415 | -1.188 | -0.407 | -1.152 | 0.499 | 0 | 0.015 | 0.015 | 0.071 | 0.57 | 0.65 | 0.74 | 0.75 | 0.61 | 0.7 |
| 5 | 161176573 | rs4921416 | 0.72 | -0.335 | -1.197 | 0.416 | -0.101 | 0.023 | 0.023 | 0 | 0.19 | 0.02 | 0.2 | 0.08 | 0.28 | 0.18 |
| 5 | 161177205 | rs10043528 | -1.556 | -0.382 | -1 | 0.517 | -0.001 | -0.004 | -0.004 | 0.125 | 0.65 | 0.68 | 0.81 | 0.78 | 0.68 | 0.75 |
| 5 | 161179799 | rs10463102 | -0.567 | -1.143 | -0.654 | 0.921 | -0.023 | -0.006 | -0.006 | 0.192 | 0.37 | 0.5 | 0.53 | 0.66 | 0.34 | 0.53 |
| 5 | 161180720 | rs4552666 | 0.163 | -1.669 | -0.865 | 1.014 | 0.024 | -0.039 | -0.039 | 0 | 0.26 | 0.12 | 0.19 | 0.31 | 0.2 | 0.35 |
| 5 | 161189729 | rs10035808 | -1.622 | -1.461 | -0.201 | 0.902 | 0.011 | 0.025 | 0.025 | 0 | 0.51 | 0.61 | 0.61 | 0.71 | 0.48 | 0.6 |
| 5 | 161198811 | rs12653458 | 0.444 | -1.504 | -0.047 | 0.674 | 0.026 | -0.023 | -0.023 | 0 | 0.15 | 0.08 | 0.13 | 0.15 | 0.1 | 0.28 |
| 5 | 161203641 | rs11746679 | 0.322 | -1.56 | 0.34 | 0.777 | -0.007 | -0.004 | -0.004 | 0 | 0.06 | 0.13 | 0.04 | 0.03 | 0.07 | 0.11 |
| 5 | 161207029 | rs13164907 | -0.169 | -1.538 | 0.132 | 0.7 | -0.002 | -0.017 | -0.017 | 0 | 0.14 | 0.09 | 0.07 | 0 | 0.13 | 0.05 |
| 5 | 161212950 | rs12658074 | -1.626 | -1.48 | -0.027 | 0.767 | -0.013 | -0.013 | -0.013 | 4.227 | 0.38 | 0.59 | 0.27 | 0.27 | 0.29 | 0.45 |
| 5 | 161221088 | rs9784608 | -0.851 | -1.444 | -0.119 | 0.767 | -0.001 | -0.074 | -0.074 | 0 | 0.22 | 0.45 | 0.14 | 0.12 | 0.2 | 0.17 |
| 5 | 161228374 | rs4317369 | -0.999 | -1.207 | -0.207 | 0.937 | -0.012 | -0.013 | -0.013 | 0 | 0.38 | 0.57 | 0.27 | 0.27 | 0.3 | 0.45 |
| 5 | 161247404 | rs4921420 | -1.38 | -1.213 | -0.403 | 0.668 | 0.022 | 0.045 | 0.045 | 0 | 0.49 | 0.37 | 0.39 | 0.73 | 0.45 | 0.66 |
| 5 | 161248654 | rs4302629 | 0.562 | -0.987 | -0.6 | 0.504 | -0.025 | 0.046 | 0.046 | 0 | 0.08 | 0.01 | 0.03 | 0.01 | 0.09 | 0.06 |
| 5 | 161259369 | rs4289596 | -0.801 | -1.463 | -0.25 | 0.735 | -0.006 | 0.018 | 0.018 | 0 | 0.27 | 0.34 | 0.24 | 0.55 | 0.3 | 0.42 |
| 5 | 161269895 | rs4596421 | -0.786 | -1.438 | -0.253 | 0.734 | -0.005 | 0.019 | 0.019 | 0 | 0.27 | 0.34 | 0.24 | 0.55 | 0.3 | 0.42 |
| 5 | 161274224 | rs11576001 | -1.216 | -1.609 | -0.001 | 1.19 | -0.051 | 0.04 | 0.04 | 0 | 0.31 | 0.5 | 0.25 | 0.37 | 0.2 | 0.41 |
| 5 | 161280643 | rs4340950 | -0.203 | -1.759 | -0.067 | 1.218 | -0.01 | 0.036 | 0.036 | 0.046 | 0.14 | 0.22 | 0.09 | 0.26 | 0.11 | 0.22 |
| 5 | 161284782 | rs4367330 | -2.243 | -1.872 | -0.009 | 1.121 | 0.027 | 0.062 | 0.062 | 0 | 0.51 | 0.65 | 0.43 | 0.74 | 0.46 | 0.62 |
| 5 | 161284829 | rs4263535 | -1.181 | -1.866 | 0.058 | 1.084 | -0.06 | 0.055 | 0.055 | 0.22 | 0.29 | 0.53 | 0.2 | 0.44 | 0.17 | 0.35 |
| 5 | 161302211 | rs4428455 | 2.487 | -1.692 | -0.478 | 0.578 | -0.068 | 0.088 | 0.088 | 0 | 0.33 | 0.08 | 0.18 | 0.16 | 0.37 | 0.18 |
| 5 | 161305071 | rs6878494 | -2.032 | -1.876 | -0.08 | 1.019 | 0.028 | 0.042 | 0.042 | 0 | 0.51 | 0.64 | 0.43 | 0.61 | 0.45 | 0.63 |
| 5 | 161310002 | rs12658807 | 0.439 | -1.852 | -0.573 | 1.006 | 0.017 | 0.008 | 0.008 | 0 | 0.12 | 0.09 | 0.1 | 0.08 | 0.06 | 0.12 |
| 5 | 161313159 | rs4554269 | -1.294 | -1.84 | -0.582 | 1.027 | -0.002 | 0.006 | 0.006 | 0 | 0.35 | 0.39 | 0.29 | 0.5 | 0.36 | 0.43 |
| 5 | 161315812 | rs17052610 | 0.444 | -1.831 | -0.666 | 1.016 | 0.039 | 0.002 | 0.002 | 0 | 0.12 | 0.03 | 0.09 | 0.08 | 0.06 | 0.12 |
| 5 | 161325934 | rs998754 | 1.228 | -1.686 | -0.546 | 1.092 | -0.003 | 0.003 | 0.003 | 0 | 0.6 | 0.58 | 0.68 | 0.5 | 0.61 | 0.57 |
| 5 | 161349821 | rs7730737 | 0.278 | -1.582 | -0.737 | 1.037 | -0.036 | -0.054 | -0.054 | 1.362 | 0.75 | 0.97 | 0.79 | 0.76 | 0.72 | 0.78 |
| 5 | 161354185 | rs1902795 | 0.278 | -1.582 | -0.735 | 1.037 | -0.032 | -0.05 | -0.05 | 0 | 0.75 | 0.96 | 0.79 | 0.76 | 0.72 | 0.78 |
| 5 | 161390744 | rs745850 | 0.594 | -1.492 | -0.715 | 0.949 | 0.005 | 0.048 | 0.048 | 0 | 0.16 | 0.07 | 0.14 | 0.03 | 0.17 | 0.16 |
| 5 | 161391350 | rs17059916 | -0.482 | -1.281 | -0.652 | 1.064 | -0.013 | 0.049 | 0.049 | 0.14 | 0.15 | 0.25 | 0.1 | 0.26 | 0.1 | 0.2 |
| 5 | 161400376 | rs11745164 | 1.404 | -1.122 | -0.682 | 1.082 | 0 | 0 | 0 | 0 | 0.51 | 0.5 | 0.58 | 0.42 | 0.55 | 0.46 |
| 5 | 161429682 | rs1895272 | 0.177 | -0.204 | -0.666 | 1.552 | -0.03 | -0.056 | -0.056 | 1.113 | 0.89 | 0.98 | 0.87 | 0.75 | 0.82 | 0.88 |
| 5 | 161434200 | rs6891725 | -0.752 | -0.509 | -0.193 | 1.203 | -0.007 | 0.088 | 0.088 | 0 | 0.1 | 0.24 | 0.07 | 0.22 | 0.08 | 0.11 |
| 5 | 161446231 | rs1477310 | -0.904 | -1.074 | -0.917 | 0.873 | 0.079 | -0.013 | -0.013 | 0 | 0.39 | 0.82 | 0.29 | 0.35 | 0.28 | 0.36 |
| 5 | 161447161 | rs7713655 | -0.561 | -1.239 | -0.133 | 0.849 | -0.049 | 0.05 | 0.05 | 0.087 | 0.14 | 0.42 | 0.11 | 0.15 | 0.06 | 0.2 |
| 5 | 161453085 | rs2422102 | 0.354 | -1.106 | -0.939 | 0.685 | 0.003 | -0.008 | -0.008 | 0 | 0.12 | 0.09 | 0.11 | 0.14 | 0.11 | 0.22 |
| 5 | 161454458 | rs6887435 | -0.345 | -0.767 | -0.853 | 1.101 | -0.033 | -0.046 | -0.046 | 0 | 0.92 | 1 | 0.88 | 0.69 | 0.88 | 0.88 |
| 5 | 161464255 | rs11956128 | -0.27 | -0.678 | -0.445 | 1.079 | -0.021 | -0.002 | -0.002 | 0 | 0.23 | 0.34 | 0.17 | 0.2 | 0.22 | 0.16 |
| 5 | 161466651 | rs17052622 | 0.736 | -0.618 | -1.547 | 1.093 | 0.013 | -0.01 | -0.01 | 0.013 | 0.17 | 0.05 | 0.12 | 0.18 | 0.17 | 0.1 |
| 5 | 161471690 | rs209333 | 0.251 | -0.322 | -1.666 | 0.787 | 0.03 | -0.026 | -0.026 | 0.003 | 0.61 | 0.27 | 0.67 | 0.49 | 0.66 | 0.71 |
| 5 | 161473771 | rs2472015 | -0.623 | 0.554 | -1.589 | 0.386 | -0.036 | 0.055 | 0.055 | 0 | 0.14 | 0.05 | 0.33 | 0.04 | 0.18 | 0.18 |
| 5 | 161484328 | rs4140810 | 0.274 | 0.113 | -1.506 | 0.961 | 0.001 | -0.053 | -0.053 | 0.167 | 0.09 | 0 | 0.01 | 0.21 | 0.05 | 0.11 |
| 5 | 161487377 | rs209341 | 0.178 | -0.39 | -1.255 | 0.788 | -0.033 | 0.005 | 0.005 | 1.096 | 0.33 | 0.54 | 0.43 | 0.36 | 0.27 | 0.37 |
| 5 | 161494852 | rs3219203 | 1.518 | -0.206 | -1.439 | 0.801 | 0.001 | 0.019 | 0.019 | 0 | 0.07 | 0.02 | 0.04 | 0.02 | 0.08 | 0.03 |
| 5 | 161495852 | rs2268583 | 0.89 | 0.215 | -1.49 | 0.925 | -0.012 | 0.012 | 0.012 | 0 | 0.05 | 0.02 | 0.24 | 0.22 | 0.07 | 0.21 |
| 5 | 161498737 | rs183294 | -0.545 | -0.332 | -1.892 | 0.392 | -0.002 | -0.001 | -0.001 | 0 | 0.55 | 0.56 | 0.64 | 0.61 | 0.6 | 0.65 |
| 5 | 161510796 | rs209350 | -0.791 | -0.36 | -1.556 | 0.513 | 0.002 | -0.004 | -0.004 | 0.04 | 0.73 | 0.76 | 0.82 | 0.65 | 0.79 | 0.78 |
| 5 | 161515215 | rs2268582 | 1.3 | -0.531 | -1.588 | 0.363 | 0.022 | 0.051 | 0.051 | 4.301 | 0.2 | 0.07 | 0.06 | 0.07 | 0.14 | 0.16 |
| 5 | 161517059 | rs209353 | -0.741 | -0.514 | -1.517 | 0.348 | 0.003 | 0.004 | 0.004 | 0.363 | 0.54 | 0.59 | 0.43 | 0.4 | 0.54 | 0.49 |
| 5 | 161517570 | rs209354 | 0.436 | 0.182 | -0.863 | 0.884 | 0.039 | -0.01 | -0.01 | 1.399 | 0.33 | 0.46 | 0.36 | 0.33 | 0.4 | 0.34 |
| 5 | 161522556 | rs11135176 | 0.927 | -0.03 | -1.269 | 1.066 | -0.025 | -0.011 | -0.011 | 0 | 0.05 | 0 | 0.23 | 0.22 | 0.07 | 0.22 |
| 5 | 161522955 | rs211017 | 1.345 | 0.482 | -1.365 | 0.608 | -0.066 | 0.009 | 0.009 | 0 | 0.15 | 0.01 | 0.1 | 0.09 | 0.2 | 0.13 |

| | | | | | | | | | | | | | | | | |
|---|---|---|---|---|---|---|---|---|---|---|---|---|---|---|---|---|
| 5 | 161528280 | rs211037 | 0.087 | 0.331 | -0.713 | 0.76 | -0.018 | 0.086 | 0.086 | 0 | 0.26 | 0.47 | 0.25 | 0.6 | 0.24 | 0.22 |
| 5 | 161530735 | rs211030 | -0.192 | 0.179 | -0.948 | 0.553 | 0.004 | 0.005 | 0.005 | 0 | 0.67 | 0.7 | 0.78 | 0.67 | 0.71 | 0.78 |
| 5 | 161536928 | rs210985 | -1.197 | 0.106 | -0.781 | 0.574 | -0.002 | -0.001 | -0.001 | 0 | 0.79 | 0.79 | 0.86 | 0.72 | 0.8 | 0.86 |
| 5 | 161537130 | rs210984 | -1.187 | 0.114 | -0.779 | 0.576 | -0.002 | -0.001 | -0.001 | 0 | 0.79 | 0.79 | 0.86 | 0.72 | 0.8 | 0.86 |
| 5 | 161541735 | rs169792 | 0.422 | 0.554 | -0.98 | 0.545 | -0.003 | -0.004 | -0.004 | 0 | 0.6 | 0.71 | 0.54 | 0.4 | 0.61 | 0.48 |
| 5 | 161542609 | rs17060096 | 0.932 | 0.79 | -0.993 | 0.127 | -0.004 | 0.008 | 0.008 | 0 | 0.11 | 0.09 | 0.08 | 0.05 | 0.1 | 0.08 |
| 5 | 161544331 | rs365054 | -0.574 | 0.43 | -0.479 | 0.003 | -0.003 | -0.044 | -0.044 | 0 | 0.37 | 0.44 | 0.5 | 0.85 | 0.35 | 0.55 |
| 5 | 161561803 | rs2422106 | -0.278 | 0.505 | -0.498 | 0.283 | -0.007 | 0.013 | 0.013 | 0 | 0.64 | 0.49 | 0.72 | 0.49 | 0.68 | 0.69 |
| 5 | 161568643 | rs13173675 | 1.884 | 0.839 | -1.011 | 0.089 | -0.035 | 0.041 | 0.041 | 0 | 0.08 | 0.01 | 0.07 | 0 | 0.13 | 0.08 |
| 5 | 161576418 | rs211014 | 0.415 | 0.698 | -0.523 | 0.316 | 0 | 0.082 | 0.082 | 0 | 0.27 | 0.43 | 0.21 | 0.52 | 0.26 | 0.28 |
| 5 | 161587753 | rs17462590 | 1.205 | 0.911 | -0.956 | 0.242 | -0.021 | 0.023 | 0.023 | 0 | 0.11 | 0.05 | 0.08 | 0.01 | 0.12 | 0.13 |
| 5 | 161600375 | rs980982 | 1.871 | 1.154 | -1.044 | 0.537 | -0.034 | -0.032 | -0.032 | 0 | 0.06 | 0.01 | 0.33 | 0.15 | 0.1 | 0.24 |
| 5 | 161624736 | rs399588 | -0.504 | -0.309 | -0.697 | -0.055 | 0.002 | 0.004 | 0.004 | 0.087 | 0.48 | 0.58 | 0.35 | 0.52 | 0.44 | 0.44 |
| 5 | 161635453 | rs446878 | -0.314 | 0.591 | -0.162 | 0.708 | -0.007 | 0.037 | 0.037 | 0 | 0.3 | 0.37 | 0.2 | 0.51 | 0.28 | 0.3 |
| 5 | 161644983 | rs141748 | -1.198 | 1.083 | 0.26 | 0.386 | 0 | 0.002 | 0.002 | 0 | 0.46 | 0.46 | 0.33 | 0.61 | 0.42 | 0.47 |
| 5 | 161645103 | rs10491327 | -0.097 | 1.065 | 0.237 | 0.369 | -0.001 | 0.074 | 0.074 | 0 | 0.22 | 0.08 | 0.09 | 0.05 | 0.15 | 0.18 |
| 5 | 161646240 | rs211001 | -0.281 | 0.8 | 0.219 | 0.465 | -0.006 | -0.023 | -0.023 | 0 | 0.22 | 0.22 | 0.22 | 0.46 | 0.27 | 0.28 |
| 5 | 161648147 | rs17462769 | 0.116 | 0.898 | -0.122 | 0.527 | -0.044 | -0.062 | -0.062 | 0 | 0.1 | 0.02 | 0.16 | 0.23 | 0.17 | 0.15 |
| 5 | 161653612 | rs6879662 | -1.21 | 0.677 | 0.123 | 0.239 | 0.006 | 0.012 | 0.012 | 0 | 0.47 | 0.62 | 0.35 | 0.61 | 0.41 | 0.47 |
| 5 | 161661324 | rs6890529 | -0.285 | 1.24 | 0.076 | 0.833 | -0.016 | -0.039 | -0.039 | 0.008 | 0.24 | 0.19 | 0.23 | 0.39 | 0.31 | 0.28 |
| 5 | 161661341 | rs6890533 | -0.431 | 1.225 | 0.104 | 0.694 | 0.008 | 0.039 | 0.039 | 0 | 0.27 | 0.37 | 0.26 | 0.51 | 0.32 | 0.29 |
| 5 | 161662347 | rs7716847 | -0.066 | 1.386 | 0.405 | 0.97 | 0.009 | 0.253 | 0.253 | 0 | 0.13 | 0.36 | 0.18 | 0.38 | 0.13 | 0.19 |
| 5 | 161666833 | rs7721992 | -1.248 | 1.642 | 0.013 | 0.502 | 0.001 | 0.008 | 0.008 | 0.682 | 0.58 | 0.64 | 0.47 | 0.62 | 0.6 | 0.46 |
| 5 | 161670428 | rs6556584 | 0.367 | 1.288 | 0.202 | 0.053 | -0.004 | -0.052 | -0.052 | 3.652 | 0.64 | 0.52 | 0.79 | 0.83 | 0.69 | 0.74 |
| 5 | 161671879 | rs7720600 | -0.36 | 1.297 | 0.233 | 0.003 | -0.004 | -0.021 | -0.021 | 0.325 | 0.75 | 0.65 | 0.88 | 0.82 | 0.78 | 0.84 |
| 5 | 161677267 | rs11135183 | 1.331 | 1.415 | 0.123 | 0.053 | -0.01 | 0.012 | 0.012 | 0.007 | 0.47 | 0.27 | 0.57 | 0.57 | 0.5 | 0.54 |
| 5 | 161681172 | rs12108728 | 0.044 | 1.005 | -0.511 | 0.153 | 0 | -0.029 | -0.029 | 0 | 0.11 | 0.13 | 0.08 | 0.01 | 0.09 | 0.11 |
| 5 | 161684398 | rs6879659 | -0.331 | 0.928 | -0.655 | 0.489 | 0 | 0.008 | 0.008 | 0 | 0.16 | 0.17 | 0.22 | 0.23 | 0.19 | 0.19 |
| 5 | 161692818 | rs5022631 | 0.224 | 0.945 | -0.68 | -0.316 | 0.003 | -0.06 | -0.06 | 0 | 0.67 | 0.6 | 0.81 | 0.87 | 0.69 | 0.76 |
| 5 | 161697175 | rs17060196 | -0.767 | 0.074 | -1.335 | -0.288 | 0.046 | 0.02 | 0.02 | 0.103 | 0.19 | 0.05 | 0.07 | 0.12 | 0.07 | 0.11 |
| 5 | 161698334 | rs4146151 | 1.058 | 0.278 | -0.747 | -0.351 | -0.002 | -0.002 | -0.002 | 0 | 0.73 | 0.76 | 0.85 | 0.81 | 0.82 | 0.84 |
| 5 | 161702316 | rs2963162 | -1.367 | 0.483 | 0.054 | -0.552 | -0.013 | -0.032 | -0.032 | 0 | 0.3 | 0.42 | 0.18 | 0.18 | 0.21 | 0.16 |
| 5 | 161708708 | rs2963185 | -1.395 | 0.322 | -0.325 | -0.569 | 0.005 | 0.012 | 0.012 | 0 | 0.28 | 0.22 | 0.17 | 0.18 | 0.22 | 0.26 |
| 5 | 161709827 | rs17060201 | 0.27 | 0.752 | -1.262 | -0.579 | -0.042 | -0.102 | -0.102 | 0.565 | 0.22 | 0.05 | 0.21 | 0.59 | 0.27 | 0.46 |
| 5 | 161710676 | rs2963172 | -0.167 | 1.073 | -1.118 | -1.153 | -0.044 | 0.057 | 0.057 | 0 | 0.1 | 0.01 | 0.06 | 0 | 0.15 | 0.1 |
| 5 | 161728497 | rs12516970 | 0.116 | 0.703 | -1.369 | -1.382 | 0.021 | 0.035 | 0.035 | 0.033 | 0.21 | 0.04 | 0.17 | 0.13 | 0.13 | 0.11 |
| 5 | 161730087 | rs4360077 | 0.16 | 0.644 | -1.079 | -1.334 | -0.023 | -0.015 | -0.015 | 0 | 0.68 | 0.6 | 0.76 | 0.81 | 0.81 | 0.81 |
| 5 | 161735710 | rs4634392 | 1.331 | 0.765 | -1.172 | -1.213 | -0.09 | 0.19 | 0.19 | 0 | 0.35 | 0.02 | 0.39 | 0.05 | 0.41 | 0.27 |
| 5 | 161738753 | rs4601080 | 0.642 | 0.412 | -0.614 | -0.985 | 0.001 | 0.01 | 0.01 | 0 | 0.58 | 0.59 | 0.67 | 0.8 | 0.62 | 0.71 |
| 5 | 161747304 | rs4376311 | -0.64 | 0.326 | -0.161 | -1.347 | -0.007 | -0.012 | -0.012 | 0 | 0.61 | 0.33 | 0.52 | 0.98 | 0.54 | 0.76 |
| 5 | 161749087 | rs13154478 | -0.394 | -0.157 | -0.301 | -1.259 | 0.007 | 0.052 | 0.052 | 0 | 0.19 | 0.05 | 0.19 | 0.08 | 0.16 | 0.09 |
| 5 | 161766576 | rs13155424 | -1.048 | -0.26 | 0.741 | -1.506 | -0.009 | 0.007 | 0.007 | 0 | 0.55 | 0.63 | 0.49 | 0.97 | 0.46 | 0.69 |
| 5 | 161767993 | rs7708929 | -0.414 | -0.373 | 1.403 | -1.137 | -0.013 | -0.008 | -0.008 | 0 | 0.21 | 0.31 | 0.28 | 0.25 | 0.17 | 0.11 |
| 5 | 161778116 | rs10462933 | -0.344 | -0.357 | 1.425 | -1.136 | 0.001 | 0.035 | 0.035 | 0 | 0.66 | 0.76 | 0.8 | 0.28 | 0.71 | 0.43 |
| 5 | 161790414 | rs6871616 | -0.56 | -0.433 | 1.393 | -1.023 | -0.008 | 0.018 | 0.018 | 0 | 0.69 | 0.82 | 0.81 | 0.32 | 0.72 | 0.45 |
| 5 | 161799865 | rs13162582 | -0.405 | -0.324 | 1.591 | -0.991 | 0.005 | 0.007 | 0.007 | 0 | 0.21 | 0.18 | 0.21 | 0.08 | 0.16 | 0.1 |
| 5 | 161814203 | rs10463107 | 0.471 | -0.405 | 1.753 | -1.007 | -0.011 | 0.018 | 0.018 | 0.106 | 0.31 | 0.15 | 0.19 | 0.67 | 0.29 | 0.55 |
| 5 | 161817847 | rs11949899 | -1.827 | -0.365 | 2.574 | 0.153 | -0.043 | 0.001 | 0.001 | 0 | 0.25 | 0.48 | 0.3 | 0.3 | 0.17 | 0.16 |
| 5 | 161818529 | rs10515838 | -1.058 | -0.162 | 1.84 | -0.167 | 0.004 | 0.01 | 0.01 | 0 | 0.19 | 0.14 | 0.2 | 0.08 | 0.16 | 0.1 |
| 5 | 161837675 | rs10069666 | -1.698 | -0.342 | 2.458 | -0.262 | -0.03 | -0.096 | -0.096 | 0 | 0.27 | 0.42 | 0.27 | 0.08 | 0.19 | 0.11 |
| 5 | 161838225 | rs4560590 | -1.341 | -0.129 | 2.385 | 0.12 | 0.003 | 0.005 | 0.005 | 0 | 0.22 | 0.16 | 0.27 | 0.26 | 0.18 | 0.12 |
| 5 | 161838281 | rs6885337 | -1.888 | -0.15 | 2.71 | 0.111 | -0.015 | -0.018 | -0.018 | 0.079 | 0.25 | 0.37 | 0.3 | 0.25 | 0.19 | 0.12 |
| 5 | 161839524 | rs10515839 | -1.207 | -0.138 | 2.352 | 0.129 | -0.01 | 0.078 | 0.078 | 0 | 0.21 | 0.04 | 0.22 | 0.09 | 0.17 | 0.1 |
| 5 | 161843588 | rs4244436 | 1.929 | -0.205 | 2.855 | 0.51 | -0.033 | 0.021 | 0.021 | 0 | 0.72 | 0.45 | 0.67 | 0.62 | 0.8 | 0.77 |
| 5 | 161848766 | rs10046030 | -1.492 | -0.005 | 2.185 | 0.476 | 0.004 | 0 | 0 | 0 | 0.22 | 0.19 | 0.3 | 0.28 | 0.19 | 0.15 |
| 5 | 161855861 | rs4632841 | 1.603 | 0.399 | 2.389 | 0.174 | 0.063 | 0.113 | 0.113 | 0 | 0.93 | 0.72 | 0.93 | 0.89 | 0.91 | 0.81 |
| 5 | 161863484 | rs10060909 | -1.523 | 0.377 | 1.912 | -0.166 | -0.016 | -0.05 | -0.05 | 0 | 0.22 | 0.4 | 0.24 | 0.11 | 0.16 | 0.11 |
| 5 | 161865725 | rs6874049 | -0.73 | 0.357 | 1.886 | -0.194 | -0.019 | -0.003 | -0.003 | 0 | 0.12 | 0.02 | 0.17 | 0.11 | 0.12 | 0.07 |
| 5 | 161866702 | rs4521525 | -1.188 | 0.451 | 1.872 | -0.125 | -0.015 | -0.05 | -0.05 | 0 | 0.21 | 0.4 | 0.24 | 0.12 | 0.16 | 0.11 |
| 5 | 161868013 | rs6556614 | 1.577 | 0.455 | 1.12 | -0.086 | -0.005 | -0.03 | -0.03 | 1.265 | 0.72 | 0.34 | 0.68 | 0.86 | 0.75 | 0.71 |
| 5 | 161875282 | rs6881035 | -1.215 | 0.468 | 0.45 | 0.016 | -0.004 | -0.02 | -0.02 | 2.246 | 0.21 | 0.3 | 0.23 | 0.1 | 0.16 | 0.11 |
| 5 | 161879758 | rs10866733 | 1.277 | 1.553 | 0.88 | -0.008 | 0.082 | -0.056 | -0.056 | 0.185 | 0.8 | 0.64 | 0.79 | 0.93 | 0.77 | 0.87 |
| 5 | 161880996 | rs9313920 | 0.944 | 1.242 | 0.513 | 0.084 | 0.011 | 0.024 | 0.024 | 0 | 0.51 | 0.4 | 0.53 | 0.34 | 0.57 | 0.32 |
| 5 | 161885547 | rs11135211 | 0.843 | 1.057 | 0.208 | -0.173 | -0.02 | -0.091 | -0.091 | 0 | 0.76 | 0.45 | 0.82 | 0.95 | 0.84 | 0.89 |
| 5 | 161893436 | rs4485963 | 0.104 | 0.973 | 0.973 | -0.235 | 0.004 | -0.038 | -0.038 | 0 | 0.86 | 0.84 | 0.89 | 0.95 | 0.87 | 0.91 |
| 5 | 161897136 | rs11135214 | 0.199 | 0.981 | 0.611 | 0.214 | -0.052 | -0.029 | -0.029 | 0 | 0.11 | 0.01 | 0.08 | 0.19 | 0.17 | 0.1 |
| 5 | 161897227 | rs12653193 | 2.229 | 0.978 | 0.806 | 0.106 | -0.025 | 0.132 | 0.132 | 0 | 0.57 | 0.17 | 0.55 | 0.7 | 0.57 | 0.6 |
| 5 | 161906784 | rs11954409 | 2.171 | 1.028 | 0.729 | 0.347 | -0.002 | 0.03 | 0.03 | 0 | 0.54 | 0.4 | 0.51 | 0.27 | 0.56 | 0.34 |
| 5 | 161912216 | rs10042997 | -0.917 | 1.002 | 0.835 | 0.358 | -0.032 | -0.072 | -0.072 | 0.008 | 0.19 | 0.06 | 0.29 | 0.63 | 0.24 | 0.55 |

| | | | | | | | | | | | | | | | | |
|---|---|---|---|---|---|---|---|---|---|---|---|---|---|---|---|---|
| 5 | 161917408 | rs2901679 | 1.287 | 0.847 | 1.321 | 0.31 | -0.02 | -0.136 | -0.136 | 0.181 | 0.73 | 0.56 | 0.8 | 0.95 | 0.8 | 0.89 |
| 5 | 161926659 | rs1484360 | 1.74 | 0.983 | 1.877 | 0.551 | -0.009 | 0.161 | 0.161 | 0 | 0.82 | 0.42 | 0.72 | 0.62 | 0.87 | 0.65 |
| 5 | 161934450 | rs1947990 | 2.079 | 0.728 | 1.762 | 0.184 | -0.004 | 0.015 | 0.015 | 0 | 0.65 | 0.44 | 0.61 | 0.52 | 0.71 | 0.56 |
| 5 | 161938435 | rs13175982 | 1.771 | 0.774 | 1.914 | 0.298 | -0.011 | 0.175 | 0.175 | 0 | 0.82 | 0.42 | 0.72 | 0.56 | 0.87 | 0.65 |
| 5 | 161944905 | rs12187749 | 1.782 | 0.774 | 1.914 | 0.298 | -0.011 | 0.172 | 0.172 | 0 | 0.82 | 0.41 | 0.72 | 0.57 | 0.87 | 0.65 |
| 5 | 161955579 | rs7356698 | 1.603 | 0.795 | 2.117 | 0.305 | 0 | 0.241 | 0.241 | 0 | 0.83 | 0.54 | 0.72 | 0.57 | 0.87 | 0.65 |
| 5 | 161968042 | rs1843120 | 0.358 | 0.745 | 2.083 | 0.276 | 0.027 | 0.005 | 0.005 | 0.659 | 0.92 | 0.51 | 0.93 | 1 | 0.98 | 0.98 |
| 5 | 161973937 | rs17060479 | 0.954 | 0.859 | 1.608 | 0.291 | 0 | 0.044 | 0.044 | 0 | 0.05 | 0.07 | 0.09 | 0.2 | 0.11 | 0.11 |
| 5 | 161994889 | rs12651722 | -1.549 | 0.731 | 1.741 | 0.013 | -0.002 | -0.003 | -0.003 | 0 | 0.25 | 0.26 | 0.34 | 0.49 | 0.28 | 0.38 |
| 5 | 161996134 | rs17561944 | -0.572 | 0.745 | 1.71 | 0.013 | 0.001 | 0.034 | 0.034 | 0 | 0.17 | 0.08 | 0.12 | 0.06 | 0.15 | 0.05 |
| 5 | 161997490 | rs10063408 | -1.678 | 0.732 | 1.758 | 0.016 | -0.015 | -0.052 | -0.052 | 5.911 | 0.26 | 0.13 | 0.33 | 0.49 | 0.28 | 0.39 |
| 5 | 161997779 | rs12655394 | -0.393 | 0.683 | 1.746 | -0.02 | -0.013 | 0.058 | 0.058 | 0 | 0.16 | 0.03 | 0.11 | 0.06 | 0.15 | 0.05 |
| 5 | 162016266 | rs973148 | 1.97 | 0.583 | 2.072 | -0.064 | -0.002 | 0.037 | 0.037 | 0 | 0.67 | 0.59 | 0.61 | 0.51 | 0.69 | 0.6 |
| 5 | 162023083 | rs1484358 | 2.142 | 0.552 | 2.225 | -0.08 | -0.001 | 0.001 | 0.001 | 0 | 0.67 | 0.38 | 0.61 | 0.51 | 0.71 | 0.59 |
| 5 | 162023980 | rs2431268 | 1.554 | 0.558 | 2.157 | -0.076 | 0.009 | 0.051 | 0.051 | 0 | 0.73 | 0.65 | 0.63 | 0.51 | 0.7 | 0.61 |
| 5 | 162030548 | rs6862669 | -1.341 | 0.556 | 2.02 | -0.103 | -0.004 | -0.029 | -0.029 | 0 | 0.23 | 0.26 | 0.16 | 0.06 | 0.19 | 0.08 |
| 5 | 162031048 | rs7708539 | -1.175 | 0.558 | 2.021 | -0.102 | -0.006 | -0.032 | -0.032 | 0 | 0.22 | 0.25 | 0.15 | 0.06 | 0.17 | 0.08 |
| 5 | 162031118 | rs12514510 | -1.28 | 0.599 | 1.862 | -0.078 | -0.011 | 0.033 | 0.033 | 0 | 0.09 | 0.08 | 0.22 | 0.42 | 0.13 | 0.33 |
| 5 | 162039311 | rs17060579 | -0.668 | 0.723 | 2.016 | -0.853 | 0 | 0.008 | 0.008 | 0 | 0.17 | 0.15 | 0.13 | 0.06 | 0.17 | 0.06 |
| 5 | 162043850 | rs172676 | 1.713 | 0.701 | 2.219 | -0.855 | -0.009 | -0.077 | -0.077 | 0 | 0.72 | 0.43 | 0.66 | 0.91 | 0.75 | 0.89 |
| 5 | 162043995 | rs178158 | 1.424 | 0.706 | 2.213 | -0.852 | 0.082 | 0.023 | 0.023 | 0.004 | 0.94 | 0.68 | 0.81 | 0.97 | 0.92 | 0.97 |
| 5 | 162044495 | rs17563003 | -0.86 | 0.707 | 1.86 | -0.856 | -0.004 | 0.013 | 0.013 | 0 | 0.18 | 0.11 | 0.13 | 0.06 | 0.18 | 0.06 |
| 5 | 162056535 | rs312326 | 1.807 | 0.449 | 1.669 | -0.847 | -0.005 | -0.044 | -0.044 | 0 | 0.71 | 0.4 | 0.65 | 0.92 | 0.75 | 0.89 |
| 5 | 162060365 | rs312312 | 1.723 | 0.385 | 1.44 | -0.815 | -0.01 | -0.061 | -0.061 | 0 | 0.71 | 0.43 | 0.66 | 0.92 | 0.75 | 0.89 |
| 5 | 162063354 | rs17497621 | -0.517 | 0.633 | 0.431 | -0.684 | -0.009 | 0.052 | 0.052 | 0 | 0.16 | 0.03 | 0.11 | 0.05 | 0.15 | 0.06 |
| 5 | 162083518 | rs7711337 | -1.253 | 0.529 | 0.142 | -1.407 | 0 | -0.066 | -0.066 | 0.104 | 0.4 | 0.57 | 0.42 | 0.02 | 0.41 | 0.21 |
| 5 | 162086807 | rs10515841 | 0.128 | 0.689 | -0.365 | -1.049 | -0.016 | 0.018 | 0.018 | 0 | 0.07 | 0.03 | 0.17 | 0.02 | 0.1 | 0.07 |
| 5 | 162088705 | rs7700528 | -0.582 | 0.63 | -0.187 | -1.076 | 0.002 | -0.041 | -0.041 | 0.752 | 0.12 | 0.18 | 0.21 | 0.02 | 0.12 | 0.09 |
| 5 | 162089602 | rs17564123 | -0.246 | 0.481 | -0.34 | -0.762 | -0.001 | -0.04 | -0.04 | 0.438 | 0.13 | 0.12 | 0.1 | 0 | 0.13 | 0.06 |
| 5 | 162095410 | rs312305 | -0.113 | 0.48 | -0.671 | -0.132 | -0.005 | 0.134 | 0.134 | 0 | 0.79 | 0.56 | 0.81 | 0.57 | 0.82 | 0.8 |
| 5 | 162098997 | rs7442737 | 0.24 | 0.58 | -1.892 | -0.459 | 0.008 | 0.063 | 0.063 | 0.003 | 0.23 | 0.1 | 0.14 | 0 | 0.23 | 0.11 |
| 5 | 162099186 | rs1818818 | 0.24 | 0.58 | -1.892 | -0.459 | 0.008 | 0.063 | 0.063 | 0.003 | 0.23 | 0.1 | 0.14 | 0 | 0.23 | 0.11 |
| 5 | 162100311 | rs699096 | 0.325 | 0.736 | -1.831 | 0.02 | -0.038 | -0.017 | -0.017 | 0.502 | 0.44 | 0.14 | 0.59 | 0.53 | 0.5 | 0.56 |
| 5 | 162107672 | rs245609 | -0.305 | -0.094 | -1.493 | 0.225 | 0.021 | -0.015 | -0.015 | 0 | 0.36 | 0.26 | 0.23 | 0.55 | 0.25 | 0.33 |
| 5 | 162124220 | rs245600 | 0.319 | -0.01 | -2.068 | -0.431 | -0.005 | 0.091 | 0.091 | 0.063 | 0.31 | 0.12 | 0.39 | 0.16 | 0.31 | 0.23 |
| 5 | 162136349 | rs245608 | 0.006 | -0.144 | -1.673 | -0.405 | 0.018 | 0.02 | 0.02 | 0 | 0.43 | 0.32 | 0.27 | 0.62 | 0.36 | 0.53 |
| 5 | 162148034 | rs20464 | 0.033 | -0.233 | -1.451 | -0.495 | 0.008 | 0.007 | 0.007 | 0 | 0.43 | 0.38 | 0.27 | 0.61 | 0.35 | 0.51 |
| 5 | 162153595 | rs6869306 | 1.419 | -0.405 | -0.807 | -0.336 | -0.054 | -0.007 | -0.007 | 1.061 | 0.2 | 0.08 | 0.29 | 0.23 | 0.33 | 0.27 |
| 5 | 162156346 | rs9313939 | 0.739 | -0.413 | -0.628 | -0.344 | 0.073 | -0.021 | -0.021 | 0 | 0.24 | 0.47 | 0.34 | 0.22 | 0.33 | 0.27 |
| 5 | 162157339 | rs2562321 | -0.863 | -0.428 | -0.617 | -0.362 | -0.016 | 0.095 | 0.095 | 0.115 | 0.69 | 0.91 | 0.61 | 0.84 | 0.69 | 0.77 |
| 5 | 162175154 | rs3914991 | -0.421 | -0.365 | -0.799 | -0.503 | 0.001 | -0.021 | -0.021 | 0.84 | 0.54 | 0.5 | 0.55 | 0.74 | 0.49 | 0.64 |
| 5 | 162185009 | rs17060763 | 0.762 | -0.535 | -1.094 | -0.492 | -0.012 | 0.003 | 0.003 | 0.158 | 0.07 | 0 | 0.03 | 0.03 | 0.07 | 0.17 |
| 5 | 162187646 | rs6892150 | 0.377 | -0.525 | -1.083 | -0.495 | -0.012 | -0.004 | -0.004 | 0.003 | 0.09 | 0.05 | 0.27 | 0.12 | 0.13 | 0.13 |
| 5 | 162188705 | rs6879531 | 0.074 | -0.586 | -0.487 | -0.498 | -0.008 | -0.009 | -0.009 | 0.014 | 0.46 | 0.54 | 0.29 | 0.62 | 0.36 | 0.48 |
| 5 | 162193227 | rs17060800 | 0.095 | -1.164 | -0.466 | -0.175 | 0.002 | -0.029 | -0.029 | 0.104 | 0.32 | 0.25 | 0.24 | 0.49 | 0.32 | 0.41 |
| 5 | 162201035 | rs6862855 | 0.308 | -0.695 | -0.362 | 0.215 | -0.008 | -0.001 | -0.001 | 0 | 0.39 | 0.41 | 0.23 | 0.58 | 0.3 | 0.32 |
| 5 | 162211967 | rs6556633 | 0.47 | -0.76 | -0.394 | 0.186 | 0.004 | -0.001 | -0.001 | 0.046 | 0.37 | 0.35 | 0.21 | 0.57 | 0.28 | 0.31 |
| 5 | 162220463 | rs1579443 | 0.148 | -0.77 | -0.448 | 0.141 | -0.006 | -0.006 | -0.006 | 0 | 0.4 | 0.43 | 0.24 | 0.61 | 0.31 | 0.32 |
| 5 | 162221666 | rs543502 | 0.003 | -0.798 | -0.387 | 0.068 | -0.001 | -0.001 | -0.001 | 0 | 0.38 | 0.39 | 0.6 | 0.35 | 0.49 | 0.58 |
| 5 | 162229592 | rs1288208 | -1.329 | -0.685 | -0.449 | -0.015 | -0.023 | 0.12 | 0.12 | 0.01 | 0.77 | 0.96 | 0.83 | 0.95 | 0.76 | 0.89 |
| 5 | 162232358 | rs1288205 | 1.627 | -0.737 | -0.509 | 0.013 | -0.133 | -0.029 | -0.029 | 0.106 | 0.21 | 0.01 | 0.27 | 0.2 | 0.32 | 0.3 |
| 5 | 162233773 | rs7715197 | 0.193 | -0.869 | -0.313 | 0.476 | -0.016 | 0.002 | 0.002 | 0 | 0.39 | 0.53 | 0.25 | 0.41 | 0.3 | 0.28 |
| 5 | 162234478 | rs17060856 | 0.46 | -0.807 | -0.304 | 0.581 | -0.005 | 0.001 | 0.001 | 0.072 | 0.06 | 0.05 | 0.02 | 0.07 | 0.03 | 0.16 |
| 5 | 162245076 | rs17060894 | 1.144 | -0.766 | -0.439 | 0.375 | 0.009 | 0.01 | 0.01 | 0 | 0.19 | 0.14 | 0.11 | 0.05 | 0.12 | 0.1 |
| 5 | 162246174 | rs7731154 | -0.01 | -0.974 | -0.47 | 0.88 | -0.037 | 0.015 | 0.015 | 0 | 0.61 | 0.77 | 0.38 | 0.34 | 0.51 | 0.38 |
| 5 | 162258547 | rs1372381 | 0.81 | -1.014 | -0.368 | 0.892 | 0.001 | 0.002 | 0.002 | 0 | 0.37 | 0.37 | 0.24 | 0.27 | 0.37 | 0.25 |
| 5 | 162265024 | rs6885189 | -0.262 | -0.521 | -0.788 | 1.02 | 0.012 | 0.007 | 0.007 | 0 | 0.65 | 0.62 | 0.39 | 0.42 | 0.57 | 0.54 |
| 5 | 162267912 | rs17060930 | 1.022 | -0.479 | -0.854 | 1.007 | -0.001 | -0.001 | -0.001 | 0.912 | 0.05 | 0.04 | 0.02 | 0.05 | 0.05 | 0.17 |
| 5 | 162297896 | rs4566832 | 0.309 | -0.141 | -0.932 | 1.228 | -0.011 | -0.003 | -0.003 | 0 | 0.61 | 0.69 | 0.46 | 0.4 | 0.55 | 0.51 |
| 5 | 162306793 | rs828718 | -0.246 | -0.063 | -1.005 | 1.374 | 0.001 | -0.039 | -0.039 | 0 | 0.31 | 0.15 | 0.38 | 0.35 | 0.28 | 0.25 |
| 5 | 162307027 | rs828720 | -1.121 | 0.146 | -0.858 | 1.252 | 0.012 | -0.003 | -0.003 | 0 | 0.77 | 0.76 | 0.77 | 0.76 | 0.73 | 0.7 |
| 5 | 162313282 | rs920604 | 0.555 | 0.069 | -0.971 | 1.293 | 0.014 | 0.016 | 0.016 | 0 | 0.48 | 0.61 | 0.36 | 0.57 | 0.41 | 0.46 |
| 5 | 162318650 | rs828732 | -1.444 | 0.105 | -1.239 | 1.364 | -0.047 | 0.008 | 0.008 | 0 | 0.79 | 0.93 | 0.78 | 0.8 | 0.73 | 0.74 |
| 5 | 162318689 | rs828733 | -1.286 | 0.112 | -1.233 | 1.369 | -0.006 | 0.001 | 0.001 | 0 | 0.78 | 0.83 | 0.76 | 0.76 | 0.73 | 0.7 |
| 5 | 162321929 | rs828736 | -0.289 | -0.239 | -1.211 | 1.315 | -0.01 | -0.005 | -0.005 | 0 | 0.68 | 0.73 | 0.47 | 0.57 | 0.56 | 0.58 |
| 5 | 162337257 | rs17569780 | 0.9 | -0.068 | -1.345 | 0.656 | -0.004 | 0.128 | 0.128 | 0 | 0.2 | 0.02 | 0.1 | 0 | 0.15 | 0.11 |
| 5 | 162365189 | rs2054228 | -1.16 | 0.041 | -1.307 | 0.802 | 0.019 | 0.054 | 0.054 | 0 | 0.81 | 0.9 | 0.93 | 1 | 0.87 | 0.91 |
| 5 | 162365528 | rs2926054 | -0.235 | 1.026 | -0.957 | 1.838 | -0.092 | -0.004 | -0.004 | 0 | 0.28 | 0.11 | 0.52 | 0.75 | 0.4 | 0.41 |
| 5 | 162371114 | rs1531263 | -1.259 | 1.051 | -1.004 | 1.89 | 0.001 | -0.038 | -0.038 | 0 | 0.43 | 0.4 | 0.59 | 0.9 | 0.47 | 0.56 |

| | | | | | | | | | | | | | | | |
|---|---|---|---|---|---|---|---|---|---|---|---|---|---|---|---|
| 5 | 162372793 | rs2963094 | 0.625 | 0.949 | -0.978 | 2.233 | -0.066 | -0.044 | -0.044 | 0 | 0.22 | 0.08 | 0.22 | 0.29 | 0.27 | 0.26 |
| 5 | 162373515 | rs7704329 | -1.11 | 1.006 | -1.077 | 2.064 | -0.001 | -0.03 | -0.03 | 0 | 0.45 | 0.41 | 0.63 | 0.89 | 0.5 | 0.58 |
| 5 | 162373967 | rs1993556 | -0.64 | 1.779 | -0.418 | 2.695 | -0.034 | 0.036 | 0.036 | 0 | 0.06 | 0 | 0.23 | 0.43 | 0.13 | 0.12 |
| 5 | 162374256 | rs1993557 | -0.347 | 1.21 | -0.279 | 1.322 | 0.041 | 0.039 | 0.039 | 0 | 0.3 | 0.39 | 0.57 | 0.62 | 0.42 | 0.45 |
| 5 | 162381246 | rs11135232 | -0.767 | 0.279 | -1.098 | 1.245 | 0.025 | -0.02 | -0.02 | 0 | 0.33 | 0.82 | 0.27 | 0.43 | 0.24 | 0.34 |
| 5 | 162384064 | rs2926053 | 0.465 | -0.054 | -1.23 | 0.509 | 0.014 | 0.026 | 0.026 | 0.406 | 0.82 | 0.86 | 0.92 | 0.95 | 0.88 | 0.93 |
| 5 | 162386462 | rs1598919 | 0.66 | -0.383 | -1.693 | 0.107 | 0.027 | 0.079 | 0.079 | 0.974 | 0.74 | 0.85 | 0.85 | 0.94 | 0.78 | 0.9 |
| 5 | 162386484 | rs1598918 | 1.194 | -0.346 | -0.718 | 0.986 | 0.001 | -0.001 | -0.001 | 0 | 0.59 | 0.44 | 0.47 | 0.41 | 0.64 | 0.57 |
| 5 | 162395073 | rs17061102 | 0.091 | -0.333 | 0.009 | 1.444 | 0.022 | 0.015 | 0.015 | 0 | 0.11 | 0.16 | 0.26 | 0.14 | 0.14 | 0.2 |
| 5 | 162411692 | rs2034943 | 0.698 | -0.58 | 0.234 | 1.312 | -0.041 | 0.04 | 0.04 | 0.715 | 0.7 | 0.54 | 0.72 | 0.57 | 0.78 | 0.78 |
| 5 | 162412044 | rs2125342 | -0.159 | -1.259 | 0.038 | 1.124 | -0.008 | 0.023 | 0.023 | 0 | 0.27 | 0.31 | 0.43 | 0.4 | 0.22 | 0.45 |
| 5 | 162412788 | rs10866738 | 1.337 | -1.046 | -0.209 | 0.817 | -0.041 | 0.151 | 0.151 | 3.322 | 0.43 | 0.13 | 0.29 | 0.16 | 0.55 | 0.33 |
| 5 | 162415684 | rs1458349 | -0.215 | -1.101 | 0.863 | 0.81 | -0.011 | 0.04 | 0.04 | 0 | 0.26 | 0.33 | 0.44 | 0.45 | 0.22 | 0.44 |
| 5 | 162416571 | rs11741192 | -0.031 | -1.082 | 0.512 | 0.279 | 0.023 | 0.039 | 0.039 | 0 | 0.67 | 0.78 | 0.81 | 0.98 | 0.72 | 0.84 |
| 5 | 162426738 | rs1870313 | 0.012 | -1.07 | 0.654 | 0.614 | 0.006 | -0.029 | -0.029 | 0 | 0.26 | 0.19 | 0.42 | 0.47 | 0.22 | 0.46 |
| 5 | 162440022 | rs1531267 | 0.792 | -1.142 | 0.587 | 0.488 | -0.026 | -0.041 | -0.041 | 0 | 0.42 | 0.76 | 0.43 | 0.54 | 0.51 | 0.45 |
| 5 | 162444591 | rs13160797 | 1.296 | -1.238 | 0.403 | 0.529 | -0.113 | 0.101 | 0.101 | 0.443 | 0.34 | 0.09 | 0.27 | 0.18 | 0.5 | 0.32 |
| 5 | 162488220 | rs11746612 | 1.054 | -1.619 | 0.875 | 0.281 | 0.001 | 0.096 | 0.096 | 0.005 | 0.4 | 0.25 | 0.34 | 0.19 | 0.6 | 0.34 |
| 5 | 162492185 | rs2963198 | -0.481 | -2.003 | 0.764 | 0.33 | 0.052 | -0.056 | -0.056 | 0.077 | 0.47 | 0.11 | 0.49 | 0.47 | 0.35 | 0.51 |
| 5 | 162499589 | rs13169504 | 1.297 | -1.892 | 1.044 | 0.38 | -0.006 | 0.046 | 0.046 | 0.049 | 0.38 | 0.3 | 0.34 | 0.19 | 0.59 | 0.34 |
| 5 | 162501105 | rs2926058 | 0.314 | -2.111 | 0.54 | 0.236 | -0.037 | -0.156 | -0.156 | 3.919 | 0.27 | 0.02 | 0.4 | 0.45 | 0.24 | 0.42 |
| 5 | 162518431 | rs6884376 | -0.521 | -2.346 | -0.128 | 0.441 | -0.002 | -0.088 | -0.088 | 0 | 0.78 | 0.93 | 0.67 | 0.64 | 0.83 | 0.64 |
| 5 | 162520134 | rs11135239 | 1.283 | -2.555 | 0.312 | 0.372 | 0 | -0.001 | -0.001 | 0 | 0.42 | 0.44 | 0.45 | 0.27 | 0.6 | 0.35 |
| 5 | 162524013 | rs4074952 | 1.821 | -2.525 | 0.018 | 0.387 | -0.01 | 0.04 | 0.04 | 0.075 | 0.38 | 0.21 | 0.42 | 0.27 | 0.6 | 0.34 |
| 5 | 162524331 | rs4254931 | -0.442 | -2.315 | -0.18 | 0.445 | -0.018 | -0.1 | -0.1 | 0 | 0.77 | 0.95 | 0.66 | 0.63 | 0.82 | 0.64 |
| 5 | 162531495 | rs9313953 | -0.686 | -2.348 | -0.071 | 0.45 | 0.006 | -0.082 | -0.082 | 3.264 | 0.79 | 0.95 | 0.67 | 0.63 | 0.84 | 0.65 |
| 5 | 162536955 | rs13179142 | 1.524 | -2.158 | 0.236 | -0.344 | -0.004 | 0.001 | 0.001 | 0 | 0.3 | 0.31 | 0.35 | 0.15 | 0.51 | 0.27 |
| 5 | 162550726 | rs4336377 | 0.005 | -2.663 | 0.457 | -0.492 | -0.054 | 0.046 | 0.046 | 0.188 | 0.23 | 0.39 | 0.17 | 0.35 | 0.13 | 0.21 |
| 5 | 162551332 | rs10515850 | 0.594 | -2.679 | 0.721 | -0.581 | -0.015 | -0.065 | -0.065 | 0 | 0.19 | 0.23 | 0.1 | 0 | 0.12 | 0.06 |
| 5 | 162554798 | rs17585996 | 2.649 | -1.45 | -0.278 | -0.878 | -0.096 | 0.002 | 0.002 | 0 | 0.16 | 0.01 | 0.23 | 0.12 | 0.32 | 0.16 |
| 5 | 162556075 | rs6556657 | -0.394 | -2.816 | -0.088 | -0.749 | 0.03 | -0.056 | -0.056 | 0 | 0.58 | 0.85 | 0.53 | 0.51 | 0.67 | 0.51 |
| 5 | 162558675 | rs13168087 | 1.383 | -2.688 | 0.077 | -0.686 | -0.067 | 0.055 | 0.055 | 0.076 | 0.29 | 0.17 | 0.34 | 0.15 | 0.51 | 0.28 |
| 5 | 162561438 | rs4242212 | 0.464 | -2.428 | 0.23 | -0.557 | -0.012 | 0.057 | 0.057 | 0.51 | 0.13 | 0.17 | 0.08 | 0.36 | 0.1 | 0.22 |
| 5 | 162571983 | rs6864537 | 1.319 | -2.37 | 0.065 | -0.805 | -0.049 | -0.088 | -0.088 | 1.217 | 0.14 | 0.32 | 0.09 | 0 | 0.06 | 0.02 |
| 5 | 162585645 | rs12187751 | 1.437 | -1.614 | -0.408 | -0.336 | -0.001 | 0.003 | 0.003 | 0 | 0.08 | 0.06 | 0.08 | 0.12 | 0.08 | 0.09 |
| 5 | 162600918 | rs4868932 | -0.194 | -1.316 | -0.264 | -0.35 | 0.022 | -0.007 | -0.007 | 0 | 0.46 | 0.67 | 0.47 | 0.5 | 0.61 | 0.49 |
| 5 | 162617260 | rs10077466 | -0.075 | -1.036 | -0.748 | -0.324 | 0.004 | -0.004 | -0.004 | 0.601 | 0.45 | 0.61 | 0.46 | 0.5 | 0.61 | 0.51 |
| 5 | 162626996 | rs2964356 | 0.266 | -0.76 | -0.903 | 0.013 | 0.001 | -0.003 | -0.003 | 0 | 0.4 | 0.38 | 0.45 | 0.52 | 0.61 | 0.48 |
| 5 | 162632615 | rs2913568 | -0.626 | -0.35 | -1.659 | -0.161 | -0.108 | 0.07 | 0.07 | 0.014 | 0.65 | 0.95 | 0.64 | 0.81 | 0.57 | 0.67 |
| 5 | 162633207 | rs12656289 | 0.456 | -0.86 | -1.618 | 0.376 | 0.013 | -0.009 | -0.009 | 0 | 0.14 | 0.1 | 0.17 | 0.5 | 0.08 | 0.27 |
| 5 | 162647465 | rs17061325 | 1.328 | -1.24 | -1.795 | -0.077 | 0.024 | 0.048 | 0.048 | 0.614 | 0.14 | 0.05 | 0.04 | 0 | 0.07 | 0.07 |
| 5 | 162647507 | rs17061327 | 0.621 | -1.107 | -1.72 | 0.138 | 0.013 | -0.01 | -0.01 | 0 | 0.14 | 0.1 | 0.17 | 0.5 | 0.08 | 0.27 |
| 5 | 162651614 | rs2114997 | 0.066 | -0.781 | -1.029 | 0.127 | 0.001 | -0.001 | -0.001 | 0 | 0.35 | 0.58 | 0.4 | 0.68 | 0.4 | 0.53 |
| 5 | 162652551 | rs4448013 | -0.601 | -1.132 | -1.061 | 0.078 | 0.121 | -0.092 | -0.092 | 0 | 0.4 | 0.89 | 0.28 | 0.48 | 0.21 | 0.42 |
| 5 | 162652827 | rs4551093 | -0.619 | -1.148 | -1.023 | 0.052 | 0.115 | -0.146 | -0.146 | 0 | 0.4 | 0.94 | 0.29 | 0.48 | 0.21 | 0.42 |
| 5 | 162657020 | rs1529437 | 1.369 | -0.071 | -0.782 | 0.423 | -0.041 | -0.015 | -0.015 | 1.974 | 0.19 | 0.09 | 0.22 | 0.25 | 0.27 | 0.24 |
| 5 | 162657404 | rs2913556 | -0.737 | -0.326 | -1.053 | 0.432 | -0.003 | 0.095 | 0.095 | 1.527 | 0.52 | 0.7 | 0.5 | 0.73 | 0.54 | 0.65 |
| 5 | 162664528 | rs2901731 | -0.687 | -0.249 | -0.991 | 0.475 | 0.006 | 0.066 | 0.066 | 1.796 | 0.52 | 0.65 | 0.49 | 0.73 | 0.54 | 0.64 |
| 5 | 162671160 | rs2913593 | 0.368 | 0.795 | -1.309 | 0.72 | -0.044 | -0.017 | -0.017 | 1.021 | 0.24 | 0.11 | 0.32 | 0.3 | 0.39 | 0.3 |
| 5 | 162677234 | rs959011 | 0.114 | 0.191 | -0.756 | 0.234 | 0.006 | 0.035 | 0.035 | 0.397 | 0.48 | 0.64 | 0.44 | 0.27 | 0.44 | 0.35 |
| 5 | 162679226 | rs12233971 | 0.442 | 0.331 | -1.109 | 0.721 | -0.036 | -0.046 | -0.046 | 0 | 0.13 | 0.01 | 0.07 | 0.15 | 0.1 | 0.15 |
| 5 | 162680419 | rs6893843 | -0.487 | 0.228 | -1.206 | 0.721 | -0.016 | -0.054 | -0.054 | 0 | 0.91 | 0.98 | 0.81 | 0.74 | 0.89 | 0.78 |
| 5 | 162683796 | rs214115 | 0.186 | 0.313 | -1.31 | -0.195 | -0.006 | 0.06 | 0.06 | 1.435 | 0.2 | 0.07 | 0.09 | 0.04 | 0.17 | 0.15 |
| 5 | 162688282 | rs300185 | 0.929 | 0.247 | -0.782 | -0.015 | -0.001 | -0.002 | -0.002 | 0 | 0.44 | 0.52 | 0.32 | 0.21 | 0.44 | 0.32 |
| 5 | 162698920 | rs300195 | 0.284 | 0.268 | -0.727 | 0.012 | 0.003 | 0.012 | 0.012 | 0.079 | 0.19 | 0.14 | 0.09 | 0.04 | 0.14 | 0.15 |
| 5 | 162702999 | rs1896618 | 0.272 | 0.433 | -0.76 | 0.124 | -0.006 | 0.011 | 0.011 | 0 | 0.65 | 0.69 | 0.43 | 0.25 | 0.59 | 0.46 |
| 5 | 162710860 | rs1862166 | 0.221 | 0.439 | -0.757 | 0.127 | -0.004 | 0.01 | 0.01 | 0 | 0.64 | 0.65 | 0.41 | 0.25 | 0.59 | 0.46 |
| 5 | 162716850 | rs1154758 | 1.244 | 0.27 | -0.504 | 0.014 | -0.01 | 0.01 | 0.01 | 0 | 0.35 | 0.25 | 0.28 | 0.24 | 0.43 | 0.35 |
| 5 | 162719547 | rs13159031 | 1.332 | 0.012 | -0.946 | -0.156 | -0.008 | 0.03 | 0.03 | 0 | 0.09 | 0.03 | 0.04 | 0 | 0.07 | 0.07 |
| 5 | 162722554 | rs2964380 | 0.009 | 0.013 | -0.791 | 0.309 | -0.006 | -0.01 | -0.01 | 0.028 | 0.67 | 0.72 | 0.62 | 0.58 | 0.62 | 0.55 |
| 5 | 162727341 | rs2913578 | -0.458 | 0.141 | -0.702 | 0.278 | 0.007 | 0.072 | 0.072 | 0 | 0.88 | 0.83 | 0.83 | 0.64 | 0.85 | 0.8 |
| 5 | 162730903 | rs2964381 | 0.515 | 0.238 | -0.8 | 0.243 | -0.006 | -0.011 | -0.011 | 0 | 0.59 | 0.63 | 0.56 | 0.54 | 0.55 | 0.62 |
| 5 | 162732021 | rs300206 | 1.067 | 0.157 | -1.399 | -0.05 | 0.006 | -0.077 | -0.077 | 0 | 0.15 | 0.04 | 0.23 | 0.42 | 0.11 | 0.25 |
| 5 | 162732792 | rs182462 | 0.429 | 0.329 | -0.895 | -0.135 | -0.004 | 0.021 | 0.021 | 0 | 0.44 | 0.39 | 0.29 | 0.18 | 0.46 | 0.45 |
| 5 | 162732988 | rs2964382 | -0.554 | 0.766 | -1.443 | 0.28 | -0.045 | -0.064 | -0.064 | 0 | 0.9 | 1 | 0.83 | 0.64 | 0.85 | 0.82 |
| 5 | 162734871 | rs300209 | 0.287 | 0.454 | -0.932 | -0.701 | -0.028 | 0.067 | 0.067 | 0 | 0.34 | 0.19 | 0.24 | 0.18 | 0.4 | 0.37 |
| 5 | 162737689 | rs10515854 | -0.638 | 0.207 | -0.288 | -0.071 | -0.019 | -0.047 | -0.047 | 0.181 | 0.09 | 0.18 | 0.04 | 0 | 0.04 | 0.01 |
| 5 | 162748171 | rs300236 | 0.153 | 0.384 | -0.555 | -0.123 | 0 | 0.014 | 0.014 | 0 | 0.52 | 0.55 | 0.57 | 0.34 | 0.55 | 0.48 |
| 5 | 162748604 | rs12186371 | 0.024 | 0.281 | -0.42 | -0.353 | -0.001 | -0.028 | -0.028 | 0.428 | 0.08 | 0.14 | 0.05 | 0.02 | 0.08 | 0.18 |

| | | | | | | | | | | | | | | | | |
|---|---|---|---|---|---|---|---|---|---|---|---|---|---|---|---|---|
| 5 | 162750370 | rs300238 | 0.136 | -0.327 | 0.03 | -1.018 | 0.002 | -0.064 | -0.064 | 0.283 | 0.44 | 0.52 | 0.47 | 0.06 | 0.51 | 0.47 |
| 5 | 162750871 | rs17061510 | 0.811 | -0.701 | -0.179 | -1.327 | -0.02 | -0.005 | -0.005 | 0.068 | 0.32 | 0.21 | 0.34 | 0.64 | 0.33 | 0.32 |
| 5 | 162751751 | rs300211 | 1.285 | -0.77 | -0.095 | -1.421 | -0.005 | -0.023 | -0.023 | 0 | 0.35 | 0.32 | 0.41 | 0.04 | 0.43 | 0.28 |
| 5 | 162753587 | rs6897401 | -0.258 | -0.809 | -0.06 | -1.317 | -0.034 | 0.003 | 0.003 | 0 | 0.24 | 0.39 | 0.21 | 0.3 | 0.2 | 0.38 |
| 5 | 162754151 | rs11742012 | 1.005 | -0.81 | -0.072 | -1.315 | -0.017 | 0.014 | 0.014 | 0.001 | 0.16 | 0.2 | 0.16 | 0.28 | 0.11 | 0.2 |
| 5 | 162759022 | rs300213 | 0.764 | -0.841 | -0.107 | -1.309 | -0.044 | -0.026 | -0.026 | 0.003 | 0.67 | 0.53 | 0.76 | 0.7 | 0.76 | 0.61 |
| 5 | 162759175 | rs300214 | -0.048 | -0.858 | -0.101 | -1.317 | -0.024 | -0.001 | -0.001 | 4.267 | 0.76 | 0.71 | 0.81 | 0.72 | 0.85 | 0.79 |
| 5 | 162759623 | rs2431212 | 0.19 | -0.84 | -0.139 | -1.274 | -0.006 | -0.033 | -0.033 | 0.167 | 0.85 | 0.8 | 0.92 | 0.98 | 0.88 | 0.8 |
| 5 | 162769175 | rs734442 | 0.773 | -0.662 | -0.106 | -1.385 | -0.001 | -0.001 | -0.001 | 0 | 0.27 | 0.28 | 0.35 | 0.64 | 0.28 | 0.31 |
| 5 | 162772259 | rs13178546 | -0.481 | -1.009 | -0.161 | -1.593 | 0 | -0.037 | -0.037 | 0 | 0.48 | 0.53 | 0.49 | 0.93 | 0.38 | 0.55 |
| 5 | 162774224 | rs1363152 | 1.208 | -1.12 | -0.236 | -1.801 | -0.012 | -0.023 | -0.023 | 0.041 | 0.21 | 0.13 | 0.31 | 0.64 | 0.24 | 0.25 |
| 5 | 162778654 | rs9313962 | 1.791 | -0.977 | -0.289 | -1.902 | -0.003 | -0.017 | -0.017 | 0 | 0.32 | 0.3 | 0.42 | 0.05 | 0.44 | 0.27 |
| 5 | 162779115 | rs4869033 | 1.043 | -1.203 | -0.248 | -1.778 | 0.025 | 0.033 | 0.033 | 2.001 | 0.57 | 0.67 | 0.77 | 0.69 | 0.71 | 0.56 |
| 5 | 162796184 | rs999581 | 0.892 | -1.188 | -0.215 | -1.68 | -0.016 | -0.018 | -0.018 | 0 | 0.61 | 0.54 | 0.75 | 0.69 | 0.73 | 0.6 |
| 5 | 162801076 | rs13159369 | 0.371 | -1.137 | -0.246 | -1.746 | 0 | -0.029 | -0.029 | 0 | 0.15 | 0.15 | 0.08 | 0.02 | 0.15 | 0.2 |
| 5 | 162803931 | rs967052 | 0.808 | -1.204 | -0.051 | -1.732 | -0.021 | -0.038 | -0.038 | 0 | 0.62 | 0.5 | 0.75 | 0.71 | 0.72 | 0.6 |
| 5 | 162811930 | rs17061586 | -0.011 | -1.129 | -0.104 | -1.729 | -0.001 | -0.037 | -0.037 | 0 | 0.17 | 0.22 | 0.09 | 0.03 | 0.16 | 0.19 |
| 5 | 162819235 | rs11952005 | 0.272 | -1.102 | -0.159 | -1.717 | -0.002 | -0.009 | -0.009 | 0.72 | 0.16 | 0.12 | 0.08 | 0.03 | 0.16 | 0.19 |
| 5 | 162826613 | rs13186063 | 0.864 | -0.668 | -0.345 | -1.54 | -0.045 | -0.059 | -0.059 | 0 | 0.23 | 0.09 | 0.33 | 0.55 | 0.28 | 0.31 |
| 5 | 162831587 | rs10040287 | 1.254 | -0.766 | -0.162 | -1.595 | -0.016 | -0.036 | -0.036 | 0.097 | 0.08 | 0.16 | 0.05 | 0 | 0.03 | 0.02 |
| 5 | 162832105 | rs9313968 | 1.768 | -0.894 | -0.128 | -1.726 | 0.001 | -0.029 | -0.029 | 0 | 0.32 | 0.29 | 0.4 | 0.06 | 0.44 | 0.22 |
| 5 | 162835160 | rs6870488 | 1.427 | -0.756 | -0.343 | -1.52 | -0.015 | 0.021 | 0.021 | 0.055 | 0.14 | 0.17 | 0.12 | 0.36 | 0.09 | 0.19 |
| 5 | 162836090 | rs13172782 | 0.54 | -0.53 | -0.48 | -1.389 | -0.021 | -0.035 | -0.035 | 0.047 | 0.25 | 0.15 | 0.34 | 0.55 | 0.28 | 0.35 |
| 5 | 162841338 | rs4307107 | 0.858 | -0.383 | -0.333 | -1.669 | 0.011 | -0.089 | -0.089 | 0.083 | 0.4 | 0.44 | 0.46 | 0.06 | 0.48 | 0.25 |
| 5 | 162847832 | rs2112770 | -0.176 | -0.489 | -0.464 | -1.617 | -0.004 | 0.01 | 0.01 | 0 | 0.83 | 0.89 | 0.92 | 0.98 | 0.84 | 0.79 |
| 5 | 162866548 | rs299322 | -0.713 | -0.833 | -0.137 | -0.541 | 0.001 | 0.041 | 0.041 | 0 | 0.32 | 0.6 | 0.33 | 0.54 | 0.3 | 0.26 |
| 5 | 162867195 | rs2069347 | 1.4 | -0.906 | -0.325 | -0.257 | -0.009 | 0.014 | 0.014 | 0 | 0.39 | 0.29 | 0.44 | 0.17 | 0.48 | 0.3 |
| 5 | 162873507 | rs422356 | -0.51 | -1.004 | -0.452 | -0.355 | 0.001 | 0.041 | 0.041 | 0 | 0.32 | 0.6 | 0.33 | 0.54 | 0.3 | 0.26 |
| 5 | 162876836 | rs299302 | -0.51 | -1.004 | -0.453 | -0.355 | 0.001 | 0.042 | 0.042 | 0 | 0.32 | 0.6 | 0.33 | 0.54 | 0.3 | 0.26 |
| 5 | 162877645 | rs6877450 | 1.238 | -1.032 | -0.293 | 0.038 | -0.015 | 0.019 | 0.019 | 0.588 | 0.08 | 0.24 | 0.06 | 0.04 | 0.04 | 0.02 |
| 5 | 162886028 | rs213505 | 0.191 | -1.083 | -0.244 | 0.013 | 0.007 | 0.025 | 0.025 | 0 | 0.23 | 0.28 | 0.25 | 0.49 | 0.26 | 0.26 |
| 5 | 162886563 | rs2287776 | 2.24 | -1.101 | -0.288 | 0.013 | 0.002 | -0.027 | -0.027 | 0 | 0.06 | 0 | 0.03 | 0.04 | 0.04 | 0.02 |
| 5 | 162887650 | rs2303076 | 0.253 | -1.156 | -0.304 | -0.003 | 0.033 | -0.021 | -0.021 | 0 | 0.28 | 0.07 | 0.23 | 0.3 | 0.21 | 0.44 |
| 5 | 162889878 | rs10076669 | 1.408 | -0.925 | -0.223 | -0.006 | -0.012 | 0.023 | 0.023 | 0.405 | 0.07 | 0.22 | 0.06 | 0.04 | 0.04 | 0.02 |
| 5 | 162895573 | rs299285 | -0.606 | -0.919 | -0.215 | -0.012 | 0.003 | 0.062 | 0.062 | 0 | 0.32 | 0.55 | 0.33 | 0.53 | 0.3 | 0.27 |
| 5 | 162896650 | rs299284 | 0.942 | -0.887 | -0.25 | -0.021 | -0.013 | 0.014 | 0.014 | 0 | 0.11 | 0.07 | 0.07 | 0.01 | 0.14 | 0.09 |
| 5 | 162901156 | rs2303078 | 2.257 | -0.884 | -0.27 | -0.002 | 0.002 | -0.027 | -0.027 | 0 | 0.06 | 0 | 0.03 | 0.04 | 0.04 | 0.02 |
| 5 | 162902516 | rs299290 | 0.088 | -0.867 | -0.226 | -0.047 | 0.007 | 0.029 | 0.029 | 0.017 | 0.24 | 0.29 | 0.26 | 0.48 | 0.26 | 0.25 |
| 5 | 162909315 | rs10866742 | 0.497 | -0.888 | -0.361 | -0.974 | 0.021 | -0.066 | -0.066 | 0.253 | 0.26 | 0.07 | 0.44 | 0.69 | 0.21 | 0.43 |
| 5 | 162909716 | rs299295 | 0.187 | -0.503 | -0.229 | -1.236 | 0.006 | -0.047 | -0.047 | 0.388 | 0.23 | 0.29 | 0.19 | 0.07 | 0.25 | 0.24 |
| 5 | 162916428 | rs299316 | -0.175 | -0.548 | -0.202 | -1.217 | -0.002 | -0.004 | -0.004 | 2.009 | 0.87 | 0.85 | 0.93 | 0.93 | 0.9 | 0.89 |
| 5 | 162917708 | rs10515860 | 2.002 | -0.392 | -0.128 | -1.348 | -0.035 | 0.032 | 0.032 | 0 | 0.1 | 0.01 | 0.1 | 0.06 | 0.11 | 0.15 |
| 5 | 162919599 | rs4599554 | -0.273 | -0.665 | 0.231 | -1.464 | 0 | 0.012 | 0.012 | 0 | 0.22 | 0.17 | 0.12 | 0.06 | 0.19 | 0.11 |
| 5 | 162922823 | rs299289 | -0.847 | -0.836 | 0.17 | -1.634 | 0 | -0.008 | -0.008 | 0 | 0.37 | 0.62 | 0.29 | 0.12 | 0.32 | 0.28 |
| 5 | 162930747 | rs10515861 | 2.179 | -0.1 | -0.695 | -2.152 | -0.1 | 0.028 | 0.028 | 0 | 0.21 | 0.02 | 0.17 | 0.08 | 0.31 | 0.25 |
| 5 | 162933960 | rs6882306 | 0.481 | 0.027 | 0.456 | -1.571 | 0.001 | 0.004 | 0.004 | 0 | 0.76 | 0.79 | 0.87 | 0.94 | 0.81 | 0.87 |
| 5 | 162941739 | rs7721639 | -0.092 | 0.567 | 0.055 | -1.48 | 0.017 | -0.086 | -0.086 | 1.88 | 0.17 | 0.33 | 0.12 | 0.01 | 0.17 | 0.12 |
| 5 | 162944408 | rs4869089 | 1.579 | 0.707 | -0.29 | -1.73 | -0.058 | 0.103 | 0.103 | 0.464 | 0.43 | 0.08 | 0.55 | 0.77 | 0.45 | 0.49 |
| 5 | 162947884 | rs7716203 | -1.19 | 0.641 | 0.04 | -1.645 | -0.009 | 0.088 | 0.088 | 0.591 | 0.45 | 0.74 | 0.34 | 0.18 | 0.45 | 0.36 |
| 5 | 162949215 | rs729352 | -0.288 | 0.757 | 0.087 | -1.332 | -0.006 | -0.025 | -0.025 | 0 | 0.28 | 0.41 | 0.22 | 0.17 | 0.28 | 0.24 |
| 5 | 162956242 | rs4400146 | -0.576 | 0.526 | -0.026 | -1.387 | -0.002 | 0.006 | 0.006 | 0 | 0.33 | 0.35 | 0.25 | 0.14 | 0.34 | 0.31 |
| 5 | 162956290 | rs4400147 | 0.143 | 0.561 | 0.35 | -1.632 | -0.001 | -0.011 | -0.011 | 0.062 | 0.37 | 0.33 | 0.35 | 0.16 | 0.43 | 0.45 |
| 5 | 162957627 | rs1496864 | -1.332 | 0.603 | 1.198 | -0.476 | 0.004 | 0 | 0 | 0 | 0.28 | 0.32 | 0.31 | 0.69 | 0.35 | 0.37 |
| 5 | 162958311 | rs7700282 | 1.076 | 0.704 | 0.708 | -0.491 | -0.029 | 0.031 | 0.031 | 0 | 0.09 | 0.03 | 0.07 | 0.01 | 0.14 | 0.08 |
| 5 | 162958461 | rs7722476 | -1.856 | 0.55 | 0.832 | -0.534 | -0.004 | 0.033 | 0.033 | 0 | 0.45 | 0.63 | 0.44 | 0.75 | 0.47 | 0.59 |
| 5 | 162966722 | rs6897573 | -0.481 | 0.235 | 0.338 | -0.068 | 0.003 | -0.006 | -0.006 | 0 | 0.92 | 0.96 | 0.95 | 0.94 | 0.93 | 0.97 |
| 5 | 162969756 | rs919258 | -0.011 | 0.402 | 0.49 | 0.033 | 0.011 | 0.107 | 0.107 | 0 | 0.09 | 0.11 | 0.18 | 0.62 | 0.14 | 0.27 |
| 5 | 162973665 | rs17061876 | 0.217 | 0.422 | 0.282 | 0.012 | -0.04 | -0.039 | -0.039 | 0.863 | 0.09 | 0.01 | 0.15 | 0.61 | 0.14 | 0.27 |
| 5 | 162978283 | rs17061883 | -0.327 | 0.182 | 0.689 | -0.179 | -0.003 | 0.021 | 0.021 | 1.157 | 0.11 | 0.42 | 0.08 | 0.06 | 0.07 | 0.03 |
| 5 | 162978954 | rs297965 | 1.087 | 0.34 | 0.009 | -0.089 | 0.015 | 0.023 | 0.023 | 0 | 0.64 | 0.29 | 0.62 | 0.28 | 0.66 | 0.49 |
| 5 | 162984955 | rs4301242 | -0.51 | 0.571 | -0.14 | 0.538 | 0.018 | 0.174 | 0.174 | 0 | 0.1 | 0.19 | 0.19 | 0.58 | 0.13 | 0.3 |
| 5 | 162989134 | rs297958 | -0.619 | 0.238 | -0.671 | -0.354 | 0.017 | 0.066 | 0.066 | 0 | 0.45 | 0.72 | 0.41 | 0.7 | 0.42 | 0.56 |
| 5 | 162989613 | rs177075 | -0.025 | 0.175 | -0.918 | -0.639 | -0.016 | 0.069 | 0.069 | 0 | 0.17 | 0.05 | 0.1 | 0.01 | 0.17 | 0.13 |
| 5 | 162989974 | rs2217265 | 1.044 | -0.367 | -0.499 | -0.661 | 0.011 | 0.087 | 0.087 | 0 | 0.47 | 0.24 | 0.53 | 0.3 | 0.56 | 0.38 |
| 5 | 162991763 | rs294598 | 0.897 | -0.737 | 0.197 | -0.907 | -0.021 | -0.135 | -0.135 | 0 | 0.81 | 0.54 | 0.86 | 0.99 | 0.87 | 0.91 |
| 5 | 162991956 | rs294597 | -0.1 | -0.447 | -0.122 | -0.822 | -0.004 | -0.011 | -0.011 | 0 | 0.33 | 0.15 | 0.33 | 0.65 | 0.32 | 0.54 |
| 5 | 162997351 | rs985254 | 0.966 | -0.44 | 0.596 | -0.853 | -0.002 | -0.01 | -0.01 | 0.084 | 0.53 | 0.53 | 0.57 | 0.29 | 0.55 | 0.38 |
| 5 | 162997442 | rs985253 | 0.472 | -0.517 | 0.351 | -0.621 | -0.003 | 0.044 | 0.044 | 0 | 0.62 | 0.72 | 0.76 | 0.9 | 0.63 | 0.63 |
| 5 | 163005105 | rs6556682 | -0.142 | -0.536 | -0.096 | -0.456 | -0.006 | 0.08 | 0.08 | 0.589 | 0.72 | 0.85 | 0.81 | 0.9 | 0.71 | 0.65 |

| | | | | | | | | | | | | | | | | |
|---|---|---|---|---|---|---|---|---|---|---|---|---|---|---|---|---|
| 5 | 163008349 | rs6862542 | 0.538 | -0.44 | 0.192 | -0.418 | -0.005 | -0.009 | -0.009 | 0 | 0.6 | 0.5 | 0.59 | 0.3 | 0.62 | 0.4 |
| 5 | 163021335 | rs6876181 | 0.781 | -0.193 | -0.732 | -0.001 | 0.015 | -0.053 | -0.053 | 0 | 0.26 | 0.16 | 0.38 | 0.35 | 0.25 | 0.2 |
| 5 | 163028221 | rs979113 | 0.383 | 0.167 | -0.194 | -0.766 | -0.006 | 0.021 | 0.021 | 0 | 0.41 | 0.39 | 0.56 | 0.84 | 0.46 | 0.51 |
| 5 | 163028782 | rs1432655 | 0.375 | 0.166 | -0.204 | -0.88 | -0.006 | 0.022 | 0.022 | 0 | 0.41 | 0.39 | 0.56 | 0.86 | 0.46 | 0.51 |
| 5 | 163028887 | rs1368287 | 0.779 | 0.209 | -0.279 | -0.653 | -0.024 | 0.042 | 0.042 | 0.001 | 0.35 | 0.19 | 0.5 | 0.77 | 0.4 | 0.46 |
| 5 | 163037864 | rs17668553 | 1.084 | 0.714 | -0.251 | -0.566 | -0.026 | -0.012 | -0.012 | 0 | 0.05 | 0 | 0.07 | 0.05 | 0.05 | 0.11 |
| 5 | 163043385 | rs292410 | 0.484 | 0.544 | -0.402 | -0.83 | -0.015 | 0.112 | 0.112 | 0.017 | 0.45 | 0.19 | 0.51 | 0.79 | 0.44 | 0.49 |
| 5 | 163045944 | rs6878378 | -0.945 | 0.525 | 0.381 | -0.977 | 0.054 | 0.024 | 0.024 | 0 | 0.14 | 0.4 | 0.16 | 0.13 | 0.16 | 0.18 |
| 5 | 163045980 | rs1368288 | -0.515 | 0.49 | 0.525 | -1.001 | 0.007 | 0.064 | 0.064 | 0.542 | 0.6 | 0.73 | 0.68 | 0.92 | 0.6 | 0.67 |
| 5 | 163046827 | rs17605858 | -0.3 | 0.514 | 0.267 | -0.975 | -0.054 | -0.037 | -0.037 | 0 | 0.11 | 0.02 | 0.12 | 0.13 | 0.16 | 0.18 |
| 5 | 163051408 | rs292418 | 0.498 | 0.515 | 0.397 | -0.774 | -0.001 | 0.049 | 0.049 | 0.004 | 0.45 | 0.3 | 0.52 | 0.76 | 0.44 | 0.49 |
| 5 | 163062738 | rs4100211 | 0.808 | 0.4 | 0.337 | -0.751 | -0.003 | 0.011 | 0.011 | 1.214 | 0.37 | 0.31 | 0.5 | 0.76 | 0.38 | 0.46 |
| 5 | 163063208 | rs4869095 | 0.176 | 0.446 | 0.603 | -0.858 | -0.012 | 0.063 | 0.063 | 2.05 | 0.46 | 0.65 | 0.59 | 0.87 | 0.49 | 0.54 |
| 5 | 163063248 | rs10070913 | -1.21 | 0.623 | 0.786 | -0.42 | 0.072 | 0.067 | 0.067 | 0 | 0.09 | 0.36 | 0.09 | 0.11 | 0.12 | 0.08 |
| 5 | 163069251 | rs158836 | -0.708 | 0.37 | -0.038 | -0.759 | -0.011 | 0.045 | 0.045 | 1.255 | 0.75 | 0.86 | 0.79 | 0.93 | 0.74 | 0.72 |
| 5 | 163070118 | rs158837 | -0.514 | 0.403 | 0.167 | -0.573 | -0.001 | 0.021 | 0.021 | 0.01 | 0.72 | 0.8 | 0.79 | 0.87 | 0.74 | 0.7 |
| 5 | 163070267 | rs6895532 | 1.366 | 0.3 | 0.214 | -0.793 | -0.011 | 0.039 | 0.039 | 0.012 | 0.41 | 0.25 | 0.54 | 0.72 | 0.44 | 0.47 |
| 5 | 163072880 | rs158842 | 0.189 | 0.385 | 0.622 | -0.604 | 0.001 | -0.004 | -0.004 | 0 | 0.66 | 0.63 | 0.73 | 0.77 | 0.63 | 0.62 |
| 5 | 163083532 | rs158850 | 0.462 | 0.3 | 0.311 | -0.972 | -0.001 | 0.026 | 0.026 | 0 | 0.26 | 0.19 | 0.17 | 0.12 | 0.24 | 0.28 |
| 5 | 163083956 | rs6893725 | -0.941 | 0.143 | 0.86 | -0.995 | -0.008 | -0.027 | -0.027 | 0.118 | 0.17 | 0.29 | 0.17 | 0.08 | 0.16 | 0.19 |
| 5 | 163084821 | rs249597 | 0.571 | 0.104 | 0.752 | -1.107 | 0.001 | -0.037 | -0.037 | 7.637 | 0.09 | 0.16 | 0.07 | 0 | 0.08 | 0.03 |
| 5 | 163085029 | rs17062147 | 0.374 | 0.117 | 0.794 | -1.071 | -0.001 | -0.019 | -0.019 | 0.116 | 0.1 | 0.11 | 0.11 | 0 | 0.08 | 0.12 |
| 5 | 163085666 | rs158851 | 0.883 | 0.17 | 0.699 | -1.069 | 0.003 | 0.021 | 0.021 | 0.036 | 0.56 | 0.41 | 0.65 | 0.8 | 0.6 | 0.53 |
| 5 | 163086195 | rs158854 | 0.923 | 0.087 | 0.671 | -0.92 | 0.008 | 0.022 | 0.022 | 0.129 | 0.55 | 0.41 | 0.65 | 0.73 | 0.6 | 0.52 |
| 5 | 163086339 | rs249598 | -0.068 | 0.141 | 0.754 | -0.935 | -0.003 | -0.011 | -0.011 | 0 | 0.68 | 0.41 | 0.76 | 0.81 | 0.71 | 0.65 |
| 5 | 163086702 | rs549297 | 1.367 | 0.198 | 0.9 | -0.893 | 0.076 | 0.082 | 0.082 | 0 | 0.92 | 0.59 | 0.92 | 0.93 | 0.91 | 0.93 |
| 5 | 163095528 | rs693516 | -1.247 | 0.244 | 0.511 | -0.856 | -0.019 | 0.106 | 0.106 | 1.028 | 0.53 | 0.78 | 0.44 | 0.24 | 0.48 | 0.51 |
| 5 | 163097842 | rs424217 | 0.889 | 0.412 | 0.457 | -0.878 | -0.025 | 0.021 | 0.021 | 0.339 | 0.24 | 0.1 | 0.17 | 0.15 | 0.24 | 0.29 |
| 5 | 163100680 | rs292404 | -0.07 | 0.414 | 0.453 | -0.908 | 0.001 | -0.018 | -0.018 | 0 | 0.21 | 0.23 | 0.19 | 0.03 | 0.17 | 0.16 |
| 5 | 163102099 | rs292406 | 0.841 | 0.537 | 0.327 | -0.902 | -0.008 | 0.013 | 0.013 | 0.065 | 0.25 | 0.2 | 0.18 | 0.15 | 0.25 | 0.29 |
| 5 | 163103426 | rs3133041 | -0.028 | 0.445 | 0.247 | -0.951 | 0.003 | 0.012 | 0.012 | 0 | 0.21 | 0.2 | 0.19 | 0.03 | 0.19 | 0.15 |
| 5 | 163119723 | rs648765 | -1.202 | 0.504 | -0.265 | -0.889 | -0.032 | 0.067 | 0.067 | 1.234 | 0.55 | 0.81 | 0.44 | 0.29 | 0.51 | 0.53 |
| 5 | 163119808 | rs10057965 | -1.438 | 0.435 | 0.404 | -0.787 | -0.006 | -0.04 | -0.04 | 0 | 0.31 | 0.61 | 0.27 | 0.1 | 0.28 | 0.22 |
| 5 | 163124620 | rs531314 | 1.034 | 0.833 | 0.29 | 0.097 | 0.033 | 0.056 | 0.056 | 0 | 0.94 | 0.83 | 0.94 | 0.93 | 0.91 | 0.93 |
| 5 | 163126801 | rs489835 | 0.448 | 0.9 | -0.481 | -0.203 | -0.013 | 0.032 | 0.032 | 0 | 0.63 | 0.24 | 0.65 | 0.79 | 0.63 | 0.84 |
| 5 | 163131540 | rs2156273 | 0.016 | 0.946 | 0.179 | -0.37 | -0.007 | 0.012 | 0.012 | 0 | 0.74 | 0.53 | 0.75 | 0.79 | 0.78 | 0.92 |
| 5 | 163135314 | rs502112 | 0.526 | 1.226 | -0.021 | -0.475 | 0.007 | -0.004 | -0.004 | 1.661 | 0.69 | 0.5 | 0.68 | 0.76 | 0.69 | 0.73 |
| 5 | 163137051 | rs4869096 | -0.843 | 1.177 | 0.057 | -0.564 | 0.012 | -0.038 | -0.038 | 0.009 | 0.11 | 0.14 | 0.08 | 0 | 0.16 | 0.09 |
| 5 | 163137635 | rs516512 | 0.008 | 1.183 | 0.084 | -0.518 | -0.014 | 0.001 | 0.001 | 0 | 0.73 | 0.52 | 0.75 | 0.79 | 0.78 | 0.93 |
| 5 | 163137979 | rs520002 | 0.736 | 1.326 | 0.04 | -0.585 | 0 | 0.002 | 0.002 | 0 | 0.64 | 0.33 | 0.65 | 0.76 | 0.66 | 0.73 |
| 5 | 163143860 | rs580015 | 1.237 | 1.139 | 0.058 | -0.687 | -0.016 | 0.033 | 0.033 | 0 | 0.57 | 0.34 | 0.6 | 0.75 | 0.52 | 0.63 |
| 5 | 163146444 | rs7702817 | -1.997 | 1.166 | 0.034 | -0.754 | 0.013 | -0.037 | -0.037 | 1.377 | 0.12 | 0.25 | 0.1 | 0.03 | 0.14 | 0.23 |
| 5 | 163150126 | rs564735 | 1.711 | 1.13 | -0.151 | -0.729 | -0.015 | 0.118 | 0.118 | 0.047 | 0.5 | 0.11 | 0.55 | 0.75 | 0.47 | 0.59 |
| 5 | 163161444 | rs2033534 | -0.341 | 1.288 | 0 | -0.685 | 0.038 | -0.045 | -0.045 | 0 | 0.08 | 0.16 | 0.06 | 0.01 | 0.15 | 0.07 |
| 5 | 163165467 | rs17610788 | -1.58 | 1.277 | -0.108 | -0.688 | -0.024 | 0.023 | 0.023 | 0 | 0.15 | 0.1 | 0.14 | 0.04 | 0.27 | 0.26 |
| 5 | 163180840 | rs1943049 | 1.118 | 0.774 | 0.053 | -0.508 | -0.014 | 0.029 | 0.029 | 0 | 0.6 | 0.26 | 0.62 | 0.75 | 0.59 | 0.69 |
| 5 | 163187708 | rs1943047 | 0.83 | 0.843 | 0.062 | -0.498 | -0.015 | 0.019 | 0.019 | 0 | 0.62 | 0.32 | 0.64 | 0.77 | 0.59 | 0.86 |
| 5 | 163213383 | rs2212797 | 1.452 | 0.839 | -0.099 | -0.485 | -0.045 | 0.064 | 0.064 | 0.021 | 0.56 | 0.19 | 0.6 | 0.74 | 0.55 | 0.66 |
| 5 | 163221209 | rs10062993 | -0.21 | 0.663 | 0.19 | -0.499 | 0.023 | -0.153 | -0.153 | 0.428 | 0.77 | 0.59 | 0.83 | 1 | 0.79 | 0.93 |
| 5 | 163222890 | rs290143 | -0.004 | 0.634 | -0.341 | -0.547 | 0.007 | 0.032 | 0.032 | 0 | 0.13 | 0 | 0.1 | 0.07 | 0.1 | 0.12 |
| 5 | 163231017 | rs4426936 | 0.112 | 0.823 | -0.02 | -0.351 | -0.006 | 0.002 | 0.002 | 0 | 0.71 | 0.41 | 0.74 | 0.78 | 0.77 | 0.92 |
| 5 | 163256746 | rs2112250 | -1.446 | 0.938 | -0.069 | -0.376 | 0.019 | -0.039 | -0.039 | 0.001 | 0.15 | 0.22 | 0.14 | 0.04 | 0.22 | 0.25 |
| 5 | 163257741 | rs1833543 | -1.625 | 0.874 | -0.151 | -0.415 | -0.047 | 0.05 | 0.05 | 0.012 | 0.45 | 0.82 | 0.41 | 0.27 | 0.46 | 0.36 |
| 5 | 163263560 | rs7713636 | -0.2 | 0.869 | -0.104 | -0.406 | 0.021 | -0.036 | -0.036 | 0 | 0.08 | 0.09 | 0.09 | 0.01 | 0.15 | 0.07 |
| 5 | 163263914 | rs7714147 | -1.181 | 0.844 | -0.104 | -0.419 | -0.019 | 0.033 | 0.033 | 0 | 0.4 | 0.73 | 0.37 | 0.24 | 0.4 | 0.17 |
| 5 | 163264549 | rs10050929 | -0.098 | 0.78 | 0.082 | -0.34 | -0.001 | 0.011 | 0.011 | 0 | 0.28 | 0.52 | 0.26 | 0.23 | 0.24 | 0.08 |
| 5 | 163265421 | rs446288 | 1.919 | 0.817 | -0.272 | -0.318 | -0.023 | -0.002 | -0.002 | 0 | 0.43 | 0.18 | 0.49 | 0.66 | 0.43 | 0.51 |
| 5 | 163273631 | rs6881878 | -0.68 | 0.911 | -0.026 | -0.306 | 0.002 | 0 | 0 | 0 | 0.34 | 0.65 | 0.3 | 0.26 | 0.24 | 0.25 |
| 5 | 163275989 | rs7712391 | 0.042 | 0.889 | 0.27 | -0.33 | 0.003 | 0.014 | 0.014 | 0 | 0.27 | 0.46 | 0.26 | 0.23 | 0.23 | 0.08 |
| 5 | 163276078 | rs290113 | 0.259 | 1.004 | 0.213 | -0.206 | -0.028 | 0.046 | 0.046 | 0 | 0.79 | 0.92 | 0.81 | 0.88 | 0.72 | 0.81 |
| 5 | 163282404 | rs1363159 | -0.443 | 0.879 | 0.245 | -0.2 | -0.001 | -0.001 | -0.001 | 0.057 | 0.32 | 0.64 | 0.28 | 0.23 | 0.24 | 0.1 |
| 5 | 163290591 | rs1489525 | -1.834 | 0.998 | 0.208 | -0.112 | -0.004 | 0.017 | 0.017 | 0.031 | 0.08 | 0.22 | 0.08 | 0.1 | 0.05 | 0.19 |
| 5 | 163294029 | rs410056 | 0.528 | 1 | 0.338 | -0.161 | -0.006 | 0.015 | 0.015 | 0 | 0.78 | 0.86 | 0.8 | 0.9 | 0.72 | 0.81 |
| 5 | 163294054 | rs371486 | 1.149 | 0.998 | 0.179 | -0.121 | 0.001 | -0.042 | -0.042 | 0 | 0.7 | 0.62 | 0.72 | 0.81 | 0.66 | 0.58 |
| 5 | 163306841 | rs9313986 | 0.568 | 0.641 | 0.433 | 0.268 | -0.016 | -0.013 | -0.013 | 0 | 0.7 | 0.49 | 0.85 | 0.83 | 0.78 | 0.89 |
| 5 | 163313040 | rs17062468 | -1.254 | 1.343 | 0.168 | 0.555 | -0.01 | -0.01 | -0.01 | 0 | 0.07 | 0.03 | 0.14 | 0.14 | 0.12 | 0.19 |
| 5 | 163313224 | rs288488 | 1.791 | 1.261 | 0.538 | 0.635 | 0.008 | -0.009 | -0.009 | 0 | 0.46 | 0.39 | 0.46 | 0.57 | 0.42 | 0.55 |
| 5 | 163318721 | rs17062481 | -1.181 | 1.265 | 0.665 | 0.656 | -0.004 | -0.004 | -0.004 | 0 | 0.08 | 0.06 | 0.16 | 0.12 | 0.11 | 0.2 |
| 5 | 163320942 | rs11953746 | -0.506 | 1.027 | 0.942 | 0.401 | 0 | -0.004 | -0.004 | 0.068 | 0.21 | 0.19 | 0.25 | 0.13 | 0.25 | 0.17 |

| | | | | | | | | | | | | | | | | |
|---|---|---|---|---|---|---|---|---|---|---|---|---|---|---|---|---|
| 5 | 163324339 | rs257688 | -0.45 | 0.89 | 1.195 | 0.433 | 0.003 | -0.006 | -0.006 | 1.492 | 0.79 | 0.87 | 0.91 | 0.85 | 0.84 | 0.96 |
| 5 | 163335429 | rs1946198 | 1.708 | 0.882 | 1.346 | 0.467 | 0.004 | 0.002 | 0.002 | 0 | 0.48 | 0.39 | 0.52 | 0.6 | 0.48 | 0.59 |
| 5 | 163338570 | rs1345605 | 0.083 | 1.061 | 1.918 | 0.298 | -0.003 | 0.052 | 0.052 | 0.556 | 0.18 | 0.36 | 0.11 | 0.15 | 0.13 | 0.04 |
| 5 | 163339463 | rs989521 | 0.302 | 1.359 | 1.831 | 0.256 | 0.012 | 0.028 | 0.028 | 1.369 | 0.78 | 0.51 | 0.86 | 0.85 | 0.83 | 0.95 |
| 5 | 163341396 | rs2546322 | -1.156 | 1.465 | 1.708 | 0.488 | -0.006 | 0.006 | 0.006 | 0 | 0.27 | 0.23 | 0.34 | 0.21 | 0.34 | 0.35 |
| 5 | 163345048 | rs2544976 | -1.2 | 1.743 | 1.709 | 0.509 | -0.025 | 0.019 | 0.019 | 0.17 | 0.28 | 0.18 | 0.35 | 0.22 | 0.37 | 0.35 |
| 5 | 163345379 | rs17062522 | 2.021 | 1.589 | 1.782 | 0.482 | 0.034 | 0.02 | 0.02 | 0 | 0.5 | 0.28 | 0.5 | 0.63 | 0.45 | 0.58 |
| 5 | 163348798 | rs2544997 | 0.415 | 1.426 | 1.854 | 0.512 | 0.006 | 0.011 | 0.011 | 1.602 | 0.78 | 0.45 | 0.85 | 0.86 | 0.82 | 0.94 |
| 5 | 163352383 | rs10037467 | 2.598 | 1.337 | 1.679 | 0.246 | 0.034 | 0.166 | 0.166 | 0 | 0.43 | 0.17 | 0.21 | 0.06 | 0.35 | 0.4 |
| 5 | 163352472 | rs10067650 | 0.041 | 1.374 | 1.907 | 0.212 | -0.001 | 0.044 | 0.044 | 1.108 | 0.19 | 0.43 | 0.12 | 0.14 | 0.14 | 0.06 |
| 5 | 163354246 | rs1421868 | 1.98 | 1.43 | 1.687 | 0.472 | 0.022 | 0.006 | 0.006 | 0 | 0.51 | 0.28 | 0.52 | 0.58 | 0.47 | 0.58 |
| 5 | 163357967 | rs1422542 | -0.946 | 1.461 | 1.509 | 0.472 | -0.03 | 0.009 | 0.009 | 0.315 | 0.27 | 0.15 | 0.33 | 0.26 | 0.35 | 0.35 |
| 5 | 163374345 | rs11954477 | -1.469 | 1.442 | 1.561 | 0.367 | -0.002 | -0.001 | -0.001 | 0.957 | 0.45 | 0.42 | 0.42 | 0.41 | 0.47 | 0.44 |
| 5 | 163375406 | rs13159590 | 2.304 | 1.417 | 1.377 | 0.469 | 0.028 | 0.184 | 0.184 | 0 | 0.43 | 0.17 | 0.22 | 0.08 | 0.36 | 0.38 |
| 5 | 163378129 | rs1105581 | 1.812 | 1.408 | 1.549 | 0.467 | 0.019 | 0.026 | 0.026 | 0 | 0.5 | 0.22 | 0.51 | 0.59 | 0.48 | 0.56 |
| 5 | 163381044 | rs4868825 | 0.14 | 1.084 | 1.699 | 0.601 | -0.014 | 0.034 | 0.034 | 0 | 0.78 | 0.64 | 0.81 | 0.75 | 0.85 | 0.77 |
| 5 | 163382675 | rs7720295 | -1.298 | 1.097 | 1.667 | 0.533 | 0.007 | -0.035 | -0.035 | 0 | 0.26 | 0.37 | 0.26 | 0.16 | 0.31 | 0.2 |
| 5 | 163393047 | rs6878709 | 1.43 | 1.132 | 1.332 | 0.485 | -0.005 | -0.003 | -0.003 | 0 | 0.67 | 0.44 | 0.74 | 0.75 | 0.74 | 0.72 |
| 5 | 163400151 | rs11135295 | 0.115 | 1.286 | 0.989 | 0.721 | -0.001 | 0.002 | 0.002 | 0 | 0.78 | 0.8 | 0.82 | 0.75 | 0.85 | 0.77 |
| 5 | 163407199 | rs6889332 | -1.382 | 1.264 | 1.078 | 0.983 | 0.022 | -0.007 | -0.007 | 0.69 | 0.23 | 0.6 | 0.29 | 0.26 | 0.28 | 0.2 |
| 5 | 163407778 | rs10515868 | 1.836 | 1.188 | 0.898 | 0.618 | -0.012 | 0.007 | 0.007 | 0 | 0.56 | 0.19 | 0.55 | 0.59 | 0.58 | 0.56 |
| 5 | 163408317 | rs7734266 | 0.278 | 0.877 | 0.904 | 0.694 | -0.002 | 0.006 | 0.006 | 0.28 | 0.78 | 0.76 | 0.82 | 0.78 | 0.84 | 0.79 |
| 5 | 163408924 | rs954667 | -1.661 | 1.264 | 1.206 | 0.26 | 0.032 | -0.057 | -0.057 | 0.184 | 0.22 | 0.49 | 0.24 | 0.19 | 0.27 | 0.17 |
| 5 | 163409675 | rs13157489 | -1.881 | 1.362 | 1.068 | 0.113 | 0.003 | -0.074 | -0.074 | 0 | 0.28 | 0.52 | 0.3 | 0.19 | 0.32 | 0.18 |
| 5 | 163410578 | rs13436169 | -1.855 | 1.375 | 1.074 | 0.115 | 0.004 | -0.074 | -0.074 | 0 | 0.28 | 0.52 | 0.3 | 0.19 | 0.32 | 0.17 |
| 5 | 163418856 | rs1363073 | -1.268 | 1.365 | 0.663 | 0.268 | 0.001 | -0.012 | -0.012 | 0 | 0.21 | 0.26 | 0.24 | 0.19 | 0.24 | 0.13 |
| 5 | 163428473 | rs10454979 | -1.894 | 1.202 | 0.883 | 0.413 | 0.052 | -0.035 | -0.035 | 0 | 0.13 | 0.37 | 0.16 | 0.04 | 0.19 | 0.14 |
| 5 | 163431998 | rs6859007 | -0.852 | 1.41 | 0.749 | -0.024 | 0.001 | -0.048 | -0.048 | 0 | 0.36 | 0.45 | 0.3 | 0.18 | 0.33 | 0.16 |
| 5 | 163433005 | rs4868730 | -1.056 | 1.493 | 0.755 | 0.025 | 0.084 | 0.006 | 0.006 | 0 | 0.08 | 0.29 | 0.11 | 0.04 | 0.14 | 0.11 |
| 5 | 163438424 | rs9313997 | -1.493 | 1.412 | 0.259 | 0.006 | -0.032 | 0.183 | 0.183 | 0.147 | 0.46 | 0.78 | 0.42 | 0.21 | 0.47 | 0.27 |
| 5 | 163447265 | rs10056554 | -0.845 | 1.433 | 0.336 | 0.071 | -0.001 | 0.005 | 0.005 | 0 | 0.37 | 0.39 | 0.3 | 0.19 | 0.35 | 0.16 |
| 5 | 163450450 | rs7443765 | -0.838 | 1.429 | 0.29 | 0.067 | -0.001 | -0.003 | -0.003 | 0 | 0.37 | 0.38 | 0.29 | 0.19 | 0.33 | 0.16 |
| 5 | 163453678 | rs12188908 | -1.049 | 1.344 | 0.341 | -0.002 | 0.075 | 0.003 | 0.003 | 0 | 0.09 | 0.35 | 0.11 | 0.04 | 0.13 | 0.11 |
| 5 | 163465470 | rs7446327 | -1.344 | 1.314 | -0.248 | 0.062 | -0.023 | 0.182 | 0.182 | 0 | 0.47 | 0.78 | 0.41 | 0.22 | 0.46 | 0.26 |
| 5 | 163472839 | rs1363072 | -1.411 | 1.271 | -0.497 | 0.409 | -0.076 | 0.153 | 0.153 | 0 | 0.48 | 0.88 | 0.42 | 0.27 | 0.48 | 0.27 |
| 5 | 163502464 | rs888640 | -1.328 | 1.4 | -0.485 | 0.938 | 0.017 | 0.022 | 0.022 | 0 | 0.5 | 0.75 | 0.57 | 0.64 | 0.52 | 0.47 |
| 5 | 163502789 | rs877353 | -0.606 | 1.395 | -0.309 | 0.79 | 0.001 | 0.009 | 0.009 | 0 | 0.34 | 0.35 | 0.27 | 0.14 | 0.29 | 0.16 |
| 5 | 163508338 | rs6893586 | -0.932 | 1.452 | -0.644 | 1.035 | 0.005 | -0.016 | -0.016 | 0 | 0.43 | 0.71 | 0.5 | 0.62 | 0.42 | 0.36 |
| 5 | 163511478 | rs7443611 | -1.226 | 1.144 | -1.395 | 0.799 | 0 | 0.109 | 0.109 | 0 | 0.59 | 0.85 | 0.66 | 0.81 | 0.59 | 0.48 |
| 5 | 163511740 | rs9313998 | -0.508 | 0.924 | -1.175 | 0.868 | -0.002 | 0.011 | 0.011 | 0 | 0.38 | 0.78 | 0.41 | 0.39 | 0.36 | 0.33 |
| 5 | 163511874 | rs13163167 | -0.073 | 0.942 | -0.951 | 0.517 | 0.001 | 0.022 | 0.022 | 0 | 0.25 | 0.27 | 0.41 | 0.53 | 0.23 | 0.3 |
| 5 | 163511991 | rs9313999 | -0.741 | 0.86 | -1.535 | 0.659 | 0.013 | -0.023 | -0.023 | 0 | 0.55 | 0.78 | 0.61 | 0.57 | 0.57 | 0.45 |
| 5 | 163517674 | rs10462943 | -0.252 | 1.133 | -1.374 | 0.803 | -0.009 | -0.043 | -0.043 | 0 | 0.34 | 0.2 | 0.43 | 0.54 | 0.36 | 0.28 |
| 5 | 163517699 | rs10462944 | -0.278 | 1.157 | -1.363 | 0.814 | -0.008 | -0.043 | -0.043 | 0 | 0.35 | 0.2 | 0.43 | 0.54 | 0.36 | 0.3 |
| 5 | 163520613 | rs12517450 | -0.49 | 1.11 | -1.619 | 0.778 | -0.014 | -0.003 | -0.003 | 0 | 0.64 | 0.85 | 0.65 | 0.66 | 0.62 | 0.55 |
| 5 | 163523259 | rs6899006 | -0.474 | 1.177 | -1.244 | 0.834 | 0.004 | 0.01 | 0.01 | 0 | 0.38 | 0.4 | 0.34 | 0.46 | 0.39 | 0.27 |
| 5 | 163526403 | rs6875996 | 0.869 | 1.326 | -1.068 | 0.753 | 0.005 | 0.001 | 0.001 | 0 | 0.66 | 0.71 | 0.69 | 0.6 | 0.63 | 0.57 |
| 5 | 163529022 | rs13188234 | 0.289 | 1.355 | -1.122 | 0.666 | 0.002 | 0.002 | 0.002 | 0 | 0.77 | 0.8 | 0.74 | 0.6 | 0.69 | 0.67 |
| 5 | 163530578 | rs1895139 | 1.206 | 1.323 | -1.134 | 0.628 | -0.016 | -0.002 | -0.002 | 0.076 | 0.64 | 0.75 | 0.69 | 0.6 | 0.63 | 0.58 |
| 5 | 163533508 | rs1421630 | 0.404 | 1.84 | -0.941 | 1.136 | -0.017 | -0.017 | -0.017 | 0 | 0.79 | 0.85 | 0.75 | 0.6 | 0.71 | 0.69 |
| 5 | 163541120 | rs7701462 | -0.755 | 2.157 | -1.191 | 1.262 | 0.034 | 0.068 | 0.068 | 0 | 0.07 | 0.12 | 0.15 | 0.22 | 0.11 | 0.2 |
| 5 | 163542018 | rs4868852 | 0.014 | 1.812 | -1.725 | 0.867 | -0.091 | -0.07 | -0.07 | 0 | 0.85 | 0.99 | 0.88 | 0.8 | 0.79 | 0.88 |
| 5 | 163549241 | rs7447133 | 0.149 | 0.958 | -1.845 | 0.34 | 0.007 | 0.03 | 0.03 | 0 | 0.25 | 0.11 | 0.21 | 0.14 | 0.25 | 0.28 |
| 5 | 163551178 | rs4868854 | 0.469 | 0.777 | -2.021 | -0.027 | 0.017 | 0.081 | 0.081 | 0 | 0.59 | 0.76 | 0.61 | 0.8 | 0.61 | 0.54 |
| 5 | 163555945 | rs2052446 | 0.057 | 0.911 | -1.275 | 0.133 | 0.003 | 0.022 | 0.022 | 0 | 0.85 | 0.89 | 0.82 | 0.94 | 0.85 | 0.81 |
| 5 | 163580334 | rs10070091 | 1.144 | 1.004 | -1.395 | 0.161 | -0.02 | 0.019 | 0.019 | 0 | 0.62 | 0.73 | 0.63 | 0.72 | 0.6 | 0.67 |
| 5 | 163586269 | rs7732882 | 0.225 | 0.973 | -1.442 | 0.137 | -0.064 | -0.019 | -0.019 | 0 | 0.75 | 0.93 | 0.83 | 0.76 | 0.72 | 0.75 |
| 5 | 163586956 | rs4868858 | -0.075 | 0.979 | -1.43 | 0.137 | -0.004 | -0.011 | -0.011 | 0 | 0.13 | 0.16 | 0.19 | 0.05 | 0.13 | 0.11 |
| 5 | 163589883 | rs2901801 | 0.296 | 0.995 | -1.421 | 0.154 | -0.029 | -0.002 | -0.002 | 0 | 0.74 | 0.86 | 0.82 | 0.78 | 0.72 | 0.75 |
| 5 | 163601262 | rs1990943 | 1.365 | 0.833 | -1.378 | 0.088 | -0.014 | 0.021 | 0.021 | 0 | 0.61 | 0.69 | 0.63 | 0.74 | 0.6 | 0.64 |
| 5 | 163602147 | rs1422767 | -1.413 | 0.999 | -1.574 | 0.195 | -0.02 | 0.028 | 0.028 | 0 | 0.94 | 1 | 0.98 | 1 | 0.94 | 0.92 |
| 5 | 163602752 | rs12188891 | 0.275 | 0.977 | -1.586 | 0.194 | -0.095 | -0.079 | -0.079 | 0.261 | 0.18 | 0.01 | 0.15 | 0.21 | 0.22 | 0.17 |
| 5 | 163617119 | rs6872031 | 0.549 | 0.782 | -1.102 | 0.067 | -0.014 | 0.003 | 0.003 | 0 | 0.68 | 0.81 | 0.68 | 0.73 | 0.67 | 0.62 |
| 5 | 163617442 | rs988498 | -0.923 | 0.749 | -1.121 | 0.034 | -0.003 | 0.003 | 0.003 | 0 | 0.36 | 0.32 | 0.4 | 0.26 | 0.38 | 0.3 |
| 5 | 163619697 | rs1159332 | -0.904 | 0.753 | -1.119 | 0.006 | -0.003 | 0.002 | 0.002 | 0 | 0.36 | 0.32 | 0.4 | 0.27 | 0.39 | 0.3 |
| 5 | 163621832 | rs2161384 | 0.334 | 0.774 | -1.323 | 0.029 | -0.092 | -0.003 | -0.003 | 0 | 0.7 | 0.94 | 0.78 | 0.79 | 0.68 | 0.72 |
| 5 | 163623015 | rs12188542 | -0.854 | 0.768 | -1.287 | 0.036 | -0.045 | 0.015 | 0.015 | 0 | 0.35 | 0.17 | 0.39 | 0.26 | 0.38 | 0.29 |
| 5 | 163630080 | rs4868871 | 1.031 | 0.79 | -1.21 | -0.098 | -0.02 | 0.044 | 0.044 | 0 | 0.59 | 0.72 | 0.67 | 0.78 | 0.56 | 0.71 |
| 5 | 163633239 | rs7446825 | 1.229 | 0.781 | -1.224 | 0.122 | 0 | -0.001 | -0.001 | 1.003 | 0.45 | 0.5 | 0.47 | 0.57 | 0.41 | 0.51 |

| | | | | | | | | | | | | | | | | |
|---|---|---|---|---|---|---|---|---|---|---|---|---|---|---|---|---|
| 5 | 163634891 | rs10042643 | -0.109 | 0.709 | -0.866 | 0.367 | -0.021 | -0.022 | -0.022 | 0 | 0.25 | 0.41 | 0.3 | 0.22 | 0.27 | 0.2 |
| 5 | 163645278 | rs17063000 | -0.433 | 1.157 | -1.37 | 0.412 | -0.065 | -0.023 | -0.023 | 0 | 0.23 | 0.05 | 0.21 | 0.21 | 0.26 | 0.2 |
| 5 | 163675157 | rs17063034 | -1.049 | 0.315 | -0.284 | -0.055 | -0.022 | -0.035 | -0.035 | 0 | 0.08 | 0.36 | 0.04 | 0 | 0.01 | 0.02 |
| 5 | 163698956 | rs17285868 | 0.035 | 0.843 | -0.83 | -0.343 | -0.056 | 0.046 | 0.046 | 0 | 0.12 | 0.01 | 0.13 | 0.03 | 0.18 | 0.05 |
| 5 | 163702195 | rs2964917 | -1.466 | 0.809 | -0.9 | -0.343 | 0.001 | -0.035 | -0.035 | 0 | 0.37 | 0.51 | 0.48 | 0.29 | 0.37 | 0.38 |
| 5 | 163713876 | rs4091221 | -0.765 | 1.107 | -1.366 | -0.472 | -0.066 | 0.101 | 0.101 | 0 | 0.23 | 0.05 | 0.29 | 0.05 | 0.27 | 0.26 |
| 5 | 163714075 | rs2964912 | -1.08 | 0.765 | -1.316 | -0.095 | -0.013 | -0.028 | -0.028 | 1.136 | 0.85 | 0.95 | 0.89 | 0.86 | 0.83 | 0.88 |
| 5 | 163731007 | rs3926173 | -0.115 | 0.711 | -0.865 | -0.047 | 0.004 | 0.018 | 0.018 | 0 | 0.16 | 0.22 | 0.16 | 0.25 | 0.15 | 0.11 |
| 5 | 163739709 | rs2915850 | -0.721 | -0.006 | -1.01 | -0.538 | 0.014 | 0.029 | 0.029 | 0.073 | 0.75 | 0.82 | 0.83 | 0.93 | 0.79 | 0.74 |
| 5 | 163742363 | rs10515871 | -1.132 | 0.129 | -0.28 | -0.09 | -0.041 | 0.041 | 0.041 | 0 | 0.2 | 0.62 | 0.32 | 0.33 | 0.14 | 0.22 |
| 5 | 163748783 | rs11745553 | 0.394 | 0.238 | -0.548 | -0.374 | -0.007 | 0.083 | 0.083 | 2.308 | 0.21 | 0.08 | 0.27 | 0.02 | 0.22 | 0.06 |
| 5 | 163749514 | rs17063165 | -0.52 | 0.095 | 0.757 | 0.346 | 0.019 | 0.132 | 0.132 | 0 | 0.08 | 0.41 | 0.15 | 0.17 | 0.05 | 0.03 |
| 5 | 163753090 | rs1030674 | -1.898 | 0.131 | 0.061 | 0.306 | -0.007 | 0.009 | 0.009 | 0 | 0.24 | 0.66 | 0.36 | 0.33 | 0.22 | 0.23 |
| 5 | 163753765 | rs11741206 | -1.412 | 0.227 | 0.217 | 0.601 | 0.008 | 0.15 | 0.15 | 0.031 | 0.15 | 0.5 | 0.31 | 0.32 | 0.15 | 0.16 |
| 5 | 163763100 | rs17063176 | -0.532 | 0.368 | -0.658 | 0.644 | 0.016 | 0.013 | 0.013 | 0.53 | 0.23 | 0.11 | 0.31 | 0.21 | 0.2 | 0.11 |
| 5 | 163767509 | rs17063187 | -0.99 | 0.644 | -0.101 | 0.802 | 0.01 | 0.067 | 0.067 | 1.054 | 0.07 | 0.16 | 0.1 | 0.14 | 0.09 | 0.21 |
| 5 | 163770824 | rs6874303 | -0.169 | -0.461 | -0.236 | 0.463 | -0.051 | 0.001 | 0.001 | 0.153 | 0.28 | 0.54 | 0.2 | 0.22 | 0.18 | 0.31 |
| 5 | 163771041 | rs17287615 | 0.075 | -0.008 | -0.321 | 0.539 | -0.005 | -0.014 | -0.014 | 0.887 | 0.42 | 0.38 | 0.52 | 0.59 | 0.4 | 0.47 |
| 5 | 163775141 | rs7719164 | 0.044 | -0.505 | 0.165 | 0.496 | -0.026 | -0.001 | -0.001 | 0 | 0.1 | 0.25 | 0.08 | 0.05 | 0.06 | 0.06 |
| 5 | 163776271 | rs4563644 | -0.843 | -0.493 | -0.346 | 0.439 | -0.011 | 0.028 | 0.028 | 0 | 0.43 | 0.75 | 0.59 | 0.64 | 0.42 | 0.58 |
| 5 | 163778421 | rs10515876 | 0.235 | -0.543 | -0.298 | 0.559 | -0.045 | -0.084 | -0.084 | 1.049 | 0.25 | 0.12 | 0.5 | 0.5 | 0.31 | 0.3 |
| 5 | 163779956 | rs7726288 | -0.817 | -0.545 | -0.372 | 0.299 | -0.015 | 0.008 | 0.008 | 0 | 0.66 | 0.75 | 0.69 | 0.72 | 0.61 | 0.7 |
| 5 | 163780571 | rs10059065 | -0.42 | -0.453 | 0.063 | 1.009 | -0.019 | 0.024 | 0.024 | 0 | 0.12 | 0.17 | 0.14 | 0.15 | 0.07 | 0.19 |
| 5 | 163786953 | rs11135310 | -0.028 | -0.23 | -0.505 | 1.037 | 0.019 | 0.087 | 0.087 | 0.127 | 0.27 | 0.53 | 0.47 | 0.54 | 0.31 | 0.27 |
| 5 | 163789191 | rs17360177 | -0.853 | -0.255 | -0.581 | 0.947 | -0.001 | 0.01 | 0.01 | 0.019 | 0.38 | 0.64 | 0.61 | 0.7 | 0.37 | 0.44 |
| 5 | 163797230 | rs6556745 | -0.486 | 0.577 | -0.47 | 1.035 | 0 | 0.055 | 0.055 | 0 | 0.33 | 0.46 | 0.46 | 0.52 | 0.31 | 0.25 |
| 5 | 163803300 | rs4868875 | 0.063 | 0.529 | -0.716 | 0.795 | 0.002 | 0.002 | 0.002 | 0 | 0.59 | 0.39 | 0.71 | 0.6 | 0.65 | 0.69 |
| 5 | 163811558 | rs3923169 | -0.701 | 0.924 | -0.539 | 1.212 | 0.006 | 0.029 | 0.029 | 0.006 | 0.27 | 0.31 | 0.4 | 0.39 | 0.3 | 0.39 |
| 5 | 163815143 | rs6556747 | -0.85 | 0.894 | -0.434 | 1.031 | -0.029 | 0.029 | 0.029 | 0 | 0.14 | 0.08 | 0.12 | 0.01 | 0.21 | 0.07 |
| 5 | 163818238 | rs9314018 | -1.864 | 0.494 | 0.059 | 1.325 | 0.002 | -0.01 | -0.01 | 0 | 0.36 | 0.57 | 0.47 | 0.66 | 0.4 | 0.52 |
| 5 | 163820512 | rs10058433 | -0.054 | 0.515 | 0.09 | 1.454 | -0.016 | 0.244 | 0.244 | 0 | 0.12 | 0.27 | 0.25 | 0.4 | 0.07 | 0.29 |
| 5 | 163836520 | rs17063302 | -0.401 | 0.554 | 0.065 | 1.682 | 0.001 | 0.005 | 0.005 | 0 | 0.09 | 0.07 | 0.08 | 0.3 | 0.1 | 0.15 |
| 5 | 163847072 | rs4374775 | -0.66 | 0.332 | -0.207 | 1.397 | -0.06 | 0.062 | 0.062 | 0 | 0.11 | 0.02 | 0.12 | 0.01 | 0.22 | 0.06 |
| 5 | 163849800 | rs10515877 | -0.09 | 0.381 | 0.001 | 1.461 | -0.007 | 0.148 | 0.148 | 0 | 0.07 | 0.18 | 0.07 | 0.3 | 0.08 | 0.15 |
| 5 | 163856389 | rs4573014 | 0.2 | -0.251 | -0.279 | 0.586 | -0.023 | -0.026 | -0.026 | 0 | 0.33 | 0.51 | 0.38 | 0.21 | 0.25 | 0.29 |
| 5 | 163862515 | rs6898157 | 0.732 | -0.193 | -0.277 | 0.692 | -0.001 | 0.022 | 0.022 | 0.779 | 0.51 | 0.4 | 0.45 | 0.45 | 0.49 | 0.54 |
| 5 | 163863245 | rs13189611 | 1.027 | 0.332 | -0.608 | 0.061 | -0.011 | 0.138 | 0.138 | 0.132 | 0.33 | 0.11 | 0.19 | 0.01 | 0.31 | 0.27 |
| 5 | 163867010 | rs4130719 | 0.724 | 0.418 | -0.263 | 0.331 | -0.021 | -0.025 | -0.025 | 0 | 0.32 | 0.45 | 0.37 | 0.21 | 0.25 | 0.29 |
| 5 | 163867990 | rs9314024 | 0.246 | 0.44 | -0.409 | 0.446 | -0.019 | -0.03 | -0.03 | 0 | 0.84 | 0.89 | 0.82 | 0.68 | 0.75 | 0.82 |
| 5 | 163870623 | rs7448596 | 0.638 | 0.365 | -0.096 | 0.368 | -0.014 | -0.019 | -0.019 | 0 | 0.32 | 0.43 | 0.38 | 0.21 | 0.26 | 0.29 |
| 5 | 163880552 | rs12516149 | 0.612 | 0.814 | -0.274 | 0.331 | -0.02 | 0.018 | 0.018 | 0 | 0.08 | 0.04 | 0.09 | 0.01 | 0.15 | 0.04 |
| 5 | 163881521 | rs17063392 | 0.651 | 0.818 | -0.272 | 0.332 | -0.02 | 0.018 | 0.018 | 0 | 0.07 | 0.04 | 0.09 | 0.01 | 0.15 | 0.04 |
| 5 | 163882988 | rs13164902 | 0.495 | 0.733 | -0.208 | 0.388 | -0.001 | 0.046 | 0.046 | 0.344 | 0.51 | 0.31 | 0.42 | 0.41 | 0.48 | 0.53 |
| 5 | 163892077 | rs6556758 | 0.547 | 0.876 | -0.207 | 0.587 | 0.003 | 0.014 | 0.014 | 0 | 0.84 | 0.82 | 0.81 | 0.68 | 0.74 | 0.82 |
| 5 | 163892479 | rs6556759 | -0.413 | 0.843 | -0.186 | 0.618 | 0.005 | 0.132 | 0.132 | 0 | 0.91 | 0.87 | 0.89 | 0.6 | 0.89 | 0.87 |
| 5 | 163894378 | rs7445653 | -0.41 | 0.848 | -0.197 | 0.545 | 0.001 | 0.068 | 0.068 | 0 | 0.91 | 0.89 | 0.9 | 0.64 | 0.89 | 0.87 |
| 5 | 163897418 | rs1352547 | 0.522 | 0.871 | -0.192 | 0.523 | 0.004 | 0.021 | 0.021 | 0 | 0.84 | 0.81 | 0.81 | 0.64 | 0.74 | 0.82 |
| 5 | 163901446 | rs1387395 | 0.405 | 0.841 | -0.238 | 0.557 | -0.015 | -0.024 | -0.024 | 0.67 | 0.84 | 0.86 | 0.82 | 0.64 | 0.75 | 0.82 |
| 5 | 163902176 | rs17063513 | 0.68 | 0.694 | -0.105 | 0.516 | -0.014 | -0.03 | -0.03 | 6.942 | 0.31 | 0.42 | 0.37 | 0.18 | 0.25 | 0.27 |
| 5 | 163912271 | rs17369872 | -0.507 | 0.759 | 0.75 | 0.557 | -0.016 | 0.023 | 0.023 | 0.367 | 0.48 | 0.72 | 0.57 | 0.54 | 0.51 | 0.45 |
| 5 | 163920917 | rs9314037 | 0.593 | 0.747 | 0.801 | 0.514 | -0.003 | -0.006 | -0.006 | 0 | 0.32 | 0.61 | 0.39 | 0.18 | 0.26 | 0.27 |
| 5 | 163922165 | rs10515882 | 0.713 | 0.777 | 0.773 | 0.499 | -0.023 | 0.022 | 0.022 | 0 | 0.08 | 0.04 | 0.08 | 0 | 0.15 | 0.04 |
| 5 | 163927797 | rs10045861 | 0.817 | 0.72 | 1.094 | 0.523 | -0.002 | -0.003 | -0.003 | 0 | 0.3 | 0.28 | 0.37 | 0.21 | 0.25 | 0.27 |
| 5 | 163928148 | rs17068611 | 0.612 | 0.739 | 0.996 | 0.535 | 0.001 | -0.016 | -0.016 | 0 | 0.08 | 0.07 | 0.09 | 0 | 0.15 | 0.05 |
| 5 | 163929870 | rs9314038 | -0.055 | 1.158 | 0.908 | 0.487 | -0.018 | 0.054 | 0.054 | 0.622 | 0.41 | 0.71 | 0.48 | 0.23 | 0.42 | 0.38 |
| 5 | 163941186 | rs357181 | 1.359 | 0.551 | 0.435 | -0.004 | -0.01 | 0.09 | 0.09 | 0.067 | 0.24 | 0.09 | 0.15 | 0.02 | 0.24 | 0.21 |
| 5 | 163948616 | rs2054923 | -0.401 | 0.927 | 0.451 | 0.454 | -0.041 | -0.043 | -0.043 | 0 | 0.9 | 0.94 | 0.87 | 0.79 | 0.82 | 0.9 |
| 5 | 163949907 | rs890781 | 0.27 | 0.651 | 0.485 | 0.267 | -0.013 | -0.026 | -0.026 | 0.393 | 0.43 | 0.25 | 0.41 | 0.57 | 0.44 | 0.5 |
| 5 | 163953132 | rs2932132 | -0.173 | 0.661 | 0.782 | 0.208 | -0.001 | 0.002 | 0.002 | 0 | 0.51 | 0.59 | 0.48 | 0.53 | 0.51 | 0.52 |
| 5 | 163954210 | rs17068422 | 1.846 | 0.754 | 0.349 | 0.209 | -0.052 | -0.065 | -0.065 | 0 | 0.1 | 0.01 | 0.21 | 0.35 | 0.15 | 0.17 |
| 5 | 163957570 | rs11958300 | 1.1 | 0.53 | 0.668 | 0.102 | 0.027 | -0.025 | -0.025 | 0 | 0.06 | 0.18 | 0.06 | 0 | 0.07 | 0.02 |
| 5 | 163967787 | rs1428076 | -0.419 | 0.523 | 0.454 | 0.106 | -0.007 | -0.026 | -0.026 | 0 | 0.56 | 0.72 | 0.58 | 0.44 | 0.56 | 0.49 |
| 5 | 163977979 | rs9314040 | -0.443 | 0.603 | 0.477 | 0.117 | -0.003 | -0.028 | -0.028 | 0 | 0.56 | 0.75 | 0.59 | 0.44 | 0.57 | 0.51 |
| 5 | 163988372 | rs10043471 | 0.929 | 0.54 | 0.569 | 0.206 | 0.017 | -0.03 | -0.03 | 2.553 | 0.06 | 0.15 | 0.06 | 0 | 0.07 | 0.02 |
| 5 | 163992315 | rs165064 | -0.589 | -0.096 | 0.31 | 0.111 | -0.018 | 0.023 | 0.023 | 0 | 0.34 | 0.49 | 0.42 | 0.35 | 0.29 | 0.38 |
| 5 | 163993043 | rs165065 | 0.116 | 0.389 | 0.884 | -0.039 | 0.14 | 0.019 | 0.019 | 0 | 0.91 | 0.65 | 0.87 | 0.91 | 0.83 | 0.9 |
| 5 | 164001035 | rs165070 | -0.834 | 0.416 | 0.707 | -0.027 | 0.036 | -0.005 | -0.005 | 0 | 0.93 | 0.89 | 0.91 | 0.93 | 0.83 | 0.91 |
| 5 | 164019936 | rs165071 | 0.979 | 0.378 | 0.848 | -0.448 | -0.02 | 0.063 | 0.063 | 0 | 0.58 | 0.73 | 0.52 | 0.72 | 0.51 | 0.56 |
| 5 | 164056594 | rs17065898 | 0.971 | 0.131 | 0.74 | -0.662 | 0.023 | -0.11 | -0.11 | 1.799 | 0.18 | 0.38 | 0.15 | 0.02 | 0.21 | 0.06 |

| | | | | | | | | | | | | | | | | |
|---|---|---|---|---|---|---|---|---|---|---|---|---|---|---|---|---|
| 5 | 164088373 | rs1345611 | 1.468 | 0.173 | 0.485 | -0.828 | 0.006 | 0.092 | 0.092 | 0.008 | 0.5 | 0.28 | 0.47 | 0.71 | 0.47 | 0.57 |
| 5 | 164103083 | rs10077775 | -0.534 | 0.1 | 0.236 | -0.554 | 0.003 | 0.008 | 0.008 | 0 | 0.8 | 0.73 | 0.66 | 0.68 | 0.75 | 0.62 |
| 5 | 164104963 | rs9918105 | 0.963 | -0.013 | 0.191 | -0.63 | -0.002 | 0.003 | 0.003 | 0.061 | 0.57 | 0.41 | 0.48 | 0.66 | 0.48 | 0.53 |
| 5 | 164114596 | rs17066799 | 0.686 | -0.254 | 0.365 | -0.536 | 0.011 | -0.032 | -0.032 | 0 | 0.15 | 0.34 | 0.11 | 0.1 | 0.13 | 0.08 |
| 5 | 164148598 | rs1299355 | -0.473 | -0.215 | 0.085 | -0.584 | -0.005 | -0.014 | -0.014 | 0.189 | 0.79 | 0.8 | 0.89 | 0.89 | 0.84 | 0.88 |
| 5 | 164150604 | rs4868886 | 2.38 | -0.203 | -1.062 | -1.259 | -0.075 | 0.194 | 0.194 | 0 | 0.29 | 0.02 | 0.19 | 0.03 | 0.33 | 0.25 |
| 5 | 164150931 | rs2547009 | 0.71 | 0.003 | -0.677 | -0.674 | -0.003 | -0.01 | -0.01 | 0 | 0.19 | 0.19 | 0.37 | 0.24 | 0.23 | 0.34 |
| 5 | 164170049 | rs11953266 | 1.066 | -0.044 | -0.641 | -0.727 | 0.023 | 0.007 | 0.007 | 0 | 0.57 | 0.25 | 0.48 | 0.6 | 0.6 | 0.48 |
| 5 | 164170398 | rs11135323 | 1.507 | -0.084 | -0.767 | -0.792 | 0.037 | 0.071 | 0.071 | 0.062 | 0.5 | 0.21 | 0.48 | 0.62 | 0.56 | 0.48 |
| 5 | 164172766 | rs10043454 | 0.997 | 0.053 | -0.627 | -0.632 | 0.017 | 0.018 | 0.018 | 0 | 0.58 | 0.32 | 0.49 | 0.66 | 0.61 | 0.54 |
| 5 | 164177109 | rs10061064 | -0.877 | 0.036 | -0.573 | -0.726 | -0.003 | -0.004 | -0.004 | 0 | 0.81 | 0.82 | 0.68 | 0.8 | 0.77 | 0.67 |
| 5 | 164179112 | rs10063667 | 0.67 | 0.108 | -0.694 | -0.644 | 0.014 | 0.03 | 0.03 | 0 | 0.59 | 0.32 | 0.54 | 0.69 | 0.61 | 0.54 |
| 5 | 164199304 | rs17068161 | 0.426 | -0.349 | -0.584 | -0.718 | -0.015 | -0.022 | -0.022 | 0 | 0.12 | 0.15 | 0.05 | 0.06 | 0.07 | 0.1 |
| 5 | 164202406 | rs11739100 | 1.637 | 0.113 | -1.055 | -0.884 | -0.106 | 0.188 | 0.188 | 0 | 0.31 | 0.02 | 0.28 | 0.05 | 0.38 | 0.28 |
| 5 | 164222067 | rs2909760 | -0.358 | 0.397 | -0.262 | 0.006 | -0.008 | -0.025 | -0.025 | 0 | 0.29 | 0.76 | 0.32 | 0.6 | 0.26 | 0.25 |
| 5 | 164234310 | rs7717101 | -0.041 | 0.292 | 0.255 | 0.111 | 0.018 | 0.099 | 0.099 | 0.545 | 0.73 | 0.49 | 0.71 | 0.43 | 0.75 | 0.75 |
| 5 | 164237749 | rs4259187 | -0.336 | 0.312 | 0.297 | 0.126 | 0.011 | 0.098 | 0.098 | 0.23 | 0.76 | 0.56 | 0.73 | 0.52 | 0.78 | 0.82 |
| 5 | 164242082 | rs6896586 | -0.344 | 0.307 | -0.387 | 0.128 | 0.019 | -0.038 | -0.038 | 0 | 0.3 | 0.21 | 0.32 | 0.41 | 0.25 | 0.46 |
| 5 | 164259172 | rs7735637 | -0.028 | 0.23 | -0.237 | 0.671 | 0.001 | 0.015 | 0.015 | 0.061 | 0.76 | 0.71 | 0.74 | 0.52 | 0.77 | 0.83 |
| 5 | 164279457 | rs10042881 | 0.294 | 0.412 | -0.282 | 0.679 | 0 | 0.015 | 0.015 | 0.174 | 0.21 | 0.24 | 0.37 | 0.49 | 0.22 | 0.15 |
| 5 | 164284810 | rs4421125 | 0.055 | 0.18 | -0.402 | 0.137 | -0.012 | -0.01 | -0.01 | 0 | 0.62 | 0.71 | 0.61 | 0.58 | 0.53 | 0.43 |
| 5 | 164286098 | rs2963134 | -0.627 | -0.248 | -0.514 | 0.263 | 0.061 | -0.007 | -0.007 | 0 | 0.57 | 0.96 | 0.7 | 0.67 | 0.67 | 0.55 |
| 5 | 164308225 | rs3923448 | -0.396 | -0.131 | -0.466 | 0.386 | 0.008 | 0.002 | 0.002 | 0 | 0.52 | 0.49 | 0.66 | 0.61 | 0.62 | 0.49 |
| 5 | 164314792 | rs2963114 | 1.218 | 0.027 | -0.697 | 0.442 | -0.001 | 0.004 | 0.004 | 0 | 0.07 | 0.03 | 0.03 | 0.07 | 0.07 | 0.11 |
| 5 | 164319435 | rs4583905 | 0.243 | 0.449 | -0.744 | 0.554 | -0.018 | -0.03 | -0.03 | 7.215 | 0.24 | 0.18 | 0.41 | 0.34 | 0.26 | 0.25 |
| 5 | 164321629 | rs6556784 | 0.108 | 0.388 | -0.44 | 0.018 | 0.014 | -0.021 | -0.021 | 0.4 | 0.31 | 0.33 | 0.28 | 0.16 | 0.4 | 0.28 |
| 5 | 164322053 | rs2962867 | -1.191 | -0.296 | -0.188 | -0.177 | -0.014 | -0.036 | -0.036 | 0 | 0.68 | 0.69 | 0.83 | 0.85 | 0.76 | 0.76 |
| 5 | 164325241 | rs2962868 | -0.109 | 0.563 | 0.031 | 0.435 | -0.006 | -0.01 | -0.01 | 0.159 | 0.29 | 0.21 | 0.26 | 0.32 | 0.34 | 0.4 |
| 5 | 164334097 | rs13171726 | 0.06 | 0.53 | 0.337 | 0.273 | 0.022 | 0.091 | 0.091 | 0.242 | 0.25 | 0.6 | 0.48 | 0.47 | 0.27 | 0.25 |
| 5 | 164337870 | rs2962872 | 0.013 | 0.522 | 0.439 | 0.225 | 0.001 | 0.002 | 0.002 | 0.144 | 0.31 | 0.28 | 0.28 | 0.32 | 0.35 | 0.37 |
| 5 | 164339317 | rs4516878 | 0.297 | 0.475 | 0.929 | 0.174 | 0.019 | 0.067 | 0.067 | 0 | 0.2 | 0.3 | 0.45 | 0.46 | 0.24 | 0.3 |
| 5 | 164341317 | rs10038674 | -0.139 | 0.378 | 1.057 | 0.123 | 0.018 | 0.065 | 0.065 | 0.082 | 0.23 | 0.39 | 0.48 | 0.47 | 0.27 | 0.24 |
| 5 | 164346536 | rs7734804 | 1.317 | 0.456 | 1.232 | 0.18 | 0.023 | 0.002 | 0.002 | 0.122 | 0.05 | 0.25 | 0.04 | 0 | 0.03 | 0.02 |
| 5 | 164368235 | rs10064772 | -1.386 | 0.343 | 1.143 | 0.065 | 0.035 | 0.101 | 0.101 | 0.345 | 0.52 | 0.71 | 0.73 | 0.78 | 0.6 | 0.65 |
| 5 | 164369360 | rs6867769 | 0.229 | 0.64 | 1.273 | 0.317 | 0.022 | 0.072 | 0.072 | 0 | 0.17 | 0.27 | 0.43 | 0.46 | 0.23 | 0.29 |
| 5 | 164370690 | rs10066478 | 1.349 | 0.737 | 1.195 | 0.529 | -0.009 | 0.006 | 0.006 | 0 | 0.13 | 0.13 | 0.4 | 0.38 | 0.2 | 0.18 |
| 5 | 164391457 | rs4429870 | 0.23 | 0.136 | 1.323 | 0.231 | -0.011 | -0.012 | -0.012 | 0 | 0.73 | 0.81 | 0.73 | 0.7 | 0.65 | 0.69 |
| 5 | 164396328 | rs11750806 | -1.129 | 0.151 | 1.064 | 0.355 | -0.011 | 0.088 | 0.088 | 0 | 0.91 | 0.87 | 0.92 | 0.75 | 0.92 | 0.79 |
| 5 | 164413592 | rs10057578 | 0.49 | 0.081 | -0.066 | 0.224 | 0.002 | 0.004 | 0.004 | 0.346 | 0.67 | 0.43 | 0.67 | 0.68 | 0.63 | 0.59 |
| 5 | 164435168 | rs13175452 | 0.493 | -0.073 | -1.153 | -0.801 | -0.033 | 0.23 | 0.23 | 0 | 0.46 | 0.22 | 0.32 | 0.1 | 0.43 | 0.35 |
| 5 | 164461215 | rs4392642 | -0.716 | 0.026 | -0.393 | 0.281 | 0.006 | 0.019 | 0.019 | 0 | 0.12 | 0.24 | 0.17 | 0.13 | 0.15 | 0.13 |
| 5 | 164490804 | rs4571494 | 0.517 | 0.312 | -0.633 | 0.295 | 0.004 | 0.017 | 0.017 | 0.087 | 0.2 | 0.27 | 0.41 | 0.61 | 0.27 | 0.36 |
| 5 | 164493042 | rs6879291 | -1.322 | -0.399 | -1.452 | -0.045 | 0.044 | 0.091 | 0.091 | 0 | 0.72 | 0.96 | 0.89 | 0.89 | 0.83 | 0.75 |
| 5 | 164507412 | rs1946187 | -0.079 | -0.067 | -0.276 | -0.254 | 0.02 | 0.011 | 0.011 | 0 | 0.33 | 0.59 | 0.53 | 0.61 | 0.39 | 0.45 |
| 5 | 164511382 | rs17072568 | 0.099 | -0.121 | -0.615 | -0.253 | 0.003 | 0.002 | 0.002 | 0 | 0.11 | 0.08 | 0.11 | 0.08 | 0.09 | 0.11 |
| 5 | 164529549 | rs7735043 | -0.765 | -0.062 | -0.686 | -0.29 | -0.026 | 0.077 | 0.077 | 0 | 0.46 | 0.8 | 0.66 | 0.68 | 0.48 | 0.56 |
| 5 | 164532052 | rs4613738 | 0.054 | -0.135 | -0.484 | -0.238 | -0.007 | -0.006 | -0.006 | 2.296 | 0.11 | 0.12 | 0.11 | 0.06 | 0.1 | 0.11 |
| 5 | 164532314 | rs10073086 | 0.076 | -0.114 | -0.073 | -0.285 | 0.038 | 0.021 | 0.021 | 0 | 0.32 | 0.51 | 0.51 | 0.61 | 0.4 | 0.44 |
| 5 | 164541466 | rs1421691 | 1.037 | 0.202 | -0.281 | -0.138 | 0.031 | 0.083 | 0.083 | 0 | 0.05 | 0.09 | 0.1 | 0.24 | 0.14 | 0.07 |
| 5 | 164561180 | rs4868751 | -0.29 | 0.091 | -0.192 | -0.192 | -0.028 | -0.02 | -0.02 | 0.692 | 0.13 | 0.21 | 0.12 | 0.08 | 0.09 | 0.11 |
| 5 | 164564797 | rs1990866 | -0.642 | 0.017 | -0.027 | -0.162 | 0.005 | 0.039 | 0.039 | 0 | 0.17 | 0.3 | 0.22 | 0.3 | 0.22 | 0.18 |
| 5 | 164570320 | rs4397144 | -0.349 | 0.029 | 0 | -0.088 | -0.001 | 0.003 | 0.003 | 0 | 0.16 | 0.18 | 0.21 | 0.32 | 0.22 | 0.18 |
| 5 | 164583556 | rs1421713 | 0.287 | -0.015 | 0.137 | -0.344 | 0.05 | 0.038 | 0.038 | 0 | 0.31 | 0.47 | 0.51 | 0.59 | 0.39 | 0.44 |
| 5 | 164591126 | rs1421699 | -1.439 | -0.626 | -0.421 | -1.107 | 0.002 | 0.156 | 0.156 | 0 | 0.6 | 0.88 | 0.75 | 0.87 | 0.65 | 0.68 |
| 5 | 164600070 | rs7735460 | 0.341 | -0.596 | -0.335 | -0.92 | -0.013 | -0.005 | -0.005 | 0.985 | 0.12 | 0.14 | 0.1 | 0.08 | 0.07 | 0.11 |
| 5 | 164616822 | rs4868917 | 1.471 | -0.14 | -0.161 | -0.693 | -0.009 | -0.007 | -0.007 | 0.18 | 0.17 | 0.15 | 0.4 | 0.49 | 0.26 | 0.35 |
| 5 | 164622936 | rs2431127 | 1.382 | 0.044 | -0.5 | -0.665 | -0.017 | 0.012 | 0.012 | 0 | 0.07 | 0.03 | 0.1 | 0.29 | 0.16 | 0.09 |
| 5 | 164629768 | rs173792 | -0.272 | -0.171 | -0.809 | -1.121 | 0 | 0.054 | 0.054 | 0 | 0.82 | 0.91 | 0.9 | 0.99 | 0.87 | 0.88 |
| 5 | 164631794 | rs12153515 | 2.101 | -0.213 | -0.499 | -1.157 | -0.007 | -0.003 | -0.003 | 0 | 0.12 | 0.11 | 0.08 | 0 | 0.12 | 0.08 |
| 5 | 164637211 | rs13178865 | 1.174 | -0.639 | -0.639 | -1.093 | 0.051 | 0.074 | 0.074 | 0 | 0.41 | 0.25 | 0.27 | 0.2 | 0.31 | 0.34 |
| 5 | 164638825 | rs1422681 | -0.278 | -0.401 | -0.557 | -0.953 | -0.022 | -0.048 | -0.048 | 0 | 0.83 | 0.91 | 0.84 | 0.71 | 0.77 | 0.87 |
| 5 | 164642505 | rs7728827 | 1.257 | -0.603 | -0.572 | -1.112 | 0.005 | 0.18 | 0.18 | 0 | 0.39 | 0.11 | 0.21 | 0.13 | 0.32 | 0.31 |
| 5 | 164646665 | rs988216 | 2.03 | -0.587 | -0.579 | -1.119 | 0.03 | 0.125 | 0.125 | 0 | 0.31 | 0.06 | 0.17 | 0.12 | 0.25 | 0.24 |
| 5 | 164650294 | rs4868753 | 0.729 | -0.595 | -0.268 | -0.837 | 0.043 | 0.115 | 0.115 | 0 | 0.51 | 0.3 | 0.33 | 0.21 | 0.39 | 0.42 |
| 5 | 164651812 | rs10069860 | 0.256 | -0.599 | -0.316 | -0.814 | -0.028 | -0.012 | -0.012 | 2.033 | 0.12 | 0.18 | 0.12 | 0.08 | 0.07 | 0.11 |
| 5 | 164663019 | rs1422679 | 0.667 | -0.539 | -0.285 | -0.964 | 0.01 | 0.017 | 0.017 | 0 | 0.52 | 0.43 | 0.35 | 0.18 | 0.39 | 0.43 |
| 5 | 164668926 | rs10064661 | 0.762 | -0.52 | 0.151 | -0.977 | 0.006 | -0.003 | -0.003 | 0 | 0.51 | 0.4 | 0.33 | 0.17 | 0.4 | 0.43 |
| 5 | 164669629 | rs10866752 | -0.206 | -0.368 | 0.021 | -0.922 | 0.006 | -0.052 | -0.052 | 0.12 | 0.38 | 0.48 | 0.6 | 0.82 | 0.54 | 0.48 |
| 5 | 164675267 | rs1347154 | 1.373 | -0.371 | -0.346 | -0.906 | -0.02 | 0.207 | 0.207 | 0 | 0.39 | 0.06 | 0.2 | 0.1 | 0.32 | 0.33 |

| | | | | | | | | | | | | | | | | |
|---|---|---|---|---|---|---|---|---|---|---|---|---|---|---|---|---|
| 5 | 164679036 | rs6556813 | -0.1 | 0.381 | 0.175 | -0.396 | 0.031 | 0.04 | 0.04 | 0 | 0.32 | 0.51 | 0.51 | 0.53 | 0.38 | 0.44 |
| 5 | 164692816 | rs7732563 | 0.627 | 0.323 | 0.165 | -0.402 | -0.001 | 0.002 | 0.002 | 0 | 0.47 | 0.29 | 0.32 | 0.43 | 0.48 | 0.4 |
| 5 | 164695671 | rs11135362 | 0.516 | 0.274 | 0.206 | -0.394 | 0.004 | 0.007 | 0.007 | 0 | 0.49 | 0.25 | 0.31 | 0.44 | 0.48 | 0.4 |
| 5 | 164696018 | rs12186829 | 0.827 | 0.345 | 0.006 | -0.479 | -0.047 | -0.131 | -0.131 | 0.017 | 0.28 | 0.08 | 0.4 | 0.47 | 0.31 | 0.46 |
| 5 | 164701201 | rs17074571 | -0.127 | 0.46 | 0.702 | -0.423 | 0.076 | 0.176 | 0.176 | 0 | 0.12 | 0.42 | 0.15 | 0.25 | 0.17 | 0.09 |
| 5 | 164703292 | rs17074579 | 1.076 | 0.493 | 0.595 | -0.437 | 0.005 | 0.034 | 0.034 | 0 | 0.09 | 0.1 | 0.11 | 0.26 | 0.17 | 0.09 |
| 5 | 164703698 | rs17074593 | 0.726 | 0.499 | 0.603 | -0.434 | 0.011 | 0.044 | 0.044 | 0 | 0.1 | 0.15 | 0.12 | 0.26 | 0.17 | 0.09 |
| 5 | 164710430 | rs1433001 | -0.473 | 0.47 | 0.72 | -0.512 | 0.008 | -0.05 | -0.05 | 0 | 0.8 | 0.69 | 0.76 | 0.91 | 0.79 | 0.85 |
| 5 | 164712235 | rs1368421 | 1.077 | 0.383 | 0.659 | -0.502 | -0.025 | -0.013 | -0.013 | 0.017 | 0.66 | 0.19 | 0.61 | 0.65 | 0.62 | 0.76 |
| 5 | 164712787 | rs13180364 | 1.16 | 0.18 | 0.558 | -0.667 | 0.042 | 0.128 | 0.128 | 0 | 0.4 | 0.11 | 0.2 | 0.18 | 0.32 | 0.3 |
| 5 | 164723605 | rs13190045 | 0.078 | 0.314 | 0.745 | -0.522 | -0.03 | -0.056 | -0.056 | 0 | 0.84 | 0.62 | 0.82 | 0.9 | 0.88 | 0.88 |
| 5 | 164751379 | rs6556817 | -0.629 | 0.296 | 0.692 | -0.446 | -0.007 | -0.017 | -0.017 | 0 | 0.9 | 0.72 | 0.85 | 0.94 | 0.91 | 0.93 |
| 5 | 164759496 | rs10515893 | 1.128 | 0.299 | 0.307 | -0.524 | 0.043 | 0.106 | 0.106 | 0 | 0.38 | 0.11 | 0.18 | 0.18 | 0.29 | 0.29 |
| 5 | 164762427 | rs10045813 | -0.451 | 0.51 | 0.604 | -0.635 | 0.011 | -0.062 | -0.062 | 0.822 | 0.78 | 0.65 | 0.76 | 0.89 | 0.79 | 0.84 |
| 5 | 164764508 | rs17075358 | 1.471 | 0.519 | 0.433 | -0.763 | 0.036 | 0.05 | 0.05 | 0 | 0.34 | 0.19 | 0.16 | 0.11 | 0.21 | 0.27 |
| 5 | 164766745 | rs17075367 | 0.675 | 0.743 | 0.508 | -0.532 | 0.021 | 0.043 | 0.043 | 0 | 0.11 | 0.17 | 0.14 | 0.24 | 0.21 | 0.11 |
| 5 | 164767433 | rs10515892 | 0.776 | 0.742 | 0.51 | -0.532 | 0.016 | 0.035 | 0.035 | 0 | 0.11 | 0.15 | 0.14 | 0.24 | 0.21 | 0.11 |
| 5 | 164768057 | rs10039383 | -0.537 | 0.753 | 0.526 | -0.578 | 0 | -0.01 | -0.01 | 0.089 | 0.46 | 0.55 | 0.62 | 0.79 | 0.58 | 0.57 |
| 5 | 164769627 | rs2217644 | -0.031 | 0.755 | 0.056 | -0.323 | -0.009 | -0.025 | -0.025 | 0 | 0.34 | 0.26 | 0.47 | 0.55 | 0.37 | 0.46 |
| 5 | 164771289 | rs1432988 | -0.415 | 0.8 | 0.187 | -0.312 | 0.016 | -0.054 | -0.054 | 0 | 0.78 | 0.66 | 0.76 | 0.89 | 0.78 | 0.85 |
| 5 | 164774447 | rs12515866 | 0.182 | 0.69 | -0.272 | 0.224 | -0.002 | -0.019 | -0.019 | 0 | 0.28 | 0.25 | 0.44 | 0.47 | 0.3 | 0.39 |
| 5 | 164786237 | rs10054506 | -0.774 | 0.159 | 0.24 | 0.052 | -0.025 | -0.011 | -0.011 | 0.003 | 0.14 | 0.39 | 0.1 | 0.13 | 0.09 | 0.07 |
| 5 | 164786295 | rs9790890 | 0.023 | 1.086 | 0.361 | 0.104 | 0.007 | 0.005 | 0.005 | 0 | 0.16 | 0.16 | 0.22 | 0.24 | 0.32 | 0.15 |
| 5 | 164788169 | rs13175231 | 0.036 | 0.271 | 0.076 | -0.46 | 0 | 0.012 | 0.012 | 0 | 0.48 | 0.31 | 0.38 | 0.4 | 0.54 | 0.43 |
| 5 | 164790544 | rs7719707 | -0.746 | 0.324 | -0.395 | -0.502 | 0.015 | 0.007 | 0.007 | 0 | 0.31 | 0.58 | 0.32 | 0.3 | 0.37 | 0.17 |
| 5 | 164792081 | rs1428506 | -0.593 | 0.618 | -0.416 | -0.092 | -0.041 | -0.053 | -0.053 | 0 | 0.64 | 0.8 | 0.47 | 0.46 | 0.59 | 0.46 |
| 5 | 164793828 | rs13360433 | 0.176 | 1.147 | 0.25 | 0.383 | 0.006 | 0.009 | 0.009 | 0 | 0.19 | 0.24 | 0.21 | 0.26 | 0.24 | 0.15 |
| 5 | 164797846 | rs6880019 | -0.846 | 1.137 | 0.712 | 0.307 | 0.001 | 0.023 | 0.023 | 0.063 | 0.28 | 0.58 | 0.27 | 0.33 | 0.26 | 0.17 |
| 5 | 164799990 | rs4562063 | 0.171 | 1.063 | 0.941 | 0.411 | 0.005 | 0.016 | 0.016 | 0 | 0.19 | 0.23 | 0.22 | 0.28 | 0.22 | 0.13 |
| 5 | 164816370 | rs9314067 | -0.401 | 1.027 | 0.829 | 0.379 | 0.008 | 0.052 | 0.052 | 1.792 | 0.26 | 0.53 | 0.27 | 0.32 | 0.24 | 0.17 |
| 5 | 164823575 | rs2431617 | -0.603 | 0.9 | -0.018 | -0.001 | -0.028 | -0.051 | -0.051 | 0 | 0.61 | 0.75 | 0.48 | 0.42 | 0.57 | 0.46 |
| 5 | 164828190 | rs7707060 | 0.409 | 0.713 | 0.429 | -0.042 | -0.011 | -0.024 | -0.024 | 0 | 0.38 | 0.38 | 0.26 | 0.18 | 0.34 | 0.27 |
| 5 | 164838018 | rs1428503 | 0.432 | 0.998 | 0.55 | 0.022 | 0.015 | 0.015 | 0.015 | 0.119 | 0.22 | 0.3 | 0.21 | 0.26 | 0.24 | 0.17 |
| 5 | 164840351 | rs7719623 | 0.345 | 1.078 | 0.405 | -0.061 | -0.015 | -0.029 | -0.029 | 0 | 0.38 | 0.39 | 0.26 | 0.17 | 0.33 | 0.27 |
| 5 | 164846823 | rs13179338 | 0.08 | 1.215 | 0.143 | -0.176 | -0.003 | -0.013 | -0.013 | 0.553 | 0.44 | 0.48 | 0.33 | 0.17 | 0.42 | 0.29 |
| 5 | 164852945 | rs1422686 | 0.195 | 1.095 | -0.052 | -0.468 | 0.001 | 0.001 | 0.001 | 0.434 | 0.3 | 0.27 | 0.23 | 0.27 | 0.28 | 0.2 |
| 5 | 164857058 | rs17422062 | 0.848 | 1.156 | 0.159 | -0.569 | -0.008 | -0.046 | -0.046 | 0.118 | 0.3 | 0.35 | 0.22 | 0.15 | 0.28 | 0.25 |
| 5 | 164858434 | rs1422690 | -0.555 | 0.886 | 0.198 | -0.506 | -0.014 | -0.007 | -0.007 | 0 | 0.61 | 0.66 | 0.46 | 0.41 | 0.56 | 0.46 |
| 5 | 164859465 | rs1469064 | 0.902 | 1.01 | 0.097 | -0.498 | 0 | -0.008 | -0.008 | 0.061 | 0.31 | 0.3 | 0.25 | 0.15 | 0.35 | 0.26 |
| 5 | 164865386 | rs10077267 | 0.414 | 0.914 | -0.189 | -0.493 | 0.006 | -0.047 | -0.047 | 0.314 | 0.37 | 0.47 | 0.33 | 0.16 | 0.43 | 0.27 |
| 5 | 164866479 | rs1422683 | 1.031 | 0.671 | -0.283 | -0.444 | 0.002 | -0.026 | -0.026 | 0.203 | 0.2 | 0.16 | 0.16 | 0.26 | 0.16 | 0.17 |
| 5 | 164877178 | rs9314069 | -0.108 | 0.891 | -0.25 | -0.453 | 0 | 0.001 | 0.001 | 0 | 0.53 | 0.52 | 0.44 | 0.43 | 0.55 | 0.44 |
| 5 | 164877481 | rs12522758 | 0.796 | 0.885 | -0.256 | -0.502 | 0.012 | -0.064 | -0.064 | 0.062 | 0.32 | 0.39 | 0.29 | 0.16 | 0.4 | 0.27 |
| 5 | 164881372 | rs4392645 | -0.761 | 0.727 | -0.29 | -0.302 | 0.01 | -0.025 | -0.025 | 0 | 0.64 | 0.61 | 0.61 | 0.71 | 0.54 | 0.68 |
| 5 | 164881560 | rs4392646 | 0.459 | 0.887 | -0.599 | -0.185 | -0.002 | -0.014 | -0.014 | 0 | 0.36 | 0.35 | 0.42 | 0.42 | 0.34 | 0.5 |
| 5 | 164883307 | rs3980160 | -0.118 | 0.88 | -0.558 | -0.359 | 0 | 0.003 | 0.003 | 0 | 0.55 | 0.52 | 0.58 | 0.68 | 0.5 | 0.62 |
| 5 | 164884538 | rs7702763 | 1.065 | 1.17 | -0.901 | -0.105 | -0.005 | -0.089 | -0.089 | 0 | 0.26 | 0.14 | 0.29 | 0.44 | 0.22 | 0.48 |
| 5 | 164900842 | rs5002678 | 0.969 | 1.235 | -0.631 | -0.14 | 0 | -0.001 | -0.001 | 0 | 0.22 | 0.2 | 0.18 | 0.25 | 0.18 | 0.15 |
| 5 | 164904060 | rs981236 | 0.117 | 1.205 | -0.653 | -0.292 | -0.007 | 0.037 | 0.037 | 0 | 0.45 | 0.39 | 0.51 | 0.7 | 0.42 | 0.61 |
| 5 | 164907971 | rs6864913 | 1.431 | 1.085 | -0.675 | -0.098 | -0.019 | -0.103 | -0.103 | 0 | 0.26 | 0.1 | 0.35 | 0.44 | 0.27 | 0.44 |
| 5 | 164909667 | rs12152953 | 0.939 | 1.149 | -0.672 | -0.108 | 0 | 0 | 0 | 0 | 0.2 | 0.18 | 0.17 | 0.24 | 0.14 | 0.15 |
| 5 | 164909862 | rs12153294 | 0.616 | 0.912 | -0.663 | -0.099 | -0.001 | 0 | 0 | 0.51 | 0.22 | 0.19 | 0.19 | 0.24 | 0.19 | 0.16 |
| 5 | 164912247 | rs17423436 | 0.995 | 0.792 | -0.563 | 0.093 | -0.017 | -0.005 | -0.005 | 0 | 0.23 | 0.13 | 0.19 | 0.25 | 0.23 | 0.13 |
| 5 | 164915687 | rs17338894 | 0.962 | 0.8 | -0.526 | 0.062 | -0.003 | 0 | 0 | 0.348 | 0.23 | 0.19 | 0.2 | 0.24 | 0.23 | 0.13 |
| 5 | 164916724 | rs17338964 | 1.264 | 0.637 | -0.641 | -0.175 | 0.013 | 0.038 | 0.038 | 0 | 0.21 | 0.11 | 0.16 | 0.14 | 0.17 | 0.27 |
| 5 | 164917641 | rs13165594 | 2 | 0.667 | -0.736 | -0.119 | -0.023 | -0.062 | -0.062 | 0 | 0.21 | 0.12 | 0.35 | 0.37 | 0.25 | 0.31 |
| 5 | 164919132 | rs10476513 | 0.271 | -0.36 | 0.257 | -0.393 | 0.001 | 0.013 | 0.013 | 0.008 | 0.23 | 0.17 | 0.18 | 0.15 | 0.2 | 0.29 |
| 5 | 164924423 | rs183158 | 0.314 | -0.497 | 0.292 | -0.337 | 0.004 | -0.021 | -0.021 | 0 | 0.8 | 0.76 | 0.85 | 0.92 | 0.74 | 0.79 |
| 5 | 164928879 | rs2431619 | -0.976 | -0.552 | 0.317 | -0.298 | -0.038 | 0.031 | 0.031 | 0 | 0.91 | 1 | 0.9 | 0.99 | 0.81 | 0.93 |
| 5 | 164931799 | rs6896868 | 0.474 | -0.485 | 0.566 | -0.182 | -0.012 | -0.047 | -0.047 | 1.851 | 0.08 | 0.2 | 0.04 | 0 | 0.03 | 0.03 |
| 5 | 164934294 | rs6882745 | 0.201 | -0.369 | -0.167 | -0.104 | -0.001 | 0.004 | 0.004 | 0 | 0.76 | 0.68 | 0.74 | 0.76 | 0.7 | 0.76 |
| 5 | 164935737 | rs966590 | 0.539 | 0.149 | -0.599 | 0.35 | 0.001 | 0.001 | 0.001 | 0 | 0.06 | 0.09 | 0.12 | 0.14 | 0.06 | 0.05 |
| 5 | 164937499 | rs17339394 | 0.514 | 0.233 | -0.714 | 0.386 | -0.041 | 0.037 | 0.037 | 0 | 0.43 | 0.21 | 0.54 | 0.63 | 0.48 | 0.44 |
| 5 | 164942881 | rs2246518 | -1.043 | 0.32 | 0.071 | 0.949 | -0.026 | 0.001 | 0.001 | 0 | 0.15 | 0.31 | 0.15 | 0.14 | 0.09 | 0.08 |
| 5 | 164953114 | rs10213969 | -1.068 | 0.095 | 0.486 | 2.222 | -0.047 | 0.242 | 0.242 | 0 | 0.1 | 0.31 | 0.12 | 0.3 | 0.03 | 0.16 |
| 5 | 164953812 | rs10064269 | -1.75 | 0.184 | 0.664 | 2.303 | -0.002 | 0.176 | 0.176 | 0 | 0.16 | 0.41 | 0.35 | 0.5 | 0.14 | 0.26 |
| 5 | 164956787 | rs6871907 | -0.375 | 0.35 | 0.753 | 2.428 | 0.005 | 0.037 | 0.037 | 0 | 0.06 | 0.1 | 0.11 | 0.26 | 0.09 | 0.13 |
| 5 | 164957355 | rs11135373 | -1.53 | 0.308 | 0.831 | 2.435 | -0.005 | 0.126 | 0.126 | 0 | 0.16 | 0.31 | 0.34 | 0.51 | 0.14 | 0.26 |
| 5 | 164961753 | rs17339906 | 0.335 | 0.014 | -0.059 | 1.716 | -0.002 | 0.026 | 0.026 | 0 | 0.2 | 0.06 | 0.1 | 0.18 | 0.18 | 0.32 |

| | | | | | | | | | | | | | | | | |
|---|---|---|---|---|---|---|---|---|---|---|---|---|---|---|---|---|
| 5 | 164966529 | rs9314073 | -1.039 | -0.458 | 0.591 | 1.498 | -0.054 | 0.185 | 0.185 | 0.041 | 0.16 | 0.39 | 0.18 | 0.32 | 0.11 | 0.2 |
| 5 | 164971497 | rs9314075 | -0.183 | -0.259 | 0.49 | 0.377 | -0.024 | -0.039 | -0.039 | 0 | 0.08 | 0.21 | 0.05 | 0 | 0.05 | 0.04 |
| 5 | 164979056 | rs17064277 | -0.566 | -0.442 | 0.078 | 0.303 | -0.001 | -0.001 | -0.001 | 5.141 | 0.39 | 0.42 | 0.46 | 0.66 | 0.34 | 0.54 |
| 5 | 164986951 | rs10079921 | -1.339 | -0.182 | 0.48 | -0.272 | 0.005 | -0.035 | -0.035 | 0 | 0.07 | 0.11 | 0.04 | 0 | 0.03 | 0.01 |
| 5 | 164990768 | rs9314079 | -0.562 | -0.212 | 0.717 | -0.292 | -0.002 | 0.012 | 0.012 | 0.072 | 0.08 | 0.12 | 0.05 | 0.1 | 0.07 | 0.08 |
| 5 | 164991553 | rs7721625 | -0.61 | 0.436 | 1.005 | -0.603 | 0.011 | -0.008 | -0.008 | 0 | 0.1 | 0.15 | 0.09 | 0.06 | 0.13 | 0.08 |
| 5 | 164992054 | rs1159244 | -1.397 | -0.002 | 0.707 | -1.659 | -0.026 | 0.111 | 0.111 | 0.512 | 0.54 | 0.78 | 0.34 | 0.22 | 0.47 | 0.39 |
| 5 | 164992281 | rs1159245 | -0.621 | -0.157 | 0.773 | -2.071 | 0.027 | 0.048 | 0.048 | 3.193 | 0.34 | 0.21 | 0.16 | 0.15 | 0.26 | 0.15 |
| 5 | 164992859 | rs12519948 | -0.356 | -0.152 | 1.252 | -2.088 | 0.054 | -0.029 | -0.029 | 0 | 0.07 | 0.28 | 0.08 | 0.01 | 0.09 | 0.16 |
| 5 | 164993708 | rs7715320 | -1.359 | 0.159 | 0.793 | -2.143 | 0.001 | 0.134 | 0.134 | 2.467 | 0.49 | 0.73 | 0.3 | 0.22 | 0.44 | 0.42 |
| 5 | 164999502 | rs10053913 | -1.033 | 0.193 | 0.6 | -2.107 | 0.013 | 0.056 | 0.056 | 2.528 | 0.46 | 0.37 | 0.25 | 0.21 | 0.4 | 0.39 |
| 5 | 165002334 | rs4421124 | -1.012 | -0.13 | 0.303 | -2.572 | 0.001 | 0.033 | 0.033 | 0 | 0.28 | 0.17 | 0.16 | 0.07 | 0.28 | 0.13 |
| 5 | 165010625 | rs6871175 | -0.808 | 0.964 | 0.552 | -2.365 | 0 | -0.026 | -0.026 | 0 | 0.16 | 0.19 | 0.09 | 0.06 | 0.14 | 0.11 |
| 5 | 165012867 | rs7703411 | -1.506 | 0.72 | 0.063 | -2.562 | -0.003 | 0.085 | 0.085 | 0.238 | 0.46 | 0.67 | 0.31 | 0.16 | 0.4 | 0.46 |
| 5 | 165015242 | rs17064971 | -1.142 | 1.427 | -0.024 | -2.455 | 0 | -0.029 | -0.029 | 0 | 0.38 | 0.63 | 0.27 | 0.15 | 0.36 | 0.45 |
| 5 | 165015268 | rs17064972 | -1.133 | 1.213 | -0.319 | -2.357 | 0.009 | -0.073 | -0.073 | 2.401 | 0.42 | 0.55 | 0.31 | 0.13 | 0.43 | 0.49 |
| 5 | 165017688 | rs7728593 | -0.717 | 1.803 | -0.948 | -1.695 | 0.004 | -0.023 | -0.023 | 0 | 0.14 | 0.17 | 0.11 | 0.05 | 0.13 | 0.15 |
| 5 | 165017751 | rs17064997 | -0.678 | 1.819 | -0.98 | -1.687 | 0.003 | -0.02 | -0.02 | 0 | 0.14 | 0.16 | 0.11 | 0.05 | 0.13 | 0.15 |
| 5 | 165024869 | rs2431171 | -0.3 | 0.542 | -1.461 | -1.274 | -0.007 | 0.141 | 0.141 | 1.556 | 0.6 | 0.85 | 0.4 | 0.18 | 0.57 | 0.56 |
| 5 | 165032300 | rs17065217 | -0.955 | 0.161 | -0.939 | -0.612 | -0.002 | -0.002 | -0.002 | 1.56 | 0.19 | 0.25 | 0.26 | 0.12 | 0.16 | 0.12 |
| 5 | 165046698 | rs262747 | -0.657 | -0.516 | -1.287 | -1.508 | 0.017 | 0.057 | 0.057 | 0 | 0.41 | 0.73 | 0.49 | 0.21 | 0.42 | 0.38 |
| 5 | 165047392 | rs262748 | 0.011 | -0.703 | -0.315 | -1.69 | -0.007 | -0.023 | -0.023 | 0.033 | 0.25 | 0.31 | 0.2 | 0.09 | 0.21 | 0.19 |
| 5 | 165047881 | rs17341948 | -0.571 | -0.653 | 0.022 | -1.688 | -0.006 | 0.002 | 0.002 | 0 | 0.14 | 0.08 | 0.26 | 0.12 | 0.19 | 0.19 |
| 5 | 165048301 | rs262749 | -0.972 | -0.975 | 0.084 | -1.177 | -0.001 | 0.005 | 0.005 | 0 | 0.27 | 0.2 | 0.38 | 0.21 | 0.33 | 0.29 |
| 5 | 165049603 | rs17065396 | 0.667 | -0.197 | -0.296 | -0.45 | -0.001 | 0 | 0 | 0 | 0.22 | 0.16 | 0.12 | 0.44 | 0.2 | 0.12 |
| 5 | 165050288 | rs10515900 | 1.191 | -0.399 | -0.889 | -0.775 | -0.009 | -0.071 | -0.071 | 0.023 | 0.11 | 0.01 | 0.13 | 0.28 | 0.09 | 0.33 |
| 5 | 165051330 | rs1030154 | 1.269 | -0.373 | -0.899 | -0.905 | 0.003 | 0.021 | 0.021 | 0.123 | 0.19 | 0.16 | 0.13 | 0.44 | 0.21 | 0.16 |
| 5 | 165052058 | rs10515901 | 1.067 | -0.272 | -1.049 | -0.833 | -0.015 | -0.044 | -0.044 | 1.546 | 0.08 | 0.01 | 0.12 | 0.29 | 0.07 | 0.31 |
| 5 | 165053875 | rs10463158 | 0.129 | -0.737 | -0.913 | -1.573 | -0.035 | 0.136 | 0.136 | 0 | 0.58 | 0.7 | 0.6 | 0.89 | 0.53 | 0.69 |
| 5 | 165054606 | rs155222 | 0.789 | -0.165 | -0.545 | -1.144 | -0.008 | -0.023 | -0.023 | 1.932 | 0.14 | 0.26 | 0.14 | 0.07 | 0.14 | 0.1 |
| 5 | 165056030 | rs155223 | -1.233 | 0.32 | -1.113 | -1.067 | -0.072 | 0.071 | 0.071 | 0 | 0.87 | 0.98 | 0.8 | 0.99 | 0.77 | 0.91 |
| 5 | 165060876 | rs262753 | -0.489 | -0.343 | -0.547 | -0.638 | -0.007 | 0.04 | 0.04 | 0.28 | 0.32 | 0.38 | 0.38 | 0.42 | 0.32 | 0.54 |
| 5 | 165065045 | rs262756 | 0.212 | 0.742 | -1.123 | -0.141 | 0.004 | 0 | 0 | 1.551 | 0.55 | 0.42 | 0.48 | 0.4 | 0.43 | 0.31 |
| 5 | 165067120 | rs1549133 | -0.049 | 0.839 | -1.409 | -0.421 | -0.005 | 0.013 | 0.013 | 0.081 | 0.74 | 0.71 | 0.71 | 0.84 | 0.71 | 0.77 |
| 5 | 165069995 | rs10515902 | 1.236 | 0.908 | -1.3 | -0.037 | 0.021 | 0.003 | 0.003 | 0 | 0.09 | 0.02 | 0.11 | 0.52 | 0.05 | 0.36 |
| 5 | 165070154 | rs256409 | 1.072 | 1.214 | -0.867 | -0.445 | 0.081 | -0.023 | -0.023 | 0 | 0.1 | 0.36 | 0.11 | 0.03 | 0.12 | 0.1 |
| 5 | 165071427 | rs262761 | 0.489 | 0.486 | -0.998 | -0.408 | 0.01 | -0.014 | -0.014 | 0 | 0.59 | 0.49 | 0.53 | 0.29 | 0.57 | 0.33 |
| 5 | 165072241 | rs196129 | 0.617 | 1.674 | -0.341 | -0.659 | -0.007 | -0.009 | -0.009 | 0.015 | 0.12 | 0.18 | 0.11 | 0.06 | 0.08 | 0.1 |
| 5 | 165072903 | rs262764 | -0.032 | 1.955 | -0.499 | -0.359 | 0.103 | -0.039 | -0.039 | 0 | 0.81 | 0.57 | 0.72 | 0.89 | 0.69 | 0.74 |
| 5 | 165079273 | rs262775 | -0.122 | 1.921 | -0.603 | -0.372 | 0.077 | -0.06 | -0.06 | 1.762 | 0.81 | 0.63 | 0.72 | 0.93 | 0.68 | 0.77 |
| 5 | 165080168 | rs12189065 | 0.026 | 0.762 | -1.392 | -0.025 | 0.036 | 0.096 | 0.096 | 1.264 | 0.52 | 0.27 | 0.49 | 0.8 | 0.41 | 0.6 |
| 5 | 165083316 | rs17065659 | -0.67 | 1.386 | -1.588 | 0.987 | -0.097 | -0.104 | -0.104 | 0.769 | 0.18 | 0.01 | 0.25 | 0.38 | 0.28 | 0.31 |
| 5 | 165083795 | rs262714 | -0.689 | 0.743 | -1.717 | 0.805 | -0.009 | 0.041 | 0.041 | 1.073 | 0.6 | 0.8 | 0.71 | 0.73 | 0.61 | 0.62 |
| 5 | 165085583 | rs116953 | -0.686 | 0.308 | -0.072 | 1.335 | 0.008 | 0.083 | 0.083 | 0 | 0.2 | 0.39 | 0.23 | 0.38 | 0.21 | 0.25 |
| 5 | 165087285 | rs10045268 | -0.865 | 0.232 | -0.261 | 1.124 | -0.009 | 0.047 | 0.047 | 0 | 0.31 | 0.59 | 0.36 | 0.58 | 0.28 | 0.43 |
| 5 | 165093364 | rs11952674 | -0.475 | -0.31 | -0.131 | 0.972 | -0.034 | -0.002 | -0.002 | 0 | 0.14 | 0.32 | 0.1 | 0.12 | 0.05 | 0.02 |
| 5 | 165093913 | rs11135378 | -1.367 | -0.456 | -0.488 | 1.76 | -0.012 | 0.032 | 0.032 | 2.6 | 0.23 | 0.33 | 0.33 | 0.58 | 0.15 | 0.22 |
| 5 | 165094688 | rs262720 | -0.178 | -0.403 | -0.595 | 0.991 | -0.006 | -0.048 | -0.048 | 0 | 0.62 | 0.45 | 0.7 | 0.86 | 0.65 | 0.64 |
| 5 | 165096435 | rs3909281 | 1.045 | -0.686 | -0.711 | 1.099 | -0.058 | 0.033 | 0.033 | 0 | 0.41 | 0.11 | 0.4 | 0.22 | 0.53 | 0.39 |
| 5 | 165096645 | rs17065681 | 0.814 | -0.631 | -0.669 | 1.254 | -0.009 | 0.025 | 0.025 | 0 | 0.06 | 0 | 0.21 | 0.49 | 0.07 | 0.18 |
| 5 | 165098394 | rs262728 | 0.645 | -0.734 | -0.713 | 1.062 | 0.001 | 0.006 | 0.006 | 0 | 0.46 | 0.42 | 0.43 | 0.22 | 0.55 | 0.46 |
| 5 | 165099402 | rs262733 | -0.661 | -0.86 | -0.636 | 0.793 | 0 | 0.015 | 0.015 | 0.01 | 0.67 | 0.76 | 0.75 | 0.86 | 0.67 | 0.65 |
| 5 | 165103181 | rs10515905 | 0.177 | -0.711 | -0.62 | 0.569 | 0.001 | -0.002 | -0.002 | 0 | 0.14 | 0.22 | 0.12 | 0.13 | 0.12 | 0.04 |
| 5 | 165106225 | rs11749716 | -0.402 | -0.689 | -0.413 | 1 | -0.019 | 0.026 | 0.026 | 4.802 | 0.26 | 0.37 | 0.35 | 0.63 | 0.15 | 0.21 |
| 5 | 165107100 | rs516732 | 0.702 | -0.548 | -0.384 | 0.85 | 0.003 | -0.009 | -0.009 | 2.176 | 0.41 | 0.47 | 0.42 | 0.24 | 0.53 | 0.44 |
| 5 | 165107162 | rs594673 | 0.129 | -0.539 | -0.481 | 0.834 | 0.024 | 0.015 | 0.015 | 0 | 0.5 | 0.42 | 0.5 | 0.37 | 0.62 | 0.48 |
| 5 | 165121041 | rs2861301 | 0.938 | -0.347 | -0.751 | 0.627 | -0.125 | 0.077 | 0.077 | 0 | 0.34 | 0.09 | 0.38 | 0.16 | 0.5 | 0.37 |
| 5 | 165132974 | rs262741 | 0.594 | -0.422 | -0.038 | 0.866 | -0.005 | 0.002 | 0.002 | 0 | 0.43 | 0.34 | 0.44 | 0.39 | 0.53 | 0.52 |
| 5 | 165136390 | rs6556822 | -0.938 | -0.23 | -0.202 | 0.44 | 0 | -0.034 | -0.034 | 0 | 0.7 | 0.72 | 0.78 | 0.93 | 0.68 | 0.7 |
| 5 | 165138708 | rs4868949 | 1.087 | -0.373 | -0.531 | 0.824 | -0.12 | -0.112 | -0.112 | 0 | 0.37 | 0.11 | 0.53 | 0.49 | 0.5 | 0.45 |
| 5 | 165138955 | rs6867953 | 0.117 | -0.262 | -0.033 | 0.673 | 0.008 | 0.051 | 0.051 | 0.007 | 0.58 | 0.36 | 0.7 | 0.85 | 0.63 | 0.61 |
| 5 | 165144196 | rs10036126 | -0.477 | -0.375 | -0.093 | 0.807 | -0.008 | -0.029 | -0.029 | 0 | 0.11 | 0.37 | 0.07 | 0.04 | 0.04 | 0.07 |
| 5 | 165145533 | rs263897 | -0.924 | -0.291 | -0.947 | 0.616 | -0.011 | 0.021 | 0.021 | 0 | 0.87 | 0.93 | 0.89 | 0.92 | 0.81 | 0.79 |
| 5 | 165148145 | rs12657428 | 0.592 | -0.376 | -0.825 | 0.961 | -0.001 | 0.246 | 0.246 | 0.197 | 0.06 | 0.22 | 0.1 | 0.33 | 0.04 | 0.15 |
| 5 | 165151403 | rs263895 | -0.909 | -0.121 | -1.83 | 0.708 | 0.016 | -0.003 | -0.003 | 0 | 0.51 | 0.9 | 0.39 | 0.46 | 0.4 | 0.49 |
| 5 | 165151524 | rs263894 | -1.732 | 0.041 | -1.91 | 0.728 | -0.094 | -0.097 | -0.097 | 0.014 | 0.64 | 0.9 | 0.5 | 0.58 | 0.54 | 0.55 |
| 5 | 165154143 | rs7714184 | 1.052 | -0.132 | -1.914 | 0.636 | -0.035 | 0.062 | 0.062 | 0 | 0.14 | 0.01 | 0.1 | 0.06 | 0.18 | 0.22 |
| 5 | 165156082 | rs263892 | -0.326 | 0.122 | -1.623 | 0.913 | 0.061 | 0.043 | 0.043 | 0 | 0.5 | 0.89 | 0.38 | 0.43 | 0.38 | 0.48 |
| 5 | 165160664 | rs13179158 | 1.994 | 0.338 | -1.456 | 0.764 | -0.039 | -0.029 | -0.029 | 0 | 0.08 | 0.01 | 0.1 | 0.1 | 0.13 | 0.04 |

| | | | | | | | | | | | | | | | | |
|---|---|---|---|---|---|---|---|---|---|---|---|---|---|---|---|---|
| 5 | 165170651 | rs263886 | -0.605 | 0.531 | -1.711 | 0.878 | -0.08 | 0.02 | 0.02 | 0 | 0.57 | 0.91 | 0.49 | 0.34 | 0.45 | 0.46 |
| 5 | 165180743 | rs6860172 | -1.01 | 0.283 | -1.65 | 0.788 | -0.13 | -0.048 | -0.048 | 0.327 | 0.66 | 0.92 | 0.49 | 0.64 | 0.52 | 0.7 |
| 5 | 165184847 | rs17065850 | 0.622 | 0.142 | -1.009 | 0.328 | 0 | 0.007 | 0.007 | 0 | 0.34 | 0.29 | 0.25 | 0.17 | 0.32 | 0.12 |
| 5 | 165186172 | rs7728710 | 0.503 | 0.276 | -0.755 | 0.592 | -0.005 | 0.034 | 0.034 | 0 | 0.43 | 0.31 | 0.32 | 0.26 | 0.45 | 0.45 |
| 5 | 165187703 | rs10866756 | -0.495 | 0.072 | -0.16 | 0.699 | -0.017 | 0.026 | 0.026 | 0.448 | 0.09 | 0.29 | 0.06 | 0.1 | 0.02 | 0.05 |
| 5 | 165199466 | rs10036921 | 0.114 | 0.227 | -0.169 | 0.4 | -0.011 | -0.056 | -0.056 | 0 | 0.44 | 0.49 | 0.23 | 0.09 | 0.31 | 0.38 |
| 5 | 165209042 | rs17065876 | -0.328 | 0.587 | -0.401 | 0.822 | 0.001 | 0.026 | 0.026 | 0 | 0.12 | 0.13 | 0.31 | 0.34 | 0.18 | 0.19 |
| 5 | 165214300 | rs2112355 | -0.753 | 0.59 | -0.538 | 0.898 | -0.04 | -0.059 | -0.059 | 3.857 | 0.16 | 0.07 | 0.42 | 0.7 | 0.25 | 0.41 |
| 5 | 165214813 | rs6897791 | 0.925 | 0.785 | -0.321 | 0.475 | 0.023 | 0.001 | 0.001 | 0 | 0.22 | 0.33 | 0.2 | 0.21 | 0.25 | 0.12 |
| 5 | 165225255 | rs1895207 | -1.401 | 0.772 | -0.707 | 0.455 | -0.051 | 0.115 | 0.115 | 0 | 0.81 | 0.99 | 0.87 | 0.97 | 0.8 | 0.87 |
| 5 | 165228923 | rs7707503 | 1.299 | 0.463 | -0.5 | 0.402 | -0.034 | -0.005 | -0.005 | 0 | 0.08 | 0.13 | 0.02 | 0.05 | 0 | 0.04 |
| 5 | 165229136 | rs10053367 | 1.334 | 0.548 | -0.327 | 0.434 | 0.004 | -0.002 | -0.002 | 0 | 0.2 | 0.23 | 0.19 | 0.18 | 0.24 | 0.06 |
| 5 | 165230027 | rs1895208 | -0.14 | 0.622 | -0.416 | 0.511 | -0.042 | -0.039 | -0.039 | 0.639 | 0.68 | 0.87 | 0.57 | 0.65 | 0.63 | 0.68 |
| 5 | 165235055 | rs7711967 | 1.042 | 0.539 | -0.168 | 0.554 | 0.008 | 0.023 | 0.023 | 0 | 0.35 | 0.29 | 0.23 | 0.17 | 0.32 | 0.42 |
| 5 | 165236250 | rs13169962 | -0.43 | 0.809 | -0.359 | 1.154 | -0.003 | 0.009 | 0.009 | 0 | 0.12 | 0.12 | 0.3 | 0.33 | 0.18 | 0.19 |
| 5 | 165243209 | rs2861622 | 0.171 | 0.745 | -0.272 | 0.913 | 0.001 | -0.037 | -0.037 | 0.354 | 0.28 | 0.42 | 0.23 | 0.15 | 0.25 | 0.1 |
| 5 | 165244680 | rs248274 | -1.029 | 0.766 | -0.336 | 0.927 | 0.002 | -0.037 | -0.037 | 0 | 0.45 | 0.54 | 0.64 | 0.84 | 0.48 | 0.49 |
| 5 | 165247018 | rs9942373 | 0.006 | 0.733 | -0.284 | 0.717 | -0.003 | -0.039 | -0.039 | 0.595 | 0.29 | 0.42 | 0.23 | 0.15 | 0.25 | 0.1 |
| 5 | 165251964 | rs409459 | -0.681 | 0.649 | -0.662 | 1.285 | -0.017 | -0.025 | -0.025 | 0 | 0.11 | 0.08 | 0.3 | 0.36 | 0.17 | 0.2 |
| 5 | 165254436 | rs10043418 | -0.947 | 0.107 | -0.191 | 0.849 | -0.046 | 0.032 | 0.032 | 5.031 | 0.19 | 0.33 | 0.06 | 0.24 | 0.05 | 0.15 |
| 5 | 165260961 | rs248272 | -0.287 | 0.74 | -0.785 | 0.246 | 0.009 | -0.075 | -0.075 | 2.21 | 0.58 | 0.46 | 0.67 | 0.92 | 0.5 | 0.79 |
| 5 | 165264791 | rs10462951 | 0.695 | 0.655 | -1.269 | 0.701 | -0.001 | 0.001 | 0.001 | 0 | 0.39 | 0.35 | 0.24 | 0.32 | 0.22 | 0.44 |
| 5 | 165265353 | rs17065942 | 1.099 | 0.601 | -1.139 | 0.678 | 0.019 | 0.007 | 0.007 | 0 | 0.33 | 0.24 | 0.23 | 0.28 | 0.21 | 0.34 |
| 5 | 165265844 | rs1643967 | -1.29 | 1.543 | -1.55 | 0.622 | -0.071 | 0.106 | 0.106 | 0.55 | 0.83 | 0.99 | 0.85 | 0.99 | 0.75 | 0.85 |
| 5 | 165266083 | rs2921533 | -1.271 | 1.147 | -0.882 | 1.281 | 0.002 | 0.011 | 0.011 | 0 | 0.2 | 0.22 | 0.43 | 0.6 | 0.28 | 0.36 |
| 5 | 165272875 | rs12654578 | 0.618 | 1.043 | -1.162 | 1.679 | -0.027 | -0.005 | -0.005 | 0 | 0.05 | 0.01 | 0.1 | 0.31 | 0.09 | 0.21 |
| 5 | 165275263 | rs17051732 | -0.311 | 1.017 | -1.097 | 1.499 | -0.007 | -0.009 | -0.009 | 0 | 0.09 | 0.09 | 0.09 | 0.2 | 0.08 | 0.08 |
| 5 | 165278640 | rs32764 | -0.761 | 0.934 | -1.127 | 1.34 | -0.048 | -0.026 | -0.026 | 0 | 0.13 | 0.04 | 0.32 | 0.14 | 0.2 | 0.13 |
| 5 | 165279647 | rs10515908 | 0.001 | 0.259 | -0.678 | 1.438 | -0.004 | 0.038 | 0.038 | 0 | 0.06 | 0.05 | 0.1 | 0.34 | 0.11 | 0.22 |
| 5 | 165279809 | rs10515910 | -0.259 | 0.253 | -0.635 | 1.435 | 0.009 | 0.083 | 0.083 | 0 | 0.07 | 0.1 | 0.1 | 0.34 | 0.11 | 0.22 |
| 5 | 165280020 | rs17065989 | -0.036 | 0.294 | -0.624 | 1.46 | 0.007 | 0.077 | 0.077 | 0 | 0.07 | 0.06 | 0.1 | 0.34 | 0.11 | 0.22 |
| 5 | 165286236 | rs6871247 | -0.845 | 0.029 | -0.42 | 1.125 | -0.008 | 0.142 | 0.142 | 0.02 | 0.18 | 0.44 | 0.16 | 0.41 | 0.15 | 0.3 |
| 5 | 165288921 | rs26438 | -0.163 | 0.249 | -0.712 | 0.447 | 0.01 | 0.012 | 0.012 | 0 | 0.49 | 0.56 | 0.56 | 0.72 | 0.38 | 0.7 |
| 5 | 165293321 | rs32720 | -0.46 | 0.755 | -0.551 | 0.702 | 0 | -0.003 | -0.003 | 0.151 | 0.4 | 0.65 | 0.37 | 0.48 | 0.34 | 0.37 |
| 5 | 165295670 | rs39978 | -1.221 | 0.658 | 0.22 | 1.197 | 0.009 | 0.176 | 0.176 | 0 | 0.08 | 0.17 | 0.21 | 0.36 | 0.09 | 0.15 |
| 5 | 165295952 | rs6869622 | -0.135 | 0.492 | 0.336 | 0.514 | -0.022 | -0.08 | -0.08 | 0 | 0.22 | 0.47 | 0.13 | 0.07 | 0.14 | 0.17 |
| 5 | 165297790 | rs40588 | 0.068 | 0.752 | -0.248 | 0.561 | -0.005 | -0.008 | -0.008 | 4.507 | 0.62 | 0.68 | 0.56 | 0.51 | 0.6 | 0.68 |
| 5 | 165300970 | rs17066036 | -0.009 | 0.721 | -0.227 | 0.952 | -0.003 | 0.087 | 0.087 | 0 | 0.05 | 0.08 | 0.1 | 0.3 | 0.06 | 0.12 |
| 5 | 165302936 | rs6898223 | 0.533 | 0.614 | -0.631 | 0.466 | -0.016 | 0.043 | 0.043 | 0 | 0.19 | 0.08 | 0.11 | 0.07 | 0.15 | 0.21 |
| 5 | 165308011 | rs10475487 | -0.689 | 1.068 | -0.491 | -0.181 | -0.01 | 0.04 | 0.04 | 0 | 0.41 | 0.38 | 0.3 | 0.1 | 0.49 | 0.28 |
| 5 | 165308407 | rs26435 | -1.178 | 0.947 | -0.472 | 0.425 | -0.003 | 0.011 | 0.011 | 0 | 0.73 | 0.69 | 0.54 | 0.69 | 0.71 | 0.69 |
| 5 | 165308685 | rs26436 | 1.998 | 0.828 | -0.367 | 0.049 | 0.031 | 0.044 | 0.044 | 0 | 0.23 | 0.11 | 0.11 | 0.13 | 0.13 | 0.25 |
| 5 | 165308760 | rs4868760 | -1.238 | 1.254 | -0.152 | 0.277 | 0.009 | 0.015 | 0.015 | 0 | 0.49 | 0.59 | 0.43 | 0.42 | 0.56 | 0.4 |
| 5 | 165309208 | rs10475761 | 0.173 | 1.399 | -0.534 | -0.324 | 0 | -0.029 | -0.029 | 0 | 0.22 | 0.28 | 0.17 | 0.04 | 0.18 | 0.2 |
| 5 | 165311430 | rs7734483 | 0.658 | 1.35 | -1.186 | -0.192 | -0.009 | 0.046 | 0.046 | 0 | 0.19 | 0.08 | 0.1 | 0.09 | 0.15 | 0.2 |
| 5 | 165312291 | rs10515911 | 0.431 | 1.742 | -1.157 | -0.166 | -0.063 | 0.053 | 0.053 | 0 | 0.16 | 0.03 | 0.11 | 0.06 | 0.27 | 0.08 |
| 5 | 165313230 | rs10060086 | -0.457 | 1.422 | -0.653 | -0.083 | -0.011 | 0.027 | 0.027 | 0 | 0.41 | 0.67 | 0.3 | 0.16 | 0.43 | 0.28 |
| 5 | 165313869 | rs396769 | 0.499 | 0.774 | -0.63 | -0.033 | 0.008 | 0.013 | 0.013 | 0 | 0.3 | 0.25 | 0.19 | 0.14 | 0.2 | 0.3 |
| 5 | 165319876 | rs2861624 | -0.596 | 1.026 | -0.551 | 0.499 | 0.006 | 0.008 | 0.008 | 0.913 | 0.5 | 0.4 | 0.42 | 0.45 | 0.45 | 0.42 |
| 5 | 165328421 | rs1982325 | -0.66 | 1.021 | -0.008 | 0.618 | -0.002 | -0.003 | -0.003 | 0.193 | 0.52 | 0.6 | 0.45 | 0.49 | 0.46 | 0.43 |
| 5 | 165330062 | rs2161185 | 0.223 | 0.853 | -0.161 | 0.697 | -0.022 | 0.001 | 0.001 | 0 | 0.27 | 0.32 | 0.13 | 0.24 | 0.15 | 0.2 |
| 5 | 165334368 | rs1116099 | -0.969 | 1.154 | -0.121 | 0.655 | -0.001 | -0.001 | -0.001 | 0.187 | 0.48 | 0.47 | 0.42 | 0.44 | 0.42 | 0.35 |
| 5 | 165335027 | rs4868761 | 0.394 | 1.318 | -0.151 | 0.579 | -0.048 | -0.039 | -0.039 | 3.214 | 0.19 | 0.06 | 0.27 | 0.24 | 0.26 | 0.15 |
| 5 | 165348080 | rs6873292 | -1.809 | 0.874 | 0.326 | 0.331 | 0.044 | 0.026 | 0.026 | 0.082 | 0.05 | 0.32 | 0.05 | 0.03 | 0.03 | 0.02 |
| 5 | 165350417 | rs6884869 | 0.011 | 1.071 | 0.356 | 0.404 | 0.009 | -0.019 | -0.019 | 0 | 0.21 | 0.31 | 0.29 | 0.16 | 0.24 | 0.14 |
| 5 | 165358649 | rs4868953 | -0.932 | 0.818 | 0.34 | 0.614 | -0.002 | -0.002 | -0.002 | 1.487 | 0.45 | 0.54 | 0.28 | 0.39 | 0.37 | 0.3 |
| 5 | 165372292 | rs1975525 | 1.262 | 0.39 | -0.212 | 1.027 | -0.031 | -0.081 | -0.081 | 0.638 | 0.13 | 0.01 | 0.1 | 0.19 | 0.16 | 0.09 |
| 5 | 165374321 | rs26443 | -0.451 | 0.655 | 0.971 | 1.042 | 0.022 | -0.078 | -0.078 | 3.509 | 0.21 | 0.42 | 0.2 | 0.08 | 0.26 | 0.15 |
| 5 | 165390453 | rs248548 | -1.387 | 0.695 | 0.888 | 1.141 | -0.001 | 0.007 | 0.007 | 0.48 | 0.42 | 0.46 | 0.29 | 0.45 | 0.41 | 0.34 |
| 5 | 165390686 | rs7737136 | -0.151 | 0.646 | 0.91 | 1.15 | -0.045 | 0.047 | 0.047 | 0 | 0.26 | 0.34 | 0.11 | 0.36 | 0.12 | 0.19 |
| 5 | 165402097 | rs830230 | -1.414 | 0.675 | 0.886 | 1.173 | 0.004 | 0.001 | 0.001 | 0.44 | 0.42 | 0.46 | 0.42 | 0.55 | 0.45 | 0.36 |
| 5 | 165405770 | rs7733351 | -0.329 | 0.663 | 0.915 | 1.084 | -0.043 | 0.048 | 0.048 | 0 | 0.27 | 0.41 | 0.13 | 0.33 | 0.14 | 0.2 |
| 5 | 165406607 | rs2591580 | 0.681 | 0.804 | 0.73 | 0.952 | -0.075 | 0.02 | 0.02 | 0 | 0.14 | 0.01 | 0.14 | 0.08 | 0.23 | 0.16 |
| 5 | 165407709 | rs2591578 | 0.313 | 0.788 | 0.906 | 0.948 | -0.004 | 0.002 | 0.002 | 0 | 0.16 | 0.16 | 0.17 | 0.09 | 0.26 | 0.18 |
| 5 | 165411778 | rs153284 | 0.446 | 0.739 | 0.778 | 0.97 | -0.082 | -0.029 | -0.029 | 0 | 0.15 | 0.01 | 0.26 | 0.18 | 0.26 | 0.19 |
| 5 | 165419506 | rs1445716 | -1.703 | 0.465 | 0.989 | 1.045 | 0.005 | 0.014 | 0.014 | 0.857 | 0.45 | 0.64 | 0.3 | 0.39 | 0.42 | 0.39 |
| 5 | 165422195 | rs13354366 | -0.462 | 0.475 | 1.026 | 1.012 | 0.009 | -0.006 | -0.006 | 0 | 0.2 | 0.22 | 0.31 | 0.18 | 0.31 | 0.21 |
| 5 | 165423817 | rs930598 | -0.407 | 0.32 | 0.967 | 0.996 | -0.038 | 0.018 | 0.018 | 0 | 0.27 | 0.4 | 0.15 | 0.3 | 0.16 | 0.21 |
| 5 | 165424767 | rs6880888 | -1.333 | 0.337 | 0.978 | 0.993 | -0.005 | 0.005 | 0.005 | 0 | 0.42 | 0.49 | 0.39 | 0.47 | 0.4 | 0.38 |

| | | | | | | | | | | | | | | | | |
|---|---|---|---|---|---|---|---|---|---|---|---|---|---|---|---|---|
| 5 | 165429598 | rs3812010 | -0.225 | 0.331 | 0.856 | 0.993 | -0.041 | 0.033 | 0.033 | 0 | 0.25 | 0.37 | 0.11 | 0.31 | 0.14 | 0.19 |
| 5 | 165431940 | rs7729971 | -1.23 | 0.418 | 0.756 | 1.007 | -0.006 | 0.007 | 0.007 | 0.378 | 0.43 | 0.49 | 0.39 | 0.5 | 0.41 | 0.39 |
| 5 | 165438394 | rs6555670 | -1.63 | 0.419 | 0.266 | 0.995 | 0.018 | -0.016 | -0.016 | 0.122 | 0.49 | 0.84 | 0.47 | 0.51 | 0.44 | 0.43 |
| 5 | 165451462 | rs31736 | -1.947 | 0.031 | 0.407 | 1.295 | 0.01 | -0.032 | -0.032 | 0 | 0.47 | 0.87 | 0.46 | 0.5 | 0.46 | 0.42 |
| 5 | 165456536 | rs10515916 | -0.656 | 0.24 | 1.391 | 1.28 | -0.026 | 0.1 | 0.1 | 0.323 | 0.15 | 0.25 | 0.08 | 0.33 | 0.1 | 0.15 |
| 5 | 165462856 | rs31733 | -2.122 | 0.358 | 1.706 | 1.212 | 0 | -0.012 | -0.012 | 0 | 0.41 | 0.68 | 0.43 | 0.52 | 0.46 | 0.38 |
| 5 | 165463582 | rs31732 | -0.451 | 0.382 | 2.004 | 1.202 | -0.003 | 0.001 | 0.001 | 0.945 | 0.21 | 0.23 | 0.16 | 0.07 | 0.25 | 0.18 |
| 5 | 165465607 | rs7712594 | -2.444 | 0.382 | 1.846 | 1.213 | -0.005 | -0.043 | -0.043 | 0 | 0.43 | 0.83 | 0.45 | 0.52 | 0.44 | 0.37 |
| 5 | 165466747 | rs10057613 | -2.254 | 0.4 | 1.856 | 1.221 | -0.001 | 0.002 | 0.002 | 1.949 | 0.42 | 0.83 | 0.31 | 0.39 | 0.4 | 0.35 |
| 5 | 165473194 | rs830383 | 1.892 | 0.313 | 1.865 | 1.026 | 0.02 | -0.034 | -0.034 | 1.75 | 0.6 | 0.17 | 0.71 | 0.59 | 0.64 | 0.69 |
| 5 | 165478262 | rs11949873 | -0.827 | 0.418 | 1.572 | 1.157 | -0.02 | 0.149 | 0.149 | 0.298 | 0.18 | 0.38 | 0.12 | 0.36 | 0.15 | 0.17 |
| 5 | 165487589 | rs41012 | 0.065 | 1.007 | 0.912 | 0.704 | -0.073 | -0.016 | -0.016 | 0 | 0.85 | 0.99 | 0.74 | 0.84 | 0.75 | 0.85 |
| 5 | 165495408 | rs31772 | 1.484 | 0.716 | 1.42 | 0.573 | 0 | -0.009 | -0.009 | 0 | 0.69 | 0.32 | 0.65 | 0.87 | 0.69 | 0.78 |
| 5 | 165511601 | rs2861667 | 1.874 | 1.498 | 2.069 | 1.061 | 0.027 | -0.046 | -0.046 | 0 | 0.78 | 0.41 | 0.66 | 0.85 | 0.68 | 0.79 |
| 5 | 165512267 | rs716710 | 1.986 | 1.406 | 2.081 | 1.056 | 0.025 | -0.053 | -0.053 | 0 | 0.78 | 0.42 | 0.68 | 0.85 | 0.69 | 0.79 |
| 5 | 165514755 | rs13184263 | 1.778 | 1.398 | 2.346 | 1.205 | 0.091 | 0.012 | 0.012 | 0.077 | 0.86 | 0.53 | 0.74 | 0.87 | 0.82 | 0.83 |
| 5 | 165518937 | rs2644731 | 1.844 | 1.517 | 2.361 | 1.38 | 0.044 | 0.013 | 0.013 | 0 | 0.86 | 0.68 | 0.74 | 0.84 | 0.79 | 0.84 |
| 5 | 165520051 | rs11741434 | -0.411 | 1.655 | 1.563 | 1.612 | -0.022 | -0.032 | -0.032 | 0 | 0.17 | 0.03 | 0.29 | 0.18 | 0.18 | 0.16 |
| 5 | 165522828 | rs17066349 | -0.746 | 1.42 | 1.799 | 1.979 | 0.069 | 0.065 | 0.065 | 0 | 0.06 | 0.18 | 0.08 | 0.19 | 0.13 | 0.12 |
| 5 | 165533735 | rs4490611 | -1.838 | 1.186 | 1.799 | 2.066 | 0.091 | 0.172 | 0.172 | 0 | 0.13 | 0.42 | 0.31 | 0.29 | 0.16 | 0.11 |
| 5 | 165536461 | rs11957197 | -2.053 | 1.125 | 1.969 | 2.332 | 0.069 | 0.139 | 0.139 | 0.004 | 0.14 | 0.38 | 0.26 | 0.3 | 0.17 | 0.15 |
| 5 | 165578886 | rs10071148 | -1.763 | 1.151 | 2.469 | 2.184 | 0.066 | 0.115 | 0.115 | 0 | 0.12 | 0.35 | 0.27 | 0.29 | 0.14 | 0.09 |
| 5 | 165579358 | rs4537081 | 2.24 | 0.737 | 2.52 | 2.223 | 0.003 | 0.057 | 0.057 | 0 | 0.75 | 0.41 | 0.63 | 0.62 | 0.79 | 0.75 |
| 5 | 165582883 | rs830664 | 2.258 | 0.831 | 2.466 | 2.261 | 0.015 | 0.075 | 0.075 | 0 | 0.78 | 0.44 | 0.56 | 0.6 | 0.79 | 0.81 |
| 5 | 165591821 | rs830672 | 2.251 | 0.79 | 2.482 | 2.216 | -0.008 | 0.043 | 0.043 | 0.954 | 0.78 | 0.33 | 0.57 | 0.6 | 0.82 | 0.8 |
| 5 | 165595428 | rs11741022 | 2.017 | 0.939 | 2.61 | 2.218 | 0.013 | 0.082 | 0.082 | 1.334 | 0.86 | 0.34 | 0.7 | 0.71 | 0.88 | 0.9 |
| 5 | 165603524 | rs13189015 | -1.732 | 0.801 | 2.066 | 2.249 | 0.026 | 0.035 | 0.035 | 0.026 | 0.13 | 0.16 | 0.22 | 0.27 | 0.19 | 0.16 |
| 5 | 165608363 | rs830694 | 2.165 | 0.247 | 2.366 | 2.457 | -0.048 | 0.067 | 0.067 | 6.004 | 0.91 | 0.24 | 0.77 | 0.81 | 1 | 0.94 |
| 5 | 165609740 | rs830677 | 0.719 | 0.58 | 1.669 | 2.288 | -0.024 | -0.02 | -0.02 | 2.771 | 0.64 | 0.22 | 0.55 | 0.53 | 0.62 | 0.6 |
| 5 | 165610244 | rs830678 | -0.201 | -0.098 | 1.36 | 2.395 | -0.002 | 0.008 | 0.008 | 0 | 0.33 | 0.35 | 0.29 | 0.37 | 0.36 | 0.32 |
| 5 | 165625456 | rs6882017 | -1.607 | 0.314 | 0.327 | 1.61 | 0.019 | 0.115 | 0.115 | 0.295 | 0.07 | 0.27 | 0.09 | 0.15 | 0.07 | 0.09 |
| 5 | 165642395 | rs2055354 | -1.341 | 0.296 | -1.106 | 1.281 | -0.023 | -0.065 | -0.065 | 0.868 | 0.13 | 0.01 | 0.12 | 0.21 | 0.16 | 0.25 |
| 5 | 165649648 | rs2173635 | 1.133 | -1.046 | -1.565 | 0.842 | 0.095 | 0.028 | 0.028 | 0.747 | 0.71 | 0.19 | 0.84 | 0.79 | 0.86 | 0.78 |
| 5 | 165650569 | rs10039913 | -0.782 | -1.145 | -1.641 | 0.518 | 0 | 0.028 | 0.028 | 0 | 0.15 | 0.12 | 0.16 | 0.04 | 0.13 | 0.13 |
| 5 | 165653331 | rs11955631 | 1.965 | -1.297 | -0.348 | 0.587 | -0.029 | -0.043 | -0.043 | 0.843 | 0.67 | 0.47 | 0.76 | 0.86 | 0.78 | 0.7 |
| 5 | 165654235 | rs17066470 | -1.781 | -1.385 | 0.164 | 0.531 | 0.01 | 0.009 | 0.009 | 0 | 0.24 | 0.21 | 0.18 | 0.05 | 0.17 | 0.18 |
| 5 | 165655853 | rs1353920 | 2.027 | -1.461 | 0.19 | 0.663 | -0.051 | 0.014 | 0.014 | 1.895 | 0.82 | 0.33 | 0.88 | 0.78 | 0.93 | 0.84 |
| 5 | 165658790 | rs12513464 | -2.014 | -1.022 | 1.223 | 0.524 | 0.04 | -0.048 | -0.048 | 0 | 0.11 | 0.51 | 0.13 | 0.03 | 0.09 | 0.06 |
| 5 | 165666696 | rs11134419 | 1.307 | -1.153 | 1.196 | 0.765 | -0.031 | -0.012 | -0.012 | 0 | 0.85 | 0.38 | 0.71 | 0.89 | 0.92 | 0.89 |
| 5 | 165669831 | rs4868970 | 0.262 | -0.347 | 0.04 | 1.178 | -0.073 | -0.108 | -0.108 | 0.485 | 0.18 | 0.04 | 0.36 | 0.43 | 0.23 | 0.37 |
| 5 | 165670128 | rs2695990 | 0.917 | -0.435 | 0.283 | 1.665 | -0.034 | 0.049 | 0.049 | 0.819 | 0.84 | 0.34 | 0.88 | 0.78 | 0.9 | 0.83 |
| 5 | 165670188 | rs4868971 | -0.176 | -0.546 | -0.318 | 1.588 | -0.098 | -0.155 | -0.155 | 0 | 0.25 | 0.04 | 0.42 | 0.49 | 0.28 | 0.39 |
| 5 | 165672653 | rs17066533 | -0.403 | -0.643 | 0.496 | 1.538 | -0.016 | 0.115 | 0.115 | 0.371 | 0.13 | 0.52 | 0.12 | 0.17 | 0.07 | 0.13 |
| 5 | 165674398 | rs9313313 | 0.28 | -0.223 | -0.358 | 1.43 | -0.141 | 0.014 | 0.014 | 0.054 | 0.27 | 0.02 | 0.2 | 0.2 | 0.32 | 0.27 |
| 5 | 165675683 | rs1392280 | -2.048 | -0.77 | -0.514 | 1.406 | 0.004 | 0.002 | 0.002 | 0 | 0.63 | 0.73 | 0.5 | 0.6 | 0.57 | 0.7 |
| 5 | 165676703 | rs17380256 | -0.162 | -0.723 | -0.674 | 1.33 | -0.084 | 0.033 | 0.033 | 0 | 0.34 | 0.04 | 0.2 | 0.25 | 0.32 | 0.32 |
| 5 | 165680168 | rs17066554 | -0.553 | -0.826 | -0.144 | 1.544 | 0.037 | 0.035 | 0.035 | 0 | 0.16 | 0.33 | 0.19 | 0.2 | 0.18 | 0.22 |
| 5 | 165688851 | rs2695985 | 0.77 | -0.686 | -0.242 | 1.433 | -0.004 | 0.035 | 0.035 | 0.022 | 0.48 | 0.35 | 0.46 | 0.68 | 0.53 | 0.51 |
| 5 | 165692655 | rs17465447 | 0.394 | -0.522 | -0.262 | 1.518 | -0.036 | -0.026 | -0.026 | 0.328 | 0.11 | 0.05 | 0.16 | 0.16 | 0.17 | 0.22 |
| 5 | 165692938 | rs2650084 | 0.449 | -0.578 | -0.283 | 1.509 | 0.006 | 0.038 | 0.038 | 0 | 0.52 | 0.35 | 0.49 | 0.7 | 0.57 | 0.52 |
| 5 | 165693202 | rs2695983 | 1.689 | -0.592 | -0.256 | 1.604 | 0.008 | 0.006 | 0.006 | 0 | 0.38 | 0.19 | 0.32 | 0.38 | 0.41 | 0.3 |
| 5 | 165694220 | rs7710884 | 0.947 | -0.553 | -0.235 | 1.628 | 0.02 | 0.025 | 0.025 | 0.181 | 0.09 | 0.12 | 0.14 | 0.17 | 0.13 | 0.21 |
| 5 | 165694928 | rs2695979 | 0.679 | -0.568 | -0.237 | 1.592 | -0.001 | 0 | 0 | 0 | 0.49 | 0.33 | 0.46 | 0.53 | 0.54 | 0.51 |
| 5 | 165695551 | rs2695978 | 0.689 | -0.567 | -0.239 | 1.547 | -0.001 | 0.034 | 0.034 | 0.168 | 0.49 | 0.35 | 0.46 | 0.65 | 0.53 | 0.51 |
| 5 | 165696562 | rs13178475 | 0.377 | -0.593 | -0.119 | 1.502 | -0.012 | 0.025 | 0.025 | 0 | 0.36 | 0.28 | 0.25 | 0.24 | 0.36 | 0.37 |
| 5 | 165699441 | rs17465517 | 0.437 | -0.673 | -0.311 | 1.491 | -0.17 | 0.06 | 0.06 | 0 | 0.34 | 0.02 | 0.23 | 0.2 | 0.38 | 0.31 |
| 5 | 165703913 | rs1320005 | 1.327 | -0.577 | -0.104 | 1.607 | -0.04 | 0.201 | 0.201 | 4.626 | 0.89 | 0.36 | 0.7 | 0.72 | 0.93 | 0.85 |
| 5 | 165704020 | rs17066578 | -1.09 | -0.575 | -0.054 | 1.602 | 0.002 | 0.254 | 0.254 | 3.09 | 0.08 | 0.53 | 0.11 | 0.18 | 0.03 | 0.11 |
| 5 | 165709922 | rs2141244 | 1.679 | -0.534 | -0.28 | 1.538 | 0.003 | 0.013 | 0.013 | 0 | 0.36 | 0.19 | 0.32 | 0.33 | 0.4 | 0.28 |
| 5 | 165712524 | rs17066591 | -0.016 | -0.465 | -0.377 | 1.463 | -0.12 | -0.071 | -0.071 | 2.311 | 0.42 | 0.05 | 0.25 | 0.41 | 0.41 | 0.37 |
| 5 | 165719650 | rs955423 | -0.836 | -0.341 | -0.125 | 1.441 | -0.008 | 0 | 0 | 0.009 | 0.56 | 0.49 | 0.41 | 0.53 | 0.54 | 0.63 |
| 5 | 165721819 | rs1696939 | -0.964 | -0.32 | -0.05 | 1.421 | -0.009 | -0.003 | -0.003 | 0 | 0.57 | 0.52 | 0.42 | 0.57 | 0.55 | 0.64 |
| 5 | 165722142 | rs1875185 | 0.695 | -0.342 | 0.03 | 1.324 | 0.024 | 0.085 | 0.085 | 0.4 | 0.94 | 0.71 | 0.73 | 0.9 | 0.94 | 0.93 |
| 5 | 165723824 | rs1527821 | -0.768 | -0.398 | -0.018 | 1.364 | -0.005 | -0.005 | -0.005 | 0 | 0.55 | 0.51 | 0.41 | 0.42 | 0.54 | 0.62 |
| 5 | 165726275 | rs951664 | -0.242 | -0.177 | 0.171 | 1.417 | 0.041 | 0.065 | 0.065 | 1.436 | 0.14 | 0.44 | 0.17 | 0.16 | 0.14 | 0.27 |
| 5 | 165728353 | rs1392282 | -0.324 | -0.148 | 0.284 | 1.425 | 0.017 | 0.021 | 0.021 | 0.049 | 0.15 | 0.33 | 0.16 | 0.17 | 0.14 | 0.27 |
| 5 | 165731079 | rs2088689 | -0.832 | -0.301 | -0.023 | 1.322 | -0.007 | -0.005 | -0.005 | 0.741 | 0.22 | 0.64 | 0.45 | 0.21 | 0.2 | 0.36 |
| 5 | 165741622 | rs6871234 | -0.938 | -0.291 | 0.081 | 1.36 | -0.027 | 0.043 | 0.043 | 0 | 0.09 | 0.24 | 0.28 | 0.14 | 0.07 | 0.09 |
| 5 | 165745076 | rs11742752 | 0.239 | -0.253 | -0.215 | 1.277 | 0.024 | -0.001 | -0.001 | 0 | 0.16 | 0.26 | 0.19 | 0.14 | 0.21 | 0.29 |

| | | | | | | | | | | | | | | | |
|---|---|---|---|---|---|---|---|---|---|---|---|---|---|---|---|
| 5 | 165746687 | rs7700439 | -0.661 | -0.456 | -0.645 | 1.109 | -0.051 | -0.009 | -0.009 | 0 | 0.12 | 0.34 | 0.23 | 0.12 | 0.05 | 0.07 |
| 5 | 165747205 | rs1540718 | 1.679 | -0.609 | -1.058 | 1.055 | -0.021 | -0.023 | -0.023 | 0 | 0.31 | 0.07 | 0.25 | 0.29 | 0.34 | 0.25 |
| 5 | 165747691 | rs1799553 | -0.443 | -0.556 | -1.125 | 1.025 | -0.128 | 0.092 | 0.092 | 0.009 | 0.58 | 0.96 | 0.76 | 0.73 | 0.6 | 0.63 |
| 5 | 165750226 | rs10516013 | 0.5 | -0.38 | -1.387 | 0.886 | -0.107 | 0.092 | 0.092 | 0.064 | 0.4 | 0.01 | 0.23 | 0.24 | 0.36 | 0.37 |
| 5 | 165752621 | rs6863659 | -1.678 | -0.783 | -1.24 | 1.18 | 0.071 | -0.123 | -0.123 | 3.424 | 0.38 | 0.93 | 0.53 | 0.47 | 0.28 | 0.44 |
| 5 | 165752915 | rs4868974 | 0.524 | -0.348 | -0.858 | 1.206 | -0.002 | -0.005 | -0.005 | 0 | 0.45 | 0.3 | 0.33 | 0.48 | 0.42 | 0.41 |
| 5 | 165752952 | rs6864006 | 0.552 | -0.345 | -0.868 | 1.064 | -0.003 | 0.055 | 0.055 | 0 | 0.45 | 0.29 | 0.33 | 0.33 | 0.42 | 0.41 |
| 5 | 165753097 | rs830818 | 0.753 | -0.436 | -0.818 | 1.198 | -0.003 | -0.002 | -0.002 | 0.007 | 0.31 | 0.32 | 0.29 | 0.29 | 0.34 | 0.24 |
| 5 | 165758259 | rs1392381 | 0.043 | -0.369 | -0.582 | 1.243 | 0.042 | 0.036 | 0.036 | 0.056 | 0.81 | 0.54 | 0.58 | 0.77 | 0.78 | 0.74 |
| 5 | 165758618 | rs1392382 | 0.854 | -0.224 | -0.577 | 1.475 | 0.004 | 0.047 | 0.047 | 0.112 | 0.75 | 0.29 | 0.63 | 0.6 | 0.74 | 0.72 |
| 5 | 165762515 | rs2861807 | -0.742 | -0.869 | -0.083 | 1.48 | -0.009 | 0.131 | 0.131 | 0.009 | 0.06 | 0.18 | 0.08 | 0.17 | 0.03 | 0.06 |
| 5 | 165763260 | rs2133726 | -0.97 | -0.872 | -0.05 | 1.478 | -0.002 | 0.17 | 0.17 | 0 | 0.06 | 0.23 | 0.08 | 0.17 | 0.03 | 0.06 |
| 5 | 165767694 | rs1527814 | 0.441 | -1.07 | -0.422 | 1.217 | 0.013 | 0.013 | 0.013 | 0 | 0.28 | 0.07 | 0.21 | 0.21 | 0.28 | 0.23 |
| 5 | 165769258 | rs1527818 | 0.482 | -1.071 | -0.437 | 1.16 | 0.008 | 0.011 | 0.011 | 0 | 0.27 | 0.06 | 0.2 | 0.21 | 0.28 | 0.22 |
| 5 | 165771482 | rs1799548 | -0.497 | -1.063 | -0.324 | 1.125 | -0.019 | -0.015 | -0.015 | 0.693 | 0.4 | 0.42 | 0.42 | 0.3 | 0.34 | 0.33 |
| 5 | 165775731 | rs1799550 | -0.645 | -1.14 | -0.457 | 1.105 | 0.024 | 0.016 | 0.016 | 0 | 0.43 | 0.35 | 0.43 | 0.34 | 0.35 | 0.34 |
| 5 | 165778894 | rs1566430 | -0.741 | -1.049 | -0.433 | 0.843 | -0.049 | -0.01 | -0.01 | 0 | 0.16 | 0.3 | 0.23 | 0.14 | 0.08 | 0.17 |
| 5 | 165779913 | rs830822 | -0.42 | -0.296 | -0.597 | 0.95 | 0.004 | -0.001 | -0.001 | 0 | 0.44 | 0.56 | 0.52 | 0.51 | 0.38 | 0.38 |
| 5 | 165788235 | rs1392383 | 0.359 | -0.091 | -0.448 | 0.934 | 0.014 | 0.011 | 0.011 | 0 | 0.56 | 0.34 | 0.65 | 0.59 | 0.57 | 0.65 |
| 5 | 165788822 | rs1527824 | -0.029 | -0.156 | 0.081 | 0.892 | 0.03 | 0.028 | 0.028 | 0.011 | 0.84 | 0.53 | 0.8 | 0.82 | 0.84 | 0.82 |
| 5 | 165794574 | rs1392384 | -0.429 | -0.176 | 0.522 | 0.892 | 0.027 | 0.015 | 0.015 | 0 | 0.87 | 0.7 | 0.82 | 0.83 | 0.83 | 0.82 |
| 5 | 165798321 | rs17066756 | 0.302 | 0.15 | -0.084 | 1.304 | -0.031 | -0.039 | -0.039 | 0 | 0.07 | 0.01 | 0.08 | 0.17 | 0.1 | 0.06 |
| 5 | 165800113 | rs1392385 | -0.163 | 0.097 | 0.049 | 1.28 | 0.003 | 0.035 | 0.035 | 0 | 0.08 | 0.07 | 0.09 | 0.18 | 0.1 | 0.06 |
| 5 | 165800248 | rs1799573 | 0.996 | 0.037 | 0.047 | 1.157 | 0.007 | 0.047 | 0.047 | 0 | 0.76 | 0.39 | 0.71 | 0.65 | 0.73 | 0.76 |
| 5 | 165811122 | rs350921 | 1.088 | 0.086 | -0.337 | 1.069 | -0.004 | 0.072 | 0.072 | 0 | 0.5 | 0.23 | 0.55 | 0.39 | 0.46 | 0.61 |
| 5 | 165820029 | rs1799577 | 0.737 | -0.087 | -0.055 | 0.985 | -0.002 | -0.022 | -0.022 | 0 | 0.76 | 0.33 | 0.73 | 0.79 | 0.77 | 0.76 |
| 5 | 165821645 | rs1392378 | 0.768 | -0.116 | -0.032 | 1.033 | -0.001 | -0.015 | -0.015 | 0.05 | 0.76 | 0.35 | 0.73 | 0.76 | 0.76 | 0.75 |
| 5 | 165825034 | rs17066814 | 0.54 | -0.291 | -0.266 | 0.987 | -0.007 | 0.001 | 0.001 | 0 | 0.09 | 0.12 | 0.12 | 0.1 | 0.1 | 0.13 |
| 5 | 165826149 | rs2861762 | 0.535 | -0.292 | -0.209 | 1.223 | 0 | 0.057 | 0.057 | 0 | 0.76 | 0.34 | 0.68 | 0.59 | 0.76 | 0.76 |
| 5 | 165827247 | rs350933 | -0.306 | -0.325 | 0.234 | 1.103 | -0.001 | -0.059 | -0.059 | 0 | 0.1 | 0.34 | 0.07 | 0.03 | 0.07 | 0.06 |
| 5 | 165827805 | rs830920 | -0.942 | -0.384 | -0.017 | 1.082 | 0.015 | -0.009 | -0.009 | 0.176 | 0.44 | 0.65 | 0.39 | 0.46 | 0.44 | 0.33 |
| 5 | 165832501 | rs1696923 | 0.805 | -0.386 | -0.188 | 0.993 | 0 | 0.012 | 0.012 | 0 | 0.25 | 0.14 | 0.14 | 0.19 | 0.26 | 0.13 |
| 5 | 165833633 | rs11134425 | 0.944 | -0.417 | -0.204 | 0.937 | -0.023 | -0.023 | -0.023 | 1.45 | 0.49 | 0.1 | 0.52 | 0.49 | 0.47 | 0.54 |
| 5 | 165836374 | rs350934 | 1.024 | -0.554 | -0.443 | 0.85 | 0.007 | -0.014 | -0.014 | 0.018 | 0.22 | 0.29 | 0.15 | 0.18 | 0.26 | 0.13 |
| 5 | 165844724 | rs1839508 | 1.175 | -0.379 | -1.277 | 0.96 | -0.113 | 0.03 | 0.03 | 0 | 0.42 | 0.03 | 0.41 | 0.3 | 0.39 | 0.47 |
| 5 | 165847422 | rs17066876 | 0.074 | -0.689 | -0.538 | 0.845 | -0.008 | -0.002 | -0.002 | 0 | 0.1 | 0.13 | 0.12 | 0.12 | 0.11 | 0.14 |
| 5 | 165856217 | rs350926 | -1.214 | -0.675 | -0.653 | 0.851 | -0.044 | 0.089 | 0.089 | 0 | 0.21 | 0.52 | 0.26 | 0.3 | 0.18 | 0.2 |
| 5 | 165859261 | rs751821 | 0.361 | -0.448 | -1.862 | 0.634 | 0.016 | 0.002 | 0.002 | 0.934 | 0.76 | 0.22 | 0.77 | 0.82 | 0.86 | 0.92 |
| 5 | 165882543 | rs2020170 | 0.158 | 0.388 | -1.934 | 0.583 | 0.029 | -0.015 | -0.015 | 0 | 0.51 | 0.19 | 0.49 | 0.56 | 0.58 | 0.78 |
| 5 | 165884072 | rs7728439 | 0.078 | 0.413 | -1.918 | 0.5 | 0.03 | 0.005 | 0.005 | 0.278 | 0.49 | 0.19 | 0.63 | 0.59 | 0.57 | 0.77 |
| 5 | 165885226 | rs2964283 | 0.403 | 1.467 | -1.592 | 0.125 | -0.035 | 0.062 | 0.062 | 0 | 0.69 | 0.84 | 0.62 | 0.87 | 0.58 | 0.59 |
| 5 | 165886820 | rs1799594 | 1.06 | 1.745 | -1.166 | 0.503 | 0.004 | 0.002 | 0.002 | 0.106 | 0.75 | 0.29 | 0.79 | 0.75 | 0.75 | 0.81 |
| 5 | 165900741 | rs6896975 | -1.3 | 1.797 | -1.295 | 0.351 | -0.038 | 0.071 | 0.071 | 0 | 0.15 | 0.03 | 0.11 | 0.01 | 0.16 | 0.12 |
| 5 | 165901316 | rs10516014 | 0.021 | 1.965 | -1.611 | 0.526 | -0.01 | -0.055 | -0.055 | 0 | 0.35 | 0.15 | 0.39 | 0.58 | 0.37 | 0.55 |
| 5 | 165901702 | rs1450613 | 0.015 | 1.884 | -1.288 | 0.544 | 0.001 | -0.004 | -0.004 | 2.178 | 0.07 | 0.06 | 0.03 | 0.12 | 0.05 | 0.07 |
| 5 | 165904364 | rs1405911 | 1.593 | 1.914 | -0.633 | 0.377 | 0.018 | 0.062 | 0.062 | 0.009 | 0.42 | 0.26 | 0.44 | 0.2 | 0.42 | 0.29 |
| 5 | 165909103 | rs10066767 | -0.646 | 1.961 | 0.379 | 0.715 | 0.004 | -0.015 | -0.015 | 0 | 0.07 | 0.09 | 0.04 | 0.02 | 0.07 | 0.07 |
| 5 | 165914041 | rs2961863 | 1.87 | 1.986 | 0.361 | 0.671 | 0.052 | -0.089 | -0.089 | 0 | 0.84 | 0.41 | 0.9 | 0.97 | 0.85 | 0.91 |
| 5 | 165919692 | rs2964287 | 1.737 | 1.967 | 0.294 | 0.684 | 0.007 | 0.002 | 0.002 | 0.969 | 0.8 | 0.27 | 0.84 | 0.79 | 0.82 | 0.88 |
| 5 | 165928002 | rs10076410 | -1.386 | 2.1 | 0.443 | 0.609 | 0.003 | -0.029 | -0.029 | 0 | 0.09 | 0.14 | 0.05 | 0.01 | 0.07 | 0.04 |
| 5 | 165929354 | rs13357804 | -0.953 | 1.748 | -0.633 | 0.445 | -0.003 | -0.006 | -0.006 | 0 | 0.08 | 0.05 | 0.04 | 0.01 | 0.07 | 0.04 |
| 5 | 165933768 | rs10516015 | -1.296 | 1.685 | -0.791 | 0.405 | -0.029 | 0.052 | 0.052 | 0 | 0.1 | 0.01 | 0.06 | 0.01 | 0.09 | 0.04 |
| 5 | 165939570 | rs10462882 | -0.113 | 1.472 | -0.694 | 0.666 | -0.004 | -0.022 | -0.022 | 0 | 0.31 | 0.17 | 0.52 | 0.66 | 0.33 | 0.5 |
| 5 | 165943379 | rs896088 | 1.43 | 0.967 | -0.13 | 0.746 | -0.004 | 0.005 | 0.005 | 1.769 | 0.74 | 0.17 | 0.8 | 0.74 | 0.82 | 0.77 |
| 5 | 165945586 | rs4868988 | -1.656 | 0.924 | 0.639 | 0.695 | 0.043 | 0.001 | 0.001 | 0 | 0.16 | 0.28 | 0.15 | 0.15 | 0.18 | 0.13 |
| 5 | 165947835 | rs2961857 | 2.602 | 0.875 | 0.652 | 0.976 | -0.006 | -0.004 | -0.004 | 0 | 0.7 | 0.17 | 0.78 | 0.7 | 0.77 | 0.72 |
| 5 | 165952411 | rs1020503 | 1.154 | 0.704 | 1.173 | 0.952 | 0 | 0.068 | 0.068 | 0.569 | 0.92 | 0.59 | 0.95 | 0.93 | 1 | 0.94 |
| 5 | 165954588 | rs1450626 | 2.171 | 0.704 | 1.115 | 0.978 | -0.045 | 0.079 | 0.079 | 0.332 | 0.88 | 0.46 | 0.91 | 0.84 | 0.97 | 0.9 |
| 5 | 165956290 | rs1450624 | 2.294 | 0.703 | 1.182 | 0.996 | -0.051 | 0.118 | 0.118 | 0.034 | 0.87 | 0.48 | 0.91 | 0.8 | 0.97 | 0.89 |
| 5 | 165973383 | rs988345 | 1.869 | 0.699 | 1.252 | 0.955 | 0.033 | -0.003 | -0.003 | 0.007 | 0.79 | 0.57 | 0.84 | 0.82 | 0.82 | 0.77 |
| 5 | 165976787 | rs1422860 | 0.485 | 0.851 | 0.267 | 1.053 | -0.035 | -0.085 | -0.085 | 0 | 0.29 | 0.03 | 0.49 | 0.42 | 0.28 | 0.39 |
| 5 | 165978860 | rs2964302 | 2.699 | 0.769 | 1.228 | 0.961 | 0.001 | 0.001 | 0.001 | 0.104 | 0.7 | 0.19 | 0.79 | 0.69 | 0.79 | 0.74 |
| 5 | 165984684 | rs2923169 | 0.939 | 0.929 | 1.247 | 0.939 | -0.004 | -0.004 | -0.004 | 0 | 0.93 | 0.93 | 0.97 | 0.88 | 0.95 | 0.89 |
| 5 | 165985356 | rs349606 | 1.04 | 0.927 | 1.246 | 0.947 | -0.004 | -0.005 | -0.005 | 2.365 | 0.93 | 0.93 | 0.97 | 0.88 | 0.95 | 0.89 |
| 5 | 165989827 | rs7446300 | -2.128 | 0.884 | 1.634 | 0.906 | 0.005 | -0.044 | -0.044 | 0.038 | 0.17 | 0.55 | 0.14 | 0.07 | 0.13 | 0.11 |
| 5 | 165992475 | rs1376158 | 1.383 | 0.941 | 1.56 | 0.91 | -0.003 | 0.001 | 0.001 | 0 | 0.9 | 0.81 | 0.94 | 0.91 | 0.93 | 0.91 |
| 5 | 165995991 | rs10071637 | -1.872 | 0.957 | 1.564 | 0.909 | 0.004 | -0.001 | -0.001 | 0 | 0.13 | 0.19 | 0.08 | 0.13 | 0.12 | 0.14 |
| 5 | 166002094 | rs6555701 | -2.547 | 0.915 | 1.681 | 0.922 | -0.03 | 0.015 | 0.015 | 0.372 | 0.16 | 0.64 | 0.14 | 0.2 | 0.09 | 0.11 |
| 5 | 166009956 | rs7724643 | -2.695 | 0.999 | 1.635 | 0.912 | 0.001 | 0.001 | 0.001 | 0 | 0.19 | 0.26 | 0.12 | 0.26 | 0.15 | 0.17 |

| | | | | | | | | | | | | | | | | |
|---|---|---|---|---|---|---|---|---|---|---|---|---|---|---|---|---|
| 5 | 166017077 | rs4868994 | -2.67 | 0.699 | 1.857 | 0.884 | 0.003 | 0.019 | 0.019 | 0 | 0.2 | 0.49 | 0.14 | 0.19 | 0.12 | 0.17 |
| 5 | 166021016 | rs9313330 | -3.102 | 0.724 | 1.902 | 0.952 | 0.001 | -0.002 | -0.002 | 0 | 0.26 | 0.79 | 0.26 | 0.33 | 0.17 | 0.22 |
| 5 | 166023384 | rs10035111 | -1.541 | 0.925 | 1.729 | 0.811 | 0.01 | 0.002 | 0.002 | 0 | 0.12 | 0.16 | 0.12 | 0.16 | 0.11 | 0.12 |
| 5 | 166025196 | rs17067250 | -2.34 | 0.607 | 1.676 | 0.734 | 0.014 | 0.035 | 0.035 | 0.103 | 0.11 | 0.53 | 0.11 | 0.07 | 0.06 | 0.07 |
| 5 | 166026376 | rs17067254 | -1.216 | 0.679 | 0.759 | 0.608 | -0.002 | -0.024 | -0.024 | 0 | 0.07 | 0.21 | 0.05 | 0.01 | 0.03 | 0.05 |
| 5 | 166040907 | rs17475596 | 0.118 | 0.727 | 0.408 | 0.452 | -0.002 | -0.013 | -0.013 | 0 | 0.13 | 0.13 | 0.07 | 0.08 | 0.12 | 0.1 |
| 5 | 166042592 | rs10516028 | -0.832 | 0.564 | 0.002 | 0.507 | 0.006 | -0.025 | -0.025 | 0 | 0.08 | 0.01 | 0.1 | 0.12 | 0.05 | 0.13 |
| 5 | 166047609 | rs10475790 | -1.041 | 0.426 | 0.558 | 0.586 | 0.008 | 0.009 | 0.009 | 0.193 | 0.2 | 0.54 | 0.18 | 0.18 | 0.14 | 0.2 |
| 5 | 166048608 | rs6555705 | -1.785 | 0.293 | 0.136 | 0.611 | 0.031 | 0.021 | 0.021 | 0.068 | 0.41 | 0.81 | 0.36 | 0.39 | 0.32 | 0.43 |
| 5 | 166049371 | rs347991 | 1.407 | 0.365 | -0.78 | 0.344 | -0.082 | -0.004 | -0.004 | 0.635 | 0.35 | 0.04 | 0.26 | 0.24 | 0.44 | 0.27 |
| 5 | 166061635 | rs6878847 | -0.245 | 0.65 | -0.062 | 0.171 | 0.003 | -0.013 | -0.013 | 0 | 0.23 | 0.21 | 0.17 | 0.12 | 0.29 | 0.17 |
| 5 | 166068292 | rs347994 | -0.59 | 0.186 | 0.145 | 0.041 | -0.004 | 0.01 | 0.01 | 2.087 | 0.66 | 0.64 | 0.57 | 0.53 | 0.67 | 0.68 |
| 5 | 166072943 | rs347989 | 1.238 | 0.278 | -0.302 | -0.241 | -0.04 | -0.089 | -0.089 | 0 | 0.25 | 0.07 | 0.19 | 0.36 | 0.28 | 0.34 |
| 5 | 166073001 | rs347988 | 1.937 | 0.275 | -0.399 | -0.226 | -0.079 | -0.121 | -0.121 | 0 | 0.2 | 0.02 | 0.17 | 0.31 | 0.25 | 0.25 |
| 5 | 166075339 | rs347984 | -0.914 | 0.316 | -0.337 | -0.322 | -0.008 | 0.01 | 0.01 | 0.935 | 0.63 | 0.61 | 0.57 | 0.53 | 0.68 | 0.64 |
| 5 | 166077206 | rs347980 | 1.27 | 0.306 | -0.43 | -0.254 | -0.06 | -0.129 | -0.129 | 0.082 | 0.24 | 0.03 | 0.18 | 0.36 | 0.27 | 0.3 |
| 5 | 166081123 | rs2961851 | 1.998 | 0.298 | -0.208 | -0.283 | -0.035 | -0.065 | -0.065 | 0.212 | 0.19 | 0.05 | 0.17 | 0.34 | 0.26 | 0.22 |
| 5 | 166081596 | rs10070707 | -0.877 | 0.242 | -0.131 | -0.355 | -0.001 | -0.05 | -0.05 | 2.559 | 0.43 | 0.55 | 0.4 | 0.19 | 0.43 | 0.42 |
| 5 | 166085869 | rs1349687 | -0.85 | 0.237 | -0.135 | -0.356 | -0.001 | -0.046 | -0.046 | 0 | 0.43 | 0.55 | 0.33 | 0.22 | 0.43 | 0.41 |
| 5 | 166086640 | rs6880340 | -0.361 | 0.233 | -0.111 | -0.366 | 0.001 | -0.054 | -0.054 | 0.115 | 0.38 | 0.51 | 0.31 | 0.18 | 0.4 | 0.33 |
| 5 | 166087904 | rs10060573 | 0.229 | 0.409 | -0.093 | -0.488 | -0.008 | -0.008 | -0.008 | 0 | 0.34 | 0.35 | 0.26 | 0.38 | 0.38 | 0.38 |
| 5 | 166089387 | rs1422859 | 0.79 | 0.491 | -0.382 | -0.456 | -0.023 | -0.019 | -0.019 | 0 | 0.05 | 0 | 0.16 | 0.08 | 0.09 | 0.11 |
| 5 | 166093550 | rs1452835 | -1.163 | 0.241 | 0.006 | -0.554 | -0.005 | -0.044 | -0.044 | 0 | 0.66 | 0.85 | 0.51 | 0.55 | 0.68 | 0.63 |
| 5 | 166094680 | rs1862403 | -0.591 | 0.201 | 0.071 | -0.603 | -0.024 | -0.013 | -0.013 | 0 | 0.6 | 0.79 | 0.49 | 0.56 | 0.59 | 0.57 |
| 5 | 166099369 | rs6892194 | -0.473 | 0.273 | -0.031 | -0.575 | 0.01 | -0.029 | -0.029 | 0 | 0.13 | 0.21 | 0.07 | 0.04 | 0.12 | 0.16 |
| 5 | 166099753 | rs347434 | 0.978 | 0.121 | -0.244 | -0.564 | -0.001 | 0.057 | 0.057 | 0 | 0.2 | 0.11 | 0.12 | 0.07 | 0.22 | 0.15 |
| 5 | 166109016 | rs2112708 | 0.006 | 0.177 | -0.034 | -0.421 | -0.004 | -0.004 | -0.004 | 0.125 | 0.41 | 0.44 | 0.38 | 0.47 | 0.45 | 0.49 |
| 5 | 166122413 | rs347423 | 0.482 | 0.33 | -0.456 | -0.316 | 0.002 | -0.013 | -0.013 | 0 | 0.38 | 0.18 | 0.27 | 0.43 | 0.34 | 0.4 |
| 5 | 166126427 | rs2161407 | -1.446 | 0.384 | -0.537 | -0.681 | -0.011 | 0.037 | 0.037 | 0.045 | 0.83 | 0.87 | 0.88 | 0.92 | 0.78 | 0.88 |
| 5 | 166127026 | rs1363634 | -0.571 | 0.49 | -0.036 | -0.88 | 0.028 | -0.044 | -0.044 | 0 | 0.17 | 0.25 | 0.37 | 0.08 | 0.2 | 0.16 |
| 5 | 166127214 | rs347431 | 0.444 | 0.463 | -0.047 | -0.867 | 0.012 | -0.059 | -0.059 | 0 | 0.66 | 0.62 | 0.52 | 0.84 | 0.59 | 0.71 |
| 5 | 166132929 | rs6883481 | -0.419 | -0.042 | -0.686 | -0.545 | 0 | 0.037 | 0.037 | 0.517 | 0.47 | 0.7 | 0.62 | 0.55 | 0.47 | 0.49 |
| 5 | 166148030 | rs13189908 | -1.401 | 0.091 | -0.984 | -0.827 | -0.052 | 0.085 | 0.085 | 0 | 0.8 | 0.96 | 0.87 | 0.92 | 0.75 | 0.85 |
| 5 | 166153356 | rs10475801 | 1.065 | -0.065 | -0.728 | -0.592 | 0.002 | -0.03 | -0.03 | 0 | 0.26 | 0.2 | 0.25 | 0.4 | 0.29 | 0.27 |
| 5 | 166159747 | rs6555706 | -0.491 | 0.278 | -0.556 | -0.698 | -0.012 | 0.006 | 0.006 | 0.32 | 0.44 | 0.7 | 0.6 | 0.52 | 0.46 | 0.42 |
| 5 | 166163013 | rs17400657 | 0.435 | 0.262 | -0.623 | -0.779 | -0.06 | 0.044 | 0.044 | 0.006 | 0.11 | 0.02 | 0.08 | 0.04 | 0.15 | 0.1 |
| 5 | 166166054 | rs6860339 | -0.397 | -0.035 | -0.48 | -0.701 | -0.009 | 0.004 | 0.004 | 0.348 | 0.44 | 0.69 | 0.59 | 0.52 | 0.45 | 0.42 |
| 5 | 166173738 | rs10056300 | -0.381 | -0.304 | -1.003 | -0.67 | 0.015 | 0.051 | 0.051 | 0 | 0.55 | 0.39 | 0.7 | 0.8 | 0.6 | 0.72 |
| 5 | 166175290 | rs4868776 | -0.283 | -0.37 | -1.161 | -0.67 | 0.033 | 0.105 | 0.105 | 0 | 0.55 | 0.27 | 0.7 | 0.8 | 0.62 | 0.73 |
| 5 | 166176112 | rs2112711 | -0.563 | 0.502 | -0.999 | -0.47 | -0.01 | -0.005 | -0.005 | 3.364 | 0.29 | 0.24 | 0.5 | 0.36 | 0.36 | 0.43 |
| 5 | 166180652 | rs6865926 | 1.606 | 0.276 | -0.986 | 0.112 | 0.007 | -0.016 | -0.016 | 0 | 0.23 | 0.2 | 0.2 | 0.33 | 0.2 | 0.23 |
| 5 | 166186881 | rs12515962 | 0.868 | 0.18 | -0.732 | -0.258 | 0.014 | 0.003 | 0.003 | 0 | 0.31 | 0.32 | 0.27 | 0.35 | 0.39 | 0.35 |
| 5 | 166193640 | rs1422821 | -0.428 | 0.221 | -1.245 | -0.335 | 0.012 | 0.015 | 0.015 | 1.024 | 0.68 | 0.73 | 0.64 | 0.82 | 0.78 | 0.69 |
| 5 | 166196443 | rs12515573 | 1.307 | -0.11 | -0.97 | 0.186 | -0.001 | 0.015 | 0.015 | 0 | 0.42 | 0.31 | 0.32 | 0.32 | 0.44 | 0.31 |
| 5 | 166197772 | rs1422824 | 0.113 | 0.066 | -0.835 | -0.456 | 0.006 | 0.047 | 0.047 | 0.002 | 0.57 | 0.66 | 0.56 | 0.81 | 0.65 | 0.59 |
| 5 | 166198761 | rs2059125 | 0.341 | -0.357 | -0.353 | 0.092 | -0.023 | -0.009 | -0.009 | 0 | 0.07 | 0.03 | 0.14 | 0.28 | 0.15 | 0.25 |
| 5 | 166199420 | rs11134434 | 0.006 | -0.338 | -0.371 | 0.078 | 0.061 | 0.168 | 0.168 | 0 | 0.11 | 0.36 | 0.16 | 0.3 | 0.16 | 0.25 |
| 5 | 166199780 | rs12520515 | 0.825 | -0.52 | -1.116 | -0.598 | 0 | -0.01 | -0.01 | 0 | 0.2 | 0.15 | 0.13 | 0.09 | 0.24 | 0.1 |
| 5 | 166200269 | rs11743567 | 0.264 | -0.941 | -1.08 | 0.137 | 0.016 | 0.015 | 0.015 | 1.709 | 0.82 | 0.87 | 0.78 | 0.93 | 0.92 | 0.87 |
| 5 | 166208165 | rs7709975 | 0.467 | -0.779 | -1.015 | 0.185 | 0.027 | 0.028 | 0.028 | 1.028 | 0.79 | 0.86 | 0.77 | 0.93 | 0.9 | 0.85 |
| 5 | 166221489 | rs12658839 | 0.633 | -0.366 | -0.424 | 0.488 | -0.038 | -0.024 | -0.024 | 0 | 0.82 | 0.47 | 0.88 | 0.89 | 0.91 | 0.93 |
| 5 | 166224576 | rs10516026 | -0.376 | -0.441 | -0.357 | 1.327 | 0.005 | 0.129 | 0.129 | 0 | 0.07 | 0.18 | 0.09 | 0.19 | 0.05 | 0.06 |
| 5 | 166232483 | rs6879854 | 0.478 | -0.489 | -0.371 | 1.225 | -0.04 | -0.028 | -0.028 | 0.437 | 0.81 | 0.48 | 0.88 | 0.89 | 0.91 | 0.93 |
| 5 | 166233056 | rs1432929 | 0.264 | -0.496 | -1.082 | 1.072 | -0.019 | 0.012 | 0.012 | 0 | 0.15 | 0 | 0.23 | 0.06 | 0.13 | 0.1 |
| 5 | 166245766 | rs4404699 | 0.641 | 1.091 | -0.78 | 1.759 | 0.019 | 0.016 | 0.016 | 0.27 | 0.73 | 0.63 | 0.61 | 0.36 | 0.66 | 0.45 |
| 5 | 166266941 | rs2116798 | -0.019 | 1.34 | -1.26 | 1.627 | -0.029 | -0.019 | -0.019 | 0 | 0.11 | 0.03 | 0.16 | 0.15 | 0.14 | 0.34 |
| 5 | 166268144 | rs1560667 | 1.748 | 0.974 | -1.464 | 1.64 | -0.032 | -0.003 | -0.003 | 0 | 0.55 | 0.16 | 0.38 | 0.41 | 0.56 | 0.44 |
| 5 | 166269027 | rs2116797 | 0.461 | 1.065 | -0.571 | 1.986 | 0.004 | 0.022 | 0.022 | 0.069 | 0.73 | 0.66 | 0.57 | 0.63 | 0.72 | 0.78 |
| 5 | 166272043 | rs1432927 | 0.217 | 1.444 | -1.442 | 2.048 | -0.004 | -0.006 | -0.006 | 0 | 0.11 | 0.1 | 0.1 | 0.15 | 0.13 | 0.1 |
| 5 | 166273624 | rs1432926 | -0.855 | 1.647 | -1.137 | 1.816 | -0.051 | -0.011 | -0.011 | 0 | 0.13 | 0.04 | 0.2 | 0.13 | 0.23 | 0.33 |
| 5 | 166277928 | rs7716178 | -0.962 | 1.717 | -1.022 | 1.906 | -0.008 | -0.009 | -0.009 | 0 | 0.05 | 0.03 | 0.16 | 0.12 | 0.15 | 0.24 |
| 5 | 166281216 | rs6882813 | 0.949 | 1.273 | -0.964 | 1.811 | -0.018 | 0.021 | 0.021 | 0 | 0.75 | 0.41 | 0.6 | 0.64 | 0.8 | 0.81 |
| 5 | 166283293 | rs6894057 | 0.885 | 1.37 | -1.251 | 1.502 | -0.041 | -0.041 | -0.041 | 0 | 0.08 | 0 | 0.13 | 0.19 | 0.12 | 0.16 |
| 5 | 166284870 | rs13157881 | 1.129 | 1.346 | -1.183 | 1.458 | 0 | 0.005 | 0.005 | 0 | 0.71 | 0.37 | 0.54 | 0.47 | 0.7 | 0.73 |
| 5 | 166289264 | rs6888143 | -0.831 | 1.27 | -1.616 | 0.774 | -0.046 | 0.015 | 0.015 | 0 | 0.19 | 0.05 | 0.16 | 0.19 | 0.21 | 0.25 |
| 5 | 166302658 | rs1432925 | -0.351 | 1.057 | -1.412 | 0.76 | -0.012 | 0.003 | 0.003 | 0 | 0.41 | 0.42 | 0.47 | 0.46 | 0.52 | 0.46 |
| 5 | 166302781 | rs17067690 | 0.509 | 0.963 | -1.539 | 1.194 | -0.003 | -0.002 | -0.002 | 0 | 0.12 | 0.14 | 0.07 | 0.21 | 0.11 | 0.2 |
| 5 | 166304508 | rs11952823 | 0.485 | 0.111 | -1.272 | 1.012 | -0.028 | 0.041 | 0.041 | 0 | 0.16 | 0.23 | 0.25 | 0.2 | 0.09 | 0.07 |
| 5 | 166309007 | rs2161425 | 0.069 | 0.328 | -1.47 | 0.614 | 0.01 | 0.007 | 0.007 | 0 | 0.38 | 0.35 | 0.63 | 0.44 | 0.47 | 0.3 |

| | | | | | | | | | | | | | | | | |
|---|---|---|---|---|---|---|---|---|---|---|---|---|---|---|---|---|
| 5 | 166310797 | rs6874869 | 1.172 | 0.398 | -1.56 | 0.552 | -0.023 | -0.033 | -0.033 | 0 | 0.12 | 0.07 | 0.07 | 0.33 | 0.12 | 0.22 |
| 5 | 166320265 | rs7706146 | 1.587 | -0.171 | -1.726 | -0.402 | -0.009 | 0.113 | 0.113 | 0 | 0.21 | 0.04 | 0.12 | 0.02 | 0.22 | 0.21 |
| 5 | 166325332 | rs7724662 | -0.117 | 0.479 | -1.062 | 0.36 | 0.014 | -0.002 | -0.002 | 1.937 | 0.3 | 0.32 | 0.55 | 0.3 | 0.38 | 0.25 |
| 5 | 166331561 | rs6893678 | 0.354 | 0.416 | -1.281 | 0.279 | 0.007 | -0.01 | -0.01 | 0 | 0.63 | 0.66 | 0.43 | 0.7 | 0.55 | 0.64 |
| 5 | 166332314 | rs3866446 | -0.191 | 0.298 | -1.316 | 0.425 | 0.001 | 0.014 | 0.014 | 0 | 0.27 | 0.25 | 0.38 | 0.36 | 0.28 | 0.19 |
| 5 | 166333068 | rs10057765 | 0.056 | 0.427 | -1.278 | 0.333 | 0.007 | 0.017 | 0.017 | 0 | 0.24 | 0.24 | 0.35 | 0.32 | 0.28 | 0.19 |
| 5 | 166333887 | rs4145651 | -0.144 | 0.272 | -0.798 | 0.199 | 0.001 | -0.012 | -0.012 | 0 | 0.92 | 0.95 | 0.97 | 0.99 | 0.92 | 0.87 |
| 5 | 166336283 | rs13187697 | 0.846 | 0.325 | -1.061 | 0.186 | -0.003 | -0.009 | -0.009 | 0.236 | 0.68 | 0.37 | 0.63 | 0.71 | 0.71 | 0.6 |
| 5 | 166337984 | rs13180091 | -0.167 | 0.829 | -1.518 | 0.611 | -0.013 | -0.006 | -0.006 | 0 | 0.15 | 0.06 | 0.26 | 0.16 | 0.19 | 0.19 |
| 5 | 166341244 | rs11960122 | -0.59 | 0.573 | -1.951 | 0.546 | -0.02 | -0.004 | -0.004 | 0.022 | 0.3 | 0.11 | 0.21 | 0.25 | 0.3 | 0.32 |
| 5 | 166341947 | rs10043614 | 0.69 | 0.431 | -1.427 | 0.708 | 0.001 | 0 | 0 | 0 | 0.62 | 0.57 | 0.56 | 0.4 | 0.58 | 0.58 |
| 5 | 166344410 | rs10070095 | 0.307 | 0.238 | -1.185 | 0.438 | -0.012 | -0.02 | -0.02 | 0 | 0.1 | 0.14 | 0.06 | 0.05 | 0.04 | 0.17 |
| 5 | 166345022 | rs2161424 | 0.172 | 0.511 | -1.387 | 0.645 | -0.057 | -0.059 | -0.059 | 0 | 0.76 | 0.89 | 0.64 | 0.39 | 0.67 | 0.75 |
| 5 | 166355313 | rs6863941 | -0.866 | 0.426 | -0.867 | 0.392 | 0.069 | 0.273 | 0.273 | 0.165 | 0.14 | 0.47 | 0.14 | 0.42 | 0.19 | 0.19 |
| 5 | 166359355 | rs10060669 | 0.328 | -0.456 | -1.292 | -0.335 | -0.036 | 0.012 | 0.012 | 1.713 | 0.38 | 0.55 | 0.39 | 0.4 | 0.29 | 0.38 |
| 5 | 166364505 | rs9313356 | -0.786 | -0.209 | -1.609 | -0.714 | 0.002 | -0.022 | -0.022 | 0 | 0.81 | 0.79 | 0.7 | 0.93 | 0.81 | 0.86 |
| 5 | 166365731 | rs12520749 | 0.402 | -0.565 | -1.725 | -0.555 | -0.003 | -0.001 | -0.001 | 0 | 0.4 | 0.4 | 0.4 | 0.38 | 0.3 | 0.42 |
| 5 | 166370461 | rs10045267 | 0.533 | -0.497 | -1.772 | -0.575 | -0.004 | 0 | 0 | 0 | 0.4 | 0.41 | 0.41 | 0.41 | 0.3 | 0.43 |
| 5 | 166379772 | rs3853241 | 1.468 | -0.557 | -2.207 | -0.854 | 0.002 | 0.023 | 0.023 | 0.163 | 0.16 | 0.01 | 0.07 | 0.05 | 0.11 | 0.16 |
| 5 | 166386727 | rs17067773 | 0.71 | -0.232 | -1.821 | -0.506 | 0.003 | 0.071 | 0.071 | 0 | 0.09 | 0.11 | 0.09 | 0.33 | 0.14 | 0.1 |
| 5 | 166389709 | rs384227 | 0.367 | 0.102 | -2.029 | -0.91 | -0.031 | 0.066 | 0.066 | 0.508 | 0.22 | 0.09 | 0.31 | 0.05 | 0.24 | 0.16 |
| 5 | 166391475 | rs445602 | -1.472 | 0.431 | -1.972 | -0.689 | -0.031 | -0.02 | -0.02 | 0.013 | 0.85 | 0.99 | 0.94 | 0.86 | 0.88 | 0.83 |
| 5 | 166391625 | rs173416 | -1.472 | 0.431 | -1.972 | -0.7 | -0.031 | -0.01 | -0.01 | 0 | 0.85 | 0.99 | 0.94 | 0.87 | 0.88 | 0.84 |
| 5 | 166401207 | rs13358998 | 1.111 | 0.671 | -1.377 | -0.529 | -0.026 | 0.058 | 0.058 | 0.004 | 0.22 | 0.39 | 0.32 | 0.38 | 0.19 | 0.26 |
| 5 | 166408664 | rs402360 | 0.646 | 1.043 | -2.299 | -0.648 | 0.007 | 0.013 | 0.013 | 0 | 0.23 | 0.21 | 0.38 | 0.34 | 0.33 | 0.19 |
| 5 | 166414576 | rs39956 | -0.068 | 0.125 | -2.486 | -1.547 | 0.102 | 0.246 | 0.246 | 0 | 0.49 | 0.15 | 0.36 | 0.16 | 0.64 | 0.42 |
| 5 | 166416817 | rs7707688 | 1.692 | 0.03 | -2.358 | -1.551 | -0.045 | 0.093 | 0.093 | 0 | 0.18 | 0.62 | 0.26 | 0.37 | 0.12 | 0.22 |
| 5 | 166416988 | rs17067797 | 1.837 | 0.061 | -2.205 | -1.341 | -0.002 | 0.031 | 0.031 | 0 | 0.07 | 0.01 | 0.2 | 0.01 | 0.07 | 0.09 |
| 5 | 166417375 | rs3853242 | 0.353 | 0.764 | -1.974 | -1.107 | 0.001 | 0.004 | 0.004 | 0.683 | 0.39 | 0.72 | 0.43 | 0.42 | 0.42 | 0.38 |
| 5 | 166423776 | rs2062696 | -0.91 | -0.545 | -1.758 | -0.789 | -0.003 | 0.064 | 0.064 | 0 | 0.91 | 0.92 | 0.89 | 0.54 | 0.9 | 0.91 |
| 5 | 166424076 | rs1501806 | -0.609 | -0.641 | -1.775 | -0.871 | 0.007 | -0.012 | -0.012 | 0 | 0.68 | 0.76 | 0.63 | 0.53 | 0.77 | 0.65 |
| 5 | 166426291 | rs17067831 | 0.924 | -0.018 | -2.119 | -0.54 | -0.026 | 0.051 | 0.051 | 0.756 | 0.23 | 0.63 | 0.31 | 0.37 | 0.17 | 0.23 |
| 5 | 166428172 | rs6878841 | -0.558 | -0.164 | -1.92 | 0.359 | 0.023 | 0.194 | 0.194 | 0.144 | 0.92 | 0.85 | 0.88 | 0.52 | 0.9 | 0.91 |
| 5 | 166428641 | rs10516031 | 0.368 | 0.042 | -1.964 | -0.084 | -0.039 | 0.095 | 0.095 | 0 | 0.18 | 0.01 | 0.12 | 0.03 | 0.18 | 0.17 |
| 5 | 166430508 | rs1501803 | -0.306 | 0.354 | -1.338 | 0.522 | 0.022 | 0.048 | 0.048 | 0.057 | 0.49 | 0.78 | 0.52 | 0.41 | 0.56 | 0.42 |
| 5 | 166436489 | rs1501799 | 0.184 | -0.059 | -0.168 | 1.022 | -0.004 | -0.006 | -0.006 | 0.155 | 0.55 | 0.52 | 0.39 | 0.34 | 0.53 | 0.38 |
| 5 | 166439275 | rs10462884 | 0.463 | 0.117 | 0.482 | 0.941 | 0.003 | 0.064 | 0.064 | 0 | 0.47 | 0.33 | 0.33 | 0.24 | 0.41 | 0.33 |
| 5 | 166439453 | rs13168143 | 0.498 | 0.095 | 0.425 | 0.986 | -0.001 | 0.099 | 0.099 | 0 | 0.46 | 0.28 | 0.32 | 0.23 | 0.41 | 0.33 |
| 5 | 166443856 | rs1027635 | 0.552 | -0.42 | 0.733 | 1.362 | 0.004 | 0.085 | 0.085 | 0 | 0.84 | 0.67 | 0.9 | 0.7 | 0.87 | 0.87 |
| 5 | 166445495 | rs17067854 | 0.2 | -0.878 | 0.134 | 1.296 | -0.014 | -0.028 | -0.028 | 0.076 | 0.37 | 0.21 | 0.58 | 0.46 | 0.46 | 0.56 |
| 5 | 166448679 | rs2902101 | 0.303 | -0.927 | -0.019 | 1.316 | -0.038 | -0.046 | -0.046 | 0 | 0.35 | 0.14 | 0.57 | 0.44 | 0.47 | 0.56 |
| 5 | 166456108 | rs13180352 | -0.009 | -0.872 | 0.038 | 1.268 | -0.001 | -0.009 | -0.009 | 0.629 | 0.4 | 0.36 | 0.6 | 0.49 | 0.47 | 0.58 |
| 5 | 166471331 | rs10070819 | 0.704 | -0.829 | -0.577 | 1.383 | -0.036 | 0.184 | 0.184 | 0.002 | 0.46 | 0.19 | 0.31 | 0.14 | 0.46 | 0.29 |
| 5 | 166473150 | rs2937576 | -1.05 | -1.037 | -0.686 | 1.418 | 0.02 | -0.022 | -0.022 | 0 | 0.25 | 0.72 | 0.14 | 0.41 | 0.11 | 0.21 |
| 5 | 166473584 | rs2910045 | -0.615 | -0.855 | -0.851 | 1.43 | -0.09 | -0.119 | -0.119 | 0.291 | 0.66 | 0.94 | 0.44 | 0.53 | 0.53 | 0.41 |
| 5 | 166477333 | rs10053470 | 0.856 | -0.818 | -0.775 | 1.322 | -0.091 | 0.083 | 0.083 | 0 | 0.39 | 0.12 | 0.3 | 0.2 | 0.43 | 0.21 |
| 5 | 166488639 | rs2163784 | -0.313 | -1.05 | -0.942 | 1.475 | -0.07 | -0.094 | -0.094 | 0.163 | 0.59 | 0.93 | 0.42 | 0.54 | 0.5 | 0.31 |
| 5 | 166489037 | rs1432950 | 0.889 | -1.01 | -0.804 | 1.563 | -0.083 | 0.136 | 0.136 | 0 | 0.4 | 0.14 | 0.3 | 0.14 | 0.42 | 0.2 |
| 5 | 166492773 | rs7727473 | -0.692 | -1.025 | -0.845 | 1.56 | -0.082 | -0.111 | -0.111 | 2.029 | 0.64 | 0.93 | 0.44 | 0.57 | 0.53 | 0.4 |
| 5 | 166496034 | rs7706460 | -0.901 | -1.012 | 0.069 | 1.59 | -0.043 | 0.089 | 0.089 | 0 | 0.22 | 0.4 | 0.1 | 0.39 | 0.11 | 0.19 |
| 5 | 166509116 | rs2915855 | 0.931 | -0.802 | -0.584 | 1.297 | -0.119 | 0.126 | 0.126 | 0 | 0.38 | 0.09 | 0.3 | 0.14 | 0.42 | 0.2 |
| 5 | 166525026 | rs2910044 | -0.029 | -0.891 | -0.214 | 1.486 | 0.021 | -0.006 | -0.006 | 0 | 0.52 | 0.65 | 0.41 | 0.49 | 0.5 | 0.31 |
| 5 | 166542621 | rs2915858 | -1.061 | -0.369 | -0.887 | 1.37 | -0.134 | 0.086 | 0.086 | 0 | 0.67 | 0.96 | 0.73 | 0.85 | 0.61 | 0.83 |
| 5 | 166548420 | rs2915860 | 0.416 | -0.731 | -0.545 | 1.844 | 0.015 | -0.073 | -0.073 | 0 | 0.27 | 0.01 | 0.31 | 0.33 | 0.23 | 0.26 |
| 5 | 166565306 | rs11952439 | -1.243 | 0.024 | 1.086 | 2.205 | -0.014 | 0.108 | 0.108 | 1.339 | 0.11 | 0.58 | 0.22 | 0.19 | 0.08 | 0.19 |
| 5 | 166568986 | rs11949431 | -1.22 | 0.009 | 1.777 | 2.436 | 0 | 0.279 | 0.279 | 0.607 | 0.11 | 0.53 | 0.24 | 0.34 | 0.08 | 0.21 |
| 5 | 166573520 | rs10041589 | -1.646 | 0.041 | 1.76 | 2.392 | -0.002 | -0.012 | -0.012 | 0 | 0.15 | 0.56 | 0.24 | 0.08 | 0.13 | 0.23 |
| 5 | 166584234 | rs10039891 | -1.51 | 0.173 | 1.861 | 2.488 | 0.026 | 0.255 | 0.255 | 0.257 | 0.14 | 0.46 | 0.24 | 0.37 | 0.13 | 0.24 |
| 5 | 166608309 | rs7703323 | -0.599 | 0.323 | 0.847 | 1.906 | -0.174 | 0.094 | 0.094 | 0 | 0.65 | 0.97 | 0.71 | 0.81 | 0.57 | 0.68 |
| 5 | 166630143 | rs918503 | -1.57 | 0.752 | 1.533 | 1.565 | 0.049 | -0.006 | -0.006 | 0 | 0.13 | 0.39 | 0.2 | 0.06 | 0.15 | 0.21 |
| 5 | 166649746 | rs10080168 | -1.594 | 0.803 | 1.052 | 1.597 | 0.051 | 0.033 | 0.033 | 0.181 | 0.14 | 0.43 | 0.2 | 0.12 | 0.15 | 0.19 |
| 5 | 166666643 | rs4869021 | 1.122 | 0.847 | 0.615 | 1.982 | 0.003 | 0.037 | 0.037 | 0.083 | 0.8 | 0.32 | 0.69 | 0.62 | 0.77 | 0.69 |
| 5 | 166670709 | rs17068169 | -0.973 | 0.949 | 0.864 | 2.278 | 0.001 | 0.014 | 0.014 | 0 | 0.06 | 0.03 | 0.19 | 0.22 | 0.08 | 0.12 |
| 5 | 166674933 | rs1158065 | -2.366 | 0.832 | 2.123 | 2.998 | 0.051 | 0.326 | 0.326 | 0 | 0.14 | 0.42 | 0.26 | 0.48 | 0.17 | 0.22 |
| 5 | 166678525 | rs888991 | 1.121 | 1.857 | 2.016 | 3.727 | 0.011 | -0.003 | -0.003 | 0 | 0.92 | 0.73 | 0.94 | 0.94 | 0.94 | 0.85 |
| 5 | 166688301 | rs4285251 | 1.723 | 2.04 | 1.297 | 2.604 | -0.016 | -0.027 | -0.027 | 0 | 0.57 | 0.27 | 0.53 | 0.47 | 0.51 | 0.39 |
| 5 | 166692411 | rs17068206 | -0.976 | 1.71 | 1.895 | 2.866 | 0.01 | 0.05 | 0.05 | 0.28 | 0.07 | 0.15 | 0.2 | 0.21 | 0.09 | 0.14 |
| 5 | 166693842 | rs1895285 | -0.185 | 1.601 | 2.118 | 2.892 | 0.013 | 0.13 | 0.13 | 0 | 0.07 | 0.15 | 0.05 | 0.27 | 0.09 | 0.1 |
| 5 | 166707713 | rs7700378 | -0.823 | 1.616 | 1.997 | 2.655 | -0.001 | 0 | 0 | 0 | 0.07 | 0.05 | 0.2 | 0.2 | 0.09 | 0.14 |

| | | | | | | | | | | | | | | | | |
|---|---|---|---|---|---|---|---|---|---|---|---|---|---|---|---|---|
| 5 | 166708644 | rs10052446 | 0.288 | 1.542 | 2.368 | 2.452 | 0.003 | -0.019 | -0.019 | 2.889 | 0.93 | 0.88 | 0.96 | 0.97 | 0.93 | 0.9 |
| 5 | 166713726 | rs6884481 | 0.366 | 1.585 | 2.497 | 2.443 | 0.046 | -0.006 | -0.006 | 0 | 0.94 | 0.75 | 0.96 | 0.98 | 0.93 | 0.9 |
| 5 | 166716981 | rs10043619 | -1.264 | 1.555 | 2.626 | 2.71 | 0.063 | 0.04 | 0.04 | 0.033 | 0.1 | 0.41 | 0.1 | 0.12 | 0.12 | 0.07 |
| 5 | 166718807 | rs7731005 | -1.628 | 1.518 | 2.575 | 3.093 | 0.036 | 0.235 | 0.235 | 0.13 | 0.12 | 0.4 | 0.09 | 0.38 | 0.12 | 0.15 |
| 5 | 166744741 | rs4242220 | 0.16 | 0.944 | 1.403 | 2.314 | 0.025 | 0.084 | 0.084 | 0 | 0.53 | 0.28 | 0.65 | 0.26 | 0.6 | 0.48 |
| 5 | 166748590 | rs4868790 | -0.236 | 0.686 | 1.615 | 2.656 | -0.003 | 0.002 | 0.002 | 0 | 0.55 | 0.54 | 0.68 | 0.48 | 0.59 | 0.59 |
| 5 | 166759856 | rs4868792 | -0.087 | 0.478 | 0.876 | 2.551 | 0.052 | -0.01 | -0.01 | 0.958 | 0.51 | 0.25 | 0.32 | 0.51 | 0.6 | 0.51 |
| 5 | 166763437 | rs13165082 | 0.062 | 0.171 | 0.841 | 2.126 | -0.008 | 0.201 | 0.201 | 0.049 | 0.45 | 0.21 | 0.3 | 0.17 | 0.54 | 0.4 |
| 5 | 166773534 | rs4869030 | -0.862 | 0.222 | 1.285 | 2.102 | -0.012 | -0.012 | -0.012 | 0 | 0.58 | 0.52 | 0.69 | 0.64 | 0.67 | 0.69 |
| 5 | 166775014 | rs17498388 | 0.604 | 0.431 | 0.841 | 1.719 | -0.102 | 0.088 | 0.088 | 0 | 0.21 | 0.08 | 0.17 | 0.03 | 0.36 | 0.27 |
| 5 | 166775284 | rs10077097 | -0.885 | 0.532 | 1.083 | 1.933 | -0.005 | -0.003 | -0.003 | 0 | 0.58 | 0.56 | 0.7 | 0.64 | 0.68 | 0.72 |
| 5 | 166778493 | rs4869032 | -0.702 | 0.658 | 1.15 | 1.789 | -0.008 | 0 | 0 | 0 | 0.56 | 0.52 | 0.68 | 0.58 | 0.64 | 0.7 |
| 5 | 166792419 | rs2902120 | -0.273 | 1.882 | 1.589 | 1.56 | 0.066 | -0.024 | -0.024 | 0 | 0.11 | 0.18 | 0.15 | 0.03 | 0.25 | 0.16 |
| 5 | 166793393 | rs1422531 | -1.321 | 1.788 | 1.438 | 1.66 | 0.018 | 0.031 | 0.031 | 0 | 0.18 | 0.22 | 0.18 | 0.4 | 0.26 | 0.34 |
| 5 | 166795002 | rs10045539 | 1.745 | 1.497 | 0.928 | 1.629 | 0.04 | 0.052 | 0.052 | 1.007 | 0.74 | 0.36 | 0.67 | 0.54 | 0.65 | 0.62 |
| 5 | 166797936 | rs11739983 | -1.805 | 0.645 | 0.554 | 0.998 | 0.006 | 0.015 | 0.015 | 0 | 0.25 | 0.58 | 0.27 | 0.3 | 0.23 | 0.35 |
| 5 | 166806876 | rs11749791 | -0.164 | 0.924 | 0.354 | 0.866 | -0.029 | 0.026 | 0.026 | 0.01 | 0.13 | 0.09 | 0.13 | 0.03 | 0.19 | 0.15 |
| 5 | 166820303 | rs1895264 | -0.737 | 0.749 | 0.161 | 0.671 | -0.07 | -0.01 | -0.01 | 0 | 0.62 | 0.81 | 0.51 | 0.55 | 0.5 | 0.61 |
| 5 | 166823660 | rs7727820 | -0.705 | 0.726 | 0.148 | 0.698 | 0.033 | 0.02 | 0.02 | 0 | 0.21 | 0.32 | 0.25 | 0.27 | 0.23 | 0.31 |
| 5 | 166824102 | rs11749309 | -0.207 | 0.956 | -0.356 | 0.934 | -0.031 | -0.027 | -0.027 | 2.416 | 0.16 | 0.09 | 0.22 | 0.25 | 0.21 | 0.29 |
| 5 | 166831694 | rs6863407 | -0.081 | -0.121 | -0.115 | 0.465 | -0.02 | -0.004 | -0.004 | 0 | 0.58 | 0.52 | 0.75 | 0.71 | 0.73 | 0.69 |
| 5 | 166837602 | rs10043275 | -0.103 | 0.147 | -0.292 | 0.365 | 0.002 | 0.001 | 0.001 | 0 | 0.6 | 0.59 | 0.76 | 0.73 | 0.73 | 0.7 |
| 5 | 166847334 | rs6879403 | -0.661 | 1.532 | -0.216 | 0.289 | 0.038 | -0.014 | -0.014 | 0 | 0.27 | 0.6 | 0.28 | 0.26 | 0.38 | 0.28 |
| 5 | 166848596 | rs9313380 | 0.268 | 1.099 | -0.315 | 0.498 | 0.038 | 0.105 | 0.105 | 0.036 | 0.81 | 0.57 | 0.84 | 0.68 | 0.79 | 0.66 |
| 5 | 166852229 | rs13170051 | -0.306 | 0.527 | -1.321 | -0.314 | -0.002 | 0.039 | 0.039 | 0 | 0.21 | 0.05 | 0.22 | 0.11 | 0.21 | 0.28 |
| 5 | 166861763 | rs13156620 | -0.386 | 0.271 | -0.622 | -0.751 | -0.029 | -0.156 | -0.156 | 0 | 0.27 | 0.44 | 0.23 | 0.02 | 0.24 | 0.12 |
| 5 | 166866436 | rs6887079 | -0.108 | 0.535 | -1.073 | -0.629 | 0.008 | 0.075 | 0.075 | 0 | 0.5 | 0.61 | 0.51 | 0.63 | 0.45 | 0.66 |
| 5 | 166868882 | rs4549550 | 0.116 | 0.687 | -1.767 | -1.051 | -0.091 | 0.029 | 0.029 | 0 | 0.26 | 0.04 | 0.22 | 0.2 | 0.36 | 0.26 |
| 5 | 166869355 | rs7724862 | 0.531 | 0.709 | -2.1 | -0.508 | -0.032 | -0.059 | -0.059 | 0.383 | 0.17 | 0.05 | 0.27 | 0.26 | 0.22 | 0.22 |
| 5 | 166870801 | rs6884191 | 0.14 | 0.46 | -1.949 | -0.647 | -0.002 | 0.018 | 0.018 | 0 | 0.66 | 0.76 | 0.63 | 0.76 | 0.67 | 0.77 |
| 5 | 166876427 | rs1030668 | -0.09 | 0.302 | -1.554 | -0.327 | 0 | 0.006 | 0.006 | 0 | 0.13 | 0.11 | 0.23 | 0.26 | 0.13 | 0.25 |
| 5 | 166881177 | rs1030669 | -0.608 | 0.564 | -1.816 | 0.471 | -0.013 | 0.003 | 0.003 | 0 | 0.38 | 0.2 | 0.43 | 0.61 | 0.46 | 0.46 |
| 5 | 166882756 | rs4869044 | 0.28 | 0.653 | -1.683 | 0.546 | 0 | 0.009 | 0.009 | 0.09 | 0.12 | 0.08 | 0.22 | 0.26 | 0.13 | 0.14 |
| 5 | 166885759 | rs6555747 | -0.33 | 0.423 | -0.924 | 0.971 | -0.005 | -0.017 | -0.017 | 0.013 | 0.41 | 0.49 | 0.58 | 0.66 | 0.41 | 0.48 |
| 5 | 166891209 | rs17499927 | 0.843 | 0.389 | 0.224 | 1.765 | 0 | 0.02 | 0.02 | 0 | 0.51 | 0.38 | 0.35 | 0.24 | 0.46 | 0.41 |
| 5 | 166893257 | rs1025482 | -0.792 | 0.393 | 0.395 | 1.865 | 0.003 | 0 | 0 | 0 | 0.45 | 0.5 | 0.47 | 0.56 | 0.53 | 0.51 |
| 5 | 166900671 | rs12515748 | 1.401 | 0.151 | 0.703 | 1.826 | -0.011 | 0.065 | 0.065 | 1.242 | 0.93 | 0.74 | 0.83 | 0.84 | 0.98 | 0.89 |
| 5 | 166901459 | rs13154921 | 0.119 | 0.322 | 0.608 | 1.978 | -0.004 | 0.006 | 0.006 | 0 | 0.71 | 0.65 | 0.67 | 0.58 | 0.74 | 0.78 |
| 5 | 166904555 | rs1432879 | 0.364 | 0.561 | 0.415 | 1.762 | -0.05 | 0.016 | 0.016 | 0 | 0.09 | 0.01 | 0.07 | 0.05 | 0.17 | 0.11 |
| 5 | 166904890 | rs6555750 | 0.696 | 0.448 | 0.941 | 2.017 | -0.013 | 0.019 | 0.019 | 0.046 | 0.66 | 0.38 | 0.5 | 0.47 | 0.7 | 0.72 |
| 5 | 166905160 | rs7719299 | 0.132 | 0.53 | 1.48 | 2.419 | -0.001 | -0.014 | -0.014 | 0.088 | 0.17 | 0.18 | 0.13 | 0.04 | 0.23 | 0.11 |
| 5 | 166909369 | rs12519007 | -0.343 | 0.387 | 1.491 | 2.724 | -0.001 | -0.007 | -0.007 | 0 | 0.18 | 0.17 | 0.13 | 0.23 | 0.21 | 0.12 |
| 5 | 166915827 | rs6555757 | 0.733 | 0.354 | 1.782 | 2.603 | -0.023 | 0.026 | 0.026 | 0.347 | 0.95 | 0.85 | 0.86 | 0.89 | 0.99 | 0.99 |
| 5 | 166918858 | rs12652771 | 1.53 | 0.331 | 1.863 | 2.764 | -0.003 | 0.17 | 0.17 | 0.892 | 0.95 | 0.73 | 0.82 | 0.83 | 0.99 | 0.9 |
| 5 | 166919900 | rs12659544 | -0.293 | 0.333 | 1.785 | 2.837 | 0 | 0.03 | 0.03 | 0 | 0.18 | 0.22 | 0.25 | 0.35 | 0.21 | 0.12 |
| 5 | 166925454 | rs1025481 | 1.22 | 0.323 | 1.745 | 2.648 | -0.031 | 0.085 | 0.085 | 0 | 0.73 | 0.41 | 0.55 | 0.48 | 0.77 | 0.76 |
| 5 | 166932455 | rs1432880 | 1.692 | 0.215 | 1.976 | 2.73 | -0.001 | 0.003 | 0.003 | 0 | 0.65 | 0.4 | 0.48 | 0.42 | 0.62 | 0.69 |
| 5 | 166937076 | rs4869051 | 1.229 | 0.22 | 1.915 | 2.582 | 0.012 | -0.039 | -0.039 | 0 | 0.7 | 0.44 | 0.75 | 0.82 | 0.64 | 0.72 |
| 5 | 166942326 | rs13181123 | 1.676 | 0.154 | 1.62 | 2.804 | 0 | -0.014 | -0.014 | 0.22 | 0.65 | 0.23 | 0.43 | 0.39 | 0.62 | 0.67 |
| 5 | 166951962 | rs1560660 | 0.42 | -0.25 | 1.796 | 2.663 | -0.023 | 0.065 | 0.065 | 0 | 0.72 | 0.87 | 0.78 | 0.87 | 0.64 | 0.78 |
| 5 | 166952417 | rs1432884 | 0.555 | -0.236 | 1.57 | 2.495 | -0.004 | 0.016 | 0.016 | 0 | 0.69 | 0.76 | 0.76 | 0.83 | 0.64 | 0.78 |
| 5 | 166952636 | rs4869052 | 0.764 | -0.219 | 1.556 | 2.503 | -0.002 | 0.002 | 0.002 | 0 | 0.68 | 0.69 | 0.75 | 0.83 | 0.64 | 0.78 |
| 5 | 166952906 | rs4309994 | 1.095 | -0.165 | 0.952 | 2.472 | 0.003 | -0.033 | -0.033 | 0 | 0.63 | 0.28 | 0.68 | 0.5 | 0.62 | 0.67 |
| 5 | 166957450 | rs12654925 | 0.114 | 0.21 | 0.63 | 2.434 | -0.001 | 0 | 0 | 0 | 0.08 | 0.08 | 0.07 | 0.16 | 0.11 | 0.22 |
| 5 | 166960247 | rs6555763 | 1.865 | 0.178 | 0.642 | 2.434 | -0.018 | 0.181 | 0.181 | 0.094 | 0.51 | 0.21 | 0.34 | 0.19 | 0.5 | 0.41 |
| 5 | 166967131 | rs6890751 | 1.46 | -0.193 | 0.543 | 2.52 | 0.023 | 0.025 | 0.025 | 0.471 | 0.57 | 0.21 | 0.4 | 0.4 | 0.61 | 0.64 |
| 5 | 166978898 | rs7709731 | -0.765 | 0.262 | 0.577 | 2.271 | 0.001 | 0 | 0 | 0 | 0.12 | 0.1 | 0.07 | 0.08 | 0.11 | 0.05 |
| 5 | 166980847 | rs1808380 | 0.918 | 0.43 | 0.523 | 2.182 | -0.008 | 0.018 | 0.018 | 0 | 0.81 | 0.86 | 0.69 | 0.87 | 0.77 | 0.75 |
| 5 | 166984125 | rs6886928 | -0.697 | 0.261 | 0.794 | 2.445 | -0.013 | 0.03 | 0.03 | 0.014 | 0.16 | 0.21 | 0.2 | 0.33 | 0.08 | 0.27 |
| 5 | 166986309 | rs6898357 | 1.48 | 0.887 | 1.64 | 2.764 | 0.023 | 0.002 | 0.002 | 0 | 0.75 | 0.43 | 0.72 | 0.75 | 0.72 | 0.73 |
| 5 | 166987039 | rs13157086 | 1.959 | 0.81 | 1.813 | 2.957 | -0.004 | -0.03 | -0.03 | 0 | 0.72 | 0.24 | 0.55 | 0.61 | 0.72 | 0.65 |
| 5 | 166987674 | rs883322 | 1.755 | 0.876 | 1.894 | 3.002 | -0.012 | -0.028 | -0.028 | 0 | 0.67 | 0.25 | 0.52 | 0.62 | 0.65 | 0.66 |
| 5 | 166988506 | rs888975 | -0.485 | 0.599 | 2.186 | 2.997 | -0.004 | 0.011 | 0.011 | 0 | 0.18 | 0.3 | 0.27 | 0.22 | 0.16 | 0.16 |
| 5 | 166992760 | rs11738110 | 1.434 | 0.551 | 2.03 | 3.002 | 0.007 | 0.002 | 0.002 | 0 | 0.71 | 0.38 | 0.58 | 0.7 | 0.67 | 0.67 |
| 5 | 166999828 | rs13153563 | 1.934 | 0.071 | 2.177 | 3.254 | 0.005 | -0.05 | -0.05 | 0 | 0.65 | 0.13 | 0.64 | 0.54 | 0.68 | 0.56 |
| 5 | 167007393 | rs9313385 | 1.477 | 0.334 | 1.54 | 3.31 | 0.007 | 0.07 | 0.07 | 0.821 | 0.72 | 0.43 | 0.73 | 0.59 | 0.71 | 0.66 |
| 5 | 167015105 | rs175874 | 1.09 | 0.37 | 1.375 | 2.902 | -0.002 | 0.017 | 0.017 | 0 | 0.9 | 0.95 | 0.92 | 0.99 | 0.88 | 0.86 |
| 5 | 167019011 | rs278016 | 1.277 | 0.781 | 0.395 | 2.45 | 0.01 | 0.032 | 0.032 | 0 | 0.74 | 0.6 | 0.72 | 0.68 | 0.71 | 0.66 |
| 5 | 167024001 | rs1903110 | -0.724 | 0.922 | 0.14 | 2.371 | 0.003 | 0.013 | 0.013 | 0.449 | 0.19 | 0.28 | 0.28 | 0.24 | 0.18 | 0.17 |

| | | | | | | | | | | | | | | | | |
|---|---|---|---|---|---|---|---|---|---|---|---|---|---|---|---|---|
| 5 | 167027636 | rs13159986 | 0.66 | 1.018 | -0.429 | 1.987 | 0.011 | 0.043 | 0.043 | 0.401 | 0.73 | 0.56 | 0.61 | 0.67 | 0.71 | 0.68 |
| 5 | 167059493 | rs2336899 | -0.042 | 0.427 | -1.49 | 0.701 | -0.094 | 0.115 | 0.115 | 0 | 0.36 | 0.13 | 0.24 | 0.13 | 0.44 | 0.3 |
| 5 | 167059744 | rs279392 | 0.631 | -0.449 | -0.225 | 2.123 | 0.002 | 0.114 | 0.114 | 0.357 | 0.84 | 0.8 | 0.58 | 0.51 | 0.77 | 0.59 |
| 5 | 167064247 | rs279389 | -1.392 | -0.951 | 0.181 | 2.732 | -0.04 | 0.169 | 0.169 | 0 | 0.15 | 0.33 | 0.14 | 0.49 | 0.07 | 0.37 |
| 5 | 167070649 | rs185763 | 0.708 | -1.038 | -0.148 | 2.324 | -0.015 | -0.005 | -0.005 | 0 | 0.85 | 0.91 | 0.63 | 0.85 | 0.8 | 0.84 |
| 5 | 167075842 | rs279396 | -1.473 | -1.528 | -0.035 | 2.762 | -0.014 | 0.102 | 0.102 | 3.314 | 0.06 | 0.09 | 0.17 | 0.46 | 0.01 | 0.28 |
| 5 | 167076076 | rs279395 | 0.066 | -1.331 | -0.064 | 2.441 | -0.004 | -0.004 | -0.004 | 0 | 0.89 | 0.94 | 0.78 | 0.92 | 0.86 | 0.89 |
| 5 | 167077450 | rs203307 | -1.666 | -1.325 | 0.17 | 2.263 | -0.007 | 0.002 | 0.002 | 2.476 | 0.18 | 0.23 | 0.12 | 0.22 | 0.12 | 0.19 |
| 5 | 167082005 | rs17068914 | -1.662 | -1.343 | 0.668 | 2.229 | -0.027 | 0.003 | 0.003 | 0 | 0.23 | 0.3 | 0.45 | 0.71 | 0.16 | 0.47 |
| 5 | 167082941 | rs10072830 | -1.947 | -1.555 | 1.101 | 2.331 | -0.007 | 0.479 | 0.479 | 4.04 | 0.08 | 0.47 | 0.2 | 0.36 | 0 | 0.28 |
| 5 | 167084107 | rs2125576 | -2.036 | -1.543 | 0.173 | 2.271 | 0.027 | -0.058 | -0.058 | 0.475 | 0.28 | 0.18 | 0.3 | 0.49 | 0.22 | 0.33 |
| 5 | 167090534 | rs17068925 | -1.477 | -1.22 | 0.559 | 2.631 | 0 | 0.017 | 0.017 | 0 | 0.18 | 0.2 | 0.27 | 0.48 | 0.18 | 0.22 |
| 5 | 167123725 | rs898172 | -1.531 | -0.973 | 0.448 | 2.327 | 0.008 | -0.018 | -0.018 | 0 | 0.22 | 0.17 | 0.27 | 0.48 | 0.18 | 0.23 |
| 5 | 167140480 | rs6555766 | -1.893 | -0.298 | 1.082 | 1.898 | -0.004 | -0.042 | -0.042 | 0 | 0.12 | 0.16 | 0.05 | 0 | 0.09 | 0.09 |
| 5 | 167141603 | rs279402 | -1.827 | 0.354 | 1.912 | 2.667 | 0.045 | 0.23 | 0.23 | 0.223 | 0.12 | 0.31 | 0.31 | 0.41 | 0.18 | 0.23 |
| 5 | 167146649 | rs7713448 | -2.225 | 0.373 | 2.136 | 2.945 | 0.066 | 0.283 | 0.283 | 0.11 | 0.14 | 0.39 | 0.34 | 0.49 | 0.19 | 0.25 |
| 5 | 167147874 | rs10057680 | -1.53 | 0.295 | 2.118 | 2.776 | -0.002 | 0.012 | 0.012 | 0 | 0.24 | 0.26 | 0.26 | 0.32 | 0.18 | 0.22 |
| 5 | 167153322 | rs2337017 | -1.52 | 0.218 | 2.25 | 2.667 | 0.02 | 0.127 | 0.127 | 0 | 0.11 | 0.17 | 0.3 | 0.42 | 0.18 | 0.23 |
| 5 | 167160432 | rs7717495 | -2.025 | 0.211 | 2.526 | 2.512 | 0.006 | 0.042 | 0.042 | 0.034 | 0.29 | 0.38 | 0.34 | 0.44 | 0.26 | 0.3 |
| 5 | 167168733 | rs10064701 | -1.858 | 0.225 | 2.964 | 2.337 | 0.004 | -0.071 | -0.071 | 0 | 0.12 | 0.25 | 0.05 | 0 | 0.09 | 0.09 |
| 5 | 167168918 | rs7736577 | -1.165 | 0.242 | 3.078 | 2.294 | 0 | -0.024 | -0.024 | 0.009 | 0.05 | 0.12 | 0.06 | 0 | 0.08 | 0.04 |
| 5 | 167169182 | rs10065043 | -2.498 | 0.249 | 3.112 | 2.75 | 0.013 | 0.089 | 0.089 | 0.11 | 0.16 | 0.2 | 0.31 | 0.44 | 0.18 | 0.27 |
| 5 | 167169471 | rs6862936 | -3.299 | 0.171 | 3.282 | 2.714 | 0.004 | 0.053 | 0.053 | 0.57 | 0.25 | 0.64 | 0.39 | 0.44 | 0.26 | 0.31 |
| 5 | 167173269 | rs9313387 | -2.702 | 0.174 | 3.01 | 2.391 | 0.002 | -0.123 | -0.123 | 0.02 | 0.18 | 0.4 | 0.11 | 0.01 | 0.17 | 0.13 |
| 5 | 167196597 | rs1023629 | -2.603 | 0.17 | 2.651 | 2.564 | -0.001 | 0.069 | 0.069 | 0.064 | 0.09 | 0.32 | 0.2 | 0.17 | 0.09 | 0.11 |
| 5 | 167209606 | rs11948139 | -2.06 | 0.125 | 2.798 | 2.467 | 0.007 | -0.086 | -0.086 | 0 | 0.13 | 0.34 | 0.06 | 0 | 0.09 | 0.1 |
| 5 | 167219268 | rs10454965 | -1.12 | -0.104 | 2.108 | 2.687 | -0.022 | -0.018 | -0.018 | 0.021 | 0.08 | 0.03 | 0.38 | 0.58 | 0.1 | 0.28 |
| 5 | 167226979 | rs1549309 | -2.222 | -0.057 | 2.21 | 2.508 | 0.006 | -0.007 | -0.007 | 0 | 0.16 | 0.23 | 0.13 | 0.12 | 0.2 | 0.17 |
| 5 | 167230303 | rs7714651 | -1.736 | -0.033 | 1.958 | 2.583 | 0.03 | 0.203 | 0.203 | 0 | 0.1 | 0.3 | 0.3 | 0.5 | 0.11 | 0.2 |
| 5 | 167240944 | rs10454960 | -3.107 | -0.029 | 1.182 | 2.498 | 0 | 0 | 0 | 0 | 0.3 | 0.71 | 0.43 | 0.56 | 0.3 | 0.33 |
| 5 | 167241065 | rs10454966 | -2.503 | -0.051 | 1.22 | 1.684 | -0.016 | -0.1 | -0.1 | 0.038 | 0.22 | 0.67 | 0.13 | 0.01 | 0.19 | 0.12 |
| 5 | 167250036 | rs10060956 | -1.584 | 0.369 | 0.282 | 2.127 | 0.011 | 0.006 | 0.006 | 0 | 0.1 | 0.14 | 0.07 | 0.1 | 0.1 | 0.12 |
| 5 | 167266430 | rs12521018 | -1.369 | 0.261 | -0.169 | 2.446 | -0.044 | -0.072 | -0.072 | 0 | 0.11 | 0.01 | 0.41 | 0.63 | 0.12 | 0.34 |
| 5 | 167269157 | rs12515610 | -1.586 | -0.231 | -0.371 | 2.669 | 0.008 | -0.059 | -0.059 | 0 | 0.24 | 0.09 | 0.16 | 0.37 | 0.21 | 0.38 |
| 5 | 167270009 | rs17434346 | 1.607 | -0.439 | -1.276 | 1.948 | -0.057 | 0.174 | 0.174 | 0 | 0.44 | 0.11 | 0.46 | 0.17 | 0.56 | 0.43 |
| 5 | 167272333 | rs12109973 | 0.256 | -0.775 | -0.142 | 1.815 | -0.016 | 0.027 | 0.027 | 0 | 0.67 | 0.44 | 0.62 | 0.53 | 0.75 | 0.79 |
| 5 | 167275532 | rs17069307 | 0.59 | -0.859 | 0.723 | 2.136 | 0.006 | 0.01 | 0.01 | 0 | 0.21 | 0.11 | 0.12 | 0.16 | 0.16 | 0.13 |
| 5 | 167276307 | rs7725644 | -1.509 | -0.916 | 0.842 | 2.105 | -0.01 | 0.013 | 0.013 | 1.171 | 0.32 | 0.62 | 0.23 | 0.38 | 0.27 | 0.39 |
| 5 | 167280392 | rs2973656 | 0.703 | -0.989 | 0.821 | 1.751 | -0.004 | 0.021 | 0.021 | 0 | 0.79 | 0.8 | 0.68 | 0.44 | 0.84 | 0.72 |
| 5 | 167285008 | rs1477284 | 1.85 | -1.027 | 0.299 | 1.664 | -0.009 | 0.163 | 0.163 | 0 | 0.47 | 0.09 | 0.48 | 0.27 | 0.6 | 0.5 |
| 5 | 167296428 | rs1035373 | 0.757 | -0.891 | 0.303 | 1.588 | 0.004 | -0.001 | -0.001 | 0 | 0.15 | 0.1 | 0.04 | 0.15 | 0.08 | 0.12 |
| 5 | 167303747 | rs2193981 | 1.979 | -0.91 | 0.303 | 1.205 | 0.008 | 0.175 | 0.175 | 0 | 0.48 | 0.15 | 0.46 | 0.23 | 0.58 | 0.51 |
| 5 | 167304609 | rs17526010 | -0.711 | -0.951 | 0.299 | 1.192 | 0.003 | 0.015 | 0.015 | 0 | 0.32 | 0.04 | 0.34 | 0.75 | 0.26 | 0.38 |
| 5 | 167310427 | rs1421985 | 1.887 | -0.933 | 0.336 | 1.204 | 0.008 | 0.2 | 0.2 | 0 | 0.49 | 0.11 | 0.48 | 0.23 | 0.6 | 0.59 |
| 5 | 167315343 | rs7449290 | 0.869 | -0.929 | 1.127 | 1.153 | 0.024 | -0.083 | -0.083 | 1.631 | 0.85 | 0.42 | 0.83 | 0.98 | 0.85 | 0.98 |
| 5 | 167317659 | rs1421982 | 1.159 | -0.982 | 0.483 | 1.749 | 0.039 | -0.069 | -0.069 | 0.149 | 0.64 | 0.11 | 0.6 | 0.55 | 0.76 | 0.79 |
| 5 | 167324809 | rs2909798 | -0.83 | -0.949 | 0.489 | 1.577 | 0.015 | 0.042 | 0.042 | 0 | 0.32 | 0.07 | 0.35 | 0.76 | 0.26 | 0.39 |
| 5 | 167337407 | rs10074148 | -2.616 | 0.066 | 0.882 | 1.996 | -0.048 | -0.04 | -0.04 | 0 | 0.41 | 0.83 | 0.41 | 0.53 | 0.42 | 0.43 |
| 5 | 167340025 | rs12109455 | -1.059 | -0.247 | 0.942 | 2.247 | -0.027 | -0.098 | -0.098 | 0.371 | 0.2 | 0.05 | 0.15 | 0.38 | 0.21 | 0.33 |
| 5 | 167341594 | rs11134480 | -2.35 | -0.279 | 1.589 | 2.213 | 0.032 | 0.088 | 0.088 | 0 | 0.11 | 0.34 | 0.13 | 0.2 | 0.12 | 0.2 |
| 5 | 167343309 | rs7708105 | -2.243 | -0.302 | 1.627 | 2.005 | -0.001 | 0.026 | 0.026 | 0.015 | 0.29 | 0.6 | 0.29 | 0.52 | 0.29 | 0.41 |
| 5 | 167346289 | rs32402 | -1.25 | -0.261 | 0.896 | 2.018 | -0.028 | -0.106 | -0.106 | 0 | 0.22 | 0.08 | 0.21 | 0.47 | 0.24 | 0.38 |
| 5 | 167351652 | rs39950 | -2.335 | -0.618 | 1.246 | 1.587 | -0.009 | -0.007 | -0.007 | 0 | 0.42 | 0.53 | 0.39 | 0.59 | 0.38 | 0.44 |
| 5 | 167358788 | rs865403 | -0.981 | -0.571 | 1.399 | 1.51 | -0.006 | -0.001 | -0.001 | 0 | 0.13 | 0.15 | 0.08 | 0.11 | 0.09 | 0.03 |
| 5 | 167365439 | rs17436436 | -0.767 | -0.6 | 1.709 | 1.563 | 0.017 | 0.003 | 0.003 | 0 | 0.07 | 0.15 | 0.09 | 0.09 | 0.12 | 0.08 |
| 5 | 167366815 | rs40583 | -1.95 | -0.671 | 1.849 | 1.546 | -0.002 | -0.008 | -0.008 | 0 | 0.19 | 0.41 | 0.18 | 0.18 | 0.19 | 0.1 |
| 5 | 167367337 | rs6882229 | -0.877 | -0.66 | 1.879 | 1.554 | 0.025 | 0.046 | 0.046 | 0 | 0.06 | 0.28 | 0.04 | 0.08 | 0.04 | 0.11 |
| 5 | 167369522 | rs32415 | -2.656 | -0.666 | 1.787 | 1.55 | 0 | 0 | 0 | 0.035 | 0.26 | 0.76 | 0.23 | 0.26 | 0.25 | 0.22 |
| 5 | 167370263 | rs9647570 | -0.52 | -0.709 | 1.159 | 1.729 | -0.018 | -0.119 | -0.119 | 0 | 0.17 | 0.01 | 0.34 | 0.55 | 0.13 | 0.25 |
| 5 | 167372668 | rs7724997 | -0.971 | -0.646 | 1.32 | 1.284 | -0.001 | 0.013 | 0.013 | 0 | 0.22 | 0.26 | 0.2 | 0.4 | 0.17 | 0.34 |
| 5 | 167373493 | rs32409 | -0.74 | -0.592 | 0.992 | 1.046 | -0.008 | -0.005 | -0.005 | 0.012 | 0.12 | 0.18 | 0.09 | 0.09 | 0.09 | 0.03 |
| 5 | 167373709 | rs2617978 | -1.539 | -0.367 | 0.834 | 1.204 | -0.002 | 0.053 | 0.053 | 0 | 0.28 | 0.45 | 0.3 | 0.49 | 0.29 | 0.4 |
| 5 | 167377392 | rs32406 | 1.562 | 0.004 | 0.805 | 0.545 | 0.002 | -0.043 | -0.043 | 0 | 0.75 | 0.45 | 0.83 | 0.81 | 0.74 | 0.75 |
| 5 | 167377835 | rs864788 | 1.104 | -0.442 | 0.582 | 0.214 | 0.002 | -0.01 | -0.01 | 0 | 0.78 | 0.7 | 0.82 | 0.83 | 0.76 | 0.77 |
| 5 | 167380478 | rs9284975 | -0.582 | -0.872 | 0.57 | 0.062 | -0.052 | -0.036 | -0.036 | 0 | 0.14 | 0.29 | 0.21 | 0.07 | 0.06 | 0.1 |
| 5 | 167389940 | rs918526 | -1.425 | -0.603 | 0.324 | 0.26 | -0.011 | 0.028 | 0.028 | 0 | 0.31 | 0.44 | 0.38 | 0.42 | 0.3 | 0.4 |
| 5 | 167392011 | rs2337033 | -1.391 | -0.348 | -0.462 | -0.073 | 0.005 | 0.006 | 0.006 | 0.201 | 0.33 | 0.38 | 0.28 | 0.38 | 0.38 | 0.35 |
| 5 | 167392547 | rs1123570 | -1.727 | -0.167 | -0.659 | 0.025 | 0.002 | 0.005 | 0.005 | 0 | 0.48 | 0.91 | 0.51 | 0.54 | 0.44 | 0.47 |
| 5 | 167394010 | rs17069454 | -0.333 | -0.092 | -0.507 | -0.103 | -0.004 | 0.011 | 0.011 | 0 | 0.1 | 0.06 | 0.06 | 0 | 0.11 | 0.05 |

| | | | | | | | | | | | | | | | | |
|---|---|---|---|---|---|---|---|---|---|---|---|---|---|---|---|---|
| 5 | 167397009 | rs1422933 | -0.677 | -0.202 | -0.394 | -0.133 | -0.004 | -0.046 | -0.046 | 0 | 0.12 | 0.17 | 0.07 | 0 | 0.11 | 0.05 |
| 5 | 167397487 | rs994890 | -1.395 | -0.177 | -0.963 | -0.875 | -0.037 | 0.036 | 0.036 | 0 | 0.72 | 0.83 | 0.78 | 0.98 | 0.62 | 0.85 |
| 5 | 167397540 | rs1422931 | 0.074 | -1.3 | -0.52 | -0.181 | -0.049 | 0.025 | 0.025 | 0 | 0.33 | 0.56 | 0.49 | 0.64 | 0.18 | 0.48 |
| 5 | 167397613 | rs994889 | -0.49 | -0.869 | -0.882 | 0.084 | 0.058 | 0.015 | 0.015 | 0.17 | 0.46 | 0.75 | 0.57 | 0.64 | 0.34 | 0.53 |
| 5 | 167400620 | rs2546930 | -0.427 | -0.763 | -1.021 | 0.235 | 0.031 | -0.008 | -0.008 | 0 | 0.44 | 0.65 | 0.42 | 0.53 | 0.32 | 0.49 |
| 5 | 167407738 | rs889068 | -0.387 | -0.989 | -1.532 | -0.602 | 0.05 | 0.212 | 0.212 | 0 | 0.55 | 0.86 | 0.69 | 0.92 | 0.34 | 0.64 |
| 5 | 167417245 | rs10516039 | 0.222 | -1.32 | -0.827 | 0.222 | -0.001 | 0.001 | 0.001 | 0 | 0.21 | 0.23 | 0.11 | 0.28 | 0.12 | 0.24 |
| 5 | 167426029 | rs12522181 | -0.853 | -1.781 | -0.377 | 0.249 | -0.001 | 0 | 0 | 0 | 0.34 | 0.35 | 0.38 | 0.37 | 0.27 | 0.35 |
| 5 | 167430057 | rs2617958 | -0.089 | -2.007 | 0.632 | 0.169 | -0.043 | 0.048 | 0.048 | 0.121 | 0.24 | 0.44 | 0.12 | 0.29 | 0.11 | 0.23 |
| 5 | 167435974 | rs17069574 | 0.302 | -1.903 | 0.442 | 0.168 | -0.006 | 0.008 | 0.008 | 0 | 0.12 | 0.08 | 0.27 | 0.09 | 0.15 | 0.11 |
| 5 | 167435994 | rs17069578 | 0.262 | -1.903 | 0.451 | 0.168 | 0 | -0.001 | -0.001 | 0 | 0.13 | 0.11 | 0.27 | 0.09 | 0.15 | 0.11 |
| 5 | 167437715 | rs12153623 | 0.571 | -1.934 | 0.532 | 0.188 | 0.004 | -0.006 | -0.006 | 0 | 0.19 | 0.19 | 0.09 | 0.28 | 0.09 | 0.23 |
| 5 | 167445006 | rs2569047 | -1.25 | -2.067 | 0.543 | 0.058 | 0.021 | 0 | 0 | 0 | 0.43 | 0.67 | 0.46 | 0.39 | 0.3 | 0.37 |
| 5 | 167456359 | rs889067 | -0.974 | -1.971 | 0.394 | 0.164 | -0.002 | 0 | 0 | 0 | 0.39 | 0.6 | 0.43 | 0.38 | 0.27 | 0.36 |
| 5 | 167465427 | rs247994 | -1.224 | -1.834 | 0.279 | 0.33 | 0.004 | -0.004 | -0.004 | 0.552 | 0.39 | 0.61 | 0.42 | 0.42 | 0.28 | 0.42 |
| 5 | 167466119 | rs7714553 | 0.742 | -2.078 | 0.442 | 0.614 | 0.007 | -0.023 | -0.023 | 0 | 0.13 | 0.09 | 0.07 | 0.25 | 0.1 | 0.2 |
| 5 | 167466419 | rs1368360 | -0.724 | -1.911 | 0.786 | 0.51 | -0.044 | 0.01 | 0.01 | 0.017 | 0.16 | 0.45 | 0.31 | 0.13 | 0.1 | 0.11 |
| 5 | 167472891 | rs10516041 | 0.339 | -1.852 | 0.682 | 0.576 | -0.019 | 0.02 | 0.02 | 0 | 0.08 | 0.18 | 0.27 | 0.11 | 0.05 | 0.09 |
| 5 | 167477752 | rs247995 | -0.223 | -3.121 | 0.169 | 0.709 | 0.004 | -0.009 | -0.009 | 0.073 | 0.79 | 0.83 | 0.77 | 0.64 | 0.9 | 0.82 |
| 5 | 167478125 | rs918527 | -0.298 | -3.053 | -0.079 | 0.245 | -0.031 | 0.006 | 0.006 | 0 | 0.34 | 0.54 | 0.34 | 0.34 | 0.22 | 0.34 |
| 5 | 167482305 | rs3860783 | 0.285 | -2.991 | 0.047 | 0.239 | 0.025 | 0.001 | 0.001 | 0 | 0.26 | 0.17 | 0.12 | 0.24 | 0.18 | 0.27 |
| 5 | 167484417 | rs1347135 | -0.618 | -2.929 | 0.082 | 0.21 | -0.02 | 0 | 0 | 0 | 0.39 | 0.56 | 0.42 | 0.37 | 0.26 | 0.37 |
| 5 | 167485170 | rs247998 | 0.67 | -2.754 | 0.2 | 0.056 | 0.024 | 0.029 | 0.029 | 0.009 | 0.46 | 0.38 | 0.37 | 0.28 | 0.64 | 0.46 |
| 5 | 167485418 | rs247999 | -0.007 | -1.295 | -0.609 | -0.127 | -0.005 | 0.009 | 0.009 | 0 | 0.37 | 0.68 | 0.43 | 0.36 | 0.35 | 0.39 |
| 5 | 167488049 | rs6555782 | 0.261 | -0.414 | -0.255 | 0.012 | 0.004 | -0.008 | -0.008 | 0 | 0.45 | 0.63 | 0.37 | 0.48 | 0.55 | 0.44 |
| 5 | 167488864 | rs7705199 | 0.821 | -0.156 | 0.006 | -0.032 | 0 | 0.08 | 0.08 | 0 | 0.27 | 0.42 | 0.16 | 0.47 | 0.2 | 0.35 |
| 5 | 167493558 | rs248002 | 1.479 | 0.077 | -0.354 | -0.192 | 0.003 | -0.015 | -0.015 | 2.96 | 0.09 | 0.11 | 0.26 | 0.02 | 0.14 | 0.07 |
| 5 | 167495972 | rs248003 | -0.238 | -0.329 | -0.512 | -0.307 | -0.004 | 0.007 | 0.007 | 0 | 0.4 | 0.39 | 0.55 | 0.15 | 0.62 | 0.43 |
| 5 | 167496863 | rs248004 | 0.22 | -0.45 | -0.8 | -0.052 | -0.012 | 0.027 | 0.027 | 0 | 0.23 | 0.13 | 0.33 | 0.14 | 0.27 | 0.34 |
| 5 | 167499769 | rs9313395 | 2.201 | -0.627 | -0.893 | 0.126 | 0.005 | -0.025 | -0.025 | 0.372 | 0.12 | 0.05 | 0.07 | 0.19 | 0.1 | 0.14 |
| 5 | 167499832 | rs9313396 | 0.278 | -0.769 | -0.648 | 0.078 | 0.008 | 0.001 | 0.001 | 0 | 0.59 | 0.62 | 0.45 | 0.56 | 0.66 | 0.51 |
| 5 | 167515847 | rs6859101 | 1.345 | -1.084 | -0.774 | 0.227 | -0.012 | 0.014 | 0.014 | 0 | 0.14 | 0.14 | 0.26 | 0.3 | 0.06 | 0.22 |
| 5 | 167517336 | rs2878892 | 0.082 | -0.876 | -0.94 | -0.21 | 0.013 | 0.034 | 0.034 | 0 | 0.42 | 0.33 | 0.19 | 0.22 | 0.3 | 0.39 |
| 5 | 167519135 | rs17531616 | 0.51 | -0.701 | -1.401 | -0.661 | -0.019 | 0.145 | 0.145 | 0 | 0.29 | 0.06 | 0.12 | 0.03 | 0.24 | 0.26 |
| 5 | 167520826 | rs7736530 | 1.252 | -0.715 | -1.406 | -0.647 | -0.019 | 0.08 | 0.08 | 0 | 0.23 | 0.07 | 0.11 | 0.03 | 0.2 | 0.23 |
| 5 | 167522265 | rs11738706 | 0.937 | -0.59 | -1.002 | -0.139 | -0.009 | 0.126 | 0.126 | 0.099 | 0.13 | 0.28 | 0.11 | 0.38 | 0.13 | 0.21 |
| 5 | 167522971 | rs2116763 | -0.661 | -0.339 | -1.238 | -1.032 | -0.015 | 0.132 | 0.132 | 0 | 0.72 | 0.93 | 0.9 | 0.98 | 0.8 | 0.73 |
| 5 | 167529727 | rs7708643 | 0.07 | 0.001 | -0.715 | -0.615 | 0.001 | -0.003 | -0.003 | 1.945 | 0.57 | 0.45 | 0.32 | 0.66 | 0.48 | 0.69 |
| 5 | 167534296 | rs13155210 | 0.365 | 0.393 | -0.608 | -0.809 | 0.064 | -0.109 | -0.109 | 0 | 0.25 | 0.41 | 0.42 | 0.04 | 0.45 | 0.16 |
| 5 | 167537605 | rs2569040 | 0.32 | 0.715 | -0.69 | -0.771 | -0.004 | -0.019 | -0.019 | 0 | 0.24 | 0.17 | 0.5 | 0.32 | 0.27 | 0.22 |
| 5 | 167543205 | rs7722031 | 1.683 | 0.831 | -0.627 | -0.795 | 0.024 | -0.013 | -0.013 | 5.114 | 0.08 | 0.12 | 0.23 | 0.04 | 0.2 | 0.07 |
| 5 | 167543292 | rs12188893 | 0.214 | 0.514 | -0.581 | -1.374 | -0.001 | 0.017 | 0.017 | 0 | 0.5 | 0.57 | 0.39 | 0.65 | 0.55 | 0.55 |
| 5 | 167543830 | rs2617972 | -1.531 | 0.439 | -0.739 | -1.397 | -0.03 | 0.106 | 0.106 | 0 | 0.76 | 0.95 | 0.91 | 0.98 | 0.82 | 0.76 |
| 5 | 167545075 | rs2569049 | 1.673 | 0.434 | -0.755 | -1.375 | 0.04 | -0.073 | -0.073 | 0 | 0.15 | 0.04 | 0.24 | 0.29 | 0.05 | 0.14 |
| 5 | 167546025 | rs1077046 | -0.233 | 0.388 | -0.722 | -1.426 | -0.003 | 0.039 | 0.039 | 0 | 0.49 | 0.36 | 0.59 | 0.34 | 0.45 | 0.46 |
| 5 | 167547671 | rs2617974 | -1.302 | 0.469 | -0.664 | -1.464 | 0.016 | -0.074 | -0.074 | 0 | 0.83 | 0.74 | 0.81 | 1 | 0.75 | 0.92 |
| 5 | 167552605 | rs1897558 | 0.604 | 0.454 | -0.672 | -1.433 | 0.001 | -0.009 | -0.009 | 0 | 0.25 | 0.21 | 0.49 | 0.33 | 0.26 | 0.22 |
| 5 | 167553880 | rs10055910 | 1.152 | 0.301 | -0.695 | -1.372 | 0.021 | -0.073 | -0.073 | 0 | 0.16 | 0.26 | 0.18 | 0 | 0.23 | 0.08 |
| 5 | 167557514 | rs4976574 | -0.423 | 0.18 | -1.099 | -1.372 | 0.035 | 0.044 | 0.044 | 0.587 | 0.59 | 0.82 | 0.66 | 0.69 | 0.77 | 0.63 |
| 5 | 167560809 | rs919739 | -0.134 | 0.365 | -1.055 | -1.289 | -0.001 | -0.055 | -0.055 | 0 | 0.59 | 0.62 | 0.71 | 0.98 | 0.57 | 0.68 |
| 5 | 167567313 | rs4976537 | 0.28 | 0.333 | -1.651 | -0.731 | -0.011 | -0.019 | -0.019 | 0 | 0.28 | 0.07 | 0.41 | 0.27 | 0.28 | 0.22 |
| 5 | 167568377 | rs7723815 | -0.158 | 0.029 | -1.291 | -0.957 | -0.029 | 0.005 | 0.005 | 0 | 0.34 | 0.48 | 0.24 | 0.66 | 0.28 | 0.43 |
| 5 | 167574792 | rs2337129 | 0.991 | 0.311 | -2.13 | -0.516 | -0.042 | -0.042 | -0.042 | 0.734 | 0.14 | 0.01 | 0.13 | 0.18 | 0.2 | 0.18 |
| 5 | 167582769 | rs7709145 | -1.244 | 0.04 | -2.556 | -1.057 | -0.041 | 0.123 | 0.123 | 0 | 0.79 | 0.99 | 0.92 | 0.98 | 0.84 | 0.8 |
| 5 | 167586356 | rs4976576 | 1.024 | -0.597 | -2.805 | -0.986 | -0.066 | -0.097 | -0.097 | 0.188 | 0.23 | 0.02 | 0.37 | 0.31 | 0.26 | 0.23 |
| 5 | 167589814 | rs9637846 | 1.037 | -0.699 | -2.131 | -0.918 | -0.016 | 0.117 | 0.117 | 0 | 0.16 | 0.25 | 0.11 | 0.51 | 0.13 | 0.22 |
| 5 | 167591402 | rs891933 | -0.183 | -0.736 | -2.574 | -1.127 | -0.136 | -0.116 | -0.116 | 0 | 0.43 | 0.95 | 0.48 | 0.53 | 0.49 | 0.38 |
| 5 | 167593445 | rs7712496 | 1.649 | -0.752 | -2.665 | -1.029 | -0.067 | 0.023 | 0.023 | 0 | 0.21 | 0.01 | 0.18 | 0.16 | 0.23 | 0.23 |
| 5 | 167593968 | rs1036193 | -0.733 | -0.741 | -2.654 | -1.086 | -0.086 | -0.007 | -0.007 | 0.137 | 0.58 | 0.96 | 0.57 | 0.66 | 0.61 | 0.57 |
| 5 | 167594563 | rs1560654 | -0.316 | -0.92 | -1.982 | -1.15 | -0.031 | 0.006 | 0.006 | 0 | 0.35 | 0.47 | 0.22 | 0.63 | 0.25 | 0.41 |
| 5 | 167595121 | rs17663412 | 1.521 | -0.669 | -1.877 | -0.438 | -0.008 | -0.017 | -0.017 | 0 | 0.14 | 0.01 | 0.08 | 0.13 | 0.11 | 0.17 |
| 5 | 167596170 | rs13176788 | 1.218 | -0.681 | -1.87 | -0.474 | -0.033 | -0.069 | -0.069 | 0 | 0.24 | 0.03 | 0.36 | 0.33 | 0.23 | 0.29 |
| 5 | 167596624 | rs7728598 | -0.272 | -0.812 | -1.672 | -0.647 | 0.006 | 0.002 | 0.002 | 0 | 0.49 | 0.44 | 0.53 | 0.96 | 0.42 | 0.57 |
| 5 | 167596923 | rs7711412 | 1.149 | -0.658 | -1.48 | -0.576 | 0.071 | -0.102 | -0.102 | 0 | 0.22 | 0.4 | 0.34 | 0.03 | 0.36 | 0.16 |
| 5 | 167598091 | rs7716939 | 0.135 | -0.697 | -1.339 | -0.204 | -0.004 | -0.003 | -0.003 | 0 | 0.31 | 0.68 | 0.42 | 0.65 | 0.4 | 0.3 |
| 5 | 167598272 | rs1560651 | -0.801 | -0.442 | -0.801 | -0.077 | -0.001 | -0.061 | -0.061 | 0 | 0.52 | 0.53 | 0.57 | 0.95 | 0.44 | 0.62 |
| 5 | 167600218 | rs7707882 | -0.353 | -0.721 | -0.53 | 0.776 | -0.039 | 0.109 | 0.109 | 0 | 0.2 | 0.38 | 0.13 | 0.58 | 0.12 | 0.29 |
| 5 | 167600914 | rs6555785 | -0.316 | -0.385 | -0.348 | 1.041 | 0.021 | 0.335 | 0.335 | 2.545 | 0.08 | 0.32 | 0.03 | 0.46 | 0.04 | 0.15 |
| 5 | 167603402 | rs9313398 | -0.43 | -0.384 | -1.214 | 0.776 | -0.031 | -0.112 | -0.112 | 0 | 0.73 | 0.96 | 0.56 | 0.6 | 0.72 | 0.63 |

| | | | | | | | | | | | | | | | |
|---|---|---|---|---|---|---|---|---|---|---|---|---|---|---|---|
| 5 | 167605733 | rs4976578 | -0.917 | -0.546 | -1.058 | 0.678 | -0.02 | 0.057 | 0.057 | 0 | 0.81 | 0.98 | 0.88 | 0.93 | 0.79 | 0.77 |
| 5 | 167607224 | rs10214212 | 0.941 | -0.848 | -0.323 | 0.662 | -0.011 | -0.01 | -0.01 | 0.189 | 0.57 | 0.54 | 0.61 | 0.31 | 0.65 | 0.55 |
| 5 | 167609373 | rs4976579 | -0.852 | -0.931 | -0.25 | 1.007 | 0.023 | -0.002 | -0.002 | 0 | 0.34 | 0.79 | 0.21 | 0.38 | 0.27 | 0.33 |
| 5 | 167611397 | rs10069488 | 0.246 | -0.658 | -0.121 | 0.999 | 0 | 0.002 | 0.002 | 0.051 | 0.7 | 0.4 | 0.8 | 0.62 | 0.71 | 0.62 |
| 5 | 167611655 | rs2132545 | 0.254 | -0.657 | -0.121 | 1 | 0 | 0.005 | 0.005 | 0.047 | 0.71 | 0.4 | 0.81 | 0.62 | 0.71 | 0.64 |
| 5 | 167612389 | rs4976582 | 1.307 | -0.633 | -0.195 | 1.096 | 0.023 | 0.087 | 0.087 | 0 | 0.49 | 0.13 | 0.54 | 0.31 | 0.59 | 0.51 |
| 5 | 167615217 | rs1472357 | -1.475 | -0.63 | -0.212 | 0.627 | -0.035 | 0.051 | 0.051 | 0.687 | 0.83 | 0.98 | 0.87 | 0.93 | 0.78 | 0.78 |
| 5 | 167616743 | rs11134493 | 0.923 | -0.606 | -0.042 | 0.725 | 0.004 | 0.002 | 0.002 | 0 | 0.55 | 0.42 | 0.72 | 0.52 | 0.6 | 0.51 |
| 5 | 167618989 | rs903851 | -0.046 | -0.583 | -0.352 | 0.498 | 0.001 | 0.037 | 0.037 | 0.128 | 0.69 | 0.81 | 0.77 | 0.83 | 0.67 | 0.68 |
| 5 | 167620250 | rs903850 | 1.515 | -0.524 | -0.585 | 0.629 | 0.02 | -0.069 | -0.069 | 0 | 0.17 | 0.05 | 0.25 | 0.3 | 0.13 | 0.35 |
| 5 | 167620270 | rs7702720 | 1.739 | -0.365 | -0.697 | 0.54 | -0.119 | 0.198 | 0.198 | 0.689 | 0.32 | 0.03 | 0.36 | 0.02 | 0.42 | 0.16 |
| 5 | 167621784 | rs1472356 | -0.869 | 0.38 | 0.154 | 1.342 | -0.054 | 0.076 | 0.076 | 0.599 | 0.83 | 0.98 | 0.86 | 0.97 | 0.76 | 0.79 |
| 5 | 167622914 | rs13180204 | -0.222 | -0.292 | 0.931 | 2.156 | 0.014 | 0.019 | 0.019 | 0 | 0.17 | 0.01 | 0.08 | 0.1 | 0.11 | 0.11 |
| 5 | 167623089 | rs17731087 | -0.124 | -0.262 | 1.21 | 2.238 | 0.012 | 0.122 | 0.122 | 0 | 0.06 | 0.12 | 0.03 | 0.35 | 0.05 | 0.14 |
| 5 | 167626142 | rs2291784 | -1.303 | -0.255 | 1.273 | 2.014 | 0.076 | 0.184 | 0.184 | 0.008 | 0.09 | 0.33 | 0.05 | 0.29 | 0.08 | 0.15 |
| 5 | 167626696 | rs3797719 | -1.893 | -0.295 | 1.191 | 2.04 | -0.014 | 0.009 | 0.009 | 0.091 | 0.32 | 0.61 | 0.21 | 0.64 | 0.2 | 0.28 |
| 5 | 167633724 | rs6898935 | -1.406 | -0.16 | 1.056 | 1.898 | 0.094 | 0.206 | 0.206 | 0.129 | 0.09 | 0.39 | 0.06 | 0.29 | 0.08 | 0.15 |
| 5 | 167636982 | rs6897788 | -0.301 | -0.162 | 0.592 | 1.921 | 0.036 | 0.149 | 0.149 | 0.204 | 0.07 | 0.25 | 0.04 | 0.29 | 0.05 | 0.14 |
| 5 | 167643545 | rs6885116 | -0.736 | -0.314 | 0.292 | 1.66 | -0.012 | -0.01 | -0.01 | 0.04 | 0.2 | 0.26 | 0.09 | 0.09 | 0.1 | 0.11 |
| 5 | 167645010 | rs3756603 | -0.275 | -0.323 | -0.082 | 1.845 | 0.011 | 0.284 | 0.284 | 0.017 | 0.05 | 0.27 | 0.08 | 0.26 | 0.02 | 0.01 |
| 5 | 167645888 | rs1363560 | -1.345 | -0.193 | -1.14 | 1.224 | -0.013 | 0.023 | 0.023 | 0.028 | 0.31 | 0.64 | 0.22 | 0.59 | 0.22 | 0.23 |
| 5 | 167648722 | rs7714057 | -0.99 | 0.257 | -0.893 | 1.587 | -0.019 | 0.18 | 0.18 | 1.847 | 0.15 | 0.68 | 0.14 | 0.48 | 0.08 | 0.13 |
| 5 | 167651603 | rs4976585 | 0.444 | 0.924 | -2.257 | 0.548 | -0.107 | 0.041 | 0.041 | 0.004 | 0.7 | 0.96 | 0.79 | 0.83 | 0.56 | 0.75 |
| 5 | 167653364 | rs3733985 | -0.325 | 0.584 | -2.654 | 0.51 | -0.082 | -0.008 | -0.008 | 0 | 0.58 | 0.92 | 0.59 | 0.64 | 0.54 | 0.53 |
| 5 | 167658698 | rs1008746 | -0.091 | 0.717 | -1.454 | 0.351 | 0.009 | -0.002 | -0.002 | 1.241 | 0.31 | 0.6 | 0.25 | 0.28 | 0.38 | 0.39 |
| 5 | 167668843 | rs11959012 | 0.078 | 0.536 | -1.263 | -0.408 | -0.001 | -0.005 | -0.005 | 0.442 | 0.45 | 0.45 | 0.41 | 0.18 | 0.46 | 0.57 |
| 5 | 167669233 | rs6887837 | 0.078 | 0.437 | -1.042 | -0.681 | 0.003 | -0.039 | -0.039 | 0.124 | 0.09 | 0.17 | 0.06 | 0 | 0.08 | 0.17 |
| 5 | 167669347 | rs6888483 | 0.347 | 0.866 | -1.179 | -0.797 | 0 | 0.016 | 0.016 | 0 | 0.38 | 0.29 | 0.38 | 0.18 | 0.42 | 0.39 |
| 5 | 167672105 | rs1960425 | -0.361 | 0.828 | -1.164 | -1.014 | 0 | 0.001 | 0.001 | 0.008 | 0.51 | 0.48 | 0.45 | 0.18 | 0.54 | 0.6 |
| 5 | 167674370 | rs3733988 | -0.493 | 0.819 | -1.237 | -0.971 | 0.002 | 0.019 | 0.019 | 0 | 0.5 | 0.44 | 0.45 | 0.17 | 0.54 | 0.58 |
| 5 | 167674472 | rs17070023 | 0.121 | 0.826 | -1.232 | -0.97 | 0.005 | -0.04 | -0.04 | 0.635 | 0.1 | 0.18 | 0.06 | 0 | 0.08 | 0.16 |
| 5 | 167675392 | rs12189266 | -0.044 | 0.407 | -1.517 | -1.142 | -0.029 | 0.164 | 0.164 | 0.444 | 0.44 | 0.16 | 0.39 | 0.14 | 0.47 | 0.56 |
| 5 | 167686974 | rs6889331 | -0.859 | 0.209 | -1.211 | -1.365 | -0.029 | 0.002 | 0.002 | 2.028 | 0.72 | 0.87 | 0.57 | 0.21 | 0.71 | 0.73 |
| 5 | 167689296 | rs17632540 | -0.014 | 0.04 | -1.489 | -1.746 | -0.028 | 0.048 | 0.048 | 0 | 0.24 | 0.15 | 0.16 | 0 | 0.3 | 0.39 |
| 5 | 167700077 | rs717036 | -0.784 | -0.3 | -1.406 | -1.382 | -0.051 | 0.03 | 0.03 | 0.505 | 0.66 | 0.85 | 0.53 | 0.25 | 0.62 | 0.65 |
| 5 | 167702041 | rs11748886 | 0.009 | -0.52 | -0.175 | -1.002 | -0.044 | -0.043 | -0.043 | 0 | 0.13 | 0.31 | 0.06 | 0.04 | 0.06 | 0.05 |
| 5 | 167708905 | rs918465 | 0.531 | 0.08 | -0.759 | -0.804 | -0.021 | 0.034 | 0.034 | 0 | 0.15 | 0.07 | 0.11 | 0 | 0.23 | 0.2 |
| 5 | 167718512 | rs12655262 | 0.04 | -0.539 | -0.304 | -0.066 | -0.015 | 0.019 | 0.019 | 0.873 | 0.23 | 0.35 | 0.3 | 0.73 | 0.17 | 0.35 |
| 5 | 167721201 | rs6875551 | -0.625 | -0.986 | -0.447 | -0.65 | -0.008 | 0.073 | 0.073 | 0.19 | 0.6 | 0.79 | 0.7 | 0.83 | 0.46 | 0.64 |
| 5 | 167727179 | rs2253028 | 0.083 | -1.16 | -0.038 | -0.655 | -0.03 | 0.015 | 0.015 | 1.635 | 0.27 | 0.6 | 0.33 | 0.73 | 0.18 | 0.33 |
| 5 | 167729035 | rs7700355 | 0.773 | -0.986 | 0.123 | -0.599 | 0.016 | 0.017 | 0.017 | 0 | 0.23 | 0.18 | 0.25 | 0.08 | 0.12 | 0.15 |
| 5 | 167731435 | rs6555796 | 0.808 | -0.369 | -0.156 | 0.239 | -0.001 | -0.006 | -0.006 | 0 | 0.41 | 0.38 | 0.29 | 0.49 | 0.37 | 0.4 |
| 5 | 167734916 | rs7703791 | 0.672 | -0.445 | -0.153 | 0.11 | 0 | -0.014 | -0.014 | 0.097 | 0.43 | 0.4 | 0.26 | 0.08 | 0.37 | 0.29 |
| 5 | 167735131 | rs7724276 | -0.279 | -0.348 | -0.1 | 0.233 | 0.04 | 0.075 | 0.075 | 0.895 | 0.73 | 0.57 | 0.67 | 0.58 | 0.66 | 0.68 |
| 5 | 167738356 | rs2112556 | -0.165 | -0.204 | -0.339 | 0.156 | 0.011 | 0.068 | 0.068 | 0 | 0.36 | 0.28 | 0.44 | 0.12 | 0.35 | 0.32 |
| 5 | 167743409 | rs749235 | 1.05 | -0.133 | -0.883 | 0.175 | 0.007 | 0.022 | 0.022 | 0 | 0.22 | 0.16 | 0.31 | 0.1 | 0.19 | 0.19 |
| 5 | 167743909 | rs17632741 | 0.201 | 0.256 | -0.498 | 0.754 | -0.039 | 0.04 | 0.04 | 0 | 0.1 | 0 | 0.11 | 0.02 | 0.16 | 0.13 |
| 5 | 167744530 | rs4976544 | -0.176 | 0.265 | -0.336 | 1.026 | -0.005 | -0.009 | -0.009 | 0 | 0.79 | 0.85 | 0.74 | 0.57 | 0.72 | 0.71 |
| 5 | 167748188 | rs11949188 | -0.614 | -0.456 | 0.403 | 1.198 | -0.02 | 0.169 | 0.169 | 0 | 0.14 | 0.32 | 0.12 | 0.4 | 0.08 | 0.18 |
| 5 | 167755986 | rs2337214 | -1.356 | 0.17 | 0.767 | 0.43 | -0.009 | -0.089 | -0.089 | 2.825 | 0.16 | 0.51 | 0.14 | 0.03 | 0.15 | 0.13 |
| 5 | 167764220 | rs10064972 | -1.626 | 0.361 | 0.679 | 0.923 | 0.072 | 0.157 | 0.157 | 0.262 | 0.05 | 0.38 | 0.09 | 0.08 | 0.04 | 0.01 |
| 5 | 167764600 | rs17731499 | -0.73 | 0.133 | -0.497 | 1.164 | -0.013 | -0.092 | -0.092 | 0 | 0.17 | 0 | 0.08 | 0.22 | 0.16 | 0.24 |
| 5 | 167768114 | rs1035421 | 0.218 | 0.767 | -0.339 | 0.673 | -0.012 | 0.013 | 0.013 | 0 | 0.76 | 0.83 | 0.82 | 0.85 | 0.73 | 0.77 |
| 5 | 167773074 | rs6555802 | 0.996 | 0.545 | 0.185 | 0.201 | -0.008 | 0.051 | 0.051 | 0 | 0.62 | 0.27 | 0.73 | 0.8 | 0.66 | 0.67 |
| 5 | 167780666 | rs10078070 | 0.307 | 0.571 | 0.592 | 0.173 | 0.007 | -0.024 | -0.024 | 0 | 0.05 | 0.13 | 0.04 | 0.03 | 0.04 | 0.04 |
| 5 | 167781775 | rs6880261 | 0.355 | 0.425 | 0.651 | 0.999 | 0.001 | 0.004 | 0.004 | 0 | 0.68 | 0.66 | 0.59 | 0.43 | 0.67 | 0.69 |
| 5 | 167784624 | rs2059090 | 0.372 | 0.655 | 0.361 | 0.943 | -0.001 | -0.001 | -0.001 | 0 | 0.06 | 0.03 | 0.06 | 0.05 | 0.08 | 0.1 |
| 5 | 167785204 | rs998864 | -0.683 | 0.682 | 0.996 | 0.876 | 0.03 | -0.042 | -0.042 | 0 | 0.1 | 0.29 | 0.06 | 0.05 | 0.06 | 0.1 |
| 5 | 167787583 | rs6873727 | 1.057 | 0.156 | 0.247 | 0.692 | -0.001 | 0.05 | 0.05 | 0 | 0.52 | 0.28 | 0.47 | 0.34 | 0.56 | 0.56 |
| 5 | 167790204 | rs10475878 | -1 | 0.144 | 0.403 | 0.66 | -0.003 | 0.004 | 0.004 | 0 | 0.35 | 0.64 | 0.43 | 0.57 | 0.3 | 0.33 |
| 5 | 167794776 | rs1030182 | 0.321 | 0.354 | 0.505 | 1.137 | -0.001 | -0.001 | -0.001 | 0 | 0.55 | 0.47 | 0.68 | 0.47 | 0.57 | 0.66 |
| 5 | 167797440 | rs6555805 | 1.115 | 0.334 | -0.496 | 0.359 | 0.059 | 0.02 | 0.02 | 0 | 0.35 | 0.11 | 0.23 | 0.23 | 0.25 | 0.46 |
| 5 | 167798135 | rs2287705 | -0.47 | 0.396 | -0.452 | 0.871 | -0.029 | 0.148 | 0.148 | 0 | 0.11 | 0.16 | 0.28 | 0.43 | 0.06 | 0.11 |
| 5 | 167798585 | rs2287703 | 0.055 | 1.016 | -1.01 | 0.022 | -0.019 | 0.058 | 0.058 | 0 | 0.61 | 0.74 | 0.7 | 0.73 | 0.58 | 0.66 |
| 5 | 167806016 | rs4976598 | 0.224 | 1.036 | -0.869 | -0.1 | 0.006 | 0.06 | 0.06 | 0 | 0.53 | 0.67 | 0.62 | 0.71 | 0.48 | 0.58 |
| 5 | 167807992 | rs13171394 | -0.446 | 0.622 | -1.062 | -0.557 | 0.021 | 0.002 | 0.002 | 0 | 0.53 | 0.7 | 0.45 | 0.37 | 0.59 | 0.61 |
| 5 | 167809259 | rs2161299 | 0.63 | 0.917 | -0.474 | -0.821 | 0.011 | -0.049 | -0.049 | 0 | 0.12 | 0.19 | 0.1 | 0 | 0.14 | 0.09 |
| 5 | 167810327 | rs2337135 | 0.964 | 0.775 | -0.355 | -1.19 | -0.001 | 0.003 | 0.003 | 0 | 0.23 | 0.19 | 0.22 | 0.31 | 0.26 | 0.41 |
| 5 | 167811567 | rs4320284 | -0.374 | 0.639 | -0.08 | -1.276 | -0.001 | -0.001 | -0.001 | 0 | 0.4 | 0.37 | 0.33 | 0.38 | 0.42 | 0.52 |

| | | | | | | | | | | | | | | | |
|---|---|---|---|---|---|---|---|---|---|---|---|---|---|---|---|
| 5 | 167812698 | rs13169010 | -1.26 | 0.573 | -0.149 | -1.366 | 0.013 | -0.004 | -0.004 | 0 | 0.55 | 0.64 | 0.45 | 0.38 | 0.58 | 0.61 |
| 5 | 167816521 | rs7737462 | -0.625 | 0.952 | -0.208 | -1.566 | 0.023 | -0.002 | -0.002 | 1.315 | 0.46 | 0.65 | 0.35 | 0.38 | 0.37 | 0.52 |
| 5 | 167817777 | rs11738934 | -2.251 | 0.563 | -1.07 | -2.203 | -0.046 | 0.093 | 0.093 | 0 | 0.78 | 0.94 | 0.84 | 1 | 0.73 | 0.79 |
| 5 | 167820982 | rs2112555 | 1.331 | 0.614 | -0.697 | -1.905 | -0.008 | -0.02 | -0.02 | 0 | 0.23 | 0.14 | 0.22 | 0.31 | 0.26 | 0.41 |
| 5 | 167822652 | rs10051783 | -0.578 | -0.368 | -0.432 | -1.6 | -0.003 | -0.01 | -0.01 | 0 | 0.18 | 0.22 | 0.11 | 0.11 | 0.16 | 0.12 |
| 5 | 167824840 | rs10064053 | -0.451 | -0.34 | -0.394 | -1.597 | -0.008 | -0.031 | -0.031 | 0 | 0.17 | 0.26 | 0.11 | 0.07 | 0.15 | 0.11 |
| 5 | 167826166 | rs6555807 | -0.327 | -0.027 | -0.444 | -1.639 | 0.002 | 0.001 | 0.001 | 0 | 0.47 | 0.53 | 0.37 | 0.35 | 0.44 | 0.56 |
| 5 | 167829061 | rs17731641 | 1.024 | -0.132 | -0.631 | -1.442 | 0.015 | -0.058 | -0.058 | 0 | 0.25 | 0.08 | 0.21 | 0.31 | 0.2 | 0.4 |
| 5 | 167829892 | rs6862868 | -0.355 | -0.143 | -0.43 | -1.37 | 0.001 | 0.001 | 0.001 | 0 | 0.44 | 0.59 | 0.31 | 0.32 | 0.4 | 0.48 |
| 5 | 167831555 | rs11134510 | 0.351 | -0.275 | -0.624 | -1.304 | -0.001 | -0.005 | -0.005 | 0.053 | 0.44 | 0.43 | 0.58 | 0.69 | 0.33 | 0.59 |
| 5 | 167832067 | rs10041542 | -0.915 | 0.083 | -0.169 | -0.947 | 0.009 | -0.072 | -0.072 | 0 | 0.23 | 0.4 | 0.18 | 0.11 | 0.26 | 0.19 |
| 5 | 167832644 | rs17551539 | -0.526 | 0.239 | -1.029 | -0.767 | -0.002 | 0.017 | 0.017 | 0 | 0.18 | 0.12 | 0.13 | 0.07 | 0.21 | 0.16 |
| 5 | 167834583 | rs17551581 | 0.352 | 0.528 | -0.825 | -0.442 | 0.003 | -0.039 | -0.039 | 0.066 | 0.12 | 0.13 | 0.1 | 0 | 0.14 | 0.09 |
| 5 | 167835539 | rs17551608 | 0.563 | 0.631 | -1.319 | -0.416 | -0.04 | 0.053 | 0.053 | 0 | 0.11 | 0.03 | 0.09 | 0 | 0.16 | 0.09 |
| 5 | 167836641 | rs1004940 | -1.013 | 0.143 | -1.397 | -0.755 | -0.062 | 0.142 | 0.142 | 0 | 0.63 | 0.86 | 0.73 | 0.99 | 0.55 | 0.7 |
| 5 | 167837919 | rs7700510 | -0.992 | -0.556 | -1.258 | 0.231 | 0.102 | 0.012 | 0.012 | 0.086 | 0.44 | 0.81 | 0.53 | 0.67 | 0.33 | 0.32 |
| 5 | 167838535 | rs3926392 | -0.456 | -0.636 | -0.751 | 0.437 | 0.008 | -0.006 | -0.006 | 1.502 | 0.32 | 0.65 | 0.44 | 0.67 | 0.22 | 0.29 |
| 5 | 167842039 | rs10077631 | -1.021 | -0.118 | -0.9 | 0.235 | 0.016 | 0.109 | 0.109 | 0.331 | 0.46 | 0.79 | 0.59 | 0.77 | 0.45 | 0.34 |
| 5 | 167844236 | rs7722673 | 1.236 | -0.053 | -0.961 | 0.188 | 0.007 | 0.034 | 0.034 | 0.527 | 0.11 | 0.04 | 0.04 | 0 | 0.08 | 0.08 |
| 5 | 167847974 | rs1422418 | 0.745 | -0.16 | -0.434 | 0.723 | -0.004 | -0.031 | -0.031 | 0 | 0.12 | 0.16 | 0.08 | 0 | 0.1 | 0.04 |
| 5 | 167853274 | rs9686714 | -0.22 | -0.427 | 0.288 | 1.057 | -0.049 | 0.092 | 0.092 | 0 | 0.12 | 0.29 | 0.12 | 0.23 | 0.08 | 0.08 |
| 5 | 167858004 | rs729149 | 0.994 | -0.653 | 0.095 | 0.827 | 0.005 | -0.005 | -0.005 | 1.864 | 0.2 | 0.16 | 0.14 | 0.22 | 0.18 | 0.38 |
| 5 | 167858372 | rs3822659 | 0.066 | -0.494 | -0.072 | 0.857 | -0.043 | 0.093 | 0.093 | 0 | 0.12 | 0.29 | 0.07 | 0.23 | 0.07 | 0.08 |
| 5 | 167859364 | rs4976608 | 0.464 | -0.068 | 0.128 | 1.162 | -0.016 | 0.022 | 0.022 | 0.278 | 0.14 | 0.26 | 0.2 | 0.17 | 0.11 | 0.07 |
| 5 | 167861846 | rs12516671 | 0.351 | 0.299 | -0.35 | 1.153 | -0.02 | 0.032 | 0.032 | 0 | 0.05 | 0.03 | 0.33 | 0.34 | 0.17 | 0.07 |
| 5 | 167863844 | rs4976609 | -0.148 | 0.15 | -0.324 | 0.975 | 0.013 | 0.018 | 0.018 | 0 | 0.28 | 0.36 | 0.48 | 0.6 | 0.37 | 0.47 |
| 5 | 167867953 | rs4976553 | -0.416 | 0.543 | -1.008 | 0.625 | 0.053 | 0.122 | 0.122 | 0 | 0.1 | 0.19 | 0.34 | 0.38 | 0.2 | 0.12 |
| 5 | 167868809 | rs3733980 | -0.877 | -0.56 | -0.835 | 0.809 | 0.07 | 0.02 | 0.02 | 0 | 0.42 | 0.76 | 0.4 | 0.61 | 0.34 | 0.52 |
| 5 | 167872381 | rs10042345 | -0.711 | 0.01 | -0.599 | 0.871 | 0.069 | 0.036 | 0.036 | 0.009 | 0.47 | 0.69 | 0.26 | 0.45 | 0.41 | 0.57 |
| 5 | 167873938 | rs1009069 | -1.051 | 0.558 | 0.255 | 1.354 | 0.021 | 0.117 | 0.117 | 0 | 0.09 | 0.26 | 0.06 | 0.19 | 0.09 | 0.07 |
| 5 | 167885622 | rs2085448 | 0.239 | 0.925 | -0.574 | 1.462 | -0.028 | -0.059 | -0.059 | 0 | 0.85 | 0.97 | 0.82 | 0.74 | 0.82 | 0.74 |
| 5 | 167891830 | rs3203960 | 1.493 | 1.241 | -0.272 | 0.299 | 0.003 | -0.006 | -0.006 | 0 | 0.54 | 0.4 | 0.33 | 0.04 | 0.45 | 0.46 |
| 5 | 167892338 | rs12515835 | -0.261 | 0.506 | 0.571 | 1.614 | -0.053 | 0.122 | 0.122 | 0 | 0.13 | 0.28 | 0.28 | 0.26 | 0.04 | 0.12 |
| 5 | 167895953 | rs17731898 | -0.101 | 0.585 | 0.442 | 1.865 | 0.002 | -0.058 | -0.058 | 0 | 0.14 | 0.05 | 0.17 | 0.45 | 0.13 | 0.24 |
| 5 | 167898299 | rs6890606 | -1.399 | 0.597 | 0.623 | 1.633 | -0.039 | -0.005 | -0.005 | 0 | 0.29 | 0.57 | 0.46 | 0.72 | 0.18 | 0.35 |
| 5 | 167902778 | rs244890 | -0.885 | 1.371 | 0.58 | 1.655 | 0.008 | 0.157 | 0.157 | 0 | 0.17 | 0.35 | 0.32 | 0.34 | 0.16 | 0.19 |
| 5 | 167905111 | rs865410 | 0.456 | 1.42 | 0.527 | 0.621 | -0.037 | -0.073 | -0.073 | 0 | 0.35 | 0.49 | 0.16 | 0.09 | 0.19 | 0.16 |
| 5 | 167906422 | rs244904 | -0.214 | 1.39 | 0.446 | 0.622 | 0.006 | 0.016 | 0.016 | 0 | 0.51 | 0.58 | 0.43 | 0.68 | 0.5 | 0.44 |
| 5 | 167907762 | rs6876713 | 0.349 | 1.467 | 0.242 | 0.729 | 0.008 | -0.069 | -0.069 | 0 | 0.15 | 0.03 | 0.18 | 0.53 | 0.1 | 0.2 |
| 5 | 167913510 | rs244903 | -0.799 | 1.553 | 0.354 | 0.81 | -0.012 | 0.082 | 0.082 | 1.757 | 0.59 | 0.76 | 0.71 | 0.89 | 0.52 | 0.5 |
| 5 | 167913603 | rs193466 | -0.415 | 1.523 | 0.434 | 1.103 | 0.026 | 0.037 | 0.037 | 1.939 | 0.55 | 0.68 | 0.62 | 0.72 | 0.34 | 0.42 |
| 5 | 167916628 | rs582906 | -0.255 | 1.49 | 0.537 | 1.144 | 0.025 | 0.007 | 0.007 | 2.879 | 0.52 | 0.62 | 0.37 | 0.62 | 0.33 | 0.39 |
| 5 | 167924928 | rs2445785 | -0.811 | 1.5 | 0.529 | 1.112 | -0.012 | 0.08 | 0.08 | 1.879 | 0.59 | 0.76 | 0.71 | 0.89 | 0.52 | 0.5 |
| 5 | 167932109 | rs244889 | -0.247 | 1.49 | 0.534 | 1.129 | 0.026 | 0.008 | 0.008 | 2.98 | 0.52 | 0.62 | 0.37 | 0.62 | 0.33 | 0.39 |
| 5 | 167936423 | rs2432645 | -0.796 | 1.379 | 0.515 | 1.094 | -0.012 | 0.085 | 0.085 | 1.733 | 0.58 | 0.76 | 0.71 | 0.89 | 0.51 | 0.5 |
| 5 | 167941990 | rs7709842 | -0.48 | 1.557 | 0.587 | 1.415 | 0.018 | 0.006 | 0.006 | 0 | 0.05 | 0.05 | 0.33 | 0.17 | 0.19 | 0.08 |
| 5 | 167945647 | rs3797711 | -1.098 | 1.555 | 0.603 | 1.432 | 0.062 | 0.125 | 0.125 | 0 | 0.06 | 0.14 | 0.34 | 0.27 | 0.19 | 0.11 |
| 5 | 167957055 | rs12654410 | -0.282 | 1.288 | 1.009 | 1.28 | -0.019 | 0.136 | 0.136 | 0 | 0.19 | 0.34 | 0.22 | 0.52 | 0.14 | 0.24 |
| 5 | 167957554 | rs3733977 | 0.02 | 1.261 | 0.999 | 1.286 | -0.003 | 0.035 | 0.035 | 0 | 0.18 | 0.18 | 0.21 | 0.53 | 0.14 | 0.23 |
| 5 | 167965753 | rs4976562 | -0.67 | 1.479 | 0.962 | 1.24 | -0.012 | -0.012 | -0.012 | 0 | 0.24 | 0.2 | 0.54 | 0.7 | 0.34 | 0.32 |
| 5 | 167980525 | rs158898 | 0.416 | 1.808 | 0.953 | 0.76 | 0.001 | -0.023 | -0.023 | 0.426 | 0.43 | 0.42 | 0.26 | 0.15 | 0.38 | 0.48 |
| 5 | 167983347 | rs158896 | -1.103 | 1.755 | 0.653 | 0.897 | 0.009 | -0.016 | -0.016 | 0 | 0.68 | 0.87 | 0.61 | 0.76 | 0.73 | 0.83 |
| 5 | 167984721 | rs13185224 | 0.468 | 1.637 | 0.711 | 1.002 | -0.005 | -0.061 | -0.061 | 2.734 | 0.18 | 0.06 | 0.23 | 0.47 | 0.16 | 0.25 |
| 5 | 167987499 | rs11952767 | 1.781 | 1.532 | 0.682 | 0.871 | -0.026 | -0.027 | -0.027 | 0 | 0.05 | 0.14 | 0.03 | 0 | 0 | 0 |
| 5 | 168006700 | rs149476 | -0.61 | 1.979 | 0.64 | 0.874 | 0.017 | 0.018 | 0.018 | 1.212 | 0.62 | 0.75 | 0.57 | 0.76 | 0.7 | 0.83 |
| 5 | 168019254 | rs292491 | 0.968 | 1.998 | 0.7 | 0.828 | 0.022 | 0.064 | 0.064 | 0.295 | 0.37 | 0.23 | 0.22 | 0.14 | 0.36 | 0.48 |
| 5 | 168033163 | rs158910 | 0.209 | 2.041 | 0.587 | 0.852 | -0.005 | 0.038 | 0.038 | 0 | 0.46 | 0.4 | 0.35 | 0.22 | 0.55 | 0.53 |
| 5 | 168039021 | rs58700 | -0.587 | 1.965 | 0.524 | 0.968 | -0.061 | -0.032 | -0.032 | 0.773 | 0.64 | 0.84 | 0.51 | 0.67 | 0.52 | 0.78 |
| 5 | 168053136 | rs188364 | -0.819 | 1.765 | 0.154 | 0.8 | -0.035 | -0.015 | -0.015 | 0 | 0.89 | 0.95 | 0.89 | 0.92 | 0.83 | 0.88 |
| 5 | 168056680 | rs292482 | -0.853 | 1.725 | 0.711 | 1.237 | 0.026 | 0.004 | 0.004 | 0.754 | 0.65 | 0.76 | 0.59 | 0.76 | 0.75 | 0.82 |
| 5 | 168080275 | rs4867894 | 0.36 | 2.163 | 0.192 | 1.491 | -0.015 | -0.024 | -0.024 | 4.047 | 0.06 | 0.01 | 0.11 | 0.08 | 0.22 | 0.04 |
| 5 | 168083311 | rs17732425 | -1.112 | 2.119 | 0.393 | 1.485 | 0.073 | 0.079 | 0.079 | 0 | 0.1 | 0.18 | 0.35 | 0.26 | 0.24 | 0.07 |
| 5 | 168093522 | rs1048307 | 0.117 | 1.848 | -0.004 | 1.702 | -0.015 | -0.079 | -0.079 | 0 | 0.18 | 0.04 | 0.21 | 0.42 | 0.17 | 0.24 |
| 5 | 168103440 | rs1870563 | -1.736 | 1.951 | 0.33 | 1.64 | 0.181 | 0.274 | 0.274 | 0.301 | 0.14 | 0.48 | 0.39 | 0.33 | 0.25 | 0.15 |
| 5 | 168109259 | rs292488 | -1.369 | 1.772 | -0.004 | 1.536 | 0.022 | 0.079 | 0.079 | 0.017 | 0.72 | 0.93 | 0.88 | 0.93 | 0.81 | 0.85 |
| 5 | 168115864 | rs898401 | -2.407 | 2.659 | 1.007 | 2.542 | 0.154 | 0.234 | 0.234 | 0.597 | 0.14 | 0.51 | 0.4 | 0.33 | 0.24 | 0.15 |
| 5 | 168124999 | rs6869359 | -0.403 | 2.865 | 0.651 | 2.626 | -0.036 | -0.078 | -0.078 | 0 | 0.17 | 0.05 | 0.21 | 0.44 | 0.19 | 0.31 |
| 5 | 168133829 | rs4868489 | -1.726 | 2.888 | 0.683 | 2.625 | 0.092 | 0.088 | 0.088 | 0 | 0.27 | 0.44 | 0.35 | 0.61 | 0.41 | 0.4 |
| 5 | 168134156 | rs11746577 | -0.487 | 2.879 | 0.705 | 2.519 | -0.01 | -0.025 | -0.025 | 0 | 0.17 | 0.11 | 0.22 | 0.45 | 0.19 | 0.31 |

| | | | | | | | | | | | | | | | | |
|---|---|---|---|---|---|---|---|---|---|---|---|---|---|---|---|---|
| 5 | 168139239 | rs1864952 | -0.75 | 2.804 | 0.317 | 2.413 | 0.013 | 0.011 | 0.011 | 0 | 0.06 | 0.18 | 0.03 | 0.03 | 0.04 | 0.02 |
| 5 | 168147329 | rs7715777 | -1.719 | 2.66 | -0.032 | 2.402 | 0.059 | 0.068 | 0.068 | 0 | 0.3 | 0.44 | 0.37 | 0.56 | 0.45 | 0.37 |
| 5 | 168155311 | rs1585399 | -0.487 | 2.494 | -0.903 | 2.423 | -0.035 | -0.081 | -0.081 | 0.709 | 0.16 | 0.03 | 0.21 | 0.44 | 0.19 | 0.31 |
| 5 | 168164744 | rs11743823 | -0.469 | 2.538 | -0.874 | 1.86 | -0.018 | -0.057 | -0.057 | 0.112 | 0.16 | 0.07 | 0.45 | 0.57 | 0.2 | 0.32 |
| 5 | 168165399 | rs10079276 | -1.19 | 2.709 | -0.654 | 1.481 | 0.046 | 0.024 | 0.024 | 0 | 0.17 | 0.24 | 0.16 | 0.23 | 0.28 | 0.13 |
| 5 | 168167292 | rs2337552 | -1.081 | 2.244 | -1.136 | 1.276 | 0.017 | 0.016 | 0.016 | 0 | 0.31 | 0.6 | 0.57 | 0.54 | 0.39 | 0.3 |
| 5 | 168167381 | rs7728435 | -0.445 | 1.526 | -2.196 | 1.158 | -0.09 | -0.134 | -0.134 | 0 | 0.36 | 0.07 | 0.54 | 0.56 | 0.36 | 0.36 |
| 5 | 168170445 | rs10462978 | -0.548 | 1.507 | -1.765 | 1.459 | -0.033 | -0.051 | -0.051 | 0 | 0.08 | 0.02 | 0.07 | 0.19 | 0.14 | 0.15 |
| 5 | 168170575 | rs7737922 | -0.978 | 0.853 | -1.161 | 1.448 | -0.01 | 0.056 | 0.056 | 0.873 | 0.32 | 0.43 | 0.5 | 0.48 | 0.35 | 0.4 |
| 5 | 168172273 | rs4519935 | -1.235 | 0.876 | -0.757 | 1.568 | -0.004 | 0.003 | 0.003 | 0.36 | 0.36 | 0.56 | 0.54 | 0.55 | 0.38 | 0.48 |
| 5 | 168175040 | rs4502845 | 1.085 | 0.9 | -0.555 | 0.349 | -0.002 | -0.165 | -0.165 | 0 | 0.23 | 0.5 | 0.21 | 0.01 | 0.18 | 0.12 |
| 5 | 168175291 | rs3822657 | -0.345 | 1.048 | -0.12 | 0.628 | -0.013 | -0.019 | -0.019 | 0 | 0.12 | 0.06 | 0.12 | 0.27 | 0.18 | 0.23 |
| 5 | 168179043 | rs10060585 | -1.136 | 1.142 | 0.172 | 0.318 | 0.046 | 0.113 | 0.113 | 0 | 0.18 | 0.46 | 0.19 | 0.23 | 0.24 | 0.25 |
| 5 | 168179549 | rs11746188 | -1.473 | 1.877 | 0.311 | 0.374 | 0.078 | 0.09 | 0.09 | 0 | 0.22 | 0.54 | 0.27 | 0.28 | 0.35 | 0.28 |
| 5 | 168180081 | rs10036727 | -1.319 | 1.736 | 0.223 | 0.152 | -0.018 | 0.009 | 0.009 | 2.289 | 0.42 | 0.71 | 0.51 | 0.53 | 0.53 | 0.53 |
| 5 | 168180589 | rs1549910 | -0.815 | 1.771 | -0.207 | 0.165 | 0 | 0 | 0 | 0.748 | 0.33 | 0.34 | 0.37 | 0.47 | 0.35 | 0.49 |
| 5 | 168180670 | rs1549909 | -1.303 | 1.68 | -0.188 | 0.163 | -0.014 | 0.004 | 0.004 | 1.734 | 0.41 | 0.35 | 0.48 | 0.53 | 0.53 | 0.54 |
| 5 | 168182992 | rs7718883 | -1.876 | 1.485 | -0.904 | -0.2 | -0.081 | 0.217 | 0.217 | 0 | 0.69 | 0.98 | 0.8 | 0.95 | 0.73 | 0.81 |
| 5 | 168183736 | rs6862713 | -0.294 | 1.345 | -0.146 | 0.459 | 0.001 | -0.082 | -0.082 | 0 | 0.26 | 0.13 | 0.33 | 0.4 | 0.28 | 0.4 |
| 5 | 168187671 | rs11742226 | -0.642 | 1.198 | 0.726 | 0.192 | 0.008 | -0.032 | -0.032 | 0.245 | 0.08 | 0.15 | 0.08 | 0 | 0.14 | 0.03 |
| 5 | 168188680 | rs17070430 | -1.142 | 1.331 | 0.965 | 0.259 | 0.069 | 0.022 | 0.022 | 0 | 0.06 | 0.15 | 0.08 | 0.07 | 0.09 | 0.02 |
| 5 | 168191066 | rs11745421 | -0.528 | 1.306 | 0.967 | 0.25 | 0 | -0.018 | -0.018 | 0 | 0.07 | 0.07 | 0.07 | 0 | 0.11 | 0.04 |
| 5 | 168193194 | rs17070438 | -0.593 | 1.349 | 1.085 | 0.254 | 0.02 | 0.015 | 0.015 | 0 | 0.08 | 0.18 | 0.09 | 0.07 | 0.09 | 0.08 |
| 5 | 168194176 | rs6860854 | -1.434 | 1.032 | 1.07 | 0.258 | 0.005 | 0.102 | 0.102 | 0 | 0.51 | 0.69 | 0.65 | 0.7 | 0.54 | 0.62 |
| 5 | 168196034 | rs9784690 | -0.129 | 1.079 | 0.675 | 0.469 | 0.027 | -0.059 | -0.059 | 0 | 0.31 | 0.13 | 0.39 | 0.56 | 0.25 | 0.48 |
| 5 | 168196959 | rs6877545 | 0.224 | 1.243 | 0.866 | 0.554 | 0.007 | 0.008 | 0.008 | 0 | 0.06 | 0.06 | 0.09 | 0.08 | 0.09 | 0.08 |
| 5 | 168197659 | rs2337556 | -1.084 | 1.24 | 1 | 0.459 | 0.004 | -0.038 | -0.038 | 0.46 | 0.41 | 0.55 | 0.58 | 0.71 | 0.42 | 0.59 |
| 5 | 168197801 | rs1875976 | -0.916 | 1.077 | 0.428 | 0.478 | 0.014 | 0.043 | 0.043 | 0.037 | 0.38 | 0.27 | 0.5 | 0.66 | 0.35 | 0.58 |
| 5 | 168197826 | rs2082400 | -0.692 | 1.084 | 0.342 | 0.438 | 0.025 | -0.01 | -0.01 | 0.241 | 0.36 | 0.16 | 0.44 | 0.59 | 0.31 | 0.54 |
| 5 | 168198305 | rs868103 | -0.886 | 0.988 | 0.597 | 1.018 | -0.013 | 0.048 | 0.048 | 0 | 0.18 | 0.35 | 0.27 | 0.28 | 0.16 | 0.29 |
| 5 | 168198967 | rs2053046 | -0.231 | 1.277 | 0.537 | 0.453 | -0.031 | 0.022 | 0.022 | 0 | 0.32 | 0.51 | 0.4 | 0.57 | 0.24 | 0.47 |
| 5 | 168199301 | rs10516050 | -1.251 | 1.934 | 0.151 | 0.793 | 0.052 | 0.027 | 0.027 | 0 | 0.08 | 0.14 | 0.15 | 0.12 | 0.17 | 0.1 |
| 5 | 168199624 | rs4867902 | -0.352 | 1.456 | -0.412 | 0.522 | 0 | 0.001 | 0.001 | 0 | 0.49 | 0.52 | 0.55 | 0.63 | 0.49 | 0.54 |
| 5 | 168202641 | rs4867903 | -1.432 | 2.104 | -0.337 | 0.911 | 0.053 | -0.002 | -0.002 | 0 | 0.09 | 0.14 | 0.12 | 0.07 | 0.18 | 0.18 |
| 5 | 168202728 | rs17665552 | -0.8 | 1.15 | -0.72 | 1.555 | 0.017 | -0.038 | -0.038 | 0.016 | 0.22 | 0.15 | 0.26 | 0.36 | 0.16 | 0.27 |
| 5 | 168203455 | rs17733068 | -1.559 | 1.411 | -0.527 | 2.058 | 0.009 | 0.057 | 0.057 | 0 | 0.13 | 0.19 | 0.38 | 0.38 | 0.17 | 0.23 |
| 5 | 168205202 | rs13185422 | -0.501 | 1.13 | -1.116 | 1.623 | 0.003 | -0.006 | -0.006 | 0 | 0.16 | 0.06 | 0.11 | 0.18 | 0.16 | 0.12 |
| 5 | 168207696 | rs17665652 | -0.311 | 0.628 | -0.713 | 1.716 | -0.01 | 0.017 | 0.017 | 0 | 0.14 | 0.27 | 0.19 | 0.15 | 0.11 | 0.07 |
| 5 | 168208107 | rs871097 | 0.992 | 0.344 | -2.196 | 0.999 | -0.063 | 0.05 | 0.05 | 0 | 0.29 | 0.01 | 0.29 | 0.16 | 0.26 | 0.22 |
| 5 | 168208381 | rs871098 | -0.272 | 1.083 | -1.842 | 0.421 | 0.006 | -0.019 | -0.019 | 0.226 | 0.24 | 0.44 | 0.31 | 0.16 | 0.25 | 0.13 |
| 5 | 168211254 | rs2163761 | -0.193 | 1.411 | -1.796 | 0.367 | -0.005 | 0.006 | 0.006 | 0 | 0.32 | 0.45 | 0.33 | 0.34 | 0.29 | 0.19 |
| 5 | 168212096 | rs10475891 | -1.217 | 1.553 | -2.258 | 0.793 | -0.127 | -0.12 | -0.12 | 0.438 | 0.56 | 0.97 | 0.53 | 0.58 | 0.54 | 0.54 |
| 5 | 168212168 | rs11134531 | -0.662 | 1.288 | -2.148 | 0.601 | -0.001 | 0 | 0 | 0.03 | 0.4 | 0.42 | 0.42 | 0.36 | 0.42 | 0.29 |
| 5 | 168212951 | rs891921 | -1.122 | 1.269 | -1.997 | 0.605 | -0.026 | 0.049 | 0.049 | 1.289 | 0.88 | 0.99 | 0.94 | 0.96 | 0.88 | 0.86 |
| 5 | 168213007 | rs2292143 | 0.659 | 1.123 | -1.685 | 0.826 | -0.014 | 0.041 | 0.041 | 0 | 0.15 | 0.35 | 0.2 | 0.17 | 0.12 | 0.06 |
| 5 | 168215850 | rs4298263 | 0.23 | 0.82 | -1.805 | 0.697 | -0.004 | 0.003 | 0.003 | 0 | 0.18 | 0.19 | 0.19 | 0.18 | 0.13 | 0.11 |
| 5 | 168216489 | rs2247645 | -0.235 | 1.035 | -2.256 | -0.08 | 0.012 | 0.047 | 0.047 | 0 | 0.38 | 0.73 | 0.21 | 0.24 | 0.29 | 0.46 |
| 5 | 168216767 | rs891928 | 0.55 | 0.599 | -1.497 | -0.124 | -0.007 | -0.059 | -0.059 | 0 | 0.25 | 0.51 | 0.13 | 0.18 | 0.18 | 0.37 |
| 5 | 168220102 | rs7728604 | -1.254 | 0.35 | -1.781 | 0.113 | -0.133 | -0.089 | -0.089 | 0.38 | 0.73 | 0.97 | 0.53 | 0.71 | 0.59 | 0.73 |
| 5 | 168223681 | rs4867939 | -0.606 | -0.679 | -1.146 | -0.04 | 0.048 | 0.022 | 0.022 | 0 | 0.68 | 0.54 | 0.48 | 0.64 | 0.56 | 0.64 |
| 5 | 168225838 | rs11134532 | -1.296 | -0.501 | -1.189 | -0.034 | -0.044 | -0.046 | -0.046 | 0.062 | 0.8 | 0.88 | 0.56 | 0.64 | 0.64 | 0.65 |
| 5 | 168231649 | rs11134533 | 0.31 | -1.416 | -0.599 | -0.225 | -0.015 | 0.023 | 0.023 | 0 | 0.31 | 0.36 | 0.3 | 0.44 | 0.23 | 0.21 |
| 5 | 168231793 | rs10071604 | 1.237 | -1.662 | -0.522 | -0.116 | -0.018 | 0.01 | 0.01 | 0 | 0.14 | 0.2 | 0.08 | 0.15 | 0.06 | 0.08 |
| 5 | 168232757 | rs17070509 | 1.641 | -1.693 | -0.221 | -0.257 | 0.015 | -0.045 | -0.045 | 0.836 | 0.08 | 0.23 | 0.07 | 0 | 0.08 | 0.01 |
| 5 | 168233413 | rs3797716 | -0.786 | -1.99 | -0.553 | -0.483 | -0.001 | -0.001 | -0.001 | 0.844 | 0.58 | 0.61 | 0.47 | 0.59 | 0.5 | 0.53 |
| 5 | 168233474 | rs3797715 | 1.996 | -2.006 | -0.573 | -0.474 | -0.001 | -0.054 | -0.054 | 0.562 | 0.15 | 0.03 | 0.2 | 0.29 | 0.17 | 0.13 |
| 5 | 168234991 | rs867477 | 1.274 | -2.047 | -0.461 | -0.526 | -0.022 | -0.044 | -0.044 | 0 | 0.16 | 0.23 | 0.09 | 0.05 | 0.1 | 0.16 |
| 5 | 168235180 | rs883242 | -1.949 | -1.993 | -0.453 | -0.494 | 0.002 | 0.002 | 0.002 | 0 | 0.74 | 0.75 | 0.55 | 0.66 | 0.61 | 0.69 |
| 5 | 168236650 | rs10038650 | 1.558 | -1.959 | -0.58 | -0.466 | -0.011 | 0.008 | 0.008 | 0 | 0.14 | 0.21 | 0.08 | 0.15 | 0.07 | 0.08 |
| 5 | 168237679 | rs4867597 | 1.043 | -1.644 | -0.982 | -0.573 | -0.025 | -0.047 | -0.047 | 1.467 | 0.16 | 0.26 | 0.09 | 0.05 | 0.1 | 0.16 |
| 5 | 168243169 | rs7708939 | -0.3 | -1.44 | -0.99 | -0.437 | 0 | -0.001 | -0.001 | 0.073 | 0.41 | 0.6 | 0.37 | 0.44 | 0.34 | 0.26 |
| 5 | 168246101 | rs17635284 | 1.645 | -1.684 | -0.99 | -0.106 | -0.031 | -0.001 | -0.001 | 0 | 0.18 | 0.02 | 0.11 | 0.17 | 0.18 | 0.31 |
| 5 | 168246880 | rs10475894 | -0.341 | -1.5 | -0.673 | -0.052 | 0.024 | 0.021 | 0.021 | 0 | 0.52 | 0.71 | 0.42 | 0.61 | 0.43 | 0.52 |
| 5 | 168252636 | rs12691283 | -0.12 | -1.098 | -0.049 | 0.288 | -0.004 | -0.002 | -0.002 | 0 | 0.28 | 0.64 | 0.14 | 0.21 | 0.17 | 0.23 |
| 5 | 168254897 | rs7726584 | 0.26 | -0.376 | 0.808 | 0.305 | 0.005 | -0.006 | -0.006 | 0 | 0.08 | 0.24 | 0.05 | 0.06 | 0.06 | 0.07 |
| 5 | 168256240 | rs4282339 | 0.318 | -0.439 | 0.569 | 0.274 | 0.011 | 0.019 | 0.019 | 0 | 0.26 | 0.17 | 0.15 | 0.19 | 0.21 | 0.32 |
| 5 | 168257350 | rs13170900 | -0.942 | -0.332 | 0.541 | 0.327 | 0.013 | -0.001 | -0.001 | 0 | 0.42 | 0.19 | 0.37 | 0.46 | 0.41 | 0.46 |
| 5 | 168258281 | rs6883428 | 0.45 | -0.123 | 0.619 | 0.157 | 0.006 | -0.005 | -0.005 | 0 | 0.08 | 0.21 | 0.05 | 0.06 | 0.06 | 0.07 |
| 5 | 168261021 | rs2974427 | -1.256 | 0.1 | 0.701 | 0.503 | 0.003 | -0.019 | -0.019 | 0 | 0.42 | 0.21 | 0.38 | 0.51 | 0.42 | 0.43 |

| | | | | | | | | | | | | | | | | |
|---|---|---|---|---|---|---|---|---|---|---|---|---|---|---|---|---|
| 5 | 168265244 | rs2974428 | -0.564 | 0.769 | 0.747 | 0.671 | -0.005 | -0.005 | -0.005 | 0 | 0.32 | 0.62 | 0.29 | 0.28 | 0.26 | 0.38 |
| 5 | 168265891 | rs1979923 | -0.085 | 0.78 | 0.07 | 0.712 | 0 | -0.003 | -0.003 | 0 | 0.27 | 0.32 | 0.27 | 0.28 | 0.26 | 0.37 |
| 5 | 168273353 | rs10068957 | -0.918 | 0.893 | -1.082 | 0.599 | 0.003 | -0.005 | -0.005 | 0.68 | 0.18 | 0.18 | 0.23 | 0.26 | 0.2 | 0.16 |
| 5 | 168279411 | rs2938763 | -1.44 | 0.948 | -1.393 | 0.649 | 0.016 | 0.004 | 0.004 | 0 | 0.48 | 0.75 | 0.51 | 0.53 | 0.45 | 0.53 |
| 5 | 168280867 | rs12516521 | -0.769 | 1.108 | -1.23 | 0.574 | -0.02 | -0.055 | -0.055 | 0 | 0.15 | 0.03 | 0.21 | 0.26 | 0.18 | 0.14 |
| 5 | 168283164 | rs6555833 | -0.827 | 1.235 | -1.216 | 0.624 | -0.038 | -0.065 | -0.065 | 0.007 | 0.15 | 0.01 | 0.21 | 0.26 | 0.18 | 0.14 |
| 5 | 168286301 | rs17635646 | -1.202 | 1.37 | -0.652 | 0.373 | 0.003 | -0.005 | -0.005 | 0 | 0.28 | 0.32 | 0.2 | 0.29 | 0.3 | 0.3 |
| 5 | 168287088 | rs7717597 | -0.714 | 1.46 | -0.7 | 0.432 | -0.007 | -0.022 | -0.022 | 0 | 0.11 | 0.08 | 0.1 | 0.23 | 0.11 | 0.1 |
| 5 | 168292739 | rs2938769 | -1.401 | 1.016 | -0.813 | 0.287 | 0.017 | 0.031 | 0.031 | 0 | 0.42 | 0.77 | 0.47 | 0.4 | 0.4 | 0.36 |
| 5 | 168297997 | rs1504955 | -0.026 | 0.699 | -0.006 | 0.23 | -0.038 | -0.06 | -0.06 | 0.204 | 0.17 | 0.37 | 0.28 | 0.06 | 0.13 | 0.12 |
| 5 | 168298658 | rs1466947 | -0.257 | 0.807 | -0.035 | 0.196 | -0.03 | -0.071 | -0.071 | 0 | 0.19 | 0.41 | 0.28 | 0.06 | 0.14 | 0.16 |
| 5 | 168305161 | rs13183458 | -0.212 | 1.04 | -0.664 | 0.486 | 0.002 | -0.009 | -0.009 | 0 | 0.25 | 0.16 | 0.33 | 0.26 | 0.24 | 0.32 |
| 5 | 168305571 | rs13188090 | 1.456 | 1.558 | -0.996 | 0.7 | 0.006 | 0.014 | 0.014 | 0.017 | 0.4 | 0.19 | 0.24 | 0.31 | 0.36 | 0.28 |
| 5 | 168306155 | rs6897408 | 1.337 | 1.478 | -1.118 | 0.688 | -0.023 | -0.018 | -0.018 | 0.053 | 0.4 | 0.09 | 0.22 | 0.31 | 0.34 | 0.28 |
| 5 | 168307047 | rs2938772 | -0.555 | 1.58 | -0.633 | 0.642 | 0.005 | 0.001 | 0.001 | 0 | 0.4 | 0.37 | 0.5 | 0.46 | 0.36 | 0.46 |
| 5 | 168309626 | rs2938774 | -1.063 | 2.35 | -0.549 | 0.474 | 0.005 | -0.016 | -0.016 | 0 | 0.35 | 0.44 | 0.46 | 0.32 | 0.42 | 0.28 |
| 5 | 168313431 | rs7707709 | 0.333 | 1.751 | -0.645 | -0.017 | -0.03 | 0.022 | 0.022 | 0 | 0.63 | 0.2 | 0.45 | 0.72 | 0.52 | 0.63 |
| 5 | 168313961 | rs4616917 | 0.783 | 1.062 | -0.268 | 0.306 | 0.051 | 0.054 | 0.054 | 0 | 0.47 | 0.18 | 0.38 | 0.59 | 0.34 | 0.46 |
| 5 | 168320056 | rs11134544 | 0.818 | 1.131 | -0.542 | 0.432 | 0.04 | 0.065 | 0.065 | 0.03 | 0.48 | 0.16 | 0.38 | 0.6 | 0.36 | 0.46 |
| 5 | 168323234 | rs7714853 | 0.633 | 0.955 | -0.137 | 0.72 | 0.056 | 0.163 | 0.163 | 0.507 | 0.93 | 0.57 | 0.93 | 0.89 | 0.96 | 0.94 |
| 5 | 168334615 | rs11134549 | 0.272 | 1.209 | -1.375 | 0.717 | -0.04 | 0.045 | 0.045 | 0 | 0.55 | 0.22 | 0.43 | 0.63 | 0.47 | 0.55 |
| 5 | 168344280 | rs12655322 | -1.211 | 1.252 | -0.89 | 1.048 | -0.02 | -0.018 | -0.018 | 0 | 0.1 | 0.06 | 0.12 | 0.11 | 0.17 | 0.14 |
| 5 | 168352808 | rs10063042 | -2.25 | 0.838 | -0.014 | 0.798 | -0.01 | -0.009 | -0.009 | 0.11 | 0.19 | 0.31 | 0.17 | 0.14 | 0.21 | 0.16 |
| 5 | 168353413 | rs891958 | -1.07 | 0.844 | -0.089 | 0.801 | -0.012 | -0.021 | -0.021 | 0 | 0.1 | 0.03 | 0.11 | 0.11 | 0.15 | 0.12 |
| 5 | 168355647 | rs1982062 | -2.202 | 0.532 | 0.081 | 1.444 | 0.006 | -0.068 | -0.068 | 0.204 | 0.36 | 0.82 | 0.49 | 0.48 | 0.35 | 0.34 |
| 5 | 168363190 | rs6879973 | -2.363 | 0.381 | -0.179 | 1.21 | 0.017 | -0.063 | -0.063 | 2.949 | 0.4 | 0.83 | 0.52 | 0.48 | 0.42 | 0.36 |
| 5 | 168363604 | rs919753 | -1.446 | 0.331 | 0.9 | 1.191 | -0.016 | 0.051 | 0.051 | 0.849 | 0.13 | 0.4 | 0.07 | 0.13 | 0.1 | 0.06 |
| 5 | 168376294 | rs6555837 | -1.928 | 0.289 | 1.095 | 1.129 | -0.007 | 0.01 | 0.01 | 0.228 | 0.3 | 0.55 | 0.46 | 0.34 | 0.33 | 0.28 |
| 5 | 168380180 | rs752498 | -2.289 | 0.271 | 2.151 | 1.084 | 0.001 | 0.011 | 0.011 | 0.163 | 0.33 | 0.47 | 0.47 | 0.37 | 0.39 | 0.32 |
| 5 | 168380438 | rs11745658 | -1.764 | 0.276 | 2.007 | 1.08 | 0.001 | 0.012 | 0.012 | 0 | 0.27 | 0.33 | 0.44 | 0.35 | 0.33 | 0.29 |
| 5 | 168385263 | rs6555841 | -2.583 | 0.265 | 1.762 | 1.479 | 0.009 | -0.038 | -0.038 | 0.024 | 0.38 | 0.78 | 0.47 | 0.53 | 0.35 | 0.33 |
| 5 | 168389938 | rs11749816 | -1.473 | 0.266 | 2.085 | 1.742 | -0.001 | -0.008 | -0.008 | 0 | 0.2 | 0.17 | 0.34 | 0.27 | 0.22 | 0.19 |
| 5 | 168391557 | rs1432910 | -2.867 | 0.563 | 1.71 | 1.452 | -0.034 | -0.038 | -0.038 | 0.23 | 0.44 | 0.88 | 0.56 | 0.55 | 0.48 | 0.33 |
| 5 | 168391942 | rs6885618 | -2.118 | 0.675 | 1.997 | 1.214 | -0.002 | 0.002 | 0.002 | 0 | 0.36 | 0.6 | 0.51 | 0.4 | 0.38 | 0.28 |
| 5 | 168395006 | rs9313435 | -0.307 | 0.92 | 1.447 | 1.313 | 0 | -0.01 | -0.01 | 0.038 | 0.05 | 0.09 | 0.03 | 0.02 | 0.06 | 0.03 |
| 5 | 168398541 | rs7724207 | -1.619 | 0.926 | 1.573 | 1.411 | 0.002 | 0.032 | 0.032 | 0 | 0.24 | 0.31 | 0.38 | 0.38 | 0.25 | 0.2 |
| 5 | 168400206 | rs13165487 | -1.914 | 0.903 | 1.58 | 1.416 | 0.012 | 0.012 | 0.012 | 0 | 0.29 | 0.38 | 0.48 | 0.4 | 0.4 | 0.23 |
| 5 | 168411374 | rs6875247 | -1.624 | 0.703 | 1.325 | 1.097 | 0.003 | -0.03 | -0.03 | 0 | 0.06 | 0.12 | 0.04 | 0 | 0.05 | 0 |
| 5 | 168423907 | rs11747642 | -0.621 | 0.458 | 0.437 | 0.784 | 0.029 | 0.177 | 0.177 | 0 | 0.49 | 0.24 | 0.34 | 0.11 | 0.43 | 0.5 |
| 5 | 168430327 | rs10055309 | -1.013 | 0.277 | 0.815 | 0.806 | -0.005 | -0.015 | -0.015 | 2.186 | 0.07 | 0.2 | 0.05 | 0.03 | 0.07 | 0.05 |
| 5 | 168437170 | rs17666792 | -1.557 | 0.289 | 1.548 | 2.084 | 0.023 | 0.246 | 0.246 | 0.335 | 0.1 | 0.19 | 0.26 | 0.53 | 0.11 | 0.12 |
| 5 | 168439476 | rs2163783 | 0.07 | 0.127 | 1.573 | 1.975 | 0.018 | 0.033 | 0.033 | 0 | 0.38 | 0.31 | 0.26 | 0.21 | 0.27 | 0.39 |
| 5 | 168440502 | rs17734379 | -0.796 | -0.04 | 1.71 | 2.49 | -0.009 | -0.016 | -0.016 | 1.288 | 0.07 | 0.01 | 0.13 | 0.17 | 0.09 | 0.09 |
| 5 | 168445489 | rs11954861 | -2.382 | -0.07 | 2.119 | 2.206 | 0.089 | 0.085 | 0.085 | 1.085 | 0.05 | 0.41 | 0.04 | 0.01 | 0.02 | 0.02 |
| 5 | 168445608 | rs2278391 | -1.42 | -0.084 | 2.099 | 2.324 | 0.003 | 0.067 | 0.067 | 0 | 0.12 | 0.11 | 0.27 | 0.53 | 0.16 | 0.16 |
| 5 | 168449182 | rs2278390 | -2.808 | -0.143 | 2.123 | 2.324 | 0.004 | 0.159 | 0.159 | 0.891 | 0.18 | 0.57 | 0.31 | 0.55 | 0.18 | 0.17 |
| 5 | 168452477 | rs6890327 | -1.356 | -0.198 | 1.659 | 1.744 | 0.006 | 0.119 | 0.119 | 0.875 | 0.22 | 0.39 | 0.34 | 0.53 | 0.22 | 0.22 |
| 5 | 168457109 | rs1432900 | -0.504 | -0.205 | 1.967 | 1.762 | 0.026 | -0.01 | -0.01 | 0 | 0.07 | 0.29 | 0.05 | 0 | 0.04 | 0.04 |
| 5 | 168465514 | rs17734491 | -0.745 | -0.161 | 1.84 | 1.98 | 0.032 | 0.238 | 0.238 | 0 | 0.06 | 0.15 | 0.12 | 0.43 | 0.08 | 0.08 |
| 5 | 168466203 | rs17734503 | -1.524 | -0.17 | 1.708 | 1.98 | 0.101 | 0.48 | 0.48 | 1.87 | 0.08 | 0.44 | 0.14 | 0.43 | 0.08 | 0.1 |
| 5 | 168468537 | rs10516056 | 0.67 | -0.28 | 0.805 | 1.456 | -0.008 | 0.012 | 0.012 | 0 | 0.32 | 0.01 | 0.19 | 0.21 | 0.24 | 0.38 |
| 5 | 168469978 | rs7717558 | -2.627 | -0.094 | 1.699 | 1.727 | 0.003 | -0.014 | -0.014 | 0 | 0.29 | 0.72 | 0.38 | 0.54 | 0.26 | 0.24 |
| 5 | 168477797 | rs11743301 | -0.634 | -0.033 | 1.529 | 1.76 | -0.015 | -0.081 | -0.081 | 0 | 0.15 | 0.29 | 0.11 | 0 | 0.11 | 0.11 |
| 5 | 168477929 | rs11134555 | -1.527 | -0.016 | 0.705 | 1.847 | -0.007 | 0.048 | 0.048 | 0.379 | 0.25 | 0.62 | 0.34 | 0.55 | 0.2 | 0.21 |
| 5 | 168483594 | rs7735291 | -1.738 | 0.095 | 0.562 | 1.866 | -0.001 | 0.044 | 0.044 | 0.037 | 0.23 | 0.65 | 0.26 | 0.5 | 0.23 | 0.19 |
| 5 | 168484494 | rs13154825 | 0.086 | 0.146 | 0.301 | 1.675 | -0.004 | 0.038 | 0.038 | 0 | 0.07 | 0 | 0.05 | 0.02 | 0.05 | 0.05 |
| 5 | 168485970 | rs10516057 | -1.272 | 0.077 | 0.998 | 1.872 | 0.036 | 0.102 | 0.102 | 0 | 0.09 | 0.2 | 0.08 | 0.26 | 0.12 | 0.07 |
| 5 | 168488958 | rs9284977 | -1.499 | 0.063 | 0.802 | 1.644 | -0.001 | 0.004 | 0.004 | 0 | 0.3 | 0.3 | 0.2 | 0.35 | 0.28 | 0.33 |
| 5 | 168489333 | rs1469683 | -0.388 | 0.148 | 0.712 | 1.927 | -0.008 | -0.005 | -0.005 | 0 | 0.05 | 0.02 | 0.11 | 0.23 | 0.06 | 0.04 |
| 5 | 168491180 | rs4868223 | 0.906 | 0.667 | 0.668 | 1.406 | 0.01 | 0.017 | 0.017 | 0 | 0.52 | 0.29 | 0.43 | 0.53 | 0.44 | 0.6 |
| 5 | 168494738 | rs12656030 | 0.302 | 0.724 | 1.033 | 1.565 | 0 | 0.014 | 0.014 | 0 | 0.06 | 0.06 | 0.11 | 0.22 | 0.05 | 0.05 |
| 5 | 168494914 | rs7701811 | -0.146 | 0.953 | 1.28 | 1.285 | 0.011 | 0.003 | 0.003 | 0 | 0.63 | 0.56 | 0.47 | 0.57 | 0.51 | 0.66 |
| 5 | 168495141 | rs7701998 | 0.726 | 1.133 | 1.1 | 1.423 | -0.017 | 0.016 | 0.016 | 0.05 | 0.07 | 0.03 | 0.05 | 0.03 | 0.09 | 0.06 |
| 5 | 168498677 | rs6864266 | 1.105 | 0.993 | 0.996 | 1.291 | 0.01 | 0.137 | 0.137 | 0 | 0.49 | 0.16 | 0.37 | 0.24 | 0.44 | 0.57 |
| 5 | 168499469 | rs6869485 | 1.062 | 1.633 | 1.006 | 1.292 | -0.003 | 0.087 | 0.087 | 0 | 0.5 | 0.2 | 0.4 | 0.28 | 0.47 | 0.59 |
| 5 | 168501451 | rs10054190 | 0.692 | 1.46 | 0.922 | 1.29 | -0.017 | 0.017 | 0.017 | 0.061 | 0.07 | 0.04 | 0.05 | 0.03 | 0.09 | 0.06 |
| 5 | 168502329 | rs188433 | -1.032 | 1.849 | 0.951 | 1.206 | -0.023 | 0.015 | 0.015 | 0 | 0.77 | 0.81 | 0.68 | 0.84 | 0.66 | 0.71 |
| 5 | 168505703 | rs295986 | -1.231 | 1.931 | 0.867 | 1.29 | -0.048 | 0.031 | 0.031 | 0 | 0.77 | 0.89 | 0.79 | 0.86 | 0.67 | 0.76 |
| 5 | 168506022 | rs1533150 | 1.081 | 1.674 | 0.797 | 1.307 | 0.001 | 0.121 | 0.121 | 0 | 0.51 | 0.17 | 0.45 | 0.25 | 0.51 | 0.58 |

| | | | | | | | | | | | | | | | | |
|---|---|---|---|---|---|---|---|---|---|---|---|---|---|---|---|---|
| 5 | 168508432 | rs2668040 | -0.945 | 1.681 | 1.106 | 1.385 | 0 | -0.001 | -0.001 | 0 | 0.76 | 0.73 | 0.72 | 0.79 | 0.72 | 0.77 |
| 5 | 168509732 | rs7735612 | 1.064 | 1.581 | -0.08 | 0.354 | 0.009 | 0.087 | 0.087 | 0 | 0.43 | 0.16 | 0.25 | 0.24 | 0.4 | 0.5 |
| 5 | 168513184 | rs296009 | 0.837 | 1.681 | 0.347 | 0.535 | 0.017 | 0.004 | 0.004 | 0.349 | 0.08 | 0.26 | 0.08 | 0.03 | 0.08 | 0.05 |
| 5 | 168513289 | rs296008 | 1.101 | 1.784 | 0.086 | 0.739 | 0.015 | 0.014 | 0.014 | 0 | 0.43 | 0.35 | 0.27 | 0.31 | 0.35 | 0.31 |
| 5 | 168513628 | rs296007 | -1.497 | 1.983 | -0.119 | 0.939 | -0.093 | -0.112 | -0.112 | 0 | 0.83 | 0.98 | 0.84 | 0.64 | 0.71 | 0.77 |
| 5 | 168514867 | rs296006 | 0.235 | 1.859 | 0.307 | 1.002 | 0 | 0.006 | 0.006 | 0 | 0.54 | 0.42 | 0.46 | 0.33 | 0.46 | 0.42 |
| 5 | 168517022 | rs167194 | 0.794 | 1.926 | -0.014 | 0.956 | -0.001 | 0.005 | 0.005 | 0 | 0.43 | 0.4 | 0.42 | 0.34 | 0.45 | 0.38 |
| 5 | 168518373 | rs296004 | 0.336 | 1.727 | -0.656 | 0.834 | 0.018 | 0.007 | 0.007 | 0 | 0.56 | 0.72 | 0.55 | 0.59 | 0.44 | 0.41 |
| 5 | 168521868 | rs296002 | -0.322 | 0.971 | -0.704 | 1.045 | -0.05 | -0.029 | -0.029 | 0 | 0.67 | 0.88 | 0.78 | 0.64 | 0.61 | 0.54 |
| 5 | 168522312 | rs11134558 | -0.572 | 0.626 | 0.208 | 1.53 | -0.029 | 0.094 | 0.094 | 0 | 0.18 | 0.34 | 0.33 | 0.31 | 0.13 | 0.13 |
| 5 | 168524685 | rs13173842 | -0.028 | 1.377 | -0.405 | 1.327 | -0.001 | -0.001 | -0.001 | 3.404 | 0.79 | 0.78 | 0.7 | 0.69 | 0.7 | 0.74 |
| 5 | 168525690 | rs10067964 | 0.674 | 1.496 | 0.324 | 1.624 | 0.081 | 0.088 | 0.088 | 0.09 | 0.8 | 0.57 | 0.68 | 0.67 | 0.68 | 0.71 |
| 5 | 168525867 | rs295999 | 0.705 | 1.1 | 0.304 | 1.259 | 0.006 | 0.023 | 0.023 | 0 | 0.68 | 0.59 | 0.67 | 0.61 | 0.73 | 0.51 |
| 5 | 168532467 | rs295996 | -0.777 | 0.888 | -0.012 | 1.133 | 0.001 | -0.002 | -0.002 | 0 | 0.1 | 0.09 | 0.13 | 0.17 | 0.06 | 0.12 |
| 5 | 168539240 | rs2569114 | -1.016 | 0.651 | -0.15 | 0.926 | -0.001 | 0 | 0 | 0 | 0.39 | 0.62 | 0.46 | 0.58 | 0.39 | 0.33 |
| 5 | 168540553 | rs12522573 | -0.032 | 1.263 | -0.724 | 0.776 | -0.018 | -0.029 | -0.029 | 0.223 | 0.2 | 0.14 | 0.27 | 0.32 | 0.29 | 0.18 |
| 5 | 168547979 | rs2668054 | -0.197 | 0.885 | -0.88 | 0.099 | -0.017 | 0.031 | 0.031 | 0 | 0.82 | 0.87 | 0.84 | 0.97 | 0.79 | 0.78 |
| 5 | 168550565 | rs297808 | 0.073 | 0.348 | -0.42 | 0.479 | 0.017 | 0.071 | 0.071 | 0.045 | 0.23 | 0.33 | 0.3 | 0.43 | 0.36 | 0.22 |
| 5 | 168554256 | rs17070834 | 1.115 | 0.349 | -1.317 | -0.662 | 0.003 | 0.057 | 0.057 | 0 | 0.26 | 0.14 | 0.18 | 0.04 | 0.25 | 0.14 |
| 5 | 168555866 | rs188431 | 0.366 | 1.323 | -0.977 | 0.059 | -0.029 | 0.028 | 0.028 | 0.151 | 0.71 | 0.77 | 0.76 | 0.93 | 0.63 | 0.7 |
| 5 | 168558803 | rs2668046 | -0.572 | 0.927 | -0.406 | 1.064 | 0.012 | 0.154 | 0.154 | 0.067 | 0.24 | 0.47 | 0.19 | 0.47 | 0.27 | 0.24 |
| 5 | 168559167 | rs876285 | 0.601 | 1.105 | -1.24 | 1.243 | -0.027 | -0.074 | -0.074 | 0.302 | 0.69 | 0.8 | 0.63 | 0.44 | 0.64 | 0.56 |
| 5 | 168560179 | rs888688 | -0.053 | 0.593 | -1.667 | 0.709 | -0.009 | 0.015 | 0.015 | 0.055 | 0.17 | 0.08 | 0.13 | 0.14 | 0.16 | 0.31 |
| 5 | 168561395 | rs1421765 | -0.168 | 1.174 | -2.104 | 0.896 | -0.073 | -0.014 | -0.014 | 0 | 0.8 | 0.9 | 0.54 | 0.75 | 0.63 | 0.81 |
| 5 | 168561511 | rs1421766 | 0.403 | 0.544 | -2.329 | 0.788 | -0.09 | 0.008 | 0.008 | 0 | 0.68 | 0.85 | 0.48 | 0.69 | 0.54 | 0.59 |
| 5 | 168563018 | rs10071654 | 0.692 | -0.335 | -1.794 | 0.289 | 0.014 | 0.157 | 0.157 | 0 | 0.47 | 0.21 | 0.31 | 0.15 | 0.42 | 0.38 |
| 5 | 168564806 | rs1421769 | 0.838 | -0.446 | -1.1 | 0.264 | 0.012 | 0.099 | 0.099 | 0 | 0.45 | 0.22 | 0.31 | 0.24 | 0.42 | 0.37 |
| 5 | 168570446 | rs12522564 | 0.323 | -0.447 | -1.052 | 0.196 | -0.004 | 0.011 | 0.011 | 0.053 | 0.21 | 0.16 | 0.17 | 0.15 | 0.21 | 0.31 |
| 5 | 168575951 | rs297832 | 0.901 | -0.86 | -0.72 | -0.045 | 0.001 | -0.016 | -0.016 | 1.168 | 0.4 | 0.42 | 0.28 | 0.16 | 0.34 | 0.11 |
| 5 | 168578995 | rs4868317 | 0.421 | -0.184 | -1.209 | -0.071 | -0.036 | 0.033 | 0.033 | 0.236 | 0.21 | 0.09 | 0.19 | 0.13 | 0.26 | 0.35 |
| 5 | 168579302 | rs1363129 | 1.636 | -0.289 | -1.18 | -0.162 | 0.012 | 0.088 | 0.088 | 0.472 | 0.29 | 0.1 | 0.15 | 0 | 0.24 | 0.07 |
| 5 | 168579843 | rs10475901 | 1.933 | -0.31 | -1.162 | -0.143 | 0.012 | 0.066 | 0.066 | 0.887 | 0.27 | 0.11 | 0.14 | 0 | 0.23 | 0.05 |
| 5 | 168582131 | rs297824 | -0.095 | -0.165 | -1.295 | 0.714 | -0.088 | -0.033 | -0.033 | 0.082 | 0.64 | 0.92 | 0.49 | 0.65 | 0.58 | 0.55 |
| 5 | 168584463 | rs190935 | -1.552 | 0.214 | -0.774 | 0.404 | -0.008 | 0.046 | 0.046 | 0 | 0.81 | 0.94 | 0.81 | 0.91 | 0.81 | 0.76 |
| 5 | 168590436 | rs297817 | 1.456 | -0.101 | -0.356 | 0.777 | 0 | -0.018 | -0.018 | 0 | 0.32 | 0.25 | 0.25 | 0.34 | 0.3 | 0.17 |
| 5 | 168592148 | rs7704526 | -0.663 | 0.006 | 0.179 | 0.729 | -0.013 | 0.086 | 0.086 | 0 | 0.29 | 0.46 | 0.44 | 0.5 | 0.29 | 0.27 |
| 5 | 168592495 | rs297819 | -1.548 | 0.06 | -0.04 | 0.634 | -0.006 | 0.032 | 0.032 | 0.021 | 0.75 | 0.86 | 0.78 | 0.89 | 0.75 | 0.72 |
| 5 | 168595003 | rs297823 | 1.877 | 0.045 | -0.016 | 0.597 | -0.032 | 0.114 | 0.114 | 0 | 0.3 | 0.1 | 0.23 | 0.09 | 0.31 | 0.08 |
| 5 | 168596883 | rs10462980 | -0.555 | 0.577 | 0.381 | 0.962 | -0.018 | 0.038 | 0.038 | 0 | 0.13 | 0.06 | 0.09 | 0.05 | 0.16 | 0.28 |
| 5 | 168598187 | rs9313446 | 0.193 | 0.465 | 0.684 | 1.118 | -0.011 | -0.036 | -0.036 | 0 | 0.2 | 0.15 | 0.36 | 0.41 | 0.23 | 0.24 |
| 5 | 168599222 | rs17070929 | -0.296 | 0.487 | 1.194 | 1.126 | -0.013 | -0.007 | -0.007 | 0 | 0.09 | 0.33 | 0.06 | 0.05 | 0.05 | 0.13 |
| 5 | 168602104 | rs297868 | -0.261 | 0.682 | 0.755 | 1.108 | 0.001 | 0 | 0 | 0 | 0.09 | 0.11 | 0.1 | 0.12 | 0.13 | 0.18 |
| 5 | 168602271 | rs297869 | -1.033 | 0.658 | 0.669 | 0.903 | -0.004 | 0.022 | 0.022 | 0 | 0.75 | 0.84 | 0.8 | 0.88 | 0.76 | 0.73 |
| 5 | 168610154 | rs11740487 | 0.01 | 0.697 | 0.453 | 0.957 | -0.002 | 0 | 0 | 0.002 | 0.06 | 0.19 | 0.02 | 0.03 | 0.03 | 0.12 |
| 5 | 168610788 | rs297879 | 0.4 | 0.915 | -0.674 | 0.944 | -0.001 | -0.003 | -0.003 | 0 | 0.58 | 0.35 | 0.64 | 0.58 | 0.6 | 0.36 |
| 5 | 168611732 | rs9313447 | -1.061 | 1.225 | 0.054 | 1.588 | 0.021 | 0.136 | 0.136 | 0.073 | 0.15 | 0.51 | 0.15 | 0.29 | 0.15 | 0.34 |
| 5 | 168616894 | rs1421763 | -0.562 | 1.133 | -0.853 | 1.352 | -0.012 | -0.03 | -0.03 | 0 | 0.19 | 0.15 | 0.26 | 0.37 | 0.22 | 0.19 |
| 5 | 168620924 | rs12515660 | -0.813 | 0.954 | -0.547 | 0.934 | -0.001 | 0.002 | 0.002 | 0 | 0.07 | 0.06 | 0.06 | 0.04 | 0.08 | 0.16 |
| 5 | 168623883 | rs2544823 | -0.048 | 1.019 | -0.326 | 0.668 | 0 | -0.001 | -0.001 | 0 | 0.39 | 0.43 | 0.42 | 0.23 | 0.42 | 0.3 |
| 5 | 168626236 | rs297850 | -0.082 | 0.903 | -0.132 | 1.015 | -0.002 | 0.052 | 0.052 | 0 | 0.15 | 0.21 | 0.23 | 0.45 | 0.13 | 0.11 |
| 5 | 168627478 | rs297852 | -0.884 | 0.869 | 0.165 | 0.856 | -0.002 | -0.013 | -0.013 | 1.68 | 0.58 | 0.62 | 0.71 | 0.77 | 0.61 | 0.45 |
| 5 | 168627798 | rs17735410 | 0.808 | 0.859 | 0.125 | 0.884 | -0.006 | -0.005 | -0.005 | 3.252 | 0.06 | 0.01 | 0.07 | 0.09 | 0.08 | 0.04 |
| 5 | 168629393 | rs297855 | -1.483 | 0.684 | 0.177 | 0.723 | -0.008 | 0.001 | 0.001 | 0.109 | 0.26 | 0.41 | 0.43 | 0.67 | 0.22 | 0.35 |
| 5 | 168630451 | rs297858 | -1.273 | 0.699 | 0.195 | 0.736 | 0.014 | 0.161 | 0.161 | 0 | 0.1 | 0.33 | 0.21 | 0.23 | 0.09 | 0.21 |
| 5 | 168630615 | rs1345591 | 0.089 | 0.697 | 0.232 | 0.735 | -0.004 | -0.006 | -0.006 | 0.562 | 0.06 | 0.13 | 0.07 | 0.04 | 0.06 | 0.15 |
| 5 | 168633029 | rs17643060 | 1.611 | 0.65 | 0.164 | 0.651 | -0.09 | 0.176 | 0.176 | 0.011 | 0.28 | 0.02 | 0.2 | 0 | 0.31 | 0.09 |
| 5 | 168633447 | rs11738097 | -0.05 | 0.639 | 0.232 | 0.768 | 0.002 | -0.048 | -0.048 | 0 | 0.16 | 0.08 | 0.23 | 0.45 | 0.13 | 0.13 |
| 5 | 168634766 | rs12514330 | -0.253 | 0.639 | 0.243 | 0.768 | -0.001 | 0.032 | 0.032 | 0 | 0.17 | 0.2 | 0.24 | 0.45 | 0.13 | 0.13 |
| 5 | 168637369 | rs10079795 | -0.047 | 0.676 | 0.234 | 0.737 | -0.006 | -0.008 | -0.008 | 1.534 | 0.07 | 0.14 | 0.07 | 0.04 | 0.06 | 0.15 |
| 5 | 168642564 | rs4868388 | -0.202 | 0.727 | 0.217 | 0.781 | 0 | 0.007 | 0.007 | 0 | 0.16 | 0.16 | 0.35 | 0.63 | 0.16 | 0.16 |
| 5 | 168644363 | rs6867453 | 0.057 | 0.732 | 0.247 | 0.701 | 0.004 | -0.012 | -0.012 | 0 | 0.43 | 0.55 | 0.49 | 0.32 | 0.47 | 0.28 |
| 5 | 168646504 | rs17643126 | 1.405 | 0.814 | 0.016 | 0.615 | -0.04 | 0.09 | 0.09 | 0.041 | 0.28 | 0.1 | 0.21 | 0 | 0.31 | 0.09 |
| 5 | 168649621 | rs10866632 | -0.05 | 0.686 | 0.273 | 0.863 | -0.001 | 0.145 | 0.145 | 0 | 0.15 | 0.32 | 0.23 | 0.44 | 0.13 | 0.1 |
| 5 | 168652877 | rs6868507 | 0.251 | 0.399 | 0.682 | 0.649 | -0.004 | 0.02 | 0.02 | 0 | 0.44 | 0.3 | 0.42 | 0.33 | 0.45 | 0.37 |
| 5 | 168653502 | rs17735629 | -0.212 | 0.368 | 0.514 | 0.573 | 0.02 | -0.054 | -0.054 | 0 | 0.16 | 0.08 | 0.22 | 0.45 | 0.1 | 0.15 |
| 5 | 168653516 | rs17668091 | 1.111 | 0.313 | 0.424 | 0.625 | 0.003 | -0.011 | -0.011 | 0.058 | 0.05 | 0.01 | 0.07 | 0.1 | 0.07 | 0.04 |
| 5 | 168654041 | rs17556758 | 1.799 | 0.365 | 0.325 | 0.475 | -0.082 | 0.141 | 0.141 | 0 | 0.26 | 0.1 | 0.21 | 0 | 0.32 | 0.1 |
| 5 | 168654113 | rs6893511 | 0.063 | 0.265 | 0.882 | 0.766 | 0.004 | 0.019 | 0.019 | 0 | 0.51 | 0.59 | 0.62 | 0.63 | 0.5 | 0.29 |
| 5 | 168658371 | rs10043015 | -0.619 | 0.642 | 0.759 | 0.84 | -0.006 | -0.001 | -0.001 | 0 | 0.58 | 0.61 | 0.63 | 0.59 | 0.7 | 0.43 |

| | | | | | | | | | | | | | | | | |
|---|---|---|---|---|---|---|---|---|---|---|---|---|---|---|---|---|
| 5 | 168663295 | rs10056028 | -0.273 | 0.807 | 0.714 | 0.806 | 0 | -0.001 | -0.001 | 0.588 | 0.56 | 0.59 | 0.59 | 0.53 | 0.59 | 0.28 |
| 5 | 168663943 | rs7719642 | -1.248 | 0.8 | 0.59 | 0.758 | 0.015 | 0.052 | 0.052 | 0 | 0.67 | 0.83 | 0.77 | 0.8 | 0.74 | 0.64 |
| 5 | 168668816 | rs745683 | 0.14 | 0.678 | 0.48 | 0.762 | 0 | -0.001 | -0.001 | 0 | 0.47 | 0.48 | 0.51 | 0.44 | 0.5 | 0.25 |
| 5 | 168677340 | rs4867775 | 0.789 | 1.053 | 0.06 | 0.687 | -0.044 | 0.214 | 0.214 | 0.015 | 0.41 | 0.08 | 0.28 | 0.13 | 0.4 | 0.31 |
| 5 | 168677793 | rs972031 | -1.356 | 0.811 | 0.268 | 0.564 | 0.006 | 0.026 | 0.026 | 0 | 0.7 | 0.74 | 0.79 | 0.8 | 0.74 | 0.65 |
| 5 | 168681309 | rs7716873 | 1.349 | 0.906 | -0.024 | 0.273 | -0.036 | 0.193 | 0.193 | 0.115 | 0.33 | 0.07 | 0.19 | 0.02 | 0.29 | 0.28 |
| 5 | 168688650 | rs11134564 | -0.319 | 1.001 | 0.302 | 0.868 | 0.005 | -0.01 | -0.01 | 0 | 0.1 | 0.07 | 0.24 | 0.15 | 0.07 | 0.07 |
| 5 | 168697807 | rs6555852 | -0.641 | 1.089 | 0.376 | 1.406 | -0.065 | -0.032 | -0.032 | 0 | 0.65 | 0.81 | 0.61 | 0.6 | 0.52 | 0.53 |
| 5 | 168699444 | rs3899063 | -1.424 | 1.02 | 0.957 | 1.591 | -0.026 | 0.113 | 0.113 | 0 | 0.22 | 0.38 | 0.33 | 0.48 | 0.16 | 0.21 |
| 5 | 168699952 | rs12521978 | 0.586 | 0.958 | 0.785 | 1.703 | -0.013 | -0.035 | -0.035 | 0.012 | 0.07 | 0.03 | 0.02 | 0.26 | 0.07 | 0.1 |
| 5 | 168700302 | rs888784 | 1.713 | 1.184 | 1.104 | 1.639 | 0.022 | 0.081 | 0.081 | 0 | 0.75 | 0.57 | 0.61 | 0.5 | 0.67 | 0.67 |
| 5 | 168702491 | rs7712111 | -0.176 | 1.187 | 1.314 | 1.576 | -0.002 | -0.004 | -0.004 | 1.878 | 0.42 | 0.5 | 0.41 | 0.68 | 0.38 | 0.45 |
| 5 | 168702931 | rs755063 | 1.358 | 1.173 | 1.314 | 1.534 | 0.083 | 0.004 | 0.004 | 0.117 | 0.83 | 0.62 | 0.65 | 0.83 | 0.74 | 0.78 |
| 5 | 168703123 | rs755062 | 1.143 | 1.193 | 0.789 | 1.449 | -0.001 | 0.184 | 0.184 | 0.095 | 0.43 | 0.17 | 0.27 | 0.13 | 0.37 | 0.33 |
| 5 | 168707562 | rs11134568 | -0.184 | 1.129 | 0.705 | 1.86 | 0.02 | 0.006 | 0.006 | 0 | 0.94 | 0.93 | 0.92 | 0.89 | 0.81 | 0.85 |
| 5 | 168709067 | rs888781 | 0.982 | 1.197 | 0.968 | 1.711 | -0.006 | 0.011 | 0.011 | 0 | 0.3 | 0.15 | 0.33 | 0.27 | 0.32 | 0.37 |
| 5 | 168710367 | rs751346 | 0.714 | 1.191 | 0.999 | 1.784 | -0.004 | -0.004 | -0.004 | 0 | 0.86 | 0.85 | 0.7 | 0.76 | 0.75 | 0.78 |
| 5 | 168710470 | rs751347 | 1.797 | 1.138 | 1.253 | 1.787 | 0.026 | 0.056 | 0.056 | 0.01 | 0.73 | 0.49 | 0.59 | 0.42 | 0.66 | 0.65 |
| 5 | 168710623 | rs751348 | 1.299 | 1.133 | 1.165 | 1.454 | 0.008 | 0.027 | 0.027 | 0.029 | 0.42 | 0.31 | 0.26 | 0.14 | 0.34 | 0.28 |
| 5 | 168712320 | rs750585 | 0.454 | 1.143 | 0.778 | 1.64 | -0.001 | 0.038 | 0.038 | 0 | 0.08 | 0.09 | 0.03 | 0.28 | 0.07 | 0.12 |
| 5 | 168715942 | rs4339366 | 1.573 | 0.805 | 0.647 | 1.368 | 0.008 | 0.043 | 0.043 | 0 | 0.69 | 0.49 | 0.62 | 0.48 | 0.64 | 0.64 |
| 5 | 168721192 | rs4432908 | -0.497 | 1.017 | -0.249 | 1.149 | -0.02 | -0.003 | -0.003 | 0 | 0.89 | 0.92 | 0.92 | 0.91 | 0.77 | 0.84 |
| 5 | 168734867 | rs6864818 | -0.619 | 1.029 | -0.434 | 1.153 | -0.022 | -0.007 | -0.007 | 0 | 0.89 | 0.9 | 0.92 | 0.9 | 0.77 | 0.84 |
| 5 | 168734887 | rs6883683 | 0.925 | 0.914 | -0.44 | 1.111 | 0.003 | 0.001 | 0.001 | 0 | 0.44 | 0.26 | 0.28 | 0.41 | 0.4 | 0.37 |
| 5 | 168749737 | rs6555855 | -0.199 | 1.433 | -0.935 | 1.117 | -0.018 | -0.009 | -0.009 | 0 | 0.89 | 0.92 | 0.89 | 0.89 | 0.77 | 0.83 |
| 5 | 168750365 | rs1422033 | -0.008 | 0.918 | -0.792 | 1.008 | 0.004 | -0.004 | -0.004 | 0 | 0.32 | 0.18 | 0.37 | 0.34 | 0.31 | 0.38 |
| 5 | 168760826 | rs1001869 | -0.644 | 1.136 | -0.715 | 0.937 | -0.051 | -0.065 | -0.065 | 0.327 | 0.29 | 0.14 | 0.48 | 0.45 | 0.44 | 0.38 |
| 5 | 168762823 | rs7444781 | -0.432 | 1.037 | -0.341 | 0.263 | 0 | -0.011 | -0.011 | 0.515 | 0.9 | 0.9 | 0.92 | 0.97 | 0.88 | 0.92 |
| 5 | 168774121 | rs1895203 | -0.39 | 0.974 | -0.211 | 0.494 | -0.023 | -0.031 | -0.031 | 0 | 0.27 | 0.17 | 0.35 | 0.58 | 0.36 | 0.25 |
| 5 | 168775141 | rs4868525 | 1.235 | 0.332 | -0.223 | 0.251 | -0.023 | 0.042 | 0.042 | 0.604 | 0.41 | 0.16 | 0.48 | 0.32 | 0.48 | 0.46 |
| 5 | 168781649 | rs6879310 | 0.016 | 0.808 | 0.582 | 0.884 | 0.002 | 0.022 | 0.022 | 0 | 0.2 | 0.17 | 0.31 | 0.48 | 0.3 | 0.2 |
| 5 | 168793246 | rs10044168 | -0.497 | -0.204 | 1.154 | 0.512 | -0.08 | -0.095 | -0.095 | 0 | 0.14 | 0.39 | 0.07 | 0 | 0.04 | 0.13 |
| 5 | 168811335 | rs9313461 | -0.56 | -0.169 | 1.115 | 0.659 | -0.086 | 0.011 | 0.011 | 0.074 | 0.15 | 0.45 | 0.09 | 0.09 | 0.05 | 0.12 |
| 5 | 168828290 | rs6555860 | 0.594 | 0.775 | -0.485 | 0.611 | 0.007 | -0.027 | -0.027 | 0 | 0.54 | 0.17 | 0.54 | 0.55 | 0.57 | 0.62 |
| 5 | 168837488 | rs6898594 | 0.289 | 0.991 | 0.076 | 0.452 | 0.004 | -0.004 | -0.004 | 0.635 | 0.58 | 0.49 | 0.5 | 0.45 | 0.55 | 0.53 |
| 5 | 168838551 | rs6864218 | -1.403 | 0.828 | 0.653 | 0.285 | 0.045 | 0.037 | 0.037 | 0.368 | 0.05 | 0.23 | 0.12 | 0.01 | 0.04 | 0.05 |
| 5 | 168839082 | rs6864664 | -1.683 | 1.095 | 0.697 | 0.257 | 0.093 | 0.026 | 0.026 | 0.001 | 0.07 | 0.28 | 0.15 | 0.01 | 0.1 | 0.06 |
| 5 | 168840498 | rs731398 | -0.628 | 0.672 | 0.208 | -0.584 | -0.006 | -0.058 | -0.058 | 0 | 0.27 | 0.28 | 0.19 | 0.03 | 0.23 | 0.24 |
| 5 | 168841357 | rs12522101 | 0.052 | 0.617 | -0.748 | -0.193 | 0.003 | -0.075 | -0.075 | 0 | 0.33 | 0.15 | 0.24 | 0.47 | 0.27 | 0.31 |
| 5 | 168841670 | rs13182979 | 0.476 | 1.233 | -0.677 | -1.229 | -0.01 | 0.004 | 0.004 | 0 | 0.26 | 0.18 | 0.17 | 0.04 | 0.32 | 0.19 |
| 5 | 168843083 | rs2112333 | -0.758 | 1.404 | -0.966 | -1.536 | -0.032 | 0.059 | 0.059 | 0 | 0.86 | 0.98 | 0.91 | 0.97 | 0.83 | 0.92 |
| 5 | 168843349 | rs929757 | -0.033 | 1.174 | -0.309 | -1.581 | 0.002 | -0.042 | -0.042 | 0 | 0.38 | 0.38 | 0.26 | 0.1 | 0.45 | 0.34 |
| 5 | 168844957 | rs7729168 | 0.882 | 1.114 | -0.005 | -1.167 | 0.011 | 0.017 | 0.017 | 0 | 0.65 | 0.43 | 0.47 | 0.6 | 0.59 | 0.57 |
| 5 | 168847968 | rs13177680 | 0.829 | 1.446 | 0.064 | -0.794 | -0.002 | -0.022 | -0.022 | 0 | 0.26 | 0.21 | 0.14 | 0.04 | 0.29 | 0.16 |
| 5 | 168853268 | rs10475909 | 0.969 | 1.373 | 0.284 | -0.85 | 0.011 | 0.001 | 0.001 | 0 | 0.45 | 0.38 | 0.34 | 0.57 | 0.29 | 0.43 |
| 5 | 168854538 | rs2338656 | 0.917 | 1.376 | 0.343 | -0.853 | -0.01 | -0.007 | -0.007 | 0 | 0.46 | 0.47 | 0.37 | 0.61 | 0.34 | 0.44 |
| 5 | 168868506 | rs4868643 | -0.786 | 1.454 | 0.24 | -0.841 | -0.008 | -0.007 | -0.007 | 0.034 | 0.85 | 0.84 | 0.76 | 0.81 | 0.77 | 0.88 |
| 5 | 168869424 | rs6865393 | 0.836 | 1.204 | 0.336 | -0.752 | -0.01 | 0.007 | 0.007 | 0 | 0.26 | 0.18 | 0.15 | 0.04 | 0.31 | 0.19 |
| 5 | 168870650 | rs12652441 | 0.318 | 1.211 | 0.351 | -0.697 | 0.013 | -0.018 | -0.018 | 1.32 | 0.29 | 0.4 | 0.35 | 0.24 | 0.36 | 0.28 |
| 5 | 168885702 | rs2431281 | -0.322 | 1.315 | 0.247 | -0.089 | -0.017 | 0.011 | 0.011 | 0 | 0.26 | 0.23 | 0.31 | 0.19 | 0.33 | 0.29 |
| 5 | 168897517 | rs255740 | -0.495 | 1.367 | -0.169 | -0.214 | 0.006 | 0.006 | 0.006 | 0 | 0.27 | 0.35 | 0.48 | 0.34 | 0.34 | 0.37 |
| 5 | 168899806 | rs11958127 | 0.232 | 0.722 | -0.475 | -0.416 | 0 | -0.001 | -0.001 | 0 | 0.26 | 0.28 | 0.22 | 0.62 | 0.25 | 0.3 |
| 5 | 168899987 | rs13156607 | -0.432 | 0.85 | -0.365 | -1.279 | 0.004 | -0.12 | -0.12 | 0 | 0.64 | 0.6 | 0.78 | 0.97 | 0.6 | 0.68 |
| 5 | 168910494 | rs10059417 | 0.282 | 0.155 | -0.636 | -0.88 | -0.023 | 0.007 | 0.007 | 0 | 0.4 | 0.33 | 0.36 | 0.57 | 0.41 | 0.35 |
| 5 | 168912305 | rs2545384 | 0.647 | 0.027 | -0.683 | -0.955 | -0.082 | -0.026 | -0.026 | 0 | 0.35 | 0.17 | 0.28 | 0.57 | 0.38 | 0.34 |
| 5 | 168922672 | rs2546707 | -0.475 | -0.258 | -0.619 | -1.068 | 0.008 | 0.01 | 0.01 | 0.015 | 0.56 | 0.45 | 0.59 | 0.77 | 0.61 | 0.58 |
| 5 | 168924485 | rs2559976 | 0.826 | -0.164 | -0.81 | -0.984 | -0.058 | -0.016 | -0.016 | 0 | 0.35 | 0.21 | 0.27 | 0.57 | 0.36 | 0.34 |
| 5 | 168930665 | rs29809 | -0.245 | -0.261 | -0.695 | -1.077 | 0.027 | 0.051 | 0.051 | 0.013 | 0.56 | 0.36 | 0.53 | 0.76 | 0.59 | 0.6 |
| 5 | 168938303 | rs2559974 | -1.263 | -0.568 | -0.206 | -1.207 | 0.015 | 0.034 | 0.034 | 0 | 0.67 | 0.75 | 0.68 | 0.8 | 0.68 | 0.71 |
| 5 | 168948878 | rs12654922 | 0.358 | -0.375 | -0.426 | -1.079 | -0.009 | 0.003 | 0.003 | 0 | 0.18 | 0.12 | 0.16 | 0.21 | 0.23 | 0.25 |
| 5 | 168963357 | rs17644016 | 0.633 | -0.369 | -0.47 | -1.082 | -0.018 | -0.009 | -0.009 | 0.025 | 0.17 | 0.05 | 0.13 | 0.2 | 0.18 | 0.24 |
| 5 | 168970505 | rs255827 | -0.163 | -0.786 | -0.07 | -1.114 | 0.026 | 0.043 | 0.043 | 0.072 | 0.57 | 0.35 | 0.55 | 0.75 | 0.63 | 0.61 |
| 5 | 168990341 | rs2546691 | 0.114 | -0.855 | -0.478 | -0.873 | 0.08 | 0.128 | 0.128 | 0 | 0.52 | 0.17 | 0.49 | 0.69 | 0.58 | 0.55 |
| 5 | 169014301 | rs255824 | -1.175 | -1.004 | -0.681 | -0.843 | -0.015 | 0.097 | 0.097 | 0.596 | 0.69 | 0.99 | 0.86 | 0.89 | 0.72 | 0.8 |
| 5 | 169017230 | rs17668965 | 1.199 | -0.944 | -0.9 | -0.172 | -0.023 | -0.047 | -0.047 | 0 | 0.08 | 0 | 0.06 | 0.18 | 0.08 | 0.13 |
| 5 | 169019631 | rs6892893 | -0.943 | -0.989 | -0.069 | -0.552 | 0.018 | 0.056 | 0.056 | 0.142 | 0.67 | 0.83 | 0.84 | 0.87 | 0.67 | 0.78 |
| 5 | 169021610 | rs4493692 | -0.579 | -0.883 | 0.155 | -0.556 | -0.022 | -0.026 | -0.026 | 0 | 0.63 | 0.53 | 0.59 | 0.73 | 0.67 | 0.7 |
| 5 | 169028481 | rs3797713 | -0.646 | -0.879 | -0.005 | -0.554 | -0.014 | -0.016 | -0.016 | 0 | 0.63 | 0.59 | 0.6 | 0.73 | 0.67 | 0.7 |
| 5 | 169031150 | rs3777084 | -0.646 | -0.879 | -0.005 | -0.554 | -0.014 | -0.016 | -0.016 | 0 | 0.63 | 0.59 | 0.6 | 0.73 | 0.67 | 0.7 |

| | | | | | | | | | | | | | | | | |
|---|---|---|---|---|---|---|---|---|---|---|---|---|---|---|---|---|
| 5 | 169031218 | rs2042248 | -0.666 | -0.878 | -0.054 | -0.554 | 0.006 | 0.007 | 0.007 | 0 | 0.63 | 0.66 | 0.6 | 0.73 | 0.67 | 0.7 |
| 5 | 169062922 | rs264866 | -0.661 | -0.864 | -0.189 | -0.547 | -0.011 | -0.013 | -0.013 | 0 | 0.63 | 0.58 | 0.59 | 0.73 | 0.67 | 0.7 |
| 5 | 169074056 | rs169082 | 0.281 | -0.336 | -0.643 | -0.176 | -0.02 | -0.011 | -0.011 | 0 | 0.4 | 0.3 | 0.47 | 0.54 | 0.48 | 0.36 |
| 5 | 169080798 | rs36101422 | -0.099 | -0.585 | -0.767 | -0.309 | 0.008 | -0.001 | -0.001 | 0 | 0.1 | 0.03 | 0.04 | 0.1 | 0.06 | 0.16 |
| 5 | 169081315 | rs12656935 | -0.157 | -0.586 | -0.768 | -0.309 | 0.008 | 0 | 0 | 0.198 | 0.1 | 0.03 | 0.04 | 0.1 | 0.06 | 0.16 |
| 5 | 169093679 | rs6883997 | 0.009 | -0.696 | -0.486 | -0.507 | 0.044 | 0.068 | 0.068 | 0 | 0.44 | 0.3 | 0.25 | 0.21 | 0.38 | 0.39 |
| 5 | 169095809 | rs12523168 | -0.192 | -0.719 | -0.406 | -0.488 | 0.01 | 0.012 | 0.012 | 0 | 0.61 | 0.43 | 0.39 | 0.49 | 0.57 | 0.6 |
| 5 | 169096774 | rs7721990 | -1.007 | -0.719 | -0.45 | -0.489 | 0.044 | 0.049 | 0.049 | 0 | 0.72 | 0.61 | 0.45 | 0.52 | 0.63 | 0.73 |
| 5 | 169099398 | rs7713640 | 0.634 | -0.694 | -0.685 | -0.48 | 0.007 | -0.029 | -0.029 | 0.1 | 0.25 | 0.11 | 0.17 | 0.29 | 0.23 | 0.31 |
| 5 | 169099623 | rs4867877 | 1.081 | -0.289 | -0.792 | -0.325 | 0.007 | 0.005 | 0.005 | 0 | 0.1 | 0.1 | 0.04 | 0.02 | 0.06 | 0.12 |
| 5 | 169109140 | rs9313467 | -0.566 | 0.243 | -0.583 | 0.058 | 0.031 | 0.035 | 0.035 | 0 | 0.72 | 0.63 | 0.45 | 0.52 | 0.63 | 0.71 |
| 5 | 169112649 | rs13183751 | -0.087 | -0.168 | -0.814 | -0.413 | 0.022 | 0.044 | 0.044 | 0 | 0.31 | 0.17 | 0.2 | 0.17 | 0.31 | 0.23 |
| 5 | 169122803 | rs3734099 | -0.855 | 0.626 | 0.123 | 0.063 | 0.095 | 0.007 | 0.007 | 0.063 | 0.1 | 0.38 | 0.08 | 0.09 | 0.12 | 0.08 |
| 5 | 169124478 | rs2112705 | -0.637 | 0.437 | -1.158 | 0.235 | -0.018 | -0.04 | -0.04 | 1.087 | 0.17 | 0.02 | 0.07 | 0.15 | 0.17 | 0.2 |
| 5 | 169126349 | rs2306562 | -0.488 | 0.641 | -1.168 | -0.01 | -0.035 | 0.008 | 0.008 | 0 | 0.16 | 0.03 | 0.09 | 0.12 | 0.19 | 0.17 |
| 5 | 169126552 | rs2112704 | -0.622 | 0.098 | -0.934 | 0.069 | -0.02 | 0.006 | 0.006 | 0.256 | 0.07 | 0.11 | 0.02 | 0.1 | 0.02 | 0.22 |
| 5 | 169127097 | rs2112703 | -0.617 | 0.083 | -1.078 | -0.037 | -0.035 | 0.018 | 0.018 | 0 | 0.18 | 0.04 | 0.09 | 0.12 | 0.2 | 0.17 |
| 5 | 169129284 | rs2306560 | -1.139 | -0.118 | -0.428 | 0.079 | -0.001 | -0.002 | -0.002 | 0 | 0.22 | 0.29 | 0.43 | 0.2 | 0.22 | 0.2 |
| 5 | 169129422 | rs2306559 | 1.371 | -0.17 | -0.744 | -1.191 | -0.011 | 0.038 | 0.038 | 0 | 0.4 | 0.2 | 0.33 | 0.68 | 0.45 | 0.31 |
| 5 | 169129842 | rs1363627 | 1.535 | -0.189 | -0.745 | -1.077 | -0.068 | 0.002 | 0.002 | 0 | 0.38 | 0.05 | 0.32 | 0.62 | 0.44 | 0.3 |
| 5 | 169129866 | rs1345723 | 0.154 | -0.129 | -0.68 | -1.129 | -0.002 | -0.052 | -0.052 | 0.217 | 0.62 | 0.44 | 0.76 | 0.88 | 0.67 | 0.51 |
| 5 | 169132375 | rs4867879 | 0.978 | -0.247 | -0.368 | -1.08 | -0.008 | 0.05 | 0.05 | 0 | 0.43 | 0.27 | 0.67 | 0.69 | 0.48 | 0.37 |
| 5 | 169138029 | rs373808 | -0.038 | -0.218 | 0.217 | -1.47 | -0.013 | -0.002 | -0.002 | 0 | 0.76 | 0.78 | 0.81 | 0.98 | 0.74 | 0.77 |
| 5 | 169146825 | rs371856 | 0.53 | -0.447 | 0.528 | -1.499 | -0.015 | 0.016 | 0.016 | 0 | 0.63 | 0.63 | 0.71 | 0.74 | 0.56 | 0.61 |
| 5 | 169148580 | rs10038749 | 1.744 | -0.574 | 0.539 | -1.847 | -0.005 | 0.03 | 0.03 | 0 | 0.35 | 0.23 | 0.58 | 0.74 | 0.4 | 0.35 |
| 5 | 169150178 | rs262883 | -0.409 | -0.522 | 1.119 | -1.668 | 0.014 | -0.09 | -0.09 | 0 | 0.8 | 0.7 | 0.82 | 0.98 | 0.78 | 0.87 |
| 5 | 169150610 | rs12652482 | 1.553 | -0.635 | 1.001 | -1.66 | -0.033 | 0.142 | 0.142 | 0 | 0.43 | 0.2 | 0.64 | 0.81 | 0.48 | 0.53 |
| 5 | 169155445 | rs262878 | -0.438 | -0.729 | 1.414 | -1.565 | -0.028 | -0.032 | -0.032 | 0.214 | 0.16 | 0.2 | 0.06 | 0.05 | 0.08 | 0.22 |
| 5 | 169165336 | rs262844 | -1.704 | -0.806 | 1.13 | -1.399 | 0.01 | 0.009 | 0.009 | 0 | 0.29 | 0.74 | 0.45 | 0.18 | 0.15 | 0.27 |
| 5 | 169173312 | rs13184159 | -0.294 | -0.704 | 1.157 | -1.269 | -0.004 | 0.02 | 0.02 | 0 | 0.06 | 0.11 | 0.34 | 0.15 | 0.04 | 0.14 |
| 5 | 169178062 | rs746468 | -0.199 | -0.661 | 1.467 | -1.294 | -0.003 | 0.023 | 0.023 | 0 | 0.06 | 0.11 | 0.34 | 0.15 | 0.04 | 0.14 |
| 5 | 169178278 | rs90213 | -1.74 | -0.86 | 1.468 | -1.418 | 0.002 | 0.002 | 0.002 | 0 | 0.31 | 0.71 | 0.45 | 0.26 | 0.15 | 0.4 |
| 5 | 169191428 | rs9918252 | -1.078 | -0.777 | 1.539 | -1.362 | 0.006 | 0.148 | 0.148 | 0 | 0.08 | 0.32 | 0.36 | 0.2 | 0.04 | 0.15 |
| 5 | 169192826 | rs12153495 | 0.97 | -0.754 | 1.268 | -1.383 | -0.054 | -0.008 | -0.008 | 0 | 0.13 | 0.01 | 0.08 | 0.11 | 0.18 | 0.14 |
| 5 | 169193066 | rs438261 | -0.026 | -0.891 | 1.639 | -1.348 | -0.022 | -0.036 | -0.036 | 0 | 0.07 | 0.15 | 0.02 | 0 | 0.02 | 0.09 |
| 5 | 169196282 | rs10055949 | 0.314 | -0.881 | 1.657 | -1.342 | -0.021 | -0.018 | -0.018 | 3.247 | 0.11 | 0.23 | 0.05 | 0.06 | 0.06 | 0.14 |
| 5 | 169198080 | rs2625384 | -1.004 | -0.827 | 1.237 | -1.284 | -0.04 | 0.014 | 0.014 | 0 | 0.17 | 0.28 | 0.38 | 0.2 | 0.08 | 0.26 |
| 5 | 169199103 | rs1013192 | 2.097 | -0.717 | 0.496 | -1.482 | -0.084 | -0.032 | -0.032 | 0 | 0.37 | 0.06 | 0.3 | 0.61 | 0.45 | 0.34 |
| 5 | 169199323 | rs1013193 | -0.596 | -0.771 | 0.791 | -1.136 | -0.015 | 0.014 | 0.014 | 0 | 0.14 | 0.2 | 0.37 | 0.2 | 0.08 | 0.24 |
| 5 | 169201727 | rs7735431 | -1.769 | -0.618 | 0.766 | -0.557 | 0.003 | 0 | 0 | 3.305 | 0.45 | 0.62 | 0.54 | 0.57 | 0.37 | 0.55 |
| 5 | 169203598 | rs966595 | -0.711 | -0.706 | 0.697 | -0.492 | 0.001 | -0.006 | -0.006 | 0 | 0.3 | 0.31 | 0.46 | 0.52 | 0.29 | 0.37 |
| 5 | 169204246 | rs17669654 | -0.525 | -0.742 | 0.601 | -0.449 | -0.021 | 0.075 | 0.075 | 0 | 0.16 | 0.25 | 0.38 | 0.44 | 0.1 | 0.23 |
| 5 | 169207603 | rs2046252 | 0.557 | -0.444 | 0.729 | -0.476 | 0.051 | 0.097 | 0.097 | 0 | 0.74 | 0.49 | 0.65 | 0.49 | 0.69 | 0.59 |
| 5 | 169209137 | rs261581 | 0.793 | -0.449 | 0.199 | -0.423 | -0.003 | 0.003 | 0.003 | 0 | 0.15 | 0.1 | 0.09 | 0.13 | 0.19 | 0.16 |
| 5 | 169209994 | rs1863992 | 1.531 | -0.499 | -0.066 | -0.503 | -0.051 | -0.084 | -0.084 | 0 | 0.59 | 0.1 | 0.58 | 0.44 | 0.61 | 0.42 |
| 5 | 169215669 | rs17559708 | -0.188 | -0.495 | -0.113 | -0.644 | -0.038 | 0.155 | 0.155 | 0 | 0.29 | 0.04 | 0.15 | 0.08 | 0.3 | 0.21 |
| 5 | 169217218 | rs10054949 | 0.156 | -0.539 | 0.043 | -0.57 | -0.004 | -0.015 | -0.015 | 0 | 0.12 | 0.33 | 0.09 | 0.06 | 0.09 | 0.14 |
| 5 | 169219999 | rs9313473 | -0.189 | -0.157 | 0.066 | -0.414 | -0.019 | 0.116 | 0.116 | 0.165 | 0.09 | 0.16 | 0.19 | 0.37 | 0.04 | 0.23 |
| 5 | 169222099 | rs261606 | -0.96 | 0 | 0.186 | -0.755 | -0.024 | -0.053 | -0.053 | 0.098 | 0.37 | 0.49 | 0.18 | 0.17 | 0.3 | 0.31 |
| 5 | 169226130 | rs261590 | 0.661 | 0.135 | -0.333 | -0.412 | 0.003 | -0.037 | -0.037 | 0.155 | 0.76 | 0.35 | 0.82 | 0.87 | 0.79 | 0.8 |
| 5 | 169238342 | rs155074 | -1.522 | 0.514 | -1.345 | 0.178 | -0.036 | -0.038 | -0.038 | 0.098 | 0.67 | 0.88 | 0.71 | 0.63 | 0.62 | 0.65 |
| 5 | 169238673 | rs261016 | -0.672 | 0.315 | -0.735 | -0.226 | -0.004 | -0.072 | -0.072 | 0 | 0.25 | 0.55 | 0.17 | 0.1 | 0.2 | 0.2 |
| 5 | 169239816 | rs155073 | -1.458 | 0.374 | -1.068 | 0.103 | -0.016 | 0.132 | 0.132 | 0 | 0.55 | 0.8 | 0.5 | 0.23 | 0.55 | 0.45 |
| 5 | 169242458 | rs261010 | -0.626 | 0.3 | -0.986 | 0.063 | 0.009 | -0.054 | -0.054 | 0.02 | 0.21 | 0.59 | 0.17 | 0.1 | 0.18 | 0.18 |
| 5 | 169245684 | rs391800 | -1.218 | 0.273 | -1.139 | 0.059 | -0.003 | 0.119 | 0.119 | 0 | 0.44 | 0.72 | 0.48 | 0.18 | 0.41 | 0.29 |
| 5 | 169253887 | rs246770 | -2.625 | 0.338 | -1.306 | 0.312 | -0.033 | -0.033 | -0.033 | 0 | 0.68 | 0.91 | 0.72 | 0.67 | 0.65 | 0.58 |
| 5 | 169255522 | rs39824 | -2.602 | 0.341 | -1.305 | 0.357 | -0.032 | -0.041 | -0.041 | 0.106 | 0.68 | 0.91 | 0.72 | 0.64 | 0.65 | 0.58 |
| 5 | 169257000 | rs2161148 | 0.407 | 0.462 | -1.107 | 0.271 | -0.019 | 0.078 | 0.078 | 0.842 | 0.22 | 0.03 | 0.23 | 0.08 | 0.23 | 0.16 |
| 5 | 169259125 | rs10054135 | -0.247 | 0.543 | -0.332 | 0.215 | 0.071 | 0.005 | 0.005 | 0 | 0.08 | 0.27 | 0.16 | 0.06 | 0.08 | 0.07 |
| 5 | 169260680 | rs396387 | -0.867 | 0.553 | -0.898 | 0.177 | 0.003 | -0.045 | -0.045 | 0 | 0.21 | 0.62 | 0.17 | 0.08 | 0.18 | 0.13 |
| 5 | 169266513 | rs261622 | -1.026 | 0.726 | -0.773 | 0.337 | -0.004 | 0.078 | 0.078 | 0 | 0.43 | 0.65 | 0.39 | 0.16 | 0.42 | 0.29 |
| 5 | 169268191 | rs261625 | -1.585 | 0.65 | -1.233 | 0.677 | -0.055 | -0.043 | -0.043 | 0 | 0.54 | 0.9 | 0.54 | 0.58 | 0.5 | 0.54 |
| 5 | 169275842 | rs30084 | -1.682 | 0.924 | -0.879 | 0.487 | -0.02 | 0.115 | 0.115 | 0 | 0.6 | 0.93 | 0.53 | 0.84 | 0.61 | 0.58 |
| 5 | 169277123 | rs152111 | 0.372 | 0.841 | -0.974 | 0.657 | 0.005 | -0.103 | -0.103 | 0.01 | 0.24 | 0.08 | 0.16 | 0.62 | 0.21 | 0.25 |
| 5 | 169278116 | rs17737597 | 1.097 | 0.87 | -0.967 | 0.64 | -0.024 | -0.022 | -0.022 | 0 | 0.08 | 0.03 | 0.17 | 0.11 | 0.14 | 0.07 |
| 5 | 169280342 | rs261077 | -0.617 | 0.792 | -0.901 | 0.381 | 0.002 | 0.031 | 0.031 | 2.256 | 0.38 | 0.13 | 0.34 | 0.73 | 0.39 | 0.33 |
| 5 | 169282375 | rs261039 | -0.776 | 0.661 | -0.614 | 0.424 | 0.015 | -0.024 | -0.024 | 0.055 | 0.2 | 0.66 | 0.17 | 0.11 | 0.21 | 0.12 |
| 5 | 169288136 | rs1482333 | 1.999 | 0.585 | -0.756 | 0.423 | -0.045 | 0.116 | 0.116 | 0 | 0.41 | 0.1 | 0.49 | 0.19 | 0.43 | 0.43 |
| 5 | 169289206 | rs10866641 | 2.036 | 0.533 | -0.759 | 0.423 | -0.053 | 0.113 | 0.113 | 0 | 0.41 | 0.08 | 0.49 | 0.19 | 0.43 | 0.43 |

| | | | | | | | | | | | | | | | | |
|---|---|---|---|---|---|---|---|---|---|---|---|---|---|---|---|---|
| 5 | 169291188 | rs261028 | -0.467 | 0.55 | -0.504 | 0.43 | 0.001 | -0.046 | -0.046 | 1.68 | 0.17 | 0.51 | 0.11 | 0.08 | 0.13 | 0.1 |
| 5 | 169299685 | rs6872720 | 1.995 | 0.413 | -0.65 | 0.458 | -0.05 | 0.159 | 0.159 | 0 | 0.41 | 0.11 | 0.46 | 0.13 | 0.43 | 0.39 |
| 5 | 169303604 | rs17670189 | 0.78 | 0.293 | -0.619 | 0.529 | -0.015 | -0.015 | -0.015 | 0 | 0.1 | 0.02 | 0.17 | 0.12 | 0.12 | 0.1 |
| 5 | 169304372 | rs261076 | -1 | 0.289 | -0.566 | 0.546 | 0.003 | -0.007 | -0.007 | 0.409 | 0.23 | 0.53 | 0.27 | 0.22 | 0.24 | 0.2 |
| 5 | 169309080 | rs261074 | -0.189 | 0.31 | -0.55 | 0.559 | 0.026 | 0.001 | 0.001 | 0 | 0.13 | 0.5 | 0.1 | 0.1 | 0.12 | 0.1 |
| 5 | 169311992 | rs261073 | -0.714 | 0.315 | -0.63 | 0.547 | -0.007 | -0.107 | -0.107 | 0 | 0.77 | 0.97 | 0.81 | 0.38 | 0.79 | 0.76 |
| 5 | 169313219 | rs4867897 | 1.884 | 0.376 | -0.592 | 0.447 | -0.029 | 0.145 | 0.145 | 0 | 0.43 | 0.2 | 0.47 | 0.13 | 0.45 | 0.41 |
| 5 | 169314869 | rs6879439 | 1.501 | 0.335 | -0.617 | 0.474 | 0.01 | 0.072 | 0.072 | 0 | 0.47 | 0.4 | 0.53 | 0.16 | 0.52 | 0.54 |
| 5 | 169315624 | rs261072 | 0.648 | 0.513 | 0.004 | 0.595 | 0.078 | 0.069 | 0.069 | 0 | 0.08 | 0.33 | 0.06 | 0.07 | 0.07 | 0.06 |
| 5 | 169323719 | rs261065 | -0.331 | 0.519 | -0.911 | 0.973 | -0.042 | -0.096 | -0.096 | 0 | 0.75 | 0.94 | 0.77 | 0.43 | 0.71 | 0.74 |
| 5 | 169324616 | rs11741998 | 1.687 | 0.43 | -0.999 | 0.542 | -0.05 | 0.122 | 0.122 | 0 | 0.4 | 0.15 | 0.53 | 0.15 | 0.44 | 0.4 |
| 5 | 169326062 | rs261060 | -0.929 | 0.285 | -0.206 | 0.734 | -0.033 | -0.054 | -0.054 | 2.006 | 0.2 | 0.52 | 0.1 | 0.1 | 0.14 | 0.11 |
| 5 | 169330699 | rs166354 | 0.583 | 0.428 | -0.417 | 1.084 | 0.002 | 0.035 | 0.035 | 0 | 0.66 | 0.59 | 0.73 | 0.48 | 0.67 | 0.65 |
| 5 | 169331239 | rs261050 | -0.978 | 0.287 | -1.284 | 0.726 | 0.008 | -0.001 | -0.001 | 0 | 0.45 | 0.74 | 0.44 | 0.54 | 0.42 | 0.37 |
| 5 | 169331450 | rs261049 | -0.638 | 0.108 | -1.07 | 0.767 | -0.009 | 0.043 | 0.043 | 0.417 | 0.29 | 0.58 | 0.35 | 0.52 | 0.26 | 0.2 |
| 5 | 169332156 | rs171862 | -0.47 | 0.293 | -1.049 | 0.749 | 0 | 0 | 0 | 0.08 | 0.71 | 0.69 | 0.78 | 0.76 | 0.72 | 0.67 |
| 5 | 169338485 | rs261007 | -0.87 | 0.568 | -0.954 | 0.839 | 0.016 | 0.077 | 0.077 | 0 | 0.28 | 0.52 | 0.36 | 0.52 | 0.31 | 0.24 |
| 5 | 169341951 | rs11741522 | 1.543 | 0.413 | -1.161 | 0.708 | -0.002 | 0.006 | 0.006 | 0 | 0.33 | 0.29 | 0.4 | 0.22 | 0.37 | 0.4 |
| 5 | 169347388 | rs33369 | -1.17 | 0.38 | -1.397 | 0.912 | -0.002 | 0.003 | 0.003 | 0 | 0.38 | 0.6 | 0.38 | 0.53 | 0.35 | 0.39 |
| 5 | 169348810 | rs978914 | 1.195 | 0.94 | -1.091 | 1.19 | -0.021 | 0.044 | 0.044 | 0 | 0.32 | 0.16 | 0.4 | 0.21 | 0.37 | 0.38 |
| 5 | 169349995 | rs12522080 | -0.262 | 0.878 | -0.493 | 2.136 | -0.014 | -0.052 | -0.052 | 1.498 | 0.13 | 0 | 0.11 | 0.2 | 0.12 | 0.12 |
| 5 | 169350967 | rs33373 | -0.992 | 0.897 | -0.323 | 1.657 | 0.025 | 0.136 | 0.136 | 0 | 0.23 | 0.49 | 0.34 | 0.54 | 0.25 | 0.22 |
| 5 | 169352662 | rs12657308 | -1.053 | 0.482 | -0.768 | 1.36 | -0.004 | 0.008 | 0.008 | 0 | 0.14 | 0.17 | 0.28 | 0.2 | 0.14 | 0.15 |
| 5 | 169355553 | rs4867577 | -0.619 | 1.052 | -1.619 | 1.437 | -0.089 | -0.188 | -0.188 | 0.44 | 0.25 | 0.02 | 0.43 | 0.46 | 0.31 | 0.37 |
| 5 | 169356966 | rs6896240 | -1.061 | 0.687 | -2.346 | 0.402 | -0.048 | -0.038 | -0.038 | 0 | 0.63 | 0.87 | 0.63 | 0.55 | 0.55 | 0.67 |
| 5 | 169358418 | rs10475927 | -0.489 | 0.335 | -1.383 | 0 | 0.001 | 0.001 | 0.001 | 0.049 | 0.38 | 0.42 | 0.51 | 0.54 | 0.41 | 0.42 |
| 5 | 169359522 | rs17071888 | 1.173 | 0.077 | -1.21 | -0.138 | -0.03 | -0.061 | -0.061 | 0 | 0.16 | 0.21 | 0.05 | 0 | 0.06 | 0.15 |
| 5 | 169361187 | rs13181166 | 0.672 | 0.224 | -1.304 | 0.183 | -0.083 | -0.154 | -0.154 | 1.047 | 0.22 | 0.02 | 0.42 | 0.41 | 0.28 | 0.35 |
| 5 | 169362010 | rs259903 | 0.171 | 0.128 | -0.349 | 0.209 | 0.049 | 0.032 | 0.032 | 0.155 | 0.1 | 0.35 | 0.07 | 0.09 | 0.09 | 0.11 |
| 5 | 169362082 | rs6895199 | 0.56 | 0.012 | -1.089 | 0.04 | 0.011 | 0.011 | 0.011 | 0 | 0.36 | 0.28 | 0.31 | 0.29 | 0.3 | 0.32 |
| 5 | 169368053 | rs4867900 | 0.286 | -0.285 | -1.201 | -0.194 | 0.016 | 0.003 | 0.003 | 0.175 | 0.45 | 0.31 | 0.48 | 0.42 | 0.41 | 0.52 |
| 5 | 169371376 | rs7716429 | 1.233 | -0.142 | -1.008 | -0.434 | 0.007 | -0.049 | -0.049 | 0.151 | 0.2 | 0.25 | 0.16 | 0.03 | 0.26 | 0.11 |
| 5 | 169372753 | rs259915 | -1.028 | -0.282 | -1.361 | -0.22 | -0.1 | -0.026 | -0.026 | 0.48 | 0.75 | 0.98 | 0.69 | 0.77 | 0.7 | 0.77 |
| 5 | 169379216 | rs1182091 | 0.076 | -0.234 | -1.259 | -0.14 | -0.001 | -0.003 | -0.003 | 0 | 0.3 | 0.66 | 0.2 | 0.21 | 0.29 | 0.21 |
| 5 | 169380639 | rs6555878 | 0.076 | -0.234 | -1.259 | -0.14 | -0.001 | -0.003 | -0.003 | 0.005 | 0.3 | 0.66 | 0.2 | 0.21 | 0.29 | 0.21 |
| 5 | 169386640 | rs11134598 | -1.099 | -0.16 | -1.497 | -0.411 | -0.1 | -0.015 | -0.015 | 0.328 | 0.75 | 0.98 | 0.68 | 0.77 | 0.69 | 0.76 |
| 5 | 169387087 | rs1422694 | 0.024 | 0.33 | -0.925 | -0.339 | 0.001 | -0.052 | -0.052 | 1.133 | 0.27 | 0.55 | 0.18 | 0.12 | 0.27 | 0.11 |
| 5 | 169391322 | rs6555879 | -0.417 | 0.027 | -1.815 | -0.19 | -0.073 | -0.119 | -0.119 | 0 | 0.75 | 0.94 | 0.67 | 0.6 | 0.67 | 0.66 |
| 5 | 169391558 | rs10067968 | 0.486 | 0.03 | -1.471 | -0.291 | -0.004 | -0.008 | -0.008 | 0 | 0.61 | 0.64 | 0.45 | 0.49 | 0.54 | 0.58 |
| 5 | 169394845 | rs6883231 | 0.094 | 0.157 | -1.371 | -0.367 | -0.013 | 0.039 | 0.039 | 0 | 0.29 | 0.2 | 0.23 | 0.12 | 0.36 | 0.13 |
| 5 | 169398287 | rs7702092 | 0.11 | 0.175 | -1.408 | -0.371 | -0.003 | 0.002 | 0.002 | 0 | 0.3 | 0.25 | 0.37 | 0.12 | 0.38 | 0.15 |
| 5 | 169398878 | rs2161369 | -1.19 | -0.047 | -1.998 | -0.393 | -0.106 | 0.005 | 0.005 | 0 | 0.7 | 0.98 | 0.73 | 0.76 | 0.67 | 0.74 |
| 5 | 169400124 | rs1363576 | -0.531 | -0.006 | -1.72 | -0.379 | 0 | 0 | 0 | 0.003 | 0.53 | 0.51 | 0.65 | 0.53 | 0.61 | 0.44 |
| 5 | 169401557 | rs13184164 | 0.395 | 0.038 | -2.706 | -0.516 | -0.017 | 0.052 | 0.052 | 0 | 0.27 | 0.13 | 0.17 | 0 | 0.32 | 0.12 |
| 5 | 169402577 | rs11134601 | 1.177 | 0.38 | -2.858 | -0.002 | -0.048 | -0.069 | -0.069 | 0 | 0.16 | 0.05 | 0.28 | 0.4 | 0.25 | 0.28 |
| 5 | 169403385 | rs2338809 | -1.374 | 0.443 | -3.153 | -0.274 | -0.085 | -0.079 | -0.079 | 0 | 0.75 | 0.99 | 0.76 | 0.75 | 0.74 | 0.76 |
| 5 | 169404088 | rs9313476 | -0.089 | 0.836 | -2.329 | -0.126 | -0.022 | 0.108 | 0.108 | 0 | 0.35 | 0.15 | 0.24 | 0.1 | 0.41 | 0.18 |
| 5 | 169410474 | rs6873329 | 0.738 | 0.857 | -2.416 | -0.196 | -0.005 | 0.112 | 0.112 | 0.01 | 0.23 | 0.04 | 0.16 | 0 | 0.23 | 0.08 |
| 5 | 169434978 | rs2042237 | -0.146 | 0.824 | -1.957 | 0.151 | 0.002 | 0.027 | 0.027 | 0 | 0.42 | 0.35 | 0.39 | 0.32 | 0.45 | 0.33 |
| 5 | 169435396 | rs2287726 | -0.186 | 0.856 | -1.893 | -0.06 | 0.003 | 0.019 | 0.019 | 0.013 | 0.43 | 0.38 | 0.39 | 0.21 | 0.42 | 0.3 |
| 5 | 169435713 | rs2287727 | 0.083 | 1.017 | -1.854 | 0.56 | -0.083 | -0.1 | -0.1 | 0 | 0.17 | 0.03 | 0.3 | 0.39 | 0.28 | 0.31 |
| 5 | 169436128 | rs10056571 | -0.693 | 0.126 | -1.073 | -0.184 | -0.003 | -0.054 | -0.054 | 10.603 | 0.1 | 0.2 | 0.04 | 0 | 0.09 | 0.06 |
| 5 | 169439742 | rs1477317 | -0.017 | -0.007 | -1.926 | -0.259 | 0.051 | 0.132 | 0.132 | 0 | 0.52 | 0.81 | 0.65 | 0.76 | 0.42 | 0.58 |
| 5 | 169440622 | rs10063533 | -0.274 | -0.11 | -2.067 | -0.701 | -0.079 | 0.16 | 0.16 | 0 | 0.61 | 0.89 | 0.7 | 0.86 | 0.53 | 0.6 |
| 5 | 169441749 | rs749965 | -0.438 | -0.836 | -2.417 | -0.748 | -0.126 | 0.126 | 0.126 | 0 | 0.63 | 0.93 | 0.71 | 0.84 | 0.54 | 0.66 |
| 5 | 169444363 | rs17738444 | -0.256 | -1.603 | -1.376 | -0.733 | -0.023 | -0.027 | -0.027 | 0 | 0.27 | 0.35 | 0.33 | 0.15 | 0.17 | 0.29 |
| 5 | 169449956 | rs6884504 | -0.749 | -1.399 | -0.332 | -0.621 | 0.05 | 0.033 | 0.033 | 1.29 | 0.09 | 0.36 | 0.19 | 0.07 | 0.1 | 0.11 |
| 5 | 169450448 | rs6859561 | -0.329 | -1.397 | -0.127 | -0.623 | 0.017 | 0.019 | 0.019 | 0.51 | 0.07 | 0.23 | 0.17 | 0.07 | 0.08 | 0.09 |
| 5 | 169450932 | rs12655995 | -0.763 | -2.1 | -0.097 | -0.16 | -0.009 | 0 | 0 | 0 | 0.44 | 0.46 | 0.32 | 0.58 | 0.35 | 0.44 |
| 5 | 169451927 | rs10044477 | -1.608 | -2.157 | -0.218 | -0.168 | -0.072 | 0.013 | 0.013 | 0 | 0.57 | 0.91 | 0.52 | 0.67 | 0.48 | 0.61 |
| 5 | 169453707 | rs6876868 | 0.634 | -2.339 | 0.414 | -0.162 | -0.017 | -0.043 | -0.043 | 0.034 | 0.22 | 0.11 | 0.26 | 0.51 | 0.29 | 0.32 |
| 5 | 169454134 | rs12653623 | -1.307 | -2.256 | 0.371 | -0.183 | 0.037 | 0.02 | 0.02 | 0 | 0.49 | 0.6 | 0.33 | 0.61 | 0.36 | 0.52 |
| 5 | 169460787 | rs10044461 | -0.962 | -2.257 | 0.261 | -0.179 | 0.034 | 0.009 | 0.009 | 0.031 | 0.46 | 0.65 | 0.32 | 0.61 | 0.33 | 0.45 |
| 5 | 169461399 | rs3763047 | -1.014 | -2.252 | 0.34 | -0.177 | 0.032 | 0.01 | 0.01 | 0.043 | 0.46 | 0.64 | 0.31 | 0.61 | 0.33 | 0.5 |
| 5 | 169461547 | rs3763048 | -1.264 | -2.184 | 0.364 | -0.117 | 0.043 | 0.02 | 0.02 | 0 | 0.49 | 0.64 | 0.33 | 0.6 | 0.35 | 0.51 |
| 5 | 169463466 | rs6863396 | -0.228 | -2.174 | 0.417 | -0.244 | 0.026 | 0.026 | 0.026 | 0 | 0.22 | 0.14 | 0.05 | 0.07 | 0.07 | 0.22 |
| 5 | 169465861 | rs7715478 | -1.212 | -1.945 | 0.334 | -0.052 | -0.01 | 0.005 | 0.005 | 0 | 0.59 | 0.6 | 0.49 | 0.64 | 0.5 | 0.64 |
| 5 | 169468243 | rs2270895 | -0.625 | -1.883 | -0.484 | -0.051 | -0.014 | 0.063 | 0.063 | 0 | 0.52 | 0.19 | 0.45 | 0.64 | 0.48 | 0.61 |
| 5 | 169471511 | rs10040662 | -1.264 | -1.626 | -0.668 | 0.132 | -0.076 | 0 | 0 | 0 | 0.62 | 0.85 | 0.51 | 0.63 | 0.5 | 0.67 |

| | | | | | | | | | | | | | | | | |
|---|---|---|---|---|---|---|---|---|---|---|---|---|---|---|---|---|
| 5 | 169471888 | rs11744647 | -0.537 | -1.245 | -0.422 | 0.224 | -0.01 | -0.04 | -0.04 | 0 | 0.36 | 0.22 | 0.34 | 0.51 | 0.38 | 0.48 |
| 5 | 169472788 | rs10074769 | -0.452 | -1.151 | 0.23 | 0.136 | -0.012 | -0.036 | -0.036 | 0 | 0.08 | 0.16 | 0.03 | 0 | 0.04 | 0.01 |
| 5 | 169474325 | rs905155 | -1.421 | -0.755 | 0.49 | 0.316 | -0.001 | 0 | 0 | 0 | 0.32 | 0.32 | 0.45 | 0.62 | 0.37 | 0.39 |
| 5 | 169476723 | rs956303 | -0.984 | -0.724 | 0.759 | 0.446 | 0.028 | -0.023 | -0.023 | 0 | 0.19 | 0.41 | 0.16 | 0.13 | 0.24 | 0.21 |
| 5 | 169477011 | rs1316638 | -0.798 | -0.751 | 0.56 | 0.476 | 0.001 | 0.001 | 0.001 | 0 | 0.15 | 0.09 | 0.29 | 0.49 | 0.12 | 0.23 |
| 5 | 169477815 | rs1010317 | -2.327 | -0.809 | 0.753 | 0.426 | -0.001 | 0.004 | 0.004 | 0.137 | 0.38 | 0.68 | 0.47 | 0.61 | 0.39 | 0.45 |
| 5 | 169479134 | rs13360445 | -1.59 | -1.226 | 0.787 | 0.774 | -0.026 | 0.131 | 0.131 | 0 | 0.19 | 0.33 | 0.3 | 0.47 | 0.13 | 0.25 |
| 5 | 169481754 | rs13358993 | -1.038 | -1.62 | 0.585 | 0.156 | -0.015 | -0.049 | -0.049 | 5.866 | 0.38 | 0.54 | 0.23 | 0.16 | 0.29 | 0.4 |
| 5 | 169485227 | rs6864400 | -1.4 | -1.874 | 0.757 | 0.401 | -0.012 | 0.005 | 0.005 | 0 | 0.4 | 0.48 | 0.46 | 0.51 | 0.32 | 0.46 |
| 5 | 169486815 | rs10045458 | -0.61 | -1.829 | 0.557 | 0.341 | 0.003 | -0.033 | -0.033 | 0.141 | 0.3 | 0.31 | 0.18 | 0.03 | 0.24 | 0.25 |
| 5 | 169488692 | rs17738770 | -1.189 | -1.825 | 0.512 | 0.458 | -0.001 | 0 | 0 | 0 | 0.38 | 0.42 | 0.44 | 0.49 | 0.3 | 0.46 |
| 5 | 169489226 | rs1553297 | -1.687 | -1.78 | 0.314 | 0.058 | 0.066 | 0.073 | 0.073 | 0.041 | 0.5 | 0.85 | 0.53 | 0.63 | 0.38 | 0.61 |
| 5 | 169491247 | rs7700353 | -0.966 | -1.119 | 0.034 | -0.384 | -0.002 | -0.061 | -0.061 | 5.118 | 0.31 | 0.38 | 0.18 | 0.03 | 0.25 | 0.27 |
| 5 | 169494844 | rs6555883 | -0.925 | -1.167 | -0.375 | 0.006 | -0.015 | -0.008 | -0.008 | 0.052 | 0.33 | 0.56 | 0.32 | 0.7 | 0.28 | 0.44 |
| 5 | 169497638 | rs10462994 | -1.161 | -1.026 | -0.551 | 0.018 | 0.004 | 0.001 | 0.001 | 0 | 0.38 | 0.64 | 0.33 | 0.67 | 0.29 | 0.45 |
| 5 | 169497800 | rs10462995 | -1.507 | -0.837 | -1.068 | -0.32 | -0.033 | 0.058 | 0.058 | 0.203 | 0.58 | 0.85 | 0.45 | 0.72 | 0.47 | 0.59 |
| 5 | 169501527 | rs6890267 | -0.264 | -0.872 | -0.702 | 0.112 | 0.061 | 0.068 | 0.068 | 1.035 | 0.09 | 0.48 | 0.09 | 0.09 | 0.07 | 0.1 |
| 5 | 169503440 | rs10866644 | -0.192 | -0.751 | -1.417 | 0.299 | -0.013 | -0.143 | -0.143 | 0.105 | 0.21 | 0.03 | 0.2 | 0.58 | 0.17 | 0.31 |
| 5 | 169504743 | rs1045168 | -0.759 | -0.623 | -1.02 | -0.633 | 0.029 | 0.044 | 0.044 | 0 | 0.46 | 0.38 | 0.26 | 0.11 | 0.33 | 0.34 |
| 5 | 169505256 | rs6868470 | -0.876 | -0.731 | -0.662 | -0.811 | -0.001 | -0.02 | -0.02 | 0 | 0.48 | 0.51 | 0.31 | 0.18 | 0.35 | 0.43 |
| 5 | 169505282 | rs6889919 | -0.724 | -0.754 | -0.222 | -0.777 | 0.026 | 0.004 | 0.004 | 0 | 0.11 | 0.23 | 0.08 | 0.07 | 0.09 | 0.07 |
| 5 | 169507067 | rs9885452 | -0.929 | -0.557 | -1.241 | -0.88 | 0.024 | 0.031 | 0.031 | 0.067 | 0.5 | 0.7 | 0.35 | 0.56 | 0.39 | 0.5 |
| 5 | 169508461 | rs13181782 | 0.931 | 0.347 | -1.805 | -1.19 | -0.079 | 0.028 | 0.028 | 0 | 0.19 | 0.02 | 0.25 | 0.11 | 0.25 | 0.16 |
| 5 | 169510630 | rs6555884 | 0.161 | 0.855 | -1.653 | -1.087 | -0.048 | 0.135 | 0.135 | 0 | 0.42 | 0.19 | 0.4 | 0.19 | 0.45 | 0.32 |
| 5 | 169512063 | rs2338833 | -0.687 | 0.699 | -1.64 | -0.983 | -0.013 | 0.05 | 0.05 | 0.086 | 0.23 | 0.68 | 0.19 | 0.5 | 0.15 | 0.26 |
| 5 | 169512748 | rs17562083 | 0.926 | 0.845 | -1.683 | -1.118 | -0.113 | 0.149 | 0.149 | 0.413 | 0.34 | 0.02 | 0.34 | 0.11 | 0.38 | 0.24 |
| 5 | 169512860 | rs10516070 | 0.135 | 0.875 | -1.608 | -1.103 | -0.024 | 0.188 | 0.188 | 0 | 0.47 | 0.19 | 0.38 | 0.17 | 0.45 | 0.32 |
| 5 | 169517852 | rs13164348 | -0.062 | 0.895 | -1.582 | -1.085 | 0.018 | 0.166 | 0.166 | 0 | 0.5 | 0.24 | 0.42 | 0.19 | 0.5 | 0.42 |
| 5 | 169522956 | rs4867913 | 0.713 | 1.286 | -1.663 | -1.07 | 0.003 | 0.093 | 0.093 | 0 | 0.36 | 0.18 | 0.36 | 0.13 | 0.35 | 0.29 |
| 5 | 169524499 | rs6555885 | -1.7 | 1.13 | -1.47 | -0.94 | -0.02 | -0.03 | -0.03 | 0.023 | 0.72 | 0.92 | 0.63 | 0.71 | 0.66 | 0.67 |
| 5 | 169526837 | rs17562153 | -1.14 | 1.137 | -1.313 | -0.948 | -0.011 | -0.005 | -0.005 | 0 | 0.65 | 0.8 | 0.6 | 0.65 | 0.59 | 0.59 |
| 5 | 169527642 | rs1501644 | 0.046 | 1.084 | -1.119 | -0.95 | 0.025 | 0.21 | 0.21 | 0.016 | 0.18 | 0.56 | 0.19 | 0.47 | 0.15 | 0.26 |
| 5 | 169529089 | rs7704953 | 0.14 | 1.125 | -1.255 | -0.984 | -0.007 | 0.159 | 0.159 | 0 | 0.44 | 0.18 | 0.39 | 0.17 | 0.41 | 0.33 |
| 5 | 169529903 | rs17738959 | -1.142 | 1.21 | -1.295 | -0.978 | 0.018 | 0.042 | 0.042 | 0.057 | 0.27 | 0.64 | 0.25 | 0.52 | 0.29 | 0.29 |
| 5 | 169529932 | rs7729998 | 0.66 | 1.308 | -1.348 | -1.023 | 0.055 | 0.103 | 0.103 | 0 | 0.39 | 0.2 | 0.32 | 0.17 | 0.3 | 0.31 |
| 5 | 169530135 | rs6555886 | -2.314 | 1.185 | -0.695 | -0.826 | 0.004 | -0.014 | -0.014 | 0 | 0.23 | 0.27 | 0.15 | 0.12 | 0.27 | 0.17 |
| 5 | 169532633 | rs10064122 | -0.207 | 1.113 | -0.636 | -0.759 | 0.013 | 0.159 | 0.159 | 0.063 | 0.19 | 0.6 | 0.19 | 0.48 | 0.15 | 0.27 |
| 5 | 169534173 | rs4315934 | -0.084 | 1.216 | -0.556 | -0.865 | -0.014 | 0.203 | 0.203 | 0.075 | 0.52 | 0.81 | 0.57 | 0.83 | 0.53 | 0.68 |
| 5 | 169535522 | rs10063424 | 0.403 | 1.328 | -0.387 | -0.749 | 0.001 | -0.01 | -0.01 | 0.529 | 0.92 | 0.9 | 0.93 | 0.99 | 0.9 | 0.94 |
| 5 | 169535734 | rs6873124 | -0.192 | 1.332 | -0.329 | -0.742 | 0.016 | 0.174 | 0.174 | 0.053 | 0.18 | 0.57 | 0.19 | 0.48 | 0.15 | 0.26 |
| 5 | 169536055 | rs6555887 | 0.63 | 1.34 | -0.366 | -0.743 | 0.004 | -0.02 | -0.02 | 0 | 0.92 | 0.85 | 0.93 | 0.99 | 0.91 | 0.94 |
| 5 | 169536150 | rs6555888 | 1.985 | 1.359 | -0.381 | -0.757 | 0.001 | -0.007 | -0.007 | 0.008 | 0.85 | 0.84 | 0.88 | 0.94 | 0.79 | 0.91 |
| 5 | 169538159 | rs17562317 | -0.053 | 1.362 | -0.437 | -0.736 | -0.028 | 0.021 | 0.021 | 0 | 0.07 | 0.02 | 0.05 | 0.05 | 0.11 | 0.03 |
| 5 | 169538372 | rs4867918 | 1.741 | 1.193 | -0.424 | -0.747 | -0.03 | -0.038 | -0.038 | 0.208 | 0.67 | 0.23 | 0.7 | 0.46 | 0.65 | 0.65 |
| 5 | 169540204 | rs1477320 | 0.488 | 1.062 | -0.32 | -0.496 | -0.012 | 0.009 | 0.009 | 0 | 0.89 | 0.94 | 0.9 | 0.94 | 0.81 | 0.91 |
| 5 | 169540547 | rs12653012 | 1.578 | 0.979 | -0.314 | -0.532 | -0.014 | -0.019 | -0.019 | 0 | 0.68 | 0.29 | 0.71 | 0.46 | 0.65 | 0.65 |
| 5 | 169543629 | rs12656959 | 0.82 | 0.928 | -0.326 | -0.53 | -0.001 | 0 | 0 | 0 | 0.07 | 0.07 | 0.03 | 0.07 | 0.07 | 0.08 |
| 5 | 169545642 | rs6880991 | 1.386 | 0.803 | -0.484 | -0.637 | 0.021 | 0.147 | 0.147 | 0.767 | 0.33 | 0.07 | 0.31 | 0.11 | 0.27 | 0.23 |
| 5 | 169551625 | rs4867922 | 1.963 | 0.948 | -0.369 | -0.543 | -0.041 | -0.032 | -0.032 | 0 | 0.65 | 0.21 | 0.68 | 0.44 | 0.61 | 0.62 |
| 5 | 169563367 | rs12515896 | 0.505 | 0.904 | -0.286 | -0.508 | 0.005 | 0.082 | 0.082 | 0.208 | 0.78 | 0.36 | 0.79 | 0.49 | 0.81 | 0.71 |
| 5 | 169566044 | rs1469066 | 1.934 | 0.259 | -0.436 | -0.576 | 0 | -0.009 | -0.009 | 0 | 0.59 | 0.24 | 0.7 | 0.43 | 0.64 | 0.63 |
| 5 | 169567312 | rs13172115 | -0.315 | 0.339 | -0.464 | -0.741 | -0.055 | 0.017 | 0.017 | 0 | 0.09 | 0.01 | 0.09 | 0.06 | 0.18 | 0.09 |
| 5 | 169569829 | rs11743915 | 1.02 | 0.241 | -0.497 | -0.476 | 0.019 | 0.091 | 0.091 | 0 | 0.34 | 0.15 | 0.32 | 0.17 | 0.3 | 0.34 |
| 5 | 169571480 | rs13169329 | 1.782 | 0.215 | -0.385 | -0.193 | -0.03 | -0.033 | -0.033 | 0.537 | 0.26 | 0.11 | 0.38 | 0.31 | 0.35 | 0.3 |
| 5 | 169572933 | rs11134616 | 0.519 | 0.169 | -0.067 | -0.068 | -0.023 | 0.051 | 0.051 | 0 | 0.75 | 0.39 | 0.81 | 0.55 | 0.84 | 0.73 |
| 5 | 169575176 | rs4867926 | -0.595 | 0.224 | 0.022 | -0.61 | -0.006 | -0.044 | -0.044 | 0 | 0.57 | 0.51 | 0.53 | 0.8 | 0.61 | 0.65 |
| 5 | 169578246 | rs889004 | 0.765 | 0.361 | 0.212 | -0.291 | 0.035 | 0.001 | 0.001 | 0 | 0.41 | 0.25 | 0.39 | 0.43 | 0.34 | 0.4 |
| 5 | 169580540 | rs1544757 | 1.237 | 0.77 | 0.75 | 0.211 | -0.004 | -0.043 | -0.043 | 1.048 | 0.73 | 0.45 | 0.81 | 0.82 | 0.78 | 0.77 |
| 5 | 169583173 | rs1035424 | -0.122 | 0.65 | 0.715 | 0.219 | 0.007 | -0.038 | -0.038 | 0 | 0.85 | 0.6 | 0.87 | 0.93 | 0.88 | 0.89 |
| 5 | 169584072 | rs4616915 | 1.335 | 0.776 | 0.495 | 0.269 | 0.012 | -0.034 | -0.034 | 0.11 | 0.69 | 0.39 | 0.74 | 0.8 | 0.67 | 0.68 |
| 5 | 169590807 | rs4634366 | -1.593 | 0.779 | 0.261 | 0.543 | 0.014 | 0.112 | 0.112 | 0 | 0.15 | 0.39 | 0.15 | 0.33 | 0.16 | 0.22 |
| 5 | 169590905 | rs4478338 | -0.986 | 0.715 | 0.251 | 0.534 | -0.033 | 0.098 | 0.098 | 0 | 0.12 | 0.29 | 0.08 | 0.33 | 0.05 | 0.15 |
| 5 | 169591321 | rs4329030 | 1.687 | 0.644 | -0.464 | 0.017 | -0.058 | 0.124 | 0.124 | 0.025 | 0.31 | 0.02 | 0.35 | 0.11 | 0.29 | 0.3 |
| 5 | 169592906 | rs10475541 | 1.063 | 0.65 | -0.153 | -0.126 | -0.003 | -0.088 | -0.088 | 4.53 | 0.66 | 0.5 | 0.78 | 0.88 | 0.72 | 0.71 |
| 5 | 169593191 | rs13359356 | -0.275 | 0.982 | 0.244 | 0.293 | -0.023 | 0.089 | 0.089 | 0 | 0.1 | 0.19 | 0.06 | 0.33 | 0.05 | 0.14 |
| 5 | 169593543 | rs5014394 | 0.366 | 1.022 | 0.102 | 0.07 | -0.048 | 0.037 | 0.037 | 0.061 | 0.11 | 0.01 | 0.05 | 0.04 | 0.12 | 0.08 |
| 5 | 169594544 | rs12651794 | -0.823 | 1.075 | 0.203 | 0.106 | 0.017 | -0.018 | -0.018 | 1.905 | 0.16 | 0.19 | 0.15 | 0.06 | 0.26 | 0.21 |
| 5 | 169595780 | rs889007 | 0.417 | 0.649 | 0.199 | 0.089 | -0.054 | -0.074 | -0.074 | 0.111 | 0.79 | 0.51 | 0.86 | 0.89 | 0.9 | 0.86 |
| 5 | 169603013 | rs10078694 | -0.724 | 0.974 | 0.018 | 0.146 | 0 | -0.003 | -0.003 | 0 | 0.15 | 0.14 | 0.1 | 0.06 | 0.19 | 0.16 |

| | | | | | | | | | | | | | | | | |
|---|---|---|---|---|---|---|---|---|---|---|---|---|---|---|---|---|
| 5 | 169606312 | rs7712919 | -0.955 | 0.913 | 0.085 | 0.13 | -0.002 | 0.006 | 0.006 | 2.899 | 0.21 | 0.16 | 0.09 | 0.1 | 0.18 | 0.16 |
| 5 | 169606878 | rs1363602 | 1.458 | 0.921 | 0.014 | 0.275 | -0.006 | -0.06 | -0.06 | 0.467 | 0.21 | 0.07 | 0.29 | 0.38 | 0.25 | 0.21 |
| 5 | 169607547 | rs12654236 | -0.488 | 0.931 | 0.032 | 0.123 | -0.001 | -0.003 | -0.003 | 0 | 0.15 | 0.13 | 0.09 | 0.06 | 0.18 | 0.14 |
| 5 | 169609479 | rs4242155 | 0.157 | 0.551 | 0.502 | 0.086 | -0.003 | -0.048 | -0.048 | 0 | 0.85 | 0.57 | 0.86 | 0.92 | 0.91 | 0.86 |
| 5 | 169611199 | rs11739905 | 1.661 | 0.641 | 0.085 | 0.011 | -0.002 | 0.164 | 0.164 | 0 | 0.39 | 0.13 | 0.38 | 0.13 | 0.33 | 0.34 |
| 5 | 169611443 | rs4077963 | 1.472 | 0.662 | 0.058 | 0.164 | -0.01 | -0.082 | -0.082 | 0.605 | 0.21 | 0.06 | 0.28 | 0.39 | 0.26 | 0.23 |
| 5 | 169615391 | rs1990942 | 0.705 | 0.564 | 0.588 | 0.011 | -0.021 | -0.052 | -0.052 | 0 | 0.79 | 0.67 | 0.84 | 0.86 | 0.86 | 0.81 |
| 5 | 169619246 | rs2287744 | 0.914 | 0.585 | 0.103 | 0.157 | 0.004 | 0.003 | 0.003 | 2.338 | 0.07 | 0.04 | 0.02 | 0.06 | 0.05 | 0.05 |
| 5 | 169622208 | rs6863241 | 0.087 | 0.848 | 0.204 | 0.29 | 0.038 | 0.082 | 0.082 | 0 | 0.09 | 0.16 | 0.11 | 0.32 | 0.19 | 0.18 |
| 5 | 169625672 | rs4867943 | 1.579 | 0.57 | -0.174 | -0.399 | -0.06 | 0.196 | 0.196 | 0.541 | 0.38 | 0.06 | 0.37 | 0.12 | 0.32 | 0.33 |
| 5 | 169625713 | rs10044209 | -0.028 | 0.749 | 0.658 | -0.14 | 0.02 | -0.035 | -0.035 | 0 | 0.35 | 0.5 | 0.41 | 0.71 | 0.47 | 0.45 |
| 5 | 169627707 | rs10447196 | 0.889 | 0.791 | 1.325 | 0.216 | 0.029 | -0.015 | -0.015 | 0 | 0.87 | 0.68 | 0.81 | 0.9 | 0.85 | 0.84 |
| 5 | 169629225 | rs4867946 | 0.94 | 0.851 | 0.897 | 0.23 | -0.028 | -0.086 | -0.086 | 0 | 0.78 | 0.5 | 0.79 | 0.91 | 0.83 | 0.83 |
| 5 | 169629403 | rs4867947 | -1.02 | 1.323 | 0.398 | 0.783 | 0.011 | 0.026 | 0.026 | 0 | 0.09 | 0.16 | 0.09 | 0.27 | 0.16 | 0.2 |
| 5 | 169632261 | rs6889648 | 1.135 | 0.39 | 0.728 | 0.114 | 0.005 | -0.056 | -0.056 | 0 | 0.9 | 0.42 | 0.89 | 1 | 0.92 | 0.98 |
| 5 | 169633156 | rs12515639 | 1.167 | 0.338 | 0.732 | 0.124 | 0.019 | -0.037 | -0.037 | 0 | 0.9 | 0.53 | 0.91 | 1 | 0.93 | 0.99 |
| 5 | 169633843 | rs4323245 | -0.382 | -0.151 | -1.086 | 0.179 | -0.004 | -0.096 | -0.096 | 0 | 0.67 | 0.98 | 0.73 | 0.6 | 0.78 | 0.52 |
| 5 | 169635536 | rs6870894 | -0.966 | 0.196 | -1.62 | -0.551 | -0.033 | 0.194 | 0.194 | 0.665 | 0.41 | 0.9 | 0.31 | 0.1 | 0.38 | 0.24 |
| 5 | 169640260 | rs11743735 | 0.997 | -0.119 | -1.094 | -0.488 | -0.107 | -0.143 | -0.143 | 0 | 0.28 | 0.06 | 0.44 | 0.49 | 0.39 | 0.28 |
| 5 | 169644032 | rs13179298 | -0.185 | 0.023 | -0.999 | -0.692 | 0.001 | 0.003 | 0.003 | 0 | 0.26 | 0.24 | 0.16 | 0.06 | 0.21 | 0.15 |
| 5 | 169644845 | rs1025003 | -1.624 | 0.023 | -1.282 | -0.642 | 0.043 | -0.063 | -0.063 | 0 | 0.7 | 0.96 | 0.75 | 0.59 | 0.77 | 0.53 |
| 5 | 169645236 | rs11743438 | 0.382 | 0.038 | -0.613 | -0.655 | -0.012 | -0.008 | -0.008 | 0 | 0.4 | 0.35 | 0.55 | 0.53 | 0.54 | 0.36 |
| 5 | 169648022 | rs4867593 | -1.524 | -0.631 | -0.238 | -0.718 | -0.032 | 0.168 | 0.168 | 0.459 | 0.39 | 0.9 | 0.31 | 0.1 | 0.35 | 0.24 |
| 5 | 169648258 | rs4867594 | 0.508 | -0.555 | 0.14 | -0.454 | 0.005 | -0.021 | -0.021 | 0.126 | 0.08 | 0.11 | 0.09 | 0 | 0.12 | 0.05 |
| 5 | 169661414 | rs918506 | -0.576 | -0.699 | 0.216 | -0.495 | 0 | 0.008 | 0.008 | 0 | 0.51 | 0.47 | 0.6 | 0.56 | 0.51 | 0.49 |
| 5 | 169663258 | rs1508759 | -1.052 | -0.699 | -0.549 | -0.493 | -0.098 | 0.007 | 0.007 | 0.102 | 0.57 | 0.92 | 0.63 | 0.61 | 0.51 | 0.5 |
| 5 | 169666462 | rs7716359 | 0.035 | -0.703 | -0.79 | -0.412 | 0.022 | -0.039 | -0.039 | 1.532 | 0.61 | 0.88 | 0.52 | 0.44 | 0.63 | 0.59 |
| 5 | 169685163 | rs315717 | -0.503 | -0.401 | -0.822 | -0.099 | -0.001 | 0.004 | 0.004 | 0 | 0.55 | 0.68 | 0.59 | 0.57 | 0.47 | 0.44 |
| 5 | 169686007 | rs432621 | -0.464 | -0.209 | -1.076 | -0.178 | 0.001 | -0.001 | -0.001 | 0 | 0.53 | 0.54 | 0.59 | 0.53 | 0.47 | 0.45 |
| 5 | 169689519 | rs2338871 | -0.575 | -0.143 | -1.612 | 0.005 | -0.027 | -0.094 | -0.094 | 0 | 0.15 | 0.03 | 0.3 | 0.3 | 0.18 | 0.19 |
| 5 | 169690247 | rs10475933 | -0.266 | -0.082 | -1.267 | -1.222 | -0.003 | 0.005 | 0.005 | 0 | 0.71 | 0.7 | 0.74 | 0.91 | 0.68 | 0.85 |
| 5 | 169692554 | rs2338872 | -0.456 | 0.022 | -1.434 | -1.037 | -0.015 | 0.173 | 0.173 | 0 | 0.59 | 0.77 | 0.68 | 0.96 | 0.57 | 0.71 |
| 5 | 169696181 | rs315751 | -0.352 | 0.767 | -0.979 | -0.404 | 0.014 | -0.035 | -0.035 | 0 | 0.41 | 0.47 | 0.54 | 0.83 | 0.43 | 0.53 |
| 5 | 169708691 | rs2338873 | -0.707 | 0.622 | -1.368 | -0.43 | -0.013 | 0.059 | 0.059 | 1.809 | 0.15 | 0.05 | 0.07 | 0.04 | 0.12 | 0.11 |
| 5 | 169715038 | rs315721 | -1.023 | 0.033 | -0.881 | -0.428 | -0.019 | -0.015 | -0.015 | 0.051 | 0.36 | 0.39 | 0.33 | 0.38 | 0.3 | 0.35 |
| 5 | 169719672 | rs10050695 | -0.37 | -0.013 | -0.706 | -0.431 | 0.008 | 0.009 | 0.009 | 0.043 | 0.16 | 0.14 | 0.07 | 0.04 | 0.11 | 0.1 |
| 5 | 169720726 | rs17072430 | -0.055 | -0.096 | -0.67 | -0.347 | -0.002 | -0.001 | -0.001 | 0 | 0.07 | 0.09 | 0.03 | 0.04 | 0.04 | 0.05 |
| 5 | 169722897 | rs3804250 | -0.606 | -0.513 | -1.275 | -0.539 | -0.002 | -0.009 | -0.009 | 1.658 | 0.16 | 0.01 | 0.14 | 0.09 | 0.12 | 0.17 |
| 5 | 169723113 | rs182618 | -0.918 | -0.202 | -1.635 | -0.944 | -0.023 | -0.059 | -0.059 | 0 | 0.75 | 0.85 | 0.68 | 0.51 | 0.72 | 0.63 |
| 5 | 169723229 | rs6866733 | -0.991 | -0.163 | -0.636 | -0.854 | -0.01 | 0.001 | 0.001 | 0 | 0.25 | 0.26 | 0.19 | 0.22 | 0.18 | 0.3 |
| 5 | 169731198 | rs315782 | 0.015 | 0.066 | -0.585 | -0.942 | -0.007 | -0.002 | -0.002 | 0.205 | 0.91 | 0.89 | 0.88 | 0.93 | 0.91 | 0.93 |
| 5 | 169731858 | rs17648477 | 0.232 | -0.271 | -1.173 | -1.635 | 0.005 | 0.055 | 0.055 | 0 | 0.2 | 0.05 | 0.1 | 0 | 0.17 | 0.16 |
| 5 | 169734070 | rs876302 | 0.387 | 0.292 | -0.655 | -1.307 | 0.019 | 0.021 | 0.021 | 0 | 0.17 | 0.19 | 0.25 | 0.33 | 0.21 | 0.29 |
| 5 | 169735463 | rs315789 | 0.99 | -0.034 | -1.229 | -1.614 | 0.009 | -0.072 | -0.072 | 0 | 0.25 | 0.14 | 0.3 | 0.52 | 0.21 | 0.27 |
| 5 | 169735920 | rs315791 | 0.589 | -0.068 | -1.23 | -1.839 | -0.012 | 0.013 | 0.013 | 0 | 0.45 | 0.38 | 0.49 | 0.53 | 0.5 | 0.37 |
| 5 | 169741340 | rs6555894 | 0.491 | 0.117 | -1.236 | -2.085 | -0.01 | -0.025 | -0.025 | 0 | 0.66 | 0.62 | 0.76 | 0.85 | 0.7 | 0.64 |
| 5 | 169742593 | rs876315 | 0.766 | 0.155 | -1.163 | -1.98 | 0.002 | 0.046 | 0.046 | 4.372 | 0.47 | 0.62 | 0.59 | 0.85 | 0.42 | 0.55 |
| 5 | 169742646 | rs906357 | 0.803 | 0.186 | -1.282 | -1.966 | 0 | -0.005 | -0.005 | 0.163 | 0.46 | 0.53 | 0.57 | 0.85 | 0.42 | 0.54 |
| 5 | 169742942 | rs925080 | -0.202 | -0.248 | -1.441 | -1.249 | 0.028 | 0.069 | 0.069 | 0 | 0.19 | 0.04 | 0.08 | 0.01 | 0.14 | 0.16 |
| 5 | 169743561 | rs315796 | -0.815 | 0.56 | -0.799 | -0.391 | -0.002 | -0.002 | -0.002 | 0.007 | 0.08 | 0.14 | 0.04 | 0.06 | 0.07 | 0.13 |
| 5 | 169746734 | rs315805 | 0.643 | 0.96 | -1.055 | -0.39 | -0.008 | 0.018 | 0.018 | 0 | 0.85 | 0.89 | 0.89 | 0.98 | 0.82 | 0.86 |
| 5 | 169750158 | rs315808 | 0.668 | 0.923 | -1.071 | -0.407 | -0.009 | 0.021 | 0.021 | 0 | 0.85 | 0.89 | 0.89 | 0.98 | 0.82 | 0.86 |
| 5 | 169755799 | rs10516073 | 0.071 | 1.42 | -1.322 | -1.323 | -0.019 | 0.029 | 0.029 | 0 | 0.3 | 0.27 | 0.2 | 0.08 | 0.34 | 0.18 |
| 5 | 169757533 | rs315780 | 0.532 | 1.123 | -1.452 | -1.519 | -0.004 | 0.009 | 0.009 | 0 | 0.34 | 0.23 | 0.6 | 0.69 | 0.34 | 0.5 |
| 5 | 169759095 | rs12513574 | 1.173 | 0.475 | -1.564 | -0.529 | -0.032 | -0.134 | -0.134 | 0.493 | 0.22 | 0.03 | 0.28 | 0.44 | 0.26 | 0.38 |
| 5 | 169759207 | rs11134622 | 0.53 | -0.05 | -1.541 | -0.974 | -0.002 | 0.008 | 0.008 | 0.637 | 0.46 | 0.56 | 0.3 | 0.27 | 0.48 | 0.29 |
| 5 | 169759990 | rs315778 | 0.481 | 0.142 | -1.733 | -1.247 | -0.023 | 0.008 | 0.008 | 0 | 0.32 | 0.18 | 0.59 | 0.7 | 0.38 | 0.46 |
| 5 | 169760934 | rs17739676 | 1.073 | -0.374 | -1.29 | -0.901 | -0.009 | 0.045 | 0.045 | 0 | 0.06 | 0.06 | 0.26 | 0.18 | 0.02 | 0.03 |
| 5 | 169761334 | rs315773 | -0.377 | -0.582 | -1.442 | -0.583 | -0.04 | -0.072 | -0.072 | 0 | 0.64 | 0.91 | 0.63 | 0.59 | 0.59 | 0.49 |
| 5 | 169765425 | rs7725699 | 1.514 | -1.024 | -0.446 | -0.482 | 0.022 | 0.009 | 0.009 | 0 | 0.51 | 0.39 | 0.49 | 0.53 | 0.58 | 0.54 |
| 5 | 169765792 | rs6555895 | 0.829 | -0.749 | -0.231 | -0.322 | -0.008 | -0.011 | -0.011 | 0 | 0.74 | 0.67 | 0.62 | 0.8 | 0.79 | 0.72 |
| 5 | 169767523 | rs315761 | -0.584 | -0.369 | 0.092 | 0.053 | -0.013 | 0.082 | 0.082 | 0 | 0.1 | 0.28 | 0.28 | 0.19 | 0.05 | 0.13 |
| 5 | 169770089 | rs315756 | -1.251 | -0.53 | 0.398 | -0.025 | -0.074 | 0.064 | 0.064 | 0.049 | 0.17 | 0.47 | 0.31 | 0.23 | 0.07 | 0.13 |
| 5 | 169774699 | rs11134624 | -0.477 | -0.335 | 0.362 | -0.05 | 0.001 | -0.005 | -0.005 | 0 | 0.14 | 0.14 | 0.09 | 0.21 | 0.15 | 0.1 |
| 5 | 169776833 | rs906362 | -0.828 | -0.373 | 0.652 | -0.072 | -0.013 | 0.041 | 0.041 | 0.101 | 0.16 | 0.37 | 0.11 | 0.21 | 0.15 | 0.11 |
| 5 | 169777436 | rs906358 | -0.534 | -0.366 | 0.579 | -0.069 | -0.001 | 0.001 | 0.001 | 0 | 0.14 | 0.18 | 0.09 | 0.21 | 0.15 | 0.1 |
| 5 | 169780757 | rs6894729 | -0.501 | -0.36 | 0.47 | -0.05 | 0.002 | -0.007 | -0.007 | 0.066 | 0.14 | 0.15 | 0.2 | 0.22 | 0.15 | 0.11 |
| 5 | 169781990 | rs6860569 | -1.718 | -0.396 | 0.301 | -0.201 | -0.002 | 0.004 | 0.004 | 0 | 0.35 | 0.61 | 0.42 | 0.41 | 0.32 | 0.35 |
| 5 | 169782920 | rs11134625 | -1.134 | -0.53 | 0.308 | -0.047 | -0.038 | 0.077 | 0.077 | 0 | 0.26 | 0.55 | 0.35 | 0.39 | 0.21 | 0.26 |

| | | | | | | | | | | | | | | | | |
|---|---|---|---|---|---|---|---|---|---|---|---|---|---|---|---|---|
| 5 | 169787576 | rs6555897 | -0.401 | 0.185 | 0.265 | -0.055 | 0.013 | -0.023 | -0.023 | 0 | 0.14 | 0.22 | 0.18 | 0.06 | 0.16 | 0.09 |
| 5 | 169787623 | rs4867600 | -0.499 | 0.261 | 0.294 | -0.037 | 0.025 | -0.021 | -0.021 | 3.964 | 0.14 | 0.23 | 0.18 | 0.06 | 0.18 | 0.09 |
| 5 | 169788967 | rs905784 | -1.612 | 0.125 | 0.483 | 0.021 | 0.028 | 0.011 | 0.011 | 0 | 0.27 | 0.59 | 0.4 | 0.27 | 0.32 | 0.27 |
| 5 | 169790212 | rs4867961 | -0.383 | 0.124 | 0.367 | -0.009 | -0.016 | 0.016 | 0.016 | 0 | 0.14 | 0.12 | 0.18 | 0.06 | 0.19 | 0.09 |
| 5 | 169790857 | rs6893608 | -0.93 | 0.089 | 0.578 | 0.025 | 0.02 | 0.105 | 0.105 | 0 | 0.12 | 0.34 | 0.21 | 0.21 | 0.13 | 0.19 |
| 5 | 169793712 | rs7710512 | -0.395 | 0.205 | 0.479 | 0.057 | 0.024 | 0.078 | 0.078 | 0 | 0.08 | 0.19 | 0.19 | 0.21 | 0.12 | 0.19 |
| 5 | 169798156 | rs11134626 | -0.374 | 0.289 | 0.355 | 0.273 | -0.032 | -0.004 | -0.004 | 0 | 0.14 | 0.07 | 0.18 | 0.13 | 0.2 | 0.1 |
| 5 | 169805123 | rs314155 | -1.421 | -0.64 | 0.085 | 0.46 | 0.002 | -0.001 | -0.001 | 0 | 0.54 | 0.65 | 0.52 | 0.51 | 0.48 | 0.57 |
| 5 | 169805411 | rs314156 | -0.232 | -1.095 | -0.003 | 0.323 | -0.032 | -0.016 | -0.016 | 2.322 | 0.21 | 0.29 | 0.13 | 0.13 | 0.1 | 0.13 |
| 5 | 169806001 | rs314157 | -1.172 | -1.092 | -0.06 | 0.323 | -0.042 | -0.013 | -0.013 | 0.118 | 0.39 | 0.52 | 0.34 | 0.37 | 0.28 | 0.47 |
| 5 | 169806216 | rs2301147 | -0.111 | -1.045 | -0.354 | 0.362 | 0.044 | 0.031 | 0.031 | 0 | 0.2 | 0.1 | 0.11 | 0.13 | 0.1 | 0.13 |
| 5 | 169809402 | rs703505 | -1.433 | -0.908 | -0.299 | 0.339 | 0.018 | 0 | 0 | 0 | 0.47 | 0.64 | 0.47 | 0.51 | 0.38 | 0.54 |
| 5 | 169810450 | rs2071161 | 0.765 | -0.712 | -0.564 | 0.492 | -0.008 | -0.018 | -0.018 | 0 | 0.1 | 0.06 | 0.15 | 0.14 | 0.1 | 0.07 |
| 5 | 169810796 | rs11739136 | 0.765 | -0.712 | -0.564 | 0.474 | -0.008 | -0.016 | -0.016 | 0 | 0.1 | 0.05 | 0.14 | 0.14 | 0.1 | 0.07 |
| 5 | 169811409 | rs950910 | -0.879 | -0.813 | -0.476 | 0.46 | -0.007 | -0.006 | -0.006 | 0 | 0.29 | 0.34 | 0.4 | 0.37 | 0.37 | 0.43 |
| 5 | 169812303 | rs2071156 | 0.575 | -0.725 | -0.316 | 0.534 | -0.002 | 0.018 | 0.018 | 0 | 0.15 | 0.28 | 0.21 | 0.22 | 0.17 | 0.31 |
| 5 | 169813844 | rs314111 | -0.354 | -0.804 | -0.462 | 0.654 | -0.005 | 0 | 0 | 0 | 0.19 | 0.2 | 0.13 | 0.2 | 0.09 | 0.14 |
| 5 | 169815484 | rs314107 | -1.1 | -0.588 | -0.703 | 0.5 | -0.001 | -0.001 | -0.001 | 0 | 0.57 | 0.63 | 0.52 | 0.55 | 0.47 | 0.55 |
| 5 | 169816565 | rs827778 | 0.132 | -1.428 | -1.299 | 0.812 | 0.068 | -0.051 | -0.051 | 0 | 0.79 | 0.94 | 0.91 | 0.69 | 0.86 | 0.79 |
| 5 | 169816977 | rs10050842 | 0.476 | -0.743 | -1.056 | 0.473 | 0.004 | 0.02 | 0.02 | 0 | 0.5 | 0.55 | 0.52 | 0.28 | 0.46 | 0.3 |
| 5 | 169817040 | rs10065969 | 0.78 | -0.329 | -0.878 | 0.113 | 0.011 | -0.016 | -0.016 | 0 | 0.26 | 0.3 | 0.24 | 0.16 | 0.27 | 0.19 |
| 5 | 169818569 | rs314127 | 0.312 | -0.501 | -1.113 | 0.396 | -0.009 | 0.026 | 0.026 | 0 | 0.6 | 0.75 | 0.76 | 0.66 | 0.6 | 0.54 |
| 5 | 169821358 | rs9313503 | -0.543 | -0.898 | -0.279 | 0.448 | -0.039 | 0.029 | 0.029 | 0 | 0.3 | 0.45 | 0.23 | 0.37 | 0.19 | 0.36 |
| 5 | 169823097 | rs314121 | -0.148 | -0.551 | -0.383 | 0.198 | 0.005 | -0.021 | -0.021 | 0 | 0.25 | 0.22 | 0.12 | 0.03 | 0.16 | 0.17 |
| 5 | 169823446 | rs314118 | -0.226 | -0.535 | -0.342 | 0.447 | -0.002 | 0 | 0 | 0 | 0.26 | 0.25 | 0.18 | 0.31 | 0.18 | 0.22 |
| 5 | 169827557 | rs1972058 | -0.699 | 0.356 | 0.113 | 0.137 | 0.038 | 0.061 | 0.061 | 0.292 | 0.21 | 0.56 | 0.25 | 0.38 | 0.26 | 0.21 |
| 5 | 169827756 | rs314137 | -0.867 | 0.407 | -0.269 | -0.3 | 0.001 | -0.007 | -0.007 | 0 | 0.54 | 0.71 | 0.6 | 0.5 | 0.55 | 0.59 |
| 5 | 169831371 | rs314143 | -0.621 | 0.476 | -0.514 | -0.359 | -0.02 | -0.008 | -0.008 | 0 | 0.32 | 0.79 | 0.48 | 0.36 | 0.37 | 0.36 |
| 5 | 169831642 | rs314144 | -1.629 | 0.415 | -0.884 | -0.607 | -0.06 | -0.132 | -0.132 | 0 | 0.71 | 0.94 | 0.69 | 0.51 | 0.67 | 0.76 |
| 5 | 169833843 | rs827782 | 0.675 | 0.797 | 0.132 | -0.466 | 0.05 | -0.038 | -0.038 | 0 | 0.09 | 0.17 | 0.16 | 0.02 | 0.15 | 0.05 |
| 5 | 169834318 | rs7736689 | -0.357 | 0.744 | 0.165 | -0.48 | -0.013 | 0.038 | 0.038 | 1.129 | 0.22 | 0.57 | 0.32 | 0.33 | 0.22 | 0.29 |
| 5 | 169834750 | rs827784 | -1.005 | 0.761 | -0.141 | -0.466 | -0.021 | -0.011 | -0.011 | 0.107 | 0.31 | 0.79 | 0.49 | 0.36 | 0.37 | 0.35 |
| 5 | 169835524 | rs17740022 | 1.175 | 0.649 | -0.078 | -0.52 | 0.002 | -0.018 | -0.018 | 4.584 | 0.05 | 0.02 | 0.02 | 0.15 | 0.06 | 0.07 |
| 5 | 169836156 | rs2656845 | -1.023 | 0.665 | 0.138 | -0.56 | 0.014 | -0.021 | -0.021 | 0 | 0.2 | 0.63 | 0.25 | 0.17 | 0.24 | 0.15 |
| 5 | 169837712 | rs2656848 | -0.5 | 0.54 | 0.39 | -0.258 | 0.037 | 0.1 | 0.1 | 0.088 | 0.09 | 0.39 | 0.08 | 0.15 | 0.08 | 0.1 |
| 5 | 169843630 | rs13175143 | 0.707 | 0.999 | 0.176 | -0.123 | 0.041 | 0.051 | 0.051 | 0.002 | 0.12 | 0.24 | 0.3 | 0.21 | 0.19 | 0.19 |
| 5 | 169847029 | rs314165 | -0.249 | 0.471 | -0.254 | -0.739 | -0.003 | -0.04 | -0.04 | 0 | 0.42 | 0.44 | 0.33 | 0.21 | 0.38 | 0.42 |
| 5 | 169850856 | rs871105 | 0.93 | 0.475 | -0.832 | -0.764 | -0.017 | 0.089 | 0.089 | 0 | 0.21 | 0.06 | 0.2 | 0.05 | 0.21 | 0.23 |
| 5 | 169854585 | rs7735377 | -1.582 | 0.285 | -1.065 | -0.764 | -0.047 | -0.139 | -0.139 | 0.015 | 0.71 | 0.95 | 0.68 | 0.5 | 0.68 | 0.79 |
| 5 | 169856568 | rs12516919 | 0.764 | 0.465 | -1.158 | -0.976 | -0.053 | 0.122 | 0.122 | 0.495 | 0.27 | 0.06 | 0.24 | 0.06 | 0.3 | 0.27 |
| 5 | 169859436 | rs13172742 | 1.132 | 0.455 | -0.714 | -0.833 | 0.017 | 0.011 | 0.011 | 0.274 | 0.13 | 0.18 | 0.19 | 0.17 | 0.2 | 0.14 |
| 5 | 169861577 | rs10052852 | -1.822 | 0.371 | -0.656 | -0.874 | -0.014 | -0.117 | -0.117 | 0 | 0.74 | 0.94 | 0.7 | 0.49 | 0.76 | 0.8 |
| 5 | 169863955 | rs6896882 | 0.057 | 0.154 | -0.434 | -1.052 | 0.008 | 0.065 | 0.065 | 0.83 | 0.41 | 0.27 | 0.28 | 0.23 | 0.39 | 0.46 |
| 5 | 169865015 | rs13156152 | -0.571 | 0.172 | -0.423 | -1.035 | 0 | 0 | 0 | 2.707 | 0.54 | 0.45 | 0.47 | 0.4 | 0.58 | 0.59 |
| 5 | 169866944 | rs13166619 | -0.535 | 0.304 | -0.246 | -0.943 | -0.001 | -0.021 | -0.021 | 0 | 0.37 | 0.44 | 0.4 | 0.25 | 0.39 | 0.34 |
| 5 | 169869515 | rs13182150 | 1.075 | 0.368 | -0.664 | -0.722 | 0.016 | 0.017 | 0.017 | 0 | 0.15 | 0.21 | 0.19 | 0.19 | 0.19 | 0.14 |
| 5 | 169869644 | rs1553532 | 0.181 | 0.244 | -1.323 | -0.755 | -0.033 | 0.038 | 0.038 | 0 | 0.37 | 0.13 | 0.37 | 0.31 | 0.38 | 0.53 |
| 5 | 169870339 | rs11134628 | -1.597 | 0.455 | -1.023 | -0.344 | -0.019 | -0.101 | -0.101 | 1.02 | 0.78 | 0.9 | 0.72 | 0.52 | 0.77 | 0.88 |
| 5 | 169871779 | rs17072667 | 0.179 | 0.452 | -0.517 | -0.303 | -0.004 | 0.023 | 0.023 | 0 | 0.14 | 0.17 | 0.08 | 0.26 | 0.15 | 0.2 |
| 5 | 169872183 | rs7712677 | -1.08 | 0.685 | -0.762 | -0.513 | -0.029 | -0.074 | -0.074 | 0 | 0.64 | 0.86 | 0.6 | 0.49 | 0.61 | 0.76 |
| 5 | 169872995 | rs6877607 | -0.447 | 0.645 | -0.622 | -0.844 | 0.012 | 0.021 | 0.021 | 1.232 | 0.5 | 0.65 | 0.43 | 0.42 | 0.5 | 0.61 |
| 5 | 169877807 | rs4867965 | -1.001 | 1.114 | -0.65 | -0.837 | 0.001 | -0.049 | -0.049 | 0 | 0.65 | 0.84 | 0.62 | 0.44 | 0.65 | 0.72 |
| 5 | 169879318 | rs17564179 | -0.492 | 1.133 | -0.82 | -1.035 | 0.036 | 0.016 | 0.016 | 1.018 | 0.34 | 0.42 | 0.38 | 0.37 | 0.48 | 0.4 |
| 5 | 169879780 | rs959461 | -0.51 | 0.619 | -0.903 | -0.953 | 0.027 | 0.101 | 0.101 | 0 | 0.71 | 0.88 | 0.79 | 0.92 | 0.82 | 0.71 |
| 5 | 169880071 | rs1504924 | -0.675 | 1.237 | -0.241 | -0.186 | 0.007 | 0.016 | 0.016 | 0.21 | 0.26 | 0.29 | 0.28 | 0.34 | 0.31 | 0.27 |
| 5 | 169891630 | rs1115192 | 0.303 | 0.591 | -0.441 | -0.585 | 0.048 | 0.084 | 0.084 | 0 | 0.37 | 0.16 | 0.25 | 0.13 | 0.25 | 0.38 |
| 5 | 169892745 | rs12656294 | 0.371 | 1.607 | 0.416 | 0.489 | 0.003 | -0.021 | -0.021 | 0 | 0.23 | 0.16 | 0.28 | 0.51 | 0.25 | 0.16 |
| 5 | 169893751 | rs10041897 | -1.456 | 2.576 | 1.447 | 1.054 | 0.072 | 0.109 | 0.109 | 0 | 0.17 | 0.35 | 0.16 | 0.33 | 0.25 | 0.23 |
| 5 | 169893895 | rs2135023 | -0.586 | 2.186 | 0.739 | 1.176 | 0.003 | 0.045 | 0.045 | 0 | 0.08 | 0.07 | 0.07 | 0.28 | 0.11 | 0.16 |
| 5 | 169908768 | rs17740227 | -0.974 | 2.177 | 1.059 | 1.065 | 0.024 | 0.143 | 0.143 | 0.247 | 0.12 | 0.28 | 0.34 | 0.38 | 0.09 | 0.32 |
| 5 | 169910062 | rs10076544 | -1.452 | 2.007 | 1.409 | 1.005 | 0.008 | 0.124 | 0.124 | 0 | 0.15 | 0.37 | 0.27 | 0.35 | 0.11 | 0.24 |
| 5 | 169913253 | rs10056890 | -0.139 | 2.304 | 0.863 | 1.153 | -0.006 | 0.004 | 0.004 | 0.137 | 0.12 | 0.1 | 0.25 | 0.33 | 0.15 | 0.22 |
| 5 | 169916008 | rs10063281 | -0.916 | 2.265 | 1.143 | 1.043 | 0.044 | 0.122 | 0.122 | 0.013 | 0.16 | 0.3 | 0.38 | 0.4 | 0.15 | 0.36 |
| 5 | 169918840 | rs7448059 | -0.819 | 2.064 | 1.03 | 0.388 | 0.108 | 0 | 0 | 0 | 0.12 | 0.32 | 0.25 | 0.12 | 0.22 | 0.18 |
| 5 | 169918872 | rs7710817 | -0.761 | 1.93 | 0.959 | 0.418 | 0.117 | 0.009 | 0.009 | 0.265 | 0.12 | 0.37 | 0.25 | 0.12 | 0.2 | 0.18 |
| 5 | 169923732 | rs888934 | 0.25 | 1.047 | -0.148 | 0.721 | 0 | 0.002 | 0.002 | 0.008 | 0.89 | 0.84 | 0.91 | 0.8 | 0.9 | 0.92 |
| 5 | 169923903 | rs888936 | 0.588 | 0.887 | -0.314 | 0.409 | 0.001 | -0.003 | -0.003 | 0.07 | 0.73 | 0.72 | 0.79 | 0.73 | 0.73 | 0.81 |
| 5 | 169924695 | rs4867606 | 0.088 | 1.141 | -1.433 | -0.14 | -0.072 | 0.13 | 0.13 | 0 | 0.25 | 0.05 | 0.18 | 0.05 | 0.31 | 0.2 |
| 5 | 169926969 | rs12521917 | 0.541 | 0.668 | -0.427 | 0.329 | -0.012 | 0.019 | 0.019 | 0 | 0.72 | 0.34 | 0.72 | 0.66 | 0.75 | 0.78 |

| | | | | | | | | | | | | | | | | |
|---|---|---|---|---|---|---|---|---|---|---|---|---|---|---|---|---|
| 5 | 169928357 | rs1978440 | 0.412 | 0.964 | -0.274 | -0.124 | 0 | -0.022 | -0.022 | 0 | 0.66 | 0.69 | 0.77 | 0.87 | 0.63 | 0.75 |
| 5 | 169933055 | rs12517302 | -0.594 | 1.583 | 0.229 | 0.713 | 0.04 | 0.085 | 0.085 | 0.034 | 0.2 | 0.33 | 0.38 | 0.4 | 0.3 | 0.37 |
| 5 | 169934120 | rs7734708 | 0.627 | 1.302 | -0.062 | 0.635 | -0.007 | -0.014 | -0.014 | 0 | 0.25 | 0.21 | 0.28 | 0.42 | 0.25 | 0.26 |
| 5 | 169934412 | rs2339091 | 0.512 | 0.848 | 0.145 | 0.481 | -0.016 | 0.039 | 0.039 | 0 | 0.24 | 0.28 | 0.18 | 0.43 | 0.16 | 0.33 |
| 5 | 169937406 | rs11740245 | 0.885 | 1.229 | -0.032 | 0.517 | -0.013 | -0.014 | -0.014 | 0 | 0.26 | 0.22 | 0.35 | 0.43 | 0.32 | 0.28 |
| 5 | 169940759 | rs10475944 | 0.537 | 0.933 | 0.29 | 0.609 | -0.027 | -0.001 | -0.001 | 0 | 0.22 | 0.26 | 0.08 | 0.16 | 0.08 | 0.2 |
| 5 | 169947870 | rs11738748 | -0.103 | 1.772 | 0.454 | 0.941 | 0.072 | 0.076 | 0.076 | 0 | 0.26 | 0.52 | 0.36 | 0.47 | 0.35 | 0.35 |
| 5 | 169948268 | rs11738868 | 0.458 | 1.55 | 0.182 | 1.128 | 0.016 | 0.036 | 0.036 | 0.951 | 0.19 | 0.26 | 0.28 | 0.46 | 0.25 | 0.27 |
| 5 | 169948896 | rs9313506 | -0.597 | 1.508 | 0.031 | 1.205 | 0.013 | 0.012 | 0.012 | 0 | 0.22 | 0.21 | 0.42 | 0.32 | 0.33 | 0.29 |
| 5 | 169951771 | rs11744046 | -0.224 | 0.626 | 0.108 | 1.233 | 0 | 0 | 0 | 0 | 0.44 | 0.45 | 0.42 | 0.46 | 0.48 | 0.46 |
| 5 | 169952910 | rs4867975 | -0.958 | 0.238 | -0.26 | 1.068 | -0.061 | -0.079 | -0.079 | 0 | 0.69 | 0.9 | 0.56 | 0.58 | 0.63 | 0.65 |
| 5 | 169954512 | rs6873409 | 0.556 | -0.015 | -0.179 | 1.314 | 0.005 | -0.007 | -0.007 | 0 | 0.1 | 0.08 | 0.03 | 0.23 | 0.06 | 0.14 |
| 5 | 169954546 | rs6873133 | -1.117 | -0.019 | -0.004 | 1.123 | 0.081 | 0.076 | 0.076 | 0.515 | 0.08 | 0.51 | 0.07 | 0.07 | 0.06 | 0.04 |
| 5 | 169956789 | rs7724503 | 0.519 | -0.193 | -0.258 | 1.346 | 0.026 | 0.027 | 0.027 | 0.66 | 0.23 | 0.07 | 0.1 | 0.19 | 0.15 | 0.19 |
| 5 | 169956981 | rs17672975 | 1.097 | -0.142 | 0.038 | 1.378 | -0.02 | 0.002 | 0.002 | 0.015 | 0.3 | 0.17 | 0.35 | 0.23 | 0.35 | 0.3 |
| 5 | 169957076 | rs6885463 | -1.115 | 0.008 | -0.073 | 1.299 | -0.064 | -0.039 | -0.039 | 0.062 | 0.71 | 0.89 | 0.56 | 0.7 | 0.63 | 0.67 |
| 5 | 169957614 | rs6555903 | 0.009 | -0.02 | 0.745 | 1.297 | -0.002 | 0.009 | 0.009 | 0 | 0.46 | 0.57 | 0.45 | 0.55 | 0.48 | 0.46 |
| 5 | 169959437 | rs4640812 | -0.529 | -0.098 | 1.253 | 1.041 | -0.012 | -0.007 | -0.007 | 0 | 0.29 | 0.67 | 0.16 | 0.24 | 0.17 | 0.2 |
| 5 | 169960814 | rs10072375 | -0.354 | -0.092 | 1.546 | 1.044 | -0.043 | -0.025 | -0.025 | 0 | 0.28 | 0.51 | 0.15 | 0.24 | 0.17 | 0.2 |
| 5 | 169960826 | rs10072376 | -0.012 | -0.105 | 1.808 | 1.038 | 0.011 | 0.015 | 0.015 | 0.952 | 0.25 | 0.17 | 0.11 | 0.11 | 0.17 | 0.2 |
| 5 | 169962828 | rs10076552 | -0.025 | -0.081 | 1.643 | 0.971 | -0.042 | -0.085 | -0.085 | 0.009 | 0.25 | 0.42 | 0.14 | 0.14 | 0.16 | 0.2 |
| 5 | 169966669 | rs2112601 | -0.382 | -0.032 | 1.199 | 0.883 | -0.012 | -0.05 | -0.05 | 0.132 | 0.28 | 0.67 | 0.16 | 0.08 | 0.16 | 0.19 |
| 5 | 169978127 | rs11958729 | -0.189 | -0.077 | 1.23 | 0.985 | -0.046 | -0.042 | -0.042 | 0 | 0.25 | 0.56 | 0.12 | 0.17 | 0.13 | 0.19 |
| 5 | 169982036 | rs9313511 | -0.763 | 0.46 | 0.985 | 1.422 | 0.015 | -0.02 | -0.02 | 0 | 0.39 | 0.73 | 0.27 | 0.46 | 0.28 | 0.37 |
| 5 | 169989879 | rs13155413 | 0.895 | 0.501 | 0.901 | 1.476 | -0.003 | -0.002 | -0.002 | 0 | 0.29 | 0.21 | 0.3 | 0.28 | 0.31 | 0.23 |
| 5 | 169992533 | rs10065828 | -0.282 | 1.203 | 1.137 | 1.3 | 0.008 | -0.014 | -0.014 | 1.044 | 0.66 | 0.52 | 0.53 | 0.66 | 0.61 | 0.61 |
| 5 | 169996594 | rs12652223 | 0.403 | 1.218 | 0.842 | 1.251 | -0.025 | 0.016 | 0.016 | 0 | 0.32 | 0.19 | 0.45 | 0.27 | 0.39 | 0.39 |
| 5 | 169997669 | rs3860759 | 0.11 | 1.252 | 0.768 | 1.053 | -0.04 | 0.023 | 0.023 | 0 | 0.34 | 0.21 | 0.47 | 0.28 | 0.45 | 0.41 |
| 5 | 169999591 | rs4867981 | 0.062 | 1.273 | 0.766 | 1.116 | -0.041 | -0.013 | -0.013 | 2.41 | 0.35 | 0.21 | 0.48 | 0.35 | 0.45 | 0.42 |
| 5 | 170004035 | rs4242158 | 0.708 | 1.081 | 0.881 | 0.457 | 0.021 | 0.018 | 0.018 | 0 | 0.51 | 0.3 | 0.38 | 0.55 | 0.44 | 0.41 |
| 5 | 170013811 | rs4867984 | 1.682 | 1.226 | 0.821 | 0.27 | 0.032 | -0.024 | -0.024 | 0 | 0.82 | 0.58 | 0.83 | 0.91 | 0.82 | 0.79 |
| 5 | 170017162 | rs10064808 | -1.779 | 0.984 | 0.675 | 0.34 | 0.02 | 0.032 | 0.032 | 0 | 0.07 | 0.19 | 0.11 | 0.08 | 0.09 | 0.09 |
| 5 | 170024636 | rs6555905 | 0.958 | 1.017 | 0.348 | 0.009 | 0.015 | -0.07 | -0.07 | 0 | 0.87 | 0.71 | 0.91 | 0.99 | 0.87 | 0.87 |
| 5 | 170027924 | rs2036561 | -0.643 | 1.286 | -0.634 | -0.514 | -0.068 | 0.129 | 0.129 | 0 | 0.34 | 0.15 | 0.21 | 0.11 | 0.44 | 0.29 |
| 5 | 170028569 | rs905809 | -0.643 | 1.286 | -0.634 | -0.514 | -0.068 | 0.129 | 0.129 | 0 | 0.34 | 0.15 | 0.21 | 0.11 | 0.44 | 0.29 |
| 5 | 170031270 | rs6877169 | -0.539 | 0.686 | -0.202 | -0.166 | -0.024 | 0.133 | 0.133 | 0 | 0.77 | 0.58 | 0.78 | 0.51 | 0.83 | 0.69 |
| 5 | 170031313 | rs905813 | -0.651 | 0.652 | 0.172 | -0.183 | -0.013 | 0.071 | 0.071 | 0 | 0.78 | 0.71 | 0.78 | 0.51 | 0.83 | 0.69 |
| 5 | 170034897 | rs12523655 | 0.704 | 0.654 | 0.268 | -0.254 | 0.005 | -0.003 | -0.003 | 0 | 0.43 | 0.4 | 0.56 | 0.4 | 0.39 | 0.37 |
| 5 | 170036025 | rs2879337 | 0.232 | 1.248 | 0.051 | -0.579 | -0.005 | 0.004 | 0.004 | 0 | 0.16 | 0.18 | 0.11 | 0.09 | 0.21 | 0.14 |
| 5 | 170037711 | rs1013922 | -0.838 | 1.178 | 0.176 | -0.487 | -0.01 | 0.053 | 0.053 | 0 | 0.78 | 0.73 | 0.78 | 0.51 | 0.83 | 0.68 |
| 5 | 170037922 | rs2089192 | 1.079 | 1.006 | -0.053 | -0.916 | -0.001 | 0.019 | 0.019 | 0 | 0.19 | 0.18 | 0.41 | 0.41 | 0.18 | 0.25 |
| 5 | 170042704 | rs11134640 | 1.246 | 0.8 | -0.092 | -0.98 | 0 | -0.022 | -0.022 | 0 | 0.23 | 0.19 | 0.13 | 0 | 0.19 | 0.12 |
| 5 | 170043173 | rs1875978 | -0.122 | 0.815 | -0.254 | -0.445 | 0.003 | -0.012 | -0.012 | 0 | 0.6 | 0.76 | 0.66 | 0.51 | 0.6 | 0.52 |
| 5 | 170047653 | rs2202437 | -0.525 | 0.902 | -0.098 | -0.333 | 0.002 | -0.011 | -0.011 | 0 | 0.89 | 0.84 | 0.94 | 0.92 | 0.89 | 0.89 |
| 5 | 170052330 | rs905823 | -0.301 | 0.966 | 0.032 | 0.015 | 0.003 | 0 | 0 | 0.081 | 0.39 | 0.54 | 0.48 | 0.66 | 0.41 | 0.53 |
| 5 | 170059227 | rs11960314 | -0.395 | 0.665 | -0.267 | 0.177 | -0.004 | 0.04 | 0.04 | 0 | 0.45 | 0.22 | 0.44 | 0.38 | 0.42 | 0.48 |
| 5 | 170066527 | rs2279873 | -0.387 | 0.566 | -0.295 | 0.488 | -0.009 | -0.025 | -0.025 | 0 | 0.44 | 0.2 | 0.43 | 0.51 | 0.43 | 0.51 |
| 5 | 170070942 | rs883848 | -0.187 | 0.737 | 0.4 | 0.425 | 0 | -0.001 | -0.001 | 0 | 0.41 | 0.36 | 0.42 | 0.46 | 0.35 | 0.46 |
| 5 | 170071480 | rs4867623 | -0.94 | 0.967 | 0.73 | 0.535 | 0.068 | 0.053 | 0.053 | 1.088 | 0.11 | 0.46 | 0.1 | 0.12 | 0.13 | 0.19 |
| 5 | 170072378 | rs9968589 | -1.48 | 0.961 | 0.713 | 0.551 | 0.092 | 0.08 | 0.08 | 1.362 | 0.16 | 0.53 | 0.14 | 0.25 | 0.22 | 0.24 |
| 5 | 170072498 | rs9968699 | 0.065 | 1.032 | 0.86 | 0.509 | 0.039 | 0.025 | 0.025 | 0.033 | 0.06 | 0.18 | 0.06 | 0.09 | 0.09 | 0.1 |
| 5 | 170095549 | rs6875696 | -0.088 | 0.546 | -0.224 | 0.342 | 0.01 | -0.054 | -0.054 | 0.384 | 0.35 | 0.14 | 0.38 | 0.45 | 0.32 | 0.42 |
| 5 | 170096737 | rs6555910 | 0.155 | -0.119 | -0.938 | -0.496 | 0.016 | -0.024 | -0.024 | 0 | 0.39 | 0.19 | 0.37 | 0.52 | 0.34 | 0.43 |
| 5 | 170099077 | rs4868011 | 1.232 | -0.227 | -1.228 | -0.48 | -0.021 | -0.103 | -0.103 | 0 | 0.21 | 0.04 | 0.29 | 0.52 | 0.21 | 0.29 |
| 5 | 170099466 | rs4868012 | 0.197 | -0.217 | -0.342 | -1.536 | 0.005 | 0.002 | 0.002 | 0 | 0.45 | 0.58 | 0.4 | 0.59 | 0.35 | 0.52 |
| 5 | 170104598 | rs6897199 | -0.104 | -0.231 | -0.405 | -1.954 | 0.01 | 0.036 | 0.036 | 0 | 0.13 | 0.05 | 0.1 | 0.08 | 0.08 | 0.13 |
| 5 | 170115523 | rs2202440 | 0.297 | 0.185 | 0.142 | -2.09 | 0.007 | -0.086 | -0.086 | 0.019 | 0.57 | 0.49 | 0.58 | 0.94 | 0.46 | 0.78 |
| 5 | 170124362 | rs875184 | -0.863 | 0.289 | 0.113 | -2.146 | -0.033 | 0.038 | 0.038 | 0.003 | 0.77 | 0.84 | 0.68 | 0.94 | 0.6 | 0.8 |
| 5 | 170127770 | rs10053768 | 0.05 | 0.173 | 0.307 | -2.079 | -0.026 | -0.095 | -0.095 | 0.01 | 0.18 | 0.3 | 0.08 | 0.01 | 0.14 | 0.04 |
| 5 | 170129532 | rs13171706 | 0.05 | 0.173 | 0.307 | -2.079 | -0.026 | -0.095 | -0.095 | 3.619 | 0.18 | 0.3 | 0.08 | 0.01 | 0.14 | 0.04 |
| 5 | 170135225 | rs102686 | 1.749 | 0.055 | 0.338 | -2.221 | 0.014 | 0.041 | 0.041 | 0 | 0.31 | 0.25 | 0.14 | 0.12 | 0.24 | 0.29 |
| 5 | 170136830 | rs172944 | -0.51 | 0.042 | 0.396 | -2.164 | -0.052 | 0.083 | 0.083 | 0 | 0.69 | 0.82 | 0.66 | 0.95 | 0.54 | 0.77 |
| 5 | 170143423 | rs7713498 | -0.686 | 0.037 | 0.406 | -2.115 | 0.011 | 0.016 | 0.016 | 0.566 | 0.48 | 0.43 | 0.76 | 0.86 | 0.59 | 0.62 |
| 5 | 170145584 | rs6555913 | -0.36 | 0.112 | 0.208 | -1.585 | 0 | -0.061 | -0.061 | 0.001 | 0.49 | 0.5 | 0.43 | 0.05 | 0.59 | 0.23 |
| 5 | 170168980 | rs2194161 | -0.611 | 0.284 | -0.026 | -1.165 | 0.008 | 0.007 | 0.007 | 0 | 0.62 | 0.6 | 0.8 | 0.85 | 0.7 | 0.66 |
| 5 | 170173510 | rs4642385 | -0.439 | 0.17 | -0.251 | 0.027 | 0.001 | -0.005 | -0.005 | 0 | 0.32 | 0.35 | 0.59 | 0.73 | 0.28 | 0.39 |
| 5 | 170178139 | rs7701794 | -0.542 | -0.048 | -0.639 | 0.061 | 0.039 | 0.027 | 0.027 | 0.236 | 0.43 | 0.76 | 0.55 | 0.64 | 0.31 | 0.42 |
| 5 | 170185337 | rs12514867 | -1.456 | -0.205 | 0.673 | -0.25 | -0.004 | 0.002 | 0.002 | 0 | 0.1 | 0.42 | 0.21 | 0.05 | 0.05 | 0.12 |
| 5 | 170185684 | rs12515465 | -1.583 | -0.198 | 0.811 | -0.247 | -0.005 | 0.001 | 0.001 | 0 | 0.1 | 0.38 | 0.21 | 0.05 | 0.05 | 0.12 |

| | | | | | | | | | | | | | | | | |
|---|---|---|---|---|---|---|---|---|---|---|---|---|---|---|---|---|
| 5 | 170187643 | rs9313524 | 0.144 | 0.609 | 0.261 | -0.511 | -0.017 | 0.007 | 0.007 | 0.425 | 0.72 | 0.78 | 0.79 | 0.84 | 0.61 | 0.87 |
| 5 | 170196638 | rs888810 | 0.076 | 0.086 | -0.014 | -0.683 | -0.059 | 0.063 | 0.063 | 3.165 | 0.63 | 0.85 | 0.75 | 0.84 | 0.55 | 0.82 |
| 5 | 170197358 | rs6555915 | -1.099 | 0.332 | 0.993 | -0.736 | 0.071 | 0.016 | 0.016 | 0 | 0.09 | 0.39 | 0.21 | 0.04 | 0.13 | 0.08 |
| 5 | 170197779 | rs10475958 | 0.551 | -0.332 | 0.957 | -0.93 | -0.002 | -0.006 | -0.006 | 0 | 0.46 | 0.45 | 0.51 | 0.8 | 0.41 | 0.61 |
| 5 | 170199228 | rs12173106 | 0.238 | -0.331 | 0.859 | -0.948 | 0.015 | 0.031 | 0.031 | 0.42 | 0.49 | 0.59 | 0.67 | 0.83 | 0.43 | 0.65 |
| 5 | 170200635 | rs1559160 | 0.632 | -0.11 | 0.169 | -0.851 | 0.013 | 0.076 | 0.076 | 0 | 0.43 | 0.19 | 0.48 | 0.81 | 0.4 | 0.62 |
| 5 | 170202984 | rs7722456 | 0.434 | 0.402 | 1.154 | -1.378 | 0.003 | -0.069 | -0.069 | 0 | 0.79 | 0.67 | 0.86 | 0.97 | 0.79 | 0.9 |
| 5 | 170207363 | rs11745599 | 1.271 | 0.399 | 1.621 | -0.831 | 0 | -0.007 | -0.007 | 0.507 | 0.71 | 0.34 | 0.66 | 0.82 | 0.7 | 0.74 |
| 5 | 170207491 | rs11746250 | 1.366 | 0.338 | 1.657 | -0.851 | -0.001 | 0.016 | 0.016 | 0.244 | 0.7 | 0.27 | 0.66 | 0.82 | 0.7 | 0.73 |
| 5 | 170209095 | rs7716934 | 0.204 | 0.299 | 1.2 | -0.81 | -0.037 | 0.067 | 0.067 | 3.909 | 0.55 | 0.82 | 0.78 | 0.75 | 0.54 | 0.71 |
| 5 | 170210237 | rs7722677 | -0.762 | 0.239 | 1.474 | -0.736 | -0.024 | -0.042 | -0.042 | 0 | 0.08 | 0.26 | 0.06 | 0 | 0.06 | 0.04 |
| 5 | 170210761 | rs929761 | -0.308 | 0.097 | 1.621 | -0.358 | 0.018 | 0.124 | 0.124 | 0 | 0.08 | 0.21 | 0.16 | 0.23 | 0.06 | 0.18 |
| 5 | 170210960 | rs2278000 | 0.315 | 0.183 | 1.529 | -0.472 | 0.012 | 0.017 | 0.017 | 0 | 0.36 | 0.25 | 0.15 | 0.21 | 0.3 | 0.25 |
| 5 | 170211251 | rs732157 | 0.536 | 0.298 | 1.35 | -0.505 | 0.014 | 0.072 | 0.072 | 1.938 | 0.48 | 0.72 | 0.71 | 0.75 | 0.44 | 0.66 |
| 5 | 170213757 | rs11134653 | -0.099 | 0.228 | 1.414 | -0.533 | 0.011 | 0.071 | 0.071 | 0 | 0.07 | 0.17 | 0.16 | 0.17 | 0.07 | 0.19 |
| 5 | 170214501 | rs1158443 | 1.086 | 0.319 | 1.497 | -0.302 | -0.008 | 0.018 | 0.018 | 0.587 | 0.4 | 0.52 | 0.55 | 0.5 | 0.36 | 0.47 |
| 5 | 170215670 | rs7725726 | -1.552 | 0.208 | 1.196 | -0.295 | -0.002 | 0.08 | 0.08 | 2.327 | 0.11 | 0.52 | 0.18 | 0.16 | 0.07 | 0.19 |
| 5 | 170218798 | rs984535 | -0.598 | 0.375 | 0.556 | -0.246 | 0.025 | 0.023 | 0.023 | 0 | 0.47 | 0.77 | 0.35 | 0.38 | 0.4 | 0.46 |
| 5 | 170221924 | rs7729540 | 0.479 | 0.518 | 0.509 | -0.007 | -0.018 | -0.019 | -0.019 | 0.013 | 0.22 | 0.14 | 0.13 | 0.25 | 0.26 | 0.13 |
| 5 | 170222885 | rs1812910 | -0.486 | 0.697 | 1.158 | -0.096 | 0.027 | 0.03 | 0.03 | 0 | 0.35 | 0.54 | 0.37 | 0.41 | 0.4 | 0.4 |
| 5 | 170223210 | rs9313525 | -1.079 | 0.466 | 0.924 | -0.214 | -0.03 | -0.027 | -0.027 | 0 | 0.46 | 0.88 | 0.4 | 0.41 | 0.42 | 0.52 |
| 5 | 170233262 | rs1862242 | -1.183 | 0.427 | 1.422 | -0.237 | -0.005 | -0.021 | -0.021 | 0.276 | 0.22 | 0.52 | 0.26 | 0.16 | 0.17 | 0.39 |
| 5 | 170234721 | rs10063254 | 0.552 | 0.285 | 1.463 | 0.021 | -0.014 | -0.017 | -0.017 | 1.398 | 0.23 | 0.16 | 0.13 | 0.25 | 0.26 | 0.13 |
| 5 | 170238904 | rs3805458 | 0.361 | 0.315 | 1.604 | 0.091 | 0.015 | 0.026 | 0.026 | 0.752 | 0.23 | 0.35 | 0.15 | 0.26 | 0.26 | 0.14 |
| 5 | 170239112 | rs1063310 | 0.467 | 0.299 | 1.624 | 0.083 | 0.007 | 0.013 | 0.013 | 0.874 | 0.22 | 0.28 | 0.14 | 0.26 | 0.26 | 0.14 |
| 5 | 170240423 | rs11134654 | 0.022 | 0.48 | 1.442 | -0.279 | 0.006 | -0.061 | -0.061 | 0 | 0.85 | 0.76 | 0.89 | 1 | 0.81 | 0.96 |
| 5 | 170240494 | rs3805455 | 0.727 | 0.447 | 1.464 | -0.223 | -0.015 | 0.018 | 0.018 | 1.007 | 0.61 | 0.26 | 0.72 | 0.73 | 0.55 | 0.82 |
| 5 | 170245846 | rs6886019 | 1.157 | 0.581 | 1.224 | -0.274 | 0 | -0.013 | -0.013 | 0.193 | 0.05 | 0.05 | 0.12 | 0 | 0.06 | 0.03 |
| 5 | 170246958 | rs1422161 | 1.616 | 0.485 | 1.178 | -0.253 | -0.03 | -0.023 | -0.023 | 0.474 | 0.47 | 0.15 | 0.45 | 0.6 | 0.48 | 0.55 |
| 5 | 170258216 | rs4867634 | 1.109 | 0.371 | 1.44 | -0.243 | 0.006 | 0.032 | 0.032 | 1.964 | 0.53 | 0.4 | 0.74 | 0.73 | 0.56 | 0.7 |
| 5 | 170263363 | rs11745293 | -0.221 | 0.586 | 1.074 | -0.266 | 0.002 | -0.034 | -0.034 | 0 | 0.88 | 0.85 | 0.9 | 1 | 0.84 | 0.96 |
| 5 | 170275387 | rs10073556 | 0.596 | 0.511 | 1.252 | -0.029 | 0.014 | 0.022 | 0.022 | 1.32 | 0.23 | 0.35 | 0.27 | 0.26 | 0.25 | 0.26 |
| 5 | 170275961 | rs11958845 | 0.276 | 0.61 | 1.246 | 0.033 | -0.007 | 0.039 | 0.039 | 0 | 0.1 | 0.26 | 0.15 | 0.14 | 0.08 | 0.13 |
| 5 | 170279080 | rs10223131 | 0.098 | 0.572 | 1.227 | 0.029 | -0.002 | 0.008 | 0.008 | 1.895 | 0.28 | 0.42 | 0.28 | 0.26 | 0.26 | 0.27 |
| 5 | 170282686 | rs6555925 | 0.812 | 0.282 | 1.342 | 0.084 | 0.002 | -0.005 | -0.005 | 0 | 0.79 | 0.7 | 0.75 | 0.85 | 0.76 | 0.84 |
| 5 | 170283091 | rs4868038 | 1.206 | 0.389 | 1.546 | 0.029 | 0.022 | -0.024 | -0.024 | 0 | 0.76 | 0.53 | 0.71 | 0.85 | 0.72 | 0.83 |
| 5 | 170319789 | rs1422155 | 0.977 | -0.418 | 0.399 | 0.297 | 0.037 | -0.151 | -0.151 | 0 | 0.29 | 0.01 | 0.17 | 0.44 | 0.17 | 0.28 |
| 5 | 170328399 | rs4868043 | 1.436 | 0.709 | 2.113 | 0.531 | 0.022 | 0.003 | 0.003 | 0.118 | 0.77 | 0.31 | 0.64 | 0.78 | 0.68 | 0.92 |
| 5 | 170369601 | rs7701805 | 0.878 | 0.685 | 2.61 | 0.524 | 0.031 | 0.014 | 0.014 | 0 | 0.81 | 0.67 | 0.68 | 0.78 | 0.69 | 0.92 |
| 5 | 170376149 | rs579535 | 0.877 | 0.686 | 2.44 | 0.514 | -0.012 | 0.007 | 0.007 | 0 | 0.06 | 0.03 | 0.1 | 0 | 0.14 | 0.03 |
| 5 | 170394572 | rs10051958 | 1.468 | 0.63 | 2.39 | 0.529 | 0.027 | 0.006 | 0.006 | 0.014 | 0.77 | 0.36 | 0.64 | 0.78 | 0.68 | 0.89 |
| 5 | 170408962 | rs10037007 | 1.635 | 0.612 | 2.543 | 0.582 | -0.015 | -0.024 | -0.024 | 1.36 | 0.75 | 0.17 | 0.57 | 0.7 | 0.68 | 0.81 |
| 5 | 170458381 | rs4242161 | 1.733 | 0.622 | 2.494 | 0.587 | -0.004 | -0.006 | -0.006 | 0 | 0.75 | 0.22 | 0.57 | 0.69 | 0.68 | 0.81 |
| 5 | 170459675 | rs7730898 | 0.488 | 0.657 | 2.415 | 0.503 | 0.045 | 0.007 | 0.007 | 0 | 0.88 | 0.82 | 0.72 | 0.89 | 0.73 | 0.92 |
| 5 | 170499222 | rs812407 | 1.402 | 0.682 | 1.978 | 0.516 | 0.026 | 0.005 | 0.005 | 0 | 0.77 | 0.36 | 0.64 | 0.78 | 0.68 | 0.89 |
| 5 | 170522906 | rs2446046 | 1.408 | 0.692 | 1.98 | 0.488 | 0.023 | 0.004 | 0.004 | 0.075 | 0.77 | 0.32 | 0.63 | 0.78 | 0.68 | 0.89 |
| 5 | 170536837 | rs17073355 | 1.759 | 0.203 | 1.433 | 0.567 | -0.001 | -0.002 | -0.002 | 0 | 0.06 | 0.08 | 0.04 | 0.11 | 0.04 | 0.01 |
| 5 | 170562476 | rs10516077 | -1.235 | 0.202 | 1.796 | 0.635 | -0.053 | 0.052 | 0.052 | 0.819 | 0.23 | 0.5 | 0.19 | 0.31 | 0.14 | 0.24 |
| 5 | 170568740 | rs4868069 | 0.71 | 1.117 | 2.003 | 0.504 | 0.051 | 0 | 0 | 0 | 0.8 | 0.63 | 0.7 | 0.83 | 0.7 | 0.85 |
| 5 | 170570675 | rs7731162 | 0.673 | 1.053 | 2.155 | 0.53 | -0.022 | 0.009 | 0.009 | 2.302 | 0.08 | 0.17 | 0.04 | 0.11 | 0.04 | 0.08 |
| 5 | 170573778 | rs2339233 | 0.821 | 1.083 | 2.1 | 0.53 | 0.035 | -0.001 | -0.001 | 2.592 | 0.8 | 0.68 | 0.71 | 0.83 | 0.71 | 0.85 |
| 5 | 170579645 | rs7723591 | 0.846 | 0.934 | 1.912 | 0.543 | 0 | -0.002 | -0.002 | 0 | 0.07 | 0.08 | 0.04 | 0.11 | 0.04 | 0.04 |
| 5 | 170600883 | rs17073388 | 0.64 | 0.62 | 2.018 | 0.566 | -0.017 | 0.003 | 0.003 | 2.887 | 0.08 | 0.13 | 0.04 | 0.11 | 0.04 | 0.04 |
| 5 | 170617859 | rs6890394 | 1.583 | 0.705 | 2.19 | 0.555 | 0.014 | -0.003 | -0.003 | 0.502 | 0.7 | 0.34 | 0.63 | 0.72 | 0.64 | 0.76 |
| 5 | 170623391 | rs11134679 | 0.69 | 0.708 | 2.204 | 0.543 | 0.015 | -0.001 | -0.001 | 2.704 | 0.81 | 0.78 | 0.7 | 0.83 | 0.68 | 0.84 |
| 5 | 170637472 | rs10074614 | 1.701 | 0.672 | 2.332 | 0.595 | -0.004 | -0.005 | -0.005 | 0.009 | 0.69 | 0.26 | 0.56 | 0.63 | 0.63 | 0.68 |
| 5 | 170640827 | rs4867648 | 0.013 | 0.75 | 1.92 | 0.478 | 0.003 | -0.062 | -0.062 | 1.523 | 0.84 | 0.8 | 0.86 | 1 | 0.82 | 0.92 |
| 5 | 170703078 | rs2914341 | 0.783 | 0.946 | 1.563 | 0.7 | 0.042 | 0.1 | 0.1 | 0.081 | 0.94 | 0.87 | 0.79 | 0.78 | 0.87 | 0.88 |
| 5 | 170714609 | rs2964519 | 1.672 | 0.47 | 1.303 | 0.547 | 0.001 | 0 | 0 | 2.436 | 0.65 | 0.27 | 0.59 | 0.68 | 0.66 | 0.77 |
| 5 | 170725989 | rs1044997 | 0.853 | 0.568 | 0.876 | 0.596 | -0.027 | -0.011 | -0.011 | 0 | 0.05 | 0.01 | 0.19 | 0.21 | 0.09 | 0.11 |
| 5 | 170726391 | rs17680600 | 0.853 | 0.568 | 0.876 | 0.596 | -0.027 | -0.011 | -0.011 | 0.032 | 0.05 | 0.01 | 0.19 | 0.21 | 0.09 | 0.11 |
| 5 | 170726798 | rs2914332 | 0.49 | 0.584 | 1.002 | 0.485 | 0.024 | -0.124 | -0.124 | 0 | 0.81 | 0.55 | 0.82 | 1 | 0.78 | 0.91 |
| 5 | 170732406 | rs2964538 | 0.274 | 0.627 | 0.672 | 0.537 | 0.035 | -0.126 | -0.126 | 0 | 0.82 | 0.51 | 0.82 | 1 | 0.78 | 0.92 |
| 5 | 170736044 | rs3806930 | 0.751 | 0.784 | -0.314 | 0.741 | -0.028 | -0.013 | -0.013 | 0 | 0.05 | 0.01 | 0.19 | 0.21 | 0.09 | 0.11 |
| 5 | 170741595 | rs2964534 | 1.498 | 0.403 | -0.36 | 0.567 | -0.011 | -0.033 | -0.033 | 0 | 0.57 | 0.08 | 0.48 | 0.6 | 0.6 | 0.65 |
| 5 | 170745820 | rs891009 | 0.987 | 0.678 | 0.232 | 0.503 | 0.001 | -0.001 | -0.001 | 0 | 0.67 | 0.34 | 0.57 | 0.69 | 0.63 | 0.74 |
| 5 | 170747271 | rs2914339 | 0.787 | 0.933 | 0.622 | 0.509 | 0.017 | -0.002 | -0.002 | 0 | 0.74 | 0.56 | 0.76 | 0.8 | 0.69 | 0.81 |
| 5 | 170749733 | rs6880753 | -0.203 | 0.969 | 0.624 | 0.504 | 0.017 | 0.034 | 0.034 | 0 | 0.05 | 0.17 | 0.21 | 0.07 | 0.06 | 0.14 |
| 5 | 170757560 | rs4868080 | -1.07 | 0.982 | 0.824 | 0.394 | 0.012 | 0.006 | 0.006 | 0 | 0.17 | 0.33 | 0.33 | 0.16 | 0.19 | 0.16 |

| | | | | | | | | | | | | | | | | |
|---|---|---|---|---|---|---|---|---|---|---|---|---|---|---|---|---|
| 5 | 170758661 | rs9686608 | -0.429 | 0.904 | 0.673 | 0.375 | 0.001 | -0.011 | -0.011 | 0 | 0.16 | 0.21 | 0.15 | 0.09 | 0.18 | 0.03 |
| 5 | 170758982 | rs11134683 | -2.058 | 1.114 | 0.841 | -0.039 | -0.001 | 0.039 | 0.039 | 0 | 0.37 | 0.71 | 0.44 | 0.17 | 0.37 | 0.21 |
| 5 | 170762656 | rs6555940 | -0.185 | 0.857 | 0.753 | 0.088 | -0.015 | 0.019 | 0.019 | 0 | 0.14 | 0.1 | 0.14 | 0.09 | 0.18 | 0.03 |
| 5 | 170778824 | rs7715167 | -1.688 | 0.558 | -0.021 | 0.266 | -0.002 | -0.024 | -0.024 | 0 | 0.63 | 0.73 | 0.45 | 0.51 | 0.62 | 0.62 |
| 5 | 170782385 | rs10073796 | -0.496 | 0.171 | 0.956 | 0.798 | 0.008 | 0.026 | 0.026 | 0 | 0.18 | 0.3 | 0.3 | 0.22 | 0.2 | 0.16 |
| 5 | 170784640 | rs7736331 | -1.777 | 0.027 | 1.092 | 0.74 | 0 | -0.004 | -0.004 | 0 | 0.36 | 0.64 | 0.45 | 0.24 | 0.38 | 0.22 |
| 5 | 170794695 | rs9313540 | -0.943 | 0.092 | 1.271 | 0.621 | 0.006 | -0.078 | -0.078 | 1.623 | 0.2 | 0.32 | 0.21 | 0.03 | 0.21 | 0.12 |
| 5 | 170795723 | rs7703205 | -0.876 | 0.131 | 1.038 | 0.64 | 0.004 | -0.054 | -0.054 | 0.08 | 0.19 | 0.28 | 0.2 | 0.04 | 0.21 | 0.12 |
| 5 | 170798093 | rs10063817 | -0.789 | 0.13 | 1.065 | 0.639 | 0.003 | -0.045 | -0.045 | 0 | 0.19 | 0.25 | 0.21 | 0.04 | 0.21 | 0.11 |
| 5 | 170801952 | rs7707008 | -0.369 | -0.324 | 0.845 | 0.504 | 0.023 | 0.013 | 0.013 | 0 | 0.23 | 0.1 | 0.09 | 0 | 0.18 | 0.07 |
| 5 | 170814023 | rs28411384 | 0.007 | -0.216 | 0.769 | 0.666 | -0.026 | -0.045 | -0.045 | 0.014 | 0.31 | 0.15 | 0.22 | 0.42 | 0.35 | 0.4 |
| 5 | 170816401 | rs11134696 | -0.435 | -0.246 | 0.728 | 0.771 | 0.001 | 0.003 | 0.003 | 0 | 0.35 | 0.43 | 0.42 | 0.53 | 0.38 | 0.52 |
| 5 | 170817415 | rs17748299 | 2.43 | -0.436 | 0.563 | 0.829 | 0.007 | -0.016 | -0.016 | 0 | 0.07 | 0.02 | 0.05 | 0.12 | 0.03 | 0.01 |
| 5 | 170823601 | rs12189191 | -0.23 | 0.332 | 0.902 | 0.846 | -0.049 | -0.074 | -0.074 | 1.856 | 0.31 | 0.08 | 0.22 | 0.42 | 0.35 | 0.4 |
| 5 | 170838791 | rs11745536 | -1.891 | 0.302 | 0.663 | 0.518 | 0.006 | 0.002 | 0.002 | 0 | 0.58 | 0.53 | 0.52 | 0.53 | 0.56 | 0.6 |
| 5 | 170840504 | rs4302608 | -1.935 | 0.314 | 0.538 | 0.524 | -0.001 | -0.001 | -0.001 | 0 | 0.59 | 0.62 | 0.53 | 0.53 | 0.56 | 0.6 |
| 5 | 170843198 | rs10463007 | -0.777 | 0.602 | -0.13 | 0.903 | 0 | 0.001 | 0.001 | 0 | 0.35 | 0.43 | 0.42 | 0.53 | 0.38 | 0.52 |
| 5 | 170851624 | rs4559013 | -1.582 | 0.51 | -0.14 | 0.873 | 0 | -0.001 | -0.001 | 0 | 0.53 | 0.55 | 0.47 | 0.52 | 0.56 | 0.58 |
| 5 | 170854389 | rs9313543 | 0.093 | 0.084 | -0.062 | 1.151 | 0.001 | -0.009 | -0.009 | 0 | 0.18 | 0.15 | 0.1 | 0.24 | 0.18 | 0.33 |
| 5 | 170863509 | rs4076077 | -1.987 | 0.067 | -0.053 | 1.032 | 0.001 | -0.001 | -0.001 | 0.063 | 0.53 | 0.58 | 0.47 | 0.52 | 0.52 | 0.56 |
| 5 | 170864021 | rs4073717 | -0.639 | -0.235 | -0.047 | 1.053 | -0.003 | -0.003 | -0.003 | 0.82 | 0.23 | 0.26 | 0.23 | 0.18 | 0.21 | 0.21 |
| 5 | 170868059 | rs10065728 | 0.218 | -0.236 | -0.075 | 0.938 | -0.004 | -0.025 | -0.025 | 0.069 | 0.17 | 0.1 | 0.09 | 0.25 | 0.18 | 0.33 |
| 5 | 170869582 | rs10077440 | -1.421 | -0.135 | -0.06 | 0.95 | -0.002 | -0.003 | -0.003 | 0.458 | 0.43 | 0.47 | 0.35 | 0.38 | 0.41 | 0.59 |
| 5 | 170875097 | rs4620037 | -0.629 | 0.353 | 0.068 | 1.548 | -0.001 | -0.001 | -0.001 | 0 | 0.22 | 0.27 | 0.24 | 0.23 | 0.21 | 0.21 |
| 5 | 170875176 | rs6891250 | -0.771 | 0.601 | -0.074 | 1.702 | -0.041 | -0.052 | -0.052 | 0 | 0.1 | 0.04 | 0.13 | 0.17 | 0.14 | 0.04 |
| 5 | 170882607 | rs2043278 | 0.098 | 0.508 | 0.034 | 1.136 | -0.031 | -0.086 | -0.086 | 1.907 | 0.79 | 0.91 | 0.82 | 0.58 | 0.74 | 0.69 |
| 5 | 170886468 | rs734840 | 1.14 | 0.67 | -0.01 | 1.016 | 0 | -0.007 | -0.007 | 0 | 0.53 | 0.46 | 0.54 | 0.34 | 0.48 | 0.47 |
| 5 | 170889081 | rs3112453 | 0.676 | 0.7 | -0.229 | 1.461 | 0.001 | 0.05 | 0.05 | 0.017 | 0.76 | 0.7 | 0.83 | 0.63 | 0.78 | 0.7 |
| 5 | 170898298 | rs3097846 | -0.924 | 0.415 | -0.517 | 1.065 | -0.014 | -0.003 | -0.003 | 1.72 | 0.2 | 0.33 | 0.25 | 0.17 | 0.16 | 0.16 |
| 5 | 170899404 | rs1545033 | 0.346 | 0.278 | -1.494 | 1.587 | -0.034 | -0.139 | -0.139 | 0 | 0.82 | 0.99 | 0.91 | 0.64 | 0.83 | 0.72 |
| 5 | 170901463 | rs10059996 | 0.81 | 0.3 | -1.659 | 0.99 | 0.088 | 0.122 | 0.122 | 0 | 0.51 | 0.22 | 0.59 | 0.32 | 0.62 | 0.34 |
| 5 | 170907977 | rs3112448 | -0.68 | 0.022 | -0.782 | 0.823 | -0.019 | -0.034 | -0.034 | 0 | 0.38 | 0.49 | 0.27 | 0.22 | 0.33 | 0.34 |
| 5 | 170909410 | rs11745375 | 0.916 | 0.578 | -0.916 | 1.149 | -0.022 | 0.015 | 0.015 | 0.06 | 0.38 | 0.27 | 0.44 | 0.31 | 0.45 | 0.22 |
| 5 | 170910612 | rs6884838 | -0.401 | 0.685 | -0.752 | 0.857 | -0.019 | 0.009 | 0.009 | 0 | 0.13 | 0.07 | 0.12 | 0.06 | 0.17 | 0.1 |
| 5 | 170912189 | rs4868092 | -0.084 | 0.549 | -1 | 0.111 | 0.01 | 0.035 | 0.035 | 1.051 | 0.35 | 0.22 | 0.43 | 0.13 | 0.3 | 0.31 |
| 5 | 170912802 | rs4868093 | -0.722 | 0.756 | -0.663 | -0.372 | 0.015 | -0.051 | -0.051 | 0 | 0.91 | 0.79 | 0.92 | 0.99 | 0.89 | 0.94 |
| 5 | 170915499 | rs751937 | -0.128 | 1.331 | -1.037 | 0.198 | -0.036 | -0.044 | -0.044 | 0 | 0.72 | 0.86 | 0.61 | 0.55 | 0.65 | 0.56 |
| 5 | 170915685 | rs751935 | 0.11 | 1.228 | -0.347 | -0.209 | -0.001 | 0.01 | 0.01 | 1.954 | 0.41 | 0.33 | 0.43 | 0.24 | 0.42 | 0.2 |
| 5 | 170916236 | rs3097838 | -0.27 | 1.019 | -0.331 | -0.309 | 0.012 | 0.063 | 0.063 | 0.773 | 0.5 | 0.62 | 0.55 | 0.28 | 0.49 | 0.35 |
| 5 | 170920703 | rs7711912 | 0.015 | 0.693 | -0.747 | -0.246 | 0.011 | 0.044 | 0.044 | 3.101 | 0.45 | 0.69 | 0.51 | 0.28 | 0.41 | 0.36 |
| 5 | 170922132 | rs17652519 | 0.902 | 0.98 | -0.866 | -0.237 | -0.037 | 0.032 | 0.032 | 0 | 0.06 | 0.01 | 0.06 | 0.01 | 0.17 | 0.04 |
| 5 | 170923152 | rs6883796 | 0.642 | 0.976 | -0.741 | -0.239 | 0.004 | -0.007 | -0.007 | 1.425 | 0.06 | 0.08 | 0.07 | 0.01 | 0.17 | 0.04 |
| 5 | 170925376 | rs891971 | -1.22 | 0.558 | -1.119 | -0.156 | 0 | -0.016 | -0.016 | 2.528 | 0.79 | 0.87 | 0.55 | 0.69 | 0.78 | 0.61 |
| 5 | 170926856 | rs7706553 | 0.645 | 0.599 | -0.91 | -0.062 | -0.004 | 0.033 | 0.033 | 2.729 | 0.48 | 0.31 | 0.36 | 0.63 | 0.49 | 0.42 |
| 5 | 170929434 | rs6882257 | -0.567 | 1.014 | -0.439 | 0.108 | 0.054 | 0.017 | 0.017 | 0.533 | 0.19 | 0.56 | 0.21 | 0.14 | 0.28 | 0.16 |
| 5 | 170936089 | rs748915 | -0.042 | 0.455 | -0.314 | 0.802 | -0.002 | 0.167 | 0.167 | 0 | 0.86 | 0.74 | 0.75 | 0.5 | 0.86 | 0.67 |
| 5 | 170936574 | rs919757 | -0.106 | 0.26 | -0.49 | -0.034 | 0.005 | 0.001 | 0.001 | 0 | 0.16 | 0.21 | 0.15 | 0.24 | 0.16 | 0.19 |
| 5 | 170937874 | rs6898626 | -0.636 | 0.063 | -1.061 | -0.265 | -0.003 | 0.037 | 0.037 | 0 | 0.49 | 0.65 | 0.46 | 0.36 | 0.47 | 0.4 |
| 5 | 170948994 | rs17681152 | 0.185 | -0.008 | -1.38 | 0.097 | -0.023 | -0.051 | -0.051 | 0 | 0.2 | 0.02 | 0.3 | 0.29 | 0.19 | 0.15 |
| 5 | 170951257 | rs891981 | -0.716 | -0.442 | -1.044 | -0.15 | 0.018 | 0.007 | 0.007 | 0 | 0.43 | 0.7 | 0.53 | 0.42 | 0.39 | 0.32 |
| 5 | 170951754 | rs10072113 | -0.793 | -0.752 | -0.916 | -0.287 | -0.006 | 0 | 0 | 0 | 0.1 | 0.1 | 0.03 | 0.14 | 0.03 | 0.06 |
| 5 | 170953973 | rs10044726 | -0.907 | -0.564 | -0.899 | -0.578 | -0.01 | 0.002 | 0.002 | 1.095 | 0.61 | 0.7 | 0.62 | 0.4 | 0.55 | 0.31 |
| 5 | 170964248 | rs891980 | -0.313 | -0.367 | -0.166 | -0.514 | -0.038 | -0.146 | -0.146 | 1.636 | 0.21 | 0.51 | 0.12 | 0.03 | 0.13 | 0.07 |
| 5 | 170967262 | rs7732542 | -0.535 | -0.143 | -0.329 | 0.055 | -0.003 | -0.005 | -0.005 | 0 | 0.62 | 0.65 | 0.65 | 0.5 | 0.59 | 0.4 |
| 5 | 170970488 | rs13158791 | -0.168 | -0.286 | -0.459 | -0.193 | -0.008 | 0.066 | 0.066 | 0 | 0.49 | 0.34 | 0.59 | 0.37 | 0.5 | 0.28 |
| 5 | 170971827 | rs7729819 | -0.48 | -0.158 | -0.222 | -0.355 | -0.002 | -0.002 | -0.002 | 2.385 | 0.55 | 0.57 | 0.61 | 0.52 | 0.5 | 0.39 |
| 5 | 170971863 | rs7729835 | -0.725 | -0.213 | -0.116 | -0.268 | -0.001 | 0.003 | 0.003 | 0 | 0.57 | 0.52 | 0.66 | 0.53 | 0.61 | 0.39 |
| 5 | 170972663 | rs11134701 | 0.194 | -0.073 | 0.416 | 0.2 | 0 | 0.06 | 0.06 | 0 | 0.14 | 0.36 | 0.14 | 0.2 | 0.12 | 0.17 |
| 5 | 170975190 | rs6867961 | 0.294 | 0.109 | -0.184 | 0.102 | -0.001 | -0.003 | -0.003 | 0 | 0.21 | 0.21 | 0.39 | 0.24 | 0.21 | 0.13 |
| 5 | 170982892 | rs10059204 | 0.524 | 0.239 | 0.197 | -0.028 | -0.003 | -0.026 | -0.026 | 4.024 | 0.07 | 0.11 | 0.03 | 0 | 0.08 | 0.07 |
| 5 | 170982902 | rs13175457 | 0.419 | 0.576 | 0.141 | 0.486 | -0.036 | -0.061 | -0.061 | 1.053 | 0.26 | 0.11 | 0.2 | 0.43 | 0.29 | 0.33 |
| 5 | 170983223 | rs1864993 | 0.907 | 0.269 | 1.506 | 0.306 | -0.016 | 0.067 | 0.067 | 0.006 | 0.86 | 0.55 | 0.87 | 0.81 | 0.9 | 0.75 |
| 5 | 170992935 | rs4868100 | 0.695 | 0.339 | 1.729 | 0.301 | 0.011 | 0.06 | 0.06 | 0.38 | 0.88 | 0.66 | 0.88 | 0.81 | 0.86 | 0.75 |
| 5 | 170993900 | rs1432963 | 0.893 | 0.333 | 2.026 | 0.119 | 0.043 | 0.024 | 0.024 | 0.086 | 0.8 | 0.47 | 0.78 | 0.77 | 0.73 | 0.67 |
| 5 | 170995850 | rs1432962 | 1.468 | 0.226 | 1.971 | 0.046 | -0.001 | -0.009 | -0.009 | 0 | 0.75 | 0.28 | 0.77 | 0.77 | 0.74 | 0.65 |
| 5 | 170998733 | rs2339319 | 1.494 | 0.315 | 2.09 | 0.06 | 0.007 | -0.012 | -0.012 | 2.479 | 0.76 | 0.35 | 0.77 | 0.77 | 0.74 | 0.65 |
| 5 | 171009918 | rs11134705 | 1.247 | 0.451 | 1.835 | 0.18 | 0.025 | 0.017 | 0.017 | 0.073 | 0.75 | 0.59 | 0.66 | 0.71 | 0.69 | 0.58 |
| 5 | 171010042 | rs11134706 | 1.673 | 0.387 | 1.932 | 0.15 | 0 | -0.001 | -0.001 | 0.84 | 0.71 | 0.29 | 0.64 | 0.71 | 0.68 | 0.58 |
| 5 | 171014336 | rs11134707 | 1.458 | 0.399 | 1.999 | 0.167 | 0.02 | 0.012 | 0.012 | 0.043 | 0.73 | 0.37 | 0.65 | 0.71 | 0.68 | 0.55 |

| | | | | | | | | | | | | | | | | |
|---|---|---|---|---|---|---|---|---|---|---|---|---|---|---|---|---|
| 5 | 171020891 | rs154367 | 0.847 | 0.45 | 1.241 | 0.214 | 0.01 | 0.006 | 0.006 | 0 | 0.81 | 0.75 | 0.8 | 0.77 | 0.73 | 0.66 |
| 5 | 171024600 | rs11134709 | 1.046 | 0.457 | 0.788 | 0.169 | 0.004 | 0.002 | 0.002 | 0.875 | 0.7 | 0.4 | 0.62 | 0.7 | 0.68 | 0.52 |
| 5 | 171036839 | rs1382901 | -0.039 | -0.229 | -0.005 | -0.561 | -0.023 | -0.106 | -0.106 | 1.294 | 0.29 | 0.3 | 0.18 | 0.01 | 0.19 | 0.07 |
| 5 | 171039622 | rs2879407 | 0.551 | 0.225 | -0.676 | 0.17 | 0.028 | -0.043 | -0.043 | 5.108 | 0.35 | 0.23 | 0.38 | 0.42 | 0.3 | 0.31 |
| 5 | 171041805 | rs745443 | 0.304 | 1.285 | -0.081 | 0.42 | -0.007 | -0.005 | -0.005 | 0.039 | 0.79 | 0.82 | 0.74 | 0.69 | 0.69 | 0.68 |
| 5 | 171047194 | rs1678787 | 1.239 | 1.603 | 0.462 | 0.642 | 0.057 | 0.045 | 0.045 | 0.338 | 0.81 | 0.4 | 0.72 | 0.68 | 0.71 | 0.64 |
| 5 | 171047866 | rs2436344 | 1.008 | 0.83 | -0.05 | 0.019 | -0.035 | 0.025 | 0.025 | 1.279 | 0.63 | 0.28 | 0.66 | 0.69 | 0.6 | 0.52 |
| 5 | 171049530 | rs10039848 | 1.042 | 0.142 | 0.022 | 0.12 | -0.001 | 0 | 0 | 0 | 0.51 | 0.56 | 0.66 | 0.52 | 0.54 | 0.47 |
| 5 | 171057231 | rs17681645 | 1.562 | 0.308 | -0.655 | 0.296 | -0.031 | -0.041 | -0.041 | 0.221 | 0.42 | 0.13 | 0.47 | 0.44 | 0.45 | 0.39 |
| 5 | 171060858 | rs257355 | 0.418 | 0.686 | -0.304 | 0.379 | 0.01 | -0.002 | -0.002 | 0 | 0.69 | 0.48 | 0.68 | 0.67 | 0.63 | 0.6 |
| 5 | 171065519 | rs257344 | -0.979 | 0.868 | -0.622 | 0.286 | 0.029 | 0.071 | 0.071 | 0 | 0.49 | 0.65 | 0.33 | 0.26 | 0.37 | 0.45 |
| 5 | 171066113 | rs33375 | 0.047 | 1.504 | -0.727 | 0.933 | -0.003 | 0.001 | 0.001 | 0 | 0.89 | 0.89 | 0.72 | 0.71 | 0.84 | 0.87 |
| 5 | 171068259 | rs254870 | 0.665 | 1.038 | -0.851 | 0.584 | -0.036 | -0.018 | -0.018 | 0.167 | 0.67 | 0.78 | 0.6 | 0.63 | 0.61 | 0.58 |
| 5 | 171069852 | rs7708413 | -0.242 | 0.369 | -0.331 | 0.128 | 0.017 | 0.039 | 0.039 | 0.057 | 0.12 | 0.24 | 0.08 | 0.16 | 0.1 | 0.16 |
| 5 | 171070387 | rs10516086 | -0.195 | 0.222 | -0.858 | -0.09 | 0.022 | 0.035 | 0.035 | 0 | 0.14 | 0.05 | 0.05 | 0 | 0.09 | 0.12 |
| 5 | 171074931 | rs254891 | 0.32 | 0.72 | -0.296 | 0.318 | 0.01 | -0.008 | -0.008 | 0.299 | 0.73 | 0.43 | 0.68 | 0.77 | 0.71 | 0.8 |
| 5 | 171080345 | rs12110228 | -0.237 | 0.745 | -0.296 | 0.575 | 0.005 | -0.01 | -0.01 | 0 | 0.24 | 0.16 | 0.13 | 0.27 | 0.25 | 0.16 |
| 5 | 171081906 | rs257352 | 0.132 | 0.424 | -0.109 | 0.118 | -0.01 | -0.072 | -0.072 | 0.026 | 0.37 | 0.51 | 0.35 | 0.14 | 0.33 | 0.26 |
| 5 | 171093383 | rs26408 | -0.634 | 0.48 | -0.074 | 0.217 | 0.013 | -0.003 | -0.003 | 0 | 0.58 | 0.47 | 0.45 | 0.4 | 0.55 | 0.41 |
| 5 | 171094151 | rs26411 | -0.709 | 0.106 | 0.307 | 0.839 | 0.028 | 0.007 | 0.007 | 0.031 | 0.47 | 0.73 | 0.53 | 0.56 | 0.4 | 0.43 |
| 5 | 171098125 | rs26413 | 0.16 | -0.049 | 0.153 | 0.443 | 0.003 | 0.041 | 0.041 | 0.023 | 0.31 | 0.28 | 0.29 | 0.11 | 0.28 | 0.23 |
| 5 | 171099903 | rs6879367 | -0.199 | 0.114 | 0.038 | 0.708 | -0.008 | 0.04 | 0.04 | 0 | 0.07 | 0.19 | 0.16 | 0.11 | 0.03 | 0.08 |
| 5 | 171101726 | rs33376 | -0.894 | 0.12 | -0.299 | 0.658 | -0.004 | 0.001 | 0.001 | 3.102 | 0.55 | 0.51 | 0.53 | 0.49 | 0.46 | 0.43 |
| 5 | 171101874 | rs12656049 | -0.584 | 0.146 | -0.64 | 1.112 | -0.031 | 0.069 | 0.069 | 0.304 | 0.18 | 0.26 | 0.23 | 0.38 | 0.16 | 0.12 |
| 5 | 171102163 | rs7722148 | -0.583 | 0.703 | -1.199 | 0.737 | 0.01 | 0.039 | 0.039 | 0.98 | 0.39 | 0.72 | 0.5 | 0.29 | 0.33 | 0.33 |
| 5 | 171103318 | rs931117 | -0.792 | 1.483 | -1.217 | 1.306 | -0.021 | -0.011 | -0.011 | 0 | 0.18 | 0.13 | 0.15 | 0.22 | 0.2 | 0.15 |
| 5 | 171104059 | rs194231 | 1.076 | 1.305 | -1.346 | 2.234 | -0.026 | -0.027 | -0.027 | 0 | 0.92 | 0.99 | 0.71 | 0.43 | 0.9 | 0.77 |
| 5 | 171105669 | rs33874 | 0.771 | 1.336 | -0.874 | 1.977 | 0.011 | 0.049 | 0.049 | 0.015 | 0.78 | 0.33 | 0.71 | 0.63 | 0.79 | 0.76 |
| 5 | 171106689 | rs11134715 | 0.588 | 1.005 | -1.624 | 0.229 | 0.029 | 0.09 | 0.09 | 0.901 | 0.36 | 0.2 | 0.17 | 0.01 | 0.24 | 0.3 |
| 5 | 171107253 | rs11134716 | 0.01 | 1.145 | -1.043 | 0.723 | -0.002 | -0.002 | -0.002 | 0.222 | 0.11 | 0.1 | 0.16 | 0.17 | 0.13 | 0.12 |
| 5 | 171112476 | rs6555968 | -0.149 | 0.852 | -0.859 | 0.086 | 0 | -0.001 | -0.001 | 0 | 0.59 | 0.65 | 0.52 | 0.34 | 0.52 | 0.54 |
| 5 | 171112535 | rs6893055 | 0.935 | 0.658 | -1.453 | -0.39 | -0.043 | -0.005 | -0.005 | 0 | 0.25 | 0.02 | 0.17 | 0.18 | 0.25 | 0.16 |
| 5 | 171115557 | rs7729995 | -0.214 | 1.095 | -1.018 | -0.475 | 0.008 | -0.053 | -0.053 | 0 | 0.35 | 0.6 | 0.35 | 0.16 | 0.3 | 0.4 |
| 5 | 171115769 | rs7710392 | -0.38 | 0.845 | -1.004 | -0.697 | 0.024 | -0.033 | -0.033 | 0 | 0.19 | 0.59 | 0.26 | 0.13 | 0.14 | 0.25 |
| 5 | 171119140 | rs2339325 | -0.084 | 0.5 | -0.634 | -0.671 | -0.026 | -0.094 | -0.094 | 2.539 | 0.25 | 0.47 | 0.14 | 0.04 | 0.17 | 0.22 |
| 5 | 171125482 | rs11134717 | -0.131 | -0.104 | -0.718 | -1.017 | -0.006 | 0.041 | 0.041 | 1.399 | 0.55 | 0.67 | 0.45 | 0.28 | 0.45 | 0.49 |
| 5 | 171125658 | rs11750546 | 0.788 | -0.105 | -0.258 | -1.151 | -0.003 | -0.043 | -0.043 | 0 | 0.32 | 0.39 | 0.2 | 0.14 | 0.28 | 0.22 |
| 5 | 171131700 | rs254907 | 0.076 | 0.123 | -0.849 | -0.5 | 0.012 | -0.056 | -0.056 | 0 | 0.67 | 0.17 | 0.6 | 0.58 | 0.68 | 0.66 |
| 5 | 171133746 | rs12653066 | 1.486 | 0.43 | -0.693 | -0.341 | -0.021 | -0.004 | -0.004 | 0 | 0.1 | 0.05 | 0.29 | 0.36 | 0.26 | 0.11 |
| 5 | 171134072 | rs1835934 | -0.189 | 0.308 | -0.77 | -0.451 | -0.005 | -0.009 | -0.009 | 0 | 0.3 | 0.26 | 0.42 | 0.41 | 0.4 | 0.26 |
| 5 | 171143166 | rs171718 | -0.107 | 0.631 | -0.876 | -0.153 | -0.057 | -0.069 | -0.069 | 0 | 0.63 | 0.16 | 0.54 | 0.5 | 0.58 | 0.65 |
| 5 | 171150423 | rs10516088 | 0.898 | 0.349 | -1.183 | -0.262 | -0.093 | 0.056 | 0.056 | 7.815 | 0.33 | 0.05 | 0.23 | 0.19 | 0.36 | 0.22 |
| 5 | 171151231 | rs10516089 | 1.139 | 0.388 | -0.513 | -0.249 | 0.007 | 0.049 | 0.049 | 0.819 | 0.27 | 0.37 | 0.37 | 0.38 | 0.32 | 0.2 |
| 5 | 171157267 | rs3101876 | -0.701 | 0.471 | -0.875 | -0.205 | 0.026 | 0.022 | 0.022 | 0 | 0.93 | 0.86 | 0.94 | 0.88 | 0.91 | 0.86 |
| 5 | 171161075 | rs7734319 | 0.237 | 0.435 | -0.998 | -0.231 | -0.005 | -0.004 | -0.004 | 0.57 | 0.59 | 0.72 | 0.58 | 0.53 | 0.6 | 0.4 |
| 5 | 171162858 | rs33858 | -1.345 | 0.675 | -1.359 | -0.285 | 0.025 | 0.008 | 0.008 | 0 | 0.94 | 0.88 | 0.94 | 0.92 | 0.92 | 0.92 |
| 5 | 171162922 | rs17568593 | 1.154 | 0.47 | -1.689 | -0.361 | -0.046 | 0.078 | 0.078 | 7.83 | 0.29 | 0.03 | 0.18 | 0.11 | 0.28 | 0.14 |
| 5 | 171168572 | rs12658131 | 0.789 | 0.398 | -1.215 | -0.173 | 0.004 | 0.024 | 0.024 | 0.845 | 0.26 | 0.29 | 0.35 | 0.38 | 0.31 | 0.19 |
| 5 | 171172825 | rs13184840 | 0.342 | 0.279 | -1.747 | -0.509 | 0.007 | 0.06 | 0.06 | 0 | 0.16 | 0.05 | 0.12 | 0.04 | 0.13 | 0.19 |
| 5 | 171174165 | rs3734102 | 0.418 | 0.505 | -1.995 | -0.544 | -0.05 | 0.118 | 0.118 | 5.542 | 0.28 | 0.03 | 0.18 | 0.08 | 0.27 | 0.16 |
| 5 | 171174351 | rs11960939 | 0.147 | -0.027 | -1.495 | -0.156 | -0.002 | 0.001 | 0.001 | 0 | 0.57 | 0.52 | 0.58 | 0.53 | 0.6 | 0.45 |
| 5 | 171175270 | rs33878 | -0.032 | -0.091 | -1.079 | 0.246 | 0.001 | -0.005 | -0.005 | 0.028 | 0.24 | 0.23 | 0.32 | 0.35 | 0.22 | 0.33 |
| 5 | 171175454 | rs32474 | -0.609 | -0.513 | -1.251 | -0.128 | -0.009 | 0.032 | 0.032 | 0 | 0.76 | 0.82 | 0.77 | 0.86 | 0.73 | 0.76 |
| 5 | 171175659 | rs26415 | 0.591 | -0.452 | -0.985 | -0.129 | 0 | -0.004 | -0.004 | 0 | 0.53 | 0.63 | 0.46 | 0.51 | 0.51 | 0.43 |
| 5 | 171176078 | rs33880 | 0.629 | -0.734 | -0.614 | -0.358 | 0.016 | 0.043 | 0.043 | 0 | 0.68 | 0.16 | 0.69 | 0.8 | 0.72 | 0.65 |
| 5 | 171177694 | rs6880586 | -0.025 | -0.703 | -0.295 | -0.037 | -0.012 | 0.027 | 0.027 | 0 | 0.05 | 0.2 | 0.04 | 0.05 | 0.01 | 0.11 |
| 5 | 171181237 | rs153750 | -0.774 | -0.701 | -1.227 | -0.082 | -0.027 | -0.029 | -0.029 | 4.872 | 0.66 | 0.91 | 0.7 | 0.58 | 0.63 | 0.75 |
| 5 | 171188178 | rs4868120 | -0.008 | -0.597 | -0.805 | -0.094 | 0.001 | -0.022 | -0.022 | 0 | 0.23 | 0.34 | 0.15 | 0.12 | 0.21 | 0.3 |
| 5 | 171189571 | rs33852 | 0.877 | -0.779 | -1.204 | -0.264 | -0.008 | -0.031 | -0.031 | 0 | 0.39 | 0.08 | 0.43 | 0.33 | 0.34 | 0.37 |
| 5 | 171193790 | rs33857 | 1.768 | -0.869 | -1.32 | -0.287 | -0.047 | -0.086 | -0.086 | 0 | 0.24 | 0.02 | 0.3 | 0.26 | 0.22 | 0.25 |
| 5 | 171194238 | rs153741 | 0.204 | -0.904 | -1.204 | -0.322 | -0.009 | 0.008 | 0.008 | 0 | 0.46 | 0.26 | 0.44 | 0.37 | 0.43 | 0.55 |
| 5 | 171195033 | rs249516 | 0.168 | -0.783 | -1.056 | -0.338 | -0.018 | -0.048 | -0.048 | 0 | 0.45 | 0.21 | 0.51 | 0.46 | 0.44 | 0.48 |
| 5 | 171197597 | rs697036 | -0.129 | 0.409 | -0.192 | 0.289 | -0.01 | 0.091 | 0.091 | 0 | 0.33 | 0.12 | 0.25 | 0.13 | 0.31 | 0.31 |
| 5 | 171202017 | rs26401 | -0.772 | 1.33 | 1.267 | 1.035 | 0.066 | 0.089 | 0.089 | 0 | 0.27 | 0.54 | 0.38 | 0.53 | 0.43 | 0.26 |
| 5 | 171203248 | rs702072 | 0.56 | 1.424 | 0.906 | 0.98 | -0.005 | 0.009 | 0.009 | 0 | 0.14 | 0.07 | 0.09 | 0.07 | 0.16 | 0.2 |
| 5 | 171203724 | rs26406 | -0.412 | 1.551 | 1.389 | 1.496 | 0.04 | 0.031 | 0.031 | 0 | 0.07 | 0.22 | 0.08 | 0.11 | 0.12 | 0.09 |
| 5 | 171204055 | rs6869153 | -1.558 | 1.53 | 1.322 | 1.507 | 0.009 | 0.036 | 0.036 | 0.084 | 0.12 | 0.26 | 0.1 | 0.17 | 0.12 | 0.24 |
| 5 | 171207593 | rs17073971 | -0.92 | 0.957 | 1.604 | 1.321 | -0.021 | 0.039 | 0.039 | 0 | 0.05 | 0.21 | 0.04 | 0.06 | 0.01 | 0.15 |
| 5 | 171210392 | rs11948648 | -0.625 | 1.082 | 0.922 | 1.39 | 0.036 | 0.062 | 0.062 | 0 | 0.19 | 0.29 | 0.31 | 0.43 | 0.32 | 0.13 |

| | | | | | | | | | | | | | | | | |
|---|---|---|---|---|---|---|---|---|---|---|---|---|---|---|---|---|
| 5 | 171213245 | rs697042 | -1.275 | 1.207 | 0.687 | 1.448 | -0.005 | -0.012 | -0.012 | 0.034 | 0.25 | 0.68 | 0.34 | 0.48 | 0.32 | 0.23 |
| 5 | 171214479 | rs702077 | -1.961 | 0.345 | 1.109 | 1.018 | -0.018 | 0.052 | 0.052 | 0.165 | 0.09 | 0.51 | 0.06 | 0.06 | 0.02 | 0.16 |
| 5 | 171239418 | rs10463010 | -0.792 | 1.292 | -0.452 | 1.338 | -0.024 | -0.04 | -0.04 | 0 | 0.12 | 0.08 | 0.23 | 0.47 | 0.13 | 0.18 |
| 5 | 171240222 | rs11948630 | -0.436 | 1.144 | 0.652 | 1.083 | 0.048 | 0.014 | 0.014 | 0.133 | 0.07 | 0.38 | 0.07 | 0.01 | 0.06 | 0.02 |
| 5 | 171241767 | rs11134724 | 0.216 | 1.204 | 0.158 | 0.213 | 0.003 | 0.106 | 0.106 | 0.345 | 0.36 | 0.16 | 0.22 | 0.06 | 0.37 | 0.18 |
| 5 | 171242214 | rs838794 | -0.619 | 1.265 | 0.475 | 0.261 | 0.005 | -0.007 | -0.007 | 0.11 | 0.49 | 0.28 | 0.47 | 0.54 | 0.53 | 0.37 |
| 5 | 171242519 | rs838792 | -0.21 | 1.157 | 0.748 | 0.035 | 0.078 | 0.037 | 0.037 | 0.386 | 0.07 | 0.46 | 0.09 | 0.01 | 0.07 | 0.02 |
| 5 | 171245547 | rs10067974 | -0.157 | 1.265 | 0.3 | -0.163 | 0.007 | 0.058 | 0.058 | 1.483 | 0.41 | 0.27 | 0.23 | 0.06 | 0.41 | 0.2 |
| 5 | 171246209 | rs11742661 | -1.174 | 1.209 | 0.523 | 0.144 | 0.005 | -0.017 | -0.017 | 0 | 0.6 | 0.71 | 0.59 | 0.54 | 0.64 | 0.42 |
| 5 | 171249781 | rs17074035 | -0.218 | 0.687 | 0.736 | -0.52 | 0.03 | -0.014 | -0.014 | 0 | 0.07 | 0.28 | 0.06 | 0.01 | 0.07 | 0.02 |
| 5 | 171250237 | rs10039348 | -0.481 | 0.729 | -0.123 | -0.592 | 0.004 | -0.047 | -0.047 | 0.059 | 0.17 | 0.19 | 0.1 | 0 | 0.19 | 0.07 |
| 5 | 171252847 | rs11740782 | -0.297 | 0.882 | -0.484 | -0.334 | -0.025 | 0.174 | 0.174 | 2.355 | 0.34 | 0.07 | 0.17 | 0.05 | 0.34 | 0.2 |
| 5 | 171260128 | rs1529702 | 1.432 | 1.093 | 0.921 | 0.649 | 0.097 | 0.102 | 0.102 | 0.009 | 0.92 | 0.63 | 0.82 | 0.87 | 0.88 | 0.97 |
| 5 | 171260919 | rs13165807 | 0.175 | -0.013 | 0.066 | 0.443 | 0.127 | 0.086 | 0.086 | 0.266 | 0.51 | 0.16 | 0.6 | 0.67 | 0.71 | 0.45 |
| 5 | 171265304 | rs702090 | 1.186 | 0.466 | 0.993 | 0.647 | 0.04 | -0.023 | -0.023 | 0 | 0.94 | 0.78 | 0.94 | 1 | 0.94 | 0.99 |
| 5 | 171267207 | rs6555975 | 1.528 | 0.403 | 0.971 | 1.051 | 0.009 | 0.029 | 0.029 | 0 | 0.91 | 0.8 | 0.81 | 0.86 | 0.88 | 0.85 |
| 5 | 171268203 | rs6891834 | 0.443 | 0.099 | 1.244 | 0.675 | 0.021 | 0.014 | 0.014 | 0 | 0.41 | 0.31 | 0.3 | 0.29 | 0.19 | 0.54 |
| 5 | 171268780 | rs6555976 | 0.174 | 0.795 | 1.277 | 0.594 | 0.009 | 0.009 | 0.009 | 0 | 0.46 | 0.49 | 0.36 | 0.3 | 0.29 | 0.61 |
| 5 | 171272773 | rs11749347 | 0.454 | 0.422 | 1.011 | 0.847 | 0.017 | -0.005 | -0.005 | 0 | 0.42 | 0.32 | 0.32 | 0.48 | 0.22 | 0.58 |
| 5 | 171273406 | rs6890298 | 0.207 | 0.424 | 0.824 | 0.847 | 0.023 | -0.003 | -0.003 | 0 | 0.44 | 0.53 | 0.34 | 0.49 | 0.22 | 0.59 |
| 5 | 171274225 | rs702097 | 0.143 | 0.803 | 0.771 | 0.972 | 0.062 | -0.005 | -0.005 | 0 | 0.52 | 0.68 | 0.39 | 0.51 | 0.3 | 0.61 |
| 5 | 171274708 | rs13153931 | 0.223 | 1.047 | 0.259 | 1.173 | 0.036 | 0.015 | 0.015 | 0.499 | 0.49 | 0.3 | 0.39 | 0.56 | 0.36 | 0.6 |
| 5 | 171274885 | rs702098 | 0.111 | 0.717 | 0.113 | 1.099 | 0.005 | 0 | 0 | 0.059 | 0.54 | 0.55 | 0.38 | 0.55 | 0.32 | 0.59 |
| 5 | 171283469 | rs4868126 | 0.387 | 1.27 | 0.195 | 1.135 | 0.015 | 0.023 | 0.023 | 0.079 | 0.71 | 0.35 | 0.66 | 0.62 | 0.64 | 0.79 |
| 5 | 171319387 | rs2569091 | 1.469 | 1.039 | 2.951 | 0.692 | 0.08 | 0.003 | 0.003 | 0.439 | 0.92 | 0.54 | 0.91 | 1 | 0.93 | 0.97 |
| 5 | 171324289 | rs17682767 | -0.533 | 0.863 | 2.56 | 0.773 | -0.005 | -0.021 | -0.021 | 0 | 0.07 | 0.01 | 0.12 | 0.09 | 0.06 | 0.08 |
| 5 | 171335293 | rs10475992 | 0.494 | 0.575 | 1.914 | 0.667 | -0.033 | 0.072 | 0.072 | 0 | 0.46 | 0.13 | 0.53 | 0.35 | 0.53 | 0.54 |
| 5 | 171341849 | rs9313564 | 0.232 | 0.342 | 2.277 | 0.732 | 0.022 | 0.014 | 0.014 | 0 | 0.44 | 0.31 | 0.35 | 0.59 | 0.4 | 0.39 |
| 5 | 171386091 | rs9313565 | 0.717 | -0.015 | 1.935 | 0.311 | -0.011 | -0.099 | -0.099 | 0.116 | 0.84 | 0.63 | 0.87 | 0.98 | 0.87 | 0.79 |
| 5 | 171401779 | rs4868133 | 0.937 | -0.081 | 2.041 | 0.429 | -0.017 | -0.112 | -0.112 | 2.987 | 0.36 | 0.05 | 0.21 | 0.52 | 0.34 | 0.31 |
| 5 | 171409661 | rs10080218 | 1.442 | 0.14 | 2.745 | 0.518 | 0.036 | -0.03 | -0.03 | 0 | 0.92 | 0.72 | 0.89 | 0.98 | 0.91 | 0.86 |
| 5 | 171419658 | rs7707358 | -1.409 | 0.233 | 2.748 | 0.589 | 0.075 | 0.007 | 0.007 | 0.222 | 0.07 | 0.38 | 0.08 | 0 | 0.07 | 0.01 |
| 5 | 171443065 | rs10038862 | -1.926 | 0.261 | 2.786 | 0.602 | 0.078 | 0.001 | 0.001 | 0.156 | 0.08 | 0.47 | 0.08 | 0 | 0.07 | 0.01 |
| 5 | 171446570 | rs10068774 | -1.179 | 0.271 | 2.692 | 0.607 | 0.07 | 0.005 | 0.005 | 0.085 | 0.06 | 0.36 | 0.07 | 0 | 0.07 | 0.01 |
| 5 | 171465421 | rs7715886 | -1.789 | 0.234 | 2.274 | 0.589 | 0.066 | -0.005 | -0.005 | 0.366 | 0.08 | 0.5 | 0.09 | 0 | 0.07 | 0.01 |
| 5 | 171468278 | rs7727588 | -1.868 | 0.082 | 1.817 | 0.609 | -0.027 | -0.035 | -0.035 | 0 | 0.14 | 0.69 | 0.11 | 0.05 | 0.08 | 0.05 |
| 5 | 171469833 | rs6555988 | -1.094 | 0.254 | 1.61 | 0.674 | 0.031 | 0.016 | 0.016 | 1.422 | 0.11 | 0.5 | 0.09 | 0.05 | 0.08 | 0.03 |
| 5 | 171472575 | rs4868136 | -0.418 | 0.67 | 0.372 | 1.112 | 0.022 | -0.005 | -0.005 | 0 | 0.48 | 0.38 | 0.56 | 0.54 | 0.61 | 0.51 |
| 5 | 171475689 | rs10035578 | 0.935 | 2.292 | 0.29 | 1.934 | -0.012 | -0.001 | -0.001 | 0.193 | 0.89 | 0.92 | 0.64 | 0.6 | 0.76 | 0.7 |
| 5 | 171479787 | rs3812024 | -0.83 | 1.376 | -0.156 | 1.02 | 0.024 | 0.075 | 0.075 | 0.059 | 0.56 | 0.92 | 0.74 | 0.73 | 0.6 | 0.42 |
| 5 | 171480533 | rs4075704 | 0.191 | 2.122 | -0.297 | 1.377 | -0.037 | -0.014 | -0.014 | 2.097 | 0.07 | 0.03 | 0.38 | 0.36 | 0.18 | 0.13 |
| 5 | 171480733 | rs4075705 | -0.408 | 1.262 | 0.652 | 0.64 | -0.001 | 0.005 | 0.005 | 0 | 0.43 | 0.39 | 0.31 | 0.4 | 0.41 | 0.29 |
| 5 | 171481606 | rs10155597 | -0.525 | 1.328 | 0.502 | 0.652 | -0.001 | -0.007 | -0.007 | 0.005 | 0.45 | 0.47 | 0.33 | 0.4 | 0.42 | 0.29 |
| 5 | 171484604 | rs17074262 | -0.996 | 0.33 | 0.47 | -0.286 | -0.025 | -0.047 | -0.047 | 0 | 0.21 | 0.29 | 0.12 | 0.04 | 0.1 | 0.23 |
| 5 | 171484631 | rs13157965 | 0.201 | 0.952 | 0.597 | -0.182 | -0.011 | 0.013 | 0.013 | 0 | 0.24 | 0.14 | 0.22 | 0.2 | 0.26 | 0.1 |
| 5 | 171485293 | rs4379217 | -1.166 | 0.534 | 0.827 | -0.19 | -0.037 | -0.098 | -0.098 | 0 | 0.17 | 0.45 | 0.11 | 0.03 | 0.1 | 0.21 |
| 5 | 171491332 | rs3812023 | -0.691 | 1.09 | 0.655 | 0.3 | 0 | 0.001 | 0.001 | 0 | 0.42 | 0.52 | 0.64 | 0.53 | 0.55 | 0.33 |
| 5 | 171491682 | rs3812021 | -0.748 | 1.027 | 1.242 | 0.648 | -0.004 | -0.025 | -0.025 | 3.4 | 0.08 | 0.24 | 0.07 | 0.03 | 0.05 | 0.13 |
| 5 | 171494680 | rs4868137 | -1.482 | 1.363 | 1.256 | 0.762 | 0.025 | 0.025 | 0.025 | 0 | 0.21 | 0.32 | 0.45 | 0.4 | 0.34 | 0.32 |
| 5 | 171503796 | rs17074274 | -1.568 | 1.23 | 1.876 | -0.045 | 0.037 | -0.096 | -0.096 | 0 | 0.15 | 0.43 | 0.14 | 0.02 | 0.14 | 0.12 |
| 5 | 171507531 | rs6866684 | -1.397 | 1.144 | 1.688 | -0.13 | 0.011 | 0.031 | 0.031 | 1.147 | 0.15 | 0.48 | 0.12 | 0.13 | 0.12 | 0.05 |
| 5 | 171509644 | rs11741056 | -2.418 | 1.153 | 1.484 | -0.136 | -0.001 | 0.014 | 0.014 | 1.329 | 0.29 | 0.73 | 0.25 | 0.16 | 0.26 | 0.17 |
| 5 | 171510180 | rs10516093 | -1.432 | 1.416 | 1.674 | 0.03 | 0.016 | -0.064 | -0.064 | 2.555 | 0.14 | 0.27 | 0.13 | 0.03 | 0.13 | 0.12 |
| 5 | 171515242 | rs4467720 | -2.149 | 1.231 | 0.967 | 0.061 | 0.001 | -0.009 | -0.009 | 0.134 | 0.23 | 0.71 | 0.21 | 0.14 | 0.21 | 0.15 |
| 5 | 171519290 | rs10043350 | -2.059 | 0.971 | 0.819 | -0.093 | 0.003 | 0.006 | 0.006 | 2.744 | 0.29 | 0.72 | 0.21 | 0.14 | 0.21 | 0.16 |
| 5 | 171522275 | rs10053012 | -1.117 | 1.226 | 0.368 | -0.07 | 0.001 | -0.009 | -0.009 | 1.746 | 0.18 | 0.18 | 0.11 | 0.04 | 0.12 | 0.11 |
| 5 | 171523436 | rs4868141 | -0.421 | 1.81 | 0.247 | 0.593 | -0.015 | -0.014 | -0.014 | 1.498 | 0.08 | 0.06 | 0.23 | 0.19 | 0.19 | 0.14 |
| 5 | 171529235 | rs11745517 | -1.521 | 1.272 | 0.458 | 0.235 | 0.001 | 0.003 | 0.003 | 0 | 0.07 | 0.07 | 0.06 | 0.09 | 0.08 | 0.06 |
| 5 | 171529721 | rs6870207 | -2.211 | 1.286 | 0.847 | 0.208 | 0.008 | 0.01 | 0.01 | 0 | 0.09 | 0.1 | 0.07 | 0.1 | 0.1 | 0.08 |
| 5 | 171529948 | rs11134728 | 2.246 | 1.519 | 1.586 | -0.128 | 0.037 | -0.036 | -0.036 | 0 | 0.77 | 0.58 | 0.76 | 0.87 | 0.75 | 0.79 |
| 5 | 171531953 | rs3103593 | 1.143 | 1.517 | 1.618 | -0.102 | 0.027 | -0.045 | -0.045 | 0 | 0.87 | 0.75 | 0.85 | 0.97 | 0.86 | 0.88 |
| 5 | 171533629 | rs2306963 | 2.011 | 1.619 | 1.892 | -0.15 | 0.03 | -0.022 | -0.022 | 0 | 0.78 | 0.68 | 0.77 | 0.85 | 0.74 | 0.78 |
| 5 | 171542877 | rs3111491 | -2.551 | 1.039 | 1.968 | -0.115 | -0.011 | 0.012 | 0.012 | 0 | 0.14 | 0.28 | 0.14 | 0.12 | 0.1 | 0.09 |
| 5 | 171547550 | rs3776763 | -1.166 | 1.206 | 1.796 | -0.077 | 0.01 | 0.027 | 0.027 | 0 | 0.05 | 0.06 | 0.09 | 0.11 | 0.06 | 0.03 |
| 5 | 171552219 | rs7706554 | -1.493 | 1.16 | 1.931 | -0.264 | 0.015 | -0.043 | -0.043 | 0 | 0.13 | 0.27 | 0.13 | 0.02 | 0.15 | 0.12 |
| 5 | 171554473 | rs2279514 | 2.097 | 1.008 | 1.69 | -0.396 | 0 | -0.001 | -0.001 | 0 | 0.71 | 0.29 | 0.71 | 0.87 | 0.73 | 0.77 |
| 5 | 171556078 | rs3776760 | -2.085 | 0.754 | 1.96 | -0.305 | 0.008 | -0.032 | -0.032 | 0 | 0.11 | 0.3 | 0.09 | 0.03 | 0.1 | 0.08 |
| 5 | 171556788 | rs3756508 | -1.675 | 1.111 | 1.722 | -0.635 | -0.004 | 0.005 | 0.005 | 0 | 0.36 | 0.35 | 0.31 | 0.25 | 0.37 | 0.27 |
| 5 | 171557137 | rs3776759 | -1.279 | 1.011 | 2.278 | -0.76 | 0.024 | -0.025 | -0.025 | 0 | 0.24 | 0.52 | 0.24 | 0.21 | 0.24 | 0.08 |

| | | | | | | | | | | | | | | | | |
|---|---|---|---|---|---|---|---|---|---|---|---|---|---|---|---|---|
| 5 | 171559860 | rs9313577 | -2.307 | 0.968 | 2.133 | -0.78 | -0.019 | 0.043 | 0.043 | 0 | 0.38 | 0.75 | 0.35 | 0.25 | 0.37 | 0.23 |
| 5 | 171560741 | rs902498 | -0.578 | 1.056 | 1.714 | -0.733 | -0.02 | 0 | 0 | 0.722 | 0.19 | 0.11 | 0.21 | 0.21 | 0.23 | 0.08 |
| 5 | 171571754 | rs6878620 | -1.53 | 0.886 | 1.728 | -0.799 | 0.002 | -0.024 | -0.024 | 0 | 0.14 | 0.19 | 0.1 | 0.04 | 0.13 | 0.11 |
| 5 | 171572577 | rs3776754 | -2.035 | 0.956 | 1.461 | -0.718 | -0.002 | 0.003 | 0.003 | 0 | 0.44 | 0.42 | 0.37 | 0.33 | 0.47 | 0.35 |
| 5 | 171575460 | rs10050886 | -1.376 | 0.27 | 1.241 | -0.531 | -0.055 | -0.074 | -0.074 | 0 | 0.2 | 0.44 | 0.12 | 0.07 | 0.12 | 0.2 |
| 5 | 171576629 | rs3797328 | -1.216 | 0.792 | -0.019 | -0.207 | -0.003 | -0.026 | -0.026 | 0 | 0.46 | 0.75 | 0.36 | 0.53 | 0.44 | 0.47 |
| 5 | 171577700 | rs6871819 | -1.263 | 0.921 | -0.464 | -0.168 | -0.044 | -0.071 | -0.071 | 0 | 0.46 | 0.87 | 0.37 | 0.55 | 0.44 | 0.47 |
| 5 | 171579380 | rs10223190 | -0.187 | 1.008 | 0.01 | -0.198 | 0.001 | -0.037 | -0.037 | 1.225 | 0.1 | 0.16 | 0.07 | 0 | 0.1 | 0.15 |
| 5 | 171579797 | rs10223207 | -0.267 | 1.002 | 0.056 | -0.2 | 0.001 | -0.042 | -0.042 | 0.277 | 0.11 | 0.17 | 0.07 | 0 | 0.1 | 0.15 |
| 5 | 171581445 | rs4868146 | -0.568 | 0.452 | 0.01 | -0.455 | 0.002 | 0.009 | 0.009 | 0 | 0.36 | 0.64 | 0.3 | 0.31 | 0.36 | 0.24 |
| 5 | 171581724 | rs3776751 | -0.539 | 0.76 | -0.305 | 0.29 | 0.001 | 0.052 | 0.052 | 0 | 0.16 | 0.23 | 0.31 | 0.36 | 0.15 | 0.18 |
| 5 | 171590162 | rs17074352 | -0.653 | 0.694 | -0.354 | 0.237 | -0.003 | 0.016 | 0.016 | 0 | 0.16 | 0.23 | 0.26 | 0.25 | 0.14 | 0.2 |
| 5 | 171590679 | rs10064789 | -0.061 | 0.964 | 0.449 | 0.133 | 0.054 | 0.01 | 0.01 | 0.094 | 0.06 | 0.28 | 0.06 | 0.03 | 0.07 | 0.1 |
| 5 | 171593223 | rs2339363 | -1.055 | 0.977 | 0.233 | 0.184 | 0 | 0.02 | 0.02 | 0 | 0.51 | 0.36 | 0.58 | 0.61 | 0.5 | 0.45 |
| 5 | 171593466 | rs11134733 | -1.963 | 0.972 | 0.471 | 0.173 | -0.019 | -0.012 | -0.012 | 0.218 | 0.6 | 0.85 | 0.69 | 0.64 | 0.57 | 0.55 |
| 5 | 171593748 | rs17074365 | -0.096 | 0.662 | 0.789 | 0.109 | 0.057 | 0.014 | 0.014 | 0 | 0.06 | 0.24 | 0.06 | 0.03 | 0.07 | 0.1 |
| 5 | 171597483 | rs3797326 | -2.204 | 0.26 | 0.349 | 0.16 | 0.036 | 0.036 | 0.036 | 0.855 | 0.44 | 0.84 | 0.53 | 0.34 | 0.35 | 0.45 |
| 5 | 171598868 | rs6863178 | -0.926 | -0.068 | 0.753 | 0.041 | -0.013 | -0.089 | -0.089 | 0.052 | 0.22 | 0.37 | 0.27 | 0.03 | 0.19 | 0.15 |
| 5 | 171600193 | rs10475571 | -0.359 | -0.075 | 0.534 | 0.1 | -0.019 | 0.065 | 0.065 | 0.036 | 0.15 | 0.28 | 0.16 | 0.29 | 0.09 | 0.21 |
| 5 | 171602932 | rs3776747 | -0.824 | -0.047 | -0.201 | 0.111 | -0.038 | 0.053 | 0.053 | 0.275 | 0.18 | 0.62 | 0.18 | 0.29 | 0.09 | 0.21 |
| 5 | 171603387 | rs7735779 | -2.251 | -0.056 | -0.233 | 0.08 | 0.046 | 0.028 | 0.028 | 0.388 | 0.39 | 0.83 | 0.44 | 0.32 | 0.28 | 0.35 |
| 5 | 171603528 | rs17700365 | -0.719 | 0.062 | -0.358 | 0.095 | 0.002 | 0.001 | 0.001 | 0 | 0.21 | 0.21 | 0.26 | 0.03 | 0.19 | 0.14 |
| 5 | 171605562 | rs2172048 | -2.066 | 0.306 | -0.715 | 0.088 | 0.031 | 0.087 | 0.087 | 1.617 | 0.48 | 0.86 | 0.52 | 0.34 | 0.41 | 0.47 |
| 5 | 171616404 | rs17700489 | -0.081 | 0.239 | -0.49 | 0.059 | 0 | -0.007 | -0.007 | 0.425 | 0.11 | 0.13 | 0.19 | 0.02 | 0.08 | 0.09 |
| 5 | 171623828 | rs6874860 | -1.927 | -0.001 | -0.776 | 0.347 | 0.058 | 0.006 | 0.006 | 0.095 | 0.41 | 0.84 | 0.49 | 0.37 | 0.29 | 0.42 |
| 5 | 171626110 | rs17074413 | -0.785 | 0.112 | -0.318 | 0.288 | -0.017 | -0.042 | -0.042 | 0.019 | 0.15 | 0.27 | 0.22 | 0.03 | 0.09 | 0.13 |
| 5 | 171627391 | rs10070923 | -0.196 | 0.13 | -1.138 | 0.057 | -0.013 | -0.006 | -0.006 | 0 | 0.63 | 0.71 | 0.69 | 0.63 | 0.63 | 0.48 |
| 5 | 171632058 | rs6890478 | 2.27 | -0.025 | -1.687 | 0.094 | -0.095 | -0.032 | -0.032 | 0 | 0.3 | 0.01 | 0.3 | 0.29 | 0.39 | 0.14 |
| 5 | 171632616 | rs6876698 | -0.287 | 0.156 | -1.536 | 0.126 | -0.007 | -0.009 | -0.009 | 0 | 0.63 | 0.78 | 0.71 | 0.64 | 0.65 | 0.44 |
| 5 | 171633449 | rs17074437 | 0.264 | 0.135 | -1.36 | 0.128 | -0.001 | 0.04 | 0.04 | 0 | 0.08 | 0.1 | 0.11 | 0.31 | 0.07 | 0.13 |
| 5 | 171633632 | rs9313585 | -0.123 | 0.077 | -1.416 | -0.02 | 0.001 | 0 | 0 | 0 | 0.61 | 0.72 | 0.71 | 0.63 | 0.65 | 0.4 |
| 5 | 171634194 | rs10491494 | 1.364 | -0.131 | -1.456 | -0.277 | 0.003 | 0.01 | 0.01 | 0 | 0.07 | 0.05 | 0.05 | 0 | 0.06 | 0.08 |
| 5 | 171634651 | rs6866600 | -0.154 | -0.329 | -0.951 | -0.327 | -0.003 | 0.012 | 0.012 | 0 | 0.19 | 0.16 | 0.25 | 0.03 | 0.18 | 0.14 |
| 5 | 171636926 | rs9043 | 0.261 | -0.174 | -1.08 | 0.034 | 0.001 | 0.009 | 0.009 | 0 | 0.27 | 0.35 | 0.3 | 0.39 | 0.3 | 0.45 |
| 5 | 171641500 | rs12520760 | 1.149 | 0.064 | -1.206 | -0.026 | -0.072 | -0.135 | -0.135 | 0 | 0.19 | 0.02 | 0.21 | 0.37 | 0.24 | 0.35 |
| 5 | 171644830 | rs12655315 | 0.452 | 0.153 | -1.03 | 0.094 | -0.004 | -0.014 | -0.014 | 1.395 | 0.26 | 0.26 | 0.29 | 0.39 | 0.31 | 0.45 |
| 5 | 171648625 | rs6876038 | -1.533 | 0.081 | -0.595 | 0.049 | -0.004 | 0.019 | 0.019 | 0 | 0.55 | 0.6 | 0.63 | 0.75 | 0.48 | 0.78 |
| 5 | 171650965 | rs13358102 | -0.643 | 0.109 | -0.533 | 0.2 | -0.003 | -0.01 | -0.01 | 0 | 0.38 | 0.37 | 0.52 | 0.49 | 0.4 | 0.61 |
| 5 | 171652920 | rs4868151 | -0.559 | 0.175 | -0.578 | 0.179 | 0.001 | -0.009 | -0.009 | 0.081 | 0.35 | 0.37 | 0.52 | 0.49 | 0.36 | 0.57 |
| 5 | 171653986 | rs6556004 | -1.341 | 0.002 | -0.688 | 0.219 | -0.008 | 0.001 | 0.001 | 0.238 | 0.54 | 0.75 | 0.6 | 0.5 | 0.52 | 0.7 |
| 5 | 171656546 | rs13165241 | 0.909 | 0.07 | -0.812 | 0.311 | -0.002 | -0.01 | -0.01 | 0.288 | 0.1 | 0.04 | 0.2 | 0.1 | 0.09 | 0.15 |
| 5 | 171656863 | rs10491493 | 0.461 | 0.151 | -0.389 | 0.365 | 0.002 | 0.012 | 0.012 | 0 | 0.26 | 0.35 | 0.3 | 0.39 | 0.29 | 0.45 |
| 5 | 171662653 | rs7734469 | -0.422 | 0.063 | -0.551 | 0.235 | 0 | 0 | 0 | 0 | 0.37 | 0.59 | 0.22 | 0.27 | 0.23 | 0.29 |
| 5 | 171666023 | rs10463015 | 1.102 | 0.015 | -0.901 | 0.344 | 0.024 | -0.025 | -0.025 | 0 | 0.18 | 0.06 | 0.11 | 0.26 | 0.11 | 0.19 |
| 5 | 171687711 | rs7736157 | -1.652 | 0.955 | -1.379 | 0.685 | -0.139 | 0 | 0 | 0 | 0.71 | 0.98 | 0.71 | 0.77 | 0.59 | 0.87 |
| 5 | 171694066 | rs10063708 | -0.522 | 1.082 | -0.434 | 1.043 | -0.01 | -0.015 | -0.015 | 0 | 0.27 | 0.26 | 0.29 | 0.38 | 0.32 | 0.46 |
| 5 | 171717713 | rs10069019 | -0.069 | 0.578 | -0.029 | 1.245 | -0.001 | 0.003 | 0.003 | 0 | 0.32 | 0.37 | 0.32 | 0.37 | 0.3 | 0.39 |
| 5 | 171733971 | rs4560562 | -0.527 | 1.204 | 0.632 | 1.39 | -0.017 | -0.01 | -0.01 | 0 | 0.76 | 0.82 | 0.64 | 0.67 | 0.64 | 0.86 |
| 5 | 171742292 | rs7715800 | -1.4 | 0.886 | 0.676 | 1.467 | -0.006 | 0.004 | 0.004 | 0 | 0.1 | 0.19 | 0.09 | 0.12 | 0.11 | 0.12 |
| 5 | 171753739 | rs2339423 | -0.157 | 0.932 | 1.07 | 1.464 | 0.041 | 0.019 | 0.019 | 0 | 0.71 | 0.52 | 0.61 | 0.68 | 0.63 | 0.86 |
| 5 | 171756937 | rs17074634 | -0.588 | 0.884 | 1.038 | 1.747 | 0.003 | 0.06 | 0.06 | 0 | 0.06 | 0.09 | 0.07 | 0.18 | 0.11 | 0.1 |
| 5 | 171762100 | rs889023 | -0.303 | 0.879 | 0.987 | 1.243 | -0.002 | -0.001 | -0.001 | 0 | 0.72 | 0.75 | 0.63 | 0.7 | 0.62 | 0.85 |
| 5 | 171762575 | rs2339428 | 0.553 | 0.67 | 1.277 | 1.248 | 0.014 | 0.01 | 0.01 | 0 | 0.61 | 0.48 | 0.54 | 0.56 | 0.52 | 0.75 |
| 5 | 171763862 | rs869366 | 0.611 | 0.6 | 1.335 | 0.769 | 0.008 | 0.005 | 0.005 | 0 | 0.6 | 0.43 | 0.53 | 0.57 | 0.52 | 0.75 |
| 5 | 171764158 | rs4424012 | -0.096 | 0.751 | 1.272 | 0.412 | 0.007 | 0.002 | 0.002 | 0 | 0.74 | 0.71 | 0.62 | 0.7 | 0.63 | 0.83 |
| 5 | 171780501 | rs4562058 | -0.395 | -0.315 | 0.216 | -0.495 | 0.026 | 0.043 | 0.043 | 0 | 0.43 | 0.63 | 0.29 | 0.12 | 0.31 | 0.48 |
| 5 | 171780628 | rs13187132 | 0.855 | -0.434 | -0.139 | -0.653 | -0.02 | 0.046 | 0.046 | 0 | 0.09 | 0.02 | 0.07 | 0.02 | 0.12 | 0.08 |
| 5 | 171785702 | rs17703000 | -0.161 | -0.588 | -0.009 | -0.63 | 0.015 | 0.007 | 0.007 | 0 | 0.09 | 0.03 | 0.07 | 0.03 | 0.05 | 0.24 |
| 5 | 171785892 | rs2569232 | -1.296 | -0.181 | -0.116 | -0.817 | -0.026 | 0.025 | 0.025 | 0 | 0.73 | 0.86 | 0.74 | 0.76 | 0.65 | 0.87 |
| 5 | 171786861 | rs13182040 | 0.234 | -0.223 | 0.613 | -0.934 | -0.011 | -0.045 | -0.045 | 0 | 0.35 | 0.47 | 0.21 | 0.09 | 0.29 | 0.25 |
| 5 | 171787120 | rs1541916 | 0.107 | -0.137 | 0.914 | -0.548 | 0.022 | 0.03 | 0.03 | 0 | 0.3 | 0.45 | 0.46 | 0.64 | 0.32 | 0.38 |
| 5 | 171800897 | rs2731717 | 0.295 | -0.159 | 1.127 | -0.698 | 0.024 | 0.024 | 0.024 | 0 | 0.28 | 0.4 | 0.45 | 0.66 | 0.31 | 0.38 |
| 5 | 171808086 | rs7721076 | -0.059 | -0.178 | 0.957 | -0.745 | -0.02 | -0.004 | -0.004 | 0 | 0.08 | 0.15 | 0.08 | 0.03 | 0.03 | 0.21 |
| 5 | 171808589 | rs7725430 | 0.015 | -0.175 | 0.882 | -0.744 | -0.017 | -0.004 | -0.004 | 0 | 0.08 | 0.12 | 0.07 | 0.03 | 0.03 | 0.21 |
| 5 | 171817636 | rs2731729 | 0.132 | -0.216 | 1.185 | -0.672 | 0.025 | 0.027 | 0.027 | 0 | 0.29 | 0.44 | 0.45 | 0.65 | 0.31 | 0.38 |
| 5 | 171820857 | rs6556019 | -0.262 | -0.511 | 1.277 | -0.705 | -0.001 | 0.001 | 0.001 | 0 | 0.33 | 0.62 | 0.48 | 0.64 | 0.32 | 0.37 |
| 5 | 171837388 | rs12657598 | -0.865 | -0.579 | 1.211 | -0.76 | 0.015 | 0.032 | 0.032 | 0 | 0.42 | 0.67 | 0.54 | 0.69 | 0.36 | 0.62 |
| 5 | 171839729 | rs10077377 | -1.357 | -0.618 | 1.463 | -0.796 | -0.034 | -0.047 | -0.047 | 0.181 | 0.15 | 0.37 | 0.12 | 0.03 | 0.09 | 0.26 |
| 5 | 171841019 | rs10476027 | 0.146 | -0.6 | 1.112 | -0.791 | 0.003 | 0.006 | 0.006 | 0 | 0.3 | 0.39 | 0.45 | 0.66 | 0.31 | 0.37 |

| | | | | | | | | | | | | | | | | |
|---|---|---|---|---|---|---|---|---|---|---|---|---|---|---|---|---|
| 5 | 171844378 | rs4446483 | -1.002 | -0.611 | 1.099 | -0.795 | 0.027 | 0.065 | 0.065 | 0 | 0.43 | 0.78 | 0.55 | 0.69 | 0.36 | 0.62 |
| 5 | 171847674 | rs17653344 | -0.398 | -0.54 | 0.904 | -0.758 | 0 | -0.001 | -0.001 | 0 | 0.36 | 0.4 | 0.51 | 0.69 | 0.35 | 0.58 |
| 5 | 171849471 | rs17074773 | 0.56 | -0.464 | 0.763 | -0.708 | -0.003 | -0.007 | -0.007 | 0 | 0.25 | 0.22 | 0.42 | 0.66 | 0.28 | 0.36 |
| 5 | 171855765 | rs4867675 | -0.027 | -0.479 | 0.774 | -0.706 | 0.005 | 0.016 | 0.016 | 0 | 0.31 | 0.47 | 0.46 | 0.66 | 0.31 | 0.37 |
| 5 | 171860077 | rs2569217 | -0.036 | -0.461 | -0.371 | -1.053 | -0.002 | 0.003 | 0.003 | 0 | 0.46 | 0.56 | 0.41 | 0.25 | 0.47 | 0.46 |
| 5 | 171865548 | rs2731688 | 1.342 | -0.539 | -1.348 | -1.071 | -0.084 | 0.045 | 0.045 | 0 | 0.28 | 0.06 | 0.27 | 0.21 | 0.36 | 0.17 |
| 5 | 171866575 | rs11747030 | 0.723 | -0.114 | -1.391 | -0.841 | -0.003 | -0.017 | -0.017 | 0 | 0.22 | 0.15 | 0.4 | 0.67 | 0.24 | 0.32 |
| 5 | 171867995 | rs919761 | -0.358 | 0.14 | -1.507 | -0.849 | 0.004 | 0.083 | 0.083 | 0 | 0.71 | 0.84 | 0.83 | 0.9 | 0.73 | 0.81 |
| 5 | 171876084 | rs2731693 | 0.114 | 0.14 | -1.457 | -0.789 | -0.04 | 0.146 | 0.146 | 0.024 | 0.39 | 0.86 | 0.33 | 0.15 | 0.37 | 0.18 |
| 5 | 171883304 | rs6865951 | -0.515 | 0.343 | -1.425 | -0.717 | -0.009 | 0.018 | 0.018 | 0 | 0.11 | 0.04 | 0.07 | 0.03 | 0.12 | 0.24 |
| 5 | 171883508 | rs1560682 | 0.62 | 0.322 | -1.251 | -0.189 | -0.029 | 0.006 | 0.006 | 0 | 0.35 | 0.28 | 0.48 | 0.31 | 0.42 | 0.25 |
| 5 | 171889524 | rs2569248 | -0.185 | 0.031 | -1.265 | -0.149 | 0.012 | 0.055 | 0.055 | 0 | 0.5 | 0.78 | 0.55 | 0.33 | 0.51 | 0.45 |
| 5 | 171897090 | rs2569235 | -0.183 | -0.206 | -1.401 | -0.017 | -0.012 | -0.026 | -0.026 | 0 | 0.67 | 0.82 | 0.77 | 0.4 | 0.64 | 0.56 |
| 5 | 171903310 | rs10042482 | -0.854 | -0.06 | -2.009 | -0.823 | -0.082 | 0.155 | 0.155 | 0 | 0.63 | 0.86 | 0.64 | 0.91 | 0.52 | 0.76 |
| 5 | 171908794 | rs10059733 | 0.203 | -0.361 | -1.898 | -0.13 | 0.008 | -0.061 | -0.061 | 0 | 0.26 | 0.18 | 0.43 | 0.51 | 0.24 | 0.28 |
| 5 | 171909024 | rs11958604 | 0.753 | -0.362 | -1.931 | 0.164 | 0.009 | 0.018 | 0.018 | 0 | 0.2 | 0.16 | 0.24 | 0.14 | 0.16 | 0.21 |
| 5 | 171910504 | rs4868171 | 0.715 | -0.429 | -1.181 | 0.468 | 0.002 | -0.001 | -0.001 | 1.096 | 0.16 | 0.1 | 0.07 | 0.31 | 0.11 | 0.14 |
| 5 | 171910863 | rs12656621 | -0.18 | -0.449 | -1.153 | 0.461 | 0.004 | -0.008 | -0.008 | 0.137 | 0.21 | 0.13 | 0.26 | 0.69 | 0.19 | 0.21 |
| 5 | 171911313 | rs11134751 | -1.175 | -0.378 | -0.989 | 0.059 | 0.017 | -0.011 | -0.011 | 0.016 | 0.37 | 0.66 | 0.2 | 0.4 | 0.27 | 0.48 |
| 5 | 171913126 | rs11747598 | 0.013 | -0.209 | -1.071 | 0.105 | -0.005 | -0.013 | -0.013 | 0 | 0.17 | 0.21 | 0.09 | 0.09 | 0.15 | 0.34 |
| 5 | 171917008 | rs4868176 | 0.1 | 0.048 | -1.273 | -0.093 | 0.001 | 0.001 | 0.001 | 3.214 | 0.16 | 0.12 | 0.08 | 0.32 | 0.11 | 0.14 |
| 5 | 171918774 | rs891998 | 0.76 | 0.841 | -1.838 | -1.678 | -0.161 | 0.135 | 0.135 | 0 | 0.25 | 0.02 | 0.2 | 0.05 | 0.39 | 0.21 |
| 5 | 171922102 | rs6898730 | -0.564 | 0.762 | -0.78 | -1.671 | 0.032 | -0.013 | -0.013 | 0 | 0.17 | 0.24 | 0.21 | 0.13 | 0.2 | 0.21 |
| 5 | 171922211 | rs6899161 | -0.502 | 0.322 | -0.994 | -2.342 | -0.002 | -0.011 | -0.011 | 0 | 0.65 | 0.61 | 0.53 | 0.23 | 0.65 | 0.55 |
| 5 | 171922637 | rs11745118 | 1.281 | 0.014 | -1.431 | -2.563 | -0.089 | 0.162 | 0.162 | 0.047 | 0.31 | 0.04 | 0.21 | 0.05 | 0.35 | 0.2 |
| 5 | 171927246 | rs11745941 | 0.227 | -0.168 | -0.994 | -2.467 | 0.035 | 0.096 | 0.096 | 1.548 | 0.5 | 0.66 | 0.47 | 0.17 | 0.56 | 0.41 |
| 5 | 171936932 | rs10078424 | -0.666 | 0.012 | -1.012 | -2.454 | -0.004 | 0.01 | 0.01 | 0 | 0.64 | 0.71 | 0.54 | 0.2 | 0.61 | 0.51 |
| 5 | 171936950 | rs1347156 | -2.176 | -0.223 | -1.281 | -2.597 | -0.02 | 0.029 | 0.029 | 0.006 | 0.85 | 0.98 | 0.93 | 0.93 | 0.87 | 0.94 |
| 5 | 171937352 | rs10071005 | -0.851 | -0.264 | -0.747 | -2.753 | 0.017 | -0.052 | -0.052 | 0 | 0.26 | 0.53 | 0.27 | 0.12 | 0.22 | 0.21 |
| 5 | 171946349 | rs6894995 | 0.03 | -0.172 | -0.801 | -2.697 | -0.005 | 0.001 | 0.001 | 0.036 | 0.17 | 0.15 | 0.23 | 0.12 | 0.19 | 0.2 |
| 5 | 171947476 | rs2163792 | 1.624 | -0.226 | -1.066 | -2.914 | -0.065 | 0.136 | 0.136 | 1.148 | 0.27 | 0.01 | 0.2 | 0.06 | 0.31 | 0.18 |
| 5 | 171952332 | rs11741321 | -0.694 | -0.407 | -0.574 | -2.778 | 0 | -0.008 | -0.008 | 0 | 0.61 | 0.42 | 0.49 | 0.24 | 0.59 | 0.48 |
| 5 | 171954233 | rs1433020 | -0.969 | -0.372 | -0.542 | -2.761 | -0.006 | 0.03 | 0.03 | 0 | 0.64 | 0.74 | 0.51 | 0.24 | 0.59 | 0.48 |
| 5 | 171960073 | rs1433021 | -0.4 | -0.369 | -0.554 | -2.809 | 0.009 | 0.036 | 0.036 | 0.708 | 0.56 | 0.4 | 0.46 | 0.18 | 0.59 | 0.44 |
| 5 | 171961443 | rs6883952 | -1.238 | -0.375 | -0.185 | -1.85 | -0.018 | 0.029 | 0.029 | 0 | 0.24 | 0.16 | 0.21 | 0.14 | 0.23 | 0.25 |
| 5 | 171962306 | rs7724869 | -1.647 | -0.143 | -0.523 | -1.687 | 0.014 | 0.061 | 0.061 | 0.351 | 0.42 | 0.72 | 0.33 | 0.18 | 0.32 | 0.35 |
| 5 | 171969572 | rs10516094 | -1.926 | 0.965 | -0.61 | -1.119 | -0.029 | 0.028 | 0.028 | 0.202 | 0.19 | 0.1 | 0.19 | 0.11 | 0.25 | 0.23 |
| 5 | 171971486 | rs1541918 | -1.682 | 0.476 | -0.545 | -1 | -0.005 | 0.017 | 0.017 | 0.114 | 0.15 | 0.07 | 0.12 | 0.1 | 0.12 | 0.14 |
| 5 | 171981503 | rs7736185 | -1.025 | 0.919 | -0.351 | -0.848 | -0.004 | 0.002 | 0.002 | 0 | 0.07 | 0.06 | 0.05 | 0.06 | 0.09 | 0.08 |
| 5 | 171991913 | rs17655319 | -0.636 | -0.03 | -0.435 | -0.969 | 0.029 | 0.06 | 0.06 | 0 | 0.21 | 0.07 | 0.08 | 0.06 | 0.12 | 0.16 |
| 5 | 171993280 | rs10059684 | -0.959 | -0.138 | 0.27 | -1.037 | -0.048 | -0.087 | -0.087 | 0.274 | 0.26 | 0.49 | 0.11 | 0.06 | 0.12 | 0.16 |
| 5 | 172003493 | rs10476041 | -1.651 | 0.948 | 1.047 | -0.412 | 0.119 | 0.085 | 0.085 | 0.082 | 0.07 | 0.28 | 0.16 | 0.08 | 0.14 | 0.09 |
| 5 | 172003564 | rs10476042 | -2.528 | 0.846 | 1.261 | -0.558 | 0.069 | -0.019 | -0.019 | 0 | 0.2 | 0.5 | 0.23 | 0.11 | 0.25 | 0.21 |
| 5 | 172004039 | rs10063826 | -2.316 | 0.925 | 1.571 | -0.496 | 0.13 | 0.095 | 0.095 | 3.4 | 0.09 | 0.49 | 0.18 | 0.08 | 0.14 | 0.09 |
| 5 | 172007558 | rs6876578 | -1.039 | 0.904 | 1.466 | -0.559 | -0.011 | 0.014 | 0.014 | 0 | 0.1 | 0.07 | 0.06 | 0.04 | 0.1 | 0.11 |
| 5 | 172010881 | rs6556031 | -2.514 | 0.868 | 1.803 | -0.558 | 0.062 | -0.024 | -0.024 | 1.04 | 0.18 | 0.56 | 0.23 | 0.1 | 0.24 | 0.21 |
| 5 | 172012595 | rs7714953 | -2.524 | 0.854 | 1.788 | -0.564 | 0.065 | -0.025 | -0.025 | 1.451 | 0.18 | 0.56 | 0.23 | 0.1 | 0.24 | 0.21 |
| 5 | 172013286 | rs714715 | -2.713 | 0.853 | 1.818 | -0.565 | 0.045 | -0.029 | -0.029 | 0 | 0.19 | 0.63 | 0.24 | 0.1 | 0.24 | 0.21 |
| 5 | 172016544 | rs11750720 | -2.049 | 0.902 | 2.017 | -0.586 | 0.139 | 0.072 | 0.072 | 2.237 | 0.08 | 0.42 | 0.16 | 0.06 | 0.14 | 0.09 |
| 5 | 172021610 | rs1876628 | 1.273 | 0.228 | 1.199 | -0.844 | 0 | -0.02 | -0.02 | 0 | 0.86 | 0.87 | 0.92 | 0.98 | 0.88 | 0.89 |
| 5 | 172023245 | rs1107775 | -2.105 | -0.06 | 1.836 | -0.889 | 0.041 | 0.032 | 0.032 | 4.065 | 0.06 | 0.51 | 0.05 | 0 | 0.01 | 0 |
| 5 | 172023436 | rs12108820 | -2.081 | -0.159 | 1.432 | -0.963 | 0.001 | -0.011 | -0.011 | 0 | 0.2 | 0.73 | 0.13 | 0.04 | 0.12 | 0.13 |
| 5 | 172031249 | rs931802 | -0.218 | 0.317 | 0.569 | -0.617 | -0.009 | 0.119 | 0.119 | 0.75 | 0.52 | 0.77 | 0.67 | 0.77 | 0.54 | 0.67 |
| 5 | 172031548 | rs17075007 | -0.959 | 0.675 | 0.515 | -0.808 | 0 | -0.023 | -0.023 | 0 | 0.16 | 0.19 | 0.17 | 0.04 | 0.2 | 0.15 |
| 5 | 172034100 | rs931804 | 0.244 | -0.086 | 0.667 | -0.187 | 0.025 | 0.032 | 0.032 | 0 | 0.22 | 0.36 | 0.36 | 0.28 | 0.27 | 0.25 |
| 5 | 172035404 | rs2002387 | -0.225 | 0.123 | 0.626 | -0.119 | -0.003 | 0 | 0 | 0.573 | 0.37 | 0.36 | 0.3 | 0.46 | 0.27 | 0.45 |
| 5 | 172036332 | rs4867681 | -1.298 | 0.598 | 0.889 | 0.012 | 0.003 | 0.016 | 0.016 | 0 | 0.4 | 0.29 | 0.47 | 0.18 | 0.38 | 0.34 |
| 5 | 172037144 | rs12520206 | -0.517 | 0.746 | 0.963 | 0.13 | -0.001 | 0.001 | 0.001 | 0 | 0.25 | 0.22 | 0.35 | 0.14 | 0.27 | 0.23 |
| 5 | 172038164 | rs17075031 | -0.626 | 0.59 | 0.694 | 0.668 | 0.009 | 0.236 | 0.236 | 0 | 0.11 | 0.33 | 0.13 | 0.41 | 0.07 | 0.29 |
| 5 | 172040365 | rs2089190 | -1.02 | 0.86 | -0.22 | 0.496 | -0.024 | -0.032 | -0.032 | 0.014 | 0.4 | 0.78 | 0.59 | 0.56 | 0.43 | 0.55 |
| 5 | 172043315 | rs3964610 | -1.071 | 0.965 | 0.296 | 0.696 | 0.001 | 0.002 | 0.002 | 0 | 0.39 | 0.56 | 0.39 | 0.51 | 0.44 | 0.53 |
| 5 | 172048029 | rs7724724 | -1.208 | 1.513 | 1.053 | 0.899 | 0.055 | 0.032 | 0.032 | 0.898 | 0.22 | 0.43 | 0.21 | 0.26 | 0.26 | 0.17 |
| 5 | 172050910 | rs949845 | -1.242 | 1.531 | 0.993 | 0.882 | 0.056 | 0.03 | 0.03 | 0.743 | 0.22 | 0.43 | 0.21 | 0.25 | 0.26 | 0.17 |
| 5 | 172051979 | rs6896660 | -1.156 | 1.469 | 0.621 | 0.803 | 0.012 | 0 | 0 | 0 | 0.3 | 0.31 | 0.45 | 0.3 | 0.39 | 0.3 |
| 5 | 172054921 | rs6894799 | -0.871 | 1.032 | 0.958 | 1.095 | 0.007 | 0.078 | 0.078 | 0 | 0.1 | 0.19 | 0.11 | 0.25 | 0.1 | 0.13 |
| 5 | 172061532 | rs13360786 | -1.539 | 0.839 | 1.587 | 1.013 | 0.008 | 0.137 | 0.137 | 0 | 0.13 | 0.39 | 0.14 | 0.25 | 0.1 | 0.15 |
| 5 | 172067768 | rs2055741 | -1.64 | 0.839 | 1.632 | 0.848 | 0.003 | 0.052 | 0.052 | 0 | 0.16 | 0.35 | 0.14 | 0.27 | 0.14 | 0.15 |
| 5 | 172071962 | rs7730108 | -0.913 | 0.086 | 0.602 | 0.133 | 0.015 | 0.102 | 0.102 | 0.801 | 0.55 | 0.84 | 0.47 | 0.76 | 0.45 | 0.67 |
| 5 | 172072365 | rs13183447 | -1.352 | 0.731 | 0.35 | -0.031 | -0.064 | 0.114 | 0.114 | 3.103 | 0.63 | 0.9 | 0.75 | 0.84 | 0.62 | 0.78 |

| | | | | | | | | | | | | | | | | |
|---|---|---|---|---|---|---|---|---|---|---|---|---|---|---|---|---|
| 5 | 172073780 | rs883150 | -1.298 | 0.598 | 0.994 | 0.009 | 0.024 | -0.038 | -0.038 | 0.009 | 0.16 | 0.37 | 0.31 | 0.1 | 0.18 | 0.11 |
| 5 | 172079889 | rs6556035 | -0.121 | 0.821 | 0.866 | -0.234 | 0.016 | 0.084 | 0.084 | 0.645 | 0.49 | 0.66 | 0.58 | 0.26 | 0.52 | 0.35 |
| 5 | 172087950 | rs905796 | 0.549 | 0.027 | 0.876 | -0.028 | 0 | -0.001 | -0.001 | 1.599 | 0.55 | 0.43 | 0.53 | 0.57 | 0.58 | 0.64 |
| 5 | 172092344 | rs11750421 | -1.246 | 0.025 | 0.908 | -0.163 | -0.024 | -0.051 | -0.051 | 0.003 | 0.17 | 0.55 | 0.22 | 0.1 | 0.13 | 0.12 |
| 5 | 172092499 | rs11738054 | -0.631 | 0.014 | 1.182 | -0.168 | 0.002 | -0.02 | -0.02 | 1.324 | 0.12 | 0.3 | 0.18 | 0.09 | 0.13 | 0.11 |
| 5 | 172092620 | rs10035202 | 0.494 | 0.009 | 0.918 | -0.055 | 0.003 | 0.001 | 0.001 | 0.077 | 0.55 | 0.43 | 0.53 | 0.57 | 0.57 | 0.62 |
| 5 | 172096366 | rs7718156 | -1.796 | -0.045 | 0.719 | -0.043 | -0.014 | -0.019 | -0.019 | 0 | 0.22 | 0.69 | 0.26 | 0.16 | 0.17 | 0.18 |
| 5 | 172099097 | rs6556038 | -0.911 | 0.087 | 0.913 | 0.039 | 0.03 | -0.014 | -0.014 | 0 | 0.13 | 0.46 | 0.19 | 0.08 | 0.12 | 0.12 |
| 5 | 172102341 | rs11741671 | 0.613 | 0.142 | -0.049 | 0.015 | 0.008 | 0.017 | 0.017 | 1.079 | 0.51 | 0.65 | 0.53 | 0.54 | 0.5 | 0.49 |
| 5 | 172102820 | rs13359269 | 0.312 | 0.352 | -0.774 | 0.09 | -0.001 | -0.01 | -0.01 | 0 | 0.09 | 0.1 | 0.05 | 0.03 | 0.08 | 0.12 |
| 5 | 172104628 | rs4867685 | -0.883 | 0.342 | -1.025 | 0.18 | -0.067 | -0.076 | -0.076 | 0 | 0.73 | 0.98 | 0.75 | 0.71 | 0.69 | 0.81 |
| 5 | 172106310 | rs1027745 | 0.916 | 0.621 | -0.559 | 0.462 | -0.036 | -0.056 | -0.056 | 0 | 0.1 | 0.03 | 0.2 | 0.29 | 0.11 | 0.11 |
| 5 | 172113235 | rs747829 | -0.02 | 0.454 | -0.208 | 0.246 | -0.003 | -0.032 | -0.032 | 0.169 | 0.09 | 0.12 | 0.06 | 0.03 | 0.09 | 0.11 |
| 5 | 172116576 | rs1064174 | -0.411 | 0.439 | -0.135 | 0.269 | -0.003 | 0.003 | 0.003 | 0.01 | 0.61 | 0.63 | 0.55 | 0.53 | 0.6 | 0.66 |
| 5 | 172117144 | rs1109845 | 0.402 | 0.457 | -0.881 | 0.292 | -0.018 | -0.005 | -0.005 | 0 | 0.44 | 0.34 | 0.43 | 0.49 | 0.44 | 0.5 |
| 5 | 172122001 | rs6894903 | 1.036 | 0.644 | -1.236 | 0.17 | -0.013 | 0.008 | 0.008 | 0.049 | 0.29 | 0.21 | 0.19 | 0.18 | 0.3 | 0.38 |
| 5 | 172127113 | rs9686679 | 0.583 | 0.607 | -1.613 | 0.549 | -0.013 | -0.067 | -0.067 | 0.924 | 0.15 | 0.05 | 0.23 | 0.31 | 0.14 | 0.12 |
| 5 | 172127304 | rs792691 | 0.168 | 0.563 | -1.232 | 0.328 | 0.002 | -0.012 | -0.012 | 0 | 0.34 | 0.18 | 0.37 | 0.38 | 0.33 | 0.26 |
| 5 | 172138212 | rs172860 | -0.451 | 0.777 | -1.846 | -0.075 | -0.078 | 0.155 | 0.155 | 1.232 | 0.46 | 0.95 | 0.47 | 0.2 | 0.49 | 0.51 |
| 5 | 172138787 | rs324337 | -0.481 | 1.159 | -0.588 | -0.28 | 0.025 | -0.036 | -0.036 | 0 | 0.17 | 0.4 | 0.18 | 0.07 | 0.17 | 0.19 |
| 5 | 172139785 | rs11748876 | 0.296 | 1.222 | -0.583 | -0.27 | 0.003 | -0.01 | -0.01 | 0 | 0.09 | 0.11 | 0.07 | 0.02 | 0.11 | 0.1 |
| 5 | 172145370 | rs7443927 | -0.114 | 1.142 | -0.963 | 0.091 | -0.005 | -0.028 | -0.028 | 0 | 0.91 | 0.98 | 0.81 | 0.76 | 0.87 | 0.91 |
| 5 | 172148182 | rs4868200 | 0.583 | 1.095 | -0.638 | -0.211 | -0.025 | 0.03 | 0.03 | 0.303 | 0.24 | 0.18 | 0.25 | 0.11 | 0.3 | 0.21 |
| 5 | 172150990 | rs4867686 | 0.488 | 1.344 | 0.037 | -0.404 | -0.012 | 0.025 | 0.025 | 0.371 | 0.25 | 0.22 | 0.24 | 0.07 | 0.27 | 0.2 |
| 5 | 172155007 | rs2662233 | 0.658 | 1.391 | 0.075 | -0.305 | -0.007 | -0.011 | -0.011 | 0.967 | 0.82 | 0.86 | 0.75 | 0.71 | 0.76 | 0.73 |
| 5 | 172156007 | rs322387 | -0.818 | 1.331 | 0.248 | -0.321 | 0.036 | -0.055 | -0.055 | 0.025 | 0.19 | 0.48 | 0.18 | 0.08 | 0.19 | 0.28 |
| 5 | 172156062 | rs322388 | 1.466 | 1.21 | 0.016 | -0.315 | 0.043 | 0.052 | 0.052 | 0.395 | 0.48 | 0.23 | 0.42 | 0.61 | 0.39 | 0.42 |
| 5 | 172157768 | rs322394 | 1.356 | 1.146 | 0.363 | -0.045 | 0.004 | -0.003 | -0.003 | 1.834 | 0.71 | 0.38 | 0.66 | 0.69 | 0.67 | 0.61 |
| 5 | 172173353 | rs322376 | 0.747 | 0.263 | -0.383 | 0.174 | 0.017 | -0.029 | -0.029 | 0 | 0.18 | 0.05 | 0.14 | 0.29 | 0.15 | 0.19 |
| 5 | 172176461 | rs322372 | 0.167 | 0.281 | 0.06 | 0.044 | 0.051 | 0.011 | 0.011 | 0.438 | 0.95 | 0.73 | 0.96 | 0.99 | 0.94 | 0.94 |
| 5 | 172176904 | rs2161375 | -0.011 | 0.567 | -0.852 | -0.304 | -0.105 | -0.012 | -0.012 | 0 | 0.31 | 0.1 | 0.36 | 0.34 | 0.47 | 0.34 |
| 5 | 172179584 | rs6872156 | -0.26 | 0.27 | -0.192 | -0.148 | -0.029 | 0.01 | 0.01 | 0 | 0.75 | 0.53 | 0.77 | 0.68 | 0.79 | 0.76 |
| 5 | 172188213 | rs322359 | 1.311 | 0.343 | -0.478 | -0.054 | 0.008 | -0.038 | -0.038 | 0 | 0.17 | 0.04 | 0.17 | 0.3 | 0.15 | 0.18 |
| 5 | 172188448 | rs322358 | -0.122 | 0.808 | -0.408 | -0.155 | 0.003 | -0.015 | -0.015 | 0 | 0.4 | 0.25 | 0.4 | 0.53 | 0.42 | 0.4 |
| 5 | 172192868 | rs322353 | -0.158 | 0.418 | -0.412 | -0.439 | -0.002 | -0.002 | -0.002 | 0 | 0.54 | 0.36 | 0.52 | 0.59 | 0.53 | 0.51 |
| 5 | 172194873 | rs322351 | 0.452 | 0.172 | -0.839 | -0.584 | -0.022 | 0.118 | 0.118 | 0 | 0.44 | 0.12 | 0.37 | 0.28 | 0.48 | 0.4 |
| 5 | 172196711 | rs2431663 | 0.763 | 0.338 | -0.105 | -0.593 | 0.047 | -0.004 | -0.004 | 0.528 | 0.92 | 0.52 | 0.95 | 1 | 0.94 | 0.99 |
| 5 | 172196997 | rs7702178 | 1.688 | 0.274 | -0.994 | -0.656 | 0.032 | -0.087 | -0.087 | 2.532 | 0.24 | 0.11 | 0.24 | 0.37 | 0.16 | 0.28 |
| 5 | 172198952 | rs322382 | 0.285 | 0.357 | -0.756 | -0.89 | 0.1 | -0.099 | -0.099 | 0 | 0.79 | 0.44 | 0.76 | 0.95 | 0.69 | 0.78 |
| 5 | 172201093 | rs11746792 | 0.689 | 0.492 | -1.817 | -0.91 | -0.027 | 0.05 | 0.05 | 0 | 0.31 | 0.14 | 0.31 | 0.23 | 0.39 | 0.27 |
| 5 | 172201974 | rs17658295 | 0.536 | 0.501 | -1.837 | -0.938 | -0.031 | 0.056 | 0.056 | 0 | 0.32 | 0.15 | 0.32 | 0.23 | 0.42 | 0.3 |
| 5 | 172204629 | rs4868204 | 1.654 | 0.178 | -1.676 | -0.789 | 0.045 | -0.085 | -0.085 | 0 | 0.25 | 0.11 | 0.28 | 0.39 | 0.15 | 0.27 |
| 5 | 172205000 | rs2249619 | 0.902 | 0.73 | -1.52 | -0.915 | -0.001 | 0.087 | 0.087 | 0 | 0.2 | 0.08 | 0.12 | 0 | 0.17 | 0.16 |
| 5 | 172213368 | rs2161373 | 1.147 | 0.733 | -1.469 | -0.836 | -0.001 | 0.11 | 0.11 | 0 | 0.22 | 0.03 | 0.14 | 0 | 0.21 | 0.19 |
| 5 | 172213577 | rs2112662 | 0.646 | 0.824 | -1.409 | -0.663 | 0.035 | 0.103 | 0.103 | 0.837 | 0.49 | 0.18 | 0.56 | 0.66 | 0.44 | 0.51 |
| 5 | 172213709 | rs2112661 | 1.718 | 0.717 | -1.485 | -0.566 | 0.024 | -0.057 | -0.057 | 0 | 0.23 | 0.09 | 0.3 | 0.41 | 0.15 | 0.26 |
| 5 | 172216133 | rs737468 | 0.68 | 0.893 | -1.463 | -0.644 | 0.001 | 0.037 | 0.037 | 0.401 | 0.29 | 0.15 | 0.27 | 0.25 | 0.29 | 0.24 |
| 5 | 172216355 | rs17715219 | 0.429 | 1.448 | -1.302 | -0.656 | 0.002 | -0.003 | -0.003 | 0 | 0.11 | 0.09 | 0.12 | 0.04 | 0.2 | 0.17 |
| 5 | 172218144 | rs12518554 | 0.637 | 1.106 | -1.403 | -0.522 | 0.007 | 0.047 | 0.047 | 0.717 | 0.31 | 0.14 | 0.26 | 0.24 | 0.29 | 0.26 |
| 5 | 172218380 | rs17075181 | 1.136 | 0.405 | -1.39 | -0.073 | 0.015 | -0.172 | -0.172 | 0 | 0.23 | 0.01 | 0.21 | 0.45 | 0.17 | 0.28 |
| 5 | 172221586 | rs749864 | 1.333 | 0.392 | -1.249 | -0.103 | 0.037 | -0.097 | -0.097 | 0.076 | 0.22 | 0.05 | 0.22 | 0.41 | 0.15 | 0.28 |
| 5 | 172230003 | rs10057387 | 0.42 | 0.721 | -0.508 | -0.242 | 0.018 | -0.001 | -0.001 | 0 | 0.49 | 0.44 | 0.41 | 0.44 | 0.41 | 0.5 |
| 5 | 172231762 | rs6877686 | -0.877 | 1.042 | -0.494 | -0.279 | 0.065 | -0.013 | -0.013 | 0.167 | 0.09 | 0.3 | 0.06 | 0.07 | 0.08 | 0.24 |
| 5 | 172233065 | rs2291045 | -0.711 | 1.035 | -0.692 | -0.3 | 0.008 | -0.033 | -0.033 | 0 | 0.09 | 0.16 | 0.06 | 0.04 | 0.09 | 0.19 |
| 5 | 172243695 | rs7715908 | 0.255 | 1.107 | -1.108 | -0.083 | -0.001 | 0.008 | 0.008 | 0 | 0.66 | 0.41 | 0.56 | 0.39 | 0.7 | 0.56 |
| 5 | 172246071 | rs17075237 | 1.139 | 1.018 | -1.247 | 0.21 | 0.024 | -0.027 | -0.027 | 0 | 0.17 | 0.04 | 0.26 | 0.25 | 0.11 | 0.14 |
| 5 | 172246156 | rs17075238 | 0.891 | 1.075 | -1.309 | -0.357 | 0.004 | -0.008 | -0.008 | 2.06 | 0.26 | 0.24 | 0.23 | 0.37 | 0.18 | 0.22 |
| 5 | 172246457 | rs2291044 | 0.082 | 1.777 | -1.072 | 0.122 | -0.056 | 0.063 | 0.063 | 0 | 0.17 | 0.04 | 0.15 | 0.02 | 0.27 | 0.15 |
| 5 | 172248631 | rs4868214 | -0.646 | 0.825 | -0.453 | 0.607 | -0.001 | 0.007 | 0.007 | 0 | 0.35 | 0.53 | 0.47 | 0.61 | 0.3 | 0.46 |
| 5 | 172250261 | rs6870663 | 0.812 | 0.591 | -0.637 | 0.747 | 0.006 | -0.009 | -0.009 | 0 | 0.65 | 0.31 | 0.5 | 0.36 | 0.71 | 0.67 |
| 5 | 172256024 | rs1564259 | 0.591 | 0.659 | 0.33 | 0.855 | 0.01 | 0.034 | 0.034 | 0 | 0.74 | 0.6 | 0.75 | 0.65 | 0.75 | 0.69 |
| 5 | 172268003 | rs751988 | -1.348 | -0.112 | -0.097 | 0.781 | 0.009 | 0.04 | 0.04 | 0 | 0.49 | 0.65 | 0.56 | 0.72 | 0.42 | 0.52 |
| 5 | 172269595 | rs792978 | -0.034 | -0.248 | 0.294 | 0.766 | -0.015 | 0.025 | 0.025 | 0 | 0.23 | 0.35 | 0.31 | 0.38 | 0.15 | 0.22 |
| 5 | 172271007 | rs792975 | -0.061 | -0.598 | 0.221 | 0.703 | 0.004 | -0.006 | -0.006 | 3.721 | 0.23 | 0.24 | 0.31 | 0.34 | 0.16 | 0.21 |
| 5 | 172272207 | rs792969 | 0.243 | -0.796 | 0.149 | 0.629 | -0.034 | 0.032 | 0.032 | 0 | 0.73 | 0.59 | 0.66 | 0.64 | 0.81 | 0.75 |
| 5 | 172272571 | rs710109 | 0.058 | -0.63 | -0.694 | 0.712 | 0.022 | -0.029 | -0.029 | 0 | 0.22 | 0.19 | 0.27 | 0.35 | 0.15 | 0.22 |
| 5 | 172276328 | rs729342 | 0.238 | -0.438 | -1.073 | 0.656 | 0 | -0.001 | -0.001 | 0 | 0.65 | 0.32 | 0.63 | 0.6 | 0.76 | 0.7 |
| 5 | 172276614 | rs729341 | -0.387 | 0.015 | -1.293 | 0.607 | -0.023 | -0.06 | -0.06 | 0 | 0.16 | 0.02 | 0.15 | 0.26 | 0.19 | 0.22 |
| 5 | 172283930 | rs13155834 | -0.869 | -0.583 | -1.026 | -0.256 | 0.035 | 0.037 | 0.037 | 0 | 0.25 | 0.14 | 0.14 | 0.01 | 0.12 | 0.23 |

| | | | | | | | | | | | | | | | | |
|---|---|---|---|---|---|---|---|---|---|---|---|---|---|---|---|---|
| 5 | 172285605 | rs13186108 | 0.831 | -0.443 | -0.539 | 0.2 | -0.012 | -0.005 | -0.005 | 0 | 0.65 | 0.45 | 0.6 | 0.65 | 0.68 | 0.72 |
| 5 | 172286508 | rs793017 | 0.503 | 0.156 | 0.23 | 0.349 | 0.095 | -0.041 | -0.041 | 0.022 | 0.92 | 0.74 | 0.92 | 1 | 0.87 | 0.98 |
| 5 | 172288907 | rs11743007 | 0.523 | -0.025 | 0.394 | 0.333 | 0.055 | -0.01 | -0.01 | 0 | 0.92 | 0.62 | 0.96 | 1 | 0.97 | 0.99 |
| 5 | 172291249 | rs13176814 | -0.572 | -0.036 | -0.465 | 0.5 | 0 | 0.01 | 0.01 | 0 | 0.13 | 0.12 | 0.28 | 0.34 | 0.12 | 0.26 |
| 5 | 172293965 | rs4868226 | 0.021 | 0.27 | -1.262 | 0.441 | -0.038 | 0.023 | 0.023 | 0.018 | 0.45 | 0.13 | 0.51 | 0.68 | 0.53 | 0.58 |
| 5 | 172294916 | rs793026 | 1.105 | 0.474 | -0.019 | 1.06 | -0.016 | 0.059 | 0.059 | 0 | 0.79 | 0.6 | 0.68 | 0.63 | 0.83 | 0.73 |
| 5 | 172298073 | rs10074697 | -0.545 | 0.571 | -0.515 | 1.253 | -0.017 | -0.029 | -0.029 | 0 | 0.14 | 0.07 | 0.3 | 0.32 | 0.24 | 0.24 |
| 5 | 172298997 | rs793019 | 0.768 | 0.473 | -0.386 | 1.218 | 0.013 | 0.11 | 0.11 | 0.104 | 0.75 | 0.47 | 0.61 | 0.55 | 0.73 | 0.72 |
| 5 | 172309853 | rs6881530 | 0.103 | 0.462 | -1.62 | 0.838 | -0.06 | -0.057 | -0.057 | 0 | 0.42 | 0.06 | 0.33 | 0.47 | 0.38 | 0.44 |
| 5 | 172314795 | rs498221 | 0.567 | -0.092 | -0.539 | 0.775 | -0.047 | -0.011 | -0.011 | 0.278 | 0.84 | 0.66 | 0.9 | 0.88 | 0.95 | 0.81 |
| 5 | 172321908 | rs490928 | 0.246 | 0.368 | -0.657 | 1.043 | 0.001 | -0.007 | -0.007 | 0 | 0.25 | 0.13 | 0.33 | 0.32 | 0.26 | 0.27 |
| 5 | 172322142 | rs6871139 | 0.498 | 0.496 | -0.66 | 1.259 | 0.036 | 0.104 | 0.104 | 0.003 | 0.17 | 0.42 | 0.21 | 0.26 | 0.22 | 0.2 |
| 5 | 172324558 | rs472959 | 0.241 | 0.719 | -0.83 | 1 | -0.004 | 0.002 | 0.002 | 0 | 0.56 | 0.44 | 0.44 | 0.4 | 0.48 | 0.51 |
| 5 | 172331057 | rs34839124 | 0.69 | 0.657 | -0.491 | 0.482 | -0.002 | 0.038 | 0.038 | 0 | 0.46 | 0.4 | 0.4 | 0.18 | 0.48 | 0.44 |
| 5 | 172338616 | rs10071392 | 0.09 | 0.648 | -0.206 | 0.914 | -0.021 | 0.002 | 0.002 | 0 | 0.63 | 0.61 | 0.59 | 0.37 | 0.75 | 0.68 |
| 5 | 172340822 | rs17660328 | -0.917 | 0.745 | -0.529 | 0.786 | -0.01 | 0.03 | 0.03 | 0 | 0.3 | 0.19 | 0.2 | 0.18 | 0.31 | 0.35 |
| 5 | 172341917 | rs17716949 | -0.388 | 0.64 | -0.791 | 0.826 | -0.009 | 0.03 | 0.03 | 0.34 | 0.23 | 0.08 | 0.16 | 0.14 | 0.22 | 0.21 |
| 5 | 172345535 | rs481221 | -0.898 | 0.637 | -1.111 | 0.817 | -0.044 | 0.015 | 0.015 | 0 | 0.61 | 0.84 | 0.55 | 0.61 | 0.54 | 0.66 |
| 5 | 172345986 | rs732677 | 0.88 | 0.004 | -1.263 | -0.189 | -0.01 | 0.06 | 0.06 | 0 | 0.1 | 0 | 0.03 | 0 | 0.1 | 0.05 |
| 5 | 172346588 | rs2161134 | 1.113 | -0.191 | -1.003 | -0.838 | -0.011 | 0.017 | 0.017 | 0 | 0.32 | 0.3 | 0.26 | 0.09 | 0.36 | 0.24 |
| 5 | 172346648 | rs888699 | -0.191 | -0.147 | -0.627 | -0.656 | 0.035 | 0.016 | 0.016 | 0 | 0.12 | 0.2 | 0.1 | 0.19 | 0.22 | 0.23 |
| 5 | 172350638 | rs485710 | 0.133 | -0.489 | -0.42 | -0.652 | 0.016 | 0.032 | 0.032 | 0.104 | 0.47 | 0.66 | 0.4 | 0.3 | 0.61 | 0.51 |
| 5 | 172355039 | rs514627 | 1.058 | -0.469 | -0.346 | -0.781 | 0.007 | -0.058 | -0.058 | 0 | 0.36 | 0.48 | 0.29 | 0.11 | 0.39 | 0.28 |
| 5 | 172360034 | rs2339745 | 0.384 | -0.871 | -0.841 | -1.161 | -0.005 | -0.008 | -0.008 | 0 | 0.22 | 0.23 | 0.2 | 0.12 | 0.16 | 0.21 |
| 5 | 172361688 | rs524160 | -0.68 | -1.107 | -1.126 | -1.012 | 0.025 | -0.034 | -0.034 | 0.023 | 0.7 | 0.91 | 0.6 | 0.4 | 0.76 | 0.72 |
| 5 | 172364312 | rs793228 | -0.452 | -1.126 | -1.098 | -1.028 | 0.036 | -0.026 | -0.026 | 0 | 0.67 | 0.9 | 0.6 | 0.4 | 0.76 | 0.71 |
| 5 | 172369769 | rs9885155 | -0.721 | -0.53 | -0.578 | -0.662 | 0 | -0.004 | -0.004 | 0 | 0.37 | 0.6 | 0.34 | 0.3 | 0.39 | 0.45 |
| 5 | 172376178 | rs544421 | -1.025 | -1.083 | -1.02 | -1.063 | -0.03 | 0.143 | 0.143 | 0 | 0.57 | 0.86 | 0.69 | 0.86 | 0.52 | 0.69 |
| 5 | 172378857 | rs539936 | -1.102 | -0.711 | -0.063 | -0.628 | -0.023 | -0.065 | -0.065 | 0 | 0.11 | 0.63 | 0.08 | 0.01 | 0.03 | 0.1 |
| 5 | 172382902 | rs2305896 | -0.292 | -0.552 | -0.9 | -0.488 | -0.084 | -0.029 | -0.029 | 0 | 0.33 | 0.13 | 0.51 | 0.66 | 0.45 | 0.37 |
| 5 | 172385569 | rs7731785 | -0.174 | -1.4 | 0.474 | -0.425 | 0.044 | 0.032 | 0.032 | 0 | 0.06 | 0.26 | 0.04 | 0.01 | 0.03 | 0.07 |
| 5 | 172388212 | rs9784719 | -1.752 | -1.624 | 0.989 | -0.543 | 0.028 | 0.003 | 0.003 | 0 | 0.07 | 0.36 | 0.05 | 0.01 | 0.02 | 0.05 |
| 5 | 172395581 | rs17075427 | -0.516 | -1.66 | 0.648 | -0.607 | 0.013 | 0.021 | 0.021 | 0 | 0.06 | 0.02 | 0.02 | 0.01 | 0.02 | 0.08 |
| 5 | 172395664 | rs11749686 | 1.136 | -2.765 | 0.181 | -1.024 | 0.126 | 0.076 | 0.076 | 0.423 | 0.58 | 0.23 | 0.73 | 0.75 | 0.81 | 0.56 |
| 5 | 172398550 | rs11739415 | 1.144 | -2.771 | 0.202 | -1.027 | 0.124 | 0.075 | 0.075 | 0.456 | 0.58 | 0.23 | 0.73 | 0.75 | 0.81 | 0.56 |
| 5 | 172398764 | rs11749083 | 1.161 | -2.774 | 0.034 | -1.028 | 0.181 | 0.085 | 0.085 | 0.504 | 0.58 | 0.12 | 0.71 | 0.75 | 0.81 | 0.56 |
| 5 | 172402686 | rs17718105 | -0.411 | -2.981 | 0.091 | -1.037 | 0.003 | 0.018 | 0.018 | 0 | 0.09 | 0.06 | 0.05 | 0.04 | 0.06 | 0.15 |
| 5 | 172403528 | rs2004997 | 0.224 | -3.08 | 0.317 | -1.059 | -0.081 | -0.019 | -0.019 | 0.79 | 0.71 | 0.59 | 0.81 | 0.8 | 0.89 | 0.73 |
| 5 | 172405033 | rs1643600 | -1.403 | -3.016 | 0.163 | -1.128 | 0.132 | 0.092 | 0.092 | 0.129 | 0.41 | 0.77 | 0.27 | 0.24 | 0.18 | 0.44 |
| 5 | 172405897 | rs17718169 | 0.064 | -2.27 | 0.368 | -1.187 | 0 | -0.001 | -0.001 | 0 | 0.05 | 0.08 | 0.04 | 0.04 | 0.05 | 0.13 |
| 5 | 172410509 | rs6895379 | -1.468 | -2.323 | 0.601 | -1.239 | -0.025 | -0.065 | -0.065 | 0 | 0.14 | 0.34 | 0.08 | 0.05 | 0.07 | 0.19 |
| 5 | 172427650 | rs2431214 | 0.045 | -2.646 | 0.595 | -1.365 | -0.076 | -0.079 | -0.079 | 4.535 | 0.25 | 0.36 | 0.07 | 0.04 | 0.11 | 0.08 |
| 5 | 172431581 | rs522133 | 0.128 | -2.643 | 0.425 | -1.364 | -0.04 | -0.042 | -0.042 | 0 | 0.24 | 0.28 | 0.07 | 0.04 | 0.1 | 0.08 |
| 5 | 172437641 | rs10053713 | 0.396 | -2.646 | 0.396 | -1.365 | -0.015 | -0.016 | -0.016 | 0 | 0.23 | 0.2 | 0.06 | 0.04 | 0.1 | 0.08 |
| 5 | 172444136 | rs499326 | -1.099 | -2.481 | -0.201 | -1.229 | 0.106 | 0.13 | 0.13 | 0 | 0.51 | 0.72 | 0.58 | 0.82 | 0.29 | 0.69 |
| 5 | 172449207 | rs4868236 | 0.794 | -2.067 | -0.212 | -0.768 | 0.009 | -0.027 | -0.027 | 0 | 0.15 | 0.07 | 0.43 | 0.73 | 0.11 | 0.4 |
| 5 | 172454998 | rs6865010 | 0.019 | -1.833 | -0.126 | -1.163 | -0.026 | -0.028 | -0.028 | 0 | 0.25 | 0.43 | 0.13 | 0.15 | 0.2 | 0.17 |
| 5 | 172461004 | rs6868972 | 0.492 | -1.789 | -0.322 | -1.156 | -0.008 | -0.015 | -0.015 | 0 | 0.21 | 0.21 | 0.1 | 0.05 | 0.17 | 0.1 |
| 5 | 172461243 | rs4868237 | 1.06 | -1.575 | -0.839 | -1.009 | 0.001 | -0.074 | -0.074 | 0.231 | 0.14 | 0.02 | 0.42 | 0.73 | 0.11 | 0.4 |
| 5 | 172469848 | rs7719172 | -1.454 | -1.191 | -1.093 | -1.646 | -0.132 | 0.196 | 0.196 | 0 | 0.61 | 0.94 | 0.71 | 0.9 | 0.48 | 0.84 |
| 5 | 172472573 | rs17729841 | -0.219 | -1.151 | -0.484 | -1.259 | -0.003 | -0.003 | -0.003 | 0 | 0.1 | 0.09 | 0.07 | 0.01 | 0.12 | 0.17 |
| 5 | 172475038 | rs10057441 | 0.177 | -1.716 | 0.085 | -1.451 | -0.04 | -0.069 | -0.069 | 0.122 | 0.22 | 0.26 | 0.06 | 0.02 | 0.07 | 0.11 |
| 5 | 172480336 | rs251253 | 0.064 | -1.458 | -0.16 | -1.761 | 0.012 | 0.067 | 0.067 | 0 | 0.44 | 0.3 | 0.39 | 0.16 | 0.65 | 0.38 |
| 5 | 172483023 | rs29775 | -1.189 | -1.113 | 0.069 | -1.459 | 0.012 | 0.011 | 0.011 | 0 | 0.63 | 0.72 | 0.52 | 0.28 | 0.72 | 0.59 |
| 5 | 172485431 | rs10054054 | 0.191 | -1.179 | 0.136 | -1.389 | -0.076 | -0.122 | -0.122 | 0 | 0.23 | 0.35 | 0.07 | 0.01 | 0.07 | 0.17 |
| 5 | 172492049 | rs13356857 | -0.163 | -1.14 | 0.17 | -1.369 | -0.056 | -0.069 | -0.069 | 1.806 | 0.25 | 0.41 | 0.09 | 0.12 | 0.1 | 0.26 |
| 5 | 172500046 | rs17075560 | 0.221 | -1.144 | 0.178 | -1.372 | -0.076 | -0.122 | -0.122 | 0.026 | 0.23 | 0.35 | 0.07 | 0.01 | 0.07 | 0.17 |
| 5 | 172511602 | rs431604 | -1.339 | -1.17 | 0.175 | -1.466 | 0.021 | 0.017 | 0.017 | 0 | 0.64 | 0.86 | 0.53 | 0.28 | 0.72 | 0.59 |
| 5 | 172515643 | rs370164 | 0.113 | -1.169 | 0.28 | -1.466 | -0.006 | 0.03 | 0.03 | 0 | 0.38 | 0.3 | 0.42 | 0.17 | 0.62 | 0.33 |
| 5 | 172526570 | rs251230 | 1.544 | -0.966 | 0.237 | -1.591 | -0.079 | 0.002 | 0.002 | 0 | 0.23 | 0.08 | 0.28 | 0.16 | 0.39 | 0.15 |
| 5 | 172534347 | rs17662639 | 0.326 | -1.046 | 0.178 | -1.572 | -0.03 | 0.046 | 0.046 | 0 | 0.1 | 0.03 | 0.07 | 0.01 | 0.12 | 0.15 |
| 5 | 172543173 | rs10067669 | 1.085 | -1.163 | 0.414 | -1.609 | 0.015 | -0.004 | -0.004 | 0.033 | 0.18 | 0.11 | 0.03 | 0 | 0.06 | 0.07 |
| 5 | 172560060 | rs10055297 | -0.111 | -1.159 | 0.415 | -1.607 | -0.053 | -0.067 | -0.067 | 0.404 | 0.25 | 0.4 | 0.09 | 0.12 | 0.1 | 0.26 |
| 5 | 172562142 | rs3131912 | -1.335 | -1.103 | 0.377 | -1.606 | 0.011 | 0.009 | 0.009 | 0 | 0.63 | 0.73 | 0.51 | 0.29 | 0.72 | 0.59 |
| 5 | 172568371 | rs17662869 | 1.064 | -1.042 | 0.099 | -1.551 | 0.015 | 0 | 0 | 3.655 | 0.18 | 0.11 | 0.03 | 0 | 0.06 | 0.07 |
| 5 | 172568979 | rs10476062 | 1.082 | -1.042 | 0.099 | -1.563 | 0.015 | -0.004 | -0.004 | 0 | 0.18 | 0.11 | 0.03 | 0 | 0.06 | 0.07 |
| 5 | 172570220 | rs171725 | -0.065 | -0.919 | 0.054 | -1.503 | -0.006 | 0.027 | 0.027 | 0 | 0.38 | 0.32 | 0.42 | 0.17 | 0.62 | 0.33 |
| 5 | 172591337 | rs8543 | 0.305 | -0.993 | -0.039 | -1.548 | -0.042 | 0.049 | 0.049 | 0 | 0.1 | 0.01 | 0.07 | 0.01 | 0.12 | 0.15 |
| 5 | 172595861 | rs39796 | 0.651 | -1.023 | 0.191 | -1.388 | 0.005 | -0.002 | -0.002 | 0 | 0.28 | 0.28 | 0.35 | 0.17 | 0.51 | 0.2 |

| | | | | | | | | | | | | | | | | |
|---|---|---|---|---|---|---|---|---|---|---|---|---|---|---|---|---|
| 5 | 172599318 | rs2250766 | -0.754 | -1.071 | 0.235 | -0.745 | -0.006 | -0.049 | -0.049 | 2.439 | 0.56 | 0.81 | 0.48 | 0.46 | 0.64 | 0.52 |
| 5 | 172606994 | rs6859142 | 1.067 | -1.109 | 0.231 | -1.327 | 0.021 | 0.008 | 0.008 | 1.526 | 0.17 | 0.1 | 0.03 | 0 | 0.05 | 0.05 |
| 5 | 172611648 | rs255313 | -0.671 | -1 | 0.298 | -1.252 | 0.004 | 0.016 | 0.016 | 0 | 0.55 | 0.64 | 0.46 | 0.33 | 0.63 | 0.52 |
| 5 | 172613028 | rs255315 | 0.958 | -1.035 | 0.463 | -1.279 | 0.001 | -0.001 | -0.001 | 0 | 0.26 | 0.25 | 0.34 | 0.22 | 0.51 | 0.2 |
| 5 | 172619253 | rs7737036 | 0.608 | -1.078 | 0.362 | -1.272 | -0.059 | -0.084 | -0.084 | 0 | 0.2 | 0.21 | 0.04 | 0.01 | 0.05 | 0.16 |
| 5 | 172621971 | rs9313618 | -0.482 | -0.349 | 0.109 | -0.882 | 0.002 | 0.002 | 0.002 | 0 | 0.07 | 0.1 | 0.06 | 0.09 | 0.07 | 0.15 |
| 5 | 172624482 | rs7701353 | -0.983 | -0.197 | 0.025 | -0.87 | -0.029 | -0.01 | -0.01 | 0 | 0.77 | 0.8 | 0.72 | 0.78 | 0.62 | 0.83 |
| 5 | 172628038 | rs12109897 | -0.266 | -0.338 | 0.114 | -0.704 | 0.034 | -0.022 | -0.022 | 0 | 0.07 | 0.19 | 0.08 | 0 | 0.12 | 0.04 |
| 5 | 172631051 | rs4868240 | 0.925 | -0.681 | -0.905 | -0.653 | 0.04 | -0.048 | -0.048 | 0.096 | 0.33 | 0.14 | 0.4 | 0.61 | 0.22 | 0.38 |
| 5 | 172632919 | rs11950713 | -0.813 | -0.691 | -0.713 | -1.029 | 0.056 | 0.165 | 0.165 | 0 | 0.45 | 0.8 | 0.31 | 0.15 | 0.37 | 0.46 |
| 5 | 172634592 | rs1123336 | -0.799 | -0.67 | -0.761 | -1.04 | 0.054 | 0.155 | 0.155 | 0 | 0.45 | 0.8 | 0.31 | 0.15 | 0.37 | 0.46 |
| 5 | 172644520 | rs7356620 | -0.071 | -0.768 | -0.84 | -1.078 | 0.017 | 0.133 | 0.133 | 0 | 0.37 | 0.17 | 0.24 | 0.07 | 0.3 | 0.38 |
| 5 | 172655663 | rs3132138 | -1.288 | -0.545 | -0.737 | -0.695 | -0.035 | -0.028 | -0.028 | 0.054 | 0.76 | 0.85 | 0.76 | 0.71 | 0.62 | 0.83 |
| 5 | 172656107 | rs12514371 | 1.341 | -0.801 | -0.747 | -0.82 | 0.037 | -0.061 | -0.061 | 0 | 0.25 | 0.16 | 0.35 | 0.55 | 0.14 | 0.21 |
| 5 | 172657021 | rs3131917 | -0.547 | -0.748 | -0.258 | -0.889 | 0.03 | 0.01 | 0.01 | 0 | 0.64 | 0.51 | 0.63 | 0.61 | 0.5 | 0.59 |
| 5 | 172659511 | rs703752 | 0.129 | -0.549 | -0.248 | -0.892 | -0.002 | 0.035 | 0.035 | 0.03 | 0.34 | 0.25 | 0.26 | 0.07 | 0.34 | 0.36 |
| 5 | 172663454 | rs3095870 | 0.177 | -0.465 | -0.266 | -0.902 | -0.005 | 0.057 | 0.057 | 0.019 | 0.34 | 0.22 | 0.26 | 0.07 | 0.34 | 0.36 |
| 5 | 172664163 | rs6882776 | -0.115 | -0.244 | -0.162 | -0.47 | 0.003 | -0.001 | -0.001 | 0.634 | 0.38 | 0.35 | 0.46 | 0.63 | 0.27 | 0.44 |
| 5 | 172673319 | rs6884881 | -1.033 | -0.087 | -1.125 | -0.705 | -0.044 | 0.025 | 0.025 | 0.004 | 0.63 | 0.86 | 0.74 | 0.75 | 0.54 | 0.73 |
| 5 | 172674820 | rs6873973 | -0.235 | -0.504 | -0.715 | -0.216 | -0.01 | -0.046 | -0.046 | 2.127 | 0.17 | 0.35 | 0.13 | 0.1 | 0.12 | 0.27 |
| 5 | 172675594 | rs2161118 | -0.347 | -0.12 | -0.764 | -0.028 | -0.011 | -0.002 | -0.002 | 0.064 | 0.43 | 0.5 | 0.58 | 0.66 | 0.33 | 0.49 |
| 5 | 172678983 | rs4867697 | -0.28 | 0.36 | -1 | -0.253 | -0.004 | -0.018 | -0.018 | 0 | 0.41 | 0.53 | 0.55 | 0.74 | 0.37 | 0.55 |
| 5 | 172679377 | rs10476070 | -0.361 | 0.283 | -1.536 | -0.535 | 0.021 | 0.131 | 0.131 | 0.723 | 0.54 | 0.76 | 0.65 | 0.82 | 0.36 | 0.69 |
| 5 | 172680870 | rs10072402 | -0.746 | 0.242 | -1.298 | -0.164 | 0.005 | 0.03 | 0.03 | 0 | 0.17 | 0.09 | 0.27 | 0.03 | 0.14 | 0.11 |
| 5 | 172682217 | rs6884482 | -0.668 | 0.062 | -0.986 | 0.32 | -0.006 | 0.003 | 0.003 | 0 | 0.31 | 0.36 | 0.22 | 0.35 | 0.28 | 0.2 |
| 5 | 172682382 | rs6884962 | -1.17 | -0.323 | -0.616 | 0.681 | -0.004 | -0.013 | -0.013 | 0 | 0.35 | 0.45 | 0.57 | 0.68 | 0.36 | 0.41 |
| 5 | 172686386 | rs10475590 | -0.765 | -0.305 | -0.649 | 1.277 | 0.007 | 0.013 | 0.013 | 0 | 0.18 | 0.19 | 0.4 | 0.22 | 0.26 | 0.17 |
| 5 | 172690831 | rs10040431 | -0.957 | -0.321 | -0.709 | 1.711 | -0.002 | -0.001 | -0.001 | 0.532 | 0.22 | 0.24 | 0.49 | 0.4 | 0.29 | 0.2 |
| 5 | 172690902 | rs3132145 | 0.633 | -0.694 | -1.101 | 1.621 | 0.028 | -0.09 | -0.09 | 0 | 0.84 | 0.96 | 0.87 | 0.64 | 0.89 | 0.72 |
| 5 | 172693710 | rs3095876 | -0.931 | 0.327 | -0.548 | 1.035 | 0.018 | -0.005 | -0.005 | 0 | 0.25 | 0.3 | 0.45 | 0.2 | 0.37 | 0.22 |
| 5 | 172699491 | rs9313626 | -0.382 | -1.998 | -1.262 | 0.689 | 0.137 | -0.1 | -0.1 | 6.202 | 0.64 | 0.91 | 0.75 | 0.49 | 0.83 | 0.74 |
| 5 | 172700938 | rs3132148 | 0.285 | -2.106 | -0.262 | 0.49 | -0.03 | 0.12 | 0.12 | 0 | 0.2 | 0.31 | 0.39 | 0.42 | 0.18 | 0.28 |
| 5 | 172705469 | rs4472273 | -0.191 | -2.173 | -0.139 | 0.131 | 0.017 | -0.058 | -0.058 | 0 | 0.24 | 0.09 | 0.41 | 0.34 | 0.24 | 0.26 |
| 5 | 172709271 | rs13162731 | -0.827 | -2.32 | 0.508 | -0.144 | -0.023 | 0.015 | 0.015 | 0 | 0.32 | 0.5 | 0.49 | 0.36 | 0.27 | 0.28 |
| 5 | 172710744 | rs4505973 | -0.007 | -1.873 | 0.524 | -0.15 | -0.006 | -0.025 | -0.025 | 0.313 | 0.11 | 0.19 | 0.27 | 0.07 | 0.09 | 0.08 |
| 5 | 172717383 | rs4365856 | -0.56 | -2.239 | 0.542 | -0.29 | -0.039 | 0.004 | 0.004 | 0.025 | 0.62 | 0.42 | 0.71 | 0.38 | 0.83 | 0.73 |
| 5 | 172719020 | rs4242165 | -1.187 | -2.281 | 0.203 | -0.322 | 0.088 | -0.075 | -0.075 | 0 | 0.69 | 0.89 | 0.76 | 0.41 | 0.85 | 0.78 |
| 5 | 172719187 | rs4868251 | -0.241 | -2.22 | 0.276 | -0.297 | 0.092 | 0.008 | 0.008 | 0.053 | 0.57 | 0.82 | 0.5 | 0.36 | 0.74 | 0.68 |
| 5 | 172722050 | rs12519191 | -0.248 | -1.848 | 0.138 | -0.086 | 0.006 | 0.002 | 0.002 | 0 | 0.24 | 0.16 | 0.37 | 0.2 | 0.24 | 0.24 |
| 5 | 172724558 | rs6898846 | -0.853 | -1.824 | -0.083 | -0.072 | 0.052 | -0.016 | -0.016 | 0.861 | 0.63 | 0.68 | 0.72 | 0.37 | 0.83 | 0.71 |
| 5 | 172725229 | rs10073580 | 0.726 | -1.729 | -0.14 | -0.122 | -0.002 | -0.022 | -0.022 | 0 | 0.38 | 0.36 | 0.33 | 0.18 | 0.58 | 0.47 |
| 5 | 172726085 | rs12519360 | 0.857 | -1.65 | -0.855 | 0.304 | 0.015 | -0.016 | -0.016 | 1.136 | 0.15 | 0.05 | 0.12 | 0.16 | 0.16 | 0.17 |
| 5 | 172726334 | rs4242166 | -0.02 | -1.092 | -0.59 | 0.747 | 0.024 | -0.013 | -0.013 | 1.081 | 0.64 | 0.64 | 0.73 | 0.39 | 0.83 | 0.73 |
| 5 | 172730408 | rs7710233 | 0.361 | 0.847 | -0.302 | 0.535 | 0.007 | 0.006 | 0.006 | 0 | 0.72 | 0.77 | 0.64 | 0.65 | 0.59 | 0.66 |
| 5 | 172736289 | rs874719 | -0.945 | 0.728 | 0.025 | 0.364 | 0.003 | -0.016 | -0.016 | 4.825 | 0.09 | 0.22 | 0.23 | 0.05 | 0.05 | 0.07 |
| 5 | 172737199 | rs6879065 | -1.273 | 0.225 | 0.288 | 0.078 | -0.004 | -0.004 | -0.004 | 0 | 0.44 | 0.56 | 0.61 | 0.43 | 0.48 | 0.45 |
| 5 | 172738233 | rs17675949 | -1.106 | 0.227 | 0.553 | -0.053 | -0.009 | -0.009 | -0.009 | 0 | 0.41 | 0.63 | 0.59 | 0.41 | 0.48 | 0.48 |
| 5 | 172740329 | rs1023617 | -0.431 | 0.459 | 0.498 | -0.009 | 0.008 | 0.013 | 0.013 | 0 | 0.28 | 0.32 | 0.29 | 0.31 | 0.36 | 0.28 |
| 5 | 172746086 | rs2053681 | -0.7 | 0.303 | -0.664 | -0.203 | 0.002 | -0.007 | -0.007 | 0 | 0.47 | 0.38 | 0.59 | 0.41 | 0.58 | 0.34 |
| 5 | 172746539 | rs3756530 | -1.282 | -0.02 | -0.582 | -0.247 | -0.025 | -0.051 | -0.051 | 0 | 0.09 | 0.34 | 0.07 | 0.02 | 0.04 | 0.05 |
| 5 | 172749522 | rs4868257 | -0.667 | 0.431 | -1.043 | -0.121 | -0.03 | -0.028 | -0.028 | 0 | 0.45 | 0.23 | 0.6 | 0.41 | 0.55 | 0.35 |
| 5 | 172754364 | rs3797456 | -0.962 | 0.445 | -1.038 | -0.239 | 0.016 | -0.008 | -0.008 | 0 | 0.25 | 0.17 | 0.14 | 0.25 | 0.2 | 0.26 |
| 5 | 172758076 | rs3756528 | -0.217 | 0.928 | -0.87 | -1.031 | -0.004 | -0.053 | -0.053 | 0 | 0.62 | 0.63 | 0.59 | 0.94 | 0.55 | 0.67 |
| 5 | 172764398 | rs895312 | -0.899 | 0.904 | -0.909 | -0.014 | -0.005 | -0.026 | -0.026 | 0 | 0.09 | 0.03 | 0.14 | 0.16 | 0.08 | 0.18 |
| 5 | 172766002 | rs7730758 | -1.067 | 1.765 | -0.26 | -0.339 | 0.08 | -0.019 | -0.019 | 0 | 0.08 | 0.2 | 0.11 | 0.06 | 0.15 | 0.12 |
| 5 | 172769501 | rs6556070 | -0.556 | 0.897 | -0.606 | -0.7 | 0.023 | 0.049 | 0.049 | 0 | 0.43 | 0.71 | 0.28 | 0.3 | 0.39 | 0.36 |
| 5 | 172772253 | rs921004 | 0.585 | 1.21 | -0.935 | -0.998 | 0.001 | 0.066 | 0.066 | 0.157 | 0.46 | 0.29 | 0.65 | 0.66 | 0.5 | 0.56 |
| 5 | 172772751 | rs11738262 | 0.822 | 0.986 | -0.821 | -1.177 | 0 | 0 | 0 | 0 | 0.08 | 0.07 | 0.04 | 0.13 | 0.1 | 0.09 |
| 5 | 172772955 | rs13154786 | -0.044 | 0.544 | -0.236 | -1.405 | -0.012 | -0.03 | -0.03 | 0 | 0.24 | 0.36 | 0.16 | 0.17 | 0.22 | 0.26 |
| 5 | 172774626 | rs6871935 | 0.007 | 0.541 | 0.061 | -1.53 | -0.008 | -0.046 | -0.046 | 0 | 0.69 | 0.61 | 0.8 | 0.83 | 0.71 | 0.82 |
| 5 | 172778834 | rs895313 | 0.878 | 0.995 | -0.416 | -0.916 | -0.024 | -0.028 | -0.028 | 0 | 0.28 | 0.16 | 0.47 | 0.61 | 0.35 | 0.48 |
| 5 | 172779475 | rs11954548 | -0.67 | 0.627 | -0.016 | -0.858 | 0.005 | -0.008 | -0.008 | 0 | 0.31 | 0.22 | 0.23 | 0.3 | 0.23 | 0.23 |
| 5 | 172780104 | rs17733311 | 0.037 | 0.725 | 0.296 | -0.72 | 0.003 | 0.026 | 0.026 | 0 | 0.19 | 0.26 | 0.2 | 0.17 | 0.16 | 0.14 |
| 5 | 172781401 | rs7717113 | -1.161 | 1.475 | 0.27 | -0.123 | 0.086 | 0.143 | 0.143 | 0 | 0.21 | 0.45 | 0.28 | 0.37 | 0.25 | 0.32 |
| 5 | 172783848 | rs1445788 | -0.08 | 0.969 | 0.241 | -0.345 | 0.023 | 0.014 | 0.014 | 0 | 0.12 | 0.26 | 0.1 | 0.19 | 0.13 | 0.22 |
| 5 | 172784614 | rs1862428 | -0.658 | 0.608 | -0.51 | -0.777 | -0.015 | 0.071 | 0.071 | 0.533 | 0.76 | 0.86 | 0.82 | 0.96 | 0.72 | 0.84 |
| 5 | 172785705 | rs10476071 | -0.119 | 0.431 | -0.62 | -0.759 | 0.009 | 0.072 | 0.072 | 0 | 0.67 | 0.8 | 0.64 | 0.85 | 0.64 | 0.76 |
| 5 | 172794876 | rs12518320 | 1.211 | 0.388 | -0.295 | 0.121 | 0.002 | -0.002 | -0.002 | 0 | 0.53 | 0.36 | 0.43 | 0.5 | 0.46 | 0.36 |
| 5 | 172797393 | rs10069570 | 0.709 | 0.741 | -0.199 | 0.124 | 0.012 | -0.045 | -0.045 | 0 | 0.14 | 0.21 | 0.08 | 0.01 | 0.15 | 0.25 |

| | | | | | | | | | | | | | | | | |
|---|---|---|---|---|---|---|---|---|---|---|---|---|---|---|---|---|
| 5 | 172797829 | rs12515371 | 1.033 | 0.659 | -0.202 | 0.263 | 0 | 0 | 0 | 0.119 | 0.07 | 0.06 | 0.17 | 0.12 | 0.08 | 0.11 |
| 5 | 172798489 | rs1000939 | -0.308 | 0.595 | -0.188 | 0.34 | -0.01 | 0.003 | 0.003 | 0.008 | 0.12 | 0.16 | 0.2 | 0.13 | 0.08 | 0.14 |
| 5 | 172801558 | rs10866663 | 0.333 | 0.708 | -0.319 | 0.372 | 0.003 | 0.01 | 0.01 | 0 | 0.77 | 0.74 | 0.7 | 0.65 | 0.7 | 0.76 |
| 5 | 172802890 | rs13165739 | -0.46 | 0.769 | -0.465 | 0.394 | 0.016 | 0.024 | 0.024 | 0.101 | 0.25 | 0.42 | 0.25 | 0.34 | 0.24 | 0.35 |
| 5 | 172804868 | rs6884147 | 1.387 | 0.787 | -0.459 | 0.391 | 0.009 | 0.001 | 0.001 | 0 | 0.62 | 0.51 | 0.62 | 0.62 | 0.55 | 0.48 |
| 5 | 172805020 | rs11134793 | -0.462 | 0.739 | -0.454 | 0.397 | 0.016 | 0.024 | 0.024 | 0.101 | 0.25 | 0.42 | 0.25 | 0.34 | 0.24 | 0.35 |
| 5 | 172806980 | rs11134794 | 0.751 | 0.786 | -0.637 | 0.284 | 0.005 | -0.021 | -0.021 | 0 | 0.71 | 0.64 | 0.77 | 0.93 | 0.64 | 0.57 |
| 5 | 172807062 | rs921006 | -0.395 | 0.791 | -0.636 | 0.283 | -0.007 | 0.007 | 0.007 | 0 | 0.87 | 0.88 | 0.86 | 0.96 | 0.79 | 0.83 |
| 5 | 172807184 | rs4385222 | 0.659 | 0.761 | -0.654 | 0.31 | 0.005 | -0.021 | -0.021 | 0 | 0.71 | 0.64 | 0.77 | 0.93 | 0.64 | 0.57 |
| 5 | 172813227 | rs17075870 | 0.447 | 0.732 | -0.778 | 0.589 | 0.001 | -0.001 | -0.001 | 0.199 | 0.09 | 0.08 | 0.13 | 0.23 | 0.08 | 0.07 |
| 5 | 172818061 | rs10061327 | 0.203 | 0.82 | -0.926 | 0.485 | -0.031 | -0.019 | -0.019 | 0 | 0.8 | 0.91 | 0.75 | 0.76 | 0.71 | 0.77 |
| 5 | 172818916 | rs6556072 | -0.499 | 0.707 | -0.909 | 0.48 | -0.001 | 0 | 0 | 0.496 | 0.26 | 0.7 | 0.32 | 0.24 | 0.23 | 0.37 |
| 5 | 172821732 | rs11744838 | 0.731 | 0.75 | -1.206 | 0.56 | 0.017 | -0.004 | -0.004 | 0 | 0.16 | 0.26 | 0.1 | 0.12 | 0.17 | 0.28 |
| 5 | 172822746 | rs7707883 | -1.158 | 0.594 | -2.032 | 0.624 | -0.003 | -0.04 | -0.04 | 0 | 0.49 | 0.88 | 0.59 | 0.48 | 0.53 | 0.62 |
| 5 | 172823738 | rs1445792 | -0.233 | 0.355 | -1.028 | 0.989 | 0.004 | -0.027 | -0.027 | 0 | 0.09 | 0.03 | 0.12 | 0.2 | 0.08 | 0.08 |
| 5 | 172824585 | rs6556074 | 0.255 | 0.299 | -0.72 | 0.938 | -0.016 | 0.016 | 0.016 | 0.008 | 0.67 | 0.5 | 0.74 | 0.69 | 0.77 | 0.55 |
| 5 | 172827954 | rs4868270 | 0 | 0.816 | -1.296 | 0.957 | -0.075 | -0.062 | -0.062 | 0 | 0.21 | 0.04 | 0.18 | 0.31 | 0.28 | 0.24 |
| 5 | 172829302 | rs7720513 | -0.198 | 1.236 | -1.172 | 0.967 | 0.007 | -0.001 | -0.001 | 0 | 0.09 | 0.09 | 0.12 | 0.09 | 0.14 | 0.12 |
| 5 | 172836544 | rs13168894 | -0.074 | 1.236 | -1.17 | 0.967 | 0.007 | 0 | 0 | 0 | 0.08 | 0.08 | 0.14 | 0.1 | 0.14 | 0.12 |
| 5 | 172838825 | rs4615311 | -1.347 | 0.548 | -0.905 | 0.842 | 0.007 | 0.008 | 0.008 | 0 | 0.58 | 0.69 | 0.6 | 0.66 | 0.64 | 0.64 |
| 5 | 172839289 | rs10780116 | -0.501 | 0.3 | -0.416 | 0.83 | -0.009 | -0.019 | -0.019 | 0 | 0.41 | 0.33 | 0.4 | 0.48 | 0.45 | 0.45 |
| 5 | 172839584 | rs17075905 | 0.299 | 0.236 | -0.421 | 0.797 | 0.001 | -0.004 | -0.004 | 0 | 0.17 | 0.11 | 0.22 | 0.17 | 0.17 | 0.22 |
| 5 | 172842872 | rs10463029 | -1.157 | 0.224 | -0.401 | 0.681 | 0.001 | 0.002 | 0.002 | 0 | 0.49 | 0.57 | 0.48 | 0.57 | 0.49 | 0.51 |
| 5 | 172844615 | rs6870544 | 0.058 | 0.113 | -0.802 | 0.434 | -0.014 | 0.019 | 0.019 | 0 | 0.16 | 0.07 | 0.15 | 0.09 | 0.23 | 0.17 |
| 5 | 172845597 | rs4867710 | 1.143 | 0.125 | -1.031 | 0.483 | -0.055 | -0.044 | -0.044 | 0.341 | 0.21 | 0.08 | 0.15 | 0.3 | 0.27 | 0.23 |
| 5 | 172852005 | rs17075931 | 0.314 | 0.211 | -0.958 | 0.521 | 0.001 | -0.006 | -0.006 | 0 | 0.17 | 0.13 | 0.23 | 0.22 | 0.17 | 0.22 |
| 5 | 172852878 | rs1445787 | -1.261 | 0.366 | -0.89 | 0.325 | -0.005 | -0.002 | -0.002 | 0.29 | 0.61 | 0.61 | 0.59 | 0.66 | 0.68 | 0.65 |
| 5 | 172856628 | rs6889586 | 0.002 | 0.106 | -0.239 | 0.365 | -0.015 | -0.021 | -0.021 | 0 | 0.15 | 0.33 | 0.1 | 0.09 | 0.12 | 0.14 |
| 5 | 172856883 | rs6889931 | -0.695 | 0.084 | -1.166 | 0.322 | 0.003 | 0.014 | 0.014 | 0.377 | 0.31 | 0.57 | 0.3 | 0.36 | 0.36 | 0.38 |
| 5 | 172860596 | rs11134797 | 0.4 | 0.063 | -1.053 | 0.712 | -0.007 | -0.018 | -0.018 | 0 | 0.06 | 0.07 | 0.04 | 0 | 0.03 | 0.01 |
| 5 | 172864110 | rs6878606 | -0.547 | 0.314 | -1.326 | 0.893 | 0.011 | -0.012 | -0.012 | 0.237 | 0.37 | 0.7 | 0.34 | 0.37 | 0.34 | 0.36 |
| 5 | 172869595 | rs11743405 | -0.099 | 0.225 | -1.587 | 0.747 | 0.059 | 0.026 | 0.026 | 0 | 0.46 | 0.19 | 0.43 | 0.32 | 0.37 | 0.41 |
| 5 | 172869675 | rs17735046 | -1.14 | 1.121 | -1.083 | 0.829 | 0.019 | 0.01 | 0.01 | 0 | 0.15 | 0.24 | 0.18 | 0.27 | 0.24 | 0.23 |
| 5 | 172870009 | rs2112618 | 0.432 | 0.306 | -1.688 | 0.471 | 0.046 | -0.005 | -0.005 | 0 | 0.75 | 0.89 | 0.65 | 0.8 | 0.82 | 0.74 |
| 5 | 172881712 | rs11744776 | -0.724 | 1.581 | -0.653 | 0.139 | 0.099 | -0.016 | -0.016 | 0 | 0.19 | 0.32 | 0.14 | 0.1 | 0.26 | 0.15 |
| 5 | 172886527 | rs750824 | -0.345 | 0.972 | -0.545 | -0.078 | -0.014 | 0.002 | 0.002 | 3.264 | 0.66 | 0.59 | 0.62 | 0.38 | 0.65 | 0.71 |
| 5 | 172892328 | rs17075980 | -1.295 | 0.947 | 0.071 | -0.219 | 0.123 | 0.011 | 0.011 | 0.207 | 0.13 | 0.63 | 0.18 | 0.1 | 0.21 | 0.09 |
| 5 | 172896796 | rs1422658 | -0.314 | 0.843 | 0.173 | -0.359 | 0.007 | 0.002 | 0.002 | 4.469 | 0.62 | 0.7 | 0.64 | 0.37 | 0.63 | 0.64 |
| 5 | 172902359 | rs750168 | 0.592 | 0.644 | 0.042 | -0.525 | 0.014 | 0.098 | 0.098 | 1.218 | 0.48 | 0.34 | 0.48 | 0.28 | 0.41 | 0.54 |
| 5 | 172908863 | rs811013 | 0.75 | 0.787 | -0.12 | -0.525 | 0.13 | 0.008 | 0.008 | 0.608 | 0.86 | 0.66 | 0.83 | 0.9 | 0.77 | 0.9 |
| 5 | 172910955 | rs254786 | 0.339 | 0.307 | -0.681 | -0.318 | 0.008 | -0.057 | -0.057 | 0 | 0.36 | 0.05 | 0.33 | 0.62 | 0.37 | 0.36 |
| 5 | 172919765 | rs254790 | 0.169 | 0.3 | -0.494 | -0.332 | 0.031 | 0.007 | 0.007 | 2.223 | 0.37 | 0.22 | 0.35 | 0.62 | 0.37 | 0.34 |
| 5 | 172924384 | rs254797 | -0.274 | 1.113 | -0.361 | -0.253 | -0.007 | 0.007 | 0.007 | 0.843 | 0.44 | 0.22 | 0.33 | 0.58 | 0.48 | 0.38 |
| 5 | 172926863 | rs13160187 | 0.165 | 1.278 | -0.416 | -0.27 | -0.036 | 0.03 | 0.03 | 0 | 0.06 | 0.01 | 0.06 | 0 | 0.12 | 0.05 |
| 5 | 172927862 | rs29656 | -0.224 | 1.074 | -0.276 | -0.269 | 0.004 | 0.01 | 0.01 | 0.901 | 0.45 | 0.27 | 0.33 | 0.57 | 0.48 | 0.41 |
| 5 | 172930329 | rs7724785 | -0.569 | 0.947 | 0.145 | -0.387 | -0.007 | -0.012 | -0.012 | 0.389 | 0.65 | 0.74 | 0.71 | 0.44 | 0.54 | 0.68 |
| 5 | 172939755 | rs6891752 | -0.612 | 0.393 | 0.407 | -0.942 | -0.003 | -0.023 | -0.023 | 0 | 0.29 | 0.4 | 0.18 | 0.17 | 0.2 | 0.29 |
| 5 | 172942459 | rs1422772 | 0.745 | 0.457 | 0.458 | -0.935 | 0 | 0.008 | 0.008 | 0 | 0.35 | 0.28 | 0.52 | 0.27 | 0.34 | 0.37 |
| 5 | 172950691 | rs11746666 | 0.502 | 0.412 | 0.529 | -0.967 | 0.002 | 0.021 | 0.021 | 0 | 0.36 | 0.3 | 0.53 | 0.28 | 0.34 | 0.39 |
| 5 | 172956807 | rs10076027 | 0.283 | 0.515 | 0.627 | -1.394 | -0.004 | -0.056 | -0.056 | 0 | 0.36 | 0.46 | 0.29 | 0.15 | 0.33 | 0.31 |
| 5 | 172958165 | rs2112620 | 1.289 | 0.236 | 0.528 | -1.353 | 0.015 | 0.022 | 0.022 | 0 | 0.24 | 0.17 | 0.19 | 0.13 | 0.21 | 0.21 |
| 5 | 172960825 | rs17686195 | -0.633 | 0.309 | 0.496 | -1.263 | 0.002 | -0.025 | -0.025 | 9.118 | 0.46 | 0.48 | 0.6 | 0.25 | 0.46 | 0.47 |
| 5 | 172962139 | rs9313635 | -1.767 | 0.116 | -0.25 | -1.302 | -0.135 | -0.112 | -0.112 | 0 | 0.7 | 0.97 | 0.75 | 0.4 | 0.54 | 0.64 |
| 5 | 172962814 | rs6870707 | -0.205 | -0.057 | -0.2 | -1.496 | -0.022 | 0.02 | 0.02 | 0 | 0.19 | 0.11 | 0.38 | 0.12 | 0.24 | 0.26 |
| 5 | 172967460 | rs11744306 | -0.588 | -0.136 | -0.109 | -1.507 | 0.125 | 0.122 | 0.122 | 0.411 | 0.51 | 0.83 | 0.37 | 0.28 | 0.31 | 0.38 |
| 5 | 172968491 | rs17052028 | -0.702 | -0.029 | 0.348 | -1.583 | 0.067 | 0.117 | 0.117 | 0.457 | 0.52 | 0.73 | 0.32 | 0.15 | 0.33 | 0.36 |
| 5 | 172969495 | rs11949667 | -0.082 | -0.135 | 0.476 | -1.643 | 0.028 | 0.034 | 0.034 | 0 | 0.25 | 0.25 | 0.08 | 0.01 | 0.12 | 0.16 |
| 5 | 172975691 | rs10866665 | -1.466 | -0.169 | 0.747 | -1.712 | 0.009 | -0.019 | -0.019 | 0.017 | 0.66 | 0.63 | 0.68 | 0.27 | 0.56 | 0.57 |
| 5 | 172978276 | rs6867566 | -1.419 | -0.183 | 0.739 | -1.718 | 0.008 | -0.019 | -0.019 | 0 | 0.65 | 0.63 | 0.68 | 0.26 | 0.56 | 0.57 |
| 5 | 172979183 | rs6891923 | -1.273 | -0.153 | 0.696 | -1.704 | -0.003 | 0.019 | 0.019 | 0 | 0.41 | 0.42 | 0.47 | 0.13 | 0.35 | 0.37 |
| 5 | 172980714 | rs12520258 | -1.235 | -0.149 | 0.626 | -1.703 | -0.006 | 0.056 | 0.056 | 0 | 0.41 | 0.37 | 0.46 | 0.13 | 0.35 | 0.37 |
| 5 | 172984114 | rs889014 | -1.454 | -0.274 | 0.669 | -1.775 | 0.002 | 0.006 | 0.006 | 0 | 0.43 | 0.6 | 0.45 | 0.13 | 0.34 | 0.36 |
| 5 | 172985611 | rs4473785 | -1.888 | -0.327 | 0.383 | -1.732 | -0.134 | -0.114 | -0.114 | 2.21 | 0.7 | 0.97 | 0.73 | 0.39 | 0.57 | 0.61 |
| 5 | 172988569 | rs12332588 | 1.172 | -0.349 | 0.578 | -1.727 | -0.018 | 0.003 | 0.003 | 0 | 0.27 | 0.39 | 0.27 | 0.27 | 0.23 | 0.25 |
| 5 | 172991573 | rs2112676 | 0.218 | -0.378 | 0.565 | -1.82 | -0.013 | 0.012 | 0.012 | 0 | 0.17 | 0.1 | 0.36 | 0.11 | 0.22 | 0.23 |
| 5 | 172995003 | rs12517954 | 1.254 | -0.393 | 0.329 | -1.795 | -0.01 | 0.001 | 0.001 | 0 | 0.27 | 0.35 | 0.26 | 0.27 | 0.23 | 0.25 |
| 5 | 172995160 | rs12517986 | -1.57 | -0.413 | 0.366 | -1.856 | -0.003 | -0.077 | -0.077 | 0 | 0.43 | 0.51 | 0.45 | 0.13 | 0.34 | 0.37 |
| 5 | 172997978 | rs17738166 | 0.367 | -0.197 | 0.048 | -1.637 | -0.01 | 0.073 | 0.073 | 0 | 0.37 | 0.2 | 0.52 | 0.24 | 0.39 | 0.38 |
| 5 | 173002143 | rs7705011 | -1.052 | -0.042 | 0.577 | -1.336 | -0.013 | -0.029 | -0.029 | 0 | 0.39 | 0.52 | 0.47 | 0.27 | 0.29 | 0.36 |

| | | | | | | | | | | | | | | | | |
|---|---|---|---|---|---|---|---|---|---|---|---|---|---|---|---|---|
| 5 | 173002891 | rs4868293 | -1.096 | -0.501 | 0.25 | -1.926 | 0.003 | -0.017 | -0.017 | 0 | 0.57 | 0.52 | 0.53 | 0.76 | 0.52 | 0.64 |
| 5 | 173003224 | rs2339962 | 0.039 | -0.746 | 0.568 | -1.62 | 0.001 | -0.032 | -0.032 | 1.514 | 0.2 | 0.18 | 0.17 | 0.08 | 0.21 | 0.14 |
| 5 | 173003451 | rs1077613 | -0.781 | -0.819 | 0.622 | -1.405 | -0.014 | -0.025 | -0.025 | 0 | 0.43 | 0.46 | 0.31 | 0.73 | 0.36 | 0.5 |
| 5 | 173004206 | rs7715674 | -0.911 | -0.872 | 0.709 | -0.331 | -0.003 | 0 | 0 | 0.917 | 0.45 | 0.53 | 0.41 | 0.53 | 0.37 | 0.55 |
| 5 | 173004277 | rs13168401 | 0.819 | -0.908 | 0.451 | -1.143 | -0.029 | 0.017 | 0.017 | 4.03 | 0.21 | 0.29 | 0.14 | 0.66 | 0.14 | 0.36 |
| 5 | 173004662 | rs17076110 | -1.409 | -0.837 | 0.636 | -1.291 | -0.028 | -0.019 | -0.019 | 0.071 | 0.13 | 0.52 | 0.17 | 0.07 | 0.05 | 0.06 |
| 5 | 173005250 | rs2112675 | -1.022 | -0.636 | -0.025 | -1.358 | 0.004 | 0.019 | 0.019 | 0 | 0.51 | 0.43 | 0.52 | 0.81 | 0.48 | 0.61 |
| 5 | 173006447 | rs10077254 | -0.772 | -0.695 | -0.069 | -1.415 | -0.008 | 0.024 | 0.024 | 0 | 0.3 | 0.23 | 0.38 | 0.16 | 0.33 | 0.28 |
| 5 | 173007836 | rs12152984 | -0.43 | -0.637 | -0.097 | -1.405 | -0.016 | 0.02 | 0.02 | 0 | 0.26 | 0.16 | 0.33 | 0.09 | 0.33 | 0.25 |
| 5 | 173009201 | rs6885014 | 1.492 | -0.734 | -0.149 | -1.45 | -0.019 | -0.029 | -0.029 | 0.098 | 0.06 | 0.17 | 0.03 | 0 | 0.01 | 0 |
| 5 | 173010992 | rs6884856 | 0.424 | -0.793 | 0.026 | -1.541 | 0 | -0.042 | -0.042 | 0 | 0.08 | 0.24 | 0.05 | 0 | 0.03 | 0.03 |
| 5 | 173011115 | rs746214 | -0.902 | -0.829 | -0.016 | -1.557 | 0.002 | 0.086 | 0.086 | 0.049 | 0.46 | 0.31 | 0.46 | 0.75 | 0.48 | 0.57 |
| 5 | 173011353 | rs746215 | 0.595 | -0.74 | -0.228 | -1.475 | -0.046 | 0.018 | 0.018 | 0.188 | 0.18 | 0.05 | 0.14 | 0.13 | 0.23 | 0.17 |
| 5 | 173016484 | rs12716293 | -1.529 | -0.777 | -0.239 | -1.445 | -0.008 | 0.027 | 0.027 | 0.942 | 0.55 | 0.6 | 0.51 | 0.75 | 0.5 | 0.61 |
| 5 | 173018372 | rs442253 | -1.828 | -1.132 | 0.013 | -0.746 | 0.007 | -0.105 | -0.105 | 0 | 0.85 | 0.98 | 0.89 | 0.39 | 0.87 | 0.68 |
| 5 | 173018537 | rs10516100 | -0.135 | -0.87 | 0.328 | -1.166 | 0 | -0.042 | -0.042 | 0.834 | 0.08 | 0.24 | 0.05 | 0 | 0.03 | 0.03 |
| 5 | 173021618 | rs6876600 | -0.323 | -0.807 | -0.245 | -1.137 | -0.024 | -0.079 | -0.079 | 1.101 | 0.12 | 0.35 | 0.06 | 0 | 0.04 | 0.04 |
| 5 | 173022413 | rs421239 | 0.715 | -0.668 | -1.206 | -0.971 | 0.057 | 0.062 | 0.062 | 1.898 | 0.57 | 0.31 | 0.52 | 0.24 | 0.72 | 0.4 |
| 5 | 173023672 | rs428826 | -0.994 | 0.33 | -1.14 | -1.399 | 0 | 0.026 | 0.026 | 1.25 | 0.39 | 0.66 | 0.39 | 0.83 | 0.34 | 0.42 |
| 5 | 173028193 | rs258868 | -0.077 | 0.611 | -0.941 | -1.403 | 0.011 | -0.019 | -0.019 | 1.05 | 0.08 | 0.13 | 0.05 | 0.01 | 0.1 | 0.05 |
| 5 | 173029602 | rs174056 | -1.312 | 0.545 | -1.596 | -1.501 | 0.018 | 0.119 | 0.119 | 1.685 | 0.49 | 0.69 | 0.4 | 0.83 | 0.42 | 0.53 |
| 5 | 173036161 | rs2279188 | 1.034 | 0.414 | -1.594 | -1.274 | 0.001 | -0.021 | -0.021 | 0 | 0.11 | 0.04 | 0.12 | 0.64 | 0.1 | 0.28 |
| 5 | 173036786 | rs10866666 | 1.47 | 1.261 | -1.924 | -1.568 | -0.061 | 0.069 | 0.069 | 0 | 0.12 | 0.02 | 0.13 | 0 | 0.22 | 0.05 |
| 5 | 173037054 | rs258862 | -0.753 | 0.438 | -1.18 | -1.285 | -0.003 | -0.008 | -0.008 | 0 | 0.43 | 0.54 | 0.29 | 0.76 | 0.32 | 0.47 |
| 5 | 173060082 | rs10476080 | -0.793 | -0.466 | -1.577 | -1.077 | 0.054 | 0.092 | 0.092 | 1.237 | 0.45 | 0.71 | 0.37 | 0.81 | 0.32 | 0.49 |
| 5 | 173062963 | rs2339995 | -0.709 | -0.063 | -1.132 | -0.663 | 0.015 | 0.023 | 0.023 | 1.31 | 0.35 | 0.7 | 0.36 | 0.81 | 0.27 | 0.39 |
| 5 | 173065757 | rs17076219 | -1.02 | 0.677 | -0.84 | -0.076 | 0.007 | 0.027 | 0.027 | 0 | 0.05 | 0.14 | 0.11 | 0.08 | 0.09 | 0.06 |
| 5 | 173073664 | rs6860785 | -0.694 | 0.764 | -1.357 | -0.404 | -0.014 | 0.02 | 0.02 | 0.216 | 0.22 | 0.19 | 0.25 | 0.14 | 0.29 | 0.3 |
| 5 | 173076223 | rs11740316 | 0.447 | 0.548 | -2.176 | -0.91 | -0.035 | 0.08 | 0.08 | 2.162 | 0.38 | 0.07 | 0.46 | 0.22 | 0.43 | 0.3 |
| 5 | 173084783 | rs9313641 | -0.476 | 0.471 | -0.952 | -0.992 | 0.018 | -0.112 | -0.112 | 0 | 0.17 | 0.5 | 0.17 | 0.02 | 0.2 | 0.19 |
| 5 | 173094151 | rs10060840 | 0.463 | 0.199 | -1.141 | -1.075 | -0.018 | 0.02 | 0.02 | 1.705 | 0.25 | 0.15 | 0.39 | 0.13 | 0.38 | 0.22 |
| 5 | 173102585 | rs10053217 | 0.091 | -0.078 | -0.342 | -1.16 | 0.021 | -0.089 | -0.089 | 3.632 | 0.29 | 0.47 | 0.38 | 0.12 | 0.38 | 0.22 |
| 5 | 173103337 | rs791645 | 0.946 | -0.5 | -1.033 | -1.157 | -0.001 | -0.114 | -0.114 | 0 | 0.19 | 0.03 | 0.15 | 0.62 | 0.14 | 0.31 |
| 5 | 173103862 | rs13166404 | 0.796 | -0.438 | -0.938 | -1.141 | 0.019 | -0.075 | -0.075 | 0 | 0.28 | 0.38 | 0.37 | 0.12 | 0.37 | 0.22 |
| 5 | 173106601 | rs17076260 | 0.476 | -0.798 | -1.109 | -1.377 | 0.014 | -0.001 | -0.001 | 0 | 0.13 | 0.06 | 0.13 | 0.1 | 0.1 | 0.15 |
| 5 | 173116066 | rs791641 | 0.098 | -0.542 | -1.009 | -1.335 | 0 | -0.002 | -0.002 | 0 | 0.48 | 0.48 | 0.53 | 0.74 | 0.52 | 0.54 |
| 5 | 173120513 | rs703724 | -0.173 | -0.604 | -1.067 | -1.736 | 0 | -0.024 | -0.024 | 0.119 | 0.56 | 0.63 | 0.5 | 0.84 | 0.33 | 0.59 |
| 5 | 173121339 | rs13183493 | 0.272 | -0.47 | -1.322 | -1.821 | 0.027 | 0.136 | 0.136 | 0.135 | 0.51 | 0.3 | 0.39 | 0.12 | 0.46 | 0.35 |
| 5 | 173122092 | rs11740647 | 0.07 | -0.006 | -1.355 | -1.514 | -0.011 | 0.1 | 0.1 | 0.356 | 0.41 | 0.23 | 0.5 | 0.19 | 0.48 | 0.31 |
| 5 | 173124502 | rs4868303 | 0.065 | -0.042 | -1.394 | -1.5 | -0.013 | 0.112 | 0.112 | 0.222 | 0.41 | 0.2 | 0.5 | 0.19 | 0.47 | 0.31 |
| 5 | 173128358 | rs7710739 | 0.648 | -0.061 | -1.151 | -1.345 | -0.035 | 0.09 | 0.09 | 0 | 0.3 | 0.12 | 0.32 | 0.13 | 0.35 | 0.19 |
| 5 | 173128421 | rs2561053 | -0.042 | 0.561 | -0.884 | -1.443 | 0.004 | 0.042 | 0.042 | 0 | 0.55 | 0.61 | 0.45 | 0.72 | 0.42 | 0.66 |
| 5 | 173129604 | rs2546725 | -0.563 | 0.82 | -0.812 | -1.294 | -0.004 | 0.004 | 0.004 | 0 | 0.92 | 0.91 | 0.91 | 0.95 | 0.81 | 0.87 |
| 5 | 173131457 | rs4868304 | 0.846 | 0.409 | -0.789 | -1.316 | 0.009 | 0.008 | 0.008 | 0.256 | 0.19 | 0.11 | 0.14 | 0.11 | 0.17 | 0.17 |
| 5 | 173132309 | rs10074473 | 0.296 | -0.031 | -0.777 | -1.197 | -0.013 | 0.023 | 0.023 | 0 | 0.54 | 0.7 | 0.67 | 0.34 | 0.51 | 0.36 |
| 5 | 173133180 | rs17690039 | 0.13 | 0.074 | -0.708 | -1.224 | 0.003 | 0.011 | 0.011 | 0.148 | 0.1 | 0.11 | 0.07 | 0 | 0.11 | 0.04 |
| 5 | 173134328 | rs746956 | 1.145 | 0.175 | -0.127 | -1.237 | 0.004 | -0.033 | -0.033 | 1.68 | 0.31 | 0.27 | 0.38 | 0.16 | 0.33 | 0.19 |
| 5 | 173134895 | rs10072660 | 1.206 | 0.201 | -0.276 | -1.224 | -0.003 | 0.008 | 0.008 | 2.783 | 0.29 | 0.19 | 0.37 | 0.16 | 0.33 | 0.19 |
| 5 | 173136182 | rs749408 | -0.404 | -0.02 | -0.67 | -1.311 | -0.02 | 0.024 | 0.024 | 1.003 | 0.59 | 0.71 | 0.61 | 0.27 | 0.51 | 0.34 |
| 5 | 173136667 | rs755643 | 1.593 | -0.173 | -0.824 | -1.25 | -0.042 | 0.02 | 0.02 | 4.171 | 0.26 | 0.04 | 0.36 | 0.18 | 0.28 | 0.17 |
| 5 | 173138434 | rs791623 | 0.704 | -0.24 | -0.625 | -0.951 | 0.022 | -0.075 | -0.075 | 0 | 0.31 | 0.12 | 0.16 | 0.6 | 0.21 | 0.5 |
| 5 | 173138522 | rs4868306 | 0.099 | -0.168 | -0.598 | -0.921 | -0.001 | 0.001 | 0.001 | 0 | 0.06 | 0.12 | 0.15 | 0.08 | 0.08 | 0.02 |
| 5 | 173138928 | rs812064 | 0.019 | -0.1 | -0.287 | -0.961 | -0.011 | -0.009 | -0.009 | 0 | 0.42 | 0.58 | 0.35 | 0.67 | 0.29 | 0.52 |
| 5 | 173141093 | rs17076316 | 1.16 | 0.205 | -0.821 | -1.11 | -0.021 | 0.011 | 0.011 | 1.104 | 0.32 | 0.12 | 0.39 | 0.25 | 0.35 | 0.21 |
| 5 | 173141268 | rs8180392 | 0.35 | -0.03 | -0.336 | -1.367 | 0.078 | -0.14 | -0.14 | 0.634 | 0.76 | 0.48 | 0.73 | 0.94 | 0.62 | 0.78 |
| 5 | 173143508 | rs17076328 | 1.556 | 0.144 | -0.899 | -1.268 | -0.009 | 0.002 | 0.002 | 1.759 | 0.3 | 0.12 | 0.38 | 0.25 | 0.32 | 0.19 |
| 5 | 173143912 | rs12651740 | 1.19 | 0.162 | -1.06 | -1.117 | -0.03 | -0.104 | -0.104 | 0.513 | 0.25 | 0.03 | 0.26 | 0.61 | 0.18 | 0.42 |
| 5 | 173144887 | rs13178761 | 0.57 | -0.175 | -0.606 | -1.254 | 0.002 | 0.018 | 0.018 | 0.863 | 0.38 | 0.39 | 0.45 | 0.25 | 0.42 | 0.24 |
| 5 | 173146043 | rs2339989 | 1.157 | -1.242 | -0.274 | -0.977 | -0.035 | -0.002 | -0.002 | 0 | 0.17 | 0.28 | 0.11 | 0.14 | 0.09 | 0.15 |
| 5 | 173146525 | rs2339990 | 0.057 | -1.24 | -0.282 | -0.926 | -0.019 | -0.014 | -0.014 | 0 | 0.43 | 0.47 | 0.22 | 0.67 | 0.25 | 0.55 |
| 5 | 173146887 | rs10064888 | -0.273 | -1.187 | -0.432 | -0.963 | 0.029 | -0.065 | -0.065 | 0 | 0.75 | 0.71 | 0.75 | 0.94 | 0.55 | 0.75 |
| 5 | 173149799 | rs10516104 | 0.328 | -1.931 | 0.263 | -0.341 | -0.065 | -0.002 | -0.002 | 0 | 0.19 | 0.39 | 0.11 | 0.14 | 0.08 | 0.16 |
| 5 | 173151171 | rs6869795 | 0.491 | -1.877 | -0.031 | -0.317 | -0.05 | -0.002 | -0.002 | 0 | 0.18 | 0.34 | 0.12 | 0.15 | 0.09 | 0.16 |
| 5 | 173151427 | rs1107406 | 0.278 | -1.889 | -0.367 | -0.325 | 0.016 | 0.011 | 0.011 | 0 | 0.33 | 0.26 | 0.55 | 0.26 | 0.31 | 0.21 |
| 5 | 173154285 | rs13179033 | -0.582 | -1.754 | -0.435 | -0.24 | 0.032 | 0 | 0 | 3.976 | 0.53 | 0.65 | 0.67 | 0.42 | 0.39 | 0.37 |
| 5 | 173156504 | rs10223202 | -0.038 | -0.951 | -0.929 | -0.067 | 0.014 | 0.002 | 0.002 | 0 | 0.3 | 0.24 | 0.53 | 0.26 | 0.29 | 0.21 |
| 5 | 173160609 | rs2879659 | -0.872 | -0.606 | -0.648 | -0.277 | -0.039 | 0.069 | 0.069 | 0.33 | 0.68 | 0.89 | 0.6 | 0.8 | 0.6 | 0.72 |
| 5 | 173161774 | rs6861562 | -0.313 | 0.192 | 0.217 | 0.411 | -0.016 | -0.025 | -0.025 | 0 | 0.29 | 0.21 | 0.34 | 0.5 | 0.32 | 0.42 |
| 5 | 173163893 | rs10078322 | -1.153 | 0.821 | 0.618 | 0.73 | -0.005 | -0.002 | -0.002 | 0 | 0.22 | 0.68 | 0.2 | 0.22 | 0.19 | 0.19 |

| | | | | | | | | | | | | | | | | |
|---|---|---|---|---|---|---|---|---|---|---|---|---|---|---|---|---|
| 5 | 173167983 | rs4868318 | -1.399 | 0.982 | 0.429 | 0.834 | -0.008 | 0.038 | 0.038 | 0.002 | 0.43 | 0.67 | 0.49 | 0.71 | 0.48 | 0.56 |
| 5 | 173168007 | rs6895993 | -0.682 | 1.016 | 0.428 | 0.795 | 0.002 | -0.009 | -0.009 | 0.027 | 0.17 | 0.21 | 0.14 | 0.14 | 0.17 | 0.17 |
| 5 | 173168142 | rs4868319 | -0.546 | 0.897 | 0.239 | 1.127 | 0.042 | 0.069 | 0.069 | 0 | 0.25 | 0.36 | 0.34 | 0.51 | 0.31 | 0.39 |
| 5 | 173180696 | rs17076407 | -0.994 | -0.139 | 0.174 | 0.814 | -0.03 | 0.062 | 0.062 | 0.273 | 0.1 | 0.25 | 0.09 | 0.19 | 0.06 | 0.15 |
| 5 | 173183624 | rs1363617 | 0.284 | -0.05 | -0.33 | 0.246 | 0.007 | 0.01 | 0.01 | 0 | 0.56 | 0.37 | 0.66 | 0.79 | 0.66 | 0.7 |
| 5 | 173190646 | rs2561746 | -0.945 | -0.086 | 0.088 | 0.312 | -0.003 | -0.005 | -0.005 | 0 | 0.15 | 0.18 | 0.18 | 0.12 | 0.14 | 0.1 |
| 5 | 173191687 | rs13164950 | -0.655 | -0.154 | -0.283 | 1.206 | 0.002 | 0.003 | 0.003 | 0.03 | 0.15 | 0.11 | 0.28 | 0.56 | 0.13 | 0.44 |
| 5 | 173195754 | rs2594830 | -1.695 | -0.202 | 0.581 | 0.786 | 0.033 | 0.014 | 0.014 | 0.05 | 0.08 | 0.31 | 0.15 | 0.07 | 0.09 | 0.11 |
| 5 | 173202901 | rs2594834 | -1.759 | -0.295 | 0.531 | 0.74 | 0.004 | -0.001 | -0.001 | 0 | 0.07 | 0.16 | 0.13 | 0.07 | 0.09 | 0.08 |
| 5 | 173203875 | rs4868324 | -0.529 | -0.401 | 0.283 | 1.328 | -0.002 | -0.029 | -0.029 | 0 | 0.08 | 0.02 | 0.26 | 0.69 | 0.07 | 0.34 |
| 5 | 173204210 | rs812618 | -0.525 | 0.246 | -0.067 | -0.071 | -0.022 | 0.023 | 0.023 | 0 | 0.79 | 0.92 | 0.85 | 1 | 0.74 | 0.93 |
| 5 | 173205058 | rs10056714 | 0.909 | 0.09 | 0.688 | -0.364 | 0.003 | -0.044 | -0.044 | 5.099 | 0.66 | 0.4 | 0.63 | 0.92 | 0.62 | 0.79 |
| 5 | 173205225 | rs791357 | -0.4 | 0.16 | 0.875 | -0.396 | 0.001 | -0.047 | -0.047 | 2.725 | 0.78 | 0.83 | 0.82 | 1 | 0.73 | 0.93 |
| 5 | 173206446 | rs7447290 | -0.614 | 0.121 | 0.776 | 0.668 | 0 | 0.016 | 0.016 | 0.262 | 0.14 | 0.13 | 0.38 | 0.57 | 0.14 | 0.39 |
| 5 | 173207311 | rs16821 | 1.474 | -0.539 | 0.643 | 0.082 | -0.006 | 0.039 | 0.039 | 0 | 0.51 | 0.23 | 0.29 | 0.4 | 0.48 | 0.35 |
| 5 | 173208162 | rs9313655 | -1.064 | -0.372 | 0.934 | 0.031 | 0.027 | -0.038 | -0.038 | 0 | 0.09 | 0.23 | 0.15 | 0.01 | 0.12 | 0.05 |
| 5 | 173208608 | rs360018 | 0.33 | -0.437 | 0.9 | 0.033 | -0.016 | -0.036 | -0.036 | 0 | 0.68 | 0.78 | 0.5 | 0.44 | 0.64 | 0.55 |
| 5 | 173213312 | rs2339992 | 1.798 | -0.618 | 0.036 | -0.409 | -0.064 | 0.057 | 0.057 | 0 | 0.34 | 0.06 | 0.22 | 0.2 | 0.4 | 0.17 |
| 5 | 173213376 | rs791363 | -0.639 | -0.685 | 0.28 | -0.462 | 0.006 | 0.018 | 0.018 | 0 | 0.68 | 0.71 | 0.83 | 0.89 | 0.73 | 0.82 |
| 5 | 173214249 | rs791365 | 1.678 | -0.691 | 0.502 | -0.412 | -0.006 | 0.017 | 0.017 | 0 | 0.37 | 0.31 | 0.25 | 0.2 | 0.4 | 0.17 |
| 5 | 173215611 | rs13160587 | 0.018 | -0.348 | 0.399 | 0.181 | 0.001 | 0.01 | 0.01 | 0.393 | 0.15 | 0.13 | 0.38 | 0.56 | 0.15 | 0.41 |
| 5 | 173216595 | rs791369 | -1.448 | -0.03 | 0.688 | 0.09 | 0.007 | -0.032 | -0.032 | 0 | 0.17 | 0.39 | 0.2 | 0.12 | 0.17 | 0.23 |
| 5 | 173217328 | rs17076500 | 0.461 | 0.175 | 0.185 | 0.765 | 0.001 | 0.022 | 0.022 | 0 | 0.17 | 0.16 | 0.35 | 0.42 | 0.16 | 0.2 |
| 5 | 173220065 | rs2042245 | 0.512 | 0.029 | 0.205 | -0.373 | 0.006 | -0.017 | -0.017 | 0 | 0.68 | 0.52 | 0.44 | 0.28 | 0.69 | 0.48 |
| 5 | 173223238 | rs791371 | -0.518 | 0.014 | -0.513 | -0.942 | 0.068 | 0.133 | 0.133 | 0.015 | 0.5 | 0.77 | 0.3 | 0.19 | 0.34 | 0.57 |
| 5 | 173226092 | rs791377 | 1.305 | 0.171 | -0.509 | -0.772 | -0.016 | 0.016 | 0.016 | 0 | 0.28 | 0.15 | 0.21 | 0.18 | 0.35 | 0.12 |
| 5 | 173232541 | rs7704485 | 0.89 | -0.248 | -0.798 | -1.203 | 0.011 | 0.038 | 0.038 | 0 | 0.21 | 0.14 | 0.1 | 0.1 | 0.19 | 0.27 |
| 5 | 173233656 | rs2546741 | 0.177 | -0.205 | -0.183 | -0.895 | 0.002 | 0.002 | 0.002 | 0 | 0.67 | 0.65 | 0.42 | 0.35 | 0.61 | 0.63 |
| 5 | 173234258 | rs2546738 | 0.203 | -0.29 | 0.052 | -1.048 | -0.008 | -0.062 | -0.062 | 1.227 | 0.11 | 0.28 | 0.06 | 0 | 0.08 | 0.05 |
| 5 | 173235253 | rs2546734 | -1.538 | -0.064 | -0.697 | -0.962 | -0.061 | 0.02 | 0.02 | 0 | 0.83 | 0.94 | 0.86 | 0.9 | 0.76 | 0.93 |
| 5 | 173235500 | rs2594852 | 1.179 | -0.688 | -0.272 | -0.83 | 0.007 | 0.01 | 0.01 | 0 | 0.11 | 0.12 | 0.04 | 0.05 | 0.05 | 0.2 |
| 5 | 173237531 | rs2546722 | -0.836 | -0.725 | -0.2 | -0.817 | -0.045 | 0.02 | 0.02 | 0 | 0.3 | 0.48 | 0.47 | 0.66 | 0.2 | 0.51 |
| 5 | 173242125 | rs2546775 | 0.344 | -0.726 | -0.645 | -1.165 | 0.025 | 0.03 | 0.03 | 0 | 0.26 | 0.17 | 0.07 | 0.18 | 0.15 | 0.34 |
| 5 | 173242579 | rs2546772 | 0.282 | -0.056 | -1.149 | -1.665 | -0.023 | 0.125 | 0.125 | 0 | 0.45 | 0.21 | 0.23 | 0.24 | 0.46 | 0.45 |
| 5 | 173249692 | rs12652980 | -1.85 | -0.095 | -1.582 | -1.878 | -0.065 | 0.092 | 0.092 | 0 | 0.79 | 0.99 | 0.85 | 0.95 | 0.76 | 0.93 |
| 5 | 173253193 | rs2659182 | 1.423 | 0.12 | -1.042 | -1.693 | -0.051 | 0.048 | 0.048 | 0 | 0.19 | 0.06 | 0.16 | 0.05 | 0.32 | 0.11 |
| 5 | 173254352 | rs1506726 | 0.317 | -0.146 | -0.168 | -1.551 | 0.06 | 0.049 | 0.049 | 0.677 | 0.94 | 0.54 | 0.88 | 1 | 0.99 | 0.97 |
| 5 | 173255480 | rs2659186 | 0.178 | -0.112 | -0.402 | -1.531 | 0 | 0.055 | 0.055 | 0 | 0.48 | 0.37 | 0.26 | 0.24 | 0.48 | 0.45 |
| 5 | 173260625 | rs7448268 | 0.033 | 0.198 | -0.347 | -1.439 | -0.005 | -0.017 | -0.017 | 0 | 0.56 | 0.47 | 0.46 | 0.19 | 0.69 | 0.35 |
| 5 | 173260645 | rs360014 | 0.082 | 0.204 | -0.356 | -1.459 | -0.011 | -0.023 | -0.023 | 0 | 0.56 | 0.48 | 0.47 | 0.22 | 0.7 | 0.35 |
| 5 | 173261584 | rs17750318 | 0.48 | 0.057 | -0.422 | -1.473 | -0.003 | 0.09 | 0.09 | 0 | 0.41 | 0.22 | 0.27 | 0.17 | 0.45 | 0.24 |
| 5 | 173262693 | rs7704517 | -0.909 | -0.342 | -0.05 | -1.372 | -0.008 | -0.012 | -0.012 | 0 | 0.27 | 0.39 | 0.53 | 0.72 | 0.3 | 0.46 |
| 5 | 173266547 | rs2594841 | 0.526 | -0.352 | -0.071 | -1.579 | -0.008 | -0.022 | -0.022 | 1.206 | 0.73 | 0.6 | 0.77 | 0.86 | 0.72 | 0.71 |
| 5 | 173267556 | rs2594850 | -0.991 | -0.311 | -0.121 | -1.313 | -0.01 | -0.041 | -0.041 | 0.028 | 0.91 | 0.82 | 0.91 | 1 | 0.96 | 0.9 |
| 5 | 173270927 | rs4867730 | -0.11 | -0.478 | -0.392 | -0.967 | 0.003 | 0 | 0 | 0.354 | 0.42 | 0.55 | 0.61 | 0.6 | 0.29 | 0.42 |
| 5 | 173272518 | rs12187599 | 2.088 | -0.564 | -0.585 | -1.024 | 0.003 | 0.057 | 0.057 | 1.464 | 0.19 | 0.06 | 0.09 | 0.04 | 0.16 | 0.03 |
| 5 | 173275241 | rs10051599 | -0.049 | -0.486 | -0.373 | -0.791 | -0.005 | 0 | 0 | 0.527 | 0.42 | 0.56 | 0.61 | 0.57 | 0.28 | 0.42 |
| 5 | 173277070 | rs165548 | -0.057 | -0.268 | -0.41 | -0.947 | -0.002 | -0.032 | -0.032 | 0.363 | 0.91 | 0.83 | 0.93 | 1 | 0.9 | 0.97 |
| 5 | 173279842 | rs359457 | 0.381 | 0.207 | -0.436 | -1.223 | -0.011 | -0.007 | -0.007 | 0 | 0.68 | 0.69 | 0.43 | 0.4 | 0.59 | 0.62 |
| 5 | 173284933 | rs2659187 | 1.071 | 0.202 | -0.466 | -1.464 | -0.018 | -0.001 | -0.001 | 0 | 0.69 | 0.51 | 0.49 | 0.33 | 0.69 | 0.57 |
| 5 | 173285204 | rs359445 | 0.773 | 0.2 | -0.372 | -1.472 | 0.032 | 0.038 | 0.038 | 0.228 | 0.11 | 0.16 | 0.08 | 0.14 | 0.19 | 0.15 |
| 5 | 173285545 | rs6894235 | 1.681 | -0.102 | -0.451 | -1.766 | -0.002 | -0.037 | -0.037 | 0 | 0.31 | 0.34 | 0.21 | 0.04 | 0.22 | 0.16 |
| 5 | 173287851 | rs875741 | 1.079 | -0.079 | -0.515 | -1.767 | 0 | -0.003 | -0.003 | 0.69 | 0.53 | 0.45 | 0.32 | 0.26 | 0.4 | 0.47 |
| 5 | 173289036 | rs2594845 | 1.742 | -0.026 | -0.577 | -1.541 | -0.015 | 0.018 | 0.018 | 0 | 0.22 | 0.06 | 0.09 | 0.21 | 0.18 | 0.31 |
| 5 | 173292099 | rs359436 | 0.651 | -0.263 | -0.223 | -1.519 | -0.03 | -0.01 | -0.01 | 0 | 0.13 | 0.49 | 0.16 | 0.05 | 0.05 | 0.19 |
| 5 | 173297883 | rs10052344 | 1.486 | -0.276 | -0.619 | -1.256 | -0.001 | 0.105 | 0.105 | 0 | 0.08 | 0.09 | 0.37 | 0.52 | 0.07 | 0.25 |
| 5 | 173298949 | rs17076646 | 2.991 | -0.303 | -0.966 | -1.54 | -0.025 | 0.091 | 0.091 | 1.914 | 0.2 | 0.01 | 0.09 | 0.04 | 0.16 | 0.03 |
| 5 | 173300687 | rs359470 | 0.697 | -0.314 | -0.801 | -1.666 | -0.014 | 0.042 | 0.042 | 0.178 | 0.71 | 0.54 | 0.49 | 0.37 | 0.69 | 0.64 |
| 5 | 173305432 | rs359477 | 1.344 | -0.133 | -1.062 | -1.718 | -0.095 | 0.152 | 0.152 | 2.737 | 0.36 | 0.11 | 0.26 | 0.1 | 0.45 | 0.13 |
| 5 | 173311438 | rs2659178 | 1.967 | -0.188 | -0.908 | -1.622 | -0.005 | 0.012 | 0.012 | 0 | 0.22 | 0.09 | 0.09 | 0.22 | 0.17 | 0.31 |
| 5 | 173314999 | rs359465 | 2.959 | -0.165 | -0.966 | -1.603 | 0.001 | 0.086 | 0.086 | 1.78 | 0.2 | 0.03 | 0.09 | 0.04 | 0.16 | 0.03 |
| 5 | 173316729 | rs359467 | 0.701 | -0.095 | -0.71 | -1.592 | -0.013 | 0.032 | 0.032 | 0.165 | 0.72 | 0.56 | 0.49 | 0.36 | 0.69 | 0.64 |
| 5 | 173334792 | rs2594867 | 1.036 | -0.09 | -0.76 | -1.656 | 0 | -0.004 | -0.004 | 0 | 0.57 | 0.52 | 0.32 | 0.29 | 0.41 | 0.54 |
| 5 | 173339222 | rs7736263 | 1.682 | 0.044 | -0.852 | -1.594 | -0.093 | 0.014 | 0.014 | 0.848 | 0.15 | 0.03 | 0.17 | 0.07 | 0.29 | 0.1 |
| 5 | 173345409 | rs17695328 | 1.693 | -0.025 | -0.942 | -1.58 | -0.042 | -0.031 | -0.031 | 0 | 0.07 | 0.01 | 0.07 | 0.11 | 0.18 | 0.08 |
| 5 | 173349116 | rs10059384 | 0.765 | -0.176 | -0.817 | -1.665 | -0.01 | -0.022 | -0.022 | 0 | 0.35 | 0.41 | 0.22 | 0.27 | 0.24 | 0.51 |
| 5 | 173352763 | rs1106693 | -1.096 | -0.228 | -0.775 | -1.693 | 0.003 | -0.005 | -0.005 | 0 | 0.93 | 0.88 | 0.95 | 1 | 0.94 | 0.98 |
| 5 | 173359051 | rs6869682 | 1.348 | -0.21 | -0.779 | -1.667 | -0.015 | -0.015 | -0.015 | 0 | 0.12 | 0.32 | 0.13 | 0.05 | 0.07 | 0.19 |
| 5 | 173376506 | rs12652907 | 1.808 | 0.154 | -0.884 | -1.697 | -0.003 | 0.114 | 0.114 | 0 | 0.08 | 0.09 | 0.35 | 0.53 | 0.05 | 0.25 |

| | | | | | | | | | | | | | | | | |
|---|---|---|---|---|---|---|---|---|---|---|---|---|---|---|---|---|
| 5 | 173380346 | rs969518 | -0.122 | 0.149 | -1.058 | -1.778 | -0.002 | -0.004 | -0.004 | 0 | 0.52 | 0.5 | 0.41 | 0.33 | 0.53 | 0.61 |
| 5 | 173383194 | rs1564823 | 1.529 | 0.428 | -1.338 | -1.5 | -0.093 | 0.014 | 0.014 | 0.848 | 0.15 | 0.03 | 0.17 | 0.07 | 0.29 | 0.1 |
| 5 | 173390382 | rs6875935 | 1.381 | 0.063 | -0.548 | -1.246 | 0.022 | 0.01 | 0.01 | 0.842 | 0.6 | 0.27 | 0.36 | 0.35 | 0.63 | 0.51 |
| 5 | 173394166 | rs1484802 | 1.05 | 0.052 | -0.519 | -0.754 | 0.007 | 0.039 | 0.039 | 0 | 0.07 | 0.04 | 0.3 | 0.49 | 0.05 | 0.24 |
| 5 | 173394487 | rs7724743 | 1.331 | -0.067 | -0.63 | -1.214 | 0.01 | -0.001 | -0.001 | 0 | 0.17 | 0.07 | 0.04 | 0.22 | 0.08 | 0.31 |
| 5 | 173401662 | rs7734214 | 0.147 | -0.058 | -0.463 | -1.416 | -0.011 | -0.042 | -0.042 | 1.495 | 0.25 | 0.3 | 0.17 | 0.16 | 0.16 | 0.37 |
| 5 | 173406806 | rs1905655 | 0.151 | 0.228 | -0.262 | -1.34 | -0.007 | 0.033 | 0.033 | 0 | 0.8 | 0.71 | 0.53 | 0.44 | 0.81 | 0.71 |
| 5 | 173409953 | rs12518748 | -0.849 | 0.8 | -0.383 | -1.777 | -0.001 | -0.011 | -0.011 | 0 | 0.35 | 0.32 | 0.25 | 0.16 | 0.32 | 0.43 |
| 5 | 173412816 | rs7726709 | 0.883 | 0.592 | -0.14 | -1.55 | -0.002 | -0.051 | -0.051 | 0 | 0.1 | 0.23 | 0.13 | 0.01 | 0.07 | 0.12 |
| 5 | 173413036 | rs7727165 | -0.127 | 0.617 | -0.154 | -1.547 | -0.012 | -0.043 | -0.043 | 0 | 0.26 | 0.31 | 0.17 | 0.16 | 0.16 | 0.4 |
| 5 | 173415407 | rs11744584 | 0.778 | 1 | -0.326 | -1.613 | -0.022 | 0.003 | 0.003 | 0.044 | 0.06 | 0.02 | 0.04 | 0.05 | 0.14 | 0.05 |
| 5 | 173418926 | rs7703555 | -0.52 | 0.892 | -0.046 | -1.81 | 0.02 | -0.077 | -0.077 | 0 | 0.24 | 0.32 | 0.23 | 0.05 | 0.28 | 0.25 |
| 5 | 173422382 | rs1422958 | -0.9 | 0.565 | 0.166 | -1.936 | -0.023 | -0.142 | -0.142 | 0 | 0.2 | 0.47 | 0.19 | 0.01 | 0.14 | 0.15 |
| 5 | 173424464 | rs7724254 | 0.665 | 0.652 | 0.159 | -1.893 | 0.012 | -0.064 | -0.064 | 0 | 0.11 | 0.31 | 0.14 | 0.01 | 0.08 | 0.12 |
| 5 | 173472820 | rs7341139 | 0.714 | 0.655 | 0.045 | -1.891 | 0.006 | -0.062 | -0.062 | 0 | 0.11 | 0.29 | 0.14 | 0.01 | 0.08 | 0.12 |
| 5 | 173481713 | rs10036246 | -0.802 | 0.616 | -0.531 | -1.901 | -0.023 | -0.119 | -0.119 | 0.122 | 0.2 | 0.54 | 0.19 | 0.01 | 0.14 | 0.16 |
| 5 | 173491300 | rs17076802 | 0.1 | 0.742 | -1.577 | -1.713 | -0.033 | -0.027 | -0.027 | 0 | 0.1 | 0 | 0.04 | 0.09 | 0.11 | 0.16 |
| 5 | 173504522 | rs4868342 | 0.456 | 0.907 | -1.589 | -1.663 | -0.043 | 0.034 | 0.034 | 0.004 | 0.77 | 0.88 | 0.82 | 0.88 | 0.69 | 0.76 |
| 5 | 173504716 | rs6891369 | 0.76 | 1.269 | -1.377 | -1.54 | -0.004 | -0.046 | -0.046 | 0 | 0.11 | 0.22 | 0.13 | 0.01 | 0.07 | 0.11 |
| 5 | 173511012 | rs17076822 | 0.904 | 1.35 | -1.958 | -1.261 | -0.031 | -0.032 | -0.032 | 0 | 0.07 | 0 | 0.04 | 0.09 | 0.09 | 0.12 |
| 5 | 173512434 | rs17708750 | 0.526 | 1.177 | -1.737 | -1.576 | -0.068 | 0.046 | 0.046 | 0 | 0.12 | 0.01 | 0.09 | 0.04 | 0.19 | 0.13 |
| 5 | 173534877 | rs4457100 | -0.775 | 0.381 | -1.285 | -1.373 | 0.005 | 0.007 | 0.007 | 0 | 0.3 | 0.71 | 0.26 | 0.09 | 0.22 | 0.24 |
| 5 | 173538655 | rs10062841 | -1.333 | 0.509 | -1.458 | -1.36 | 0.008 | 0.096 | 0.096 | 0 | 0.45 | 0.74 | 0.37 | 0.15 | 0.39 | 0.38 |
| 5 | 173539058 | rs6893176 | 0.731 | 0.134 | -1.119 | -1.485 | -0.016 | -0.039 | -0.039 | 0 | 0.7 | 0.47 | 0.76 | 0.88 | 0.8 | 0.7 |
| 5 | 173539502 | rs4296822 | 1.158 | 0.478 | -2.006 | -1.843 | 0.015 | 0.167 | 0.167 | 0 | 0.36 | 0.15 | 0.26 | 0.07 | 0.34 | 0.2 |
| 5 | 173547383 | rs11739519 | -0.715 | 0.917 | -1.38 | -1.495 | -0.008 | 0.021 | 0.021 | 0 | 0.09 | 0.04 | 0.06 | 0.01 | 0.11 | 0.04 |
| 5 | 173553251 | rs7712944 | 0.428 | 1.139 | -1.295 | -1.435 | -0.004 | 0.003 | 0.003 | 0 | 0.07 | 0.05 | 0.05 | 0.03 | 0.07 | 0.04 |
| 5 | 173553760 | rs6877384 | -0.719 | 0.975 | -1.27 | -1.361 | -0.002 | -0.013 | -0.013 | 0 | 0.23 | 0.18 | 0.36 | 0.75 | 0.23 | 0.5 |
| 5 | 173553879 | rs6877562 | -1.362 | 0.791 | -0.942 | -1.419 | 0 | -0.012 | -0.012 | 1.414 | 0.38 | 0.42 | 0.46 | 0.78 | 0.36 | 0.56 |
| 5 | 173554239 | rs6878270 | -0.633 | 1.153 | -0.833 | -0.616 | -0.02 | -0.047 | -0.047 | 0.586 | 0.14 | 0.05 | 0.3 | 0.71 | 0.17 | 0.37 |
| 5 | 173555467 | rs10516110 | -0.201 | 1.122 | -0.644 | -0.681 | -0.038 | -0.037 | -0.037 | 0 | 0.08 | 0.05 | 0.18 | 0.16 | 0.16 | 0.13 |
| 5 | 173583051 | rs1368272 | -0.683 | 0.497 | -0.173 | -1.126 | 0.012 | -0.03 | -0.03 | 0 | 0.33 | 0.49 | 0.53 | 0.75 | 0.36 | 0.38 |
| 5 | 173584572 | rs2431744 | -0.539 | 0.551 | -0.318 | -0.97 | 0.001 | 0.004 | 0.004 | 0 | 0.14 | 0.13 | 0.15 | 0.1 | 0.14 | 0.15 |
| 5 | 173586469 | rs13167802 | -1.368 | 0.187 | -0.73 | -1.722 | -0.052 | 0.194 | 0.194 | 0 | 0.66 | 0.88 | 0.82 | 0.98 | 0.63 | 0.76 |
| 5 | 173587528 | rs10452507 | 0.151 | -0.055 | 0.506 | -1.551 | -0.037 | -0.093 | -0.093 | 0 | 0.14 | 0.29 | 0.07 | 0.03 | 0.08 | 0.09 |
| 5 | 173587641 | rs10866671 | 0.504 | 0.079 | 0.406 | -1.461 | 0.011 | 0.01 | 0.01 | 0 | 0.2 | 0.21 | 0.42 | 0.74 | 0.24 | 0.3 |
| 5 | 173589734 | rs10476096 | -0.049 | -0.083 | 0.368 | -1.175 | -0.034 | -0.111 | -0.111 | 0.953 | 0.15 | 0.41 | 0.09 | 0.03 | 0.08 | 0.09 |
| 5 | 173592360 | rs10516111 | -0.302 | -0.149 | 0.242 | -1.172 | -0.07 | -0.141 | -0.141 | 0 | 0.19 | 0.47 | 0.09 | 0.03 | 0.09 | 0.12 |
| 5 | 173594220 | rs7716878 | -0.705 | 0.442 | -0.271 | -0.904 | -0.006 | -0.033 | -0.033 | 0 | 0.42 | 0.46 | 0.37 | 0.25 | 0.38 | 0.4 |
| 5 | 173601704 | rs871503 | -0.586 | 0.363 | -0.952 | -0.655 | -0.041 | 0.104 | 0.104 | 0 | 0.55 | 0.84 | 0.64 | 0.75 | 0.46 | 0.6 |
| 5 | 173602922 | rs6878578 | 0.204 | -0.5 | -0.268 | -1.107 | -0.045 | -0.061 | -0.061 | 0.003 | 0.19 | 0.26 | 0.07 | 0.06 | 0.06 | 0.18 |
| 5 | 173608806 | rs884343 | 0.344 | 0.007 | -0.842 | -1.153 | -0.017 | 0.018 | 0.018 | 0 | 0.35 | 0.37 | 0.57 | 0.8 | 0.49 | 0.44 |
| 5 | 173609274 | rs1560639 | -0.244 | 0.893 | -1.326 | -0.645 | -0.034 | 0.152 | 0.152 | 0 | 0.59 | 0.84 | 0.68 | 0.86 | 0.55 | 0.66 |
| 5 | 173611384 | rs17076923 | -0.577 | 1.021 | -0.74 | 0.436 | -0.01 | -0.026 | -0.026 | 0 | 0.1 | 0.04 | 0.07 | 0.18 | 0.12 | 0.14 |
| 5 | 173613616 | rs10476097 | 0.158 | 0.95 | -0.594 | 0.09 | 0.001 | -0.016 | -0.016 | 0 | 0.55 | 0.55 | 0.54 | 0.29 | 0.54 | 0.45 |
| 5 | 173614754 | rs17076933 | -1.433 | 1.094 | -0.261 | 0.116 | 0.026 | -0.035 | -0.035 | 0 | 0.07 | 0.16 | 0.09 | 0.02 | 0.1 | 0.04 |
| 5 | 173616645 | rs6887234 | -0.792 | 0.845 | -0.108 | 0.256 | -0.003 | 0.005 | 0.005 | 0.051 | 0.18 | 0.3 | 0.18 | 0.14 | 0.18 | 0.22 |
| 5 | 173616699 | rs6556120 | 0.247 | 0.324 | -0.514 | 0.156 | 0.022 | -0.052 | -0.052 | 0.248 | 0.33 | 0.2 | 0.27 | 0.54 | 0.25 | 0.37 |
| 5 | 173616773 | rs6866159 | 0.306 | 0.218 | -0.384 | 0.057 | 0.001 | -0.002 | -0.002 | 0 | 0.13 | 0.15 | 0.15 | 0.08 | 0.15 | 0.11 |
| 5 | 173620286 | rs7705993 | 0.892 | 0.724 | -0.104 | -0.037 | -0.003 | 0.012 | 0.012 | 0 | 0.45 | 0.53 | 0.43 | 0.31 | 0.47 | 0.37 |
| 5 | 173622863 | rs7711162 | 0.025 | 0.043 | -0.676 | -0.787 | 0.017 | -0.149 | -0.149 | 0.471 | 0.66 | 0.53 | 0.78 | 0.96 | 0.65 | 0.81 |
| 5 | 173625154 | rs6863418 | 0.153 | -0.037 | -0.695 | -0.687 | 0.006 | -0.064 | -0.064 | 0 | 0.66 | 0.44 | 0.73 | 0.89 | 0.65 | 0.78 |
| 5 | 173628743 | rs10035757 | -0.189 | -0.245 | -0.75 | -0.689 | -0.007 | 0.108 | 0.108 | 0 | 0.17 | 0.27 | 0.32 | 0.55 | 0.17 | 0.34 |
| 5 | 173630504 | rs11952072 | 0.53 | -0.36 | -0.961 | -0.895 | -0.002 | 0.011 | 0.011 | 0 | 0.17 | 0.13 | 0.16 | 0.09 | 0.19 | 0.12 |
| 5 | 173631215 | rs10044426 | -0.14 | -0.338 | -0.962 | -0.859 | -0.008 | 0.107 | 0.107 | 0 | 0.17 | 0.28 | 0.32 | 0.55 | 0.17 | 0.34 |
| 5 | 173633945 | rs4867745 | 0.324 | -0.273 | -0.942 | -0.894 | -0.005 | 0.124 | 0.124 | 0.039 | 0.14 | 0.26 | 0.34 | 0.61 | 0.14 | 0.33 |
| 5 | 173635263 | rs1864924 | 1.667 | -0.248 | -1.273 | -0.953 | -0.081 | 0.036 | 0.036 | 3.039 | 0.34 | 0.02 | 0.28 | 0.22 | 0.32 | 0.29 |
| 5 | 173637293 | rs2890803 | 0.864 | -0.148 | -0.533 | -0.904 | 0.039 | -0.054 | -0.054 | 0 | 0.1 | 0.23 | 0.08 | 0 | 0.12 | 0.04 |
| 5 | 173637603 | rs11952064 | 0.189 | -0.084 | -0.497 | -0.674 | 0.001 | 0.015 | 0.015 | 0 | 0.65 | 0.5 | 0.54 | 0.36 | 0.65 | 0.55 |
| 5 | 173640709 | rs1864926 | -0.96 | 0.119 | -1.016 | -0.76 | -0.021 | 0.065 | 0.065 | 0 | 0.8 | 0.89 | 0.88 | 0.97 | 0.79 | 0.88 |
| 5 | 173641418 | rs6556123 | 0.246 | 0.08 | -0.912 | -0.64 | -0.006 | 0.081 | 0.081 | 0.629 | 0.59 | 0.71 | 0.69 | 0.87 | 0.57 | 0.73 |
| 5 | 173642435 | rs4867746 | 0.17 | -0.508 | -0.54 | -0.092 | 0.009 | 0.011 | 0.011 | 0 | 0.51 | 0.37 | 0.57 | 0.38 | 0.54 | 0.44 |
| 5 | 173647712 | rs4295404 | -0.484 | -1.094 | -0.306 | 0.478 | 0.005 | -0.02 | -0.02 | 0 | 0.33 | 0.22 | 0.35 | 0.5 | 0.28 | 0.26 |
| 5 | 173648438 | rs10475598 | -1.215 | -1.404 | -0.174 | 0.425 | 0.057 | 0.109 | 0.109 | 0 | 0.44 | 0.8 | 0.48 | 0.77 | 0.32 | 0.45 |
| 5 | 173650528 | rs10476105 | -0.376 | -1.402 | 0.018 | 0.854 | -0.006 | 0.138 | 0.138 | 0 | 0.13 | 0.2 | 0.15 | 0.4 | 0.11 | 0.12 |
| 5 | 173650876 | rs187162 | 0.267 | -1.01 | -0.457 | -0.074 | -0.003 | -0.011 | -0.011 | 0 | 0.27 | 0.41 | 0.23 | 0.27 | 0.27 | 0.32 |
| 5 | 173652714 | rs356067 | 0.156 | -1.271 | -0.663 | 0.012 | -0.012 | 0.064 | 0.064 | 0 | 0.39 | 0.22 | 0.35 | 0.23 | 0.39 | 0.26 |
| 5 | 173654128 | rs4868363 | 1.086 | -1.226 | -1.085 | -0.308 | -0.037 | -0.031 | -0.031 | 0.502 | 0.2 | 0.14 | 0.3 | 0.39 | 0.37 | 0.42 |
| 5 | 173655405 | rs356065 | -0.233 | -1.455 | -1.255 | -0.484 | 0.116 | 0.026 | 0.026 | 0.049 | 0.55 | 0.82 | 0.6 | 0.41 | 0.71 | 0.63 |

| | | | | | | | | | | | | | | | | |
|---|---|---|---|---|---|---|---|---|---|---|---|---|---|---|---|---|
| 5 | 173660347 | rs4868364 | 1.83 | -0.869 | -0.999 | -0.216 | -0.093 | -0.015 | -0.015 | 0 | 0.14 | 0.03 | 0.23 | 0.13 | 0.33 | 0.23 |
| 5 | 173673921 | rs242258 | 0.039 | -1.513 | -0.9 | -0.453 | -0.011 | 0.012 | 0.012 | 0 | 0.3 | 0.33 | 0.27 | 0.54 | 0.21 | 0.21 |
| 5 | 173675031 | rs4868367 | -0.284 | -1.948 | -1.215 | -0.589 | -0.009 | 0 | 0 | 0 | 0.59 | 0.59 | 0.66 | 0.61 | 0.74 | 0.62 |
| 5 | 173679727 | rs2082381 | 0.429 | -1.896 | -1.109 | -0.688 | 0 | 0 | 0 | 0 | 0.48 | 0.49 | 0.48 | 0.35 | 0.59 | 0.38 |
| 5 | 173682973 | rs1432621 | 0.379 | -1.897 | -1.161 | -0.673 | -0.023 | 0.154 | 0.154 | 0 | 0.16 | 0.39 | 0.17 | 0.33 | 0.11 | 0.17 |
| 5 | 173683048 | rs17773992 | 2.722 | -1.852 | -1.314 | -0.781 | -0.026 | 0.058 | 0.058 | 0 | 0.15 | 0.07 | 0.11 | 0.01 | 0.18 | 0.1 |
| 5 | 173683416 | rs4868371 | 2.414 | -1.733 | -1.378 | -0.722 | -0.093 | -0.015 | -0.015 | 0.84 | 0.14 | 0.03 | 0.22 | 0.12 | 0.32 | 0.2 |
| 5 | 173683914 | rs1392391 | -2.031 | -1.771 | -1.391 | -0.734 | 0.08 | 0.11 | 0.11 | 0.923 | 0.75 | 0.98 | 0.83 | 0.94 | 0.85 | 0.78 |
| 5 | 173684268 | rs6556125 | 2.206 | -1.34 | -1.104 | -0.548 | -0.093 | -0.015 | -0.015 | 0.112 | 0.14 | 0.03 | 0.22 | 0.12 | 0.32 | 0.2 |
| 5 | 173684581 | rs7720654 | 1.417 | -1.397 | -0.916 | -0.532 | -0.005 | -0.148 | -0.148 | 0.657 | 0.21 | 0.46 | 0.16 | 0.01 | 0.18 | 0.1 |
| 5 | 173685519 | rs17077133 | 1.695 | -1.285 | -0.842 | -0.242 | -0.03 | -0.035 | -0.035 | 0.527 | 0.08 | 0.04 | 0.17 | 0.26 | 0.15 | 0.21 |
| 5 | 173685646 | rs891891 | 1.715 | -1.337 | -0.772 | -0.27 | 0.005 | -0.016 | -0.016 | 0 | 0.07 | 0.27 | 0.03 | 0.01 | 0.02 | 0.07 |
| 5 | 173685728 | rs17077140 | 1.609 | -1.123 | -1.206 | -0.154 | -0.031 | -0.036 | -0.036 | 0 | 0.09 | 0.04 | 0.17 | 0.26 | 0.15 | 0.21 |
| 5 | 173685962 | rs17077141 | 0.628 | -1.324 | -1.15 | -0.101 | -0.001 | -0.013 | -0.013 | 5.55 | 0.09 | 0.06 | 0.14 | 0.32 | 0.09 | 0.09 |
| 5 | 173686539 | rs1432619 | 0.165 | -0.922 | -0.908 | -0.281 | -0.008 | -0.01 | -0.01 | 0 | 0.41 | 0.51 | 0.55 | 0.4 | 0.6 | 0.51 |
| 5 | 173687247 | rs6893633 | 1.511 | -0.921 | -1.557 | -0.554 | 0.012 | 0.104 | 0.104 | 0 | 0.24 | 0.05 | 0.19 | 0.05 | 0.2 | 0.18 |
| 5 | 173687472 | rs17077144 | 1.145 | -0.03 | -0.962 | -0.144 | 0.024 | 0.041 | 0.041 | 0 | 0.11 | 0.21 | 0.21 | 0.26 | 0.16 | 0.28 |
| 5 | 173687757 | rs4867750 | 1.328 | 0.091 | -1.227 | -0.059 | -0.078 | -0.022 | -0.022 | 0.04 | 0.14 | 0.04 | 0.21 | 0.14 | 0.28 | 0.2 |
| 5 | 173688699 | rs1835904 | 0.39 | -1.437 | -0.432 | 0.234 | -0.006 | -0.007 | -0.007 | 0 | 0.65 | 0.7 | 0.79 | 0.78 | 0.75 | 0.83 |
| 5 | 173690809 | rs181451 | -0.351 | -1.175 | -1.051 | 0.544 | 0.032 | -0.021 | -0.021 | 0 | 0.81 | 0.93 | 0.91 | 0.79 | 0.9 | 0.93 |
| 5 | 173699270 | rs1432616 | 0.128 | -1.113 | -0.915 | 0.615 | 0.005 | -0.025 | -0.025 | 0 | 0.72 | 0.83 | 0.73 | 0.64 | 0.75 | 0.78 |
| 5 | 173699598 | rs17077184 | 0.853 | -1.373 | -0.269 | 0.622 | -0.028 | 0.028 | 0.028 | 0 | 0.13 | 0.23 | 0.09 | 0.23 | 0.05 | 0.08 |
| 5 | 173701609 | rs189592 | 0.347 | -1.325 | 0.063 | 0.607 | -0.001 | 0.025 | 0.025 | 0 | 0.86 | 0.86 | 0.82 | 0.67 | 0.89 | 0.89 |
| 5 | 173709075 | rs7711316 | -1.16 | -1.37 | 0.664 | 0.663 | 0.034 | 0.343 | 0.343 | 0 | 0.09 | 0.32 | 0.2 | 0.38 | 0.08 | 0.11 |
| 5 | 173710328 | rs10064200 | -0.516 | -1.313 | 0.715 | 0.764 | 0.013 | 0.205 | 0.205 | 0 | 0.06 | 0.21 | 0.16 | 0.31 | 0.06 | 0.09 |
| 5 | 173711503 | rs10516115 | -0.319 | -1.383 | 0.748 | 0.733 | 0 | 0.136 | 0.136 | 0 | 0.06 | 0.11 | 0.16 | 0.31 | 0.06 | 0.09 |
| 5 | 173711724 | rs1909706 | 0.412 | -1.713 | 0.637 | 0.486 | -0.008 | -0.007 | -0.007 | 0.575 | 0.26 | 0.33 | 0.24 | 0.25 | 0.26 | 0.34 |
| 5 | 173715191 | rs6894337 | 1.431 | -1.821 | 0.495 | 0.486 | 0.008 | -0.011 | -0.011 | 0 | 0.12 | 0.1 | 0.08 | 0.23 | 0.06 | 0.05 |
| 5 | 173717971 | rs17077288 | -0.743 | -1.849 | 0.399 | 0.381 | 0.004 | -0.003 | -0.003 | 0 | 0.38 | 0.62 | 0.42 | 0.56 | 0.33 | 0.53 |
| 5 | 173720227 | rs12658280 | 0.44 | -1.512 | 0.023 | 0.05 | -0.016 | -0.009 | -0.009 | 0 | 0.26 | 0.33 | 0.23 | 0.25 | 0.25 | 0.34 |
| 5 | 173720675 | rs2936995 | 0.996 | -1.459 | -0.981 | -0.139 | -0.133 | 0.03 | 0.03 | 0 | 0.33 | 0.07 | 0.45 | 0.21 | 0.52 | 0.39 |
| 5 | 173722032 | rs12656362 | 0.417 | -1.543 | -0.476 | 0.009 | -0.008 | -0.004 | -0.004 | 0.14 | 0.23 | 0.25 | 0.2 | 0.23 | 0.2 | 0.31 |
| 5 | 173728190 | rs2936997 | -0.479 | -1.548 | -0.66 | 0.21 | -0.04 | 0.033 | 0.033 | 0 | 0.69 | 0.67 | 0.72 | 0.49 | 0.8 | 0.81 |
| 5 | 173730624 | rs11741853 | 1.216 | 0.418 | -1.661 | -0.202 | -0.104 | -0.059 | -0.059 | 2.117 | 0.15 | 0.03 | 0.22 | 0.2 | 0.3 | 0.26 |
| 5 | 173730678 | rs2964078 | 0.163 | 0.092 | -1.088 | -0.506 | -0.006 | 0.092 | 0.092 | 0 | 0.3 | 0.16 | 0.24 | 0.05 | 0.27 | 0.26 |
| 5 | 173734349 | rs2937000 | -0.134 | 0.14 | -0.769 | -0.117 | 0 | 0.004 | 0.004 | 0 | 0.78 | 0.83 | 0.71 | 0.94 | 0.79 | 0.83 |
| 5 | 173734366 | rs11738411 | 0.361 | 0.12 | -0.958 | 0.194 | 0.006 | 0.015 | 0.015 | 0 | 0.54 | 0.4 | 0.41 | 0.33 | 0.62 | 0.48 |
| 5 | 173738577 | rs2964083 | -0.514 | 0.095 | -0.962 | 0.386 | 0.011 | -0.009 | -0.009 | 0.194 | 0.3 | 0.75 | 0.45 | 0.37 | 0.23 | 0.4 |
| 5 | 173743117 | rs1560049 | -0.452 | 0.148 | -0.789 | 0.571 | 0.005 | 0.018 | 0.018 | 0 | 0.09 | 0.05 | 0.17 | 0.39 | 0.09 | 0.13 |
| 5 | 173749575 | rs6861937 | 0.044 | 0.746 | -0.652 | 0.058 | 0.001 | -0.015 | -0.015 | 0 | 0.2 | 0.24 | 0.24 | 0.06 | 0.3 | 0.31 |
| 5 | 173750017 | rs10058766 | 1.585 | 0.369 | -0.565 | -0.486 | 0.006 | -0.008 | -0.008 | 0 | 0.35 | 0.24 | 0.35 | 0.45 | 0.29 | 0.39 |
| 5 | 173750173 | rs10866673 | 1.429 | 0.387 | -0.784 | -0.358 | 0.001 | -0.001 | -0.001 | 0.254 | 0.38 | 0.34 | 0.39 | 0.45 | 0.34 | 0.4 |
| 5 | 173750565 | rs2162728 | 1.179 | -0.419 | 0.102 | 0.354 | 0.001 | -0.005 | -0.005 | 0 | 0.62 | 0.64 | 0.62 | 0.51 | 0.6 | 0.71 |
| 5 | 173753330 | rs2936985 | 0.848 | -0.462 | -0.21 | 0.182 | -0.003 | -0.051 | -0.051 | 0 | 0.72 | 0.7 | 0.79 | 0.95 | 0.71 | 0.89 |
| 5 | 173753932 | rs2913474 | -1.179 | -0.247 | -1.392 | 0.414 | -0.008 | 0.043 | 0.043 | 0 | 0.18 | 0.21 | 0.2 | 0.45 | 0.12 | 0.18 |
| 5 | 173755438 | rs17077332 | -0.605 | -0.184 | -1.344 | 0.349 | -0.041 | -0.051 | -0.051 | 0.07 | 0.09 | 0.2 | 0.05 | 0 | 0.02 | 0.01 |
| 5 | 173756503 | rs2964098 | -1.207 | -0.188 | -1.428 | 0.363 | -0.045 | -0.047 | -0.047 | 0 | 0.17 | 0.29 | 0.07 | 0.04 | 0.05 | 0.04 |
| 5 | 173756569 | rs17077334 | -0.391 | 0.328 | -2.013 | 0.651 | -0.007 | -0.007 | -0.007 | 0 | 0.08 | 0.09 | 0.04 | 0 | 0.02 | 0.01 |
| 5 | 173758070 | rs904720 | 1.009 | 1.171 | -1.999 | 1.067 | -0.088 | -0.165 | -0.165 | 0 | 0.74 | 0.97 | 0.67 | 0.54 | 0.64 | 0.74 |
| 5 | 173760983 | rs2936976 | 0.55 | 1.316 | -1.448 | 0.326 | -0.054 | 0.026 | 0.026 | 0 | 0.87 | 0.98 | 0.88 | 0.96 | 0.8 | 0.92 |
| 5 | 173763567 | rs2964102 | 0.647 | 1.168 | -1.037 | 0.599 | 0 | 0.001 | 0.001 | 0 | 0.78 | 0.79 | 0.84 | 0.79 | 0.73 | 0.91 |
| 5 | 173767508 | rs2913466 | -0.281 | 1.25 | -0.651 | -0.601 | 0.004 | -0.124 | -0.124 | 0.009 | 0.25 | 0.4 | 0.23 | 0.02 | 0.25 | 0.13 |
| 5 | 173767646 | rs2964106 | 0.931 | 2.058 | -0.304 | -1.517 | -0.002 | -0.037 | -0.037 | 0 | 0.66 | 0.65 | 0.69 | 0.96 | 0.57 | 0.84 |
| 5 | 173770556 | rs753829 | 1.233 | 3.106 | -0.14 | -1.976 | -0.029 | 0.064 | 0.064 | 0 | 0.6 | 0.72 | 0.62 | 0.93 | 0.49 | 0.77 |
| 5 | 173772185 | rs4544836 | -0.062 | 2.25 | 0.056 | -1.265 | 0.009 | 0.087 | 0.087 | 0 | 0.17 | 0.39 | 0.12 | 0.36 | 0.16 | 0.15 |
| 5 | 173772593 | rs4242178 | 0.787 | 2.127 | 0.002 | -1.454 | 0.003 | 0.035 | 0.035 | 0.005 | 0.17 | 0.12 | 0.16 | 0.01 | 0.14 | 0.13 |
| 5 | 173776218 | rs1863952 | 0.197 | 2.393 | 0.024 | -1.43 | -0.023 | 0.031 | 0.031 | 0 | 0.77 | 0.86 | 0.8 | 0.94 | 0.65 | 0.89 |
| 5 | 173777372 | rs2909728 | 2.064 | 2.189 | 0.03 | -1.403 | 0.017 | 0.012 | 0.012 | 0 | 0.41 | 0.19 | 0.5 | 0.59 | 0.36 | 0.6 |
| 5 | 173777520 | rs12659144 | 0.015 | 2.134 | 0.572 | -1.377 | 0.015 | 0.05 | 0.05 | 0 | 0.15 | 0.29 | 0.11 | 0.34 | 0.16 | 0.16 |
| 5 | 173781009 | rs2964109 | 0.378 | 1.638 | 0.309 | -1.53 | -0.011 | -0.029 | -0.029 | 0 | 0.08 | 0.16 | 0.05 | 0.03 | 0.06 | 0.04 |
| 5 | 173781916 | rs2909726 | -0.918 | 1.602 | 0.408 | -1.518 | -0.004 | 0.027 | 0.027 | 0 | 0.23 | 0.38 | 0.15 | 0.36 | 0.21 | 0.2 |
| 5 | 173782172 | rs1863954 | -0.068 | 1.643 | 0.28 | -1.524 | -0.013 | 0.014 | 0.014 | 0 | 0.77 | 0.84 | 0.79 | 0.94 | 0.66 | 0.89 |
| 5 | 173782309 | rs17077406 | 0.69 | 1.62 | 0.261 | -1.525 | -0.001 | -0.035 | -0.035 | 0 | 0.07 | 0.17 | 0.04 | 0 | 0.05 | 0.01 |
| 5 | 173782331 | rs12521800 | 2.242 | 1.621 | 0.181 | -1.43 | 0.015 | -0.029 | -0.029 | 0.011 | 0.35 | 0.16 | 0.46 | 0.54 | 0.28 | 0.56 |
| 5 | 173788582 | rs989062 | -0.426 | 2.029 | 0.207 | -1.413 | 0.012 | -0.021 | -0.021 | 0.116 | 0.91 | 0.91 | 0.86 | 1 | 0.78 | 0.96 |
| 5 | 173791376 | rs962880 | 1.658 | 1.786 | 0.363 | -1.378 | -0.004 | 0.008 | 0.008 | 0 | 0.55 | 0.43 | 0.66 | 0.55 | 0.49 | 0.7 |
| 5 | 173793906 | rs2936959 | -1.264 | 1.467 | 0.229 | -1.253 | 0 | 0 | 0 | 0.369 | 0.37 | 0.37 | 0.33 | 0.44 | 0.42 | 0.28 |
| 5 | 173794156 | rs2964115 | -1.765 | 0.949 | -1.042 | -1.798 | -0.008 | 0.019 | 0.019 | 0 | 0.85 | 0.9 | 0.85 | 0.95 | 0.85 | 0.77 |
| 5 | 173795792 | rs1875189 | -2.009 | 1.352 | -0.945 | -1.612 | 0.022 | -0.068 | -0.068 | 0 | 0.65 | 0.84 | 0.6 | 0.53 | 0.71 | 0.49 |

| | | | | | | | | | | | | | | | | |
|---|---|---|---|---|---|---|---|---|---|---|---|---|---|---|---|---|
| 5 | 173795901 | rs1875190 | 2.231 | 1.476 | -0.975 | -1.613 | -0.007 | 0.023 | 0.023 | 0 | 0.07 | 0.03 | 0.11 | 0.38 | 0.08 | 0.16 |
| 5 | 173799591 | rs2936957 | -2.069 | 1.46 | -1.022 | -1.628 | 0.022 | -0.068 | -0.068 | 0 | 0.65 | 0.84 | 0.6 | 0.52 | 0.71 | 0.49 |
| 5 | 173800923 | rs1818061 | -1.379 | 1.448 | -0.801 | -1.962 | 0.01 | 0.08 | 0.08 | 0 | 0.57 | 0.73 | 0.46 | 0.14 | 0.61 | 0.32 |
| 5 | 173801412 | rs723320 | 1.061 | 1.087 | -1.046 | -1.58 | -0.072 | 0.042 | 0.042 | 0 | 0.17 | 0.03 | 0.18 | 0.05 | 0.21 | 0.17 |
| 5 | 173806760 | rs2936952 | -0.068 | 0.824 | -0.187 | -1.225 | 0.005 | 0.006 | 0.006 | 0 | 0.19 | 0.14 | 0.09 | 0.07 | 0.15 | 0.11 |
| 5 | 173810054 | rs13185171 | 0.061 | 1.207 | 0.27 | -0.967 | 0.035 | -0.048 | -0.048 | 0 | 0.09 | 0.22 | 0.11 | 0 | 0.13 | 0.03 |
| 5 | 173812410 | rs4868392 | -0.191 | 0.696 | 0.345 | -0.015 | 0.001 | 0.001 | 0.001 | 0 | 0.41 | 0.43 | 0.49 | 0.43 | 0.39 | 0.54 |
| 5 | 173812533 | rs2909716 | -0.704 | -0.379 | 0.75 | 0.129 | -0.014 | 0.106 | 0.106 | 0.092 | 0.09 | 0.39 | 0.06 | 0.08 | 0.03 | 0.03 |
| 5 | 173815831 | rs2936948 | -0.622 | -0.314 | 0.431 | 0.072 | 0.012 | 0.016 | 0.016 | 0 | 0.34 | 0.51 | 0.34 | 0.43 | 0.45 | 0.29 |
| 5 | 173816715 | rs2913496 | -0.602 | -0.296 | 0.44 | 0.081 | 0.013 | 0.017 | 0.017 | 0 | 0.34 | 0.51 | 0.34 | 0.43 | 0.44 | 0.26 |
| 5 | 173818378 | rs1895140 | 0.564 | -0.18 | 0.46 | 0.14 | -0.013 | 0.012 | 0.012 | 0 | 0.37 | 0.44 | 0.42 | 0.41 | 0.27 | 0.54 |
| 5 | 173823541 | rs1862134 | -0.962 | -0.156 | 0.475 | 0.065 | -0.005 | -0.013 | -0.013 | 0 | 0.62 | 0.55 | 0.7 | 0.78 | 0.68 | 0.76 |
| 5 | 173827151 | rs2936940 | -1.347 | -0.385 | 0.723 | 0.093 | -0.026 | -0.045 | -0.045 | 0 | 0.67 | 0.49 | 0.73 | 0.82 | 0.76 | 0.83 |
| 5 | 173828150 | rs6876769 | 0.615 | -0.455 | 0.737 | 0.165 | -0.003 | -0.025 | -0.025 | 0 | 0.37 | 0.22 | 0.42 | 0.44 | 0.32 | 0.57 |
| 5 | 173833752 | rs2936934 | 0.007 | -0.445 | 0.591 | 0.137 | 0 | -0.001 | -0.001 | 0 | 0.43 | 0.38 | 0.45 | 0.5 | 0.35 | 0.6 |
| 5 | 173835016 | rs4867763 | -0.853 | -0.362 | 0.388 | 0.088 | -0.006 | -0.006 | -0.006 | 0 | 0.62 | 0.37 | 0.69 | 0.76 | 0.73 | 0.79 |
| 5 | 173838067 | rs7721965 | -0.044 | -0.294 | -0.231 | 0.162 | -0.01 | 0.005 | 0.005 | 0 | 0.41 | 0.48 | 0.43 | 0.42 | 0.31 | 0.56 |
| 5 | 173848018 | rs6556132 | 0.073 | 0.123 | -1.362 | 0.408 | -0.001 | -0.043 | -0.043 | 0 | 0.34 | 0.18 | 0.41 | 0.42 | 0.31 | 0.52 |
| 5 | 173851524 | rs2909707 | 0.006 | 0.148 | -1.267 | -0.15 | -0.01 | 0.069 | 0.069 | 0 | 0.54 | 0.69 | 0.47 | 0.33 | 0.5 | 0.47 |
| 5 | 173852472 | rs11745617 | 0.937 | 0.264 | -1.506 | -0.349 | -0.008 | 0.022 | 0.022 | 0 | 0.17 | 0.1 | 0.18 | 0.15 | 0.17 | 0.31 |
| 5 | 173853681 | rs4446484 | 0.27 | 0.033 | -1.721 | -0.247 | -0.046 | 0.031 | 0.031 | 0 | 0.42 | 0.21 | 0.5 | 0.62 | 0.49 | 0.48 |
| 5 | 173854793 | rs2909705 | 1.312 | -0.193 | -1.738 | -0.41 | -0.007 | 0.061 | 0.061 | 0 | 0.28 | 0.08 | 0.21 | 0.15 | 0.24 | 0.17 |
| 5 | 173854937 | rs2964057 | 0.31 | -0.411 | -1.235 | -0.316 | 0.006 | 0.015 | 0.015 | 0 | 0.7 | 0.29 | 0.71 | 0.77 | 0.72 | 0.65 |
| 5 | 173859366 | rs1863957 | 0.251 | -0.258 | -1.385 | -0.392 | -0.016 | 0.015 | 0.015 | 0.217 | 0.3 | 0.18 | 0.27 | 0.23 | 0.35 | 0.17 |
| 5 | 173859454 | rs12514199 | 0.443 | -0.558 | -0.997 | -0.943 | -0.017 | 0.031 | 0.031 | 0.02 | 0.38 | 0.34 | 0.37 | 0.22 | 0.44 | 0.42 |
| 5 | 173863445 | rs2913530 | -0.212 | -0.759 | -0.869 | -0.824 | 0.003 | -0.024 | -0.024 | 0 | 0.56 | 0.41 | 0.57 | 0.74 | 0.53 | 0.58 |
| 5 | 173872772 | rs11134841 | 0.731 | -1.216 | -1.109 | -0.266 | 0.034 | 0.006 | 0.006 | 0 | 0.22 | 0.13 | 0.22 | 0.13 | 0.1 | 0.2 |
| 5 | 173874986 | rs794733 | -0.202 | -0.956 | -0.739 | -1.252 | 0.007 | -0.165 | -0.165 | 0 | 0.74 | 0.48 | 0.81 | 1 | 0.75 | 0.89 |
| 5 | 173875005 | rs17723581 | 1.124 | -0.738 | -1.058 | -0.996 | -0.048 | -0.017 | -0.017 | 0 | 0.27 | 0.05 | 0.23 | 0.26 | 0.32 | 0.17 |
| 5 | 173875557 | rs890754 | 1.581 | -0.75 | -0.94 | -0.9 | -0.023 | 0.033 | 0.033 | 0 | 0.2 | 0.12 | 0.19 | 0.14 | 0.21 | 0.28 |
| 5 | 173880643 | rs171371 | 0.837 | -0.895 | -0.735 | -0.72 | -0.011 | 0.011 | 0.011 | 0 | 0.27 | 0.26 | 0.36 | 0.56 | 0.19 | 0.4 |
| 5 | 173885707 | rs1963576 | 1.343 | -0.457 | -1.287 | -0.991 | -0.013 | 0.06 | 0.06 | 0 | 0.21 | 0.04 | 0.17 | 0.06 | 0.23 | 0.16 |
| 5 | 173888281 | rs890755 | 0.541 | -0.859 | -1.065 | -0.931 | -0.045 | 0.094 | 0.094 | 0 | 0.42 | 0.18 | 0.38 | 0.24 | 0.47 | 0.46 |
| 5 | 173891894 | rs12109471 | 0.187 | -0.574 | -0.593 | -0.914 | 0.039 | -0.144 | -0.144 | 0.059 | 0.22 | 0.35 | 0.16 | 0 | 0.26 | 0.08 |
| 5 | 173901938 | rs11742276 | 0.878 | -0.386 | -1.295 | -0.711 | -0.074 | 0.082 | 0.082 | 0.224 | 0.35 | 0.12 | 0.34 | 0.21 | 0.43 | 0.4 |
| 5 | 173905800 | rs2636047 | -0.967 | -0.446 | -0.889 | -0.733 | 0.062 | -0.078 | -0.078 | 0 | 0.82 | 0.59 | 0.86 | 0.96 | 0.8 | 0.87 |
| 5 | 173908150 | rs794740 | 0.034 | -0.892 | -1.952 | -0.736 | 0.074 | 0.119 | 0.119 | 0.331 | 0.45 | 0.2 | 0.48 | 0.75 | 0.37 | 0.47 |
| 5 | 173911948 | rs2560332 | 0.429 | -0.379 | -1.471 | -0.211 | -0.024 | 0.058 | 0.058 | 0.739 | 0.64 | 0.24 | 0.66 | 0.79 | 0.62 | 0.81 |
| 5 | 173913221 | rs1366137 | 0.435 | -0.376 | -1.464 | -0.209 | -0.019 | 0.052 | 0.052 | 0.809 | 0.64 | 0.26 | 0.66 | 0.79 | 0.62 | 0.81 |
| 5 | 173914450 | rs2560337 | 0.435 | -0.376 | -1.464 | -0.209 | -0.019 | 0.052 | 0.052 | 0.809 | 0.64 | 0.25 | 0.66 | 0.79 | 0.62 | 0.81 |
| 5 | 173924667 | rs6876247 | 1.13 | -0.098 | -1.695 | 0.226 | -0.063 | -0.014 | -0.014 | 0 | 0.23 | 0.06 | 0.23 | 0.23 | 0.31 | 0.24 |
| 5 | 173928775 | rs354993 | 0.068 | -1.708 | -1.045 | 0.659 | 0.002 | 0.015 | 0.015 | 0.357 | 0.78 | 0.74 | 0.78 | 0.71 | 0.77 | 0.87 |
| 5 | 173928808 | rs354992 | 0.542 | -2.157 | -1.057 | 0.806 | 0 | -0.001 | -0.001 | 0.168 | 0.11 | 0.13 | 0.15 | 0.14 | 0.1 | 0.29 |
| 5 | 173933655 | rs10463037 | 1.239 | -2.147 | -1.084 | 0.848 | 0.015 | 0.116 | 0.116 | 0.024 | 0.08 | 0.17 | 0.14 | 0.25 | 0.12 | 0.06 |
| 5 | 173935749 | rs354986 | 0.372 | -2.453 | -0.453 | 0.663 | -0.026 | -0.024 | -0.024 | 2.448 | 0.78 | 0.6 | 0.78 | 0.87 | 0.87 | 0.82 |
| 5 | 173938767 | rs10060172 | 1.867 | -2.935 | -0.829 | 0.471 | 0.057 | 0.091 | 0.091 | 0.025 | 0.53 | 0.23 | 0.42 | 0.3 | 0.63 | 0.45 |
| 5 | 173939850 | rs17077686 | 1.293 | -2.653 | -0.826 | 0.509 | 0.001 | -0.003 | -0.003 | 0.196 | 0.1 | 0.08 | 0.17 | 0.27 | 0.03 | 0.13 |
| 5 | 173941032 | rs354983 | -1.204 | -2.663 | -0.879 | 0.281 | -0.025 | 0.05 | 0.05 | 0.087 | 0.89 | 0.97 | 0.93 | 1 | 0.89 | 0.98 |
| 5 | 173942065 | rs10516117 | 1.231 | -2.736 | -0.774 | 0.572 | 0.001 | 0 | 0 | 0.183 | 0.1 | 0.08 | 0.17 | 0.3 | 0.03 | 0.14 |
| 5 | 173944093 | rs6865356 | -0.343 | -2.857 | -0.509 | 0.662 | -0.036 | 0.105 | 0.105 | 0 | 0.21 | 0.32 | 0.33 | 0.53 | 0.15 | 0.28 |
| 5 | 173944166 | rs703739 | 0.636 | -3.163 | -0.891 | 0.312 | 0.056 | 0.074 | 0.074 | 0 | 0.45 | 0.24 | 0.39 | 0.32 | 0.62 | 0.43 |
| 5 | 173945466 | rs10475601 | 1.175 | -2.899 | -0.781 | 0.236 | 0.005 | 0.077 | 0.077 | 0 | 0.1 | 0.19 | 0.14 | 0.17 | 0.12 | 0.06 |
| 5 | 173950287 | rs17063641 | 1.315 | -2.958 | -0.997 | 0.121 | 0.001 | 0.003 | 0.003 | 0 | 0.11 | 0.09 | 0.17 | 0.37 | 0.03 | 0.15 |
| 5 | 173951029 | rs13185484 | 2.661 | -2.63 | -1.247 | -0.267 | -0.137 | -0.057 | -0.057 | 0 | 0.2 | 0.02 | 0.21 | 0.22 | 0.33 | 0.21 |
| 5 | 173951543 | rs10065187 | -0.279 | -2.717 | -0.876 | -0.113 | -0.006 | 0.03 | 0.03 | 0 | 0.3 | 0.35 | 0.39 | 0.53 | 0.28 | 0.26 |
| 5 | 173952640 | rs6556141 | -1.042 | -2.513 | -0.576 | -0.571 | -0.037 | -0.059 | -0.059 | 0 | 0.73 | 0.65 | 0.84 | 0.89 | 0.85 | 0.78 |
| 5 | 173955389 | rs11747479 | 1.207 | -2.18 | -0.929 | -0.451 | 0.011 | 0.048 | 0.048 | 0 | 0.23 | 0.13 | 0.15 | 0.09 | 0.2 | 0.2 |
| 5 | 173956047 | rs1490046 | 0.865 | -1.726 | -1.049 | -0.287 | 0.001 | -0.013 | -0.013 | 1.251 | 0.09 | 0.09 | 0.06 | 0.01 | 0.09 | 0.09 |
| 5 | 173957118 | rs13179988 | 1.202 | -1.369 | -1.182 | -0.016 | -0.093 | 0.054 | 0.054 | 0 | 0.33 | 0.14 | 0.3 | 0.21 | 0.43 | 0.25 |
| 5 | 173970989 | rs11134853 | -0.619 | -1.26 | -0.287 | 0.605 | -0.017 | 0.054 | 0.054 | 0.148 | 0.24 | 0.28 | 0.35 | 0.47 | 0.15 | 0.34 |
| 5 | 173972852 | rs7718237 | -1.075 | -1.59 | -0.066 | 0.271 | 0.003 | 0.006 | 0.006 | 0 | 0.47 | 0.49 | 0.59 | 0.74 | 0.4 | 0.64 |
| 5 | 173974707 | rs10039242 | 1.322 | -1.86 | -0.379 | -0.282 | 0.006 | 0 | 0 | 0.36 | 0.49 | 0.56 | 0.48 | 0.43 | 0.57 | 0.29 |
| 5 | 173975145 | rs923520 | -0.71 | -1.848 | -0.366 | -0.269 | -0.002 | 0 | 0 | 0.045 | 0.4 | 0.39 | 0.46 | 0.55 | 0.28 | 0.57 |
| 5 | 173978470 | rs10516124 | 0.218 | -1.764 | -0.814 | -0.256 | 0.003 | 0.021 | 0.021 | 0 | 0.87 | 0.91 | 0.93 | 0.98 | 0.84 | 0.86 |
| 5 | 173978606 | rs13164416 | 1.122 | -1.777 | -0.612 | -0.197 | 0.024 | -0.01 | -0.01 | 0 | 0.52 | 0.72 | 0.51 | 0.43 | 0.57 | 0.29 |
| 5 | 173980686 | rs10063092 | -0.391 | -1.534 | -0.171 | -0.167 | -0.007 | -0.056 | -0.056 | 0 | 0.1 | 0.24 | 0.05 | 0.01 | 0.1 | 0.11 |
| 5 | 173981106 | rs11134856 | -0.798 | -1.664 | -0.173 | 0.137 | -0.018 | 0.016 | 0.016 | 0 | 0.13 | 0.25 | 0.15 | 0.14 | 0.11 | 0.26 |
| 5 | 173984116 | rs750229 | -1.553 | -1.592 | -0.555 | 0.3 | -0.007 | 0.012 | 0.012 | 0 | 0.39 | 0.48 | 0.41 | 0.51 | 0.37 | 0.54 |
| 5 | 173985281 | rs7736798 | -1.109 | -1.566 | -0.54 | 0.143 | -0.003 | -0.008 | -0.008 | 0 | 0.82 | 0.79 | 0.81 | 0.89 | 0.87 | 0.81 |

| | | | | | | | | | | | | | | | | |
|---|---|---|---|---|---|---|---|---|---|---|---|---|---|---|---|---|
| 5 | 173986118 | rs4868424 | -1.109 | -1.566 | -0.54 | 0.143 | -0.003 | -0.008 | -0.008 | 2.103 | 0.82 | 0.79 | 0.81 | 0.89 | 0.87 | 0.81 |
| 5 | 173993166 | rs1387768 | -1.101 | -1.82 | -0.607 | 0.081 | -0.028 | 0.036 | 0.036 | 0.245 | 0.31 | 0.41 | 0.35 | 0.5 | 0.28 | 0.49 |
| 5 | 173993649 | rs6889902 | 1.976 | -1.741 | -0.662 | 0.027 | -0.005 | 0.017 | 0.017 | 0 | 0.45 | 0.31 | 0.4 | 0.38 | 0.51 | 0.26 |
| 5 | 173995084 | rs4868428 | 1.491 | -1.731 | -0.669 | 0.016 | 0.001 | 0.029 | 0.029 | 0 | 0.16 | 0.04 | 0.16 | 0.1 | 0.12 | 0.2 |
| 5 | 173996446 | rs7707148 | -2.114 | -1.679 | -0.568 | 0.05 | -0.01 | 0.004 | 0.004 | 0 | 0.58 | 0.68 | 0.61 | 0.62 | 0.51 | 0.74 |
| 5 | 173996545 | rs17063736 | 0.377 | -1.318 | 0.041 | 0.432 | -0.012 | 0.01 | 0.01 | 0 | 0.09 | 0.06 | 0.06 | 0 | 0.09 | 0.05 |
| 5 | 173996604 | rs7725112 | -0.547 | -1.364 | -0.026 | 0.487 | -0.046 | 0.095 | 0.095 | 0 | 0.19 | 0.4 | 0.21 | 0.34 | 0.17 | 0.22 |
| 5 | 173999598 | rs4312897 | 0.154 | -0.601 | -0.554 | 0.233 | 0.022 | -0.056 | -0.056 | 1.437 | 0.89 | 0.77 | 0.92 | 0.99 | 0.91 | 0.94 |
| 5 | 174004543 | rs12516352 | -0.655 | -0.893 | -0.926 | 0.578 | 0.001 | 0.002 | 0.002 | 0 | 0.07 | 0.07 | 0.1 | 0.15 | 0.03 | 0.17 |
| 5 | 174008419 | rs6883108 | 0.009 | -1.244 | -0.738 | 0.574 | -0.045 | 0.038 | 0.038 | 0 | 0.27 | 0.48 | 0.31 | 0.39 | 0.19 | 0.37 |
| 5 | 174010424 | rs12514122 | 0.577 | -1.248 | -1.455 | 0.365 | -0.005 | -0.017 | -0.017 | 0 | 0.07 | 0.08 | 0.04 | 0 | 0.06 | 0.03 |
| 5 | 174011260 | rs13179274 | 0.891 | -0.793 | -1.736 | 0.402 | -0.104 | -0.102 | -0.102 | 0.147 | 0.4 | 0.09 | 0.36 | 0.46 | 0.49 | 0.27 |
| 5 | 174020823 | rs17727501 | -0.384 | -0.77 | -0.575 | 0.18 | 0 | -0.012 | -0.012 | 2.608 | 0.07 | 0.05 | 0.08 | 0 | 0.1 | 0.02 |
| 5 | 174025065 | rs12515393 | 1.006 | -1.578 | -1.076 | 0.219 | 0.091 | 0.058 | 0.058 | 0.228 | 0.52 | 0.22 | 0.57 | 0.67 | 0.67 | 0.75 |
| 5 | 174026875 | rs11748851 | -0.311 | -1.805 | -0.738 | 0.143 | 0.001 | -0.048 | -0.048 | 0 | 0.12 | 0.14 | 0.1 | 0 | 0.11 | 0.02 |
| 5 | 174027661 | rs4868431 | 0.067 | -1.778 | -0.713 | 0.106 | -0.042 | 0.011 | 0.011 | 0 | 0.68 | 0.55 | 0.69 | 0.68 | 0.77 | 0.81 |
| 5 | 174032728 | rs4868432 | 1.475 | -1.746 | -0.957 | 0.12 | 0.003 | 0.014 | 0.014 | 1.559 | 0.75 | 0.28 | 0.76 | 0.91 | 0.79 | 0.9 |
| 5 | 174036190 | rs12186684 | 1.579 | -1.289 | -1.212 | 0.521 | -0.159 | -0.128 | -0.128 | 1.545 | 0.4 | 0.04 | 0.41 | 0.55 | 0.51 | 0.56 |
| 5 | 174036233 | rs10866682 | 0.955 | -1.42 | -0.089 | 0.523 | 0.028 | 0.038 | 0.038 | 2.976 | 0.88 | 0.51 | 0.86 | 0.92 | 0.9 | 0.97 |
| 5 | 174038768 | rs7723208 | -0.565 | -1.696 | -0.799 | 0.409 | 0.046 | 0.028 | 0.028 | 0.009 | 0.33 | 0.2 | 0.24 | 0.22 | 0.19 | 0.2 |
| 5 | 174050806 | rs11746728 | -0.709 | -1.543 | -0.065 | 0.417 | -0.013 | -0.009 | -0.009 | 1.396 | 0.41 | 0.48 | 0.34 | 0.32 | 0.35 | 0.27 |
| 5 | 174051117 | rs17063801 | -0.48 | -1.396 | -0.206 | 0.516 | -0.005 | 0.008 | 0.008 | 0 | 0.36 | 0.42 | 0.33 | 0.36 | 0.33 | 0.31 |
| 5 | 174060774 | rs6878215 | -1.475 | -1.736 | 0.368 | 0.211 | -0.025 | -0.009 | -0.009 | 0 | 0.28 | 0.66 | 0.18 | 0.24 | 0.13 | 0.24 |
| 5 | 174062145 | rs4280892 | -1.374 | -1.685 | 0.528 | 0.241 | -0.076 | -0.026 | -0.026 | 0.015 | 0.27 | 0.48 | 0.16 | 0.24 | 0.13 | 0.25 |
| 5 | 174069193 | rs7723988 | -1.493 | -1.099 | 0.776 | 0.146 | 0.024 | 0.004 | 0.004 | 0 | 0.06 | 0.18 | 0.08 | 0.05 | 0.09 | 0.04 |
| 5 | 174069668 | rs11749255 | -0.219 | -1.428 | 0.395 | -0.258 | 0.017 | 0.058 | 0.058 | 0 | 0.37 | 0.17 | 0.3 | 0.24 | 0.3 | 0.28 |
| 5 | 174069956 | rs4867771 | 0.76 | -1.543 | 0.884 | -0.316 | 0.017 | 0.071 | 0.071 | 0 | 0.88 | 0.6 | 0.88 | 0.8 | 0.9 | 0.8 |
| 5 | 174070820 | rs6864763 | -0.238 | -1.63 | 0.816 | -0.312 | 0.007 | 0.136 | 0.136 | 0.1 | 0.65 | 0.93 | 0.79 | 0.85 | 0.73 | 0.79 |
| 5 | 174071838 | rs13163544 | 1.123 | -1.81 | 0.842 | -0.386 | 0.013 | 0.036 | 0.036 | 0.143 | 0.46 | 0.3 | 0.61 | 0.59 | 0.58 | 0.52 |
| 5 | 174074391 | rs4868438 | 1.513 | -1.819 | 0.589 | -0.41 | -0.099 | -0.04 | -0.04 | 1.704 | 0.39 | 0.1 | 0.53 | 0.51 | 0.49 | 0.48 |
| 5 | 174076669 | rs11744357 | -0.126 | -1.942 | 0.647 | -0.649 | 0.04 | 0.07 | 0.07 | 0 | 0.37 | 0.16 | 0.21 | 0.2 | 0.28 | 0.26 |
| 5 | 174076982 | rs7720077 | -0.056 | -1.923 | 0.826 | -0.659 | 0.018 | 0.024 | 0.024 | 0 | 0.36 | 0.24 | 0.21 | 0.11 | 0.25 | 0.19 |
| 5 | 174082558 | rs1507808 | 0.868 | -1.733 | 0.879 | -0.536 | 0.001 | 0.014 | 0.014 | 0 | 0.47 | 0.45 | 0.58 | 0.55 | 0.52 | 0.54 |
| 5 | 174101621 | rs10051893 | 0.694 | -1.404 | 0.323 | -1.164 | 0.005 | 0.05 | 0.05 | 0.093 | 0.57 | 0.32 | 0.63 | 0.91 | 0.55 | 0.82 |
| 5 | 174120131 | rs17796183 | 0.091 | -1.557 | -0.005 | -1.124 | 0.032 | 0.091 | 0.091 | 1.668 | 0.28 | 0.11 | 0.19 | 0.05 | 0.22 | 0.17 |
| 5 | 174121496 | rs1567207 | 0.292 | -1.6 | 0.95 | -1.289 | -0.009 | -0.071 | -0.071 | 0 | 0.64 | 0.61 | 0.73 | 0.94 | 0.67 | 0.82 |
| 5 | 174123135 | rs884823 | 1.735 | -2.125 | 0.897 | -0.821 | -0.01 | 0.016 | 0.016 | 1.063 | 0.75 | 0.33 | 0.66 | 0.66 | 0.77 | 0.72 |
| 5 | 174124562 | rs12659443 | -1.109 | -2.085 | 1.03 | -0.772 | 0.05 | 0.038 | 0.038 | 0 | 0.5 | 0.71 | 0.52 | 0.39 | 0.45 | 0.47 |
| 5 | 174132754 | rs10061074 | -0.57 | -2.066 | 1.592 | -0.692 | 0.022 | 0.005 | 0.005 | 0 | 0.05 | 0.25 | 0.04 | 0.01 | 0.02 | 0.09 |
| 5 | 174134301 | rs6556154 | 0.864 | -2.129 | 1.278 | -0.685 | 0.067 | 0.015 | 0.015 | 0 | 0.52 | 0.25 | 0.52 | 0.6 | 0.61 | 0.53 |
| 5 | 174135863 | rs11952298 | -0.434 | -1.953 | 0.831 | -1.445 | 0.023 | 0.055 | 0.055 | 0 | 0.38 | 0.3 | 0.27 | 0.05 | 0.34 | 0.22 |
| 5 | 174145260 | rs1567202 | 0.037 | -1.009 | 0.481 | -1.234 | 0.055 | 0.147 | 0.147 | 0.678 | 0.5 | 0.32 | 0.66 | 0.82 | 0.63 | 0.74 |
| 5 | 174153037 | rs4868442 | 0.276 | -0.868 | 0.087 | -1.195 | 0.007 | 0.113 | 0.113 | 1.19 | 0.45 | 0.24 | 0.65 | 0.81 | 0.59 | 0.68 |
| 5 | 174156168 | rs4242182 | 1.871 | -0.476 | 1.233 | -0.687 | -0.018 | -0.086 | -0.086 | 0.543 | 0.85 | 0.36 | 0.85 | 0.98 | 0.88 | 0.92 |
| 5 | 174157711 | rs14459 | 1.762 | -0.418 | 1.003 | -0.692 | -0.013 | -0.115 | -0.115 | 0 | 0.82 | 0.45 | 0.84 | 0.98 | 0.83 | 0.9 |
| 5 | 174164216 | rs7733337 | 1.442 | 0.188 | 0.125 | -0.406 | 0.002 | -0.056 | -0.056 | 0 | 0.87 | 0.81 | 0.91 | 1 | 0.86 | 0.93 |
| 5 | 174168311 | rs17063894 | -0.944 | 0.463 | 0.142 | 0.051 | 0.009 | 0.003 | 0.003 | 2.372 | 0.27 | 0.48 | 0.36 | 0.27 | 0.28 | 0.2 |
| 5 | 174173364 | rs12515514 | 0.018 | 0.458 | -0.763 | 0.157 | -0.002 | -0.019 | -0.019 | 0 | 0.3 | 0.15 | 0.27 | 0.34 | 0.29 | 0.23 |
| 5 | 174180561 | rs7728861 | 0.324 | 0.347 | 0 | -0.034 | -0.017 | -0.018 | -0.018 | 0 | 0.65 | 0.46 | 0.7 | 0.67 | 0.73 | 0.78 |
| 5 | 174186729 | rs7705461 | 0.448 | 0.382 | -0.131 | -0.325 | -0.003 | -0.007 | -0.007 | 0.21 | 0.69 | 0.73 | 0.66 | 0.63 | 0.72 | 0.87 |
| 5 | 174187518 | rs1125813 | 1.352 | -0.582 | -0.319 | -1.566 | -0.018 | 0.059 | 0.059 | 3.507 | 0.37 | 0.21 | 0.56 | 0.79 | 0.42 | 0.68 |
| 5 | 174189823 | rs1507806 | -0.246 | -0.903 | -0.024 | -1.805 | -0.007 | -0.019 | -0.019 | 0 | 0.24 | 0.28 | 0.16 | 0.17 | 0.15 | 0.15 |
| 5 | 174193309 | rs1507805 | 0.294 | -1.07 | 0.264 | -1.9 | -0.022 | -0.104 | -0.104 | 0.06 | 0.71 | 0.6 | 0.78 | 0.94 | 0.77 | 0.84 |
| 5 | 174196626 | rs17796794 | 0.072 | -1.119 | -0.554 | -1.946 | -0.009 | 0.052 | 0.052 | 0 | 0.21 | 0.11 | 0.15 | 0.05 | 0.24 | 0.16 |
| 5 | 174199638 | rs3935249 | 0.097 | -0.894 | -0.2 | -1.648 | 0.004 | 0.003 | 0.003 | 3.457 | 0.64 | 0.61 | 0.83 | 0.85 | 0.71 | 0.9 |
| 5 | 174201046 | rs10070886 | 0.773 | -0.753 | -0.09 | -1.45 | 0.001 | -0.006 | -0.006 | 0 | 0.39 | 0.38 | 0.62 | 0.79 | 0.43 | 0.69 |
| 5 | 174204662 | rs6556160 | 0.975 | -0.235 | -0.571 | -1.297 | -0.003 | 0.003 | 0.003 | 1.041 | 0.36 | 0.34 | 0.62 | 0.73 | 0.47 | 0.67 |
| 5 | 174207972 | rs13166147 | -0.071 | -0.368 | -1.086 | -1.146 | 0 | 0.009 | 0.009 | 0 | 0.73 | 0.8 | 0.8 | 0.95 | 0.73 | 0.81 |
| 5 | 174210668 | rs1605522 | -0.073 | -0.365 | -1.084 | -1.145 | 0 | 0.009 | 0.009 | 0 | 0.73 | 0.82 | 0.8 | 0.95 | 0.73 | 0.81 |
| 5 | 174212397 | rs12152810 | -0.014 | -0.364 | -1.122 | -0.352 | 0.012 | 0.006 | 0.006 | 0 | 0.19 | 0.02 | 0.05 | 0.16 | 0.11 | 0.07 |
| 5 | 174221703 | rs7731308 | -0.771 | -0.285 | -0.634 | -0.047 | -0.018 | 0.009 | 0.009 | 0 | 0.31 | 0.35 | 0.23 | 0.38 | 0.19 | 0.27 |
| 5 | 174223234 | rs12660057 | -0.641 | 0.335 | -0.868 | -0.311 | -0.008 | 0.026 | 0.026 | 0 | 0.19 | 0.12 | 0.15 | 0.05 | 0.22 | 0.11 |
| 5 | 174225001 | rs11949641 | -1.122 | 1.249 | -0.637 | -0.494 | -0.005 | 0 | 0 | 0.059 | 0.23 | 0.19 | 0.22 | 0.05 | 0.36 | 0.13 |
| 5 | 174241185 | rs7721551 | -0.492 | 1.177 | -0.794 | -0.621 | 0.008 | 0.01 | 0.01 | 1.275 | 0.7 | 0.68 | 0.47 | 0.44 | 0.46 | 0.52 |
| 5 | 174242357 | rs10053044 | 0.122 | 0.559 | -1.118 | -0.805 | 0.007 | 0.018 | 0.018 | 0.054 | 0.57 | 0.67 | 0.42 | 0.32 | 0.4 | 0.47 |
| 5 | 174244471 | rs11134879 | 0.627 | 0.327 | -0.75 | -0.974 | 0.012 | -0.001 | -0.001 | 0 | 0.48 | 0.51 | 0.31 | 0.12 | 0.35 | 0.27 |
| 5 | 174247900 | rs12108679 | 1.607 | 0.203 | -0.754 | -0.825 | 0.009 | 0.093 | 0.093 | 0 | 0.09 | 0.18 | 0.1 | 0.2 | 0.05 | 0.2 |
| 5 | 174249930 | rs6871270 | 0.341 | 0.504 | -0.921 | -0.897 | 0 | -0.028 | -0.028 | 0 | 0.52 | 0.58 | 0.38 | 0.11 | 0.47 | 0.28 |
| 5 | 174250895 | rs7726244 | -1.44 | 0.285 | -1.379 | -0.595 | -0.018 | -0.025 | -0.025 | 0 | 0.8 | 0.85 | 0.63 | 0.44 | 0.7 | 0.57 |

| | | | | | | | | | | | | | | | | |
|---|---|---|---|---|---|---|---|---|---|---|---|---|---|---|---|---|
| 5 | 174261760 | rs4263513 | 0.764 | -0.29 | -1.361 | -0.47 | 0.065 | 0.038 | 0.038 | 0 | 0.32 | 0.1 | 0.28 | 0.26 | 0.23 | 0.16 |
| 5 | 174272876 | rs6556164 | -0.39 | -0.041 | -1.378 | -0.013 | -0.05 | -0.08 | -0.08 | 0 | 0.67 | 0.85 | 0.64 | 0.49 | 0.61 | 0.52 |
| 5 | 174274276 | rs6556165 | -0.187 | -0.078 | -1.025 | -0.716 | 0.002 | 0.059 | 0.059 | 0.028 | 0.44 | 0.69 | 0.44 | 0.27 | 0.46 | 0.37 |
| 5 | 174279701 | rs10062555 | -0.72 | -0.007 | -0.386 | -0.796 | -0.011 | -0.067 | -0.067 | 0 | 0.3 | 0.43 | 0.27 | 0.15 | 0.29 | 0.2 |
| 5 | 174279808 | rs10062615 | -0.72 | -0.007 | -0.386 | -0.796 | -0.011 | -0.067 | -0.067 | 0 | 0.3 | 0.43 | 0.27 | 0.15 | 0.29 | 0.2 |
| 5 | 174284101 | rs11738763 | -0.089 | 0.54 | -0.499 | -0.558 | -0.022 | 0.029 | 0.029 | 0 | 0.33 | 0.27 | 0.3 | 0.15 | 0.47 | 0.26 |
| 5 | 174294991 | rs17064054 | -0.496 | 0.298 | -0.095 | -0.591 | -0.015 | -0.059 | -0.059 | 3.404 | 0.14 | 0.19 | 0.12 | 0 | 0.11 | 0.05 |
| 5 | 174295209 | rs7444161 | 0.195 | 0.441 | 0.303 | -0.586 | 0.026 | 0.017 | 0.017 | 0 | 0.07 | 0.15 | 0.16 | 0.1 | 0.13 | 0.06 |
| 5 | 174298465 | rs12517380 | -0.765 | 0.236 | 0.133 | -0.809 | -0.001 | -0.025 | -0.025 | 0 | 0.48 | 0.5 | 0.41 | 0.17 | 0.58 | 0.29 |
| 5 | 174303400 | rs961412 | -0.038 | 0.234 | 0.165 | -0.743 | -0.001 | -0.024 | -0.024 | 0.065 | 0.11 | 0.1 | 0.08 | 0 | 0.11 | 0.05 |
| 5 | 174320627 | rs6884951 | -0.671 | 0.157 | 0.296 | -0.857 | 0 | -0.02 | -0.02 | 0 | 0.43 | 0.52 | 0.57 | 0.25 | 0.59 | 0.31 |
| 5 | 174330323 | rs255626 | 0.456 | 0.271 | 0.066 | -0.77 | 0.019 | 0.011 | 0.011 | 0 | 0.07 | 0.14 | 0.24 | 0.1 | 0.13 | 0.06 |
| 5 | 174330780 | rs255627 | 0.336 | 0.363 | 0.061 | -0.828 | -0.018 | 0.018 | 0.018 | 0 | 0.3 | 0.24 | 0.28 | 0.15 | 0.45 | 0.24 |
| 5 | 174337156 | rs10075815 | -1.248 | 0.243 | -0.346 | -0.823 | 0.033 | 0.099 | 0.099 | 0.006 | 0.56 | 0.88 | 0.69 | 0.26 | 0.7 | 0.36 |
| 5 | 174338352 | rs6872302 | 1.343 | 0.005 | -0.382 | 0.118 | 0.075 | -0.056 | -0.056 | 0 | 0.41 | 0.11 | 0.16 | 0.49 | 0.27 | 0.6 |
| 5 | 174340216 | rs2431160 | -0.682 | 0.059 | -0.141 | -0.197 | -0.002 | 0.056 | 0.056 | 0 | 0.41 | 0.64 | 0.42 | 0.15 | 0.58 | 0.26 |
| 5 | 174341887 | rs2434195 | -0.11 | 0.533 | -0.135 | 0.084 | -0.041 | 0.037 | 0.037 | 0 | 0.29 | 0.18 | 0.31 | 0.13 | 0.51 | 0.24 |
| 5 | 174347952 | rs30945 | -0.796 | 0.699 | 0.212 | 0.814 | 0.016 | 0.092 | 0.092 | 0 | 0.09 | 0.18 | 0.35 | 0.32 | 0.15 | 0.14 |
| 5 | 174360217 | rs414031 | -1.515 | 1.161 | 0.363 | 0.215 | 0.036 | -0.02 | -0.02 | 0 | 0.17 | 0.71 | 0.46 | 0.11 | 0.29 | 0.2 |
| 5 | 174370490 | rs17797954 | -0.229 | 0.616 | -1.014 | -0.405 | -0.012 | 0.084 | 0.084 | 0.045 | 0.21 | 0.04 | 0.1 | 0 | 0.18 | 0.1 |
| 5 | 174370845 | rs13189951 | 0.024 | 1.025 | -0.927 | -0.089 | -0.035 | -0.025 | -0.025 | 0 | 0.06 | 0.02 | 0.23 | 0.1 | 0.19 | 0.13 |
| 5 | 174375709 | rs17064145 | -1.076 | 0.575 | -0.709 | -0.169 | -0.024 | -0.022 | -0.022 | 3.498 | 0.09 | 0.67 | 0.11 | 0.03 | 0.03 | 0.06 |
| 5 | 174379880 | rs17798338 | 0.502 | 1.424 | -1.572 | 0.394 | -0.042 | -0.052 | -0.052 | 0.032 | 0.11 | 0.04 | 0.18 | 0.34 | 0.2 | 0.08 |
| 5 | 174390710 | rs929748 | -1.141 | 0.538 | -1.93 | -0.09 | 0.006 | -0.058 | -0.058 | 0.004 | 0.48 | 0.9 | 0.56 | 0.5 | 0.43 | 0.4 |
| 5 | 174399898 | rs11134893 | -0.658 | -0.023 | -2.244 | -0.243 | -0.034 | 0.099 | 0.099 | 0 | 0.43 | 0.14 | 0.26 | 0.26 | 0.46 | 0.38 |
| 5 | 174402211 | rs6897655 | 1.356 | 0.35 | -1.828 | 0.14 | 0.103 | 0.121 | 0.121 | 0 | 0.39 | 0.15 | 0.12 | 0.17 | 0.22 | 0.41 |
| 5 | 174402671 | rs1421780 | -0.169 | 0.478 | -1.957 | 0.178 | -0.039 | 0.061 | 0.061 | 0 | 0.31 | 0.13 | 0.22 | 0.21 | 0.38 | 0.29 |
| 5 | 174403916 | rs7726898 | 0.453 | 0.681 | -1.547 | -0.299 | 0.013 | -0.008 | -0.008 | 0 | 0.22 | 0.23 | 0.35 | 0.14 | 0.33 | 0.24 |
| 5 | 174404365 | rs17064191 | 1.387 | 0.638 | -1.526 | -0.241 | 0.09 | 0.1 | 0.1 | 0 | 0.4 | 0.22 | 0.13 | 0.16 | 0.23 | 0.41 |
| 5 | 174404780 | rs4868473 | 0.121 | 1.427 | -1.46 | 0.349 | 0.007 | -0.006 | -0.006 | 0 | 0.76 | 0.76 | 0.61 | 0.85 | 0.64 | 0.74 |
| 5 | 174405982 | rs6898657 | 0.626 | 0.787 | -1.442 | 1.144 | -0.007 | -0.045 | -0.045 | 0.057 | 0.75 | 0.9 | 0.61 | 0.38 | 0.73 | 0.76 |
| 5 | 174410456 | rs13169879 | -0.667 | 0.708 | -0.932 | 0.879 | -0.002 | 0.001 | 0.001 | 0 | 0.08 | 0.08 | 0.16 | 0.06 | 0.08 | 0.11 |
| 5 | 174413569 | rs12520371 | -0.613 | 1.337 | -0.991 | 1.463 | -0.006 | -0.002 | -0.002 | 0 | 0.17 | 0.16 | 0.22 | 0.59 | 0.2 | 0.24 |
| 5 | 174414273 | rs1979926 | -0.744 | 0.418 | -1.084 | 1.129 | 0.02 | 0.016 | 0.016 | 0.033 | 0.44 | 0.64 | 0.61 | 0.64 | 0.39 | 0.45 |
| 5 | 174414636 | rs6859877 | -1.022 | 0.675 | -1.451 | 0.695 | -0.09 | 0.04 | 0.04 | 1.126 | 0.56 | 0.91 | 0.75 | 0.65 | 0.51 | 0.62 |
| 5 | 174416168 | rs748522 | -0.216 | -0.023 | -1.279 | 0.364 | -0.006 | 0.002 | 0.002 | 0 | 0.33 | 0.61 | 0.57 | 0.63 | 0.31 | 0.42 |
| 5 | 174417049 | rs17064225 | 0.345 | -0.255 | -0.431 | 0.2 | 0.03 | 0.09 | 0.09 | 0 | 0.16 | 0.22 | 0.37 | 0.58 | 0.28 | 0.27 |
| 5 | 174417633 | rs17064226 | 0.699 | -0.254 | -0.251 | 0.166 | -0.014 | -0.007 | -0.007 | 0 | 0.13 | 0.08 | 0.23 | 0.14 | 0.21 | 0.18 |
| 5 | 174417644 | rs17064227 | 1.276 | -0.344 | -0.33 | 0.083 | -0.001 | -0.017 | -0.017 | 0 | 0.07 | 0.01 | 0.17 | 0.06 | 0.08 | 0.11 |
| 5 | 174420134 | rs10053632 | -2.056 | -0.534 | -0.462 | -0.048 | -0.051 | 0.042 | 0.042 | 1.103 | 0.58 | 0.81 | 0.73 | 0.65 | 0.52 | 0.62 |
| 5 | 174423122 | rs11740726 | -1.072 | -0.636 | 0.066 | -0.024 | -0.004 | -0.094 | -0.094 | 0 | 0.35 | 0.43 | 0.32 | 0.08 | 0.33 | 0.31 |
| 5 | 174424824 | rs10052925 | 1.276 | -0.557 | 0.37 | 0.122 | -0.023 | -0.047 | -0.047 | 10.176 | 0.1 | 0.18 | 0.04 | 0 | 0.04 | 0.07 |
| 5 | 174427085 | rs6877848 | -0.423 | -0.518 | 0.453 | 0.311 | -0.046 | 0.247 | 0.247 | 0.009 | 0.14 | 0.41 | 0.31 | 0.55 | 0.09 | 0.22 |
| 5 | 174427832 | rs1862158 | -0.924 | -0.466 | -0.343 | 0.182 | 0.01 | -0.031 | -0.031 | 0.379 | 0.39 | 0.8 | 0.61 | 0.55 | 0.39 | 0.44 |
| 5 | 174428604 | rs9313702 | -0.453 | -0.133 | -0.128 | 0.419 | 0 | 0 | 0 | 0 | 0.35 | 0.39 | 0.4 | 0.47 | 0.31 | 0.33 |
| 5 | 174428848 | rs10516123 | 0.01 | 0.357 | -0.37 | 0.443 | 0.023 | 0.004 | 0.004 | 0 | 0.1 | 0.16 | 0.14 | 0.09 | 0.13 | 0.07 |
| 5 | 174429085 | rs7728260 | -1.245 | 0.399 | -1.325 | 0.123 | -0.053 | -0.094 | -0.094 | 0 | 0.8 | 0.98 | 0.9 | 0.71 | 0.8 | 0.74 |
| 5 | 174429248 | rs7728402 | -0.449 | -0.499 | -0.65 | 0.159 | 0 | -0.002 | -0.002 | 0 | 0.42 | 0.62 | 0.47 | 0.5 | 0.23 | 0.42 |
| 5 | 174434329 | rs11134896 | -0.701 | 0.293 | -0.858 | -0.153 | -0.001 | 0.001 | 0.001 | 0 | 0.44 | 0.59 | 0.49 | 0.5 | 0.52 | 0.38 |
| 5 | 174434677 | rs12522738 | 0.942 | 0.258 | -0.852 | -0.135 | -0.008 | 0.002 | 0.002 | 0 | 0.06 | 0.13 | 0.14 | 0.08 | 0.05 | 0.04 |
| 5 | 174435132 | rs981359 | -0.424 | 0.443 | -0.883 | -0.36 | 0 | 0.006 | 0.006 | 0 | 0.45 | 0.49 | 0.59 | 0.37 | 0.55 | 0.41 |
| 5 | 174436976 | rs17799508 | -0.578 | 1.608 | -0.888 | 0.149 | 0.001 | -0.022 | -0.022 | 0 | 0.3 | 0.35 | 0.37 | 0.09 | 0.41 | 0.18 |
| 5 | 174437991 | rs6869560 | 0.019 | 1.65 | -0.88 | 0.156 | 0.017 | -0.039 | -0.039 | 0.276 | 0.25 | 0.34 | 0.33 | 0.09 | 0.32 | 0.16 |
| 5 | 174438151 | rs6869749 | -1.549 | 1.004 | -0.878 | 0.185 | 0.003 | -0.001 | -0.001 | 0 | 0.47 | 0.59 | 0.51 | 0.5 | 0.56 | 0.45 |
| 5 | 174438353 | rs2216554 | 1.556 | 1.017 | -0.894 | 0.165 | 0.006 | -0.002 | -0.002 | 0 | 0.52 | 0.39 | 0.39 | 0.5 | 0.44 | 0.54 |
| 5 | 174441565 | rs17064274 | 0.408 | 1.063 | -0.967 | 0.319 | -0.004 | 0.005 | 0.005 | 0 | 0.07 | 0.15 | 0.13 | 0.09 | 0.05 | 0.05 |
| 5 | 174443233 | rs12522449 | -0.412 | 1.07 | -0.974 | 0.314 | 0 | -0.001 | -0.001 | 0.006 | 0.11 | 0.15 | 0.15 | 0.09 | 0.11 | 0.06 |
| 5 | 174454025 | rs884252 | 0.351 | 1.134 | -1.143 | 0.355 | -0.008 | 0.054 | 0.054 | 0.268 | 0.8 | 0.75 | 0.76 | 0.6 | 0.83 | 0.71 |
| 5 | 174465510 | rs2135053 | -0.395 | 0.682 | -0.912 | 0.749 | 0 | 0.003 | 0.003 | 0 | 0.42 | 0.57 | 0.59 | 0.5 | 0.5 | 0.39 |
| 5 | 174468355 | rs2036563 | 0.296 | 1.123 | -1.286 | 1.283 | 0.001 | 0.002 | 0.002 | 0.346 | 0.56 | 0.63 | 0.51 | 0.59 | 0.44 | 0.57 |
| 5 | 174470714 | rs10079096 | -0.326 | 1.388 | -1.698 | 0.968 | -0.026 | -0.045 | -0.045 | 2.626 | 0.31 | 0.12 | 0.42 | 0.36 | 0.32 | 0.34 |
| 5 | 174470775 | rs10072136 | -0.385 | 1.234 | -2.487 | 0.244 | -0.051 | -0.014 | -0.014 | 0.945 | 0.65 | 0.91 | 0.79 | 0.64 | 0.63 | 0.67 |
| 5 | 174472998 | rs10053511 | -0.123 | 1.035 | -1.773 | -0.009 | -0.006 | -0.004 | -0.004 | 0.033 | 0.69 | 0.74 | 0.72 | 0.63 | 0.63 | 0.68 |
| 5 | 174474826 | rs6859889 | 0.102 | 1.023 | -0.83 | -0.033 | 0.015 | 0.006 | 0.006 | 0 | 0.66 | 0.55 | 0.67 | 0.58 | 0.59 | 0.66 |
| 5 | 174474862 | rs1566890 | 0.489 | 0.857 | -1.106 | -0.1 | -0.008 | -0.061 | -0.061 | 0.153 | 0.29 | 0.11 | 0.46 | 0.53 | 0.31 | 0.3 |
| 5 | 174475568 | rs7723710 | 0.421 | 0.718 | -0.647 | -0.166 | 0.006 | -0.018 | -0.018 | 3.194 | 0.3 | 0.25 | 0.47 | 0.52 | 0.31 | 0.3 |
| 5 | 174476356 | rs9313705 | 0.275 | 0.511 | -0.418 | -0.492 | -0.015 | -0.111 | -0.111 | 0 | 0.4 | 0.48 | 0.23 | 0.11 | 0.29 | 0.38 |
| 5 | 174476464 | rs10214323 | 0.409 | 0.55 | -0.419 | -0.434 | -0.011 | -0.081 | -0.081 | 0 | 0.4 | 0.5 | 0.23 | 0.12 | 0.3 | 0.36 |
| 5 | 174477636 | rs11953826 | -1.169 | 0.911 | -0.427 | 0.554 | -0.069 | -0.124 | -0.124 | 0 | 0.34 | 0.89 | 0.45 | 0.53 | 0.35 | 0.3 |

| | | | | | | | | | | | | | | | | |
|---|---|---|---|---|---|---|---|---|---|---|---|---|---|---|---|---|
| 5 | 174481433 | rs10516126 | -0.942 | 1.086 | 0.295 | 0.637 | 0.016 | 0.034 | 0.034 | 0 | 0.3 | 0.56 | 0.41 | 0.52 | 0.35 | 0.32 |
| 5 | 174485054 | rs1394612 | -0.929 | 1.112 | -0.181 | 0.645 | -0.025 | -0.046 | -0.046 | 0.176 | 0.31 | 0.72 | 0.44 | 0.54 | 0.35 | 0.32 |
| 5 | 174485598 | rs871234 | -1.252 | 0.209 | 0.565 | 0.179 | 0.061 | 0.08 | 0.08 | 3.39 | 0.05 | 0.48 | 0.06 | 0.04 | 0 | 0.01 |
| 5 | 174489788 | rs6894193 | 0.437 | 0.534 | -0.313 | -0.271 | -0.003 | 0.009 | 0.009 | 0 | 0.77 | 0.82 | 0.8 | 0.96 | 0.76 | 0.94 |
| 5 | 174492784 | rs6893314 | -0.296 | -0.278 | 0.078 | 0.184 | -0.035 | 0.043 | 0.043 | 0 | 0.16 | 0.31 | 0.1 | 0.27 | 0.1 | 0.19 |
| 5 | 174493130 | rs6893965 | -0.843 | -0.506 | 0.234 | 0.356 | -0.073 | 0.089 | 0.089 | 0 | 0.2 | 0.59 | 0.13 | 0.3 | 0.1 | 0.21 |
| 5 | 174496946 | rs1394613 | -0.195 | -0.358 | -0.208 | 0.111 | -0.049 | 0.105 | 0.105 | 0.007 | 0.82 | 0.98 | 0.89 | 0.97 | 0.8 | 0.96 |
| 5 | 174498941 | rs13179042 | 1.39 | -1.508 | -0.354 | 0.551 | 0.054 | 0.017 | 0.017 | 0 | 0.49 | 0.29 | 0.59 | 0.42 | 0.64 | 0.64 |
| 5 | 174507231 | rs10041184 | -0.812 | -0.993 | -0.329 | 0.612 | -0.002 | -0.002 | -0.002 | 0 | 0.33 | 0.72 | 0.31 | 0.58 | 0.2 | 0.33 |
| 5 | 174515327 | rs949637 | 1.103 | -0.78 | -0.716 | 0.73 | -0.021 | -0.044 | -0.044 | 4.833 | 0.07 | 0 | 0.11 | 0.17 | 0.09 | 0.16 |
| 5 | 174515632 | rs1394615 | 1.446 | -0.858 | -0.63 | 0.585 | 0.01 | 0.009 | 0.009 | 0 | 0.46 | 0.26 | 0.56 | 0.41 | 0.56 | 0.62 |
| 5 | 174519247 | rs962492 | -0.891 | -0.628 | -0.247 | 0.926 | -0.02 | 0.014 | 0.014 | 0.002 | 0.33 | 0.63 | 0.31 | 0.57 | 0.22 | 0.32 |
| 5 | 174525366 | rs12153326 | -0.365 | -0.301 | -0.578 | 0.718 | -0.039 | 0.074 | 0.074 | 0.032 | 0.14 | 0.02 | 0.1 | 0.01 | 0.16 | 0.04 |
| 5 | 174526733 | rs10071330 | 0.839 | -0.57 | -0.431 | 1.043 | 0 | 0 | 0 | 0 | 0.11 | 0.05 | 0.16 | 0.28 | 0.04 | 0.1 |
| 5 | 174535069 | rs4008125 | -0.743 | -0.502 | -0.536 | 0.58 | -0.008 | 0.018 | 0.018 | 0.552 | 0.32 | 0.59 | 0.34 | 0.62 | 0.31 | 0.31 |
| 5 | 174543678 | rs6871049 | -1.343 | -0.576 | -1.216 | 0.515 | -0.001 | -0.012 | -0.012 | 0 | 0.43 | 0.89 | 0.49 | 0.64 | 0.39 | 0.35 |
| 5 | 174545220 | rs6877847 | -1.379 | -0.317 | -0.579 | 0.329 | 0.004 | -0.033 | -0.033 | 0 | 0.14 | 0.22 | 0.15 | 0.05 | 0.15 | 0.09 |
| 5 | 174550294 | rs3105418 | -1.305 | 0.258 | -1.481 | 0.046 | -0.034 | 0.042 | 0.042 | 0.424 | 0.57 | 0.93 | 0.75 | 0.74 | 0.63 | 0.48 |
| 5 | 174550786 | rs6556187 | 0.169 | 0.697 | -0.78 | 0.613 | 0.064 | 0.172 | 0.172 | 0.245 | 0.11 | 0.29 | 0.13 | 0.29 | 0.15 | 0.17 |
| 5 | 174550930 | rs6556188 | -0.521 | 0.606 | -0.564 | 0.568 | 0.074 | 0.143 | 0.143 | 0.55 | 0.15 | 0.39 | 0.16 | 0.29 | 0.2 | 0.18 |
| 5 | 174551497 | rs17064566 | 1.164 | 0.406 | -1.001 | 0.307 | -0.011 | -0.067 | -0.067 | 0 | 0.13 | 0 | 0.07 | 0.18 | 0.09 | 0.18 |
| 5 | 174555605 | rs2249160 | 0.027 | 0.837 | -0.492 | 0.463 | -0.003 | 0.001 | 0.001 | 0 | 0.32 | 0.33 | 0.46 | 0.39 | 0.33 | 0.22 |
| 5 | 174567556 | rs7707903 | -0.95 | 0.787 | -0.487 | 0.289 | 0.004 | 0.003 | 0.003 | 0.505 | 0.34 | 0.6 | 0.41 | 0.61 | 0.37 | 0.32 |
| 5 | 174569103 | rs13186454 | 0.963 | 0.731 | -0.534 | 0.364 | -0.02 | 0.063 | 0.063 | 0 | 0.15 | 0.21 | 0.22 | 0.31 | 0.11 | 0.13 |
| 5 | 174577067 | rs11742698 | 1.36 | 0.854 | -1.15 | 0.221 | 0.005 | 0.1 | 0.1 | 0 | 0.46 | 0.13 | 0.27 | 0.24 | 0.4 | 0.55 |
| 5 | 174577240 | rs6889467 | -0.827 | 0.912 | -0.856 | 0.207 | -0.013 | 0.031 | 0.031 | 0.321 | 0.84 | 0.92 | 0.74 | 0.89 | 0.79 | 0.9 |
| 5 | 174577496 | rs6893566 | -0.198 | 0.918 | -0.853 | 0.222 | 0.001 | -0.021 | -0.021 | 0 | 0.8 | 0.76 | 0.68 | 0.85 | 0.77 | 0.87 |
| 5 | 174585025 | rs6864341 | -0.093 | 1.239 | -0.886 | 0.687 | 0.009 | 0.016 | 0.016 | 0.018 | 0.09 | 0.12 | 0.08 | 0.17 | 0.14 | 0.13 |
| 5 | 174586909 | rs750494 | 0.713 | 1.102 | -0.837 | 0.547 | 0 | 0.017 | 0.017 | 0.018 | 0.52 | 0.42 | 0.32 | 0.37 | 0.51 | 0.55 |
| 5 | 174588311 | rs11134913 | 0.431 | 0.747 | -0.878 | 0.509 | 0 | 0 | 0 | 0 | 0.55 | 0.58 | 0.36 | 0.53 | 0.52 | 0.62 |
| 5 | 174594583 | rs10213884 | 0.202 | 0.7 | -0.852 | 0.517 | 0 | 0 | 0 | 0.092 | 0.58 | 0.59 | 0.39 | 0.52 | 0.57 | 0.6 |
| 5 | 174595046 | rs9686689 | -0.261 | 0.677 | -1.472 | 0.367 | -0.029 | 0.036 | 0.036 | 0 | 0.73 | 0.8 | 0.6 | 0.84 | 0.62 | 0.74 |
| 5 | 174597494 | rs1422902 | -0.288 | 0.681 | -1.47 | 0.369 | -0.029 | 0.034 | 0.034 | 0 | 0.74 | 0.8 | 0.6 | 0.84 | 0.62 | 0.75 |
| 5 | 174597961 | rs11747833 | 0.68 | 0.77 | -1.898 | 0.515 | -0.012 | 0.017 | 0.017 | 1.555 | 0.38 | 0.12 | 0.23 | 0.31 | 0.33 | 0.46 |
| 5 | 174600635 | rs7706583 | -0.246 | 0.753 | -1.608 | 0.546 | -0.024 | -0.007 | -0.007 | 0 | 0.42 | 0.28 | 0.35 | 0.53 | 0.48 | 0.56 |
| 5 | 174600839 | rs4868490 | -0.474 | 0.578 | -1.478 | 0.489 | -0.005 | -0.013 | -0.013 | 0 | 0.73 | 0.83 | 0.79 | 0.69 | 0.72 | 0.55 |
| 5 | 174601205 | rs11739264 | -0.02 | 0.706 | -1.146 | 0.429 | 0.001 | -0.003 | -0.003 | 0 | 0.12 | 0.16 | 0.08 | 0.05 | 0.13 | 0.27 |
| 5 | 174602438 | rs13169756 | 0.051 | 1.135 | -1.634 | 0.872 | -0.031 | -0.058 | -0.058 | 0 | 0.2 | 0.08 | 0.26 | 0.45 | 0.22 | 0.22 |
| 5 | 174603338 | rs7718738 | 0.045 | 0.458 | -1.218 | 0.203 | -0.014 | -0.057 | -0.057 | 0.352 | 0.35 | 0.42 | 0.21 | 0.12 | 0.29 | 0.34 |
| 5 | 174605024 | rs11956268 | 0.045 | 0.436 | -1.218 | 0.205 | -0.015 | -0.059 | -0.059 | 0.01 | 0.35 | 0.42 | 0.21 | 0.12 | 0.29 | 0.34 |
| 5 | 174605242 | rs11952967 | 0.045 | 0.436 | -1.218 | 0.204 | -0.015 | -0.057 | -0.057 | 0 | 0.35 | 0.42 | 0.21 | 0.12 | 0.29 | 0.34 |
| 5 | 174605736 | rs6894781 | -1.662 | 0.321 | -1.648 | 0.417 | 0.001 | -0.005 | -0.005 | 0.061 | 0.86 | 0.86 | 0.69 | 0.84 | 0.86 | 0.9 |
| 5 | 174606536 | rs10463049 | 1.162 | 0.522 | -1.785 | 0.577 | -0.022 | -0.046 | -0.046 | 0.009 | 0.1 | 0.02 | 0.16 | 0.24 | 0.08 | 0.1 |
| 5 | 174607841 | rs695221 | -0.236 | 0.429 | -1.581 | 0.529 | 0.026 | 0.005 | 0.005 | 0.061 | 0.49 | 0.62 | 0.41 | 0.56 | 0.39 | 0.62 |
| 5 | 174609937 | rs5005160 | 0.625 | 0.838 | -1.581 | 0.425 | -0.031 | 0.027 | 0.027 | 0.201 | 0.33 | 0.17 | 0.23 | 0.27 | 0.41 | 0.26 |
| 5 | 174611340 | rs4336372 | -0.166 | 0.153 | -1.545 | 0.172 | 0.023 | -0.004 | -0.004 | 0 | 0.56 | 0.39 | 0.42 | 0.55 | 0.68 | 0.66 |
| 5 | 174611803 | rs4868495 | 1.068 | -0.21 | -1.409 | 0.137 | -0.091 | -0.1 | -0.1 | 0 | 0.2 | 0.41 | 0.11 | 0.05 | 0.1 | 0.12 |
| 5 | 174611949 | rs1549225 | 0.125 | -0.124 | -1.45 | 0.281 | 0.05 | 0.059 | 0.059 | 1.299 | 0.51 | 0.29 | 0.37 | 0.36 | 0.63 | 0.48 |
| 5 | 174613013 | rs1549223 | 0.233 | 0.037 | -1.62 | 0.171 | -0.038 | -0.006 | -0.006 | 0.064 | 0.28 | 0.19 | 0.22 | 0.27 | 0.34 | 0.35 |
| 5 | 174613184 | rs11738870 | 0.395 | -0.111 | -1.601 | 0.16 | 0.003 | 0.003 | 0.003 | 0.22 | 0.4 | 0.54 | 0.24 | 0.3 | 0.35 | 0.26 |
| 5 | 174614408 | rs7711234 | 1.638 | 0.052 | -1.653 | 0.181 | -0.004 | -0.007 | -0.007 | 0.965 | 0.13 | 0.09 | 0.11 | 0.2 | 0.15 | 0.14 |
| 5 | 174614480 | rs11747115 | 2.052 | -0.033 | -1.67 | 0.162 | -0.021 | 0.074 | 0.074 | 0.096 | 0.2 | 0.08 | 0.12 | 0.07 | 0.25 | 0.12 |
| 5 | 174615982 | rs11748167 | 2.076 | -0.061 | -1.667 | 0.165 | -0.017 | 0.076 | 0.076 | 0.07 | 0.2 | 0.08 | 0.12 | 0.07 | 0.25 | 0.12 |
| 5 | 174616919 | rs6886962 | -0.93 | 0.501 | -1.419 | 0.548 | -0.023 | -0.055 | -0.055 | 0.391 | 0.72 | 0.83 | 0.49 | 0.6 | 0.72 | 0.76 |
| 5 | 174618053 | rs7732078 | 0.699 | 0.457 | -0.961 | 0.519 | -0.022 | -0.021 | -0.021 | 0.908 | 0.4 | 0.45 | 0.23 | 0.31 | 0.34 | 0.28 |
| 5 | 174619417 | rs6863550 | -0.285 | 1.031 | -0.803 | 0.897 | 0 | -0.001 | -0.001 | 0 | 0.26 | 0.28 | 0.23 | 0.29 | 0.34 | 0.34 |
| 5 | 174619797 | rs17064682 | 0.642 | 1.009 | -0.663 | 1.027 | -0.008 | -0.024 | -0.024 | 0.652 | 0.09 | 0.06 | 0.16 | 0.23 | 0.07 | 0.11 |
| 5 | 174620910 | rs17808557 | 0.336 | 1.181 | -1.205 | 0.727 | -0.048 | 0.02 | 0.02 | 0 | 0.14 | 0.06 | 0.11 | 0.06 | 0.2 | 0.17 |
| 5 | 174628231 | rs1833755 | -0.689 | 0.837 | -0.709 | 0.798 | 0.026 | 0.051 | 0.051 | 0 | 0.31 | 0.48 | 0.39 | 0.47 | 0.31 | 0.46 |
| 5 | 174629627 | rs6556193 | -0.085 | 0.186 | -1.01 | 0.517 | 0.003 | 0.005 | 0.005 | 0.361 | 0.44 | 0.67 | 0.44 | 0.47 | 0.46 | 0.31 |
| 5 | 174638216 | rs4868506 | -0.567 | 0.119 | -1.351 | 0.246 | -0.028 | 0.071 | 0.071 | 0.015 | 0.6 | 0.82 | 0.6 | 0.78 | 0.59 | 0.49 |
| 5 | 174640738 | rs17064692 | 0.093 | -0.3 | -0.874 | 0.589 | 0.003 | -0.004 | -0.004 | 2.189 | 0.21 | 0.19 | 0.15 | 0.21 | 0.19 | 0.07 |
| 5 | 174642940 | rs17752592 | 0.943 | -0.308 | -0.944 | 0.234 | -0.033 | 0.061 | 0.061 | 0.226 | 0.35 | 0.23 | 0.26 | 0.14 | 0.39 | 0.4 |
| 5 | 174645778 | rs7721011 | 0.923 | -0.153 | -0.738 | 0.49 | 0.007 | 0.019 | 0.019 | 0 | 0.2 | 0.25 | 0.3 | 0.46 | 0.27 | 0.37 |
| 5 | 174646311 | rs1422919 | -0.984 | -0.58 | -0.995 | 0.175 | -0.021 | -0.042 | -0.042 | 0 | 0.91 | 0.99 | 0.84 | 0.81 | 0.9 | 0.91 |
| 5 | 174646985 | rs1345732 | -0.435 | -0.618 | -0.559 | 0.126 | -0.006 | 0.01 | 0.01 | 0 | 0.87 | 0.8 | 0.82 | 0.78 | 0.89 | 0.86 |
| 5 | 174648663 | rs6556195 | -0.765 | -0.485 | -0.86 | 0.323 | -0.001 | -0.005 | -0.005 | 0 | 0.51 | 0.49 | 0.56 | 0.65 | 0.51 | 0.49 |
| 5 | 174648786 | rs6898421 | -0.615 | 0.053 | -1.113 | 0.802 | -0.007 | 0.011 | 0.011 | 0.005 | 0.39 | 0.33 | 0.52 | 0.65 | 0.45 | 0.43 |
| 5 | 174649989 | rs1363684 | -0.417 | -0.691 | -1.706 | 1.182 | 0.015 | -0.04 | -0.04 | 0 | 0.18 | 0.05 | 0.23 | 0.19 | 0.19 | 0.06 |

| | | | | | | | | | | | | | | | | |
|---|---|---|---|---|---|---|---|---|---|---|---|---|---|---|---|---|
| 5 | 174651783 | rs1122550 | -0.577 | -0.829 | -2.355 | 0.774 | -0.047 | 0.066 | 0.066 | 0.019 | 0.59 | 0.94 | 0.64 | 0.68 | 0.62 | 0.53 |
| 5 | 174652883 | rs889063 | -0.439 | -0.742 | -2.299 | 0.77 | -0.044 | 0.033 | 0.033 | 0.118 | 0.56 | 0.94 | 0.6 | 0.6 | 0.58 | 0.53 |
| 5 | 174659054 | rs10038738 | 0.273 | -0.167 | -2.159 | 0.887 | -0.029 | 0.015 | 0.015 | 0.588 | 0.31 | 0.81 | 0.25 | 0.25 | 0.38 | 0.32 |
| 5 | 174659422 | rs1549221 | 0.175 | -0.125 | -2.157 | 0.75 | -0.024 | -0.007 | -0.007 | 0 | 0.32 | 0.81 | 0.25 | 0.3 | 0.38 | 0.33 |
| 5 | 174661472 | rs1007387 | 0.268 | -0.033 | -2.412 | 0.775 | -0.009 | -0.003 | -0.003 | 0.163 | 0.31 | 0.73 | 0.23 | 0.3 | 0.38 | 0.33 |
| 5 | 174663244 | rs1549229 | 0.232 | -0.093 | -2.782 | 0.289 | -0.017 | 0.022 | 0.022 | 0 | 0.49 | 0.83 | 0.55 | 0.54 | 0.5 | 0.5 |
| 5 | 174663604 | rs1549228 | 0.322 | 0.693 | -2.32 | 0.369 | -0.032 | -0.009 | -0.009 | 0.404 | 0.7 | 0.93 | 0.61 | 0.68 | 0.69 | 0.66 |
| 5 | 174675119 | rs12657640 | -0.408 | 0.797 | -1.618 | 0.209 | 0.006 | 0.01 | 0.01 | 0.007 | 0.05 | 0.04 | 0.02 | 0.01 | 0.04 | 0.05 |
| 5 | 174677671 | rs11749820 | -0.225 | 1.212 | -2.54 | -0.219 | -0.138 | 0.052 | 0.052 | 0 | 0.29 | 0.02 | 0.22 | 0.11 | 0.41 | 0.27 |
| 5 | 174684808 | rs12656156 | -1.034 | 0.661 | -0.648 | 0.01 | -0.009 | -0.042 | -0.042 | 0 | 0.09 | 0.28 | 0.05 | 0.01 | 0.06 | 0.05 |
| 5 | 174687482 | rs10057481 | -0.108 | 0.709 | -0.619 | 0.196 | -0.002 | 0.009 | 0.009 | 0 | 0.46 | 0.62 | 0.42 | 0.57 | 0.44 | 0.54 |
| 5 | 174691429 | rs13175086 | -0.718 | 0.437 | 0.249 | 0.596 | -0.009 | 0.032 | 0.032 | 4.139 | 0.09 | 0.4 | 0.04 | 0.11 | 0.04 | 0.06 |
| 5 | 174696248 | rs13160120 | -0.941 | 0.425 | 0.089 | 0.761 | 0.003 | 0.038 | 0.038 | 0.012 | 0.27 | 0.56 | 0.21 | 0.36 | 0.25 | 0.23 |
| 5 | 174706296 | rs11953534 | -0.478 | 0.384 | 0.427 | 0.691 | 0.002 | 0.047 | 0.047 | 2.929 | 0.09 | 0.4 | 0.05 | 0.11 | 0.05 | 0.06 |
| 5 | 174708042 | rs3955072 | 0.625 | -0.011 | -0.032 | 0.698 | 0.01 | -0.01 | -0.01 | 0.203 | 0.19 | 0.11 | 0.19 | 0.21 | 0.14 | 0.25 |
| 5 | 174708482 | rs17754498 | 0.472 | 0.289 | -0.125 | 0.852 | -0.01 | 0.023 | 0.023 | 0 | 0.15 | 0.06 | 0.12 | 0.1 | 0.15 | 0.14 |
| 5 | 174709899 | rs1471107 | -0.826 | 0.282 | -0.446 | 0.651 | -0.002 | -0.009 | -0.009 | 0.101 | 0.66 | 0.66 | 0.75 | 0.87 | 0.55 | 0.64 |
| 5 | 174729526 | rs6888523 | -0.869 | 0.496 | -0.701 | 0.651 | -0.096 | 0.075 | 0.075 | 0.017 | 0.73 | 0.93 | 0.78 | 0.87 | 0.57 | 0.69 |
| 5 | 174743514 | rs1902602 | -0.424 | 0.317 | -0.479 | 0.673 | 0.007 | 0.01 | 0.01 | 0 | 0.45 | 0.59 | 0.4 | 0.63 | 0.37 | 0.48 |
| 5 | 174745357 | rs12189407 | 0.388 | 0.786 | -0.451 | 1.173 | 0.004 | -0.057 | -0.057 | 0 | 0.22 | 0.1 | 0.19 | 0.38 | 0.18 | 0.32 |
| 5 | 174747627 | rs4867791 | 0.751 | 1.207 | -0.631 | 1.344 | -0.111 | 0.062 | 0.062 | 0 | 0.23 | 0.03 | 0.19 | 0.09 | 0.37 | 0.23 |
| 5 | 174749051 | rs11960551 | -0.329 | 0.978 | -0.473 | 1.338 | -0.019 | 0.078 | 0.078 | 0 | 0.28 | 0.44 | 0.23 | 0.48 | 0.21 | 0.35 |
| 5 | 174756703 | rs2471015 | 0.373 | 0.923 | -0.988 | 0.832 | 0.035 | 0.038 | 0.038 | 0.425 | 0.49 | 0.71 | 0.56 | 0.44 | 0.4 | 0.38 |
| 5 | 174759912 | rs930875 | -0.16 | 1.797 | -1.297 | 0.434 | -0.11 | 0.13 | 0.13 | 0 | 0.75 | 0.97 | 0.79 | 0.96 | 0.61 | 0.75 |
| 5 | 174760060 | rs11956185 | -1.676 | 1.808 | -0.028 | 1.35 | 0.182 | 0.276 | 0.276 | 0.31 | 0.08 | 0.44 | 0.08 | 0.21 | 0.13 | 0.05 |
| 5 | 174761095 | rs12652255 | -0.459 | 1.067 | -0.966 | 1.48 | 0.006 | -0.069 | -0.069 | 0.087 | 0.22 | 0.04 | 0.21 | 0.29 | 0.17 | 0.29 |
| 5 | 174761461 | rs897748 | -1.138 | 0.904 | -0.282 | 1.694 | -0.021 | 0.015 | 0.015 | 0 | 0.31 | 0.48 | 0.3 | 0.67 | 0.24 | 0.38 |
| 5 | 174761977 | rs6891429 | -0.742 | 0.874 | -0.197 | 1.888 | 0.031 | 0.279 | 0.279 | 0 | 0.09 | 0.36 | 0.08 | 0.33 | 0.07 | 0.06 |
| 5 | 174762119 | rs2471014 | 1.198 | 1.139 | -0.647 | 1.233 | 0.019 | 0.052 | 0.052 | 0 | 0.43 | 0.33 | 0.47 | 0.21 | 0.38 | 0.26 |
| 5 | 174763322 | rs966775 | -0.641 | 0.711 | -0.281 | 1.462 | -0.014 | 0.027 | 0.027 | 0 | 0.23 | 0.31 | 0.24 | 0.34 | 0.17 | 0.33 |
| 5 | 174765845 | rs2445806 | 1.11 | 1.397 | -0.782 | 1.434 | -0.044 | -0.033 | -0.033 | 0 | 0.88 | 0.97 | 0.81 | 0.83 | 0.84 | 0.76 |
| 5 | 174766716 | rs4424010 | 0.138 | 1.016 | -0.907 | 0.763 | -0.007 | 0.041 | 0.041 | 0 | 0.43 | 0.31 | 0.43 | 0.23 | 0.56 | 0.43 |
| 5 | 174768731 | rs4242185 | 0.7 | 1.183 | -0.972 | 0.656 | -0.005 | 0.009 | 0.009 | 0 | 0.34 | 0.3 | 0.36 | 0.13 | 0.41 | 0.29 |
| 5 | 174769272 | rs10516130 | 0.975 | 0.337 | -0.88 | 0.638 | -0.05 | -0.013 | -0.013 | 0 | 0.22 | 0.49 | 0.27 | 0.16 | 0.14 | 0.2 |
| 5 | 174769302 | rs2471010 | 0.631 | 0.35 | -1.004 | 0.65 | -0.025 | 0.039 | 0.039 | 0 | 0.57 | 0.79 | 0.63 | 0.29 | 0.55 | 0.5 |
| 5 | 174773989 | rs2445803 | -0.158 | 0.246 | -1.068 | 0.558 | -0.001 | -0.046 | -0.046 | 0 | 0.68 | 0.86 | 0.74 | 0.36 | 0.73 | 0.64 |
| 5 | 174774285 | rs17064931 | -0.037 | 0.182 | -0.972 | 0.6 | -0.049 | -0.032 | -0.032 | 0 | 0.09 | 0 | 0.08 | 0.1 | 0.16 | 0.14 |
| 5 | 174777691 | rs1379273 | 0.847 | -0.214 | -0.706 | 0.593 | -0.055 | -0.036 | -0.036 | 0 | 0.23 | 0.52 | 0.28 | 0.13 | 0.14 | 0.2 |
| 5 | 174782119 | rs251925 | 0.532 | 0.443 | -0.793 | 1.027 | -0.015 | -0.054 | -0.054 | 0 | 0.65 | 0.86 | 0.69 | 0.37 | 0.69 | 0.58 |
| 5 | 174785110 | rs4868515 | -0.991 | -0.051 | -0.068 | 0.811 | 0.03 | -0.019 | -0.019 | 0 | 0.15 | 0.34 | 0.13 | 0.1 | 0.16 | 0.09 |
| 5 | 174785408 | rs4868517 | -1.732 | -0.39 | -0.415 | 1.137 | 0.001 | 0.031 | 0.031 | 0 | 0.33 | 0.48 | 0.29 | 0.47 | 0.34 | 0.28 |
| 5 | 174788181 | rs12515797 | -0.204 | -0.31 | -0.265 | 0.449 | 0.001 | -0.006 | -0.006 | 0 | 0.13 | 0.11 | 0.08 | 0 | 0.14 | 0.02 |
| 5 | 174788968 | rs265941 | -1.106 | -0.352 | -0.244 | 0.472 | 0.006 | -0.063 | -0.063 | 1.434 | 0.21 | 0.4 | 0.15 | 0.1 | 0.21 | 0.1 |
| 5 | 174794364 | rs703745 | 0.837 | -0.188 | -0.645 | 1.233 | -0.028 | -0.042 | -0.042 | 0.019 | 0.69 | 0.87 | 0.59 | 0.29 | 0.7 | 0.57 |
| 5 | 174800946 | rs2644656 | 0.719 | -0.579 | -0.782 | 1.446 | -0.016 | -0.056 | -0.056 | 0 | 0.76 | 0.97 | 0.72 | 0.78 | 0.82 | 0.63 |
| 5 | 174803794 | rs265945 | 1.917 | -0.678 | -0.228 | 1.348 | 0.014 | 0.048 | 0.048 | 0 | 0.56 | 0.28 | 0.53 | 0.3 | 0.61 | 0.47 |
| 5 | 174804903 | rs265949 | 1.88 | -0.717 | 0.196 | 1.325 | 0.004 | 0.014 | 0.014 | 0 | 0.56 | 0.42 | 0.54 | 0.3 | 0.6 | 0.47 |
| 5 | 174808210 | rs2544727 | 0.887 | -0.424 | 0.45 | 1.64 | -0.001 | 0.065 | 0.065 | 0.303 | 0.75 | 0.51 | 0.65 | 0.65 | 0.8 | 0.63 |
| 5 | 174809308 | rs2431135 | 0.769 | -0.428 | 0.867 | 1.638 | 0 | 0.02 | 0.02 | 0.062 | 0.77 | 0.75 | 0.67 | 0.65 | 0.8 | 0.63 |
| 5 | 174810090 | rs7445117 | 0.132 | -0.047 | 0.528 | 1.841 | -0.024 | -0.069 | -0.069 | 0 | 0.82 | 0.98 | 0.72 | 0.77 | 0.85 | 0.67 |
| 5 | 174810955 | rs2607180 | 0.333 | -0.098 | 0.564 | 1.591 | -0.03 | -0.078 | -0.078 | 0.176 | 0.8 | 0.95 | 0.7 | 0.65 | 0.8 | 0.63 |
| 5 | 174816228 | rs2471007 | -0.83 | -0.092 | 1.004 | 1.344 | -0.006 | 0.078 | 0.078 | 0 | 0.18 | 0.52 | 0.15 | 0.34 | 0.18 | 0.13 |
| 5 | 174817026 | rs2544732 | -0.398 | 0.017 | -0.171 | 1.401 | -0.004 | 0.024 | 0.024 | 0 | 0.16 | 0.24 | 0.12 | 0.34 | 0.18 | 0.13 |
| 5 | 174832617 | rs2644630 | 0.351 | 0.176 | -1.484 | 1.022 | -0.073 | -0.049 | -0.049 | 0.021 | 0.74 | 0.99 | 0.72 | 0.77 | 0.77 | 0.65 |
| 5 | 174836280 | rs265984 | 0.775 | -0.233 | -1.314 | 1.087 | 0.004 | 0.037 | 0.037 | 0 | 0.49 | 0.23 | 0.45 | 0.45 | 0.54 | 0.41 |
| 5 | 174837538 | rs1121582 | -0.674 | -0.867 | -0.682 | 0.9 | -0.029 | 0.004 | 0.004 | 0 | 0.24 | 0.35 | 0.21 | 0.24 | 0.15 | 0.18 |
| 5 | 174842248 | rs265987 | -0.577 | -0.593 | -1.233 | 0.864 | 0.045 | -0.03 | -0.03 | 0 | 0.4 | 0.71 | 0.27 | 0.48 | 0.29 | 0.26 |
| 5 | 174844038 | rs2544738 | 0.083 | 0.009 | -0.872 | 0.636 | -0.002 | 0.003 | 0.003 | 0.048 | 0.58 | 0.57 | 0.59 | 0.55 | 0.6 | 0.53 |
| 5 | 174846909 | rs265993 | 0.212 | 0.379 | -0.633 | 0.005 | 0.004 | -0.043 | -0.043 | 0 | 0.31 | 0.39 | 0.21 | 0.13 | 0.34 | 0.27 |
| 5 | 174847021 | rs11958400 | 0.521 | 0.431 | -0.699 | 0.044 | -0.006 | 0.007 | 0.007 | 0 | 0.27 | 0.26 | 0.2 | 0.12 | 0.34 | 0.26 |
| 5 | 174849907 | rs11954565 | -0.063 | 0.112 | -0.644 | 0.122 | -0.003 | 0.004 | 0.004 | 0 | 0.4 | 0.42 | 0.49 | 0.53 | 0.35 | 0.27 |
| 5 | 174850778 | rs2644635 | 0.011 | 0.112 | -0.664 | 0.114 | -0.005 | 0.012 | 0.012 | 0 | 0.39 | 0.47 | 0.49 | 0.52 | 0.35 | 0.26 |
| 5 | 174851190 | rs1587176 | 0.024 | 0.513 | -0.925 | 0.266 | -0.008 | -0.011 | -0.011 | 0.006 | 0.18 | 0.25 | 0.1 | 0.12 | 0.19 | 0.25 |
| 5 | 174854625 | rs266003 | -0.777 | 0.846 | -1.382 | 0.369 | -0.081 | -0.043 | -0.043 | 0.772 | 0.77 | 0.94 | 0.67 | 0.75 | 0.68 | 0.64 |
| 5 | 174855149 | rs266004 | 0.074 | 0.609 | -1.079 | 0.417 | -0.012 | -0.002 | -0.002 | 0 | 0.61 | 0.66 | 0.54 | 0.62 | 0.51 | 0.4 |
| 5 | 174855411 | rs2644640 | 0.68 | 0.584 | -1.642 | 0.664 | -0.055 | -0.052 | -0.052 | 0 | 0.13 | 0.01 | 0.17 | 0.17 | 0.17 | 0.08 |
| 5 | 174858052 | rs17065021 | 0.461 | 0.66 | -1.174 | 1.073 | -0.001 | 0.007 | 0.007 | 0 | 0.15 | 0.18 | 0.14 | 0.22 | 0.16 | 0.07 |
| 5 | 174858902 | rs2453737 | 0.105 | 0.182 | -1.135 | 0.703 | 0.001 | 0 | 0 | 0 | 0.43 | 0.57 | 0.43 | 0.4 | 0.36 | 0.42 |
| 5 | 174860411 | rs265971 | -1.488 | 0.535 | -1.152 | 0.822 | -0.024 | 0.018 | 0.018 | 0.144 | 0.78 | 0.88 | 0.76 | 0.81 | 0.73 | 0.6 |

| | | | | | | | | | | | | | | | | |
|---|---|---|---|---|---|---|---|---|---|---|---|---|---|---|---|---|
| 5 | 174860516 | rs265972 | -0.378 | 0.344 | -1.221 | 0.785 | -0.018 | 0.014 | 0.014 | 0.099 | 0.59 | 0.74 | 0.58 | 0.64 | 0.53 | 0.5 |
| 5 | 174860699 | rs265973 | -0.491 | 0.208 | -0.786 | 0.777 | -0.004 | -0.002 | -0.002 | 1.843 | 0.6 | 0.61 | 0.6 | 0.55 | 0.55 | 0.4 |
| 5 | 174861240 | rs265974 | -0.843 | -0.463 | -0.807 | 0.718 | 0.031 | -0.035 | -0.035 | 0.54 | 0.43 | 0.83 | 0.4 | 0.44 | 0.35 | 0.3 |
| 5 | 174861613 | rs10039221 | 1.175 | -0.324 | -0.69 | 0.694 | 0.001 | -0.003 | -0.003 | 0 | 0.17 | 0.13 | 0.18 | 0.18 | 0.16 | 0.09 |
| 5 | 174862420 | rs265976 | 0.238 | -0.511 | -0.179 | 0.704 | -0.007 | -0.011 | -0.011 | 0 | 0.21 | 0.57 | 0.18 | 0.19 | 0.19 | 0.2 |
| 5 | 174863502 | rs11747728 | 0.996 | -0.458 | -0.442 | 0.988 | 0.004 | -0.007 | -0.007 | 0.004 | 0.17 | 0.13 | 0.15 | 0.21 | 0.14 | 0.08 |
| 5 | 174863576 | rs835541 | -0.04 | -0.657 | -0.71 | 0.731 | -0.006 | 0.035 | 0.035 | 0 | 0.46 | 0.29 | 0.4 | 0.41 | 0.45 | 0.39 |
| 5 | 174863690 | rs835616 | -1.618 | -0.163 | -0.67 | 0.555 | -0.127 | -0.028 | -0.028 | 0.226 | 0.78 | 0.98 | 0.75 | 0.8 | 0.65 | 0.61 |
| 5 | 174863891 | rs835540 | 0.84 | -0.793 | -0.766 | 0.365 | 0.017 | -0.082 | -0.082 | 0 | 0.29 | 0.13 | 0.35 | 0.47 | 0.23 | 0.27 |
| 5 | 174863905 | rs265978 | -1.168 | -0.387 | 0.156 | 0.248 | 0.03 | -0.02 | -0.02 | 0.084 | 0.74 | 0.69 | 0.67 | 0.84 | 0.59 | 0.58 |
| 5 | 174866091 | rs11746641 | 1.082 | -0.787 | 0.073 | 0.296 | -0.005 | 0 | 0 | 0 | 0.2 | 0.21 | 0.17 | 0.17 | 0.17 | 0.14 |
| 5 | 174866920 | rs251937 | 0.573 | -0.933 | 0.219 | 0.253 | 0.02 | -0.015 | -0.015 | 0.018 | 0.37 | 0.25 | 0.4 | 0.41 | 0.28 | 0.3 |
| 5 | 174867899 | rs4867798 | 0.149 | -0.992 | 0.128 | 0.212 | 0.007 | -0.012 | -0.012 | 0.209 | 0.35 | 0.29 | 0.27 | 0.47 | 0.32 | 0.3 |
| 5 | 174868700 | rs686 | -0.841 | -0.718 | 0.324 | 0.201 | 0.033 | -0.045 | -0.045 | 0 | 0.7 | 0.41 | 0.65 | 0.85 | 0.59 | 0.61 |
| 5 | 174870150 | rs4532 | -1.544 | -0.512 | -0.403 | 0.297 | -0.102 | 0.039 | 0.039 | 0 | 0.77 | 0.94 | 0.72 | 0.85 | 0.6 | 0.6 |
| 5 | 174870196 | rs5326 | 0.714 | -0.55 | -0.581 | 0.307 | 0 | 0 | 0 | 2.522 | 0.16 | 0.16 | 0.1 | 0.27 | 0.14 | 0.16 |
| 5 | 174870902 | rs265981 | -1.418 | -0.455 | -0.458 | 0.352 | -0.159 | 0.015 | 0.015 | 0 | 0.77 | 0.98 | 0.72 | 0.85 | 0.6 | 0.61 |
| 5 | 174878467 | rs10063286 | -0.08 | -0.895 | 0.112 | 0.49 | -0.037 | 0.079 | 0.079 | 0 | 0.08 | 0.2 | 0.06 | 0.2 | 0.02 | 0.13 |
| 5 | 174890313 | rs1355077 | 0.497 | -1.014 | -0.485 | 0.164 | -0.004 | -0.028 | -0.028 | 0 | 0.4 | 0.23 | 0.44 | 0.46 | 0.41 | 0.33 |
| 5 | 174891621 | rs2453741 | -1.322 | -0.979 | -1.121 | 0.083 | -0.122 | 0.063 | 0.063 | 0 | 0.73 | 0.94 | 0.71 | 0.85 | 0.57 | 0.57 |
| 5 | 174892501 | rs1039089 | 1.275 | -0.633 | -0.646 | 0.233 | 0.001 | -0.001 | -0.001 | 0.805 | 0.17 | 0.17 | 0.17 | 0.16 | 0.14 | 0.1 |
| 5 | 174893280 | rs1496133 | -0.431 | -0.758 | -0.349 | 0.186 | -0.004 | 0.004 | 0.004 | 0 | 0.59 | 0.65 | 0.65 | 0.62 | 0.54 | 0.43 |
| 5 | 174893557 | rs1508763 | -0.241 | -0.638 | -0.208 | 0.378 | -0.002 | 0.003 | 0.003 | 0 | 0.42 | 0.48 | 0.48 | 0.46 | 0.41 | 0.33 |
| 5 | 174896514 | rs1022040 | -0.8 | -0.616 | -0.167 | 0.248 | -0.006 | -0.001 | -0.001 | 0 | 0.29 | 0.6 | 0.22 | 0.35 | 0.22 | 0.12 |
| 5 | 174898012 | rs2644648 | -0.075 | -0.15 | 0.157 | -0.184 | 0.012 | -0.028 | -0.028 | 0 | 0.35 | 0.41 | 0.35 | 0.17 | 0.43 | 0.24 |
| 5 | 174901052 | rs6861032 | -0.539 | -0.858 | 0.289 | 0.208 | -0.004 | 0.003 | 0.003 | 0 | 0.14 | 0.17 | 0.17 | 0.18 | 0.12 | 0.12 |
| 5 | 174909302 | rs267368 | -0.012 | -1.229 | 0.341 | -0.09 | 0.001 | 0.002 | 0.002 | 0 | 0.48 | 0.53 | 0.57 | 0.32 | 0.62 | 0.29 |
| 5 | 174910713 | rs17065085 | 0.789 | -1.14 | 0.104 | -0.078 | -0.004 | 0.032 | 0.032 | 0 | 0.07 | 0.01 | 0.09 | 0.02 | 0.07 | 0.08 |
| 5 | 174911675 | rs267388 | -0.56 | -1.426 | 0.262 | -0.202 | 0.023 | 0.011 | 0.011 | 0 | 0.58 | 0.77 | 0.61 | 0.35 | 0.65 | 0.42 |
| 5 | 174916655 | rs267376 | -0.456 | -1.415 | 0.517 | -0.196 | 0.025 | 0.015 | 0.015 | 0 | 0.57 | 0.72 | 0.61 | 0.35 | 0.65 | 0.42 |
| 5 | 174918392 | rs267382 | 0.143 | -1.414 | 0.635 | -0.143 | -0.001 | -0.001 | -0.001 | 0 | 0.47 | 0.51 | 0.49 | 0.31 | 0.58 | 0.34 |
| 5 | 174919450 | rs267384 | 1.181 | -1.231 | 0.495 | -0.059 | -0.05 | 0.026 | 0.026 | 0 | 0.3 | 0.14 | 0.37 | 0.16 | 0.49 | 0.18 |
| 5 | 174921431 | rs7718943 | -0.098 | -1.732 | 0.536 | 0.152 | -0.041 | -0.031 | -0.031 | 1.51 | 0.31 | 0.43 | 0.28 | 0.29 | 0.21 | 0.51 |
| 5 | 174930452 | rs11134927 | -0.913 | -1.894 | 0.546 | 0.173 | 0.032 | 0.013 | 0.013 | 0 | 0.43 | 0.59 | 0.37 | 0.45 | 0.33 | 0.57 |
| 5 | 174934482 | rs6882734 | 0.26 | -1.865 | 0.757 | 0.203 | 0.01 | 0.025 | 0.025 | 0 | 0.12 | 0.18 | 0.1 | 0.15 | 0.12 | 0.07 |
| 5 | 174935571 | rs2937463 | 0.429 | -1.989 | 0.975 | 0.177 | -0.05 | -0.016 | -0.016 | 0 | 0.25 | 0.57 | 0.2 | 0.26 | 0.14 | 0.4 |
| 5 | 174937193 | rs2662170 | 0.737 | -2.013 | 1.177 | 0.164 | -0.045 | -0.007 | -0.007 | 0 | 0.23 | 0.44 | 0.18 | 0.26 | 0.14 | 0.4 |
| 5 | 174943264 | rs283013 | 1.061 | -2.019 | 1.154 | 0.162 | -0.002 | 0 | 0 | 0 | 0.21 | 0.21 | 0.16 | 0.26 | 0.14 | 0.41 |
| 5 | 174948557 | rs157474 | -0.068 | -2.042 | 0.926 | 0.148 | 0.057 | 0.026 | 0.026 | 0 | 0.3 | 0.76 | 0.3 | 0.3 | 0.16 | 0.49 |
| 5 | 174950674 | rs157477 | 0.455 | -2.032 | 0.978 | 0.153 | -0.022 | -0.006 | -0.006 | 2.448 | 0.25 | 0.62 | 0.21 | 0.28 | 0.14 | 0.41 |
| 5 | 174953743 | rs198039 | 0.608 | -2.032 | 1.129 | 0.153 | -0.054 | -0.012 | -0.012 | 0 | 0.24 | 0.48 | 0.19 | 0.26 | 0.14 | 0.41 |
| 5 | 174954861 | rs156373 | -0.205 | -2.075 | 0.941 | 0.173 | 0.018 | 0.01 | 0.01 | 0 | 0.31 | 0.7 | 0.32 | 0.3 | 0.17 | 0.5 |
| 5 | 174955584 | rs273078 | 0.603 | -2.016 | 0.929 | 0.17 | -0.05 | -0.009 | -0.009 | 0 | 0.24 | 0.52 | 0.2 | 0.27 | 0.14 | 0.39 |
| 5 | 174956467 | rs267399 | -0.126 | -2.004 | 0.588 | 0.174 | -0.016 | -0.006 | -0.006 | 0.44 | 0.3 | 0.64 | 0.29 | 0.29 | 0.15 | 0.47 |
| 5 | 174957482 | rs2471017 | -0.476 | -1.636 | -0.13 | 0.215 | -0.042 | -0.031 | -0.031 | 0 | 0.86 | 0.99 | 0.9 | 0.85 | 0.83 | 0.91 |
| 5 | 174963151 | rs182064 | 1.236 | -1.72 | -0.031 | 0.289 | 0.028 | -0.031 | -0.031 | 0 | 0.36 | 0.5 | 0.39 | 0.21 | 0.51 | 0.19 |
| 5 | 174965974 | rs2662176 | 0.306 | -1.407 | -0.483 | 0.512 | -0.029 | -0.002 | -0.002 | 0 | 0.23 | 0.38 | 0.18 | 0.27 | 0.15 | 0.41 |
| 5 | 174966057 | rs2453746 | 0.856 | -1.278 | -0.481 | 0.579 | -0.003 | -0.001 | -0.001 | 0.055 | 0.6 | 0.57 | 0.64 | 0.6 | 0.68 | 0.44 |
| 5 | 174968560 | rs2457087 | 0.242 | -0.503 | -1.044 | 0.62 | -0.037 | -0.013 | -0.013 | 0.573 | 0.86 | 0.99 | 0.9 | 0.9 | 0.84 | 0.94 |
| 5 | 174969467 | rs267393 | -0.323 | -0.976 | -0.769 | 0.686 | 0.052 | 0.069 | 0.069 | 0 | 0.47 | 0.63 | 0.52 | 0.73 | 0.29 | 0.65 |
| 5 | 174969611 | rs12109728 | -0.213 | -1.047 | -0.081 | 1.326 | -0.003 | 0.026 | 0.026 | 0 | 0.3 | 0.34 | 0.28 | 0.51 | 0.23 | 0.41 |
| 5 | 174969879 | rs13164796 | -0.588 | -1.241 | 0.057 | 1.115 | -0.008 | 0.008 | 0.008 | 0 | 0.14 | 0.2 | 0.15 | 0.18 | 0.14 | 0.12 |
| 5 | 174974576 | rs10476160 | -1.17 | -1.137 | -0.093 | 0.839 | -0.038 | 0.011 | 0.011 | 0.038 | 0.26 | 0.61 | 0.22 | 0.25 | 0.2 | 0.18 |
| 5 | 174982432 | rs10038582 | -1.108 | -1.14 | -0.182 | 0.927 | -0.042 | 0.014 | 0.014 | 0.798 | 0.24 | 0.63 | 0.21 | 0.23 | 0.17 | 0.14 |
| 5 | 174990352 | rs10044281 | -0.691 | -0.917 | -0.615 | 0.439 | -0.059 | 0.017 | 0.017 | 3.178 | 0.16 | 0.56 | 0.21 | 0.17 | 0.06 | 0.17 |
| 5 | 174998993 | rs200461 | -0.533 | -0.172 | -1.513 | 0.182 | -0.01 | 0.052 | 0.052 | 0 | 0.56 | 0.72 | 0.65 | 0.73 | 0.51 | 0.5 |
| 5 | 175000507 | rs9313721 | -0.641 | -0.123 | -1.71 | 0.571 | 0.01 | -0.024 | -0.024 | 0 | 0.18 | 0.16 | 0.18 | 0.38 | 0.16 | 0.19 |
| 5 | 175002424 | rs2940479 | -0.196 | 0.599 | -2.001 | -0.878 | -0.068 | 0.137 | 0.137 | 0.141 | 0.38 | 0.79 | 0.43 | 0.01 | 0.5 | 0.16 |
| 5 | 175006696 | rs2937457 | -0.818 | 0.484 | -1.806 | -0.329 | 0.005 | 0.022 | 0.022 | 0.082 | 0.13 | 0.07 | 0.08 | 0.09 | 0.1 | 0.21 |
| 5 | 175007689 | rs2937456 | -0.273 | 0.404 | -1.508 | -0.011 | 0.003 | 0.019 | 0.019 | 1.235 | 0.35 | 0.55 | 0.54 | 0.59 | 0.39 | 0.33 |
| 5 | 175010396 | rs10042293 | -0.282 | 0.151 | -1.684 | 0.672 | 0.023 | -0.053 | -0.053 | 0 | 0.17 | 0.08 | 0.2 | 0.38 | 0.1 | 0.17 |
| 5 | 175012469 | rs7735518 | -0.35 | -0.194 | -1.317 | 0.071 | 0.003 | -0.028 | -0.028 | 0 | 0.34 | 0.42 | 0.31 | 0.14 | 0.34 | 0.41 |
| 5 | 175012488 | rs867339 | 0.35 | -0.305 | -1.428 | 0.295 | 0.022 | -0.058 | -0.058 | 0 | 0.16 | 0.04 | 0.18 | 0.34 | 0.1 | 0.17 |
| 5 | 175015522 | rs6895025 | -1.197 | -0.329 | -0.979 | 0.409 | -0.037 | 0.002 | 0.002 | 0.125 | 0.58 | 0.82 | 0.59 | 0.61 | 0.48 | 0.66 |
| 5 | 175018135 | rs1472606 | -0.574 | 0.111 | -1.084 | 0.039 | 0.013 | -0.004 | -0.004 | 0 | 0.76 | 0.75 | 0.76 | 0.78 | 0.67 | 0.82 |
| 5 | 175019755 | rs2937451 | -0.49 | 0.183 | -1.073 | -0.036 | 0.011 | -0.005 | -0.005 | 0 | 0.74 | 0.75 | 0.75 | 0.78 | 0.64 | 0.8 |
| 5 | 175022459 | rs606095 | -0.738 | -0.017 | -0.615 | 0.169 | 0.006 | 0.002 | 0.002 | 0.386 | 0.52 | 0.69 | 0.65 | 0.61 | 0.44 | 0.68 |
| 5 | 175023656 | rs2134176 | -0.841 | 0.032 | -0.137 | 0.285 | -0.012 | 0.002 | 0.002 | 0.241 | 0.3 | 0.39 | 0.29 | 0.36 | 0.26 | 0.31 |
| 5 | 175024726 | rs2115512 | -0.095 | -0.248 | 0.271 | 0.416 | -0.055 | 0.045 | 0.045 | 0.047 | 0.2 | 0.34 | 0.2 | 0.34 | 0.09 | 0.18 |

| | | | | | | | | | | | | | | | | |
|---|---|---|---|---|---|---|---|---|---|---|---|---|---|---|---|---|
| 5 | 175025995 | rs6861000 | -0.186 | -0.15 | 0.106 | 0.498 | -0.015 | 0.004 | 0.004 | 0 | 0.23 | 0.24 | 0.4 | 0.23 | 0.33 | 0.37 |
| 5 | 175029274 | rs870258 | -1.594 | -0.107 | 0.212 | 0.571 | 0.004 | 0.001 | 0.001 | 0 | 0.47 | 0.61 | 0.61 | 0.55 | 0.44 | 0.62 |
| 5 | 175030363 | rs2913217 | 0.571 | 0.344 | 0.236 | 1.137 | 0.024 | -0.074 | -0.074 | 0.508 | 0.76 | 0.89 | 0.76 | 0.55 | 0.84 | 0.78 |
| 5 | 175031628 | rs1501745 | -1.422 | 1.103 | 1.088 | 0.642 | 0.038 | -0.005 | -0.005 | 0 | 0.22 | 0.61 | 0.38 | 0.23 | 0.26 | 0.35 |
| 5 | 175032264 | rs2913216 | 0.015 | 1.152 | 1.163 | 0.675 | 0.037 | -0.003 | -0.003 | 1.238 | 0.05 | 0.21 | 0.03 | 0.02 | 0.05 | 0.07 |
| 5 | 175037358 | rs1019808 | -0.144 | 1.158 | 0.842 | 0.714 | 0.059 | 0.038 | 0.038 | 0 | 0.14 | 0.33 | 0.33 | 0.21 | 0.2 | 0.26 |
| 5 | 175044569 | rs13160891 | 0.074 | 1.181 | 0.502 | 0.721 | 0.028 | 0.016 | 0.016 | 0 | 0.13 | 0.24 | 0.32 | 0.2 | 0.21 | 0.28 |
| 5 | 175052459 | rs610101 | -1.545 | 0.919 | -0.166 | 0.829 | -0.003 | -0.001 | -0.001 | 0.124 | 0.36 | 0.77 | 0.48 | 0.35 | 0.38 | 0.43 |
| 5 | 175052567 | rs2913229 | -1.05 | 1.488 | -0.159 | 0.846 | 0.042 | 0.011 | 0.011 | 0 | 0.25 | 0.54 | 0.4 | 0.26 | 0.32 | 0.38 |
| 5 | 175055515 | rs890707 | 0.467 | 1.38 | -0.554 | 0.801 | 0 | -0.001 | -0.001 | 0 | 0.57 | 0.43 | 0.65 | 0.38 | 0.52 | 0.45 |
| 5 | 175059713 | rs17572757 | 1.55 | 0.936 | -0.454 | 0.466 | 0.005 | -0.02 | -0.02 | 0 | 0.05 | 0.08 | 0.03 | 0.01 | 0.07 | 0.02 |
| 5 | 175065368 | rs2937448 | 0.225 | 1.028 | -1.004 | 0.017 | 0.02 | 0.094 | 0.094 | 0 | 0.5 | 0.35 | 0.4 | 0.08 | 0.53 | 0.44 |
| 5 | 175068588 | rs4868527 | 0.525 | 0.953 | -1.423 | 0.392 | 0.033 | 0.096 | 0.096 | 0 | 0.5 | 0.2 | 0.38 | 0.31 | 0.44 | 0.46 |
| 5 | 175070333 | rs590774 | -0.934 | -0.022 | -0.528 | 0.597 | -0.004 | 0.145 | 0.145 | 0.545 | 0.49 | 0.85 | 0.33 | 0.24 | 0.43 | 0.28 |
| 5 | 175083754 | rs2963854 | -0.273 | 0.006 | 0.228 | 0.727 | -0.012 | 0.004 | 0.004 | 3.002 | 0.36 | 0.59 | 0.19 | 0.37 | 0.22 | 0.16 |
| 5 | 175088435 | rs659562 | -0.694 | 1.157 | 0.73 | 0.806 | -0.017 | 0.018 | 0.018 | 0 | 0.16 | 0.12 | 0.14 | 0.09 | 0.24 | 0.12 |
| 5 | 175093375 | rs6864183 | -0.868 | -0.032 | 1.882 | 1.135 | 0.001 | -0.004 | -0.004 | 0 | 0.41 | 0.54 | 0.46 | 0.58 | 0.52 | 0.33 |
| 5 | 175107463 | rs645574 | -2.867 | -0.791 | 4.06 | 1.273 | -0.024 | 0.103 | 0.103 | 0.42 | 0.11 | 0.55 | 0.09 | 0.14 | 0.04 | 0.12 |
| 5 | 175109219 | rs2067474 | -2.314 | -0.787 | 3.983 | 1.251 | -0.015 | 0.075 | 0.075 | 0 | 0.07 | 0.2 | 0.05 | 0.14 | 0.04 | 0.12 |
| 5 | 175110779 | rs1800689 | -1.269 | -0.784 | 3.286 | 1.22 | -0.017 | 0.005 | 0.005 | 0 | 0.08 | 0 | 0.04 | 0.05 | 0.07 | 0.05 |
| 5 | 175120476 | rs13162969 | -2.14 | -0.777 | 3.417 | 1.224 | -0.005 | -0.032 | -0.032 | 0 | 0.11 | 0.34 | 0.08 | 0.05 | 0.07 | 0.05 |
| 5 | 175120898 | rs632994 | -2.513 | -0.768 | 2.533 | 1.26 | 0.004 | -0.002 | -0.002 | 0 | 0.22 | 0.77 | 0.15 | 0.2 | 0.12 | 0.2 |
| 5 | 175127131 | rs13356830 | -3.38 | -0.126 | 3.333 | 1.109 | 0.006 | 0.046 | 0.046 | 1.797 | 0.14 | 0.59 | 0.14 | 0.14 | 0.1 | 0.06 |
| 5 | 175131097 | rs10079693 | -3.414 | -0.175 | 3.212 | 0.877 | 0.008 | 0.014 | 0.014 | 0.905 | 0.14 | 0.54 | 0.13 | 0.09 | 0.1 | 0.05 |
| 5 | 175136782 | rs735051 | -3.055 | -0.184 | 2.555 | 0.532 | 0.01 | -0.026 | -0.026 | 0.075 | 0.1 | 0.2 | 0.08 | 0.04 | 0.1 | 0.05 |
| 5 | 175138430 | rs10035429 | -2.925 | -0.195 | 2.548 | 0.525 | 0.012 | -0.027 | -0.027 | 0.022 | 0.1 | 0.2 | 0.08 | 0.04 | 0.11 | 0.05 |
| 5 | 175143632 | rs718907 | -2.516 | -0.317 | 2.547 | 0.598 | 0.007 | -0.008 | -0.008 | 0.929 | 0.07 | 0.13 | 0.06 | 0.04 | 0.08 | 0.06 |
| 5 | 175155256 | rs10039854 | -2.459 | -0.322 | 2.246 | 0.594 | 0.027 | -0.002 | -0.002 | 0.117 | 0.11 | 0.34 | 0.07 | 0.05 | 0.09 | 0.06 |
| 5 | 175158653 | rs6875659 | -2.659 | -0.505 | 0.943 | 0.535 | 0.04 | 0.062 | 0.062 | 2.516 | 0.19 | 0.93 | 0.14 | 0.05 | 0.09 | 0.09 |
| 5 | 175164610 | rs10866687 | 2.456 | -0.501 | 1.03 | 0.497 | 0.022 | 0.029 | 0.029 | 2.214 | 0.8 | 0.17 | 0.87 | 0.95 | 0.9 | 0.91 |
| 5 | 175174011 | rs970282 | 1.433 | -0.431 | 0.567 | 0.556 | 0.001 | -0.019 | -0.019 | 0 | 0.93 | 0.89 | 0.96 | 1 | 0.93 | 0.95 |
| 5 | 175177445 | rs12658312 | 1.673 | -0.381 | -0.615 | 0.59 | 0.043 | 0.064 | 0.064 | 2.485 | 0.8 | 0.16 | 0.88 | 0.95 | 0.9 | 0.91 |
| 5 | 175202064 | rs11749986 | 1.037 | -0.247 | -0.343 | 1.547 | -0.006 | 0.023 | 0.023 | 0 | 0.84 | 0.8 | 0.9 | 0.69 | 0.9 | 0.81 |
| 5 | 175202905 | rs4141517 | 1.237 | 0.486 | -0.455 | 1.757 | -0.001 | -0.004 | -0.004 | 1.299 | 0.77 | 0.76 | 0.77 | 0.68 | 0.76 | 0.76 |
| 5 | 175204022 | rs2382072 | 0.778 | 0.303 | -0.863 | 1.267 | -0.019 | 0.02 | 0.02 | 0 | 0.45 | 0.21 | 0.49 | 0.44 | 0.47 | 0.55 |
| 5 | 175204783 | rs4868534 | -0.605 | 0.226 | -1.61 | 0.307 | -0.008 | 0.029 | 0.029 | 0 | 0.23 | 0.07 | 0.2 | 0.16 | 0.28 | 0.25 |
| 5 | 175206596 | rs2084763 | -0.394 | 0.699 | -1.991 | -1.486 | 0.001 | -0.018 | -0.018 | 0 | 0.55 | 0.52 | 0.78 | 0.85 | 0.65 | 0.58 |
| 5 | 175214075 | rs3899033 | 1.299 | 0.087 | -1.037 | -2.812 | -0.022 | -0.044 | -0.044 | 0.105 | 0.17 | 0.31 | 0.09 | 0.1 | 0.1 | 0.24 |
| 5 | 175242385 | rs17065522 | 0.68 | -0.343 | -0.192 | -2.055 | -0.033 | -0.028 | -0.028 | 0 | 0.28 | 0.35 | 0.13 | 0.14 | 0.14 | 0.3 |
| 5 | 175244097 | rs4242187 | -0.715 | -0.09 | -0.372 | -2.022 | -0.011 | 0.013 | 0.013 | 0 | 0.72 | 0.75 | 0.78 | 0.91 | 0.65 | 0.71 |
| 5 | 175245365 | rs17065524 | 0.309 | -1.317 | -0.766 | -1.952 | 0.007 | 0.007 | 0.007 | 0.47 | 0.3 | 0.3 | 0.14 | 0.17 | 0.19 | 0.34 |
| 5 | 175245433 | rs17065525 | 0.655 | -1.31 | -0.752 | -1.939 | -0.01 | -0.013 | -0.013 | 0 | 0.27 | 0.29 | 0.12 | 0.12 | 0.15 | 0.3 |
| 5 | 175249549 | rs1001120 | 0.937 | -1.095 | -0.977 | -1.501 | 0.025 | -0.012 | -0.012 | 0 | 0.19 | 0.13 | 0.12 | 0.27 | 0.09 | 0.31 |
| 5 | 175268492 | rs1544923 | 0.393 | -0.699 | -0.927 | -0.726 | -0.008 | 0.003 | 0.003 | 0 | 0.2 | 0.26 | 0.15 | 0.28 | 0.12 | 0.3 |
| 5 | 175948316 | rs9313736 | 0.174 | -1.067 | -0.524 | 0.207 | -0.001 | 0.004 | 0.004 | 0 | 0.4 | 0.35 | 0.33 | 0.31 | 0.39 | 0.27 |
| 5 | 175980113 | rs4868648 | 0.436 | -1.353 | -1.91 | -0.201 | 0.101 | 0.069 | 0.069 | 0.567 | 0.51 | 0.05 | 0.61 | 0.31 | 0.68 | 0.49 |
| 5 | 175989028 | rs4868651 | -0.171 | -1.024 | -1.303 | -0.515 | 0.016 | 0.105 | 0.105 | 0 | 0.48 | 0.33 | 0.52 | 0.2 | 0.53 | 0.37 |
| 5 | 175991998 | rs6892681 | 0.715 | -0.777 | -1.338 | -0.586 | -0.116 | 0.136 | 0.136 | 0.043 | 0.29 | 0.08 | 0.41 | 0.02 | 0.45 | 0.21 |
| 5 | 175995392 | rs11739773 | 0.54 | -1.653 | -0.379 | -0.285 | 0.042 | 0.024 | 0.024 | 0 | 0.56 | 0.34 | 0.65 | 0.36 | 0.7 | 0.49 |
| 5 | 175998246 | rs17078320 | 0.415 | -1.739 | 0.159 | -0.233 | -0.012 | 0.021 | 0.021 | 0 | 0.27 | 0.3 | 0.23 | 0.48 | 0.22 | 0.35 |
| 5 | 175998397 | rs10075586 | 0.048 | -1.766 | 0.528 | -0.246 | -0.037 | 0.054 | 0.054 | 0 | 0.3 | 0.46 | 0.23 | 0.48 | 0.21 | 0.35 |
| 5 | 176005184 | rs1565151 | -0.342 | -1.498 | 0.836 | -0.111 | -0.025 | 0.035 | 0.035 | 0.366 | 0.7 | 0.59 | 0.75 | 0.55 | 0.76 | 0.66 |
| 5 | 176005477 | rs2358747 | -0.336 | -1.536 | 0.833 | -0.109 | -0.026 | 0.036 | 0.036 | 0.085 | 0.7 | 0.59 | 0.77 | 0.55 | 0.75 | 0.66 |
| 5 | 176006611 | rs1545976 | 0.36 | -1.513 | 0.681 | -0.121 | 0.011 | -0.023 | -0.023 | 0 | 0.28 | 0.19 | 0.23 | 0.45 | 0.24 | 0.33 |
| 5 | 176008238 | rs4867840 | 0.553 | -1.459 | 0.356 | -0.192 | -0.042 | 0.015 | 0.015 | 2.06 | 0.81 | 0.42 | 0.86 | 0.82 | 0.89 | 0.8 |
| 5 | 176011264 | rs752139 | 0.291 | -1.544 | 0.291 | -0.265 | 0 | 0 | 0 | 0 | 0.61 | 0.6 | 0.66 | 0.35 | 0.7 | 0.49 |
| 5 | 176011889 | rs4868660 | 1.567 | -1.473 | 0.109 | -0.539 | -0.078 | 0.07 | 0.07 | 0 | 0.24 | 0.1 | 0.4 | 0.03 | 0.41 | 0.17 |
| 5 | 176017455 | rs2291442 | 0.909 | -1.474 | 0.167 | -0.487 | 0.001 | -0.001 | -0.001 | 0.056 | 0.31 | 0.3 | 0.23 | 0.32 | 0.24 | 0.29 |
| 5 | 176017639 | rs17078347 | 0.993 | -1.457 | -0.006 | -0.434 | -0.024 | -0.158 | -0.158 | 0 | 0.25 | 0.02 | 0.22 | 0.44 | 0.24 | 0.35 |
| 5 | 176020203 | rs34425002 | 0.883 | -1.386 | 0.052 | -0.405 | -0.003 | 0 | 0 | 0 | 0.32 | 0.32 | 0.23 | 0.32 | 0.25 | 0.29 |
| 5 | 176021956 | rs11134985 | 0.827 | -1.389 | 0.135 | -0.441 | 0.006 | -0.061 | -0.061 | 0.017 | 0.3 | 0.31 | 0.43 | 0.03 | 0.45 | 0.2 |
| 5 | 176024880 | rs4868663 | -0.504 | -1.414 | 0.158 | 0.226 | -0.024 | 0.057 | 0.057 | 0.941 | 0.88 | 0.74 | 0.9 | 0.79 | 0.93 | 0.83 |
| 5 | 176030090 | rs6878704 | 0.265 | -1.048 | 0.019 | -0.114 | -0.019 | -0.01 | -0.01 | 0 | 0.6 | 0.54 | 0.66 | 0.35 | 0.69 | 0.5 |
| 5 | 176040488 | rs7356786 | -0.533 | -0.949 | -0.305 | -0.05 | -0.007 | -0.006 | -0.006 | 0 | 0.45 | 0.45 | 0.35 | 0.71 | 0.34 | 0.53 |
| 5 | 176043333 | rs13184620 | 0.211 | -1.838 | -0.436 | 0.059 | -0.016 | 0.009 | 0.009 | 0.227 | 0.62 | 0.43 | 0.59 | 0.42 | 0.73 | 0.58 |
| 5 | 176045442 | rs11739413 | 0.096 | -1.915 | -0.37 | 0.089 | -0.007 | -0.023 | -0.023 | 0.353 | 0.29 | 0.36 | 0.22 | 0.2 | 0.24 | 0.29 |
| 5 | 176046567 | rs744307 | 0.8 | -1.763 | -1.021 | 0.211 | 0.024 | -0.065 | -0.065 | 0 | 0.11 | 0.04 | 0.19 | 0.45 | 0.07 | 0.14 |
| 5 | 176047761 | rs2075667 | 0.827 | -1.761 | -1.038 | 0.212 | 0.019 | -0.069 | -0.069 | 0.478 | 0.11 | 0.03 | 0.19 | 0.45 | 0.07 | 0.14 |
| 5 | 176049708 | rs11739753 | 1.056 | -1.471 | -1.537 | -0.614 | -0.168 | 0.214 | 0.214 | 0 | 0.32 | 0.05 | 0.43 | 0.02 | 0.46 | 0.23 |

| | | | | | | | | | | | | | | | | |
|---|---|---|---|---|---|---|---|---|---|---|---|---|---|---|---|---|
| 5 | 176050715 | rs13160179 | 0.21 | -1.512 | -1.385 | -0.443 | -0.013 | -0.087 | -0.087 | 0 | 0.22 | 0.06 | 0.1 | 0.33 | 0.23 | 0.34 |
| 5 | 176056020 | rs4242202 | 0.147 | -1.44 | -1.285 | -0.391 | 0.002 | -0.08 | -0.08 | 0.806 | 0.23 | 0.08 | 0.11 | 0.35 | 0.21 | 0.32 |
| 5 | 176060846 | rs1352303 | 0.637 | -0.955 | -0.831 | -0.902 | -0.044 | 0.041 | 0.041 | 0 | 0.33 | 0.18 | 0.43 | 0.02 | 0.45 | 0.22 |
| 5 | 176069103 | rs11134988 | 0.836 | -1.145 | -0.868 | -0.778 | -0.041 | 0.083 | 0.083 | 0 | 0.35 | 0.18 | 0.5 | 0.1 | 0.56 | 0.28 |
| 5 | 176076234 | rs4867841 | 0.081 | -1.964 | -0.293 | -0.387 | 0.019 | 0.043 | 0.043 | 0 | 0.58 | 0.66 | 0.61 | 0.24 | 0.66 | 0.47 |
| 5 | 176079574 | rs11739733 | 0.884 | -1.864 | 0.164 | -0.71 | -0.002 | -0.02 | -0.02 | 0 | 0.42 | 0.33 | 0.54 | 0.02 | 0.58 | 0.28 |
| 5 | 176080142 | rs3797465 | -0.059 | -2.361 | 0.024 | -0.389 | -0.037 | 0.07 | 0.07 | 0 | 0.71 | 0.42 | 0.69 | 0.47 | 0.84 | 0.6 |
| 5 | 176082381 | rs4867842 | 1.406 | -2.219 | -0.351 | -0.607 | -0.023 | 0.138 | 0.138 | 0 | 0.41 | 0.16 | 0.52 | 0.02 | 0.58 | 0.28 |
| 5 | 176084160 | rs4868678 | 0.276 | -2.335 | -0.3 | -0.282 | 0.045 | -0.089 | -0.089 | 0 | 0.66 | 0.83 | 0.84 | 0.53 | 0.74 | 0.71 |
| 5 | 176084894 | rs748742 | 0.308 | -2.337 | -0.277 | -0.271 | 0.044 | -0.09 | -0.09 | 0 | 0.66 | 0.82 | 0.83 | 0.54 | 0.74 | 0.71 |
| 5 | 176085168 | rs748743 | 0.28 | -2.273 | -0.273 | -0.267 | 0.042 | -0.09 | -0.09 | 0.078 | 0.66 | 0.82 | 0.84 | 0.54 | 0.74 | 0.71 |
| 5 | 176085391 | rs1063266 | 0.898 | -2.516 | 0.363 | -0.45 | -0.001 | -0.001 | -0.001 | 0 | 0.57 | 0.42 | 0.78 | 0.51 | 0.72 | 0.7 |
| 5 | 176085522 | rs2087210 | -0.496 | -2.853 | 0.539 | -0.403 | 0.011 | -0.012 | -0.012 | 0 | 0.35 | 0.31 | 0.18 | 0.49 | 0.26 | 0.29 |
| 5 | 176091411 | rs4868683 | 0.266 | -2.29 | 0.593 | -0.797 | -0.025 | -0.046 | -0.046 | 0 | 0.9 | 0.81 | 0.96 | 1 | 0.97 | 0.97 |
| 5 | 176091730 | rs4868684 | 1.286 | -2.669 | 0.184 | -1.267 | 0.015 | 0.153 | 0.153 | 0 | 0.44 | 0.21 | 0.55 | 0.01 | 0.62 | 0.31 |
| 5 | 176092285 | rs7730457 | 0.467 | -2.713 | 0.295 | -0.766 | -0.012 | 0.125 | 0.125 | 0 | 0.19 | 0.29 | 0.17 | 0.53 | 0.13 | 0.4 |
| 5 | 176096188 | rs6556280 | 0.544 | -2.819 | 0.187 | -0.838 | 0.014 | -0.015 | -0.015 | 0.081 | 0.64 | 0.63 | 0.73 | 0.54 | 0.75 | 0.7 |
| 5 | 176096236 | rs7733892 | 0.566 | -2.838 | 0.342 | -0.857 | -0.01 | 0.115 | 0.115 | 0 | 0.19 | 0.26 | 0.17 | 0.53 | 0.13 | 0.38 |
| 5 | 176096428 | rs4867848 | 0.188 | -2.582 | -0.33 | -0.632 | 0.012 | -0.097 | -0.097 | 0 | 0.25 | 0.09 | 0.22 | 0.45 | 0.21 | 0.27 |
| 5 | 176098533 | rs4868689 | 0.463 | -2.096 | -0.101 | -0.594 | -0.035 | -0.059 | -0.059 | 0 | 0.89 | 0.73 | 0.95 | 1 | 0.97 | 0.97 |
| 5 | 176101496 | rs12653525 | -0.433 | -2.183 | -0.742 | -0.292 | 0.003 | -0.006 | -0.006 | 0 | 0.24 | 0.21 | 0.17 | 0.47 | 0.14 | 0.31 |
| 5 | 176101823 | rs4867851 | 0.114 | -2.525 | -1.724 | -0.717 | 0.014 | -0.114 | -0.114 | 0 | 0.3 | 0.06 | 0.23 | 0.48 | 0.25 | 0.43 |
| 5 | 176105090 | rs1982803 | 0.869 | -2.062 | -1.533 | -1.179 | -0.083 | 0.273 | 0.273 | 0.015 | 0.38 | 0.06 | 0.53 | 0.02 | 0.6 | 0.26 |
| 5 | 176109874 | rs1487802 | -0.674 | -0.928 | -0.912 | -0.42 | 0.017 | 0.019 | 0.019 | 0 | 0.53 | 0.71 | 0.36 | 0.54 | 0.43 | 0.46 |
| 5 | 176110659 | rs17078413 | 0.708 | -1.336 | -0.852 | -0.121 | 0.01 | -0.009 | -0.009 | 0 | 0.13 | 0.11 | 0.08 | 0.29 | 0.03 | 0.2 |
| 5 | 176118380 | rs7737146 | -0.327 | -1.139 | -1.267 | -0.203 | -0.012 | 0.013 | 0.013 | 0 | 0.3 | 0.63 | 0.24 | 0.53 | 0.23 | 0.36 |
| 5 | 176123629 | rs6860786 | 0.142 | -0.058 | -1.383 | 0.173 | 0.001 | -0.037 | -0.037 | 0 | 0.23 | 0.18 | 0.12 | 0 | 0.18 | 0.09 |
| 5 | 176124970 | rs12186673 | -0.206 | 1.02 | -1.427 | 1.157 | 0.001 | 0.012 | 0.012 | 0 | 0.1 | 0.09 | 0.1 | 0.25 | 0.14 | 0.13 |
| 5 | 176127377 | rs4868696 | -0.073 | 1.574 | -1.631 | 0.871 | -0.105 | -0.089 | -0.089 | 0.156 | 0.22 | 0.07 | 0.52 | 0.41 | 0.39 | 0.26 |
| 5 | 176131444 | rs11134992 | 0.033 | 0.874 | -1.007 | 0.614 | 0.004 | 0.007 | 0.007 | 1.805 | 0.51 | 0.46 | 0.53 | 0.34 | 0.59 | 0.46 |
| 5 | 176132977 | rs7722011 | -0.075 | 0.372 | -0.79 | 1.484 | -0.001 | 0.077 | 0.077 | 0 | 0.14 | 0.14 | 0.12 | 0.42 | 0.13 | 0.14 |
| 5 | 176133254 | rs11741836 | 0.411 | 1.184 | -0.993 | 0.903 | -0.037 | 0.032 | 0.032 | 0 | 0.25 | 0.16 | 0.35 | 0.14 | 0.34 | 0.26 |
| 5 | 176133680 | rs9686813 | -0.978 | 0.384 | -0.667 | 0.655 | 0.016 | 0.05 | 0.05 | 0 | 0.48 | 0.33 | 0.51 | 0.66 | 0.52 | 0.5 |
| 5 | 176134572 | rs4868700 | -0.351 | -0.054 | -0.893 | 0.142 | -0.041 | -0.026 | -0.026 | 1.193 | 0.61 | 0.73 | 0.45 | 0.51 | 0.54 | 0.56 |
| 5 | 176135176 | rs17078434 | 1.473 | -0.251 | -0.558 | -0.455 | 0 | -0.007 | -0.007 | 0 | 0.25 | 0.26 | 0.25 | 0.15 | 0.25 | 0.2 |
| 5 | 176138039 | rs10476204 | 0.978 | -0.288 | -0.325 | -0.536 | 0.004 | -0.028 | -0.028 | 0.198 | 0.28 | 0.36 | 0.28 | 0.15 | 0.3 | 0.21 |
| 5 | 176140452 | rs2220368 | -1.663 | -0.331 | -0.562 | -0.51 | -0.005 | -0.008 | -0.008 | 0 | 0.86 | 0.81 | 0.86 | 0.92 | 0.88 | 0.84 |
| 5 | 176140801 | rs1386713 | -1.762 | -0.316 | -0.793 | -0.461 | 0.009 | 0.006 | 0.006 | 0 | 0.87 | 0.91 | 0.87 | 0.9 | 0.88 | 0.84 |
| 5 | 176142940 | rs4867853 | -0.807 | -0.393 | -0.538 | -0.519 | -0.007 | -0.034 | -0.034 | 0.016 | 0.64 | 0.58 | 0.62 | 0.77 | 0.64 | 0.63 |
| 5 | 176144471 | rs4867856 | 1.248 | -0.124 | -0.639 | -0.287 | -0.042 | -0.006 | -0.006 | 0 | 0.27 | 0.15 | 0.44 | 0.32 | 0.34 | 0.28 |
| 5 | 176144839 | rs11741326 | 0.258 | -0.154 | -0.681 | -0.282 | 0.008 | -0.041 | -0.041 | 0.865 | 0.23 | 0.15 | 0.17 | 0.44 | 0.19 | 0.23 |
| 5 | 176148768 | rs10035561 | 0.367 | -0.112 | -0.853 | -0.312 | 0.001 | 0.002 | 0.002 | 0.011 | 0.47 | 0.39 | 0.41 | 0.54 | 0.47 | 0.46 |
| 5 | 176153641 | rs7701834 | -0.137 | 0.331 | -1.477 | -0.036 | -0.044 | 0 | 0 | 0 | 0.61 | 0.77 | 0.5 | 0.57 | 0.54 | 0.56 |
| 5 | 176156228 | rs11134998 | 0.2 | 0.235 | -1.208 | 0.474 | 0.014 | -0.06 | -0.06 | 0 | 0.73 | 0.86 | 0.76 | 0.47 | 0.75 | 0.7 |
| 5 | 176158047 | rs4867858 | -0.612 | 0.607 | -0.928 | 0.272 | -0.001 | 0.005 | 0.005 | 0 | 0.86 | 0.9 | 0.86 | 0.89 | 0.86 | 0.83 |
| 5 | 176160097 | rs9313742 | -0.171 | 0.57 | -0.997 | 0.298 | -0.001 | -0.001 | -0.001 | 0 | 0.52 | 0.45 | 0.43 | 0.67 | 0.55 | 0.58 |
| 5 | 176160463 | rs11949733 | -0.134 | 0.446 | -1.104 | 0.393 | -0.017 | 0.007 | 0.007 | 0 | 0.23 | 0.17 | 0.3 | 0.23 | 0.3 | 0.21 |
| 5 | 176161032 | rs10062831 | -0.804 | -1.153 | -0.153 | 0.681 | -0.035 | 0.032 | 0.032 | 0 | 0.24 | 0.3 | 0.16 | 0.32 | 0.14 | 0.32 |
| 5 | 176162950 | rs13174602 | 0.662 | -0.441 | 0.114 | 0.845 | -0.008 | -0.004 | -0.004 | 0 | 0.38 | 0.34 | 0.53 | 0.51 | 0.45 | 0.44 |
| 5 | 176165086 | rs490332 | 0.261 | -0.495 | -0.189 | 0.653 | -0.001 | -0.002 | -0.002 | 0 | 0.47 | 0.48 | 0.58 | 0.67 | 0.5 | 0.57 |
| 5 | 176168507 | rs729823 | 0.756 | -1.219 | -0.723 | 0.599 | -0.005 | 0.017 | 0.017 | 0 | 0.2 | 0.06 | 0.17 | 0.15 | 0.21 | 0.19 |
| 5 | 176170762 | rs2306826 | 0.512 | -0.711 | -0.112 | 0.776 | -0.023 | -0.012 | -0.012 | 1.315 | 0.16 | 0.02 | 0.08 | 0.14 | 0.14 | 0.11 |
| 5 | 176176383 | rs2962783 | 0.378 | -1.539 | 0.898 | 1.184 | 0.011 | 0.027 | 0.027 | 0 | 0.47 | 0.35 | 0.55 | 0.4 | 0.54 | 0.43 |
| 5 | 176188633 | rs577557 | -2.615 | -1.497 | 2.516 | 1.451 | 0.002 | 0.294 | 0.294 | 4.994 | 0.07 | 0.55 | 0.06 | 0.25 | 0 | 0 |
| 5 | 176202730 | rs529322 | 2.519 | -1.535 | 2.653 | 1.105 | -0.017 | 0.206 | 0.206 | 5.43 | 0.91 | 0.46 | 0.94 | 0.83 | 1 | 1 |
| 5 | 176209414 | rs551357 | -2.867 | -1.592 | 2.829 | 0.896 | -0.046 | 0.056 | 0.056 | 1.631 | 0.13 | 0.57 | 0.1 | 0.16 | 0.06 | 0.06 |
| 5 | 176211401 | rs501666 | -2.423 | -1.583 | 3.214 | 0.904 | -0.022 | 0.046 | 0.046 | 0 | 0.09 | 0.27 | 0.07 | 0.11 | 0.05 | 0.06 |
| 5 | 176212290 | rs2940527 | 2.522 | -1.81 | 2.992 | 0.955 | -0.049 | 0.087 | 0.087 | 0.213 | 0.81 | 0.42 | 0.8 | 0.72 | 0.87 | 0.79 |
| 5 | 176212812 | rs691889 | -2.41 | -1.796 | 2.825 | 0.866 | -0.043 | 0.07 | 0.07 | 1.145 | 0.13 | 0.51 | 0.1 | 0.15 | 0.07 | 0.06 |
| 5 | 176221957 | rs511567 | 1.977 | -1.883 | 2.054 | 0.541 | 0.007 | 0.003 | 0.003 | 0.109 | 0.75 | 0.25 | 0.82 | 0.81 | 0.88 | 0.81 |
| 5 | 176235075 | rs692403 | 1.854 | -1.939 | 1.987 | 0.301 | 0.018 | 0.015 | 0.015 | 0.132 | 0.73 | 0.25 | 0.81 | 0.85 | 0.88 | 0.81 |
| 5 | 176242895 | rs691799 | 1.873 | -1.873 | 2.001 | 0.016 | 0.002 | 0.004 | 0.004 | 1.258 | 0.77 | 0.25 | 0.85 | 0.92 | 0.93 | 0.91 |
| 5 | 176255153 | rs2962830 | 0.944 | -2.032 | 2.425 | 0.025 | -0.012 | -0.031 | -0.031 | 0 | 0.94 | 0.77 | 0.96 | 1 | 0.98 | 0.99 |
| 5 | 176255646 | rs692843 | 2.12 | -2.102 | 1.829 | 0.062 | 0.07 | 0.055 | 0.055 | 1.66 | 0.76 | 0.17 | 0.86 | 0.91 | 0.94 | 0.91 |
| 5 | 176255657 | rs692825 | -1.357 | -1.726 | 1.903 | 0.269 | -0.061 | -0.001 | -0.001 | 0 | 0.12 | 0.29 | 0.07 | 0.09 | 0.03 | 0.07 |
| 5 | 176273370 | rs480782 | 1.529 | -1.642 | 1.522 | -0.134 | -0.04 | -0.098 | -0.098 | 0 | 0.84 | 0.4 | 0.77 | 1 | 0.93 | 0.9 |
| 5 | 176284713 | rs1000144 | 0.634 | -1.707 | 0.949 | 0.737 | 0.015 | -0.027 | -0.027 | 0.155 | 0.82 | 0.88 | 0.84 | 0.66 | 0.89 | 0.82 |
| 5 | 176285598 | rs11135004 | 1.412 | -2.123 | 0.914 | 0.395 | 0.047 | -0.014 | -0.014 | 0 | 0.64 | 0.3 | 0.61 | 0.63 | 0.81 | 0.71 |
| 5 | 176294895 | rs11954962 | -0.647 | -1.775 | 0.702 | 0.311 | -0.003 | -0.022 | -0.022 | 0 | 0.08 | 0.1 | 0.18 | 0 | 0.06 | 0.07 |

| | | | | | | | | | | | | | | | | |
|---|---|---|---|---|---|---|---|---|---|---|---|---|---|---|---|---|
| 5 | 176305678 | rs478253 | 0.985 | -1.121 | -0.421 | -0.091 | -0.136 | 0.01 | 0.01 | 0 | 0.27 | 0.02 | 0.22 | 0.18 | 0.4 | 0.19 |
| 5 | 176305953 | rs11953674 | -0.728 | -1.244 | 0.793 | 0.198 | -0.022 | -0.061 | -0.061 | 0 | 0.1 | 0.22 | 0.19 | 0.01 | 0.06 | 0.06 |
| 5 | 176307504 | rs3736432 | 0.664 | -1.13 | 0.5 | 0.45 | -0.029 | 0.002 | 0.002 | 0 | 0.68 | 0.47 | 0.62 | 0.68 | 0.84 | 0.74 |
| 5 | 176314434 | rs2278493 | -0.117 | 0.646 | -0.675 | 0.919 | -0.056 | -0.08 | -0.08 | 0.729 | 0.25 | 0.07 | 0.43 | 0.4 | 0.33 | 0.41 |
| 5 | 176314459 | rs2278492 | -0.243 | 0.505 | -0.075 | 0.876 | -0.005 | -0.008 | -0.008 | 0 | 0.26 | 0.18 | 0.45 | 0.39 | 0.33 | 0.4 |
| 5 | 176316201 | rs691057 | -0.892 | 0.077 | -0.346 | 0.172 | -0.001 | -0.011 | -0.011 | 0 | 0.58 | 0.73 | 0.64 | 0.58 | 0.55 | 0.63 |
| 5 | 176324980 | rs691541 | 0.342 | -0.439 | 0.318 | -0.27 | -0.055 | -0.072 | -0.072 | 0 | 0.16 | 0.28 | 0.08 | 0.01 | 0.03 | 0.11 |
| 5 | 176328336 | rs3762974 | 0.452 | -0.201 | 0.131 | 0.092 | -0.067 | -0.121 | -0.121 | 0.008 | 0.3 | 0.08 | 0.49 | 0.49 | 0.35 | 0.42 |
| 5 | 176330236 | rs9313744 | -0.704 | 0.468 | -0.3 | -0.314 | -0.021 | 0.026 | 0.026 | 0 | 0.91 | 0.97 | 0.92 | 0.99 | 0.84 | 0.92 |
| 5 | 176331832 | rs2962835 | 0.668 | 0.19 | -0.08 | -0.275 | -0.065 | -0.083 | -0.083 | 0 | 0.17 | 0.31 | 0.09 | 0.01 | 0.03 | 0.12 |
| 5 | 176334601 | rs1472920 | 1.562 | 0.12 | 0.186 | -0.201 | 0.009 | -0.002 | -0.002 | 1.251 | 0.31 | 0.28 | 0.24 | 0.28 | 0.28 | 0.28 |
| 5 | 176338955 | rs2962842 | 0.94 | -0.061 | 0.475 | -0.277 | -0.009 | -0.01 | -0.01 | 0 | 0.4 | 0.36 | 0.33 | 0.48 | 0.45 | 0.36 |
| 5 | 176343061 | rs2860534 | 1.527 | -0.326 | 0.425 | -0.315 | 0.009 | -0.002 | -0.002 | 0 | 0.31 | 0.28 | 0.24 | 0.28 | 0.28 | 0.28 |
| 5 | 176350336 | rs10052228 | 0.515 | -0.269 | 0.331 | -0.284 | -0.065 | -0.083 | -0.083 | 0 | 0.17 | 0.31 | 0.08 | 0.01 | 0.03 | 0.12 |
| 5 | 176366896 | rs10516137 | 0.032 | -0.283 | 0.307 | -0.309 | -0.052 | -0.082 | -0.082 | 0 | 0.2 | 0.31 | 0.09 | 0.01 | 0.05 | 0.14 |
| 5 | 176370402 | rs13360277 | -0.072 | -0.335 | 0.702 | -0.334 | -0.106 | -0.149 | -0.149 | 1.088 | 0.21 | 0.43 | 0.1 | 0.01 | 0.05 | 0.14 |
| 5 | 176378574 | rs365132 | -0.36 | -0.07 | 0.435 | -0.271 | -0.08 | -0.073 | -0.073 | 1.061 | 0.61 | 0.88 | 0.44 | 0.49 | 0.49 | 0.5 |
| 5 | 176382995 | rs3733876 | 1.211 | -0.01 | 0.662 | -0.264 | -0.017 | -0.017 | -0.017 | 0.368 | 0.07 | 0.02 | 0.09 | 0.19 | 0.16 | 0.08 |
| 5 | 176402401 | rs1700490 | -0.349 | -0.036 | 0.683 | -0.264 | -0.061 | -0.055 | -0.055 | 0.196 | 0.61 | 0.84 | 0.44 | 0.49 | 0.49 | 0.5 |
| 5 | 176445698 | rs353467 | -0.362 | 0.063 | 0.627 | -0.272 | -0.079 | -0.07 | -0.07 | 0.776 | 0.61 | 0.88 | 0.44 | 0.49 | 0.49 | 0.5 |
| 5 | 176470431 | rs6870999 | 1.593 | 0.541 | 0.19 | -0.603 | -0.026 | -0.149 | -0.149 | 0 | 0.18 | 0.01 | 0.38 | 0.51 | 0.17 | 0.35 |
| 5 | 176475355 | rs9313747 | 1.023 | 0.263 | 0.412 | -0.697 | -0.062 | -0.08 | -0.08 | 0.272 | 0.17 | 0.3 | 0.09 | 0.01 | 0.04 | 0.12 |
| 5 | 176492557 | rs251848 | 0.127 | 0.236 | 0.214 | -0.814 | -0.071 | -0.051 | -0.051 | 0 | 0.58 | 0.88 | 0.44 | 0.47 | 0.49 | 0.49 |
| 5 | 176493683 | rs6893089 | 0.271 | -0.078 | 0.401 | -0.82 | -0.106 | -0.115 | -0.115 | 0 | 0.19 | 0.42 | 0.11 | 0.02 | 0.04 | 0.12 |
| 5 | 176498882 | rs353482 | 2.042 | -0.043 | 0.599 | -0.844 | 0.012 | -0.002 | -0.002 | 0.429 | 0.31 | 0.23 | 0.24 | 0.27 | 0.28 | 0.28 |
| 5 | 176501583 | rs6867614 | -0.613 | -0.233 | 0.57 | -1.133 | -0.082 | -0.14 | -0.14 | 0 | 0.21 | 0.43 | 0.11 | 0.01 | 0.07 | 0.16 |
| 5 | 176511432 | rs451643 | -0.346 | 0.162 | -0.62 | -0.897 | -0.014 | -0.083 | -0.083 | 4.607 | 0.29 | 0.1 | 0.31 | 0.47 | 0.29 | 0.39 |
| 5 | 176516631 | rs1966265 | 0.769 | 0.455 | -0.946 | -1.177 | -0.046 | -0.046 | -0.046 | 0 | 0.11 | 0.02 | 0.32 | 0.51 | 0.18 | 0.21 |
| 5 | 176517797 | rs376618 | -2.139 | -0.149 | -1.101 | -1.954 | 0.002 | -0.089 | -0.089 | 2.238 | 0.76 | 0.63 | 0.83 | 1 | 0.74 | 0.9 |
| 5 | 176520243 | rs351855 | -0.005 | -0.373 | -0.937 | -1.612 | -0.011 | -0.077 | -0.077 | 1.278 | 0.29 | 0.11 | 0.31 | 0.46 | 0.29 | 0.39 |
| 5 | 176521456 | rs2011077 | 0.49 | -0.048 | -0.598 | -1.036 | -0.047 | -0.054 | -0.054 | 0.046 | 0.11 | 0.02 | 0.32 | 0.51 | 0.18 | 0.21 |
| 5 | 176527577 | rs6556301 | 0.077 | -0.501 | -0.197 | -1.141 | 0 | 0.001 | 0.001 | 0.068 | 0.28 | 0.21 | 0.4 | 0.52 | 0.34 | 0.36 |
| 5 | 176531772 | rs7708357 | -0.89 | -1.027 | -0.28 | -1.5 | 0 | 0.021 | 0.021 | 0.124 | 0.59 | 0.38 | 0.75 | 0.95 | 0.6 | 0.82 |
| 5 | 176541937 | rs351861 | -0.873 | -1.733 | -0.268 | -0.964 | 0.036 | -0.005 | -0.005 | 0.929 | 0.39 | 0.72 | 0.33 | 0.53 | 0.2 | 0.49 |
| 5 | 176569188 | rs9885210 | -0.852 | -1.712 | -0.184 | -1.033 | 0.046 | -0.012 | -0.012 | 0 | 0.38 | 0.77 | 0.33 | 0.53 | 0.2 | 0.37 |
| 5 | 176578072 | rs10039241 | -0.027 | -1.998 | 0.333 | -1.196 | -0.04 | 0.171 | 0.171 | 0.025 | 0.2 | 0.52 | 0.26 | 0.51 | 0.14 | 0.3 |
| 5 | 176589585 | rs244708 | -1.413 | -1.865 | -0.044 | -1.559 | -0.021 | -0.013 | -0.013 | 0.311 | 0.61 | 0.79 | 0.48 | 0.53 | 0.44 | 0.55 |
| 5 | 176598502 | rs10035180 | -1.022 | -1.928 | -0.046 | -1.562 | 0.046 | -0.012 | -0.012 | 0.081 | 0.38 | 0.77 | 0.33 | 0.53 | 0.2 | 0.37 |
| 5 | 176610012 | rs6872021 | -1.483 | -1.889 | -0.048 | -1.557 | -0.064 | 0.14 | 0.14 | 0 | 0.77 | 0.98 | 0.85 | 1 | 0.75 | 0.82 |
| 5 | 176636882 | rs1363405 | -1.052 | -1.915 | -0.033 | -1.547 | 0.048 | -0.012 | -0.012 | 0 | 0.38 | 0.77 | 0.33 | 0.53 | 0.2 | 0.37 |
| 5 | 176637240 | rs3733875 | 1.346 | -1.92 | 0.093 | -1.562 | 0.013 | -0.035 | -0.035 | 0 | 0.13 | 0.07 | 0.21 | 0.51 | 0.12 | 0.27 |
| 5 | 176637576 | rs28932178 | 0.813 | -1.93 | 0.146 | -1.566 | 0.008 | -0.042 | -0.042 | 0 | 0.16 | 0.11 | 0.22 | 0.51 | 0.14 | 0.3 |
| 5 | 176683794 | rs6890368 | 1.058 | -1.966 | 0.264 | -1.591 | -0.051 | -0.068 | -0.068 | 3.745 | 0.15 | 0.25 | 0.05 | 0 | 0.04 | 0.02 |
| 5 | 176689166 | rs7719125 | 1.46 | -2.01 | 0.296 | -1.632 | -0.038 | -0.024 | -0.024 | 0 | 0.38 | 0.17 | 0.51 | 0.47 | 0.55 | 0.45 |
| 5 | 176691279 | rs10037055 | 0.606 | -2.093 | 0.735 | -1.182 | -0.03 | 0.072 | 0.072 | 0.234 | 0.76 | 0.43 | 0.71 | 0.47 | 0.84 | 0.65 |
| 5 | 176700885 | rs12660023 | 1.283 | -2.012 | 0.723 | -1.142 | 0.07 | -0.019 | -0.019 | 0 | 0.62 | 0.22 | 0.66 | 0.47 | 0.8 | 0.63 |
| 5 | 176718361 | rs6886255 | 0.52 | -1.879 | 0.766 | -1.064 | -0.032 | 0.105 | 0.105 | 0.08 | 0.77 | 0.46 | 0.72 | 0.47 | 0.84 | 0.65 |
| 5 | 176719355 | rs11953271 | -0.438 | -1.834 | 0.84 | -1.057 | 0 | -0.011 | -0.011 | 0 | 0.07 | 0.07 | 0.03 | 0.02 | 0.04 | 0.08 |
| 5 | 176726635 | rs3088050 | 0.827 | -1.917 | 0.783 | -1.012 | -0.056 | -0.076 | -0.076 | 0 | 0.12 | 0.01 | 0.31 | 0.43 | 0.17 | 0.33 |
| 5 | 176728458 | rs1128287 | -0.755 | -1.636 | 1.384 | -0.486 | -0.013 | -0.031 | -0.031 | 0 | 0.86 | 0.9 | 0.75 | 0.47 | 0.8 | 0.73 |
| 5 | 176728817 | rs28710014 | -1.886 | -1.666 | 1.513 | -0.536 | 0.012 | -0.021 | -0.021 | 0 | 0.09 | 0.46 | 0.06 | 0.02 | 0.02 | 0.07 |
| 5 | 176731452 | rs28362590 | 0.015 | -1.516 | 1.163 | -0.444 | -0.012 | -0.044 | -0.044 | 2.349 | 0.82 | 0.87 | 0.73 | 0.45 | 0.79 | 0.67 |
| 5 | 176735187 | rs10051822 | 1.24 | -1.6 | 1.152 | -0.331 | 0.035 | -0.023 | -0.023 | 0 | 0.63 | 0.25 | 0.68 | 0.51 | 0.81 | 0.62 |
| 5 | 176737309 | rs7732626 | -0.533 | -1.597 | 1.281 | -0.632 | -0.08 | -0.136 | -0.136 | 0 | 0.21 | 0.53 | 0.09 | 0.02 | 0.07 | 0.08 |
| 5 | 176740244 | rs11949767 | 0.264 | -1.514 | 1.024 | -0.588 | 0.006 | 0.08 | 0.08 | 0 | 0.77 | 0.63 | 0.72 | 0.44 | 0.76 | 0.63 |
| 5 | 176740704 | rs11952102 | 0.251 | -1.512 | 1.017 | -0.679 | 0.007 | 0.072 | 0.072 | 0 | 0.77 | 0.63 | 0.72 | 0.43 | 0.76 | 0.63 |
| 5 | 176740996 | rs6885410 | 0.828 | -1.217 | 0.047 | -0.562 | 0.044 | 0 | 0 | 0 | 0.62 | 0.2 | 0.64 | 0.42 | 0.73 | 0.61 |
| 5 | 176755342 | rs6876677 | -0.85 | -0.929 | 0.09 | -1.127 | 0.055 | 0.162 | 0.162 | 0 | 0.48 | 0.73 | 0.32 | 0.08 | 0.35 | 0.29 |
| 5 | 176765989 | rs11949401 | -0.805 | -0.938 | 0.128 | -1.131 | 0.046 | 0.128 | 0.128 | 0 | 0.47 | 0.7 | 0.32 | 0.08 | 0.35 | 0.27 |
| 5 | 176773091 | rs9313758 | -0.835 | -0.916 | 0.025 | -1.116 | 0.054 | 0.16 | 0.16 | 0 | 0.47 | 0.72 | 0.32 | 0.08 | 0.35 | 0.29 |
| 5 | 176774403 | rs4532376 | -0.789 | -0.804 | -0.354 | -1.081 | -0.009 | -0.072 | -0.072 | 0.048 | 0.45 | 0.48 | 0.3 | 0.08 | 0.34 | 0.27 |
| 5 | 176779857 | rs10064229 | -0.772 | -1.238 | -0.312 | -0.829 | -0.076 | -0.073 | -0.073 | 0 | 0.24 | 0.55 | 0.17 | 0.08 | 0.09 | 0.12 |
| 5 | 176784512 | rs4075958 | -0.123 | -0.894 | -0.823 | -0.658 | -0.048 | 0.023 | 0.023 | 4.184 | 0.25 | 0.07 | 0.24 | 0.14 | 0.28 | 0.24 |
| 5 | 176788570 | rs4976646 | -1.089 | -1.513 | -0.205 | -0.697 | 0.013 | 0.022 | 0.022 | 0.147 | 0.47 | 0.62 | 0.34 | 0.24 | 0.35 | 0.39 |
| 5 | 176789719 | rs6556312 | -0.129 | -1.139 | -0.656 | -0.057 | 0.037 | 0.084 | 0.084 | 0.126 | 0.15 | 0.33 | 0.3 | 0.25 | 0.16 | 0.22 |
| 5 | 176794191 | rs12654812 | -0.57 | -1.311 | -0.836 | -0.124 | 0.01 | 0.024 | 0.024 | 0 | 0.41 | 0.32 | 0.33 | 0.23 | 0.35 | 0.39 |
| 5 | 176797343 | rs4074995 | 0.114 | -0.628 | -1.071 | -0.081 | -0.038 | 0.023 | 0.023 | 4.672 | 0.25 | 0.08 | 0.23 | 0.14 | 0.29 | 0.24 |
| 5 | 176798306 | rs11746443 | 0.082 | -0.529 | -1.151 | -0.056 | -0.046 | 0.014 | 0.014 | 4.357 | 0.25 | 0.05 | 0.23 | 0.14 | 0.28 | 0.24 |
| 5 | 176801131 | rs10866705 | -0.166 | -0.456 | -1.084 | 0.363 | -0.005 | -0.013 | -0.013 | 0 | 0.76 | 0.81 | 0.71 | 0.7 | 0.72 | 0.73 |

| | | | | | | | | | | | | | | | | |
|---|---|---|---|---|---|---|---|---|---|---|---|---|---|---|---|---|
| 5 | 176809618 | rs7708314 | -0.56 | -0.431 | -1.937 | -0.236 | -0.029 | -0.007 | -0.007 | 0 | 0.14 | 0.02 | 0.21 | 0.11 | 0.11 | 0.17 |
| 5 | 176814656 | rs33921462 | -0.039 | -0.35 | -1.793 | -0.469 | -0.089 | -0.038 | -0.038 | 0 | 0.26 | 0.03 | 0.23 | 0.19 | 0.29 | 0.23 |
| 5 | 176817143 | rs3812035 | -0.498 | -0.342 | -1.581 | -0.434 | -0.018 | 0.026 | 0.026 | 1.011 | 0.35 | 0.18 | 0.28 | 0.21 | 0.33 | 0.3 |
| 5 | 176830627 | rs17876032 | 0.296 | -0.995 | -1.708 | -0.264 | 0.096 | 0.105 | 0.105 | 2.692 | 0.51 | 0.06 | 0.46 | 0.21 | 0.68 | 0.33 |
| 5 | 176850425 | rs6556324 | -1.066 | -0.385 | 0.419 | 0.01 | 0 | -0.023 | -0.023 | 0 | 0.08 | 0.39 | 0.04 | 0 | 0.02 | 0.02 |
| 5 | 176857270 | rs2731665 | 0.456 | 0.029 | -0.089 | -0.047 | -0.003 | 0.074 | 0.074 | 0.265 | 0.48 | 0.26 | 0.35 | 0.25 | 0.47 | 0.27 |
| 5 | 176864269 | rs2545796 | 0.61 | -0.007 | -0.079 | 0.048 | -0.012 | 0.074 | 0.074 | 0.164 | 0.47 | 0.24 | 0.35 | 0.26 | 0.5 | 0.28 |
| 5 | 176867276 | rs335434 | 0.589 | -0.017 | -0.071 | 0.044 | -0.009 | 0.064 | 0.064 | 0.173 | 0.47 | 0.26 | 0.35 | 0.26 | 0.5 | 0.28 |
| 5 | 176872494 | rs2731662 | 0.029 | -0.021 | -0.006 | 0.038 | 0 | -0.013 | -0.013 | 0.202 | 0.54 | 0.51 | 0.38 | 0.26 | 0.53 | 0.3 |
| 5 | 176877110 | rs461264 | -0.178 | 0.008 | -0.054 | 0.053 | -0.012 | -0.03 | -0.03 | 0.149 | 0.06 | 0.23 | 0.03 | 0 | 0.02 | 0.02 |
| 5 | 176877624 | rs465670 | 0.644 | 0.01 | -0.062 | 0.052 | -0.011 | 0.063 | 0.063 | 0.239 | 0.47 | 0.24 | 0.34 | 0.26 | 0.5 | 0.28 |
| 5 | 176885499 | rs2544809 | 0.112 | -0.021 | -0.098 | 0.045 | -0.001 | -0.01 | -0.01 | 0 | 0.53 | 0.49 | 0.37 | 0.26 | 0.52 | 0.29 |
| 5 | 176890067 | rs163199 | 0.407 | 0.173 | -0.606 | 0.194 | -0.006 | 0.056 | 0.056 | 0.195 | 0.47 | 0.26 | 0.35 | 0.26 | 0.5 | 0.27 |
| 5 | 176896412 | rs2029164 | -0.075 | 0.324 | -1.094 | 0.41 | 0.004 | -0.018 | -0.018 | 0 | 0.08 | 0.02 | 0.12 | 0.2 | 0.06 | 0.27 |
| 5 | 176898619 | rs335420 | 0.356 | 0.386 | -0.982 | 0.376 | -0.018 | 0.056 | 0.056 | 2.991 | 0.47 | 0.16 | 0.34 | 0.33 | 0.5 | 0.28 |
| 5 | 176916516 | rs335462 | -0.017 | 0.61 | -0.691 | 0.527 | 0.001 | 0.014 | 0.014 | 0.008 | 0.53 | 0.39 | 0.38 | 0.39 | 0.53 | 0.5 |
| 5 | 176929130 | rs1422268 | 0.76 | 0.951 | -0.217 | 1.546 | 0.032 | 0.259 | 0.259 | 0 | 0.95 | 0.78 | 0.86 | 0.69 | 0.94 | 0.92 |
| 5 | 176929974 | rs2544814 | -0.015 | 0.093 | 0.227 | 1.172 | 0 | 0.026 | 0.026 | 0 | 0.48 | 0.4 | 0.53 | 0.35 | 0.49 | 0.4 |
| 5 | 176938343 | rs2630758 | 0.069 | -0.16 | 0.073 | 0.852 | 0.002 | 0.054 | 0.054 | 0.355 | 0.48 | 0.25 | 0.58 | 0.63 | 0.48 | 0.44 |
| 5 | 176940384 | rs335438 | 0.632 | -0.52 | 0.17 | 0.738 | 0.015 | -0.057 | -0.057 | 0.015 | 0.41 | 0.45 | 0.32 | 0.21 | 0.47 | 0.44 |
| 5 | 176948121 | rs3805550 | -0.166 | -0.56 | 0.315 | 1.096 | -0.037 | 0.055 | 0.055 | 0 | 0.11 | 0.25 | 0.1 | 0.17 | 0.05 | 0.12 |
| 5 | 176970435 | rs163197 | 0.42 | -0.404 | 0.343 | 0.932 | -0.055 | 0.111 | 0.111 | 0.358 | 0.87 | 0.52 | 0.88 | 0.83 | 0.95 | 0.88 |
| 5 | 176973151 | rs335454 | 0.091 | -0.404 | 0.405 | 0.932 | -0.042 | 0.114 | 0.114 | 0 | 0.88 | 0.6 | 0.89 | 0.83 | 0.95 | 0.88 |
| 5 | 176975693 | rs335423 | 0.435 | -0.406 | 0.498 | 0.93 | -0.055 | 0.107 | 0.107 | 0.358 | 0.87 | 0.48 | 0.88 | 0.83 | 0.95 | 0.88 |
| 5 | 176977611 | rs28425483 | 0.891 | -0.69 | -0.139 | 0.615 | 0.014 | -0.051 | -0.051 | 0 | 0.2 | 0.02 | 0.26 | 0.29 | 0.15 | 0.16 |
| 5 | 176984163 | rs335472 | -0.053 | -0.653 | 0.221 | 0.571 | -0.04 | 0.141 | 0.141 | 0 | 0.12 | 0.45 | 0.11 | 0.17 | 0.05 | 0.12 |
| 5 | 176985515 | rs1001530 | -0.031 | -0.652 | 0.226 | 0.571 | -0.044 | 0.109 | 0.109 | 0 | 0.12 | 0.37 | 0.11 | 0.17 | 0.04 | 0.12 |
| 5 | 176990151 | rs335482 | 0.507 | -0.062 | -0.2 | -0.079 | 0.006 | -0.005 | -0.005 | 0.012 | 0.27 | 0.29 | 0.24 | 0.24 | 0.37 | 0.35 |
| 5 | 177005613 | rs168080 | 0.368 | -0.931 | -0.189 | 0.265 | 0 | 0.005 | 0.005 | 0 | 0.49 | 0.59 | 0.49 | 0.45 | 0.52 | 0.47 |
| 5 | 177011863 | rs430797 | 0.368 | -0.932 | -0.19 | 0.265 | 0 | 0.005 | 0.005 | 0 | 0.49 | 0.59 | 0.49 | 0.45 | 0.51 | 0.47 |
| 5 | 177020032 | rs34582406 | 0.5 | -1.027 | -0.316 | 0.258 | -0.07 | 0.04 | 0.04 | 0 | 0.3 | 0.15 | 0.27 | 0.22 | 0.36 | 0.43 |
| 5 | 177025763 | rs28375402 | 0.929 | -1.084 | -0.155 | 0.389 | 0.001 | -0.001 | -0.001 | 0.267 | 0.4 | 0.48 | 0.41 | 0.42 | 0.44 | 0.38 |
| 5 | 177027405 | rs2306760 | 0.49 | -0.997 | -0.477 | 0.525 | 0.049 | -0.112 | -0.112 | 0 | 0.18 | 0.01 | 0.17 | 0.33 | 0.09 | 0.13 |
| 5 | 177041772 | rs13169820 | 0.51 | -1.141 | -0.392 | 0.406 | -0.101 | 0.029 | 0.029 | 0 | 0.29 | 0.09 | 0.26 | 0.22 | 0.35 | 0.43 |
| 5 | 177041793 | rs7720519 | 0.561 | -1.169 | -0.197 | 0.478 | 0.002 | -0.001 | -0.001 | 0.022 | 0.09 | 0.15 | 0.1 | 0.05 | 0.11 | 0.08 |
| 5 | 177043366 | rs4458584 | 0.918 | -1.257 | -0.148 | 0.603 | -0.005 | -0.005 | -0.005 | 11.91 | 0.41 | 0.47 | 0.39 | 0.4 | 0.41 | 0.38 |
| 5 | 177044707 | rs6600989 | -0.063 | -1.388 | -0.545 | 0.645 | -0.004 | 0.005 | 0.005 | 0 | 0.6 | 0.59 | 0.62 | 0.73 | 0.54 | 0.49 |
| 5 | 177047009 | rs7727897 | 0.233 | -1.195 | -0.895 | 0.752 | 0.04 | -0.047 | -0.047 | 0 | 0.18 | 0.07 | 0.18 | 0.33 | 0.09 | 0.1 |
| 5 | 177120697 | rs10060053 | -1.014 | -0.884 | -0.884 | 0.612 | 0 | 0 | 0 | 0 | 0.6 | 0.65 | 0.57 | 0.59 | 0.55 | 0.61 |
| 5 | 177135730 | rs17052274 | -0.331 | -1.386 | -0.737 | 1.216 | 0.017 | -0.081 | -0.081 | 0 | 0.17 | 0.06 | 0.21 | 0.35 | 0.12 | 0.1 |
| 5 | 177138298 | rs6601075 | -0.634 | -1.146 | -0.177 | 0.427 | -0.014 | -0.047 | -0.047 | 0 | 0.13 | 0.29 | 0.12 | 0.04 | 0.11 | 0.12 |
| 5 | 177260129 | rs4958955 | 0.195 | -0.989 | -0.873 | 0.936 | -0.049 | -0.043 | -0.043 | 0 | 0.4 | 0.06 | 0.37 | 0.57 | 0.39 | 0.53 |
| 5 | 177373622 | rs7718261 | 1.178 | -1.269 | 0.869 | 1.147 | -0.084 | 0.047 | 0.047 | 3.515 | 0.75 | 0.42 | 0.76 | 0.68 | 0.9 | 0.85 |
| 5 | 177377648 | rs4958945 | 1.178 | -1.267 | 0.867 | 1.149 | -0.084 | 0.047 | 0.047 | 3.515 | 0.75 | 0.43 | 0.76 | 0.68 | 0.9 | 0.85 |
| 5 | 177378845 | rs7705931 | 1.178 | -1.267 | 0.867 | 1.149 | -0.084 | 0.047 | 0.047 | 3.515 | 0.75 | 0.42 | 0.76 | 0.68 | 0.9 | 0.85 |
| 5 | 177386313 | rs11739986 | 0.755 | -0.985 | -0.054 | 1.216 | -0.031 | 0.013 | 0.013 | 3.426 | 0.79 | 0.75 | 0.83 | 0.69 | 0.93 | 0.92 |
| 5 | 177391446 | rs6864151 | -1.455 | 0.263 | 0.148 | 0.677 | 0.042 | -0.017 | -0.017 | 3.855 | 0.06 | 0.2 | 0.02 | 0 | 0.06 | 0.01 |
| 5 | 177400550 | rs7709539 | 0.724 | 0.12 | -0.297 | 0.991 | 0.002 | -0.009 | -0.009 | 0 | 0.85 | 0.85 | 0.9 | 0.78 | 0.86 | 0.92 |
| 5 | 177401509 | rs9329125 | 1.355 | -0.632 | -0.217 | 0.636 | 0.01 | 0.002 | 0.002 | 0.911 | 0.76 | 0.78 | 0.89 | 0.78 | 0.86 | 0.91 |
| 5 | 177409826 | rs4308500 | -0.794 | -0.764 | -0.307 | 0.801 | 0.005 | -0.015 | -0.015 | 2.085 | 0.1 | 0.08 | 0.07 | 0.21 | 0.09 | 0.07 |
| 5 | 177410416 | rs7708103 | 0.552 | -1.769 | -0.474 | 0.709 | -0.001 | -0.001 | -0.001 | 0 | 0.33 | 0.29 | 0.37 | 0.38 | 0.27 | 0.33 |
| 5 | 177413659 | rs4413548 | -0.522 | -1.54 | -0.253 | 0.501 | -0.014 | -0.003 | -0.003 | 0 | 0.56 | 0.52 | 0.57 | 0.44 | 0.65 | 0.53 |
| 5 | 177418754 | rs6883364 | 0.028 | -1.48 | -0.033 | 0.105 | -0.002 | -0.002 | -0.002 | 0 | 0.38 | 0.36 | 0.46 | 0.44 | 0.53 | 0.46 |
| 5 | 177419356 | rs4604209 | 0.063 | 0.035 | 0.073 | -0.102 | 0.002 | 0.005 | 0.005 | 0 | 0.2 | 0.2 | 0.16 | 0.34 | 0.26 | 0.16 |
| 5 | 177419967 | rs1800197 | 0.71 | 0.241 | 0.065 | 0.001 | -0.022 | 0.019 | 0.019 | 0 | 0.18 | 0.12 | 0.3 | 0.1 | 0.27 | 0.3 |
| 5 | 177422823 | rs4072924 | -0.721 | 0.034 | 0.579 | 0.014 | -0.001 | -0.001 | -0.001 | 0 | 0.39 | 0.58 | 0.5 | 0.44 | 0.53 | 0.46 |
| 5 | 177441189 | rs4389708 | -0.024 | -0.065 | 0.383 | 0.569 | -0.029 | -0.05 | -0.05 | 0 | 0.11 | 0.03 | 0.07 | 0.18 | 0.17 | 0.11 |
| 5 | 177483118 | rs11953798 | 1.081 | -0.816 | 0.676 | 0.404 | -0.002 | 0 | 0 | 0.445 | 0.53 | 0.52 | 0.51 | 0.53 | 0.46 | 0.49 |
| 5 | 177484841 | rs4246820 | 0.075 | -0.81 | 0.796 | 0.466 | -0.025 | -0.055 | -0.055 | 0 | 0.13 | 0.02 | 0.08 | 0.29 | 0.19 | 0.15 |
| 5 | 177485273 | rs7449056 | -1.505 | -2.256 | 0.35 | -0.281 | -0.011 | -0.031 | -0.031 | 0.158 | 0.57 | 0.74 | 0.53 | 0.47 | 0.58 | 0.6 |
| 5 | 177487613 | rs10038486 | -0.41 | -2.009 | 0.733 | -0.433 | 0.021 | -0.052 | -0.052 | 0 | 0.33 | 0.44 | 0.39 | 0.18 | 0.36 | 0.35 |
| 5 | 177492159 | rs10066995 | 0.239 | -2.109 | 0.896 | -0.551 | -0.042 | -0.055 | -0.055 | 0.002 | 0.19 | 0.36 | 0.09 | 0.06 | 0.12 | 0.14 |
| 5 | 177493867 | rs4246823 | 0.2 | -2.079 | 0.479 | -0.601 | 0.004 | -0.023 | -0.023 | 0 | 0.72 | 0.71 | 0.65 | 0.88 | 0.72 | 0.69 |
| 5 | 177494159 | rs7443165 | 0.463 | -2.002 | 0.043 | -0.612 | -0.027 | -0.04 | -0.04 | 0 | 0.67 | 0.37 | 0.82 | 0.93 | 0.83 | 0.81 |
| 5 | 177500500 | rs11745142 | -0.093 | -1.357 | -0.128 | -0.268 | 0.001 | -0.003 | -0.003 | 0.03 | 0.19 | 0.17 | 0.28 | 0.11 | 0.24 | 0.3 |
| 5 | 177505021 | rs10066407 | 0.356 | -1.145 | -0.154 | 0.124 | 0.008 | -0.008 | -0.008 | 3.627 | 0.74 | 0.77 | 0.83 | 0.71 | 0.75 | 0.8 |
| 5 | 177505652 | rs12515321 | 1.652 | -0.527 | -0.063 | 0.039 | -0.003 | -0.002 | -0.002 | 0 | 0.41 | 0.39 | 0.48 | 0.46 | 0.44 | 0.4 |
| 5 | 177507767 | rs4976806 | 0.138 | -0.02 | -0.166 | 0.046 | 0.005 | -0.016 | -0.016 | 0.012 | 0.71 | 0.64 | 0.66 | 0.82 | 0.69 | 0.61 |
| 5 | 177511743 | rs4380683 | -1.25 | -0.36 | -0.014 | 0.255 | -0.014 | 0.022 | 0.022 | 0 | 0.28 | 0.57 | 0.21 | 0.33 | 0.27 | 0.25 |

| | | | | | | | | | | | | | | | | |
|---|---|---|---|---|---|---|---|---|---|---|---|---|---|---|---|---|
| 5 | 177518612 | rs6881402 | 0.028 | -0.543 | -0.177 | -0.223 | 0.003 | -0.019 | -0.019 | 0 | 0.14 | 0.14 | 0.07 | 0 | 0.06 | 0.04 |
| 5 | 177535370 | rs6601211 | -0.081 | 0.754 | 0.123 | -0.177 | 0.036 | -0.014 | -0.014 | 0 | 0.71 | 0.62 | 0.67 | 0.79 | 0.66 | 0.59 |
| 5 | 177540223 | rs7721220 | 1.446 | 0.391 | 0.095 | -0.159 | 0.02 | 0.004 | 0.004 | 0 | 0.42 | 0.35 | 0.39 | 0.44 | 0.28 | 0.32 |
| 5 | 177546161 | rs10903275 | -0.059 | 0.917 | 0.123 | -0.19 | -0.061 | 0.05 | 0.05 | 1.242 | 0.32 | 0.17 | 0.41 | 0.13 | 0.45 | 0.43 |
| 5 | 177546281 | rs3935147 | 1.409 | 0.9 | 0.514 | -0.27 | 0.017 | 0.005 | 0.005 | 0 | 0.44 | 0.39 | 0.4 | 0.44 | 0.28 | 0.34 |
| 5 | 177549491 | rs3812080 | 0.945 | 0.975 | 0.554 | -0.206 | -0.005 | -0.009 | -0.009 | 0.903 | 0.16 | 0.12 | 0.15 | 0.27 | 0.23 | 0.21 |
| 5 | 177551169 | rs3812075 | -0.067 | 0.979 | 0.476 | -0.199 | 0.015 | 0.048 | 0.048 | 0.745 | 0.78 | 0.63 | 0.82 | 0.71 | 0.74 | 0.78 |
| 5 | 177556429 | rs13436876 | -0.011 | 0.865 | 0.52 | -0.199 | -0.002 | 0.034 | 0.034 | 0.717 | 0.21 | 0.3 | 0.17 | 0.29 | 0.24 | 0.22 |
| 5 | 177558929 | rs10036388 | -0.304 | 0.821 | 0.317 | -0.183 | -0.028 | 0.051 | 0.051 | 0 | 0.36 | 0.21 | 0.43 | 0.16 | 0.5 | 0.45 |
| 5 | 177562815 | rs7379278 | 0.096 | 0.818 | 0.334 | -0.1 | -0.003 | 0.059 | 0.059 | 1.368 | 0.77 | 0.63 | 0.82 | 0.65 | 0.75 | 0.76 |
| 5 | 177574421 | rs6601221 | 0.064 | 1.752 | -0.162 | 0.178 | -0.05 | -0.037 | -0.037 | 0 | 0.67 | 0.87 | 0.66 | 0.44 | 0.58 | 0.54 |
| 5 | 177577138 | rs7446073 | 0.347 | 0.956 | -0.177 | -0.198 | -0.019 | 0.055 | 0.055 | 0.485 | 0.62 | 0.76 | 0.56 | 0.86 | 0.48 | 0.55 |
| 5 | 177591495 | rs4976741 | 0.461 | -0.262 | -0.506 | 0.094 | -0.06 | 0.038 | 0.038 | 0 | 0.28 | 0.41 | 0.29 | 0.46 | 0.12 | 0.27 |
| 5 | 177604268 | rs17080239 | 0.577 | 0.826 | -0.955 | 0.081 | -0.034 | -0.001 | -0.001 | 0 | 0.2 | 0.08 | 0.24 | 0.13 | 0.23 | 0.26 |
| 5 | 177609226 | rs6601223 | 0.829 | 0.703 | 0.101 | 0.147 | 0.154 | 0.029 | 0.029 | 0.085 | 0.93 | 0.55 | 0.91 | 1 | 0.92 | 0.99 |
| 5 | 177609540 | rs9329133 | 0.376 | -0.13 | -0.021 | 0.202 | -0.013 | 0.011 | 0.011 | 0.362 | 0.3 | 0.34 | 0.3 | 0.5 | 0.15 | 0.29 |
| 5 | 177612293 | rs7719717 | 0.597 | 0.006 | -0.188 | 0.086 | 0.032 | -0.092 | -0.092 | 0.155 | 0.86 | 0.53 | 0.86 | 0.98 | 0.87 | 0.93 |
| 5 | 177612634 | rs7715618 | 0.523 | 0.043 | -0.16 | 0.104 | 0.042 | -0.087 | -0.087 | 0 | 0.87 | 0.53 | 0.85 | 0.98 | 0.87 | 0.93 |
| 5 | 177612983 | rs13169133 | 0.498 | 0.037 | -0.342 | 0.176 | 0.058 | -0.01 | -0.01 | 1.959 | 0.56 | 0.17 | 0.55 | 0.49 | 0.72 | 0.63 |
| 5 | 177614931 | rs9784730 | -0.284 | 0.702 | -1.085 | 0.319 | 0.016 | 0.032 | 0.032 | 0.063 | 0.22 | 0.32 | 0.23 | 0.32 | 0.28 | 0.21 |
| 5 | 177616112 | rs2913797 | 1.122 | 0.67 | -1.686 | -0.086 | -0.001 | 0.006 | 0.006 | 0 | 0.08 | 0.07 | 0.06 | 0.02 | 0.07 | 0.07 |
| 5 | 177616669 | rs2961669 | 0.377 | 0.579 | -1.946 | -0.089 | -0.073 | 0.101 | 0.101 | 0.434 | 0.43 | 0.13 | 0.41 | 0.19 | 0.52 | 0.5 |
| 5 | 177620189 | rs2961671 | 0.961 | 0.861 | -1.483 | 0.261 | -0.011 | 0.009 | 0.009 | 0 | 0.07 | 0.02 | 0.06 | 0.08 | 0.07 | 0.08 |
| 5 | 177623076 | rs2913815 | 0.413 | 0.347 | -2.07 | 0.22 | -0.029 | 0.129 | 0.129 | 0.383 | 0.47 | 0.17 | 0.3 | 0.22 | 0.45 | 0.46 |
| 5 | 177626134 | rs7733159 | -0.605 | -0.446 | -1.465 | 0.782 | 0.002 | -0.003 | -0.003 | 9.198 | 0.27 | 0.27 | 0.33 | 0.54 | 0.17 | 0.49 |
| 5 | 177627170 | rs2913816 | -0.564 | -0.147 | -0.885 | 0.297 | 0 | -0.008 | -0.008 | 0 | 0.35 | 0.39 | 0.5 | 0.7 | 0.32 | 0.42 |
| 5 | 177630040 | rs2913817 | 0.624 | 0.193 | -1.085 | 1.051 | -0.044 | -0.078 | -0.078 | 0 | 0.13 | 0.02 | 0.11 | 0.34 | 0.19 | 0.23 |
| 5 | 177632527 | rs2961674 | 0.697 | -0.026 | -0.948 | -0.293 | -0.006 | -0.033 | -0.033 | 0 | 0.25 | 0.26 | 0.13 | 0.07 | 0.17 | 0.22 |
| 5 | 177636199 | rs1059367 | 1.6 | 0.053 | -1.08 | -0.029 | -0.022 | -0.026 | -0.026 | 0 | 0.06 | 0.01 | 0.13 | 0.17 | 0.08 | 0.07 |
| 5 | 177636571 | rs2913752 | 1.026 | -0.029 | -1.012 | -0.154 | 0.01 | 0.052 | 0.052 | 0 | 0.23 | 0.11 | 0.11 | 0.07 | 0.17 | 0.22 |
| 5 | 177637447 | rs2913751 | -0.353 | 0.229 | -0.835 | -0.421 | -0.011 | -0.008 | -0.008 | 0 | 0.63 | 0.71 | 0.61 | 0.46 | 0.52 | 0.61 |
| 5 | 177637882 | rs17759 | 1.064 | 0.541 | -1.004 | -0.513 | -0.04 | -0.051 | -0.051 | 0 | 0.29 | 0.05 | 0.37 | 0.37 | 0.27 | 0.31 |
| 5 | 177640394 | rs17577590 | 0.199 | 0.721 | -0.507 | -0.375 | 0.001 | -0.009 | -0.009 | 0 | 0.38 | 0.43 | 0.48 | 0.39 | 0.34 | 0.37 |
| 5 | 177640813 | rs11747964 | 0.349 | 0.878 | -0.735 | -0.419 | 0.068 | -0.017 | -0.017 | 0 | 0.09 | 0.37 | 0.11 | 0.03 | 0.08 | 0.06 |
| 5 | 177647696 | rs2961680 | -0.592 | 0.938 | -1.316 | -0.332 | -0.003 | -0.005 | -0.005 | 0.053 | 0.77 | 0.78 | 0.73 | 0.53 | 0.74 | 0.71 |
| 5 | 177648488 | rs1366223 | -1.22 | 0.682 | -2.071 | -0.271 | -0.046 | -0.082 | -0.082 | 2.51 | 0.82 | 0.91 | 0.85 | 0.64 | 0.76 | 0.75 |
| 5 | 177649986 | rs3749813 | 1.518 | 0.599 | -2.207 | -0.394 | -0.067 | -0.066 | -0.066 | 0.908 | 0.14 | 0.02 | 0.14 | 0.27 | 0.22 | 0.19 |
| 5 | 177651040 | rs6887384 | 0.192 | 0.941 | -1.538 | -0.576 | 0.109 | -0.014 | -0.014 | 0 | 0.1 | 0.41 | 0.12 | 0.03 | 0.1 | 0.06 |
| 5 | 177651852 | rs13355310 | 1.674 | 1.013 | -2.158 | -0.391 | -0.001 | -0.058 | -0.058 | 0 | 0.14 | 0.2 | 0.13 | 0 | 0.2 | 0.09 |
| 5 | 177651980 | rs2961683 | -0.524 | 1.142 | -2.567 | 0.432 | -0.036 | -0.061 | -0.061 | 3.071 | 0.83 | 0.9 | 0.86 | 0.64 | 0.77 | 0.74 |
| 5 | 177652086 | rs2961684 | -0.916 | 0.302 | -1.967 | -0.191 | -0.003 | 0.065 | 0.065 | 0.007 | 0.34 | 0.17 | 0.32 | 0.18 | 0.29 | 0.34 |
| 5 | 177652884 | rs10062421 | -0.857 | 0.674 | -0.902 | 0.633 | 0 | -0.003 | -0.003 | 0 | 0.08 | 0.09 | 0.14 | 0.11 | 0.06 | 0.13 |
| 5 | 177661902 | rs7719352 | -1.06 | 0.717 | -0.855 | 0.219 | 0 | 0.001 | 0.001 | 0.586 | 0.21 | 0.7 | 0.17 | 0.09 | 0.17 | 0.1 |
| 5 | 177664297 | rs7715762 | -0.634 | 0.678 | 0.1 | 0.156 | -0.004 | -0.043 | -0.043 | 0.045 | 0.1 | 0.23 | 0.09 | 0.02 | 0.1 | 0.01 |
| 5 | 177665342 | rs1544926 | 1.386 | 0.866 | -1.07 | -0.187 | -0.029 | 0.21 | 0.21 | 0.001 | 0.47 | 0.1 | 0.42 | 0.18 | 0.44 | 0.49 |
| 5 | 177667225 | rs17052412 | -0.169 | 0.887 | -0.953 | -0.124 | 0.079 | 0.019 | 0.019 | 0.015 | 0.1 | 0.48 | 0.08 | 0.07 | 0.07 | 0.07 |
| 5 | 177667642 | rs11746584 | -0.364 | 0.863 | -0.975 | -0.051 | 0.047 | 0.012 | 0.012 | 0.07 | 0.11 | 0.51 | 0.08 | 0.08 | 0.07 | 0.09 |
| 5 | 177668965 | rs2241580 | -0.256 | 0.808 | -0.679 | 0.443 | 0.118 | 0.062 | 0.062 | 0.01 | 0.16 | 0.48 | 0.25 | 0.22 | 0.23 | 0.19 |
| 5 | 177669413 | rs2241579 | -0.72 | 0.62 | -0.918 | 1.096 | 0.002 | 0.001 | 0.001 | 0 | 0.37 | 0.57 | 0.36 | 0.62 | 0.41 | 0.42 |
| 5 | 177675217 | rs2913851 | 1.267 | 0.76 | -1.381 | 0.952 | -0.046 | 0.202 | 0.202 | 1.049 | 0.5 | 0.1 | 0.44 | 0.22 | 0.48 | 0.39 |
| 5 | 177675465 | rs6894255 | 1.723 | 0.375 | -1.59 | 0.543 | -0.015 | 0.131 | 0.131 | 0.224 | 0.35 | 0.14 | 0.29 | 0.14 | 0.35 | 0.29 |
| 5 | 177678526 | rs6422346 | -0.175 | 0.619 | -0.318 | 1.059 | 0.113 | 0.071 | 0.071 | 0 | 0.86 | 0.64 | 0.75 | 0.79 | 0.77 | 0.81 |
| 5 | 177679654 | rs2033200 | 1.498 | 0.594 | -1.031 | 0.94 | -0.073 | 0.17 | 0.17 | 0.041 | 0.47 | 0.05 | 0.42 | 0.21 | 0.45 | 0.47 |
| 5 | 177679728 | rs12652833 | -0.239 | 0.294 | -0.772 | 1.237 | 0.017 | -0.022 | -0.022 | 0 | 0.31 | 0.15 | 0.16 | 0.37 | 0.25 | 0.27 |
| 5 | 177682664 | rs752283 | 1.118 | -0.548 | -0.567 | 1.016 | 0.004 | 0.059 | 0.059 | 0 | 0.42 | 0.09 | 0.38 | 0.28 | 0.41 | 0.46 |
| 5 | 177683377 | rs890802 | 0.275 | -0.337 | -0.269 | 1.002 | 0.003 | -0.005 | -0.005 | 0 | 0.55 | 0.6 | 0.58 | 0.43 | 0.49 | 0.5 |
| 5 | 177686791 | rs2973748 | -1.343 | -0.604 | 0.079 | 1.488 | -0.011 | 0.04 | 0.04 | 0.178 | 0.11 | 0.22 | 0.16 | 0.22 | 0.08 | 0.04 |
| 5 | 177690539 | rs2278230 | -1.289 | -1.28 | 0.421 | 1.15 | -0.035 | 0.011 | 0.011 | 0 | 0.34 | 0.41 | 0.29 | 0.56 | 0.24 | 0.38 |
| 5 | 177690978 | rs7713983 | -1.698 | -1.08 | 0.843 | 1.003 | 0.032 | 0.019 | 0.019 | 0.089 | 0.42 | 0.67 | 0.41 | 0.63 | 0.32 | 0.44 |
| 5 | 177692247 | rs2913844 | 1.347 | -1.453 | 1.468 | 1.318 | -0.012 | 0.016 | 0.016 | 0 | 0.72 | 0.63 | 0.6 | 0.32 | 0.75 | 0.55 |
| 5 | 177701071 | rs2973758 | 2.173 | -1.528 | 1.259 | 1.125 | 0.06 | 0.128 | 0.128 | 0.41 | 0.58 | 0.23 | 0.53 | 0.2 | 0.69 | 0.49 |
| 5 | 177704027 | rs2973762 | 1.218 | -1.422 | 1.307 | 1.335 | 0.016 | 0.047 | 0.047 | 0 | 0.89 | 0.82 | 0.8 | 0.74 | 0.88 | 0.84 |
| 5 | 177704937 | rs752572 | 1.906 | -1.482 | 1.426 | 1.31 | 0.003 | 0 | 0 | 0 | 0.68 | 0.73 | 0.57 | 0.32 | 0.7 | 0.52 |
| 5 | 177705831 | rs2913838 | 1.876 | -1.665 | 2.005 | 1.287 | -0.005 | 0.074 | 0.074 | 0.023 | 0.77 | 0.64 | 0.74 | 0.63 | 0.82 | 0.79 |
| 5 | 177707204 | rs2033204 | -0.952 | -1.665 | 2.239 | 1.311 | -0.001 | 0.009 | 0.009 | 0 | 0.21 | 0.18 | 0.25 | 0.43 | 0.18 | 0.33 |
| 5 | 177707274 | rs11744096 | -0.375 | -1.766 | 2.235 | 1.334 | -0.021 | 0.029 | 0.029 | 0 | 0.11 | 0.19 | 0.06 | 0.12 | 0.06 | 0.05 |
| 5 | 177707427 | rs2973765 | 0.772 | -1.707 | 1.913 | 1.363 | 0.003 | 0.129 | 0.129 | 0 | 0.94 | 0.84 | 0.89 | 0.77 | 0.94 | 0.86 |
| 5 | 177711520 | rs10464068 | -0.128 | -1.696 | 1.606 | 1.335 | 0.03 | -0.07 | -0.07 | 0.185 | 0.15 | 0.05 | 0.19 | 0.42 | 0.14 | 0.3 |
| 5 | 177713191 | rs2913836 | 2.477 | -1.642 | 1.637 | 1.064 | 0.068 | 0.235 | 0.235 | 0.263 | 0.53 | 0.17 | 0.49 | 0.17 | 0.64 | 0.44 |

| | | | | | | | | | | | | | | | | |
|---|---|---|---|---|---|---|---|---|---|---|---|---|---|---|---|---|
| 5 | 177713480 | rs2913835 | 2.076 | -1.597 | 1.606 | 1.077 | 0.036 | 0.041 | 0.041 | 0.017 | 0.59 | 0.34 | 0.59 | 0.25 | 0.7 | 0.45 |
| 5 | 177715291 | rs890801 | 2.029 | -1.572 | 1.601 | 1.091 | 0.016 | 0.018 | 0.018 | 0 | 0.6 | 0.42 | 0.59 | 0.25 | 0.7 | 0.46 |
| 5 | 177715452 | rs6892747 | -0.88 | -1.534 | 1.62 | 1.244 | 0.015 | -0.034 | -0.034 | 0 | 0.21 | 0.13 | 0.24 | 0.44 | 0.18 | 0.36 |
| 5 | 177717640 | rs2913826 | 0.59 | -1.951 | 1.565 | 0.842 | -0.053 | 0.011 | 0.011 | 0 | 0.84 | 0.65 | 0.85 | 0.85 | 0.92 | 0.83 |
| 5 | 177717723 | rs2973775 | 1.654 | -1.961 | 1.481 | 0.851 | -0.045 | 0 | 0 | 0 | 0.7 | 0.47 | 0.78 | 0.72 | 0.84 | 0.74 |
| 5 | 177718389 | rs890807 | 1.758 | -1.816 | 1.463 | 0.846 | -0.015 | 0.004 | 0.004 | 0 | 0.68 | 0.51 | 0.7 | 0.66 | 0.78 | 0.71 |
| 5 | 177720154 | rs7380466 | 1.439 | -1.674 | 0.856 | 0.891 | -0.007 | -0.004 | -0.004 | 1.928 | 0.76 | 0.71 | 0.75 | 0.76 | 0.83 | 0.74 |
| 5 | 177727491 | rs7704172 | 1.788 | -1.839 | 0.778 | 0.413 | 0.009 | 0.006 | 0.006 | 0 | 0.6 | 0.68 | 0.56 | 0.32 | 0.72 | 0.43 |
| 5 | 177731143 | rs2913761 | 1.132 | -1.777 | 0.641 | 0.528 | 0.009 | -0.008 | -0.008 | 2.145 | 0.72 | 0.78 | 0.68 | 0.37 | 0.79 | 0.54 |
| 5 | 177731221 | rs2913762 | 1.185 | -1.783 | 0.614 | 0.524 | 0.01 | -0.009 | -0.009 | 0 | 0.72 | 0.79 | 0.68 | 0.37 | 0.79 | 0.52 |
| 5 | 177735777 | rs2913766 | -0.18 | -1.667 | 0.396 | 0.613 | 0.01 | -0.031 | -0.031 | 0 | 0.87 | 0.94 | 0.7 | 0.42 | 0.87 | 0.69 |
| 5 | 177743408 | rs6883861 | -0.579 | -1.864 | 0.595 | 0.419 | -0.028 | -0.07 | -0.07 | 3.788 | 0.13 | 0.23 | 0.06 | 0 | 0.06 | 0.1 |
| 5 | 177744380 | rs2162765 | -0.346 | -1.527 | 0.078 | 0.654 | 0.005 | -0.04 | -0.04 | 0 | 0.15 | 0.07 | 0.31 | 0.58 | 0.15 | 0.32 |
| 5 | 177744496 | rs2973701 | 0.498 | -1.478 | 0.148 | 0.486 | 0.004 | -0.005 | -0.005 | 0 | 0.86 | 0.88 | 0.94 | 0.85 | 0.89 | 0.89 |
| 5 | 177746863 | rs11745722 | -1.697 | -1.673 | 0.131 | 0.133 | -0.003 | -0.017 | -0.017 | 0.162 | 0.52 | 0.52 | 0.49 | 0.79 | 0.37 | 0.66 |
| 5 | 177747691 | rs17081072 | -0.256 | -1.567 | 0.033 | 0.735 | 0.012 | -0.039 | -0.039 | 0.28 | 0.13 | 0.05 | 0.27 | 0.55 | 0.12 | 0.34 |
| 5 | 177747796 | rs17081075 | -0.322 | -0.496 | 0.149 | -0.346 | 0.064 | -0.037 | -0.037 | 0.285 | 0.25 | 0.46 | 0.33 | 0.15 | 0.32 | 0.23 |
| 5 | 177748169 | rs2973695 | -0.628 | -0.712 | -0.005 | -0.288 | 0.063 | -0.048 | -0.048 | 0 | 0.29 | 0.52 | 0.35 | 0.2 | 0.34 | 0.24 |
| 5 | 177749643 | rs2913784 | -0.636 | -0.538 | -0.007 | -0.394 | 0.048 | -0.061 | -0.061 | 0.69 | 0.29 | 0.52 | 0.35 | 0.15 | 0.32 | 0.24 |
| 5 | 177751488 | rs890813 | -1.398 | -0.79 | 0.04 | -0.51 | -0.003 | 0.006 | 0.006 | 0 | 0.43 | 0.63 | 0.41 | 0.21 | 0.37 | 0.35 |
| 5 | 177751751 | rs749425 | -0.536 | -1.004 | -0.286 | -0.135 | 0.003 | 0.004 | 0.004 | 0 | 0.13 | 0.11 | 0.1 | 0.07 | 0.08 | 0.12 |
| 5 | 177754580 | rs2973689 | 1.158 | -1.293 | -0.267 | 0.287 | 0.007 | -0.003 | -0.003 | 0 | 0.61 | 0.63 | 0.67 | 0.45 | 0.77 | 0.6 |
| 5 | 177755179 | rs11958296 | -0.948 | -0.897 | -0.139 | -0.176 | -0.004 | -0.024 | -0.024 | 0 | 0.13 | 0.11 | 0.05 | 0 | 0.07 | 0.12 |
| 5 | 177766183 | rs4976761 | -0.331 | -0.506 | -0.128 | 0.194 | -0.002 | 0.018 | 0.018 | 0.51 | 0.07 | 0.07 | 0.13 | 0.23 | 0.11 | 0.21 |
| 5 | 177766580 | rs2973808 | -1.553 | -0.176 | -0.051 | -0.644 | -0.004 | 0.116 | 0.116 | 2.31 | 0.47 | 0.73 | 0.64 | 0.8 | 0.49 | 0.66 |
| 5 | 177767548 | rs17081090 | -0.476 | 0.157 | 0.614 | -0.507 | 0.003 | -0.027 | -0.027 | 0 | 0.18 | 0.13 | 0.08 | 0 | 0.12 | 0.17 |
| 5 | 177772890 | rs6601232 | -1.023 | 0.275 | 0.764 | -0.549 | -0.011 | -0.066 | -0.066 | 3.192 | 0.26 | 0.34 | 0.11 | 0.1 | 0.2 | 0.25 |
| 5 | 177775092 | rs7730805 | -0.384 | 0.38 | 0.514 | -0.181 | 0.035 | 0.073 | 0.073 | 0.175 | 0.06 | 0.21 | 0.02 | 0.1 | 0.05 | 0.06 |
| 5 | 177776148 | rs751181 | -1.429 | 0.338 | -0.136 | -0.24 | 0.032 | 0.129 | 0.129 | 4.392 | 0.5 | 0.71 | 0.66 | 0.84 | 0.53 | 0.67 |
| 5 | 177779566 | rs7715128 | 1.025 | 0.673 | -0.259 | 0.399 | 0.008 | -0.027 | -0.027 | 0 | 0.58 | 0.53 | 0.44 | 0.26 | 0.54 | 0.42 |
| 5 | 177779994 | rs2913810 | 0.597 | -0.145 | -0.593 | 0.673 | 0.002 | 0.001 | 0.001 | 0.052 | 0.73 | 0.78 | 0.83 | 0.76 | 0.79 | 0.64 |
| 5 | 177780183 | rs7727323 | -0.433 | -0.159 | -0.664 | 0.588 | -0.01 | 0.017 | 0.017 | 0.452 | 0.05 | 0.12 | 0.02 | 0.11 | 0.02 | 0.06 |
| 5 | 177787515 | rs7729589 | -0.356 | -0.318 | -1.346 | 0.091 | 0.023 | 0.07 | 0.07 | 0.089 | 0.3 | 0.11 | 0.17 | 0.13 | 0.25 | 0.27 |
| 5 | 177788462 | rs10440797 | 0.518 | 0.332 | -1.635 | 0.179 | -0.095 | 0.053 | 0.053 | 0 | 0.27 | 0.12 | 0.25 | 0.15 | 0.45 | 0.25 |
| 5 | 177790952 | rs10464069 | -0.694 | -0.037 | -0.926 | 1.214 | 0.003 | 0.099 | 0.099 | 0 | 0.09 | 0.09 | 0.35 | 0.51 | 0.11 | 0.27 |
| 5 | 177793761 | rs2910111 | -0.881 | -0.548 | -0.083 | 1.343 | -0.038 | 0.214 | 0.214 | 0 | 0.13 | 0.32 | 0.17 | 0.43 | 0.08 | 0.31 |
| 5 | 177796121 | rs2973713 | -1.206 | -1.001 | -0.03 | 0.857 | 0.039 | 0.099 | 0.099 | 0.034 | 0.45 | 0.72 | 0.59 | 0.82 | 0.32 | 0.58 |
| 5 | 177798295 | rs2910119 | -0.956 | -1.229 | 0.463 | 0.887 | -0.031 | 0.068 | 0.068 | 0 | 0.23 | 0.55 | 0.37 | 0.27 | 0.17 | 0.24 |
| 5 | 177798812 | rs2973717 | -1.538 | -1.216 | 0.362 | 0.856 | 0.04 | 0.084 | 0.084 | 0.021 | 0.45 | 0.73 | 0.6 | 0.78 | 0.32 | 0.59 |
| 5 | 177802161 | rs2973720 | -1.214 | -1.14 | 0.394 | 0.465 | -0.015 | 0.002 | 0.002 | 0 | 0.41 | 0.53 | 0.47 | 0.38 | 0.29 | 0.41 |
| 5 | 177802373 | rs12519834 | 0.403 | -0.714 | 0.012 | 0.22 | -0.008 | 0.086 | 0.086 | 0 | 0.18 | 0.03 | 0.11 | 0.08 | 0.13 | 0.16 |
| 5 | 177811754 | rs13178731 | -1.414 | -0.509 | 0.872 | 0.497 | 0.026 | -0.03 | -0.03 | 0 | 0.07 | 0.25 | 0.06 | 0.01 | 0.07 | 0.04 |
| 5 | 177815313 | rs2973737 | -0.489 | -0.291 | -0.157 | 0.441 | -0.009 | 0.006 | 0.006 | 0 | 0.54 | 0.7 | 0.54 | 0.42 | 0.45 | 0.55 |
| 5 | 177820134 | rs2973739 | -0.472 | -0.399 | -0.606 | 0.428 | -0.065 | 0.046 | 0.046 | 0 | 0.74 | 0.87 | 0.76 | 0.87 | 0.56 | 0.81 |
| 5 | 177826469 | rs7722612 | 0.228 | -0.588 | -0.311 | 0.515 | 0.011 | -0.006 | -0.006 | 0 | 0.4 | 0.63 | 0.47 | 0.41 | 0.3 | 0.48 |
| 5 | 177830605 | rs4976774 | 0.384 | -0.602 | -0.84 | 0.7 | 0.005 | -0.063 | -0.063 | 0 | 0.37 | 0.11 | 0.39 | 0.4 | 0.3 | 0.48 |
| 5 | 177830769 | rs2973742 | -0.21 | -1.257 | -0.203 | 0.901 | -0.023 | 0.009 | 0.009 | 0 | 0.37 | 0.38 | 0.32 | 0.47 | 0.25 | 0.35 |
| 5 | 177840382 | rs10070303 | -1.218 | -1.683 | 0.208 | 1.309 | -0.072 | 0.119 | 0.119 | 0.005 | 0.24 | 0.46 | 0.24 | 0.37 | 0.11 | 0.25 |
| 5 | 177841886 | rs1110326 | -0.81 | -1.731 | 0.514 | 1.444 | -0.03 | 0.134 | 0.134 | 0 | 0.12 | 0.22 | 0.19 | 0.4 | 0.05 | 0.13 |
| 5 | 177842788 | rs7716953 | -0.518 | -1.8 | -0.386 | 0.933 | 0.095 | 0.109 | 0.109 | 0 | 0.5 | 0.79 | 0.54 | 0.7 | 0.35 | 0.52 |
| 5 | 177843645 | rs11960640 | 0.235 | -1.749 | 0.33 | 0.599 | -0.002 | -0.004 | -0.004 | 0 | 0.38 | 0.36 | 0.32 | 0.34 | 0.32 | 0.39 |
| 5 | 177845760 | rs4349734 | -1.048 | -1.145 | -0.088 | 0.819 | -0.077 | -0.062 | -0.062 | 0 | 0.69 | 0.84 | 0.39 | 0.51 | 0.52 | 0.61 |
| 5 | 177848109 | rs11956401 | -1.046 | -1.602 | 0.926 | 0.516 | -0.037 | -0.075 | -0.075 | 0 | 0.26 | 0.57 | 0.15 | 0.07 | 0.15 | 0.15 |
| 5 | 177848429 | rs17081205 | -0.797 | -1.635 | 1.198 | 0.521 | 0.01 | -0.005 | -0.005 | 0 | 0.13 | 0.46 | 0.11 | 0.07 | 0.08 | 0.05 |
| 5 | 177851623 | rs4361533 | -0.397 | -1.89 | 1.182 | 0.688 | -0.003 | -0.03 | -0.03 | 7.224 | 0.15 | 0.15 | 0.05 | 0 | 0.07 | 0.1 |
| 5 | 177851690 | rs17648414 | 0.936 | -2.483 | 0.668 | 0.293 | -0.015 | 0.163 | 0.163 | 0 | 0.37 | 0.1 | 0.22 | 0.12 | 0.36 | 0.31 |
| 5 | 177854235 | rs10063441 | 0.31 | -2.529 | 1.1 | 0.368 | 0.067 | -0.02 | -0.02 | 0 | 0.07 | 0.31 | 0.09 | 0.01 | 0.07 | 0.02 |
| 5 | 177856605 | rs10037451 | -1.704 | -2.506 | 0.808 | 0.401 | -0.197 | -0.055 | -0.055 | 0.316 | 0.69 | 0.98 | 0.5 | 0.68 | 0.52 | 0.68 |
| 5 | 177857534 | rs7728076 | -0.393 | -2.474 | 0.995 | 0.484 | -0.055 | 0.193 | 0.193 | 0.178 | 0.1 | 0.27 | 0.13 | 0.55 | 0.01 | 0.25 |
| 5 | 177863091 | rs6422348 | -1.598 | -2.493 | 0.655 | 0.238 | 0.082 | -0.047 | -0.047 | 0 | 0.32 | 0.86 | 0.21 | 0.38 | 0.16 | 0.34 |
| 5 | 177866785 | rs4383738 | 0.818 | -2.485 | 0.648 | 0.247 | 0 | 0.003 | 0.003 | 0 | 0.38 | 0.12 | 0.29 | 0.3 | 0.37 | 0.34 |
| 5 | 177867175 | rs4618447 | -0.938 | -2.203 | 1.196 | 0.423 | -0.061 | 0.242 | 0.242 | 0.271 | 0.11 | 0.35 | 0.07 | 0.37 | 0.01 | 0.2 |
| 5 | 177875446 | rs11956256 | -0.939 | -1.736 | 0.513 | -0.661 | -0.018 | -0.112 | -0.112 | 0 | 0.23 | 0.35 | 0.14 | 0.01 | 0.16 | 0.14 |
| 5 | 177876309 | rs4976725 | -0.116 | -1.186 | 0.192 | -0.361 | 0.001 | -0.013 | -0.013 | 0.864 | 0.94 | 0.91 | 0.95 | 1 | 0.91 | 0.98 |
| 5 | 177880278 | rs12652427 | -0.321 | -2.001 | 0.001 | -0.409 | 0.059 | 0.014 | 0.014 | 0.865 | 0.43 | 0.8 | 0.33 | 0.6 | 0.33 | 0.48 |
| 5 | 177881763 | rs6868991 | -0.251 | -2.164 | 0.853 | -0.323 | -0.002 | 0.464 | 0.464 | 0 | 0.07 | 0.34 | 0.1 | 0.51 | 0.01 | 0.23 |
| 5 | 177882547 | rs2647688 | 0.35 | -2.146 | 1.071 | -0.417 | -0.014 | -0.086 | -0.086 | 0.435 | 0.36 | 0.45 | 0.22 | 0.09 | 0.32 | 0.25 |
| 5 | 177883242 | rs2546629 | 0.29 | -2.146 | 1.104 | -0.417 | -0.018 | -0.103 | -0.103 | 0 | 0.36 | 0.54 | 0.23 | 0.09 | 0.32 | 0.25 |
| 5 | 177888966 | rs1846371 | 1.1 | -2.072 | 0.448 | -0.759 | -0.032 | -0.016 | -0.016 | 0 | 0.33 | 0.22 | 0.57 | 0.4 | 0.49 | 0.3 |

| | | | | | | | | | | | | | | | | |
|---|---|---|---|---|---|---|---|---|---|---|---|---|---|---|---|---|
| 5 | 177891489 | rs10039491 | 0.927 | -2.233 | 0.46 | -0.886 | -0.005 | 0.098 | 0.098 | 0 | 0.3 | 0.14 | 0.23 | 0.09 | 0.32 | 0.25 |
| 5 | 177892956 | rs10069282 | 0.898 | -2.465 | 0.257 | -1.057 | -0.003 | 0.023 | 0.023 | 0 | 0.25 | 0.2 | 0.22 | 0.09 | 0.29 | 0.23 |
| 5 | 177893248 | rs7706759 | 0.455 | -2.734 | 0.622 | -0.999 | -0.081 | 0.065 | 0.065 | 0 | 0.7 | 0.46 | 0.81 | 0.48 | 0.88 | 0.57 |
| 5 | 177895383 | rs17081346 | -0.223 | -2.759 | 0.539 | -1.08 | -0.033 | 0.244 | 0.244 | 0 | 0.07 | 0.19 | 0.09 | 0.52 | 0.01 | 0.26 |
| 5 | 177897992 | rs2546636 | 0.21 | -2.78 | 0.689 | -1.124 | -0.034 | 0.032 | 0.032 | 0 | 0.73 | 0.65 | 0.84 | 0.48 | 0.9 | 0.61 |
| 5 | 177899270 | rs13159176 | 0.668 | -2.773 | 0.467 | -1.209 | 0.051 | 0.04 | 0.04 | 0 | 0.49 | 0.27 | 0.61 | 0.39 | 0.61 | 0.4 |
| 5 | 177902122 | rs13187323 | -0.709 | -2.925 | 0.354 | -1.444 | -0.018 | 0.005 | 0.005 | 0 | 0.31 | 0.38 | 0.23 | 0.64 | 0.21 | 0.38 |
| 5 | 177905384 | rs603852 | 1.861 | -2.549 | -0.097 | -1.413 | -0.032 | -0.001 | -0.001 | 0.654 | 0.29 | 0.21 | 0.52 | 0.31 | 0.46 | 0.31 |
| 5 | 177906089 | rs13165074 | 2.265 | -2.474 | -0.563 | -1.372 | -0.14 | -0.047 | -0.047 | 0 | 0.26 | 0.05 | 0.49 | 0.31 | 0.46 | 0.31 |
| 5 | 177912033 | rs1106072 | 2.143 | -2.406 | -0.467 | -1.335 | -0.113 | -0.028 | -0.028 | 0.084 | 0.27 | 0.09 | 0.5 | 0.3 | 0.46 | 0.29 |
| 5 | 177913591 | rs684612 | -0.769 | -2.688 | -0.046 | -1.196 | -0.035 | 0.045 | 0.045 | 0 | 0.31 | 0.5 | 0.24 | 0.57 | 0.21 | 0.35 |
| 5 | 177916502 | rs1996223 | 2.1 | -2.438 | -0.821 | -1.119 | -0.111 | -0.027 | -0.027 | 0 | 0.27 | 0.09 | 0.51 | 0.3 | 0.5 | 0.29 |
| 5 | 177918571 | rs11249547 | 0.368 | -2.404 | -0.767 | -1.433 | -0.002 | 0.056 | 0.056 | 0 | 0.22 | 0.11 | 0.2 | 0.06 | 0.26 | 0.22 |
| 5 | 177932674 | rs2672818 | 0.13 | -2.158 | -1.089 | -1.611 | 0.033 | 0.139 | 0.139 | 0 | 0.73 | 0.96 | 0.92 | 0.93 | 0.83 | 0.79 |
| 5 | 177935694 | rs536664 | 1.375 | -1.985 | -0.59 | -1.898 | 0.001 | -0.017 | -0.017 | 0 | 0.45 | 0.47 | 0.69 | 0.87 | 0.63 | 0.64 |
| 5 | 177936898 | rs585219 | 1.085 | -1.897 | -0.393 | -1.872 | 0.043 | 0.065 | 0.065 | 0.376 | 0.51 | 0.43 | 0.82 | 0.87 | 0.73 | 0.67 |
| 5 | 177939207 | rs2546646 | -0.024 | -1.667 | -0.819 | -1.744 | 0.005 | -0.011 | -0.011 | 0.241 | 0.73 | 0.8 | 0.9 | 0.93 | 0.84 | 0.8 |
| 5 | 177942798 | rs6868242 | 2.038 | -1.01 | -1.285 | -1.112 | -0.119 | -0.028 | -0.028 | 0.61 | 0.28 | 0.08 | 0.49 | 0.28 | 0.45 | 0.3 |
| 5 | 177948971 | rs577450 | -0.703 | -1.849 | -0.772 | -0.947 | 0.027 | 0.018 | 0.018 | 4.465 | 0.45 | 0.6 | 0.33 | 0.64 | 0.27 | 0.44 |
| 5 | 177949429 | rs2546644 | -0.36 | -1.854 | -0.927 | -0.668 | 0.008 | 0.049 | 0.049 | 0 | 0.85 | 0.95 | 0.95 | 0.94 | 0.88 | 0.84 |
| 5 | 177954516 | rs2672826 | -0.494 | -1.832 | -0.867 | -0.645 | 0.003 | 0.038 | 0.038 | 0 | 0.87 | 0.93 | 0.95 | 0.97 | 0.89 | 0.88 |
| 5 | 177956908 | rs7713018 | -0.417 | -2.651 | -0.41 | -0.528 | -0.035 | 0.046 | 0.046 | 0.411 | 0.24 | 0.34 | 0.22 | 0.57 | 0.13 | 0.31 |
| 5 | 177960131 | rs517338 | 1.472 | -2.49 | -0.396 | -0.898 | 0 | 0 | 0 | 0 | 0.42 | 0.4 | 0.55 | 0.37 | 0.58 | 0.37 |
| 5 | 177960432 | rs4976788 | 1.848 | -2.429 | -0.174 | -1.042 | -0.039 | 0 | 0 | 0.225 | 0.36 | 0.24 | 0.49 | 0.37 | 0.51 | 0.36 |
| 5 | 177960872 | rs489777 | 0.049 | -2.564 | -0.091 | -1.15 | 0.002 | -0.057 | -0.057 | 0 | 0.13 | 0.22 | 0.08 | 0.04 | 0.09 | 0.11 |
| 5 | 177961860 | rs553780 | 1.304 | -2.442 | 0.297 | -1.074 | 0.067 | -0.011 | -0.011 | 2.058 | 0.05 | 0.19 | 0.06 | 0 | 0.07 | 0.02 |
| 5 | 177963524 | rs515242 | 1.587 | -3.212 | 0.323 | -0.75 | 0 | -0.002 | -0.002 | 0 | 0.41 | 0.4 | 0.55 | 0.37 | 0.58 | 0.37 |
| 5 | 177964020 | rs694350 | 0.401 | -3.189 | 0.513 | -0.748 | 0.048 | -0.025 | -0.025 | 0 | 0.14 | 0.26 | 0.19 | 0.07 | 0.19 | 0.16 |
| 5 | 177964444 | rs2546643 | -0.44 | -3.844 | 0.261 | -0.703 | 0.018 | 0.009 | 0.009 | 0 | 0.85 | 0.88 | 0.95 | 0.9 | 0.94 | 0.83 |
| 5 | 177964789 | rs6888802 | -0.215 | -3.94 | 0.317 | -0.776 | -0.007 | -0.033 | -0.033 | 0 | 0.12 | 0.15 | 0.15 | 0 | 0.1 | 0.1 |
| 5 | 177965633 | rs564195 | 0.14 | -3.806 | 0.444 | -0.716 | 0.022 | 0.002 | 0.002 | 0 | 0.19 | 0.29 | 0.21 | 0.17 | 0.26 | 0.21 |
| 5 | 177965696 | rs688074 | -0.884 | -3.87 | 0.354 | -0.726 | -0.006 | -0.029 | -0.029 | 0 | 0.41 | 0.53 | 0.39 | 0.17 | 0.38 | 0.31 |
| 5 | 177966921 | rs596279 | -0.866 | -3.879 | 0.323 | -0.725 | -0.014 | -0.041 | -0.041 | 9.18 | 0.4 | 0.51 | 0.38 | 0.17 | 0.34 | 0.31 |
| 5 | 177968251 | rs4976733 | -2.114 | -3.736 | 0.06 | -0.649 | -0.042 | -0.005 | -0.005 | 0.119 | 0.66 | 0.87 | 0.59 | 0.7 | 0.44 | 0.69 |
| 5 | 177968768 | rs472238 | -1.82 | -4.026 | 0.304 | -0.826 | 0.032 | 0.11 | 0.11 | 0 | 0.58 | 0.83 | 0.36 | 0.19 | 0.34 | 0.51 |
| 5 | 177968915 | rs4976790 | -0.695 | -4.492 | 0.503 | -0.915 | -0.033 | -0.05 | -0.05 | 0 | 0.35 | 0.38 | 0.11 | 0.05 | 0.1 | 0.24 |
| 5 | 177969010 | rs642624 | 0.017 | -4.324 | 0.51 | -0.838 | 0.002 | -0.018 | -0.018 | 0 | 0.21 | 0.33 | 0.22 | 0.15 | 0.23 | 0.25 |
| 5 | 177971352 | rs535858 | -0.843 | -4.345 | 0.559 | -0.881 | -0.04 | -0.063 | -0.063 | 0.568 | 0.35 | 0.35 | 0.17 | 0.06 | 0.13 | 0.24 |
| 5 | 177974127 | rs504596 | -0.144 | -4.044 | 0.228 | -0.67 | -0.011 | -0.027 | -0.027 | 0 | 0.8 | 0.63 | 0.87 | 0.87 | 0.82 | 0.74 |
| 5 | 177977679 | rs564487 | 1.007 | -3.956 | -0.28 | -0.595 | -0.03 | -0.058 | -0.058 | 0 | 0.6 | 0.49 | 0.74 | 0.84 | 0.73 | 0.59 |
| 5 | 177980792 | rs1545751 | -0.488 | -3.235 | -0.849 | 0.174 | -0.004 | 0.003 | 0.003 | 0 | 0.18 | 0.2 | 0.24 | 0.41 | 0.05 | 0.32 |
| 5 | 177982013 | rs262012 | -1.336 | -2.544 | -1.306 | -0.032 | -0.043 | 0.005 | 0.005 | 0.833 | 0.65 | 0.91 | 0.59 | 0.72 | 0.38 | 0.66 |
| 5 | 177990886 | rs262037 | -0.075 | -3.067 | -1.012 | -0.163 | -0.089 | -0.098 | -0.098 | 0 | 0.32 | 0.53 | 0.17 | 0.05 | 0.07 | 0.18 |
| 5 | 177991258 | rs185493 | -1.04 | -2.139 | -1.851 | 0.275 | -0.046 | -0.021 | -0.021 | 0 | 0.64 | 0.97 | 0.59 | 0.72 | 0.37 | 0.65 |
| 5 | 177995909 | rs625820 | -0.299 | -1.454 | -0.54 | 0.534 | -0.003 | 0.011 | 0.011 | 0 | 0.22 | 0.24 | 0.27 | 0.29 | 0.27 | 0.23 |
| 5 | 177999447 | rs262065 | -1.2 | -1.69 | 0.175 | 0.258 | -0.083 | -0.047 | -0.047 | 1.519 | 0.2 | 0.5 | 0.26 | 0.12 | 0.06 | 0.18 |
| 5 | 178000014 | rs7716274 | -0.658 | -1.397 | 0.277 | 0.298 | -0.036 | -0.012 | -0.012 | 0.046 | 0.08 | 0.24 | 0.04 | 0.02 | 0.01 | 0.06 |
| 5 | 178010145 | rs555417 | 0.573 | -0.939 | -0.627 | 0.008 | 0.004 | 0.04 | 0.04 | 0 | 0.36 | 0.16 | 0.31 | 0.26 | 0.39 | 0.2 |
| 5 | 178011820 | rs6600941 | -0.476 | -2.068 | -0.404 | 0.347 | -0.035 | 0.001 | 0.001 | 0 | 0.35 | 0.6 | 0.25 | 0.41 | 0.12 | 0.4 |
| 5 | 178014123 | rs7732660 | 1.037 | -1.446 | -0.506 | 0.139 | -0.008 | 0.044 | 0.044 | 0.114 | 0.42 | 0.28 | 0.37 | 0.26 | 0.48 | 0.21 |
| 5 | 178015835 | rs262028 | -0.538 | -1.906 | -0.396 | 0.017 | 0.006 | 0.002 | 0.002 | 0.776 | 0.31 | 0.75 | 0.27 | 0.34 | 0.1 | 0.3 |
| 5 | 178016218 | rs262029 | -0.227 | -1.951 | -0.121 | 0.014 | -0.065 | -0.043 | -0.043 | 0 | 0.7 | 0.82 | 0.61 | 0.64 | 0.59 | 0.55 |
| 5 | 178017695 | rs1967419 | -0.242 | -2.297 | 0.512 | -0.063 | -0.087 | 0.036 | 0.036 | 0.549 | 0.23 | 0.46 | 0.23 | 0.35 | 0.08 | 0.26 |
| 5 | 178019031 | rs721143 | -0.195 | -2.285 | 0.508 | -0.082 | -0.086 | 0.03 | 0.03 | 0.433 | 0.23 | 0.45 | 0.23 | 0.34 | 0.08 | 0.27 |
| 5 | 178034999 | rs775700 | 0.015 | -1.681 | 0.169 | -0.149 | -0.006 | 0.016 | 0.016 | 0.599 | 0.89 | 0.94 | 0.94 | 1 | 0.87 | 0.91 |
| 5 | 178038291 | rs10051137 | 0.472 | -1.729 | 0.409 | -0.098 | 0.026 | -0.021 | -0.021 | 3.29 | 0.18 | 0.12 | 0.18 | 0.34 | 0.08 | 0.28 |
| 5 | 178054930 | rs13360296 | 0.303 | -1.74 | 0.518 | -0.096 | -0.032 | 0.015 | 0.015 | 0 | 0.19 | 0.24 | 0.19 | 0.35 | 0.09 | 0.24 |
| 5 | 178062743 | rs3925516 | 0.189 | -1.557 | 0.235 | -0.08 | 0.002 | -0.017 | -0.017 | 0 | 0.91 | 0.84 | 0.93 | 0.98 | 0.9 | 0.86 |
| 5 | 178068976 | rs12719855 | 0.96 | -1.69 | 0.435 | -0.112 | -0.037 | -0.009 | -0.009 | 0 | 0.66 | 0.41 | 0.69 | 0.64 | 0.81 | 0.63 |
| 5 | 178072229 | rs3925518 | -1.104 | -1.405 | 0.133 | -0.09 | 0.001 | 0.026 | 0.026 | 0 | 0.55 | 0.65 | 0.51 | 0.68 | 0.46 | 0.7 |
| 5 | 178072635 | rs4413550 | 0.117 | -1.439 | 0.17 | -0.09 | -0.104 | 0.038 | 0.038 | 0.008 | 0.22 | 0.47 | 0.22 | 0.34 | 0.09 | 0.23 |
| 5 | 178073319 | rs12189095 | -0.058 | -1.336 | -0.073 | -0.099 | -0.021 | -0.026 | -0.026 | 0 | 0.33 | 0.18 | 0.29 | 0.34 | 0.37 | 0.47 |
| 5 | 178083799 | rs1023916 | 1.088 | -1.143 | -0.482 | -0.051 | -0.082 | -0.11 | -0.11 | 0 | 0.21 | 0.04 | 0.25 | 0.34 | 0.31 | 0.35 |
| 5 | 178084166 | rs7723662 | 0.123 | -0.846 | -0.009 | 0.271 | -0.072 | 0.023 | 0.023 | 0 | 0.21 | 0.36 | 0.13 | 0.33 | 0.08 | 0.25 |
| 5 | 178085216 | rs12523265 | 0.967 | -0.199 | -0.292 | 0.218 | -0.048 | 0.005 | 0.005 | 0.587 | 0.34 | 0.16 | 0.49 | 0.31 | 0.45 | 0.17 |
| 5 | 178089232 | rs6897878 | -0.586 | -0.09 | -0.042 | 0.279 | -0.002 | -0.003 | -0.003 | 2.061 | 0.38 | 0.43 | 0.25 | 0.29 | 0.3 | 0.48 |
| 5 | 178090633 | rs6884557 | -1.214 | 0.185 | 0.487 | 0.787 | -0.013 | 0.038 | 0.038 | 0 | 0.24 | 0.51 | 0.28 | 0.39 | 0.21 | 0.3 |
| 5 | 178094484 | rs10516147 | -0.536 | 0.589 | -0.004 | 1.247 | -0.041 | -0.09 | -0.09 | 0 | 0.19 | 0.05 | 0.25 | 0.38 | 0.25 | 0.36 |
| 5 | 178096908 | rs11249561 | -0.239 | 0.55 | 0.717 | 1.228 | 0.027 | 0.109 | 0.109 | 0 | 0.08 | 0.21 | 0.09 | 0.21 | 0.07 | 0.18 |

| | | | | | | | | | | | | | | | | |
|---|---|---|---|---|---|---|---|---|---|---|---|---|---|---|---|---|
| 5 | 178097049 | rs4419608 | -0.585 | 0.536 | 0.542 | 1.264 | -0.015 | -0.047 | -0.047 | 0 | 0.2 | 0.09 | 0.25 | 0.37 | 0.25 | 0.35 |
| 5 | 178100927 | rs10071738 | 0.361 | 0.836 | -0.202 | 0.254 | -0.007 | -0.039 | -0.039 | 0 | 0.28 | 0.21 | 0.45 | 0.39 | 0.3 | 0.27 |
| 5 | 178103405 | rs10076182 | -0.778 | 1.151 | 0.667 | 0.406 | 0.003 | 0.013 | 0.013 | 0 | 0.25 | 0.32 | 0.26 | 0.38 | 0.33 | 0.37 |
| 5 | 178117620 | rs7719405 | 0.477 | 1.507 | 0.621 | 0.522 | 0.03 | 0.075 | 0.075 | 0 | 0.74 | 0.63 | 0.69 | 0.39 | 0.68 | 0.47 |
| 5 | 178120334 | rs7715476 | 0.719 | 1.441 | -0.486 | -0.041 | 0.014 | 0.001 | 0.001 | 1.906 | 0.5 | 0.27 | 0.55 | 0.41 | 0.46 | 0.35 |
| 5 | 178121599 | rs6866344 | 0.148 | 1.216 | -0.577 | 0.17 | -0.052 | -0.062 | -0.062 | 0 | 0.2 | 0.07 | 0.22 | 0.38 | 0.28 | 0.37 |
| 5 | 178125921 | rs13173933 | -0.164 | 0.817 | -0.35 | -0.052 | 0.003 | 0.005 | 0.005 | 0.644 | 0.67 | 0.37 | 0.76 | 0.75 | 0.73 | 0.73 |
| 5 | 178125974 | rs6859031 | -0.164 | 0.817 | -0.371 | -0.052 | 0.004 | 0.008 | 0.008 | 0.644 | 0.67 | 0.36 | 0.75 | 0.75 | 0.73 | 0.73 |
| 5 | 178136005 | rs1079487 | 0.096 | 0.885 | -0.858 | 0.027 | -0.031 | -0.026 | -0.026 | 0 | 0.24 | 0.11 | 0.25 | 0.34 | 0.28 | 0.42 |
| 5 | 178136440 | rs11740283 | 0.085 | 0.884 | -0.859 | 0.026 | -0.031 | -0.026 | -0.026 | 0.026 | 0.24 | 0.11 | 0.25 | 0.34 | 0.28 | 0.42 |
| 5 | 178138449 | rs6898275 | 0.702 | 0.924 | -0.788 | -0.04 | -0.022 | -0.05 | -0.05 | 0 | 0.42 | 0.19 | 0.5 | 0.41 | 0.45 | 0.31 |
| 5 | 178162596 | rs6873580 | 0.13 | 0.323 | -0.076 | -0.183 | -0.009 | -0.006 | -0.006 | 0 | 0.31 | 0.61 | 0.26 | 0.27 | 0.27 | 0.32 |
| 5 | 178165521 | rs10043919 | -0.146 | 0.181 | -0.424 | -0.494 | 0.013 | 0.044 | 0.044 | 0.11 | 0.55 | 0.64 | 0.36 | 0.3 | 0.43 | 0.41 |
| 5 | 178166903 | rs10055721 | 0.241 | 0.344 | -0.228 | -0.264 | -0.007 | -0.004 | -0.004 | 0 | 0.3 | 0.62 | 0.26 | 0.28 | 0.27 | 0.32 |
| 5 | 178172559 | rs10066888 | 0.75 | 0.776 | 0.501 | 0.454 | 0.039 | 0.012 | 0.012 | 2.229 | 0.51 | 0.34 | 0.4 | 0.44 | 0.37 | 0.44 |
| 5 | 178179729 | rs10062847 | 0.024 | 1.297 | 0.884 | 0.565 | 0.032 | 0.102 | 0.102 | 0 | 0.75 | 0.61 | 0.65 | 0.57 | 0.71 | 0.76 |
| 5 | 178180299 | rs4280887 | -0.364 | 1.408 | 0.761 | 0.47 | 0.011 | -0.001 | -0.001 | 0 | 0.23 | 0.26 | 0.3 | 0.26 | 0.29 | 0.35 |
| 5 | 178184042 | rs13157698 | 0.168 | 1.479 | 0.648 | 0.239 | -0.006 | -0.011 | -0.011 | 0.031 | 0.19 | 0.15 | 0.22 | 0.34 | 0.21 | 0.35 |
| 5 | 178187961 | rs7710807 | 0.165 | 1.336 | 0.875 | -0.75 | 0.002 | -0.037 | -0.037 | 0 | 0.58 | 0.57 | 0.34 | 0.23 | 0.51 | 0.41 |
| 5 | 178188257 | rs7711283 | 0.371 | 1.199 | 0.531 | -0.719 | -0.005 | -0.031 | -0.031 | 0 | 0.54 | 0.57 | 0.3 | 0.22 | 0.42 | 0.38 |
| 5 | 178193081 | rs4410655 | 0.165 | 1.173 | -0.01 | -0.758 | 0.002 | -0.077 | -0.077 | 0.641 | 0.54 | 0.49 | 0.36 | 0.07 | 0.55 | 0.23 |
| 5 | 178195673 | rs11949276 | -1.227 | 1.532 | 0.149 | 0.289 | 0.033 | -0.002 | -0.002 | 2.714 | 0.27 | 0.47 | 0.32 | 0.3 | 0.34 | 0.32 |
| 5 | 178196024 | rs11949338 | -1.192 | 1.482 | -0.036 | 0.287 | 0.021 | -0.003 | -0.003 | 0 | 0.27 | 0.42 | 0.31 | 0.3 | 0.33 | 0.31 |
| 5 | 178203585 | rs9717043 | -0.929 | 1.682 | -0.998 | 0.128 | 0 | 0.001 | 0.001 | 0 | 0.06 | 0.06 | 0.11 | 0.07 | 0.09 | 0.06 |
| 5 | 178204459 | rs4074670 | -0.683 | 1.876 | -1.168 | 0.154 | -0.063 | 0.014 | 0.014 | 0 | 0.18 | 0.04 | 0.17 | 0.16 | 0.24 | 0.25 |
| 5 | 178207609 | rs12655129 | -1.079 | 1.741 | -0.821 | 0.136 | 0.001 | -0.001 | -0.001 | 0.065 | 0.13 | 0.2 | 0.14 | 0.09 | 0.16 | 0.12 |
| 5 | 178208644 | rs12516527 | -0.825 | 2.084 | -1.2 | -0.642 | -0.035 | 0.083 | 0.083 | 0 | 0.31 | 0.21 | 0.33 | 0.05 | 0.41 | 0.29 |
| 5 | 178213713 | rs10455076 | -0.509 | 1.565 | -0.649 | 0.054 | -0.012 | 0.07 | 0.07 | 0 | 0.15 | 0.27 | 0.11 | 0.35 | 0.1 | 0.25 |
| 5 | 178214780 | rs7444303 | 1.047 | 1.146 | -1.204 | -0.548 | 0.005 | 0.04 | 0.04 | 0 | 0.22 | 0.11 | 0.15 | 0.08 | 0.24 | 0.12 |
| 5 | 178217756 | rs11955306 | 0.199 | 1.366 | -0.915 | -0.909 | 0.002 | 0.015 | 0.015 | 0 | 0.25 | 0.16 | 0.16 | 0.01 | 0.23 | 0.25 |
| 5 | 178220184 | rs7733056 | -0.109 | 1.436 | -0.609 | -0.402 | 0.001 | -0.008 | -0.008 | 0 | 0.28 | 0.29 | 0.25 | 0.15 | 0.29 | 0.28 |
| 5 | 178224547 | rs6877160 | 0.702 | 1.288 | -0.699 | -0.112 | 0.008 | 0.037 | 0.037 | 0 | 0.31 | 0.16 | 0.2 | 0.27 | 0.31 | 0.33 |
| 5 | 178239210 | rs6865763 | -0.38 | 1.117 | -0.648 | -0.52 | 0.007 | 0.037 | 0.037 | 0 | 0.52 | 0.64 | 0.43 | 0.33 | 0.49 | 0.39 |
| 5 | 178239484 | rs7706887 | 0.378 | 1.021 | -0.076 | -0.705 | -0.002 | -0.016 | -0.016 | 0 | 0.23 | 0.28 | 0.19 | 0.15 | 0.23 | 0.15 |
| 5 | 178241495 | rs10794696 | 0.649 | 0.909 | -0.023 | -0.747 | -0.003 | -0.07 | -0.07 | 0 | 0.26 | 0.28 | 0.18 | 0.01 | 0.24 | 0.2 |
| 5 | 178246618 | rs10055695 | 1.355 | 0.987 | -0.201 | -0.285 | -0.033 | -0.07 | -0.07 | 0 | 0.13 | 0.03 | 0.26 | 0.32 | 0.14 | 0.23 |
| 5 | 178252366 | rs10068395 | 0.789 | 0.78 | -0.001 | -0.587 | -0.002 | -0.046 | -0.046 | 0 | 0.2 | 0.19 | 0.1 | 0.01 | 0.17 | 0.11 |
| 5 | 178254864 | rs6600976 | -0.002 | 0.875 | 0.049 | -0.639 | 0.001 | -0.001 | -0.001 | 0 | 0.55 | 0.47 | 0.43 | 0.58 | 0.56 | 0.51 |
| 5 | 178263416 | rs6877165 | 0.254 | 0.724 | 0.021 | -0.63 | -0.004 | 0.041 | 0.041 | 0.008 | 0.24 | 0.34 | 0.23 | 0.31 | 0.24 | 0.14 |
| 5 | 178264000 | rs12055051 | 1.087 | 0.76 | -0.127 | -0.533 | -0.005 | 0.028 | 0.028 | 0 | 0.29 | 0.17 | 0.2 | 0.28 | 0.32 | 0.35 |
| 5 | 178266604 | rs11952214 | 1.087 | 0.76 | -0.127 | -0.533 | -0.005 | 0.028 | 0.028 | 0 | 0.29 | 0.17 | 0.2 | 0.28 | 0.32 | 0.35 |
| 5 | 178267705 | rs12515789 | -0.856 | 0.769 | -0.304 | -0.628 | 0.006 | 0.026 | 0.026 | 0 | 0.8 | 0.75 | 0.62 | 0.59 | 0.79 | 0.71 |
| 5 | 178268306 | rs13174784 | -0.214 | 0.608 | -0.357 | -1.069 | 0.039 | 0.155 | 0.155 | 0 | 0.5 | 0.7 | 0.38 | 0.15 | 0.45 | 0.37 |
| 5 | 178269682 | rs13157175 | -0.123 | 0.799 | -0.262 | -1.077 | 0.027 | -0.01 | -0.01 | 0 | 0.75 | 0.65 | 0.48 | 0.29 | 0.71 | 0.65 |
| 5 | 178270810 | rs6871932 | 0.274 | 0.766 | -0.46 | -0.863 | 0.011 | 0.048 | 0.048 | 0.216 | 0.47 | 0.34 | 0.3 | 0.28 | 0.47 | 0.42 |
| 5 | 178274144 | rs7708351 | -0.219 | 0.784 | -0.573 | -0.979 | -0.003 | 0.012 | 0.012 | 0 | 0.44 | 0.41 | 0.36 | 0.16 | 0.46 | 0.26 |
| 5 | 178279114 | rs7447547 | -0.288 | 0.328 | -0.414 | -0.947 | -0.012 | -0.023 | -0.023 | 0 | 0.57 | 0.49 | 0.44 | 0.24 | 0.63 | 0.35 |
| 5 | 178286356 | rs7445694 | 0.139 | 0.68 | -0.541 | -1.003 | -0.013 | 0.036 | 0.036 | 0 | 0.41 | 0.32 | 0.33 | 0.24 | 0.48 | 0.3 |
| 5 | 178293562 | rs6877197 | 0.392 | 0.917 | -0.777 | -1.148 | 0.003 | 0.036 | 0.036 | 0.915 | 0.23 | 0.12 | 0.12 | 0.09 | 0.23 | 0.14 |
| 5 | 178296936 | rs10479516 | -0.884 | 0.944 | -0.38 | -1.504 | 0.009 | 0.019 | 0.019 | 0 | 0.52 | 0.52 | 0.42 | 0.25 | 0.62 | 0.36 |
| 5 | 178299188 | rs10040956 | -0.863 | 0.933 | -0.308 | -1.509 | 0 | -0.004 | -0.004 | 0.915 | 0.52 | 0.47 | 0.41 | 0.25 | 0.61 | 0.36 |
| 5 | 178300191 | rs6870926 | 0.11 | 0.995 | -0.161 | -1.499 | 0.054 | -0.041 | -0.041 | 0 | 0.08 | 0.27 | 0.11 | 0 | 0.12 | 0.05 |
| 5 | 178303868 | rs7448421 | 0.811 | 1.023 | -0.383 | -1.482 | -0.039 | 0.025 | 0.025 | 0 | 0.21 | 0.11 | 0.19 | 0.15 | 0.27 | 0.17 |
| 5 | 178315686 | rs1046724 | 0.827 | 1.014 | -0.386 | -1.495 | -0.043 | 0.067 | 0.067 | 0 | 0.21 | 0.09 | 0.15 | 0.03 | 0.27 | 0.17 |
| 5 | 178325515 | rs4444956 | -0.336 | 0.931 | -0.367 | -1.513 | -0.034 | 0.133 | 0.133 | 0.019 | 0.43 | 0.19 | 0.28 | 0.12 | 0.5 | 0.31 |
| 5 | 178327082 | rs10044051 | -1.04 | 0.858 | -0.051 | -0.844 | -0.023 | 0.014 | 0.014 | 0 | 0.59 | 0.5 | 0.5 | 0.5 | 0.71 | 0.55 |
| 5 | 178329316 | rs10055777 | -0.388 | 0.888 | -0.13 | -0.851 | 0.021 | -0.003 | -0.003 | 0.083 | 0.52 | 0.39 | 0.42 | 0.5 | 0.59 | 0.49 |
| 5 | 178329846 | rs953741 | 0.907 | 0.948 | -0.181 | -0.893 | -0.09 | 0.089 | 0.089 | 0 | 0.22 | 0.06 | 0.16 | 0.04 | 0.34 | 0.18 |
| 5 | 178330030 | rs10035887 | -1.03 | 0.822 | -0.123 | -0.895 | -0.006 | 0.004 | 0.004 | 0 | 0.58 | 0.47 | 0.49 | 0.5 | 0.64 | 0.52 |
| 5 | 178335516 | rs13171869 | 1.077 | 0.808 | -0.207 | -0.896 | -0.074 | 0.087 | 0.087 | 0 | 0.2 | 0.04 | 0.14 | 0.04 | 0.27 | 0.17 |
| 5 | 178337122 | rs2004503 | -0.353 | 0.809 | -0.216 | -0.904 | 0.009 | 0.004 | 0.004 | 0 | 0.5 | 0.38 | 0.4 | 0.5 | 0.52 | 0.48 |
| 5 | 178337215 | rs732531 | 1.006 | 0.796 | -0.332 | -0.812 | -0.04 | 0.018 | 0.018 | 0 | 0.21 | 0.11 | 0.19 | 0.17 | 0.27 | 0.17 |
| 5 | 178340304 | rs7723421 | -0.206 | 0.732 | -0.176 | -0.827 | -0.011 | -0.012 | -0.012 | 3.798 | 0.37 | 0.45 | 0.3 | 0.33 | 0.37 | 0.43 |
| 5 | 178341750 | rs1995079 | 0.674 | 0.572 | -0.085 | -0.581 | 0.009 | -0.013 | -0.013 | 0 | 0.41 | 0.33 | 0.5 | 0.5 | 0.35 | 0.39 |
| 5 | 178353321 | rs11750407 | 0.935 | 0.238 | -0.308 | -0.297 | -0.006 | -0.066 | -0.066 | 0.097 | 0.29 | 0.12 | 0.38 | 0.37 | 0.28 | 0.37 |
| 5 | 178357931 | rs17081834 | 0.857 | 0.141 | -0.137 | -0.343 | -0.017 | 0.002 | 0.002 | 0 | 0.12 | 0.21 | 0.06 | 0.12 | 0.08 | 0.03 |
| 5 | 178359822 | rs12187167 | 1.192 | 0.311 | -0.297 | -0.397 | -0.008 | 0.012 | 0.012 | 0 | 0.06 | 0.02 | 0.06 | 0 | 0.11 | 0.04 |
| 5 | 178359989 | rs12186577 | 0.347 | 0.261 | -0.034 | -0.347 | 0.027 | 0.043 | 0.043 | 0 | 0.07 | 0.2 | 0.12 | 0.12 | 0.11 | 0.04 |
| 5 | 178360056 | rs12186584 | 1.197 | 0.295 | -0.311 | -0.367 | -0.008 | 0.012 | 0.012 | 0 | 0.06 | 0.02 | 0.06 | 0 | 0.11 | 0.04 |

| | | | | | | | | | | | | | | | | |
|---|---|---|---|---|---|---|---|---|---|---|---|---|---|---|---|---|
| 5 | 178363254 | rs13362387 | -0.121 | -0.069 | -0.425 | -0.497 | 0.006 | -0.011 | -0.011 | 0 | 0.61 | 0.33 | 0.53 | 0.51 | 0.67 | 0.64 |
| 5 | 178368096 | rs17664029 | -0.134 | 0.163 | -0.638 | -0.701 | -0.011 | 0.031 | 0.031 | 0 | 0.33 | 0.21 | 0.28 | 0.17 | 0.38 | 0.27 |
| 5 | 178369673 | rs10060764 | 0.082 | 0.548 | -0.261 | -0.508 | 0.008 | 0.006 | 0.006 | 0 | 0.5 | 0.55 | 0.62 | 0.53 | 0.55 | 0.55 |
| 5 | 178379681 | rs4700746 | 1.039 | 0.389 | -0.221 | -0.087 | -0.003 | -0.011 | -0.011 | 0.593 | 0.3 | 0.24 | 0.42 | 0.36 | 0.29 | 0.35 |
| 5 | 178384832 | rs6867653 | 0.892 | 0.216 | -0.252 | -0.14 | -0.029 | 0 | 0 | 1.195 | 0.14 | 0.26 | 0.06 | 0.12 | 0.06 | 0.04 |
| 5 | 178385585 | rs10070619 | 0.384 | 0.31 | -0.26 | -0.226 | 0.027 | -0.064 | -0.064 | 0.451 | 0.18 | 0.26 | 0.11 | 0.05 | 0.2 | 0.23 |
| 5 | 178390237 | rs903981 | 0.207 | 0.152 | -0.351 | -0.103 | 0.012 | -0.005 | -0.005 | 0 | 0.47 | 0.35 | 0.59 | 0.46 | 0.57 | 0.52 |
| 5 | 178391860 | rs6867221 | 0.925 | 0.18 | -0.497 | -0.111 | -0.012 | -0.031 | -0.031 | 0 | 0.3 | 0.11 | 0.28 | 0.34 | 0.31 | 0.37 |
| 5 | 178391902 | rs12719860 | -0.915 | 0.187 | -0.441 | -0.107 | 0.002 | -0.001 | -0.001 | 10.552 | 0.68 | 0.64 | 0.69 | 0.66 | 0.69 | 0.63 |
| 5 | 178400475 | rs7443667 | -0.011 | 0.046 | -0.496 | -0.171 | 0.011 | 0.012 | 0.012 | 0 | 0.53 | 0.37 | 0.66 | 0.63 | 0.6 | 0.74 |
| 5 | 178403949 | rs9637932 | 0.939 | 0.023 | -0.87 | -0.158 | -0.025 | -0.008 | -0.008 | 0 | 0.14 | 0.25 | 0.07 | 0.12 | 0.08 | 0.04 |
| 5 | 178405251 | rs4700765 | 1.478 | 0.22 | -1.271 | -0.15 | -0.05 | -0.087 | -0.087 | 0.17 | 0.28 | 0.14 | 0.42 | 0.37 | 0.3 | 0.32 |
| 5 | 178405729 | rs2856354 | 1.359 | -0.041 | -0.971 | 0.2 | 0.005 | -0.028 | -0.028 | 0 | 0.06 | 0 | 0.07 | 0.23 | 0.03 | 0.22 |
| 5 | 178406388 | rs7725272 | 1.052 | 0.1 | -0.936 | 0.158 | -0.036 | 0.058 | 0.058 | 0.187 | 0.24 | 0.1 | 0.22 | 0.11 | 0.31 | 0.15 |
| 5 | 178408602 | rs17078853 | 0.932 | -0.393 | -0.298 | 0.745 | 0.013 | -0.036 | -0.036 | 0.023 | 0.06 | 0 | 0.05 | 0.14 | 0.04 | 0.08 |
| 5 | 178409285 | rs10464073 | -0.319 | -0.39 | -0.292 | 0.735 | 0.003 | -0.038 | -0.038 | 0.957 | 0.12 | 0.04 | 0.14 | 0.23 | 0.13 | 0.25 |
| 5 | 178410151 | rs2071247 | -1.256 | -0.949 | -0.101 | 0.787 | -0.026 | 0.055 | 0.055 | 0 | 0.2 | 0.22 | 0.2 | 0.37 | 0.17 | 0.29 |
| 5 | 178410213 | rs17078877 | 0.571 | -0.557 | -0.698 | 0.563 | 0.013 | -0.036 | -0.036 | 0 | 0.06 | 0 | 0.05 | 0.14 | 0.04 | 0.08 |
| 5 | 178410538 | rs1475950 | -0.502 | -0.217 | -0.338 | 0.278 | -0.018 | -0.007 | -0.007 | 0 | 0.14 | 0.22 | 0.07 | 0.12 | 0.08 | 0.04 |
| 5 | 178413947 | rs11746675 | -0.292 | -0.501 | -0.672 | 0.405 | 0.006 | 0.004 | 0.004 | 0.742 | 0.46 | 0.4 | 0.61 | 0.59 | 0.64 | 0.51 |
| 5 | 178415937 | rs2067011 | -0.132 | 0.082 | -0.662 | 0.384 | 0 | -0.002 | -0.002 | 0 | 0.41 | 0.39 | 0.53 | 0.59 | 0.55 | 0.51 |
| 5 | 178416288 | rs2071246 | -0.799 | -0.827 | -0.813 | -0.074 | 0.03 | 0.028 | 0.028 | 0.788 | 0.66 | 0.78 | 0.68 | 0.7 | 0.75 | 0.68 |
| 5 | 178424544 | rs2078183 | -0.001 | -1.143 | -0.582 | 0.227 | 0.011 | -0.038 | -0.038 | 0.196 | 0.38 | 0.49 | 0.52 | 0.2 | 0.54 | 0.28 |
| 5 | 178426150 | rs4700775 | 0.016 | -0.713 | -0.619 | -0.314 | -0.032 | 0.015 | 0.015 | 0 | 0.63 | 0.46 | 0.64 | 0.47 | 0.69 | 0.6 |
| 5 | 178430000 | rs11249609 | 1.164 | -0.749 | -0.77 | -0.348 | -0.072 | -0.001 | -0.001 | 0.213 | 0.32 | 0.01 | 0.29 | 0.29 | 0.31 | 0.44 |
| 5 | 178430375 | rs2173096 | 0.569 | -0.669 | -0.617 | -0.342 | 0.029 | -0.032 | -0.032 | 0 | 0.3 | 0.35 | 0.33 | 0.17 | 0.39 | 0.16 |
| 5 | 178434806 | rs6601002 | -0.066 | -0.895 | -0.765 | -0.433 | 0 | -0.01 | -0.01 | 0 | 0.47 | 0.48 | 0.39 | 0.24 | 0.45 | 0.26 |
| 5 | 178436190 | rs12658310 | -0.06 | -0.891 | -0.755 | -0.432 | 0 | -0.01 | -0.01 | 0.323 | 0.47 | 0.48 | 0.39 | 0.24 | 0.45 | 0.26 |
| 5 | 178436617 | rs7700958 | 0.297 | -0.659 | -0.657 | -0.243 | 0.001 | 0.093 | 0.093 | 0 | 0.2 | 0.31 | 0.29 | 0.47 | 0.24 | 0.3 |
| 5 | 178447321 | rs6873398 | -0.557 | -0.543 | -0.628 | -0.646 | 0.024 | -0.034 | -0.034 | 0.058 | 0.69 | 0.62 | 0.67 | 0.85 | 0.61 | 0.84 |
| 5 | 178452569 | rs11738515 | 1.37 | -0.774 | -0.845 | -0.555 | 0.041 | -0.014 | -0.014 | 0 | 0.14 | 0.04 | 0.05 | 0.1 | 0.06 | 0.09 |
| 5 | 178453904 | rs11249610 | 1.238 | -1.081 | -1.169 | -0.639 | -0.057 | -0.037 | -0.037 | 0 | 0.34 | 0.01 | 0.36 | 0.33 | 0.31 | 0.43 |
| 5 | 178454737 | rs7717952 | -0.876 | -1.063 | -0.94 | -0.798 | 0.003 | -0.002 | -0.002 | 0 | 0.77 | 0.76 | 0.68 | 0.87 | 0.61 | 0.85 |
| 5 | 178455198 | rs10464075 | 0.819 | -1.017 | -1.108 | -0.76 | 0.028 | -0.024 | -0.024 | 0.478 | 0.47 | 0.07 | 0.41 | 0.43 | 0.37 | 0.52 |
| 5 | 178455954 | rs7732931 | 0.819 | -1.017 | -1.108 | -0.76 | 0.028 | -0.024 | -0.024 | 4.445 | 0.47 | 0.07 | 0.41 | 0.43 | 0.37 | 0.52 |
| 5 | 178461594 | rs11741906 | 0.819 | -1.017 | -1.145 | -0.76 | 0.013 | -0.041 | -0.041 | 0 | 0.47 | 0.05 | 0.41 | 0.43 | 0.37 | 0.52 |
| 5 | 178482994 | rs7711420 | 0.104 | -0.91 | -0.211 | -0.598 | -0.006 | 0.074 | 0.074 | 0 | 0.22 | 0.37 | 0.2 | 0.43 | 0.24 | 0.31 |
| 5 | 178483806 | rs4701028 | 0.729 | -0.487 | -0.434 | -0.743 | 0.011 | 0.046 | 0.046 | 0.107 | 0.43 | 0.23 | 0.44 | 0.34 | 0.38 | 0.47 |
| 5 | 178487452 | rs12522073 | 1.016 | -0.853 | -0.516 | -0.794 | -0.108 | 0.035 | 0.035 | 0 | 0.16 | 0.02 | 0.23 | 0.1 | 0.33 | 0.13 |
| 5 | 178491000 | rs6886245 | 1.495 | -1.048 | -0.352 | -0.754 | -0.017 | -0.026 | -0.026 | 0 | 0.06 | 0.15 | 0.01 | 0.01 | 0.02 | 0.02 |
| 5 | 178495923 | rs2411912 | 1.663 | -1.364 | -0.657 | -0.992 | 0.031 | -0.016 | -0.016 | 0 | 0.24 | 0.11 | 0.3 | 0.3 | 0.17 | 0.36 |
| 5 | 178503640 | rs4700780 | 0.27 | -0.894 | -0.098 | -0.775 | 0.008 | 0.036 | 0.036 | 0 | 0.23 | 0.31 | 0.25 | 0.43 | 0.29 | 0.35 |
| 5 | 178507069 | rs1445846 | -0.214 | -0.619 | -0.19 | -0.824 | 0.014 | -0.019 | -0.019 | 0 | 0.77 | 0.74 | 0.74 | 0.89 | 0.67 | 0.85 |
| 5 | 178507090 | rs1445845 | -0.239 | -0.612 | -0.187 | -0.821 | 0.015 | -0.019 | -0.019 | 1.628 | 0.77 | 0.74 | 0.74 | 0.89 | 0.67 | 0.85 |
| 5 | 178507270 | rs1445844 | -0.295 | -0.625 | -0.198 | -0.827 | 0.005 | -0.009 | -0.009 | 0 | 0.78 | 0.77 | 0.74 | 0.89 | 0.67 | 0.85 |
| 5 | 178510048 | rs1049532 | -0.386 | -0.758 | -0.189 | -0.862 | -0.004 | -0.001 | -0.001 | 0 | 0.43 | 0.54 | 0.34 | 0.55 | 0.35 | 0.45 |
| 5 | 178513141 | rs1136377 | -0.206 | -0.733 | -0.24 | -0.934 | 0.049 | -0.049 | -0.049 | 0 | 0.77 | 0.67 | 0.73 | 0.89 | 0.67 | 0.85 |
| 5 | 178516483 | rs1562234 | -0.431 | -0.755 | -0.231 | -0.868 | 0.004 | 0.002 | 0.002 | 0 | 0.43 | 0.59 | 0.34 | 0.57 | 0.37 | 0.45 |
| 5 | 178539382 | rs1863918 | 1.454 | -0.701 | -0.742 | -0.845 | 0.001 | -0.001 | -0.001 | 0 | 0.34 | 0.05 | 0.39 | 0.31 | 0.3 | 0.4 |
| 5 | 178540656 | rs10479525 | -0.429 | -0.701 | -0.663 | -0.863 | 0.056 | 0.032 | 0.032 | 0.164 | 0.46 | 0.74 | 0.35 | 0.58 | 0.37 | 0.45 |
| 5 | 178540975 | rs1054480 | 1.326 | -0.823 | -0.793 | -0.701 | -0.014 | -0.016 | -0.016 | 0 | 0.33 | 0.04 | 0.38 | 0.31 | 0.3 | 0.4 |
| 5 | 178547567 | rs2288382 | 0.263 | -0.716 | -0.01 | -0.211 | -0.035 | 0.119 | 0.119 | 0 | 0.14 | 0.29 | 0.09 | 0.38 | 0.07 | 0.21 |
| 5 | 178552518 | rs6860046 | -0.644 | 0.416 | -0.543 | -0.541 | -0.058 | 0.016 | 0.016 | 1.36 | 0.43 | 0.83 | 0.44 | 0.57 | 0.46 | 0.48 |
| 5 | 178553341 | rs6601018 | -0.561 | 0.039 | 0.11 | -0.238 | 0.002 | 0.006 | 0.006 | 0 | 0.72 | 0.75 | 0.69 | 0.57 | 0.62 | 0.62 |
| 5 | 178555045 | rs2303644 | 0.786 | 0.349 | -0.861 | -0.199 | -0.078 | -0.006 | -0.006 | 5.001 | 0.12 | 0.01 | 0.21 | 0.1 | 0.26 | 0.14 |
| 5 | 178557142 | rs2251105 | 0.201 | 0.115 | -0.84 | -0.335 | 0.002 | 0.024 | 0.024 | 0.264 | 0.47 | 0.4 | 0.53 | 0.43 | 0.51 | 0.48 |
| 5 | 178562967 | rs1972715 | 0.311 | 0.21 | -0.903 | -0.313 | 0.048 | -0.014 | -0.014 | 0.083 | 0.14 | 0.25 | 0.23 | 0.1 | 0.26 | 0.14 |
| 5 | 178567054 | rs2303638 | 1.22 | 0.025 | -1.18 | -0.307 | 0.011 | -0.031 | -0.031 | 0 | 0.32 | 0.05 | 0.3 | 0.33 | 0.25 | 0.34 |
| 5 | 178572268 | rs11746863 | 0.613 | 0.095 | -0.962 | -0.345 | 0.002 | 0.003 | 0.003 | 0 | 0.23 | 0.2 | 0.13 | 0.14 | 0.11 | 0.15 |
| 5 | 178573564 | rs7736323 | -0.115 | 1.67 | -0.655 | -0.102 | 0 | 0.002 | 0.002 | 0 | 0.13 | 0.14 | 0.22 | 0.1 | 0.26 | 0.13 |
| 5 | 178577392 | rs2411906 | -0.066 | 1.717 | -0.979 | 0.046 | 0.027 | 0.049 | 0.049 | 0 | 0.69 | 0.53 | 0.66 | 0.57 | 0.62 | 0.62 |
| 5 | 178579914 | rs3776818 | 0.342 | 0.255 | -0.881 | -0.037 | 0.005 | -0.001 | -0.001 | 0.567 | 0.55 | 0.58 | 0.45 | 0.47 | 0.35 | 0.44 |
| 5 | 178595870 | rs7735461 | -0.062 | 0.711 | -1.178 | -0.659 | -0.019 | 0.077 | 0.077 | 0 | 0.56 | 0.68 | 0.48 | 0.18 | 0.54 | 0.39 |
| 5 | 178604377 | rs4701077 | 0.133 | 0.367 | -1.117 | -0.722 | 0.014 | 0.044 | 0.044 | 0 | 0.4 | 0.26 | 0.54 | 0.77 | 0.37 | 0.54 |
| 5 | 178604465 | rs6862563 | 0.488 | 0.634 | -1.698 | -0.924 | -0.009 | 0.022 | 0.022 | 0 | 0.1 | 0.03 | 0.09 | 0 | 0.14 | 0.08 |
| 5 | 178607770 | rs888763 | -0.138 | 0.495 | -1.552 | -1.151 | 0.023 | 0.182 | 0.182 | 0 | 0.51 | 0.8 | 0.65 | 0.86 | 0.45 | 0.63 |
| 5 | 178609060 | rs3797600 | -0.102 | 0.193 | -1.549 | -1.143 | 0.032 | 0.078 | 0.078 | 0 | 0.48 | 0.7 | 0.61 | 0.8 | 0.4 | 0.6 |
| 5 | 178610028 | rs3776819 | -0.624 | 0.405 | -1.735 | -1.012 | -0.088 | 0.152 | 0.152 | 0 | 0.59 | 0.9 | 0.71 | 0.85 | 0.54 | 0.72 |
| 5 | 178614693 | rs252070 | 0.36 | 0.519 | -1.561 | -0.975 | -0.02 | 0.074 | 0.074 | 0 | 0.42 | 0.25 | 0.31 | 0.15 | 0.45 | 0.27 |

| | | | | | | | | | | | | | | | | |
|---|---|---|---|---|---|---|---|---|---|---|---|---|---|---|---|---|
| 5 | 178614867 | rs171667 | -0.277 | 0.731 | -1.99 | -0.012 | -0.071 | -0.123 | -0.123 | 0 | 0.76 | 0.98 | 0.63 | 0.66 | 0.73 | 0.79 |
| 5 | 178616135 | rs252071 | 0.138 | 0.857 | -1.424 | -0.668 | -0.033 | 0.121 | 0.121 | 0 | 0.33 | 0.13 | 0.21 | 0.05 | 0.37 | 0.19 |
| 5 | 178617862 | rs9329070 | -0.916 | 0.712 | -0.153 | -0.462 | 0.018 | -0.116 | -0.116 | 0 | 0.26 | 0.43 | 0.27 | 0.09 | 0.25 | 0.2 |
| 5 | 178617938 | rs252072 | 1.392 | 0.473 | -0.276 | 0.315 | -0.001 | 0.021 | 0.021 | 0 | 0.58 | 0.34 | 0.62 | 0.68 | 0.61 | 0.69 |
| 5 | 178618707 | rs6895999 | -0.355 | 0.474 | 0.134 | 0.229 | 0.004 | -0.021 | -0.021 | 2.637 | 0.06 | 0.07 | 0.05 | 0 | 0.06 | 0.03 |
| 5 | 178619190 | rs6860197 | -0.077 | 0.265 | 0.309 | 0.428 | -0.008 | -0.002 | -0.002 | 0.315 | 0.1 | 0.07 | 0.11 | 0 | 0.14 | 0.12 |
| 5 | 178619774 | rs10074081 | -1.439 | -0.177 | 1.27 | 0.928 | -0.035 | -0.02 | -0.02 | 0.7 | 0.12 | 0.37 | 0.15 | 0.08 | 0.05 | 0.05 |
| 5 | 178620555 | rs4700788 | -1.906 | -0.394 | 0.915 | 1.527 | -0.071 | 0.104 | 0.104 | 0 | 0.2 | 0.56 | 0.33 | 0.48 | 0.1 | 0.35 |
| 5 | 178621576 | rs3822601 | -1.894 | -0.382 | 0.848 | 1.541 | -0.075 | 0.118 | 0.118 | 0 | 0.2 | 0.57 | 0.33 | 0.48 | 0.1 | 0.35 |
| 5 | 178624101 | rs2279979 | -1.173 | 0.044 | 0.021 | 1.655 | -0.062 | 0.246 | 0.246 | 0 | 0.12 | 0.44 | 0.21 | 0.4 | 0.05 | 0.32 |
| 5 | 178627708 | rs3797610 | -0.705 | 1.346 | -1.437 | 1.229 | -0.099 | -0.048 | -0.048 | 5.127 | 0.21 | 0.03 | 0.16 | 0.23 | 0.26 | 0.17 |
| 5 | 178630445 | rs252061 | -0.65 | 1.104 | -1.406 | 2.01 | -0.028 | -0.082 | -0.082 | 0.01 | 0.12 | 0.01 | 0.19 | 0.44 | 0.12 | 0.31 |
| 5 | 178632482 | rs252064 | -1.647 | 1.276 | -0.83 | 1.945 | 0.004 | 0.032 | 0.032 | 0 | 0.2 | 0.22 | 0.34 | 0.56 | 0.25 | 0.34 |
| 5 | 178633183 | rs252066 | -0.853 | 1.317 | -1.178 | 1.898 | -0.036 | -0.08 | -0.08 | 0 | 0.12 | 0.02 | 0.21 | 0.44 | 0.15 | 0.31 |
| 5 | 178634619 | rs423552 | 0.354 | 0.411 | -1.234 | 0.791 | 0.008 | -0.006 | -0.006 | 0 | 0.85 | 0.88 | 0.94 | 0.99 | 0.91 | 0.92 |
| 5 | 178634672 | rs398829 | -0.67 | 0.452 | -1.232 | 0.83 | -0.001 | 0 | 0 | 0.238 | 0.4 | 0.41 | 0.26 | 0.63 | 0.28 | 0.54 |
| 5 | 178634683 | rs11750821 | -1.373 | 1.078 | -1.47 | 0.585 | -0.036 | -0.021 | -0.021 | 0 | 0.07 | 0.01 | 0.09 | 0.11 | 0.1 | 0.07 |
| 5 | 178635005 | rs416646 | -0.425 | 0.914 | -1.229 | 0.516 | 0 | -0.002 | -0.002 | 0.514 | 0.4 | 0.56 | 0.33 | 0.51 | 0.35 | 0.31 |
| 5 | 178636202 | rs337807 | -0.574 | 1.149 | -1.548 | 0.448 | -0.023 | -0.066 | -0.066 | 0 | 0.2 | 0.1 | 0.24 | 0.35 | 0.2 | 0.22 |
| 5 | 178637124 | rs337806 | -0.721 | 1.611 | -1.461 | -0.213 | -0.001 | 0 | 0 | 0 | 0.82 | 0.84 | 0.83 | 0.86 | 0.72 | 0.89 |
| 5 | 178640760 | rs466750 | -0.591 | 1.88 | -1.269 | -0.675 | -0.031 | 0.042 | 0.042 | 0 | 0.29 | 0.19 | 0.29 | 0.1 | 0.36 | 0.26 |
| 5 | 178640982 | rs442406 | -0.192 | 1.641 | -1.017 | -0.905 | 0.001 | -0.004 | -0.004 | 2.595 | 0.22 | 0.25 | 0.23 | 0.12 | 0.28 | 0.3 |
| 5 | 178642524 | rs464156 | -0.564 | 1.186 | -0.967 | -0.691 | -0.002 | 0.018 | 0.018 | 0 | 0.41 | 0.36 | 0.36 | 0.13 | 0.49 | 0.46 |
| 5 | 178645310 | rs461898 | -0.671 | 1.02 | -1.201 | -0.394 | -0.001 | 0.003 | 0.003 | 0 | 0.53 | 0.56 | 0.55 | 0.63 | 0.45 | 0.66 |
| 5 | 178647388 | rs411804 | 0.242 | 1.164 | -1.476 | -0.636 | -0.025 | 0.053 | 0.053 | 0 | 0.86 | 0.95 | 0.92 | 0.96 | 0.84 | 0.76 |
| 5 | 178652282 | rs458752 | 0.419 | 0.989 | -1.414 | -0.851 | -0.012 | 0.043 | 0.043 | 0.064 | 0.78 | 0.85 | 0.87 | 0.99 | 0.75 | 0.87 |
| 5 | 178659691 | rs456909 | 0.848 | 1.1 | -0.734 | -0.248 | 0.022 | 0.009 | 0.009 | 0 | 0.62 | 0.57 | 0.63 | 0.64 | 0.56 | 0.7 |
| 5 | 178663338 | rs469090 | 0.787 | 0.261 | -1.542 | -0.933 | -0.003 | -0.029 | -0.029 | 1.465 | 0.15 | 0.02 | 0.25 | 0.18 | 0.11 | 0.21 |
| 5 | 178663408 | rs469568 | -0.043 | -0.285 | -1.112 | -1.688 | 0.002 | -0.003 | -0.003 | 0 | 0.53 | 0.6 | 0.36 | 0.06 | 0.57 | 0.32 |
| 5 | 178665185 | rs11746802 | 0.731 | -0.193 | -0.98 | -1.774 | 0.008 | -0.141 | -0.141 | 0 | 0.33 | 0.47 | 0.25 | 0.05 | 0.36 | 0.2 |
| 5 | 178665240 | rs469099 | 1.026 | -0.2 | -1.078 | -1.759 | -0.046 | 0.112 | 0.112 | 0 | 0.18 | 0.01 | 0.09 | 0.01 | 0.21 | 0.12 |
| 5 | 178665824 | rs469262 | -0.34 | -0.351 | -0.893 | -1.854 | 0.004 | 0.058 | 0.058 | 0.015 | 0.58 | 0.73 | 0.4 | 0.06 | 0.62 | 0.35 |
| 5 | 178667984 | rs467604 | -0.322 | -0.369 | -0.854 | -1.842 | 0.007 | 0.037 | 0.037 | 0.015 | 0.58 | 0.68 | 0.51 | 0.1 | 0.63 | 0.35 |
| 5 | 178671164 | rs4700790 | 0.246 | -0.226 | -0.214 | -1.666 | 0.009 | -0.112 | -0.112 | 0 | 0.39 | 0.52 | 0.4 | 0.08 | 0.42 | 0.25 |
| 5 | 178674076 | rs4701085 | 1.522 | -0.46 | -0.613 | -1.752 | -0.105 | 0.119 | 0.119 | 0.229 | 0.25 | 0.03 | 0.3 | 0.08 | 0.34 | 0.19 |
| 5 | 178675065 | rs7716112 | -0.954 | -0.708 | -0.166 | -1.475 | -0.037 | -0.091 | -0.091 | 0.018 | 0.14 | 0.62 | 0.1 | 0 | 0.06 | 0.03 |
| 5 | 178681856 | rs6875299 | -0.671 | -0.682 | -0.001 | -1.383 | -0.031 | -0.044 | -0.044 | 0 | 0.09 | 0.63 | 0.06 | 0 | 0.01 | 0.01 |
| 5 | 178686332 | rs17607156 | 1.156 | -0.541 | -0.317 | -1.689 | -0.04 | 0.097 | 0.097 | 1.284 | 0.3 | 0.12 | 0.33 | 0.08 | 0.38 | 0.25 |
| 5 | 178686438 | rs340125 | 0.091 | -0.782 | 0.08 | -1.361 | -0.013 | -0.145 | -0.145 | 0 | 0.79 | 0.51 | 0.87 | 0.98 | 0.8 | 0.92 |
| 5 | 178687683 | rs340122 | -0.354 | -0.975 | -0.962 | -0.661 | 0.004 | -0.004 | -0.004 | 0 | 0.2 | 0.2 | 0.22 | 0.28 | 0.15 | 0.43 |
| 5 | 178688093 | rs340121 | 0.772 | -0.874 | -1.628 | -0.935 | 0.003 | 0.037 | 0.037 | 0.83 | 0.69 | 0.21 | 0.72 | 0.79 | 0.69 | 0.65 |
| 5 | 178689856 | rs9885424 | -0.017 | -0.791 | -1.465 | -0.382 | 0 | 0.011 | 0.011 | 0.279 | 0.07 | 0.07 | 0.08 | 0.2 | 0.04 | 0.19 |
| 5 | 178690776 | rs42875 | -0.029 | -0.865 | -1.662 | -0.103 | 0.02 | -0.018 | -0.018 | 0 | 0.15 | 0.08 | 0.1 | 0.2 | 0.11 | 0.15 |
| 5 | 178698709 | rs901253 | 0.286 | -0.605 | -2.299 | 0.199 | -0.018 | -0.018 | -0.018 | 0 | 0.48 | 0.14 | 0.53 | 0.44 | 0.51 | 0.44 |
| 5 | 178700899 | rs2013658 | 0.989 | -0.3 | -2.193 | 0.306 | -0.069 | 0.014 | 0.014 | 0 | 0.27 | 0.09 | 0.4 | 0.22 | 0.37 | 0.2 |
| 5 | 178707077 | rs459770 | -0.401 | -0.599 | -1.515 | 0.483 | 0.004 | -0.013 | -0.013 | 0 | 0.34 | 0.29 | 0.37 | 0.47 | 0.32 | 0.48 |
| 5 | 178712059 | rs6888613 | -0.451 | -1.128 | -1.505 | 0.093 | 0.002 | 0.015 | 0.015 | 0.359 | 0.45 | 0.41 | 0.37 | 0.25 | 0.43 | 0.39 |
| 5 | 178713712 | rs338881 | 0.4 | -0.955 | -1.267 | 0.177 | -0.021 | -0.071 | -0.071 | 3.734 | 0.19 | 0.31 | 0.12 | 0.03 | 0.15 | 0.1 |
| 5 | 178716389 | rs181797 | 0.275 | -0.595 | -1.165 | 0.447 | 0 | 0 | 0 | 0 | 0.22 | 0.2 | 0.32 | 0.22 | 0.24 | 0.28 |
| 5 | 178719364 | rs4700798 | 1.103 | -0.561 | -1.985 | 0.579 | -0.121 | -0.015 | -0.015 | 0 | 0.3 | 0.07 | 0.39 | 0.28 | 0.37 | 0.23 |
| 5 | 178719705 | rs1862266 | -0.721 | -0.446 | -0.692 | 1.277 | -0.017 | 0.048 | 0.048 | 0 | 0.13 | 0.39 | 0.1 | 0.14 | 0.1 | 0.09 |
| 5 | 178720968 | rs166031 | -0.405 | -0.116 | -1.509 | 1.36 | -0.057 | -0.068 | -0.068 | 0 | 0.28 | 0.06 | 0.34 | 0.29 | 0.33 | 0.36 |
| 5 | 178724727 | rs467053 | -0.41 | 0.335 | -0.895 | 1.183 | 0.022 | 0.008 | 0.008 | 0 | 0.45 | 0.37 | 0.45 | 0.56 | 0.42 | 0.6 |
| 5 | 178730062 | rs2431423 | -0.933 | 1.167 | -0.813 | 1.709 | -0.025 | -0.046 | -0.046 | 0 | 0.22 | 0.13 | 0.37 | 0.32 | 0.28 | 0.19 |
| 5 | 178731921 | rs1961716 | 0.837 | 0.845 | -0.106 | 1.68 | -0.009 | 0.072 | 0.072 | 0 | 0.84 | 0.7 | 0.88 | 0.78 | 0.85 | 0.84 |
| 5 | 178733902 | rs6868803 | -1.013 | 1.077 | -0.93 | 1.644 | -0.005 | -0.03 | -0.03 | 0.111 | 0.13 | 0.05 | 0.09 | 0.18 | 0.14 | 0.18 |
| 5 | 178737752 | rs878932 | 0.895 | 1.389 | -0.719 | 1.617 | -0.003 | -0.012 | -0.012 | 1.312 | 0.74 | 0.76 | 0.64 | 0.6 | 0.71 | 0.87 |
| 5 | 178739016 | rs878933 | -0.571 | 1.72 | -1.02 | 1.625 | -0.055 | -0.075 | -0.075 | 0 | 0.14 | 0.04 | 0.29 | 0.27 | 0.2 | 0.11 |
| 5 | 178742614 | rs340113 | 1.17 | 1.082 | 0.012 | 0.844 | -0.014 | -0.055 | -0.055 | 0.021 | 0.9 | 0.7 | 0.95 | 1 | 0.94 | 0.98 |
| 5 | 178746447 | rs163321 | -0.649 | 1.273 | -0.706 | 0.813 | -0.023 | 0.043 | 0.043 | 0 | 0.39 | 0.24 | 0.38 | 0.72 | 0.41 | 0.59 |
| 5 | 178747939 | rs163497 | 1.041 | 1.269 | -0.066 | 1.812 | 0.017 | 0.182 | 0.182 | 0 | 0.84 | 0.7 | 0.86 | 0.62 | 0.82 | 0.87 |
| 5 | 178749829 | rs455437 | 0.493 | 0.696 | -0.107 | 1.351 | -0.01 | 0 | 0 | 0 | 0.82 | 0.78 | 0.91 | 0.83 | 0.87 | 0.86 |
| 5 | 178750377 | rs422172 | 0.954 | -0.149 | -1.137 | 1.322 | 0.052 | 0.047 | 0.047 | 0 | 0.56 | 0.74 | 0.6 | 0.35 | 0.69 | 0.48 |
| 5 | 178752095 | rs2450191 | 1.009 | -0.271 | -1.082 | 0.231 | -0.001 | -0.022 | -0.022 | 0 | 0.24 | 0.25 | 0.13 | 0.05 | 0.22 | 0.05 |
| 5 | 178752189 | rs4701095 | 1.075 | 1.124 | -1.385 | 0.18 | -0.047 | -0.029 | -0.029 | 0 | 0.08 | 0.01 | 0.16 | 0.13 | 0.18 | 0.13 |
| 5 | 178752351 | rs4701096 | -0.558 | 0.654 | -1.31 | 0.134 | -0.003 | -0.009 | -0.009 | 0.012 | 0.58 | 0.59 | 0.79 | 0.76 | 0.64 | 0.8 |
| 5 | 178754468 | rs11956034 | 0.433 | 0.515 | -0.485 | 0.291 | -0.006 | 0.009 | 0.009 | 0 | 0.28 | 0.24 | 0.33 | 0.51 | 0.2 | 0.4 |
| 5 | 178756557 | rs443941 | -0.703 | 0.191 | -0.266 | 0.4 | 0.001 | -0.007 | -0.007 | 0.136 | 0.55 | 0.55 | 0.5 | 0.67 | 0.54 | 0.64 |
| 5 | 178758909 | rs340419 | -1.115 | 0.451 | -0.247 | 0.277 | -0.004 | 0.007 | 0.007 | 0.013 | 0.64 | 0.69 | 0.65 | 0.75 | 0.59 | 0.83 |

| | | | | | | | | | | | | | | | | |
|---|---|---|---|---|---|---|---|---|---|---|---|---|---|---|---|---|
| 5 | 178759900 | rs11745920 | 1.099 | 0.519 | -0.754 | 0.707 | 0.023 | -0.09 | -0.09 | 0.23 | 0.23 | 0.03 | 0.16 | 0.31 | 0.16 | 0.37 |
| 5 | 178761358 | rs7734278 | -0.642 | 0.414 | -0.436 | 0.666 | -0.01 | -0.008 | -0.008 | 0 | 0.46 | 0.26 | 0.49 | 0.54 | 0.47 | 0.67 |
| 5 | 178762092 | rs3846697 | -0.306 | 1.149 | -0.16 | 0.942 | -0.005 | -0.002 | -0.002 | 0 | 0.2 | 0.16 | 0.29 | 0.29 | 0.28 | 0.29 |
| 5 | 178766209 | rs26813 | 0.769 | 0.865 | 0.137 | 0.893 | -0.02 | -0.009 | -0.009 | 0 | 0.18 | 0.27 | 0.14 | 0.14 | 0.14 | 0.17 |
| 5 | 178766925 | rs7733548 | 0.328 | 0.693 | -0.388 | 0.883 | -0.001 | -0.015 | -0.015 | 0 | 0.18 | 0.13 | 0.2 | 0.35 | 0.18 | 0.3 |
| 5 | 178770100 | rs7736370 | 1.062 | 0.713 | -0.56 | 0.911 | -0.005 | -0.051 | -0.051 | 0 | 0.13 | 0.06 | 0.14 | 0.31 | 0.14 | 0.27 |
| 5 | 178770485 | rs12513486 | 0.342 | 0.904 | -0.648 | 0.519 | -0.039 | -0.014 | -0.014 | 1.612 | 0.16 | 0.08 | 0.28 | 0.2 | 0.24 | 0.26 |
| 5 | 178773041 | rs153813 | -0.5 | 0.698 | -0.181 | 0.424 | -0.005 | 0.009 | 0.009 | 0 | 0.39 | 0.55 | 0.37 | 0.5 | 0.36 | 0.51 |
| 5 | 178780789 | rs255022 | -0.683 | 0.238 | -0.246 | 0.228 | -0.003 | 0.008 | 0.008 | 0.112 | 0.48 | 0.59 | 0.52 | 0.37 | 0.48 | 0.53 |
| 5 | 178782813 | rs340413 | -1.53 | 0.091 | -0.326 | -0.5 | -0.01 | -0.058 | -0.058 | 0 | 0.69 | 0.73 | 0.84 | 0.97 | 0.82 | 0.89 |
| 5 | 178785054 | rs255025 | 0.245 | 0.311 | 0.013 | 0.044 | -0.022 | -0.058 | -0.058 | 0 | 0.24 | 0.42 | 0.17 | 0.08 | 0.18 | 0.2 |
| 5 | 178790953 | rs340434 | -0.881 | 0.593 | -0.264 | 0.129 | -0.002 | -0.023 | -0.023 | 0.449 | 0.53 | 0.57 | 0.63 | 0.75 | 0.57 | 0.7 |
| 5 | 178796142 | rs368080 | 0.139 | 0.952 | -0.33 | 0.938 | -0.006 | 0.008 | 0.008 | 0.051 | 0.9 | 0.93 | 0.89 | 0.61 | 0.89 | 0.79 |
| 5 | 178805612 | rs1026985 | 0.561 | 0.741 | -0.006 | 0.637 | 0.001 | -0.004 | -0.004 | 0 | 0.62 | 0.6 | 0.66 | 0.75 | 0.57 | 0.66 |
| 5 | 178807044 | rs560137 | -0.836 | 0.802 | -0.044 | 0.618 | -0.012 | 0.03 | 0.03 | 0 | 0.39 | 0.27 | 0.39 | 0.66 | 0.44 | 0.53 |
| 5 | 178807114 | rs1032150 | 0.064 | 0.762 | -0.172 | 0.599 | -0.007 | -0.005 | -0.005 | 0 | 0.29 | 0.16 | 0.26 | 0.25 | 0.33 | 0.29 |
| 5 | 178820635 | rs6883577 | 1.029 | 0.577 | -0.253 | 0.511 | 0.001 | -0.01 | -0.01 | 0.059 | 0.55 | 0.45 | 0.56 | 0.32 | 0.55 | 0.46 |
| 5 | 178826910 | rs7731436 | -0.075 | 0.252 | -1.094 | 0.825 | 0.027 | -0.121 | -0.121 | 0 | 0.26 | 0.06 | 0.23 | 0.49 | 0.21 | 0.4 |
| 5 | 178827859 | rs7736755 | 0.383 | 0.173 | -0.967 | 0.122 | 0.036 | 0.043 | 0.043 | 0.009 | 0.21 | 0.06 | 0.14 | 0.1 | 0.16 | 0.17 |
| 5 | 178840519 | rs11249625 | 0.211 | 0.095 | -0.902 | -0.536 | -0.007 | -0.028 | -0.028 | 0 | 0.66 | 0.62 | 0.66 | 0.83 | 0.65 | 0.71 |
| 5 | 178841876 | rs12652161 | 0.211 | 0.335 | -1.055 | -0.872 | 0 | -0.004 | -0.004 | 0 | 0.4 | 0.44 | 0.46 | 0.65 | 0.34 | 0.51 |
| 5 | 178842930 | rs11743636 | 1.338 | 0.767 | -1.323 | -0.541 | -0.047 | -0.043 | -0.043 | 0.038 | 0.07 | 0 | 0.13 | 0.15 | 0.17 | 0.1 |
| 5 | 178846679 | rs7718190 | -0.072 | -0.194 | -1.246 | -1.033 | -0.027 | 0.001 | 0.001 | 0.174 | 0.35 | 0.47 | 0.42 | 0.59 | 0.29 | 0.51 |
| 5 | 178849450 | rs1317022 | -0.181 | -0.25 | -1.34 | -1.191 | -0.008 | -0.006 | -0.006 | 0 | 0.42 | 0.48 | 0.48 | 0.59 | 0.36 | 0.54 |
| 5 | 178850328 | rs7735477 | 1.144 | -0.198 | -1.478 | -1.283 | 0.006 | -0.005 | -0.005 | 0 | 0.24 | 0.27 | 0.2 | 0.18 | 0.27 | 0.17 |
| 5 | 178851394 | rs2411390 | 0.017 | 0.187 | -1.477 | -0.932 | -0.008 | 0.061 | 0.061 | 0 | 0.59 | 0.73 | 0.62 | 0.78 | 0.56 | 0.68 |
| 5 | 178856589 | rs2009877 | 1.357 | 0.364 | -1.902 | -0.772 | -0.097 | 0.006 | 0.006 | 0.461 | 0.18 | 0.01 | 0.15 | 0.16 | 0.27 | 0.15 |
| 5 | 178858393 | rs11249626 | 0.144 | 0.01 | -1.452 | -0.481 | 0 | -0.011 | -0.011 | 0 | 0.47 | 0.48 | 0.5 | 0.69 | 0.4 | 0.53 |
| 5 | 178864168 | rs4701113 | 0.145 | 0.547 | -1.789 | 0.114 | -0.033 | 0.061 | 0.061 | 0.062 | 0.46 | 0.17 | 0.53 | 0.64 | 0.54 | 0.6 |
| 5 | 178884087 | rs6601036 | 0.587 | 0.439 | -0.883 | 0.983 | 0.003 | -0.008 | -0.008 | 1.064 | 0.82 | 0.88 | 0.59 | 0.46 | 0.88 | 0.66 |
| 5 | 178886655 | rs4701151 | -1.557 | 0.505 | -0.22 | 0.963 | -0.003 | -0.015 | -0.015 | 0 | 0.33 | 0.49 | 0.59 | 0.67 | 0.31 | 0.51 |
| 5 | 178887382 | rs11743487 | 0.271 | 0.515 | -0.585 | 0.863 | 0.002 | -0.001 | -0.001 | 0 | 0.83 | 0.9 | 0.59 | 0.46 | 0.88 | 0.65 |
| 5 | 178894709 | rs2411385 | 1.324 | 0.451 | -0.225 | 0.738 | -0.003 | 0.005 | 0.005 | 0 | 0.69 | 0.62 | 0.43 | 0.36 | 0.69 | 0.5 |
| 5 | 178897465 | rs4701169 | 0.341 | 0.52 | -0.724 | 0.626 | -0.001 | -0.003 | -0.003 | 0.789 | 0.88 | 0.88 | 0.92 | 0.83 | 0.87 | 0.9 |
| 5 | 178897590 | rs4701170 | -0.407 | 0.494 | -0.527 | 1.076 | -0.01 | -0.031 | -0.031 | 0 | 0.16 | 0.06 | 0.47 | 0.49 | 0.16 | 0.34 |
| 5 | 178899320 | rs10464087 | -0.29 | 0.17 | -0.554 | 0.891 | 0.011 | -0.021 | -0.021 | 0 | 0.14 | 0.06 | 0.42 | 0.53 | 0.09 | 0.32 |
| 5 | 178899878 | rs10464090 | 0.183 | 0.163 | -0.545 | 0.866 | 0.008 | -0.006 | -0.006 | 4.801 | 0.85 | 0.93 | 0.59 | 0.47 | 0.91 | 0.68 |
| 5 | 178901458 | rs12516534 | 0.695 | -0.204 | -0.66 | 0.69 | 0.002 | -0.001 | -0.001 | 0 | 0.79 | 0.84 | 0.56 | 0.45 | 0.87 | 0.62 |
| 5 | 178903149 | rs7726821 | 1.116 | 0.275 | -0.438 | 0.202 | -0.001 | 0.001 | 0.001 | 0 | 0.66 | 0.62 | 0.39 | 0.36 | 0.64 | 0.52 |
| 5 | 178904395 | rs6897650 | 0.109 | 0.543 | -0.472 | 0.193 | -0.026 | 0.016 | 0.016 | 0 | 0.83 | 0.9 | 0.85 | 0.9 | 0.76 | 0.83 |
| 5 | 178912074 | rs6601042 | -0.642 | 0.318 | -0.306 | 0.441 | 0.002 | 0.016 | 0.016 | 0 | 0.09 | 0.11 | 0.06 | 0.17 | 0.1 | 0.08 |
| 5 | 178912694 | rs17616436 | 0.341 | 0.138 | -0.99 | -0.434 | -0.026 | 0.087 | 0.087 | 0.202 | 0.44 | 0.18 | 0.29 | 0.27 | 0.49 | 0.47 |
| 5 | 178918122 | rs11748822 | -0.329 | 0.251 | -0.207 | -0.061 | -0.002 | 0.083 | 0.083 | 0 | 0.71 | 0.55 | 0.56 | 0.52 | 0.69 | 0.68 |
| 5 | 178921861 | rs11249631 | 0.075 | 0.138 | -0.179 | -0.162 | -0.003 | -0.009 | -0.009 | 0 | 0.62 | 0.53 | 0.51 | 0.35 | 0.6 | 0.58 |
| 5 | 178921964 | rs11249632 | 0.339 | 0.459 | -0.064 | 0.159 | 0.028 | 0.032 | 0.032 | 0 | 0.1 | 0.15 | 0.07 | 0.17 | 0.1 | 0.06 |
| 5 | 178931310 | rs11249638 | 0.931 | 0.241 | 0.076 | -0.069 | 0.012 | 0.042 | 0.042 | 0 | 0.78 | 0.68 | 0.6 | 0.35 | 0.8 | 0.72 |
| 5 | 178935665 | rs6864599 | -0.024 | -0.739 | -0.025 | -0.755 | -0.031 | -0.076 | -0.076 | 0.019 | 0.2 | 0.32 | 0.08 | 0.02 | 0.11 | 0.16 |
| 5 | 178936722 | rs9329079 | -0.991 | -0.72 | 0.038 | -0.666 | -0.021 | 0.052 | 0.052 | 0 | 0.23 | 0.43 | 0.52 | 0.62 | 0.17 | 0.39 |
| 5 | 178944407 | rs936817 | -0.081 | -0.317 | -0.11 | -1.701 | 0.001 | -0.057 | -0.057 | 0 | 0.42 | 0.39 | 0.31 | 0.07 | 0.39 | 0.44 |
| 5 | 178947661 | rs6862387 | 0.737 | -0.026 | -0.245 | -2.211 | -0.007 | 0.005 | 0.005 | 0.032 | 0.32 | 0.28 | 0.3 | 0.25 | 0.42 | 0.25 |
| 5 | 178951106 | rs4701187 | -0.456 | 0.02 | -0.338 | -1.768 | 0 | 0.054 | 0.054 | 0 | 0.94 | 0.9 | 0.73 | 0.34 | 0.95 | 0.82 |
| 5 | 178961710 | rs6876210 | 1.184 | -0.313 | -0.657 | -2.277 | 0.011 | 0.103 | 0.103 | 0 | 0.24 | 0.06 | 0.1 | 0.01 | 0.18 | 0.21 |
| 5 | 178974997 | rs7733389 | -0.04 | -0.468 | -0.152 | -2.373 | 0.008 | 0.022 | 0.022 | 0.148 | 0.47 | 0.41 | 0.52 | 0.7 | 0.38 | 0.54 |
| 5 | 178976966 | rs10074573 | -0.141 | -0.495 | 0.014 | -2.376 | -0.004 | -0.013 | -0.013 | 0.077 | 0.56 | 0.55 | 0.51 | 0.28 | 0.64 | 0.47 |
| 5 | 178982446 | rs4701131 | 0.033 | -0.436 | 0.007 | -2.381 | 0.01 | 0.035 | 0.035 | 1.141 | 0.45 | 0.36 | 0.49 | 0.72 | 0.37 | 0.53 |
| 5 | 178986436 | rs7702303 | 0.13 | -0.561 | 0.088 | -2.381 | 0.009 | 0.023 | 0.023 | 0.883 | 0.43 | 0.38 | 0.47 | 0.71 | 0.35 | 0.52 |
| 5 | 178990305 | rs7706275 | 0.557 | -0.585 | 0.068 | -2.372 | 0.004 | 0.001 | 0.001 | 0 | 0.33 | 0.26 | 0.42 | 0.65 | 0.26 | 0.41 |
| 5 | 178991794 | rs4700810 | -0.04 | -0.503 | -0.147 | -2.579 | 0.021 | 0.048 | 0.048 | 0 | 0.46 | 0.37 | 0.51 | 0.73 | 0.37 | 0.47 |
| 5 | 178997692 | rs4701133 | 1.262 | -0.16 | -0.132 | -2.801 | 0.039 | -0.023 | -0.023 | 3.71 | 0.27 | 0.36 | 0.29 | 0.18 | 0.38 | 0.18 |
| 5 | 179000230 | rs10076298 | 1.904 | -0.624 | -0.412 | -2.86 | 0.026 | 0.077 | 0.077 | 0 | 0.23 | 0.06 | 0.08 | 0.02 | 0.15 | 0.18 |
| 5 | 179020622 | rs4701135 | 0.98 | -0.507 | -0.537 | -2.751 | 0.004 | 0.02 | 0.02 | 0.585 | 0.12 | 0.04 | 0.08 | 0.09 | 0.11 | 0.07 |
| 5 | 179025550 | rs4701136 | -0.353 | -0.502 | -0.429 | -2.812 | 0.002 | -0.007 | -0.007 | 0 | 0.73 | 0.77 | 0.8 | 0.91 | 0.74 | 0.65 |
| 5 | 179026192 | rs4701137 | -0.293 | -0.509 | -0.314 | -2.797 | -0.003 | -0.038 | -0.038 | 0.11 | 0.73 | 0.7 | 0.86 | 0.91 | 0.74 | 0.66 |
| 5 | 179034069 | rs4700812 | -0.576 | -0.676 | -0.124 | -2.908 | 0.006 | -0.053 | -0.053 | 0.436 | 0.58 | 0.45 | 0.72 | 0.94 | 0.69 | 0.58 |
| 5 | 179034260 | rs4701140 | -0.405 | -0.249 | 0.126 | -2.698 | -0.01 | 0.011 | 0.011 | 2.097 | 0.62 | 0.66 | 0.67 | 0.78 | 0.52 | 0.72 |
| 5 | 179035183 | rs35654688 | 1.88 | -0.311 | -0.077 | -2.453 | 0.003 | 0.024 | 0.024 | 0 | 0.1 | 0.06 | 0.05 | 0.04 | 0.09 | 0.11 |
| 5 | 179040113 | rs12518041 | 0.963 | -0.162 | -0.454 | -2.443 | 0.017 | -0.012 | -0.012 | 0 | 0.28 | 0.31 | 0.31 | 0.17 | 0.43 | 0.19 |
| 5 | 179044292 | rs4131080 | 0.657 | -0.193 | -0.719 | -2.467 | 0.032 | 0.042 | 0.042 | 0.308 | 0.4 | 0.27 | 0.44 | 0.69 | 0.3 | 0.51 |
| 5 | 179045142 | rs4701143 | 1.375 | -0.203 | -0.713 | -2.466 | 0.034 | -0.008 | -0.008 | 0 | 0.3 | 0.14 | 0.38 | 0.65 | 0.2 | 0.4 |

| | | | | | | | | | | | | | | | | |
|---|---|---|---|---|---|---|---|---|---|---|---|---|---|---|---|---|
| 5 | 179048449 | rs4535419 | 0.781 | 0.041 | -0.535 | -2.326 | 0.017 | -0.012 | -0.012 | 2.078 | 0.29 | 0.31 | 0.31 | 0.17 | 0.43 | 0.18 |
| 5 | 179049329 | rs4563584 | 0.985 | 0.03 | -1.278 | -2.107 | 0.031 | 0.018 | 0.018 | 0.003 | 0.35 | 0.12 | 0.48 | 0.69 | 0.25 | 0.58 |
| 5 | 179050188 | rs1136267 | 0.734 | 0.343 | -0.9 | -1.892 | 0.04 | 0.078 | 0.078 | 0.617 | 0.46 | 0.29 | 0.54 | 0.73 | 0.35 | 0.69 |
| 5 | 179051538 | rs4078141 | 0.734 | 0.343 | -0.9 | -1.892 | 0.04 | 0.078 | 0.078 | 0.617 | 0.46 | 0.29 | 0.54 | 0.73 | 0.35 | 0.69 |
| 5 | 179060995 | rs13162707 | 0.793 | 0.305 | -1.503 | -1.823 | 0.017 | 0.12 | 0.12 | 1.181 | 0.46 | 0.16 | 0.56 | 0.73 | 0.36 | 0.69 |
| 5 | 179066006 | rs6875168 | 0.275 | 0.858 | -1.4 | -1.177 | -0.019 | -0.033 | -0.033 | 0 | 0.13 | 0.02 | 0.06 | 0.12 | 0.14 | 0.07 |
| 5 | 179078068 | rs28564876 | -0.073 | 1.241 | -0.754 | -0.975 | 0.002 | -0.001 | -0.001 | 0.84 | 0.26 | 0.24 | 0.27 | 0.18 | 0.41 | 0.19 |
| 5 | 179093431 | rs7718486 | -0.151 | 0.445 | -0.32 | -0.656 | -0.004 | 0 | 0 | 0.714 | 0.68 | 0.63 | 0.48 | 0.29 | 0.72 | 0.48 |
| 5 | 179094108 | rs4331859 | -0.803 | 0.568 | 0.27 | -0.279 | 0.003 | -0.03 | -0.03 | 0 | 0.16 | 0.23 | 0.16 | 0.09 | 0.16 | 0.14 |
| 5 | 179108502 | rs12515399 | 0.268 | 0.361 | -0.232 | 0.392 | -0.001 | -0.002 | -0.002 | 0.088 | 0.43 | 0.5 | 0.45 | 0.46 | 0.39 | 0.6 |
| 5 | 179121263 | rs7700649 | 0.479 | -0.561 | 0.059 | 0.51 | -0.028 | 0.009 | 0.009 | 0 | 0.73 | 0.36 | 0.8 | 0.74 | 0.82 | 0.82 |
| 5 | 179131143 | rs12374446 | 1.145 | -0.419 | 0.28 | 0.19 | -0.012 | -0.012 | -0.012 | 0 | 0.48 | 0.17 | 0.56 | 0.62 | 0.51 | 0.66 |
| 5 | 179135815 | rs6893300 | 0.203 | -0.558 | 0.401 | 0.124 | -0.023 | 0.01 | 0.01 | 0 | 0.23 | 0.12 | 0.23 | 0.15 | 0.32 | 0.16 |
| 5 | 179153797 | rs7734102 | -0.276 | -0.593 | 0.554 | 0.1 | -0.016 | 0.016 | 0.016 | 0.168 | 0.27 | 0.18 | 0.26 | 0.16 | 0.35 | 0.18 |
| 5 | 179154998 | rs11738720 | -0.287 | -0.597 | 0.552 | 0.098 | -0.016 | 0.017 | 0.017 | 0.144 | 0.27 | 0.18 | 0.26 | 0.16 | 0.35 | 0.18 |
| 5 | 179156449 | rs7566 | 0.168 | -0.665 | 1.188 | 0.089 | -0.049 | 0.026 | 0.026 | 0 | 0.77 | 0.56 | 0.83 | 0.77 | 0.86 | 0.83 |
| 5 | 179156782 | rs6639 | -0.232 | -0.626 | 1.111 | 0.08 | -0.014 | 0.015 | 0.015 | 0.147 | 0.26 | 0.18 | 0.25 | 0.16 | 0.34 | 0.18 |
| 5 | 179159781 | rs7723450 | -0.012 | -0.65 | 1.202 | 0.049 | 0.02 | -0.029 | -0.029 | 0 | 0.19 | 0.14 | 0.15 | 0.23 | 0.14 | 0.17 |
| 5 | 179166974 | rs10516146 | 1.415 | -0.617 | 1.279 | 0.074 | -0.004 | 0.004 | 0.004 | 2.269 | 0.49 | 0.22 | 0.56 | 0.62 | 0.5 | 0.65 |
| 5 | 179168097 | rs17617068 | 0.266 | -0.612 | 1.369 | 0.073 | -0.071 | 0.005 | 0.005 | 0 | 0.23 | 0.07 | 0.23 | 0.15 | 0.32 | 0.17 |
| 5 | 179177271 | rs6601073 | 1.374 | -0.657 | 1.394 | 0.057 | -0.004 | 0.007 | 0.007 | 2.269 | 0.49 | 0.22 | 0.56 | 0.62 | 0.5 | 0.64 |
| 5 | 179186197 | rs6865507 | -0.077 | -0.683 | 1.385 | -0.019 | -0.001 | -0.025 | -0.025 | 0 | 0.05 | 0.1 | 0.03 | 0 | 0.03 | 0.02 |
| 5 | 179201847 | rs6895902 | -0.491 | -0.635 | 0.969 | -0.015 | -0.011 | 0.018 | 0.018 | 0.017 | 0.29 | 0.2 | 0.26 | 0.15 | 0.36 | 0.2 |
| 5 | 179202688 | rs3797762 | -0.508 | -0.635 | 0.969 | -0.015 | -0.011 | 0.018 | 0.018 | 0.013 | 0.29 | 0.2 | 0.26 | 0.15 | 0.36 | 0.2 |
| 5 | 179207547 | rs1019443 | 1.358 | -0.692 | 0.284 | 0.1 | 0.006 | -0.01 | -0.01 | 0 | 0.51 | 0.22 | 0.59 | 0.61 | 0.54 | 0.65 |
| 5 | 179214818 | rs12522926 | 1.468 | -0.603 | 0.263 | 0.142 | -0.018 | -0.028 | -0.028 | 0 | 0.48 | 0.19 | 0.56 | 0.6 | 0.51 | 0.65 |
| 5 | 179220638 | rs730012 | 0.063 | -0.374 | 0.269 | 0.202 | -0.099 | 0.004 | 0.004 | 0 | 0.23 | 0.04 | 0.22 | 0.15 | 0.31 | 0.18 |
| 5 | 179227286 | rs166624 | -0.789 | -0.653 | 0.57 | 0.332 | -0.038 | 0.013 | 0.013 | 0.232 | 0.24 | 0.41 | 0.18 | 0.23 | 0.16 | 0.13 |
| 5 | 179231248 | rs12652934 | 1.155 | -0.429 | -0.42 | 0.366 | 0.007 | -0.018 | -0.018 | 0.093 | 0.49 | 0.17 | 0.57 | 0.6 | 0.52 | 0.66 |
| 5 | 179240910 | rs10516140 | 0.673 | -0.723 | 0.117 | 0.631 | -0.07 | 0.04 | 0.04 | 0 | 0.75 | 0.52 | 0.83 | 0.75 | 0.85 | 0.85 |
| 5 | 179248774 | rs502729 | -0.051 | -0.057 | -1.159 | -0.143 | 0.015 | 0.125 | 0.125 | 1.196 | 0.45 | 0.22 | 0.6 | 0.8 | 0.42 | 0.55 |
| 5 | 179249073 | rs513165 | -0.5 | -0.932 | -1.922 | 0.15 | 0.061 | -0.07 | -0.07 | 0 | 0.34 | 0.81 | 0.52 | 0.53 | 0.25 | 0.5 |
| 5 | 179250130 | rs7711505 | 0.016 | -0.9 | -1.73 | -0.202 | 0.006 | 0.012 | 0.012 | 0 | 0.1 | 0.08 | 0.05 | 0 | 0.05 | 0.01 |
| 5 | 179254247 | rs10056257 | -0.321 | -0.55 | -1.202 | -0.667 | -0.004 | -0.026 | -0.026 | 0 | 0.45 | 0.48 | 0.65 | 0.77 | 0.45 | 0.6 |
| 5 | 179260009 | rs2241349 | 0.018 | -0.742 | -1.259 | 0.048 | -0.009 | -0.019 | -0.019 | 0 | 0.23 | 0.21 | 0.46 | 0.61 | 0.3 | 0.51 |
| 5 | 179260153 | rs4935 | -1.534 | -1.174 | -0.78 | -0.54 | -0.022 | 0.058 | 0.058 | 0 | 0.6 | 0.73 | 0.73 | 0.84 | 0.51 | 0.65 |
| 5 | 179260528 | rs155788 | -0.455 | -1.416 | -0.114 | -0.16 | 0.013 | -0.015 | -0.015 | 0 | 0.16 | 0.46 | 0.16 | 0.11 | 0.09 | 0.04 |
| 5 | 179264731 | rs10277 | -1.672 | -1.18 | -0.124 | -0.326 | -0.026 | 0.059 | 0.059 | 0 | 0.6 | 0.78 | 0.73 | 0.76 | 0.52 | 0.63 |
| 5 | 179264915 | rs1065154 | -1.626 | -1.159 | -0.668 | -0.319 | -0.093 | 0.126 | 0.126 | 0 | 0.6 | 0.92 | 0.74 | 0.76 | 0.51 | 0.62 |
| 5 | 179267783 | rs248247 | 0.431 | -1.117 | -0.689 | 0.443 | -0.024 | -0.05 | -0.05 | 0.097 | 0.25 | 0.16 | 0.23 | 0.47 | 0.33 | 0.42 |
| 5 | 179267949 | rs248248 | -2.026 | -0.453 | -1.868 | -0.643 | -0.062 | 0.071 | 0.071 | 0 | 0.76 | 0.95 | 0.85 | 0.9 | 0.71 | 0.79 |
| 5 | 179273820 | rs748197 | 0.267 | -0.528 | -1.368 | -0.252 | -0.009 | 0.078 | 0.078 | 0 | 0.36 | 0.17 | 0.61 | 0.76 | 0.4 | 0.59 |
| 5 | 179285752 | rs10060182 | 0.241 | -0.764 | -1.372 | -0.235 | 0.051 | 0.073 | 0.073 | 0 | 0.35 | 0.17 | 0.57 | 0.76 | 0.3 | 0.57 |
| 5 | 179290845 | rs30386 | -0.568 | 0.214 | -0.829 | -0.471 | 0.003 | 0.024 | 0.024 | 0.919 | 0.54 | 0.43 | 0.67 | 0.77 | 0.45 | 0.64 |
| 5 | 179295060 | rs30384 | 0.707 | -0.538 | -1.019 | -0.188 | 0.035 | 0.011 | 0.011 | 0.011 | 0.22 | 0.12 | 0.15 | 0.14 | 0.13 | 0.16 |
| 5 | 179297262 | rs407437 | 1.382 | -0.499 | 0.093 | -0.299 | 0.033 | 0.029 | 0.029 | 6.726 | 0.06 | 0.44 | 0.04 | 0 | 0.01 | 0 |
| 5 | 179313656 | rs88183 | -0.546 | -0.407 | -1.033 | -0.436 | 0.045 | 0.109 | 0.109 | 0.117 | 0.52 | 0.28 | 0.63 | 0.77 | 0.41 | 0.59 |
| 5 | 179327513 | rs32458 | -0.074 | -0.366 | -1.882 | -0.195 | 0.007 | -0.026 | -0.026 | 0 | 0.63 | 0.29 | 0.4 | 0.46 | 0.67 | 0.53 |
| 5 | 179327827 | rs32461 | -0.97 | -0.116 | -2.773 | -0.844 | -0.178 | -0.036 | -0.036 | 0 | 0.76 | 0.97 | 0.77 | 0.78 | 0.56 | 0.74 |
| 5 | 179328228 | rs32462 | -0.084 | -1.088 | -2.341 | -0.449 | 0.076 | 0.065 | 0.065 | 0.677 | 0.35 | 0.81 | 0.18 | 0.15 | 0.15 | 0.2 |
| 5 | 179333740 | rs155796 | -0.219 | -0.263 | -2.33 | -0.416 | 0.054 | 0.12 | 0.12 | 0.167 | 0.51 | 0.26 | 0.63 | 0.77 | 0.4 | 0.59 |
| 5 | 179345996 | rs26074 | -0.163 | -0.11 | -2.782 | -0.358 | 0.068 | 0.147 | 0.147 | 0.186 | 0.5 | 0.2 | 0.63 | 0.75 | 0.4 | 0.6 |
| 5 | 179362813 | rs17079808 | 1.205 | -0.128 | -2.789 | -0.204 | 0.009 | 0.007 | 0.007 | 0 | 0.18 | 0.11 | 0.19 | 0.11 | 0.1 | 0.03 |
| 5 | 179362964 | rs518075 | -0.476 | -0.162 | -2.94 | 0.002 | 0.048 | 0.054 | 0.054 | 0.18 | 0.5 | 0.23 | 0.63 | 0.62 | 0.41 | 0.57 |
| 5 | 179365949 | rs13164817 | 1.611 | 0.234 | -3.236 | -0.063 | -0.024 | 0.02 | 0.02 | 0 | 0.11 | 0.03 | 0.07 | 0.07 | 0.16 | 0.12 |
| 5 | 179368372 | rs17616317 | 1.8 | 0.103 | -3.259 | -0.202 | 0.01 | 0.02 | 0.02 | 0 | 0.1 | 0.03 | 0.06 | 0.05 | 0.08 | 0.12 |
| 5 | 179368681 | rs42414 | -0.883 | 0.312 | -3.329 | -0.163 | -0.078 | 0.012 | 0.012 | 1.848 | 0.67 | 0.84 | 0.73 | 0.72 | 0.57 | 0.62 |
| 5 | 179369288 | rs10491478 | 0.356 | 0.658 | -2.533 | 0.696 | -0.016 | -0.095 | -0.095 | 0.045 | 0.23 | 0.08 | 0.41 | 0.48 | 0.25 | 0.4 |
| 5 | 179371900 | rs248321 | -0.703 | 1.411 | -2.174 | 0.808 | -0.019 | -0.016 | -0.016 | 0.133 | 0.85 | 0.87 | 0.85 | 0.79 | 0.7 | 0.82 |
| 5 | 179377650 | rs248327 | -0.622 | 1.235 | -2.232 | 0.14 | -0.044 | -0.05 | -0.05 | 0 | 0.82 | 0.87 | 0.58 | 0.38 | 0.65 | 0.57 |
| 5 | 179391811 | rs2162760 | 0.335 | 0.238 | -2.008 | 0.181 | -0.009 | -0.048 | -0.048 | 0.195 | 0.09 | 0 | 0.05 | 0.16 | 0.07 | 0.1 |
| 5 | 179396502 | rs248221 | -0.409 | 0.377 | -2.218 | -0.423 | 0.016 | 0.056 | 0.056 | 0 | 0.27 | 0.11 | 0.25 | 0.11 | 0.2 | 0.08 |
| 5 | 179400847 | rs3756616 | -0.16 | 0.536 | -2.199 | -0.665 | 0.02 | 0.176 | 0.176 | 0 | 0.47 | 0.84 | 0.39 | 0.15 | 0.36 | 0.31 |
| 5 | 179401333 | rs17079868 | 0.207 | 0.298 | -1.992 | -0.375 | 0.008 | -0.005 | -0.005 | 0 | 0.07 | 0.04 | 0.11 | 0.09 | 0.03 | 0.02 |
| 5 | 179401490 | rs11739898 | 0.115 | 0.842 | -1.656 | -0.572 | -0.001 | 0 | 0 | 0.051 | 0.16 | 0.16 | 0.17 | 0.15 | 0.17 | 0.03 |
| 5 | 179404375 | rs27017 | -0.75 | 0.834 | -2.506 | -0.575 | -0.034 | -0.064 | -0.064 | 0.113 | 0.56 | 0.91 | 0.55 | 0.46 | 0.56 | 0.33 |
| 5 | 179407229 | rs28011 | -1.239 | 0.6 | -3.069 | -0.882 | -0.047 | -0.137 | -0.137 | 0 | 0.78 | 0.96 | 0.68 | 0.52 | 0.73 | 0.58 |
| 5 | 179430530 | rs248318 | 1.031 | 0.623 | -3.399 | -1.223 | -0.012 | -0.065 | -0.065 | 0.848 | 0.16 | 0.04 | 0.12 | 0.29 | 0.15 | 0.17 |
| 5 | 179436278 | rs6873317 | 1.376 | 0.658 | -3.336 | -1.412 | -0.001 | 0.001 | 0.001 | 0.408 | 0.18 | 0.17 | 0.19 | 0.15 | 0.2 | 0.05 |

| | | | | | | | | | | | | | | | | |
|---|---|---|---|---|---|---|---|---|---|---|---|---|---|---|---|---|
| 5 | 179454428 | rs10036834 | -0.439 | 0.586 | -3.441 | -1.35 | -0.007 | 0.01 | 0.01 | 2.51 | 0.69 | 0.79 | 0.72 | 0.73 | 0.69 | 0.83 |
| 5 | 179454508 | rs10045947 | 0.862 | 0.45 | -2.875 | -1.522 | -0.007 | -0.002 | -0.002 | 0 | 0.19 | 0.14 | 0.2 | 0.19 | 0.21 | 0.14 |
| 5 | 179454656 | rs13166286 | 1.718 | 0.426 | -2.867 | -1.575 | -0.013 | -0.004 | -0.004 | 0.047 | 0.12 | 0.04 | 0.09 | 0.12 | 0.11 | 0.12 |
| 5 | 179457898 | rs10479453 | 0.068 | 0.463 | -2.782 | -2.041 | 0.037 | 0.107 | 0.107 | 0 | 0.52 | 0.72 | 0.5 | 0.73 | 0.41 | 0.72 |
| 5 | 179470930 | rs10061480 | 0.371 | 0.42 | -1.246 | -1.603 | -0.003 | 0.002 | 0.002 | 1.187 | 0.6 | 0.44 | 0.64 | 0.58 | 0.62 | 0.76 |
| 5 | 179484838 | rs7734329 | -0.735 | 0.275 | -1.138 | -1.51 | -0.002 | -0.048 | -0.048 | 0 | 0.77 | 0.48 | 0.76 | 0.85 | 0.77 | 0.93 |
| 5 | 179495532 | rs1348523 | 0.71 | 0.296 | -1.167 | -1.52 | 0 | -0.009 | -0.009 | 0.01 | 0.18 | 0.17 | 0.19 | 0.06 | 0.19 | 0.1 |
| 5 | 179497249 | rs4700858 | 0.172 | 0.387 | -1.166 | -1.302 | -0.011 | 0.008 | 0.008 | 0.593 | 0.64 | 0.47 | 0.71 | 0.62 | 0.66 | 0.75 |
| 5 | 179500722 | rs4700862 | 0.155 | 0.434 | -0.683 | -1.153 | -0.002 | 0.011 | 0.011 | 0.317 | 0.63 | 0.42 | 0.69 | 0.6 | 0.64 | 0.78 |
| 5 | 179503416 | rs6898203 | -1.123 | 0.369 | -0.903 | -0.984 | -0.023 | -0.007 | -0.007 | 0 | 0.83 | 0.91 | 0.8 | 0.83 | 0.77 | 0.97 |
| 5 | 179510642 | rs33916112 | 0.489 | 0.174 | -1.413 | -0.988 | 0.015 | 0.09 | 0.09 | 0 | 0.22 | 0.08 | 0.12 | 0.06 | 0.17 | 0.25 |
| 5 | 179524425 | rs7728341 | 0.293 | 0.349 | -1.402 | -0.708 | -0.023 | -0.041 | -0.041 | 0 | 0.48 | 0.11 | 0.51 | 0.56 | 0.47 | 0.63 |
| 5 | 179527895 | rs2278662 | -1.082 | 0.469 | -1.204 | -0.451 | -0.034 | -0.008 | -0.008 | 0 | 0.84 | 0.92 | 0.8 | 0.84 | 0.77 | 0.96 |
| 5 | 179529124 | rs7725201 | -0.77 | 0.43 | -0.861 | -0.431 | 0.001 | 0.001 | 0.001 | 0 | 0.82 | 0.81 | 0.79 | 0.8 | 0.77 | 0.96 |
| 5 | 179529895 | rs4700708 | 0.685 | 0.249 | -1.38 | -0.21 | -0.027 | -0.137 | -0.137 | 2.158 | 0.35 | 0.06 | 0.41 | 0.55 | 0.35 | 0.47 |
| 5 | 179532944 | rs6894296 | -1.221 | 0.5 | -0.715 | -0.004 | -0.027 | -0.038 | -0.038 | 0.515 | 0.84 | 0.91 | 0.82 | 0.8 | 0.79 | 0.96 |
| 5 | 179537115 | rs4282280 | -0.026 | 0.337 | -0.096 | -0.063 | -0.002 | 0.008 | 0.008 | 0.631 | 0.53 | 0.39 | 0.39 | 0.31 | 0.49 | 0.63 |
| 5 | 179537727 | rs12655994 | 0.684 | 0.113 | 0.097 | -0.041 | -0.004 | 0.028 | 0.028 | 0 | 0.3 | 0.45 | 0.42 | 0.48 | 0.3 | 0.33 |
| 5 | 179543280 | rs17080045 | 0.597 | 0.157 | -0.026 | -0.021 | -0.003 | 0.034 | 0.034 | 0 | 0.3 | 0.43 | 0.4 | 0.48 | 0.3 | 0.32 |
| 5 | 179543546 | rs11741189 | 0.352 | 0.115 | -0.603 | -0.471 | 0.016 | 0.052 | 0.052 | 0 | 0.25 | 0.12 | 0.14 | 0.06 | 0.19 | 0.26 |
| 5 | 179547579 | rs4700874 | -0.288 | 0.513 | -0.369 | 0.033 | 0 | -0.008 | -0.008 | 10.347 | 0.6 | 0.42 | 0.65 | 0.77 | 0.6 | 0.73 |
| 5 | 179555654 | rs6895192 | 0.168 | 0.579 | -0.691 | 0.085 | 0 | -0.013 | -0.013 | 0 | 0.46 | 0.52 | 0.43 | 0.25 | 0.46 | 0.34 |
| 5 | 179556518 | rs1867455 | -0.289 | 1.25 | -1.196 | 0.288 | -0.004 | -0.024 | -0.024 | 0 | 0.7 | 0.82 | 0.6 | 0.53 | 0.67 | 0.7 |
| 5 | 179563435 | rs11546322 | -0.321 | 0.693 | -1.335 | 0.047 | 0.011 | 0.038 | 0.038 | 0 | 0.41 | 0.26 | 0.54 | 0.71 | 0.39 | 0.52 |
| 5 | 179564141 | rs4624755 | -0.364 | 0.707 | -1.047 | 0.719 | -0.017 | 0.022 | 0.022 | 0 | 0.55 | 0.68 | 0.48 | 0.52 | 0.52 | 0.56 |
| 5 | 179570737 | rs12516993 | -0.669 | 0.763 | -0.857 | 1.067 | 0 | 0 | 0 | 0 | 0.19 | 0.16 | 0.21 | 0.26 | 0.18 | 0.2 |
| 5 | 179584778 | rs4389652 | 0.067 | 0.987 | -1.616 | 0.358 | -0.071 | -0.007 | -0.007 | 1.287 | 0.21 | 0.03 | 0.13 | 0.09 | 0.23 | 0.24 |
| 5 | 179585309 | rs3940643 | 0.375 | 0.54 | -1.293 | 0.391 | -0.017 | 0.002 | 0.002 | 0 | 0.45 | 0.45 | 0.61 | 0.52 | 0.32 | 0.37 |
| 5 | 179585577 | rs6894856 | -0.783 | 1.505 | -1.176 | 0.426 | -0.008 | 0.003 | 0.003 | 0.362 | 0.83 | 0.89 | 0.9 | 0.88 | 0.75 | 0.88 |
| 5 | 179585792 | rs6897922 | 0.455 | 0.333 | -0.733 | -0.124 | -0.022 | 0.005 | 0.005 | 0.566 | 0.42 | 0.46 | 0.52 | 0.6 | 0.23 | 0.43 |
| 5 | 179586456 | rs7720463 | -0.771 | 0.33 | -0.674 | -0.245 | 0.023 | -0.056 | -0.056 | 0.763 | 0.69 | 0.79 | 0.53 | 0.45 | 0.78 | 0.7 |
| 5 | 179594023 | rs13356323 | -0.216 | 0.184 | 0.059 | -0.261 | 0.007 | -0.004 | -0.004 | 0 | 0.56 | 0.56 | 0.5 | 0.43 | 0.69 | 0.63 |
| 5 | 179597647 | rs1867459 | 0.591 | 0.404 | -0.375 | -0.321 | -0.042 | 0.09 | 0.09 | 0 | 0.29 | 0.11 | 0.26 | 0.07 | 0.3 | 0.26 |
| 5 | 179598578 | rs882702 | -0.354 | 0.265 | 0.034 | -0.243 | 0.001 | 0.006 | 0.006 | 0 | 0.49 | 0.6 | 0.45 | 0.34 | 0.55 | 0.58 |
| 5 | 179603337 | rs6601089 | 0.75 | 0.01 | 0.141 | 0.03 | -0.032 | -0.019 | -0.019 | 0.194 | 0.66 | 0.55 | 0.75 | 0.65 | 0.73 | 0.68 |
| 5 | 179604217 | rs1867457 | 0.18 | 0.175 | 0.278 | 0.062 | 0 | -0.004 | -0.004 | 0.322 | 0.13 | 0.15 | 0.1 | 0.12 | 0.13 | 0.13 |
| 5 | 179607388 | rs13153506 | 0.908 | 0.238 | -0.22 | 0.064 | -0.001 | 0 | 0 | 0 | 0.62 | 0.37 | 0.56 | 0.63 | 0.5 | 0.62 |
| 5 | 179608360 | rs11749727 | 1.45 | 0.29 | -0.829 | 0.132 | 0.006 | -0.056 | -0.056 | 0 | 0.46 | 0.1 | 0.42 | 0.41 | 0.38 | 0.47 |
| 5 | 179608656 | rs7380160 | -0.178 | 0.467 | -0.476 | 0.231 | 0.079 | 0.022 | 0.022 | 0 | 0.22 | 0.39 | 0.32 | 0.25 | 0.37 | 0.25 |
| 5 | 179611713 | rs4367258 | -0.141 | 0.338 | -1.189 | 0.796 | -0.002 | -0.001 | -0.001 | 0 | 0.8 | 0.82 | 0.91 | 0.77 | 0.88 | 0.82 |
| 5 | 179611897 | rs4367257 | -0.141 | 0.338 | -1.189 | 0.796 | -0.002 | -0.001 | -0.001 | 0 | 0.8 | 0.82 | 0.91 | 0.77 | 0.88 | 0.82 |
| 5 | 179613262 | rs13168609 | 1.074 | 0.58 | -1.374 | 0.774 | -0.011 | -0.032 | -0.032 | 0.292 | 0.48 | 0.1 | 0.44 | 0.4 | 0.43 | 0.48 |
| 5 | 179614645 | rs4700720 | 0.005 | 0.4 | -1.207 | 0.768 | -0.009 | 0 | 0 | 0.018 | 0.78 | 0.78 | 0.91 | 0.77 | 0.87 | 0.81 |
| 5 | 179615478 | rs6601095 | -0.721 | 1.52 | -0.637 | 0.81 | 0.004 | 0 | 0 | 0.079 | 0.3 | 0.68 | 0.47 | 0.37 | 0.43 | 0.32 |
| 5 | 179617799 | rs4400089 | 0.94 | 1.525 | -0.237 | 1.232 | 0.024 | 0.091 | 0.091 | 0 | 0.82 | 0.68 | 0.86 | 0.62 | 0.79 | 0.74 |
| 5 | 179619222 | rs7737438 | 0.241 | 1.31 | -0.636 | 0.348 | -0.003 | -0.031 | -0.031 | 0 | 0.65 | 0.39 | 0.61 | 0.91 | 0.66 | 0.76 |
| 5 | 179621485 | rs4700914 | 0.43 | 1.221 | -0.909 | 0.675 | 0.001 | 0.017 | 0.017 | 0.478 | 0.52 | 0.36 | 0.52 | 0.65 | 0.51 | 0.56 |
| 5 | 179623637 | rs7720553 | -0.093 | 1.073 | -1.259 | 0.166 | -0.02 | 0.059 | 0.059 | 0 | 0.12 | 0.02 | 0.13 | 0 | 0.13 | 0.11 |
| 5 | 179639885 | rs4273568 | 0.792 | 0.916 | -0.502 | 0.743 | 0.017 | 0.144 | 0.144 | 0.028 | 0.95 | 0.81 | 0.95 | 0.87 | 0.96 | 0.82 |
| 5 | 179641129 | rs1465982 | 0.659 | 0.615 | -0.667 | 0.382 | 0.061 | -0.002 | -0.002 | 0 | 0.45 | 0.28 | 0.45 | 0.56 | 0.31 | 0.35 |
| 5 | 179641662 | rs3111532 | 0.658 | 1.026 | -0.257 | -0.092 | 0.005 | 0 | 0 | 0 | 0.62 | 0.5 | 0.59 | 0.62 | 0.49 | 0.58 |
| 5 | 179643073 | rs6893212 | 1.837 | 0.551 | -0.634 | -0.184 | 0.027 | -0.036 | -0.036 | 0.128 | 0.36 | 0.26 | 0.4 | 0.45 | 0.29 | 0.23 |
| 5 | 179648989 | rs6882940 | -1.098 | 1.416 | -0.762 | -0.304 | 0.047 | 0.006 | 0.006 | 0 | 0.88 | 0.81 | 0.86 | 0.89 | 0.75 | 0.85 |
| 5 | 179649862 | rs3099065 | 1.012 | 1.458 | -0.753 | -0.374 | -0.021 | -0.024 | -0.024 | 0.185 | 0.11 | 0.04 | 0.08 | 0.22 | 0.15 | 0.08 |
| 5 | 179653428 | rs12656368 | 1.781 | 1.172 | -0.762 | -0.535 | 0.018 | 0 | 0 | 0.012 | 0.48 | 0.4 | 0.53 | 0.48 | 0.45 | 0.41 |
| 5 | 179653647 | rs3111521 | -0.419 | 1.205 | -0.758 | -0.652 | 0.017 | -0.012 | -0.012 | 0.27 | 0.24 | 0.32 | 0.26 | 0.23 | 0.25 | 0.18 |
| 5 | 179657422 | rs6601103 | 1.795 | 0.965 | -0.596 | -0.647 | -0.001 | 0.066 | 0.066 | 0 | 0.44 | 0.29 | 0.48 | 0.76 | 0.43 | 0.33 |
| 5 | 179662384 | rs1127575 | -1.36 | 0.975 | -0.345 | -0.723 | -0.004 | 0 | 0 | 0 | 0.42 | 0.55 | 0.45 | 0.15 | 0.49 | 0.5 |
| 5 | 179663251 | rs9605 | -0.626 | 1.105 | -0.357 | -0.749 | 0.025 | -0.036 | -0.036 | 0.26 | 0.29 | 0.52 | 0.29 | 0.15 | 0.4 | 0.41 |
| 5 | 179667806 | rs6894217 | 0.446 | 1.099 | -0.107 | -0.747 | 0.076 | -0.012 | -0.012 | 0.02 | 0.17 | 0.33 | 0.17 | 0.1 | 0.27 | 0.18 |
| 5 | 179670516 | rs4601008 | 1.144 | 1.093 | -0.077 | -0.751 | 0 | -0.004 | -0.004 | 0 | 0.11 | 0.13 | 0.12 | 0.05 | 0.13 | 0.23 |
| 5 | 179672825 | rs4362908 | -1.474 | 1.079 | -0.104 | -0.748 | -0.003 | -0.003 | -0.003 | 0 | 0.42 | 0.54 | 0.45 | 0.15 | 0.49 | 0.5 |
| 5 | 179676661 | rs11743623 | 0.531 | 1.013 | -0.097 | -0.703 | 0.003 | -0.053 | -0.053 | 0 | 0.13 | 0.32 | 0.16 | 0.03 | 0.1 | 0.11 |
| 5 | 179677517 | rs4147385 | 0.747 | 1.196 | -0.168 | -0.701 | -0.004 | -0.003 | -0.003 | 0 | 0.16 | 0.14 | 0.18 | 0.22 | 0.2 | 0.22 |
| 5 | 179679760 | rs6863088 | 2.757 | 0.859 | -0.286 | -0.719 | 0.022 | -0.017 | -0.017 | 0 | 0.36 | 0.27 | 0.4 | 0.55 | 0.29 | 0.26 |
| 5 | 179680233 | rs11249687 | -0.326 | 0.625 | 0.016 | -0.715 | 0.009 | 0.01 | 0.01 | 0 | 0.86 | 0.86 | 0.93 | 0.91 | 0.92 | 0.83 |
| 5 | 179681514 | rs11956696 | 0.93 | 0.634 | 0.05 | -0.726 | 0.042 | -0.028 | -0.028 | 0 | 0.11 | 0.35 | 0.14 | 0.03 | 0.13 | 0.16 |
| 5 | 179682832 | rs17627593 | -0.977 | 0.568 | 0.067 | -0.671 | 0.011 | -0.023 | -0.023 | 0 | 0.32 | 0.55 | 0.33 | 0.17 | 0.4 | 0.4 |
| 5 | 179687013 | rs17080136 | -0.264 | 0.599 | 0.016 | -0.642 | 0.061 | -0.051 | -0.051 | 0 | 0.25 | 0.37 | 0.27 | 0.11 | 0.39 | 0.31 |

| | | | | | | | | | | | | | | | | |
|---|---|---|---|---|---|---|---|---|---|---|---|---|---|---|---|---|
| 5 | 179687772 | rs6868333 | 1.339 | 0.694 | -0.362 | -0.513 | 0 | 0.072 | 0.072 | 0.189 | 0.54 | 0.3 | 0.6 | 0.74 | 0.52 | 0.42 |
| 5 | 179694680 | rs7713083 | 0.666 | 0.801 | -0.241 | -0.635 | 0.015 | -0.051 | -0.051 | 0 | 0.69 | 0.57 | 0.45 | 0.86 | 0.58 | 0.73 |
| 5 | 179695430 | rs4700730 | -0.427 | 0.037 | -0.968 | -0.25 | -0.008 | 0.046 | 0.046 | 0 | 0.34 | 0.25 | 0.29 | 0.23 | 0.38 | 0.28 |
| 5 | 179698554 | rs6601105 | -1.158 | -0.473 | -1.128 | -0.307 | -0.007 | 0.037 | 0.037 | 0.003 | 0.55 | 0.36 | 0.48 | 0.37 | 0.58 | 0.55 |
| 5 | 179700603 | rs2112593 | -0.218 | -0.799 | -0.651 | 0.427 | -0.021 | 0.005 | 0.005 | 0.018 | 0.3 | 0.64 | 0.28 | 0.36 | 0.16 | 0.26 |
| 5 | 179712060 | rs6867398 | 0.711 | 0.173 | -0.859 | 0.357 | 0.01 | -0.003 | -0.003 | 0 | 0.14 | 0.2 | 0.18 | 0.09 | 0.25 | 0.16 |
| 5 | 179721688 | rs3749791 | 0.613 | -0.556 | -0.797 | 0.501 | -0.002 | 0.003 | 0.003 | 0 | 0.42 | 0.56 | 0.52 | 0.62 | 0.4 | 0.48 |
| 5 | 179722533 | rs12520876 | -0.907 | -0.95 | -1.265 | 0.351 | 0.024 | -0.006 | -0.006 | 1.36 | 0.45 | 0.39 | 0.42 | 0.52 | 0.34 | 0.6 |
| 5 | 179727957 | rs7725 | 0.596 | -0.407 | -1.043 | 0.759 | -0.019 | -0.051 | -0.051 | 0 | 0.18 | 0.09 | 0.19 | 0.37 | 0.19 | 0.26 |
| 5 | 179728707 | rs4700931 | -0.66 | -0.326 | -1.172 | 0.596 | -0.04 | 0.002 | 0.002 | 0 | 0.38 | 0.26 | 0.41 | 0.65 | 0.48 | 0.48 |
| 5 | 179729519 | rs1035411 | -0.585 | -0.477 | -1.138 | 0.454 | 0.023 | 0.069 | 0.069 | 0 | 0.6 | 0.7 | 0.65 | 0.9 | 0.66 | 0.63 |
| 5 | 179729625 | rs2303005 | 0.494 | -0.098 | -1.518 | 0.646 | -0.006 | -0.018 | -0.018 | 0 | 0.2 | 0.19 | 0.29 | 0.47 | 0.23 | 0.29 |
| 5 | 179730478 | rs2892152 | 1.53 | -0.09 | -1.637 | 0.705 | -0.068 | -0.055 | -0.055 | 0.17 | 0.17 | 0.02 | 0.12 | 0.18 | 0.25 | 0.2 |
| 5 | 179731695 | rs729652 | 1.187 | -0.153 | -1.349 | 0.676 | -0.035 | -0.034 | -0.034 | 0 | 0.35 | 0.17 | 0.32 | 0.37 | 0.43 | 0.29 |
| 5 | 179736367 | rs2386854 | 0.531 | -0.148 | -1.394 | 0.743 | -0.026 | -0.072 | -0.072 | 0 | 0.19 | 0.1 | 0.28 | 0.47 | 0.23 | 0.29 |
| 5 | 179738237 | rs3805705 | 1.375 | -0.093 | -1.43 | 0.702 | -0.036 | -0.026 | -0.026 | 0 | 0.17 | 0.06 | 0.11 | 0.18 | 0.25 | 0.19 |
| 5 | 179738638 | rs11744256 | -0.779 | -0.921 | -0.888 | 0.742 | 0 | -0.001 | -0.001 | 0 | 0.31 | 0.34 | 0.34 | 0.38 | 0.28 | 0.42 |
| 5 | 179740827 | rs2303007 | 0.063 | -0.666 | -1.03 | 0.93 | -0.029 | -0.056 | -0.056 | 0.082 | 0.18 | 0.06 | 0.22 | 0.4 | 0.23 | 0.31 |
| 5 | 179741534 | rs3805703 | 0.921 | -0.581 | -0.988 | 0.855 | -0.006 | -0.008 | -0.008 | 2.011 | 0.3 | 0.33 | 0.32 | 0.19 | 0.24 | 0.26 |
| 5 | 179743731 | rs2303008 | -0.199 | -0.267 | -1.021 | 1.277 | -0.045 | -0.068 | -0.068 | 0 | 0.17 | 0.06 | 0.23 | 0.38 | 0.22 | 0.28 |
| 5 | 179745607 | rs2303009 | 0.332 | -0.011 | -1.076 | 1.372 | -0.017 | -0.015 | -0.015 | 0 | 0.09 | 0.04 | 0.09 | 0.37 | 0.13 | 0.21 |
| 5 | 179748631 | rs7707082 | -0.072 | -0.254 | -0.845 | 1.009 | -0.003 | -0.015 | -0.015 | 0 | 0.2 | 0.15 | 0.18 | 0.38 | 0.19 | 0.26 |
| 5 | 179751111 | rs3828686 | -1.09 | -1.392 | -0.487 | 0.044 | -0.007 | -0.006 | -0.006 | 0 | 0.35 | 0.61 | 0.3 | 0.16 | 0.19 | 0.41 |
| 5 | 179751441 | rs3749788 | 0.011 | -0.615 | -0.066 | 0.145 | 0.006 | -0.057 | -0.057 | 0 | 0.11 | 0.29 | 0.16 | 0.01 | 0.1 | 0.09 |
| 5 | 179761769 | rs6859159 | 0.781 | -0.168 | -0.805 | 0.496 | -0.01 | -0.017 | -0.017 | 2.541 | 0.13 | 0.07 | 0.11 | 0.19 | 0.17 | 0.13 |
| 5 | 179763018 | rs13179839 | 0.143 | -1.02 | -0.628 | -0.085 | -0.051 | -0.079 | -0.079 | 0 | 0.16 | 0.24 | 0.04 | 0 | 0.05 | 0.09 |
| 5 | 179763789 | rs2303011 | 1.575 | -0.592 | -1.582 | 0.094 | -0.047 | -0.043 | -0.043 | 0.143 | 0.18 | 0 | 0.19 | 0.17 | 0.18 | 0.11 |
| 5 | 179769964 | rs4432844 | -0.2 | -0.363 | -1.13 | 0.103 | -0.003 | -0.016 | -0.016 | 0 | 0.37 | 0.36 | 0.37 | 0.29 | 0.34 | 0.35 |
| 5 | 179781898 | rs1422572 | -1.027 | -0.556 | -0.806 | 0.438 | -0.017 | 0.003 | 0.003 | 0 | 0.33 | 0.38 | 0.25 | 0.36 | 0.28 | 0.32 |
| 5 | 179783770 | rs10041598 | -0.351 | -0.336 | -0.422 | 0.537 | 0.003 | 0.031 | 0.031 | 0 | 0.07 | 0.16 | 0.09 | 0.13 | 0.07 | 0.08 |
| 5 | 179789391 | rs12652232 | 0.673 | -0.283 | -0.966 | 0.793 | 0.011 | 0.074 | 0.074 | 0.07 | 0.08 | 0.11 | 0.06 | 0.23 | 0.11 | 0.12 |
| 5 | 179790552 | rs918481 | 1.106 | -0.056 | -1.175 | 0.479 | -0.053 | 0.107 | 0.107 | 0 | 0.26 | 0.08 | 0.34 | 0.1 | 0.33 | 0.2 |
| 5 | 179793256 | rs1862326 | -0.765 | -0.448 | -0.193 | 1.269 | -0.02 | 0.056 | 0.056 | 0.671 | 0.2 | 0.3 | 0.16 | 0.39 | 0.15 | 0.25 |
| 5 | 179794208 | rs10464059 | -0.793 | -0.435 | -0.21 | 1.064 | -0.022 | 0.044 | 0.044 | 0.904 | 0.21 | 0.32 | 0.13 | 0.32 | 0.15 | 0.26 |
| 5 | 179797036 | rs2386861 | 0.579 | -0.077 | -0.302 | 0.264 | 0.001 | 0.004 | 0.004 | 0 | 0.55 | 0.41 | 0.58 | 0.42 | 0.55 | 0.44 |
| 5 | 179802263 | rs10053999 | -1.227 | -0.14 | 0.635 | 0.423 | 0.01 | 0.038 | 0.038 | 0 | 0.19 | 0.55 | 0.14 | 0.17 | 0.2 | 0.28 |
| 5 | 179802420 | rs7712252 | 0.129 | -0.136 | 0.381 | 0.158 | -0.002 | 0.014 | 0.014 | 0 | 0.11 | 0.04 | 0.03 | 0 | 0.09 | 0.06 |
| 5 | 179804005 | rs7722078 | -0.978 | -0.204 | 0.781 | 0.155 | -0.017 | 0.023 | 0.023 | 0 | 0.15 | 0.26 | 0.07 | 0.15 | 0.1 | 0.12 |
| 5 | 179805370 | rs6601116 | -1.482 | -0.308 | 0.515 | 0.189 | 0.008 | 0.018 | 0.018 | 0 | 0.34 | 0.71 | 0.33 | 0.21 | 0.32 | 0.39 |
| 5 | 179806356 | rs7380895 | -1.065 | 0.013 | 0.165 | -0.093 | 0.003 | -0.053 | -0.053 | 0 | 0.29 | 0.36 | 0.29 | 0.07 | 0.3 | 0.32 |
| 5 | 179812850 | rs7715370 | 0.217 | -0.731 | 0.091 | 0.474 | -0.01 | -0.003 | -0.003 | 0.37 | 0.65 | 0.61 | 0.7 | 0.64 | 0.76 | 0.75 |
| 5 | 179818868 | rs7733582 | 0.587 | -0.772 | 0.325 | 0.598 | -0.012 | 0.101 | 0.101 | 0 | 0.81 | 0.52 | 0.8 | 0.69 | 0.85 | 0.83 |
| 5 | 179821445 | rs7726800 | -0.045 | -0.806 | 0.251 | 0.513 | -0.003 | -0.01 | -0.01 | 0 | 0.64 | 0.63 | 0.81 | 0.77 | 0.71 | 0.75 |
| 5 | 179823603 | rs10479572 | 0.637 | -0.833 | -0.01 | 0.491 | -0.012 | 0.021 | 0.021 | 0 | 0.75 | 0.43 | 0.77 | 0.7 | 0.75 | 0.77 |
| 5 | 179827609 | rs6601123 | 1.331 | -0.931 | -0.612 | 0.423 | -0.107 | -0.109 | -0.109 | 0 | 0.42 | 0.07 | 0.61 | 0.47 | 0.53 | 0.55 |
| 5 | 179828925 | rs9329110 | -0.321 | -0.927 | -0.306 | 0.147 | 0.006 | 0.024 | 0.024 | 0 | 0.36 | 0.33 | 0.19 | 0.23 | 0.34 | 0.28 |
| 5 | 179831373 | rs13168292 | -0.05 | -1.003 | -0.21 | 0.446 | 0.01 | 0.014 | 0.014 | 0 | 0.28 | 0.2 | 0.14 | 0.23 | 0.22 | 0.23 |
| 5 | 179837009 | rs10039851 | -0.747 | -0.898 | 0.428 | 0.747 | -0.023 | 0.022 | 0.022 | 0.01 | 0.18 | 0.31 | 0.21 | 0.22 | 0.13 | 0.13 |
| 5 | 179840538 | rs34108387 | -0.537 | -1.094 | -0.25 | 0.552 | 0.032 | 0.041 | 0.041 | 0 | 0.37 | 0.19 | 0.17 | 0.28 | 0.26 | 0.28 |
| 5 | 179850802 | rs4700952 | -1.246 | -1.117 | 0.498 | 0.961 | -0.034 | 0.028 | 0.028 | 0.608 | 0.26 | 0.58 | 0.24 | 0.38 | 0.16 | 0.17 |
| 5 | 179856266 | rs1828172 | 1.324 | -0.924 | 0.116 | 0.807 | 0.021 | 0.022 | 0.022 | 0 | 0.56 | 0.36 | 0.63 | 0.39 | 0.62 | 0.69 |
| 5 | 179862734 | rs11958726 | 0.559 | -0.885 | -0.119 | 0.776 | 0.012 | -0.001 | -0.001 | 3.691 | 0.14 | 0.09 | 0.06 | 0.14 | 0.1 | 0.06 |
| 5 | 179863366 | rs951921 | 1.107 | -0.362 | 0.023 | 0.575 | 0.027 | 0.015 | 0.015 | 0.011 | 0.53 | 0.28 | 0.7 | 0.43 | 0.6 | 0.71 |
| 5 | 179863807 | rs1848480 | 1.072 | -0.318 | -0.016 | 0.596 | 0.016 | 0.01 | 0.01 | 0 | 0.54 | 0.36 | 0.71 | 0.43 | 0.6 | 0.71 |
| 5 | 179867194 | rs13185599 | 0.162 | -0.402 | -0.617 | 1.168 | 0.019 | -0.026 | -0.026 | 0 | 0.73 | 0.82 | 0.84 | 0.43 | 0.83 | 0.83 |
| 5 | 179877022 | rs6893735 | 0.289 | 0.495 | -0.325 | 0.67 | 0 | 0.005 | 0.005 | 0 | 0.54 | 0.51 | 0.58 | 0.38 | 0.59 | 0.57 |
| 5 | 179881433 | rs6886960 | -1.126 | 1.129 | 0.4 | 0.574 | 0.04 | 0.023 | 0.023 | 0 | 0.12 | 0.3 | 0.12 | 0.11 | 0.15 | 0.23 |
| 5 | 179888268 | rs7729820 | -0.052 | 1.186 | -0.012 | 0.455 | 0.023 | 0.041 | 0.041 | 0.073 | 0.48 | 0.68 | 0.37 | 0.36 | 0.53 | 0.58 |
| 5 | 179890300 | rs7726530 | -0.805 | 1.124 | 0.225 | 0.897 | 0.044 | -0.057 | -0.057 | 0.097 | 0.67 | 0.83 | 0.51 | 0.44 | 0.76 | 0.68 |
| 5 | 179895013 | rs7700744 | -0.772 | 1.127 | 0.25 | 0.679 | 0.027 | -0.022 | -0.022 | 0 | 0.67 | 0.78 | 0.5 | 0.4 | 0.75 | 0.61 |
| 5 | 179899660 | rs11739992 | -1.265 | 1.907 | 0.625 | 0.942 | 0.024 | -0.017 | -0.017 | 0.042 | 0.08 | 0.11 | 0.08 | 0.01 | 0.17 | 0.1 |
| 5 | 179905742 | rs7730828 | -1.616 | 1.756 | 0.819 | 1.006 | 0.031 | 0.102 | 0.102 | 1.347 | 0.14 | 0.6 | 0.16 | 0.28 | 0.13 | 0.23 |
| 5 | 179918722 | rs6875637 | -1.938 | 2.008 | 1.385 | 0.959 | 0.105 | 0.134 | 0.134 | 0 | 0.11 | 0.28 | 0.14 | 0.27 | 0.17 | 0.12 |
| 5 | 179925212 | rs10464103 | -0.053 | 2.058 | 0.908 | 0.87 | -0.004 | 0.099 | 0.099 | 0.011 | 0.54 | 0.25 | 0.38 | 0.24 | 0.55 | 0.43 |
| 5 | 179925350 | rs4538575 | -1.205 | 2.138 | 1.121 | 0.866 | 0.043 | -0.071 | -0.071 | 0 | 0.67 | 0.88 | 0.56 | 0.49 | 0.73 | 0.56 |
| 5 | 179938884 | rs937919 | 0.811 | 2.084 | 1.201 | 0.09 | 0.006 | 0.026 | 0.026 | 0.017 | 0.37 | 0.26 | 0.52 | 0.76 | 0.34 | 0.46 |
| 5 | 179943955 | rs17630822 | -0.693 | 2.494 | 0.548 | 0.203 | 0.006 | 0.003 | 0.003 | 0 | 0.05 | 0.06 | 0.08 | 0.1 | 0.08 | 0.05 |
| 5 | 179980527 | rs11738129 | -0.247 | 1.803 | 0.523 | 0.217 | 0.005 | 0.094 | 0.094 | 1.154 | 0.52 | 0.29 | 0.39 | 0.21 | 0.55 | 0.42 |
| 5 | 180003085 | rs10069 | -1.288 | 1.217 | 0.42 | 0.624 | 0.002 | -0.001 | -0.001 | 5.119 | 0.11 | 0.12 | 0.14 | 0.12 | 0.18 | 0.15 |

| | | | | | | | | | | | | | | | | |
|---|---|---|---|---|---|---|---|---|---|---|---|---|---|---|---|---|
| 5 | 180010639 | rs11957360 | 0.042 | 0.388 | -0.244 | 0.767 | 0 | 0.01 | 0.01 | 0.243 | 0.5 | 0.5 | 0.51 | 0.78 | 0.5 | 0.64 |
| 5 | 180014616 | rs13186854 | 0.726 | 0.185 | -1.249 | 1.018 | -0.107 | -0.015 | -0.015 | 0.314 | 0.33 | 0.06 | 0.39 | 0.31 | 0.42 | 0.46 |
| 5 | 180015141 | rs2447736 | -0.397 | 0.125 | -1.298 | 1.027 | -0.014 | -0.05 | -0.05 | 0 | 0.27 | 0.2 | 0.39 | 0.44 | 0.34 | 0.22 |
| 5 | 180016644 | rs368686 | 0.309 | -0.152 | -1.34 | 0.598 | -0.07 | 0.136 | 0.136 | 0 | 0.3 | 0.06 | 0.19 | 0.07 | 0.31 | 0.2 |
| 5 | 180019395 | rs307803 | -0.389 | -0.203 | -0.722 | 0.83 | -0.005 | 0.034 | 0.034 | 0 | 0.38 | 0.24 | 0.25 | 0.27 | 0.37 | 0.34 |
| 5 | 180028743 | rs2279622 | -1.741 | -0.284 | 0.432 | 1.585 | 0.007 | 0.19 | 0.19 | 0.124 | 0.1 | 0.35 | 0.07 | 0.2 | 0.06 | 0.15 |
| 5 | 180032263 | rs11747066 | 0.38 | -0.466 | -0.232 | 1.382 | -0.007 | -0.024 | -0.024 | 0 | 0.3 | 0.09 | 0.17 | 0.29 | 0.26 | 0.32 |
| 5 | 180033292 | rs10058772 | 0.33 | -0.529 | -0.262 | 1.363 | -0.004 | 0.015 | 0.015 | 0 | 0.31 | 0.08 | 0.17 | 0.23 | 0.26 | 0.32 |
| 5 | 180040918 | rs2242217 | 1.474 | -0.636 | -0.009 | 1.498 | -0.053 | 0.042 | 0.042 | 0.597 | 0.41 | 0.21 | 0.42 | 0.3 | 0.5 | 0.48 |
| 5 | 180041662 | rs400330 | 0.4 | -0.66 | 0.318 | 1.557 | -0.012 | 0.007 | 0.007 | 0.264 | 0.6 | 0.53 | 0.76 | 0.48 | 0.68 | 0.63 |
| 5 | 180047412 | rs382266 | 0.844 | -0.79 | 0.451 | 2.111 | -0.028 | 0.184 | 0.184 | 0 | 0.88 | 0.76 | 0.95 | 0.63 | 0.94 | 0.84 |
| 5 | 180051987 | rs3797104 | -0.77 | -0.874 | 0.221 | 1.384 | -0.012 | 0.006 | 0.006 | 0 | 0.18 | 0.26 | 0.1 | 0.21 | 0.14 | 0.09 |
| 5 | 180052108 | rs307823 | 1.02 | -0.844 | 0.291 | 1.117 | -0.032 | 0.054 | 0.054 | 0 | 0.74 | 0.42 | 0.83 | 0.49 | 0.8 | 0.79 |
| 5 | 180055188 | rs3797102 | 0.815 | -0.022 | -0.965 | 0.306 | -0.179 | -0.079 | -0.079 | 0 | 0.31 | 0.03 | 0.49 | 0.35 | 0.45 | 0.41 |
| 5 | 180058963 | rs2290983 | 0.775 | 0.518 | 0.381 | -0.176 | 0.007 | -0.006 | -0.006 | 0 | 0.58 | 0.39 | 0.65 | 0.47 | 0.6 | 0.55 |
| 5 | 180060790 | rs10085025 | 0.471 | 0.532 | 0.797 | -0.421 | 0.081 | -0.01 | -0.01 | 0.085 | 0.92 | 0.67 | 0.92 | 0.98 | 0.9 | 0.89 |
| 5 | 180065455 | rs4700745 | 1.186 | 0.467 | 0.664 | -0.443 | 0.009 | 0.052 | 0.052 | 0 | 0.53 | 0.33 | 0.61 | 0.36 | 0.54 | 0.52 |
| 5 | 180067675 | rs307806 | -1.198 | 0.755 | 0.132 | -0.963 | 0.012 | -0.038 | -0.038 | 0.024 | 0.12 | 0.22 | 0.11 | 0.04 | 0.13 | 0.24 |
| 5 | 180068741 | rs11748431 | 0.162 | 0.798 | -0.096 | -1.113 | -0.002 | 0.039 | 0.039 | 0 | 0.25 | 0.13 | 0.15 | 0.02 | 0.27 | 0.15 |
| 5 | 180071276 | rs11960332 | -0.193 | 1.087 | -0.276 | -0.834 | 0 | 0.024 | 0.024 | 0.011 | 0.15 | 0.15 | 0.28 | 0.53 | 0.14 | 0.19 |
| 5 | 180074248 | rs307814 | -0.784 | 0.803 | -0.223 | -0.621 | 0.031 | 0.037 | 0.037 | 0.089 | 0.53 | 0.29 | 0.55 | 0.62 | 0.6 | 0.61 |
| 5 | 180076999 | rs1382302 | -0.589 | 0.775 | -0.478 | -0.72 | -0.013 | 0.013 | 0.013 | 0 | 0.48 | 0.18 | 0.47 | 0.57 | 0.51 | 0.49 |
| 5 | 180077487 | rs307805 | -0.593 | 1.211 | -0.068 | -0.066 | 0 | -0.056 | -0.056 | 0 | 0.13 | 0.26 | 0.09 | 0.02 | 0.11 | 0.25 |
| 5 | 180081766 | rs6890771 | -0.527 | 1.248 | -0.393 | 0.295 | 0.001 | 0 | 0 | 0.848 | 0.55 | 0.58 | 0.51 | 0.57 | 0.53 | 0.61 |
| 5 | 180094042 | rs7705017 | -1.047 | 0.939 | -0.277 | 0.583 | 0.02 | -0.009 | -0.009 | 0.061 | 0.06 | 0.16 | 0.04 | 0.03 | 0.09 | 0.03 |
| 5 | 180096972 | rs11959164 | -0.429 | 1.041 | -0.625 | 0.565 | 0.002 | -0.001 | -0.001 | 1.123 | 0.47 | 0.54 | 0.47 | 0.55 | 0.46 | 0.46 |
| 5 | 180109638 | rs10042049 | -0.233 | 1 | -0.771 | 0.572 | 0.001 | -0.028 | -0.028 | 0 | 0.33 | 0.22 | 0.41 | 0.52 | 0.35 | 0.28 |
| 5 | 180115479 | rs11740857 | -0.115 | 0.571 | -1.658 | 0.398 | -0.046 | 0.076 | 0.076 | 0 | 0.37 | 0.13 | 0.44 | 0.22 | 0.58 | 0.26 |
| 5 | 180115817 | rs11249743 | 0.041 | -0.872 | -0.926 | 0.878 | -0.003 | 0.011 | 0.011 | 0 | 0.22 | 0.29 | 0.26 | 0.28 | 0.2 | 0.34 |
| 5 | 180116840 | rs3852150 | 0.215 | -1.512 | -0.102 | 1.663 | -0.017 | 0.024 | 0.024 | 0 | 0.32 | 0.36 | 0.2 | 0.47 | 0.22 | 0.39 |
| 5 | 180116874 | rs4074596 | 0.991 | -1.597 | -0.315 | 1.634 | -0.006 | 0.082 | 0.082 | 0 | 0.41 | 0.13 | 0.57 | 0.25 | 0.6 | 0.34 |
| 5 | 180118945 | rs17080484 | 0.64 | -1.959 | -0.089 | 1.93 | 0.013 | 0.036 | 0.036 | 0 | 0.22 | 0.08 | 0.07 | 0.14 | 0.16 | 0.27 |
| 5 | 180119918 | rs10055973 | 0.952 | -2.361 | 0.428 | 2.032 | 0.038 | 0.052 | 0.052 | 0 | 0.59 | 0.33 | 0.62 | 0.29 | 0.72 | 0.45 |
| 5 | 180120192 | rs7731780 | -0.603 | -2.345 | 0.893 | 2.504 | 0.009 | 0.253 | 0.253 | 0.5 | 0.08 | 0.29 | 0.21 | 0.3 | 0.06 | 0.14 |
| 5 | 180120528 | rs6601136 | 0.138 | -2.342 | 1.012 | 2.398 | 0.007 | -0.01 | -0.01 | 0 | 0.7 | 0.76 | 0.84 | 0.59 | 0.79 | 0.6 |
| 5 | 180128788 | rs6601138 | 0.011 | -2.398 | 1.142 | 2.522 | -0.011 | 0.104 | 0.104 | 0.217 | 0.92 | 0.84 | 0.91 | 0.72 | 0.95 | 0.87 |
| 5 | 180131124 | rs4700754 | -1.405 | -2.533 | 1.462 | 2.66 | -0.018 | 0.215 | 0.215 | 0.358 | 0.12 | 0.35 | 0.27 | 0.48 | 0.08 | 0.17 |
| 5 | 180131514 | rs13161948 | 1.596 | -2.607 | 1.24 | 1.946 | 0.002 | 0.194 | 0.194 | 0 | 0.44 | 0.16 | 0.48 | 0.16 | 0.58 | 0.25 |
| 5 | 180135005 | rs9791144 | 1.301 | -2.681 | 1.24 | 1.93 | 0.069 | 0.131 | 0.131 | 0 | 0.56 | 0.17 | 0.59 | 0.3 | 0.72 | 0.43 |
| 5 | 180154779 | rs10479580 | 0.487 | -2.719 | 1.489 | 2.08 | 0.001 | -0.002 | -0.002 | 0 | 0.67 | 0.67 | 0.69 | 0.42 | 0.75 | 0.49 |
| 5 | 180157191 | rs12719876 | 1.197 | -2.715 | 1.498 | 2.053 | 0.062 | 0.099 | 0.099 | 0 | 0.57 | 0.19 | 0.6 | 0.31 | 0.73 | 0.44 |
| 5 | 180165367 | rs10479581 | 1.026 | -2.653 | 1.257 | 1.912 | 0.03 | 0.045 | 0.045 | 0 | 0.6 | 0.25 | 0.61 | 0.32 | 0.73 | 0.45 |
| 5 | 180165553 | rs17080555 | -0.19 | -2.508 | 1.088 | 1.899 | -0.011 | 0.018 | 0.018 | 0.002 | 0.07 | 0.14 | 0.03 | 0.12 | 0.04 | 0.07 |
| 5 | 180168265 | rs17702447 | 0.478 | -2.235 | 0.847 | 1.875 | -0.034 | -0.014 | -0.014 | 0 | 0.13 | 0.01 | 0.12 | 0.11 | 0.15 | 0.17 |
| 5 | 180170852 | rs11743612 | 1.453 | -2.698 | 0.914 | 1.58 | 0.01 | 0.032 | 0.032 | 0.017 | 0.48 | 0.37 | 0.49 | 0.34 | 0.59 | 0.38 |
| 5 | 180171716 | rs10903267 | 0.276 | -2.748 | 1.011 | 1.562 | -0.011 | -0.034 | -0.034 | 0 | 0.79 | 0.7 | 0.83 | 0.89 | 0.8 | 0.81 |
| 5 | 180176497 | rs6886113 | -0.319 | -2.729 | 1.096 | 1.574 | 0.006 | -0.037 | -0.037 | 0.316 | 0.28 | 0.23 | 0.33 | 0.55 | 0.22 | 0.41 |
| 5 | 180179514 | rs7713468 | 1.067 | -2.525 | 1.177 | 1.313 | 0.001 | 0.003 | 0.003 | 0 | 0.49 | 0.46 | 0.5 | 0.33 | 0.59 | 0.4 |
| 5 | 180199253 | rs7702121 | 0.082 | -2.348 | 1.014 | 1.399 | -0.036 | -0.036 | -0.036 | 0 | 0.16 | 0.01 | 0.09 | 0.17 | 0.13 | 0.15 |
| 5 | 180202156 | rs2387293 | -1.212 | -2.298 | 1.166 | 1.277 | -0.018 | -0.024 | -0.024 | 0.022 | 0.31 | 0.37 | 0.23 | 0.18 | 0.2 | 0.36 |
| 5 | 180202977 | rs17080585 | -0.782 | -2.303 | 1.075 | 1.323 | -0.008 | 0.076 | 0.076 | 0 | 0.06 | 0.15 | 0.06 | 0.16 | 0.03 | 0.15 |
| 5 | 180206079 | rs6881079 | -1.547 | -2.134 | 0.655 | 1.119 | -0.005 | -0.001 | -0.001 | 0 | 0.38 | 0.61 | 0.3 | 0.34 | 0.23 | 0.51 |
| 5 | 180207020 | rs676699 | -1.665 | -1.972 | -0.227 | 0.959 | 0.087 | -0.013 | -0.013 | 0.172 | 0.42 | 0.84 | 0.32 | 0.4 | 0.24 | 0.54 |
| 5 | 180209454 | rs655437 | -1.615 | -2.188 | -0.187 | 0.948 | 0.095 | -0.009 | -0.009 | 0 | 0.42 | 0.83 | 0.31 | 0.4 | 0.22 | 0.53 |
| 5 | 180211280 | rs3733754 | -1.013 | -1.436 | 0.11 | 0.502 | -0.053 | -0.059 | -0.059 | 0.004 | 0.16 | 0.3 | 0.16 | 0.05 | 0.07 | 0.22 |
| 5 | 180218831 | rs2070924 | -0.624 | -0.909 | -0.233 | 1.038 | -0.006 | 0.02 | 0.02 | 0.367 | 0.08 | 0.1 | 0.04 | 0.21 | 0.05 | 0.16 |
| 5 | 180226571 | rs4700986 | -1.193 | -2.06 | -0.206 | 0.463 | 0.045 | 0.032 | 0.032 | 1.262 | 0.47 | 0.68 | 0.33 | 0.39 | 0.29 | 0.52 |
| 5 | 180226673 | rs655601 | -0.707 | -2.453 | 0.157 | 0.426 | -0.059 | -0.04 | -0.04 | 2.717 | 0.26 | 0.63 | 0.22 | 0.1 | 0.11 | 0.23 |
| 5 | 180240026 | rs624335 | -1.231 | -1.726 | 0.399 | 0.606 | -0.01 | 0.026 | 0.026 | 0 | 0.16 | 0.34 | 0.1 | 0.24 | 0.14 | 0.19 |
| 5 | 180245926 | rs1872183 | -1.001 | -1.767 | -0.341 | 1.477 | 0.013 | -0.065 | -0.065 | 0 | 0.12 | 0.03 | 0.18 | 0.42 | 0.07 | 0.13 |
| 5 | 180255963 | rs4700988 | 1.314 | -1.996 | -0.203 | 0.438 | 0.001 | -0.02 | -0.02 | 0.057 | 0.79 | 0.28 | 0.84 | 0.96 | 0.8 | 0.85 |
| 5 | 180280901 | rs2619764 | 0.894 | -2.216 | 0.39 | 0.328 | -0.018 | -0.048 | -0.048 | 0 | 0.9 | 0.82 | 0.93 | 1 | 0.91 | 0.98 |
| 5 | 180285668 | rs2619756 | 2.334 | -2.083 | 0.391 | 0.137 | -0.002 | -0.047 | -0.047 | 0.956 | 0.72 | 0.36 | 0.83 | 0.96 | 0.76 | 0.79 |
| 5 | 180301648 | rs34362037 | -1.785 | -2.566 | -0.15 | 0.792 | 0.042 | -0.065 | -0.065 | 0 | 0.24 | 0.11 | 0.31 | 0.64 | 0.16 | 0.4 |
| 5 | 180302462 | rs7715533 | -0.548 | -1.918 | -0.834 | -0.9 | 0 | -0.003 | -0.003 | 0 | 0.43 | 0.59 | 0.37 | 0.19 | 0.38 | 0.36 |
| 5 | 180326502 | rs2619739 | -0.262 | -1.968 | -0.764 | -1.003 | 0.018 | 0.04 | 0.04 | 0 | 0.13 | 0.07 | 0.05 | 0.04 | 0.1 | 0.2 |
| 5 | 180341845 | rs4700772 | 0.779 | -0.456 | -0.909 | -0.912 | -0.072 | -0.066 | -0.066 | 0 | 0.18 | 0.08 | 0.33 | 0.25 | 0.29 | 0.11 |
| 5 | 180342600 | rs4701005 | -0.404 | -0.827 | -0.463 | -1.134 | -0.004 | -0.004 | -0.004 | 0 | 0.32 | 0.67 | 0.42 | 0.35 | 0.38 | 0.27 |
| 5 | 180433439 | rs7713324 | 0.211 | -1.081 | -0.149 | -1.368 | -0.052 | -0.08 | -0.08 | 0 | 0.14 | 0.28 | 0.08 | 0.02 | 0.05 | 0.1 |

| | | | | | | | | | | | | | | | | |
|---|---|---|---|---|---|---|---|---|---|---|---|---|---|---|---|---|
| 5 | 180433451 | rs7717283 | 0.211 | -1.081 | -0.149 | -1.368 | -0.052 | -0.08 | -0.08 | 0.012 | 0.14 | 0.28 | 0.08 | 0.02 | 0.05 | 0.1 |
| 5 | 180433806 | rs10057843 | 0.211 | -1.112 | 0.353 | -1.383 | -0.052 | -0.08 | -0.08 | 0.21 | 0.14 | 0.28 | 0.08 | 0.02 | 0.05 | 0.1 |
| 5 | 180433880 | rs10072450 | 0.211 | -1.112 | 0.353 | -1.383 | -0.052 | -0.08 | -0.08 | 1.529 | 0.14 | 0.28 | 0.08 | 0.02 | 0.05 | 0.1 |
| 5 | 180433966 | rs10067377 | 0.232 | -1.112 | 0.353 | -1.383 | -0.052 | -0.079 | -0.079 | 0 | 0.14 | 0.28 | 0.08 | 0.02 | 0.05 | 0.1 |
| 5 | 180439488 | rs7721042 | 0.891 | -0.949 | 0.272 | -1.368 | 0.065 | 0.055 | 0.055 | 0 | 0.19 | 0.33 | 0.35 | 0.25 | 0.3 | 0.1 |
| 5 | 180449895 | rs4337818 | -0.924 | -1.151 | -0.142 | -1.469 | -0.01 | 0.019 | 0.019 | 0.006 | 0.47 | 0.77 | 0.5 | 0.36 | 0.44 | 0.37 |
| 5 | 180452560 | rs4701015 | -1.516 | -1.29 | -0.298 | -1.548 | -0.028 | -0.022 | -0.022 | 1.046 | 0.6 | 0.78 | 0.53 | 0.4 | 0.53 | 0.49 |
| 5 | 180455372 | rs11249756 | 1.305 | -1.213 | -0.178 | -1.537 | -0.009 | -0.009 | -0.009 | 0 | 0.18 | 0.17 | 0.34 | 0.26 | 0.31 | 0.11 |
| 5 | 180457135 | rs6601163 | -1.829 | -1.267 | -0.56 | -1.688 | -0.026 | -0.036 | -0.036 | 0 | 0.67 | 0.79 | 0.55 | 0.4 | 0.6 | 0.57 |
| 5 | 180458813 | rs6868418 | 1.482 | -0.769 | -0.229 | -1.178 | -0.051 | -0.03 | -0.03 | 1.828 | 0.14 | 0.07 | 0.28 | 0.15 | 0.28 | 0.08 |
| 5 | 180467482 | rs6873621 | -0.122 | -1.006 | 0.098 | -1.241 | 0.041 | 0.025 | 0.025 | 0 | 0.33 | 0.51 | 0.43 | 0.35 | 0.41 | 0.26 |
| 5 | 180474080 | rs7709346 | -1.325 | -1.191 | -0.225 | -1.284 | -0.03 | -0.028 | -0.028 | 0.02 | 0.6 | 0.78 | 0.54 | 0.4 | 0.55 | 0.47 |
| 5 | 180485353 | rs3806869 | 0.842 | -1.48 | -0.114 | -1.302 | -0.016 | -0.024 | -0.024 | 0 | 0.13 | 0.14 | 0.04 | 0.02 | 0.03 | 0.09 |
| 5 | 180485720 | rs1467 | -0.847 | -1.535 | 0.06 | -1.357 | -0.003 | 0.009 | 0.009 | 0 | 0.48 | 0.64 | 0.49 | 0.38 | 0.49 | 0.39 |
| 5 | 180485949 | rs2277935 | -0.282 | -1.039 | 0.182 | -1.116 | 0.009 | 0.006 | 0.006 | 0 | 0.35 | 0.38 | 0.44 | 0.37 | 0.45 | 0.29 |
| 5 | 180487085 | rs10434808 | 0.968 | -0.769 | -0.316 | -1.004 | -0.013 | -0.011 | -0.011 | 0 | 0.41 | 0.34 | 0.49 | 0.59 | 0.47 | 0.52 |
| 5 | 180489103 | rs12654960 | 1.404 | -1.825 | -0.542 | -0.656 | -0.031 | -0.143 | -0.143 | 2.989 | 0.23 | 0.03 | 0.25 | 0.54 | 0.25 | 0.35 |
| 5 | 180492405 | rs888949 | 0.828 | -2.347 | 0.645 | -1.01 | -0.061 | -0.057 | -0.057 | 0 | 0.71 | 0.6 | 0.84 | 0.84 | 0.86 | 0.76 |
| 5 | 180493946 | rs1862333 | -0.458 | -1.691 | 0.032 | -0.845 | -0.017 | -0.061 | -0.061 | 0 | 0.34 | 0.34 | 0.12 | 0.04 | 0.15 | 0.26 |
| 5 | 180494088 | rs962352 | 0.281 | -1.443 | -1.082 | -0.665 | -0.001 | -0.005 | -0.005 | 0.834 | 0.92 | 0.95 | 0.91 | 0.89 | 0.92 | 0.93 |
| 5 | 180495340 | rs1579303 | 0.463 | -1.158 | -1.062 | -0.496 | 0.062 | 0.067 | 0.067 | 0 | 0.55 | 0.34 | 0.76 | 0.75 | 0.75 | 0.61 |
| 5 | 180501193 | rs6601167 | -0.581 | -0.932 | -1.406 | -0.537 | 0.042 | 0.076 | 0.076 | 0 | 0.32 | 0.83 | 0.24 | 0.17 | 0.23 | 0.31 |
| 5 | 180502321 | rs7379313 | -1.022 | -0.883 | -1.147 | -0.253 | 0.012 | -0.002 | -0.002 | 0.278 | 0.14 | 0.09 | 0.13 | 0.16 | 0.1 | 0.23 |
| 5 | 180508615 | rs13167249 | -0.267 | -0.375 | -0.571 | -0.532 | -0.001 | -0.004 | -0.004 | 0.258 | 0.62 | 0.67 | 0.74 | 0.87 | 0.55 | 0.58 |
| 5 | 180511976 | rs729129 | -0.415 | -0.965 | -0.725 | -0.468 | 0.022 | 0.012 | 0.012 | 1.321 | 0.3 | 0.29 | 0.31 | 0.22 | 0.18 | 0.28 |
| 5 | 180512315 | rs729128 | -0.394 | -1.058 | -0.604 | -0.302 | 0.007 | -0.002 | -0.002 | 0 | 0.21 | 0.21 | 0.26 | 0.22 | 0.11 | 0.23 |
| 5 | 180513467 | rs730545 | 0.098 | -0.484 | -0.528 | -0.164 | -0.031 | -0.032 | -0.032 | 1.406 | 0.32 | 0.22 | 0.44 | 0.52 | 0.38 | 0.23 |
| 5 | 180514652 | rs13358985 | -0.845 | -1.213 | 0.106 | -0.541 | -0.07 | -0.122 | -0.122 | 3.1 | 0.17 | 0.45 | 0.16 | 0.02 | 0.09 | 0.08 |
| 5 | 180527455 | rs4339306 | 0.389 | -0.278 | -1.157 | -0.429 | -0.11 | 0.104 | 0.104 | 0 | 0.43 | 0.07 | 0.3 | 0.22 | 0.47 | 0.52 |
| 5 | 180535802 | rs17705959 | 0.234 | -0.558 | -0.898 | -0.18 | -0.044 | -0.017 | -0.017 | 0.014 | 0.5 | 0.12 | 0.41 | 0.4 | 0.5 | 0.65 |
| 5 | 180538233 | rs7380392 | 0.076 | -0.812 | -0.334 | -0.338 | 0.001 | -0.004 | -0.004 | 0.023 | 0.59 | 0.49 | 0.42 | 0.26 | 0.56 | 0.55 |
| 5 | 180541251 | rs7379517 | -0.135 | -0.882 | -0.402 | -0.296 | -0.011 | 0 | 0 | 0 | 0.07 | 0.07 | 0.09 | 0.15 | 0.03 | 0.16 |
| 5 | 180541464 | rs10069087 | 0.093 | -1.136 | -0.164 | -0.436 | -0.091 | -0.138 | -0.138 | 0.022 | 0.17 | 0.37 | 0.07 | 0 | 0.07 | 0.03 |
| 5 | 180543544 | rs7380326 | -0.227 | -0.978 | -0.894 | -0.317 | -0.003 | -0.002 | -0.002 | 0 | 0.07 | 0.08 | 0.1 | 0.15 | 0.03 | 0.16 |
| 5 | 180548624 | rs888943 | 0.884 | -0.773 | -1.157 | -0.227 | -0.094 | 0.026 | 0.026 | 0.02 | 0.41 | 0.09 | 0.34 | 0.27 | 0.48 | 0.47 |
| 5 | 180549211 | rs2052499 | 0.891 | -0.743 | -1.097 | -0.219 | -0.051 | 0.045 | 0.045 | 0 | 0.41 | 0.21 | 0.35 | 0.26 | 0.48 | 0.48 |
| 5 | 180554812 | rs1363537 | 0.002 | 0.488 | -1.05 | 0.166 | -0.041 | -0.053 | -0.053 | 0 | 0.71 | 0.84 | 0.54 | 0.42 | 0.62 | 0.69 |
| 5 | 180563838 | rs10479482 | -0.176 | 0.499 | -1.544 | -0.709 | 0.014 | 0.185 | 0.185 | 3.199 | 0.55 | 0.83 | 0.56 | 0.84 | 0.44 | 0.61 |
| 5 | 180567719 | rs11249769 | -0.024 | 1.208 | -0.765 | -0.978 | -0.001 | 0.001 | 0.001 | 0.042 | 0.58 | 0.54 | 0.59 | 0.82 | 0.55 | 0.67 |
| 5 | 180578750 | rs2892345 | -0.131 | -0.735 | 0.222 | -1.74 | -0.047 | -0.082 | -0.082 | 0.728 | 0.29 | 0.52 | 0.23 | 0.09 | 0.22 | 0.16 |
| 6 | 240484 | rs12213916 | -0.924 | -0.331 | 1.269 | 0.554 | -0.014 | 0.012 | 0.012 | 0 | 0.07 | 0.04 | 0.07 | 0.01 | 0.11 | 0.06 |
| 6 | 246178 | rs13213850 | -1.937 | 0.783 | 2.257 | 1.391 | 0.039 | -0.031 | -0.031 | 0 | 0.08 | 0.24 | 0.09 | 0.01 | 0.08 | 0.04 |
| 6 | 253428 | rs6596865 | -2.154 | 0.761 | 1.565 | 1.383 | 0.008 | 0.118 | 0.118 | 0 | 0.16 | 0.47 | 0.25 | 0.26 | 0.14 | 0.14 |
| 6 | 274847 | rs12212311 | -1.528 | 1.313 | -0.141 | 0.843 | -0.029 | 0.037 | 0.037 | 0 | 0.08 | 0.03 | 0.04 | 0 | 0.1 | 0.06 |
| 6 | 312247 | rs17133064 | -1.607 | 0.767 | 0.474 | 1.768 | -0.016 | 0.17 | 0.17 | 0 | 0.06 | 0.17 | 0.1 | 0.27 | 0.03 | 0.1 |
| 6 | 378970 | rs9501958 | -1.638 | 0.926 | 0.976 | 0.914 | 0.054 | -0.013 | -0.013 | 0 | 0.08 | 0.27 | 0.08 | 0.02 | 0.08 | 0.04 |
| 6 | 380341 | rs6900267 | 1.82 | 0.073 | 1.65 | 1.261 | -0.007 | 0.068 | 0.068 | 0.876 | 0.87 | 0.6 | 0.81 | 0.85 | 0.93 | 0.87 |
| 6 | 382897 | rs9378774 | 0.328 | -0.173 | 0.971 | 0.877 | 0.043 | 0.037 | 0.037 | 0.144 | 0.59 | 0.3 | 0.6 | 0.3 | 0.67 | 0.46 |
| 6 | 389319 | rs9503555 | -2.175 | 0.409 | 1.611 | 1.775 | 0.026 | 0.088 | 0.088 | 0.278 | 0.1 | 0.24 | 0.14 | 0.31 | 0.11 | 0.12 |
| 6 | 390105 | rs6899334 | -1.808 | 0.408 | 1.821 | 1.625 | 0.006 | -0.008 | -0.008 | 0 | 0.08 | 0.07 | 0.07 | 0.03 | 0.11 | 0.03 |
| 6 | 395634 | rs6930635 | -2.281 | 0.535 | 2.444 | 1.68 | 0.048 | -0.031 | -0.031 | 0 | 0.09 | 0.2 | 0.07 | 0.03 | 0.12 | 0.04 |
| 6 | 397290 | rs1775589 | -2.486 | 0.981 | 2.284 | 2.223 | -0.02 | 0.097 | 0.097 | 0 | 0.21 | 0.33 | 0.3 | 0.52 | 0.15 | 0.3 |
| 6 | 398775 | rs2671422 | -2.391 | 0.982 | 2.389 | 2.221 | -0.017 | 0.158 | 0.158 | 0 | 0.19 | 0.39 | 0.3 | 0.51 | 0.13 | 0.3 |
| 6 | 403604 | rs1877179 | 1.972 | 1.05 | 2.652 | 2.325 | 0.042 | 0.229 | 0.229 | 0 | 0.93 | 0.7 | 0.87 | 0.71 | 0.91 | 0.91 |
| 6 | 403799 | rs3778607 | 0.995 | 0.899 | 1.657 | 1.39 | -0.023 | 0.185 | 0.185 | 0.804 | 0.5 | 0.2 | 0.47 | 0.26 | 0.49 | 0.42 |
| 6 | 404632 | rs2001508 | -2.334 | 0.68 | 2.16 | 1.504 | 0.068 | 0.223 | 0.223 | 0.006 | 0.11 | 0.34 | 0.23 | 0.38 | 0.18 | 0.12 |
| 6 | 405873 | rs17825664 | -0.892 | 0.693 | 1.821 | 1.298 | -0.009 | 0.009 | 0.009 | 0 | 0.05 | 0.03 | 0.05 | 0.02 | 0.08 | 0.03 |
| 6 | 405900 | rs3800262 | 0.045 | 0.11 | 1.571 | 1.172 | 0.008 | -0.005 | -0.005 | 0 | 0.74 | 0.73 | 0.75 | 0.75 | 0.82 | 0.72 |
| 6 | 408079 | rs1050976 | 1.967 | 0.264 | 0.846 | 1.042 | -0.146 | 0.094 | 0.094 | 1.417 | 0.47 | 0.03 | 0.44 | 0.31 | 0.48 | 0.44 |
| 6 | 408246 | rs7768807 | -0.558 | 0.223 | 1.272 | 1.01 | -0.006 | 0.009 | 0.009 | 0 | 0.3 | 0.34 | 0.28 | 0.27 | 0.27 | 0.31 |
| 6 | 410493 | rs1877175 | 1.727 | 0.305 | 0.912 | 1.039 | 0.002 | -0.001 | -0.001 | 0.419 | 0.8 | 0.77 | 0.84 | 0.82 | 0.77 | 0.81 |
| 6 | 411064 | rs872071 | 1.98 | 0.195 | 0.607 | 0.868 | -0.138 | 0.099 | 0.099 | 2.442 | 0.48 | 0.03 | 0.44 | 0.31 | 0.47 | 0.43 |
| 6 | 411554 | rs6906608 | -0.593 | 0.331 | 0.724 | 0.87 | -0.021 | -0.016 | -0.016 | 0.836 | 0.12 | 0.06 | 0.09 | 0.14 | 0.15 | 0.16 |
| 6 | 412741 | rs7757906 | 1.493 | 0.321 | 0.803 | 0.805 | 0.044 | 0.074 | 0.074 | 0.167 | 0.8 | 0.65 | 0.74 | 0.57 | 0.71 | 0.73 |
| 6 | 417727 | rs9378805 | 1.65 | 0.436 | -0.136 | 0.679 | -0.079 | 0.151 | 0.151 | 2.046 | 0.5 | 0.05 | 0.44 | 0.31 | 0.47 | 0.42 |
| 6 | 421317 | rs9378808 | 0.414 | 0.015 | 0.09 | 0.766 | 0.004 | -0.048 | -0.048 | 0.293 | 0.2 | 0.1 | 0.22 | 0.25 | 0.18 | 0.25 |
| 6 | 422408 | rs9378374 | 2.124 | -0.521 | -0.22 | 0.561 | 0.003 | -0.034 | -0.034 | 0 | 0.74 | 0.19 | 0.7 | 0.57 | 0.74 | 0.74 |
| 6 | 427152 | rs7748534 | 1.06 | -0.935 | -0.239 | 0.402 | 0.006 | -0.009 | -0.009 | 0 | 0.68 | 0.29 | 0.64 | 0.47 | 0.7 | 0.62 |
| 6 | 428983 | rs9502066 | 0.313 | -0.83 | 0.18 | 0.48 | -0.003 | 0.031 | 0.031 | 0.052 | 0.17 | 0.22 | 0.2 | 0.27 | 0.17 | 0.27 |

| | | | | | | | | | | | | | | | | |
|---|---|---|---|---|---|---|---|---|---|---|---|---|---|---|---|---|
| 6 | 431891 | rs9405666 | 0.997 | -1.071 | -0.189 | 0.083 | -0.008 | 0 | 0 | 0 | 0.6 | 0.44 | 0.6 | 0.48 | 0.59 | 0.56 |
| 6 | 439985 | rs872468 | -0.055 | -1.329 | -0.425 | 0.159 | 0.001 | -0.006 | -0.006 | 0 | 0.12 | 0.05 | 0.09 | 0.13 | 0.12 | 0.14 |
| 6 | 441883 | rs6920655 | 1.273 | -1.702 | -0.367 | -0.098 | 0.036 | 0.011 | 0.011 | 0 | 0.54 | 0.3 | 0.58 | 0.48 | 0.62 | 0.54 |
| 6 | 442206 | rs4959880 | 1.033 | -1.621 | 0.245 | 0.222 | 0.041 | 0.208 | 0.208 | 1.095 | 0.83 | 0.51 | 0.8 | 0.62 | 0.81 | 0.77 |
| 6 | 442357 | rs9392017 | 0.055 | -1.53 | -0.4 | -0.233 | 0.01 | -0.037 | -0.037 | 0 | 0.29 | 0.16 | 0.27 | 0.39 | 0.25 | 0.47 |
| 6 | 442410 | rs7749710 | -0.029 | -1.77 | -0.475 | -0.251 | -0.019 | 0.056 | 0.056 | 0 | 0.11 | 0.01 | 0.07 | 0.02 | 0.13 | 0.07 |
| 6 | 444600 | rs9405675 | 0.955 | -1.878 | -0.311 | -0.217 | 0.056 | 0.071 | 0.071 | 0 | 0.56 | 0.21 | 0.56 | 0.36 | 0.63 | 0.47 |
| 6 | 449358 | rs9405681 | 1.156 | -1.744 | -0.013 | -0.064 | -0.031 | 0.01 | 0.01 | 0.435 | 0.71 | 0.47 | 0.76 | 0.74 | 0.75 | 0.87 |
| 6 | 449451 | rs9405683 | 0.051 | -1.754 | 0.001 | -0.02 | -0.011 | 0.024 | 0.024 | 0 | 0.28 | 0.34 | 0.29 | 0.39 | 0.24 | 0.45 |
| 6 | 451925 | rs4311550 | 1.189 | -1.689 | -0.191 | -0.063 | -0.059 | 0.076 | 0.076 | 1.141 | 0.44 | 0.1 | 0.49 | 0.34 | 0.51 | 0.3 |
| 6 | 455419 | rs908026 | 1.146 | -1.726 | 0.084 | -0.093 | -0.002 | 0.007 | 0.007 | 0 | 0.43 | 0.36 | 0.5 | 0.34 | 0.5 | 0.29 |
| 6 | 457748 | rs4959270 | 1.406 | -1.615 | -0.083 | -0.054 | -0.046 | 0.035 | 0.035 | 0 | 0.38 | 0.2 | 0.47 | 0.34 | 0.48 | 0.28 |
| 6 | 460732 | rs2316795 | 0.542 | -1.989 | -0.104 | 0.032 | -0.029 | 0.01 | 0.01 | 0.64 | 0.72 | 0.54 | 0.78 | 0.74 | 0.76 | 0.87 |
| 6 | 462299 | rs962517 | 1.384 | -1.889 | -0.864 | 0.058 | -0.133 | 0.007 | 0.007 | 0.165 | 0.35 | 0.07 | 0.48 | 0.27 | 0.49 | 0.31 |
| 6 | 463753 | rs908024 | -0.224 | -1.879 | -0.64 | 0.169 | 0.011 | 0 | 0 | 0 | 0.4 | 0.31 | 0.3 | 0.44 | 0.29 | 0.58 |
| 6 | 464093 | rs908023 | 0.4 | -1.826 | -0.761 | 0.142 | -0.034 | 0.016 | 0.016 | 0.99 | 0.72 | 0.52 | 0.77 | 0.73 | 0.77 | 0.89 |
| 6 | 464426 | rs868094 | 0.7 | -0.866 | -1.321 | 0.053 | -0.076 | 0.037 | 0.037 | 0 | 0.36 | 0.13 | 0.48 | 0.28 | 0.49 | 0.31 |
| 6 | 466033 | rs1540771 | 0.387 | -0.905 | -1.436 | 0.084 | -0.081 | 0.069 | 0.069 | 0.232 | 0.39 | 0.11 | 0.49 | 0.28 | 0.51 | 0.34 |
| 6 | 466368 | rs1540767 | -0.951 | -1.242 | -0.416 | 0.701 | -0.016 | 0.016 | 0.016 | 0 | 0.34 | 0.56 | 0.26 | 0.63 | 0.3 | 0.63 |
| 6 | 467414 | rs4275100 | 1.139 | -1.532 | -0.3 | 0.949 | -0.041 | 0.025 | 0.025 | 0 | 0.66 | 0.43 | 0.65 | 0.55 | 0.8 | 0.69 |
| 6 | 467533 | rs4339504 | 0.671 | -0.95 | -0.553 | 0.972 | -0.024 | 0.007 | 0.007 | 0 | 0.85 | 0.8 | 0.81 | 0.8 | 0.93 | 0.79 |
| 6 | 467773 | rs4959923 | 0.803 | -0.864 | -0.575 | 0.973 | -0.012 | 0.001 | 0.001 | 0 | 0.83 | 0.8 | 0.8 | 0.81 | 0.91 | 0.78 |
| 6 | 470611 | rs4959924 | -0.036 | -0.168 | -1.429 | 0.987 | -0.151 | -0.074 | -0.074 | 0 | 0.37 | 0.04 | 0.44 | 0.38 | 0.51 | 0.32 |
| 6 | 471435 | rs9503944 | -1.02 | -0.746 | 0.137 | 1.278 | -0.005 | 0.066 | 0.066 | 0.017 | 0.11 | 0.25 | 0.14 | 0.15 | 0.07 | 0.28 |
| 6 | 473248 | rs6597037 | 1.131 | -1.119 | 0.5 | 1.037 | -0.039 | -0.008 | -0.008 | 0 | 0.79 | 0.64 | 0.8 | 0.83 | 0.86 | 0.66 |
| 6 | 474252 | rs9392029 | 1.284 | -1.187 | 0.648 | 1.01 | -0.052 | -0.016 | -0.016 | 0 | 0.77 | 0.54 | 0.78 | 0.83 | 0.85 | 0.67 |
| 6 | 475154 | rs9503977 | 1.375 | -1.072 | 0.554 | 0.97 | -0.038 | -0.016 | -0.016 | 0 | 0.77 | 0.57 | 0.77 | 0.83 | 0.82 | 0.66 |
| 6 | 477738 | rs7750350 | 1.338 | -0.997 | 0.425 | 1.591 | -0.017 | 0.05 | 0.05 | 0.71 | 0.69 | 0.53 | 0.53 | 0.5 | 0.72 | 0.53 |
| 6 | 480628 | rs9392573 | 1.239 | -1.165 | 0.52 | 1.232 | -0.032 | 0.038 | 0.038 | 1.495 | 0.71 | 0.46 | 0.56 | 0.62 | 0.78 | 0.62 |
| 6 | 483817 | rs10900954 | 0.981 | -1.297 | -0.121 | 0.908 | 0.071 | -0.016 | -0.016 | 0 | 0.59 | 0.14 | 0.5 | 0.63 | 0.71 | 0.55 |
| 6 | 485078 | rs974455 | 0.921 | -1.333 | 0.305 | 0.92 | 0.015 | 0.002 | 0.002 | 0.202 | 0.62 | 0.34 | 0.57 | 0.69 | 0.73 | 0.6 |
| 6 | 485586 | rs11242914 | 1.122 | -1.368 | -0.294 | 1.043 | 0.101 | -0.025 | -0.025 | 1.215 | 0.57 | 0.11 | 0.54 | 0.65 | 0.72 | 0.5 |
| 6 | 485620 | rs6597054 | -0.32 | -1.353 | 0.106 | 0.497 | -0.041 | -0.022 | -0.022 | 0.131 | 0.17 | 0.25 | 0.12 | 0.07 | 0.08 | 0.28 |
| 6 | 486193 | rs12952 | 0.511 | -0.899 | -0.747 | 0.751 | 0.06 | 0.041 | 0.041 | 0.98 | 0.6 | 0.21 | 0.7 | 0.75 | 0.71 | 0.56 |
| 6 | 496555 | rs9405242 | 0.064 | -0.337 | -0.926 | 0.611 | -0.019 | 0.005 | 0.005 | 0 | 0.38 | 0.31 | 0.39 | 0.71 | 0.56 | 0.42 |
| 6 | 511080 | rs9392615 | -0.634 | -0.852 | -0.479 | 1.219 | -0.021 | 0.101 | 0.101 | 0.049 | 0.2 | 0.4 | 0.23 | 0.44 | 0.14 | 0.14 |
| 6 | 516350 | rs2493013 | 1.066 | -0.904 | -0.367 | 1.547 | -0.018 | 0.016 | 0.016 | 0.868 | 0.7 | 0.51 | 0.77 | 0.42 | 0.78 | 0.48 |
| 6 | 518059 | rs9504180 | -0.294 | -0.851 | -0.285 | 1.503 | -0.004 | -0.005 | -0.005 | 0 | 0.18 | 0.27 | 0.13 | 0.14 | 0.13 | 0.1 |
| 6 | 518078 | rs9392056 | 1.041 | -1.572 | -1.006 | 0.643 | 0.046 | -0.042 | -0.042 | 0 | 0.52 | 0.12 | 0.5 | 0.48 | 0.63 | 0.71 |
| 6 | 526744 | rs2473475 | 0.127 | -1.537 | -0.901 | 0.623 | -0.036 | -0.154 | -0.154 | 0.511 | 0.24 | 0.41 | 0.31 | 0.05 | 0.19 | 0.12 |
| 6 | 530459 | rs2473479 | -0.176 | -1.557 | -0.914 | 0.677 | -0.032 | 0.029 | 0.029 | 0 | 0.27 | 0.54 | 0.33 | 0.38 | 0.19 | 0.15 |
| 6 | 530476 | rs6938671 | 0.202 | -1.553 | -0.922 | 0.67 | -0.018 | 0.016 | 0.016 | 0 | 0.7 | 0.45 | 0.64 | 0.62 | 0.76 | 0.82 |
| 6 | 533866 | rs2493037 | 0.567 | -1.813 | -1.116 | 0.45 | -0.036 | -0.116 | -0.116 | 0 | 0.16 | 0.42 | 0.22 | 0.02 | 0.12 | 0.06 |
| 6 | 535839 | rs2493040 | -1.116 | -1.819 | -1.435 | 0.476 | 0.065 | -0.068 | -0.068 | 0 | 0.45 | 0.88 | 0.47 | 0.52 | 0.32 | 0.25 |
| 6 | 537157 | rs11242982 | 1.541 | -1.39 | -2.37 | -0.188 | -0.14 | -0.075 | -0.075 | 0 | 0.25 | 0.02 | 0.25 | 0.3 | 0.39 | 0.18 |
| 6 | 541668 | rs6904638 | -0.04 | -1.104 | -1.72 | -0.134 | 0.001 | 0.008 | 0.008 | 0 | 0.26 | 0.24 | 0.21 | 0.18 | 0.2 | 0.15 |
| 6 | 542159 | rs6918152 | 0.705 | -0.848 | -1.967 | 0.048 | 0.044 | -0.043 | -0.043 | 0 | 0.52 | 0.12 | 0.5 | 0.48 | 0.63 | 0.72 |
| 6 | 549364 | rs2473490 | -0.297 | 0.366 | -0.975 | -0.185 | 0.001 | -0.019 | -0.019 | 0 | 0.39 | 0.61 | 0.27 | 0.16 | 0.4 | 0.22 |
| 6 | 566509 | rs17134857 | 0.79 | 0.465 | -1.586 | -0.174 | -0.024 | -0.002 | -0.002 | 0 | 0.36 | 0.1 | 0.52 | 0.33 | 0.4 | 0.4 |
| 6 | 566741 | rs2476842 | -0.8 | 0.901 | -0.779 | -0.14 | 0.012 | 0.095 | 0.095 | 0 | 0.49 | 0.71 | 0.51 | 0.7 | 0.5 | 0.44 |
| 6 | 570556 | rs846167 | 0.625 | 1.003 | -0.215 | 0.501 | 0.014 | 0.201 | 0.201 | 0 | 0.94 | 0.87 | 0.92 | 0.59 | 0.91 | 0.94 |
| 6 | 572480 | rs2073009 | 0.817 | 1.104 | -0.004 | 0.066 | -0.006 | 0.015 | 0.015 | 0 | 0.59 | 0.36 | 0.67 | 0.35 | 0.58 | 0.63 |
| 6 | 576902 | rs708038 | 0.111 | 1.32 | 0.448 | 0.187 | 0.014 | 0.007 | 0.007 | 0 | 0.2 | 0.28 | 0.16 | 0.19 | 0.26 | 0.17 |
| 6 | 577820 | rs9504361 | 1.404 | 1.199 | 0.124 | -0.168 | 0.004 | 0.077 | 0.077 | 0 | 0.48 | 0.31 | 0.48 | 0.25 | 0.5 | 0.37 |
| 6 | 580362 | rs1150824 | 1.228 | 0.585 | 0.587 | 0.337 | -0.001 | 0 | 0 | 0 | 0.6 | 0.42 | 0.67 | 0.38 | 0.6 | 0.6 |
| 6 | 586348 | rs3757120 | -0.18 | 0.198 | 0.399 | -0.424 | -0.009 | 0.039 | 0.039 | 0 | 0.17 | 0.07 | 0.07 | 0.05 | 0.15 | 0.1 |
| 6 | 594078 | rs3823133 | -0.477 | 0.042 | 1.059 | -0.479 | -0.025 | -0.02 | -0.02 | 0.495 | 0.09 | 0.25 | 0.04 | 0.09 | 0.04 | 0.14 |
| 6 | 594407 | rs3799311 | 2.093 | 0.116 | 1.45 | -0.404 | -0.002 | 0.003 | 0.003 | 0 | 0.7 | 0.34 | 0.66 | 0.64 | 0.74 | 0.48 |
| 6 | 597871 | rs4960092 | 0.65 | 0.168 | 1.677 | -0.374 | 0.025 | 0.019 | 0.019 | 0 | 0.85 | 0.79 | 0.86 | 0.86 | 0.8 | 0.85 |
| 6 | 602983 | rs17135357 | 0.3 | 0.201 | 1.57 | -0.373 | -0.02 | -0.022 | -0.022 | 0.016 | 0.08 | 0.04 | 0.11 | 0.13 | 0.13 | 0.11 |
| 6 | 604158 | rs1747593 | -0.294 | 0.158 | 1.619 | -0.392 | -0.024 | -0.02 | -0.02 | 0.005 | 0.09 | 0.23 | 0.04 | 0.09 | 0.04 | 0.14 |
| 6 | 605003 | rs6911258 | 1.882 | 0.148 | 1.768 | -0.417 | 0.007 | 0 | 0 | 0 | 0.76 | 0.49 | 0.81 | 0.78 | 0.76 | 0.7 |
| 6 | 605217 | rs3863219 | -0.299 | 0.177 | 1.72 | -0.442 | -0.021 | -0.02 | -0.02 | 0 | 0.09 | 0.23 | 0.04 | 0.09 | 0.04 | 0.14 |
| 6 | 606568 | rs4960096 | 2.097 | 0.039 | 1.457 | -0.616 | -0.009 | 0.046 | 0.046 | 0 | 0.53 | 0.24 | 0.57 | 0.64 | 0.57 | 0.38 |
| 6 | 610088 | rs1737556 | -0.15 | 0.09 | 1.433 | -0.396 | 0.018 | 0.015 | 0.015 | 0 | 0.08 | 0.06 | 0.03 | 0.09 | 0.04 | 0.14 |
| 6 | 612788 | rs9378432 | -0.226 | 0.124 | 1.459 | -0.39 | 0.001 | -0.008 | -0.008 | 0 | 0.95 | 0.96 | 0.98 | 1 | 0.93 | 0.95 |
| 6 | 623284 | rs1150859 | -1.688 | 0.109 | 1.401 | -0.342 | 0.001 | 0.102 | 0.102 | 0 | 0.22 | 0.48 | 0.31 | 0.36 | 0.19 | 0.48 |
| 6 | 625268 | rs9405865 | 0.767 | -0.055 | 0.823 | -0.703 | -0.013 | 0 | 0 | 0 | 0.79 | 0.88 | 0.81 | 0.86 | 0.74 | 0.84 |
| 6 | 625835 | rs2294670 | -0.8 | -0.063 | 1.255 | -0.624 | -0.028 | -0.023 | -0.023 | 0 | 0.1 | 0.32 | 0.05 | 0.09 | 0.04 | 0.14 |

| | | | | | | | | | | | | | | | | |
|---|---|---|---|---|---|---|---|---|---|---|---|---|---|---|---|---|
| 6 | 627382 | rs7772761 | -0.473 | 0.426 | 1.009 | -0.527 | 0.004 | -0.063 | -0.063 | 0 | 0.2 | 0.19 | 0.1 | 0 | 0.17 | 0.09 |
| 6 | 642289 | rs9378445 | 1.279 | 0.499 | 1.128 | 0.061 | 0.053 | 0.135 | 0.135 | 0 | 0.83 | 0.56 | 0.7 | 0.72 | 0.79 | 0.66 |
| 6 | 650071 | rs1612811 | 1.653 | -0.543 | 1.173 | -0.642 | -0.024 | -0.025 | -0.025 | 0 | 0.81 | 0.59 | 0.79 | 0.86 | 0.86 | 0.75 |
| 6 | 654482 | rs9502388 | -1.163 | -0.572 | 1.218 | -0.654 | -0.008 | 0 | 0 | 0 | 0.13 | 0.21 | 0.09 | 0.13 | 0.08 | 0.21 |
| 6 | 654985 | rs9405889 | 1.812 | -0.971 | 1.211 | -1.282 | 0.02 | 0.083 | 0.083 | 0 | 0.55 | 0.31 | 0.65 | 0.79 | 0.65 | 0.52 |
| 6 | 656143 | rs1211554 | 1.984 | -1.009 | 1.477 | -1.195 | -0.054 | -0.035 | -0.035 | 0 | 0.79 | 0.43 | 0.87 | 0.86 | 0.9 | 0.78 |
| 6 | 656343 | rs17136239 | -1.116 | -1.168 | 1.02 | -1.267 | 0.025 | 0.002 | 0.002 | 0 | 0.16 | 0.08 | 0.08 | 0.14 | 0.08 | 0.21 |
| 6 | 656555 | rs1766848 | -0.384 | -0.874 | 1.139 | -1.153 | -0.022 | 0.011 | 0.011 | 0 | 0.07 | 0.01 | 0.13 | 0.07 | 0.1 | 0.06 |
| 6 | 665448 | rs1747579 | -0.186 | -0.733 | 1.321 | -1.063 | 0.001 | -0.003 | -0.003 | 0 | 0.07 | 0.07 | 0.14 | 0.07 | 0.09 | 0.06 |
| 6 | 668627 | rs7454457 | 1.268 | -0.811 | 1.261 | -1.432 | 0.003 | -0.051 | -0.051 | 0.115 | 0.6 | 0.53 | 0.65 | 0.8 | 0.63 | 0.53 |
| 6 | 672798 | rs12333170 | -1.138 | -0.839 | 1.285 | -1.401 | -0.015 | -0.045 | -0.045 | 0 | 0.13 | 0.26 | 0.17 | 0.07 | 0.12 | 0.07 |
| 6 | 680678 | rs1890361 | 1.837 | -0.879 | 1.308 | -1.479 | 0.01 | 0.105 | 0.105 | 0 | 0.53 | 0.35 | 0.62 | 0.8 | 0.6 | 0.52 |
| 6 | 682543 | rs4959379 | -0.924 | -0.852 | 1.321 | -1.479 | -0.012 | -0.043 | -0.043 | 0 | 0.13 | 0.24 | 0.17 | 0.07 | 0.12 | 0.07 |
| 6 | 687753 | rs2039713 | 1.041 | -0.89 | 1.192 | -1.371 | 0.001 | -0.035 | -0.035 | 1.699 | 0.66 | 0.63 | 0.79 | 0.86 | 0.72 | 0.58 |
| 6 | 689565 | rs2317079 | 0.752 | -0.888 | 1.24 | -1.444 | -0.005 | -0.123 | -0.123 | 0 | 0.71 | 0.65 | 0.84 | 1 | 0.77 | 0.78 |
| 6 | 696058 | rs4959381 | -1.577 | -1.422 | 1.449 | -1.014 | -0.028 | -0.031 | -0.031 | 0 | 0.3 | 0.4 | 0.16 | 0.19 | 0.19 | 0.35 |
| 6 | 697754 | rs1890366 | 1.682 | -1.268 | 1.544 | -0.956 | -0.02 | -0.036 | -0.036 | 0 | 0.69 | 0.43 | 0.81 | 0.83 | 0.78 | 0.68 |
| 6 | 701536 | rs11243093 | -0.117 | -1.113 | 1.428 | -0.99 | -0.021 | 0.009 | 0.009 | 0 | 0.07 | 0.02 | 0.18 | 0.03 | 0.1 | 0.06 |
| 6 | 702398 | rs2804731 | 1.912 | -1.135 | 1.597 | -1 | -0.01 | -0.035 | -0.035 | 0 | 0.63 | 0.47 | 0.63 | 0.8 | 0.68 | 0.62 |
| 6 | 708461 | rs1890364 | 1.79 | -1.256 | 1.689 | -1.341 | -0.024 | -0.103 | -0.103 | 0 | 0.68 | 0.51 | 0.66 | 0.9 | 0.73 | 0.73 |
| 6 | 709555 | rs2804736 | 0.343 | -0.854 | 1.806 | -1.144 | -0.003 | -0.02 | -0.02 | 0 | 0.86 | 0.83 | 0.9 | 0.93 | 0.88 | 0.81 |
| 6 | 709765 | rs11968829 | -1.851 | -0.799 | 1.618 | -1.074 | -0.001 | -0.001 | -0.001 | 0 | 0.15 | 0.17 | 0.22 | 0.13 | 0.14 | 0.17 |
| 6 | 711150 | rs2244443 | 1.643 | -1.078 | 1.643 | -1.369 | -0.013 | -0.039 | -0.039 | 1.464 | 0.64 | 0.43 | 0.49 | 0.8 | 0.71 | 0.69 |
| 6 | 711792 | rs17238124 | -2.067 | -1.151 | 1.322 | -1.407 | 0.003 | 0.006 | 0.006 | 0 | 0.2 | 0.18 | 0.39 | 0.13 | 0.19 | 0.24 |
| 6 | 715282 | rs2788222 | 2.078 | -1.094 | 1.455 | -1.942 | 0.006 | -0.008 | -0.008 | 0.773 | 0.58 | 0.42 | 0.47 | 0.87 | 0.66 | 0.67 |
| 6 | 720465 | rs2788223 | 1.562 | -1.097 | 1.543 | -1.872 | -0.027 | -0.082 | -0.082 | 0 | 0.7 | 0.45 | 0.81 | 0.9 | 0.8 | 0.77 |
| 6 | 721999 | rs6900976 | -1.63 | -0.754 | 1.571 | -1.599 | -0.01 | -0.064 | -0.064 | 0 | 0.11 | 0.22 | 0.07 | 0 | 0.08 | 0.1 |
| 6 | 722716 | rs7739861 | -1.687 | -0.755 | 1.593 | -1.599 | -0.007 | -0.068 | -0.068 | 0 | 0.11 | 0.3 | 0.07 | 0 | 0.08 | 0.1 |
| 6 | 722957 | rs7770942 | -1.643 | -0.749 | 1.464 | -1.593 | -0.01 | -0.066 | -0.066 | 0.499 | 0.11 | 0.26 | 0.07 | 0 | 0.08 | 0.1 |
| 6 | 728418 | rs9328385 | -2.008 | -0.798 | 1.083 | -1.687 | -0.002 | -0.083 | -0.083 | 2.604 | 0.2 | 0.29 | 0.42 | 0.01 | 0.22 | 0.2 |
| 6 | 732846 | rs17137319 | -1.597 | -0.62 | 1.279 | -1.472 | 0.019 | -0.063 | -0.063 | 0.649 | 0.13 | 0.25 | 0.38 | 0.01 | 0.16 | 0.11 |
| 6 | 733136 | rs4959408 | -0.664 | -0.594 | 1.005 | -1.458 | 0.007 | -0.025 | -0.025 | 0 | 0.09 | 0.1 | 0.34 | 0.01 | 0.12 | 0.1 |
| 6 | 733163 | rs2804744 | 1.632 | -0.847 | 0.691 | -1.685 | -0.002 | -0.032 | -0.032 | 1.665 | 0.65 | 0.41 | 0.5 | 0.9 | 0.71 | 0.75 |
| 6 | 742311 | rs2788234 | 0.207 | -0.517 | 0.54 | -1.333 | 0.013 | 0.029 | 0.029 | 0.427 | 0.85 | 0.91 | 0.92 | 0.9 | 0.89 | 0.89 |
| 6 | 743488 | rs2788235 | 0.167 | -0.675 | 0.505 | -1.329 | 0.025 | 0.031 | 0.031 | 0.392 | 0.85 | 0.92 | 0.91 | 0.9 | 0.9 | 0.89 |
| 6 | 744573 | rs2788236 | 1.015 | -0.712 | 0.554 | -1.452 | 0.019 | 0.029 | 0.029 | 0 | 0.78 | 0.84 | 0.89 | 0.9 | 0.85 | 0.85 |
| 6 | 753246 | rs2788245 | 1.648 | -0.675 | 0.546 | -1.529 | -0.008 | -0.043 | -0.043 | 0 | 0.69 | 0.41 | 0.67 | 0.89 | 0.74 | 0.78 |
| 6 | 756884 | rs2788197 | 0.226 | -0.543 | -0.026 | -1.155 | 0.025 | 0.037 | 0.037 | 0.404 | 0.85 | 0.95 | 0.92 | 0.9 | 0.91 | 0.89 |
| 6 | 758665 | rs9505014 | -1.245 | 0.179 | 0.551 | -0.782 | 0.013 | -0.035 | -0.035 | 0.117 | 0.19 | 0.27 | 0.28 | 0.07 | 0.23 | 0.27 |
| 6 | 764818 | rs9392857 | -0.851 | 0.08 | 0.165 | -0.615 | 0.005 | 0.017 | 0.017 | 0 | 0.29 | 0.23 | 0.36 | 0.77 | 0.29 | 0.48 |
| 6 | 767471 | rs2804765 | 0.845 | -0.376 | 0.942 | -1.242 | 0.004 | -0.045 | -0.045 | 0.123 | 0.77 | 0.55 | 0.75 | 0.85 | 0.81 | 0.75 |
| 6 | 770769 | rs13211623 | -0.464 | -0.349 | 1.201 | -1.252 | 0.018 | -0.024 | -0.024 | 0 | 0.08 | 0.14 | 0.09 | 0.02 | 0.12 | 0.05 |
| 6 | 771504 | rs7762873 | 1.665 | -0.514 | 1.095 | -1.347 | -0.007 | -0.009 | -0.009 | 0 | 0.84 | 0.32 | 0.86 | 0.86 | 0.88 | 0.91 |
| 6 | 775505 | rs6937708 | -1.754 | -0.506 | 0.914 | -1.693 | -0.005 | 0.021 | 0.021 | 0 | 0.58 | 0.69 | 0.65 | 0.89 | 0.55 | 0.75 |
| 6 | 776262 | rs9406006 | 2.265 | -0.667 | 1.212 | -1.007 | 0.016 | 0.036 | 0.036 | 0.72 | 0.64 | 0.24 | 0.66 | 0.77 | 0.72 | 0.66 |
| 6 | 780441 | rs13219492 | 1.655 | -0.82 | 1.366 | -1.044 | -0.02 | -0.039 | -0.039 | 0.271 | 0.8 | 0.5 | 0.85 | 0.9 | 0.85 | 0.86 |
| 6 | 787110 | rs4960299 | 0.726 | -0.736 | 1.515 | -0.966 | -0.02 | -0.037 | -0.037 | 0.021 | 0.83 | 0.53 | 0.79 | 0.92 | 0.89 | 0.73 |
| 6 | 788169 | rs9328417 | -1.34 | -0.715 | 1.397 | -0.939 | 0.003 | -0.065 | -0.065 | 0 | 0.14 | 0.34 | 0.11 | 0.01 | 0.12 | 0.04 |
| 6 | 791318 | rs9392150 | 0.5 | -0.755 | 1.42 | -1.026 | -0.02 | -0.032 | -0.032 | 0.33 | 0.82 | 0.72 | 0.81 | 0.92 | 0.89 | 0.73 |
| 6 | 792032 | rs6932682 | 1.773 | -0.691 | 1.26 | -1.043 | 0.016 | 0.069 | 0.069 | 0 | 0.65 | 0.25 | 0.68 | 0.9 | 0.72 | 0.68 |
| 6 | 792837 | rs9392155 | 0.016 | -0.717 | 0.821 | 0.032 | -0.011 | 0.05 | 0.05 | 2.994 | 0.79 | 0.61 | 0.86 | 0.67 | 0.82 | 0.92 |
| 6 | 792983 | rs9505192 | 1.678 | -0.822 | 0.67 | 0.028 | -0.014 | -0.004 | -0.004 | 0.419 | 0.49 | 0.17 | 0.57 | 0.58 | 0.54 | 0.6 |
| 6 | 795414 | rs6597267 | 0.044 | -1.017 | 1.114 | 0.179 | -0.021 | 0.089 | 0.089 | 3.251 | 0.78 | 0.52 | 0.84 | 0.67 | 0.82 | 0.87 |
| 6 | 796416 | rs7751939 | 0.938 | -1.088 | 1.11 | 0.117 | 0.003 | 0.001 | 0.001 | 0.084 | 0.64 | 0.32 | 0.71 | 0.67 | 0.67 | 0.61 |
| 6 | 797780 | rs17138546 | -0.298 | -1.037 | 1.125 | 0.027 | 0.009 | -0.072 | -0.072 | 0 | 0.15 | 0.25 | 0.09 | 0 | 0.14 | 0.26 |
| 6 | 798331 | rs10223528 | 1.779 | -1.047 | 0.992 | 0.05 | 0.005 | 0.016 | 0.016 | 1.846 | 0.48 | 0.23 | 0.66 | 0.62 | 0.54 | 0.51 |
| 6 | 798559 | rs7775255 | -0.002 | -1.148 | 0.874 | 0.552 | -0.008 | 0.044 | 0.044 | 2.114 | 0.8 | 0.69 | 0.85 | 0.7 | 0.82 | 0.87 |
| 6 | 799618 | rs7753332 | -0.732 | -1.012 | 1.416 | 0.881 | 0.014 | -0.092 | -0.092 | 0.265 | 0.16 | 0.31 | 0.09 | 0 | 0.14 | 0.25 |
| 6 | 800299 | rs7752216 | -1.854 | -1.006 | 1.229 | 0.878 | -0.019 | -0.078 | -0.078 | 0 | 0.26 | 0.47 | 0.15 | 0.1 | 0.2 | 0.33 |
| 6 | 802769 | rs7756344 | -1.326 | -0.886 | 1.016 | 0.929 | -0.001 | -0.05 | -0.05 | 0 | 0.2 | 0.35 | 0.12 | 0.09 | 0.16 | 0.29 |
| 6 | 802938 | rs7768872 | -1.001 | -0.818 | 0.682 | 0.934 | -0.034 | -0.063 | -0.063 | 0 | 0.23 | 0.37 | 0.13 | 0.09 | 0.14 | 0.34 |
| 6 | 803882 | rs9405338 | 1.731 | -0.893 | 0.503 | 1.054 | 0.038 | 0.068 | 0.068 | 0.128 | 0.64 | 0.12 | 0.66 | 0.75 | 0.73 | 0.54 |
| 6 | 805207 | rs11758860 | -1.147 | -0.865 | 0.486 | 1.039 | -0.007 | -0.059 | -0.059 | 3.189 | 0.24 | 0.35 | 0.14 | 0.09 | 0.21 | 0.36 |
| 6 | 811952 | rs7761470 | 1.53 | -0.78 | 0.39 | 1.301 | 0.045 | -0.046 | -0.046 | 0.483 | 0.58 | 0.09 | 0.52 | 0.36 | 0.69 | 0.4 |
| 6 | 812922 | rs4360172 | 0.936 | -0.626 | 0.591 | 0.255 | -0.021 | 0.042 | 0.042 | 0.831 | 0.81 | 0.47 | 0.76 | 0.76 | 0.85 | 0.83 |
| 6 | 821327 | rs9328435 | -0.908 | -0.184 | 0.304 | 0.165 | -0.03 | -0.148 | -0.148 | 0 | 0.21 | 0.43 | 0.12 | 0 | 0.14 | 0.31 |
| 6 | 827388 | rs4959455 | -1.031 | 0.372 | -0.157 | 1.616 | 0.002 | 0.17 | 0.17 | 0.024 | 0.05 | 0.09 | 0.13 | 0.45 | 0.05 | 0.17 |
| 6 | 829223 | rs4959456 | -1.059 | 0.517 | 0.08 | 1.187 | -0.006 | 0.112 | 0.112 | 0.721 | 0.16 | 0.28 | 0.19 | 0.38 | 0.13 | 0.21 |
| 6 | 830293 | rs4960355 | -0.814 | 0.406 | -1.181 | 0.539 | -0.139 | -0.093 | -0.093 | 0 | 0.68 | 0.95 | 0.51 | 0.69 | 0.57 | 0.68 |

| | | | | | | | | | | | | | | | | |
|---|---|---|---|---|---|---|---|---|---|---|---|---|---|---|---|---|
| 6 | 831356 | rs7757643 | -1.011 | -0.214 | -0.15 | 0.408 | -0.003 | 0.005 | 0.005 | 1.582 | 0.38 | 0.59 | 0.3 | 0.44 | 0.32 | 0.33 |
| 6 | 833579 | rs2317222 | -0.88 | 0.165 | -0.174 | 0.697 | 0.011 | 0.098 | 0.098 | 2.751 | 0.2 | 0.38 | 0.22 | 0.38 | 0.21 | 0.26 |
| 6 | 840475 | rs7770224 | -0.282 | -0.483 | -0.634 | 0.008 | 0.033 | 0.032 | 0.032 | 3.171 | 0.42 | 0.66 | 0.36 | 0.28 | 0.32 | 0.35 |
| 6 | 841861 | rs9406064 | -0.132 | -0.435 | -0.217 | -0.273 | 0.001 | -0.01 | -0.01 | 0 | 0.33 | 0.38 | 0.19 | 0.3 | 0.28 | 0.29 |
| 6 | 858212 | rs9378515 | -0.482 | -1.113 | 0.799 | -0.448 | 0.009 | 0.015 | 0.015 | 0 | 0.45 | 0.44 | 0.3 | 0.42 | 0.36 | 0.43 |
| 6 | 858970 | rs1572438 | -0.286 | -1.099 | 0.868 | -0.459 | -0.001 | 0.014 | 0.014 | 0 | 0.4 | 0.3 | 0.27 | 0.4 | 0.33 | 0.41 |
| 6 | 859115 | rs3799325 | -0.565 | -1.081 | 0.86 | -0.45 | 0.008 | 0.015 | 0.015 | 0 | 0.45 | 0.44 | 0.3 | 0.42 | 0.36 | 0.43 |
| 6 | 866857 | rs17140186 | -0.409 | -2.157 | 0.468 | -1.177 | -0.039 | -0.041 | -0.041 | 4.438 | 0.23 | 0.3 | 0.1 | 0.08 | 0.13 | 0.23 |
| 6 | 866963 | rs11753072 | 0.599 | -2.521 | 0.15 | -1.387 | -0.113 | -0.058 | -0.058 | 0 | 0.38 | 0.08 | 0.6 | 0.54 | 0.53 | 0.3 |
| 6 | 870244 | rs884126 | -1.058 | -2.384 | 0.192 | -1.306 | -0.032 | -0.017 | -0.017 | 0 | 0.67 | 0.74 | 0.39 | 0.47 | 0.48 | 0.71 |
| 6 | 870776 | rs11754896 | 0.463 | -2.455 | 0.29 | -1.269 | 0.031 | 0.033 | 0.033 | 2.152 | 0.18 | 0.1 | 0.07 | 0.06 | 0.1 | 0.21 |
| 6 | 877124 | rs9505394 | -0.945 | -2.152 | 0.604 | -1.048 | 0.036 | 0.037 | 0.037 | 2.191 | 0.07 | 0.38 | 0.05 | 0.02 | 0.04 | 0.06 |
| 6 | 879853 | rs9379179 | -0.009 | -1.709 | -0.314 | -1.043 | -0.013 | 0.052 | 0.052 | 0 | 0.44 | 0.21 | 0.27 | 0.38 | 0.38 | 0.45 |
| 6 | 883074 | rs12207278 | -0.43 | -1.741 | -0.238 | -0.938 | -0.001 | -0.001 | -0.001 | 1.584 | 0.1 | 0.12 | 0.1 | 0.08 | 0.04 | 0.11 |
| 6 | 887977 | rs4960397 | 0.369 | -1.376 | -0.137 | -1.039 | 0.016 | -0.017 | -0.017 | 0.032 | 0.33 | 0.42 | 0.21 | 0.3 | 0.33 | 0.25 |
| 6 | 888109 | rs11754408 | -0.14 | -0.869 | -0.486 | -1.224 | 0.055 | 0.081 | 0.081 | 0 | 0.44 | 0.32 | 0.24 | 0.22 | 0.34 | 0.46 |
| 6 | 888239 | rs13203409 | 0.763 | -0.147 | -0.417 | -0.555 | -0.082 | -0.106 | -0.106 | 0.021 | 0.17 | 0.04 | 0.51 | 0.49 | 0.29 | 0.2 |
| 6 | 888734 | rs6922858 | 0.676 | -0.912 | 0.241 | -1.145 | -0.038 | -0.081 | -0.081 | 0.065 | 0.8 | 0.39 | 0.9 | 0.99 | 0.9 | 0.88 |
| 6 | 889708 | rs9328467 | 1.215 | -0.214 | 0.317 | -0.828 | 0.028 | -0.073 | -0.073 | 0.702 | 0.88 | 0.59 | 0.94 | 1 | 0.92 | 0.92 |
| 6 | 890387 | rs9393002 | 0.27 | -0.142 | -0.721 | -1.124 | 0.019 | 0.103 | 0.103 | 0.396 | 0.46 | 0.31 | 0.69 | 0.79 | 0.54 | 0.48 |
| 6 | 890831 | rs12210966 | 0.404 | 0.066 | -1.248 | -0.094 | 0.021 | -0.072 | -0.072 | 0 | 0.26 | 0.09 | 0.17 | 0.36 | 0.17 | 0.26 |
| 6 | 891191 | rs7760887 | -0.452 | -0.092 | -0.126 | -0.322 | 0.034 | 0.04 | 0.04 | 0 | 0.09 | 0.4 | 0.12 | 0.08 | 0.07 | 0.19 |
| 6 | 891210 | rs2756318 | -0.458 | -0.832 | -0.501 | -1.032 | 0.012 | -0.005 | -0.005 | 0.706 | 0.52 | 0.67 | 0.38 | 0.57 | 0.42 | 0.53 |
| 6 | 892309 | rs9379189 | 0.991 | -0.937 | -0.444 | -0.998 | 0.013 | -0.024 | -0.024 | 0 | 0.26 | 0.2 | 0.19 | 0.39 | 0.18 | 0.25 |
| 6 | 893800 | rs2756315 | -0.447 | -1.031 | -0.52 | -1.118 | 0.011 | 0.004 | 0.004 | 0 | 0.49 | 0.61 | 0.38 | 0.47 | 0.4 | 0.52 |
| 6 | 895605 | rs4960405 | 1.118 | -0.874 | -0.748 | -1.161 | -0.046 | 0.074 | 0.074 | 0 | 0.13 | 0.03 | 0.09 | 0 | 0.16 | 0.08 |
| 6 | 896838 | rs9406137 | 1.017 | -0.692 | -0.601 | -0.989 | 0.006 | -0.042 | -0.042 | 0.168 | 0.25 | 0.15 | 0.19 | 0.39 | 0.2 | 0.26 |
| 6 | 903764 | rs2784968 | -0.502 | -0.477 | -0.369 | -1.314 | 0.002 | -0.002 | -0.002 | 0 | 0.48 | 0.53 | 0.37 | 0.23 | 0.43 | 0.38 |
| 6 | 904145 | rs2756313 | -0.111 | -0.634 | -0.465 | -1.584 | 0 | -0.02 | -0.02 | 0 | 0.29 | 0.29 | 0.26 | 0.15 | 0.25 | 0.18 |
| 6 | 907550 | rs1055368 | 0.159 | -0.706 | -0.37 | -1.235 | -0.019 | -0.002 | -0.002 | 2.743 | 0.45 | 0.28 | 0.56 | 0.37 | 0.49 | 0.31 |
| 6 | 911118 | rs9505480 | -0.601 | -0.58 | 0.221 | -1.222 | 0.022 | 0.088 | 0.088 | 0.059 | 0.08 | 0.32 | 0.04 | 0.15 | 0.05 | 0.06 |
| 6 | 911832 | rs4997129 | -0.854 | -0.775 | -0.052 | -1.203 | 0.011 | 0.029 | 0.029 | 0 | 0.7 | 0.51 | 0.68 | 0.63 | 0.67 | 0.54 |
| 6 | 915792 | rs1414773 | 0.134 | -0.854 | 0.035 | -1.178 | -0.029 | -0.031 | -0.031 | 0 | 0.21 | 0.32 | 0.11 | 0.15 | 0.16 | 0.2 |
| 6 | 920240 | rs2784973 | -1.697 | -0.663 | -0.239 | -1.112 | -0.037 | -0.088 | -0.088 | 2.178 | 0.81 | 0.93 | 0.76 | 0.63 | 0.79 | 0.74 |
| 6 | 921867 | rs9502739 | 0.251 | -0.661 | -0.06 | -1.141 | 0.063 | 0.004 | 0.004 | 0 | 0.05 | 0.24 | 0.04 | 0 | 0.06 | 0.16 |
| 6 | 921899 | rs2145915 | -1.705 | -0.662 | -0.131 | -1.14 | -0.037 | -0.088 | -0.088 | 0 | 0.81 | 0.93 | 0.76 | 0.63 | 0.79 | 0.74 |
| 6 | 925581 | rs9505521 | 0.03 | -0.503 | -0.24 | -1.302 | 0.11 | 0.031 | 0.031 | 0 | 0.06 | 0.32 | 0.05 | 0 | 0.07 | 0.15 |
| 6 | 927700 | rs7740771 | -0.654 | -0.62 | -0.753 | -1.313 | -0.002 | -0.034 | -0.034 | 0 | 0.65 | 0.46 | 0.55 | 0.57 | 0.64 | 0.18 |
| 6 | 930792 | rs12202545 | 0.483 | -0.184 | -0.939 | -1.364 | -0.05 | -0.055 | -0.055 | 0.202 | 0.3 | 0.13 | 0.49 | 0.34 | 0.37 | 0.27 |
| 6 | 931640 | rs12200073 | 0.172 | -0.194 | -0.943 | -1.403 | -0.043 | -0.029 | -0.029 | 0.046 | 0.34 | 0.21 | 0.52 | 0.34 | 0.43 | 0.31 |
| 6 | 938900 | rs1764109 | -0.741 | -0.146 | -0.306 | -1.13 | -0.003 | -0.022 | -0.022 | 0 | 0.27 | 0.33 | 0.14 | 0.12 | 0.23 | 0.2 |
| 6 | 942941 | rs9505552 | -0.334 | -0.09 | -0.353 | -1.124 | 0.067 | -0.001 | -0.001 | 0 | 0.08 | 0.27 | 0.05 | 0.07 | 0.08 | 0.13 |
| 6 | 948181 | rs11752902 | 0.387 | -0.108 | -0.626 | -0.895 | 0.011 | 0.054 | 0.054 | 0 | 0.18 | 0.04 | 0.09 | 0.06 | 0.15 | 0.13 |
| 6 | 951385 | rs1336302 | 1.153 | -0.085 | -0.706 | -0.743 | -0.026 | -0.022 | -0.022 | 0 | 0.34 | 0.21 | 0.54 | 0.35 | 0.36 | 0.26 |
| 6 | 955499 | rs716923 | 0.176 | -0.078 | -0.624 | -0.653 | 0.011 | 0.022 | 0.022 | 0 | 0.2 | 0.15 | 0.11 | 0.06 | 0.16 | 0.13 |
| 6 | 955559 | rs723106 | 0.158 | 0.128 | -0.275 | -0.385 | 0 | 0.003 | 0.003 | 0 | 0.59 | 0.48 | 0.67 | 0.41 | 0.58 | 0.46 |
| 6 | 958905 | rs6925446 | -0.762 | -0.294 | -0.155 | -0.395 | -0.019 | 0.03 | 0.03 | 0.093 | 0.18 | 0.4 | 0.27 | 0.21 | 0.15 | 0.22 |
| 6 | 965576 | rs2756293 | -0.865 | -0.185 | -0.433 | -0.228 | 0 | 0 | 0 | 0 | 0.77 | 0.8 | 0.8 | 0.7 | 0.77 | 0.72 |
| 6 | 966237 | rs945542 | -0.824 | -0.249 | -0.589 | -0.327 | -0.008 | -0.007 | -0.007 | 0 | 0.75 | 0.87 | 0.8 | 0.7 | 0.72 | 0.72 |
| 6 | 967985 | rs9379282 | 0.352 | -0.538 | -0.269 | -0.383 | 0.005 | -0.003 | -0.003 | 0.906 | 0.39 | 0.32 | 0.26 | 0.39 | 0.36 | 0.39 |
| 6 | 971454 | rs9505651 | -1.689 | -0.959 | 0.459 | -0.787 | -0.026 | -0.039 | -0.039 | 0.006 | 0.21 | 0.6 | 0.13 | 0.14 | 0.17 | 0.14 |
| 6 | 972155 | rs11755910 | -0.548 | -0.878 | 0.05 | -0.776 | -0.013 | -0.013 | -0.013 | 0 | 0.1 | 0.05 | 0.33 | 0.13 | 0.15 | 0.09 |
| 6 | 972898 | rs2025122 | -0.777 | -0.905 | 0.075 | -0.793 | -0.005 | 0.029 | 0.029 | 0 | 0.13 | 0.08 | 0.07 | 0.07 | 0.15 | 0.09 |
| 6 | 979803 | rs2784956 | -2.855 | -0.991 | 0.019 | -0.859 | 0.017 | 0.05 | 0.05 | 0.652 | 0.41 | 0.88 | 0.51 | 0.31 | 0.39 | 0.34 |
| 6 | 980237 | rs6597398 | -2.233 | -0.603 | 0.569 | -0.694 | -0.016 | -0.01 | -0.01 | 4.662 | 0.17 | 0.72 | 0.1 | 0.12 | 0.09 | 0.17 |
| 6 | 984646 | rs2095580 | -0.916 | -0.53 | 0.486 | -0.782 | -0.016 | -0.007 | -0.007 | 0 | 0.14 | 0.05 | 0.35 | 0.13 | 0.18 | 0.11 |
| 6 | 994790 | rs7738087 | 2.639 | -0.665 | 0.441 | -0.837 | 0.032 | 0.08 | 0.08 | 0 | 0.61 | 0.13 | 0.49 | 0.75 | 0.63 | 0.71 |
| 6 | 997118 | rs7753372 | -1.167 | -0.529 | 0.921 | -0.664 | 0.021 | 0.063 | 0.063 | 0.248 | 0.08 | 0.46 | 0.05 | 0.08 | 0.02 | 0.03 |
| 6 | 998062 | rs845869 | 1.637 | -0.438 | 0.543 | -0.621 | -0.004 | -0.003 | -0.003 | 1.537 | 0.74 | 0.29 | 0.56 | 0.76 | 0.77 | 0.82 |
| 6 | 1004455 | rs7357076 | -1.592 | -0.433 | 0.663 | -0.71 | -0.021 | -0.018 | -0.018 | 0 | 0.25 | 0.55 | 0.26 | 0.21 | 0.2 | 0.16 |
| 6 | 1008959 | rs845896 | 1.12 | -0.169 | -0.13 | -0.864 | -0.017 | -0.07 | -0.07 | 0 | 0.77 | 0.67 | 0.69 | 0.96 | 0.78 | 0.78 |
| 6 | 1016984 | rs10793858 | 1.151 | 0.046 | -0.919 | -0.715 | 0.006 | 0.085 | 0.085 | 0.022 | 0.63 | 0.11 | 0.47 | 0.79 | 0.61 | 0.72 |
| 6 | 1017009 | rs845890 | -0.771 | 0.011 | 0.395 | -0.798 | 0.028 | 0.053 | 0.053 | 0 | 0.14 | 0.36 | 0.2 | 0.18 | 0.18 | 0.06 |
| 6 | 1019362 | rs845885 | 1.16 | -0.069 | 1.602 | -0.93 | -0.023 | -0.142 | -0.142 | 0 | 0.8 | 0.55 | 0.7 | 0.99 | 0.83 | 0.85 |
| 6 | 1022191 | rs1737701 | 0.706 | -0.043 | 1.419 | -0.954 | -0.015 | 0.077 | 0.077 | 0 | 0.87 | 0.99 | 0.76 | 1 | 0.87 | 0.86 |
| 6 | 1022228 | rs909989 | 0.706 | -0.043 | 1.419 | -0.954 | -0.015 | 0.077 | 0.077 | 0 | 0.87 | 0.99 | 0.76 | 1 | 0.87 | 0.86 |
| 6 | 1022562 | rs845880 | -0.63 | 0.558 | 0.989 | -1.693 | 0.01 | -0.009 | -0.009 | 0 | 0.28 | 0.36 | 0.29 | 0.26 | 0.39 | 0.25 |
| 6 | 1023154 | rs707777 | 1.387 | 0.834 | 1.054 | -1.808 | 0.018 | -0.147 | -0.147 | 0 | 0.8 | 0.58 | 0.67 | 1 | 0.78 | 0.83 |
| 6 | 1028789 | rs1764123 | 0.497 | 0.397 | -0.105 | -2.246 | -0.026 | 0.171 | 0.171 | 0.141 | 0.66 | 0.93 | 0.62 | 0.93 | 0.68 | 0.67 |

| | | | | | | | | | | | | | | | | |
|---|---|---|---|---|---|---|---|---|---|---|---|---|---|---|---|---|
| 6 | 1034050 | rs6934943 | 0.774 | 0.448 | 0.229 | -2.248 | -0.011 | -0.08 | -0.08 | 0 | 0.62 | 0.58 | 0.59 | 0.93 | 0.68 | 0.66 |
| 6 | 1038522 | rs2317453 | 0.201 | 0.705 | -0.023 | -2.099 | -0.069 | 0.072 | 0.072 | 0 | 0.88 | 0.99 | 0.7 | 1 | 0.81 | 0.86 |
| 6 | 1041861 | rs9393100 | 0.376 | 0.407 | -0.013 | -1.84 | 0.023 | -0.006 | -0.006 | 0 | 0.21 | 0.17 | 0.1 | 0.18 | 0.12 | 0.27 |
| 6 | 1045503 | rs1331484 | 1.89 | 0.775 | -0.658 | -2.156 | -0.082 | -0.11 | -0.11 | 2.709 | 0.28 | 0.03 | 0.26 | 0.63 | 0.27 | 0.25 |
| 6 | 1047230 | rs11759510 | 0.883 | 0.425 | -0.502 | -2.044 | 0.045 | -0.025 | -0.025 | 0 | 0.18 | 0.06 | 0.05 | 0.18 | 0.07 | 0.24 |
| 6 | 1049017 | rs6925338 | 0.785 | 0.483 | -0.001 | -2.108 | 0.007 | 0.018 | 0.018 | 0.342 | 0.62 | 0.36 | 0.55 | 0.82 | 0.68 | 0.59 |
| 6 | 1049358 | rs6597402 | 0.053 | 0.083 | 0.409 | -1.855 | -0.082 | -0.002 | -0.002 | 0.818 | 0.25 | 0.47 | 0.13 | 0.18 | 0.12 | 0.27 |
| 6 | 1050172 | rs6899610 | 0.452 | 0.253 | -0.136 | -1.768 | -0.001 | -0.001 | -0.001 | 0 | 0.22 | 0.19 | 0.1 | 0.18 | 0.12 | 0.27 |
| 6 | 1053816 | rs12194400 | 0.495 | 0.783 | -0.515 | -1.48 | -0.021 | 0.03 | 0.03 | 0 | 0.41 | 0.28 | 0.36 | 0.74 | 0.42 | 0.41 |
| 6 | 1055655 | rs9505667 | -0.775 | 0.634 | -0.298 | -0.802 | -0.032 | -0.07 | -0.07 | 1.302 | 0.62 | 0.9 | 0.77 | 0.49 | 0.62 | 0.75 |
| 6 | 1056155 | rs11966212 | -2.338 | 0.096 | 1.321 | -0.728 | -0.019 | -0.027 | -0.027 | 1.802 | 0.31 | 0.63 | 0.22 | 0.2 | 0.24 | 0.34 |
| 6 | 1056944 | rs9393101 | -1.045 | 0.227 | 1.762 | -0.394 | -0.007 | -0.006 | -0.006 | 0.064 | 0.11 | 0.18 | 0.07 | 0.13 | 0.1 | 0.15 |
| 6 | 1057018 | rs9393103 | -1.381 | 0.226 | 1.725 | -0.395 | -0.024 | -0.02 | -0.02 | 0.469 | 0.13 | 0.33 | 0.09 | 0.13 | 0.1 | 0.16 |
| 6 | 1057990 | rs6917581 | -1.871 | 0.249 | 1.058 | -0.397 | -0.044 | -0.027 | -0.027 | 0 | 0.22 | 0.6 | 0.15 | 0.17 | 0.14 | 0.23 |
| 6 | 1063883 | rs2143396 | -2.305 | 0.357 | 1.054 | -0.414 | -0.008 | -0.012 | -0.012 | 0 | 0.28 | 0.67 | 0.21 | 0.22 | 0.24 | 0.3 |
| 6 | 1067284 | rs9501682 | -1.389 | 0.362 | 2.013 | -0.379 | -0.002 | 0.024 | 0.024 | 0.35 | 0.07 | 0.28 | 0.05 | 0.04 | 0.01 | 0.06 |
| 6 | 1074922 | rs6905782 | -2.489 | 0.433 | 1.914 | -0.401 | 0.074 | -0.017 | -0.017 | 0 | 0.09 | 0.52 | 0.11 | 0.02 | 0.11 | 0.06 |
| 6 | 1075055 | rs6929735 | -2.413 | 0.437 | 1.947 | -0.402 | 0.105 | -0.007 | -0.007 | 0 | 0.08 | 0.48 | 0.1 | 0 | 0.11 | 0.04 |
| 6 | 1078564 | rs2073019 | -2.09 | 0.286 | 1.848 | -0.283 | -0.042 | -0.019 | -0.019 | 0 | 0.2 | 0.51 | 0.11 | 0.21 | 0.13 | 0.19 |
| 6 | 1079216 | rs6596781 | -2.917 | 0.152 | 1.879 | -0.236 | -0.014 | -0.014 | -0.014 | 0 | 0.27 | 0.64 | 0.2 | 0.24 | 0.22 | 0.27 |
| 6 | 1080098 | rs7750356 | -1.935 | 0.358 | 2.242 | -0.117 | -0.02 | 0.012 | 0.012 | 0 | 0.18 | 0.31 | 0.11 | 0.24 | 0.13 | 0.21 |
| 6 | 1080950 | rs7747742 | -2.232 | 0.52 | 2.192 | -0.038 | 0 | -0.029 | -0.029 | 0 | 0.17 | 0.45 | 0.15 | 0.17 | 0.15 | 0.21 |
| 6 | 1088964 | rs7770094 | -2.449 | 0.673 | 2.03 | 0.023 | 0.009 | 0.01 | 0.01 | 1.604 | 0.2 | 0.51 | 0.16 | 0.24 | 0.18 | 0.21 |
| 6 | 1098413 | rs7356952 | -1.987 | 1.281 | 1.698 | -0.012 | 0.061 | -0.031 | -0.031 | 0 | 0.06 | 0.11 | 0.08 | 0 | 0.15 | 0.03 |
| 6 | 1099806 | rs7745182 | -1.712 | 0.997 | 1.767 | 0.242 | -0.001 | 0.022 | 0.022 | 0 | 0.08 | 0.08 | 0.07 | 0.25 | 0.07 | 0.15 |
| 6 | 1100777 | rs2050158 | -0.636 | 0.803 | 0.94 | -0.114 | -0.084 | -0.059 | -0.059 | 0 | 0.63 | 0.94 | 0.66 | 0.46 | 0.57 | 0.67 |
| 6 | 1107689 | rs7738538 | -1.907 | 0.446 | 2.41 | 0.172 | 0.002 | 0.008 | 0.008 | 0.01 | 0.08 | 0.36 | 0.05 | 0.04 | 0.03 | 0.06 |
| 6 | 1115234 | rs9392255 | -2.966 | 0.42 | 2.438 | 0.123 | -0.018 | 0.003 | 0.003 | 0.903 | 0.19 | 0.52 | 0.1 | 0.22 | 0.13 | 0.15 |
| 6 | 1115338 | rs9378276 | -3.031 | 0.42 | 2.473 | 0.119 | -0.01 | -0.001 | -0.001 | 0 | 0.21 | 0.68 | 0.14 | 0.28 | 0.16 | 0.23 |
| 6 | 1115897 | rs1997773 | -1.391 | 0.448 | 2.113 | 0.175 | 0.022 | 0.067 | 0.067 | 0 | 0.07 | 0.18 | 0.06 | 0.18 | 0.07 | 0.09 |
| 6 | 1123382 | rs7754106 | -2.269 | 0.457 | 1.94 | 0.31 | -0.029 | 0.055 | 0.055 | 1.616 | 0.17 | 0.62 | 0.13 | 0.3 | 0.11 | 0.21 |
| 6 | 1125882 | rs9502831 | -2.003 | 0.61 | 1.133 | 0.245 | -0.016 | 0.073 | 0.073 | 1.681 | 0.16 | 0.49 | 0.1 | 0.24 | 0.11 | 0.15 |
| 6 | 1128724 | rs9392256 | -1.31 | 0.911 | 0.537 | 0.328 | 0.01 | 0.074 | 0.074 | 0.902 | 0.1 | 0.3 | 0.07 | 0.2 | 0.09 | 0.09 |
| 6 | 1132076 | rs13215872 | -1.386 | 1.05 | -0.383 | -0.024 | -0.024 | 0.041 | 0.041 | 0 | 0.07 | 0.01 | 0.04 | 0 | 0.09 | 0.04 |
| 6 | 1135502 | rs6915235 | -0.912 | 1.183 | -0.379 | -0.207 | -0.001 | -0.002 | -0.002 | 0 | 0.17 | 0.17 | 0.17 | 0.08 | 0.23 | 0.15 |
| 6 | 1136474 | rs879243 | -0.755 | 0.346 | -1.26 | -1.052 | -0.005 | 0.143 | 0.143 | 0 | 0.22 | 0.01 | 0.09 | 0 | 0.16 | 0.08 |
| 6 | 1137160 | rs240685 | 0.202 | 0.334 | -1.38 | -0.76 | 0.002 | 0.003 | 0.003 | 0 | 0.43 | 0.41 | 0.39 | 0.57 | 0.45 | 0.6 |
| 6 | 1139117 | rs1331489 | 0.308 | 0.055 | -1.869 | -0.303 | -0.024 | -0.045 | -0.045 | 0 | 0.24 | 0.07 | 0.27 | 0.29 | 0.25 | 0.23 |
| 6 | 1144109 | rs9502839 | -0.232 | -0.328 | -0.938 | -0.327 | 0.008 | 0.001 | 0.001 | 0 | 0.38 | 0.31 | 0.2 | 0.33 | 0.35 | 0.45 |
| 6 | 1145265 | rs12215602 | 0.35 | -0.129 | -1.185 | -0.34 | -0.058 | -0.08 | -0.08 | 0 | 0.24 | 0.03 | 0.27 | 0.28 | 0.25 | 0.23 |
| 6 | 1146443 | rs3817936 | 0.676 | 0.485 | -0.783 | -0.332 | 0.01 | -0.004 | -0.004 | 0 | 0.2 | 0.27 | 0.32 | 0.2 | 0.25 | 0.18 |
| 6 | 1149483 | rs1331487 | -0.514 | 0.212 | -0.788 | -0.765 | -0.001 | 0.003 | 0.003 | 0 | 0.64 | 0.65 | 0.7 | 0.49 | 0.68 | 0.55 |
| 6 | 1149700 | rs7454978 | -0.085 | 0.458 | -0.774 | -0.889 | 0 | -0.008 | -0.008 | 0 | 0.32 | 0.32 | 0.37 | 0.2 | 0.35 | 0.21 |
| 6 | 1149856 | rs9392258 | 1.247 | 0.944 | -1.029 | -0.632 | -0.004 | -0.002 | -0.002 | 0 | 0.17 | 0.11 | 0.18 | 0.19 | 0.15 | 0.16 |
| 6 | 1152071 | rs11751734 | 0.606 | 0.846 | -1.084 | -0.527 | -0.072 | 0.046 | 0.046 | 0.345 | 0.17 | 0.01 | 0.26 | 0.06 | 0.23 | 0.15 |
| 6 | 1161892 | rs9391907 | 0.849 | 1.01 | -0.937 | -0.273 | -0.029 | -0.004 | -0.004 | 0 | 0.2 | 0.09 | 0.2 | 0.2 | 0.23 | 0.18 |
| 6 | 1163451 | rs7748810 | -0.074 | 1.139 | -0.132 | -0.451 | 0.001 | -0.063 | -0.063 | 1.59 | 0.17 | 0.34 | 0.26 | 0.05 | 0.13 | 0.13 |
| 6 | 1163541 | rs410173 | 0.199 | 1.409 | -0.409 | 0.014 | 0.013 | 0.019 | 0.019 | 0 | 0.82 | 0.73 | 0.77 | 0.7 | 0.75 | 0.8 |
| 6 | 1167422 | rs1161761 | -0.687 | 1.309 | -0.429 | 0.24 | 0.007 | 0.027 | 0.027 | 0 | 0.64 | 0.78 | 0.72 | 0.72 | 0.7 | 0.72 |
| 6 | 1167794 | rs6940789 | 0.458 | 1.524 | -0.176 | 0.287 | 0.007 | -0.033 | -0.033 | 0.622 | 0.39 | 0.24 | 0.29 | 0.44 | 0.4 | 0.49 |
| 6 | 1169047 | rs9405444 | -0.679 | 1.446 | 0.479 | -0.17 | 0.019 | 0.009 | 0.009 | 0 | 0.17 | 0.23 | 0.36 | 0.27 | 0.2 | 0.21 |
| 6 | 1169198 | rs9391909 | -0.08 | 1.498 | 0.272 | -0.029 | 0.005 | 0.006 | 0.006 | 0 | 0.07 | 0.07 | 0.14 | 0.21 | 0.1 | 0.12 |
| 6 | 1185373 | rs238013 | -1.633 | 1.058 | 0.482 | -0.366 | -0.003 | -0.024 | -0.024 | 0 | 0.24 | 0.69 | 0.25 | 0.21 | 0.23 | 0.27 |
| 6 | 1185475 | rs238428 | -1.302 | 1 | 1.141 | -0.392 | 0.011 | -0.075 | -0.075 | 0 | 0.21 | 0.44 | 0.21 | 0.12 | 0.23 | 0.2 |
| 6 | 1189589 | rs9378280 | 0.805 | 1.241 | 0.815 | -0.648 | -0.005 | 0.01 | 0.01 | 0 | 0.37 | 0.23 | 0.29 | 0.6 | 0.41 | 0.42 |
| 6 | 1193067 | rs164257 | -0.834 | 1.22 | 0.692 | -0.627 | 0.004 | 0.033 | 0.033 | 0 | 0.58 | 0.67 | 0.5 | 0.72 | 0.63 | 0.62 |
| 6 | 1193415 | rs4959523 | -0.128 | 1.235 | 0.421 | -0.703 | -0.007 | 0.021 | 0.021 | 0 | 0.13 | 0.07 | 0.14 | 0.12 | 0.12 | 0.18 |
| 6 | 1193467 | rs238016 | -1.802 | 1.192 | 0.208 | -0.794 | -0.079 | 0.078 | 0.078 | 0 | 0.7 | 0.98 | 0.73 | 0.86 | 0.7 | 0.78 |
| 6 | 1193988 | rs238018 | -0.934 | 1.193 | 0.218 | -0.787 | -0.058 | 0.122 | 0.122 | 0 | 0.61 | 0.92 | 0.53 | 0.8 | 0.63 | 0.69 |
| 6 | 1199956 | rs460922 | -0.926 | 1.245 | 0.608 | -0.664 | -0.003 | -0.038 | -0.038 | 2.78 | 0.23 | 0.35 | 0.34 | 0.18 | 0.19 | 0.26 |
| 6 | 1202082 | rs238026 | -1.495 | 1.211 | 0.724 | -0.671 | 0.005 | -0.071 | -0.071 | 0 | 0.29 | 0.47 | 0.41 | 0.18 | 0.3 | 0.29 |
| 6 | 1209774 | rs458399 | -1.572 | 1.339 | -0.14 | -0.595 | -0.015 | 0.051 | 0.051 | 0 | 0.68 | 0.85 | 0.7 | 0.78 | 0.68 | 0.71 |
| 6 | 1211523 | rs7750580 | 0.462 | 1.362 | 0.079 | -0.489 | 0 | 0.014 | 0.014 | 0 | 0.39 | 0.26 | 0.28 | 0.6 | 0.39 | 0.41 |
| 6 | 1214888 | rs369656 | -1.376 | 1.316 | -0.071 | -0.643 | -0.009 | 0.126 | 0.126 | 0 | 0.65 | 0.86 | 0.7 | 0.87 | 0.66 | 0.78 |
| 6 | 1216376 | rs459162 | -1.664 | 1.629 | -0.003 | -0.709 | 0.007 | -0.02 | -0.02 | 0.194 | 0.3 | 0.62 | 0.47 | 0.25 | 0.34 | 0.4 |
| 6 | 1216558 | rs4959529 | 0.645 | 1.578 | 0.076 | -0.545 | 0.005 | 0.01 | 0.01 | 0 | 0.35 | 0.24 | 0.23 | 0.61 | 0.32 | 0.38 |
| 6 | 1221088 | rs238031 | -0.935 | 1.501 | 0.328 | -0.429 | 0.047 | 0.037 | 0.037 | 0 | 0.16 | 0.5 | 0.25 | 0.24 | 0.19 | 0.2 |
| 6 | 1221215 | rs171126 | -0.769 | 1.494 | 0.232 | -0.606 | -0.018 | 0.199 | 0.199 | 0 | 0.52 | 0.78 | 0.52 | 0.85 | 0.54 | 0.65 |
| 6 | 1222588 | rs238035 | -0.647 | 1.5 | 0.239 | -0.609 | 0.015 | 0.201 | 0.201 | 0 | 0.5 | 0.76 | 0.45 | 0.85 | 0.46 | 0.57 |

| | | | | | | | | | | | | | | | | |
|---|---|---|---|---|---|---|---|---|---|---|---|---|---|---|---|---|
| 6 | 1223054 | rs238037 | -1.197 | 1.565 | 0.526 | -0.592 | 0.059 | -0.033 | -0.033 | 0 | 0.18 | 0.42 | 0.28 | 0.16 | 0.26 | 0.26 |
| 6 | 1223075 | rs1866018 | 0.56 | 1.525 | 0.522 | -0.374 | -0.009 | -0.002 | -0.002 | 0 | 0.34 | 0.36 | 0.24 | 0.6 | 0.28 | 0.38 |
| 6 | 1223093 | rs238038 | -0.79 | 1.551 | 0.336 | -0.34 | 0.039 | 0.047 | 0.047 | 0 | 0.16 | 0.4 | 0.22 | 0.24 | 0.19 | 0.19 |
| 6 | 1224584 | rs9405449 | 0.484 | 1.19 | -0.415 | -0.635 | 0.005 | 0.015 | 0.015 | 0 | 0.37 | 0.18 | 0.24 | 0.59 | 0.36 | 0.39 |
| 6 | 1226045 | rs9405450 | 0.282 | 0.823 | -0.345 | -0.623 | -0.005 | 0.029 | 0.029 | 0 | 0.38 | 0.27 | 0.33 | 0.64 | 0.44 | 0.48 |
| 6 | 1228065 | rs12526976 | 0.111 | 1 | -0.576 | -0.668 | -0.011 | 0.048 | 0.048 | 0 | 0.11 | 0.02 | 0.15 | 0.05 | 0.08 | 0.13 |
| 6 | 1228644 | rs238048 | -1.124 | 1.162 | -0.643 | -0.685 | -0.082 | 0.16 | 0.16 | 0 | 0.62 | 0.97 | 0.68 | 0.87 | 0.61 | 0.75 |
| 6 | 1229712 | rs238049 | -0.41 | 1.091 | 0.333 | -0.162 | 0.003 | 0.001 | 0.001 | 0 | 0.12 | 0.13 | 0.17 | 0.21 | 0.13 | 0.17 |
| 6 | 1232766 | rs238051 | -0.659 | 1.155 | -0.295 | -0.17 | 0.043 | 0.085 | 0.085 | 0 | 0.14 | 0.39 | 0.22 | 0.29 | 0.13 | 0.19 |
| 6 | 1233229 | rs238053 | -0.858 | 1.189 | -0.725 | -0.015 | -0.002 | -0.014 | -0.014 | 0 | 0.27 | 0.53 | 0.38 | 0.34 | 0.23 | 0.33 |
| 6 | 1239940 | rs238064 | -0.616 | 1.148 | -1.356 | 0.006 | -0.033 | 0.111 | 0.111 | 0 | 0.7 | 0.92 | 0.8 | 0.92 | 0.71 | 0.9 |
| 6 | 1252894 | rs2118263 | -0.036 | 1.337 | -0.84 | 0.821 | -0.033 | -0.014 | -0.014 | 0 | 0.3 | 0.19 | 0.3 | 0.6 | 0.42 | 0.48 |
| 6 | 1254509 | rs12176547 | -0.599 | 0.549 | -0.564 | 1.314 | 0.002 | -0.007 | -0.007 | 0.038 | 0.19 | 0.18 | 0.15 | 0.34 | 0.19 | 0.37 |
| 6 | 1256449 | rs2008503 | -0.567 | 0.513 | -0.638 | 1.301 | 0.003 | -0.057 | -0.057 | 0 | 0.19 | 0.08 | 0.14 | 0.35 | 0.18 | 0.37 |
| 6 | 1257128 | rs2438072 | -1.143 | 0.509 | -0.569 | 0.608 | 0.002 | 0.105 | 0.105 | 0.316 | 0.63 | 0.84 | 0.53 | 0.9 | 0.67 | 0.84 |
| 6 | 1261007 | rs2496305 | -0.095 | 0.145 | -0.047 | 1.215 | -0.029 | -0.016 | -0.016 | 2.306 | 0.36 | 0.2 | 0.28 | 0.43 | 0.45 | 0.43 |
| 6 | 1261609 | rs2459601 | -0.805 | 0.165 | 0.563 | 1.377 | 0.002 | 0.008 | 0.008 | 0 | 0.5 | 0.55 | 0.39 | 0.66 | 0.55 | 0.78 |
| 6 | 1261815 | rs9378605 | -0.319 | -0.569 | 0.579 | 1.846 | -0.017 | 0.064 | 0.064 | 0 | 0.16 | 0.2 | 0.07 | 0.5 | 0.07 | 0.28 |
| 6 | 1264403 | rs2459602 | 0.034 | -0.15 | 0.173 | 1.423 | 0.001 | -0.027 | -0.027 | 2.371 | 0.27 | 0.16 | 0.26 | 0.39 | 0.28 | 0.39 |
| 6 | 1270046 | rs2496281 | -0.189 | -0.507 | 0.549 | 1.385 | -0.016 | 0.021 | 0.021 | 0 | 0.29 | 0.42 | 0.26 | 0.4 | 0.23 | 0.38 |
| 6 | 1274536 | rs2496291 | -0.208 | -0.492 | 0.538 | 1.393 | -0.024 | 0.033 | 0.033 | 0 | 0.29 | 0.5 | 0.26 | 0.4 | 0.23 | 0.39 |
| 6 | 1275006 | rs9392268 | 0.1 | -0.479 | 0.051 | 1.379 | 0.007 | -0.084 | -0.084 | 3.765 | 0.13 | 0 | 0.05 | 0.35 | 0.07 | 0.26 |
| 6 | 1277151 | rs2252693 | -0.794 | -0.308 | 0.377 | 0.819 | -0.013 | 0.007 | 0.007 | 0.46 | 0.38 | 0.65 | 0.34 | 0.47 | 0.31 | 0.55 |
| 6 | 1277197 | rs2496301 | 0 | 0.098 | -0.559 | 0.201 | 0.012 | 0.006 | 0.006 | 0.012 | 0.25 | 0.14 | 0.24 | 0.26 | 0.22 | 0.39 |
| 6 | 1281814 | rs2496302 | -0.428 | 0.443 | -0.378 | -0.414 | 0.005 | 0.071 | 0.071 | 0 | 0.55 | 0.65 | 0.62 | 0.88 | 0.42 | 0.82 |
| 6 | 1282564 | rs1437282 | -1.113 | 0.508 | 0.193 | -0.106 | 0.018 | -0.068 | -0.068 | 0 | 0.14 | 0.35 | 0.1 | 0.02 | 0.14 | 0.08 |
| 6 | 1284697 | rs1550198 | -0.086 | 0.074 | -0.85 | 0.214 | 0.044 | -0.019 | -0.019 | 0.012 | 0.58 | 0.73 | 0.55 | 0.61 | 0.71 | 0.47 |
| 6 | 1286450 | rs2459596 | -0.806 | 0.679 | -0.595 | -0.325 | 0.003 | -0.034 | -0.034 | 0 | 0.14 | 0.17 | 0.09 | 0.01 | 0.16 | 0.07 |
| 6 | 1289462 | rs12209110 | -0.915 | 0.526 | -1.03 | -0.715 | -0.002 | 0.041 | 0.041 | 1.885 | 0.13 | 0.05 | 0.04 | 0.02 | 0.11 | 0.1 |
| 6 | 1289815 | rs6937290 | 0.109 | -0.536 | -1.079 | -2.353 | -0.009 | -0.094 | -0.094 | 0 | 0.33 | 0.33 | 0.17 | 0.01 | 0.25 | 0.19 |
| 6 | 1289926 | rs1437288 | -0.694 | 0.53 | -0.758 | -1.227 | 0.034 | 0.142 | 0.142 | 0.022 | 0.83 | 0.69 | 0.67 | 0.52 | 0.8 | 0.66 |
| 6 | 1295151 | rs6596805 | 0.668 | 0.498 | -0.866 | -1.312 | 0 | 0.009 | 0.009 | 0 | 0.59 | 0.47 | 0.47 | 0.48 | 0.6 | 0.51 |
| 6 | 1295625 | rs9391920 | 0.28 | 0.476 | -0.794 | -1.152 | 0.003 | 0.044 | 0.044 | 0 | 0.71 | 0.62 | 0.52 | 0.51 | 0.71 | 0.61 |
| 6 | 1296166 | rs11753974 | 0.969 | 0.066 | -1.207 | -1.463 | -0.019 | 0.089 | 0.089 | 0 | 0.15 | 0.01 | 0.08 | 0.01 | 0.14 | 0.08 |
| 6 | 1301766 | rs6925983 | -0.437 | 0.357 | -0.626 | -0.775 | 0.025 | 0.09 | 0.09 | 0 | 0.81 | 0.71 | 0.62 | 0.38 | 0.77 | 0.66 |
| 6 | 1302607 | rs4959539 | 0.566 | 0.146 | -0.268 | -1.159 | -0.002 | 0.011 | 0.011 | 0.022 | 0.54 | 0.31 | 0.4 | 0.36 | 0.57 | 0.5 |
| 6 | 1307692 | rs4959540 | -0.907 | 0.689 | 0.307 | -1.417 | -0.002 | -0.025 | -0.025 | 0 | 0.24 | 0.21 | 0.39 | 0 | 0.28 | 0.2 |
| 6 | 1307846 | rs11242674 | 1.301 | 0.773 | 0.884 | -0.465 | 0 | -0.006 | -0.006 | 0 | 0.66 | 0.44 | 0.4 | 0.69 | 0.63 | 0.61 |
| 6 | 1309171 | rs882003 | -0.128 | 0.762 | 0.744 | -0.614 | -0.007 | -0.05 | -0.05 | 0 | 0.17 | 0.2 | 0.21 | 0 | 0.16 | 0.12 |
| 6 | 1309918 | rs893235 | -0.099 | 0.752 | 0.807 | -0.615 | -0.001 | -0.029 | -0.029 | 0 | 0.17 | 0.18 | 0.22 | 0 | 0.17 | 0.12 |
| 6 | 1311644 | rs11755376 | -0.155 | 0.767 | 0.749 | -0.62 | -0.004 | -0.047 | -0.047 | 0 | 0.17 | 0.2 | 0.22 | 0 | 0.18 | 0.12 |
| 6 | 1316451 | rs2006798 | 1.136 | 0.813 | 0.925 | -0.244 | 0.011 | 0.052 | 0.052 | 0 | 0.82 | 0.68 | 0.62 | 0.64 | 0.81 | 0.73 |
| 6 | 1318878 | rs11242675 | 1.397 | 1.307 | 1.157 | 0.17 | 0.026 | 0.057 | 0.057 | 0 | 0.71 | 0.48 | 0.53 | 0.42 | 0.66 | 0.59 |
| 6 | 1331850 | rs6903955 | 1.82 | 1.209 | 0.474 | 0.297 | 0.038 | 0.044 | 0.044 | 0 | 0.5 | 0.3 | 0.3 | 0.39 | 0.43 | 0.37 |
| 6 | 1332197 | rs7746577 | 0.792 | 1.425 | 0.45 | 0.216 | -0.002 | -0.003 | -0.003 | 0 | 0.66 | 0.7 | 0.52 | 0.41 | 0.61 | 0.54 |
| 6 | 1334355 | rs7739076 | 1.62 | 1.439 | 0.327 | 0.343 | 0 | -0.001 | -0.001 | 0.492 | 0.55 | 0.58 | 0.33 | 0.44 | 0.43 | 0.46 |
| 6 | 1334448 | rs7739254 | 0.789 | 1.4 | 0.445 | 0.465 | 0.012 | 0.034 | 0.034 | 0 | 0.67 | 0.6 | 0.51 | 0.45 | 0.62 | 0.62 |
| 6 | 1335969 | rs9689020 | 1.421 | 1.291 | 0.367 | 0.323 | -0.002 | -0.001 | -0.001 | 0.867 | 0.67 | 0.35 | 0.4 | 0.63 | 0.63 | 0.62 |
| 6 | 1336867 | rs7750826 | -0.964 | 1.634 | 0.464 | 0.554 | 0.016 | 0.023 | 0.023 | 0.549 | 0.16 | 0.17 | 0.37 | 0.35 | 0.22 | 0.3 |
| 6 | 1337153 | rs12204462 | 0.562 | 1.743 | 0.446 | 0.555 | -0.021 | 0.018 | 0.018 | 0.017 | 0.05 | 0 | 0.03 | 0.01 | 0.08 | 0.08 |
| 6 | 1337311 | rs11242679 | -0.6 | 1.755 | 0.473 | 0.54 | 0.009 | 0.091 | 0.091 | 0.242 | 0.11 | 0.17 | 0.21 | 0.35 | 0.11 | 0.22 |
| 6 | 1337981 | rs11966550 | -1.024 | 1.75 | 0.786 | 0.456 | 0.035 | 0.067 | 0.067 | 0 | 0.18 | 0.3 | 0.39 | 0.34 | 0.22 | 0.24 |
| 6 | 1340189 | rs9502893 | 1.192 | 1.162 | 0.556 | -0.047 | -0.003 | 0.012 | 0.012 | 0 | 0.56 | 0.41 | 0.37 | 0.61 | 0.57 | 0.49 |
| 6 | 1340548 | rs12200309 | -0.616 | 1.345 | 0.315 | -0.174 | 0.007 | -0.021 | -0.021 | 0 | 0.11 | 0.1 | 0.19 | 0 | 0.19 | 0.16 |
| 6 | 1348230 | rs9405136 | -0.886 | 1.156 | 0.388 | -0.108 | -0.009 | -0.012 | -0.012 | 0 | 0.35 | 0.4 | 0.46 | 0.34 | 0.31 | 0.34 |
| 6 | 1349828 | rs9392279 | -0.205 | 1.26 | -0.456 | -0.141 | -0.002 | -0.052 | -0.052 | 0 | 0.44 | 0.19 | 0.59 | 0.6 | 0.58 | 0.56 |
| 6 | 1351928 | rs938721 | -0.007 | 1.798 | -0.238 | -0.126 | -0.003 | 0.002 | 0.002 | 0.45 | 0.59 | 0.45 | 0.59 | 0.66 | 0.5 | 0.56 |
| 6 | 1359690 | rs7759626 | -0.084 | 1.902 | -0.637 | 0.187 | 0.02 | -0.006 | -0.006 | 0 | 0.65 | 0.54 | 0.62 | 0.66 | 0.57 | 0.65 |
| 6 | 1361844 | rs1737766 | -0.481 | 2.124 | -1.744 | 0.375 | -0.017 | -0.028 | -0.028 | 0 | 0.76 | 0.86 | 0.83 | 0.66 | 0.71 | 0.72 |
| 6 | 1361996 | rs9392284 | -0.341 | 2.156 | -1.451 | 0.188 | -0.033 | -0.045 | -0.045 | 0 | 0.11 | 0.02 | 0.24 | 0.28 | 0.15 | 0.18 |
| 6 | 1368415 | rs1737753 | -0.297 | 1.458 | -1.154 | -0.119 | -0.009 | -0.037 | -0.037 | 1.857 | 0.87 | 0.96 | 0.9 | 0.87 | 0.85 | 0.92 |
| 6 | 1369271 | rs12190545 | 0.442 | 1.205 | -1.143 | -0.081 | 0.004 | -0.018 | -0.018 | 0 | 0.11 | 0 | 0.03 | 0.11 | 0.07 | 0.1 |
| 6 | 1373145 | rs2317906 | 0.067 | 1.859 | -0.886 | -0.298 | -0.02 | 0.023 | 0.023 | 0 | 0.05 | 0.02 | 0.04 | 0 | 0.06 | 0.07 |
| 6 | 1374277 | rs10498654 | 0.156 | 1.751 | -0.539 | -0.386 | -0.003 | -0.022 | -0.022 | 0 | 0.28 | 0.28 | 0.12 | 0.06 | 0.23 | 0.21 |
| 6 | 1374892 | rs6899796 | 0.053 | 1.698 | -0.436 | -0.249 | -0.003 | -0.005 | -0.005 | 0 | 0.35 | 0.31 | 0.23 | 0.06 | 0.39 | 0.24 |
| 6 | 1377218 | rs10214673 | 0.261 | 1.773 | -0.732 | 0.09 | 0.01 | 0.03 | 0.03 | 0.08 | 0.4 | 0.24 | 0.56 | 0.78 | 0.35 | 0.57 |
| 6 | 1380746 | rs1711969 | -0.375 | 1.966 | -0.243 | 0.1 | 0.008 | -0.039 | -0.039 | 0 | 0.34 | 0.42 | 0.28 | 0.19 | 0.4 | 0.24 |
| 6 | 1385259 | rs9405137 | -1.131 | 2.652 | 0.081 | 0.581 | 0.097 | -0.012 | -0.012 | 2.654 | 0.12 | 0.29 | 0.18 | 0.11 | 0.25 | 0.13 |
| 6 | 1386697 | rs1878475 | -0.821 | 1.894 | -0.281 | 1.363 | 0 | 0.014 | 0.014 | 0 | 0.08 | 0.09 | 0.06 | 0.18 | 0.08 | 0.18 |

| | | | | | | | | | | | | | | | | |
|---|---|---|---|---|---|---|---|---|---|---|---|---|---|---|---|---|
| 6 | 1386737 | rs1711970 | 0.09 | 1.856 | -0.301 | 1.143 | 0.013 | 0.008 | 0.008 | 0 | 0.37 | 0.51 | 0.3 | 0.4 | 0.37 | 0.51 |
| 6 | 1392108 | rs1711968 | 0.345 | 1.51 | -1.1 | 0.591 | -0.02 | 0.048 | 0.048 | 0 | 0.6 | 0.79 | 0.69 | 0.74 | 0.56 | 0.7 |
| 6 | 1395043 | rs2293783 | -0.297 | 1.399 | -1.214 | 0.394 | 0.016 | -0.003 | -0.003 | 0 | 0.32 | 0.16 | 0.23 | 0.27 | 0.23 | 0.21 |
| 6 | 1397839 | rs4959557 | 0.752 | 1.596 | -0.627 | -0.044 | 0.031 | -0.069 | -0.069 | 0.031 | 0.86 | 0.74 | 0.77 | 1 | 0.76 | 0.89 |
| 6 | 1402537 | rs1711973 | 0.794 | 1.525 | -0.765 | -0.246 | -0.013 | 0 | 0 | 0.055 | 0.85 | 0.88 | 0.85 | 1 | 0.75 | 0.9 |
| 6 | 1404628 | rs746095 | 0.257 | 1.316 | -1.607 | -0.784 | -0.009 | 0.157 | 0.157 | 0 | 0.51 | 0.75 | 0.68 | 0.8 | 0.49 | 0.6 |
| 6 | 1405284 | rs731394 | 0.409 | 1.148 | -1.579 | -0.697 | 0.025 | 0.082 | 0.082 | 0 | 0.34 | 0.13 | 0.17 | 0.2 | 0.26 | 0.3 |
| 6 | 1406885 | rs1711953 | 0.206 | 0.996 | -1.388 | -0.239 | 0.002 | 0.007 | 0.007 | 0.008 | 0.39 | 0.6 | 0.51 | 0.62 | 0.33 | 0.54 |
| 6 | 1407099 | rs9501718 | 0.563 | 0.495 | -1.214 | -0.041 | -0.013 | 0.095 | 0.095 | 0 | 0.22 | 0.37 | 0.19 | 0.39 | 0.15 | 0.36 |
| 6 | 1408267 | rs9328051 | -0.025 | 0.717 | -1.372 | -0.803 | 0.005 | -0.06 | -0.06 | 0 | 0.7 | 0.68 | 0.61 | 0.99 | 0.51 | 0.82 |
| 6 | 1408465 | rs17133898 | -0.648 | 0.077 | -1.655 | -0.661 | -0.026 | -0.007 | -0.007 | 0 | 0.25 | 0.37 | 0.32 | 0.22 | 0.16 | 0.13 |
| 6 | 1408677 | rs11753773 | 0.648 | 0.364 | -1.694 | -0.264 | 0.063 | 0 | 0 | 0 | 0.5 | 0.6 | 0.4 | 0.55 | 0.3 | 0.66 |
| 6 | 1412674 | rs9378623 | 0.054 | 0.323 | -1.317 | 0.189 | 0.006 | -0.013 | -0.013 | 0 | 0.17 | 0.17 | 0.2 | 0.23 | 0.15 | 0.16 |
| 6 | 1416849 | rs1711959 | -0.473 | 0.529 | -1.296 | 0.39 | -0.002 | 0.019 | 0.019 | 0 | 0.13 | 0.19 | 0.19 | 0.24 | 0.12 | 0.18 |
| 6 | 1418887 | rs6596819 | -0.289 | 0.449 | -1.444 | -0.362 | 0.025 | 0.008 | 0.008 | 0 | 0.3 | 0.19 | 0.27 | 0.22 | 0.24 | 0.29 |
| 6 | 1424362 | rs4317453 | -0.378 | 1.644 | -1.278 | -0.7 | -0.053 | 0.042 | 0.042 | 0 | 0.14 | 0.05 | 0.16 | 0 | 0.32 | 0.1 |
| 6 | 1435220 | rs9502924 | -0.758 | 1.043 | -1.021 | -1.314 | -0.035 | 0.058 | 0.058 | 0 | 0.23 | 0.1 | 0.18 | 0 | 0.32 | 0.13 |
| 6 | 1439976 | rs11242687 | -0.616 | 0.572 | -1.435 | -1.907 | -0.075 | 0.104 | 0.104 | 0 | 0.24 | 0.09 | 0.28 | 0 | 0.38 | 0.12 |
| 6 | 1441282 | rs9405475 | -0.024 | -0.12 | -1.112 | -1.623 | -0.019 | 0.012 | 0.012 | 0.021 | 0.68 | 0.4 | 0.59 | 0.4 | 0.83 | 0.53 |
| 6 | 1441494 | rs2493158 | 0.795 | -0.221 | -0.852 | -1.681 | -0.01 | 0.081 | 0.081 | 0.243 | 0.19 | 0.28 | 0.35 | 0.6 | 0.13 | 0.45 |
| 6 | 1443099 | rs10458124 | 0.424 | -0.145 | -1.004 | -1.649 | -0.001 | -0.001 | -0.001 | 0 | 0.58 | 0.42 | 0.45 | 0.41 | 0.54 | 0.47 |
| 6 | 1443413 | rs11242692 | 0.746 | -0.169 | -1.518 | -1.996 | 0.006 | 0.01 | 0.01 | 0 | 0.48 | 0.46 | 0.35 | 0.27 | 0.37 | 0.41 |
| 6 | 1443760 | rs9328052 | 0.606 | -0.021 | -1.352 | -1.96 | -0.002 | 0.006 | 0.006 | 0 | 0.1 | 0.1 | 0.09 | 0.11 | 0.15 | 0.06 |
| 6 | 1443801 | rs17202895 | 0.908 | -0.013 | -1.358 | -1.989 | -0.038 | 0.029 | 0.029 | 0 | 0.12 | 0.06 | 0.15 | 0 | 0.28 | 0.08 |
| 6 | 1445115 | rs6937443 | 0.309 | -0.163 | -1.356 | -1.944 | 0 | 0.002 | 0.002 | 0 | 0.52 | 0.5 | 0.33 | 0.36 | 0.45 | 0.39 |
| 6 | 1446047 | rs1922938 | -0.205 | -0.309 | -1.331 | -1.875 | -0.016 | 0.002 | 0.002 | 0 | 0.63 | 0.63 | 0.49 | 0.36 | 0.75 | 0.45 |
| 6 | 1447308 | rs975513 | 1.079 | -0.281 | -1.344 | -1.658 | -0.001 | -0.001 | -0.001 | 0 | 0.32 | 0.32 | 0.21 | 0.25 | 0.28 | 0.31 |
| 6 | 1449121 | rs2745636 | -0.005 | -0.471 | -1.449 | -1.576 | 0.011 | -0.005 | -0.005 | 0.018 | 0.39 | 0.45 | 0.69 | 0.68 | 0.51 | 0.62 |
| 6 | 1450581 | rs12213362 | 0.433 | -0.189 | -1.339 | -1.272 | 0 | -0.009 | -0.009 | 0 | 0.41 | 0.46 | 0.27 | 0.32 | 0.42 | 0.34 |
| 6 | 1451437 | rs932408 | -0.252 | 0.013 | -1.339 | -0.942 | -0.003 | 0.026 | 0.026 | 0 | 0.46 | 0.62 | 0.57 | 0.68 | 0.46 | 0.59 |
| 6 | 1451582 | rs932410 | 0.147 | -0.146 | -1.639 | -0.262 | -0.001 | 0.052 | 0.052 | 0.182 | 0.46 | 0.27 | 0.52 | 0.71 | 0.45 | 0.67 |
| 6 | 1452004 | rs9405481 | -0.615 | -0.283 | -1.252 | 1.313 | 0.008 | 0.023 | 0.023 | 0.009 | 0.1 | 0.08 | 0.09 | 0.4 | 0.04 | 0.08 |
| 6 | 1454972 | rs2816252 | 0.145 | 0.346 | -1.6 | 1.156 | -0.059 | -0.063 | -0.063 | 0.028 | 0.73 | 0.94 | 0.62 | 0.66 | 0.77 | 0.45 |
| 6 | 1455570 | rs1474721 | -0.478 | 0.147 | -0.887 | 0.607 | -0.017 | -0.038 | -0.038 | 0 | 0.26 | 0.49 | 0.17 | 0.15 | 0.23 | 0.19 |
| 6 | 1464671 | rs17133981 | -0.397 | 0.177 | -0.826 | 0.909 | -0.001 | -0.002 | -0.002 | 0.009 | 0.09 | 0.07 | 0.03 | 0.1 | 0.1 | 0.05 |
| 6 | 1464679 | rs2816272 | 0.29 | 0.017 | -0.561 | 1.246 | -0.001 | -0.001 | -0.001 | 0.695 | 0.6 | 0.58 | 0.52 | 0.59 | 0.68 | 0.35 |
| 6 | 1479768 | rs6927891 | 0.081 | 0.083 | -0.54 | 0.514 | 0.021 | 0.029 | 0.029 | 0 | 0.49 | 0.62 | 0.43 | 0.65 | 0.53 | 0.37 |
| 6 | 1480008 | rs13192008 | -0.266 | -0.174 | -0.703 | 0.061 | -0.007 | -0.042 | -0.042 | 0 | 0.88 | 0.87 | 0.94 | 0.98 | 0.89 | 0.92 |
| 6 | 1486425 | rs4434508 | -0.181 | 0.423 | -0.972 | -0.168 | -0.001 | -0.04 | -0.04 | 0 | 0.85 | 0.86 | 0.9 | 0.98 | 0.82 | 0.91 |
| 6 | 1488725 | rs7762812 | 0.209 | 0.47 | -0.975 | -0.164 | 0.001 | -0.013 | -0.013 | 0 | 0.82 | 0.85 | 0.85 | 0.96 | 0.7 | 0.88 |
| 6 | 1490104 | rs9502950 | 0.838 | 0.039 | -0.056 | -0.262 | -0.004 | 0.002 | 0.002 | 0.029 | 0.42 | 0.36 | 0.37 | 0.64 | 0.45 | 0.35 |
| 6 | 1490582 | rs6906804 | -0.408 | -0.771 | -0.323 | -0.584 | -0.001 | 0 | 0 | 2.737 | 0.28 | 0.31 | 0.43 | 0.32 | 0.22 | 0.37 |
| 6 | 1490941 | rs4131572 | 0.621 | -0.437 | -0.82 | -0.54 | 0 | -0.02 | -0.02 | 0 | 0.1 | 0.03 | 0.11 | 0.64 | 0.07 | 0.21 |
| 6 | 1492662 | rs4618572 | -0.471 | -0.244 | -0.601 | -0.803 | 0.002 | -0.045 | -0.045 | 0 | 0.88 | 0.86 | 0.92 | 0.98 | 0.85 | 0.92 |
| 6 | 1494957 | rs11752453 | -0.101 | -0.445 | -0.864 | -0.778 | 0.003 | 0.03 | 0.03 | 0 | 0.19 | 0.12 | 0.21 | 0.08 | 0.2 | 0.14 |
| 6 | 1495954 | rs4146690 | -0.089 | -0.743 | -0.379 | -0.937 | -0.004 | -0.027 | -0.027 | 0 | 0.87 | 0.64 | 0.9 | 0.95 | 0.93 | 0.94 |
| 6 | 1498457 | rs9405484 | 0.302 | -0.995 | -0.758 | -0.598 | 0.02 | -0.088 | -0.088 | 0 | 0.25 | 0.1 | 0.18 | 0.61 | 0.19 | 0.37 |
| 6 | 1500565 | rs12660207 | -0.631 | -1.219 | -0.746 | -0.759 | -0.001 | -0.001 | -0.001 | 0 | 0.29 | 0.31 | 0.42 | 0.3 | 0.22 | 0.37 |
| 6 | 1502398 | rs13211750 | 0.32 | -0.215 | -1.111 | -0.851 | 0.014 | -0.011 | -0.011 | 0 | 0.29 | 0.38 | 0.42 | 0.23 | 0.35 | 0.38 |
| 6 | 1504122 | rs10900909 | -0.217 | -0.151 | -1.592 | -0.903 | -0.048 | 0.101 | 0.101 | 0 | 0.33 | 0.11 | 0.3 | 0.11 | 0.4 | 0.19 |
| 6 | 1505511 | rs4443540 | -0.219 | 0.024 | -1.41 | -0.638 | -0.012 | 0.026 | 0.026 | 0 | 0.18 | 0.07 | 0.22 | 0.2 | 0.2 | 0.26 |
| 6 | 1506180 | rs7763653 | -0.375 | 0.28 | -1.294 | -0.507 | -0.022 | 0.002 | 0.002 | 0 | 0.26 | 0.16 | 0.47 | 0.3 | 0.33 | 0.37 |
| 6 | 1509347 | rs9392301 | 0.469 | 0.455 | -1.595 | 0.532 | 0.007 | -0.072 | -0.072 | 0 | 0.13 | 0.02 | 0.04 | 0.35 | 0.11 | 0.19 |
| 6 | 1510231 | rs6905465 | -0.898 | 0.655 | -0.89 | -0.462 | 0.011 | -0.087 | -0.087 | 0 | 0.78 | 0.74 | 0.83 | 1 | 0.73 | 0.86 |
| 6 | 1513383 | rs7755969 | -0.837 | 0.629 | -0.768 | -0.475 | 0.018 | -0.113 | -0.113 | 1.334 | 0.77 | 0.7 | 0.82 | 1 | 0.73 | 0.86 |
| 6 | 1514367 | rs7761242 | -0.916 | 0.669 | -0.977 | -0.454 | -0.002 | -0.04 | -0.04 | 1.368 | 0.78 | 0.81 | 0.83 | 1 | 0.73 | 0.86 |
| 6 | 1515374 | rs3935776 | 0.087 | 0.279 | -0.747 | 0.194 | -0.001 | 0 | 0 | 0 | 0.32 | 0.38 | 0.26 | 0.33 | 0.29 | 0.3 |
| 6 | 1528282 | rs11242703 | 0.19 | 0.433 | -0.838 | 0.283 | -0.002 | -0.002 | -0.002 | 0.803 | 0.51 | 0.45 | 0.61 | 0.54 | 0.49 | 0.57 |
| 6 | 1529787 | rs2569865 | 0.35 | 0.688 | -1.557 | 0.057 | -0.04 | -0.026 | -0.026 | 0 | 0.46 | 0.19 | 0.58 | 0.5 | 0.53 | 0.59 |
| 6 | 1529872 | rs2816283 | -0.272 | 0.42 | -1.128 | 0.061 | -0.017 | -0.007 | -0.007 | 0 | 0.65 | 0.51 | 0.82 | 0.67 | 0.73 | 0.75 |
| 6 | 1530985 | rs6937266 | 0.378 | 0.551 | -1.577 | -0.316 | -0.081 | 0.12 | 0.12 | 2.249 | 0.24 | 0.09 | 0.26 | 0.04 | 0.34 | 0.16 |
| 6 | 1532165 | rs11962452 | -1.029 | 0.52 | -1.336 | -0.008 | -0.037 | -0.033 | -0.033 | 0 | 0.46 | 0.74 | 0.49 | 0.47 | 0.47 | 0.46 |
| 6 | 1533447 | rs4959569 | -0.665 | 0.329 | -1.291 | 0.043 | 0.019 | -0.064 | -0.064 | 0.621 | 0.38 | 0.74 | 0.33 | 0.47 | 0.3 | 0.36 |
| 6 | 1539528 | rs2569829 | 0.493 | 0.455 | -0.682 | 0.185 | 0.037 | 0.042 | 0.042 | 0.017 | 0.16 | 0.29 | 0.23 | 0.17 | 0.19 | 0.15 |
| 6 | 1545513 | rs2317965 | -0.96 | 0.6 | -1.142 | 0.232 | -0.049 | -0.073 | -0.073 | 0.028 | 0.31 | 0.76 | 0.47 | 0.47 | 0.38 | 0.3 |
| 6 | 1550886 | rs2317966 | 0.773 | 0.214 | -0.741 | 0.397 | 0.001 | -0.002 | -0.002 | 0.144 | 0.62 | 0.34 | 0.55 | 0.59 | 0.64 | 0.44 |
| 6 | 1558449 | rs1535011 | 0.92 | 0.019 | -0.168 | 0.496 | -0.015 | 0.068 | 0.068 | 0 | 0.21 | 0.28 | 0.11 | 0.38 | 0.17 | 0.16 |
| 6 | 1564371 | rs2317967 | 1.171 | 0.577 | -0.327 | 0.122 | -0.003 | -0.003 | -0.003 | 0 | 0.1 | 0.1 | 0.2 | 0 | 0.24 | 0.12 |
| 6 | 1568046 | rs2206602 | -0.303 | 0.446 | -0.402 | 0.127 | 0 | 0.002 | 0.002 | 0 | 0.6 | 0.71 | 0.77 | 0.62 | 0.59 | 0.66 |

| | | | | | | | | | | | | | | | | |
|---|---|---|---|---|---|---|---|---|---|---|---|---|---|---|---|---|
| 6 | 1569619 | rs12530463 | 1.08 | 0.711 | -0.672 | -0.109 | -0.039 | 0.035 | 0.035 | 0.088 | 0.1 | 0.05 | 0.19 | 0 | 0.24 | 0.11 |
| 6 | 1573613 | rs2338 | 0.779 | 0.344 | -0.604 | -0.049 | 0.026 | 0.023 | 0.023 | 4.218 | 0.37 | 0.24 | 0.51 | 0.31 | 0.31 | 0.31 |
| 6 | 1573756 | rs2206603 | 0.776 | 0.344 | -0.604 | -0.049 | 0.026 | 0.023 | 0.023 | 4.148 | 0.37 | 0.23 | 0.51 | 0.31 | 0.31 | 0.31 |
| 6 | 1577436 | rs6934209 | -0.014 | 1.039 | -0.269 | 0.167 | 0.075 | 0.056 | 0.056 | 0.395 | 0.17 | 0.31 | 0.23 | 0.25 | 0.28 | 0.25 |
| 6 | 1577662 | rs2569838 | -0.86 | 0.541 | -0.951 | -0.736 | -0.006 | 0.017 | 0.017 | 0 | 0.79 | 0.86 | 0.86 | 1 | 0.78 | 0.8 |
| 6 | 1579190 | rs9378638 | 0.035 | -0.009 | -0.843 | -0.476 | 0.049 | 0.019 | 0.019 | 5.736 | 0.36 | 0.15 | 0.5 | 0.36 | 0.28 | 0.26 |
| 6 | 1580403 | rs9378300 | -0.567 | 0.366 | -0.329 | -0.076 | 0 | 0.001 | 0.001 | 0 | 0.06 | 0.04 | 0.02 | 0.08 | 0.03 | 0.08 |
| 6 | 1580604 | rs2816221 | -0.529 | 0.021 | -0.054 | -0.174 | -0.022 | -0.037 | -0.037 | 1.922 | 0.12 | 0.26 | 0.06 | 0.05 | 0.07 | 0.13 |
| 6 | 1580838 | rs9405141 | -0.433 | 0.306 | 0.05 | 0.316 | -0.001 | 0.015 | 0.015 | 0.245 | 0.13 | 0.16 | 0.28 | 0.29 | 0.11 | 0.25 |
| 6 | 1580957 | rs2816222 | -0.679 | 0.459 | -0.108 | -0.595 | 0 | -0.03 | -0.03 | 0 | 0.43 | 0.42 | 0.62 | 0.92 | 0.42 | 0.57 |
| 6 | 1581064 | rs6925535 | -0.798 | 1.952 | 0.148 | 0.352 | 0.037 | -0.05 | -0.05 | 0 | 0.17 | 0.23 | 0.21 | 0.06 | 0.24 | 0.18 |
| 6 | 1581803 | rs2816223 | 1.053 | 1.124 | -0.199 | 1.167 | 0.002 | 0.025 | 0.025 | 0.995 | 0.83 | 0.84 | 0.57 | 0.66 | 0.76 | 0.65 |
| 6 | 1582162 | rs910016 | 0.497 | 0.962 | -0.607 | 0.5 | -0.021 | 0.034 | 0.034 | 0.885 | 0.94 | 0.99 | 0.86 | 1 | 0.89 | 0.87 |
| 6 | 1582559 | rs6916316 | -1 | 0.802 | -0.258 | 0.902 | -0.007 | 0.008 | 0.008 | 0.684 | 0.27 | 0.31 | 0.25 | 0.33 | 0.22 | 0.21 |
| 6 | 1583600 | rs7772118 | -0.88 | 0.815 | -0.481 | 0.88 | 0.001 | -0.002 | -0.002 | 0 | 0.24 | 0.22 | 0.23 | 0.28 | 0.2 | 0.17 |
| 6 | 1584384 | rs2816226 | -1.053 | 1.981 | -0.142 | 0.332 | 0.056 | -0.05 | -0.05 | 0 | 0.28 | 0.44 | 0.38 | 0.15 | 0.39 | 0.21 |
| 6 | 1584451 | rs2745575 | -0.167 | 1.72 | -0.674 | 0.536 | -0.012 | 0.002 | 0.002 | 0 | 0.45 | 0.72 | 0.58 | 0.42 | 0.51 | 0.42 |
| 6 | 1584974 | rs4959580 | -0.259 | 1.953 | -0.693 | 0.093 | -0.008 | 0.001 | 0.001 | 0 | 0.2 | 0.15 | 0.18 | 0.02 | 0.26 | 0.09 |
| 6 | 1589002 | rs742554 | -0.67 | 1.47 | -0.389 | 0.274 | -0.001 | 0.037 | 0.037 | 0 | 0.44 | 0.63 | 0.44 | 0.87 | 0.41 | 0.62 |
| 6 | 1590396 | rs2281272 | -0.274 | 0.934 | -0.318 | 0.981 | -0.002 | 0.027 | 0.027 | 0 | 0.13 | 0.16 | 0.08 | 0.26 | 0.1 | 0.27 |
| 6 | 1591011 | rs2076513 | -0.326 | 0.922 | -0.305 | 0.759 | -0.003 | -0.004 | -0.004 | 0 | 0.4 | 0.54 | 0.34 | 0.62 | 0.34 | 0.51 |
| 6 | 1593851 | rs4591901 | -1.247 | 0.702 | 0.007 | 0.909 | 0.004 | 0.051 | 0.051 | 0 | 0.11 | 0.31 | 0.11 | 0.18 | 0.11 | 0.15 |
| 6 | 1595216 | rs2249437 | 0.209 | 0.67 | -1.215 | 0.391 | -0.053 | 0.062 | 0.062 | 0 | 0.58 | 0.8 | 0.63 | 0.73 | 0.54 | 0.49 |
| 6 | 1598157 | rs2745580 | -0.652 | -0.108 | -0.329 | -0.059 | 0.007 | -0.037 | -0.037 | 0 | 0.14 | 0.5 | 0.13 | 0.05 | 0.11 | 0.2 |
| 6 | 1598227 | rs2745581 | -0.794 | -0.2 | -1.726 | -0.821 | -0.076 | 0.132 | 0.132 | 0 | 0.67 | 0.89 | 0.73 | 0.95 | 0.59 | 0.75 |
| 6 | 1598942 | rs2235715 | 0.721 | -1.25 | -2.022 | -0.532 | 0.049 | -0.006 | -0.006 | 0 | 0.25 | 0.03 | 0.35 | 0.2 | 0.17 | 0.15 |
| 6 | 1603402 | rs2745593 | -0.291 | -0.555 | -1.273 | -0.364 | -0.01 | 0.062 | 0.062 | 0 | 0.54 | 0.68 | 0.65 | 0.87 | 0.54 | 0.74 |
| 6 | 1606031 | rs2745596 | -0.74 | -0.137 | -1.614 | -0.207 | -0.102 | 0.11 | 0.11 | 0 | 0.67 | 0.89 | 0.7 | 0.92 | 0.57 | 0.76 |
| 6 | 1607603 | rs2235718 | 0.037 | -0.236 | -1.233 | 0.468 | -0.006 | -0.033 | -0.033 | 0 | 0.1 | 0.05 | 0.07 | 0.24 | 0.13 | 0.26 |
| 6 | 1613686 | rs2745599 | -0.404 | -0.586 | -1.306 | -0.399 | -0.04 | 0.116 | 0.116 | 0 | 0.61 | 0.88 | 0.61 | 0.85 | 0.44 | 0.71 |
| 6 | 1617143 | rs9501744 | -0.448 | -0.291 | -0.039 | -0.245 | -0.015 | -0.088 | -0.088 | 0.016 | 0.22 | 0.25 | 0.12 | 0 | 0.14 | 0.04 |
| 6 | 1620037 | rs2569881 | -0.5 | -0.318 | 0.143 | -0.236 | -0.035 | -0.157 | -0.157 | 0.846 | 0.22 | 0.39 | 0.13 | 0 | 0.13 | 0.03 |
| 6 | 1621404 | rs9405497 | -1.344 | 0.027 | 0.39 | 0.677 | 0.051 | 0.188 | 0.188 | 0.739 | 0.12 | 0.41 | 0.31 | 0.39 | 0.12 | 0.14 |
| 6 | 1622957 | rs9502981 | -0.44 | -0.071 | -0.037 | 0.378 | -0.003 | -0.021 | -0.021 | 0.523 | 0.09 | 0.19 | 0.09 | 0.04 | 0.1 | 0.17 |
| 6 | 1623990 | rs9501746 | -1.065 | -0.458 | -0.402 | 0.444 | -0.037 | -0.029 | -0.029 | 0 | 0.21 | 0.63 | 0.2 | 0.1 | 0.15 | 0.17 |
| 6 | 1626544 | rs2012025 | -0.926 | -0.124 | -1.639 | 0.591 | 0.014 | -0.05 | -0.05 | 0 | 0.48 | 0.89 | 0.59 | 0.54 | 0.4 | 0.4 |
| 6 | 1627632 | rs9392312 | -0.872 | 0.024 | -1.396 | 0.864 | -0.035 | -0.076 | -0.076 | 0.208 | 0.13 | 0.03 | 0.31 | 0.43 | 0.14 | 0.16 |
| 6 | 1628359 | rs1006883 | -0.036 | 0.404 | -1.978 | 0.745 | -0.057 | -0.026 | -0.026 | 0 | 0.42 | 0.08 | 0.43 | 0.74 | 0.39 | 0.61 |
| 6 | 1633682 | rs742556 | -0.913 | 0.987 | -0.883 | 1.654 | 0.008 | 0.019 | 0.019 | 0 | 0.23 | 0.26 | 0.28 | 0.51 | 0.29 | 0.37 |
| 6 | 1639642 | rs873709 | 0.932 | 0.655 | -1.309 | 0.78 | -0.008 | 0.09 | 0.09 | 0 | 0.17 | 0.04 | 0.1 | 0 | 0.15 | 0.05 |
| 6 | 1639800 | rs2569861 | 0.113 | 0.733 | -1.234 | 0.727 | -0.005 | 0.033 | 0.033 | 0 | 0.22 | 0.13 | 0.16 | 0.01 | 0.23 | 0.12 |
| 6 | 1640551 | rs2569862 | -1.327 | 0.4 | -0.422 | 0.423 | -0.001 | -0.024 | -0.024 | 0.537 | 0.52 | 0.56 | 0.71 | 0.8 | 0.52 | 0.56 |
| 6 | 1641007 | rs2007297 | -1.287 | 0.43 | 0.327 | 0.484 | -0.001 | 0.149 | 0.149 | 0 | 0.51 | 0.76 | 0.71 | 0.8 | 0.5 | 0.55 |
| 6 | 1648955 | rs12527698 | 0.307 | 0.514 | 0.634 | 0.651 | -0.025 | -0.037 | -0.037 | 0 | 0.07 | 0.01 | 0.26 | 0.17 | 0.1 | 0.1 |
| 6 | 1649347 | rs2569844 | -1.171 | 0.34 | 0.823 | 0.709 | 0.014 | -0.001 | -0.001 | 0 | 0.29 | 0.42 | 0.36 | 0.64 | 0.33 | 0.48 |
| 6 | 1656487 | rs9392317 | -0.017 | 0.331 | 0.764 | 0.806 | -0.006 | -0.003 | -0.003 | 1.543 | 0.91 | 0.85 | 0.91 | 0.93 | 0.94 | 0.86 |
| 6 | 1660002 | rs2814827 | -0.201 | 0.381 | 1.082 | 1.023 | 0.047 | 0.14 | 0.14 | 0.103 | 0.16 | 0.38 | 0.24 | 0.54 | 0.2 | 0.26 |
| 6 | 1661570 | rs2814807 | -0.572 | 0.399 | 0.801 | 0.615 | -0.007 | 0.05 | 0.05 | 0 | 0.43 | 0.25 | 0.42 | 0.62 | 0.45 | 0.45 |
| 6 | 1663631 | rs7759035 | 1.515 | 0.401 | 0.93 | 0.507 | -0.003 | 0.045 | 0.045 | 0.317 | 0.41 | 0.26 | 0.23 | 0.18 | 0.38 | 0.37 |
| 6 | 1664675 | rs2569885 | 0.391 | 0.334 | 0.988 | 0.517 | 0.076 | 0.025 | 0.025 | 0.011 | 0.92 | 0.54 | 0.92 | 0.97 | 0.94 | 0.92 |
| 6 | 1670717 | rs2246345 | 0.182 | 0.412 | 0.821 | 0.703 | -0.019 | -0.024 | -0.024 | 0 | 0.14 | 0.1 | 0.21 | 0.51 | 0.19 | 0.25 |
| 6 | 1670985 | rs2814811 | 0.626 | 0.386 | 0.881 | 0.677 | 0.012 | 0.019 | 0.019 | 0 | 0.55 | 0.37 | 0.44 | 0.69 | 0.58 | 0.63 |
| 6 | 1679526 | rs1962132 | -0.111 | 0.423 | 0.779 | 0.733 | 0 | 0.019 | 0.019 | 0 | 0.93 | 0.87 | 0.74 | 0.79 | 0.91 | 0.9 |
| 6 | 1681040 | rs4060649 | 0.021 | 0.452 | 0.829 | 0.738 | 0.025 | 0.123 | 0.123 | 0 | 0.16 | 0.27 | 0.22 | 0.51 | 0.19 | 0.25 |
| 6 | 1682567 | rs2814818 | -0.116 | 0.406 | 0.831 | 0.643 | 0.009 | 0.019 | 0.019 | 0 | 0.35 | 0.32 | 0.28 | 0.1 | 0.32 | 0.27 |
| 6 | 1682704 | rs9766727 | 0.116 | 0.379 | 0.794 | 0.617 | -0.005 | -0.014 | -0.014 | 0 | 0.15 | 0.3 | 0.16 | 0.1 | 0.13 | 0.22 |
| 6 | 1683192 | rs7756485 | -0.248 | 0.408 | 0.365 | 0.603 | -0.011 | -0.052 | -0.052 | 0 | 0.36 | 0.42 | 0.28 | 0.1 | 0.33 | 0.27 |
| 6 | 1683711 | rs2814819 | -0.077 | 0.442 | 0.147 | 0.629 | 0.001 | 0.028 | 0.028 | 0 | 0.92 | 0.84 | 0.74 | 0.79 | 0.91 | 0.9 |
| 6 | 1684571 | rs9405144 | 0.156 | 0.528 | 0.011 | 0.546 | 0.001 | 0.036 | 0.036 | 0 | 0.14 | 0.17 | 0.14 | 0.51 | 0.17 | 0.23 |
| 6 | 1686599 | rs2814820 | -1.076 | 0.486 | -0.316 | 0.574 | -0.025 | 0.044 | 0.044 | 1.3 | 0.53 | 0.76 | 0.5 | 0.59 | 0.51 | 0.52 |
| 6 | 1688391 | rs4461777 | 0.458 | 0.499 | -0.385 | 0.541 | -0.001 | -0.003 | -0.003 | 0.759 | 0.57 | 0.57 | 0.4 | 0.26 | 0.52 | 0.6 |
| 6 | 1688525 | rs4357177 | -0.074 | 0.467 | -1.533 | 0.514 | 0 | 0.034 | 0.034 | 0 | 0.93 | 0.88 | 0.74 | 0.77 | 0.91 | 0.9 |
| 6 | 1689999 | rs11242715 | 0.373 | 0.833 | -1.164 | 0.545 | -0.032 | 0.13 | 0.13 | 1.645 | 0.4 | 0.13 | 0.24 | 0.18 | 0.39 | 0.37 |
| 6 | 1695439 | rs9392322 | 0.147 | 0.728 | -0.679 | 1.495 | 0.01 | -0.015 | -0.015 | 0 | 0.61 | 0.29 | 0.61 | 0.46 | 0.63 | 0.59 |
| 6 | 1696841 | rs11968464 | -0.568 | 0.62 | -0.822 | 1.81 | -0.001 | 0.001 | 0.001 | 0 | 0.08 | 0.09 | 0.26 | 0.24 | 0.1 | 0.07 |
| 6 | 1698198 | rs4538755 | 0.202 | 0.207 | -0.395 | 1.296 | 0.024 | 0.005 | 0.005 | 0 | 0.53 | 0.33 | 0.43 | 0.47 | 0.6 | 0.56 |
| 6 | 1701839 | rs9503009 | 0.12 | 0.079 | -0.628 | 1.791 | 0.009 | 0 | 0 | 0 | 0.09 | 0.07 | 0.28 | 0.33 | 0.05 | 0.06 |
| 6 | 1703290 | rs9378305 | 0.12 | 0.079 | -0.628 | 1.945 | 0.009 | 0.015 | 0.015 | 0.317 | 0.09 | 0.07 | 0.29 | 0.42 | 0.05 | 0.06 |
| 6 | 1707254 | rs9503012 | 0.238 | -0.71 | -1.349 | 0.389 | 0.026 | 0.063 | 0.063 | 0 | 0.64 | 0.26 | 0.76 | 0.91 | 0.71 | 0.79 |

| | | | | | | | | | | | | | | | | |
|---|---|---|---|---|---|---|---|---|---|---|---|---|---|---|---|---|
| 6 | 1707399 | rs9503013 | 0.125 | -1.109 | -1.265 | 0.461 | -0.026 | -0.015 | -0.015 | 0 | 0.81 | 0.3 | 0.87 | 0.91 | 0.92 | 0.88 |
| 6 | 1708871 | rs9405506 | 0.059 | -2.098 | -1.078 | 0.164 | -0.014 | -0.003 | -0.003 | 0 | 0.42 | 0.57 | 0.31 | 0.47 | 0.24 | 0.42 |
| 6 | 1709037 | rs12197419 | -0.232 | -2.75 | -1.234 | 0.13 | 0.07 | -0.002 | -0.002 | 0.238 | 0.6 | 0.27 | 0.71 | 0.55 | 0.75 | 0.67 |
| 6 | 1709291 | rs9392324 | -0.398 | -1.958 | 0.186 | 0.074 | -0.007 | -0.015 | -0.015 | 0.016 | 0.37 | 0.39 | 0.3 | 0.21 | 0.27 | 0.16 |
| 6 | 1709703 | rs12206258 | -0.253 | -0.68 | 0.665 | 0.995 | 0.006 | 0.011 | 0.011 | 0 | 0.34 | 0.28 | 0.25 | 0.2 | 0.33 | 0.17 |
| 6 | 1712223 | rs7746883 | 0.441 | -0.681 | 0.607 | 0.927 | -0.021 | -0.139 | -0.139 | 0.359 | 0.26 | 0.39 | 0.17 | 0 | 0.23 | 0.08 |
| 6 | 1712283 | rs7765344 | 1.367 | -0.799 | 0.367 | 1.034 | 0.06 | -0.027 | -0.027 | 0.009 | 0.48 | 0.14 | 0.53 | 0.47 | 0.61 | 0.71 |
| 6 | 1712306 | rs7765461 | 0.884 | -0.785 | 0.415 | 0.714 | -0.012 | -0.021 | -0.021 | 0.209 | 0.89 | 0.9 | 0.93 | 0.84 | 0.86 | 0.86 |
| 6 | 1713390 | rs12203614 | 0.215 | -0.75 | 0.435 | 0.736 | -0.011 | 0.002 | 0.002 | 0 | 0.05 | 0.03 | 0.04 | 0.15 | 0.1 | 0.09 |
| 6 | 1713692 | rs6936881 | 1.329 | -0.762 | 0.427 | 0.725 | 0.06 | 0.004 | 0.004 | 0.361 | 0.48 | 0.14 | 0.55 | 0.57 | 0.61 | 0.72 |
| 6 | 1714220 | rs1022547 | -0.22 | -0.079 | 1.105 | 0.372 | -0.026 | -0.174 | -0.174 | 0.117 | 0.74 | 0.56 | 0.83 | 1 | 0.77 | 0.92 |
| 6 | 1714815 | rs1022550 | -0.395 | -0.032 | 1.029 | 0.379 | -0.004 | -0.068 | -0.068 | 0 | 0.75 | 0.74 | 0.83 | 1 | 0.77 | 0.92 |
| 6 | 1716354 | rs12200877 | -0.525 | 0.399 | 0.786 | 0.72 | 0.032 | 0.134 | 0.134 | 0.017 | 0.09 | 0.16 | 0.22 | 0.32 | 0.16 | 0.23 |
| 6 | 1716710 | rs761147 | 0.8 | 0.458 | 0.792 | 0.486 | 0.021 | 0.065 | 0.065 | 0 | 0.61 | 0.29 | 0.71 | 0.89 | 0.69 | 0.83 |
| 6 | 1721914 | rs4959149 | -1.067 | 0.971 | 0.774 | 0.832 | 0.07 | 0.187 | 0.187 | 1.888 | 0.15 | 0.44 | 0.3 | 0.32 | 0.19 | 0.26 |
| 6 | 1726162 | rs9378310 | 0.882 | 1.047 | 0.285 | 0.801 | -0.015 | -0.011 | -0.011 | 0 | 0.57 | 0.64 | 0.5 | 0.57 | 0.5 | 0.56 |
| 6 | 1729592 | rs3800030 | -1.166 | 1.133 | -0.158 | 0.826 | 0.003 | 0.023 | 0.023 | 1.807 | 0.17 | 0.2 | 0.27 | 0.38 | 0.24 | 0.32 |
| 6 | 1730861 | rs9392330 | -0.476 | 1.254 | -0.378 | 0.529 | -0.07 | 0.15 | 0.15 | 0 | 0.77 | 0.98 | 0.84 | 1 | 0.73 | 0.92 |
| 6 | 1735174 | rs12189843 | 1.023 | 0.775 | 0.132 | 0.97 | -0.014 | -0.026 | -0.026 | 0 | 0.84 | 0.3 | 0.84 | 0.89 | 0.9 | 0.92 |
| 6 | 1742513 | rs12200248 | 0.755 | 1.151 | 0.083 | 1.103 | 0.005 | 0.043 | 0.043 | 0 | 0.52 | 0.29 | 0.47 | 0.66 | 0.49 | 0.58 |
| 6 | 1742794 | rs3734739 | 0.755 | 1.151 | 0.083 | 1.111 | 0.005 | 0.041 | 0.041 | 0 | 0.52 | 0.29 | 0.47 | 0.66 | 0.49 | 0.58 |
| 6 | 1757229 | rs7740069 | 0.51 | 1.16 | 0.172 | 1.062 | -0.001 | -0.013 | -0.013 | 0 | 0.1 | 0.17 | 0.06 | 0.07 | 0.08 | 0.08 |
| 6 | 1759740 | rs10484696 | 1.273 | 1.175 | -0.147 | 1.196 | -0.015 | -0.038 | -0.038 | 0 | 0.07 | 0 | 0.06 | 0.14 | 0.1 | 0.09 |
| 6 | 1760556 | rs3800036 | 0.248 | 1.184 | -0.089 | 1.21 | 0.003 | -0.002 | -0.002 | 1.467 | 0.61 | 0.45 | 0.53 | 0.67 | 0.57 | 0.63 |
| 6 | 1763561 | rs3800042 | 1.064 | 1.279 | -0.989 | 1.258 | 0.009 | -0.006 | -0.006 | 0 | 0.43 | 0.22 | 0.4 | 0.46 | 0.37 | 0.44 |
| 6 | 1764324 | rs3800043 | -0.032 | 1.715 | -1.495 | 1.054 | -0.023 | -0.028 | -0.028 | 0 | 0.3 | 0.15 | 0.38 | 0.33 | 0.35 | 0.36 |
| 6 | 1764507 | rs6926560 | 0.377 | 1.504 | -1.299 | 0.89 | -0.003 | -0.004 | -0.004 | 0 | 0.95 | 0.96 | 0.94 | 1 | 0.91 | 0.99 |
| 6 | 1766973 | rs3800046 | -0.057 | 1.549 | -1.263 | 1.258 | -0.029 | 0.064 | 0.064 | 0 | 0.56 | 0.26 | 0.46 | 0.69 | 0.55 | 0.65 |
| 6 | 1769245 | rs12196461 | -1.794 | 1.695 | -0.649 | 1.898 | -0.003 | -0.007 | -0.007 | 0 | 0.26 | 0.27 | 0.34 | 0.43 | 0.31 | 0.47 |
| 6 | 1773297 | rs3800052 | -0.9 | 1.909 | -0.965 | 2.165 | -0.026 | -0.018 | -0.018 | 0.024 | 0.05 | 0 | 0.12 | 0.21 | 0.09 | 0.13 |
| 6 | 1775436 | rs2318089 | -1.42 | 1.993 | -0.972 | 2.077 | -0.051 | -0.065 | -0.065 | 0.309 | 0.17 | 0.1 | 0.3 | 0.34 | 0.28 | 0.47 |
| 6 | 1775863 | rs722585 | -1.459 | 1.981 | -0.966 | 1.831 | -0.043 | -0.048 | -0.048 | 2.173 | 0.18 | 0.11 | 0.3 | 0.31 | 0.28 | 0.47 |
| 6 | 1780691 | rs9503023 | -1.158 | 1.243 | -0.338 | 1.261 | 0.002 | 0.004 | 0.004 | 0 | 0.08 | 0.19 | 0.18 | 0.09 | 0.1 | 0.19 |
| 6 | 1780915 | rs3800061 | -0.403 | 1.545 | -0.205 | 2.221 | 0.007 | 0.066 | 0.066 | 0.413 | 0.08 | 0.09 | 0.04 | 0.25 | 0.08 | 0.14 |
| 6 | 1781167 | rs3823269 | -1.18 | 1.668 | -0.426 | 2.592 | -0.01 | -0.024 | -0.024 | 0 | 0.1 | 0.04 | 0.12 | 0.25 | 0.08 | 0.2 |
| 6 | 1781381 | rs689035 | -0.823 | 1.743 | -0.296 | 2.651 | 0.003 | 0.04 | 0.04 | 0 | 0.07 | 0.08 | 0.06 | 0.19 | 0.11 | 0.06 |
| 6 | 1782040 | rs3823271 | -1.242 | 1.597 | -0.107 | 1.956 | -0.001 | -0.002 | -0.002 | 0 | 0.28 | 0.38 | 0.17 | 0.3 | 0.29 | 0.38 |
| 6 | 1783630 | rs3800065 | -0.792 | 0.583 | 0.025 | 1.542 | -0.032 | 0.035 | 0.035 | 0 | 0.31 | 0.52 | 0.24 | 0.52 | 0.22 | 0.37 |
| 6 | 1784096 | rs6919059 | -1.149 | 0.784 | -0.236 | 1.17 | -0.001 | -0.013 | -0.013 | 0 | 0.44 | 0.51 | 0.33 | 0.72 | 0.41 | 0.49 |
| 6 | 1785725 | rs226443 | -0.946 | 1.051 | -0.156 | 1.399 | -0.006 | -0.01 | -0.01 | 0.515 | 0.14 | 0.1 | 0.09 | 0.21 | 0.13 | 0.15 |
| 6 | 1786032 | rs3800070 | -0.477 | 1.084 | -0.059 | 1.362 | -0.009 | -0.014 | -0.014 | 0 | 0.12 | 0.04 | 0.06 | 0.18 | 0.11 | 0.11 |
| 6 | 1786340 | rs226445 | -0.135 | 0.787 | -0.176 | 0.31 | 0.005 | 0.005 | 0.005 | 1.985 | 0.48 | 0.56 | 0.22 | 0.21 | 0.36 | 0.33 |
| 6 | 1797160 | rs226456 | 0.574 | 0.823 | 0.083 | 0.509 | 0.006 | 0.01 | 0.01 | 0 | 0.38 | 0.36 | 0.34 | 0.22 | 0.29 | 0.31 |
| 6 | 1801390 | rs3778526 | -0.381 | 1.165 | 0.187 | 1.298 | 0.051 | 0.091 | 0.091 | 0 | 0.1 | 0.21 | 0.12 | 0.24 | 0.19 | 0.11 |
| 6 | 1807658 | rs226462 | -0.031 | 1.229 | -0.393 | 1.148 | -0.006 | 0.028 | 0.028 | 1.3 | 0.55 | 0.77 | 0.44 | 0.4 | 0.42 | 0.42 |
| 6 | 1807959 | rs226463 | -0.503 | 1.031 | -0.874 | 1.013 | -0.045 | 0.078 | 0.078 | 0.373 | 0.64 | 0.95 | 0.64 | 0.79 | 0.61 | 0.54 |
| 6 | 1815691 | rs226468 | -0.242 | 0.674 | -0.612 | 1.031 | -0.027 | -0.023 | -0.023 | 0 | 0.58 | 0.81 | 0.55 | 0.49 | 0.44 | 0.42 |
| 6 | 1816632 | rs9378314 | -0.683 | 0.502 | 0.21 | 1.227 | -0.016 | 0.054 | 0.054 | 0.854 | 0.18 | 0.37 | 0.2 | 0.29 | 0.11 | 0.09 |
| 6 | 1823732 | rs3778530 | -0.113 | 0.896 | -0.403 | 1.143 | -0.021 | -0.024 | -0.024 | 0 | 0.68 | 0.78 | 0.56 | 0.59 | 0.51 | 0.6 |
| 6 | 1824441 | rs1157673 | -0.451 | 1.156 | -0.427 | 1.282 | -0.002 | -0.029 | -0.029 | 0 | 0.74 | 0.46 | 0.7 | 0.84 | 0.73 | 0.7 |
| 6 | 1824483 | rs1157674 | 0.314 | 0.03 | -0.594 | 0.834 | 0 | -0.001 | -0.001 | 0 | 0.53 | 0.46 | 0.52 | 0.45 | 0.41 | 0.51 |
| 6 | 1825105 | rs6596856 | 0.866 | -0.194 | -0.818 | 0.662 | -0.034 | 0.001 | 0.001 | 0 | 0.23 | 0.41 | 0.24 | 0.25 | 0.19 | 0.25 |
| 6 | 1825466 | rs9328071 | -0.072 | -0.027 | -1.145 | 0.676 | 0.004 | 0.032 | 0.032 | 0 | 0.43 | 0.15 | 0.33 | 0.34 | 0.3 | 0.35 |
| 6 | 1828165 | rs9328072 | -0.347 | -0.004 | -0.912 | 0.954 | 0.001 | -0.002 | -0.002 | 0.078 | 0.37 | 0.34 | 0.3 | 0.47 | 0.31 | 0.36 |
| 6 | 1830831 | rs3823276 | -0.3 | 0.366 | -1.099 | 0.91 | -0.022 | -0.044 | -0.044 | 0.088 | 0.12 | 0.01 | 0.05 | 0.19 | 0.1 | 0.11 |
| 6 | 1831050 | rs3800083 | -0.005 | 0.282 | -1.127 | 0.664 | 0.008 | 0.031 | 0.031 | 0 | 0.25 | 0.13 | 0.25 | 0.18 | 0.17 | 0.24 |
| 6 | 1832936 | rs3778533 | -1.04 | 0.06 | -0.253 | 0.962 | -0.024 | 0.027 | 0.027 | 0 | 0.19 | 0.37 | 0.07 | 0.22 | 0.11 | 0.12 |
| 6 | 1834156 | rs3800086 | -1.02 | 0.272 | -0.85 | 0.901 | -0.006 | -0.006 | -0.006 | 0 | 0.11 | 0.05 | 0.04 | 0.07 | 0.09 | 0.12 |
| 6 | 1837828 | rs3800091 | 0.29 | 0.207 | -1.839 | 0.588 | 0.023 | 0.174 | 0.174 | 1.373 | 0.49 | 0.11 | 0.6 | 0.25 | 0.62 | 0.37 |
| 6 | 1842762 | rs12192948 | 0.502 | -0.312 | -0.761 | 1.421 | 0.001 | -0.004 | -0.004 | 0.383 | 0.31 | 0.3 | 0.27 | 0.24 | 0.31 | 0.34 |
| 6 | 1856514 | rs912310 | 0.283 | 0.378 | -1.204 | 1.068 | -0.056 | -0.028 | -0.028 | 0 | 0.12 | 0.02 | 0.33 | 0.12 | 0.26 | 0.14 |
| 6 | 1859793 | rs6940806 | -0.09 | 0.199 | -0.508 | 1.848 | 0.019 | 0.024 | 0.024 | 1.112 | 0.46 | 0.68 | 0.33 | 0.35 | 0.37 | 0.51 |
| 6 | 1863854 | rs3800105 | -0.232 | 0.239 | -0.301 | 1.759 | 0.001 | -0.001 | -0.001 | 0 | 0.13 | 0.13 | 0.05 | 0.11 | 0.06 | 0.17 |
| 6 | 1866779 | rs12190459 | 0.275 | 0.393 | -0.359 | 1.771 | -0.044 | 0.034 | 0.034 | 0 | 0.1 | 0.02 | 0.25 | 0.02 | 0.21 | 0.1 |
| 6 | 1871901 | rs6596858 | -0.312 | 0.223 | -0.065 | 1.913 | 0.002 | -0.002 | -0.002 | 1.477 | 0.13 | 0.09 | 0.04 | 0.11 | 0.06 | 0.17 |
| 6 | 1877958 | rs9378661 | 0.436 | 0.199 | -0.227 | 1.869 | -0.014 | 0.171 | 0.171 | 1.393 | 0.49 | 0.08 | 0.51 | 0.23 | 0.61 | 0.36 |
| 6 | 1880964 | rs3800116 | 0.744 | 0.179 | 0.211 | 1.879 | 0.004 | 0.002 | 0.002 | 0.059 | 0.32 | 0.37 | 0.34 | 0.36 | 0.32 | 0.35 |
| 6 | 1896442 | rs1998519 | -0.363 | 0.271 | 0.107 | 1.778 | -0.009 | 0.012 | 0.012 | 0.037 | 0.13 | 0.16 | 0.12 | 0.11 | 0.07 | 0.17 |
| 6 | 1899671 | rs3800122 | 0.254 | 0.313 | -0.215 | 1.495 | 0.018 | 0.094 | 0.094 | 0.702 | 0.51 | 0.27 | 0.58 | 0.33 | 0.6 | 0.4 |

| | | | | | | | | | | | | | | | | |
|---|---|---|---|---|---|---|---|---|---|---|---|---|---|---|---|---|
| 6 | 1907155 | rs3800128 | -0.498 | -0.093 | -0.993 | 1.221 | -0.128 | -0.033 | -0.033 | 0 | 0.61 | 0.93 | 0.49 | 0.66 | 0.48 | 0.63 |
| 6 | 1908210 | rs2875711 | 0.708 | -0.046 | -0.252 | 1.572 | 0.011 | 0.027 | 0.027 | 0.336 | 0.27 | 0.38 | 0.39 | 0.38 | 0.29 | 0.32 |
| 6 | 1910108 | rs10484693 | -0.235 | -0.141 | -0.494 | 1.222 | 0.01 | -0.008 | -0.008 | 0 | 0.14 | 0.01 | 0.04 | 0.07 | 0.06 | 0.15 |
| 6 | 1913138 | rs1570535 | -0.662 | -0.12 | -0.477 | 0.959 | 0.018 | 0.221 | 0.221 | 1.351 | 0.48 | 0.9 | 0.47 | 0.85 | 0.36 | 0.63 |
| 6 | 1915129 | rs3778552 | 0.383 | 0.422 | 0.172 | 1.175 | 0.004 | -0.028 | -0.028 | 0 | 0.26 | 0.38 | 0.2 | 0.21 | 0.28 | 0.32 |
| 6 | 1918262 | rs3800131 | -0.728 | 0.449 | 0.232 | 0.662 | -0.003 | -0.053 | -0.053 | 3.963 | 0.55 | 0.52 | 0.67 | 0.83 | 0.63 | 0.75 |
| 6 | 1921848 | rs3800136 | -0.429 | 0.956 | 0.451 | 1.188 | 0.045 | 0.101 | 0.101 | 0 | 0.11 | 0.31 | 0.17 | 0.18 | 0.15 | 0.14 |
| 6 | 1935155 | rs912308 | -0.15 | 0.927 | -0.01 | 1.341 | 0.007 | -0.021 | -0.021 | 0 | 0.16 | 0.06 | 0.11 | 0.17 | 0.13 | 0.17 |
| 6 | 1942538 | rs3800143 | 0.311 | 0.813 | 0.086 | 1.432 | -0.008 | 0.004 | 0.004 | 0.79 | 0.3 | 0.2 | 0.38 | 0.3 | 0.35 | 0.28 |
| 6 | 1943154 | rs3800144 | -0.164 | 0.924 | -0.179 | 1.466 | 0.013 | -0.021 | -0.021 | 0 | 0.16 | 0.06 | 0.11 | 0.16 | 0.12 | 0.17 |
| 6 | 1954508 | rs7759192 | -0.381 | 1.221 | 0.225 | 1.191 | 0.021 | 0.053 | 0.053 | 0.194 | 0.1 | 0.19 | 0.16 | 0.18 | 0.15 | 0.14 |
| 6 | 1955398 | rs9378664 | 0.223 | 1.19 | 0.144 | 1.118 | 0.006 | -0.016 | -0.016 | 0 | 0.41 | 0.3 | 0.46 | 0.46 | 0.41 | 0.44 |
| 6 | 1957861 | rs2761239 | -0.263 | 0.811 | -0.173 | 0.504 | 0.004 | 0.032 | 0.032 | 1.651 | 0.52 | 0.66 | 0.63 | 0.64 | 0.56 | 0.58 |
| 6 | 1976501 | rs1408002 | 0.082 | 0.752 | -0.146 | 0.492 | 0.006 | 0.027 | 0.027 | 3.237 | 0.48 | 0.39 | 0.34 | 0.33 | 0.4 | 0.41 |
| 6 | 1988385 | rs10223785 | -0.217 | 0.778 | 0.186 | 1.126 | -0.004 | 0.066 | 0.066 | 0 | 0.05 | 0.13 | 0.03 | 0.17 | 0.03 | 0.23 |
| 6 | 1988782 | rs7755470 | -1.48 | 0.793 | 0.989 | 0.744 | 0.026 | 0.022 | 0.022 | 0.11 | 0.1 | 0.39 | 0.12 | 0.12 | 0.09 | 0.17 |
| 6 | 1988924 | rs9405157 | -0.806 | 1.698 | 1.557 | 0.942 | 0.115 | 0.127 | 0.127 | 0 | 0.11 | 0.29 | 0.21 | 0.18 | 0.22 | 0.16 |
| 6 | 1996943 | rs4959620 | -0.349 | 1.405 | 0.683 | 0.452 | -0.006 | -0.037 | -0.037 | 0 | 0.67 | 0.75 | 0.59 | 0.48 | 0.64 | 0.72 |
| 6 | 1997212 | rs4959621 | 0.607 | 1.149 | 0.942 | 0.184 | 0.007 | 0.024 | 0.024 | 0 | 0.5 | 0.54 | 0.38 | 0.3 | 0.41 | 0.43 |
| 6 | 2012739 | rs6596867 | 0.534 | 1.041 | 0.942 | 0.16 | -0.005 | 0.011 | 0.011 | 0.217 | 0.57 | 0.58 | 0.41 | 0.29 | 0.45 | 0.42 |
| 6 | 2028237 | rs9392354 | -0.376 | 1.08 | 0.853 | 0.341 | -0.028 | -0.036 | -0.036 | 0.615 | 0.14 | 0.08 | 0.21 | 0.2 | 0.25 | 0.21 |
| 6 | 2032259 | rs9392356 | -1.201 | 1.252 | 1.013 | 0.19 | 0.011 | 0.01 | 0.01 | 0.016 | 0.19 | 0.25 | 0.23 | 0.19 | 0.27 | 0.32 |
| 6 | 2037632 | rs6910252 | -0.707 | 1.143 | 0.979 | 0.179 | -0.002 | -0.015 | -0.015 | 0.077 | 0.74 | 0.77 | 0.63 | 0.49 | 0.71 | 0.73 |
| 6 | 2039027 | rs9503076 | -0.595 | 1.107 | 1.262 | 0.158 | 0.004 | 0.054 | 0.054 | 0.125 | 0.73 | 0.65 | 0.62 | 0.49 | 0.71 | 0.73 |
| 6 | 2048584 | rs4959630 | 0.637 | 1.109 | 1.285 | 0.109 | -0.001 | 0.018 | 0.018 | 0.486 | 0.55 | 0.42 | 0.39 | 0.29 | 0.44 | 0.4 |
| 6 | 2052811 | rs9405541 | 0.049 | 1.257 | 1.369 | 0.109 | -0.004 | -0.003 | -0.003 | 0.036 | 0.11 | 0.11 | 0.2 | 0.18 | 0.22 | 0.2 |
| 6 | 2059110 | rs476594 | -0.977 | 1.136 | 1.493 | 0.04 | 0.002 | 0.068 | 0.068 | 0.898 | 0.47 | 0.69 | 0.61 | 0.71 | 0.56 | 0.6 |
| 6 | 2062447 | rs635234 | 0.455 | 1.114 | 1.4 | 0.039 | -0.004 | -0.01 | -0.01 | 1.04 | 0.27 | 0.25 | 0.38 | 0.52 | 0.32 | 0.27 |
| 6 | 2064648 | rs676015 | -0.693 | 1.414 | 1.456 | -0.136 | 0.005 | 0.01 | 0.01 | 0.525 | 0.46 | 0.57 | 0.63 | 0.72 | 0.63 | 0.59 |
| 6 | 2077339 | rs9391943 | -0.002 | 1.353 | 0.765 | 0.038 | -0.034 | -0.034 | -0.034 | 0 | 0.12 | 0.07 | 0.2 | 0.18 | 0.25 | 0.2 |
| 6 | 2093432 | rs1004779 | -0.544 | 0.849 | 1.031 | -0.344 | 0.001 | 0.037 | 0.037 | 0.834 | 0.47 | 0.64 | 0.61 | 0.71 | 0.56 | 0.6 |
| 6 | 2097654 | rs9503096 | -0.132 | 0.365 | 0.434 | -0.844 | -0.013 | 0.029 | 0.029 | 0 | 0.31 | 0.23 | 0.29 | 0.09 | 0.36 | 0.28 |
| 6 | 2097686 | rs4959631 | -0.188 | 0.53 | 0.567 | -0.253 | 0 | 0.002 | 0.002 | 0 | 0.5 | 0.48 | 0.61 | 0.63 | 0.57 | 0.48 |
| 6 | 2122454 | rs7768483 | -0.607 | -0.512 | 0.397 | -0.156 | -0.066 | -0.029 | -0.029 | 0 | 0.25 | 0.55 | 0.13 | 0.16 | 0.1 | 0.26 |
| 6 | 2123744 | rs3800156 | 0.38 | 1.054 | -0.142 | 0.52 | -0.098 | -0.129 | -0.129 | 0.597 | 0.2 | 0.03 | 0.36 | 0.51 | 0.35 | 0.27 |
| 6 | 2139450 | rs11970276 | -0.244 | 0.477 | 0.358 | 0.53 | -0.038 | -0.033 | -0.033 | 0 | 0.12 | 0.03 | 0.12 | 0.12 | 0.21 | 0.21 |
| 6 | 2145136 | rs4959167 | -1.336 | 0.28 | 0.915 | 0.423 | -0.012 | 0.048 | 0.048 | 12.853 | 0.09 | 0.6 | 0.15 | 0.12 | 0.02 | 0.16 |
| 6 | 2190846 | rs9392371 | -0.502 | 0.71 | -0.138 | 0.37 | -0.045 | -0.05 | -0.05 | 2.155 | 0.13 | 0.03 | 0.21 | 0.17 | 0.23 | 0.24 |
| 6 | 2199634 | rs12528102 | -0.498 | 0.627 | -0.104 | 0.377 | -0.02 | -0.027 | -0.027 | 0 | 0.14 | 0.07 | 0.21 | 0.18 | 0.22 | 0.24 |
| 6 | 2205806 | rs6908269 | -0.856 | 0.279 | 0.29 | 0.201 | -0.007 | 0.032 | 0.032 | 1.842 | 0.19 | 0.54 | 0.16 | 0.21 | 0.17 | 0.21 |
| 6 | 2209577 | rs234915 | 0.222 | 0.939 | -0.489 | -0.065 | -0.052 | 0.026 | 0.026 | 0.71 | 0.74 | 0.85 | 0.56 | 0.81 | 0.46 | 0.69 |
| 6 | 2209680 | rs234914 | 0.387 | 0.856 | 0.292 | -0.442 | 0.035 | -0.021 | -0.021 | 0 | 0.73 | 0.65 | 0.56 | 0.8 | 0.45 | 0.69 |
| 6 | 2209860 | rs9378324 | 0.831 | -0.272 | -0.125 | -0.532 | 0.01 | 0.008 | 0.008 | 1.976 | 0.53 | 0.42 | 0.21 | 0.4 | 0.32 | 0.48 |
| 6 | 2210734 | rs234912 | -0.487 | -0.12 | -0.563 | -0.958 | 0.024 | 0.078 | 0.078 | 0 | 0.42 | 0.74 | 0.55 | 0.76 | 0.37 | 0.52 |
| 6 | 2223002 | rs9392374 | 0.932 | -0.295 | -0.364 | -0.845 | 0.014 | 0.006 | 0.006 | 0 | 0.18 | 0.12 | 0.09 | 0.16 | 0.14 | 0.19 |
| 6 | 2236926 | rs3800191 | 0.932 | -0.374 | -0.441 | -0.831 | 0.032 | 0.028 | 0.028 | 0.025 | 0.35 | 0.27 | 0.12 | 0.22 | 0.18 | 0.29 |
| 6 | 2247049 | rs12198593 | 0.624 | -0.169 | -0.697 | -0.702 | 0.01 | -0.074 | -0.074 | 0 | 0.15 | 0.02 | 0.31 | 0.42 | 0.11 | 0.19 |
| 6 | 2248825 | rs335963 | -0.109 | 0.051 | -0.714 | -0.823 | -0.096 | 0.009 | 0.009 | 0.01 | 0.76 | 0.89 | 0.6 | 0.79 | 0.51 | 0.69 |
| 6 | 2276503 | rs13202573 | 0.902 | 0.009 | -0.717 | -0.817 | 0.019 | -0.079 | -0.079 | 0.096 | 0.16 | 0.03 | 0.32 | 0.41 | 0.12 | 0.19 |
| 6 | 2277237 | rs2479017 | -0.694 | 0.286 | -0.821 | -0.923 | -0.119 | 0.099 | 0.099 | 0 | 0.82 | 0.95 | 0.82 | 0.98 | 0.61 | 0.9 |
| 6 | 2293020 | rs12193414 | 1.125 | -0.113 | -0.222 | -0.909 | 0.005 | 0.005 | 0.005 | 0.048 | 0.35 | 0.32 | 0.13 | 0.22 | 0.19 | 0.3 |
| 6 | 2329626 | rs2505658 | 0.999 | 0.009 | 0.138 | -1.014 | 0.001 | 0.007 | 0.007 | 0 | 0.19 | 0.06 | 0.11 | 0.17 | 0.19 | 0.2 |
| 6 | 2337811 | rs12210306 | 1.355 | -0.35 | 0.072 | -1.267 | -0.009 | -0.063 | -0.063 | 0 | 0.23 | 0.24 | 0.08 | 0 | 0.14 | 0.13 |
| 6 | 2354697 | rs1475411 | -0.251 | 0.39 | -0.293 | -0.843 | -0.087 | 0.012 | 0.012 | 0.011 | 0.77 | 0.89 | 0.61 | 0.79 | 0.53 | 0.68 |
| 6 | 2362742 | rs4959171 | 0.305 | 1.013 | -0.035 | -0.552 | -0.035 | -0.01 | -0.01 | 0 | 0.16 | 0.11 | 0.3 | 0.21 | 0.3 | 0.28 |
| 6 | 2366199 | rs11242760 | 0.217 | -0.37 | 0.526 | -0.065 | -0.004 | -0.027 | -0.027 | 0.13 | 0.17 | 0.08 | 0.34 | 0.41 | 0.18 | 0.25 |
| 6 | 2374647 | rs12525582 | 0.294 | -0.468 | 1.089 | -0.111 | -0.018 | -0.083 | -0.083 | 0 | 0.17 | 0.05 | 0.33 | 0.42 | 0.18 | 0.25 |
| 6 | 2374821 | rs1883587 | -0.111 | -0.764 | 0.902 | -0.412 | 0.054 | -0.068 | -0.068 | 6.017 | 0.71 | 0.92 | 0.85 | 0.58 | 0.79 | 0.71 |
| 6 | 2375660 | rs4959177 | 0.069 | -0.627 | 0.845 | -0.503 | -0.005 | 0.001 | 0.001 | 1.08 | 0.11 | 0.08 | 0.2 | 0.14 | 0.15 | 0.21 |
| 6 | 2379182 | rs11242762 | 1.493 | -0.602 | 0.949 | -0.589 | -0.093 | 0.194 | 0.194 | 0 | 0.38 | 0.09 | 0.28 | 0 | 0.45 | 0.25 |
| 6 | 2382395 | rs9503168 | 0.941 | -0.706 | 0.94 | -0.556 | -0.002 | 0.155 | 0.155 | 0 | 0.49 | 0.15 | 0.34 | 0.26 | 0.53 | 0.39 |
| 6 | 2386913 | rs4959652 | -1.192 | -0.599 | 1.223 | -0.36 | -0.007 | -0.054 | -0.054 | 0 | 0.42 | 0.47 | 0.6 | 0.84 | 0.4 | 0.6 |
| 6 | 2390151 | rs9405166 | 0.609 | -0.583 | 0.982 | -0.262 | 0.017 | -0.064 | -0.064 | 0 | 0.1 | 0.01 | 0.05 | 0.23 | 0.06 | 0.13 |
| 6 | 2399099 | rs12210286 | 0.173 | -0.42 | 1.533 | -0.116 | -0.001 | 0.015 | 0.015 | 0.496 | 0.21 | 0.19 | 0.33 | 0.43 | 0.19 | 0.26 |
| 6 | 2403974 | rs6920321 | -1.014 | -0.39 | 1.546 | -0.176 | 0.003 | 0.009 | 0.009 | 2.142 | 0.33 | 0.59 | 0.56 | 0.58 | 0.35 | 0.48 |
| 6 | 2412228 | rs4959663 | -0.215 | -0.368 | 1.494 | -0.165 | 0.046 | 0.053 | 0.053 | 0.014 | 0.11 | 0.35 | 0.22 | 0.17 | 0.14 | 0.21 |
| 6 | 2412255 | rs4959664 | -1.127 | -0.363 | 1.183 | -0.16 | -0.001 | -0.001 | -0.001 | 0 | 0.34 | 0.65 | 0.56 | 0.58 | 0.35 | 0.48 |
| 6 | 2413588 | rs3734505 | 0.073 | -0.267 | 0.415 | -0.31 | 0.01 | 0.012 | 0.012 | 0 | 0.09 | 0.11 | 0.19 | 0.17 | 0.14 | 0.21 |
| 6 | 2428540 | rs2325876 | 1.443 | -0.35 | 0.006 | -0.633 | -0.038 | 0.077 | 0.077 | 0 | 0.35 | 0.18 | 0.27 | 0 | 0.42 | 0.22 |

| | | | | | | | | | | | | | | | | |
|---|---|---|---|---|---|---|---|---|---|---|---|---|---|---|---|---|
| 6 | 2429573 | rs13196905 | -0.91 | -0.059 | -0.416 | -0.169 | 0.006 | 0.001 | 0.001 | 1.797 | 0.34 | 0.42 | 0.47 | 0.61 | 0.37 | 0.49 |
| 6 | 2431429 | rs13194198 | -0.155 | -0.014 | -0.393 | -0.083 | 0.031 | 0.039 | 0.039 | 0 | 0.11 | 0.27 | 0.14 | 0.19 | 0.15 | 0.23 |
| 6 | 2437996 | rs6940091 | -0.79 | -0.39 | -0.565 | -0.17 | 0.022 | -0.011 | -0.011 | 0.101 | 0.81 | 0.97 | 0.88 | 0.8 | 0.85 | 0.85 |
| 6 | 2438231 | rs11758482 | -0.817 | 0.346 | 0.258 | 0.702 | -0.001 | 0.022 | 0.022 | 0 | 0.23 | 0.28 | 0.34 | 0.41 | 0.22 | 0.26 |
| 6 | 2444222 | rs11242772 | -0.288 | 0.317 | -0.062 | 0.638 | -0.068 | -0.139 | -0.139 | 3.425 | 0.19 | 0.01 | 0.19 | 0.37 | 0.22 | 0.26 |
| 6 | 2445280 | rs9503194 | -1.796 | 0.225 | 0.084 | 0.577 | 0 | -0.009 | -0.009 | 0.046 | 0.44 | 0.49 | 0.59 | 0.8 | 0.42 | 0.64 |
| 6 | 2445903 | rs9392388 | -0.007 | 0.202 | 0.037 | 0.7 | 0.017 | -0.02 | -0.02 | 0 | 0.1 | 0.08 | 0.12 | 0.24 | 0.05 | 0.14 |
| 6 | 2447602 | rs17609498 | -0.268 | 0.225 | 0.069 | 0.709 | -0.004 | -0.004 | -0.004 | 0 | 0.1 | 0.09 | 0.24 | 0.19 | 0.14 | 0.23 |
| 6 | 2447699 | rs742487 | -0.339 | 0.173 | -0.06 | 0.642 | -0.024 | 0.036 | 0.036 | 0.162 | 0.12 | 0.2 | 0.13 | 0.24 | 0.05 | 0.14 |
| 6 | 2449186 | rs761098 | -1.158 | 0.256 | -0.203 | 0.585 | -0.015 | -0.016 | -0.016 | 0 | 0.33 | 0.24 | 0.47 | 0.54 | 0.39 | 0.49 |
| 6 | 2449872 | rs9501843 | -1.152 | 0.319 | -0.108 | -0.405 | 0.001 | -0.028 | -0.028 | 0 | 0.43 | 0.55 | 0.6 | 0.77 | 0.43 | 0.52 |
| 6 | 2452024 | rs9391949 | -0.491 | 0.757 | -1.249 | -0.169 | -0.002 | -0.007 | -0.007 | 7.632 | 0.32 | 0.16 | 0.51 | 0.65 | 0.29 | 0.37 |
| 6 | 2453470 | rs4959669 | -0.332 | 0.754 | -1.072 | -0.019 | -0.002 | -0.003 | -0.003 | 0 | 0.09 | 0.07 | 0.08 | 0.12 | 0.13 | 0.15 |
| 6 | 2455853 | rs6919576 | -0.815 | 0.322 | -1.063 | 0.278 | -0.004 | -0.013 | -0.013 | 1.823 | 0.29 | 0.16 | 0.51 | 0.66 | 0.28 | 0.36 |
| 6 | 2461567 | rs17135071 | -0.886 | -0.087 | -0.616 | -0.171 | -0.006 | -0.032 | -0.032 | 0.147 | 0.12 | 0.13 | 0.05 | 0 | 0.05 | 0.04 |
| 6 | 2462807 | rs9391950 | 0.27 | -0.124 | -1.133 | -0.254 | -0.014 | -0.025 | -0.025 | 0.776 | 0.3 | 0.21 | 0.26 | 0.67 | 0.33 | 0.41 |
| 6 | 2464385 | rs7772576 | -0.953 | -0.592 | -0.402 | -0.903 | -0.05 | -0.143 | -0.143 | 0 | 0.28 | 0.46 | 0.14 | 0.05 | 0.18 | 0.1 |
| 6 | 2466482 | rs4634509 | -0.609 | -0.671 | 0.065 | -0.935 | -0.001 | 0 | 0 | 0 | 0.56 | 0.6 | 0.38 | 0.72 | 0.51 | 0.51 |
| 6 | 2467372 | rs17135104 | 0.527 | -0.744 | 0.306 | -0.963 | -0.014 | -0.009 | -0.009 | 0 | 0.07 | 0.08 | 0.02 | 0.05 | 0.03 | 0.03 |
| 6 | 2467475 | rs9405561 | 0.774 | -0.605 | 0.389 | -0.859 | -0.004 | -0.01 | -0.01 | 0 | 0.31 | 0.27 | 0.28 | 0.68 | 0.33 | 0.41 |
| 6 | 2470570 | rs6899721 | 0.426 | -0.598 | 0.077 | -0.976 | -0.01 | 0.026 | 0.026 | 0 | 0.4 | 0.3 | 0.47 | 0.16 | 0.44 | 0.41 |
| 6 | 2477490 | rs7751264 | 0.382 | -0.721 | 0.267 | -0.727 | -0.003 | 0.108 | 0.108 | 0 | 0.46 | 0.24 | 0.51 | 0.18 | 0.52 | 0.45 |
| 6 | 2477763 | rs4959186 | -0.374 | -0.829 | 0.421 | -0.866 | -0.004 | 0.03 | 0.03 | 0.045 | 0.6 | 0.64 | 0.55 | 0.17 | 0.57 | 0.48 |
| 6 | 2479379 | rs9501848 | -0.136 | -0.8 | 0.378 | -0.805 | -0.038 | -0.073 | -0.073 | 0 | 0.12 | 0.25 | 0.04 | 0 | 0.04 | 0.03 |
| 6 | 2486804 | rs10046356 | 0.091 | -0.704 | -0.521 | -0.756 | -0.024 | -0.051 | -0.051 | 0.002 | 0.11 | 0.18 | 0.04 | 0 | 0.04 | 0.03 |
| 6 | 2501118 | rs35374935 | 0.056 | -0.476 | -1.257 | -0.529 | -0.018 | -0.043 | -0.043 | 0.734 | 0.11 | 0.01 | 0.07 | 0.16 | 0.09 | 0.11 |
| 6 | 2507448 | rs879098 | 0.275 | -0.129 | -1.23 | -1.075 | -0.088 | 0.181 | 0.181 | 0 | 0.34 | 0.08 | 0.4 | 0.01 | 0.43 | 0.31 |
| 6 | 2509443 | rs1994124 | -0.225 | 0.1 | -0.869 | -0.273 | -0.005 | 0.066 | 0.066 | 0 | 0.58 | 0.7 | 0.55 | 0.2 | 0.56 | 0.49 |
| 6 | 2511716 | rs12212791 | -0.727 | 0.186 | -0.802 | 0.796 | 0 | -0.014 | -0.014 | 0 | 0.18 | 0.16 | 0.34 | 0.51 | 0.22 | 0.32 |
| 6 | 2512075 | rs1772987 | 0.381 | -0.223 | -0.399 | 0.268 | -0.031 | 0.022 | 0.022 | 0 | 0.73 | 0.59 | 0.77 | 0.72 | 0.82 | 0.75 |
| 6 | 2514312 | rs967430 | 0.627 | -0.036 | -0.814 | -0.089 | 0 | -0.001 | -0.001 | 2.006 | 0.72 | 0.73 | 0.51 | 0.33 | 0.63 | 0.56 |
| 6 | 2514911 | rs764450 | 0.083 | -0.113 | -0.259 | -0.653 | -0.007 | -0.04 | -0.04 | 0 | 0.27 | 0.32 | 0.2 | 0.09 | 0.25 | 0.2 |
| 6 | 2520336 | rs7764946 | -0.009 | 0.277 | -0.103 | -0.302 | -0.006 | 0.029 | 0.029 | 0 | 0.08 | 0.01 | 0.04 | 0 | 0.09 | 0.08 |
| 6 | 2523195 | rs7766538 | -0.405 | 0.265 | 0.053 | -0.412 | 0.029 | -0.047 | -0.047 | 1.362 | 0.7 | 0.81 | 0.74 | 0.53 | 0.78 | 0.69 |
| 6 | 2523683 | rs7742406 | 0.295 | 0.311 | 0.044 | -0.407 | -0.008 | 0.029 | 0.029 | 0 | 0.08 | 0.01 | 0.04 | 0 | 0.1 | 0.08 |
| 6 | 2524003 | rs1772982 | 0.363 | 0.296 | 0.238 | -0.391 | 0.017 | -0.011 | -0.011 | 1.038 | 0.6 | 0.66 | 0.69 | 0.52 | 0.68 | 0.61 |
| 6 | 2531100 | rs1531815 | 0.427 | 0.316 | 0.487 | -0.8 | -0.012 | -0.058 | -0.058 | 0 | 0.14 | 0.23 | 0.07 | 0 | 0.08 | 0.02 |
| 6 | 2532384 | rs1564633 | 0.534 | 0.858 | 0.823 | -0.97 | 0.007 | -0.011 | -0.011 | 0 | 0.5 | 0.5 | 0.32 | 0.12 | 0.4 | 0.32 |
| 6 | 2536050 | rs931242 | 0.602 | 0.863 | 0.528 | -0.863 | 0.007 | 0.001 | 0.001 | 0.087 | 0.13 | 0.1 | 0.06 | 0 | 0.08 | 0.01 |
| 6 | 2542395 | rs1773001 | -0.018 | 0.937 | 0.745 | -0.933 | -0.022 | 0.044 | 0.044 | 0 | 0.6 | 0.67 | 0.37 | 0.12 | 0.49 | 0.4 |
| 6 | 2546384 | rs1614248 | 0.102 | 0.931 | 0.649 | -0.936 | -0.042 | 0.097 | 0.097 | 0.036 | 0.6 | 0.73 | 0.37 | 0.12 | 0.49 | 0.4 |
| 6 | 2555246 | rs2449447 | -0.409 | 1.159 | 0.686 | -0.713 | 0.01 | -0.017 | -0.017 | 2.31 | 0.67 | 0.68 | 0.65 | 0.52 | 0.73 | 0.66 |
| 6 | 2557509 | rs2170616 | 0.347 | 1.218 | 0.645 | -0.703 | -0.025 | 0.029 | 0.029 | 0 | 0.07 | 0.03 | 0.29 | 0.41 | 0.24 | 0.26 |
| 6 | 2558291 | rs1680870 | 0.071 | 1.174 | 0.713 | -0.744 | -0.041 | 0.112 | 0.112 | 0.111 | 0.6 | 0.74 | 0.37 | 0.12 | 0.49 | 0.4 |
| 6 | 2560663 | rs1680875 | -0.253 | 1.131 | 1.006 | -0.729 | -0.011 | 0.014 | 0.014 | 0 | 0.65 | 0.55 | 0.65 | 0.52 | 0.73 | 0.66 |
| 6 | 2560712 | rs1773013 | -0.408 | 1.057 | 0.85 | -0.538 | -0.015 | -0.032 | -0.032 | 0 | 0.94 | 0.99 | 0.89 | 0.79 | 0.92 | 0.88 |
| 6 | 2561352 | rs4959190 | 0.228 | 1.091 | 1.128 | -0.784 | 0.002 | -0.001 | -0.001 | 0 | 0.56 | 0.39 | 0.59 | 0.52 | 0.63 | 0.58 |
| 6 | 2563775 | rs17208835 | 1.36 | 0.985 | 1.182 | -1.051 | 0.006 | 0.123 | 0.123 | 0 | 0.35 | 0.14 | 0.23 | 0.11 | 0.31 | 0.3 |
| 6 | 2565752 | rs12195826 | -0.09 | 0.996 | 1.268 | -0.985 | -0.003 | 0 | 0 | 0 | 0.6 | 0.54 | 0.61 | 0.53 | 0.68 | 0.59 |
| 6 | 2567677 | rs1772990 | -0.666 | 0.969 | 1.225 | -0.999 | 0.016 | -0.047 | -0.047 | 0.134 | 0.68 | 0.83 | 0.67 | 0.53 | 0.73 | 0.64 |
| 6 | 2573436 | rs1772994 | 0.721 | 0.99 | 1.3 | -1.034 | 0.006 | -0.002 | -0.002 | 0 | 0.43 | 0.37 | 0.28 | 0.12 | 0.36 | 0.35 |
| 6 | 2573748 | rs2794553 | 0.122 | 1.016 | 1.303 | -0.955 | -0.009 | -0.043 | -0.043 | 0 | 0.08 | 0.15 | 0.05 | 0 | 0.04 | 0.05 |
| 6 | 2575480 | rs9405566 | -0.519 | 1.208 | 1.184 | -0.895 | 0.055 | 0.132 | 0.132 | 0.001 | 0.11 | 0.17 | 0.32 | 0.42 | 0.29 | 0.28 |
| 6 | 2576644 | rs2624921 | 0.065 | 1.004 | 0.619 | -0.931 | 0.02 | 0.008 | 0.008 | 0 | 0.55 | 0.68 | 0.6 | 0.54 | 0.64 | 0.63 |
| 6 | 2579405 | rs1254722 | 0.442 | 0.784 | 0.502 | -0.913 | 0 | -0.045 | -0.045 | 0.067 | 0.51 | 0.43 | 0.3 | 0.12 | 0.38 | 0.31 |
| 6 | 2585649 | rs9392400 | -0.404 | 1.1 | 0.051 | 0.16 | -0.05 | -0.031 | -0.031 | 0 | 0.09 | 0.02 | 0.29 | 0.39 | 0.16 | 0.23 |
| 6 | 2589139 | rs4959684 | -1.01 | 1.219 | -0.02 | -0.359 | -0.001 | 0.127 | 0.127 | 0 | 0.59 | 0.81 | 0.72 | 0.87 | 0.61 | 0.61 |
| 6 | 2591489 | rs6934075 | 0.504 | 1.294 | -0.157 | -0.307 | 0.004 | 0.001 | 0.001 | 0 | 0.07 | 0.06 | 0.05 | 0 | 0.05 | 0.06 |
| 6 | 2591603 | rs9391956 | -0.403 | 1.356 | 0.04 | -0.322 | 0.016 | -0.049 | -0.049 | 0 | 0.35 | 0.39 | 0.57 | 0.87 | 0.44 | 0.52 |
| 6 | 2595478 | rs1001729 | 0.247 | 1.169 | -0.6 | 0.034 | 0.001 | 0.012 | 0.012 | 0 | 0.25 | 0.26 | 0.28 | 0.5 | 0.27 | 0.29 |
| 6 | 2599592 | rs17209317 | 0.236 | 1.076 | -1.142 | -0.673 | -0.01 | 0.14 | 0.14 | 0 | 0.41 | 0.18 | 0.29 | 0.09 | 0.41 | 0.39 |
| 6 | 2602357 | rs9392407 | -0.342 | 1.179 | -0.588 | 0.468 | 0.007 | 0.034 | 0.034 | 0 | 0.35 | 0.52 | 0.38 | 0.49 | 0.38 | 0.35 |
| 6 | 2604645 | rs9503226 | -0.738 | 1.079 | -0.837 | 0.141 | -0.001 | 0.001 | 0.001 | 0 | 0.07 | 0.08 | 0.08 | 0.04 | 0.09 | 0.1 |
| 6 | 2608995 | rs6918808 | -0.094 | 1.106 | -1.838 | -0.334 | -0.068 | 0.092 | 0.092 | 0 | 0.42 | 0.12 | 0.51 | 0.76 | 0.48 | 0.64 |
| 6 | 2609373 | rs9503230 | 0.441 | 0.942 | -1.244 | -1.161 | 0.003 | -0.076 | -0.076 | 0.983 | 0.24 | 0.21 | 0.15 | 0 | 0.23 | 0.14 |
| 6 | 2613944 | rs7755154 | 0.049 | 1.245 | -1.004 | -0.89 | 0.006 | 0.053 | 0.053 | 0.008 | 0.2 | 0.27 | 0.33 | 0.47 | 0.26 | 0.22 |
| 6 | 2617122 | rs4959689 | 0.163 | 0.916 | -0.599 | -1.136 | -0.014 | 0.059 | 0.059 | 0 | 0.44 | 0.26 | 0.25 | 0.03 | 0.45 | 0.26 |
| 6 | 2619850 | rs1384537 | 0.185 | 0.973 | -0.659 | -1.149 | -0.005 | -0.021 | -0.021 | 0.193 | 0.44 | 0.3 | 0.25 | 0.03 | 0.43 | 0.26 |
| 6 | 2621887 | rs9503232 | 1.023 | 0.948 | -0.48 | -1.178 | 0.005 | -0.097 | -0.097 | 0.165 | 0.25 | 0.25 | 0.15 | 0 | 0.23 | 0.15 |

| | | | | | | | | | | | | | | | | |
|---|---|---|---|---|---|---|---|---|---|---|---|---|---|---|---|---|
| 6 | 2622911 | rs1545849 | -0.568 | 0.932 | -0.344 | -1.155 | 0.03 | 0.067 | 0.067 | 0.143 | 0.85 | 0.6 | 0.86 | 0.73 | 0.81 | 0.87 |
| 6 | 2623423 | rs9501863 | 0.823 | 0.952 | -0.439 | -1.18 | -0.012 | 0.057 | 0.057 | 0 | 0.19 | 0.08 | 0.1 | 0.03 | 0.21 | 0.12 |
| 6 | 2623820 | rs6902511 | 1.162 | 0.904 | -0.471 | -1.183 | -0.027 | -0.167 | -0.167 | 0.018 | 0.31 | 0.04 | 0.39 | 0.49 | 0.29 | 0.51 |
| 6 | 2623930 | rs9503233 | 0.85 | 0.891 | -0.469 | -1.182 | -0.013 | 0.054 | 0.054 | 0 | 0.19 | 0.08 | 0.1 | 0.03 | 0.21 | 0.12 |
| 6 | 2624053 | rs17135340 | 0.948 | 0.932 | -0.384 | -0.995 | -0.025 | -0.03 | -0.03 | 0 | 0.11 | 0.06 | 0.12 | 0.27 | 0.19 | 0.14 |
| 6 | 2624262 | rs1150656 | 0.975 | 0.698 | -0.136 | -0.901 | -0.006 | 0.06 | 0.06 | 1.265 | 0.08 | 0.15 | 0.21 | 0.21 | 0.07 | 0.08 |
| 6 | 2624374 | rs9378720 | 0.91 | 0.703 | -0.163 | -0.901 | -0.012 | -0.157 | -0.157 | 0.06 | 0.32 | 0.06 | 0.39 | 0.49 | 0.29 | 0.52 |
| 6 | 2626824 | rs12526097 | 0.93 | 0.734 | -0.057 | -0.819 | -0.003 | 0.098 | 0.098 | 1.1 | 0.08 | 0.17 | 0.21 | 0.21 | 0.07 | 0.09 |
| 6 | 2632108 | rs1038335 | -0.172 | 0.818 | -0.03 | -0.944 | 0.003 | 0.005 | 0.005 | 0 | 0.56 | 0.37 | 0.36 | 0.3 | 0.62 | 0.4 |
| 6 | 2635495 | rs9328110 | -0.294 | 0.925 | -0.551 | -0.992 | -0.048 | -0.036 | -0.036 | 0 | 0.23 | 0.12 | 0.34 | 0.68 | 0.35 | 0.36 |
| 6 | 2640997 | rs6922032 | -0.416 | 0.241 | -0.283 | -1.88 | 0.002 | -0.097 | -0.097 | 0 | 0.81 | 0.75 | 0.87 | 1 | 0.83 | 0.9 |
| 6 | 2644095 | rs718604 | 0.668 | 0.097 | -0.805 | -1.972 | -0.025 | -0.081 | -0.081 | 0 | 0.33 | 0.06 | 0.43 | 0.53 | 0.31 | 0.48 |
| 6 | 2644630 | rs2170614 | 0.052 | 0.337 | -0.404 | -2.021 | -0.007 | 0.046 | 0.046 | 0 | 0.24 | 0.46 | 0.32 | 0.43 | 0.23 | 0.24 |
| 6 | 2644839 | rs6903273 | 0.708 | 0.582 | -0.542 | -2.228 | 0 | -0.044 | -0.044 | 0 | 0.18 | 0.15 | 0.12 | 0 | 0.17 | 0.11 |
| 6 | 2661853 | rs9503244 | 0.158 | 0.35 | -0.382 | -2.266 | 0.013 | -0.106 | -0.106 | 0.839 | 0.37 | 0.35 | 0.23 | 0.04 | 0.44 | 0.26 |
| 6 | 2662079 | rs12525061 | 0.877 | 0.157 | -0.681 | -1.968 | -0.031 | -0.03 | -0.03 | 0.028 | 0.1 | 0.01 | 0.2 | 0.15 | 0.12 | 0.09 |
| 6 | 2665938 | rs34857522 | 1.085 | 0.124 | -0.443 | -2.048 | 0.001 | -0.06 | -0.06 | 0 | 0.16 | 0.14 | 0.11 | 0 | 0.15 | 0.09 |
| 6 | 2666428 | rs17135412 | 1.352 | 0.232 | -0.411 | -1.614 | -0.007 | -0.021 | -0.021 | 0.111 | 0.1 | 0.04 | 0.07 | 0.31 | 0.11 | 0.12 |
| 6 | 2666478 | rs12203065 | 0.323 | 0.239 | 0.068 | -1.889 | 0.014 | -0.031 | -0.031 | 0 | 0.31 | 0.41 | 0.35 | 0.18 | 0.39 | 0.26 |
| 6 | 2666549 | rs12203132 | 0.292 | 0.232 | 0.085 | -1.893 | 0.015 | -0.034 | -0.034 | 0.269 | 0.31 | 0.42 | 0.35 | 0.18 | 0.39 | 0.26 |
| 6 | 2666848 | rs9503245 | 0.316 | 0.226 | 0.077 | -1.896 | 0.015 | -0.032 | -0.032 | 0.999 | 0.31 | 0.41 | 0.35 | 0.18 | 0.39 | 0.26 |
| 6 | 2666932 | rs733985 | 0.204 | 0.075 | -0.095 | -2.19 | -0.023 | -0.054 | -0.054 | 1.145 | 0.19 | 0.11 | 0.3 | 0.44 | 0.23 | 0.31 |
| 6 | 2667031 | rs6904200 | -0.006 | 0.077 | -0.19 | -2.144 | -0.002 | 0.006 | 0.006 | 0 | 0.41 | 0.43 | 0.64 | 0.62 | 0.56 | 0.5 |
| 6 | 2667689 | rs9501873 | 0.581 | 0.104 | -0.095 | -2.14 | 0.018 | -0.036 | -0.036 | 1.245 | 0.3 | 0.4 | 0.35 | 0.18 | 0.39 | 0.26 |
| 6 | 2667963 | rs7762519 | 1.051 | -0.128 | -0.349 | -2.162 | 0.055 | 0.092 | 0.092 | 0 | 0.35 | 0.14 | 0.25 | 0.07 | 0.25 | 0.29 |
| 6 | 2668872 | rs9405575 | 0.917 | 0.228 | -0.432 | -1.8 | -0.035 | -0.048 | -0.048 | 0 | 0.11 | 0.02 | 0.24 | 0.43 | 0.16 | 0.23 |
| 6 | 2670186 | rs12200414 | 1.017 | 0.259 | -0.367 | -2.063 | -0.027 | 0.041 | 0.041 | 0 | 0.08 | 0.04 | 0.06 | 0 | 0.08 | 0.08 |
| 6 | 2670633 | rs4959698 | 0.867 | 0.186 | -0.168 | -2.028 | -0.005 | 0.003 | 0.003 | 0 | 0.12 | 0.18 | 0.22 | 0.14 | 0.11 | 0.13 |
| 6 | 2671994 | rs9328113 | 1.164 | 0.358 | -0.276 | -2.008 | -0.025 | 0.027 | 0.027 | 0 | 0.19 | 0.12 | 0.12 | 0.07 | 0.28 | 0.16 |
| 6 | 2675271 | rs9328114 | 0.431 | 0.168 | -0.142 | -2.025 | -0.021 | 0.02 | 0.02 | 0.009 | 0.29 | 0.22 | 0.18 | 0.07 | 0.36 | 0.24 |
| 6 | 2678750 | rs4959699 | -0.384 | -0.26 | -0.028 | -2.184 | -0.003 | -0.077 | -0.077 | 0 | 0.39 | 0.54 | 0.56 | 0.88 | 0.42 | 0.49 |
| 6 | 2679676 | rs12523992 | 0.952 | -0.217 | -0.303 | -1.837 | -0.009 | 0.011 | 0.011 | 0 | 0.11 | 0.06 | 0.26 | 0.42 | 0.17 | 0.28 |
| 6 | 2680732 | rs2038760 | 1.425 | -0.89 | -0.26 | -2.156 | 0.01 | 0.015 | 0.015 | 0 | 0.21 | 0.09 | 0.18 | 0.05 | 0.15 | 0.19 |
| 6 | 2680767 | rs2038761 | 0.968 | -0.934 | -0.451 | -2.073 | -0.021 | -0.086 | -0.086 | 0 | 0.29 | 0.07 | 0.37 | 0.42 | 0.31 | 0.34 |
| 6 | 2680805 | rs6939996 | 1.056 | -0.695 | -0.348 | -2.007 | -0.068 | 0.097 | 0.097 | 0.137 | 0.28 | 0.16 | 0.17 | 0.07 | 0.34 | 0.26 |
| 6 | 2681511 | rs7761747 | 0.547 | -0.593 | 0.037 | -1.626 | -0.001 | 0.001 | 0.001 | 0.079 | 0.23 | 0.24 | 0.29 | 0.46 | 0.24 | 0.2 |
| 6 | 2686488 | rs9392414 | -1.566 | -0.394 | -0.092 | -1.986 | 0 | -0.006 | -0.006 | 0 | 0.78 | 0.77 | 0.87 | 0.9 | 0.72 | 0.81 |
| 6 | 2686681 | rs9405578 | -0.35 | -0.404 | -0.015 | -1.752 | -0.005 | -0.012 | -0.012 | 3.196 | 0.37 | 0.52 | 0.37 | 0.43 | 0.34 | 0.3 |
| 6 | 2687067 | rs9503252 | 0.813 | -0.427 | -0.312 | -1.436 | 0.001 | 0.01 | 0.01 | 0.077 | 0.11 | 0.17 | 0.08 | 0.3 | 0.12 | 0.08 |
| 6 | 2687729 | rs2126243 | 0.58 | -0.404 | -0.457 | -1.403 | -0.013 | -0.074 | -0.074 | 0 | 0.3 | 0.08 | 0.38 | 0.44 | 0.31 | 0.36 |
| 6 | 2689064 | rs6935206 | -0.526 | -0.383 | 0.467 | -1.066 | 0.001 | 0.037 | 0.037 | 0 | 0.13 | 0.33 | 0.09 | 0.3 | 0.12 | 0.11 |
| 6 | 2693442 | rs9391962 | -0.569 | -0.36 | 0.008 | -1.3 | -0.004 | -0.012 | -0.012 | 0 | 0.26 | 0.29 | 0.27 | 0.16 | 0.23 | 0.25 |
| 6 | 2705016 | rs9503260 | -0.067 | -0.398 | 0.06 | -1.09 | 0 | 0.013 | 0.013 | 0.121 | 0.19 | 0.16 | 0.26 | 0.15 | 0.2 | 0.15 |
| 6 | 2706569 | rs9405180 | 1.094 | -1 | 0.494 | -1.615 | -0.025 | 0.014 | 0.014 | 0.609 | 0.38 | 0.26 | 0.47 | 0.57 | 0.49 | 0.56 |
| 6 | 2711123 | rs1599908 | 0.202 | -1.386 | 0.585 | -1.852 | -0.012 | -0.053 | -0.053 | 0 | 0.27 | 0.29 | 0.18 | 0.02 | 0.16 | 0.18 |
| 6 | 2711622 | rs6930454 | 0.516 | -1.364 | 0.53 | -1.837 | -0.011 | -0.013 | -0.013 | 0 | 0.65 | 0.54 | 0.65 | 0.6 | 0.66 | 0.74 |
| 6 | 2711880 | rs942490 | 0.159 | -1.347 | 0.528 | -1.821 | -0.005 | 0.005 | 0.005 | 0 | 0.85 | 0.85 | 0.92 | 0.74 | 0.85 | 0.89 |
| 6 | 2716860 | rs1461656 | 0.25 | -1.369 | 0.601 | -1.837 | 0 | -0.029 | -0.029 | 0.874 | 0.27 | 0.23 | 0.18 | 0.02 | 0.17 | 0.17 |
| 6 | 2717090 | rs4959710 | 0.287 | -1.2 | 0.763 | -1.626 | -0.007 | 0.008 | 0.008 | 0 | 0.85 | 0.82 | 0.92 | 0.74 | 0.85 | 0.89 |
| 6 | 2719315 | rs9503281 | 0.628 | -1.152 | 0.615 | -1.589 | -0.001 | 0.011 | 0.011 | 0.041 | 0.19 | 0.16 | 0.26 | 0.15 | 0.2 | 0.15 |
| 6 | 2722312 | rs7771292 | -0.075 | -1.115 | 0.631 | -1.624 | -0.009 | -0.066 | -0.066 | 0.319 | 0.12 | 0.25 | 0.08 | 0 | 0.12 | 0.04 |
| 6 | 2726653 | rs9285964 | -0.85 | -1.197 | 0.649 | -1.563 | -0.005 | 0.001 | 0.001 | 0 | 0.34 | 0.56 | 0.36 | 0.44 | 0.35 | 0.26 |
| 6 | 2728600 | rs4246059 | 0.774 | -1.567 | 0.109 | -1.827 | 0.047 | 0.117 | 0.117 | 0 | 0.54 | 0.29 | 0.6 | 0.83 | 0.59 | 0.65 |
| 6 | 2728935 | rs912699 | -0.153 | -1.539 | 0.489 | -1.574 | -0.068 | -0.073 | -0.073 | 0 | 0.83 | 0.58 | 0.83 | 0.95 | 0.92 | 0.84 |
| 6 | 2736688 | rs9503288 | -0.506 | -1.255 | 0.036 | -1.704 | 0.005 | 0.018 | 0.018 | 0.574 | 0.39 | 0.35 | 0.24 | 0.4 | 0.38 | 0.26 |
| 6 | 2740501 | rs9501884 | 0.433 | -1.072 | -0.332 | -1.727 | -0.002 | 0.016 | 0.016 | 1.019 | 0.26 | 0.24 | 0.19 | 0.11 | 0.25 | 0.16 |
| 6 | 2741655 | rs4382335 | -0.456 | -1.225 | -0.422 | -1.471 | 0.041 | -0.036 | -0.036 | 0 | 0.25 | 0.13 | 0.19 | 0.35 | 0.21 | 0.26 |
| 6 | 2743245 | rs7775946 | 0.424 | -1.004 | -0.472 | -1.502 | -0.033 | -0.014 | -0.014 | 0 | 0.67 | 0.6 | 0.75 | 0.67 | 0.74 | 0.7 |
| 6 | 2743674 | rs17135660 | -0.076 | -0.207 | -0.946 | -1.614 | -0.001 | -0.017 | -0.017 | 0 | 0.07 | 0.05 | 0.03 | 0 | 0.07 | 0.01 |
| 6 | 2744554 | rs17135661 | -0.542 | 0.17 | -0.958 | -1.485 | -0.007 | 0.03 | 0.03 | 2.774 | 0.3 | 0.26 | 0.21 | 0.12 | 0.3 | 0.19 |
| 6 | 2745372 | rs11759003 | 0.085 | 0.097 | -1.134 | -1.35 | 0.037 | 0.092 | 0.092 | 0 | 0.55 | 0.26 | 0.73 | 0.98 | 0.64 | 0.67 |
| 6 | 2749381 | rs2296356 | -0.696 | 0.289 | -1.102 | 0.158 | -0.004 | -0.044 | -0.044 | 0 | 0.11 | 0.03 | 0.27 | 0.32 | 0.12 | 0.12 |
| 6 | 2749661 | rs2296355 | -1.465 | -0.173 | -0.847 | -0.335 | -0.008 | 0.005 | 0.005 | 0 | 0.34 | 0.35 | 0.38 | 0.39 | 0.3 | 0.39 |
| 6 | 2753152 | rs17135673 | -0.192 | -0.233 | -0.474 | -0.336 | -0.007 | -0.051 | -0.051 | 0.847 | 0.11 | 0.03 | 0.27 | 0.32 | 0.12 | 0.12 |
| 6 | 2759201 | rs12202342 | 0.642 | -0.192 | -0.211 | -0.268 | -0.031 | -0.086 | -0.086 | 0.384 | 0.76 | 0.58 | 0.87 | 0.93 | 0.82 | 0.73 |
| 6 | 2760373 | rs10080269 | 0.409 | -0.21 | -0.138 | -0.295 | -0.007 | -0.025 | -0.025 | 0 | 0.79 | 0.76 | 0.89 | 0.93 | 0.83 | 0.74 |
| 6 | 2760872 | rs1381326 | -0.283 | -0.197 | -0.27 | -0.248 | -0.002 | 0.003 | 0.003 | 0.074 | 0.12 | 0.08 | 0.1 | 0.07 | 0.15 | 0.2 |
| 6 | 2774052 | rs160666 | 0.819 | -0.047 | -1.11 | 0.243 | -0.028 | -0.081 | -0.081 | 0 | 0.26 | 0.08 | 0.3 | 0.36 | 0.3 | 0.4 |
| 6 | 2775431 | rs9328128 | 1.076 | -0.272 | -0.56 | 0.164 | 0.006 | -0.019 | -0.019 | 0 | 0.4 | 0.42 | 0.34 | 0.24 | 0.42 | 0.31 |

| | | | | | | | | | | | | | | | | |
|---|---|---|---|---|---|---|---|---|---|---|---|---|---|---|---|---|
| 6 | 2776303 | rs160699 | 0.066 | -0.265 | -0.994 | 0.21 | -0.002 | -0.019 | -0.019 | 0 | 0.39 | 0.28 | 0.39 | 0.43 | 0.44 | 0.53 |
| 6 | 2779882 | rs898768 | -0.637 | -0.266 | -1.076 | 0.487 | 0.011 | -0.072 | -0.072 | 0 | 0.2 | 0.04 | 0.24 | 0.33 | 0.14 | 0.16 |
| 6 | 2783985 | rs3823104 | -0.083 | 0.27 | -0.896 | 0.226 | 0.016 | -0.026 | -0.026 | 0 | 0.65 | 0.51 | 0.67 | 0.78 | 0.6 | 0.56 |
| 6 | 2784571 | rs160703 | 0.801 | -1.465 | -1.225 | 0.293 | 0.018 | -0.025 | -0.025 | 0 | 0.19 | 0.1 | 0.21 | 0.26 | 0.2 | 0.33 |
| 6 | 2784797 | rs3734499 | 0.064 | -1.666 | -1.527 | 0.485 | -0.008 | -0.066 | -0.066 | 0 | 0.18 | 0.07 | 0.3 | 0.4 | 0.19 | 0.14 |
| 6 | 2785055 | rs160702 | 0.199 | -1.783 | -1.68 | 0.296 | 0.027 | -0.016 | -0.016 | 0 | 0.3 | 0.12 | 0.26 | 0.31 | 0.26 | 0.36 |
| 6 | 2785112 | rs160701 | 1.344 | -1.587 | -1.658 | 0.456 | 0.008 | -0.053 | -0.053 | 0 | 0.17 | 0.06 | 0.22 | 0.31 | 0.19 | 0.3 |
| 6 | 2787925 | rs460677 | -0.393 | -1.604 | -1.712 | 0.395 | -0.014 | 0.043 | 0.043 | 0.195 | 0.44 | 0.07 | 0.54 | 0.71 | 0.44 | 0.51 |
| 6 | 2789371 | rs315002 | -0.906 | -1.717 | -1.297 | 0.184 | -0.021 | 0.033 | 0.033 | 0.995 | 0.62 | 0.31 | 0.68 | 0.81 | 0.58 | 0.58 |
| 6 | 2793954 | rs6942083 | -0.769 | -1.925 | -0.943 | 0.583 | 0.002 | -0.048 | -0.048 | 0 | 0.29 | 0.11 | 0.33 | 0.38 | 0.24 | 0.17 |
| 6 | 2798619 | rs9885826 | 0.033 | -1.705 | -0.583 | 0.744 | -0.004 | 0.005 | 0.005 | 1.063 | 0.11 | 0.09 | 0.1 | 0.14 | 0.11 | 0.07 |
| 6 | 2799060 | rs160690 | 0.49 | -1.59 | -0.431 | 0.46 | -0.019 | 0.007 | 0.007 | 0.073 | 0.73 | 0.68 | 0.8 | 0.71 | 0.8 | 0.71 |
| 6 | 2807359 | rs7766569 | -1.086 | -1.076 | -0.614 | 0.216 | 0.008 | 0.036 | 0.036 | 0 | 0.32 | 0.25 | 0.28 | 0.22 | 0.28 | 0.21 |
| 6 | 2807760 | rs1350849 | -0.537 | -0.862 | -0.861 | 0.324 | -0.031 | -0.015 | -0.015 | 2.843 | 0.19 | 0.04 | 0.23 | 0.22 | 0.22 | 0.17 |
| 6 | 2812175 | rs9405587 | -0.945 | -0.948 | -0.307 | 0.296 | -0.005 | -0.002 | -0.002 | 0 | 0.29 | 0.33 | 0.28 | 0.29 | 0.23 | 0.16 |
| 6 | 2814466 | rs4351304 | -0.726 | -0.934 | -0.278 | 0.27 | -0.004 | 0.004 | 0.004 | 0.034 | 0.25 | 0.3 | 0.25 | 0.29 | 0.19 | 0.16 |
| 6 | 2814525 | rs10900929 | 0.718 | -1.311 | -0.696 | 0.221 | 0.008 | -0.002 | -0.002 | 0 | 0.21 | 0.11 | 0.2 | 0.19 | 0.2 | 0.28 |
| 6 | 2818875 | rs2118940 | -0.77 | -1.345 | -0.53 | 0.201 | 0 | -0.001 | -0.001 | 0 | 0.47 | 0.48 | 0.46 | 0.48 | 0.39 | 0.44 |
| 6 | 2820878 | rs9391970 | -0.385 | -1.146 | -0.522 | 0.307 | -0.01 | 0.017 | 0.017 | 0.19 | 0.25 | 0.42 | 0.26 | 0.29 | 0.19 | 0.16 |
| 6 | 2826677 | rs6940098 | -1.025 | -1.139 | -0.714 | 0.299 | 0.001 | -0.009 | -0.009 | 0 | 0.55 | 0.62 | 0.5 | 0.48 | 0.45 | 0.47 |
| 6 | 2826717 | rs11242799 | 0.512 | -0.966 | -0.827 | 0.698 | 0.015 | -0.001 | -0.001 | 0.183 | 0.21 | 0.1 | 0.19 | 0.18 | 0.16 | 0.26 |
| 6 | 2828026 | rs6912537 | -0.189 | -0.121 | -0.735 | 1.22 | -0.001 | 0.003 | 0.003 | 0 | 0.16 | 0.15 | 0.2 | 0.29 | 0.14 | 0.14 |
| 6 | 2830523 | rs316365 | -0.13 | 0.451 | -0.879 | 1.31 | -0.017 | -0.039 | -0.039 | 0.027 | 0.07 | 0 | 0.19 | 0.25 | 0.1 | 0.05 |
| 6 | 2831191 | rs10793840 | -0.221 | 0.319 | -0.701 | 0.782 | -0.01 | -0.014 | -0.014 | 0 | 0.61 | 0.71 | 0.53 | 0.54 | 0.46 | 0.53 |
| 6 | 2831909 | rs9503309 | -0.777 | 0.044 | 0.119 | 0.335 | -0.006 | -0.058 | -0.058 | 0 | 0.14 | 0.36 | 0.08 | 0.03 | 0.07 | 0.05 |
| 6 | 2832055 | rs316364 | -0.816 | 0.292 | -0.276 | 0.684 | -0.008 | 0.014 | 0.014 | 0 | 0.38 | 0.47 | 0.44 | 0.49 | 0.35 | 0.36 |
| 6 | 2832078 | rs316363 | -0.303 | 0.451 | -0.62 | 0.138 | 0.009 | 0.019 | 0.019 | 0 | 0.5 | 0.58 | 0.64 | 0.73 | 0.48 | 0.65 |
| 6 | 2832331 | rs316362 | -0.263 | 0.761 | -1.311 | 0.569 | -0.035 | -0.04 | -0.04 | 0 | 0.32 | 0.11 | 0.43 | 0.36 | 0.34 | 0.25 |
| 6 | 2833739 | rs15286 | 0.227 | 1.18 | -0.178 | 0.661 | 0.014 | 0.012 | 0.012 | 0 | 0.82 | 0.75 | 0.82 | 0.75 | 0.72 | 0.73 |
| 6 | 2836600 | rs2293771 | 0.033 | 1.073 | -0.323 | 0.445 | -0.002 | 0.021 | 0.021 | 0.118 | 0.18 | 0.08 | 0.13 | 0.17 | 0.13 | 0.19 |
| 6 | 2837846 | rs316338 | 0.709 | 0.968 | 0.312 | 0.439 | 0.012 | 0.014 | 0.014 | 0.464 | 0.74 | 0.39 | 0.76 | 0.7 | 0.68 | 0.74 |
| 6 | 2838046 | rs316339 | 0.024 | 1.085 | 0.201 | 0.514 | 0.042 | 0.042 | 0.042 | 0 | 0.82 | 0.67 | 0.8 | 0.74 | 0.72 | 0.75 |
| 6 | 2838102 | rs316340 | 1.201 | 0.446 | -0.66 | 0.344 | -0.109 | -0.046 | -0.046 | 0 | 0.46 | 0.09 | 0.58 | 0.52 | 0.52 | 0.53 |
| 6 | 2841556 | rs316346 | 0.577 | 0.802 | 0.235 | 0.721 | 0.014 | 0.017 | 0.017 | 0 | 0.66 | 0.56 | 0.68 | 0.52 | 0.61 | 0.58 |
| 6 | 2842830 | rs409346 | 0.196 | 1.102 | 0.68 | 0.523 | 0.07 | 0.037 | 0.037 | 0.019 | 0.83 | 0.73 | 0.78 | 0.75 | 0.68 | 0.73 |
| 6 | 2843335 | rs405616 | 0.776 | 1.122 | 0.678 | 0.434 | 0.007 | 0.002 | 0.002 | 0.277 | 0.58 | 0.43 | 0.52 | 0.61 | 0.47 | 0.65 |
| 6 | 2847576 | rs9503317 | -0.019 | 1.094 | 0.307 | 0.417 | -0.001 | 0 | 0 | 0 | 0.22 | 0.24 | 0.28 | 0.28 | 0.2 | 0.24 |
| 6 | 2854367 | rs412303 | 1.436 | 0.956 | 0.035 | 0.459 | 0.011 | -0.058 | -0.058 | 0.043 | 0.4 | 0.12 | 0.39 | 0.47 | 0.36 | 0.45 |
| 6 | 2856192 | rs386595 | 0.351 | 1.067 | -0.099 | 0.344 | -0.027 | -0.001 | -0.001 | 0 | 0.65 | 0.77 | 0.65 | 0.6 | 0.58 | 0.57 |
| 6 | 2862053 | rs414861 | 1.407 | 0.982 | -0.383 | 0.468 | 0.036 | -0.065 | -0.065 | 0.163 | 0.34 | 0.1 | 0.38 | 0.41 | 0.28 | 0.43 |
| 6 | 2863203 | rs392120 | -0.193 | 1.355 | -0.323 | 0.365 | -0.032 | -0.025 | -0.025 | 0 | 0.81 | 0.85 | 0.75 | 0.68 | 0.63 | 0.71 |
| 6 | 2864295 | rs380779 | 0.863 | 0.779 | -0.311 | 0.438 | 0.038 | -0.057 | -0.057 | 0.007 | 0.39 | 0.14 | 0.4 | 0.45 | 0.3 | 0.45 |
| 6 | 2869031 | rs6932052 | -1.139 | 0.681 | 0.045 | 0.495 | 0.008 | -0.005 | -0.005 | 0 | 0.64 | 0.73 | 0.58 | 0.6 | 0.71 | 0.56 |
| 6 | 2871085 | rs9392436 | -1.252 | 0.737 | -0.495 | 0.322 | 0.025 | -0.09 | -0.09 | 0.007 | 0.66 | 0.93 | 0.59 | 0.58 | 0.7 | 0.56 |
| 6 | 2872759 | rs11753600 | -0.86 | 1.219 | 0.189 | 0.532 | 0.012 | 0.044 | 0.044 | 0 | 0.06 | 0.13 | 0.13 | 0.11 | 0.11 | 0.08 |
| 6 | 2876581 | rs2252076 | 0.534 | 0.75 | 0.217 | 0.025 | -0.025 | -0.075 | -0.075 | 0 | 0.71 | 0.62 | 0.84 | 0.87 | 0.84 | 0.81 |
| 6 | 2885791 | rs318470 | 0.067 | 0.72 | -0.129 | 0.08 | -0.002 | -0.019 | -0.019 | 0.075 | 0.91 | 0.94 | 0.95 | 1 | 0.96 | 0.98 |
| 6 | 2886067 | rs12530071 | -0.363 | 1.805 | -0.023 | 0.168 | 0.017 | 0.004 | 0.004 | 0.019 | 0.11 | 0.12 | 0.16 | 0.11 | 0.25 | 0.11 |
| 6 | 2886795 | rs318473 | 0.667 | 1.571 | -0.308 | 0.099 | -0.013 | 0.093 | 0.093 | 0 | 0.53 | 0.33 | 0.64 | 0.74 | 0.54 | 0.63 |
| 6 | 2887652 | rs318477 | 0.714 | 1.623 | -0.583 | 0.142 | -0.006 | 0.084 | 0.084 | 0.007 | 0.52 | 0.33 | 0.64 | 0.75 | 0.55 | 0.63 |
| 6 | 2888213 | rs318481 | 0.751 | 1.646 | -0.926 | 0.466 | -0.009 | 0.056 | 0.056 | 0 | 0.5 | 0.24 | 0.63 | 0.64 | 0.53 | 0.61 |
| 6 | 2889071 | rs13196459 | -0.543 | 1.861 | -0.456 | 0.94 | 0.004 | 0.024 | 0.024 | 0 | 0.13 | 0.17 | 0.17 | 0.32 | 0.18 | 0.23 |
| 6 | 2889616 | rs9392442 | -0.469 | 1.665 | -0.566 | 0.514 | -0.002 | -0.016 | -0.016 | 0 | 0.11 | 0.15 | 0.1 | 0.1 | 0.1 | 0.17 |
| 6 | 2890190 | rs1052886 | -0.928 | 2.062 | -0.889 | 0.674 | -0.005 | -0.006 | -0.006 | 0.884 | 0.1 | 0.06 | 0.12 | 0.12 | 0.25 | 0.1 |
| 6 | 2890283 | rs318486 | 0.334 | 1.1 | -1.154 | 0.122 | 0.012 | 0.016 | 0.016 | 0 | 0.56 | 0.7 | 0.62 | 0.56 | 0.67 | 0.55 |
| 6 | 2893008 | rs318489 | 0.204 | 1.069 | -1.54 | 0.004 | -0.044 | 0.086 | 0.086 | 0 | 0.67 | 0.93 | 0.71 | 0.8 | 0.64 | 0.59 |
| 6 | 2899392 | rs9503330 | -0.743 | 1.082 | -0.567 | 0.425 | -0.003 | -0.018 | -0.018 | 0 | 0.09 | 0.12 | 0.05 | 0.01 | 0.05 | 0.07 |
| 6 | 2902393 | rs3799229 | -0.844 | 1.396 | 0.056 | 0.579 | 0.091 | 0.042 | 0.042 | 0 | 0.06 | 0.24 | 0.06 | 0.07 | 0.13 | 0.11 |
| 6 | 2906147 | rs11242807 | -0.978 | 1.38 | 0.019 | 0.593 | 0.044 | 0.079 | 0.079 | 0 | 0.06 | 0.19 | 0.11 | 0.19 | 0.13 | 0.18 |
| 6 | 2910998 | rs318496 | -0.184 | 0.963 | -0.717 | 0.432 | 0.014 | -0.024 | -0.024 | 0.054 | 0.78 | 0.93 | 0.85 | 0.78 | 0.82 | 0.63 |
| 6 | 2912277 | rs318497 | 0.699 | 0.761 | -0.17 | -0.167 | -0.001 | -0.001 | -0.001 | 0.432 | 0.44 | 0.38 | 0.55 | 0.58 | 0.46 | 0.29 |
| 6 | 2914609 | rs6940344 | -1.185 | 0.848 | -0.598 | -0.806 | -0.044 | 0.178 | 0.178 | 0 | 0.39 | 0.84 | 0.28 | 0.17 | 0.42 | 0.25 |
| 6 | 2915664 | rs7759176 | -1.022 | 0.675 | 0.34 | -0.628 | -0.005 | -0.052 | -0.052 | 0.204 | 0.3 | 0.56 | 0.24 | 0.13 | 0.27 | 0.24 |
| 6 | 2931879 | rs7751676 | 0.049 | 0.698 | 0.252 | -0.574 | -0.008 | -0.031 | -0.031 | 0.325 | 0.08 | 0.1 | 0.05 | 0.01 | 0.07 | 0.06 |
| 6 | 2934641 | rs2165125 | -0.169 | 0.695 | 0.53 | -0.586 | 0.021 | -0.141 | -0.141 | 0.058 | 0.2 | 0.54 | 0.14 | 0 | 0.2 | 0.08 |
| 6 | 2937347 | rs11242810 | -0.541 | 0.739 | 0.468 | -0.467 | 0.015 | -0.049 | -0.049 | 0 | 0.23 | 0.63 | 0.15 | 0.17 | 0.25 | 0.1 |
| 6 | 2945331 | rs318427 | 0.961 | 0.892 | 0.241 | 0.139 | 0.014 | 0.006 | 0.006 | 0 | 0.85 | 0.78 | 0.84 | 0.8 | 0.79 | 0.87 |
| 6 | 2947783 | rs13218926 | 0.888 | 1.048 | -0.175 | 0.186 | -0.013 | -0.033 | -0.033 | 0 | 0.34 | 0.25 | 0.28 | 0.38 | 0.35 | 0.4 |
| 6 | 2951849 | rs7763055 | -0.595 | 0.721 | 0.176 | -0.315 | -0.053 | -0.075 | -0.075 | 0 | 0.25 | 0.38 | 0.09 | 0.17 | 0.15 | 0.19 |

| | | | | | | | | | | | | | | | | |
|---|---|---|---|---|---|---|---|---|---|---|---|---|---|---|---|---|
| 6 | 2952289 | rs2756071 | 1.003 | 0.719 | 0.172 | -0.367 | -0.006 | -0.023 | -0.023 | 0 | 0.68 | 0.46 | 0.76 | 0.79 | 0.7 | 0.78 |
| 6 | 2953445 | rs3818276 | -0.124 | 0.804 | 0.135 | -0.328 | 0.005 | -0.003 | -0.003 | 0 | 0.91 | 0.88 | 0.85 | 0.97 | 0.85 | 0.93 |
| 6 | 2955802 | rs2295769 | 1.596 | 0.86 | -0.093 | -0.262 | -0.083 | -0.058 | -0.058 | 0.206 | 0.3 | 0.03 | 0.25 | 0.27 | 0.3 | 0.29 |
| 6 | 2956443 | rs10900931 | -0.594 | 0.81 | 0.048 | -0.295 | -0.044 | -0.079 | -0.079 | 0 | 0.25 | 0.38 | 0.1 | 0.17 | 0.19 | 0.19 |
| 6 | 2958095 | rs3799200 | 0.343 | 0.847 | 0.1 | -0.312 | 0.017 | -0.008 | -0.008 | 0.178 | 0.86 | 0.63 | 0.81 | 0.93 | 0.83 | 0.86 |
| 6 | 2958548 | rs3778253 | -0.58 | 0.851 | 0.017 | -0.317 | -0.044 | -0.079 | -0.079 | 0.12 | 0.25 | 0.38 | 0.1 | 0.17 | 0.19 | 0.19 |
| 6 | 2959513 | rs2236277 | 0.963 | 0.945 | -0.311 | -0.223 | -0.018 | -0.063 | -0.063 | 0 | 0.3 | 0.15 | 0.45 | 0.48 | 0.34 | 0.37 |
| 6 | 2963370 | rs9378350 | 0.836 | 0.855 | -0.19 | -0.124 | 0.005 | 0.003 | 0.003 | 0 | 0.66 | 0.34 | 0.73 | 0.91 | 0.74 | 0.74 |
| 6 | 2966681 | rs7751512 | 0.612 | 1.155 | -0.079 | 0.126 | -0.025 | -0.06 | -0.06 | 1.965 | 0.3 | 0.14 | 0.46 | 0.45 | 0.34 | 0.37 |
| 6 | 2970568 | rs7743193 | 1.592 | 0.933 | 0.157 | 0.202 | 0.007 | 0.028 | 0.028 | 0 | 0.66 | 0.26 | 0.74 | 0.74 | 0.66 | 0.69 |
| 6 | 2981772 | rs4389803 | 0.754 | 0.937 | -0.015 | 0.149 | 0 | 0.001 | 0.001 | 0.235 | 0.49 | 0.57 | 0.4 | 0.51 | 0.48 | 0.41 |
| 6 | 2986088 | rs11242815 | 0.096 | 0.95 | -0.483 | 0.56 | -0.01 | -0.021 | -0.021 | 0 | 0.3 | 0.26 | 0.45 | 0.55 | 0.38 | 0.38 |
| 6 | 2989582 | rs2326106 | 0.763 | 1.062 | -0.43 | 0.598 | 0.019 | 0.042 | 0.042 | 0 | 0.86 | 0.72 | 0.82 | 0.71 | 0.79 | 0.86 |
| 6 | 2990373 | rs6920013 | 0.465 | 0.883 | -0.61 | 0.561 | -0.03 | 0.01 | 0.01 | 0.539 | 0.59 | 0.68 | 0.49 | 0.36 | 0.48 | 0.53 |
| 6 | 2990406 | rs6920145 | 1.789 | 0.815 | -0.662 | 0.542 | -0.018 | -0.009 | -0.009 | 0.184 | 0.4 | 0.07 | 0.36 | 0.34 | 0.35 | 0.39 |
| 6 | 2990894 | rs1054132 | -0.417 | 0.76 | -0.367 | 0.507 | -0.001 | 0.001 | 0.001 | 0 | 0.42 | 0.57 | 0.44 | 0.37 | 0.43 | 0.41 |
| 6 | 2991195 | rs8583 | 1.521 | 0.721 | -0.765 | 0.521 | -0.037 | -0.032 | -0.032 | 0 | 0.39 | 0.05 | 0.36 | 0.34 | 0.35 | 0.39 |
| 6 | 2997544 | rs9405188 | -0.783 | 0.041 | 0.005 | 0.599 | -0.003 | -0.003 | -0.003 | 0 | 0.18 | 0.24 | 0.21 | 0.14 | 0.18 | 0.13 |
| 6 | 2999729 | rs2070999 | -1.666 | 0.155 | -0.389 | 0.6 | -0.101 | 0.013 | 0.013 | 0 | 0.61 | 0.96 | 0.65 | 0.72 | 0.6 | 0.61 |
| 6 | 3000303 | rs2071002 | -1.585 | 0.614 | 0.494 | 0.808 | 0.015 | 0.021 | 0.021 | 0 | 0.25 | 0.39 | 0.32 | 0.34 | 0.29 | 0.26 |
| 6 | 3002238 | rs4149352 | -0.519 | 0.129 | 0.521 | 0.811 | -0.001 | -0.004 | -0.004 | 0 | 0.77 | 0.8 | 0.72 | 0.68 | 0.76 | 0.72 |
| 6 | 3006807 | rs4149360 | -0.398 | 0.239 | 0.204 | 0.7 | -0.023 | -0.021 | -0.021 | 0.315 | 0.14 | 0.03 | 0.21 | 0.12 | 0.19 | 0.1 |
| 6 | 3010390 | rs1143684 | -0.737 | 0.195 | 0.293 | 0.818 | -0.045 | -0.096 | -0.096 | 0.848 | 0.79 | 0.98 | 0.77 | 0.66 | 0.79 | 0.72 |
| 6 | 3011149 | rs9378352 | 0.64 | 0.095 | 1.006 | 0.753 | -0.001 | 0.005 | 0.005 | 0 | 0.62 | 0.43 | 0.6 | 0.5 | 0.6 | 0.57 |
| 6 | 3014483 | rs3823096 | 1.715 | 0.038 | 1.02 | 0.656 | -0.011 | 0.019 | 0.019 | 0 | 0.46 | 0.24 | 0.37 | 0.38 | 0.4 | 0.47 |
| 6 | 3015790 | rs1049115 | -0.171 | -0.765 | 0.864 | 0.696 | -0.005 | -0.033 | -0.033 | 0.006 | 0.71 | 0.93 | 0.73 | 0.65 | 0.71 | 0.69 |
| 6 | 3015865 | rs4149367 | -0.749 | -0.745 | 1.147 | 0.708 | 0.043 | 0.128 | 0.128 | 0 | 0.07 | 0.36 | 0.1 | 0.16 | 0.07 | 0.12 |
| 6 | 3016107 | rs11242817 | -0.369 | -0.574 | 0.336 | 0.643 | -0.005 | -0.005 | -0.005 | 0 | 0.14 | 0.08 | 0.21 | 0.12 | 0.19 | 0.1 |
| 6 | 3016130 | rs4149369 | -0.106 | -0.56 | 0.328 | 0.65 | -0.006 | -0.034 | -0.034 | 0 | 0.72 | 0.93 | 0.73 | 0.65 | 0.71 | 0.69 |
| 6 | 3017580 | rs1885298 | -0.401 | -0.531 | 0.984 | 0.817 | 0 | 0 | 0 | 0 | 0.15 | 0.15 | 0.22 | 0.12 | 0.19 | 0.1 |
| 6 | 3019927 | rs6955 | -0.966 | -0.527 | 0.974 | 0.819 | -0.001 | -0.001 | -0.001 | 0.151 | 0.16 | 0.2 | 0.22 | 0.12 | 0.19 | 0.1 |
| 6 | 3027131 | rs911536 | -1.062 | -0.864 | 1.599 | 1.239 | 0.025 | 0.162 | 0.162 | 0 | 0.13 | 0.28 | 0.22 | 0.3 | 0.15 | 0.17 |
| 6 | 3029127 | rs1885300 | 0.488 | -1.18 | 1.613 | 1.118 | 0.022 | 0.008 | 0.008 | 0 | 0.7 | 0.85 | 0.72 | 0.68 | 0.75 | 0.71 |
| 6 | 3032804 | rs6918464 | 1.404 | -1.201 | 1.599 | 1.089 | -0.002 | -0.001 | -0.001 | 0.396 | 0.56 | 0.52 | 0.46 | 0.38 | 0.58 | 0.43 |
| 6 | 3035613 | rs6920099 | 1.399 | -1.387 | 1.495 | 0.985 | -0.007 | -0.003 | -0.003 | 0 | 0.54 | 0.57 | 0.49 | 0.38 | 0.59 | 0.5 |
| 6 | 3036661 | rs7762140 | -2.215 | -1.638 | 1.165 | 0.913 | 0.066 | -0.028 | -0.028 | 0 | 0.35 | 0.84 | 0.35 | 0.33 | 0.26 | 0.38 |
| 6 | 3039609 | rs1040521 | 1.786 | -1.536 | 1.363 | 0.92 | 0.068 | 0.059 | 0.059 | 0 | 0.5 | 0.28 | 0.43 | 0.36 | 0.57 | 0.36 |
| 6 | 3041200 | rs7774283 | 1.694 | -1.555 | 1.357 | 0.816 | 0.02 | 0.009 | 0.009 | 0 | 0.52 | 0.44 | 0.44 | 0.39 | 0.57 | 0.38 |
| 6 | 3044887 | rs4959762 | -0.302 | -1.232 | 0.924 | 1.059 | -0.026 | -0.081 | -0.081 | 0 | 0.82 | 0.98 | 0.73 | 0.69 | 0.82 | 0.74 |
| 6 | 3046537 | rs9503375 | -0.815 | -1.194 | 1.526 | 0.966 | 0.018 | 0.147 | 0.147 | 0 | 0.12 | 0.27 | 0.21 | 0.32 | 0.15 | 0.17 |
| 6 | 3049982 | rs7761969 | 1.021 | -1.153 | 1.412 | 0.726 | 0.009 | -0.006 | -0.006 | 1.268 | 0.58 | 0.68 | 0.46 | 0.4 | 0.56 | 0.4 |
| 6 | 3055752 | rs11242820 | 1.854 | -1.138 | 0.707 | 0.694 | -0.041 | -0.074 | -0.074 | 0 | 0.47 | 0.03 | 0.4 | 0.39 | 0.54 | 0.36 |
| 6 | 3055877 | rs7765538 | -1.337 | -1.14 | 1.037 | 0.762 | -0.012 | 0.005 | 0.005 | 0.15 | 0.28 | 0.56 | 0.31 | 0.29 | 0.28 | 0.33 |
| 6 | 3057413 | rs7775816 | 0.545 | -0.869 | 1.157 | 0.948 | -0.024 | 0.008 | 0.008 | 0 | 0.76 | 0.64 | 0.67 | 0.68 | 0.83 | 0.69 |
| 6 | 3064983 | rs9328140 | -0.147 | -0.63 | 0.577 | 1.067 | 0.003 | -0.008 | -0.008 | 0 | 0.82 | 0.85 | 0.69 | 0.68 | 0.84 | 0.73 |
| 6 | 3067074 | rs9503383 | 0.829 | -0.486 | 0.543 | 1.136 | -0.008 | 0.003 | 0.003 | 0 | 0.62 | 0.53 | 0.54 | 0.53 | 0.67 | 0.52 |
| 6 | 3071913 | rs6596945 | -1.187 | -0.049 | 0.663 | 1.05 | -0.007 | 0.037 | 0.037 | 0 | 0.23 | 0.43 | 0.26 | 0.28 | 0.21 | 0.23 |
| 6 | 3075553 | rs2326173 | 0.097 | -0.281 | 0.399 | 0.771 | 0 | -0.001 | -0.001 | 0 | 0.6 | 0.46 | 0.63 | 0.41 | 0.64 | 0.58 |
| 6 | 3075698 | rs3757082 | 0.098 | -0.323 | 0.443 | 0.751 | 0.008 | -0.008 | -0.008 | 1.761 | 0.6 | 0.43 | 0.63 | 0.41 | 0.64 | 0.58 |
| 6 | 3077141 | rs2272990 | 0.912 | -0.608 | 1.625 | 0.994 | 0.048 | 0.223 | 0.223 | 0 | 0.95 | 0.79 | 0.9 | 0.73 | 0.93 | 0.93 |
| 6 | 3080324 | rs2064310 | 1.261 | -0.607 | 1.819 | 0.987 | 0.064 | 0.256 | 0.256 | 0.112 | 0.94 | 0.71 | 0.89 | 0.72 | 0.93 | 0.93 |
| 6 | 3085477 | rs909966 | -1.766 | -0.818 | 1.65 | 0.145 | -0.023 | -0.043 | -0.043 | 0 | 0.08 | 0.21 | 0.04 | 0 | 0.02 | 0.12 |
| 6 | 3095388 | rs11967255 | -2.161 | -0.775 | 1.811 | 0.113 | -0.006 | -0.043 | -0.043 | 0 | 0.1 | 0.52 | 0.06 | 0 | 0.03 | 0.08 |
| 6 | 3099418 | rs7745598 | -2.099 | -0.757 | 1.822 | 0.122 | 0.004 | -0.034 | -0.034 | 0 | 0.09 | 0.53 | 0.06 | 0 | 0.03 | 0.08 |
| 6 | 3104696 | rs10498658 | 0.31 | -0.321 | 0.994 | -0.459 | -0.071 | 0.087 | 0.087 | 12.226 | 0.16 | 0.02 | 0.16 | 0 | 0.26 | 0.04 |
| 6 | 3108907 | rs9503400 | -2.316 | -1.289 | 2.012 | 0.279 | -0.039 | -0.019 | -0.019 | 0 | 0.11 | 0.66 | 0.13 | 0.06 | 0.03 | 0.1 |
| 6 | 3112530 | rs7739011 | 2.275 | -1.508 | 2.097 | 0.22 | -0.042 | -0.042 | -0.042 | 0 | 0.84 | 0.37 | 0.85 | 0.93 | 0.93 | 0.86 |
| 6 | 3130428 | rs1009263 | -1.804 | -1.345 | 2.177 | 0.38 | -0.021 | 0.023 | 0.023 | 0 | 0.05 | 0.15 | 0.09 | 0.07 | 0.01 | 0.04 |
| 6 | 3133459 | rs9632529 | -2.154 | -1.359 | 2.342 | 0.468 | 0 | 0.092 | 0.092 | 0 | 0.07 | 0.33 | 0.1 | 0.08 | 0.01 | 0.11 |
| 6 | 3143765 | rs6933330 | 0.333 | -1.733 | 1.109 | 0.818 | -0.045 | -0.129 | -0.129 | 0 | 0.72 | 0.92 | 0.54 | 0.52 | 0.74 | 0.56 |
| 6 | 3144113 | rs9405612 | 0.823 | -2.691 | 1.296 | 0.215 | 0.051 | -0.008 | -0.008 | 0.087 | 0.53 | 0.76 | 0.49 | 0.52 | 0.69 | 0.42 |
| 6 | 3145127 | rs9392464 | -0.109 | -2.642 | 1.282 | 0.288 | -0.068 | -0.135 | -0.135 | 0 | 0.73 | 0.95 | 0.61 | 0.6 | 0.74 | 0.58 |
| 6 | 3149271 | rs13206421 | 0.27 | -2.553 | 1.378 | 0.204 | -0.023 | -0.05 | -0.05 | 5.737 | 0.17 | 0.17 | 0.05 | 0.01 | 0.05 | 0.11 |
| 6 | 3153383 | rs3088000 | 0.253 | -2.621 | 1.856 | 0.171 | -0.03 | -0.059 | -0.059 | 0 | 0.17 | 0.19 | 0.05 | 0.01 | 0.05 | 0.11 |
| 6 | 3157399 | rs909962 | 0.649 | -2.751 | 2.361 | 0.172 | -0.014 | 0.023 | 0.023 | 2.116 | 0.63 | 0.47 | 0.52 | 0.52 | 0.73 | 0.47 |
| 6 | 3162378 | rs9392465 | 1.119 | -2.605 | 2.303 | 0.343 | 0.034 | -0.016 | -0.016 | 0.526 | 0.46 | 0.35 | 0.48 | 0.54 | 0.69 | 0.36 |
| 6 | 3174947 | rs9501935 | 0.665 | -3.216 | 2.48 | 0.667 | 0.07 | 0.016 | 0.016 | 1.64 | 0.7 | 0.96 | 0.88 | 0.79 | 0.87 | 0.73 |
| 6 | 3176888 | rs11970635 | -2.396 | -2.634 | 3.468 | 1.06 | 0.038 | 0.344 | 0.344 | 4.905 | 0.07 | 0.5 | 0.04 | 0.28 | 0.01 | 0.11 |
| 6 | 3185712 | rs7754077 | -2.896 | -2.642 | 3.143 | 1.056 | -0.051 | 0.097 | 0.097 | 7.42 | 0.09 | 0.71 | 0.07 | 0.28 | 0.02 | 0.15 |

| | | | | | | | | | | | | | | | | |
|---|---|---|---|---|---|---|---|---|---|---|---|---|---|---|---|---|
| 6 | 3193440 | rs9503437 | -2.63 | -2.619 | 3.208 | 1.068 | -0.001 | 0.284 | 0.284 | 2.804 | 0.08 | 0.52 | 0.06 | 0.28 | 0.01 | 0.14 |
| 6 | 3200962 | rs9503448 | -2.164 | -2.418 | 2.2 | 0.18 | 0.032 | 0.028 | 0.028 | 3.814 | 0.06 | 0.52 | 0.03 | 0 | 0 | 0.05 |
| 6 | 3205325 | rs6929344 | 0.842 | -2.966 | 1.7 | 0.132 | -0.015 | -0.007 | -0.007 | 0 | 0.58 | 0.44 | 0.71 | 0.66 | 0.72 | 0.51 |
| 6 | 3205434 | rs11242827 | 0.857 | -3.017 | 1.972 | 0.089 | -0.027 | -0.014 | -0.014 | 0 | 0.58 | 0.5 | 0.7 | 0.66 | 0.71 | 0.51 |
| 6 | 3207258 | rs9405622 | 0.939 | -3.397 | 2.314 | 0.208 | -0.042 | -0.009 | -0.009 | 1.701 | 0.62 | 0.44 | 0.77 | 0.69 | 0.81 | 0.57 |
| 6 | 3209588 | rs7769489 | 2.019 | -2.626 | 2.849 | 0.414 | -0.02 | 0.011 | 0.011 | 0.939 | 0.81 | 0.27 | 0.8 | 0.78 | 0.9 | 0.79 |
| 6 | 3212382 | rs6920128 | -2.414 | -2.442 | 3.105 | 0.103 | -0.03 | -0.028 | -0.028 | 0 | 0.07 | 0.7 | 0.05 | 0 | 0.01 | 0 |
| 6 | 3224209 | rs7772006 | -1.94 | -2.311 | 3.184 | 0.17 | 0.059 | 0.06 | 0.06 | 5.688 | 0.06 | 0.48 | 0.04 | 0 | 0.01 | 0.02 |
| 6 | 3229477 | rs7747852 | -1.715 | -2.002 | 2.915 | 0.372 | -0.018 | 0.003 | 0.003 | 6.765 | 0.08 | 0.64 | 0.06 | 0.03 | 0.01 | 0.03 |
| 6 | 3250324 | rs6596957 | -0.896 | -2.43 | 0.65 | 0.867 | 0.034 | -0.064 | -0.064 | 0 | 0.31 | 0.12 | 0.3 | 0.4 | 0.26 | 0.28 |
| 6 | 3255857 | rs9503499 | 1.109 | -3.287 | 0.619 | 0.788 | 0.013 | -0.015 | -0.015 | 0 | 0.82 | 0.89 | 0.83 | 0.69 | 0.91 | 0.77 |
| 6 | 3256895 | rs4445096 | 1.93 | -3.967 | 1.27 | 0.431 | 0.009 | -0.001 | -0.001 | 0.045 | 0.69 | 0.35 | 0.78 | 0.69 | 0.85 | 0.69 |
| 6 | 3258353 | rs7756623 | 0.992 | -3.986 | 1.234 | 0.418 | 0.047 | -0.069 | -0.069 | 1.505 | 0.82 | 0.96 | 0.83 | 0.69 | 0.91 | 0.78 |
| 6 | 3264101 | rs4959785 | 2.229 | -4.285 | 1.172 | 0.397 | 0.086 | 0.005 | 0.005 | 0 | 0.62 | 0.3 | 0.77 | 0.66 | 0.84 | 0.65 |
| 6 | 3264671 | rs13193012 | 1.293 | -4.286 | 1.361 | -0.225 | 0.018 | 0.003 | 0.003 | 1.987 | 0.62 | 0.72 | 0.74 | 0.66 | 0.77 | 0.72 |
| 6 | 3270006 | rs1127485 | 1.996 | -4.204 | 2.287 | 0.424 | -0.051 | 0.003 | 0.003 | 0 | 0.67 | 0.63 | 0.79 | 0.66 | 0.89 | 0.69 |
| 6 | 3270121 | rs3813485 | 1.597 | -4.202 | 1.897 | 0.379 | -0.021 | 0.007 | 0.007 | 0 | 0.75 | 0.73 | 0.8 | 0.67 | 0.9 | 0.75 |
| 6 | 3270578 | rs1127473 | -1.228 | -4.252 | 1.481 | 0.33 | -0.002 | 0 | 0 | 0 | 0.14 | 0.13 | 0.08 | 0.12 | 0.06 | 0.1 |
| 6 | 3272807 | rs9501975 | -1.292 | -4.251 | 1.481 | 0.331 | -0.001 | 0.001 | 0.001 | 0 | 0.15 | 0.14 | 0.08 | 0.12 | 0.06 | 0.1 |
| 6 | 3277298 | rs7775219 | 2.395 | -4.319 | 1.173 | 0.691 | 0.208 | -0.09 | -0.09 | 0 | 0.61 | 0.11 | 0.75 | 0.55 | 0.89 | 0.61 |
| 6 | 3287118 | rs6910086 | 0.713 | -4.078 | 1.165 | 0.773 | 0.035 | -0.075 | -0.075 | 0 | 0.88 | 0.98 | 0.88 | 0.68 | 0.95 | 0.78 |
| 6 | 3287486 | rs4959796 | -2.15 | -4.179 | 1.563 | 0.627 | -0.024 | -0.01 | -0.01 | 0 | 0.24 | 0.68 | 0.18 | 0.12 | 0.06 | 0.18 |
| 6 | 3288384 | rs9405623 | -2.161 | -4.274 | 1.64 | 0.648 | -0.026 | 0.027 | 0.027 | 0 | 0.28 | 0.32 | 0.28 | 0.47 | 0.13 | 0.33 |
| 6 | 3290726 | rs4076221 | -2.273 | -4.241 | 1.154 | 0.773 | 0.03 | -0.026 | -0.026 | 1.172 | 0.33 | 0.72 | 0.31 | 0.46 | 0.11 | 0.39 |
| 6 | 3290819 | rs4587211 | -0.38 | -4.173 | 0.829 | 0.79 | -0.048 | -0.056 | -0.056 | 0.281 | 0.1 | 0.31 | 0.03 | 0 | 0.01 | 0.08 |
| 6 | 3291336 | rs9378356 | -1.897 | -3.98 | -0.422 | 0.896 | 0.198 | -0.109 | -0.109 | 0.543 | 0.39 | 0.9 | 0.33 | 0.47 | 0.11 | 0.41 |
| 6 | 3294824 | rs9378357 | -1.452 | -3.09 | 0.018 | 1.08 | -0.105 | 0.068 | 0.068 | 0 | 0.24 | 0.48 | 0.16 | 0.35 | 0.06 | 0.31 |
| 6 | 3296997 | rs4305775 | 1.006 | -2.522 | -1.157 | 0.983 | 0.004 | -0.075 | -0.075 | 0.029 | 0.78 | 0.99 | 0.77 | 0.74 | 0.85 | 0.76 |
| 6 | 3297021 | rs9378359 | -1.37 | -2.882 | -1.105 | 0.742 | 0.07 | 0.01 | 0.01 | 0 | 0.26 | 0.1 | 0.22 | 0.23 | 0.13 | 0.29 |
| 6 | 3299656 | rs9501981 | -0.156 | -2.775 | -0.663 | 0.515 | -0.017 | -0.029 | -0.029 | 0 | 0.06 | 0.09 | 0.02 | 0 | 0.02 | 0.02 |
| 6 | 3300964 | rs9501984 | 0.171 | -1.997 | -1.096 | 0.526 | -0.004 | 0.006 | 0.006 | 0.005 | 0.46 | 0.63 | 0.43 | 0.42 | 0.5 | 0.43 |
| 6 | 3302217 | rs4959232 | 0.338 | -1.144 | -1.6 | 0.815 | -0.08 | -0.104 | -0.104 | 0.213 | 0.24 | 0.02 | 0.27 | 0.36 | 0.29 | 0.23 |
| 6 | 3312592 | rs9405629 | -0.536 | -0.996 | -0.878 | 1.026 | -0.014 | 0.059 | 0.059 | 0 | 0.14 | 0.23 | 0.17 | 0.24 | 0.1 | 0.11 |
| 6 | 3316746 | rs9378784 | 0.36 | -0.742 | -1.059 | 1.04 | -0.013 | -0.023 | -0.023 | 1.675 | 0.28 | 0.17 | 0.4 | 0.6 | 0.35 | 0.31 |
| 6 | 3317016 | rs4959804 | -0.398 | -0.738 | -0.457 | 1.184 | -0.004 | -0.001 | -0.001 | 0 | 0.76 | 0.69 | 0.72 | 0.78 | 0.82 | 0.71 |
| 6 | 3334247 | rs11968806 | -0.647 | -1.299 | -0.488 | 0.993 | -0.063 | -0.022 | -0.022 | 0 | 0.29 | 0.48 | 0.17 | 0.23 | 0.18 | 0.26 |
| 6 | 3336800 | rs9392482 | -0.914 | -1.289 | -0.456 | 1.116 | -0.051 | 0.019 | 0.019 | 0 | 0.34 | 0.48 | 0.28 | 0.45 | 0.25 | 0.33 |
| 6 | 3340717 | rs9503551 | 0.422 | -0.827 | -0.899 | 0.962 | 0.017 | -0.019 | -0.019 | 1.547 | 0.15 | 0.09 | 0.1 | 0.23 | 0.12 | 0.24 |
| 6 | 3341675 | rs6917161 | 0.331 | 0.016 | -1.241 | 0.521 | -0.061 | -0.043 | -0.043 | 0 | 0.85 | 0.99 | 0.63 | 0.86 | 0.81 | 0.76 |
| 6 | 3343544 | rs10498660 | 0.836 | 0.843 | -1.242 | 0.222 | -0.003 | -0.004 | -0.004 | 0 | 0.4 | 0.25 | 0.29 | 0.35 | 0.4 | 0.34 |
| 6 | 3349908 | rs744375 | 0.373 | 1.129 | -1.155 | 0.072 | 0.009 | -0.012 | -0.012 | 0 | 0.35 | 0.66 | 0.33 | 0.52 | 0.29 | 0.4 |
| 6 | 3354261 | rs1887843 | 0.782 | 1.25 | -1.326 | 0.031 | -0.039 | 0.007 | 0.007 | 1.339 | 0.46 | 0.11 | 0.32 | 0.36 | 0.46 | 0.35 |
| 6 | 3368057 | rs1475474 | 0.775 | 1.035 | -1.382 | -0.116 | 0.015 | -0.133 | -0.133 | 0 | 0.32 | 0.08 | 0.29 | 0.52 | 0.29 | 0.42 |
| 6 | 3369235 | rs9503566 | 1.064 | 1.093 | -0.545 | -0.219 | 0 | 0.001 | 0.001 | 0.511 | 0.4 | 0.34 | 0.27 | 0.33 | 0.37 | 0.32 |
| 6 | 3369270 | rs9378365 | 0.581 | 1.322 | -0.003 | -0.111 | 0.02 | -0.012 | -0.012 | 0.006 | 0.78 | 0.64 | 0.61 | 0.85 | 0.71 | 0.75 |
| 6 | 3371291 | rs981946 | 0.62 | 1.084 | 0.223 | -0.172 | -0.001 | -0.004 | -0.004 | 0 | 0.39 | 0.49 | 0.27 | 0.33 | 0.37 | 0.29 |
| 6 | 3374575 | rs6919934 | -0.103 | 1.032 | 0.037 | -0.261 | 0 | 0.001 | 0.001 | 0.512 | 0.52 | 0.49 | 0.38 | 0.33 | 0.52 | 0.33 |
| 6 | 3375518 | rs3734217 | -0.837 | 0.974 | 0.257 | 0.323 | 0.011 | 0.078 | 0.078 | 0 | 0.1 | 0.31 | 0.09 | 0.16 | 0.09 | 0.06 |
| 6 | 3380220 | rs9503572 | 0.72 | 1.078 | -0.424 | 0.239 | -0.089 | -0.007 | -0.007 | 3.226 | 0.8 | 0.98 | 0.67 | 0.87 | 0.74 | 0.75 |
| 6 | 3384053 | rs9405636 | 0.093 | 1.182 | -0.283 | 0.429 | -0.049 | -0.045 | -0.045 | 0 | 0.91 | 0.99 | 0.71 | 0.86 | 0.86 | 0.85 |
| 6 | 3398515 | rs3813479 | 0.669 | 1.293 | -0.277 | 0.067 | 0.008 | 0.07 | 0.07 | 3.147 | 0.36 | 0.2 | 0.26 | 0.18 | 0.34 | 0.24 |
| 6 | 3398623 | rs3813477 | 0.403 | 1.052 | -0.253 | 0.133 | 0.018 | -0.07 | -0.07 | 0 | 0.39 | 0.14 | 0.3 | 0.52 | 0.36 | 0.48 |
| 6 | 3400610 | rs6923610 | 0.438 | 1.105 | -0.107 | 0.118 | 0.006 | -0.007 | -0.007 | 0.28 | 0.41 | 0.35 | 0.32 | 0.53 | 0.36 | 0.5 |
| 6 | 3400769 | rs12204791 | -0.7 | 1.295 | -0.494 | -0.316 | 0.01 | -0.034 | -0.034 | 1.541 | 0.09 | 0.13 | 0.08 | 0 | 0.13 | 0.03 |
| 6 | 3402520 | rs9392489 | 0.3 | 0.706 | -0.754 | 0.029 | 0.003 | 0.011 | 0.011 | 1.894 | 0.5 | 0.43 | 0.38 | 0.67 | 0.52 | 0.61 |
| 6 | 3403063 | rs6936313 | 0.425 | 1.228 | -1.196 | 0.366 | -0.07 | -0.029 | -0.029 | 0.15 | 0.85 | 0.99 | 0.61 | 0.87 | 0.79 | 0.85 |
| 6 | 3407646 | rs6596966 | -0.734 | 0.377 | -0.546 | 0.076 | -0.052 | -0.036 | -0.036 | 0.484 | 0.16 | 0.47 | 0.11 | 0.06 | 0.1 | 0.18 |
| 6 | 3409112 | rs4959824 | -0.904 | 0.644 | -0.548 | 0.039 | -0.027 | -0.046 | -0.046 | 0 | 0.18 | 0.46 | 0.18 | 0.06 | 0.14 | 0.19 |
| 6 | 3410247 | rs2277076 | 0.451 | 0.741 | -1.171 | 0.329 | -0.02 | 0.015 | 0.015 | 0 | 0.41 | 0.26 | 0.35 | 0.42 | 0.46 | 0.41 |
| 6 | 3411111 | rs10498662 | 0.334 | 0.647 | -1.296 | 0.683 | 0.018 | -0.044 | -0.044 | 0.595 | 0.35 | 0.12 | 0.25 | 0.36 | 0.32 | 0.37 |
| 6 | 3411158 | rs1410661 | 0.507 | 0.775 | -0.555 | 0.888 | 0.029 | 0.029 | 0.029 | 0.35 | 0.8 | 0.46 | 0.63 | 0.79 | 0.8 | 0.81 |
| 6 | 3412240 | rs4959825 | 0.186 | 0.426 | -0.525 | 0.922 | -0.006 | 0.008 | 0.008 | 0.279 | 0.69 | 0.63 | 0.55 | 0.57 | 0.71 | 0.78 |
| 6 | 3415302 | rs4452682 | 0.343 | 0.433 | -0.384 | 0.554 | -0.002 | 0.022 | 0.022 | 0 | 0.49 | 0.33 | 0.34 | 0.3 | 0.41 | 0.6 |
| 6 | 3417643 | rs6923735 | -1.042 | 0.299 | 0.248 | 1.028 | -0.001 | -0.02 | -0.02 | 0.215 | 0.33 | 0.3 | 0.38 | 0.44 | 0.38 | 0.18 |
| 6 | 3420406 | rs13204048 | -0.875 | 1.049 | 0.754 | 1.373 | 0.002 | 0.01 | 0.01 | 0.385 | 0.21 | 0.26 | 0.33 | 0.45 | 0.39 | 0.16 |
| 6 | 3433318 | rs17309827 | -1.187 | 0.96 | 0.562 | 1.292 | -0.059 | -0.098 | -0.098 | 0.184 | 0.24 | 0.09 | 0.32 | 0.46 | 0.38 | 0.17 |
| 6 | 3435112 | rs4959832 | -1.554 | 0.906 | 1.043 | 1.227 | 0.024 | 0.052 | 0.052 | 0 | 0.28 | 0.46 | 0.38 | 0.46 | 0.4 | 0.18 |
| 6 | 3436881 | rs4959835 | -0.698 | 1.093 | 0.921 | 1.352 | -0.002 | -0.009 | -0.009 | 2.63 | 0.18 | 0.17 | 0.24 | 0.33 | 0.21 | 0.1 |
| 6 | 3438738 | rs12197689 | 1.07 | 1.184 | 1.349 | 1.307 | 0.008 | 0.059 | 0.059 | 0.008 | 0.75 | 0.38 | 0.53 | 0.65 | 0.74 | 0.85 |

| | | | | | | | | | | | | | | | | |
|---|---|---|---|---|---|---|---|---|---|---|---|---|---|---|---|---|
| 6 | 3440314 | rs9328172 | 1.567 | 1.172 | 1.157 | 1.066 | 0.022 | 0.021 | 0.021 | 0 | 0.68 | 0.5 | 0.44 | 0.58 | 0.56 | 0.78 |
| 6 | 3442900 | rs6921583 | 1.524 | 1.207 | 1.141 | 1.07 | 0.021 | 0.02 | 0.02 | 0 | 0.68 | 0.52 | 0.44 | 0.57 | 0.55 | 0.77 |
| 6 | 3442988 | rs17250853 | -0.839 | 1.251 | 0.536 | 1.094 | -0.001 | -0.001 | -0.001 | 0 | 0.09 | 0.09 | 0.13 | 0.11 | 0.19 | 0.08 |
| 6 | 3444970 | rs729131 | 0.415 | 1.117 | 0.226 | 1.053 | -0.055 | -0.098 | -0.098 | 0 | 0.83 | 0.98 | 0.68 | 0.71 | 0.79 | 0.9 |
| 6 | 3446263 | rs9503598 | 1.601 | 0.951 | 0.43 | 0.703 | -0.036 | -0.011 | -0.011 | 0.036 | 0.55 | 0.24 | 0.36 | 0.54 | 0.42 | 0.68 |
| 6 | 3447277 | rs9502008 | 0.896 | 1.11 | 1.391 | 0.671 | 0.003 | 0.003 | 0.003 | 0 | 0.72 | 0.69 | 0.61 | 0.7 | 0.68 | 0.8 |
| 6 | 3452974 | rs9378798 | -0.117 | 1.321 | 1.255 | 0.689 | -0.05 | -0.08 | -0.08 | 0 | 0.84 | 0.97 | 0.68 | 0.7 | 0.78 | 0.9 |
| 6 | 3454032 | rs11242859 | 1.695 | 1.248 | 2.174 | 0.683 | -0.004 | -0.003 | -0.003 | 0.204 | 0.58 | 0.38 | 0.38 | 0.52 | 0.45 | 0.68 |
| 6 | 3462885 | rs4959838 | -0.392 | 1.184 | 1.855 | 0.646 | 0.005 | -0.045 | -0.045 | 0 | 0.09 | 0.17 | 0.06 | 0.01 | 0.08 | 0.09 |
| 6 | 3463536 | rs11242860 | 1.008 | 1.197 | 1.9 | 0.705 | -0.028 | -0.041 | -0.041 | 0 | 0.72 | 0.82 | 0.47 | 0.52 | 0.55 | 0.79 |
| 6 | 3468224 | rs7775474 | 1.393 | 1.202 | 1.996 | 0.71 | -0.018 | -0.026 | -0.026 | 0 | 0.67 | 0.76 | 0.44 | 0.52 | 0.52 | 0.77 |
| 6 | 3470203 | rs7772129 | -1.559 | 1.018 | 2.262 | 0.88 | 0.026 | -0.018 | -0.018 | 10.58 | 0.14 | 0.34 | 0.16 | 0.09 | 0.13 | 0.12 |
| 6 | 3470242 | rs9328178 | 1.203 | 0.992 | 2.199 | 0.934 | 0.011 | 0.024 | 0.024 | 0 | 0.68 | 0.57 | 0.52 | 0.59 | 0.65 | 0.77 |
| 6 | 3471259 | rs9328180 | 1.411 | 1.117 | 1.9 | 1.006 | -0.001 | -0.009 | -0.009 | 0 | 0.62 | 0.35 | 0.45 | 0.59 | 0.63 | 0.71 |
| 6 | 3472175 | rs7356995 | 1.064 | 1.166 | 1.746 | 1.004 | 0.011 | 0.016 | 0.016 | 0.354 | 0.67 | 0.5 | 0.58 | 0.64 | 0.67 | 0.77 |
| 6 | 3474807 | rs12211852 | 1.318 | 1.104 | 1.32 | 1.043 | -0.015 | 0.006 | 0.006 | 0.643 | 0.63 | 0.23 | 0.51 | 0.63 | 0.61 | 0.7 |
| 6 | 3476276 | rs4959241 | -1.347 | 1.352 | 1.068 | 0.953 | 0 | -0.003 | -0.003 | 0.148 | 0.51 | 0.63 | 0.54 | 0.49 | 0.47 | 0.38 |
| 6 | 3481313 | rs10900943 | -0.808 | 1.648 | 1.297 | 0.69 | 0.004 | -0.01 | -0.01 | 0 | 0.27 | 0.28 | 0.28 | 0.01 | 0.23 | 0.19 |
| 6 | 3481662 | rs913534 | -0.652 | 1.692 | 1.811 | 0.696 | 0.019 | 0.107 | 0.107 | 0 | 0.23 | 0.39 | 0.25 | 0.49 | 0.24 | 0.19 |
| 6 | 3485134 | rs1205032 | -1.893 | 1.656 | 1.804 | 0.689 | -0.013 | -0.015 | -0.015 | 0 | 0.52 | 0.76 | 0.63 | 0.5 | 0.47 | 0.38 |
| 6 | 3487423 | rs748842 | -1.914 | 1.654 | 1.745 | 0.688 | -0.02 | -0.022 | -0.022 | 0 | 0.52 | 0.79 | 0.63 | 0.51 | 0.47 | 0.38 |
| 6 | 3488625 | rs7763703 | -0.932 | 1.689 | 1.721 | 0.923 | 0.018 | 0.129 | 0.129 | 0 | 0.24 | 0.46 | 0.37 | 0.5 | 0.24 | 0.19 |
| 6 | 3496602 | rs7760989 | -1.578 | 1.38 | 0.694 | 0.766 | 0.005 | 0.007 | 0.007 | 0 | 0.48 | 0.62 | 0.6 | 0.47 | 0.45 | 0.36 |
| 6 | 3499963 | rs4959247 | -0.83 | 1.678 | 0.358 | 0.217 | -0.007 | 0.061 | 0.061 | 2.552 | 0.25 | 0.07 | 0.27 | 0.01 | 0.26 | 0.17 |
| 6 | 3503685 | rs4373399 | -1.143 | 1.553 | 0.516 | 1.048 | 0.005 | 0.022 | 0.022 | 0 | 0.3 | 0.38 | 0.31 | 0.37 | 0.26 | 0.33 |
| 6 | 3513246 | rs880287 | -0.669 | 1.307 | 0.502 | 1.115 | 0 | -0.024 | -0.024 | 0 | 0.13 | 0.07 | 0.11 | 0.31 | 0.11 | 0.15 |
| 6 | 3516504 | rs4959847 | -1.556 | 0.918 | 0.448 | 0.8 | 0.022 | -0.016 | -0.016 | 0 | 0.39 | 0.77 | 0.3 | 0.6 | 0.29 | 0.43 |
| 6 | 3527310 | rs4959849 | -0.993 | 0.882 | 1.406 | -0.016 | -0.017 | 0.005 | 0.005 | 0.716 | 0.22 | 0.65 | 0.18 | 0.29 | 0.15 | 0.15 |
| 6 | 3531912 | rs9502014 | -0.614 | 1.329 | 1.446 | -0.189 | 0.094 | 0.06 | 0.06 | 0 | 0.05 | 0.28 | 0.08 | 0.13 | 0.11 | 0.06 |
| 6 | 3533221 | rs9502015 | -1.216 | 1.263 | 1.248 | -0.223 | 0.089 | 0.05 | 0.05 | 3.468 | 0.09 | 0.54 | 0.1 | 0.13 | 0.11 | 0.06 |
| 6 | 3535405 | rs1010210 | -1.261 | 1.445 | 0.75 | -0.314 | -0.001 | -0.006 | -0.006 | 0 | 0.27 | 0.69 | 0.26 | 0.27 | 0.27 | 0.18 |
| 6 | 3543701 | rs9503618 | -1.262 | 1.715 | -0.026 | -0.365 | -0.08 | 0.035 | 0.035 | 1.375 | 0.38 | 0.89 | 0.46 | 0.31 | 0.44 | 0.51 |
| 6 | 3547303 | rs7740160 | -1.016 | 1.849 | 0.302 | -0.526 | -0.032 | 0.074 | 0.074 | 0.646 | 0.32 | 0.8 | 0.31 | 0.04 | 0.36 | 0.39 |
| 6 | 3548999 | rs6596983 | -1.098 | 2.074 | 0.27 | -0.642 | -0.049 | 0.099 | 0.099 | 3.088 | 0.35 | 0.8 | 0.34 | 0.04 | 0.41 | 0.4 |
| 6 | 3549292 | rs13200960 | -0.069 | 2.183 | 0.507 | -0.553 | -0.004 | 0.004 | 0.004 | 0.184 | 0.19 | 0.17 | 0.18 | 0.05 | 0.2 | 0.19 |
| 6 | 3567555 | rs6905808 | 0.424 | 1.84 | -0.211 | -0.467 | 0 | -0.021 | -0.021 | 0 | 0.64 | 0.41 | 0.65 | 0.93 | 0.55 | 0.59 |
| 6 | 3569274 | rs9405647 | 0.752 | 1.602 | -1.336 | -0.361 | -0.019 | 0.086 | 0.086 | 0.496 | 0.53 | 0.17 | 0.54 | 0.66 | 0.45 | 0.44 |
| 6 | 3572371 | rs875085 | -0.998 | 1.451 | -0.788 | 0.541 | -0.034 | -0.052 | -0.052 | 0 | 0.2 | 0.07 | 0.2 | 0.33 | 0.23 | 0.39 |
| 6 | 3573222 | rs878803 | -1.456 | 1.497 | 0.201 | 0.216 | 0.072 | -0.018 | -0.018 | 0 | 0.1 | 0.26 | 0.11 | 0.04 | 0.13 | 0.1 |
| 6 | 3573808 | rs1883612 | 0.391 | 1.104 | -0.079 | 0.295 | 0.001 | 0.001 | 0.001 | 0 | 0.58 | 0.36 | 0.67 | 0.65 | 0.58 | 0.47 |
| 6 | 3574454 | rs9638039 | -0.4 | 0.506 | 0.402 | 0.58 | -0.001 | 0.001 | 0.001 | 0 | 0.35 | 0.33 | 0.39 | 0.45 | 0.32 | 0.47 |
| 6 | 3579754 | rs1205017 | -0.983 | 0.064 | 0.141 | 0.574 | 0.004 | 0.01 | 0.01 | 0.005 | 0.62 | 0.72 | 0.67 | 0.62 | 0.64 | 0.62 |
| 6 | 3580235 | rs1205019 | -0.932 | -0.06 | 0.555 | 0.511 | -0.002 | -0.004 | -0.004 | 0 | 0.59 | 0.57 | 0.66 | 0.62 | 0.64 | 0.48 |
| 6 | 3580339 | rs1205020 | -0.944 | -0.066 | 0.552 | 0.508 | -0.002 | -0.004 | -0.004 | 0 | 0.6 | 0.57 | 0.66 | 0.62 | 0.64 | 0.49 |
| 6 | 3582177 | rs1205023 | 0.048 | -0.044 | -0.244 | 0.515 | -0.185 | -0.044 | -0.044 | 0.708 | 0.34 | 0.04 | 0.34 | 0.28 | 0.42 | 0.21 |
| 6 | 3584259 | rs6902488 | -0.961 | 0.221 | 0.994 | 0.852 | -0.011 | 0.119 | 0.119 | 0 | 0.06 | 0.24 | 0.18 | 0.16 | 0.03 | 0.12 |
| 6 | 3589672 | rs6917333 | -1.41 | 0.301 | 1.627 | 0.702 | 0.05 | 0.129 | 0.129 | 0.518 | 0.08 | 0.5 | 0.21 | 0.15 | 0.04 | 0.09 |
| 6 | 3591250 | rs9503635 | -0.72 | 0.261 | 1.339 | 0.535 | 0 | -0.01 | -0.01 | 0 | 0.08 | 0.08 | 0.05 | 0.01 | 0.06 | 0.12 |
| 6 | 3603720 | rs13215178 | -1.539 | 0.091 | 1.453 | 0.441 | -0.01 | -0.006 | -0.006 | 0 | 0.18 | 0.24 | 0.28 | 0.17 | 0.18 | 0.22 |
| 6 | 3603809 | rs1205026 | 0.343 | 0.586 | 0.605 | 0.427 | -0.139 | -0.096 | -0.096 | 0 | 0.37 | 0.03 | 0.24 | 0.38 | 0.37 | 0.23 |
| 6 | 3605434 | rs1205028 | -1.499 | 0.694 | 0.747 | 0.442 | -0.015 | -0.05 | -0.05 | 0 | 0.64 | 0.8 | 0.59 | 0.56 | 0.6 | 0.58 |
| 6 | 3607159 | rs9502026 | -2.046 | 0.545 | 1.228 | 0.517 | -0.01 | -0.017 | -0.017 | 0 | 0.23 | 0.62 | 0.23 | 0.18 | 0.22 | 0.22 |
| 6 | 3609712 | rs7766993 | -1.655 | 0.688 | 0.534 | 0.624 | -0.002 | -0.003 | -0.003 | 0 | 0.19 | 0.23 | 0.19 | 0.18 | 0.2 | 0.22 |
| 6 | 3619044 | rs13219314 | -0.569 | 0.768 | 0.199 | 0.636 | -0.003 | 0.004 | 0.004 | 0 | 0.09 | 0.05 | 0.1 | 0.06 | 0.13 | 0.08 |
| 6 | 3626372 | rs7738922 | -0.354 | 0.594 | 0.121 | 0.663 | -0.008 | -0.007 | -0.007 | 0 | 0.06 | 0.03 | 0.08 | 0.06 | 0.1 | 0.06 |
| 6 | 3626708 | rs4959855 | 1.134 | 0.651 | 0.217 | 0.642 | 0 | 0.003 | 0.003 | 1.07 | 0.36 | 0.36 | 0.53 | 0.45 | 0.39 | 0.42 |
| 6 | 3627623 | rs7743738 | -1.452 | 0.498 | 0.098 | 0.472 | -0.004 | -0.009 | -0.009 | 0 | 0.21 | 0.26 | 0.16 | 0.18 | 0.2 | 0.35 |
| 6 | 3629689 | rs6596989 | -0.617 | 0.605 | -0.629 | 0.539 | -0.105 | -0.026 | -0.026 | 0.014 | 0.69 | 0.99 | 0.8 | 0.78 | 0.71 | 0.83 |
| 6 | 3636411 | rs9392512 | -0.133 | 0.642 | -0.431 | 0.547 | -0.051 | -0.032 | -0.032 | 0 | 0.63 | 0.88 | 0.76 | 0.61 | 0.6 | 0.79 |
| 6 | 3639205 | rs9378806 | 1.623 | 0.598 | -0.13 | 1.07 | -0.008 | 0.01 | 0.01 | 1.009 | 0.35 | 0.26 | 0.54 | 0.32 | 0.37 | 0.41 |
| 6 | 3639598 | rs6904497 | -0.563 | 0 | 0.468 | 1.46 | 0.041 | 0.238 | 0.238 | 0.028 | 0.1 | 0.36 | 0.13 | 0.28 | 0.12 | 0.08 |
| 6 | 3642866 | rs9503656 | -1.777 | -0.142 | -0.603 | 1.343 | 0.033 | -0.023 | -0.023 | 0 | 0.25 | 0.87 | 0.23 | 0.23 | 0.19 | 0.3 |
| 6 | 3643787 | rs6940109 | -1.829 | -0.113 | -0.672 | 1.276 | 0.055 | -0.038 | -0.038 | 0.73 | 0.29 | 0.88 | 0.26 | 0.31 | 0.22 | 0.38 |
| 6 | 3644751 | rs9392004 | -0.058 | 0.22 | -0.915 | 1.325 | -0.048 | -0.12 | -0.12 | 0.05 | 0.74 | 0.98 | 0.85 | 0.62 | 0.77 | 0.88 |
| 6 | 3651622 | rs13202552 | -1.26 | 0.327 | -0.156 | 1.265 | -0.022 | -0.074 | -0.074 | 0 | 0.23 | 0.34 | 0.19 | 0.06 | 0.23 | 0.31 |
| 6 | 3653224 | rs9503662 | -0.659 | 0.297 | -0.061 | 1.457 | 0.098 | 0.482 | 0.482 | 3.201 | 0.05 | 0.49 | 0.07 | 0.25 | 0.01 | 0.05 |
| 6 | 3655479 | rs880645 | 0.311 | 0.306 | -0.399 | 1.192 | -0.021 | -0.013 | -0.013 | 0 | 0.07 | 0.01 | 0.08 | 0.05 | 0.11 | 0.06 |
| 6 | 3657174 | rs13200516 | 0.311 | 0.306 | -0.394 | 1.192 | -0.017 | -0.01 | -0.01 | 1.077 | 0.07 | 0 | 0.08 | 0.05 | 0.11 | 0.06 |
| 6 | 3660425 | rs10458075 | -0.652 | 0.281 | -0.267 | 1.196 | -0.029 | -0.082 | -0.082 | 0 | 0.16 | 0.31 | 0.11 | 0.02 | 0.11 | 0.25 |

| | | | | | | | | | | | | | | | | |
|---|---|---|---|---|---|---|---|---|---|---|---|---|---|---|---|---|
| 6 | 3661014 | rs226913 | 0.502 | 0.308 | -0.966 | 1.329 | -0.017 | -0.069 | -0.069 | 0 | 0.19 | 0.01 | 0.12 | 0.21 | 0.15 | 0.08 |
| 6 | 3665573 | rs9502039 | -1.883 | 0.334 | -0.587 | 1.541 | 0.038 | -0.033 | -0.033 | 0 | 0.28 | 0.85 | 0.26 | 0.31 | 0.24 | 0.35 |
| 6 | 3680949 | rs12212080 | -0.635 | 0.632 | -0.404 | 1.436 | 0.003 | -0.005 | -0.005 | 0 | 0.09 | 0.1 | 0.1 | 0.05 | 0.15 | 0.21 |
| 6 | 3684936 | rs6901819 | 0.098 | 0.516 | -0.555 | 1.561 | -0.029 | -0.049 | -0.049 | 0 | 0.09 | 0.01 | 0.04 | 0.18 | 0.1 | 0.03 |
| 6 | 3695754 | rs226933 | 0.015 | 0.854 | -0.645 | 1.425 | -0.064 | -0.048 | -0.048 | 0.759 | 0.2 | 0.02 | 0.19 | 0.2 | 0.22 | 0.12 |
| 6 | 3697149 | rs9328186 | -0.385 | 0.819 | -0.089 | 1.504 | -0.025 | 0.092 | 0.092 | 0.013 | 0.24 | 0.43 | 0.16 | 0.43 | 0.18 | 0.21 |
| 6 | 3698134 | rs6909042 | -0.858 | 0.907 | 0.125 | 1.635 | 0.155 | 0.424 | 0.424 | 0.608 | 0.06 | 0.46 | 0.08 | 0.26 | 0.07 | 0.2 |
| 6 | 3699703 | rs226930 | -1.533 | 0.798 | -0.205 | 1.545 | -0.01 | 0.08 | 0.08 | 1.315 | 0.49 | 0.91 | 0.4 | 0.65 | 0.43 | 0.47 |
| 6 | 3701232 | rs226927 | -1.247 | 0.887 | 0.063 | 1.664 | 0.007 | 0.025 | 0.025 | 0.523 | 0.47 | 0.63 | 0.37 | 0.65 | 0.43 | 0.34 |
| 6 | 3701286 | rs226926 | -0.128 | 0.863 | -0.288 | 1.422 | -0.002 | 0.001 | 0.001 | 0.001 | 0.22 | 0.2 | 0.21 | 0.22 | 0.23 | 0.12 |
| 6 | 3701644 | rs226924 | -0.374 | 0.78 | -0.484 | 1.606 | -0.007 | 0.007 | 0.007 | 0 | 0.45 | 0.36 | 0.36 | 0.38 | 0.47 | 0.46 |
| 6 | 3708478 | rs226940 | -0.51 | 0.926 | -1.376 | 0.911 | -0.054 | 0.074 | 0.074 | 3.887 | 0.56 | 0.91 | 0.53 | 0.68 | 0.58 | 0.51 |
| 6 | 3712111 | rs9392005 | 0.434 | 0.964 | -1.436 | 0.939 | -0.065 | 0.139 | 0.139 | 0.015 | 0.5 | 0.11 | 0.28 | 0.28 | 0.56 | 0.45 |
| 6 | 3712882 | rs13219323 | 0.327 | 0.954 | -0.736 | 1.084 | 0.005 | 0.014 | 0.014 | 0 | 0.54 | 0.26 | 0.32 | 0.39 | 0.62 | 0.54 |
| 6 | 3713527 | rs1535005 | 0.679 | 0.957 | -0.739 | 1.085 | -0.07 | 0.138 | 0.138 | 0 | 0.5 | 0.11 | 0.28 | 0.28 | 0.56 | 0.45 |
| 6 | 3714512 | rs9378811 | 0.319 | 0.906 | -0.531 | 1.058 | 0.007 | 0.014 | 0.014 | 0 | 0.54 | 0.29 | 0.32 | 0.38 | 0.62 | 0.54 |
| 6 | 3717124 | rs226947 | -0.757 | 0.527 | 0.452 | 1.53 | -0.039 | 0.108 | 0.108 | 2.75 | 0.21 | 0.42 | 0.28 | 0.33 | 0.11 | 0.29 |
| 6 | 3723332 | rs742541 | 0.278 | 0.737 | 0.78 | 1.709 | 0.015 | 0.008 | 0.008 | 0 | 0.13 | 0.22 | 0.13 | 0.14 | 0.14 | 0.07 |
| 6 | 3730290 | rs226956 | -0.116 | 0.537 | 1.255 | 1.693 | -0.039 | 0.041 | 0.041 | 0.218 | 0.16 | 0.29 | 0.25 | 0.24 | 0.05 | 0.19 |
| 6 | 3731723 | rs192021 | -0.18 | 0.543 | 1.421 | 1.701 | -0.04 | 0.038 | 0.038 | 0.478 | 0.16 | 0.28 | 0.25 | 0.24 | 0.05 | 0.19 |
| 6 | 3733519 | rs9503724 | 1.676 | 0.206 | 0.751 | 1.315 | -0.102 | 0.123 | 0.123 | 0.092 | 0.44 | 0.1 | 0.28 | 0.27 | 0.54 | 0.42 |
| 6 | 3737700 | rs11242868 | -0.924 | 0.195 | 0.912 | 1.217 | -0.003 | 0 | 0 | 0 | 0.36 | 0.29 | 0.43 | 0.39 | 0.35 | 0.32 |
| 6 | 3739193 | rs4959871 | -0.859 | 0.576 | 0.771 | 1.108 | -0.009 | 0.007 | 0.007 | 0 | 0.4 | 0.35 | 0.46 | 0.4 | 0.41 | 0.33 |
| 6 | 3739835 | rs12193236 | -0.18 | 0.467 | 0.674 | 0.815 | -0.006 | 0.006 | 0.006 | 0 | 0.24 | 0.16 | 0.3 | 0.19 | 0.24 | 0.14 |
| 6 | 3740307 | rs11242871 | -0.816 | 0.455 | 0.598 | 0.825 | 0.005 | 0.009 | 0.009 | 0 | 0.41 | 0.48 | 0.47 | 0.53 | 0.41 | 0.39 |
| 6 | 3742857 | rs9503725 | 1.459 | 0.622 | 0.219 | 0.861 | -0.076 | 0.107 | 0.107 | 0.016 | 0.36 | 0.12 | 0.24 | 0.19 | 0.5 | 0.41 |
| 6 | 3749106 | rs6930339 | 0.472 | 0.705 | 0.317 | 1.096 | 0.003 | -0.006 | -0.006 | 0.003 | 0.54 | 0.61 | 0.42 | 0.54 | 0.65 | 0.57 |
| 6 | 3756223 | rs6596992 | 0.814 | 0.503 | 0.388 | 1.031 | -0.002 | 0.037 | 0.037 | 0.337 | 0.79 | 0.68 | 0.62 | 0.38 | 0.8 | 0.75 |
| 6 | 3762272 | rs9503728 | 0.093 | 0.296 | 0.455 | 0.606 | -0.029 | -0.033 | -0.033 | 0 | 0.31 | 0.58 | 0.18 | 0.19 | 0.18 | 0.25 |
| 6 | 3764125 | rs6596996 | 1.249 | 0.635 | 0.029 | 0.133 | -0.041 | 0.096 | 0.096 | 0.099 | 0.38 | 0.17 | 0.28 | 0.17 | 0.5 | 0.39 |
| 6 | 3780061 | rs2002521 | -0.011 | 0.397 | 0.298 | 0.383 | -0.004 | -0.017 | -0.017 | 0 | 0.07 | 0.09 | 0.03 | 0 | 0.04 | 0.04 |
| 6 | 3780451 | rs6596998 | -0.283 | 0.578 | 0.627 | 1.044 | -0.019 | 0.005 | 0.005 | 0 | 0.42 | 0.51 | 0.27 | 0.46 | 0.29 | 0.32 |
| 6 | 3781719 | rs6596999 | -0.266 | 0.564 | 0.095 | 0.996 | -0.008 | -0.001 | -0.001 | 0.057 | 0.41 | 0.52 | 0.26 | 0.41 | 0.28 | 0.31 |
| 6 | 3783763 | rs12208688 | -0.627 | 0.669 | -0.049 | 1.048 | 0.016 | 0.004 | 0.004 | 0 | 0.48 | 0.58 | 0.48 | 0.54 | 0.4 | 0.45 |
| 6 | 3789181 | rs9502072 | -0.499 | 1.02 | -0.185 | 1.207 | 0.003 | 0.005 | 0.005 | 0.347 | 0.17 | 0.19 | 0.26 | 0.22 | 0.22 | 0.2 |
| 6 | 3797198 | rs12197988 | 0.233 | 0.818 | -0.311 | 0.946 | 0.024 | 0.027 | 0.027 | 0 | 0.42 | 0.27 | 0.26 | 0.35 | 0.36 | 0.35 |
| 6 | 3797857 | rs6925590 | -0.326 | 0.848 | 0.259 | 0.935 | 0.004 | 0 | 0 | 0.02 | 0.49 | 0.46 | 0.37 | 0.5 | 0.45 | 0.42 |
| 6 | 3801605 | rs11753146 | 0.762 | 0.372 | -0.405 | 0.661 | -0.046 | 0.102 | 0.102 | 0 | 0.28 | 0.02 | 0.15 | 0.1 | 0.25 | 0.28 |
| 6 | 3804556 | rs4275102 | 0.598 | 0.456 | 0.035 | 0.994 | 0.001 | 0.005 | 0.005 | 0 | 0.37 | 0.38 | 0.39 | 0.43 | 0.39 | 0.32 |
| 6 | 3808632 | rs9392011 | 0.999 | 0.451 | -0.054 | 0.99 | -0.035 | -0.04 | -0.04 | 0 | 0.34 | 0.15 | 0.37 | 0.42 | 0.42 | 0.32 |
| 6 | 3808855 | rs9378384 | -0.422 | 0.446 | 0.343 | 0.897 | 0.014 | 0.015 | 0.015 | 0.365 | 0.78 | 0.64 | 0.63 | 0.71 | 0.77 | 0.69 |
| 6 | 3809510 | rs12197082 | 1.21 | 0.296 | 0.477 | 0.796 | 0.015 | 0.09 | 0.09 | 0 | 0.32 | 0.15 | 0.17 | 0.1 | 0.24 | 0.3 |
| 6 | 3810805 | rs6940448 | 0.356 | 0.328 | 0.629 | 0.844 | 0.005 | 0.009 | 0.009 | 0.728 | 0.4 | 0.34 | 0.24 | 0.3 | 0.34 | 0.36 |
| 6 | 3811329 | rs2326359 | -0.149 | 0.429 | 0.607 | 1.05 | 0.003 | 0.04 | 0.04 | 0 | 0.52 | 0.36 | 0.4 | 0.34 | 0.48 | 0.61 |
| 6 | 3812036 | rs12214729 | 1.237 | 0.528 | 0.246 | 1.055 | -0.044 | 0.02 | 0.02 | 1.961 | 0.11 | 0.01 | 0.17 | 0.04 | 0.14 | 0.17 |
| 6 | 3816394 | rs9378385 | 0.241 | 0.397 | 0.675 | 1.255 | 0.001 | 0.002 | 0.002 | 0 | 0.37 | 0.36 | 0.39 | 0.41 | 0.43 | 0.33 |
| 6 | 3817299 | rs6916614 | 0.276 | 0.238 | 0.425 | 1.139 | -0.009 | 0.046 | 0.046 | 0 | 0.09 | 0.16 | 0.07 | 0.21 | 0.09 | 0.06 |
| 6 | 3820229 | rs2002837 | -0.041 | 0.072 | -0.174 | 0.72 | 0.035 | 0.094 | 0.094 | 0 | 0.51 | 0.79 | 0.39 | 0.28 | 0.39 | 0.53 |
| 6 | 3824776 | rs9503750 | -0.811 | 0.124 | -0.095 | 1.295 | -0.043 | -0.004 | -0.004 | 0 | 0.72 | 0.89 | 0.55 | 0.67 | 0.65 | 0.74 |
| 6 | 3826298 | rs4370404 | -0.287 | 0.016 | -0.345 | 1.203 | -0.005 | -0.065 | -0.065 | 0 | 0.18 | 0.06 | 0.14 | 0.38 | 0.24 | 0.2 |
| 6 | 3830849 | rs2326364 | 0.573 | -0.183 | 0.08 | 1.168 | -0.056 | 0.011 | 0.011 | 0 | 0.27 | 0.08 | 0.31 | 0.25 | 0.35 | 0.25 |
| 6 | 3833922 | rs6597005 | -0.407 | -0.291 | 0.554 | 1.173 | -0.001 | -0.001 | -0.001 | 0 | 0.39 | 0.59 | 0.39 | 0.43 | 0.4 | 0.36 |
| 6 | 3836353 | rs9328196 | -0.193 | -0.214 | 0.097 | 1.183 | -0.001 | -0.001 | -0.001 | 0 | 0.37 | 0.41 | 0.37 | 0.37 | 0.39 | 0.34 |
| 6 | 3837198 | rs17318866 | 0.317 | 0.147 | -0.505 | 0.899 | -0.047 | 0.016 | 0.016 | 0 | 0.28 | 0.11 | 0.31 | 0.22 | 0.35 | 0.26 |
| 6 | 3839422 | rs9405672 | -1.451 | 0.239 | 0.119 | 1.031 | 0.023 | 0.023 | 0.023 | 0.011 | 0.24 | 0.37 | 0.18 | 0.3 | 0.28 | 0.26 |
| 6 | 3839513 | rs9392527 | -1.247 | -0.395 | -0.092 | 1.217 | -0.001 | -0.001 | -0.001 | 0 | 0.23 | 0.21 | 0.13 | 0.23 | 0.21 | 0.29 |
| 6 | 3839544 | rs6927199 | 0.803 | -0.257 | -0.078 | 1.293 | 0.002 | 0 | 0 | 0 | 0.84 | 0.78 | 0.75 | 0.8 | 0.81 | 0.87 |
| 6 | 3839792 | rs9392528 | -0.435 | -0.405 | -0.698 | 1.599 | -0.06 | -0.054 | -0.054 | 0 | 0.85 | 0.97 | 0.71 | 0.77 | 0.78 | 0.77 |
| 6 | 3842494 | rs9503755 | 0.87 | -0.709 | -0.396 | 1.525 | 0.001 | -0.004 | -0.004 | 1.227 | 0.42 | 0.46 | 0.38 | 0.38 | 0.49 | 0.34 |
| 6 | 3844275 | rs10498664 | 0.885 | -0.714 | -0.44 | 1.513 | 0.002 | -0.006 | -0.006 | 0 | 0.42 | 0.45 | 0.38 | 0.38 | 0.49 | 0.34 |
| 6 | 3845399 | rs9392016 | -0.365 | -0.811 | -0.166 | 1.62 | 0.016 | 0.166 | 0.166 | 0.007 | 0.08 | 0.41 | 0.07 | 0.14 | 0.02 | 0.16 |
| 6 | 3848633 | rs7740880 | -0.365 | -0.811 | -0.184 | 1.62 | 0.018 | 0.171 | 0.171 | 0.007 | 0.08 | 0.41 | 0.07 | 0.14 | 0.01 | 0.16 |
| 6 | 3849305 | rs2239713 | -0.333 | -0.737 | -1.057 | 1.444 | -0.032 | -0.013 | -0.013 | 0.578 | 0.73 | 0.89 | 0.61 | 0.72 | 0.69 | 0.69 |
| 6 | 3852692 | rs4959877 | 0.003 | -1.782 | -0.725 | 1.241 | 0.004 | 0.02 | 0.02 | 0.106 | 0.11 | 0.21 | 0.07 | 0.16 | 0.12 | 0.09 |
| 6 | 3852794 | rs2490036 | 0.386 | -1.745 | -1.771 | 1.072 | -0.036 | -0.078 | -0.078 | 0 | 0.19 | 0.01 | 0.15 | 0.21 | 0.2 | 0.15 |
| 6 | 3853906 | rs6901207 | 0.806 | -1.251 | -1.243 | 0.412 | 0.004 | -0.009 | -0.009 | 0 | 0.33 | 0.4 | 0.35 | 0.16 | 0.44 | 0.26 |
| 6 | 3854438 | rs6910945 | 0.896 | -1.211 | -1.822 | 0.432 | -0.031 | 0.033 | 0.033 | 0.028 | 0.32 | 0.28 | 0.34 | 0.16 | 0.44 | 0.26 |
| 6 | 3857624 | rs2496388 | 1.102 | -1.703 | -2.118 | 0.779 | 0.103 | -0.064 | -0.064 | 0.558 | 0.66 | 0.13 | 0.68 | 0.59 | 0.84 | 0.7 |
| 6 | 3859096 | rs2252854 | 0.837 | -0.923 | -1.49 | 0.896 | 0.006 | -0.013 | -0.013 | 0.547 | 0.77 | 0.86 | 0.72 | 0.61 | 0.83 | 0.85 |

| | | | | | | | | | | | | | | | | |
|---|---|---|---|---|---|---|---|---|---|---|---|---|---|---|---|---|
| 6 | 3860010 | rs2490045 | 0.857 | -0.926 | -1.49 | 0.903 | 0.006 | -0.012 | -0.012 | 0.831 | 0.77 | 0.86 | 0.72 | 0.61 | 0.83 | 0.84 |
| 6 | 3861834 | rs2490049 | 1.296 | -0.783 | -1.684 | 0.742 | 0.05 | 0.022 | 0.022 | 0 | 0.51 | 0.2 | 0.49 | 0.42 | 0.65 | 0.54 |
| 6 | 3862593 | rs2490050 | -0.229 | -0.77 | -1.784 | 0.748 | -0.01 | -0.029 | -0.029 | 0 | 0.8 | 0.94 | 0.68 | 0.73 | 0.83 | 0.69 |
| 6 | 3864693 | rs9918376 | -0.26 | -0.444 | -1.791 | 0.41 | -0.02 | 0.008 | 0.008 | 0.051 | 0.11 | 0.1 | 0.04 | 0.09 | 0.05 | 0.05 |
| 6 | 3865850 | rs2490051 | 0.179 | 0.241 | -2.159 | 0.061 | 0.041 | 0.011 | 0.011 | 0 | 0.51 | 0.14 | 0.49 | 0.41 | 0.65 | 0.54 |
| 6 | 3885152 | rs4145940 | -0.092 | 0.826 | -1.251 | -1.216 | 0.003 | 0.019 | 0.019 | 0 | 0.46 | 0.57 | 0.4 | 0.17 | 0.45 | 0.4 |
| 6 | 3885376 | rs11242890 | -1.27 | 1.879 | -1.595 | -2.039 | -0.064 | 0.102 | 0.102 | 0.014 | 0.19 | 0.03 | 0.25 | 0.01 | 0.26 | 0.24 |
| 6 | 3887022 | rs6914431 | -1.547 | 2.321 | -1.446 | -2.164 | -0.076 | 0.086 | 0.086 | 0 | 0.2 | 0.07 | 0.24 | 0.03 | 0.28 | 0.22 |
| 6 | 3895914 | rs4959883 | -1.857 | 1.449 | -0.217 | -2.289 | 0.015 | -0.153 | -0.153 | 0 | 0.23 | 0.52 | 0.24 | 0.03 | 0.22 | 0.24 |
| 6 | 3896005 | rs4959884 | -1.444 | 1.612 | 0.034 | -2.206 | 0.01 | -0.062 | -0.062 | 0 | 0.18 | 0.3 | 0.23 | 0.03 | 0.23 | 0.22 |
| 6 | 3907174 | rs13213674 | -1.227 | 1.445 | 0.435 | -2.603 | 0.019 | -0.126 | -0.126 | 0 | 0.23 | 0.6 | 0.18 | 0.01 | 0.3 | 0.3 |
| 6 | 3908474 | rs4296932 | -1.683 | 1.085 | -0.148 | -3.125 | -0.047 | 0.276 | 0.276 | 0.153 | 0.45 | 0.91 | 0.48 | 0.11 | 0.48 | 0.34 |
| 6 | 3912275 | rs13196874 | -1.394 | 0.871 | 0.881 | -3.227 | -0.001 | -0.069 | -0.069 | 0 | 0.18 | 0.44 | 0.3 | 0.09 | 0.17 | 0.17 |
| 6 | 3913345 | rs4959894 | -2.044 | 0.977 | 0.725 | -3.23 | -0.075 | 0.286 | 0.286 | 0.071 | 0.46 | 0.92 | 0.5 | 0.12 | 0.51 | 0.35 |
| 6 | 3913630 | rs4074617 | -1.863 | 0.977 | 0.897 | -3.23 | -0.01 | 0.011 | 0.011 | 0.021 | 0.42 | 0.67 | 0.46 | 0.1 | 0.5 | 0.35 |
| 6 | 3919215 | rs6925458 | -1.219 | 0.763 | 1.154 | -2.829 | 0.023 | 0.015 | 0.015 | 0 | 0.11 | 0.37 | 0.26 | 0.09 | 0.13 | 0.15 |
| 6 | 3934615 | rs7768618 | 0.956 | 0.368 | 1.136 | -3.164 | 0.005 | -0.064 | -0.064 | 0 | 0.59 | 0.46 | 0.55 | 0.91 | 0.57 | 0.65 |
| 6 | 3940533 | rs9392536 | 1.188 | 0.346 | 0.953 | -3.155 | -0.055 | 0.259 | 0.259 | 0 | 0.54 | 0.1 | 0.51 | 0.88 | 0.55 | 0.65 |
| 6 | 3942134 | rs6925554 | -1.723 | 0.395 | 1.185 | -3.044 | -0.011 | -0.063 | -0.063 | 0 | 0.11 | 0.41 | 0.07 | 0.02 | 0.05 | 0 |
| 6 | 3944424 | rs4959901 | -0.163 | 0.538 | 1.178 | -3.025 | -0.037 | -0.031 | -0.031 | 0.622 | 0.06 | 0.01 | 0.24 | 0.09 | 0.14 | 0.16 |
| 6 | 3944564 | rs9328207 | -1.554 | 0.45 | 1.278 | -3.09 | 0.006 | -0.045 | -0.045 | 0 | 0.1 | 0.4 | 0.07 | 0.02 | 0.05 | 0 |
| 6 | 3946972 | rs6597013 | 0.119 | 0.475 | 1.283 | -3.172 | -0.001 | -0.176 | -0.176 | 2.242 | 0.71 | 0.52 | 0.83 | 1 | 0.73 | 0.82 |
| 6 | 3947354 | rs4959902 | -1.788 | 0.492 | 1.379 | -2.943 | 0.024 | -0.042 | -0.042 | 0.063 | 0.17 | 0.47 | 0.32 | 0.11 | 0.18 | 0.17 |
| 6 | 3947984 | rs9405676 | 0.862 | 0.62 | 0.757 | -2.737 | -0.003 | 0.067 | 0.067 | 0 | 0.51 | 0.38 | 0.54 | 0.76 | 0.5 | 0.56 |
| 6 | 3949423 | rs9378389 | 1.017 | 0.429 | -0.151 | -2.744 | -0.035 | 0.199 | 0.199 | 0 | 0.48 | 0.22 | 0.5 | 0.81 | 0.5 | 0.57 |
| 6 | 3950674 | rs4959264 | 0.958 | 0.511 | -0.362 | -2.67 | -0.001 | 0.076 | 0.076 | 0 | 0.49 | 0.38 | 0.52 | 0.81 | 0.52 | 0.57 |
| 6 | 3951007 | rs6932988 | 0.046 | 0.795 | -1.458 | -2.406 | -0.024 | 0.126 | 0.126 | 0 | 0.69 | 0.91 | 0.86 | 1 | 0.71 | 0.8 |
| 6 | 3956160 | rs11755366 | -0.748 | 0.784 | -0.585 | -1.067 | 0.002 | 0.005 | 0.005 | 0 | 0.12 | 0.16 | 0.27 | 0.16 | 0.15 | 0.22 |
| 6 | 3957271 | rs6597017 | 0.317 | 0.634 | -0.883 | -0.018 | -0.001 | 0.001 | 0.001 | 0 | 0.7 | 0.66 | 0.61 | 0.4 | 0.68 | 0.6 |
| 6 | 3958190 | rs6597018 | -0.232 | 0.286 | -1.314 | 0.215 | 0.013 | 0.018 | 0.018 | 0 | 0.43 | 0.6 | 0.3 | 0.21 | 0.4 | 0.3 |
| 6 | 3958481 | rs9502099 | -0.288 | 0.614 | -2.341 | 0.658 | -0.113 | -0.136 | -0.136 | 0.198 | 0.57 | 0.97 | 0.38 | 0.42 | 0.43 | 0.52 |
| 6 | 3960888 | rs9405684 | -0.472 | 0.608 | -2.107 | 0.395 | -0.021 | 0.025 | 0.025 | 0.168 | 0.1 | 0.01 | 0.09 | 0.04 | 0.13 | 0.02 |
| 6 | 3963696 | rs9405685 | -0.044 | 0.789 | -2.124 | 0.369 | -0.024 | 0.019 | 0.019 | 0.108 | 0.07 | 0.01 | 0.06 | 0.03 | 0.1 | 0.02 |
| 6 | 3963841 | rs9405214 | -0.237 | 0.66 | -3.3 | 0.556 | -0.067 | -0.106 | -0.106 | 0 | 0.59 | 0.94 | 0.6 | 0.47 | 0.64 | 0.61 |
| 6 | 3965523 | rs7747045 | -0.071 | -0.072 | -2.591 | -0.181 | -0.036 | -0.095 | -0.095 | 3.113 | 0.24 | 0.61 | 0.3 | 0.04 | 0.18 | 0.05 |
| 6 | 3965812 | rs4959911 | -0.681 | 0.116 | -2.779 | 0.166 | -0.074 | -0.038 | -0.038 | 0.058 | 0.59 | 0.95 | 0.59 | 0.37 | 0.64 | 0.6 |
| 6 | 3972060 | rs11242894 | -0.536 | -0.202 | -2.069 | 0.592 | -0.01 | 0.094 | 0.094 | 0 | 0.48 | 0.82 | 0.62 | 0.68 | 0.54 | 0.59 |
| 6 | 3977149 | rs6925753 | 0.4 | -0.255 | -1.908 | 0.655 | -0.003 | 0.094 | 0.094 | 0 | 0.52 | 0.2 | 0.39 | 0.32 | 0.46 | 0.41 |
| 6 | 3978257 | rs2326366 | -0.216 | -0.335 | -0.855 | 0.212 | -0.017 | -0.011 | -0.011 | 0 | 0.66 | 0.75 | 0.67 | 0.58 | 0.56 | 0.45 |
| 6 | 3979677 | rs2326368 | 0.901 | -0.24 | -0.198 | -0.235 | -0.017 | 0.009 | 0.009 | 2.195 | 0.66 | 0.41 | 0.73 | 0.58 | 0.7 | 0.72 |
| 6 | 3979751 | rs9392025 | 1.161 | 0.315 | 0.311 | -0.428 | 0 | 0.001 | 0.001 | 0 | 0.51 | 0.48 | 0.47 | 0.51 | 0.47 | 0.42 |
| 6 | 3981940 | rs7757311 | 1.841 | 0.301 | 0.31 | -0.899 | -0.014 | 0.019 | 0.019 | 1.063 | 0.4 | 0.28 | 0.42 | 0.67 | 0.39 | 0.49 |
| 6 | 3982898 | rs4330570 | 1.136 | 0.431 | 0.38 | -0.935 | -0.007 | 0.01 | 0.01 | 0.036 | 0.61 | 0.35 | 0.55 | 0.67 | 0.59 | 0.59 |
| 6 | 3988311 | rs11242895 | 0.049 | 0.722 | 0.077 | -0.984 | -0.023 | 0.054 | 0.054 | 0 | 0.16 | 0.07 | 0.11 | 0.01 | 0.16 | 0.19 |
| 6 | 3988620 | rs6597022 | 1.125 | 0.743 | 0.304 | -0.887 | 0.002 | 0.002 | 0.002 | 0.005 | 0.57 | 0.38 | 0.51 | 0.62 | 0.54 | 0.65 |
| 6 | 3989106 | rs9378836 | 1.357 | 0.753 | -0.074 | -0.864 | 0.022 | 0.023 | 0.023 | 0.102 | 0.55 | 0.21 | 0.49 | 0.62 | 0.54 | 0.66 |
| 6 | 3989654 | rs9328211 | 0.311 | 1.191 | 0.316 | -0.778 | 0.015 | 0.023 | 0.023 | 0 | 0.83 | 0.76 | 0.82 | 0.74 | 0.75 | 0.79 |
| 6 | 3990096 | rs9392544 | -0.754 | 0.959 | 0.106 | -1.32 | -0.026 | 0.07 | 0.07 | 0 | 0.21 | 0.08 | 0.18 | 0.01 | 0.25 | 0.23 |
| 6 | 3996978 | rs6597023 | 0.234 | 1.182 | 0.342 | -0.888 | -0.005 | -0.014 | -0.014 | 0 | 0.89 | 0.92 | 0.88 | 0.78 | 0.89 | 0.87 |
| 6 | 4008166 | rs6912179 | -1.38 | 1.266 | 0.723 | -1.159 | 0.019 | -0.042 | -0.042 | 0.003 | 0.28 | 0.44 | 0.37 | 0.18 | 0.35 | 0.36 |
| 6 | 4009504 | rs6938648 | -0.982 | 1.332 | 0.842 | -1.178 | -0.003 | -0.02 | -0.02 | 0 | 0.24 | 0.35 | 0.32 | 0.18 | 0.25 | 0.35 |
| 6 | 4012929 | rs10458166 | -0.161 | 1.306 | 0.588 | -1.247 | 0.01 | 0.068 | 0.068 | 0 | 0.18 | 0.06 | 0.11 | 0.01 | 0.17 | 0.26 |
| 6 | 4013741 | rs2147374 | 2.428 | 0.799 | 0.33 | -1.551 | -0.06 | -0.029 | -0.029 | 0 | 0.39 | 0.12 | 0.42 | 0.6 | 0.43 | 0.44 |
| 6 | 4016348 | rs12110567 | -0.923 | 0.721 | 0.924 | -1.305 | -0.021 | -0.081 | -0.081 | 0.034 | 0.25 | 0.34 | 0.14 | 0.01 | 0.18 | 0.13 |
| 6 | 4018568 | rs17137491 | -0.4 | 0.738 | 0.945 | -1.299 | -0.046 | -0.107 | -0.107 | 0 | 0.21 | 0.33 | 0.09 | 0 | 0.11 | 0.05 |
| 6 | 4022611 | rs2181894 | 1.042 | 0.926 | 0.494 | -1.182 | 0.008 | 0.041 | 0.041 | 0.791 | 0.68 | 0.8 | 0.77 | 0.79 | 0.75 | 0.72 |
| 6 | 4030913 | rs1408457 | 1.304 | 0.98 | 0.614 | -1.246 | -0.009 | 0.017 | 0.017 | 0 | 0.67 | 0.74 | 0.74 | 0.79 | 0.64 | 0.69 |
| 6 | 4031148 | rs723796 | 0.22 | 1.089 | 0.988 | -1.31 | -0.007 | 0.019 | 0.019 | 0 | 0.14 | 0.3 | 0.21 | 0.16 | 0.13 | 0.06 |
| 6 | 4031998 | rs9503893 | 1.278 | 1.11 | 0.939 | -1.292 | -0.007 | 0.028 | 0.028 | 0.15 | 0.65 | 0.74 | 0.73 | 0.79 | 0.64 | 0.69 |
| 6 | 4033065 | rs6924389 | 0.151 | 1.142 | 0.816 | -1.272 | -0.005 | -0.008 | -0.008 | 0 | 0.9 | 0.93 | 0.9 | 0.79 | 0.91 | 0.83 |
| 6 | 4039475 | rs3761787 | 0.868 | 1.131 | 1.004 | -1.199 | 0.003 | -0.001 | -0.001 | 0 | 0.85 | 0.86 | 0.81 | 0.79 | 0.75 | 0.74 |
| 6 | 4043847 | rs707980 | 1.517 | 0.992 | 1.008 | -1.245 | -0.001 | -0.001 | -0.001 | 0 | 0.63 | 0.33 | 0.7 | 0.79 | 0.65 | 0.69 |
| 6 | 4054749 | rs853401 | 0.356 | 0.818 | 1.009 | -1.156 | -0.012 | -0.015 | -0.015 | 0 | 0.88 | 0.93 | 0.83 | 0.72 | 0.86 | 0.77 |
| 6 | 4057542 | rs9503907 | 0.678 | 0.715 | 0.376 | -1.14 | 0.006 | 0.02 | 0.02 | 0 | 0.12 | 0.06 | 0.05 | 0.01 | 0.11 | 0.04 |
| 6 | 4058782 | rs12198921 | 1.832 | 0.762 | 0.131 | -1.148 | -0.11 | 0.011 | 0.011 | 2.367 | 0.49 | 0.03 | 0.49 | 0.63 | 0.52 | 0.62 |
| 6 | 4063681 | rs853411 | 1.39 | 1.188 | 1.035 | -0.63 | -0.007 | 0.022 | 0.022 | 0.013 | 0.67 | 0.74 | 0.73 | 0.78 | 0.65 | 0.69 |
| 6 | 4067046 | rs7357049 | -1.632 | 1.389 | 1.862 | -0.326 | 0.051 | 0.083 | 0.083 | 1.089 | 0.15 | 0.5 | 0.15 | 0.25 | 0.14 | 0.22 |
| 6 | 4068335 | rs623861 | -1.438 | 1.581 | 2.042 | -0.125 | 0.135 | 0.12 | 0.12 | 0 | 0.05 | 0.22 | 0.24 | 0.1 | 0.13 | 0.04 |
| 6 | 4068716 | rs593291 | 2.187 | 1.436 | 2.237 | -0.308 | 0.048 | 0.072 | 0.072 | 0 | 0.78 | 0.46 | 0.65 | 0.65 | 0.74 | 0.74 |

| | | | | | | | | | | | | | | | | |
|---|---|---|---|---|---|---|---|---|---|---|---|---|---|---|---|---|
| 6 | 4069685 | rs639905 | -1.711 | 1.463 | 2.128 | -0.154 | 0.084 | 0.18 | 0.18 | 0.008 | 0.15 | 0.41 | 0.3 | 0.35 | 0.19 | 0.19 |
| 6 | 4079875 | rs665728 | -1.833 | 1.137 | 1.185 | -0.423 | -0.001 | -0.004 | -0.004 | 0 | 0.38 | 0.84 | 0.4 | 0.37 | 0.35 | 0.33 |
| 6 | 4096678 | rs608195 | -1.724 | 1.426 | 1.725 | -0.455 | 0 | -0.001 | -0.001 | 1.048 | 0.37 | 0.58 | 0.39 | 0.36 | 0.36 | 0.27 |
| 6 | 4097315 | rs1414102 | -1.162 | 1.04 | 1.802 | -0.573 | -0.034 | -0.124 | -0.124 | 0 | 0.25 | 0.38 | 0.12 | 0 | 0.17 | 0.13 |
| 6 | 4101567 | rs626080 | -1.775 | 1.584 | 1.61 | -0.529 | 0.001 | -0.001 | -0.001 | 0.579 | 0.4 | 0.52 | 0.43 | 0.36 | 0.49 | 0.29 |
| 6 | 4104567 | rs582008 | -1.146 | 1.648 | 1.882 | -0.413 | 0.019 | 0.116 | 0.116 | 0 | 0.13 | 0.35 | 0.24 | 0.34 | 0.15 | 0.16 |
| 6 | 4109593 | rs649725 | -1.425 | 1.657 | 1.579 | -0.441 | 0.01 | 0.005 | 0.005 | 0 | 0.31 | 0.37 | 0.36 | 0.36 | 0.33 | 0.22 |
| 6 | 4111429 | rs678285 | 0.132 | 1.686 | 1.519 | -0.431 | -0.016 | -0.007 | -0.007 | 1.752 | 0.07 | 0.02 | 0.17 | 0.34 | 0.06 | 0.1 |
| 6 | 4114788 | rs633290 | -1.44 | 1.675 | 1.63 | -0.441 | 0.017 | 0.008 | 0.008 | 0 | 0.31 | 0.4 | 0.36 | 0.36 | 0.33 | 0.22 |
| 6 | 4116262 | rs7166 | -1.536 | 1.683 | 1.507 | -0.437 | 0.002 | 0.001 | 0.001 | 0 | 0.3 | 0.29 | 0.36 | 0.36 | 0.33 | 0.22 |
| 6 | 4122249 | rs643232 | -1.38 | 1.682 | 1.526 | -0.437 | 0.006 | 0.003 | 0.003 | 0 | 0.3 | 0.34 | 0.36 | 0.36 | 0.33 | 0.22 |
| 6 | 4125474 | rs11751668 | 0.319 | 1.697 | 1.412 | -0.365 | -0.016 | -0.025 | -0.025 | 0 | 0.06 | 0.01 | 0.03 | 0.27 | 0.05 | 0.09 |
| 6 | 4125655 | rs659305 | -1.362 | 1.697 | 1.437 | -0.372 | 0.003 | 0.001 | 0.001 | 0 | 0.3 | 0.3 | 0.36 | 0.36 | 0.33 | 0.22 |
| 6 | 4133855 | rs3177253 | -0.564 | 1.653 | 1.369 | -0.403 | -0.017 | 0.026 | 0.026 | 0 | 0.22 | 0.11 | 0.17 | 0.02 | 0.27 | 0.12 |
| 6 | 4134031 | rs853422 | -1.333 | 1.547 | 1.686 | -0.316 | 0.064 | 0.075 | 0.075 | 0.647 | 0.09 | 0.55 | 0.22 | 0.07 | 0.09 | 0.07 |
| 6 | 4145534 | rs9503936 | -1.652 | 1.67 | 1.167 | -0.591 | 0.009 | -0.012 | -0.012 | 0 | 0.34 | 0.37 | 0.23 | 0.29 | 0.36 | 0.21 |
| 6 | 4146704 | rs9405702 | -0.59 | 1.645 | 0.677 | -0.47 | -0.001 | 0.013 | 0.013 | 0.705 | 0.09 | 0.12 | 0.05 | 0.27 | 0.07 | 0.09 |
| 6 | 4148182 | rs11755258 | -1.106 | 1.425 | 0.569 | -0.883 | -0.007 | -0.01 | -0.01 | 0 | 0.3 | 0.22 | 0.2 | 0.02 | 0.36 | 0.19 |
| 6 | 4151518 | rs4605929 | -1.678 | 1.479 | -0.103 | -0.539 | -0.079 | 0.085 | 0.085 | 1.418 | 0.41 | 0.89 | 0.34 | 0.25 | 0.4 | 0.28 |
| 6 | 4153137 | rs653674 | -0.094 | 2.439 | 0.312 | -0.48 | 0.015 | 0 | 0 | 0 | 0.06 | 0.09 | 0.24 | 0.07 | 0.09 | 0.21 |
| 6 | 4153297 | rs6935622 | -0.889 | 2.1 | 0.754 | -0.591 | 0.004 | -0.045 | -0.045 | 0 | 0.09 | 0.23 | 0.05 | 0 | 0.09 | 0.06 |
| 6 | 4154018 | rs13206717 | -0.679 | 2.393 | 0.537 | -0.349 | 0.143 | 0.011 | 0.011 | 0 | 0.07 | 0.2 | 0.13 | 0.03 | 0.19 | 0.05 |
| 6 | 4158024 | rs618997 | -0.915 | 2.13 | -0.267 | 0.052 | -0.02 | -0.007 | -0.007 | 0 | 0.38 | 0.29 | 0.52 | 0.54 | 0.42 | 0.52 |
| 6 | 4158369 | rs6923069 | 0.86 | 2.076 | -0.026 | -0.155 | 0.001 | -0.003 | -0.003 | 2.677 | 0.56 | 0.47 | 0.45 | 0.54 | 0.53 | 0.41 |
| 6 | 4158522 | rs2025267 | -0.064 | 1.759 | -0.612 | -0.331 | -0.058 | 0.034 | 0.034 | 0 | 0.8 | 0.95 | 0.8 | 0.87 | 0.75 | 0.89 |
| 6 | 4164620 | rs650359 | -0.152 | 1.668 | -0.394 | -0.097 | -0.027 | -0.006 | -0.006 | 0 | 0.09 | 0.03 | 0.2 | 0.06 | 0.11 | 0.22 |
| 6 | 4172642 | rs622774 | -0.226 | 1.656 | -0.284 | -0.187 | -0.013 | 0.007 | 0.007 | 0 | 0.09 | 0.05 | 0.21 | 0.06 | 0.11 | 0.22 |
| 6 | 4177527 | rs9378848 | -1.386 | 1.38 | 0.579 | -0.511 | -0.001 | -0.014 | -0.014 | 0 | 0.28 | 0.67 | 0.34 | 0.15 | 0.29 | 0.37 |
| 6 | 4179398 | rs9405708 | -0.217 | 1.328 | 0.431 | -0.154 | -0.002 | 0.035 | 0.035 | 0 | 0.09 | 0.13 | 0.11 | 0.26 | 0.06 | 0.16 |
| 6 | 4187067 | rs666105 | -0.293 | 1.396 | 0.579 | -0.206 | -0.002 | -0.002 | -0.002 | 0 | 0.11 | 0.11 | 0.21 | 0.14 | 0.14 | 0.28 |
| 6 | 4189651 | rs600996 | 0.191 | 1.323 | 0.449 | -0.281 | -0.031 | 0.006 | 0.006 | 0 | 0.1 | 0.03 | 0.2 | 0.06 | 0.11 | 0.22 |
| 6 | 4189964 | rs12212264 | 1.722 | 1.23 | 0.289 | -0.578 | 0.003 | -0.057 | -0.057 | 0 | 0.37 | 0.18 | 0.34 | 0.48 | 0.39 | 0.32 |
| 6 | 4190450 | rs12663111 | 0.25 | 1.242 | 0.299 | -0.525 | -0.002 | 0.036 | 0.036 | 0 | 0.08 | 0.12 | 0.11 | 0.27 | 0.06 | 0.15 |
| 6 | 4190681 | rs615712 | 0.29 | 1.248 | 0.378 | -0.556 | -0.006 | -0.004 | -0.004 | 0 | 0.1 | 0.1 | 0.21 | 0.14 | 0.14 | 0.29 |
| 6 | 4207455 | rs853190 | -0.479 | 1.271 | 0.265 | -0.609 | -0.085 | 0.06 | 0.06 | 0 | 0.76 | 0.98 | 0.79 | 0.89 | 0.74 | 0.84 |
| 6 | 4207930 | rs685716 | 0.026 | 1.186 | 0.398 | -0.586 | -0.036 | -0.026 | -0.026 | 0.013 | 0.67 | 0.86 | 0.68 | 0.63 | 0.68 | 0.69 |
| 6 | 4209535 | rs6933147 | 0.042 | 1.162 | 0.719 | -0.616 | -0.01 | -0.076 | -0.076 | 1.652 | 0.16 | 0.29 | 0.12 | 0 | 0.14 | 0.08 |
| 6 | 4210658 | rs17538963 | 0.649 | 1.221 | 0.226 | -0.601 | 0 | 0.004 | 0.004 | 0 | 0.1 | 0.05 | 0.06 | 0 | 0.1 | 0.03 |
| 6 | 4210746 | rs621445 | 0.703 | 1.181 | 0.421 | -0.421 | 0 | 0.039 | 0.039 | 0 | 0.49 | 0.35 | 0.55 | 0.63 | 0.53 | 0.6 |
| 6 | 4210858 | rs12213573 | 1.379 | 0.985 | 0.183 | -0.402 | 0.006 | -0.033 | -0.033 | 0.042 | 0.37 | 0.23 | 0.34 | 0.48 | 0.39 | 0.32 |
| 6 | 4228049 | rs2810793 | 0.356 | 0.587 | 0.025 | -0.31 | 0.007 | 0.013 | 0.013 | 0 | 0.1 | 0.06 | 0.18 | 0.05 | 0.08 | 0.2 |
| 6 | 4229688 | rs2105042 | 1.065 | 0.57 | -0.126 | -0.262 | -0.013 | -0.069 | -0.069 | 0.063 | 0.36 | 0.16 | 0.38 | 0.47 | 0.43 | 0.32 |
| 6 | 4245958 | rs7755249 | 0.311 | 0.326 | 0.152 | -0.129 | 0.005 | -0.011 | -0.011 | 2.54 | 0.17 | 0.14 | 0.15 | 0.27 | 0.12 | 0.34 |
| 6 | 4249646 | rs4259288 | 0.38 | 0.348 | 0.053 | -0.673 | 0.042 | 0.057 | 0.057 | 0 | 0.61 | 0.29 | 0.67 | 0.85 | 0.75 | 0.77 |
| 6 | 4255387 | rs12204780 | 0.368 | 0.319 | 0.056 | -0.692 | 0.044 | 0.056 | 0.056 | 0 | 0.61 | 0.29 | 0.67 | 0.86 | 0.76 | 0.78 |
| 6 | 4255708 | rs9504008 | 0.027 | 0.257 | 0.747 | -0.767 | 0.001 | -0.083 | -0.083 | 0.897 | 0.13 | 0.27 | 0.09 | 0 | 0.12 | 0.03 |
| 6 | 4260231 | rs6597048 | 0.027 | 0.257 | 0.751 | -0.767 | 0 | -0.082 | -0.082 | 0 | 0.13 | 0.27 | 0.09 | 0 | 0.12 | 0.03 |
| 6 | 4260723 | rs9328233 | 0.373 | 0.244 | 0.821 | -0.774 | 0 | -0.029 | -0.029 | 0 | 0.12 | 0.1 | 0.07 | 0 | 0.12 | 0.03 |
| 6 | 4263349 | rs6937085 | -0.551 | 0.168 | 1.085 | -0.798 | -0.008 | -0.016 | -0.016 | 0 | 0.37 | 0.59 | 0.33 | 0.15 | 0.25 | 0.22 |
| 6 | 4266832 | rs9504021 | -0.379 | 0.348 | 0.571 | -0.777 | 0.024 | 0.035 | 0.035 | 0 | 0.34 | 0.24 | 0.18 | 0.12 | 0.24 | 0.18 |
| 6 | 4283916 | rs9378854 | 0.581 | 0.434 | 0.445 | -0.428 | 0.012 | -0.039 | -0.039 | 0.457 | 0.17 | 0.08 | 0.14 | 0.34 | 0.11 | 0.36 |
| 6 | 4284640 | rs10223671 | -0.466 | 0.436 | 0.482 | -0.468 | -0.004 | 0.003 | 0.003 | 0 | 0.29 | 0.32 | 0.22 | 0.34 | 0.23 | 0.39 |
| 6 | 4292043 | rs7740637 | -0.3 | 0.3 | 0.393 | -0.517 | 0.027 | -0.011 | -0.011 | 0 | 0.4 | 0.31 | 0.25 | 0.46 | 0.23 | 0.51 |
| 6 | 4296918 | rs17565917 | 0.704 | 0.282 | 0.204 | -0.525 | 0.057 | 0.027 | 0.027 | 0 | 0.21 | 0.06 | 0.1 | 0.12 | 0.12 | 0.15 |
| 6 | 4297286 | rs1999022 | -1.235 | 0.293 | 0.348 | -0.555 | 0.049 | -0.035 | -0.035 | 0.007 | 0.54 | 0.8 | 0.48 | 0.49 | 0.36 | 0.56 |
| 6 | 4298538 | rs17500205 | 0.641 | 0.196 | 0.295 | -0.758 | 0 | -0.023 | -0.023 | 0 | 0.1 | 0.1 | 0.07 | 0 | 0.12 | 0.03 |
| 6 | 4299380 | rs6597051 | -1.226 | 0.341 | -0.154 | -0.727 | -0.052 | -0.023 | -0.023 | 0.37 | 0.6 | 0.87 | 0.53 | 0.59 | 0.43 | 0.64 |
| 6 | 4301551 | rs1327993 | -1.126 | 0.458 | 0.244 | -1.238 | -0.002 | -0.007 | -0.007 | 2.852 | 0.37 | 0.62 | 0.35 | 0.15 | 0.25 | 0.22 |
| 6 | 4304046 | rs7759565 | -0.76 | 0.816 | 0.532 | -0.939 | 0.017 | -0.068 | -0.068 | 0.853 | 0.15 | 0.45 | 0.23 | 0.04 | 0.12 | 0.06 |
| 6 | 4304633 | rs6597052 | 1.156 | 1.008 | -0.013 | -0.44 | 0 | -0.041 | -0.041 | 0 | 0.7 | 0.53 | 0.62 | 0.85 | 0.7 | 0.84 |
| 6 | 4309521 | rs932847 | 0.931 | 2.074 | -0.536 | -0.244 | -0.055 | 0.053 | 0.053 | 0 | 0.91 | 0.97 | 0.86 | 1 | 0.8 | 0.93 |
| 6 | 4310125 | rs1327992 | 1.558 | 2.104 | -0.562 | -0.282 | 0 | -0.018 | -0.018 | 0.176 | 0.87 | 0.86 | 0.83 | 1 | 0.78 | 0.93 |
| 6 | 4313727 | rs9378858 | 1.004 | 2.115 | -0.983 | -0.101 | -0.056 | 0.054 | 0.054 | 0 | 0.9 | 0.98 | 0.86 | 1 | 0.8 | 0.93 |
| 6 | 4317398 | rs1887574 | -0.384 | 1.683 | -0.886 | -0.038 | -0.009 | 0.015 | 0.015 | 1.65 | 0.36 | 0.7 | 0.39 | 0.26 | 0.41 | 0.21 |
| 6 | 4319573 | rs13209318 | -1.314 | 2.331 | 0.091 | -0.223 | 0.083 | -0.044 | -0.044 | 0.533 | 0.13 | 0.4 | 0.11 | 0.04 | 0.19 | 0.11 |
| 6 | 4320190 | rs2326477 | 0.27 | 1.842 | -0.538 | -0.011 | 0.015 | -0.023 | -0.023 | 0 | 0.4 | 0.22 | 0.4 | 0.49 | 0.33 | 0.32 |
| 6 | 4323299 | rs10900955 | -0.345 | 1.63 | -0.444 | -0.116 | 0.002 | -0.006 | -0.006 | 0 | 0.56 | 0.73 | 0.53 | 0.51 | 0.58 | 0.64 |
| 6 | 4338841 | rs9392582 | 0.157 | 1.14 | -0.613 | -0.008 | -0.005 | -0.014 | -0.014 | 0 | 0.21 | 0.14 | 0.16 | 0.22 | 0.21 | 0.34 |
| 6 | 4349650 | rs1265208 | 0.808 | 0.839 | -0.369 | -0.173 | -0.003 | -0.013 | -0.013 | 1.248 | 0.48 | 0.26 | 0.49 | 0.5 | 0.45 | 0.37 |

| | | | | | | | | | | | | | | | | |
|---|---|---|---|---|---|---|---|---|---|---|---|---|---|---|---|---|
| 6 | 4349814 | rs1265207 | 0.889 | 0.825 | -0.392 | -0.165 | -0.024 | -0.037 | -0.037 | 0.011 | 0.47 | 0.14 | 0.48 | 0.5 | 0.45 | 0.37 |
| 6 | 4351746 | rs557994 | -0.162 | 0.549 | -0.028 | -0.318 | -0.002 | -0.001 | -0.001 | 0 | 0.76 | 0.72 | 0.68 | 0.74 | 0.7 | 0.73 |
| 6 | 4352851 | rs10900956 | 0.134 | 0.528 | 0.275 | -0.333 | 0.002 | 0 | 0 | 0 | 0.23 | 0.22 | 0.31 | 0.26 | 0.3 | 0.27 |
| 6 | 4354439 | rs4959277 | 0.404 | 0.397 | 0.376 | -0.302 | -0.001 | -0.006 | -0.006 | 0 | 0.2 | 0.16 | 0.16 | 0.22 | 0.22 | 0.34 |
| 6 | 4355314 | rs4959279 | -0.027 | 0.352 | 0.601 | -0.37 | -0.007 | -0.004 | -0.004 | 4.686 | 0.24 | 0.31 | 0.18 | 0.23 | 0.23 | 0.35 |
| 6 | 4359684 | rs811226 | 0.684 | 0.689 | 0.256 | -0.303 | 0.006 | 0.004 | 0.004 | 0 | 0.52 | 0.29 | 0.53 | 0.52 | 0.49 | 0.46 |
| 6 | 4362258 | rs170948 | -0.346 | 0.272 | -0.086 | 0.151 | -0.006 | 0.002 | 0.002 | 0 | 0.86 | 0.8 | 0.89 | 0.84 | 0.91 | 0.92 |
| 6 | 4363482 | rs9328246 | 0.005 | 0.312 | -0.004 | 0.439 | -0.002 | -0.018 | -0.018 | 0 | 0.18 | 0.09 | 0.13 | 0.23 | 0.18 | 0.33 |
| 6 | 4364141 | rs17138098 | 0.07 | 0.022 | -0.404 | 0.619 | 0.009 | -0.006 | -0.006 | 0.896 | 0.12 | 0.1 | 0.18 | 0.11 | 0.09 | 0.06 |
| 6 | 4371153 | rs9392592 | 0.855 | -0.149 | -0.488 | 0.681 | -0.005 | -0.035 | -0.035 | 0 | 0.09 | 0 | 0.02 | 0.13 | 0.08 | 0.14 |
| 6 | 4373213 | rs9405232 | 0.605 | -0.1 | -0.14 | 0.995 | -0.004 | -0.003 | -0.003 | 0 | 0.6 | 0.65 | 0.49 | 0.61 | 0.55 | 0.69 |
| 6 | 4376376 | rs541051 | -0.025 | -0.141 | -0.136 | 0.582 | -0.046 | -0.044 | -0.044 | 1.009 | 0.26 | 0.16 | 0.32 | 0.36 | 0.34 | 0.29 |
| 6 | 4378534 | rs7760710 | 1.112 | -0.14 | -0.282 | 0.558 | 0.023 | 0.024 | 0.024 | 0 | 0.06 | 0.01 | 0.03 | 0 | 0.03 | 0.01 |
| 6 | 4378997 | rs1265203 | -0.439 | -0.065 | -0.211 | 0.608 | 0.001 | 0.001 | 0.001 | 0 | 0.29 | 0.32 | 0.34 | 0.36 | 0.37 | 0.29 |
| 6 | 4379511 | rs17138114 | 0.171 | -0.074 | -0.259 | 0.627 | 0.015 | -0.015 | -0.015 | 0 | 0.11 | 0.08 | 0.19 | 0.15 | 0.08 | 0.06 |
| 6 | 4380902 | rs1265200 | 0.034 | -0.049 | -0.33 | 0.625 | -0.092 | -0.086 | -0.086 | 0.336 | 0.24 | 0.05 | 0.31 | 0.35 | 0.34 | 0.28 |
| 6 | 4383332 | rs4142082 | 1.112 | 0.142 | -0.222 | 0.343 | -0.003 | 0.036 | 0.036 | 0 | 0.39 | 0.23 | 0.34 | 0.27 | 0.37 | 0.39 |
| 6 | 4384960 | rs4959949 | 0.741 | 0.497 | -0.181 | 0.754 | 0 | 0.002 | 0.002 | 0 | 0.59 | 0.52 | 0.46 | 0.5 | 0.55 | 0.65 |
| 6 | 4391872 | rs1023017 | -0.454 | 0.668 | -0.213 | 0.354 | -0.022 | 0.018 | 0.018 | 0 | 0.07 | 0.02 | 0.06 | 0 | 0.11 | 0.02 |
| 6 | 4394906 | rs2326488 | -1.107 | -0.522 | 0.434 | 0.558 | -0.091 | 0.007 | 0.007 | 0.674 | 0.17 | 0.44 | 0.21 | 0.13 | 0.08 | 0.05 |
| 6 | 4395451 | rs168126 | 1.728 | -0.118 | 1.483 | 0.364 | 0.078 | 0.009 | 0.009 | 0.208 | 0.93 | 0.59 | 0.92 | 1 | 0.92 | 0.99 |
| 6 | 4395504 | rs339294 | 0.298 | -0.381 | 0.763 | 0.189 | 0.016 | 0.037 | 0.037 | 0 | 0.46 | 0.31 | 0.57 | 0.67 | 0.57 | 0.56 |
| 6 | 4398169 | rs446293 | -0.075 | -0.039 | 1.275 | 0.547 | 0.019 | 0.026 | 0.026 | 0 | 0.51 | 0.52 | 0.62 | 0.75 | 0.62 | 0.53 |
| 6 | 4400709 | rs885185 | 0.395 | -0.692 | 0.447 | 0.272 | 0.089 | 0.027 | 0.027 | 2.046 | 0.52 | 0.14 | 0.61 | 0.62 | 0.64 | 0.61 |
| 6 | 4401627 | rs11242924 | 0.925 | -0.472 | 0.549 | 0.381 | -0.094 | -0.086 | -0.086 | 0 | 0.36 | 0.08 | 0.48 | 0.52 | 0.46 | 0.4 |
| 6 | 4402634 | rs9502175 | 0.267 | -0.614 | 1.026 | 0.406 | -0.012 | 0.16 | 0.16 | 1.274 | 0.93 | 0.7 | 0.93 | 0.84 | 0.97 | 0.99 |
| 6 | 4403802 | rs6921285 | 0.428 | -0.543 | 1.086 | 0.387 | 0.029 | -0.03 | -0.03 | 0 | 0.42 | 0.19 | 0.54 | 0.53 | 0.59 | 0.48 |
| 6 | 4415975 | rs10900961 | 0.881 | -0.524 | 1.012 | 0.1 | -0.063 | 0.029 | 0.029 | 0.024 | 0.36 | 0.13 | 0.44 | 0.68 | 0.45 | 0.4 |
| 6 | 4425733 | rs13208005 | -0.059 | -0.375 | 1.213 | 0.079 | 0.014 | -0.008 | -0.008 | 1.503 | 0.06 | 0.08 | 0.08 | 0 | 0.16 | 0.09 |
| 6 | 4427554 | rs17138253 | 1.241 | -0.229 | 1.185 | -0.118 | 0.039 | -0.087 | -0.087 | 0 | 0.83 | 0.65 | 0.84 | 1 | 0.76 | 0.9 |
| 6 | 4431113 | rs4403297 | 0.872 | -0.55 | 1.161 | 0.058 | -0.067 | 0.032 | 0.032 | 0 | 0.37 | 0.14 | 0.45 | 0.67 | 0.46 | 0.4 |
| 6 | 4433060 | rs9392046 | -0.519 | -0.571 | 1.186 | 0.046 | -0.005 | 0.007 | 0.007 | 0 | 0.58 | 0.62 | 0.51 | 0.32 | 0.52 | 0.59 |
| 6 | 4436299 | rs953323 | 1.017 | -0.575 | 0.95 | 0.166 | -0.187 | -0.137 | -0.137 | 0.004 | 0.36 | 0.02 | 0.43 | 0.56 | 0.45 | 0.4 |
| 6 | 4445953 | rs4959976 | -0.25 | -0.583 | 1.219 | 0.113 | -0.007 | -0.012 | -0.012 | 0 | 0.53 | 0.51 | 0.44 | 0.32 | 0.38 | 0.55 |
| 6 | 4448061 | rs12207060 | 0.993 | -0.452 | 0.965 | 0.181 | -0.166 | -0.105 | -0.105 | 0 | 0.36 | 0.03 | 0.43 | 0.58 | 0.45 | 0.39 |
| 6 | 4448377 | rs17570071 | 0.993 | -0.452 | 0.965 | 0.223 | -0.166 | -0.129 | -0.129 | 0 | 0.36 | 0.03 | 0.43 | 0.56 | 0.45 | 0.39 |
| 6 | 4452666 | rs9378878 | 0.439 | -0.752 | 0.851 | -0.087 | -0.017 | 0.045 | 0.045 | 0.311 | 0.2 | 0.03 | 0.14 | 0.11 | 0.21 | 0.27 |
| 6 | 4456168 | rs4959980 | 0.38 | -0.962 | 1.059 | -0.24 | -0.054 | -0.042 | -0.042 | 1.988 | 0.29 | 0.47 | 0.31 | 0.2 | 0.16 | 0.28 |
| 6 | 4460769 | rs1265296 | 0.575 | -1.088 | 1.193 | 0.217 | -0.001 | -0.002 | -0.002 | 2.711 | 0.64 | 0.49 | 0.66 | 0.64 | 0.6 | 0.66 |
| 6 | 4462171 | rs7762514 | -0.439 | -1.04 | 1.172 | 0.235 | -0.031 | -0.016 | -0.016 | 0.471 | 0.3 | 0.36 | 0.24 | 0.28 | 0.27 | 0.29 |
| 6 | 4464320 | rs4959982 | -0.792 | -0.788 | 1.164 | 0.407 | 0.009 | 0.011 | 0.011 | 0 | 0.59 | 0.5 | 0.67 | 0.57 | 0.54 | 0.55 |
| 6 | 4464630 | rs9405746 | -0.428 | -0.926 | 0.797 | 0.41 | 0.068 | -0.023 | -0.023 | 0 | 0.54 | 0.24 | 0.59 | 0.56 | 0.39 | 0.5 |
| 6 | 4471472 | rs4959985 | 0.04 | -1.051 | 0.967 | 0.105 | -0.02 | -0.013 | -0.013 | 0 | 0.33 | 0.46 | 0.47 | 0.3 | 0.26 | 0.33 |
| 6 | 4472587 | rs7746329 | -0.092 | -0.893 | 1.067 | 0.073 | 0.013 | -0.008 | -0.008 | 0.803 | 0.24 | 0.51 | 0.23 | 0.28 | 0.27 | 0.23 |
| 6 | 4476202 | rs911566 | 0.106 | -1.411 | 0.895 | -0.199 | 0.005 | 0.005 | 0.005 | 0.648 | 0.34 | 0.62 | 0.43 | 0.25 | 0.19 | 0.35 |
| 6 | 4481421 | rs1011088 | -0.299 | -1.358 | 0.809 | -0.279 | 0.033 | 0.038 | 0.038 | 1.538 | 0.45 | 0.6 | 0.48 | 0.26 | 0.3 | 0.3 |
| 6 | 4484855 | rs1265256 | 0.436 | -1.559 | -0.133 | -0.522 | 0.095 | 0.184 | 0.184 | 2.234 | 0.52 | 0.15 | 0.54 | 0.74 | 0.66 | 0.65 |
| 6 | 4490506 | rs2150232 | 0.224 | -1.521 | 0.49 | -0.531 | -0.013 | -0.092 | -0.092 | 0 | 0.84 | 0.63 | 0.91 | 0.99 | 0.88 | 0.97 |
| 6 | 4492079 | rs4959299 | 0.146 | -1.765 | 0.396 | -0.574 | 0.017 | 0.017 | 0.017 | 0.321 | 0.36 | 0.25 | 0.39 | 0.26 | 0.23 | 0.31 |
| 6 | 4495016 | rs1265242 | 0.683 | -1.022 | -0.302 | -0.583 | -0.084 | -0.071 | -0.071 | 0 | 0.28 | 0.12 | 0.33 | 0.5 | 0.41 | 0.44 |
| 6 | 4497145 | rs1265239 | -0.788 | -1.717 | -0.151 | -0.667 | 0.002 | 0 | 0 | 0 | 0.75 | 0.8 | 0.78 | 0.73 | 0.74 | 0.83 |
| 6 | 4499029 | rs1265238 | 0.235 | -1.887 | -0.069 | -0.596 | -0.019 | -0.014 | -0.014 | 0 | 0.66 | 0.56 | 0.72 | 0.7 | 0.69 | 0.76 |
| 6 | 4499772 | rs1265236 | -0.178 | -1.85 | -0.348 | -0.579 | -0.005 | -0.006 | -0.006 | 1.904 | 0.69 | 0.67 | 0.73 | 0.73 | 0.69 | 0.82 |
| 6 | 4500682 | rs1265235 | 0.984 | -1.668 | -1.125 | -0.531 | -0.163 | -0.181 | -0.181 | 1.948 | 0.29 | 0.03 | 0.33 | 0.43 | 0.44 | 0.44 |
| 6 | 4501489 | rs2181015 | 0.827 | -1.594 | -1.292 | -0.312 | 0.004 | 0.004 | 0.004 | 0.01 | 0.31 | 0.23 | 0.36 | 0.25 | 0.23 | 0.26 |
| 6 | 4504891 | rs9378403 | -0.303 | -0.671 | -1.095 | 0.058 | 0.025 | 0.031 | 0.031 | 0 | 0.27 | 0.15 | 0.22 | 0.19 | 0.25 | 0.16 |
| 6 | 4508326 | rs9328255 | -0.657 | -0.573 | -1.011 | 0.167 | 0.035 | 0.013 | 0.013 | 0 | 0.32 | 0.25 | 0.27 | 0.32 | 0.29 | 0.21 |
| 6 | 4512439 | rs17138372 | 1.12 | 0.458 | -1.619 | 0.583 | -0.132 | -0.131 | -0.131 | 0.037 | 0.32 | 0.04 | 0.37 | 0.38 | 0.37 | 0.35 |
| 6 | 4514319 | rs1265224 | 0.119 | 0.162 | -1.104 | 0.124 | 0.002 | -0.002 | -0.002 | 0 | 0.24 | 0.21 | 0.25 | 0.11 | 0.23 | 0.26 |
| 6 | 4515551 | rs11242955 | 1.163 | 0.37 | -0.594 | 0.295 | -0.022 | 0.065 | 0.065 | 0 | 0.44 | 0.28 | 0.46 | 0.7 | 0.47 | 0.46 |
| 6 | 4515915 | rs6932387 | -0.133 | 0.451 | -0.503 | 0.208 | -0.061 | 0.051 | 0.051 | 0 | 0.28 | 0.07 | 0.26 | 0.17 | 0.32 | 0.23 |
| 6 | 4517448 | rs9378885 | 1.389 | 0.455 | -0.369 | 0.408 | -0.05 | -0.01 | -0.01 | 0 | 0.4 | 0.17 | 0.42 | 0.57 | 0.43 | 0.37 |
| 6 | 4519370 | rs1265216 | -1.312 | 0.301 | 0.231 | 0.479 | -0.012 | 0.007 | 0.007 | 0 | 0.57 | 0.63 | 0.55 | 0.43 | 0.55 | 0.62 |
| 6 | 4520511 | rs1265214 | 0.229 | 0.22 | 0.131 | 0.431 | 0 | -0.004 | -0.004 | 0 | 0.24 | 0.22 | 0.25 | 0.12 | 0.24 | 0.31 |
| 6 | 4524187 | rs4959304 | 0.092 | 0.278 | -0.356 | 0.414 | -0.025 | -0.024 | -0.024 | 0 | 0.13 | 0.04 | 0.21 | 0.18 | 0.14 | 0.14 |
| 6 | 4530500 | rs11968164 | 1.847 | 0.098 | -0.073 | 0.597 | -0.027 | -0.019 | -0.019 | 0 | 0.38 | 0.15 | 0.37 | 0.36 | 0.36 | 0.35 |
| 6 | 4534174 | rs1923322 | -0.509 | 0.076 | 0.021 | 0.556 | 0.011 | 0.066 | 0.066 | 0 | 0.77 | 0.91 | 0.76 | 0.89 | 0.77 | 0.71 |
| 6 | 4539480 | rs6907555 | -1.334 | 0.025 | 0.484 | 0.825 | -0.006 | -0.033 | -0.033 | 0.266 | 0.39 | 0.76 | 0.39 | 0.53 | 0.41 | 0.35 |
| 6 | 4548562 | rs6908122 | -0.275 | 0.074 | 0.077 | 1.094 | -0.005 | 0.036 | 0.036 | 0 | 0.09 | 0.18 | 0.06 | 0.17 | 0.07 | 0.1 |

| | | | | | | | | | | | | | | | | |
|---|---|---|---|---|---|---|---|---|---|---|---|---|---|---|---|---|
| 6 | 4551432 | rs403026 | -1.258 | -0.011 | -0.646 | 0.904 | -0.061 | -0.112 | -0.112 | 0.405 | 0.45 | 0.96 | 0.46 | 0.54 | 0.48 | 0.35 |
| 6 | 4560611 | rs440354 | -0.681 | -0.279 | -0.073 | 0.966 | 0.01 | -0.004 | -0.004 | 0 | 0.22 | 0.17 | 0.17 | 0.24 | 0.2 | 0.16 |
| 6 | 4563856 | rs437035 | -1.099 | -0.343 | 0.844 | 0.805 | -0.023 | 0.013 | 0.013 | 0 | 0.27 | 0.68 | 0.21 | 0.37 | 0.21 | 0.18 |
| 6 | 4569182 | rs407626 | -0.841 | 0.399 | 0.19 | 0.578 | -0.082 | -0.019 | -0.019 | 0 | 0.86 | 0.98 | 0.74 | 0.88 | 0.8 | 0.71 |
| 6 | 4572377 | rs17138501 | -0.735 | 0.016 | 0.606 | 0.663 | -0.089 | 0.115 | 0.115 | 0.087 | 0.19 | 0.47 | 0.12 | 0.34 | 0.09 | 0.14 |
| 6 | 4574440 | rs11242959 | 1.872 | 0.003 | -0.523 | 0.342 | -0.009 | -0.098 | -0.098 | 0.089 | 0.36 | 0.03 | 0.26 | 0.38 | 0.29 | 0.4 |
| 6 | 4588126 | rs9504156 | 0.496 | 0.104 | 0.294 | 0.461 | 0.005 | -0.031 | -0.031 | 0.12 | 0.07 | 0.11 | 0.09 | 0.01 | 0.12 | 0.04 |
| 6 | 4588847 | rs17138527 | -0.605 | -0.049 | 0.248 | 0.383 | -0.017 | -0.05 | -0.05 | 0.163 | 0.16 | 0.18 | 0.12 | 0.01 | 0.15 | 0.05 |
| 6 | 4590547 | rs4960005 | -0.358 | 0.583 | 0.527 | 0.707 | -0.005 | -0.051 | -0.051 | 0 | 0.11 | 0.19 | 0.11 | 0.01 | 0.14 | 0.04 |
| 6 | 4599442 | rs9504162 | -0.644 | 0.785 | 0.018 | 0.763 | -0.025 | 0.008 | 0.008 | 0 | 0.22 | 0.13 | 0.25 | 0.16 | 0.29 | 0.14 |
| 6 | 4600610 | rs7772028 | 0.282 | 0.444 | 0.156 | 0.916 | 0.01 | 0.027 | 0.027 | 0 | 0.08 | 0.11 | 0.19 | 0.17 | 0.09 | 0.09 |
| 6 | 4600758 | rs11965481 | -1.012 | 0.154 | 0.181 | 0.979 | -0.009 | 0.008 | 0.008 | 0.273 | 0.2 | 0.73 | 0.2 | 0.33 | 0.17 | 0.16 |
| 6 | 4604939 | rs13204938 | -1.11 | 0.451 | 0.115 | 0.905 | -0.006 | -0.021 | -0.021 | 0 | 0.39 | 0.69 | 0.51 | 0.49 | 0.41 | 0.27 |
| 6 | 4607325 | rs9502208 | -0.734 | 0.678 | -0.359 | 0.555 | 0.025 | -0.073 | -0.073 | 0 | 0.13 | 0.36 | 0.16 | 0.02 | 0.16 | 0.04 |
| 6 | 4608253 | rs2007513 | 0.002 | 0.801 | -1.401 | 0.769 | 0.024 | 0.07 | 0.07 | 0 | 0.22 | 0.38 | 0.28 | 0.33 | 0.28 | 0.18 |
| 6 | 4611829 | rs9405753 | -0.064 | 0.819 | -2.609 | 0.189 | -0.082 | 0.009 | 0.009 | 1.995 | 0.48 | 0.96 | 0.54 | 0.58 | 0.46 | 0.31 |
| 6 | 4615655 | rs12206701 | -0.258 | 0.285 | -1.826 | 0.126 | 0.008 | 0.028 | 0.028 | 0 | 0.26 | 0.12 | 0.12 | 0.2 | 0.22 | 0.23 |
| 6 | 4617817 | rs714112 | -0.378 | 0.466 | -1.865 | 0.113 | -0.01 | 0.022 | 0.022 | 1.177 | 0.32 | 0.12 | 0.28 | 0.25 | 0.28 | 0.33 |
| 6 | 4629654 | rs234427 | -1.006 | 0.202 | -1.438 | 0.382 | 0.011 | -0.003 | -0.003 | 0 | 0.19 | 0.83 | 0.17 | 0.15 | 0.13 | 0.11 |
| 6 | 4644601 | rs12204701 | -0.979 | 0.216 | -0.251 | 0.275 | 0.027 | -0.04 | -0.04 | 0.801 | 0.08 | 0.28 | 0.04 | 0 | 0.07 | 0.01 |
| 6 | 4646578 | rs1765828 | -1.502 | 0.138 | 0.746 | 0.316 | 0.004 | 0.04 | 0.04 | 0.39 | 0.11 | 0.6 | 0.13 | 0.11 | 0.07 | 0.01 |
| 6 | 4648010 | rs4960011 | -1.056 | 0.652 | 0.317 | 0.255 | -0.013 | -0.017 | -0.017 | 0 | 0.12 | 0.14 | 0.42 | 0.21 | 0.18 | 0.19 |
| 6 | 4651958 | rs9504181 | -0.296 | 0.735 | 0.547 | -0.063 | 0.056 | -0.034 | -0.034 | 0 | 0.09 | 0.3 | 0.1 | 0 | 0.1 | 0.01 |
| 6 | 4655223 | rs12215365 | -1.611 | 0.72 | 0.781 | 0.215 | -0.018 | 0.018 | 0.018 | 0.419 | 0.17 | 0.43 | 0.09 | 0.15 | 0.13 | 0.09 |
| 6 | 4655282 | rs1022249 | -0.469 | 0.864 | 0.763 | 0.199 | -0.039 | -0.032 | -0.032 | 0 | 0.09 | 0.04 | 0.42 | 0.17 | 0.21 | 0.15 |
| 6 | 4655316 | rs400004 | -2.772 | 0.833 | 0.808 | 0.184 | -0.018 | -0.024 | -0.024 | 0 | 0.3 | 0.84 | 0.54 | 0.32 | 0.34 | 0.25 |
| 6 | 4655490 | rs371778 | -0.833 | 0.961 | 0.306 | 0.248 | 0.001 | -0.001 | -0.001 | 0 | 0.11 | 0.16 | 0.43 | 0.17 | 0.21 | 0.16 |
| 6 | 4655992 | rs393587 | -2.232 | 0.932 | 0.301 | 0.182 | 0.025 | 0.034 | 0.034 | 0.017 | 0.28 | 0.6 | 0.52 | 0.32 | 0.34 | 0.25 |
| 6 | 4656073 | rs408603 | -1.465 | 1.212 | -0.361 | 0.395 | -0.003 | -0.003 | -0.003 | 0.112 | 0.18 | 0.21 | 0.46 | 0.27 | 0.27 | 0.21 |
| 6 | 4658893 | rs396367 | -0.666 | 1.046 | -0.564 | 0.399 | -0.008 | 0.001 | 0.001 | 0 | 0.22 | 0.14 | 0.13 | 0.21 | 0.21 | 0.29 |
| 6 | 4661844 | rs9378408 | -0.401 | 0.479 | -0.529 | 0.44 | 0.005 | -0.012 | -0.012 | 0 | 0.24 | 0.09 | 0.14 | 0.24 | 0.23 | 0.33 |
| 6 | 4662666 | rs13191293 | 0.456 | 0.353 | -0.428 | 0.699 | 0 | 0.012 | 0.012 | 0.093 | 0.15 | 0.18 | 0.25 | 0.25 | 0.16 | 0.16 |
| 6 | 4664262 | rs1971444 | -0.16 | 0.337 | -0.241 | 0.699 | 0.01 | -0.003 | -0.003 | 0 | 0.24 | 0.14 | 0.14 | 0.24 | 0.23 | 0.33 |
| 6 | 4665135 | rs429909 | 0.427 | 0.541 | -0.444 | 0.241 | -0.017 | -0.006 | -0.006 | 0.175 | 0.3 | 0.22 | 0.34 | 0.3 | 0.36 | 0.39 |
| 6 | 4666878 | rs437875 | -0.756 | 0.511 | 0.265 | 0.253 | -0.001 | -0.001 | -0.001 | 0 | 0.57 | 0.56 | 0.51 | 0.54 | 0.59 | 0.72 |
| 6 | 4668941 | rs761181 | 0.252 | 0.477 | 0.3 | -0.009 | -0.001 | 0.014 | 0.014 | 0.305 | 0.52 | 0.43 | 0.56 | 0.63 | 0.5 | 0.42 |
| 6 | 4670710 | rs9502219 | 1.592 | 0.437 | 0.139 | -0.033 | -0.016 | 0.003 | 0.003 | 0.41 | 0.15 | 0.08 | 0.23 | 0.14 | 0.21 | 0.24 |
| 6 | 4671240 | rs408884 | -0.863 | 0.423 | 0.195 | -0.046 | -0.001 | -0.014 | -0.014 | 0 | 0.94 | 0.93 | 0.96 | 1 | 0.94 | 0.98 |
| 6 | 4674242 | rs9392622 | 1.768 | 0.184 | -0.102 | -0.204 | -0.023 | -0.064 | -0.064 | 0.347 | 0.13 | 0.03 | 0.17 | 0.35 | 0.14 | 0.17 |
| 6 | 4674712 | rs432695 | 2.103 | 0.095 | 0.097 | -1.321 | -0.01 | 0.003 | 0.003 | 0.359 | 0.15 | 0.11 | 0.23 | 0.14 | 0.21 | 0.22 |
| 6 | 4676000 | rs392108 | 0.002 | -0.044 | 0.197 | -1.343 | -0.004 | -0.019 | -0.019 | 0.035 | 0.69 | 0.59 | 0.8 | 0.77 | 0.7 | 0.66 |
| 6 | 4676555 | rs7768553 | 0.204 | -0.063 | 0.167 | -1.349 | -0.008 | -0.032 | -0.032 | 0 | 0.67 | 0.53 | 0.8 | 0.77 | 0.7 | 0.66 |
| 6 | 4678260 | rs6918242 | 0.39 | -0.021 | 0.172 | -1.419 | -0.005 | 0.011 | 0.011 | 0 | 0.24 | 0.33 | 0.25 | 0.32 | 0.23 | 0.33 |
| 6 | 4679459 | rs9504197 | -0.02 | 0.055 | 0.017 | -1.424 | -0.001 | 0.002 | 0.002 | 0.29 | 0.54 | 0.58 | 0.48 | 0.54 | 0.5 | 0.41 |
| 6 | 4682543 | rs9502221 | 0.023 | -0.455 | 0.019 | -1.207 | 0.001 | 0 | 0 | 0.024 | 0.37 | 0.37 | 0.44 | 0.44 | 0.38 | 0.48 |
| 6 | 4683536 | rs6939368 | 0.927 | -0.489 | -0.53 | -1.275 | 0.005 | 0.017 | 0.017 | 1.544 | 0.18 | 0.14 | 0.11 | 0.03 | 0.17 | 0.05 |
| 6 | 4684896 | rs9504202 | 1.319 | 0.054 | -0.662 | -0.728 | -0.019 | -0.002 | -0.002 | 0 | 0.14 | 0.08 | 0.23 | 0.14 | 0.21 | 0.23 |
| 6 | 4688031 | rs6903613 | 0.344 | -0.134 | -0.67 | -0.924 | -0.001 | -0.006 | -0.006 | 0 | 0.57 | 0.53 | 0.52 | 0.55 | 0.6 | 0.46 |
| 6 | 4688861 | rs4960016 | 0.561 | 0.224 | -1.349 | -0.753 | -0.021 | -0.082 | -0.082 | 1.366 | 0.16 | 0.01 | 0.19 | 0.3 | 0.13 | 0.26 |
| 6 | 4690760 | rs2875955 | -0.669 | 0.661 | -1.144 | -0.528 | -0.011 | 0.059 | 0.059 | 1.547 | 0.61 | 0.76 | 0.56 | 0.24 | 0.59 | 0.47 |
| 6 | 4691632 | rs1513664 | -1.246 | 0.643 | -0.715 | -0.228 | -0.001 | -0.018 | -0.018 | 0 | 0.74 | 0.77 | 0.75 | 0.56 | 0.71 | 0.69 |
| 6 | 4694078 | rs5002346 | 0.898 | 1.036 | -1.114 | 0.085 | -0.072 | -0.027 | -0.027 | 0 | 0.15 | 0.01 | 0.23 | 0.15 | 0.22 | 0.25 |
| 6 | 4694899 | rs7747158 | -0.488 | 0.435 | -0.491 | 0.387 | -0.003 | 0.036 | 0.036 | 0 | 0.65 | 0.51 | 0.7 | 0.56 | 0.66 | 0.67 |
| 6 | 4701144 | rs4959312 | -0.479 | 0.794 | -1.336 | 0.439 | 0.041 | 0.025 | 0.025 | 0 | 0.5 | 0.35 | 0.61 | 0.47 | 0.57 | 0.62 |
| 6 | 4705849 | rs13197190 | -0.161 | 1.436 | -1.966 | 0.414 | -0.079 | 0.043 | 0.043 | 0 | 0.25 | 0.1 | 0.28 | 0.14 | 0.35 | 0.38 |
| 6 | 4711062 | rs1849564 | -0.472 | 1.047 | -1.963 | 0.782 | -0.064 | -0.082 | -0.082 | 0.409 | 0.23 | 0.02 | 0.31 | 0.33 | 0.22 | 0.25 |
| 6 | 4715297 | rs9378905 | 0.028 | 0.541 | -1.725 | 0.702 | -0.003 | -0.002 | -0.002 | 0 | 0.32 | 0.19 | 0.46 | 0.7 | 0.34 | 0.44 |
| 6 | 4715325 | rs6928621 | -0.336 | 0.347 | -1.035 | 0.806 | -0.006 | -0.002 | -0.002 | 0.013 | 0.07 | 0.1 | 0.05 | 0.05 | 0.05 | 0.03 |
| 6 | 4715882 | rs3812183 | -0.573 | 0.259 | -0.263 | 1.044 | 0 | 0.008 | 0.008 | 0.769 | 0.39 | 0.54 | 0.43 | 0.45 | 0.38 | 0.31 |
| 6 | 4715980 | rs3812180 | -0.039 | 0.098 | 0.3 | 1.112 | 0.013 | 0.013 | 0.013 | 0.508 | 0.22 | 0.27 | 0.32 | 0.27 | 0.22 | 0.05 |
| 6 | 4716016 | rs3812179 | -0.065 | 0.09 | 0.382 | 1.107 | 0.017 | 0.017 | 0.017 | 0.494 | 0.22 | 0.31 | 0.33 | 0.27 | 0.22 | 0.05 |
| 6 | 4716037 | rs3812178 | -0.065 | 0.09 | 0.382 | 1.107 | 0.017 | 0.017 | 0.017 | 0.494 | 0.22 | 0.31 | 0.33 | 0.27 | 0.22 | 0.05 |
| 6 | 4717213 | rs7760489 | -0.47 | -0.045 | 0.7 | 1.057 | 0.007 | 0.005 | 0.005 | 0.581 | 0.25 | 0.35 | 0.34 | 0.27 | 0.23 | 0.06 |
| 6 | 4721046 | rs7767658 | -0.373 | -0.056 | 0.946 | 1.237 | 0.004 | 0.057 | 0.057 | 0 | 0.09 | 0.25 | 0.06 | 0.15 | 0.08 | 0.16 |
| 6 | 4731340 | rs12203138 | -1.083 | 0.053 | 1.152 | 1.183 | 0.001 | 0.022 | 0.022 | 0 | 0.29 | 0.4 | 0.34 | 0.42 | 0.29 | 0.2 |
| 6 | 4731576 | rs6917691 | -0.432 | -0.088 | 1.142 | 1.167 | 0.004 | 0.06 | 0.06 | 0 | 0.09 | 0.32 | 0.07 | 0.15 | 0.08 | 0.15 |
| 6 | 4733025 | rs4960037 | -0.878 | -0.188 | 1.393 | 1.122 | 0.001 | -0.002 | -0.002 | 0 | 0.31 | 0.32 | 0.34 | 0.3 | 0.29 | 0.16 |
| 6 | 4735035 | rs13196069 | -0.904 | -0.268 | 1.418 | 1.097 | 0.003 | -0.001 | -0.001 | 1.798 | 0.31 | 0.31 | 0.33 | 0.29 | 0.27 | 0.16 |
| 6 | 4736583 | rs4960042 | 0.637 | -0.254 | 1.501 | 1.002 | 0 | 0.059 | 0.059 | 0 | 0.9 | 0.67 | 0.93 | 0.85 | 0.91 | 0.84 |

| | | | | | | | | | | | | | | | | |
|---|---|---|---|---|---|---|---|---|---|---|---|---|---|---|---|---|
| 6 | 4737179 | rs12665403 | -0.168 | -0.186 | 1.009 | 1.061 | -0.006 | -0.012 | -0.012 | 0 | 0.21 | 0.16 | 0.21 | 0.21 | 0.2 | 0.06 |
| 6 | 4738346 | rs9405775 | 1.085 | -0.175 | 0.992 | 1.05 | 0.007 | 0 | 0 | 1.241 | 0.59 | 0.36 | 0.66 | 0.62 | 0.64 | 0.68 |
| 6 | 4742265 | rs9328275 | 1.156 | -0.209 | 1.007 | 0.891 | 0.018 | 0.013 | 0.013 | 0.919 | 0.58 | 0.31 | 0.55 | 0.38 | 0.64 | 0.64 |
| 6 | 4744063 | rs6920511 | 0.912 | -0.161 | 1.208 | 0.211 | -0.042 | -0.012 | -0.012 | 0 | 0.81 | 0.56 | 0.87 | 0.82 | 0.84 | 0.74 |
| 6 | 4747159 | rs1040045 | 1.372 | 0.692 | 0.428 | -0.187 | -0.017 | 0.051 | 0.051 | 2.673 | 0.76 | 0.14 | 0.8 | 0.93 | 0.72 | 0.87 |
| 6 | 4762144 | rs4502986 | -1.722 | -0.132 | 1.146 | -0.153 | -0.035 | -0.02 | -0.02 | 0 | 0.2 | 0.34 | 0.1 | 0.11 | 0.11 | 0.27 |
| 6 | 4769767 | rs110284 | -1.211 | -0.062 | 2.123 | 0.045 | 0.058 | 0.007 | 0.007 | 0.205 | 0.06 | 0.32 | 0.07 | 0 | 0.06 | 0.02 |
| 6 | 4773086 | rs6928581 | -1.837 | -0.06 | 1.779 | 0.216 | 0.011 | 0.052 | 0.052 | 0 | 0.05 | 0.33 | 0.03 | 0.04 | 0.01 | 0.12 |
| 6 | 4787500 | rs6918380 | -1.808 | -0.035 | 0.783 | 0.08 | 0.012 | 0.01 | 0.01 | 0.022 | 0.15 | 0.12 | 0.05 | 0.08 | 0.08 | 0.28 |
| 6 | 4789259 | rs2038069 | -2.314 | 0.742 | 1.786 | 0.02 | 0.035 | -0.021 | -0.021 | 0.221 | 0.17 | 0.53 | 0.12 | 0.08 | 0.18 | 0.27 |
| 6 | 4799647 | rs11967994 | -2.084 | 1.028 | 2.472 | 0.207 | 0.026 | 0.04 | 0.04 | 0 | 0.11 | 0.23 | 0.14 | 0.11 | 0.11 | 0.19 |
| 6 | 4810410 | rs6918626 | -1.618 | 0.966 | 2.791 | 0.175 | -0.015 | 0.019 | 0.019 | 0 | 0.08 | 0.04 | 0.05 | 0.03 | 0.1 | 0.16 |
| 6 | 4810841 | rs9504259 | -2.325 | 0.844 | 2.956 | 0.044 | -0.003 | 0.006 | 0.006 | 0 | 0.11 | 0.08 | 0.08 | 0.05 | 0.13 | 0.16 |
| 6 | 4821146 | rs7747591 | -3.978 | 0.768 | 3.567 | 0.193 | -0.001 | 0.007 | 0.007 | 0.354 | 0.25 | 0.73 | 0.25 | 0.2 | 0.23 | 0.34 |
| 6 | 4824198 | rs7763686 | 2.144 | 0.661 | 2.528 | -0.923 | -0.049 | 0.115 | 0.115 | 0.077 | 0.55 | 0.07 | 0.64 | 0.82 | 0.59 | 0.5 |
| 6 | 4864003 | rs6926191 | -1.543 | 0.705 | 2.615 | -0.81 | -0.004 | -0.004 | -0.004 | 0 | 0.07 | 0.04 | 0.04 | 0.05 | 0.07 | 0.15 |
| 6 | 4864183 | rs12198622 | -1.658 | 0.467 | 2.422 | -1.05 | -0.017 | 0.037 | 0.037 | 1.07 | 0.11 | 0.05 | 0.11 | 0.01 | 0.16 | 0.09 |
| 6 | 4895977 | rs9405783 | 2.839 | 0.496 | 2.767 | -0.864 | 0 | -0.003 | -0.003 | 0 | 0.75 | 0.23 | 0.73 | 0.82 | 0.75 | 0.65 |
| 6 | 4896184 | rs6916304 | 1.836 | 0.506 | 2.856 | -0.79 | 0.064 | 0.21 | 0.21 | 4.569 | 0.95 | 0.38 | 0.88 | 0.91 | 0.99 | 0.97 |
| 6 | 4904380 | rs1417657 | 1.96 | 0.178 | 2.426 | -0.906 | 0.019 | 0.077 | 0.077 | 0 | 0.57 | 0.17 | 0.66 | 0.75 | 0.62 | 0.52 |
| 6 | 4910650 | rs1891047 | 2.084 | 0.612 | 2.421 | -0.563 | 0.015 | 0.056 | 0.056 | 0.347 | 0.83 | 0.32 | 0.78 | 0.78 | 0.81 | 0.81 |
| 6 | 4915028 | rs9405785 | 1.901 | 0.651 | 2.211 | -0.58 | -0.009 | 0.011 | 0.011 | 0 | 0.58 | 0.22 | 0.61 | 0.64 | 0.51 | 0.51 |
| 6 | 4915203 | rs4144546 | 1.9 | 0.549 | 2.635 | -0.253 | 0.007 | 0.007 | 0.007 | 0 | 0.87 | 0.74 | 0.91 | 0.86 | 0.86 | 0.83 |
| 6 | 4927010 | rs6905856 | 1.854 | 0.602 | 1.523 | -0.806 | -0.001 | 0.011 | 0.011 | 0 | 0.74 | 0.26 | 0.75 | 0.86 | 0.71 | 0.69 |
| 6 | 4929389 | rs7755300 | -2.094 | 0.783 | 2.08 | -0.747 | 0.031 | -0.011 | -0.011 | 0 | 0.12 | 0.16 | 0.11 | 0.04 | 0.2 | 0.15 |
| 6 | 4936863 | rs3804547 | 2.971 | 0.52 | 3.023 | -0.371 | -0.002 | -0.005 | -0.005 | 0 | 0.77 | 0.2 | 0.71 | 0.73 | 0.73 | 0.7 |
| 6 | 4939281 | rs7738117 | 2.432 | 0.764 | 2.923 | -0.645 | 0.037 | 0.048 | 0.048 | 0 | 0.74 | 0.52 | 0.72 | 0.73 | 0.67 | 0.69 |
| 6 | 4940123 | rs17284610 | -0.734 | 0.894 | 2.618 | -0.734 | -0.049 | -0.031 | -0.031 | 0.001 | 0.1 | 0.01 | 0.09 | 0.11 | 0.15 | 0.19 |
| 6 | 4940912 | rs4960053 | 2.673 | 0.844 | 3.044 | -0.758 | -0.012 | 0.004 | 0.004 | 0 | 0.8 | 0.4 | 0.77 | 0.83 | 0.81 | 0.87 |
| 6 | 4941853 | rs947154 | 2.003 | 0.78 | 3.26 | -0.768 | 0.005 | 0.13 | 0.13 | 0 | 0.92 | 0.58 | 0.86 | 0.9 | 0.98 | 0.96 |
| 6 | 4950275 | rs812134 | 2.369 | 0.794 | 2.242 | -0.904 | -0.001 | 0.004 | 0.004 | 0.453 | 0.62 | 0.18 | 0.62 | 0.67 | 0.62 | 0.61 |
| 6 | 4953879 | rs7776245 | -2.702 | 0.946 | 2.438 | -1.283 | 0.024 | -0.052 | -0.052 | 0 | 0.13 | 0.64 | 0.13 | 0.02 | 0.16 | 0.06 |
| 6 | 4956003 | rs2793207 | -0.337 | 0.883 | 1.781 | -1.464 | 0.003 | 0.003 | 0.003 | 0.059 | 0.05 | 0.04 | 0.07 | 0 | 0.04 | 0.01 |
| 6 | 4956221 | rs6597101 | 2.326 | 0.632 | 1.984 | -1.618 | -0.008 | -0.032 | -0.032 | 0 | 0.83 | 0.76 | 0.76 | 0.94 | 0.8 | 0.87 |
| 6 | 4958175 | rs795816 | 2.271 | 0.62 | 2.121 | -1.623 | -0.022 | -0.043 | -0.043 | 0 | 0.85 | 0.69 | 0.82 | 0.94 | 0.84 | 0.87 |
| 6 | 4958482 | rs6907963 | 2.044 | 0.364 | 1.545 | -2.098 | -0.016 | -0.051 | -0.051 | 0 | 0.62 | 0.5 | 0.53 | 0.82 | 0.63 | 0.65 |
| 6 | 4959847 | rs1246934 | -1.444 | 0.433 | 1.669 | -2.087 | 0.006 | -0.031 | -0.031 | 0.437 | 0.26 | 0.3 | 0.29 | 0.12 | 0.29 | 0.3 |
| 6 | 4960420 | rs2985678 | 1.574 | 0.396 | 1.728 | -2.063 | -0.011 | -0.023 | -0.023 | 0 | 0.89 | 0.75 | 0.83 | 0.94 | 0.86 | 0.88 |
| 6 | 4961165 | rs1246936 | 2.037 | 0.399 | 1.611 | -2.078 | 0 | -0.009 | -0.009 | 0.015 | 0.62 | 0.45 | 0.53 | 0.82 | 0.55 | 0.57 |
| 6 | 4961322 | rs2764158 | -1.566 | 0.3 | 1.507 | -2.028 | -0.008 | -0.055 | -0.055 | 0.018 | 0.27 | 0.37 | 0.32 | 0.12 | 0.24 | 0.22 |
| 6 | 4962641 | rs1246938 | -0.567 | 0.045 | 0.837 | -2.325 | -0.008 | -0.059 | -0.059 | 0.393 | 0.18 | 0.19 | 0.12 | 0 | 0.12 | 0.08 |
| 6 | 4967378 | rs2793208 | 0.821 | 0.001 | 0.719 | -2.525 | 0.002 | -0.066 | -0.066 | 0.662 | 0.72 | 0.7 | 0.72 | 0.98 | 0.68 | 0.75 |
| 6 | 4973879 | rs9405788 | 0.363 | -0.191 | 0.384 | -2.659 | -0.015 | 0.034 | 0.034 | 0 | 0.11 | 0.03 | 0.06 | 0.03 | 0.11 | 0.07 |
| 6 | 4978235 | rs2764176 | 0.686 | 0.002 | 0.669 | -2.481 | 0.004 | -0.083 | -0.083 | 0 | 0.72 | 0.68 | 0.71 | 0.98 | 0.68 | 0.75 |
| 6 | 4984231 | rs808734 | 0.938 | 0.015 | 0.575 | -2.466 | 0.011 | -0.042 | -0.042 | 0 | 0.91 | 0.77 | 0.85 | 0.98 | 0.84 | 0.84 |
| 6 | 4985005 | rs808733 | 0.14 | -0.275 | 0.671 | -2.671 | -0.007 | -0.03 | -0.03 | 0 | 0.26 | 0.29 | 0.26 | 0.12 | 0.21 | 0.2 |
| 6 | 4991024 | rs275259 | -0.681 | -0.552 | 1.452 | -1.953 | -0.028 | -0.087 | -0.087 | 0 | 0.3 | 0.46 | 0.29 | 0.12 | 0.24 | 0.26 |
| 6 | 4995017 | rs1246985 | 1.113 | -0.601 | 1.469 | -1.966 | -0.003 | -0.015 | -0.015 | 0 | 0.6 | 0.42 | 0.55 | 0.82 | 0.61 | 0.59 |
| 6 | 4998963 | rs1749144 | -1.147 | -0.613 | 1.534 | -1.978 | -0.021 | -0.051 | -0.051 | 0 | 0.37 | 0.52 | 0.32 | 0.17 | 0.28 | 0.36 |
| 6 | 5002737 | rs1749147 | -0.218 | -0.738 | 1.383 | -1.946 | -0.008 | -0.022 | -0.022 | 0 | 0.25 | 0.28 | 0.26 | 0.12 | 0.21 | 0.2 |
| 6 | 5004596 | rs431060 | 1.471 | -0.665 | 1.28 | -1.873 | 0.043 | 0.111 | 0.111 | 0.04 | 0.55 | 0.26 | 0.52 | 0.81 | 0.57 | 0.54 |
| 6 | 5006839 | rs707966 | -1.03 | -0.534 | 1.209 | -1.892 | -0.004 | -0.017 | -0.017 | 1.662 | 0.34 | 0.37 | 0.31 | 0.17 | 0.28 | 0.35 |
| 6 | 5024363 | rs2764142 | 1.687 | -0.215 | 1.573 | -1.202 | 0.031 | -0.027 | -0.027 | 0.649 | 0.92 | 0.76 | 0.84 | 0.99 | 0.85 | 0.89 |
| 6 | 5026772 | rs1108041 | -1.734 | -0.237 | 1.64 | -1.21 | 0.024 | -0.004 | -0.004 | 0 | 0.09 | 0.29 | 0.18 | 0.05 | 0.15 | 0.08 |
| 6 | 5029168 | rs877229 | 2.778 | -0.234 | 1.52 | -1.198 | -0.009 | -0.081 | -0.081 | 0 | 0.82 | 0.52 | 0.78 | 0.94 | 0.78 | 0.74 |
| 6 | 5030523 | rs17139235 | -1.517 | -0.153 | 1.24 | -1.149 | -0.01 | -0.021 | -0.021 | 0 | 0.09 | 0.2 | 0.06 | 0.05 | 0.07 | 0.15 |
| 6 | 5043326 | rs12200195 | -2.09 | -0.148 | 1.194 | -1.148 | -0.019 | -0.041 | -0.041 | 8.665 | 0.13 | 0.25 | 0.08 | 0.05 | 0.1 | 0.21 |
| 6 | 5044769 | rs2793259 | 2.229 | -0.091 | 1.312 | -1.022 | -0.005 | -0.064 | -0.064 | 3.395 | 0.84 | 0.53 | 0.79 | 0.94 | 0.81 | 0.79 |
| 6 | 5051556 | rs11964945 | -1.686 | -0.24 | 0.183 | -0.606 | -0.041 | -0.002 | -0.002 | 0 | 0.18 | 0.39 | 0.11 | 0.14 | 0.11 | 0.19 |
| 6 | 5054524 | rs11967105 | -1.06 | -0.071 | -0.937 | -0.349 | -0.001 | -0.001 | -0.001 | 0.012 | 0.13 | 0.15 | 0.08 | 0.14 | 0.07 | 0.08 |
| 6 | 5058794 | rs1246994 | -0.057 | 0.718 | -1.936 | -0.341 | -0.171 | -0.048 | -0.048 | 0.344 | 0.35 | 0.03 | 0.51 | 0.72 | 0.47 | 0.39 |
| 6 | 5059245 | rs1246996 | 0.276 | -0.079 | -1.265 | -1.214 | -0.014 | -0.043 | -0.043 | 0 | 0.41 | 0.38 | 0.31 | 0.09 | 0.27 | 0.23 |
| 6 | 5061580 | rs9405253 | 0.512 | -0.616 | -1.631 | -1.56 | 0.015 | 0.025 | 0.025 | 0 | 0.17 | 0.08 | 0.2 | 0.04 | 0.13 | 0.07 |
| 6 | 5064140 | rs11963503 | 0.01 | -0.906 | -1.007 | -1.857 | -0.001 | -0.021 | -0.021 | 5.247 | 0.25 | 0.23 | 0.1 | 0.03 | 0.16 | 0.2 |
| 6 | 5064920 | rs9405254 | -0.829 | -1.016 | -0.631 | -1.903 | -0.002 | -0.02 | -0.02 | 0 | 0.44 | 0.38 | 0.33 | 0.11 | 0.32 | 0.28 |
| 6 | 5067324 | rs4560685 | -1.531 | 0.157 | -1.273 | -2.319 | -0.05 | -0.029 | -0.029 | 0 | 0.67 | 0.98 | 0.6 | 0.26 | 0.65 | 0.65 |
| 6 | 5071492 | rs9392653 | -0.562 | -0.244 | -0.69 | -2.221 | 0.006 | -0.008 | -0.008 | 0.721 | 0.41 | 0.34 | 0.29 | 0.09 | 0.29 | 0.27 |
| 6 | 5075864 | rs666518 | -1.232 | -0.28 | -0.804 | -2.103 | -0.001 | 0.014 | 0.014 | 0 | 0.89 | 0.99 | 0.85 | 0.94 | 0.9 | 0.94 |
| 6 | 5080264 | rs376210 | 0.612 | -0.386 | -0.197 | -2.267 | -0.009 | -0.028 | -0.028 | 0.547 | 0.64 | 0.48 | 0.72 | 0.9 | 0.75 | 0.76 |

| | | | | | | | | | | | | | | | | |
|---|---|---|---|---|---|---|---|---|---|---|---|---|---|---|---|---|
| 6 | 5080904 | rs1265351 | 0.446 | -0.397 | -0.222 | -2.267 | -0.03 | -0.081 | -0.081 | 0 | 0.67 | 0.62 | 0.74 | 0.92 | 0.76 | 0.76 |
| 6 | 5082246 | rs200853 | 0.565 | -0.438 | -0.137 | -2.16 | -0.024 | -0.039 | -0.039 | 0 | 0.71 | 0.45 | 0.74 | 0.9 | 0.81 | 0.83 |
| 6 | 5084139 | rs6597110 | 0.655 | -0.175 | -0.151 | -1.911 | -0.015 | -0.038 | -0.038 | 0 | 0.12 | 0.19 | 0.06 | 0.02 | 0.07 | 0.1 |
| 6 | 5089278 | rs463951 | 0.746 | 0.526 | -0.799 | -2.168 | -0.017 | 0.15 | 0.15 | 0.486 | 0.47 | 0.18 | 0.54 | 0.77 | 0.52 | 0.45 |
| 6 | 5094746 | rs17139342 | -0.073 | 0.235 | 0.186 | -2.276 | -0.022 | -0.14 | -0.14 | 0.297 | 0.19 | 0.43 | 0.13 | 0.01 | 0.13 | 0.06 |
| 6 | 5095147 | rs462347 | -0.755 | 0.092 | -1.056 | -1.862 | 0.031 | -0.027 | -0.027 | 1.889 | 0.66 | 0.91 | 0.54 | 0.64 | 0.67 | 0.66 |
| 6 | 5096756 | rs6597112 | 0.472 | 0.6 | -0.772 | -1.973 | 0.002 | -0.005 | -0.005 | 3.596 | 0.11 | 0.14 | 0.04 | 0.07 | 0.1 | 0.28 |
| 6 | 5096972 | rs6911123 | 0.669 | 0.886 | -0.315 | -2.198 | 0.034 | -0.048 | -0.048 | 0 | 0.11 | 0.2 | 0.18 | 0 | 0.17 | 0.06 |
| 6 | 5100903 | rs17139385 | -0.005 | 0.717 | -0.368 | -2.104 | 0 | -0.002 | -0.002 | 0 | 0.13 | 0.17 | 0.05 | 0.13 | 0.11 | 0.38 |
| 6 | 5101520 | rs421473 | 0.525 | 0.536 | -0.439 | -2.279 | 0.008 | 0.014 | 0.014 | 0.105 | 0.58 | 0.4 | 0.66 | 0.79 | 0.69 | 0.49 |
| 6 | 5106588 | rs6923992 | 0.036 | 0.874 | -0.115 | -2.411 | -0.001 | -0.099 | -0.099 | 0 | 0.25 | 0.39 | 0.14 | 0.07 | 0.2 | 0.33 |
| 6 | 5106807 | rs685187 | -0.956 | 0.981 | 0.041 | -2.481 | 0.031 | -0.091 | -0.091 | 0 | 0.3 | 0.56 | 0.26 | 0.13 | 0.31 | 0.45 |
| 6 | 5107943 | rs11242990 | 0.796 | 1.066 | 0.064 | -2.384 | 0.052 | -0.053 | -0.053 | 2.293 | 0.11 | 0.27 | 0.18 | 0 | 0.17 | 0.06 |
| 6 | 5108965 | rs467373 | 0.624 | 1.04 | 0.082 | -2.4 | 0.073 | -0.061 | -0.061 | 0 | 0.12 | 0.35 | 0.19 | 0.01 | 0.17 | 0.06 |
| 6 | 5109119 | rs393628 | -1.184 | 0.948 | -0.137 | -2.413 | 0.004 | -0.034 | -0.034 | 0.059 | 0.89 | 0.98 | 0.83 | 0.88 | 0.92 | 0.82 |
| 6 | 5109592 | rs9606 | 1.354 | 0.901 | -0.147 | -2.477 | -0.027 | 0.171 | 0.171 | 0.276 | 0.47 | 0.12 | 0.47 | 0.8 | 0.52 | 0.43 |
| 6 | 5112814 | rs441482 | 0.293 | 0.879 | 0.129 | -2.502 | 0.007 | -0.085 | -0.085 | 0 | 0.64 | 0.5 | 0.54 | 0.91 | 0.63 | 0.81 |
| 6 | 5115596 | rs457759 | 1.038 | 0.892 | 0.051 | -2.546 | -0.021 | 0.173 | 0.173 | 2.349 | 0.52 | 0.19 | 0.49 | 0.84 | 0.53 | 0.61 |
| 6 | 5116366 | rs464681 | 1.15 | 0.922 | 0.003 | -2.521 | 0 | 0.186 | 0.186 | 0.721 | 0.49 | 0.17 | 0.48 | 0.8 | 0.52 | 0.45 |
| 6 | 5118251 | rs375064 | 0.574 | 0.986 | 0.047 | -2.516 | 0 | -0.037 | -0.037 | 0 | 0.61 | 0.47 | 0.54 | 0.86 | 0.62 | 0.71 |
| 6 | 5119062 | rs458108 | -1.482 | 1.14 | -0.128 | -2.42 | -0.008 | -0.021 | -0.021 | 0 | 0.92 | 1 | 0.84 | 0.93 | 0.93 | 0.92 |
| 6 | 5119696 | rs467304 | 1.312 | 1.15 | -0.398 | -2.343 | -0.107 | -0.138 | -0.138 | 0.001 | 0.21 | 0.01 | 0.22 | 0.62 | 0.29 | 0.29 |
| 6 | 5120585 | rs7754867 | 0.45 | 1.362 | 0.085 | -2.256 | 0.002 | -0.007 | -0.007 | 0 | 0.1 | 0.14 | 0.05 | 0.07 | 0.11 | 0.26 |
| 6 | 5124416 | rs373696 | -0.507 | 1.343 | 0.233 | -1.841 | 0.036 | -0.037 | -0.037 | 0.021 | 0.81 | 0.62 | 0.64 | 0.92 | 0.75 | 0.88 |
| 6 | 5130064 | rs11242992 | -0.661 | 1.362 | 0.1 | -1.851 | 0.043 | -0.046 | -0.046 | 0.016 | 0.23 | 0.42 | 0.24 | 0.12 | 0.29 | 0.41 |
| 6 | 5130155 | rs465481 | -1.468 | 1.341 | 0.098 | -1.848 | -0.003 | 0 | 0 | 0 | 0.91 | 0.89 | 0.83 | 0.92 | 0.93 | 0.92 |
| 6 | 5130429 | rs464606 | 0.829 | 1.364 | -0.019 | -1.83 | 0.032 | -0.04 | -0.04 | 0 | 0.67 | 0.46 | 0.59 | 0.8 | 0.64 | 0.51 |
| 6 | 5142842 | rs6597115 | 0.844 | 1.574 | 0.205 | -1.616 | 0.103 | -0.056 | -0.056 | 1.283 | 0.11 | 0.36 | 0.19 | 0 | 0.18 | 0.05 |
| 6 | 5144192 | rs399120 | -0.942 | 1.382 | 0.164 | -1.837 | -0.002 | -0.01 | -0.01 | 0 | 0.74 | 0.78 | 0.73 | 0.91 | 0.8 | 0.87 |
| 6 | 5147040 | rs10484845 | 0.158 | 1.477 | 0.206 | -1.525 | 0 | -0.001 | -0.001 | 0 | 0.12 | 0.15 | 0.05 | 0.14 | 0.11 | 0.33 |
| 6 | 5148115 | rs4960068 | 0.743 | 1.179 | 0.291 | -1.522 | 0.098 | -0.057 | -0.057 | 0 | 0.11 | 0.38 | 0.19 | 0 | 0.16 | 0.04 |
| 6 | 5148918 | rs6922265 | 0.344 | 0.907 | -0.045 | -1.377 | -0.004 | -0.005 | -0.005 | 0 | 0.1 | 0.14 | 0.04 | 0.08 | 0.07 | 0.27 |
| 6 | 5152786 | rs9378939 | 0.326 | 0.895 | -0.329 | -1.372 | 0.026 | 0.032 | 0.032 | 0 | 0.5 | 0.6 | 0.43 | 0.77 | 0.38 | 0.47 |
| 6 | 5154108 | rs4959332 | 0.466 | 0.943 | -0.206 | -1.367 | 0.004 | 0.032 | 0.032 | 0.3 | 0.53 | 0.4 | 0.47 | 0.76 | 0.49 | 0.52 |
| 6 | 5155210 | rs7766717 | -0.797 | 1.321 | -0.059 | -1.07 | 0.002 | -0.013 | -0.013 | 0 | 0.75 | 0.75 | 0.73 | 0.86 | 0.74 | 0.8 |
| 6 | 5157471 | rs6917225 | 0.701 | 1.069 | -0.673 | -1.161 | 0.022 | 0.038 | 0.038 | 0 | 0.35 | 0.28 | 0.31 | 0.15 | 0.31 | 0.26 |
| 6 | 5159064 | rs119408 | -0.169 | 1.404 | -0.76 | -0.494 | -0.003 | 0.002 | 0.002 | 0.941 | 0.39 | 0.4 | 0.4 | 0.7 | 0.4 | 0.52 |
| 6 | 5159417 | rs1016598 | 0.158 | 1.473 | -1.495 | -0.411 | -0.027 | -0.053 | -0.053 | 0 | 0.33 | 0.12 | 0.25 | 0.59 | 0.34 | 0.46 |
| 6 | 5162330 | rs11242995 | 0.22 | 1.385 | -1.446 | 0.136 | -0.096 | -0.114 | -0.114 | 0.421 | 0.3 | 0.05 | 0.35 | 0.52 | 0.39 | 0.48 |
| 6 | 5172009 | rs12661523 | 0.657 | 1.408 | -1.481 | 0.245 | -0.057 | -0.096 | -0.096 | 1.392 | 0.23 | 0.06 | 0.21 | 0.43 | 0.29 | 0.24 |
| 6 | 5172917 | rs1970010 | 0.226 | 0.396 | -0.876 | -0.357 | -0.004 | 0.004 | 0.004 | 0.475 | 0.28 | 0.49 | 0.31 | 0.26 | 0.26 | 0.25 |
| 6 | 5177985 | rs10458086 | 0.507 | 0.204 | -1.003 | -0.521 | -0.009 | -0.003 | -0.003 | 0 | 0.41 | 0.37 | 0.44 | 0.61 | 0.5 | 0.57 |
| 6 | 5180888 | rs445711 | 0.298 | 0.319 | -0.809 | -0.393 | -0.003 | -0.002 | -0.002 | 0 | 0.29 | 0.61 | 0.32 | 0.25 | 0.27 | 0.25 |
| 6 | 5199067 | rs2746228 | -0.064 | -0.197 | -0.749 | -0.437 | 0.035 | 0.121 | 0.121 | 2.402 | 0.62 | 0.86 | 0.69 | 0.86 | 0.69 | 0.8 |
| 6 | 5200311 | rs444607 | -0.046 | -0.196 | -0.727 | -0.436 | 0.036 | 0.102 | 0.102 | 2.256 | 0.62 | 0.83 | 0.69 | 0.86 | 0.69 | 0.8 |
| 6 | 5220965 | rs4607474 | 0.344 | -0.311 | -0.849 | -0.338 | -0.005 | 0.044 | 0.044 | 0.702 | 0.29 | 0.01 | 0.24 | 0.14 | 0.24 | 0.18 |
| 6 | 5253063 | rs1570085 | 0.138 | -0.264 | -0.296 | -0.042 | -0.033 | 0.026 | 0.026 | 0.629 | 0.25 | 0.56 | 0.26 | 0.26 | 0.19 | 0.15 |
| 6 | 5257610 | rs2753244 | 0.134 | -0.298 | -0.131 | -0.059 | -0.032 | 0.027 | 0.027 | 0.817 | 0.25 | 0.55 | 0.26 | 0.25 | 0.19 | 0.13 |
| 6 | 5261601 | rs9504370 | -0.446 | -0.322 | -0.301 | -0.246 | -0.005 | -0.002 | -0.002 | 0.171 | 0.33 | 0.66 | 0.32 | 0.25 | 0.25 | 0.16 |
| 6 | 5265853 | rs730594 | -0.623 | -0.38 | -0.983 | -1.076 | -0.002 | 0.066 | 0.066 | 0.899 | 0.71 | 0.96 | 0.76 | 0.86 | 0.75 | 0.78 |
| 6 | 5272087 | rs7768590 | 1.381 | -0.348 | -0.753 | -0.97 | -0.004 | -0.045 | -0.045 | 0 | 0.08 | 0.17 | 0.07 | 0 | 0.07 | 0.03 |
| 6 | 5281762 | rs9328293 | -0.067 | -0.311 | -0.943 | -1.353 | 0.002 | 0.126 | 0.126 | 0.116 | 0.58 | 0.8 | 0.7 | 0.89 | 0.61 | 0.74 |
| 6 | 5282415 | rs17140075 | -0.204 | -0.621 | -0.737 | -1.29 | -0.016 | -0.14 | -0.14 | 0 | 0.28 | 0.39 | 0.18 | 0 | 0.2 | 0.15 |
| 6 | 5282950 | rs1535298 | 1.063 | -0.484 | -1.2 | -1.168 | -0.081 | 0.096 | 0.096 | 1.959 | 0.39 | 0.14 | 0.44 | 0.81 | 0.5 | 0.65 |
| 6 | 5286306 | rs17140092 | 1.453 | -0.553 | -1.404 | -1.33 | -0.006 | -0.042 | -0.042 | 0.156 | 0.08 | 0.15 | 0.05 | 0 | 0.05 | 0.03 |
| 6 | 5291511 | rs17736905 | 1.279 | -0.425 | -1.896 | -1.25 | -0.028 | 0.076 | 0.076 | 0.003 | 0.14 | 0.03 | 0.1 | 0 | 0.13 | 0.07 |
| 6 | 5294170 | rs9392669 | 0.92 | -0.155 | -1.872 | -1.094 | -0.169 | -0.034 | -0.034 | 0 | 0.37 | 0.03 | 0.45 | 0.71 | 0.49 | 0.58 |
| 6 | 5294886 | rs7755157 | 0.914 | -0.147 | -1.869 | -1.092 | -0.168 | -0.034 | -0.034 | 0 | 0.37 | 0.03 | 0.45 | 0.72 | 0.49 | 0.58 |
| 6 | 5299689 | rs6905457 | -0.126 | 0.281 | -1.595 | -0.804 | -0.103 | -0.025 | -0.025 | 0 | 0.74 | 0.98 | 0.64 | 0.79 | 0.68 | 0.7 |
| 6 | 5304085 | rs6597125 | 0.335 | -0.054 | -0.88 | -0.939 | -0.005 | -0.028 | -0.028 | 0 | 0.66 | 0.57 | 0.57 | 0.8 | 0.65 | 0.83 |
| 6 | 5304851 | rs6924717 | 0.528 | 0.059 | -0.544 | -1.004 | 0 | -0.019 | -0.019 | 0 | 0.17 | 0.27 | 0.11 | 0.1 | 0.13 | 0.2 |
| 6 | 5306457 | rs9502289 | 0.054 | -0.177 | -0.207 | -0.917 | -0.037 | -0.065 | -0.065 | 0 | 0.14 | 0.42 | 0.09 | 0.03 | 0.07 | 0.03 |
| 6 | 5306757 | rs9405818 | -1.068 | 0.346 | -0.559 | -0.567 | -0.002 | 0 | 0 | 4.212 | 0.35 | 0.69 | 0.35 | 0.33 | 0.39 | 0.25 |
| 6 | 5307508 | rs6926122 | -0.982 | -0.457 | -0.425 | -0.74 | -0.016 | -0.059 | -0.059 | 0 | 0.26 | 0.68 | 0.17 | 0.03 | 0.17 | 0.1 |
| 6 | 5312080 | rs9405821 | 0.486 | 0.124 | -0.547 | -0.629 | -0.104 | -0.1 | -0.1 | 0 | 0.16 | 0.03 | 0.26 | 0.3 | 0.29 | 0.16 |
| 6 | 5320622 | rs708036 | -0.72 | 0.018 | -0.329 | -0.651 | 0.02 | 0.01 | 0.01 | 0 | 0.31 | 0.58 | 0.38 | 0.33 | 0.39 | 0.22 |
| 6 | 5337783 | rs2552298 | -0.735 | -0.024 | -0.697 | -0.776 | 0.005 | 0.016 | 0.016 | 0.504 | 0.51 | 0.56 | 0.69 | 0.8 | 0.59 | 0.53 |
| 6 | 5343889 | rs811275 | 0.667 | 0.01 | -0.467 | -0.753 | -0.017 | -0.048 | -0.048 | 0 | 0.11 | 0.29 | 0.09 | 0.03 | 0.1 | 0.02 |
| 6 | 5360749 | rs797125 | -1.569 | -0.02 | -1.307 | -0.817 | -0.021 | 0.07 | 0.07 | 0 | 0.64 | 0.85 | 0.75 | 0.79 | 0.66 | 0.62 |

| | | | | | | | | | | | | | | | | |
|---|---|---|---|---|---|---|---|---|---|---|---|---|---|---|---|---|
| 6 | 5362054 | rs797126 | -0.867 | -0.008 | -1.406 | -0.796 | 0.093 | -0.011 | -0.011 | 0 | 0.48 | 0.79 | 0.48 | 0.49 | 0.36 | 0.45 |
| 6 | 5364290 | rs9405828 | 0.778 | 0.312 | -1.649 | -0.639 | -0.104 | -0.098 | -0.098 | 0.013 | 0.16 | 0.03 | 0.27 | 0.29 | 0.29 | 0.15 |
| 6 | 5366604 | rs708037 | -0.137 | -0.749 | -0.821 | -0.585 | 0.005 | -0.067 | -0.067 | 0 | 0.13 | 0.44 | 0.13 | 0.03 | 0.13 | 0.03 |
| 6 | 5367273 | rs797147 | -0.959 | -1.426 | -0.767 | -0.549 | -0.064 | -0.127 | -0.127 | 0 | 0.24 | 0.53 | 0.15 | 0.03 | 0.15 | 0.05 |
| 6 | 5371349 | rs2432808 | -0.24 | -0.482 | -1.552 | 0.069 | 0.002 | -0.082 | -0.082 | 0 | 0.23 | 0.09 | 0.32 | 0.47 | 0.21 | 0.37 |
| 6 | 5374744 | rs17140294 | -0.476 | -0.206 | -0.904 | -0.379 | -0.001 | -0.022 | -0.022 | 0 | 0.11 | 0.11 | 0.05 | 0 | 0.07 | 0.03 |
| 6 | 5379058 | rs11243007 | -0.41 | -0.328 | -1.362 | -0.458 | 0.016 | 0.091 | 0.091 | 0.702 | 0.24 | 0.06 | 0.19 | 0.08 | 0.2 | 0.21 |
| 6 | 5379228 | rs2432749 | -0.715 | -0.388 | -1.122 | -0.187 | 0.011 | 0.007 | 0.007 | 0 | 0.7 | 0.69 | 0.53 | 0.63 | 0.53 | 0.72 |
| 6 | 5379525 | rs6932040 | 0.763 | -1.269 | -1.189 | -0.364 | 0.02 | 0.028 | 0.028 | 0 | 0.22 | 0.13 | 0.22 | 0.13 | 0.16 | 0.17 |
| 6 | 5384079 | rs2326627 | 0.291 | -1.496 | -0.311 | -0.434 | -0.034 | 0.112 | 0.112 | 0.612 | 0.21 | 0.43 | 0.11 | 0.42 | 0.14 | 0.34 |
| 6 | 5385623 | rs4295527 | -0.192 | -1.504 | -0.336 | -0.474 | -0.008 | 0.003 | 0.003 | 0 | 0.42 | 0.51 | 0.33 | 0.55 | 0.3 | 0.51 |
| 6 | 5386086 | rs4960084 | 1.091 | -1.463 | -0.371 | -0.459 | 0.015 | 0.031 | 0.031 | 0 | 0.21 | 0.08 | 0.22 | 0.13 | 0.16 | 0.17 |
| 6 | 5387513 | rs2432755 | -1.312 | -1.082 | -0.21 | -0.695 | -0.018 | -0.017 | -0.017 | 0 | 0.74 | 0.79 | 0.57 | 0.66 | 0.59 | 0.75 |
| 6 | 5388057 | rs6910655 | -0.14 | -1.071 | -0.145 | -0.595 | -0.003 | 0 | 0 | 0 | 0.43 | 0.53 | 0.34 | 0.58 | 0.3 | 0.52 |
| 6 | 5389729 | rs1895646 | -0.403 | -1.747 | -0.046 | -0.365 | 0.086 | 0 | 0 | 0.198 | 0.46 | 0.71 | 0.42 | 0.62 | 0.27 | 0.49 |
| 6 | 5389764 | rs17140384 | 0.772 | -1.808 | 0.513 | -0.364 | -0.028 | -0.038 | -0.038 | 0.286 | 0.07 | 0.23 | 0.03 | 0.02 | 0.03 | 0.08 |
| 6 | 5389883 | rs1895645 | -1.431 | -1.831 | 0.455 | -0.483 | -0.041 | -0.021 | -0.021 | 0 | 0.71 | 0.79 | 0.63 | 0.7 | 0.55 | 0.71 |
| 6 | 5390860 | rs2503814 | 0.481 | -2.075 | 0.399 | -0.441 | 0.028 | -0.045 | -0.045 | 0 | 0.27 | 0.17 | 0.27 | 0.49 | 0.19 | 0.34 |
| 6 | 5391419 | rs2432794 | 0.036 | -2.056 | 0.5 | -0.493 | -0.036 | -0.071 | -0.071 | 0 | 0.16 | 0.35 | 0.09 | 0.05 | 0.08 | 0.11 |
| 6 | 5395285 | rs2032989 | -0.643 | -2.002 | 0.125 | -0.484 | 0.073 | 0.152 | 0.152 | 0 | 0.48 | 0.76 | 0.39 | 0.21 | 0.37 | 0.4 |
| 6 | 5397254 | rs2432785 | 0.377 | -1.978 | 0.095 | -0.421 | -0.04 | -0.051 | -0.051 | 0 | 0.09 | 0.25 | 0.04 | 0.02 | 0.03 | 0.08 |
| 6 | 5398329 | rs2326606 | 0.756 | -1.669 | -0.509 | -0.249 | 0.006 | -0.16 | -0.16 | 1.509 | 0.23 | 0.01 | 0.25 | 0.49 | 0.17 | 0.32 |
| 6 | 5417143 | rs6904616 | 0.929 | -1.629 | -0.32 | -0.259 | -0.002 | 0 | 0 | 0 | 0.1 | 0.11 | 0.09 | 0.11 | 0.05 | 0.09 |
| 6 | 5419000 | rs713513 | -0.537 | -1.65 | -0.256 | -0.475 | 0 | 0.002 | 0.002 | 0 | 0.6 | 0.57 | 0.52 | 0.58 | 0.5 | 0.62 |
| 6 | 5424762 | rs11752854 | 0.771 | -1.36 | -0.72 | -0.079 | 0.002 | -0.155 | -0.155 | 0.789 | 0.21 | 0.01 | 0.23 | 0.48 | 0.16 | 0.3 |
| 6 | 5435274 | rs9405267 | 0.043 | -0.8 | -0.235 | -0.523 | 0.005 | -0.003 | -0.003 | 0 | 0.54 | 0.47 | 0.67 | 0.86 | 0.59 | 0.61 |
| 6 | 5435456 | rs2432761 | -0.055 | -0.938 | -0.318 | -0.527 | 0.007 | -0.013 | -0.013 | 0.117 | 0.62 | 0.76 | 0.6 | 0.49 | 0.66 | 0.53 |
| 6 | 5441653 | rs9405833 | 0.832 | -0.687 | -0.552 | -0.342 | -0.024 | -0.025 | -0.025 | 0 | 0.31 | 0.26 | 0.44 | 0.41 | 0.48 | 0.31 |
| 6 | 5444001 | rs200198 | -0.224 | -0.924 | -0.489 | -0.494 | 0.049 | -0.004 | -0.004 | 0.032 | 0.49 | 0.64 | 0.43 | 0.56 | 0.33 | 0.54 |
| 6 | 5446551 | rs2472866 | 0.015 | -0.729 | -0.456 | 0.08 | -0.028 | -0.059 | -0.059 | 0 | 0.23 | 0.1 | 0.26 | 0.32 | 0.25 | 0.31 |
| 6 | 5448113 | rs183221 | -0.654 | -0.789 | -0.515 | -0.061 | 0.015 | 0.052 | 0.052 | 0 | 0.8 | 0.9 | 0.86 | 0.97 | 0.8 | 0.83 |
| 6 | 5449240 | rs17140468 | 0.585 | -0.791 | -0.567 | 0.003 | -0.049 | -0.082 | -0.082 | 0 | 0.15 | 0.01 | 0.16 | 0.23 | 0.16 | 0.17 |
| 6 | 5452658 | rs414830 | 0.073 | -0.701 | -0.468 | -0.16 | -0.003 | -0.032 | -0.032 | 1.983 | 0.69 | 0.61 | 0.77 | 0.92 | 0.7 | 0.73 |
| 6 | 5454013 | rs2503838 | 0.074 | -0.611 | -1.332 | 0 | -0.027 | -0.075 | -0.075 | 0 | 0.23 | 0.1 | 0.28 | 0.38 | 0.26 | 0.32 |
| 6 | 5455161 | rs2791458 | -0.723 | -0.345 | -1.372 | 0.172 | -0.002 | 0.017 | 0.017 | 0.971 | 0.86 | 0.92 | 0.88 | 0.96 | 0.83 | 0.91 |
| 6 | 5457827 | rs9405834 | 0.417 | -0.215 | -1.364 | 0.427 | -0.033 | -0.032 | -0.032 | 0.521 | 0.39 | 0.14 | 0.43 | 0.43 | 0.4 | 0.34 |
| 6 | 5460662 | rs9405270 | 1.02 | 0.295 | -0.644 | 0.173 | -0.012 | 0.012 | 0.012 | 1.985 | 0.08 | 0.11 | 0.03 | 0.09 | 0.03 | 0.07 |
| 6 | 5461970 | rs2432798 | -0.109 | 1.019 | -1.148 | 0.112 | -0.047 | -0.024 | -0.024 | 0 | 0.09 | 0 | 0.11 | 0.09 | 0.16 | 0.16 |
| 6 | 5468259 | rs2503800 | -0.39 | 1.172 | -0.957 | 0.098 | -0.007 | -0.007 | -0.007 | 0 | 0.18 | 0.14 | 0.27 | 0.22 | 0.26 | 0.31 |
| 6 | 5472657 | rs760612 | -0.578 | 0.515 | -0.836 | -0.31 | 0.003 | -0.028 | -0.028 | 0 | 0.07 | 0.19 | 0.06 | 0.02 | 0.07 | 0.03 |
| 6 | 5475485 | rs433295 | 0.336 | 0.443 | -1.33 | -0.207 | -0.001 | 0.008 | 0.008 | 0 | 0.84 | 0.91 | 0.89 | 1 | 0.83 | 0.9 |
| 6 | 5480096 | rs200215 | -0.11 | 0.665 | -1.732 | 0.214 | -0.06 | 0.019 | 0.019 | 0.245 | 0.57 | 0.85 | 0.5 | 0.37 | 0.49 | 0.6 |
| 6 | 5481989 | rs2503801 | 0.26 | 0.024 | -0.675 | 0.452 | -0.01 | -0.001 | -0.001 | 0 | 0.62 | 0.51 | 0.66 | 0.32 | 0.68 | 0.57 |
| 6 | 5482597 | rs191833 | -0.485 | 0.242 | -0.164 | 0.483 | 0 | 0.001 | 0.001 | 0 | 0.49 | 0.51 | 0.52 | 0.45 | 0.47 | 0.63 |
| 6 | 5486254 | rs7774359 | -0.619 | 0.286 | 0.171 | 0.228 | 0 | -0.029 | -0.029 | 0 | 0.27 | 0.57 | 0.19 | 0.21 | 0.26 | 0.33 |
| 6 | 5486756 | rs7740897 | -0.939 | 0.247 | 0.677 | 0.273 | 0.036 | 0.089 | 0.089 | 0.971 | 0.11 | 0.52 | 0.1 | 0.21 | 0.07 | 0.2 |
| 6 | 5487630 | rs386501 | -0.764 | 0.185 | 0.375 | 0.675 | 0.001 | -0.002 | -0.002 | 0 | 0.37 | 0.63 | 0.57 | 0.33 | 0.34 | 0.38 |
| 6 | 5488980 | rs9504412 | 0.359 | 0.267 | 0.386 | 0.136 | 0 | -0.023 | -0.023 | 0 | 0.16 | 0.14 | 0.09 | 0.04 | 0.18 | 0.12 |
| 6 | 5489201 | rs9392082 | -0.222 | 0.103 | -0.384 | 0.378 | 0.004 | -0.008 | -0.008 | 0 | 0.22 | 0.15 | 0.13 | 0.35 | 0.22 | 0.25 |
| 6 | 5491723 | rs574716 | -0.533 | 0.549 | -0.369 | 0.217 | -0.006 | 0 | 0 | 0 | 0.33 | 0.24 | 0.3 | 0.39 | 0.35 | 0.35 |
| 6 | 5492019 | rs12204105 | -0.361 | 0.504 | -0.184 | -0.019 | -0.001 | -0.018 | -0.018 | 0 | 0.29 | 0.32 | 0.33 | 0.12 | 0.31 | 0.28 |
| 6 | 5496553 | rs2472867 | -0.177 | 0.127 | -0.451 | 0.284 | -0.002 | 0 | 0 | 0 | 0.72 | 0.72 | 0.86 | 0.67 | 0.77 | 0.85 |
| 6 | 5501389 | rs9405272 | -0.207 | 0.586 | -0.722 | 0.437 | -0.021 | -0.06 | -0.06 | 0 | 0.09 | 0.01 | 0.13 | 0.21 | 0.1 | 0.19 |
| 6 | 5507182 | rs9405840 | -0.636 | 0.514 | -0.259 | -0.26 | -0.002 | -0.002 | -0.002 | 0.091 | 0.44 | 0.56 | 0.49 | 0.36 | 0.58 | 0.55 |
| 6 | 5508618 | rs1015139 | 0.116 | 1.546 | 0.004 | -0.315 | 0.07 | 0.014 | 0.014 | 0 | 0.1 | 0.24 | 0.16 | 0.13 | 0.13 | 0.2 |
| 6 | 5510272 | rs9502309 | 0.413 | 1.657 | -0.176 | -0.294 | 0.005 | -0.001 | -0.001 | 0.321 | 0.09 | 0.12 | 0.06 | 0.13 | 0.13 | 0.19 |
| 6 | 5511420 | rs10900970 | 0.23 | 1.52 | -0.019 | -0.331 | 0 | 0 | 0 | 0 | 0.11 | 0.08 | 0.15 | 0.23 | 0.12 | 0.19 |
| 6 | 5513550 | rs12204218 | 0.46 | 1.359 | -0.291 | -0.706 | -0.081 | 0.116 | 0.116 | 0.041 | 0.19 | 0.03 | 0.16 | 0 | 0.31 | 0.13 |
| 6 | 5518218 | rs17140739 | 0.356 | 1.404 | -0.177 | -0.671 | 0.004 | -0.001 | -0.001 | 0 | 0.09 | 0.12 | 0.14 | 0.14 | 0.13 | 0.21 |
| 6 | 5520850 | rs6910841 | 0.2 | 0.553 | -0.073 | -1.171 | 0.003 | 0.003 | 0.003 | 0 | 0.14 | 0.11 | 0.18 | 0.05 | 0.1 | 0.07 |
| 6 | 5523184 | rs9328309 | 0.252 | 0.516 | 0.092 | -1.151 | -0.006 | 0.041 | 0.041 | 0.226 | 0.12 | 0.17 | 0.15 | 0.21 | 0.12 | 0.19 |
| 6 | 5523888 | rs7763423 | -1.363 | 0.378 | 0.096 | -1.26 | -0.004 | 0.105 | 0.105 | 0 | 0.5 | 0.75 | 0.54 | 0.26 | 0.54 | 0.41 |
| 6 | 5524393 | rs9405273 | -1.484 | 0.379 | 0.083 | -1.257 | -0.004 | 0.106 | 0.106 | 0 | 0.5 | 0.75 | 0.54 | 0.26 | 0.54 | 0.4 |
| 6 | 5531253 | rs6923467 | -0.524 | 0.316 | 0.299 | -0.595 | -0.014 | -0.049 | -0.049 | 0.058 | 0.18 | 0.33 | 0.22 | 0.05 | 0.13 | 0.08 |
| 6 | 5534271 | rs9405274 | -0.214 | 0.479 | -0.285 | -0.312 | 0.002 | 0.091 | 0.091 | 0.093 | 0.12 | 0.24 | 0.16 | 0.21 | 0.11 | 0.19 |
| 6 | 5541761 | rs4499976 | 0.562 | 0.774 | -0.776 | -0.73 | -0.003 | -0.043 | -0.043 | 0.287 | 0.85 | 0.87 | 0.79 | 0.98 | 0.87 | 0.9 |
| 6 | 5545046 | rs2326613 | 0.686 | 1.129 | -0.979 | -0.906 | -0.019 | 0.013 | 0.013 | 0 | 0.74 | 0.87 | 0.71 | 0.98 | 0.67 | 0.76 |
| 6 | 5545654 | rs7749599 | 0.455 | 0.363 | -0.495 | -0.279 | -0.002 | -0.042 | -0.042 | 0 | 0.88 | 0.89 | 0.84 | 0.98 | 0.92 | 0.9 |
| 6 | 5547614 | rs4960098 | -0.669 | -0.237 | -0.74 | 0.54 | 0.075 | -0.006 | -0.006 | 0 | 0.43 | 0.74 | 0.5 | 0.48 | 0.28 | 0.44 |

| | | | | | | | | | | | | | | | | |
|---|---|---|---|---|---|---|---|---|---|---|---|---|---|---|---|---|
| 6 | 5549885 | rs9504433 | -0.641 | -0.839 | 0.097 | 1.159 | -0.037 | 0.007 | 0.007 | 1.676 | 0.26 | 0.37 | 0.3 | 0.33 | 0.14 | 0.29 |
| 6 | 5551232 | rs6937827 | 0.756 | 0.008 | 0.236 | 1.955 | 0.055 | 0.128 | 0.128 | 0 | 0.82 | 0.65 | 0.72 | 0.63 | 0.77 | 0.65 |
| 6 | 5555588 | rs10484311 | -0.213 | -0.276 | -0.222 | 1.773 | -0.008 | 0.001 | 0.001 | 3.152 | 0.2 | 0.24 | 0.28 | 0.3 | 0.1 | 0.23 |
| 6 | 5560506 | rs7746509 | -1.362 | -0.305 | -0.012 | 2.006 | -0.021 | 0.019 | 0.019 | 0.299 | 0.32 | 0.39 | 0.39 | 0.52 | 0.22 | 0.47 |
| 6 | 5560766 | rs2326646 | -0.316 | -0.315 | -0.028 | 2.284 | 0.022 | -0.022 | -0.022 | 1.915 | 0.2 | 0.16 | 0.31 | 0.33 | 0.09 | 0.2 |
| 6 | 5561678 | rs4960100 | -0.896 | 0.525 | 0.049 | 2.08 | 0.004 | 0.036 | 0.036 | 0 | 0.06 | 0.09 | 0.04 | 0.16 | 0.07 | 0.19 |
| 6 | 5562546 | rs9378429 | 0.102 | 0.843 | -0.217 | 1.767 | -0.062 | -0.043 | -0.043 | 5.921 | 0.9 | 0.99 | 0.82 | 0.85 | 0.8 | 0.84 |
| 6 | 5567978 | rs11752390 | -0.826 | -0.019 | -0.293 | 2.05 | 0.043 | -0.029 | -0.029 | 0 | 0.2 | 0.1 | 0.21 | 0.21 | 0.14 | 0.23 |
| 6 | 5568083 | rs9392698 | 0.428 | 0.334 | 0.194 | 2.035 | 0.004 | -0.001 | -0.001 | 0 | 0.85 | 0.78 | 0.8 | 0.88 | 0.79 | 0.88 |
| 6 | 5571672 | rs17140870 | -0.986 | 0.037 | 0.622 | 2.427 | 0.039 | 0.252 | 0.252 | 1.36 | 0.1 | 0.51 | 0.23 | 0.27 | 0.06 | 0.17 |
| 6 | 5574319 | rs9378430 | -1.162 | 0.208 | 0.748 | 2.085 | 0.017 | -0.028 | -0.028 | 0 | 0.26 | 0.24 | 0.27 | 0.4 | 0.22 | 0.28 |
| 6 | 5575967 | rs4960104 | 1.46 | 0.245 | 0.749 | 2.058 | 0.006 | -0.014 | -0.014 | 0.073 | 0.6 | 0.62 | 0.53 | 0.49 | 0.58 | 0.6 |
| 6 | 5581224 | rs9378968 | -1.088 | 0.214 | 0.748 | 2.131 | 0.034 | -0.08 | -0.08 | 0 | 0.24 | 0.14 | 0.31 | 0.46 | 0.19 | 0.29 |
| 6 | 5584422 | rs6597147 | 0.321 | 0.204 | 0.74 | 1.707 | -0.005 | 0 | 0 | 0 | 0.85 | 0.84 | 0.8 | 0.88 | 0.8 | 0.87 |
| 6 | 5585770 | rs7767977 | -1.11 | 0.187 | 0.607 | 1.75 | 0.036 | -0.095 | -0.095 | 6.717 | 0.24 | 0.08 | 0.22 | 0.41 | 0.2 | 0.27 |
| 6 | 5592246 | rs2142736 | 0.236 | 0.267 | 0.628 | 1.424 | -0.026 | -0.01 | -0.01 | 0 | 0.86 | 0.93 | 0.84 | 0.87 | 0.8 | 0.87 |
| 6 | 5592955 | rs4960110 | 1.574 | 0.158 | 1.011 | 1.449 | -0.012 | 0.022 | 0.022 | 0.616 | 0.63 | 0.5 | 0.55 | 0.54 | 0.64 | 0.62 |
| 6 | 5593550 | rs2013289 | 0.314 | 0.543 | 0.623 | 1.653 | -0.028 | -0.002 | -0.002 | 0 | 0.87 | 0.93 | 0.84 | 0.88 | 0.8 | 0.87 |
| 6 | 5594199 | rs9504439 | 1.679 | 0.759 | 0.818 | 1.529 | 0.002 | -0.032 | -0.032 | 0 | 0.58 | 0.19 | 0.6 | 0.56 | 0.61 | 0.54 |
| 6 | 5594488 | rs9378433 | -0.355 | 0.761 | 0.649 | 1.579 | -0.018 | -0.106 | -0.106 | 0 | 0.16 | 0.01 | 0.15 | 0.32 | 0.14 | 0.26 |
| 6 | 5595194 | rs9504440 | -0.861 | 0.814 | 1.58 | 1.391 | 0.043 | -0.022 | -0.022 | 0 | 0.08 | 0.39 | 0.05 | 0 | 0.05 | 0.03 |
| 6 | 5596984 | rs9378969 | 1.722 | 0.899 | 1.498 | 1.589 | -0.011 | 0.03 | 0.03 | 0.05 | 0.59 | 0.26 | 0.59 | 0.7 | 0.59 | 0.61 |
| 6 | 5597160 | rs7744153 | 0.348 | 0.984 | 1.126 | 1.15 | -0.028 | -0.012 | -0.012 | 0 | 0.86 | 0.93 | 0.84 | 0.87 | 0.8 | 0.85 |
| 6 | 5598143 | rs9378434 | -1.327 | 0.944 | 1.452 | 1.207 | -0.004 | -0.004 | -0.004 | 0 | 0.24 | 0.27 | 0.31 | 0.21 | 0.22 | 0.26 |
| 6 | 5598984 | rs4323358 | -1.391 | 1.03 | 1.493 | 0.388 | 0 | -0.068 | -0.068 | 2.627 | 0.24 | 0.29 | 0.22 | 0.04 | 0.23 | 0.21 |
| 6 | 5599207 | rs9378971 | 1.867 | 1.106 | 1.353 | 0.428 | 0.004 | -0.045 | -0.045 | 0.076 | 0.61 | 0.49 | 0.63 | 0.86 | 0.59 | 0.66 |
| 6 | 5599657 | rs9378972 | 1.709 | 1.095 | 1.345 | 0.419 | 0.005 | -0.027 | -0.027 | 1.327 | 0.6 | 0.49 | 0.6 | 0.83 | 0.56 | 0.62 |
| 6 | 5604113 | rs9504449 | -1.565 | 0.974 | 0.983 | 0.393 | -0.005 | -0.115 | -0.115 | 0.357 | 0.26 | 0.46 | 0.22 | 0.04 | 0.22 | 0.23 |
| 6 | 5604836 | rs11755552 | -0.388 | 1.039 | 0.039 | 0.465 | -0.029 | 0.063 | 0.063 | 1.347 | 0.15 | 0.03 | 0.15 | 0.04 | 0.17 | 0.16 |
| 6 | 5607234 | rs941427 | -0.531 | 1.034 | 0.151 | 0.49 | 0.004 | -0.041 | -0.041 | 3 | 0.07 | 0.18 | 0.04 | 0 | 0.05 | 0.03 |
| 6 | 5608766 | rs7738833 | -1.208 | 1.051 | 0.133 | 0.49 | 0 | -0.005 | -0.005 | 0 | 0.23 | 0.21 | 0.2 | 0.03 | 0.21 | 0.17 |
| 6 | 5611577 | rs2092141 | 0.489 | 1.586 | -0.229 | 0.785 | -0.072 | -0.01 | -0.01 | 0 | 0.83 | 0.98 | 0.82 | 0.86 | 0.75 | 0.86 |
| 6 | 5618987 | rs11962015 | -0.985 | 1.361 | 0.02 | 0.627 | -0.013 | 0.031 | 0.031 | 3.042 | 0.16 | 0.14 | 0.18 | 0.04 | 0.18 | 0.23 |
| 6 | 5620993 | rs7746914 | -0.976 | 1.309 | 0.277 | 1.459 | 0.005 | 0.031 | 0.031 | 3.481 | 0.18 | 0.21 | 0.26 | 0.35 | 0.22 | 0.27 |
| 6 | 5622232 | rs4959341 | -0.862 | 1.14 | -0.177 | 0.757 | -0.017 | 0.006 | 0.006 | 0.599 | 0.46 | 0.27 | 0.57 | 0.59 | 0.49 | 0.47 |
| 6 | 5623410 | rs7743714 | -1.121 | 1.263 | 0.352 | 0.652 | 0.005 | 0.029 | 0.029 | 0.998 | 0.52 | 0.71 | 0.63 | 0.63 | 0.57 | 0.51 |
| 6 | 5625287 | rs2326632 | -0.046 | 0.57 | 0.411 | 0.683 | -0.012 | -0.026 | -0.026 | 0.23 | 0.31 | 0.4 | 0.35 | 0.22 | 0.29 | 0.14 |
| 6 | 5630468 | rs4960116 | -1.362 | 0.524 | -0.378 | 0.493 | -0.079 | 0.245 | 0.245 | 2.123 | 0.57 | 0.9 | 0.72 | 0.94 | 0.56 | 0.67 |
| 6 | 5630682 | rs4960117 | -0.036 | 0.867 | -0.472 | 0.963 | -0.002 | 0.008 | 0.008 | 0 | 0.17 | 0.17 | 0.25 | 0.4 | 0.24 | 0.29 |
| 6 | 5632356 | rs6597153 | -0.989 | 0.985 | -0.509 | 0.685 | 0.008 | 0.003 | 0.003 | 0.952 | 0.47 | 0.42 | 0.65 | 0.93 | 0.47 | 0.63 |
| 6 | 5636137 | rs9405858 | -0.012 | 1.252 | -1.155 | 0.93 | -0.001 | 0.011 | 0.011 | 0 | 0.17 | 0.17 | 0.25 | 0.41 | 0.25 | 0.3 |
| 6 | 5637740 | rs12525101 | -0.309 | 0.733 | -1.226 | 0.358 | -0.045 | -0.006 | -0.006 | 0 | 0.25 | 0.02 | 0.31 | 0.21 | 0.26 | 0.14 |
| 6 | 5644736 | rs7741352 | -0.677 | 0.907 | -0.334 | 0.123 | -0.005 | -0.03 | -0.03 | 0 | 0.29 | 0.59 | 0.3 | 0.18 | 0.26 | 0.21 |
| 6 | 5646316 | rs4960120 | 0.536 | 1.221 | -0.13 | -0.352 | 0.017 | -0.087 | -0.087 | 0 | 0.76 | 0.37 | 0.8 | 0.99 | 0.75 | 0.82 |
| 6 | 5648087 | rs10484314 | 0.754 | 1.166 | -0.776 | 0.154 | 0.092 | -0.007 | -0.007 | 0 | 0.49 | 0.17 | 0.43 | 0.58 | 0.37 | 0.52 |
| 6 | 5649266 | rs4960121 | 0.027 | 1.342 | -0.084 | 0.142 | 0.075 | -0.1 | -0.1 | 0 | 0.82 | 0.5 | 0.8 | 0.98 | 0.77 | 0.86 |
| 6 | 5653994 | rs4960124 | 0.622 | 1.193 | -0.57 | 0.245 | 0.07 | 0.006 | 0.006 | 1.881 | 0.51 | 0.29 | 0.44 | 0.57 | 0.38 | 0.56 |
| 6 | 5656275 | rs765878 | 0.675 | 1.172 | -0.611 | 0.774 | 0.039 | 0.059 | 0.059 | 3.211 | 0.51 | 0.37 | 0.46 | 0.3 | 0.39 | 0.47 |
| 6 | 5659478 | rs6597156 | 0.199 | 1.3 | -0.15 | 0.795 | 0.032 | 0.05 | 0.05 | 0.022 | 0.78 | 0.47 | 0.79 | 0.69 | 0.75 | 0.8 |
| 6 | 5661403 | rs1977057 | 0.435 | 1.285 | -0.839 | 0.798 | 0.002 | 0.003 | 0.003 | 0.279 | 0.76 | 0.28 | 0.77 | 0.67 | 0.73 | 0.78 |
| 6 | 5663598 | rs11243027 | -0.079 | 1.579 | -1.031 | 0.998 | -0.053 | -0.081 | -0.081 | 3.923 | 0.26 | 0.09 | 0.33 | 0.41 | 0.34 | 0.3 |
| 6 | 5664388 | rs1008198 | -0.383 | 1.684 | -1.01 | 0.786 | -0.003 | 0.032 | 0.032 | 0.24 | 0.43 | 0.26 | 0.44 | 0.7 | 0.54 | 0.41 |
| 6 | 5664464 | rs2064108 | 0.061 | 2.08 | -1.005 | 0.725 | -0.059 | 0.002 | 0.002 | 2.564 | 0.32 | 0.14 | 0.32 | 0.29 | 0.45 | 0.3 |
| 6 | 5665387 | rs12190311 | 0.408 | 1.971 | -0.66 | 0.332 | -0.005 | 0.081 | 0.081 | 4.114 | 0.51 | 0.24 | 0.42 | 0.65 | 0.45 | 0.59 |
| 6 | 5666064 | rs3828857 | 0.091 | 1.932 | 0.189 | 0.394 | 0.005 | -0.009 | -0.009 | 0 | 0.4 | 0.44 | 0.37 | 0.35 | 0.46 | 0.35 |
| 6 | 5676459 | rs9405859 | -0.13 | 1.693 | 0.625 | 0.083 | -0.023 | 0.066 | 0.066 | 0.233 | 0.44 | 0.2 | 0.43 | 0.29 | 0.51 | 0.41 |
| 6 | 5686407 | rs4959347 | -0.102 | 1.602 | 1.121 | 0.032 | 0 | 0.012 | 0.012 | 0 | 0.44 | 0.54 | 0.46 | 0.29 | 0.49 | 0.37 |
| 6 | 5694713 | rs12215495 | 0.482 | 1.26 | 0.065 | 0.053 | -0.023 | -0.059 | -0.059 | 1.509 | 0.13 | 0.01 | 0.06 | 0.15 | 0.11 | 0.22 |
| 6 | 5695126 | rs11752777 | -0.065 | 1.049 | 0.795 | -0.657 | -0.001 | 0.003 | 0.003 | 0.05 | 0.46 | 0.51 | 0.47 | 0.29 | 0.53 | 0.42 |
| 6 | 5701541 | rs9405281 | 0.404 | 0.912 | 0.036 | -0.602 | -0.08 | 0.048 | 0.048 | 0.275 | 0.4 | 0.07 | 0.4 | 0.27 | 0.51 | 0.38 |
| 6 | 5704014 | rs1005521 | 0.394 | 0.077 | 0.376 | -0.861 | 0.005 | 0.023 | 0.023 | 1.904 | 0.4 | 0.23 | 0.34 | 0.68 | 0.34 | 0.51 |
| 6 | 5710228 | rs13210778 | -0.111 | -0.34 | 0.674 | -1.511 | 0.001 | 0.001 | 0.001 | 0.376 | 0.52 | 0.49 | 0.47 | 0.18 | 0.56 | 0.45 |
| 6 | 5710354 | rs13191290 | 0.046 | -0.331 | 0.446 | -1.507 | 0.014 | 0.118 | 0.118 | 0.521 | 0.49 | 0.25 | 0.45 | 0.18 | 0.56 | 0.43 |
| 6 | 5712081 | rs1977058 | 0.111 | -0.374 | -0.342 | -1.637 | -0.028 | 0.228 | 0.228 | 1.644 | 0.48 | 0.1 | 0.44 | 0.15 | 0.55 | 0.45 |
| 6 | 5717397 | rs6918658 | 0.487 | -0.923 | 0.19 | -1.829 | -0.008 | 0.058 | 0.058 | 0.954 | 0.39 | 0.31 | 0.42 | 0.8 | 0.37 | 0.52 |
| 6 | 5722306 | rs11243032 | 0.704 | -1.089 | 0.104 | -1.876 | -0.035 | 0.122 | 0.122 | 0.787 | 0.37 | 0.16 | 0.41 | 0.78 | 0.37 | 0.52 |
| 6 | 5722398 | rs13206042 | -0.579 | -1.124 | 0.173 | -1.876 | -0.016 | 0.153 | 0.153 | 1.453 | 0.58 | 0.81 | 0.49 | 0.22 | 0.59 | 0.44 |
| 6 | 5723118 | rs7741438 | 0.132 | -1.661 | 0.457 | -2.453 | -0.004 | 0.021 | 0.021 | 0.131 | 0.55 | 0.64 | 0.59 | 0.84 | 0.5 | 0.72 |
| 6 | 5723227 | rs6924018 | -0.442 | -0.499 | 1.088 | -1.777 | -0.016 | -0.075 | -0.075 | 0 | 0.21 | 0.26 | 0.11 | 0.04 | 0.17 | 0.21 |

| | | | | | | | | | | | | | | | | |
|---|---|---|---|---|---|---|---|---|---|---|---|---|---|---|---|---|
| 6 | 5723400 | rs1569464 | 0.104 | -0.324 | 0.445 | -1.68 | -0.094 | 0.105 | 0.105 | 0 | 0.24 | 0.08 | 0.28 | 0.12 | 0.37 | 0.2 |
| 6 | 5732820 | rs6926774 | 1.277 | -0.428 | 0.548 | -1.533 | -0.058 | 0.267 | 0.267 | 1.244 | 0.46 | 0.08 | 0.52 | 0.83 | 0.38 | 0.54 |
| 6 | 5735207 | rs6597161 | 0.379 | 0.223 | 1.221 | -0.716 | -0.009 | 0.005 | 0.005 | 0.169 | 0.69 | 0.64 | 0.65 | 0.84 | 0.55 | 0.72 |
| 6 | 5737122 | rs2145980 | 0.444 | 0.138 | 1.174 | -0.639 | 0.011 | -0.031 | -0.031 | 0.081 | 0.69 | 0.55 | 0.64 | 0.8 | 0.55 | 0.72 |
| 6 | 5741436 | rs9504502 | -2.189 | -0.701 | 2.107 | 0.328 | -0.033 | 0.151 | 0.151 | 0 | 0.09 | 0.61 | 0.21 | 0.28 | 0.04 | 0.17 |
| 6 | 5744992 | rs7740019 | -1.847 | -0.764 | 2.615 | 0.1 | -0.02 | -0.059 | -0.059 | 0 | 0.18 | 0.27 | 0.11 | 0.06 | 0.11 | 0.18 |
| 6 | 5746943 | rs9504505 | -2.586 | -0.855 | 2.374 | 0.181 | 0.02 | -0.007 | -0.007 | 0 | 0.24 | 0.78 | 0.31 | 0.35 | 0.16 | 0.3 |
| 6 | 5773821 | rs7766672 | -2.411 | -0.884 | 2.358 | 0.213 | -0.004 | 0.001 | 0.001 | 0.041 | 0.23 | 0.71 | 0.32 | 0.35 | 0.14 | 0.3 |
| 6 | 5773955 | rs17133302 | -1.237 | -0.868 | 2.842 | 0.153 | -0.003 | -0.098 | -0.098 | 4.301 | 0.15 | 0.32 | 0.12 | 0.01 | 0.1 | 0.14 |
| 6 | 5777131 | rs1125513 | -1.965 | -0.793 | 2.246 | 0.087 | 0.008 | 0.1 | 0.1 | 0 | 0.11 | 0.33 | 0.23 | 0.25 | 0.11 | 0.13 |
| 6 | 5777514 | rs1535586 | -0.604 | -0.71 | 1.612 | 0.058 | 0.018 | -0.032 | -0.032 | 0 | 0.06 | 0.15 | 0.05 | 0 | 0.09 | 0.03 |
| 6 | 5785789 | rs1108304 | -2.06 | -0.732 | 1.58 | 0.318 | -0.005 | 0 | 0 | 0 | 0.22 | 0.71 | 0.32 | 0.31 | 0.13 | 0.29 |
| 6 | 5786114 | rs1815271 | -1.513 | -0.488 | 1.257 | 0.163 | -0.026 | 0.049 | 0.049 | 0 | 0.19 | 0.46 | 0.3 | 0.25 | 0.12 | 0.24 |
| 6 | 5788014 | rs4960137 | -1.48 | -0.299 | 1.077 | 0.199 | -0.031 | 0.066 | 0.066 | 0 | 0.19 | 0.47 | 0.3 | 0.27 | 0.12 | 0.22 |
| 6 | 5794948 | rs9502339 | -0.913 | -0.198 | 0.275 | -0.698 | 0.01 | 0.09 | 0.09 | 0.008 | 0.33 | 0.78 | 0.22 | 0.05 | 0.28 | 0.2 |
| 6 | 5800202 | rs9502340 | -0.765 | -0.658 | 1.224 | -0.57 | -0.035 | -0.09 | -0.09 | 0.018 | 0.25 | 0.41 | 0.21 | 0.06 | 0.18 | 0.25 |
| 6 | 5808446 | rs7758823 | -0.967 | -0.496 | 0.91 | -0.563 | 0.022 | 0.037 | 0.037 | 0.164 | 0.28 | 0.78 | 0.22 | 0.05 | 0.19 | 0.21 |
| 6 | 5822192 | rs1407795 | 0.502 | -0.12 | 0.647 | -0.62 | 0.011 | 0.064 | 0.064 | 0 | 0.4 | 0.24 | 0.37 | 0.13 | 0.34 | 0.41 |
| 6 | 5831567 | rs9392718 | 0.362 | -0.104 | 0.582 | -0.611 | 0.012 | 0.043 | 0.043 | 0 | 0.42 | 0.28 | 0.38 | 0.13 | 0.34 | 0.37 |
| 6 | 5842924 | rs9504549 | -0.32 | 0.294 | 0.305 | 0.179 | 0.005 | 0.01 | 0.01 | 0 | 0.68 | 0.61 | 0.63 | 0.6 | 0.62 | 0.72 |
| 6 | 5855453 | rs10900977 | 0.222 | 0.007 | 0.059 | -0.289 | -0.001 | -0.001 | -0.001 | 0 | 0.5 | 0.5 | 0.54 | 0.44 | 0.46 | 0.49 |
| 6 | 5855762 | rs4959359 | -0.608 | -0.048 | -0.599 | -0.2 | -0.062 | -0.06 | -0.06 | 0 | 0.69 | 0.84 | 0.69 | 0.47 | 0.6 | 0.61 |
| 6 | 5863343 | rs748903 | -0.256 | -0.54 | -0.535 | -0.268 | -0.014 | -0.066 | -0.066 | 0 | 0.14 | 0.04 | 0.14 | 0.34 | 0.14 | 0.13 |
| 6 | 5869552 | rs6914546 | -1.296 | -0.705 | -0.622 | -0.348 | 0.007 | -0.011 | -0.011 | 0.004 | 0.22 | 0.8 | 0.27 | 0.4 | 0.15 | 0.21 |
| 6 | 5877163 | rs12663663 | -0.327 | -0.657 | -0.78 | -0.351 | 0.001 | -0.009 | -0.009 | 0 | 0.16 | 0.17 | 0.15 | 0.34 | 0.14 | 0.14 |
| 6 | 5877846 | rs12528005 | -0.231 | -0.636 | -1.447 | -0.34 | -0.025 | -0.082 | -0.082 | 0 | 0.14 | 0.03 | 0.14 | 0.34 | 0.14 | 0.14 |
| 6 | 5878720 | rs6937541 | -0.067 | -0.539 | -1.533 | -0.875 | 0.016 | 0.04 | 0.04 | 0 | 0.19 | 0.07 | 0.1 | 0 | 0.17 | 0.16 |
| 6 | 5879417 | rs2146453 | -0.349 | 0.073 | -1.566 | -0.515 | 0.008 | -0.032 | -0.032 | 0 | 0.06 | 0.13 | 0.04 | 0 | 0.06 | 0.02 |
| 6 | 5884859 | rs1407793 | -0.455 | -0.002 | -1.385 | -0.198 | 0.048 | 0.044 | 0.044 | 0 | 0.41 | 0.76 | 0.36 | 0.33 | 0.31 | 0.36 |
| 6 | 5896220 | rs2181351 | -0.345 | 0.361 | -1.842 | -0.884 | 0.003 | 0.178 | 0.178 | 0.385 | 0.42 | 0.83 | 0.31 | 0.14 | 0.37 | 0.31 |
| 6 | 5896461 | rs12195664 | -0.226 | 1.438 | -2.171 | -0.672 | -0.032 | -0.014 | -0.014 | 0.806 | 0.11 | 0.06 | 0.18 | 0.15 | 0.21 | 0.19 |
| 6 | 5898375 | rs12193326 | -0.57 | 1.164 | -1.867 | -0.81 | -0.023 | -0.004 | -0.004 | 2.059 | 0.12 | 0.09 | 0.2 | 0.14 | 0.21 | 0.2 |
| 6 | 5900268 | rs1590328 | -0.777 | 1.164 | -1.867 | -0.81 | -0.005 | 0 | 0 | 0.006 | 0.13 | 0.13 | 0.2 | 0.14 | 0.21 | 0.2 |
| 6 | 5905209 | rs6597169 | -0.744 | 1.227 | -1.842 | -0.813 | -0.003 | 0 | 0 | 0 | 0.13 | 0.13 | 0.2 | 0.13 | 0.21 | 0.2 |
| 6 | 5906900 | rs9504579 | -1.612 | 0.431 | -0.97 | -1.507 | 0.011 | -0.064 | -0.064 | 0 | 0.28 | 0.44 | 0.27 | 0.13 | 0.29 | 0.26 |
| 6 | 5909164 | rs7775306 | -1.582 | 0.472 | -0.551 | -1.513 | 0.015 | -0.074 | -0.074 | 0.324 | 0.27 | 0.47 | 0.27 | 0.13 | 0.29 | 0.26 |
| 6 | 5910329 | rs1001290 | -1.69 | -0.346 | -0.426 | -2.041 | -0.002 | -0.023 | -0.023 | 0 | 0.37 | 0.61 | 0.32 | 0.15 | 0.35 | 0.37 |
| 6 | 5919551 | rs9328336 | 0.027 | -0.184 | -0.95 | -2.426 | 0.015 | 0.022 | 0.022 | 0 | 0.48 | 0.55 | 0.45 | 0.29 | 0.58 | 0.47 |
| 6 | 5923152 | rs998360 | 0.288 | -0.246 | -0.92 | -2.385 | -0.001 | -0.027 | -0.027 | 0.017 | 0.45 | 0.52 | 0.29 | 0.15 | 0.42 | 0.36 |
| 6 | 5927559 | rs13197916 | 0.693 | -0.353 | -0.507 | -2.591 | 0.005 | -0.034 | -0.034 | 0 | 0.34 | 0.36 | 0.26 | 0.15 | 0.36 | 0.23 |
| 6 | 5928289 | rs12193231 | 0.251 | -0.06 | -0.758 | -2.232 | -0.043 | -0.014 | -0.014 | 1.296 | 0.15 | 0.06 | 0.19 | 0.16 | 0.2 | 0.19 |
| 6 | 5934718 | rs6939701 | -0.59 | -0.175 | -0.704 | -2.264 | 0.006 | 0.005 | 0.005 | 0 | 0.65 | 0.72 | 0.51 | 0.32 | 0.65 | 0.57 |
| 6 | 5938368 | rs3018 | 0.001 | -0.297 | -0.444 | -2.328 | 0.003 | 0.059 | 0.059 | 1.536 | 0.5 | 0.59 | 0.32 | 0.16 | 0.46 | 0.37 |
| 6 | 5945951 | rs10458090 | -0.306 | -0.123 | -0.243 | -1.487 | 0.011 | 0 | 0 | 0 | 0.4 | 0.52 | 0.52 | 0.41 | 0.48 | 0.47 |
| 6 | 5946029 | rs9504599 | -0.263 | -0.533 | -0.454 | -1.875 | 0.019 | 0.012 | 0.012 | 1.402 | 0.21 | 0.12 | 0.07 | 0.02 | 0.13 | 0.11 |
| 6 | 5947139 | rs4959364 | -0.449 | -0.841 | -0.156 | -1.978 | 0.003 | 0.027 | 0.027 | 0 | 0.51 | 0.68 | 0.55 | 0.41 | 0.5 | 0.54 |
| 6 | 5947883 | rs9405884 | 0.712 | -0.615 | -0.001 | -1.998 | 0.004 | -0.017 | -0.017 | 0 | 0.26 | 0.32 | 0.23 | 0.17 | 0.29 | 0.23 |
| 6 | 5954189 | rs4492236 | -1.463 | -0.712 | -0.464 | -1.56 | -0.076 | -0.122 | -0.122 | 1.902 | 0.64 | 0.92 | 0.61 | 0.49 | 0.6 | 0.58 |
| 6 | 5958615 | rs11759185 | 0.444 | -0.629 | -0.518 | -1.523 | -0.067 | -0.059 | -0.059 | 3.171 | 0.12 | 0.03 | 0.27 | 0.24 | 0.18 | 0.24 |
| 6 | 5960053 | rs9504609 | -0.589 | -0.66 | -0.356 | -1.542 | 0.02 | 0.025 | 0.025 | 0.439 | 0.51 | 0.61 | 0.56 | 0.42 | 0.57 | 0.52 |
| 6 | 5961601 | rs1555536 | -0.086 | -0.352 | -0.211 | -1.32 | -0.01 | -0.02 | -0.02 | 0 | 0.23 | 0.17 | 0.29 | 0.3 | 0.21 | 0.31 |
| 6 | 5964831 | rs4960150 | 0.243 | -0.205 | 0.022 | -0.955 | 0.008 | -0.033 | -0.033 | 0.262 | 0.27 | 0.39 | 0.24 | 0.17 | 0.3 | 0.23 |
| 6 | 5969617 | rs6935001 | -0.611 | -0.017 | -0.064 | -0.644 | -0.001 | 0 | 0 | 0.112 | 0.65 | 0.69 | 0.65 | 0.49 | 0.66 | 0.61 |
| 6 | 5976788 | rs7745233 | -1.059 | -0.752 | -0.013 | 0.323 | -0.031 | -0.001 | -0.001 | 0 | 0.14 | 0.24 | 0.12 | 0.17 | 0.04 | 0.27 |
| 6 | 5977833 | rs1885728 | 0.24 | -1.032 | -0.05 | -0.109 | 0.008 | 0.014 | 0.014 | 0 | 0.55 | 0.55 | 0.49 | 0.17 | 0.67 | 0.42 |
| 6 | 5980711 | rs6942389 | 0.122 | -0.994 | 0.222 | 0.309 | -0.007 | -0.012 | -0.012 | 0 | 0.57 | 0.48 | 0.5 | 0.34 | 0.65 | 0.47 |
| 6 | 5983363 | rs11243046 | -1.054 | -0.789 | 0.355 | 0.519 | -0.048 | -0.006 | -0.006 | 0 | 0.12 | 0.36 | 0.05 | 0.09 | 0.03 | 0.25 |
| 6 | 5992490 | rs2208870 | -0.584 | -1.031 | -0.445 | 0.629 | -0.035 | -0.059 | -0.059 | 9.079 | 0.67 | 0.88 | 0.54 | 0.44 | 0.66 | 0.71 |
| 6 | 5994992 | rs12333117 | 0.601 | -0.815 | -0.323 | -0.121 | 0 | -0.003 | -0.003 | 0.137 | 0.35 | 0.33 | 0.36 | 0.31 | 0.4 | 0.35 |
| 6 | 5995651 | rs7748174 | -0.667 | -0.771 | -0.204 | 0.232 | -0.004 | -0.012 | -0.012 | 0 | 0.66 | 0.71 | 0.53 | 0.39 | 0.68 | 0.54 |
| 6 | 5997619 | rs1887131 | -0.969 | -0.612 | -0.452 | 0.477 | -0.02 | -0.084 | -0.084 | 0 | 0.68 | 0.89 | 0.57 | 0.52 | 0.69 | 0.73 |
| 6 | 6001381 | rs582186 | -0.223 | -0.697 | -0.274 | 0.113 | 0.001 | 0.003 | 0.003 | 1.461 | 0.45 | 0.55 | 0.56 | 0.73 | 0.38 | 0.53 |
| 6 | 6008062 | rs582677 | 0.537 | -0.57 | -0.635 | 0.764 | 0.013 | -0.04 | -0.04 | 0 | 0.31 | 0.15 | 0.32 | 0.39 | 0.26 | 0.23 |
| 6 | 6009848 | rs3763180 | -0.425 | 0.081 | -0.796 | 0.146 | -0.009 | -0.019 | -0.019 | 0 | 0.56 | 0.78 | 0.56 | 0.5 | 0.54 | 0.54 |
| 6 | 6010539 | rs4960155 | -0.45 | -0.004 | -0.867 | 0.02 | -0.005 | -0.01 | -0.01 | 0 | 0.52 | 0.78 | 0.52 | 0.5 | 0.51 | 0.47 |
| 6 | 6012391 | rs9379002 | -0.071 | 0.507 | -0.331 | -0.312 | -0.015 | 0.025 | 0.025 | 0 | 0.38 | 0.49 | 0.36 | 0.5 | 0.29 | 0.4 |
| 6 | 6013003 | rs9405284 | -0.752 | 0.535 | -0.238 | -0.259 | 0.027 | 0.011 | 0.011 | 0.283 | 0.25 | 0.52 | 0.32 | 0.28 | 0.31 | 0.38 |
| 6 | 6017169 | rs1475157 | 0.456 | 0.337 | -0.301 | -0.288 | 0.014 | 0.006 | 0.006 | 0 | 0.26 | 0.14 | 0.18 | 0.22 | 0.17 | 0.12 |
| 6 | 6019942 | rs667875 | 0.59 | 0.439 | -0.472 | -0.428 | 0.017 | 0.073 | 0.073 | 0.681 | 0.52 | 0.24 | 0.47 | 0.59 | 0.41 | 0.38 |

| | | | | | | | | | | | | | | | | |
|---|---|---|---|---|---|---|---|---|---|---|---|---|---|---|---|---|
| 6 | 6020072 | rs17141559 | 0.744 | 0.502 | -0.386 | -0.301 | 0.005 | -0.003 | -0.003 | 0 | 0.24 | 0.09 | 0.17 | 0.21 | 0.17 | 0.12 |
| 6 | 6027082 | rs9328341 | -0.09 | 0.692 | 0.123 | -0.124 | -0.002 | 0.071 | 0.071 | 0.137 | 0.13 | 0.28 | 0.18 | 0.29 | 0.12 | 0.36 |
| 6 | 6027104 | rs4959366 | 0.182 | 0.726 | -0.027 | -0.237 | -0.002 | 0.012 | 0.012 | 0 | 0.37 | 0.38 | 0.32 | 0.22 | 0.36 | 0.18 |
| 6 | 6034281 | rs17298948 | 0.933 | 0.707 | -0.213 | -0.409 | 0 | -0.042 | -0.042 | 0 | 0.21 | 0.23 | 0.15 | 0 | 0.2 | 0.07 |
| 6 | 6037120 | rs629744 | -1.35 | 0.777 | 0.012 | -0.292 | -0.007 | 0.008 | 0.008 | 0.355 | 0.8 | 0.85 | 0.81 | 0.88 | 0.75 | 0.76 |
| 6 | 6037327 | rs640887 | 1.355 | 0.707 | 0.049 | -0.21 | 0.006 | -0.014 | -0.014 | 0.045 | 0.29 | 0.22 | 0.3 | 0.37 | 0.25 | 0.2 |
| 6 | 6040482 | rs9504631 | 0.046 | 0.746 | 0.092 | -0.201 | 0.008 | 0.083 | 0.083 | 0.108 | 0.13 | 0.28 | 0.2 | 0.3 | 0.12 | 0.37 |
| 6 | 6047249 | rs9502396 | 0.173 | 0.739 | 0.142 | -0.201 | -0.002 | 0.027 | 0.027 | 0 | 0.18 | 0.19 | 0.19 | 0.33 | 0.18 | 0.16 |
| 6 | 6048115 | rs1819282 | 0.003 | 0.556 | 0.253 | -0.172 | 0.003 | 0.002 | 0.002 | 0 | 0.38 | 0.36 | 0.38 | 0.45 | 0.43 | 0.4 |
| 6 | 6048156 | rs1831016 | 0.247 | 0.556 | 0.228 | -0.202 | 0 | -0.019 | -0.019 | 0.049 | 0.33 | 0.42 | 0.32 | 0.17 | 0.32 | 0.4 |
| 6 | 6051117 | rs4959367 | 0.802 | 0.454 | 0.238 | -0.199 | -0.003 | -0.016 | -0.016 | 0 | 0.27 | 0.24 | 0.18 | 0 | 0.27 | 0.12 |
| 6 | 6051522 | rs1854005 | -1.495 | 0.431 | 0.057 | -0.156 | 0.007 | -0.016 | -0.016 | 0.822 | 0.71 | 0.77 | 0.7 | 0.63 | 0.75 | 0.8 |
| 6 | 6052055 | rs9405897 | 0.128 | 0.5 | 0.405 | -0.223 | 0.004 | -0.083 | -0.083 | 0 | 0.32 | 0.52 | 0.31 | 0.18 | 0.29 | 0.39 |
| 6 | 6054401 | rs13210587 | -0.042 | 0.576 | -0.043 | -0.426 | 0.001 | -0.027 | -0.027 | 0 | 0.35 | 0.36 | 0.25 | 0.12 | 0.36 | 0.22 |
| 6 | 6058587 | rs13212236 | 0.328 | 0.693 | 0.07 | -0.058 | 0.005 | -0.003 | -0.003 | 0.005 | 0.08 | 0.09 | 0.05 | 0.05 | 0.09 | 0.05 |
| 6 | 6058775 | rs6928498 | -0.676 | 0.664 | -0.2 | -0.225 | 0.012 | 0.089 | 0.089 | 0.023 | 0.48 | 0.84 | 0.33 | 0.31 | 0.4 | 0.22 |
| 6 | 6059613 | rs1482873 | 0.803 | 0.757 | -0.045 | 0.103 | -0.02 | -0.017 | -0.017 | 0 | 0.22 | 0.22 | 0.35 | 0.38 | 0.33 | 0.53 |
| 6 | 6063348 | rs17141663 | 1.096 | 0.601 | 0.163 | -0.002 | 0.005 | -0.014 | -0.014 | 0 | 0.11 | 0.08 | 0.1 | 0.18 | 0.09 | 0.06 |
| 6 | 6063424 | rs9502404 | 0.837 | 0.623 | 0.156 | -0.15 | 0 | -0.035 | -0.035 | 0 | 0.26 | 0.23 | 0.14 | 0 | 0.23 | 0.13 |
| 6 | 6064369 | rs2054641 | 0.859 | 0.632 | 0.16 | -0.145 | 0 | -0.036 | -0.036 | 1.62 | 0.26 | 0.23 | 0.14 | 0 | 0.23 | 0.13 |
| 6 | 6074202 | rs1905214 | -0.482 | 0.811 | -0.279 | -0.475 | 0.006 | -0.005 | -0.005 | 0 | 0.43 | 0.49 | 0.51 | 0.66 | 0.51 | 0.68 |
| 6 | 6080068 | rs922868 | 0.049 | 0.653 | -0.651 | -0.79 | -0.001 | -0.074 | -0.074 | 0 | 0.39 | 0.4 | 0.2 | 0.03 | 0.32 | 0.18 |
| 6 | 6080436 | rs6910246 | -0.885 | 0.751 | -0.765 | -0.682 | 0.003 | 0.015 | 0.015 | 0.667 | 0.66 | 0.72 | 0.47 | 0.26 | 0.66 | 0.54 |
| 6 | 6081071 | rs11757860 | 1.497 | 0.848 | -0.896 | -0.651 | -0.087 | 0.042 | 0.042 | 0 | 0.16 | 0.01 | 0.18 | 0.11 | 0.25 | 0.26 |
| 6 | 6082485 | rs4142291 | -1.323 | 0.665 | -0.852 | -0.509 | -0.024 | 0.024 | 0.024 | 0.124 | 0.72 | 0.62 | 0.62 | 0.65 | 0.76 | 0.82 |
| 6 | 6084559 | rs9392745 | 0.104 | 0.664 | -0.834 | -0.545 | -0.045 | -0.032 | -0.032 | 0 | 0.31 | 0.18 | 0.4 | 0.62 | 0.43 | 0.63 |
| 6 | 6087974 | rs4960164 | 0.933 | 0.814 | -0.676 | -1.006 | 0.015 | 0.077 | 0.077 | 0.385 | 0.26 | 0.14 | 0.12 | 0 | 0.17 | 0.13 |
| 6 | 6088055 | rs17141697 | 0.4 | 1.102 | -0.301 | -0.386 | 0.016 | -0.078 | -0.078 | 0 | 0.15 | 0.3 | 0.1 | 0.02 | 0.17 | 0.06 |
| 6 | 6091039 | rs1905210 | 0.312 | 1.029 | -0.53 | 0.078 | 0 | 0.014 | 0.014 | 0 | 0.08 | 0.09 | 0.06 | 0.12 | 0.09 | 0.1 |
| 6 | 6092876 | rs9504664 | -0.116 | 1.184 | -0.554 | -0.131 | 0.008 | -0.01 | -0.01 | 1.901 | 0.27 | 0.34 | 0.26 | 0.23 | 0.34 | 0.35 |
| 6 | 6093220 | rs9405900 | 0.289 | 1.408 | -0.805 | 0.437 | -0.024 | 0.026 | 0.026 | 0 | 0.06 | 0.01 | 0.16 | 0.39 | 0.09 | 0.28 |
| 6 | 6093449 | rs6915313 | 0.349 | 1.207 | -1.016 | -0.754 | 0.001 | 0.035 | 0.035 | 0 | 0.34 | 0.29 | 0.19 | 0.12 | 0.29 | 0.24 |
| 6 | 6094026 | rs1552195 | -0.171 | 1.152 | -1.03 | -0.769 | 0.036 | 0.139 | 0.139 | 0 | 0.51 | 0.3 | 0.37 | 0.24 | 0.54 | 0.49 |
| 6 | 6097536 | rs7755737 | 0.334 | 1.353 | -0.615 | -0.783 | 0.008 | -0.066 | -0.066 | 0 | 0.13 | 0.3 | 0.09 | 0.02 | 0.13 | 0.06 |
| 6 | 6114382 | rs2876006 | 0.671 | 1.514 | -1.196 | -0.337 | -0.085 | 0.042 | 0.042 | 0 | 0.16 | 0.02 | 0.18 | 0.11 | 0.25 | 0.25 |
| 6 | 6129217 | rs2220053 | 0.384 | 1.558 | -1.099 | -0.293 | -0.05 | 0.04 | 0.04 | 0 | 0.18 | 0.1 | 0.22 | 0.13 | 0.27 | 0.27 |
| 6 | 6129505 | rs1482871 | -0.967 | 0.994 | -1.426 | -0.237 | 0.012 | -0.038 | -0.038 | 0 | 0.74 | 0.82 | 0.65 | 0.52 | 0.75 | 0.79 |
| 6 | 6130218 | rs2326699 | -0.503 | 0.7 | -1.523 | -0.641 | 0.002 | 0.014 | 0.014 | 0.036 | 0.24 | 0.62 | 0.26 | 0.32 | 0.23 | 0.3 |
| 6 | 6131324 | rs1351656 | 0.306 | 0.321 | -1.381 | -0.799 | 0.05 | 0.129 | 0.129 | 0.209 | 0.38 | 0.18 | 0.19 | 0.1 | 0.26 | 0.24 |
| 6 | 6132436 | rs9502416 | -0.443 | 0.658 | -1.366 | -0.509 | 0.003 | -0.02 | -0.02 | 0 | 0.67 | 0.72 | 0.56 | 0.53 | 0.68 | 0.72 |
| 6 | 6132722 | rs6917409 | -0.291 | 1.037 | -1.531 | -0.639 | -0.009 | -0.005 | -0.005 | 0 | 0.53 | 0.67 | 0.39 | 0.44 | 0.49 | 0.48 |
| 6 | 6132752 | rs11756027 | 0.867 | 1.004 | -1.74 | -0.942 | -0.038 | 0.057 | 0.057 | 0 | 0.19 | 0.06 | 0.19 | 0.13 | 0.23 | 0.29 |
| 6 | 6133434 | rs13197045 | 0.556 | 0.954 | -1.099 | -0.973 | 0.011 | 0.053 | 0.053 | 0 | 0.37 | 0.29 | 0.25 | 0.11 | 0.31 | 0.22 |
| 6 | 6134186 | rs13212444 | 0.392 | 0.634 | -1.139 | -1.028 | -0.016 | 0.013 | 0.013 | 0 | 0.1 | 0.26 | 0.07 | 0.09 | 0.07 | 0.12 |
| 6 | 6137108 | rs2170618 | 0.646 | 0.634 | -1.329 | -1.204 | 0.014 | 0.052 | 0.052 | 0.788 | 0.3 | 0.21 | 0.2 | 0.09 | 0.23 | 0.19 |
| 6 | 6139569 | rs7772158 | -0.604 | 0.679 | -1.181 | -1.092 | -0.011 | 0.042 | 0.042 | 0 | 0.52 | 0.77 | 0.4 | 0.35 | 0.46 | 0.47 |
| 6 | 6143944 | rs9504688 | 0.728 | 0.731 | -1.067 | -1.09 | 0.013 | 0.038 | 0.038 | 0.788 | 0.3 | 0.22 | 0.2 | 0.09 | 0.23 | 0.19 |
| 6 | 6143961 | rs9502421 | 0.347 | 0.734 | -1.102 | -1.117 | 0.017 | 0.079 | 0.079 | 0 | 0.36 | 0.23 | 0.25 | 0.11 | 0.3 | 0.22 |
| 6 | 6144857 | rs3024466 | 1.766 | 0.689 | -1.225 | -1.16 | 0.004 | 0.054 | 0.054 | 4.687 | 0.22 | 0.1 | 0.15 | 0 | 0.16 | 0.08 |
| 6 | 6145409 | rs3024486 | 0.611 | 0.678 | -0.976 | -1.165 | -0.017 | 0.026 | 0.026 | 0 | 0.06 | 0 | 0.04 | 0.01 | 0.07 | 0.02 |
| 6 | 6145692 | rs1050783 | 1.54 | 0.678 | -0.966 | -1.158 | -0.007 | -0.084 | -0.084 | 0.076 | 0.23 | 0.27 | 0.16 | 0 | 0.16 | 0.08 |
| 6 | 6145726 | rs1050782 | -0.817 | 0.613 | -0.992 | -1.131 | -0.021 | 0.01 | 0.01 | 0 | 0.54 | 0.78 | 0.4 | 0.37 | 0.47 | 0.48 |
| 6 | 6148217 | rs1032045 | -0.114 | 0.616 | -1.06 | -1.183 | 0.005 | 0.003 | 0.003 | 0 | 0.46 | 0.57 | 0.33 | 0.36 | 0.37 | 0.44 |
| 6 | 6150122 | rs3778361 | 0.959 | 0.591 | -1.082 | -1.176 | 0.006 | -0.024 | -0.024 | 0 | 0.09 | 0 | 0.05 | 0.09 | 0.07 | 0.12 |
| 6 | 6151819 | rs3024459 | 1.172 | 0.495 | -1.346 | -1.492 | 0 | -0.021 | -0.021 | 0 | 0.11 | 0.1 | 0.07 | 0 | 0.11 | 0.03 |
| 6 | 6152555 | rs13213508 | 0.846 | 0.593 | -0.943 | -1.199 | 0.005 | -0.001 | -0.001 | 0.555 | 0.1 | 0.07 | 0.05 | 0.09 | 0.08 | 0.12 |
| 6 | 6153002 | rs13206518 | 0.879 | 0.621 | -0.966 | -1.211 | 0.001 | -0.018 | -0.018 | 0 | 0.1 | 0.01 | 0.05 | 0.09 | 0.08 | 0.12 |
| 6 | 6156004 | rs4960169 | 0.881 | 0.651 | -0.94 | -1.216 | 0.024 | 0.075 | 0.075 | 0.438 | 0.31 | 0.17 | 0.2 | 0.09 | 0.24 | 0.2 |
| 6 | 6157881 | rs3024455 | 0.935 | 0.687 | -0.913 | -1.198 | 0.02 | 0.065 | 0.065 | 0.494 | 0.31 | 0.19 | 0.2 | 0.09 | 0.24 | 0.2 |
| 6 | 6157957 | rs2326708 | -0.559 | 0.778 | -0.829 | -1.165 | 0.018 | 0.038 | 0.038 | 0.009 | 0.22 | 0.42 | 0.19 | 0.25 | 0.23 | 0.28 |
| 6 | 6158149 | rs381061 | -1.282 | 1.038 | -1.039 | -0.938 | -0.029 | -0.099 | -0.099 | 0 | 0.74 | 0.92 | 0.6 | 0.52 | 0.7 | 0.77 |
| 6 | 6159049 | rs4960171 | -0.651 | 0.519 | -0.67 | -0.958 | 0 | 0.017 | 0.017 | 0 | 0.53 | 0.62 | 0.37 | 0.34 | 0.45 | 0.48 |
| 6 | 6161109 | rs6927469 | 0.791 | 0.533 | -1.101 | -0.884 | 0.004 | 0.1 | 0.1 | 2.027 | 0.27 | 0.09 | 0.18 | 0.02 | 0.23 | 0.1 |
| 6 | 6161432 | rs9392752 | -0.202 | 0.858 | -0.335 | -0.074 | 0.001 | 0.06 | 0.06 | 0 | 0.13 | 0.23 | 0.11 | 0.26 | 0.13 | 0.24 |
| 6 | 6162060 | rs6928884 | -0.114 | 1.047 | -0.918 | -0.14 | -0.052 | -0.015 | -0.015 | 0.064 | 0.6 | 0.78 | 0.43 | 0.37 | 0.48 | 0.47 |
| 6 | 6163088 | rs6911594 | -0.677 | 0.987 | -0.328 | -0.34 | 0.009 | -0.018 | -0.018 | 0 | 0.38 | 0.52 | 0.36 | 0.3 | 0.51 | 0.56 |
| 6 | 6163492 | rs13202850 | 0.296 | 1.183 | -0.267 | -0.438 | -0.004 | -0.004 | -0.004 | 0 | 0.12 | 0.11 | 0.1 | 0.14 | 0.14 | 0.26 |
| 6 | 6164170 | rs4960173 | 0.966 | 0.669 | -0.249 | -0.717 | 0.063 | 0.083 | 0.083 | 2.061 | 0.39 | 0.23 | 0.22 | 0.08 | 0.22 | 0.2 |
| 6 | 6165468 | rs434602 | 0.038 | 0.818 | -0.463 | -0.487 | -0.014 | -0.01 | -0.01 | 1.171 | 0.36 | 0.2 | 0.34 | 0.62 | 0.38 | 0.4 |

| | | | | | | | | | | | | | | | |
|---|---|---|---|---|---|---|---|---|---|---|---|---|---|---|---|
| 6 | 6165834 | rs922866 | 1.43 | 0.632 | 0.227 | -0.803 | -0.053 | -0.155 | -0.155 | 1.875 | 0.3 | 0.43 | 0.2 | 0 | 0.16 | 0.06 |
| 6 | 6168194 | rs900401 | 1.295 | 0.71 | -0.142 | -0.297 | -0.01 | -0.086 | -0.086 | 0 | 0.19 | 0.06 | 0.23 | 0.53 | 0.22 | 0.22 |
| 6 | 6169218 | rs3823193 | -0.48 | 0.705 | 0.167 | -0.755 | -0.012 | 0.001 | 0.001 | 0 | 0.24 | 0.34 | 0.15 | 0.21 | 0.2 | 0.39 |
| 6 | 6169656 | rs7769202 | -0.099 | 0.737 | -0.139 | -1.056 | 0.007 | -0.003 | -0.003 | 0 | 0.29 | 0.32 | 0.27 | 0.26 | 0.36 | 0.42 |
| 6 | 6170449 | rs1609584 | 0.182 | 0.101 | -0.253 | -1.247 | -0.014 | -0.105 | -0.105 | 0 | 0.35 | 0.39 | 0.23 | 0.02 | 0.22 | 0.09 |
| 6 | 6171672 | rs755656 | 0 | 0.324 | -0.284 | -0.975 | -0.001 | -0.016 | -0.016 | 0 | 0.54 | 0.48 | 0.33 | 0.14 | 0.47 | 0.53 |
| 6 | 6172302 | rs406238 | -0.447 | 0.5 | -0.478 | 0.021 | -0.028 | -0.035 | -0.035 | 0.31 | 0.75 | 0.84 | 0.45 | 0.62 | 0.63 | 0.73 |
| 6 | 6172680 | rs387528 | -0.456 | 0.482 | -0.478 | 0.029 | -0.028 | -0.034 | -0.034 | 0.035 | 0.75 | 0.84 | 0.44 | 0.62 | 0.64 | 0.73 |
| 6 | 6173370 | rs17141840 | -0.072 | -0.061 | -0.372 | -0.124 | -0.024 | 0.024 | 0.024 | 0.009 | 0.15 | 0.29 | 0.11 | 0.16 | 0.11 | 0.15 |
| 6 | 6173987 | rs786735 | 0.478 | 0.18 | -0.747 | -0.555 | -0.001 | -0.007 | -0.007 | 0 | 0.2 | 0.21 | 0.15 | 0.1 | 0.2 | 0.17 |
| 6 | 6174856 | rs5986 | 0.405 | 0.407 | -1.128 | -0.482 | -0.006 | 0.033 | 0.033 | 0 | 0.11 | 0.04 | 0.07 | 0 | 0.11 | 0.03 |
| 6 | 6174866 | rs5982 | 0.211 | 0.264 | -0.869 | -0.106 | 0.008 | 0.006 | 0.006 | 3.549 | 0.29 | 0.15 | 0.1 | 0.36 | 0.21 | 0.38 |
| 6 | 6175591 | rs407447 | -0.101 | 0.076 | -0.631 | 0.199 | -0.011 | -0.003 | -0.003 | 0 | 0.32 | 0.43 | 0.28 | 0.27 | 0.31 | 0.31 |
| 6 | 6175975 | rs11969912 | -0.329 | 0.614 | -0.442 | 0.297 | 0.006 | -0.013 | -0.013 | 0 | 0.34 | 0.36 | 0.27 | 0.34 | 0.34 | 0.38 |
| 6 | 6176580 | rs7740009 | 0.399 | -0.032 | -0.48 | 0.854 | -0.011 | 0 | 0 | 0.496 | 0.58 | 0.44 | 0.58 | 0.4 | 0.67 | 0.48 |
| 6 | 6176625 | rs7774391 | 0.353 | -0.063 | -0.407 | 0.903 | -0.011 | 0.001 | 0.001 | 3.212 | 0.58 | 0.43 | 0.59 | 0.4 | 0.69 | 0.47 |
| 6 | 6177044 | rs3778354 | 0.725 | 0.291 | -0.533 | 0.946 | -0.001 | -0.001 | -0.001 | 0 | 0.51 | 0.47 | 0.49 | 0.38 | 0.51 | 0.43 |
| 6 | 6178328 | rs3116567 | 1.005 | 0.246 | 0.078 | 1.414 | 0.012 | 0.164 | 0.164 | 0 | 0.82 | 0.6 | 0.77 | 0.57 | 0.84 | 0.65 |
| 6 | 6179882 | rs3778353 | -0.523 | 0.144 | -0.205 | 0.841 | -0.02 | 0.024 | 0.024 | 0 | 0.28 | 0.23 | 0.24 | 0.17 | 0.32 | 0.21 |
| 6 | 6181035 | rs3799559 | -0.68 | -0.431 | 0.099 | 1.39 | -0.034 | 0.082 | 0.082 | 0 | 0.23 | 0.35 | 0.28 | 0.52 | 0.13 | 0.37 |
| 6 | 6181526 | rs3024443 | -0.59 | -0.486 | -0.118 | 1.395 | -0.024 | 0.044 | 0.044 | 0 | 0.23 | 0.3 | 0.27 | 0.52 | 0.13 | 0.37 |
| 6 | 6182605 | rs2274393 | -0.782 | -0.454 | -0.196 | 1.149 | -0.022 | 0.014 | 0.014 | 0 | 0.35 | 0.51 | 0.34 | 0.56 | 0.25 | 0.44 |
| 6 | 6184819 | rs3799558 | -0.935 | 0.038 | -0.583 | 0.202 | -0.039 | 0.024 | 0.024 | 0 | 0.64 | 0.76 | 0.63 | 0.75 | 0.54 | 0.66 |
| 6 | 6185017 | rs3799557 | 0.438 | -0.414 | -0.521 | 0.456 | 0.012 | -0.007 | -0.007 | 0 | 0.22 | 0.15 | 0.26 | 0.25 | 0.14 | 0.35 |
| 6 | 6191521 | rs10085324 | 0.361 | -0.326 | -0.26 | 0.435 | -0.005 | 0.204 | 0.204 | 0 | 0.05 | 0.14 | 0.08 | 0.47 | 0.04 | 0.09 |
| 6 | 6196604 | rs3024436 | -1.077 | -0.882 | -0.487 | -0.018 | -0.057 | -0.004 | -0.004 | 0 | 0.6 | 0.85 | 0.45 | 0.68 | 0.54 | 0.67 |
| 6 | 6197412 | rs3024481 | 0.96 | -0.9 | -0.482 | 0.024 | 0.006 | 0.001 | 0.001 | 0 | 0.06 | 0.03 | 0 | 0 | 0.01 | 0 |
| 6 | 6198195 | rs3024429 | 0.267 | -0.832 | -0.476 | 0.091 | -0.002 | 0.098 | 0.098 | 0 | 0.06 | 0.08 | 0.05 | 0.45 | 0.06 | 0.11 |
| 6 | 6201515 | rs7746985 | 0.798 | -1.013 | -0.356 | -0.28 | 0 | -0.025 | -0.025 | 0.23 | 0.15 | 0.16 | 0.12 | 0 | 0.16 | 0.08 |
| 6 | 6202734 | rs1482868 | -0.373 | -1.028 | -0.514 | -0.333 | 0.027 | 0.067 | 0.067 | 0 | 0.42 | 0.25 | 0.26 | 0.24 | 0.34 | 0.48 |
| 6 | 6204341 | rs9392757 | 0.735 | -0.846 | -0.851 | -0.138 | -0.001 | 0.032 | 0.032 | 0 | 0.05 | 0 | 0.03 | 0.44 | 0.04 | 0.1 |
| 6 | 6205842 | rs2085575 | -1.941 | -0.893 | -0.809 | -0.301 | -0.071 | -0.034 | -0.034 | 0 | 0.68 | 0.86 | 0.45 | 0.69 | 0.57 | 0.69 |
| 6 | 6211419 | rs3024422 | 0.564 | -0.801 | -0.449 | -0.357 | -0.002 | -0.047 | -0.047 | 0 | 0.15 | 0.19 | 0.12 | 0 | 0.15 | 0.06 |
| 6 | 6218703 | rs3799552 | 0.678 | -0.936 | -0.915 | -0.611 | 0.007 | 0.002 | 0.002 | 0 | 0.21 | 0.16 | 0.13 | 0.22 | 0.14 | 0.34 |
| 6 | 6220029 | rs1993552 | 0.088 | -0.772 | -0.887 | -0.626 | -0.011 | 0.021 | 0.021 | 0 | 0.16 | 0.11 | 0.13 | 0.02 | 0.2 | 0.18 |
| 6 | 6222737 | rs3024415 | -1.556 | -0.728 | -1.207 | -0.551 | -0.048 | -0.029 | -0.029 | 0 | 0.69 | 0.84 | 0.46 | 0.69 | 0.58 | 0.68 |
| 6 | 6223294 | rs3024409 | -0.122 | -0.609 | -1.61 | -0.516 | 0.023 | 0.067 | 0.067 | 0 | 0.35 | 0.17 | 0.24 | 0.21 | 0.3 | 0.37 |
| 6 | 6226147 | rs9502425 | -0.772 | 0.077 | -0.759 | -0.149 | 0 | 0.074 | 0.074 | 0 | 0.07 | 0.11 | 0.05 | 0.45 | 0.06 | 0.11 |
| 6 | 6226951 | rs1742928 | 0.53 | 0.019 | -0.884 | -0.648 | 0.026 | -0.005 | -0.005 | 0 | 0.59 | 0.36 | 0.65 | 0.52 | 0.66 | 0.67 |
| 6 | 6228025 | rs1742929 | -0.753 | 1.397 | -0.902 | -1.028 | 0.006 | 0 | 0 | 0 | 0.11 | 0.12 | 0.07 | 0.16 | 0.13 | 0.15 |
| 6 | 6229348 | rs1781785 | -0.696 | 1.724 | -1.168 | -1.347 | 0.002 | -0.021 | -0.021 | 0 | 0.45 | 0.52 | 0.46 | 0.22 | 0.45 | 0.39 |
| 6 | 6241281 | rs1781795 | -0.787 | 1.412 | -1.807 | -0.783 | -0.019 | -0.01 | -0.01 | 0.284 | 0.09 | 0.05 | 0.07 | 0.15 | 0.13 | 0.15 |
| 6 | 6243797 | rs1742925 | -0.405 | 0.796 | -1.8 | -0.874 | 0.059 | 0.135 | 0.135 | 1.08 | 0.55 | 0.76 | 0.55 | 0.22 | 0.66 | 0.48 |
| 6 | 6246984 | rs1267843 | -0.203 | 1.044 | -1.54 | -0.668 | -0.006 | 0.097 | 0.097 | 0 | 0.43 | 0.75 | 0.44 | 0.16 | 0.53 | 0.33 |
| 6 | 6249276 | rs3024391 | -0.093 | 0.978 | -2.019 | 0.067 | -0.057 | -0.1 | -0.1 | 0 | 0.66 | 0.94 | 0.63 | 0.57 | 0.65 | 0.67 |
| 6 | 6249645 | rs3024388 | 0.241 | 0.81 | -2.332 | -0.342 | 0.004 | -0.027 | -0.027 | 1.215 | 0.31 | 0.18 | 0.25 | 0.45 | 0.26 | 0.45 |
| 6 | 6249914 | rs3024384 | 0.133 | 0.803 | -1.962 | -0.628 | 0 | -0.016 | -0.016 | 0 | 0.07 | 0.07 | 0.04 | 0 | 0.06 | 0.07 |
| 6 | 6250501 | rs3024377 | 0.393 | 0.683 | -1.222 | -0.907 | -0.008 | -0.052 | -0.052 | 0 | 0.12 | 0.16 | 0.16 | 0 | 0.13 | 0.04 |
| 6 | 6251504 | rs3024367 | -0.76 | 0.619 | -0.725 | -0.982 | 0.016 | -0.149 | -0.149 | 0 | 0.23 | 0.51 | 0.23 | 0 | 0.25 | 0.15 |
| 6 | 6253017 | rs6597197 | -1.301 | 0.614 | -0.71 | -0.868 | 0.001 | 0.041 | 0.041 | 0 | 0.34 | 0.73 | 0.38 | 0.12 | 0.38 | 0.21 |
| 6 | 6258630 | rs9504738 | -0.221 | 0.757 | 0.141 | -0.833 | 0.063 | 0.001 | 0.001 | 2.245 | 0.07 | 0.29 | 0.12 | 0 | 0.07 | 0.04 |
| 6 | 6260749 | rs3901123 | 0.364 | 0.415 | -0.372 | -0.775 | -0.001 | 0.006 | 0.006 | 0 | 0.12 | 0.11 | 0.05 | 0.26 | 0.08 | 0.29 |
| 6 | 6263242 | rs13204446 | -0.166 | 0.184 | -0.335 | -0.907 | -0.002 | 0.012 | 0.012 | 0 | 0.17 | 0.11 | 0.18 | 0.09 | 0.2 | 0.14 |
| 6 | 6267304 | rs3024358 | -0.319 | 0.21 | -0.4 | -0.952 | 0.003 | 0.008 | 0.008 | 0 | 0.19 | 0.15 | 0.19 | 0 | 0.18 | 0.11 |
| 6 | 6274084 | rs3863222 | -0.747 | 0.176 | -0.436 | -0.991 | 0.004 | 0.008 | 0.008 | 0 | 0.34 | 0.29 | 0.26 | 0.28 | 0.3 | 0.41 |
| 6 | 6274161 | rs3851513 | 0.577 | 0.444 | -1.028 | -0.911 | -0.008 | 0.021 | 0.021 | 0 | 0.2 | 0.16 | 0.2 | 0.18 | 0.2 | 0.17 |
| 6 | 6274570 | rs3851514 | -1.348 | 0.339 | -1.237 | -0.645 | -0.035 | -0.094 | -0.094 | 0 | 0.72 | 0.9 | 0.66 | 0.57 | 0.7 | 0.66 |
| 6 | 6275932 | rs9328348 | -0.038 | 0.138 | -0.687 | -0.681 | -0.004 | 0.001 | 0.001 | 0 | 0.32 | 0.36 | 0.19 | 0.35 | 0.33 | 0.44 |
| 6 | 6280910 | rs7770172 | -0.216 | 0.312 | -1.287 | -0.413 | -0.013 | -0.018 | -0.018 | 0 | 0.54 | 0.74 | 0.44 | 0.48 | 0.55 | 0.59 |
| 6 | 6282778 | rs9405911 | -0.17 | 0.203 | -1.297 | -0.422 | -0.021 | -0.021 | -0.021 | 0 | 0.53 | 0.78 | 0.43 | 0.49 | 0.52 | 0.57 |
| 6 | 6283666 | rs749005 | -0.273 | 0.288 | -1.534 | -0.608 | 0 | -0.018 | -0.018 | 0 | 0.16 | 0.18 | 0.2 | 0.03 | 0.17 | 0.1 |
| 6 | 6284125 | rs9502429 | -0.199 | 0.212 | -1.601 | -0.406 | -0.047 | -0.011 | -0.011 | 0.678 | 0.55 | 0.81 | 0.52 | 0.44 | 0.51 | 0.57 |
| 6 | 6289857 | rs1267914 | 0.3 | -0.859 | -1.937 | 0.049 | 0.033 | -0.088 | -0.088 | 0 | 0.3 | 0.07 | 0.19 | 0.44 | 0.23 | 0.26 |
| 6 | 6290003 | rs11243069 | -0.122 | -0.904 | -1.202 | -0.061 | -0.025 | 0.068 | 0.068 | 0 | 0.11 | 0.01 | 0.15 | 0 | 0.13 | 0.06 |
| 6 | 6292430 | rs3024354 | -1.352 | -1.093 | -1.534 | 0.158 | 0.082 | -0.001 | -0.001 | 0 | 0.35 | 0.89 | 0.35 | 0.32 | 0.22 | 0.35 |
| 6 | 6303246 | rs3024346 | 0.042 | -0.239 | -0.831 | 0.224 | 0.002 | 0.004 | 0.004 | 0 | 0.09 | 0.06 | 0.06 | 0.01 | 0.09 | 0.03 |
| 6 | 6305477 | rs2295753 | -0.903 | -0.514 | -0.284 | 0.523 | -0.045 | 0.071 | 0.071 | 0.736 | 0.26 | 0.42 | 0.25 | 0.33 | 0.16 | 0.32 |
| 6 | 6306405 | rs714408 | 0.726 | -0.457 | -0.52 | 0.09 | -0.001 | -0.017 | -0.017 | 0 | 0.35 | 0.27 | 0.28 | 0.48 | 0.37 | 0.31 |
| 6 | 6307443 | rs17378819 | -0.705 | 0.169 | -0.679 | -0.087 | -0.017 | 0.009 | 0.009 | 0 | 0.09 | 0.06 | 0.17 | 0.01 | 0.2 | 0.08 |

| | | | | | | | | | | | | | | | | |
|---|---|---|---|---|---|---|---|---|---|---|---|---|---|---|---|---|
| 6 | 6308750 | rs1742934 | 0.912 | 0.05 | -0.62 | -0.025 | -0.003 | -0.019 | -0.019 | 0.076 | 0.33 | 0.27 | 0.28 | 0.48 | 0.36 | 0.3 |
| 6 | 6309698 | rs1742937 | -0.922 | 0.461 | -0.35 | -0.427 | 0.019 | 0.024 | 0.024 | 1.391 | 0.24 | 0.41 | 0.39 | 0.25 | 0.32 | 0.32 |
| 6 | 6310536 | rs2755413 | -0.52 | 0.444 | -1.135 | -1.535 | -0.087 | 0.099 | 0.099 | 0 | 0.71 | 0.95 | 0.79 | 0.88 | 0.72 | 0.75 |
| 6 | 6311188 | rs1674045 | -0.463 | 0.401 | -1.113 | -1.528 | -0.078 | 0.095 | 0.095 | 0 | 0.71 | 0.93 | 0.78 | 0.87 | 0.71 | 0.74 |
| 6 | 6314893 | rs3024339 | -0.302 | -0.163 | -0.247 | -1.013 | -0.003 | -0.033 | -0.033 | 0.646 | 0.08 | 0.11 | 0.11 | 0 | 0.06 | 0.05 |
| 6 | 6316802 | rs11243078 | -1.505 | -0.284 | -0.201 | -1.093 | -0.033 | -0.101 | -0.101 | 0 | 0.19 | 0.28 | 0.14 | 0 | 0.1 | 0.19 |
| 6 | 6318126 | rs3024321 | -1.714 | -0.221 | -0.426 | -1.075 | 0.003 | -0.078 | -0.078 | 0 | 0.25 | 0.27 | 0.3 | 0 | 0.31 | 0.23 |
| 6 | 6318795 | rs5985 | -0.896 | 0.175 | -0.644 | -0.576 | 0.005 | -0.037 | -0.037 | 0.709 | 0.14 | 0.16 | 0.27 | 0 | 0.24 | 0.09 |
| 6 | 6320808 | rs2815822 | 0.244 | -0.773 | -0.864 | -0.45 | 0.024 | 0.03 | 0.03 | 0 | 0.81 | 0.88 | 0.89 | 0.95 | 0.89 | 0.8 |
| 6 | 6322875 | rs3024304 | 0.493 | -1.49 | -0.66 | 0.052 | -0.004 | -0.001 | -0.001 | 0 | 0.54 | 0.52 | 0.56 | 0.49 | 0.76 | 0.43 |
| 6 | 6322978 | rs1267858 | -0.046 | -1.932 | -0.933 | 0.484 | 0.038 | -0.048 | -0.048 | 0 | 0.18 | 0.11 | 0.24 | 0.31 | 0.07 | 0.2 |
| 6 | 6325739 | rs3844196 | 0.384 | -0.976 | -1.448 | -0.275 | -0.056 | -0.092 | -0.092 | 1.369 | 0.24 | 0.08 | 0.22 | 0.48 | 0.31 | 0.24 |
| 6 | 6326858 | rs1781789 | -0.412 | -1.06 | -0.949 | -0.523 | -0.007 | -0.017 | -0.017 | 0.744 | 0.8 | 0.83 | 0.89 | 0.93 | 0.89 | 0.67 |
| 6 | 6331873 | rs927332 | -0.187 | -1.057 | -0.516 | -0.311 | -0.001 | 0.004 | 0.004 | 2.518 | 0.47 | 0.45 | 0.51 | 0.65 | 0.45 | 0.52 |
| 6 | 6331936 | rs1318606 | 0.112 | -1.146 | -0.585 | -0.455 | -0.015 | -0.025 | -0.025 | 0 | 0.39 | 0.22 | 0.46 | 0.65 | 0.39 | 0.43 |
| 6 | 6336584 | rs3851518 | 1.403 | -1.593 | -0.951 | -0.853 | -0.13 | 0.163 | 0.163 | 0.141 | 0.25 | 0.04 | 0.26 | 0.01 | 0.42 | 0.13 |
| 6 | 6336922 | rs3863223 | 0.092 | -2.158 | -0.602 | -0.94 | 0.003 | 0.002 | 0.002 | 0 | 0.35 | 0.56 | 0.28 | 0.44 | 0.37 | 0.37 |
| 6 | 6337506 | rs11243081 | 1.046 | -2.13 | -0.626 | -0.794 | -0.102 | -0.121 | -0.121 | 0 | 0.23 | 0.02 | 0.19 | 0.41 | 0.3 | 0.27 |
| 6 | 6338541 | rs1674070 | -1.011 | -2.254 | -0.341 | -0.629 | 0.01 | 0.006 | 0.006 | 5.236 | 0.29 | 0.71 | 0.36 | 0.24 | 0.17 | 0.27 |
| 6 | 6339905 | rs1674065 | 0.798 | -2.073 | -0.58 | -0.497 | -0.01 | 0.009 | 0.009 | 15.232 | 0.12 | 0.16 | 0.22 | 0.23 | 0.04 | 0.13 |
| 6 | 6341754 | rs9502437 | 0.726 | -1.911 | -0.609 | -0.537 | 0.135 | 0.065 | 0.065 | 0 | 0.5 | 0.2 | 0.45 | 0.41 | 0.72 | 0.4 |
| 6 | 6342180 | rs17379732 | 0.849 | -1.649 | -0.089 | -0.591 | 0.006 | -0.026 | -0.026 | 0 | 0.06 | 0.1 | 0.04 | 0.01 | 0.06 | 0.09 |
| 6 | 6342377 | rs17142071 | -0.089 | -1.754 | 0.379 | -0.666 | 0.022 | -0.045 | -0.045 | 1.539 | 0.09 | 0.24 | 0.06 | 0 | 0.07 | 0.1 |
| 6 | 6343177 | rs6597199 | -0.438 | -1.446 | -0.273 | -0.351 | 0.038 | 0.048 | 0.048 | 0 | 0.5 | 0.63 | 0.44 | 0.63 | 0.33 | 0.57 |
| 6 | 6344007 | rs9328354 | 1.156 | -1.15 | -0.57 | -0.581 | 0.011 | -0.072 | -0.072 | 1.419 | 0.27 | 0.32 | 0.31 | 0 | 0.39 | 0.14 |
| 6 | 6348954 | rs2038553 | 0.061 | -1.333 | -0.792 | 0.197 | -0.021 | 0.041 | 0.041 | 0 | 0.29 | 0.3 | 0.23 | 0.39 | 0.2 | 0.41 |
| 6 | 6350381 | rs9392770 | -0.255 | -0.499 | -1.14 | 0.129 | -0.051 | -0.105 | -0.105 | 4.773 | 0.79 | 0.96 | 0.76 | 0.62 | 0.74 | 0.73 |
| 6 | 6351730 | rs13206706 | 0.268 | 0.109 | -1.065 | -0.956 | -0.032 | 0.02 | 0.02 | 0.937 | 0.29 | 0.2 | 0.34 | 0 | 0.42 | 0.18 |
| 6 | 6355433 | rs4960192 | -0.056 | -0.241 | -0.883 | -0.279 | 0 | -0.001 | -0.001 | 0.003 | 0.67 | 0.75 | 0.73 | 0.58 | 0.74 | 0.64 |
| 6 | 6355602 | rs6923580 | 0.361 | -0.488 | -0.658 | -0.339 | 0.003 | 0.021 | 0.021 | 3.361 | 0.12 | 0.44 | 0.18 | 0.13 | 0.07 | 0.08 |
| 6 | 6355950 | rs6924158 | -0.167 | -0.563 | -0.711 | -0.376 | 0.002 | -0.002 | -0.002 | 0 | 0.66 | 0.75 | 0.73 | 0.57 | 0.74 | 0.64 |
| 6 | 6356142 | rs9405917 | 0.145 | -0.339 | -0.489 | -0.184 | -0.001 | 0.116 | 0.116 | 0 | 0.1 | 0.23 | 0.11 | 0.21 | 0.09 | 0.09 |
| 6 | 6356482 | rs13198304 | 0.849 | -0.303 | -0.496 | -0.202 | -0.005 | 0.002 | 0.002 | 1.375 | 0.1 | 0.19 | 0.17 | 0.13 | 0.07 | 0.08 |
| 6 | 6358001 | rs6939615 | 0.76 | -0.303 | -0.551 | -0.201 | -0.005 | 0.001 | 0.001 | 1.439 | 0.11 | 0.26 | 0.17 | 0.13 | 0.07 | 0.08 |
| 6 | 6361816 | rs7774436 | 0.929 | -0.441 | -0.72 | -0.388 | -0.018 | -0.098 | -0.098 | 0.4 | 0.2 | 0.25 | 0.1 | 0 | 0.16 | 0.14 |
| 6 | 6366427 | rs988256 | -0.482 | -0.084 | -1.877 | -0.344 | -0.044 | -0.099 | -0.099 | 0 | 0.69 | 0.94 | 0.69 | 0.42 | 0.66 | 0.45 |
| 6 | 6366541 | rs1058242 | 1.254 | 0.386 | -1.42 | -0.888 | -0.039 | 0.063 | 0.063 | 1.68 | 0.26 | 0.14 | 0.31 | 0.07 | 0.33 | 0.15 |
| 6 | 6368578 | rs1902946 | -0.035 | 0.266 | -1.097 | -0.406 | -0.016 | -0.018 | -0.018 | 0 | 0.17 | 0.34 | 0.22 | 0.13 | 0.12 | 0.17 |
| 6 | 6369055 | rs17393473 | 0.203 | 0.456 | -1.318 | 0.074 | -0.027 | -0.004 | -0.004 | 0 | 0.18 | 0.05 | 0.13 | 0.16 | 0.24 | 0.34 |
| 6 | 6376547 | rs13206136 | 0.337 | 0.481 | -1.045 | -0.365 | -0.009 | 0.017 | 0.017 | 0.31 | 0.36 | 0.38 | 0.49 | 0.2 | 0.4 | 0.21 |
| 6 | 6381420 | rs4267999 | 0.026 | -0.389 | -1.072 | -0.191 | 0.019 | -0.022 | -0.022 | 0.183 | 0.38 | 0.15 | 0.22 | 0.39 | 0.33 | 0.41 |
| 6 | 6384148 | rs4543420 | 0.177 | -1.182 | -0.418 | -0.543 | -0.074 | 0.104 | 0.104 | 0 | 0.73 | 0.5 | 0.59 | 0.47 | 0.84 | 0.69 |
| 6 | 6385850 | rs6935596 | -0.649 | -1.288 | -0.131 | -0.387 | -0.019 | 0 | 0 | 0.373 | 0.89 | 0.87 | 0.82 | 0.88 | 0.96 | 0.93 |
| 6 | 6386396 | rs2326728 | 0.211 | -1.118 | -0.017 | -0.464 | 0.008 | 0.001 | 0.001 | 0 | 0.06 | 0.14 | 0.06 | 0.05 | 0.09 | 0.11 |
| 6 | 6389848 | rs12196470 | 0.797 | -1.47 | -0.418 | -0.934 | 0.096 | -0.002 | -0.002 | 0 | 0.49 | 0.13 | 0.44 | 0.42 | 0.61 | 0.46 |
| 6 | 6393064 | rs2326766 | 1.606 | -1.31 | -0.599 | -0.981 | -0.155 | 0.157 | 0.157 | 0.969 | 0.27 | 0.05 | 0.3 | 0.03 | 0.42 | 0.17 |
| 6 | 6397716 | rs9405920 | 0.738 | -1.374 | -0.431 | -0.719 | -0.01 | -0.044 | -0.044 | 0.022 | 0.09 | 0 | 0.17 | 0.43 | 0.08 | 0.19 |
| 6 | 6400475 | rs9328361 | 0.88 | -1.38 | -0.217 | -1.255 | -0.003 | -0.043 | -0.043 | 0 | 0.18 | 0.14 | 0.09 | 0 | 0.15 | 0.13 |
| 6 | 6407229 | rs1929089 | 0.458 | -1.498 | -0.018 | -1.074 | -0.006 | -0.002 | -0.002 | 0 | 0.78 | 0.74 | 0.73 | 0.83 | 0.84 | 0.82 |
| 6 | 6409308 | rs2815816 | -0.987 | -1.202 | -0.452 | -0.936 | -0.053 | -0.012 | -0.012 | 0.214 | 0.79 | 0.97 | 0.83 | 0.78 | 0.73 | 0.75 |
| 6 | 6410663 | rs4455715 | 0.776 | -1.763 | -0.007 | -1.202 | -0.001 | 0 | 0 | 0 | 0.45 | 0.57 | 0.55 | 0.59 | 0.52 | 0.51 |
| 6 | 6420874 | rs7768676 | -0.046 | -1.777 | 0.162 | -1.309 | -0.035 | -0.057 | -0.057 | 0 | 0.23 | 0.3 | 0.12 | 0.06 | 0.14 | 0.15 |
| 6 | 6423728 | rs10900987 | -0.186 | -1.919 | -0.253 | -1.497 | -0.041 | -0.018 | -0.018 | 0.223 | 0.65 | 0.88 | 0.65 | 0.65 | 0.64 | 0.63 |
| 6 | 6425334 | rs2876023 | 1.512 | -1.856 | -0.155 | -1.414 | 0.013 | -0.057 | -0.057 | 0 | 0.3 | 0.32 | 0.34 | 0.03 | 0.43 | 0.22 |
| 6 | 6432541 | rs9405926 | 0.813 | -1.791 | -0.172 | -1.258 | 0.002 | -0.041 | -0.041 | 0.311 | 0.09 | 0 | 0.18 | 0.54 | 0.06 | 0.26 |
| 6 | 6432964 | rs7749518 | 1.715 | -1.785 | -0.148 | -1.302 | -0.058 | 0.076 | 0.076 | 0 | 0.29 | 0.17 | 0.33 | 0.03 | 0.42 | 0.21 |
| 6 | 6433911 | rs2774511 | 0.637 | -1.677 | -0.162 | -1.18 | 0.008 | 0.002 | 0.002 | 0 | 0.12 | 0.11 | 0.16 | 0.1 | 0.06 | 0.11 |
| 6 | 6435068 | rs2774512 | 0.243 | -1.696 | -0.118 | -1.17 | -0.005 | -0.014 | -0.014 | 0 | 0.55 | 0.51 | 0.69 | 0.69 | 0.58 | 0.62 |
| 6 | 6445863 | rs1761928 | 0.347 | -1.093 | -0.203 | -1.306 | 0.003 | 0.002 | 0.002 | 0 | 0.12 | 0.1 | 0.16 | 0.12 | 0.09 | 0.11 |
| 6 | 6448063 | rs7767612 | -0.545 | -0.868 | 0.01 | -1.336 | -0.007 | -0.026 | -0.026 | 0 | 0.24 | 0.25 | 0.13 | 0.06 | 0.18 | 0.15 |
| 6 | 6448219 | rs7772062 | 0.041 | -0.56 | 0.237 | -0.836 | -0.012 | -0.016 | -0.016 | 0 | 0.1 | 0.17 | 0.07 | 0.04 | 0.08 | 0.07 |
| 6 | 6451982 | rs7760746 | 0.851 | 0.071 | -0.553 | -0.907 | -0.025 | 0.021 | 0.021 | 0 | 0.25 | 0.16 | 0.33 | 0.03 | 0.38 | 0.2 |
| 6 | 6455153 | rs9379036 | -0.541 | -0.537 | -0.208 | -0.008 | -0.031 | -0.011 | -0.011 | 1.97 | 0.29 | 0.4 | 0.18 | 0.26 | 0.22 | 0.3 |
| 6 | 6467682 | rs4960199 | -0.266 | -0.914 | -0.522 | 0.923 | 0.032 | -0.078 | -0.078 | 0 | 0.12 | 0.02 | 0.2 | 0.34 | 0.03 | 0.08 |
| 6 | 6467872 | rs2027118 | -1.004 | -1.123 | -0.557 | 0.813 | 0.025 | -0.03 | -0.03 | 0 | 0.32 | 0.2 | 0.3 | 0.59 | 0.22 | 0.33 |
| 6 | 6470892 | rs12198111 | 1.627 | -0.834 | -0.987 | 0.158 | -0.019 | 0.056 | 0.056 | 0.263 | 0.28 | 0.17 | 0.29 | 0.01 | 0.34 | 0.25 |
| 6 | 6474600 | rs1537624 | 0.817 | -0.931 | -0.983 | 0.204 | 0.011 | 0.017 | 0.017 | 0 | 0.4 | 0.2 | 0.5 | 0.35 | 0.37 | 0.33 |
| 6 | 6476278 | rs945144 | -0.583 | -0.671 | -0.89 | 0.149 | -0.046 | -0.038 | -0.038 | 0 | 0.69 | 0.81 | 0.64 | 0.6 | 0.6 | 0.67 |
| 6 | 6477849 | rs9405302 | -0.673 | -1.075 | -1.127 | 0.097 | -0.016 | -0.002 | -0.002 | 0.198 | 0.39 | 0.6 | 0.33 | 0.6 | 0.21 | 0.34 |

| | | | | | | | | | | | | | | | | |
|---|---|---|---|---|---|---|---|---|---|---|---|---|---|---|---|---|
| 6 | 6479161 | rs12192707 | 1.339 | -0.593 | -2.082 | -0.145 | -0.005 | 0.107 | 0.107 | 0.077 | 0.33 | 0.18 | 0.32 | 0.01 | 0.38 | 0.28 |
| 6 | 6486785 | rs11243096 | 0.482 | -0.039 | -2.108 | 1.093 | -0.045 | -0.07 | -0.07 | 0.31 | 0.17 | 0.01 | 0.23 | 0.2 | 0.18 | 0.11 |
| 6 | 6487306 | rs6929885 | 0.565 | 0.239 | -1.364 | 1.876 | -0.087 | -0.033 | -0.033 | 0 | 0.7 | 0.87 | 0.67 | 0.26 | 0.63 | 0.6 |
| 6 | 6487931 | rs708039 | -0.025 | 0.882 | -1.054 | 1.924 | -0.045 | -0.044 | -0.044 | 0.066 | 0.08 | 0.02 | 0.09 | 0.26 | 0.16 | 0.15 |
| 6 | 6491347 | rs1535705 | 0.525 | 0.768 | -1.04 | 1.653 | -0.033 | -0.054 | -0.054 | 0.412 | 0.81 | 0.88 | 0.79 | 0.51 | 0.74 | 0.76 |
| 6 | 6495031 | rs11965794 | -0.429 | 0.076 | -0.926 | 1.275 | -0.005 | -0.019 | -0.019 | 0 | 0.16 | 0.09 | 0.18 | 0.24 | 0.14 | 0.3 |
| 6 | 6495522 | rs6919894 | -1.204 | -0.346 | -0.313 | 1.046 | -0.021 | -0.002 | -0.002 | 0 | 0.29 | 0.56 | 0.29 | 0.28 | 0.2 | 0.36 |
| 6 | 6505406 | rs849057 | 0.893 | -0.22 | -0.606 | 0.993 | -0.005 | -0.008 | -0.008 | 0.909 | 0.64 | 0.26 | 0.66 | 0.6 | 0.67 | 0.69 |
| 6 | 6512190 | rs708041 | 0.165 | -0.11 | -0.709 | 1.143 | -0.068 | 0.024 | 0.024 | 0.196 | 0.83 | 0.96 | 0.85 | 0.89 | 0.77 | 0.83 |
| 6 | 6516047 | rs849050 | 0.298 | -0.081 | -0.126 | 0.809 | 0.001 | -0.001 | -0.001 | 0.24 | 0.85 | 0.8 | 0.84 | 0.89 | 0.77 | 0.81 |
| 6 | 6517089 | rs708052 | 1.155 | -0.117 | -0.827 | 0.372 | -0.011 | 0.092 | 0.092 | 0.248 | 0.33 | 0.19 | 0.27 | 0.06 | 0.37 | 0.28 |
| 6 | 6520931 | rs1535708 | 0.474 | -0.027 | -1.035 | 0.977 | -0.047 | -0.143 | -0.143 | 0.123 | 0.27 | 0.06 | 0.38 | 0.49 | 0.31 | 0.36 |
| 6 | 6523310 | rs708057 | 1.458 | -0.329 | -1.254 | 0.75 | -0.066 | 0.124 | 0.124 | 0.301 | 0.33 | 0.04 | 0.28 | 0.13 | 0.36 | 0.23 |
| 6 | 6524922 | rs708059 | 0.178 | -0.402 | -0.952 | 0.966 | -0.012 | 0.011 | 0.011 | 0.253 | 0.78 | 0.87 | 0.83 | 0.81 | 0.75 | 0.79 |
| 6 | 6527344 | rs9405937 | 0.517 | -0.297 | -0.743 | 1.402 | -0.016 | -0.042 | -0.042 | 0 | 0.28 | 0.12 | 0.23 | 0.34 | 0.3 | 0.35 |
| 6 | 6531684 | rs1761937 | -0.743 | -0.263 | 0.149 | 1.067 | -0.011 | -0.05 | -0.05 | 0 | 0.14 | 0.38 | 0.11 | 0.02 | 0.07 | 0.15 |
| 6 | 6535234 | rs1889508 | 0.558 | -0.18 | -0.314 | 1.339 | -0.035 | -0.068 | -0.068 | 0 | 0.27 | 0.11 | 0.4 | 0.39 | 0.32 | 0.36 |
| 6 | 6535323 | rs1761938 | 1.184 | -0.06 | 0.23 | 1.173 | -0.002 | -0.019 | -0.019 | 0.018 | 0.68 | 0.46 | 0.72 | 0.79 | 0.71 | 0.66 |
| 6 | 6541883 | rs742341 | -0.919 | 0.022 | 0.44 | 1.432 | -0.008 | 0.133 | 0.133 | 0 | 0.09 | 0.18 | 0.09 | 0.27 | 0.06 | 0.05 |
| 6 | 6542167 | rs742342 | -0.919 | 0.022 | 0.44 | 1.432 | -0.008 | 0.133 | 0.133 | 0 | 0.09 | 0.18 | 0.09 | 0.27 | 0.06 | 0.05 |
| 6 | 6544702 | rs1294468 | -0.587 | -0.051 | 0.367 | 1.128 | 0 | 0.002 | 0.002 | 0 | 0.44 | 0.5 | 0.55 | 0.7 | 0.42 | 0.5 |
| 6 | 6550156 | rs2774504 | -0.364 | 0.153 | 0.394 | 1.35 | 0 | -0.001 | -0.001 | 0 | 0.39 | 0.6 | 0.49 | 0.45 | 0.39 | 0.48 |
| 6 | 6552994 | rs594074 | 0.018 | 0.481 | -0.203 | 1.265 | -0.021 | -0.017 | -0.017 | 0.091 | 0.88 | 0.93 | 0.85 | 0.79 | 0.8 | 0.81 |
| 6 | 6558761 | rs681505 | 0.081 | 0.495 | -0.203 | 1.188 | -0.016 | -0.017 | -0.017 | 0 | 0.88 | 0.94 | 0.85 | 0.78 | 0.82 | 0.81 |
| 6 | 6559568 | rs4441992 | 0.121 | 0.237 | 0.162 | 1.162 | 0.002 | 0.037 | 0.037 | 0 | 0.31 | 0.4 | 0.45 | 0.49 | 0.35 | 0.43 |
| 6 | 6566290 | rs9504848 | -0.126 | 0.283 | -0.342 | 0.72 | -0.001 | -0.001 | -0.001 | 0 | 0.43 | 0.43 | 0.49 | 0.45 | 0.48 | 0.47 |
| 6 | 6566395 | rs9328369 | -0.135 | 0.461 | -0.212 | 0.594 | 0.016 | 0.02 | 0.02 | 0 | 0.79 | 0.67 | 0.81 | 0.75 | 0.79 | 0.75 |
| 6 | 6574384 | rs1999652 | 0.638 | 0.66 | -0.262 | 0.551 | 0.018 | 0.026 | 0.026 | 0 | 0.72 | 0.57 | 0.68 | 0.68 | 0.72 | 0.7 |
| 6 | 6578128 | rs9405942 | -0.089 | 0.683 | -0.259 | 0.533 | 0.025 | -0.013 | -0.013 | 0.084 | 0.78 | 0.67 | 0.79 | 0.86 | 0.77 | 0.74 |
| 6 | 6582123 | rs4959388 | 1.138 | 0.622 | -0.38 | 0.42 | 0.007 | 0.066 | 0.066 | 1.535 | 0.33 | 0.18 | 0.26 | 0.13 | 0.3 | 0.26 |
| 6 | 6584787 | rs9392810 | 0.742 | 0.682 | -0.357 | 0.425 | -0.006 | 0.078 | 0.078 | 0.772 | 0.37 | 0.22 | 0.31 | 0.13 | 0.38 | 0.3 |
| 6 | 6588881 | rs977785 | 0.242 | 0.644 | -0.238 | 0.599 | 0.023 | 0.048 | 0.048 | 0 | 0.77 | 0.63 | 0.72 | 0.67 | 0.76 | 0.73 |
| 6 | 6591304 | rs3804465 | 0.515 | 0.454 | -0.226 | 0.454 | -0.002 | -0.003 | -0.003 | 0 | 0.34 | 0.34 | 0.42 | 0.55 | 0.36 | 0.37 |
| 6 | 6592023 | rs9328375 | 0.38 | 0.734 | -0.22 | 0.459 | 0.01 | 0.025 | 0.025 | 0 | 0.7 | 0.54 | 0.69 | 0.66 | 0.71 | 0.66 |
| 6 | 6592095 | rs10214456 | 0.434 | 1.067 | -0.038 | 0.617 | 0.006 | 0.021 | 0.021 | 0.167 | 0.37 | 0.21 | 0.35 | 0.32 | 0.39 | 0.35 |
| 6 | 6592191 | rs9328376 | 0.087 | 1.019 | -0.176 | -0.076 | 0.016 | 0.072 | 0.072 | 0.243 | 0.39 | 0.23 | 0.31 | 0.22 | 0.4 | 0.36 |
| 6 | 6593049 | rs4246076 | 0.475 | 0.797 | -0.15 | -0.225 | -0.018 | 0.129 | 0.129 | 0.107 | 0.36 | 0.13 | 0.3 | 0.13 | 0.41 | 0.34 |
| 6 | 6593324 | rs9405309 | 0.025 | 0.915 | 0.236 | -0.125 | -0.002 | 0.002 | 0.002 | 0 | 0.26 | 0.2 | 0.23 | 0.1 | 0.35 | 0.27 |
| 6 | 6593695 | rs9502478 | 2.313 | 0.569 | 1.245 | 0.844 | -0.041 | -0.02 | -0.02 | 0 | 0.77 | 0.51 | 0.83 | 0.79 | 0.84 | 0.8 |
| 6 | 6594031 | rs3761985 | 1.321 | 0.566 | 1.056 | 0.994 | -0.011 | -0.009 | -0.009 | 0 | 0.91 | 0.92 | 0.85 | 0.82 | 0.84 | 0.83 |
| 6 | 6594321 | rs9328377 | 1.978 | 0.553 | 1.197 | 0.831 | -0.004 | -0.011 | -0.011 | 1.268 | 0.8 | 0.45 | 0.82 | 0.79 | 0.83 | 0.78 |
| 6 | 6601050 | rs17142362 | -0.764 | 0.008 | 0.799 | 0.694 | 0.031 | 0.033 | 0.033 | 0 | 0.06 | 0.36 | 0.02 | 0 | 0.01 | 0.03 |
| 6 | 6610609 | rs7740529 | -1.521 | -0.151 | 0.375 | 1.034 | -0.056 | 0.053 | 0.053 | 0.982 | 0.19 | 0.6 | 0.16 | 0.31 | 0.09 | 0.22 |
| 6 | 6612197 | rs909790 | -0.019 | 0.599 | -0.04 | 0.111 | -0.005 | -0.006 | -0.006 | 0 | 0.69 | 0.75 | 0.71 | 0.93 | 0.58 | 0.83 |
| 6 | 6614433 | rs2294460 | -0.002 | 0.831 | -0.022 | 0.669 | 0.005 | -0.007 | -0.007 | 0 | 0.71 | 0.73 | 0.7 | 0.8 | 0.62 | 0.82 |
| 6 | 6614501 | rs2294461 | -0.371 | 0.76 | 0.079 | 1.327 | -0.001 | 0.001 | 0.001 | 0.52 | 0.12 | 0.11 | 0.16 | 0.18 | 0.09 | 0.09 |
| 6 | 6618373 | rs3789766 | -0.781 | 0.959 | 0.049 | 1.624 | 0.023 | 0.091 | 0.091 | 0 | 0.22 | 0.42 | 0.28 | 0.51 | 0.23 | 0.4 |
| 6 | 6618549 | rs6934234 | 1.113 | 0.557 | -0.248 | 0.94 | 0.024 | 0.061 | 0.061 | 0.117 | 0.42 | 0.29 | 0.4 | 0.28 | 0.34 | 0.36 |
| 6 | 6619698 | rs763405 | 0.033 | 0.89 | -0.423 | 1.322 | 0.026 | 0.016 | 0.016 | 0 | 0.13 | 0.25 | 0.13 | 0.14 | 0.16 | 0.16 |
| 6 | 6621962 | rs7770797 | 0.013 | 0.714 | -1.179 | 1.099 | -0.098 | -0.089 | -0.089 | 0.048 | 0.69 | 0.93 | 0.71 | 0.62 | 0.59 | 0.66 |
| 6 | 6623677 | rs9504873 | -0.259 | 0.83 | -1.504 | 0.675 | -0.053 | -0.038 | -0.038 | 0 | 0.27 | 0.05 | 0.32 | 0.27 | 0.32 | 0.22 |
| 6 | 6625704 | rs3804476 | -0.741 | 0.688 | -1.629 | 0.312 | -0.06 | 0.036 | 0.036 | 0.001 | 0.37 | 0.08 | 0.38 | 0.27 | 0.44 | 0.29 |
| 6 | 6626263 | rs6922002 | -0.031 | -0.205 | -0.592 | 0.18 | 0.003 | -0.004 | -0.004 | 0.46 | 0.16 | 0.4 | 0.1 | 0.1 | 0.11 | 0.22 |
| 6 | 6626983 | rs17142392 | -0.644 | -0.718 | -0.43 | -0.519 | -0.034 | -0.027 | -0.027 | 0.528 | 0.25 | 0.36 | 0.11 | 0.15 | 0.15 | 0.33 |
| 6 | 6627102 | rs1076672 | 0.464 | -0.657 | -0.461 | -0.46 | -0.015 | 0.007 | 0.007 | 0.006 | 0.11 | 0.15 | 0.06 | 0.1 | 0.07 | 0.19 |
| 6 | 6627184 | rs1076673 | -1.155 | -0.788 | -0.629 | -0.536 | 0.048 | 0.096 | 0.096 | 4.083 | 0.4 | 0.76 | 0.2 | 0.16 | 0.25 | 0.43 |
| 6 | 6629987 | rs3827783 | 0.702 | -0.334 | -0.737 | -0.451 | 0.006 | -0.051 | -0.051 | 0 | 0.19 | 0.04 | 0.27 | 0.27 | 0.17 | 0.18 |
| 6 | 6630337 | rs3804479 | -0.49 | 0.085 | -0.324 | -0.418 | 0.009 | 0.004 | 0.004 | 0.021 | 0.21 | 0.15 | 0.07 | 0.04 | 0.15 | 0.18 |
| 6 | 6631674 | rs11961398 | -0.859 | 0.323 | -0.013 | -0.22 | 0.009 | -0.086 | -0.086 | 0.069 | 0.19 | 0.38 | 0.1 | 0.04 | 0.15 | 0.15 |
| 6 | 6635363 | rs9502483 | 0.415 | 0.2 | -0.474 | 0.115 | 0.005 | -0.028 | -0.028 | 0.021 | 0.23 | 0.04 | 0.28 | 0.23 | 0.19 | 0.23 |
| 6 | 6637861 | rs3804485 | 0.191 | 0.383 | -0.358 | 0.195 | 0.019 | 0.033 | 0.033 | 0 | 0.45 | 0.66 | 0.43 | 0.34 | 0.36 | 0.43 |
| 6 | 6637995 | rs3804486 | 0.739 | 0.227 | -0.247 | 0.326 | 0.009 | 0.004 | 0.004 | 0 | 0.36 | 0.25 | 0.35 | 0.32 | 0.3 | 0.32 |
| 6 | 6639425 | rs2064236 | 0.53 | 0.267 | -0.291 | 0.373 | -0.011 | -0.006 | -0.006 | 0.101 | 0.38 | 0.38 | 0.37 | 0.34 | 0.3 | 0.36 |
| 6 | 6642058 | rs7774142 | 0.16 | 0.266 | -0.516 | 0.309 | -0.01 | 0.036 | 0.036 | 0 | 0.34 | 0.5 | 0.5 | 0.48 | 0.26 | 0.38 |
| 6 | 6642595 | rs3804488 | -0.203 | 0.881 | -0.77 | 0.753 | -0.031 | -0.028 | -0.028 | 0 | 0.71 | 0.8 | 0.71 | 0.61 | 0.56 | 0.71 |
| 6 | 6653139 | rs3804490 | 0.398 | 0.555 | -0.241 | 0.71 | 0.019 | 0.005 | 0.005 | 0.597 | 0.25 | 0.4 | 0.19 | 0.22 | 0.25 | 0.26 |
| 6 | 6654517 | rs3765282 | -0.183 | -0.142 | -0.461 | 0.599 | -0.028 | -0.012 | -0.012 | 0 | 0.28 | 0.33 | 0.26 | 0.27 | 0.2 | 0.22 |
| 6 | 6655522 | rs760893 | -0.486 | 0.018 | -0.725 | 0.807 | -0.003 | 0 | 0 | 0.304 | 0.43 | 0.41 | 0.49 | 0.48 | 0.34 | 0.46 |
| 6 | 6655893 | rs3765289 | 0.112 | -0.342 | -0.576 | 0.757 | -0.032 | -0.002 | -0.002 | 0 | 0.23 | 0.33 | 0.25 | 0.28 | 0.18 | 0.2 |

| | | | | | | | | | | | | | | | | |
|---|---|---|---|---|---|---|---|---|---|---|---|---|---|---|---|---|
| 6 | 6656198 | rs760894 | -0.383 | -0.053 | -0.761 | 0.844 | 0.002 | 0 | 0 | 0 | 0.47 | 0.45 | 0.49 | 0.5 | 0.32 | 0.48 |
| 6 | 6656487 | rs7745381 | -0.775 | 0.48 | -1.155 | 1.156 | -0.048 | -0.012 | -0.012 | 0 | 0.72 | 0.84 | 0.69 | 0.7 | 0.57 | 0.74 |
| 6 | 6660214 | rs12529598 | 0.205 | 0.148 | -1.148 | 0.88 | 0.014 | -0.064 | -0.064 | 0 | 0.12 | 0.01 | 0.19 | 0.26 | 0.08 | 0.08 |
| 6 | 6662286 | rs201010 | -0.805 | 0.589 | -0.531 | 0.326 | 0.003 | -0.025 | -0.025 | 0 | 0.74 | 0.75 | 0.72 | 0.89 | 0.65 | 0.78 |
| 6 | 6664030 | rs9379053 | 0.158 | 0.595 | -0.599 | 1.144 | 0 | -0.023 | -0.023 | 0 | 0.15 | 0.09 | 0.19 | 0.19 | 0.1 | 0.23 |
| 6 | 6665771 | rs201016 | -0.117 | 0.783 | -0.679 | 0.979 | -0.01 | -0.023 | -0.023 | 0 | 0.3 | 0.24 | 0.36 | 0.67 | 0.33 | 0.41 |
| 6 | 6666708 | rs12660371 | -0.179 | 0.621 | -0.106 | 1.499 | 0 | 0.016 | 0.016 | 0 | 0.15 | 0.17 | 0.17 | 0.23 | 0.11 | 0.24 |
| 6 | 6668181 | rs9405313 | -0.175 | 0.609 | -0.096 | 1.498 | -0.001 | 0.006 | 0.006 | 0 | 0.15 | 0.15 | 0.18 | 0.23 | 0.11 | 0.24 |
| 6 | 6669148 | rs12663300 | -0.032 | 0.604 | -0.096 | 1.495 | 0 | 0.015 | 0.015 | 0.025 | 0.15 | 0.17 | 0.18 | 0.23 | 0.11 | 0.24 |
| 6 | 6675339 | rs2050056 | -0.837 | 0.685 | 0.67 | 2.015 | 0.031 | 0.219 | 0.219 | 0.381 | 0.16 | 0.36 | 0.27 | 0.45 | 0.14 | 0.21 |
| 6 | 6675654 | rs12529068 | -0.37 | 0.577 | 0.665 | 2.073 | -0.002 | 0.013 | 0.013 | 0.692 | 0.12 | 0.11 | 0.24 | 0.29 | 0.09 | 0.08 |
| 6 | 6677443 | rs9502486 | -0.635 | 0.54 | 0.838 | 2.144 | -0.017 | 0.111 | 0.111 | 0.536 | 0.13 | 0.27 | 0.25 | 0.31 | 0.09 | 0.08 |
| 6 | 6679166 | rs6924103 | -1.109 | 0.422 | 0.328 | 1.131 | 0 | -0.006 | -0.006 | 0 | 0.51 | 0.55 | 0.44 | 0.29 | 0.44 | 0.55 |
| 6 | 6680510 | rs3761986 | -0.562 | 0.294 | 0.336 | 1.309 | 0.002 | -0.001 | -0.001 | 0.147 | 0.4 | 0.44 | 0.36 | 0.4 | 0.38 | 0.53 |
| 6 | 6682321 | rs9405967 | 0.046 | 0.255 | -0.38 | -0.014 | -0.079 | 0.042 | 0.042 | 2.256 | 0.24 | 0.05 | 0.28 | 0.19 | 0.26 | 0.34 |
| 6 | 6684357 | rs7741565 | -0.856 | 0.247 | -0.23 | -0.066 | 0.007 | 0.035 | 0.035 | 0 | 0.43 | 0.67 | 0.4 | 0.22 | 0.34 | 0.4 |
| 6 | 6692142 | rs115560 | 0.151 | 0.574 | -0.71 | 0.206 | -0.056 | 0.057 | 0.057 | 2.84 | 0.7 | 0.87 | 0.7 | 0.78 | 0.68 | 0.62 |
| 6 | 6693320 | rs2235493 | 0.001 | 0.464 | -0.351 | 0.731 | -0.011 | -0.024 | -0.024 | 0 | 0.09 | 0.04 | 0.11 | 0.42 | 0.08 | 0.15 |
| 6 | 6703516 | rs12207082 | -0.573 | 0.223 | 0.035 | 0.305 | -0.022 | 0.161 | 0.161 | 0 | 0.23 | 0.46 | 0.18 | 0.51 | 0.15 | 0.26 |
| 6 | 6709118 | rs201036 | 0.797 | 0.123 | -0.288 | -0.139 | 0.022 | -0.022 | -0.022 | 0 | 0.62 | 0.33 | 0.77 | 0.49 | 0.74 | 0.67 |
| 6 | 6710351 | rs201037 | -0.517 | 0.101 | -0.086 | -0.199 | -0.001 | -0.033 | -0.033 | 0.053 | 0.86 | 0.85 | 0.95 | 1 | 0.88 | 0.93 |
| 6 | 6711680 | rs9405314 | -0.028 | 0.479 | -0.877 | -0.691 | -0.002 | 0.015 | 0.015 | 0.053 | 0.33 | 0.29 | 0.27 | 0.19 | 0.31 | 0.36 |
| 6 | 6712175 | rs9405969 | 0.946 | 0.023 | -0.419 | -0.355 | -0.011 | -0.018 | -0.018 | 0 | 0.12 | 0.03 | 0.11 | 0.17 | 0.09 | 0.24 |
| 6 | 6712345 | rs201039 | -0.291 | 0.08 | 0.239 | -0.207 | 0 | -0.053 | -0.053 | 0 | 0.86 | 0.83 | 0.95 | 1 | 0.88 | 0.93 |
| 6 | 6714550 | rs201043 | 0.611 | 0.347 | -0.037 | 0.047 | 0.05 | -0.045 | -0.045 | 0 | 0.61 | 0.25 | 0.76 | 0.49 | 0.74 | 0.67 |
| 6 | 6720219 | rs201050 | 0.712 | 0.928 | 0.455 | -0.078 | 0.021 | 0.002 | 0.002 | 0.045 | 0.51 | 0.33 | 0.67 | 0.5 | 0.62 | 0.64 |
| 6 | 6721713 | rs201051 | 0.326 | 0.057 | 0.751 | -0.051 | -0.002 | -0.001 | -0.001 | 0 | 0.57 | 0.49 | 0.76 | 0.55 | 0.75 | 0.73 |
| 6 | 6722492 | rs201053 | 0.656 | 0.16 | 0.316 | 0.009 | 0.084 | 0 | 0 | 0.228 | 0.5 | 0.25 | 0.69 | 0.55 | 0.68 | 0.65 |
| 6 | 6726927 | rs12662158 | -0.205 | 0.352 | 1.159 | 0.613 | -0.008 | 0.094 | 0.094 | 0 | 0.11 | 0.18 | 0.11 | 0.43 | 0.04 | 0.23 |
| 6 | 6728616 | rs9504905 | -1.547 | 0.444 | 1.989 | -0.581 | 0.063 | -0.001 | -0.001 | 0.152 | 0.07 | 0.36 | 0.08 | 0 | 0.06 | 0.02 |
| 6 | 6730512 | rs201061 | -0.491 | 0.493 | 1.511 | -0.915 | -0.008 | 0.026 | 0.026 | 0 | 0.59 | 0.69 | 0.65 | 0.73 | 0.53 | 0.73 |
| 6 | 6736109 | rs9504909 | -0.758 | 0.463 | 0.979 | -0.911 | 0.001 | -0.066 | -0.066 | 0 | 0.13 | 0.19 | 0.13 | 0 | 0.13 | 0.06 |
| 6 | 6738355 | rs1294409 | -0.233 | 0.762 | 0.69 | -0.539 | 0.001 | 0.074 | 0.074 | 0 | 0.57 | 0.73 | 0.51 | 0.26 | 0.62 | 0.4 |
| 6 | 6738752 | rs1294410 | -0.416 | 0.767 | 0.422 | -0.532 | -0.012 | 0.072 | 0.072 | 0 | 0.59 | 0.79 | 0.51 | 0.26 | 0.63 | 0.4 |
| 6 | 6742752 | rs1294420 | 0.091 | 0.729 | 0.179 | -0.403 | 0.002 | 0.042 | 0.042 | 0 | 0.51 | 0.36 | 0.41 | 0.26 | 0.56 | 0.35 |
| 6 | 6745163 | rs1294433 | 0.169 | 0.806 | -0.265 | -0.364 | 0.007 | 0.104 | 0.104 | 0 | 0.5 | 0.27 | 0.4 | 0.26 | 0.55 | 0.35 |
| 6 | 6752059 | rs1294438 | 0.134 | 0.724 | -0.551 | -0.262 | 0.014 | 0.038 | 0.038 | 0 | 0.41 | 0.21 | 0.5 | 0.7 | 0.35 | 0.59 |
| 6 | 6755091 | rs2181398 | 0.105 | 0.901 | -0.742 | -0.492 | -0.014 | 0.022 | 0.022 | 0.11 | 0.37 | 0.18 | 0.41 | 0.72 | 0.4 | 0.54 |
| 6 | 6755865 | rs6938520 | 0.509 | 0.838 | -0.008 | -0.729 | 0.011 | -0.065 | -0.065 | 0.163 | 0.78 | 0.64 | 0.68 | 0.94 | 0.77 | 0.83 |
| 6 | 6761642 | rs2876028 | -0.499 | 0.304 | -0.42 | -0.55 | 0.007 | 0.004 | 0.004 | 0 | 0.37 | 0.61 | 0.3 | 0.33 | 0.28 | 0.38 |
| 6 | 6761715 | rs2876029 | 0.364 | 0.449 | -0.62 | -0.821 | -0.014 | -0.013 | -0.013 | 1.487 | 0.09 | 0.07 | 0.12 | 0.15 | 0.15 | 0.06 |
| 6 | 6762167 | rs2326778 | -0.083 | 0.408 | -0.289 | -0.742 | 0.01 | 0.015 | 0.015 | 0 | 0.25 | 0.19 | 0.15 | 0.19 | 0.21 | 0.23 |
| 6 | 6779888 | rs4960242 | -0.244 | 0.4 | -0.088 | -0.917 | -0.009 | -0.061 | -0.061 | 0 | 0.3 | 0.46 | 0.21 | 0.16 | 0.27 | 0.25 |
| 6 | 6781726 | rs10485084 | 0.312 | 0.851 | -1.294 | -0.711 | -0.122 | 0.05 | 0.05 | 0 | 0.25 | 0.01 | 0.23 | 0.13 | 0.35 | 0.3 |
| 6 | 6784416 | rs4959407 | 0.051 | 0.687 | -0.702 | -0.853 | -0.001 | -0.009 | -0.009 | 0 | 0.57 | 0.49 | 0.43 | 0.25 | 0.52 | 0.5 |
| 6 | 6786253 | rs9502503 | -0.376 | 1.195 | -0.25 | -0.296 | 0.038 | 0.024 | 0.024 | 0 | 0.16 | 0.36 | 0.16 | 0.18 | 0.18 | 0.22 |
| 6 | 6786425 | rs1573948 | 0.133 | 0.988 | -0.865 | -0.133 | -0.017 | -0.031 | -0.031 | 0.056 | 0.12 | 0.01 | 0.11 | 0.14 | 0.12 | 0.15 |
| 6 | 6788490 | rs17142568 | 0.137 | 0.633 | -0.515 | -0.944 | -0.011 | -0.052 | -0.052 | 0.242 | 0.18 | 0.25 | 0.12 | 0.04 | 0.15 | 0.11 |
| 6 | 6788525 | rs9504932 | -0.854 | 0.749 | -0.262 | -0.978 | 0.002 | -0.021 | -0.021 | 0 | 0.3 | 0.44 | 0.25 | 0.22 | 0.29 | 0.2 |
| 6 | 6790060 | rs1892462 | -0.611 | 0.741 | -0.51 | -1.46 | -0.006 | 0.03 | 0.03 | 0 | 0.6 | 0.73 | 0.51 | 0.23 | 0.57 | 0.52 |
| 6 | 6793702 | rs7740821 | 0.173 | 0.736 | -0.518 | -1.221 | -0.003 | -0.028 | -0.028 | 0 | 0.42 | 0.5 | 0.35 | 0.15 | 0.4 | 0.4 |
| 6 | 6794340 | rs12213204 | 0.572 | 1.105 | -0.597 | -0.406 | -0.001 | 0 | 0 | 0.618 | 0.1 | 0.1 | 0.08 | 0.14 | 0.13 | 0.08 |
| 6 | 6803704 | rs2876031 | -0.197 | 0.8 | -0.678 | 0.117 | 0.003 | -0.008 | -0.008 | 0.237 | 0.19 | 0.15 | 0.14 | 0.58 | 0.15 | 0.29 |
| 6 | 6804795 | rs1892460 | 0.197 | 0.98 | -0.536 | -0.899 | 0.005 | -0.011 | -0.011 | 0.357 | 0.7 | 0.65 | 0.59 | 0.83 | 0.61 | 0.57 |
| 6 | 6804900 | rs1317392 | -0.212 | 0.567 | -0.883 | -1.049 | -0.041 | 0.005 | 0.005 | 0 | 0.86 | 0.93 | 0.74 | 0.85 | 0.79 | 0.79 |
| 6 | 6805787 | rs1892459 | -0.197 | 0.144 | -0.869 | -1.517 | 0.007 | 0.092 | 0.092 | 1.748 | 0.52 | 0.71 | 0.51 | 0.9 | 0.5 | 0.48 |
| 6 | 6806171 | rs2326825 | 0.007 | 0.265 | -1.317 | -1.551 | -0.076 | 0.056 | 0.056 | 0 | 0.84 | 0.95 | 0.79 | 0.93 | 0.74 | 0.76 |
| 6 | 6806247 | rs6927306 | -0.115 | -0.326 | -1.079 | -1.118 | -0.024 | 0.069 | 0.069 | 0.878 | 0.57 | 0.69 | 0.56 | 0.73 | 0.54 | 0.44 |
| 6 | 6806970 | rs11969257 | 0.177 | -0.076 | -0.927 | -0.706 | -0.005 | -0.009 | -0.009 | 0.135 | 0.6 | 0.65 | 0.54 | 0.42 | 0.47 | 0.49 |
| 6 | 6807071 | rs6911009 | -0.119 | 0.367 | -0.564 | -0.862 | -0.01 | -0.002 | -0.002 | 0 | 0.64 | 0.65 | 0.62 | 0.41 | 0.56 | 0.34 |
| 6 | 6807206 | rs9378479 | 0.033 | 0.769 | -1.068 | -0.478 | -0.025 | -0.02 | -0.02 | 0.029 | 0.29 | 0.21 | 0.34 | 0.29 | 0.34 | 0.19 |
| 6 | 6808208 | rs4960248 | -0.205 | 0.276 | -0.515 | 0.017 | 0.004 | 0 | 0 | 0 | 0.51 | 0.47 | 0.42 | 0.5 | 0.43 | 0.24 |
| 6 | 6808762 | rs11755624 | -0.382 | 0.864 | -0.311 | -0.168 | 0.015 | 0.002 | 0.002 | 0 | 0.06 | 0.16 | 0.06 | 0.08 | 0.05 | 0.09 |
| 6 | 6811806 | rs4959411 | -0.094 | 0.823 | -0.712 | -0.981 | -0.01 | 0.001 | 0.001 | 0.213 | 0.58 | 0.68 | 0.53 | 0.94 | 0.52 | 0.64 |
| 6 | 6815016 | rs2326827 | -0.515 | 0.314 | -1.251 | -0.552 | 0.033 | -0.003 | -0.003 | 0 | 0.53 | 0.75 | 0.29 | 0.55 | 0.39 | 0.48 |
| 6 | 6816332 | rs9502509 | -0.16 | -0.416 | -0.695 | -0.795 | 0.011 | -0.082 | -0.082 | 0 | 0.28 | 0.44 | 0.19 | 0.11 | 0.26 | 0.32 |
| 6 | 6817791 | rs4530912 | -0.778 | -0.804 | -0.951 | -0.662 | 0.02 | -0.003 | -0.003 | 0 | 0.38 | 0.69 | 0.23 | 0.61 | 0.3 | 0.45 |
| 6 | 6818562 | rs9379064 | -0.078 | -0.356 | -1.326 | -0.815 | -0.111 | -0.159 | -0.159 | 0 | 0.21 | 0.01 | 0.21 | 0.52 | 0.27 | 0.24 |
| 6 | 6819029 | rs7766251 | 1.496 | -0.893 | -1.819 | -1.783 | -0.055 | 0.206 | 0.206 | 0 | 0.39 | 0.13 | 0.37 | 0.07 | 0.47 | 0.3 |

| | | | | | | | | | | | | | | | | |
|---|---|---|---|---|---|---|---|---|---|---|---|---|---|---|---|---|
| 6 | 6823042 | rs17566008 | 0.76 | -1.28 | -2.089 | -2.192 | 0.006 | 0.006 | 0.006 | 0 | 0.17 | 0.13 | 0.07 | 0.01 | 0.13 | 0.09 |
| 6 | 6823174 | rs9504940 | -0.158 | -1.571 | -1.379 | -2.246 | -0.023 | -0.079 | -0.079 | 0.027 | 0.31 | 0.42 | 0.14 | 0.06 | 0.2 | 0.17 |
| 6 | 6824153 | rs9502510 | -1.437 | -1.385 | -1.578 | -1.598 | -0.006 | 0.006 | 0.006 | 0.436 | 0.63 | 0.66 | 0.43 | 0.76 | 0.55 | 0.44 |
| 6 | 6825878 | rs1892457 | -0.557 | -1.393 | -1.642 | -1.329 | -0.08 | -0.172 | -0.172 | 1.723 | 0.75 | 0.99 | 0.77 | 0.47 | 0.72 | 0.8 |
| 6 | 6826108 | rs1892458 | 1.39 | -0.933 | -1.302 | -1.508 | 0.001 | -0.002 | -0.002 | 0 | 0.34 | 0.32 | 0.48 | 0.24 | 0.39 | 0.49 |
| 6 | 6829312 | rs1286053 | -0.818 | -0.996 | -1.391 | -1.549 | -0.09 | 0.125 | 0.125 | 0.123 | 0.78 | 0.97 | 0.68 | 1 | 0.66 | 0.76 |
| 6 | 6833987 | rs13215298 | 1.582 | -0.796 | -1.188 | -1.259 | -0.02 | 0.037 | 0.037 | 0 | 0.08 | 0.01 | 0.05 | 0.01 | 0.08 | 0.01 |
| 6 | 6834674 | rs1286046 | 1.222 | -0.899 | -1.031 | -1.299 | -0.009 | 0.01 | 0.01 | 2.341 | 0.36 | 0.27 | 0.55 | 0.22 | 0.45 | 0.38 |
| 6 | 6836159 | rs1286044 | 1.038 | -1.175 | -0.835 | -1.267 | -0.014 | 0.026 | 0.026 | 0.98 | 0.39 | 0.27 | 0.57 | 0.23 | 0.47 | 0.52 |
| 6 | 6841023 | rs6929727 | 0.828 | -1.168 | -0.557 | -1.408 | 0.002 | -0.027 | -0.027 | 0 | 0.61 | 0.6 | 0.46 | 0.81 | 0.6 | 0.44 |
| 6 | 6843733 | rs1286037 | -0.42 | -0.866 | -0.211 | -1.15 | -0.044 | -0.011 | -0.011 | 0 | 0.36 | 0.48 | 0.39 | 0.27 | 0.24 | 0.28 |
| 6 | 6846862 | rs1927273 | -0.606 | -0.655 | -0.25 | -1.026 | -0.014 | -0.008 | -0.008 | 0 | 0.43 | 0.48 | 0.43 | 0.28 | 0.31 | 0.33 |
| 6 | 6848153 | rs943608 | -0.667 | -0.579 | -0.914 | -0.983 | 0.02 | 0.014 | 0.014 | 0 | 0.44 | 0.6 | 0.43 | 0.28 | 0.31 | 0.33 |
| 6 | 6848537 | rs1286024 | -0.684 | -0.516 | -1.044 | -1.092 | -0.096 | 0.142 | 0.142 | 0 | 0.78 | 0.97 | 0.78 | 1 | 0.7 | 0.78 |
| 6 | 6848893 | rs10900995 | 1.242 | -0.836 | -0.78 | -0.763 | 0.004 | -0.001 | -0.001 | 1.138 | 0.34 | 0.38 | 0.35 | 0.72 | 0.39 | 0.45 |
| 6 | 6851165 | rs2326828 | -0.578 | -1.414 | -1.624 | -1.121 | -0.13 | 0.171 | 0.171 | 2.111 | 0.75 | 0.98 | 0.59 | 1 | 0.63 | 0.74 |
| 6 | 6854396 | rs4960254 | 1.526 | -1.446 | -1.48 | -0.783 | -0.107 | -0.042 | -0.042 | 0 | 0.28 | 0.04 | 0.37 | 0.75 | 0.36 | 0.26 |
| 6 | 6855722 | rs7751407 | -0.207 | -1.702 | -0.977 | -0.847 | -0.062 | -0.016 | -0.016 | 0 | 0.3 | 0.52 | 0.14 | 0.21 | 0.13 | 0.23 |
| 6 | 6857640 | rs17142630 | -0.111 | -1.626 | -1.131 | -0.809 | -0.041 | -0.01 | -0.01 | 0.069 | 0.31 | 0.57 | 0.14 | 0.22 | 0.13 | 0.26 |
| 6 | 6858879 | rs6916313 | 0.325 | -0.912 | -0.771 | -1.007 | -0.004 | -0.039 | -0.039 | 0 | 0.59 | 0.62 | 0.51 | 0.96 | 0.47 | 0.53 |
| 6 | 6860330 | rs9405978 | -0.638 | -0.791 | -0.603 | -0.007 | 0 | 0.024 | 0.024 | 0 | 0.16 | 0.16 | 0.17 | 0.48 | 0.16 | 0.13 |
| 6 | 6860677 | rs6597232 | 0.007 | -1.142 | -0.687 | -0.598 | -0.029 | 0.029 | 0.029 | 2.841 | 0.65 | 0.77 | 0.52 | 0.76 | 0.54 | 0.41 |
| 6 | 6868584 | rs10900997 | 1.419 | -1.149 | -1.067 | -0.524 | -0.105 | 0.031 | 0.031 | 1.075 | 0.36 | 0.02 | 0.3 | 0.24 | 0.38 | 0.36 |
| 6 | 6869872 | rs9405979 | -0.574 | -0.774 | -0.108 | 0.233 | -0.003 | 0.077 | 0.077 | 0 | 0.13 | 0.19 | 0.16 | 0.54 | 0.12 | 0.11 |
| 6 | 6873602 | rs6908267 | 0.261 | -0.282 | 0.082 | 0.224 | -0.023 | 0.009 | 0.009 | 7.791 | 0.72 | 0.84 | 0.6 | 0.75 | 0.65 | 0.55 |
| 6 | 6873698 | rs6929018 | 0.45 | -0.321 | 0.154 | 0.055 | -0.001 | 0 | 0 | 0 | 0.69 | 0.72 | 0.5 | 0.21 | 0.61 | 0.5 |
| 6 | 6877411 | rs6910011 | 0.446 | -0.291 | 0.144 | 0.054 | -0.01 | 0.008 | 0.008 | 6.97 | 0.69 | 0.76 | 0.52 | 0.23 | 0.61 | 0.49 |
| 6 | 6879520 | rs1887510 | 1.042 | -0.427 | 0.235 | 0.047 | 0 | -0.007 | -0.007 | 0 | 0.6 | 0.6 | 0.48 | 0.22 | 0.56 | 0.43 |
| 6 | 6880482 | rs6922731 | 0.162 | -0.435 | 0.092 | -0.187 | -0.016 | -0.001 | -0.001 | 0 | 0.76 | 0.86 | 0.64 | 0.77 | 0.67 | 0.6 |
| 6 | 6882117 | rs1887509 | 1.108 | -0.769 | 0.299 | -0.505 | -0.023 | 0.01 | 0.01 | 4.69 | 0.44 | 0.23 | 0.4 | 0.57 | 0.5 | 0.4 |
| 6 | 6883618 | rs2182802 | 0.318 | -0.661 | 0.749 | -1.048 | -0.023 | -0.095 | -0.095 | 0.116 | 0.69 | 0.56 | 0.86 | 0.92 | 0.78 | 0.64 |
| 6 | 6884733 | rs4960263 | 0.348 | -0.609 | 0.662 | -0.927 | 0.001 | -0.039 | -0.039 | 0 | 0.87 | 0.87 | 0.89 | 1 | 0.86 | 0.94 |
| 6 | 6886326 | rs11967438 | 0.633 | -0.511 | 0.437 | -0.862 | -0.012 | -0.054 | -0.054 | 0 | 0.63 | 0.38 | 0.83 | 0.91 | 0.77 | 0.58 |
| 6 | 6887373 | rs9504956 | -0.147 | -0.546 | 0.497 | -0.767 | -0.054 | -0.085 | -0.085 | 0 | 0.26 | 0.59 | 0.08 | 0.09 | 0.1 | 0.37 |
| 6 | 6887857 | rs1887507 | 1.047 | 0.595 | -0.003 | 0.195 | 0.006 | -0.023 | -0.023 | 0.018 | 0.64 | 0.45 | 0.43 | 0.71 | 0.58 | 0.7 |
| 6 | 6893991 | rs17142659 | -0.686 | 0.528 | -0.624 | 1.118 | -0.003 | -0.016 | -0.016 | 0 | 0.17 | 0.1 | 0.41 | 0.28 | 0.21 | 0.21 |
| 6 | 6901007 | rs3748060 | 0.538 | 0.365 | -0.154 | 1.369 | -0.008 | 0.124 | 0.124 | 0.068 | 0.78 | 0.65 | 0.7 | 0.45 | 0.77 | 0.74 |
| 6 | 6904825 | rs11965947 | -0.323 | 0.436 | 0.072 | 0.792 | 0.002 | -0.016 | -0.016 | 0.15 | 0.08 | 0.18 | 0.05 | 0.03 | 0.08 | 0.03 |
| 6 | 6905830 | rs9502516 | 0.488 | 0.54 | -0.479 | -0.049 | -0.008 | 0.008 | 0.008 | 0 | 0.29 | 0.23 | 0.31 | 0.16 | 0.37 | 0.44 |
| 6 | 6906508 | rs2148976 | 0.025 | 0.428 | -0.514 | -0.044 | 0.008 | -0.014 | -0.014 | 0 | 0.37 | 0.4 | 0.36 | 0.19 | 0.45 | 0.47 |
| 6 | 6906545 | rs12198471 | 0.07 | -0.2 | -1.03 | -0.588 | 0.014 | 0.175 | 0.175 | 0.062 | 0.34 | 0.11 | 0.17 | 0.04 | 0.29 | 0.23 |
| 6 | 6908368 | rs7738221 | -0.51 | -1.056 | -0.389 | -0.308 | -0.003 | -0.001 | -0.001 | 0.279 | 0.45 | 0.53 | 0.44 | 0.66 | 0.36 | 0.36 |
| 6 | 6912695 | rs6908529 | -0.605 | -0.455 | -0.005 | -0.171 | 0 | -0.001 | -0.001 | 0 | 0.45 | 0.54 | 0.59 | 0.75 | 0.42 | 0.3 |
| 6 | 6913661 | rs2802454 | 0.768 | -0.519 | -0.011 | -0.212 | 0.029 | 0.115 | 0.115 | 0 | 0.51 | 0.29 | 0.36 | 0.16 | 0.54 | 0.67 |
| 6 | 6918126 | rs2768999 | 1.055 | -0.315 | 0.352 | 0.578 | 0.015 | 0.246 | 0.246 | 0 | 0.93 | 0.82 | 0.86 | 0.61 | 0.92 | 0.89 |
| 6 | 6926410 | rs2876048 | -0.382 | -0.591 | -0.019 | -0.666 | -0.005 | -0.071 | -0.071 | 0 | 0.24 | 0.29 | 0.25 | 0 | 0.21 | 0.15 |
| 6 | 6929780 | rs7757080 | -1.084 | -0.241 | 0.028 | -0.213 | -0.01 | -0.022 | -0.022 | 1.737 | 0.06 | 0.12 | 0.04 | 0.01 | 0.04 | 0.11 |
| 6 | 6933182 | rs17142699 | -1.403 | -0.116 | 0.354 | 0.039 | 0 | 0.024 | 0.024 | 0 | 0.06 | 0.16 | 0.06 | 0.07 | 0.05 | 0.14 |
| 6 | 6936571 | rs2768990 | 0.1 | 0.041 | -0.021 | -0.961 | -0.001 | 0.004 | 0.004 | 0 | 0.34 | 0.29 | 0.24 | 0.06 | 0.32 | 0.24 |
| 6 | 6942514 | rs2094440 | -0.487 | 0.281 | -0.015 | -1.436 | 0 | -0.055 | -0.055 | 0 | 0.53 | 0.59 | 0.38 | 0.08 | 0.45 | 0.37 |
| 6 | 6942564 | rs2094442 | -0.203 | 0.196 | -0.049 | -1.515 | 0.002 | 0.002 | 0.002 | 2.519 | 0.2 | 0.3 | 0.14 | 0.08 | 0.13 | 0.16 |
| 6 | 6942918 | rs2490294 | -0.158 | 0.724 | 0.307 | -0.877 | 0 | -0.055 | -0.055 | 0 | 0.53 | 0.59 | 0.38 | 0.08 | 0.45 | 0.37 |
| 6 | 6952323 | rs2802426 | -0.285 | 0.583 | 0.394 | -0.066 | -0.003 | -0.001 | -0.001 | 0 | 0.6 | 0.7 | 0.44 | 0.36 | 0.48 | 0.48 |
| 6 | 6954493 | rs2802422 | 0.306 | 0.615 | 0.426 | -0.533 | 0.001 | 0.025 | 0.025 | 4.392 | 0.49 | 0.45 | 0.34 | 0.14 | 0.43 | 0.38 |
| 6 | 6956966 | rs6918634 | -1.379 | 0.522 | 1.047 | 0.346 | 0.014 | 0.071 | 0.071 | 0 | 0.08 | 0.3 | 0.1 | 0.13 | 0.07 | 0.16 |
| 6 | 6957239 | rs6907319 | -2.186 | 0.44 | 1.002 | 0.388 | -0.025 | 0.056 | 0.056 | 0 | 0.17 | 0.48 | 0.15 | 0.2 | 0.12 | 0.24 |
| 6 | 6959468 | rs2765353 | 1.284 | 0.748 | 0.982 | 0.373 | -0.005 | 0.002 | 0.002 | 0 | 0.92 | 0.79 | 0.88 | 0.94 | 0.95 | 0.88 |
| 6 | 6959748 | rs7770144 | 1.155 | 1.453 | 1.083 | 0.438 | -0.001 | 0.011 | 0.011 | 0.198 | 0.54 | 0.56 | 0.58 | 0.63 | 0.45 | 0.47 |
| 6 | 6959879 | rs7771496 | 1.595 | 1.519 | 1.356 | 0.473 | 0.025 | -0.002 | -0.002 | 3.775 | 0.48 | 0.23 | 0.49 | 0.54 | 0.38 | 0.4 |
| 6 | 6960310 | rs1536241 | -1.033 | 1.532 | 1.366 | 0.357 | -0.004 | 0.005 | 0.005 | 0.089 | 0.4 | 0.39 | 0.31 | 0.3 | 0.5 | 0.41 |
| 6 | 6963959 | rs2765357 | -0.106 | 1.942 | 1.154 | 0.384 | -0.04 | 0.018 | 0.018 | 0.159 | 0.18 | 0.04 | 0.19 | 0.17 | 0.27 | 0.14 |
| 6 | 6964649 | rs4142382 | 0.964 | 1.784 | 1.167 | 0.377 | -0.042 | 0.012 | 0.012 | 0.225 | 0.61 | 0.81 | 0.65 | 0.64 | 0.49 | 0.57 |
| 6 | 6973110 | rs1980810 | 0.635 | 1.115 | 1.236 | 0.488 | -0.015 | -0.002 | -0.002 | 0 | 0.67 | 0.84 | 0.72 | 0.65 | 0.61 | 0.62 |
| 6 | 6973630 | rs9379070 | 1.327 | 1.083 | 1.351 | 0.529 | -0.01 | -0.001 | -0.001 | 0 | 0.57 | 0.24 | 0.56 | 0.56 | 0.51 | 0.44 |
| 6 | 6973651 | rs4960278 | -1.544 | 1.033 | 1.438 | 0.452 | 0.051 | 0.142 | 0.142 | 0.003 | 0.07 | 0.47 | 0.09 | 0.09 | 0.03 | 0.15 |
| 6 | 6978633 | rs1892512 | 1.157 | 1.154 | 0.883 | 0.458 | -0.007 | 0.014 | 0.014 | 0.192 | 0.57 | 0.35 | 0.58 | 0.64 | 0.5 | 0.5 |
| 6 | 6981639 | rs7759019 | -1.805 | 0.643 | 1.074 | 0.603 | 0.058 | 0.057 | 0.057 | 2.672 | 0.11 | 0.48 | 0.12 | 0.1 | 0.11 | 0.07 |
| 6 | 6982279 | rs6597241 | -2.457 | 0.67 | 1.056 | 0.573 | -0.003 | -0.012 | -0.012 | 0 | 0.2 | 0.64 | 0.19 | 0.14 | 0.18 | 0.2 |
| 6 | 6985715 | rs7341301 | -1.856 | 0.245 | 0.225 | 0.873 | -0.014 | 0.011 | 0.011 | 0 | 0.12 | 0.26 | 0.11 | 0.12 | 0.08 | 0.18 |

| | | | | | | | | | | | | | | | | |
|---|---|---|---|---|---|---|---|---|---|---|---|---|---|---|---|---|
| 6 | 6985996 | rs9502527 | -1.893 | 0.245 | 0.226 | 0.873 | -0.018 | 0.015 | 0.015 | 0 | 0.13 | 0.32 | 0.12 | 0.12 | 0.08 | 0.18 |
| 6 | 6986617 | rs2765366 | 0.523 | 0.519 | -0.422 | 1.129 | -0.007 | -0.048 | -0.048 | 3.921 | 0.91 | 1 | 0.94 | 0.84 | 0.94 | 0.8 |
| 6 | 6988005 | rs7761644 | -1.461 | 0.746 | -0.022 | 1.004 | 0 | -0.001 | -0.001 | 0 | 0.09 | 0.11 | 0.07 | 0.13 | 0.06 | 0.09 |
| 6 | 6991304 | rs12206270 | -0.742 | 0.823 | 0.018 | 0.925 | 0 | 0.001 | 0.001 | 0 | 0.06 | 0.07 | 0.03 | 0.05 | 0.06 | 0.02 |
| 6 | 6994388 | rs4960285 | -1.012 | 0.914 | 0.322 | 1.517 | 0.022 | 0.11 | 0.11 | 0 | 0.07 | 0.18 | 0.15 | 0.31 | 0.11 | 0.26 |
| 6 | 6996479 | rs10458204 | -0.74 | 0.557 | -0.395 | 0.561 | 0.007 | 0.004 | 0.004 | 0 | 0.16 | 0.05 | 0.15 | 0.14 | 0.16 | 0.11 |
| 6 | 7005642 | rs12196079 | -0.774 | 0.53 | -0.408 | 0.429 | 0.008 | 0.021 | 0.021 | 0.064 | 0.16 | 0.05 | 0.14 | 0.1 | 0.15 | 0.11 |
| 6 | 7015106 | rs12213148 | -0.367 | 0.512 | -0.224 | 0.272 | 0.001 | -0.006 | -0.006 | 0 | 0.06 | 0.03 | 0.02 | 0.05 | 0.05 | 0.02 |
| 6 | 7015958 | rs1750242 | 0.821 | 0.865 | 0.383 | 0.237 | 0.033 | 0.032 | 0.032 | 0.192 | 0.89 | 0.72 | 0.88 | 0.87 | 0.88 | 0.76 |
| 6 | 7020114 | rs490213 | 1.596 | 0.927 | 0.325 | 0.27 | -0.002 | -0.002 | -0.002 | 0 | 0.87 | 0.86 | 0.9 | 0.82 | 0.83 | 0.76 |
| 6 | 7023672 | rs480774 | -1.963 | -0.101 | 0.767 | 0.018 | -0.021 | -0.052 | -0.052 | 0.628 | 0.29 | 0.45 | 0.37 | 0.86 | 0.28 | 0.43 |
| 6 | 7026277 | rs12201780 | -0.045 | -0.052 | 0.063 | 0.201 | -0.01 | -0.021 | -0.021 | 0.805 | 0.05 | 0 | 0.02 | 0.05 | 0.04 | 0.02 |
| 6 | 7026562 | rs12201938 | -0.628 | -0.07 | 0.127 | 0.196 | 0.006 | 0.054 | 0.054 | 0.379 | 0.13 | 0.02 | 0.07 | 0.03 | 0.13 | 0.07 |
| 6 | 7034459 | rs1615495 | 0.298 | -0.066 | 0.034 | -0.383 | -0.001 | -0.003 | -0.003 | 0 | 0.79 | 0.8 | 0.83 | 0.86 | 0.8 | 0.63 |
| 6 | 7037477 | rs473437 | 0.094 | -0.043 | -0.048 | -0.333 | 0.002 | 0.002 | 0.002 | 0 | 0.84 | 0.89 | 0.91 | 0.87 | 0.86 | 0.72 |
| 6 | 7047957 | rs12215917 | -1.117 | -0.159 | -0.115 | -0.221 | 0.017 | -0.013 | -0.013 | 0 | 0.13 | 0.07 | 0.05 | 0.12 | 0.13 | 0.07 |
| 6 | 7071510 | rs9405997 | 1.76 | -0.138 | 0 | -0.279 | 0 | -0.001 | -0.001 | 0 | 0.84 | 0.83 | 0.86 | 0.86 | 0.85 | 0.7 |
| 6 | 7071873 | rs9505022 | -0.499 | -0.099 | 0.02 | -0.277 | 0.007 | -0.019 | -0.019 | 0 | 0.05 | 0.08 | 0.06 | 0 | 0.09 | 0.04 |
| 6 | 7074654 | rs6908957 | -0.804 | -0.179 | -0.168 | -0.321 | 0.019 | 0.005 | 0.005 | 4.154 | 0.11 | 0.07 | 0.05 | 0.06 | 0.04 | 0.02 |
| 6 | 7075782 | rs611615 | 0.39 | -0.124 | 0.318 | -0.286 | 0.03 | 0.066 | 0.066 | 0.005 | 0.95 | 0.83 | 0.93 | 0.92 | 0.96 | 0.76 |
| 6 | 7080522 | rs6912908 | -0.869 | -0.602 | 0.456 | -0.79 | -0.037 | -0.023 | -0.023 | 0 | 0.29 | 0.52 | 0.36 | 0.82 | 0.24 | 0.44 |
| 6 | 7086433 | rs486655 | -1.376 | -0.215 | 0.698 | -0.09 | -0.035 | -0.021 | -0.021 | 0.472 | 0.15 | 0.29 | 0.1 | 0.07 | 0.09 | 0.06 |
| 6 | 7090193 | rs4960288 | 1.533 | -0.189 | 0.602 | -0.049 | -0.033 | 0.006 | 0.006 | 0.045 | 0.79 | 0.65 | 0.85 | 0.83 | 0.88 | 0.69 |
| 6 | 7131524 | rs508362 | -1.179 | 0.194 | 0.646 | -0.02 | 0.01 | -0.019 | -0.019 | 0.154 | 0.07 | 0.21 | 0.03 | 0.03 | 0.07 | 0.06 |
| 6 | 7156901 | rs1285882 | -2.01 | 0.637 | 1.454 | -0.495 | 0.011 | -0.051 | -0.051 | 0.042 | 0.2 | 0.53 | 0.13 | 0.06 | 0.21 | 0.11 |
| 6 | 7166238 | rs9405328 | -2.369 | 0.323 | 1.373 | 0.27 | -0.006 | 0.007 | 0.007 | 0.012 | 0.24 | 0.55 | 0.14 | 0.32 | 0.22 | 0.17 |
| 6 | 7176744 | rs2326880 | -1.297 | 0.399 | 1.358 | 0.269 | 0.019 | 0.018 | 0.018 | 0 | 0.11 | 0.18 | 0.07 | 0.24 | 0.15 | 0.11 |
| 6 | 7181993 | rs9392862 | -2.07 | 0.356 | 1.434 | 0.246 | 0.004 | -0.001 | -0.001 | 0 | 0.16 | 0.28 | 0.09 | 0.23 | 0.16 | 0.15 |
| 6 | 7197346 | rs4959431 | 0.07 | -0.034 | 0.393 | 0.03 | 0.001 | 0.001 | 0.001 | 0 | 0.43 | 0.59 | 0.34 | 0.44 | 0.42 | 0.22 |
| 6 | 7207990 | rs6597255 | -0.536 | -0.132 | 0.56 | -0.222 | -0.001 | 0.059 | 0.059 | 0.021 | 0.55 | 0.7 | 0.52 | 0.79 | 0.52 | 0.76 |
| 6 | 7211751 | rs1334577 | 0.307 | -0.044 | 0.197 | -0.425 | 0.012 | -0.079 | -0.079 | 0.464 | 0.27 | 0.14 | 0.34 | 0.5 | 0.22 | 0.57 |
| 6 | 7211818 | rs1334576 | -0.334 | 0.147 | 0.268 | -0.488 | -0.008 | -0.004 | -0.004 | 0.219 | 0.35 | 0.32 | 0.43 | 0.53 | 0.41 | 0.64 |
| 6 | 7213279 | rs6906505 | -0.994 | 0.297 | 0.274 | -0.463 | -0.009 | 0.078 | 0.078 | 0.019 | 0.52 | 0.71 | 0.49 | 0.79 | 0.47 | 0.72 |
| 6 | 7217517 | rs6931262 | -0.324 | 0.44 | 0.394 | -0.212 | -0.01 | 0.12 | 0.12 | 0.043 | 0.06 | 0.29 | 0.03 | 0.23 | 0.03 | 0.05 |
| 6 | 7225995 | rs2714357 | -1.555 | 0.489 | -0.107 | -0.587 | 0 | 0.001 | 0.001 | 1.037 | 0.26 | 0.72 | 0.21 | 0.26 | 0.34 | 0.21 |
| 6 | 7232186 | rs9505086 | -0.287 | 0.244 | -0.28 | -0.133 | -0.001 | -0.001 | -0.001 | 0 | 0.41 | 0.55 | 0.43 | 0.4 | 0.4 | 0.41 |
| 6 | 7238069 | rs2714335 | -1.453 | 0.271 | -0.854 | -0.104 | -0.064 | -0.002 | -0.002 | 0 | 0.72 | 0.95 | 0.6 | 0.79 | 0.62 | 0.79 |
| 6 | 7247248 | rs2256596 | -1.717 | 0.328 | -0.88 | -0.141 | -0.056 | -0.017 | -0.017 | 0 | 0.75 | 0.95 | 0.63 | 0.79 | 0.66 | 0.78 |
| 6 | 7250960 | rs2714338 | -0.867 | 0.585 | 0.019 | 0.815 | 0.071 | 0.089 | 0.089 | 0.27 | 0.22 | 0.39 | 0.27 | 0.41 | 0.32 | 0.2 |
| 6 | 7251451 | rs1061637 | -0.275 | 0.39 | -0.081 | 0.845 | -0.002 | 0.043 | 0.043 | 0.05 | 0.17 | 0.27 | 0.2 | 0.37 | 0.16 | 0.16 |
| 6 | 7253731 | rs2714341 | -0.844 | 0.256 | -0.227 | 0.749 | 0.012 | 0.009 | 0.009 | 0.11 | 0.46 | 0.65 | 0.49 | 0.61 | 0.54 | 0.56 |
| 6 | 7254641 | rs2714342 | 0.006 | 0.325 | 0.129 | 0.995 | 0.008 | 0.029 | 0.029 | 0.008 | 0.13 | 0.19 | 0.19 | 0.2 | 0.15 | 0.08 |
| 6 | 7257272 | rs11965592 | -0.101 | 0.199 | -0.059 | 0.844 | -0.005 | 0.036 | 0.036 | 0 | 0.09 | 0.1 | 0.06 | 0.18 | 0.05 | 0.18 |
| 6 | 7260776 | rs2714344 | -0.522 | 0.208 | -0.269 | 0.673 | -0.022 | -0.021 | -0.021 | 0.011 | 0.39 | 0.31 | 0.43 | 0.41 | 0.42 | 0.39 |
| 6 | 7260883 | rs4960298 | 1.533 | 0.047 | -0.508 | 0.512 | -0.012 | 0.032 | 0.032 | 0 | 0.15 | 0.04 | 0.08 | 0.13 | 0.14 | 0.24 |
| 6 | 7261985 | rs2764082 | -0.508 | 0.499 | 0.042 | 0.635 | 0.001 | -0.004 | -0.004 | 1.141 | 0.41 | 0.59 | 0.39 | 0.31 | 0.45 | 0.44 |
| 6 | 7266055 | rs2842362 | -0.278 | 0.406 | 0.242 | 0.614 | -0.006 | -0.014 | -0.014 | 0 | 0.38 | 0.46 | 0.33 | 0.29 | 0.29 | 0.34 |
| 6 | 7267002 | rs9968917 | -0.49 | 0.392 | 0.37 | 0.785 | -0.032 | 0.097 | 0.097 | 0 | 0.09 | 0.24 | 0.03 | 0.23 | 0.03 | 0.08 |
| 6 | 7267582 | rs2842363 | -1.12 | 0.427 | 0.004 | 0.845 | -0.013 | 0.018 | 0.018 | 0.503 | 0.64 | 0.77 | 0.5 | 0.76 | 0.4 | 0.69 |
| 6 | 7268031 | rs11243147 | -0.744 | 0.391 | 0.283 | 0.705 | -0.003 | -0.001 | -0.001 | 0 | 0.59 | 0.64 | 0.55 | 0.56 | 0.49 | 0.68 |
| 6 | 7274354 | rs9505109 | 0.336 | 0.287 | 0.206 | 0.864 | 0 | 0.002 | 0.002 | 0.09 | 0.3 | 0.33 | 0.29 | 0.34 | 0.32 | 0.4 |
| 6 | 7278329 | rs2764085 | 0.402 | 0.117 | 0.252 | 0.349 | -0.005 | -0.03 | -0.03 | 0 | 0.3 | 0.33 | 0.19 | 0.16 | 0.22 | 0.35 |
| 6 | 7281654 | rs1050226 | 0.59 | 0.129 | -0.309 | 0.374 | -0.039 | -0.065 | -0.065 | 0 | 0.34 | 0.2 | 0.32 | 0.41 | 0.41 | 0.42 |
| 6 | 7282297 | rs9406015 | 0.276 | 0.114 | -0.212 | 0.452 | 0.01 | 0.016 | 0.016 | 0.509 | 0.1 | 0.19 | 0.06 | 0.15 | 0.08 | 0.05 |
| 6 | 7288613 | rs11719 | 1.102 | 0.059 | -0.177 | 0.361 | 0 | -0.035 | -0.035 | 0 | 0.26 | 0.3 | 0.18 | 0.16 | 0.23 | 0.33 |
| 6 | 7289166 | rs3087986 | 0.083 | 0.121 | -0.215 | 0.444 | -0.006 | -0.028 | -0.028 | 0.566 | 0.4 | 0.29 | 0.33 | 0.4 | 0.41 | 0.45 |
| 6 | 7290437 | rs9505118 | 0.083 | 0.121 | -0.215 | 0.444 | -0.006 | -0.028 | -0.028 | 0.566 | 0.4 | 0.29 | 0.33 | 0.4 | 0.42 | 0.45 |
| 6 | 7291732 | rs9505122 | 0.083 | 0.121 | -0.215 | 0.444 | -0.006 | -0.028 | -0.028 | 0 | 0.4 | 0.29 | 0.33 | 0.4 | 0.41 | 0.45 |
| 6 | 7295605 | rs9502587 | 0.083 | 0.152 | -0.225 | 0.307 | -0.005 | -0.027 | -0.027 | 0.417 | 0.4 | 0.29 | 0.33 | 0.41 | 0.41 | 0.45 |
| 6 | 7302035 | rs7748736 | -0.73 | 0.061 | -0.269 | 0.283 | 0 | -0.002 | -0.002 | 0 | 0.48 | 0.47 | 0.38 | 0.56 | 0.49 | 0.5 |
| 6 | 7302507 | rs2764092 | -1.776 | 0.289 | -0.226 | 0.422 | -0.005 | -0.031 | -0.031 | 0 | 0.76 | 0.86 | 0.58 | 0.72 | 0.72 | 0.82 |
| 6 | 7309150 | rs3778333 | -0.98 | 0.496 | -0.145 | 0.301 | 0.044 | 0.005 | 0.005 | 0.562 | 0.22 | 0.51 | 0.15 | 0.26 | 0.25 | 0.18 |
| 6 | 7312637 | rs2714354 | 0.584 | 0.671 | -0.423 | 0.538 | 0.014 | -0.006 | -0.006 | 0 | 0.34 | 0.26 | 0.28 | 0.31 | 0.28 | 0.44 |
| 6 | 7317608 | rs9328413 | -0.499 | 1.156 | -0.488 | 0.869 | 0.049 | 0.012 | 0.012 | 0.113 | 0.15 | 0.36 | 0.1 | 0.14 | 0.18 | 0.13 |
| 6 | 7323740 | rs9505150 | 0.088 | 1.281 | -0.743 | 0.844 | 0.047 | 0.001 | 0.001 | 0.03 | 0.11 | 0.24 | 0.09 | 0.08 | 0.16 | 0.11 |
| 6 | 7327846 | rs9505168 | -0.454 | 0.811 | -0.77 | 0.625 | 0.001 | 0.002 | 0.002 | 0 | 0.57 | 0.52 | 0.54 | 0.74 | 0.54 | 0.66 |
| 6 | 7328580 | rs9406022 | -0.368 | 0.481 | -1.091 | 1.124 | -0.021 | -0.051 | -0.051 | 0 | 0.34 | 0.2 | 0.27 | 0.49 | 0.33 | 0.43 |
| 6 | 7330169 | rs9379089 | -1.186 | -0.213 | -0.37 | 1.287 | -0.008 | 0.062 | 0.062 | 0 | 0.08 | 0.11 | 0.07 | 0.21 | 0.06 | 0.13 |
| 6 | 7332556 | rs1325031 | -1.14 | -0.49 | -0.216 | 0.601 | 0.006 | 0.039 | 0.039 | 0 | 0.3 | 0.4 | 0.42 | 0.44 | 0.3 | 0.3 |

| | | | | | | | | | | | | | | | | |
|---|---|---|---|---|---|---|---|---|---|---|---|---|---|---|---|---|
| 6 | 7336518 | rs2038828 | -0.932 | -1.06 | -0.03 | -0.09 | 0.023 | 0.002 | 0.002 | 2.732 | 0.48 | 0.33 | 0.46 | 0.44 | 0.4 | 0.4 |
| 6 | 7357849 | rs2764077 | -1.746 | 0.026 | 0.604 | 0.444 | 0.002 | 0.008 | 0.008 | 0 | 0.39 | 0.57 | 0.47 | 0.44 | 0.39 | 0.37 |
| 6 | 7366898 | rs7769554 | -1.983 | 0.003 | 0.59 | 0.417 | -0.052 | -0.052 | -0.052 | 0 | 0.43 | 0.9 | 0.51 | 0.55 | 0.41 | 0.38 |
| 6 | 7368917 | rs1324502 | -1.99 | 0.002 | 0.589 | 0.417 | -0.052 | -0.051 | -0.051 | 0 | 0.43 | 0.89 | 0.5 | 0.55 | 0.41 | 0.38 |
| 6 | 7374207 | rs2876098 | 0.219 | 0.039 | 0.582 | 0.455 | -0.011 | -0.01 | -0.01 | 0 | 0.09 | 0.07 | 0.15 | 0.11 | 0.15 | 0.06 |
| 6 | 7380267 | rs11243161 | 2.159 | 0.101 | 0.33 | 0.225 | -0.116 | -0.098 | -0.098 | 0 | 0.4 | 0.03 | 0.37 | 0.42 | 0.41 | 0.52 |
| 6 | 7390077 | rs6597266 | -2.257 | 0.837 | 1.532 | 1.022 | 0.01 | 0.028 | 0.028 | 0 | 0.34 | 0.45 | 0.34 | 0.44 | 0.35 | 0.29 |
| 6 | 7391921 | rs4959432 | 1.343 | 0.9 | 1.513 | 0.997 | 0.018 | 0.019 | 0.019 | 0.525 | 0.91 | 0.68 | 0.94 | 0.9 | 0.94 | 0.97 |
| 6 | 7393452 | rs2274212 | 2.162 | 0.897 | 1.349 | 1.015 | -0.032 | -0.015 | -0.015 | 0 | 0.58 | 0.26 | 0.48 | 0.46 | 0.56 | 0.62 |
| 6 | 7393786 | rs9505195 | -0.401 | 0.897 | 1.366 | 0.99 | 0.038 | -0.005 | -0.005 | 0 | 0.05 | 0.23 | 0.03 | 0 | 0.02 | 0.02 |
| 6 | 7399889 | rs11243164 | 2.417 | 0.897 | 1.361 | 0.991 | -0.03 | -0.005 | -0.005 | 6.342 | 0.56 | 0.24 | 0.48 | 0.46 | 0.56 | 0.62 |
| 6 | 7408678 | rs6927113 | 2.106 | 0.898 | 1.362 | 0.992 | -0.006 | -0.006 | -0.006 | 0 | 0.63 | 0.38 | 0.51 | 0.46 | 0.58 | 0.64 |
| 6 | 7411831 | rs12530477 | -1.312 | 0.887 | 1.367 | 1.001 | 0.006 | 0.001 | 0.001 | 0.132 | 0.18 | 0.18 | 0.2 | 0.14 | 0.26 | 0.21 |
| 6 | 7414739 | rs2296258 | -0.485 | 0.888 | 1.327 | 1.006 | 0.011 | -0.057 | -0.057 | 0 | 0.13 | 0.05 | 0.22 | 0.31 | 0.11 | 0.12 |
| 6 | 7419922 | rs11243168 | 1.884 | 0.898 | 1.37 | 0.872 | -0.039 | -0.009 | -0.009 | 0 | 0.55 | 0.19 | 0.47 | 0.46 | 0.55 | 0.61 |
| 6 | 7424510 | rs11243169 | 1.092 | 0.919 | 1.386 | 0.883 | 0.074 | 0.07 | 0.07 | 0 | 0.92 | 0.63 | 0.92 | 0.91 | 0.93 | 0.97 |
| 6 | 7427198 | rs10447470 | 1.618 | 0.945 | 0.748 | 0.892 | -0.091 | -0.056 | -0.056 | 0.016 | 0.58 | 0.1 | 0.47 | 0.45 | 0.56 | 0.61 |
| 6 | 7437357 | rs12191747 | 1.481 | 1.014 | 0.641 | 0.976 | -0.046 | -0.041 | -0.041 | 1.93 | 0.6 | 0.2 | 0.49 | 0.46 | 0.58 | 0.63 |
| 6 | 7444223 | rs12199530 | 1.693 | 1.23 | 0.774 | 1.086 | -0.049 | -0.045 | -0.045 | 0 | 0.61 | 0.19 | 0.49 | 0.46 | 0.57 | 0.62 |
| 6 | 7470321 | rs11756237 | 1.662 | 1.248 | 0.669 | 0.959 | -0.067 | -0.076 | -0.076 | 2.773 | 0.58 | 0.14 | 0.49 | 0.49 | 0.57 | 0.62 |
| 6 | 7476908 | rs2806192 | 1.77 | 1.595 | 0.728 | 1.171 | -0.039 | -0.043 | -0.043 | 0 | 0.61 | 0.21 | 0.49 | 0.49 | 0.58 | 0.62 |
| 6 | 7481928 | rs2763122 | 1.687 | 1.84 | 1.107 | 0.898 | 0.099 | 0.071 | 0.071 | 0.025 | 0.85 | 0.54 | 0.87 | 0.82 | 0.8 | 0.92 |
| 6 | 7497110 | rs823695 | -1.019 | 1.741 | 0.053 | 0.969 | 0.041 | 0.018 | 0.018 | 0 | 0.1 | 0.27 | 0.08 | 0.08 | 0.15 | 0.06 |
| 6 | 7500570 | rs6939137 | -1.279 | 1.228 | -0.251 | 0.758 | 0.005 | -0.006 | -0.006 | 0 | 0.09 | 0.13 | 0.09 | 0.06 | 0.09 | 0.09 |
| 6 | 7500699 | rs4959442 | -1.576 | 1.118 | -0.62 | 1.189 | -0.006 | -0.033 | -0.033 | 0 | 0.19 | 0.12 | 0.28 | 0.36 | 0.17 | 0.16 |
| 6 | 7501676 | rs9406032 | -1.009 | 1.222 | -0.088 | 0.827 | 0.025 | 0.008 | 0.008 | 0 | 0.31 | 0.52 | 0.42 | 0.36 | 0.34 | 0.4 |
| 6 | 7505244 | rs12214176 | -1.209 | 1.111 | 1.256 | 0.586 | 0.022 | -0.026 | -0.026 | 0 | 0.31 | 0.56 | 0.39 | 0.23 | 0.35 | 0.4 |
| 6 | 7506490 | rs2806181 | -1.437 | 0.141 | -0.07 | -0.074 | -0.119 | -0.033 | -0.033 | 0 | 0.6 | 0.95 | 0.56 | 0.35 | 0.56 | 0.5 |
| 6 | 7515692 | rs2051088 | -0.248 | 0.344 | 0.155 | -0.011 | -0.035 | 0.018 | 0.018 | 0 | 0.26 | 0.16 | 0.36 | 0.18 | 0.35 | 0.38 |
| 6 | 7517239 | rs6597269 | -0.477 | -0.057 | 0.493 | -0.201 | -0.05 | -0.051 | -0.051 | 0 | 0.23 | 0.45 | 0.17 | 0.13 | 0.18 | 0.11 |
| 6 | 7519529 | rs2798164 | -0.426 | 0.038 | 0.17 | -0.181 | -0.047 | 0.025 | 0.025 | 0.001 | 0.28 | 0.16 | 0.4 | 0.18 | 0.39 | 0.38 |
| 6 | 7524824 | rs13191992 | -0.613 | -0.385 | 0.464 | -0.334 | -0.057 | -0.016 | -0.016 | 0 | 0.17 | 0.21 | 0.04 | 0.11 | 0.05 | 0.09 |
| 6 | 7527538 | rs6597270 | -0.573 | 0.35 | 0.259 | -0.366 | -0.007 | -0.006 | -0.006 | 0 | 0.12 | 0.1 | 0.1 | 0 | 0.16 | 0.2 |
| 6 | 7528113 | rs9406039 | -0.18 | 0.557 | 0.116 | -0.15 | -0.029 | -0.034 | -0.034 | 0 | 0.13 | 0.07 | 0.3 | 0.26 | 0.23 | 0.15 |
| 6 | 7539954 | rs1322219 | -1.547 | 0.298 | 0.283 | -0.346 | -0.005 | 0.005 | 0.005 | 0 | 0.45 | 0.57 | 0.52 | 0.41 | 0.52 | 0.5 |
| 6 | 7542274 | rs36087964 | -1.525 | 0.247 | 0.469 | -0.499 | -0.003 | -0.018 | -0.018 | 0 | 0.08 | 0.12 | 0.04 | 0.02 | 0.06 | 0.04 |
| 6 | 7543783 | rs2806167 | -0.614 | 0.53 | 0.086 | -0.365 | 0.001 | 0 | 0 | 0.383 | 0.38 | 0.58 | 0.4 | 0.38 | 0.38 | 0.32 |
| 6 | 7546404 | rs1998323 | -1.275 | 0.578 | -0.008 | -0.499 | 0.02 | 0.006 | 0.006 | 0 | 0.58 | 0.75 | 0.54 | 0.39 | 0.62 | 0.51 |
| 6 | 7552020 | rs2744365 | -0.743 | 0.4 | 0.478 | -0.949 | -0.01 | -0.08 | -0.08 | 0 | 0.3 | 0.36 | 0.17 | 0.06 | 0.25 | 0.25 |
| 6 | 7552162 | rs2237103 | -0.782 | 0.404 | 0.438 | -0.947 | -0.014 | -0.105 | -0.105 | 0 | 0.31 | 0.43 | 0.18 | 0.06 | 0.25 | 0.25 |
| 6 | 7557048 | rs10484325 | -0.582 | 0.662 | 0.402 | -0.751 | 0.044 | -0.05 | -0.05 | 0 | 0.09 | 0.21 | 0.09 | 0 | 0.14 | 0.05 |
| 6 | 7558318 | rs10484326 | -0.369 | 0.225 | 0.048 | -1.242 | 0.017 | 0.045 | 0.045 | 0 | 0.29 | 0.16 | 0.15 | 0.05 | 0.26 | 0.15 |
| 6 | 7561661 | rs2757632 | -1.002 | 0.357 | 0.341 | -1.326 | -0.003 | -0.099 | -0.099 | 0.312 | 0.4 | 0.49 | 0.24 | 0.05 | 0.42 | 0.27 |
| 6 | 7563232 | rs2076295 | -0.884 | 1.128 | 0.135 | -0.372 | 0.04 | 0.045 | 0.045 | 0 | 0.33 | 0.51 | 0.42 | 0.47 | 0.41 | 0.31 |
| 6 | 7564461 | rs1358904 | -0.164 | 1.08 | 0.208 | -0.116 | 0.025 | 0.074 | 0.074 | 0 | 0.26 | 0.39 | 0.35 | 0.47 | 0.28 | 0.25 |
| 6 | 7566386 | rs2842683 | -0.269 | 0.935 | -0.36 | -0.119 | 0.01 | 0.049 | 0.049 | 0 | 0.53 | 0.73 | 0.45 | 0.33 | 0.49 | 0.43 |
| 6 | 7570963 | rs2076302 | -1.335 | 0.853 | -0.107 | -0.144 | 0 | 0.001 | 0.001 | 0 | 0.75 | 0.82 | 0.79 | 0.8 | 0.77 | 0.69 |
| 6 | 7573945 | rs9791321 | -0.251 | 0.631 | -0.694 | 0.03 | -0.055 | -0.04 | -0.04 | 0 | 0.37 | 0.21 | 0.41 | 0.47 | 0.42 | 0.31 |
| 6 | 7573956 | rs4960330 | -0.556 | 0.733 | 0.047 | 0.164 | 0 | -0.002 | -0.002 | 2.406 | 0.56 | 0.52 | 0.55 | 0.59 | 0.52 | 0.53 |
| 6 | 7574536 | rs7741957 | 0.274 | 0.565 | 0.205 | 0.584 | -0.025 | 0.187 | 0.187 | 0 | 0.05 | 0.16 | 0.15 | 0.29 | 0.01 | 0.09 |
| 6 | 7574805 | rs2259208 | -0.335 | 1.282 | -0.118 | 0.266 | -0.05 | -0.062 | -0.062 | 0 | 0.72 | 0.87 | 0.65 | 0.56 | 0.64 | 0.68 |
| 6 | 7574852 | rs2295859 | 0.604 | 1.346 | -0.148 | 0.319 | -0.001 | -0.002 | -0.002 | 0 | 0.06 | 0.08 | 0.05 | 0.08 | 0.06 | 0.04 |
| 6 | 7576527 | rs1016835 | -0.814 | 0.942 | 0.238 | 0.065 | 0.001 | 0.005 | 0.005 | 0 | 0.79 | 0.61 | 0.79 | 0.81 | 0.8 | 0.73 |
| 6 | 7579412 | rs2806225 | -0.298 | 1.214 | 0.125 | -0.045 | 0.017 | -0.024 | -0.024 | 0.019 | 0.73 | 0.6 | 0.74 | 0.8 | 0.71 | 0.72 |
| 6 | 7580958 | rs2076299 | -0.424 | 0.605 | 0.232 | -0.043 | -0.017 | 0.197 | 0.197 | 0 | 0.09 | 0.31 | 0.18 | 0.29 | 0.04 | 0.16 |
| 6 | 7581636 | rs6929069 | 0.77 | 0.639 | 0.276 | -0.131 | -0.046 | 0.017 | 0.017 | 0 | 0.2 | 0.38 | 0.19 | 0.19 | 0.14 | 0.25 |
| 6 | 7584617 | rs2076300 | 1.542 | 0.72 | 0.242 | -0.154 | -0.009 | 0.173 | 0.173 | 0 | 0.37 | 0.14 | 0.24 | 0.07 | 0.38 | 0.26 |
| 6 | 7586308 | rs12250 | 0.281 | 0.656 | 0.406 | 0.046 | -0.017 | -0.021 | -0.021 | 0 | 0.25 | 0.39 | 0.23 | 0.2 | 0.22 | 0.27 |
| 6 | 7590180 | rs6931192 | 0.271 | 0.665 | 0.411 | 0.057 | -0.017 | -0.019 | -0.019 | 0 | 0.26 | 0.4 | 0.24 | 0.2 | 0.22 | 0.27 |
| 6 | 7590828 | rs9328431 | 0.136 | 0.581 | 0.422 | 0.224 | -0.013 | 0.087 | 0.087 | 0 | 0.08 | 0.14 | 0.17 | 0.28 | 0.04 | 0.15 |
| 6 | 7595126 | rs12664272 | 0.22 | 0.623 | -0.022 | -0.44 | -0.023 | 0.04 | 0.04 | 0 | 0.26 | 0.34 | 0.35 | 0.48 | 0.17 | 0.4 |
| 6 | 7596192 | rs2806222 | 0.704 | 0.828 | -0.172 | -0.686 | 0 | -0.045 | -0.045 | 0 | 0.47 | 0.49 | 0.31 | 0.08 | 0.47 | 0.28 |
| 6 | 7600469 | rs6597271 | 0.629 | 1.02 | -0.141 | -0.595 | 0.102 | -0.029 | -0.029 | 0.065 | 0.09 | 0.35 | 0.07 | 0 | 0.1 | 0.01 |
| 6 | 7601720 | rs3734586 | 0.329 | 0.932 | -0.197 | -0.562 | 0.025 | -0.048 | -0.048 | 0 | 0.25 | 0.17 | 0.34 | 0.48 | 0.17 | 0.41 |
| 6 | 7602857 | rs9502622 | 0.233 | 1.455 | 0.308 | -0.335 | 0.094 | -0.036 | -0.036 | 0 | 0.09 | 0.35 | 0.07 | 0 | 0.1 | 0.02 |
| 6 | 7606255 | rs2757597 | -0.412 | 1.506 | 0.289 | -0.213 | -0.05 | -0.042 | -0.042 | 0 | 0.73 | 0.86 | 0.66 | 0.6 | 0.65 | 0.69 |
| 6 | 7610036 | rs2757598 | -0.4 | 1.539 | 0.307 | -0.211 | -0.046 | -0.038 | -0.038 | 0.549 | 0.73 | 0.86 | 0.66 | 0.6 | 0.64 | 0.69 |
| 6 | 7610486 | rs2806218 | -0.376 | 1.48 | 0.257 | -0.222 | -0.048 | -0.041 | -0.041 | 3.318 | 0.73 | 0.85 | 0.66 | 0.6 | 0.65 | 0.69 |
| 6 | 7610817 | rs1044764 | 0.391 | 1.475 | 0.217 | -0.242 | 0.102 | -0.029 | -0.029 | 0.027 | 0.09 | 0.35 | 0.07 | 0 | 0.1 | 0.01 |

| | | | | | | | | | | | | | | | | |
|---|---|---|---|---|---|---|---|---|---|---|---|---|---|---|---|---|
| 6 | 7612038 | rs1044761 | 0.391 | 1.475 | 0.214 | -0.242 | 0.101 | -0.03 | -0.03 | 0.932 | 0.09 | 0.35 | 0.07 | 0 | 0.1 | 0.01 |
| 6 | 7612268 | rs2757599 | -0.419 | 1.516 | 0.158 | -0.229 | -0.065 | -0.054 | -0.054 | 0 | 0.73 | 0.89 | 0.66 | 0.61 | 0.65 | 0.69 |
| 6 | 7614629 | rs7767437 | 0.027 | 1.245 | 0.673 | -0.476 | 0.102 | -0.031 | -0.031 | 0 | 0.09 | 0.36 | 0.07 | 0 | 0.1 | 0.01 |
| 6 | 7618813 | rs9378496 | -0.547 | 1.043 | -0.389 | -0.188 | -0.004 | -0.066 | -0.066 | 0.63 | 0.3 | 0.16 | 0.43 | 0.54 | 0.28 | 0.38 |
| 6 | 7623854 | rs9406042 | -0.655 | 1.078 | -0.284 | -0.266 | 0.001 | -0.008 | -0.008 | 0 | 0.37 | 0.37 | 0.48 | 0.56 | 0.35 | 0.44 |
| 6 | 7624941 | rs2181111 | 0.709 | 1.535 | 0.225 | -0.688 | 0.004 | -0.047 | -0.047 | 0 | 0.5 | 0.48 | 0.35 | 0.05 | 0.44 | 0.27 |
| 6 | 7626819 | rs9379120 | 0.479 | 1.486 | 0.182 | -0.436 | -0.002 | 0.003 | 0.003 | 0 | 0.59 | 0.33 | 0.69 | 0.65 | 0.61 | 0.53 |
| 6 | 7629966 | rs13212018 | 0.623 | 1.555 | 0.572 | -0.414 | -0.004 | 0.001 | 0.001 | 0.061 | 0.62 | 0.39 | 0.59 | 0.7 | 0.49 | 0.54 |
| 6 | 7633658 | rs9502630 | 0.898 | 1.605 | 0.72 | -0.777 | 0.008 | -0.009 | -0.009 | 0 | 0.51 | 0.45 | 0.33 | 0.04 | 0.48 | 0.29 |
| 6 | 7639693 | rs7453895 | -0.299 | 1.711 | 0.782 | -0.649 | 0.043 | -0.034 | -0.034 | 0 | 0.08 | 0.18 | 0.13 | 0 | 0.21 | 0.08 |
| 6 | 7640927 | rs11969065 | -0.231 | 1.718 | 0.719 | -0.644 | 0.068 | -0.039 | -0.039 | 0 | 0.08 | 0.22 | 0.13 | 0 | 0.21 | 0.08 |
| 6 | 7641836 | rs9406044 | 0.028 | 1.358 | 0.643 | -0.249 | 0.016 | 0.055 | 0.055 | 0 | 0.73 | 0.58 | 0.71 | 0.69 | 0.71 | 0.61 |
| 6 | 7643033 | rs9392164 | -0.482 | 1.414 | 0.018 | -0.278 | -0.019 | -0.042 | -0.042 | 0 | 0.22 | 0.14 | 0.38 | 0.66 | 0.23 | 0.29 |
| 6 | 7644889 | rs7767637 | -0.1 | 1.43 | 0.06 | -0.202 | -0.007 | -0.016 | -0.016 | 2.216 | 0.75 | 0.79 | 0.72 | 0.71 | 0.71 | 0.61 |
| 6 | 7646108 | rs9406045 | -0.673 | 1.663 | -0.212 | 0.244 | -0.029 | -0.061 | -0.061 | 1.492 | 0.2 | 0.1 | 0.38 | 0.65 | 0.22 | 0.29 |
| 6 | 7649849 | rs9392168 | -0.603 | 1.694 | -0.273 | 0.26 | -0.04 | -0.076 | -0.076 | 0.024 | 0.19 | 0.08 | 0.38 | 0.65 | 0.22 | 0.29 |
| 6 | 7652173 | rs2327016 | -0.188 | 1.947 | 0 | 0.061 | -0.003 | 0.001 | 0.001 | 0 | 0.13 | 0.13 | 0.37 | 0.71 | 0.21 | 0.25 |
| 6 | 7654391 | rs1555479 | -0.096 | 1.964 | 0.237 | -0.217 | 0.004 | -0.005 | -0.005 | 0 | 0.78 | 0.79 | 0.77 | 0.85 | 0.75 | 0.65 |
| 6 | 7667453 | rs9378503 | -0.628 | 1.655 | 0.013 | 0.16 | 0.008 | 0.019 | 0.019 | 0 | 0.17 | 0.22 | 0.39 | 0.68 | 0.24 | 0.26 |
| 6 | 7668709 | rs9505252 | -0.517 | 0.398 | 0.115 | -0.816 | -0.07 | -0.096 | -0.096 | 0.074 | 0.18 | 0.37 | 0.06 | 0.02 | 0.07 | 0.09 |
| 6 | 7668923 | rs1885487 | 0.246 | 1.718 | -0.074 | -0.518 | 0 | -0.002 | -0.002 | 0 | 0.77 | 0.73 | 0.74 | 0.86 | 0.71 | 0.66 |
| 6 | 7669953 | rs9328434 | -0.987 | 1.925 | -0.219 | -0.113 | 0.007 | -0.037 | -0.037 | 0 | 0.09 | 0.11 | 0.15 | 0.01 | 0.11 | 0.05 |
| 6 | 7673719 | rs1322237 | 0.712 | 2.203 | -0.601 | 0.562 | -0.004 | -0.003 | -0.003 | 0.022 | 0.85 | 0.89 | 0.61 | 0.32 | 0.79 | 0.6 |
| 6 | 7676267 | rs12214941 | -1.069 | 1.948 | -0.652 | 0.149 | -0.034 | 0.032 | 0.032 | 0 | 0.07 | 0.03 | 0.08 | 0 | 0.2 | 0.1 |
| 6 | 7680228 | rs6920279 | -0.331 | 0.683 | -0.522 | 0.088 | -0.017 | -0.017 | -0.017 | 0.107 | 0.08 | 0.08 | 0.01 | 0 | 0 | 0.03 |
| 6 | 7685326 | rs4959451 | -0.029 | 1.147 | -1.426 | 0.626 | -0.091 | 0.184 | 0.184 | 0.668 | 0.61 | 0.95 | 0.67 | 0.87 | 0.56 | 0.86 |
| 6 | 7704133 | rs2146064 | -0.06 | 1.25 | -0.469 | 0.479 | -0.016 | 0.191 | 0.191 | 0.937 | 0.59 | 0.85 | 0.68 | 0.87 | 0.58 | 0.86 |
| 6 | 7706004 | rs6927500 | 0.287 | 1.005 | 0.49 | 0.885 | -0.002 | 0.054 | 0.054 | 0.031 | 0.49 | 0.35 | 0.34 | 0.25 | 0.46 | 0.54 |
| 6 | 7718590 | rs13216054 | 0.453 | 0.925 | 0.514 | 0.816 | -0.03 | 0.113 | 0.113 | 0 | 0.46 | 0.2 | 0.32 | 0.23 | 0.46 | 0.53 |
| 6 | 7720059 | rs12198986 | 0.47 | 0.926 | 0.514 | 0.816 | -0.031 | 0.112 | 0.112 | 0 | 0.46 | 0.2 | 0.32 | 0.23 | 0.46 | 0.53 |
| 6 | 7726057 | rs1107495 | -0.737 | 1.287 | 0.531 | 0.966 | 0.066 | 0.223 | 0.223 | 0.028 | 0.07 | 0.16 | 0.36 | 0.45 | 0.18 | 0.24 |
| 6 | 7729614 | rs270417 | -0.48 | 1.06 | -0.204 | 0.517 | -0.004 | 0.191 | 0.191 | 4.451 | 0.66 | 0.93 | 0.8 | 0.97 | 0.69 | 0.88 |
| 6 | 7730944 | rs7753111 | -0.601 | 0.952 | -0.098 | 0.436 | 0.008 | 0.008 | 0.008 | 0.628 | 0.44 | 0.48 | 0.3 | 0.27 | 0.39 | 0.3 |
| 6 | 7733862 | rs6910759 | -1.047 | 1.139 | -0.949 | 0.341 | -0.028 | 0.119 | 0.119 | 0 | 0.52 | 0.88 | 0.68 | 0.73 | 0.57 | 0.5 |
| 6 | 7734059 | rs6900447 | 0.496 | 1.009 | -0.981 | 0.209 | 0.051 | 0.131 | 0.131 | 6.318 | 0.23 | 0.05 | 0.1 | 0 | 0.18 | 0.09 |
| 6 | 7735788 | rs932659 | -0.938 | 1.735 | -0.704 | 1.123 | 0 | 0.05 | 0.05 | 0.793 | 0.07 | 0.04 | 0.19 | 0.35 | 0.19 | 0.22 |
| 6 | 7736743 | rs2068361 | -0.133 | 0.719 | -0.486 | 0.434 | 0.028 | 0.028 | 0.028 | 4.038 | 0.34 | 0.21 | 0.3 | 0.25 | 0.25 | 0.13 |
| 6 | 7737808 | rs270406 | 0.258 | 0.884 | -0.593 | 0.426 | 0.06 | 0.051 | 0.051 | 1.226 | 0.5 | 0.68 | 0.58 | 0.59 | 0.59 | 0.72 |
| 6 | 7738931 | rs2876117 | 0.335 | 1.309 | -0.761 | 0.191 | -0.054 | 0.079 | 0.079 | 0 | 0.36 | 0.17 | 0.34 | 0.17 | 0.5 | 0.45 |
| 6 | 7738959 | rs911749 | -0.603 | 1.141 | -0.729 | 0.314 | 0.003 | -0.006 | -0.006 | 0 | 0.21 | 0.17 | 0.3 | 0.31 | 0.18 | 0.11 |
| 6 | 7739175 | rs98899 | 0.386 | 0.767 | -0.264 | 0.031 | 0.064 | -0.028 | -0.028 | 0.007 | 0.91 | 0.54 | 0.9 | 1 | 0.92 | 0.96 |
| 6 | 7741401 | rs4143048 | 0.049 | 0.721 | -0.771 | 0.17 | 0.011 | -0.047 | -0.047 | 0.365 | 0.13 | 0.04 | 0.15 | 0.28 | 0.08 | 0.05 |
| 6 | 7742590 | rs12214906 | 0.776 | 0.698 | -0.764 | 0.071 | -0.035 | 0.055 | 0.055 | 0 | 0.36 | 0.2 | 0.41 | 0.19 | 0.51 | 0.45 |
| 6 | 7743663 | rs9505270 | -1.147 | 0.764 | -0.28 | 0.07 | -0.001 | -0.001 | -0.001 | 0.517 | 0.27 | 0.3 | 0.35 | 0.3 | 0.24 | 0.15 |
| 6 | 7749643 | rs270413 | 0.752 | 0.468 | -0.714 | -0.359 | -0.024 | 0.075 | 0.075 | 0 | 0.39 | 0.19 | 0.41 | 0.19 | 0.52 | 0.45 |
| 6 | 7749972 | rs17685208 | 0.289 | 0.003 | -0.781 | -0.361 | 0.003 | -0.048 | -0.048 | 0 | 0.15 | 0.04 | 0.17 | 0.27 | 0.12 | 0.06 |
| 6 | 7757374 | rs270404 | 0.689 | -0.017 | -0.716 | -0.418 | 0.027 | 0.076 | 0.076 | 0 | 0.46 | 0.31 | 0.48 | 0.19 | 0.59 | 0.48 |
| 6 | 7759341 | rs10498670 | 0.172 | 0.023 | -0.776 | -0.374 | -0.009 | -0.065 | -0.065 | 0 | 0.15 | 0.02 | 0.16 | 0.28 | 0.11 | 0.06 |
| 6 | 7762715 | rs270378 | -0.063 | 0.071 | -0.403 | -0.219 | 0.004 | -0.003 | -0.003 | 0.462 | 0.61 | 0.39 | 0.65 | 0.47 | 0.7 | 0.55 |
| 6 | 7763858 | rs17673852 | 0.594 | 0.286 | -0.75 | -0.136 | -0.002 | -0.009 | -0.009 | 0 | 0.07 | 0.05 | 0.11 | 0 | 0.1 | 0.06 |
| 6 | 7765840 | rs270398 | -0.919 | -0.027 | -0.625 | -0.029 | 0.019 | -0.03 | -0.03 | 0.544 | 0.74 | 0.82 | 0.81 | 0.49 | 0.81 | 0.63 |
| 6 | 7780436 | rs11243205 | 0.228 | 1.824 | -0.466 | 0.163 | 0.057 | 0.022 | 0.022 | 0.514 | 0.06 | 0.07 | 0.26 | 0.17 | 0.23 | 0.27 |
| 6 | 7785459 | rs9328444 | -0.388 | 1.625 | -0.72 | 0.069 | -0.045 | -0.073 | -0.073 | 0.578 | 0.18 | 0.08 | 0.4 | 0.52 | 0.3 | 0.31 |
| 6 | 7788602 | rs9379137 | -0.659 | 1.202 | -0.527 | -0.161 | -0.009 | -0.02 | -0.02 | 0 | 0.31 | 0.28 | 0.49 | 0.55 | 0.37 | 0.37 |
| 6 | 7789943 | rs13216391 | 0.109 | 0.824 | -0.371 | -0.607 | -0.01 | -0.094 | -0.094 | 1.664 | 0.19 | 0.31 | 0.15 | 0 | 0.16 | 0.09 |
| 6 | 7793274 | rs911751 | -0.972 | 0.79 | -0.532 | -0.177 | 0.003 | 0 | 0 | 0.12 | 0.5 | 0.61 | 0.64 | 0.55 | 0.52 | 0.46 |
| 6 | 7807436 | rs927400 | 0.808 | 0.744 | -0.608 | -0.292 | 0.012 | 0.061 | 0.061 | 0.033 | 0.5 | 0.33 | 0.29 | 0.26 | 0.45 | 0.42 |
| 6 | 7810021 | rs2326994 | -0.606 | 0.007 | -0.405 | -0.984 | 0.001 | 0.068 | 0.068 | 2.102 | 0.67 | 0.81 | 0.84 | 0.99 | 0.71 | 0.84 |
| 6 | 7824236 | rs6913143 | -0.685 | 0.344 | 0.63 | -0.243 | 0.019 | -0.04 | -0.04 | 0.218 | 0.08 | 0.17 | 0.05 | 0 | 0.1 | 0.08 |
| 6 | 7825257 | rs10498673 | -0.962 | 0.321 | 0.81 | -0.255 | 0.055 | -0.035 | -0.035 | 0 | 0.09 | 0.27 | 0.06 | 0 | 0.1 | 0.08 |
| 6 | 7830141 | rs267180 | -0.271 | 0.086 | 0.692 | 0.287 | 0.002 | 0.061 | 0.061 | 0.541 | 0.23 | 0.36 | 0.41 | 0.6 | 0.25 | 0.32 |
| 6 | 7834019 | rs267186 | -0.895 | 0.227 | 0.511 | 0.003 | -0.002 | 0.002 | 0.002 | 2.268 | 0.36 | 0.36 | 0.56 | 0.74 | 0.38 | 0.47 |
| 6 | 7838510 | rs6940057 | 0.229 | 0.145 | 0.782 | -0.124 | -0.002 | -0.017 | -0.017 | 0 | 0.06 | 0.22 | 0.05 | 0 | 0.04 | 0 |
| 6 | 7838618 | rs267187 | -0.588 | 0.086 | 0.693 | -0.105 | 0.009 | -0.008 | -0.008 | 0 | 0.31 | 0.4 | 0.56 | 0.79 | 0.34 | 0.45 |
| 6 | 7842307 | rs267191 | -1.301 | 0.054 | 0.735 | -0.094 | 0.001 | 0.023 | 0.023 | 1.05 | 0.43 | 0.64 | 0.62 | 0.79 | 0.43 | 0.49 |
| 6 | 7846500 | rs267193 | -0.953 | 0.242 | 0.838 | 0.053 | 0.003 | -0.014 | -0.014 | 0.352 | 0.37 | 0.44 | 0.57 | 0.79 | 0.39 | 0.49 |
| 6 | 7857732 | rs267203 | -0.895 | -0.274 | 0.631 | -0.176 | 0.002 | -0.026 | -0.026 | 0.31 | 0.76 | 0.82 | 0.84 | 1 | 0.76 | 0.94 |
| 6 | 7860317 | rs267205 | -0.615 | -0.178 | 1.212 | 0.557 | -0.007 | 0.045 | 0.045 | 0.476 | 0.25 | 0.36 | 0.47 | 0.68 | 0.24 | 0.35 |
| 6 | 7868866 | rs1237087 | -0.369 | -0.063 | 1.446 | 0.633 | 0.002 | 0.014 | 0.014 | 1.456 | 0.23 | 0.28 | 0.48 | 0.61 | 0.27 | 0.34 |

| | | | | | | | | | | | | | | | | |
|---|---|---|---|---|---|---|---|---|---|---|---|---|---|---|---|---|
| 6 | 7870593 | rs9505293 | -0.674 | -0.064 | 1.416 | 0.47 | -0.004 | -0.011 | -0.011 | 0 | 0.14 | 0.22 | 0.11 | 0.11 | 0.13 | 0.11 |
| 6 | 7872469 | rs592849 | -1.111 | -0.031 | 1.547 | 0.589 | -0.011 | 0.013 | 0.013 | 1.807 | 0.33 | 0.55 | 0.44 | 0.6 | 0.29 | 0.36 |
| 6 | 7875033 | rs1225930 | -0.278 | -0.136 | 1.512 | 0.108 | 0.009 | 0.052 | 0.052 | 2.771 | 0.26 | 0.48 | 0.5 | 0.66 | 0.28 | 0.35 |
| 6 | 7881931 | rs1043784 | -0.862 | 0.207 | 0.979 | 0.2 | 0.005 | -0.019 | -0.019 | 0 | 0.15 | 0.52 | 0.13 | 0.1 | 0.13 | 0.12 |
| 6 | 7887223 | rs372578 | -0.468 | -0.05 | 0.44 | -0.573 | -0.024 | 0.028 | 0.028 | 0.008 | 0.29 | 0.49 | 0.49 | 0.67 | 0.25 | 0.37 |
| 6 | 7889465 | rs1225946 | -0.457 | 0.075 | 0.478 | -0.455 | -0.01 | 0.009 | 0.009 | 0.799 | 0.31 | 0.6 | 0.52 | 0.67 | 0.3 | 0.35 |
| 6 | 7890272 | rs1225948 | -0.05 | 0.243 | -0.084 | -0.348 | 0.001 | 0.015 | 0.015 | 0 | 0.25 | 0.33 | 0.49 | 0.67 | 0.27 | 0.34 |
| 6 | 7891160 | rs13873 | -0.013 | 0.204 | -0.153 | -0.323 | 0.001 | 0.037 | 0.037 | 0.07 | 0.25 | 0.42 | 0.49 | 0.65 | 0.27 | 0.34 |
| 6 | 7897400 | rs420901 | -0.801 | 0.469 | -1.195 | -0.682 | -0.066 | 0.194 | 0.194 | 2.493 | 0.68 | 0.95 | 0.82 | 0.97 | 0.67 | 0.81 |
| 6 | 7907139 | rs632177 | -0.666 | 0.41 | -0.235 | 0.392 | 0.001 | -0.003 | -0.003 | 0 | 0.29 | 0.29 | 0.5 | 0.67 | 0.3 | 0.37 |
| 6 | 7911702 | rs3812162 | 0.826 | 0.611 | -0.817 | 0.081 | -0.036 | 0.165 | 0.165 | 0.101 | 0.46 | 0.19 | 0.4 | 0.19 | 0.48 | 0.42 |
| 6 | 7912390 | rs1743635 | -0.19 | 0.341 | -0.287 | 0.149 | 0 | -0.001 | -0.001 | 0.463 | 0.78 | 0.78 | 0.85 | 0.87 | 0.77 | 0.77 |
| 6 | 7913546 | rs1632346 | -0.166 | 0.168 | -0.654 | 0.798 | 0.026 | -0.025 | -0.025 | 0.469 | 0.15 | 0.08 | 0.19 | 0.32 | 0.11 | 0.15 |
| 6 | 7913556 | rs1620656 | 0.192 | 0.713 | 0.072 | 0.522 | 0.002 | -0.018 | -0.018 | 1.028 | 0.76 | 0.77 | 0.84 | 0.87 | 0.76 | 0.77 |
| 6 | 7917086 | rs1653098 | -0.902 | 0.724 | 0.35 | 0.964 | 0.006 | 0.043 | 0.043 | 0.321 | 0.2 | 0.37 | 0.27 | 0.34 | 0.22 | 0.15 |
| 6 | 7917528 | rs9505309 | -0.698 | 0.827 | 0.243 | 0.943 | -0.001 | -0.006 | -0.006 | 0.033 | 0.18 | 0.19 | 0.21 | 0.28 | 0.2 | 0.15 |
| 6 | 7918311 | rs6922018 | -0.919 | 1.089 | 0.325 | 0.973 | 0.004 | 0.004 | 0.004 | 0 | 0.21 | 0.26 | 0.27 | 0.28 | 0.28 | 0.21 |
| 6 | 7920361 | rs1594467 | 0.741 | 0.916 | -0.014 | -0.291 | 0.002 | -0.026 | -0.026 | 0.568 | 0.46 | 0.46 | 0.26 | 0.08 | 0.35 | 0.44 |
| 6 | 7920808 | rs419588 | -0.469 | 1.032 | -0.364 | 0.259 | -0.101 | -0.137 | -0.137 | 1.38 | 0.71 | 0.98 | 0.6 | 0.36 | 0.64 | 0.7 |
| 6 | 7920904 | rs365936 | -0.203 | 0.457 | -0.287 | -0.137 | -0.006 | -0.007 | -0.007 | 0 | 0.45 | 0.68 | 0.5 | 0.42 | 0.41 | 0.63 |
| 6 | 7924658 | rs400299 | -0.098 | -0.009 | 0.156 | -0.859 | -0.001 | -0.003 | -0.003 | 0 | 0.68 | 0.37 | 0.69 | 0.74 | 0.73 | 0.5 |
| 6 | 7938773 | rs155487 | 0.894 | 0.137 | 0.566 | -1.018 | 0.135 | 0.122 | 0.122 | 0.04 | 0.89 | 0.6 | 0.77 | 0.81 | 0.83 | 0.78 |
| 6 | 7939671 | rs7759627 | 0.312 | -0.234 | 0.708 | -1.189 | -0.003 | -0.007 | -0.007 | 0 | 0.25 | 0.28 | 0.09 | 0.08 | 0.14 | 0.3 |
| 6 | 7943101 | rs2748359 | 0.082 | -0.271 | 0.526 | -1.725 | 0 | -0.01 | -0.01 | 0 | 0.63 | 0.41 | 0.62 | 0.79 | 0.64 | 0.7 |
| 6 | 7949806 | rs155502 | -0.939 | -0.388 | 0.6 | -1.663 | -0.025 | -0.006 | -0.006 | 0 | 0.79 | 0.64 | 0.88 | 0.81 | 0.89 | 0.69 |
| 6 | 7951721 | rs155504 | 0.631 | -0.159 | 0.832 | -1.504 | 0.033 | 0.04 | 0.04 | 0.019 | 0.93 | 0.77 | 0.89 | 0.88 | 0.91 | 0.89 |
| 6 | 7960625 | rs2748363 | 1.19 | -0.394 | 0.249 | -1.904 | -0.156 | -0.058 | -0.058 | 0 | 0.29 | 0.02 | 0.47 | 0.71 | 0.4 | 0.36 |
| 6 | 7960742 | rs9505329 | 1.105 | -0.447 | 0.622 | -2.015 | 0.012 | 0.009 | 0.009 | 0.026 | 0.25 | 0.13 | 0.15 | 0.02 | 0.2 | 0.09 |
| 6 | 7961189 | rs155476 | 0.285 | -0.463 | 0.751 | -2.027 | -0.012 | -0.112 | -0.112 | 0 | 0.32 | 0.48 | 0.2 | 0.02 | 0.27 | 0.15 |
| 6 | 7967946 | rs1150893 | 0.14 | -0.532 | 0.472 | -1.713 | 0.005 | -0.028 | -0.028 | 0.031 | 0.4 | 0.42 | 0.63 | 0.79 | 0.51 | 0.52 |
| 6 | 7973380 | rs7763447 | -0.482 | -0.366 | -0.22 | -1.694 | -0.002 | -0.003 | -0.003 | 0.103 | 0.63 | 0.62 | 0.42 | 0.17 | 0.51 | 0.53 |
| 6 | 7973917 | rs6928832 | -0.145 | -0.341 | -0.24 | -1.625 | 0.003 | 0.012 | 0.012 | 5.166 | 0.55 | 0.56 | 0.36 | 0.17 | 0.41 | 0.48 |
| 6 | 7975135 | rs4959462 | 0.436 | -0.525 | -0.763 | -1.514 | 0.066 | 0.167 | 0.167 | 0 | 0.51 | 0.18 | 0.67 | 0.77 | 0.59 | 0.43 |
| 6 | 7975259 | rs6597292 | 0.058 | -0.304 | -0.109 | -1.357 | -0.007 | -0.03 | -0.03 | 0 | 0.46 | 0.5 | 0.26 | 0.16 | 0.36 | 0.51 |
| 6 | 7979533 | rs7775460 | -0.71 | 0.299 | 0.31 | -0.893 | 0.08 | 0.003 | 0.003 | 0 | 0.15 | 0.45 | 0.22 | 0.11 | 0.21 | 0.14 |
| 6 | 7981671 | rs197123 | -0.256 | 0.123 | -0.477 | -1.572 | -0.005 | 0.012 | 0.012 | 0.038 | 0.63 | 0.66 | 0.7 | 0.8 | 0.63 | 0.57 |
| 6 | 7985800 | rs197127 | -0.803 | 0.605 | -0.475 | -1.639 | 0.027 | 0.007 | 0.007 | 0 | 0.9 | 0.86 | 0.8 | 0.87 | 0.79 | 0.89 |
| 6 | 7988189 | rs4959464 | 1.338 | 0.044 | -0.935 | -1.892 | -0.09 | -0.089 | -0.089 | 0.345 | 0.21 | 0.03 | 0.38 | 0.71 | 0.31 | 0.33 |
| 6 | 7994253 | rs13195836 | 0.867 | -0.239 | -0.897 | -2.067 | 0.013 | 0.081 | 0.081 | 0 | 0.26 | 0.09 | 0.1 | 0.04 | 0.17 | 0.31 |
| 6 | 7999843 | rs2180165 | -1.811 | -0.42 | -0.801 | -2.224 | 0.085 | 0.093 | 0.093 | 0.329 | 0.7 | 0.88 | 0.86 | 1 | 0.82 | 0.86 |
| 6 | 8001117 | rs9328448 | 0.752 | -0.149 | 0.13 | -2.131 | 0.067 | -0.002 | -0.002 | 0 | 0.11 | 0.28 | 0.18 | 0.08 | 0.18 | 0.09 |
| 6 | 8001278 | rs910552 | 1.327 | -0.12 | 0.005 | -2.014 | -0.043 | -0.043 | -0.043 | 1.163 | 0.22 | 0.11 | 0.37 | 0.71 | 0.31 | 0.33 |
| 6 | 8002142 | rs9505333 | 0.083 | -0.126 | 0.559 | -2.17 | -0.005 | -0.021 | -0.021 | 1.501 | 0.35 | 0.29 | 0.21 | 0.03 | 0.35 | 0.35 |
| 6 | 8002705 | rs12204273 | 0.186 | 0.074 | 0.664 | -1.719 | 0.102 | 0.067 | 0.067 | 0 | 0.07 | 0.28 | 0.18 | 0.14 | 0.16 | 0.07 |
| 6 | 8003485 | rs11243217 | 0.707 | 0.098 | 0.621 | -1.666 | 0.09 | 0.06 | 0.06 | 0 | 0.06 | 0.16 | 0.2 | 0.14 | 0.16 | 0.06 |
| 6 | 8005381 | rs9406074 | -0.451 | -0.242 | 0.85 | -1.86 | 0.003 | 0.001 | 0.001 | 1.517 | 0.59 | 0.4 | 0.61 | 0.83 | 0.64 | 0.73 |
| 6 | 8009354 | rs9392186 | -0.451 | -0.242 | 0.85 | -1.86 | 0.003 | 0.001 | 0.001 | 1.517 | 0.59 | 0.4 | 0.61 | 0.83 | 0.64 | 0.73 |
| 6 | 8009760 | rs7775656 | -1.057 | -0.268 | 0.897 | -1.872 | -0.056 | -0.102 | -0.102 | 0.03 | 0.64 | 0.55 | 0.81 | 0.97 | 0.8 | 0.78 |
| 6 | 8013121 | rs2057184 | -1.083 | -0.269 | 0.739 | -1.872 | -0.024 | -0.058 | -0.058 | 2.837 | 0.65 | 0.61 | 0.82 | 0.97 | 0.8 | 0.79 |
| 6 | 8013793 | rs12215188 | 0.851 | -0.244 | 0.753 | -1.87 | 0.083 | 0.054 | 0.054 | 0 | 0.06 | 0.15 | 0.2 | 0.14 | 0.16 | 0.06 |
| 6 | 8014990 | rs2815151 | -0.457 | -0.247 | 0.752 | -1.872 | -0.014 | -0.038 | -0.038 | 0 | 0.59 | 0.46 | 0.61 | 0.83 | 0.64 | 0.73 |
| 6 | 8015129 | rs2748376 | -0.465 | -0.244 | 0.753 | -1.87 | -0.014 | -0.038 | -0.038 | 0 | 0.59 | 0.46 | 0.61 | 0.83 | 0.64 | 0.73 |
| 6 | 8015275 | rs10458217 | 0.794 | -0.076 | 0.698 | -1.747 | 0.083 | 0.054 | 0.054 | 0 | 0.06 | 0.15 | 0.2 | 0.14 | 0.16 | 0.06 |
| 6 | 8015720 | rs3734590 | 0.794 | -0.076 | 0.698 | -1.747 | 0.083 | 0.054 | 0.054 | 0 | 0.06 | 0.15 | 0.2 | 0.14 | 0.16 | 0.06 |
| 6 | 8015821 | rs3734591 | 0.794 | -0.076 | 0.698 | -1.747 | 0.083 | 0.054 | 0.054 | 0 | 0.06 | 0.15 | 0.2 | 0.14 | 0.16 | 0.06 |
| 6 | 8019175 | rs9505336 | 0.018 | -0.067 | 0.71 | -1.85 | 0.069 | -0.009 | -0.009 | 0 | 0.14 | 0.45 | 0.17 | 0.08 | 0.2 | 0.11 |
| 6 | 8020467 | rs10458106 | 0.871 | -0.032 | 0.708 | -1.842 | 0.083 | 0.054 | 0.054 | 0 | 0.06 | 0.15 | 0.2 | 0.14 | 0.16 | 0.06 |
| 6 | 8026969 | rs2744009 | -0.415 | -0.206 | 0.732 | -1.965 | -0.007 | -0.062 | -0.062 | 0.163 | 0.56 | 0.53 | 0.67 | 0.94 | 0.62 | 0.76 |
| 6 | 8028980 | rs2744008 | 0.189 | -0.224 | 0.49 | -1.945 | -0.032 | 0.14 | 0.14 | 0.551 | 0.48 | 0.17 | 0.45 | 0.78 | 0.45 | 0.63 |
| 6 | 8038317 | rs2142958 | 0.128 | -0.227 | 0.22 | -1.782 | 0.064 | -0.029 | -0.029 | 0 | 0.13 | 0.31 | 0.16 | 0.06 | 0.19 | 0.09 |
| 6 | 8046602 | rs2206098 | -1.563 | -0.402 | 0.017 | -1.887 | 0.104 | 0.167 | 0.167 | 0.928 | 0.72 | 0.95 | 0.85 | 1 | 0.83 | 0.87 |
| 6 | 8054562 | rs9505348 | -0.108 | 0.611 | 0.704 | -1.037 | 0.07 | -0.018 | -0.018 | 0 | 0.13 | 0.31 | 0.16 | 0.07 | 0.2 | 0.11 |
| 6 | 8060534 | rs12663430 | 0.037 | 0.875 | 0.24 | 0.328 | -0.005 | -0.035 | -0.035 | 0 | 0.07 | 0.02 | 0.17 | 0.18 | 0.07 | 0.16 |
| 6 | 8062532 | rs12207627 | -0.094 | 1.3 | 0.842 | 0.637 | 0.083 | 0.06 | 0.06 | 0 | 0.06 | 0.16 | 0.2 | 0.16 | 0.16 | 0.06 |
| 6 | 8066286 | rs9505351 | 0.429 | 0.599 | 0.7 | 0.779 | 0.042 | 0.112 | 0.112 | 0 | 0.16 | 0.27 | 0.27 | 0.58 | 0.25 | 0.22 |
| 6 | 8071562 | rs9392958 | -0.428 | -0.292 | 0.345 | 0.276 | -0.002 | 0.002 | 0.002 | 0 | 0.34 | 0.29 | 0.37 | 0.19 | 0.36 | 0.4 |
| 6 | 8073368 | rs2744013 | -0.149 | -0.327 | 0.532 | 0.519 | 0.017 | 0.018 | 0.018 | 0.474 | 0.31 | 0.39 | 0.38 | 0.66 | 0.39 | 0.32 |
| 6 | 8078267 | rs2815162 | 0.395 | -0.333 | 0.107 | 0.93 | -0.07 | -0.085 | -0.085 | 0 | 0.22 | 0.08 | 0.23 | 0.46 | 0.32 | 0.22 |
| 6 | 8084877 | rs9379163 | -0.62 | -0.295 | 0.124 | 0.926 | -0.07 | -0.084 | -0.084 | 0 | 0.78 | 0.92 | 0.77 | 0.54 | 0.68 | 0.78 |

| | | | | | | | | | | | | | | | | |
|---|---|---|---|---|---|---|---|---|---|---|---|---|---|---|---|---|
| 6 | 8086500 | rs12199241 | 0.05 | -0.746 | 0.221 | 0.915 | -0.034 | 0.069 | 0.069 | 1.096 | 0.25 | 0.08 | 0.14 | 0.09 | 0.26 | 0.22 |
| 6 | 8101635 | rs7768487 | 0.595 | -0.832 | 0.485 | 1.006 | -0.008 | 0.103 | 0.103 | 0.27 | 0.06 | 0.16 | 0.11 | 0.21 | 0.03 | 0.04 |
| 6 | 8103048 | rs3757096 | 0.131 | -0.825 | 0.423 | 0.979 | -0.04 | 0.076 | 0.076 | 0.803 | 0.25 | 0.07 | 0.14 | 0.08 | 0.26 | 0.23 |
| 6 | 8106028 | rs1316319 | -0.832 | -0.886 | 0.726 | 0.952 | -0.002 | -0.01 | -0.01 | 0.767 | 0.34 | 0.4 | 0.27 | 0.3 | 0.29 | 0.28 |
| 6 | 8110994 | rs2039520 | -1.712 | -0.921 | 0.879 | 0.932 | 0.008 | -0.001 | -0.001 | 0 | 0.5 | 0.62 | 0.54 | 0.48 | 0.47 | 0.53 |
| 6 | 8111791 | rs11243232 | -1.072 | -0.793 | 0.833 | 0.958 | 0.001 | -0.001 | -0.001 | 0 | 0.42 | 0.39 | 0.46 | 0.43 | 0.38 | 0.5 |
| 6 | 8112440 | rs6597300 | -0.864 | -0.811 | 0.868 | 1.004 | -0.026 | 0.084 | 0.084 | 0 | 0.18 | 0.36 | 0.33 | 0.34 | 0.14 | 0.29 |
| 6 | 8113343 | rs6930629 | 0.445 | -0.789 | 0.89 | 1 | -0.005 | 0.129 | 0.129 | 0.812 | 0.07 | 0.24 | 0.12 | 0.23 | 0.03 | 0.04 |
| 6 | 8114669 | rs10458218 | 0.093 | -0.625 | 0.445 | 0.576 | -0.061 | 0.086 | 0.086 | 1.893 | 0.25 | 0.03 | 0.14 | 0.09 | 0.26 | 0.22 |
| 6 | 8115492 | rs6900034 | -0.401 | -0.1 | 1.017 | 1.12 | -0.002 | 0.001 | 0.001 | 0 | 0.11 | 0.12 | 0.22 | 0.12 | 0.11 | 0.25 |
| 6 | 8125408 | rs9378524 | -0.838 | -0.038 | 1.048 | 1.069 | -0.009 | 0.014 | 0.014 | 0 | 0.4 | 0.31 | 0.42 | 0.29 | 0.45 | 0.55 |
| 6 | 8129573 | rs12200068 | 0.107 | 0.068 | 0.787 | 1.014 | -0.049 | 0.081 | 0.081 | 0 | 0.25 | 0.08 | 0.2 | 0.06 | 0.33 | 0.26 |
| 6 | 8134765 | rs12213485 | -0.839 | 0.078 | 0.882 | 1.114 | -0.032 | 0.02 | 0.02 | 0 | 0.39 | 0.2 | 0.42 | 0.29 | 0.45 | 0.45 |
| 6 | 8141269 | rs9392965 | 0.484 | 0.003 | 1.155 | 1.45 | -0.006 | 0.062 | 0.062 | 1.014 | 0.05 | 0.08 | 0.08 | 0.2 | 0.04 | 0.04 |
| 6 | 8156183 | rs1334905 | -0.056 | 0 | 1.007 | 1.436 | 0 | -0.02 | -0.02 | 0 | 0.09 | 0.04 | 0.2 | 0.15 | 0.09 | 0.16 |
| 6 | 8158396 | rs9379168 | 0.134 | 0.119 | 1.586 | 1.442 | 0.041 | 0.198 | 0.198 | 1.33 | 0.06 | 0.24 | 0.1 | 0.2 | 0.04 | 0.04 |
| 6 | 8170017 | rs9406106 | -0.038 | 0.026 | 1.253 | 1.381 | 0.032 | 0.286 | 0.286 | 0.96 | 0.07 | 0.35 | 0.1 | 0.25 | 0.04 | 0.05 |
| 6 | 8177933 | rs11243239 | 1.295 | 0.126 | 0.509 | 0.403 | 0.011 | 0.159 | 0.159 | 1.169 | 0.35 | 0.07 | 0.22 | 0.09 | 0.24 | 0.25 |
| 6 | 8178303 | rs2327034 | 0.121 | -0.032 | 0.648 | 0.529 | -0.039 | 0.191 | 0.191 | 0.064 | 0.12 | 0.29 | 0.26 | 0.38 | 0.05 | 0.15 |
| 6 | 8178956 | rs10901021 | 1.285 | 0.29 | 0.18 | 0.397 | 0.012 | 0.161 | 0.161 | 1.136 | 0.35 | 0.07 | 0.22 | 0.09 | 0.25 | 0.25 |
| 6 | 8180286 | rs515077 | -0.454 | 0.592 | 0.516 | 0.687 | 0.001 | -0.003 | -0.003 | 0.215 | 0.37 | 0.63 | 0.47 | 0.49 | 0.37 | 0.47 |
| 6 | 8181299 | rs12719578 | 0.363 | 1.256 | 0.652 | 1.232 | 0.05 | 0.112 | 0.112 | 0.79 | 0.79 | 0.42 | 0.72 | 0.56 | 0.7 | 0.72 |
| 6 | 8184485 | rs7768905 | -0.346 | 0.921 | 0.202 | 0.699 | -0.001 | -0.009 | -0.009 | 0 | 0.08 | 0.08 | 0.06 | 0 | 0.11 | 0.1 |
| 6 | 8193128 | rs491511 | 0.427 | 0.871 | 0.617 | 0.29 | 0.005 | -0.03 | -0.03 | 0 | 0.63 | 0.56 | 0.42 | 0.11 | 0.52 | 0.5 |
| 6 | 8195366 | rs12214024 | -0.23 | 0.984 | 0.673 | 0.271 | 0.001 | -0.021 | -0.021 | 1.531 | 0.09 | 0.13 | 0.07 | 0 | 0.12 | 0.1 |
| 6 | 8200369 | rs9379173 | 1.518 | 0.74 | 0.8 | -0.102 | 0.034 | 0.088 | 0.088 | 0 | 0.41 | 0.25 | 0.23 | 0.06 | 0.25 | 0.27 |
| 6 | 8205113 | rs9378526 | 0.112 | 0.579 | 1.163 | 0.564 | 0.042 | 0.063 | 0.063 | 0.085 | 0.71 | 0.49 | 0.48 | 0.44 | 0.57 | 0.59 |
| 6 | 8207135 | rs7760150 | -1.208 | 0.589 | 1.168 | 0.555 | 0.005 | -0.114 | -0.114 | 7.046 | 0.24 | 0.41 | 0.18 | 0.05 | 0.26 | 0.23 |
| 6 | 8211191 | rs2327036 | -1.477 | 0.591 | 1.182 | 0.579 | -0.01 | -0.001 | -0.001 | 0 | 0.34 | 0.43 | 0.26 | 0.38 | 0.31 | 0.3 |
| 6 | 8211654 | rs12216170 | 0.197 | 0.596 | 0.947 | 0.56 | -0.023 | 0.056 | 0.056 | 0.229 | 0.1 | 0.02 | 0.05 | 0 | 0.11 | 0.11 |
| 6 | 8214304 | rs533632 | -0.401 | 0.565 | 1.085 | 0.953 | -0.048 | -0.053 | -0.053 | 0 | 0.77 | 0.89 | 0.66 | 0.58 | 0.56 | 0.71 |
| 6 | 8222599 | rs12200008 | -0.738 | 0.501 | 1.381 | 0.924 | 0.004 | -0.006 | -0.006 | 0 | 0.22 | 0.24 | 0.14 | 0.33 | 0.17 | 0.19 |
| 6 | 8222774 | rs9379174 | -1.101 | 0.523 | 1.377 | 0.916 | 0.003 | -0.006 | -0.006 | 0 | 0.25 | 0.25 | 0.18 | 0.35 | 0.22 | 0.2 |
| 6 | 8227374 | rs6597304 | -1.689 | 0.566 | 1.194 | 0.934 | -0.009 | -0.003 | -0.003 | 0 | 0.35 | 0.58 | 0.27 | 0.38 | 0.31 | 0.3 |
| 6 | 8230547 | rs13194449 | -1.556 | 0.615 | 0.948 | 0.973 | -0.003 | -0.001 | -0.001 | 0.341 | 0.34 | 0.39 | 0.25 | 0.38 | 0.31 | 0.3 |
| 6 | 8233002 | rs2876124 | 0.301 | 1.17 | 0.586 | 0.886 | 0.026 | 0.029 | 0.029 | 0 | 0.17 | 0.23 | 0.3 | 0.19 | 0.27 | 0.26 |
| 6 | 8235538 | rs12662470 | -0.527 | 0.79 | 0.277 | 0.794 | -0.016 | 0.017 | 0.017 | 0.195 | 0.09 | 0.06 | 0.1 | 0.05 | 0.14 | 0.11 |
| 6 | 8237195 | rs9406110 | -1.552 | 0.433 | 0.37 | 1.162 | -0.021 | 0.019 | 0.019 | 0 | 0.28 | 0.47 | 0.21 | 0.31 | 0.23 | 0.18 |
| 6 | 8240327 | rs9392980 | -0.807 | 0.19 | 0.284 | 1.468 | -0.033 | 0.068 | 0.068 | 0.005 | 0.14 | 0.29 | 0.12 | 0.25 | 0.06 | 0.07 |
| 6 | 8245115 | rs9505400 | 0.048 | 0.17 | -0.655 | 1.025 | 0.018 | 0.03 | 0.03 | 0 | 0.49 | 0.29 | 0.58 | 0.45 | 0.59 | 0.51 |
| 6 | 8247159 | rs9379177 | -0.296 | 0.085 | -0.353 | 1.147 | -0.012 | -0.019 | -0.019 | 0.396 | 0.7 | 0.39 | 0.68 | 0.77 | 0.75 | 0.79 |
| 6 | 8256964 | rs1413904 | -0.094 | 0.24 | -0.804 | 0.895 | -0.004 | 0.033 | 0.033 | 0 | 0.44 | 0.25 | 0.28 | 0.31 | 0.44 | 0.44 |
| 6 | 8260336 | rs1199414 | -0.057 | 0.164 | -0.367 | 1.069 | -0.023 | 0.029 | 0.029 | 0.496 | 0.64 | 0.56 | 0.66 | 0.55 | 0.7 | 0.59 |
| 6 | 8266473 | rs685655 | -0.223 | 0.669 | -0.16 | 0.936 | 0.001 | 0.002 | 0.002 | 0 | 0.61 | 0.61 | 0.42 | 0.35 | 0.62 | 0.58 |
| 6 | 8266826 | rs9392985 | -0.174 | 0.734 | 0.109 | 0.806 | 0.009 | -0.025 | -0.025 | 0 | 0.16 | 0.16 | 0.13 | 0.29 | 0.14 | 0.25 |
| 6 | 8269415 | rs621896 | -0.886 | 0.915 | 0.746 | 0.738 | -0.003 | 0.001 | 0.001 | 0 | 0.45 | 0.65 | 0.38 | 0.38 | 0.48 | 0.52 |
| 6 | 8273873 | rs591705 | -0.212 | 1.53 | 1.002 | 0.798 | 0.042 | -0.05 | -0.05 | 0 | 0.17 | 0.28 | 0.17 | 0.03 | 0.28 | 0.18 |
| 6 | 8278455 | rs663716 | -0.642 | 1.467 | 0.854 | 0.788 | 0.059 | 0.043 | 0.043 | 0 | 0.21 | 0.44 | 0.24 | 0.23 | 0.3 | 0.29 |
| 6 | 8291520 | rs9502704 | -0.992 | 1.467 | 0.931 | 0.544 | 0.054 | 0.096 | 0.096 | 0 | 0.06 | 0.2 | 0.1 | 0.1 | 0.09 | 0.01 |
| 6 | 8293936 | rs17143507 | -0.245 | 1.31 | 0.648 | 0.537 | -0.055 | -0.061 | -0.061 | 0 | 0.1 | 0 | 0.12 | 0.22 | 0.16 | 0.17 |
| 6 | 8304628 | rs9392991 | -0.328 | 1.389 | 0.62 | -0.186 | -0.026 | 0.092 | 0.092 | 0 | 0.3 | 0.13 | 0.23 | 0.02 | 0.33 | 0.27 |
| 6 | 8312174 | rs9406114 | -0.024 | 1.458 | 0.665 | -0.24 | -0.006 | 0.026 | 0.026 | 0 | 0.26 | 0.16 | 0.19 | 0.02 | 0.27 | 0.25 |
| 6 | 8318647 | rs4959477 | 0.074 | 1.459 | 0.618 | -0.238 | -0.013 | 0.053 | 0.053 | 0 | 0.25 | 0.12 | 0.18 | 0.01 | 0.27 | 0.25 |
| 6 | 8318698 | rs4959478 | -0.901 | 1.473 | 0.719 | -0.144 | 0.006 | -0.01 | -0.01 | 0 | 0.41 | 0.45 | 0.34 | 0.28 | 0.45 | 0.48 |
| 6 | 8326905 | rs2817084 | 0.857 | 1.064 | 0.279 | 0.082 | -0.019 | 0.057 | 0.057 | 0 | 0.53 | 0.18 | 0.61 | 0.69 | 0.53 | 0.49 |
| 6 | 8334960 | rs2327076 | -0.927 | 1.301 | 0.215 | 0.925 | 0.021 | 0.067 | 0.067 | 0.068 | 0.16 | 0.31 | 0.17 | 0.29 | 0.21 | 0.26 |
| 6 | 8344007 | rs2327042 | -0.357 | 0.972 | -0.34 | 0.151 | -0.021 | 0.05 | 0.05 | 1.35 | 0.61 | 0.73 | 0.39 | 0.22 | 0.57 | 0.51 |
| 6 | 8346462 | rs1591996 | -0.143 | 0.317 | -0.796 | 0.019 | 0.011 | 0.075 | 0.075 | 0.151 | 0.44 | 0.67 | 0.31 | 0.11 | 0.42 | 0.4 |
| 6 | 8348773 | rs1538889 | 0.813 | 0.273 | -0.566 | 0.173 | -0.061 | -0.032 | -0.032 | 0.116 | 0.35 | 0.15 | 0.62 | 0.63 | 0.44 | 0.46 |
| 6 | 8349511 | rs2185503 | -0.223 | 0.003 | -0.051 | -0.188 | -0.002 | -0.007 | -0.007 | 0.071 | 0.39 | 0.56 | 0.24 | 0.15 | 0.29 | 0.36 |
| 6 | 8350917 | rs9328464 | -0.759 | 0.221 | 0.152 | -0.317 | 0 | -0.008 | -0.008 | 0 | 0.53 | 0.44 | 0.35 | 0.26 | 0.49 | 0.49 |
| 6 | 8351628 | rs6900613 | -0.083 | 0.119 | 0.11 | -0.387 | -0.001 | -0.04 | -0.04 | 0 | 0.35 | 0.35 | 0.28 | 0.03 | 0.36 | 0.28 |
| 6 | 8352004 | rs7453084 | 0.031 | 0.22 | -0.131 | -0.299 | -0.037 | 0.027 | 0.027 | 0 | 0.21 | 0.34 | 0.09 | 0.23 | 0.14 | 0.21 |
| 6 | 8354119 | rs9393000 | -0.292 | 0.738 | -0.762 | -0.564 | -0.008 | 0.031 | 0.031 | 0.131 | 0.33 | 0.23 | 0.27 | 0.03 | 0.36 | 0.28 |
| 6 | 8357237 | rs1572852 | -0.087 | 1.178 | -0.898 | -0.272 | -0.024 | 0.017 | 0.017 | 0 | 0.17 | 0.1 | 0.18 | 0 | 0.26 | 0.19 |
| 6 | 8362964 | rs9379188 | -0.224 | 0.817 | -0.924 | 0.299 | -0.011 | -0.006 | -0.006 | 0 | 0.12 | 0.03 | 0.09 | 0.12 | 0.14 | 0.11 |
| 6 | 8362985 | rs9406122 | -0.763 | 0.235 | -1.42 | 0.758 | 0.024 | -0.054 | -0.054 | 0 | 0.28 | 0.15 | 0.19 | 0.43 | 0.19 | 0.54 |
| 6 | 8364433 | rs9406123 | 0.437 | 0.144 | -1.224 | 0.566 | 0.005 | -0.004 | -0.004 | 0.141 | 0.92 | 0.94 | 0.93 | 0.93 | 0.94 | 0.93 |
| 6 | 8367406 | rs12208331 | -0.49 | -0.034 | -1.049 | 0.357 | 0.006 | 0.011 | 0.011 | 0.28 | 0.37 | 0.28 | 0.28 | 0.35 | 0.34 | 0.38 |

| | | | | | | | | | | | | | | | | |
|---|---|---|---|---|---|---|---|---|---|---|---|---|---|---|---|---|
| 6 | 8369286 | rs12198143 | -0.01 | -0.159 | -1.012 | 0.335 | -0.028 | -0.012 | -0.012 | 0 | 0.11 | 0.26 | 0.07 | 0.11 | 0.08 | 0.16 |
| 6 | 8371230 | rs6926953 | 0.297 | 0.408 | -1.129 | -0.07 | -0.013 | 0.011 | 0.011 | 0 | 0.18 | 0.19 | 0.26 | 0.1 | 0.26 | 0.22 |
| 6 | 8376617 | rs9393004 | 1.16 | 0.095 | -1.211 | 0.133 | 0.011 | 0.04 | 0.04 | 1.436 | 0.48 | 0.21 | 0.51 | 0.39 | 0.55 | 0.43 |
| 6 | 8377927 | rs11966093 | 0.929 | 0.2 | -1.174 | 0.14 | 0.01 | -0.022 | -0.022 | 0 | 0.1 | 0.13 | 0.14 | 0 | 0.19 | 0.19 |
| 6 | 8378609 | rs10484750 | -0.395 | 0.097 | -0.876 | 0.145 | -0.054 | 0.045 | 0.045 | 1.871 | 0.36 | 0.25 | 0.36 | 0.26 | 0.46 | 0.41 |
| 6 | 8379684 | rs4959484 | 0.521 | 0.137 | -0.79 | 0.209 | 0.003 | -0.003 | -0.003 | 0 | 0.12 | 0.15 | 0.15 | 0.06 | 0.19 | 0.22 |
| 6 | 8379808 | rs6597313 | 0.255 | 0.014 | 0.099 | 0.202 | -0.021 | 0.112 | 0.112 | 0 | 0.85 | 0.6 | 0.79 | 0.74 | 0.89 | 0.79 |
| 6 | 8380193 | rs11243253 | -0.33 | 0.064 | -0.444 | 0.126 | -0.06 | 0.035 | 0.035 | 8.727 | 0.3 | 0.16 | 0.24 | 0.21 | 0.4 | 0.42 |
| 6 | 8381005 | rs10484749 | 1.702 | 0.101 | -0.447 | 0.204 | -0.02 | -0.012 | -0.012 | 0 | 0.08 | 0.01 | 0.12 | 0.1 | 0.07 | 0.02 |
| 6 | 8388757 | rs12201401 | 0.762 | 0.141 | -0.179 | 0.321 | -0.054 | -0.029 | -0.029 | 0.131 | 0.18 | 0.01 | 0.1 | 0.16 | 0.2 | 0.2 |
| 6 | 8390129 | rs1934789 | 1.163 | 0.101 | -0.15 | 0.323 | 0.003 | 0.012 | 0.012 | 0.679 | 0.14 | 0.1 | 0.16 | 0.11 | 0.12 | 0.05 |
| 6 | 8391116 | rs1199389 | -1.33 | 0.069 | -0.03 | 0.36 | -0.006 | -0.009 | -0.009 | 0 | 0.29 | 0.75 | 0.37 | 0.31 | 0.31 | 0.44 |
| 6 | 8392223 | rs9379190 | 1.152 | -0.264 | 0.023 | 0.44 | -0.015 | -0.012 | -0.012 | 0.325 | 0.52 | 0.18 | 0.51 | 0.49 | 0.48 | 0.26 |
| 6 | 8392434 | rs1199391 | -0.592 | -0.235 | 0.343 | 0.417 | -0.021 | 0.12 | 0.12 | 0.005 | 0.16 | 0.53 | 0.22 | 0.25 | 0.11 | 0.22 |
| 6 | 8400901 | rs6901239 | 0.85 | -0.154 | -0.033 | 0.177 | -0.038 | -0.027 | -0.027 | 0 | 0.43 | 0.22 | 0.48 | 0.48 | 0.48 | 0.53 |
| 6 | 8412911 | rs4960404 | 1.448 | -0.093 | 0.015 | 0.162 | 0.039 | -0.036 | -0.036 | 2.169 | 0.07 | 0.12 | 0.11 | 0.01 | 0.14 | 0.19 |
| 6 | 8414341 | rs3823108 | 0.52 | -0.166 | -0.106 | 0.246 | 0.003 | -0.003 | -0.003 | 0.365 | 0.08 | 0.06 | 0.08 | 0.07 | 0.07 | 0.07 |
| 6 | 8419561 | rs6902448 | -0.889 | -0.157 | -0.272 | 0.229 | -0.079 | -0.012 | -0.012 | 0 | 0.8 | 0.98 | 0.88 | 0.85 | 0.74 | 0.79 |
| 6 | 8420464 | rs6935430 | 1.511 | -0.18 | -0.243 | 0.247 | -0.026 | -0.058 | -0.058 | 0.734 | 0.33 | 0.06 | 0.33 | 0.38 | 0.33 | 0.22 |
| 6 | 8424284 | rs2184580 | -0.849 | -0.118 | 0.027 | 0.288 | 0.005 | 0.005 | 0.005 | 0 | 0.44 | 0.75 | 0.52 | 0.42 | 0.42 | 0.54 |
| 6 | 8428853 | rs9406138 | 0.45 | -0.128 | 0.265 | 0.343 | 0.008 | -0.007 | -0.007 | 0 | 0.25 | 0.18 | 0.3 | 0.3 | 0.19 | 0.19 |
| 6 | 8447793 | rs2184582 | -0.11 | -0.069 | 0.23 | 0.249 | 0.03 | -0.051 | -0.051 | 0 | 0.7 | 0.56 | 0.7 | 0.84 | 0.63 | 0.58 |
| 6 | 8468978 | rs7774752 | -0.11 | -0.069 | 0.23 | 0.249 | 0.03 | -0.051 | -0.051 | 0.062 | 0.7 | 0.56 | 0.7 | 0.84 | 0.63 | 0.58 |
| 6 | 8476862 | rs6913560 | 0.15 | -0.199 | 0.386 | 0.29 | -0.01 | 0.009 | 0.009 | 0 | 0.09 | 0.17 | 0.08 | 0.07 | 0.07 | 0.07 |
| 6 | 8483819 | rs1335639 | 0.658 | -0.256 | 0.354 | 0.317 | -0.004 | 0.014 | 0.014 | 6.51 | 0.58 | 0.33 | 0.61 | 0.71 | 0.56 | 0.48 |
| 6 | 8516062 | rs9406173 | -0.119 | -0.453 | 0.101 | 0.447 | 0.008 | -0.012 | -0.012 | 0 | 0.3 | 0.3 | 0.24 | 0.42 | 0.22 | 0.29 |
| 6 | 8528235 | rs17144047 | 0.89 | -0.324 | -0.099 | 0.449 | 0.002 | -0.039 | -0.039 | 0.813 | 0.19 | 0.07 | 0.15 | 0.29 | 0.15 | 0.19 |
| 6 | 8533784 | rs230244 | -0.735 | -0.117 | -0.193 | 0.226 | -0.068 | -0.022 | -0.022 | 1.923 | 0.8 | 0.98 | 0.84 | 0.84 | 0.77 | 0.8 |
| 6 | 8534460 | rs6942177 | 0.084 | -0.077 | 0.062 | 0.166 | 0.02 | -0.031 | -0.031 | 0 | 0.7 | 0.65 | 0.7 | 0.84 | 0.63 | 0.58 |
| 6 | 8542393 | rs1796684 | 0.408 | -0.046 | -0.069 | 0.162 | 0.032 | -0.037 | -0.037 | 0 | 0.11 | 0.18 | 0.14 | 0.03 | 0.19 | 0.29 |
| 6 | 8545975 | rs1854412 | 1.68 | -0.557 | -0.557 | 0.158 | -0.061 | -0.111 | -0.111 | 0 | 0.25 | 0.03 | 0.3 | 0.38 | 0.28 | 0.2 |
| 6 | 8546104 | rs330109 | 0.381 | -0.72 | -0.131 | 0.416 | 0.002 | -0.013 | -0.013 | 0.03 | 0.52 | 0.76 | 0.6 | 0.49 | 0.56 | 0.54 |
| 6 | 8555544 | rs330126 | 0.875 | -0.845 | -0.147 | 0.427 | 0.008 | -0.011 | -0.011 | 0 | 0.2 | 0.14 | 0.16 | 0.28 | 0.15 | 0.19 |
| 6 | 8561815 | rs850188 | -0.15 | -0.773 | -0.214 | 0.47 | -0.022 | -0.021 | -0.021 | 1.434 | 0.72 | 0.89 | 0.76 | 0.77 | 0.71 | 0.73 |
| 6 | 8565164 | rs850180 | -0.41 | -0.675 | -0.063 | 0.466 | -0.008 | -0.001 | -0.001 | 0.022 | 0.77 | 0.88 | 0.83 | 0.8 | 0.78 | 0.76 |
| 6 | 8573749 | rs850173 | -0.403 | -0.728 | 0.4 | 0.617 | -0.003 | -0.007 | -0.007 | 0 | 0.36 | 0.57 | 0.31 | 0.32 | 0.3 | 0.48 |
| 6 | 8575284 | rs850170 | 0.088 | -0.736 | 0.518 | 0.606 | -0.015 | 0.015 | 0.015 | 0 | 0.29 | 0.16 | 0.27 | 0.28 | 0.3 | 0.45 |
| 6 | 8575472 | rs17144157 | -0.755 | -0.711 | 0.594 | 0.648 | -0.001 | 0.136 | 0.136 | 0 | 0.05 | 0.14 | 0.07 | 0.19 | 0.01 | 0.1 |
| 6 | 8584311 | rs850154 | -0.525 | -0.74 | 0.602 | 0.58 | -0.004 | -0.018 | -0.018 | 1.434 | 0.35 | 0.49 | 0.3 | 0.33 | 0.3 | 0.48 |
| 6 | 8609514 | rs1410771 | -0.245 | -0.781 | 0.501 | 0.566 | -0.004 | 0.022 | 0.022 | 0 | 0.68 | 0.81 | 0.73 | 0.72 | 0.7 | 0.54 |
| 6 | 8620503 | rs6597343 | 0.73 | -0.87 | 0.497 | 0.506 | 0.002 | 0.002 | 0.002 | 0 | 0.46 | 0.41 | 0.54 | 0.52 | 0.49 | 0.35 |
| 6 | 8621243 | rs1328874 | 0.66 | -0.84 | 0.596 | 0.838 | 0 | 0 | 0 | 0.017 | 0.46 | 0.53 | 0.45 | 0.41 | 0.48 | 0.35 |
| 6 | 8621262 | rs2984091 | 1.055 | -0.887 | 0.559 | 0.84 | -0.071 | -0.084 | -0.084 | 0 | 0.18 | 0.02 | 0.26 | 0.26 | 0.22 | 0.18 |
| 6 | 8632173 | rs12192533 | 1.655 | -1.002 | 0.81 | 0.782 | -0.011 | -0.009 | -0.009 | 0 | 0.06 | 0.1 | 0.04 | 0.05 | 0.04 | 0.07 |
| 6 | 8639771 | rs9406183 | 0.385 | -0.988 | 0.555 | 0.822 | 0.009 | -0.003 | -0.003 | 0 | 0.49 | 0.6 | 0.47 | 0.46 | 0.48 | 0.37 |
| 6 | 8645541 | rs4446612 | 1.139 | -1.046 | 0.498 | 0.904 | -0.001 | 0.003 | 0.003 | 0.986 | 0.38 | 0.34 | 0.37 | 0.33 | 0.4 | 0.29 |
| 6 | 8668039 | rs9379238 | 0.665 | -1.105 | 0.494 | 0.88 | 0 | 0 | 0 | 0 | 0.45 | 0.41 | 0.44 | 0.41 | 0.47 | 0.35 |
| 6 | 8669722 | rs2987579 | -0.202 | -1.072 | 0.448 | 0.879 | 0.002 | 0.007 | 0.007 | 0 | 0.66 | 0.76 | 0.72 | 0.67 | 0.69 | 0.52 |
| 6 | 8690217 | rs1576263 | 0.917 | -0.992 | 0.228 | 1.027 | 0 | 0.013 | 0.013 | 0.583 | 0.45 | 0.4 | 0.48 | 0.31 | 0.54 | 0.5 |
| 6 | 8690438 | rs1590802 | 0.173 | -1.001 | 0.218 | 1.031 | 0.008 | -0.002 | -0.002 | 0 | 0.56 | 0.63 | 0.57 | 0.42 | 0.61 | 0.57 |
| 6 | 8698414 | rs9406195 | 0.175 | -0.998 | -0.303 | 1.07 | -0.005 | -0.005 | -0.005 | 0 | 0.54 | 0.49 | 0.59 | 0.54 | 0.62 | 0.58 |
| 6 | 8702472 | rs2987574 | -0.605 | -0.921 | -0.477 | 1.018 | 0.023 | 0.033 | 0.033 | 0 | 0.76 | 0.89 | 0.87 | 0.8 | 0.84 | 0.75 |
| 6 | 8713230 | rs2987586 | -0.259 | -1.028 | -0.71 | 1.108 | -0.01 | 0.058 | 0.058 | 2.532 | 0.31 | 0.49 | 0.39 | 0.46 | 0.3 | 0.24 |
| 6 | 8717287 | rs2987583 | -0.259 | -1.028 | -0.71 | 1.108 | -0.01 | 0.058 | 0.058 | 2.532 | 0.31 | 0.49 | 0.39 | 0.46 | 0.3 | 0.24 |
| 6 | 8727059 | rs2208732 | 0.101 | -0.775 | -0.694 | 0.868 | 0.016 | -0.014 | -0.014 | 0 | 0.24 | 0.51 | 0.36 | 0.18 | 0.24 | 0.19 |
| 6 | 8730524 | rs9406197 | 1.428 | -0.822 | -1.28 | 0.828 | -0.121 | -0.089 | -0.089 | 0 | 0.28 | 0.02 | 0.25 | 0.32 | 0.35 | 0.2 |
| 6 | 8733517 | rs3005141 | -0.539 | -0.804 | -0.243 | 0.968 | 0.018 | -0.019 | -0.019 | 0 | 0.25 | 0.54 | 0.36 | 0.18 | 0.25 | 0.19 |
| 6 | 8747817 | rs12210654 | -0.091 | -0.744 | -0.744 | 1.007 | -0.017 | 0.002 | 0.002 | 0.933 | 0.79 | 0.87 | 0.67 | 0.82 | 0.75 | 0.81 |
| 6 | 8749360 | rs6934483 | 0.732 | -0.364 | -0.586 | 0.938 | 0.022 | 0.007 | 0.007 | 0.054 | 0.5 | 0.36 | 0.46 | 0.49 | 0.58 | 0.54 |
| 6 | 8776850 | rs4478454 | 0.652 | -0.33 | -0.403 | 1.524 | -0.023 | 0.032 | 0.032 | 2.939 | 0.66 | 0.49 | 0.65 | 0.55 | 0.73 | 0.59 |
| 6 | 8794058 | rs9406199 | -0.929 | -0.616 | -0.89 | 1.931 | 0.04 | -0.062 | -0.062 | 0.771 | 0.15 | 0.05 | 0.18 | 0.49 | 0.06 | 0.24 |
| 6 | 8796609 | rs9378571 | -0.513 | -0.7 | -0.775 | 1.258 | 0.001 | 0.001 | 0.001 | 0 | 0.14 | 0.14 | 0.17 | 0.07 | 0.11 | 0.13 |
| 6 | 8803042 | rs7765012 | -0.496 | -0.841 | -0.483 | 1.278 | 0.001 | -0.001 | -0.001 | 0 | 0.13 | 0.14 | 0.18 | 0.05 | 0.11 | 0.2 |
| 6 | 8808011 | rs4959502 | -0.142 | -1.625 | -0.847 | 1.127 | 0.027 | 0.012 | 0.012 | 0.215 | 0.6 | 0.67 | 0.84 | 0.63 | 0.77 | 0.57 |
| 6 | 8813921 | rs4621671 | 0.537 | -1.387 | -0.277 | 1.489 | 0.032 | -0.021 | -0.021 | 0 | 0.43 | 0.21 | 0.47 | 0.44 | 0.62 | 0.33 |
| 6 | 8832274 | rs9379246 | -0.063 | -0.45 | -0.34 | 0.899 | -0.003 | 0.005 | 0.005 | 0 | 0.13 | 0.12 | 0.18 | 0.04 | 0.15 | 0.18 |
| 6 | 8835912 | rs6901566 | -0.005 | -0.91 | -0.598 | 0.668 | 0.011 | 0.004 | 0.004 | 0 | 0.24 | 0.18 | 0.3 | 0.21 | 0.18 | 0.2 |
| 6 | 8838259 | rs4588725 | 0.723 | -1.303 | -0.63 | 1.057 | 0.02 | -0.019 | -0.019 | 0 | 0.14 | 0.1 | 0.05 | 0.26 | 0.08 | 0.28 |
| 6 | 8840051 | rs6920189 | -0.034 | -1.22 | -0.434 | 1.037 | -0.006 | 0.001 | 0.001 | 0 | 0.17 | 0.17 | 0.35 | 0.15 | 0.21 | 0.22 |

| | | | | | | | | | | | | | | | | |
|---|---|---|---|---|---|---|---|---|---|---|---|---|---|---|---|---|
| 6 | 8840357 | rs6597356 | -1.085 | -1.048 | -0.17 | 0.992 | -0.003 | 0 | 0 | 0.729 | 0.54 | 0.59 | 0.55 | 0.51 | 0.41 | 0.67 |
| 6 | 8843399 | rs13219016 | -0.063 | -0.318 | -0.631 | 0.654 | 0.023 | -0.028 | -0.028 | 0 | 0.23 | 0.14 | 0.22 | 0.33 | 0.2 | 0.33 |
| 6 | 8844901 | rs4421242 | -1.143 | -0.32 | -0.614 | 0.517 | 0.025 | 0.023 | 0.023 | 0.982 | 0.63 | 0.61 | 0.61 | 0.52 | 0.52 | 0.7 |
| 6 | 8845576 | rs4326271 | -0.259 | -0.134 | -1.479 | 0.626 | 0.005 | -0.044 | -0.044 | 0.034 | 0.26 | 0.15 | 0.27 | 0.38 | 0.26 | 0.37 |
| 6 | 8846671 | rs3922759 | 0.614 | 0.432 | -1.462 | 0.734 | -0.039 | 0.036 | 0.036 | 0 | 0.08 | 0 | 0.05 | 0.01 | 0.11 | 0.04 |
| 6 | 8849750 | rs9328507 | 0.571 | 0.364 | -1.176 | 0.988 | -0.099 | -0.106 | -0.106 | 0 | 0.3 | 0.04 | 0.28 | 0.42 | 0.43 | 0.28 |
| 6 | 8853850 | rs7741799 | 0.098 | 0.312 | -0.889 | 0.738 | 0 | 0 | 0 | 0 | 0.79 | 0.81 | 0.81 | 0.77 | 0.79 | 0.89 |
| 6 | 8855948 | rs9393069 | 0.75 | 0.073 | -1.024 | 0.346 | 0.003 | 0.003 | 0.003 | 0.059 | 0.61 | 0.68 | 0.7 | 0.63 | 0.64 | 0.75 |
| 6 | 8857222 | rs4246082 | 0.048 | 0.209 | -0.946 | -0.136 | 0.018 | 0.069 | 0.069 | 0 | 0.53 | 0.35 | 0.59 | 0.23 | 0.57 | 0.52 |
| 6 | 8860590 | rs12205102 | 0.608 | 0.619 | -1.038 | -0.255 | -0.069 | 0.145 | 0.145 | 0 | 0.37 | 0.12 | 0.42 | 0.11 | 0.45 | 0.36 |
| 6 | 8863330 | rs4960464 | 0.532 | 0.433 | -0.864 | -0.076 | -0.025 | 0.099 | 0.099 | 0 | 0.42 | 0.28 | 0.47 | 0.12 | 0.52 | 0.37 |
| 6 | 8869571 | rs4959505 | -0.29 | 0.358 | -0.571 | 0.461 | 0.004 | 0.006 | 0.006 | 0 | 0.39 | 0.57 | 0.45 | 0.58 | 0.44 | 0.47 |
| 6 | 8870738 | rs4555971 | -0.116 | 0.831 | -0.552 | 0.797 | 0.004 | 0.006 | 0.006 | 0 | 0.43 | 0.49 | 0.55 | 0.41 | 0.5 | 0.39 |
| 6 | 8870769 | rs4636072 | 0.173 | 0.98 | -1.162 | 1.035 | -0.033 | -0.04 | -0.04 | 0 | 0.34 | 0.21 | 0.42 | 0.38 | 0.42 | 0.34 |
| 6 | 8871624 | rs4529337 | -0.181 | 0.846 | -1.466 | 0.913 | -0.039 | -0.089 | -0.089 | 0 | 0.38 | 0.08 | 0.4 | 0.48 | 0.36 | 0.53 |
| 6 | 8871713 | rs4566922 | -0.151 | 0.707 | -1.203 | 0.988 | -0.074 | -0.07 | -0.07 | 1.577 | 0.42 | 0.1 | 0.46 | 0.5 | 0.49 | 0.55 |
| 6 | 8877784 | rs9393071 | 0.866 | 0.163 | 0.188 | 1.019 | -0.023 | 0.053 | 0.053 | 0.089 | 0.92 | 0.68 | 0.96 | 0.9 | 0.97 | 0.94 |
| 6 | 8880099 | rs4276549 | 1.067 | 0.027 | 0.132 | 0.797 | -0.03 | -0.02 | -0.02 | 0.062 | 0.84 | 0.27 | 0.91 | 0.9 | 0.91 | 0.93 |
| 6 | 8894137 | rs4285366 | 1.587 | 0.307 | -0.107 | 1.326 | -0.004 | 0.005 | 0.005 | 0.112 | 0.76 | 0.29 | 0.67 | 0.69 | 0.8 | 0.85 |
| 6 | 8898039 | rs6930767 | 1.186 | 0.185 | 0.323 | 1.449 | -0.014 | 0.125 | 0.125 | 0.001 | 0.85 | 0.43 | 0.83 | 0.69 | 0.88 | 0.87 |
| 6 | 8899270 | rs11752885 | 0.714 | 0.157 | -0.746 | 0.781 | -0.119 | 0.165 | 0.165 | 0.008 | 0.36 | 0.06 | 0.33 | 0.13 | 0.46 | 0.42 |
| 6 | 8901524 | rs9378578 | 0.786 | -0.042 | -0.372 | 0.885 | -0.005 | 0.157 | 0.157 | 0.001 | 0.85 | 0.47 | 0.83 | 0.69 | 0.88 | 0.87 |
| 6 | 8904861 | rs12215694 | 0.382 | 0.096 | -1.087 | 0.971 | -0.062 | 0.118 | 0.118 | 0 | 0.42 | 0.1 | 0.53 | 0.25 | 0.55 | 0.53 |
| 6 | 8905429 | rs4475363 | -0.569 | -0.089 | 0.291 | 1.551 | 0.023 | 0.101 | 0.101 | 0 | 0.06 | 0.13 | 0.13 | 0.2 | 0.09 | 0.07 |
| 6 | 8908960 | rs4246086 | 0.95 | -0.774 | 0.278 | 1.638 | -0.069 | 0.037 | 0.037 | 0.195 | 0.74 | 0.57 | 0.7 | 0.74 | 0.87 | 0.74 |
| 6 | 8917897 | rs4351309 | 1.161 | -0.447 | -0.657 | 1.775 | 0.017 | 0.005 | 0.005 | 0.719 | 0.57 | 0.07 | 0.47 | 0.41 | 0.66 | 0.63 |
| 6 | 8923203 | rs9406223 | 0.528 | -0.631 | 0.109 | 1.943 | 0.019 | -0.013 | -0.013 | 0 | 0.77 | 0.48 | 0.75 | 0.8 | 0.77 | 0.89 |
| 6 | 8927894 | rs4521641 | 0.709 | -0.579 | 0.158 | 1.934 | 0.012 | -0.019 | -0.019 | 0 | 0.76 | 0.44 | 0.74 | 0.81 | 0.77 | 0.87 |
| 6 | 8929421 | rs4449671 | 1.152 | -0.639 | -0.039 | 1.913 | -0.02 | -0.002 | -0.002 | 2.48 | 0.67 | 0.12 | 0.69 | 0.68 | 0.71 | 0.85 |
| 6 | 8936505 | rs9379263 | -0.388 | -0.61 | 0.102 | 1.944 | -0.017 | 0.045 | 0.045 | 0 | 0.12 | 0.12 | 0.23 | 0.27 | 0.06 | 0.22 |
| 6 | 8944188 | rs9379265 | 1.567 | -0.766 | -0.084 | 2.09 | 0.026 | 0.137 | 0.137 | 0.705 | 0.55 | 0.08 | 0.41 | 0.23 | 0.65 | 0.5 |
| 6 | 8944529 | rs7739228 | -0.006 | -0.749 | -0.085 | 2.322 | 0.028 | -0.068 | -0.068 | 3.241 | 0.1 | 0.02 | 0.26 | 0.22 | 0.06 | 0.21 |
| 6 | 8945690 | rs9406226 | 1.07 | -0.759 | 0.139 | 2.304 | 0 | -0.01 | -0.01 | 1.33 | 0.67 | 0.25 | 0.68 | 0.5 | 0.71 | 0.72 |
| 6 | 8950441 | rs13211231 | 1.567 | -0.761 | 0.098 | 2.218 | 0.032 | 0.124 | 0.124 | 0.165 | 0.56 | 0.12 | 0.41 | 0.23 | 0.65 | 0.49 |
| 6 | 8951083 | rs4316043 | -0.181 | -0.428 | 0.528 | 2.243 | -0.008 | 0.035 | 0.035 | 0 | 0.1 | 0.27 | 0.11 | 0.18 | 0.06 | 0.15 |
| 6 | 8952927 | rs9505568 | 0.649 | -0.332 | 0.485 | 2.211 | 0.029 | 0.083 | 0.083 | 0.022 | 0.77 | 0.5 | 0.79 | 0.63 | 0.77 | 0.86 |
| 6 | 8967879 | rs9505573 | 0.118 | -0.173 | -0.219 | 1.766 | 0.004 | 0 | 0 | 1.039 | 0.5 | 0.45 | 0.4 | 0.57 | 0.45 | 0.55 |
| 6 | 8973552 | rs9328513 | -0.93 | -0.181 | -0.326 | 1.835 | -0.036 | 0.032 | 0.032 | 0.849 | 0.67 | 0.94 | 0.59 | 0.8 | 0.64 | 0.66 |
| 6 | 8975508 | rs11751589 | 0.32 | -0.15 | -0.094 | 1.939 | 0.001 | 0.017 | 0.017 | 3.218 | 0.4 | 0.33 | 0.29 | 0.25 | 0.38 | 0.41 |
| 6 | 8976039 | rs9502754 | 0.394 | -0.153 | -0.124 | 1.939 | -0.002 | -0.001 | -0.001 | 0.037 | 0.41 | 0.44 | 0.3 | 0.4 | 0.38 | 0.42 |
| 6 | 8976234 | rs17403006 | 0.515 | -0.114 | -0.587 | 1.84 | -0.045 | 0.06 | 0.06 | 4.004 | 0.38 | 0.09 | 0.28 | 0.24 | 0.38 | 0.4 |
| 6 | 8982655 | rs6907965 | 0.032 | 0.001 | -0.566 | 1.883 | 0.006 | -0.001 | -0.001 | 0 | 0.48 | 0.27 | 0.39 | 0.58 | 0.44 | 0.56 |
| 6 | 8983001 | rs1997702 | 0.142 | 0.064 | -0.554 | 1.786 | -0.01 | -0.008 | -0.008 | 3.355 | 0.46 | 0.2 | 0.38 | 0.46 | 0.45 | 0.54 |
| 6 | 8985287 | rs4960473 | -0.108 | 0.163 | -0.429 | 1.696 | -0.005 | 0.019 | 0.019 | 0.133 | 0.52 | 0.36 | 0.4 | 0.6 | 0.45 | 0.58 |
| 6 | 8990207 | rs9379274 | 0.447 | 0.238 | -0.52 | 1.826 | 0.003 | -0.007 | -0.007 | 0.223 | 0.39 | 0.19 | 0.32 | 0.37 | 0.34 | 0.45 |
| 6 | 8990652 | rs742301 | 0.443 | 0.261 | -0.543 | 1.849 | -0.003 | -0.027 | -0.027 | 0 | 0.1 | 0.02 | 0.06 | 0.21 | 0.09 | 0.13 |
| 6 | 8991886 | rs6597377 | -0.054 | 0.31 | -0.277 | 1.876 | 0.012 | 0.019 | 0.019 | 0.164 | 0.5 | 0.32 | 0.39 | 0.59 | 0.43 | 0.6 |
| 6 | 8995233 | rs10484331 | 0.839 | 0.386 | -0.402 | 1.773 | -0.025 | -0.046 | -0.046 | 0 | 0.31 | 0.1 | 0.28 | 0.32 | 0.3 | 0.37 |
| 6 | 8998811 | rs2064197 | -0.266 | 0.86 | 0.193 | 1.659 | 0.039 | 0.043 | 0.043 | 0 | 0.14 | 0.22 | 0.2 | 0.27 | 0.21 | 0.12 |
| 6 | 8999646 | rs2327112 | -0.038 | 0.536 | 0.089 | 1.537 | -0.005 | 0.031 | 0.031 | 0.159 | 0.52 | 0.4 | 0.4 | 0.41 | 0.49 | 0.6 |
| 6 | 9008032 | rs2050037 | -0.241 | 0.808 | -0.151 | 1.328 | 0.033 | 0.23 | 0.23 | 0 | 0.15 | 0.34 | 0.32 | 0.47 | 0.16 | 0.16 |
| 6 | 9015569 | rs382854 | -0.613 | 0.861 | -1.533 | 1.096 | 0.016 | 0.095 | 0.095 | 0.028 | 0.51 | 0.77 | 0.7 | 0.64 | 0.52 | 0.52 |
| 6 | 9017266 | rs17403999 | -0.049 | 1.133 | -1.818 | 0.54 | -0.064 | 0.096 | 0.096 | 0 | 0.33 | 0.1 | 0.36 | 0.14 | 0.36 | 0.34 |
| 6 | 9020424 | rs384323 | 0.088 | 1.322 | -1.867 | 0.861 | -0.072 | -0.019 | -0.019 | 0 | 0.37 | 0.1 | 0.3 | 0.36 | 0.49 | 0.55 |
| 6 | 9022100 | rs421424 | -0.313 | 1.181 | -1.628 | 1.144 | -0.007 | -0.014 | -0.014 | 0 | 0.24 | 0.2 | 0.25 | 0.29 | 0.25 | 0.18 |
| 6 | 9026671 | rs11243295 | -0.49 | 1.279 | -1.501 | 1.425 | -0.003 | -0.007 | -0.007 | 0 | 0.25 | 0.26 | 0.22 | 0.38 | 0.28 | 0.21 |
| 6 | 9026712 | rs11243296 | -0.34 | 1.075 | -2.038 | 1.004 | 0.007 | 0.008 | 0.008 | 0 | 0.64 | 0.33 | 0.48 | 0.68 | 0.69 | 0.68 |
| 6 | 9031654 | rs380915 | -0.102 | 0.577 | -1.943 | 1.017 | -0.007 | -0.012 | -0.012 | 0 | 0.23 | 0.18 | 0.2 | 0.3 | 0.24 | 0.21 |
| 6 | 9031961 | rs9505593 | -0.004 | 0.069 | -1.186 | 0.678 | 0.035 | 0.031 | 0.031 | 2.95 | 0.07 | 0.48 | 0.05 | 0 | 0 | 0 |
| 6 | 9042512 | rs11752691 | 0.969 | 0.062 | -1.529 | 0.651 | -0.066 | 0.073 | 0.073 | 0 | 0.44 | 0.1 | 0.31 | 0.29 | 0.51 | 0.46 |
| 6 | 9044593 | rs9505597 | 0.366 | 0.024 | -1.491 | 0.707 | 0.006 | -0.014 | -0.014 | 0 | 0.26 | 0.17 | 0.43 | 0.32 | 0.24 | 0.34 |
| 6 | 9048002 | rs7738581 | 0.973 | 0.068 | -1.474 | 0.693 | -0.071 | 0.069 | 0.069 | 0 | 0.44 | 0.1 | 0.3 | 0.29 | 0.51 | 0.45 |
| 6 | 9055743 | rs6597378 | -0.201 | -0.057 | -1.24 | 0.375 | 0.005 | 0.015 | 0.015 | 0.723 | 0.8 | 0.76 | 0.81 | 0.7 | 0.79 | 0.81 |
| 6 | 9060163 | rs12210482 | -0.168 | -0.196 | -0.221 | 0.531 | -0.014 | 0.021 | 0.021 | 0 | 0.34 | 0.43 | 0.51 | 0.46 | 0.28 | 0.39 |
| 6 | 9065677 | rs2224791 | 1.111 | -0.198 | -0.278 | 0.538 | -0.076 | 0.126 | 0.126 | 0 | 0.43 | 0.1 | 0.27 | 0.24 | 0.5 | 0.42 |
| 6 | 9066118 | rs2057512 | 0.925 | -0.168 | -0.243 | 0.565 | -0.005 | 0.05 | 0.05 | 0 | 0.46 | 0.33 | 0.3 | 0.24 | 0.5 | 0.42 |
| 6 | 9068966 | rs2209126 | -0.029 | -0.28 | -0.048 | 0.396 | 0.011 | -0.029 | -0.029 | 0.21 | 0.31 | 0.26 | 0.49 | 0.46 | 0.28 | 0.39 |
| 6 | 9076749 | rs13208208 | -0.423 | -0.288 | -0.17 | 0.379 | -0.02 | 0.022 | 0.022 | 0 | 0.35 | 0.47 | 0.51 | 0.46 | 0.28 | 0.39 |
| 6 | 9077515 | rs11243301 | -0.38 | -0.255 | 0.037 | 0.401 | -0.026 | 0.036 | 0.036 | 0 | 0.34 | 0.5 | 0.52 | 0.46 | 0.28 | 0.4 |

| | | | | | | | | | | | | | | | | |
|---|---|---|---|---|---|---|---|---|---|---|---|---|---|---|---|---|
| 6 | 9079324 | rs9406232 | 0.618 | -0.253 | 0.501 | 0.37 | -0.005 | 0.008 | 0.008 | 0.046 | 0.38 | 0.5 | 0.29 | 0.42 | 0.35 | 0.37 |
| 6 | 9102386 | rs4626470 | -0.325 | -0.488 | -0.065 | -0.068 | -0.081 | 0.157 | 0.157 | 0.13 | 0.63 | 0.88 | 0.69 | 0.91 | 0.55 | 0.6 |
| 6 | 9103538 | rs9406234 | -0.565 | -0.71 | 0.429 | -0.037 | -0.028 | 0.25 | 0.25 | 1.119 | 0.13 | 0.52 | 0.05 | 0.35 | 0.06 | 0.03 |
| 6 | 9103828 | rs9379278 | -0.432 | -0.712 | 0.538 | -0.013 | -0.017 | 0.238 | 0.238 | 0.906 | 0.13 | 0.5 | 0.05 | 0.26 | 0.05 | 0.02 |
| 6 | 9105062 | rs7756210 | 0.288 | -0.841 | 0.026 | -0.102 | 0.021 | -0.056 | -0.056 | 0 | 0.44 | 0.81 | 0.31 | 0.54 | 0.38 | 0.37 |
| 6 | 9116357 | rs2146959 | -0.104 | -0.72 | 0.173 | -0.256 | -0.007 | 0.058 | 0.058 | 0.742 | 0.4 | 0.25 | 0.26 | 0.05 | 0.42 | 0.29 |
| 6 | 9123728 | rs11760199 | 0.891 | -0.46 | 0.808 | 0.414 | -0.042 | 0.075 | 0.075 | 0 | 0.29 | 0.07 | 0.25 | 0.19 | 0.32 | 0.34 |
| 6 | 9131436 | rs1555601 | -0.435 | -0.341 | 0.773 | 0.202 | -0.016 | -0.068 | -0.068 | 1.331 | 0.11 | 0.28 | 0.03 | 0 | 0.05 | 0.02 |
| 6 | 9131684 | rs1555605 | 0.216 | -0.342 | 0.211 | 0.605 | 0.015 | -0.044 | -0.044 | 0.291 | 0.43 | 0.8 | 0.31 | 0.53 | 0.38 | 0.37 |
| 6 | 9138264 | rs2327138 | 0.485 | -0.714 | 0.305 | 0.432 | 0.046 | 0.134 | 0.134 | 0 | 0.5 | 0.3 | 0.33 | 0.25 | 0.56 | 0.51 |
| 6 | 9139583 | rs13209110 | -0.738 | -1.727 | 1.053 | 0.997 | -0.036 | 0.011 | 0.011 | 0.01 | 0.42 | 0.52 | 0.52 | 0.47 | 0.28 | 0.43 |
| 6 | 9139653 | rs767022 | 0.793 | -1.764 | 1.15 | 1.147 | 0.112 | -0.007 | -0.007 | 0 | 0.56 | 0.19 | 0.47 | 0.44 | 0.72 | 0.56 |
| 6 | 9142017 | rs12205137 | -0.349 | -1.704 | 0.522 | 0.778 | 0.031 | -0.016 | -0.016 | 0.667 | 0.34 | 0.22 | 0.49 | 0.36 | 0.24 | 0.38 |
| 6 | 9159899 | rs35343570 | -0.069 | -1.765 | 0.104 | 0.921 | 0.012 | 0.007 | 0.007 | 0 | 0.06 | 0.01 | 0.29 | 0.27 | 0.04 | 0.09 |
| 6 | 9170896 | rs2143361 | -0.706 | -1.621 | 0.365 | 0.665 | 0.058 | -0.018 | -0.018 | 0 | 0.43 | 0.62 | 0.53 | 0.46 | 0.28 | 0.43 |
| 6 | 9178954 | rs16892001 | -0.457 | -1.212 | -0.382 | 0.702 | 0.056 | -0.03 | -0.03 | 0 | 0.35 | 0.18 | 0.49 | 0.36 | 0.26 | 0.38 |
| 6 | 9181572 | rs1324546 | -0.473 | -1.14 | -0.606 | 0.74 | -0.018 | 0.003 | 0.003 | 0 | 0.37 | 0.42 | 0.5 | 0.36 | 0.26 | 0.38 |
| 6 | 9249495 | rs4464773 | 0.358 | -0.376 | -1.586 | 0.118 | 0.01 | 0.194 | 0.194 | 0 | 0.49 | 0.1 | 0.45 | 0.28 | 0.6 | 0.42 |
| 6 | 9255500 | rs1830947 | 0.17 | -0.371 | -1.603 | 0.923 | -0.015 | -0.034 | -0.034 | 0.469 | 0.09 | 0.03 | 0.08 | 0.26 | 0.1 | 0.16 |
| 6 | 9275355 | rs9379412 | 0.759 | -0.4 | -1.407 | 0.585 | 0.003 | 0.169 | 0.169 | 0 | 0.47 | 0.19 | 0.4 | 0.28 | 0.55 | 0.4 |
| 6 | 9275594 | rs7768303 | 0.147 | -0.865 | -0.683 | 0.272 | 0.002 | -0.008 | -0.008 | 0.001 | 0.72 | 0.74 | 0.57 | 0.54 | 0.73 | 0.62 |
| 6 | 9282211 | rs9358619 | -0.098 | -1.032 | -0.815 | 0.139 | -0.004 | -0.059 | -0.059 | 0.658 | 0.84 | 0.91 | 0.66 | 0.55 | 0.8 | 0.79 |
| 6 | 9283127 | rs7756320 | 0.135 | -1.073 | -0.734 | 0.07 | -0.001 | -0.014 | -0.014 | 0.7 | 0.11 | 0.1 | 0.08 | 0 | 0.06 | 0.17 |
| 6 | 9283312 | rs9393394 | 1.517 | -1.03 | -0.51 | 0.11 | 0.013 | 0.15 | 0.15 | 0 | 0.47 | 0.24 | 0.4 | 0.28 | 0.55 | 0.4 |
| 6 | 9285085 | rs9393418 | 0.103 | -0.859 | -0.493 | 0.28 | 0.001 | -0.022 | -0.022 | 0 | 0.83 | 0.86 | 0.65 | 0.54 | 0.79 | 0.79 |
| 6 | 9289146 | rs9379615 | 0.769 | -0.978 | 0.413 | 0.212 | -0.014 | 0.004 | 0.004 | 0 | 0.58 | 0.54 | 0.49 | 0.54 | 0.65 | 0.56 |
| 6 | 9296868 | rs1359638 | 0.193 | -0.652 | 0.629 | 0.775 | 0 | 0.013 | 0.013 | 0 | 0.09 | 0.09 | 0.06 | 0.27 | 0.11 | 0.16 |
| 6 | 9298042 | rs9467302 | 1.031 | -0.5 | -0.04 | -0.097 | -0.031 | 0.074 | 0.074 | 0 | 0.41 | 0.24 | 0.3 | 0.28 | 0.43 | 0.42 |
| 6 | 9298640 | rs16892041 | -0.032 | -0.514 | 0.722 | -0.184 | 0.025 | 0.061 | 0.061 | 0 | 0.15 | 0.34 | 0.1 | 0.27 | 0.17 | 0.23 |
| 6 | 9299057 | rs7748490 | 0.34 | -0.522 | 0.728 | -0.221 | 0.027 | -0.013 | -0.013 | 1.291 | 0.06 | 0.25 | 0.04 | 0 | 0.06 | 0.07 |
| 6 | 9300312 | rs9358817 | 0.451 | -0.568 | 0.559 | 0.01 | 0.043 | 0.01 | 0.01 | 0 | 0.5 | 0.25 | 0.35 | 0.58 | 0.54 | 0.58 |
| 6 | 9304007 | rs7746692 | 0.002 | -0.631 | 0.793 | -0.497 | -0.003 | -0.04 | -0.04 | 1.118 | 0.26 | 0.34 | 0.25 | 0.06 | 0.26 | 0.17 |
| 6 | 9305544 | rs9348682 | 1.444 | -0.641 | 0.674 | -0.435 | -0.036 | 0.077 | 0.077 | 0 | 0.4 | 0.2 | 0.3 | 0.28 | 0.43 | 0.31 |
| 6 | 9308085 | rs9295671 | 1.278 | -0.441 | 0.634 | -0.43 | -0.008 | 0.036 | 0.036 | 0 | 0.42 | 0.32 | 0.31 | 0.29 | 0.43 | 0.34 |
| 6 | 9308123 | rs7768472 | -0.341 | -0.501 | 0.895 | -0.305 | 0.001 | -0.019 | -0.019 | 0 | 0.79 | 0.84 | 0.61 | 0.62 | 0.79 | 0.77 |
| 6 | 9312345 | rs1409096 | -0.59 | -0.365 | 0.734 | -0.235 | 0 | 0.004 | 0.004 | 0 | 0.35 | 0.35 | 0.3 | 0.33 | 0.36 | 0.43 |
| 6 | 9313277 | rs16892087 | 0.822 | -0.097 | 0.646 | -0.143 | -0.001 | 0.011 | 0.011 | 0 | 0.1 | 0.09 | 0.06 | 0.28 | 0.1 | 0.26 |
| 6 | 9325123 | rs9379986 | 0.361 | -0.081 | 0.075 | -0.78 | 0 | -0.011 | -0.011 | 0 | 0.64 | 0.43 | 0.52 | 0.34 | 0.64 | 0.44 |
| 6 | 9334307 | rs7762096 | 0.34 | 0.021 | -0.501 | -0.804 | -0.009 | 0.039 | 0.039 | 0 | 0.17 | 0.1 | 0.19 | 0.05 | 0.19 | 0.08 |
| 6 | 9337268 | rs17213073 | 0.12 | 0.09 | -0.789 | -0.937 | -0.054 | 0.119 | 0.119 | 0 | 0.2 | 0.05 | 0.19 | 0.04 | 0.23 | 0.11 |
| 6 | 9340964 | rs9393943 | 1.157 | -0.122 | -1.06 | -1.349 | -0.028 | 0.083 | 0.083 | 0 | 0.36 | 0.19 | 0.21 | 0.22 | 0.38 | 0.29 |
| 6 | 9344751 | rs10946994 | 1.141 | -0.19 | -1.106 | -1.388 | -0.044 | 0.086 | 0.086 | 0 | 0.34 | 0.14 | 0.21 | 0.22 | 0.38 | 0.28 |
| 6 | 9350263 | rs1409093 | 0.041 | -0.491 | -0.653 | -1.429 | 0.008 | 0.069 | 0.069 | 0.443 | 0.63 | 0.82 | 0.5 | 0.26 | 0.65 | 0.51 |
| 6 | 9368299 | rs1325814 | 0.211 | -0.485 | -1.132 | -1.474 | 0.002 | 0.046 | 0.046 | 0.289 | 0.66 | 0.81 | 0.45 | 0.26 | 0.66 | 0.58 |
| 6 | 9380976 | rs1952110 | 1.426 | -0.284 | -0.842 | -1.565 | -0.028 | 0.064 | 0.064 | 0.003 | 0.34 | 0.22 | 0.21 | 0.19 | 0.38 | 0.28 |
| 6 | 9384993 | rs2209543 | 0.234 | -0.37 | -0.35 | -1.21 | 0 | -0.003 | -0.003 | 1.238 | 0.68 | 0.7 | 0.46 | 0.25 | 0.69 | 0.53 |
| 6 | 9389312 | rs2294736 | 1.165 | -0.254 | -0.322 | -0.822 | -0.013 | 0.003 | 0.003 | 0.014 | 0.1 | 0.18 | 0.04 | 0.1 | 0.05 | 0.02 |
| 6 | 9390592 | rs214432 | -0.508 | -0.805 | -0.095 | -0.35 | -0.012 | 0.026 | 0.026 | 1.994 | 0.28 | 0.49 | 0.41 | 0.27 | 0.26 | 0.27 |
| 6 | 9391468 | rs214434 | 0.406 | -0.874 | -0.14 | -0.306 | 0 | -0.009 | -0.009 | 0 | 0.2 | 0.14 | 0.34 | 0.23 | 0.19 | 0.19 |
| 6 | 9393712 | rs214438 | -0.892 | -1.011 | -0.6 | -0.538 | -0.005 | 0.111 | 0.111 | 0 | 0.72 | 0.89 | 0.79 | 0.89 | 0.72 | 0.77 |
| 6 | 9394089 | rs214439 | -0.523 | -1.136 | -0.069 | 0.002 | -0.019 | 0.098 | 0.098 | 1.49 | 0.27 | 0.44 | 0.47 | 0.48 | 0.24 | 0.31 |
| 6 | 9398094 | rs214443 | 0.63 | -1.158 | -0.354 | 0.009 | 0 | 0.002 | 0.002 | 3.792 | 0.17 | 0.11 | 0.32 | 0.23 | 0.14 | 0.16 |
| 6 | 9399038 | rs214445 | -1.371 | -0.762 | -0.826 | -0.373 | -0.023 | 0.049 | 0.049 | 0 | 0.52 | 0.87 | 0.6 | 0.58 | 0.44 | 0.42 |
| 6 | 9401979 | rs214447 | 0.855 | -0.755 | -0.373 | -0.217 | 0.022 | 0.051 | 0.051 | 0 | 0.51 | 0.28 | 0.59 | 0.64 | 0.52 | 0.5 |
| 6 | 9402068 | rs6457811 | -0.261 | -0.751 | -0.142 | -0.157 | -0.012 | 0.093 | 0.093 | 0.031 | 0.14 | 0.37 | 0.16 | 0.26 | 0.12 | 0.19 |
| 6 | 9404980 | rs214450 | -0.088 | -0.674 | -0.68 | -0.171 | -0.004 | 0.014 | 0.014 | 2.826 | 0.21 | 0.23 | 0.37 | 0.27 | 0.19 | 0.2 |
| 6 | 9405449 | rs13212842 | 1.008 | -0.508 | -0.868 | 0.035 | -0.046 | -0.005 | -0.005 | 0.269 | 0.37 | 0.15 | 0.27 | 0.4 | 0.4 | 0.39 |
| 6 | 9405559 | rs16892185 | -0.082 | -0.582 | -0.33 | 0.304 | 0 | -0.007 | -0.007 | 0 | 0.07 | 0.1 | 0.05 | 0 | 0.07 | 0.08 |
| 6 | 9410202 | rs7754614 | -0.099 | -0.874 | 0.228 | 0.002 | -0.021 | -0.035 | -0.035 | 0 | 0.15 | 0.5 | 0.08 | 0.06 | 0.1 | 0.2 |
| 6 | 9411973 | rs2327167 | 0.983 | -0.966 | -0.425 | 0.05 | -0.016 | -0.018 | -0.018 | 0 | 0.33 | 0.24 | 0.35 | 0.52 | 0.4 | 0.44 |
| 6 | 9413201 | rs2017505 | -1.292 | -1.177 | -1.058 | -0.599 | -0.059 | 0.063 | 0.063 | 0.012 | 0.82 | 0.93 | 0.77 | 0.97 | 0.72 | 0.81 |
| 6 | 9413488 | rs957122 | -0.186 | -1.611 | -1.329 | -0.754 | -0.006 | 0.017 | 0.017 | 0.204 | 0.62 | 0.8 | 0.52 | 0.62 | 0.64 | 0.7 |
| 6 | 9418280 | rs9380620 | -0.269 | -1.249 | -0.532 | -0.545 | 0.006 | 0 | 0 | 0 | 0.38 | 0.36 | 0.25 | 0.39 | 0.26 | 0.29 |
| 6 | 9422071 | rs10947666 | 1.872 | -1.249 | -0.591 | -0.311 | -0.012 | -0.013 | -0.013 | 1.203 | 0.22 | 0.16 | 0.32 | 0.29 | 0.28 | 0.31 |
| 6 | 9423799 | rs11756620 | 0.81 | -1.312 | -0.591 | -0.347 | 0.037 | -0.057 | -0.057 | 0 | 0.26 | 0.08 | 0.19 | 0.36 | 0.16 | 0.13 |
| 6 | 9433183 | rs6458053 | -0.502 | -1.16 | -0.33 | -0.475 | 0.001 | -0.068 | -0.068 | 0 | 0.67 | 0.65 | 0.67 | 0.94 | 0.63 | 0.75 |
| 6 | 9433574 | rs12203857 | 1.336 | -1.203 | -0.294 | -0.465 | -0.012 | -0.021 | -0.021 | 2.452 | 0.28 | 0.22 | 0.41 | 0.52 | 0.34 | 0.45 |
| 6 | 9436244 | rs1015137 | -0.191 | -1.138 | -0.2 | -0.463 | -0.004 | -0.001 | -0.001 | 0 | 0.39 | 0.4 | 0.25 | 0.4 | 0.29 | 0.3 |
| 6 | 9440886 | rs9380830 | 1.402 | -1.113 | -0.212 | -0.391 | 0.001 | 0.026 | 0.026 | 0 | 0.06 | 0.06 | 0.09 | 0.25 | 0.06 | 0.14 |

| | | | | | | | | | | | | | | | | |
|---|---|---|---|---|---|---|---|---|---|---|---|---|---|---|---|---|
| 6 | 9442330 | rs7771515 | 0.168 | -1.173 | -0.176 | -0.621 | 0.005 | -0.003 | -0.003 | 4.904 | 0.32 | 0.3 | 0.22 | 0.39 | 0.22 | 0.2 |
| 6 | 9455253 | rs9357334 | 1.288 | -1.117 | -0.09 | -0.586 | -0.013 | -0.022 | -0.022 | 0 | 0.28 | 0.22 | 0.41 | 0.54 | 0.34 | 0.46 |
| 6 | 9457140 | rs742513 | 0.143 | -1.069 | -0.033 | -0.563 | 0 | -0.005 | -0.005 | 0 | 0.07 | 0.2 | 0.04 | 0.03 | 0.06 | 0.09 |
| 6 | 9458329 | rs761116 | -0.475 | -1.103 | -0.069 | -0.614 | 0.002 | -0.068 | -0.068 | 0 | 0.67 | 0.65 | 0.67 | 0.94 | 0.63 | 0.75 |
| 6 | 9466991 | rs10755718 | 0.249 | -1.091 | -0.181 | -0.571 | -0.007 | 0.059 | 0.059 | 0.507 | 0.54 | 0.33 | 0.6 | 0.9 | 0.5 | 0.57 |
| 6 | 9469238 | rs9381107 | 0.707 | -1.225 | -0.254 | -0.575 | 0.04 | -0.024 | -0.024 | 0.016 | 0.27 | 0.16 | 0.19 | 0.36 | 0.16 | 0.15 |
| 6 | 9472085 | rs9369355 | 0.24 | -1.21 | -0.315 | -0.755 | -0.007 | 0.069 | 0.069 | 1.027 | 0.54 | 0.33 | 0.6 | 0.9 | 0.5 | 0.57 |
| 6 | 9479015 | rs12192884 | 2.112 | -1.073 | -0.334 | -0.338 | -0.005 | -0.005 | -0.005 | 0 | 0.21 | 0.16 | 0.22 | 0.29 | 0.28 | 0.3 |
| 6 | 9479371 | rs2206547 | -0.713 | -1.094 | -0.25 | -0.661 | -0.015 | 0.063 | 0.063 | 1.941 | 0.69 | 0.81 | 0.58 | 0.92 | 0.63 | 0.75 |
| 6 | 9485556 | rs1001384 | 0.224 | -1.039 | -0.229 | -0.444 | 0.017 | -0.02 | -0.02 | 0.905 | 0.33 | 0.1 | 0.21 | 0.32 | 0.24 | 0.22 |
| 6 | 9485571 | rs1001383 | -0.387 | -0.933 | -0.163 | -0.403 | -0.007 | -0.004 | -0.004 | 0 | 0.42 | 0.58 | 0.27 | 0.35 | 0.3 | 0.31 |
| 6 | 9489930 | rs7739703 | 1.551 | -0.316 | -0.49 | -0.342 | 0.005 | -0.024 | -0.024 | 1.491 | 0.07 | 0.1 | 0.03 | 0 | 0.08 | 0.08 |
| 6 | 9489952 | rs7739844 | -0.028 | -0.543 | -0.618 | -0.469 | 0.01 | -0.007 | -0.007 | 0.064 | 0.68 | 0.61 | 0.51 | 0.66 | 0.62 | 0.61 |
| 6 | 9490667 | rs9472218 | 0.09 | -0.608 | -0.55 | -0.269 | -0.003 | -0.003 | -0.003 | 1.665 | 0.6 | 0.6 | 0.48 | 0.6 | 0.62 | 0.58 |
| 6 | 9493302 | rs12194169 | 1.021 | -0.497 | -0.5 | -0.103 | -0.006 | 0.068 | 0.068 | 0 | 0.07 | 0.1 | 0.09 | 0.25 | 0.06 | 0.14 |
| 6 | 9500269 | rs7765744 | 0.81 | -0.482 | -0.521 | -0.579 | -0.009 | -0.009 | -0.009 | 0 | 0.07 | 0.11 | 0.02 | 0.04 | 0.06 | 0.06 |
| 6 | 9500813 | rs17213227 | 1.764 | -0.301 | -0.644 | -0.431 | -0.023 | -0.023 | -0.023 | 0 | 0.21 | 0.12 | 0.23 | 0.31 | 0.27 | 0.33 |
| 6 | 9501534 | rs9381374 | -0.635 | -0.14 | -0.484 | -0.423 | 0.003 | 0.003 | 0.003 | 0 | 0.51 | 0.53 | 0.37 | 0.54 | 0.46 | 0.42 |
| 6 | 9501859 | rs9381380 | -0.883 | -0.122 | -0.622 | -1.289 | 0.001 | -0.012 | -0.012 | 0 | 0.76 | 0.78 | 0.63 | 0.84 | 0.75 | 0.78 |
| 6 | 9502395 | rs9395094 | 0.046 | 0.123 | -1.348 | -0.985 | -0.007 | 0.04 | 0.04 | 0.045 | 0.47 | 0.41 | 0.47 | 0.73 | 0.5 | 0.62 |
| 6 | 9512346 | rs9357504 | 0.242 | 0.36 | -1.967 | -0.403 | 0.016 | -0.059 | -0.059 | 1.833 | 0.14 | 0.04 | 0.1 | 0.34 | 0.11 | 0.17 |
| 6 | 9514315 | rs12196647 | 1.237 | 0.382 | -2.013 | -0.616 | -0.017 | -0.037 | -0.037 | 0 | 0.25 | 0.14 | 0.31 | 0.41 | 0.3 | 0.35 |
| 6 | 9517257 | rs2294732 | 0.823 | 0.154 | -1.634 | -0.046 | -0.011 | 0.021 | 0.021 | 0.101 | 0.12 | 0.04 | 0.07 | 0.07 | 0.12 | 0.07 |
| 6 | 9517502 | rs2294729 | -0.262 | 0.263 | -1.407 | 0.263 | 0.013 | -0.058 | -0.058 | 0.16 | 0.14 | 0.04 | 0.1 | 0.33 | 0.11 | 0.17 |
| 6 | 9519288 | rs7747168 | -0.023 | 0.646 | -1.103 | -0.234 | 0.011 | -0.003 | -0.003 | 2.895 | 0.51 | 0.34 | 0.47 | 0.51 | 0.57 | 0.56 |
| 6 | 9521502 | rs6921813 | -0.316 | 1.2 | -1.302 | -1.089 | -0.005 | 0.183 | 0.183 | 0 | 0.39 | 0.15 | 0.22 | 0.03 | 0.38 | 0.3 |
| 6 | 9524194 | rs9296557 | 1.011 | 1.972 | 0.492 | -0.385 | 0.228 | 0.04 | 0.04 | 0 | 0.93 | 0.67 | 0.91 | 0.94 | 0.82 | 0.92 |
| 6 | 9537611 | rs4715102 | 0.32 | 1.321 | 0.085 | -1.492 | -0.002 | 0.015 | 0.015 | 0 | 0.59 | 0.37 | 0.68 | 0.88 | 0.6 | 0.64 |
| 6 | 9544144 | rs7757300 | 0.645 | 1.724 | -0.224 | -1.848 | -0.017 | 0.244 | 0.244 | 0 | 0.51 | 0.17 | 0.63 | 0.88 | 0.52 | 0.63 |
| 6 | 9545130 | rs10948513 | 0.531 | 1.976 | -0.941 | -1.229 | -0.069 | -0.095 | -0.095 | 0 | 0.17 | 0.01 | 0.32 | 0.29 | 0.23 | 0.19 |
| 6 | 9558460 | rs980414 | -0.156 | 1.438 | -0.405 | -2.42 | 0.006 | -0.079 | -0.079 | 0 | 0.74 | 0.64 | 0.75 | 0.96 | 0.69 | 0.76 |
| 6 | 9561817 | rs9395670 | 0.297 | 0.877 | -0.122 | -2.473 | 0.008 | 0.048 | 0.048 | 0 | 0.51 | 0.37 | 0.66 | 0.87 | 0.56 | 0.64 |
| 6 | 9565483 | rs9349592 | 0.021 | 0.342 | -0.115 | -2.612 | 0.002 | -0.071 | -0.071 | 0 | 0.58 | 0.48 | 0.69 | 0.87 | 0.58 | 0.66 |
| 6 | 9576203 | rs4715321 | 0.487 | 0.764 | -0.447 | -2.547 | 0.003 | 0.051 | 0.051 | 0 | 0.46 | 0.34 | 0.6 | 0.87 | 0.47 | 0.63 |
| 6 | 9582038 | rs17797675 | 0.037 | 0.515 | -0.859 | -2.42 | -0.006 | 0.051 | 0.051 | 0.116 | 0.14 | 0.04 | 0.19 | 0.06 | 0.16 | 0.1 |
| 6 | 9582818 | rs3918445 | 0.109 | 0.334 | -0.621 | -2.419 | -0.001 | 0.113 | 0.113 | 0.47 | 0.49 | 0.31 | 0.51 | 0.85 | 0.51 | 0.67 |
| 6 | 9583208 | rs16892381 | 1.312 | 0.233 | -0.812 | -2.273 | -0.024 | -0.101 | -0.101 | 0.936 | 0.23 | 0.06 | 0.29 | 0.5 | 0.24 | 0.24 |
| 6 | 9583352 | rs992003 | 0.307 | 0.303 | -0.465 | -2.902 | -0.001 | 0.111 | 0.111 | 0.352 | 0.49 | 0.31 | 0.51 | 0.85 | 0.51 | 0.67 |
| 6 | 9597209 | rs9349739 | 0.347 | -0.047 | -0.631 | -2.984 | -0.004 | 0.087 | 0.087 | 0.394 | 0.47 | 0.31 | 0.47 | 0.87 | 0.49 | 0.61 |
| 6 | 9598025 | rs7765001 | -0.604 | 0.051 | -1.086 | -2.948 | -0.071 | 0.164 | 0.164 | 0 | 0.74 | 0.98 | 0.82 | 0.94 | 0.74 | 0.78 |
| 6 | 9608887 | rs855250 | -0.402 | -0.404 | -0.421 | -2.769 | -0.012 | -0.076 | -0.076 | 0 | 0.22 | 0.28 | 0.13 | 0.03 | 0.16 | 0.13 |
| 6 | 9618701 | rs864111 | -0.298 | -0.524 | -0.068 | -3.01 | -0.039 | -0.09 | -0.09 | 1.197 | 0.22 | 0.35 | 0.17 | 0.01 | 0.12 | 0.08 |
| 6 | 9620646 | rs12207681 | 0.458 | -0.44 | -0.135 | -3.002 | 0.004 | -0.022 | -0.022 | 0 | 0.1 | 0.1 | 0.1 | 0.03 | 0.13 | 0.09 |
| 6 | 9662073 | rs9464676 | 1.357 | -0.453 | -0.298 | -3.471 | 0.004 | -0.027 | -0.027 | 0.955 | 0.2 | 0.19 | 0.19 | 0.06 | 0.26 | 0.27 |
| 6 | 9665839 | rs12203770 | 0.5 | -0.311 | -0.739 | -3.242 | -0.008 | 0.035 | 0.035 | 0.384 | 0.1 | 0.05 | 0.08 | 0.03 | 0.1 | 0.09 |
| 6 | 9669475 | rs1328140 | 0.755 | -0.735 | -0.704 | -3.257 | -0.008 | 0.062 | 0.062 | 0 | 0.41 | 0.27 | 0.52 | 0.89 | 0.47 | 0.55 |
| 6 | 9709940 | rs1328134 | -0.068 | -0.758 | -1.233 | -3.196 | 0.005 | 0.09 | 0.09 | 0 | 0.39 | 0.25 | 0.47 | 0.91 | 0.42 | 0.49 |
| 6 | 9715511 | rs9464727 | 0.121 | -0.519 | -1.293 | -3.271 | 0.001 | -0.11 | -0.11 | 0.063 | 0.41 | 0.44 | 0.35 | 0.06 | 0.38 | 0.34 |
| 6 | 9729119 | rs1410364 | 0.781 | -0.521 | -1.192 | -3.149 | -0.004 | -0.006 | -0.006 | 0.132 | 0.23 | 0.15 | 0.19 | 0.03 | 0.24 | 0.17 |
| 6 | 9730967 | rs10484244 | 0.809 | -0.469 | -1.177 | -3.139 | -0.014 | 0.113 | 0.113 | 0.023 | 0.23 | 0.04 | 0.17 | 0.03 | 0.21 | 0.17 |
| 6 | 9736138 | rs7739767 | 0.987 | -0.635 | -1.531 | -3.136 | -0.007 | 0.012 | 0.012 | 0.011 | 0.12 | 0.01 | 0.24 | 0.07 | 0.13 | 0.05 |
| 6 | 9736976 | rs2327200 | 1.148 | -0.219 | -1.138 | -2.915 | 0.078 | -0.123 | -0.123 | 0 | 0.23 | 0.42 | 0.28 | 0.06 | 0.3 | 0.3 |
| 6 | 9740771 | rs10949272 | -0.155 | -0.214 | -1.535 | -1.942 | -0.006 | 0.054 | 0.054 | 0.637 | 0.24 | 0.11 | 0.19 | 0.03 | 0.23 | 0.17 |
| 6 | 9745037 | rs1410366 | -0.181 | -0.284 | -1.532 | -1.943 | -0.003 | -0.009 | -0.009 | 0 | 0.24 | 0.16 | 0.19 | 0.03 | 0.24 | 0.17 |
| 6 | 9748363 | rs6941642 | -0.115 | -0.28 | -1.612 | -1.941 | -0.013 | 0.057 | 0.057 | 0.038 | 0.23 | 0.1 | 0.19 | 0.03 | 0.24 | 0.17 |
| 6 | 9750430 | rs10949284 | -0.794 | -0.291 | -1.64 | -1.949 | 0.015 | 0.082 | 0.082 | 0 | 0.32 | 0.11 | 0.21 | 0.03 | 0.26 | 0.2 |
| 6 | 9758853 | rs12204248 | -0.391 | 0.478 | -1.19 | -1.46 | -0.005 | -0.001 | -0.001 | 0 | 0.23 | 0.13 | 0.19 | 0.03 | 0.24 | 0.17 |
| 6 | 9765893 | rs9383052 | 0.51 | 0.103 | -1.239 | -0.341 | 0.009 | -0.013 | -0.013 | 0 | 0.77 | 0.89 | 0.61 | 0.72 | 0.76 | 0.78 |
| 6 | 9767013 | rs1927594 | -0.049 | 0.014 | -0.927 | 0.579 | 0 | 0.115 | 0.115 | 0 | 0.06 | 0.11 | 0.05 | 0.28 | 0.05 | 0.06 |
| 6 | 9770871 | rs1206915 | 0.028 | -0.106 | -0.926 | 0.796 | -0.012 | 0.137 | 0.137 | 1.902 | 0.05 | 0.1 | 0.04 | 0.32 | 0.02 | 0.08 |
| 6 | 9773164 | rs1206917 | 0.017 | -0.179 | -0.975 | 0.081 | -0.001 | -0.002 | -0.002 | 0 | 0.44 | 0.61 | 0.52 | 0.44 | 0.44 | 0.49 |
| 6 | 9774694 | rs12208751 | 0.622 | -0.006 | -0.939 | 0.088 | 0.001 | -0.001 | -0.001 | 0 | 0.08 | 0.08 | 0.15 | 0.05 | 0.14 | 0.18 |
| 6 | 9776194 | rs9370818 | -0.139 | -0.233 | -1.43 | -0.365 | 0.059 | -0.026 | -0.026 | 0.002 | 0.31 | 0.13 | 0.16 | 0.34 | 0.17 | 0.21 |
| 6 | 9780097 | rs2223236 | 0.09 | 0.172 | -1.226 | -0.071 | -0.001 | -0.003 | -0.003 | 0 | 0.6 | 0.64 | 0.62 | 0.46 | 0.57 | 0.59 |
| 6 | 9784181 | rs2049926 | 0.446 | 0.216 | -0.831 | -0.435 | -0.003 | 0.034 | 0.034 | 0 | 0.55 | 0.29 | 0.54 | 0.35 | 0.54 | 0.45 |
| 6 | 9788600 | rs12662884 | 0.444 | 0.2 | -0.837 | -0.468 | -0.001 | 0.062 | 0.062 | 0 | 0.54 | 0.24 | 0.52 | 0.35 | 0.52 | 0.45 |
| 6 | 9789093 | rs7767721 | -0.232 | 0.229 | -0.14 | -0.072 | 0.041 | 0.369 | 0.369 | 0 | 0.06 | 0.25 | 0.1 | 0.45 | 0.1 | 0.12 |
| 6 | 9801457 | rs9296990 | 0.469 | 0.111 | -0.606 | -1.518 | 0.005 | -0.024 | -0.024 | 0.152 | 0.32 | 0.41 | 0.45 | 0.76 | 0.39 | 0.35 |
| 6 | 9801994 | rs6917809 | -0.303 | 0.042 | -0.639 | -1.594 | -0.012 | 0.089 | 0.089 | 0 | 0.63 | 0.86 | 0.67 | 0.79 | 0.65 | 0.57 |

| | | | | | | | | | | | | | | | | |
|---|---|---|---|---|---|---|---|---|---|---|---|---|---|---|---|---|
| 6 | 9804986 | rs9476916 | 1.344 | 0.147 | -0.539 | -1.429 | -0.018 | 0.058 | 0.058 | 0 | 0.22 | 0.09 | 0.19 | 0.03 | 0.23 | 0.18 |
| 6 | 9805059 | rs6905506 | -0.958 | 0.095 | -0.848 | -1.668 | 0.003 | 0.074 | 0.074 | 0 | 0.85 | 0.98 | 0.92 | 1 | 0.86 | 0.94 |
| 6 | 9806230 | rs17613393 | 1.061 | 0.498 | -0.359 | -1.012 | -0.062 | 0.123 | 0.123 | 0 | 0.22 | 0.02 | 0.19 | 0.03 | 0.22 | 0.18 |
| 6 | 9810702 | rs4716001 | -0.361 | 0.795 | -0.296 | 0.108 | 0 | -0.001 | -0.001 | 0 | 0.29 | 0.28 | 0.36 | 0.72 | 0.3 | 0.32 |
| 6 | 9822157 | rs855375 | -0.92 | 1.378 | 0.096 | 0.369 | 0.064 | 0.013 | 0.013 | 0 | 0.19 | 0.41 | 0.33 | 0.2 | 0.28 | 0.37 |
| 6 | 9822482 | rs1555035 | -0.063 | 1.182 | -0.228 | 0.271 | 0 | 0.005 | 0.005 | 0 | 0.36 | 0.34 | 0.45 | 0.34 | 0.41 | 0.29 |
| 6 | 9822908 | rs1555037 | 0.575 | 1.03 | -0.425 | 0.31 | 0.002 | 0.003 | 0.003 | 0 | 0.19 | 0.19 | 0.17 | 0.14 | 0.19 | 0.19 |
| 6 | 9822953 | rs1555038 | 0.522 | 1.14 | -0.462 | 0.309 | -0.016 | 0.008 | 0.008 | 0 | 0.21 | 0.18 | 0.23 | 0.13 | 0.3 | 0.2 |
| 6 | 9823155 | rs1557007 | 0.437 | 1.125 | -0.724 | 0.009 | -0.024 | 0.042 | 0.042 | 0.01 | 0.08 | 0.01 | 0.04 | 0 | 0.08 | 0.02 |
| 6 | 9823784 | rs6459434 | -0.103 | 0.824 | -0.357 | -0.162 | 0.003 | -0.008 | -0.008 | 0.237 | 0.39 | 0.44 | 0.44 | 0.3 | 0.48 | 0.32 |
| 6 | 9825495 | rs9296996 | 0.498 | 1.054 | -0.527 | -0.306 | 0 | -0.034 | -0.034 | 0 | 0.24 | 0.26 | 0.2 | 0.04 | 0.35 | 0.23 |
| 6 | 9831530 | rs1934775 | -0.471 | 0.681 | -1.148 | 0.054 | -0.008 | -0.022 | -0.022 | 0.302 | 0.76 | 0.81 | 0.77 | 0.67 | 0.71 | 0.77 |
| 6 | 9835194 | rs855367 | -0.13 | 0.538 | -1.316 | -0.062 | -0.011 | -0.018 | -0.018 | 0.377 | 0.59 | 0.81 | 0.61 | 0.43 | 0.63 | 0.51 |
| 6 | 9835428 | rs13213170 | 1.244 | 0.619 | -1.4 | -0.121 | -0.087 | 0.053 | 0.053 | 0.049 | 0.19 | 0.01 | 0.12 | 0.08 | 0.25 | 0.2 |
| 6 | 9842214 | rs760618 | 0.304 | 0.529 | -1.402 | 0.07 | -0.024 | -0.047 | -0.047 | 0 | 0.23 | 0.12 | 0.26 | 0.28 | 0.22 | 0.31 |
| 6 | 9842319 | rs707782 | -0.858 | -0.008 | -1.01 | 0.21 | 0.009 | -0.022 | -0.022 | 1.726 | 0.8 | 0.86 | 0.86 | 0.7 | 0.84 | 0.82 |
| 6 | 9846872 | rs1206973 | 1.033 | -0.218 | -0.802 | 0.09 | -0.002 | 0.005 | 0.005 | 0 | 0.21 | 0.23 | 0.22 | 0.13 | 0.27 | 0.19 |
| 6 | 9848666 | rs12524134 | 1.125 | -0.402 | -1.045 | -0.051 | -0.006 | 0.063 | 0.063 | 0 | 0.07 | 0.09 | 0.06 | 0.29 | 0.04 | 0.13 |
| 6 | 9850927 | rs2185489 | -1.266 | -0.199 | -0.924 | 0.088 | -0.012 | 0.017 | 0.017 | 0 | 0.79 | 0.85 | 0.78 | 0.87 | 0.73 | 0.81 |
| 6 | 9856368 | rs11752484 | 0.61 | -0.318 | -1.054 | 0.054 | -0.107 | -0.021 | -0.021 | 0 | 0.4 | 0.13 | 0.37 | 0.36 | 0.45 | 0.5 |
| 6 | 9869358 | rs9396671 | 0.345 | -0.611 | -0.452 | 0.149 | -0.039 | 0.017 | 0.017 | 0.007 | 0.18 | 0.46 | 0.26 | 0.21 | 0.12 | 0.12 |
| 6 | 9875386 | rs9367915 | 0.947 | -0.28 | -0.681 | 0.245 | 0.01 | 0.006 | 0.006 | 0 | 0.17 | 0.08 | 0.14 | 0.12 | 0.15 | 0.15 |
| 6 | 9895130 | rs7739081 | 0.166 | -0.436 | 0.108 | 0.426 | -0.037 | 0.043 | 0.043 | 0 | 0.23 | 0.39 | 0.17 | 0.31 | 0.15 | 0.18 |
| 6 | 9906609 | rs6459485 | 0.335 | -0.689 | 0.6 | 0.067 | -0.026 | -0.049 | -0.049 | 0 | 0.16 | 0.31 | 0.11 | 0.05 | 0.12 | 0.14 |
| 6 | 9906751 | rs1925768 | -0.593 | -0.929 | 0.609 | 0.109 | -0.045 | 0.059 | 0.059 | 0 | 0.75 | 0.58 | 0.83 | 0.68 | 0.84 | 0.83 |
| 6 | 9908321 | rs6939240 | 0.727 | -0.93 | 0.754 | 0.109 | 0.01 | -0.014 | -0.014 | 0 | 0.23 | 0.13 | 0.14 | 0.32 | 0.16 | 0.17 |
| 6 | 9912324 | rs12176510 | 1.271 | -0.917 | 0.252 | -0.091 | 0.004 | 0.022 | 0.022 | 0 | 0.19 | 0.1 | 0.16 | 0.12 | 0.18 | 0.2 |
| 6 | 9914294 | rs9396715 | 0.072 | -0.855 | -0.146 | -0.126 | 0.001 | -0.002 | -0.002 | 0 | 0.43 | 0.41 | 0.59 | 0.48 | 0.56 | 0.53 |
| 6 | 9941223 | rs9396752 | 1.497 | -0.214 | -1.146 | 0.059 | -0.117 | -0.077 | -0.077 | 0 | 0.18 | 0.01 | 0.29 | 0.28 | 0.29 | 0.21 |
| 6 | 9941445 | rs9297028 | 1.106 | -0.37 | -1.128 | 0.183 | 0.019 | -0.032 | -0.032 | 0.354 | 0.22 | 0.11 | 0.14 | 0.31 | 0.17 | 0.18 |
| 6 | 9943593 | rs9383254 | 1.105 | -0.383 | -1.035 | 0.029 | -0.017 | 0.005 | 0.005 | 0.008 | 0.19 | 0.05 | 0.15 | 0.13 | 0.17 | 0.19 |
| 6 | 9944367 | rs4716126 | 0.002 | -0.343 | -0.65 | 0.093 | -0.045 | -0.041 | -0.041 | 0 | 0.08 | 0.02 | 0.2 | 0.14 | 0.15 | 0.19 |
| 6 | 9949384 | rs729254 | -0.012 | -0.639 | -0.018 | -0.07 | -0.02 | -0.025 | -0.025 | 0.022 | 0.86 | 0.73 | 0.91 | 0.91 | 0.9 | 0.9 |
| 6 | 9952059 | rs9396763 | 0.595 | -0.684 | 0.212 | -0.106 | -0.045 | -0.055 | -0.055 | 0.009 | 0.72 | 0.46 | 0.82 | 0.87 | 0.79 | 0.79 |
| 6 | 9952313 | rs9370969 | 0.05 | 0.032 | -0.044 | 0.342 | -0.009 | -0.017 | -0.017 | 0.052 | 0.88 | 0.73 | 0.92 | 0.91 | 0.9 | 0.9 |
| 6 | 9955326 | rs2327219 | -0.016 | 0.225 | -0.696 | 0.809 | -0.021 | -0.059 | -0.059 | 0.969 | 0.72 | 0.85 | 0.62 | 0.57 | 0.67 | 0.55 |
| 6 | 9956562 | rs13217591 | 0.656 | 0.55 | -1.319 | -0.093 | -0.09 | 0.096 | 0.096 | 0 | 0.19 | 0.01 | 0.24 | 0.04 | 0.23 | 0.19 |
| 6 | 9959207 | rs11965246 | 0.095 | 0.346 | -0.52 | -0.006 | -0.005 | 0.01 | 0.01 | 0.163 | 0.1 | 0.29 | 0.05 | 0.1 | 0.07 | 0.15 |
| 6 | 9976215 | rs9396778 | 0.821 | 0.057 | -1.287 | -0.304 | -0.06 | 0.088 | 0.088 | 0.014 | 0.32 | 0.03 | 0.26 | 0.12 | 0.33 | 0.25 |
| 6 | 9977413 | rs201265 | 0.726 | 0.19 | -1.278 | 0.262 | -0.014 | 0.048 | 0.048 | 0 | 0.07 | 0.03 | 0.21 | 0.51 | 0.08 | 0.23 |
| 6 | 9982500 | rs3765276 | 1.232 | 0.148 | -1.159 | 0.136 | 0 | -0.012 | -0.012 | 0.06 | 0.08 | 0.11 | 0.04 | 0.04 | 0.08 | 0.06 |
| 6 | 9986985 | rs969527 | -0.43 | 0.028 | -0.499 | 0.028 | -0.001 | 0 | 0 | 0 | 0.65 | 0.68 | 0.6 | 0.27 | 0.65 | 0.6 |
| 6 | 9988811 | rs10484260 | 0.624 | 0.073 | -0.283 | 0.099 | -0.005 | 0.006 | 0.006 | 0.004 | 0.1 | 0.28 | 0.05 | 0.1 | 0.07 | 0.15 |
| 6 | 9994389 | rs201260 | -0.152 | 0.226 | -0.555 | 0.178 | 0.002 | 0.006 | 0.006 | 0 | 0.42 | 0.37 | 0.5 | 0.63 | 0.42 | 0.5 |
| 6 | 9994519 | rs17621880 | 0.929 | 0.493 | -0.87 | 0.021 | -0.03 | 0.089 | 0.089 | 0 | 0.15 | 0.01 | 0.09 | 0.01 | 0.17 | 0.06 |
| 6 | 10005707 | rs201255 | -0.018 | 0.288 | -0.431 | 0.317 | 0.004 | 0.015 | 0.015 | 0.2 | 0.24 | 0.36 | 0.44 | 0.65 | 0.26 | 0.44 |
| 6 | 10008957 | rs9396808 | 1.087 | 0.317 | -0.426 | 0.57 | -0.004 | -0.005 | -0.005 | 0.394 | 0.15 | 0.13 | 0.18 | 0.14 | 0.17 | 0.19 |
| 6 | 10010063 | rs2179366 | 1.121 | 0.316 | -0.432 | 0.569 | -0.007 | -0.009 | -0.009 | 0.115 | 0.15 | 0.11 | 0.18 | 0.14 | 0.17 | 0.19 |
| 6 | 10011084 | rs201252 | -0.825 | -0.063 | -0.353 | 0.209 | -0.004 | -0.022 | -0.022 | 0 | 0.39 | 0.49 | 0.51 | 0.74 | 0.37 | 0.62 |
| 6 | 10012023 | rs201251 | -0.777 | -0.113 | -0.553 | -0.148 | -0.006 | -0.003 | -0.003 | 0.45 | 0.58 | 0.48 | 0.64 | 0.67 | 0.62 | 0.67 |
| 6 | 10013636 | rs201250 | -0.661 | -0.066 | -0.5 | -0.04 | -0.008 | 0.066 | 0.066 | 0 | 0.26 | 0.41 | 0.34 | 0.6 | 0.22 | 0.43 |
| 6 | 10019083 | rs9371028 | 1.361 | 0.239 | -0.727 | -0.11 | -0.036 | -0.044 | -0.044 | 0 | 0.14 | 0.01 | 0.16 | 0.13 | 0.15 | 0.18 |
| 6 | 10020243 | rs201237 | 0.902 | 0.241 | -0.638 | -0.256 | -0.038 | 0.092 | 0.092 | 0 | 0.34 | 0.19 | 0.32 | 0.07 | 0.4 | 0.25 |
| 6 | 10022285 | rs201233 | 0.685 | 0.213 | -0.539 | -0.27 | -0.012 | 0.02 | 0.02 | 0 | 0.35 | 0.33 | 0.33 | 0.07 | 0.4 | 0.25 |
| 6 | 10032432 | rs201224 | -0.554 | 0.177 | -0.447 | -0.116 | -0.007 | -0.017 | -0.017 | 0 | 0.66 | 0.56 | 0.56 | 0.28 | 0.67 | 0.62 |
| 6 | 10035866 | rs12529014 | 0.357 | 0.22 | -0.481 | 0.375 | -0.004 | 0.052 | 0.052 | 0 | 0.08 | 0.08 | 0.16 | 0.48 | 0.1 | 0.24 |
| 6 | 10035875 | rs2327221 | 0.308 | 0.063 | -0.558 | 0.081 | 0.004 | 0.01 | 0.01 | 0 | 0.27 | 0.18 | 0.21 | 0.21 | 0.24 | 0.35 |
| 6 | 10036770 | rs1883408 | -0.038 | 0.077 | -0.533 | 0.09 | 0 | -0.026 | -0.026 | 0.758 | 0.51 | 0.51 | 0.41 | 0.15 | 0.52 | 0.44 |
| 6 | 10038006 | rs7341284 | 0.135 | 0.398 | -0.501 | 0.175 | -0.005 | 0.005 | 0.005 | 0 | 0.39 | 0.43 | 0.62 | 0.71 | 0.47 | 0.64 |
| 6 | 10041138 | rs6459596 | 0.713 | 0.485 | -0.711 | 0.373 | 0.033 | 0.068 | 0.068 | 0 | 0.25 | 0.39 | 0.39 | 0.53 | 0.33 | 0.38 |
| 6 | 10041880 | rs9297055 | 0.969 | -0.133 | -0.725 | 0.773 | -0.038 | -0.061 | -0.061 | 0.087 | 0.14 | 0.02 | 0.23 | 0.18 | 0.16 | 0.27 |
| 6 | 10042928 | rs855404 | -0.845 | -0.425 | -0.703 | 0.159 | -0.021 | -0.013 | -0.013 | 0 | 0.72 | 0.71 | 0.82 | 0.81 | 0.78 | 0.87 |
| 6 | 10044051 | rs855402 | -0.506 | -0.275 | -0.8 | 0.246 | -0.036 | -0.029 | -0.029 | 0.416 | 0.67 | 0.61 | 0.8 | 0.81 | 0.76 | 0.85 |
| 6 | 10047472 | rs855399 | -0.288 | -0.103 | -0.824 | 0.316 | -0.019 | -0.018 | -0.018 | 0 | 0.62 | 0.61 | 0.79 | 0.77 | 0.74 | 0.75 |
| 6 | 10047914 | rs855398 | 0.253 | 0.29 | -0.726 | 0.76 | -0.003 | -0.004 | -0.004 | 0 | 0.46 | 0.55 | 0.62 | 0.39 | 0.58 | 0.48 |
| 6 | 10048422 | rs855397 | 0.174 | -0.188 | -0.699 | 0.75 | 0.005 | -0.003 | -0.003 | 0.152 | 0.62 | 0.42 | 0.76 | 0.58 | 0.7 | 0.74 |
| 6 | 10048851 | rs13219920 | 0.219 | 0.058 | -0.531 | -0.194 | -0.001 | -0.01 | -0.01 | 0.055 | 0.55 | 0.47 | 0.38 | 0.78 | 0.52 | 0.53 |
| 6 | 10049137 | rs855395 | -0.499 | -0.177 | -0.177 | 0.201 | 0.007 | 0.009 | 0.009 | 0.876 | 0.38 | 0.6 | 0.57 | 0.47 | 0.43 | 0.57 |
| 6 | 10049717 | rs855394 | -0.451 | -0.119 | 0.011 | 0.265 | 0.043 | 0.1 | 0.1 | 0.039 | 0.21 | 0.58 | 0.24 | 0.38 | 0.29 | 0.34 |

| | | | | | | | | | | | | | | | | |
|---|---|---|---|---|---|---|---|---|---|---|---|---|---|---|---|---|
| 6 | 10058811 | rs760892 | -0.31 | -0.361 | 0.073 | 0.059 | 0.001 | -0.015 | -0.015 | 4.833 | 0.55 | 0.47 | 0.58 | 0.22 | 0.49 | 0.48 |
| 6 | 10065843 | rs13192915 | 0.887 | -0.588 | 0.013 | -0.045 | 0.001 | 0.024 | 0.024 | 0 | 0.24 | 0.14 | 0.15 | 0.15 | 0.26 | 0.19 |
| 6 | 10067722 | rs9383382 | 1.446 | -0.426 | 0.128 | 0.172 | -0.007 | -0.032 | -0.032 | 0 | 0.26 | 0.17 | 0.2 | 0.41 | 0.24 | 0.19 |
| 6 | 10069734 | rs6922037 | 0.829 | -0.422 | 0.182 | -0.198 | -0.01 | -0.019 | -0.019 | 0 | 0.32 | 0.44 | 0.25 | 0.77 | 0.25 | 0.35 |
| 6 | 10079333 | rs1322825 | -0.081 | -0.597 | 0.23 | -0.147 | 0.007 | 0.011 | 0.011 | 0 | 0.38 | 0.34 | 0.28 | 0.18 | 0.36 | 0.25 |
| 6 | 10081139 | rs6924077 | -0.932 | -0.58 | -0.311 | -0.018 | -0.082 | -0.014 | -0.014 | 0.013 | 0.61 | 0.86 | 0.68 | 0.59 | 0.52 | 0.63 |
| 6 | 10084375 | rs6459626 | 0.493 | -0.401 | -0.439 | 0.094 | 0.011 | 0.179 | 0.179 | 1.32 | 0.21 | 0.45 | 0.4 | 0.42 | 0.17 | 0.39 |
| 6 | 10086919 | rs7751450 | -0.026 | -0.327 | -1.098 | 0.099 | 0.021 | 0.045 | 0.045 | 0 | 0.37 | 0.27 | 0.26 | 0.17 | 0.35 | 0.25 |
| 6 | 10088295 | rs9371078 | 1.152 | 0.158 | -1.29 | -0.089 | -0.032 | -0.054 | -0.054 | 0 | 0.28 | 0.11 | 0.2 | 0.4 | 0.3 | 0.22 |
| 6 | 10091216 | rs7747662 | 0.066 | 0.307 | -0.948 | -0.294 | -0.005 | 0.015 | 0.015 | 0 | 0.25 | 0.18 | 0.21 | 0.07 | 0.31 | 0.16 |
| 6 | 10091425 | rs17624038 | 0.25 | 0.585 | -0.803 | 0.225 | -0.045 | 0.028 | 0.028 | 0 | 0.15 | 0.02 | 0.33 | 0.05 | 0.14 | 0.19 |
| 6 | 10094268 | rs11751970 | 0.408 | 0.643 | -0.827 | 0.256 | -0.055 | 0.017 | 0.017 | 0.095 | 0.14 | 0.01 | 0.26 | 0.05 | 0.14 | 0.19 |
| 6 | 10095619 | rs11756005 | -0.966 | 0.33 | -0.632 | 0.394 | -0.004 | -0.012 | -0.012 | 0 | 0.13 | 0.12 | 0.1 | 0.13 | 0.14 | 0.15 |
| 6 | 10097016 | rs13209414 | 0.138 | 0.642 | -0.447 | 0.543 | -0.011 | -0.016 | -0.016 | 0 | 0.18 | 0.12 | 0.35 | 0.25 | 0.2 | 0.27 |
| 6 | 10099016 | rs957610 | -1.155 | -0.175 | 0.563 | -0.078 | -0.018 | 0.069 | 0.069 | 0 | 0.15 | 0.4 | 0.14 | 0.24 | 0.12 | 0.2 |
| 6 | 10100647 | rs11759305 | -1.009 | -0.1 | 0.584 | -0.399 | -0.026 | 0.01 | 0.01 | 0 | 0.16 | 0.28 | 0.08 | 0.15 | 0.09 | 0.14 |
| 6 | 10102217 | rs2876189 | -0.4 | 0.092 | 0.552 | -0.492 | 0.032 | 0.029 | 0.029 | 0.064 | 0.36 | 0.75 | 0.26 | 0.22 | 0.3 | 0.3 |
| 6 | 10103275 | rs7755414 | -0.466 | 0.168 | 0.592 | -0.442 | 0.035 | 0.045 | 0.045 | 0.04 | 0.4 | 0.75 | 0.26 | 0.22 | 0.33 | 0.31 |
| 6 | 10105007 | rs2224660 | -1.708 | 0.585 | 1.659 | 0.262 | 0.05 | 0.194 | 0.194 | 1.299 | 0.06 | 0.55 | 0.05 | 0.12 | 0.02 | 0.07 |
| 6 | 10106467 | rs4716306 | 0.795 | 1.057 | 0.737 | -0.296 | 0.06 | -0.121 | -0.121 | 0 | 0.84 | 0.57 | 0.84 | 1 | 0.8 | 0.85 |
| 6 | 10109357 | rs1207005 | 0.629 | 1.264 | -0.929 | -0.161 | -0.003 | -0.033 | -0.033 | 0 | 0.78 | 0.79 | 0.81 | 1 | 0.75 | 0.84 |
| 6 | 10111700 | rs4716310 | -0.788 | 1.406 | -1.45 | 1.046 | -0.053 | -0.112 | -0.112 | 0.242 | 0.22 | 0.04 | 0.36 | 0.57 | 0.27 | 0.38 |
| 6 | 10112297 | rs1889211 | -1.393 | 1.181 | -0.989 | 1.589 | -0.002 | 0.021 | 0.021 | 0 | 0.13 | 0.16 | 0.17 | 0.29 | 0.11 | 0.18 |
| 6 | 10112415 | rs1207009 | 0.058 | 1.285 | -1.928 | 0.915 | -0.051 | -0.017 | -0.017 | 0 | 0.57 | 0.88 | 0.41 | 0.59 | 0.52 | 0.58 |
| 6 | 10112580 | rs1209347 | -0.07 | 1.071 | -0.877 | 0.455 | 0.041 | -0.005 | -0.005 | 0 | 0.11 | 0.37 | 0.1 | 0.08 | 0.12 | 0.13 |
| 6 | 10112607 | rs4557511 | 0.583 | 0.827 | -0.851 | 0.889 | 0 | -0.004 | -0.004 | 0 | 0.33 | 0.21 | 0.23 | 0.3 | 0.3 | 0.42 |
| 6 | 10113579 | rs1212597 | -0.129 | 0.969 | -1.005 | 0.75 | -0.002 | 0.002 | 0.002 | 0 | 0.6 | 0.6 | 0.59 | 0.33 | 0.56 | 0.52 |
| 6 | 10116849 | rs4712358 | 0.552 | 0.901 | -1.717 | 0.097 | -0.057 | 0.171 | 0.171 | 0 | 0.28 | 0.05 | 0.36 | 0.01 | 0.28 | 0.1 |
| 6 | 10126207 | rs942012 | 0.26 | 1.442 | -0.93 | 0.712 | 0.013 | 0.013 | 0.013 | 0 | 0.69 | 0.57 | 0.75 | 0.67 | 0.67 | 0.58 |
| 6 | 10127684 | rs12527216 | 0.53 | 1.106 | -1.054 | 1.06 | -0.014 | -0.006 | -0.006 | 0 | 0.4 | 0.53 | 0.45 | 0.33 | 0.33 | 0.3 |
| 6 | 10128271 | rs10456848 | -0.222 | 1.235 | -1.235 | 1.736 | -0.014 | -0.077 | -0.077 | 0 | 0.18 | 0.1 | 0.13 | 0.32 | 0.19 | 0.28 |
| 6 | 10131177 | rs10456221 | -0.411 | 1.473 | -1.075 | 1.653 | -0.025 | -0.055 | -0.055 | 0 | 0.2 | 0.09 | 0.18 | 0.5 | 0.28 | 0.42 |
| 6 | 10141017 | rs6650997 | 0.109 | 1.326 | -0.209 | 1.138 | 0.012 | -0.044 | -0.044 | 0 | 0.1 | 0.32 | 0.05 | 0 | 0.07 | 0.03 |
| 6 | 10142268 | rs2327251 | 0.654 | 1.088 | -0.294 | 0.712 | 0.048 | 0.089 | 0.089 | 2.96 | 0.52 | 0.69 | 0.39 | 0.01 | 0.34 | 0.15 |
| 6 | 10148486 | rs6922273 | 0.03 | 0.918 | -0.722 | 1.213 | -0.027 | -0.069 | -0.069 | 0.286 | 0.2 | 0.09 | 0.19 | 0.57 | 0.28 | 0.42 |
| 6 | 10150762 | rs9358202 | 1.263 | 0.907 | -0.785 | 0.956 | -0.057 | 0.02 | 0.02 | 0.319 | 0.41 | 0.09 | 0.4 | 0.26 | 0.38 | 0.28 |
| 6 | 10151450 | rs12193757 | -0.158 | 0.713 | -0.497 | 0.674 | -0.002 | -0.011 | -0.011 | 0 | 0.06 | 0.05 | 0.03 | 0.01 | 0.04 | 0.09 |
| 6 | 10152592 | rs914932 | 0.869 | 0.825 | -0.239 | 0.769 | 0.003 | 0.056 | 0.056 | 1.883 | 0.5 | 0.42 | 0.46 | 0.18 | 0.47 | 0.28 |
| 6 | 10153009 | rs6914993 | -0.039 | 0.963 | -0.189 | 1.192 | -0.002 | -0.006 | -0.006 | 0 | 0.79 | 0.79 | 0.85 | 0.63 | 0.75 | 0.56 |
| 6 | 10153944 | rs7761624 | 0.69 | 0.728 | 0.035 | 0.78 | -0.009 | 0.047 | 0.047 | 0.185 | 0.47 | 0.38 | 0.45 | 0.22 | 0.46 | 0.29 |
| 6 | 10158948 | rs9465334 | 0.434 | 0.932 | 0.084 | 0.573 | 0.042 | 0.165 | 0.165 | 0.085 | 0.45 | 0.75 | 0.37 | 0.01 | 0.35 | 0.14 |
| 6 | 10163721 | rs9358212 | -0.188 | 0.893 | -0.042 | 0.887 | -0.002 | -0.03 | -0.03 | 2.071 | 0.53 | 0.83 | 0.56 | 0.53 | 0.53 | 0.41 |
| 6 | 10173148 | rs11751873 | 0.703 | 0.851 | 0.108 | 0.695 | 0.02 | 0.019 | 0.019 | 0 | 0.4 | 0.27 | 0.35 | 0.31 | 0.32 | 0.24 |
| 6 | 10173476 | rs6908117 | 0.727 | 1.002 | 0.185 | 0.838 | 0.027 | 0.027 | 0.027 | 0 | 0.42 | 0.28 | 0.35 | 0.31 | 0.32 | 0.25 |
| 6 | 10173613 | rs6935000 | -0.013 | 0.747 | 0.165 | 1.105 | -0.003 | 0.001 | 0.001 | 0 | 0.53 | 0.52 | 0.53 | 0.49 | 0.53 | 0.47 |
| 6 | 10174570 | rs9358225 | -1.269 | 0.825 | 0.488 | 1.922 | -0.024 | 0.117 | 0.117 | 0.304 | 0.07 | 0.13 | 0.07 | 0.31 | 0.03 | 0.12 |
| 6 | 10177613 | rs2327252 | -1.674 | 1.118 | 0.385 | 1.285 | 0.078 | 0.243 | 0.243 | 1.519 | 0.15 | 0.36 | 0.24 | 0.49 | 0.2 | 0.34 |
| 6 | 10178094 | rs9465426 | -0.625 | 0.791 | -0.042 | 0.531 | 0 | 0.016 | 0.016 | 0 | 0.18 | 0.14 | 0.07 | 0.06 | 0.2 | 0.3 |
| 6 | 10179035 | rs1029123 | 0.6 | 0.628 | 0.091 | 0.788 | 0.01 | -0.015 | -0.015 | 0 | 0.67 | 0.75 | 0.61 | 0.58 | 0.71 | 0.67 |
| 6 | 10179366 | rs1029124 | 0.471 | 0.639 | 0.238 | 0.962 | 0 | 0.087 | 0.087 | 0.086 | 0.9 | 0.77 | 0.91 | 0.72 | 0.93 | 0.91 |
| 6 | 10179707 | rs1537488 | 0.319 | 0.617 | 0.457 | 0.951 | 0.017 | 0.144 | 0.144 | 0 | 0.9 | 0.69 | 0.9 | 0.72 | 0.93 | 0.91 |
| 6 | 10180565 | rs4712416 | -0.845 | 1.166 | 0.408 | 1.248 | 0.034 | 0.114 | 0.114 | 0 | 0.07 | 0.08 | 0.16 | 0.39 | 0.17 | 0.28 |
| 6 | 10180750 | rs1999565 | 1.209 | 0.819 | 0.501 | 0.772 | -0.001 | -0.002 | -0.002 | 0 | 0.57 | 0.45 | 0.44 | 0.21 | 0.53 | 0.39 |
| 6 | 10186289 | rs17631613 | -0.38 | -0.12 | 0.133 | 0.225 | 0.008 | 0.058 | 0.058 | 0 | 0.23 | 0.07 | 0.29 | 0.11 | 0.23 | 0.21 |
| 6 | 10190058 | rs6919094 | 0.528 | -0.3 | -0.239 | -0.221 | -0.035 | 0.008 | 0.008 | 0.525 | 0.61 | 0.11 | 0.66 | 0.64 | 0.64 | 0.74 |
| 6 | 10196666 | rs1931023 | -0.222 | -0.286 | 0.482 | -0.228 | 0 | 0 | 0 | 1.676 | 0.74 | 0.33 | 0.79 | 0.68 | 0.75 | 0.78 |
| 6 | 10197494 | rs17556163 | 0.725 | -0.425 | 0.398 | -0.317 | -0.034 | 0.062 | 0.062 | 1.877 | 0.19 | 0.08 | 0.16 | 0.01 | 0.23 | 0.15 |
| 6 | 10200678 | rs9368113 | 1.035 | -0.692 | 0.152 | -0.49 | -0.045 | 0.044 | 0.044 | 0.149 | 0.47 | 0.15 | 0.42 | 0.59 | 0.48 | 0.57 |
| 6 | 10201975 | rs6899608 | 0.017 | -0.689 | 0.34 | -0.487 | 0.008 | 0.04 | 0.04 | 0 | 0.74 | 0.85 | 0.62 | 0.91 | 0.73 | 0.78 |
| 6 | 10203618 | rs4712428 | 1.327 | -0.674 | -0.129 | -0.47 | -0.131 | -0.087 | -0.087 | 0.114 | 0.4 | 0.04 | 0.35 | 0.55 | 0.44 | 0.52 |
| 6 | 10211332 | rs9460420 | 0.112 | -0.717 | 0.38 | -0.506 | 0.001 | -0.01 | -0.01 | 0.529 | 0.69 | 0.33 | 0.73 | 0.95 | 0.73 | 0.82 |
| 6 | 10211788 | rs9465546 | 1.35 | -0.748 | 0.795 | -0.485 | 0.005 | 0.017 | 0.017 | 0 | 0.91 | 0.54 | 0.9 | 0.95 | 0.96 | 0.95 |
| 6 | 10214451 | rs9368130 | 0.49 | -0.859 | 0.216 | -0.381 | -0.001 | -0.035 | -0.035 | 0 | 0.11 | 0.07 | 0.27 | 0.31 | 0.13 | 0.12 |
| 6 | 10218897 | rs1157210 | 0.805 | -0.808 | 0.201 | -0.454 | -0.036 | 0.07 | 0.07 | 2.158 | 0.19 | 0.06 | 0.16 | 0.01 | 0.23 | 0.13 |
| 6 | 10219563 | rs10806909 | 0.739 | -0.815 | 0.445 | -0.101 | -0.026 | 0.009 | 0.009 | 0 | 0.45 | 0.28 | 0.41 | 0.56 | 0.49 | 0.55 |
| 6 | 10222874 | rs9368137 | 0.886 | -0.724 | 0.072 | -0.053 | -0.091 | -0.04 | -0.04 | 1.052 | 0.42 | 0.1 | 0.38 | 0.55 | 0.49 | 0.55 |
| 6 | 10231925 | rs6900553 | 0.791 | -0.959 | 0.571 | 0.184 | -0.052 | 0.064 | 0.064 | 0.026 | 0.79 | 0.41 | 0.6 | 0.63 | 0.85 | 0.84 |
| 6 | 10235805 | rs4712449 | 1.003 | -0.994 | -0.087 | 0.027 | -0.08 | -0.079 | -0.079 | 1.695 | 0.41 | 0.09 | 0.37 | 0.54 | 0.5 | 0.5 |
| 6 | 10236008 | rs4712450 | 0.875 | -0.986 | 0.078 | 0.056 | 0.003 | 0.088 | 0.088 | 0.543 | 0.92 | 0.56 | 0.89 | 0.92 | 0.99 | 0.95 |

| | | | | | | | | | | | | | | | | |
|---|---|---|---|---|---|---|---|---|---|---|---|---|---|---|---|---|
| 6 | 10240416 | rs9358303 | 0.822 | -0.64 | -1.169 | 0.068 | -0.105 | -0.103 | -0.103 | 0 | 0.41 | 0.06 | 0.36 | 0.53 | 0.5 | 0.51 |
| 6 | 10241822 | rs9358305 | 0.26 | -0.043 | -0.471 | -0.344 | -0.002 | -0.049 | -0.049 | 0.111 | 0.6 | 0.57 | 0.75 | 0.91 | 0.63 | 0.69 |
| 6 | 10243211 | rs12526330 | -0.803 | -0.49 | -1.039 | 0.625 | 0.038 | -0.062 | -0.062 | 1.077 | 0.26 | 0.12 | 0.21 | 0.41 | 0.18 | 0.25 |
| 6 | 10245760 | rs1023276 | 0.163 | 0.241 | -1.528 | 0.66 | -0.076 | -0.082 | -0.082 | 0 | 0.23 | 0.1 | 0.28 | 0.61 | 0.36 | 0.27 |
| 6 | 10246691 | rs955506 | 0.727 | -1.066 | -1.138 | 0.233 | 0.003 | -0.004 | -0.004 | 0 | 0.68 | 0.32 | 0.79 | 0.66 | 0.81 | 0.61 |
| 6 | 10256675 | rs9350235 | 0.631 | -1.103 | -0.713 | 0.216 | -0.034 | 0.01 | 0.01 | 0.139 | 0.69 | 0.39 | 0.8 | 0.66 | 0.81 | 0.6 |
| 6 | 10256743 | rs9368175 | 0.209 | -1.38 | -0.243 | 0.283 | -0.005 | 0 | 0 | 0 | 0.89 | 0.89 | 0.96 | 0.89 | 0.91 | 0.87 |
| 6 | 10259637 | rs6456333 | 0.011 | -1.458 | 0.424 | 0.271 | -0.067 | -0.005 | -0.005 | 0 | 0.12 | 0.3 | 0.06 | 0.1 | 0.02 | 0.13 |
| 6 | 10260935 | rs1931017 | 1.487 | -1.654 | 0.588 | 0.231 | -0.027 | 0.004 | 0.004 | 0.365 | 0.69 | 0.36 | 0.81 | 0.66 | 0.84 | 0.62 |
| 6 | 10261644 | rs11963800 | 0.245 | -1.586 | 0.791 | 0.255 | -0.053 | 0.002 | 0.002 | 0 | 0.11 | 0.21 | 0.06 | 0.12 | 0.02 | 0.13 |
| 6 | 10267089 | rs4710928 | 1.216 | -1.601 | 0.895 | 0.235 | -0.044 | 0.046 | 0.046 | 0.466 | 0.8 | 0.48 | 0.86 | 0.78 | 0.85 | 0.83 |
| 6 | 10267228 | rs4710929 | 1.54 | -1.652 | 0.794 | 0.219 | 0.052 | -0.019 | -0.019 | 0 | 0.68 | 0.25 | 0.8 | 0.66 | 0.84 | 0.62 |
| 6 | 10268828 | rs2184176 | 1.235 | -1.288 | 0.62 | -0.213 | -0.028 | 0.148 | 0.148 | 0 | 0.44 | 0.24 | 0.52 | 0.06 | 0.48 | 0.34 |
| 6 | 10269064 | rs6940110 | 0.589 | -1.341 | 0.692 | -0.182 | -0.003 | 0.001 | 0.001 | 0 | 0.56 | 0.43 | 0.58 | 0.17 | 0.5 | 0.47 |
| 6 | 10270377 | rs1556547 | 0.324 | -1.262 | 0.454 | -0.059 | -0.001 | -0.001 | -0.001 | 0 | 0.61 | 0.41 | 0.65 | 0.27 | 0.63 | 0.51 |
| 6 | 10271276 | rs4512211 | -0.106 | -1.281 | 0.557 | -0.036 | -0.086 | 0.007 | 0.007 | 0.042 | 0.86 | 0.6 | 0.93 | 0.87 | 0.98 | 0.87 |
| 6 | 10273963 | rs1333652 | -0.043 | -1.006 | 0.511 | -0.102 | 0.004 | -0.01 | -0.01 | 0 | 0.62 | 0.51 | 0.66 | 0.27 | 0.64 | 0.58 |
| 6 | 10282741 | rs6456357 | -1.006 | -1.007 | 0.569 | -0.075 | -0.027 | 0.046 | 0.046 | 0 | 0.17 | 0.31 | 0.13 | 0.21 | 0.14 | 0.24 |
| 6 | 10293348 | rs7756736 | 0.851 | -1.043 | 0.517 | -0.21 | 0.017 | 0.085 | 0.085 | 0 | 0.49 | 0.34 | 0.56 | 0.07 | 0.57 | 0.35 |
| 6 | 10293566 | rs7755897 | -0.828 | -0.921 | 0.295 | 0.196 | -0.019 | 0.007 | 0.007 | 0 | 0.12 | 0.16 | 0.1 | 0.18 | 0.07 | 0.15 |
| 6 | 10295743 | rs7753334 | -0.906 | -0.616 | -0.123 | -0.009 | -0.006 | 0.083 | 0.083 | 0 | 0.55 | 0.84 | 0.56 | 0.72 | 0.55 | 0.58 |
| 6 | 10295815 | rs7757564 | -0.89 | -0.09 | 0.437 | -0.048 | 0.159 | 0.03 | 0.03 | 0 | 0.06 | 0.3 | 0.16 | 0.01 | 0.15 | 0.03 |
| 6 | 10295896 | rs13217528 | 0.399 | -0.277 | -0.023 | -0.056 | -0.081 | -0.109 | -0.109 | 0 | 0.33 | 0.03 | 0.28 | 0.66 | 0.33 | 0.41 |
| 6 | 10296336 | rs1333657 | 0.311 | -0.245 | 0.38 | 0.277 | -0.027 | -0.062 | -0.062 | 0 | 0.36 | 0.16 | 0.27 | 0.46 | 0.36 | 0.42 |
| 6 | 10297906 | rs7764232 | -0.107 | -0.174 | 0.856 | 0.277 | 0.002 | 0.003 | 0.003 | 0 | 0.42 | 0.45 | 0.44 | 0.47 | 0.51 | 0.54 |
| 6 | 10298095 | rs9465867 | -1.508 | -0.423 | 0.809 | 0.098 | 0.002 | 0.071 | 0.071 | 0 | 0.62 | 0.78 | 0.63 | 0.96 | 0.62 | 0.75 |
| 6 | 10298301 | rs9465868 | -1.526 | -0.422 | 0.799 | 0.098 | 0.001 | 0.078 | 0.078 | 0 | 0.62 | 0.81 | 0.63 | 0.96 | 0.62 | 0.75 |
| 6 | 10300656 | rs7756830 | -1 | -0.37 | 0.653 | 0.391 | -0.017 | 0.054 | 0.054 | 0 | 0.19 | 0.24 | 0.18 | 0.5 | 0.12 | 0.21 |
| 6 | 10303945 | rs9348442 | -0.671 | -0.307 | 0.343 | 0.449 | 0.014 | -0.006 | -0.006 | 0 | 0.13 | 0.08 | 0.14 | 0.47 | 0.09 | 0.2 |
| 6 | 10305925 | rs4710948 | 0.528 | -0.191 | 0.261 | 0.477 | -0.023 | -0.019 | -0.019 | 0.989 | 0.26 | 0.21 | 0.21 | 0.35 | 0.32 | 0.29 |
| 6 | 10308594 | rs7752724 | 0.513 | 0.061 | 0.104 | 0.474 | 0.003 | 0.008 | 0.008 | 0.129 | 0.26 | 0.27 | 0.17 | 0.35 | 0.27 | 0.27 |
| 6 | 10313354 | rs4712557 | -0.795 | -0.067 | -0.639 | 0.052 | 0.036 | 0.105 | 0.105 | 0 | 0.61 | 0.8 | 0.66 | 0.96 | 0.67 | 0.72 |
| 6 | 10313664 | rs12209059 | -0.788 | 0.423 | -0.73 | 0.794 | -0.029 | -0.069 | -0.069 | 3.674 | 0.24 | 0.11 | 0.37 | 0.48 | 0.32 | 0.3 |
| 6 | 10316035 | rs9368249 | -0.446 | 0.372 | -0.702 | 0.978 | -0.02 | -0.071 | -0.071 | 0 | 0.21 | 0.11 | 0.33 | 0.49 | 0.28 | 0.23 |
| 6 | 10316981 | rs10946418 | -0.56 | -0.07 | -0.385 | 0.343 | 0.002 | 0 | 0 | 0 | 0.55 | 0.45 | 0.58 | 0.79 | 0.62 | 0.6 |
| 6 | 10324497 | rs10456009 | -0.891 | -0.079 | -0.919 | 0.168 | -0.002 | 0.108 | 0.108 | 0 | 0.62 | 0.81 | 0.52 | 0.91 | 0.59 | 0.68 |
| 6 | 10324944 | rs12199168 | -0.715 | -0.19 | -0.867 | 0.76 | -0.001 | -0.071 | -0.071 | 0 | 0.23 | 0.12 | 0.2 | 0.46 | 0.22 | 0.35 |
| 6 | 10331755 | rs6940204 | -1.096 | -0.84 | -0.184 | 0.587 | -0.06 | 0.09 | 0.09 | 0.56 | 0.22 | 0.65 | 0.15 | 0.38 | 0.09 | 0.08 |
| 6 | 10337251 | rs6908920 | -0.607 | -0.679 | -0.947 | 0.112 | 0.071 | -0.05 | -0.05 | 0.271 | 0.46 | 0.91 | 0.3 | 0.52 | 0.32 | 0.33 |
| 6 | 10342168 | rs9465973 | 0.269 | -0.013 | -0.68 | 0.024 | 0.001 | 0 | 0 | 0 | 0.26 | 0.24 | 0.4 | 0.44 | 0.34 | 0.43 |
| 6 | 10342349 | rs11755338 | 0.325 | -0.006 | -1.093 | 0.028 | -0.094 | -0.083 | -0.083 | 0 | 0.24 | 0.09 | 0.39 | 0.44 | 0.34 | 0.43 |
| 6 | 10343701 | rs12204307 | 0.957 | -0.098 | -1.242 | -0.493 | -0.082 | 0.15 | 0.15 | 0 | 0.23 | 0.01 | 0.14 | 0.03 | 0.26 | 0.11 |
| 6 | 10345571 | rs6456402 | 0.003 | -0.045 | -0.243 | -0.184 | 0.023 | 0.057 | 0.057 | 0 | 0.52 | 0.62 | 0.49 | 0.81 | 0.57 | 0.53 |
| 6 | 10349672 | rs4448079 | -0.292 | 0.292 | 0.006 | 0.138 | -0.002 | -0.004 | -0.004 | 0 | 0.78 | 0.81 | 0.55 | 0.43 | 0.73 | 0.74 |
| 6 | 10356484 | rs667342 | 0.196 | 0.431 | 0.8 | -0.229 | 0 | -0.056 | -0.056 | 0 | 0.46 | 0.53 | 0.47 | 0.94 | 0.48 | 0.64 |
| 6 | 10358695 | rs606699 | -0.775 | 0.347 | 0.036 | -0.302 | 0.004 | 0.173 | 0.173 | 0 | 0.69 | 0.96 | 0.62 | 0.97 | 0.73 | 0.75 |
| 6 | 10362596 | rs9295506 | -0.196 | 0.405 | 0.264 | -0.254 | 0.003 | 0.008 | 0.008 | 0 | 0.3 | 0.69 | 0.28 | 0.52 | 0.31 | 0.3 |
| 6 | 10365025 | rs492696 | 0.631 | 0.561 | -0.18 | -0.167 | -0.026 | -0.009 | -0.009 | 0 | 0.36 | 0.15 | 0.33 | 0.45 | 0.4 | 0.42 |
| 6 | 10368708 | rs484730 | 0.029 | 0.093 | 0.111 | -0.163 | 0 | 0 | 0 | 0 | 0.57 | 0.41 | 0.45 | 0.49 | 0.65 | 0.53 |
| 6 | 10369675 | rs10484886 | -0.19 | 0.191 | -0.501 | 0.153 | 0 | 0.014 | 0.014 | 0 | 0.06 | 0.11 | 0.03 | 0.1 | 0.05 | 0.19 |
| 6 | 10374314 | rs16893301 | -0.35 | 0.095 | -0.591 | 0.072 | 0 | -0.001 | -0.001 | 0.022 | 0.1 | 0.11 | 0.03 | 0.1 | 0.06 | 0.19 |
| 6 | 10375395 | rs809951 | 0.524 | 0.714 | -0.68 | 0.211 | 0 | -0.022 | -0.022 | 0 | 0.32 | 0.15 | 0.26 | 0.45 | 0.33 | 0.4 |
| 6 | 10380456 | rs638501 | 0.049 | -0.075 | -0.293 | -0.47 | 0 | -0.011 | -0.011 | 0 | 0.57 | 0.47 | 0.56 | 0.88 | 0.68 | 0.56 |
| 6 | 10387251 | rs480965 | -0.139 | 0.085 | -0.259 | -0.542 | -0.005 | -0.046 | -0.046 | 0 | 0.2 | 0.26 | 0.1 | 0.03 | 0.14 | 0.08 |
| 6 | 10388026 | rs999436 | -0.052 | 0.433 | -1.064 | -0.74 | -0.06 | -0.028 | -0.028 | 0 | 0.17 | 0.04 | 0.28 | 0.14 | 0.21 | 0.27 |
| 6 | 10388653 | rs3818408 | -0.591 | 0.431 | -1.103 | -0.413 | -0.018 | -0.011 | -0.011 | 0 | 0.42 | 0.37 | 0.58 | 0.47 | 0.46 | 0.51 |
| 6 | 10388968 | rs533746 | -1.38 | 0.292 | -1.29 | -0.444 | -0.005 | -0.049 | -0.049 | 0 | 0.63 | 0.82 | 0.7 | 0.5 | 0.6 | 0.58 |
| 6 | 10389971 | rs503532 | -0.639 | 0.031 | -1.191 | -0.122 | 0.034 | 0.034 | 0.034 | 0 | 0.44 | 0.79 | 0.4 | 0.39 | 0.35 | 0.31 |
| 6 | 10392845 | rs618337 | -0.42 | -0.063 | -0.984 | -0.244 | -0.003 | -0.003 | -0.003 | 0 | 0.26 | 0.7 | 0.33 | 0.26 | 0.22 | 0.22 |
| 6 | 10395206 | rs537112 | -0.356 | 0.002 | -0.45 | -0.216 | 0.087 | 0.224 | 0.224 | 0 | 0.07 | 0.36 | 0.13 | 0.25 | 0.07 | 0.02 |
| 6 | 10395572 | rs533558 | -0.827 | -0.168 | -1.67 | -0.296 | 0.023 | 0.085 | 0.085 | 0 | 0.43 | 0.78 | 0.43 | 0.29 | 0.37 | 0.31 |
| 6 | 10403183 | rs303050 | -0.687 | -0.261 | -1.379 | 0.011 | 0.01 | 0.003 | 0.003 | 0.057 | 0.27 | 0.74 | 0.23 | 0.27 | 0.15 | 0.09 |
| 6 | 10403650 | rs3798696 | 0.595 | 0.054 | -1.537 | -0.359 | 0.012 | 0.018 | 0.018 | 0 | 0.15 | 0.09 | 0.09 | 0.03 | 0.11 | 0.07 |
| 6 | 10412188 | rs1675414 | -0.424 | 0.419 | -1.867 | -0.072 | -0.14 | 0.065 | 0.065 | 0.059 | 0.39 | 0.08 | 0.47 | 0.19 | 0.44 | 0.5 |
| 6 | 10419476 | rs303055 | -0.688 | -0.17 | -1.586 | 0.32 | -0.046 | 0.015 | 0.015 | 0 | 0.6 | 0.84 | 0.64 | 0.64 | 0.51 | 0.64 |
| 6 | 10420098 | rs1621700 | -0.616 | 0.264 | -1.642 | 0.521 | -0.089 | -0.011 | -0.011 | 0 | 0.13 | 0.01 | 0.34 | 0.07 | 0.23 | 0.18 |
| 6 | 10424468 | rs6924202 | -0.499 | -0.562 | -1.206 | 0.643 | 0.034 | 0.023 | 0.023 | 0 | 0.48 | 0.8 | 0.54 | 0.35 | 0.38 | 0.57 |
| 6 | 10428481 | rs16870286 | -0.866 | -0.777 | -0.973 | 0.77 | -0.036 | -0.068 | -0.068 | 0 | 0.13 | 0.03 | 0.3 | 0.34 | 0.18 | 0.16 |
| 6 | 10430171 | rs303061 | 0.223 | -1.361 | -0.312 | 0.157 | 0.008 | -0.004 | -0.004 | 0.008 | 0.29 | 0.24 | 0.18 | 0.3 | 0.21 | 0.43 |

| | | | | | | | | | | | | | | | | |
|---|---|---|---|---|---|---|---|---|---|---|---|---|---|---|---|---|
| 6 | 10430978 | rs303063 | 0.505 | -1.45 | -0.302 | 0.14 | 0.017 | -0.011 | -0.011 | 0 | 0.27 | 0.19 | 0.16 | 0.3 | 0.18 | 0.41 |
| 6 | 10433825 | rs4710998 | 0.468 | -1.45 | -0.378 | 0.123 | 0.026 | -0.017 | -0.017 | 1.036 | 0.27 | 0.15 | 0.17 | 0.3 | 0.19 | 0.41 |
| 6 | 10444626 | rs534660 | -1.028 | -1.835 | 0.113 | -0.465 | -0.01 | -0.095 | -0.095 | 0 | 0.27 | 0.45 | 0.38 | 0.06 | 0.2 | 0.22 |
| 6 | 10449258 | rs1348 | 0.558 | -1.598 | 0.263 | -0.108 | 0.022 | 0.01 | 0.01 | 0 | 0.28 | 0.15 | 0.16 | 0.15 | 0.18 | 0.41 |
| 6 | 10452107 | rs303074 | -1.434 | -1.626 | -0.359 | 0.1 | -0.07 | -0.057 | -0.057 | 0 | 0.6 | 0.91 | 0.52 | 0.59 | 0.46 | 0.69 |
| 6 | 10453884 | rs4711005 | -0.222 | -1.152 | -0.14 | -0.396 | 0.004 | -0.009 | -0.009 | 0.405 | 0.1 | 0.13 | 0.21 | 0.04 | 0.12 | 0.15 |
| 6 | 10456706 | rs9348512 | -0.221 | -1.096 | -0.004 | -0.296 | -0.001 | -0.002 | -0.002 | 0 | 0.4 | 0.35 | 0.4 | 0.19 | 0.32 | 0.58 |
| 6 | 10458854 | rs7758450 | -0.042 | -1.266 | -0.059 | -0.372 | -0.004 | -0.028 | -0.028 | 0 | 0.21 | 0.21 | 0.06 | 0.03 | 0.09 | 0.08 |
| 6 | 10458926 | rs9358532 | -0.048 | -0.743 | -0.104 | -0.113 | 0.04 | 0.072 | 0.072 | 1.716 | 0.41 | 0.18 | 0.22 | 0.23 | 0.33 | 0.52 |
| 6 | 10460373 | rs9348513 | -0.086 | -0.634 | -0.11 | -0.111 | 0.06 | 0.097 | 0.097 | 0.838 | 0.43 | 0.18 | 0.35 | 0.18 | 0.31 | 0.5 |
| 6 | 10460560 | rs470790 | -0.843 | -0.267 | 0.049 | 0.246 | -0.019 | -0.023 | -0.023 | 0 | 0.59 | 0.76 | 0.52 | 0.49 | 0.47 | 0.67 |
| 6 | 10461119 | rs470764 | -0.451 | 0.118 | -0.007 | 0.506 | 0.038 | 0.065 | 0.065 | 0 | 0.09 | 0.28 | 0.13 | 0.22 | 0.13 | 0.14 |
| 6 | 10464025 | rs501436 | -0.574 | -0.176 | -0.232 | 0.286 | -0.005 | -0.006 | -0.006 | 0.288 | 0.59 | 0.63 | 0.52 | 0.49 | 0.47 | 0.67 |
| 6 | 10466038 | rs3889361 | 0.039 | 0.452 | -0.937 | 0.287 | 0.012 | 0.023 | 0.023 | 0 | 0.39 | 0.16 | 0.43 | 0.33 | 0.36 | 0.53 |
| 6 | 10467695 | rs796102 | -0.141 | 0.669 | -0.901 | 0.722 | -0.028 | -0.035 | -0.035 | 0 | 0.18 | 0.1 | 0.15 | 0.23 | 0.22 | 0.17 |
| 6 | 10473602 | rs796294 | -0.519 | 0.477 | -1.069 | 0.782 | -0.056 | -0.079 | -0.079 | 0.563 | 0.82 | 0.99 | 0.67 | 0.74 | 0.81 | 0.64 |
| 6 | 10474840 | rs611542 | 0.14 | 0.115 | -0.361 | 0.391 | 0.012 | -0.024 | -0.024 | 0 | 0.31 | 0.2 | 0.33 | 0.46 | 0.28 | 0.22 |
| 6 | 10481390 | rs502431 | 0.289 | 0.27 | -0.479 | -0.067 | 0.015 | 0.008 | 0.008 | 0.092 | 0.25 | 0.14 | 0.29 | 0.23 | 0.23 | 0.21 |
| 6 | 10482087 | rs9689588 | -0.787 | 0.25 | 0.337 | 0.014 | -0.004 | 0.015 | 0.015 | 0 | 0.06 | 0.15 | 0.05 | 0.13 | 0.05 | 0.05 |
| 6 | 10493771 | rs6905006 | -0.832 | 0.516 | 0.731 | -0.5 | 0.018 | -0.059 | -0.059 | 0 | 0.21 | 0.47 | 0.14 | 0.16 | 0.19 | 0.25 |
| 6 | 10496883 | rs496862 | -0.998 | -0.038 | 0.553 | -0.975 | 0.017 | 0.022 | 0.022 | 2.639 | 0.29 | 0.78 | 0.18 | 0.17 | 0.18 | 0.17 |
| 6 | 10501254 | rs512750 | 0.185 | -0.131 | 0.959 | -1.069 | -0.001 | -0.013 | -0.013 | 0 | 0.14 | 0.12 | 0.05 | 0.01 | 0.11 | 0.06 |
| 6 | 10503010 | rs796101 | -1.267 | -0.056 | 1.245 | -1.066 | -0.01 | -0.021 | -0.021 | 0.086 | 0.28 | 0.7 | 0.18 | 0.17 | 0.18 | 0.17 |
| 6 | 10513955 | rs3798745 | 0.37 | 0.333 | 0.976 | -1.161 | 0.003 | 0.049 | 0.049 | 0 | 0.44 | 0.21 | 0.46 | 0.63 | 0.39 | 0.58 |
| 6 | 10514226 | rs634103 | -0.455 | 0.323 | 1.338 | -1.2 | 0.009 | -0.012 | -0.012 | 0.02 | 0.62 | 0.57 | 0.55 | 0.65 | 0.51 | 0.7 |
| 6 | 10527046 | rs1226066 | 0.321 | 0.289 | 1.271 | -0.947 | -0.006 | -0.036 | -0.036 | 0 | 0.11 | 0.12 | 0.04 | 0.01 | 0.08 | 0.05 |
| 6 | 10528794 | rs560194 | -0.36 | 0.803 | 1.338 | -0.79 | 0.016 | -0.008 | -0.008 | 0 | 0.64 | 0.58 | 0.56 | 0.61 | 0.5 | 0.72 |
| 6 | 10539289 | rs1226013 | -0.416 | 0.995 | 1.123 | -0.722 | 0.008 | -0.004 | -0.004 | 0 | 0.64 | 0.61 | 0.52 | 0.6 | 0.52 | 0.73 |
| 6 | 10541213 | rs504083 | -1.693 | 1.55 | 1.567 | -0.529 | 0.004 | -0.011 | -0.011 | 0 | 0.34 | 0.62 | 0.24 | 0.25 | 0.32 | 0.3 |
| 6 | 10542527 | rs581151 | -0.643 | 1.645 | 1.356 | -0.485 | 0.001 | -0.083 | -0.083 | 2.758 | 0.15 | 0.35 | 0.07 | 0.01 | 0.1 | 0.03 |
| 6 | 10545728 | rs16870458 | -0.711 | 1.815 | 0.266 | -0.328 | -0.004 | -0.003 | -0.003 | 5.709 | 0.15 | 0.13 | 0.15 | 0.25 | 0.2 | 0.22 |
| 6 | 10555289 | rs3756948 | -0.7 | 1.769 | -0.021 | -0.33 | -0.016 | -0.019 | -0.019 | 0 | 0.17 | 0.11 | 0.25 | 0.3 | 0.22 | 0.24 |
| 6 | 10556872 | rs2230906 | -0.847 | 1.279 | 0.125 | -0.531 | -0.011 | 0.044 | 0.044 | 0.222 | 0.14 | 0.03 | 0.05 | 0.01 | 0.13 | 0.05 |
| 6 | 10566078 | rs9466768 | -1.549 | 1.214 | 1.06 | -0.414 | 0.06 | -0.001 | -0.001 | 0 | 0.07 | 0.32 | 0.04 | 0 | 0.04 | 0.07 |
| 6 | 10571260 | rs9295586 | 1.045 | 1.615 | 0.92 | -0.51 | -0.001 | -0.006 | -0.006 | 0.09 | 0.56 | 0.44 | 0.66 | 0.65 | 0.51 | 0.61 |
| 6 | 10571992 | rs566298 | 0.635 | 2.019 | 1.145 | -0.606 | 0.008 | 0.005 | 0.005 | 0 | 0.71 | 0.61 | 0.73 | 0.65 | 0.67 | 0.66 |
| 6 | 10573910 | rs1225999 | 0.766 | 1.954 | 1.651 | -0.652 | 0.02 | 0.011 | 0.011 | 0 | 0.7 | 0.46 | 0.72 | 0.66 | 0.67 | 0.66 |
| 6 | 10574118 | rs624678 | 1.266 | 2.018 | 1.438 | -0.659 | -0.005 | -0.007 | -0.007 | 0 | 0.57 | 0.49 | 0.64 | 0.64 | 0.44 | 0.59 |
| 6 | 10580668 | rs596145 | -1.495 | 1.355 | 1.699 | -0.892 | -0.099 | -0.146 | -0.146 | 0 | 0.21 | 0.44 | 0.08 | 0.01 | 0.06 | 0.03 |
| 6 | 10580910 | rs597152 | -1.574 | 1.457 | 1.702 | -0.888 | 0 | -0.148 | -0.148 | 1.048 | 0.22 | 0.45 | 0.1 | 0.01 | 0.15 | 0.05 |
| 6 | 10586006 | rs569693 | -1.354 | 1.632 | 1.714 | -0.801 | 0.019 | -0.124 | -0.124 | 0 | 0.19 | 0.49 | 0.1 | 0.01 | 0.15 | 0.05 |
| 6 | 10594260 | rs13204330 | -0.492 | 1.68 | 1.194 | -0.795 | -0.031 | 0.045 | 0.045 | 4.168 | 0.13 | 0.04 | 0.07 | 0 | 0.14 | 0.06 |
| 6 | 10598086 | rs4712775 | 1.397 | 1.958 | 1.539 | -0.619 | -0.007 | 0.013 | 0.013 | 0.361 | 0.57 | 0.33 | 0.73 | 0.63 | 0.52 | 0.47 |
| 6 | 10599279 | rs9466910 | -0.307 | 1.939 | 1.018 | -0.595 | -0.031 | 0.046 | 0.046 | 4.467 | 0.13 | 0.04 | 0.06 | 0 | 0.14 | 0.06 |
| 6 | 10599534 | rs17576994 | 1.163 | 1.892 | 1.338 | -0.458 | 0 | -0.007 | -0.007 | 0.256 | 0.57 | 0.48 | 0.69 | 0.61 | 0.47 | 0.42 |
| 6 | 10600880 | rs7755748 | 1.363 | 1.455 | 0.63 | -0.053 | -0.047 | -0.02 | -0.02 | 0 | 0.59 | 0.08 | 0.53 | 0.61 | 0.56 | 0.44 |
| 6 | 10602483 | rs9393456 | 1.015 | 1.219 | 1.119 | -0.043 | 0.005 | 0.005 | 0.005 | 0 | 0.69 | 0.35 | 0.7 | 0.65 | 0.62 | 0.49 |
| 6 | 10606611 | rs9466918 | 1.65 | 0.829 | 0.617 | -0.204 | 0.008 | 0.03 | 0.03 | 0 | 0.55 | 0.16 | 0.46 | 0.64 | 0.57 | 0.39 |
| 6 | 10613457 | rs7764197 | -1.047 | 1.047 | 1.193 | 0.049 | 0 | 0.114 | 0.114 | 1.094 | 0.12 | 0.37 | 0.11 | 0.21 | 0.07 | 0.25 |
| 6 | 10615078 | rs942631 | 0.497 | 1.224 | 0.795 | -0.148 | -0.016 | 0.012 | 0.012 | 0 | 0.81 | 0.84 | 0.7 | 0.85 | 0.68 | 0.71 |
| 6 | 10621547 | rs9460944 | -1.398 | 0.545 | 1.08 | -0.013 | -0.03 | -0.047 | -0.047 | 0 | 0.1 | 0.31 | 0.12 | 0 | 0.04 | 0.03 |
| 6 | 10624071 | rs4449626 | -1.169 | 0.614 | 0.753 | 0.022 | -0.028 | -0.042 | -0.042 | 0.551 | 0.09 | 0.24 | 0.11 | 0 | 0.04 | 0.03 |
| 6 | 10628014 | rs13210512 | -0.898 | 1.221 | 0.148 | 0.596 | -0.029 | -0.028 | -0.028 | 0 | 0.06 | 0.01 | 0.11 | 0.18 | 0.09 | 0.19 |
| 6 | 10642259 | rs4712802 | 0.962 | 1.751 | 0.512 | 0.117 | -0.039 | -0.005 | -0.005 | 0 | 0.72 | 0.81 | 0.56 | 0.71 | 0.55 | 0.49 |
| 6 | 10643835 | rs6456591 | -1.73 | 0.558 | 0.448 | -0.289 | 0.052 | 0.06 | 0.06 | 2.248 | 0.19 | 0.91 | 0.18 | 0.05 | 0.07 | 0.14 |
| 6 | 10647272 | rs9885719 | -1.796 | 0.59 | 1.592 | -0.174 | -0.021 | 0.013 | 0.013 | 1.863 | 0.1 | 0.52 | 0.05 | 0.04 | 0.02 | 0.11 |
| 6 | 10652375 | rs6919481 | -0.911 | 0.63 | 1.896 | -0.133 | 0.015 | 0.008 | 0.008 | 0.004 | 0.06 | 0.23 | 0.03 | 0 | 0.02 | 0.05 |
| 6 | 10686061 | rs4712827 | 0.56 | 0.512 | 1.369 | -0.653 | 0.01 | -0.039 | -0.039 | 0 | 0.66 | 0.56 | 0.6 | 0.8 | 0.55 | 0.55 |
| 6 | 10687746 | rs7749306 | -2.142 | 0.022 | 2.152 | -0.291 | -0.042 | -0.021 | -0.021 | 0.443 | 0.11 | 0.78 | 0.06 | 0.04 | 0.02 | 0.08 |
| 6 | 10709075 | rs6912993 | -1.88 | 0.174 | 1.033 | -0.198 | -0.036 | -0.009 | -0.009 | 0 | 0.08 | 0.8 | 0.06 | 0.04 | 0.01 | 0.06 |
| 6 | 10723449 | rs1045911 | -1.212 | 0.113 | 1.085 | -0.417 | -0.041 | -0.048 | -0.048 | 0.057 | 0.19 | 0.45 | 0.11 | 0.08 | 0.14 | 0.11 |
| 6 | 10724560 | rs3756954 | -0.792 | -0.079 | -0.359 | -0.077 | -0.092 | -0.047 | -0.047 | 0 | 0.65 | 0.98 | 0.68 | 0.69 | 0.68 | 0.67 |
| 6 | 10729020 | rs2153073 | 0.013 | -0.054 | 0.019 | 0.201 | 0.018 | -0.013 | -0.013 | 0 | 0.52 | 0.21 | 0.53 | 0.53 | 0.61 | 0.65 |
| 6 | 10733296 | rs506569 | -0.232 | -0.08 | 0.004 | 0.183 | -0.005 | -0.031 | -0.031 | 0.459 | 0.48 | 0.91 | 0.48 | 0.44 | 0.39 | 0.35 |
| 6 | 10738813 | rs529124 | -0.153 | -0.218 | -0.003 | 0.075 | 0.014 | -0.015 | -0.015 | 0 | 0.47 | 0.85 | 0.47 | 0.44 | 0.37 | 0.35 |
| 6 | 10739432 | rs1233887 | 0.72 | -0.214 | -0.126 | 0.098 | -0.089 | -0.04 | -0.04 | 0 | 0.35 | 0.02 | 0.32 | 0.29 | 0.32 | 0.33 |
| 6 | 10743972 | rs521417 | -0.221 | -0.203 | 0.251 | 0.125 | 0.005 | -0.004 | -0.004 | 0 | 0.44 | 0.71 | 0.48 | 0.44 | 0.38 | 0.36 |
| 6 | 10745066 | rs545019 | 0.491 | -0.175 | 0.257 | 0.077 | -0.018 | 0.002 | 0.002 | 0.279 | 0.36 | 0.25 | 0.44 | 0.32 | 0.34 | 0.35 |
| 6 | 10746227 | rs494723 | -0.122 | -0.545 | -0.111 | -0.012 | 0.042 | -0.016 | -0.016 | 0 | 0.41 | 0.75 | 0.41 | 0.44 | 0.27 | 0.31 |

| | | | | | | | | | | | | | | | | |
|---|---|---|---|---|---|---|---|---|---|---|---|---|---|---|---|---|
| 6 | 10758957 | rs2018035 | 0.16 | -0.906 | -0.928 | 0.015 | 0.07 | -0.094 | -0.094 | 0 | 0.25 | 0.01 | 0.19 | 0.36 | 0.11 | 0.21 |
| 6 | 10762997 | rs501948 | -0.687 | -0.866 | -1.204 | -0.468 | 0.012 | 0.076 | 0.076 | 0.057 | 0.78 | 0.97 | 0.79 | 0.92 | 0.79 | 0.81 |
| 6 | 10763312 | rs526495 | -0.306 | -1.029 | -0.443 | -0.552 | -0.012 | -0.032 | -0.032 | 0 | 0.74 | 0.74 | 0.78 | 0.91 | 0.79 | 0.81 |
| 6 | 10764727 | rs126405 | -0.696 | -0.933 | -0.853 | -0.503 | 0.012 | 0.076 | 0.076 | 0.057 | 0.78 | 0.97 | 0.79 | 0.92 | 0.8 | 0.8 |
| 6 | 10775446 | rs478791 | -0.683 | -0.734 | -0.87 | -0.445 | -0.006 | 0.053 | 0.053 | 0.053 | 0.77 | 0.96 | 0.76 | 0.87 | 0.77 | 0.81 |
| 6 | 10791859 | rs518954 | 0.553 | -0.949 | -0.272 | -0.529 | 0.023 | -0.011 | -0.011 | 0 | 0.33 | 0.24 | 0.25 | 0.33 | 0.22 | 0.24 |
| 6 | 10792197 | rs521802 | -0.459 | -0.766 | 0.183 | -0.574 | -0.012 | -0.024 | -0.024 | 0 | 0.73 | 0.74 | 0.78 | 0.85 | 0.79 | 0.81 |
| 6 | 10792965 | rs517546 | -0.997 | -0.767 | -0.063 | -0.582 | 0.002 | 0.049 | 0.049 | 0.041 | 0.78 | 0.99 | 0.8 | 0.85 | 0.81 | 0.85 |
| 6 | 10794536 | rs474519 | -0.247 | -0.36 | 0.427 | -0.592 | -0.032 | -0.07 | -0.07 | 0.174 | 0.72 | 0.5 | 0.77 | 0.85 | 0.77 | 0.81 |
| 6 | 10809432 | rs9366631 | 0.735 | -0.323 | -0.113 | -0.58 | 0.008 | 0.008 | 0.008 | 0 | 0.21 | 0.15 | 0.12 | 0.1 | 0.15 | 0.12 |
| 6 | 10811678 | rs4379275 | -0.035 | -0.336 | -0.555 | -0.556 | -0.002 | 0 | 0 | 0.027 | 0.43 | 0.45 | 0.53 | 0.52 | 0.59 | 0.56 |
| 6 | 10817505 | rs2009118 | -0.714 | -0.139 | -1.344 | -0.394 | -0.007 | -0.112 | -0.112 | 0 | 0.67 | 0.99 | 0.67 | 0.62 | 0.75 | 0.74 |
| 6 | 10829319 | rs518204 | -0.721 | 0.508 | -1.187 | -0.265 | -0.118 | -0.106 | -0.106 | 0 | 0.64 | 0.99 | 0.68 | 0.61 | 0.65 | 0.54 |
| 6 | 10836938 | rs3846839 | -0.37 | 0.856 | -0.462 | 0.127 | -0.015 | -0.03 | -0.03 | 0 | 0.34 | 0.26 | 0.4 | 0.5 | 0.41 | 0.33 |
| 6 | 10852772 | rs6905789 | -0.991 | 0.859 | -0.073 | -0.096 | 0.016 | -0.044 | -0.044 | 0.044 | 0.08 | 0.18 | 0.09 | 0 | 0.14 | 0.07 |
| 6 | 10863325 | rs6456719 | -0.891 | 0.983 | -1.026 | -0.146 | -0.124 | -0.042 | -0.042 | 0.313 | 0.64 | 0.97 | 0.54 | 0.69 | 0.61 | 0.55 |
| 6 | 10864932 | rs4712983 | 0.368 | 0.671 | -1.073 | 0.099 | 0.009 | -0.09 | -0.09 | 0 | 0.2 | 0.07 | 0.13 | 0.46 | 0.14 | 0.24 |
| 6 | 10872061 | rs6929329 | 1.103 | 0.794 | -0.503 | -0.024 | -0.001 | 0.017 | 0.017 | 0 | 0.23 | 0.12 | 0.13 | 0.23 | 0.22 | 0.14 |
| 6 | 10875903 | rs12201787 | -0.52 | 0.777 | -0.077 | -0.122 | -0.003 | -0.003 | -0.003 | 0.041 | 0.09 | 0.07 | 0.1 | 0 | 0.14 | 0.1 |
| 6 | 10876036 | rs2076258 | -0.098 | 0.745 | -0.072 | -0.066 | 0.004 | 0.008 | 0.008 | 0 | 0.44 | 0.37 | 0.25 | 0.69 | 0.37 | 0.36 |
| 6 | 10877867 | rs9379881 | -0.139 | 0.649 | -0.479 | -0.023 | -0.048 | 0.079 | 0.079 | 0 | 0.47 | 0.15 | 0.31 | 0.69 | 0.5 | 0.43 |
| 6 | 10879996 | rs3756957 | 0.992 | 0.904 | -0.355 | 0.152 | -0.019 | 0.029 | 0.029 | 0 | 0.39 | 0.09 | 0.44 | 0.29 | 0.35 | 0.5 |
| 6 | 10882070 | rs16870746 | -0.926 | 0.853 | 0.476 | 0.446 | 0.074 | 0.087 | 0.087 | 0.167 | 0.08 | 0.49 | 0.17 | 0.06 | 0.08 | 0.02 |
| 6 | 10882175 | rs2275387 | -1.62 | 0.826 | 0.457 | 0.43 | -0.018 | -0.013 | -0.013 | 0 | 0.13 | 0.56 | 0.2 | 0.06 | 0.08 | 0.06 |
| 6 | 10885971 | rs6456737 | 0.322 | 0.972 | -0.25 | 0.488 | 0.001 | 0.047 | 0.047 | 0 | 0.28 | 0.11 | 0.17 | 0.24 | 0.26 | 0.21 |
| 6 | 10886585 | rs12211124 | 1.172 | 0.976 | -0.439 | 0.532 | -0.035 | 0.006 | 0.006 | 0 | 0.39 | 0.08 | 0.55 | 0.28 | 0.35 | 0.52 |
| 6 | 10887251 | rs9366663 | 0.203 | 1.318 | 0.496 | 0.898 | 0.015 | 0.031 | 0.031 | 0 | 0.73 | 0.65 | 0.78 | 0.66 | 0.68 | 0.75 |
| 6 | 10892220 | rs9368446 | 0.471 | 1.398 | 0.054 | 0.93 | -0.076 | 0.046 | 0.046 | 0 | 0.27 | 0.05 | 0.21 | 0.19 | 0.38 | 0.22 |
| 6 | 10892499 | rs9366664 | 0.014 | 1.396 | 0.616 | 1.042 | 0.011 | 0 | 0 | 0 | 0.32 | 0.34 | 0.26 | 0.37 | 0.44 | 0.23 |
| 6 | 10893120 | rs946351 | -0.141 | 1.306 | 0.661 | 1.037 | 0 | -0.019 | -0.019 | 0.132 | 0.24 | 0.19 | 0.16 | 0.33 | 0.19 | 0.24 |
| 6 | 10894255 | rs9368453 | 0.323 | 1.582 | 0.449 | 1.096 | -0.073 | 0.03 | 0.03 | 0 | 0.27 | 0.05 | 0.21 | 0.23 | 0.38 | 0.23 |
| 6 | 10896053 | rs13194859 | -0.295 | 1.461 | 0.684 | 1.185 | 0.034 | 0.142 | 0.142 | 0 | 0.23 | 0.45 | 0.19 | 0.44 | 0.24 | 0.2 |
| 6 | 10897488 | rs2153157 | -0.702 | 1.427 | -0.191 | 0.964 | 0.001 | 0.077 | 0.077 | 0 | 0.51 | 0.78 | 0.39 | 0.68 | 0.51 | 0.42 |
| 6 | 10899254 | rs9467852 | -0.233 | 1.846 | -0.413 | 1.079 | -0.027 | -0.052 | -0.052 | 0 | 0.22 | 0.15 | 0.51 | 0.5 | 0.27 | 0.37 |
| 6 | 10904964 | rs1890486 | -0.546 | 1.579 | -0.167 | 0.93 | 0 | 0.003 | 0.003 | 0 | 0.77 | 0.76 | 0.84 | 0.68 | 0.75 | 0.83 |
| 6 | 10906046 | rs10484890 | -0.073 | 2.078 | -0.401 | 0.752 | 0.049 | -0.023 | -0.023 | 0 | 0.05 | 0.1 | 0.08 | 0.01 | 0.14 | 0.04 |
| 6 | 10906154 | rs9357021 | 0.926 | 1.271 | -1.101 | 0.552 | 0.046 | 0.073 | 0.073 | 0 | 0.35 | 0.12 | 0.26 | 0.21 | 0.26 | 0.3 |
| 6 | 10906256 | rs4713039 | 0.022 | 1.342 | -0.898 | 0.924 | -0.038 | -0.12 | -0.12 | 0.126 | 0.23 | 0.03 | 0.11 | 0.38 | 0.19 | 0.3 |
| 6 | 10909689 | rs4711127 | 0.027 | 1.362 | -0.991 | 0.572 | -0.026 | -0.029 | -0.029 | 0 | 0.49 | 0.11 | 0.54 | 0.48 | 0.46 | 0.51 |
| 6 | 10911282 | rs4713044 | 0.965 | 1.744 | 0.041 | 0.325 | 0.106 | -0.091 | -0.091 | 0.615 | 0.85 | 0.59 | 0.82 | 1 | 0.71 | 0.81 |
| 6 | 10922638 | rs12190237 | -0.5 | 1.792 | 0.151 | 0.336 | 0.056 | -0.044 | -0.044 | 0.241 | 0.11 | 0.19 | 0.14 | 0 | 0.27 | 0.17 |
| 6 | 10923522 | rs4713064 | -0.417 | 1.554 | 0.033 | 0.462 | -0.003 | -0.002 | -0.002 | 3.803 | 0.36 | 0.41 | 0.54 | 0.4 | 0.5 | 0.55 |
| 6 | 10925803 | rs1767777 | 1 | 1.387 | 0.167 | 0.17 | 0.006 | 0.011 | 0.011 | 0 | 0.26 | 0.15 | 0.17 | 0.06 | 0.24 | 0.13 |
| 6 | 10926942 | rs12213249 | 0.943 | 1.055 | -0.208 | 0.698 | -0.004 | -0.056 | -0.056 | 1.59 | 0.31 | 0.03 | 0.25 | 0.32 | 0.24 | 0.28 |
| 6 | 10932569 | rs12199131 | 0.845 | 1.144 | -0.189 | 0.626 | -0.017 | -0.006 | -0.006 | 0 | 0.32 | 0.03 | 0.27 | 0.26 | 0.26 | 0.28 |
| 6 | 10933378 | rs12214825 | 0.864 | 1.143 | -0.197 | 0.626 | -0.027 | -0.016 | -0.016 | 0 | 0.32 | 0.01 | 0.27 | 0.26 | 0.27 | 0.28 |
| 6 | 10947182 | rs2788872 | -0.082 | 1.177 | 0.633 | 0.793 | -0.009 | 0.031 | 0.031 | 0 | 0.72 | 0.6 | 0.6 | 0.54 | 0.78 | 0.62 |
| 6 | 10952243 | rs1225741 | 0.406 | 1.182 | 0.977 | 0.43 | -0.001 | -0.018 | -0.018 | 0.184 | 0.35 | 0.35 | 0.28 | 0.09 | 0.47 | 0.25 |
| 6 | 10955408 | rs1225746 | 0.971 | 0.97 | 0.95 | 0.747 | -0.007 | 0.157 | 0.157 | 0.645 | 0.94 | 0.78 | 0.94 | 0.8 | 0.98 | 0.97 |
| 6 | 10956649 | rs7759825 | 0.635 | 1.057 | 0.738 | 0.602 | 0.005 | 0.012 | 0.012 | 0 | 0.33 | 0.2 | 0.29 | 0.24 | 0.29 | 0.34 |
| 6 | 10961454 | rs2295599 | 0.294 | 0.828 | 0.352 | 0.555 | 0.002 | -0.081 | -0.081 | 0.035 | 0.2 | 0.1 | 0.33 | 0.43 | 0.17 | 0.39 |
| 6 | 10981053 | rs8523 | -0.342 | 0.602 | 0.324 | -0.366 | -0.002 | 0.029 | 0.029 | 0 | 0.54 | 0.63 | 0.67 | 0.91 | 0.44 | 0.73 |
| 6 | 10982359 | rs17606561 | 1.101 | 0.597 | -0.058 | -0.127 | -0.028 | -0.005 | -0.005 | 0 | 0.3 | 0.01 | 0.27 | 0.26 | 0.23 | 0.22 |
| 6 | 10982652 | rs1225737 | -0.639 | 0.414 | 0.073 | -0.341 | -0.051 | 0.163 | 0.163 | 0 | 0.6 | 0.85 | 0.7 | 0.91 | 0.44 | 0.66 |
| 6 | 10982848 | rs3734397 | -1.792 | 0.96 | -0.355 | -0.547 | -0.032 | 0.118 | 0.118 | 0.051 | 0.73 | 0.99 | 0.84 | 0.94 | 0.75 | 0.89 |
| 6 | 10982973 | rs3734398 | -0.161 | 0.925 | 0.179 | -0.547 | 0.003 | 0.147 | 0.147 | 0 | 0.54 | 0.25 | 0.64 | 0.91 | 0.43 | 0.66 |
| 6 | 10990493 | rs2281591 | 0.84 | 1.29 | 0.421 | -0.526 | -0.031 | -0.029 | -0.029 | 0 | 0.34 | 0.51 | 0.3 | 0.27 | 0.23 | 0.24 |
| 6 | 10990751 | rs911196 | 0.755 | 1.29 | 0.133 | -0.231 | -0.04 | -0.146 | -0.146 | 0 | 0.19 | 0.02 | 0.33 | 0.46 | 0.18 | 0.37 |
| 6 | 11001276 | rs3798709 | 0.755 | 1.31 | -0.176 | -0.22 | -0.04 | -0.146 | -0.146 | 0 | 0.19 | 0.02 | 0.33 | 0.46 | 0.18 | 0.37 |
| 6 | 11003875 | rs976081 | -1.756 | 1.355 | -0.196 | -0.261 | -0.03 | 0.121 | 0.121 | 0.297 | 0.73 | 0.99 | 0.84 | 0.94 | 0.75 | 0.89 |
| 6 | 11005842 | rs2295602 | -0.547 | 1.125 | 0.228 | -0.021 | -0.009 | 0.05 | 0.05 | 0 | 0.57 | 0.71 | 0.68 | 0.9 | 0.44 | 0.67 |
| 6 | 11022154 | rs3756963 | 0.525 | 0.802 | -0.29 | 0.216 | 0.01 | 0.009 | 0.009 | 0 | 0.32 | 0.25 | 0.28 | 0.29 | 0.24 | 0.24 |
| 6 | 11042165 | rs9468304 | 0.069 | 0.586 | -1.184 | 0.405 | -0.022 | -0.099 | -0.099 | 0 | 0.2 | 0.05 | 0.35 | 0.63 | 0.18 | 0.4 |
| 6 | 11074302 | rs4713165 | -0.089 | 0.816 | -0.991 | 0.013 | 0.016 | 0.127 | 0.127 | 0.092 | 0.51 | 0.24 | 0.61 | 0.74 | 0.42 | 0.61 |
| 6 | 11080825 | rs12665478 | -0.098 | 0.82 | -0.988 | 0.015 | 0.015 | 0.125 | 0.125 | 0.113 | 0.51 | 0.24 | 0.61 | 0.74 | 0.42 | 0.61 |
| 6 | 11102871 | rs1099 | 0.415 | 0.67 | -0.994 | 0.665 | -0.027 | -0.112 | -0.112 | 1.987 | 0.16 | 0.01 | 0.29 | 0.34 | 0.16 | 0.32 |
| 6 | 11103537 | rs13214666 | 0.703 | 0.545 | -0.695 | -0.047 | -0.008 | -0.063 | -0.063 | 0 | 0.26 | 0.31 | 0.15 | 0.02 | 0.18 | 0.19 |
| 6 | 11103888 | rs9393931 | 0.556 | 0.478 | -0.758 | -0.095 | -0.001 | 0.019 | 0.019 | 0 | 0.18 | 0.19 | 0.2 | 0.39 | 0.16 | 0.21 |
| 6 | 11109831 | rs12527972 | 1.059 | 0.601 | -0.92 | 0.078 | -0.026 | -0.116 | -0.116 | 0.007 | 0.16 | 0.01 | 0.3 | 0.36 | 0.16 | 0.33 |

| | | | | | | | | | | | | | | | | |
|---|---|---|---|---|---|---|---|---|---|---|---|---|---|---|---|---|
| 6 | 11122023 | rs2224653 | -1.126 | 0.746 | -0.635 | -0.05 | -0.016 | 0.148 | 0.148 | 0 | 0.67 | 0.89 | 0.79 | 0.9 | 0.68 | 0.82 |
| 6 | 11138038 | rs3734399 | 0.86 | 0.733 | -0.673 | 0.409 | -0.027 | -0.116 | -0.116 | 0 | 0.16 | 0.01 | 0.29 | 0.36 | 0.16 | 0.33 |
| 6 | 11145688 | rs11970707 | -0.186 | 0.703 | -0.002 | -0.026 | -0.008 | 0.052 | 0.052 | 0 | 0.21 | 0.32 | 0.2 | 0.35 | 0.18 | 0.26 |
| 6 | 11146221 | rs17559298 | 1.045 | 0.531 | 0.116 | -0.715 | -0.003 | -0.035 | -0.035 | 0 | 0.25 | 0.23 | 0.17 | 0.02 | 0.19 | 0.13 |
| 6 | 11146875 | rs4713206 | 0.082 | 0.569 | 0.245 | -0.916 | 0.009 | 0.03 | 0.03 | 0 | 0.53 | 0.4 | 0.64 | 0.68 | 0.61 | 0.66 |
| 6 | 11148438 | rs9393950 | 0.583 | 0.715 | 0.305 | -0.799 | 0.008 | -0.028 | -0.028 | 0 | 0.3 | 0.16 | 0.44 | 0.63 | 0.31 | 0.47 |
| 6 | 11149203 | rs9368594 | 0.306 | 0.72 | 0.567 | -0.839 | -0.001 | -0.001 | -0.001 | 0 | 0.34 | 0.37 | 0.46 | 0.65 | 0.33 | 0.53 |
| 6 | 11153633 | rs7752029 | 0.958 | 0.539 | 0.638 | -0.752 | -0.002 | 0.011 | 0.011 | 0 | 0.16 | 0.17 | 0.18 | 0.28 | 0.14 | 0.18 |
| 6 | 11158816 | rs10946984 | 1.156 | 0.656 | 0.103 | -0.999 | -0.061 | 0.091 | 0.091 | 0 | 0.28 | 0.12 | 0.24 | 0.05 | 0.39 | 0.2 |
| 6 | 11161863 | rs9468530 | 0.094 | 0.59 | 0.231 | -0.882 | 0.008 | 0.044 | 0.044 | 0 | 0.46 | 0.68 | 0.45 | 0.35 | 0.53 | 0.4 |
| 6 | 11174591 | rs4552715 | 0.775 | 0.519 | 0.039 | -0.563 | 0.019 | -0.099 | -0.099 | 0 | 0.28 | 0.1 | 0.39 | 0.57 | 0.26 | 0.45 |
| 6 | 11176704 | rs7743913 | -0.036 | 0.41 | 0.126 | -1.138 | 0.014 | 0.032 | 0.032 | 0.034 | 0.52 | 0.64 | 0.43 | 0.35 | 0.55 | 0.42 |
| 6 | 11178462 | rs9295806 | -0.969 | 0.388 | 0.088 | -1.163 | -0.005 | 0.005 | 0.005 | 0 | 0.82 | 0.88 | 0.88 | 1 | 0.83 | 0.89 |
| 6 | 11182995 | rs2327338 | -0.92 | 0.377 | 0.082 | -1.169 | -0.004 | 0.007 | 0.007 | 0 | 0.81 | 0.88 | 0.87 | 1 | 0.83 | 0.89 |
| 6 | 11183587 | rs1044417 | -0.311 | 0.543 | 0.194 | -0.9 | 0.019 | 0.049 | 0.049 | 0 | 0.52 | 0.69 | 0.43 | 0.35 | 0.55 | 0.44 |
| 6 | 11184090 | rs1570372 | -0.311 | 0.543 | 0.194 | -0.9 | 0.019 | 0.049 | 0.049 | 0 | 0.52 | 0.69 | 0.43 | 0.35 | 0.55 | 0.44 |
| 6 | 11185184 | rs1911 | 0.603 | 0.607 | 0.315 | -0.977 | -0.012 | 0.025 | 0.025 | 0.632 | 0.21 | 0.3 | 0.18 | 0.3 | 0.16 | 0.2 |
| 6 | 11185533 | rs1050775 | -0.235 | 0.403 | -0.078 | -0.881 | 0.017 | 0.044 | 0.044 | 0.049 | 0.52 | 0.65 | 0.42 | 0.35 | 0.55 | 0.42 |
| 6 | 11187925 | rs17561071 | 0.744 | 0.458 | -0.422 | -0.877 | 0.004 | 0.027 | 0.027 | 0 | 0.23 | 0.16 | 0.16 | 0.02 | 0.19 | 0.1 |
| 6 | 11190626 | rs941982 | -0.794 | 0.605 | -0.234 | -0.526 | -0.011 | 0.011 | 0.011 | 0 | 0.88 | 0.95 | 0.92 | 1 | 0.86 | 0.97 |
| 6 | 11192575 | rs1465131 | -0.38 | 0.529 | -0.063 | -0.565 | -0.001 | -0.026 | -0.026 | 0 | 0.86 | 0.85 | 0.9 | 1 | 0.86 | 0.97 |
| 6 | 11194511 | rs3798727 | 0.659 | 0.303 | -0.422 | -0.318 | -0.002 | 0.034 | 0.034 | 0.191 | 0.4 | 0.2 | 0.34 | 0.34 | 0.34 | 0.32 |
| 6 | 11198498 | rs10947009 | 0.029 | 0.23 | 0.128 | 0.108 | -0.033 | -0.042 | -0.042 | 0.124 | 0.41 | 0.23 | 0.5 | 0.51 | 0.5 | 0.63 |
| 6 | 11199234 | rs3798731 | 0.3 | 0.322 | 0.522 | 0.032 | 0.029 | -0.085 | -0.085 | 1.593 | 0.33 | 0.13 | 0.41 | 0.51 | 0.3 | 0.55 |
| 6 | 11200574 | rs726230 | -0.652 | 0.217 | 0.508 | -0.014 | 0.111 | 0.087 | 0.087 | 0.068 | 0.13 | 0.54 | 0.17 | 0.13 | 0.23 | 0.09 |
| 6 | 11201157 | rs3798734 | -0.609 | 0.223 | 0.474 | -0.011 | 0.125 | 0.099 | 0.099 | 0 | 0.13 | 0.5 | 0.17 | 0.13 | 0.23 | 0.09 |
| 6 | 11201468 | rs3798735 | 0.53 | 0.215 | 0.098 | 0.11 | 0.021 | -0.08 | -0.08 | 0 | 0.28 | 0.05 | 0.32 | 0.37 | 0.26 | 0.46 |
| 6 | 11202357 | rs2146342 | 0.551 | 0.222 | 0.101 | 0.117 | 0.02 | -0.078 | -0.078 | 0 | 0.28 | 0.05 | 0.32 | 0.36 | 0.26 | 0.46 |
| 6 | 11208880 | rs3102 | -0.11 | 0.508 | 0.71 | 0.242 | 0.085 | -0.043 | -0.043 | 0 | 0.1 | 0.3 | 0.11 | 0.01 | 0.22 | 0.09 |
| 6 | 11212959 | rs3798743 | -0.648 | 0.501 | 0.511 | 0.293 | 0.122 | 0.101 | 0.101 | 0.004 | 0.13 | 0.5 | 0.17 | 0.13 | 0.23 | 0.09 |
| 6 | 11213832 | rs3734404 | -0.44 | 0.264 | -0.005 | -0.024 | 0.015 | 0.018 | 0.018 | 0 | 0.34 | 0.6 | 0.38 | 0.41 | 0.43 | 0.31 |
| 6 | 11214216 | rs9295823 | -0.291 | 0.273 | -0.251 | -0.712 | -0.003 | 0.004 | 0.004 | 0 | 0.85 | 0.9 | 0.83 | 0.86 | 0.86 | 0.88 |
| 6 | 11217843 | rs2018334 | 0.091 | 0.094 | -0.122 | -1.08 | -0.001 | 0.039 | 0.039 | 0 | 0.52 | 0.35 | 0.59 | 0.76 | 0.49 | 0.64 |
| 6 | 11217897 | rs2018336 | 0.381 | -0.884 | -0.2 | -1.073 | -0.05 | 0.105 | 0.105 | 0 | 0.25 | 0.02 | 0.16 | 0.08 | 0.23 | 0.21 |
| 6 | 11220417 | rs6457107 | -0.765 | -0.75 | 0.037 | -1.098 | -0.001 | -0.026 | -0.026 | 0.076 | 0.88 | 0.91 | 0.92 | 1 | 0.89 | 0.97 |
| 6 | 11223208 | rs7739306 | 0.058 | -0.777 | 0.125 | -1.081 | -0.003 | 0.058 | 0.058 | 0 | 0.05 | 0.24 | 0.09 | 0.13 | 0.04 | 0.08 |
| 6 | 11224111 | rs2025677 | -1.069 | -0.701 | -0.297 | -1.078 | -0.007 | 0.018 | 0.018 | 0 | 0.89 | 0.96 | 0.92 | 1 | 0.89 | 0.97 |
| 6 | 11224748 | rs7769173 | 0.481 | -0.604 | -0.268 | -0.741 | -0.006 | 0.001 | 0.001 | 0 | 0.44 | 0.27 | 0.56 | 0.6 | 0.55 | 0.51 |
| 6 | 11227328 | rs6457131 | 0.277 | -0.707 | -0.289 | -0.732 | 0.016 | 0.022 | 0.022 | 0 | 0.3 | 0.17 | 0.2 | 0.21 | 0.24 | 0.2 |
| 6 | 11227385 | rs2182336 | -0.728 | -0.341 | -0.362 | -0.775 | 0.002 | 0.003 | 0.003 | 1.999 | 0.77 | 0.81 | 0.85 | 0.94 | 0.81 | 0.88 |
| 6 | 11227987 | rs9348836 | 0.216 | 0.05 | -0.359 | -0.277 | -0.014 | 0.002 | 0.002 | 0.127 | 0.42 | 0.32 | 0.54 | 0.64 | 0.53 | 0.6 |
| 6 | 11229953 | rs11970705 | -0.335 | -0.286 | -0.143 | 0.105 | -0.022 | 0.013 | 0.013 | 0 | 0.23 | 0.28 | 0.13 | 0.22 | 0.17 | 0.18 |
| 6 | 11235682 | rs2182337 | -0.617 | 0.376 | -0.521 | 0.019 | 0.002 | 0.002 | 0.002 | 0 | 0.19 | 0.16 | 0.13 | 0.13 | 0.14 | 0.1 |
| 6 | 11236678 | rs9368621 | 0.394 | 0.324 | -0.551 | -0.313 | 0.001 | -0.008 | -0.008 | 0 | 0.8 | 0.81 | 0.88 | 0.94 | 0.85 | 0.9 |
| 6 | 11239119 | rs9468690 | 0.107 | 0.462 | -0.511 | -0.699 | -0.001 | -0.014 | -0.014 | 0 | 0.41 | 0.38 | 0.45 | 0.2 | 0.36 | 0.52 |
| 6 | 11239518 | rs9461574 | 0.07 | 0.506 | -0.425 | -0.378 | -0.004 | -0.014 | -0.014 | 0 | 0.41 | 0.44 | 0.47 | 0.28 | 0.39 | 0.53 |
| 6 | 11239943 | rs9393992 | -0.041 | 0.594 | -0.436 | -0.29 | -0.001 | 0.053 | 0.053 | 0 | 0.23 | 0.33 | 0.25 | 0.61 | 0.25 | 0.3 |
| 6 | 11242203 | rs12209631 | -0.035 | 1.139 | -0.714 | -0.725 | -0.039 | 0.088 | 0.088 | 0 | 0.25 | 0.12 | 0.16 | 0.01 | 0.27 | 0.09 |
| 6 | 11243136 | rs767231 | -0.921 | 0.833 | -0.278 | -0.814 | 0.004 | 0.038 | 0.038 | 0 | 0.43 | 0.68 | 0.38 | 0.77 | 0.43 | 0.38 |
| 6 | 11243857 | rs744970 | -0.705 | 0.742 | -0.183 | -1.008 | -0.008 | -0.12 | -0.12 | 0 | 0.35 | 0.49 | 0.21 | 0.04 | 0.29 | 0.12 |
| 6 | 11247387 | rs6908326 | -0.904 | 0.275 | -0.009 | 0.087 | -0.022 | 0.059 | 0.059 | 0.304 | 0.24 | 0.62 | 0.22 | 0.58 | 0.23 | 0.26 |
| 6 | 11248362 | rs16871157 | -0.204 | 0.102 | -0.733 | -0.168 | 0.018 | -0.007 | -0.007 | 0 | 0.17 | 0.1 | 0.1 | 0.17 | 0.1 | 0.26 |
| 6 | 11255358 | rs6457200 | -0.47 | 0.603 | -0.482 | -0.507 | 0.004 | 0.018 | 0.018 | 0 | 0.54 | 0.63 | 0.32 | 0.15 | 0.38 | 0.31 |
| 6 | 11256000 | rs7766626 | -0.559 | 0.371 | 0.121 | -0.556 | -0.001 | -0.012 | -0.012 | 1.236 | 0.4 | 0.47 | 0.33 | 0.76 | 0.43 | 0.53 |
| 6 | 11266942 | rs9468726 | -1.373 | -0.576 | 0.498 | -0.49 | -0.048 | -0.004 | -0.004 | 0 | 0.28 | 0.49 | 0.23 | 0.28 | 0.19 | 0.17 |
| 6 | 11268188 | rs11964334 | -1.223 | -0.904 | 0.776 | -0.828 | -0.038 | -0.065 | -0.065 | 0 | 0.23 | 0.36 | 0.15 | 0.05 | 0.14 | 0.07 |
| 6 | 11270637 | rs4713332 | -0.02 | -0.776 | 0.576 | -0.735 | 0.024 | 0.014 | 0.014 | 0 | 0.11 | 0.06 | 0.08 | 0.05 | 0.05 | 0.05 |
| 6 | 11272532 | rs6920739 | -0.912 | -0.831 | 0.929 | -0.771 | -0.022 | -0.099 | -0.099 | 1.456 | 0.27 | 0.43 | 0.19 | 0.07 | 0.23 | 0.13 |
| 6 | 11274179 | rs11967893 | -0.405 | -0.699 | 0.711 | -0.593 | -0.016 | -0.007 | -0.007 | 0.039 | 0.12 | 0.17 | 0.09 | 0.05 | 0.05 | 0.05 |
| 6 | 11274670 | rs11967989 | -0.384 | -0.697 | 0.505 | -0.592 | 0.014 | 0.009 | 0.009 | 0 | 0.12 | 0.15 | 0.09 | 0.05 | 0.05 | 0.05 |
| 6 | 11286280 | rs7748486 | -1.566 | -0.704 | 0.448 | -0.656 | 0.034 | 0.027 | 0.027 | 0.857 | 0.41 | 0.72 | 0.3 | 0.29 | 0.29 | 0.2 |
| 6 | 11287746 | rs4713341 | -1.067 | 0.126 | 0.636 | -0.582 | -0.004 | 0.007 | 0.007 | 0.146 | 0.19 | 0.13 | 0.16 | 0.14 | 0.17 | 0.16 |
| 6 | 11288206 | rs2327394 | -2.252 | 0.061 | 0.795 | -0.363 | -0.003 | -0.004 | -0.004 | 0 | 0.34 | 0.6 | 0.38 | 0.26 | 0.27 | 0.24 |
| 6 | 11288980 | rs16871207 | -1.101 | 0.075 | 0.922 | -0.627 | -0.014 | -0.016 | -0.016 | 0.965 | 0.13 | 0.2 | 0.21 | 0.11 | 0.09 | 0.09 |
| 6 | 11295283 | rs12663655 | 0.086 | 0.307 | 0.648 | -0.601 | -0.01 | -0.051 | -0.051 | 0 | 0.21 | 0.12 | 0.24 | 0.64 | 0.24 | 0.38 |
| 6 | 11296124 | rs2142742 | -0.839 | 0.304 | 0.739 | -0.733 | -0.006 | -0.046 | -0.046 | 0.147 | 0.13 | 0.18 | 0.15 | 0.03 | 0.1 | 0.08 |
| 6 | 11305230 | rs4713379 | -1.015 | 0.426 | 0.77 | -0.75 | 0.004 | -0.049 | -0.049 | 0.267 | 0.19 | 0.4 | 0.18 | 0.07 | 0.16 | 0.14 |
| 6 | 11312130 | rs4713396 | -1.736 | 0.701 | 0.528 | -0.728 | 0.001 | 0.042 | 0.042 | 0 | 0.49 | 0.62 | 0.59 | 0.79 | 0.49 | 0.58 |
| 6 | 11317315 | rs10484451 | -0.377 | 0.057 | 0.092 | -0.539 | -0.001 | -0.007 | -0.007 | 0 | 0.08 | 0.06 | 0.15 | 0 | 0.08 | 0.03 |

| | | | | | | | | | | | | | | | | |
|---|---|---|---|---|---|---|---|---|---|---|---|---|---|---|---|---|
| 6 | 11322090 | rs12110566 | -1.097 | 0.102 | 0.087 | -0.529 | -0.001 | 0.009 | 0.009 | 0 | 0.15 | 0.13 | 0.25 | 0.05 | 0.14 | 0.06 |
| 6 | 11322636 | rs16871253 | 0.114 | 0.124 | 0.099 | -0.492 | 0 | 0 | 0 | 0 | 0.07 | 0.08 | 0.11 | 0.05 | 0.05 | 0.04 |
| 6 | 11323865 | rs9791189 | 0.055 | -0.057 | -0.264 | -0.399 | 0.007 | -0.041 | -0.041 | 1.128 | 0.2 | 0.12 | 0.19 | 0.64 | 0.23 | 0.37 |
| 6 | 11325777 | rs11755688 | -1.128 | -0.009 | -0.416 | -0.873 | -0.007 | -0.016 | -0.016 | 0 | 0.39 | 0.52 | 0.32 | 0.73 | 0.38 | 0.56 |
| 6 | 11327021 | rs742115 | -1.727 | 0.05 | -0.595 | -0.915 | -0.018 | 0.093 | 0.093 | 0 | 0.54 | 0.75 | 0.61 | 0.77 | 0.51 | 0.62 |
| 6 | 11329423 | rs967473 | -0.535 | 0.112 | -0.544 | -0.796 | -0.006 | 0.019 | 0.019 | 0 | 0.15 | 0.12 | 0.29 | 0.07 | 0.14 | 0.06 |
| 6 | 11329600 | rs1883239 | -1.723 | 0.027 | -0.153 | -0.413 | -0.024 | -0.093 | -0.093 | 0.022 | 0.18 | 0.33 | 0.1 | 0.04 | 0.13 | 0.17 |
| 6 | 11330169 | rs11757904 | -0.907 | 0.662 | -1.316 | -0.049 | -0.022 | 0.048 | 0.048 | 0 | 0.09 | 0.02 | 0.07 | 0 | 0.11 | 0.08 |
| 6 | 11332132 | rs7738900 | -0.367 | 0.256 | -1.353 | 0.101 | 0.053 | 0.051 | 0.051 | 2.737 | 0.4 | 0.75 | 0.34 | 0.25 | 0.25 | 0.43 |
| 6 | 11333684 | rs4713432 | 0.045 | 0.865 | -1.688 | 0.328 | -0.067 | -0.009 | -0.009 | 0 | 0.67 | 0.9 | 0.77 | 0.74 | 0.62 | 0.76 |
| 6 | 11335478 | rs6940151 | 0.088 | 0.885 | -1.725 | 0.311 | -0.082 | -0.007 | -0.007 | 0.925 | 0.64 | 0.89 | 0.76 | 0.72 | 0.6 | 0.74 |
| 6 | 11339769 | rs2064112 | 0.677 | 1.336 | -1.242 | -0.073 | 0.032 | 0.188 | 0.188 | 0 | 0.48 | 0.26 | 0.45 | 0.08 | 0.39 | 0.34 |
| 6 | 11346572 | rs2179179 | -0.961 | 1.533 | 0.399 | 1.311 | 0.064 | 0.202 | 0.202 | 0 | 0.13 | 0.26 | 0.25 | 0.55 | 0.22 | 0.28 |
| 6 | 11356849 | rs6935281 | 0.017 | 0.955 | -0.493 | 0.203 | -0.012 | 0.069 | 0.069 | 0.233 | 0.47 | 0.69 | 0.53 | 0.3 | 0.53 | 0.5 |
| 6 | 11357449 | rs1883235 | -0.015 | 1.469 | -0.365 | 0.459 | 0.001 | 0 | 0 | 0 | 0.29 | 0.38 | 0.25 | 0.27 | 0.28 | 0.18 |
| 6 | 11357978 | rs7775768 | 0.865 | 0.801 | 0.16 | -0.06 | -0.002 | 0.046 | 0.046 | 0 | 0.37 | 0.27 | 0.39 | 0.18 | 0.4 | 0.36 |
| 6 | 11360163 | rs3798752 | 0.258 | 0.697 | 0.271 | 0.267 | -0.029 | -0.002 | -0.002 | 0 | 0.65 | 0.79 | 0.61 | 0.38 | 0.61 | 0.51 |
| 6 | 11367261 | rs11756278 | 0.705 | 0.579 | 0.366 | 0.215 | 0.001 | 0 | 0 | 0.63 | 0.26 | 0.27 | 0.21 | 0.17 | 0.2 | 0.15 |
| 6 | 11371615 | rs7741863 | 0.415 | 0.428 | 0.324 | 0.093 | -0.008 | 0.062 | 0.062 | 0 | 0.44 | 0.68 | 0.46 | 0.25 | 0.49 | 0.38 |
| 6 | 11378141 | rs7750329 | -1.24 | 0.36 | 0.599 | 0.013 | -0.004 | 0.045 | 0.045 | 0 | 0.65 | 0.74 | 0.62 | 0.83 | 0.6 | 0.64 |
| 6 | 11379146 | rs9469022 | -0.662 | 0.359 | 0.657 | 0.012 | 0 | -0.006 | -0.006 | 0 | 0.58 | 0.56 | 0.56 | 0.78 | 0.51 | 0.62 |
| 6 | 11383976 | rs9296000 | -0.704 | 0.353 | 0.567 | 0.033 | 0.015 | -0.018 | -0.018 | 1.247 | 0.38 | 0.47 | 0.41 | 0.66 | 0.41 | 0.5 |
| 6 | 11390177 | rs4074864 | 0.687 | 0.309 | 0.532 | 0.019 | 0.01 | 0.017 | 0.017 | 0 | 0.27 | 0.22 | 0.21 | 0.17 | 0.2 | 0.15 |
| 6 | 11390934 | rs9461714 | -0.489 | 0.336 | 0.524 | 0.071 | -0.001 | -0.001 | -0.001 | 0 | 0.35 | 0.37 | 0.36 | 0.61 | 0.33 | 0.47 |
| 6 | 11403701 | rs9469055 | -0.307 | 0.452 | 0.759 | -0.222 | 0.003 | 0.021 | 0.021 | 0.003 | 0.39 | 0.64 | 0.3 | 0.24 | 0.34 | 0.18 |
| 6 | 11412916 | rs7772068 | 1.244 | 0.176 | 0.632 | -0.444 | -0.018 | 0.063 | 0.063 | 0 | 0.36 | 0.23 | 0.36 | 0.09 | 0.43 | 0.28 |
| 6 | 11415271 | rs4713491 | 0.145 | -0.074 | 0.752 | -0.512 | -0.001 | -0.002 | -0.002 | 0 | 0.67 | 0.68 | 0.61 | 0.29 | 0.69 | 0.4 |
| 6 | 11422923 | rs9394080 | 0.474 | 0.021 | 0.863 | -0.526 | 0 | -0.016 | -0.016 | 0 | 0.45 | 0.45 | 0.43 | 0.13 | 0.55 | 0.31 |
| 6 | 11424279 | rs4076156 | 0.371 | -0.375 | 0.853 | -0.386 | -0.035 | -0.021 | -0.021 | 0 | 0.27 | 0.37 | 0.2 | 0.19 | 0.16 | 0.12 |
| 6 | 11429737 | rs4713501 | 0.703 | 0.143 | 0.908 | -0.456 | -0.011 | 0.051 | 0.051 | 0 | 0.4 | 0.29 | 0.4 | 0.12 | 0.46 | 0.27 |
| 6 | 11432812 | rs4476816 | -0.485 | 0.178 | 0.951 | 0.07 | -0.03 | -0.077 | -0.077 | 0 | 0.74 | 0.89 | 0.67 | 0.47 | 0.71 | 0.45 |
| 6 | 11436564 | rs11961030 | -0.185 | 0.007 | 1.088 | 0.033 | -0.031 | -0.013 | -0.013 | 0 | 0.34 | 0.53 | 0.26 | 0.28 | 0.22 | 0.16 |
| 6 | 11437537 | rs4711289 | -0.171 | 0.018 | 0.98 | 0.039 | -0.029 | -0.012 | -0.012 | 0 | 0.34 | 0.47 | 0.25 | 0.28 | 0.22 | 0.16 |
| 6 | 11445854 | rs9461735 | 0.794 | 0.296 | 0.459 | -0.614 | -0.019 | 0.032 | 0.032 | 0.066 | 0.31 | 0.21 | 0.39 | 0.08 | 0.43 | 0.25 |
| 6 | 11458263 | rs10947259 | -0.859 | -0.315 | 0.795 | -0.374 | -0.002 | -0.009 | -0.009 | 0 | 0.12 | 0.14 | 0.03 | 0.04 | 0.05 | 0.1 |
| 6 | 11461202 | rs4713527 | 0.807 | -0.826 | 1.114 | -0.62 | -0.035 | -0.055 | -0.055 | 0 | 0.64 | 0.44 | 0.83 | 0.78 | 0.78 | 0.79 |
| 6 | 11463219 | rs16871366 | -0.111 | -0.931 | 1.198 | -0.7 | -0.002 | -0.029 | -0.029 | 0.073 | 0.07 | 0.11 | 0.04 | 0 | 0.05 | 0 |
| 6 | 11467864 | rs7760983 | 1.262 | -0.96 | 1.514 | -0.469 | 0.029 | 0.024 | 0.024 | 0 | 0.62 | 0.32 | 0.82 | 0.71 | 0.78 | 0.77 |
| 6 | 11468610 | rs9380307 | 1.262 | -0.96 | 1.52 | -0.469 | 0.027 | 0.022 | 0.022 | 0 | 0.62 | 0.32 | 0.82 | 0.71 | 0.77 | 0.77 |
| 6 | 11470114 | rs10947294 | -0.108 | -0.551 | 1.469 | -0.252 | 0.001 | -0.001 | -0.001 | 0 | 0.06 | 0.06 | 0.04 | 0.16 | 0.05 | 0.09 |
| 6 | 11488926 | rs9469223 | -0.047 | -0.801 | 1.573 | -0.538 | -0.003 | -0.025 | -0.025 | 0 | 0.15 | 0.19 | 0.08 | 0.02 | 0.1 | 0.02 |
| 6 | 11490909 | rs6914420 | -0.332 | -0.832 | 1.035 | -0.446 | -0.001 | -0.002 | -0.002 | 0 | 0.36 | 0.38 | 0.7 | 0.58 | 0.55 | 0.39 |
| 6 | 11494660 | rs6900740 | -0.166 | -0.902 | 1.001 | -0.507 | -0.022 | -0.028 | -0.028 | 0.019 | 0.09 | 0.22 | 0.03 | 0.03 | 0.04 | 0.05 |
| 6 | 11494940 | rs12198774 | -0.605 | -0.924 | 1.048 | -0.512 | -0.001 | 0.003 | 0.003 | 0 | 0.38 | 0.46 | 0.7 | 0.58 | 0.55 | 0.38 |
| 6 | 11495216 | rs12198950 | -0.605 | -0.924 | 1.048 | -0.512 | -0.001 | 0.003 | 0.003 | 0 | 0.38 | 0.46 | 0.7 | 0.58 | 0.55 | 0.38 |
| 6 | 11495818 | rs17676117 | 0.79 | -0.887 | 0.864 | -0.517 | 0.006 | 0.009 | 0.009 | 0 | 0.14 | 0.11 | 0.08 | 0 | 0.11 | 0.02 |
| 6 | 11499392 | rs566568 | 0.637 | -0.952 | 0.734 | -0.519 | -0.034 | -0.039 | -0.039 | 0.757 | 0.75 | 0.46 | 0.85 | 0.82 | 0.83 | 0.88 |
| 6 | 11500818 | rs594164 | 0.541 | -0.884 | 0.366 | -0.418 | 0.071 | 0.024 | 0.024 | 0 | 0.53 | 0.32 | 0.23 | 0.43 | 0.38 | 0.65 |
| 6 | 11513834 | rs555034 | 1.745 | -0.901 | 0.065 | -0.432 | 0.079 | 0.052 | 0.052 | 0 | 0.38 | 0.05 | 0.16 | 0.24 | 0.28 | 0.5 |
| 6 | 11516046 | rs562508 | 1.302 | -0.894 | 0.105 | -0.418 | 0.107 | 0.004 | 0.004 | 0 | 0.44 | 0.09 | 0.2 | 0.4 | 0.33 | 0.6 |
| 6 | 11518303 | rs9461825 | -0.824 | -0.837 | 0.663 | -0.492 | -0.047 | -0.117 | -0.117 | 0.002 | 0.25 | 0.47 | 0.12 | 0.05 | 0.15 | 0.07 |
| 6 | 11540409 | rs6907207 | -0.793 | -0.735 | 0.597 | -0.346 | -0.014 | 0.017 | 0.017 | 0 | 0.11 | 0.21 | 0.06 | 0.2 | 0.08 | 0.14 |
| 6 | 11550364 | rs4713618 | -0.172 | -0.83 | 0.783 | -0.419 | -0.008 | -0.011 | -0.011 | 0 | 0.33 | 0.31 | 0.68 | 0.55 | 0.52 | 0.33 |
| 6 | 11551050 | rs17397079 | 0.683 | -0.895 | 0.63 | -0.475 | -0.006 | -0.05 | -0.05 | 0.143 | 0.15 | 0.17 | 0.08 | 0 | 0.11 | 0.02 |
| 6 | 11577466 | rs12212790 | 0.668 | -0.787 | 0.503 | -0.39 | 0.002 | -0.003 | -0.003 | 0 | 0.06 | 0.04 | 0.04 | 0.16 | 0.05 | 0.09 |
| 6 | 11579093 | rs595877 | 0.226 | -0.825 | 0.587 | -0.42 | 0.006 | 0.017 | 0.017 | 0 | 0.66 | 0.47 | 0.29 | 0.45 | 0.46 | 0.67 |
| 6 | 11581990 | rs10947419 | 0.547 | -0.682 | -0.091 | -0.173 | 0.002 | -0.003 | -0.003 | 0 | 0.06 | 0.04 | 0.04 | 0.16 | 0.05 | 0.09 |
| 6 | 11586435 | rs598907 | 1.608 | -0.84 | -0.195 | -0.279 | 0.084 | 0.049 | 0.049 | 0 | 0.38 | 0.06 | 0.16 | 0.28 | 0.29 | 0.53 |
| 6 | 11589656 | rs6900578 | 0.753 | 0.573 | -0.014 | -0.454 | 0.016 | -0.006 | -0.006 | 0.006 | 0.06 | 0.11 | 0.15 | 0.04 | 0.17 | 0.07 |
| 6 | 11590140 | rs9469574 | 0.099 | 0.538 | -0.048 | -0.399 | 0.007 | 0.002 | 0.002 | 0 | 0.12 | 0.11 | 0.18 | 0.17 | 0.22 | 0.15 |
| 6 | 11600883 | rs576862 | 0.035 | 1.112 | 0.022 | -0.477 | -0.008 | 0.001 | 0.001 | 0 | 0.26 | 0.23 | 0.46 | 0.69 | 0.28 | 0.32 |
| 6 | 11601134 | rs12215876 | 0.089 | 1.229 | -0.166 | -0.29 | 0.01 | 0.004 | 0.004 | 0 | 0.12 | 0.12 | 0.19 | 0.17 | 0.22 | 0.16 |
| 6 | 11610425 | rs576100 | -1.144 | 1.004 | -0.601 | 0.196 | 0.014 | -0.006 | -0.006 | 0 | 0.53 | 0.72 | 0.73 | 0.54 | 0.55 | 0.47 |
| 6 | 11610581 | rs561332 | -0.409 | -0.193 | -0.475 | -0.119 | 0.001 | 0.038 | 0.038 | 0 | 0.24 | 0.37 | 0.49 | 0.36 | 0.23 | 0.22 |
| 6 | 11614620 | rs563605 | -1.093 | -0.079 | -0.889 | -0.132 | 0.072 | 0.102 | 0.102 | 0 | 0.67 | 0.87 | 0.85 | 0.88 | 0.76 | 0.62 |
| 6 | 11616089 | rs707811 | -1.172 | 0.632 | -0.356 | 0.819 | 0.013 | 0.04 | 0.04 | 0 | 0.16 | 0.22 | 0.42 | 0.31 | 0.2 | 0.2 |
| 6 | 11616687 | rs11753744 | -1.021 | 0.715 | -0.471 | 0.559 | 0.006 | -0.003 | -0.003 | 0 | 0.07 | 0.1 | 0.08 | 0.05 | 0.1 | 0.09 |
| 6 | 11618167 | rs479526 | 0.257 | 0.278 | -0.637 | 0.38 | 0.008 | -0.001 | -0.001 | 0.276 | 0.51 | 0.56 | 0.6 | 0.52 | 0.47 | 0.58 |
| 6 | 11618187 | rs4574630 | -0.459 | 0.378 | -0.761 | 0.193 | -0.002 | -0.006 | -0.006 | 0.3 | 0.67 | 0.7 | 0.77 | 0.58 | 0.66 | 0.7 |

| | | | | | | | | | | | | | | | | |
|---|---|---|---|---|---|---|---|---|---|---|---|---|---|---|---|---|
| 6 | 11620846 | rs471693 | 0.426 | -0.097 | -1.221 | 0.252 | 0.003 | -0.028 | -0.028 | 22.085 | 0.1 | 0.04 | 0.32 | 0.33 | 0.09 | 0.17 |
| 6 | 11623628 | rs546173 | -0.634 | -0.188 | -0.615 | 0.181 | 0 | 0.032 | 0.032 | 0 | 0.66 | 0.55 | 0.76 | 0.55 | 0.67 | 0.7 |
| 6 | 11625376 | rs9461953 | -0.462 | -0.606 | -1.62 | -0.428 | 0.006 | 0.056 | 0.056 | 0 | 0.23 | 0.12 | 0.14 | 0.05 | 0.19 | 0.12 |
| 6 | 11625711 | rs9469705 | -0.21 | -0.9 | -1.617 | -0.571 | 0.021 | 0.171 | 0.171 | 0 | 0.52 | 0.23 | 0.42 | 0.22 | 0.57 | 0.52 |
| 6 | 11627539 | rs9469713 | 1.185 | -1.078 | -1.609 | -0.413 | -0.081 | 0.075 | 0.075 | 0.052 | 0.27 | 0.09 | 0.28 | 0.16 | 0.38 | 0.39 |
| 6 | 11631630 | rs539317 | 0.21 | -1.057 | -1.003 | 0.126 | 0.002 | -0.036 | -0.036 | 0 | 0.1 | 0.04 | 0.22 | 0.26 | 0.09 | 0.18 |
| 6 | 11632868 | rs915006 | -1.183 | -1.517 | -1.094 | 0.033 | 0.008 | 0 | 0 | 0 | 0.37 | 0.64 | 0.42 | 0.39 | 0.32 | 0.39 |
| 6 | 11634722 | rs821437 | 0.87 | -0.939 | -1.061 | 0.232 | -0.012 | 0 | 0 | 0 | 0.06 | 0.13 | 0.03 | 0.03 | 0.05 | 0.02 |
| 6 | 11636481 | rs9469732 | -0.222 | -0.706 | -1.396 | 0.318 | 0.006 | 0.006 | 0.006 | 0.768 | 0.12 | 0.09 | 0.05 | 0.04 | 0.05 | 0.05 |
| 6 | 11637483 | rs548304 | -0.693 | 0.06 | -1.41 | 0.615 | -0.071 | -0.066 | -0.066 | 0 | 0.19 | 0.02 | 0.3 | 0.24 | 0.22 | 0.26 |
| 6 | 11639524 | rs6915482 | -0.916 | 0.053 | -1.067 | 0.473 | -0.005 | 0.01 | 0.01 | 0 | 0.08 | 0.04 | 0.05 | 0.07 | 0.1 | 0.16 |
| 6 | 11640004 | rs549655 | -0.651 | -0.064 | -1.386 | 0.801 | -0.057 | -0.045 | -0.045 | 0 | 0.37 | 0.21 | 0.4 | 0.48 | 0.49 | 0.57 |
| 6 | 11641800 | rs821438 | 0.163 | 0.089 | -1.334 | 0.768 | -0.026 | -0.031 | -0.031 | 0.322 | 0.23 | 0.18 | 0.26 | 0.4 | 0.34 | 0.37 |
| 6 | 11643081 | rs530313 | 0.184 | 0.087 | -1.342 | 0.767 | -0.035 | -0.042 | -0.042 | 0.309 | 0.23 | 0.14 | 0.26 | 0.39 | 0.34 | 0.38 |
| 6 | 11647007 | rs9348938 | 0.345 | 0.086 | -1.357 | 0.778 | -0.06 | -0.068 | -0.068 | 0 | 0.22 | 0.1 | 0.25 | 0.39 | 0.34 | 0.29 |
| 6 | 11648113 | rs12213930 | 0.354 | 0.085 | -1.357 | 0.928 | -0.06 | -0.006 | -0.006 | 0 | 0.21 | 0.1 | 0.25 | 0.27 | 0.34 | 0.29 |
| 6 | 11650140 | rs13204757 | 0.334 | -0.721 | -1.339 | 0.54 | 0.003 | 0.024 | 0.024 | 0 | 0.19 | 0.11 | 0.12 | 0.08 | 0.17 | 0.12 |
| 6 | 11654095 | rs863699 | -1.252 | -0.751 | -0.461 | 0.858 | 0.005 | 0.008 | 0.008 | 0 | 0.58 | 0.65 | 0.49 | 0.65 | 0.63 | 0.69 |
| 6 | 11656361 | rs1150582 | -0.913 | -0.852 | -0.241 | 0.741 | 0.006 | 0.002 | 0.002 | 0.333 | 0.36 | 0.63 | 0.2 | 0.3 | 0.26 | 0.38 |
| 6 | 11657162 | rs7764199 | 0.731 | -0.771 | -0.329 | 0.895 | -0.053 | -0.017 | -0.017 | 0.261 | 0.23 | 0.09 | 0.27 | 0.31 | 0.37 | 0.32 |
| 6 | 11658168 | rs1150584 | -1.35 | -0.929 | 0.014 | 0.905 | 0.002 | 0.026 | 0.026 | 0 | 0.6 | 0.74 | 0.5 | 0.71 | 0.64 | 0.7 |
| 6 | 11660514 | rs1206001 | 0.191 | -0.741 | 0.24 | 0.988 | 0.018 | 0.018 | 0.018 | 0 | 0.27 | 0.39 | 0.33 | 0.41 | 0.38 | 0.32 |
| 6 | 11661269 | rs1150593 | -0.67 | -0.834 | -0.095 | 0.544 | -0.032 | -0.062 | -0.062 | 2.422 | 0.8 | 0.71 | 0.89 | 0.93 | 0.86 | 0.92 |
| 6 | 11664422 | rs1206002 | -0.286 | -0.063 | 0.727 | 1.132 | 0.03 | 0.047 | 0.047 | 0 | 0.3 | 0.52 | 0.35 | 0.43 | 0.38 | 0.32 |
| 6 | 11677538 | rs2022016 | -0.184 | -0.053 | 0.494 | 1.137 | 0.026 | 0.039 | 0.039 | 0 | 0.29 | 0.47 | 0.34 | 0.43 | 0.38 | 0.32 |
| 6 | 11683950 | rs2142895 | -0.219 | 0.008 | 0.398 | 1.164 | 0.025 | 0.039 | 0.039 | 0 | 0.29 | 0.47 | 0.34 | 0.43 | 0.38 | 0.33 |
| 6 | 11684342 | rs6934072 | 0.172 | -0.004 | 0.431 | 1.205 | -0.005 | -0.002 | -0.002 | 0 | 0.26 | 0.21 | 0.28 | 0.33 | 0.37 | 0.32 |
| 6 | 11690213 | rs11753959 | 0.198 | 0.076 | -0.095 | 0.925 | -0.005 | -0.002 | -0.002 | 1.347 | 0.26 | 0.21 | 0.28 | 0.33 | 0.37 | 0.32 |
| 6 | 11693147 | rs6934095 | -1.314 | 0.061 | -0.181 | 0.939 | 0.011 | 0.051 | 0.051 | 0.003 | 0.49 | 0.7 | 0.45 | 0.65 | 0.56 | 0.61 |
| 6 | 11705700 | rs742223 | 0.79 | 0.767 | -0.656 | 0.538 | -0.026 | 0.032 | 0.032 | 0 | 0.06 | 0.01 | 0.05 | 0.01 | 0.09 | 0.06 |
| 6 | 11707809 | rs9366872 | -0.49 | 0.558 | -0.393 | 0.824 | -0.01 | -0.002 | -0.002 | 0.466 | 0.28 | 0.21 | 0.38 | 0.33 | 0.38 | 0.32 |
| 6 | 11709097 | rs9366874 | -0.228 | 0.685 | -0.811 | 0.678 | -0.044 | -0.002 | -0.002 | 1.79 | 0.24 | 0.1 | 0.37 | 0.3 | 0.35 | 0.29 |
| 6 | 11710405 | rs9348953 | -0.783 | 0.023 | -0.118 | 0.263 | -0.047 | -0.075 | -0.075 | 0 | 0.24 | 0.49 | 0.11 | 0.16 | 0.19 | 0.45 |
| 6 | 11710554 | rs210920 | -1.197 | 0.073 | -0.293 | 0.275 | -0.001 | -0.003 | -0.003 | 0.82 | 0.17 | 0.19 | 0.16 | 0.13 | 0.16 | 0.23 |
| 6 | 11710560 | rs210919 | -0.479 | 0.606 | -0.359 | 0.723 | 0.003 | 0.006 | 0.006 | 0 | 0.16 | 0.15 | 0.28 | 0.33 | 0.29 | 0.22 |
| 6 | 11711968 | rs210917 | 0.182 | -0.588 | -0.476 | -0.349 | -0.021 | 0.041 | 0.041 | 0.146 | 0.83 | 0.94 | 0.89 | 1 | 0.79 | 0.89 |
| 6 | 11712586 | rs9380473 | -0.257 | -0.628 | -0.395 | 0.21 | 0 | 0.123 | 0.123 | 0 | 0.06 | 0.07 | 0.04 | 0.48 | 0.04 | 0.14 |
| 6 | 11714806 | rs3734273 | 1.659 | -1.296 | -0.371 | -0.325 | 0.029 | 0.127 | 0.127 | 0 | 0.48 | 0.19 | 0.57 | 0.27 | 0.56 | 0.44 |
| 6 | 11716759 | rs1413346 | 0.248 | -1.399 | -0.315 | -0.4 | 0.002 | 0.016 | 0.016 | 0 | 0.08 | 0.04 | 0.04 | 0 | 0.06 | 0.04 |
| 6 | 11717934 | rs6933892 | 1.25 | -1.38 | 0.051 | -0.353 | 0.024 | 0.045 | 0.045 | 0 | 0.56 | 0.33 | 0.62 | 0.27 | 0.62 | 0.46 |
| 6 | 11718127 | rs210910 | -0.368 | -1.389 | 0.042 | -0.375 | -0.023 | 0.043 | 0.043 | 0 | 0.83 | 0.94 | 0.89 | 1 | 0.79 | 0.89 |
| 6 | 11718181 | rs2235427 | 0.913 | -1.375 | 0.055 | -0.344 | 0.005 | 0.007 | 0.007 | 0.48 | 0.62 | 0.37 | 0.65 | 0.76 | 0.66 | 0.6 |
| 6 | 11721129 | rs2235422 | 0.848 | -1.599 | 0.219 | -0.358 | -0.002 | -0.021 | -0.021 | 0 | 0.67 | 0.72 | 0.72 | 0.44 | 0.66 | 0.68 |
| 6 | 11721301 | rs9394270 | 0.229 | -1.598 | 0.213 | -0.401 | -0.005 | 0.037 | 0.037 | 0.087 | 0.74 | 0.8 | 0.77 | 0.93 | 0.69 | 0.82 |
| 6 | 11723636 | rs2076185 | 0.122 | -1.521 | 0.242 | -0.274 | 0 | 0.124 | 0.124 | 0 | 0.06 | 0.07 | 0.04 | 0.48 | 0.04 | 0.14 |
| 6 | 11723832 | rs2076184 | -1.207 | -1.635 | 0.615 | -0.317 | -0.069 | 0.122 | 0.122 | 0 | 0.17 | 0.5 | 0.15 | 0.65 | 0.07 | 0.34 |
| 6 | 11724542 | rs210903 | 0.638 | -1.619 | 0.832 | -0.343 | -0.024 | -0.052 | -0.052 | 0 | 0.72 | 0.58 | 0.79 | 0.84 | 0.76 | 0.69 |
| 6 | 11726626 | rs4713831 | 1.29 | -1.613 | 0.715 | -0.349 | -0.003 | -0.001 | -0.001 | 0 | 0.65 | 0.39 | 0.74 | 0.35 | 0.73 | 0.55 |
| 6 | 11730612 | rs9348963 | 1.812 | -1.888 | 0.183 | -0.782 | -0.066 | 0.051 | 0.051 | 0.622 | 0.39 | 0.15 | 0.41 | 0.73 | 0.47 | 0.39 |
| 6 | 11732667 | rs9296142 | -0.661 | -1.976 | 0.179 | -0.842 | -0.036 | 0.008 | 0.008 | 0 | 0.15 | 0.26 | 0.11 | 0.16 | 0.08 | 0.26 |
| 6 | 11732823 | rs2235418 | 2.5 | -1.841 | -0.233 | -0.842 | -0.167 | 0.011 | 0.011 | 0.48 | 0.32 | 0.02 | 0.37 | 0.25 | 0.43 | 0.25 |
| 6 | 11733777 | rs6908760 | -0.26 | -1.902 | 0.143 | -0.738 | -0.006 | -0.026 | -0.026 | 0.101 | 0.19 | 0.26 | 0.23 | 0.05 | 0.15 | 0.17 |
| 6 | 11737370 | rs12189748 | -0.854 | -1.901 | 0.098 | -0.732 | 0 | -0.014 | -0.014 | 0 | 0.24 | 0.28 | 0.24 | 0.05 | 0.19 | 0.21 |
| 6 | 11739127 | rs6911562 | 0.225 | -1.776 | 0.013 | -0.658 | -0.001 | -0.007 | -0.007 | 0 | 0.15 | 0.17 | 0.2 | 0.05 | 0.14 | 0.17 |
| 6 | 11740036 | rs210890 | -0.576 | -1.729 | 0.044 | -0.739 | 0.009 | -0.059 | -0.059 | 0.011 | 0.84 | 0.81 | 0.88 | 1 | 0.8 | 0.89 |
| 6 | 11741114 | rs12529085 | -0.112 | -1.683 | -0.106 | -0.457 | -0.021 | -0.031 | -0.031 | 0 | 0.12 | 0.07 | 0.12 | 0.49 | 0.14 | 0.18 |
| 6 | 11745652 | rs2294425 | 2.327 | -1.936 | -0.285 | -0.681 | -0.119 | 0.03 | 0.03 | 0.345 | 0.31 | 0.05 | 0.36 | 0.25 | 0.42 | 0.25 |
| 6 | 11754206 | rs2235409 | -1.392 | -2.038 | 0.01 | -0.662 | -0.023 | 0.009 | 0.009 | 0.046 | 0.3 | 0.55 | 0.31 | 0.68 | 0.22 | 0.38 |
| 6 | 11759109 | rs6938865 | -0.125 | -1.976 | 0.02 | -0.846 | 0.001 | -0.009 | -0.009 | 0 | 0.76 | 0.77 | 0.77 | 0.94 | 0.79 | 0.71 |
| 6 | 11759376 | rs2294418 | -0.053 | -1.891 | -0.018 | -0.557 | -0.024 | -0.045 | -0.045 | 0 | 0.13 | 0.08 | 0.12 | 0.49 | 0.14 | 0.2 |
| 6 | 11759662 | rs9368866 | 0.56 | -2.11 | 0.105 | -0.958 | -0.01 | -0.058 | -0.058 | 0.597 | 0.62 | 0.57 | 0.68 | 0.92 | 0.64 | 0.64 |
| 6 | 11760259 | rs9394295 | 0.182 | -2.019 | -0.192 | -0.932 | 0.003 | -0.004 | -0.004 | 0.018 | 0.71 | 0.73 | 0.74 | 0.95 | 0.76 | 0.68 |
| 6 | 11760807 | rs9368867 | -1.281 | -2.101 | -0.194 | -0.883 | -0.006 | -0.009 | -0.009 | 0 | 0.37 | 0.56 | 0.37 | 0.72 | 0.32 | 0.42 |
| 6 | 11761177 | rs4713856 | -0.229 | -2.137 | -0.144 | -0.692 | -0.06 | 0.032 | 0.032 | 1.06 | 0.2 | 0.37 | 0.22 | 0.24 | 0.16 | 0.22 |
| 6 | 11762311 | rs12211506 | 0.641 | -1.944 | -0.112 | -0.795 | -0.015 | -0.096 | -0.096 | 0 | 0.62 | 0.52 | 0.71 | 0.94 | 0.7 | 0.62 |
| 6 | 11762961 | rs2235396 | 0.659 | -1.924 | -0.108 | -0.786 | -0.011 | -0.083 | -0.083 | 0 | 0.62 | 0.54 | 0.71 | 0.94 | 0.7 | 0.62 |
| 6 | 11764205 | rs2235391 | -0.107 | -1.52 | -0.291 | -0.238 | -0.015 | -0.026 | -0.026 | 0 | 0.12 | 0.09 | 0.17 | 0.52 | 0.13 | 0.17 |
| 6 | 11767620 | rs6457825 | -0.353 | -1.756 | 0.037 | -0.481 | -0.035 | -0.064 | -0.064 | 0 | 0.23 | 0.4 | 0.21 | 0.08 | 0.18 | 0.28 |
| 6 | 11768932 | rs11753766 | 0.394 | -1.717 | 0.012 | -0.528 | -0.002 | -0.04 | -0.04 | 0.037 | 0.07 | 0.15 | 0.04 | 0 | 0.05 | 0.06 |

| | | | | | | | | | | | | | | | | |
|---|---|---|---|---|---|---|---|---|---|---|---|---|---|---|---|---|
| 6 | 11769280 | rs9470042 | -1.707 | -1.935 | -0.077 | -0.557 | 0.029 | 0.022 | 0.022 | 0.243 | 0.48 | 0.7 | 0.42 | 0.6 | 0.38 | 0.53 |
| 6 | 11769395 | rs16871788 | -0.122 | -1.456 | 0.115 | -0.35 | -0.01 | -0.075 | -0.075 | 0 | 0.17 | 0.32 | 0.11 | 0.03 | 0.16 | 0.13 |
| 6 | 11770114 | rs9394302 | -0.379 | -1.134 | -0.079 | 0.15 | -0.008 | -0.001 | -0.001 | 0 | 0.12 | 0.13 | 0.12 | 0.48 | 0.13 | 0.15 |
| 6 | 11770705 | rs760788 | 0.755 | -1.434 | -0.016 | -0.173 | 0.002 | 0.007 | 0.007 | 0 | 0.55 | 0.45 | 0.61 | 0.32 | 0.55 | 0.53 |
| 6 | 11770777 | rs760787 | 0.724 | -1.363 | 0.052 | -0.682 | -0.006 | -0.036 | -0.036 | 0 | 0.6 | 0.52 | 0.65 | 0.81 | 0.59 | 0.68 |
| 6 | 11770803 | rs742225 | 0.653 | -1.266 | 0.394 | -0.414 | -0.015 | -0.033 | -0.033 | 0 | 0.62 | 0.52 | 0.61 | 0.77 | 0.67 | 0.63 |
| 6 | 11770814 | rs742224 | 0.833 | -1.091 | -0.341 | -0.345 | 0.031 | 0.081 | 0.081 | 0.167 | 0.55 | 0.28 | 0.57 | 0.77 | 0.63 | 0.55 |
| 6 | 11771259 | rs9470047 | -0.772 | -0.765 | 0.119 | 0.216 | -0.026 | 0.024 | 0.024 | 1.167 | 0.15 | 0.55 | 0.17 | 0.16 | 0.1 | 0.17 |
| 6 | 11774238 | rs4713896 | -0.987 | -0.851 | -0.404 | 0.062 | -0.012 | -0.004 | -0.004 | 0 | 0.34 | 0.59 | 0.25 | 0.67 | 0.21 | 0.45 |
| 6 | 11774583 | rs6903956 | -0.299 | -0.672 | -1.154 | -0.743 | 0.001 | -0.05 | -0.05 | 0.088 | 0.68 | 0.63 | 0.65 | 0.93 | 0.62 | 0.82 |
| 6 | 11774626 | rs12204492 | 0.984 | -0.551 | -1.6 | -0.547 | -0.121 | 0.021 | 0.021 | 1.254 | 0.34 | 0.06 | 0.4 | 0.27 | 0.41 | 0.38 |
| 6 | 11774792 | rs6929400 | -0.107 | -0.566 | -1.52 | -0.821 | -0.054 | 0.122 | 0.122 | 0.374 | 0.69 | 0.85 | 0.67 | 0.92 | 0.61 | 0.83 |
| 6 | 11774859 | rs9380521 | -0.378 | -0.907 | -0.841 | 0.169 | -0.014 | 0.039 | 0.039 | 0.006 | 0.21 | 0.26 | 0.17 | 0.47 | 0.13 | 0.33 |
| 6 | 11775274 | rs2235386 | 0.432 | -0.061 | -1.612 | -0.161 | -0.056 | 0.016 | 0.016 | 0.079 | 0.35 | 0.12 | 0.38 | 0.28 | 0.4 | 0.37 |
| 6 | 11775721 | rs2235385 | 0.575 | -0.517 | -0.936 | -0.116 | 0 | -0.001 | -0.001 | 0 | 0.56 | 0.39 | 0.71 | 0.47 | 0.72 | 0.58 |
| 6 | 11776631 | rs2235384 | 0.078 | -0.746 | -0.563 | -0.092 | -0.047 | 0.054 | 0.054 | 0 | 0.71 | 0.57 | 0.81 | 0.54 | 0.84 | 0.67 |
| 6 | 11777985 | rs10947559 | 0.079 | -0.744 | -0.718 | -0.068 | -0.05 | 0.06 | 0.06 | 0 | 0.71 | 0.55 | 0.81 | 0.54 | 0.83 | 0.65 |
| 6 | 11779088 | rs2076189 | -0.191 | -0.119 | -1.429 | -0.248 | 0.025 | -0.017 | -0.017 | 0 | 0.31 | 0.43 | 0.4 | 0.23 | 0.4 | 0.28 |
| 6 | 11779499 | rs2076188 | -0.164 | 0.144 | -1.703 | -0.309 | 0.027 | 0 | 0 | 0.036 | 0.59 | 0.31 | 0.75 | 0.4 | 0.75 | 0.54 |
| 6 | 11786165 | rs9368883 | 0.378 | -0.203 | -1.929 | -0.021 | -0.095 | 0.015 | 0.015 | 0.51 | 0.39 | 0.08 | 0.39 | 0.32 | 0.43 | 0.35 |
| 6 | 11790493 | rs12199311 | 0.21 | -0.501 | -1.492 | 0.302 | -0.023 | -0.054 | -0.054 | 0.052 | 0.36 | 0.2 | 0.43 | 0.51 | 0.41 | 0.48 |
| 6 | 11792273 | rs9296163 | -0.662 | -2.413 | -1.833 | -0.596 | -0.002 | -0.089 | -0.089 | 0.611 | 0.68 | 0.95 | 0.6 | 0.41 | 0.7 | 0.46 |
| 6 | 11793197 | rs4713922 | 0.983 | -2.837 | -1.194 | -0.946 | 0.001 | 0.009 | 0.009 | 0.736 | 0.38 | 0.5 | 0.37 | 0.4 | 0.56 | 0.34 |
| 6 | 11795990 | rs9296165 | -0.505 | -2.828 | -0.949 | -0.964 | -0.06 | -0.126 | -0.126 | 0 | 0.31 | 0.44 | 0.23 | 0.02 | 0.16 | 0.13 |
| 6 | 11798414 | rs4713925 | 0.982 | -2.547 | -0.85 | -0.786 | -0.059 | -0.076 | -0.076 | 1.186 | 0.35 | 0.07 | 0.34 | 0.4 | 0.56 | 0.34 |
| 6 | 11799148 | rs4711432 | -0.117 | -2.509 | -0.584 | -0.982 | 0.052 | -0.015 | -0.015 | 0 | 0.53 | 0.78 | 0.54 | 0.52 | 0.59 | 0.4 |
| 6 | 11803437 | rs9368892 | -0.057 | -2.174 | -0.274 | -1.026 | 0.026 | -0.002 | -0.002 | 0.709 | 0.49 | 0.67 | 0.52 | 0.51 | 0.56 | 0.38 |
| 6 | 11806433 | rs1883324 | -0.773 | -1.818 | 0.462 | -0.399 | 0.03 | -0.019 | -0.019 | 0 | 0.41 | 0.73 | 0.6 | 0.5 | 0.35 | 0.63 |
| 6 | 11808064 | rs7743245 | 0.17 | -1.829 | 0.429 | -0.405 | 0 | -0.037 | -0.037 | 0 | 0.28 | 0.17 | 0.42 | 0.48 | 0.3 | 0.53 |
| 6 | 11814975 | rs9470184 | -1.065 | -1.787 | 0.465 | -0.549 | 0.076 | -0.029 | -0.029 | 0 | 0.44 | 0.86 | 0.56 | 0.48 | 0.35 | 0.62 |
| 6 | 11818316 | rs1883326 | 0.238 | -1.754 | 0.467 | -0.532 | -0.001 | -0.029 | -0.029 | 1.657 | 0.27 | 0.17 | 0.36 | 0.42 | 0.29 | 0.52 |
| 6 | 11820508 | rs9470202 | -0.647 | -1.751 | 0.474 | -0.61 | -0.081 | -0.08 | -0.08 | 0 | 0.16 | 0.65 | 0.19 | 0.05 | 0.05 | 0.1 |
| 6 | 11821628 | rs9296173 | -0.282 | -1.726 | 0.456 | -0.599 | -0.06 | -0.083 | -0.083 | 0.018 | 0.14 | 0.44 | 0.19 | 0.02 | 0.05 | 0.09 |
| 6 | 11825414 | rs1569731 | -0.092 | -1.564 | 0.07 | -0.505 | -0.003 | 0.023 | 0.023 | 4.789 | 0.29 | 0.32 | 0.37 | 0.47 | 0.29 | 0.52 |
| 6 | 11827597 | rs6936286 | -0.646 | -1.308 | -0.387 | -0.413 | 0.051 | -0.047 | -0.047 | 0 | 0.41 | 0.87 | 0.57 | 0.49 | 0.34 | 0.59 |
| 6 | 11829240 | rs13194405 | -0.194 | -0.498 | -0.073 | 0.114 | 0 | -0.009 | -0.009 | 0 | 0.3 | 0.25 | 0.38 | 0.44 | 0.33 | 0.55 |
| 6 | 11834477 | rs13208185 | 0.946 | -0.714 | 0.725 | -0.23 | 0.002 | 0.003 | 0.003 | 0.022 | 0.61 | 0.31 | 0.62 | 0.61 | 0.67 | 0.72 |
| 6 | 11834552 | rs10485419 | -0.9 | -0.606 | 1.099 | -1.034 | 0.05 | 0.029 | 0.029 | 0 | 0.12 | 0.31 | 0.15 | 0.23 | 0.18 | 0.15 |
| 6 | 11834883 | rs9462160 | 0.978 | -0.948 | 1.01 | -1.287 | -0.004 | 0.005 | 0.005 | 0 | 0.74 | 0.74 | 0.72 | 0.75 | 0.73 | 0.75 |
| 6 | 11835432 | rs10485420 | 0.177 | -1.027 | 1.349 | -1.765 | -0.039 | -0.077 | -0.077 | 0 | 0.14 | 0.26 | 0.06 | 0 | 0.05 | 0.04 |
| 6 | 11835787 | rs9380549 | -0.64 | -1.036 | 0.973 | -1.791 | 0.012 | -0.038 | -0.038 | 0.182 | 0.18 | 0.35 | 0.12 | 0.17 | 0.18 | 0.19 |
| 6 | 11835876 | rs10947590 | 0.779 | -1.115 | 1.054 | -1.774 | -0.053 | -0.16 | -0.16 | 0.047 | 0.73 | 0.46 | 0.75 | 0.99 | 0.86 | 0.89 |
| 6 | 11844684 | rs9368918 | -0.992 | -0.96 | 0.486 | -1.7 | 0.013 | 0.03 | 0.03 | 0.566 | 0.34 | 0.73 | 0.19 | 0.17 | 0.24 | 0.23 |
| 6 | 11846840 | rs4711449 | 0.779 | -1.167 | 0.858 | -1.291 | 0 | -0.023 | -0.023 | 0 | 0.88 | 0.87 | 0.85 | 0.99 | 0.91 | 0.93 |
| 6 | 11852967 | rs910153 | 0.758 | -1.163 | 0.838 | -1.289 | 0 | -0.021 | -0.021 | 0.001 | 0.88 | 0.87 | 0.85 | 0.99 | 0.91 | 0.93 |
| 6 | 11870256 | rs4713981 | 0.718 | -1.123 | 0.653 | -1.366 | -0.02 | -0.116 | -0.116 | 0 | 0.65 | 0.45 | 0.75 | 0.99 | 0.81 | 0.87 |
| 6 | 11871008 | rs6457927 | 1.737 | -0.31 | 0.938 | -0.421 | 0.028 | -0.084 | -0.084 | 0.379 | 0.88 | 0.52 | 0.94 | 1 | 0.92 | 0.94 |
| 6 | 11874298 | rs7740058 | 1.126 | -0.194 | -0.095 | 0.159 | 0.003 | -0.005 | -0.005 | 0 | 0.86 | 0.88 | 0.7 | 0.81 | 0.87 | 0.61 |
| 6 | 11881672 | rs13192449 | -0.185 | 0.111 | -0.88 | 0.257 | -0.087 | -0.014 | -0.014 | 2.845 | 0.29 | 0.05 | 0.4 | 0.24 | 0.43 | 0.35 |
| 6 | 11884990 | rs9357221 | 1.004 | -0.57 | -0.511 | 0.614 | 0.004 | 0.026 | 0.026 | 0 | 0.51 | 0.21 | 0.36 | 0.42 | 0.49 | 0.27 |
| 6 | 11887007 | rs12211229 | 0.045 | -0.201 | -0.683 | 0.717 | -0.049 | -0.011 | -0.011 | 0 | 0.29 | 0.05 | 0.39 | 0.24 | 0.39 | 0.35 |
| 6 | 11898806 | rs6457952 | 0.657 | -0.315 | 0.324 | 0.672 | -0.013 | 0.027 | 0.027 | 0 | 0.84 | 0.77 | 0.79 | 0.66 | 0.92 | 0.63 |
| 6 | 11902591 | rs6937349 | 0.516 | -0.201 | -0.132 | 0.729 | 0.013 | -0.032 | -0.032 | 0.046 | 0.87 | 0.9 | 0.8 | 0.66 | 0.92 | 0.63 |
| 6 | 11908166 | rs17406952 | 0.492 | 0.001 | 0 | 0.453 | 0.005 | 0.014 | 0.014 | 0 | 0.51 | 0.34 | 0.36 | 0.42 | 0.49 | 0.27 |
| 6 | 11912213 | rs9366921 | -0.679 | -0.69 | -1.063 | -0.152 | 0.024 | 0.082 | 0.082 | 0 | 0.23 | 0.04 | 0.08 | 0.12 | 0.15 | 0.19 |
| 6 | 11913828 | rs209773 | -0.882 | -2.305 | -1.863 | -0.34 | 0.172 | 0.133 | 0.133 | 0.009 | 0.51 | 0.84 | 0.21 | 0.31 | 0.25 | 0.34 |
| 6 | 11918582 | rs876829 | 0.202 | -3.051 | -0.521 | -0.617 | -0.028 | -0.119 | -0.119 | 0.165 | 0.17 | 0.38 | 0.08 | 0 | 0.08 | 0.03 |
| 6 | 11920517 | rs169713 | -1.143 | -3.2 | 0.117 | -0.683 | 0.06 | 0.066 | 0.066 | 0.027 | 0.48 | 0.61 | 0.2 | 0.17 | 0.24 | 0.33 |
| 6 | 11921997 | rs17533974 | -0.299 | -3.588 | 0.325 | -0.87 | 0.024 | 0.012 | 0.012 | 1.3 | 0.14 | 0.06 | 0.04 | 0.11 | 0.06 | 0.22 |
| 6 | 11922362 | rs23805 | -1.514 | -3.554 | 0.416 | -0.845 | 0.099 | 0.083 | 0.083 | 0.682 | 0.5 | 0.68 | 0.21 | 0.31 | 0.25 | 0.33 |
| 6 | 11923165 | rs209779 | -1.215 | -3.799 | 0.641 | -0.982 | -0.06 | -0.042 | -0.042 | 3.341 | 0.38 | 0.47 | 0.15 | 0.25 | 0.17 | 0.25 |
| 6 | 11926263 | rs2327461 | -2.015 | -3.207 | -1.084 | -0.403 | -0.163 | -0.091 | -0.091 | 0 | 0.71 | 0.94 | 0.49 | 0.63 | 0.52 | 0.53 |
| 6 | 11930487 | rs9380643 | -1.838 | -3.676 | -1.032 | -0.658 | 0.076 | 0.009 | 0.009 | 1.89 | 0.58 | 0.94 | 0.35 | 0.34 | 0.26 | 0.44 |
| 6 | 11934964 | rs10947661 | -1.196 | -3.714 | -1 | -0.611 | 0.193 | 0.236 | 0.236 | 0.775 | 0.51 | 0.88 | 0.21 | 0.18 | 0.22 | 0.32 |
| 6 | 11936514 | rs2327462 | 1.976 | -3.569 | -1.112 | -0.74 | 0.023 | -0.038 | -0.038 | 0 | 0.4 | 0.05 | 0.63 | 0.64 | 0.71 | 0.54 |
| 6 | 11938672 | rs6904014 | -0.133 | -3.485 | -0.595 | -0.918 | -0.071 | -0.09 | -0.09 | 0 | 0.13 | 0.53 | 0.07 | 0 | 0.02 | 0 |
| 6 | 11938797 | rs169715 | -0.001 | -3.371 | -0.648 | -0.836 | 0.002 | 0.008 | 0.008 | 0 | 0.93 | 0.96 | 0.84 | 0.98 | 0.95 | 0.87 |
| 6 | 11942371 | rs1205852 | -0.473 | -3.543 | -0.724 | -0.958 | 0.101 | 0.145 | 0.145 | 4.026 | 0.37 | 0.82 | 0.18 | 0.06 | 0.17 | 0.09 |
| 6 | 11945778 | rs16872095 | 0.659 | -3.447 | -0.622 | -0.881 | -0.023 | -0.031 | -0.031 | 0 | 0.12 | 0.26 | 0.06 | 0.06 | 0.08 | 0.08 |

| | | | | | | | | | | | | | | | | |
|---|---|---|---|---|---|---|---|---|---|---|---|---|---|---|---|---|
| 6 | 11948078 | rs1205886 | 0.435 | -3.206 | -1.061 | -0.726 | 0.005 | 0.01 | 0.01 | 0 | 0.92 | 0.94 | 0.84 | 0.98 | 0.94 | 0.87 |
| 6 | 11951031 | rs209816 | -0.405 | -3.292 | -1.062 | -0.802 | 0.116 | 0.18 | 0.18 | 4.293 | 0.37 | 0.85 | 0.18 | 0.06 | 0.17 | 0.09 |
| 6 | 11952845 | rs10947670 | 1.729 | -3.229 | -1.197 | -0.571 | 0.025 | -0.032 | -0.032 | 0 | 0.4 | 0.05 | 0.63 | 0.65 | 0.72 | 0.54 |
| 6 | 11959888 | rs6941983 | 0.364 | -3.283 | -0.535 | -0.776 | 0.004 | -0.02 | -0.02 | 0 | 0.58 | 0.35 | 0.69 | 0.92 | 0.78 | 0.77 |
| 6 | 11960383 | rs209803 | -0.042 | -3.465 | -0.244 | -0.868 | -0.053 | -0.073 | -0.073 | 0 | 0.12 | 0.35 | 0.04 | 0 | 0.02 | 0 |
| 6 | 11961620 | rs209799 | -0.383 | -3.766 | 0.238 | -0.975 | -0.073 | -0.028 | -0.028 | 0 | 0.32 | 0.46 | 0.11 | 0.27 | 0.13 | 0.25 |
| 6 | 11961951 | rs169716 | -0.475 | -3.758 | 0.297 | -0.97 | -0.08 | 0.026 | 0.026 | 0 | 0.2 | 0.38 | 0.06 | 0.27 | 0.06 | 0.25 |
| 6 | 11961970 | rs6921943 | 2.188 | -3.818 | 0.214 | -1.007 | 0.054 | 0.028 | 0.028 | 1.168 | 0.4 | 0.11 | 0.64 | 0.65 | 0.71 | 0.53 |
| 6 | 11963222 | rs186017 | -0.042 | -3.645 | 0.292 | -0.933 | -0.022 | -0.043 | -0.043 | 0 | 0.81 | 0.7 | 0.79 | 0.92 | 0.87 | 0.78 |
| 6 | 11965698 | rs7775818 | -0.088 | -3.56 | 0.213 | -0.831 | -0.034 | -0.015 | -0.015 | 0.309 | 0.17 | 0.2 | 0.05 | 0.12 | 0.05 | 0.24 |
| 6 | 11969231 | rs11963539 | -0.267 | -3.536 | 0.405 | -0.843 | -0.028 | -0.041 | -0.041 | 0 | 0.2 | 0.28 | 0.21 | 0.08 | 0.12 | 0.2 |
| 6 | 11969531 | rs1543573 | -1.561 | -2.431 | -0.115 | -0.726 | 0.051 | 0.22 | 0.22 | 0 | 0.83 | 0.58 | 0.79 | 0.49 | 0.75 | 0.69 |
| 6 | 11972512 | rs6907381 | -0.455 | -2.535 | -0.019 | -0.752 | -0.065 | -0.085 | -0.085 | 0 | 0.24 | 0.49 | 0.23 | 0.08 | 0.12 | 0.2 |
| 6 | 11972709 | rs9296224 | 0.612 | -2.799 | -0.171 | -0.886 | 0.051 | 0.039 | 0.039 | 0 | 0.21 | 0.12 | 0.07 | 0.12 | 0.09 | 0.26 |
| 6 | 11972742 | rs913021 | 0.146 | -2.816 | -0.256 | -0.941 | 0.01 | 0.002 | 0.002 | 0 | 0.51 | 0.41 | 0.66 | 0.52 | 0.6 | 0.43 |
| 6 | 11974582 | rs6457997 | -1.681 | -2.74 | -0.337 | -0.868 | -0.005 | -0.009 | -0.009 | 0 | 0.74 | 0.74 | 0.75 | 0.64 | 0.69 | 0.69 |
| 6 | 11976739 | rs9470631 | 0.632 | -2.664 | -0.209 | -0.823 | -0.009 | -0.014 | -0.014 | 0.821 | 0.2 | 0.22 | 0.21 | 0.08 | 0.12 | 0.2 |
| 6 | 11980411 | rs13216585 | 2.07 | -2.317 | -0.488 | -1.001 | -0.02 | -0.023 | -0.023 | 1.848 | 0.14 | 0.08 | 0.14 | 0.26 | 0.25 | 0.09 |
| 6 | 11982461 | rs4711507 | -2.055 | -2.445 | -0.363 | -0.937 | 0.015 | 0.043 | 0.043 | 0 | 0.86 | 0.78 | 0.8 | 0.64 | 0.75 | 0.7 |
| 6 | 11985333 | rs11752244 | 0.739 | -2.535 | -0.193 | -1.025 | 0.003 | 0.002 | 0.002 | 0 | 0.18 | 0.16 | 0.2 | 0.08 | 0.12 | 0.2 |
| 6 | 11986572 | rs12528610 | 1.665 | -2.638 | -0.322 | -1.094 | -0.11 | -0.035 | -0.035 | 0 | 0.29 | 0.09 | 0.45 | 0.28 | 0.47 | 0.23 |
| 6 | 11986665 | rs11760074 | 0.683 | -2.599 | -0.239 | -1.049 | -0.02 | -0.028 | -0.028 | 0 | 0.19 | 0.23 | 0.2 | 0.08 | 0.12 | 0.2 |
| 6 | 11999268 | rs6905791 | -0.243 | -2.039 | -0.322 | -1.018 | 0 | -0.021 | -0.021 | 0 | 0.65 | 0.69 | 0.61 | 0.52 | 0.65 | 0.54 |
| 6 | 12020241 | rs6925772 | -1.817 | -1.509 | -0.836 | -0.498 | 0.013 | 0.02 | 0.02 | 0 | 0.86 | 0.76 | 0.82 | 0.83 | 0.81 | 0.8 |
| 6 | 12029312 | rs9349054 | 2.119 | -1.465 | -1.36 | -0.476 | -0.024 | -0.029 | -0.029 | 6.22 | 0.11 | 0.04 | 0.17 | 0.18 | 0.19 | 0.2 |
| 6 | 12029424 | rs2327506 | 1.413 | -1.496 | -1.463 | -0.521 | -0.064 | -0.088 | -0.088 | 0 | 0.32 | 0.12 | 0.49 | 0.47 | 0.54 | 0.38 |
| 6 | 12033251 | rs9296237 | 0.771 | -1.769 | -1.429 | -0.777 | 0.043 | 0.023 | 0.023 | 0 | 0.17 | 0.09 | 0.04 | 0.12 | 0.06 | 0.18 |
| 6 | 12035329 | rs12525800 | 1.298 | -1.302 | -1.301 | -0.569 | -0.07 | -0.015 | -0.015 | 0.749 | 0.32 | 0.12 | 0.33 | 0.29 | 0.45 | 0.35 |
| 6 | 12042205 | rs12234132 | 0.893 | -1.837 | -1.239 | -0.594 | 0.038 | 0.061 | 0.061 | 0 | 0.26 | 0.12 | 0.1 | 0.13 | 0.14 | 0.16 |
| 6 | 12044054 | rs9470843 | -0.409 | -0.631 | -1.322 | -0.761 | 0.017 | 0.045 | 0.045 | 0.093 | 0.29 | 0.18 | 0.26 | 0.13 | 0.25 | 0.24 |
| 6 | 12048827 | rs1570989 | -2.146 | -0.124 | -0.942 | -0.265 | 0.005 | 0.003 | 0.003 | 1.075 | 0.85 | 0.82 | 0.81 | 0.81 | 0.78 | 0.79 |
| 6 | 12059318 | rs6933280 | 0.602 | -0.412 | -0.899 | -0.283 | -0.005 | 0.002 | 0.002 | 0 | 0.08 | 0.1 | 0.16 | 0.08 | 0.06 | 0.09 |
| 6 | 12059954 | rs13218440 | 0.074 | -0.295 | -0.891 | -0.316 | -0.001 | 0.006 | 0.006 | 0 | 0.39 | 0.41 | 0.39 | 0.42 | 0.41 | 0.42 |
| 6 | 12063868 | rs6930779 | 0.844 | 0.046 | -1.257 | -1.041 | -0.004 | 0.002 | 0.002 | 0 | 0.08 | 0.1 | 0.16 | 0.08 | 0.06 | 0.1 |
| 6 | 12069697 | rs4714170 | 1.817 | 0.026 | -1.424 | -1.292 | -0.006 | -0.014 | -0.014 | 0.155 | 0.21 | 0.15 | 0.35 | 0.28 | 0.26 | 0.27 |
| 6 | 12071284 | rs6916070 | 2.04 | -0.099 | -1.742 | -1.626 | -0.08 | 0.127 | 0.127 | 0 | 0.24 | 0.06 | 0.17 | 0 | 0.27 | 0.13 |
| 6 | 12071388 | rs16872237 | 0.989 | -0.214 | -1.237 | -1.554 | -0.004 | -0.032 | -0.032 | 0 | 0.13 | 0.13 | 0.08 | 0 | 0.12 | 0.06 |
| 6 | 12076171 | rs9470903 | 0.729 | -0.099 | -0.897 | -2.391 | -0.002 | -0.017 | -0.017 | 0 | 0.27 | 0.24 | 0.35 | 0.47 | 0.29 | 0.37 |
| 6 | 12078712 | rs2327508 | -1.08 | -0.303 | -0.826 | -2.508 | 0.001 | -0.002 | -0.002 | 0 | 0.68 | 0.63 | 0.69 | 0.71 | 0.62 | 0.66 |
| 6 | 12080585 | rs9462447 | 0.912 | -0.512 | -1.23 | -2.362 | 0.003 | 0.003 | 0.003 | 0 | 0.29 | 0.25 | 0.22 | 0.25 | 0.15 | 0.19 |
| 6 | 12080649 | rs9394517 | 1.11 | 0.078 | -1.062 | -2.234 | 0 | -0.004 | -0.004 | 1.414 | 0.07 | 0.06 | 0.05 | 0.08 | 0.06 | 0.11 |
| 6 | 12083440 | rs9394520 | 1.115 | 0.083 | -1.059 | -2.238 | -0.001 | -0.002 | -0.002 | 0 | 0.07 | 0.06 | 0.05 | 0.07 | 0.06 | 0.11 |
| 6 | 12087892 | rs10947756 | 0.741 | -0.306 | -1.404 | -2.558 | -0.01 | 0.011 | 0.011 | 0 | 0.39 | 0.21 | 0.4 | 0.66 | 0.4 | 0.37 |
| 6 | 12092498 | rs9296262 | 0.421 | -0.094 | -0.587 | -2.158 | 0.015 | 0.009 | 0.009 | 0.04 | 0.17 | 0.15 | 0.08 | 0.12 | 0.09 | 0.14 |
| 6 | 12093331 | rs6910367 | -0.466 | 0.098 | -0.137 | -2.211 | 0.048 | 0.281 | 0.281 | 0.222 | 0.84 | 0.54 | 0.7 | 0.53 | 0.82 | 0.71 |
| 6 | 12099278 | rs9380781 | 1.338 | 0.028 | -0.674 | -2.447 | -0.021 | 0.022 | 0.022 | 0.643 | 0.28 | 0.11 | 0.25 | 0.22 | 0.34 | 0.29 |
| 6 | 12102615 | rs7768491 | 2.078 | -0.002 | -0.871 | -2.517 | 0.027 | 0.028 | 0.028 | 0 | 0.21 | 0.05 | 0.19 | 0.18 | 0.12 | 0.11 |
| 6 | 12120445 | rs2228215 | 1.599 | -0.599 | -0.708 | -2.343 | -0.049 | 0.004 | 0.004 | 1.077 | 0.26 | 0.06 | 0.25 | 0.22 | 0.34 | 0.28 |
| 6 | 12122174 | rs2228210 | 1.622 | -0.604 | -0.716 | -2.345 | -0.084 | -0.021 | -0.021 | 0.192 | 0.26 | 0.02 | 0.25 | 0.22 | 0.34 | 0.29 |
| 6 | 12123249 | rs2228220 | 0.25 | -0.752 | -0.616 | -2.464 | -0.001 | -0.001 | -0.001 | 0 | 0.17 | 0.18 | 0.1 | 0.13 | 0.12 | 0.19 |
| 6 | 12124855 | rs2228213 | 1.764 | -0.729 | -0.646 | -2.461 | -0.048 | 0.002 | 0.002 | 1.044 | 0.26 | 0.06 | 0.25 | 0.22 | 0.33 | 0.28 |
| 6 | 12129047 | rs7741589 | 1.416 | -0.739 | -0.597 | -2.523 | -0.018 | -0.013 | -0.013 | 0 | 0.12 | 0.08 | 0.28 | 0.47 | 0.18 | 0.29 |
| 6 | 12129596 | rs9357299 | 1.767 | -0.744 | -0.601 | -2.525 | -0.068 | -0.008 | -0.008 | 0 | 0.26 | 0.04 | 0.25 | 0.22 | 0.34 | 0.28 |
| 6 | 12129936 | rs3734271 | 0.707 | -0.807 | -0.555 | -2.542 | 0.015 | 0.012 | 0.012 | 0.05 | 0.16 | 0.12 | 0.08 | 0.1 | 0.09 | 0.15 |
| 6 | 12149138 | rs220018 | 0.524 | -0.79 | -0.478 | -2.548 | 0.003 | -0.001 | -0.001 | 3.946 | 0.66 | 0.68 | 0.6 | 0.4 | 0.68 | 0.52 |
| 6 | 12150359 | rs3777763 | 1.139 | -0.816 | -0.602 | -2.586 | -0.034 | 0.12 | 0.12 | 0 | 0.35 | 0.1 | 0.31 | 0.18 | 0.34 | 0.24 |
| 6 | 12151610 | rs1998528 | 1.719 | -0.631 | -0.712 | -2.458 | -0.043 | 0.005 | 0.005 | 1.063 | 0.26 | 0.07 | 0.25 | 0.22 | 0.33 | 0.28 |
| 6 | 12155649 | rs2157494 | -0.012 | -0.392 | -0.636 | -2.217 | -0.008 | -0.03 | -0.03 | 0 | 0.24 | 0.17 | 0.33 | 0.57 | 0.26 | 0.44 |
| 6 | 12159112 | rs7747677 | -0.967 | 0.16 | -0.123 | -1.84 | -0.002 | 0.028 | 0.028 | 0 | 0.75 | 0.72 | 0.66 | 0.55 | 0.74 | 0.64 |
| 6 | 12159699 | rs3777755 | 1.482 | 0.399 | -0.688 | -1.775 | -0.072 | -0.03 | -0.03 | 0 | 0.2 | 0.03 | 0.22 | 0.22 | 0.3 | 0.24 |
| 6 | 12170927 | rs219963 | -0.486 | -0.59 | 0.541 | -1.168 | -0.033 | 0.009 | 0.009 | 0.334 | 0.15 | 0.37 | 0.09 | 0.13 | 0.07 | 0.14 |
| 6 | 12173757 | rs10947819 | 1.092 | -0.582 | -0.168 | -1.278 | 0.018 | 0.035 | 0.035 | 0.436 | 0.19 | 0.08 | 0.17 | 0.16 | 0.12 | 0.12 |
| 6 | 12174132 | rs9367018 | -0.093 | -0.525 | -0.695 | -1.236 | 0 | 0 | 0 | 0 | 0.13 | 0.15 | 0.08 | 0.11 | 0.09 | 0.15 |
| 6 | 12174911 | rs12189595 | 0.426 | -0.305 | -0.997 | -1.246 | 0.01 | 0.033 | 0.033 | 0.024 | 0.24 | 0.14 | 0.21 | 0.16 | 0.18 | 0.16 |
| 6 | 12175433 | rs12209865 | 0.637 | -0.282 | -1.319 | -0.938 | -0.052 | -0.009 | -0.009 | 0 | 0.23 | 0.07 | 0.21 | 0.24 | 0.31 | 0.26 |
| 6 | 12175447 | rs1111404 | 0.226 | -0.248 | -1.304 | -0.92 | -0.03 | 0.012 | 0.012 | 1.27 | 0.27 | 0.15 | 0.25 | 0.26 | 0.36 | 0.32 |
| 6 | 12176183 | rs10755704 | 0.096 | -0.225 | -1.335 | -0.908 | -0.027 | 0.018 | 0.018 | 0.309 | 0.28 | 0.16 | 0.25 | 0.26 | 0.36 | 0.32 |
| 6 | 12179747 | rs12192252 | 0.672 | 0.039 | -1.54 | -0.58 | 0.004 | 0.024 | 0.024 | 0 | 0.19 | 0.07 | 0.17 | 0.19 | 0.14 | 0.13 |
| 6 | 12181293 | rs16872375 | -0.126 | -0.082 | -1.451 | -0.526 | 0.007 | 0.003 | 0.003 | 0 | 0.12 | 0.08 | 0.07 | 0.07 | 0.08 | 0.15 |

| | | | | | | | | | | | | | | | | |
|---|---|---|---|---|---|---|---|---|---|---|---|---|---|---|---|---|
| 6 | 12183050 | rs10947826 | 0.246 | 0.462 | -2.077 | -0.335 | -0.044 | -0.012 | -0.012 | 0 | 0.21 | 0.06 | 0.2 | 0.24 | 0.29 | 0.27 |
| 6 | 12183952 | rs11755555 | 0.01 | -0.039 | -1.469 | -0.347 | 0.014 | 0.023 | 0.023 | 0 | 0.18 | 0.07 | 0.17 | 0.18 | 0.12 | 0.1 |
| 6 | 12196910 | rs1040992 | -0.115 | 0.015 | -1.659 | -0.665 | -0.021 | 0.137 | 0.137 | 0 | 0.23 | 0.02 | 0.11 | 0.04 | 0.2 | 0.17 |
| 6 | 12199517 | rs219976 | 0.314 | -0.482 | -1.094 | -1.008 | -0.024 | -0.041 | -0.041 | 0 | 0.87 | 0.78 | 0.95 | 0.95 | 0.94 | 0.79 |
| 6 | 12200317 | rs219978 | -0.35 | -0.154 | -0.844 | -0.303 | 0.001 | 0 | 0 | 0 | 0.47 | 0.48 | 0.49 | 0.53 | 0.46 | 0.42 |
| 6 | 12200946 | rs1023483 | -0.266 | -0.157 | -0.808 | -0.305 | 0.004 | 0 | 0 | 0 | 0.45 | 0.31 | 0.49 | 0.54 | 0.46 | 0.42 |
| 6 | 12201764 | rs9380902 | 0.052 | 0.136 | -0.369 | 0.449 | -0.007 | -0.032 | -0.032 | 0.811 | 0.21 | 0.16 | 0.36 | 0.49 | 0.26 | 0.24 |
| 6 | 12204493 | rs1034451 | 0.992 | -0.065 | 0.324 | 0.808 | -0.032 | 0.073 | 0.073 | 1.47 | 0.75 | 0.47 | 0.88 | 0.62 | 0.85 | 0.77 |
| 6 | 12205124 | rs9471195 | -1.54 | 0.194 | 1.182 | 1.119 | 0.108 | 0.339 | 0.339 | 0.961 | 0.05 | 0.34 | 0.04 | 0.19 | 0.04 | 0.02 |
| 6 | 12205259 | rs219982 | 1.271 | 0.262 | 1.297 | 0.876 | 0.035 | 0.092 | 0.092 | 0.627 | 0.83 | 0.58 | 0.89 | 0.71 | 0.86 | 0.91 |
| 6 | 12208819 | rs219986 | 0.382 | 0.094 | 0.51 | 0.788 | 0.022 | 0.028 | 0.028 | 0.831 | 0.53 | 0.4 | 0.38 | 0.7 | 0.43 | 0.39 |
| 6 | 12209692 | rs219988 | -0.084 | 0.037 | 0.549 | 0.742 | -0.018 | 0.029 | 0.029 | 0 | 0.61 | 0.76 | 0.41 | 0.74 | 0.46 | 0.45 |
| 6 | 12211902 | rs219999 | 0.505 | -0.409 | 0.17 | 0.43 | 0 | 0.002 | 0.002 | 0 | 0.24 | 0.27 | 0.24 | 0.35 | 0.24 | 0.12 |
| 6 | 12213473 | rs220007 | 0.099 | -0.316 | -0.097 | 0.579 | 0.037 | 0.013 | 0.013 | 3.397 | 0.45 | 0.66 | 0.24 | 0.57 | 0.28 | 0.36 |
| 6 | 12214326 | rs9367034 | 0.732 | -0.068 | -0.234 | 0.38 | -0.049 | 0.038 | 0.038 | 0.043 | 0.21 | 0.06 | 0.24 | 0.2 | 0.27 | 0.15 |
| 6 | 12216503 | rs1794854 | 0.088 | -0.215 | -0.081 | 0.54 | -0.011 | 0.018 | 0.018 | 1.409 | 0.88 | 0.62 | 0.94 | 0.86 | 0.94 | 0.98 |
| 6 | 12216646 | rs1794856 | 1.431 | -0.278 | -0.387 | 0.503 | -0.152 | -0.107 | -0.107 | 0 | 0.23 | 0.02 | 0.46 | 0.24 | 0.4 | 0.46 |
| 6 | 12216966 | rs10755709 | 0.185 | -0.208 | -0.222 | 0.279 | 0.033 | 0.008 | 0.008 | 2.698 | 0.46 | 0.37 | 0.31 | 0.41 | 0.3 | 0.43 |
| 6 | 12217422 | rs7356945 | 0.174 | -0.228 | -0.2 | 0.283 | 0.029 | 0.018 | 0.018 | 0.79 | 0.46 | 0.36 | 0.31 | 0.32 | 0.3 | 0.43 |
| 6 | 12217861 | rs4714277 | 0.026 | -0.482 | -0.23 | 0.237 | 0.003 | -0.05 | -0.05 | 1.571 | 0.32 | 0.45 | 0.31 | 0.3 | 0.33 | 0.17 |
| 6 | 12217999 | rs6908010 | -0.514 | -0.487 | -0.258 | 0.251 | 0.016 | 0.016 | 0.016 | 1.726 | 0.66 | 0.6 | 0.47 | 0.59 | 0.54 | 0.51 |
| 6 | 12219785 | rs12662518 | 0.586 | -0.418 | -0.35 | 0.218 | 0.048 | 0.016 | 0.016 | 0.76 | 0.35 | 0.24 | 0.16 | 0.27 | 0.21 | 0.34 |
| 6 | 12220894 | rs10456102 | 0.958 | -0.329 | -0.054 | 0.654 | -0.071 | -0.006 | -0.006 | 0 | 0.2 | 0.03 | 0.16 | 0.26 | 0.26 | 0.09 |
| 6 | 12223922 | rs17592421 | 1.218 | -0.811 | -0.096 | 0.446 | -0.049 | -0.014 | -0.014 | 0 | 0.29 | 0.12 | 0.38 | 0.24 | 0.41 | 0.52 |
| 6 | 12224328 | rs9471238 | 0.851 | -0.78 | -0.599 | 0.464 | 0.038 | -0.021 | -0.021 | 0 | 0.24 | 0.18 | 0.12 | 0.27 | 0.13 | 0.26 |
| 6 | 12224531 | rs2327538 | 0.815 | -0.14 | -0.423 | 0.428 | 0 | -0.001 | -0.001 | 0.044 | 0.55 | 0.56 | 0.52 | 0.51 | 0.55 | 0.78 |
| 6 | 12224716 | rs10947850 | -0.565 | -0.068 | -0.782 | 0.45 | -0.03 | -0.059 | -0.059 | 0.361 | 0.14 | 0.28 | 0.15 | 0.05 | 0.08 | 0.07 |
| 6 | 12229938 | rs9380934 | -0.475 | 0.005 | -0.771 | 0.5 | -0.003 | -0.003 | -0.003 | 0.158 | 0.37 | 0.34 | 0.44 | 0.5 | 0.39 | 0.21 |
| 6 | 12236036 | rs565744 | 0.349 | -0.055 | -0.535 | 0.043 | -0.021 | -0.08 | -0.08 | 0 | 0.8 | 0.68 | 0.8 | 0.95 | 0.87 | 0.92 |
| 6 | 12240805 | rs7749495 | 0.34 | 0.668 | -1.325 | 0.401 | -0.108 | 0.003 | 0.003 | 0.009 | 0.23 | 0.03 | 0.28 | 0.25 | 0.33 | 0.13 |
| 6 | 12245416 | rs9349146 | 0.503 | -0.143 | -0.74 | 0.809 | 0.031 | 0.001 | 0.001 | 2.048 | 0.29 | 0.34 | 0.4 | 0.26 | 0.4 | 0.53 |
| 6 | 12248805 | rs557421 | -0.357 | -0.476 | -1.003 | 0.617 | -0.026 | 0.039 | 0.039 | 0 | 0.62 | 0.67 | 0.53 | 0.71 | 0.55 | 0.79 |
| 6 | 12253021 | rs9380949 | 0.814 | -0.421 | -1.204 | 0.676 | -0.139 | -0.071 | -0.071 | 0 | 0.27 | 0.02 | 0.37 | 0.26 | 0.4 | 0.52 |
| 6 | 12255290 | rs4714340 | 0.512 | -0.476 | -1.072 | 0.424 | 0.022 | 0.046 | 0.046 | 1.85 | 0.32 | 0.43 | 0.4 | 0.49 | 0.4 | 0.52 |
| 6 | 12258294 | rs7744944 | 0.059 | -0.613 | -1.834 | 0.212 | 0.082 | 0 | 0 | 0 | 0.29 | 0.11 | 0.12 | 0.23 | 0.15 | 0.3 |
| 6 | 12260470 | rs9367057 | -0.065 | 0.335 | -1.691 | 0.52 | -0.085 | -0.005 | -0.005 | 0 | 0.41 | 0.12 | 0.61 | 0.63 | 0.47 | 0.69 |
| 6 | 12261921 | rs6458155 | -0.707 | 0.242 | -0.942 | 0.223 | -0.01 | 0.005 | 0.005 | 0 | 0.36 | 0.51 | 0.36 | 0.56 | 0.36 | 0.44 |
| 6 | 12262514 | rs9394696 | -0.333 | 0.633 | -1.571 | 0.468 | -0.026 | -0.075 | -0.075 | 1.307 | 0.2 | 0.07 | 0.26 | 0.53 | 0.27 | 0.37 |
| 6 | 12263693 | rs13192093 | 0.15 | 1.122 | -1.348 | -0.104 | -0.035 | 0.061 | 0.061 | 0 | 0.37 | 0.18 | 0.23 | 0.27 | 0.4 | 0.37 |
| 6 | 12273540 | rs521824 | -0.072 | 0.99 | -1.227 | -0.208 | -0.007 | 0.085 | 0.085 | 0 | 0.44 | 0.25 | 0.42 | 0.21 | 0.55 | 0.43 |
| 6 | 12277622 | rs6931867 | -0.746 | 0.533 | -0.307 | -0.366 | -0.007 | -0.043 | -0.043 | 0 | 0.21 | 0.39 | 0.14 | 0.16 | 0.19 | 0.34 |
| 6 | 12286838 | rs9380978 | -0.678 | 0.436 | -0.344 | -0.189 | -0.002 | -0.008 | -0.008 | 0.036 | 0.2 | 0.25 | 0.12 | 0.21 | 0.19 | 0.35 |
| 6 | 12292528 | rs1800542 | -1.134 | 0.22 | 0.355 | -0.499 | 0.016 | -0.028 | -0.028 | 7.554 | 0.08 | 0.32 | 0.04 | 0 | 0.04 | 0.06 |
| 6 | 12292772 | rs2070699 | 0.729 | 0.291 | -0.947 | -0.156 | -0.158 | -0.12 | -0.12 | 0.328 | 0.37 | 0.04 | 0.5 | 0.54 | 0.47 | 0.37 |
| 6 | 12295503 | rs1626492 | -0.424 | -0.031 | -0.832 | -0.631 | 0.025 | 0.032 | 0.032 | 0 | 0.64 | 0.29 | 0.75 | 0.96 | 0.71 | 0.82 |
| 6 | 12295814 | rs2071943 | -0.581 | 0.089 | -0.422 | 0.121 | 0.002 | -0.01 | -0.01 | 0 | 0.24 | 0.15 | 0.14 | 0.29 | 0.21 | 0.43 |
| 6 | 12296255 | rs5370 | -0.61 | 0.078 | -0.386 | 0.118 | 0.003 | -0.008 | -0.008 | 0.023 | 0.25 | 0.16 | 0.15 | 0.29 | 0.21 | 0.43 |
| 6 | 12297028 | rs9296344 | -1.327 | 0.155 | 0.645 | -0.039 | 0.074 | 0.028 | 0.028 | 0.111 | 0.08 | 0.38 | 0.04 | 0.04 | 0.06 | 0.09 |
| 6 | 12297853 | rs4714384 | -1.131 | -0.253 | 0.121 | -0.273 | 0.022 | 0.006 | 0.006 | 0 | 0.43 | 0.7 | 0.39 | 0.64 | 0.37 | 0.32 |
| 6 | 12298333 | rs9296345 | 0.024 | -0.441 | 0.52 | -0.44 | -0.013 | -0.058 | -0.058 | 0 | 0.11 | 0.23 | 0.08 | 0.01 | 0.12 | 0.09 |
| 6 | 12298984 | rs2859338 | 1.233 | -0.441 | -0.192 | -0.946 | -0.147 | 0.175 | 0.175 | 0.821 | 0.31 | 0.03 | 0.3 | 0.03 | 0.43 | 0.21 |
| 6 | 12302569 | rs11755836 | -0.148 | -0.312 | 0.704 | -0.073 | -0.042 | -0.115 | -0.115 | 0 | 0.16 | 0.02 | 0.23 | 0.52 | 0.14 | 0.13 |
| 6 | 12304829 | rs16872602 | -0.688 | -0.359 | 0.982 | -0.094 | -0.03 | -0.006 | -0.006 | 0.17 | 0.14 | 0.4 | 0.1 | 0.1 | 0.13 | 0.13 |
| 6 | 12306232 | rs17594577 | 0.894 | -0.353 | 0.941 | -0.075 | -0.002 | -0.003 | -0.003 | 0.841 | 0.06 | 0.1 | 0.04 | 0.04 | 0.06 | 0.04 |
| 6 | 12307264 | rs9471438 | -2.062 | -0.359 | 1.28 | -0.047 | -0.002 | 0.008 | 0.008 | 0 | 0.39 | 0.51 | 0.42 | 0.6 | 0.4 | 0.37 |
| 6 | 12315910 | rs9381015 | -2.422 | -0.259 | 1.505 | 0.365 | 0.017 | 0.002 | 0.002 | 0.383 | 0.42 | 0.77 | 0.41 | 0.65 | 0.38 | 0.38 |
| 6 | 12319527 | rs17765653 | -1.813 | -0.235 | 1.506 | 0.376 | 0.002 | 0.015 | 0.015 | 0 | 0.33 | 0.4 | 0.38 | 0.61 | 0.35 | 0.33 |
| 6 | 12325761 | rs16872657 | -1.046 | -0.2 | 1.432 | 0.279 | 0.012 | -0.062 | -0.062 | 0 | 0.17 | 0.36 | 0.15 | 0.07 | 0.21 | 0.22 |
| 6 | 12327232 | rs10947946 | -0.352 | -0.126 | 0.83 | 0.403 | -0.032 | -0.106 | -0.106 | 0.609 | 0.16 | 0.03 | 0.24 | 0.52 | 0.14 | 0.11 |
| 6 | 12329292 | rs4379286 | 1.955 | -0.085 | 0.914 | 0.351 | 0.015 | 0.019 | 0.019 | 0.045 | 0.55 | 0.2 | 0.57 | 0.36 | 0.6 | 0.59 |
| 6 | 12331807 | rs4714429 | -0.396 | 0.129 | 0.809 | 0.585 | -0.046 | -0.094 | -0.094 | 0 | 0.16 | 0.04 | 0.27 | 0.52 | 0.18 | 0.18 |
| 6 | 12341753 | rs9471517 | -1.291 | -0.182 | 1.287 | 0.291 | -0.055 | -0.077 | -0.077 | 1.017 | 0.23 | 0.44 | 0.11 | 0.1 | 0.16 | 0.2 |
| 6 | 12342789 | rs1317584 | -0.664 | 0.287 | 0.992 | 0.516 | -0.01 | -0.01 | -0.01 | 0.049 | 0.19 | 0.13 | 0.31 | 0.46 | 0.24 | 0.19 |
| 6 | 12344262 | rs10807258 | -2.026 | 0.158 | 0.961 | 0.417 | -0.004 | -0.002 | -0.002 | 0.025 | 0.43 | 0.64 | 0.43 | 0.57 | 0.42 | 0.4 |
| 6 | 12349737 | rs6899475 | -2.033 | 0.347 | 0.924 | 0.45 | -0.009 | 0.003 | 0.003 | 0 | 0.45 | 0.69 | 0.43 | 0.56 | 0.45 | 0.43 |
| 6 | 12352001 | rs2095342 | -0.389 | 0.468 | 0.377 | 0.634 | -0.069 | -0.058 | -0.058 | 0 | 0.18 | 0.04 | 0.26 | 0.32 | 0.21 | 0.18 |
| 6 | 12354387 | rs2050288 | -0.998 | 0.456 | 0.577 | 0.566 | 0.032 | 0.027 | 0.027 | 0 | 0.26 | 0.36 | 0.3 | 0.37 | 0.25 | 0.21 |
| 6 | 12354681 | rs12332935 | -1.266 | 0.425 | 0.072 | 0.392 | 0.004 | -0.009 | -0.009 | 0 | 0.19 | 0.09 | 0.13 | 0.17 | 0.2 | 0.22 |
| 6 | 12357003 | rs4714456 | -1.633 | -0.19 | 0.34 | 0.361 | -0.01 | 0.014 | 0.014 | 0.509 | 0.33 | 0.57 | 0.31 | 0.49 | 0.29 | 0.29 |

| | | | | | | | | | | | | | | | | |
|---|---|---|---|---|---|---|---|---|---|---|---|---|---|---|---|---|
| 6 | 12359302 | rs6925906 | -0.958 | -0.002 | -0.115 | -0.367 | -0.011 | -0.04 | -0.04 | 0 | 0.15 | 0.17 | 0.06 | 0.02 | 0.11 | 0.14 |
| 6 | 12366819 | rs4714469 | 0.26 | 0.208 | -0.344 | -0.5 | 0.011 | 0.017 | 0.017 | 0.025 | 0.56 | 0.61 | 0.65 | 0.88 | 0.63 | 0.74 |
| 6 | 12367517 | rs9381072 | 0.248 | 0.165 | -0.278 | -0.572 | 0.011 | 0.007 | 0.007 | 0.019 | 0.57 | 0.62 | 0.66 | 0.93 | 0.63 | 0.77 |
| 6 | 12368221 | rs9462715 | -0.687 | 0.229 | 0.305 | 0.458 | 0.026 | 0.122 | 0.122 | 0 | 0.08 | 0.17 | 0.13 | 0.21 | 0.05 | 0.21 |
| 6 | 12368551 | rs9394804 | 0.508 | -0.382 | 0.146 | -0.335 | -0.025 | -0.025 | -0.025 | 0 | 0.72 | 0.4 | 0.83 | 0.88 | 0.8 | 0.86 |
| 6 | 12369148 | rs6934829 | -0.896 | 0.557 | 0.207 | 0.404 | 0.01 | 0.025 | 0.025 | 0.115 | 0.38 | 0.47 | 0.39 | 0.51 | 0.39 | 0.41 |
| 6 | 12376259 | rs6458229 | 1.28 | 0.308 | -0.072 | 0.44 | -0.065 | 0.024 | 0.024 | 0.012 | 0.43 | 0.08 | 0.44 | 0.35 | 0.38 | 0.45 |
| 6 | 12376658 | rs4714488 | 0.601 | 0.333 | 0.451 | 0.378 | 0 | 0 | 0 | 0 | 0.6 | 0.57 | 0.62 | 0.47 | 0.51 | 0.61 |
| 6 | 12377393 | rs9462724 | 0.342 | 0.341 | 0.515 | 0.373 | 0.004 | 0.001 | 0.001 | 0 | 0.05 | 0.05 | 0.1 | 0.09 | 0.05 | 0.12 |
| 6 | 12384564 | rs9471621 | 0.503 | 0.333 | 0.561 | 0.367 | 0.002 | 0.001 | 0.001 | 0 | 0.59 | 0.51 | 0.61 | 0.47 | 0.5 | 0.6 |
| 6 | 12384972 | rs10947982 | 1.011 | 0.3 | 0.636 | 0.359 | -0.001 | 0.003 | 0.003 | 0 | 0.52 | 0.44 | 0.51 | 0.37 | 0.45 | 0.47 |
| 6 | 12385692 | rs10947983 | 1.102 | 0.469 | 0.066 | 0.438 | -0.034 | 0.052 | 0.052 | 0.005 | 0.49 | 0.15 | 0.54 | 0.45 | 0.43 | 0.57 |
| 6 | 12388165 | rs994565 | -0.012 | 1.33 | 0.424 | 0.992 | -0.081 | -0.128 | -0.128 | 0 | 0.21 | 0.06 | 0.26 | 0.47 | 0.37 | 0.31 |
| 6 | 12395762 | rs1034237 | 2.097 | 0.845 | 1.18 | 0.748 | 0.005 | 0.008 | 0.008 | 0 | 0.85 | 0.64 | 0.84 | 0.85 | 0.88 | 0.82 |
| 6 | 12396138 | rs16872754 | -1.027 | 0.899 | 1.056 | 0.749 | 0.001 | -0.006 | -0.006 | 0 | 0.08 | 0.13 | 0.1 | 0.08 | 0.05 | 0.16 |
| 6 | 12396682 | rs1334305 | -2.045 | 0.883 | 1.104 | 0.808 | 0 | 0.009 | 0.009 | 0 | 0.14 | 0.28 | 0.16 | 0.18 | 0.12 | 0.19 |
| 6 | 12397466 | rs6922970 | -1.545 | 0.896 | 0.895 | 1.045 | 0 | 0.003 | 0.003 | 0 | 0.11 | 0.09 | 0.15 | 0.24 | 0.14 | 0.19 |
| 6 | 12401910 | rs2050289 | -1.555 | 0.769 | 1.216 | 1.43 | -0.01 | 0.011 | 0.011 | 0.355 | 0.24 | 0.35 | 0.24 | 0.54 | 0.21 | 0.25 |
| 6 | 12403680 | rs9369310 | 1.201 | 0.999 | 1.53 | 1.428 | 0.001 | 0.032 | 0.032 | 1.026 | 0.75 | 0.63 | 0.64 | 0.51 | 0.71 | 0.71 |
| 6 | 12406400 | rs9367114 | 1.508 | 0.372 | 1.516 | 1.056 | -0.007 | 0.026 | 0.026 | 1.245 | 0.47 | 0.31 | 0.43 | 0.48 | 0.4 | 0.54 |
| 6 | 12406520 | rs9471684 | 0.025 | 0.378 | 1.597 | 1.073 | -0.015 | 0.031 | 0.031 | 0 | 0.25 | 0.17 | 0.21 | 0.07 | 0.33 | 0.16 |
| 6 | 12409416 | rs12216318 | 0.263 | 0.398 | 1.543 | 1.054 | -0.037 | 0.062 | 0.062 | 0 | 0.23 | 0.11 | 0.2 | 0.02 | 0.33 | 0.15 |
| 6 | 12417093 | rs756250 | -0.313 | 0.523 | 1.685 | 1.207 | 0.004 | 0.005 | 0.005 | 0 | 0.07 | 0.05 | 0.08 | 0.18 | 0.09 | 0.16 |
| 6 | 12417263 | rs9471712 | -0.152 | 0.494 | 1.657 | 1.178 | -0.027 | 0.072 | 0.072 | 0 | 0.29 | 0.16 | 0.25 | 0.04 | 0.39 | 0.18 |
| 6 | 12423025 | rs7740766 | 1.977 | 0.467 | 1.238 | 1.231 | 0.006 | 0.041 | 0.041 | 0.767 | 0.41 | 0.27 | 0.36 | 0.3 | 0.32 | 0.38 |
| 6 | 12426180 | rs9394861 | 0.879 | 0.487 | 1.177 | 1.612 | -0.009 | 0.047 | 0.047 | 0 | 0.78 | 0.74 | 0.71 | 0.57 | 0.8 | 0.73 |
| 6 | 12427049 | rs9471757 | -0.881 | 0.731 | 0.518 | 1.764 | -0.027 | -0.07 | -0.07 | 0 | 0.21 | 0.07 | 0.25 | 0.48 | 0.27 | 0.35 |
| 6 | 12429371 | rs9381141 | 0.663 | 0.833 | 0.579 | 1.001 | -0.001 | 0.009 | 0.009 | 0 | 0.53 | 0.62 | 0.43 | 0.52 | 0.46 | 0.52 |
| 6 | 12429862 | rs13203590 | 0.911 | 0.817 | 0.615 | 0.986 | 0.005 | 0.013 | 0.013 | 0.129 | 0.49 | 0.56 | 0.39 | 0.37 | 0.39 | 0.4 |
| 6 | 12446475 | rs6901053 | -1.169 | 0.394 | 0.607 | 1.299 | 0 | -0.013 | -0.013 | 0 | 0.23 | 0.2 | 0.31 | 0.53 | 0.22 | 0.3 |
| 6 | 12452295 | rs9471843 | -1.644 | 0.202 | 1.264 | 0.63 | -0.02 | -0.052 | -0.052 | 0 | 0.15 | 0.62 | 0.16 | 0.06 | 0.09 | 0.17 |
| 6 | 12458242 | rs4714604 | -1.585 | 0.413 | 1.234 | 1.155 | 0.001 | 0.058 | 0.058 | 0.109 | 0.15 | 0.51 | 0.17 | 0.24 | 0.12 | 0.22 |
| 6 | 12458796 | rs7760491 | -1.467 | 0.425 | 1.694 | 1.161 | 0.012 | 0.063 | 0.063 | 0.008 | 0.13 | 0.32 | 0.15 | 0.24 | 0.11 | 0.22 |
| 6 | 12463796 | rs9296384 | -0.607 | 0.127 | 1.649 | 1.455 | -0.011 | 0.036 | 0.036 | 2.109 | 0.25 | 0.32 | 0.31 | 0.31 | 0.22 | 0.29 |
| 6 | 12471230 | rs9381187 | -0.321 | 0.091 | 1.71 | 1.845 | -0.001 | -0.001 | -0.001 | 0.507 | 0.23 | 0.2 | 0.3 | 0.22 | 0.21 | 0.27 |
| 6 | 12472256 | rs6458287 | -1.668 | -0.183 | 1.638 | 1.623 | 0.086 | 0.043 | 0.043 | 3.56 | 0.45 | 0.81 | 0.49 | 0.58 | 0.31 | 0.45 |
| 6 | 12478460 | rs9369371 | -0.704 | 0.391 | 2.114 | 1.591 | -0.01 | 0.005 | 0.005 | 0.143 | 0.27 | 0.35 | 0.33 | 0.23 | 0.25 | 0.35 |
| 6 | 12483980 | rs9369375 | -1.795 | 0.051 | 2.192 | 1.562 | 0.058 | 0.038 | 0.038 | 1.913 | 0.46 | 0.77 | 0.49 | 0.57 | 0.37 | 0.47 |
| 6 | 12488729 | rs1999622 | -0.418 | 0.062 | 2.215 | 1.558 | -0.006 | 0.002 | 0.002 | 2.034 | 0.24 | 0.26 | 0.3 | 0.23 | 0.2 | 0.29 |
| 6 | 12494502 | rs9394922 | -1.02 | 0.044 | 2.591 | 1.595 | -0.006 | 0.003 | 0.003 | 1.741 | 0.29 | 0.6 | 0.33 | 0.33 | 0.21 | 0.37 |
| 6 | 12504967 | rs1413513 | -0.65 | -0.113 | 1.996 | 0.991 | -0.024 | -0.013 | -0.013 | 5.872 | 0.38 | 0.4 | 0.4 | 0.29 | 0.28 | 0.39 |
| 6 | 12507307 | rs16872856 | -0.87 | -0.02 | 1.802 | 0.881 | 0.016 | 0.007 | 0.007 | 0.039 | 0.06 | 0.16 | 0.06 | 0.06 | 0.04 | 0.1 |
| 6 | 12507954 | rs9471946 | -0.718 | 0.002 | 1.772 | 0.892 | 0.015 | 0.007 | 0.007 | 0 | 0.05 | 0.12 | 0.05 | 0.06 | 0.04 | 0.1 |
| 6 | 12511030 | rs12199205 | -1.327 | -0.004 | 1.561 | 0.922 | -0.008 | 0.033 | 0.033 | 0 | 0.19 | 0.35 | 0.16 | 0.26 | 0.17 | 0.22 |
| 6 | 12512344 | rs9369394 | -1.549 | -0.052 | 1.034 | 1.156 | 0.03 | 0.012 | 0.012 | 2.124 | 0.53 | 0.69 | 0.53 | 0.59 | 0.41 | 0.45 |
| 6 | 12514092 | rs9349250 | -0.385 | -0.066 | 0.901 | 1.171 | 0.032 | -0.016 | -0.016 | 0 | 0.34 | 0.17 | 0.37 | 0.38 | 0.27 | 0.32 |
| 6 | 12516431 | rs10948062 | -1.02 | 0.549 | 0.646 | 0.963 | 0.002 | 0.007 | 0.007 | 0.505 | 0.55 | 0.67 | 0.54 | 0.6 | 0.48 | 0.49 |
| 6 | 12516585 | rs12200432 | -0.79 | 0.402 | 0.657 | 0.933 | 0.011 | 0.013 | 0.013 | 0 | 0.5 | 0.57 | 0.49 | 0.56 | 0.46 | 0.43 |
| 6 | 12517124 | rs2095343 | 0.908 | 0.992 | 0.637 | 0.535 | 0.029 | -0.018 | -0.018 | 0 | 0.77 | 0.69 | 0.65 | 0.83 | 0.63 | 0.78 |
| 6 | 12520288 | rs6914768 | 1.188 | 1.103 | 0.841 | 0.275 | 0.007 | -0.028 | -0.028 | 0 | 0.72 | 0.4 | 0.57 | 0.84 | 0.63 | 0.73 |
| 6 | 12521039 | rs6920298 | 1.219 | 1.032 | 1.264 | 0.192 | 0.019 | 0.127 | 0.127 | 0.279 | 0.87 | 0.53 | 0.88 | 0.77 | 0.89 | 0.8 |
| 6 | 12521061 | rs9369399 | 1.132 | 0.527 | -0.369 | 0.563 | -0.113 | -0.068 | -0.068 | 0.252 | 0.4 | 0.08 | 0.33 | 0.45 | 0.44 | 0.39 |
| 6 | 12523257 | rs4714657 | 1.193 | 0.303 | -0.134 | 0.716 | 0.076 | -0.077 | -0.077 | 0.018 | 0.23 | 0.07 | 0.17 | 0.28 | 0.12 | 0.17 |
| 6 | 12525129 | rs9472002 | -0.034 | 0.428 | 0.138 | 0.482 | 0.001 | 0.004 | 0.004 | 0 | 0.08 | 0.11 | 0.05 | 0.1 | 0.07 | 0.17 |
| 6 | 12531551 | rs2504899 | -0.108 | 0.407 | 0.536 | 0.608 | -0.023 | 0.003 | 0.003 | 0 | 0.63 | 0.59 | 0.7 | 0.41 | 0.77 | 0.62 |
| 6 | 12532019 | rs9472015 | -1.808 | 1.822 | 0.994 | 0.377 | 0.076 | -0.049 | -0.049 | 0.317 | 0.09 | 0.3 | 0.11 | 0 | 0.12 | 0.06 |
| 6 | 12532323 | rs12665596 | -1.075 | 1.434 | 0.393 | 0.538 | -0.026 | 0.001 | 0.001 | 0.789 | 0.22 | 0.1 | 0.28 | 0.19 | 0.29 | 0.41 |
| 6 | 12536342 | rs2504857 | -2.234 | 1.399 | 1.006 | 0.554 | 0.011 | -0.026 | -0.026 | 0 | 0.24 | 0.56 | 0.3 | 0.15 | 0.23 | 0.33 |
| 6 | 12537326 | rs17698884 | -0.786 | 1.404 | 1.042 | 0.551 | -0.01 | -0.005 | -0.005 | 0.003 | 0.14 | 0.11 | 0.22 | 0.15 | 0.18 | 0.32 |
| 6 | 12540017 | rs9296412 | -1.921 | 1.39 | 1.243 | 0.847 | 0.028 | 0.036 | 0.036 | 0 | 0.21 | 0.35 | 0.28 | 0.25 | 0.24 | 0.35 |
| 6 | 12540827 | rs7745473 | -0.883 | 1.389 | 0.882 | 0.85 | -0.01 | -0.015 | -0.015 | 0.105 | 0.15 | 0.11 | 0.22 | 0.25 | 0.18 | 0.34 |
| 6 | 12544251 | rs6907466 | -0.397 | 1.428 | 0.838 | 0.816 | 0.013 | -0.025 | -0.025 | 3.554 | 0.05 | 0.15 | 0.05 | 0 | 0.04 | 0 |
| 6 | 12544672 | rs2490445 | -2.347 | 1.314 | 0.445 | 0.91 | -0.05 | -0.107 | -0.107 | 0 | 0.32 | 0.92 | 0.46 | 0.39 | 0.29 | 0.39 |
| 6 | 12558084 | rs17766596 | -0.601 | 1.435 | 1.236 | 0.858 | -0.017 | -0.009 | -0.009 | 0 | 0.13 | 0.1 | 0.22 | 0.15 | 0.2 | 0.28 |
| 6 | 12558110 | rs2504890 | -1.656 | 1.359 | 1.462 | 0.885 | -0.006 | 0.091 | 0.091 | 2.144 | 0.09 | 0.34 | 0.06 | 0.12 | 0.02 | 0.02 |
| 6 | 12558421 | rs2327567 | -0.642 | 1.361 | 1.069 | 0.885 | -0.001 | -0.002 | -0.002 | 0 | 0.14 | 0.15 | 0.24 | 0.15 | 0.21 | 0.28 |
| 6 | 12559239 | rs2504892 | -2.111 | 1.381 | 0.956 | 0.723 | 0.001 | 0.009 | 0.009 | 0.329 | 0.24 | 0.63 | 0.31 | 0.27 | 0.24 | 0.3 |
| 6 | 12569231 | rs9369414 | 0.589 | 0.998 | 0.336 | 0.777 | 0.001 | -0.073 | -0.073 | 0 | 0.15 | 0.01 | 0.21 | 0.28 | 0.12 | 0.24 |
| 6 | 12575449 | rs17616713 | 0.616 | 1.299 | 0.324 | -0.045 | -0.06 | 0.07 | 0.07 | 0 | 0.12 | 0.01 | 0.1 | 0 | 0.17 | 0.13 |

| | | | | | | | | | | | | | | | | |
|---|---|---|---|---|---|---|---|---|---|---|---|---|---|---|---|---|
| 6 | 12575720 | rs7770994 | -1.638 | 0.852 | 1.267 | 0.716 | 0.024 | 0.044 | 0.044 | 0 | 0.2 | 0.33 | 0.28 | 0.27 | 0.23 | 0.3 |
| 6 | 12580740 | rs9357418 | -0.024 | 0.989 | 0.545 | 0.864 | -0.048 | -0.144 | -0.144 | 0 | 0.22 | 0.02 | 0.31 | 0.44 | 0.25 | 0.36 |
| 6 | 12581864 | rs7744769 | 1.761 | 1.058 | 1.969 | 0.696 | 0.068 | 0.101 | 0.101 | 0.193 | 0.86 | 0.54 | 0.81 | 0.82 | 0.82 | 0.84 |
| 6 | 12581921 | rs7744797 | -1.321 | 0.319 | 2.253 | 0.019 | -0.005 | 0.005 | 0.005 | 0 | 0.08 | 0.14 | 0.12 | 0.09 | 0.05 | 0.14 |
| 6 | 12595275 | rs6927364 | 1.91 | 0.029 | 0.461 | 0.04 | 0.024 | 0.032 | 0.032 | 5.092 | 0.79 | 0.18 | 0.76 | 0.89 | 0.88 | 0.82 |
| 6 | 12596998 | rs542119 | 1.181 | 0.159 | 0.53 | 0.102 | -0.002 | -0.01 | -0.01 | 0 | 0.88 | 0.82 | 0.84 | 0.91 | 0.9 | 0.85 |
| 6 | 12604081 | rs6905435 | -1.123 | 0.037 | 0.667 | 0.027 | -0.002 | -0.015 | -0.015 | 0 | 0.07 | 0.3 | 0.05 | 0 | 0.01 | 0 |
| 6 | 12604455 | rs2516225 | -1.906 | 0.038 | -0.52 | -0.043 | 0.027 | 0.057 | 0.057 | 3.99 | 0.28 | 0.88 | 0.29 | 0.17 | 0.19 | 0.28 |
| 6 | 12607041 | rs2504914 | -1.692 | -0.009 | -0.494 | 0.041 | 0.031 | 0.033 | 0.033 | 3.468 | 0.25 | 0.91 | 0.27 | 0.18 | 0.14 | 0.29 |
| 6 | 12613231 | rs519167 | 1.27 | 0.129 | -0.348 | -0.378 | 0.009 | 0.039 | 0.039 | 0 | 0.66 | 0.23 | 0.69 | 0.84 | 0.68 | 0.53 |
| 6 | 12616984 | rs9349266 | -0.673 | 0.071 | 0.654 | -0.428 | 0.004 | 0.026 | 0.026 | 0 | 0.11 | 0.2 | 0.14 | 0.15 | 0.1 | 0.23 |
| 6 | 12616996 | rs531602 | 1.615 | 0.528 | -0.682 | -0.312 | -0.088 | -0.048 | -0.048 | 1.018 | 0.42 | 0.02 | 0.33 | 0.35 | 0.44 | 0.28 |
| 6 | 12618970 | rs471603 | 0.33 | 0.34 | 0.34 | -0.584 | -0.003 | -0.109 | -0.109 | 0.258 | 0.8 | 0.66 | 0.84 | 0.99 | 0.79 | 0.76 |
| 6 | 12627866 | rs497757 | 0.586 | 0.493 | 0.062 | -0.54 | 0.011 | 0.054 | 0.054 | 1.137 | 0.69 | 0.21 | 0.72 | 0.91 | 0.71 | 0.8 |
| 6 | 12636430 | rs7754760 | 0.58 | 0.465 | 0.309 | -0.547 | 0.01 | 0.035 | 0.035 | 1.385 | 0.69 | 0.26 | 0.72 | 0.91 | 0.71 | 0.83 |
| 6 | 12637286 | rs472072 | 1.164 | 0.553 | 0.081 | -0.573 | -0.021 | 0.173 | 0.173 | 2.081 | 0.6 | 0.09 | 0.59 | 0.85 | 0.61 | 0.6 |
| 6 | 12645681 | rs12665303 | -1.576 | 0.627 | 0.846 | -0.006 | 0.004 | 0.016 | 0.016 | 0 | 0.25 | 0.38 | 0.35 | 0.67 | 0.22 | 0.6 |
| 6 | 12653770 | rs7768223 | -0.683 | 0.726 | 0.685 | 0.151 | -0.002 | -0.023 | -0.023 | 0.012 | 0.15 | 0.09 | 0.21 | 0.52 | 0.12 | 0.36 |
| 6 | 12661620 | rs9369484 | -0.904 | 0.685 | 1.048 | 0.096 | 0.083 | 0.112 | 0.112 | 0.053 | 0.08 | 0.33 | 0.14 | 0.15 | 0.1 | 0.24 |
| 6 | 12664559 | rs2327588 | -0.956 | 0.685 | 1.045 | 0.095 | 0.076 | 0.104 | 0.104 | 0.026 | 0.09 | 0.33 | 0.14 | 0.15 | 0.1 | 0.25 |
| 6 | 12693454 | rs558134 | 0.809 | 0.59 | -0.062 | -0.762 | -0.017 | 0.027 | 0.027 | 0.549 | 0.39 | 0.1 | 0.37 | 0.32 | 0.42 | 0.27 |
| 6 | 12693886 | rs9357455 | -0.311 | 0.775 | -0.005 | -0.418 | -0.006 | 0.003 | 0.003 | 0 | 0.12 | 0.09 | 0.21 | 0.57 | 0.13 | 0.37 |
| 6 | 12695971 | rs471799 | -0.515 | 0.776 | 0.266 | -0.89 | 0.009 | 0.086 | 0.086 | 2.571 | 0.66 | 0.8 | 0.77 | 0.91 | 0.69 | 0.85 |
| 6 | 12712516 | rs570678 | 1.022 | 0.631 | 0.119 | -0.271 | -0.007 | -0.026 | -0.026 | 0 | 0.43 | 0.11 | 0.38 | 0.39 | 0.45 | 0.34 |
| 6 | 12715593 | rs16873301 | -0.953 | 0.651 | 0.498 | -0.686 | 0.017 | -0.078 | -0.078 | 0 | 0.13 | 0.3 | 0.15 | 0.02 | 0.12 | 0.21 |
| 6 | 12716522 | rs511574 | -0.605 | 0.837 | -0.117 | -0.715 | -0.064 | 0.142 | 0.142 | 2.111 | 0.71 | 0.94 | 0.78 | 0.96 | 0.68 | 0.87 |
| 6 | 12718156 | rs475543 | 0.204 | 0.738 | 0.365 | -0.682 | -0.009 | 0.015 | 0.015 | 0 | 0.58 | 0.38 | 0.57 | 0.88 | 0.57 | 0.81 |
| 6 | 12720399 | rs522923 | -0.683 | 0.794 | -0.161 | -0.686 | -0.067 | 0.133 | 0.133 | 0 | 0.71 | 0.93 | 0.75 | 0.96 | 0.67 | 0.88 |
| 6 | 12722987 | rs560810 | 0.131 | 0.792 | 0.031 | -0.547 | -0.031 | 0.109 | 0.109 | 0 | 0.58 | 0.27 | 0.57 | 0.88 | 0.57 | 0.81 |
| 6 | 12728818 | rs491451 | 0.071 | 0.988 | 0.092 | 0.109 | 0.005 | 0.1 | 0.1 | 0 | 0.09 | 0.08 | 0.24 | 0.55 | 0.12 | 0.29 |
| 6 | 12732095 | rs570295 | -0.525 | 0.833 | -0.042 | -0.035 | -0.031 | 0.11 | 0.11 | 0 | 0.63 | 0.82 | 0.72 | 0.96 | 0.62 | 0.85 |
| 6 | 12734891 | rs9472446 | 0.644 | 1.006 | -0.592 | 0.161 | 0.034 | 0.019 | 0.019 | 0.073 | 0.43 | 0.16 | 0.3 | 0.31 | 0.41 | 0.41 |
| 6 | 12750933 | rs6458439 | -0.297 | 0.945 | -0.365 | -0.624 | 0 | 0.117 | 0.117 | 0.054 | 0.45 | 0.73 | 0.32 | 0.14 | 0.4 | 0.26 |
| 6 | 12768218 | rs12202891 | 0.816 | 1.075 | -0.509 | 0.35 | -0.033 | -0.071 | -0.071 | 0 | 0.13 | 0.01 | 0.29 | 0.34 | 0.16 | 0.17 |
| 6 | 12786272 | rs9472551 | -0.029 | 0.103 | 0.439 | -0.973 | -0.001 | -0.074 | -0.074 | 0.055 | 0.44 | 0.57 | 0.37 | 0.03 | 0.42 | 0.29 |
| 6 | 12789191 | rs6458461 | -0.029 | 0.103 | 0.439 | -0.973 | -0.001 | -0.074 | -0.074 | 0.273 | 0.44 | 0.57 | 0.37 | 0.03 | 0.42 | 0.29 |
| 6 | 12797055 | rs4714870 | 0.197 | 0.093 | 0.475 | -0.976 | -0.007 | -0.137 | -0.137 | 0 | 0.41 | 0.52 | 0.26 | 0.03 | 0.35 | 0.25 |
| 6 | 12800332 | rs17617207 | -0.177 | 0.09 | 0.469 | -0.979 | 0.001 | -0.012 | -0.012 | 2.869 | 0.45 | 0.61 | 0.38 | 0.03 | 0.42 | 0.29 |
| 6 | 12803314 | rs12215445 | 0.462 | -0.136 | -0.028 | -0.849 | -0.019 | -0.094 | -0.094 | 0 | 0.19 | 0.01 | 0.17 | 0.35 | 0.13 | 0.23 |
| 6 | 12804619 | rs1953089 | 0.078 | -0.047 | 0.511 | -1.556 | -0.007 | -0.137 | -0.137 | 0 | 0.41 | 0.52 | 0.26 | 0.04 | 0.35 | 0.25 |
| 6 | 12817608 | rs13215300 | -0.465 | -0.204 | 0.4 | -1.581 | -0.013 | -0.037 | -0.037 | 0.006 | 0.1 | 0.1 | 0.05 | 0 | 0.08 | 0.05 |
| 6 | 12822566 | rs10434854 | -1.264 | -0.293 | 0.822 | -1.53 | -0.016 | 0.008 | 0.008 | 0.64 | 0.16 | 0.3 | 0.19 | 0.17 | 0.15 | 0.17 |
| 6 | 12833077 | rs12528956 | 0.605 | -0.213 | 0.735 | -1.606 | -0.016 | 0.106 | 0.106 | 0 | 0.31 | 0.13 | 0.18 | 0 | 0.26 | 0.14 |
| 6 | 12841306 | rs3945322 | -1.209 | -0.172 | 0.784 | -1.109 | -0.007 | 0.02 | 0.02 | 0.369 | 0.14 | 0.29 | 0.17 | 0.17 | 0.14 | 0.17 |
| 6 | 12859886 | rs9296494 | 0.181 | -0.17 | 0.524 | -1.571 | 0.025 | 0.057 | 0.057 | 0 | 0.55 | 0.35 | 0.73 | 1 | 0.66 | 0.82 |
| 6 | 12864134 | rs9472752 | 0.579 | -0.248 | 0.433 | -1.464 | 0.017 | 0.191 | 0.191 | 0 | 0.5 | 0.21 | 0.61 | 0.82 | 0.59 | 0.69 |
| 6 | 12871736 | rs1360579 | 0.236 | -0.249 | 0.697 | -1.743 | -0.003 | -0.085 | -0.085 | 0.022 | 0.34 | 0.39 | 0.18 | 0 | 0.26 | 0.17 |
| 6 | 12874309 | rs1571997 | 0.729 | 0.017 | 0.761 | -1.799 | -0.011 | 0.17 | 0.17 | 0 | 0.48 | 0.24 | 0.58 | 0.86 | 0.52 | 0.74 |
| 6 | 12875054 | rs1332841 | 0.065 | -0.394 | 0.529 | -1.841 | -0.002 | -0.08 | -0.08 | 0.044 | 0.35 | 0.39 | 0.19 | 0 | 0.26 | 0.17 |
| 6 | 12889004 | rs1332844 | 0.333 | -0.323 | 0.81 | -1.739 | 0.002 | -0.074 | -0.074 | 0 | 0.52 | 0.54 | 0.65 | 0.93 | 0.62 | 0.68 |
| 6 | 12892236 | rs6902050 | 1.076 | -0.544 | 1.005 | -1.88 | -0.022 | -0.062 | -0.062 | 0 | 0.88 | 0.7 | 0.84 | 0.94 | 0.92 | 0.84 |
| 6 | 12898292 | rs13219256 | 0.359 | -0.502 | 0.555 | -1.881 | -0.003 | 0.145 | 0.145 | 0.011 | 0.35 | 0.16 | 0.18 | 0.01 | 0.3 | 0.15 |
| 6 | 12901441 | rs9369640 | 0.39 | -0.571 | 0.868 | -1.852 | 0.033 | 0.099 | 0.099 | 0.088 | 0.52 | 0.29 | 0.63 | 0.97 | 0.62 | 0.81 |
| 6 | 12903957 | rs9349379 | 1.214 | -0.155 | 0.367 | -1.555 | -0.071 | 0.076 | 0.076 | 0.086 | 0.35 | 0.03 | 0.38 | 0.69 | 0.4 | 0.51 |
| 6 | 12909738 | rs6916096 | -1.065 | -0.693 | 1.606 | -1.678 | 0 | -0.004 | -0.004 | 0 | 0.09 | 0.47 | 0.05 | 0.02 | 0.03 | 0.02 |
| 6 | 12909874 | rs7454157 | -0.641 | -0.729 | 1.748 | -1.765 | 0.012 | -0.014 | -0.014 | 0.011 | 0.48 | 0.66 | 0.36 | 0.08 | 0.39 | 0.33 |
| 6 | 12914690 | rs11754198 | -0.652 | -0.667 | 1.756 | -1.594 | 0.002 | 0.004 | 0.004 | 0.124 | 0.07 | 0.35 | 0.03 | 0.02 | 0.03 | 0.02 |
| 6 | 12927845 | rs4714990 | -0.855 | -0.66 | 1.282 | -1.053 | 0.036 | 0.129 | 0.129 | 0.06 | 0.52 | 0.85 | 0.39 | 0.26 | 0.39 | 0.34 |
| 6 | 12927883 | rs12176425 | -1.331 | -0.697 | 1.536 | -1.02 | -0.026 | 0.226 | 0.226 | 3.251 | 0.1 | 0.58 | 0.07 | 0.2 | 0.04 | 0.03 |
| 6 | 12929033 | rs9296526 | -1.578 | -0.683 | 1.513 | -1.012 | -0.045 | 0.173 | 0.173 | 4.849 | 0.11 | 0.65 | 0.08 | 0.2 | 0.04 | 0.03 |
| 6 | 12930061 | rs2210514 | -1.142 | -0.628 | 1.354 | -0.984 | -0.001 | 0.298 | 0.298 | 1.488 | 0.09 | 0.53 | 0.07 | 0.2 | 0.04 | 0.03 |
| 6 | 12934687 | rs7739181 | 0.297 | -0.599 | 0.966 | -1.081 | -0.038 | 0.205 | 0.205 | 0.434 | 0.37 | 0.09 | 0.29 | 0 | 0.34 | 0.18 |
| 6 | 12941017 | rs6941877 | -1.2 | -0.262 | 1.624 | -0.483 | 0.001 | 0.279 | 0.279 | 1.077 | 0.08 | 0.46 | 0.05 | 0.2 | 0.03 | 0.03 |
| 6 | 12948388 | rs6922782 | -0.282 | -0.328 | 1.345 | -1.156 | -0.002 | -0.093 | -0.093 | 0.004 | 0.36 | 0.52 | 0.23 | 0.13 | 0.31 | 0.29 |
| 6 | 12959815 | rs13194360 | -0.134 | -0.672 | 1.305 | -1.261 | -0.046 | -0.08 | -0.08 | 0.297 | 0.26 | 0.47 | 0.13 | 0.11 | 0.17 | 0.23 |
| 6 | 12961672 | rs6458568 | -0.217 | -0.687 | 0.004 | -1.916 | 0.011 | 0.16 | 0.16 | 0 | 0.58 | 0.84 | 0.34 | 0.14 | 0.41 | 0.49 |
| 6 | 12962995 | rs13211739 | 1.147 | -0.771 | -0.041 | -1.985 | 0.023 | 0.128 | 0.128 | 0 | 0.29 | 0.11 | 0.17 | 0.03 | 0.24 | 0.26 |
| 6 | 12963962 | rs4715016 | -0.079 | -0.419 | 0.032 | -1.717 | 0.007 | 0.176 | 0.176 | 0 | 0.58 | 0.86 | 0.34 | 0.14 | 0.41 | 0.49 |
| 6 | 12974653 | rs12203322 | 0.588 | -0.479 | -0.156 | -1.063 | -0.022 | -0.063 | -0.063 | 0 | 0.1 | 0.01 | 0.19 | 0.32 | 0.1 | 0.15 |

| | | | | | | | | | | | | | | | | |
|---|---|---|---|---|---|---|---|---|---|---|---|---|---|---|---|---|
| 6 | 13007933 | rs4349811 | -0.044 | -0.751 | 0.125 | -1.135 | 0.036 | 0.139 | 0.139 | 0 | 0.58 | 0.84 | 0.34 | 0.17 | 0.39 | 0.49 |
| 6 | 13022728 | rs9381634 | 0.977 | -0.17 | 0.097 | -0.71 | -0.044 | -0.119 | -0.119 | 0 | 0.2 | 0.03 | 0.29 | 0.51 | 0.26 | 0.23 |
| 6 | 13023788 | rs4129937 | -0.241 | -0.831 | 0.814 | -1.163 | -0.035 | -0.04 | -0.04 | 2.023 | 0.73 | 0.45 | 0.77 | 0.87 | 0.82 | 0.76 |
| 6 | 13030256 | rs9369762 | -0.147 | -0.794 | 1.192 | -1.177 | -0.042 | -0.059 | -0.059 | 0 | 0.73 | 0.49 | 0.77 | 0.87 | 0.8 | 0.73 |
| 6 | 13033576 | rs9463377 | 0.77 | -0.884 | 1.154 | -1.254 | -0.011 | -0.066 | -0.066 | 0.217 | 0.3 | 0.33 | 0.19 | 0.05 | 0.22 | 0.26 |
| 6 | 13048051 | rs7743585 | 0.484 | -0.952 | 1.534 | -1.238 | -0.037 | -0.058 | -0.058 | 0 | 0.32 | 0.47 | 0.21 | 0.14 | 0.23 | 0.27 |
| 6 | 13055406 | rs9296592 | 0.756 | -0.956 | 1.456 | -1.247 | 0.012 | -0.014 | -0.014 | 0 | 0.3 | 0.3 | 0.19 | 0.05 | 0.22 | 0.27 |
| 6 | 13060568 | rs7738252 | 0.747 | -0.978 | 1.349 | -1.232 | -0.015 | -0.041 | -0.041 | 0 | 0.3 | 0.34 | 0.19 | 0.09 | 0.22 | 0.27 |
| 6 | 13064501 | rs4715077 | -0.707 | -1.004 | 1.163 | -1.414 | 0.005 | 0.015 | 0.015 | 0 | 0.77 | 0.83 | 0.8 | 0.93 | 0.8 | 0.89 |
| 6 | 13078644 | rs9473427 | 0.536 | -1.052 | 1.23 | -1.424 | -0.028 | -0.069 | -0.069 | 0 | 0.36 | 0.59 | 0.22 | 0.11 | 0.22 | 0.43 |
| 6 | 13100821 | rs1937774 | -0.11 | -0.787 | 1.264 | -1.094 | -0.016 | -0.046 | -0.046 | 0.157 | 0.75 | 0.65 | 0.78 | 0.93 | 0.8 | 0.9 |
| 6 | 13109416 | rs11751831 | 0.159 | -0.529 | 0.102 | -0.947 | -0.003 | -0.012 | -0.012 | 1.371 | 0.23 | 0.24 | 0.2 | 0.07 | 0.19 | 0.1 |
| 6 | 13111241 | rs9349459 | 0.854 | -0.115 | -0.284 | -0.52 | -0.065 | -0.123 | -0.123 | 2.578 | 0.2 | 0.03 | 0.35 | 0.5 | 0.31 | 0.23 |
| 6 | 13111431 | rs11755240 | 0.106 | -0.291 | 0.08 | -0.831 | -0.002 | -0.008 | -0.008 | 1.876 | 0.23 | 0.24 | 0.2 | 0.07 | 0.19 | 0.1 |
| 6 | 13116222 | rs6921616 | 0.468 | -0.142 | 0.162 | -0.814 | 0.024 | 0.027 | 0.027 | 0 | 0.33 | 0.24 | 0.21 | 0.11 | 0.25 | 0.51 |
| 6 | 13117477 | rs7740162 | 0.427 | -0.151 | 0.185 | -0.813 | 0.018 | 0.015 | 0.015 | 0 | 0.34 | 0.26 | 0.22 | 0.11 | 0.25 | 0.51 |
| 6 | 13135340 | rs6925345 | 0.379 | 0.182 | -0.348 | -0.095 | -0.148 | -0.065 | -0.065 | 0 | 0.23 | 0.01 | 0.21 | 0.24 | 0.32 | 0.24 |
| 6 | 13137765 | rs10485363 | 0.903 | -0.56 | -0.025 | -0.369 | -0.013 | 0.04 | 0.04 | 0 | 0.15 | 0.07 | 0.17 | 0 | 0.16 | 0.03 |
| 6 | 13137859 | rs1223521 | 0.853 | -0.745 | -0.252 | -0.449 | 0.003 | 0.133 | 0.133 | 0.129 | 0.27 | 0.06 | 0.24 | 0.1 | 0.22 | 0.48 |
| 6 | 13145926 | rs11966433 | -0.147 | -0.726 | 0.502 | -0.27 | -0.026 | -0.047 | -0.047 | 0 | 0.23 | 0.41 | 0.21 | 0.15 | 0.19 | 0.08 |
| 6 | 13149492 | rs9296614 | 0.45 | -0.94 | 0.423 | -0.306 | 0.028 | 0.08 | 0.08 | 1.094 | 0.46 | 0.17 | 0.51 | 0.64 | 0.58 | 0.43 |
| 6 | 13157870 | rs16873638 | -0.151 | -0.947 | 0.41 | -0.325 | -0.03 | -0.049 | -0.049 | 0 | 0.23 | 0.42 | 0.21 | 0.15 | 0.19 | 0.08 |
| 6 | 13161879 | rs1223545 | 0.783 | -1.158 | 0.388 | -0.34 | 0.011 | 0.012 | 0.012 | 0.036 | 0.28 | 0.25 | 0.25 | 0.08 | 0.21 | 0.49 |
| 6 | 13164648 | rs12192359 | 0.07 | -0.958 | 0.708 | -0.492 | 0.002 | -0.054 | -0.054 | 0.012 | 0.29 | 0.32 | 0.3 | 0.1 | 0.31 | 0.09 |
| 6 | 13165689 | rs10948504 | 0.746 | -1.047 | 0.579 | -0.526 | -0.016 | 0.015 | 0.015 | 0 | 0.23 | 0.11 | 0.27 | 0.01 | 0.29 | 0.05 |
| 6 | 13171036 | rs1223549 | 0.91 | -1.08 | 0.423 | -0.361 | 0.019 | 0.027 | 0.027 | 0 | 0.27 | 0.23 | 0.25 | 0.14 | 0.2 | 0.49 |
| 6 | 13171279 | rs10456623 | 1.16 | -0.731 | 0.037 | -0.211 | -0.059 | -0.04 | -0.04 | 0.829 | 0.18 | 0.01 | 0.15 | 0.22 | 0.16 | 0.2 |
| 6 | 13171426 | rs1223550 | 0.698 | -0.725 | 0.052 | -0.208 | -0.034 | -0.021 | -0.021 | 0 | 0.3 | 0.46 | 0.27 | 0.26 | 0.21 | 0.52 |
| 6 | 13173147 | rs9349494 | 1.313 | -0.768 | 0.031 | -0.173 | -0.08 | -0.144 | -0.144 | 0 | 0.21 | 0.03 | 0.15 | 0.42 | 0.31 | 0.15 |
| 6 | 13173833 | rs1223552 | 0.27 | -0.68 | 0.089 | -0.616 | 0.001 | 0.02 | 0.02 | 0 | 0.5 | 0.49 | 0.42 | 0.68 | 0.51 | 0.67 |
| 6 | 13174712 | rs7771293 | 0.241 | -0.634 | 0.084 | -0.631 | 0.002 | -0.114 | -0.114 | 0 | 0.31 | 0.45 | 0.42 | 0.09 | 0.33 | 0.12 |
| 6 | 13180371 | rs1150613 | -0.391 | -0.136 | 0.347 | -0.255 | 0.005 | -0.059 | -0.059 | 0.128 | 0.44 | 0.52 | 0.36 | 0.84 | 0.48 | 0.39 |
| 6 | 13181124 | rs1150611 | 0.709 | 0.06 | -0.731 | -0.175 | -0.005 | -0.002 | -0.002 | 0 | 0.19 | 0.13 | 0.16 | 0.22 | 0.16 | 0.2 |
| 6 | 13182995 | rs3823445 | 0.613 | 0.518 | -0.7 | 0.252 | -0.082 | -0.117 | -0.117 | 0 | 0.21 | 0.04 | 0.16 | 0.54 | 0.31 | 0.19 |
| 6 | 13195458 | rs1150600 | -0.333 | -0.201 | 0.126 | 0.22 | 0.007 | 0.038 | 0.038 | 2.482 | 0.28 | 0.43 | 0.31 | 0.61 | 0.31 | 0.15 |
| 6 | 13201744 | rs10807412 | 1.038 | -0.35 | -2.184 | -1.016 | -0.153 | 0.126 | 0.126 | 6.393 | 0.36 | 0.05 | 0.42 | 0.16 | 0.44 | 0.32 |
| 6 | 13202091 | rs1150594 | -0.502 | 0.326 | -1.835 | -1.261 | 0.038 | 0.119 | 0.119 | 0.339 | 0.72 | 0.91 | 0.76 | 0.95 | 0.77 | 0.52 |
| 6 | 13203570 | rs12193156 | -0.434 | 0.831 | -1.173 | -0.382 | -0.036 | 0.086 | 0.086 | 1.23 | 0.23 | 0.07 | 0.23 | 0 | 0.27 | 0.05 |
| 6 | 13209794 | rs1150616 | 0.265 | 0.894 | -0.73 | 0.55 | -0.001 | 0.002 | 0.002 | 1.303 | 0.53 | 0.32 | 0.53 | 0.57 | 0.58 | 0.45 |
| 6 | 13210958 | rs4715157 | -1.03 | 0.707 | 0.235 | 0.974 | -0.011 | 0.018 | 0.018 | 0 | 0.09 | 0.2 | 0.25 | 0.13 | 0.06 | 0.11 |
| 6 | 13211953 | rs4715158 | -1.546 | 0.555 | 0.794 | 0.94 | -0.02 | 0.022 | 0.022 | 0 | 0.11 | 0.29 | 0.26 | 0.14 | 0.06 | 0.11 |
| 6 | 13216058 | rs4715166 | 0.474 | 0.285 | 0.65 | 0.766 | 0.003 | 0.003 | 0.003 | 0.243 | 0.64 | 0.35 | 0.7 | 0.29 | 0.7 | 0.46 |
| 6 | 13223996 | rs12205644 | 0.786 | 0.209 | 0.609 | 0.752 | 0.011 | 0.018 | 0.018 | 0.498 | 0.61 | 0.32 | 0.69 | 0.27 | 0.69 | 0.44 |
| 6 | 13225077 | rs6458720 | 0.351 | -0.094 | 1.397 | 0.626 | -0.003 | 0 | 0 | 0.258 | 0.69 | 0.64 | 0.75 | 0.28 | 0.73 | 0.5 |
| 6 | 13226792 | rs1232381 | 0.635 | 0.051 | 1.544 | 1.465 | 0.005 | 0.126 | 0.126 | 0 | 0.94 | 0.95 | 0.92 | 0.58 | 0.9 | 0.88 |
| 6 | 13231980 | rs12660341 | -0.496 | -0.078 | 1.74 | 1.297 | -0.009 | 0.017 | 0.017 | 0 | 0.09 | 0.21 | 0.26 | 0.13 | 0.06 | 0.1 |
| 6 | 13238239 | rs384366 | -0.96 | -0.169 | 2.109 | 1.266 | -0.005 | 0.002 | 0.002 | 0.786 | 0.31 | 0.37 | 0.43 | 0.43 | 0.23 | 0.49 |
| 6 | 13240272 | rs202062 | -0.75 | -0.193 | 2.339 | 1.459 | -0.005 | 0.036 | 0.036 | 0 | 0.09 | 0.28 | 0.27 | 0.13 | 0.06 | 0.1 |
| 6 | 13261256 | rs393032 | -1.467 | -0.191 | 2.557 | 1.426 | -0.012 | -0.02 | -0.02 | 0 | 0.14 | 0.25 | 0.29 | 0.1 | 0.09 | 0.16 |
| 6 | 13262589 | rs426518 | -1.767 | -0.187 | 2.555 | 1.44 | -0.023 | -0.015 | -0.015 | 0 | 0.16 | 0.48 | 0.31 | 0.12 | 0.1 | 0.16 |
| 6 | 13264026 | rs202064 | 1.024 | 0.115 | 1.771 | 1.788 | 0.004 | 0.102 | 0.102 | 0 | 0.93 | 0.94 | 0.9 | 0.45 | 0.89 | 0.87 |
| 6 | 13267681 | rs366904 | 1.803 | 0.027 | 2.568 | 1.764 | 0.064 | 0.509 | 0.509 | 0 | 0.9 | 0.59 | 0.86 | 0.53 | 0.91 | 0.87 |
| 6 | 13268211 | rs202072 | 2.272 | -0.082 | 2.734 | 1.653 | 0.009 | 0.19 | 0.19 | 0 | 0.84 | 0.46 | 0.83 | 0.52 | 0.86 | 0.81 |
| 6 | 13273318 | rs202054 | 1.464 | -0.338 | 2.514 | 1.565 | 0.003 | 0.002 | 0.002 | 0.003 | 0.62 | 0.41 | 0.7 | 0.32 | 0.67 | 0.47 |
| 6 | 13274123 | rs202051 | 1.611 | -0.28 | 2.648 | 1.809 | -0.003 | 0.081 | 0.081 | 0 | 0.9 | 0.87 | 0.88 | 0.64 | 0.9 | 0.83 |
| 6 | 13275950 | rs202044 | 2.769 | -0.328 | 2.984 | 1.708 | 0.003 | 0.235 | 0.235 | 0 | 0.81 | 0.51 | 0.82 | 0.61 | 0.84 | 0.78 |
| 6 | 13276415 | rs1512414 | 2.894 | -0.396 | 2.844 | 1.643 | -0.007 | 0.036 | 0.036 | 0.085 | 0.73 | 0.37 | 0.55 | 0.57 | 0.78 | 0.67 |
| 6 | 13280641 | rs413120 | 2.515 | -0.398 | 2.849 | 1.69 | 0.007 | 0.194 | 0.194 | 0 | 0.83 | 0.64 | 0.83 | 0.58 | 0.84 | 0.77 |
| 6 | 13284204 | rs435466 | -2.653 | -0.467 | 2.684 | 1.743 | -0.044 | -0.041 | -0.041 | 0 | 0.19 | 0.65 | 0.33 | 0.16 | 0.11 | 0.18 |
| 6 | 13287400 | rs693758 | -2.404 | -0.467 | 2.706 | 1.748 | -0.043 | 0.022 | 0.022 | 0.33 | 0.11 | 0.62 | 0.06 | 0.14 | 0.03 | 0.07 |
| 6 | 13288850 | rs3734878 | 0.061 | -0.523 | 2.357 | 1.709 | 0.012 | -0.008 | -0.008 | 0 | 0.2 | 0.03 | 0.13 | 0.21 | 0.18 | 0.3 |
| 6 | 13292076 | rs202032 | -0.686 | -0.464 | 2.413 | 1.759 | 0.016 | 0.046 | 0.046 | 0.239 | 0.29 | 0.07 | 0.39 | 0.23 | 0.25 | 0.4 |
| 6 | 13294866 | rs6910418 | 1.944 | -0.533 | 2.512 | 1.935 | 0.062 | 0.126 | 0.126 | 0.234 | 0.51 | 0.22 | 0.41 | 0.26 | 0.59 | 0.34 |
| 6 | 13295502 | rs4602719 | 1.437 | -0.817 | 2.435 | 2.093 | -0.005 | 0.042 | 0.042 | 0 | 0.91 | 0.94 | 0.89 | 0.67 | 0.88 | 0.83 |
| 6 | 13295515 | rs1550526 | 1.582 | -0.819 | 2.445 | 2.092 | -0.001 | 0.055 | 0.055 | 0.01 | 0.91 | 0.92 | 0.89 | 0.67 | 0.88 | 0.84 |
| 6 | 13295536 | rs454563 | 1.437 | -0.817 | 2.438 | 2.093 | -0.005 | 0.042 | 0.042 | 0 | 0.91 | 0.94 | 0.89 | 0.66 | 0.88 | 0.83 |
| 6 | 13296206 | rs7774863 | -1.842 | -0.845 | 2.493 | 1.981 | 0.044 | 0.045 | 0.045 | 0.093 | 0.46 | 0.69 | 0.55 | 0.69 | 0.38 | 0.64 |
| 6 | 13296310 | rs449727 | -0.928 | -0.865 | 2.781 | 1.985 | -0.035 | -0.028 | -0.028 | 0.922 | 0.33 | 0.42 | 0.44 | 0.32 | 0.26 | 0.48 |
| 6 | 13296439 | rs402705 | -0.887 | -0.864 | 2.867 | 1.985 | -0.018 | -0.014 | -0.014 | 0 | 0.33 | 0.37 | 0.43 | 0.32 | 0.26 | 0.48 |

| | | | | | | | | | | | | | | | | |
|---|---|---|---|---|---|---|---|---|---|---|---|---|---|---|---|---|
| 6 | 13298011 | rs9296641 | -2.041 | -0.825 | 2.494 | 2.014 | 0.005 | 0.149 | 0.149 | 0 | 0.13 | 0.25 | 0.12 | 0.37 | 0.12 | 0.17 |
| 6 | 13304658 | rs202023 | -1.013 | -1.08 | 2.01 | 1.551 | 0.025 | 0.021 | 0.021 | 0.459 | 0.33 | 0.25 | 0.42 | 0.32 | 0.26 | 0.48 |
| 6 | 13311823 | rs2439542 | -1.16 | -1.067 | 1.966 | 1.559 | -0.006 | -0.004 | -0.004 | 0 | 0.35 | 0.35 | 0.43 | 0.32 | 0.26 | 0.47 |
| 6 | 13317084 | rs480122 | -1.06 | -1.2 | 1.505 | 2.458 | 0.014 | -0.024 | -0.024 | 0 | 0.26 | 0.16 | 0.25 | 0.42 | 0.2 | 0.41 |
| 6 | 13317408 | rs2496130 | -1.365 | -1.422 | 2.166 | 1.967 | -0.014 | -0.031 | -0.031 | 0.658 | 0.4 | 0.4 | 0.36 | 0.22 | 0.25 | 0.43 |
| 6 | 13318030 | rs3823441 | -1.384 | -1.297 | 2.082 | 2.47 | -0.006 | 0.055 | 0.055 | 0 | 0.17 | 0.22 | 0.1 | 0.4 | 0.13 | 0.2 |
| 6 | 13318609 | rs555100 | -0.413 | -1.455 | 2.038 | 2.185 | 0.026 | 0.061 | 0.061 | 1.519 | 0.28 | 0.16 | 0.32 | 0.12 | 0.22 | 0.41 |
| 6 | 13321097 | rs2496132 | -0.547 | -1.468 | 2.243 | 2.178 | 0 | -0.008 | -0.008 | 0.104 | 0.3 | 0.3 | 0.33 | 0.12 | 0.22 | 0.41 |
| 6 | 13323895 | rs2458304 | -1.744 | -1.357 | 2.302 | 2.363 | 0.077 | 0.032 | 0.032 | 0.061 | 0.51 | 0.74 | 0.46 | 0.6 | 0.36 | 0.61 |
| 6 | 13331097 | rs1512411 | -1.16 | -1.287 | 2.182 | 2.262 | -0.007 | 0.004 | 0.004 | 2.495 | 0.39 | 0.39 | 0.41 | 0.4 | 0.33 | 0.57 |
| 6 | 13332467 | rs499818 | -0.371 | -1.234 | 1.76 | 2.113 | 0.021 | 0.103 | 0.103 | 0.484 | 0.29 | 0.1 | 0.33 | 0.12 | 0.24 | 0.41 |
| 6 | 13336815 | rs1866863 | -1.411 | -0.937 | 1.952 | 2.034 | 0.007 | 0.012 | 0.012 | 0.317 | 0.49 | 0.49 | 0.54 | 0.7 | 0.39 | 0.61 |
| 6 | 13338569 | rs573668 | -0.204 | -0.968 | 1.852 | 1.99 | 0.012 | -0.005 | -0.005 | 0 | 0.32 | 0.17 | 0.44 | 0.31 | 0.28 | 0.43 |
| 6 | 13340967 | rs9370020 | -0.251 | -0.942 | 1.706 | 2.005 | 0.006 | -0.001 | -0.001 | 0 | 0.32 | 0.22 | 0.45 | 0.31 | 0.28 | 0.43 |
| 6 | 13342122 | rs2496167 | -1.062 | -0.958 | 1.843 | 2.014 | 0.005 | 0.002 | 0.002 | 0 | 0.43 | 0.54 | 0.49 | 0.4 | 0.31 | 0.46 |
| 6 | 13342557 | rs539629 | -0.854 | -0.897 | 1.436 | 2.048 | 0 | 0 | 0 | 0 | 0.42 | 0.35 | 0.47 | 0.4 | 0.31 | 0.46 |
| 6 | 13343870 | rs3001963 | -1.51 | -0.852 | 1.434 | 2.175 | 0.068 | 0.28 | 0.28 | 0.603 | 0.1 | 0.41 | 0.09 | 0.28 | 0.09 | 0.17 |
| 6 | 13344616 | rs569038 | -1.596 | -0.852 | 1.392 | 2.184 | 0.065 | 0.283 | 0.283 | 0.775 | 0.1 | 0.43 | 0.09 | 0.28 | 0.09 | 0.17 |
| 6 | 13346701 | rs2254292 | -0.963 | -0.794 | 0.355 | 2.207 | -0.007 | -0.001 | -0.001 | 0.868 | 0.43 | 0.53 | 0.49 | 0.4 | 0.31 | 0.45 |
| 6 | 13346867 | rs2292477 | -1.291 | -0.421 | 0.041 | 2.577 | 0.022 | 0.148 | 0.148 | 0.056 | 0.07 | 0.14 | 0.07 | 0.27 | 0.09 | 0.17 |
| 6 | 13349274 | rs6919374 | -0.624 | -0.436 | 0.035 | 2.536 | 0.002 | 0.002 | 0.002 | 0 | 0.09 | 0.07 | 0.03 | 0.09 | 0.02 | 0.03 |
| 6 | 13350427 | rs6926971 | -1.471 | -0.396 | 0.009 | 2.461 | -0.014 | 0.072 | 0.072 | 0 | 0.16 | 0.21 | 0.09 | 0.37 | 0.11 | 0.19 |
| 6 | 13353734 | rs9381968 | -1.451 | 0.11 | -0.252 | 1.968 | -0.001 | 0.049 | 0.049 | 0.213 | 0.12 | 0.15 | 0.15 | 0.51 | 0.12 | 0.17 |
| 6 | 13355485 | rs497505 | 0.383 | 0.313 | -0.984 | 0.475 | -0.06 | 0.075 | 0.075 | 0.871 | 0.53 | 0.1 | 0.62 | 0.65 | 0.56 | 0.49 |
| 6 | 13361927 | rs2496160 | 0.232 | 0.263 | 0.162 | 0.249 | -0.009 | 0.027 | 0.027 | 0 | 0.57 | 0.37 | 0.65 | 0.7 | 0.55 | 0.56 |
| 6 | 13373569 | rs517803 | 0.321 | 0.224 | 0.214 | 0.264 | 0.003 | -0.017 | -0.017 | 0.681 | 0.57 | 0.54 | 0.65 | 0.68 | 0.53 | 0.53 |
| 6 | 13384746 | rs531256 | 0.229 | 0.547 | -0.451 | 0.425 | 0.02 | -0.032 | -0.032 | 0 | 0.6 | 0.56 | 0.67 | 0.71 | 0.55 | 0.56 |
| 6 | 13391155 | rs821292 | 0.296 | 0.497 | -1.274 | 0.685 | -0.035 | -0.04 | -0.04 | 0.052 | 0.42 | 0.19 | 0.61 | 0.56 | 0.52 | 0.54 |
| 6 | 13391347 | rs17700842 | -0.444 | -0.257 | -1.791 | 0.67 | -0.031 | -0.091 | -0.091 | 0.34 | 0.29 | 0.04 | 0.2 | 0.3 | 0.28 | 0.19 |
| 6 | 13395558 | rs536916 | 0.77 | -0.779 | -1.036 | 0.448 | 0.042 | 0.014 | 0.014 | 0 | 0.57 | 0.39 | 0.75 | 0.63 | 0.71 | 0.76 |
| 6 | 13400667 | rs523221 | 0.397 | -1.19 | -0.506 | 0.233 | -0.009 | 0.06 | 0.06 | 0 | 0.74 | 0.53 | 0.82 | 0.65 | 0.77 | 0.87 |
| 6 | 13421241 | rs16874142 | -0.593 | -1.186 | -0.358 | 0.019 | -0.009 | -0.013 | -0.013 | 0 | 0.17 | 0.13 | 0.07 | 0.03 | 0.05 | 0.11 |
| 6 | 13425532 | rs9296672 | -0.621 | -0.99 | -0.025 | 0.314 | -0.001 | 0.002 | 0.002 | 0 | 0.24 | 0.25 | 0.16 | 0.31 | 0.22 | 0.13 |
| 6 | 13441424 | rs4715295 | 1.226 | -0.342 | -0.844 | 0.219 | 0.011 | 0.021 | 0.021 | 0 | 0.55 | 0.13 | 0.7 | 0.61 | 0.63 | 0.71 |
| 6 | 13444738 | rs9395787 | 0.68 | 0.064 | -0.056 | 0.473 | 0.025 | 0.021 | 0.021 | 0 | 0.74 | 0.6 | 0.79 | 0.72 | 0.69 | 0.83 |
| 6 | 13451226 | rs551499 | 0.861 | 0.507 | -1.312 | 0.563 | -0.105 | -0.107 | -0.107 | 0 | 0.58 | 0.1 | 0.55 | 0.59 | 0.57 | 0.57 |
| 6 | 13452404 | rs13213519 | -0.098 | 0.621 | -0.862 | 0.652 | -0.003 | 0.002 | 0.002 | 0 | 0.19 | 0.19 | 0.25 | 0.19 | 0.18 | 0.28 |
| 6 | 13455437 | rs2434396 | 1.901 | 0.139 | -1.993 | -0.003 | -0.118 | -0.148 | -0.148 | 0 | 0.41 | 0.06 | 0.47 | 0.51 | 0.44 | 0.45 |
| 6 | 13456354 | rs2841565 | 0.078 | 0.218 | -1.347 | 0.137 | 0.016 | 0.005 | 0.005 | 0 | 0.84 | 0.82 | 0.85 | 0.82 | 0.75 | 0.85 |
| 6 | 13464827 | rs13196546 | 0.332 | 0.309 | -1.505 | 0.264 | -0.002 | -0.046 | -0.046 | 0 | 0.18 | 0.08 | 0.24 | 0.24 | 0.18 | 0.24 |
| 6 | 13466830 | rs2841555 | 0.971 | 0.456 | -1.568 | 0.193 | -0.083 | -0.151 | -0.151 | 1.604 | 0.61 | 0.08 | 0.57 | 0.55 | 0.63 | 0.57 |
| 6 | 13485083 | rs2115063 | 0.891 | -0.4 | -1.481 | 0.035 | 0.083 | 0.027 | 0.027 | 0 | 0.63 | 0.16 | 0.77 | 0.73 | 0.78 | 0.76 |
| 6 | 13489400 | rs2059804 | -0.101 | -0.375 | -1.383 | -0.075 | 0.048 | 0.066 | 0.066 | 0 | 0.21 | 0.07 | 0.1 | 0.1 | 0.13 | 0.12 |
| 6 | 13490904 | rs2494774 | 1.389 | -0.125 | -1.618 | 0.069 | -0.056 | -0.119 | -0.119 | 1.029 | 0.43 | 0.08 | 0.61 | 0.54 | 0.61 | 0.48 |
| 6 | 13506872 | rs2560790 | 0.677 | -0.633 | -0.366 | -0.34 | -0.045 | -0.04 | -0.04 | 1.123 | 0.71 | 0.49 | 0.8 | 0.79 | 0.81 | 0.82 |
| 6 | 13519372 | rs2560776 | 1.138 | -0.65 | -0.402 | -0.458 | 0.009 | 0.008 | 0.008 | 0 | 0.58 | 0.43 | 0.75 | 0.78 | 0.73 | 0.73 |
| 6 | 13524036 | rs9474418 | -0.894 | -0.126 | -0.436 | -0.212 | -0.023 | -0.06 | -0.06 | 1.742 | 0.15 | 0.19 | 0.07 | 0.01 | 0.08 | 0.08 |
| 6 | 13554043 | rs4712032 | -0.244 | 0.228 | -1.321 | -0.202 | 0.001 | 0.074 | 0.074 | 0 | 0.38 | 0.19 | 0.4 | 0.26 | 0.36 | 0.35 |
| 6 | 13561042 | rs12216101 | -0.016 | 0.208 | -1.453 | -0.103 | -0.013 | 0.018 | 0.018 | 0.337 | 0.33 | 0.06 | 0.22 | 0.27 | 0.31 | 0.28 |
| 6 | 13568642 | rs9382221 | 0.2 | 0.493 | -0.821 | 0.158 | 0.025 | -0.002 | -0.002 | 0 | 0.34 | 0.46 | 0.29 | 0.64 | 0.37 | 0.35 |
| 6 | 13572686 | rs2804922 | 0.09 | 0.152 | -0.918 | -0.222 | -0.001 | -0.012 | -0.012 | 0.505 | 0.3 | 0.36 | 0.32 | 0.12 | 0.3 | 0.34 |
| 6 | 13576708 | rs2328676 | 0.229 | 0.03 | -0.769 | -0.285 | -0.002 | 0.022 | 0.022 | 0 | 0.28 | 0.22 | 0.31 | 0.12 | 0.3 | 0.34 |
| 6 | 13584583 | rs2804918 | -0.747 | 0.189 | -0.605 | -0.272 | 0.044 | 0 | 0 | 0 | 0.67 | 0.57 | 0.71 | 0.36 | 0.61 | 0.65 |
| 6 | 13587980 | rs2841512 | -0.806 | 0.138 | -0.49 | -0.731 | 0 | -0.012 | -0.012 | 0 | 0.88 | 0.92 | 0.93 | 1 | 0.89 | 0.96 |
| 6 | 13590185 | rs4712047 | 0.359 | 0.369 | -0.778 | -0.237 | 0.015 | -0.001 | -0.001 | 1.556 | 0.51 | 0.61 | 0.39 | 0.54 | 0.41 | 0.47 |
| 6 | 13594666 | rs2804921 | 1.033 | 0.489 | -0.844 | -0.323 | 0.001 | -0.019 | -0.019 | 0 | 0.2 | 0.19 | 0.25 | 0.06 | 0.26 | 0.18 |
| 6 | 13598122 | rs7759452 | 1.372 | 0.231 | -1.138 | -0.363 | -0.024 | 0.064 | 0.064 | 0.494 | 0.3 | 0.05 | 0.18 | 0.21 | 0.26 | 0.23 |
| 6 | 13599303 | rs3757261 | 1.384 | 0.351 | -1.173 | -0.402 | -0.007 | 0.061 | 0.061 | 0.583 | 0.3 | 0.05 | 0.18 | 0.21 | 0.24 | 0.23 |
| 6 | 13601528 | rs2841514 | 0.198 | 0.258 | -0.845 | -0.456 | 0.002 | -0.012 | -0.012 | 0 | 0.6 | 0.46 | 0.63 | 0.26 | 0.58 | 0.48 |
| 6 | 13607990 | rs2841515 | 0.298 | 0.243 | -0.825 | -0.464 | -0.009 | 0.026 | 0.026 | 0 | 0.59 | 0.34 | 0.66 | 0.27 | 0.58 | 0.48 |
| 6 | 13610188 | rs2841517 | 0.201 | 0.238 | -0.827 | -0.466 | 0.001 | -0.009 | -0.009 | 0 | 0.6 | 0.45 | 0.67 | 0.27 | 0.58 | 0.48 |
| 6 | 13613783 | rs2841523 | 0.29 | 0.235 | -0.851 | -0.382 | -0.007 | 0.02 | 0.02 | 0 | 0.59 | 0.35 | 0.63 | 0.3 | 0.58 | 0.49 |
| 6 | 13623324 | rs12209803 | 0.354 | 0.232 | -0.802 | 0.072 | -0.007 | 0.016 | 0.016 | 0 | 0.58 | 0.32 | 0.65 | 0.37 | 0.59 | 0.48 |
| 6 | 13625413 | rs10948800 | 1.661 | -0.223 | -1.289 | -0.328 | -0.034 | -0.05 | -0.05 | 0.181 | 0.2 | 0.03 | 0.12 | 0.3 | 0.19 | 0.18 |
| 6 | 13627151 | rs9382303 | 1.784 | 0.282 | -1.318 | 0.709 | -0.031 | 0.028 | 0.028 | 0 | 0.06 | 0 | 0.1 | 0.4 | 0.1 | 0.05 |
| 6 | 13628357 | rs7742342 | -0.646 | 0.323 | -1.041 | 0.629 | -0.004 | -0.005 | -0.005 | 0 | 0.82 | 0.87 | 0.81 | 0.58 | 0.78 | 0.9 |
| 6 | 13630864 | rs10807467 | 0.308 | 0.259 | -0.997 | 0.474 | 0.003 | -0.016 | -0.016 | 0 | 0.59 | 0.46 | 0.64 | 0.28 | 0.58 | 0.49 |
| 6 | 13634982 | rs707816 | -0.272 | 0.923 | -0.094 | 0.896 | 0.018 | -0.006 | -0.006 | 0 | 0.34 | 0.52 | 0.55 | 0.29 | 0.38 | 0.41 |
| 6 | 13638451 | rs2071835 | 1.726 | 1.151 | -0.302 | 1.133 | -0.036 | 0.001 | 0.001 | 0 | 0.06 | 0.01 | 0.1 | 0.32 | 0.12 | 0.06 |

| | | | | | | | | | | | | | | | | |
|---|---|---|---|---|---|---|---|---|---|---|---|---|---|---|---|---|
| 6 | 13638984 | rs4715451 | -1.247 | 0.557 | -0.048 | 0.649 | -0.031 | -0.062 | -0.062 | 0 | 0.63 | 0.86 | 0.72 | 0.46 | 0.6 | 0.7 |
| 6 | 13643604 | rs10484251 | 1.175 | 0.506 | -0.147 | -0.208 | -0.004 | 0.031 | 0.031 | 0 | 0.25 | 0.12 | 0.27 | 0.13 | 0.27 | 0.24 |
| 6 | 13648864 | rs16874275 | 0.608 | 0.403 | -0.114 | -0.189 | 0.002 | -0.002 | -0.002 | 0 | 0.07 | 0.06 | 0.06 | 0.06 | 0.04 | 0.14 |
| 6 | 13657040 | rs3846912 | 0.026 | 0.782 | 0.046 | -0.476 | -0.006 | -0.035 | -0.035 | 0 | 0.4 | 0.41 | 0.53 | 0.16 | 0.38 | 0.3 |
| 6 | 13659726 | rs204231 | 0.971 | 0.842 | 0.015 | -0.418 | 0.04 | 0.056 | 0.056 | 0 | 0.15 | 0.29 | 0.13 | 0.23 | 0.18 | 0.26 |
| 6 | 13661270 | rs2299027 | 0.892 | 0.524 | -0.232 | -0.91 | -0.001 | 0.022 | 0.022 | 0 | 0.25 | 0.14 | 0.27 | 0.13 | 0.27 | 0.24 |
| 6 | 13685551 | rs11965800 | 1.242 | 0.519 | -0.381 | -0.908 | 0.019 | 0.046 | 0.046 | 0.067 | 0.11 | 0.03 | 0.06 | 0.03 | 0.07 | 0.06 |
| 6 | 13700419 | rs11759518 | 0.995 | 0.623 | -0.323 | -0.835 | -0.01 | 0.05 | 0.05 | 0 | 0.24 | 0.08 | 0.27 | 0.13 | 0.27 | 0.22 |
| 6 | 13709180 | rs2237092 | -0.045 | 0.559 | -0.001 | -0.512 | -0.013 | -0.034 | -0.034 | 0.021 | 0.41 | 0.48 | 0.57 | 0.25 | 0.4 | 0.32 |
| 6 | 13715303 | rs24023 | -0.843 | 0.762 | -0.26 | -0.28 | -0.007 | 0.013 | 0.013 | 0 | 0.55 | 0.69 | 0.55 | 0.39 | 0.58 | 0.58 |
| 6 | 13719673 | rs412970 | -0.169 | 0.783 | -0.442 | -0.288 | -0.002 | 0.007 | 0.007 | 0 | 0.45 | 0.6 | 0.46 | 0.28 | 0.49 | 0.51 |
| 6 | 13727788 | rs6905662 | 1.257 | 0.894 | -0.476 | -0.089 | 0.017 | 0.008 | 0.008 | 0 | 0.1 | 0.03 | 0.09 | 0.11 | 0.08 | 0.08 |
| 6 | 13733565 | rs204236 | 0.615 | 1.458 | -0.062 | 0.027 | 0.038 | 0.023 | 0.023 | 0 | 0.15 | 0.28 | 0.13 | 0.16 | 0.2 | 0.26 |
| 6 | 13739490 | rs10456703 | -0.018 | 1.461 | -0.249 | 0.034 | 0.041 | 0.018 | 0.018 | 1.895 | 0.19 | 0.41 | 0.17 | 0.16 | 0.23 | 0.26 |
| 6 | 13741615 | rs4712100 | 0.039 | 0.983 | -0.838 | -0.06 | 0.008 | 0.047 | 0.047 | 0 | 0.29 | 0.14 | 0.29 | 0.13 | 0.28 | 0.28 |
| 6 | 13755516 | rs1573701 | 0.238 | 1.403 | -0.606 | 0.244 | 0.009 | -0.007 | -0.007 | 0 | 0.21 | 0.26 | 0.16 | 0.14 | 0.23 | 0.24 |
| 6 | 13755585 | rs1990665 | 0.27 | 0.517 | -0.682 | -0.046 | -0.016 | 0.046 | 0.046 | 0.037 | 0.24 | 0.04 | 0.26 | 0.15 | 0.21 | 0.22 |
| 6 | 13763788 | rs4512225 | 0.296 | 0.144 | 0.328 | 0.488 | -0.001 | 0.001 | 0.001 | 0 | 0.36 | 0.49 | 0.31 | 0.43 | 0.32 | 0.47 |
| 6 | 13769838 | rs9396115 | -0.452 | 0.129 | 0.347 | 0.549 | 0.001 | 0.001 | 0.001 | 0 | 0.54 | 0.5 | 0.49 | 0.52 | 0.52 | 0.57 |
| 6 | 13777209 | rs4715562 | -0.686 | 0.035 | 0.38 | 0.577 | -0.007 | 0.02 | 0.02 | 0 | 0.3 | 0.4 | 0.27 | 0.36 | 0.33 | 0.29 |
| 6 | 13780517 | rs9396146 | 0.799 | 0.102 | 0.194 | 0.623 | 0.007 | -0.013 | -0.013 | 0 | 0.31 | 0.21 | 0.26 | 0.39 | 0.24 | 0.44 |
| 6 | 13782620 | rs1062066 | -0.474 | -0.278 | 0.209 | 0.445 | -0.003 | -0.035 | -0.035 | 0 | 0.27 | 0.15 | 0.25 | 0.36 | 0.33 | 0.27 |
| 6 | 13787578 | rs3734672 | 1.109 | -0.15 | 0.123 | 0.374 | -0.022 | 0.033 | 0.033 | 0 | 0.25 | 0.1 | 0.23 | 0.15 | 0.21 | 0.3 |
| 6 | 13789235 | rs10223600 | -0.186 | -0.462 | 0.15 | 0.318 | 0.008 | 0.034 | 0.034 | 0 | 0.13 | 0.07 | 0.11 | 0 | 0.13 | 0.07 |
| 6 | 13792600 | rs6926691 | -1.242 | -0.336 | 0.99 | 0.799 | 0.008 | 0.056 | 0.056 | 0 | 0.23 | 0.38 | 0.21 | 0.38 | 0.29 | 0.25 |
| 6 | 13794110 | rs4715568 | -0.511 | -0.153 | 0.601 | 0.736 | -0.006 | -0.033 | -0.033 | 0 | 0.59 | 0.73 | 0.54 | 0.53 | 0.6 | 0.62 |
| 6 | 13794521 | rs3823246 | -0.271 | -0.095 | 0.68 | 1.17 | -0.001 | 0.012 | 0.012 | 0 | 0.59 | 0.36 | 0.54 | 0.69 | 0.65 | 0.65 |
| 6 | 13801615 | rs3180196 | -0.235 | 0.365 | 0.413 | 0.905 | -0.05 | -0.049 | -0.049 | 1.009 | 0.08 | 0.01 | 0.09 | 0.3 | 0.18 | 0.07 |
| 6 | 13805189 | rs9396165 | 0.542 | -0.488 | 0.685 | 0.646 | -0.057 | -0.005 | -0.005 | 0 | 0.46 | 0.13 | 0.39 | 0.46 | 0.49 | 0.55 |
| 6 | 13824087 | rs1204156 | -0.152 | -0.033 | 0.552 | 0.708 | -0.003 | 0.002 | 0.002 | 0.091 | 0.61 | 0.69 | 0.5 | 0.71 | 0.59 | 0.68 |
| 6 | 13827708 | rs13202002 | -0.525 | 0.183 | -0.179 | 0.726 | -0.019 | 0.035 | 0.035 | 0.015 | 0.15 | 0.37 | 0.1 | 0.21 | 0.11 | 0.19 |
| 6 | 13830987 | rs1204159 | 0.205 | 0.656 | -0.876 | 0.535 | -0.029 | -0.014 | -0.014 | 0.215 | 0.62 | 0.87 | 0.53 | 0.7 | 0.59 | 0.68 |
| 6 | 13836166 | rs1204166 | -1.052 | 0.295 | 0.194 | 0.173 | -0.032 | -0.1 | -0.1 | 0 | 0.14 | 0.4 | 0.1 | 0 | 0.11 | 0.06 |
| 6 | 13836672 | rs1204168 | -0.963 | 0.946 | -0.39 | 0.432 | 0.001 | 0.005 | 0.005 | 0.147 | 0.44 | 0.5 | 0.61 | 0.4 | 0.52 | 0.33 |
| 6 | 13836706 | rs1204169 | -0.983 | 0.934 | -0.402 | 0.419 | 0 | 0.004 | 0.004 | 0.202 | 0.45 | 0.5 | 0.62 | 0.4 | 0.53 | 0.33 |
| 6 | 13847687 | rs6899735 | -0.271 | 1.631 | -0.587 | 1.119 | -0.044 | -0.064 | -0.064 | 0 | 0.26 | 0.13 | 0.49 | 0.36 | 0.32 | 0.47 |
| 6 | 13854317 | rs12662217 | -0.328 | 1.469 | -0.851 | 0.875 | -0.08 | -0.079 | -0.079 | 0.099 | 0.33 | 0.11 | 0.53 | 0.33 | 0.41 | 0.5 |
| 6 | 13857781 | rs9382641 | -0.343 | 1.445 | -0.841 | 0.863 | -0.077 | -0.077 | -0.077 | 0 | 0.33 | 0.1 | 0.53 | 0.33 | 0.41 | 0.5 |
| 6 | 13861949 | rs9475717 | -0.304 | 1.708 | -0.533 | 1.003 | 0.041 | 0.041 | 0.041 | 0.583 | 0.06 | 0.11 | 0.08 | 0.15 | 0.11 | 0.03 |
| 6 | 13866326 | rs1204224 | -0.785 | 0.546 | -0.398 | 0.304 | 0.001 | 0.009 | 0.009 | 0 | 0.49 | 0.37 | 0.59 | 0.32 | 0.49 | 0.61 |
| 6 | 13870749 | rs723584 | -0.124 | 0.267 | -0.474 | 0.128 | -0.048 | -0.079 | -0.079 | 0 | 0.17 | 0.29 | 0.06 | 0 | 0.08 | 0.11 |
| 6 | 13877123 | rs1204250 | -0.853 | 0.299 | -0.677 | 0.337 | 0.012 | 0.013 | 0.013 | 0 | 0.59 | 0.75 | 0.71 | 0.52 | 0.61 | 0.67 |
| 6 | 13889438 | rs6938515 | -0.103 | 1.22 | -1.378 | 0.305 | -0.143 | -0.052 | -0.052 | 0 | 0.33 | 0.04 | 0.52 | 0.23 | 0.45 | 0.34 |
| 6 | 13897337 | rs4715640 | -0.114 | 1.182 | -1.414 | 0.108 | -0.019 | -0.006 | -0.006 | 0 | 0.16 | 0.09 | 0.11 | 0.15 | 0.18 | 0.14 |
| 6 | 13897986 | rs2274088 | 1.042 | 1.369 | -1.838 | -0.045 | -0.082 | -0.055 | -0.055 | 0.169 | 0.27 | 0.02 | 0.42 | 0.18 | 0.3 | 0.21 |
| 6 | 13921016 | rs1330118 | 0.493 | 0.595 | -0.624 | -0.015 | -0.009 | 0.002 | 0.002 | 0.453 | 0.42 | 0.41 | 0.29 | 0.66 | 0.34 | 0.51 |
| 6 | 13929502 | rs280143 | 0.707 | 0.624 | -0.022 | -0.123 | -0.153 | -0.015 | -0.015 | 0 | 0.35 | 0.03 | 0.46 | 0.19 | 0.44 | 0.28 |
| 6 | 13933410 | rs2057536 | -0.721 | 0.628 | 0.335 | -0.129 | 0.003 | -0.005 | -0.005 | 0.639 | 0.63 | 0.63 | 0.51 | 0.81 | 0.56 | 0.71 |
| 6 | 13934527 | rs9475806 | 0.939 | 0.316 | -0.442 | -0.12 | 0.002 | -0.048 | -0.048 | 0 | 0.33 | 0.18 | 0.3 | 0.6 | 0.32 | 0.51 |
| 6 | 13935066 | rs7745956 | -0.585 | 0.518 | -0.517 | 0.147 | -0.007 | 0.01 | 0.01 | 0 | 0.25 | 0.12 | 0.17 | 0.18 | 0.24 | 0.18 |
| 6 | 13948389 | rs1192057 | 0.784 | 0.388 | 0.173 | 0.982 | 0.069 | 0.129 | 0.129 | 0 | 0.89 | 0.56 | 0.81 | 0.84 | 0.89 | 0.77 |
| 6 | 13952482 | rs6917710 | 0.526 | -0.076 | -0.064 | 0.88 | -0.015 | 0.02 | 0.02 | 3.177 | 0.24 | 0.28 | 0.14 | 0.52 | 0.16 | 0.27 |
| 6 | 13952853 | rs9475975 | 0.708 | -0.119 | -0.013 | 0.892 | -0.045 | 0.071 | 0.071 | 4.865 | 0.23 | 0.35 | 0.14 | 0.49 | 0.15 | 0.25 |
| 6 | 13953227 | rs280169 | 1.246 | -0.054 | 0.049 | 1.024 | 0.027 | 0.089 | 0.089 | 0.052 | 0.87 | 0.4 | 0.78 | 0.81 | 0.89 | 0.75 |
| 6 | 13958654 | rs4715723 | 0.712 | 0.165 | 0.094 | 1.125 | 0.11 | 0.043 | 0.043 | 0 | 0.89 | 0.5 | 0.81 | 0.89 | 0.89 | 0.75 |
| 6 | 13962457 | rs280182 | 1.183 | 0.328 | -0.926 | 0.95 | -0.037 | 0.058 | 0.058 | 0.276 | 0.38 | 0.03 | 0.27 | 0.18 | 0.37 | 0.27 |
| 6 | 13963283 | rs280184 | 0.521 | 0.953 | -0.377 | 1.072 | -0.022 | -0.058 | -0.058 | 0 | 0.54 | 0.15 | 0.39 | 0.39 | 0.49 | 0.42 |
| 6 | 13978275 | rs280162 | 0.531 | 0.953 | -0.37 | 1.069 | -0.02 | -0.059 | -0.059 | 0 | 0.54 | 0.15 | 0.39 | 0.38 | 0.5 | 0.42 |
| 6 | 13979526 | rs2147201 | -0.08 | 0.765 | -0.392 | 1.12 | -0.022 | -0.018 | -0.018 | 0.21 | 0.31 | 0.21 | 0.41 | 0.49 | 0.39 | 0.31 |
| 6 | 13995112 | rs1003024 | -0.143 | 0.766 | -0.523 | 1.104 | -0.015 | -0.014 | -0.014 | 0 | 0.32 | 0.26 | 0.42 | 0.52 | 0.39 | 0.33 |
| 6 | 14006021 | rs1555051 | -0.062 | 0.798 | -0.573 | 1.106 | -0.024 | -0.011 | -0.011 | 0.044 | 0.31 | 0.22 | 0.42 | 0.43 | 0.4 | 0.3 |
| 6 | 14007050 | rs9396422 | -0.322 | 0.822 | -0.584 | 1.085 | -0.006 | -0.001 | -0.001 | 0 | 0.36 | 0.32 | 0.43 | 0.46 | 0.41 | 0.33 |
| 6 | 14009231 | rs10484348 | -0.584 | 0.804 | -0.763 | 1.143 | -0.036 | 0.045 | 0.045 | 0 | 0.41 | 0.26 | 0.56 | 0.73 | 0.47 | 0.5 |
| 6 | 14009291 | rs16874548 | -0.842 | 0.618 | -0.661 | 1.735 | -0.013 | -0.039 | -0.039 | 1.412 | 0.35 | 0.32 | 0.45 | 0.49 | 0.35 | 0.59 |
| 6 | 14009619 | rs4715861 | -0.996 | 0.181 | 0.098 | 1.73 | -0.031 | 0.035 | 0.035 | 0 | 0.17 | 0.28 | 0.11 | 0.17 | 0.08 | 0.24 |
| 6 | 14013760 | rs10484347 | -0.197 | 0.962 | -0.378 | 1.83 | -0.045 | -0.037 | -0.037 | 0 | 0.07 | 0.01 | 0.17 | 0.16 | 0.11 | 0.22 |
| 6 | 14014793 | rs9382889 | -1.664 | 0.579 | -0.295 | 1.315 | 0.003 | 0.017 | 0.017 | 0.073 | 0.73 | 0.78 | 0.8 | 0.88 | 0.71 | 0.87 |
| 6 | 14014915 | rs6939259 | -1.121 | 0.599 | 0.012 | 1.461 | 0.004 | 0.025 | 0.025 | 0.268 | 0.3 | 0.54 | 0.36 | 0.41 | 0.28 | 0.47 |
| 6 | 14019157 | rs9367805 | -1.01 | 0.621 | 0.025 | 1.387 | 0.009 | 0.043 | 0.043 | 0.376 | 0.29 | 0.37 | 0.34 | 0.46 | 0.28 | 0.47 |

| | | | | | | | | | | | | | | | | |
|---|---|---|---|---|---|---|---|---|---|---|---|---|---|---|---|---|
| 6 | 14020445 | rs35772589 | 0.134 | 0.014 | -0.043 | 1.059 | -0.015 | -0.029 | -0.029 | 0 | 0.06 | 0.1 | 0.03 | 0 | 0.03 | 0.03 |
| 6 | 14030470 | rs1023107 | 0.561 | 0.031 | -0.566 | 1.015 | -0.029 | -0.004 | -0.004 | 0 | 0.31 | 0.12 | 0.38 | 0.36 | 0.34 | 0.33 |
| 6 | 14036774 | rs6459332 | -0.982 | 0.476 | -0.615 | 0.129 | -0.018 | 0.084 | 0.084 | 1.884 | 0.63 | 0.78 | 0.72 | 0.93 | 0.57 | 0.82 |
| 6 | 14040480 | rs9382934 | -0.894 | 0.444 | -0.273 | 0.918 | 0 | 0.02 | 0.02 | 0 | 0.21 | 0.28 | 0.27 | 0.29 | 0.18 | 0.39 |
| 6 | 14044820 | rs12197169 | -0.134 | 0.523 | -0.419 | 0.837 | -0.001 | -0.002 | -0.002 | 0 | 0.17 | 0.2 | 0.17 | 0.17 | 0.17 | 0.16 |
| 6 | 14045170 | rs6924356 | -1.144 | 0.492 | -1.565 | -0.183 | -0.098 | 0.15 | 0.15 | 0.504 | 0.71 | 0.98 | 0.78 | 0.95 | 0.64 | 0.81 |
| 6 | 14046411 | rs6915882 | -0.545 | 0.255 | -1.26 | -0.41 | -0.013 | 0.205 | 0.205 | 1.552 | 0.5 | 0.8 | 0.58 | 0.83 | 0.49 | 0.45 |
| 6 | 14046490 | rs10949221 | 0.812 | 0.257 | -1.007 | -0.246 | -0.001 | -0.004 | -0.004 | 0.267 | 0.17 | 0.17 | 0.16 | 0.19 | 0.16 | 0.17 |
| 6 | 14053646 | rs9396467 | 0.379 | -0.046 | -0.025 | -0.117 | 0.002 | 0.007 | 0.007 | 0 | 0.28 | 0.29 | 0.38 | 0.41 | 0.33 | 0.23 |
| 6 | 14054510 | rs6905903 | -0.877 | -0.012 | -0.092 | -0.296 | -0.002 | -0.001 | -0.001 | 0 | 0.83 | 0.82 | 0.82 | 0.58 | 0.83 | 0.79 |
| 6 | 14056150 | rs16874628 | -0.512 | -0.224 | -0.01 | -0.442 | -0.029 | -0.052 | -0.052 | 0 | 0.1 | 0.34 | 0.05 | 0 | 0.04 | 0.02 |
| 6 | 14062176 | rs10949222 | 0.461 | -0.103 | 0.051 | -0.4 | -0.006 | -0.019 | -0.019 | 0 | 0.27 | 0.19 | 0.36 | 0.41 | 0.32 | 0.22 |
| 6 | 14064273 | rs9476476 | 0.227 | -0.131 | 0.139 | -0.462 | -0.003 | 0.007 | 0.007 | 0.472 | 0.53 | 0.42 | 0.45 | 0.18 | 0.5 | 0.55 |
| 6 | 14065118 | rs9382936 | -0.51 | 0 | 0.084 | -0.307 | 0 | -0.027 | -0.027 | 0 | 0.74 | 0.78 | 0.81 | 0.54 | 0.78 | 0.73 |
| 6 | 14065237 | rs9476478 | 1.025 | -0.314 | -0.296 | -0.571 | -0.039 | 0.083 | 0.083 | 1.394 | 0.33 | 0.16 | 0.38 | 0.13 | 0.39 | 0.47 |
| 6 | 14065782 | rs2142856 | -0.517 | -0.462 | 0.029 | -0.677 | -0.014 | 0.005 | 0.005 | 0 | 0.33 | 0.45 | 0.38 | 0.38 | 0.32 | 0.2 |
| 6 | 14067170 | rs9476481 | 0.628 | -0.461 | -0.581 | -0.665 | -0.023 | 0 | 0 | 0 | 0.4 | 0.2 | 0.31 | 0.4 | 0.49 | 0.38 |
| 6 | 14073320 | rs9296918 | 0.017 | -0.273 | -0.697 | -0.574 | 0.048 | 0.071 | 0.071 | 0 | 0.49 | 0.74 | 0.33 | 0.4 | 0.44 | 0.37 |
| 6 | 14085309 | rs9382938 | -0.728 | -0.953 | 0.083 | -0.447 | -0.062 | 0.038 | 0.038 | 1.178 | 0.25 | 0.54 | 0.11 | 0.39 | 0.12 | 0.27 |
| 6 | 14085980 | rs17430541 | 0.302 | -0.378 | -0.202 | -0.669 | -0.019 | 0.047 | 0.047 | 0 | 0.14 | 0.09 | 0.29 | 0.05 | 0.22 | 0.07 |
| 6 | 14086264 | rs1028322 | -0.461 | 0.245 | 0.201 | -0.892 | 0.004 | 0 | 0 | 0 | 0.63 | 0.64 | 0.7 | 0.4 | 0.64 | 0.62 |
| 6 | 14086393 | rs1028323 | 0.944 | 0.088 | 0.192 | -1.051 | 0.01 | 0.064 | 0.064 | 0.278 | 0.36 | 0.15 | 0.2 | 0.21 | 0.36 | 0.25 |
| 6 | 14088091 | rs11756591 | 0.371 | -0.146 | 0.49 | -1.254 | 0 | 0.057 | 0.057 | 1.686 | 0.46 | 0.65 | 0.49 | 0.05 | 0.53 | 0.2 |
| 6 | 14088302 | rs2013807 | 0.483 | -0.089 | 0.648 | -1.223 | -0.001 | 0.007 | 0.007 | 0.92 | 0.45 | 0.39 | 0.48 | 0.05 | 0.53 | 0.2 |
| 6 | 14090808 | rs16874682 | 0.703 | 0.058 | 0.186 | -1.087 | 0 | -0.02 | -0.02 | 0 | 0.17 | 0.24 | 0.3 | 0.14 | 0.21 | 0.09 |
| 6 | 14093556 | rs2092204 | -0.289 | 0.272 | -0.272 | -0.93 | -0.008 | 0.016 | 0.016 | 0 | 0.58 | 0.6 | 0.53 | 0.36 | 0.57 | 0.39 |
| 6 | 14096658 | rs12529514 | 0.732 | 0.38 | -0.628 | -0.711 | -0.003 | -0.039 | -0.039 | 0.158 | 0.07 | 0.01 | 0.03 | 0.2 | 0.06 | 0.14 |
| 6 | 14105507 | rs9464654 | -0.208 | 0.308 | -0.599 | -0.831 | 0.004 | 0.097 | 0.097 | 1.033 | 0.54 | 0.72 | 0.51 | 0.34 | 0.53 | 0.34 |
| 6 | 14105768 | rs12524473 | 1.422 | 0.3 | -0.954 | -0.774 | -0.011 | 0.07 | 0.07 | 1.07 | 0.21 | 0.05 | 0.13 | 0 | 0.24 | 0.09 |
| 6 | 14109729 | rs6934409 | -0.965 | 0.736 | -1.25 | -0.081 | -0.091 | -0.069 | -0.069 | 0 | 0.83 | 0.99 | 0.87 | 0.81 | 0.76 | 0.88 |
| 6 | 14126422 | rs6929821 | 0.642 | 0.151 | -0.716 | 0.009 | 0.005 | 0.026 | 0.026 | 0 | 0.3 | 0.18 | 0.17 | 0.2 | 0.31 | 0.22 |
| 6 | 14131804 | rs7743206 | 0.144 | -0.024 | -0.467 | 0.409 | 0.004 | 0.008 | 0.008 | 0 | 0.08 | 0.15 | 0.09 | 0.18 | 0.13 | 0.09 |
| 6 | 14134083 | rs9296925 | 0.471 | -0.854 | -0.408 | 0.377 | -0.006 | -0.003 | -0.003 | 0.051 | 0.4 | 0.35 | 0.43 | 0.39 | 0.5 | 0.52 |
| 6 | 14134592 | rs853360 | -0.391 | -0.641 | -0.75 | 0.132 | 0 | -0.006 | -0.006 | 0 | 0.73 | 0.73 | 0.69 | 0.83 | 0.7 | 0.86 |
| 6 | 14136428 | rs1050650 | -0.438 | -0.052 | -0.964 | 0.735 | -0.029 | -0.019 | -0.019 | 0.016 | 0.62 | 0.77 | 0.61 | 0.61 | 0.51 | 0.48 |
| 6 | 14138292 | rs9370729 | 0.43 | -0.323 | -1.281 | 0.431 | 0.014 | 0.132 | 0.132 | 0 | 0.44 | 0.09 | 0.45 | 0.18 | 0.56 | 0.43 |
| 6 | 14143961 | rs853372 | -0.057 | -0.781 | -0.285 | 0.102 | -0.014 | -0.036 | -0.036 | 0.079 | 0.23 | 0.32 | 0.28 | 0.12 | 0.22 | 0.11 |
| 6 | 14144207 | rs745247 | 0.235 | -0.262 | 0.149 | 0.073 | 0 | 0.012 | 0.012 | 0 | 0.61 | 0.67 | 0.68 | 0.19 | 0.63 | 0.49 |
| 6 | 14146403 | rs7745829 | -1.266 | -0.435 | 0.405 | 0.917 | -0.006 | 0.25 | 0.25 | 0.317 | 0.13 | 0.47 | 0.19 | 0.33 | 0.09 | 0.24 |
| 6 | 14146661 | rs1537149 | -0.632 | -0.15 | 0.477 | 0.25 | 0.054 | 0 | 0 | 0 | 0.06 | 0.23 | 0.11 | 0 | 0.06 | 0.03 |
| 6 | 14147085 | rs7754804 | -0.532 | -0.138 | 0.454 | 0.257 | 0.056 | 0.002 | 0.002 | 0 | 0.05 | 0.22 | 0.1 | 0 | 0.06 | 0.03 |
| 6 | 14149263 | rs1547616 | 0.021 | -0.431 | 0.142 | 0.35 | 0.03 | 0.003 | 0.003 | 0 | 0.62 | 0.7 | 0.79 | 0.38 | 0.77 | 0.53 |
| 6 | 14150230 | rs6459336 | 0.86 | 0.016 | 0.193 | -0.272 | 0.005 | -0.108 | -0.108 | 0.078 | 0.36 | 0.4 | 0.33 | 0.01 | 0.34 | 0.34 |
| 6 | 14162229 | rs1535359 | 0.397 | -0.214 | 0.08 | 0.262 | 0 | -0.009 | -0.009 | 0 | 0.62 | 0.48 | 0.57 | 0.71 | 0.61 | 0.67 |
| 6 | 14162949 | rs853340 | 0.148 | -0.546 | 0.158 | 0.257 | 0.006 | -0.013 | -0.013 | 0.056 | 0.12 | 0.05 | 0.06 | 0.22 | 0.09 | 0.2 |
| 6 | 14167198 | rs853346 | -0.901 | -0.901 | 0.521 | 0.643 | -0.008 | 0.002 | 0.002 | 0 | 0.34 | 0.57 | 0.24 | 0.6 | 0.23 | 0.5 |
| 6 | 14170358 | rs853354 | -1.151 | -0.848 | 0.329 | 0.559 | 0.042 | -0.016 | -0.016 | 0.518 | 0.4 | 0.74 | 0.35 | 0.6 | 0.31 | 0.54 |
| 6 | 14175404 | rs853377 | -0.853 | -0.68 | 0.116 | 0.439 | -0.052 | 0.066 | 0.066 | 2.135 | 0.79 | 0.97 | 0.72 | 0.93 | 0.71 | 0.93 |
| 6 | 14183028 | rs853386 | -1.539 | -0.916 | 0.85 | 1.039 | 0.042 | -0.019 | -0.019 | 0 | 0.4 | 0.76 | 0.35 | 0.61 | 0.31 | 0.54 |
| 6 | 14187776 | rs9370732 | 0.013 | -0.316 | 0.864 | 1.204 | 0.082 | 0.151 | 0.151 | 0 | 0.08 | 0.2 | 0.16 | 0.22 | 0.16 | 0.04 |
| 6 | 14190496 | rs9396476 | -0.234 | -0.595 | 1.036 | 1.567 | -0.006 | 0.013 | 0.013 | 0 | 0.21 | 0.24 | 0.15 | 0.56 | 0.14 | 0.36 |
| 6 | 14190706 | rs853388 | -1.355 | -0.525 | 0.914 | 0.983 | 0.056 | 0.034 | 0.034 | 0.865 | 0.42 | 0.73 | 0.34 | 0.75 | 0.3 | 0.58 |
| 6 | 14201634 | rs1556388 | 0.731 | -0.592 | 0.872 | 1.131 | -0.027 | 0.02 | 0.02 | 0 | 0.71 | 0.63 | 0.79 | 0.32 | 0.8 | 0.51 |
| 6 | 14203502 | rs1929320 | 0.429 | -0.446 | 0.629 | 0.688 | -0.038 | -0.028 | -0.028 | 0 | 0.06 | 0.01 | 0.14 | 0.24 | 0.15 | 0.04 |
| 6 | 14203872 | rs6459338 | 0.859 | -1.06 | 1 | 0.714 | -0.014 | 0.006 | 0.006 | 0 | 0.69 | 0.66 | 0.77 | 0.34 | 0.77 | 0.45 |
| 6 | 14210463 | rs1537147 | 0.725 | -1.076 | 0.967 | 0.791 | -0.012 | 0.01 | 0.01 | 0 | 0.72 | 0.73 | 0.81 | 0.34 | 0.8 | 0.48 |
| 6 | 14215829 | rs6911584 | -0.834 | -1.026 | 1.216 | 0.466 | -0.021 | -0.045 | -0.045 | 0 | 0.14 | 0.38 | 0.13 | 0.08 | 0.1 | 0.07 |
| 6 | 14238900 | rs9370734 | 0.287 | -1.121 | 1.164 | 0.589 | -0.002 | 0.025 | 0.025 | 0 | 0.95 | 0.87 | 0.92 | 0.86 | 0.96 | 0.89 |
| 6 | 14240594 | rs1537151 | 0.811 | -1.109 | 1.107 | 0.567 | -0.002 | -0.002 | -0.002 | 0 | 0.84 | 0.79 | 0.8 | 0.77 | 0.86 | 0.82 |
| 6 | 14244274 | rs12207359 | 0.104 | -1.102 | 0.851 | 0.566 | -0.026 | 0.001 | 0.001 | 0.111 | 0.1 | 0.01 | 0.11 | 0.09 | 0.1 | 0.07 |
| 6 | 14244517 | rs9370735 | 1.158 | -1.125 | 1.023 | 0.556 | -0.024 | -0.023 | -0.023 | 0 | 0.79 | 0.39 | 0.77 | 0.77 | 0.86 | 0.82 |
| 6 | 14249646 | rs7739748 | 1.729 | -1.524 | 0.157 | 0.125 | -0.059 | 0.216 | 0.216 | 0.017 | 0.44 | 0.04 | 0.56 | 0.1 | 0.58 | 0.43 |
| 6 | 14260486 | rs9476529 | 1.503 | -1.272 | 0.21 | 0.629 | 0.009 | 0.007 | 0.007 | 0 | 0.6 | 0.39 | 0.65 | 0.63 | 0.69 | 0.76 |
| 6 | 14265882 | rs4712230 | 0.671 | -1.253 | 0.35 | 0.579 | -0.002 | 0.102 | 0.102 | 0.121 | 0.91 | 0.48 | 0.89 | 0.85 | 0.96 | 0.89 |
| 6 | 14268073 | rs9464673 | -1.188 | -1.373 | 0.227 | 0.474 | 0.019 | 0.05 | 0.05 | 0 | 0.42 | 0.27 | 0.32 | 0.74 | 0.37 | 0.43 |
| 6 | 14271921 | rs944486 | 0.857 | -0.656 | -0.317 | 0.549 | -0.016 | 0.014 | 0.014 | 0 | 0.73 | 0.43 | 0.78 | 0.71 | 0.79 | 0.85 |
| 6 | 14275069 | rs9367830 | -0.563 | 0.032 | -0.852 | 0.27 | -0.042 | -0.007 | -0.007 | 0 | 0.22 | 0.07 | 0.22 | 0.2 | 0.26 | 0.12 |
| 6 | 14278169 | rs6917540 | 0.846 | -0.378 | -1.43 | -0.218 | -0.008 | 0.202 | 0.202 | 0 | 0.48 | 0.06 | 0.56 | 0.2 | 0.58 | 0.44 |
| 6 | 14278200 | rs2183674 | 0.989 | -0.199 | -0.911 | -0.271 | 0.001 | 0.133 | 0.133 | 0 | 0.45 | 0.19 | 0.55 | 0.18 | 0.56 | 0.42 |

| | | | | | | | | | | | | | | | | |
|---|---|---|---|---|---|---|---|---|---|---|---|---|---|---|---|---|
| 6 | 14289082 | rs2094923 | 0.805 | -0.128 | -0.681 | -0.455 | -0.025 | 0.145 | 0.145 | 0 | 0.45 | 0.16 | 0.58 | 0.18 | 0.55 | 0.36 |
| 6 | 14291089 | rs2274822 | -0.434 | -0.425 | -0.412 | -0.358 | 0.026 | 0.043 | 0.043 | 0 | 0.35 | 0.19 | 0.23 | 0.23 | 0.24 | 0.22 |
| 6 | 14291231 | rs3818671 | -0.415 | -0.402 | 0.036 | -0.568 | -0.003 | -0.009 | -0.009 | 0.1 | 0.37 | 0.36 | 0.26 | 0.19 | 0.24 | 0.25 |
| 6 | 14295178 | rs11759411 | 1.138 | -0.319 | -0.433 | -0.777 | -0.106 | 0.161 | 0.161 | 0 | 0.43 | 0.03 | 0.53 | 0.14 | 0.55 | 0.35 |
| 6 | 14304370 | rs4715885 | -0.378 | -0.687 | 0.207 | -0.976 | 0.001 | -0.005 | -0.005 | 0.078 | 0.34 | 0.32 | 0.37 | 0.16 | 0.28 | 0.29 |
| 6 | 14310129 | rs6933937 | 0.483 | -0.847 | 0.291 | -1.084 | -0.007 | -0.042 | -0.042 | 0 | 0.22 | 0.24 | 0.22 | 0.09 | 0.19 | 0.11 |
| 6 | 14310886 | rs4712235 | 0.404 | -0.99 | 0.615 | -0.89 | -0.014 | 0.02 | 0.02 | 0 | 0.68 | 0.49 | 0.67 | 0.34 | 0.72 | 0.52 |
| 6 | 14311488 | rs10456759 | 0.112 | -0.972 | 0.839 | -0.868 | -0.011 | -0.021 | -0.021 | 0 | 0.16 | 0.26 | 0.16 | 0.06 | 0.08 | 0.16 |
| 6 | 14312081 | rs12526130 | 1.082 | -0.677 | 0.532 | -0.636 | -0.059 | 0.022 | 0.022 | 0.189 | 0.36 | 0.15 | 0.4 | 0.24 | 0.49 | 0.37 |
| 6 | 14314883 | rs6905081 | 0.323 | -0.68 | 0.321 | -0.691 | -0.001 | -0.015 | -0.015 | 0 | 0.37 | 0.36 | 0.36 | 0.16 | 0.27 | 0.23 |
| 6 | 14318451 | rs11751940 | 0.412 | -0.509 | 0.151 | -0.601 | -0.007 | 0.025 | 0.025 | 0.011 | 0.45 | 0.31 | 0.44 | 0.26 | 0.52 | 0.4 |
| 6 | 14321020 | rs6920737 | 0.041 | -0.008 | -0.32 | -0.378 | -0.018 | -0.053 | -0.053 | 0.01 | 0.45 | 0.54 | 0.45 | 0.15 | 0.29 | 0.25 |
| 6 | 14322665 | rs10949239 | 0.315 | 0.478 | -0.133 | 0.27 | 0.025 | 0.148 | 0.148 | 0 | 0.81 | 0.63 | 0.84 | 0.41 | 0.78 | 0.62 |
| 6 | 14327412 | rs6930492 | -0.176 | -0.151 | -0.347 | -0.678 | -0.01 | -0.038 | -0.038 | 0 | 0.42 | 0.43 | 0.44 | 0.15 | 0.3 | 0.23 |
| 6 | 14328155 | rs6936230 | -0.381 | -0.358 | -0.309 | -0.899 | -0.001 | -0.021 | -0.021 | 0 | 0.44 | 0.52 | 0.31 | 0.2 | 0.4 | 0.3 |
| 6 | 14328324 | rs6459348 | -0.339 | -0.506 | -0.433 | -1.007 | 0.022 | 0.026 | 0.026 | 0 | 0.42 | 0.66 | 0.23 | 0.29 | 0.31 | 0.25 |
| 6 | 14329647 | rs7752155 | -0.744 | -0.542 | -0.286 | -1.208 | 0.016 | -0.057 | -0.057 | 0.392 | 0.32 | 0.46 | 0.29 | 0.18 | 0.32 | 0.24 |
| 6 | 14331711 | rs7764117 | -0.362 | -0.86 | -0.164 | -1.055 | -0.044 | -0.055 | -0.055 | 0.558 | 0.24 | 0.39 | 0.12 | 0.11 | 0.14 | 0.14 |
| 6 | 14336818 | rs9476537 | -0.068 | -0.677 | -0.955 | -0.527 | 0.028 | -0.027 | -0.027 | 0 | 0.28 | 0.1 | 0.15 | 0.33 | 0.26 | 0.22 |
| 6 | 14337193 | rs719343 | -0.026 | -0.746 | -0.95 | -0.522 | 0.04 | -0.028 | -0.028 | 0.784 | 0.27 | 0.1 | 0.15 | 0.32 | 0.25 | 0.21 |
| 6 | 14345024 | rs737548 | -0.423 | -0.734 | -0.579 | -0.559 | -0.008 | -0.002 | -0.002 | 0 | 0.57 | 0.48 | 0.3 | 0.4 | 0.4 | 0.33 |
| 6 | 14349114 | rs6899596 | -0.564 | -0.775 | -0.845 | -0.504 | 0.037 | -0.04 | -0.04 | 0.016 | 0.55 | 0.8 | 0.3 | 0.55 | 0.38 | 0.4 |
| 6 | 14349627 | rs9464677 | 0.784 | 0.758 | -0.52 | -0.276 | 0.001 | 0.001 | 0.001 | 0 | 0.62 | 0.58 | 0.63 | 0.46 | 0.51 | 0.34 |
| 6 | 14350345 | rs12205119 | 0.436 | 0.963 | -0.691 | -0.002 | 0.006 | 0.088 | 0.088 | 0 | 0.48 | 0.27 | 0.66 | 0.29 | 0.49 | 0.4 |
| 6 | 14352513 | rs6913034 | 1.108 | 2.104 | 0.101 | 0.878 | 0.025 | 0.053 | 0.053 | 0 | 0.82 | 0.73 | 0.81 | 0.47 | 0.71 | 0.5 |
| 6 | 14353238 | rs10949240 | 0.221 | 2.058 | 0.112 | 0.951 | 0.02 | 0.057 | 0.057 | 0 | 0.9 | 0.87 | 0.86 | 0.62 | 0.78 | 0.68 |
| 6 | 14355492 | rs9476540 | 1.166 | 1.397 | 0.284 | 0.392 | -0.001 | 0 | 0 | 0.12 | 0.61 | 0.59 | 0.5 | 0.29 | 0.49 | 0.25 |
| 6 | 14358168 | rs4715890 | 1.041 | 1.368 | 0.38 | 0.377 | 0 | -0.004 | -0.004 | 0 | 0.6 | 0.56 | 0.5 | 0.29 | 0.49 | 0.25 |
| 6 | 14361203 | rs6940613 | -0.581 | 1.106 | 0.062 | 0.072 | -0.001 | -0.022 | -0.022 | 0 | 0.15 | 0.16 | 0.06 | 0.01 | 0.08 | 0.03 |
| 6 | 14364727 | rs9464681 | 1.066 | 1.536 | 0.235 | 0.382 | -0.059 | -0.004 | -0.004 | 0 | 0.66 | 0.84 | 0.55 | 0.32 | 0.51 | 0.25 |
| 6 | 14367567 | rs9476543 | 1.776 | 1.152 | 1.011 | 0.494 | 0.002 | 0.044 | 0.044 | 1.315 | 0.52 | 0.32 | 0.48 | 0.3 | 0.51 | 0.22 |
| 6 | 14371336 | rs10949241 | 1.136 | 1.248 | 0.582 | 0.601 | -0.062 | -0.003 | -0.003 | 0 | 0.66 | 0.84 | 0.56 | 0.29 | 0.57 | 0.25 |
| 6 | 14371419 | rs7762317 | 0.558 | 1.475 | 0.595 | 1.089 | -0.013 | 0.041 | 0.041 | 0 | 0.94 | 0.98 | 0.94 | 0.56 | 0.91 | 0.65 |
| 6 | 14382803 | rs12208770 | 1.514 | 1.567 | 0.83 | 0.774 | 0.011 | -0.005 | -0.005 | 0.484 | 0.62 | 0.56 | 0.45 | 0.29 | 0.59 | 0.27 |
| 6 | 14394504 | rs9476551 | 0.899 | 1.439 | 0.963 | 1.661 | -0.016 | 0.058 | 0.058 | 0 | 0.94 | 0.97 | 0.82 | 0.55 | 0.89 | 0.67 |
| 6 | 14395097 | rs9986461 | -1.931 | 1.199 | 1.448 | 1.409 | 0.007 | -0.03 | -0.03 | 0 | 0.38 | 0.45 | 0.41 | 0.22 | 0.37 | 0.44 |
| 6 | 14395373 | rs11969660 | -0.819 | 1.357 | 1.398 | 1.279 | 0.009 | -0.019 | -0.019 | 0 | 0.1 | 0.15 | 0.09 | 0.02 | 0.09 | 0.1 |
| 6 | 14396349 | rs4715891 | -2.394 | 1.324 | 1.976 | 1.694 | 0.013 | -0.011 | -0.011 | 0 | 0.39 | 0.48 | 0.53 | 0.69 | 0.42 | 0.75 |
| 6 | 14407713 | rs2073270 | -1.144 | 1.485 | 1.671 | 1.89 | -0.032 | -0.056 | -0.056 | 1.801 | 0.14 | 0.05 | 0.39 | 0.53 | 0.21 | 0.51 |
| 6 | 14408104 | rs9476556 | -1.734 | 1.467 | 2.029 | 1.839 | 0.101 | -0.026 | -0.026 | 0.246 | 0.11 | 0.43 | 0.09 | 0.02 | 0.09 | 0.12 |
| 6 | 14410699 | rs4715895 | -1.466 | 1.557 | 1.286 | 1.926 | -0.072 | -0.135 | -0.135 | 1.098 | 0.19 | 0.03 | 0.41 | 0.53 | 0.27 | 0.52 |
| 6 | 14429562 | rs12662871 | -1.826 | 1.371 | 1.499 | 1.486 | 0.017 | 0.011 | 0.011 | 0 | 0.15 | 0.26 | 0.1 | 0.2 | 0.14 | 0.22 |
| 6 | 14441229 | rs2223220 | -1.509 | 1.421 | 1.444 | 1.692 | -0.051 | -0.05 | -0.05 | 0 | 0.26 | 0.17 | 0.48 | 0.64 | 0.38 | 0.6 |
| 6 | 14446670 | rs10949246 | 2.099 | 1.107 | 1.494 | 1.636 | -0.023 | 0.007 | 0.007 | 2.688 | 0.62 | 0.65 | 0.46 | 0.28 | 0.53 | 0.28 |
| 6 | 14451276 | rs9370742 | 1.213 | -0.185 | 1.517 | 1.414 | 0.006 | 0.014 | 0.014 | 0.039 | 0.86 | 0.92 | 0.94 | 0.92 | 0.87 | 0.85 |
| 6 | 14456739 | rs9296932 | 1.395 | -0.211 | 1.88 | 1.401 | -0.015 | -0.031 | -0.031 | 0 | 0.84 | 0.59 | 0.91 | 0.92 | 0.87 | 0.85 |
| 6 | 14465299 | rs4715902 | -1.418 | -0.124 | 1.569 | 1.883 | -0.026 | -0.042 | -0.042 | 0 | 0.38 | 0.18 | 0.48 | 0.6 | 0.41 | 0.64 |
| 6 | 14470099 | rs9349942 | -1.948 | -0.262 | 2.169 | 1.879 | 0.001 | 0.018 | 0.018 | 2.582 | 0.45 | 0.68 | 0.57 | 0.72 | 0.44 | 0.75 |
| 6 | 14486933 | rs1883155 | -1.5 | -0.198 | 2.356 | 1.84 | -0.006 | 0.011 | 0.011 | 3.164 | 0.4 | 0.36 | 0.53 | 0.69 | 0.43 | 0.68 |
| 6 | 14490011 | rs9396498 | -1.563 | -0.194 | 2.435 | 1.855 | -0.004 | -0.003 | -0.003 | 0 | 0.39 | 0.37 | 0.53 | 0.62 | 0.43 | 0.66 |
| 6 | 14492975 | rs4715908 | -1.393 | -0.191 | 2.278 | 1.857 | -0.039 | -0.033 | -0.033 | 0 | 0.38 | 0.19 | 0.51 | 0.62 | 0.43 | 0.66 |
| 6 | 14493643 | rs6941351 | -2.772 | -0.159 | 2.406 | 1.897 | 0.02 | 0.117 | 0.117 | 4.245 | 0.07 | 0.48 | 0.06 | 0.09 | 0.01 | 0.08 |
| 6 | 14498175 | rs9476574 | -1.385 | -0.063 | 1.798 | 1.894 | -0.055 | -0.053 | -0.053 | 0 | 0.37 | 0.16 | 0.5 | 0.61 | 0.43 | 0.64 |
| 6 | 14507185 | rs13437378 | -2.007 | -0.079 | 2.245 | 1.902 | -0.007 | 0.014 | 0.014 | 1.797 | 0.18 | 0.57 | 0.15 | 0.17 | 0.14 | 0.4 |
| 6 | 14509537 | rs7760753 | -2.517 | -0.091 | 2.354 | 2.094 | 0.025 | 0.116 | 0.116 | 7.601 | 0.07 | 0.59 | 0.06 | 0.08 | 0.01 | 0.1 |
| 6 | 14517404 | rs7751475 | -2.442 | -0.107 | 2.189 | 2.501 | -0.017 | 0.199 | 0.199 | 6.115 | 0.11 | 0.66 | 0.2 | 0.31 | 0.06 | 0.12 |
| 6 | 14533220 | rs6903810 | -2.312 | -0.201 | 2.712 | 2.416 | 0.043 | 0.039 | 0.039 | 4.086 | 0.05 | 0.33 | 0.04 | 0 | 0 | 0 |
| 6 | 14541122 | rs1883151 | -0.367 | 0.135 | 2.426 | 2.867 | -0.034 | -0.011 | -0.011 | 0 | 0.05 | 0 | 0.18 | 0.24 | 0.12 | 0.13 |
| 6 | 14545779 | rs9396504 | -0.748 | 0.076 | 2.265 | 2.767 | -0.025 | -0.022 | -0.022 | 0 | 0.08 | 0.01 | 0.05 | 0.1 | 0.08 | 0.34 |
| 6 | 14554627 | rs6937863 | -1.611 | 0.05 | 2.804 | 2.811 | 0.025 | 0.358 | 0.358 | 0.664 | 0.07 | 0.29 | 0.17 | 0.31 | 0.05 | 0.1 |
| 6 | 14558978 | rs1267480 | -1.726 | -0.009 | 2.774 | 2.627 | 0.013 | 0.021 | 0.021 | 0 | 0.12 | 0.18 | 0.25 | 0.25 | 0.21 | 0.15 |
| 6 | 14560180 | rs9396510 | -2.063 | -0.194 | 2.951 | 2.643 | 0.002 | 0.319 | 0.319 | 0.111 | 0.09 | 0.3 | 0.17 | 0.32 | 0.06 | 0.11 |
| 6 | 14570894 | rs2327835 | 1.598 | -0.009 | 2.261 | 2.57 | 0.004 | 0.013 | 0.013 | 0 | 0.89 | 0.86 | 0.77 | 0.76 | 0.83 | 0.86 |
| 6 | 14577131 | rs2876366 | 2.261 | -0.686 | 2.069 | 2.233 | 0.013 | -0.012 | -0.012 | 0 | 0.61 | 0.21 | 0.48 | 0.49 | 0.65 | 0.65 |
| 6 | 14579439 | rs9476591 | -1.759 | -0.659 | 2.204 | 2.179 | 0.005 | -0.004 | -0.004 | 0 | 0.28 | 0.67 | 0.3 | 0.27 | 0.17 | 0.21 |
| 6 | 14581154 | rs9464699 | -0.242 | -0.683 | 2.537 | 2.17 | -0.003 | -0.024 | -0.024 | 0.056 | 0.18 | 0.16 | 0.1 | 0.02 | 0.12 | 0.09 |
| 6 | 14586786 | rs9382987 | -1.366 | -0.72 | 2.78 | 2.174 | -0.031 | 0.028 | 0.028 | 0 | 0.25 | 0.32 | 0.26 | 0.27 | 0.17 | 0.18 |
| 6 | 14593016 | rs6899842 | -1.381 | -0.761 | 2.849 | 2.166 | -0.043 | 0.042 | 0.042 | 0 | 0.25 | 0.32 | 0.24 | 0.27 | 0.17 | 0.19 |
| 6 | 14596044 | rs6932984 | -1.52 | -0.726 | 2.829 | 2.152 | -0.039 | 0.022 | 0.022 | 0.07 | 0.26 | 0.53 | 0.28 | 0.27 | 0.17 | 0.19 |

| | | | | | | | | | | | | | | | | |
|---|---|---|---|---|---|---|---|---|---|---|---|---|---|---|---|---|
| 6 | 14597572 | rs6940298 | -2.246 | -0.6 | 2.587 | 2.224 | 0.003 | 0.307 | 0.307 | 3.95 | 0.1 | 0.48 | 0.19 | 0.23 | 0.05 | 0.12 |
| 6 | 14598308 | rs10484339 | -0.657 | -0.542 | 2.194 | 2.042 | -0.016 | -0.023 | -0.023 | 0.227 | 0.08 | 0.01 | 0.05 | 0.16 | 0.08 | 0.38 |
| 6 | 14598820 | rs7770731 | -1.577 | -0.591 | 2.375 | 1.985 | -0.036 | -0.065 | -0.065 | 0.91 | 0.28 | 0.5 | 0.18 | 0.19 | 0.2 | 0.48 |
| 6 | 14599167 | rs857475 | 1.849 | -0.581 | 2.3 | 1.988 | 0 | -0.008 | -0.008 | 0 | 0.68 | 0.34 | 0.77 | 0.81 | 0.76 | 0.51 |
| 6 | 14599431 | rs857473 | -1.672 | -0.691 | 1.965 | 2.05 | 0.001 | 0.031 | 0.031 | 0 | 0.14 | 0.14 | 0.34 | 0.47 | 0.18 | 0.24 |
| 6 | 14601073 | rs12204995 | 2.623 | -0.719 | 1.546 | 1.996 | 0.01 | 0.111 | 0.111 | 1.87 | 0.52 | 0.14 | 0.42 | 0.34 | 0.58 | 0.27 |
| 6 | 14606414 | rs9476611 | -0.399 | -0.47 | 1.834 | 1.136 | -0.003 | -0.036 | -0.036 | 0.085 | 0.18 | 0.18 | 0.12 | 0.02 | 0.15 | 0.1 |
| 6 | 14607135 | rs12197964 | 1.743 | -0.385 | 1.973 | 1.338 | 0.046 | 0.11 | 0.11 | 0.288 | 0.89 | 0.71 | 0.78 | 0.75 | 0.87 | 0.86 |
| 6 | 14608299 | rs6932589 | 1.838 | -0.51 | 1.369 | 1.027 | 0.008 | 0.009 | 0.009 | 1.07 | 0.58 | 0.14 | 0.57 | 0.61 | 0.65 | 0.38 |
| 6 | 14613673 | rs6910689 | 1.78 | -0.45 | 1.374 | 1.107 | 0.015 | 0.022 | 0.022 | 0.254 | 0.62 | 0.29 | 0.61 | 0.66 | 0.69 | 0.41 |
| 6 | 14614112 | rs6920206 | -0.784 | -0.077 | 0.786 | 1.131 | 0.009 | 0.03 | 0.03 | 0.498 | 0.26 | 0.15 | 0.19 | 0.18 | 0.23 | 0.45 |
| 6 | 14615489 | rs857445 | -0.657 | -0.112 | 0.738 | 1.366 | -0.027 | -0.013 | -0.013 | 0.29 | 0.06 | 0.02 | 0.16 | 0.21 | 0.11 | 0.13 |
| 6 | 14621353 | rs11759651 | -0.91 | -1.453 | 0.215 | 0.052 | 0.02 | 0.02 | 0.02 | 0.031 | 0.39 | 0.34 | 0.25 | 0.15 | 0.23 | 0.37 |
| 6 | 14624215 | rs9370770 | 1.234 | -1.102 | 0.078 | 0.247 | 0.076 | 0.069 | 0.069 | 1.551 | 0.62 | 0.14 | 0.62 | 0.76 | 0.76 | 0.55 |
| 6 | 14624986 | rs9382990 | 0.786 | -1.128 | 0.72 | 0.244 | -0.005 | 0.031 | 0.031 | 0 | 0.86 | 0.84 | 0.79 | 0.76 | 0.86 | 0.75 |
| 6 | 14626729 | rs857477 | -1.633 | -1.237 | 1.028 | 0.176 | 0.003 | 0.003 | 0.003 | 0 | 0.3 | 0.27 | 0.3 | 0.23 | 0.24 | 0.39 |
| 6 | 14628648 | rs11964010 | 1.523 | -1.114 | 1.217 | 0.297 | -0.025 | 0.039 | 0.039 | 0.133 | 0.82 | 0.4 | 0.78 | 0.78 | 0.86 | 0.76 |
| 6 | 14631611 | rs857484 | 1.173 | -1.203 | 0.965 | 0.081 | 0.038 | 0.039 | 0.039 | 4.666 | 0.77 | 0.19 | 0.82 | 1 | 0.9 | 0.84 |
| 6 | 14636963 | rs707824 | 1.096 | -1.121 | 1.7 | 0.132 | 0.011 | -0.014 | -0.014 | 0 | 0.78 | 0.26 | 0.84 | 1 | 0.9 | 0.84 |
| 6 | 14639232 | rs17385119 | -0.359 | -1.017 | 1.563 | 0.181 | 0.017 | 0.092 | 0.092 | 1.248 | 0.18 | 0.06 | 0.14 | 0 | 0.14 | 0.16 |
| 6 | 14642101 | rs857424 | -0.852 | -0.345 | 1.613 | 0.636 | 0.002 | 0.051 | 0.051 | 0 | 0.3 | 0.46 | 0.33 | 0.57 | 0.31 | 0.43 |
| 6 | 14644319 | rs713460 | -0.402 | -0.158 | 1.16 | 1.083 | -0.014 | 0.001 | 0.001 | 0 | 0.33 | 0.1 | 0.22 | 0.28 | 0.29 | 0.33 |
| 6 | 14647764 | rs7747013 | 1.138 | 1.629 | 2.132 | 1.868 | -0.009 | -0.011 | -0.011 | 0 | 0.65 | 0.7 | 0.54 | 0.51 | 0.47 | 0.49 |
| 6 | 14653049 | rs2027491 | -2.188 | 1.573 | 2.438 | 1.767 | -0.013 | -0.017 | -0.017 | 3.532 | 0.08 | 0.53 | 0.04 | 0 | 0 | 0.02 |
| 6 | 14657742 | rs10484340 | -0.154 | 1.714 | 1.85 | 1.899 | -0.031 | -0.01 | -0.01 | 0 | 0.21 | 0.15 | 0.32 | 0.23 | 0.39 | 0.31 |
| 6 | 14659151 | rs857416 | 1.812 | 1.668 | 1.943 | 2.015 | -0.01 | -0.014 | -0.014 | 0 | 0.92 | 0.58 | 0.93 | 1 | 0.99 | 0.97 |
| 6 | 14659783 | rs857415 | 1.538 | 1.684 | 1.028 | 2.247 | -0.061 | -0.042 | -0.042 | 0 | 0.57 | 0.11 | 0.5 | 0.46 | 0.46 | 0.47 |
| 6 | 14661538 | rs761833 | -0.859 | 1.609 | 1.072 | 2.112 | 0.026 | 0.008 | 0.008 | 0.134 | 0.29 | 0.58 | 0.36 | 0.3 | 0.41 | 0.42 |
| 6 | 14664322 | rs857413 | 1.653 | 1.692 | 0.351 | 2.199 | -0.006 | 0.009 | 0.009 | 4.318 | 0.8 | 0.27 | 0.84 | 0.72 | 0.87 | 0.82 |
| 6 | 14668457 | rs7764728 | -0.976 | 1.747 | 0.13 | 2.049 | 0.016 | 0.008 | 0.008 | 0 | 0.35 | 0.4 | 0.41 | 0.58 | 0.53 | 0.58 |
| 6 | 14670887 | rs707821 | -1.405 | 1.679 | -0.088 | 2.429 | 0 | 0.026 | 0.026 | 0 | 0.07 | 0.1 | 0.15 | 0.25 | 0.12 | 0.17 |
| 6 | 14671614 | rs857410 | -1.356 | 1.143 | -0.028 | 2.02 | 0.004 | 0.035 | 0.035 | 0.916 | 0.14 | 0.12 | 0.43 | 0.59 | 0.24 | 0.3 |
| 6 | 14686471 | rs6459369 | -1.365 | 0.291 | 0.176 | 1.16 | -0.013 | 0.004 | 0.004 | 0.276 | 0.22 | 0.7 | 0.14 | 0.27 | 0.14 | 0.17 |
| 6 | 14691612 | rs1267514 | 1.461 | -0.215 | 1.136 | 0.991 | -0.047 | 0.039 | 0.039 | 0 | 0.77 | 0.49 | 0.88 | 0.67 | 0.87 | 0.81 |
| 6 | 14698808 | rs857433 | 1.814 | -0.347 | 1.026 | 0.922 | 0.068 | -0.036 | -0.036 | 0 | 0.73 | 0.12 | 0.84 | 0.67 | 0.86 | 0.81 |
| 6 | 14699918 | rs16875557 | -1.107 | -0.416 | 1.816 | 0.888 | -0.011 | 0.043 | 0.043 | 0 | 0.16 | 0.24 | 0.09 | 0.33 | 0.12 | 0.16 |
| 6 | 14702667 | rs6937972 | 0.376 | -0.703 | 0.846 | -0.482 | -0.015 | 0.093 | 0.093 | 0 | 0.25 | 0.07 | 0.14 | 0.06 | 0.28 | 0.18 |
| 6 | 14706641 | rs864495 | 1.289 | -0.621 | 1.242 | -0.327 | -0.021 | -0.037 | -0.037 | 0 | 0.9 | 0.55 | 0.96 | 1 | 1 | 0.97 |
| 6 | 14712493 | rs764650 | 1.125 | 0.137 | 0.97 | -0.005 | 0.039 | 0.039 | 0.039 | 0 | 0.8 | 0.5 | 0.75 | 0.73 | 0.78 | 0.7 |
| 6 | 14714195 | rs1267505 | 0.593 | -0.699 | 0.095 | -0.732 | 0.092 | 0.233 | 0.233 | 0.128 | 0.61 | 0.07 | 0.66 | 0.92 | 0.77 | 0.77 |
| 6 | 14715882 | rs1267499 | 0.538 | -0.587 | 0.16 | -0.753 | 0.091 | 0.22 | 0.22 | 0.085 | 0.61 | 0.09 | 0.66 | 0.93 | 0.77 | 0.76 |
| 6 | 14719496 | rs17119 | 0.206 | -0.337 | 0.483 | -0.583 | -0.043 | -0.109 | -0.109 | 0.009 | 0.65 | 0.55 | 0.68 | 0.92 | 0.76 | 0.76 |
| 6 | 14737509 | rs2038016 | 1.374 | 0.362 | 0.812 | -0.229 | 0.025 | -0.038 | -0.038 | 0 | 0.9 | 0.7 | 0.85 | 0.98 | 0.89 | 0.9 |
| 6 | 14737615 | rs9370778 | 1.374 | 0.362 | 0.812 | -0.229 | 0.025 | -0.038 | -0.038 | 0 | 0.9 | 0.7 | 0.85 | 0.98 | 0.89 | 0.9 |
| 6 | 14743783 | rs9476648 | 2.004 | -0.229 | 1.269 | -0.726 | 0.001 | -0.019 | -0.019 | 0.569 | 0.79 | 0.22 | 0.79 | 0.98 | 0.86 | 0.83 |
| 6 | 14745960 | rs9464714 | 1.398 | -0.25 | 2.086 | -0.731 | 0.053 | -0.041 | -0.041 | 0.387 | 0.89 | 0.51 | 0.84 | 0.98 | 0.89 | 0.9 |
| 6 | 14748675 | rs2472793 | 1.327 | -0.258 | 2.207 | -0.586 | 0.045 | -0.011 | -0.011 | 0 | 0.89 | 0.58 | 0.85 | 0.96 | 0.89 | 0.9 |
| 6 | 14749648 | rs1474561 | 0.552 | -0.108 | 0.84 | -0.613 | -0.055 | -0.031 | -0.031 | 0 | 0.39 | 0.09 | 0.43 | 0.37 | 0.39 | 0.42 |
| 6 | 14756166 | rs10484344 | -0.933 | -0.388 | 1.59 | -0.701 | -0.024 | -0.049 | -0.049 | 0 | 0.09 | 0.23 | 0.06 | 0 | 0.03 | 0.06 |
| 6 | 14756493 | rs7741301 | -1.152 | -0.389 | 1.493 | -0.717 | -0.026 | -0.061 | -0.061 | 0.209 | 0.11 | 0.36 | 0.08 | 0 | 0.03 | 0.06 |
| 6 | 14762428 | rs926242 | 0.588 | -0.325 | 0.951 | -0.748 | -0.007 | 0.004 | 0.004 | 0.156 | 0.92 | 0.92 | 0.87 | 0.96 | 0.89 | 0.9 |
| 6 | 14762768 | rs7773235 | -1.374 | -0.385 | 1.412 | -0.827 | -0.033 | -0.075 | -0.075 | 0 | 0.12 | 0.5 | 0.09 | 0 | 0.05 | 0.06 |
| 6 | 14763148 | rs2472797 | 2.151 | -0.438 | 0.999 | -0.837 | 0.021 | 0.046 | 0.046 | 1.373 | 0.78 | 0.19 | 0.77 | 0.96 | 0.84 | 0.85 |
| 6 | 14767419 | rs2472802 | 2.25 | -0.378 | 0.945 | -0.88 | 0.02 | 0.064 | 0.064 | 0.628 | 0.78 | 0.14 | 0.77 | 0.96 | 0.84 | 0.8 |
| 6 | 14772897 | rs2500081 | -2.459 | -0.696 | 1.605 | -0.964 | -0.034 | -0.075 | -0.075 | 0 | 0.21 | 0.64 | 0.25 | 0.03 | 0.12 | 0.18 |
| 6 | 14781867 | rs1014610 | -1.874 | -0.757 | 1.902 | -0.981 | 0.006 | -0.024 | -0.024 | 0.039 | 0.12 | 0.4 | 0.15 | 0.03 | 0.07 | 0.15 |
| 6 | 14783210 | rs2472811 | -2.881 | -0.686 | 1.628 | -1.109 | -0.026 | -0.067 | -0.067 | 0.092 | 0.26 | 0.62 | 0.29 | 0.05 | 0.17 | 0.23 |
| 6 | 14787728 | rs2500090 | -2.749 | -0.834 | 0.957 | -1.127 | 0.074 | 0.111 | 0.111 | 4.821 | 0.25 | 0.92 | 0.27 | 0.04 | 0.12 | 0.23 |
| 6 | 14790216 | rs2472814 | -2.028 | -0.491 | 1.485 | -0.897 | 0.013 | -0.007 | -0.007 | 0 | 0.11 | 0.33 | 0.16 | 0.04 | 0.08 | 0.21 |
| 6 | 14792476 | rs768543 | -0.584 | -0.205 | 0.846 | -0.948 | -0.025 | 0.03 | 0.03 | 0 | 0.06 | 0.02 | 0.04 | 0 | 0.07 | 0.12 |
| 6 | 14795962 | rs233500 | -1.926 | -0.353 | 1.394 | -0.93 | 0.003 | -0.019 | -0.019 | 0 | 0.1 | 0.28 | 0.17 | 0.03 | 0.07 | 0.16 |
| 6 | 14797073 | rs233504 | -1.122 | -0.32 | 0.194 | -0.465 | -0.014 | 0 | 0 | 0 | 0.55 | 0.92 | 0.48 | 0.39 | 0.4 | 0.61 |
| 6 | 14802687 | rs233492 | -1.721 | -0.241 | 1.194 | -0.407 | 0.023 | 0.005 | 0.005 | 1.889 | 0.08 | 0.33 | 0.1 | 0.02 | 0.03 | 0.1 |
| 6 | 14805514 | rs233494 | 1.402 | 0.752 | 0.796 | -0.493 | -0.027 | -0.003 | -0.003 | 0 | 0.95 | 0.96 | 0.88 | 0.96 | 0.88 | 0.96 |
| 6 | 14806317 | rs233495 | 1.459 | 0.236 | 0.743 | -0.029 | -0.029 | 0.049 | 0.049 | 0.447 | 0.46 | 0.2 | 0.5 | 0.61 | 0.57 | 0.34 |
| 6 | 14807697 | rs4715925 | -0.449 | -0.013 | 1.096 | -0.129 | 0.037 | 0.017 | 0.017 | 0.018 | 0.37 | 0.2 | 0.3 | 0.35 | 0.29 | 0.56 |
| 6 | 14823968 | rs11970548 | -2.327 | 0.545 | 1.595 | 0.099 | 0.163 | 0.023 | 0.023 | 0.744 | 0.1 | 0.43 | 0.15 | 0.05 | 0.13 | 0.08 |
| 6 | 14839778 | rs6939905 | -2.099 | 0.486 | 1.766 | 0.104 | 0.062 | 0.034 | 0.034 | 0.447 | 0.08 | 0.35 | 0.12 | 0.05 | 0.06 | 0.08 |
| 6 | 14840050 | rs2031298 | -2.039 | 0.492 | 1.72 | 0.107 | 0.042 | 0.019 | 0.019 | 0.179 | 0.08 | 0.29 | 0.12 | 0.05 | 0.06 | 0.08 |

| | | | | | | | | | | | | | | | | |
|---|---|---|---|---|---|---|---|---|---|---|---|---|---|---|---|---|
| 6 | 14845799 | rs1259003 | 2.452 | 0.334 | 1.606 | -0.037 | 0.064 | -0.032 | -0.032 | 1.444 | 0.86 | 0.6 | 0.83 | 0.93 | 0.85 | 0.9 |
| 6 | 14848468 | rs404238 | 1.708 | 0.011 | 1.054 | 0.02 | -0.138 | -0.058 | -0.058 | 0.202 | 0.44 | 0.03 | 0.48 | 0.59 | 0.55 | 0.34 |
| 6 | 14864129 | rs1258993 | 2.627 | 0.179 | 2.405 | 0.109 | 0.096 | 0.02 | 0.02 | 0.924 | 0.89 | 0.46 | 0.83 | 0.93 | 0.89 | 0.89 |
| 6 | 14879389 | rs394849 | 1.698 | 0.205 | 2.338 | 0.063 | 0.061 | 0.006 | 0.006 | 0 | 0.93 | 0.78 | 0.85 | 0.94 | 0.9 | 0.95 |
| 6 | 14884524 | rs373633 | 2.46 | 0.179 | 2.299 | 0.056 | 0.094 | 0.004 | 0.004 | 0.158 | 0.89 | 0.56 | 0.83 | 0.94 | 0.89 | 0.9 |
| 6 | 14887524 | rs374136 | 1.306 | -0.135 | 1.772 | 0.131 | 0.003 | 0.002 | 0.002 | 0.034 | 0.61 | 0.34 | 0.58 | 0.62 | 0.67 | 0.49 |
| 6 | 14894523 | rs94860 | 0.934 | -0.297 | 1.486 | 0.119 | -0.005 | -0.001 | -0.001 | 1.616 | 0.65 | 0.31 | 0.68 | 0.67 | 0.75 | 0.5 |
| 6 | 14906120 | rs10485309 | 0.075 | 0.152 | 0.77 | 0.284 | -0.019 | -0.029 | -0.029 | 2.39 | 0.05 | 0.01 | 0.14 | 0.12 | 0.09 | 0.1 |
| 6 | 14926320 | rs6908102 | -1.548 | 0.2 | 1.235 | -0.016 | 0.04 | -0.101 | -0.101 | 0.269 | 0.21 | 0.44 | 0.2 | 0.06 | 0.2 | 0.26 |
| 6 | 14944991 | rs16875934 | -1.449 | 0.267 | 1.195 | 0.22 | 0.049 | -0.001 | -0.001 | 0 | 0.06 | 0.24 | 0.1 | 0.06 | 0.07 | 0.1 |
| 6 | 14950033 | rs9383005 | 1.359 | 0.15 | 0.759 | 0.367 | -0.014 | -0.013 | -0.013 | 0 | 0.4 | 0.29 | 0.49 | 0.38 | 0.53 | 0.22 |
| 6 | 14950520 | rs441370 | -0.692 | -0.322 | 0.523 | 0.202 | 0.014 | -0.015 | -0.015 | 0 | 0.23 | 0.22 | 0.13 | 0.41 | 0.16 | 0.35 |
| 6 | 14956881 | rs432518 | 1.925 | -0.481 | 0.692 | 0.304 | -0.039 | -0.036 | -0.036 | 0.024 | 0.37 | 0.28 | 0.42 | 0.38 | 0.52 | 0.2 |
| 6 | 14956927 | rs7741153 | -1.185 | -0.584 | 0.92 | 0.229 | 0.008 | -0.017 | -0.017 | 0 | 0.28 | 0.27 | 0.3 | 0.48 | 0.24 | 0.47 |
| 6 | 14956983 | rs404305 | 0.238 | -0.631 | 0.835 | 0.046 | -0.018 | -0.076 | -0.076 | 0.409 | 0.66 | 0.55 | 0.72 | 0.86 | 0.76 | 0.67 |
| 6 | 14957708 | rs4715947 | -0.269 | -0.605 | 0.7 | 0.202 | 0.055 | -0.079 | -0.079 | 0 | 0.22 | 0.06 | 0.2 | 0.39 | 0.17 | 0.36 |
| 6 | 14966295 | rs2327868 | 0.58 | -0.557 | 1.281 | -0.289 | 0.042 | -0.012 | -0.012 | 0 | 0.93 | 0.7 | 0.87 | 1 | 0.96 | 0.83 |
| 6 | 14970341 | rs6908722 | 0.099 | -0.53 | 1.16 | -0.296 | -0.001 | -0.007 | -0.007 | 0 | 0.28 | 0.26 | 0.15 | 0.15 | 0.21 | 0.16 |
| 6 | 14977206 | rs6902553 | 2.205 | -0.622 | 0.902 | -0.166 | -0.112 | -0.035 | -0.035 | 0 | 0.37 | 0.13 | 0.47 | 0.53 | 0.52 | 0.2 |
| 6 | 14977563 | rs837170 | -0.283 | -0.661 | 1.249 | -0.2 | 0 | -0.006 | -0.006 | 2.24 | 0.72 | 0.73 | 0.84 | 0.85 | 0.79 | 0.83 |
| 6 | 14984975 | rs2661095 | 1.298 | -0.651 | 1.43 | -0.213 | 0.003 | -0.001 | -0.001 | 0 | 0.5 | 0.58 | 0.73 | 0.7 | 0.65 | 0.5 |
| 6 | 14988767 | rs2327869 | -1.382 | -0.53 | 1.302 | -0.247 | 0.009 | 0.01 | 0.01 | 1.106 | 0.36 | 0.67 | 0.37 | 0.34 | 0.26 | 0.64 |
| 6 | 14996169 | rs9476703 | -1.021 | -0.613 | 1.408 | -0.252 | -0.056 | -0.015 | -0.015 | 4.126 | 0.3 | 0.38 | 0.25 | 0.32 | 0.17 | 0.58 |
| 6 | 14997706 | rs6459385 | 0.52 | -0.442 | 1.372 | -0.269 | 0.01 | -0.042 | -0.042 | 7.721 | 0.67 | 0.6 | 0.65 | 0.84 | 0.69 | 0.77 |
| 6 | 14999452 | rs1259061 | 0.461 | -0.13 | 0.974 | -0.133 | 0.038 | -0.008 | -0.008 | 0 | 0.94 | 0.87 | 0.8 | 0.96 | 0.88 | 0.93 |
| 6 | 15001506 | rs1259065 | 1.554 | -0.135 | 0.869 | -0.17 | -0.012 | 0.013 | 0.013 | 0 | 0.43 | 0.34 | 0.5 | 0.57 | 0.53 | 0.3 |
| 6 | 15001664 | rs1259066 | 1.696 | -0.072 | 0.839 | -0.144 | -0.021 | 0.014 | 0.014 | 0 | 0.41 | 0.3 | 0.42 | 0.57 | 0.53 | 0.22 |
| 6 | 15009137 | rs724626 | 0.369 | 0.075 | 0.36 | -0.267 | 0.018 | -0.002 | -0.002 | 0.556 | 0.94 | 0.93 | 0.81 | 0.96 | 0.87 | 0.92 |
| 6 | 15013875 | rs1259074 | 0.286 | 0.08 | 0.341 | -0.109 | 0.057 | 0.012 | 0.012 | 0 | 0.94 | 0.9 | 0.8 | 0.91 | 0.87 | 0.93 |
| 6 | 15015307 | rs1259078 | 1.517 | 0.404 | -0.297 | -0.454 | 0.001 | -0.002 | -0.002 | 2.319 | 0.37 | 0.34 | 0.39 | 0.57 | 0.47 | 0.22 |
| 6 | 15016472 | rs12211166 | 0.064 | 0.389 | -1.154 | -0.813 | -0.013 | 0.045 | 0.045 | 0 | 0.23 | 0.13 | 0.13 | 0.12 | 0.19 | 0.13 |
| 6 | 15022437 | rs4715958 | -0.769 | -0.05 | -1.338 | -1.055 | 0.061 | 0.157 | 0.157 | 0.031 | 0.38 | 0.8 | 0.24 | 0.06 | 0.21 | 0.24 |
| 6 | 15025298 | rs4580854 | 0.727 | 0.706 | -0.804 | -0.279 | -0.021 | 0.038 | 0.038 | 0.732 | 0.61 | 0.32 | 0.51 | 0.71 | 0.61 | 0.37 |
| 6 | 15032722 | rs11754347 | 0.324 | -0.124 | -0.799 | -0.208 | -0.013 | 0.025 | 0.025 | 0 | 0.46 | 0.26 | 0.57 | 0.58 | 0.46 | 0.55 |
| 6 | 15034289 | rs2179990 | 0.108 | -0.251 | -0.496 | -0.3 | -0.001 | 0.021 | 0.021 | 0 | 0.5 | 0.41 | 0.58 | 0.56 | 0.47 | 0.57 |
| 6 | 15041943 | rs2038288 | 0.471 | -0.23 | -1.194 | -0.259 | -0.013 | 0.006 | 0.006 | 0 | 0.17 | 0.12 | 0.28 | 0.11 | 0.21 | 0.29 |
| 6 | 15044804 | rs1080472 | 0.615 | -0.057 | -0.462 | -0.19 | -0.019 | 0.071 | 0.071 | 0 | 0.48 | 0.26 | 0.53 | 0.59 | 0.47 | 0.6 |
| 6 | 15049404 | rs6929546 | -0.147 | -0.451 | -0.693 | -0.312 | -0.004 | 0.053 | 0.053 | 0.427 | 0.09 | 0.01 | 0.05 | 0 | 0.09 | 0.05 |
| 6 | 15059299 | rs6914891 | 0.216 | -0.043 | -0.747 | -0.08 | 0.002 | 0.021 | 0.021 | 0 | 0.54 | 0.68 | 0.52 | 0.54 | 0.42 | 0.58 |
| 6 | 15062971 | rs7744623 | -0.884 | -0.78 | -0.56 | -0.34 | -0.022 | -0.044 | -0.044 | 0.301 | 0.26 | 0.66 | 0.17 | 0.04 | 0.18 | 0.17 |
| 6 | 15066724 | rs3846947 | -0.532 | 0.055 | -0.688 | 0.417 | -0.001 | -0.011 | -0.011 | 0.806 | 0.37 | 0.3 | 0.45 | 0.5 | 0.44 | 0.49 |
| 6 | 15067436 | rs4715959 | 0.832 | -0.154 | -0.865 | 0.467 | 0.011 | -0.007 | -0.007 | 0 | 0.76 | 0.81 | 0.74 | 0.75 | 0.83 | 0.72 |
| 6 | 15067521 | rs4715960 | 0.548 | -0.022 | -1.049 | -0.14 | 0.011 | 0.063 | 0.063 | 0 | 0.37 | 0.27 | 0.35 | 0.16 | 0.3 | 0.27 |
| 6 | 15072641 | rs7751222 | 0.805 | -0.413 | -0.869 | -0.71 | 0.002 | -0.015 | -0.015 | 0 | 0.14 | 0.09 | 0.06 | 0 | 0.11 | 0.02 |
| 6 | 15072927 | rs7755465 | 1.042 | 0.014 | -0.161 | 0.116 | -0.003 | -0.004 | -0.004 | 0 | 0.34 | 0.36 | 0.33 | 0.45 | 0.43 | 0.43 |
| 6 | 15081252 | rs1739091 | 0.354 | -0.197 | 0.183 | 0.182 | 0.026 | -0.028 | -0.028 | 1.413 | 0.19 | 0.09 | 0.24 | 0.26 | 0.11 | 0.19 |
| 6 | 15089151 | rs6941421 | 1.088 | -0.164 | 0.426 | 0.61 | -0.011 | -0.001 | -0.001 | 0 | 0.33 | 0.28 | 0.28 | 0.37 | 0.44 | 0.42 |
| 6 | 15089309 | rs1770342 | -0.533 | -0.303 | 0.302 | 0.808 | 0.008 | 0.035 | 0.035 | 1.456 | 0.67 | 0.77 | 0.66 | 0.75 | 0.74 | 0.61 |
| 6 | 15089680 | rs1770341 | -0.404 | -0.304 | 0.446 | 0.807 | 0.008 | 0.021 | 0.021 | 1.461 | 0.66 | 0.7 | 0.65 | 0.76 | 0.74 | 0.61 |
| 6 | 15092665 | rs2747507 | 0.544 | -0.248 | 0.269 | 0.847 | 0.001 | -0.001 | -0.001 | 0.557 | 0.1 | 0.08 | 0.19 | 0.18 | 0.05 | 0.06 |
| 6 | 15093021 | rs6906466 | 0.921 | -0.225 | 0.315 | 0.874 | -0.003 | -0.009 | -0.009 | 0 | 0.17 | 0.09 | 0.11 | 0.21 | 0.2 | 0.07 |
| 6 | 15109703 | rs9383023 | 0.333 | -0.261 | 0.257 | 0.895 | -0.013 | -0.008 | -0.008 | 0 | 0.21 | 0.1 | 0.16 | 0.2 | 0.25 | 0.13 |
| 6 | 15121215 | rs2791609 | 0.317 | -0.165 | 0.542 | 1.091 | -0.005 | 0.012 | 0.012 | 0.047 | 0.1 | 0.13 | 0.19 | 0.17 | 0.05 | 0.06 |
| 6 | 15133865 | rs1884121 | -0.181 | -0.149 | 0.686 | 0.968 | 0.005 | 0.032 | 0.032 | 1.38 | 0.66 | 0.7 | 0.65 | 0.78 | 0.73 | 0.61 |
| 6 | 15134641 | rs9476746 | 0.863 | -0.183 | 0.735 | 0.831 | 0.001 | 0.012 | 0.012 | 0 | 0.48 | 0.41 | 0.4 | 0.38 | 0.57 | 0.47 |
| 6 | 15135210 | rs7766425 | 1.013 | -0.235 | 0.644 | 0.859 | -0.016 | 0.027 | 0.027 | 0 | 0.46 | 0.23 | 0.41 | 0.43 | 0.55 | 0.48 |
| 6 | 15136808 | rs2747498 | 0.608 | -0.307 | 1.026 | 1.159 | 0.009 | -0.012 | -0.012 | 0 | 0.56 | 0.27 | 0.54 | 0.45 | 0.61 | 0.5 |
| 6 | 15137727 | rs2144159 | -0.125 | -0.342 | 1.26 | 1.135 | 0.001 | -0.003 | -0.003 | 0.211 | 0.65 | 0.59 | 0.75 | 0.57 | 0.74 | 0.56 |
| 6 | 15138938 | rs9396561 | -0.843 | -0.278 | 1.127 | 1.253 | 0.012 | 0.024 | 0.024 | 0 | 0.17 | 0.29 | 0.33 | 0.19 | 0.18 | 0.1 |
| 6 | 15143556 | rs9370795 | -1.235 | -0.268 | 1.123 | 1.252 | 0.111 | 0.111 | 0.111 | 0.018 | 0.06 | 0.28 | 0.19 | 0.12 | 0.09 | 0.05 |
| 6 | 15145910 | rs2253602 | 0.98 | -0.277 | 1.134 | 1.161 | 0.003 | 0.037 | 0.037 | 0.476 | 0.49 | 0.35 | 0.43 | 0.37 | 0.56 | 0.49 |
| 6 | 15157289 | rs9476757 | -1.012 | -0.205 | 1.084 | 1.073 | -0.014 | -0.025 | -0.025 | 0 | 0.17 | 0.27 | 0.27 | 0.08 | 0.13 | 0.07 |
| 6 | 15163801 | rs9383032 | -0.88 | -0.169 | 1.128 | 1.103 | 0.009 | 0.032 | 0.032 | 0 | 0.17 | 0.31 | 0.32 | 0.19 | 0.15 | 0.09 |
| 6 | 15170802 | rs6935115 | -1.882 | -0.02 | 1.156 | 1.086 | 0.086 | 0.067 | 0.067 | 0.865 | 0.08 | 0.3 | 0.21 | 0.12 | 0.12 | 0.06 |
| 6 | 15208761 | rs12524343 | -0.817 | -0.175 | 0.805 | 1.065 | 0.003 | -0.051 | -0.051 | 0 | 0.17 | 0.36 | 0.28 | 0.06 | 0.13 | 0.1 |
| 6 | 15215150 | rs12664987 | 1.189 | 0.44 | -0.599 | 0.986 | -0.057 | -0.125 | -0.125 | 0 | 0.25 | 0.02 | 0.25 | 0.36 | 0.3 | 0.26 |
| 6 | 15218671 | rs12198482 | 0.131 | 0.972 | -0.067 | 1.145 | -0.016 | 0.003 | 0.003 | 0.065 | 0.23 | 0.08 | 0.2 | 0.27 | 0.21 | 0.37 |
| 6 | 15222515 | rs9349973 | -0.845 | 1.13 | 0.087 | 1.117 | -0.001 | -0.001 | -0.001 | 0.003 | 0.2 | 0.19 | 0.34 | 0.2 | 0.25 | 0.13 |
| 6 | 15240444 | rs6915344 | -1.378 | 0.783 | 0.148 | 1.031 | 0.006 | 0.028 | 0.028 | 0.28 | 0.51 | 0.72 | 0.52 | 0.41 | 0.41 | 0.54 |

| | | | | | | | | | | | | | | | | |
|---|---|---|---|---|---|---|---|---|---|---|---|---|---|---|---|---|
| 6 | 15247393 | rs12530202 | 0.155 | 0.842 | -0.079 | 1.057 | -0.002 | 0.002 | 0.002 | 0.012 | 0.12 | 0.11 | 0.25 | 0.06 | 0.12 | 0.04 |
| 6 | 15250052 | rs2295954 | 0.605 | 0.616 | -0.177 | 1.191 | -0.003 | 0.005 | 0.005 | 0.219 | 0.1 | 0.05 | 0.12 | 0.06 | 0.08 | 0.02 |
| 6 | 15256571 | rs10949286 | -0.669 | 0.433 | 0.068 | 1.157 | 0.012 | -0.002 | -0.002 | 0 | 0.34 | 0.2 | 0.25 | 0.33 | 0.26 | 0.46 |
| 6 | 15257883 | rs17619730 | 0.649 | 0.417 | -0.084 | 1.054 | 0.001 | 0.015 | 0.015 | 0 | 0.09 | 0 | 0.04 | 0.06 | 0.05 | 0.16 |
| 6 | 15259116 | rs11966818 | 0.22 | 0.431 | 0.343 | 1.235 | 0.003 | 0.037 | 0.037 | 0 | 0.11 | 0.34 | 0.13 | 0.09 | 0.08 | 0.01 |
| 6 | 15268051 | rs7766973 | -1.995 | 0.432 | 0.293 | 1.224 | 0.023 | 0.006 | 0.006 | 0.059 | 0.47 | 0.63 | 0.38 | 0.41 | 0.37 | 0.48 |
| 6 | 15271696 | rs7453429 | -0.081 | 0.404 | 0.239 | 1.271 | -0.018 | -0.009 | -0.009 | 0 | 0.23 | 0.07 | 0.23 | 0.26 | 0.21 | 0.35 |
| 6 | 15273252 | rs752861 | -0.625 | 0.436 | 0.246 | 1.143 | 0.008 | -0.011 | -0.011 | 1.041 | 0.31 | 0.15 | 0.26 | 0.33 | 0.25 | 0.42 |
| 6 | 15275305 | rs6921502 | -2.255 | 0.566 | 0.201 | 1.237 | 0.001 | 0.01 | 0.01 | 0.488 | 0.54 | 0.66 | 0.49 | 0.55 | 0.46 | 0.55 |
| 6 | 15276772 | rs12175318 | 0.693 | 0.724 | -0.143 | 1.307 | -0.019 | -0.033 | -0.033 | 0.698 | 0.07 | 0.01 | 0.07 | 0.14 | 0.09 | 0.07 |
| 6 | 15279490 | rs1322753 | -1.867 | 0.696 | -0.06 | 1.107 | 0.015 | 0.03 | 0.03 | 1.038 | 0.5 | 0.27 | 0.46 | 0.55 | 0.46 | 0.55 |
| 6 | 15280207 | rs9396578 | 0.063 | 0.703 | -0.233 | 1.202 | -0.008 | -0.043 | -0.043 | 0 | 0.16 | 0.06 | 0.18 | 0.25 | 0.17 | 0.08 |
| 6 | 15281420 | rs6459404 | -2.047 | 0.585 | 0.146 | 1.184 | 0.001 | 0.004 | 0.004 | 0.652 | 0.53 | 0.54 | 0.48 | 0.55 | 0.46 | 0.55 |
| 6 | 15281454 | rs9370809 | -1.341 | 0.936 | -0.288 | 1.226 | -0.041 | -0.041 | -0.041 | 0 | 0.29 | 0.08 | 0.36 | 0.31 | 0.32 | 0.29 |
| 6 | 15284527 | rs7757613 | -0.925 | 0.572 | 0.464 | 1.17 | 0.003 | -0.014 | -0.014 | 0 | 0.12 | 0.41 | 0.06 | 0.05 | 0.07 | 0.09 |
| 6 | 15300590 | rs794781 | -1.709 | 0.539 | 0.432 | 1.516 | -0.012 | 0.009 | 0.009 | 0.236 | 0.25 | 0.42 | 0.17 | 0.25 | 0.2 | 0.26 |
| 6 | 15305142 | rs794785 | 0.221 | 0.556 | 0.642 | 1.377 | 0.022 | 0.011 | 0.011 | 0 | 0.08 | 0.15 | 0.1 | 0.1 | 0.14 | 0.07 |
| 6 | 15308120 | rs12215596 | 0.638 | 0.55 | 0.62 | 1.379 | 0.005 | 0.012 | 0.012 | 0 | 0.08 | 0.02 | 0.04 | 0.05 | 0.04 | 0.12 |
| 6 | 15312085 | rs3954 | -1.314 | 0.448 | 0.697 | 1.385 | 0 | -0.014 | -0.014 | 0.006 | 0.19 | 0.3 | 0.15 | 0.15 | 0.18 | 0.19 |
| 6 | 15313789 | rs794791 | -2.526 | 0.376 | 0.583 | 1.554 | 0.008 | 0.002 | 0.002 | 0 | 0.45 | 0.62 | 0.3 | 0.52 | 0.37 | 0.47 |
| 6 | 15316052 | rs12199895 | 0.596 | 0.579 | 0.576 | 1.616 | -0.009 | -0.01 | -0.01 | 0 | 0.08 | 0.1 | 0.04 | 0.05 | 0.03 | 0.12 |
| 6 | 15321859 | rs957387 | 0.344 | 0.588 | 0.458 | 1.73 | -0.001 | -0.018 | -0.018 | 0 | 0.09 | 0.04 | 0.05 | 0.15 | 0.09 | 0.15 |
| 6 | 15323484 | rs16876223 | 0.508 | 0.525 | 0.567 | 1.528 | -0.012 | 0.003 | 0.003 | 0 | 0.05 | 0.1 | 0.01 | 0.05 | 0.01 | 0.07 |
| 6 | 15328073 | rs7765425 | -0.158 | 0.871 | 0.555 | 1.432 | -0.017 | -0.027 | -0.027 | 0 | 0.12 | 0.17 | 0.06 | 0.05 | 0.04 | 0.12 |
| 6 | 15334789 | rs707831 | -0.217 | 0.932 | 0.403 | 1.466 | -0.041 | -0.031 | -0.031 | 0 | 0.93 | 1 | 0.91 | 0.9 | 0.87 | 0.93 |
| 6 | 15346005 | rs13193457 | 0.409 | 0.921 | 0.456 | 1.532 | -0.001 | 0.036 | 0.036 | 0 | 0.07 | 0.12 | 0.07 | 0.19 | 0.08 | 0.01 |
| 6 | 15349589 | rs707836 | 0.145 | 0.915 | 0.398 | 1.091 | 0.015 | 0.007 | 0.007 | 0 | 0.92 | 0.87 | 0.9 | 0.9 | 0.86 | 0.93 |
| 6 | 15352243 | rs707838 | 0.676 | 0.902 | 0.262 | 1.097 | 0.002 | -0.017 | -0.017 | 0 | 0.8 | 0.71 | 0.85 | 0.85 | 0.82 | 0.81 |
| 6 | 15357299 | rs794788 | 0.833 | 0.935 | 0.209 | 1.114 | 0.003 | -0.015 | -0.015 | 0 | 0.8 | 0.72 | 0.85 | 0.84 | 0.82 | 0.81 |
| 6 | 15364430 | rs13195345 | -1.003 | 1.492 | 0.004 | 1.515 | 0.009 | 0.036 | 0.036 | 0 | 0.2 | 0.32 | 0.2 | 0.38 | 0.24 | 0.23 |
| 6 | 15365977 | rs12195286 | -0.267 | 1.615 | -0.121 | 1.684 | -0.006 | -0.007 | -0.007 | 0 | 0.09 | 0.06 | 0.08 | 0.15 | 0.13 | 0.16 |
| 6 | 15369898 | rs13205612 | 0.09 | 1.28 | -0.398 | 1.022 | 0.018 | 0.038 | 0.038 | 0.488 | 0.18 | 0.07 | 0.09 | 0.09 | 0.12 | 0.16 |
| 6 | 15371243 | rs3846949 | -0.135 | 0.797 | -0.298 | 0.81 | 0 | 0.003 | 0.003 | 0 | 0.36 | 0.49 | 0.31 | 0.33 | 0.32 | 0.31 |
| 6 | 15374962 | rs2072819 | 1.309 | 1.111 | -0.411 | 0.95 | 0.001 | -0.005 | -0.005 | 0 | 0.35 | 0.4 | 0.5 | 0.38 | 0.4 | 0.42 |
| 6 | 15385281 | rs6924733 | 0.496 | 1.112 | -0.44 | 1.06 | -0.001 | -0.004 | -0.004 | 0 | 0.53 | 0.45 | 0.58 | 0.46 | 0.5 | 0.58 |
| 6 | 15389246 | rs9396584 | -0.011 | 1.014 | -0.385 | 0.661 | 0.017 | 0.074 | 0.074 | 0.073 | 0.85 | 0.64 | 0.86 | 0.78 | 0.85 | 0.84 |
| 6 | 15394340 | rs10484355 | 1.191 | 0.877 | -0.575 | 0.519 | -0.004 | 0.048 | 0.048 | 0 | 0.15 | 0.03 | 0.08 | 0.09 | 0.11 | 0.15 |
| 6 | 15397441 | rs11756544 | 0.141 | 0.981 | -0.523 | 0.367 | -0.058 | 0.111 | 0.111 | 0.236 | 0.24 | 0.04 | 0.15 | 0.09 | 0.24 | 0.25 |
| 6 | 15397635 | rs9383054 | 1.386 | 0.978 | -0.466 | 0.439 | 0 | 0.02 | 0.02 | 0.08 | 0.37 | 0.45 | 0.53 | 0.55 | 0.4 | 0.5 |
| 6 | 15400114 | rs2237154 | -0.118 | 1.021 | -0.418 | 0.461 | -0.006 | -0.019 | -0.019 | 0 | 0.65 | 0.68 | 0.69 | 0.64 | 0.65 | 0.76 |
| 6 | 15408528 | rs2237150 | 1.38 | 1.055 | -0.264 | 0.655 | 0.001 | 0.018 | 0.018 | 0 | 0.37 | 0.45 | 0.53 | 0.48 | 0.41 | 0.51 |
| 6 | 15434412 | rs2299049 | 1.235 | 1.032 | -0.45 | 0.745 | -0.006 | -0.037 | -0.037 | 6.011 | 0.22 | 0.08 | 0.17 | 0.25 | 0.19 | 0.09 |
| 6 | 15437390 | rs2282827 | 0.475 | 1.502 | -0.488 | 0.712 | -0.01 | -0.026 | -0.026 | 0 | 0.57 | 0.65 | 0.61 | 0.57 | 0.48 | 0.64 |
| 6 | 15440339 | rs2237143 | -0.497 | 1.502 | -0.579 | 0.781 | -0.016 | -0.014 | -0.014 | 0 | 0.88 | 0.87 | 0.86 | 0.71 | 0.79 | 0.82 |
| 6 | 15445799 | rs2237142 | 1.765 | 1.289 | -0.613 | 0.742 | 0.008 | 0.005 | 0.005 | 0.36 | 0.17 | 0.03 | 0.14 | 0.14 | 0.14 | 0.03 |
| 6 | 15446447 | rs2237140 | 0.512 | 1.28 | -0.471 | 0.746 | -0.014 | 0.001 | 0.001 | 0 | 0.58 | 0.68 | 0.61 | 0.64 | 0.48 | 0.65 |
| 6 | 15449441 | rs4712251 | 0.808 | 1.581 | -0.102 | 1.014 | 0.038 | 0.002 | 0.002 | 0.033 | 0.36 | 0.23 | 0.24 | 0.35 | 0.26 | 0.16 |
| 6 | 15451898 | rs4715976 | 0.762 | 1.59 | -0.084 | 0.997 | -0.017 | -0.011 | -0.011 | 0.062 | 0.74 | 0.71 | 0.75 | 0.71 | 0.62 | 0.68 |
| 6 | 15453583 | rs2299045 | 1.108 | 1.598 | -0.14 | 0.988 | -0.01 | 0.018 | 0.018 | 0 | 0.37 | 0.45 | 0.51 | 0.47 | 0.36 | 0.52 |
| 6 | 15459934 | rs9476827 | 1.148 | 1.6 | -0.168 | 1.099 | -0.01 | 0.003 | 0.003 | 0 | 0.37 | 0.44 | 0.45 | 0.42 | 0.35 | 0.5 |
| 6 | 15462871 | rs11759978 | 0.564 | 1.625 | -0.628 | 0.783 | 0.001 | -0.015 | -0.015 | 1.899 | 0.08 | 0.08 | 0.08 | 0 | 0.13 | 0.08 |
| 6 | 15465765 | rs17455404 | 0.729 | 1.411 | -0.682 | 0.867 | 0.03 | 0.023 | 0.023 | 0.113 | 0.34 | 0.07 | 0.29 | 0.28 | 0.26 | 0.16 |
| 6 | 15476619 | rs2064097 | 0.771 | 1.789 | -0.39 | 0.628 | 0 | 0 | 0 | 0 | 0.74 | 0.7 | 0.75 | 0.81 | 0.61 | 0.68 |
| 6 | 15479224 | rs2049950 | 0.376 | 1.854 | -0.423 | 0.682 | 0.002 | 0 | 0 | 0 | 0.82 | 0.79 | 0.83 | 0.81 | 0.75 | 0.76 |
| 6 | 15489578 | rs742100 | 0.692 | 1.479 | -0.645 | 0.715 | 0.042 | 0.035 | 0.035 | 0.037 | 0.34 | 0.09 | 0.22 | 0.3 | 0.25 | 0.16 |
| 6 | 15490202 | rs2327897 | 0.44 | 1.723 | -0.516 | 0.569 | 0.005 | -0.025 | -0.025 | 0 | 0.09 | 0.1 | 0.08 | 0 | 0.15 | 0.08 |
| 6 | 15490359 | rs2237124 | 0.971 | 1.615 | 0.388 | 0.576 | -0.002 | 0.003 | 0.003 | 0 | 0.37 | 0.39 | 0.51 | 0.51 | 0.35 | 0.52 |
| 6 | 15494177 | rs2237120 | 0.956 | 1.099 | 0.051 | 0.15 | -0.01 | -0.041 | -0.041 | 0 | 0.22 | 0.13 | 0.36 | 0.44 | 0.28 | 0.39 |
| 6 | 15499419 | rs7769291 | 1.434 | 1.128 | -0.089 | 0.109 | 0.043 | 0.029 | 0.029 | 0.099 | 0.34 | 0.09 | 0.22 | 0.3 | 0.25 | 0.16 |
| 6 | 15500687 | rs2237113 | -0.398 | 1.154 | 0.251 | 0.07 | 0.001 | 0 | 0 | 0.621 | 0.49 | 0.58 | 0.6 | 0.51 | 0.5 | 0.6 |
| 6 | 15502083 | rs2237112 | 0.627 | 1.212 | 0.466 | 0.148 | 0.028 | -0.011 | -0.011 | 1.101 | 0.73 | 0.66 | 0.75 | 0.83 | 0.6 | 0.67 |
| 6 | 15503575 | rs2282816 | -0.751 | 1.269 | 0.401 | 0.4 | -0.012 | -0.006 | -0.006 | 0 | 0.92 | 0.93 | 0.89 | 0.92 | 0.87 | 0.88 |
| 6 | 15503919 | rs2282815 | 0.6 | 1.137 | 0.616 | 0.322 | 0.002 | -0.013 | -0.013 | 0.588 | 0.66 | 0.54 | 0.68 | 0.72 | 0.68 | 0.58 |
| 6 | 15504589 | rs760659 | 0.1 | 0.768 | 0.467 | 0.072 | 0.009 | -0.076 | -0.076 | 0 | 0.74 | 0.53 | 0.74 | 0.91 | 0.76 | 0.75 |
| 6 | 15506637 | rs16876555 | 0.731 | 0.626 | 0.474 | 0.45 | -0.055 | 0.063 | 0.063 | 0 | 0.17 | 0.44 | 0.17 | 0.25 | 0.07 | 0.13 |
| 6 | 15508110 | rs2072824 | -0.001 | 0.946 | -0.111 | 0.317 | -0.005 | 0.004 | 0.004 | 1.556 | 0.82 | 0.83 | 0.85 | 0.9 | 0.79 | 0.77 |
| 6 | 15510332 | rs3800496 | 0.763 | 1.294 | -0.611 | 0.209 | -0.038 | -0.099 | -0.099 | 0.732 | 0.19 | 0.06 | 0.3 | 0.48 | 0.28 | 0.36 |
| 6 | 15513482 | rs2235258 | 0.324 | 1.081 | -0.41 | 0.047 | 0.001 | -0.001 | -0.001 | 0 | 0.66 | 0.3 | 0.65 | 0.67 | 0.75 | 0.67 |
| 6 | 15517659 | rs2072822 | 0.759 | 1.262 | -0.851 | 0.155 | -0.044 | -0.104 | -0.104 | 0 | 0.18 | 0.05 | 0.29 | 0.44 | 0.27 | 0.36 |

| | | | | | | | | | | | | | | | | |
|---|---|---|---|---|---|---|---|---|---|---|---|---|---|---|---|---|
| 6 | 15517684 | rs909626 | 1.671 | 0.854 | -1.18 | -0.722 | 0.056 | 0.071 | 0.071 | 0.086 | 0.28 | 0.1 | 0.18 | 0.14 | 0.14 | 0.12 |
| 6 | 15519885 | rs13198512 | 0.309 | 0.982 | -1.13 | -0.758 | -0.008 | 0.156 | 0.156 | 0 | 0.5 | 0.22 | 0.36 | 0.22 | 0.48 | 0.31 |
| 6 | 15519914 | rs13198533 | 1.256 | 1.72 | -0.997 | -0.713 | -0.02 | 0.011 | 0.011 | 0 | 0.16 | 0.11 | 0.14 | 0 | 0.23 | 0.11 |
| 6 | 15523101 | rs1047631 | 0.206 | 1.608 | -0.705 | -0.53 | 0.021 | -0.037 | -0.037 | 3.392 | 0.11 | 0.2 | 0.1 | 0.02 | 0.15 | 0.15 |
| 6 | 15524480 | rs742106 | -0.09 | 1.83 | -1.056 | -0.036 | -0.022 | -0.029 | -0.029 | 0 | 0.26 | 0.15 | 0.44 | 0.58 | 0.36 | 0.49 |
| 6 | 15525157 | rs16876575 | 1.346 | 0.608 | -1.385 | -0.701 | 0.057 | 0.013 | 0.013 | 0 | 0.27 | 0.07 | 0.18 | 0.28 | 0.16 | 0.15 |
| 6 | 15525863 | rs9464793 | 1.667 | 0.237 | -0.99 | -0.865 | -0.033 | -0.029 | -0.029 | 0 | 0.17 | 0.23 | 0.16 | 0.08 | 0.07 | 0.11 |
| 6 | 15525997 | rs6937379 | -1.994 | 0.498 | -1.034 | -1.056 | -0.02 | 0.027 | 0.027 | 0 | 0.83 | 0.9 | 0.85 | 1 | 0.78 | 0.92 |
| 6 | 15526417 | rs4712253 | 0.271 | 0.61 | -0.716 | -0.785 | 0.001 | 0 | 0 | 0 | 0.47 | 0.44 | 0.4 | 0.47 | 0.43 | 0.46 |
| 6 | 15528249 | rs7762368 | 1.615 | 0.385 | -0.74 | -0.897 | 0.047 | 0.011 | 0.011 | 0 | 0.26 | 0.07 | 0.17 | 0.27 | 0.16 | 0.15 |
| 6 | 15530784 | rs9296979 | 0.842 | 0.556 | -0.451 | -0.673 | -0.063 | -0.053 | -0.053 | 0 | 0.19 | 0.38 | 0.17 | 0.07 | 0.08 | 0.14 |
| 6 | 15532594 | rs9296980 | 1.467 | 0.587 | -0.596 | -0.652 | -0.034 | -0.024 | -0.024 | 0 | 0.16 | 0.23 | 0.15 | 0.08 | 0.07 | 0.13 |
| 6 | 15538436 | rs875462 | -1.904 | 0.851 | -0.689 | -0.777 | -0.025 | 0.028 | 0.028 | 0 | 0.84 | 0.91 | 0.86 | 1 | 0.76 | 0.91 |
| 6 | 15538685 | rs875463 | 0.923 | 0.683 | -0.495 | -0.425 | -0.05 | -0.053 | -0.053 | 0 | 0.2 | 0.32 | 0.17 | 0.07 | 0.08 | 0.14 |
| 6 | 15539010 | rs9396592 | 0.232 | 0.936 | -0.722 | -0.186 | -0.014 | -0.016 | -0.016 | 0 | 0.26 | 0.22 | 0.42 | 0.59 | 0.37 | 0.47 |
| 6 | 15547476 | rs1040410 | 1.169 | 0.347 | -0.678 | -0.323 | -0.031 | -0.025 | -0.025 | 0 | 0.18 | 0.23 | 0.16 | 0.07 | 0.08 | 0.13 |
| 6 | 15550658 | rs9370823 | 0.719 | 0.316 | -0.686 | -0.183 | -0.015 | -0.03 | -0.03 | 2.231 | 0.19 | 0.11 | 0.3 | 0.45 | 0.28 | 0.35 |
| 6 | 15560583 | rs9476849 | 1.025 | -0.004 | -0.662 | -0.776 | -0.031 | -0.027 | -0.027 | 5.26 | 0.18 | 0.23 | 0.16 | 0.07 | 0.08 | 0.13 |
| 6 | 15565031 | rs9358063 | -0.531 | 0.639 | -1.039 | -0.329 | -0.01 | 0.037 | 0.037 | 2.761 | 0.42 | 0.37 | 0.52 | 0.88 | 0.52 | 0.67 |
| 6 | 15598880 | rs12199640 | 0.783 | 0.481 | -1.147 | 0.395 | -0.001 | -0.057 | -0.057 | 1.226 | 0.09 | 0.01 | 0.04 | 0.27 | 0.04 | 0.1 |
| 6 | 15604919 | rs7752070 | 0.424 | -0.12 | -0.879 | -0.302 | -0.055 | -0.1 | -0.1 | 0 | 0.25 | 0.41 | 0.19 | 0.07 | 0.13 | 0.15 |
| 6 | 15615637 | rs3829893 | 1.233 | -0.432 | -1.207 | -0.704 | 0.045 | -0.024 | -0.024 | 0 | 0.26 | 0.07 | 0.19 | 0.32 | 0.17 | 0.15 |
| 6 | 15616897 | rs12524690 | 1.198 | -0.402 | -1.194 | -0.664 | 0.044 | -0.024 | -0.024 | 0 | 0.26 | 0.07 | 0.19 | 0.32 | 0.17 | 0.15 |
| 6 | 15625808 | rs2743868 | -0.499 | -0.001 | -1.186 | -0.891 | -0.01 | -0.002 | -0.002 | 0 | 0.68 | 0.69 | 0.53 | 0.39 | 0.52 | 0.43 |
| 6 | 15633432 | rs1011313 | -1.424 | 0.458 | -1.198 | -0.589 | -0.024 | -0.037 | -0.037 | 0.234 | 0.92 | 0.96 | 0.87 | 0.83 | 0.9 | 0.87 |
| 6 | 15638140 | rs16876746 | 1.642 | 0.35 | -0.811 | -0.702 | 0.026 | -0.02 | -0.02 | 0.086 | 0.05 | 0.13 | 0.03 | 0 | 0.05 | 0.06 |
| 6 | 15653649 | rs2619522 | 0.483 | 0.419 | -0.944 | -0.721 | -0.033 | -0.075 | -0.075 | 0 | 0.26 | 0.42 | 0.23 | 0.09 | 0.19 | 0.22 |
| 6 | 15656155 | rs2619520 | -1.242 | -0.002 | -1.001 | -0.435 | -0.025 | -0.041 | -0.041 | 0 | 0.92 | 0.96 | 0.87 | 0.82 | 0.89 | 0.87 |
| 6 | 15658905 | rs1997679 | 0.362 | -0.24 | -0.827 | -0.921 | -0.011 | -0.079 | -0.079 | 0 | 0.36 | 0.45 | 0.31 | 0.07 | 0.31 | 0.22 |
| 6 | 15661461 | rs9476886 | 0.027 | -1.078 | -0.881 | -0.511 | 0.01 | 0.009 | 0.009 | 4.534 | 0.49 | 0.44 | 0.36 | 0.39 | 0.28 | 0.32 |
| 6 | 15663847 | rs2619536 | 0.859 | -1.151 | -0.31 | -0.923 | -0.042 | -0.101 | -0.101 | 0 | 0.17 | 0.32 | 0.16 | 0 | 0.09 | 0.05 |
| 6 | 15668661 | rs742208 | 0.46 | -1.29 | -0.235 | -0.993 | -0.054 | -0.118 | -0.118 | 0.697 | 0.2 | 0.4 | 0.17 | 0 | 0.1 | 0.06 |
| 6 | 15671731 | rs2769563 | 0.987 | -1.27 | -0.253 | -0.981 | -0.038 | -0.096 | -0.096 | 0.069 | 0.17 | 0.31 | 0.16 | 0 | 0.09 | 0.05 |
| 6 | 15674176 | rs760760 | 0.601 | -0.798 | -0.312 | -0.565 | -0.001 | 0.041 | 0.041 | 0.031 | 0.12 | 0.12 | 0.38 | 0.46 | 0.18 | 0.24 |
| 6 | 15675641 | rs7755766 | 0.954 | -0.46 | -0.326 | -0.784 | 0.014 | -0.02 | -0.02 | 0 | 0.11 | 0.13 | 0.09 | 0.02 | 0.14 | 0.15 |
| 6 | 15677343 | rs977746 | 1.61 | -0.568 | -0.67 | -1.023 | -0.033 | 0.051 | 0.051 | 0.014 | 0.23 | 0.12 | 0.17 | 0 | 0.32 | 0.11 |
| 6 | 15678724 | rs6910708 | 0.794 | -0.668 | -0.623 | -0.892 | -0.002 | 0.037 | 0.037 | 0.028 | 0.12 | 0.12 | 0.37 | 0.46 | 0.18 | 0.24 |
| 6 | 15684445 | rs6914388 | -0.117 | -1.044 | -0.444 | -1.443 | -0.014 | -0.136 | -0.136 | 0.549 | 0.31 | 0.54 | 0.26 | 0.03 | 0.23 | 0.22 |
| 6 | 15684656 | rs17473769 | 1.725 | -1.002 | -0.608 | -1.396 | -0.056 | 0.073 | 0.073 | 0 | 0.2 | 0.06 | 0.16 | 0 | 0.31 | 0.1 |
| 6 | 15686225 | rs11963843 | 1.239 | -1.027 | -0.491 | -1.412 | 0.001 | -0.051 | -0.051 | 0.344 | 0.23 | 0.23 | 0.19 | 0 | 0.31 | 0.1 |
| 6 | 15686699 | rs9296989 | -0.236 | -0.855 | -0.542 | -1.323 | 0.013 | 0.064 | 0.064 | 0 | 0.52 | 0.67 | 0.43 | 0.03 | 0.55 | 0.32 |
| 6 | 15690314 | rs6906528 | 1.046 | -0.752 | -0.738 | -0.844 | -0.015 | 0.012 | 0.012 | 0 | 0.24 | 0.13 | 0.18 | 0 | 0.32 | 0.1 |
| 6 | 15697116 | rs2031819 | -0.725 | -1.269 | -0.862 | -0.514 | 0.003 | 0.003 | 0.003 | 0 | 0.66 | 0.68 | 0.49 | 0.34 | 0.67 | 0.58 |
| 6 | 15697582 | rs2031818 | 0.47 | -1.384 | -1.273 | -0.584 | -0.047 | 0.102 | 0.102 | 0.013 | 0.4 | 0.12 | 0.32 | 0.26 | 0.52 | 0.45 |
| 6 | 15698548 | rs7450621 | 0.463 | -1.38 | -1.323 | -0.559 | -0.029 | 0.084 | 0.084 | 0.017 | 0.4 | 0.16 | 0.32 | 0.27 | 0.52 | 0.45 |
| 6 | 15712067 | rs2022348 | 0.4 | -1.364 | -1.4 | -0.542 | -0.065 | 0.08 | 0.08 | 0.156 | 0.39 | 0.1 | 0.31 | 0.27 | 0.51 | 0.45 |
| 6 | 15713873 | rs13200381 | -0.234 | -1.449 | -1.245 | -0.555 | 0.038 | 0.083 | 0.083 | 0.172 | 0.5 | 0.26 | 0.35 | 0.34 | 0.57 | 0.53 |
| 6 | 15719924 | rs4282401 | -1.151 | -1.475 | -1.424 | -0.322 | -0.007 | -0.038 | -0.038 | 0 | 0.91 | 0.96 | 0.94 | 0.85 | 0.9 | 0.86 |
| 6 | 15721214 | rs17409155 | 0.969 | -1.426 | -1.514 | -0.384 | -0.115 | 0.161 | 0.161 | 0 | 0.32 | 0.09 | 0.26 | 0.05 | 0.46 | 0.25 |
| 6 | 15722099 | rs6930152 | 0.969 | -1.29 | -0.971 | 0.093 | -0.021 | 0.01 | 0.01 | 0 | 0.07 | 0.13 | 0.03 | 0.07 | 0.05 | 0.06 |
| 6 | 15722501 | rs9464819 | 1.264 | -1.199 | -1.196 | -0.01 | 0.004 | -0.092 | -0.092 | 0.01 | 0.33 | 0.32 | 0.28 | 0 | 0.41 | 0.15 |
| 6 | 15722834 | rs9476895 | 0.71 | -1.44 | -0.35 | 0.852 | 0.02 | 0.03 | 0.03 | 0 | 0.53 | 0.38 | 0.44 | 0.28 | 0.61 | 0.51 |
| 6 | 15723361 | rs2022345 | 0.2 | -0.479 | -0.88 | 1.301 | -0.041 | -0.164 | -0.164 | 0 | 0.24 | 0.04 | 0.44 | 0.5 | 0.23 | 0.28 |
| 6 | 15736008 | rs2619514 | -0.014 | -0.55 | -0.578 | 1.053 | -0.01 | 0.021 | 0.021 | 0.821 | 0.9 | 0.81 | 0.94 | 0.86 | 0.91 | 0.85 |
| 6 | 15736512 | rs12530267 | 0.526 | -0.38 | -0.97 | 1.293 | -0.011 | -0.027 | -0.027 | 0 | 0.2 | 0.06 | 0.31 | 0.25 | 0.2 | 0.15 |
| 6 | 15739843 | rs2619513 | -0.508 | -0.2 | -0.896 | 1.021 | -0.019 | -0.015 | -0.015 | 0.034 | 0.89 | 0.95 | 0.83 | 0.85 | 0.81 | 0.82 |
| 6 | 15741773 | rs9396600 | 0.301 | -0.759 | -0.868 | 1.206 | 0.05 | -0.077 | -0.077 | 0 | 0.2 | 0.12 | 0.27 | 0.46 | 0.12 | 0.23 |
| 6 | 15746084 | rs1887605 | -0.289 | -0.471 | -0.433 | 0.056 | 0.001 | -0.049 | -0.049 | 0 | 0.27 | 0.34 | 0.25 | 0.07 | 0.26 | 0.24 |
| 6 | 15746398 | rs1927582 | 0.894 | -0.396 | -0.407 | -0.053 | 0.004 | -0.028 | -0.028 | 0 | 0.06 | 0.08 | 0.04 | 0 | 0.04 | 0.05 |
| 6 | 15746779 | rs990062 | 0.26 | -0.56 | -0.352 | -0.143 | -0.005 | 0.01 | 0.01 | 0 | 0.18 | 0.13 | 0.21 | 0.07 | 0.22 | 0.18 |
| 6 | 15747582 | rs7760793 | 0.766 | -0.645 | -0.262 | -0.791 | 0.009 | -0.07 | -0.07 | 0 | 0.27 | 0.24 | 0.22 | 0 | 0.36 | 0.15 |
| 6 | 15748472 | rs12192181 | 0.925 | -0.86 | -0.21 | -0.327 | -0.014 | 0.004 | 0.004 | 0.722 | 0.38 | 0.43 | 0.36 | 0.49 | 0.21 | 0.34 |
| 6 | 15748726 | rs9476903 | -0.115 | -0.825 | -0.17 | -0.371 | -0.004 | -0.073 | -0.073 | 0.086 | 0.45 | 0.43 | 0.42 | 0.07 | 0.58 | 0.33 |
| 6 | 15750432 | rs17475195 | 1.673 | -0.774 | -0.201 | -0.343 | -0.008 | -0.023 | -0.023 | 0.044 | 0.21 | 0.15 | 0.18 | 0 | 0.32 | 0.1 |
| 6 | 15751845 | rs7738190 | 0.848 | -0.778 | -0.216 | -0.332 | 0.028 | -0.114 | -0.114 | 0 | 0.27 | 0.3 | 0.21 | 0 | 0.36 | 0.15 |
| 6 | 15758316 | rs4715996 | -1.438 | -0.72 | -0.287 | -0.11 | -0.061 | -0.064 | -0.064 | 0 | 0.9 | 0.99 | 0.84 | 0.74 | 0.8 | 0.82 |
| 6 | 15760644 | rs1109287 | -0.731 | -0.998 | -0.325 | -0.356 | 0.001 | 0.004 | 0.004 | 0 | 0.59 | 0.56 | 0.49 | 0.25 | 0.61 | 0.52 |
| 6 | 15764638 | rs441539 | 0.165 | -0.954 | -0.359 | -0.361 | -0.002 | 0.001 | 0.001 | 0.734 | 0.38 | 0.33 | 0.28 | 0.18 | 0.39 | 0.33 |
| 6 | 15771988 | rs6913914 | 1.631 | -0.874 | -0.279 | -0.339 | 0.007 | -0.025 | -0.025 | 0.384 | 0.05 | 0.06 | 0.03 | 0 | 0.04 | 0.05 |

| | | | | | | | | | | | | | | | | |
|---|---|---|---|---|---|---|---|---|---|---|---|---|---|---|---|---|
| 6 | 15777670 | rs7739977 | 0.561 | -0.944 | -0.373 | -0.278 | -0.011 | 0.005 | 0.005 | 1.186 | 0.38 | 0.59 | 0.37 | 0.49 | 0.23 | 0.35 |
| 6 | 15786602 | rs74111 | 0.267 | -0.309 | -0.146 | -0.399 | -0.016 | 0.011 | 0.011 | 0.321 | 0.37 | 0.25 | 0.38 | 0.07 | 0.53 | 0.28 |
| 6 | 15789191 | rs9464828 | 0.516 | -0.375 | -0.093 | -0.09 | 0.001 | -0.008 | -0.008 | 0 | 0.16 | 0.12 | 0.21 | 0.06 | 0.22 | 0.18 |
| 6 | 15792723 | rs4712257 | 0.563 | -0.673 | -0.012 | 0.39 | 0.006 | -0.004 | -0.004 | 0 | 0.37 | 0.37 | 0.34 | 0.42 | 0.21 | 0.34 |
| 6 | 15802026 | rs1883450 | -0.509 | 0.355 | -1.036 | 0.86 | -0.06 | -0.062 | -0.062 | 0 | 0.9 | 0.99 | 0.83 | 0.67 | 0.8 | 0.81 |
| 6 | 15802466 | rs6914658 | -0.212 | 0.028 | -0.955 | 0.483 | 0.002 | 0.002 | 0.002 | 0 | 0.48 | 0.54 | 0.47 | 0.22 | 0.59 | 0.42 |
| 6 | 15803232 | rs387816 | -0.212 | 0.028 | -0.955 | 0.494 | 0.002 | 0.003 | 0.003 | 0 | 0.48 | 0.54 | 0.47 | 0.23 | 0.59 | 0.42 |
| 6 | 15813357 | rs4715998 | 0.612 | 0.063 | -1.001 | 0.167 | -0.01 | 0.001 | 0.001 | 2.276 | 0.41 | 0.43 | 0.36 | 0.46 | 0.21 | 0.41 |
| 6 | 15817628 | rs373949 | -0.313 | 0.375 | -0.645 | -0.41 | -0.001 | -0.047 | -0.047 | 0 | 0.45 | 0.49 | 0.43 | 0.06 | 0.58 | 0.31 |
| 6 | 15817878 | rs13216265 | -0.656 | 0.492 | -0.917 | 0.565 | -0.027 | 0.038 | 0.038 | 0.391 | 0.19 | 0.08 | 0.21 | 0.07 | 0.23 | 0.2 |
| 6 | 15819270 | rs13437271 | -1 | 0.058 | -0.4 | 0.99 | -0.014 | 0.024 | 0.024 | 0.032 | 0.17 | 0.34 | 0.19 | 0.23 | 0.14 | 0.27 |
| 6 | 15819385 | rs382182 | 0.464 | 0.392 | -0.696 | 0.848 | 0.006 | -0.011 | -0.011 | 0 | 0.58 | 0.46 | 0.61 | 0.63 | 0.57 | 0.42 |
| 6 | 15819455 | rs4712258 | 0.303 | 0.109 | -1.628 | 0.667 | -0.105 | -0.009 | -0.009 | 0.319 | 0.37 | 0.08 | 0.27 | 0.29 | 0.42 | 0.28 |
| 6 | 15820425 | rs522264 | -0.753 | -1.064 | 0.082 | 1.277 | -0.041 | 0.025 | 0.025 | 0 | 0.35 | 0.53 | 0.46 | 0.42 | 0.22 | 0.45 |
| 6 | 15825826 | rs9476952 | 0.144 | -1.13 | 0.491 | 1.401 | 0.006 | -0.021 | -0.021 | 0 | 0.25 | 0.2 | 0.41 | 0.35 | 0.26 | 0.33 |
| 6 | 15828029 | rs220884 | -0.733 | -0.664 | 1.253 | 1.47 | 0.04 | 0 | 0 | 1.947 | 0.47 | 0.72 | 0.54 | 0.44 | 0.41 | 0.55 |
| 6 | 15830025 | rs523848 | 0.181 | -0.738 | 1.7 | 1.395 | -0.004 | 0.001 | 0.001 | 0.024 | 0.33 | 0.36 | 0.46 | 0.35 | 0.3 | 0.33 |
| 6 | 15837290 | rs13211010 | -1.206 | -0.557 | 2.089 | 1.5 | -0.001 | -0.004 | -0.004 | 3.533 | 0.13 | 0.33 | 0.07 | 0.09 | 0.11 | 0.21 |
| 6 | 15838552 | rs9476953 | 0.349 | -0.604 | 2.156 | 1.496 | 0.006 | -0.002 | -0.002 | 0 | 0.36 | 0.32 | 0.47 | 0.35 | 0.32 | 0.38 |
| 6 | 15840433 | rs16877105 | -0.884 | -0.557 | 2.275 | 1.477 | 0 | -0.002 | -0.002 | 0 | 0.11 | 0.22 | 0.06 | 0.09 | 0.11 | 0.21 |
| 6 | 15840845 | rs220938 | -0.702 | -0.469 | 2.3 | 1.787 | 0.042 | 0.028 | 0.028 | 0 | 0.51 | 0.76 | 0.56 | 0.55 | 0.44 | 0.63 |
| 6 | 15843042 | rs10484351 | -1.184 | -0.486 | 2.264 | 1.747 | -0.001 | -0.004 | -0.004 | 0 | 0.13 | 0.29 | 0.07 | 0.09 | 0.11 | 0.21 |
| 6 | 15843551 | rs117348 | -0.708 | -0.478 | 1.938 | 1.764 | 0.041 | 0.005 | 0.005 | 0 | 0.5 | 0.7 | 0.55 | 0.45 | 0.43 | 0.59 |
| 6 | 15845797 | rs220933 | -0.66 | -0.471 | 1.802 | 1.762 | 0.056 | 0.004 | 0.004 | 0 | 0.51 | 0.77 | 0.56 | 0.45 | 0.43 | 0.59 |
| 6 | 15852965 | rs433285 | 1.576 | -0.495 | 1.718 | 1.95 | -0.012 | 0.078 | 0.078 | 0 | 0.77 | 0.65 | 0.78 | 0.5 | 0.81 | 0.63 |
| 6 | 15856619 | rs9370836 | 0.261 | -0.264 | 1.884 | 2.037 | 0.001 | 0.002 | 0.002 | 0.061 | 0.39 | 0.37 | 0.45 | 0.35 | 0.39 | 0.39 |
| 6 | 15858996 | rs6459417 | 1.055 | -0.295 | 1.771 | 1.699 | -0.021 | 0.073 | 0.073 | 0 | 0.85 | 0.77 | 0.82 | 0.59 | 0.9 | 0.81 |
| 6 | 15860342 | rs6459418 | 2.053 | -0.272 | 1.734 | 1.667 | 0.002 | 0.037 | 0.037 | 0 | 0.72 | 0.62 | 0.73 | 0.5 | 0.72 | 0.59 |
| 6 | 15862865 | rs17419416 | -1.11 | -0.17 | 1.69 | 1.472 | 0.015 | -0.008 | -0.008 | 0 | 0.13 | 0.2 | 0.09 | 0.09 | 0.18 | 0.23 |
| 6 | 15864284 | rs10080664 | 0.551 | -0.192 | 1.382 | 1.542 | 0.008 | -0.023 | -0.023 | 0 | 0.34 | 0.3 | 0.41 | 0.51 | 0.32 | 0.39 |
| 6 | 15873122 | rs6459420 | 1.319 | -0.512 | 1.75 | 1.946 | -0.018 | 0.06 | 0.06 | 0 | 0.92 | 0.83 | 0.88 | 0.76 | 0.96 | 0.94 |
| 6 | 15882899 | rs6903604 | 2.604 | -0.595 | 1.932 | 1.89 | -0.006 | 0.006 | 0.006 | 0.287 | 0.72 | 0.31 | 0.73 | 0.53 | 0.82 | 0.64 |
| 6 | 15883689 | rs881018 | 2.151 | -0.618 | 2.152 | 1.913 | -0.012 | 0.076 | 0.076 | 0.002 | 0.78 | 0.46 | 0.75 | 0.62 | 0.83 | 0.79 |
| 6 | 15886077 | rs1013050 | 2.758 | -0.607 | 2.168 | 1.938 | -0.02 | 0.028 | 0.028 | 0.28 | 0.72 | 0.36 | 0.73 | 0.53 | 0.82 | 0.64 |
| 6 | 15886775 | rs9476971 | -1.016 | -0.485 | 2.053 | 1.871 | 0.119 | 0.001 | 0.001 | 0.182 | 0.06 | 0.32 | 0.06 | 0 | 0.08 | 0.03 |
| 6 | 15900124 | rs4712263 | 1.601 | -0.554 | 1.421 | 1.973 | -0.007 | 0.014 | 0.014 | 0 | 0.85 | 0.83 | 0.81 | 0.62 | 0.91 | 0.82 |
| 6 | 15916396 | rs707840 | -0.462 | -0.537 | 1.36 | 1.626 | -0.006 | 0.001 | 0.001 | 0 | 0.05 | 0.05 | 0.01 | 0.07 | 0.01 | 0.16 |
| 6 | 15923183 | rs6932531 | -0.823 | -0.653 | 1.299 | 1.554 | -0.044 | 0.01 | 0.01 | 0.397 | 0.61 | 0.84 | 0.64 | 0.6 | 0.53 | 0.76 |
| 6 | 15923323 | rs6932716 | -1.051 | -0.677 | 1.831 | 1.611 | -0.011 | 0.05 | 0.05 | 0 | 0.1 | 0.32 | 0.1 | 0.17 | 0.08 | 0.14 |
| 6 | 15927034 | rs2096212 | -1.657 | -0.706 | 1.981 | 1.6 | 0 | -0.009 | -0.009 | 0 | 0.13 | 0.27 | 0.07 | 0.08 | 0.09 | 0.19 |
| 6 | 15931538 | rs9296994 | 0.687 | -0.734 | 1.637 | 1.617 | 0.007 | -0.016 | -0.016 | 0.581 | 0.38 | 0.26 | 0.47 | 0.35 | 0.35 | 0.43 |
| 6 | 15933483 | rs11966220 | -0.901 | -0.601 | 1.653 | 1.307 | -0.015 | 0.003 | 0.003 | 0 | 0.07 | 0.07 | 0.02 | 0.08 | 0.01 | 0.16 |
| 6 | 15936579 | rs16877356 | -1.783 | -0.547 | 1.698 | 1.24 | 0.05 | 0.007 | 0.007 | 0.004 | 0.16 | 0.51 | 0.16 | 0.18 | 0.18 | 0.16 |
| 6 | 15943821 | rs9383094 | -0.452 | -0.521 | 1.224 | 1.229 | -0.017 | 0.01 | 0.01 | 0 | 0.05 | 0.06 | 0.01 | 0.1 | 0.01 | 0.16 |
| 6 | 15945111 | rs7745832 | -0.951 | -0.58 | 1.204 | 1.27 | -0.028 | 0.012 | 0.012 | 0 | 0.07 | 0.11 | 0.06 | 0.16 | 0.03 | 0.09 |
| 6 | 15945195 | rs7745980 | -0.379 | -0.25 | 1.136 | 1.271 | -0.003 | 0.005 | 0.005 | 0.015 | 0.51 | 0.76 | 0.55 | 0.42 | 0.45 | 0.61 |
| 6 | 15947495 | rs6941696 | 0.468 | -0.188 | 0.558 | 0.898 | 0.005 | 0.007 | 0.007 | 0.575 | 0.38 | 0.28 | 0.42 | 0.32 | 0.36 | 0.42 |
| 6 | 15950027 | rs1539422 | 0.317 | 0.209 | -0.066 | 0.151 | -0.041 | 0.165 | 0.165 | 0.979 | 0.4 | 0.11 | 0.29 | 0.04 | 0.44 | 0.22 |
| 6 | 15958117 | rs9358075 | -0.066 | 0.036 | 0.137 | 1.089 | -0.027 | 0.017 | 0.017 | 2.037 | 0.47 | 0.2 | 0.36 | 0.39 | 0.51 | 0.49 |
| 6 | 15962288 | rs9370850 | 1.368 | 0.151 | 0.152 | 0.677 | 0.007 | -0.036 | -0.036 | 0 | 0.93 | 0.83 | 0.96 | 1 | 0.94 | 0.96 |
| 6 | 15966535 | rs12204956 | 0.481 | 0.013 | -0.361 | 0.704 | -0.144 | 0.175 | 0.175 | 1.655 | 0.37 | 0.03 | 0.29 | 0.11 | 0.43 | 0.22 |
| 6 | 15970805 | rs1891518 | 0.271 | -0.162 | 0.962 | 1.337 | -0.006 | -0.005 | -0.005 | 0.439 | 0.43 | 0.47 | 0.5 | 0.46 | 0.39 | 0.34 |
| 6 | 15985534 | rs1935787 | 0.562 | -0.218 | 1 | 1.554 | -0.038 | 0.069 | 0.069 | 1.79 | 0.41 | 0.15 | 0.3 | 0.26 | 0.42 | 0.3 |
| 6 | 15986282 | rs4716020 | 1.107 | -0.171 | 1.134 | 1.729 | 0.004 | 0.084 | 0.084 | 1.176 | 0.89 | 0.82 | 0.74 | 0.71 | 0.87 | 0.7 |
| 6 | 15990916 | rs1544214 | 0.51 | -0.174 | 1.436 | 1.837 | 0 | 0.001 | 0.001 | 1.535 | 0.42 | 0.38 | 0.41 | 0.45 | 0.4 | 0.36 |
| 6 | 15992308 | rs6924791 | -0.027 | -0.169 | 1.435 | 1.338 | -0.001 | -0.005 | -0.005 | 0 | 0.51 | 0.47 | 0.65 | 0.73 | 0.52 | 0.65 |
| 6 | 15995565 | rs6459440 | 0.028 | -0.201 | 1.148 | 1.336 | 0.006 | 0.047 | 0.047 | 1.382 | 0.5 | 0.43 | 0.33 | 0.26 | 0.47 | 0.34 |
| 6 | 15996478 | rs1891517 | 0.226 | -0.173 | 0.885 | 0.911 | -0.003 | 0.103 | 0.103 | 0.809 | 0.46 | 0.32 | 0.32 | 0.12 | 0.46 | 0.23 |
| 6 | 15999438 | rs1891519 | 1.546 | 0.553 | 1.272 | 1.443 | 0.058 | 0.045 | 0.045 | 0.369 | 0.84 | 0.59 | 0.71 | 0.83 | 0.83 | 0.72 |
| 6 | 15999893 | rs6928380 | 1.779 | 0.455 | 1.398 | 1.563 | 0.044 | 0.074 | 0.074 | 1.504 | 0.87 | 0.63 | 0.74 | 0.82 | 0.87 | 0.7 |
| 6 | 16006964 | rs12215509 | 0.693 | 0.691 | 1.029 | 1.059 | -0.009 | 0.086 | 0.086 | 0 | 0.34 | 0.13 | 0.29 | 0.16 | 0.36 | 0.15 |
| 6 | 16007935 | rs10498686 | -0.865 | 0.397 | 1.075 | 1.281 | 0.041 | 0.085 | 0.085 | 0.019 | 0.16 | 0.38 | 0.28 | 0.27 | 0.17 | 0.26 |
| 6 | 16017042 | rs10456781 | 0.655 | 0.352 | 1.047 | 1.9 | -0.008 | 0.007 | 0.007 | 0 | 0.39 | 0.21 | 0.37 | 0.36 | 0.39 | 0.4 |
| 6 | 16019115 | rs6921953 | 0.712 | 0.663 | 0.925 | 1.896 | 0 | -0.023 | -0.023 | 0 | 0.93 | 0.87 | 0.96 | 0.99 | 0.94 | 0.95 |
| 6 | 16025120 | rs16877437 | -0.822 | 0.611 | 0.941 | 2.013 | 0.006 | 0.015 | 0.015 | 0 | 0.14 | 0.18 | 0.25 | 0.27 | 0.17 | 0.26 |
| 6 | 16032433 | rs1935784 | 0.256 | 0.291 | 0.885 | 2.324 | 0 | -0.002 | -0.002 | 0 | 0.47 | 0.5 | 0.45 | 0.41 | 0.46 | 0.45 |
| 6 | 16036551 | rs4712274 | -0.079 | 0.307 | 0.904 | 2.385 | 0.005 | 0.012 | 0.012 | 0 | 0.51 | 0.64 | 0.45 | 0.54 | 0.47 | 0.57 |
| 6 | 16047221 | rs7772440 | -0.792 | 0.332 | 0.89 | 2.372 | 0.031 | 0.029 | 0.029 | 0 | 0.15 | 0.27 | 0.26 | 0.21 | 0.17 | 0.28 |
| 6 | 16047734 | rs10498685 | 1.138 | 0.242 | 0.725 | 2.371 | -0.077 | 0.021 | 0.021 | 0 | 0.33 | 0.05 | 0.28 | 0.2 | 0.37 | 0.15 |

| | | | | | | | | | | | | | | | | |
|---|---|---|---|---|---|---|---|---|---|---|---|---|---|---|---|---|
| 6 | 16051027 | rs2327948 | -0.133 | 0.613 | 0.753 | 2.384 | 0.003 | 0.002 | 0.002 | 0 | 0.49 | 0.54 | 0.39 | 0.6 | 0.41 | 0.5 |
| 6 | 16056026 | rs13192674 | -0.813 | 0.768 | 0.723 | 2.408 | 0.017 | 0.012 | 0.012 | 0 | 0.14 | 0.2 | 0.26 | 0.2 | 0.18 | 0.29 |
| 6 | 16059645 | rs2275757 | -0.087 | 0.529 | 0.252 | 1.769 | -0.008 | -0.047 | -0.047 | 0 | 0.4 | 0.49 | 0.32 | 0.21 | 0.36 | 0.36 |
| 6 | 16060085 | rs1267719 | -1.2 | 0.834 | -0.222 | 1.845 | -0.125 | 0.097 | 0.097 | 2.131 | 0.59 | 0.96 | 0.64 | 0.81 | 0.56 | 0.79 |
| 6 | 16060209 | rs1891514 | -0.409 | 0.635 | 0.123 | 2.271 | 0.018 | 0.006 | 0.006 | 0 | 0.45 | 0.68 | 0.4 | 0.59 | 0.43 | 0.5 |
| 6 | 16062613 | rs9370863 | -0.265 | 0.603 | -0.498 | 1.771 | -0.029 | 0.007 | 0.007 | 0.067 | 0.82 | 0.96 | 0.85 | 0.87 | 0.84 | 0.88 |
| 6 | 16068024 | rs7772752 | 0.857 | 0.819 | -0.389 | 2.005 | 0 | -0.007 | -0.007 | 0.099 | 0.55 | 0.52 | 0.65 | 0.63 | 0.53 | 0.57 |
| 6 | 16071974 | rs909560 | -0.074 | 0.893 | -0.362 | 2.031 | -0.003 | 0.029 | 0.029 | 0 | 0.78 | 0.67 | 0.76 | 0.68 | 0.81 | 0.81 |
| 6 | 16075936 | rs7767544 | 1.104 | 0.846 | -0.363 | 1.903 | 0.006 | 0.064 | 0.064 | 0 | 0.51 | 0.32 | 0.57 | 0.29 | 0.51 | 0.42 |
| 6 | 16077203 | rs9396641 | 1.128 | 0.849 | -0.4 | 1.905 | 0.008 | 0.082 | 0.082 | 0 | 0.51 | 0.27 | 0.56 | 0.29 | 0.51 | 0.42 |
| 6 | 16080109 | rs7775095 | -0.177 | 0.804 | -0.373 | 1.962 | -0.001 | 0.06 | 0.06 | 0 | 0.29 | 0.5 | 0.28 | 0.58 | 0.33 | 0.47 |
| 6 | 16081474 | rs9396642 | 1.997 | 0.905 | -0.996 | 1.317 | -0.059 | 0.101 | 0.101 | 0.118 | 0.34 | 0.03 | 0.3 | 0.14 | 0.32 | 0.27 |
| 6 | 16087234 | rs2294270 | -0.343 | 0.72 | -0.287 | 1.53 | -0.002 | -0.009 | -0.009 | 0.283 | 0.79 | 0.77 | 0.84 | 0.88 | 0.82 | 0.89 |
| 6 | 16089085 | rs16877617 | -0.519 | 0.85 | 0.013 | 2.033 | 0.004 | 0.025 | 0.025 | 0 | 0.28 | 0.4 | 0.21 | 0.38 | 0.33 | 0.39 |
| 6 | 16098290 | rs6459448 | 0.789 | 0.259 | 0.415 | 2.351 | -0.001 | -0.003 | -0.003 | 0 | 0.54 | 0.59 | 0.42 | 0.45 | 0.48 | 0.42 |
| 6 | 16098990 | rs11966052 | -1.037 | -1.085 | 0.747 | 2.819 | -0.01 | 0.09 | 0.09 | 0 | 0.14 | 0.19 | 0.26 | 0.32 | 0.14 | 0.22 |
| 6 | 16109163 | rs2294261 | -1.521 | -1.427 | 0.202 | 2.244 | -0.164 | 0.26 | 0.26 | 0.118 | 0.59 | 0.96 | 0.65 | 0.92 | 0.51 | 0.68 |
| 6 | 16110383 | rs9367895 | -1.5 | -0.858 | 1.606 | 3.06 | 0.068 | 0.517 | 0.517 | 3.462 | 0.06 | 0.48 | 0.08 | 0.34 | 0.01 | 0.04 |
| 6 | 16110445 | rs2327949 | -1.397 | -0.896 | 1.079 | 3.055 | -0.008 | -0.02 | -0.02 | 0 | 0.17 | 0.12 | 0.29 | 0.27 | 0.23 | 0.29 |
| 6 | 16113733 | rs2480 | -1.655 | -1.014 | 0.795 | 2.402 | -0.076 | 0.108 | 0.108 | 0 | 0.5 | 0.94 | 0.64 | 0.69 | 0.48 | 0.61 |
| 6 | 16114706 | rs11969250 | -0.717 | -0.959 | 1.276 | 2.8 | 0.055 | 0.18 | 0.18 | 0 | 0.08 | 0.18 | 0.2 | 0.4 | 0.12 | 0.16 |
| 6 | 16116748 | rs10484357 | -0.195 | -1.191 | 1.282 | 2.428 | 0.02 | -0.01 | -0.01 | 1.491 | 0.22 | 0.18 | 0.12 | 0.22 | 0.16 | 0.25 |
| 6 | 16120819 | rs9464867 | -1.331 | -0.614 | 1.981 | 2.635 | 0 | 0.087 | 0.087 | 0.163 | 0.21 | 0.34 | 0.31 | 0.41 | 0.22 | 0.33 |
| 6 | 16127407 | rs3757354 | -1.328 | -0.561 | 1.994 | 2.634 | -0.003 | 0.077 | 0.077 | 0.04 | 0.21 | 0.36 | 0.31 | 0.4 | 0.21 | 0.33 |
| 6 | 16130333 | rs909562 | -1.752 | 0.226 | 2.105 | 2.839 | 0.101 | 0.31 | 0.31 | 0.068 | 0.1 | 0.37 | 0.27 | 0.39 | 0.12 | 0.16 |
| 6 | 16132409 | rs1011616 | -1.24 | 0.312 | 1.356 | 2.884 | 0.048 | 0.172 | 0.172 | 0 | 0.08 | 0.19 | 0.25 | 0.38 | 0.12 | 0.16 |
| 6 | 16133986 | rs2237106 | -1.061 | -0.854 | 1.505 | 2.929 | -0.01 | 0.244 | 0.244 | 0 | 0.05 | 0.18 | 0.17 | 0.34 | 0.02 | 0.15 |
| 6 | 16144614 | rs2056937 | -1.241 | -1.016 | 1.485 | 2.281 | 0.015 | -0.022 | -0.022 | 0 | 0.36 | 0.65 | 0.41 | 0.46 | 0.29 | 0.43 |
| 6 | 16147349 | rs2072781 | -1.496 | -0.895 | 1.726 | 2.549 | 0.038 | 0.224 | 0.224 | 0 | 0.07 | 0.26 | 0.06 | 0.26 | 0.08 | 0.12 |
| 6 | 16148058 | rs2205795 | -0.753 | -1.085 | 1.489 | 2.153 | 0.016 | -0.01 | -0.01 | 0 | 0.26 | 0.13 | 0.32 | 0.2 | 0.21 | 0.29 |
| 6 | 16153745 | rs2072779 | -1.084 | -0.606 | 1.757 | 2.32 | -0.014 | -0.01 | -0.01 | 0.085 | 0.28 | 0.39 | 0.35 | 0.22 | 0.24 | 0.39 |
| 6 | 16179113 | rs11966292 | -1.801 | -0.151 | 1.843 | 2.479 | -0.003 | 0.161 | 0.161 | 0.298 | 0.08 | 0.24 | 0.16 | 0.32 | 0.05 | 0.07 |
| 6 | 16182921 | rs7773751 | -2.073 | -0.161 | 1.942 | 2.473 | 0.009 | 0.244 | 0.244 | 0.827 | 0.09 | 0.31 | 0.16 | 0.32 | 0.05 | 0.07 |
| 6 | 16186284 | rs11965222 | -1.817 | -0.052 | 1.221 | 1.587 | 0.001 | -0.007 | -0.007 | 0 | 0.08 | 0.15 | 0.04 | 0.07 | 0.07 | 0.05 |
| 6 | 16203561 | rs2092107 | -1.671 | -0.383 | 1.614 | 1.79 | -0.022 | 0.022 | 0.022 | 0 | 0.16 | 0.28 | 0.18 | 0.15 | 0.11 | 0.16 |
| 6 | 16209140 | rs6926350 | -1.157 | -0.808 | 1.824 | 1.826 | -0.022 | 0.007 | 0.007 | 0 | 0.08 | 0.16 | 0.06 | 0.13 | 0.02 | 0.04 |
| 6 | 16209635 | rs4712277 | -2.11 | -0.603 | 1.894 | 1.717 | -0.008 | 0.001 | 0.001 | 0 | 0.21 | 0.4 | 0.33 | 0.17 | 0.17 | 0.24 |
| 6 | 16219422 | rs7750030 | -1.479 | -0.58 | 1.93 | 1.75 | -0.006 | 0.002 | 0.002 | 0 | 0.14 | 0.16 | 0.17 | 0.15 | 0.1 | 0.16 |
| 6 | 16235577 | rs9370880 | -2.092 | -0.658 | 2.008 | 1.767 | -0.038 | 0.038 | 0.038 | 0.22 | 0.19 | 0.31 | 0.22 | 0.28 | 0.11 | 0.2 |
| 6 | 16246785 | rs3818407 | -1.81 | -0.633 | 2.258 | 2.255 | -0.019 | 0.074 | 0.074 | 0 | 0.15 | 0.34 | 0.14 | 0.22 | 0.12 | 0.22 |
| 6 | 16256043 | rs10484358 | -1.023 | -0.517 | 2.063 | 1.716 | 0.007 | 0.019 | 0.019 | 0 | 0.09 | 0.05 | 0.04 | 0 | 0.08 | 0.07 |
| 6 | 16268059 | rs9367907 | -1.474 | -0.613 | 2.414 | 1.733 | 0.01 | 0.025 | 0.025 | 0 | 0.06 | 0.21 | 0.02 | 0.07 | 0.02 | 0.05 |
| 6 | 16279097 | rs739473 | -1.419 | -0.612 | 2.41 | 1.733 | 0.016 | 0.032 | 0.032 | 0 | 0.06 | 0.22 | 0.02 | 0.07 | 0.02 | 0.05 |
| 6 | 16279176 | rs739472 | -1.917 | -0.653 | 2.357 | 1.893 | 0.001 | 0.172 | 0.172 | 0 | 0.09 | 0.33 | 0.13 | 0.22 | 0.04 | 0.18 |
| 6 | 16280056 | rs2157871 | -1.1 | -0.432 | 1.528 | 1.38 | 0.007 | 0.024 | 0.024 | 0 | 0.09 | 0.04 | 0.04 | 0 | 0.08 | 0.06 |
| 6 | 16289328 | rs11969354 | -2.355 | -0.549 | 1.405 | 1.176 | -0.003 | -0.043 | -0.043 | 0 | 0.23 | 0.25 | 0.21 | 0.03 | 0.21 | 0.28 |
| 6 | 16292760 | rs4716059 | -1.646 | -0.253 | 1.156 | 1.458 | 0 | -0.013 | -0.013 | 0.112 | 0.13 | 0.16 | 0.16 | 0.03 | 0.12 | 0.2 |
| 6 | 16300824 | rs16877817 | -1.729 | 0.4 | 1.206 | 2.374 | 0.018 | 0.104 | 0.104 | 0 | 0.11 | 0.19 | 0.1 | 0.36 | 0.17 | 0.16 |
| 6 | 16318811 | rs179996 | -0.654 | -0.004 | 0.106 | 1.395 | 0.075 | 0.066 | 0.066 | 0.21 | 0.51 | 0.9 | 0.64 | 0.62 | 0.59 | 0.65 |
| 6 | 16320320 | rs1886349 | -0.598 | -0.326 | -0.197 | 1.151 | 0 | 0.032 | 0.032 | 0 | 0.09 | 0.06 | 0.16 | 0.37 | 0.05 | 0.17 |
| 6 | 16321792 | rs179993 | 0.264 | 0.353 | -0.682 | 1.2 | -0.139 | -0.016 | -0.016 | 0 | 0.2 | 0.02 | 0.38 | 0.15 | 0.38 | 0.22 |
| 6 | 16321827 | rs2237224 | -0.299 | 0.054 | 0.563 | 1.727 | 0.029 | -0.021 | -0.021 | 0 | 0.15 | 0.25 | 0.15 | 0.07 | 0.18 | 0.2 |
| 6 | 16322055 | rs909522 | -0.427 | -0.047 | 0.88 | 1.666 | 0.057 | -0.04 | -0.04 | 0 | 0.17 | 0.38 | 0.16 | 0.07 | 0.18 | 0.21 |
| 6 | 16323392 | rs2073519 | -0.793 | -0.102 | 1.032 | 1.643 | 0.031 | 0.182 | 0.182 | 0 | 0.21 | 0.44 | 0.21 | 0.46 | 0.2 | 0.38 |
| 6 | 16332730 | rs2073518 | -1.183 | -0.633 | 1.437 | 1.886 | -0.039 | 0.203 | 0.203 | 0 | 0.12 | 0.26 | 0.18 | 0.41 | 0.05 | 0.19 |
| 6 | 16343056 | rs9383153 | -1.138 | -0.505 | 1.528 | 1.908 | -0.043 | 0.141 | 0.141 | 0 | 0.13 | 0.27 | 0.19 | 0.39 | 0.06 | 0.19 |
| 6 | 16362246 | rs179974 | -0.421 | -0.389 | 0.873 | 1.087 | 0.064 | 0.046 | 0.046 | 0 | 0.63 | 0.89 | 0.87 | 0.73 | 0.75 | 0.71 |
| 6 | 16362388 | rs179973 | -0.24 | -0.369 | 0.965 | 1.004 | 0.001 | 0.001 | 0.001 | 0 | 0.58 | 0.42 | 0.83 | 0.67 | 0.75 | 0.69 |
| 6 | 16387803 | rs9464886 | -0.897 | 0.906 | 2.456 | 2.063 | 0.039 | -0.014 | -0.014 | 1.475 | 0.09 | 0.19 | 0.13 | 0.04 | 0.17 | 0.07 |
| 6 | 16388451 | rs179957 | -0.049 | 0.855 | 1.652 | 0.825 | 0.055 | 0.003 | 0.003 | 0.106 | 0.51 | 0.31 | 0.7 | 0.53 | 0.58 | 0.63 |
| 6 | 16391928 | rs10949354 | -1.005 | 1.149 | 1.926 | 1.609 | -0.001 | 0.09 | 0.09 | 0 | 0.08 | 0.07 | 0.05 | 0.45 | 0.08 | 0.24 |
| 6 | 16393833 | rs7762645 | -1.103 | 1.174 | 1.858 | 1.458 | 0.209 | 0.09 | 0.09 | 0 | 0.06 | 0.39 | 0.11 | 0.03 | 0.13 | 0.04 |
| 6 | 16394282 | rs9383156 | -0.573 | 1.138 | 0.783 | 1.719 | -0.021 | -0.025 | -0.025 | 0 | 0.05 | 0.03 | 0.13 | 0.11 | 0.07 | 0.07 |
| 6 | 16398138 | rs179944 | -1.596 | 1.134 | 0.905 | 1.97 | 0.015 | 0.233 | 0.233 | 0 | 0.05 | 0.15 | 0.18 | 0.38 | 0.07 | 0.16 |
| 6 | 16398318 | rs179943 | -1.599 | 1.129 | 0.903 | 1.95 | 0.034 | 0.32 | 0.32 | 0 | 0.05 | 0.18 | 0.18 | 0.39 | 0.07 | 0.15 |
| 6 | 16399407 | rs9396669 | -1.872 | 0.996 | 1.033 | 1.83 | 0.018 | 0.255 | 0.255 | 0.077 | 0.08 | 0.24 | 0.19 | 0.39 | 0.08 | 0.18 |
| 6 | 16399557 | rs3819405 | -0.471 | 0.705 | 0.679 | 0.777 | -0.012 | -0.008 | -0.008 | 0.327 | 0.38 | 0.5 | 0.47 | 0.31 | 0.34 | 0.5 |
| 6 | 16399647 | rs4565296 | -0.578 | 1.107 | 0.633 | 0.534 | 0.012 | 0.007 | 0.007 | 0 | 0.06 | 0.18 | 0.05 | 0.04 | 0.07 | 0.02 |
| 6 | 16399680 | rs4431439 | -0.735 | 1.634 | 0.285 | -0.08 | -0.012 | 0.012 | 0.012 | 0 | 0.31 | 0.29 | 0.45 | 0.04 | 0.34 | 0.21 |

| | | | | | | | | | | | | | | | | |
|---|---|---|---|---|---|---|---|---|---|---|---|---|---|---|---|---|
| 6 | 16399841 | rs3819403 | -1.269 | 1.58 | 0.796 | 0.412 | 0.119 | 0.08 | 0.08 | 0 | 0.05 | 0.31 | 0.07 | 0.04 | 0.08 | 0.01 |
| 6 | 16400090 | rs3819400 | -1.408 | 1.545 | 0.44 | 0.149 | 0.02 | 0.07 | 0.07 | 1.195 | 0.29 | 0.44 | 0.28 | 0.49 | 0.36 | 0.47 |
| 6 | 16407345 | rs9464904 | -0.54 | 1.517 | -0.358 | -1.455 | -0.015 | -0.025 | -0.025 | 0.333 | 0.39 | 0.58 | 0.49 | 0.04 | 0.49 | 0.32 |
| 6 | 16410186 | rs404143 | -0.757 | 1.378 | -0.779 | -1.44 | -0.008 | 0.029 | 0.029 | 0 | 0.3 | 0.25 | 0.39 | 0.93 | 0.32 | 0.52 |
| 6 | 16415864 | rs4716068 | 0.243 | 1.114 | -1.12 | -1.441 | -0.009 | 0.047 | 0.047 | 0 | 0.16 | 0.06 | 0.08 | 0.05 | 0.15 | 0.21 |
| 6 | 16420309 | rs2237213 | -0.089 | 0.973 | -0.891 | -0.452 | 0.003 | -0.005 | -0.005 | 0.123 | 0.19 | 0.15 | 0.25 | 0.56 | 0.13 | 0.28 |
| 6 | 16420489 | rs16878121 | 0.37 | 1.397 | -0.423 | -0.776 | 0.024 | -0.065 | -0.065 | 0 | 0.11 | 0.27 | 0.2 | 0 | 0.16 | 0.04 |
| 6 | 16420840 | rs2237209 | -0.217 | 1.452 | -0.468 | -0.771 | 0.007 | 0.061 | 0.061 | 0 | 0.11 | 0.14 | 0.13 | 0.35 | 0.14 | 0.23 |
| 6 | 16421663 | rs1016222 | -2.195 | 1.417 | -1.091 | -1.406 | -0.045 | 0.147 | 0.147 | 0.548 | 0.75 | 0.98 | 0.87 | 0.98 | 0.77 | 0.87 |
| 6 | 16423412 | rs2282841 | -0.544 | 1.596 | -0.097 | -1.176 | 0.064 | 0.019 | 0.019 | 0 | 0.08 | 0.29 | 0.06 | 0.04 | 0.09 | 0.19 |
| 6 | 16424289 | rs10484361 | 0.559 | 1.598 | -0.321 | -1.441 | -0.001 | 0.013 | 0.013 | 0.053 | 0.24 | 0.14 | 0.23 | 0.01 | 0.25 | 0.12 |
| 6 | 16424956 | rs2237204 | 0.395 | 1.451 | -0.325 | -0.731 | 0.007 | -0.014 | -0.014 | 0.133 | 0.19 | 0.18 | 0.2 | 0.39 | 0.11 | 0.24 |
| 6 | 16425007 | rs2237203 | -0.304 | 1.534 | -0.184 | -0.788 | 0.023 | 0.015 | 0.015 | 0 | 0.19 | 0.35 | 0.26 | 0.22 | 0.26 | 0.29 |
| 6 | 16425304 | rs932411 | -0.227 | 1.429 | -0.147 | -0.924 | -0.002 | 0.006 | 0.006 | 0.081 | 0.32 | 0.42 | 0.42 | 0.58 | 0.3 | 0.36 |
| 6 | 16425403 | rs932412 | -0.819 | 1.293 | 0.064 | -0.407 | -0.001 | -0.003 | -0.003 | 0 | 0.62 | 0.66 | 0.69 | 0.63 | 0.63 | 0.69 |
| 6 | 16432284 | rs1015568 | -0.365 | 1.183 | -0.348 | -0.643 | 0.006 | 0.054 | 0.054 | 0 | 0.49 | 0.36 | 0.48 | 0.1 | 0.52 | 0.49 |
| 6 | 16433223 | rs4716071 | 0.212 | 1.221 | -0.85 | -0.745 | -0.035 | 0.16 | 0.16 | 0 | 0.38 | 0.15 | 0.39 | 0.01 | 0.39 | 0.24 |
| 6 | 16437642 | rs2237198 | -0.41 | 1.395 | -0.83 | -0.25 | 0.012 | 0.002 | 0.002 | 0 | 0.08 | 0.1 | 0.07 | 0.07 | 0.12 | 0.24 |
| 6 | 16441033 | rs1709172 | -0.598 | 1.091 | -1.093 | 0.265 | 0.012 | -0.068 | -0.068 | 0.007 | 0.6 | 0.85 | 0.72 | 0.56 | 0.63 | 0.68 |
| 6 | 16449304 | rs10484363 | 0.583 | 0.915 | -1.041 | 0.117 | 0.009 | 0.111 | 0.111 | 0.175 | 0.34 | 0.14 | 0.27 | 0.18 | 0.33 | 0.24 |
| 6 | 16466893 | rs2876420 | 0.08 | 0.952 | -0.783 | 0.394 | 0.011 | 0.059 | 0.059 | 0 | 0.43 | 0.31 | 0.46 | 0.77 | 0.45 | 0.62 |
| 6 | 16468304 | rs17593746 | 0.399 | 0.717 | -0.992 | 0.394 | 0.001 | 0.059 | 0.059 | 0.125 | 0.29 | 0.16 | 0.27 | 0.18 | 0.31 | 0.23 |
| 6 | 16474309 | rs1514328 | 0.039 | 0.813 | -0.549 | 0.409 | -0.008 | -0.05 | -0.05 | 0.462 | 0.28 | 0.38 | 0.31 | 0.15 | 0.24 | 0.19 |
| 6 | 16476928 | rs7750263 | 0.328 | 0.88 | -0.64 | 0.902 | -0.01 | 0.031 | 0.031 | 0 | 0.45 | 0.23 | 0.28 | 0.43 | 0.49 | 0.49 |
| 6 | 16477220 | rs9477141 | 0.59 | 1.069 | -0.45 | 0.445 | -0.005 | -0.023 | -0.023 | 0.266 | 0.23 | 0.27 | 0.34 | 0.15 | 0.21 | 0.24 |
| 6 | 16478072 | rs2237195 | 0.015 | 1.248 | -0.573 | 0.79 | -0.009 | 0.039 | 0.039 | 0 | 0.43 | 0.23 | 0.27 | 0.4 | 0.47 | 0.48 |
| 6 | 16479778 | rs7766262 | 0.344 | 1.308 | -1.125 | 0.692 | -0.092 | 0.192 | 0.192 | 0.046 | 0.38 | 0.04 | 0.23 | 0.15 | 0.41 | 0.31 |
| 6 | 16481618 | rs981459 | 0.041 | 1.339 | -0.782 | 0.752 | -0.001 | 0.001 | 0.001 | 0 | 0.44 | 0.38 | 0.3 | 0.41 | 0.5 | 0.51 |
| 6 | 16487244 | rs235146 | 0 | 1.03 | -0.863 | 0.738 | -0.005 | 0.022 | 0.022 | 0.531 | 0.43 | 0.37 | 0.47 | 0.26 | 0.42 | 0.42 |
| 6 | 16487406 | rs235147 | -0.256 | 0.847 | -0.834 | 0.968 | -0.002 | 0.002 | 0.002 | 0.007 | 0.57 | 0.49 | 0.71 | 0.63 | 0.53 | 0.8 |
| 6 | 16488572 | rs235148 | 0.464 | 0.693 | -0.321 | 0.522 | 0.046 | -0.078 | -0.078 | 0 | 0.88 | 0.66 | 0.92 | 1 | 0.88 | 0.97 |
| 6 | 16493362 | rs12204397 | 0.297 | 0.928 | -0.957 | 0.664 | -0.028 | 0.012 | 0.012 | 0 | 0.46 | 0.2 | 0.33 | 0.44 | 0.48 | 0.34 |
| 6 | 16493863 | rs9350008 | 0.679 | 0.551 | -0.658 | 0.599 | 0 | 0 | 0 | 0 | 0.64 | 0.4 | 0.6 | 0.64 | 0.65 | 0.67 |
| 6 | 16495008 | rs2237188 | -0.138 | -0.078 | -0.585 | 0.297 | -0.031 | -0.025 | -0.025 | 0.112 | 0.3 | 0.17 | 0.25 | 0.37 | 0.36 | 0.43 |
| 6 | 16497812 | rs1399219 | -0.032 | -0.398 | -0.491 | 0.301 | -0.008 | -0.015 | -0.015 | 0 | 0.29 | 0.21 | 0.23 | 0.37 | 0.32 | 0.41 |
| 6 | 16500860 | rs235158 | 0.386 | -0.574 | -0.478 | 0.373 | 0.011 | -0.008 | -0.008 | 0 | 0.08 | 0.08 | 0.16 | 0.15 | 0.02 | 0.21 |
| 6 | 16503887 | rs4716075 | -0.874 | -0.645 | 0.029 | 0.301 | 0.006 | -0.009 | -0.009 | 0 | 0.42 | 0.62 | 0.44 | 0.52 | 0.37 | 0.64 |
| 6 | 16508469 | rs4716077 | -0.132 | -0.865 | 0.225 | -0.047 | -0.009 | -0.044 | -0.044 | 0 | 0.25 | 0.44 | 0.37 | 0.15 | 0.18 | 0.37 |
| 6 | 16513846 | rs2237187 | 0.626 | -0.38 | -0.18 | 0.212 | -0.063 | -0.066 | -0.066 | 0 | 0.12 | 0.03 | 0.13 | 0.23 | 0.18 | 0.12 |
| 6 | 16514408 | rs16878314 | 0.61 | -1.309 | -0.287 | -0.427 | -0.015 | -0.041 | -0.041 | 0 | 0.17 | 0.16 | 0.13 | 0 | 0.11 | 0.05 |
| 6 | 16516192 | rs2299088 | 0.545 | -1.081 | -0.6 | -0.151 | -0.028 | -0.104 | -0.104 | 0.663 | 0.14 | 0.02 | 0.1 | 0.39 | 0.15 | 0.25 |
| 6 | 16519398 | rs3806137 | 0.279 | -0.793 | -0.304 | -0.629 | -0.022 | -0.004 | -0.004 | 0.181 | 0.26 | 0.19 | 0.3 | 0.25 | 0.33 | 0.28 |
| 6 | 16519707 | rs4716079 | 1.117 | -0.753 | -0.3 | -0.649 | 0.006 | -0.038 | -0.038 | 0 | 0.14 | 0.16 | 0.18 | 0.01 | 0.16 | 0.17 |
| 6 | 16519724 | rs3817837 | -0.156 | -1.189 | 0.1 | -0.447 | -0.003 | 0.02 | 0.02 | 0 | 0.47 | 0.45 | 0.56 | 0.77 | 0.5 | 0.71 |
| 6 | 16522720 | rs6459473 | 0.468 | -1.13 | -0.018 | -0.296 | 0.014 | -0.001 | -0.001 | 1.003 | 0.28 | 0.3 | 0.31 | 0.25 | 0.35 | 0.3 |
| 6 | 16522826 | rs9477153 | 0.663 | -0.643 | -0.745 | -0.024 | 0.011 | -0.047 | -0.047 | 0 | 0.22 | 0.18 | 0.35 | 0.53 | 0.23 | 0.47 |
| 6 | 16528671 | rs2237185 | -1.034 | 0.243 | -0.876 | 0.072 | -0.074 | 0.066 | 0.066 | 0 | 0.72 | 0.94 | 0.79 | 0.85 | 0.66 | 0.82 |
| 6 | 16536821 | rs7764112 | -0.761 | 0.803 | 0.046 | 0.54 | -0.015 | 0.024 | 0.024 | 0 | 0.06 | 0.21 | 0.08 | 0.07 | 0.03 | 0.05 |
| 6 | 16542106 | rs3765273 | -0.681 | 1.152 | -0.591 | 0.403 | -0.011 | 0.008 | 0.008 | 0 | 0.3 | 0.24 | 0.31 | 0.21 | 0.37 | 0.3 |
| 6 | 16543263 | rs9350009 | -1.244 | 0.844 | -0.37 | 0.919 | -0.024 | -0.043 | -0.043 | 0 | 0.07 | 0 | 0.11 | 0.21 | 0.08 | 0.08 |
| 6 | 16553926 | rs2237182 | -0.983 | 1.088 | -0.318 | 0.303 | -0.006 | 0 | 0 | 0 | 0.19 | 0.09 | 0.19 | 0.19 | 0.26 | 0.2 |
| 6 | 16560141 | rs11756866 | -1.23 | 0.95 | 0.457 | 0.111 | 0.043 | -0.061 | -0.061 | 0.305 | 0.1 | 0.21 | 0.12 | 0 | 0.13 | 0.02 |
| 6 | 16565192 | rs9477166 | -1.507 | 0.161 | 0.697 | 0.359 | -0.033 | 0.013 | 0.013 | 0.128 | 0.15 | 0.59 | 0.15 | 0.14 | 0.11 | 0.16 |
| 6 | 16570089 | rs236954 | 1.032 | 0.466 | 0.434 | 0.047 | 0.01 | -0.012 | -0.012 | 0 | 0.69 | 0.6 | 0.71 | 0.73 | 0.58 | 0.73 |
| 6 | 16572720 | rs721010 | 1.545 | 0.237 | 0.415 | -0.206 | -0.043 | 0.088 | 0.088 | 0.99 | 0.6 | 0.12 | 0.65 | 0.73 | 0.53 | 0.72 |
| 6 | 16573829 | rs7770397 | -0.648 | -0.632 | 1.032 | 0.183 | 0.062 | 0.014 | 0.014 | 0 | 0.05 | 0.28 | 0.04 | 0 | 0.04 | 0.01 |
| 6 | 16578284 | rs236965 | 0.955 | -0.653 | 0.555 | 0.058 | -0.002 | -0.028 | -0.028 | 0 | 0.82 | 0.67 | 0.83 | 0.89 | 0.81 | 0.85 |
| 6 | 16580575 | rs3806139 | 2.132 | -0.761 | 0.526 | 0.137 | 0.048 | 0.036 | 0.036 | 2.363 | 0.67 | 0.11 | 0.73 | 0.72 | 0.73 | 0.8 |
| 6 | 16583147 | rs236968 | -1.951 | -0.549 | 0.757 | 0.279 | -0.039 | 0.016 | 0.016 | 0.906 | 0.15 | 0.55 | 0.1 | 0.17 | 0.09 | 0.06 |
| 6 | 16588243 | rs236974 | -1.732 | -0.481 | 0.81 | 0.485 | -0.019 | 0.072 | 0.072 | 0.84 | 0.13 | 0.5 | 0.08 | 0.2 | 0.07 | 0.06 |
| 6 | 16589157 | rs236976 | -1.345 | -0.643 | 0.61 | 0.41 | -0.048 | 0.081 | 0.081 | 2.879 | 0.12 | 0.53 | 0.07 | 0.17 | 0.04 | 0.05 |
| 6 | 16591760 | rs236981 | 0.925 | -0.847 | 0.152 | 0.883 | -0.012 | 0.017 | 0.017 | 0 | 0.71 | 0.65 | 0.74 | 0.41 | 0.76 | 0.67 |
| 6 | 16598133 | rs6931635 | 1.563 | -0.767 | 0.108 | 0.632 | 0.05 | 0.054 | 0.054 | 0.67 | 0.57 | 0.24 | 0.66 | 0.27 | 0.68 | 0.61 |
| 6 | 16600679 | rs1997716 | 1.063 | -0.34 | 0.219 | 0.634 | -0.008 | 0.025 | 0.025 | 0 | 0.72 | 0.68 | 0.82 | 0.54 | 0.76 | 0.78 |
| 6 | 16601587 | rs2237172 | 1.287 | 0.458 | -0.311 | 0.877 | -0.001 | -0.012 | -0.012 | 2.978 | 0.74 | 0.24 | 0.76 | 0.53 | 0.75 | 0.76 |
| 6 | 16605411 | rs4716085 | 0.597 | 0.57 | -0.382 | 0.847 | 0.006 | -0.063 | -0.063 | 0.006 | 0.77 | 0.35 | 0.71 | 0.95 | 0.75 | 0.86 |
| 6 | 16607777 | rs3806141 | 0.583 | 0.535 | -0.415 | 1.083 | 0.001 | -0.018 | -0.018 | 4.949 | 0.69 | 0.31 | 0.67 | 0.88 | 0.68 | 0.76 |
| 6 | 16609060 | rs742328 | 0.68 | 0.605 | -0.933 | 0.811 | -0.005 | -0.041 | -0.041 | 0 | 0.63 | 0.21 | 0.59 | 0.51 | 0.64 | 0.71 |
| 6 | 16626066 | rs12665284 | -0.638 | 0.738 | -0.83 | 0.224 | -0.003 | -0.001 | -0.001 | 0.102 | 0.12 | 0.12 | 0.2 | 0.02 | 0.16 | 0.08 |

| | | | | | | | | | | | | | | | | |
|---|---|---|---|---|---|---|---|---|---|---|---|---|---|---|---|---|
| 6 | 16627255 | rs2299070 | -0.761 | -0.51 | -0.94 | -0.024 | 0.024 | 0.009 | 0.009 | 0 | 0.38 | 0.31 | 0.27 | 0.34 | 0.19 | 0.28 |
| 6 | 16628317 | rs1003384 | -1.012 | -1.135 | -1.032 | -0.389 | 0.17 | 0.078 | 0.078 | 0 | 0.5 | 0.79 | 0.35 | 0.4 | 0.23 | 0.35 |
| 6 | 16628526 | rs16878682 | 0.002 | -1.67 | -1.296 | -0.701 | 0.012 | -0.063 | -0.063 | 0 | 0.2 | 0.05 | 0.23 | 0.32 | 0.16 | 0.23 |
| 6 | 16630898 | rs17603856 | -0.186 | -0.666 | -1.855 | -1.076 | -0.099 | -0.015 | -0.015 | 0.039 | 0.2 | 0.09 | 0.43 | 0.18 | 0.35 | 0.21 |
| 6 | 16631418 | rs7745276 | -0.148 | -1.146 | -1.509 | -1.143 | -0.051 | -0.075 | -0.075 | 0.027 | 0.21 | 0.29 | 0.07 | 0.02 | 0.05 | 0.05 |
| 6 | 16633968 | rs2299068 | -2.101 | -1.581 | -1.683 | -1.216 | -0.042 | -0.081 | -0.081 | 1.796 | 0.71 | 0.91 | 0.67 | 0.57 | 0.59 | 0.58 |
| 6 | 16634454 | rs16878696 | 0.255 | -1.611 | -1.601 | -1.265 | -0.025 | -0.076 | -0.076 | 0.138 | 0.16 | 0.48 | 0.09 | 0.05 | 0.07 | 0.07 |
| 6 | 16635641 | rs6899788 | -0.429 | -1.746 | -1.577 | -1.335 | -0.032 | -0.046 | -0.046 | 0 | 0.3 | 0.64 | 0.12 | 0.06 | 0.07 | 0.12 |
| 6 | 16639120 | rs729770 | -0.795 | -1.588 | -1.655 | -1.331 | 0.031 | -0.072 | -0.072 | 0 | 0.29 | 0.59 | 0.36 | 0.08 | 0.36 | 0.26 |
| 6 | 16647158 | rs2237166 | -0.844 | -1.619 | -1.89 | -1.276 | -0.075 | -0.049 | -0.049 | 0.541 | 0.39 | 0.19 | 0.52 | 0.51 | 0.5 | 0.5 |
| 6 | 16652519 | rs2143067 | 0.039 | -1.12 | -1.795 | -0.974 | -0.069 | 0.048 | 0.048 | 0 | 0.13 | 0.07 | 0.25 | 0.03 | 0.27 | 0.15 |
| 6 | 16655201 | rs2281152 | -1.148 | -1.446 | -1.842 | -1.117 | -0.016 | -0.042 | -0.042 | 0.044 | 0.59 | 0.79 | 0.55 | 0.56 | 0.52 | 0.61 |
| 6 | 16659095 | rs909786 | -1.923 | -1.619 | -1.678 | -0.754 | -0.076 | 0.053 | 0.053 | 0.004 | 0.73 | 0.95 | 0.72 | 0.86 | 0.57 | 0.72 |
| 6 | 16659539 | rs2073514 | -0.976 | -1.687 | -1.618 | -0.71 | -0.055 | 0.091 | 0.091 | 0 | 0.61 | 0.88 | 0.67 | 0.8 | 0.51 | 0.63 |
| 6 | 16662064 | rs9477201 | 0.036 | -1.65 | -1.383 | -0.598 | 0.018 | -0.003 | -0.003 | 0.212 | 0.38 | 0.62 | 0.25 | 0.48 | 0.2 | 0.39 |
| 6 | 16664565 | rs3812189 | 0.786 | -1.954 | -1.59 | -0.82 | 0.084 | 0.026 | 0.026 | 0 | 0.36 | 0.2 | 0.21 | 0.32 | 0.2 | 0.37 |
| 6 | 16670279 | rs1014984 | -1 | -1.609 | -1.208 | -0.609 | -0.004 | 0.002 | 0.002 | 0 | 0.7 | 0.75 | 0.71 | 0.73 | 0.6 | 0.7 |
| 6 | 16670901 | rs723054 | 0.083 | -1.671 | -0.857 | -0.659 | -0.015 | 0.048 | 0.048 | 0.029 | 0.55 | 0.37 | 0.62 | 0.67 | 0.53 | 0.62 |
| 6 | 16672173 | rs7758530 | 0.056 | -1.729 | -0.802 | -0.629 | 0.039 | -0.022 | -0.022 | 0 | 0.1 | 0.39 | 0.08 | 0.05 | 0.06 | 0.07 |
| 6 | 16673598 | rs7739556 | 0.691 | -1.878 | -0.951 | -0.682 | -0.033 | -0.011 | -0.011 | 0.169 | 0.38 | 0.56 | 0.26 | 0.36 | 0.23 | 0.42 |
| 6 | 16673629 | rs7760476 | 1.496 | -1.865 | -0.99 | -0.67 | 0.071 | -0.008 | -0.008 | 0 | 0.3 | 0.16 | 0.19 | 0.32 | 0.19 | 0.35 |
| 6 | 16675733 | rs9464933 | -0.545 | -1.675 | -0.863 | -0.655 | 0.045 | 0.03 | 0.03 | 0 | 0.57 | 0.69 | 0.63 | 0.65 | 0.71 | 0.59 |
| 6 | 16677571 | rs3812192 | 0.965 | -1.519 | -0.927 | -0.651 | 0 | 0 | 0 | 6.337 | 0.3 | 0.24 | 0.31 | 0.28 | 0.48 | 0.3 |
| 6 | 16681323 | rs3793103 | -0.423 | -1.463 | -0.399 | -0.721 | 0.041 | 0.085 | 0.085 | 0 | 0.54 | 0.72 | 0.58 | 0.68 | 0.59 | 0.6 |
| 6 | 16683672 | rs10484365 | -0.074 | -1.306 | -0.267 | -0.701 | 0 | -0.014 | -0.014 | 0.975 | 0.07 | 0.09 | 0.16 | 0 | 0.08 | 0.04 |
| 6 | 16686066 | rs6903850 | 0.021 | -1.305 | -0.3 | -0.706 | -0.012 | 0.023 | 0.023 | 0.025 | 0.15 | 0.1 | 0.22 | 0.02 | 0.21 | 0.06 |
| 6 | 16687493 | rs17669102 | 0.422 | -1.456 | -0.314 | -0.697 | -0.029 | 0.045 | 0.045 | 0 | 0.18 | 0.09 | 0.32 | 0.01 | 0.22 | 0.17 |
| 6 | 16689882 | rs9464935 | -0.528 | -1.299 | -0.179 | -0.617 | 0.018 | -0.087 | -0.087 | 0.018 | 0.2 | 0.44 | 0.14 | 0.06 | 0.2 | 0.09 |
| 6 | 16692186 | rs9370920 | -0.58 | -1.192 | -0.889 | -0.437 | -0.028 | -0.027 | -0.027 | 0 | 0.35 | 0.22 | 0.47 | 0.36 | 0.42 | 0.29 |
| 6 | 16694177 | rs12204969 | 0.662 | -0.99 | -1.107 | -0.555 | -0.01 | 0.017 | 0.017 | 0.805 | 0.1 | 0.06 | 0.06 | 0.01 | 0.14 | 0.02 |
| 6 | 16694386 | rs1407830 | -0.974 | -1.126 | -1.197 | -0.342 | 0.064 | 0.026 | 0.026 | 0 | 0.7 | 0.85 | 0.8 | 0.73 | 0.81 | 0.67 |
| 6 | 16695473 | rs3812199 | -0.092 | 0.521 | -0.288 | 0.427 | -0.001 | -0.005 | -0.005 | 0.199 | 0.41 | 0.45 | 0.35 | 0.34 | 0.42 | 0.4 |
| 6 | 16696724 | rs1359067 | -0.887 | 0.499 | -0.379 | 0.507 | 0.019 | 0.081 | 0.081 | 0 | 0.59 | 0.78 | 0.7 | 0.75 | 0.61 | 0.62 |
| 6 | 16698022 | rs3812204 | -0.459 | 0.489 | -0.279 | 0.481 | -0.001 | -0.001 | -0.001 | 0 | 0.41 | 0.57 | 0.46 | 0.63 | 0.31 | 0.38 |
| 6 | 16699330 | rs9383201 | 0.266 | 0.62 | -0.285 | 0.585 | 0.009 | 0.049 | 0.049 | 0 | 0.25 | 0.35 | 0.4 | 0.45 | 0.24 | 0.34 |
| 6 | 16699949 | rs3812205 | -0.039 | 0.957 | -0.246 | 0.531 | 0 | -0.001 | -0.001 | 0 | 0.18 | 0.16 | 0.07 | 0.19 | 0.07 | 0.07 |
| 6 | 16700833 | rs3812207 | -0.184 | 1.386 | -0.686 | 0.679 | -0.005 | 0 | 0 | 0 | 0.33 | 0.25 | 0.29 | 0.32 | 0.37 | 0.31 |
| 6 | 16706041 | rs531089 | -0.035 | 1.547 | -0.595 | 0.713 | -0.01 | -0.007 | -0.007 | 0.006 | 0.54 | 0.31 | 0.39 | 0.51 | 0.57 | 0.36 |
| 6 | 16712658 | rs707848 | 0.096 | 1.085 | -0.102 | 0.837 | 0.019 | 0 | 0 | 0 | 0.66 | 0.48 | 0.61 | 0.64 | 0.54 | 0.66 |
| 6 | 16714617 | rs1262918 | 0.148 | 0.728 | -0.511 | 0.925 | -0.003 | -0.002 | -0.002 | 0.881 | 0.32 | 0.33 | 0.39 | 0.26 | 0.26 | 0.29 |
| 6 | 16717736 | rs3812216 | 0.648 | 0.574 | -0.162 | 0.859 | 0.004 | -0.002 | -0.002 | 0.001 | 0.24 | 0.18 | 0.18 | 0.21 | 0.16 | 0.12 |
| 6 | 16722046 | rs13216386 | 0.848 | 0.099 | -0.6 | 0.869 | -0.04 | 0.098 | 0.098 | 0.115 | 0.24 | 0.06 | 0.2 | 0.07 | 0.25 | 0.25 |
| 6 | 16722436 | rs3828867 | -0.26 | 0.039 | -0.286 | 0.751 | 0.008 | -0.001 | -0.001 | 0 | 0.44 | 0.41 | 0.3 | 0.59 | 0.35 | 0.45 |
| 6 | 16724881 | rs3828868 | 0.248 | 0.047 | -0.162 | 0.742 | 0.006 | 0 | 0 | 0.084 | 0.37 | 0.33 | 0.25 | 0.38 | 0.29 | 0.38 |
| 6 | 16728780 | rs1001210 | -0.673 | 0.02 | -0.233 | 0.804 | 0.019 | -0.007 | -0.007 | 0 | 0.23 | 0.16 | 0.22 | 0.76 | 0.15 | 0.28 |
| 6 | 16729739 | rs1150635 | 0.287 | -0.025 | -0.082 | 0.703 | 0.001 | -0.001 | -0.001 | 0 | 0.36 | 0.39 | 0.26 | 0.38 | 0.29 | 0.39 |
| 6 | 16731200 | rs6459484 | 0.257 | 0.027 | -0.334 | 0.795 | -0.009 | 0.039 | 0.039 | 0 | 0.09 | 0.17 | 0.06 | 0.22 | 0.06 | 0.07 |
| 6 | 16731311 | rs10484366 | 1.111 | 0.114 | -0.539 | 0.706 | 0.024 | -0.046 | -0.046 | 0 | 0.1 | 0.02 | 0.07 | 0.31 | 0.03 | 0.12 |
| 6 | 16736206 | rs4716092 | 0.239 | 0.097 | -0.559 | 0.665 | -0.012 | -0.048 | -0.048 | 0 | 0.35 | 0.08 | 0.23 | 0.38 | 0.29 | 0.39 |
| 6 | 16737108 | rs3793119 | 0.934 | 0.161 | -0.327 | 0.961 | 0.024 | -0.046 | -0.046 | 0 | 0.1 | 0.02 | 0.07 | 0.32 | 0.03 | 0.13 |
| 6 | 16740750 | rs598908 | -0.929 | 0.438 | -0.318 | 0.355 | 0.006 | -0.007 | -0.007 | 0 | 0.77 | 0.75 | 0.68 | 0.85 | 0.59 | 0.71 |
| 6 | 16741886 | rs3812227 | -0.639 | -0.061 | -0.522 | 0.519 | 0.027 | -0.065 | -0.065 | 0 | 0.27 | 0.17 | 0.3 | 0.59 | 0.17 | 0.32 |
| 6 | 16742033 | rs697739 | -0.662 | -0.263 | -0.396 | 0.177 | 0.038 | 0.072 | 0.072 | 0 | 0.51 | 0.66 | 0.51 | 0.78 | 0.32 | 0.43 |
| 6 | 16742765 | rs1144695 | -0.752 | -0.344 | -0.462 | 0.038 | 0.049 | 0.152 | 0.152 | 0 | 0.53 | 0.74 | 0.52 | 0.86 | 0.33 | 0.44 |
| 6 | 16742925 | rs1144696 | 0.86 | -0.459 | -0.146 | 0.071 | -0.029 | 0.035 | 0.035 | 0.267 | 0.24 | 0.41 | 0.21 | 0.28 | 0.16 | 0.12 |
| 6 | 16742952 | rs570917 | 1.003 | -0.257 | -0.296 | -0.1 | -0.028 | 0.073 | 0.073 | 0 | 0.27 | 0.13 | 0.31 | 0.06 | 0.31 | 0.34 |
| 6 | 16743334 | rs1144698 | 0.306 | -0.321 | -0.275 | 0.084 | -0.026 | 0.026 | 0.026 | 0 | 0.29 | 0.42 | 0.24 | 0.36 | 0.19 | 0.2 |
| 6 | 16743776 | rs1150639 | 0.903 | -0.314 | -0.254 | 0.116 | -0.028 | 0.027 | 0.027 | 0.041 | 0.24 | 0.39 | 0.22 | 0.28 | 0.16 | 0.13 |
| 6 | 16743909 | rs16878970 | 0.861 | -0.366 | -0.288 | 0.291 | -0.001 | 0.006 | 0.006 | 0 | 0.07 | 0.08 | 0.08 | 0.15 | 0.05 | 0.06 |
| 6 | 16744169 | rs493352 | -0.095 | -0.346 | -0.373 | 0.293 | -0.004 | -0.011 | -0.011 | 0 | 0.59 | 0.62 | 0.61 | 0.5 | 0.54 | 0.51 |
| 6 | 16744616 | rs651702 | -0.625 | -0.363 | -0.352 | 0.285 | 0.004 | -0.015 | -0.015 | 0.021 | 0.72 | 0.68 | 0.7 | 0.84 | 0.57 | 0.65 |
| 6 | 16744687 | rs1144700 | 0.239 | -0.362 | -0.531 | 0.249 | -0.042 | -0.011 | -0.011 | 1.203 | 0.37 | 0.45 | 0.3 | 0.62 | 0.19 | 0.27 |
| 6 | 16745530 | rs547491 | -1.176 | -0.243 | -1.159 | 0.286 | -0.034 | 0.043 | 0.043 | 0 | 0.77 | 0.86 | 0.72 | 0.94 | 0.59 | 0.72 |
| 6 | 16746624 | rs3806152 | 1.006 | -0.319 | -1.139 | 0.434 | 0.022 | -0.025 | -0.025 | 1.069 | 0.12 | 0.07 | 0.09 | 0.35 | 0.03 | 0.13 |
| 6 | 16754366 | rs2301594 | 0.965 | 0.001 | -0.806 | 0.112 | 0.043 | -0.014 | -0.014 | 0 | 0.32 | 0.12 | 0.26 | 0.61 | 0.18 | 0.27 |
| 6 | 16757006 | rs631661 | 0.805 | 0.423 | -0.929 | -0.381 | -0.086 | 0.081 | 0.081 | 0 | 0.24 | 0.05 | 0.16 | 0.08 | 0.28 | 0.25 |
| 6 | 16758112 | rs2927 | -0.664 | 0.573 | -0.546 | -0.253 | -0.028 | 0.031 | 0.031 | 0 | 0.78 | 0.85 | 0.72 | 0.92 | 0.59 | 0.71 |
| 6 | 16763241 | rs471716 | -0.057 | 0.312 | -0.39 | -0.356 | -0.003 | 0.021 | 0.021 | 0.001 | 0.34 | 0.27 | 0.21 | 0.15 | 0.33 | 0.3 |
| 6 | 16769798 | rs623897 | -0.339 | 0.849 | -0.993 | -0.05 | -0.037 | -0.068 | -0.068 | 0 | 0.76 | 0.86 | 0.66 | 0.51 | 0.62 | 0.75 |

| | | | | | | | | | | | | | | | |
|---|---|---|---|---|---|---|---|---|---|---|---|---|---|---|---|
| 6 | 16769840 | rs16879127 | 0.302 | 0.359 | -0.622 | 0.252 | 0.013 | 0.004 | 0.004 | 1.386 | 0.38 | 0.27 | 0.43 | 0.35 | 0.3 | 0.42 |
| 6 | 16771465 | rs12664358 | -0.509 | 0.673 | -0.424 | 0.315 | 0.001 | -0.005 | -0.005 | 0 | 0.05 | 0.06 | 0.02 | 0.12 | 0.04 | 0.08 |
| 6 | 16772207 | rs9396710 | -0.984 | 0.26 | 0.188 | 0.533 | -0.032 | 0.142 | 0.142 | 0 | 0.14 | 0.35 | 0.2 | 0.62 | 0.08 | 0.24 |
| 6 | 16772225 | rs628059 | 0.042 | 0.672 | -0.228 | -0.5 | 0.016 | -0.021 | -0.021 | 0 | 0.65 | 0.6 | 0.55 | 0.73 | 0.58 | 0.66 |
| 6 | 16772252 | rs628021 | -0.174 | 0.8 | -0.201 | -0.644 | -0.012 | 0.021 | 0.021 | 0.012 | 0.7 | 0.87 | 0.64 | 0.76 | 0.68 | 0.66 |
| 6 | 16773382 | rs602206 | -0.722 | 0.704 | 0.142 | -0.954 | -0.01 | -0.02 | -0.02 | 0.651 | 0.39 | 0.57 | 0.33 | 0.73 | 0.31 | 0.43 |
| 6 | 16774161 | rs598948 | 0.44 | 0.719 | -0.302 | -1.025 | -0.066 | 0.075 | 0.075 | 0.343 | 0.23 | 0.04 | 0.1 | 0.08 | 0.24 | 0.2 |
| 6 | 16777131 | rs528785 | -0.537 | 0.854 | 0.034 | -1.131 | 0.005 | 0.019 | 0.019 | 0.376 | 0.41 | 0.66 | 0.33 | 0.73 | 0.35 | 0.43 |
| 6 | 16793870 | rs765528 | 0.022 | 1.39 | -0.065 | -0.976 | 0 | -0.001 | -0.001 | 0.151 | 0.18 | 0.16 | 0.37 | 0.22 | 0.29 | 0.18 |
| 6 | 16795923 | rs4618530 | 0.114 | 1.297 | -0.255 | -1.191 | -0.006 | -0.022 | -0.022 | 0 | 0.27 | 0.16 | 0.23 | 0.61 | 0.28 | 0.4 |
| 6 | 16796614 | rs642859 | 1.225 | 1.126 | -0.205 | -1.365 | -0.001 | -0.048 | -0.048 | 0.022 | 0.43 | 0.43 | 0.34 | 0.07 | 0.4 | 0.28 |
| 6 | 16799604 | rs12205787 | 0.411 | 1.321 | -1.013 | -1.187 | -0.036 | 0.056 | 0.056 | 0.126 | 0.2 | 0.04 | 0.12 | 0.12 | 0.22 | 0.26 |
| 6 | 16801767 | rs10949381 | -0.605 | 1.149 | -0.884 | -1.111 | -0.034 | 0.013 | 0.013 | 0 | 0.39 | 0.24 | 0.49 | 0.44 | 0.51 | 0.49 |
| 6 | 16801866 | rs607341 | 0.264 | 0.937 | -0.614 | -0.631 | -0.027 | 0.095 | 0.095 | 0.053 | 0.83 | 0.73 | 0.84 | 0.51 | 0.9 | 0.78 |
| 6 | 16804245 | rs6905556 | 0.593 | 1.57 | -1.232 | -0.855 | -0.035 | -0.025 | -0.025 | 1.47 | 0.07 | 0 | 0.07 | 0.09 | 0.1 | 0.05 |
| 6 | 16806220 | rs11752823 | 0.266 | 1.56 | -1.173 | -0.754 | 0.002 | -0.024 | -0.024 | 0.022 | 0.24 | 0.18 | 0.21 | 0.57 | 0.23 | 0.39 |
| 6 | 16806341 | rs685183 | -0.536 | 1.242 | -1.251 | -0.943 | -0.012 | 0.046 | 0.046 | 0 | 0.79 | 0.95 | 0.87 | 0.86 | 0.82 | 0.87 |
| 6 | 16806573 | rs7740575 | -0.295 | 1.747 | -0.515 | 0.098 | -0.003 | -0.017 | -0.017 | 0 | 0.25 | 0.2 | 0.23 | 0.61 | 0.26 | 0.43 |
| 6 | 16807092 | rs6902696 | -0.232 | 1.857 | -0.392 | 0.205 | 0.045 | 0.032 | 0.032 | 0 | 0.1 | 0.16 | 0.32 | 0.15 | 0.23 | 0.13 |
| 6 | 16807562 | rs13193662 | -0.228 | 0.893 | -0.833 | 0.508 | -0.003 | -0.016 | -0.016 | 0 | 0.21 | 0.17 | 0.19 | 0.55 | 0.22 | 0.37 |
| 6 | 16807715 | rs604217 | -0.566 | 0.375 | -0.775 | -0.129 | 0.006 | 0.038 | 0.038 | 0 | 0.28 | 0.17 | 0.26 | 0.09 | 0.25 | 0.23 |
| 6 | 16814304 | rs1150644 | -0.721 | -0.189 | -0.054 | 1.118 | 0.003 | -0.002 | -0.002 | 0 | 0.37 | 0.36 | 0.35 | 0.59 | 0.28 | 0.23 |
| 6 | 16814885 | rs1407536 | -0.688 | -0.395 | -0.037 | 1.976 | 0.038 | -0.029 | -0.029 | 0 | 0.34 | 0.07 | 0.23 | 0.34 | 0.27 | 0.17 |
| 6 | 16817368 | rs1240808 | -1.294 | -0.264 | 0.55 | 2.254 | -0.008 | -0.029 | -0.029 | 0 | 0.44 | 0.51 | 0.46 | 0.8 | 0.36 | 0.43 |
| 6 | 16818625 | rs9396712 | -0.548 | -0.351 | 0.658 | 2.28 | 0.041 | -0.063 | -0.063 | 2.965 | 0.35 | 0.15 | 0.33 | 0.59 | 0.28 | 0.23 |
| 6 | 16819108 | rs12525399 | -0.892 | -0.435 | 0.776 | 2.262 | 0.053 | -0.03 | -0.03 | 0 | 0.38 | 0.24 | 0.34 | 0.59 | 0.27 | 0.23 |
| 6 | 16821679 | rs9383211 | -0.219 | -0.63 | 0.6 | 2.371 | 0.026 | -0.088 | -0.088 | 0 | 0.26 | 0.01 | 0.17 | 0.34 | 0.18 | 0.13 |
| 6 | 16823746 | rs4716098 | -0.24 | -0.587 | 0.639 | 2.038 | 0 | 0 | 0 | 0 | 0.06 | 0.05 | 0.05 | 0 | 0.07 | 0.04 |
| 6 | 16825137 | rs1240820 | -1.178 | -0.586 | 0.776 | 2.123 | -0.015 | -0.024 | -0.024 | 0 | 0.4 | 0.43 | 0.44 | 0.79 | 0.31 | 0.36 |
| 6 | 16841508 | rs12204538 | -2.248 | -0.376 | 1.394 | 2.058 | 0.05 | 0.235 | 0.235 | 0.868 | 0.07 | 0.46 | 0.15 | 0.19 | 0.06 | 0.2 |
| 6 | 16856088 | rs9370929 | -0.209 | -0.487 | 0.684 | 2.257 | -0.004 | -0.065 | -0.065 | 0 | 0.2 | 0.04 | 0.18 | 0.3 | 0.18 | 0.1 |
| 6 | 16862985 | rs1240825 | -1.383 | -0.529 | 1.211 | 2.172 | 0.046 | 0.201 | 0.201 | 0.129 | 0.05 | 0.28 | 0.13 | 0.18 | 0.06 | 0.17 |
| 6 | 16869419 | rs770938 | -1.707 | -0.65 | 1.201 | 2.066 | -0.006 | 0.091 | 0.091 | 0.012 | 0.11 | 0.43 | 0.15 | 0.18 | 0.07 | 0.18 |
| 6 | 16882702 | rs3006076 | -1.612 | -0.665 | 0.98 | 2.116 | -0.022 | 0.02 | 0.02 | 0 | 0.33 | 0.59 | 0.34 | 0.46 | 0.25 | 0.28 |
| 6 | 16892412 | rs9367931 | -0.209 | -0.333 | -0.215 | 1.852 | -0.009 | -0.078 | -0.078 | 0.18 | 0.17 | 0.01 | 0.11 | 0.24 | 0.13 | 0.06 |
| 6 | 16899443 | rs4145201 | -0.927 | -0.018 | 0.262 | 1.735 | -0.008 | 0.006 | 0.006 | 0 | 0.32 | 0.42 | 0.37 | 0.62 | 0.25 | 0.43 |
| 6 | 16899778 | rs6909376 | -1.348 | 1.624 | 0.731 | 1.417 | 0.024 | -0.013 | -0.013 | 0.054 | 0.21 | 0.27 | 0.35 | 0.14 | 0.33 | 0.29 |
| 6 | 16901138 | rs4074661 | -1.347 | 2.396 | 0.899 | 1.39 | 0.088 | -0.021 | -0.021 | 0.334 | 0.21 | 0.34 | 0.33 | 0.15 | 0.42 | 0.29 |
| 6 | 16910129 | rs9350018 | -1.533 | 2.241 | 0.963 | 2.081 | 0.111 | 0.108 | 0.108 | 0.884 | 0.19 | 0.4 | 0.34 | 0.33 | 0.34 | 0.34 |
| 6 | 16916543 | rs9396724 | -1.625 | 1.532 | 0.868 | 2.261 | 0.051 | 0.283 | 0.283 | 0.573 | 0.06 | 0.39 | 0.05 | 0.19 | 0.03 | 0.11 |
| 6 | 16918550 | rs1473730 | -0.981 | 1.481 | -0.244 | 1.886 | -0.02 | -0.058 | -0.058 | 0.011 | 0.13 | 0.01 | 0.06 | 0.19 | 0.1 | 0.05 |
| 6 | 16920208 | rs9383214 | -1.544 | 2.33 | 0.216 | 1.862 | 0.073 | 0.11 | 0.11 | 0.063 | 0.26 | 0.48 | 0.38 | 0.47 | 0.38 | 0.23 |
| 6 | 16921131 | rs9297020 | 0.033 | 2.036 | -0.542 | 0.963 | -0.001 | 0.021 | 0.021 | 0 | 0.74 | 0.78 | 0.83 | 0.82 | 0.78 | 0.63 |
| 6 | 16921147 | rs6910676 | 0.476 | 2.399 | -0.325 | 1.188 | -0.004 | -0.013 | -0.013 | 0 | 0.4 | 0.4 | 0.26 | 0.28 | 0.35 | 0.28 |
| 6 | 16927529 | rs927366 | 0.446 | 2.478 | -0.186 | 1.537 | -0.021 | -0.02 | -0.02 | 0.504 | 0.38 | 0.5 | 0.2 | 0.3 | 0.23 | 0.29 |
| 6 | 16929712 | rs7763768 | 0.865 | 2.953 | -0.153 | 1.914 | 0.02 | 0.016 | 0.016 | 0 | 0.76 | 0.76 | 0.42 | 0.5 | 0.49 | 0.64 |
| 6 | 16933420 | rs9477259 | -0.437 | 3.27 | -0.3 | 2.185 | 0.017 | 0 | 0 | 0.627 | 0.08 | 0.11 | 0.41 | 0.1 | 0.24 | 0.21 |
| 6 | 16944576 | rs736778 | 0.566 | 2.701 | -0.218 | 2.049 | 0 | -0.001 | -0.001 | 0 | 0.41 | 0.55 | 0.28 | 0.39 | 0.31 | 0.38 |
| 6 | 16946758 | rs4461735 | 1.24 | 2.586 | -0.311 | 1.991 | 0.018 | 0.114 | 0.114 | 0.304 | 0.33 | 0.05 | 0.17 | 0.15 | 0.24 | 0.34 |
| 6 | 16948451 | rs17770697 | 1.131 | 2.628 | -0.248 | 2.013 | 0.04 | 0.109 | 0.109 | 0 | 0.34 | 0.13 | 0.18 | 0.15 | 0.24 | 0.34 |
| 6 | 16948558 | rs1033762 | 0.129 | 2.525 | -0.233 | 2.034 | -0.055 | -0.077 | -0.077 | 0.828 | 0.85 | 0.96 | 0.84 | 0.59 | 0.72 | 0.86 |
| 6 | 16949332 | rs7767720 | 0.233 | 2.416 | -0.162 | 1.897 | 0.004 | 0.036 | 0.036 | 0 | 0.44 | 0.4 | 0.56 | 0.2 | 0.41 | 0.48 |
| 6 | 16956041 | rs9477268 | -0.973 | 2.693 | 0.044 | 2.061 | 0.046 | -0.034 | -0.034 | 0 | 0.11 | 0.27 | 0.38 | 0.05 | 0.18 | 0.15 |
| 6 | 16966702 | rs10456797 | 0.289 | 2.677 | 0.036 | 2.365 | 0.011 | 0.056 | 0.056 | 0.409 | 0.85 | 0.82 | 0.81 | 0.54 | 0.75 | 0.88 |
| 6 | 16969155 | rs7740716 | 0.351 | 2.681 | -0.056 | 2.361 | -0.011 | -0.006 | -0.006 | 0 | 0.85 | 0.91 | 0.84 | 0.57 | 0.75 | 0.88 |
| 6 | 16970970 | rs7382170 | 0.481 | 2.393 | 0.187 | 1.831 | -0.003 | -0.005 | -0.005 | 0 | 0.42 | 0.38 | 0.26 | 0.31 | 0.34 | 0.42 |
| 6 | 16972975 | rs10807603 | -0.349 | 2.456 | 0.147 | 1.841 | 0.006 | 0.032 | 0.032 | 0 | 0.53 | 0.62 | 0.55 | 0.36 | 0.51 | 0.57 |
| 6 | 16973835 | rs911594 | 0.06 | 2.63 | 0.006 | 2.002 | -0.039 | -0.054 | -0.054 | 0.155 | 0.85 | 0.95 | 0.82 | 0.5 | 0.72 | 0.88 |
| 6 | 16974703 | rs9297025 | 0.858 | 2.076 | 0.332 | 1.727 | -0.045 | -0.023 | -0.023 | 0.869 | 0.35 | 0.47 | 0.29 | 0.26 | 0.22 | 0.34 |
| 6 | 16975862 | rs9396726 | -0.35 | 2.41 | 0.198 | 1.671 | 0.009 | 0.001 | 0.001 | 0.042 | 0.08 | 0.1 | 0.26 | 0.11 | 0.17 | 0.16 |
| 6 | 16984459 | rs9383219 | 0.133 | 2.268 | 0.189 | 1.646 | 0.005 | 0.017 | 0.017 | 0 | 0.44 | 0.4 | 0.53 | 0.27 | 0.4 | 0.5 |
| 6 | 16989325 | rs9358118 | -0.765 | 2.418 | 0.062 | 1.722 | 0.029 | 0.01 | 0.01 | 0.224 | 0.1 | 0.15 | 0.29 | 0.13 | 0.19 | 0.17 |
| 6 | 17005239 | rs13215385 | 1.191 | 2.007 | -0.246 | 1.567 | 0.051 | 0.093 | 0.093 | 0 | 0.31 | 0.05 | 0.24 | 0.16 | 0.22 | 0.34 |
| 6 | 17012670 | rs7755677 | 0.414 | 2.197 | 0.198 | 1.905 | 0.001 | 0.001 | 0.001 | 0 | 0.42 | 0.5 | 0.27 | 0.3 | 0.29 | 0.38 |
| 6 | 17013429 | rs10498687 | 1.452 | 2.389 | -0.079 | 2.02 | 0.032 | 0.045 | 0.045 | 0 | 0.31 | 0.05 | 0.27 | 0.21 | 0.24 | 0.34 |
| 6 | 17013648 | rs9297026 | 0.722 | 2.589 | -0.061 | 2.136 | -0.031 | 0.02 | 0.02 | 0.094 | 0.4 | 0.14 | 0.53 | 0.34 | 0.44 | 0.51 |
| 6 | 17014527 | rs9370949 | -0.336 | 2.503 | -0.01 | 2.167 | -0.014 | -0.011 | -0.011 | 0 | 0.09 | 0.07 | 0.25 | 0.13 | 0.2 | 0.17 |
| 6 | 17015485 | rs11753264 | 0.509 | 2.661 | -0.028 | 1.899 | -0.036 | -0.07 | -0.07 | 0 | 0.12 | 0.01 | 0.11 | 0.23 | 0.13 | 0.05 |
| 6 | 17021077 | rs10949393 | 0.46 | 2.445 | 0.185 | 1.826 | 0.001 | 0.002 | 0.002 | 0 | 0.39 | 0.33 | 0.24 | 0.27 | 0.29 | 0.39 |

| | | | | | | | | | | | | | | | | |
|---|---|---|---|---|---|---|---|---|---|---|---|---|---|---|---|---|
| 6 | 17022069 | rs11963899 | 0.547 | 2.434 | 0.307 | 1.27 | 0 | -0.004 | -0.004 | 0 | 0.13 | 0.14 | 0.16 | 0.08 | 0.14 | 0.11 |
| 6 | 17023108 | rs9396740 | 1.111 | 2.465 | 0.286 | 1.275 | 0.024 | -0.018 | -0.018 | 0 | 0.33 | 0.09 | 0.28 | 0.33 | 0.27 | 0.37 |
| 6 | 17031611 | rs4434476 | -0.275 | 2.111 | 0.633 | 1.129 | 0 | 0 | 0 | 0 | 0.52 | 0.46 | 0.34 | 0.48 | 0.41 | 0.43 |
| 6 | 17034315 | rs7740186 | 0.814 | 1.816 | 0.97 | 1.182 | 0 | -0.025 | -0.025 | 0 | 0.44 | 0.18 | 0.52 | 0.45 | 0.43 | 0.4 |
| 6 | 17043546 | rs9396742 | 0.81 | 1.859 | 1.028 | 1.142 | 0.01 | -0.015 | -0.015 | 0 | 0.45 | 0.24 | 0.52 | 0.47 | 0.43 | 0.39 |
| 6 | 17048176 | rs10949397 | -1.153 | 1.787 | 1.669 | 0.443 | 0.014 | -0.052 | -0.052 | 0 | 0.1 | 0.39 | 0.14 | 0.01 | 0.1 | 0.07 |
| 6 | 17048225 | rs10949399 | -0.937 | 1.828 | 1.604 | 0.456 | -0.002 | -0.051 | -0.051 | 0 | 0.09 | 0.18 | 0.12 | 0 | 0.1 | 0.07 |
| 6 | 17054240 | rs4142714 | 0.154 | 1.791 | 1.336 | 0.408 | 0.043 | 0.004 | 0.004 | 0 | 0.78 | 0.69 | 0.81 | 0.78 | 0.73 | 0.8 |
| 6 | 17061546 | rs7745845 | 1.085 | 1.533 | 1.435 | 0.716 | 0.037 | -0.039 | -0.039 | 0.021 | 0.91 | 0.67 | 0.87 | 1 | 0.89 | 0.91 |
| 6 | 17064764 | rs4613810 | 1.338 | 1.516 | 1.137 | 0.707 | 0.03 | -0.057 | -0.057 | 0 | 0.89 | 0.56 | 0.86 | 1 | 0.89 | 0.91 |
| 6 | 17066353 | rs4360123 | 1.058 | 1.633 | 1.191 | 0.767 | 0.048 | -0.041 | -0.041 | 0 | 0.9 | 0.56 | 0.86 | 1 | 0.89 | 0.91 |
| 6 | 17081711 | rs4236169 | -0.876 | 2.119 | 1.227 | 1.159 | 0.018 | 0.006 | 0.006 | 0 | 0.33 | 0.61 | 0.35 | 0.37 | 0.4 | 0.32 |
| 6 | 17083832 | rs9370958 | 0.134 | 1.752 | -0.019 | 0.915 | -0.029 | 0.009 | 0.009 | 0 | 0.25 | 0.02 | 0.16 | 0.22 | 0.22 | 0.29 |
| 6 | 17087510 | rs7741340 | -0.572 | 1.221 | 0.58 | 1.463 | -0.04 | 0.049 | 0.049 | 0 | 0.41 | 0.8 | 0.4 | 0.68 | 0.44 | 0.39 |
| 6 | 17096354 | rs6926443 | -0.388 | 1.216 | 0.943 | 1.575 | -0.018 | 0.152 | 0.152 | 0 | 0.09 | 0.18 | 0.08 | 0.42 | 0.05 | 0.14 |
| 6 | 17102858 | rs1473746 | -0.633 | 1.253 | 0.579 | 1.506 | -0.039 | 0.049 | 0.049 | 0 | 0.4 | 0.8 | 0.4 | 0.68 | 0.44 | 0.4 |
| 6 | 17103517 | rs927629 | 0.432 | 1.409 | 0.488 | 1.277 | -0.035 | -0.009 | -0.009 | 0 | 0.2 | 0.09 | 0.16 | 0.26 | 0.27 | 0.17 |
| 6 | 17106718 | rs4716108 | 0.408 | 1.151 | 0.387 | 1.128 | 0.016 | 0.01 | 0.01 | 0 | 0.63 | 0.53 | 0.55 | 0.44 | 0.59 | 0.56 |
| 6 | 17107168 | rs4716110 | 1.18 | 1.068 | -0.017 | 0.536 | 0 | -0.026 | -0.026 | 0 | 0.33 | 0.37 | 0.32 | 0.07 | 0.36 | 0.33 |
| 6 | 17108786 | rs13210795 | 0.406 | 1.168 | -0.247 | 0.701 | -0.005 | -0.035 | -0.035 | 1.632 | 0.25 | 0.03 | 0.19 | 0.21 | 0.18 | 0.29 |
| 6 | 17115648 | rs6914177 | 1.186 | 1.116 | 0.024 | 0.523 | -0.005 | 0.01 | 0.01 | 0 | 0.33 | 0.27 | 0.31 | 0.07 | 0.37 | 0.34 |
| 6 | 17122836 | rs9350030 | 0.05 | 0.739 | -0.222 | 0.895 | 0.047 | -0.04 | -0.04 | 1.482 | 0.29 | 0.11 | 0.22 | 0.39 | 0.23 | 0.23 |
| 6 | 17150480 | rs4716123 | -0.68 | 1.327 | -0.154 | 0.917 | -0.004 | 0.03 | 0.03 | 0 | 0.55 | 0.25 | 0.45 | 0.63 | 0.46 | 0.58 |
| 6 | 17151765 | rs9350031 | 0.55 | 1.301 | -0.41 | 1.254 | 0.013 | 0.026 | 0.026 | 0 | 0.23 | 0.09 | 0.17 | 0.21 | 0.18 | 0.17 |
| 6 | 17152561 | rs17621654 | -0.256 | 1.326 | -0.4 | 1.543 | 0.018 | -0.024 | -0.024 | 0.156 | 0.3 | 0.11 | 0.22 | 0.38 | 0.27 | 0.26 |
| 6 | 17154680 | rs1408299 | -0.113 | 1.515 | 0.318 | 1.751 | 0.035 | 0.157 | 0.157 | 0.45 | 0.93 | 0.74 | 0.82 | 0.8 | 0.92 | 0.93 |
| 6 | 17154856 | rs1408300 | 1.269 | 1.53 | 0.267 | 1.679 | 0.014 | -0.042 | -0.042 | 0 | 0.38 | 0.49 | 0.38 | 0.18 | 0.46 | 0.36 |
| 6 | 17158078 | rs9477339 | 0.511 | 1.45 | 0.705 | 1.858 | 0.004 | -0.015 | -0.015 | 0 | 0.23 | 0.13 | 0.22 | 0.24 | 0.19 | 0.32 |
| 6 | 17176032 | rs1535546 | 0.081 | 1.733 | 0.978 | 1.621 | 0.033 | 0.189 | 0.189 | 0.317 | 0.07 | 0.22 | 0.17 | 0.25 | 0.09 | 0.08 |
| 6 | 17177658 | rs12528226 | 0.051 | 1.719 | 0.933 | 1.614 | 0.028 | 0.178 | 0.178 | 0.361 | 0.07 | 0.21 | 0.17 | 0.25 | 0.08 | 0.08 |
| 6 | 17181845 | rs4716127 | 1.472 | 1.551 | -0.011 | 0.977 | 0.004 | 0.036 | 0.036 | 0 | 0.22 | 0.17 | 0.19 | 0.07 | 0.24 | 0.26 |
| 6 | 17192800 | rs9350032 | 0.777 | 1.405 | -0.252 | 0.877 | -0.008 | 0.092 | 0.092 | 0.12 | 0.32 | 0.1 | 0.26 | 0.22 | 0.29 | 0.2 |
| 6 | 17200210 | rs9465001 | -0.339 | 1.312 | -0.036 | 0.918 | -0.039 | -0.024 | -0.024 | 0 | 0.58 | 0.82 | 0.62 | 0.49 | 0.52 | 0.66 |
| 6 | 17210051 | rs1547103 | -0.095 | 1.252 | 0.102 | 1.082 | -0.006 | 0.023 | 0.023 | 0 | 0.32 | 0.41 | 0.4 | 0.42 | 0.26 | 0.38 |
| 6 | 17213063 | rs9465003 | 0.567 | 1.272 | -0.718 | 0.61 | 0 | 0.017 | 0.017 | 0 | 0.24 | 0.08 | 0.22 | 0.17 | 0.19 | 0.31 |
| 6 | 17226021 | rs9370968 | 0.567 | 1.156 | -0.484 | 1.012 | -0.02 | -0.031 | -0.031 | 0 | 0.06 | 0.01 | 0.03 | 0.15 | 0.07 | 0.09 |
| 6 | 17226085 | rs1923747 | -0.346 | 0.915 | -0.606 | 0.745 | -0.038 | -0.074 | -0.074 | 0.235 | 0.66 | 0.84 | 0.48 | 0.39 | 0.59 | 0.67 |
| 6 | 17234722 | rs6923624 | -0.308 | 0.901 | -0.455 | 0.76 | -0.017 | -0.052 | -0.052 | 0.53 | 0.65 | 0.78 | 0.48 | 0.4 | 0.6 | 0.67 |
| 6 | 17237051 | rs6940891 | 1.671 | 1.134 | -0.506 | 0.49 | -0.009 | -0.027 | -0.027 | 0 | 0.22 | 0.28 | 0.16 | 0.08 | 0.2 | 0.23 |
| 6 | 17241154 | rs1322846 | -0.002 | 1.22 | -1.016 | 0.652 | -0.016 | 0.006 | 0.006 | 0 | 0.4 | 0.28 | 0.29 | 0.35 | 0.41 | 0.44 |
| 6 | 17248307 | rs6459529 | 0.198 | 1.231 | -0.938 | 0.744 | -0.012 | -0.006 | -0.006 | 0 | 0.57 | 0.66 | 0.41 | 0.4 | 0.48 | 0.62 |
| 6 | 17249496 | rs9477367 | -0.273 | 1.183 | -1.154 | 1.273 | -0.002 | 0.009 | 0.009 | 0 | 0.91 | 0.94 | 0.75 | 0.67 | 0.89 | 0.89 |
| 6 | 17250282 | rs1175988 | -0.259 | 1.171 | -1.143 | 1.275 | 0 | 0.013 | 0.013 | 0 | 0.91 | 0.94 | 0.75 | 0.67 | 0.89 | 0.89 |
| 6 | 17252342 | rs13217920 | 0.559 | 1.09 | -1.243 | 1.226 | -0.009 | -0.013 | -0.013 | 0 | 0.07 | 0.03 | 0.05 | 0.16 | 0.08 | 0.1 |
| 6 | 17261189 | rs6921352 | 0.1 | 1.21 | -1.3 | 1.018 | -0.019 | -0.017 | -0.017 | 0 | 0.6 | 0.7 | 0.44 | 0.4 | 0.54 | 0.67 |
| 6 | 17268236 | rs3886378 | 1.159 | 1.002 | -1.355 | 0.589 | -0.003 | 0.028 | 0.028 | 0 | 0.27 | 0.14 | 0.21 | 0.17 | 0.26 | 0.37 |
| 6 | 17271896 | rs823727 | 0.021 | 1.241 | -1.104 | 0.622 | -0.005 | -0.007 | -0.007 | 0 | 0.4 | 0.53 | 0.46 | 0.59 | 0.36 | 0.38 |
| 6 | 17277198 | rs1359006 | 1.066 | 1.147 | -1.335 | 0.645 | -0.014 | 0.035 | 0.035 | 0 | 0.32 | 0.51 | 0.41 | 0.44 | 0.28 | 0.34 |
| 6 | 17277353 | rs1359007 | -0.1 | 1.315 | -1.227 | 0.623 | -0.011 | 0.029 | 0.029 | 0 | 0.68 | 0.69 | 0.67 | 0.77 | 0.61 | 0.76 |
| 6 | 17277820 | rs9396764 | -0.291 | 1.347 | -1.43 | 0.64 | -0.067 | 0.1 | 0.1 | 0 | 0.69 | 0.9 | 0.69 | 0.77 | 0.62 | 0.76 |
| 6 | 17283321 | rs3749855 | 0.242 | 0.483 | -1.194 | 0.968 | -0.016 | -0.03 | -0.03 | 0 | 0.08 | 0.02 | 0.04 | 0.17 | 0.07 | 0.06 |
| 6 | 17284382 | rs12210130 | 0.704 | 0.414 | -1.282 | 0.548 | 0.008 | 0.02 | 0.02 | 0 | 0.18 | 0.16 | 0.15 | 0.19 | 0.16 | 0.09 |
| 6 | 17289545 | rs4716132 | 0.432 | 0.451 | -1.414 | 0.574 | -0.01 | 0.06 | 0.06 | 0.122 | 0.23 | 0.06 | 0.16 | 0.19 | 0.2 | 0.09 |
| 6 | 17290619 | rs864280 | -0.997 | 0.394 | -1.75 | 0.387 | -0.047 | 0.059 | 0.059 | 0 | 0.68 | 0.88 | 0.74 | 0.81 | 0.65 | 0.61 |
| 6 | 17296172 | rs9370972 | 0.374 | 0.272 | -1.24 | 0.574 | 0 | -0.007 | -0.007 | 0.038 | 0.42 | 0.63 | 0.36 | 0.43 | 0.42 | 0.42 |
| 6 | 17301686 | rs12199579 | 0.57 | 0.29 | -1.327 | 0.478 | 0.005 | 0.065 | 0.065 | 0.041 | 0.23 | 0.1 | 0.17 | 0.18 | 0.2 | 0.1 |
| 6 | 17314356 | rs1745072 | 0.848 | 0.233 | -1.034 | 0.744 | -0.023 | -0.064 | -0.064 | 1.149 | 0.27 | 0.34 | 0.17 | 0.09 | 0.23 | 0.21 |
| 6 | 17314690 | rs1772127 | 0.54 | 0.174 | -0.928 | 0.97 | -0.001 | -0.036 | -0.036 | 1.388 | 0.36 | 0.46 | 0.25 | 0.2 | 0.34 | 0.27 |
| 6 | 17315838 | rs1745074 | 0.163 | 0.267 | -1.178 | 0.942 | 0.005 | -0.02 | -0.02 | 0 | 0.44 | 0.76 | 0.38 | 0.44 | 0.42 | 0.4 |
| 6 | 17318116 | rs9477384 | -0.38 | 0.24 | -0.945 | 0.897 | 0.029 | -0.009 | -0.009 | 0 | 0.55 | 0.24 | 0.62 | 0.6 | 0.6 | 0.6 |
| 6 | 17325605 | rs12216482 | -0.481 | 0.478 | -0.906 | 0.786 | -0.018 | -0.007 | -0.007 | 0.014 | 0.18 | 0.06 | 0.12 | 0.23 | 0.18 | 0.1 |
| 6 | 17330001 | rs12203365 | -0.889 | 0.316 | -0.71 | 0.501 | 0.001 | 0.015 | 0.015 | 0.55 | 0.28 | 0.24 | 0.21 | 0.15 | 0.23 | 0.17 |
| 6 | 17330855 | rs12526082 | -0.718 | 0.394 | -1.068 | 0.641 | 0.023 | -0.005 | -0.005 | 0.144 | 0.59 | 0.35 | 0.5 | 0.41 | 0.64 | 0.58 |
| 6 | 17333338 | rs4440469 | 0.646 | 0.431 | -0.69 | 0.967 | -0.017 | -0.014 | -0.014 | 0 | 0.33 | 0.29 | 0.3 | 0.37 | 0.41 | 0.43 |
| 6 | 17333372 | rs4587177 | 0.938 | 0.554 | -0.911 | 0.944 | -0.102 | -0.031 | -0.031 | 0 | 0.3 | 0.05 | 0.25 | 0.26 | 0.36 | 0.41 |
| 6 | 17343655 | rs13197759 | -0.657 | 0.176 | -0.706 | 0.916 | -0.016 | 0.01 | 0.01 | 0 | 0.61 | 0.54 | 0.53 | 0.52 | 0.67 | 0.62 |
| 6 | 17351791 | rs6930531 | 0.071 | 0.236 | -0.533 | 0.628 | 0.001 | 0.018 | 0.018 | 1.509 | 0.17 | 0.11 | 0.12 | 0.07 | 0.13 | 0.12 |
| 6 | 17356569 | rs9383276 | 0.776 | 0.118 | -0.638 | 0.778 | -0.053 | -0.025 | -0.025 | 0 | 0.43 | 0.08 | 0.44 | 0.38 | 0.43 | 0.38 |
| 6 | 17358636 | rs6925396 | -0.631 | 0.453 | -0.574 | 0.323 | 0.007 | -0.001 | -0.001 | 0 | 0.52 | 0.5 | 0.44 | 0.46 | 0.55 | 0.59 |

| | | | | | | | | | | | | | | | | |
|---|---|---|---|---|---|---|---|---|---|---|---|---|---|---|---|---|
| 6 | 17364146 | rs6459534 | -0.702 | 0.507 | -0.574 | -0.104 | 0.004 | 0.003 | 0.003 | 0 | 0.52 | 0.51 | 0.44 | 0.4 | 0.55 | 0.6 |
| 6 | 17368972 | rs4716138 | -0.038 | 0.738 | -0.32 | 0.38 | -0.054 | -0.034 | -0.034 | 0.291 | 0.67 | 0.82 | 0.69 | 0.67 | 0.59 | 0.52 |
| 6 | 17376666 | rs9350040 | -0.006 | 0.881 | -0.507 | 0.454 | 0.002 | 0.001 | 0.001 | 0 | 0.64 | 0.58 | 0.67 | 0.67 | 0.58 | 0.52 |
| 6 | 17386895 | rs7773313 | -0.737 | 0.784 | 0.015 | -0.07 | 0.015 | -0.049 | -0.049 | 0 | 0.18 | 0.42 | 0.13 | 0.09 | 0.17 | 0.11 |
| 6 | 17395812 | rs1555111 | -0.135 | 0.864 | -0.767 | -0.708 | -0.016 | 0.052 | 0.052 | 0 | 0.28 | 0.18 | 0.23 | 0.15 | 0.3 | 0.35 |
| 6 | 17395933 | rs10080250 | -0.609 | 0.377 | -0.459 | -0.812 | 0.01 | 0.081 | 0.081 | 0 | 0.35 | 0.79 | 0.31 | 0.18 | 0.31 | 0.36 |
| 6 | 17398530 | rs7748415 | -0.2 | 0.455 | -1.113 | -1.466 | -0.008 | 0.047 | 0.047 | 0 | 0.81 | 0.91 | 0.81 | 0.97 | 0.8 | 0.87 |
| 6 | 17408833 | rs4716142 | -0.129 | 0.1 | -0.336 | -1.423 | -0.002 | -0.015 | -0.015 | 0 | 0.34 | 0.63 | 0.28 | 0.19 | 0.3 | 0.36 |
| 6 | 17417227 | rs7755120 | 0.079 | 0.369 | -0.665 | -1.426 | -0.038 | 0.09 | 0.09 | 0 | 0.3 | 0.1 | 0.37 | 0.14 | 0.34 | 0.32 |
| 6 | 17422536 | rs10485376 | -0.028 | 0.361 | -0.762 | -1.682 | -0.004 | -0.031 | -0.031 | 5.432 | 0.11 | 0.17 | 0.09 | 0.01 | 0.09 | 0.09 |
| 6 | 17427133 | rs1555112 | 0.177 | 0.538 | -0.934 | -1.422 | -0.035 | 0.069 | 0.069 | 0 | 0.29 | 0.07 | 0.36 | 0.15 | 0.32 | 0.31 |
| 6 | 17441419 | rs6918762 | -0.202 | 0.626 | -0.883 | -1.574 | -0.07 | 0.158 | 0.158 | 0 | 0.61 | 0.91 | 0.52 | 0.86 | 0.54 | 0.56 |
| 6 | 17442326 | rs3777686 | -0.15 | 0.61 | -0.761 | -1.454 | -0.037 | 0.115 | 0.115 | 0 | 0.61 | 0.85 | 0.52 | 0.83 | 0.54 | 0.56 |
| 6 | 17444328 | rs1320998 | -0.103 | 0.599 | -0.682 | -1.46 | -0.03 | 0.111 | 0.111 | 0 | 0.6 | 0.83 | 0.51 | 0.83 | 0.54 | 0.56 |
| 6 | 17446523 | rs910042 | 0.115 | 0.437 | -0.317 | -1.543 | 0.001 | 0.135 | 0.135 | 0 | 0.55 | 0.28 | 0.46 | 0.84 | 0.54 | 0.56 |
| 6 | 17457057 | rs910043 | -0.891 | 0.387 | 1.099 | -1.298 | 0.04 | 0.075 | 0.075 | 5.136 | 0.05 | 0.52 | 0.05 | 0.02 | 0 | 0 |
| 6 | 17464687 | rs9383287 | -0.258 | 0.616 | 0.602 | -1.625 | -0.051 | 0.108 | 0.108 | 0 | 0.62 | 0.82 | 0.52 | 0.85 | 0.54 | 0.59 |
| 6 | 17470693 | rs1320994 | -0.231 | 0.659 | 0.396 | -1.319 | -0.034 | 0.091 | 0.091 | 0 | 0.59 | 0.8 | 0.51 | 0.8 | 0.54 | 0.59 |
| 6 | 17473927 | rs1320995 | 0.208 | 0.712 | 0.009 | -1.291 | -0.038 | 0.185 | 0.185 | 0 | 0.52 | 0.17 | 0.45 | 0.78 | 0.53 | 0.59 |
| 6 | 17481372 | rs16879756 | 0.034 | 1.08 | -0.125 | -0.252 | -0.07 | -0.066 | -0.066 | 1.387 | 0.1 | 0.01 | 0.17 | 0.34 | 0.23 | 0.24 |
| 6 | 17483875 | rs3777700 | -0.021 | 0.517 | 0.352 | -0.873 | 0.009 | -0.018 | -0.018 | 0 | 0.66 | 0.33 | 0.71 | 0.97 | 0.73 | 0.83 |
| 6 | 17494348 | rs10807611 | -0.816 | 1.146 | 0.358 | 0.624 | 0.009 | -0.003 | -0.003 | 0 | 0.48 | 0.41 | 0.6 | 0.48 | 0.55 | 0.61 |
| 6 | 17494463 | rs823433 | -0.593 | 1.322 | 0.662 | 0.986 | 0.059 | 0.112 | 0.112 | 0 | 0.07 | 0.19 | 0.1 | 0.26 | 0.14 | 0.13 |
| 6 | 17502936 | rs707860 | -1.177 | 0.676 | 0.769 | 0.87 | 0.067 | 0.067 | 0.067 | 0.428 | 0.57 | 0.75 | 0.74 | 0.76 | 0.71 | 0.72 |
| 6 | 17507670 | rs2273011 | -1.161 | 0.844 | 0.872 | 0.88 | 0.072 | 0.084 | 0.084 | 0.71 | 0.53 | 0.72 | 0.74 | 0.75 | 0.7 | 0.71 |
| 6 | 17510429 | rs9297036 | 0.052 | 0.79 | 0.339 | 1.104 | 0.009 | -0.037 | -0.037 | 0 | 0.1 | 0.03 | 0.06 | 0.12 | 0.07 | 0.1 |
| 6 | 17511487 | rs926583 | -0.374 | 0.955 | 0.796 | 1.161 | 0.034 | 0.177 | 0.177 | 0 | 0.06 | 0.21 | 0.07 | 0.25 | 0.07 | 0.12 |
| 6 | 17514637 | rs9367952 | -0.002 | 1.192 | 0.46 | 0.995 | -0.122 | -0.066 | -0.066 | 0 | 0.26 | 0.07 | 0.37 | 0.36 | 0.46 | 0.35 |
| 6 | 17529407 | rs7755827 | -0.272 | 1.139 | 0.458 | 0.758 | 0.004 | 0.092 | 0.092 | 0 | 0.06 | 0.1 | 0.06 | 0.24 | 0.07 | 0.12 |
| 6 | 17535126 | rs16879781 | -0.521 | 0.951 | 0.511 | 0.631 | -0.001 | -0.021 | -0.021 | 0 | 0.12 | 0.1 | 0.19 | 0.02 | 0.08 | 0.14 |
| 6 | 17535454 | rs6939935 | -1.994 | 0.853 | 0.704 | 0.571 | -0.015 | -0.013 | -0.013 | 0 | 0.29 | 0.58 | 0.37 | 0.27 | 0.25 | 0.31 |
| 6 | 17537797 | rs9465047 | -1.412 | 0.851 | 0.694 | 0.399 | -0.035 | -0.147 | -0.147 | 0.015 | 0.24 | 0.49 | 0.3 | 0.02 | 0.18 | 0.2 |
| 6 | 17542304 | rs3777708 | -0.743 | 0.984 | 0.515 | 0.457 | -0.025 | 0.007 | 0.007 | 0 | 0.33 | 0.25 | 0.45 | 0.73 | 0.53 | 0.52 |
| 6 | 17543036 | rs2296249 | -1.011 | 0.907 | 0.217 | 0.591 | 0.009 | 0.019 | 0.019 | 0 | 0.45 | 0.62 | 0.56 | 0.73 | 0.63 | 0.57 |
| 6 | 17543946 | rs6914958 | -1.235 | 1.084 | -0.112 | 0.802 | -0.008 | 0.001 | 0.001 | 0 | 0.17 | 0.2 | 0.25 | 0.25 | 0.15 | 0.26 |
| 6 | 17547788 | rs4712301 | -0.883 | 1.948 | -0.774 | 1.217 | -0.144 | -0.095 | -0.095 | 1.428 | 0.28 | 0.03 | 0.33 | 0.37 | 0.4 | 0.32 |
| 6 | 17550158 | rs6925145 | 1.29 | 1.566 | -0.562 | 1.244 | 0.058 | 0.055 | 0.055 | 0.344 | 0.49 | 0.2 | 0.36 | 0.39 | 0.38 | 0.53 |
| 6 | 17551377 | rs3777709 | -0.896 | 2.495 | 0.12 | 1.576 | -0.029 | -0.01 | -0.01 | 0.113 | 0.34 | 0.21 | 0.5 | 0.37 | 0.43 | 0.35 |
| 6 | 17554899 | rs1984515 | 1.391 | 2.311 | 0.412 | 1.53 | 0.02 | 0.023 | 0.023 | 0.453 | 0.55 | 0.3 | 0.38 | 0.39 | 0.39 | 0.53 |
| 6 | 17571000 | rs2876453 | -0.496 | 2.614 | 0.104 | 1.434 | -0.017 | -0.087 | -0.087 | 0 | 0.22 | 0.07 | 0.24 | 0.35 | 0.22 | 0.25 |
| 6 | 17572764 | rs6923384 | -1.159 | 2.728 | 0.941 | 1.049 | 0.06 | -0.025 | -0.025 | 0 | 0.06 | 0.16 | 0.07 | 0.01 | 0.13 | 0.03 |
| 6 | 17603200 | rs4142551 | -0.499 | 2.956 | 0.854 | 1.079 | -0.014 | 0.012 | 0.012 | 0.05 | 0.12 | 0.08 | 0.26 | 0.05 | 0.22 | 0.1 |
| 6 | 17610090 | rs1055111 | -0.499 | 2.956 | 0.878 | 1.079 | -0.012 | 0.009 | 0.009 | 0.05 | 0.12 | 0.08 | 0.26 | 0.05 | 0.22 | 0.1 |
| 6 | 17614952 | rs9350051 | 1.721 | 2.787 | 1.683 | 1.441 | 0.055 | 0.056 | 0.056 | 0.12 | 0.7 | 0.47 | 0.54 | 0.55 | 0.54 | 0.71 |
| 6 | 17616556 | rs3777712 | -1.65 | 2.991 | 2.06 | 1.56 | 0.002 | -0.004 | -0.004 | 0 | 0.36 | 0.63 | 0.4 | 0.7 | 0.39 | 0.44 |
| 6 | 17624060 | rs1830818 | 0.463 | 2.747 | 2.117 | 1.423 | 0.033 | -0.009 | -0.009 | 0.067 | 0.93 | 0.83 | 0.94 | 0.96 | 0.89 | 0.96 |
| 6 | 17627494 | rs12524105 | -0.112 | 2.749 | 2.116 | 1.401 | -0.008 | 0.008 | 0.008 | 0.019 | 0.12 | 0.09 | 0.25 | 0.05 | 0.22 | 0.1 |
| 6 | 17633061 | rs2274136 | 0.794 | 2.722 | 2.175 | 1.431 | 0.008 | -0.022 | -0.022 | 0.057 | 0.85 | 0.74 | 0.8 | 0.88 | 0.8 | 0.87 |
| 6 | 17635050 | rs7770657 | 2.264 | 2.674 | 2.11 | 1.451 | 0.029 | 0.133 | 0.133 | 0 | 0.51 | 0.16 | 0.33 | 0.26 | 0.41 | 0.45 |
| 6 | 17661619 | rs10447411 | -0.498 | 2.767 | 1.933 | 1.413 | 0.037 | -0.027 | -0.027 | 0.581 | 0.13 | 0.19 | 0.27 | 0.04 | 0.25 | 0.1 |
| 6 | 17675246 | rs2228375 | -0.107 | 2.81 | 1.616 | 1.415 | -0.078 | 0.028 | 0.028 | 0.1 | 0.12 | 0.02 | 0.28 | 0.05 | 0.24 | 0.1 |
| 6 | 17676470 | rs9396788 | 1.925 | 2.808 | 1.693 | 1.421 | -0.003 | -0.003 | -0.003 | 0.018 | 0.68 | 0.32 | 0.51 | 0.55 | 0.53 | 0.7 |
| 6 | 17695858 | rs9297042 | -0.145 | 2.591 | 0.791 | 1.68 | 0.006 | -0.055 | -0.055 | 0.266 | 0.16 | 0.05 | 0.18 | 0.31 | 0.13 | 0.25 |
| 6 | 17697963 | rs12211005 | -1.047 | 2.887 | 0.965 | 1.595 | 0.065 | 0.107 | 0.107 | 0.046 | 0.14 | 0.25 | 0.27 | 0.33 | 0.26 | 0.15 |
| 6 | 17699322 | rs12199222 | -0.992 | 2.817 | 0.975 | 1.606 | 0.063 | 0.111 | 0.111 | 0.046 | 0.14 | 0.25 | 0.27 | 0.33 | 0.25 | 0.15 |
| 6 | 17706404 | rs13219815 | -0.123 | 2.795 | 0.425 | 1.084 | 0.001 | 0.004 | 0.004 | 0 | 0.09 | 0.09 | 0.13 | 0.08 | 0.1 | 0.09 |
| 6 | 17711461 | rs10949437 | -0.631 | 2.815 | 0.418 | 1.238 | -0.01 | 0.008 | 0.008 | 0.019 | 0.12 | 0.09 | 0.24 | 0.05 | 0.22 | 0.1 |
| 6 | 17721741 | rs2038743 | -0.944 | 1.87 | -0.054 | 1.594 | -0.005 | 0.061 | 0.061 | 0 | 0.21 | 0.27 | 0.15 | 0.5 | 0.19 | 0.18 |
| 6 | 17725183 | rs7743695 | -1.103 | 1.856 | -1.094 | 1.334 | 0.007 | 0.051 | 0.051 | 0 | 0.59 | 0.92 | 0.72 | 0.76 | 0.66 | 0.56 |
| 6 | 17744282 | rs10456805 | -0.772 | 2.391 | -0.628 | 1.647 | 0.144 | 0.284 | 0.284 | 0.05 | 0.06 | 0.29 | 0.21 | 0.26 | 0.16 | 0.07 |
| 6 | 17744751 | rs9396801 | -0.161 | 2.144 | -1.744 | 1.31 | -0.066 | -0.042 | -0.042 | 0 | 0.13 | 0.02 | 0.18 | 0.19 | 0.21 | 0.14 |
| 6 | 17744780 | rs1969914 | 0.64 | 0.866 | -1.435 | 0.897 | 0.049 | -0.012 | -0.012 | 0 | 0.4 | 0.12 | 0.25 | 0.3 | 0.28 | 0.4 |
| 6 | 17754961 | rs17714754 | 0.424 | 0.821 | -1.587 | 0.765 | 0.039 | -0.078 | -0.078 | 0.853 | 0.39 | 0.07 | 0.21 | 0.42 | 0.27 | 0.39 |
| 6 | 17755884 | rs9396802 | 0.63 | 0.502 | -1.107 | -0.332 | -0.004 | -0.029 | -0.029 | 0 | 0.6 | 0.51 | 0.53 | 0.8 | 0.6 | 0.63 |
| 6 | 17755901 | rs11756169 | -1.272 | 0.056 | -1.077 | -0.883 | -0.004 | -0.04 | -0.04 | 0 | 0.18 | 0.17 | 0.09 | 0.01 | 0.13 | 0.09 |
| 6 | 17764279 | rs1044654 | -0.213 | 0.193 | -1.321 | -0.238 | 0.009 | -0.016 | -0.016 | 0 | 0.2 | 0.06 | 0.31 | 0.19 | 0.24 | 0.14 |
| 6 | 17764755 | rs2296197 | -0.1 | 0.184 | -1.326 | -0.242 | 0.01 | -0.016 | -0.016 | 0.387 | 0.2 | 0.06 | 0.31 | 0.19 | 0.24 | 0.14 |
| 6 | 17772764 | rs1924466 | 0.792 | 0.096 | -0.966 | -0.762 | -0.001 | 0.016 | 0.016 | 0.53 | 0.47 | 0.66 | 0.48 | 0.59 | 0.44 | 0.44 |
| 6 | 17778716 | rs4712311 | -0.397 | -0.306 | -0.943 | -0.928 | 0.002 | -0.044 | -0.044 | 0.385 | 0.91 | 0.88 | 0.94 | 0.99 | 0.93 | 0.93 |

| | | | | | | | | | | | | | | | | |
|---|---|---|---|---|---|---|---|---|---|---|---|---|---|---|---|---|
| 6 | 17781450 | rs942376 | 0.622 | -0.255 | -1.088 | -0.927 | 0.03 | -0.025 | -0.025 | 0.108 | 0.19 | 0.06 | 0.3 | 0.2 | 0.21 | 0.12 |
| 6 | 17794494 | rs3734234 | 1.373 | -0.411 | -0.93 | -1.296 | -0.084 | -0.014 | -0.014 | 0.47 | 0.39 | 0.15 | 0.42 | 0.58 | 0.43 | 0.38 |
| 6 | 17795185 | rs6459571 | -0.588 | -0.404 | 0.119 | -1.323 | -0.003 | -0.037 | -0.037 | 0.05 | 0.29 | 0.35 | 0.18 | 0.2 | 0.25 | 0.42 |
| 6 | 17799575 | rs3734235 | 0.117 | -0.348 | 0.162 | -1.295 | -0.015 | 0.027 | 0.027 | 0.06 | 0.23 | 0.1 | 0.16 | 0.2 | 0.24 | 0.37 |
| 6 | 17808321 | rs733697 | 1.333 | -0.378 | 0.367 | -1.17 | -0.04 | 0.041 | 0.041 | 0 | 0.46 | 0.23 | 0.45 | 0.59 | 0.46 | 0.4 |
| 6 | 17813825 | rs10456809 | -0.629 | -0.38 | 0.927 | -1.192 | 0 | -0.001 | -0.001 | 0.007 | 0.39 | 0.34 | 0.46 | 0.4 | 0.45 | 0.47 |
| 6 | 17813893 | rs10456810 | 0.075 | -0.354 | 0.936 | -1.188 | -0.001 | 0 | 0 | 0 | 0.18 | 0.14 | 0.13 | 0.2 | 0.21 | 0.35 |
| 6 | 17821673 | rs7749814 | -1.241 | -0.386 | 0.985 | -1.137 | -0.001 | 0.004 | 0.004 | 0.068 | 0.52 | 0.49 | 0.56 | 0.41 | 0.54 | 0.6 |
| 6 | 17822774 | rs10498688 | -0.1 | -0.385 | 0.942 | -1.152 | 0.011 | 0.007 | 0.007 | 0 | 0.29 | 0.21 | 0.39 | 0.21 | 0.29 | 0.19 |
| 6 | 17823124 | rs7770400 | -1.429 | -0.365 | 1.011 | -1.119 | -0.005 | 0.012 | 0.012 | 0 | 0.53 | 0.6 | 0.57 | 0.41 | 0.54 | 0.6 |
| 6 | 17831419 | rs2277080 | -0.666 | -0.397 | 0.895 | -1.151 | -0.009 | 0.038 | 0.038 | 0 | 0.3 | 0.16 | 0.18 | 0.2 | 0.28 | 0.39 |
| 6 | 17834198 | rs4716189 | -1.001 | -0.645 | 0.827 | -0.976 | 0.01 | 0.029 | 0.029 | 0 | 0.52 | 0.3 | 0.51 | 0.4 | 0.5 | 0.54 |
| 6 | 17834942 | rs17689501 | -0.017 | -0.59 | 0.558 | -0.955 | -0.015 | 0.022 | 0.022 | 0 | 0.21 | 0.08 | 0.12 | 0.19 | 0.21 | 0.33 |
| 6 | 17847860 | rs11754145 | 0.728 | -0.644 | 0.577 | -0.786 | 0.011 | -0.023 | -0.023 | 0 | 0.22 | 0.09 | 0.39 | 0.21 | 0.26 | 0.15 |
| 6 | 17856704 | rs665966 | 1.336 | -0.721 | 0.72 | -0.876 | -0.038 | -0.002 | -0.002 | 0 | 0.41 | 0.25 | 0.38 | 0.58 | 0.46 | 0.4 |
| 6 | 17871072 | rs12201755 | -0.085 | -0.889 | 0.764 | -0.824 | -0.003 | 0.01 | 0.01 | 0 | 0.2 | 0.12 | 0.13 | 0.2 | 0.2 | 0.33 |
| 6 | 17881217 | rs9367972 | 0.395 | -0.966 | 1.094 | -0.873 | -0.004 | -0.001 | -0.001 | 0 | 0.58 | 0.6 | 0.49 | 0.59 | 0.58 | 0.53 |
| 6 | 17881428 | rs9383330 | 1.401 | -0.994 | 0.938 | -0.913 | -0.034 | -0.008 | -0.008 | 2.441 | 0.42 | 0.22 | 0.38 | 0.57 | 0.46 | 0.4 |
| 6 | 17884879 | rs760849 | 1.345 | -0.992 | 0.968 | -0.91 | -0.009 | -0.003 | -0.003 | 0 | 0.42 | 0.23 | 0.4 | 0.57 | 0.49 | 0.41 |
| 6 | 17885402 | rs724916 | -0.723 | -1.161 | 1.154 | -0.667 | -0.019 | 0.009 | 0.009 | 0 | 0.78 | 0.72 | 0.62 | 0.79 | 0.77 | 0.85 |
| 6 | 17890002 | rs9477550 | 0.684 | -1.141 | 1.012 | -0.644 | 0.036 | 0.034 | 0.034 | 0 | 0.6 | 0.3 | 0.51 | 0.77 | 0.65 | 0.72 |
| 6 | 17905008 | rs12199169 | -0.212 | -0.986 | 0.988 | -0.618 | -0.009 | 0.024 | 0.024 | 0 | 0.1 | 0.22 | 0.05 | 0.19 | 0.06 | 0.17 |
| 6 | 17907735 | rs9371024 | -0.794 | -1.22 | 1.155 | -0.711 | -0.034 | 0.025 | 0.025 | 0 | 0.78 | 0.65 | 0.62 | 0.78 | 0.77 | 0.85 |
| 6 | 17913077 | rs1014146 | 0.314 | -1.121 | 1.089 | -0.719 | -0.014 | 0.014 | 0.014 | 0 | 0.67 | 0.43 | 0.56 | 0.59 | 0.68 | 0.68 |
| 6 | 17925426 | rs7765428 | 0.897 | -1.28 | 1.06 | -0.866 | 0.055 | 0.019 | 0.019 | 0 | 0.54 | 0.25 | 0.49 | 0.58 | 0.6 | 0.61 |
| 6 | 17931276 | rs6923237 | -0.23 | -1.31 | 1.105 | -0.871 | -0.009 | 0.011 | 0.011 | 0 | 0.11 | 0.18 | 0.04 | 0.19 | 0.06 | 0.17 |
| 6 | 17934110 | rs2876471 | -0.421 | -1.184 | 0.999 | -0.809 | -0.01 | -0.013 | -0.013 | 0 | 0.22 | 0.27 | 0.1 | 0.21 | 0.13 | 0.24 |
| 6 | 17935418 | rs4712321 | -0.587 | -1.243 | 0.973 | -0.787 | -0.05 | 0.03 | 0.03 | 0 | 0.77 | 0.55 | 0.6 | 0.78 | 0.76 | 0.85 |
| 6 | 17979139 | rs12201624 | -0.724 | -1.113 | 0.768 | -1.117 | -0.015 | -0.035 | -0.035 | 0.378 | 0.24 | 0.39 | 0.12 | 0.21 | 0.17 | 0.24 |
| 6 | 17992528 | rs2050029 | -1.092 | -1.145 | 0.522 | -0.904 | 0.017 | 0.009 | 0.009 | 0 | 0.8 | 0.9 | 0.63 | 0.84 | 0.76 | 0.85 |
| 6 | 17999212 | rs1569542 | -0.606 | -1.123 | 0.43 | -0.855 | -0.01 | -0.013 | -0.013 | 0 | 0.23 | 0.29 | 0.11 | 0.21 | 0.16 | 0.25 |
| 6 | 18003488 | rs2820221 | 1.346 | -0.557 | -0.374 | -0.473 | -0.1 | -0.077 | -0.077 | 0.276 | 0.33 | 0.12 | 0.3 | 0.47 | 0.41 | 0.35 |
| 6 | 18005589 | rs760848 | 0.513 | -0.64 | 0.102 | -0.227 | 0.018 | 0.01 | 0.01 | 0 | 0.53 | 0.36 | 0.45 | 0.55 | 0.58 | 0.6 |
| 6 | 18021075 | rs7766167 | -0.726 | -0.98 | -0.04 | -0.184 | 0.092 | 0.031 | 0.031 | 0 | 0.45 | 0.78 | 0.54 | 0.43 | 0.34 | 0.38 |
| 6 | 18025864 | rs11751607 | -0.677 | -1.027 | 0.364 | -0.191 | -0.002 | -0.001 | -0.001 | 0 | 0.43 | 0.46 | 0.52 | 0.43 | 0.34 | 0.37 |
| 6 | 18030079 | rs9383343 | 0.196 | -1.085 | 0.057 | -0.009 | 0.011 | -0.029 | -0.029 | 2.065 | 0.06 | 0.01 | 0.01 | 0.16 | 0.03 | 0.1 |
| 6 | 18030178 | rs4507599 | -0.131 | -1.084 | 0.398 | -0.486 | 0.01 | 0.005 | 0.005 | 0 | 0.28 | 0.25 | 0.45 | 0.24 | 0.24 | 0.25 |
| 6 | 18033349 | rs10807621 | -0.598 | -0.642 | -0.126 | -0.928 | 0.035 | 0.087 | 0.087 | 0.015 | 0.47 | 0.28 | 0.52 | 0.27 | 0.41 | 0.34 |
| 6 | 18038081 | rs6459587 | -1.203 | -0.805 | 0.13 | -0.816 | 0.025 | 0.078 | 0.078 | 0.505 | 0.44 | 0.73 | 0.49 | 0.18 | 0.38 | 0.37 |
| 6 | 18044772 | rs20510 | 0.485 | -0.82 | 0.103 | -0.787 | 0.01 | -0.012 | -0.012 | 0 | 0.14 | 0.1 | 0.3 | 0.18 | 0.13 | 0.17 |
| 6 | 18045178 | rs5008499 | 0.463 | -0.844 | 0.089 | -0.802 | 0.011 | -0.011 | -0.011 | 0 | 0.15 | 0.1 | 0.3 | 0.18 | 0.13 | 0.17 |
| 6 | 18055908 | rs9477599 | -0.973 | -0.813 | 0.404 | -0.929 | -0.033 | -0.099 | -0.099 | 0.08 | 0.15 | 0.52 | 0.09 | 0.02 | 0.09 | 0.09 |
| 6 | 18072657 | rs4380756 | -1.19 | -0.721 | 0.552 | -1.05 | -0.009 | -0.141 | -0.141 | 0.125 | 0.29 | 0.57 | 0.18 | 0.03 | 0.25 | 0.2 |
| 6 | 18084765 | rs6939758 | -0.906 | -0.732 | 0.5 | -0.948 | 0.002 | 0.004 | 0.004 | 0.279 | 0.36 | 0.66 | 0.38 | 0.19 | 0.3 | 0.34 |
| 6 | 18085650 | rs6459591 | -1.034 | -0.712 | 0.348 | -0.938 | -0.006 | -0.028 | -0.028 | 2.244 | 0.35 | 0.59 | 0.37 | 0.19 | 0.3 | 0.34 |
| 6 | 18088988 | rs12201541 | 0.578 | -0.458 | -0.385 | -0.847 | -0.059 | 0.074 | 0.074 | 0.022 | 0.12 | 0.01 | 0.07 | 0 | 0.17 | 0.11 |
| 6 | 18099594 | rs4716220 | -0.743 | -0.505 | -0.623 | -0.412 | 0.004 | -0.004 | -0.004 | 0 | 0.79 | 0.83 | 0.82 | 0.75 | 0.8 | 0.8 |
| 6 | 18100374 | rs5029614 | -0.524 | -0.528 | -0.6 | -0.527 | -0.015 | 0.079 | 0.079 | 0 | 0.35 | 0.2 | 0.37 | 0.17 | 0.35 | 0.36 |
| 6 | 18100865 | rs9358148 | 0.641 | -0.471 | 0.032 | 0.363 | -0.004 | 0.002 | 0.002 | 0.303 | 0.73 | 0.3 | 0.62 | 0.68 | 0.8 | 0.72 |
| 6 | 18102370 | rs6908777 | -0.84 | -0.08 | 0.805 | 0.008 | 0.053 | -0.032 | -0.032 | 0 | 0.1 | 0.35 | 0.13 | 0.03 | 0.11 | 0.1 |
| 6 | 18103281 | rs12194394 | 0.154 | -0.443 | 0.387 | -0.07 | -0.018 | 0.002 | 0.002 | 0 | 0.46 | 0.33 | 0.59 | 0.41 | 0.49 | 0.41 |
| 6 | 18106076 | rs4294009 | 0.653 | -0.453 | 0.34 | -0.063 | -0.043 | -0.052 | -0.052 | 0 | 0.39 | 0.18 | 0.49 | 0.39 | 0.4 | 0.32 |
| 6 | 18110443 | rs6899431 | -0.269 | -0.573 | 0.637 | 0.028 | 0.005 | -0.001 | -0.001 | 0.042 | 0.52 | 0.65 | 0.69 | 0.45 | 0.54 | 0.48 |
| 6 | 18115120 | rs12524744 | 0.638 | -0.671 | 0.495 | 0.094 | 0.004 | -0.033 | -0.033 | 0 | 0.06 | 0.01 | 0.17 | 0.19 | 0.05 | 0.04 |
| 6 | 18115715 | rs11967409 | 0.282 | -0.649 | 0.672 | 0.059 | 0.005 | -0.038 | -0.038 | 0 | 0.08 | 0.2 | 0.11 | 0.02 | 0.09 | 0.09 |
| 6 | 18117368 | rs6909725 | 0.192 | -0.651 | 0.705 | 0.058 | 0.009 | -0.04 | -0.04 | 0 | 0.08 | 0.23 | 0.11 | 0.02 | 0.09 | 0.09 |
| 6 | 18118098 | rs9477630 | 0.349 | -0.673 | 0.889 | 0.049 | -0.006 | -0.007 | -0.007 | 0 | 0.4 | 0.31 | 0.5 | 0.39 | 0.4 | 0.32 |
| 6 | 18119112 | rs10456213 | -0.289 | -0.697 | 0.971 | 0.035 | 0.012 | 0.015 | 0.015 | 0 | 0.51 | 0.66 | 0.66 | 0.41 | 0.54 | 0.44 |
| 6 | 18119477 | rs4588678 | 0.359 | -0.66 | 0.772 | 0.074 | -0.049 | -0.041 | -0.041 | 0.143 | 0.4 | 0.18 | 0.51 | 0.38 | 0.43 | 0.32 |
| 6 | 18122506 | rs10949483 | 0.54 | -0.563 | 0.726 | 0.091 | -0.031 | -0.034 | -0.034 | 0.798 | 0.39 | 0.2 | 0.49 | 0.39 | 0.4 | 0.32 |
| 6 | 18127067 | rs34889843 | 0.374 | -0.644 | 0.801 | 0.144 | 0.01 | -0.043 | -0.043 | 0 | 0.07 | 0.01 | 0.17 | 0.19 | 0.05 | 0.04 |
| 6 | 18131464 | rs9367980 | 0.188 | -0.812 | 0.543 | -0.038 | 0.024 | 0.027 | 0.027 | 0 | 0.65 | 0.3 | 0.65 | 0.74 | 0.73 | 0.6 |
| 6 | 18132822 | rs2842947 | -0.223 | -0.916 | 0.762 | -0.085 | 0.007 | 0.005 | 0.005 | 0 | 0.71 | 0.78 | 0.77 | 0.76 | 0.78 | 0.62 |
| 6 | 18135178 | rs2842950 | -0.39 | -0.914 | 0.766 | -0.09 | 0.007 | 0.009 | 0.009 | 0 | 0.72 | 0.81 | 0.81 | 0.81 | 0.78 | 0.65 |
| 6 | 18135683 | rs2842951 | 0.17 | -0.918 | 0.801 | -0.094 | 0.005 | 0.004 | 0.004 | 0 | 0.67 | 0.36 | 0.74 | 0.81 | 0.76 | 0.65 |
| 6 | 18137714 | rs4712327 | 0.17 | -0.909 | 0.672 | -0.089 | 0.005 | 0.004 | 0.004 | 0 | 0.67 | 0.36 | 0.74 | 0.81 | 0.76 | 0.65 |
| 6 | 18139214 | rs2842934 | -0.276 | -0.889 | 0.526 | -0.064 | 0.009 | 0.006 | 0.006 | 0 | 0.71 | 0.81 | 0.78 | 0.76 | 0.78 | 0.63 |
| 6 | 18140175 | rs2842935 | 0.194 | -0.795 | 0.204 | -0.179 | 0.021 | 0.035 | 0.035 | 0 | 0.66 | 0.3 | 0.69 | 0.78 | 0.73 | 0.63 |
| 6 | 18140250 | rs2842936 | 0.471 | -0.891 | -0.157 | -0.26 | 0.062 | 0.138 | 0.138 | 0 | 0.58 | 0.22 | 0.66 | 0.78 | 0.67 | 0.54 |

| | | | | | | | | | | | | | | | |
|---|---|---|---|---|---|---|---|---|---|---|---|---|---|---|---|
| 6 | 18141308 | rs12663332 | -0.349 | -0.546 | -0.088 | -0.178 | 0.001 | -0.009 | -0.009 | 0 | 0.14 | 0.14 | 0.25 | 0.2 | 0.15 | 0.08 |
| 6 | 18141330 | rs2842938 | -0.273 | -0.7 | 0.26 | -0.424 | 0.002 | -0.017 | -0.017 | 0 | 0.87 | 0.76 | 0.85 | 0.91 | 0.84 | 0.76 |
| 6 | 18143769 | rs2518463 | 0.587 | -0.611 | -0.022 | -0.36 | 0.019 | 0.047 | 0.047 | 0 | 0.53 | 0.72 | 0.56 | 0.39 | 0.54 | 0.42 |
| 6 | 18145411 | rs9297056 | -0.357 | -0.615 | -0.05 | -0.3 | -0.017 | -0.067 | -0.067 | 0 | 0.38 | 0.15 | 0.36 | 0.52 | 0.4 | 0.33 |
| 6 | 18147780 | rs3898137 | 1.683 | -0.631 | 0.064 | -0.411 | -0.039 | 0.022 | 0.022 | 0 | 0.34 | 0.16 | 0.34 | 0.25 | 0.4 | 0.23 |
| 6 | 18148019 | rs4449636 | 0.641 | -0.509 | -0.144 | -0.261 | 0.02 | 0.05 | 0.05 | 0 | 0.53 | 0.72 | 0.56 | 0.38 | 0.54 | 0.42 |
| 6 | 18148513 | rs2518471 | -0.46 | -0.111 | 0.176 | 0.162 | 0.007 | -0.006 | -0.006 | 0 | 0.92 | 0.94 | 0.93 | 0.9 | 0.93 | 0.75 |
| 6 | 18157968 | rs2876476 | -0.276 | -0.08 | 0.273 | 0.149 | -0.003 | 0 | 0 | 0 | 0.91 | 0.9 | 0.9 | 0.91 | 0.91 | 0.75 |
| 6 | 18161563 | rs4716224 | -0.108 | -0.208 | 0.105 | 0.02 | -0.017 | -0.079 | -0.079 | 0 | 0.32 | 0.15 | 0.35 | 0.53 | 0.34 | 0.27 |
| 6 | 18164073 | rs9396834 | -0.242 | -0.212 | 0.196 | 0.018 | 0 | -0.027 | -0.027 | 1.131 | 0.33 | 0.27 | 0.36 | 0.53 | 0.34 | 0.27 |
| 6 | 18170524 | rs7744164 | 0.61 | -0.2 | 0.225 | -0.01 | 0.004 | 0.016 | 0.016 | 0 | 0.57 | 0.64 | 0.55 | 0.39 | 0.58 | 0.48 |
| 6 | 18171477 | rs1011620 | 0.022 | -0.301 | 0.469 | 0.025 | -0.012 | -0.004 | -0.004 | 0.914 | 0.88 | 0.84 | 0.89 | 0.91 | 0.9 | 0.74 |
| 6 | 18181374 | rs12665749 | 0.535 | -0.259 | 0.151 | -0.079 | -0.001 | 0 | 0 | 0 | 0.14 | 0.12 | 0.14 | 0.14 | 0.12 | 0.19 |
| 6 | 18185410 | rs6932667 | -0.288 | -0.264 | 0.266 | 0.142 | 0.002 | 0 | 0 | 0 | 0.91 | 0.93 | 0.92 | 0.93 | 0.93 | 0.75 |
| 6 | 18186547 | rs440163 | 0.339 | -0.255 | 0.322 | 0.138 | -0.006 | -0.002 | -0.002 | 0.02 | 0.1 | 0.13 | 0.1 | 0.07 | 0.09 | 0.25 |
| 6 | 18187584 | rs764099 | 0.325 | -0.332 | -0.106 | 0.368 | -0.011 | 0.04 | 0.04 | 0 | 0.07 | 0.11 | 0.16 | 0.19 | 0.03 | 0.04 |
| 6 | 18197475 | rs214621 | -0.676 | 0.003 | 0.246 | -0.17 | -0.042 | -0.038 | -0.038 | 0 | 0.16 | 0.53 | 0.16 | 0.07 | 0.1 | 0.26 |
| 6 | 18203126 | rs214609 | -0.673 | 0.568 | -0.792 | 0.16 | -0.041 | -0.084 | -0.084 | 0 | 0.83 | 0.98 | 0.9 | 0.66 | 0.83 | 0.82 |
| 6 | 18208319 | rs214596 | -0.705 | 0.501 | -0.237 | -0.124 | -0.001 | -0.03 | -0.03 | 0 | 0.33 | 0.37 | 0.32 | 0.22 | 0.32 | 0.5 |
| 6 | 18209582 | rs214590 | 0.356 | 1.251 | -0.617 | 0.148 | 0.013 | 0.004 | 0.004 | 0 | 0.51 | 0.24 | 0.55 | 0.48 | 0.43 | 0.53 |
| 6 | 18209763 | rs6911008 | -1.149 | 1.659 | -0.137 | 0.059 | 0.02 | -0.006 | -0.006 | 0 | 0.23 | 0.57 | 0.29 | 0.26 | 0.25 | 0.29 |
| 6 | 18213988 | rs6903583 | -0.657 | 1.437 | -0.305 | -0.481 | 0.022 | -0.057 | -0.057 | 3.1 | 0.16 | 0.21 | 0.17 | 0.01 | 0.21 | 0.09 |
| 6 | 18223409 | rs214505 | -0.157 | 0.786 | -0.882 | -1.574 | 0.015 | 0.014 | 0.014 | 0.022 | 0.36 | 0.63 | 0.41 | 0.1 | 0.22 | 0.39 |
| 6 | 18225173 | rs3476 | 1.224 | 0.502 | -1.152 | -1.333 | -0.013 | -0.05 | -0.05 | 0 | 0.2 | 0.03 | 0.11 | 0.25 | 0.21 | 0.25 |
| 6 | 18235776 | rs4712329 | 1.312 | 0.324 | -1.289 | -0.845 | -0.017 | -0.062 | -0.062 | 0 | 0.18 | 0.13 | 0.18 | 0.44 | 0.25 | 0.14 |
| 6 | 18237661 | rs1013062 | 1.322 | 0.29 | -1.166 | -0.954 | -0.013 | -0.053 | -0.053 | 2.766 | 0.2 | 0.03 | 0.11 | 0.25 | 0.21 | 0.25 |
| 6 | 18256625 | rs17336208 | 1.235 | -0.212 | -1.334 | -1.201 | 0 | 0.033 | 0.033 | 0.316 | 0.07 | 0.01 | 0.19 | 0.01 | 0.04 | 0.05 |
| 6 | 18258516 | rs172520 | 0.043 | -0.173 | -1.296 | -1.178 | 0.017 | 0.016 | 0.016 | 0 | 0.36 | 0.63 | 0.41 | 0.1 | 0.22 | 0.39 |
| 6 | 18265698 | rs214575 | -0.715 | -0.164 | -1.29 | -1.169 | -0.027 | -0.032 | -0.032 | 0 | 0.69 | 0.84 | 0.69 | 0.35 | 0.64 | 0.73 |
| 6 | 18267095 | rs214568 | 0.015 | -0.33 | -1.003 | -1.234 | 0.012 | 0.009 | 0.009 | 0.06 | 0.35 | 0.61 | 0.41 | 0.1 | 0.22 | 0.39 |
| 6 | 18280885 | rs214553 | 0.004 | -0.341 | -1.007 | -1.283 | 0.013 | 0.004 | 0.004 | 0.057 | 0.35 | 0.61 | 0.37 | 0.08 | 0.21 | 0.39 |
| 6 | 18284716 | rs214544 | 0.094 | -0.658 | -0.483 | -1.453 | -0.032 | -0.037 | -0.037 | 0 | 0.16 | 0.26 | 0.27 | 0.07 | 0.08 | 0.33 |
| 6 | 18294663 | rs445149 | 0.27 | -0.716 | -0.464 | -1.536 | -0.035 | -0.086 | -0.086 | 0.31 | 0.67 | 0.58 | 0.66 | 0.93 | 0.8 | 0.62 |
| 6 | 18295504 | rs7753514 | -0.871 | -0.913 | -0.023 | -1.612 | 0.004 | -0.06 | -0.06 | 0.274 | 0.89 | 0.81 | 0.93 | 1 | 0.92 | 0.97 |
| 6 | 18300782 | rs1324432 | 0.431 | -1.279 | -0.186 | -1.818 | -0.048 | -0.1 | -0.1 | 0 | 0.74 | 0.49 | 0.84 | 0.94 | 0.85 | 0.68 |
| 6 | 18301503 | rs7747741 | 0.835 | -1.051 | -0.32 | -1.73 | -0.024 | 0.015 | 0.015 | 0.09 | 0.33 | 0.25 | 0.44 | 0.25 | 0.43 | 0.31 |
| 6 | 18312515 | rs9477695 | 1.1 | -0.744 | -0.489 | -1.789 | -0.042 | 0.038 | 0.038 | 0 | 0.13 | 0.06 | 0.16 | 0.01 | 0.22 | 0.11 |
| 6 | 18321125 | rs728471 | -0.768 | -0.771 | -0.4 | -1.874 | 0 | -0.026 | -0.026 | 0.138 | 0.87 | 0.89 | 0.93 | 1 | 0.87 | 0.94 |
| 6 | 18323876 | rs912530 | 0.552 | -1.075 | -0.636 | -1.85 | -0.027 | -0.029 | -0.029 | 0 | 0.38 | 0.28 | 0.42 | 0.46 | 0.52 | 0.31 |
| 6 | 18326873 | rs13210382 | 1.161 | -1.124 | -0.566 | -1.84 | 0.005 | -0.002 | -0.002 | 0 | 0.11 | 0.1 | 0.1 | 0.01 | 0.05 | 0.06 |
| 6 | 18330781 | rs9477700 | -0.344 | -1.048 | -0.067 | -1.905 | -0.005 | 0.005 | 0.005 | 0 | 0.34 | 0.4 | 0.36 | 0.28 | 0.36 | 0.56 |
| 6 | 18334298 | rs2328243 | -0.651 | -1.015 | 0.109 | -1.934 | 0.041 | -0.097 | -0.097 | 0 | 0.86 | 0.68 | 0.91 | 1 | 0.87 | 0.94 |
| 6 | 18336777 | rs725550 | 1.384 | -0.877 | -0.145 | -1.938 | -0.026 | 0.02 | 0.02 | 0 | 0.11 | 0.07 | 0.15 | 0.01 | 0.21 | 0.11 |
| 6 | 18338338 | rs9358158 | 2.04 | -1.032 | -0.304 | -1.921 | 0.004 | -0.064 | -0.064 | 0 | 0.16 | 0.05 | 0.26 | 0.27 | 0.17 | 0.18 |
| 6 | 18340287 | rs1408442 | 1.286 | -1.12 | -0.267 | -1.928 | 0.002 | -0.053 | -0.053 | 0.89 | 0.29 | 0.18 | 0.28 | 0.45 | 0.31 | 0.2 |
| 6 | 18340298 | rs11967050 | 0.703 | -1.132 | -0.001 | -1.973 | -0.01 | 0.009 | 0.009 | 3.894 | 0.15 | 0.23 | 0.17 | 0.22 | 0.12 | 0.17 |
| 6 | 18346644 | rs2328225 | -0.183 | -1.096 | -0.103 | -1.959 | 0 | -0.037 | -0.037 | 0.22 | 0.72 | 0.59 | 0.76 | 0.78 | 0.75 | 0.77 |
| 6 | 18347266 | rs7753875 | 0.282 | -1.051 | -0.079 | -1.678 | -0.002 | -0.002 | -0.002 | 0 | 0.36 | 0.35 | 0.47 | 0.37 | 0.41 | 0.55 |
| 6 | 18348630 | rs16880565 | 0.279 | -1.069 | -0.025 | -1.686 | -0.016 | -0.004 | -0.004 | 0 | 0.08 | 0.12 | 0.06 | 0.06 | 0.03 | 0.28 |
| 6 | 18349366 | rs1408445 | -0.826 | -1.072 | -0.053 | -1.835 | 0 | -0.037 | -0.037 | 0 | 0.87 | 0.85 | 0.91 | 1 | 0.88 | 0.95 |
| 6 | 18351530 | rs9383372 | 0.833 | -1.09 | -0.324 | -1.819 | 0.026 | -0.047 | -0.047 | 0 | 0.24 | 0.14 | 0.36 | 0.35 | 0.2 | 0.43 |
| 6 | 18352976 | rs1951924 | -0.324 | -1.178 | -0.237 | -1.872 | -0.001 | -0.041 | -0.041 | 0 | 0.72 | 0.57 | 0.74 | 0.78 | 0.75 | 0.76 |
| 6 | 18353562 | rs9297068 | 0.22 | -0.959 | -0.18 | -1.824 | 0.001 | 0 | 0 | 0 | 0.36 | 0.39 | 0.51 | 0.36 | 0.4 | 0.54 |
| 6 | 18365591 | rs1815296 | 1.097 | -0.668 | -0.152 | -1.675 | 0.086 | -0.061 | -0.061 | 0 | 0.11 | 0.23 | 0.15 | 0.01 | 0.21 | 0.11 |
| 6 | 18374600 | rs9396857 | 1.066 | -1.047 | -0.633 | -1.755 | 0 | -0.015 | -0.015 | 0 | 0.25 | 0.19 | 0.2 | 0.42 | 0.29 | 0.22 |
| 6 | 18385889 | rs7748189 | 0.304 | -0.879 | -0.626 | -1.798 | -0.005 | -0.023 | -0.023 | 0.616 | 0.62 | 0.42 | 0.72 | 0.78 | 0.7 | 0.73 |
| 6 | 18387265 | rs9371051 | 1.075 | -0.695 | -0.952 | -1.546 | -0.016 | -0.074 | -0.074 | 0 | 0.21 | 0.1 | 0.17 | 0.42 | 0.26 | 0.16 |
| 6 | 18387692 | rs9396858 | 0.546 | -0.776 | -0.825 | -1.594 | 0.002 | -0.021 | -0.021 | 0 | 0.26 | 0.24 | 0.2 | 0.42 | 0.28 | 0.18 |
| 6 | 18387815 | rs764885 | 0.85 | -0.712 | -0.761 | -1.547 | -0.002 | 0 | 0 | 4.228 | 0.17 | 0.17 | 0.15 | 0.21 | 0.15 | 0.18 |
| 6 | 18394727 | rs11754093 | 1.451 | -0.115 | -0.941 | -0.951 | -0.026 | -0.065 | -0.065 | 0 | 0.16 | 0.08 | 0.14 | 0.42 | 0.23 | 0.15 |
| 6 | 18396689 | rs6941991 | -0.562 | -0.05 | -0.384 | -0.882 | 0.001 | 0 | 0 | 0 | 0.36 | 0.39 | 0.54 | 0.36 | 0.4 | 0.54 |
| 6 | 18403259 | rs12199402 | 0.521 | -0.514 | -0.632 | -0.441 | -0.008 | -0.079 | -0.079 | 0.77 | 0.16 | 0.03 | 0.33 | 0.3 | 0.17 | 0.16 |
| 6 | 18404020 | rs9350070 | -0.172 | -0.564 | -0.389 | -0.513 | -0.01 | 0.006 | 0.006 | 0 | 0.22 | 0.28 | 0.36 | 0.26 | 0.19 | 0.24 |
| 6 | 18406854 | rs9367995 | 0.72 | -0.268 | -0.239 | -0.8 | -0.001 | -0.063 | -0.063 | 0 | 0.88 | 0.72 | 0.9 | 1 | 0.89 | 0.96 |
| 6 | 18407590 | rs2147215 | -0.026 | 0.182 | -0.978 | -1.03 | -0.034 | 0.081 | 0.081 | 0 | 0.21 | 0.04 | 0.17 | 0.1 | 0.22 | 0.16 |
| 6 | 18407743 | rs1408263 | 0.612 | -0.175 | -0.482 | -0.509 | -0.01 | 0 | 0 | 0 | 0.64 | 0.54 | 0.55 | 0.67 | 0.69 | 0.58 |
| 6 | 18418708 | rs2038745 | 0.283 | -0.31 | -0.014 | -0.555 | -0.043 | -0.027 | -0.027 | 0 | 0.75 | 0.46 | 0.79 | 0.83 | 0.83 | 0.87 |
| 6 | 18424299 | rs516487 | 0.5 | -0.196 | 0.188 | -0.605 | -0.03 | -0.041 | -0.041 | 0 | 0.72 | 0.5 | 0.77 | 0.83 | 0.78 | 0.84 |
| 6 | 18426608 | rs500720 | 0.994 | 0.535 | 0.067 | -0.917 | -0.001 | -0.014 | -0.014 | 0 | 0.64 | 0.42 | 0.55 | 0.82 | 0.63 | 0.77 |

| | | | | | | | | | | | | | | | | |
|---|---|---|---|---|---|---|---|---|---|---|---|---|---|---|---|---|
| 6 | 18431695 | rs478281 | -0.682 | 0.476 | -0.001 | -0.878 | -0.011 | 0.002 | 0.002 | 0 | 0.4 | 0.5 | 0.39 | 0.4 | 0.34 | 0.34 |
| 6 | 18434676 | rs4716253 | 0.573 | 0.585 | -0.384 | -0.799 | 0.008 | -0.015 | -0.015 | 0 | 0.08 | 0.07 | 0.22 | 0.01 | 0.14 | 0.07 |
| 6 | 18438876 | rs6927687 | 1.885 | 0.507 | -1.273 | -1.311 | -0.049 | -0.144 | -0.144 | 0 | 0.25 | 0.04 | 0.13 | 0.45 | 0.25 | 0.24 |
| 6 | 18439027 | rs4712335 | 0.845 | 0.519 | -1.005 | -1.183 | -0.002 | -0.003 | -0.003 | 0 | 0.08 | 0.06 | 0.22 | 0.01 | 0.14 | 0.07 |
| 6 | 18441067 | rs7774742 | -0.109 | 0.372 | -1.01 | -1.263 | -0.003 | 0.002 | 0.002 | 0 | 0.13 | 0.13 | 0.27 | 0.08 | 0.15 | 0.25 |
| 6 | 18441336 | rs9477745 | 1.255 | -0.078 | -0.775 | -1.288 | -0.022 | 0.005 | 0.005 | 0.013 | 0.13 | 0.14 | 0.09 | 0.17 | 0.04 | 0.1 |
| 6 | 18441624 | rs10949509 | 0.942 | 0.038 | -1.426 | -1.42 | -0.042 | -0.056 | -0.056 | 0.005 | 0.43 | 0.12 | 0.46 | 0.53 | 0.52 | 0.44 |
| 6 | 18446747 | rs496519 | 0.88 | 0.371 | -1.167 | -1.13 | -0.028 | -0.047 | -0.047 | 0 | 0.18 | 0.06 | 0.24 | 0.23 | 0.21 | 0.19 |
| 6 | 18458536 | rs13196052 | 0.791 | 0.397 | -1.036 | -1.152 | -0.01 | 0.015 | 0.015 | 0 | 0.07 | 0.04 | 0.05 | 0 | 0.1 | 0.04 |
| 6 | 18458637 | rs474465 | 0.608 | 0.332 | -0.762 | -1.098 | 0 | -0.001 | -0.001 | 3.253 | 0.2 | 0.19 | 0.25 | 0.24 | 0.21 | 0.19 |
| 6 | 18462492 | rs10807623 | 1.059 | 0.201 | -1.043 | -1.244 | -0.03 | -0.049 | -0.049 | 0 | 0.41 | 0.19 | 0.26 | 0.52 | 0.42 | 0.41 |
| 6 | 18465710 | rs480936 | -0.14 | -0.09 | -0.511 | -1.411 | -0.014 | 0 | 0 | 0 | 0.94 | 0.89 | 0.95 | 0.93 | 0.99 | 0.82 |
| 6 | 18466750 | rs527984 | 0.482 | -0.105 | -0.613 | -1.476 | -0.012 | -0.036 | -0.036 | 0 | 0.19 | 0.06 | 0.22 | 0.22 | 0.2 | 0.19 |
| 6 | 18467582 | rs531972 | 0.154 | 0.116 | -0.355 | -1.512 | -0.009 | -0.008 | -0.008 | 0.153 | 0.92 | 0.89 | 0.94 | 0.93 | 0.96 | 0.81 |
| 6 | 18471865 | rs12198115 | 0.242 | 0.408 | -0.436 | -1.663 | -0.045 | 0.036 | 0.036 | 0 | 0.18 | 0.07 | 0.21 | 0.08 | 0.26 | 0.16 |
| 6 | 18476660 | rs532347 | 0.946 | 0.131 | -0.111 | -1.772 | -0.005 | -0.03 | -0.03 | 0 | 0.92 | 0.83 | 0.92 | 0.97 | 0.95 | 0.91 |
| 6 | 18481019 | rs567621 | 0.541 | 0.14 | -0.105 | -1.759 | -0.002 | -0.014 | -0.014 | 1.499 | 0.93 | 0.89 | 0.92 | 0.96 | 0.95 | 0.9 |
| 6 | 18487593 | rs4716267 | 0.66 | 0.108 | 0.127 | -1.896 | 0.014 | -0.05 | -0.05 | 0 | 0.75 | 0.64 | 0.72 | 0.85 | 0.72 | 0.6 |
| 6 | 18493389 | rs4716269 | -0.572 | 0.194 | 0.355 | -1.991 | 0.01 | -0.032 | -0.032 | 6.913 | 0.24 | 0.31 | 0.37 | 0.13 | 0.26 | 0.26 |
| 6 | 18496595 | rs1408498 | 0.77 | 0.213 | 0.265 | -2.007 | 0.011 | -0.034 | -0.034 | 0 | 0.62 | 0.6 | 0.47 | 0.8 | 0.51 | 0.51 |
| 6 | 18497778 | rs4716273 | -0.836 | 0.233 | 0.243 | -2.015 | 0.009 | -0.012 | -0.012 | 0.712 | 0.86 | 0.84 | 0.81 | 0.9 | 0.79 | 0.71 |
| 6 | 18499926 | rs7755288 | 0.286 | 0.249 | 0.155 | -1.993 | 0.004 | -0.013 | -0.013 | 0.171 | 0.18 | 0.21 | 0.29 | 0.09 | 0.21 | 0.15 |
| 6 | 18501982 | rs1359341 | 0.458 | 0.203 | 0.149 | -2.011 | -0.037 | 0.052 | 0.052 | 0.211 | 0.66 | 0.76 | 0.52 | 0.81 | 0.56 | 0.54 |
| 6 | 18502289 | rs6942330 | 0.779 | -0.321 | 0.207 | -2.489 | -0.047 | -0.135 | -0.135 | 0.154 | 0.2 | 0.42 | 0.14 | 0.01 | 0.09 | 0.08 |
| 6 | 18508270 | rs12662585 | 1.499 | -0.4 | -0.086 | -2.446 | -0.054 | 0.104 | 0.104 | 0.339 | 0.37 | 0.08 | 0.3 | 0.76 | 0.41 | 0.37 |
| 6 | 18510324 | rs912552 | 1.286 | -0.384 | -0.16 | -2.418 | -0.005 | 0.086 | 0.086 | 0.646 | 0.41 | 0.24 | 0.34 | 0.76 | 0.46 | 0.4 |
| 6 | 18510379 | rs12529334 | 0.804 | -0.275 | -0.072 | -2.311 | -0.03 | 0.032 | 0.032 | 0 | 0.16 | 0.06 | 0.27 | 0.09 | 0.21 | 0.15 |
| 6 | 18515976 | rs6459621 | -0.417 | -0.283 | -0.155 | -2.268 | -0.002 | -0.061 | -0.061 | 0 | 0.38 | 0.56 | 0.42 | 0.1 | 0.31 | 0.23 |
| 6 | 18525867 | rs932355 | 1.543 | -0.146 | -0.674 | -1.827 | -0.029 | -0.075 | -0.075 | 0 | 0.31 | 0.09 | 0.27 | 0.64 | 0.34 | 0.3 |
| 6 | 18531115 | rs4645427 | 0.448 | -0.211 | -0.409 | -2.016 | 0.004 | 0.098 | 0.098 | 0.072 | 0.57 | 0.27 | 0.59 | 0.85 | 0.64 | 0.51 |
| 6 | 18538045 | rs9368002 | 0.196 | -0.209 | -0.433 | -1.999 | -0.008 | 0.005 | 0.005 | 0 | 0.25 | 0.14 | 0.32 | 0.21 | 0.3 | 0.21 |
| 6 | 18540282 | rs1016115 | -0.622 | -0.315 | 0.091 | -2.132 | 0.044 | -0.03 | -0.03 | 0 | 0.8 | 0.6 | 0.73 | 0.86 | 0.77 | 0.61 |
| 6 | 18541514 | rs1408496 | -0.803 | -0.329 | 0.078 | -2.146 | -0.001 | -0.009 | -0.009 | 0.388 | 0.48 | 0.52 | 0.47 | 0.22 | 0.43 | 0.31 |
| 6 | 18544463 | rs9350079 | 0.52 | -0.295 | 0.068 | -2.099 | 0.002 | 0.019 | 0.019 | 0 | 0.54 | 0.44 | 0.41 | 0.64 | 0.47 | 0.39 |
| 6 | 18559445 | rs9396878 | 0.555 | -0.261 | 0.087 | -2.079 | 0.002 | 0.071 | 0.071 | 0 | 0.57 | 0.32 | 0.6 | 0.85 | 0.64 | 0.51 |
| 6 | 18559687 | rs6927679 | -0.076 | -0.287 | 0.32 | -2.143 | 0.002 | -0.066 | -0.066 | 0.162 | 0.67 | 0.45 | 0.69 | 0.88 | 0.72 | 0.62 |
| 6 | 18561460 | rs12215669 | -0.013 | -0.344 | 0.427 | -1.984 | -0.006 | -0.025 | -0.025 | 0 | 0.1 | 0.15 | 0.1 | 0.04 | 0.08 | 0.11 |
| 6 | 18578387 | rs10484370 | -0.204 | -0.337 | 0.419 | -1.957 | 0.024 | 0.002 | 0.002 | 0.341 | 0.08 | 0.33 | 0.11 | 0.04 | 0.05 | 0.1 |
| 6 | 18579752 | rs6925505 | -1.291 | -0.213 | -0.272 | -2.013 | -0.031 | 0.073 | 0.073 | 0 | 0.57 | 0.74 | 0.58 | 0.26 | 0.51 | 0.42 |
| 6 | 18581172 | rs2064175 | -0.526 | -0.335 | -0.251 | -2.042 | -0.019 | -0.078 | -0.078 | 0.745 | 0.3 | 0.61 | 0.24 | 0.04 | 0.17 | 0.2 |
| 6 | 18585348 | rs9477792 | -0.964 | -0.004 | -0.389 | -1.776 | -0.019 | 0.053 | 0.053 | 0 | 0.56 | 0.67 | 0.56 | 0.26 | 0.49 | 0.42 |
| 6 | 18596034 | rs7770329 | 0.145 | -0.311 | -0.237 | -1.893 | -0.001 | -0.01 | -0.01 | 0 | 0.23 | 0.2 | 0.2 | 0.06 | 0.15 | 0.15 |
| 6 | 18602913 | rs9350086 | -0.623 | 0.035 | -0.24 | -1.915 | -0.002 | -0.001 | -0.001 | 0.076 | 0.81 | 0.8 | 0.82 | 0.91 | 0.74 | 0.7 |
| 6 | 18609828 | rs6930066 | 0.244 | -0.173 | -0.138 | -2.03 | 0.004 | -0.063 | -0.063 | 0 | 0.63 | 0.55 | 0.69 | 0.87 | 0.64 | 0.62 |
| 6 | 18610789 | rs4716291 | 2.126 | -0.654 | -0.349 | -2.262 | -0.001 | -0.022 | -0.022 | 0 | 0.25 | 0.23 | 0.22 | 0.65 | 0.28 | 0.28 |
| 6 | 18614099 | rs17642987 | -0.05 | -0.619 | -0.238 | -2.233 | 0.007 | 0.019 | 0.019 | 0.278 | 0.33 | 0.22 | 0.4 | 0.22 | 0.31 | 0.3 |
| 6 | 18624174 | rs2223382 | -1.278 | -0.44 | -0.122 | -2.088 | 0.004 | -0.007 | -0.007 | 0 | 0.83 | 0.79 | 0.83 | 0.9 | 0.77 | 0.7 |
| 6 | 18630338 | rs17643301 | 2.104 | -0.511 | -0.326 | -2.079 | -0.004 | 0.032 | 0.032 | 1.705 | 0.07 | 0 | 0.07 | 0.01 | 0.04 | 0.05 |
| 6 | 18637193 | rs11966801 | 1.295 | -0.522 | -0.248 | -2.095 | 0.011 | 0.064 | 0.064 | 0 | 0.06 | 0.12 | 0.04 | 0.08 | 0.04 | 0.07 |
| 6 | 18655570 | rs1474618 | -0.668 | -0.53 | -0.355 | -2.212 | -0.021 | 0.053 | 0.053 | 3.496 | 0.73 | 0.81 | 0.76 | 0.87 | 0.7 | 0.64 |
| 6 | 18656883 | rs9477809 | -0.943 | -0.813 | -0.448 | -2.436 | 0.01 | 0.069 | 0.069 | 0.276 | 0.48 | 0.55 | 0.53 | 0.22 | 0.42 | 0.35 |
| 6 | 18663606 | rs9465181 | 1.3 | -0.924 | -0.24 | -2.318 | -0.001 | -0.005 | -0.005 | 1.001 | 0.4 | 0.55 | 0.34 | 0.66 | 0.39 | 0.34 |
| 6 | 18666001 | rs9465185 | 2.002 | -0.993 | -0.26 | -2.286 | -0.001 | -0.019 | -0.019 | 10.718 | 0.3 | 0.27 | 0.26 | 0.64 | 0.32 | 0.32 |
| 6 | 18666712 | rs1534927 | -0.339 | -0.974 | -0.384 | -2.278 | 0.009 | 0.024 | 0.024 | 0.929 | 0.32 | 0.18 | 0.4 | 0.22 | 0.31 | 0.3 |
| 6 | 18668020 | rs17644348 | -0.646 | -1.179 | -0.497 | -2.367 | 0.023 | 0.074 | 0.074 | 0.692 | 0.42 | 0.2 | 0.45 | 0.22 | 0.37 | 0.31 |
| 6 | 18668726 | rs6904170 | 2.442 | -1.028 | -0.422 | -2.287 | -0.007 | 0.026 | 0.026 | 0 | 0.07 | 0.03 | 0.04 | 0 | 0.06 | 0.01 |
| 6 | 18669821 | rs2206087 | -0.97 | -0.88 | -0.114 | -2.032 | 0.015 | 0.032 | 0.032 | 0.066 | 0.43 | 0.28 | 0.49 | 0.23 | 0.37 | 0.36 |
| 6 | 18681772 | rs4498360 | 1.027 | -0.7 | -0.398 | -1.886 | 0.003 | 0.009 | 0.009 | 0 | 0.49 | 0.47 | 0.41 | 0.65 | 0.45 | 0.39 |
| 6 | 18688972 | rs9371088 | 0.959 | -0.82 | -0.377 | -1.861 | -0.032 | 0.045 | 0.045 | 0 | 0.17 | 0.02 | 0.28 | 0.1 | 0.19 | 0.13 |
| 6 | 18691010 | rs9368017 | -0.208 | -0.738 | -0.275 | -1.812 | 0.015 | 0.015 | 0.015 | 0.474 | 0.3 | 0.17 | 0.39 | 0.22 | 0.27 | 0.28 |
| 6 | 18692084 | rs9383431 | -0.179 | -0.194 | 0.045 | -1.681 | 0.014 | -0.029 | -0.029 | 0 | 0.74 | 0.64 | 0.73 | 0.86 | 0.71 | 0.64 |
| 6 | 18699142 | rs2208857 | -0.77 | 0.218 | -0.16 | -1.383 | 0.008 | -0.006 | -0.006 | 0.463 | 0.83 | 0.76 | 0.82 | 0.89 | 0.76 | 0.7 |
| 6 | 18701589 | rs9383432 | -0.012 | 0.214 | -0.31 | -1.374 | 0 | 0.002 | 0.002 | 0.581 | 0.23 | 0.14 | 0.31 | 0.19 | 0.23 | 0.2 |
| 6 | 18708372 | rs1570394 | -0.285 | 0.583 | -0.103 | -1.154 | -0.004 | 0.003 | 0.003 | 0.036 | 0.83 | 0.8 | 0.82 | 0.86 | 0.76 | 0.7 |
| 6 | 18712544 | rs16881089 | -1.436 | 0.246 | 0.358 | -1.108 | 0.003 | 0.006 | 0.006 | 1.101 | 0.12 | 0.26 | 0.15 | 0.13 | 0.09 | 0.12 |
| 6 | 18713640 | rs9477835 | 0.313 | -0.046 | 0.371 | -1.511 | 0 | -0.009 | -0.009 | 0.499 | 0.46 | 0.5 | 0.43 | 0.73 | 0.49 | 0.61 |
| 6 | 18716821 | rs9477837 | -0.497 | -0.337 | 0.414 | -1.853 | -0.028 | 0.057 | 0.057 | 0 | 0.65 | 0.85 | 0.61 | 0.76 | 0.64 | 0.74 |
| 6 | 18716933 | rs16881104 | -0.07 | -0.393 | 0.609 | -1.905 | 0.002 | -0.008 | -0.008 | 0 | 0.1 | 0.2 | 0.12 | 0.09 | 0.07 | 0.09 |
| 6 | 18733216 | rs980959 | 0.044 | -0.468 | 0.966 | -1.979 | -0.012 | -0.03 | -0.03 | 1.102 | 0.1 | 0.16 | 0.05 | 0.02 | 0.09 | 0.07 |

| | | | | | | | | | | | | | | | | |
|---|---|---|---|---|---|---|---|---|---|---|---|---|---|---|---|---|
| 6 | 18740159 | rs1998416 | 0.571 | -0.429 | 0.964 | -1.976 | 0.001 | -0.015 | -0.015 | 0.964 | 0.45 | 0.51 | 0.43 | 0.73 | 0.49 | 0.62 |
| 6 | 18750940 | rs9477861 | 1.332 | -0.462 | 0.919 | -1.991 | -0.002 | 0.017 | 0.017 | 0.862 | 0.37 | 0.34 | 0.37 | 0.73 | 0.41 | 0.6 |
| 6 | 18751370 | rs1033819 | 0.449 | -0.438 | 0.939 | -1.986 | -0.003 | 0.036 | 0.036 | 0.066 | 0.47 | 0.63 | 0.44 | 0.73 | 0.48 | 0.62 |
| 6 | 18760100 | rs16881153 | 0.006 | -0.215 | 0.958 | -1.599 | 0.003 | -0.031 | -0.031 | 0 | 0.08 | 0.15 | 0.06 | 0 | 0.07 | 0.02 |
| 6 | 18762620 | rs7767234 | 1.324 | -0.185 | 0.973 | -1.639 | -0.014 | 0.027 | 0.027 | 0.219 | 0.42 | 0.28 | 0.44 | 0.64 | 0.5 | 0.58 |
| 6 | 18766010 | rs9477872 | 0.952 | -0.865 | 0.952 | -1.301 | 0 | 0.024 | 0.024 | 0 | 0.47 | 0.42 | 0.49 | 0.64 | 0.54 | 0.59 |
| 6 | 18769716 | rs16881169 | -0.764 | -0.787 | 1.008 | -1.144 | -0.03 | -0.007 | -0.007 | 0 | 0.12 | 0.21 | 0.09 | 0.11 | 0.06 | 0.14 |
| 6 | 18773098 | rs1418968 | -1.366 | -0.785 | 0.867 | -1.091 | -0.056 | -0.026 | -0.026 | 0 | 0.18 | 0.4 | 0.12 | 0.13 | 0.09 | 0.15 |
| 6 | 18775937 | rs7774998 | 0.17 | -0.461 | 0.1 | -1.253 | 0 | -0.003 | -0.003 | 1.028 | 0.71 | 0.71 | 0.64 | 0.75 | 0.71 | 0.74 |
| 6 | 18779046 | rs13216194 | 1.629 | -0.354 | -0.532 | -1.175 | -0.1 | -0.078 | -0.078 | 0.061 | 0.36 | 0.04 | 0.28 | 0.62 | 0.42 | 0.49 |
| 6 | 18781294 | rs6928682 | 1.089 | -0.272 | 0.529 | -1.211 | 0.004 | 0.137 | 0.137 | 0.59 | 0.52 | 0.21 | 0.43 | 0.72 | 0.56 | 0.67 |
| 6 | 18793472 | rs9297093 | 0.621 | -0.563 | 1.198 | -1.201 | -0.004 | -0.034 | -0.034 | 0.078 | 0.6 | 0.53 | 0.56 | 0.73 | 0.62 | 0.59 |
| 6 | 18798316 | rs9477887 | 0.1 | -0.611 | 1.236 | -1.225 | -0.008 | -0.026 | -0.026 | 0 | 0.08 | 0.07 | 0.04 | 0.01 | 0.06 | 0.11 |
| 6 | 18815501 | rs1343132 | -0.378 | -0.635 | 0.98 | -1.205 | -0.017 | -0.035 | -0.035 | 0 | 0.1 | 0.21 | 0.05 | 0.01 | 0.06 | 0.19 |
| 6 | 18816027 | rs17654008 | 1.174 | -0.638 | 0.407 | -1.248 | -0.01 | 0.059 | 0.059 | 0 | 0.1 | 0.01 | 0.06 | 0 | 0.08 | 0.01 |
| 6 | 18827886 | rs2798142 | 0.116 | -0.597 | 0.496 | -1.187 | -0.013 | 0.019 | 0.019 | 0 | 0.7 | 0.81 | 0.62 | 0.74 | 0.67 | 0.76 |
| 6 | 18831438 | rs4142204 | 0.952 | -0.462 | -0.092 | -1.104 | 0.016 | 0.122 | 0.122 | 0.108 | 0.53 | 0.22 | 0.52 | 0.73 | 0.59 | 0.56 |
| 6 | 18842559 | rs1334768 | 0.501 | -0.469 | 0.286 | -0.64 | -0.009 | -0.02 | -0.02 | 0.006 | 0.92 | 0.88 | 0.95 | 0.99 | 0.94 | 0.81 |
| 6 | 18862782 | rs6911193 | 0.89 | -0.47 | 0.129 | -0.599 | -0.015 | 0.005 | 0.005 | 2.199 | 0.4 | 0.25 | 0.36 | 0.66 | 0.48 | 0.37 |
| 6 | 18864497 | rs9297095 | -0.043 | -0.599 | 0.03 | -0.668 | 0.002 | -0.009 | -0.009 | 0 | 0.07 | 0.07 | 0.07 | 0.1 | 0.05 | 0.07 |
| 6 | 18867178 | rs1891657 | 1.562 | -0.778 | -0.036 | -0.772 | -0.052 | -0.053 | -0.053 | 2.425 | 0.33 | 0.12 | 0.32 | 0.63 | 0.45 | 0.33 |
| 6 | 18872816 | rs4712357 | 0.051 | -0.713 | -0.04 | -0.729 | 0 | 0.007 | 0.007 | 0 | 0.07 | 0.07 | 0.07 | 0.04 | 0.05 | 0.08 |
| 6 | 18879615 | rs1891659 | 0.156 | -1.033 | 0.342 | -1.052 | -0.007 | 0.008 | 0.008 | 0.033 | 0.33 | 0.22 | 0.41 | 0.27 | 0.34 | 0.31 |
| 6 | 18882998 | rs7744327 | 0.149 | -1.053 | 0.684 | -1.062 | -0.009 | -0.015 | -0.015 | 0 | 0.9 | 0.87 | 0.95 | 0.98 | 0.94 | 0.8 |
| 6 | 18884991 | rs1343138 | 0.321 | -1.083 | 0.815 | -1.077 | -0.001 | -0.014 | -0.014 | 0 | 0.84 | 0.82 | 0.86 | 0.98 | 0.87 | 0.75 |
| 6 | 18897387 | rs9358183 | -0.679 | -1.14 | 0.824 | -1.071 | 0.003 | 0.015 | 0.015 | 0.445 | 0.45 | 0.59 | 0.5 | 0.32 | 0.39 | 0.38 |
| 6 | 18914005 | rs9383446 | -0.078 | -1.018 | 0.89 | -1.008 | 0.005 | 0.006 | 0.006 | 0 | 0.08 | 0.06 | 0.04 | 0.04 | 0.04 | 0.09 |
| 6 | 18914624 | rs988458 | 0.378 | -1.048 | 1.068 | -1.007 | 0.005 | 0.021 | 0.021 | 0.189 | 0.57 | 0.67 | 0.54 | 0.71 | 0.61 | 0.51 |
| 6 | 18938869 | rs989496 | 0.347 | -1.029 | 0.856 | -1.063 | 0.005 | 0.004 | 0.004 | 0.554 | 0.06 | 0.04 | 0.04 | 0.04 | 0.05 | 0.06 |
| 6 | 18960583 | rs16881537 | 0.808 | -1.096 | 0.768 | -1.192 | -0.027 | -0.023 | -0.023 | 0 | 0.17 | 0.09 | 0.3 | 0.23 | 0.21 | 0.25 |
| 6 | 18966110 | rs9465237 | 0.444 | -1.287 | 0.983 | -1.126 | 0.009 | 0.011 | 0.011 | 0.006 | 0.58 | 0.63 | 0.57 | 0.73 | 0.67 | 0.58 |
| 6 | 18972484 | rs17658803 | 0.107 | -1.306 | 0.85 | -1.126 | -0.013 | 0.03 | 0.03 | 0 | 0.11 | 0.05 | 0.11 | 0.04 | 0.09 | 0.1 |
| 6 | 18975071 | rs13196368 | 1.682 | -1.312 | 1.12 | -1.119 | 0.004 | 0 | 0 | 0 | 0.38 | 0.39 | 0.38 | 0.64 | 0.5 | 0.41 |
| 6 | 18976445 | rs834362 | -0.298 | -1.315 | 1.043 | -1.227 | -0.006 | 0.014 | 0.014 | 0.015 | 0.69 | 0.81 | 0.6 | 0.74 | 0.7 | 0.79 |
| 6 | 18980752 | rs12529004 | 0.809 | -1.403 | 1.016 | -1.213 | -0.001 | -0.006 | -0.006 | 0 | 0.05 | 0.02 | 0.02 | 0.03 | 0.03 | 0.04 |
| 6 | 18982296 | rs986759 | 0.758 | -1.441 | 1.12 | -1.137 | 0.02 | 0.036 | 0.036 | 0 | 0.53 | 0.67 | 0.52 | 0.7 | 0.61 | 0.55 |
| 6 | 18984299 | rs834372 | -1.774 | -1.443 | 1.135 | -1.157 | -0.018 | -0.038 | -0.038 | 0 | 0.31 | 0.4 | 0.23 | 0.12 | 0.21 | 0.39 |
| 6 | 18986548 | rs16881553 | 0.744 | -1.257 | 0.724 | -1.02 | -0.001 | -0.006 | -0.006 | 1.954 | 0.05 | 0.02 | 0.02 | 0.03 | 0.03 | 0.04 |
| 6 | 18987143 | rs7743567 | 0.372 | -1.255 | 0.78 | -1.025 | 0.011 | 0.022 | 0.022 | 0 | 0.59 | 0.7 | 0.55 | 0.72 | 0.64 | 0.6 |
| 6 | 18989461 | rs634162 | -0.043 | -1.028 | 0.595 | -0.902 | -0.009 | 0.003 | 0.003 | 0 | 0.07 | 0.04 | 0.08 | 0.04 | 0.09 | 0.14 |
| 6 | 18999707 | rs472659 | -0.065 | -1.032 | 0.702 | -0.929 | -0.005 | 0.003 | 0.003 | 0 | 0.07 | 0.05 | 0.08 | 0.04 | 0.09 | 0.14 |
| 6 | 19002293 | rs834367 | -0.31 | -1.295 | 0.877 | -1.079 | 0.001 | 0.001 | 0.001 | 0.136 | 0.72 | 0.75 | 0.66 | 0.78 | 0.77 | 0.82 |
| 6 | 19002681 | rs834366 | -0.393 | -1.301 | 0.829 | -1.082 | 0.002 | 0.003 | 0.003 | 0.179 | 0.73 | 0.77 | 0.67 | 0.78 | 0.77 | 0.82 |
| 6 | 19008948 | rs1977368 | 0.676 | -1.321 | 1.162 | -1.148 | 0 | -0.013 | -0.013 | 0 | 0.58 | 0.55 | 0.49 | 0.74 | 0.62 | 0.64 |
| 6 | 19013033 | rs1407500 | -1.147 | -1.41 | 1.223 | -1.123 | -0.044 | -0.046 | -0.046 | 0.089 | 0.24 | 0.39 | 0.18 | 0.13 | 0.15 | 0.29 |
| 6 | 19015964 | rs524559 | -0.561 | -0.976 | 0.978 | -1.002 | -0.027 | 0.032 | 0.032 | 0 | 0.13 | 0.07 | 0.16 | 0.04 | 0.15 | 0.2 |
| 6 | 19017212 | rs17716154 | 1.094 | -0.945 | 1.035 | -0.937 | -0.013 | 0.017 | 0.017 | 0 | 0.06 | 0.04 | 0.09 | 0 | 0.07 | 0.07 |
| 6 | 19021288 | rs1983876 | 0.021 | -1.04 | 1.339 | -0.935 | -0.019 | -0.016 | -0.016 | 0 | 0.69 | 0.54 | 0.63 | 0.78 | 0.77 | 0.75 |
| 6 | 19024910 | rs7757753 | -0.005 | -1.007 | 1.137 | -0.92 | -0.001 | -0.001 | -0.001 | 0 | 0.7 | 0.71 | 0.65 | 0.77 | 0.76 | 0.75 |
| 6 | 19028788 | rs975303 | -0.144 | -1.049 | 0.833 | -0.393 | -0.006 | -0.017 | -0.017 | 0 | 0.15 | 0.05 | 0.11 | 0.15 | 0.15 | 0.15 |
| 6 | 19034411 | rs10484380 | -0.402 | -1.043 | 1.271 | -0.444 | 0.039 | -0.038 | -0.038 | 0 | 0.11 | 0.37 | 0.17 | 0.03 | 0.1 | 0.11 |
| 6 | 19038132 | rs9368028 | -0.388 | -1.057 | 1.327 | -0.427 | 0.005 | 0.013 | 0.013 | 0 | 0.18 | 0.32 | 0.18 | 0.17 | 0.18 | 0.19 |
| 6 | 19051695 | rs926274 | 0.983 | -1.142 | 1.261 | -0.503 | 0.005 | 0.008 | 0.008 | 0 | 0.61 | 0.5 | 0.47 | 0.6 | 0.62 | 0.61 |
| 6 | 19055259 | rs10949576 | 2.043 | -1.347 | 1.074 | -0.626 | -0.093 | -0.118 | -0.118 | 0 | 0.36 | 0.08 | 0.31 | 0.53 | 0.47 | 0.37 |
| 6 | 19059547 | rs9477957 | -0.441 | -1.37 | 1.228 | -0.584 | -0.062 | -0.113 | -0.113 | 0 | 0.25 | 0.44 | 0.16 | 0.08 | 0.14 | 0.24 |
| 6 | 19066605 | rs763687 | -0.881 | -1.188 | 0.66 | -0.444 | -0.009 | -0.032 | -0.032 | 0.045 | 0.85 | 0.92 | 0.75 | 0.78 | 0.86 | 0.86 |
| 6 | 19071311 | rs2064081 | -0.751 | -1.182 | 0.66 | -0.445 | -0.002 | -0.007 | -0.007 | 0.323 | 0.84 | 0.88 | 0.74 | 0.78 | 0.85 | 0.86 |
| 6 | 19078368 | rs2842386 | -0.967 | -1.213 | 0.821 | -0.866 | 0.015 | 0.024 | 0.024 | 0 | 0.42 | 0.66 | 0.33 | 0.23 | 0.32 | 0.43 |
| 6 | 19094442 | rs2743000 | -0.949 | -1.159 | 0.417 | -0.776 | 0.053 | 0.085 | 0.085 | 0.043 | 0.41 | 0.78 | 0.34 | 0.23 | 0.29 | 0.38 |
| 6 | 19098413 | rs2743005 | -0.728 | -1.185 | 0.314 | -0.709 | -0.019 | -0.028 | -0.028 | 3.095 | 0.84 | 0.92 | 0.75 | 0.82 | 0.85 | 0.81 |
| 6 | 19100507 | rs6931466 | -0.368 | -1.431 | 0.589 | -0.947 | 0.003 | 0.006 | 0.006 | 0 | 0.29 | 0.22 | 0.2 | 0.16 | 0.25 | 0.25 |
| 6 | 19112000 | rs6939229 | 1.72 | -1.119 | 0.138 | -0.816 | -0.067 | -0.05 | -0.05 | 0.213 | 0.37 | 0.14 | 0.36 | 0.6 | 0.49 | 0.37 |
| 6 | 19144144 | rs10498691 | 0.712 | -1.063 | 0.47 | -0.739 | 0.001 | -0.025 | -0.025 | 0 | 0.07 | 0.09 | 0.08 | 0 | 0.05 | 0.07 |
| 6 | 19147207 | rs9350102 | -0.087 | -1.203 | 0.625 | -0.731 | -0.012 | 0.009 | 0.009 | 0 | 0.07 | 0.12 | 0.09 | 0.09 | 0.03 | 0.18 |
| 6 | 19152514 | rs12530087 | 0.304 | -1.366 | 0.778 | -0.864 | -0.002 | -0.005 | -0.005 | 0 | 0.18 | 0.28 | 0.22 | 0.14 | 0.15 | 0.19 |
| 6 | 19164109 | rs9368036 | -1.19 | -1.459 | 0.944 | -0.9 | -0.032 | -0.023 | -0.023 | 0 | 0.13 | 0.36 | 0.17 | 0.09 | 0.06 | 0.13 |
| 6 | 19167996 | rs4075048 | -1.288 | -1.603 | 0.925 | -1.038 | -0.013 | -0.019 | -0.019 | 0 | 0.29 | 0.66 | 0.26 | 0.13 | 0.15 | 0.24 |
| 6 | 19169348 | rs10498693 | 0.953 | -1.596 | 0.901 | -1.037 | 0.011 | 0.038 | 0.038 | 0 | 0.13 | 0.06 | 0.08 | 0 | 0.09 | 0.02 |
| 6 | 19173487 | rs9358193 | -1.142 | -1.603 | 0.817 | -1.031 | -0.04 | -0.05 | -0.05 | 0.176 | 0.29 | 0.59 | 0.25 | 0.13 | 0.15 | 0.24 |

| | | | | | | | | | | | | | | | | |
|---|---|---|---|---|---|---|---|---|---|---|---|---|---|---|---|---|
| 6 | 19177636 | rs6939219 | 0.695 | -1.564 | 0.713 | -1.021 | -0.005 | -0.025 | -0.025 | 0 | 0.62 | 0.56 | 0.63 | 0.83 | 0.7 | 0.62 |
| 6 | 19190261 | rs9368038 | -1.213 | -1.223 | 0.867 | -0.843 | 0.011 | 0.012 | 0.012 | 0.208 | 0.35 | 0.68 | 0.34 | 0.13 | 0.21 | 0.31 |
| 6 | 19193980 | rs9784816 | 0.471 | -1.154 | 0.809 | -0.787 | -0.009 | 0.001 | 0.001 | 0 | 0.06 | 0.12 | 0.09 | 0.07 | 0.03 | 0.08 |
| 6 | 19195066 | rs9350106 | -0.55 | -1.173 | 0.882 | -0.815 | -0.016 | -0.025 | -0.025 | 0.5 | 0.25 | 0.36 | 0.23 | 0.12 | 0.16 | 0.24 |
| 6 | 19195308 | rs6919994 | -0.202 | -0.79 | 0.814 | -0.377 | 0.015 | -0.053 | -0.053 | 0.008 | 0.1 | 0.27 | 0.1 | 0 | 0.06 | 0.07 |
| 6 | 19196491 | rs9366275 | -0.262 | -0.595 | -0.079 | -0.03 | -0.012 | 0.013 | 0.013 | 0 | 0.06 | 0.13 | 0.09 | 0.1 | 0.03 | 0.17 |
| 6 | 19203108 | rs10946296 | 0.497 | -0.097 | -0.485 | 0.085 | -0.039 | -0.044 | -0.044 | 0.624 | 0.37 | 0.15 | 0.31 | 0.59 | 0.46 | 0.36 |
| 6 | 19211776 | rs9460306 | -1.263 | -0.294 | 0.658 | 0.316 | 0.012 | 0.142 | 0.142 | 0.512 | 0.12 | 0.47 | 0.3 | 0.23 | 0.1 | 0.25 |
| 6 | 19212088 | rs9465275 | -0.701 | -0.314 | 1.126 | 0.404 | 0.004 | 0.093 | 0.093 | 0 | 0.06 | 0.22 | 0.12 | 0.15 | 0.02 | 0.08 |
| 6 | 19214889 | rs1858504 | -0.661 | -0.983 | 0.211 | -0.779 | 0.008 | 0.009 | 0.009 | 0 | 0.47 | 0.58 | 0.44 | 0.27 | 0.35 | 0.41 |
| 6 | 19216822 | rs9368039 | -0.861 | -1.2 | 0.117 | -1.004 | -0.037 | 0.026 | 0.026 | 0 | 0.23 | 0.49 | 0.36 | 0.26 | 0.19 | 0.34 |
| 6 | 19218841 | rs9295421 | -0.134 | -1.161 | 0.063 | -1.178 | 0.003 | 0.001 | 0.001 | 0.186 | 0.32 | 0.3 | 0.32 | 0.1 | 0.26 | 0.31 |
| 6 | 19227858 | rs9465305 | -0.322 | -1.203 | 0.217 | -1.198 | 0.004 | 0.001 | 0.001 | 0.279 | 0.33 | 0.29 | 0.32 | 0.12 | 0.26 | 0.32 |
| 6 | 19228938 | rs1858513 | -0.972 | -1.154 | 0.253 | -1.158 | -0.001 | 0.006 | 0.006 | 0 | 0.11 | 0.21 | 0.24 | 0.12 | 0.1 | 0.26 |
| 6 | 19235450 | rs7762189 | -0.234 | -1.182 | 0.222 | -1.161 | 0.002 | 0.013 | 0.013 | 0 | 0.05 | 0.13 | 0.17 | 0.1 | 0.07 | 0.17 |
| 6 | 19244975 | rs4712373 | -1.207 | -1.352 | 0.849 | -1.197 | 0.047 | 0.072 | 0.072 | 0 | 0.51 | 0.68 | 0.52 | 0.27 | 0.38 | 0.49 |
| 6 | 19245298 | rs9460319 | 0.051 | -1.312 | 0.816 | -1.177 | -0.009 | -0.024 | -0.024 | 0 | 0.06 | 0.09 | 0.08 | 0.02 | 0.03 | 0.08 |
| 6 | 19253687 | rs9465317 | -1.706 | -0.59 | 0.627 | -1.08 | -0.037 | -0.066 | -0.066 | 0 | 0.74 | 0.89 | 0.73 | 0.51 | 0.66 | 0.68 |
| 6 | 19256851 | rs4712374 | -0.074 | -0.614 | 0.564 | -1.207 | -0.002 | -0.02 | -0.02 | 0 | 0.39 | 0.39 | 0.35 | 0.12 | 0.3 | 0.36 |
| 6 | 19261762 | rs10214554 | 0.484 | -0.549 | 0.588 | -0.975 | -0.005 | -0.017 | -0.017 | 0 | 0.06 | 0.07 | 0.08 | 0.02 | 0.03 | 0.08 |
| 6 | 19268860 | rs941991 | -0.137 | -0.692 | 0.69 | -1.025 | 0.014 | 0.017 | 0.017 | 0 | 0.45 | 0.45 | 0.43 | 0.27 | 0.35 | 0.48 |
| 6 | 19281736 | rs12195637 | 1.502 | -0.583 | 0.67 | -0.931 | -0.027 | -0.045 | -0.045 | 2.65 | 0.26 | 0.12 | 0.27 | 0.5 | 0.35 | 0.25 |
| 6 | 19287493 | rs9295427 | -0.737 | -0.987 | 1.165 | -0.902 | -0.004 | 0.049 | 0.049 | 0 | 0.08 | 0.21 | 0.19 | 0.16 | 0.05 | 0.3 |
| 6 | 19289788 | rs12202040 | 0.155 | -1.146 | 1.207 | -1.086 | 0.015 | -0.009 | -0.009 | 0 | 0.31 | 0.25 | 0.17 | 0.02 | 0.2 | 0.15 |
| 6 | 19293110 | rs9350120 | -1.083 | -1.044 | 1.023 | -1.053 | 0.037 | 0.065 | 0.065 | 0.053 | 0.52 | 0.67 | 0.5 | 0.25 | 0.39 | 0.57 |
| 6 | 19295872 | rs12189951 | 0.853 | -0.96 | 0.654 | -1.078 | 0.026 | 0.067 | 0.067 | 1.55 | 0.24 | 0.11 | 0.09 | 0.01 | 0.17 | 0.07 |
| 6 | 19297457 | rs12205836 | 0.088 | -0.866 | 0.753 | -1.063 | 0.026 | 0.047 | 0.047 | 1.246 | 0.3 | 0.17 | 0.17 | 0.02 | 0.2 | 0.14 |
| 6 | 19298425 | rs1495978 | -1.002 | -0.946 | 0.904 | -1.021 | 0.016 | 0.026 | 0.026 | 0 | 0.51 | 0.59 | 0.49 | 0.26 | 0.39 | 0.56 |
| 6 | 19299263 | rs10155664 | -0.283 | -0.782 | 1.14 | -0.822 | -0.029 | -0.022 | -0.022 | 0 | 0.19 | 0.38 | 0.2 | 0.14 | 0.14 | 0.31 |
| 6 | 19303176 | rs1390459 | -0.276 | -0.8 | 1.231 | -0.824 | -0.035 | 0.021 | 0.021 | 0 | 0.83 | 0.62 | 0.83 | 0.83 | 0.89 | 0.76 |
| 6 | 19303601 | rs12528086 | 0.064 | -0.924 | 1 | -0.933 | -0.004 | -0.023 | -0.023 | 0.344 | 0.25 | 0.28 | 0.28 | 0.11 | 0.23 | 0.24 |
| 6 | 19303981 | rs7754477 | -0.458 | -0.983 | 0.909 | -0.982 | 0.007 | 0.017 | 0.017 | 0 | 0.31 | 0.28 | 0.36 | 0.13 | 0.27 | 0.34 |
| 6 | 19304567 | rs11967991 | 0.619 | -0.885 | 0.898 | -0.945 | 0.004 | -0.019 | -0.019 | 0.784 | 0.2 | 0.2 | 0.11 | 0.01 | 0.15 | 0.06 |
| 6 | 19316742 | rs9350122 | -0.003 | -0.678 | 1.481 | -0.331 | -0.026 | 0.002 | 0.002 | 0 | 0.16 | 0.29 | 0.16 | 0.16 | 0.11 | 0.24 |
| 6 | 19323345 | rs16882031 | -0.888 | -0.515 | 1.448 | -0.444 | 0.024 | 0.074 | 0.074 | 0 | 0.06 | 0.2 | 0.23 | 0.12 | 0.08 | 0.17 |
| 6 | 19328118 | rs7750685 | -1.683 | -0.705 | 1.535 | -0.549 | -0.033 | 0.01 | 0.01 | 0.14 | 0.29 | 0.55 | 0.44 | 0.29 | 0.23 | 0.51 |
| 6 | 19334386 | rs10081157 | -0.898 | -0.431 | 1.67 | -0.632 | -0.012 | -0.02 | -0.02 | 2.456 | 0.2 | 0.39 | 0.37 | 0.12 | 0.18 | 0.28 |
| 6 | 19339645 | rs9465339 | -0.509 | -0.448 | 1.574 | -0.64 | -0.01 | -0.005 | -0.005 | 0.9 | 0.17 | 0.29 | 0.25 | 0.13 | 0.16 | 0.2 |
| 6 | 19349880 | rs9465351 | -0.213 | -0.396 | 1.558 | -0.687 | -0.022 | -0.068 | -0.068 | 0 | 0.15 | 0.23 | 0.15 | 0 | 0.11 | 0.11 |
| 6 | 19353011 | rs7773088 | -0.417 | -0.427 | 1.841 | -0.619 | 0.005 | 0.007 | 0.007 | 0 | 0.17 | 0.38 | 0.3 | 0.15 | 0.18 | 0.22 |
| 6 | 19359154 | rs6911832 | -0.838 | -0.429 | 1.866 | -0.623 | -0.01 | -0.013 | -0.013 | 0 | 0.19 | 0.34 | 0.34 | 0.15 | 0.18 | 0.31 |
| 6 | 19360281 | rs2132404 | -0.186 | -0.371 | 1.666 | -0.574 | -0.008 | 0.001 | 0.001 | 0 | 0.16 | 0.25 | 0.3 | 0.14 | 0.16 | 0.26 |
| 6 | 19362058 | rs4712392 | -0.648 | -0.008 | 1.315 | -0.764 | -0.002 | -0.001 | -0.001 | 2.115 | 0.35 | 0.58 | 0.32 | 0.34 | 0.3 | 0.21 |
| 6 | 19363073 | rs1463780 | -1.378 | -0.093 | 1.206 | -0.777 | -0.03 | -0.073 | -0.073 | 1.272 | 0.45 | 0.85 | 0.66 | 0.5 | 0.44 | 0.54 |
| 6 | 19368281 | rs12193482 | 0.965 | -0.074 | 1.264 | -0.621 | 0.013 | 0.009 | 0.009 | 0.739 | 0.11 | 0.11 | 0.05 | 0 | 0.08 | 0.02 |
| 6 | 19373325 | rs1580731 | -0.498 | -0.098 | 1.504 | -0.623 | 0.002 | 0.013 | 0.013 | 0 | 0.3 | 0.42 | 0.3 | 0.34 | 0.3 | 0.2 |
| 6 | 19377333 | rs9368060 | -0.916 | 0.306 | 1.379 | -0.734 | -0.009 | -0.012 | -0.012 | 0 | 0.4 | 0.63 | 0.58 | 0.5 | 0.45 | 0.47 |
| 6 | 19380131 | rs9368062 | -0.008 | 0.613 | 1.46 | -0.44 | 0.008 | 0.026 | 0.026 | 0 | 0.21 | 0.27 | 0.24 | 0.34 | 0.22 | 0.18 |
| 6 | 19381355 | rs11755216 | 0.05 | 0.694 | 1.305 | -0.397 | 0.021 | 0.053 | 0.053 | 0 | 0.2 | 0.25 | 0.23 | 0.34 | 0.22 | 0.18 |
| 6 | 19381386 | rs10946313 | -0.957 | 0.702 | 1.051 | -0.419 | 0.038 | 0.068 | 0.068 | 0 | 0.3 | 0.49 | 0.52 | 0.49 | 0.37 | 0.45 |
| 6 | 19382736 | rs9460343 | -0.065 | 0.666 | 0.624 | -0.321 | 0.004 | 0.008 | 0.008 | 0 | 0.07 | 0.06 | 0.26 | 0.15 | 0.15 | 0.26 |
| 6 | 19387225 | rs6919702 | 0.057 | -0.107 | 0.623 | -1.144 | 0.018 | -0.003 | -0.003 | 0 | 0.39 | 0.44 | 0.51 | 0.61 | 0.45 | 0.49 |
| 6 | 19387399 | rs4144187 | 0.009 | -0.336 | 0.145 | -1.391 | 0.006 | 0.051 | 0.051 | 0 | 0.41 | 0.3 | 0.34 | 0.77 | 0.35 | 0.53 |
| 6 | 19390126 | rs9350132 | 1.137 | -0.409 | -0.021 | -1.335 | 0.017 | -0.034 | -0.034 | 0.005 | 0.17 | 0.12 | 0.15 | 0.35 | 0.09 | 0.21 |
| 6 | 19390189 | rs1495982 | -0.487 | -0.479 | 0.381 | -1.442 | -0.007 | 0.003 | 0.003 | 0 | 0.43 | 0.49 | 0.52 | 0.53 | 0.33 | 0.62 |
| 6 | 19390622 | rs9366292 | 1.278 | -0.301 | 0.036 | -1.464 | -0.074 | -0.139 | -0.139 | 0 | 0.2 | 0.04 | 0.17 | 0.41 | 0.26 | 0.22 |
| 6 | 19392417 | rs9368065 | -0.318 | -0.32 | 0.124 | -1.474 | -0.005 | 0.004 | 0.004 | 0 | 0.42 | 0.43 | 0.52 | 0.53 | 0.33 | 0.62 |
| 6 | 19400053 | rs2101874 | 0.918 | -0.26 | -0.063 | -1.379 | -0.036 | -0.1 | -0.1 | 0 | 0.22 | 0.14 | 0.17 | 0.45 | 0.24 | 0.22 |
| 6 | 19401418 | rs9460351 | 1.15 | -0.265 | 0.029 | -1.373 | -0.011 | -0.036 | -0.036 | 0.231 | 0.2 | 0.17 | 0.17 | 0.44 | 0.23 | 0.2 |
| 6 | 19407446 | rs9295432 | 0.353 | -0.37 | 0.103 | -1.466 | 0.008 | -0.022 | -0.022 | 0 | 0.23 | 0.29 | 0.37 | 0.17 | 0.23 | 0.4 |
| 6 | 19413687 | rs9348375 | 0.202 | -0.499 | 0.193 | -1.479 | -0.012 | 0.014 | 0.014 | 0 | 0.24 | 0.29 | 0.21 | 0.36 | 0.15 | 0.34 |
| 6 | 19414172 | rs9358218 | 1.002 | -0.492 | 0.172 | -1.472 | -0.01 | 0.027 | 0.027 | 0 | 0.18 | 0.21 | 0.17 | 0.35 | 0.11 | 0.21 |
| 6 | 19416421 | rs9368069 | 0.765 | -0.486 | 0.158 | -1.469 | -0.014 | 0.028 | 0.028 | 0 | 0.19 | 0.24 | 0.17 | 0.35 | 0.11 | 0.22 |
| 6 | 19418123 | rs6905315 | 0.003 | -0.58 | 0.1 | -1.555 | 0 | 0.002 | 0.002 | 0 | 0.26 | 0.2 | 0.38 | 0.17 | 0.25 | 0.43 |
| 6 | 19419004 | rs9465383 | 0.847 | -0.57 | -0.042 | -1.542 | -0.023 | 0.018 | 0.018 | 0 | 0.2 | 0.06 | 0.34 | 0.17 | 0.22 | 0.31 |
| 6 | 19425275 | rs6915939 | 0.444 | -0.592 | 0.051 | -1.505 | -0.034 | 0.047 | 0.047 | 0 | 0.21 | 0.29 | 0.19 | 0.36 | 0.11 | 0.32 |
| 6 | 19427108 | rs1463779 | -1.385 | -0.658 | -0.149 | -1.852 | -0.002 | -0.075 | -0.075 | 0 | 0.7 | 0.69 | 0.81 | 1 | 0.71 | 0.91 |
| 6 | 19427710 | rs9368070 | 0.782 | -0.591 | -0.72 | -1.576 | -0.101 | -0.142 | -0.142 | 0.273 | 0.22 | 0.06 | 0.21 | 0.43 | 0.33 | 0.25 |
| 6 | 19427830 | rs16882175 | 0.458 | -1.025 | -0.312 | -1.304 | -0.01 | -0.023 | -0.023 | 0 | 0.06 | 0.1 | 0.04 | 0 | 0.03 | 0.12 |

| | | | | | | | | | | | | | | | | |
|---|---|---|---|---|---|---|---|---|---|---|---|---|---|---|---|---|
| 6 | 19428772 | rs16882179 | 0.989 | -0.945 | -0.39 | -1.19 | -0.036 | 0.102 | 0.102 | 0.07 | 0.16 | 0.28 | 0.16 | 0.35 | 0.08 | 0.2 |
| 6 | 19434157 | rs9368072 | 0.442 | -0.085 | -1.292 | -0.882 | -0.048 | -0.098 | -0.098 | 0 | 0.21 | 0.1 | 0.16 | 0.46 | 0.27 | 0.23 |
| 6 | 19436035 | rs12211780 | 0.929 | -0.906 | -1.041 | -1.212 | -0.027 | 0.012 | 0.012 | 0.063 | 0.26 | 0.31 | 0.27 | 0.32 | 0.15 | 0.22 |
| 6 | 19436935 | rs1565992 | -0.17 | -0.924 | -1.049 | -1.296 | -0.016 | -0.003 | -0.003 | 0.25 | 0.42 | 0.45 | 0.34 | 0.34 | 0.25 | 0.38 |
| 6 | 19437583 | rs2172963 | -0.711 | -0.376 | -0.931 | -1.107 | 0.053 | -0.118 | -0.118 | 0 | 0.79 | 0.59 | 0.79 | 0.96 | 0.7 | 0.9 |
| 6 | 19443752 | rs4144260 | 0.201 | -0.438 | -1.168 | -0.817 | -0.028 | 0.006 | 0.006 | 0 | 0.32 | 0.39 | 0.28 | 0.33 | 0.18 | 0.22 |
| 6 | 19444511 | rs4560636 | -0.029 | 0.443 | -1.556 | 0.205 | -0.071 | 0 | 0 | 0 | 0.44 | 0.12 | 0.47 | 0.63 | 0.49 | 0.54 |
| 6 | 19452488 | rs9350145 | 0.041 | 0.063 | -0.863 | 0.451 | 0.013 | 0.002 | 0.002 | 0 | 0.52 | 0.54 | 0.69 | 0.47 | 0.63 | 0.76 |
| 6 | 19452510 | rs10946317 | -0.211 | -0.349 | -0.894 | 0.416 | 0.057 | -0.012 | -0.012 | 0.362 | 0.48 | 0.34 | 0.28 | 0.5 | 0.32 | 0.26 |
| 6 | 19452664 | rs9295438 | 1.365 | -1.52 | 0.156 | -0.175 | -0.034 | 0.072 | 0.072 | 0 | 0.83 | 0.43 | 0.73 | 0.72 | 0.9 | 0.68 |
| 6 | 19453594 | rs4348303 | 0.806 | -2.072 | 0.084 | -1.491 | -0.016 | -0.04 | -0.04 | 0 | 0.72 | 0.68 | 0.77 | 0.87 | 0.82 | 0.64 |
| 6 | 19453993 | rs4388279 | 1.241 | -2.764 | 0.567 | -1.542 | -0.065 | -0.056 | -0.056 | 0 | 0.65 | 0.49 | 0.84 | 0.76 | 0.82 | 0.67 |
| 6 | 19455877 | rs4235993 | 1.511 | -2.807 | 0.127 | -2.282 | -0.043 | 0.058 | 0.058 | 0.218 | 0.37 | 0.19 | 0.56 | 0.7 | 0.49 | 0.44 |
| 6 | 19456474 | rs4507577 | 0.767 | -3.147 | 0.481 | -2.41 | -0.067 | -0.07 | -0.07 | 0 | 0.7 | 0.55 | 0.87 | 0.93 | 0.89 | 0.85 |
| 6 | 19457709 | rs13220516 | -0.786 | -3.028 | 0.274 | -2.417 | 0.01 | 0.007 | 0.007 | 0.006 | 0.25 | 0.21 | 0.1 | 0.07 | 0.12 | 0.16 |
| 6 | 19468120 | rs4235996 | -0.259 | -2.611 | 0.055 | -2.238 | 0.002 | 0.011 | 0.011 | 1.048 | 0.3 | 0.23 | 0.27 | 0.21 | 0.34 | 0.33 |
| 6 | 19468243 | rs9283867 | 1.655 | -2.857 | -0.135 | -2.393 | -0.059 | 0.055 | 0.055 | 0.142 | 0.36 | 0.14 | 0.54 | 0.7 | 0.52 | 0.5 |
| 6 | 19471680 | rs4452634 | -0.263 | -2.628 | 0.232 | -2.291 | -0.012 | -0.068 | -0.068 | 0.084 | 0.31 | 0.5 | 0.29 | 0.2 | 0.35 | 0.33 |
| 6 | 19478455 | rs16882265 | -0.378 | -2.617 | 0.176 | -2.269 | -0.006 | -0.015 | -0.015 | 0.504 | 0.26 | 0.23 | 0.13 | 0.08 | 0.12 | 0.21 |
| 6 | 19492203 | rs11757407 | -1.141 | -2.359 | 0.065 | -2.173 | -0.017 | 0.074 | 0.074 | 0.109 | 0.61 | 0.73 | 0.44 | 0.29 | 0.48 | 0.47 |
| 6 | 19492819 | rs6921277 | 0.466 | -2.415 | 0.136 | -2.269 | -0.018 | -0.052 | -0.052 | 0 | 0.19 | 0.31 | 0.2 | 0.08 | 0.17 | 0.14 |
| 6 | 19512065 | rs2223288 | 1.284 | -2.39 | -0.038 | -2.242 | -0.045 | 0.071 | 0.071 | 0.374 | 0.38 | 0.15 | 0.55 | 0.71 | 0.52 | 0.53 |
| 6 | 19525075 | rs4712415 | 1.164 | -2.401 | -0.032 | -2.247 | -0.033 | 0.077 | 0.077 | 0.237 | 0.39 | 0.17 | 0.55 | 0.71 | 0.52 | 0.53 |
| 6 | 19526053 | rs2743582 | -0.217 | -2.3 | -0.007 | -2.227 | -0.022 | -0.022 | -0.022 | 0 | 0.94 | 0.98 | 0.94 | 0.83 | 0.86 | 0.87 |
| 6 | 19529446 | rs2205861 | 0.107 | -2.368 | 0.172 | -2.27 | 0.005 | -0.024 | -0.024 | 0 | 0.73 | 0.53 | 0.71 | 0.75 | 0.69 | 0.71 |
| 6 | 19534362 | rs9460378 | 0.613 | -2.344 | -0.093 | -2.265 | 0.019 | 0.023 | 0.023 | 0.008 | 0.91 | 0.79 | 0.9 | 0.83 | 0.86 | 0.84 |
| 6 | 19537814 | rs760668 | 0.541 | -2.371 | -0.525 | -2.272 | -0.003 | -0.016 | -0.016 | 0 | 0.69 | 0.42 | 0.67 | 0.7 | 0.65 | 0.68 |
| 6 | 19538647 | rs12215504 | 0.442 | -1.943 | -0.527 | -2.242 | -0.032 | -0.084 | -0.084 | 0 | 0.21 | 0.31 | 0.12 | 0.01 | 0.11 | 0.13 |
| 6 | 19548810 | rs9348385 | 0.083 | -1.066 | -0.749 | -1.742 | 0.088 | 0.168 | 0.168 | 1.013 | 0.55 | 0.26 | 0.73 | 0.81 | 0.68 | 0.64 |
| 6 | 19549868 | rs4710911 | 1.177 | 0.383 | -0.314 | -0.668 | 0.088 | 0.083 | 0.083 | 0.152 | 0.91 | 0.68 | 0.89 | 0.83 | 0.85 | 0.84 |
| 6 | 19556168 | rs2743595 | -0.099 | 0.095 | -1.536 | -0.94 | -0.028 | 0.177 | 0.177 | 0.576 | 0.61 | 0.92 | 0.46 | 0.13 | 0.41 | 0.36 |
| 6 | 19558904 | rs9465438 | -1.029 | 0.303 | -0.649 | -0.137 | -0.002 | -0.006 | -0.006 | 0.072 | 0.06 | 0.14 | 0.04 | 0.04 | 0.05 | 0.02 |
| 6 | 19560024 | rs10484720 | -0.839 | 0.607 | -0.478 | 0.01 | 0 | 0.003 | 0.003 | 0 | 0.22 | 0.29 | 0.18 | 0.25 | 0.23 | 0.27 |
| 6 | 19560177 | rs2743596 | -1.306 | 1.051 | -0.334 | -0.283 | 0.003 | -0.017 | -0.017 | 0 | 0.18 | 0.22 | 0.16 | 0.11 | 0.23 | 0.09 |
| 6 | 19561987 | rs760669 | 0.376 | 1.046 | -0.71 | 0.199 | -0.04 | 0.014 | 0.014 | 0 | 0.82 | 0.94 | 0.86 | 0.9 | 0.73 | 0.86 |
| 6 | 19563321 | rs2743597 | 0.435 | 1.023 | -0.771 | 0.132 | -0.042 | 0.022 | 0.022 | 0.033 | 0.81 | 0.94 | 0.86 | 0.9 | 0.73 | 0.85 |
| 6 | 19564461 | rs9460382 | -0.713 | 0.628 | -0.11 | 0.349 | -0.002 | 0.005 | 0.005 | 0 | 0.13 | 0.21 | 0.09 | 0.17 | 0.09 | 0.12 |
| 6 | 19566284 | rs1883233 | -1.034 | 0.019 | 0.022 | 0.174 | -0.016 | -0.029 | -0.029 | 0 | 0.28 | 0.41 | 0.24 | 0.16 | 0.2 | 0.22 |
| 6 | 19567942 | rs7745130 | -0.458 | 0.399 | 0.237 | 0.083 | 0.011 | 0.103 | 0.103 | 0.464 | 0.52 | 0.21 | 0.45 | 0.29 | 0.51 | 0.46 |
| 6 | 19570634 | rs2179176 | -0.996 | 0.291 | 1.013 | 0.207 | 0.004 | 0.051 | 0.051 | 0.023 | 0.59 | 0.77 | 0.54 | 0.23 | 0.6 | 0.45 |
| 6 | 19570818 | rs2817282 | -1.598 | 0.349 | 1.331 | 0.227 | 0.057 | 0.048 | 0.048 | 3.771 | 0.05 | 0.42 | 0.04 | 0 | 0 | 0.01 |
| 6 | 19589595 | rs2743610 | -2.068 | 0.353 | 1.277 | 0.231 | 0.058 | 0.094 | 0.094 | 2.968 | 0.08 | 0.48 | 0.09 | 0.06 | 0.09 | 0.04 |
| 6 | 19599795 | rs2817297 | -0.963 | 0.416 | 0.894 | 0.259 | 0.004 | 0.009 | 0.009 | 0.406 | 0.59 | 0.78 | 0.54 | 0.34 | 0.6 | 0.51 |
| 6 | 19604412 | rs2817283 | -0.756 | 0.339 | 0.904 | 0.287 | 0.002 | 0.004 | 0.004 | 0 | 0.57 | 0.65 | 0.53 | 0.34 | 0.59 | 0.49 |
| 6 | 19610858 | rs6933602 | -0.935 | 0.414 | 1.09 | 0.153 | 0.007 | 0.008 | 0.008 | 0.786 | 0.57 | 0.73 | 0.53 | 0.36 | 0.6 | 0.46 |
| 6 | 19613078 | rs1936905 | -0.623 | 0.281 | 1.221 | -0.007 | 0.011 | 0.047 | 0.047 | 0.257 | 0.51 | 0.39 | 0.46 | 0.29 | 0.51 | 0.42 |
| 6 | 19613858 | rs9366302 | -0.469 | 0.267 | 0.902 | -0.014 | -0.017 | 0.196 | 0.196 | 0.724 | 0.48 | 0.14 | 0.44 | 0.18 | 0.51 | 0.35 |
| 6 | 19617841 | rs1120462 | -0.853 | 0.365 | 0.934 | -0.015 | 0.002 | 0.026 | 0.026 | 1.009 | 0.53 | 0.61 | 0.46 | 0.29 | 0.49 | 0.43 |
| 6 | 19628340 | rs2328483 | -1.016 | 0.897 | 0.836 | 0.041 | 0.001 | -0.006 | -0.006 | 0 | 0.37 | 0.59 | 0.36 | 0.26 | 0.32 | 0.35 |
| 6 | 19628633 | rs6930954 | -0.565 | 0.933 | 0.688 | 0.041 | -0.051 | 0.06 | 0.06 | 1.902 | 0.32 | 0.12 | 0.31 | 0.16 | 0.32 | 0.27 |
| 6 | 19642528 | rs9460391 | 0.858 | 1.387 | 0.986 | 0.043 | 0.015 | -0.01 | -0.01 | 0.24 | 0.67 | 0.54 | 0.66 | 0.74 | 0.68 | 0.69 |
| 6 | 19643688 | rs7743224 | -0.211 | 0.63 | 0.68 | 0.032 | -0.024 | -0.017 | -0.017 | 0.315 | 0.87 | 0.97 | 0.78 | 0.85 | 0.84 | 0.81 |
| 6 | 19644251 | rs1936896 | 0.218 | -0.462 | 0.701 | -0.131 | 0.002 | -0.04 | -0.04 | 0 | 0.14 | 0.18 | 0.14 | 0.02 | 0.18 | 0.08 |
| 6 | 19647981 | rs9348389 | 0.698 | -0.65 | 0.464 | 0.207 | 0.011 | 0.021 | 0.021 | 0 | 0.62 | 0.32 | 0.65 | 0.77 | 0.65 | 0.67 |
| 6 | 19650665 | rs1892187 | -0.634 | -0.761 | 0.382 | 0.184 | -0.033 | 0.018 | 0.018 | 0 | 0.77 | 0.89 | 0.71 | 0.82 | 0.72 | 0.75 |
| 6 | 19653999 | rs2876555 | -0.133 | -0.673 | 0.474 | 0.271 | 0.002 | 0.01 | 0.01 | 0 | 0.69 | 0.71 | 0.68 | 0.82 | 0.69 | 0.73 |
| 6 | 19655292 | rs9358258 | 0.194 | -0.506 | 0.041 | 0.317 | -0.012 | -0.028 | -0.028 | 0 | 0.65 | 0.44 | 0.67 | 0.78 | 0.69 | 0.69 |
| 6 | 19659057 | rs9348390 | 1.34 | -0.369 | -0.657 | 0.545 | -0.162 | -0.048 | -0.048 | 0 | 0.42 | 0.03 | 0.46 | 0.62 | 0.44 | 0.46 |
| 6 | 19660872 | rs4712423 | 0.49 | -0.053 | -0.368 | 0.817 | 0.015 | 0.013 | 0.013 | 0 | 0.6 | 0.31 | 0.61 | 0.67 | 0.6 | 0.62 |
| 6 | 19664481 | rs12197609 | -0.053 | 0.196 | -0.779 | 0.61 | -0.048 | 0.04 | 0.04 | 1.074 | 0.13 | 0.02 | 0.12 | 0.06 | 0.17 | 0.17 |
| 6 | 19665957 | rs9460394 | -0.507 | -0.366 | -0.075 | 1.23 | -0.007 | 0.018 | 0.018 | 0 | 0.07 | 0.09 | 0.11 | 0.16 | 0.08 | 0.04 |
| 6 | 19667419 | rs9358259 | 0.468 | -0.351 | -0.115 | 0.857 | -0.007 | -0.037 | -0.037 | 0.914 | 0.65 | 0.46 | 0.65 | 0.75 | 0.64 | 0.68 |
| 6 | 19668213 | rs9358260 | 0.596 | -0.238 | -0.77 | 1.115 | 0.042 | -0.049 | -0.049 | 0.266 | 0.54 | 0.15 | 0.49 | 0.51 | 0.54 | 0.61 |
| 6 | 19669842 | rs9368104 | 0.376 | -0.169 | -0.389 | 0.717 | 0.003 | 0.032 | 0.032 | 0 | 0.65 | 0.71 | 0.65 | 0.79 | 0.63 | 0.71 |
| 6 | 19670986 | rs2031255 | 0.6 | -0.27 | -0.204 | 1.074 | 0.002 | -0.002 | -0.002 | 0 | 0.56 | 0.43 | 0.51 | 0.49 | 0.55 | 0.65 |
| 6 | 19672138 | rs9465476 | -0.404 | -0.375 | -0.378 | 0.585 | -0.001 | -0.022 | -0.022 | 0 | 0.14 | 0.15 | 0.13 | 0.02 | 0.18 | 0.07 |
| 6 | 19672799 | rs9356704 | 0.575 | 0.056 | -1.323 | 0.75 | -0.056 | 0.002 | 0.002 | 0 | 0.39 | 0.14 | 0.39 | 0.4 | 0.37 | 0.53 |
| 6 | 19672913 | rs16882448 | -0.821 | 0.099 | -0.531 | 1.136 | 0.02 | 0.049 | 0.049 | 0 | 0.06 | 0.16 | 0.07 | 0.14 | 0.08 | 0.1 |
| 6 | 19673765 | rs1033872 | -0.344 | -0.632 | -1.498 | 1.022 | 0.103 | -0.035 | -0.035 | 0.3 | 0.42 | 0.88 | 0.32 | 0.41 | 0.26 | 0.39 |

| | | | | | | | | | | | | | | | | |
|---|---|---|---|---|---|---|---|---|---|---|---|---|---|---|---|---|
| 6 | 19677864 | rs9356705 | -0.379 | -0.658 | -0.721 | 1.457 | 0.01 | 0.19 | 0.19 | 0 | 0.06 | 0.22 | 0.03 | 0.23 | 0.04 | 0.06 |
| 6 | 19677912 | rs17547028 | -0.092 | -1.024 | -1.307 | 1.01 | -0.004 | 0.037 | 0.037 | 0.087 | 0.14 | 0.03 | 0.07 | 0.07 | 0.12 | 0.12 |
| 6 | 19679772 | rs9358262 | -0.93 | -1.13 | -0.624 | 0.825 | 0.032 | -0.025 | -0.025 | 1.91 | 0.55 | 0.76 | 0.37 | 0.48 | 0.39 | 0.5 |
| 6 | 19687502 | rs16882470 | 0.622 | -1.676 | -0.129 | 0.77 | 0.005 | 0.005 | 0.005 | 0 | 0.16 | 0.12 | 0.05 | 0.07 | 0.09 | 0.06 |
| 6 | 19688331 | rs9366310 | -0.027 | -1.724 | 0.486 | 0.641 | -0.003 | 0.186 | 0.186 | 1.094 | 0.19 | 0.44 | 0.2 | 0.41 | 0.14 | 0.32 |
| 6 | 19691473 | rs13218434 | 0.821 | -1.727 | 0.542 | 0.619 | 0.002 | 0.017 | 0.017 | 0.689 | 0.13 | 0.18 | 0.16 | 0.15 | 0.1 | 0.24 |
| 6 | 19692306 | rs9358263 | 0.053 | -1.715 | 0.611 | 1.039 | 0.02 | 0.306 | 0.306 | 0 | 0.06 | 0.25 | 0.07 | 0.33 | 0.04 | 0.07 |
| 6 | 19693784 | rs6923485 | 0.765 | -1.577 | 0.402 | 0.47 | 0.001 | 0.013 | 0.013 | 0.722 | 0.13 | 0.18 | 0.16 | 0.15 | 0.1 | 0.24 |
| 6 | 19697189 | rs9368109 | -0.251 | -1.376 | 0.432 | 0.264 | -0.004 | -0.03 | -0.03 | 0 | 0.94 | 0.86 | 0.96 | 1 | 0.96 | 0.99 |
| 6 | 19698464 | rs1936899 | 0.153 | -1.4 | 0.457 | 0.244 | -0.042 | -0.113 | -0.113 | 0 | 0.78 | 0.63 | 0.88 | 1 | 0.87 | 0.93 |
| 6 | 19700166 | rs6940685 | -0.496 | -0.31 | 0.158 | 0.546 | -0.001 | -0.004 | -0.004 | 0.89 | 0.45 | 0.52 | 0.4 | 0.42 | 0.42 | 0.45 |
| 6 | 19703618 | rs9465496 | -0.46 | -0.228 | 0.544 | 0.641 | 0.016 | 0.058 | 0.058 | 0 | 0.26 | 0.45 | 0.33 | 0.42 | 0.28 | 0.31 |
| 6 | 19706407 | rs9358265 | -1.076 | -0.469 | 0.187 | 0.547 | 0.04 | 0.064 | 0.064 | 0 | 0.43 | 0.74 | 0.23 | 0.31 | 0.34 | 0.25 |
| 6 | 19706553 | rs9465498 | 0.896 | -0.492 | 0.286 | 0.523 | -0.001 | 0.012 | 0.012 | 0 | 0.12 | 0.12 | 0.08 | 0.07 | 0.11 | 0.09 |
| 6 | 19706799 | rs1016252 | 1.275 | -0.599 | 0.273 | 0.425 | -0.009 | 0.058 | 0.058 | 0 | 0.16 | 0.03 | 0.07 | 0 | 0.15 | 0.09 |
| 6 | 19708229 | rs17547572 | 1.025 | -0.603 | 0.451 | 0.574 | -0.012 | -0.009 | -0.009 | 0 | 0.11 | 0.06 | 0.11 | 0.34 | 0.16 | 0.22 |
| 6 | 19712293 | rs6926312 | -1.746 | -0.63 | 0.429 | 0.522 | -0.035 | 0.024 | 0.024 | 0.161 | 0.54 | 0.8 | 0.35 | 0.65 | 0.5 | 0.47 |
| 6 | 19717442 | rs2147044 | -1.061 | -0.603 | 0.536 | 0.623 | 0.004 | -0.017 | -0.017 | 3.075 | 0.27 | 0.76 | 0.22 | 0.58 | 0.24 | 0.3 |
| 6 | 19721889 | rs6911293 | -0.172 | -0.601 | 0.872 | 0.64 | 0.023 | 0.125 | 0.125 | 0 | 0.19 | 0.34 | 0.19 | 0.58 | 0.23 | 0.3 |
| 6 | 19726320 | rs9366313 | 0.878 | -0.714 | 0.744 | 0.621 | -0.005 | 0.067 | 0.067 | 0 | 0.07 | 0.13 | 0.04 | 0.24 | 0.07 | 0.07 |
| 6 | 19729003 | rs9356708 | -1.307 | -0.935 | 0.569 | 0.417 | 0.064 | 0.096 | 0.096 | 0.011 | 0.46 | 0.81 | 0.26 | 0.32 | 0.33 | 0.25 |
| 6 | 19729381 | rs7769632 | -0.286 | -0.99 | 0.696 | 0.44 | -0.03 | -0.117 | -0.117 | 0 | 0.27 | 0.62 | 0.14 | 0 | 0.16 | 0.09 |
| 6 | 19732492 | rs6920825 | -0.128 | -1.004 | 0.881 | 0.467 | -0.01 | -0.008 | -0.008 | 0.128 | 0.27 | 0.36 | 0.16 | 0.25 | 0.22 | 0.16 |
| 6 | 19739943 | rs9460408 | -1.053 | -1.052 | 0.695 | 0.765 | 0.055 | 0.014 | 0.014 | 0.523 | 0.35 | 0.82 | 0.36 | 0.3 | 0.23 | 0.17 |
| 6 | 19742243 | rs1209816 | -0.851 | -1.268 | 0.415 | 0.564 | 0.003 | -0.002 | -0.002 | 0 | 0.92 | 0.94 | 0.8 | 0.95 | 0.95 | 0.97 |
| 6 | 19743486 | rs9358267 | 0.699 | -1.097 | 0.553 | 0.69 | -0.003 | 0.106 | 0.106 | 0 | 0.08 | 0.24 | 0.07 | 0.25 | 0.07 | 0.07 |
| 6 | 19748549 | rs11759345 | 1.145 | -0.48 | 0.216 | 0.764 | -0.061 | -0.065 | -0.065 | 0.11 | 0.11 | 0.01 | 0.13 | 0.32 | 0.2 | 0.23 |
| 6 | 19751668 | rs12192729 | -0.52 | -0.578 | 0.348 | 0.618 | 0.012 | 0.013 | 0.013 | 0 | 0.33 | 0.72 | 0.18 | 0.2 | 0.22 | 0.16 |
| 6 | 19752263 | rs12197522 | 0.863 | -0.591 | -0.159 | 0.714 | -0.008 | 0.004 | 0.004 | 0 | 0.12 | 0.14 | 0.04 | 0.17 | 0.07 | 0.09 |
| 6 | 19753061 | rs7766034 | -1.364 | -0.751 | -0.363 | 0.656 | -0.028 | 0.053 | 0.053 | 0.848 | 0.53 | 0.87 | 0.49 | 0.65 | 0.44 | 0.46 |
| 6 | 19756008 | rs9350175 | 0.446 | -0.728 | -0.139 | 0.599 | -0.027 | 0.048 | 0.048 | 0 | 0.14 | 0.26 | 0.05 | 0.24 | 0.07 | 0.09 |
| 6 | 19759362 | rs9295447 | -1.853 | -0.342 | -0.847 | 0.254 | -0.042 | -0.03 | -0.03 | 0 | 0.88 | 0.99 | 0.85 | 0.9 | 0.89 | 0.82 |
| 6 | 19759587 | rs9295448 | 0.162 | -0.087 | -0.63 | 0.478 | -0.008 | -0.027 | -0.027 | 0.002 | 0.23 | 0.16 | 0.18 | 0.59 | 0.25 | 0.32 |
| 6 | 19760008 | rs11758935 | 0.998 | -0.085 | -0.636 | 0.528 | -0.041 | -0.05 | -0.05 | 0 | 0.11 | 0.04 | 0.14 | 0.31 | 0.18 | 0.23 |
| 6 | 19761215 | rs6916251 | -1.368 | -0.221 | -0.484 | 0.295 | 0.007 | 0.062 | 0.062 | 0.365 | 0.53 | 0.79 | 0.34 | 0.69 | 0.44 | 0.45 |
| 6 | 19761718 | rs6456259 | -0.081 | -0.262 | -0.377 | 0.231 | -0.039 | -0.145 | -0.145 | 1.378 | 0.27 | 0.6 | 0.16 | 0 | 0.17 | 0.07 |
| 6 | 19776659 | rs6901079 | -0.615 | -0.206 | -0.491 | 0.043 | 0.006 | 0.008 | 0.008 | 1.862 | 0.35 | 0.66 | 0.23 | 0.15 | 0.27 | 0.26 |
| 6 | 19780956 | rs9465523 | -0.218 | -0.292 | -0.352 | -0.046 | 0.022 | -0.067 | -0.067 | 0 | 0.16 | 0.49 | 0.14 | 0.05 | 0.13 | 0.17 |
| 6 | 19785588 | rs7739264 | -1.719 | -0.231 | -0.66 | 0.017 | -0.073 | 0.049 | 0.049 | 0.626 | 0.6 | 0.9 | 0.42 | 0.75 | 0.53 | 0.58 |
| 6 | 19790487 | rs760795 | 0.19 | -0.304 | -0.292 | 0.132 | -0.044 | 0.048 | 0.048 | 0 | 0.2 | 0.46 | 0.1 | 0.28 | 0.12 | 0.12 |
| 6 | 19790560 | rs760794 | -1.512 | -0.307 | -0.329 | 0.113 | -0.012 | 0.07 | 0.07 | 0.442 | 0.53 | 0.85 | 0.35 | 0.7 | 0.44 | 0.45 |
| 6 | 19790571 | rs17483277 | -0.31 | -0.313 | -0.325 | 0.11 | -0.001 | -0.003 | -0.003 | 0 | 0.33 | 0.35 | 0.26 | 0.42 | 0.33 | 0.33 |
| 6 | 19790809 | rs2223361 | -1.471 | -0.277 | -0.346 | 0.13 | -0.012 | 0.07 | 0.07 | 0.442 | 0.53 | 0.85 | 0.35 | 0.7 | 0.44 | 0.45 |
| 6 | 19792518 | rs1997688 | 0.606 | 0.009 | -0.298 | 0.187 | 0.023 | 0.057 | 0.057 | 0 | 0.15 | 0.25 | 0.2 | 0.45 | 0.24 | 0.29 |
| 6 | 19793162 | rs11752817 | 1.05 | 0.008 | -0.329 | 0.225 | -0.08 | -0.08 | -0.08 | 0 | 0.12 | 0.01 | 0.16 | 0.34 | 0.23 | 0.23 |
| 6 | 19793829 | rs6921866 | -0.234 | 0.233 | -0.329 | 0.247 | 0.006 | 0.05 | 0.05 | 0.089 | 0.21 | 0.34 | 0.2 | 0.42 | 0.23 | 0.29 |
| 6 | 19794585 | rs7759616 | -1.595 | 0.181 | -0.34 | 0.184 | -0.02 | 0.08 | 0.08 | 0.278 | 0.53 | 0.83 | 0.38 | 0.73 | 0.49 | 0.45 |
| 6 | 19795968 | rs7769163 | 1.052 | 0.117 | -0.308 | 0.173 | 0.009 | 0.039 | 0.039 | 0 | 0.11 | 0.2 | 0.15 | 0.28 | 0.14 | 0.22 |
| 6 | 19796863 | rs2206034 | -1.518 | 0.089 | -0.334 | 0.166 | -0.014 | 0.045 | 0.045 | 0.381 | 0.53 | 0.85 | 0.37 | 0.66 | 0.44 | 0.44 |
| 6 | 19797837 | rs6456269 | -0.498 | 0.107 | -0.323 | 0.154 | -0.001 | -0.002 | -0.002 | 0.112 | 0.32 | 0.35 | 0.27 | 0.38 | 0.32 | 0.32 |
| 6 | 19798141 | rs6903595 | -1.274 | -0.004 | -0.362 | 0.158 | 0.039 | 0.047 | 0.047 | 0.454 | 0.52 | 0.78 | 0.34 | 0.66 | 0.39 | 0.44 |
| 6 | 19798169 | rs6926381 | 0.544 | -0.003 | -0.561 | 0.158 | 0.002 | 0.059 | 0.059 | 0 | 0.14 | 0.24 | 0.16 | 0.38 | 0.15 | 0.28 |
| 6 | 19798682 | rs9368123 | 0.655 | -0.04 | -0.592 | 0.142 | -0.01 | 0.015 | 0.015 | 0 | 0.17 | 0.22 | 0.07 | 0.27 | 0.11 | 0.12 |
| 6 | 19798704 | rs6904518 | -1.314 | 0.009 | -0.556 | 0.175 | 0.039 | 0.05 | 0.05 | 0.639 | 0.52 | 0.85 | 0.34 | 0.66 | 0.39 | 0.44 |
| 6 | 19799441 | rs2223360 | 0.114 | 0.038 | -0.551 | 0.194 | 0.005 | 0.001 | 0.001 | 0 | 0.27 | 0.22 | 0.13 | 0.28 | 0.22 | 0.16 |
| 6 | 19803768 | rs6907340 | -1.033 | 0.079 | -0.605 | 0.197 | 0.046 | 0.055 | 0.055 | 0.248 | 0.51 | 0.69 | 0.31 | 0.67 | 0.39 | 0.44 |
| 6 | 19808270 | rs760793 | 0.374 | 0.32 | -0.638 | 0.239 | 0.01 | 0.014 | 0.014 | 0 | 0.1 | 0.13 | 0.09 | 0.18 | 0.15 | 0.2 |
| 6 | 19809033 | rs6938418 | 0.041 | 0.173 | -0.927 | 0.234 | -0.001 | 0.005 | 0.005 | 0.709 | 0.26 | 0.33 | 0.22 | 0.38 | 0.26 | 0.29 |
| 6 | 19811991 | rs9368127 | 0.29 | 0.328 | -1.225 | 0.241 | 0.001 | 0.014 | 0.014 | 0.061 | 0.22 | 0.26 | 0.13 | 0.31 | 0.2 | 0.27 |
| 6 | 19818707 | rs4712433 | 2.129 | 0.022 | -2.365 | 0.118 | -0.067 | 0.002 | 0.002 | 0 | 0.2 | 0.03 | 0.4 | 0.13 | 0.31 | 0.21 |
| 6 | 19820658 | rs6910773 | 0.047 | 0.438 | -2.576 | 0.497 | 0.013 | 0.007 | 0.007 | 0.019 | 0.23 | 0.27 | 0.16 | 0.29 | 0.24 | 0.28 |
| 6 | 19821270 | rs7771384 | -0.48 | -0.375 | -2.074 | 0.709 | -0.012 | 0.037 | 0.037 | 3.242 | 0.27 | 0.65 | 0.22 | 0.44 | 0.23 | 0.28 |
| 6 | 19821781 | rs12527834 | 0.461 | 0.726 | -2.298 | 0.267 | -0.013 | 0.053 | 0.053 | 0.385 | 0.11 | 0.01 | 0.06 | 0 | 0.1 | 0.03 |
| 6 | 19824971 | rs9358278 | -0.256 | 0.696 | -1.899 | 0.879 | -0.001 | -0.038 | -0.038 | 0 | 0.22 | 0.13 | 0.16 | 0.55 | 0.21 | 0.28 |
| 6 | 19826630 | rs973244 | 0.191 | 0.791 | -2.109 | 0.304 | -0.008 | 0.037 | 0.037 | 0 | 0.69 | 0.87 | 0.55 | 0.82 | 0.69 | 0.65 |
| 6 | 19827905 | rs9348399 | -0.189 | 0.653 | -2.059 | 0.55 | 0.007 | -0.042 | -0.042 | 0 | 0.16 | 0.03 | 0.27 | 0.23 | 0.15 | 0.32 |
| 6 | 19827919 | rs971144 | 0.169 | 0.663 | -2.777 | -0.157 | 0.053 | 0.127 | 0.127 | 0.692 | 0.45 | 0.81 | 0.39 | 0.21 | 0.4 | 0.37 |
| 6 | 19828213 | rs6926229 | 0.126 | 1.572 | -2.527 | -0.38 | -0.005 | 0.132 | 0.132 | 1.549 | 0.55 | 0.74 | 0.45 | 0.14 | 0.54 | 0.39 |
| 6 | 19831560 | rs6922714 | 0.375 | 0.872 | -2.318 | -0.884 | 0.001 | -0.086 | -0.086 | 0 | 0.33 | 0.62 | 0.26 | 0.02 | 0.3 | 0.19 |

| | | | | | | | | | | | | | | | | |
|---|---|---|---|---|---|---|---|---|---|---|---|---|---|---|---|---|
| 6 | 19840492 | rs1047033 | 0.1 | 1.34 | -2.219 | -0.237 | -0.066 | 0 | 0 | 2.405 | 0.22 | 0.06 | 0.26 | 0.18 | 0.34 | 0.17 |
| 6 | 19842497 | rs6917605 | -0.213 | 1.072 | -1.971 | -0.03 | 0.02 | 0.08 | 0.08 | 1.484 | 0.5 | 0.33 | 0.47 | 0.73 | 0.56 | 0.42 |
| 6 | 19844355 | rs6906699 | -0.368 | 1.246 | -1.654 | 0.077 | -0.005 | -0.004 | -0.004 | 0 | 0.19 | 0.14 | 0.23 | 0.2 | 0.25 | 0.13 |
| 6 | 19848673 | rs7749020 | -0.882 | 1.69 | -0.768 | 0.285 | 0.039 | 0.07 | 0.07 | 2.392 | 0.19 | 0.59 | 0.3 | 0.31 | 0.23 | 0.09 |
| 6 | 19850875 | rs6905466 | -1.573 | 1.35 | -0.316 | -0.189 | 0.001 | 0 | 0 | 0.267 | 0.29 | 0.66 | 0.37 | 0.31 | 0.33 | 0.26 |
| 6 | 19855933 | rs742228 | -0.402 | 1.391 | 0.682 | -0.193 | 0.085 | 0.111 | 0.111 | 0 | 0.16 | 0.35 | 0.28 | 0.28 | 0.24 | 0.09 |
| 6 | 19856058 | rs742227 | 1.058 | 1.567 | -0.092 | -0.497 | -0.032 | 0.007 | 0.007 | 1.903 | 0.12 | 0.02 | 0.09 | 0.07 | 0.16 | 0.16 |
| 6 | 19858468 | rs13203119 | 0.116 | 1.333 | 0.055 | -0.601 | 0 | -0.016 | -0.016 | 0 | 0.12 | 0.1 | 0.08 | 0.01 | 0.09 | 0.18 |
| 6 | 19859897 | rs9366320 | -1.624 | 1.297 | 0.169 | -0.809 | 0.052 | 0.121 | 0.121 | 0 | 0.54 | 0.73 | 0.66 | 0.88 | 0.64 | 0.64 |
| 6 | 19860298 | rs4618518 | 1.139 | 1.712 | 0.364 | -0.505 | -0.016 | -0.002 | -0.002 | 0 | 0.13 | 0.12 | 0.19 | 0.51 | 0.19 | 0.21 |
| 6 | 19860447 | rs932346 | -1.441 | 1.037 | -0.015 | -1.174 | 0.01 | 0.004 | 0.004 | 0 | 0.78 | 0.82 | 0.86 | 0.99 | 0.8 | 0.87 |
| 6 | 19868502 | rs6902539 | 0.767 | 1.347 | 0.303 | -1.055 | 0.084 | -0.061 | -0.061 | 0.039 | 0.11 | 0.21 | 0.18 | 0 | 0.23 | 0.03 |
| 6 | 19870333 | rs10484639 | 1.786 | 0.891 | -0.009 | -1.045 | 0.006 | -0.004 | -0.004 | 0 | 0.11 | 0.02 | 0.03 | 0.07 | 0.08 | 0.14 |
| 6 | 19871076 | rs9465549 | 0.072 | 1.014 | 0.109 | -1.062 | -0.001 | 0 | 0 | 0.056 | 0.39 | 0.54 | 0.51 | 0.4 | 0.39 | 0.32 |
| 6 | 19873584 | rs1534923 | 0.273 | 1.039 | 0.247 | -1.04 | 0.125 | 0.218 | 0.218 | 0 | 0.13 | 0.43 | 0.3 | 0.29 | 0.22 | 0.08 |
| 6 | 19876533 | rs12528660 | 1.408 | 0.958 | -0.034 | -1.021 | 0.033 | 0.075 | 0.075 | 0.015 | 0.25 | 0.08 | 0.2 | 0.11 | 0.16 | 0.22 |
| 6 | 19878461 | rs2328423 | -0.011 | 1.29 | 0.042 | -1.083 | -0.008 | -0.018 | -0.018 | 0 | 0.38 | 0.25 | 0.32 | 0.59 | 0.37 | 0.55 |
| 6 | 19879568 | rs7772056 | -0.155 | 0.919 | 0.113 | -0.77 | -0.001 | 0 | 0 | 0 | 0.39 | 0.55 | 0.52 | 0.39 | 0.4 | 0.34 |
| 6 | 19880730 | rs2876542 | 0.884 | 1.445 | 0.073 | -0.555 | -0.019 | 0.006 | 0.006 | 1.811 | 0.12 | 0.02 | 0.05 | 0.06 | 0.12 | 0.15 |
| 6 | 19884082 | rs7769600 | -0.736 | 1.488 | 0.673 | -0.275 | 0.058 | -0.076 | -0.076 | 0 | 0.12 | 0.29 | 0.16 | 0 | 0.2 | 0.04 |
| 6 | 19887307 | rs10080611 | -0.638 | 1.57 | 0.682 | -0.233 | 0.105 | -0.073 | -0.073 | 0.229 | 0.12 | 0.38 | 0.17 | 0 | 0.19 | 0.04 |
| 6 | 19891340 | rs7769434 | -0.491 | 1.387 | 0.055 | 0.059 | -0.001 | -0.002 | -0.002 | 0 | 0.33 | 0.27 | 0.42 | 0.54 | 0.38 | 0.38 |
| 6 | 19893110 | rs926367 | -0.908 | 1.384 | 0.801 | -0.233 | 0.053 | 0.026 | 0.026 | 1.123 | 0.14 | 0.47 | 0.25 | 0.13 | 0.15 | 0.18 |
| 6 | 19893932 | rs12203172 | 1.092 | 0.699 | -0.052 | -0.498 | -0.007 | 0.05 | 0.05 | 0 | 0.2 | 0.01 | 0.1 | 0.07 | 0.13 | 0.09 |
| 6 | 19894725 | rs7739799 | -1.043 | 0.894 | 0.11 | -0.726 | -0.008 | 0.013 | 0.013 | 0 | 0.82 | 0.83 | 0.87 | 1 | 0.79 | 0.89 |
| 6 | 19898770 | rs2328421 | -0.784 | 0.901 | 0.505 | -0.723 | 0.008 | -0.085 | -0.085 | 0 | 0.8 | 0.69 | 0.86 | 1 | 0.79 | 0.89 |
| 6 | 19899654 | rs5010585 | -0.456 | 0.92 | 0.787 | -0.706 | 0.004 | -0.063 | -0.063 | 1.12 | 0.77 | 0.51 | 0.73 | 0.9 | 0.78 | 0.88 |
| 6 | 19902810 | rs10946347 | 0.476 | 1.063 | 0.193 | -0.574 | 0.007 | -0.088 | -0.088 | 0 | 0.44 | 0.11 | 0.34 | 0.47 | 0.39 | 0.46 |
| 6 | 19906838 | rs1997691 | -0.831 | 0.969 | 0.777 | -0.632 | 0.01 | -0.134 | -0.134 | 0.111 | 0.79 | 0.56 | 0.84 | 1 | 0.79 | 0.89 |
| 6 | 19908495 | rs1883328 | -1.097 | 0.857 | 0.578 | -0.591 | -0.012 | -0.091 | -0.091 | 0 | 0.82 | 0.69 | 0.87 | 1 | 0.84 | 0.9 |
| 6 | 19909241 | rs6936399 | 0.997 | 0.502 | 0.405 | -0.219 | -0.018 | 0.025 | 0.025 | 1.096 | 0.12 | 0.14 | 0.11 | 0.26 | 0.05 | 0.2 |
| 6 | 19911852 | rs16882922 | 0.91 | 0.482 | 0.396 | -0.228 | -0.018 | 0.022 | 0.022 | 1.157 | 0.12 | 0.14 | 0.11 | 0.26 | 0.05 | 0.2 |
| 6 | 19916257 | rs1202312 | -0.261 | 0.63 | 0.431 | -0.184 | 0.011 | -0.009 | -0.009 | 0 | 0.66 | 0.57 | 0.63 | 0.68 | 0.6 | 0.58 |
| 6 | 19919986 | rs1202309 | -0.121 | 0.593 | 0.341 | -0.222 | -0.018 | 0.008 | 0.008 | 0.398 | 0.39 | 0.45 | 0.4 | 0.52 | 0.31 | 0.38 |
| 6 | 19920339 | rs1202308 | -1.045 | 0.967 | 0.146 | -0.288 | 0.003 | -0.011 | -0.011 | 0.907 | 0.94 | 0.95 | 0.93 | 0.98 | 0.88 | 0.84 |
| 6 | 19920904 | rs7768737 | 2.006 | 0.915 | 0.132 | -0.268 | 0.021 | -0.03 | -0.03 | 0 | 0.09 | 0.03 | 0.1 | 0.26 | 0.05 | 0.2 |
| 6 | 19921581 | rs10456227 | 1.413 | 0.968 | 0.098 | -0.298 | -0.006 | -0.011 | -0.011 | 0 | 0.07 | 0 | 0.02 | 0.06 | 0.05 | 0.06 |
| 6 | 19922354 | rs1202305 | 1.356 | 1.047 | 0.464 | -0.007 | -0.001 | -0.023 | -0.023 | 0 | 0.26 | 0.26 | 0.16 | 0.06 | 0.23 | 0.16 |
| 6 | 19923212 | rs9350196 | 1.393 | 1.207 | 0.216 | -0.154 | -0.016 | -0.039 | -0.039 | 0 | 0.16 | 0.05 | 0.16 | 0.28 | 0.15 | 0.21 |
| 6 | 19924318 | rs1931802 | 0.892 | 1.182 | 0.166 | -0.084 | -0.027 | 0.044 | 0.044 | 0 | 0.26 | 0.11 | 0.24 | 0.16 | 0.3 | 0.21 |
| 6 | 19928775 | rs1202297 | -0.737 | 0.098 | 0.11 | 0.089 | 0.003 | -0.011 | -0.011 | 1.708 | 0.94 | 0.95 | 0.93 | 0.99 | 0.89 | 0.85 |
| 6 | 19929121 | rs1202296 | 0.021 | 0.145 | 0.033 | 0.059 | -0.007 | 0.004 | 0.004 | 0 | 0.81 | 0.84 | 0.76 | 0.9 | 0.7 | 0.63 |
| 6 | 19931781 | rs1202293 | 0.003 | 0.162 | -0.064 | 0.063 | 0.005 | -0.005 | -0.005 | 0 | 0.81 | 0.83 | 0.8 | 0.86 | 0.7 | 0.61 |
| 6 | 19933160 | rs1202292 | 1.5 | -0.015 | -0.025 | -0.005 | 0.016 | 0.013 | 0.013 | 0.023 | 0.19 | 0.15 | 0.09 | 0.1 | 0.09 | 0.08 |
| 6 | 19945285 | rs1079801 | 1.353 | 0.035 | -0.418 | 0.081 | 0.012 | -0.011 | -0.011 | 0 | 0.11 | 0.1 | 0.11 | 0.26 | 0.06 | 0.2 |
| 6 | 19946730 | rs2184393 | 0.218 | 0.074 | -0.491 | 0.038 | -0.005 | 0 | 0 | 0.377 | 0.56 | 0.54 | 0.36 | 0.39 | 0.37 | 0.44 |
| 6 | 19956843 | rs2782185 | -0.236 | 0.561 | -0.006 | 0.439 | -0.022 | -0.008 | -0.008 | 1.25 | 0.38 | 0.47 | 0.28 | 0.31 | 0.29 | 0.36 |
| 6 | 19959784 | rs2782187 | 0.515 | 1.029 | -1.147 | 0.666 | -0.002 | -0.002 | -0.002 | 0.177 | 0.79 | 0.82 | 0.64 | 0.65 | 0.67 | 0.6 |
| 6 | 19966110 | rs2782195 | 0.564 | 1.205 | -1.601 | 0.713 | -0.002 | -0.002 | -0.002 | 0 | 0.78 | 0.78 | 0.63 | 0.65 | 0.67 | 0.6 |
| 6 | 19966473 | rs1931786 | 0.079 | 1.076 | -2.102 | 0.28 | -0.049 | -0.018 | -0.018 | 0 | 0.08 | 0.01 | 0.09 | 0.12 | 0.17 | 0.13 |
| 6 | 19966649 | rs12529614 | -0.865 | -0.012 | -1.722 | 0.337 | 0.026 | -0.007 | -0.007 | 0 | 0.34 | 0.2 | 0.26 | 0.31 | 0.28 | 0.35 |
| 6 | 19969480 | rs16883016 | 0.307 | 0.039 | -1.629 | 0.448 | -0.009 | 0.027 | 0.027 | 0.992 | 0.1 | 0.15 | 0.11 | 0.26 | 0.05 | 0.18 |
| 6 | 19972938 | rs12202520 | -0.506 | 0.79 | -1.638 | -0.439 | -0.009 | 0.096 | 0.096 | 0.008 | 0.24 | 0.05 | 0.14 | 0.06 | 0.24 | 0.16 |
| 6 | 19975946 | rs517374 | -0.673 | 0.859 | -1.462 | -0.155 | -0.007 | 0.023 | 0.023 | 0 | 0.2 | 0.13 | 0.22 | 0.13 | 0.26 | 0.26 |
| 6 | 19987971 | rs17560881 | -0.14 | 0.515 | -1.556 | -0.547 | 0.002 | 0.008 | 0.008 | 0 | 0.12 | 0.06 | 0.04 | 0.03 | 0.09 | 0.07 |
| 6 | 19988653 | rs9348404 | 0.62 | 0.125 | -2.563 | -0.227 | -0.011 | -0.105 | -0.105 | 0 | 0.24 | 0.02 | 0.2 | 0.33 | 0.2 | 0.28 |
| 6 | 19989673 | rs9465606 | 0.332 | 0.34 | -2.382 | -0.626 | 0.011 | 0.066 | 0.066 | 0 | 0.53 | 0.67 | 0.61 | 0.78 | 0.52 | 0.51 |
| 6 | 19990972 | rs965037 | -0.355 | 0.152 | -2.476 | -0.707 | -0.019 | 0.037 | 0.037 | 0 | 0.68 | 0.84 | 0.68 | 0.81 | 0.62 | 0.58 |
| 6 | 19992265 | rs520188 | -0.879 | 0.293 | -2.474 | -0.725 | -0.01 | 0.017 | 0.017 | 0.235 | 0.9 | 0.97 | 0.94 | 1 | 0.91 | 0.97 |
| 6 | 19996808 | rs9350204 | 1.184 | 0.144 | -2.364 | -0.557 | -0.033 | -0.133 | -0.133 | 0 | 0.23 | 0.01 | 0.19 | 0.32 | 0.2 | 0.29 |
| 6 | 19997176 | rs9465608 | 0.233 | -0.284 | -1.277 | -0.831 | -0.007 | 0 | 0 | 2.191 | 0.4 | 0.38 | 0.45 | 0.48 | 0.44 | 0.34 |
| 6 | 20001747 | rs499899 | -0.873 | -0.328 | -1.678 | -0.864 | -0.097 | -0.086 | -0.086 | 0.306 | 0.75 | 0.94 | 0.64 | 0.55 | 0.67 | 0.73 |
| 6 | 20003157 | rs7775315 | 0.656 | -0.318 | -0.73 | -0.633 | -0.041 | 0.027 | 0.027 | 0.357 | 0.31 | 0.48 | 0.26 | 0.32 | 0.21 | 0.29 |
| 6 | 20004051 | rs992893 | -0.054 | 0.175 | -0.969 | -0.421 | -0.021 | 0.022 | 0.022 | 0 | 0.71 | 0.83 | 0.68 | 0.81 | 0.64 | 0.63 |
| 6 | 20015205 | rs11753077 | -0.178 | 0.282 | -1.33 | 0.244 | -0.091 | -0.096 | -0.096 | 1.405 | 0.34 | 0.12 | 0.33 | 0.38 | 0.37 | 0.45 |
| 6 | 20016967 | rs625964 | -0.678 | 0.19 | -0.188 | 0.002 | 0.004 | 0.007 | 0.007 | 0.425 | 0.43 | 0.51 | 0.46 | 0.51 | 0.4 | 0.51 |
| 6 | 20017533 | rs623329 | -0.979 | 0.025 | -0.046 | -0.147 | -0.016 | -0.006 | -0.006 | 0 | 0.5 | 0.86 | 0.52 | 0.51 | 0.43 | 0.51 |
| 6 | 20017642 | rs622890 | -0.707 | 0.057 | 0.073 | -0.131 | -0.004 | -0.001 | -0.001 | 0 | 0.48 | 0.66 | 0.5 | 0.51 | 0.43 | 0.51 |
| 6 | 20018376 | rs492591 | -0.338 | 0.059 | 0.458 | -0.113 | -0.002 | 0.034 | 0.034 | 1.426 | 0.09 | 0.21 | 0.12 | 0.15 | 0.04 | 0.06 |

| | | | | | | | | | | | | | | | | |
|---|---|---|---|---|---|---|---|---|---|---|---|---|---|---|---|---|
| 6 | 20021842 | rs7763815 | -0.604 | 0.136 | 0.016 | -0.119 | -0.004 | -0.001 | -0.001 | 0 | 0.48 | 0.66 | 0.5 | 0.51 | 0.43 | 0.48 |
| 6 | 20028644 | rs9348408 | 0.616 | 0.955 | -0.928 | 0.422 | -0.096 | -0.059 | -0.059 | 1.252 | 0.19 | 0.01 | 0.2 | 0.21 | 0.25 | 0.19 |
| 6 | 20035068 | rs495768 | -0.469 | 0.858 | -0.158 | 0.508 | 0.001 | -0.033 | -0.033 | 0 | 0.27 | 0.66 | 0.27 | 0.17 | 0.22 | 0.18 |
| 6 | 20038974 | rs687734 | -0.461 | 0.819 | -0.081 | 0.621 | -0.001 | -0.02 | -0.02 | 0 | 0.27 | 0.65 | 0.27 | 0.22 | 0.23 | 0.2 |
| 6 | 20039645 | rs2472779 | 0.429 | 0.76 | 0.004 | 0.583 | -0.001 | 0.001 | 0.001 | 0 | 0.2 | 0.19 | 0.2 | 0.11 | 0.19 | 0.18 |
| 6 | 20042503 | rs490655 | -0.252 | 0.868 | -0.623 | 0.595 | -0.004 | -0.006 | -0.006 | 0.135 | 0.26 | 0.74 | 0.26 | 0.2 | 0.2 | 0.2 |
| 6 | 20046786 | rs558825 | -0.855 | 1.083 | -1.052 | 0.657 | -0.013 | -0.04 | -0.04 | 0 | 0.45 | 0.89 | 0.44 | 0.46 | 0.4 | 0.38 |
| 6 | 20050161 | rs9350206 | 0.673 | 0.606 | -1.168 | 0.184 | -0.096 | -0.093 | -0.093 | 0 | 0.17 | 0.01 | 0.28 | 0.31 | 0.27 | 0.22 |
| 6 | 20057330 | rs602640 | -0.359 | -0.361 | -0.335 | -0.352 | -0.002 | -0.004 | -0.004 | 0 | 0.05 | 0.07 | 0.01 | 0.04 | 0.03 | 0.04 |
| 6 | 20057889 | rs2472771 | -0.461 | -0.673 | 0.107 | -0.624 | -0.007 | -0.007 | -0.007 | 0.063 | 0.41 | 0.53 | 0.39 | 0.28 | 0.3 | 0.35 |
| 6 | 20062601 | rs4712446 | -1.045 | -0.847 | 0.446 | -0.79 | -0.052 | 0.004 | 0.004 | 0 | 0.28 | 0.49 | 0.31 | 0.28 | 0.18 | 0.22 |
| 6 | 20062778 | rs4712447 | -0.429 | -1.159 | 0.265 | -1.224 | -0.01 | -0.12 | -0.12 | 0.284 | 0.74 | 0.59 | 0.82 | 0.98 | 0.78 | 0.79 |
| 6 | 20063881 | rs592265 | -0.829 | -1.173 | -0.282 | -1.242 | -0.015 | 0.107 | 0.107 | 0.004 | 0.81 | 0.98 | 0.88 | 0.98 | 0.82 | 0.81 |
| 6 | 20064497 | rs998633 | -0.515 | -1.221 | 0.223 | -1.091 | -0.015 | 0.002 | 0.002 | 0 | 0.22 | 0.3 | 0.26 | 0.25 | 0.13 | 0.19 |
| 6 | 20066657 | rs13212039 | 0.376 | -1.257 | -0.207 | -1.032 | 0.035 | -0.02 | -0.02 | 0 | 0.16 | 0.07 | 0.13 | 0.21 | 0.07 | 0.14 |
| 6 | 20072580 | rs670210 | 0.093 | -1.153 | -0.291 | -1.189 | -0.052 | 0.151 | 0.151 | 0 | 0.7 | 0.93 | 0.74 | 0.93 | 0.68 | 0.71 |
| 6 | 20072776 | rs1931774 | 0.404 | -1.662 | -0.167 | -1.043 | 0.004 | -0.009 | -0.009 | 0.102 | 0.14 | 0.11 | 0.11 | 0.24 | 0.05 | 0.09 |
| 6 | 20082771 | rs2205618 | 0.079 | -1.395 | -0.334 | -1.132 | -0.035 | 0.146 | 0.146 | 0 | 0.71 | 0.93 | 0.75 | 0.93 | 0.72 | 0.7 |
| 6 | 20084591 | rs2457350 | -0.297 | -1.449 | -0.303 | -1.1 | -0.019 | 0.11 | 0.11 | 0.009 | 0.75 | 0.94 | 0.77 | 0.93 | 0.76 | 0.71 |
| 6 | 20089434 | rs2457346 | 0.038 | -1.625 | 0.652 | -0.945 | -0.094 | 0.083 | 0.083 | 1.293 | 0.16 | 0.44 | 0.14 | 0.24 | 0.05 | 0.09 |
| 6 | 20089466 | rs2457345 | 0.165 | -1.629 | 0.653 | -0.944 | -0.092 | 0.096 | 0.096 | 1.705 | 0.15 | 0.44 | 0.14 | 0.24 | 0.04 | 0.08 |
| 6 | 20089551 | rs6919270 | -0.328 | -0.94 | 0.704 | -0.756 | 0.007 | -0.031 | -0.031 | 0.03 | 0.06 | 0.12 | 0.04 | 0 | 0.05 | 0.02 |
| 6 | 20091293 | rs2472763 | -0.417 | -1.179 | 0.792 | -0.885 | 0.009 | 0.012 | 0.012 | 0.005 | 0.66 | 0.75 | 0.72 | 0.75 | 0.71 | 0.66 |
| 6 | 20092949 | rs2472762 | -0.432 | -1.594 | 0.351 | -1.252 | -0.067 | -0.144 | -0.144 | 0 | 0.24 | 0.6 | 0.14 | 0 | 0.1 | 0.07 |
| 6 | 20096334 | rs2457340 | -1.123 | -1.538 | 0.151 | -1.27 | -0.01 | -0.082 | -0.082 | 1.053 | 0.16 | 0.69 | 0.14 | 0 | 0.12 | 0.03 |
| 6 | 20097305 | rs9368149 | 0.392 | -1.306 | -0.236 | -0.735 | 0.023 | -0.041 | -0.041 | 0.398 | 0.21 | 0.03 | 0.25 | 0.19 | 0.15 | 0.21 |
| 6 | 20097963 | rs9348410 | 0.556 | -1.261 | -0.233 | -1.275 | -0.019 | 0.038 | 0.038 | 0.838 | 0.37 | 0.22 | 0.52 | 0.7 | 0.53 | 0.51 |
| 6 | 20099162 | rs1812333 | -0.449 | -0.912 | -0.669 | -1.05 | 0.017 | 0.041 | 0.041 | 0.435 | 0.59 | 0.71 | 0.64 | 0.71 | 0.65 | 0.57 |
| 6 | 20100222 | rs2235142 | 0.076 | -0.855 | -1.082 | -0.707 | -0.003 | -0.003 | -0.003 | 0 | 0.12 | 0.03 | 0.08 | 0.08 | 0.11 | 0.14 |
| 6 | 20101640 | rs16883345 | 0.357 | -0.77 | -0.727 | -0.653 | 0.051 | 0.065 | 0.065 | 0 | 0.14 | 0.02 | 0.04 | 0.01 | 0.06 | 0.06 |
| 6 | 20107427 | rs2457336 | 0.14 | -0.013 | -0.731 | 0.086 | 0.014 | 0.022 | 0.022 | 0.341 | 0.61 | 0.69 | 0.56 | 0.68 | 0.66 | 0.6 |
| 6 | 20109696 | rs2457334 | -0.354 | 0.468 | -0.897 | 0.508 | -0.011 | -0.029 | -0.029 | 0.034 | 0.37 | 0.33 | 0.36 | 0.43 | 0.44 | 0.51 |
| 6 | 20111100 | rs1555004 | 0.133 | 0.615 | -0.646 | 0.447 | -0.014 | -0.008 | -0.008 | 0.089 | 0.31 | 0.26 | 0.32 | 0.34 | 0.37 | 0.26 |
| 6 | 20113516 | rs1980430 | 0.155 | 0.362 | -0.151 | 0.219 | -0.024 | -0.015 | -0.015 | 0.289 | 0.76 | 0.55 | 0.71 | 0.84 | 0.81 | 0.75 |
| 6 | 20115177 | rs9368151 | 0.207 | 0.459 | -0.155 | 0.117 | 0.001 | -0.001 | -0.001 | 0.701 | 0.48 | 0.47 | 0.41 | 0.53 | 0.44 | 0.49 |
| 6 | 20116687 | rs2472768 | -0.546 | 0.657 | -0.151 | 0.186 | 0.029 | 0.061 | 0.061 | 0 | 0.62 | 0.8 | 0.79 | 0.87 | 0.71 | 0.74 |
| 6 | 20119035 | rs16883364 | -0.095 | 1.13 | -0.031 | 0.825 | -0.034 | -0.02 | -0.02 | 0 | 0.09 | 0.07 | 0.22 | 0.4 | 0.19 | 0.21 |
| 6 | 20120469 | rs2472756 | -0.384 | 1.042 | 0.234 | 0.745 | -0.004 | -0.027 | -0.027 | 0.131 | 0.17 | 0.16 | 0.08 | 0.01 | 0.09 | 0.17 |
| 6 | 20121957 | rs2457330 | -1.027 | 1.209 | -0.157 | 0.847 | -0.038 | -0.065 | -0.065 | 0.11 | 0.23 | 0.12 | 0.27 | 0.41 | 0.28 | 0.34 |
| 6 | 20124860 | rs949550 | -0.797 | 1.625 | 0.139 | 1.198 | 0.015 | 0.051 | 0.051 | 0 | 0.11 | 0.12 | 0.35 | 0.48 | 0.24 | 0.25 |
| 6 | 20126656 | rs2483766 | -0.608 | 1.542 | -0.155 | 1.189 | -0.035 | -0.029 | -0.029 | 1.383 | 0.1 | 0.05 | 0.21 | 0.39 | 0.18 | 0.19 |
| 6 | 20128557 | rs2483767 | -0.627 | 0.548 | 0.225 | 0.356 | -0.025 | -0.034 | -0.034 | 1.488 | 0.14 | 0.2 | 0.09 | 0.02 | 0.07 | 0.15 |
| 6 | 20135427 | rs843318 | 0.028 | 0.947 | -0.032 | 0.372 | -0.003 | 0.02 | 0.02 | 0 | 0.07 | 0.02 | 0.05 | 0 | 0.06 | 0.06 |
| 6 | 20144790 | rs1202196 | 0.932 | 0.968 | 0.533 | 0.562 | 0.001 | 0.005 | 0.005 | 0.008 | 0.67 | 0.4 | 0.59 | 0.52 | 0.63 | 0.66 |
| 6 | 20146507 | rs1212547 | 0.3 | 0.948 | 0.705 | 0.459 | -0.04 | -0.106 | -0.106 | 0.01 | 0.82 | 0.7 | 0.92 | 1 | 0.9 | 0.89 |
| 6 | 20146536 | rs9350215 | -0.424 | 1.225 | -0.011 | 0.677 | -0.06 | -0.071 | -0.071 | 9.054 | 0.14 | 0.03 | 0.29 | 0.48 | 0.27 | 0.23 |
| 6 | 20152095 | rs2457322 | 0.5 | 1.081 | -0.371 | 0.809 | -0.002 | -0.002 | -0.002 | 0 | 0.58 | 0.59 | 0.57 | 0.53 | 0.55 | 0.55 |
| 6 | 20163330 | rs10946363 | -0.491 | 1.328 | -0.896 | 1.317 | -0.022 | -0.065 | -0.065 | 0.033 | 0.22 | 0.11 | 0.4 | 0.53 | 0.32 | 0.38 |
| 6 | 20164196 | rs7750646 | -0.369 | 1.251 | -0.384 | 0.601 | -0.001 | -0.061 | -0.061 | 0 | 0.16 | 0.33 | 0.15 | 0.03 | 0.16 | 0.07 |
| 6 | 20167622 | rs6932299 | -0.574 | 1.036 | -1.116 | -0.019 | -0.005 | -0.043 | -0.043 | 0 | 0.38 | 0.19 | 0.44 | 0.55 | 0.41 | 0.51 |
| 6 | 20168333 | rs9348414 | 0.58 | 1.048 | -1.019 | 0.366 | 0.009 | -0.017 | -0.017 | 0 | 0.19 | 0.12 | 0.25 | 0.37 | 0.18 | 0.31 |
| 6 | 20169881 | rs943571 | -0.334 | 0.79 | -1.354 | -0.359 | -0.008 | 0.102 | 0.102 | 0 | 0.55 | 0.75 | 0.35 | 0.16 | 0.44 | 0.43 |
| 6 | 20171056 | rs9465643 | 0.202 | 0.602 | -1.389 | -0.356 | 0.007 | 0.002 | 0.002 | 0 | 0.11 | 0.11 | 0.07 | 0.01 | 0.09 | 0.12 |
| 6 | 20171926 | rs586124 | 0.731 | 0.89 | -1.256 | 0.118 | -0.052 | -0.072 | -0.072 | 0 | 0.26 | 0.12 | 0.4 | 0.45 | 0.37 | 0.24 |
| 6 | 20178992 | rs9460465 | 0.153 | -0.139 | -0.377 | -0.183 | -0.06 | -0.008 | -0.008 | 0 | 0.18 | 0.39 | 0.09 | 0.15 | 0.06 | 0.17 |
| 6 | 20180235 | rs1202209 | -0.357 | -0.1 | -0.607 | -0.229 | -0.002 | 0.003 | 0.003 | 0.015 | 0.57 | 0.61 | 0.68 | 0.7 | 0.51 | 0.54 |
| 6 | 20181505 | rs10946366 | -0.847 | -0.173 | -0.609 | -0.384 | -0.028 | 0.02 | 0.02 | 0 | 0.67 | 0.76 | 0.75 | 0.7 | 0.59 | 0.6 |
| 6 | 20182185 | rs4712460 | 0.499 | 0.283 | -0.945 | -0.125 | -0.05 | -0.042 | -0.042 | 0 | 0.37 | 0.15 | 0.58 | 0.55 | 0.44 | 0.36 |
| 6 | 20185051 | rs555017 | -0.457 | 0.323 | -0.728 | -0.035 | 0.005 | 0.009 | 0.009 | 0 | 0.53 | 0.4 | 0.67 | 0.55 | 0.61 | 0.45 |
| 6 | 20186760 | rs2328395 | -0.207 | 0.349 | -0.824 | -0.014 | -0.001 | 0.02 | 0.02 | 0 | 0.49 | 0.38 | 0.65 | 0.54 | 0.52 | 0.4 |
| 6 | 20190950 | rs9465648 | -0.454 | -0.172 | -0.571 | 0.059 | -0.047 | 0.038 | 0.038 | 0.03 | 0.11 | 0.24 | 0.04 | 0.25 | 0.04 | 0.06 |
| 6 | 20191838 | rs865226 | -0.02 | 0.251 | -1.001 | -0.025 | 0.005 | -0.014 | -0.014 | 0.393 | 0.44 | 0.32 | 0.57 | 0.46 | 0.53 | 0.37 |
| 6 | 20192190 | rs1536578 | 0.259 | 0.36 | -1.155 | 0.031 | -0.006 | -0.035 | -0.035 | 0.358 | 0.41 | 0.22 | 0.56 | 0.46 | 0.53 | 0.37 |
| 6 | 20193747 | rs7766133 | -0.032 | 0.325 | -1.151 | 0.035 | 0.045 | -0.022 | -0.022 | 0.154 | 0.46 | 0.22 | 0.59 | 0.47 | 0.61 | 0.42 |
| 6 | 20196423 | rs9358309 | 0.061 | 0.386 | -1.2 | -0.301 | -0.006 | -0.026 | -0.026 | 0 | 0.42 | 0.21 | 0.56 | 0.47 | 0.52 | 0.38 |
| 6 | 20198845 | rs943573 | 0.093 | -0.059 | -1.293 | -0.646 | -0.004 | -0.013 | -0.013 | 0.064 | 0.42 | 0.31 | 0.6 | 0.47 | 0.57 | 0.38 |
| 6 | 20200420 | rs6907076 | -0.854 | -0.424 | -1.055 | -0.801 | -0.028 | -0.074 | -0.074 | 2.015 | 0.3 | 0.45 | 0.22 | 0.14 | 0.2 | 0.24 |
| 6 | 20200664 | rs9465656 | 0.037 | -0.188 | -1.584 | -0.711 | -0.011 | -0.022 | -0.022 | 0 | 0.21 | 0.23 | 0.16 | 0.13 | 0.14 | 0.2 |
| 6 | 20201181 | rs11752473 | -0.131 | 0.654 | -2.164 | -0.239 | -0.011 | -0.03 | -0.03 | 0.124 | 0.4 | 0.21 | 0.59 | 0.47 | 0.57 | 0.38 |

| | | | | | | | | | | | | | | | | |
|---|---|---|---|---|---|---|---|---|---|---|---|---|---|---|---|---|
| 6 | 20202770 | rs10447379 | -0.335 | 0.344 | -2.312 | -0.06 | 0.035 | -0.026 | -0.026 | 0.011 | 0.45 | 0.18 | 0.63 | 0.47 | 0.63 | 0.43 |
| 6 | 20204711 | rs4712463 | -0.672 | 0.352 | -1.61 | 0.455 | 0 | -0.027 | -0.027 | 0.014 | 0.22 | 0.15 | 0.22 | 0.46 | 0.27 | 0.25 |
| 6 | 20209344 | rs1536581 | 0.477 | -0.329 | -1.471 | -0.431 | -0.008 | -0.05 | -0.05 | 0 | 0.75 | 0.35 | 0.68 | 0.88 | 0.76 | 0.83 |
| 6 | 20210374 | rs1327665 | 0.93 | -0.326 | -2.111 | -0.434 | -0.1 | 0.002 | 0.002 | 0 | 0.43 | 0.08 | 0.36 | 0.37 | 0.47 | 0.49 |
| 6 | 20213334 | rs1077919 | 0.236 | -0.109 | -1.501 | -0.008 | -0.018 | -0.05 | -0.05 | 0 | 0.78 | 0.35 | 0.73 | 0.88 | 0.83 | 0.86 |
| 6 | 20226668 | rs11751661 | -0.453 | 0 | -1.916 | 0.573 | -0.051 | -0.163 | -0.163 | 0 | 0.26 | 0.02 | 0.27 | 0.51 | 0.25 | 0.31 |
| 6 | 20237043 | rs717208 | -0.564 | -0.226 | -1.04 | -0.167 | 0.001 | -0.006 | -0.006 | 0 | 0.39 | 0.38 | 0.31 | 0.16 | 0.4 | 0.55 |
| 6 | 20237388 | rs6940079 | -1.033 | -0.793 | -0.88 | 0.578 | -0.004 | 0.025 | 0.025 | 0 | 0.06 | 0.13 | 0.04 | 0.14 | 0.08 | 0.05 |
| 6 | 20243096 | rs754344 | -1.158 | -1.011 | -0.412 | 0.227 | -0.012 | -0.01 | -0.01 | 1.809 | 0.34 | 0.66 | 0.21 | 0.27 | 0.28 | 0.46 |
| 6 | 20246018 | rs7746765 | -0.813 | -1.005 | -0.467 | 0.506 | -0.031 | 0.002 | 0.002 | 1.24 | 0.16 | 0.39 | 0.07 | 0.2 | 0.09 | 0.24 |
| 6 | 20248007 | rs4712465 | -0.248 | -0.758 | -1.216 | 1.35 | 0.016 | -0.058 | -0.058 | 0.02 | 0.1 | 0.01 | 0.13 | 0.43 | 0.05 | 0.18 |
| 6 | 20250826 | rs9465673 | -0.645 | -0.748 | -0.954 | 0.404 | 0.01 | 0.012 | 0.012 | 0 | 0.22 | 0.18 | 0.14 | 0.09 | 0.19 | 0.29 |
| 6 | 20251486 | rs9348417 | -0.294 | -0.306 | -1.622 | 0.837 | -0.046 | -0.093 | -0.093 | 0.005 | 0.24 | 0.03 | 0.33 | 0.32 | 0.28 | 0.19 |
| 6 | 20252093 | rs9366337 | -0.384 | -0.343 | -1.765 | 0.905 | -0.047 | -0.079 | -0.079 | 0 | 0.26 | 0.02 | 0.28 | 0.28 | 0.28 | 0.2 |
| 6 | 20253392 | rs11753226 | 0.782 | -0.792 | -1.652 | 0.632 | 0.02 | 0.001 | 0.001 | 0.573 | 0.68 | 0.89 | 0.79 | 0.73 | 0.72 | 0.71 |
| 6 | 20265965 | rs1887476 | 0.919 | -0.962 | -0.806 | 0.557 | 0.007 | 0.002 | 0.002 | 0.352 | 0.67 | 0.73 | 0.77 | 0.73 | 0.72 | 0.7 |
| 6 | 20274321 | rs4712469 | 0.524 | -0.951 | -0.826 | 0.563 | 0.003 | 0.004 | 0.004 | 0 | 0.91 | 0.93 | 0.94 | 0.95 | 0.94 | 0.89 |
| 6 | 20279412 | rs2328380 | 0.706 | -0.969 | -0.641 | 0.553 | -0.012 | -0.015 | -0.015 | 0 | 0.89 | 0.81 | 0.93 | 0.95 | 0.94 | 0.88 |
| 6 | 20285619 | rs12207062 | -0.201 | -1.18 | -0.326 | 0.446 | 0.013 | -0.018 | -0.018 | 1.681 | 0.23 | 0.06 | 0.15 | 0.23 | 0.23 | 0.19 |
| 6 | 20292031 | rs4712476 | 0.785 | -1.187 | -0.332 | 0.442 | 0.024 | 0 | 0 | 0 | 0.68 | 0.87 | 0.79 | 0.72 | 0.71 | 0.69 |
| 6 | 20296072 | rs6902984 | -0.741 | -0.543 | -0.554 | 0.261 | -0.036 | 0.016 | 0.016 | 0 | 0.12 | 0.01 | 0.09 | 0.05 | 0.11 | 0.12 |
| 6 | 20299481 | rs4712480 | 0.586 | -0.495 | -0.258 | 0.232 | 0.009 | 0.015 | 0.015 | 0.112 | 0.64 | 0.85 | 0.79 | 0.72 | 0.68 | 0.69 |
| 6 | 20301011 | rs4712482 | 0.836 | -0.755 | 0.459 | 0.113 | -0.012 | -0.006 | -0.006 | 0.324 | 0.61 | 0.38 | 0.7 | 0.72 | 0.72 | 0.68 |
| 6 | 20313120 | rs9465693 | -0.808 | -0.892 | 0.513 | 0.393 | -0.004 | -0.003 | -0.003 | 0.052 | 0.41 | 0.45 | 0.33 | 0.31 | 0.33 | 0.33 |
| 6 | 20324128 | rs7748774 | 0.59 | -1.166 | 1.098 | 1.078 | -0.029 | 0.015 | 0.015 | 0 | 0.61 | 0.46 | 0.69 | 0.47 | 0.73 | 0.6 |
| 6 | 20325205 | rs6917385 | -0.649 | -1.155 | 0.962 | 1.075 | -0.021 | 0.009 | 0.009 | 0.348 | 0.4 | 0.54 | 0.31 | 0.53 | 0.27 | 0.41 |
| 6 | 20335060 | rs11751348 | -0.451 | -1.138 | 0.57 | 1.12 | 0.045 | -0.069 | -0.069 | 0.436 | 0.35 | 0.11 | 0.26 | 0.45 | 0.26 | 0.33 |
| 6 | 20344428 | rs4712488 | -1.083 | -0.132 | 0.243 | 1.009 | 0.013 | -0.02 | -0.02 | 0 | 0.62 | 0.56 | 0.58 | 0.71 | 0.59 | 0.69 |
| 6 | 20344996 | rs9358331 | -1.042 | -0.174 | 0.241 | 0.987 | 0.004 | -0.008 | -0.008 | 0 | 0.62 | 0.51 | 0.58 | 0.71 | 0.6 | 0.68 |
| 6 | 20346195 | rs2208438 | 0.172 | -0.066 | 0.015 | 1.022 | -0.055 | -0.018 | -0.018 | 0 | 0.23 | 0.07 | 0.29 | 0.23 | 0.33 | 0.3 |
| 6 | 20359969 | rs9350245 | -0.878 | 0.108 | 0.124 | 0.786 | 0.002 | -0.004 | -0.004 | 0 | 0.61 | 0.62 | 0.55 | 0.71 | 0.54 | 0.67 |
| 6 | 20366062 | rs9366347 | -0.379 | -0.222 | -0.27 | 1.082 | 0.025 | -0.016 | -0.016 | 0 | 0.37 | 0.3 | 0.32 | 0.47 | 0.29 | 0.38 |
| 6 | 20366580 | rs16901546 | -0.167 | -0.24 | -0.314 | 1.043 | -0.009 | 0.006 | 0.006 | 0.356 | 0.32 | 0.35 | 0.34 | 0.58 | 0.26 | 0.3 |
| 6 | 20374040 | rs10946380 | -0.214 | 0.948 | -0.62 | -0.032 | 0.02 | -0.057 | -0.057 | 0 | 0.16 | 0.23 | 0.14 | 0.01 | 0.2 | 0.09 |
| 6 | 20382600 | rs1570155 | -0.991 | 0.714 | -1.615 | 0.149 | 0.026 | -0.064 | -0.064 | 0.229 | 0.51 | 0.89 | 0.47 | 0.43 | 0.57 | 0.49 |
| 6 | 20395257 | rs6915426 | 0.157 | 0.839 | -1.556 | 0.026 | 0.018 | -0.057 | -0.057 | 1.719 | 0.15 | 0.23 | 0.12 | 0.01 | 0.17 | 0.06 |
| 6 | 20396953 | rs10806918 | 0.24 | 0.574 | -1.852 | 0.164 | -0.153 | -0.123 | -0.123 | 0 | 0.28 | 0.02 | 0.27 | 0.33 | 0.39 | 0.31 |
| 6 | 20401909 | rs749215 | 0.268 | 0.575 | -1.377 | -0.062 | 0.027 | -0.072 | -0.072 | 0 | 0.15 | 0.27 | 0.13 | 0.01 | 0.17 | 0.06 |
| 6 | 20413721 | rs9368188 | -0.927 | 0.127 | -2.08 | 0.075 | 0.005 | 0.011 | 0.011 | 0 | 0.6 | 0.84 | 0.68 | 0.64 | 0.65 | 0.69 |
| 6 | 20414340 | rs12527393 | 0.556 | 0.476 | -2.344 | 0.127 | -0.048 | 0.023 | 0.023 | 0.349 | 0.23 | 0.04 | 0.34 | 0.18 | 0.28 | 0.29 |
| 6 | 20414798 | rs17569598 | 0.061 | 0.588 | -2.257 | 0.404 | -0.124 | -0.174 | -0.174 | 0 | 0.28 | 0.02 | 0.25 | 0.43 | 0.35 | 0.4 |
| 6 | 20415053 | rs911361 | -0.388 | 0.149 | -1.661 | 0.286 | -0.016 | 0.034 | 0.034 | 0 | 0.56 | 0.78 | 0.66 | 0.65 | 0.57 | 0.68 |
| 6 | 20416572 | rs9295464 | -0.617 | -0.976 | -0.586 | 0.559 | -0.056 | 0.071 | 0.071 | 0 | 0.28 | 0.44 | 0.2 | 0.46 | 0.17 | 0.26 |
| 6 | 20417976 | rs746720 | 1.128 | -1.003 | -1.16 | 0.127 | -0.022 | 0.038 | 0.038 | 0.597 | 0.38 | 0.19 | 0.44 | 0.21 | 0.49 | 0.3 |
| 6 | 20418329 | rs2328488 | 0.561 | -1.034 | -1.171 | 0.087 | -0.05 | 0.049 | 0.049 | 0 | 0.25 | 0.03 | 0.25 | 0.23 | 0.25 | 0.35 |
| 6 | 20425458 | rs7752992 | -0.506 | -0.713 | -0.548 | 0.293 | 0.026 | 0.023 | 0.023 | 0 | 0.53 | 0.66 | 0.47 | 0.71 | 0.37 | 0.57 |
| 6 | 20433077 | rs9465742 | 0.645 | -0.39 | -0.191 | 0.174 | -0.009 | 0.009 | 0.009 | 0.346 | 0.93 | 0.84 | 0.94 | 0.93 | 0.92 | 0.92 |
| 6 | 20442354 | rs7760528 | 0.427 | -0.673 | -0.476 | 0.012 | 0.043 | 0.08 | 0.08 | 1.135 | 0.58 | 0.23 | 0.62 | 0.71 | 0.69 | 0.53 |
| 6 | 20452735 | rs9295468 | 1.572 | -0.277 | 0.028 | 0.553 | -0.007 | -0.01 | -0.01 | 0 | 0.9 | 0.82 | 0.93 | 0.94 | 0.89 | 0.88 |
| 6 | 20462138 | rs6938550 | 1.47 | -0.259 | 0.124 | 0.551 | -0.005 | -0.007 | -0.007 | 0 | 0.91 | 0.79 | 0.93 | 0.94 | 0.9 | 0.9 |
| 6 | 20467578 | rs6926668 | 0.437 | -0.383 | -0.137 | 0.982 | 0.005 | -0.019 | -0.019 | 0.226 | 0.78 | 0.8 | 0.81 | 0.59 | 0.77 | 0.78 |
| 6 | 20470908 | rs12205236 | 1.196 | -0.414 | 0.288 | 0.905 | -0.042 | 0.083 | 0.083 | 0 | 0.7 | 0.44 | 0.77 | 0.53 | 0.76 | 0.67 |
| 6 | 20480586 | rs3828858 | -0.184 | -0.065 | -0.585 | 1.174 | 0.014 | -0.112 | -0.112 | 0 | 0.17 | 0.04 | 0.22 | 0.44 | 0.14 | 0.2 |
| 6 | 20482411 | rs7747931 | 0.633 | 0.79 | -0.174 | 0.608 | 0.005 | -0.009 | -0.009 | 0 | 0.62 | 0.63 | 0.39 | 0.31 | 0.49 | 0.53 |
| 6 | 20485898 | rs11964747 | 1.193 | 0.526 | -0.337 | -0.155 | 0.005 | 0.02 | 0.02 | 6.044 | 0.24 | 0.16 | 0.16 | 0.07 | 0.2 | 0.06 |
| 6 | 20486506 | rs4134958 | 0.468 | 1.042 | 0.274 | 0.063 | 0.06 | 0.097 | 0.097 | 1.352 | 0.76 | 0.59 | 0.53 | 0.65 | 0.64 | 0.72 |
| 6 | 20488234 | rs2328524 | 0.52 | 0.761 | -0.227 | -0.263 | -0.001 | -0.084 | -0.084 | 0.721 | 0.42 | 0.11 | 0.31 | 0.58 | 0.38 | 0.59 |
| 6 | 20492995 | rs1042317 | -0.307 | 0.821 | -0.091 | -0.081 | -0.02 | -0.022 | -0.022 | 0 | 0.83 | 0.94 | 0.7 | 0.77 | 0.74 | 0.8 |
| 6 | 20495546 | rs10946384 | 1.101 | 0.668 | -0.154 | -0.423 | -0.013 | 0.01 | 0.01 | 0 | 0.25 | 0.06 | 0.17 | 0.22 | 0.2 | 0.36 |
| 6 | 20504851 | rs7775526 | 0.856 | 0.601 | 0.204 | -0.458 | -0.009 | -0.029 | -0.029 | 0 | 0.32 | 0.45 | 0.29 | 0.2 | 0.28 | 0.16 |
| 6 | 20506815 | rs12206516 | 1.465 | 0.638 | -0.728 | -0.361 | -0.012 | -0.012 | -0.012 | 0.055 | 0.25 | 0.04 | 0.23 | 0.18 | 0.2 | 0.13 |
| 6 | 20509339 | rs942041 | -0.354 | 1.168 | -0.115 | -0.339 | -0.002 | 0.007 | 0.007 | 0 | 0.4 | 0.58 | 0.31 | 0.25 | 0.37 | 0.45 |
| 6 | 20520353 | rs17512008 | 0.702 | 0.983 | -0.927 | -0.181 | -0.013 | 0.015 | 0.015 | 0 | 0.23 | 0.05 | 0.16 | 0.2 | 0.2 | 0.33 |
| 6 | 20524382 | rs761154 | 0.006 | 1.493 | -0.857 | -0.084 | -0.029 | 0.021 | 0.021 | 0.013 | 0.61 | 0.86 | 0.59 | 0.65 | 0.57 | 0.46 |
| 6 | 20527017 | rs4429936 | 1.339 | 1.556 | -0.149 | -0.284 | -0.032 | 0.028 | 0.028 | 1.658 | 0.66 | 0.2 | 0.53 | 0.8 | 0.55 | 0.57 |
| 6 | 20533162 | rs6909467 | 0.669 | 2.007 | 0.426 | -0.331 | 0.009 | 0 | 0 | 1.745 | 0.81 | 0.81 | 0.7 | 0.82 | 0.72 | 0.68 |
| 6 | 20533383 | rs9368197 | 0.43 | 1.537 | -0.265 | -0.428 | 0.036 | -0.13 | -0.13 | 0.349 | 0.34 | 0.05 | 0.33 | 0.55 | 0.27 | 0.42 |
| 6 | 20535970 | rs6904566 | -0.213 | 1.354 | 0.884 | -0.464 | 0.069 | -0.05 | -0.05 | 0 | 0.1 | 0.31 | 0.1 | 0.01 | 0.15 | 0.04 |
| 6 | 20541106 | rs12200871 | -0.93 | 1.447 | 1.156 | -0.522 | 0.022 | -0.099 | -0.099 | 0 | 0.14 | 0.41 | 0.12 | 0 | 0.12 | 0.06 |

| | | | | | | | | | | | | | | | | |
|---|---|---|---|---|---|---|---|---|---|---|---|---|---|---|---|---|
| 6 | 20541180 | rs9348432 | 0.017 | 1.421 | 0.933 | -0.536 | 0.024 | -0.039 | -0.039 | 0 | 0.39 | 0.24 | 0.38 | 0.55 | 0.34 | 0.45 |
| 6 | 20545468 | rs9366354 | -0.592 | 1.446 | 1.128 | -0.518 | 0.002 | -0.001 | -0.001 | 0 | 0.48 | 0.53 | 0.44 | 0.55 | 0.4 | 0.48 |
| 6 | 20554951 | rs9350257 | 0.42 | 1.416 | 0.724 | -0.498 | 0.061 | -0.116 | -0.116 | 1.929 | 0.34 | 0.06 | 0.32 | 0.55 | 0.24 | 0.42 |
| 6 | 20556905 | rs9350258 | 0.385 | 1.429 | 0.726 | -0.492 | 0.073 | -0.109 | -0.109 | 0.102 | 0.36 | 0.08 | 0.32 | 0.55 | 0.24 | 0.44 |
| 6 | 20562618 | rs7772137 | 0.542 | 1.407 | 0.775 | -0.447 | 0.001 | -0.033 | -0.033 | 0.054 | 0.07 | 0.13 | 0.04 | 0 | 0.06 | 0.03 |
| 6 | 20570142 | rs2064320 | -0.464 | 1.502 | 0.588 | -0.377 | -0.073 | -0.064 | -0.064 | 0.001 | 0.88 | 0.99 | 0.76 | 0.82 | 0.78 | 0.72 |
| 6 | 20580893 | rs9368204 | 0.114 | 1.367 | 0.729 | -0.466 | 0.013 | -0.028 | -0.028 | 0 | 0.38 | 0.28 | 0.39 | 0.56 | 0.33 | 0.44 |
| 6 | 20581828 | rs7744833 | 1.572 | 1.376 | 0.765 | -0.457 | 0.007 | 0.014 | 0.014 | 0 | 0.34 | 0.21 | 0.25 | 0.26 | 0.33 | 0.2 |
| 6 | 20589341 | rs6907767 | 0.487 | 1.459 | 0.802 | -0.44 | -0.006 | -0.001 | -0.001 | 0 | 0.8 | 0.79 | 0.72 | 0.81 | 0.72 | 0.68 |
| 6 | 20601785 | rs12209806 | 1.635 | 1.362 | 0.638 | -0.418 | 0.005 | 0.02 | 0.02 | 0.053 | 0.34 | 0.17 | 0.24 | 0.26 | 0.33 | 0.19 |
| 6 | 20609241 | rs7758129 | 0.563 | 1.686 | 0.659 | -0.345 | 0.065 | 0.017 | 0.017 | 1.75 | 0.83 | 0.67 | 0.69 | 0.81 | 0.68 | 0.7 |
| 6 | 20611926 | rs9465831 | 0.132 | 1.662 | 0.471 | -0.357 | -0.006 | -0.006 | -0.006 | 0 | 0.86 | 0.87 | 0.77 | 0.81 | 0.8 | 0.71 |
| 6 | 20612311 | rs13212501 | 0.527 | 1.574 | 0.503 | -0.379 | 0.001 | -0.03 | -0.03 | 0 | 0.07 | 0.12 | 0.04 | 0 | 0.06 | 0.03 |
| 6 | 20620003 | rs9350268 | 0.123 | 1.291 | 0.408 | -0.396 | 0.045 | -0.07 | -0.07 | 0.094 | 0.35 | 0.14 | 0.36 | 0.55 | 0.28 | 0.42 |
| 6 | 20622753 | rs1569660 | 1.511 | 1.414 | 0.964 | -0.503 | 0.074 | -0.031 | -0.031 | 0 | 0.71 | 0.54 | 0.59 | 0.81 | 0.57 | 0.65 |
| 6 | 20625491 | rs9465838 | -0.321 | 1.588 | 0.903 | -0.484 | -0.044 | -0.055 | -0.055 | 0 | 0.91 | 1 | 0.85 | 0.81 | 0.87 | 0.76 |
| 6 | 20626409 | rs6917599 | 1.147 | 1.146 | 0.659 | -0.434 | 0 | -0.011 | -0.011 | 0 | 0.56 | 0.46 | 0.42 | 0.63 | 0.5 | 0.47 |
| 6 | 20630397 | rs13190734 | 0.138 | 1.261 | 0.579 | -0.418 | 0.001 | -0.002 | -0.002 | 0 | 0.08 | 0.09 | 0.12 | 0.01 | 0.19 | 0.05 |
| 6 | 20631011 | rs10484633 | 1.076 | 1.162 | 0.453 | -0.406 | -0.021 | 0.003 | 0.003 | 0 | 0.17 | 0.01 | 0.23 | 0.18 | 0.14 | 0.22 |
| 6 | 20637306 | rs9368215 | 1.514 | 1.444 | 1.315 | -0.007 | 0.049 | -0.012 | -0.012 | 0 | 0.75 | 0.61 | 0.66 | 0.81 | 0.64 | 0.7 |
| 6 | 20639409 | rs9465850 | 0.129 | 1.09 | 1.099 | 0.138 | -0.011 | 0.031 | 0.031 | 0 | 0.19 | 0.18 | 0.16 | 0.38 | 0.16 | 0.23 |
| 6 | 20640316 | rs9465851 | 0.513 | 1.141 | 1.068 | 0.069 | 0.003 | -0.011 | -0.011 | 0 | 0.69 | 0.79 | 0.68 | 0.43 | 0.7 | 0.5 |
| 6 | 20647660 | rs10484632 | -1.098 | 1.367 | 1.305 | 0.157 | 0.047 | -0.068 | -0.068 | 1.105 | 0.14 | 0.29 | 0.18 | 0.01 | 0.24 | 0.07 |
| 6 | 20648762 | rs9460540 | 1.791 | 1.179 | 1.27 | 0.051 | -0.017 | 0.048 | 0.048 | 0.369 | 0.49 | 0.12 | 0.45 | 0.41 | 0.44 | 0.4 |
| 6 | 20652765 | rs9295475 | -0.491 | 1.116 | 1.496 | 0.077 | -0.042 | 0.109 | 0.109 | 0.011 | 0.23 | 0.42 | 0.22 | 0.39 | 0.16 | 0.25 |
| 6 | 20655503 | rs16884072 | -0.517 | 1.251 | 1.388 | 0.09 | 0.054 | -0.049 | -0.049 | 0.171 | 0.1 | 0.23 | 0.13 | 0.01 | 0.19 | 0.08 |
| 6 | 20657564 | rs4712523 | -0.928 | 1.204 | 0.806 | 0.112 | 0.003 | 0.012 | 0.012 | 0.402 | 0.28 | 0.65 | 0.32 | 0.41 | 0.32 | 0.29 |
| 6 | 20660365 | rs6456369 | -1.251 | 1.193 | 0.745 | 0.1 | -0.008 | -0.018 | -0.018 | 0.684 | 0.33 | 0.75 | 0.36 | 0.4 | 0.36 | 0.28 |
| 6 | 20663335 | rs4712527 | -0.063 | 0.767 | 0.644 | 0.049 | -0.006 | -0.014 | -0.014 | 0 | 0.84 | 0.92 | 0.83 | 0.82 | 0.8 | 0.74 |
| 6 | 20679310 | rs1569699 | -1.287 | 0.62 | 0.967 | 0.281 | -0.001 | 0.013 | 0.013 | 0.281 | 0.34 | 0.63 | 0.33 | 0.47 | 0.33 | 0.29 |
| 6 | 20679709 | rs7756992 | -0.992 | 0.578 | 0.903 | 0.259 | -0.001 | 0.047 | 0.047 | 0.064 | 0.29 | 0.63 | 0.3 | 0.47 | 0.28 | 0.27 |
| 6 | 20689945 | rs7748720 | 0.13 | -0.186 | 1.273 | 0.352 | -0.001 | -0.002 | -0.002 | 0 | 0.81 | 0.86 | 0.81 | 0.82 | 0.79 | 0.73 |
| 6 | 20694884 | rs2206734 | -0.473 | -0.059 | 1.413 | 0.474 | -0.005 | 0.009 | 0.009 | 0 | 0.23 | 0.25 | 0.18 | 0.38 | 0.18 | 0.26 |
| 6 | 20713706 | rs1040558 | -0.38 | -0.066 | 1.405 | -0.004 | 0.001 | 0.003 | 0.003 | 0 | 0.23 | 0.21 | 0.2 | 0.47 | 0.18 | 0.25 |
| 6 | 20716253 | rs9295478 | -1.201 | -0.053 | 1.161 | -0.013 | -0.006 | 0.008 | 0.008 | 0.205 | 0.32 | 0.69 | 0.28 | 0.47 | 0.22 | 0.28 |
| 6 | 20717095 | rs6935599 | -1.008 | -0.118 | 1.213 | 0.138 | -0.024 | 0.042 | 0.042 | 1.092 | 0.27 | 0.66 | 0.26 | 0.47 | 0.18 | 0.26 |
| 6 | 20717404 | rs10946403 | -0.973 | -0.107 | 1.11 | 0.144 | -0.038 | 0.076 | 0.076 | 1.016 | 0.27 | 0.6 | 0.25 | 0.47 | 0.18 | 0.26 |
| 6 | 20719145 | rs9358357 | -0.874 | -0.098 | 0.969 | 0.154 | -0.034 | 0.074 | 0.074 | 1.462 | 0.27 | 0.61 | 0.25 | 0.47 | 0.18 | 0.25 |
| 6 | 20720279 | rs9356746 | 0.292 | -0.025 | 0.632 | 0.209 | 0 | 0.022 | 0.022 | 0 | 0.17 | 0.16 | 0.17 | 0.47 | 0.13 | 0.21 |
| 6 | 20728513 | rs6928012 | 1.265 | 0.048 | 0.076 | 0.146 | 0.039 | 0.066 | 0.066 | 1.023 | 0.55 | 0.24 | 0.54 | 0.35 | 0.61 | 0.46 |
| 6 | 20728731 | rs6908425 | 0.459 | 0.35 | -0.071 | 0.317 | 0 | 0.001 | 0.001 | 0 | 0.82 | 0.81 | 0.79 | 0.81 | 0.8 | 0.7 |
| 6 | 20731524 | rs7741604 | 0.211 | -0.115 | -0.486 | 0.628 | 0.003 | -0.129 | -0.129 | 0 | 0.16 | 0.01 | 0.16 | 0.41 | 0.13 | 0.19 |
| 6 | 20733389 | rs9465873 | -0.216 | -0.22 | -0.323 | -0.203 | -0.003 | 0.004 | 0.004 | 0 | 0.58 | 0.57 | 0.57 | 0.42 | 0.62 | 0.53 |
| 6 | 20734994 | rs13217519 | 0.723 | -0.327 | -0.265 | -0.329 | -0.006 | 0 | 0 | 0 | 0.26 | 0.22 | 0.28 | 0.35 | 0.27 | 0.42 |
| 6 | 20737354 | rs9465874 | 1.487 | -0.614 | -0.321 | -0.609 | -0.094 | 0.114 | 0.114 | 0.71 | 0.3 | 0.09 | 0.3 | 0.08 | 0.4 | 0.24 |
| 6 | 20737757 | rs9368228 | 0.76 | -0.581 | -0.254 | -0.65 | -0.005 | 0.001 | 0.001 | 0 | 0.27 | 0.23 | 0.28 | 0.35 | 0.27 | 0.42 |
| 6 | 20740296 | rs9350276 | 0.151 | -0.646 | -0.065 | -0.705 | 0.006 | 0.037 | 0.037 | 0 | 0.43 | 0.27 | 0.45 | 0.68 | 0.4 | 0.6 |
| 6 | 20743952 | rs9358363 | -0.648 | -0.739 | 0.107 | -0.663 | 0.001 | 0.002 | 0.002 | 0 | 0.85 | 0.88 | 0.82 | 0.88 | 0.85 | 0.87 |
| 6 | 20744727 | rs4710944 | 1.938 | -0.585 | -0.193 | -0.849 | -0.077 | 0.105 | 0.105 | 0.311 | 0.22 | 0.06 | 0.26 | 0.01 | 0.33 | 0.2 |
| 6 | 20746236 | rs7738382 | 0.261 | -0.675 | 0.275 | -0.797 | -0.031 | 0.029 | 0.029 | 0.31 | 0.65 | 0.45 | 0.61 | 0.54 | 0.74 | 0.61 |
| 6 | 20751315 | rs7747724 | 0.589 | -0.42 | 0.661 | -0.653 | 0.011 | -0.023 | -0.023 | 0 | 0.36 | 0.43 | 0.37 | 0.2 | 0.45 | 0.32 |
| 6 | 20755932 | rs2819996 | 0.109 | -0.578 | 0.586 | -0.686 | -0.006 | 0.024 | 0.024 | 0 | 0.19 | 0.21 | 0.19 | 0.34 | 0.14 | 0.26 |
| 6 | 20758760 | rs10946406 | 0.766 | -0.497 | 0.586 | -0.713 | 0.004 | -0.011 | -0.011 | 0 | 0.35 | 0.38 | 0.36 | 0.2 | 0.45 | 0.25 |
| 6 | 20766103 | rs4585542 | 0.166 | -0.467 | 0.517 | -0.726 | -0.001 | 0.02 | 0.02 | 0 | 0.47 | 0.41 | 0.45 | 0.47 | 0.42 | 0.51 |
| 6 | 20791712 | rs12208378 | 0.475 | -1.171 | 0.601 | -0.7 | -0.011 | -0.018 | -0.018 | 0 | 0.12 | 0.15 | 0.14 | 0.09 | 0.08 | 0.12 |
| 6 | 20797851 | rs12211211 | 0.825 | -1.166 | 0.569 | -0.697 | -0.001 | -0.002 | -0.002 | 0 | 0.1 | 0.11 | 0.14 | 0.09 | 0.08 | 0.12 |
| 6 | 20803463 | rs9465894 | 0.766 | -1.153 | 0.449 | -0.772 | -0.01 | 0.015 | 0.015 | 1.012 | 0.08 | 0.05 | 0.03 | 0.01 | 0.07 | 0.02 |
| 6 | 20803687 | rs6925153 | 0.809 | -0.941 | 0.598 | -0.59 | -0.022 | 0.015 | 0.015 | 0.05 | 0.37 | 0.3 | 0.36 | 0.38 | 0.43 | 0.4 |
| 6 | 20803812 | rs6456374 | 0.144 | -0.92 | 0.548 | -0.611 | -0.023 | -0.064 | -0.064 | 0.285 | 0.83 | 0.64 | 0.79 | 0.91 | 0.87 | 0.86 |
| 6 | 20804563 | rs9465895 | 1.535 | -0.791 | 0.216 | -0.456 | -0.001 | -0.003 | -0.003 | 0.002 | 0.23 | 0.16 | 0.22 | 0.19 | 0.24 | 0.19 |
| 6 | 20819976 | rs10806925 | 0.709 | -0.796 | 0.161 | -0.507 | 0.013 | -0.009 | -0.009 | 0 | 0.35 | 0.16 | 0.36 | 0.52 | 0.33 | 0.34 |
| 6 | 20821489 | rs7753109 | -0.345 | -0.851 | 0.572 | -0.623 | -0.022 | -0.079 | -0.079 | 3.58 | 0.22 | 0.33 | 0.2 | 0.09 | 0.19 | 0.17 |
| 6 | 20826281 | rs4395717 | -0.576 | -0.871 | 0.638 | -0.647 | 0.013 | 0.014 | 0.014 | 1.047 | 0.6 | 0.66 | 0.57 | 0.47 | 0.63 | 0.57 |
| 6 | 20829772 | rs9356755 | 0.752 | -0.727 | 0.575 | -0.543 | -0.018 | 0.011 | 0.011 | 0 | 0.37 | 0.3 | 0.37 | 0.38 | 0.43 | 0.39 |
| 6 | 20841593 | rs11967068 | 0.45 | -0.785 | 0.52 | -0.31 | -0.008 | 0.039 | 0.039 | 0 | 0.14 | 0.19 | 0.16 | 0.33 | 0.09 | 0.22 |
| 6 | 20849412 | rs7753271 | -0.17 | -0.855 | 0.607 | -0.403 | -0.003 | 0.012 | 0.012 | 0 | 0.22 | 0.25 | 0.22 | 0.33 | 0.2 | 0.25 |
| 6 | 20856703 | rs4712556 | 1.042 | -0.992 | 0.736 | -0.567 | 0.023 | -0.008 | -0.008 | 0.097 | 0.58 | 0.35 | 0.57 | 0.58 | 0.66 | 0.58 |
| 6 | 20858852 | rs9366363 | 0.621 | -1.039 | 1.092 | -0.591 | -0.038 | 0.043 | 0.043 | 0 | 0.69 | 0.54 | 0.71 | 0.67 | 0.74 | 0.71 |
| 6 | 20861152 | rs16901585 | -0.603 | -1.056 | 1.127 | -0.614 | -0.054 | 0.084 | 0.084 | 0.139 | 0.27 | 0.45 | 0.24 | 0.33 | 0.2 | 0.27 |

| | | | | | | | | | | | | | | | | |
|---|---|---|---|---|---|---|---|---|---|---|---|---|---|---|---|---|
| 6 | 20865629 | rs12191898 | 0.881 | -1.032 | 1.091 | -0.587 | -0.003 | -0.005 | -0.005 | 0.141 | 0.1 | 0.12 | 0.14 | 0.1 | 0.08 | 0.13 |
| 6 | 20866048 | rs7749397 | -0.059 | -1.03 | 1.07 | -0.589 | 0.002 | -0.032 | -0.032 | 0.224 | 0.93 | 0.75 | 0.92 | 1 | 0.95 | 0.91 |
| 6 | 20870093 | rs9350294 | -0.019 | -1.058 | 0.965 | -0.681 | 0.024 | 0.036 | 0.036 | 0.953 | 0.56 | 0.33 | 0.55 | 0.71 | 0.65 | 0.59 |
| 6 | 20876683 | rs4077405 | 0.099 | -0.99 | 0.671 | -0.653 | 0.018 | 0.044 | 0.044 | 0.421 | 0.51 | 0.61 | 0.52 | 0.47 | 0.52 | 0.54 |
| 6 | 20881386 | rs10946415 | 1.67 | -0.892 | 0.629 | -0.66 | -0.008 | -0.019 | -0.019 | 0 | 0.22 | 0.09 | 0.21 | 0.2 | 0.22 | 0.19 |
| 6 | 20884080 | rs4546483 | -1.025 | -0.86 | 0.628 | -0.63 | -0.001 | -0.003 | -0.003 | 0 | 0.71 | 0.67 | 0.71 | 0.81 | 0.73 | 0.72 |
| 6 | 20887954 | rs6938256 | -0.123 | -0.759 | 0.428 | -0.569 | 0.003 | 0.004 | 0.004 | 0.614 | 0.57 | 0.39 | 0.56 | 0.71 | 0.65 | 0.58 |
| 6 | 20901267 | rs7758467 | 1.092 | -0.485 | 0.059 | -0.546 | -0.026 | 0.039 | 0.039 | 0 | 0.07 | 0 | 0.06 | 0 | 0.13 | 0.03 |
| 6 | 20901311 | rs6456382 | -0.568 | -0.756 | 0.332 | -0.224 | -0.022 | 0.016 | 0.016 | 0.095 | 0.34 | 0.49 | 0.33 | 0.36 | 0.3 | 0.3 |
| 6 | 20903971 | rs9358377 | 0.93 | -0.454 | -0.32 | -0.242 | -0.034 | 0.004 | 0.004 | 0 | 0.29 | 0.21 | 0.33 | 0.39 | 0.36 | 0.46 |
| 6 | 20910936 | rs9350298 | 1.211 | -0.395 | -0.887 | -0.22 | -0.139 | -0.047 | -0.047 | 0 | 0.27 | 0.03 | 0.3 | 0.37 | 0.36 | 0.38 |
| 6 | 20911003 | rs4710958 | 0.574 | -0.26 | -0.495 | -0.169 | -0.018 | -0.031 | -0.031 | 0 | 0.3 | 0.14 | 0.29 | 0.26 | 0.29 | 0.21 |
| 6 | 20940379 | rs12196595 | 0.377 | -0.079 | -0.284 | -0.019 | 0 | -0.002 | -0.002 | 0 | 0.1 | 0.07 | 0.08 | 0.12 | 0.09 | 0.14 |
| 6 | 20943232 | rs9465941 | -0.717 | -0.22 | -0.433 | -0.31 | -0.017 | 0.025 | 0.025 | 0 | 0.9 | 0.95 | 0.94 | 1 | 0.88 | 0.97 |
| 6 | 20952163 | rs9465945 | 0.564 | -0.252 | -0.319 | -0.326 | -0.003 | 0 | 0 | 0 | 0.1 | 0.1 | 0.08 | 0.12 | 0.07 | 0.14 |
| 6 | 20954826 | rs1004172 | 1.417 | -0.57 | -0.495 | -0.288 | -0.014 | -0.05 | -0.05 | 0 | 0.23 | 0.06 | 0.3 | 0.23 | 0.23 | 0.19 |
| 6 | 20955332 | rs7775523 | 0.894 | -0.611 | -0.369 | -0.452 | -0.008 | -0.012 | -0.012 | 0.054 | 0.3 | 0.14 | 0.37 | 0.23 | 0.35 | 0.22 |
| 6 | 20966438 | rs4712573 | 0.498 | -0.829 | -0.46 | -0.778 | -0.041 | -0.007 | -0.007 | 0 | 0.46 | 0.18 | 0.46 | 0.64 | 0.54 | 0.64 |
| 6 | 20967716 | rs9460583 | 0.723 | -0.767 | 0.113 | -0.621 | -0.015 | 0.007 | 0.007 | 0.014 | 0.11 | 0.2 | 0.09 | 0.12 | 0.07 | 0.13 |
| 6 | 20971985 | rs6913817 | 0.413 | -0.733 | 0.462 | -0.658 | 0.016 | -0.026 | -0.026 | 0.336 | 0.07 | 0.26 | 0.06 | 0.01 | 0.04 | 0.01 |
| 6 | 20989286 | rs201346 | -0.339 | -0.66 | 0.008 | -0.589 | 0.002 | -0.001 | -0.001 | 0.08 | 0.58 | 0.45 | 0.54 | 0.64 | 0.66 | 0.67 |
| 6 | 20990153 | rs6456389 | 0.932 | -0.568 | -0.655 | -0.215 | -0.089 | -0.038 | -0.038 | 0 | 0.27 | 0.07 | 0.3 | 0.44 | 0.34 | 0.47 |
| 6 | 20999387 | rs9358383 | 0.212 | -0.484 | -0.243 | -0.23 | 0.014 | 0.009 | 0.009 | 0.018 | 0.35 | 0.68 | 0.41 | 0.23 | 0.27 | 0.28 |
| 6 | 21005346 | rs6932393 | 0.226 | -0.119 | -0.461 | -0.997 | -0.011 | 0.123 | 0.123 | 1.141 | 0.44 | 0.3 | 0.61 | 0.86 | 0.54 | 0.6 |
| 6 | 21008153 | rs1498426 | -0.055 | -0.198 | 0.159 | -0.919 | -0.003 | -0.036 | -0.036 | 0 | 0.52 | 0.54 | 0.65 | 0.85 | 0.58 | 0.63 |
| 6 | 21009108 | rs374654 | 0.42 | 0.037 | -0.282 | -1.126 | -0.017 | 0.015 | 0.015 | 0 | 0.17 | 0.12 | 0.15 | 0 | 0.24 | 0.07 |
| 6 | 21009888 | rs9465946 | 0.488 | -0.589 | 0.203 | -0.935 | -0.022 | -0.016 | -0.016 | 0.024 | 0.26 | 0.31 | 0.22 | 0.2 | 0.14 | 0.31 |
| 6 | 21012904 | rs6905258 | 0.259 | -0.523 | 0.483 | -1.066 | -0.06 | -0.083 | -0.083 | 0 | 0.29 | 0.47 | 0.19 | 0.15 | 0.17 | 0.32 |
| 6 | 21016532 | rs201311 | 0.039 | 0.36 | 0.256 | -0.311 | 0.022 | 0.072 | 0.072 | 0 | 0.92 | 0.92 | 0.76 | 0.41 | 0.76 | 0.7 |
| 6 | 21018655 | rs10946425 | 0.989 | 0.245 | -0.106 | -0.883 | 0.029 | 0.072 | 0.072 | 0 | 0.41 | 0.2 | 0.42 | 0.2 | 0.41 | 0.3 |
| 6 | 21025421 | rs169293 | 1.011 | 0.314 | 0.017 | -0.726 | -0.008 | -0.003 | -0.003 | 0.104 | 0.11 | 0.09 | 0.07 | 0 | 0.13 | 0.03 |
| 6 | 21028059 | rs6921781 | 0.581 | 0.222 | 0.017 | -0.672 | -0.017 | -0.044 | -0.044 | 0 | 0.09 | 0.11 | 0.05 | 0 | 0.04 | 0.03 |
| 6 | 21030153 | rs201301 | 0.281 | -0.124 | -0.6 | -0.848 | -0.008 | -0.032 | -0.032 | 0 | 0.22 | 0.09 | 0.19 | 0.19 | 0.19 | 0.14 |
| 6 | 21031079 | rs4710961 | 0.969 | -0.038 | -0.379 | -0.334 | -0.001 | 0.004 | 0.004 | 0.279 | 0.33 | 0.28 | 0.41 | 0.26 | 0.37 | 0.24 |
| 6 | 21032682 | rs415446 | 0.112 | 0.448 | -0.424 | -0.303 | -0.001 | 0 | 0 | 0 | 0.6 | 0.46 | 0.53 | 0.34 | 0.49 | 0.5 |
| 6 | 21033965 | rs201300 | 0.632 | 0.163 | -0.868 | -0.665 | 0.004 | 0.062 | 0.062 | 0.019 | 0.44 | 0.32 | 0.29 | 0.24 | 0.4 | 0.38 |
| 6 | 21034656 | rs13210277 | 1.207 | 0.304 | -0.794 | -0.571 | 0.014 | 0.046 | 0.046 | 0.166 | 0.33 | 0.21 | 0.32 | 0.13 | 0.27 | 0.34 |
| 6 | 21037940 | rs989969 | 0.72 | 0.578 | -0.626 | -0.185 | 0.004 | -0.005 | -0.005 | 0 | 0.32 | 0.23 | 0.38 | 0.29 | 0.33 | 0.2 |
| 6 | 21038922 | rs201313 | 1.291 | 0.508 | -0.956 | -0.491 | -0.035 | 0.062 | 0.062 | 0 | 0.13 | 0.05 | 0.07 | 0.01 | 0.12 | 0.08 |
| 6 | 21039951 | rs9295489 | 0.698 | 0.442 | -0.812 | -0.411 | 0.027 | 0.044 | 0.044 | 0.749 | 0.43 | 0.29 | 0.38 | 0.32 | 0.35 | 0.44 |
| 6 | 21045315 | rs201322 | 0.336 | 0.492 | -1.088 | -0.357 | -0.028 | 0.015 | 0.015 | 0.074 | 0.22 | 0.08 | 0.12 | 0.16 | 0.2 | 0.17 |
| 6 | 21046035 | rs201327 | 0.208 | 0.472 | -0.88 | -0.367 | -0.005 | 0.018 | 0.018 | 0.091 | 0.22 | 0.18 | 0.13 | 0.16 | 0.2 | 0.17 |
| 6 | 21046090 | rs201328 | 1.344 | 0.472 | -0.896 | -0.374 | -0.045 | 0.06 | 0.06 | 0 | 0.12 | 0.04 | 0.07 | 0.01 | 0.14 | 0.08 |
| 6 | 21046168 | rs201329 | 0.361 | 0.533 | -0.731 | -0.306 | -0.002 | 0.011 | 0.011 | 0.091 | 0.22 | 0.21 | 0.13 | 0.16 | 0.2 | 0.17 |
| 6 | 21062454 | rs9465956 | 1.108 | 0.399 | -0.456 | -0.456 | -0.005 | -0.036 | -0.036 | 0 | 0.34 | 0.34 | 0.35 | 0.14 | 0.27 | 0.34 |
| 6 | 21065787 | rs1001310 | 0.552 | 0.779 | -0.404 | 0.165 | -0.003 | -0.004 | -0.004 | 0 | 0.7 | 0.71 | 0.75 | 0.48 | 0.63 | 0.56 |
| 6 | 21065826 | rs4712580 | -0.275 | 0.916 | -0.404 | 0.561 | -0.006 | 0.046 | 0.046 | 0 | 0.93 | 0.98 | 0.89 | 0.64 | 0.83 | 0.73 |
| 6 | 21073029 | rs11757677 | -0.399 | 0.7 | -0.427 | 0.224 | 0.001 | -0.001 | -0.001 | 0 | 0.31 | 0.36 | 0.49 | 0.63 | 0.37 | 0.37 |
| 6 | 21073431 | rs6918997 | -0.586 | 0.543 | -0.212 | -0.116 | -0.003 | -0.052 | -0.052 | 0 | 0.91 | 0.83 | 0.9 | 1 | 0.93 | 0.92 |
| 6 | 21074193 | rs9295491 | -0.433 | 0.492 | -0.187 | 0.319 | 0.01 | 0.079 | 0.079 | 0 | 0.14 | 0.3 | 0.23 | 0.31 | 0.18 | 0.26 |
| 6 | 21074363 | rs4283868 | 0.952 | 0.458 | -0.33 | 0.046 | 0.001 | -0.019 | -0.019 | 0 | 0.59 | 0.47 | 0.41 | 0.33 | 0.55 | 0.55 |
| 6 | 21076048 | rs16884481 | 1.633 | -0.168 | -0.618 | -0.043 | 0.04 | -0.04 | -0.04 | 0 | 0.08 | 0 | 0.03 | 0.2 | 0.02 | 0.1 |
| 6 | 21076770 | rs2068072 | -0.727 | 0.084 | -0.37 | -0.005 | 0.008 | -0.021 | -0.021 | 0 | 0.33 | 0.52 | 0.51 | 0.66 | 0.38 | 0.38 |
| 6 | 21078233 | rs9358388 | -0.134 | 0.139 | -0.084 | 0.294 | 0.015 | 0.103 | 0.103 | 0 | 0.13 | 0.31 | 0.23 | 0.29 | 0.16 | 0.27 |
| 6 | 21079084 | rs10080639 | 0.682 | 0.34 | -0.42 | -0.19 | 0.015 | -0.021 | -0.021 | 0 | 0.64 | 0.54 | 0.59 | 0.69 | 0.61 | 0.55 |
| 6 | 21079573 | rs9366375 | 0.418 | 0.41 | -0.419 | -0.093 | 0.001 | 0.017 | 0.017 | 0 | 0.11 | 0.16 | 0.22 | 0.31 | 0.18 | 0.25 |
| 6 | 21079709 | rs7750461 | 3.058 | 0.095 | -0.969 | -0.841 | -0.039 | 0.108 | 0.108 | 0.093 | 0.17 | 0.02 | 0.14 | 0.02 | 0.18 | 0.12 |
| 6 | 21081206 | rs4323302 | -0.416 | 0.231 | -0.136 | -0.485 | -0.001 | -0.001 | -0.001 | 0 | 0.47 | 0.49 | 0.51 | 0.52 | 0.54 | 0.52 |
| 6 | 21081650 | rs9366376 | 0.485 | 0.461 | -0.205 | -0.534 | 0 | 0 | 0 | 0 | 0.14 | 0.17 | 0.25 | 0.27 | 0.22 | 0.26 |
| 6 | 21084430 | rs4421186 | -0.241 | -0.003 | -0.124 | -0.407 | 0.001 | 0.002 | 0.002 | 0.569 | 0.46 | 0.44 | 0.49 | 0.56 | 0.45 | 0.44 |
| 6 | 21086047 | rs4130033 | 2.088 | -0.019 | -0.165 | -0.63 | 0.018 | 0.044 | 0.044 | 1.354 | 0.28 | 0.17 | 0.19 | 0.17 | 0.23 | 0.24 |
| 6 | 21086753 | rs11964144 | -0.185 | 0.135 | -0.094 | -0.592 | -0.006 | -0.035 | -0.035 | 0 | 0.39 | 0.52 | 0.31 | 0.26 | 0.38 | 0.29 |
| 6 | 21086785 | rs10946428 | 0.292 | 0.291 | -0.487 | -0.503 | -0.003 | 0.03 | 0.03 | 0 | 0.35 | 0.26 | 0.29 | 0.26 | 0.38 | 0.29 |
| 6 | 21090622 | rs6904348 | 0.346 | 0.312 | -0.511 | -0.486 | -0.007 | 0.036 | 0.036 | 0 | 0.35 | 0.21 | 0.29 | 0.26 | 0.38 | 0.3 |
| 6 | 21100044 | rs9350319 | -0.541 | 0.571 | -0.052 | -0.21 | 0.004 | 0.016 | 0.016 | 0.032 | 0.21 | 0.31 | 0.32 | 0.27 | 0.24 | 0.36 |
| 6 | 21103457 | rs11968036 | 1.099 | 0.599 | -0.375 | -0.261 | -0.002 | -0.023 | -0.023 | 0.006 | 0.08 | 0.09 | 0.09 | 0.01 | 0.07 | 0.1 |
| 6 | 21107109 | rs9358391 | -0.524 | 0.62 | -0.372 | -0.216 | -0.004 | -0.008 | -0.008 | 0 | 0.19 | 0.19 | 0.31 | 0.26 | 0.25 | 0.36 |
| 6 | 21108114 | rs4712585 | -0.551 | 0.758 | -0.629 | -0.649 | -0.057 | 0.069 | 0.069 | 0.47 | 0.72 | 0.84 | 0.76 | 0.94 | 0.62 | 0.81 |
| 6 | 21115316 | rs9356764 | 0.005 | 0.559 | -0.568 | -0.583 | -0.043 | 0.012 | 0.012 | 4.764 | 0.64 | 0.83 | 0.54 | 0.68 | 0.46 | 0.57 |

| | | | | | | | | | | | | | | | | |
|---|---|---|---|---|---|---|---|---|---|---|---|---|---|---|---|---|
| 6 | 21116900 | rs3935207 | 0.81 | 0.258 | -0.31 | -0.917 | -0.005 | -0.074 | -0.074 | 1.304 | 0.17 | 0.28 | 0.11 | 0 | 0.16 | 0.14 |
| 6 | 21117606 | rs7764365 | 1.862 | 0.24 | -0.218 | -0.879 | 0.002 | -0.045 | -0.045 | 0 | 0.11 | 0.12 | 0.08 | 0 | 0.12 | 0.1 |
| 6 | 21119130 | rs6456397 | 1.288 | 0.202 | -0.28 | -0.425 | 0.037 | 0.031 | 0.031 | 0.014 | 0.38 | 0.23 | 0.24 | 0.35 | 0.26 | 0.35 |
| 6 | 21120754 | rs4074910 | 1.299 | 0.241 | -0.267 | -0.406 | 0.034 | 0.027 | 0.027 | 0 | 0.37 | 0.26 | 0.23 | 0.35 | 0.26 | 0.34 |
| 6 | 21120769 | rs4076112 | 0.652 | 0.626 | -0.338 | -0.545 | 0.015 | 0.068 | 0.068 | 0 | 0.5 | 0.35 | 0.32 | 0.36 | 0.4 | 0.42 |
| 6 | 21124521 | rs10946430 | 0.225 | 0.436 | -0.288 | -0.392 | -0.014 | 0.019 | 0.019 | 0.01 | 0.61 | 0.73 | 0.49 | 0.68 | 0.45 | 0.57 |
| 6 | 21130870 | rs6937439 | 0.848 | 0.351 | -0.24 | -0.302 | -0.014 | 0.083 | 0.083 | 0 | 0.23 | 0.38 | 0.24 | 0.32 | 0.2 | 0.23 |
| 6 | 21136120 | rs2446489 | 0.217 | 0.853 | -0.596 | -0.509 | -0.038 | -0.03 | -0.03 | 0 | 0.23 | 0.15 | 0.4 | 0.32 | 0.37 | 0.34 |
| 6 | 21138100 | rs9358393 | 0.436 | 0.604 | -0.538 | -0.422 | -0.003 | 0.011 | 0.011 | 0 | 0.59 | 0.63 | 0.47 | 0.67 | 0.45 | 0.55 |
| 6 | 21142660 | rs6932702 | 0.598 | 0.559 | -0.862 | -0.613 | -0.014 | 0.041 | 0.041 | 0.68 | 0.17 | 0.06 | 0.16 | 0.06 | 0.2 | 0.08 |
| 6 | 21144599 | rs10498701 | 1.142 | 0.604 | -0.813 | -0.944 | -0.001 | -0.04 | -0.04 | 0.465 | 0.23 | 0.17 | 0.16 | 0.01 | 0.25 | 0.2 |
| 6 | 21146266 | rs1466340 | -0.241 | 0.641 | -1.005 | -0.891 | 0 | -0.028 | -0.028 | 0.005 | 0.44 | 0.53 | 0.34 | 0.06 | 0.43 | 0.29 |
| 6 | 21148742 | rs9368280 | 0.693 | 0.424 | -1.187 | -0.224 | 0.015 | -0.028 | -0.028 | 0 | 0.1 | 0.06 | 0.05 | 0.26 | 0.07 | 0.18 |
| 6 | 21149064 | rs9295496 | -0.276 | 0.378 | -1.182 | -0.394 | -0.007 | -0.014 | -0.014 | 0 | 0.39 | 0.53 | 0.25 | 0.27 | 0.33 | 0.39 |
| 6 | 21152274 | rs10946433 | 0.652 | 0.567 | -1.613 | -0.198 | 0.013 | -0.022 | -0.022 | 0 | 0.1 | 0.09 | 0.05 | 0.26 | 0.07 | 0.17 |
| 6 | 21155343 | rs6456401 | -1.204 | 1.048 | -1.781 | 0.155 | -0.1 | -0.124 | -0.124 | 0 | 0.67 | 0.95 | 0.75 | 0.48 | 0.66 | 0.69 |
| 6 | 21157038 | rs4712586 | -1.362 | 1.25 | -1.316 | 0.722 | -0.051 | -0.073 | -0.073 | 0.649 | 0.89 | 0.98 | 0.93 | 0.71 | 0.85 | 0.9 |
| 6 | 21157173 | rs9368283 | -0.241 | 0.403 | -0.538 | -0.235 | -0.012 | -0.025 | -0.025 | 0 | 0.35 | 0.46 | 0.24 | 0.27 | 0.31 | 0.37 |
| 6 | 21175492 | rs12525940 | -0.042 | 1.029 | -0.38 | 0.311 | 0.003 | -0.007 | -0.007 | 0 | 0.07 | 0.06 | 0.05 | 0.1 | 0.05 | 0.04 |
| 6 | 21176124 | rs12526391 | -0.496 | 1.282 | -0.607 | 0.028 | -0.037 | 0.016 | 0.016 | 0 | 0.13 | 0.03 | 0.13 | 0.06 | 0.18 | 0.1 |
| 6 | 21191928 | rs9465991 | -0.837 | 1.599 | 0.28 | 0.169 | 0.06 | 0.048 | 0.048 | 0.302 | 0.13 | 0.4 | 0.32 | 0.21 | 0.17 | 0.18 |
| 6 | 21200282 | rs2305955 | -1.807 | 1.189 | 0.272 | -0.134 | -0.005 | 0.005 | 0.005 | 0.608 | 0.36 | 0.69 | 0.55 | 0.35 | 0.39 | 0.32 |
| 6 | 21205388 | rs10456242 | -1.097 | 0.873 | 0.631 | -0.239 | 0.01 | -0.02 | -0.02 | 0 | 0.21 | 0.47 | 0.35 | 0.21 | 0.21 | 0.17 |
| 6 | 21207550 | rs4291091 | -1.696 | 0.634 | 0.516 | -0.269 | 0.021 | -0.006 | -0.006 | 0 | 0.53 | 0.66 | 0.62 | 0.6 | 0.57 | 0.55 |
| 6 | 21208429 | rs898165 | -0.871 | 0.985 | 0.875 | 0.104 | 0.01 | 0.022 | 0.022 | 0.964 | 0.1 | 0.17 | 0.14 | 0.22 | 0.15 | 0.06 |
| 6 | 21211797 | rs12201217 | -1.386 | 0.887 | 1.332 | -0.014 | 0.003 | -0.011 | -0.011 | 0 | 0.34 | 0.38 | 0.45 | 0.29 | 0.37 | 0.45 |
| 6 | 21212926 | rs9358398 | -1.653 | 0.816 | 1.293 | -0.026 | 0.002 | 0.009 | 0.009 | 0 | 0.29 | 0.36 | 0.45 | 0.5 | 0.33 | 0.42 |
| 6 | 21213307 | rs9358400 | -0.247 | 0.843 | 1.262 | -0.027 | -0.008 | 0.05 | 0.05 | 1.555 | 0.08 | 0.16 | 0.08 | 0.17 | 0.06 | 0.04 |
| 6 | 21215485 | rs13200422 | -0.429 | 0.711 | 0.705 | 0 | -0.001 | -0.002 | -0.002 | 0 | 0.18 | 0.19 | 0.29 | 0.28 | 0.17 | 0.35 |
| 6 | 21216746 | rs10946438 | -1.322 | 0.619 | 0.422 | 0.017 | 0 | 0.001 | 0.001 | 0.178 | 0.38 | 0.64 | 0.47 | 0.36 | 0.36 | 0.47 |
| 6 | 21217185 | rs9358401 | -1.452 | 0.383 | 0.222 | -0.142 | -0.003 | -0.01 | -0.01 | 2.771 | 0.48 | 0.77 | 0.62 | 0.59 | 0.51 | 0.53 |
| 6 | 21222751 | rs16884693 | -0.26 | 0.174 | 0.595 | -0.109 | -0.062 | -0.065 | -0.065 | 0.529 | 0.11 | 0.24 | 0.06 | 0 | 0.01 | 0.01 |
| 6 | 21223140 | rs2328574 | -0.605 | 0.481 | 0.446 | -0.079 | -0.017 | -0.04 | -0.04 | 0.215 | 0.26 | 0.22 | 0.39 | 0.51 | 0.33 | 0.46 |
| 6 | 21224517 | rs7770752 | -1.212 | 0.916 | 0.726 | -0.073 | 0.113 | -0.025 | -0.025 | 0.461 | 0.13 | 0.37 | 0.17 | 0.08 | 0.18 | 0.1 |
| 6 | 21225250 | rs1870421 | -0.665 | 0.926 | 0.706 | -0.047 | 0.081 | 0.083 | 0.083 | 0 | 0.06 | 0.36 | 0.06 | 0.07 | 0.04 | 0.02 |
| 6 | 21225639 | rs9466006 | -0.891 | 0.86 | 0.66 | -0.083 | 0.031 | -0.038 | -0.038 | 0.218 | 0.1 | 0.14 | 0.14 | 0.01 | 0.15 | 0.09 |
| 6 | 21227803 | rs13202305 | -0.417 | 0.926 | 0.346 | -0.042 | -0.003 | 0 | 0 | 0 | 0.09 | 0.06 | 0.14 | 0.01 | 0.15 | 0.09 |
| 6 | 21228888 | rs4710966 | -0.988 | 0.112 | 0.089 | 0.061 | -0.005 | 0.001 | 0.001 | 1.687 | 0.44 | 0.48 | 0.47 | 0.54 | 0.4 | 0.48 |
| 6 | 21231007 | rs9460612 | -1.015 | -0.109 | -0.269 | 0.192 | 0.037 | 0.04 | 0.04 | 0.189 | 0.33 | 0.74 | 0.39 | 0.24 | 0.22 | 0.33 |
| 6 | 21233412 | rs6937811 | 0.37 | 0.283 | 0.025 | 0.425 | -0.004 | 0 | 0 | 0 | 0.76 | 0.72 | 0.79 | 0.77 | 0.78 | 0.68 |
| 6 | 21233667 | rs6915429 | 1.092 | 0.13 | 0.111 | 0.852 | 0.01 | -0.004 | -0.004 | 0 | 0.59 | 0.36 | 0.54 | 0.38 | 0.63 | 0.44 |
| 6 | 21233903 | rs10755611 | 1.042 | 0.172 | 0.536 | 1.086 | -0.014 | 0.093 | 0.093 | 0.147 | 0.81 | 0.49 | 0.85 | 0.69 | 0.89 | 0.78 |
| 6 | 21234034 | rs10946439 | -0.315 | 0.102 | 0.152 | 1.155 | 0.001 | -0.026 | -0.026 | 3.309 | 0.17 | 0.07 | 0.24 | 0.32 | 0.18 | 0.31 |
| 6 | 21234424 | rs10946440 | 1.08 | 0.234 | 0.547 | 0.961 | -0.003 | 0.089 | 0.089 | 0.063 | 0.79 | 0.49 | 0.84 | 0.7 | 0.86 | 0.79 |
| 6 | 21234711 | rs10806933 | 1.135 | 0.245 | 0.541 | 0.966 | -0.006 | 0.086 | 0.086 | 0.045 | 0.79 | 0.5 | 0.85 | 0.7 | 0.86 | 0.77 |
| 6 | 21236547 | rs1459042 | 0.987 | 0.275 | 0.523 | 0.963 | -0.031 | 0.065 | 0.065 | 0.934 | 0.79 | 0.51 | 0.83 | 0.74 | 0.87 | 0.78 |
| 6 | 21237606 | rs12202180 | 1.38 | 0.394 | 0.179 | 0.92 | 0.018 | 0.091 | 0.091 | 0.376 | 0.53 | 0.31 | 0.49 | 0.22 | 0.59 | 0.38 |
| 6 | 21242620 | rs9350336 | 0.741 | 0.16 | 0.312 | 0.778 | -0.017 | 0.01 | 0.01 | 0 | 0.79 | 0.62 | 0.83 | 0.81 | 0.81 | 0.8 |
| 6 | 21245429 | rs6925942 | 0.118 | 0.139 | 0.351 | 0.73 | -0.023 | 0.024 | 0.024 | 0 | 0.86 | 0.68 | 0.88 | 0.84 | 0.91 | 0.84 |
| 6 | 21245969 | rs2045546 | 1.741 | 0.299 | 0.409 | 0.656 | -0.004 | 0.05 | 0.05 | 0.566 | 0.47 | 0.27 | 0.43 | 0.24 | 0.49 | 0.36 |
| 6 | 21250287 | rs1459044 | -0.118 | 0.291 | 0.306 | 0.767 | -0.017 | 0.047 | 0.047 | 0 | 0.86 | 0.69 | 0.78 | 0.83 | 0.91 | 0.84 |
| 6 | 21253160 | rs12209954 | 1.939 | 0.333 | 0.132 | 0.7 | -0.07 | 0.039 | 0.039 | 0.458 | 0.43 | 0.1 | 0.4 | 0.25 | 0.48 | 0.36 |
| 6 | 21254378 | rs12527560 | 0.315 | 0.52 | 0.117 | 0.716 | -0.018 | 0.007 | 0.007 | 0 | 0.06 | 0.04 | 0.09 | 0.09 | 0.13 | 0.05 |
| 6 | 21254597 | rs2125569 | 0.013 | 0.451 | 0.137 | 0.714 | -0.012 | -0.042 | -0.042 | 0 | 0.16 | 0.05 | 0.14 | 0.44 | 0.16 | 0.37 |
| 6 | 21256119 | rs1459046 | -0.33 | 0.464 | 0.202 | 0.755 | -0.008 | -0.037 | -0.037 | 0 | 0.21 | 0.09 | 0.16 | 0.44 | 0.21 | 0.38 |
| 6 | 21256468 | rs12111545 | -0.03 | 0.483 | 0.201 | 0.769 | 0.026 | -0.012 | -0.012 | 0 | 0.07 | 0.14 | 0.1 | 0.09 | 0.13 | 0.05 |
| 6 | 21259328 | rs6935795 | 0.429 | 0.2 | 0.062 | 0.758 | -0.039 | 0.017 | 0.017 | 0.081 | 0.76 | 0.63 | 0.72 | 0.77 | 0.87 | 0.79 |
| 6 | 21260472 | rs7775958 | 1.317 | 0.402 | 0.066 | 0.621 | 0.008 | 0.06 | 0.06 | 0.633 | 0.49 | 0.13 | 0.49 | 0.33 | 0.62 | 0.41 |
| 6 | 21264120 | rs807816 | -0.178 | 0.341 | 0.15 | 0.614 | 0.008 | 0.057 | 0.057 | 0.465 | 0.14 | 0.33 | 0.23 | 0.17 | 0.1 | 0.18 |
| 6 | 21271965 | rs10946444 | 1.485 | 0.037 | -0.712 | 0.504 | -0.079 | 0.053 | 0.053 | 0.395 | 0.44 | 0.1 | 0.43 | 0.23 | 0.54 | 0.37 |
| 6 | 21277763 | rs807835 | 0.851 | 0.34 | -0.354 | 0.781 | -0.002 | -0.008 | -0.008 | 0 | 0.64 | 0.59 | 0.67 | 0.44 | 0.66 | 0.53 |
| 6 | 21281316 | rs809917 | -0.434 | -0.444 | -0.195 | 0.999 | -0.029 | 0.003 | 0.003 | 0 | 0.23 | 0.39 | 0.34 | 0.21 | 0.19 | 0.21 |
| 6 | 21282089 | rs9348464 | -1.014 | -0.026 | -0.674 | 1.036 | 0.004 | -0.096 | -0.096 | 0 | 0.26 | 0.06 | 0.2 | 0.47 | 0.18 | 0.42 |
| 6 | 21286718 | rs9466023 | -0.518 | 0.335 | -0.483 | 1.104 | -0.003 | 0.001 | 0.001 | 0 | 0.07 | 0.06 | 0.11 | 0.24 | 0.11 | 0.25 |
| 6 | 21287305 | rs9358407 | -1.34 | 0.01 | -0.372 | 1.112 | 0.005 | -0.01 | -0.01 | 0.445 | 0.34 | 0.29 | 0.27 | 0.51 | 0.24 | 0.49 |
| 6 | 21287993 | rs717512 | -0.901 | 0.649 | -0.296 | 0.761 | -0.012 | -0.015 | -0.015 | 0 | 0.08 | 0.02 | 0.2 | 0.25 | 0.12 | 0.26 |
| 6 | 21288170 | rs7764474 | -1.152 | 0.602 | -0.2 | 0.743 | 0.006 | 0.032 | 0.032 | 0 | 0.11 | 0.12 | 0.22 | 0.3 | 0.12 | 0.28 |
| 6 | 21291682 | rs807842 | 0.05 | 0.952 | -0.304 | -0.44 | -0.006 | 0.003 | 0.003 | 0 | 0.72 | 0.77 | 0.68 | 0.85 | 0.64 | 0.85 |
| 6 | 21293307 | rs2817308 | -0.134 | 1.041 | -0.324 | -0.458 | -0.002 | -0.002 | -0.002 | 0 | 0.74 | 0.77 | 0.69 | 0.86 | 0.67 | 0.87 |

| | | | | | | | | | | | | | | | | |
|---|---|---|---|---|---|---|---|---|---|---|---|---|---|---|---|---|
| 6 | 21293420 | rs12524269 | -0.051 | 1.179 | -0.359 | -0.638 | -0.004 | 0.003 | 0.003 | 0.199 | 0.09 | 0.04 | 0.11 | 0.09 | 0.13 | 0.05 |
| 6 | 21298076 | rs807846 | 0.432 | 1.05 | -0.13 | -0.739 | -0.001 | -0.009 | -0.009 | 0 | 0.65 | 0.71 | 0.6 | 0.79 | 0.57 | 0.81 |
| 6 | 21298295 | rs807848 | 0.801 | 0.643 | 0.706 | -1.067 | -0.021 | 0.074 | 0.074 | 0 | 0.57 | 0.35 | 0.5 | 0.78 | 0.51 | 0.67 |
| 6 | 21298458 | rs9466027 | -0.696 | 0.502 | 1.064 | -1.064 | 0.01 | -0.01 | -0.01 | 0 | 0.14 | 0.33 | 0.14 | 0.13 | 0.11 | 0.06 |
| 6 | 21299738 | rs807850 | 0.594 | 0.449 | 0.966 | -1.206 | 0.009 | -0.081 | -0.081 | 1.083 | 0.58 | 0.51 | 0.57 | 0.88 | 0.52 | 0.69 |
| 6 | 21301005 | rs9466028 | -0.829 | 0.479 | 0.994 | -1.215 | 0.001 | -0.045 | -0.045 | 0 | 0.14 | 0.3 | 0.14 | 0.04 | 0.14 | 0.03 |
| 6 | 21304468 | rs807853 | 0.062 | 0.456 | 0.917 | -1.232 | 0.019 | -0.119 | -0.119 | 0 | 0.67 | 0.53 | 0.6 | 0.86 | 0.61 | 0.69 |
| 6 | 21308972 | rs807860 | 0.427 | 0.658 | 0.193 | -1.144 | -0.017 | 0.03 | 0.03 | 0 | 0.62 | 0.75 | 0.64 | 0.87 | 0.56 | 0.69 |
| 6 | 21310031 | rs9466030 | -0.169 | 0.77 | 0.149 | -1.014 | -0.014 | 0.007 | 0.007 | 0.185 | 0.71 | 0.8 | 0.69 | 0.91 | 0.63 | 0.72 |
| 6 | 21312967 | rs1535604 | -0.745 | -0.786 | -0.163 | -0.969 | 0.118 | 0.203 | 0.203 | 6.83 | 0.5 | 0.8 | 0.53 | 0.78 | 0.34 | 0.5 |
| 6 | 21313844 | rs719098 | -0.743 | -0.729 | 0.237 | -0.707 | 0.061 | 0.088 | 0.088 | 5.881 | 0.5 | 0.67 | 0.31 | 0.72 | 0.32 | 0.5 |
| 6 | 21315175 | rs9358410 | -0.317 | -1.152 | 0.482 | -0.818 | -0.027 | 0.017 | 0.017 | 0 | 0.23 | 0.37 | 0.21 | 0.66 | 0.16 | 0.38 |
| 6 | 21316633 | rs12528501 | 0.477 | -1.216 | 0.195 | -0.92 | -0.002 | 0.048 | 0.048 | 7.115 | 0.14 | 0.03 | 0.06 | 0.04 | 0.11 | 0.03 |
| 6 | 21317871 | rs9358411 | -0.604 | -1.195 | 0.519 | -0.785 | -0.003 | -0.003 | -0.003 | 0 | 0.36 | 0.39 | 0.27 | 0.66 | 0.27 | 0.39 |
| 6 | 21327621 | rs12192730 | 0.9 | -1.343 | 0.462 | -0.841 | 0.043 | 0.034 | 0.034 | 2.077 | 0.53 | 0.32 | 0.68 | 0.35 | 0.72 | 0.51 |
| 6 | 21330768 | rs9368296 | -0.334 | -1.64 | 0.754 | -0.936 | -0.001 | 0.004 | 0.004 | 0 | 0.2 | 0.24 | 0.23 | 0.64 | 0.21 | 0.38 |
| 6 | 21331329 | rs10946450 | 1.783 | -1.491 | 0.636 | -1.267 | 0.028 | 0.1 | 0.1 | 2.153 | 0.46 | 0.13 | 0.59 | 0.34 | 0.58 | 0.44 |
| 6 | 21331475 | rs10946451 | 1.783 | -1.491 | 0.636 | -1.267 | 0.028 | 0.1 | 0.1 | 2.153 | 0.46 | 0.13 | 0.59 | 0.34 | 0.58 | 0.44 |
| 6 | 21331640 | rs4712606 | 0.028 | -1.533 | 1.173 | -1.333 | -0.032 | -0.068 | -0.068 | 0 | 0.74 | 0.71 | 0.9 | 0.99 | 0.91 | 0.86 |
| 6 | 21332460 | rs6907202 | -0.259 | -1.611 | 1.064 | -1.344 | 0.002 | -0.022 | -0.022 | 0 | 0.77 | 0.83 | 0.91 | 0.99 | 0.91 | 0.86 |
| 6 | 21334986 | rs9348466 | -0.219 | -1.545 | 1.197 | -0.989 | 0.001 | 0.061 | 0.061 | 0 | 0.2 | 0.31 | 0.23 | 0.62 | 0.2 | 0.38 |
| 6 | 21337024 | rs9368297 | 0.237 | -1.566 | 1.231 | -1.029 | -0.01 | -0.071 | -0.071 | 0.06 | 0.72 | 0.7 | 0.85 | 0.99 | 0.8 | 0.83 |
| 6 | 21337783 | rs12203023 | 1.833 | -1.632 | 1.045 | -1.062 | 0.02 | 0.058 | 0.058 | 2.821 | 0.47 | 0.08 | 0.59 | 0.37 | 0.61 | 0.43 |
| 6 | 21338609 | rs9348467 | 0.009 | -1.603 | 1.24 | -0.895 | -0.004 | 0.052 | 0.052 | 0 | 0.17 | 0.27 | 0.19 | 0.62 | 0.16 | 0.39 |
| 6 | 21339460 | rs7744616 | -0.921 | -1.718 | 1.223 | 0.038 | 0.022 | 0.238 | 0.238 | 0.79 | 0.1 | 0.33 | 0.15 | 0.37 | 0.08 | 0.19 |
| 6 | 21340086 | rs7749539 | 0.857 | -1.567 | 0.586 | -0.16 | 0.009 | 0.003 | 0.003 | 0.508 | 0.61 | 0.7 | 0.76 | 0.71 | 0.69 | 0.67 |
| 6 | 21342418 | rs12191693 | 2.1 | -1.699 | 0.494 | -0.261 | -0.024 | 0.077 | 0.077 | 4.083 | 0.45 | 0.12 | 0.57 | 0.34 | 0.59 | 0.48 |
| 6 | 21342509 | rs2064301 | -0.481 | -1.94 | 0.606 | -0.249 | -0.024 | -0.051 | -0.051 | 0.09 | 0.79 | 0.78 | 0.93 | 0.99 | 0.95 | 0.94 |
| 6 | 21347410 | rs1555083 | -0.602 | -1.573 | 0.448 | -0.03 | -0.039 | -0.059 | -0.059 | 0 | 0.19 | 0.06 | 0.17 | 0.28 | 0.27 | 0.26 |
| 6 | 21350628 | rs6911394 | -0.934 | -1.872 | 0.749 | -0.857 | -0.017 | 0.036 | 0.036 | 0 | 0.23 | 0.44 | 0.26 | 0.69 | 0.18 | 0.44 |
| 6 | 21357289 | rs9466040 | -1.552 | -1.838 | 0.723 | -1.043 | 0.005 | -0.071 | -0.071 | 0.597 | 0.13 | 0.36 | 0.09 | 0.01 | 0.11 | 0.03 |
| 6 | 21359292 | rs2179507 | -0.366 | -2.077 | 0.411 | -1.074 | -0.022 | -0.041 | -0.041 | 0.974 | 0.81 | 0.81 | 0.93 | 0.99 | 0.96 | 0.95 |
| 6 | 21368617 | rs6456419 | -0.417 | -1.487 | 0.729 | -0.59 | -0.003 | 0.076 | 0.076 | 0 | 0.18 | 0.36 | 0.24 | 0.63 | 0.16 | 0.45 |
| 6 | 21368678 | rs6456420 | -1.665 | -1.8 | -0.272 | -1.103 | 0.043 | 0.047 | 0.047 | 4.9 | 0.42 | 0.8 | 0.38 | 0.63 | 0.33 | 0.5 |
| 6 | 21369139 | rs17623091 | 0.042 | -1.449 | -0.1 | -0.952 | -0.011 | 0.054 | 0.054 | 5.36 | 0.15 | 0.03 | 0.11 | 0.01 | 0.17 | 0.05 |
| 6 | 21372123 | rs11754000 | -1.455 | -1.433 | 0.373 | -1.107 | -0.01 | -0.078 | -0.078 | 0 | 0.25 | 0.6 | 0.23 | 0.01 | 0.26 | 0.11 |
| 6 | 21372634 | rs2328601 | -0.242 | -0.981 | 0.189 | -1.254 | 0.004 | 0 | 0 | 0.114 | 0.34 | 0.36 | 0.35 | 0.6 | 0.35 | 0.49 |
| 6 | 21375263 | rs9348475 | 0.028 | -0.921 | -0.021 | -1.239 | -0.041 | -0.026 | -0.026 | 0.002 | 0.3 | 0.17 | 0.33 | 0.59 | 0.35 | 0.49 |
| 6 | 21386879 | rs6920208 | -1.025 | -1.005 | 0.113 | -1.684 | 0.017 | 0.05 | 0.05 | 0.185 | 0.52 | 0.69 | 0.55 | 0.62 | 0.6 | 0.59 |
| 6 | 21387232 | rs7757787 | -0.534 | -0.952 | 0.581 | -1.469 | 0.001 | -0.033 | -0.033 | 0.187 | 0.18 | 0.18 | 0.18 | 0.02 | 0.25 | 0.1 |
| 6 | 21390844 | rs6456421 | -1.088 | -0.681 | 0.74 | -1.075 | 0.007 | 0.005 | 0.005 | 1.04 | 0.06 | 0.26 | 0.02 | 0 | 0.01 | 0 |
| 6 | 21392864 | rs9295508 | 0.133 | -0.574 | 0.138 | -1.151 | -0.031 | -0.025 | -0.025 | 0 | 0.31 | 0.21 | 0.37 | 0.56 | 0.34 | 0.53 |
| 6 | 21393645 | rs9350354 | 0.93 | -0.888 | -0.136 | -1.28 | -0.02 | 0.014 | 0.014 | 0 | 0.37 | 0.22 | 0.39 | 0.41 | 0.4 | 0.34 |
| 6 | 21397637 | rs6907305 | 0.14 | -0.793 | -0.371 | -1.47 | 0.007 | -0.005 | -0.005 | 0 | 0.73 | 0.79 | 0.84 | 0.96 | 0.82 | 0.89 |
| 6 | 21398413 | rs9368309 | 0.213 | -0.842 | -0.53 | -1.51 | 0.076 | 0.113 | 0.113 | 0.9 | 0.72 | 0.88 | 0.83 | 0.96 | 0.83 | 0.91 |
| 6 | 21400062 | rs6924841 | 0.117 | -0.599 | -0.486 | -1.281 | -0.031 | -0.009 | -0.009 | 0 | 0.3 | 0.23 | 0.42 | 0.64 | 0.41 | 0.52 |
| 6 | 21402434 | rs1534993 | -0.131 | -0.374 | -0.7 | -1.279 | 0.06 | 0.119 | 0.119 | 1.05 | 0.76 | 0.96 | 0.85 | 0.97 | 0.84 | 0.91 |
| 6 | 21420553 | rs727843 | -0.925 | -0.028 | -0.298 | -0.977 | 0.009 | 0.051 | 0.051 | 0 | 0.54 | 0.31 | 0.56 | 0.68 | 0.59 | 0.61 |
| 6 | 21427143 | rs9368310 | 0.536 | -1.017 | -0.379 | -0.81 | 0.05 | 0.082 | 0.082 | 0 | 0.51 | 0.77 | 0.54 | 0.33 | 0.56 | 0.42 |
| 6 | 21439343 | rs6904446 | -0.868 | -0.904 | -0.013 | -0.661 | -0.002 | -0.01 | -0.01 | 0 | 0.55 | 0.5 | 0.48 | 0.65 | 0.48 | 0.62 |
| 6 | 21451301 | rs9466087 | -0.334 | -0.832 | 0.083 | -0.161 | -0.011 | 0.058 | 0.058 | 0.97 | 0.3 | 0.53 | 0.34 | 0.59 | 0.31 | 0.53 |
| 6 | 21452964 | rs7772439 | -0.25 | -0.75 | -0.703 | -0.134 | -0.003 | 0.037 | 0.037 | 0.623 | 0.28 | 0.38 | 0.32 | 0.59 | 0.31 | 0.53 |
| 6 | 21462277 | rs9466097 | -0.711 | -0.214 | -1.785 | 0.068 | -0.011 | 0.031 | 0.031 | 0.492 | 0.57 | 0.83 | 0.5 | 0.66 | 0.48 | 0.56 |
| 6 | 21468077 | rs9466100 | 0.258 | -0.299 | -2.33 | 0.078 | 0.03 | -0.053 | -0.053 | 0.425 | 0.36 | 0.12 | 0.29 | 0.61 | 0.31 | 0.48 |
| 6 | 21470419 | rs7759649 | -0.531 | 0.014 | -2.25 | 0.136 | -0.005 | 0.046 | 0.046 | 0.914 | 0.55 | 0.91 | 0.48 | 0.68 | 0.43 | 0.56 |
| 6 | 21476990 | rs4283883 | -0.252 | 0.21 | -1.941 | 0.389 | 0.023 | -0.017 | -0.017 | 0.009 | 0.37 | 0.13 | 0.31 | 0.67 | 0.33 | 0.5 |
| 6 | 21482225 | rs6456430 | 0.027 | 0.818 | -2.341 | 0.142 | -0.106 | -0.004 | -0.004 | 0 | 0.79 | 0.98 | 0.62 | 0.86 | 0.67 | 0.82 |
| 6 | 21487753 | rs9356777 | 0.289 | 0.156 | -2.276 | -0.637 | 0.034 | 0.079 | 0.079 | 0.292 | 0.45 | 0.33 | 0.39 | 0.79 | 0.4 | 0.47 |
| 6 | 21489747 | rs4236005 | -0.024 | 0.078 | -1.245 | -0.857 | 0.006 | -0.007 | -0.007 | 0 | 0.49 | 0.54 | 0.55 | 0.9 | 0.42 | 0.51 |
| 6 | 21497929 | rs9358435 | 0.036 | -0.086 | -0.778 | -1.049 | -0.012 | -0.07 | -0.07 | 3.807 | 0.41 | 0.5 | 0.49 | 0.94 | 0.32 | 0.33 |
| 6 | 21498491 | rs4310042 | 0.047 | -0.129 | -0.528 | -1.038 | 0 | -0.02 | -0.02 | 0 | 0.39 | 0.42 | 0.48 | 0.91 | 0.3 | 0.3 |
| 6 | 21502677 | rs13209771 | 1.141 | -0.485 | -0.995 | -0.76 | -0.021 | -0.041 | -0.041 | 0.708 | 0.16 | 0.01 | 0.07 | 0.21 | 0.15 | 0.11 |
| 6 | 21503218 | rs6456431 | -0.646 | -0.492 | -0.323 | -1.239 | 0.018 | 0.024 | 0.024 | 0.927 | 0.43 | 0.64 | 0.51 | 0.94 | 0.33 | 0.34 |
| 6 | 21517321 | rs4712619 | -1.109 | -0.292 | -0.293 | -1.227 | -0.078 | 0.122 | 0.122 | 0 | 0.74 | 0.95 | 0.7 | 0.96 | 0.62 | 0.78 |
| 6 | 21525076 | rs7770014 | -0.937 | -0.87 | 0.415 | 0.061 | -0.014 | -0.001 | -0.001 | 0 | 0.34 | 0.62 | 0.27 | 0.73 | 0.17 | 0.17 |
| 6 | 21526583 | rs6456433 | -0.957 | -0.87 | 0.415 | 0.075 | -0.014 | -0.002 | -0.002 | 0 | 0.34 | 0.62 | 0.27 | 0.75 | 0.17 | 0.17 |
| 6 | 21531793 | rs4129098 | -0.124 | -0.683 | 0.297 | 0.131 | 0.043 | -0.082 | -0.082 | 0 | 0.24 | 0.08 | 0.2 | 0.67 | 0.17 | 0.17 |
| 6 | 21539993 | rs9295512 | -0.046 | -0.573 | 0.294 | 0.269 | -0.02 | -0.142 | -0.142 | 0 | 0.25 | 0.04 | 0.21 | 0.65 | 0.21 | 0.13 |
| 6 | 21540627 | rs12660679 | -0.948 | -0.606 | 0.576 | 0.227 | -0.024 | 0.002 | 0.002 | 5.023 | 0.34 | 0.44 | 0.28 | 0.69 | 0.25 | 0.19 |

| | | | | | | | | | | | | | | | | |
|---|---|---|---|---|---|---|---|---|---|---|---|---|---|---|---|---|
| 6 | 21555608 | rs9350367 | -0.925 | -0.575 | 0.809 | 0.214 | -0.033 | 0.007 | 0.007 | 5.313 | 0.36 | 0.49 | 0.28 | 0.67 | 0.25 | 0.2 |
| 6 | 21573637 | rs9358442 | -0.907 | -0.514 | 0.974 | 0.259 | -0.042 | 0.027 | 0.027 | 5.487 | 0.32 | 0.46 | 0.27 | 0.63 | 0.21 | 0.17 |
| 6 | 21590104 | rs9368326 | -0.262 | -0.416 | 0.607 | 0.302 | -0.015 | -0.158 | -0.158 | 0.788 | 0.25 | 0.03 | 0.22 | 0.6 | 0.2 | 0.16 |
| 6 | 21602552 | rs1620996 | 0.005 | -0.293 | 0.686 | 0.356 | -0.036 | -0.037 | -0.037 | 0 | 0.93 | 0.99 | 0.79 | 0.73 | 0.89 | 0.85 |
| 6 | 21609285 | rs1622043 | 0.666 | -0.537 | 0.948 | -0.809 | 0.011 | 0.05 | 0.05 | 1.662 | 0.66 | 0.79 | 0.57 | 0.13 | 0.7 | 0.66 |
| 6 | 21621850 | rs7751661 | 0.686 | -0.579 | 1.47 | -1.022 | -0.008 | -0.035 | -0.035 | 2.568 | 0.64 | 0.57 | 0.55 | 0.12 | 0.71 | 0.67 |
| 6 | 21623715 | rs1744855 | 0.792 | -0.609 | 1.148 | -1.033 | 0.018 | 0.037 | 0.037 | 0.925 | 0.6 | 0.36 | 0.53 | 0.11 | 0.71 | 0.66 |
| 6 | 21634171 | rs1744867 | 1.124 | 0.265 | 1.9 | -0.081 | 0.012 | 0.074 | 0.074 | 0 | 0.8 | 0.67 | 0.68 | 0.32 | 0.81 | 0.73 |
| 6 | 21658881 | rs941990 | 1.095 | 0.412 | 0.612 | -0.194 | -0.008 | 0.021 | 0.021 | 0 | 0.74 | 0.41 | 0.63 | 0.31 | 0.79 | 0.73 |
| 6 | 21670816 | rs1744885 | 0.414 | 1.24 | -0.053 | 0.048 | 0.022 | -0.014 | -0.014 | 0 | 0.66 | 0.59 | 0.57 | 0.23 | 0.55 | 0.46 |
| 6 | 21690172 | rs2180418 | -0.843 | 1.285 | 0.055 | -0.16 | 0.002 | -0.054 | -0.054 | 0 | 0.17 | 0.24 | 0.12 | 0 | 0.16 | 0.08 |
| 6 | 21691009 | rs6918842 | -1.196 | 1.436 | 0.354 | -0.411 | -0.006 | 0.033 | 0.033 | 0 | 0.46 | 0.71 | 0.71 | 0.81 | 0.53 | 0.53 |
| 6 | 21691704 | rs1555529 | -0.488 | 1.871 | 0.546 | 0.554 | -0.045 | -0.053 | -0.053 | 0 | 0.14 | 0.1 | 0.34 | 0.62 | 0.24 | 0.29 |
| 6 | 21693152 | rs4712624 | -0.971 | 1.095 | 1.334 | 0.514 | 0.017 | 0.088 | 0.088 | 0.083 | 0.07 | 0.23 | 0.22 | 0.2 | 0.11 | 0.15 |
| 6 | 21701841 | rs4712626 | -1.355 | 1.28 | 2.045 | 0.115 | 0.009 | -0.029 | -0.029 | 0 | 0.27 | 0.36 | 0.38 | 0.19 | 0.33 | 0.34 |
| 6 | 21710896 | rs4712631 | -1.585 | 1.192 | 1.894 | 0.12 | 0 | 0.006 | 0.006 | 0 | 0.3 | 0.72 | 0.41 | 0.19 | 0.33 | 0.34 |
| 6 | 21716554 | rs16885170 | -1.227 | 1.241 | 1.612 | 0.328 | 0.008 | 0.02 | 0.02 | 0.506 | 0.06 | 0.08 | 0.25 | 0.19 | 0.15 | 0.22 |
| 6 | 21726770 | rs16885175 | -1.63 | 0.95 | 1.581 | 0.453 | -0.004 | 0.023 | 0.023 | 0.324 | 0.08 | 0.11 | 0.23 | 0.21 | 0.12 | 0.22 |
| 6 | 21727148 | rs7739131 | -1.674 | 1.032 | 1.882 | 1.421 | -0.009 | 0.029 | 0.029 | 0.186 | 0.28 | 0.61 | 0.38 | 0.37 | 0.22 | 0.27 |
| 6 | 21728317 | rs2180419 | -2.048 | 1.037 | 1.885 | 1.435 | -0.003 | 0.006 | 0.006 | 0.529 | 0.34 | 0.62 | 0.61 | 0.56 | 0.33 | 0.5 |
| 6 | 21729682 | rs16885189 | -1.306 | 1.286 | 1.432 | 1.649 | 0.001 | 0.024 | 0.024 | 0.309 | 0.08 | 0.12 | 0.24 | 0.22 | 0.11 | 0.22 |
| 6 | 21731736 | rs9358448 | -0.693 | 1.279 | 1.43 | 1.62 | 0.002 | 0.182 | 0.182 | 0 | 0.05 | 0.09 | 0.23 | 0.36 | 0.05 | 0.15 |
| 6 | 21734750 | rs9466168 | -0.909 | 1.095 | 1.513 | 0.727 | 0.005 | -0.1 | -0.1 | 0 | 0.18 | 0.3 | 0.16 | 0 | 0.17 | 0.11 |
| 6 | 21753645 | rs1322884 | 0.013 | 1.173 | 0.762 | 0.78 | -0.051 | 0.024 | 0.024 | 0.361 | 0.65 | 0.82 | 0.62 | 0.64 | 0.55 | 0.42 |
| 6 | 21766998 | rs9466179 | -0.866 | 0.619 | 1.416 | 0.539 | -0.013 | 0.181 | 0.181 | 2.231 | 0.21 | 0.51 | 0.39 | 0.47 | 0.16 | 0.21 |
| 6 | 21774036 | rs13197267 | -0.167 | 0.81 | 1.52 | 0.48 | -0.001 | 0 | 0 | 0 | 0.63 | 0.67 | 0.6 | 0.64 | 0.56 | 0.39 |
| 6 | 21780789 | rs3789763 | 1.077 | 0.837 | 1.227 | 0.308 | 0.022 | 0.108 | 0.108 | 1.404 | 0.41 | 0.19 | 0.22 | 0.2 | 0.36 | 0.18 |
| 6 | 21781952 | rs7761394 | -0.326 | 0.88 | 1.007 | 0.271 | -0.025 | 0.024 | 0.024 | 0.426 | 0.65 | 0.78 | 0.64 | 0.68 | 0.57 | 0.41 |
| 6 | 21786171 | rs9466182 | -0.494 | 0.929 | 1.305 | 0.139 | 0.004 | -0.058 | -0.058 | 0 | 0.13 | 0.25 | 0.13 | 0 | 0.17 | 0.06 |
| 6 | 21796241 | rs6915524 | -0.213 | 0.943 | 1.019 | 0.392 | -0.02 | 0.045 | 0.045 | 0.894 | 0.63 | 0.78 | 0.67 | 0.74 | 0.59 | 0.35 |
| 6 | 21805714 | rs6916032 | -1.028 | 0.854 | 1.067 | 0.886 | 0.002 | 0.151 | 0.151 | 0.17 | 0.16 | 0.36 | 0.34 | 0.46 | 0.18 | 0.14 |
| 6 | 21813153 | rs16901621 | -0.159 | 0.646 | -0.081 | 0.339 | 0.004 | -0.036 | -0.036 | 0 | 0.15 | 0.05 | 0.34 | 0.74 | 0.14 | 0.25 |
| 6 | 21813634 | rs12213384 | 1.529 | -0.047 | -0.808 | -0.603 | -0.008 | 0.127 | 0.127 | 2.957 | 0.21 | 0.03 | 0.15 | 0.01 | 0.17 | 0.1 |
| 6 | 21814497 | rs17198823 | 0.768 | 0.567 | -1.099 | -1.07 | -0.022 | 0.016 | 0.016 | 0 | 0.1 | 0.03 | 0.06 | 0.06 | 0.16 | 0.07 |
| 6 | 21815916 | rs4712634 | 0.73 | 0.495 | -0.467 | -1.645 | 0.004 | -0.017 | -0.017 | 0 | 0.39 | 0.32 | 0.35 | 0.01 | 0.38 | 0.18 |
| 6 | 21816316 | rs17198949 | 1.845 | 0.498 | -0.711 | -1.597 | -0.022 | 0.162 | 0.162 | 2.709 | 0.26 | 0.03 | 0.19 | 0.01 | 0.21 | 0.1 |
| 6 | 21818196 | rs2181359 | -0.887 | 0.454 | -0.066 | -1.63 | -0.007 | 0.017 | 0.017 | 0 | 0.7 | 0.76 | 0.83 | 0.88 | 0.69 | 0.55 |
| 6 | 21824770 | rs11961985 | 0.023 | 0.46 | 0.761 | -1.338 | 0.003 | -0.039 | -0.039 | 0 | 0.11 | 0.19 | 0.16 | 0 | 0.15 | 0.07 |
| 6 | 21827104 | rs912026 | 0.737 | 0.82 | 0.176 | -1.273 | -0.002 | -0.007 | -0.007 | 0 | 0.64 | 0.45 | 0.71 | 0.16 | 0.7 | 0.31 |
| 6 | 21827710 | rs4710988 | -0.176 | 0.97 | -0.015 | -1.12 | 0.002 | -0.033 | -0.033 | 0 | 0.82 | 0.77 | 0.89 | 0.89 | 0.86 | 0.61 |
| 6 | 21831292 | rs10946475 | 0.154 | 1.096 | -0.021 | -1.07 | -0.001 | -0.019 | -0.019 | 0 | 0.12 | 0.11 | 0.16 | 0 | 0.18 | 0.08 |
| 6 | 21832546 | rs16885318 | 0.986 | 1.165 | -0.36 | -0.971 | -0.008 | -0.017 | -0.017 | 0 | 0.09 | 0.05 | 0.24 | 0.14 | 0.13 | 0.06 |
| 6 | 21834059 | rs966812 | 0.802 | 1.095 | -0.318 | -0.854 | -0.002 | -0.057 | -0.057 | 1.068 | 0.14 | 0.05 | 0.15 | 0.64 | 0.15 | 0.26 |
| 6 | 21835652 | rs12208206 | 0.102 | 1.143 | -0.242 | -0.996 | 0.026 | -0.028 | -0.028 | 0 | 0.91 | 0.81 | 0.95 | 0.97 | 0.94 | 0.92 |
| 6 | 21844709 | rs11962022 | -0.81 | 1.006 | -0.334 | -0.992 | 0.007 | -0.093 | -0.093 | 0 | 0.29 | 0.44 | 0.27 | 0.11 | 0.31 | 0.43 |
| 6 | 21847532 | rs2094571 | -0.871 | 1.022 | -0.57 | -0.848 | 0 | -0.007 | -0.007 | 0.044 | 0.44 | 0.48 | 0.43 | 0.75 | 0.47 | 0.69 |
| 6 | 21847916 | rs1410524 | 0.418 | 0.983 | -0.664 | -0.056 | -0.032 | -0.053 | -0.053 | 0.399 | 0.09 | 0.01 | 0.24 | 0.14 | 0.13 | 0.06 |
| 6 | 21856821 | rs9368359 | 0.495 | 1.06 | -0.376 | -0.942 | 0.013 | -0.111 | -0.111 | 0.689 | 0.66 | 0.54 | 0.72 | 0.96 | 0.61 | 0.78 |
| 6 | 21874359 | rs2182997 | 1.127 | 0.74 | -0.588 | -0.631 | 0.006 | 0.018 | 0.018 | 0.7 | 0.47 | 0.3 | 0.54 | 0.34 | 0.46 | 0.51 |
| 6 | 21883677 | rs4712642 | 0.139 | 0.536 | -0.317 | -0.835 | 0.005 | 0.091 | 0.091 | 0 | 0.87 | 0.64 | 0.73 | 0.82 | 0.89 | 0.93 |
| 6 | 21884440 | rs952579 | 0.21 | 0.956 | -0.628 | -1.095 | 0.002 | -0.01 | -0.01 | 0 | 0.08 | 0.09 | 0.11 | 0 | 0.17 | 0.06 |
| 6 | 21884519 | rs952578 | 1.747 | 0.299 | -0.78 | -0.939 | 0.002 | 0.129 | 0.129 | 3.281 | 0.37 | 0.1 | 0.29 | 0.14 | 0.35 | 0.44 |
| 6 | 21887276 | rs883173 | 1.293 | 0.484 | -0.546 | 0.604 | -0.045 | -0.049 | -0.049 | 0 | 0.08 | 0.01 | 0.11 | 0.2 | 0.12 | 0.08 |
| 6 | 21891605 | rs12201134 | -1.146 | 0.292 | -0.379 | -0.246 | 0 | -0.01 | -0.01 | 0 | 0.24 | 0.28 | 0.39 | 0.84 | 0.25 | 0.31 |
| 6 | 21894448 | rs12210225 | 0.353 | 0.335 | -0.583 | -0.131 | -0.009 | -0.01 | -0.01 | 0 | 0.11 | 0.06 | 0.14 | 0.65 | 0.14 | 0.25 |
| 6 | 21903533 | rs7760611 | -0.799 | -0.038 | -0.56 | -0.783 | -0.006 | 0.017 | 0.017 | 0.88 | 0.39 | 0.69 | 0.39 | 0.17 | 0.51 | 0.37 |
| 6 | 21905726 | rs9350395 | -1.321 | -0.273 | -0.846 | -0.821 | 0.051 | 0.068 | 0.068 | 0.268 | 0.49 | 0.74 | 0.48 | 0.73 | 0.57 | 0.59 |
| 6 | 21908093 | rs1111737 | -1.386 | -0.447 | -0.079 | -0.585 | -0.032 | 0.12 | 0.12 | 0 | 0.16 | 0.26 | 0.12 | 0.56 | 0.07 | 0.27 |
| 6 | 21911129 | rs10946483 | 1.037 | -0.704 | 0.098 | -0.87 | -0.039 | 0 | 0 | 0 | 0.79 | 0.64 | 0.67 | 0.27 | 0.89 | 0.71 |
| 6 | 21911616 | rs7745461 | 1.17 | -0.722 | 0.162 | -0.89 | -0.035 | -0.005 | -0.005 | 0.08 | 0.78 | 0.65 | 0.65 | 0.26 | 0.88 | 0.7 |
| 6 | 21911852 | rs6456457 | -1.795 | -0.654 | 0.044 | -1.144 | -0.015 | -0.049 | -0.049 | 0.989 | 0.34 | 0.39 | 0.44 | 0.89 | 0.24 | 0.45 |
| 6 | 21912632 | rs9368368 | -1.585 | -0.797 | -0.123 | -1.376 | -0.004 | -0.01 | -0.01 | 4.575 | 0.31 | 0.32 | 0.42 | 0.82 | 0.2 | 0.46 |
| 6 | 21913741 | rs9348495 | -0.654 | -0.598 | -0.039 | -1.273 | 0.012 | 0.012 | 0.012 | 0 | 0.21 | 0.18 | 0.36 | 0.81 | 0.16 | 0.45 |
| 6 | 21921026 | rs6934973 | -1.239 | -0.458 | 0.027 | -1.129 | -0.005 | 0.004 | 0.004 | 0 | 0.34 | 0.32 | 0.42 | 0.45 | 0.29 | 0.34 |
| 6 | 21922708 | rs12213632 | -0.716 | -0.545 | -0.237 | -1.667 | 0.027 | 0.022 | 0.022 | 0 | 0.66 | 0.52 | 0.66 | 0.45 | 0.61 | 0.48 |
| 6 | 21923512 | rs9358465 | 0.105 | -0.615 | -0.118 | -1.578 | -0.006 | 0.061 | 0.061 | 0 | 0.8 | 0.68 | 0.82 | 0.32 | 0.85 | 0.61 |
| 6 | 21925491 | rs9283872 | 0.512 | -0.471 | -0.339 | -1.527 | 0.039 | 0.049 | 0.049 | 0 | 0.46 | 0.22 | 0.52 | 0.29 | 0.45 | 0.33 |
| 6 | 21929886 | rs12207699 | 0.738 | -0.609 | -0.492 | -1.495 | 0.004 | 0.084 | 0.084 | 0 | 0.42 | 0.31 | 0.32 | 0.17 | 0.44 | 0.27 |
| 6 | 21930988 | rs2039109 | -0.134 | -0.092 | -0.562 | -1.449 | -0.007 | -0.026 | -0.026 | 0 | 0.2 | 0.11 | 0.35 | 0.29 | 0.24 | 0.22 |

| | | | | | | | | | | | | | | | | |
|---|---|---|---|---|---|---|---|---|---|---|---|---|---|---|---|---|
| 6 | 21933134 | rs9358470 | 0.489 | -0.002 | -0.63 | -1.501 | 0 | -0.002 | -0.002 | 0 | 0.15 | 0.18 | 0.11 | 0.16 | 0.14 | 0.14 |
| 6 | 21934671 | rs7745964 | 0.492 | -0.013 | -0.645 | -1.506 | -0.001 | 0 | 0 | 0 | 0.15 | 0.16 | 0.11 | 0.16 | 0.14 | 0.14 |
| 6 | 21936868 | rs12210951 | 1.561 | 0.011 | -0.782 | -1.498 | 0.008 | 0.082 | 0.082 | 0 | 0.28 | 0.14 | 0.21 | 0.01 | 0.3 | 0.13 |
| 6 | 21943942 | rs12524124 | 0.252 | 0.066 | -0.382 | -0.93 | 0.004 | 0.013 | 0.013 | 0 | 0.36 | 0.45 | 0.45 | 0.55 | 0.33 | 0.51 |
| 6 | 21947118 | rs913568 | -0.051 | 0.421 | -0.433 | -1.141 | 0.001 | 0.01 | 0.01 | 0 | 0.5 | 0.46 | 0.46 | 0.24 | 0.5 | 0.38 |
| 6 | 21950971 | rs6927510 | 0.427 | 0.246 | -0.725 | -1.53 | -0.009 | 0.07 | 0.07 | 0 | 0.4 | 0.25 | 0.39 | 0.05 | 0.43 | 0.34 |
| 6 | 21955964 | rs11751362 | 0.813 | 0.542 | -0.58 | -1.414 | -0.005 | 0.015 | 0.015 | 0 | 0.26 | 0.19 | 0.14 | 0.02 | 0.25 | 0.25 |
| 6 | 21956404 | rs6903716 | -0.653 | 0.823 | -0.479 | -1.006 | 0.018 | 0.014 | 0.014 | 0.153 | 0.23 | 0.33 | 0.22 | 0.23 | 0.29 | 0.29 |
| 6 | 21967698 | rs868949 | 0.52 | 0.336 | -1.104 | -0.901 | 0.043 | -0.04 | -0.04 | 0.517 | 0.4 | 0.12 | 0.49 | 0.54 | 0.33 | 0.36 |
| 6 | 21968899 | rs1928176 | -0.703 | 0.347 | -1.138 | -1.447 | -0.021 | 0.121 | 0.121 | 1.378 | 0.4 | 0.26 | 0.62 | 0.97 | 0.47 | 0.65 |
| 6 | 21969495 | rs13212670 | -0.383 | 0.064 | -0.256 | -0.713 | -0.009 | -0.002 | -0.002 | 0 | 0.55 | 0.53 | 0.47 | 0.49 | 0.64 | 0.62 |
| 6 | 21972800 | rs11755535 | -0.053 | 0.436 | -0.122 | -1.36 | -0.004 | 0.002 | 0.002 | 0 | 0.34 | 0.33 | 0.28 | 0.16 | 0.38 | 0.46 |
| 6 | 21982253 | rs6922362 | -0.712 | -0.141 | -0.239 | -2.03 | -0.01 | 0.01 | 0.01 | 0 | 0.38 | 0.63 | 0.35 | 0.14 | 0.45 | 0.52 |
| 6 | 21986847 | rs6909060 | -0.011 | -0.021 | -0.345 | -1.801 | -0.073 | 0.057 | 0.057 | 2.626 | 0.21 | 0.1 | 0.21 | 0.09 | 0.32 | 0.45 |
| 6 | 21986895 | rs2026708 | 0.575 | -1.049 | -0.188 | -1.711 | 0.019 | 0.028 | 0.028 | 0 | 0.14 | 0.11 | 0.2 | 0 | 0.13 | 0.03 |
| 6 | 21994744 | rs9350404 | -0.037 | -0.433 | -0.34 | -1.561 | -0.103 | 0.132 | 0.132 | 0.015 | 0.37 | 0.1 | 0.36 | 0.18 | 0.5 | 0.53 |
| 6 | 21996130 | rs1074871 | 0.335 | -0.678 | 0.042 | -0.755 | -0.007 | 0.142 | 0.142 | 0 | 0.1 | 0.17 | 0.27 | 0.47 | 0.08 | 0.2 |
| 6 | 21996860 | rs2078543 | -1.185 | -1.179 | 0.358 | -1.106 | -0.002 | -0.084 | -0.084 | 0.35 | 0.66 | 0.66 | 0.75 | 0.98 | 0.66 | 0.84 |
| 6 | 21998490 | rs4712648 | 0.305 | -0.901 | 0.458 | -0.691 | -0.008 | 0.123 | 0.123 | 0 | 0.1 | 0.19 | 0.28 | 0.53 | 0.08 | 0.2 |
| 6 | 22002897 | rs9366420 | 0.683 | -1.029 | 0.431 | -0.76 | -0.05 | 0.055 | 0.055 | 0 | 0.17 | 0.28 | 0.11 | 0.31 | 0.06 | 0.11 |
| 6 | 22012566 | rs9358481 | -0.587 | -1.067 | 0.283 | -0.904 | -0.004 | 0 | 0 | 0 | 0.3 | 0.67 | 0.39 | 0.78 | 0.15 | 0.3 |
| 6 | 22018581 | rs9356791 | -0.55 | -0.827 | -0.097 | -0.852 | -0.002 | -0.005 | -0.005 | 0 | 0.42 | 0.35 | 0.43 | 0.48 | 0.51 | 0.63 |
| 6 | 22021230 | rs9350407 | 0.604 | -0.912 | 0.128 | -0.897 | -0.046 | 0.051 | 0.051 | 1.206 | 0.18 | 0.27 | 0.1 | 0.32 | 0.07 | 0.1 |
| 6 | 22021373 | rs9350408 | -0.731 | -0.849 | -0.162 | -0.887 | -0.001 | -0.001 | -0.001 | 0 | 0.45 | 0.47 | 0.44 | 0.48 | 0.51 | 0.61 |
| 6 | 22029017 | rs7744710 | 0.409 | -0.076 | -0.41 | -1.257 | -0.015 | 0.019 | 0.019 | 0.896 | 0.16 | 0.14 | 0.17 | 0.08 | 0.23 | 0.22 |
| 6 | 22031444 | rs11963286 | 0.923 | -0.137 | -0.494 | -1.235 | 0.006 | -0.008 | -0.008 | 3.761 | 0.16 | 0.13 | 0.11 | 0.29 | 0.08 | 0.1 |
| 6 | 22033121 | rs9368395 | 1.722 | -0.156 | -1.433 | -1.632 | 0.003 | 0.077 | 0.077 | 0.196 | 0.23 | 0.07 | 0.2 | 0.03 | 0.24 | 0.04 |
| 6 | 22033222 | rs11750961 | -0.164 | -0.213 | -1.36 | -1.444 | -0.004 | -0.009 | -0.009 | 0.057 | 0.29 | 0.17 | 0.26 | 0.32 | 0.3 | 0.31 |
| 6 | 22034440 | rs867863 | 0.46 | -0.014 | -1.374 | -1.281 | 0.029 | 0.034 | 0.034 | 0 | 0.4 | 0.16 | 0.29 | 0.32 | 0.36 | 0.32 |
| 6 | 22034957 | rs17261417 | 0.157 | 0.237 | -0.574 | -0.553 | 0 | 0 | 0 | 0 | 0.49 | 0.5 | 0.36 | 0.31 | 0.39 | 0.24 |
| 6 | 22036867 | rs1928174 | -0.176 | 0.969 | -1.065 | 0.106 | -0.085 | -0.046 | -0.046 | 0 | 0.3 | 0.17 | 0.31 | 0.39 | 0.4 | 0.53 |
| 6 | 22037128 | rs9356793 | -0.471 | 1.156 | -0.404 | -0.136 | 0.003 | 0.039 | 0.039 | 0 | 0.34 | 0.2 | 0.26 | 0.12 | 0.35 | 0.18 |
| 6 | 22038033 | rs9358484 | -0.041 | 0.895 | -0.537 | 0.13 | 0.004 | 0.014 | 0.014 | 0 | 0.25 | 0.12 | 0.2 | 0.17 | 0.27 | 0.09 |
| 6 | 22038380 | rs400211 | 0.158 | 1.343 | -0.412 | 0.155 | -0.039 | 0.116 | 0.116 | 0 | 0.57 | 0.83 | 0.63 | 0.8 | 0.5 | 0.56 |
| 6 | 22038526 | rs1207774 | -1.473 | 1.179 | 1.171 | 1.293 | 0.046 | 0.121 | 0.121 | 1.352 | 0.11 | 0.43 | 0.13 | 0.16 | 0.1 | 0.03 |
| 6 | 22039241 | rs1952452 | -1.288 | 1.168 | 0.767 | 1.615 | 0.012 | 0.112 | 0.112 | 0 | 0.07 | 0.13 | 0.19 | 0.19 | 0.08 | 0.14 |
| 6 | 22041133 | rs2328662 | -0.854 | 1.461 | 0.618 | 0.959 | 0.001 | -0.007 | -0.007 | 0 | 0.13 | 0.15 | 0.13 | 0.06 | 0.15 | 0.07 |
| 6 | 22042807 | rs1928172 | -1.312 | 1.033 | 1.016 | 1.499 | 0.022 | 0.095 | 0.095 | 0.202 | 0.07 | 0.17 | 0.23 | 0.19 | 0.09 | 0.15 |
| 6 | 22043393 | rs196032 | -1.822 | 0.867 | 1.203 | 1.514 | 0.003 | 0.115 | 0.115 | 0 | 0.2 | 0.43 | 0.42 | 0.33 | 0.21 | 0.31 |
| 6 | 22045644 | rs993600 | -1.539 | 1.038 | 0.733 | 1.215 | 0.011 | 0.032 | 0.032 | 0.016 | 0.13 | 0.17 | 0.2 | 0.21 | 0.16 | 0.23 |
| 6 | 22059967 | rs1207782 | 0.368 | 0.499 | 0.921 | 0.048 | -0.001 | -0.009 | -0.009 | 0.264 | 0.61 | 0.66 | 0.41 | 0.19 | 0.6 | 0.41 |
| 6 | 22060799 | rs492044 | 0.23 | 0.371 | 0.882 | 0.626 | 0 | -0.02 | -0.02 | 0 | 0.63 | 0.66 | 0.59 | 0.84 | 0.62 | 0.54 |
| 6 | 22064639 | rs9366426 | 0.623 | 0.239 | 1.391 | 0.32 | 0.015 | 0.12 | 0.12 | 0 | 0.54 | 0.28 | 0.35 | 0.15 | 0.56 | 0.41 |
| 6 | 22068174 | rs4145443 | 0.773 | 0.275 | 1.358 | 0.304 | 0.013 | 0.141 | 0.141 | 0 | 0.52 | 0.27 | 0.34 | 0.14 | 0.54 | 0.38 |
| 6 | 22069970 | rs196038 | -0.327 | 0.202 | 1.293 | 0.134 | -0.071 | 0.129 | 0.129 | 0 | 0.7 | 0.9 | 0.71 | 0.89 | 0.63 | 0.75 |
| 6 | 22070076 | rs9295533 | -0.603 | 0.186 | 1.682 | 0.245 | 0.018 | 0.043 | 0.043 | 0 | 0.07 | 0.24 | 0.13 | 0.05 | 0.03 | 0.2 |
| 6 | 22075262 | rs10484389 | -1.188 | 0.011 | 1.608 | 0.11 | -0.013 | -0.018 | -0.018 | 0.005 | 0.07 | 0.18 | 0.03 | 0.02 | 0.02 | 0.17 |
| 6 | 22077277 | rs1207778 | -0.225 | -0.389 | 1.243 | -0.127 | -0.048 | 0.119 | 0.119 | 1.432 | 0.67 | 0.9 | 0.68 | 0.86 | 0.64 | 0.73 |
| 6 | 22099060 | rs9466264 | -1.04 | -0.798 | 1.463 | 0.007 | -0.014 | -0.015 | -0.015 | 0 | 0.06 | 0.12 | 0.03 | 0.03 | 0.02 | 0.18 |
| 6 | 22100075 | rs1207775 | -0.14 | -0.471 | 1.021 | -0.466 | -0.074 | 0.139 | 0.139 | 1.619 | 0.61 | 0.85 | 0.63 | 0.86 | 0.55 | 0.72 |
| 6 | 22100367 | rs10946507 | 0.842 | -0.314 | 1.137 | -0.474 | 0.001 | 0.016 | 0.016 | 0.44 | 0.44 | 0.35 | 0.31 | 0.14 | 0.46 | 0.38 |
| 6 | 22112043 | rs9393228 | -0.119 | -0.917 | 0.488 | -0.154 | 0.049 | 0.103 | 0.103 | 1.266 | 0.47 | 0.8 | 0.57 | 0.65 | 0.41 | 0.44 |
| 6 | 22115130 | rs11759745 | 0.563 | -1.323 | 0.686 | -0.476 | 0.007 | -0.073 | -0.073 | 0 | 0.29 | 0.48 | 0.32 | 0.13 | 0.33 | 0.22 |
| 6 | 22115940 | rs10484386 | -0.242 | -1.371 | 0.675 | -0.508 | -0.039 | -0.062 | -0.062 | 0 | 0.13 | 0.22 | 0.04 | 0 | 0.04 | 0.07 |
| 6 | 22116151 | rs9283873 | 0.158 | -1.24 | 0.451 | -0.494 | -0.007 | -0.081 | -0.081 | 0 | 0.33 | 0.48 | 0.33 | 0.13 | 0.35 | 0.29 |
| 6 | 22122568 | rs7764461 | -0.066 | -0.922 | -0.225 | 0.079 | 0.022 | 0.054 | 0.054 | 0 | 0.46 | 0.67 | 0.42 | 0.31 | 0.39 | 0.55 |
| 6 | 22123695 | rs9393231 | -0.409 | -1.074 | -0.393 | -0.112 | -0.007 | 0.028 | 0.028 | 0 | 0.56 | 0.65 | 0.44 | 0.2 | 0.52 | 0.51 |
| 6 | 22125964 | rs4712653 | -0.657 | -1.4 | -0.472 | 0.535 | -0.009 | 0.134 | 0.134 | 0.014 | 0.52 | 0.9 | 0.66 | 0.73 | 0.46 | 0.67 |
| 6 | 22128438 | rs6922033 | -0.432 | -1.447 | -0.466 | 0.289 | 0.013 | 0.138 | 0.138 | 0.044 | 0.49 | 0.9 | 0.65 | 0.73 | 0.44 | 0.61 |
| 6 | 22131929 | rs9295536 | -0.408 | -1.434 | -0.46 | 0.296 | 0.023 | 0.135 | 0.135 | 0.015 | 0.48 | 0.9 | 0.65 | 0.73 | 0.43 | 0.6 |
| 6 | 22136160 | rs6915322 | -0.659 | -1.178 | -0.489 | 0.598 | -0.071 | 0.012 | 0.012 | 1.506 | 0.53 | 0.98 | 0.66 | 0.66 | 0.45 | 0.66 |
| 6 | 22136409 | rs9466269 | 0.215 | -1.143 | -0.211 | 0.328 | -0.008 | -0.031 | -0.031 | 0 | 0.41 | 0.55 | 0.38 | 0.2 | 0.38 | 0.48 |
| 6 | 22139729 | rs6934891 | 0.102 | -0.987 | -0.278 | 0.426 | 0.008 | 0.019 | 0.019 | 0 | 0.46 | 0.63 | 0.46 | 0.34 | 0.43 | 0.58 |
| 6 | 22140004 | rs6939340 | -0.458 | -0.876 | -1.024 | -0.362 | -0.032 | 0.079 | 0.079 | 0.27 | 0.53 | 0.9 | 0.66 | 0.68 | 0.48 | 0.66 |
| 6 | 22140441 | rs7774801 | -0.575 | -0.799 | -1.762 | -0.27 | 0.013 | -0.028 | -0.028 | 0 | 0.84 | 0.99 | 0.88 | 0.86 | 0.88 | 0.86 |
| 6 | 22141596 | rs9358492 | -0.48 | -0.356 | -1.019 | -0.004 | 0.023 | 0.077 | 0.077 | 0 | 0.48 | 0.74 | 0.63 | 0.68 | 0.45 | 0.66 |
| 6 | 22142054 | rs9466272 | -0.488 | -0.35 | -1.015 | -0.106 | 0.021 | 0.053 | 0.053 | 0 | 0.48 | 0.73 | 0.63 | 0.65 | 0.45 | 0.66 |
| 6 | 22150358 | rs10498705 | -0.612 | 0.196 | -0.579 | -0.366 | -0.018 | 0.02 | 0.02 | 0.673 | 0.08 | 0.03 | 0.06 | 0.01 | 0.14 | 0.09 |
| 6 | 22153416 | rs11751377 | -0.994 | -0.608 | -0.257 | -0.089 | -0.033 | -0.009 | -0.009 | 0 | 0.16 | 0.24 | 0.18 | 0.19 | 0.09 | 0.2 |

| | | | | | | | | | | | | | | | | |
|---|---|---|---|---|---|---|---|---|---|---|---|---|---|---|---|---|
| 6 | 22153591 | rs9393240 | -0.787 | -0.466 | -0.514 | -0.026 | -0.005 | -0.001 | -0.001 | 0 | 0.15 | 0.17 | 0.17 | 0.19 | 0.09 | 0.21 |
| 6 | 22155441 | rs16885715 | -0.843 | -0.495 | -0.682 | -0.658 | 0.011 | 0.004 | 0.004 | 0 | 0.23 | 0.17 | 0.13 | 0.02 | 0.17 | 0.22 |
| 6 | 22155622 | rs4571555 | -0.828 | -0.164 | -1.042 | -1.784 | -0.014 | 0.024 | 0.024 | 0 | 0.67 | 0.72 | 0.75 | 0.98 | 0.6 | 0.79 |
| 6 | 22155662 | rs4583970 | 0.991 | -1.217 | 0.148 | -1.276 | 0.013 | 0.011 | 0.011 | 0.227 | 0.64 | 0.34 | 0.78 | 0.87 | 0.81 | 0.65 |
| 6 | 22160802 | rs4289633 | -1.62 | -0.897 | 0.97 | -0.167 | 0.028 | 0.216 | 0.216 | 0 | 0.14 | 0.35 | 0.28 | 0.57 | 0.12 | 0.2 |
| 6 | 22161904 | rs9358493 | 1.652 | -1.141 | 0.385 | -0.327 | 0.036 | -0.035 | -0.035 | 1.067 | 0.6 | 0.15 | 0.55 | 0.42 | 0.73 | 0.55 |
| 6 | 22162129 | rs9358494 | 1.461 | -1.09 | 0.682 | -0.299 | 0.04 | -0.003 | -0.003 | 0 | 0.57 | 0.27 | 0.56 | 0.42 | 0.71 | 0.53 |
| 6 | 22166552 | rs9393243 | 1.215 | -1.044 | 0.842 | -0.233 | -0.003 | -0.01 | -0.01 | 0.465 | 0.65 | 0.34 | 0.77 | 0.84 | 0.79 | 0.63 |
| 6 | 22182491 | rs6940059 | 0.742 | -1.053 | 0.925 | -0.235 | -0.038 | -0.113 | -0.113 | 0.218 | 0.69 | 0.46 | 0.79 | 0.97 | 0.82 | 0.68 |
| 6 | 22188924 | rs7451606 | 0.736 | -1.038 | 0.75 | -0.267 | -0.042 | -0.121 | -0.121 | 0 | 0.69 | 0.48 | 0.79 | 0.97 | 0.82 | 0.67 |
| 6 | 22190338 | rs4712661 | -0.405 | -0.997 | 0.067 | -0.038 | -0.024 | -0.012 | -0.012 | 0.193 | 0.08 | 0.02 | 0.2 | 0.41 | 0.08 | 0.07 |
| 6 | 22194163 | rs9358509 | 1.474 | -0.974 | 0.034 | -0.175 | 0.066 | 0.006 | 0.006 | 1.739 | 0.55 | 0.13 | 0.54 | 0.38 | 0.71 | 0.53 |
| 6 | 22194664 | rs1129570 | -0.258 | -1.07 | 0.446 | -0.32 | 0.002 | -0.014 | -0.014 | 0.943 | 0.21 | 0.18 | 0.14 | 0.01 | 0.17 | 0.25 |
| 6 | 22195500 | rs7739417 | 1.238 | -0.885 | 0.778 | 0.413 | -0.048 | 0.26 | 0.26 | 0 | 0.82 | 0.56 | 0.72 | 0.52 | 0.9 | 0.85 |
| 6 | 22202275 | rs1539032 | -0.099 | -0.594 | 0.041 | -0.8 | -0.001 | -0.036 | -0.036 | 0.079 | 0.24 | 0.22 | 0.14 | 0.01 | 0.23 | 0.2 |
| 6 | 22202679 | rs7749084 | -0.229 | -1.047 | -0.144 | -1.348 | 0.001 | -0.01 | -0.01 | 0 | 0.16 | 0.14 | 0.16 | 0.04 | 0.14 | 0.13 |
| 6 | 22202725 | rs7767024 | -0.42 | -1.354 | -0.22 | -1.098 | -0.024 | 0.007 | 0.007 | 0 | 0.22 | 0.33 | 0.33 | 0.76 | 0.16 | 0.4 |
| 6 | 22207220 | rs16901634 | -0.085 | -1.488 | -0.871 | -0.063 | 0.022 | -0.042 | -0.042 | 0 | 0.11 | 0.06 | 0.18 | 0.27 | 0.06 | 0.25 |
| 6 | 22207494 | rs17262993 | 1.458 | -1.201 | -0.839 | -0.383 | -0.009 | 0.008 | 0.008 | 0 | 0.3 | 0.24 | 0.35 | 0.23 | 0.37 | 0.27 |
| 6 | 22207573 | rs17263042 | -0.054 | -0.875 | -0.769 | 0.219 | -0.016 | -0.005 | -0.005 | 0 | 0.08 | 0.05 | 0.12 | 0.06 | 0.13 | 0.07 |
| 6 | 22208508 | rs17263098 | 0.63 | -1.667 | -0.806 | 0.242 | -0.01 | 0.13 | 0.13 | 0.143 | 0.22 | 0.03 | 0.11 | 0.01 | 0.19 | 0.2 |
| 6 | 22209610 | rs13198653 | 0.644 | -1.655 | -0.816 | 0.248 | -0.014 | 0.133 | 0.133 | 0.116 | 0.22 | 0.02 | 0.11 | 0.01 | 0.19 | 0.2 |
| 6 | 22220860 | rs9393266 | -1.413 | -1.209 | 0.268 | 0.836 | -0.043 | 0.233 | 0.233 | 0.18 | 0.19 | 0.52 | 0.21 | 0.5 | 0.11 | 0.18 |
| 6 | 22222129 | rs1891086 | 0.31 | -1.439 | -0.585 | 0.168 | 0.016 | 0.015 | 0.015 | 0 | 0.59 | 0.65 | 0.59 | 0.71 | 0.75 | 0.55 |
| 6 | 22224378 | rs6937402 | -0.121 | -1.066 | -0.936 | 0.203 | 0 | -0.012 | -0.012 | 0.414 | 0.07 | 0.05 | 0.02 | 0 | 0.06 | 0.04 |
| 6 | 22231659 | rs12194364 | -0.043 | -0.819 | -1.188 | 0.063 | -0.039 | 0.035 | 0.035 | 0 | 0.09 | 0.01 | 0.09 | 0.01 | 0.16 | 0.08 |
| 6 | 22236978 | rs17204453 | 1.241 | -1.13 | -1.699 | 0.553 | -0.128 | -0.002 | -0.002 | 0.24 | 0.37 | 0.06 | 0.34 | 0.32 | 0.5 | 0.3 |
| 6 | 22238088 | rs6924146 | -0.441 | -1.395 | -1.364 | 0.796 | 0.004 | 0.006 | 0.006 | 0 | 0.17 | 0.15 | 0.11 | 0.04 | 0.15 | 0.19 |
| 6 | 22248775 | rs13196524 | -0.126 | -1.117 | -1.302 | 0.57 | 0.004 | -0.006 | -0.006 | 0 | 0.14 | 0.12 | 0.31 | 0.27 | 0.07 | 0.23 |
| 6 | 22254839 | rs9460718 | -0.302 | -1.089 | -1.483 | -0.055 | 0.004 | -0.006 | -0.006 | 0 | 0.14 | 0.12 | 0.06 | 0.01 | 0.12 | 0.17 |
| 6 | 22255514 | rs2066267 | -0.521 | -0.996 | -1.494 | 0.331 | 0.004 | 0.002 | 0.002 | 0 | 0.27 | 0.31 | 0.27 | 0.24 | 0.22 | 0.38 |
| 6 | 22263249 | rs12191980 | -0.162 | -0.497 | -1.924 | 0.221 | -0.013 | 0.051 | 0.051 | 0 | 0.09 | 0 | 0.06 | 0 | 0.07 | 0.04 |
| 6 | 22265981 | rs2455931 | 0.89 | 0.205 | -2.365 | 0.411 | -0.162 | -0.033 | -0.033 | 0 | 0.32 | 0.04 | 0.29 | 0.33 | 0.46 | 0.26 |
| 6 | 22266169 | rs16885931 | -0.415 | -0.005 | -2.337 | 0.657 | -0.065 | -0.143 | -0.143 | 0.062 | 0.19 | 0.01 | 0.28 | 0.39 | 0.23 | 0.29 |
| 6 | 22270120 | rs2066266 | 0.564 | -0.243 | -1.729 | -0.573 | 0.004 | 0.031 | 0.031 | 0 | 0.46 | 0.37 | 0.66 | 0.92 | 0.57 | 0.55 |
| 6 | 22270403 | rs6906383 | 0 | -0.706 | -1.935 | -1.063 | 0.01 | 0.112 | 0.112 | 0 | 0.28 | 0.09 | 0.17 | 0 | 0.25 | 0.16 |
| 6 | 22272753 | rs17205146 | -0.842 | -0.828 | -1.248 | -1.096 | 0.03 | 0.09 | 0.09 | 0 | 0.45 | 0.29 | 0.26 | 0.1 | 0.34 | 0.36 |
| 6 | 22274766 | rs6903568 | 0.124 | -0.653 | -1.273 | -1.024 | 0.013 | 0.058 | 0.058 | 0 | 0.27 | 0.12 | 0.16 | 0 | 0.23 | 0.16 |
| 6 | 22280214 | rs12524841 | -1.131 | -0.681 | -0.416 | 0.284 | 0.045 | 0.251 | 0.251 | 0 | 0.06 | 0.26 | 0.21 | 0.28 | 0.07 | 0.15 |
| 6 | 22282731 | rs2744119 | 0.153 | -0.085 | -1.131 | 0.752 | -0.038 | -0.055 | -0.055 | 0 | 0.09 | 0.01 | 0.18 | 0.27 | 0.11 | 0.14 |
| 6 | 22282905 | rs849874 | -1.193 | -0.462 | -0.733 | 0.108 | -0.02 | -0.001 | -0.001 | 0 | 0.3 | 0.37 | 0.29 | 0.31 | 0.16 | 0.35 |
| 6 | 22286012 | rs1205961 | 0.572 | -0.438 | -0.971 | -0.945 | 0.019 | 0.089 | 0.089 | 0 | 0.59 | 0.26 | 0.68 | 0.94 | 0.64 | 0.68 |
| 6 | 22288160 | rs1205960 | 0.585 | -0.276 | -0.517 | -0.996 | -0.006 | -0.049 | -0.049 | 0 | 0.72 | 0.65 | 0.8 | 0.89 | 0.73 | 0.7 |
| 6 | 22291367 | rs849886 | 1.21 | -0.297 | -1.089 | -0.608 | -0.051 | 0.049 | 0.049 | 0.115 | 0.45 | 0.14 | 0.45 | 0.63 | 0.49 | 0.42 |
| 6 | 22299046 | rs12210179 | -0.1 | -0.289 | -0.806 | -0.838 | -0.018 | 0.089 | 0.089 | 0 | 0.22 | 0.06 | 0.15 | 0.01 | 0.22 | 0.18 |
| 6 | 22299638 | rs849876 | 0.393 | -0.155 | -0.247 | -0.69 | -0.002 | -0.019 | -0.019 | 0.547 | 0.9 | 0.84 | 0.94 | 0.96 | 0.9 | 0.9 |
| 6 | 22307725 | rs2655426 | 0.991 | -0.529 | 0.112 | -0.723 | -0.006 | -0.031 | -0.031 | 0 | 0.59 | 0.51 | 0.63 | 0.77 | 0.63 | 0.55 |
| 6 | 22312287 | rs2473122 | 1.119 | -0.47 | 0.492 | -0.509 | 0.02 | 0.132 | 0.132 | 0 | 0.92 | 0.73 | 0.77 | 0.78 | 0.92 | 0.75 |
| 6 | 22321128 | rs2655417 | 1.565 | -0.483 | 0.671 | -0.502 | 0.06 | 0.015 | 0.015 | 0 | 0.94 | 0.74 | 0.89 | 0.95 | 0.93 | 0.87 |
| 6 | 22322004 | rs9358533 | -0.442 | -0.511 | 0.907 | -0.008 | -0.014 | 0.093 | 0.093 | 0 | 0.15 | 0.37 | 0.19 | 0.25 | 0.09 | 0.18 |
| 6 | 22323312 | rs2655420 | 1.539 | -0.425 | 0.695 | 0.104 | 0.024 | 0.156 | 0.156 | 0 | 0.9 | 0.66 | 0.76 | 0.75 | 0.91 | 0.75 |
| 6 | 22341469 | rs1205896 | 0.926 | -0.745 | 0.097 | -0.593 | 0.003 | 0.03 | 0.03 | 0 | 0.46 | 0.37 | 0.48 | 0.66 | 0.52 | 0.41 |
| 6 | 22343301 | rs1935007 | 1.875 | -0.376 | -0.126 | -0.396 | -0.151 | -0.139 | -0.139 | 0.018 | 0.31 | 0.02 | 0.28 | 0.39 | 0.4 | 0.22 |
| 6 | 22344213 | rs1033440 | -0.309 | -0.124 | 0.474 | -0.704 | -0.017 | -0.053 | -0.053 | 0 | 0.78 | 0.67 | 0.84 | 0.88 | 0.8 | 0.8 |
| 6 | 22352142 | rs1737231 | 0.855 | -0.258 | 0.389 | -0.772 | 0.001 | 0.007 | 0.007 | 0 | 0.55 | 0.45 | 0.5 | 0.66 | 0.57 | 0.54 |
| 6 | 22359969 | rs714589 | 0.752 | -0.231 | -0.209 | -0.702 | 0.001 | 0.008 | 0.008 | 0 | 0.57 | 0.62 | 0.52 | 0.66 | 0.59 | 0.54 |
| 6 | 22362945 | rs2744143 | 0.16 | -0.391 | -0.291 | -0.8 | 0.004 | 0.003 | 0.003 | 0 | 0.2 | 0.19 | 0.12 | 0.1 | 0.15 | 0.15 |
| 6 | 22380084 | rs7767991 | -0.369 | -0.138 | -0.58 | -1.229 | -0.001 | -0.02 | -0.02 | 0 | 0.39 | 0.31 | 0.31 | 0.07 | 0.39 | 0.34 |
| 6 | 22382907 | rs719332 | -0.806 | -0.281 | -0.35 | -0.6 | 0.003 | 0.003 | 0.003 | 0.656 | 0.48 | 0.55 | 0.56 | 0.53 | 0.45 | 0.63 |
| 6 | 22388887 | rs10946553 | 0.941 | -0.175 | -1.045 | -0.736 | -0.001 | -0.004 | -0.004 | 0 | 0.11 | 0.11 | 0.13 | 0.06 | 0.09 | 0.1 |
| 6 | 22389227 | rs12200760 | 0.941 | -0.118 | -1.045 | -0.736 | -0.001 | -0.004 | -0.004 | 0 | 0.11 | 0.1 | 0.13 | 0.06 | 0.09 | 0.1 |
| 6 | 22390618 | rs760853 | 1.114 | -0.108 | -1.12 | -0.826 | -0.013 | 0.034 | 0.034 | 0 | 0.16 | 0.07 | 0.1 | 0 | 0.19 | 0.14 |
| 6 | 22391406 | rs742286 | -2.082 | 0.283 | -1.471 | -1.005 | -0.055 | -0.095 | -0.095 | 1.367 | 0.76 | 0.93 | 0.75 | 0.63 | 0.69 | 0.78 |
| 6 | 22394070 | rs2064193 | 0.504 | 0.092 | -1.302 | -0.712 | 0.006 | -0.018 | -0.018 | 0 | 0.14 | 0.1 | 0.29 | 0.2 | 0.12 | 0.23 |
| 6 | 22399487 | rs6941503 | 0.744 | 0.021 | -1.1 | -0.821 | 0.015 | -0.01 | -0.01 | 0 | 0.13 | 0.42 | 0.1 | 0.09 | 0.09 | 0.11 |
| 6 | 22403373 | rs6933317 | 0.886 | -0.014 | -1.325 | -1.124 | -0.006 | 0.04 | 0.04 | 0 | 0.48 | 0.24 | 0.31 | 0.37 | 0.48 | 0.36 |
| 6 | 22403467 | rs2655439 | -1.082 | 0.007 | -1.32 | -1.146 | 0.012 | -0.041 | -0.041 | 0 | 0.84 | 0.6 | 0.84 | 0.9 | 0.86 | 0.85 |
| 6 | 22403592 | rs10484722 | 0.625 | -0.087 | -1.359 | -1.196 | -0.04 | -0.086 | -0.086 | 0 | 0.13 | 0.35 | 0.06 | 0 | 0.05 | 0.05 |
| 6 | 22404476 | rs6940071 | 0.232 | -0.05 | -1.417 | -1.158 | 0.004 | -0.019 | -0.019 | 0.011 | 0.56 | 0.5 | 0.42 | 0.69 | 0.55 | 0.52 |

| | | | | | | | | | | | | | | | | |
|---|---|---|---|---|---|---|---|---|---|---|---|---|---|---|---|---|
| 6 | 22405131 | rs12527298 | 1.45 | 0.032 | -1.567 | -1.14 | -0.097 | -0.062 | -0.062 | 0.042 | 0.37 | 0.04 | 0.27 | 0.37 | 0.44 | 0.31 |
| 6 | 22405396 | rs13193011 | 2.048 | 0.23 | -1.407 | -0.724 | -0.061 | -0.124 | -0.124 | 0 | 0.21 | 0.01 | 0.17 | 0.36 | 0.25 | 0.18 |
| 6 | 22417829 | rs2655436 | 1.106 | 0.128 | -1.32 | -0.762 | -0.067 | 0.089 | 0.089 | 0 | 0.15 | 0.01 | 0.13 | 0.01 | 0.19 | 0.1 |
| 6 | 22426161 | rs9356821 | -0.655 | -0.114 | -1.258 | -0.599 | -0.001 | -0.001 | -0.001 | 0.163 | 0.34 | 0.4 | 0.28 | 0.33 | 0.3 | 0.31 |
| 6 | 22431621 | rs7768473 | -0.209 | -0.024 | -1.25 | -0.506 | -0.037 | 0.019 | 0.019 | 1.196 | 0.72 | 0.91 | 0.59 | 0.78 | 0.7 | 0.67 |
| 6 | 22433910 | rs7773201 | 0.527 | 0.288 | -0.894 | -0.103 | 0.017 | 0.273 | 0.273 | 0 | 0.05 | 0.19 | 0.08 | 0.32 | 0.04 | 0.16 |
| 6 | 22441270 | rs6938736 | 0.24 | 0.3 | -1.375 | -1.021 | 0.008 | 0.026 | 0.026 | 0 | 0.59 | 0.58 | 0.52 | 0.78 | 0.65 | 0.64 |
| 6 | 22445521 | rs9379400 | 0.161 | 0.353 | -1.335 | -0.78 | 0.003 | 0.041 | 0.041 | 0 | 0.4 | 0.29 | 0.39 | 0.75 | 0.4 | 0.46 |
| 6 | 22446285 | rs10456256 | 0.33 | 0.364 | -1.322 | -0.749 | -0.01 | -0.015 | -0.015 | 0 | 0.12 | 0.17 | 0.13 | 0.06 | 0.08 | 0.1 |
| 6 | 22450262 | rs13212916 | 0.636 | 0.761 | -1.624 | -0.288 | -0.023 | -0.017 | -0.017 | 0 | 0.28 | 0.12 | 0.26 | 0.69 | 0.32 | 0.36 |
| 6 | 22451820 | rs4560628 | 0.156 | 0.513 | -1.269 | -0.288 | 0.001 | 0.026 | 0.026 | 0 | 0.73 | 0.76 | 0.66 | 0.85 | 0.75 | 0.74 |
| 6 | 22454221 | rs9466363 | 0.468 | 0.435 | -1.434 | 0.392 | -0.006 | -0.112 | -0.112 | 0 | 0.22 | 0.04 | 0.3 | 0.39 | 0.18 | 0.23 |
| 6 | 22457726 | rs12662946 | -0.194 | 0.412 | -1.13 | 0.664 | -0.001 | 0.046 | 0.046 | 0 | 0.09 | 0.14 | 0.1 | 0.32 | 0.1 | 0.14 |
| 6 | 22480823 | rs9356824 | -0.543 | 0.954 | -1.035 | 0.105 | -0.003 | 0.003 | 0.003 | 0 | 0.2 | 0.21 | 0.29 | 0.53 | 0.32 | 0.32 |
| 6 | 22484377 | rs9393300 | 0.797 | 0.599 | -1.365 | -0.636 | -0.011 | -0.033 | -0.033 | 0 | 0.45 | 0.18 | 0.46 | 0.42 | 0.39 | 0.47 |
| 6 | 22490056 | rs9379408 | -0.313 | 0.34 | -0.573 | -0.873 | -0.002 | 0.001 | 0.001 | 0 | 0.46 | 0.65 | 0.45 | 0.58 | 0.44 | 0.48 |
| 6 | 22491270 | rs2328715 | 0.035 | 0.173 | -0.315 | -1.007 | -0.001 | -0.103 | -0.103 | 0 | 0.38 | 0.58 | 0.28 | 0.05 | 0.34 | 0.26 |
| 6 | 22491609 | rs2504817 | -0.178 | 0.066 | 0.001 | -0.993 | -0.015 | -0.014 | -0.014 | 0 | 0.4 | 0.29 | 0.44 | 0.53 | 0.52 | 0.47 |
| 6 | 22491695 | rs2504818 | -1.749 | 0.009 | 0.044 | -0.975 | 0.025 | -0.053 | -0.053 | 0 | 0.77 | 0.9 | 0.72 | 0.6 | 0.85 | 0.72 |
| 6 | 22491997 | rs2185732 | -0.659 | 0.104 | 0.28 | -1.007 | 0.038 | -0.001 | -0.001 | 0 | 0.56 | 0.68 | 0.57 | 0.59 | 0.67 | 0.59 |
| 6 | 22495180 | rs9460748 | 1.165 | 0.093 | 0.332 | -1.11 | 0 | -0.038 | -0.038 | 0 | 0.24 | 0.26 | 0.19 | 0.02 | 0.24 | 0.19 |
| 6 | 22501407 | rs17410062 | 1.45 | 0.093 | 0.271 | -1.11 | -0.01 | 0.013 | 0.013 | 0 | 0.22 | 0.17 | 0.17 | 0.02 | 0.23 | 0.18 |
| 6 | 22501435 | rs13197009 | 0.979 | 0.13 | 0.092 | -1.084 | 0.046 | -0.065 | -0.065 | 0.917 | 0.41 | 0.06 | 0.42 | 0.41 | 0.33 | 0.41 |
| 6 | 22501947 | rs7742591 | -0.743 | 0.262 | 0.364 | -0.889 | 0.039 | 0.048 | 0.048 | 0 | 0.32 | 0.45 | 0.38 | 0.58 | 0.43 | 0.41 |
| 6 | 22502606 | rs6456494 | -0.743 | 0.262 | 0.364 | -0.889 | 0.039 | 0.048 | 0.048 | 0 | 0.32 | 0.45 | 0.38 | 0.58 | 0.43 | 0.41 |
| 6 | 22504508 | rs2000190 | -0.568 | 0.283 | 0.503 | -0.858 | 0.002 | 0.003 | 0.003 | 0 | 0.3 | 0.32 | 0.36 | 0.58 | 0.41 | 0.41 |
| 6 | 22504748 | rs2000191 | 0.716 | 0.108 | 0.196 | -0.823 | -0.01 | 0.026 | 0.026 | 0 | 0.09 | 0.02 | 0.07 | 0.02 | 0.07 | 0.07 |
| 6 | 22504878 | rs7758817 | 0.987 | -0.02 | 0.126 | -0.743 | 0.003 | -0.037 | -0.037 | 0.311 | 0.2 | 0.06 | 0.23 | 0.22 | 0.2 | 0.2 |
| 6 | 22505677 | rs1418310 | -1.866 | -0.126 | 0.107 | -0.745 | 0.03 | -0.07 | -0.07 | 0 | 0.78 | 0.91 | 0.72 | 0.6 | 0.86 | 0.73 |
| 6 | 22507985 | rs12195428 | -0.284 | 0.24 | 0.287 | -0.655 | 0 | -0.001 | -0.001 | 1.681 | 0.23 | 0.28 | 0.21 | 0.15 | 0.32 | 0.19 |
| 6 | 22508420 | rs1891347 | -0.545 | 0.198 | 0.272 | -0.662 | -0.022 | 0.013 | 0.013 | 0 | 0.43 | 0.34 | 0.43 | 0.37 | 0.51 | 0.39 |
| 6 | 22513002 | rs9356826 | 1.106 | 0.309 | 0.223 | -0.614 | -0.017 | 0.008 | 0.008 | 0.057 | 0.12 | 0.05 | 0.13 | 0.09 | 0.25 | 0.12 |
| 6 | 22513809 | rs16886349 | 0.989 | 0.283 | 0.135 | -0.567 | 0.01 | -0.039 | -0.039 | 0 | 0.2 | 0.05 | 0.21 | 0.22 | 0.19 | 0.21 |
| 6 | 22514356 | rs9466384 | -1.147 | 0.119 | 0.403 | -0.713 | 0.011 | 0.031 | 0.031 | 1.396 | 0.54 | 0.7 | 0.48 | 0.39 | 0.63 | 0.48 |
| 6 | 22515654 | rs7765404 | -0.643 | 0.152 | 0.409 | -0.644 | 0.011 | 0.081 | 0.081 | 0 | 0.89 | 0.68 | 0.9 | 0.77 | 0.89 | 0.85 |
| 6 | 22516569 | rs7752528 | -0.091 | 0.1 | 0.343 | -0.723 | 0.004 | -0.001 | -0.001 | 0.616 | 0.47 | 0.57 | 0.41 | 0.17 | 0.56 | 0.37 |
| 6 | 22517187 | rs9460753 | 1.603 | 0.106 | 0.35 | -0.706 | 0.032 | -0.057 | -0.057 | 0 | 0.23 | 0.09 | 0.35 | 0.59 | 0.15 | 0.35 |
| 6 | 22518463 | rs6904957 | -0.728 | 0.13 | 0.351 | -0.769 | -0.011 | -0.03 | -0.03 | 0.223 | 0.66 | 0.6 | 0.55 | 0.17 | 0.74 | 0.5 |
| 6 | 22525146 | rs9393307 | 0.234 | 0.16 | 0.502 | -0.736 | 0.005 | -0.007 | -0.007 | 0 | 0.42 | 0.53 | 0.42 | 0.37 | 0.5 | 0.37 |
| 6 | 22526294 | rs7757274 | -0.339 | 0.163 | 0.512 | -1.083 | -0.006 | -0.041 | -0.041 | 0 | 0.6 | 0.55 | 0.49 | 0.16 | 0.67 | 0.43 |
| 6 | 22527784 | rs9466387 | 0.925 | 0.362 | 0.657 | -0.866 | 0.001 | -0.003 | -0.003 | 0 | 0.06 | 0.08 | 0.08 | 0.02 | 0.08 | 0.07 |
| 6 | 22538612 | rs10946563 | 0.185 | 0.328 | 0.557 | -0.852 | 0.005 | -0.049 | -0.049 | 0 | 0.41 | 0.54 | 0.35 | 0.16 | 0.49 | 0.3 |
| 6 | 22540765 | rs9295552 | 0.185 | 0.326 | 0.484 | -0.844 | 0.005 | -0.047 | -0.047 | 0 | 0.41 | 0.55 | 0.35 | 0.16 | 0.49 | 0.3 |
| 6 | 22542906 | rs9295553 | 0.344 | 0.475 | 0.42 | -0.263 | 0.021 | -0.011 | -0.011 | 0.028 | 0.14 | 0.2 | 0.14 | 0.09 | 0.24 | 0.12 |
| 6 | 22550777 | rs9393310 | -0.489 | -0.074 | -0.006 | 0.036 | -0.008 | 0.001 | 0.001 | 1.535 | 0.7 | 0.69 | 0.62 | 0.4 | 0.82 | 0.59 |
| 6 | 22556207 | rs9358572 | -0.22 | -0.096 | -0.076 | 0.001 | 0.014 | 0.006 | 0.006 | 0.955 | 0.35 | 0.29 | 0.44 | 0.62 | 0.25 | 0.47 |
| 6 | 22556225 | rs13214499 | -0.395 | -0.182 | -0.143 | -0.046 | -0.013 | 0.024 | 0.024 | 0 | 0.11 | 0.04 | 0.09 | 0.02 | 0.13 | 0.14 |
| 6 | 22556254 | rs9379421 | 0.224 | -0.18 | -0.08 | 0.268 | 0.013 | -0.013 | -0.013 | 0 | 0.29 | 0.24 | 0.31 | 0.39 | 0.16 | 0.32 |
| 6 | 22556583 | rs4712694 | -0.368 | 0.034 | -0.071 | 0.362 | 0.003 | 0.059 | 0.059 | 0.288 | 0.08 | 0.12 | 0.14 | 0.23 | 0.09 | 0.12 |
| 6 | 22556791 | rs6938780 | 0.625 | -0.011 | 0.098 | -0.234 | -0.004 | -0.066 | -0.066 | 0.764 | 0.18 | 0.24 | 0.18 | 0 | 0.18 | 0.06 |
| 6 | 22565152 | rs12193939 | 0.522 | -0.064 | -0.186 | -0.16 | 0.043 | -0.056 | -0.056 | 0 | 0.27 | 0.15 | 0.29 | 0.39 | 0.17 | 0.32 |
| 6 | 22568534 | rs1361697 | -0.952 | 0.454 | -0.217 | -0.266 | -0.012 | 0.005 | 0.005 | 0 | 0.89 | 0.94 | 0.87 | 0.91 | 0.77 | 0.88 |
| 6 | 22570064 | rs2076506 | 1.346 | 0.209 | -0.283 | -0.216 | 0.058 | -0.086 | -0.086 | 0 | 0.2 | 0.06 | 0.25 | 0.39 | 0.09 | 0.23 |
| 6 | 22570189 | rs6922960 | 0.242 | 0.287 | -0.164 | -0.116 | 0 | -0.001 | -0.001 | 0.35 | 0.26 | 0.26 | 0.28 | 0.17 | 0.28 | 0.21 |
| 6 | 22570245 | rs6900627 | 0.384 | 0.288 | -0.164 | -0.158 | 0 | -0.027 | -0.027 | 0 | 0.25 | 0.24 | 0.21 | 0 | 0.27 | 0.14 |
| 6 | 22571037 | rs11751669 | 0.494 | 0.26 | -0.189 | -0.19 | -0.012 | 0.073 | 0.073 | 0 | 0.23 | 0.09 | 0.21 | 0.01 | 0.26 | 0.16 |
| 6 | 22571323 | rs3734215 | 0.413 | 0.217 | -0.197 | -0.142 | -0.001 | 0.01 | 0.01 | 0 | 0.24 | 0.11 | 0.25 | 0.17 | 0.27 | 0.21 |
| 6 | 22578106 | rs12111199 | 0.424 | 0.215 | -0.2 | -0.143 | -0.002 | 0.01 | 0.01 | 0 | 0.24 | 0.11 | 0.25 | 0.17 | 0.26 | 0.21 |
| 6 | 22581038 | rs6934462 | 1.131 | 0.182 | 0.065 | -0.209 | 0 | 0.004 | 0.004 | 0 | 0.35 | 0.32 | 0.27 | 0.31 | 0.32 | 0.39 |
| 6 | 22597820 | rs2294825 | -0.67 | 0.143 | -0.122 | -0.201 | 0.007 | 0.009 | 0.009 | 0 | 0.53 | 0.61 | 0.61 | 0.58 | 0.45 | 0.5 |
| 6 | 22597840 | rs2294827 | 0.841 | 0.093 | -0.025 | -0.149 | -0.058 | 0.047 | 0.047 | 0.208 | 0.25 | 0.44 | 0.3 | 0.39 | 0.1 | 0.24 |
| 6 | 22600813 | rs7743899 | -1.164 | 0.213 | -0.035 | -0.178 | -0.065 | -0.03 | -0.03 | 1.535 | 0.89 | 0.98 | 0.88 | 0.88 | 0.77 | 0.88 |
| 6 | 22606773 | rs6909574 | -0.421 | 0.039 | 0.106 | -0.119 | 0 | -0.014 | -0.014 | 0.961 | 0.32 | 0.37 | 0.35 | 0.19 | 0.36 | 0.26 |
| 6 | 22607026 | rs10485012 | 0.149 | 0.045 | 0.05 | -0.142 | -0.029 | 0.094 | 0.094 | 0.014 | 0.27 | 0.1 | 0.25 | 0.02 | 0.33 | 0.18 |
| 6 | 22608225 | rs9356829 | 1.173 | 0.006 | -0.038 | 0.018 | -0.009 | 0.005 | 0.005 | 0 | 0.22 | 0.24 | 0.26 | 0.36 | 0.08 | 0.26 |
| 6 | 22612757 | rs13217022 | 0.015 | 0.076 | 0.014 | -0.023 | -0.006 | -0.021 | -0.021 | 0 | 0.28 | 0.36 | 0.29 | 0.17 | 0.29 | 0.22 |
| 6 | 22615484 | rs728544 | 0.14 | 0.093 | -0.136 | 0.321 | -0.009 | 0.009 | 0.009 | 0 | 0.27 | 0.33 | 0.33 | 0.38 | 0.16 | 0.3 |
| 6 | 22623239 | rs2328723 | -0.161 | 0.557 | -0.462 | 0.393 | -0.025 | -0.02 | -0.02 | 0 | 0.89 | 0.94 | 0.81 | 0.69 | 0.75 | 0.8 |
| 6 | 22623471 | rs10456261 | 0.987 | 0.086 | -0.494 | 0.475 | 0.049 | -0.06 | -0.06 | 0.51 | 0.19 | 0.08 | 0.22 | 0.33 | 0.07 | 0.17 |

| | | | | | | | | | | | | | | | | |
|---|---|---|---|---|---|---|---|---|---|---|---|---|---|---|---|---|
| 6 | 22624817 | rs12193608 | 0.458 | 0.125 | -0.52 | 0.437 | 0.031 | -0.068 | -0.068 | 0 | 0.23 | 0.09 | 0.28 | 0.34 | 0.13 | 0.25 |
| 6 | 22631696 | rs1883653 | 0.184 | -0.02 | -0.519 | 0.314 | 0 | -0.002 | -0.002 | 0.043 | 0.67 | 0.63 | 0.6 | 0.31 | 0.68 | 0.61 |
| 6 | 22633741 | rs16886628 | -0.23 | 0.036 | -0.334 | 0.326 | 0.002 | 0.006 | 0.006 | 0 | 0.05 | 0.09 | 0.05 | 0.09 | 0.06 | 0.08 |
| 6 | 22639787 | rs6927934 | -0.182 | -0.073 | -0.408 | 0.726 | -0.033 | -0.032 | -0.032 | 0.068 | 0.88 | 0.92 | 0.79 | 0.61 | 0.76 | 0.82 |
| 6 | 22642328 | rs6921871 | -0.271 | -0.07 | -0.412 | 0.727 | -0.038 | -0.038 | -0.038 | 0.057 | 0.89 | 0.95 | 0.8 | 0.61 | 0.76 | 0.82 |
| 6 | 22643749 | rs10806959 | 0.228 | -0.33 | -0.31 | 0.045 | -0.003 | -0.002 | -0.002 | 0.479 | 0.61 | 0.63 | 0.55 | 0.58 | 0.5 | 0.69 |
| 6 | 22647683 | rs1040535 | 1.672 | -0.326 | -0.318 | 0.181 | -0.027 | 0.063 | 0.063 | 0 | 0.15 | 0.25 | 0.19 | 0.29 | 0.08 | 0.13 |
| 6 | 22656328 | rs2057016 | 0.177 | -0.428 | -0.335 | 0.005 | 0 | 0.003 | 0.003 | 0 | 0.5 | 0.48 | 0.44 | 0.32 | 0.44 | 0.55 |
| 6 | 22657821 | rs2057018 | -0.132 | -0.378 | -0.305 | 0.064 | -0.017 | -0.025 | -0.025 | 0 | 0.85 | 0.9 | 0.85 | 0.61 | 0.76 | 0.76 |
| 6 | 22661439 | rs4712701 | 0.398 | -0.439 | -0.313 | -0.018 | 0.006 | 0.007 | 0.007 | 0 | 0.59 | 0.5 | 0.6 | 0.51 | 0.48 | 0.58 |
| 6 | 22665236 | rs2223628 | -0.114 | -0.393 | -0.449 | 0.033 | -0.02 | -0.033 | -0.033 | 0 | 0.84 | 0.89 | 0.85 | 0.61 | 0.76 | 0.76 |
| 6 | 22672562 | rs4712703 | 0.902 | -0.17 | -0.961 | -0.446 | -0.035 | -0.053 | -0.053 | 0 | 0.12 | 0.01 | 0.22 | 0.22 | 0.12 | 0.27 |
| 6 | 22673453 | rs926571 | 0.984 | -0.308 | -0.874 | -0.45 | 0.003 | 0 | 0 | 0 | 0.34 | 0.3 | 0.39 | 0.28 | 0.31 | 0.21 |
| 6 | 22673972 | rs16886741 | 1.508 | -0.071 | -0.691 | -0.758 | -0.011 | -0.06 | -0.06 | 0 | 0.23 | 0.28 | 0.1 | 0.01 | 0.15 | 0.17 |
| 6 | 22677768 | rs9466435 | -0.489 | 0.143 | -1.024 | -0.455 | -0.042 | 0.035 | 0.035 | 0 | 0.69 | 0.88 | 0.61 | 0.26 | 0.63 | 0.59 |
| 6 | 22678728 | rs9366470 | 1.077 | 0.106 | -0.882 | -0.35 | 0.018 | -0.051 | -0.051 | 0 | 0.21 | 0.12 | 0.26 | 0.41 | 0.15 | 0.27 |
| 6 | 22686841 | rs2057019 | -0.676 | 0.197 | -0.745 | -0.591 | -0.005 | -0.006 | -0.006 | 0 | 0.72 | 0.77 | 0.65 | 0.36 | 0.67 | 0.68 |
| 6 | 22687498 | rs11752967 | 0.706 | 0.452 | -0.658 | -0.654 | 0 | -0.005 | -0.005 | 0 | 0.24 | 0.23 | 0.25 | 0.09 | 0.28 | 0.17 |
| 6 | 22693005 | rs9366472 | 1.019 | 0.985 | -0.852 | -0.186 | -0.04 | -0.032 | -0.032 | 0 | 0.07 | 0 | 0.08 | 0.11 | 0.14 | 0.1 |
| 6 | 22694829 | rs1555108 | -0.148 | 0.829 | -0.809 | 0.016 | -0.02 | -0.022 | -0.022 | 0 | 0.88 | 0.94 | 0.89 | 0.65 | 0.79 | 0.8 |
| 6 | 22695992 | rs9466444 | 0.707 | 0.421 | -0.62 | -0.853 | 0.031 | 0.049 | 0.049 | 0 | 0.51 | 0.64 | 0.35 | 0.01 | 0.41 | 0.28 |
| 6 | 22698186 | rs17508873 | 1.697 | 0.257 | -0.719 | -0.998 | -0.015 | 0.076 | 0.076 | 0 | 0.18 | 0.05 | 0.19 | 0 | 0.22 | 0.08 |
| 6 | 22698259 | rs4711022 | 0.86 | 0.18 | -0.787 | -0.725 | -0.039 | -0.053 | -0.053 | 0 | 0.11 | 0.02 | 0.21 | 0.22 | 0.14 | 0.26 |
| 6 | 22698796 | rs13215525 | 0.561 | -0.188 | -0.063 | -0.943 | -0.012 | -0.043 | -0.043 | 0 | 0.1 | 0.34 | 0.06 | 0 | 0.05 | 0.05 |
| 6 | 22699395 | rs9460776 | 0.788 | 0.082 | -0.142 | -0.606 | -0.005 | -0.001 | -0.001 | 0 | 0.43 | 0.58 | 0.35 | 0.31 | 0.27 | 0.33 |
| 6 | 22701495 | rs12204845 | 0.415 | 0.296 | 0.004 | -0.786 | -0.001 | -0.001 | -0.001 | 0 | 0.06 | 0.07 | 0.08 | 0.07 | 0.07 | 0.09 |
| 6 | 22704405 | rs6933929 | 0.331 | 0.362 | 0.14 | -0.931 | 0.002 | -0.022 | -0.022 | 2.016 | 0.37 | 0.66 | 0.19 | 0.01 | 0.29 | 0.22 |
| 6 | 22706635 | rs6903545 | 0.737 | -0.166 | 0.198 | -0.77 | -0.035 | -0.139 | -0.139 | 0 | 0.29 | 0.54 | 0.15 | 0.01 | 0.19 | 0.2 |
| 6 | 22708228 | rs7751689 | 0.484 | 0.131 | 0.222 | -0.404 | -0.024 | -0.014 | -0.014 | 0.017 | 0.37 | 0.47 | 0.22 | 0.27 | 0.28 | 0.34 |
| 6 | 22709430 | rs9393328 | -0.462 | -0.018 | 0.15 | -0.728 | -0.004 | -0.014 | -0.014 | 0 | 0.46 | 0.51 | 0.61 | 0.74 | 0.53 | 0.59 |
| 6 | 22709628 | rs4711023 | 1.286 | 0.132 | -0.082 | -0.583 | -0.015 | -0.056 | -0.056 | 0 | 0.09 | 0.01 | 0.19 | 0.28 | 0.09 | 0.23 |
| 6 | 22710041 | rs4712710 | -0.684 | 0.1 | 0.413 | -0.599 | 0.001 | 0.003 | 0.003 | 0 | 0.46 | 0.58 | 0.63 | 0.74 | 0.53 | 0.59 |
| 6 | 22710500 | rs16886790 | 1.444 | -0.002 | 0.402 | -0.624 | 0.018 | 0.006 | 0.006 | 0 | 0.29 | 0.19 | 0.1 | 0.25 | 0.18 | 0.25 |
| 6 | 22711708 | rs9358580 | 0.228 | 0.067 | 0.473 | -0.567 | -0.018 | 0.011 | 0.011 | 0 | 0.2 | 0.11 | 0.18 | 0.14 | 0.35 | 0.16 |
| 6 | 22711807 | rs9356834 | -0.015 | 0.019 | 0.542 | -0.564 | 0.011 | 0.011 | 0.011 | 0 | 0.35 | 0.41 | 0.41 | 0.47 | 0.44 | 0.35 |
| 6 | 22713440 | rs1024216 | 0.322 | -0.626 | 0.633 | -0.415 | 0.015 | 0.018 | 0.018 | 0.388 | 0.49 | 0.36 | 0.3 | 0.38 | 0.55 | 0.41 |
| 6 | 22713651 | rs9295562 | -0.037 | -0.354 | 0.247 | -0.304 | -0.003 | -0.024 | -0.024 | 0 | 0.31 | 0.28 | 0.4 | 0.47 | 0.34 | 0.37 |
| 6 | 22714582 | rs9295563 | -0.179 | -0.326 | 0.233 | -0.31 | -0.006 | -0.007 | -0.007 | 2.114 | 0.22 | 0.18 | 0.3 | 0.4 | 0.29 | 0.37 |
| 6 | 22720838 | rs16886814 | 1.77 | -0.32 | -0.099 | -0.27 | 0.006 | -0.084 | -0.084 | 0.278 | 0.13 | 0.02 | 0.21 | 0.33 | 0.08 | 0.17 |
| 6 | 22722608 | rs4286772 | -1.308 | -0.195 | 0.065 | -0.493 | -0.001 | -0.011 | -0.011 | 0.574 | 0.49 | 0.54 | 0.64 | 0.75 | 0.48 | 0.66 |
| 6 | 22725908 | rs12208226 | 0.06 | -0.107 | 0.139 | -0.423 | -0.005 | -0.014 | -0.014 | 0.095 | 0.12 | 0.15 | 0.12 | 0.03 | 0.11 | 0.12 |
| 6 | 22728788 | rs9295564 | -0.181 | 0.077 | -0.127 | -0.425 | -0.001 | -0.002 | -0.002 | 1.087 | 0.44 | 0.45 | 0.52 | 0.46 | 0.47 | 0.34 |
| 6 | 22729300 | rs10498713 | 0.614 | 0.207 | -0.373 | -0.409 | -0.029 | -0.007 | -0.007 | 0 | 0.11 | 0.05 | 0.1 | 0.13 | 0.16 | 0.1 |
| 6 | 22734908 | rs17206256 | 1.467 | 0.161 | -0.404 | -0.536 | -0.013 | 0.016 | 0.016 | 0 | 0.16 | 0.08 | 0.18 | 0 | 0.22 | 0.07 |
| 6 | 22735375 | rs9466453 | -0.276 | -0.037 | -0.232 | -0.261 | -0.001 | -0.001 | -0.001 | 0.102 | 0.46 | 0.48 | 0.59 | 0.46 | 0.53 | 0.41 |
| 6 | 22738731 | rs7752736 | -0.257 | -0.103 | -0.422 | -0.208 | 0.029 | -0.04 | -0.04 | 0 | 0.67 | 0.83 | 0.54 | 0.42 | 0.74 | 0.53 |
| 6 | 22739169 | rs11963356 | 1.205 | -0.099 | -0.228 | -0.158 | -0.001 | 0.004 | 0.004 | 0 | 0.34 | 0.29 | 0.16 | 0.27 | 0.31 | 0.3 |
| 6 | 22739365 | rs7739303 | 0.309 | 0.211 | -0.04 | -0.062 | -0.002 | 0.01 | 0.01 | 0.19 | 0.05 | 0.18 | 0.08 | 0.03 | 0.06 | 0.07 |
| 6 | 22741910 | rs9348532 | 0.59 | 0.194 | -0.257 | -0.092 | 0.002 | -0.007 | -0.007 | 0.186 | 0.54 | 0.44 | 0.43 | 0.58 | 0.45 | 0.53 |
| 6 | 22746117 | rs10946578 | -0.646 | 0.198 | -0.531 | 0.4 | 0.006 | -0.008 | -0.008 | 0 | 0.17 | 0.18 | 0.28 | 0.29 | 0.17 | 0.29 |
| 6 | 22746343 | rs12663096 | 0.233 | 0.36 | -0.747 | 0.453 | 0.008 | -0.031 | -0.031 | 0 | 0.12 | 0.04 | 0.19 | 0.26 | 0.11 | 0.23 |
| 6 | 22747334 | rs10456021 | -0.316 | 0.621 | -0.522 | 0.541 | 0.001 | 0.009 | 0.009 | 0 | 0.3 | 0.3 | 0.42 | 0.46 | 0.32 | 0.27 |
| 6 | 22748186 | rs9466456 | 0.111 | 0.565 | -0.629 | 0.208 | -0.001 | 0 | 0 | 0.841 | 0.3 | 0.34 | 0.27 | 0.29 | 0.3 | 0.36 |
| 6 | 22749157 | rs9466459 | 0.013 | 0.172 | -1.213 | -0.538 | 0.029 | 0.086 | 0.086 | 2.438 | 0.43 | 0.74 | 0.41 | 0.19 | 0.37 | 0.36 |
| 6 | 22749250 | rs9466460 | 0.383 | 0.856 | -1.029 | -0.812 | 0 | -0.039 | -0.039 | 0.225 | 0.19 | 0.21 | 0.13 | 0.01 | 0.25 | 0.12 |
| 6 | 22749679 | rs9466461 | -0.812 | 0.403 | -0.468 | -0.003 | 0.004 | 0.108 | 0.108 | 0 | 0.24 | 0.41 | 0.31 | 0.55 | 0.25 | 0.45 |
| 6 | 22749901 | rs4077636 | 0.584 | 0.092 | -0.765 | -1.098 | -0.025 | 0.063 | 0.063 | 1.019 | 0.45 | 0.22 | 0.51 | 0.71 | 0.47 | 0.53 |
| 6 | 22752606 | rs4305713 | -0.301 | 0.013 | -0.286 | -0.96 | -0.002 | 0.038 | 0.038 | 0 | 0.87 | 0.76 | 0.87 | 0.76 | 0.87 | 0.84 |
| 6 | 22755139 | rs4466239 | 0.25 | 0.016 | -0.719 | -0.995 | 0.024 | 0.068 | 0.068 | 3.229 | 0.55 | 0.26 | 0.6 | 0.76 | 0.6 | 0.69 |
| 6 | 22755319 | rs4510658 | 0.123 | 0.025 | -0.625 | -0.988 | 0.013 | 0.041 | 0.041 | 1.927 | 0.57 | 0.31 | 0.6 | 0.76 | 0.6 | 0.7 |
| 6 | 22757260 | rs4607423 | 1.361 | 0.25 | -0.85 | -1.193 | -0.027 | 0.037 | 0.037 | 0 | 0.07 | 0.01 | 0.04 | 0 | 0.08 | 0.01 |
| 6 | 22760170 | rs4432972 | 0.393 | 0.099 | -0.494 | -1.473 | 0.001 | 0.042 | 0.042 | 1.631 | 0.35 | 0.21 | 0.33 | 0.18 | 0.34 | 0.24 |
| 6 | 22765885 | rs9393332 | -0.496 | 0.003 | -0.164 | -1.365 | -0.007 | 0.007 | 0.007 | 2.952 | 0.53 | 0.66 | 0.55 | 0.5 | 0.51 | 0.55 |
| 6 | 22766715 | rs10498709 | 0.04 | 0.155 | 0.33 | -1.251 | 0.006 | 0.051 | 0.051 | 0 | 0.12 | 0.26 | 0.2 | 0.32 | 0.15 | 0.3 |
| 6 | 22774547 | rs13199412 | 0.372 | 0.188 | 0.223 | -1.433 | 0 | -0.028 | -0.028 | 0 | 0.1 | 0.13 | 0.09 | 0.01 | 0.12 | 0.1 |
| 6 | 22775611 | rs4711031 | 0.064 | 0.142 | 0.064 | -1.353 | -0.006 | -0.03 | -0.03 | 0 | 0.12 | 0.08 | 0.17 | 0.31 | 0.14 | 0.25 |
| 6 | 22776030 | rs4711032 | -1.251 | 0.215 | -0.007 | -1.61 | -0.001 | -0.001 | -0.001 | 0 | 0.39 | 0.62 | 0.36 | 0.33 | 0.39 | 0.49 |
| 6 | 22776453 | rs6456505 | 0.621 | 0.315 | -0.067 | -1.483 | -0.006 | -0.013 | -0.013 | 0.145 | 0.09 | 0.07 | 0.16 | 0.31 | 0.12 | 0.25 |
| 6 | 22798634 | rs9466471 | -0.689 | 0.185 | 0.034 | -1.602 | -0.005 | -0.131 | -0.131 | 0 | 0.29 | 0.53 | 0.2 | 0.02 | 0.29 | 0.23 |

| | | | | | | | | | | | | | | | | |
|---|---|---|---|---|---|---|---|---|---|---|---|---|---|---|---|---|
| 6 | 22799700 | rs6933879 | 0.396 | 0.5 | -0.37 | -1.274 | -0.001 | -0.015 | -0.015 | 0 | 0.1 | 0.09 | 0.15 | 0.3 | 0.11 | 0.26 |
| 6 | 22809110 | rs9460788 | -0.161 | 0.464 | -0.076 | -1.335 | 0.01 | -0.097 | -0.097 | 0 | 0.22 | 0.33 | 0.17 | 0.02 | 0.25 | 0.2 |
| 6 | 22818550 | rs9393337 | 0.038 | 0.661 | -0.621 | -1.241 | -0.005 | 0.005 | 0.005 | 0.323 | 0.3 | 0.3 | 0.26 | 0.29 | 0.31 | 0.44 |
| 6 | 22825009 | rs12192582 | 0.135 | 0.65 | -0.477 | -1.226 | 0.01 | 0.023 | 0.023 | 0 | 0.1 | 0.23 | 0.17 | 0.11 | 0.12 | 0.17 |
| 6 | 22827583 | rs9393338 | 1.006 | 0.349 | -0.883 | -1.349 | 0.01 | -0.078 | -0.078 | 0.614 | 0.11 | 0 | 0.21 | 0.53 | 0.07 | 0.13 |
| 6 | 22829432 | rs4712717 | -0.034 | 0.405 | -0.642 | -1.478 | -0.012 | 0.006 | 0.006 | 0 | 0.22 | 0.3 | 0.38 | 0.69 | 0.19 | 0.31 |
| 6 | 22831647 | rs1205495 | 0.026 | 0.295 | -0.63 | -1.931 | 0.011 | 0.06 | 0.06 | 0.141 | 0.47 | 0.33 | 0.41 | 0.34 | 0.54 | 0.46 |
| 6 | 22838984 | rs1074349 | -0.493 | 0.331 | -0.738 | -1.985 | -0.009 | -0.131 | -0.131 | 0 | 0.58 | 0.54 | 0.71 | 0.99 | 0.62 | 0.77 |
| 6 | 22846052 | rs1018401 | 0.282 | 0.67 | -1.082 | -1.546 | -0.078 | -0.046 | -0.046 | 0.761 | 0.27 | 0.06 | 0.26 | 0.36 | 0.32 | 0.37 |
| 6 | 22846683 | rs1308832 | 0.927 | 0.69 | -1.022 | -1.758 | -0.011 | 0.011 | 0.011 | 0.012 | 0.17 | 0.1 | 0.15 | 0 | 0.22 | 0.08 |
| 6 | 22848706 | rs6456508 | 0.315 | 0.521 | -0.602 | -1.613 | 0.012 | 0.034 | 0.034 | 0 | 0.09 | 0.29 | 0.15 | 0.11 | 0.09 | 0.16 |
| 6 | 22851846 | rs1107271 | -0.022 | 0.575 | -0.875 | -1.758 | -0.04 | 0.069 | 0.069 | 0.446 | 0.43 | 0.21 | 0.36 | 0.34 | 0.5 | 0.45 |
| 6 | 22856407 | rs1205498 | 0.117 | 0.631 | -0.863 | -1.755 | 0.034 | 0.005 | 0.005 | 0 | 0.12 | 0.26 | 0.16 | 0.11 | 0.15 | 0.18 |
| 6 | 22860722 | rs16886943 | 0.984 | 0.539 | -1.36 | -1.551 | -0.015 | 0.098 | 0.098 | 0 | 0.17 | 0.03 | 0.12 | 0 | 0.17 | 0.07 |
| 6 | 22862571 | rs9366479 | 0.126 | 0.359 | -1.486 | -1.669 | -0.03 | -0.086 | -0.086 | 0 | 0.35 | 0.07 | 0.41 | 0.54 | 0.36 | 0.3 |
| 6 | 22865377 | rs9460790 | -0.642 | 0.752 | -0.764 | -1.159 | -0.007 | 0.071 | 0.071 | 1.046 | 0.46 | 0.31 | 0.56 | 0.89 | 0.51 | 0.6 |
| 6 | 22868767 | rs10440820 | 0.958 | 1.014 | -0.994 | -1.286 | -0.016 | 0.023 | 0.023 | 0.337 | 0.17 | 0.08 | 0.1 | 0.01 | 0.19 | 0.14 |
| 6 | 22869813 | rs12664088 | 0.272 | 1.046 | -1.16 | -1.159 | -0.027 | -0.057 | -0.057 | 0 | 0.1 | 0.02 | 0.16 | 0.34 | 0.14 | 0.24 |
| 6 | 22873320 | rs9393344 | 0.53 | 0.929 | -1.01 | -1.064 | -0.008 | -0.074 | -0.074 | 0.296 | 0.19 | 0.08 | 0.29 | 0.53 | 0.19 | 0.23 |
| 6 | 22875263 | rs9379449 | 0.521 | 0.897 | -1.006 | -1.045 | -0.035 | -0.109 | -0.109 | 0.616 | 0.21 | 0.04 | 0.32 | 0.52 | 0.23 | 0.24 |
| 6 | 22876597 | rs9379450 | -0.366 | 0.476 | -0.622 | -1.637 | 0.014 | 0.113 | 0.113 | 3.449 | 0.48 | 0.27 | 0.59 | 0.88 | 0.55 | 0.61 |
| 6 | 22879014 | rs9393346 | -0.27 | 0.674 | -0.465 | -1.465 | -0.011 | 0.037 | 0.037 | 3.916 | 0.41 | 0.33 | 0.56 | 0.87 | 0.47 | 0.62 |
| 6 | 22881915 | rs9466503 | -0.97 | 0.304 | -0.058 | -1.427 | -0.011 | -0.042 | -0.042 | 0 | 0.08 | 0.23 | 0.06 | 0 | 0.05 | 0.03 |
| 6 | 22885675 | rs2254032 | -0.728 | 0.129 | -1.058 | -1.745 | -0.07 | 0.029 | 0.029 | 0 | 0.68 | 0.94 | 0.64 | 0.8 | 0.65 | 0.64 |
| 6 | 22896343 | rs13200281 | 0.41 | -0.079 | -1.131 | -2.149 | -0.008 | 0.102 | 0.102 | 0 | 0.29 | 0.11 | 0.18 | 0.01 | 0.3 | 0.17 |
| 6 | 22901378 | rs1928078 | -1.107 | -0.244 | -0.68 | -2.211 | -0.003 | 0.018 | 0.018 | 0 | 0.68 | 0.72 | 0.61 | 0.84 | 0.66 | 0.62 |
| 6 | 22901620 | rs1928079 | 0.594 | -0.315 | -0.53 | -2.26 | -0.001 | -0.001 | -0.001 | 0 | 0.3 | 0.28 | 0.38 | 0.83 | 0.35 | 0.38 |
| 6 | 22903884 | rs16887037 | 0.912 | -0.21 | -1.323 | -2.131 | -0.042 | 0.158 | 0.158 | 0 | 0.25 | 0.03 | 0.15 | 0 | 0.25 | 0.14 |
| 6 | 22905037 | rs1928082 | -0.81 | -0.153 | -1.008 | -1.583 | -0.01 | 0.019 | 0.019 | 0 | 0.68 | 0.81 | 0.62 | 0.8 | 0.66 | 0.59 |
| 6 | 22905353 | rs1928083 | 0.276 | 0.228 | -1.158 | -1.494 | -0.048 | -0.015 | -0.015 | 0 | 0.27 | 0.11 | 0.37 | 0.77 | 0.34 | 0.34 |
| 6 | 22905803 | rs9366481 | 0.233 | -0.028 | -0.539 | -1.696 | -0.003 | -0.082 | -0.082 | 0 | 0.35 | 0.41 | 0.26 | 0.07 | 0.28 | 0.33 |
| 6 | 22907216 | rs2498283 | -1.017 | 0.306 | -0.98 | -1.394 | -0.04 | 0.069 | 0.069 | 0.407 | 0.75 | 0.89 | 0.76 | 0.89 | 0.69 | 0.67 |
| 6 | 22907994 | rs4711037 | 0.999 | -0.131 | -1.506 | -0.784 | -0.003 | -0.125 | -0.125 | 0.527 | 0.17 | 0.03 | 0.25 | 0.56 | 0.13 | 0.17 |
| 6 | 22911301 | rs9366483 | 1.075 | -0.231 | -1.499 | -1.046 | -0.008 | 0.009 | 0.009 | 0 | 0.2 | 0.09 | 0.2 | 0.11 | 0.21 | 0.2 |
| 6 | 22913739 | rs2480014 | -1.618 | -0.145 | -1.375 | -0.938 | -0.04 | 0.068 | 0.068 | 0.314 | 0.75 | 0.89 | 0.76 | 0.89 | 0.69 | 0.67 |
| 6 | 22914924 | rs11963363 | 0.292 | -0.226 | -1.22 | -0.632 | -0.008 | -0.015 | -0.015 | 0 | 0.24 | 0.13 | 0.19 | 0.17 | 0.25 | 0.22 |
| 6 | 22920671 | rs11964627 | -0.428 | -0.239 | -1.239 | -0.69 | -0.011 | -0.018 | -0.018 | 0.765 | 0.33 | 0.22 | 0.32 | 0.28 | 0.37 | 0.37 |
| 6 | 22922389 | rs9358603 | 0.594 | -0.17 | -0.92 | -0.343 | 0.007 | 0.009 | 0.009 | 0.513 | 0.09 | 0.09 | 0.13 | 0.11 | 0.11 | 0.15 |
| 6 | 22923951 | rs9393356 | 0.228 | -0.616 | -0.773 | -0.685 | 0.017 | 0.041 | 0.041 | 0 | 0.47 | 0.31 | 0.62 | 0.74 | 0.54 | 0.5 |
| 6 | 22925531 | rs9358604 | 0.891 | -0.581 | -0.777 | -0.6 | -0.096 | -0.101 | -0.101 | 0 | 0.38 | 0.09 | 0.42 | 0.63 | 0.43 | 0.37 |
| 6 | 22926459 | rs9379455 | -1.034 | -0.957 | -0.251 | -0.765 | 0.002 | 0.015 | 0.015 | 0 | 0.69 | 0.76 | 0.78 | 0.94 | 0.69 | 0.78 |
| 6 | 22929284 | rs12529562 | -0.068 | -1.265 | 0.015 | -0.525 | -0.052 | 0.076 | 0.076 | 0.209 | 0.16 | 0.42 | 0.11 | 0.2 | 0.07 | 0.26 |
| 6 | 22933778 | rs4588682 | -0.323 | -1.251 | -0.036 | -0.531 | -0.045 | 0.032 | 0.032 | 0 | 0.22 | 0.45 | 0.15 | 0.21 | 0.15 | 0.28 |
| 6 | 22941834 | rs13201577 | 0 | -1.283 | 0.016 | -0.566 | -0.002 | 0 | 0 | 0 | 0.19 | 0.23 | 0.13 | 0.18 | 0.14 | 0.14 |
| 6 | 22942213 | rs4712728 | 0.565 | -1.381 | 0.151 | -0.667 | 0.012 | 0.013 | 0.013 | 0 | 0.51 | 0.39 | 0.68 | 0.75 | 0.63 | 0.54 |
| 6 | 22947966 | rs12527826 | 0.839 | -1.419 | 0.137 | -0.673 | 0.014 | -0.02 | -0.02 | 0 | 0.11 | 0.13 | 0.07 | 0.18 | 0.05 | 0.12 |
| 6 | 22953253 | rs9460805 | 1.495 | -1.404 | 0.144 | -0.666 | -0.005 | -0.029 | -0.029 | 0 | 0.08 | 0.1 | 0.06 | 0 | 0.08 | 0.02 |
| 6 | 22956252 | rs4280956 | 0.018 | -1.415 | 0.177 | -0.669 | -0.033 | 0.031 | 0.031 | 0 | 0.78 | 0.6 | 0.84 | 0.76 | 0.85 | 0.85 |
| 6 | 22957076 | rs7766780 | 0.483 | -1.282 | 0.06 | -0.602 | 0.048 | 0.049 | 0.049 | 0 | 0.51 | 0.31 | 0.67 | 0.74 | 0.63 | 0.54 |
| 6 | 22973286 | rs9466534 | 0.353 | -1.057 | -0.688 | -0.699 | 0.002 | -0.036 | -0.036 | 0 | 0.34 | 0.3 | 0.22 | 0.01 | 0.3 | 0.21 |
| 6 | 22974226 | rs9393360 | 1.279 | -0.836 | -1.089 | -0.26 | 0.008 | -0.113 | -0.113 | 0 | 0.15 | 0.01 | 0.14 | 0.51 | 0.12 | 0.18 |
| 6 | 22977272 | rs2473681 | 0.476 | -0.831 | -0.695 | -0.524 | -0.003 | -0.024 | -0.024 | 0.276 | 0.29 | 0.33 | 0.17 | 0.06 | 0.22 | 0.22 |
| 6 | 22979684 | rs2473685 | -0.305 | -0.016 | -1.559 | 0.18 | -0.133 | -0.103 | -0.103 | 0 | 0.58 | 0.93 | 0.49 | 0.36 | 0.52 | 0.59 |
| 6 | 22997063 | rs7774154 | -0.21 | 0.098 | -0.695 | 0.168 | -0.007 | -0.005 | -0.005 | 0 | 0.65 | 0.66 | 0.48 | 0.34 | 0.55 | 0.64 |
| 6 | 22999042 | rs9466547 | -0.153 | 0.04 | -0.473 | 0.138 | 0.007 | 0.004 | 0.004 | 0.659 | 0.64 | 0.6 | 0.47 | 0.34 | 0.55 | 0.63 |
| 6 | 23006527 | rs1997805 | 0.838 | -0.801 | -0.57 | -0.368 | 0.017 | 0.005 | 0.005 | 0 | 0.25 | 0.22 | 0.13 | 0.17 | 0.19 | 0.13 |
| 6 | 23006763 | rs9466553 | 0.259 | -1.212 | -0.241 | -0.646 | 0.005 | 0.024 | 0.024 | 0 | 0.33 | 0.22 | 0.31 | 0.17 | 0.35 | 0.48 |
| 6 | 23009496 | rs3950153 | 0.764 | -1.36 | -0.553 | -0.833 | -0.017 | -0.007 | -0.007 | 0 | 0.23 | 0.07 | 0.24 | 0.15 | 0.25 | 0.35 |
| 6 | 23011580 | rs9358611 | 1.331 | -0.242 | -0.856 | -0.595 | -0.039 | -0.043 | -0.043 | 0 | 0.07 | 0.02 | 0.19 | 0.15 | 0.21 | 0.18 |
| 6 | 23012043 | rs2498238 | 0.52 | -1.36 | -0.163 | -0.62 | -0.051 | -0.037 | -0.037 | 0.197 | 0.33 | 0.53 | 0.18 | 0.18 | 0.21 | 0.15 |
| 6 | 23012251 | rs2498239 | 0.657 | -1.404 | -0.092 | -0.619 | -0.044 | -0.042 | -0.042 | 0 | 0.31 | 0.47 | 0.17 | 0.13 | 0.19 | 0.15 |
| 6 | 23013575 | rs973720 | -1.662 | -1.146 | -0.533 | -0.434 | -0.01 | -0.056 | -0.056 | 0 | 0.9 | 0.99 | 0.81 | 0.51 | 0.92 | 0.88 |
| 6 | 23021257 | rs6912766 | 1.235 | -1.313 | -0.242 | -0.898 | 0.011 | 0.009 | 0.009 | 0 | 0.23 | 0.19 | 0.11 | 0.13 | 0.19 | 0.12 |
| 6 | 23025825 | rs2498253 | 0.064 | -1.455 | -0.251 | -1.053 | 0.021 | 0.021 | 0.021 | 0 | 0.51 | 0.58 | 0.36 | 0.31 | 0.46 | 0.49 |
| 6 | 23027242 | rs13219106 | 0.456 | -1.13 | -0.496 | -1.26 | -0.028 | -0.004 | -0.004 | 0.043 | 0.24 | 0.05 | 0.19 | 0.15 | 0.26 | 0.37 |
| 6 | 23029696 | rs12526121 | 0.182 | -1.112 | -0.361 | -1.257 | 0.001 | 0.002 | 0.002 | 0 | 0.27 | 0.21 | 0.22 | 0.18 | 0.27 | 0.37 |
| 6 | 23031210 | rs17669952 | 0.939 | -0.681 | -0.452 | -1.103 | -0.003 | 0.028 | 0.028 | 0 | 0.08 | 0.02 | 0.1 | 0.02 | 0.09 | 0.12 |
| 6 | 23034461 | rs2498258 | 0.775 | -0.791 | -0.494 | -1.323 | 0.022 | 0.027 | 0.027 | 0 | 0.14 | 0.09 | 0.08 | 0.01 | 0.11 | 0.14 |
| 6 | 23034890 | rs17670031 | 0.784 | -1.178 | -1.182 | -1.764 | -0.055 | 0.067 | 0.067 | 0 | 0.33 | 0.06 | 0.24 | 0.15 | 0.35 | 0.33 |

| | | | | | | | | | | | | | | | | |
|---|---|---|---|---|---|---|---|---|---|---|---|---|---|---|---|---|
| 6 | 23034984 | rs13214549 | 1.322 | -1.183 | -0.498 | -1.828 | -0.016 | -0.065 | -0.065 | 0.218 | 0.14 | 0.22 | 0.1 | 0.01 | 0.11 | 0.11 |
| 6 | 23035333 | rs2498259 | -0.134 | -1.364 | -0.096 | -1.852 | -0.001 | -0.013 | -0.013 | 0 | 0.56 | 0.57 | 0.37 | 0.15 | 0.48 | 0.49 |
| 6 | 23039736 | rs2498264 | 0.323 | -1.369 | -0.188 | -1.877 | 0.025 | 0.041 | 0.041 | 0 | 0.43 | 0.36 | 0.25 | 0.08 | 0.35 | 0.27 |
| 6 | 23039835 | rs2473729 | 0.734 | -1.34 | -0.39 | -1.861 | 0.027 | 0.149 | 0.149 | 0.311 | 0.37 | 0.15 | 0.22 | 0.08 | 0.34 | 0.25 |
| 6 | 23054938 | rs2498280 | 1.327 | -1.308 | -0.196 | -1.789 | 0.004 | -0.004 | -0.004 | 0 | 0.13 | 0.11 | 0.07 | 0.01 | 0.11 | 0.14 |
| 6 | 23057637 | rs9466588 | 0.708 | -1.297 | -0.201 | -1.782 | -0.005 | -0.036 | -0.036 | 0 | 0.17 | 0.2 | 0.09 | 0.01 | 0.13 | 0.17 |
| 6 | 23069364 | rs9466602 | 0.427 | -1.367 | -0.23 | -1.86 | 0.017 | 0.025 | 0.025 | 0 | 0.42 | 0.36 | 0.24 | 0.08 | 0.35 | 0.27 |
| 6 | 23073785 | rs4523075 | 0.139 | -0.525 | 0.134 | -1.448 | -0.1 | -0.022 | -0.022 | 0 | 0.26 | 0.11 | 0.45 | 0.76 | 0.37 | 0.3 |
| 6 | 23077311 | rs9393366 | 0.423 | -0.33 | 0.071 | -1.346 | -0.074 | -0.009 | -0.009 | 0 | 0.17 | 0.09 | 0.28 | 0.15 | 0.31 | 0.19 |
| 6 | 23078326 | rs7745131 | 0.371 | -0.676 | 0.425 | -1.515 | -0.006 | -0.019 | -0.019 | 0 | 0.3 | 0.33 | 0.18 | 0.13 | 0.24 | 0.14 |
| 6 | 23078899 | rs11759524 | 1.084 | -0.645 | 0.67 | -1.662 | 0.019 | 0.017 | 0.017 | 0 | 0.28 | 0.21 | 0.16 | 0.02 | 0.21 | 0.25 |
| 6 | 23081110 | rs12110710 | -0.074 | -0.336 | 0.858 | -0.949 | -0.003 | -0.017 | -0.017 | 0.117 | 0.05 | 0.07 | 0.02 | 0 | 0.02 | 0.03 |
| 6 | 23088578 | rs12524966 | 0.477 | -0.335 | 0.965 | -0.921 | -0.014 | -0.047 | -0.047 | 0 | 0.26 | 0.32 | 0.16 | 0.07 | 0.22 | 0.09 |
| 6 | 23112754 | rs4464782 | -0.375 | 0.204 | 1.142 | -0.474 | 0.019 | 0.001 | 0.001 | 0.169 | 0.3 | 0.4 | 0.45 | 0.72 | 0.37 | 0.29 |
| 6 | 23115750 | rs9466625 | -0.78 | 0.08 | 1.005 | -0.528 | 0.024 | -0.009 | -0.009 | 0.045 | 0.34 | 0.57 | 0.48 | 0.72 | 0.39 | 0.31 |
| 6 | 23126759 | rs4566882 | -0.563 | -0.096 | 0.869 | -0.696 | -0.014 | -0.053 | -0.053 | 0 | 0.72 | 0.67 | 0.82 | 0.92 | 0.76 | 0.88 |
| 6 | 23129600 | rs4557514 | 0.741 | -0.1 | 0.084 | -0.667 | 0.02 | 0.091 | 0.091 | 0.174 | 0.25 | 0.07 | 0.15 | 0.08 | 0.22 | 0.12 |
| 6 | 23134539 | rs9466637 | 0.682 | -0.089 | 0.062 | -0.661 | 0.016 | 0.091 | 0.091 | 0.161 | 0.25 | 0.05 | 0.14 | 0.08 | 0.22 | 0.12 |
| 6 | 23154902 | rs13200541 | 1.066 | -0.066 | -0.032 | -0.633 | -0.046 | 0.043 | 0.043 | 3.677 | 0.35 | 0.05 | 0.34 | 0.2 | 0.37 | 0.56 |
| 6 | 23155974 | rs7341287 | -0.502 | 0.023 | 0.777 | -0.593 | 0.04 | 0.002 | 0.002 | 0.218 | 0.32 | 0.46 | 0.46 | 0.71 | 0.39 | 0.3 |
| 6 | 23158950 | rs4712739 | 0.791 | 0.216 | 0.437 | -0.103 | 0.022 | -0.032 | -0.032 | 0 | 0.13 | 0.08 | 0.16 | 0.47 | 0.06 | 0.09 |
| 6 | 23160288 | rs9348547 | 0.211 | 0.463 | 0.543 | -0.23 | -0.005 | -0.008 | -0.008 | 0 | 0.11 | 0.09 | 0.16 | 0.18 | 0.27 | 0.12 |
| 6 | 23171814 | rs4712742 | 0.032 | 0.132 | 0.859 | 0.059 | -0.021 | 0.043 | 0.043 | 0 | 0.19 | 0.25 | 0.18 | 0.43 | 0.07 | 0.1 |
| 6 | 23175293 | rs12198618 | 0.062 | 0.432 | -0.496 | 0.05 | -0.084 | 0.005 | 0.005 | 0 | 0.35 | 0.12 | 0.3 | 0.28 | 0.47 | 0.2 |
| 6 | 23179698 | rs4438950 | 0.817 | -0.009 | -0.446 | 0.105 | 0.019 | -0.042 | -0.042 | 0 | 0.15 | 0.09 | 0.17 | 0.43 | 0.08 | 0.08 |
| 6 | 23181312 | rs12663119 | 1.144 | 0.381 | -0.798 | -0.174 | -0.062 | 0 | 0 | 0.144 | 0.35 | 0.05 | 0.34 | 0.24 | 0.39 | 0.6 |
| 6 | 23183971 | rs7745485 | 0.477 | 0.391 | -0.712 | -0.346 | 0.016 | 0.092 | 0.092 | 0.063 | 0.3 | 0.12 | 0.25 | 0.16 | 0.26 | 0.2 |
| 6 | 23184419 | rs4354144 | -0.219 | 0.508 | -0.579 | -0.26 | -0.026 | 0.035 | 0.035 | 0 | 0.39 | 0.26 | 0.41 | 0.32 | 0.48 | 0.28 |
| 6 | 23184825 | rs12524736 | 0.2 | 0.607 | -0.062 | 0.591 | 0.025 | 0.012 | 0.012 | 0 | 0.07 | 0.07 | 0.15 | 0.16 | 0.21 | 0.08 |
| 6 | 23188714 | rs10946595 | -0.692 | 0.521 | -0.224 | 0.521 | -0.023 | -0.023 | -0.023 | 0 | 0.14 | 0.07 | 0.2 | 0.25 | 0.26 | 0.15 |
| 6 | 23189857 | rs9466669 | -0.421 | -0.641 | 0.367 | 0.493 | 0.032 | -0.005 | -0.005 | 0.287 | 0.06 | 0.26 | 0.04 | 0 | 0.02 | 0.02 |
| 6 | 23190068 | rs9348549 | 0.938 | -0.554 | -0.846 | 0.668 | 0.062 | -0.042 | -0.042 | 0 | 0.63 | 0.2 | 0.61 | 0.44 | 0.81 | 0.74 |
| 6 | 23191842 | rs9379479 | 0.698 | -0.279 | -0.634 | 0.672 | 0.019 | -0.02 | -0.02 | 3.81 | 0.71 | 0.21 | 0.67 | 0.53 | 0.86 | 0.82 |
| 6 | 23194139 | rs4712746 | -0.828 | 0.475 | -0.669 | 0.065 | 0.003 | 0.042 | 0.042 | 0 | 0.37 | 0.29 | 0.58 | 0.85 | 0.32 | 0.59 |
| 6 | 23195237 | rs6899463 | -0.736 | 0.519 | -0.491 | 0.621 | 0.001 | -0.008 | -0.008 | 0 | 0.27 | 0.25 | 0.39 | 0.67 | 0.21 | 0.24 |
| 6 | 23198104 | rs9393377 | -0.736 | 0.519 | -0.36 | 0.612 | -0.001 | 0.004 | 0.004 | 0 | 0.27 | 0.29 | 0.39 | 0.67 | 0.21 | 0.24 |
| 6 | 23201234 | rs7771103 | -0.923 | 0.728 | -0.748 | 0.61 | 0 | -0.026 | -0.026 | 0 | 0.28 | 0.24 | 0.34 | 0.62 | 0.29 | 0.19 |
| 6 | 23202774 | rs9379481 | -1.014 | 0.084 | -0.583 | -0.644 | 0.002 | -0.012 | -0.012 | 0 | 0.47 | 0.46 | 0.63 | 0.87 | 0.44 | 0.53 |
| 6 | 23202962 | rs4639331 | -0.914 | 0.691 | -0.856 | -0.993 | 0.003 | 0.048 | 0.048 | 0 | 0.51 | 0.4 | 0.66 | 0.87 | 0.51 | 0.51 |
| 6 | 23205329 | rs7763770 | -0.637 | 0.729 | -0.74 | -0.795 | -0.001 | -0.021 | -0.021 | 0 | 0.38 | 0.35 | 0.57 | 0.87 | 0.37 | 0.48 |
| 6 | 23207033 | rs4712751 | 0.22 | 1.419 | -0.739 | -0.419 | 0.012 | 0.003 | 0.003 | 0 | 0.07 | 0.13 | 0.09 | 0.07 | 0.11 | 0.21 |
| 6 | 23208904 | rs9379482 | 0.539 | 1.23 | -1.038 | -1.135 | -0.002 | 0.042 | 0.042 | 1.093 | 0.71 | 0.26 | 0.75 | 0.92 | 0.68 | 0.77 |
| 6 | 23210222 | rs9460847 | 0.252 | 1.127 | -1.459 | -0.649 | -0.01 | -0.068 | -0.068 | 0 | 0.14 | 0.05 | 0.12 | 0.25 | 0.14 | 0.32 |
| 6 | 23212391 | rs9393380 | 0.017 | 0.947 | -2.015 | -0.858 | -0.001 | 0.076 | 0.076 | 0 | 0.34 | 0.13 | 0.43 | 0.76 | 0.31 | 0.48 |
| 6 | 23214201 | rs4247256 | -0.563 | 1.005 | -1.267 | -0.881 | 0.021 | -0.15 | -0.15 | 0 | 0.79 | 0.57 | 0.83 | 0.98 | 0.75 | 0.86 |
| 6 | 23214230 | rs4502928 | 0.035 | 0.909 | -1.723 | -0.762 | 0.009 | 0.076 | 0.076 | 0 | 0.35 | 0.18 | 0.44 | 0.76 | 0.32 | 0.49 |
| 6 | 23216134 | rs13205089 | 1.351 | 1.043 | -1.624 | -0.492 | -0.049 | 0.155 | 0.155 | 0.048 | 0.36 | 0.11 | 0.31 | 0.15 | 0.36 | 0.29 |
| 6 | 23218671 | rs9393382 | 0.515 | 0.933 | -1.191 | 0.296 | -0.002 | -0.077 | -0.077 | 0.248 | 0.2 | 0.07 | 0.31 | 0.51 | 0.17 | 0.15 |
| 6 | 23231296 | rs6456536 | 0.6 | 1.376 | -0.62 | 0.607 | -0.001 | 0.009 | 0.009 | 0.929 | 0.63 | 0.46 | 0.55 | 0.55 | 0.6 | 0.43 |
| 6 | 23236282 | rs4354145 | 1.379 | 1.775 | -1.006 | 0.728 | 0.044 | 0.039 | 0.039 | 0.221 | 0.46 | 0.2 | 0.44 | 0.6 | 0.38 | 0.32 |
| 6 | 23237776 | rs7740800 | 0.266 | 1.613 | -0.263 | 0.779 | -0.004 | 0.009 | 0.009 | 0.272 | 0.67 | 0.74 | 0.72 | 0.7 | 0.63 | 0.44 |
| 6 | 23241944 | rs9358623 | 0.475 | 1.67 | -0.423 | 0.845 | -0.023 | -0.032 | -0.032 | 0 | 0.05 | 0.01 | 0.15 | 0.15 | 0.09 | 0.1 |
| 6 | 23242514 | rs4452636 | 0.312 | 1.625 | -0.308 | 0.671 | -0.001 | 0.018 | 0.018 | 0.16 | 0.23 | 0.17 | 0.17 | 0.15 | 0.25 | 0.39 |
| 6 | 23243562 | rs4236026 | 0.686 | 1.752 | -0.301 | 0.542 | 0 | 0 | 0 | 0.103 | 0.83 | 0.84 | 0.78 | 0.78 | 0.74 | 0.73 |
| 6 | 23245281 | rs9356854 | 1.106 | 1.156 | -1.08 | 0.554 | 0.059 | -0.029 | -0.029 | 0 | 0.43 | 0.12 | 0.47 | 0.44 | 0.32 | 0.28 |
| 6 | 23264573 | rs7770298 | -0.108 | 1.539 | -0.612 | 0.179 | -0.006 | -0.006 | -0.006 | 0.448 | 0.09 | 0.07 | 0.13 | 0.2 | 0.14 | 0.12 |
| 6 | 23264838 | rs7774358 | -1.472 | 1.33 | -0.43 | -0.239 | 0.002 | -0.024 | -0.024 | 0.018 | 0.29 | 0.58 | 0.22 | 0.26 | 0.27 | 0.31 |
| 6 | 23268407 | rs9466719 | -0.647 | 1.378 | -0.881 | -0.404 | -0.001 | 0.007 | 0.007 | 0 | 0.57 | 0.71 | 0.46 | 0.4 | 0.56 | 0.51 |
| 6 | 23272919 | rs11970088 | -0.728 | 1.388 | -0.811 | -0.964 | 0.016 | 0.02 | 0.02 | 0 | 0.6 | 0.69 | 0.44 | 0.31 | 0.63 | 0.44 |
| 6 | 23278093 | rs9348557 | -0.318 | 1.238 | 0.191 | -1.763 | -0.007 | -0.052 | -0.052 | 0 | 0.68 | 0.59 | 0.81 | 0.88 | 0.72 | 0.79 |
| 6 | 23288100 | rs7453438 | 0.763 | 1.335 | 0.156 | -1.744 | 0.011 | 0.12 | 0.12 | 0 | 0.48 | 0.33 | 0.66 | 0.85 | 0.49 | 0.66 |
| 6 | 23288544 | rs4489142 | -0.231 | 1.423 | 0.264 | -1.853 | -0.032 | -0.103 | -0.103 | 0 | 0.66 | 0.51 | 0.8 | 0.94 | 0.78 | 0.89 |
| 6 | 23288843 | rs9393387 | 0.822 | 1.422 | 0.295 | -1.828 | 0.002 | 0.07 | 0.07 | 0 | 0.49 | 0.4 | 0.67 | 0.85 | 0.51 | 0.66 |
| 6 | 23294310 | rs1925437 | -0.58 | 0.583 | 0.435 | -1.74 | -0.074 | -0.105 | -0.105 | 0 | 0.74 | 0.55 | 0.87 | 0.94 | 0.87 | 0.91 |
| 6 | 23301437 | rs1925459 | -0.198 | 1 | 0.539 | -1.209 | 0.006 | -0.017 | -0.017 | 0 | 0.89 | 0.85 | 0.91 | 0.94 | 0.9 | 0.98 |
| 6 | 23310674 | rs7751597 | -0.486 | 0.76 | 0.81 | -1.235 | 0.015 | -0.026 | -0.026 | 0.407 | 0.11 | 0.35 | 0.08 | 0.02 | 0.09 | 0.12 |
| 6 | 23318780 | rs9379494 | -0.441 | 1.007 | 0.793 | -1.24 | 0.016 | -0.016 | -0.016 | 0 | 0.9 | 0.87 | 0.92 | 0.94 | 0.89 | 0.98 |
| 6 | 23320045 | rs2147855 | 1.524 | 1.142 | 0.726 | -1.319 | 0.023 | -0.033 | -0.033 | 0 | 0.7 | 0.47 | 0.77 | 0.85 | 0.64 | 0.71 |
| 6 | 23320162 | rs11968448 | -0.658 | 1.197 | 0.874 | -1.524 | 0.025 | 0 | 0 | 0 | 0.08 | 0.13 | 0.08 | 0.08 | 0.16 | 0.14 |

| | | | | | | | | | | | | | | | | |
|---|---|---|---|---|---|---|---|---|---|---|---|---|---|---|---|---|
| 6 | 23320971 | rs9358627 | 0.698 | 1.006 | 0.382 | -1.694 | -0.053 | -0.034 | -0.034 | 0 | 0.18 | 0.02 | 0.21 | 0.19 | 0.25 | 0.26 |
| 6 | 23321626 | rs6906134 | -1.663 | 1.331 | 1.234 | -1.442 | 0.043 | -0.043 | -0.043 | 0.453 | 0.2 | 0.4 | 0.15 | 0.1 | 0.24 | 0.27 |
| 6 | 23323255 | rs9356858 | 0.657 | 1.391 | 0.991 | -1.163 | 0.06 | -0.028 | -0.028 | 2.467 | 0.83 | 0.75 | 0.84 | 0.91 | 0.74 | 0.84 |
| 6 | 23335471 | rs9460868 | -0.809 | 1.114 | 0.251 | -1.288 | -0.004 | 0.025 | 0.025 | 0 | 0.47 | 0.59 | 0.29 | 0.09 | 0.42 | 0.33 |
| 6 | 23338399 | rs17274747 | 0.192 | 1.571 | 0.053 | -1.223 | 0.007 | -0.024 | -0.024 | 0 | 0.11 | 0.13 | 0.1 | 0.01 | 0.12 | 0.07 |
| 6 | 23347873 | rs9466754 | -0.038 | 1.359 | -0.117 | -1.28 | -0.031 | 0.018 | 0.018 | 0 | 0.15 | 0.1 | 0.09 | 0.08 | 0.2 | 0.2 |
| 6 | 23352409 | rs9466755 | -1.398 | 1.139 | -0.041 | -1.488 | -0.003 | -0.013 | -0.013 | 0 | 0.61 | 0.58 | 0.48 | 0.27 | 0.58 | 0.53 |
| 6 | 23355882 | rs9466756 | 0.45 | 1.089 | 0.146 | -1.507 | -0.003 | -0.025 | -0.025 | 0 | 0.54 | 0.51 | 0.62 | 0.81 | 0.63 | 0.66 |
| 6 | 23358040 | rs7763712 | 1.327 | 1.191 | -0.152 | -1.425 | 0.002 | 0.03 | 0.03 | 0 | 0.38 | 0.27 | 0.51 | 0.73 | 0.42 | 0.47 |
| 6 | 23366140 | rs9356861 | 1.255 | 0.263 | -0.117 | -1.377 | -0.026 | 0.036 | 0.036 | 0 | 0.39 | 0.27 | 0.54 | 0.73 | 0.51 | 0.48 |
| 6 | 23368008 | rs12214706 | 0.231 | 0.205 | 0.21 | -1.049 | 0.01 | 0.001 | 0.001 | 0.269 | 0.18 | 0.15 | 0.22 | 0.18 | 0.19 | 0.26 |
| 6 | 23374214 | rs6918629 | 0.469 | 0.708 | -0.111 | -0.717 | 0.016 | 0.056 | 0.056 | 0 | 0.53 | 0.42 | 0.61 | 0.8 | 0.63 | 0.63 |
| 6 | 23375531 | rs6904862 | -0.322 | -0.33 | -0.492 | -0.298 | -0.012 | -0.017 | -0.017 | 0 | 0.36 | 0.37 | 0.28 | 0.19 | 0.26 | 0.33 |
| 6 | 23378951 | rs1925458 | -1.078 | 0.324 | -0.745 | 0.597 | -0.001 | -0.009 | -0.009 | 0 | 0.13 | 0.12 | 0.24 | 0.24 | 0.12 | 0.16 |
| 6 | 23380034 | rs9379506 | -0.28 | 0.115 | -0.948 | 0.175 | 0.004 | 0.006 | 0.006 | 0.205 | 0.09 | 0 | 0.05 | 0.06 | 0.06 | 0.11 |
| 6 | 23380783 | rs942767 | -0.254 | -0.305 | -1.131 | 0.569 | 0.016 | -0.064 | -0.064 | 0 | 0.78 | 0.92 | 0.61 | 0.49 | 0.81 | 0.62 |
| 6 | 23383429 | rs9393409 | 0.263 | -0.923 | -0.709 | -0.326 | -0.009 | 0.002 | 0.002 | 0.282 | 0.55 | 0.56 | 0.46 | 0.49 | 0.64 | 0.54 |
| 6 | 23387883 | rs6456548 | -0.489 | -1.161 | -0.599 | -0.301 | 0.007 | -0.014 | -0.014 | 0 | 0.76 | 0.81 | 0.59 | 0.49 | 0.8 | 0.61 |
| 6 | 23390147 | rs1998740 | -0.131 | -1.045 | -0.296 | -0.519 | 0 | -0.001 | -0.001 | 0 | 0.58 | 0.46 | 0.49 | 0.44 | 0.7 | 0.54 |
| 6 | 23395165 | rs913011 | 0.147 | -0.135 | -0.749 | -1.09 | -0.063 | 0.133 | 0.133 | 0 | 0.26 | 0.09 | 0.19 | 0 | 0.32 | 0.15 |
| 6 | 23396997 | rs2764717 | -0.838 | 0.403 | -0.061 | -0.256 | 0.006 | 0 | 0 | 0 | 0.56 | 0.61 | 0.51 | 0.46 | 0.66 | 0.53 |
| 6 | 23401282 | rs6932335 | -0.097 | 0.014 | -0.005 | 0.14 | 0.01 | -0.008 | -0.008 | 1.224 | 0.31 | 0.39 | 0.29 | 0.29 | 0.38 | 0.36 |
| 6 | 23404276 | rs7775425 | -0.693 | -0.271 | 0.008 | 0.374 | -0.007 | -0.005 | -0.005 | 0 | 0.17 | 0.22 | 0.1 | 0.12 | 0.11 | 0.06 |
| 6 | 23408674 | rs913009 | -1.089 | -0.228 | 0.329 | 0.365 | -0.02 | -0.017 | -0.017 | 0 | 0.15 | 0.34 | 0.11 | 0.09 | 0.14 | 0.09 |
| 6 | 23413786 | rs1291389 | -1.029 | -0.474 | 0.739 | 0.951 | -0.032 | 0.075 | 0.075 | 0.094 | 0.23 | 0.42 | 0.38 | 0.3 | 0.18 | 0.26 |
| 6 | 23416375 | rs1291393 | -1.209 | -0.675 | 1.052 | 0.975 | -0.042 | 0.081 | 0.081 | 0.357 | 0.24 | 0.49 | 0.44 | 0.33 | 0.18 | 0.33 |
| 6 | 23419643 | rs943068 | -1.008 | -0.692 | 1.329 | 1.242 | -0.023 | 0.071 | 0.071 | 0.057 | 0.22 | 0.34 | 0.37 | 0.33 | 0.17 | 0.31 |
| 6 | 23421034 | rs4521580 | 1.095 | -0.28 | 0.267 | 0.597 | 0.026 | -0.053 | -0.053 | 0 | 0.44 | 0.17 | 0.36 | 0.59 | 0.4 | 0.57 |
| 6 | 23425966 | rs1291409 | -0.307 | -0.237 | 0.513 | 0.68 | -0.02 | 0.067 | 0.067 | 0 | 0.21 | 0.32 | 0.39 | 0.33 | 0.18 | 0.31 |
| 6 | 23426620 | rs411288 | -0.377 | -0.556 | 0.473 | 0.387 | 0.002 | -0.018 | -0.018 | 0.799 | 0.36 | 0.38 | 0.25 | 0.1 | 0.44 | 0.18 |
| 6 | 23433756 | rs2223494 | -0.204 | -0.455 | 0.562 | 0.172 | 0.002 | -0.01 | -0.01 | 0 | 0.24 | 0.21 | 0.3 | 0.08 | 0.38 | 0.2 |
| 6 | 23434397 | rs4601122 | 0.255 | -0.213 | 0.474 | 0.239 | 0.046 | -0.023 | -0.023 | 0.406 | 0.16 | 0.19 | 0.24 | 0.08 | 0.3 | 0.18 |
| 6 | 23435415 | rs199035 | -1.101 | -0.702 | 0.615 | 0.524 | 0 | 0 | 0 | 0.837 | 0.44 | 0.44 | 0.54 | 0.47 | 0.51 | 0.47 |
| 6 | 23437273 | rs6936349 | 0.203 | -0.647 | 0.077 | 0.678 | 0.014 | -0.036 | -0.036 | 0 | 0.15 | 0.04 | 0.14 | 0.18 | 0.1 | 0.16 |
| 6 | 23438445 | rs199032 | -1.542 | -0.775 | 0.222 | 0.251 | -0.03 | 0.018 | 0.018 | 0.042 | 0.45 | 0.8 | 0.52 | 0.29 | 0.51 | 0.33 |
| 6 | 23440413 | rs440115 | -1.069 | -0.633 | 0.739 | 0.165 | 0.012 | 0.059 | 0.059 | 0.126 | 0.08 | 0.27 | 0.11 | 0.1 | 0.06 | 0.06 |
| 6 | 23441842 | rs734749 | -0.08 | -0.942 | -0.476 | -0.197 | -0.05 | -0.043 | -0.043 | 0.058 | 0.32 | 0.07 | 0.22 | 0.3 | 0.3 | 0.31 |
| 6 | 23442229 | rs926453 | 1.122 | -0.835 | -0.446 | -0.778 | 0.002 | -0.001 | -0.001 | 0.685 | 0.37 | 0.35 | 0.44 | 0.61 | 0.29 | 0.47 |
| 6 | 23442310 | rs199028 | -0.511 | -0.376 | -0.475 | -0.853 | 0.016 | 0.004 | 0.004 | 0.015 | 0.94 | 0.97 | 0.89 | 0.94 | 0.76 | 0.94 |
| 6 | 23443422 | rs199027 | 0.331 | -0.729 | -0.255 | -0.935 | -0.006 | 0.004 | 0.004 | 0 | 0.79 | 0.76 | 0.7 | 0.89 | 0.62 | 0.8 |
| 6 | 23443962 | rs199025 | 0.289 | -1.276 | -0.277 | -1.179 | -0.008 | 0.092 | 0.092 | 1.863 | 0.59 | 0.71 | 0.59 | 0.87 | 0.42 | 0.66 |
| 6 | 23446234 | rs955978 | 0.493 | -0.858 | -0.237 | -1.153 | 0.007 | 0.012 | 0.012 | 0 | 0.49 | 0.53 | 0.57 | 0.72 | 0.42 | 0.6 |
| 6 | 23446691 | rs767943 | 0.891 | -0.738 | -0.186 | -1.103 | 0.041 | 0.054 | 0.054 | 0.613 | 0.46 | 0.33 | 0.45 | 0.78 | 0.29 | 0.56 |
| 6 | 23450286 | rs10223406 | -0.54 | -0.533 | 0.044 | -0.765 | -0.014 | -0.028 | -0.028 | 0 | 0.13 | 0.19 | 0.19 | 0.04 | 0.13 | 0.1 |
| 6 | 23454721 | rs199018 | -0.017 | 0.376 | -0.644 | -0.419 | -0.051 | -0.036 | -0.036 | 0 | 0.73 | 0.89 | 0.84 | 0.66 | 0.7 | 0.76 |
| 6 | 23454783 | rs199017 | -1.133 | 0.171 | -0.549 | -0.746 | -0.001 | -0.009 | -0.009 | 0 | 0.39 | 0.55 | 0.42 | 0.15 | 0.39 | 0.33 |
| 6 | 23465010 | rs191815 | -1.094 | 0.197 | -1.247 | -0.614 | -0.088 | 0.015 | 0.015 | 0 | 0.85 | 0.97 | 0.87 | 0.89 | 0.72 | 0.87 |
| 6 | 23466015 | rs2056986 | 1.866 | 0.289 | -1.235 | -0.482 | 0.015 | -0.069 | -0.069 | 0 | 0.3 | 0.16 | 0.39 | 0.51 | 0.26 | 0.39 |
| 6 | 23466670 | rs199004 | -0.263 | 0.13 | -1.067 | -0.519 | -0.015 | 0.007 | 0.007 | 0.693 | 0.24 | 0.29 | 0.22 | 0.28 | 0.13 | 0.19 |
| 6 | 23469680 | rs10456274 | -1.263 | 0.212 | -0.876 | -0.509 | 0.003 | 0.006 | 0.006 | 0 | 0.49 | 0.5 | 0.44 | 0.35 | 0.43 | 0.41 |
| 6 | 23475473 | rs199072 | -1.035 | 0.138 | -0.929 | -0.551 | 0.057 | -0.048 | -0.048 | 0 | 0.8 | 0.68 | 0.84 | 0.89 | 0.69 | 0.83 |
| 6 | 23479628 | rs1007959 | 1.62 | 0.152 | -1.534 | -0.174 | 0.014 | -0.04 | -0.04 | 0 | 0.31 | 0.21 | 0.39 | 0.53 | 0.26 | 0.41 |
| 6 | 23487567 | rs9466796 | -0.163 | 0.179 | -1.36 | 0.251 | 0.01 | 0.006 | 0.006 | 0 | 0.18 | 0.13 | 0.1 | 0.01 | 0.14 | 0.11 |
| 6 | 23490117 | rs199055 | 0.429 | 0.187 | -1.396 | 0.565 | -0.009 | 0.008 | 0.008 | 1.422 | 0.12 | 0.12 | 0.04 | 0.23 | 0.01 | 0.1 |
| 6 | 23493761 | rs17412056 | -0.729 | 1.106 | -1.507 | 0.036 | -0.036 | 0.06 | 0.06 | 0 | 0.27 | 0.14 | 0.21 | 0.11 | 0.31 | 0.26 |
| 6 | 23497282 | rs10946610 | -0.692 | 0.448 | -1.447 | 0.9 | 0.013 | -0.037 | -0.037 | 0 | 0.16 | 0.12 | 0.09 | 0.26 | 0.1 | 0.09 |
| 6 | 23497530 | rs199048 | -0.041 | 1.189 | -1.735 | 0.679 | -0.03 | 0.02 | 0.02 | 0 | 0.19 | 0.1 | 0.24 | 0.12 | 0.26 | 0.16 |
| 6 | 23498851 | rs4711049 | 0.147 | -0.769 | -1.777 | 1.173 | 0.009 | -0.013 | -0.013 | 0.319 | 0.23 | 0.2 | 0.27 | 0.41 | 0.12 | 0.3 |
| 6 | 23505577 | rs16887636 | -0.15 | -1.071 | -1.606 | 0.348 | 0.006 | 0.03 | 0.03 | 0 | 0.09 | 0.03 | 0.04 | 0.01 | 0.06 | 0.09 |
| 6 | 23507631 | rs199105 | -0.535 | -1.423 | -2.052 | 0.03 | 0.055 | 0.035 | 0.035 | 0 | 0.48 | 0.66 | 0.32 | 0.33 | 0.35 | 0.36 |
| 6 | 23518743 | rs9466803 | -0.691 | -1.822 | -1.128 | -0.777 | -0.01 | -0.013 | -0.013 | 0 | 0.38 | 0.44 | 0.23 | 0.27 | 0.25 | 0.28 |
| 6 | 23519714 | rs2206333 | -0.808 | -1.874 | -1.016 | -0.813 | -0.011 | -0.016 | -0.016 | 0 | 0.39 | 0.45 | 0.23 | 0.28 | 0.26 | 0.28 |
| 6 | 23520707 | rs2056985 | -0.601 | -1.686 | -1.191 | -1.132 | 0 | -0.003 | -0.003 | 0 | 0.61 | 0.57 | 0.39 | 0.32 | 0.53 | 0.48 |
| 6 | 23524218 | rs199097 | -1.506 | -1.551 | -1.281 | -1.026 | -0.077 | -0.182 | -0.182 | 0 | 0.73 | 0.96 | 0.55 | 0.5 | 0.63 | 0.63 |
| 6 | 23526341 | rs12189677 | 1.11 | -1.337 | -1.141 | -1.084 | -0.081 | 0.095 | 0.095 | 0.088 | 0.23 | 0.04 | 0.16 | 0.05 | 0.28 | 0.2 |
| 6 | 23533562 | rs199091 | -0.673 | -1.767 | -0.44 | -1.171 | 0.002 | 0.018 | 0.018 | 0.158 | 0.43 | 0.58 | 0.26 | 0.28 | 0.32 | 0.34 |
| 6 | 23542172 | rs199086 | -0.416 | -2.163 | 0.46 | -1.173 | 0.005 | -0.034 | -0.034 | 0 | 0.08 | 0.16 | 0.05 | 0.01 | 0.07 | 0.08 |
| 6 | 23551341 | rs199083 | -1.392 | -1.856 | 0.806 | -1.186 | -0.032 | -0.002 | -0.002 | 0 | 0.68 | 0.84 | 0.41 | 0.3 | 0.53 | 0.53 |
| 6 | 23554642 | rs199079 | 0.614 | -1.889 | 0.766 | -1.282 | -0.007 | -0.11 | -0.11 | 1.086 | 0.3 | 0.46 | 0.21 | 0.04 | 0.28 | 0.21 |

| | | | | | | | | | | | | | | | | |
|---|---|---|---|---|---|---|---|---|---|---|---|---|---|---|---|---|
| 6 | 23556224 | rs7749787 | -1.724 | -1.796 | 0.44 | -1.144 | -0.099 | -0.108 | -0.108 | 0 | 0.72 | 0.93 | 0.5 | 0.45 | 0.55 | 0.63 |
| 6 | 23557000 | rs6918377 | 0.329 | -1.915 | 0.573 | -1.413 | 0.015 | 0.064 | 0.064 | 0.689 | 0.12 | 0.19 | 0.1 | 0.23 | 0.1 | 0.14 |
| 6 | 23559581 | rs9466816 | -1.142 | -1.911 | 0.633 | -1.451 | -0.016 | -0.017 | -0.017 | 0 | 0.39 | 0.43 | 0.2 | 0.26 | 0.24 | 0.3 |
| 6 | 23562269 | rs199124 | 0.506 | -1.85 | 0.604 | -1.315 | -0.009 | -0.099 | -0.099 | 1.037 | 0.31 | 0.58 | 0.22 | 0.05 | 0.28 | 0.21 |
| 6 | 23565104 | rs7769725 | -0.811 | -1.753 | 0.494 | -1.177 | -0.005 | 0.01 | 0.01 | 0.395 | 0.24 | 0.26 | 0.15 | 0.25 | 0.16 | 0.21 |
| 6 | 23566016 | rs197948 | 0.311 | -1.861 | 0.186 | -1.492 | 0.007 | 0.001 | 0.001 | 0 | 0.12 | 0.12 | 0.05 | 0.01 | 0.06 | 0.09 |
| 6 | 23566794 | rs75522 | -0.932 | -2.008 | 0.363 | -1.683 | -0.004 | 0.07 | 0.07 | 0.033 | 0.58 | 0.77 | 0.32 | 0.06 | 0.42 | 0.4 |
| 6 | 23569915 | rs197951 | 0.057 | -2.078 | 0.263 | -1.73 | 0.006 | 0.003 | 0.003 | 0 | 0.39 | 0.39 | 0.23 | 0.05 | 0.34 | 0.29 |
| 6 | 23573947 | rs197956 | -0.083 | -1.698 | -0.338 | -1.459 | 0.029 | 0.075 | 0.075 | 0.056 | 0.24 | 0.13 | 0.08 | 0.02 | 0.14 | 0.18 |
| 6 | 23576589 | rs197962 | -0.658 | -1.927 | -0.564 | -1.477 | 0.001 | -0.037 | -0.037 | 0 | 0.55 | 0.61 | 0.28 | 0.06 | 0.4 | 0.39 |
| 6 | 23576753 | rs197963 | -0.71 | -1.798 | -1.568 | -1.411 | -0.002 | 0.145 | 0.145 | 0.088 | 0.58 | 0.8 | 0.31 | 0.07 | 0.41 | 0.39 |
| 6 | 23577577 | rs197965 | -0.187 | -1.46 | -1.899 | -1.092 | 0.047 | 0.123 | 0.123 | 0.598 | 0.23 | 0.05 | 0.07 | 0.02 | 0.12 | 0.17 |
| 6 | 23585283 | rs197969 | -1.12 | -1.032 | -1.764 | -0.586 | -0.112 | -0.092 | -0.092 | 0.171 | 0.7 | 0.94 | 0.48 | 0.44 | 0.54 | 0.63 |
| 6 | 23590536 | rs517002 | -0.396 | -0.868 | -1.423 | -1.109 | 0.046 | 0.107 | 0.107 | 0.682 | 0.21 | 0.04 | 0.06 | 0.01 | 0.1 | 0.18 |
| 6 | 23591084 | rs197976 | -0.309 | -0.948 | -1.31 | -1.154 | -0.008 | 0.151 | 0.151 | 0 | 0.41 | 0.19 | 0.22 | 0.06 | 0.4 | 0.36 |
| 6 | 23592348 | rs197979 | -0.741 | -0.258 | -0.98 | -0.322 | -0.025 | 0.031 | 0.031 | 0 | 0.59 | 0.74 | 0.44 | 0.32 | 0.49 | 0.44 |
| 6 | 23593375 | rs197988 | -0.736 | -0.231 | -1.099 | -0.153 | -0.03 | -0.007 | -0.007 | 0 | 0.59 | 0.75 | 0.45 | 0.39 | 0.5 | 0.44 |
| 6 | 23593767 | rs2025503 | -0.443 | -0.182 | -0.704 | -0.243 | 0.009 | 0.011 | 0.011 | 0 | 0.3 | 0.33 | 0.24 | 0.25 | 0.21 | 0.25 |
| 6 | 23595139 | rs6934968 | -0.83 | 0.402 | -0.285 | -0.16 | 0.006 | -0.031 | -0.031 | 0 | 0.22 | 0.38 | 0.2 | 0.13 | 0.18 | 0.16 |
| 6 | 23597059 | rs9379522 | -0.245 | 0.597 | -1.349 | -0.1 | -0.084 | -0.053 | -0.053 | 0 | 0.35 | 0.08 | 0.44 | 0.62 | 0.37 | 0.36 |
| 6 | 23600234 | rs197999 | 0.131 | 0.184 | -0.801 | -1.006 | 0.019 | -0.015 | -0.015 | 0.001 | 0.31 | 0.44 | 0.39 | 0.28 | 0.37 | 0.39 |
| 6 | 23602548 | rs506398 | 0.263 | 0.106 | -0.444 | -1.043 | -0.007 | -0.031 | -0.031 | 0 | 0.36 | 0.6 | 0.2 | 0.09 | 0.26 | 0.26 |
| 6 | 23604717 | rs537774 | -0.699 | 0.596 | -0.332 | -0.907 | -0.001 | 0 | 0 | 0 | 0.79 | 0.81 | 0.71 | 0.82 | 0.75 | 0.66 |
| 6 | 23612047 | rs565366 | 0.154 | 0.249 | -0.011 | -1.375 | 0.002 | 0.006 | 0.006 | 0 | 0.56 | 0.6 | 0.53 | 0.65 | 0.44 | 0.35 |
| 6 | 23614714 | rs9348578 | 0.913 | 0.007 | -0.299 | -1.556 | -0.013 | -0.045 | -0.045 | 1.337 | 0.33 | 0.11 | 0.42 | 0.62 | 0.33 | 0.33 |
| 6 | 23615399 | rs9466869 | 0.772 | -0.187 | -0.028 | -1.607 | 0.017 | 0.015 | 0.015 | 1.734 | 0.2 | 0.16 | 0.08 | 0.03 | 0.12 | 0.03 |
| 6 | 23621288 | rs541250 | -0.822 | -0.134 | -0.667 | -1.745 | -0.05 | 0.024 | 0.024 | 0 | 0.73 | 0.84 | 0.67 | 0.81 | 0.62 | 0.53 |
| 6 | 23630633 | rs9358642 | 0.961 | -0.13 | -0.765 | -1.742 | -0.087 | -0.058 | -0.058 | 0 | 0.35 | 0.06 | 0.43 | 0.62 | 0.37 | 0.36 |
| 6 | 23631118 | rs17211240 | -0.226 | 0.145 | -0.735 | -1.53 | -0.001 | 0.006 | 0.006 | 0 | 0.22 | 0.19 | 0.15 | 0.17 | 0.23 | 0.21 |
| 6 | 23640863 | rs536104 | -0.708 | 0.568 | -0.542 | -1.188 | -0.02 | 0.012 | 0.012 | 0 | 0.74 | 0.81 | 0.69 | 0.81 | 0.67 | 0.55 |
| 6 | 23645529 | rs2147454 | 0.115 | 0.227 | -0.47 | -1.107 | 0.012 | 0.007 | 0.007 | 0 | 0.14 | 0.2 | 0.13 | 0.19 | 0.17 | 0.16 |
| 6 | 23646875 | rs542063 | 0.414 | 0.212 | -0.652 | -1.201 | -0.001 | -0.002 | -0.002 | 0 | 0.55 | 0.53 | 0.53 | 0.65 | 0.43 | 0.34 |
| 6 | 23648347 | rs576816 | -0.052 | 0.683 | -1.341 | -1.009 | -0.06 | 0.027 | 0.027 | 0 | 0.71 | 0.86 | 0.67 | 0.81 | 0.6 | 0.49 |
| 6 | 23649645 | rs1321913 | -0.064 | 0.496 | -1.369 | -0.437 | -0.051 | -0.141 | -0.141 | 0 | 0.24 | 0.05 | 0.37 | 0.53 | 0.27 | 0.29 |
| 6 | 23650537 | rs528782 | -0.337 | 0.47 | -1.062 | -1.603 | -0.001 | 0.014 | 0.014 | 0 | 0.34 | 0.28 | 0.5 | 0.78 | 0.35 | 0.35 |
| 6 | 23652193 | rs9358645 | -0.189 | 1.247 | -1.361 | -1.323 | -0.021 | 0.025 | 0.025 | 0 | 0.12 | 0.06 | 0.09 | 0.04 | 0.14 | 0.11 |
| 6 | 23655168 | rs566071 | -0.095 | 1.262 | -1.48 | -1.23 | -0.041 | 0.056 | 0.056 | 0 | 0.83 | 0.98 | 0.87 | 0.95 | 0.78 | 0.69 |
| 6 | 23655229 | rs7742637 | 0.005 | 1.109 | -0.642 | -1.045 | -0.006 | 0.014 | 0.014 | 0 | 0.38 | 0.35 | 0.31 | 0.16 | 0.41 | 0.26 |
| 6 | 23656880 | rs9393425 | -0.532 | 1.263 | -0.277 | -0.191 | 0 | -0.001 | -0.001 | 0 | 0.1 | 0.14 | 0.09 | 0.11 | 0.11 | 0.17 |
| 6 | 23657168 | rs550050 | -0.571 | 1.442 | -0.333 | -0.268 | 0.033 | 0.076 | 0.076 | 0 | 0.49 | 0.66 | 0.44 | 0.27 | 0.54 | 0.48 |
| 6 | 23659465 | rs503082 | -1.378 | 1.72 | 0.032 | 0.291 | 0.025 | -0.013 | -0.013 | 0.817 | 0.18 | 0.28 | 0.24 | 0.13 | 0.24 | 0.19 |
| 6 | 23663679 | rs9356870 | 1.088 | 1.629 | 0.067 | 0.457 | 0.035 | -0.003 | -0.003 | 0.63 | 0.87 | 0.74 | 0.79 | 0.9 | 0.8 | 0.84 |
| 6 | 23667178 | rs9356871 | 0.599 | 1.205 | 0.506 | 0.549 | 0.015 | 0.016 | 0.016 | 0 | 0.8 | 0.54 | 0.85 | 0.76 | 0.8 | 0.75 |
| 6 | 23667206 | rs9379536 | 0.599 | 0.135 | -0.208 | -0.173 | 0.003 | 0.004 | 0.004 | 0 | 0.59 | 0.61 | 0.77 | 0.76 | 0.67 | 0.69 |
| 6 | 23671926 | rs501223 | 1.262 | 0.126 | -0.033 | 0.15 | -0.006 | 0.026 | 0.026 | 0 | 0.95 | 0.89 | 0.98 | 0.87 | 0.97 | 0.89 |
| 6 | 23680115 | rs766978 | 0.747 | 0.495 | 0.086 | 0.455 | 0.016 | -0.006 | -0.006 | 0 | 0.42 | 0.3 | 0.53 | 0.45 | 0.41 | 0.35 |
| 6 | 23680187 | rs543017 | -0.077 | 0.638 | 0.263 | 0.474 | 0.027 | 0.05 | 0.05 | 0 | 0.74 | 0.57 | 0.73 | 0.63 | 0.72 | 0.56 |
| 6 | 23680434 | rs791984 | 0.172 | 0.547 | 0.593 | 0.412 | 0 | 0.04 | 0.04 | 0 | 0.32 | 0.21 | 0.2 | 0.16 | 0.31 | 0.21 |
| 6 | 23680952 | rs9466899 | 0.41 | 0.586 | 0.574 | 0.42 | 0.013 | 0.011 | 0.011 | 0.003 | 0.22 | 0.29 | 0.25 | 0.25 | 0.25 | 0.35 |
| 6 | 23682287 | rs12212408 | 0.16 | 0.454 | 0.687 | 0.414 | -0.003 | 0.063 | 0.063 | 0 | 0.32 | 0.17 | 0.2 | 0.19 | 0.31 | 0.21 |
| 6 | 23686383 | rs9466901 | 0.42 | 0.861 | 0.952 | 0.417 | 0.009 | 0.006 | 0.006 | 0 | 0.21 | 0.26 | 0.25 | 0.24 | 0.25 | 0.34 |
| 6 | 23699268 | rs12207724 | 0.257 | 0.783 | 0.882 | 0.47 | -0.012 | 0.056 | 0.056 | 0 | 0.31 | 0.15 | 0.24 | 0.18 | 0.31 | 0.21 |
| 6 | 23699770 | rs13204659 | 1.017 | 0.675 | 1.044 | 0.426 | 0.017 | -0.009 | -0.009 | 0 | 0.41 | 0.19 | 0.46 | 0.42 | 0.41 | 0.35 |
| 6 | 23712752 | rs1324674 | 0.357 | -0.029 | 1.153 | 0.232 | 0 | 0.004 | 0.004 | 0.069 | 0.67 | 0.36 | 0.68 | 0.55 | 0.69 | 0.51 |
| 6 | 23716153 | rs2181938 | -0.523 | 0 | 1.204 | 0.222 | 0.015 | 0.01 | 0.01 | 0 | 0.76 | 0.67 | 0.73 | 0.75 | 0.75 | 0.66 |
| 6 | 23721032 | rs12526936 | 0.346 | 0.009 | 1.126 | 0.226 | 0 | 0.006 | 0.006 | 0.045 | 0.67 | 0.36 | 0.68 | 0.55 | 0.69 | 0.51 |
| 6 | 23723537 | rs11753322 | 0.929 | -0.002 | 0.794 | 0.255 | -0.023 | -0.045 | -0.045 | 1.511 | 0.34 | 0.09 | 0.43 | 0.37 | 0.38 | 0.31 |
| 6 | 23737077 | rs2328795 | 0.445 | -0.407 | 0.298 | 0.563 | 0.016 | -0.055 | -0.055 | 1.223 | 0.29 | 0.21 | 0.45 | 0.53 | 0.27 | 0.32 |
| 6 | 23737301 | rs6930955 | 0.19 | -0.711 | 0.294 | 0.235 | -0.002 | -0.029 | -0.029 | 0.427 | 0.33 | 0.38 | 0.28 | 0.19 | 0.32 | 0.25 |
| 6 | 23738614 | rs9358664 | 0.402 | -0.743 | 0.35 | 0.219 | -0.016 | 0.043 | 0.043 | 0 | 0.3 | 0.16 | 0.25 | 0.19 | 0.31 | 0.25 |
| 6 | 23741735 | rs13199891 | -0.038 | -0.756 | 0.339 | 0.335 | 0.001 | 0.02 | 0.02 | 0.038 | 0.08 | 0.12 | 0.15 | 0.26 | 0.05 | 0.13 |
| 6 | 23741814 | rs1408746 | -0.832 | -0.753 | 0.336 | 0.247 | 0.012 | -0.049 | -0.049 | 0 | 0.69 | 0.65 | 0.76 | 0.85 | 0.64 | 0.67 |
| 6 | 23741911 | rs3887159 | -0.832 | -0.753 | 0.336 | 0.247 | 0.012 | -0.049 | -0.049 | 0 | 0.69 | 0.65 | 0.76 | 0.85 | 0.64 | 0.67 |
| 6 | 23749404 | rs12661549 | 0.693 | -0.717 | 0.321 | 0.328 | -0.003 | -0.012 | -0.012 | 0 | 0.07 | 0.04 | 0.03 | 0.12 | 0.05 | 0.1 |
| 6 | 23751890 | rs9366524 | 0.264 | -0.903 | 0.313 | 0.222 | 0.001 | 0.042 | 0.042 | 0 | 0.33 | 0.21 | 0.27 | 0.16 | 0.31 | 0.27 |
| 6 | 23752689 | rs16888104 | 0.214 | -0.948 | 0.28 | 0.275 | 0 | -0.011 | -0.011 | 0.05 | 0.08 | 0.06 | 0.03 | 0.14 | 0.06 | 0.11 |
| 6 | 23766170 | rs10456038 | -0.797 | -0.853 | 0.283 | 0.224 | 0.018 | -0.057 | -0.057 | 0.03 | 0.72 | 0.63 | 0.76 | 0.86 | 0.67 | 0.72 |
| 6 | 23774487 | rs6911487 | -0.509 | -0.94 | 0.27 | 0.252 | -0.002 | 0 | 0 | 0 | 0.46 | 0.56 | 0.44 | 0.51 | 0.41 | 0.43 |
| 6 | 23775595 | rs9379577 | 1.048 | -0.952 | 0.038 | 0.089 | -0.034 | 0.024 | 0.024 | 0 | 0.23 | 0.1 | 0.19 | 0.13 | 0.25 | 0.16 |

| | | | | | | | | | | | | | | | | |
|---|---|---|---|---|---|---|---|---|---|---|---|---|---|---|---|---|
| 6 | 23775868 | rs9356895 | 1.455 | -0.924 | 0.038 | 0.102 | -0.007 | -0.065 | -0.065 | 0.015 | 0.26 | 0.07 | 0.33 | 0.36 | 0.27 | 0.29 |
| 6 | 23782517 | rs9393469 | 1.468 | -0.923 | 0.038 | 0.156 | -0.007 | -0.067 | -0.067 | 0.755 | 0.26 | 0.07 | 0.33 | 0.38 | 0.27 | 0.29 |
| 6 | 23783391 | rs7768453 | -0.142 | -0.927 | 0.059 | 0.13 | 0.002 | -0.008 | -0.008 | 0 | 0.1 | 0.14 | 0.17 | 0.2 | 0.09 | 0.16 |
| 6 | 23785834 | rs1325024 | -0.702 | -0.912 | 0.098 | 0.088 | 0.004 | -0.011 | -0.011 | 0 | 0.44 | 0.42 | 0.43 | 0.49 | 0.4 | 0.43 |
| 6 | 23790115 | rs1408744 | -0.31 | -0.875 | 0.276 | 0.164 | 0.001 | -0.005 | -0.005 | 0 | 0.64 | 0.62 | 0.75 | 0.73 | 0.62 | 0.61 |
| 6 | 23790159 | rs11967333 | 0.935 | -1.512 | -0.185 | -0.108 | -0.002 | 0.003 | 0.003 | 0 | 0.09 | 0.07 | 0.03 | 0.06 | 0.02 | 0.13 |
| 6 | 23790212 | rs9393479 | 1.093 | -1.557 | -0.158 | 0.09 | -0.003 | -0.007 | -0.007 | 2.096 | 0.23 | 0.16 | 0.3 | 0.35 | 0.25 | 0.2 |
| 6 | 23790996 | rs7772969 | -0.364 | -1.537 | -0.22 | 0.027 | -0.009 | -0.011 | -0.011 | 0 | 0.05 | 0.2 | 0.05 | 0 | 0.02 | 0 |
| 6 | 23792256 | rs12055681 | 0.794 | -1.543 | -0.492 | 0.108 | -0.068 | -0.078 | -0.078 | 0 | 0.26 | 0.09 | 0.36 | 0.32 | 0.32 | 0.28 |
| 6 | 23796356 | rs9356902 | 1.036 | -1.445 | -0.346 | -0.232 | -0.014 | 0.012 | 0.012 | 0 | 0.22 | 0.11 | 0.19 | 0.18 | 0.25 | 0.17 |
| 6 | 23798295 | rs9348604 | 1.05 | -1.445 | -0.346 | -0.232 | -0.014 | 0.011 | 0.011 | 0 | 0.22 | 0.11 | 0.19 | 0.18 | 0.25 | 0.17 |
| 6 | 23804221 | rs12213691 | 1.159 | -1.514 | -0.451 | -0.274 | -0.06 | 0.014 | 0.014 | 0 | 0.21 | 0.03 | 0.18 | 0.13 | 0.25 | 0.17 |
| 6 | 23812126 | rs12211578 | 1.318 | -1.522 | -0.45 | -0.282 | -0.063 | 0.011 | 0.011 | 0 | 0.2 | 0.03 | 0.17 | 0.13 | 0.24 | 0.16 |
| 6 | 23816088 | rs12192500 | 0.347 | -1.557 | -0.34 | -0.194 | 0 | -0.013 | -0.013 | 0.062 | 0.08 | 0.05 | 0.03 | 0.24 | 0.06 | 0.11 |
| 6 | 23824730 | rs9393495 | -0.534 | -1.633 | -0.2 | -0.267 | 0.008 | -0.024 | -0.024 | 0.349 | 0.63 | 0.58 | 0.75 | 0.73 | 0.61 | 0.6 |
| 6 | 23826607 | rs1886561 | 0.134 | -1.383 | -0.42 | -0.023 | -0.008 | -0.049 | -0.049 | 0 | 0.3 | 0.22 | 0.48 | 0.5 | 0.34 | 0.32 |
| 6 | 23835343 | rs1925191 | -0.919 | -0.985 | -0.331 | 0.153 | 0.001 | 0.043 | 0.043 | 0 | 0.8 | 0.75 | 0.62 | 0.52 | 0.78 | 0.79 |
| 6 | 23837322 | rs6928476 | -1.137 | -1.021 | -0.322 | 0.134 | 0.005 | 0.046 | 0.046 | 0 | 0.8 | 0.74 | 0.61 | 0.52 | 0.74 | 0.8 |
| 6 | 23844762 | rs12207777 | 0.356 | -0.96 | -0.424 | 0.078 | -0.011 | -0.02 | -0.02 | 1.05 | 0.08 | 0.04 | 0.03 | 0.25 | 0.07 | 0.1 |
| 6 | 23850593 | rs4712791 | 0.982 | -1.224 | -0.742 | -0.22 | -0.037 | 0.043 | 0.043 | 0 | 0.25 | 0.11 | 0.27 | 0.13 | 0.3 | 0.2 |
| 6 | 23850705 | rs9466936 | 0.244 | -1.334 | -0.248 | -0.333 | 0.005 | -0.007 | -0.007 | 1.274 | 0.37 | 0.35 | 0.27 | 0.31 | 0.39 | 0.39 |
| 6 | 23858310 | rs7756011 | -1.356 | -1.13 | -0.206 | 0.145 | -0.003 | -0.005 | -0.005 | 3.73 | 0.79 | 0.79 | 0.62 | 0.6 | 0.74 | 0.79 |
| 6 | 23858932 | rs9466937 | 0.562 | -1.166 | 0.055 | -0.111 | 0.003 | -0.058 | -0.058 | 0.05 | 0.3 | 0.34 | 0.24 | 0.08 | 0.31 | 0.35 |
| 6 | 23860377 | rs9393496 | 0.343 | -1.177 | 0.043 | -0.079 | 0.002 | -0.039 | -0.039 | 0 | 0.31 | 0.4 | 0.25 | 0.18 | 0.31 | 0.37 |
| 6 | 23877973 | rs12213583 | 0.337 | -0.921 | -0.359 | 0.232 | -0.024 | -0.022 | -0.022 | 0.065 | 0.07 | 0.02 | 0.04 | 0.26 | 0.08 | 0.13 |
| 6 | 23880358 | rs16888247 | 1.625 | -0.549 | -0.432 | 0.223 | -0.024 | 0.002 | 0.002 | 0.007 | 0.08 | 0.04 | 0.34 | 0.4 | 0.18 | 0.16 |
| 6 | 23881151 | rs6908183 | 1.557 | -0.84 | -0.247 | -0.132 | -0.02 | 0.043 | 0.043 | 0 | 0.25 | 0.15 | 0.21 | 0.05 | 0.3 | 0.24 |
| 6 | 23886075 | rs1397836 | -0.025 | -0.987 | 0.049 | -0.199 | 0.014 | 0.006 | 0.006 | 0.91 | 0.06 | 0.33 | 0.06 | 0 | 0.02 | 0 |
| 6 | 23890877 | rs4327688 | 1.039 | -0.866 | -0.191 | -0.147 | 0.009 | -0.057 | -0.057 | 0 | 0.27 | 0.34 | 0.24 | 0.05 | 0.31 | 0.24 |
| 6 | 23893378 | rs1397837 | -0.98 | -1.158 | -0.998 | -0.293 | -0.027 | -0.037 | -0.037 | 0 | 0.64 | 0.73 | 0.68 | 0.51 | 0.59 | 0.62 |
| 6 | 23895019 | rs6908544 | 1.202 | -0.736 | -0.717 | 0.195 | -0.014 | 0.026 | 0.026 | 0 | 0.25 | 0.17 | 0.21 | 0.05 | 0.3 | 0.24 |
| 6 | 23895830 | rs1588549 | 1.212 | -0.678 | -0.869 | 0.209 | -0.004 | 0.003 | 0.003 | 0 | 0.26 | 0.23 | 0.22 | 0.05 | 0.31 | 0.24 |
| 6 | 23900335 | rs11757520 | 1.175 | 0.274 | -1.513 | 0.765 | -0.024 | 0 | 0 | 0.587 | 0.08 | 0.04 | 0.34 | 0.39 | 0.18 | 0.16 |
| 6 | 23904420 | rs9466975 | 0.155 | 0.04 | -1.352 | 0.545 | 0.006 | 0.04 | 0.04 | 0.02 | 0.51 | 0.71 | 0.65 | 0.59 | 0.53 | 0.51 |
| 6 | 23905915 | rs9466976 | 0.249 | 0.109 | -1.456 | 0.397 | 0.002 | -0.005 | -0.005 | 0 | 0.48 | 0.63 | 0.65 | 0.48 | 0.53 | 0.5 |
| 6 | 23908557 | rs12194004 | 0.352 | 0.5 | -2.205 | -0.058 | -0.095 | 0.11 | 0.11 | 0.294 | 0.36 | 0.03 | 0.28 | 0.13 | 0.36 | 0.29 |
| 6 | 23909381 | rs507059 | 0.844 | 0.867 | -1.419 | 0.563 | -0.024 | -0.028 | -0.028 | 0 | 0.88 | 0.96 | 0.94 | 0.81 | 0.86 | 0.82 |
| 6 | 23909811 | rs481314 | 0.07 | 0.993 | -1.504 | 0.431 | -0.008 | 0.007 | 0.007 | 0.021 | 0.46 | 0.26 | 0.57 | 0.47 | 0.45 | 0.56 |
| 6 | 23913528 | rs806547 | -0.719 | 0.92 | -0.389 | -0.27 | -0.016 | -0.081 | -0.081 | 0 | 0.31 | 0.54 | 0.16 | 0.12 | 0.27 | 0.2 |
| 6 | 23915870 | rs474833 | -1.208 | 2.054 | -0.109 | 0.039 | -0.005 | 0.012 | 0.012 | 0.016 | 0.39 | 0.68 | 0.33 | 0.25 | 0.53 | 0.24 |
| 6 | 23915899 | rs474923 | -0.838 | 1.964 | -0.158 | -0.075 | -0.023 | 0.019 | 0.019 | 1.224 | 0.15 | 0.62 | 0.13 | 0.11 | 0.1 | 0.07 |
| 6 | 23918178 | rs13203985 | 0.675 | 2.001 | -0.428 | -0.213 | -0.047 | 0.019 | 0.019 | 0.015 | 0.16 | 0.02 | 0.05 | 0.1 | 0.17 | 0.14 |
| 6 | 23918873 | rs1511467 | 0.026 | 2.163 | -0.307 | -0.134 | -0.004 | 0.008 | 0.008 | 0 | 0.06 | 0.03 | 0.15 | 0.03 | 0.25 | 0.04 |
| 6 | 23920719 | rs12206829 | -0.053 | 2.156 | -0.308 | -0.134 | -0.005 | 0.009 | 0.009 | 0 | 0.07 | 0.03 | 0.15 | 0.03 | 0.26 | 0.04 |
| 6 | 23922624 | rs12200455 | -0.036 | 2.155 | -0.288 | -0.135 | 0 | 0.009 | 0.009 | 0 | 0.07 | 0.03 | 0.15 | 0.03 | 0.26 | 0.04 |
| 6 | 23924231 | rs9393510 | 0.271 | 0.913 | -0.09 | 0.013 | -0.002 | 0.007 | 0.007 | 0 | 0.07 | 0.08 | 0.11 | 0.1 | 0.05 | 0.07 |
| 6 | 23924306 | rs545967 | -0.251 | 0.824 | 0.082 | -0.062 | 0.01 | -0.052 | -0.052 | 0 | 0.1 | 0.29 | 0.08 | 0.01 | 0.09 | 0.05 |
| 6 | 23931502 | rs12190667 | -0.292 | 1.06 | -1.116 | 0.042 | -0.117 | 0.055 | 0.055 | 0.323 | 0.25 | 0.09 | 0.22 | 0.14 | 0.43 | 0.17 |
| 6 | 23931801 | rs12208958 | -0.182 | 1.028 | -1.116 | 0.042 | -0.119 | 0.056 | 0.056 | 0.336 | 0.25 | 0.09 | 0.22 | 0.14 | 0.43 | 0.17 |
| 6 | 23934445 | rs518587 | -0.739 | 0.363 | -0.951 | 0.194 | 0.026 | 0.037 | 0.037 | 0 | 0.47 | 0.77 | 0.43 | 0.35 | 0.57 | 0.33 |
| 6 | 23936906 | rs9358724 | -0.547 | -0.992 | -0.695 | -0.008 | -0.04 | -0.023 | -0.023 | 0 | 0.35 | 0.4 | 0.31 | 0.24 | 0.26 | 0.17 |
| 6 | 23941746 | rs17586672 | 0.368 | -0.208 | -1.438 | -0.598 | -0.043 | 0.09 | 0.09 | 0 | 0.26 | 0.07 | 0.2 | 0.09 | 0.33 | 0.18 |
| 6 | 23944129 | rs528715 | 0.388 | -0.579 | -0.854 | 1.049 | 0 | 0.083 | 0.083 | 0.316 | 0.92 | 0.91 | 0.78 | 0.58 | 0.92 | 0.73 |
| 6 | 23945104 | rs7750211 | -0.484 | -1.249 | -0.402 | 0.858 | -0.061 | -0.022 | -0.022 | 0 | 0.23 | 0.48 | 0.13 | 0.14 | 0.12 | 0.15 |
| 6 | 23946867 | rs550969 | 0.853 | -1.151 | -1.2 | 0.365 | 0.037 | 0.07 | 0.07 | 0 | 0.6 | 0.27 | 0.5 | 0.24 | 0.7 | 0.48 |
| 6 | 23951514 | rs6456588 | -0.823 | -0.455 | -1.004 | -0.062 | -0.031 | -0.035 | -0.035 | 0 | 0.12 | 0.16 | 0.05 | 0.01 | 0.02 | 0.03 |
| 6 | 23954438 | rs9348623 | -0.447 | 0.511 | -1.243 | 0.374 | -0.043 | -0.044 | -0.044 | 0.771 | 0.13 | 0.01 | 0.17 | 0.17 | 0.16 | 0.11 |
| 6 | 23964481 | rs483792 | -0.796 | 0.84 | -0.773 | -0.335 | 0.008 | -0.011 | -0.011 | 1.514 | 0.24 | 0.27 | 0.24 | 0.14 | 0.31 | 0.24 |
| 6 | 23968351 | rs474019 | 0.092 | 0.286 | -0.743 | -1.584 | 0.006 | -0.004 | -0.004 | 0 | 0.42 | 0.54 | 0.32 | 0.14 | 0.32 | 0.29 |
| 6 | 23972121 | rs1397842 | 0.236 | -0.256 | -0.321 | -1.977 | -0.002 | -0.023 | -0.023 | 0.015 | 0.25 | 0.27 | 0.27 | 0.13 | 0.27 | 0.22 |
| 6 | 23972386 | rs573863 | 0.875 | -0.378 | -0.332 | -1.991 | 0.039 | 0.041 | 0.041 | 0 | 0.14 | 0.07 | 0.03 | 0.01 | 0.04 | 0.07 |
| 6 | 23972951 | rs544840 | -0.97 | -0.206 | -0.59 | -2.471 | -0.005 | 0.033 | 0.033 | 0 | 0.64 | 0.68 | 0.52 | 0.22 | 0.66 | 0.56 |
| 6 | 23975629 | rs9467001 | -0.631 | -0.285 | -0.815 | -2.592 | 0.006 | -0.033 | -0.033 | 0.638 | 0.41 | 0.49 | 0.28 | 0.19 | 0.42 | 0.5 |
| 6 | 23975968 | rs13204056 | -0.426 | 0.137 | -0.993 | -2.687 | 0.012 | -0.012 | -0.012 | 0 | 0.24 | 0.3 | 0.17 | 0.17 | 0.23 | 0.46 |
| 6 | 23977847 | rs17528254 | 0.907 | 0.197 | -1.162 | -2.622 | -0.001 | -0.01 | -0.01 | 0 | 0.11 | 0.08 | 0.08 | 0.01 | 0.17 | 0.04 |
| 6 | 23978218 | rs1397840 | -0.912 | 0.039 | -1.457 | -2.502 | 0.012 | 0.016 | 0.016 | 0 | 0.63 | 0.78 | 0.56 | 0.29 | 0.68 | 0.68 |
| 6 | 23978607 | rs16888397 | 1.257 | 0.243 | -1.331 | -2.399 | 0.002 | -0.002 | -0.002 | 0 | 0.08 | 0.04 | 0.03 | 0.08 | 0.06 | 0.2 |
| 6 | 23979222 | rs9295609 | 0.5 | 0.364 | -1.249 | -2.341 | 0.024 | -0.001 | -0.001 | 0 | 0.12 | 0.17 | 0.13 | 0.07 | 0.16 | 0.22 |
| 6 | 23981466 | rs518886 | -1.342 | -0.058 | -1.745 | -2.025 | -0.025 | -0.036 | -0.036 | 0 | 0.78 | 0.84 | 0.58 | 0.3 | 0.73 | 0.76 |

| | | | | | | | | | | | | | | | | |
|---|---|---|---|---|---|---|---|---|---|---|---|---|---|---|---|---|
| 6 | 23984030 | rs508584 | 0.496 | -0.231 | -1.727 | -2.249 | 0.054 | 0.067 | 0.067 | 0 | 0.23 | 0.11 | 0.05 | 0.1 | 0.1 | 0.3 |
| 6 | 23986903 | rs12206379 | 1.048 | 0.666 | -1.49 | -1.631 | -0.023 | 0.034 | 0.034 | 0.014 | 0.19 | 0.12 | 0.15 | 0.1 | 0.23 | 0.13 |
| 6 | 23990308 | rs10484661 | 1.348 | 0.657 | -1.467 | -1.581 | -0.018 | 0.022 | 0.022 | 0 | 0.16 | 0.14 | 0.15 | 0.09 | 0.22 | 0.09 |
| 6 | 23998157 | rs554400 | 0.582 | 0.175 | -1.307 | -1.659 | -0.005 | -0.028 | -0.028 | 0 | 0.14 | 0.26 | 0.09 | 0.08 | 0.1 | 0.26 |
| 6 | 24000499 | rs1511479 | -0.113 | 0.064 | -1.978 | -2.112 | -0.004 | 0.029 | 0.029 | 0 | 0.45 | 0.34 | 0.42 | 0.16 | 0.56 | 0.39 |
| 6 | 24002463 | rs34014604 | 1.233 | -0.054 | -2.128 | -2.116 | 0.056 | 0.089 | 0.089 | 0 | 0.18 | 0.05 | 0.04 | 0.04 | 0.06 | 0.15 |
| 6 | 24005370 | rs12207326 | 0.356 | 1.189 | -1.864 | -1.639 | -0.126 | 0.018 | 0.018 | 0.397 | 0.2 | 0.01 | 0.19 | 0.08 | 0.34 | 0.25 |
| 6 | 24006893 | rs1397844 | -0.729 | 0.771 | -0.7 | -0.85 | -0.022 | -0.054 | -0.054 | 0 | 0.24 | 0.32 | 0.22 | 0.09 | 0.2 | 0.12 |
| 6 | 24009794 | rs10484660 | 0.049 | 1.393 | -1.78 | -0.558 | -0.034 | 0.031 | 0.031 | 0 | 0.16 | 0.01 | 0.17 | 0.07 | 0.18 | 0.09 |
| 6 | 24013681 | rs7765395 | 0.195 | 1.958 | -1.505 | -0.426 | -0.045 | 0.044 | 0.044 | 0 | 0.17 | 0.12 | 0.15 | 0.09 | 0.26 | 0.1 |
| 6 | 24013982 | rs1511481 | 0.613 | 1.385 | -1.051 | 0.417 | 0.002 | 0.009 | 0.009 | 0 | 0.8 | 0.83 | 0.74 | 0.9 | 0.79 | 0.9 |
| 6 | 24017908 | rs17529740 | -0.939 | 2.149 | -0.94 | 1.367 | -0.028 | -0.023 | -0.023 | 0 | 0.21 | 0.12 | 0.29 | 0.26 | 0.35 | 0.24 |
| 6 | 24019087 | rs6922630 | -1.511 | 2.091 | -0.012 | 1.151 | 0.045 | 0.026 | 0.026 | 0 | 0.26 | 0.42 | 0.42 | 0.32 | 0.39 | 0.31 |
| 6 | 24022898 | rs16888503 | -0.173 | 2.023 | 0.145 | 1.665 | 0.007 | 0.102 | 0.102 | 0 | 0.06 | 0.11 | 0.1 | 0.33 | 0.09 | 0.12 |
| 6 | 24026030 | rs9356919 | -0.44 | 1.928 | -0.249 | 1.043 | -0.008 | -0.007 | -0.007 | 3.69 | 0.35 | 0.25 | 0.5 | 0.6 | 0.43 | 0.39 |
| 6 | 24028403 | rs7761213 | -1.524 | 1.708 | 0.161 | 0.789 | 0.029 | 0.057 | 0.057 | 0.018 | 0.52 | 0.64 | 0.75 | 0.7 | 0.66 | 0.61 |
| 6 | 24034089 | rs12529384 | 1.269 | 1.788 | -0.13 | 0.778 | -0.045 | 0.04 | 0.04 | 0 | 0.18 | 0.01 | 0.24 | 0.04 | 0.21 | 0.17 |
| 6 | 24042393 | rs10946659 | 0.256 | 1.7 | 0.022 | 0.889 | 0 | 0.015 | 0.015 | 0 | 0.11 | 0.1 | 0.17 | 0.23 | 0.14 | 0.11 |
| 6 | 24046037 | rs1520968 | -0.094 | 1.799 | -0.102 | 0.768 | -0.006 | -0.003 | -0.003 | 0 | 0.1 | 0.07 | 0.2 | 0.09 | 0.2 | 0.12 |
| 6 | 24046561 | rs9393517 | -0.76 | 1.479 | 0.015 | 0.769 | 0.005 | 0.006 | 0.006 | 0.015 | 0.41 | 0.48 | 0.61 | 0.47 | 0.54 | 0.51 |
| 6 | 24052526 | rs9348626 | -0.515 | 1.546 | -0.222 | 0.889 | -0.016 | -0.014 | -0.014 | 0 | 0.36 | 0.36 | 0.57 | 0.47 | 0.52 | 0.43 |
| 6 | 24053169 | rs9358744 | -0.802 | 1.629 | -0.247 | 1.411 | 0.001 | 0.061 | 0.061 | 0 | 0.07 | 0.14 | 0.12 | 0.34 | 0.1 | 0.15 |
| 6 | 24053389 | rs9358745 | -0.404 | 1.555 | -0.352 | 1.438 | -0.006 | 0.045 | 0.045 | 0 | 0.06 | 0.06 | 0.1 | 0.33 | 0.08 | 0.15 |
| 6 | 24059407 | rs9467027 | -0.038 | 1.007 | -0.678 | 0.692 | 0.011 | -0.015 | -0.015 | 0 | 0.37 | 0.19 | 0.28 | 0.42 | 0.3 | 0.43 |
| 6 | 24059779 | rs10456299 | 0.604 | 0.989 | -0.868 | 0.75 | -0.003 | -0.014 | -0.014 | 0 | 0.24 | 0.03 | 0.12 | 0.19 | 0.17 | 0.34 |
| 6 | 24060381 | rs1277145 | 0.854 | 0.804 | 0.025 | 0.455 | -0.003 | 0.149 | 0.149 | 0 | 0.85 | 0.69 | 0.85 | 0.59 | 0.87 | 0.86 |
| 6 | 24068430 | rs1277130 | -0.084 | 1.286 | 0.253 | 0.343 | 0.032 | 0.021 | 0.021 | 0 | 0.09 | 0.17 | 0.13 | 0.11 | 0.19 | 0.14 |
| 6 | 24070837 | rs1520966 | 1.561 | 1.231 | 0.204 | 0.254 | 0.046 | 0.093 | 0.093 | 0 | 0.74 | 0.45 | 0.64 | 0.46 | 0.66 | 0.68 |
| 6 | 24073404 | rs1915463 | 0.81 | 1.289 | 0.113 | 0.317 | 0.042 | 0.286 | 0.286 | 0 | 0.87 | 0.57 | 0.83 | 0.57 | 0.87 | 0.87 |
| 6 | 24077431 | rs12210651 | 0.524 | 1.164 | 0.058 | 0.257 | 0.01 | -0.007 | -0.007 | 1.448 | 0.37 | 0.24 | 0.29 | 0.42 | 0.31 | 0.46 |
| 6 | 24078057 | rs9358749 | 0.33 | 1.236 | -0.089 | 0.473 | 0.007 | 0.033 | 0.033 | 0 | 0.93 | 0.73 | 0.94 | 0.9 | 0.97 | 0.99 |
| 6 | 24078258 | rs9460965 | 0.078 | 1.366 | -0.511 | 0.473 | -0.003 | -0.003 | -0.003 | 0.797 | 0.45 | 0.45 | 0.39 | 0.42 | 0.35 | 0.51 |
| 6 | 24080842 | rs6903992 | -0.36 | 0.665 | -0.217 | 0.18 | -0.004 | 0 | 0 | 0 | 0.54 | 0.51 | 0.51 | 0.52 | 0.54 | 0.65 |
| 6 | 24090497 | rs1402405 | -0.279 | 0.746 | -0.386 | 0.243 | 0.001 | -0.002 | -0.002 | 0 | 0.55 | 0.59 | 0.52 | 0.52 | 0.55 | 0.65 |
| 6 | 24098627 | rs1277152 | -0.291 | 0.607 | -0.548 | 0.141 | -0.002 | -0.003 | -0.003 | 0 | 0.53 | 0.55 | 0.62 | 0.63 | 0.52 | 0.61 |
| 6 | 24100190 | rs9467041 | 0.216 | 0.616 | -1.174 | -0.829 | 0.005 | 0.061 | 0.061 | 0 | 0.45 | 0.31 | 0.48 | 0.05 | 0.44 | 0.3 |
| 6 | 24101448 | rs11757735 | 0.878 | 0.491 | -1.837 | -0.708 | -0.069 | 0.211 | 0.211 | 1.113 | 0.35 | 0.03 | 0.3 | 0.03 | 0.42 | 0.23 |
| 6 | 24102723 | rs16888641 | -0.138 | 0.793 | -1.358 | 0.827 | -0.021 | 0.012 | 0.012 | 0 | 0.07 | 0.02 | 0.14 | 0.4 | 0.11 | 0.19 |
| 6 | 24105315 | rs6933683 | -0.51 | 0.478 | -1.257 | 0.706 | 0 | -0.001 | -0.001 | 0 | 0.1 | 0.11 | 0.11 | 0.14 | 0.08 | 0.18 |
| 6 | 24105330 | rs2328816 | 0.426 | -0.166 | -0.785 | -0.122 | -0.051 | -0.128 | -0.128 | 0.111 | 0.77 | 0.51 | 0.86 | 0.99 | 0.87 | 0.92 |
| 6 | 24105447 | rs9379643 | -0.238 | 0.498 | -0.978 | 0.127 | -0.003 | 0.003 | 0.003 | 0 | 0.18 | 0.18 | 0.28 | 0.41 | 0.23 | 0.26 |
| 6 | 24107093 | rs6922632 | -0.381 | -0.221 | -0.371 | -0.705 | -0.026 | -0.076 | -0.076 | 0.29 | 0.16 | 0.31 | 0.1 | 0.01 | 0.12 | 0.06 |
| 6 | 24112537 | rs1419229 | 1.271 | -0.034 | -0.057 | -0.393 | -0.005 | 0.028 | 0.028 | 0 | 0.54 | 0.31 | 0.51 | 0.3 | 0.55 | 0.49 |
| 6 | 24114594 | rs13215261 | 0.562 | -0.04 | -0.008 | -0.364 | 0.009 | 0 | 0 | 0 | 0.06 | 0.07 | 0.05 | 0.05 | 0.07 | 0.06 |
| 6 | 24116423 | rs9467044 | 2.19 | -0.141 | -0.104 | -0.612 | 0.003 | 0.157 | 0.157 | 0 | 0.42 | 0.18 | 0.33 | 0.13 | 0.43 | 0.33 |
| 6 | 24120269 | rs9467046 | 2.257 | -0.169 | -0.168 | -0.197 | -0.015 | 0.211 | 0.211 | 1.337 | 0.47 | 0.11 | 0.36 | 0.14 | 0.51 | 0.38 |
| 6 | 24121221 | rs11754578 | -0.312 | -0.085 | -0.293 | 0.188 | -0.005 | -0.005 | -0.005 | 0 | 0.2 | 0.15 | 0.23 | 0.53 | 0.23 | 0.32 |
| 6 | 24123314 | rs10946673 | -0.737 | -0.137 | -0.317 | 0.077 | 0.002 | 0.01 | 0.01 | 0.082 | 0.26 | 0.23 | 0.28 | 0.59 | 0.29 | 0.4 |
| 6 | 24126285 | rs4455659 | 0.849 | -0.132 | -0.338 | 0.062 | 0 | 0.001 | 0.001 | 0 | 0.88 | 0.84 | 0.95 | 0.92 | 0.96 | 0.95 |
| 6 | 24130821 | rs4449613 | -1.122 | 0.255 | -0.471 | 0.37 | -0.002 | -0.011 | -0.011 | 0 | 0.33 | 0.28 | 0.31 | 0.58 | 0.31 | 0.38 |
| 6 | 24133472 | rs12210008 | -0.352 | 0.282 | -0.428 | 0.688 | -0.027 | -0.033 | -0.033 | 0 | 0.06 | 0.01 | 0.12 | 0.16 | 0.08 | 0.12 |
| 6 | 24139942 | rs12202381 | -0.931 | 0.059 | 0.017 | 0.173 | -0.005 | -0.038 | -0.038 | 0.325 | 0.16 | 0.29 | 0.11 | 0.03 | 0.11 | 0.08 |
| 6 | 24140690 | rs12196541 | -0.817 | -0.007 | 0.007 | 0.13 | 0 | 0.019 | 0.019 | 0 | 0.17 | 0.14 | 0.16 | 0.05 | 0.13 | 0.08 |
| 6 | 24141920 | rs10946675 | 1.956 | -0.155 | -0.686 | 0.291 | -0.104 | 0 | 0 | 0.477 | 0.39 | 0.13 | 0.48 | 0.27 | 0.52 | 0.38 |
| 6 | 24142938 | rs12215160 | 0.474 | -0.443 | -0.066 | 0.709 | -0.019 | 0.095 | 0.095 | 0 | 0.74 | 0.59 | 0.7 | 0.45 | 0.79 | 0.7 |
| 6 | 24144867 | rs10946676 | 0.2 | -0.534 | 0.35 | 0.59 | 0.027 | -0.023 | -0.023 | 0 | 0.92 | 0.7 | 0.94 | 1 | 0.95 | 0.97 |
| 6 | 24146672 | rs3829810 | -0.989 | -0.619 | 0.267 | 0.598 | -0.007 | -0.003 | -0.003 | 0 | 0.31 | 0.14 | 0.27 | 0.27 | 0.26 | 0.32 |
| 6 | 24147483 | rs1053047 | 2.374 | -1.221 | 0.24 | 0.5 | -0.102 | 0.003 | 0.003 | 0 | 0.39 | 0.13 | 0.48 | 0.28 | 0.52 | 0.38 |
| 6 | 24147572 | rs3178 | 2.38 | -1.22 | 0.24 | 0.5 | -0.102 | 0.003 | 0.003 | 0 | 0.39 | 0.13 | 0.48 | 0.28 | 0.52 | 0.38 |
| 6 | 24157375 | rs2209544 | 0.286 | -1.166 | 0.116 | 0.206 | -0.01 | -0.054 | -0.054 | 0 | 0.12 | 0.01 | 0.13 | 0.18 | 0.11 | 0.16 |
| 6 | 24161869 | rs12211300 | 2.341 | -1.313 | 0.093 | 0.052 | -0.078 | 0.11 | 0.11 | 0.122 | 0.39 | 0.12 | 0.46 | 0.2 | 0.54 | 0.34 |
| 6 | 24163221 | rs4712804 | 1.65 | -1.294 | 0.33 | 0.111 | 0.049 | 0.093 | 0.093 | 0 | 0.55 | 0.26 | 0.55 | 0.24 | 0.65 | 0.46 |
| 6 | 24170975 | rs16888748 | 0.015 | -1.113 | 0.205 | 0.395 | -0.041 | -0.107 | -0.107 | 0.147 | 0.14 | 0.01 | 0.18 | 0.57 | 0.14 | 0.26 |
| 6 | 24173269 | rs6938529 | -0.183 | -1.114 | 0.373 | 0.364 | -0.009 | -0.037 | -0.037 | 0 | 0.16 | 0.07 | 0.24 | 0.57 | 0.16 | 0.26 |
| 6 | 24173382 | rs9467062 | -0.674 | -1.139 | 0.606 | 0.337 | 0.054 | 0.027 | 0.027 | 3.931 | 0.07 | 0.5 | 0.05 | 0 | 0.02 | 0 |
| 6 | 24173585 | rs7773379 | -0.674 | -1.139 | 0.606 | 0.337 | 0.054 | 0.027 | 0.027 | 3.931 | 0.07 | 0.49 | 0.05 | 0 | 0.02 | 0 |
| 6 | 24174186 | rs6901696 | -1.12 | -1.187 | 0.585 | 0.314 | 0.032 | 0.005 | 0.005 | 1.654 | 0.07 | 0.49 | 0.05 | 0 | 0.02 | 0 |
| 6 | 24174333 | rs6456594 | -1.345 | -1.159 | 0.624 | 0.335 | -0.006 | 0.03 | 0.03 | 0.243 | 0.25 | 0.64 | 0.3 | 0.58 | 0.19 | 0.27 |
| 6 | 24174803 | rs7744665 | -0.835 | -1.029 | 0.777 | 0.092 | 0.054 | 0.027 | 0.027 | 3.931 | 0.07 | 0.49 | 0.05 | 0 | 0.02 | 0 |

| | | | | | | | | | | | | | | | | |
|---|---|---|---|---|---|---|---|---|---|---|---|---|---|---|---|---|
| 6 | 24178306 | rs1419228 | 0.893 | -1 | 0.354 | 0.049 | -0.023 | -0.103 | -0.103 | 0.621 | 0.73 | 0.42 | 0.82 | 0.97 | 0.81 | 0.88 |
| 6 | 24185221 | rs2791972 | 1.117 | -1.039 | -0.277 | 0.191 | 0.029 | 0.041 | 0.041 | 0.56 | 0.59 | 0.29 | 0.68 | 0.78 | 0.71 | 0.66 |
| 6 | 24186590 | rs1277190 | 2.092 | -0.846 | -1.045 | 0.112 | -0.163 | 0.081 | 0.081 | 0.437 | 0.36 | 0.04 | 0.4 | 0.17 | 0.52 | 0.34 |
| 6 | 24188947 | rs9460974 | -0.662 | -0.846 | -0.362 | 0.066 | -0.001 | -0.008 | -0.008 | 0.971 | 0.23 | 0.26 | 0.16 | 0.05 | 0.19 | 0.18 |
| 6 | 24191459 | rs2027584 | -0.151 | -0.98 | -0.421 | 0.654 | -0.011 | -0.014 | -0.014 | 0 | 0.09 | 0.04 | 0.1 | 0.36 | 0.11 | 0.16 |
| 6 | 24193869 | rs1856839 | -0.086 | -1.022 | -0.393 | 0.552 | 0 | 0.025 | 0.025 | 0 | 0.18 | 0.11 | 0.11 | 0.04 | 0.15 | 0.14 |
| 6 | 24195759 | rs2799373 | 1.356 | -1.052 | -0.274 | 0.669 | -0.002 | 0.003 | 0.003 | 0 | 0.67 | 0.64 | 0.74 | 0.66 | 0.7 | 0.62 |
| 6 | 24210084 | rs9379648 | 0.386 | -1.323 | -0.48 | 0.888 | 0 | -0.067 | -0.067 | 0 | 0.1 | 0.01 | 0.14 | 0.24 | 0.06 | 0.1 |
| 6 | 24215665 | rs11755011 | 0.024 | -1.029 | -0.256 | 0.72 | -0.009 | 0.011 | 0.011 | 0 | 0.06 | 0.03 | 0.07 | 0.36 | 0.06 | 0.15 |
| 6 | 24218593 | rs1417215 | 1.991 | -1.264 | 0.555 | 0.562 | -0.019 | 0.047 | 0.047 | 0 | 0.8 | 0.41 | 0.8 | 0.63 | 0.86 | 0.77 |
| 6 | 24220393 | rs1340697 | 1.858 | -1.375 | 0.388 | 0.575 | -0.011 | 0.029 | 0.029 | 0 | 0.68 | 0.4 | 0.71 | 0.46 | 0.74 | 0.63 |
| 6 | 24221523 | rs11962116 | -0.432 | -1.35 | 0.907 | 0.575 | 0.017 | -0.015 | -0.015 | 0 | 0.05 | 0.2 | 0.08 | 0 | 0.04 | 0.01 |
| 6 | 24222452 | rs1770909 | 1.652 | -1.335 | 0.786 | 0.606 | -0.005 | 0.094 | 0.094 | 0 | 0.85 | 0.6 | 0.88 | 0.63 | 0.9 | 0.78 |
| 6 | 24228685 | rs793839 | 2.096 | -1.364 | 0.354 | 0.467 | 0.021 | -0.043 | -0.043 | 0 | 0.51 | 0.13 | 0.6 | 0.43 | 0.6 | 0.45 |
| 6 | 24232697 | rs793836 | 2.192 | -1.41 | 0.146 | 0.443 | -0.028 | -0.104 | -0.104 | 0 | 0.49 | 0.05 | 0.6 | 0.43 | 0.6 | 0.46 |
| 6 | 24233032 | rs9295616 | 0.019 | -1.5 | 0.518 | 0.393 | -0.006 | -0.026 | -0.026 | 0.904 | 0.16 | 0.15 | 0.09 | 0.03 | 0.11 | 0.12 |
| 6 | 24234933 | rs793834 | 1.504 | -1.474 | 0.379 | 0.449 | 0.023 | 0.005 | 0.005 | 0 | 0.67 | 0.26 | 0.72 | 0.61 | 0.77 | 0.72 |
| 6 | 24261060 | rs870601 | 1.547 | -1.4 | 0.19 | 0.196 | 0.04 | 0.111 | 0.111 | 1.373 | 0.58 | 0.13 | 0.68 | 0.79 | 0.69 | 0.67 |
| 6 | 24272384 | rs2792682 | 1.199 | -1.481 | 1.196 | 0.096 | -0.028 | -0.124 | -0.124 | 0.123 | 0.72 | 0.45 | 0.81 | 0.96 | 0.8 | 0.75 |
| 6 | 24273791 | rs807701 | 1.863 | -1.479 | 1.189 | 0.124 | 0.039 | 0.111 | 0.111 | 0.521 | 0.56 | 0.2 | 0.68 | 0.78 | 0.66 | 0.61 |
| 6 | 24275383 | rs807699 | 0.932 | -1.567 | 1.501 | 0.097 | -0.037 | -0.124 | -0.124 | 0 | 0.77 | 0.48 | 0.84 | 0.96 | 0.85 | 0.77 |
| 6 | 24280101 | rs3789224 | -0.488 | -1.499 | 1.5 | 0.156 | -0.011 | 0.022 | 0.022 | 0 | 0.16 | 0.25 | 0.13 | 0.18 | 0.12 | 0.14 |
| 6 | 24285571 | rs2328208 | 1.855 | -1.524 | 1.595 | 0.085 | -0.005 | -0.007 | -0.007 | 0 | 0.72 | 0.32 | 0.77 | 0.81 | 0.78 | 0.79 |
| 6 | 24286502 | rs3789226 | 0.034 | -1.573 | 1.369 | 0.051 | 0.001 | -0.006 | -0.006 | 0 | 0.15 | 0.13 | 0.09 | 0.03 | 0.11 | 0.18 |
| 6 | 24288724 | rs17302729 | -0.358 | -1.533 | 1.043 | 0.076 | -0.001 | 0.002 | 0.002 | 0 | 0.15 | 0.11 | 0.12 | 0.18 | 0.14 | 0.14 |
| 6 | 24291203 | rs2274305 | 2.154 | -1.519 | 0.714 | 0.084 | 0.014 | 0.132 | 0.132 | 1.236 | 0.54 | 0.05 | 0.67 | 0.78 | 0.67 | 0.61 |
| 6 | 24292185 | rs3804326 | 0.488 | -1.689 | 0.593 | 0.411 | 0.019 | -0.088 | -0.088 | 0 | 0.13 | 0.01 | 0.17 | 0.26 | 0.07 | 0.1 |
| 6 | 24292282 | rs3804327 | -0.843 | -1.051 | 1.115 | 0.409 | 0 | 0 | 0 | 0 | 0.17 | 0.17 | 0.15 | 0.18 | 0.18 | 0.16 |
| 6 | 24303195 | rs3846829 | -0.306 | -0.977 | 0.924 | 0.54 | -0.052 | -0.132 | -0.132 | 0 | 0.15 | 0.01 | 0.15 | 0.52 | 0.18 | 0.31 |
| 6 | 24304751 | rs707862 | -1.097 | -1.184 | 1.325 | -0.243 | -0.006 | -0.046 | -0.046 | 1.431 | 0.09 | 0.24 | 0.06 | 0 | 0.05 | 0.07 |
| 6 | 24305848 | rs707864 | 2.118 | -1.215 | 1.423 | -0.272 | -0.012 | -0.12 | -0.12 | 1.251 | 0.84 | 0.52 | 0.87 | 1 | 0.89 | 0.91 |
| 6 | 24312954 | rs16889066 | -1.021 | -1.033 | 1.163 | -0.088 | 0.005 | -0.033 | -0.033 | 0 | 0.07 | 0.17 | 0.07 | 0 | 0.06 | 0.02 |
| 6 | 24313747 | rs12196880 | -0.548 | -1.069 | 1.042 | -0.078 | -0.012 | -0.07 | -0.07 | 2.234 | 0.14 | 0.26 | 0.09 | 0.02 | 0.08 | 0.08 |
| 6 | 24318912 | rs12207932 | -0.959 | -0.948 | 0.934 | 0.034 | -0.018 | -0.005 | -0.005 | 0.017 | 0.24 | 0.24 | 0.15 | 0.18 | 0.16 | 0.25 |
| 6 | 24322287 | rs11966126 | -1.135 | -0.939 | 0.815 | -0.17 | 0.009 | -0.054 | -0.054 | 0 | 0.14 | 0.2 | 0.14 | 0.01 | 0.17 | 0.11 |
| 6 | 24323858 | rs9366567 | 0.893 | -0.987 | 0.837 | 0.087 | -0.047 | -0.017 | -0.017 | 0 | 0.76 | 0.64 | 0.84 | 0.82 | 0.84 | 0.75 |
| 6 | 24339448 | rs4052662 | 0.63 | -0.334 | 0.045 | 0.034 | -0.006 | 0.023 | 0.023 | 0 | 0.07 | 0.02 | 0.05 | 0 | 0.05 | 0.03 |
| 6 | 24342371 | rs12192603 | -1.202 | -0.6 | 0.577 | 0.167 | -0.018 | -0.013 | -0.013 | 0 | 0.27 | 0.59 | 0.25 | 0.2 | 0.23 | 0.22 |
| 6 | 24342612 | rs1535331 | 0.285 | -0.193 | 0.204 | 0.776 | -0.03 | 0.005 | 0.005 | 0 | 0.05 | 0.01 | 0.08 | 0.34 | 0.08 | 0.16 |
| 6 | 24346056 | rs1277351 | 1.995 | -0.384 | 0.11 | 0.47 | -0.009 | -0.022 | -0.022 | 1.005 | 0.53 | 0.16 | 0.59 | 0.45 | 0.57 | 0.49 |
| 6 | 24347624 | rs2753912 | 2.215 | -0.486 | 0.021 | 0.402 | -0.021 | -0.033 | -0.033 | 0 | 0.5 | 0.1 | 0.58 | 0.46 | 0.57 | 0.49 |
| 6 | 24347883 | rs793674 | 2.158 | -0.31 | 0.519 | 0.488 | 0.018 | 0.017 | 0.017 | 0 | 0.73 | 0.47 | 0.73 | 0.64 | 0.74 | 0.68 |
| 6 | 24347972 | rs6921810 | -1.132 | -0.393 | 0.679 | 0.303 | 0.061 | -0.021 | -0.021 | 2.15 | 0.08 | 0.39 | 0.12 | 0.01 | 0.07 | 0.03 |
| 6 | 24353742 | rs813227 | 0.984 | -0.278 | -0.593 | 0.298 | -0.015 | 0.022 | 0.022 | 0 | 0.48 | 0.22 | 0.47 | 0.41 | 0.51 | 0.44 |
| 6 | 24354045 | rs3765502 | -0.136 | -0.214 | -0.71 | 0.482 | -0.04 | -0.051 | -0.051 | 0.106 | 0.09 | 0 | 0.22 | 0.39 | 0.11 | 0.24 |
| 6 | 24377143 | rs793697 | 1.409 | -0.442 | -0.744 | 0.413 | -0.036 | 0.005 | 0.005 | 0.215 | 0.45 | 0.16 | 0.45 | 0.41 | 0.5 | 0.44 |
| 6 | 24380957 | rs9467129 | -0.371 | -0.694 | -0.2 | 0.076 | -0.002 | -0.026 | -0.026 | 0 | 0.38 | 0.51 | 0.28 | 0.19 | 0.38 | 0.27 |
| 6 | 24382897 | rs811103 | 0.586 | -0.519 | -0.895 | 0.231 | 0.013 | 0.092 | 0.092 | 0.36 | 0.56 | 0.22 | 0.68 | 0.8 | 0.59 | 0.68 |
| 6 | 24391075 | rs793676 | 0.762 | -0.169 | -1.63 | 0.446 | -0.114 | -0.125 | -0.125 | 0.715 | 0.4 | 0.03 | 0.52 | 0.62 | 0.55 | 0.54 |
| 6 | 24405120 | rs12055519 | -0.55 | -0.258 | -0.732 | 1.119 | -0.027 | -0.038 | -0.038 | 2.957 | 0.09 | 0.01 | 0.28 | 0.47 | 0.1 | 0.22 |
| 6 | 24413457 | rs6920147 | -1.446 | -0.681 | -0.358 | 0.534 | 0.003 | 0.004 | 0.004 | 0 | 0.38 | 0.4 | 0.48 | 0.75 | 0.35 | 0.53 |
| 6 | 24418348 | rs2793422 | 2.3 | -0.105 | -0.691 | 0.02 | -0.057 | 0.046 | 0.046 | 0 | 0.35 | 0.09 | 0.25 | 0.23 | 0.36 | 0.26 |
| 6 | 24420113 | rs1321820 | -0.789 | -0.324 | -0.246 | -0.071 | 0.009 | 0.071 | 0.071 | 0 | 0.16 | 0.25 | 0.39 | 0.48 | 0.21 | 0.27 |
| 6 | 24420234 | rs1772253 | 2.583 | -0.345 | -0.297 | -0.094 | -0.058 | 0.043 | 0.043 | 0 | 0.35 | 0.09 | 0.25 | 0.23 | 0.36 | 0.26 |
| 6 | 24420376 | rs1056283 | -1.567 | -0.181 | 0.18 | 0.102 | 0 | -0.008 | -0.008 | 0 | 0.39 | 0.51 | 0.53 | 0.75 | 0.43 | 0.53 |
| 6 | 24426959 | rs2298059 | 0.587 | -0.468 | -0.046 | 0.147 | -0.031 | -0.035 | -0.035 | 0 | 0.16 | 0.02 | 0.1 | 0.15 | 0.19 | 0.15 |
| 6 | 24429249 | rs6942189 | -0.831 | -0.477 | 0.215 | 0.188 | -0.008 | 0.12 | 0.12 | 0 | 0.16 | 0.28 | 0.35 | 0.48 | 0.14 | 0.24 |
| 6 | 24430292 | rs793683 | 1.604 | -0.552 | 0.3 | 0.142 | -0.037 | 0.097 | 0.097 | 0 | 0.77 | 0.54 | 0.6 | 0.4 | 0.84 | 0.65 |
| 6 | 24430746 | rs16889242 | 0.656 | -0.518 | 0.207 | 0.175 | -0.042 | -0.04 | -0.04 | 0 | 0.13 | 0.01 | 0.09 | 0.14 | 0.19 | 0.13 |
| 6 | 24433483 | rs9461006 | 0.364 | -0.632 | 0.502 | 0.118 | 0.012 | 0.011 | 0.011 | 0 | 0.16 | 0.21 | 0.11 | 0.15 | 0.2 | 0.14 |
| 6 | 24436760 | rs2273605 | -0.784 | -0.729 | 0.682 | 0.179 | -0.015 | 0.126 | 0.126 | 0 | 0.15 | 0.25 | 0.35 | 0.49 | 0.13 | 0.24 |
| 6 | 24438662 | rs793671 | 1.971 | -0.819 | 0.282 | -0.006 | 0.004 | -0.067 | -0.067 | 2.277 | 0.58 | 0.13 | 0.42 | 0.41 | 0.6 | 0.47 |
| 6 | 24439600 | rs9295624 | -0.328 | -0.841 | 0.375 | -0.029 | 0 | -0.008 | -0.008 | 0 | 0.16 | 0.18 | 0.22 | 0.02 | 0.24 | 0.19 |
| 6 | 24440351 | rs9461011 | -0.227 | -0.855 | 0.369 | -0.036 | 0 | -0.008 | -0.008 | 0 | 0.16 | 0.17 | 0.21 | 0.02 | 0.24 | 0.19 |
| 6 | 24441746 | rs9467160 | -0.929 | -0.857 | 0.445 | -0.038 | 0.044 | -0.098 | -0.098 | 0 | 0.2 | 0.38 | 0.23 | 0.02 | 0.25 | 0.19 |
| 6 | 24441758 | rs1742273 | -1.533 | -0.857 | 0.427 | -0.01 | -0.052 | 0.151 | 0.151 | 0 | 0.22 | 0.46 | 0.35 | 0.57 | 0.15 | 0.35 |
| 6 | 24444534 | rs793717 | 1.459 | -0.984 | 0.327 | -0.003 | 0.002 | -0.014 | -0.014 | 0 | 0.75 | 0.66 | 0.74 | 0.87 | 0.73 | 0.65 |
| 6 | 24445131 | rs793715 | 0.881 | -1.225 | 0.18 | -0.026 | 0.018 | 0.018 | 0.018 | 0 | 0.83 | 0.86 | 0.95 | 0.89 | 0.95 | 0.87 |
| 6 | 24447849 | rs9393556 | -1.6 | -1 | 0.402 | 0.095 | 0.019 | 0.077 | 0.077 | 0 | 0.28 | 0.48 | 0.47 | 0.47 | 0.34 | 0.39 |

| | | | | | | | | | | | | | | | | |
|---|---|---|---|---|---|---|---|---|---|---|---|---|---|---|---|---|
| 6 | 24448687 | rs13205780 | -1.203 | -1.001 | 0.339 | 0.029 | -0.005 | 0.01 | 0.01 | 0 | 0.24 | 0.23 | 0.24 | 0.02 | 0.28 | 0.2 |
| 6 | 24451256 | rs6456614 | -1.038 | -1.093 | 0.247 | 0.027 | -0.003 | 0.003 | 0.003 | 0.022 | 0.29 | 0.58 | 0.38 | 0.6 | 0.28 | 0.35 |
| 6 | 24454196 | rs9393558 | -1.089 | -0.838 | 0.34 | 0.175 | -0.002 | -0.001 | -0.001 | 0 | 0.24 | 0.29 | 0.46 | 0.48 | 0.33 | 0.4 |
| 6 | 24460679 | rs2793434 | 1.442 | -0.892 | 0.329 | 0.121 | -0.002 | 0.012 | 0.012 | 0 | 0.47 | 0.36 | 0.39 | 0.37 | 0.43 | 0.43 |
| 6 | 24464907 | rs9467176 | -0.488 | -0.941 | 0.624 | -0.082 | -0.039 | -0.052 | -0.052 | 0 | 0.31 | 0.52 | 0.15 | 0.11 | 0.16 | 0.19 |
| 6 | 24466721 | rs2244216 | -0.992 | -0.96 | 0.565 | 0.03 | -0.013 | 0.009 | 0.009 | 0 | 0.22 | 0.23 | 0.21 | 0.16 | 0.34 | 0.27 |
| 6 | 24467366 | rs6928435 | -0.366 | -0.964 | 0.563 | 0.016 | -0.039 | -0.052 | -0.052 | 0 | 0.3 | 0.52 | 0.15 | 0.11 | 0.16 | 0.19 |
| 6 | 24471393 | rs9358768 | -0.079 | -0.909 | 0.534 | 0.036 | -0.062 | 0.008 | 0.008 | 0 | 0.13 | 0.04 | 0.15 | 0.09 | 0.3 | 0.16 |
| 6 | 24472336 | rs2817246 | -1.128 | -0.915 | 0.53 | 0.041 | -0.014 | 0.01 | 0.01 | 0 | 0.22 | 0.23 | 0.21 | 0.16 | 0.34 | 0.27 |
| 6 | 24472423 | rs2817247 | 0.691 | -1.029 | 0.517 | 0.224 | -0.015 | 0.085 | 0.085 | 0 | 0.89 | 0.85 | 0.69 | 0.58 | 0.92 | 0.79 |
| 6 | 24476553 | rs760904 | 0.756 | -1.018 | 1.035 | 0.217 | -0.014 | 0.007 | 0.007 | 0 | 0.63 | 0.6 | 0.56 | 0.47 | 0.76 | 0.6 |
| 6 | 24476694 | rs2817204 | -0.206 | -0.989 | 0.953 | 0.247 | 0.002 | 0.019 | 0.019 | 0 | 0.93 | 0.95 | 0.7 | 0.58 | 0.9 | 0.79 |
| 6 | 24478813 | rs2760145 | -1.735 | -0.97 | 0.965 | 0.101 | -0.001 | 0.015 | 0.015 | 0 | 0.53 | 0.63 | 0.35 | 0.26 | 0.51 | 0.46 |
| 6 | 24479197 | rs2760144 | -1.069 | -0.97 | 0.986 | 0.088 | -0.005 | 0.004 | 0.004 | 0 | 0.22 | 0.25 | 0.21 | 0.16 | 0.34 | 0.27 |
| 6 | 24479845 | rs760906 | -1.749 | -0.961 | 0.969 | 0.094 | -0.001 | 0.015 | 0.015 | 0 | 0.53 | 0.63 | 0.35 | 0.26 | 0.51 | 0.46 |
| 6 | 24480295 | rs6911965 | -0.101 | -0.978 | 0.953 | 0.089 | -0.019 | -0.026 | -0.026 | 0 | 0.28 | 0.32 | 0.14 | 0.1 | 0.17 | 0.16 |
| 6 | 24481039 | rs2760140 | -1.395 | -0.921 | 1.015 | 0.114 | 0.006 | -0.01 | -0.01 | 0.789 | 0.25 | 0.31 | 0.22 | 0.17 | 0.34 | 0.3 |
| 6 | 24481299 | rs2760139 | -1.73 | -0.711 | 0.97 | 0.14 | -0.005 | 0.035 | 0.035 | 0 | 0.55 | 0.66 | 0.36 | 0.27 | 0.53 | 0.5 |
| 6 | 24487434 | rs2744570 | 0.029 | -0.583 | 0.932 | 0.219 | 0.008 | 0 | 0 | 0 | 0.12 | 0.11 | 0.15 | 0.1 | 0.29 | 0.16 |
| 6 | 24489652 | rs1126617 | -1.535 | -0.619 | 0.932 | 0.21 | 0.006 | -0.008 | -0.008 | 0 | 0.24 | 0.31 | 0.21 | 0.17 | 0.33 | 0.29 |
| 6 | 24489741 | rs2076316 | -2.045 | -0.706 | 0.957 | 0.161 | 0.027 | -0.047 | -0.047 | 0.334 | 0.3 | 0.62 | 0.26 | 0.17 | 0.37 | 0.33 |
| 6 | 24490147 | rs2076317 | -2.062 | -0.714 | 0.953 | 0.157 | 0.026 | -0.047 | -0.047 | 0.145 | 0.31 | 0.62 | 0.26 | 0.17 | 0.37 | 0.33 |
| 6 | 24490511 | rs2744572 | 0.071 | -0.761 | 1.002 | 0.284 | -0.004 | 0.081 | 0.081 | 0 | 0.92 | 0.9 | 0.69 | 0.58 | 0.92 | 0.79 |
| 6 | 24491475 | rs1883415 | -1.686 | -0.73 | 0.905 | 0.255 | 0.021 | -0.034 | -0.034 | 0 | 0.26 | 0.39 | 0.21 | 0.17 | 0.33 | 0.29 |
| 6 | 24493922 | rs3761793 | 0.298 | -0.831 | 0.812 | 0.227 | 0.011 | 0.011 | 0.011 | 0 | 0.25 | 0.17 | 0.11 | 0.1 | 0.15 | 0.17 |
| 6 | 24494463 | rs16889393 | 0.246 | -0.815 | 0.794 | 0.23 | 0.011 | 0.006 | 0.006 | 0 | 0.07 | 0.02 | 0.03 | 0.07 | 0.03 | 0.11 |
| 6 | 24494517 | rs11759284 | 2.609 | -0.812 | 0.775 | 0.281 | -0.134 | -0.072 | -0.072 | 0 | 0.34 | 0.04 | 0.32 | 0.31 | 0.4 | 0.29 |
| 6 | 24494924 | rs4646828 | 2.609 | -0.812 | 0.775 | 0.281 | -0.134 | -0.072 | -0.072 | 0.932 | 0.34 | 0.04 | 0.32 | 0.31 | 0.4 | 0.29 |
| 6 | 24496054 | rs2817213 | -1.9 | -0.748 | 0.888 | 0.215 | 0.028 | -0.036 | -0.036 | 0 | 0.28 | 0.4 | 0.22 | 0.17 | 0.35 | 0.33 |
| 6 | 24499551 | rs7775073 | -1.807 | -0.59 | 0.951 | 0.278 | 0.035 | -0.037 | -0.037 | 0.398 | 0.26 | 0.4 | 0.21 | 0.17 | 0.32 | 0.33 |
| 6 | 24501129 | rs4646834 | 0.287 | -0.614 | 0.95 | 0.274 | 0.019 | 0.022 | 0.022 | 0 | 0.27 | 0.17 | 0.12 | 0.11 | 0.19 | 0.18 |
| 6 | 24501421 | rs2245674 | -1.841 | -0.573 | 0.955 | 0.286 | 0.034 | -0.037 | -0.037 | 0 | 0.26 | 0.4 | 0.21 | 0.17 | 0.32 | 0.33 |
| 6 | 24503590 | rs2760118 | -2.222 | -0.561 | 1.032 | 0.293 | 0.039 | -0.056 | -0.056 | 0 | 0.28 | 0.47 | 0.22 | 0.17 | 0.32 | 0.32 |
| 6 | 24503898 | rs2817219 | -1.395 | -0.503 | 0.962 | 0.324 | 0.026 | -0.014 | -0.014 | 0 | 0.21 | 0.31 | 0.2 | 0.16 | 0.31 | 0.3 |
| 6 | 24504189 | rs2760117 | -1.395 | -0.503 | 0.978 | 0.324 | 0.011 | -0.006 | -0.006 | 0 | 0.21 | 0.29 | 0.2 | 0.16 | 0.31 | 0.3 |
| 6 | 24504238 | rs2760116 | -1.395 | -0.503 | 0.977 | 0.324 | 0.026 | -0.014 | -0.014 | 1.329 | 0.21 | 0.31 | 0.2 | 0.16 | 0.31 | 0.3 |
| 6 | 24505030 | rs2744583 | -1.417 | -0.56 | 0.954 | 0.303 | 0 | 0.013 | 0.013 | 0 | 0.5 | 0.55 | 0.33 | 0.27 | 0.51 | 0.47 |
| 6 | 24505221 | rs2817220 | -0.548 | -0.277 | 0.772 | 0.306 | 0.06 | -0.013 | -0.013 | 0 | 0.12 | 0.27 | 0.17 | 0.09 | 0.26 | 0.15 |
| 6 | 24507761 | rs2328824 | 1.709 | -0.331 | 0.759 | 0.403 | -0.011 | -0.015 | -0.015 | 3.993 | 0.72 | 0.67 | 0.5 | 0.42 | 0.62 | 0.5 |
| 6 | 24513636 | rs1569579 | 0.818 | -0.371 | 1.059 | 0.592 | -0.003 | 0.214 | 0.214 | 0 | 0.91 | 0.79 | 0.67 | 0.53 | 0.91 | 0.8 |
| 6 | 24514569 | rs2252525 | 2.119 | -0.517 | 1.139 | 0.598 | -0.032 | 0.078 | 0.078 | 0 | 0.78 | 0.6 | 0.64 | 0.53 | 0.86 | 0.63 |
| 6 | 24518321 | rs807513 | -0.073 | -0.443 | 1.081 | 0.592 | -0.002 | 0.041 | 0.041 | 0 | 0.93 | 0.96 | 0.69 | 0.54 | 0.92 | 0.8 |
| 6 | 24519886 | rs807515 | 1.296 | -0.509 | 1.066 | 0.509 | -0.02 | 0.15 | 0.15 | 0.018 | 0.85 | 0.63 | 0.65 | 0.54 | 0.88 | 0.8 |
| 6 | 24522794 | rs11751325 | 2.295 | -0.441 | 0.513 | 0.378 | -0.139 | -0.098 | -0.098 | 0 | 0.35 | 0.04 | 0.32 | 0.32 | 0.41 | 0.3 |
| 6 | 24524704 | rs807517 | -0.27 | -0.301 | 0.861 | -0.046 | 0.073 | -0.009 | -0.009 | 0 | 0.11 | 0.2 | 0.16 | 0.09 | 0.24 | 0.14 |
| 6 | 24526243 | rs807518 | -0.306 | -0.633 | 0.623 | 0.305 | -0.026 | 0.041 | 0.041 | 0 | 0.94 | 0.99 | 0.68 | 0.5 | 0.92 | 0.84 |
| 6 | 24526990 | rs809419 | 1.355 | -0.785 | 0.599 | 0.004 | 0.195 | 0.075 | 0.075 | 0.513 | 0.5 | 0.16 | 0.5 | 0.33 | 0.73 | 0.53 |
| 6 | 24530873 | rs2247845 | 1.056 | -0.777 | 0.593 | 0.045 | 0 | 0.03 | 0.03 | 0 | 0.86 | 0.83 | 0.63 | 0.43 | 0.9 | 0.73 |
| 6 | 24530934 | rs2267539 | -0.379 | -0.692 | 0.672 | -0.029 | 0.005 | 0.001 | 0.001 | 0 | 0.35 | 0.59 | 0.18 | 0.11 | 0.21 | 0.19 |
| 6 | 24531450 | rs12199955 | 2.139 | -0.384 | 0.543 | -0.046 | -0.104 | 0.022 | 0.022 | 0.028 | 0.4 | 0.05 | 0.35 | 0.26 | 0.54 | 0.42 |
| 6 | 24533074 | rs2744597 | -0.366 | -0.401 | 0.565 | -0.009 | -0.026 | 0.041 | 0.041 | 0 | 0.94 | 0.99 | 0.68 | 0.46 | 0.92 | 0.84 |
| 6 | 24533432 | rs2744600 | -0.74 | -0.409 | 0.822 | -0.059 | 0.001 | -0.003 | -0.003 | 0 | 0.42 | 0.55 | 0.26 | 0.17 | 0.34 | 0.32 |
| 6 | 24534193 | rs1054899 | -1.006 | -0.409 | 0.783 | -0.053 | 0.053 | 0.104 | 0.104 | 0 | 0.44 | 0.67 | 0.27 | 0.17 | 0.35 | 0.31 |
| 6 | 24537865 | rs2744603 | -0.407 | -0.441 | 1.019 | -0.067 | -0.021 | -0.035 | -0.035 | 0 | 0.36 | 0.39 | 0.21 | 0.16 | 0.28 | 0.27 |
| 6 | 24538113 | rs2817232 | 0.887 | -0.408 | 0.851 | 0.021 | -0.009 | 0.052 | 0.052 | 0 | 0.85 | 0.79 | 0.63 | 0.42 | 0.9 | 0.73 |
| 6 | 24539139 | rs1091031 | -1.378 | -0.452 | 0.832 | -0.025 | 0.014 | 0.159 | 0.159 | 0 | 0.48 | 0.75 | 0.33 | 0.17 | 0.46 | 0.43 |
| 6 | 24540539 | rs2744604 | 1.238 | -0.408 | 0.925 | 0.007 | 0.006 | 0.013 | 0.013 | 0 | 0.75 | 0.67 | 0.49 | 0.34 | 0.69 | 0.57 |
| 6 | 24540715 | rs807511 | -0.773 | -0.399 | 0.899 | -0.03 | -0.009 | 0.006 | 0.006 | 0 | 0.16 | 0.17 | 0.18 | 0.12 | 0.23 | 0.29 |
| 6 | 24541368 | rs2817239 | 1.548 | -0.442 | 0.943 | -0.067 | 0.01 | 0.079 | 0.079 | 0 | 0.45 | 0.3 | 0.38 | 0.24 | 0.54 | 0.4 |
| 6 | 24544332 | rs2744605 | -0.425 | -0.409 | 0.935 | -0.053 | 0.022 | -0.017 | -0.017 | 1.567 | 0.06 | 0.18 | 0.06 | 0.01 | 0.07 | 0.03 |
| 6 | 24544475 | rs807527 | 0.159 | -0.428 | 0.966 | -0.023 | -0.002 | 0.135 | 0.135 | 0 | 0.91 | 0.85 | 0.68 | 0.46 | 0.92 | 0.87 |
| 6 | 24544903 | rs699463 | -1.236 | -0.449 | 1.102 | -0.057 | 0.019 | -0.025 | -0.025 | 0 | 0.19 | 0.32 | 0.19 | 0.12 | 0.24 | 0.29 |
| 6 | 24544967 | rs16889440 | 0.206 | -0.518 | 1.056 | -0.095 | -0.033 | -0.025 | -0.025 | 0 | 0.27 | 0.27 | 0.11 | 0.1 | 0.14 | 0.17 |
| 6 | 24545307 | rs12196913 | 2.18 | -0.262 | 0.899 | -0.023 | -0.128 | 0.041 | 0.041 | 0 | 0.37 | 0.08 | 0.3 | 0.24 | 0.45 | 0.32 |
| 6 | 24546604 | rs2760182 | -0.288 | -0.264 | 0.918 | 0.108 | 0.028 | -0.013 | -0.013 | 0.216 | 0.05 | 0.17 | 0.05 | 0.01 | 0.07 | 0.03 |
| 6 | 24548561 | rs2760181 | -0.517 | -0.238 | 0.809 | 0.123 | 0.014 | -0.023 | -0.023 | 0 | 0.06 | 0.17 | 0.06 | 0.01 | 0.07 | 0.03 |
| 6 | 24549668 | rs807532 | 0.287 | -0.223 | 0.511 | 0.209 | -0.002 | 0.128 | 0.128 | 0 | 0.91 | 0.85 | 0.67 | 0.46 | 0.92 | 0.86 |
| 6 | 24550964 | rs9467220 | -0.499 | 0.099 | 0.528 | -0.166 | -0.013 | -0.043 | -0.043 | 0 | 0.35 | 0.42 | 0.18 | 0.11 | 0.29 | 0.23 |
| 6 | 24551664 | rs807534 | 0.333 | 0.112 | -0.116 | -0.098 | 0.002 | 0 | 0 | 0.042 | 0.08 | 0.08 | 0.09 | 0.08 | 0.13 | 0.11 |

| | | | | | | | | | | | | | | | | |
|---|---|---|---|---|---|---|---|---|---|---|---|---|---|---|---|---|
| 6 | 24551729 | rs807535 | 0.273 | 0.113 | -0.115 | -0.097 | 0.001 | 0 | 0 | 0.142 | 0.08 | 0.08 | 0.1 | 0.08 | 0.13 | 0.11 |
| 6 | 24554762 | rs2243831 | -0.485 | 0.007 | -0.148 | -0.169 | -0.022 | -0.04 | -0.04 | 0 | 0.37 | 0.53 | 0.17 | 0.11 | 0.24 | 0.23 |
| 6 | 24559281 | rs807541 | -0.685 | 0.21 | -0.785 | -0.306 | -0.011 | 0.001 | 0.001 | 0 | 0.13 | 0.14 | 0.17 | 0.11 | 0.22 | 0.24 |
| 6 | 24559433 | rs3903801 | 1.86 | 0.104 | -0.855 | -0.271 | -0.143 | 0.045 | 0.045 | 0 | 0.37 | 0.04 | 0.31 | 0.22 | 0.43 | 0.32 |
| 6 | 24564129 | rs807545 | 0.234 | 0.249 | -0.494 | -0.182 | -0.001 | 0 | 0 | 0.307 | 0.08 | 0.11 | 0.08 | 0.07 | 0.07 | 0.1 |
| 6 | 24568393 | rs12193738 | 1.839 | 0.12 | -0.786 | -0.334 | -0.098 | 0.124 | 0.124 | 0 | 0.4 | 0.1 | 0.32 | 0.19 | 0.47 | 0.3 |
| 6 | 24568452 | rs12213545 | 0.098 | 0.491 | -0.871 | 0.077 | -0.022 | -0.027 | -0.027 | 0 | 0.06 | 0.04 | 0.09 | 0.1 | 0.1 | 0.16 |
| 6 | 24575094 | rs2817198 | -0.423 | 0.131 | -0.473 | 0.044 | -0.016 | -0.044 | -0.044 | 0 | 0.29 | 0.29 | 0.12 | 0.09 | 0.18 | 0.2 |
| 6 | 24578272 | rs2760157 | -1.057 | 0.834 | 0.091 | 0.873 | 0.038 | 0.253 | 0.253 | 0 | 0.17 | 0.45 | 0.4 | 0.54 | 0.2 | 0.18 |
| 6 | 24581283 | rs807508 | -0.835 | 0.771 | -0.653 | 0.15 | 0.002 | -0.001 | -0.001 | 0 | 0.08 | 0.12 | 0.08 | 0.07 | 0.07 | 0.1 |
| 6 | 24584181 | rs6902039 | -0.396 | 0.755 | -1.656 | -0.901 | -0.011 | 0.2 | 0.2 | 0 | 0.38 | 0.11 | 0.25 | 0.04 | 0.35 | 0.31 |
| 6 | 24584366 | rs2817200 | -0.554 | 1.686 | -1.657 | -0.549 | 0.004 | 0 | 0 | 0.019 | 0.23 | 0.25 | 0.31 | 0.25 | 0.42 | 0.29 |
| 6 | 24590547 | rs5026394 | 0.713 | 0.894 | -1.307 | -0.609 | 0.014 | 0.009 | 0.009 | 0 | 0.5 | 0.39 | 0.38 | 0.41 | 0.38 | 0.42 |
| 6 | 24596478 | rs4576240 | -0.744 | 0.874 | -1.384 | -0.728 | 0.006 | 0.039 | 0.039 | 0 | 0.87 | 0.94 | 0.94 | 0.99 | 0.9 | 0.93 |
| 6 | 24605024 | rs17307478 | 1.828 | 1.564 | -1.454 | -0.791 | -0.026 | 0.025 | 0.025 | 2.027 | 0.05 | 0.01 | 0.04 | 0.01 | 0.11 | 0.03 |
| 6 | 24605424 | rs4236031 | -0.392 | 0.973 | -1.097 | -0.809 | 0.002 | -0.002 | -0.002 | 0 | 0.83 | 0.83 | 0.91 | 0.98 | 0.87 | 0.93 |
| 6 | 24612902 | rs6915373 | -0.154 | 0.961 | -0.454 | -0.534 | 0.036 | 0.42 | 0.42 | 0.143 | 0.06 | 0.26 | 0.28 | 0.47 | 0.03 | 0.15 |
| 6 | 24614392 | rs9379671 | 0.867 | 0.296 | -0.811 | -1.129 | -0.011 | -0.039 | -0.039 | 0 | 0.19 | 0.12 | 0.22 | 0.26 | 0.19 | 0.14 |
| 6 | 24615063 | rs7763790 | 0.707 | 0.004 | -0.491 | -1.358 | -0.056 | -0.074 | -0.074 | 0.287 | 0.31 | 0.38 | 0.15 | 0.13 | 0.18 | 0.31 |
| 6 | 24628472 | rs9366573 | -0.232 | 1.7 | -0.578 | -0.848 | 0.005 | -0.001 | -0.001 | 0 | 0.82 | 0.84 | 0.79 | 0.89 | 0.62 | 0.7 |
| 6 | 24632642 | rs761100 | 0.443 | 1.44 | -0.403 | -1.213 | 0.012 | -0.018 | -0.018 | 0.023 | 0.72 | 0.69 | 0.72 | 0.88 | 0.54 | 0.64 |
| 6 | 24633429 | rs7766230 | 1.129 | 0.502 | -0.707 | -1.119 | 0.035 | -0.044 | -0.044 | 0.115 | 0.31 | 0.2 | 0.38 | 0.66 | 0.19 | 0.44 |
| 6 | 24640196 | rs9356940 | -0.784 | 0.571 | -0.702 | -1.127 | 0.011 | -0.005 | -0.005 | 0 | 0.81 | 0.83 | 0.79 | 0.89 | 0.63 | 0.7 |
| 6 | 24641328 | rs9366577 | 0.956 | 0.222 | -0.631 | -0.834 | 0.004 | 0.022 | 0.022 | 0 | 0.19 | 0.13 | 0.1 | 0.06 | 0.16 | 0.11 |
| 6 | 24641605 | rs16889556 | 1.685 | 0.105 | -0.593 | -0.816 | -0.013 | -0.03 | -0.03 | 0 | 0.15 | 0.05 | 0.16 | 0.13 | 0.17 | 0.08 |
| 6 | 24645967 | rs2038135 | -0.843 | 0.236 | -0.31 | -0.645 | 0.007 | -0.002 | -0.002 | 0 | 0.81 | 0.83 | 0.79 | 0.89 | 0.63 | 0.7 |
| 6 | 24646821 | rs3756821 | 0.002 | 0.185 | -0.307 | -0.389 | -0.002 | -0.002 | -0.002 | 0 | 0.47 | 0.49 | 0.37 | 0.26 | 0.4 | 0.24 |
| 6 | 24649061 | rs9461045 | 0.908 | -0.046 | -0.345 | -0.061 | 0.033 | -0.058 | -0.058 | 0 | 0.31 | 0.2 | 0.39 | 0.63 | 0.19 | 0.44 |
| 6 | 24650237 | rs12393 | -0.288 | 0.283 | 0.271 | -0.021 | 0.032 | 0.044 | 0.044 | 1.197 | 0.06 | 0.3 | 0.07 | 0.02 | 0.02 | 0.01 |
| 6 | 24650671 | rs707887 | -0.496 | 1.043 | 0.495 | -0.132 | 0.067 | -0.014 | -0.014 | 0 | 0.84 | 0.7 | 0.82 | 0.88 | 0.68 | 0.72 |
| 6 | 24650761 | rs3087943 | -0.265 | 0.691 | 0.179 | -0.421 | 0.014 | -0.059 | -0.059 | 0.118 | 0.17 | 0.27 | 0.18 | 0.03 | 0.18 | 0.1 |
| 6 | 24651604 | rs9379679 | -0.572 | 0.198 | 0.355 | 0.54 | -0.004 | 0.219 | 0.219 | 0 | 0.07 | 0.23 | 0.29 | 0.46 | 0.03 | 0.17 |
| 6 | 24653273 | rs2294689 | -0.786 | 0.212 | 0.479 | 0.538 | 0 | 0.216 | 0.216 | 0 | 0.07 | 0.26 | 0.3 | 0.46 | 0.04 | 0.17 |
| 6 | 24653376 | rs1129644 | -0.776 | 0.49 | -0.083 | 0.054 | -0.056 | 0.032 | 0.032 | 0 | 0.77 | 0.93 | 0.84 | 0.87 | 0.66 | 0.7 |
| 6 | 24656571 | rs7768291 | 1.147 | -0.067 | 0.141 | -0.238 | 0.002 | 0.002 | 0.002 | 0.016 | 0.24 | 0.23 | 0.12 | 0.15 | 0.16 | 0.24 |
| 6 | 24657168 | rs9467251 | -0.278 | -0.076 | 0.18 | -0.2 | -0.001 | 0.206 | 0.206 | 0 | 0.07 | 0.25 | 0.3 | 0.46 | 0.04 | 0.17 |
| 6 | 24659071 | rs2143340 | 1.359 | -0.091 | 0.25 | -0.207 | 0.005 | 0.008 | 0.008 | 0.593 | 0.23 | 0.18 | 0.12 | 0.14 | 0.16 | 0.24 |
| 6 | 24659482 | rs3181230 | -0.275 | -0.085 | 0.24 | -0.197 | 0 | 0.219 | 0.219 | 0 | 0.07 | 0.26 | 0.3 | 0.46 | 0.04 | 0.17 |
| 6 | 24661599 | rs2056998 | -0.461 | -0.129 | 0.22 | -0.22 | -0.005 | 0.199 | 0.199 | 0 | 0.08 | 0.26 | 0.3 | 0.46 | 0.04 | 0.17 |
| 6 | 24665340 | rs3756819 | 1.362 | -0.372 | 0.16 | -0.257 | 0.006 | 0.01 | 0.01 | 0.586 | 0.23 | 0.18 | 0.12 | 0.14 | 0.16 | 0.24 |
| 6 | 24666007 | rs1061925 | 1.616 | -0.518 | 0.304 | -0.367 | -0.023 | -0.039 | -0.039 | 0 | 0.19 | 0.23 | 0.1 | 0.08 | 0.12 | 0.23 |
| 6 | 24667323 | rs17243157 | 1.771 | -0.517 | -0.084 | -0.369 | 0.026 | 0.046 | 0.046 | 1.09 | 0.19 | 0.05 | 0.09 | 0.08 | 0.12 | 0.24 |
| 6 | 24667662 | rs3181228 | 1.631 | -0.474 | -0.47 | -0.55 | -0.007 | 0.074 | 0.074 | 0 | 0.24 | 0.07 | 0.15 | 0.06 | 0.25 | 0.14 |
| 6 | 24667799 | rs3181227 | 1.536 | -0.49 | -0.485 | -0.575 | -0.046 | -0.054 | -0.054 | 0 | 0.17 | 0.05 | 0.21 | 0.18 | 0.21 | 0.12 |
| 6 | 24669945 | rs2223588 | -1.27 | -0.534 | -0.435 | -0.567 | -0.056 | 0.032 | 0.032 | 0 | 0.77 | 0.93 | 0.84 | 0.87 | 0.66 | 0.7 |
| 6 | 24680290 | rs6928074 | 0.732 | -0.483 | -0.433 | -0.576 | -0.04 | 0.02 | 0.02 | 0 | 0.42 | 0.18 | 0.36 | 0.24 | 0.46 | 0.26 |
| 6 | 24690449 | rs12207801 | 1.971 | -0.32 | -0.405 | -0.492 | -0.015 | 0.047 | 0.047 | 0.24 | 0.08 | 0 | 0.05 | 0.01 | 0.06 | 0.04 |
| 6 | 24692179 | rs1885211 | 0.23 | -0.349 | -0.192 | -0.468 | 0.003 | 0.007 | 0.007 | 0 | 0.61 | 0.55 | 0.47 | 0.39 | 0.57 | 0.47 |
| 6 | 24693523 | rs3777665 | 1.418 | -0.375 | -0.529 | -0.609 | 0.007 | 0.01 | 0.01 | 0.532 | 0.23 | 0.17 | 0.11 | 0.13 | 0.15 | 0.24 |
| 6 | 24693846 | rs3777664 | -1.151 | -0.261 | -0.885 | -0.533 | -0.041 | 0.025 | 0.025 | 0 | 0.78 | 0.85 | 0.85 | 0.85 | 0.67 | 0.72 |
| 6 | 24698215 | rs7765904 | 1.39 | -0.278 | -1.116 | -0.571 | 0.009 | 0.065 | 0.065 | 0 | 0.25 | 0.13 | 0.16 | 0.06 | 0.25 | 0.14 |
| 6 | 24698500 | rs12193165 | 1.203 | -0.084 | -1.068 | -0.397 | -0.063 | -0.059 | -0.059 | 0 | 0.17 | 0.02 | 0.21 | 0.16 | 0.21 | 0.12 |
| 6 | 24700235 | rs3777663 | 1.211 | -0.134 | -1.104 | -0.417 | -0.07 | -0.049 | -0.049 | 0 | 0.17 | 0.01 | 0.21 | 0.15 | 0.21 | 0.12 |
| 6 | 24704538 | rs1053598 | -0.808 | -0.05 | -0.885 | -0.289 | -0.018 | -0.043 | -0.043 | 0 | 0.74 | 0.78 | 0.59 | 0.36 | 0.67 | 0.57 |
| 6 | 24706770 | rs6931809 | 0.868 | -0.318 | -0.245 | -0.571 | -0.005 | -0.023 | -0.023 | 0 | 0.31 | 0.33 | 0.22 | 0.09 | 0.26 | 0.16 |
| 6 | 24706852 | rs6909099 | 0.868 | -0.318 | -0.245 | -0.577 | -0.005 | -0.024 | -0.024 | 1.334 | 0.31 | 0.33 | 0.22 | 0.08 | 0.26 | 0.16 |
| 6 | 24709202 | rs2294686 | 0.03 | -0.283 | -0.994 | -0.467 | -0.005 | 0.233 | 0.233 | 0 | 0.05 | 0.19 | 0.27 | 0.49 | 0.03 | 0.16 |
| 6 | 24709830 | rs9356943 | 0.958 | -0.144 | -1.38 | -0.621 | 0.009 | 0.079 | 0.079 | 0 | 0.28 | 0.13 | 0.16 | 0.06 | 0.26 | 0.16 |
| 6 | 24719978 | rs3813687 | 1.459 | -0.114 | -0.826 | -0.535 | -0.047 | -0.05 | -0.05 | 0 | 0.08 | 0 | 0.15 | 0.14 | 0.15 | 0.09 |
| 6 | 24721787 | rs9379687 | 0.126 | -0.621 | -0.796 | -0.588 | 0.013 | -0.021 | -0.021 | 0.073 | 0.14 | 0.11 | 0.3 | 0.55 | 0.09 | 0.24 |
| 6 | 24726408 | rs2145753 | 0.192 | -0.559 | -0.358 | -0.971 | -0.006 | 0.021 | 0.021 | 0.341 | 0.37 | 0.47 | 0.52 | 0.58 | 0.32 | 0.29 |
| 6 | 24729115 | rs2328830 | -1.592 | -0.55 | -0.275 | -0.397 | -0.058 | 0.097 | 0.097 | 0 | 0.6 | 0.84 | 0.74 | 0.82 | 0.48 | 0.61 |
| 6 | 24735308 | rs10946724 | -0.747 | -0.571 | -0.164 | -0.152 | 0.026 | 0.003 | 0.003 | 0 | 0.39 | 0.22 | 0.5 | 0.68 | 0.31 | 0.47 |
| 6 | 24738595 | rs2876673 | -0.413 | -0.013 | 0.234 | 0.33 | -0.005 | -0.048 | -0.048 | 0 | 0.18 | 0.1 | 0.39 | 0.56 | 0.17 | 0.21 |
| 6 | 24758824 | rs9358796 | -0.884 | -0.396 | 0.594 | 0.037 | -0.003 | 0.303 | 0.303 | 0 | 0.1 | 0.34 | 0.33 | 0.6 | 0.05 | 0.12 |
| 6 | 24760958 | rs2817739 | -1.324 | -0.413 | -0.378 | -0.393 | -0.116 | 0.188 | 0.188 | 0.904 | 0.58 | 0.98 | 0.72 | 0.86 | 0.49 | 0.67 |
| 6 | 24771469 | rs6904263 | 0.463 | 0.022 | 0.117 | 0.125 | -0.021 | -0.023 | -0.023 | 0 | 0.12 | 0.03 | 0.15 | 0.11 | 0.19 | 0.12 |
| 6 | 24781025 | rs2245409 | -0.152 | -1.05 | 0.608 | 0.416 | -0.009 | -0.01 | -0.01 | 0.045 | 0.2 | 0.26 | 0.12 | 0.1 | 0.12 | 0.26 |
| 6 | 24781735 | rs2307306 | -0.134 | -0.965 | 0.218 | -0.121 | -0.029 | -0.009 | -0.009 | 0 | 0.07 | 0.25 | 0.03 | 0.03 | 0 | 0.04 |

| | | | | | | | | | | | | | | | | |
|---|---|---|---|---|---|---|---|---|---|---|---|---|---|---|---|---|
| 6 | 24791100 | rs2817738 | -0.563 | -0.23 | -0.33 | 0.224 | -0.035 | 0.032 | 0.032 | 0 | 0.26 | 0.44 | 0.43 | 0.7 | 0.16 | 0.4 |
| 6 | 24803825 | rs4074236 | 0.224 | 0.344 | -1.236 | -0.508 | -0.029 | 0.15 | 0.15 | 0 | 0.39 | 0.14 | 0.22 | 0.09 | 0.38 | 0.28 |
| 6 | 24804860 | rs1803005 | -0.055 | 0.588 | -1.316 | 0.515 | -0.03 | -0.059 | -0.059 | 0.044 | 0.16 | 0.07 | 0.32 | 0.53 | 0.22 | 0.37 |
| 6 | 24806070 | rs1132352 | 0.363 | 0.081 | -1.05 | -0.338 | -0.002 | 0.072 | 0.072 | 0 | 0.54 | 0.27 | 0.43 | 0.19 | 0.53 | 0.43 |
| 6 | 24808771 | rs7756233 | -0.072 | -0.33 | -0.627 | -0.434 | -0.015 | -0.027 | -0.027 | 0 | 0.23 | 0.19 | 0.41 | 0.69 | 0.3 | 0.44 |
| 6 | 24809771 | rs9379692 | -0.289 | -0.332 | -0.605 | -0.435 | -0.033 | -0.046 | -0.046 | 1.058 | 0.76 | 0.42 | 0.83 | 0.88 | 0.82 | 0.85 |
| 6 | 24819447 | rs9358803 | 0.772 | -0.372 | -0.653 | -0.467 | 0.004 | 0.106 | 0.106 | 0 | 0.52 | 0.22 | 0.42 | 0.19 | 0.52 | 0.41 |
| 6 | 24822304 | rs4712856 | -0.216 | -0.315 | -0.376 | -0.575 | -0.046 | -0.133 | -0.133 | 2.734 | 0.76 | 0.45 | 0.84 | 0.96 | 0.83 | 0.81 |
| 6 | 24827691 | rs4610554 | -0.12 | -0.306 | -0.404 | -0.504 | 0.006 | 0.013 | 0.013 | 0.145 | 0.24 | 0.25 | 0.45 | 0.69 | 0.31 | 0.45 |
| 6 | 24836161 | rs6901565 | -0.487 | -0.214 | 0.134 | -0.796 | -0.024 | -0.072 | -0.072 | 0 | 0.79 | 0.66 | 0.87 | 0.95 | 0.84 | 0.81 |
| 6 | 24837206 | rs4712860 | -0.251 | -0.292 | -0.04 | -1.052 | -0.003 | 0.009 | 0.009 | 0 | 0.43 | 0.37 | 0.61 | 0.86 | 0.46 | 0.56 |
| 6 | 24844443 | rs10946731 | -0.445 | -0.112 | -0.469 | -0.914 | -0.025 | -0.084 | -0.084 | 0 | 0.79 | 0.65 | 0.86 | 0.96 | 0.84 | 0.81 |
| 6 | 24849666 | rs4451128 | -0.939 | 0.454 | -0.824 | -0.361 | -0.027 | 0.061 | 0.061 | 0 | 0.69 | 0.87 | 0.78 | 0.9 | 0.64 | 0.78 |
| 6 | 24851787 | rs9393592 | 0.324 | 0.336 | -1.202 | -0.141 | -0.075 | 0.084 | 0.084 | 0 | 0.43 | 0.11 | 0.36 | 0.2 | 0.5 | 0.31 |
| 6 | 24852751 | rs9467333 | -0.751 | 0.531 | -0.699 | 0.68 | 0.003 | 0.024 | 0.024 | 0 | 0.14 | 0.16 | 0.2 | 0.23 | 0.15 | 0.23 |
| 6 | 24856323 | rs4437459 | 0.764 | 0.48 | -0.572 | 0.727 | 0.005 | 0.049 | 0.049 | 0 | 0.87 | 0.82 | 0.8 | 0.76 | 0.85 | 0.79 |
| 6 | 24859663 | rs4364477 | 0.115 | 0.065 | -1.562 | 0.975 | 0.003 | -0.036 | -0.036 | 0 | 0.25 | 0.07 | 0.27 | 0.32 | 0.27 | 0.24 |
| 6 | 24863075 | rs4256430 | 1.133 | 0.859 | -0.942 | 0.598 | 0.001 | 0.015 | 0.015 | 0 | 0.48 | 0.44 | 0.42 | 0.34 | 0.43 | 0.4 |
| 6 | 24868080 | rs6906206 | 1.155 | 1.021 | -0.516 | 0.76 | 0.017 | 0.006 | 0.006 | 0 | 0.71 | 0.41 | 0.65 | 0.71 | 0.66 | 0.6 |
| 6 | 24870448 | rs1884159 | -1.36 | 1.021 | -0.516 | 0.752 | 0.017 | 0.017 | 0.017 | 0 | 0.29 | 0.57 | 0.34 | 0.35 | 0.34 | 0.41 |
| 6 | 24871357 | rs2747666 | -0.063 | 1.235 | -0.436 | 0.607 | 0.022 | 0.051 | 0.051 | 0 | 0.13 | 0.21 | 0.21 | 0.19 | 0.2 | 0.21 |
| 6 | 24872939 | rs10946735 | -0.126 | 0.331 | -0.745 | -0.091 | -0.006 | -0.022 | -0.022 | 0 | 0.16 | 0.16 | 0.07 | 0.02 | 0.08 | 0.11 |
| 6 | 24881498 | rs6930916 | 1.232 | 0.119 | -1.066 | -0.025 | 0.013 | -0.051 | -0.051 | 0 | 0.21 | 0.08 | 0.25 | 0.32 | 0.22 | 0.18 |
| 6 | 24882122 | rs13194040 | -0.725 | 0.48 | -0.016 | -0.25 | 0.033 | -0.011 | -0.011 | 0 | 0.11 | 0.2 | 0.11 | 0.08 | 0.18 | 0.19 |
| 6 | 24885300 | rs13192978 | -0.433 | 0.283 | 0.236 | -0.231 | -0.035 | -0.073 | -0.073 | 0 | 0.16 | 0.3 | 0.09 | 0.01 | 0.08 | 0.09 |
| 6 | 24887485 | rs2747680 | -0.044 | 0.346 | -0.101 | -0.14 | 0.01 | 0.06 | 0.06 | 0 | 0.17 | 0.36 | 0.22 | 0.17 | 0.16 | 0.19 |
| 6 | 24898856 | rs17671350 | -0.101 | 0.099 | -0.078 | -0.26 | 0.009 | 0.012 | 0.012 | 0 | 0.12 | 0.07 | 0.06 | 0.01 | 0.07 | 0.09 |
| 6 | 24899069 | rs2817722 | -0.775 | 0.256 | -0.28 | -0.257 | 0.043 | 0.028 | 0.028 | 0 | 0.25 | 0.42 | 0.3 | 0.25 | 0.31 | 0.35 |
| 6 | 24901350 | rs2817725 | -1.572 | 0.485 | -0.652 | -0.211 | -0.052 | 0.054 | 0.054 | 0 | 0.45 | 0.88 | 0.41 | 0.26 | 0.42 | 0.48 |
| 6 | 24904676 | rs2747677 | -1.379 | 0.553 | -0.584 | -0.102 | -0.067 | 0.014 | 0.014 | 0 | 0.42 | 0.9 | 0.4 | 0.27 | 0.39 | 0.44 |
| 6 | 24913789 | rs2747676 | -1.03 | 0.724 | -0.135 | 0.079 | 0.005 | -0.002 | -0.002 | 0.268 | 0.34 | 0.38 | 0.33 | 0.26 | 0.35 | 0.42 |
| 6 | 24926730 | rs606162 | -0.763 | 1.132 | 0.424 | 0.029 | 0.06 | -0.06 | -0.06 | 0.218 | 0.1 | 0.3 | 0.11 | 0.01 | 0.14 | 0.08 |
| 6 | 24931083 | rs216259 | -0.843 | 1.032 | -0.282 | 0.293 | -0.01 | -0.042 | -0.042 | 0 | 0.77 | 0.93 | 0.73 | 0.71 | 0.72 | 0.79 |
| 6 | 24931287 | rs216258 | -0.107 | 0.891 | -0.36 | 0.176 | -0.001 | 0 | 0 | 0.283 | 0.29 | 0.29 | 0.27 | 0.4 | 0.32 | 0.35 |
| 6 | 24938277 | rs13199679 | -0.914 | 0.579 | -0.305 | 0.496 | 0.004 | 0.03 | 0.03 | 0 | 0.06 | 0.11 | 0.05 | 0.13 | 0.06 | 0.08 |
| 6 | 24939867 | rs529648 | -0.291 | 0.68 | -0.333 | 0.49 | 0.004 | 0.038 | 0.038 | 0 | 0.19 | 0.2 | 0.2 | 0.49 | 0.19 | 0.27 |
| 6 | 24940849 | rs9467354 | -0.19 | 0.645 | -0.094 | 0.442 | -0.001 | -0.003 | -0.003 | 0 | 0.63 | 0.68 | 0.56 | 0.34 | 0.58 | 0.6 |
| 6 | 24941436 | rs419288 | 0.428 | 0.642 | -0.071 | 0.455 | 0.009 | 0.059 | 0.059 | 0 | 0.14 | 0.25 | 0.17 | 0.37 | 0.14 | 0.2 |
| 6 | 24941502 | rs13212828 | 1.013 | 0.426 | -0.939 | 0.166 | 0.041 | 0.066 | 0.066 | 0 | 0.31 | 0.12 | 0.18 | 0.14 | 0.23 | 0.2 |
| 6 | 24942222 | rs414243 | -0.624 | 0.844 | -0.578 | 0.891 | -0.015 | -0.06 | -0.06 | 0 | 0.75 | 0.9 | 0.69 | 0.59 | 0.68 | 0.74 |
| 6 | 24943775 | rs644748 | -0.043 | 0.679 | -0.402 | 0.9 | 0 | 0.005 | 0.005 | 0 | 0.15 | 0.17 | 0.17 | 0.38 | 0.16 | 0.23 |
| 6 | 24948508 | rs487700 | -0.571 | 0.637 | -0.534 | 0.8 | 0.007 | -0.006 | -0.006 | 1.922 | 0.76 | 0.87 | 0.71 | 0.74 | 0.73 | 0.78 |
| 6 | 24950599 | rs7759489 | 0.008 | 0.712 | -0.322 | 1.056 | 0.004 | 0.043 | 0.043 | 0 | 0.16 | 0.23 | 0.19 | 0.52 | 0.16 | 0.23 |
| 6 | 24950829 | rs680009 | 0.247 | 0.547 | 0.048 | 0.505 | -0.005 | -0.011 | -0.011 | 0 | 0.57 | 0.5 | 0.47 | 0.23 | 0.56 | 0.54 |
| 6 | 24951199 | rs647297 | 0.733 | 0.705 | 0.336 | 0.092 | 0.028 | -0.039 | -0.039 | 0 | 0.8 | 0.58 | 0.85 | 0.88 | 0.78 | 0.81 |
| 6 | 24951809 | rs387628 | -0.166 | -0.23 | 0.137 | -0.183 | -0.001 | -0.003 | -0.003 | 0 | 0.32 | 0.3 | 0.41 | 0.25 | 0.32 | 0.41 |
| 6 | 24952847 | rs127485 | 1.474 | -1.201 | 0.385 | -0.613 | -0.027 | -0.06 | -0.06 | 0 | 0.78 | 0.35 | 0.82 | 0.97 | 0.9 | 0.91 |
| 6 | 24957511 | rs6913176 | -0.638 | -1.251 | 0.689 | -0.565 | -0.029 | -0.024 | -0.024 | 0.004 | 0.09 | 0.2 | 0.05 | 0.03 | 0.02 | 0.05 |
| 6 | 24958507 | rs678784 | 1.691 | -1.287 | 0.539 | -0.568 | -0.047 | -0.06 | -0.06 | 0 | 0.8 | 0.35 | 0.84 | 0.95 | 0.93 | 0.92 |
| 6 | 24959844 | rs396599 | 1.64 | -1.283 | 0.55 | -0.571 | -0.047 | -0.067 | -0.067 | 2.045 | 0.8 | 0.36 | 0.84 | 0.97 | 0.93 | 0.93 |
| 6 | 24962428 | rs393228 | 1.64 | -1.252 | 0.549 | -0.571 | -0.043 | -0.065 | -0.065 | 0 | 0.8 | 0.36 | 0.84 | 0.97 | 0.93 | 0.93 |
| 6 | 24963024 | rs449796 | 1.64 | -1.275 | 0.549 | -0.571 | -0.045 | -0.065 | -0.065 | 0 | 0.8 | 0.36 | 0.84 | 0.97 | 0.93 | 0.93 |
| 6 | 24966689 | rs9295638 | 0.326 | -1.607 | 0.184 | -0.346 | 0.008 | -0.052 | -0.052 | 0.045 | 0.22 | 0.13 | 0.33 | 0.57 | 0.19 | 0.33 |
| 6 | 24967240 | rs10946737 | 1.208 | -1.656 | -0.298 | -0.644 | -0.123 | -0.015 | -0.015 | 0 | 0.23 | 0.01 | 0.28 | 0.14 | 0.34 | 0.35 |
| 6 | 24967549 | rs127489 | 0.805 | -1.557 | -0.048 | -0.531 | 0.02 | -0.009 | -0.009 | 0 | 0.6 | 0.35 | 0.56 | 0.38 | 0.76 | 0.59 |
| 6 | 24971403 | rs384245 | 1.011 | -1.466 | -0.044 | -0.524 | -0.062 | -0.08 | -0.08 | 0.039 | 0.81 | 0.43 | 0.87 | 0.97 | 0.94 | 0.93 |
| 6 | 24971721 | rs9461088 | -0.432 | -1.315 | -0.114 | -0.331 | -0.032 | -0.027 | -0.027 | 0 | 0.1 | 0.26 | 0.06 | 0.03 | 0.03 | 0.05 |
| 6 | 24975533 | rs9379701 | 0.57 | -1.218 | -0.423 | -0.219 | 0 | -0.122 | -0.122 | 0 | 0.21 | 0.05 | 0.31 | 0.57 | 0.18 | 0.33 |
| 6 | 24976191 | rs9393600 | -0.125 | -1.242 | -0.122 | -0.413 | 0 | -0.003 | -0.003 | 0.655 | 0.65 | 0.65 | 0.66 | 0.82 | 0.63 | 0.61 |
| 6 | 24990786 | rs4712872 | -0.051 | -1.287 | -0.1 | -0.324 | 0 | -0.003 | -0.003 | 0.045 | 0.35 | 0.33 | 0.33 | 0.18 | 0.38 | 0.38 |
| 6 | 24992226 | rs9379703 | 0.46 | -1.364 | -0.331 | -0.34 | -0.033 | -0.127 | -0.127 | 0 | 0.22 | 0.05 | 0.34 | 0.57 | 0.27 | 0.36 |
| 6 | 24993127 | rs10946739 | 0.841 | -1.132 | -0.142 | -0.616 | -0.121 | 0.024 | 0.024 | 0 | 0.25 | 0.03 | 0.26 | 0.15 | 0.36 | 0.34 |
| 6 | 24996338 | rs6935286 | -0.033 | -0.974 | -0.168 | -0.543 | -0.001 | 0 | 0 | 0.445 | 0.35 | 0.32 | 0.33 | 0.18 | 0.38 | 0.38 |
| 6 | 25006785 | rs1410689 | -0.047 | -1.703 | 0.041 | -0.089 | 0.007 | 0.042 | 0.042 | 0.296 | 0.48 | 0.6 | 0.47 | 0.25 | 0.56 | 0.47 |
| 6 | 25010999 | rs10946741 | 0.187 | -1.651 | -0.001 | -0.103 | 0 | -0.009 | -0.009 | 0 | 0.44 | 0.41 | 0.42 | 0.16 | 0.54 | 0.42 |
| 6 | 25012167 | rs2166681 | 0.19 | -1.391 | -0.901 | 0.26 | 0.009 | -0.008 | -0.008 | 0 | 0.14 | 0.12 | 0.24 | 0.53 | 0.11 | 0.26 |
| 6 | 25015681 | rs2183067 | 0.75 | -0.934 | -1.23 | -0.232 | -0.162 | 0.083 | 0.083 | 0 | 0.33 | 0.04 | 0.35 | 0.14 | 0.52 | 0.38 |
| 6 | 25027095 | rs9366597 | -0.165 | -0.797 | -0.414 | 0.486 | -0.015 | 0.033 | 0.033 | 0 | 0.23 | 0.27 | 0.28 | 0.61 | 0.14 | 0.31 |
| 6 | 25027288 | rs6918101 | 0.752 | -0.656 | -1.338 | 0.086 | -0.033 | 0.069 | 0.069 | 0.441 | 0.36 | 0.21 | 0.42 | 0.14 | 0.56 | 0.38 |

| | | | | | | | | | | | | | | | | |
|---|---|---|---|---|---|---|---|---|---|---|---|---|---|---|---|---|
| 6 | 25028663 | rs7743187 | 0.142 | -0.072 | -1.186 | 0.31 | 0.013 | 0.047 | 0.047 | 2.872 | 0.66 | 0.89 | 0.77 | 0.82 | 0.72 | 0.74 |
| 6 | 25035920 | rs303890 | -0.901 | 0.186 | -0.471 | 0.844 | -0.001 | 0.02 | 0.02 | 0 | 0.3 | 0.4 | 0.39 | 0.35 | 0.3 | 0.29 |
| 6 | 25036980 | rs303031 | -1.036 | 0.924 | -0.192 | 0.634 | -0.006 | -0.006 | -0.006 | 0 | 0.43 | 0.4 | 0.48 | 0.46 | 0.48 | 0.36 |
| 6 | 25039602 | rs303898 | 0.98 | 0.043 | -0.654 | 0.194 | -0.001 | -0.001 | -0.001 | 0 | 0.45 | 0.52 | 0.43 | 0.51 | 0.51 | 0.56 |
| 6 | 25050631 | rs12525405 | 0.015 | -0.003 | -0.945 | 0.377 | -0.061 | -0.065 | -0.065 | 2.162 | 0.13 | 0.04 | 0.24 | 0.28 | 0.21 | 0.17 |
| 6 | 25055165 | rs458664 | 0.99 | -0.304 | -0.233 | 0.2 | 0.064 | 0.08 | 0.08 | 1.184 | 0.5 | 0.3 | 0.58 | 0.75 | 0.64 | 0.68 |
| 6 | 25057499 | rs303000 | -0.6 | -0.57 | -0.105 | 0.075 | -0.007 | 0.01 | 0.01 | 0 | 0.85 | 0.91 | 0.87 | 0.87 | 0.81 | 0.88 |
| 6 | 25060346 | rs9461093 | -0.422 | -0.728 | 0.193 | -0.007 | -0.005 | -0.031 | -0.031 | 1.242 | 0.28 | 0.14 | 0.39 | 0.35 | 0.26 | 0.3 |
| 6 | 25060863 | rs12214056 | 1.843 | -0.977 | 0.172 | -0.143 | -0.014 | -0.014 | -0.014 | 0.346 | 0.39 | 0.26 | 0.35 | 0.47 | 0.43 | 0.51 |
| 6 | 25062826 | rs302992 | -0.144 | -0.968 | 0.182 | -0.123 | 0.044 | 0.103 | 0.103 | 0 | 0.15 | 0.23 | 0.28 | 0.3 | 0.19 | 0.16 |
| 6 | 25065669 | rs303878 | 1.136 | -1.001 | 0.124 | -0.268 | 0.014 | 0.022 | 0.022 | 0.7 | 0.53 | 0.53 | 0.64 | 0.77 | 0.62 | 0.67 |
| 6 | 25072522 | rs11757374 | -0.49 | -0.845 | -0.115 | -0.05 | -0.054 | -0.074 | -0.074 | 0 | 0.18 | 0.09 | 0.28 | 0.29 | 0.24 | 0.21 |
| 6 | 25077179 | rs7740534 | -0.34 | -0.86 | 0.104 | -0.097 | -0.018 | -0.038 | -0.038 | 0 | 0.1 | 0.22 | 0.05 | 0.01 | 0.05 | 0.02 |
| 6 | 25078882 | rs9295646 | 0.64 | -1.006 | 0.394 | -0.181 | -0.014 | -0.074 | -0.074 | 0 | 0.74 | 0.54 | 0.8 | 0.85 | 0.77 | 0.85 |
| 6 | 25080685 | rs10214692 | 1.837 | -1.126 | 0.254 | -0.207 | -0.069 | -0.041 | -0.041 | 0.131 | 0.53 | 0.09 | 0.48 | 0.58 | 0.53 | 0.62 |
| 6 | 25084718 | rs303006 | -0.345 | -1.004 | 0.268 | -0.157 | 0.008 | -0.014 | -0.014 | 2.133 | 0.85 | 0.79 | 0.86 | 0.87 | 0.82 | 0.89 |
| 6 | 25086062 | rs9467386 | -0.86 | -0.865 | 0.084 | -0.106 | 0.026 | 0.041 | 0.041 | 0 | 0.18 | 0.28 | 0.31 | 0.28 | 0.24 | 0.23 |
| 6 | 25094953 | rs715426 | 0.711 | -0.937 | 0.085 | -0.202 | -0.013 | -0.048 | -0.048 | 0 | 0.73 | 0.44 | 0.79 | 0.86 | 0.77 | 0.85 |
| 6 | 25106400 | rs7773802 | 1.263 | -0.929 | -0.045 | -0.205 | 0.019 | 0.063 | 0.063 | 0.055 | 0.66 | 0.26 | 0.73 | 0.84 | 0.71 | 0.81 |
| 6 | 25106783 | rs1923195 | 1.263 | -0.929 | -0.045 | -0.205 | 0.019 | 0.063 | 0.063 | 0.055 | 0.66 | 0.26 | 0.73 | 0.84 | 0.71 | 0.81 |
| 6 | 25111621 | rs3805845 | 0.76 | -1.205 | -0.264 | -0.129 | 0.044 | 0.001 | 0.001 | 0.086 | 0.15 | 0.01 | 0.12 | 0.08 | 0.06 | 0.12 |
| 6 | 25113207 | rs188567 | 0.668 | -1.013 | 0.01 | -0.036 | 0.003 | -0.018 | -0.018 | 0 | 0.05 | 0.1 | 0.03 | 0.01 | 0.04 | 0.02 |
| 6 | 25115296 | rs2273671 | 0.308 | -0.933 | -0.26 | 0.079 | -0.055 | -0.057 | -0.057 | 2.447 | 0.1 | 0.02 | 0.25 | 0.27 | 0.18 | 0.18 |
| 6 | 25118110 | rs302978 | -1.365 | -1.143 | -0.187 | -0.165 | 0.031 | 0.061 | 0.061 | 0.131 | 0.32 | 0.74 | 0.25 | 0.15 | 0.24 | 0.17 |
| 6 | 25118116 | rs302977 | -1.365 | -1.143 | -0.187 | -0.165 | 0.031 | 0.061 | 0.061 | 0.131 | 0.32 | 0.74 | 0.25 | 0.16 | 0.24 | 0.17 |
| 6 | 25122715 | rs916539 | 0.203 | -1.129 | -0.136 | -0.162 | 0.006 | -0.015 | -0.015 | 0 | 0.07 | 0.18 | 0.05 | 0.02 | 0.06 | 0.05 |
| 6 | 25122737 | rs302971 | -0.495 | -1.226 | -0.644 | -0.196 | 0.031 | 0.031 | 0.031 | 0.038 | 0.8 | 0.99 | 0.84 | 0.91 | 0.87 | 0.85 |
| 6 | 25127176 | rs302968 | -1.783 | -0.611 | -0.213 | 0.097 | 0.057 | 0.151 | 0.151 | 0.871 | 0.36 | 0.9 | 0.27 | 0.15 | 0.24 | 0.17 |
| 6 | 25138627 | rs2028973 | 0.079 | -0.529 | 0.191 | 0.164 | 0.001 | -0.012 | -0.012 | 0 | 0.13 | 0.21 | 0.13 | 0.12 | 0.13 | 0.08 |
| 6 | 25138663 | rs6908916 | -0.086 | -0.555 | 0.052 | 0.142 | -0.024 | -0.022 | -0.022 | 0 | 0.06 | 0.19 | 0.03 | 0.01 | 0.02 | 0.02 |
| 6 | 25138723 | rs303019 | -1.678 | -0.488 | -0.198 | 0.257 | -0.066 | -0.082 | -0.082 | 0 | 0.46 | 0.92 | 0.52 | 0.44 | 0.41 | 0.35 |
| 6 | 25140405 | rs9654578 | -0.026 | -0.227 | 0.507 | 0.364 | -0.018 | -0.011 | -0.011 | 0.083 | 0.05 | 0.17 | 0.03 | 0.01 | 0.02 | 0.02 |
| 6 | 25147518 | rs4712888 | 1.425 | 0.036 | 0.436 | 0.565 | 0.026 | -0.04 | -0.04 | 0 | 0.92 | 0.72 | 0.92 | 1 | 0.94 | 1 |
| 6 | 25152674 | rs6941417 | -0.514 | 0.267 | 0.401 | 0.877 | 0.018 | 0.016 | 0.016 | 0 | 0.15 | 0.21 | 0.23 | 0.21 | 0.19 | 0.12 |
| 6 | 25156197 | rs7739830 | 1.326 | 0.266 | 0.362 | 0.394 | 0.045 | -0.009 | -0.009 | 0 | 0.92 | 0.68 | 0.91 | 1 | 0.94 | 0.96 |
| 6 | 25172641 | rs9467434 | 0.117 | -0.178 | -0.923 | 0.144 | 0.042 | 0.073 | 0.073 | 0 | 0.58 | 0.84 | 0.72 | 0.77 | 0.65 | 0.71 |
| 6 | 25174070 | rs1986731 | -0.745 | 0.15 | -0.277 | -0.238 | 0.007 | -0.073 | -0.073 | 0.897 | 0.15 | 0.35 | 0.11 | 0.01 | 0.11 | 0.05 |
| 6 | 25174148 | rs1815094 | -0.384 | 0.232 | -0.374 | 0.213 | 0.002 | -0.008 | -0.008 | 0.007 | 0.83 | 0.86 | 0.85 | 0.69 | 0.85 | 0.67 |
| 6 | 25175151 | rs1880274 | -0.175 | 0.205 | -0.253 | 0.219 | -0.03 | -0.024 | -0.024 | 0 | 0.06 | 0.23 | 0.03 | 0.02 | 0 | 0.02 |
| 6 | 25175316 | rs2328848 | 1.061 | 0.702 | -0.905 | 0.308 | -0.035 | -0.086 | -0.086 | 0 | 0.26 | 0.07 | 0.43 | 0.41 | 0.3 | 0.34 |
| 6 | 25176145 | rs7752388 | 0.763 | 0.387 | -0.598 | 0.138 | -0.013 | -0.015 | -0.015 | 0 | 0.63 | 0.41 | 0.7 | 0.66 | 0.7 | 0.59 |
| 6 | 25177722 | rs6899989 | 0.263 | 1 | -1.241 | -0.056 | -0.055 | 0.028 | 0.028 | 0 | 0.17 | 0.06 | 0.16 | 0.1 | 0.26 | 0.14 |
| 6 | 25178990 | rs13198274 | 1.017 | 0.78 | -1.655 | -0.205 | -0.067 | 0.035 | 0.035 | 0.761 | 0.27 | 0.05 | 0.2 | 0.16 | 0.3 | 0.16 |
| 6 | 25179093 | rs7748632 | -0.217 | 0.336 | -0.802 | -0.192 | 0.004 | 0.011 | 0.011 | 0.537 | 0.49 | 0.51 | 0.4 | 0.31 | 0.55 | 0.31 |
| 6 | 25181447 | rs9358839 | -0.192 | 0.269 | -0.869 | -0.029 | -0.008 | 0 | 0 | 0 | 0.47 | 0.66 | 0.6 | 0.54 | 0.48 | 0.57 |
| 6 | 25181627 | rs9379729 | 0.846 | 0.465 | -1.05 | 0.188 | -0.033 | -0.113 | -0.113 | 0.632 | 0.22 | 0.07 | 0.35 | 0.38 | 0.18 | 0.41 |
| 6 | 25186512 | rs10498722 | -1.101 | 0.108 | -0.7 | -0.088 | 0.001 | -0.011 | -0.011 | 0 | 0.08 | 0.16 | 0.06 | 0.02 | 0.07 | 0.02 |
| 6 | 25192394 | rs9358840 | 0.366 | -0.385 | -0.62 | -0.394 | -0.001 | -0.001 | -0.001 | 0 | 0.31 | 0.29 | 0.3 | 0.37 | 0.36 | 0.34 |
| 6 | 25193064 | rs4144129 | -1.018 | -0.776 | -0.681 | -0.677 | 0.004 | -0.042 | -0.042 | 0 | 0.7 | 0.62 | 0.78 | 0.85 | 0.71 | 0.87 |
| 6 | 25198663 | rs9393623 | 0.398 | -0.738 | -0.895 | -0.549 | -0.001 | -0.09 | -0.09 | 0 | 0.34 | 0.14 | 0.45 | 0.43 | 0.29 | 0.51 |
| 6 | 25199042 | rs9393624 | -0.329 | -0.392 | -0.893 | -0.579 | 0.01 | -0.051 | -0.051 | 0 | 0.64 | 0.48 | 0.74 | 0.83 | 0.64 | 0.84 |
| 6 | 25200414 | rs7748938 | 1.063 | -0.516 | -0.824 | -0.132 | -0.034 | -0.049 | -0.049 | 3.095 | 0.26 | 0.07 | 0.29 | 0.29 | 0.27 | 0.38 |
| 6 | 25200842 | rs6936465 | 0.829 | -0.844 | -0.581 | -0.19 | 0.036 | -0.033 | -0.033 | 0 | 0.47 | 0.14 | 0.45 | 0.44 | 0.55 | 0.51 |
| 6 | 25212558 | rs2876687 | -0.187 | -0.874 | -0.497 | -0.13 | -0.015 | 0.004 | 0.004 | 0.043 | 0.43 | 0.46 | 0.46 | 0.54 | 0.38 | 0.46 |
| 6 | 25216159 | rs2328869 | 1.283 | -0.814 | 0.043 | -0.458 | 0.023 | 0.03 | 0.03 | 0.021 | 0.94 | 0.66 | 0.96 | 0.99 | 0.99 | 0.98 |
| 6 | 25225513 | rs1880269 | 0.579 | -0.919 | -0.707 | -0.582 | 0.075 | 0.088 | 0.088 | 0 | 0.48 | 0.15 | 0.51 | 0.7 | 0.64 | 0.63 |
| 6 | 25229450 | rs4712901 | 0.59 | -0.995 | -0.885 | -0.508 | -0.064 | -0.095 | -0.095 | 1.064 | 0.3 | 0.08 | 0.35 | 0.55 | 0.37 | 0.5 |
| 6 | 25237288 | rs11755567 | 1.104 | -1.227 | -0.483 | -0.657 | -0.023 | -0.01 | -0.01 | 1.354 | 0.22 | 0.31 | 0.16 | 0.18 | 0.19 | 0.16 |
| 6 | 25239809 | rs2064207 | 1.212 | -1.223 | -0.49 | -0.655 | -0.02 | -0.008 | -0.008 | 0 | 0.21 | 0.3 | 0.16 | 0.18 | 0.19 | 0.16 |
| 6 | 25245680 | rs769108 | -0.155 | -1.537 | -0.107 | -0.619 | -0.003 | -0.004 | -0.004 | 0 | 0.45 | 0.39 | 0.55 | 0.65 | 0.39 | 0.64 |
| 6 | 25247204 | rs7774664 | 0.727 | -1.562 | -0.339 | -0.59 | -0.068 | -0.118 | -0.118 | 0.084 | 0.29 | 0.06 | 0.34 | 0.54 | 0.35 | 0.5 |
| 6 | 25249623 | rs10080837 | 0.484 | -1.47 | -0.134 | -0.598 | -0.038 | -0.007 | -0.007 | 0.03 | 0.11 | 0.16 | 0.19 | 0.11 | 0.03 | 0.13 |
| 6 | 25251377 | rs7758340 | 0.557 | -1.457 | -0.126 | -0.591 | -0.038 | -0.007 | -0.007 | 1.585 | 0.11 | 0.16 | 0.19 | 0.12 | 0.03 | 0.13 |
| 6 | 25257580 | rs2690106 | -0.692 | -1.257 | -0.092 | -0.59 | 0.005 | 0.027 | 0.027 | 0 | 0.5 | 0.63 | 0.49 | 0.63 | 0.46 | 0.61 |
| 6 | 25257966 | rs9356970 | 0.725 | -1.224 | -0.165 | -0.538 | -0.081 | -0.116 | -0.116 | 0 | 0.28 | 0.05 | 0.33 | 0.6 | 0.35 | 0.49 |
| 6 | 25259137 | rs2744249 | -0.443 | -1.187 | 0.021 | -0.63 | -0.001 | -0.007 | -0.007 | 0 | 0.46 | 0.5 | 0.45 | 0.63 | 0.45 | 0.59 |
| 6 | 25260434 | rs10456319 | 0.393 | -0.927 | 0.024 | -0.371 | -0.01 | -0.019 | -0.019 | 0 | 0.3 | 0.18 | 0.34 | 0.61 | 0.35 | 0.51 |
| 6 | 25266147 | rs9379750 | 0.569 | -0.835 | 0.887 | -0.417 | -0.01 | -0.04 | -0.04 | 0 | 0.61 | 0.49 | 0.57 | 0.68 | 0.67 | 0.75 |
| 6 | 25266803 | rs2744254 | 0.211 | -0.752 | 1.033 | -0.453 | 0.002 | 0.007 | 0.007 | 0 | 0.65 | 0.69 | 0.63 | 0.69 | 0.71 | 0.76 |

| | | | | | | | | | | | | | | | | |
|---|---|---|---|---|---|---|---|---|---|---|---|---|---|---|---|---|
| 6 | 25274612 | rs721037 | -0.846 | -0.602 | 0.756 | -0.404 | 0.009 | -0.005 | -0.005 | 0 | 0.81 | 0.84 | 0.69 | 0.81 | 0.74 | 0.9 |
| 6 | 25274645 | rs2078527 | 0.482 | -0.694 | 0.726 | -0.448 | -0.003 | -0.026 | -0.026 | 2.719 | 0.64 | 0.47 | 0.59 | 0.68 | 0.68 | 0.74 |
| 6 | 25276282 | rs2744268 | 0.512 | -0.738 | 0.727 | -0.469 | -0.006 | -0.018 | -0.018 | 0 | 0.63 | 0.61 | 0.63 | 0.69 | 0.71 | 0.72 |
| 6 | 25277419 | rs1016855 | -0.168 | -0.244 | 0.565 | -0.316 | 0.052 | -0.047 | -0.047 | 0.259 | 0.76 | 0.55 | 0.62 | 0.81 | 0.69 | 0.89 |
| 6 | 25289097 | rs2690134 | -0.305 | -0.023 | -0.904 | -0.335 | -0.053 | 0.02 | 0.02 | 0 | 0.75 | 0.87 | 0.66 | 0.78 | 0.64 | 0.8 |
| 6 | 25292214 | rs9379751 | 1.277 | -0.316 | -0.956 | -0.088 | -0.019 | -0.021 | -0.021 | 0 | 0.47 | 0.22 | 0.46 | 0.37 | 0.51 | 0.49 |
| 6 | 25292401 | rs10456320 | -0.057 | -0.211 | -1.054 | -0.134 | 0.022 | 0.064 | 0.064 | 0 | 0.17 | 0.08 | 0.06 | 0.02 | 0.12 | 0.15 |
| 6 | 25296606 | rs2744294 | -0.354 | 0.009 | -0.949 | -0.004 | -0.05 | 0.014 | 0.014 | 0 | 0.76 | 0.88 | 0.66 | 0.79 | 0.64 | 0.8 |
| 6 | 25310585 | rs7752524 | 0.455 | -0.075 | -0.903 | -0.022 | -0.004 | 0.003 | 0.003 | 0 | 0.07 | 0.07 | 0.05 | 0.01 | 0.1 | 0.1 |
| 6 | 25311461 | rs6921343 | -0.245 | -0.221 | -0.655 | 0.101 | -0.05 | 0.067 | 0.067 | 0 | 0.22 | 0.41 | 0.16 | 0.42 | 0.12 | 0.32 |
| 6 | 25312755 | rs9295654 | -0.245 | -0.221 | -0.655 | 0.101 | -0.05 | 0.067 | 0.067 | 0 | 0.22 | 0.41 | 0.16 | 0.42 | 0.12 | 0.32 |
| 6 | 25312915 | rs9467466 | -0.245 | -0.221 | -0.655 | 0.101 | -0.05 | 0.067 | 0.067 | 0 | 0.22 | 0.41 | 0.16 | 0.42 | 0.12 | 0.32 |
| 6 | 25319626 | rs999039 | -0.366 | -0.179 | -0.643 | 0.118 | -0.051 | 0.02 | 0.02 | 0 | 0.28 | 0.41 | 0.19 | 0.43 | 0.18 | 0.34 |
| 6 | 25320920 | rs4712908 | 1.323 | -0.046 | -0.902 | 0.111 | -0.005 | -0.026 | -0.026 | 0 | 0.37 | 0.21 | 0.39 | 0.34 | 0.38 | 0.4 |
| 6 | 25322814 | rs7764258 | 0.232 | 0.346 | -1.136 | 0.302 | -0.008 | 0.032 | 0.032 | 0 | 0.12 | 0.07 | 0.09 | 0.02 | 0.15 | 0.15 |
| 6 | 25327149 | rs2744303 | 0.789 | 0.144 | -0.843 | 0.425 | 0 | 0 | 0 | 0 | 0.44 | 0.47 | 0.45 | 0.34 | 0.42 | 0.4 |
| 6 | 25328153 | rs9295655 | -0.203 | 0.57 | -0.778 | 0.393 | -0.013 | 0.009 | 0.009 | 0 | 0.26 | 0.43 | 0.2 | 0.32 | 0.22 | 0.24 |
| 6 | 25328567 | rs2690110 | -0.224 | 0.701 | -1.193 | 0.265 | -0.037 | 0.028 | 0.028 | 0 | 0.71 | 0.9 | 0.63 | 0.78 | 0.64 | 0.74 |
| 6 | 25330538 | rs2690107 | -0.913 | 0.499 | -0.27 | 0.654 | 0.003 | 0.036 | 0.036 | 0 | 0.29 | 0.48 | 0.23 | 0.46 | 0.27 | 0.35 |
| 6 | 25331237 | rs7769912 | -0.556 | 0.331 | -0.417 | 0.144 | 0.001 | -0.008 | -0.008 | 3.175 | 0.14 | 0.15 | 0.07 | 0.01 | 0.1 | 0.07 |
| 6 | 25333071 | rs5009712 | -0.852 | 0.308 | -0.52 | 0.007 | 0 | -0.01 | -0.01 | 0 | 0.12 | 0.13 | 0.11 | 0.02 | 0.16 | 0.12 |
| 6 | 25334967 | rs7762537 | -0.46 | 0.228 | -0.315 | -0.379 | 0.014 | -0.028 | -0.028 | 0.193 | 0.15 | 0.19 | 0.12 | 0.02 | 0.2 | 0.14 |
| 6 | 25335121 | rs7762836 | -1.124 | 0.193 | -0.335 | -0.386 | 0.004 | -0.047 | -0.047 | 0 | 0.25 | 0.31 | 0.2 | 0.04 | 0.28 | 0.25 |
| 6 | 25336402 | rs2254974 | -0.271 | 0.577 | -0.705 | 0.221 | -0.013 | -0.022 | -0.022 | 0 | 0.9 | 0.95 | 0.73 | 0.79 | 0.87 | 0.92 |
| 6 | 25340482 | rs2206194 | -0.837 | 0.109 | -0.568 | 0.155 | -0.001 | -0.001 | -0.001 | 0 | 0.58 | 0.66 | 0.46 | 0.59 | 0.52 | 0.64 |
| 6 | 25341553 | rs16890409 | 0.392 | -0.092 | -0.695 | 0.503 | -0.007 | -0.031 | -0.031 | 0 | 0.14 | 0.08 | 0.12 | 0.29 | 0.16 | 0.23 |
| 6 | 25341580 | rs2744229 | -0.859 | -0.575 | -0.624 | 0.286 | -0.005 | 0.005 | 0.005 | 0 | 0.38 | 0.53 | 0.38 | 0.43 | 0.39 | 0.41 |
| 6 | 25342417 | rs9467471 | 2.056 | -0.744 | -0.436 | 0.372 | -0.001 | 0 | 0 | 0 | 0.3 | 0.22 | 0.24 | 0.2 | 0.35 | 0.25 |
| 6 | 25342432 | rs1569558 | -0.442 | -0.983 | -0.206 | 0.244 | 0.004 | -0.004 | -0.004 | 0 | 0.26 | 0.29 | 0.2 | 0.32 | 0.24 | 0.34 |
| 6 | 25343805 | rs1076668 | 0.236 | -1.08 | -0.335 | 0.121 | -0.009 | -0.018 | -0.018 | 0 | 0.22 | 0.27 | 0.2 | 0.13 | 0.19 | 0.14 |
| 6 | 25346334 | rs2690095 | 0.581 | -1.051 | -0.359 | 0.138 | 0.016 | 0.021 | 0.021 | 0 | 0.57 | 0.69 | 0.51 | 0.31 | 0.61 | 0.4 |
| 6 | 25346955 | rs2690098 | 1.186 | -0.955 | -0.103 | 0.162 | 0.02 | 0.007 | 0.007 | 0 | 0.07 | 0.14 | 0.17 | 0.12 | 0.15 | 0.08 |
| 6 | 25347733 | rs6456672 | 0.813 | -1.431 | 0.364 | 0.159 | 0.028 | 0.131 | 0.131 | 0 | 0.05 | 0.24 | 0.03 | 0.14 | 0.01 | 0.12 |
| 6 | 25348189 | rs2328853 | -0.164 | -1.433 | 0.108 | 0.192 | 0.009 | -0.033 | -0.033 | 0 | 0.26 | 0.13 | 0.18 | 0.33 | 0.23 | 0.35 |
| 6 | 25351244 | rs2744235 | 1.57 | -1.363 | 0.574 | 0.256 | -0.023 | 0.005 | 0.005 | 0 | 0.36 | 0.28 | 0.41 | 0.3 | 0.51 | 0.33 |
| 6 | 25352637 | rs11967836 | -0.079 | -1.057 | 0.643 | 0.431 | 0.033 | 0.045 | 0.045 | 0 | 0.07 | 0.18 | 0.06 | 0.14 | 0.08 | 0.13 |
| 6 | 25353197 | rs2690059 | -0.732 | -0.921 | 0.347 | 0.362 | 0 | 0.005 | 0.005 | 0 | 0.88 | 0.87 | 0.7 | 0.79 | 0.85 | 0.89 |
| 6 | 25361055 | rs10946765 | 0.73 | -0.895 | 0.322 | 0.185 | 0.004 | 0.024 | 0.024 | 0 | 0.48 | 0.37 | 0.46 | 0.33 | 0.57 | 0.44 |
| 6 | 25364481 | rs13215347 | 0.412 | -0.783 | 0.3 | 0.175 | 0.003 | 0.004 | 0.004 | 0.154 | 0.55 | 0.4 | 0.49 | 0.34 | 0.64 | 0.54 |
| 6 | 25368978 | rs9467476 | -0.161 | -0.958 | 0.507 | 0.247 | -0.031 | -0.014 | -0.014 | 0 | 0.12 | 0.22 | 0.07 | 0.05 | 0.06 | 0.1 |
| 6 | 25370970 | rs9467478 | -0.465 | -0.907 | 0.707 | 0.229 | 0.001 | -0.014 | -0.014 | 0.13 | 0.14 | 0.22 | 0.11 | 0.06 | 0.13 | 0.12 |
| 6 | 25374754 | rs7760906 | -1.04 | -0.817 | 0.301 | 0.41 | -0.049 | -0.05 | -0.05 | 0.223 | 0.92 | 0.99 | 0.72 | 0.79 | 0.85 | 0.89 |
| 6 | 25379926 | rs4711082 | 0.175 | -1.256 | 0.517 | 0.161 | 0.016 | 0.016 | 0.016 | 0 | 0.56 | 0.75 | 0.52 | 0.34 | 0.62 | 0.44 |
| 6 | 25380135 | rs11757140 | 1.271 | -1.134 | 0.188 | 0 | -0.025 | 0.016 | 0.016 | 0.037 | 0.36 | 0.27 | 0.38 | 0.28 | 0.47 | 0.31 |
| 6 | 25383814 | rs215012 | 0.99 | -1.143 | 0.125 | 0.004 | 0.001 | -0.003 | -0.003 | 0 | 0.4 | 0.39 | 0.4 | 0.28 | 0.48 | 0.34 |
| 6 | 25385131 | rs16890508 | 0.295 | -0.945 | 0.011 | 0.147 | 0.021 | 0.006 | 0.006 | 0 | 0.09 | 0.04 | 0.02 | 0.05 | 0.05 | 0.1 |
| 6 | 25392345 | rs10498723 | -0.024 | -0.924 | 0.055 | 0.5 | -0.002 | -0.012 | -0.012 | 2.082 | 0.11 | 0.08 | 0.27 | 0.34 | 0.13 | 0.16 |
| 6 | 25393306 | rs716797 | -0.704 | -1.287 | -0.36 | 0.112 | 0.008 | -0.002 | -0.002 | 0 | 0.68 | 0.66 | 0.63 | 0.72 | 0.54 | 0.69 |
| 6 | 25399005 | rs9366619 | 1.573 | -1.036 | -1.001 | 0.245 | -0.041 | -0.005 | -0.005 | 0 | 0.2 | 0.09 | 0.18 | 0.17 | 0.29 | 0.19 |
| 6 | 25400323 | rs9461157 | 0.037 | -1.412 | -1.026 | -0.166 | 0.002 | 0.002 | 0.002 | 0 | 0.54 | 0.44 | 0.36 | 0.39 | 0.43 | 0.48 |
| 6 | 25400396 | rs4712929 | 0.858 | -0.677 | -1.557 | -0.555 | -0.037 | -0.029 | -0.029 | 0 | 0.08 | 0.01 | 0.13 | 0.11 | 0.1 | 0.12 |
| 6 | 25401131 | rs16890551 | 0.534 | -0.982 | -1.211 | -0.781 | 0.012 | 0.004 | 0.004 | 0.024 | 0.08 | 0.05 | 0.02 | 0.04 | 0.03 | 0.11 |
| 6 | 25402622 | rs9467496 | 0.771 | -0.792 | -1.165 | -1.013 | -0.001 | -0.009 | -0.009 | 0 | 0.1 | 0.11 | 0.06 | 0.01 | 0.12 | 0.09 |
| 6 | 25407482 | rs12201737 | -0.29 | -0.93 | -1.215 | -1.065 | 0.012 | 0.013 | 0.013 | 0 | 0.3 | 0.24 | 0.17 | 0.04 | 0.22 | 0.2 |
| 6 | 25409785 | rs6912426 | 0.095 | -0.578 | -1.078 | -0.563 | 0 | 0.032 | 0.032 | 0 | 0.37 | 0.21 | 0.29 | 0.25 | 0.38 | 0.32 |
| 6 | 25409846 | rs4440453 | 0.733 | 0.055 | -0.494 | -0.582 | -0.006 | 0.002 | 0.002 | 0 | 0.15 | 0.11 | 0.16 | 0.11 | 0.24 | 0.14 |
| 6 | 25410077 | rs4368803 | -0.1 | 0.072 | -0.472 | -0.522 | -0.004 | -0.018 | -0.018 | 0.084 | 0.38 | 0.41 | 0.27 | 0.13 | 0.33 | 0.31 |
| 6 | 25411464 | rs9358854 | -0.121 | 0.564 | -0.498 | -0.213 | -0.011 | -0.049 | -0.049 | 0 | 0.43 | 0.49 | 0.28 | 0.15 | 0.34 | 0.31 |
| 6 | 25412987 | rs927984 | 0.339 | 1.498 | -0.318 | 0.769 | 0.046 | 0.045 | 0.045 | 0 | 0.94 | 0.88 | 0.9 | 0.84 | 0.87 | 0.92 |
| 6 | 25413828 | rs1995675 | -0.374 | 1.015 | -0.587 | 0.393 | 0.011 | 0.015 | 0.015 | 0 | 0.09 | 0.04 | 0.06 | 0.06 | 0.07 | 0.13 |
| 6 | 25413922 | rs1543603 | 0.393 | 0.372 | -0.198 | 0.231 | -0.007 | -0.021 | -0.021 | 0 | 0.73 | 0.5 | 0.82 | 0.81 | 0.76 | 0.8 |
| 6 | 25414325 | rs9467504 | 0.197 | 0.662 | -0.223 | 0.753 | 0 | 0.022 | 0.022 | 0 | 0.16 | 0.2 | 0.19 | 0.25 | 0.17 | 0.17 |
| 6 | 25419318 | rs9295657 | 0.289 | 0.741 | 0.197 | 0.381 | 0.007 | -0.041 | -0.041 | 1.51 | 0.86 | 0.75 | 0.93 | 0.96 | 0.87 | 0.89 |
| 6 | 25420744 | rs2076890 | 0.276 | 1.035 | 0.058 | -0.005 | 0 | -0.005 | -0.005 | 0 | 0.13 | 0.15 | 0.12 | 0.05 | 0.15 | 0.11 |
| 6 | 25421015 | rs12191318 | 0.467 | 0.959 | -0.22 | 0.294 | -0.014 | -0.044 | -0.044 | 0 | 0.12 | 0.03 | 0.08 | 0.19 | 0.13 | 0.12 |
| 6 | 25424256 | rs12213948 | 0.697 | 1.19 | -0.432 | 0.388 | -0.005 | -0.009 | -0.009 | 0 | 0.62 | 0.62 | 0.5 | 0.46 | 0.57 | 0.62 |
| 6 | 25428954 | rs722086 | -0.378 | 1.253 | -0.287 | 0.234 | 0.027 | -0.054 | -0.054 | 0.146 | 0.85 | 0.73 | 0.9 | 0.97 | 0.82 | 0.89 |
| 6 | 25432769 | rs12524673 | 0.771 | 1.097 | -0.503 | 0.522 | 0.001 | -0.047 | -0.047 | 1.016 | 0.11 | 0.03 | 0.3 | 0.31 | 0.1 | 0.13 |
| 6 | 25435758 | rs9467515 | 0.142 | 0.887 | -0.374 | -0.196 | 0.01 | 0.016 | 0.016 | 0.177 | 0.51 | 0.53 | 0.42 | 0.32 | 0.42 | 0.45 |

| | | | | | | | | | | | | | | | | |
|---|---|---|---|---|---|---|---|---|---|---|---|---|---|---|---|---|
| 6 | 25436325 | rs9461170 | 0.109 | 0.443 | -0.379 | -0.585 | -0.051 | -0.078 | -0.078 | 1.538 | 0.32 | 0.44 | 0.22 | 0.08 | 0.17 | 0.19 |
| 6 | 25437986 | rs6900224 | -1.499 | 1.054 | -0.629 | -0.28 | -0.005 | 0 | 0 | 2.332 | 0.87 | 0.87 | 0.87 | 0.87 | 0.77 | 0.83 |
| 6 | 25442464 | rs1876372 | 0.175 | -0.017 | -0.633 | -0.372 | -0.008 | -0.018 | -0.018 | 0 | 0.16 | 0.21 | 0.12 | 0.05 | 0.09 | 0.15 |
| 6 | 25446308 | rs9358858 | 0.825 | 0.571 | -0.704 | -0.001 | -0.018 | -0.018 | -0.018 | 0 | 0.18 | 0.08 | 0.19 | 0.24 | 0.25 | 0.24 |
| 6 | 25455015 | rs10498725 | 0.555 | -0.23 | -0.733 | -0.402 | -0.003 | 0.003 | 0.003 | 0 | 0.15 | 0.15 | 0.15 | 0.09 | 0.16 | 0.14 |
| 6 | 25457251 | rs6456683 | 0.589 | -0.299 | -0.473 | -0.538 | -0.031 | -0.055 | -0.055 | 0.057 | 0.15 | 0.24 | 0.06 | 0.03 | 0.08 | 0.11 |
| 6 | 25460678 | rs12525342 | 0.826 | 0.776 | -0.964 | 0.026 | -0.068 | -0.029 | -0.029 | 0 | 0.16 | 0.02 | 0.26 | 0.15 | 0.21 | 0.24 |
| 6 | 25462824 | rs4320355 | 0.078 | 0.966 | -0.767 | 0.437 | 0.014 | 0.014 | 0.014 | 0.123 | 0.82 | 0.8 | 0.85 | 0.66 | 0.72 | 0.88 |
| 6 | 25466889 | rs301396 | 0.545 | 0.93 | -0.894 | 0.517 | 0.003 | 0.004 | 0.004 | 1.211 | 0.58 | 0.61 | 0.66 | 0.52 | 0.55 | 0.65 |
| 6 | 25468825 | rs2769703 | -0.768 | 1.204 | -1.293 | 0.572 | 0.03 | 0.039 | 0.039 | 0.581 | 0.89 | 0.88 | 0.88 | 0.66 | 0.76 | 0.9 |
| 6 | 25472787 | rs9393646 | 0.864 | 1.253 | -1.42 | 0.746 | 0.005 | 0.003 | 0.003 | 0 | 0.17 | 0.06 | 0.16 | 0.14 | 0.16 | 0.23 |
| 6 | 25475131 | rs463121 | -0.302 | 1.23 | -1.212 | 0.782 | 0.005 | 0.005 | 0.005 | 1.272 | 0.6 | 0.63 | 0.68 | 0.51 | 0.55 | 0.65 |
| 6 | 25481991 | rs10498726 | -0.238 | 1.77 | -1.383 | 0.867 | -0.067 | -0.028 | -0.028 | 0 | 0.15 | 0.02 | 0.26 | 0.15 | 0.2 | 0.25 |
| 6 | 25486154 | rs10498727 | 0.025 | 1.725 | -1.146 | 1.034 | -0.035 | -0.029 | -0.029 | 0 | 0.08 | 0.02 | 0.18 | 0.12 | 0.16 | 0.12 |
| 6 | 25486626 | rs301395 | -0.076 | 1.46 | -0.716 | 1.056 | 0.002 | -0.006 | -0.006 | 0.439 | 0.65 | 0.59 | 0.53 | 0.63 | 0.63 | 0.52 |
| 6 | 25486800 | rs9295663 | 0.612 | 1.235 | -1.119 | 0.98 | -0.017 | -0.027 | -0.027 | 0.128 | 0.2 | 0.12 | 0.12 | 0.3 | 0.24 | 0.13 |
| 6 | 25488265 | rs4712944 | 0.088 | 1.06 | -0.03 | 0.912 | 0.005 | 0.006 | 0.006 | 0 | 0.36 | 0.43 | 0.5 | 0.44 | 0.38 | 0.47 |
| 6 | 25492364 | rs1034050 | -0.24 | 1.082 | -0.021 | 0.868 | -0.002 | 0.001 | 0.001 | 0 | 0.66 | 0.7 | 0.54 | 0.63 | 0.64 | 0.52 |
| 6 | 25499895 | rs7760799 | 0.732 | 1.189 | -0.44 | 1.069 | -0.027 | -0.056 | -0.056 | 0 | 0.1 | 0.02 | 0.16 | 0.29 | 0.13 | 0.18 |
| 6 | 25504562 | rs407934 | 0.504 | 0.499 | -0.532 | 0.365 | 0 | 0.039 | 0.039 | 0.049 | 0.35 | 0.24 | 0.35 | 0.21 | 0.35 | 0.28 |
| 6 | 25516126 | rs301379 | 0.059 | -0.033 | -0.61 | -0.714 | -0.002 | -0.025 | -0.025 | 2.796 | 0.51 | 0.51 | 0.39 | 0.15 | 0.42 | 0.4 |
| 6 | 25516795 | rs17266491 | 0.266 | -0.108 | -0.821 | -0.993 | -0.007 | 0.033 | 0.033 | 0 | 0.23 | 0.13 | 0.21 | 0.1 | 0.25 | 0.25 |
| 6 | 25519272 | rs9393648 | 1.572 | -0.101 | -1.003 | -0.628 | -0.032 | -0.035 | -0.035 | 0 | 0.09 | 0.02 | 0.17 | 0.41 | 0.14 | 0.22 |
| 6 | 25519515 | rs301371 | -0.014 | -0.171 | -0.879 | -0.694 | 0.013 | 0.033 | 0.033 | 0 | 0.51 | 0.44 | 0.38 | 0.15 | 0.41 | 0.4 |
| 6 | 25520038 | rs441472 | 0.893 | -0.123 | -0.833 | -0.652 | -0.024 | -0.063 | -0.063 | 1.349 | 0.28 | 0.37 | 0.17 | 0.05 | 0.17 | 0.16 |
| 6 | 25520094 | rs16890706 | 1.612 | 0.016 | -0.989 | -0.581 | -0.043 | -0.056 | -0.056 | 0 | 0.09 | 0 | 0.07 | 0.22 | 0.14 | 0.11 |
| 6 | 25522713 | rs212939 | 1.695 | -0.755 | -1.113 | -1.03 | 0.007 | 0.008 | 0.008 | 0 | 0.17 | 0.11 | 0.11 | 0.04 | 0.13 | 0.15 |
| 6 | 25533534 | rs12526480 | 0.441 | -1.023 | -0.694 | -1.185 | 0.012 | 0.046 | 0.046 | 0 | 0.37 | 0.28 | 0.37 | 0.26 | 0.32 | 0.37 |
| 6 | 25535612 | rs150533 | 1.042 | -0.955 | -0.727 | -1.177 | -0.007 | -0.029 | -0.029 | 0 | 0.25 | 0.23 | 0.13 | 0.05 | 0.16 | 0.16 |
| 6 | 25540718 | rs214064 | -1.631 | -0.862 | -0.844 | -1.141 | -0.036 | 0.013 | 0.013 | 0 | 0.92 | 0.99 | 0.85 | 0.95 | 0.86 | 0.87 |
| 6 | 25541836 | rs190045 | 0.96 | -1.204 | -0.439 | -0.989 | -0.012 | 0.092 | 0.092 | 0 | 0.13 | 0.19 | 0.07 | 0.31 | 0.12 | 0.1 |
| 6 | 25548288 | rs441460 | -0.153 | -1.061 | -0.382 | -0.952 | 0.008 | 0.003 | 0.003 | 0 | 0.49 | 0.46 | 0.44 | 0.49 | 0.44 | 0.47 |
| 6 | 25550718 | rs10456323 | 1.035 | -1.06 | -0.469 | -1.029 | 0.018 | 0.032 | 0.032 | 0 | 0.23 | 0.15 | 0.12 | 0.05 | 0.16 | 0.16 |
| 6 | 25556249 | rs214050 | -1.634 | -0.885 | -0.47 | -0.976 | -0.012 | 0.011 | 0.011 | 0 | 0.85 | 0.89 | 0.8 | 0.94 | 0.77 | 0.86 |
| 6 | 25576377 | rs3804123 | 0.386 | -0.984 | -0.359 | -0.901 | -0.002 | 0.017 | 0.017 | 0 | 0.36 | 0.41 | 0.36 | 0.45 | 0.33 | 0.39 |
| 6 | 25582771 | rs13203202 | -0.345 | -1.247 | -0.553 | -1.216 | -0.003 | -0.001 | -0.001 | 0 | 0.54 | 0.51 | 0.44 | 0.53 | 0.42 | 0.51 |
| 6 | 25587836 | rs3788994 | 2.184 | -1.216 | -0.585 | -1.21 | -0.056 | 0.025 | 0.025 | 0 | 0.11 | 0.01 | 0.23 | 0.08 | 0.18 | 0.22 |
| 6 | 25589165 | rs7766874 | -0.384 | -1.432 | -0.396 | -1.236 | -0.002 | -0.008 | -0.008 | 0 | 0.54 | 0.52 | 0.47 | 0.62 | 0.43 | 0.53 |
| 6 | 25590224 | rs1028318 | 1.048 | -1.364 | -0.365 | -1.205 | 0.006 | 0.013 | 0.013 | 0 | 0.24 | 0.21 | 0.3 | 0.4 | 0.31 | 0.32 |
| 6 | 25594379 | rs9295665 | 1.068 | -1.134 | -0.307 | -0.998 | 0.008 | 0.075 | 0.075 | 0 | 0.13 | 0.17 | 0.24 | 0.42 | 0.16 | 0.24 |
| 6 | 25599234 | rs3788991 | -0.312 | -1.268 | -0.365 | -1.098 | 0.003 | 0.006 | 0.006 | 0 | 0.64 | 0.57 | 0.55 | 0.5 | 0.58 | 0.51 |
| 6 | 25600602 | rs9885888 | 1.362 | -1.163 | -0.578 | -1.042 | 0.007 | 0.008 | 0.008 | 0 | 0.07 | 0.06 | 0.02 | 0.02 | 0.02 | 0.1 |
| 6 | 25600968 | rs10456324 | 0.561 | -1.114 | -0.469 | -1.112 | 0.003 | 0.017 | 0.017 | 0 | 0.47 | 0.34 | 0.42 | 0.37 | 0.46 | 0.46 |
| 6 | 25603100 | rs7746950 | 0.695 | -1.229 | -0.459 | -1.042 | -0.001 | 0.03 | 0.03 | 0 | 0.46 | 0.29 | 0.41 | 0.3 | 0.47 | 0.46 |
| 6 | 25605512 | rs7740510 | -0.049 | -0.995 | -0.456 | -0.824 | 0.001 | 0.011 | 0.011 | 0 | 0.23 | 0.28 | 0.32 | 0.54 | 0.25 | 0.28 |
| 6 | 25610615 | rs2272824 | 0.891 | -0.936 | -0.411 | -0.686 | 0.016 | 0.112 | 0.112 | 0 | 0.13 | 0.22 | 0.24 | 0.42 | 0.17 | 0.24 |
| 6 | 25614046 | rs2024874 | -0.881 | -1.312 | -0.52 | -1.119 | -0.004 | 0.04 | 0.04 | 0 | 0.62 | 0.69 | 0.67 | 0.82 | 0.62 | 0.71 |
| 6 | 25617068 | rs10946785 | 0.841 | -1.206 | -0.525 | -1.064 | 0.01 | -0.005 | -0.005 | 0.001 | 0.26 | 0.29 | 0.3 | 0.25 | 0.31 | 0.32 |
| 6 | 25622212 | rs2281071 | -0.106 | -1.209 | -0.669 | -1.284 | -0.003 | -0.011 | -0.011 | 0 | 0.4 | 0.42 | 0.34 | 0.27 | 0.36 | 0.42 |
| 6 | 25622233 | rs941428 | 1.756 | -1.179 | -0.673 | -1.205 | -0.024 | -0.009 | -0.009 | 0.422 | 0.1 | 0.05 | 0.21 | 0.43 | 0.17 | 0.24 |
| 6 | 25624395 | rs7765813 | 1.753 | -1.145 | -0.733 | -1.114 | -0.026 | -0.011 | -0.011 | 2.746 | 0.05 | 0.01 | 0.06 | 0.04 | 0.08 | 0.03 |
| 6 | 25624416 | rs12183240 | 0.815 | -1.061 | -0.687 | -1.193 | 0.008 | -0.005 | -0.005 | 0 | 0.25 | 0.25 | 0.28 | 0.24 | 0.3 | 0.3 |
| 6 | 25626549 | rs6456685 | -1.888 | -1.143 | -0.708 | -1.303 | -0.01 | 0.007 | 0.007 | 0 | 0.84 | 0.89 | 0.8 | 0.87 | 0.78 | 0.87 |
| 6 | 25629658 | rs1980449 | -0.51 | -1.096 | -0.701 | -1.304 | -0.029 | 0.011 | 0.011 | 0 | 0.71 | 0.86 | 0.71 | 0.75 | 0.67 | 0.8 |
| 6 | 25629747 | rs1980450 | 0.534 | -0.843 | -0.397 | -0.98 | 0.006 | -0.009 | -0.009 | 1.432 | 0.26 | 0.31 | 0.3 | 0.25 | 0.3 | 0.31 |
| 6 | 25631230 | rs2072843 | 1.502 | -0.748 | -0.406 | -0.882 | -0.003 | 0.016 | 0.016 | 0 | 0.11 | 0.11 | 0.05 | 0.17 | 0.12 | 0.09 |
| 6 | 25641200 | rs932316 | -0.095 | -0.851 | -0.641 | -0.959 | 0.033 | 0.028 | 0.028 | 0.19 | 0.28 | 0.2 | 0.15 | 0.14 | 0.18 | 0.18 |
| 6 | 25642577 | rs10447389 | 0.601 | -0.674 | -0.491 | -0.959 | 0.009 | -0.009 | -0.009 | 0 | 0.25 | 0.29 | 0.29 | 0.25 | 0.29 | 0.31 |
| 6 | 25642991 | rs9358865 | 1.15 | -0.633 | -0.527 | -0.904 | 0.016 | 0.121 | 0.121 | 0 | 0.13 | 0.23 | 0.24 | 0.42 | 0.17 | 0.24 |
| 6 | 25646145 | rs909665 | 0.589 | -0.561 | -0.649 | -1.136 | 0.01 | -0.01 | -0.01 | 1.166 | 0.25 | 0.29 | 0.29 | 0.24 | 0.29 | 0.3 |
| 6 | 25652225 | rs2294344 | 2.289 | -0.398 | -1.073 | -1.135 | -0.051 | 0.033 | 0.033 | 0 | 0.1 | 0.01 | 0.22 | 0.08 | 0.17 | 0.21 |
| 6 | 25657463 | rs2011472 | -1.99 | -0.429 | -0.875 | -0.988 | -0.008 | 0.003 | 0.003 | 0 | 0.85 | 0.89 | 0.81 | 0.86 | 0.78 | 0.89 |
| 6 | 25663360 | rs9461190 | 1.84 | -0.645 | -0.939 | -0.985 | 0.01 | 0.052 | 0.052 | 0 | 0.17 | 0.08 | 0.11 | 0.04 | 0.14 | 0.15 |
| 6 | 25669525 | rs2876691 | -0.033 | -0.415 | -0.346 | -0.315 | 0.014 | 0.018 | 0.018 | 0 | 0.21 | 0.13 | 0.12 | 0.11 | 0.15 | 0.15 |
| 6 | 25670946 | rs12211682 | 0.922 | -0.408 | 0.264 | 0.283 | -0.002 | 0.035 | 0.035 | 0 | 0.11 | 0.15 | 0.06 | 0.17 | 0.12 | 0.08 |
| 6 | 25673267 | rs4711090 | -1.157 | -0.235 | 0.066 | 0.239 | -0.007 | 0.004 | 0.004 | 0 | 0.85 | 0.87 | 0.81 | 0.86 | 0.78 | 0.88 |
| 6 | 25674509 | rs4711091 | 1.077 | -0.335 | -0.072 | 0.168 | -0.027 | 0.139 | 0.139 | 0 | 0.46 | 0.2 | 0.44 | 0.21 | 0.46 | 0.45 |
| 6 | 25677954 | rs9467561 | 0.525 | -0.327 | -0.284 | 0.269 | 0 | -0.002 | -0.002 | 0 | 0.58 | 0.39 | 0.51 | 0.38 | 0.58 | 0.54 |
| 6 | 25679960 | rs2205935 | 0.496 | -0.006 | -0.858 | 0.439 | 0.008 | 0.102 | 0.102 | 0 | 0.14 | 0.22 | 0.25 | 0.46 | 0.17 | 0.24 |

| | | | | | | | | | | | | | | | | |
|---|---|---|---|---|---|---|---|---|---|---|---|---|---|---|---|---|
| 6 | 25682262 | rs3765236 | 0.643 | 0.178 | -1.43 | 0.018 | -0.079 | 0.151 | 0.151 | 0.036 | 0.44 | 0.13 | 0.45 | 0.22 | 0.45 | 0.45 |
| 6 | 25683280 | rs1883259 | -0.38 | 0.704 | -0.627 | 0.543 | 0.002 | -0.002 | -0.002 | 0 | 0.85 | 0.84 | 0.81 | 0.85 | 0.78 | 0.87 |
| 6 | 25683954 | rs6942120 | -0.454 | 0.224 | -0.603 | 1.277 | -0.004 | 0.02 | 0.02 | 0 | 0.19 | 0.21 | 0.22 | 0.35 | 0.15 | 0.31 |
| 6 | 25684279 | rs4712955 | 0.628 | 0.659 | -0.76 | 1.485 | -0.025 | -0.014 | -0.014 | 0.172 | 0.71 | 0.79 | 0.61 | 0.35 | 0.57 | 0.62 |
| 6 | 25684405 | rs4711092 | 0.291 | -0.01 | -0.656 | 0.958 | -0.001 | 0.002 | 0.002 | 0.005 | 0.57 | 0.44 | 0.62 | 0.59 | 0.56 | 0.63 |
| 6 | 25684653 | rs9348689 | -0.356 | -0.012 | -0.718 | 0.354 | -0.006 | 0.012 | 0.012 | 0 | 0.46 | 0.4 | 0.41 | 0.56 | 0.55 | 0.53 |
| 6 | 25686405 | rs9366627 | -0.327 | 0.168 | -0.653 | 0.183 | 0.004 | 0.008 | 0.008 | 0.221 | 0.21 | 0.23 | 0.41 | 0.66 | 0.3 | 0.34 |
| 6 | 25694491 | rs4409177 | 1.606 | -0.162 | -0.99 | -0.572 | -0.043 | 0.078 | 0.078 | 0 | 0.29 | 0.15 | 0.19 | 0.14 | 0.28 | 0.27 |
| 6 | 25699999 | rs10946793 | 0.402 | -0.114 | -0.891 | -0.549 | -0.034 | -0.045 | -0.045 | 0.093 | 0.18 | 0.1 | 0.39 | 0.66 | 0.29 | 0.33 |
| 6 | 25701718 | rs11794 | 0.408 | -0.118 | -0.893 | -0.707 | -0.034 | -0.04 | -0.04 | 0 | 0.18 | 0.1 | 0.44 | 0.68 | 0.29 | 0.33 |
| 6 | 25701903 | rs10105 | 0.246 | -0.658 | -0.377 | -1.296 | -0.001 | -0.041 | -0.041 | 0.048 | 0.25 | 0.35 | 0.22 | 0.1 | 0.26 | 0.21 |
| 6 | 25707171 | rs9379780 | 0.256 | -1.079 | 0.03 | -1.497 | 0.005 | -0.007 | -0.007 | 0.342 | 0.52 | 0.53 | 0.68 | 0.86 | 0.6 | 0.65 |
| 6 | 25708960 | rs6456688 | 0.566 | -0.755 | -0.127 | -1.494 | 0 | -0.004 | -0.004 | 6.399 | 0.52 | 0.54 | 0.39 | 0.24 | 0.49 | 0.5 |
| 6 | 25714739 | rs10946795 | 1.007 | -0.756 | -0.24 | -0.91 | 0.003 | -0.036 | -0.036 | 0 | 0.1 | 0.02 | 0.25 | 0.51 | 0.06 | 0.24 |
| 6 | 25725506 | rs9379784 | 0.356 | -0.817 | -0.095 | -1.644 | -0.017 | -0.024 | -0.024 | 0 | 0.2 | 0.15 | 0.44 | 0.66 | 0.29 | 0.33 |
| 6 | 25726343 | rs6940348 | -1.441 | -0.919 | -0.08 | -1.659 | -0.021 | -0.003 | -0.003 | 0.208 | 0.95 | 0.84 | 0.98 | 0.98 | 0.98 | 0.96 |
| 6 | 25726774 | rs4711095 | 0.314 | -0.93 | -0.063 | -1.701 | -0.032 | -0.031 | -0.031 | 0 | 0.2 | 0.1 | 0.44 | 0.68 | 0.29 | 0.33 |
| 6 | 25727054 | rs4711096 | 0.27 | -0.937 | 0.033 | -1.704 | -0.023 | -0.023 | -0.023 | 0 | 0.2 | 0.11 | 0.44 | 0.69 | 0.29 | 0.33 |
| 6 | 25727265 | rs4712960 | 0.315 | -0.935 | -0.072 | -1.703 | -0.032 | -0.031 | -0.031 | 0 | 0.2 | 0.1 | 0.44 | 0.68 | 0.29 | 0.33 |
| 6 | 25727334 | rs4712961 | 0.315 | -0.935 | -0.072 | -1.703 | -0.032 | -0.031 | -0.031 | 0.987 | 0.2 | 0.1 | 0.44 | 0.68 | 0.29 | 0.33 |
| 6 | 25727517 | rs9358872 | 0.315 | -0.942 | -0.002 | -1.716 | -0.032 | -0.031 | -0.031 | 0 | 0.2 | 0.1 | 0.44 | 0.68 | 0.29 | 0.33 |
| 6 | 25732314 | rs4711097 | 0.325 | -0.938 | -0.008 | -1.731 | -0.032 | -0.041 | -0.041 | 2.184 | 0.2 | 0.1 | 0.44 | 0.67 | 0.29 | 0.33 |
| 6 | 25732331 | rs4712962 | -1.471 | -0.934 | -0.003 | -1.821 | -0.021 | -0.003 | -0.003 | 0 | 0.95 | 0.84 | 0.98 | 0.98 | 0.98 | 0.96 |
| 6 | 25736678 | rs3922841 | 1.207 | -1.045 | -0.255 | -2.014 | -0.026 | 0.238 | 0.238 | 0.496 | 0.49 | 0.19 | 0.3 | 0.18 | 0.43 | 0.42 |
| 6 | 25738956 | rs9358876 | 0.382 | -1.005 | -0.358 | -1.939 | -0.032 | -0.03 | -0.03 | 0 | 0.2 | 0.1 | 0.44 | 0.69 | 0.29 | 0.33 |
| 6 | 25740502 | rs4712963 | 0.382 | -1.005 | -0.358 | -1.939 | -0.032 | -0.03 | -0.03 | 0 | 0.2 | 0.1 | 0.44 | 0.69 | 0.29 | 0.33 |
| 6 | 25745742 | rs7776065 | 0.086 | -1.012 | -0.187 | -1.953 | -0.003 | -0.049 | -0.049 | 0 | 0.25 | 0.39 | 0.22 | 0.11 | 0.25 | 0.21 |
| 6 | 25751721 | rs12207270 | 2.126 | -1.004 | -0.597 | -2.031 | -0.067 | 0.073 | 0.073 | 0 | 0.25 | 0.09 | 0.18 | 0.13 | 0.28 | 0.26 |
| 6 | 25754271 | rs1892256 | 2.126 | -1.004 | -0.597 | -2.031 | -0.067 | 0.073 | 0.073 | 0 | 0.25 | 0.09 | 0.18 | 0.13 | 0.28 | 0.26 |
| 6 | 25755515 | rs9358883 | 0.312 | -0.804 | -0.61 | -1.886 | -0.032 | -0.03 | -0.03 | 0 | 0.2 | 0.1 | 0.44 | 0.69 | 0.29 | 0.34 |
| 6 | 25758212 | rs9295669 | -0.451 | -0.822 | -0.601 | -2.057 | -0.005 | 0.001 | 0.001 | 0.961 | 0.41 | 0.39 | 0.62 | 0.8 | 0.54 | 0.56 |
| 6 | 25758448 | rs17268697 | 0.934 | -0.725 | -0.617 | -1.639 | 0.001 | 0.104 | 0.104 | 0.41 | 0.2 | 0.01 | 0.1 | 0.05 | 0.15 | 0.13 |
| 6 | 25769349 | rs1892251 | 0.231 | -0.677 | 0.003 | -1.658 | 0.014 | -0.052 | -0.052 | 0 | 0.88 | 0.69 | 0.9 | 0.99 | 0.91 | 0.97 |
| 6 | 25770239 | rs2328893 | 0.964 | -1.159 | -0.328 | -1.876 | 0.087 | 0.22 | 0.22 | 1.031 | 0.53 | 0.13 | 0.7 | 0.84 | 0.7 | 0.78 |
| 6 | 25772047 | rs4712972 | 0.723 | -1.18 | 0.078 | -1.877 | -0.038 | -0.107 | -0.107 | 0.04 | 0.79 | 0.47 | 0.85 | 0.97 | 0.88 | 0.95 |
| 6 | 25776949 | rs11754288 | 1.763 | -0.832 | -0.353 | -1.786 | -0.132 | 0.089 | 0.089 | 0 | 0.34 | 0.05 | 0.28 | 0.17 | 0.42 | 0.45 |
| 6 | 25780332 | rs1141034 | 0 | -0.334 | 0.051 | -1.137 | -0.04 | -0.067 | -0.067 | 0 | 0.2 | 0.09 | 0.43 | 0.67 | 0.29 | 0.33 |
| 6 | 25780811 | rs6910549 | 0.282 | -0.376 | 0.144 | -1.199 | 0.014 | -0.052 | -0.052 | 0 | 0.88 | 0.69 | 0.9 | 0.99 | 0.91 | 0.97 |
| 6 | 25781139 | rs7749149 | -0.13 | -0.373 | 0.166 | -1.196 | 0 | 0 | 0 | 0 | 0.22 | 0.21 | 0.44 | 0.74 | 0.31 | 0.34 |
| 6 | 25783315 | rs3923 | 0.77 | -0.506 | -0.337 | -1.157 | 0.005 | -0.001 | -0.001 | 0 | 0.68 | 0.59 | 0.47 | 0.31 | 0.61 | 0.64 |
| 6 | 25787817 | rs2096386 | 0.488 | -0.138 | -0.321 | -0.956 | 0.015 | 0.005 | 0.005 | 0 | 0.72 | 0.6 | 0.48 | 0.3 | 0.62 | 0.66 |
| 6 | 25794431 | rs2762353 | -1.31 | -0.34 | -1.088 | -1.195 | -0.128 | 0.103 | 0.103 | 0 | 0.63 | 0.92 | 0.7 | 0.82 | 0.56 | 0.52 |
| 6 | 25800197 | rs3823151 | -0.082 | -0.809 | -0.399 | -1.346 | -0.055 | -0.003 | -0.003 | 0 | 0.14 | 0.32 | 0.06 | 0.09 | 0.04 | 0.04 |
| 6 | 25813150 | rs1165196 | -1.487 | -0.678 | -0.586 | -1.368 | -0.129 | 0.097 | 0.097 | 0 | 0.63 | 0.92 | 0.7 | 0.82 | 0.56 | 0.52 |
| 6 | 25830298 | rs1165176 | -1.516 | -0.826 | -0.599 | -1.466 | -0.146 | 0.124 | 0.124 | 0 | 0.6 | 0.93 | 0.69 | 0.82 | 0.54 | 0.49 |
| 6 | 25841408 | rs1324087 | -0.301 | -0.769 | -0.131 | -1.468 | -0.003 | -0.005 | -0.005 | 1.231 | 0.16 | 0.29 | 0.1 | 0.11 | 0.11 | 0.05 |
| 6 | 25842203 | rs4712976 | 0.189 | -0.968 | -0.16 | -1.681 | -0.014 | -0.037 | -0.037 | 0 | 0.19 | 0.12 | 0.44 | 0.67 | 0.26 | 0.3 |
| 6 | 25842605 | rs9393672 | -1.473 | -1.057 | -0.213 | -1.763 | -0.003 | 0.026 | 0.026 | 0 | 0.52 | 0.44 | 0.63 | 0.8 | 0.52 | 0.47 |
| 6 | 25843746 | rs3887266 | 0.95 | -0.998 | -0.469 | -1.735 | 0.005 | 0.081 | 0.081 | 0.097 | 0.16 | 0.01 | 0.08 | 0.02 | 0.13 | 0.12 |
| 6 | 25849620 | rs942379 | -1.913 | -0.998 | -0.477 | -1.715 | -0.136 | 0.139 | 0.139 | 0 | 0.59 | 0.92 | 0.69 | 0.81 | 0.54 | 0.48 |
| 6 | 25855624 | rs717551 | 0.874 | -0.818 | -0.286 | -1.151 | -0.046 | -0.042 | -0.042 | 0.023 | 0.07 | 0.02 | 0.12 | 0.17 | 0.22 | 0.11 |
| 6 | 25862466 | rs1165165 | -0.652 | -1.024 | -0.184 | -1.278 | -0.007 | -0.022 | -0.022 | 0 | 0.29 | 0.33 | 0.18 | 0.13 | 0.23 | 0.17 |
| 6 | 25864625 | rs1165159 | -0.767 | -1.008 | -0.17 | -1.265 | -0.005 | -0.015 | -0.015 | 0.082 | 0.32 | 0.33 | 0.18 | 0.14 | 0.25 | 0.17 |
| 6 | 25873025 | rs501220 | -0.628 | -0.721 | 0.231 | -1.334 | -0.004 | -0.01 | -0.01 | 0 | 0.18 | 0.46 | 0.12 | 0.13 | 0.12 | 0.06 |
| 6 | 25875212 | rs1165200 | 0.431 | -0.698 | 0.047 | -1.27 | 0 | -0.007 | -0.007 | 0 | 0.8 | 0.76 | 0.84 | 0.81 | 0.76 | 0.87 |
| 6 | 25876089 | rs12664474 | -0.106 | -1.001 | -0.142 | -1.313 | 0.044 | -0.086 | -0.086 | 0 | 0.26 | 0.12 | 0.34 | 0.52 | 0.18 | 0.3 |
| 6 | 25878506 | rs11969868 | 0.462 | -0.896 | -0.058 | -1.621 | -0.018 | -0.039 | -0.039 | 0 | 0.2 | 0.26 | 0.15 | 0.06 | 0.17 | 0.23 |
| 6 | 25879539 | rs487624 | -0.594 | -0.73 | -0.236 | -1.566 | -0.001 | -0.006 | -0.006 | 0 | 0.37 | 0.37 | 0.31 | 0.28 | 0.43 | 0.38 |
| 6 | 25880494 | rs645279 | -2.173 | -0.444 | -0.825 | -1.742 | -0.077 | 0.101 | 0.101 | 0 | 0.72 | 0.94 | 0.74 | 0.88 | 0.72 | 0.67 |
| 6 | 25886921 | rs17271767 | -0.058 | -0.595 | -0.503 | -1.737 | -0.001 | 0 | 0 | 0 | 0.17 | 0.16 | 0.15 | 0.17 | 0.25 | 0.12 |
| 6 | 25887217 | rs7750960 | 0.805 | -0.563 | -0.388 | -1.806 | 0.043 | 0.002 | 0.002 | 0 | 0.06 | 0.17 | 0.07 | 0.04 | 0.09 | 0.05 |
| 6 | 25887776 | rs12209218 | 0.084 | -0.575 | -0.462 | -1.806 | -0.011 | 0.004 | 0.004 | 0 | 0.17 | 0.13 | 0.15 | 0.17 | 0.25 | 0.12 |
| 6 | 25887959 | rs670011 | -1.168 | -0.604 | -0.49 | -1.823 | -0.031 | 0.123 | 0.123 | 0 | 0.55 | 0.81 | 0.6 | 0.71 | 0.48 | 0.56 |
| 6 | 25889530 | rs9393674 | -0.344 | -0.803 | -0.201 | -1.775 | -0.012 | -0.003 | -0.003 | 0 | 0.42 | 0.43 | 0.51 | 0.6 | 0.35 | 0.5 |
| 6 | 25890834 | rs9348697 | -0.672 | -0.662 | -0.13 | -1.853 | 0 | 0.013 | 0.013 | 0 | 0.36 | 0.23 | 0.46 | 0.69 | 0.39 | 0.38 |
| 6 | 25895575 | rs6910741 | 0.176 | -0.587 | 0.07 | -1.57 | 0.03 | -0.064 | -0.064 | 0 | 0.23 | 0.09 | 0.33 | 0.63 | 0.18 | 0.29 |
| 6 | 25901133 | rs13207673 | 0.49 | -0.283 | 0.279 | -1.531 | -0.004 | 0.004 | 0.004 | 0 | 0.06 | 0.05 | 0.06 | 0.01 | 0.08 | 0.03 |
| 6 | 25901758 | rs9356991 | -0.727 | -0.242 | 0.065 | -1.792 | -0.034 | 0.011 | 0.011 | 0 | 0.35 | 0.14 | 0.46 | 0.7 | 0.4 | 0.4 |
| 6 | 25903308 | rs16891146 | 0.475 | -0.326 | 0.058 | -2.06 | -0.004 | -0.032 | -0.032 | 0 | 0.24 | 0.24 | 0.19 | 0.09 | 0.22 | 0.25 |

| | | | | | | | | | | | | | | | | |
|---|---|---|---|---|---|---|---|---|---|---|---|---|---|---|---|---|
| 6 | 25904414 | rs6910174 | -0.593 | -0.269 | -0.426 | -2.022 | -0.031 | 0.012 | 0.012 | 0 | 0.35 | 0.14 | 0.46 | 0.7 | 0.4 | 0.4 |
| 6 | 25911775 | rs3846838 | -0.622 | -0.279 | -0.369 | -2.021 | -0.023 | 0.016 | 0.016 | 0 | 0.35 | 0.15 | 0.46 | 0.71 | 0.4 | 0.4 |
| 6 | 25912103 | rs6905887 | -0.591 | -0.279 | -0.369 | -2.021 | -0.031 | 0.012 | 0.012 | 0 | 0.35 | 0.14 | 0.46 | 0.7 | 0.4 | 0.4 |
| 6 | 25912634 | rs3799369 | 1.279 | -0.4 | -0.488 | -2.069 | -0.011 | 0.045 | 0.045 | 0.509 | 0.12 | 0.01 | 0.07 | 0.03 | 0.13 | 0.09 |
| 6 | 25912816 | rs3799371 | -0.71 | -0.354 | -0.421 | -2.031 | -0.023 | 0.016 | 0.016 | 0 | 0.35 | 0.15 | 0.46 | 0.71 | 0.4 | 0.4 |
| 6 | 25913098 | rs6932113 | -0.679 | -0.354 | -0.424 | -2.03 | -0.031 | 0.012 | 0.012 | 0 | 0.35 | 0.14 | 0.46 | 0.7 | 0.4 | 0.4 |
| 6 | 25914801 | rs2071299 | -0.679 | -0.354 | -0.424 | -2.03 | -0.031 | 0.012 | 0.012 | 0 | 0.35 | 0.14 | 0.46 | 0.7 | 0.4 | 0.4 |
| 6 | 25914901 | rs2071298 | -0.679 | -0.354 | -0.424 | -2.03 | -0.031 | 0.012 | 0.012 | 0 | 0.35 | 0.14 | 0.46 | 0.7 | 0.4 | 0.4 |
| 6 | 25916979 | rs1865760 | -0.679 | -0.354 | -0.424 | -2.03 | -0.031 | 0.012 | 0.012 | 0 | 0.35 | 0.14 | 0.46 | 0.7 | 0.4 | 0.4 |
| 6 | 25918688 | rs3752421 | -0.71 | -0.354 | -0.421 | -2.031 | -0.023 | 0.016 | 0.016 | 0 | 0.35 | 0.15 | 0.46 | 0.71 | 0.4 | 0.4 |
| 6 | 25919480 | rs7450342 | -1.059 | -0.35 | -0.404 | -2.037 | -0.019 | 0.029 | 0.029 | 0 | 0.4 | 0.32 | 0.53 | 0.72 | 0.49 | 0.43 |
| 6 | 25920265 | rs7749342 | -1.047 | -0.349 | -0.403 | -2.037 | -0.019 | 0.028 | 0.028 | 0 | 0.4 | 0.33 | 0.53 | 0.72 | 0.49 | 0.43 |
| 6 | 25924158 | rs1540273 | -0.683 | -0.347 | -0.439 | -2.035 | -0.031 | 0.012 | 0.012 | 0 | 0.35 | 0.14 | 0.46 | 0.7 | 0.4 | 0.4 |
| 6 | 25926674 | rs10484431 | 1.02 | -0.435 | -0.563 | -1.993 | -0.05 | 0.064 | 0.064 | 0.025 | 0.19 | 0.03 | 0.15 | 0.06 | 0.22 | 0.23 |
| 6 | 25928376 | rs9295676 | 0.946 | -0.298 | -0.404 | -1.721 | 0.003 | 0.042 | 0.042 | 0.636 | 0.09 | 0.07 | 0.26 | 0.59 | 0.04 | 0.19 |
| 6 | 25931577 | rs199741 | -2.316 | -0.276 | -0.542 | -1.949 | -0.05 | 0.096 | 0.096 | 0 | 0.73 | 0.95 | 0.76 | 0.87 | 0.76 | 0.69 |
| 6 | 25935765 | rs2328903 | -0.878 | -0.408 | -0.306 | -1.752 | -0.014 | 0.021 | 0.021 | 0 | 0.35 | 0.18 | 0.47 | 0.72 | 0.41 | 0.4 |
| 6 | 25936787 | rs199736 | -1.982 | -0.478 | -0.209 | -1.898 | 0.006 | 0.09 | 0.09 | 0 | 0.63 | 0.76 | 0.71 | 0.86 | 0.69 | 0.63 |
| 6 | 25936944 | rs9393676 | -0.993 | -0.393 | -0.155 | -1.809 | -0.016 | 0.025 | 0.025 | 0 | 0.4 | 0.31 | 0.54 | 0.75 | 0.49 | 0.43 |
| 6 | 25937694 | rs9467638 | 0.775 | -0.437 | -0.222 | -1.751 | -0.017 | 0.08 | 0.08 | 0.241 | 0.11 | 0.12 | 0.27 | 0.62 | 0.05 | 0.21 |
| 6 | 25939053 | rs17586946 | 0.93 | -0.375 | -0.273 | -1.792 | -0.021 | 0.001 | 0.001 | 0 | 0.07 | 0.05 | 0.12 | 0.07 | 0.22 | 0.09 |
| 6 | 25939723 | rs1436307 | -0.737 | -0.464 | -0.256 | -1.808 | -0.013 | 0.021 | 0.021 | 0 | 0.35 | 0.19 | 0.47 | 0.72 | 0.41 | 0.4 |
| 6 | 25943197 | rs9358896 | -0.747 | -0.466 | -0.257 | -1.809 | -0.013 | 0.022 | 0.022 | 0 | 0.35 | 0.19 | 0.47 | 0.72 | 0.41 | 0.4 |
| 6 | 25948421 | rs1436306 | -0.737 | -0.405 | -0.229 | -1.827 | -0.018 | 0.048 | 0.048 | 0 | 0.35 | 0.19 | 0.47 | 0.76 | 0.43 | 0.4 |
| 6 | 25950382 | rs7740793 | 0.737 | -0.477 | -0.206 | -1.804 | -0.018 | 0.081 | 0.081 | 0.172 | 0.11 | 0.13 | 0.27 | 0.62 | 0.05 | 0.2 |
| 6 | 25953684 | rs199725 | -2.033 | -0.568 | -0.226 | -1.912 | 0.005 | 0.092 | 0.092 | 0 | 0.64 | 0.77 | 0.71 | 0.86 | 0.69 | 0.63 |
| 6 | 25956501 | rs6906576 | 0.782 | -0.526 | -0.224 | -1.849 | -0.017 | 0.08 | 0.08 | 0.153 | 0.11 | 0.12 | 0.27 | 0.62 | 0.05 | 0.2 |
| 6 | 25957392 | rs169219 | -1.55 | -0.428 | -0.138 | -1.92 | -0.007 | 0.045 | 0.045 | 0 | 0.58 | 0.63 | 0.64 | 0.83 | 0.61 | 0.6 |
| 6 | 25957773 | rs10946800 | -0.817 | -0.394 | -0.178 | -1.897 | -0.021 | 0.049 | 0.049 | 0 | 0.35 | 0.18 | 0.47 | 0.76 | 0.43 | 0.4 |
| 6 | 25963456 | rs199738 | -0.022 | -0.231 | 0.009 | -1.673 | -0.039 | -0.099 | -0.099 | 0 | 0.23 | 0.45 | 0.17 | 0.07 | 0.17 | 0.2 |
| 6 | 25967068 | rs2009610 | 0.8 | -0.086 | -0.529 | -1.471 | -0.043 | -0.012 | -0.012 | 0 | 0.07 | 0.02 | 0.12 | 0.07 | 0.22 | 0.09 |
| 6 | 25974204 | rs13196552 | 1.156 | -0.325 | -0.454 | -1.603 | 0.009 | -0.021 | -0.021 | 0 | 0.08 | 0.15 | 0.07 | 0.02 | 0.09 | 0.03 |
| 6 | 25974450 | rs12192649 | 0.719 | -0.171 | -0.455 | -1.603 | -0.022 | 0.001 | 0.001 | 0 | 0.07 | 0.05 | 0.12 | 0.07 | 0.21 | 0.08 |
| 6 | 25976927 | rs6925912 | 0.432 | -0.468 | 0.194 | -1.799 | -0.042 | -0.063 | -0.063 | 0 | 0.21 | 0.33 | 0.16 | 0.07 | 0.13 | 0.2 |
| 6 | 25977361 | rs6906460 | 0.432 | -0.526 | 0.005 | -1.798 | -0.052 | -0.07 | -0.07 | 0 | 0.21 | 0.35 | 0.16 | 0.07 | 0.13 | 0.2 |
| 6 | 25985396 | rs1130000 | -0.713 | -0.769 | -0.274 | -2.025 | -0.041 | 0.037 | 0.037 | 0 | 0.37 | 0.15 | 0.48 | 0.72 | 0.44 | 0.4 |
| 6 | 25990367 | rs199755 | -2.597 | -0.809 | -0.221 | -2.084 | -0.04 | 0.094 | 0.094 | 0 | 0.7 | 0.89 | 0.74 | 0.86 | 0.7 | 0.64 |
| 6 | 25991620 | rs2051538 | 0.643 | -0.848 | -0.274 | -2.106 | -0.048 | -0.067 | -0.067 | 0.014 | 0.2 | 0.35 | 0.16 | 0.07 | 0.13 | 0.21 |
| 6 | 25992309 | rs199754 | -2.715 | -0.862 | -0.361 | -2.111 | -0.094 | 0.09 | 0.09 | 0 | 0.72 | 0.95 | 0.74 | 0.86 | 0.7 | 0.64 |
| 6 | 25997458 | rs12216125 | 0.433 | -1.008 | -0.457 | -2.086 | -0.082 | 0.045 | 0.045 | 0 | 0.2 | 0.06 | 0.2 | 0.11 | 0.36 | 0.17 |
| 6 | 26000885 | rs6901039 | -2.275 | -0.996 | -0.371 | -2.132 | -0.001 | -0.012 | -0.012 | 0 | 0.67 | 0.69 | 0.71 | 0.86 | 0.69 | 0.64 |
| 6 | 26008260 | rs9393681 | -0.963 | -1.027 | -0.529 | -2.083 | -0.03 | 0.082 | 0.082 | 0 | 0.39 | 0.2 | 0.48 | 0.76 | 0.46 | 0.4 |
| 6 | 26008876 | rs10484432 | 0.893 | -0.948 | -0.531 | -1.87 | -0.025 | -0.023 | -0.023 | 0 | 0.08 | 0.05 | 0.12 | 0.16 | 0.21 | 0.09 |
| 6 | 26016462 | rs199750 | -2.859 | -1.175 | -0.584 | -1.804 | -0.098 | 0.073 | 0.073 | 0 | 0.74 | 0.95 | 0.73 | 0.86 | 0.69 | 0.65 |
| 6 | 26017171 | rs2237231 | 0.841 | -1.218 | -0.448 | -1.591 | 0.014 | -0.022 | -0.022 | 0.156 | 0.13 | 0.07 | 0.27 | 0.51 | 0.07 | 0.21 |
| 6 | 26017542 | rs16891235 | 1.415 | -1.231 | -0.33 | -1.63 | 0.001 | -0.008 | -0.008 | 0 | 0.08 | 0.13 | 0.07 | 0.01 | 0.09 | 0.04 |
| 6 | 26020431 | rs2157050 | -0.597 | -1.217 | -0.493 | -1.66 | -0.018 | 0.029 | 0.029 | 0.028 | 0.37 | 0.2 | 0.47 | 0.71 | 0.4 | 0.39 |
| 6 | 26022244 | rs3734528 | 1.418 | -1.176 | -0.473 | -1.641 | 0.001 | -0.008 | -0.008 | 0 | 0.08 | 0.13 | 0.07 | 0.01 | 0.09 | 0.03 |
| 6 | 26023206 | rs9379818 | -0.235 | -1.199 | -0.497 | -1.654 | -0.04 | 0.038 | 0.038 | 0 | 0.32 | 0.18 | 0.46 | 0.76 | 0.42 | 0.38 |
| 6 | 26027135 | rs3752420 | -0.038 | -1.238 | -0.54 | -1.669 | -0.052 | -0.013 | -0.013 | 0 | 0.31 | 0.13 | 0.45 | 0.71 | 0.39 | 0.37 |
| 6 | 26027433 | rs3752419 | -0.184 | -1.223 | -0.533 | -1.665 | -0.049 | -0.005 | -0.005 | 0.037 | 0.32 | 0.13 | 0.45 | 0.71 | 0.4 | 0.38 |
| 6 | 26030492 | rs10484433 | 1.425 | -1.196 | -0.534 | -1.689 | 0.002 | -0.009 | -0.009 | 4.794 | 0.08 | 0.13 | 0.07 | 0.01 | 0.09 | 0.03 |
| 6 | 26031613 | rs10484434 | 1.244 | -1.374 | -0.647 | -1.794 | 0.005 | 0.037 | 0.037 | 0.094 | 0.14 | 0.02 | 0.11 | 0.06 | 0.12 | 0.19 |
| 6 | 26031868 | rs2213284 | -2.601 | -1.445 | -0.699 | -1.799 | -0.107 | 0.094 | 0.094 | 0 | 0.71 | 0.94 | 0.72 | 0.86 | 0.67 | 0.64 |
| 6 | 26033506 | rs2230655 | -2.633 | -1.445 | -0.699 | -1.827 | -0.102 | 0.096 | 0.096 | 0 | 0.71 | 0.94 | 0.72 | 0.86 | 0.67 | 0.64 |
| 6 | 26033641 | rs2230654 | 0.471 | -1.494 | -0.484 | -1.844 | -0.048 | -0.064 | -0.064 | 0 | 0.2 | 0.35 | 0.16 | 0.07 | 0.14 | 0.2 |
| 6 | 26035208 | rs4401650 | -2.126 | -1.476 | -0.546 | -1.835 | -0.076 | 0.137 | 0.137 | 0 | 0.66 | 0.93 | 0.71 | 0.86 | 0.66 | 0.62 |
| 6 | 26036476 | rs1540275 | -2.603 | -1.446 | -0.543 | -1.821 | -0.107 | 0.097 | 0.097 | 0 | 0.71 | 0.94 | 0.72 | 0.86 | 0.67 | 0.63 |
| 6 | 26044064 | rs11751732 | 0.418 | -1.318 | -0.379 | -1.728 | -0.048 | -0.064 | -0.064 | 0 | 0.2 | 0.35 | 0.16 | 0.07 | 0.13 | 0.2 |
| 6 | 26044369 | rs2032447 | -1.621 | -1.349 | -0.392 | -1.764 | 0 | -0.002 | -0.002 | 1.906 | 0.61 | 0.65 | 0.69 | 0.84 | 0.65 | 0.62 |
| 6 | 26046158 | rs11755492 | 1.305 | -1.303 | -0.457 | -1.732 | 0.015 | 0.03 | 0.03 | 0 | 0.15 | 0.03 | 0.12 | 0.07 | 0.12 | 0.2 |
| 6 | 26046565 | rs7756117 | -1.715 | -1.225 | -0.367 | -1.695 | -0.003 | 0.049 | 0.049 | 0 | 0.62 | 0.72 | 0.69 | 0.86 | 0.65 | 0.62 |
| 6 | 26048931 | rs10807007 | -0.304 | -1.188 | -0.434 | -1.672 | -0.04 | 0.037 | 0.037 | 0 | 0.32 | 0.18 | 0.46 | 0.75 | 0.42 | 0.39 |
| 6 | 26050845 | rs9689245 | 0.522 | -1.205 | -0.369 | -1.663 | -0.051 | -0.061 | -0.061 | 0.702 | 0.19 | 0.34 | 0.12 | 0.07 | 0.11 | 0.2 |
| 6 | 26056549 | rs10425 | -1.924 | -0.916 | -0.909 | -1.552 | -0.075 | 0.14 | 0.14 | 0 | 0.65 | 0.93 | 0.67 | 0.86 | 0.64 | 0.63 |
| 6 | 26066198 | rs9393683 | -0.423 | -0.49 | -0.445 | -1.11 | -0.042 | 0.029 | 0.029 | 0 | 0.31 | 0.18 | 0.46 | 0.75 | 0.42 | 0.38 |
| 6 | 26069669 | rs9295684 | -0.423 | -0.49 | -0.445 | -1.11 | -0.042 | 0.029 | 0.029 | 0 | 0.31 | 0.18 | 0.46 | 0.75 | 0.42 | 0.38 |
| 6 | 26071093 | rs2051544 | -0.386 | -0.536 | -0.437 | -1.124 | -0.052 | -0.011 | -0.011 | 0 | 0.31 | 0.13 | 0.45 | 0.71 | 0.4 | 0.38 |
| 6 | 26072804 | rs6942196 | -0.427 | -0.538 | -0.448 | -1.13 | -0.034 | -0.002 | -0.002 | 0 | 0.31 | 0.18 | 0.46 | 0.71 | 0.4 | 0.38 |

| | | | | | | | | | | | | | | | | |
|---|---|---|---|---|---|---|---|---|---|---|---|---|---|---|---|---|
| 6 | 26082710 | rs9295687 | -0.411 | -0.483 | -0.822 | -1.077 | -0.035 | 0 | 0 | 0.054 | 0.31 | 0.18 | 0.46 | 0.71 | 0.4 | 0.37 |
| 6 | 26088890 | rs2794719 | -0.403 | -0.482 | -0.822 | -1.076 | -0.035 | -0.001 | -0.001 | 0 | 0.31 | 0.18 | 0.46 | 0.72 | 0.4 | 0.37 |
| 6 | 26089098 | rs9366637 | 0.658 | -0.641 | -0.781 | -0.886 | 0.013 | -0.005 | -0.005 | 0 | 0.12 | 0.07 | 0.26 | 0.51 | 0.06 | 0.19 |
| 6 | 26091179 | rs1799945 | 0.663 | -0.559 | -0.888 | -1.413 | -0.024 | 0.061 | 0.061 | 0.303 | 0.14 | 0.01 | 0.12 | 0.03 | 0.17 | 0.07 |
| 6 | 26091336 | rs2071303 | -1.233 | -0.944 | -0.674 | -1.691 | 0.022 | 0.023 | 0.023 | 0.348 | 0.45 | 0.38 | 0.42 | 0.63 | 0.37 | 0.35 |
| 6 | 26093303 | rs1800708 | 1.165 | -0.948 | -0.783 | -1.676 | 0.002 | 0.019 | 0.019 | 0 | 0.11 | 0.08 | 0.21 | 0.55 | 0.09 | 0.21 |
| 6 | 26094367 | rs1572982 | -1.92 | -1.259 | -0.554 | -1.958 | -0.006 | 0.006 | 0.006 | 0 | 0.64 | 0.64 | 0.58 | 0.7 | 0.47 | 0.55 |
| 6 | 26095931 | rs707889 | 0.325 | -1.12 | -0.567 | -2.009 | 0.109 | 0.07 | 0.07 | 0 | 0.09 | 0.26 | 0.14 | 0.17 | 0.2 | 0.07 |
| 6 | 26097194 | rs17596719 | 1.461 | -1.183 | -0.599 | -2.041 | 0.021 | 0.033 | 0.033 | 0.604 | 0.15 | 0.02 | 0.12 | 0.07 | 0.1 | 0.2 |
| 6 | 26097384 | rs6918586 | -1.407 | -1.076 | -0.438 | -1.975 | 0.031 | 0.078 | 0.078 | 0 | 0.54 | 0.66 | 0.38 | 0.14 | 0.39 | 0.34 |
| 6 | 26098755 | rs1150658 | 0.264 | -0.972 | -0.417 | -1.925 | 0.109 | 0.07 | 0.07 | 0 | 0.09 | 0.26 | 0.14 | 0.17 | 0.2 | 0.07 |
| 6 | 26102708 | rs198856 | 1.682 | -0.995 | -0.441 | -1.945 | -0.006 | 0.006 | 0.006 | 0 | 0.36 | 0.36 | 0.42 | 0.3 | 0.53 | 0.45 |
| 6 | 26104057 | rs198854 | 1.691 | -0.995 | -0.441 | -1.945 | -0.006 | 0.006 | 0.006 | 0 | 0.36 | 0.36 | 0.42 | 0.31 | 0.53 | 0.45 |
| 6 | 26104096 | rs198853 | 2.566 | -0.996 | -0.479 | -1.943 | -0.071 | 0.082 | 0.082 | 0 | 0.28 | 0.1 | 0.28 | 0.14 | 0.33 | 0.38 |
| 6 | 26104280 | rs2229768 | 0.247 | -0.943 | -0.387 | -1.889 | 0.109 | 0.07 | 0.07 | 0 | 0.09 | 0.26 | 0.14 | 0.17 | 0.2 | 0.07 |
| 6 | 26104448 | rs198852 | 2.572 | -1.338 | -0.463 | -1.944 | -0.095 | 0.082 | 0.082 | 0 | 0.28 | 0.11 | 0.31 | 0.14 | 0.37 | 0.38 |
| 6 | 26106237 | rs707891 | 0.426 | -1.34 | -0.433 | -1.945 | 0.05 | 0.05 | 0.05 | 0 | 0.08 | 0.19 | 0.11 | 0.17 | 0.15 | 0.07 |
| 6 | 26107463 | rs198846 | -0.993 | -1.455 | -0.018 | -2.029 | -0.002 | -0.037 | -0.037 | 0 | 0.84 | 0.88 | 0.86 | 0.97 | 0.82 | 0.93 |
| 6 | 26107790 | rs198845 | 2.631 | -1.386 | -0.349 | -1.98 | -0.095 | 0.085 | 0.085 | 0 | 0.28 | 0.11 | 0.3 | 0.13 | 0.37 | 0.38 |
| 6 | 26108168 | rs2051542 | 1.269 | -1.442 | -0.352 | -1.996 | 0.002 | 0.019 | 0.019 | 0 | 0.11 | 0.07 | 0.21 | 0.55 | 0.09 | 0.2 |
| 6 | 26109233 | rs707892 | 0.242 | -1.443 | -0.347 | -1.996 | 0.071 | 0.07 | 0.07 | 0 | 0.09 | 0.26 | 0.12 | 0.17 | 0.16 | 0.07 |
| 6 | 26110961 | rs6927689 | 1.081 | -1.463 | -0.362 | -2.008 | 0.003 | 0.007 | 0.007 | 0 | 0.11 | 0.07 | 0.21 | 0.55 | 0.09 | 0.21 |
| 6 | 26111756 | rs16891315 | 1.044 | -1.582 | -0.272 | -2.077 | 0.002 | -0.031 | -0.031 | 0.001 | 0.11 | 0.19 | 0.08 | 0.01 | 0.1 | 0.03 |
| 6 | 26112893 | rs933199 | 1.247 | -1.566 | -0.294 | -2.053 | 0.003 | 0.016 | 0.016 | 0 | 0.11 | 0.07 | 0.21 | 0.55 | 0.09 | 0.21 |
| 6 | 26113340 | rs198838 | 1.782 | -1.495 | -0.306 | -2.033 | -0.006 | 0.006 | 0.006 | 0 | 0.36 | 0.36 | 0.42 | 0.31 | 0.53 | 0.45 |
| 6 | 26114372 | rs198834 | 2.704 | -1.498 | -0.305 | -2.035 | -0.137 | 0.075 | 0.075 | 0 | 0.28 | 0.06 | 0.3 | 0.14 | 0.37 | 0.38 |
| 6 | 26114702 | rs13161 | 1.775 | -1.494 | -0.296 | -2.03 | -0.005 | 0.005 | 0.005 | 0 | 0.36 | 0.36 | 0.42 | 0.31 | 0.53 | 0.45 |
| 6 | 26116424 | rs707896 | 0.255 | -1.473 | -0.275 | -2.012 | 0.075 | 0.074 | 0.074 | 0 | 0.08 | 0.26 | 0.12 | 0.17 | 0.16 | 0.07 |
| 6 | 26116996 | rs198831 | 1.592 | -1.479 | -0.313 | -2.024 | -0.007 | 0.012 | 0.012 | 0 | 0.38 | 0.36 | 0.43 | 0.31 | 0.53 | 0.43 |
| 6 | 26117220 | rs707898 | 0.188 | -1.455 | -0.294 | -1.996 | 0.027 | 0.027 | 0.027 | 0 | 0.1 | 0.19 | 0.11 | 0.17 | 0.15 | 0.06 |
| 6 | 26118667 | rs16891325 | 1.12 | -1.464 | -0.322 | -2.001 | -0.042 | -0.04 | -0.04 | 0.019 | 0.07 | 0.01 | 0.1 | 0.17 | 0.15 | 0.06 |
| 6 | 26118893 | rs198829 | 1.695 | -1.512 | -0.34 | -2.041 | -0.016 | 0.023 | 0.023 | 0 | 0.37 | 0.3 | 0.43 | 0.31 | 0.53 | 0.43 |
| 6 | 26119459 | rs198828 | 1.66 | -1.481 | -0.319 | -2.016 | -0.015 | 0.025 | 0.025 | 0 | 0.38 | 0.3 | 0.43 | 0.31 | 0.53 | 0.43 |
| 6 | 26121153 | rs198826 | 1.653 | -1.474 | -0.315 | -2.012 | -0.015 | 0.025 | 0.025 | 0 | 0.38 | 0.3 | 0.42 | 0.31 | 0.52 | 0.43 |
| 6 | 26123629 | rs198821 | 1.605 | -1.475 | -0.305 | -2.013 | -0.007 | 0.012 | 0.012 | 0 | 0.38 | 0.36 | 0.43 | 0.31 | 0.53 | 0.43 |
| 6 | 26124243 | rs198820 | 2.66 | -1.46 | -0.382 | -2.008 | -0.114 | 0.082 | 0.082 | 0 | 0.28 | 0.08 | 0.31 | 0.14 | 0.37 | 0.38 |
| 6 | 26124303 | rs16891334 | 0.864 | -1.514 | -0.347 | -2.015 | 0 | -0.033 | -0.033 | 0.429 | 0.11 | 0.19 | 0.08 | 0.01 | 0.09 | 0.03 |
| 6 | 26124430 | rs198819 | 1.492 | -1.331 | -0.504 | -1.896 | -0.015 | 0.025 | 0.025 | 0 | 0.38 | 0.3 | 0.43 | 0.31 | 0.53 | 0.43 |
| 6 | 26125342 | rs129128 | -1.108 | -1.343 | -0.545 | -1.914 | -0.019 | 0.059 | 0.059 | 0.29 | 0.86 | 0.99 | 0.89 | 0.97 | 0.84 | 0.91 |
| 6 | 26127184 | rs198816 | 1.448 | -1.246 | -0.457 | -1.844 | -0.015 | 0.025 | 0.025 | 0 | 0.38 | 0.3 | 0.42 | 0.31 | 0.52 | 0.43 |
| 6 | 26128446 | rs198811 | 0.832 | -1.297 | -0.43 | -1.865 | 0.023 | 0.038 | 0.038 | 0 | 0.49 | 0.44 | 0.63 | 0.86 | 0.62 | 0.64 |
| 6 | 26128618 | rs198810 | 2.474 | -1.261 | -0.475 | -1.871 | -0.095 | 0.082 | 0.082 | 0 | 0.28 | 0.11 | 0.3 | 0.14 | 0.37 | 0.37 |
| 6 | 26129375 | rs6908263 | 0.806 | -1.141 | -0.018 | -1.288 | -0.004 | 0.071 | 0.071 | 0 | 0.11 | 0.14 | 0.22 | 0.55 | 0.09 | 0.21 |
| 6 | 26129392 | rs17531978 | 0.816 | -0.978 | -0.078 | -1.586 | 0.009 | 0.033 | 0.033 | 0.818 | 0.15 | 0.02 | 0.12 | 0.07 | 0.12 | 0.2 |
| 6 | 26132664 | rs707901 | -0.225 | -0.682 | 0.203 | -1.35 | 0.03 | 0.029 | 0.029 | 0 | 0.09 | 0.18 | 0.11 | 0.16 | 0.15 | 0.06 |
| 6 | 26135498 | rs2856646 | 0.282 | -1.164 | 0.411 | -1.474 | -0.001 | -0.017 | -0.017 | 0 | 0.67 | 0.73 | 0.73 | 0.91 | 0.72 | 0.71 |
| 6 | 26147367 | rs1183694 | -0.15 | -0.692 | 0.571 | -0.871 | 0.006 | 0.002 | 0.002 | 6.132 | 0.08 | 0.37 | 0.07 | 0 | 0.01 | 0 |
| 6 | 26148311 | rs806972 | -0.121 | -0.69 | 0.606 | -0.869 | 0.01 | 0.006 | 0.006 | 5.864 | 0.08 | 0.37 | 0.07 | 0 | 0.01 | 0 |
| 6 | 26148326 | rs806973 | -1.403 | -0.631 | 0.083 | -1.257 | -0.003 | 0.012 | 0.012 | 4.392 | 0.35 | 0.65 | 0.35 | 0.22 | 0.35 | 0.16 |
| 6 | 26151535 | rs9358908 | 1.235 | -1.519 | -0.722 | -1.548 | 0.023 | -0.119 | -0.119 | 0 | 0.19 | 0.02 | 0.28 | 0.65 | 0.12 | 0.37 |
| 6 | 26153335 | rs9379827 | 0.328 | -1.322 | -0.606 | -1.571 | -0.04 | -0.017 | -0.017 | 0 | 0.15 | 0.06 | 0.15 | 0.17 | 0.22 | 0.12 |
| 6 | 26157762 | rs3857546 | 0.113 | -1.53 | -0.523 | -1.693 | -0.005 | -0.03 | -0.03 | 0 | 0.12 | 0.16 | 0.12 | 0.03 | 0.12 | 0.04 |
| 6 | 26158079 | rs7749823 | 0.711 | -1.529 | -0.514 | -1.693 | 0.002 | -0.026 | -0.026 | 0 | 0.1 | 0.12 | 0.11 | 0.03 | 0.11 | 0.04 |
| 6 | 26158993 | rs1009181 | -1.329 | -1.525 | -0.499 | -1.694 | 0.01 | -0.013 | -0.013 | 0.034 | 0.3 | 0.38 | 0.29 | 0.22 | 0.34 | 0.16 |
| 6 | 26159356 | rs2183947 | 0.067 | -1.637 | -0.351 | -1.557 | -0.036 | -0.018 | -0.018 | 0.234 | 0.16 | 0.09 | 0.15 | 0.19 | 0.22 | 0.12 |
| 6 | 26159495 | rs806966 | 0.706 | -1.616 | -0.327 | -1.54 | 0.013 | 0.009 | 0.009 | 5.349 | 0.08 | 0.34 | 0.07 | 0 | 0.02 | 0 |
| 6 | 26162679 | rs9467684 | -0.018 | -1.737 | -0.618 | -1.599 | -0.031 | -0.029 | -0.029 | 0 | 0.06 | 0.15 | 0.03 | 0.02 | 0.01 | 0.04 |
| 6 | 26167710 | rs4486004 | -0.139 | -1.738 | -0.629 | -1.606 | 0.003 | 0 | 0 | 0.918 | 0.17 | 0.2 | 0.16 | 0.19 | 0.22 | 0.12 |
| 6 | 26169632 | rs9461237 | 0.445 | -1.724 | -0.644 | -1.598 | -0.026 | -0.027 | -0.027 | 0 | 0.05 | 0.14 | 0.03 | 0.01 | 0.01 | 0.04 |
| 6 | 26172219 | rs9379829 | 0.067 | -1.59 | -0.679 | -1.51 | -0.039 | -0.019 | -0.019 | 0 | 0.15 | 0.06 | 0.15 | 0.17 | 0.21 | 0.12 |
| 6 | 26175852 | rs9379831 | -0.634 | -1.909 | -1.084 | -1.808 | -0.009 | 0.008 | 0.008 | 1.438 | 0.28 | 0.21 | 0.2 | 0.24 | 0.33 | 0.25 |
| 6 | 26183816 | rs17533076 | 0.509 | -1.869 | -0.9 | -2.02 | 0.01 | 0.008 | 0.008 | 1.406 | 0.1 | 0.05 | 0.05 | 0.06 | 0.04 | 0.07 |
| 6 | 26184041 | rs16891375 | 0.784 | -1.975 | -0.983 | -2.09 | 0.018 | 0.029 | 0.029 | 0 | 0.11 | 0.01 | 0.04 | 0.05 | 0.11 | 0.13 |
| 6 | 26184617 | rs4496782 | -1.563 | -2.308 | -0.81 | -2.314 | -0.061 | -0.057 | -0.057 | 1.248 | 0.39 | 0.51 | 0.23 | 0.16 | 0.24 | 0.26 |
| 6 | 26187125 | rs6906367 | -1.25 | -2.449 | -0.76 | -2.324 | -0.028 | -0.022 | -0.022 | 0.023 | 0.36 | 0.39 | 0.21 | 0.16 | 0.21 | 0.24 |
| 6 | 26187334 | rs11751812 | 0.858 | -2.273 | -0.697 | -2.253 | 0.018 | 0.029 | 0.029 | 0 | 0.11 | 0.01 | 0.04 | 0.05 | 0.11 | 0.13 |
| 6 | 26187783 | rs1543681 | -1.875 | -2.467 | -0.943 | -2.444 | -0.145 | 0.093 | 0.093 | 0.323 | 0.73 | 0.94 | 0.68 | 0.92 | 0.52 | 0.71 |
| 6 | 26188864 | rs16891378 | -0.54 | -2.443 | -0.876 | -2.432 | 0.015 | 0.015 | 0.015 | 2.894 | 0.24 | 0.2 | 0.11 | 0.11 | 0.16 | 0.21 |
| 6 | 26189615 | rs806977 | -1.743 | -2.474 | -0.866 | -2.455 | -0.117 | 0.105 | 0.105 | 0.459 | 0.72 | 0.9 | 0.68 | 0.92 | 0.52 | 0.71 |

| | | | | | | | | | | | | | | | | |
|---|---|---|---|---|---|---|---|---|---|---|---|---|---|---|---|---|
| 6 | 26190747 | rs11753837 | 0.387 | -2.469 | -0.832 | -2.416 | -0.005 | -0.005 | -0.005 | 0 | 0.14 | 0.15 | 0.06 | 0.06 | 0.12 | 0.14 |
| 6 | 26192502 | rs10452600 | -0.536 | -2.471 | -0.833 | -2.417 | 0.014 | 0.014 | 0.014 | 2.93 | 0.24 | 0.2 | 0.11 | 0.12 | 0.16 | 0.21 |
| 6 | 26194113 | rs806978 | 0.873 | -2.57 | -0.847 | -2.463 | 0.004 | 0 | 0 | 5.505 | 0.08 | 0.36 | 0.07 | 0 | 0.02 | 0.01 |
| 6 | 26195502 | rs1341759 | -0.509 | -2.571 | -0.859 | -2.465 | 0.014 | 0.014 | 0.014 | 2.93 | 0.24 | 0.2 | 0.11 | 0.12 | 0.16 | 0.21 |
| 6 | 26198046 | rs4145878 | -0.83 | -2.173 | -0.888 | -2.501 | 0.002 | -0.028 | -0.028 | 1.274 | 0.61 | 0.42 | 0.6 | 0.92 | 0.54 | 0.68 |
| 6 | 26198916 | rs806792 | 0.775 | -2.437 | -0.719 | -2.271 | 0.004 | 0 | 0 | 5.505 | 0.08 | 0.36 | 0.07 | 0 | 0.02 | 0 |
| 6 | 26199014 | rs34550794 | 0.625 | -2.082 | -1.098 | -2.087 | 0.01 | 0.008 | 0.008 | 1.419 | 0.1 | 0.05 | 0.05 | 0.06 | 0.04 | 0.07 |
| 6 | 26200707 | rs3757135 | -0.644 | -2.101 | -1.08 | -2.063 | 0.014 | 0.014 | 0.014 | 2.93 | 0.24 | 0.2 | 0.11 | 0.12 | 0.16 | 0.21 |
| 6 | 26202337 | rs16891401 | -0.68 | -2.103 | -1.074 | -2.064 | 0.008 | 0.008 | 0.008 | 2.736 | 0.25 | 0.23 | 0.11 | 0.12 | 0.16 | 0.21 |
| 6 | 26203910 | rs11758351 | -0.714 | -2.107 | -0.947 | -2.064 | -0.005 | -0.004 | -0.004 | 0 | 0.25 | 0.27 | 0.12 | 0.13 | 0.16 | 0.21 |
| 6 | 26205293 | rs806795 | -1.603 | -2.067 | -1.141 | -2.195 | -0.126 | 0.096 | 0.096 | 0.453 | 0.73 | 0.9 | 0.67 | 0.92 | 0.53 | 0.74 |
| 6 | 26205718 | rs2069018 | -0.648 | -1.811 | -0.969 | -1.958 | 0.005 | 0.005 | 0.005 | 2.334 | 0.25 | 0.23 | 0.12 | 0.12 | 0.17 | 0.22 |
| 6 | 26206074 | rs989710 | 0.373 | -1.806 | -0.974 | -2.201 | 0.002 | 0.002 | 0.002 | 0.591 | 0.11 | 0.08 | 0.05 | 0.06 | 0.05 | 0.08 |
| 6 | 26207928 | rs2022448 | -0.696 | -1.836 | -0.976 | -2.22 | 0.005 | 0.005 | 0.005 | 2.334 | 0.25 | 0.23 | 0.12 | 0.12 | 0.17 | 0.22 |
| 6 | 26211368 | rs11755943 | -0.743 | -1.871 | -0.837 | -2.276 | -0.003 | -0.005 | -0.005 | 0 | 0.25 | 0.26 | 0.12 | 0.12 | 0.16 | 0.21 |
| 6 | 26215802 | rs11755618 | 0.225 | -1.889 | -0.845 | -2.287 | 0 | -0.004 | -0.004 | 0.219 | 0.14 | 0.15 | 0.06 | 0.04 | 0.11 | 0.13 |
| 6 | 26219384 | rs11753131 | -0.855 | -1.899 | -0.704 | -2.292 | 0.006 | 0.005 | 0.005 | 2.54 | 0.25 | 0.22 | 0.11 | 0.1 | 0.16 | 0.21 |
| 6 | 26221173 | rs11754368 | 0.225 | -1.898 | -0.704 | -2.291 | 0 | -0.004 | -0.004 | 0.071 | 0.14 | 0.15 | 0.05 | 0.04 | 0.11 | 0.13 |
| 6 | 26227104 | rs3818947 | 0.383 | -1.892 | -0.813 | -2.288 | 0.002 | 0.002 | 0.002 | 0.591 | 0.11 | 0.08 | 0.05 | 0.06 | 0.05 | 0.08 |
| 6 | 26231443 | rs11759904 | -0.743 | -1.897 | -0.814 | -2.291 | 0.005 | 0.003 | 0.003 | 2.472 | 0.25 | 0.23 | 0.11 | 0.1 | 0.16 | 0.21 |
| 6 | 26231608 | rs11757549 | 0.225 | -1.895 | -0.813 | -2.29 | -0.003 | -0.007 | -0.007 | 1.204 | 0.14 | 0.15 | 0.06 | 0.04 | 0.11 | 0.12 |
| 6 | 26233387 | rs10946808 | 1.337 | -2.153 | -1.28 | -2.506 | -0.086 | -0.047 | -0.047 | 0.215 | 0.25 | 0.02 | 0.38 | 0.77 | 0.3 | 0.48 |
| 6 | 26234476 | rs16891458 | 0.462 | -2.08 | -1.197 | -2.434 | 0.002 | 0.002 | 0.002 | 0.591 | 0.11 | 0.07 | 0.05 | 0.06 | 0.05 | 0.08 |
| 6 | 26235300 | rs16891464 | 0.281 | -2.063 | -1.191 | -2.424 | -0.002 | -0.007 | -0.007 | 0 | 0.14 | 0.15 | 0.06 | 0.04 | 0.11 | 0.13 |
| 6 | 26237155 | rs10484436 | -0.753 | -2.138 | -1.228 | -2.464 | 0.008 | 0.008 | 0.008 | 2.792 | 0.26 | 0.23 | 0.11 | 0.1 | 0.16 | 0.21 |
| 6 | 26237457 | rs10484437 | 0.229 | -1.951 | -1.175 | -2.339 | 0.004 | 0.004 | 0.004 | 0.954 | 0.12 | 0.07 | 0.05 | 0.06 | 0.05 | 0.08 |
| 6 | 26239404 | rs9358913 | 1.298 | -1.993 | -1.277 | -2.377 | -0.085 | -0.053 | -0.053 | 0 | 0.25 | 0.02 | 0.37 | 0.76 | 0.29 | 0.47 |
| 6 | 26240624 | rs3734533 | -1.101 | -1.776 | -0.862 | -2.128 | 0.005 | 0.004 | 0.004 | 1.198 | 0.28 | 0.27 | 0.15 | 0.12 | 0.19 | 0.2 |
| 6 | 26240649 | rs3734534 | 0.176 | -1.722 | -0.953 | -2.118 | 0.002 | -0.006 | -0.006 | 0 | 0.16 | 0.16 | 0.11 | 0.04 | 0.15 | 0.13 |
| 6 | 26244580 | rs1000308 | -0.006 | -1.705 | -0.807 | -2.074 | -0.011 | -0.007 | -0.007 | 0 | 0.12 | 0.11 | 0.05 | 0.07 | 0.05 | 0.06 |
| 6 | 26245888 | rs16891501 | -0.006 | -1.597 | -1 | -2.019 | -0.011 | -0.007 | -0.007 | 0 | 0.12 | 0.11 | 0.05 | 0.07 | 0.04 | 0.07 |
| 6 | 26246664 | rs6913828 | -0.299 | -1.6 | -1.083 | -2.021 | -0.027 | -0.012 | -0.012 | 0.087 | 0.14 | 0.28 | 0.07 | 0.07 | 0.05 | 0.06 |
| 6 | 26251601 | rs11753610 | 0.054 | -1.133 | -1.548 | -1.839 | 0.035 | 0.067 | 0.067 | 0 | 0.14 | 0.02 | 0.09 | 0.03 | 0.15 | 0.13 |
| 6 | 26255451 | rs6935954 | -1.093 | -0.39 | -1.501 | -1.738 | -0.166 | 0.075 | 0.075 | 0.055 | 0.77 | 0.96 | 0.72 | 0.92 | 0.61 | 0.75 |
| 6 | 26262306 | rs7761779 | 0.186 | -1.059 | -1.19 | -1.806 | 0.018 | 0.019 | 0.019 | 0 | 0.18 | 0.09 | 0.12 | 0.04 | 0.16 | 0.08 |
| 6 | 26267755 | rs7773004 | -0.012 | -0.685 | -1.534 | -2.022 | -0.071 | 0.139 | 0.139 | 0 | 0.6 | 0.88 | 0.64 | 0.84 | 0.49 | 0.68 |
| 6 | 26267800 | rs7773163 | -0.088 | -1.009 | -1.166 | -2.158 | -0.006 | -0.019 | -0.019 | 0 | 0.22 | 0.17 | 0.13 | 0.06 | 0.17 | 0.12 |
| 6 | 26270961 | rs11759682 | 0.451 | -1.006 | -1.169 | -2.156 | 0.016 | 0.018 | 0.018 | 0 | 0.18 | 0.1 | 0.12 | 0.06 | 0.16 | 0.08 |
| 6 | 26271713 | rs9393690 | -0.489 | -1.43 | -0.91 | -2.537 | 0.022 | -0.046 | -0.046 | 0 | 0.65 | 0.35 | 0.64 | 0.87 | 0.55 | 0.74 |
| 6 | 26272039 | rs3823157 | 0.14 | -1.424 | -0.905 | -2.516 | -0.006 | -0.019 | -0.019 | 0 | 0.22 | 0.17 | 0.13 | 0.06 | 0.17 | 0.12 |
| 6 | 26274498 | rs6901243 | 0.14 | -1.424 | -0.905 | -2.516 | -0.006 | -0.019 | -0.019 | 0 | 0.22 | 0.17 | 0.13 | 0.06 | 0.17 | 0.12 |
| 6 | 26276650 | rs9393692 | -1.306 | -1.461 | -0.553 | -2.562 | -0.07 | 0.046 | 0.046 | 0.025 | 0.76 | 0.8 | 0.72 | 0.89 | 0.58 | 0.76 |
| 6 | 26282322 | rs6940237 | -0.447 | -1.493 | -0.549 | -2.578 | 0.022 | -0.046 | -0.046 | 0.072 | 0.65 | 0.34 | 0.64 | 0.87 | 0.55 | 0.74 |
| 6 | 26285683 | rs2393593 | -1.581 | -1.498 | -0.541 | -2.58 | -0.157 | 0.053 | 0.053 | 0.055 | 0.78 | 0.93 | 0.73 | 0.89 | 0.58 | 0.76 |
| 6 | 26285744 | rs16891517 | 0.377 | -1.896 | -0.727 | -2.739 | -0.025 | 0 | 0 | 0 | 0.08 | 0.11 | 0.04 | 0.06 | 0.04 | 0.08 |
| 6 | 26285867 | rs3999544 | -1.751 | -1.896 | -0.726 | -2.739 | -0.157 | 0.053 | 0.053 | 0.055 | 0.78 | 0.93 | 0.73 | 0.89 | 0.58 | 0.76 |
| 6 | 26285908 | rs16891518 | 0.426 | -1.884 | -0.735 | -2.731 | -0.024 | 0.001 | 0.001 | 0 | 0.08 | 0.11 | 0.04 | 0.06 | 0.04 | 0.08 |
| 6 | 26286744 | rs9358918 | 0.392 | -1.882 | -0.738 | -2.733 | 0.013 | 0.128 | 0.128 | 0.16 | 0.45 | 0.17 | 0.51 | 0.81 | 0.38 | 0.62 |
| 6 | 26287043 | rs1124999 | -0.768 | -1.904 | -0.673 | -2.744 | 0.041 | -0.058 | -0.058 | 0 | 0.67 | 0.39 | 0.64 | 0.87 | 0.55 | 0.74 |
| 6 | 26289435 | rs9379843 | -0.576 | -1.898 | -0.673 | -2.741 | 0.022 | -0.046 | -0.046 | 0.058 | 0.65 | 0.34 | 0.64 | 0.87 | 0.55 | 0.74 |
| 6 | 26291601 | rs9379845 | 1.743 | -2.301 | -0.892 | -2.924 | -0.127 | -0.033 | -0.033 | 0 | 0.28 | 0.02 | 0.4 | 0.76 | 0.32 | 0.49 |
| 6 | 26293338 | rs7746807 | -0.592 | -2.186 | -0.506 | -2.872 | 0.037 | -0.067 | -0.067 | 0.247 | 0.65 | 0.43 | 0.65 | 0.87 | 0.56 | 0.74 |
| 6 | 26293841 | rs9393697 | 0.46 | -2.181 | -0.539 | -2.866 | 0.005 | 0.116 | 0.116 | 0.728 | 0.43 | 0.17 | 0.51 | 0.82 | 0.38 | 0.62 |
| 6 | 26295686 | rs2893820 | 1.037 | -2.184 | -0.55 | -2.867 | -0.061 | 0.055 | 0.055 | 0.384 | 0.36 | 0.07 | 0.47 | 0.77 | 0.34 | 0.55 |
| 6 | 26300438 | rs9467700 | 0.906 | -2.194 | -0.491 | -2.872 | -0.015 | -0.004 | -0.004 | 0.071 | 0.08 | 0.33 | 0.06 | 0.02 | 0.02 | 0.02 |
| 6 | 26303319 | rs9357004 | -0.329 | -2.257 | -0.518 | -2.91 | 0.022 | -0.064 | -0.064 | 0 | 0.61 | 0.44 | 0.59 | 0.86 | 0.54 | 0.69 |
| 6 | 26305407 | rs994379 | -0.013 | -2.042 | -0.522 | -2.836 | 0.003 | 0.101 | 0.101 | 0.047 | 0.5 | 0.72 | 0.59 | 0.8 | 0.43 | 0.57 |
| 6 | 26306282 | rs9366648 | 1.051 | -2.017 | -0.522 | -2.871 | -0.063 | 0.052 | 0.052 | 0.549 | 0.36 | 0.07 | 0.47 | 0.77 | 0.34 | 0.55 |
| 6 | 26306684 | rs1883214 | 0.022 | -2.02 | -0.509 | -2.863 | -0.018 | -0.023 | -0.023 | 0 | 0.25 | 0.2 | 0.16 | 0.1 | 0.2 | 0.15 |
| 6 | 26308287 | rs10484438 | 0.022 | -2.024 | -0.474 | -2.868 | -0.018 | -0.023 | -0.023 | 0 | 0.25 | 0.2 | 0.16 | 0.1 | 0.2 | 0.15 |
| 6 | 26313348 | rs6923139 | -0.415 | -1.977 | -0.62 | -2.835 | -0.007 | -0.066 | -0.066 | 0 | 0.14 | 0.65 | 0.12 | 0.02 | 0.09 | 0.02 |
| 6 | 26316982 | rs11966360 | -0.889 | -1.94 | -0.636 | -2.834 | 0 | -0.003 | -0.003 | 0.646 | 0.21 | 0.76 | 0.16 | 0.06 | 0.13 | 0.08 |
| 6 | 26325463 | rs3778279 | -0.08 | -1.62 | -0.74 | -2.821 | 0.003 | 0.092 | 0.092 | 0.047 | 0.5 | 0.71 | 0.59 | 0.8 | 0.43 | 0.57 |
| 6 | 26325888 | rs2179152 | -1.306 | -1.231 | -1.003 | -2.61 | -0.116 | 0.072 | 0.072 | 0.236 | 0.75 | 0.92 | 0.75 | 0.89 | 0.64 | 0.72 |
| 6 | 26327953 | rs6939048 | -1.306 | -1.231 | -1.003 | -2.61 | -0.116 | 0.072 | 0.072 | 0.236 | 0.75 | 0.92 | 0.74 | 0.89 | 0.63 | 0.72 |
| 6 | 26330492 | rs9358927 | -0.012 | -1.228 | -0.489 | -2.255 | 0.006 | 0.114 | 0.114 | 1.018 | 0.43 | 0.18 | 0.51 | 0.82 | 0.38 | 0.61 |
| 6 | 26331493 | rs11962660 | 0.55 | -1.258 | -0.292 | -2.259 | -0.015 | -0.004 | -0.004 | 0 | 0.08 | 0.33 | 0.06 | 0.02 | 0.02 | 0.02 |
| 6 | 26337264 | rs7755741 | 0.701 | -0.634 | -1.213 | -1.929 | -0.117 | -0.018 | -0.018 | 0 | 0.3 | 0.03 | 0.4 | 0.76 | 0.32 | 0.49 |
| 6 | 26338056 | rs4412193 | -0.258 | -0.612 | -1.195 | -1.92 | 0.005 | 0.113 | 0.113 | 0.946 | 0.43 | 0.17 | 0.51 | 0.82 | 0.39 | 0.61 |

| | | | | | | | | | | | | | | | | |
|---|---|---|---|---|---|---|---|---|---|---|---|---|---|---|---|---|
| 6 | 26338434 | rs2893842 | 0.498 | 0.257 | -1.433 | -1.458 | -0.116 | -0.016 | -0.016 | 0 | 0.3 | 0.03 | 0.4 | 0.76 | 0.33 | 0.51 |
| 6 | 26341030 | rs9358928 | -0.314 | 0.404 | -0.624 | -1.357 | 0.003 | -0.03 | -0.03 | 0 | 0.6 | 0.6 | 0.47 | 0.21 | 0.58 | 0.39 |
| 6 | 26345141 | rs10946813 | -1.081 | 0.318 | -0.548 | -1.107 | 0.015 | -0.017 | -0.017 | 0 | 0.32 | 0.4 | 0.3 | 0.22 | 0.41 | 0.35 |
| 6 | 26351597 | rs6456716 | -0.062 | 0.465 | -0.477 | -1.138 | 0.002 | 0.004 | 0.004 | 0 | 0.14 | 0.13 | 0.11 | 0.02 | 0.12 | 0.15 |
| 6 | 26351937 | rs6456718 | -0.224 | 0.445 | -0.361 | -1.132 | -0.006 | -0.034 | -0.034 | 0 | 0.15 | 0.22 | 0.13 | 0.02 | 0.12 | 0.15 |
| 6 | 26354247 | rs9366653 | -0.161 | 0.403 | -0.497 | -1.208 | -0.03 | 0 | 0 | 0 | 0.08 | 0.02 | 0.04 | 0.1 | 0.11 | 0.13 |
| 6 | 26354572 | rs9461245 | 0.227 | 0.384 | -0.499 | -1.224 | 0.001 | 0.003 | 0.003 | 0 | 0.13 | 0.11 | 0.11 | 0.02 | 0.11 | 0.15 |
| 6 | 26355283 | rs6933583 | -2.12 | 0.321 | -0.234 | -1.298 | 0.001 | -0.05 | -0.05 | 1.781 | 0.3 | 0.39 | 0.21 | 0.13 | 0.28 | 0.32 |
| 6 | 26355758 | rs4712980 | -2.12 | 0.321 | -0.234 | -1.298 | 0.001 | -0.05 | -0.05 | 0 | 0.3 | 0.39 | 0.21 | 0.13 | 0.28 | 0.32 |
| 6 | 26361430 | rs4712981 | -2.076 | 0.321 | -0.228 | -1.298 | 0 | -0.031 | -0.031 | 0 | 0.29 | 0.34 | 0.21 | 0.13 | 0.28 | 0.32 |
| 6 | 26365147 | rs9393709 | -2.286 | 0.365 | -0.22 | -1.288 | 0.021 | -0.025 | -0.025 | 0 | 0.32 | 0.43 | 0.3 | 0.22 | 0.4 | 0.33 |
| 6 | 26365595 | rs12203176 | 0.005 | 0.364 | -0.22 | -1.288 | 0.003 | 0.007 | 0.007 | 0 | 0.14 | 0.12 | 0.11 | 0.02 | 0.12 | 0.15 |
| 6 | 26366836 | rs9379856 | 0.095 | 0.364 | -0.236 | -1.289 | -0.023 | 0.001 | 0.001 | 0 | 0.08 | 0.03 | 0.04 | 0.1 | 0.11 | 0.13 |
| 6 | 26367218 | rs12199613 | -2.288 | 0.358 | -0.238 | -1.292 | 0.02 | -0.025 | -0.025 | 0.798 | 0.32 | 0.43 | 0.3 | 0.22 | 0.4 | 0.33 |
| 6 | 26367471 | rs9688949 | -0.144 | 0.342 | -0.25 | -1.294 | -0.004 | -0.026 | -0.026 | 0 | 0.15 | 0.21 | 0.13 | 0.02 | 0.12 | 0.15 |
| 6 | 26369549 | rs9379859 | -0.033 | 0.352 | -0.305 | -1.286 | -0.024 | 0.002 | 0.002 | 0 | 0.08 | 0.03 | 0.04 | 0.1 | 0.11 | 0.13 |
| 6 | 26372349 | rs7765566 | 1.12 | 0.397 | -0.294 | -1.288 | -0.006 | 0.012 | 0.012 | 0.853 | 0.05 | 0.01 | 0.01 | 0.01 | 0.04 | 0.05 |
| 6 | 26372786 | rs12176317 | -0.041 | 0.41 | -0.291 | -1.262 | -0.024 | 0.002 | 0.002 | 0 | 0.08 | 0.03 | 0.04 | 0.1 | 0.11 | 0.13 |
| 6 | 26373678 | rs9393713 | -0.041 | 0.41 | -0.291 | -1.262 | -0.024 | 0.002 | 0.002 | 0 | 0.08 | 0.03 | 0.04 | 0.1 | 0.11 | 0.13 |
| 6 | 26373740 | rs9393714 | -0.041 | 0.41 | -0.291 | -1.262 | -0.024 | 0.002 | 0.002 | 0 | 0.08 | 0.03 | 0.04 | 0.1 | 0.11 | 0.13 |
| 6 | 26374410 | rs9379870 | -1.956 | 0.387 | -0.297 | -1.273 | 0.001 | -0.05 | -0.05 | 1.628 | 0.3 | 0.39 | 0.21 | 0.13 | 0.28 | 0.32 |
| 6 | 26374549 | rs11758089 | 0.544 | -0.182 | -0.67 | -1.831 | 0.012 | 0.023 | 0.023 | 0.042 | 0.19 | 0.08 | 0.1 | 0.07 | 0.18 | 0.06 |
| 6 | 26374658 | rs2073526 | -2.191 | -0.604 | -0.946 | -1.669 | -0.114 | -0.021 | -0.021 | 1.138 | 0.61 | 0.95 | 0.47 | 0.33 | 0.59 | 0.42 |
| 6 | 26375159 | rs2073529 | 0.228 | -0.046 | -0.609 | -1.335 | -0.024 | -0.001 | -0.001 | 0 | 0.08 | 0.03 | 0.04 | 0.1 | 0.11 | 0.13 |
| 6 | 26376161 | rs1985732 | -1.62 | -0.059 | -0.572 | -1.391 | 0.002 | -0.031 | -0.031 | 0 | 0.3 | 0.33 | 0.22 | 0.13 | 0.29 | 0.33 |
| 6 | 26376628 | rs12214031 | -1.786 | -0.122 | -0.665 | -1.331 | 0.023 | -0.019 | -0.019 | 0.883 | 0.32 | 0.41 | 0.31 | 0.23 | 0.41 | 0.34 |
| 6 | 26377591 | rs1979 | 0.133 | 0.093 | -1.027 | -1.068 | -0.024 | -0.022 | -0.022 | 0 | 0.08 | 0.03 | 0.04 | 0.14 | 0.11 | 0.14 |
| 6 | 26378681 | rs1624064 | -1.491 | -0.41 | -1.202 | -1.254 | 0.019 | 0.161 | 0.161 | 0.74 | 0.51 | 0.85 | 0.45 | 0.22 | 0.47 | 0.29 |
| 6 | 26380545 | rs3757142 | -0.906 | -0.397 | -0.884 | -1.317 | -0.001 | -0.028 | -0.028 | 0.628 | 0.21 | 0.66 | 0.23 | 0.11 | 0.2 | 0.09 |
| 6 | 26382765 | rs12189870 | -0.906 | -0.397 | -0.884 | -1.317 | -0.001 | -0.028 | -0.028 | 0.541 | 0.21 | 0.66 | 0.23 | 0.11 | 0.2 | 0.09 |
| 6 | 26383386 | rs9461249 | 0.406 | -0.413 | -0.594 | -1.35 | 0.067 | 0.067 | 0.067 | 2.041 | 0.11 | 0.55 | 0.14 | 0.1 | 0.12 | 0.03 |
| 6 | 26383459 | rs9467741 | 0.242 | -0.415 | -0.608 | -1.351 | 0.057 | 0.056 | 0.056 | 2.369 | 0.11 | 0.57 | 0.14 | 0.1 | 0.12 | 0.03 |
| 6 | 26386762 | rs9467745 | -1.058 | -0.443 | -0.705 | -1.344 | -0.002 | -0.028 | -0.028 | 0 | 0.21 | 0.66 | 0.23 | 0.11 | 0.2 | 0.09 |
| 6 | 26387546 | rs9358938 | 0.759 | -0.396 | -0.798 | -1.271 | -0.023 | -0.022 | -0.022 | 0 | 0.06 | 0.03 | 0.04 | 0.12 | 0.12 | 0.14 |
| 6 | 26388672 | rs1796518 | -1.032 | -0.718 | -0.981 | -1.467 | -0.001 | 0.035 | 0.035 | 0 | 0.34 | 0.22 | 0.27 | 0.22 | 0.38 | 0.34 |
| 6 | 26389926 | rs7756881 | -1.23 | -0.596 | -0.861 | -1.368 | -0.006 | -0.03 | -0.03 | 0.708 | 0.22 | 0.68 | 0.23 | 0.11 | 0.2 | 0.09 |
| 6 | 26390039 | rs7757048 | -1.23 | -0.596 | -0.861 | -1.368 | -0.006 | -0.03 | -0.03 | 0.623 | 0.22 | 0.68 | 0.23 | 0.11 | 0.2 | 0.09 |
| 6 | 26390830 | rs2076029 | -1.844 | -0.629 | -0.941 | -1.416 | -0.011 | 0.059 | 0.059 | 0.356 | 0.42 | 0.81 | 0.4 | 0.26 | 0.44 | 0.38 |
| 6 | 26396727 | rs7356988 | -1.868 | -0.765 | -1.048 | -1.407 | -0.012 | 0.055 | 0.055 | 0.306 | 0.43 | 0.84 | 0.49 | 0.27 | 0.44 | 0.38 |
| 6 | 26396739 | rs7356849 | -0.669 | -0.79 | -0.699 | -1.357 | -0.049 | -0.07 | -0.07 | 0 | 0.25 | 0.65 | 0.22 | 0.07 | 0.14 | 0.18 |
| 6 | 26398914 | rs9467750 | -0.669 | -0.79 | -0.699 | -1.357 | -0.049 | -0.07 | -0.07 | 0 | 0.25 | 0.65 | 0.22 | 0.07 | 0.14 | 0.18 |
| 6 | 26401252 | rs1741738 | 0.583 | -0.6 | -1.023 | -1.303 | -0.004 | 0.026 | 0.026 | 0.644 | 0.14 | 0.01 | 0.1 | 0.07 | 0.16 | 0.06 |
| 6 | 26403703 | rs3902051 | 0.101 | -0.547 | -0.872 | -1.287 | -0.001 | -0.009 | -0.009 | 0 | 0.15 | 0.13 | 0.14 | 0.03 | 0.12 | 0.15 |
| 6 | 26404680 | rs1796519 | 0.767 | -0.597 | -0.877 | -1.311 | 0.011 | 0.024 | 0.024 | 0.89 | 0.16 | 0.07 | 0.1 | 0.07 | 0.14 | 0.05 |
| 6 | 26404958 | rs10456045 | -1.191 | -0.218 | -0.732 | -1.224 | 0.013 | 0.002 | 0.002 | 0 | 0.14 | 0.13 | 0.18 | 0.19 | 0.32 | 0.21 |
| 6 | 26405264 | rs10807008 | 0.292 | -0.367 | -0.723 | -1.264 | 0.002 | 0.007 | 0.007 | 0 | 0.14 | 0.11 | 0.13 | 0.03 | 0.12 | 0.15 |
| 6 | 26406053 | rs3857549 | -0.378 | -0.518 | -0.301 | -1.359 | -0.003 | -0.073 | -0.073 | 0 | 0.12 | 0.52 | 0.11 | 0.02 | 0.07 | 0.05 |
| 6 | 26407408 | rs4609015 | -0.536 | -0.474 | -0.608 | -1.357 | 0 | -0.014 | -0.014 | 1.069 | 0.21 | 0.16 | 0.18 | 0.03 | 0.17 | 0.2 |
| 6 | 26410227 | rs7770214 | -0.523 | -0.47 | -0.606 | -1.355 | 0 | -0.014 | -0.014 | 0.533 | 0.21 | 0.16 | 0.18 | 0.03 | 0.17 | 0.2 |
| 6 | 26410800 | rs1796520 | -2.218 | -0.623 | -0.775 | -1.462 | -0.023 | 0.051 | 0.051 | 0.703 | 0.54 | 0.91 | 0.48 | 0.34 | 0.56 | 0.44 |
| 6 | 26413088 | rs6900725 | -0.385 | -0.641 | -0.768 | -1.473 | 0 | -0.014 | -0.014 | 0 | 0.21 | 0.16 | 0.18 | 0.03 | 0.17 | 0.2 |
| 6 | 26413744 | rs4712990 | 0.463 | -0.637 | -0.74 | -1.472 | 0.001 | 0.006 | 0.006 | 0 | 0.14 | 0.11 | 0.15 | 0.03 | 0.12 | 0.14 |
| 6 | 26416399 | rs4712993 | -0.394 | -0.647 | -0.745 | -1.478 | 0 | -0.015 | -0.015 | 0 | 0.21 | 0.16 | 0.18 | 0.03 | 0.17 | 0.19 |
| 6 | 26421030 | rs7744254 | -0.435 | -0.651 | -0.747 | -1.479 | 0 | -0.014 | -0.014 | 0 | 0.21 | 0.16 | 0.18 | 0.03 | 0.17 | 0.19 |
| 6 | 26421207 | rs6456721 | -0.394 | -0.649 | -0.746 | -1.478 | 0 | -0.015 | -0.015 | 0 | 0.21 | 0.16 | 0.18 | 0.03 | 0.17 | 0.19 |
| 6 | 26421392 | rs1796521 | -2.283 | -0.699 | -0.755 | -1.461 | -0.074 | 0.01 | 0.01 | 0.719 | 0.54 | 0.94 | 0.49 | 0.34 | 0.56 | 0.43 |
| 6 | 26423090 | rs12191917 | -0.393 | -0.651 | -0.722 | -1.437 | 0 | -0.015 | -0.015 | 0.375 | 0.21 | 0.16 | 0.18 | 0.03 | 0.17 | 0.19 |
| 6 | 26423560 | rs4320356 | -2.271 | -0.65 | -0.719 | -1.451 | -0.026 | 0.044 | 0.044 | 0.592 | 0.4 | 0.84 | 0.42 | 0.27 | 0.45 | 0.38 |
| 6 | 26423856 | rs2393652 | -0.384 | -0.644 | -0.722 | -1.448 | 0 | -0.015 | -0.015 | 0 | 0.21 | 0.16 | 0.18 | 0.03 | 0.17 | 0.19 |
| 6 | 26423867 | rs2893845 | -0.384 | -0.644 | -0.722 | -1.448 | 0 | -0.015 | -0.015 | 0 | 0.21 | 0.16 | 0.18 | 0.03 | 0.17 | 0.19 |
| 6 | 26424729 | rs7751280 | -0.384 | -0.644 | -0.722 | -1.448 | 0 | -0.015 | -0.015 | 0 | 0.21 | 0.16 | 0.18 | 0.03 | 0.17 | 0.19 |
| 6 | 26426487 | rs10946829 | -0.384 | -0.644 | -0.722 | -1.448 | 0 | -0.015 | -0.015 | 0 | 0.21 | 0.16 | 0.18 | 0.03 | 0.17 | 0.19 |
| 6 | 26427063 | rs3846843 | -0.384 | -0.566 | -0.726 | -1.434 | 0 | -0.015 | -0.015 | 0 | 0.21 | 0.17 | 0.18 | 0.03 | 0.17 | 0.19 |
| 6 | 26428842 | rs1741740 | -2.156 | -0.132 | -1.218 | -1.177 | 0.01 | -0.035 | -0.035 | 0 | 0.77 | 0.98 | 0.76 | 0.75 | 0.82 | 0.78 |
| 6 | 26430848 | rs2893847 | -0.52 | 0.578 | -0.101 | -1.538 | 0.002 | 0.007 | 0.007 | 0 | 0.2 | 0.14 | 0.18 | 0.03 | 0.17 | 0.19 |
| 6 | 26430981 | rs2893848 | -0.52 | 0.578 | -0.101 | -1.538 | 0.002 | 0.007 | 0.007 | 0 | 0.2 | 0.14 | 0.18 | 0.03 | 0.17 | 0.19 |
| 6 | 26433329 | rs1624440 | -2.395 | 0.586 | -0.111 | -1.533 | -0.027 | 0.045 | 0.045 | 0.656 | 0.53 | 0.91 | 0.48 | 0.33 | 0.56 | 0.43 |
| 6 | 26440542 | rs17539219 | -0.523 | 0.443 | -0.097 | -1.606 | 0 | -0.014 | -0.014 | 0 | 0.21 | 0.16 | 0.18 | 0.03 | 0.17 | 0.19 |
| 6 | 26440652 | rs12214444 | 0.305 | 0.444 | -0.098 | -1.602 | 0.001 | 0.007 | 0.007 | 0 | 0.14 | 0.11 | 0.15 | 0.03 | 0.12 | 0.14 |

| | | | | | | | | | | | | | | | | |
|---|---|---|---|---|---|---|---|---|---|---|---|---|---|---|---|---|
| 6 | 26441548 | rs4712996 | -0.53 | 0.445 | -0.096 | -1.606 | 0 | -0.014 | -0.014 | 0 | 0.21 | 0.16 | 0.18 | 0.03 | 0.17 | 0.19 |
| 6 | 26442786 | rs7768108 | 0.267 | 0.441 | -0.095 | -1.609 | 0.002 | 0.008 | 0.008 | 0 | 0.14 | 0.11 | 0.14 | 0.03 | 0.12 | 0.14 |
| 6 | 26446209 | rs3846848 | -0.3 | 0.469 | -0.064 | -1.561 | 0 | -0.013 | -0.013 | 0.948 | 0.21 | 0.16 | 0.18 | 0.03 | 0.17 | 0.19 |
| 6 | 26446705 | rs2281027 | -0.323 | 0.299 | -0.042 | -1.497 | -0.06 | -0.02 | -0.02 | 0 | 0.12 | 0.59 | 0.09 | 0.04 | 0.02 | 0.03 |
| 6 | 26449219 | rs1796524 | 0.616 | 0.552 | -0.503 | -1.393 | 0.007 | 0.017 | 0.017 | 1.129 | 0.13 | 0.06 | 0.06 | 0.06 | 0.11 | 0.05 |
| 6 | 26451553 | rs2237236 | -1.492 | 0.754 | -0.506 | -1.215 | -0.025 | 0.026 | 0.026 | 0.299 | 0.4 | 0.84 | 0.42 | 0.29 | 0.45 | 0.38 |
| 6 | 26453742 | rs10484440 | -0.151 | 1.045 | 0.155 | -0.945 | 0.061 | 0.033 | 0.033 | 0 | 0.05 | 0.12 | 0.12 | 0.08 | 0.15 | 0.03 |
| 6 | 26454454 | rs2893849 | 0.366 | 0.836 | -0.074 | -1.13 | -0.006 | 0.041 | 0.041 | 0 | 0.85 | 0.96 | 0.85 | 0.95 | 0.85 | 0.82 |
| 6 | 26455811 | rs9295689 | 2.155 | 0.618 | -0.263 | -1.413 | -0.02 | 0.089 | 0.089 | 0.575 | 0.49 | 0.09 | 0.53 | 0.7 | 0.45 | 0.61 |
| 6 | 26456069 | rs9295690 | 1.595 | 0.471 | 0.439 | -1.406 | -0.033 | -0.038 | -0.038 | 0.871 | 0.73 | 0.67 | 0.77 | 0.85 | 0.79 | 0.7 |
| 6 | 26456443 | rs6456724 | 0.282 | 0.581 | 0.305 | -1.391 | -0.01 | 0.043 | 0.043 | 0 | 0.84 | 0.96 | 0.84 | 0.95 | 0.85 | 0.82 |
| 6 | 26458265 | rs6929846 | 1.67 | 0.571 | 0.715 | -1.397 | -0.043 | -0.094 | -0.094 | 0.044 | 0.74 | 0.43 | 0.71 | 0.92 | 0.82 | 0.81 |
| 6 | 26459244 | rs6912934 | 0.273 | 0.709 | 0.455 | -1.325 | -0.012 | 0.04 | 0.04 | 0 | 0.85 | 0.96 | 0.84 | 0.95 | 0.85 | 0.82 |
| 6 | 26460721 | rs2893856 | 0.273 | 0.709 | 0.455 | -1.325 | -0.012 | 0.04 | 0.04 | 0 | 0.85 | 0.96 | 0.84 | 0.95 | 0.85 | 0.82 |
| 6 | 26464472 | rs9467759 | -0.894 | 0.789 | 0.67 | -1.342 | 0.037 | -0.016 | -0.016 | 0 | 0.12 | 0.43 | 0.23 | 0.07 | 0.16 | 0.18 |
| 6 | 26464789 | rs6916321 | -0.566 | 0.925 | 0.505 | -1.299 | -0.006 | 0 | 0 | 0.068 | 0.85 | 0.86 | 0.88 | 0.88 | 0.76 | 0.93 |
| 6 | 26466225 | rs1977198 | 2.081 | 0.843 | 0.436 | -1.362 | -0.03 | 0.1 | 0.1 | 0.771 | 0.49 | 0.1 | 0.55 | 0.7 | 0.49 | 0.61 |
| 6 | 26467182 | rs3799380 | -1.823 | 0.667 | 0.74 | -1.384 | -0.029 | -0.034 | -0.034 | 0.304 | 0.25 | 0.64 | 0.19 | 0.16 | 0.19 | 0.2 |
| 6 | 26469445 | rs9461254 | -0.066 | 0.647 | 0.883 | -1.385 | -0.012 | -0.005 | -0.005 | 0 | 0.07 | 0.38 | 0.07 | 0.02 | 0.01 | 0 |
| 6 | 26469866 | rs9366658 | -0.787 | 0.648 | 0.936 | -1.398 | 0.007 | -0.016 | -0.016 | 0 | 0.15 | 0.21 | 0.11 | 0.12 | 0.18 | 0.17 |
| 6 | 26469875 | rs9358944 | -0.787 | 0.648 | 0.936 | -1.398 | 0.007 | -0.016 | -0.016 | 0 | 0.15 | 0.21 | 0.11 | 0.12 | 0.18 | 0.17 |
| 6 | 26470862 | rs4711112 | -0.28 | 0.691 | 0.934 | -1.442 | 0.02 | -0.005 | -0.005 | 0.073 | 0.83 | 0.77 | 0.87 | 0.88 | 0.76 | 0.93 |
| 6 | 26471596 | rs6911470 | -0.022 | 0.672 | 0.947 | -1.448 | -0.012 | -0.005 | -0.005 | 0.26 | 0.07 | 0.38 | 0.07 | 0.02 | 0.01 | 0 |
| 6 | 26471992 | rs6927012 | -0.766 | 0.686 | 0.997 | -1.45 | 0.004 | 0.014 | 0.014 | 1.811 | 0.87 | 0.93 | 0.94 | 0.94 | 0.89 | 0.95 |
| 6 | 26472655 | rs7763910 | 1.892 | 0.644 | 0.975 | -1.456 | -0.013 | 0.063 | 0.063 | 0.728 | 0.57 | 0.12 | 0.67 | 0.72 | 0.56 | 0.74 |
| 6 | 26473983 | rs7773913 | -0.82 | 0.657 | 0.981 | -1.448 | -0.045 | -0.007 | -0.007 | 0 | 0.1 | 0.43 | 0.08 | 0.04 | 0.02 | 0.03 |
| 6 | 26475784 | rs13437351 | -0.82 | 0.657 | 0.981 | -1.448 | -0.045 | -0.007 | -0.007 | 0 | 0.1 | 0.43 | 0.08 | 0.04 | 0.02 | 0.03 |
| 6 | 26477779 | rs6456728 | -2.068 | 0.663 | 0.983 | -1.449 | -0.027 | -0.036 | -0.036 | 0 | 0.25 | 0.61 | 0.19 | 0.16 | 0.2 | 0.19 |
| 6 | 26478909 | rs6906521 | -0.266 | 0.641 | 0.981 | -1.45 | 0.02 | -0.005 | -0.005 | 0 | 0.83 | 0.77 | 0.87 | 0.88 | 0.76 | 0.93 |
| 6 | 26478927 | rs9358946 | -0.739 | 0.641 | 0.979 | -1.45 | 0.007 | -0.016 | -0.016 | 2.164 | 0.15 | 0.21 | 0.11 | 0.12 | 0.18 | 0.17 |
| 6 | 26479150 | rs16891725 | -0.224 | 0.637 | 1.032 | -1.46 | 0.018 | 0.01 | 0.01 | 0 | 0.08 | 0.14 | 0.05 | 0.11 | 0.12 | 0.12 |
| 6 | 26479891 | rs9357008 | -0.267 | 0.673 | 0.965 | -1.454 | 0.02 | -0.005 | -0.005 | 0 | 0.83 | 0.78 | 0.87 | 0.88 | 0.75 | 0.94 |
| 6 | 26481642 | rs7756567 | -0.817 | 0.618 | 0.917 | -1.438 | 0.007 | -0.016 | -0.016 | 1.152 | 0.15 | 0.21 | 0.11 | 0.12 | 0.18 | 0.17 |
| 6 | 26481714 | rs6940691 | -0.049 | 0.615 | 0.911 | -1.431 | -0.012 | -0.005 | -0.005 | 0 | 0.07 | 0.38 | 0.07 | 0.02 | 0.01 | 0 |
| 6 | 26484376 | rs6456730 | -0.717 | 0.625 | 0.866 | -1.426 | -0.045 | -0.007 | -0.007 | 0.923 | 0.1 | 0.43 | 0.08 | 0.04 | 0.02 | 0.03 |
| 6 | 26487310 | rs4713001 | 0.234 | 0.615 | 0.789 | -1.438 | -0.017 | 0.019 | 0.019 | 0 | 0.09 | 0.02 | 0.14 | 0.03 | 0.11 | 0.11 |
| 6 | 26494187 | rs6910993 | -0.646 | 0.609 | 0.951 | -1.398 | -0.042 | -0.011 | -0.011 | 2.037 | 0.1 | 0.41 | 0.08 | 0.04 | 0.01 | 0.03 |
| 6 | 26495099 | rs2093169 | -0.868 | 0.831 | 0.844 | -1.315 | 0.007 | -0.014 | -0.014 | 0 | 0.15 | 0.21 | 0.11 | 0.12 | 0.18 | 0.17 |
| 6 | 26499903 | rs6926629 | 1.788 | 0.49 | 0.643 | -1.508 | 0 | 0.026 | 0.026 | 0.143 | 0.52 | 0.51 | 0.59 | 0.73 | 0.44 | 0.58 |
| 6 | 26505135 | rs3736780 | -0.856 | 0.338 | 0.764 | -1.615 | 0.004 | 0.02 | 0.02 | 1.518 | 0.86 | 0.94 | 0.94 | 0.96 | 0.89 | 0.95 |
| 6 | 26505362 | rs3736781 | 1.835 | 0.341 | 0.77 | -1.6 | 0 | 0.026 | 0.026 | 0.145 | 0.52 | 0.51 | 0.59 | 0.75 | 0.44 | 0.58 |
| 6 | 26510564 | rs1056667 | 1.827 | 0.341 | 0.79 | -1.6 | 0 | 0.023 | 0.023 | 0.163 | 0.52 | 0.52 | 0.59 | 0.75 | 0.44 | 0.57 |
| 6 | 26510748 | rs3799383 | -0.423 | 0.369 | 0.885 | -1.586 | 0.045 | 0.031 | 0.031 | 0 | 0.09 | 0.21 | 0.06 | 0.12 | 0.12 | 0.16 |
| 6 | 26514166 | rs9393729 | 1.841 | 0.357 | 0.805 | -1.61 | 0 | 0.023 | 0.023 | 0.163 | 0.52 | 0.52 | 0.59 | 0.75 | 0.44 | 0.57 |
| 6 | 26520941 | rs1535276 | 1.849 | 0.357 | 0.806 | -1.61 | 0 | 0.026 | 0.026 | 0.145 | 0.52 | 0.51 | 0.59 | 0.75 | 0.44 | 0.57 |
| 6 | 26521353 | rs6924865 | 1.841 | 0.357 | 0.805 | -1.61 | 0 | 0.023 | 0.023 | 0.163 | 0.52 | 0.52 | 0.59 | 0.75 | 0.44 | 0.58 |
| 6 | 26524313 | rs6911330 | -0.869 | 0.353 | 0.809 | -1.627 | 0.004 | 0.02 | 0.02 | 1.518 | 0.86 | 0.94 | 0.94 | 0.96 | 0.89 | 0.95 |
| 6 | 26526994 | rs9379888 | 0.256 | 0.349 | 0.891 | -1.634 | -0.018 | 0.033 | 0.033 | 0 | 0.06 | 0.14 | 0.02 | 0.11 | 0.04 | 0.12 |
| 6 | 26528250 | rs9295695 | 1.888 | 0.355 | 0.891 | -1.632 | 0 | 0.026 | 0.026 | 0.168 | 0.52 | 0.51 | 0.59 | 0.75 | 0.44 | 0.58 |
| 6 | 26533140 | rs1884945 | 0.425 | 0.378 | 0.88 | -1.646 | -0.013 | 0.016 | 0.016 | 0 | 0.08 | 0.03 | 0.14 | 0.03 | 0.12 | 0.11 |
| 6 | 26533757 | rs10946835 | 1.928 | 0.378 | 0.88 | -1.646 | 0 | 0.023 | 0.023 | 0.163 | 0.52 | 0.52 | 0.59 | 0.75 | 0.44 | 0.58 |
| 6 | 26535541 | rs2393670 | 1.928 | 0.378 | 0.88 | -1.646 | 0 | 0.023 | 0.023 | 0.163 | 0.52 | 0.52 | 0.59 | 0.75 | 0.44 | 0.58 |
| 6 | 26537801 | rs6920256 | -0.115 | 0.387 | 0.902 | -1.646 | 0.04 | 0.027 | 0.027 | 0 | 0.08 | 0.19 | 0.05 | 0.12 | 0.12 | 0.15 |
| 6 | 26537813 | rs10456047 | 0.413 | 0.403 | 0.823 | -1.627 | -0.013 | 0.016 | 0.016 | 0 | 0.08 | 0.03 | 0.14 | 0.03 | 0.12 | 0.11 |
| 6 | 26542773 | rs9467782 | 1.839 | 0.394 | 0.819 | -1.617 | 0 | 0.024 | 0.024 | 0.145 | 0.52 | 0.51 | 0.59 | 0.75 | 0.44 | 0.58 |
| 6 | 26544110 | rs9461270 | 1.85 | 0.372 | 0.809 | -1.63 | 0 | 0.025 | 0.025 | 0.051 | 0.52 | 0.52 | 0.59 | 0.75 | 0.44 | 0.58 |
| 6 | 26545632 | rs4871 | 1.829 | 0.43 | 0.793 | -1.598 | 0 | 0.024 | 0.024 | 0.145 | 0.52 | 0.52 | 0.59 | 0.75 | 0.44 | 0.58 |
| 6 | 26546048 | rs15580 | -0.842 | 0.43 | 0.764 | -1.588 | 0.004 | 0.02 | 0.02 | 1.518 | 0.86 | 0.94 | 0.94 | 0.96 | 0.89 | 0.95 |
| 6 | 26550619 | rs9986382 | 1.788 | 0.397 | 0.751 | -1.326 | 0 | 0.014 | 0.014 | 0.145 | 0.52 | 0.52 | 0.54 | 0.68 | 0.44 | 0.57 |
| 6 | 26553943 | rs2224380 | 1.788 | 0.397 | 0.752 | -1.326 | 0 | 0.015 | 0.015 | 0.145 | 0.52 | 0.51 | 0.54 | 0.68 | 0.44 | 0.57 |
| 6 | 26554968 | rs9461271 | 1.788 | 0.397 | 0.752 | -1.326 | 0 | 0.014 | 0.014 | 0.145 | 0.52 | 0.52 | 0.54 | 0.68 | 0.44 | 0.57 |
| 6 | 26555289 | rs6929908 | -0.613 | 0.438 | 0.772 | -1.339 | -0.009 | -0.007 | -0.007 | 4.825 | 0.85 | 0.91 | 0.84 | 0.84 | 0.75 | 0.92 |
| 6 | 26555879 | rs10484442 | 1.814 | 0.36 | 0.76 | -1.331 | 0 | 0.015 | 0.015 | 0.129 | 0.52 | 0.51 | 0.54 | 0.68 | 0.44 | 0.57 |
| 6 | 26557854 | rs767471 | 1.806 | 0.361 | 0.76 | -1.33 | 0 | 0.015 | 0.015 | 0.145 | 0.52 | 0.51 | 0.54 | 0.68 | 0.43 | 0.57 |
| 6 | 26565560 | rs12208211 | 0.292 | 0.314 | 0.7 | -1.359 | -0.014 | 0.02 | 0.02 | 0 | 0.09 | 0.03 | 0.14 | 0.03 | 0.12 | 0.11 |
| 6 | 26566804 | rs6456733 | 1.817 | 0.335 | 0.733 | -1.33 | 0.001 | 0.015 | 0.015 | 0.034 | 0.51 | 0.52 | 0.54 | 0.68 | 0.44 | 0.57 |
| 6 | 26568741 | rs1078679 | 1.817 | 0.335 | 0.734 | -1.362 | 0.001 | 0.014 | 0.014 | 0.034 | 0.51 | 0.52 | 0.54 | 0.68 | 0.44 | 0.57 |
| 6 | 26572165 | rs6925895 | 1.809 | 0.334 | 0.733 | -1.39 | 0.001 | 0.014 | 0.014 | 0.036 | 0.52 | 0.52 | 0.54 | 0.68 | 0.44 | 0.57 |
| 6 | 26574149 | rs6456735 | 1.785 | 0.223 | 0.649 | -1.451 | 0.003 | 0.018 | 0.018 | 0.057 | 0.49 | 0.52 | 0.53 | 0.68 | 0.42 | 0.56 |

| | | | | | | | | | | | | | | | | |
|---|---|---|---|---|---|---|---|---|---|---|---|---|---|---|---|---|
| 6 | 26575154 | rs1321482 | 1.835 | 0.226 | 0.648 | -1.449 | 0.002 | 0.015 | 0.015 | 0.008 | 0.49 | 0.52 | 0.53 | 0.68 | 0.42 | 0.56 |
| 6 | 26575986 | rs1570061 | 1.835 | 0.226 | 0.648 | -1.449 | 0.002 | 0.015 | 0.015 | 0.008 | 0.49 | 0.52 | 0.53 | 0.68 | 0.43 | 0.56 |
| 6 | 26577924 | rs6918854 | 1.835 | 0.226 | 0.648 | -1.449 | 0.002 | 0.015 | 0.015 | 0.008 | 0.49 | 0.52 | 0.53 | 0.68 | 0.43 | 0.56 |
| 6 | 26581874 | rs9467803 | 0.294 | 0.063 | 0.919 | -1.366 | -0.016 | -0.01 | -0.01 | 0.022 | 0.06 | 0.33 | 0.06 | 0.02 | 0.01 | 0 |
| 6 | 26582414 | rs9379892 | -0.714 | 0.145 | 0.616 | -1.27 | -0.01 | -0.008 | -0.008 | 0 | 0.85 | 0.91 | 0.83 | 0.83 | 0.76 | 0.92 |
| 6 | 26584668 | rs9379893 | -0.529 | 0.081 | 0.765 | -1.205 | -0.008 | -0.021 | -0.021 | 0 | 0.2 | 0.23 | 0.13 | 0.13 | 0.19 | 0.24 |
| 6 | 26590941 | rs6922256 | -0.571 | 0.057 | 0.727 | -1.335 | 0 | 0 | 0 | 0.002 | 0.8 | 0.88 | 0.89 | 0.88 | 0.87 | 0.89 |
| 6 | 26593471 | rs4368798 | 1.784 | 0.312 | 0.652 | -1.436 | 0.005 | 0.005 | 0.005 | 0.871 | 0.45 | 0.49 | 0.51 | 0.67 | 0.38 | 0.54 |
| 6 | 26595267 | rs6915555 | 0.546 | 0.315 | 0.659 | -1.421 | -0.007 | 0.002 | 0.002 | 0.28 | 0.65 | 0.73 | 0.64 | 0.79 | 0.57 | 0.78 |
| 6 | 26597333 | rs3800302 | 0.547 | 0.314 | 0.658 | -1.422 | -0.007 | 0.002 | 0.002 | 0.29 | 0.65 | 0.73 | 0.64 | 0.79 | 0.57 | 0.78 |
| 6 | 26598188 | rs3800303 | 1.764 | 0.326 | 0.66 | -1.413 | 0.005 | 0.005 | 0.005 | 0.882 | 0.45 | 0.49 | 0.51 | 0.67 | 0.38 | 0.54 |
| 6 | 26599043 | rs13978 | 0.276 | 0.25 | 0.746 | -1.441 | -0.011 | -0.004 | -0.004 | 0 | 0.06 | 0.34 | 0.07 | 0.02 | 0.01 | 0 |
| 6 | 26604140 | rs6456739 | 1.16 | 0.25 | 0.767 | -1.465 | -0.002 | 0.005 | 0.005 | 0.153 | 0.53 | 0.64 | 0.57 | 0.77 | 0.5 | 0.69 |
| 6 | 26605864 | rs12205310 | 0.616 | 0.344 | 0.557 | -1.385 | -0.035 | 0.021 | 0.021 | 0 | 0.11 | 0.02 | 0.17 | 0.02 | 0.17 | 0.09 |
| 6 | 26606512 | rs9379902 | 0.24 | 0.348 | 0.612 | -1.285 | -0.008 | 0.003 | 0.003 | 0.412 | 0.65 | 0.73 | 0.64 | 0.79 | 0.57 | 0.77 |
| 6 | 26608261 | rs9467810 | 1.516 | 0.353 | 0.625 | -1.284 | 0.005 | 0.005 | 0.005 | 0.871 | 0.45 | 0.49 | 0.51 | 0.67 | 0.38 | 0.54 |
| 6 | 26608325 | rs6919391 | 0.251 | 0.366 | 0.62 | -1.272 | -0.008 | 0.003 | 0.003 | 0.411 | 0.65 | 0.73 | 0.64 | 0.79 | 0.57 | 0.77 |
| 6 | 26612363 | rs6911476 | 0.143 | 0.388 | 0.647 | -1.238 | -0.011 | -0.004 | -0.004 | 0 | 0.07 | 0.35 | 0.07 | 0.02 | 0.01 | 0 |
| 6 | 26620312 | rs7754159 | -0.139 | 0.386 | 0.655 | -1.253 | -0.008 | -0.024 | -0.024 | 0 | 0.19 | 0.22 | 0.12 | 0.12 | 0.19 | 0.21 |
| 6 | 26622055 | rs2494697 | 0.24 | 0.391 | 0.648 | -1.265 | -0.008 | 0.004 | 0.004 | 0.229 | 0.65 | 0.73 | 0.64 | 0.79 | 0.57 | 0.75 |
| 6 | 26624822 | rs2451731 | 1.436 | 0.324 | 0.642 | -1.251 | 0.006 | 0.005 | 0.005 | 0.778 | 0.46 | 0.5 | 0.51 | 0.67 | 0.38 | 0.55 |
| 6 | 26629404 | rs2451741 | 1.436 | 0.327 | 0.576 | -1.25 | 0.005 | 0.005 | 0.005 | 0.778 | 0.46 | 0.5 | 0.51 | 0.67 | 0.38 | 0.55 |
| 6 | 26639613 | rs1027204 | 1.435 | 0.331 | 0.578 | -1.248 | 0.005 | 0.005 | 0.005 | 0.842 | 0.46 | 0.5 | 0.51 | 0.67 | 0.38 | 0.55 |
| 6 | 26639777 | rs2172007 | 1.84 | 0.334 | 0.557 | -1.246 | -0.011 | -0.024 | -0.024 | 0.19 | 0.39 | 0.15 | 0.45 | 0.65 | 0.37 | 0.55 |
| 6 | 26641060 | rs2504599 | 1.468 | 0.663 | 0.441 | -1.096 | 0.005 | 0.005 | 0.005 | 0.842 | 0.46 | 0.5 | 0.51 | 0.67 | 0.38 | 0.55 |
| 6 | 26641172 | rs2494705 | 0.288 | 0.697 | 0.411 | -1.076 | -0.008 | 0.004 | 0.004 | 0.272 | 0.65 | 0.73 | 0.64 | 0.79 | 0.57 | 0.75 |
| 6 | 26643435 | rs2504571 | 1.434 | 0.667 | 0.27 | -1.076 | 0.006 | 0.005 | 0.005 | 0.842 | 0.46 | 0.5 | 0.51 | 0.67 | 0.38 | 0.55 |
| 6 | 26648334 | rs2504600 | 1.427 | 0.666 | 0.27 | -1.076 | 0.006 | 0.006 | 0.006 | 0.816 | 0.46 | 0.5 | 0.51 | 0.67 | 0.38 | 0.55 |
| 6 | 26649018 | rs2451753 | 0.248 | 0.667 | 0.352 | -1.107 | -0.022 | 0.011 | 0.011 | 0.248 | 0.65 | 0.77 | 0.64 | 0.79 | 0.57 | 0.75 |
| 6 | 26649421 | rs2494711 | 0.248 | 0.667 | 0.352 | -1.107 | -0.022 | 0.011 | 0.011 | 0.248 | 0.65 | 0.77 | 0.64 | 0.79 | 0.57 | 0.75 |
| 6 | 26654318 | rs9379907 | -0.171 | 0.66 | 0.351 | -1.11 | -0.014 | -0.038 | -0.038 | 0 | 0.2 | 0.27 | 0.13 | 0.12 | 0.19 | 0.21 |
| 6 | 26655821 | rs2451715 | 0.261 | 0.745 | 0.394 | -1.067 | -0.022 | 0.012 | 0.012 | 0.205 | 0.65 | 0.77 | 0.64 | 0.79 | 0.57 | 0.75 |
| 6 | 26656890 | rs2504565 | 1.504 | 0.748 | 0.394 | -1.055 | 0.006 | 0.008 | 0.008 | 0.792 | 0.46 | 0.5 | 0.51 | 0.71 | 0.38 | 0.55 |
| 6 | 26663478 | rs7740197 | -0.229 | 0.846 | 0.286 | -1.036 | -0.011 | -0.03 | -0.03 | 0.749 | 0.19 | 0.27 | 0.13 | 0.12 | 0.19 | 0.21 |
| 6 | 26665835 | rs868190 | 0.477 | 1.099 | -0.293 | -0.784 | -0.022 | 0.014 | 0.014 | 0.266 | 0.65 | 0.77 | 0.64 | 0.79 | 0.57 | 0.75 |
| 6 | 26758479 | rs4711122 | -0.299 | 1.22 | -0.431 | -0.735 | -0.001 | 0 | 0 | 0 | 0.15 | 0.14 | 0.24 | 0.09 | 0.3 | 0.14 |
| 6 | 26922906 | rs9348738 | 0.281 | 0.135 | -0.683 | -0.785 | 0.005 | 0.01 | 0.01 | 0.52 | 0.2 | 0.01 | 0.12 | 0.11 | 0.13 | 0.13 |
| 6 | 26951678 | rs10946888 | 1.352 | -0.038 | -0.797 | -1.063 | 0.029 | 0.06 | 0.06 | 1.72 | 0.48 | 0.18 | 0.62 | 0.68 | 0.56 | 0.63 |
| 6 | 26988417 | rs6920115 | -0.057 | -0.212 | 0.13 | -1.222 | 0.01 | 0.003 | 0.003 | 0 | 0.06 | 0.18 | 0.08 | 0.05 | 0.11 | 0.02 |
| 6 | 26988992 | rs7741445 | 0.133 | -0.468 | 0.7 | -1.379 | 0.017 | 0.012 | 0.012 | 1.063 | 0.27 | 0.13 | 0.07 | 0.06 | 0.12 | 0.15 |
| 6 | 26992023 | rs4583981 | 1.584 | -0.317 | 0.578 | -1.326 | 0.023 | 0.063 | 0.063 | 1.562 | 0.47 | 0.17 | 0.62 | 0.69 | 0.56 | 0.63 |
| 6 | 26998567 | rs4358615 | 1.011 | -0.318 | 0.581 | -1.324 | 0.08 | 0.09 | 0.09 | 1.883 | 0.53 | 0.35 | 0.7 | 0.74 | 0.67 | 0.65 |
| 6 | 27002406 | rs9348747 | -0.654 | -0.3 | 0.923 | -1.337 | -0.001 | 0.004 | 0.004 | 1.553 | 0.41 | 0.29 | 0.21 | 0.24 | 0.22 | 0.33 |
| 6 | 27006475 | rs12527111 | 1.481 | -0.177 | 0.383 | -1.297 | 0.021 | 0.061 | 0.061 | 1.641 | 0.47 | 0.17 | 0.62 | 0.68 | 0.56 | 0.63 |
| 6 | 27024687 | rs12190473 | -0.711 | 0.31 | 0.759 | -1.031 | 0.174 | 0.041 | 0.041 | 0 | 0.05 | 0.2 | 0.1 | 0.05 | 0.21 | 0.04 |
| 6 | 27025281 | rs12199218 | -0.845 | 0.297 | 0.751 | -1.037 | 0.159 | 0.034 | 0.034 | 0 | 0.05 | 0.2 | 0.1 | 0.05 | 0.21 | 0.04 |
| 6 | 27028454 | rs7761889 | -0.433 | -0.081 | 1.025 | -1.143 | -0.051 | 0.031 | 0.031 | 0 | 0.18 | 0.41 | 0.19 | 0.2 | 0.11 | 0.19 |
| 6 | 27030924 | rs3922717 | -0.734 | 0.032 | 1 | -1.117 | 0.166 | 0.037 | 0.037 | 0 | 0.05 | 0.2 | 0.1 | 0.05 | 0.21 | 0.04 |
| 6 | 27034253 | rs7747454 | -1.666 | -0.389 | 0.769 | -1.348 | 0.029 | 0.058 | 0.058 | 1.771 | 0.52 | 0.82 | 0.38 | 0.31 | 0.44 | 0.38 |
| 6 | 27036456 | rs4424065 | -1.605 | -0.417 | 0.757 | -1.358 | 0.037 | 0.07 | 0.07 | 1.909 | 0.52 | 0.81 | 0.38 | 0.3 | 0.43 | 0.37 |
| 6 | 27039476 | rs12661273 | 0.114 | -0.479 | 0.803 | -1.405 | -0.003 | -0.013 | -0.013 | 0 | 0.27 | 0.17 | 0.07 | 0.05 | 0.11 | 0.15 |
| 6 | 27047256 | rs6904071 | 0.143 | -0.381 | 0.993 | -1.366 | 0.014 | -0.033 | -0.033 | 0.861 | 0.11 | 0.13 | 0.1 | 0.01 | 0.14 | 0.08 |
| 6 | 27057945 | rs9348750 | -1.821 | -0.447 | 1.066 | -1.379 | 0.017 | 0.015 | 0.015 | 0 | 0.26 | 0.66 | 0.3 | 0.25 | 0.32 | 0.24 |
| 6 | 27070543 | rs6456767 | -1.881 | -0.588 | 1.044 | -1.482 | 0.028 | 0.058 | 0.058 | 1.463 | 0.53 | 0.83 | 0.38 | 0.3 | 0.44 | 0.38 |
| 6 | 27080205 | rs6921388 | -1.776 | -0.603 | 1.063 | -1.489 | 0.017 | 0.014 | 0.014 | 0 | 0.26 | 0.66 | 0.3 | 0.25 | 0.32 | 0.24 |
| 6 | 27080460 | rs2393915 | -2.466 | -0.572 | 0.737 | -1.812 | -0.167 | -0.072 | -0.072 | 2.016 | 0.64 | 0.96 | 0.45 | 0.32 | 0.53 | 0.45 |
| 6 | 27090404 | rs7745603 | -0.764 | -0.484 | 0.948 | -1.741 | 0.008 | 0.032 | 0.032 | 0 | 0.2 | 0.53 | 0.23 | 0.19 | 0.21 | 0.21 |
| 6 | 27091425 | rs6913660 | 0.358 | -0.475 | 0.972 | -1.75 | 0.014 | -0.033 | -0.033 | 0 | 0.11 | 0.13 | 0.1 | 0.01 | 0.14 | 0.08 |
| 6 | 27094019 | rs6921711 | 0.417 | -0.529 | 0.919 | -1.77 | -0.006 | -0.016 | -0.016 | 0.572 | 0.26 | 0.17 | 0.07 | 0.05 | 0.11 | 0.15 |
| 6 | 27103580 | rs2022272 | -2.442 | -0.392 | 1.252 | -1.381 | 0.01 | 0.007 | 0.007 | 0.309 | 0.34 | 0.83 | 0.33 | 0.27 | 0.34 | 0.27 |
| 6 | 27114052 | rs9393790 | -1.231 | 0.157 | 1.101 | -1.394 | 0.078 | 0.053 | 0.053 | 0 | 0.21 | 0.55 | 0.28 | 0.23 | 0.31 | 0.22 |
| 6 | 27114883 | rs6919033 | 0.6 | 0.157 | 1.046 | -1.386 | -0.045 | -0.014 | -0.014 | 1.286 | 0.08 | 0.16 | 0.03 | 0.03 | 0.02 | 0.04 |
| 6 | 27119826 | rs7744759 | -1.897 | 0.235 | 1.045 | -1.387 | 0.02 | 0.008 | 0.008 | 0.004 | 0.26 | 0.67 | 0.3 | 0.24 | 0.32 | 0.23 |
| 6 | 27121224 | rs16894368 | 0.577 | 0.174 | 1.048 | -1.391 | -0.046 | -0.015 | -0.015 | 0 | 0.09 | 0.16 | 0.03 | 0.03 | 0.02 | 0.04 |
| 6 | 27121325 | rs9379960 | -1.904 | 0.226 | 0.942 | -1.37 | 0.02 | 0.008 | 0.008 | 0.004 | 0.26 | 0.67 | 0.31 | 0.24 | 0.32 | 0.23 |
| 6 | 27128307 | rs4713069 | -2.665 | -0.347 | 0.809 | -1.433 | -0.055 | 0 | 0 | 0 | 0.45 | 0.96 | 0.46 | 0.3 | 0.44 | 0.35 |
| 6 | 27128806 | rs9295730 | -1.894 | -0.135 | 1.025 | -1.275 | 0.024 | 0.011 | 0.011 | 0 | 0.25 | 0.66 | 0.3 | 0.24 | 0.32 | 0.24 |
| 6 | 27129757 | rs1028308 | -1.91 | -0.129 | 1.014 | -1.272 | 0.021 | 0.009 | 0.009 | 0.019 | 0.25 | 0.67 | 0.3 | 0.24 | 0.32 | 0.23 |
| 6 | 27138529 | rs9461335 | -1.938 | -0.128 | 1.028 | -1.272 | 0.019 | 0.008 | 0.008 | 0.02 | 0.26 | 0.67 | 0.31 | 0.24 | 0.32 | 0.23 |

| | | | | | | | | | | | | | | | | |
|---|---|---|---|---|---|---|---|---|---|---|---|---|---|---|---|---|
| 6 | 27143883 | rs13194053 | -0.026 | -0.112 | 0.817 | -1.335 | 0.013 | -0.033 | -0.033 | 0 | 0.11 | 0.14 | 0.11 | 0.01 | 0.14 | 0.08 |
| 6 | 27150599 | rs911186 | -1.404 | -0.246 | 1.237 | -1.403 | -0.012 | -0.088 | -0.088 | 1.785 | 0.21 | 0.37 | 0.18 | 0.03 | 0.18 | 0.18 |
| 6 | 27154361 | rs858972 | 0.825 | -0.118 | 1.421 | -1.206 | 0.057 | 0.075 | 0.075 | 0.22 | 0.82 | 0.49 | 0.77 | 0.78 | 0.75 | 0.86 |
| 6 | 27160095 | rs1102563 | 1.818 | -0.307 | 1.204 | -1.29 | 0.035 | 0.008 | 0.008 | 0.195 | 0.66 | 0.19 | 0.7 | 0.75 | 0.7 | 0.73 |
| 6 | 27164757 | rs6901448 | -0.92 | -0.294 | 1.363 | -1.354 | -0.051 | -0.028 | -0.028 | 0 | 0.23 | 0.53 | 0.2 | 0.15 | 0.15 | 0.16 |
| 6 | 27171437 | rs7755577 | -0.92 | -0.294 | 1.363 | -1.354 | -0.051 | -0.028 | -0.028 | 0 | 0.23 | 0.53 | 0.25 | 0.15 | 0.15 | 0.16 |
| 6 | 27186172 | rs6902268 | -1.812 | -0.406 | 0.704 | -1.647 | 0.047 | 0.047 | 0.047 | 0.111 | 0.38 | 0.82 | 0.31 | 0.23 | 0.31 | 0.28 |
| 6 | 27187355 | rs1156086 | -1.812 | -0.406 | 0.704 | -1.659 | 0.047 | 0.049 | 0.049 | 0.111 | 0.38 | 0.82 | 0.31 | 0.23 | 0.31 | 0.28 |
| 6 | 27192629 | rs4713074 | -1.82 | -0.422 | 0.696 | -1.667 | 0.048 | 0.051 | 0.051 | 0.092 | 0.38 | 0.82 | 0.36 | 0.23 | 0.31 | 0.29 |
| 6 | 27193095 | rs996247 | -1.82 | -0.422 | 0.696 | -1.667 | 0.048 | 0.051 | 0.051 | 0.092 | 0.38 | 0.82 | 0.36 | 0.23 | 0.31 | 0.29 |
| 6 | 27194072 | rs6918423 | -1.467 | -0.435 | 0.874 | -1.677 | -0.017 | -0.012 | -0.012 | 0.182 | 0.32 | 0.6 | 0.3 | 0.17 | 0.19 | 0.27 |
| 6 | 27198718 | rs9468015 | -1.903 | -0.435 | 0.854 | -1.676 | 0.048 | 0.051 | 0.051 | 0.092 | 0.38 | 0.82 | 0.36 | 0.23 | 0.31 | 0.29 |
| 6 | 27201262 | rs3800308 | -1.512 | -0.553 | 0.86 | -1.676 | -0.01 | -0.006 | -0.006 | 1.491 | 0.33 | 0.62 | 0.24 | 0.17 | 0.19 | 0.27 |
| 6 | 27207734 | rs9468018 | 0.452 | -0.637 | 1.012 | -1.664 | -0.023 | -0.027 | -0.027 | 0.144 | 0.07 | 0.12 | 0.05 | 0.01 | 0.03 | 0.08 |
| 6 | 27208165 | rs4711144 | -1.426 | -0.634 | 0.876 | -1.666 | -0.018 | -0.012 | -0.012 | 0 | 0.32 | 0.59 | 0.29 | 0.17 | 0.19 | 0.27 |
| 6 | 27210614 | rs3800312 | -1.426 | -0.626 | 0.86 | -1.659 | -0.018 | -0.012 | -0.012 | 0 | 0.32 | 0.59 | 0.29 | 0.17 | 0.19 | 0.27 |
| 6 | 27217236 | rs10946904 | -1.762 | -1.165 | 0.295 | -1.948 | 0.064 | 0.054 | 0.054 | 0.553 | 0.41 | 0.85 | 0.35 | 0.26 | 0.32 | 0.28 |
| 6 | 27220890 | rs2295603 | -2.126 | -1.389 | -0.03 | -2.231 | -0.058 | 0.082 | 0.082 | 0.83 | 0.54 | 0.91 | 0.44 | 0.27 | 0.48 | 0.38 |
| 6 | 27226965 | rs9393796 | 0.367 | -1.339 | 0.483 | -2.16 | 0.004 | 0.009 | 0.009 | 0 | 0.11 | 0.14 | 0.07 | 0.1 | 0.12 | 0.06 |
| 6 | 27229561 | rs11758636 | 0.101 | -1.452 | 0.58 | -2.184 | -0.026 | -0.028 | -0.028 | 0 | 0.09 | 0.19 | 0.06 | 0.02 | 0.04 | 0.09 |
| 6 | 27230815 | rs9295732 | 0.398 | -1.406 | 0.438 | -2.146 | -0.057 | -0.019 | -0.019 | 0.9 | 0.11 | 0.16 | 0.04 | 0.04 | 0.03 | 0.04 |
| 6 | 27235866 | rs6915678 | -1.766 | -1.502 | 0.412 | -2.255 | 0.001 | -0.001 | -0.001 | 2.343 | 0.39 | 0.57 | 0.27 | 0.13 | 0.34 | 0.25 |
| 6 | 27236048 | rs3800314 | -2.219 | -1.506 | 0.29 | -2.27 | -0.009 | 0.104 | 0.104 | 0.3 | 0.53 | 0.85 | 0.41 | 0.26 | 0.47 | 0.37 |
| 6 | 27238000 | rs17687319 | 0.84 | -1.448 | 0.292 | -2.215 | -0.028 | 0.017 | 0.017 | 0 | 0.1 | 0.01 | 0.06 | 0.03 | 0.1 | 0.07 |
| 6 | 27243480 | rs6917419 | 0.241 | -1.4 | 0.4 | -2.188 | 0.005 | 0.01 | 0.01 | 0 | 0.11 | 0.16 | 0.08 | 0.09 | 0.13 | 0.06 |
| 6 | 27244590 | rs9379968 | -2.056 | -1.273 | 0.181 | -2.268 | -0.032 | 0.102 | 0.102 | 0.126 | 0.53 | 0.85 | 0.46 | 0.26 | 0.51 | 0.37 |
| 6 | 27248931 | rs6932590 | -0.929 | -1.183 | 0.389 | -2.156 | -0.008 | -0.09 | -0.09 | 0 | 0.26 | 0.38 | 0.2 | 0.02 | 0.24 | 0.15 |
| 6 | 27256102 | rs3800316 | -1.057 | -1.067 | -0.096 | -2.085 | -0.002 | -0.04 | -0.04 | 0 | 0.35 | 0.58 | 0.24 | 0.03 | 0.25 | 0.16 |
| 6 | 27261324 | rs7746199 | 0.231 | -1.131 | 0 | -2.052 | -0.013 | -0.083 | -0.083 | 0.923 | 0.18 | 0.25 | 0.15 | 0.03 | 0.17 | 0.11 |
| 6 | 27267008 | rs4559087 | -1.781 | -1.408 | -0.935 | -2.107 | -0.068 | 0.032 | 0.032 | 0.1 | 0.55 | 0.96 | 0.49 | 0.3 | 0.47 | 0.38 |
| 6 | 27271343 | rs7771953 | -1.286 | -1.412 | -0.812 | -2.088 | 0.025 | 0.094 | 0.094 | 0.019 | 0.49 | 0.63 | 0.36 | 0.17 | 0.44 | 0.33 |
| 6 | 27277402 | rs6456773 | -1.268 | -1.655 | -0.929 | -2.197 | 0.011 | 0.027 | 0.027 | 0 | 0.95 | 1 | 0.99 | 0.99 | 0.98 | 0.92 |
| 6 | 27279852 | rs2235233 | 1.342 | -0.709 | -1.427 | -1.094 | -0.064 | -0.125 | -0.125 | 0.264 | 0.2 | 0.01 | 0.26 | 0.38 | 0.2 | 0.31 |
| 6 | 27289776 | rs6923811 | -0.934 | -1.13 | -0.782 | -0.889 | 0.016 | 0.015 | 0.015 | 0.207 | 0.44 | 0.56 | 0.29 | 0.11 | 0.33 | 0.29 |
| 6 | 27291822 | rs4145206 | 0.263 | -0.298 | -0.624 | -0.176 | 0.006 | 0.021 | 0.021 | 0 | 0.17 | 0.03 | 0.09 | 0.08 | 0.13 | 0.14 |
| 6 | 27296512 | rs9461357 | -0.168 | -0.031 | -0.928 | -0.113 | 0.019 | 0.047 | 0.047 | 0 | 0.21 | 0.06 | 0.11 | 0.08 | 0.16 | 0.16 |
| 6 | 27298200 | rs9379976 | -0.339 | 0.174 | -0.864 | -0.036 | -0.024 | 0.014 | 0.014 | 0 | 0.22 | 0.09 | 0.16 | 0.14 | 0.24 | 0.19 |
| 6 | 27298279 | rs9379977 | -0.374 | 0.187 | -0.849 | -0.026 | -0.022 | 0.015 | 0.015 | 0 | 0.22 | 0.09 | 0.17 | 0.14 | 0.24 | 0.19 |
| 6 | 27298629 | rs9379978 | -0.368 | 0.182 | -0.87 | 0.055 | -0.025 | 0.01 | 0.01 | 0 | 0.22 | 0.08 | 0.17 | 0.15 | 0.26 | 0.19 |
| 6 | 27298905 | rs721600 | -0.368 | 0.182 | -0.87 | 0.055 | -0.025 | 0.01 | 0.01 | 0 | 0.22 | 0.08 | 0.17 | 0.15 | 0.26 | 0.19 |
| 6 | 27300580 | rs1156457 | 0.332 | 0.359 | -0.992 | 0.55 | -0.015 | -0.017 | -0.017 | 0 | 0.08 | 0.03 | 0.12 | 0.28 | 0.16 | 0.21 |
| 6 | 27302825 | rs9348761 | -0.27 | 0.357 | -0.955 | 0.199 | -0.024 | 0.01 | 0.01 | 0 | 0.22 | 0.09 | 0.17 | 0.15 | 0.26 | 0.19 |
| 6 | 27303927 | rs9461362 | -0.081 | 0.277 | -0.969 | 0.001 | 0.016 | 0.034 | 0.034 | 0 | 0.21 | 0.08 | 0.13 | 0.1 | 0.18 | 0.17 |
| 6 | 27308175 | rs9461363 | -0.487 | 0.374 | -1.113 | -0.104 | -0.049 | 0.061 | 0.061 | 1.156 | 0.28 | 0.09 | 0.19 | 0.12 | 0.32 | 0.2 |
| 6 | 27310533 | rs9461366 | -0.665 | 0.611 | -0.959 | -0.143 | -0.051 | 0.055 | 0.055 | 0.952 | 0.31 | 0.2 | 0.24 | 0.12 | 0.39 | 0.2 |
| 6 | 27312078 | rs764284 | -1.281 | 0.07 | -0.484 | 0.07 | -0.005 | -0.003 | -0.003 | 0.884 | 0.4 | 0.37 | 0.41 | 0.51 | 0.49 | 0.41 |
| 6 | 27321848 | rs9368508 | 0.201 | 0.2 | -0.474 | 0.153 | -0.017 | -0.031 | -0.031 | 0.118 | 0.14 | 0.08 | 0.22 | 0.41 | 0.23 | 0.25 |
| 6 | 27326863 | rs2235232 | -1.632 | 0.031 | 0.009 | -0.036 | -0.002 | 0.003 | 0.003 | 0.008 | 0.44 | 0.69 | 0.47 | 0.53 | 0.57 | 0.4 |
| 6 | 27328577 | rs17688516 | 0.835 | 0.136 | -0.435 | -0.025 | -0.002 | -0.005 | -0.005 | 0 | 0.15 | 0.03 | 0.08 | 0.11 | 0.14 | 0.11 |
| 6 | 27329788 | rs17739727 | 0.085 | 0.113 | -0.379 | -0.012 | -0.021 | 0.021 | 0.021 | 0 | 0.2 | 0.09 | 0.13 | 0.11 | 0.23 | 0.15 |
| 6 | 27332298 | rs1011665 | -1.383 | 0.036 | -0.327 | -0.042 | -0.004 | -0.002 | -0.002 | 0 | 0.4 | 0.39 | 0.4 | 0.53 | 0.48 | 0.41 |
| 6 | 27336489 | rs6914864 | -2.161 | -0.606 | -1.035 | -0.402 | -0.128 | -0.115 | -0.115 | 0.527 | 0.7 | 0.97 | 0.61 | 0.58 | 0.68 | 0.67 |
| 6 | 27338890 | rs9468076 | -0.146 | 0.593 | -0.034 | 0.437 | -0.023 | 0.018 | 0.018 | 0 | 0.19 | 0.09 | 0.13 | 0.11 | 0.23 | 0.15 |
| 6 | 27341409 | rs7759001 | -0.148 | 0.592 | 0.066 | 0.557 | -0.006 | -0.006 | -0.006 | 0 | 0.85 | 0.88 | 0.77 | 0.59 | 0.77 | 0.74 |
| 6 | 27350329 | rs1883403 | 1.225 | 0.525 | 0.161 | 0.401 | -0.003 | -0.002 | -0.002 | 0 | 0.61 | 0.61 | 0.59 | 0.47 | 0.51 | 0.58 |
| 6 | 27351239 | rs6920362 | 0.515 | 0.526 | 0.296 | 0.404 | 0.007 | 0.093 | 0.093 | 0 | 0.79 | 0.64 | 0.72 | 0.58 | 0.74 | 0.73 |
| 6 | 27351931 | rs2393929 | 0.515 | 0.526 | 0.296 | 0.404 | 0.007 | 0.093 | 0.093 | 0 | 0.79 | 0.64 | 0.72 | 0.58 | 0.74 | 0.73 |
| 6 | 27353114 | rs2393931 | 0.64 | 0.478 | 0.274 | 0.379 | 0.004 | 0.085 | 0.085 | 0 | 0.78 | 0.64 | 0.72 | 0.58 | 0.74 | 0.73 |
| 6 | 27353852 | rs2143062 | -0.236 | 0.513 | 0.322 | 0.402 | -0.016 | -0.041 | -0.041 | 0.642 | 0.84 | 0.92 | 0.78 | 0.59 | 0.77 | 0.73 |
| 6 | 27355460 | rs6913166 | 1.391 | 0.526 | 0.324 | 0.403 | 0.002 | 0.001 | 0.001 | 0 | 0.61 | 0.58 | 0.6 | 0.47 | 0.51 | 0.58 |
| 6 | 27355842 | rs2206250 | 0.564 | 0.527 | 0.326 | 0.406 | 0.007 | 0.093 | 0.093 | 0 | 0.79 | 0.64 | 0.72 | 0.58 | 0.74 | 0.73 |
| 6 | 27356216 | rs2223464 | 0.438 | 0.525 | 0.35 | 0.405 | 0.008 | 0.084 | 0.084 | 0 | 0.8 | 0.66 | 0.73 | 0.58 | 0.74 | 0.73 |
| 6 | 27356527 | rs13220817 | 1.429 | 0.498 | 0.334 | 0.389 | 0.002 | 0 | 0 | 0 | 0.6 | 0.58 | 0.59 | 0.47 | 0.51 | 0.58 |
| 6 | 27356902 | rs10807020 | 0.623 | 0.498 | 0.435 | 0.452 | 0.007 | 0.093 | 0.093 | 0 | 0.79 | 0.64 | 0.72 | 0.58 | 0.74 | 0.73 |
| 6 | 27357308 | rs6904883 | 0.623 | 0.498 | 0.435 | 0.452 | 0.007 | 0.093 | 0.093 | 0 | 0.79 | 0.64 | 0.72 | 0.58 | 0.74 | 0.73 |
| 6 | 27358392 | rs2206251 | 0.623 | 0.498 | 0.435 | 0.452 | 0.007 | 0.093 | 0.093 | 0 | 0.79 | 0.64 | 0.72 | 0.58 | 0.74 | 0.73 |
| 6 | 27360690 | rs4711148 | 0.616 | 0.498 | 0.435 | 0.453 | 0.007 | 0.091 | 0.091 | 0 | 0.79 | 0.64 | 0.72 | 0.58 | 0.74 | 0.73 |
| 6 | 27362109 | rs6923108 | 0.511 | 0.506 | 0.434 | 0.457 | 0.009 | 0.089 | 0.089 | 0 | 0.79 | 0.65 | 0.73 | 0.58 | 0.74 | 0.73 |
| 6 | 27363137 | rs12662076 | 1.019 | 0.561 | 0.421 | 0.44 | 0.033 | 0.069 | 0.069 | 0 | 0.76 | 0.62 | 0.67 | 0.58 | 0.65 | 0.69 |

| | | | | | | | | | | | | | | | | |
|---|---|---|---|---|---|---|---|---|---|---|---|---|---|---|---|---|
| 6 | 27363898 | rs4713092 | 0.48 | 0.568 | 0.419 | 0.452 | 0.008 | 0.083 | 0.083 | 0 | 0.8 | 0.67 | 0.73 | 0.58 | 0.74 | 0.73 |
| 6 | 27365994 | rs6456777 | 0.48 | 0.568 | 0.419 | 0.452 | 0.008 | 0.083 | 0.083 | 0 | 0.8 | 0.67 | 0.73 | 0.58 | 0.74 | 0.73 |
| 6 | 27367824 | rs9969094 | 0.238 | 0.565 | 0.424 | 0.451 | 0.019 | 0.016 | 0.016 | 0.386 | 0.05 | 0.25 | 0.05 | 0.01 | 0.02 | 0 |
| 6 | 27367887 | rs6915934 | 0.446 | 0.575 | 0.4 | 0.454 | 0.008 | 0.083 | 0.083 | 0 | 0.8 | 0.67 | 0.73 | 0.58 | 0.74 | 0.73 |
| 6 | 27368031 | rs766092 | 0.446 | 0.545 | 0.403 | 0.455 | 0.008 | 0.083 | 0.083 | 0 | 0.8 | 0.67 | 0.73 | 0.58 | 0.74 | 0.73 |
| 6 | 27370046 | rs6456778 | 0.446 | 0.497 | 0.691 | 0.423 | 0.008 | 0.083 | 0.083 | 0 | 0.8 | 0.66 | 0.73 | 0.58 | 0.74 | 0.73 |
| 6 | 27371011 | rs3734575 | 0.445 | 0.497 | 0.691 | 0.423 | 0.008 | 0.083 | 0.083 | 0 | 0.8 | 0.66 | 0.73 | 0.58 | 0.74 | 0.73 |
| 6 | 27371193 | rs3734574 | 0.445 | 0.497 | 0.691 | 0.423 | 0.008 | 0.083 | 0.083 | 0 | 0.8 | 0.66 | 0.73 | 0.58 | 0.74 | 0.73 |
| 6 | 27372242 | rs4504471 | 0.445 | 0.497 | 0.691 | 0.423 | 0.008 | 0.083 | 0.083 | 0 | 0.8 | 0.66 | 0.73 | 0.58 | 0.74 | 0.73 |
| 6 | 27375611 | rs2076305 | 0.834 | 0.62 | 1.059 | 0.417 | 0.052 | 0.126 | 0.126 | 0.001 | 0.77 | 0.51 | 0.68 | 0.58 | 0.67 | 0.73 |
| 6 | 27376910 | rs6938943 | 1.361 | 0.568 | 1.09 | 0.382 | 0.009 | 0.026 | 0.026 | 0.033 | 0.71 | 0.35 | 0.64 | 0.58 | 0.64 | 0.71 |
| 6 | 27379547 | rs4713097 | 1.061 | 0.637 | 1.094 | 0.257 | 0.043 | 0.101 | 0.101 | 0 | 0.74 | 0.47 | 0.66 | 0.58 | 0.65 | 0.71 |
| 6 | 27384520 | rs6929812 | 1.478 | 0.671 | 1.113 | 0.256 | 0.03 | 0.037 | 0.037 | 0.09 | 0.7 | 0.4 | 0.62 | 0.58 | 0.57 | 0.7 |
| 6 | 27385770 | rs7754977 | 0.856 | 0.667 | 1.147 | 0.265 | 0.004 | 0.062 | 0.062 | 0 | 0.76 | 0.63 | 0.71 | 0.58 | 0.71 | 0.71 |
| 6 | 27386057 | rs6456781 | 0.31 | 0.66 | 1.179 | 0.249 | 0.008 | 0.073 | 0.073 | 0.093 | 0.81 | 0.74 | 0.75 | 0.59 | 0.76 | 0.74 |
| 6 | 27388739 | rs6456782 | 1.055 | 0.65 | 1.132 | 0.25 | 0.039 | 0.091 | 0.091 | 0.003 | 0.74 | 0.51 | 0.67 | 0.58 | 0.64 | 0.71 |
| 6 | 27389258 | rs7771598 | 1.096 | 0.651 | 1.133 | 0.251 | 0.037 | 0.089 | 0.089 | 0 | 0.73 | 0.51 | 0.67 | 0.58 | 0.64 | 0.71 |
| 6 | 27390399 | rs6456785 | 1.237 | 0.65 | 1.133 | 0.251 | 0.07 | 0.081 | 0.081 | 0.032 | 0.73 | 0.51 | 0.63 | 0.58 | 0.57 | 0.7 |
| 6 | 27395764 | rs6926142 | 1.662 | 0.64 | 1.18 | 0.246 | 0.014 | 0.017 | 0.017 | 0.155 | 0.7 | 0.34 | 0.61 | 0.58 | 0.56 | 0.7 |
| 6 | 27396066 | rs2206253 | 1.095 | 0.634 | 1.177 | 0.249 | 0.037 | 0.089 | 0.089 | 0 | 0.73 | 0.51 | 0.67 | 0.58 | 0.64 | 0.71 |
| 6 | 27398133 | rs3918438 | 1.177 | 0.616 | 1.155 | 0.24 | 0.036 | 0.089 | 0.089 | 0 | 0.73 | 0.46 | 0.66 | 0.58 | 0.64 | 0.71 |
| 6 | 27400031 | rs1569566 | 1.157 | 0.629 | 1.17 | 0.246 | 0.034 | 0.083 | 0.083 | 0.025 | 0.73 | 0.46 | 0.66 | 0.58 | 0.64 | 0.71 |
| 6 | 27402023 | rs6908086 | 1.139 | 0.629 | 1.169 | 0.247 | 0.038 | 0.092 | 0.092 | 0.007 | 0.73 | 0.46 | 0.66 | 0.58 | 0.64 | 0.71 |
| 6 | 27403605 | rs4713098 | 1.811 | 0.564 | 1.036 | 0.189 | -0.005 | 0.008 | 0.008 | 0.016 | 0.55 | 0.27 | 0.53 | 0.47 | 0.44 | 0.58 |
| 6 | 27405202 | rs4713099 | 1.812 | 0.564 | 1.013 | 0.189 | -0.004 | 0.008 | 0.008 | 0.016 | 0.55 | 0.28 | 0.53 | 0.47 | 0.44 | 0.58 |
| 6 | 27412386 | rs7772160 | 1.452 | 0.673 | 1.08 | 0.21 | 0.025 | 0.027 | 0.027 | 0.138 | 0.7 | 0.36 | 0.6 | 0.58 | 0.56 | 0.7 |
| 6 | 27418847 | rs7509 | 1.253 | 0.692 | 1.106 | 0.21 | 0.075 | 0.083 | 0.083 | 0.033 | 0.72 | 0.48 | 0.63 | 0.57 | 0.57 | 0.7 |
| 6 | 27422493 | rs2235252 | 0.988 | 0.689 | 1.089 | 0.21 | 0.04 | 0.081 | 0.081 | 0 | 0.74 | 0.53 | 0.67 | 0.58 | 0.65 | 0.71 |
| 6 | 27422794 | rs2235254 | 0.988 | 0.689 | 1.089 | 0.21 | 0.04 | 0.081 | 0.081 | 0 | 0.74 | 0.53 | 0.67 | 0.58 | 0.64 | 0.71 |
| 6 | 27424184 | rs2092121 | 0.986 | 0.689 | 1.089 | 0.21 | 0.04 | 0.081 | 0.081 | 0 | 0.74 | 0.53 | 0.67 | 0.58 | 0.65 | 0.71 |
| 6 | 27425185 | rs1883216 | 0.986 | 0.689 | 1.089 | 0.21 | 0.04 | 0.081 | 0.081 | 0 | 0.74 | 0.53 | 0.67 | 0.58 | 0.65 | 0.71 |
| 6 | 27427542 | rs7748366 | 0.973 | 0.688 | 1.089 | 0.21 | 0.041 | 0.082 | 0.082 | 0 | 0.74 | 0.54 | 0.67 | 0.58 | 0.65 | 0.71 |
| 6 | 27432848 | rs4358616 | 0.961 | 0.699 | 1.095 | 0.209 | 0.005 | 0.081 | 0.081 | 0 | 0.74 | 0.54 | 0.7 | 0.58 | 0.71 | 0.71 |
| 6 | 27434220 | rs6920408 | 0.972 | 0.684 | 1.09 | 0.209 | 0.002 | 0.082 | 0.082 | 0 | 0.74 | 0.54 | 0.7 | 0.58 | 0.71 | 0.71 |
| 6 | 27435024 | rs12211424 | 0.929 | 0.76 | 1.056 | 0.18 | -0.035 | 0.046 | 0.046 | 2.484 | 0.09 | 0.01 | 0.05 | 0 | 0.15 | 0.02 |
| 6 | 27435707 | rs2272814 | 1.197 | 0.747 | 1.065 | 0.187 | 0.002 | 0.082 | 0.082 | 0 | 0.74 | 0.54 | 0.7 | 0.58 | 0.71 | 0.71 |
| 6 | 27436245 | rs12213598 | 1.192 | 0.747 | 1.065 | 0.187 | 0.002 | 0.082 | 0.082 | 0.006 | 0.74 | 0.53 | 0.7 | 0.58 | 0.71 | 0.71 |
| 6 | 27439125 | rs2142705 | 1.196 | 0.749 | 1.066 | 0.188 | 0.002 | 0.081 | 0.081 | 0 | 0.74 | 0.53 | 0.7 | 0.58 | 0.71 | 0.71 |
| 6 | 27440004 | rs7750795 | 1.221 | 0.734 | 1.059 | 0.18 | 0.001 | 0.08 | 0.08 | 0 | 0.74 | 0.53 | 0.7 | 0.58 | 0.71 | 0.71 |
| 6 | 27443957 | rs1014227 | 1.227 | 0.745 | 1.064 | 0.186 | 0.001 | 0.08 | 0.08 | 0.014 | 0.74 | 0.53 | 0.7 | 0.58 | 0.71 | 0.71 |
| 6 | 27444714 | rs4713106 | 0.238 | 0.751 | 1.059 | 0.191 | 0.003 | 0.03 | 0.03 | 0.181 | 0.83 | 0.82 | 0.76 | 0.59 | 0.76 | 0.74 |
| 6 | 27455625 | rs6456794 | 0.252 | 0.741 | 1.072 | 0.225 | 0.002 | 0.024 | 0.024 | 0.193 | 0.83 | 0.83 | 0.77 | 0.59 | 0.76 | 0.74 |
| 6 | 27462824 | rs2393962 | 1.947 | 0.806 | 0.982 | 0.213 | -0.004 | 0.005 | 0.005 | 0.014 | 0.56 | 0.3 | 0.54 | 0.47 | 0.45 | 0.58 |
| 6 | 27463684 | rs16867808 | 0.628 | 0.561 | 0.925 | 0.221 | -0.001 | -0.01 | -0.01 | 4.352 | 0.14 | 0.02 | 0.07 | 0.11 | 0.12 | 0.12 |
| 6 | 27466712 | rs6903282 | 1.476 | 0.568 | 0.907 | 0.231 | 0.029 | 0.073 | 0.073 | 0.186 | 0.72 | 0.5 | 0.66 | 0.58 | 0.64 | 0.7 |
| 6 | 27471098 | rs10946933 | 0.904 | 0.596 | 0.749 | 0.243 | -0.001 | 0.001 | 0.001 | 0 | 0.77 | 0.76 | 0.73 | 0.58 | 0.72 | 0.71 |
| 6 | 27472223 | rs6929563 | -0.087 | 0.616 | 0.69 | 0.261 | -0.03 | -0.066 | -0.066 | 0 | 0.84 | 0.95 | 0.78 | 0.59 | 0.76 | 0.7 |
| 6 | 27480227 | rs2205829 | 1.809 | 0.707 | 0.739 | 0.233 | -0.009 | 0.006 | 0.006 | 0.041 | 0.55 | 0.31 | 0.57 | 0.47 | 0.51 | 0.58 |
| 6 | 27480917 | rs6918131 | 0.94 | 1.004 | 0.824 | 0.509 | -0.001 | 0.001 | 0.001 | 0 | 0.77 | 0.76 | 0.73 | 0.58 | 0.71 | 0.71 |
| 6 | 27483260 | rs2294308 | 1.346 | 0.957 | 0.634 | 0.514 | 0.033 | 0.064 | 0.064 | 0.188 | 0.72 | 0.53 | 0.66 | 0.57 | 0.64 | 0.7 |
| 6 | 27488602 | rs970668 | 0.536 | 0.954 | 0.48 | 0.63 | -0.005 | -0.005 | -0.005 | 0 | 0.91 | 0.89 | 0.9 | 0.71 | 0.9 | 0.83 |
| 6 | 27498955 | rs6456795 | 1.439 | 0.948 | 0.722 | 0.496 | -0.004 | 0 | 0 | 0 | 0.84 | 0.79 | 0.76 | 0.64 | 0.81 | 0.68 |
| 6 | 27501345 | rs10946935 | 1.951 | 0.449 | 0.707 | 0.297 | -0.002 | -0.014 | -0.014 | 0 | 0.65 | 0.3 | 0.61 | 0.53 | 0.64 | 0.53 |
| 6 | 27511371 | rs2064092 | -0.731 | 0.388 | 0.72 | 0.23 | -0.03 | -0.011 | -0.011 | 2.192 | 0.18 | 0.25 | 0.1 | 0.11 | 0.09 | 0.15 |
| 6 | 27515776 | rs6915987 | 2.03 | -0.035 | 0.666 | 0.086 | 0.017 | -0.026 | -0.026 | 0 | 0.57 | 0.21 | 0.58 | 0.5 | 0.62 | 0.45 |
| 6 | 27516738 | rs4711157 | -0.291 | 0.095 | 0.658 | 0.254 | -0.029 | -0.028 | -0.028 | 0.055 | 0.93 | 0.99 | 0.92 | 0.73 | 0.9 | 0.83 |
| 6 | 27519345 | rs6934794 | 1.661 | -0.015 | 0.859 | 0.204 | -0.02 | 0.024 | 0.024 | 0 | 0.69 | 0.6 | 0.7 | 0.53 | 0.8 | 0.55 |
| 6 | 27526897 | rs6908024 | 0.359 | 0.034 | 0.758 | 0.214 | -0.004 | -0.003 | -0.003 | 0 | 0.91 | 0.89 | 0.89 | 0.73 | 0.9 | 0.83 |
| 6 | 27528643 | rs6915266 | 1.984 | 0.06 | 0.832 | 0.21 | -0.002 | -0.018 | -0.018 | 0 | 0.64 | 0.29 | 0.6 | 0.52 | 0.64 | 0.54 |
| 6 | 27530562 | rs2142704 | -0.302 | 0.096 | 0.763 | 0.236 | -0.029 | -0.028 | -0.028 | 0.411 | 0.93 | 0.99 | 0.92 | 0.73 | 0.9 | 0.83 |
| 6 | 27545744 | rs6903982 | -0.648 | -0.129 | 1.27 | 0.125 | -0.074 | -0.025 | -0.025 | 0 | 0.18 | 0.4 | 0.11 | 0.11 | 0.09 | 0.13 |
| 6 | 27547792 | rs7774939 | 1.855 | -0.127 | 1.197 | 0.161 | -0.033 | 0.037 | 0.037 | 0 | 0.67 | 0.43 | 0.69 | 0.52 | 0.79 | 0.57 |
| 6 | 27548092 | rs7775817 | 1.888 | -0.128 | 1.21 | 0.161 | 0.005 | -0.007 | -0.007 | 0.111 | 0.67 | 0.33 | 0.68 | 0.52 | 0.79 | 0.57 |
| 6 | 27553784 | rs12197514 | 2.422 | -0.466 | 0.497 | -0.446 | -0.099 | -0.145 | -0.145 | 1.166 | 0.33 | 0.03 | 0.42 | 0.48 | 0.32 | 0.39 |
| 6 | 27555126 | rs4713113 | 1.988 | -0.027 | 1.312 | 0.175 | -0.006 | 0.005 | 0.005 | 0 | 0.67 | 0.36 | 0.68 | 0.52 | 0.79 | 0.57 |
| 6 | 27560587 | rs10946940 | 2.061 | 0.058 | 1.24 | 0.166 | -0.003 | -0.019 | -0.019 | 0 | 0.64 | 0.29 | 0.6 | 0.51 | 0.64 | 0.54 |
| 6 | 27566169 | rs9461392 | -0.105 | 0.043 | 1.218 | 0.21 | -0.029 | -0.028 | -0.028 | 0 | 0.93 | 0.99 | 0.92 | 0.71 | 0.9 | 0.83 |
| 6 | 27573536 | rs6456799 | 2.118 | 0.022 | 1.26 | 0.19 | -0.003 | -0.011 | -0.011 | 0 | 0.66 | 0.31 | 0.61 | 0.52 | 0.64 | 0.54 |
| 6 | 27585038 | rs9468174 | -0.235 | 0.005 | 1.357 | 0.037 | -0.03 | -0.009 | -0.009 | 0 | 0.15 | 0.21 | 0.09 | 0.11 | 0.08 | 0.13 |

| | | | | | | | | | | | | | | | | |
|---|---|---|---|---|---|---|---|---|---|---|---|---|---|---|---|---|
| 6 | 27587518 | rs6921256 | -2.374 | 0.01 | 1.356 | 0.04 | -0.008 | -0.052 | -0.052 | 0 | 0.36 | 0.78 | 0.41 | 0.46 | 0.36 | 0.46 |
| 6 | 27588727 | rs9393831 | -1.993 | -0.141 | 1.349 | 0.074 | -0.046 | 0.063 | 0.063 | 0.388 | 0.29 | 0.53 | 0.3 | 0.45 | 0.2 | 0.41 |
| 6 | 27601587 | rs2143082 | -1.16 | 0.054 | 1.276 | 0.191 | -0.006 | 0.124 | 0.124 | 0.002 | 0.14 | 0.33 | 0.21 | 0.35 | 0.12 | 0.28 |
| 6 | 27623511 | rs9366694 | -2.065 | 0.126 | 1.279 | 0.158 | 0.026 | 0.119 | 0.119 | 0 | 0.18 | 0.46 | 0.27 | 0.35 | 0.19 | 0.33 |
| 6 | 27632780 | rs16867886 | -0.559 | 0.149 | 1.374 | 0.23 | 0 | 0.019 | 0.019 | 0 | 0.1 | 0.16 | 0.09 | 0.28 | 0.08 | 0.19 |
| 6 | 27634640 | rs6917366 | -2.147 | 0.15 | 1.366 | 0.23 | 0.031 | 0.137 | 0.137 | 0 | 0.17 | 0.46 | 0.24 | 0.35 | 0.17 | 0.32 |
| 6 | 27634927 | rs9380001 | -0.514 | 0.148 | 1.334 | 0.23 | 0 | -0.002 | -0.002 | 0 | 0.1 | 0.12 | 0.09 | 0.28 | 0.08 | 0.19 |
| 6 | 27641958 | rs6903016 | -2.071 | 0.156 | 1.341 | 0.224 | 0.031 | 0.127 | 0.127 | 0 | 0.17 | 0.43 | 0.23 | 0.35 | 0.17 | 0.32 |
| 6 | 27645278 | rs9368521 | -0.602 | 0.128 | 1.269 | 0.187 | 0 | 0.011 | 0.011 | 0 | 0.1 | 0.15 | 0.1 | 0.28 | 0.08 | 0.19 |
| 6 | 27646100 | rs9357043 | -0.288 | 0.125 | 1.27 | 0.186 | -0.002 | 0.005 | 0.005 | 0 | 0.08 | 0.11 | 0.08 | 0.28 | 0.08 | 0.16 |
| 6 | 27656090 | rs9348772 | -2.251 | 0.089 | 1.571 | 0.102 | 0 | -0.001 | -0.001 | 0 | 0.32 | 0.68 | 0.36 | 0.45 | 0.31 | 0.44 |
| 6 | 27656255 | rs10946943 | -0.54 | 0.1 | 1.515 | 0.108 | 0 | 0.013 | 0.013 | 0 | 0.1 | 0.16 | 0.1 | 0.28 | 0.08 | 0.18 |
| 6 | 27656499 | rs9380006 | -1.89 | 0.108 | 1.532 | 0.107 | 0.032 | 0.146 | 0.146 | 0 | 0.16 | 0.46 | 0.23 | 0.35 | 0.16 | 0.31 |
| 6 | 27657647 | rs6920044 | -0.562 | 0.115 | 1.451 | 0.099 | 0 | 0.012 | 0.012 | 0 | 0.1 | 0.16 | 0.1 | 0.28 | 0.08 | 0.18 |
| 6 | 27660508 | rs9380007 | -2.556 | -0.108 | 1.398 | 0.023 | 0.021 | -0.06 | -0.06 | 0 | 0.42 | 0.84 | 0.4 | 0.48 | 0.35 | 0.53 |
| 6 | 27668013 | rs12663707 | -0.578 | 0.041 | 1.097 | 0.065 | 0 | 0.011 | 0.011 | 0 | 0.1 | 0.16 | 0.1 | 0.28 | 0.08 | 0.18 |
| 6 | 27670697 | rs2393997 | -1.044 | 0.125 | 0.964 | -0.046 | -0.03 | 0.033 | 0.033 | 1.337 | 0.93 | 0.99 | 0.96 | 1 | 0.86 | 0.98 |
| 6 | 27670917 | rs10484400 | -0.564 | 0.092 | 1.04 | 0.048 | 0 | 0.01 | 0.01 | 0 | 0.1 | 0.16 | 0.1 | 0.28 | 0.08 | 0.18 |
| 6 | 27676794 | rs4713118 | -1.503 | 0.156 | 1.227 | -0.011 | 0.02 | 0.014 | 0.014 | 0.006 | 0.22 | 0.48 | 0.27 | 0.18 | 0.25 | 0.3 |
| 6 | 27683572 | rs9380010 | -1.365 | 0.216 | 1.17 | 0.019 | 0.02 | 0.012 | 0.012 | 0.046 | 0.21 | 0.52 | 0.26 | 0.17 | 0.24 | 0.25 |
| 6 | 27689502 | rs9295740 | -1.314 | 0.162 | 1.155 | -0.001 | -0.025 | 0.011 | 0.011 | 0.009 | 0.21 | 0.52 | 0.24 | 0.17 | 0.18 | 0.23 |
| 6 | 27689939 | rs2056923 | -0.462 | 0.145 | 1.388 | -0.002 | -0.001 | -0.012 | -0.012 | 0 | 0.16 | 0.26 | 0.13 | 0.11 | 0.16 | 0.15 |
| 6 | 27690174 | rs2056924 | -2.114 | 0.159 | 1.415 | 0.056 | -0.003 | -0.014 | -0.014 | 0 | 0.31 | 0.73 | 0.37 | 0.45 | 0.33 | 0.4 |
| 6 | 27690905 | rs2056925 | -1.62 | 0.167 | 1.456 | 0.047 | 0.002 | -0.002 | -0.002 | 0 | 0.24 | 0.59 | 0.28 | 0.18 | 0.25 | 0.25 |
| 6 | 27693003 | rs6922574 | -1.565 | 0.184 | 1.52 | 0.058 | 0.007 | 0.002 | 0.002 | 0.21 | 0.24 | 0.54 | 0.28 | 0.18 | 0.25 | 0.25 |
| 6 | 27694421 | rs9468209 | -2.494 | 0.047 | 1.501 | 0.001 | 0.015 | -0.067 | -0.067 | 0.273 | 0.4 | 0.83 | 0.4 | 0.48 | 0.35 | 0.51 |
| 6 | 27695329 | rs9348775 | -1.574 | 0.133 | 1.51 | 0.036 | 0.006 | 0 | 0 | 0 | 0.24 | 0.54 | 0.28 | 0.18 | 0.25 | 0.25 |
| 6 | 27706180 | rs9468213 | -2.338 | 0.129 | 1.469 | 0.035 | -0.002 | -0.025 | -0.025 | 0 | 0.33 | 0.74 | 0.37 | 0.46 | 0.33 | 0.43 |
| 6 | 27709441 | rs6902689 | -1.578 | 0.089 | 1.423 | -0.024 | 0.006 | 0 | 0 | 0 | 0.23 | 0.57 | 0.29 | 0.18 | 0.25 | 0.25 |
| 6 | 27712825 | rs4713119 | -0.923 | 0.153 | 1.076 | -0.088 | -0.032 | 0.038 | 0.038 | 1.425 | 0.92 | 0.99 | 0.96 | 1 | 0.86 | 0.98 |
| 6 | 27714575 | rs6901520 | -1.743 | -0.005 | 1.457 | 0.252 | 0.003 | -0.002 | -0.002 | 0 | 0.24 | 0.62 | 0.29 | 0.18 | 0.25 | 0.25 |
| 6 | 27715243 | rs9283880 | -1.743 | -0.005 | 1.457 | 0.252 | 0.003 | -0.002 | -0.002 | 0 | 0.24 | 0.62 | 0.29 | 0.18 | 0.25 | 0.25 |
| 6 | 27719764 | rs9461406 | -1.095 | 0.013 | 1.527 | 0.261 | 0.022 | 0.017 | 0.017 | 0 | 0.18 | 0.34 | 0.26 | 0.17 | 0.22 | 0.25 |
| 6 | 27721835 | rs4713120 | -2.806 | -0.094 | 1.506 | 0.256 | 0.02 | -0.067 | -0.067 | 0 | 0.4 | 0.82 | 0.4 | 0.48 | 0.35 | 0.51 |
| 6 | 27726731 | rs7776351 | -1.942 | 0.285 | 1.595 | 0.37 | 0.003 | 0.001 | 0.001 | 0.218 | 0.24 | 0.62 | 0.29 | 0.18 | 0.25 | 0.25 |
| 6 | 27727051 | rs7742752 | -0.315 | 0.279 | 1.645 | 0.369 | 0.015 | 0.013 | 0.013 | 0 | 0.05 | 0.21 | 0.03 | 0.01 | 0.02 | 0 |
| 6 | 27730064 | rs9295746 | -1.194 | 0.302 | 1.6 | 0.378 | 0.023 | 0.021 | 0.021 | 0 | 0.18 | 0.34 | 0.26 | 0.18 | 0.22 | 0.25 |
| 6 | 27730463 | rs7759217 | -2.007 | 0.313 | 1.646 | 0.391 | 0.007 | 0.009 | 0.009 | 0 | 0.23 | 0.54 | 0.28 | 0.19 | 0.24 | 0.27 |
| 6 | 27734470 | rs9366702 | -2.007 | 0.313 | 1.668 | 0.391 | 0.006 | 0.008 | 0.008 | 0 | 0.23 | 0.55 | 0.28 | 0.19 | 0.24 | 0.27 |
| 6 | 27739380 | rs742047 | -0.984 | 0.332 | 1.651 | 0.4 | 0.023 | 0.024 | 0.024 | 0 | 0.16 | 0.27 | 0.24 | 0.18 | 0.22 | 0.25 |
| 6 | 27740438 | rs12192568 | 0.842 | 0.413 | 1.208 | 0.064 | -0.036 | 0.04 | 0.04 | 0.303 | 0.1 | 0.01 | 0.05 | 0.03 | 0.11 | 0.06 |
| 6 | 27741682 | rs6915101 | -0.523 | 0.408 | 1.714 | 0.458 | 0.037 | 0.034 | 0.034 | 0 | 0.05 | 0.27 | 0.03 | 0.01 | 0.02 | 0 |
| 6 | 27742186 | rs6901437 | -0.48 | 0.409 | 1.704 | 0.458 | 0.02 | 0.018 | 0.018 | 0 | 0.05 | 0.23 | 0.03 | 0.01 | 0.02 | 0 |
| 6 | 27746342 | rs9468227 | -0.547 | 0.396 | 1.757 | 0.461 | 0.026 | 0.024 | 0.024 | 0 | 0.05 | 0.26 | 0.03 | 0.01 | 0.02 | 0 |
| 6 | 27747042 | rs9357048 | -0.169 | 0.435 | 1.52 | 0.529 | -0.023 | -0.021 | -0.021 | 4.503 | 0.06 | 0.01 | 0.06 | 0.27 | 0.08 | 0.16 |
| 6 | 27757960 | rs200469 | -1.32 | 0.119 | 1.773 | 0.484 | -0.024 | 0.063 | 0.063 | 0 | 0.09 | 0.34 | 0.12 | 0.07 | 0.03 | 0.1 |
| 6 | 27767395 | rs9295749 | -0.402 | 0.129 | 1.559 | 0.473 | -0.018 | 0 | 0 | 0 | 0.11 | 0.14 | 0.08 | 0.11 | 0.07 | 0.12 |
| 6 | 27771684 | rs200479 | -1.096 | 0.227 | 1.799 | 0.444 | 0.003 | 0.106 | 0.106 | 0 | 0.08 | 0.33 | 0.12 | 0.07 | 0.03 | 0.08 |
| 6 | 27773904 | rs200482 | -1.005 | 0.469 | 1.927 | 0.44 | 0.046 | 0.054 | 0.054 | 0 | 0.07 | 0.23 | 0.15 | 0.07 | 0.09 | 0.1 |
| 6 | 27779506 | rs200486 | -2.198 | 0.336 | 2.119 | 0.519 | 0.002 | 0.017 | 0.017 | 0 | 0.18 | 0.43 | 0.25 | 0.14 | 0.17 | 0.22 |
| 6 | 27782031 | rs370155 | -0.953 | 0.339 | 2.115 | 0.52 | 0.046 | 0.054 | 0.054 | 0 | 0.07 | 0.23 | 0.15 | 0.07 | 0.09 | 0.1 |
| 6 | 27784428 | rs9295750 | -0.894 | 0.337 | 2.122 | 0.519 | -0.016 | -0.013 | -0.013 | 0 | 0.11 | 0.18 | 0.1 | 0.07 | 0.08 | 0.12 |
| 6 | 27788942 | rs200501 | -0.987 | 0.341 | 2.098 | 0.516 | 0.052 | 0.061 | 0.061 | 0 | 0.06 | 0.21 | 0.14 | 0.07 | 0.09 | 0.08 |
| 6 | 27796935 | rs200490 | -0.908 | 0.323 | 2.115 | 0.506 | 0.035 | 0.042 | 0.042 | 0 | 0.07 | 0.2 | 0.14 | 0.07 | 0.09 | 0.07 |
| 6 | 27804934 | rs201004 | -2.207 | 0.315 | 2.135 | 0.502 | 0 | 0.014 | 0.014 | 0 | 0.18 | 0.4 | 0.24 | 0.14 | 0.17 | 0.22 |
| 6 | 27808192 | rs201002 | -0.943 | 0.333 | 2.109 | 0.508 | 0.034 | 0.041 | 0.041 | 0 | 0.07 | 0.2 | 0.14 | 0.07 | 0.09 | 0.1 |
| 6 | 27811815 | rs200997 | -3.398 | 0.421 | 2.225 | 0.451 | 0.001 | -0.008 | -0.008 | 0 | 0.3 | 0.72 | 0.37 | 0.42 | 0.28 | 0.36 |
| 6 | 27813813 | rs200994 | -2.784 | 0.36 | 2.404 | 0.563 | 0.03 | 0.016 | 0.016 | 0 | 0.2 | 0.49 | 0.26 | 0.15 | 0.22 | 0.22 |
| 6 | 27815494 | rs200991 | -2.187 | 0.349 | 2.421 | 0.565 | -0.006 | 0.033 | 0.033 | 0 | 0.16 | 0.4 | 0.22 | 0.14 | 0.15 | 0.2 |
| 6 | 27815823 | rs200990 | -0.912 | 0.398 | 2.386 | 0.599 | 0.035 | 0.042 | 0.042 | 0 | 0.07 | 0.2 | 0.14 | 0.07 | 0.09 | 0.1 |
| 6 | 27825860 | rs9461416 | -0.896 | 0.419 | 2.371 | 0.597 | -0.025 | -0.002 | -0.002 | 0 | 0.09 | 0.17 | 0.07 | 0.07 | 0.06 | 0.09 |
| 6 | 27835930 | rs200951 | -3.308 | 0.429 | 2.258 | 0.597 | -0.001 | 0.004 | 0.004 | 0.927 | 0.28 | 0.71 | 0.35 | 0.43 | 0.26 | 0.34 |
| 6 | 27837267 | rs200953 | -0.978 | 0.601 | 2.12 | 0.459 | 0.035 | 0.042 | 0.042 | 0 | 0.07 | 0.2 | 0.14 | 0.07 | 0.09 | 0.1 |
| 6 | 27849676 | rs573179 | -2.553 | 0.549 | 2.166 | 0.442 | 0.042 | 0.017 | 0.017 | 0 | 0.18 | 0.48 | 0.25 | 0.14 | 0.21 | 0.22 |
| 6 | 27852357 | rs200979 | -0.775 | 0.757 | 1.879 | 0.527 | 0.124 | 0.058 | 0.058 | 0 | 0.05 | 0.2 | 0.15 | 0.07 | 0.14 | 0.1 |
| 6 | 27859568 | rs200968 | -0.765 | 0.757 | 1.879 | 0.527 | 0.124 | 0.058 | 0.058 | 0 | 0.05 | 0.2 | 0.15 | 0.07 | 0.14 | 0.1 |
| 6 | 27869321 | rs1873213 | -0.411 | 0.442 | 1.555 | 0.35 | -0.017 | 0.004 | 0.004 | 0 | 0.06 | 0.13 | 0.06 | 0.08 | 0.03 | 0.05 |
| 6 | 27870772 | rs9468252 | -1.075 | 0.405 | 1.606 | 0.35 | -0.024 | 0 | 0 | 0 | 0.08 | 0.18 | 0.09 | 0.09 | 0.05 | 0.1 |
| 6 | 27878738 | rs149878 | -3.252 | 0.288 | 1.798 | 0.479 | -0.004 | 0.013 | 0.013 | 0.4 | 0.29 | 0.67 | 0.37 | 0.46 | 0.26 | 0.35 |

| | | | | | | | | | | | | | | | | |
|---|---|---|---|---|---|---|---|---|---|---|---|---|---|---|---|---|
| 6 | 27881406 | rs9885649 | -1.074 | 0.331 | 1.809 | 0.469 | -0.019 | 0 | 0 | 0 | 0.08 | 0.18 | 0.09 | 0.09 | 0.05 | 0.1 |
| 6 | 27883201 | rs1497526 | -1.351 | 0.255 | 2.03 | 0.453 | -0.043 | -0.018 | -0.018 | 0 | 0.12 | 0.29 | 0.09 | 0.08 | 0.05 | 0.08 |
| 6 | 27884916 | rs9468254 | -0.499 | 0.307 | 2.038 | 0.455 | -0.017 | 0.004 | 0.004 | 0 | 0.07 | 0.14 | 0.07 | 0.08 | 0.04 | 0.06 |
| 6 | 27885356 | rs731132 | 1.843 | 0.453 | 1.146 | -0.055 | -0.15 | -0.157 | -0.157 | 0.479 | 0.34 | 0.02 | 0.37 | 0.44 | 0.36 | 0.38 |
| 6 | 27886830 | rs2130357 | -2.467 | 0.747 | 2.127 | 0.657 | 0.092 | -0.005 | -0.005 | 0 | 0.4 | 0.83 | 0.34 | 0.46 | 0.26 | 0.44 |
| 6 | 27890631 | rs6933825 | -1.255 | 0.736 | 2.265 | 0.651 | -0.086 | 0.029 | 0.029 | 0 | 0.25 | 0.44 | 0.21 | 0.36 | 0.14 | 0.3 |
| 6 | 27895213 | rs156737 | -2.948 | 0.755 | 2.26 | 0.655 | 0.064 | -0.119 | -0.119 | 1.139 | 0.46 | 0.95 | 0.5 | 0.54 | 0.36 | 0.54 |
| 6 | 27895504 | rs9468256 | -1.281 | 0.723 | 2.289 | 0.657 | -0.101 | 0.036 | 0.036 | 0 | 0.26 | 0.5 | 0.22 | 0.36 | 0.15 | 0.3 |
| 6 | 27901041 | rs7760871 | -0.473 | 0.725 | 2.293 | 0.655 | -0.015 | 0.011 | 0.011 | 0 | 0.07 | 0.18 | 0.08 | 0.08 | 0.04 | 0.06 |
| 6 | 27902992 | rs7742858 | -0.193 | 0.716 | 2.143 | 0.659 | -0.048 | -0.03 | -0.03 | 0 | 0.17 | 0.25 | 0.12 | 0.08 | 0.1 | 0.17 |
| 6 | 27904181 | rs6910968 | -0.195 | 0.793 | 2.147 | 0.676 | -0.048 | -0.03 | -0.03 | 0 | 0.17 | 0.25 | 0.12 | 0.08 | 0.1 | 0.17 |
| 6 | 27915087 | rs276370 | -1.954 | 0.782 | 2.115 | 0.68 | -0.021 | 0.001 | 0.001 | 0.436 | 0.32 | 0.64 | 0.33 | 0.45 | 0.17 | 0.39 |
| 6 | 27919688 | rs9461425 | -1.097 | 0.803 | 2.129 | 0.679 | 0.028 | -0.008 | -0.008 | 0 | 0.1 | 0.31 | 0.12 | 0.08 | 0.1 | 0.08 |
| 6 | 27925367 | rs7767176 | 0.262 | 0.752 | 2.202 | 0.66 | -0.022 | -0.045 | -0.045 | 0 | 0.1 | 0.19 | 0.1 | 0.03 | 0.08 | 0.12 |
| 6 | 27928445 | rs9295753 | -1.212 | 0.786 | 2.143 | 0.627 | -0.086 | 0.027 | 0.027 | 0 | 0.26 | 0.44 | 0.2 | 0.36 | 0.14 | 0.3 |
| 6 | 27932762 | rs2116984 | -1.261 | 0.791 | 2.125 | 0.631 | -0.102 | 0.033 | 0.033 | 0 | 0.26 | 0.5 | 0.21 | 0.36 | 0.14 | 0.3 |
| 6 | 27948574 | rs7769416 | -0.09 | 0.012 | 2.128 | 0.671 | -0.048 | -0.03 | -0.03 | 0.645 | 0.17 | 0.25 | 0.11 | 0.08 | 0.1 | 0.17 |
| 6 | 27955115 | rs12332927 | -1.806 | 0.141 | 1.791 | 0.724 | -0.002 | -0.025 | -0.025 | 0 | 0.14 | 0.32 | 0.13 | 0.09 | 0.12 | 0.15 |
| 6 | 27957079 | rs9357054 | -0.097 | 0.144 | 1.633 | 0.742 | 0.011 | -0.004 | -0.004 | 0 | 0.05 | 0.05 | 0.07 | 0.27 | 0.03 | 0.12 |
| 6 | 27965503 | rs149901 | -2.916 | 0.161 | 1.717 | 0.897 | 0.026 | 0.033 | 0.033 | 0 | 0.2 | 0.42 | 0.29 | 0.21 | 0.21 | 0.24 |
| 6 | 27969004 | rs12333142 | -0.557 | 0.165 | 1.768 | 0.9 | -0.016 | 0.001 | 0.001 | 0 | 0.07 | 0.13 | 0.07 | 0.08 | 0.04 | 0.06 |
| 6 | 27970031 | rs149946 | -2.092 | 0.164 | 1.64 | 0.904 | 0.06 | 0.061 | 0.061 | 0.017 | 0.16 | 0.41 | 0.28 | 0.2 | 0.2 | 0.17 |
| 6 | 27979625 | rs10484402 | -0.112 | 0.153 | 1.347 | 0.946 | 0.011 | -0.002 | -0.002 | 0 | 0.05 | 0.05 | 0.07 | 0.27 | 0.03 | 0.12 |
| 6 | 27981673 | rs9368540 | -0.648 | 0.168 | 1.331 | 0.955 | 0.006 | -0.022 | -0.022 | 0 | 0.06 | 0.01 | 0.06 | 0.1 | 0.05 | 0.17 |
| 6 | 27982152 | rs149971 | -2.638 | 0.081 | 1.213 | 0.92 | 0.061 | -0.084 | -0.084 | 0 | 0.47 | 0.95 | 0.5 | 0.57 | 0.37 | 0.54 |
| 6 | 27986529 | rs9368544 | -0.611 | 0.249 | 1.293 | 0.912 | -0.002 | -0.022 | -0.022 | 0 | 0.06 | 0.01 | 0.07 | 0.1 | 0.06 | 0.17 |
| 6 | 27987776 | rs149976 | -2.623 | 0.236 | 1.283 | 1.051 | 0.055 | -0.07 | -0.07 | 0.141 | 0.46 | 0.95 | 0.51 | 0.58 | 0.37 | 0.54 |
| 6 | 27998258 | rs149990 | -0.922 | 0.412 | 1.579 | 1.034 | 0.035 | 0.066 | 0.066 | 0 | 0.07 | 0.14 | 0.16 | 0.12 | 0.1 | 0.09 |
| 6 | 28000361 | rs1529749 | -0.654 | 0.412 | 1.579 | 1.034 | -0.002 | -0.022 | -0.022 | 2.022 | 0.06 | 0.01 | 0.07 | 0.1 | 0.06 | 0.17 |
| 6 | 28001610 | rs149942 | -3.121 | 0.392 | 1.763 | 1.028 | 0.028 | 0.049 | 0.049 | 0 | 0.22 | 0.41 | 0.35 | 0.3 | 0.26 | 0.34 |
| 6 | 28002388 | rs149943 | -0.931 | 0.428 | 1.728 | 1.049 | 0.035 | 0.066 | 0.066 | 0 | 0.07 | 0.14 | 0.16 | 0.12 | 0.1 | 0.09 |
| 6 | 28008331 | rs2394044 | -0.671 | -0.094 | 1.649 | 1.052 | -0.002 | -0.022 | -0.022 | 0 | 0.06 | 0.01 | 0.07 | 0.1 | 0.06 | 0.17 |
| 6 | 28014597 | rs149900 | -1.97 | -0.074 | 1.75 | 1.099 | 0.06 | 0.061 | 0.061 | 0.111 | 0.16 | 0.39 | 0.27 | 0.2 | 0.19 | 0.17 |
| 6 | 28021589 | rs203888 | -2.11 | -0.055 | 1.747 | 1.099 | 0.056 | 0.053 | 0.053 | 0.036 | 0.17 | 0.39 | 0.27 | 0.2 | 0.2 | 0.17 |
| 6 | 28022248 | rs203890 | -2.11 | -0.138 | 1.747 | 1.099 | 0.053 | 0.053 | 0.053 | 0.036 | 0.17 | 0.39 | 0.28 | 0.2 | 0.2 | 0.17 |
| 6 | 28026077 | rs9468274 | -2.11 | -0.133 | 1.748 | 1.096 | 0.053 | 0.053 | 0.053 | 0.036 | 0.17 | 0.39 | 0.28 | 0.2 | 0.2 | 0.17 |
| 6 | 28027688 | rs9468276 | -2.11 | -0.133 | 1.748 | 1.096 | 0.053 | 0.053 | 0.053 | 0.036 | 0.17 | 0.39 | 0.28 | 0.2 | 0.2 | 0.17 |
| 6 | 28028390 | rs9468277 | -2.11 | -0.133 | 1.748 | 1.096 | 0.053 | 0.053 | 0.053 | 0.036 | 0.17 | 0.39 | 0.28 | 0.2 | 0.2 | 0.17 |
| 6 | 28030499 | rs9380046 | -2.11 | -0.133 | 1.748 | 1.096 | 0.053 | 0.053 | 0.053 | 0.036 | 0.17 | 0.39 | 0.28 | 0.2 | 0.2 | 0.17 |
| 6 | 28033087 | rs149951 | -2.712 | -0.139 | 1.743 | 1.096 | 0.03 | 0.048 | 0.048 | 0 | 0.21 | 0.4 | 0.34 | 0.3 | 0.26 | 0.34 |
| 6 | 28033523 | rs10484403 | -1.991 | -0.139 | 1.919 | 1.104 | 0.027 | 0.029 | 0.029 | 0 | 0.16 | 0.3 | 0.27 | 0.2 | 0.19 | 0.17 |
| 6 | 28037893 | rs9380047 | -1.844 | -0.465 | 2.015 | 1.133 | 0.046 | 0.03 | 0.03 | 0 | 0.15 | 0.31 | 0.27 | 0.19 | 0.2 | 0.14 |
| 6 | 28039586 | rs4713135 | -1.863 | -0.466 | 2.015 | 1.133 | 0.045 | 0.03 | 0.03 | 0 | 0.15 | 0.31 | 0.27 | 0.19 | 0.2 | 0.14 |
| 6 | 28046673 | rs203876 | -2.57 | -0.468 | 1.856 | 1.166 | 0.059 | 0.078 | 0.078 | 0.003 | 0.19 | 0.42 | 0.34 | 0.3 | 0.25 | 0.31 |
| 6 | 28048624 | rs203877 | -2.638 | -0.476 | 1.853 | 1.162 | 0.043 | 0.061 | 0.061 | 0 | 0.21 | 0.42 | 0.34 | 0.3 | 0.25 | 0.31 |
| 6 | 28048996 | rs203878 | -2.633 | -0.6 | 1.786 | 1.202 | 0.033 | -0.075 | -0.075 | 0 | 0.45 | 0.95 | 0.5 | 0.59 | 0.37 | 0.52 |
| 6 | 28055495 | rs10484404 | -0.195 | -0.603 | 1.839 | 1.219 | -0.056 | -0.041 | -0.041 | 0 | 0.17 | 0.32 | 0.12 | 0.08 | 0.11 | 0.18 |
| 6 | 28058635 | rs1853097 | -1.849 | -0.559 | 1.833 | 1.236 | 0.047 | 0.031 | 0.031 | 0 | 0.15 | 0.31 | 0.27 | 0.19 | 0.2 | 0.14 |
| 6 | 28061744 | rs3823180 | -1.849 | -0.485 | 1.851 | 1.285 | 0.047 | 0.031 | 0.031 | 0 | 0.15 | 0.31 | 0.27 | 0.19 | 0.2 | 0.14 |
| 6 | 28062066 | rs203893 | -2.55 | -0.485 | 1.851 | 1.285 | 0.044 | 0.062 | 0.062 | 0 | 0.21 | 0.41 | 0.33 | 0.3 | 0.25 | 0.31 |
| 6 | 28077374 | rs203884 | -2.55 | -0.479 | 1.834 | 1.288 | 0.044 | 0.062 | 0.062 | 0 | 0.21 | 0.41 | 0.33 | 0.3 | 0.25 | 0.31 |
| 6 | 28082711 | rs9368554 | -1.851 | -0.467 | 1.812 | 1.259 | 0.047 | 0.031 | 0.031 | 0 | 0.15 | 0.31 | 0.27 | 0.19 | 0.2 | 0.14 |
| 6 | 28089699 | rs1631552 | -2.526 | -0.473 | 1.817 | 1.262 | 0.044 | 0.062 | 0.062 | 0 | 0.21 | 0.41 | 0.33 | 0.3 | 0.25 | 0.31 |
| 6 | 28092685 | rs4713139 | -0.539 | -0.458 | 1.778 | 1.276 | 0.009 | 0.015 | 0.015 | 0 | 0.06 | 0.08 | 0.16 | 0.12 | 0.1 | 0.07 |
| 6 | 28094731 | rs2275508 | -2.289 | -0.458 | 1.782 | 1.276 | 0.033 | 0.021 | 0.021 | 0 | 0.16 | 0.31 | 0.27 | 0.19 | 0.2 | 0.17 |
| 6 | 28096280 | rs1654774 | -2.782 | -0.478 | 1.715 | 1.263 | -0.004 | -0.082 | -0.082 | 0 | 0.46 | 0.97 | 0.5 | 0.59 | 0.44 | 0.55 |
| 6 | 28097193 | rs4713140 | -0.537 | -0.778 | 1.764 | 1.287 | 0.009 | 0.015 | 0.015 | 0 | 0.06 | 0.08 | 0.16 | 0.12 | 0.1 | 0.07 |
| 6 | 28100640 | rs1225710 | -2.715 | -0.779 | 1.765 | 1.287 | 0.033 | -0.075 | -0.075 | 0.091 | 0.45 | 0.95 | 0.49 | 0.59 | 0.37 | 0.52 |
| 6 | 28104634 | rs9380057 | -2.202 | -0.749 | 1.789 | 1.282 | 0.04 | 0.025 | 0.025 | 0 | 0.16 | 0.31 | 0.27 | 0.19 | 0.2 | 0.14 |
| 6 | 28105196 | rs1947862 | -2.202 | -0.749 | 1.789 | 1.282 | 0.04 | 0.025 | 0.025 | 0 | 0.16 | 0.31 | 0.26 | 0.19 | 0.2 | 0.14 |
| 6 | 28106347 | rs4713142 | -2.202 | -0.743 | 1.792 | 1.259 | 0.04 | 0.025 | 0.025 | 0 | 0.16 | 0.31 | 0.27 | 0.19 | 0.2 | 0.14 |
| 6 | 28106827 | rs4713145 | -2.202 | -0.743 | 1.792 | 1.259 | 0.04 | 0.025 | 0.025 | 0 | 0.16 | 0.31 | 0.26 | 0.19 | 0.2 | 0.14 |
| 6 | 28107357 | rs3757188 | -0.464 | -0.738 | 1.775 | 1.263 | 0.011 | 0.017 | 0.017 | 0 | 0.06 | 0.08 | 0.16 | 0.12 | 0.1 | 0.07 |
| 6 | 28107662 | rs3757186 | -1.225 | -0.804 | 1.804 | 1.232 | -0.014 | -0.004 | -0.004 | 0.948 | 0.07 | 0.08 | 0.05 | 0.06 | 0.03 | 0.05 |
| 6 | 28108232 | rs1904840 | -2.068 | -0.705 | 1.83 | 1.283 | 0.051 | 0.036 | 0.036 | 0 | 0.14 | 0.3 | 0.26 | 0.19 | 0.2 | 0.14 |
| 6 | 28108967 | rs9368555 | -2.068 | -0.705 | 1.83 | 1.283 | 0.051 | 0.036 | 0.036 | 0 | 0.14 | 0.3 | 0.26 | 0.19 | 0.2 | 0.14 |
| 6 | 28109633 | rs2277103 | -0.71 | -0.709 | 1.859 | 1.281 | -0.017 | 0.004 | 0.004 | 0 | 0.06 | 0.13 | 0.06 | 0.06 | 0.03 | 0.05 |
| 6 | 28110148 | rs868987 | -2.058 | -0.704 | 1.855 | 1.282 | 0.051 | 0.036 | 0.036 | 0 | 0.14 | 0.3 | 0.26 | 0.19 | 0.2 | 0.14 |
| 6 | 28110478 | rs1890809 | -0.536 | -0.753 | 1.873 | 1.288 | -0.064 | 0.02 | 0.02 | 0 | 0.2 | 0.37 | 0.14 | 0.28 | 0.11 | 0.2 |

| | | | | | | | | | | | | | | | | |
|---|---|---|---|---|---|---|---|---|---|---|---|---|---|---|---|---|
| 6 | 28111114 | rs2791333 | -2.774 | -0.755 | 1.782 | 1.28 | 0.039 | -0.066 | -0.066 | 3.703 | 0.46 | 0.95 | 0.49 | 0.59 | 0.37 | 0.52 |
| 6 | 28113373 | rs1225715 | -2.724 | -0.748 | 1.788 | 1.284 | 0.044 | -0.058 | -0.058 | 0 | 0.45 | 0.92 | 0.49 | 0.59 | 0.37 | 0.52 |
| 6 | 28123928 | rs1150666 | -2.93 | -0.74 | 2.025 | 1.295 | 0.046 | 0.067 | 0.067 | 0 | 0.2 | 0.4 | 0.33 | 0.3 | 0.25 | 0.31 |
| 6 | 28124469 | rs9295759 | -2.046 | -0.734 | 2.011 | 1.289 | 0.051 | 0.036 | 0.036 | 0 | 0.14 | 0.3 | 0.26 | 0.19 | 0.2 | 0.14 |
| 6 | 28126202 | rs9348796 | -2.046 | -0.734 | 2.011 | 1.289 | 0.051 | 0.036 | 0.036 | 0 | 0.14 | 0.3 | 0.26 | 0.19 | 0.2 | 0.14 |
| 6 | 28129713 | rs1225618 | -2.759 | -0.752 | 2.002 | 1.29 | 0.044 | -0.058 | -0.058 | 0 | 0.45 | 0.92 | 0.49 | 0.59 | 0.37 | 0.52 |
| 6 | 28130194 | rs16893731 | -0.704 | -0.73 | 2.025 | 1.261 | -0.017 | 0.004 | 0.004 | 0 | 0.06 | 0.13 | 0.06 | 0.06 | 0.03 | 0.05 |
| 6 | 28132358 | rs9380059 | -2.002 | -0.707 | 1.914 | 1.269 | 0.051 | 0.036 | 0.036 | 0 | 0.14 | 0.3 | 0.26 | 0.19 | 0.2 | 0.14 |
| 6 | 28134106 | rs6929559 | -0.115 | -0.717 | 1.937 | 1.271 | -0.052 | -0.037 | -0.037 | 0 | 0.17 | 0.28 | 0.12 | 0.08 | 0.11 | 0.18 |
| 6 | 28137027 | rs3173443 | -2.019 | -0.637 | 1.863 | 1.277 | 0.051 | 0.036 | 0.036 | 0 | 0.14 | 0.3 | 0.26 | 0.19 | 0.2 | 0.14 |
| 6 | 28137454 | rs4713152 | -2.019 | -0.637 | 1.863 | 1.277 | 0.051 | 0.036 | 0.036 | 0 | 0.14 | 0.3 | 0.26 | 0.19 | 0.2 | 0.14 |
| 6 | 28137730 | rs9393902 | -0.486 | -0.641 | 1.83 | 1.28 | -0.007 | 0.017 | 0.017 | 0 | 0.06 | 0.07 | 0.13 | 0.12 | 0.03 | 0.07 |
| 6 | 28143118 | rs9380064 | -0.487 | -0.63 | 1.829 | 1.288 | 0.011 | 0.017 | 0.017 | 0 | 0.06 | 0.08 | 0.16 | 0.12 | 0.1 | 0.07 |
| 6 | 28147987 | rs9295761 | -1.925 | -0.637 | 1.838 | 1.285 | 0.051 | 0.036 | 0.036 | 0 | 0.14 | 0.29 | 0.26 | 0.19 | 0.2 | 0.14 |
| 6 | 28148752 | rs1225591 | -2.752 | -0.644 | 1.856 | 1.289 | 0.049 | 0.07 | 0.07 | 0 | 0.2 | 0.4 | 0.32 | 0.3 | 0.24 | 0.31 |
| 6 | 28152567 | rs2622315 | -0.773 | -0.691 | 1.805 | 1.313 | -0.068 | 0.005 | 0.005 | 0 | 0.22 | 0.4 | 0.15 | 0.27 | 0.11 | 0.2 |
| 6 | 28158954 | rs1237873 | -1.068 | -0.466 | 1.792 | 1.331 | 0 | 0.05 | 0.05 | 0 | 0.06 | 0.11 | 0.09 | 0.16 | 0.08 | 0.15 |
| 6 | 28165099 | rs1150689 | -2.176 | -0.443 | 1.838 | 1.314 | 0.06 | 0.046 | 0.046 | 0.056 | 0.17 | 0.44 | 0.28 | 0.19 | 0.21 | 0.15 |
| 6 | 28166923 | rs1233704 | -2.176 | -0.443 | 1.838 | 1.314 | 0.06 | 0.046 | 0.046 | 0.056 | 0.17 | 0.44 | 0.28 | 0.19 | 0.21 | 0.15 |
| 6 | 28170297 | rs735765 | -0.956 | -0.431 | 1.889 | 1.302 | 0.079 | 0.104 | 0.104 | 0 | 0.07 | 0.23 | 0.18 | 0.13 | 0.12 | 0.06 |
| 6 | 28173219 | rs1233708 | -2.501 | -0.446 | 1.69 | 1.287 | 0.021 | 0.04 | 0.04 | 0 | 0.23 | 0.52 | 0.34 | 0.31 | 0.27 | 0.28 |
| 6 | 28176415 | rs1233666 | -1.324 | -0.606 | 1.646 | 1.21 | -0.083 | 0.036 | 0.036 | 0.02 | 0.26 | 0.56 | 0.28 | 0.39 | 0.16 | 0.31 |
| 6 | 28177492 | rs988084 | -0.836 | -0.605 | 1.718 | 1.213 | -0.031 | 0.112 | 0.112 | 0 | 0.06 | 0.22 | 0.1 | 0.22 | 0.01 | 0.03 |
| 6 | 28177588 | rs988083 | -0.836 | -0.605 | 1.718 | 1.213 | -0.031 | 0.112 | 0.112 | 0 | 0.06 | 0.22 | 0.1 | 0.22 | 0.01 | 0.03 |
| 6 | 28181675 | rs1150699 | -1.542 | -0.664 | 1.744 | 1.19 | -0.082 | 0.026 | 0.026 | 0 | 0.27 | 0.57 | 0.28 | 0.38 | 0.17 | 0.32 |
| 6 | 28184260 | rs1150703 | -0.377 | -0.652 | 1.757 | 1.185 | 0.001 | 0.01 | 0.01 | 0 | 0.05 | 0.11 | 0.09 | 0.21 | 0.01 | 0.03 |
| 6 | 28184371 | rs1150704 | -1.455 | -0.645 | 1.75 | 1.181 | -0.082 | 0.024 | 0.024 | 0 | 0.27 | 0.57 | 0.28 | 0.38 | 0.17 | 0.32 |
| 6 | 28185726 | rs1233664 | -1.455 | -0.645 | 1.75 | 1.181 | -0.082 | 0.024 | 0.024 | 0 | 0.27 | 0.57 | 0.28 | 0.38 | 0.17 | 0.32 |
| 6 | 28186554 | rs1736894 | -1.455 | -0.645 | 1.75 | 1.181 | -0.082 | 0.024 | 0.024 | 0 | 0.27 | 0.57 | 0.28 | 0.38 | 0.17 | 0.32 |
| 6 | 28187101 | rs1736891 | -2.599 | -0.637 | 1.598 | 1.291 | 0.039 | -0.058 | -0.058 | 0.112 | 0.46 | 0.95 | 0.48 | 0.58 | 0.38 | 0.49 |
| 6 | 28189389 | rs1736890 | -1.449 | -0.641 | 1.647 | 1.297 | -0.082 | 0.023 | 0.023 | 0 | 0.27 | 0.57 | 0.28 | 0.38 | 0.17 | 0.32 |
| 6 | 28191057 | rs9366717 | -0.242 | -0.564 | 1.634 | 1.309 | 0.012 | 0.018 | 0.018 | 0 | 0.05 | 0.07 | 0.09 | 0.11 | 0.1 | 0.08 |
| 6 | 28197469 | rs16893827 | -0.7 | -0.622 | 1.852 | 1.29 | -0.018 | 0.011 | 0.011 | 0 | 0.09 | 0.14 | 0.06 | 0.09 | 0.05 | 0.08 |
| 6 | 28197605 | rs1150707 | -2.653 | -0.642 | 1.798 | 1.291 | 0.038 | -0.058 | -0.058 | 0 | 0.46 | 0.95 | 0.48 | 0.58 | 0.38 | 0.49 |
| 6 | 28201138 | rs7206 | -2.653 | -0.649 | 1.867 | 1.289 | 0.038 | -0.058 | -0.058 | 0 | 0.46 | 0.95 | 0.48 | 0.58 | 0.38 | 0.49 |
| 6 | 28203300 | rs9380069 | -0.929 | -0.655 | 1.953 | 1.305 | 0.015 | 0.035 | 0.035 | 0 | 0.07 | 0.13 | 0.15 | 0.15 | 0.11 | 0.08 |
| 6 | 28204793 | rs11752073 | 0.172 | -0.664 | 1.977 | 1.197 | -0.042 | -0.062 | -0.062 | 0.332 | 0.15 | 0.27 | 0.11 | 0.03 | 0.1 | 0.15 |
| 6 | 28207737 | rs11758148 | -0.571 | -0.655 | 1.885 | 1.203 | -0.05 | -0.056 | -0.056 | 0.559 | 0.19 | 0.34 | 0.18 | 0.1 | 0.15 | 0.26 |
| 6 | 28208535 | rs1150711 | -2.669 | -0.65 | 1.794 | 1.204 | 0.108 | -0.027 | -0.027 | 0 | 0.43 | 0.87 | 0.44 | 0.57 | 0.32 | 0.47 |
| 6 | 28209102 | rs9295768 | -1.91 | -0.641 | 1.745 | 1.215 | 0.006 | 0.052 | 0.052 | 0 | 0.16 | 0.31 | 0.17 | 0.21 | 0.15 | 0.16 |
| 6 | 28210688 | rs967005 | -1.91 | -0.641 | 1.745 | 1.215 | 0.006 | 0.052 | 0.052 | 0 | 0.16 | 0.31 | 0.17 | 0.21 | 0.15 | 0.16 |
| 6 | 28212489 | rs7757215 | -0.648 | -0.632 | 1.656 | 1.213 | -0.017 | 0.01 | 0.01 | 0 | 0.09 | 0.16 | 0.07 | 0.09 | 0.05 | 0.08 |
| 6 | 28212748 | rs13408 | -2.81 | -0.617 | 1.385 | 1.165 | 0.038 | -0.042 | -0.042 | 2.458 | 0.48 | 0.95 | 0.51 | 0.58 | 0.4 | 0.5 |
| 6 | 28213239 | rs1150712 | -0.356 | -0.532 | 1.483 | 1.196 | 0.008 | 0.002 | 0.002 | 0 | 0.05 | 0.1 | 0.09 | 0.22 | 0.01 | 0.02 |
| 6 | 28215446 | rs1233710 | -1.583 | -0.574 | 1.649 | 1.215 | -0.041 | 0.048 | 0.048 | 0 | 0.15 | 0.28 | 0.15 | 0.35 | 0.06 | 0.13 |
| 6 | 28215641 | rs2235359 | -0.76 | -0.636 | 1.745 | 1.128 | -0.055 | -0.067 | -0.067 | 0 | 0.21 | 0.38 | 0.19 | 0.09 | 0.16 | 0.26 |
| 6 | 28216027 | rs2235358 | -0.3 | -0.629 | 1.658 | 1.131 | 0.002 | -0.007 | -0.007 | 0 | 0.05 | 0 | 0.06 | 0.06 | 0.04 | 0.11 |
| 6 | 28219826 | rs1736895 | -2.804 | -0.614 | 1.638 | 1.153 | 0.037 | -0.042 | -0.042 | 0.332 | 0.48 | 0.95 | 0.51 | 0.58 | 0.4 | 0.5 |
| 6 | 28221816 | rs10456362 | 1.32 | -0.617 | 1.739 | 1.156 | 0.023 | 0.041 | 0.041 | 0 | 0.9 | 0.79 | 0.88 | 0.87 | 0.88 | 0.92 |
| 6 | 28222980 | rs2185955 | -0.711 | -0.63 | 1.783 | 1.151 | -0.054 | -0.067 | -0.067 | 0 | 0.21 | 0.38 | 0.19 | 0.09 | 0.16 | 0.26 |
| 6 | 28223680 | rs7750106 | -1.585 | -0.631 | 1.788 | 1.154 | -0.06 | 0.078 | 0.078 | 0 | 0.16 | 0.32 | 0.16 | 0.35 | 0.06 | 0.13 |
| 6 | 28223731 | rs12214383 | 2.679 | -0.626 | 1.775 | 1.153 | 0.037 | -0.042 | -0.042 | 0.005 | 0.52 | 0.05 | 0.49 | 0.42 | 0.6 | 0.5 |
| 6 | 28226878 | rs1531681 | 2.679 | -0.626 | 1.775 | 1.153 | 0.037 | -0.042 | -0.042 | 0 | 0.52 | 0.05 | 0.49 | 0.42 | 0.6 | 0.5 |
| 6 | 28227436 | rs12000 | -2.2 | -0.729 | 1.795 | 1.161 | 0.027 | -0.007 | -0.007 | 0.62 | 0.37 | 0.7 | 0.35 | 0.44 | 0.22 | 0.39 |
| 6 | 28227604 | rs1635 | -1.594 | -0.713 | 1.815 | 1.172 | -0.06 | 0.078 | 0.078 | 0 | 0.16 | 0.32 | 0.16 | 0.35 | 0.06 | 0.13 |
| 6 | 28227633 | rs9461446 | -0.253 | -0.654 | 1.724 | 1.153 | -0.004 | 0.011 | 0.011 | 0 | 0.06 | 0.06 | 0.08 | 0.12 | 0.03 | 0.08 |
| 6 | 28228342 | rs1679709 | 1.402 | -0.653 | 1.725 | 1.153 | 0.035 | 0.058 | 0.058 | 0 | 0.9 | 0.75 | 0.87 | 0.87 | 0.88 | 0.92 |
| 6 | 28229881 | rs1778508 | 1.317 | -0.654 | 1.649 | 1.153 | 0.035 | 0.057 | 0.057 | 0 | 0.9 | 0.74 | 0.87 | 0.87 | 0.88 | 0.92 |
| 6 | 28231243 | rs9468322 | -0.317 | -0.676 | 1.613 | 1.153 | -0.005 | 0.012 | 0.012 | 0 | 0.06 | 0.07 | 0.08 | 0.12 | 0.03 | 0.08 |
| 6 | 28238969 | rs2179174 | -2.046 | -0.699 | 1.611 | 1.179 | -0.062 | 0.004 | 0.004 | 0 | 0.35 | 0.57 | 0.34 | 0.4 | 0.21 | 0.36 |
| 6 | 28239915 | rs11965538 | -1.63 | -0.734 | 1.583 | 1.206 | 0.019 | 0.042 | 0.042 | 0 | 0.11 | 0.26 | 0.13 | 0.13 | 0.12 | 0.08 |
| 6 | 28240798 | rs1778484 | -2.712 | -0.68 | 1.565 | 1.239 | 0.124 | -0.036 | -0.036 | 0 | 0.48 | 0.95 | 0.48 | 0.58 | 0.33 | 0.48 |
| 6 | 28242794 | rs1419183 | -1.258 | -0.644 | 1.612 | 1.183 | 0.034 | 0.059 | 0.059 | 0 | 0.1 | 0.26 | 0.13 | 0.13 | 0.12 | 0.08 |
| 6 | 28250236 | rs1150724 | -2.702 | -0.672 | 1.464 | 1.249 | 0.125 | -0.017 | -0.017 | 0 | 0.48 | 0.95 | 0.48 | 0.6 | 0.33 | 0.48 |
| 6 | 28250984 | rs6901575 | -1.265 | -0.6 | 1.606 | 1.14 | 0.034 | 0.059 | 0.059 | 0 | 0.1 | 0.26 | 0.13 | 0.13 | 0.12 | 0.08 |
| 6 | 28252859 | rs11756111 | -1.153 | -0.603 | 1.623 | 1.094 | -0.081 | -0.096 | -0.096 | 0 | 0.24 | 0.47 | 0.21 | 0.09 | 0.16 | 0.26 |
| 6 | 28254988 | rs16901846 | -0.029 | -0.536 | 1.932 | 1.185 | -0.022 | -0.035 | -0.035 | 0 | 0.14 | 0.19 | 0.1 | 0.03 | 0.09 | 0.15 |
| 6 | 28263721 | rs9461448 | -0.666 | -0.512 | 1.932 | 1.277 | -0.02 | 0.055 | 0.055 | 0 | 0.07 | 0.17 | 0.09 | 0.12 | 0.03 | 0.08 |
| 6 | 28264681 | rs3800324 | -0.887 | -0.583 | 1.902 | 1.338 | 0.023 | -0.025 | -0.025 | 0 | 0.11 | 0.12 | 0.14 | 0.33 | 0.05 | 0.11 |

| | | | | | | | | | | | | | | | | |
|---|---|---|---|---|---|---|---|---|---|---|---|---|---|---|---|---|
| 6 | 28264692 | rs3800325 | -1.031 | -0.656 | 1.98 | 1.306 | -0.087 | -0.052 | -0.052 | 0 | 0.23 | 0.47 | 0.18 | 0.14 | 0.14 | 0.29 |
| 6 | 28265640 | rs2743554 | -1.394 | -0.656 | 1.875 | 1.307 | -0.039 | 0.067 | 0.067 | 0 | 0.12 | 0.31 | 0.11 | 0.13 | 0.05 | 0.08 |
| 6 | 28266149 | rs2142730 | -2.764 | -0.534 | 1.816 | 1.3 | 0.055 | -0.009 | -0.009 | 0 | 0.46 | 0.93 | 0.49 | 0.6 | 0.38 | 0.5 |
| 6 | 28269407 | rs16893917 | -0.865 | -0.566 | 1.81 | 1.313 | 0.023 | -0.025 | -0.025 | 1.267 | 0.11 | 0.12 | 0.14 | 0.33 | 0.05 | 0.11 |
| 6 | 28269663 | rs1997660 | -2.727 | -0.571 | 1.792 | 1.305 | 0.126 | -0.028 | -0.028 | 0 | 0.46 | 0.95 | 0.46 | 0.6 | 0.31 | 0.48 |
| 6 | 28270251 | rs1062169 | -1.023 | -0.574 | 1.851 | 1.303 | -0.087 | -0.052 | -0.052 | 0 | 0.23 | 0.47 | 0.18 | 0.14 | 0.14 | 0.29 |
| 6 | 28271198 | rs9468333 | -2.716 | -0.569 | 1.829 | 1.301 | 0.126 | -0.028 | -0.028 | 0 | 0.46 | 0.95 | 0.46 | 0.6 | 0.31 | 0.48 |
| 6 | 28271602 | rs1339899 | -0.972 | -0.562 | 1.872 | 1.302 | -0.031 | 0.034 | 0.034 | 0 | 0.09 | 0.21 | 0.1 | 0.13 | 0.04 | 0.08 |
| 6 | 28282648 | rs853693 | -2.708 | -0.566 | 1.712 | 1.297 | 0.126 | -0.028 | -0.028 | 0 | 0.46 | 0.95 | 0.46 | 0.6 | 0.31 | 0.48 |
| 6 | 28283772 | rs4713159 | -0.857 | -0.576 | 1.734 | 1.306 | 0.023 | -0.025 | -0.025 | 0.252 | 0.11 | 0.12 | 0.14 | 0.33 | 0.05 | 0.11 |
| 6 | 28284721 | rs16893947 | -1.851 | -0.575 | 1.784 | 1.303 | -0.056 | 0.016 | 0.016 | 0 | 0.31 | 0.47 | 0.31 | 0.42 | 0.18 | 0.37 |
| 6 | 28285482 | rs853690 | -2.732 | -0.57 | 1.75 | 1.3 | 0.126 | -0.028 | -0.028 | 0.517 | 0.46 | 0.95 | 0.46 | 0.6 | 0.31 | 0.48 |
| 6 | 28288785 | rs853685 | -1.351 | -0.556 | 1.782 | 1.3 | 0.046 | 0.084 | 0.084 | 0 | 0.12 | 0.36 | 0.14 | 0.13 | 0.12 | 0.08 |
| 6 | 28291240 | rs707907 | -2.726 | -0.568 | 1.738 | 1.299 | 0.126 | -0.028 | -0.028 | 0 | 0.46 | 0.95 | 0.46 | 0.6 | 0.31 | 0.48 |
| 6 | 28292706 | rs9301 | -2.782 | -0.532 | 1.737 | 1.298 | 0.041 | -0.025 | -0.025 | 0 | 0.46 | 0.95 | 0.49 | 0.61 | 0.38 | 0.5 |
| 6 | 28294550 | rs853684 | -2.782 | -0.532 | 1.737 | 1.298 | 0.041 | -0.025 | -0.025 | 2.844 | 0.46 | 0.95 | 0.49 | 0.6 | 0.38 | 0.5 |
| 6 | 28294888 | rs4711167 | -0.868 | -0.688 | 1.798 | 1.319 | 0.015 | -0.014 | -0.014 | 0 | 0.11 | 0.12 | 0.14 | 0.33 | 0.05 | 0.11 |
| 6 | 28296863 | rs853679 | -1.348 | -0.654 | 1.808 | 1.29 | 0.047 | 0.085 | 0.085 | 0 | 0.12 | 0.36 | 0.14 | 0.13 | 0.12 | 0.08 |
| 6 | 28299295 | rs16893970 | -1.825 | -0.687 | 1.779 | 1.287 | -0.06 | 0.015 | 0.015 | 0.001 | 0.32 | 0.49 | 0.31 | 0.42 | 0.18 | 0.37 |
| 6 | 28299687 | rs853676 | -1.312 | -0.565 | 1.875 | 1.256 | 0.051 | 0.09 | 0.09 | 0 | 0.11 | 0.34 | 0.14 | 0.13 | 0.12 | 0.08 |
| 6 | 28300336 | rs17312661 | 0.188 | -0.509 | 1.75 | 1.158 | 0.01 | -0.047 | -0.047 | 0 | 0.12 | 0.21 | 0.12 | 0.03 | 0.16 | 0.13 |
| 6 | 28301143 | rs7772827 | -2.174 | -0.582 | 1.74 | 1.234 | -0.028 | 0.012 | 0.012 | 0.022 | 0.34 | 0.62 | 0.36 | 0.43 | 0.26 | 0.38 |
| 6 | 28302113 | rs16893973 | -0.982 | -0.6 | 1.755 | 1.253 | 0.006 | -0.004 | -0.004 | 0 | 0.12 | 0.17 | 0.15 | 0.33 | 0.05 | 0.11 |
| 6 | 28302278 | rs16893975 | -0.982 | -0.6 | 1.755 | 1.253 | 0.006 | -0.004 | -0.004 | 1.66 | 0.12 | 0.17 | 0.15 | 0.33 | 0.05 | 0.11 |
| 6 | 28302784 | rs1416920 | -2.279 | -0.614 | 1.819 | 1.253 | 0.055 | -0.034 | -0.034 | 0 | 0.37 | 0.79 | 0.37 | 0.48 | 0.27 | 0.41 |
| 6 | 28302884 | rs1416919 | -1.003 | -0.615 | 1.816 | 1.254 | 0.006 | -0.005 | -0.005 | 2.864 | 0.12 | 0.17 | 0.15 | 0.33 | 0.05 | 0.11 |
| 6 | 28305104 | rs723476 | -0.975 | -0.608 | 1.835 | 1.258 | 0.006 | -0.004 | -0.004 | 0.35 | 0.12 | 0.17 | 0.15 | 0.33 | 0.05 | 0.11 |
| 6 | 28308671 | rs13210258 | -1.301 | -0.6 | 1.806 | 1.248 | -0.008 | -0.013 | -0.013 | 0.019 | 0.25 | 0.62 | 0.22 | 0.16 | 0.22 | 0.3 |
| 6 | 28309569 | rs9468344 | -2.268 | -0.605 | 1.795 | 1.255 | 0.055 | -0.034 | -0.034 | 0 | 0.37 | 0.79 | 0.37 | 0.48 | 0.27 | 0.41 |
| 6 | 28309590 | rs6912584 | -1.005 | -0.606 | 1.828 | 1.253 | -0.017 | -0.07 | -0.07 | 0 | 0.22 | 0.45 | 0.21 | 0.1 | 0.21 | 0.27 |
| 6 | 28309753 | rs10484542 | -0.982 | -0.607 | 1.8 | 1.267 | 0.006 | -0.004 | -0.004 | 0 | 0.12 | 0.17 | 0.15 | 0.33 | 0.05 | 0.11 |
| 6 | 28315875 | rs213240 | -2.712 | -0.439 | 1.453 | 1.233 | 0.037 | -0.037 | -0.037 | 0 | 0.46 | 0.96 | 0.48 | 0.61 | 0.38 | 0.51 |
| 6 | 28316478 | rs12180820 | -2.23 | -0.482 | 1.524 | 1.207 | 0.057 | -0.033 | -0.033 | 0.268 | 0.37 | 0.79 | 0.36 | 0.48 | 0.27 | 0.41 |
| 6 | 28321544 | rs1474589 | -0.726 | -0.312 | 1.544 | 1.058 | -0.043 | -0.05 | -0.05 | 0 | 0.19 | 0.32 | 0.18 | 0.09 | 0.13 | 0.25 |
| 6 | 28325308 | rs6922111 | -0.73 | -0.258 | 1.558 | 1.058 | -0.003 | -0.06 | -0.06 | 0 | 0.19 | 0.36 | 0.2 | 0.1 | 0.2 | 0.25 |
| 6 | 28330264 | rs213230 | -1.449 | -0.244 | 1.599 | 1.124 | -0.023 | -0.045 | -0.045 | 0 | 0.27 | 0.46 | 0.24 | 0.16 | 0.24 | 0.32 |
| 6 | 28331252 | rs213228 | -2.316 | -0.332 | 1.655 | 1.248 | 0.036 | 0.008 | 0.008 | 0.058 | 0.38 | 0.71 | 0.35 | 0.31 | 0.3 | 0.4 |
| 6 | 28333396 | rs17313059 | -0.544 | -0.167 | 1.598 | 1.219 | 0.01 | -0.008 | -0.008 | 0 | 0.09 | 0.08 | 0.07 | 0.13 | 0.05 | 0.17 |
| 6 | 28337769 | rs12661831 | 0.865 | -0.214 | 1.044 | 0.71 | -0.034 | 0.057 | 0.057 | 0.601 | 0.11 | 0 | 0.05 | 0.01 | 0.11 | 0.05 |
| 6 | 28342921 | rs6916243 | 1.037 | -0.039 | 1.547 | 1.096 | -0.024 | -0.006 | -0.006 | 0 | 0.93 | 0.87 | 0.97 | 0.99 | 0.98 | 0.98 |
| 6 | 28346769 | rs7774788 | 0.651 | 0.03 | 1.55 | 1.105 | 0.013 | -0.006 | -0.006 | 0 | 0.89 | 0.9 | 0.92 | 0.86 | 0.95 | 0.82 |
| 6 | 28346910 | rs7774981 | -1.134 | -0.031 | 1.602 | 1.115 | -0.012 | -0.035 | -0.035 | 0 | 0.23 | 0.48 | 0.25 | 0.16 | 0.24 | 0.21 |
| 6 | 28348158 | rs1052215 | 2.572 | -0.05 | 1.548 | 1.243 | 0.025 | -0.048 | -0.048 | 0 | 0.54 | 0.04 | 0.51 | 0.41 | 0.61 | 0.51 |
| 6 | 28348660 | rs2072319 | -0.411 | 0.085 | 1.576 | 1.227 | 0.01 | -0.008 | -0.008 | 2.149 | 0.09 | 0.08 | 0.08 | 0.14 | 0.05 | 0.18 |
| 6 | 28349725 | rs3734563 | -2.037 | 0.066 | 1.577 | 1.227 | -0.022 | -0.008 | -0.008 | 0 | 0.33 | 0.56 | 0.33 | 0.3 | 0.29 | 0.39 |
| 6 | 28349926 | rs1556957 | -0.416 | 0.07 | 1.55 | 1.229 | 0.01 | -0.008 | -0.008 | 0.586 | 0.09 | 0.08 | 0.08 | 0.14 | 0.05 | 0.18 |
| 6 | 28354250 | rs3799499 | -1.125 | 0.072 | 1.577 | 1.223 | -0.012 | -0.035 | -0.035 | 0 | 0.23 | 0.48 | 0.25 | 0.16 | 0.24 | 0.21 |
| 6 | 28358195 | rs2859379 | -1.088 | 0.044 | 1.753 | 1.211 | -0.023 | -0.014 | -0.014 | 0 | 0.1 | 0.29 | 0.09 | 0.02 | 0.06 | 0.03 |
| 6 | 28358320 | rs1361385 | -2.081 | 0.074 | 1.683 | 1.224 | -0.022 | -0.008 | -0.008 | 0 | 0.33 | 0.56 | 0.33 | 0.3 | 0.29 | 0.39 |
| 6 | 28358893 | rs2232434 | -0.395 | 0.074 | 1.662 | 1.225 | 0.01 | -0.008 | -0.008 | 0.304 | 0.09 | 0.08 | 0.08 | 0.14 | 0.05 | 0.18 |
| 6 | 28359700 | rs2232428 | -0.227 | 0.054 | 1.721 | 1.17 | -0.037 | -0.07 | -0.07 | 0 | 0.17 | 0.32 | 0.15 | 0.03 | 0.13 | 0.15 |
| 6 | 28359709 | rs2232427 | -1.189 | 0.069 | 1.652 | 1.182 | -0.013 | -0.036 | -0.036 | 0.167 | 0.24 | 0.48 | 0.25 | 0.16 | 0.24 | 0.21 |
| 6 | 28362471 | rs16894060 | -1.984 | 0.053 | 1.738 | 1.191 | -0.027 | -0.008 | -0.008 | 0 | 0.32 | 0.48 | 0.32 | 0.3 | 0.29 | 0.39 |
| 6 | 28366262 | rs2232422 | 0.286 | 0.049 | 1.623 | 1.186 | -0.014 | -0.029 | -0.029 | 2.377 | 0.13 | 0.15 | 0.09 | 0.03 | 0.09 | 0.14 |
| 6 | 28367683 | rs4357130 | -2.039 | 0.019 | 1.611 | 1.189 | -0.022 | -0.008 | -0.008 | 0 | 0.33 | 0.56 | 0.33 | 0.3 | 0.29 | 0.39 |
| 6 | 28370246 | rs6907950 | -2.029 | 0.018 | 1.614 | 1.172 | -0.022 | -0.007 | -0.007 | 0 | 0.33 | 0.56 | 0.33 | 0.3 | 0.29 | 0.38 |
| 6 | 28370393 | rs6908137 | -2.044 | 0.018 | 1.616 | 1.171 | -0.023 | -0.008 | -0.008 | 0.721 | 0.33 | 0.57 | 0.34 | 0.3 | 0.29 | 0.38 |
| 6 | 28371402 | rs4254981 | -1.891 | 0.025 | 1.621 | 1.175 | -0.026 | -0.006 | -0.006 | 0 | 0.31 | 0.53 | 0.32 | 0.3 | 0.29 | 0.38 |
| 6 | 28380248 | rs1124131 | -1.889 | 0.014 | 1.618 | 1.17 | -0.026 | -0.006 | -0.006 | 0 | 0.31 | 0.53 | 0.32 | 0.3 | 0.29 | 0.39 |
| 6 | 28381140 | rs6899389 | -1.889 | 0.014 | 1.618 | 1.17 | -0.026 | -0.006 | -0.006 | 0 | 0.31 | 0.53 | 0.32 | 0.3 | 0.29 | 0.38 |
| 6 | 28382262 | rs1558205 | -2.367 | 0.005 | 1.664 | 1.173 | 0.02 | 0.005 | 0.005 | 0 | 0.38 | 0.67 | 0.35 | 0.31 | 0.3 | 0.4 |
| 6 | 28382932 | rs6922169 | -1.88 | -0.004 | 1.651 | 1.174 | -0.026 | -0.006 | -0.006 | 0 | 0.31 | 0.53 | 0.32 | 0.3 | 0.29 | 0.38 |
| 6 | 28383038 | rs17314991 | -0.374 | -0.007 | 1.628 | 1.179 | 0.01 | -0.008 | -0.008 | 0 | 0.09 | 0.08 | 0.08 | 0.15 | 0.05 | 0.18 |
| 6 | 28387443 | rs10484540 | -0.301 | -0.007 | 1.616 | 1.179 | 0.025 | -0.026 | -0.026 | 0 | 0.09 | 0.04 | 0.08 | 0.15 | 0.05 | 0.18 |
| 6 | 28390843 | rs3800328 | -1.889 | -0.012 | 1.63 | 1.171 | -0.026 | -0.006 | -0.006 | 0.175 | 0.32 | 0.52 | 0.32 | 0.3 | 0.29 | 0.38 |
| 6 | 28391465 | rs2859365 | -2.584 | 0.021 | 1.627 | 1.228 | 0.031 | -0.021 | -0.021 | 0 | 0.42 | 0.77 | 0.47 | 0.54 | 0.38 | 0.46 |
| 6 | 28395506 | rs1339898 | -2.366 | -0.008 | 1.628 | 1.178 | 0.018 | 0.005 | 0.005 | 0 | 0.38 | 0.66 | 0.35 | 0.31 | 0.3 | 0.4 |
| 6 | 28399412 | rs7740351 | -1.894 | 0.014 | 1.596 | 1.178 | -0.023 | -0.005 | -0.005 | 0 | 0.32 | 0.54 | 0.32 | 0.3 | 0.29 | 0.38 |
| 6 | 28400339 | rs16894106 | -1.136 | 0.021 | 1.598 | 1.164 | -0.008 | -0.033 | -0.033 | 0.22 | 0.23 | 0.45 | 0.25 | 0.15 | 0.24 | 0.21 |

| | | | | | | | | | | | | | | | | |
|---|---|---|---|---|---|---|---|---|---|---|---|---|---|---|---|---|
| 6 | 28400538 | rs7766356 | 0.248 | 0.102 | 1.554 | 1.208 | 0.003 | -0.027 | -0.027 | 0.137 | 0.13 | 0.16 | 0.11 | 0.03 | 0.16 | 0.14 |
| 6 | 28403225 | rs2071965 | -1.798 | 0.096 | 1.476 | 1.23 | -0.023 | -0.005 | -0.005 | 0.718 | 0.31 | 0.54 | 0.32 | 0.3 | 0.29 | 0.38 |
| 6 | 28404353 | rs719370 | -0.38 | 0.01 | 1.51 | 1.227 | -0.019 | 0 | 0 | 0 | 0.06 | 0.14 | 0.03 | 0.01 | 0.02 | 0.02 |
| 6 | 28408258 | rs916403 | -2.532 | 0.958 | 1.454 | 1.273 | -0.012 | -0.02 | -0.02 | 0 | 0.43 | 0.77 | 0.49 | 0.54 | 0.45 | 0.46 |
| 6 | 28411244 | rs13190937 | -1.01 | 0.942 | 1.495 | 1.221 | 0.048 | -0.032 | -0.032 | 0.248 | 0.23 | 0.43 | 0.27 | 0.15 | 0.31 | 0.21 |
| 6 | 28411303 | rs2531804 | -2.285 | 0.955 | 1.453 | 1.224 | 0.003 | 0.007 | 0.007 | 0 | 0.38 | 0.66 | 0.38 | 0.31 | 0.37 | 0.41 |
| 6 | 28414166 | rs2859367 | -0.189 | 0.948 | 1.46 | 1.221 | -0.018 | -0.003 | -0.003 | 0 | 0.06 | 0.12 | 0.03 | 0.01 | 0.02 | 0.02 |
| 6 | 28417222 | rs6903535 | -1.832 | 0.966 | 1.456 | 1.223 | 0.008 | -0.009 | -0.009 | 0 | 0.32 | 0.52 | 0.35 | 0.31 | 0.35 | 0.38 |
| 6 | 28423603 | rs2531811 | -1.75 | 0.875 | 1.618 | 1.426 | 0.006 | 0.157 | 0.157 | 0 | 0.13 | 0.43 | 0.16 | 0.29 | 0.1 | 0.08 |
| 6 | 28425755 | rs2531814 | -2.387 | 0.807 | 1.585 | 1.413 | -0.051 | 0.116 | 0.116 | 0 | 0.23 | 0.48 | 0.25 | 0.44 | 0.15 | 0.28 |
| 6 | 28426986 | rs2021745 | -2.398 | 0.765 | 1.576 | 1.418 | -0.048 | 0.112 | 0.112 | 0 | 0.23 | 0.53 | 0.25 | 0.44 | 0.15 | 0.28 |
| 6 | 28427130 | rs2859372 | -2.398 | 0.765 | 1.576 | 1.418 | -0.048 | 0.112 | 0.112 | 0 | 0.23 | 0.53 | 0.25 | 0.44 | 0.15 | 0.28 |
| 6 | 28428145 | rs2859374 | -2.398 | 0.765 | 1.576 | 1.418 | -0.048 | 0.112 | 0.112 | 0 | 0.23 | 0.53 | 0.25 | 0.44 | 0.15 | 0.28 |
| 6 | 28436060 | rs2531815 | -3.143 | 0.849 | 1.503 | 1.252 | 0.011 | -0.036 | -0.036 | 0 | 0.34 | 0.8 | 0.4 | 0.57 | 0.3 | 0.35 |
| 6 | 28441634 | rs12663002 | -1.433 | 0.876 | 1.474 | 1.309 | 0 | 0.011 | 0.011 | 0 | 0.15 | 0.19 | 0.21 | 0.38 | 0.13 | 0.24 |
| 6 | 28444165 | rs16894128 | -0.639 | 1.061 | 1.648 | 1.14 | 0.041 | 0.038 | 0.038 | 0 | 0.07 | 0.11 | 0.09 | 0.13 | 0.11 | 0.07 |
| 6 | 28447915 | rs1029328 | -1.355 | 1.044 | 1.735 | 1.14 | 0.071 | 0.099 | 0.099 | 0 | 0.1 | 0.29 | 0.11 | 0.18 | 0.12 | 0.1 |
| 6 | 28461221 | rs2859355 | -2.778 | 1.055 | 1.523 | 1.226 | -0.005 | -0.028 | -0.028 | 0 | 0.35 | 0.76 | 0.4 | 0.51 | 0.33 | 0.35 |
| 6 | 28463576 | rs2227228 | -2.778 | 1.055 | 1.523 | 1.226 | -0.005 | -0.028 | -0.028 | 0.772 | 0.35 | 0.76 | 0.4 | 0.51 | 0.33 | 0.35 |
| 6 | 28465355 | rs2859356 | -2.078 | 1.037 | 1.593 | 1.23 | -0.044 | 0.1 | 0.1 | 0 | 0.27 | 0.48 | 0.29 | 0.49 | 0.17 | 0.34 |
| 6 | 28466442 | rs12661782 | -0.467 | 1.018 | 1.59 | 1.241 | 0.03 | -0.087 | -0.087 | 0 | 0.14 | 0.03 | 0.19 | 0.42 | 0.09 | 0.25 |
| 6 | 28466988 | rs2191037 | -2.206 | 1.026 | 1.593 | 1.233 | -0.044 | 0.1 | 0.1 | 0 | 0.27 | 0.48 | 0.29 | 0.49 | 0.17 | 0.34 |
| 6 | 28471612 | rs9468385 | -1.309 | 1.015 | 1.605 | 1.238 | -0.043 | 0.085 | 0.085 | 0 | 0.2 | 0.29 | 0.23 | 0.48 | 0.1 | 0.31 |
| 6 | 28473325 | rs2071966 | -0.682 | 1.022 | 1.562 | 1.24 | 0.031 | -0.053 | -0.053 | 0 | 0.16 | 0.08 | 0.21 | 0.43 | 0.09 | 0.27 |
| 6 | 28480833 | rs434112 | -2.945 | 1.058 | 1.601 | 1.222 | -0.005 | -0.027 | -0.027 | 0 | 0.35 | 0.75 | 0.4 | 0.51 | 0.33 | 0.36 |
| 6 | 28483150 | rs722788 | -0.926 | 1.065 | 1.651 | 1.22 | 0.137 | -0.019 | -0.019 | 0.266 | 0.07 | 0.27 | 0.11 | 0.02 | 0.16 | 0.02 |
| 6 | 28483482 | rs406113 | -2.947 | 1.061 | 1.643 | 1.221 | -0.005 | -0.027 | -0.027 | 0.767 | 0.35 | 0.75 | 0.4 | 0.51 | 0.33 | 0.35 |
| 6 | 28486822 | rs1003359 | -2.511 | 1.065 | 1.639 | 1.226 | 0.006 | 0.024 | 0.024 | 0 | 0.31 | 0.66 | 0.39 | 0.51 | 0.31 | 0.35 |
| 6 | 28488034 | rs2078035 | -1.312 | 1.078 | 1.599 | 1.239 | -0.044 | 0.087 | 0.087 | 0 | 0.2 | 0.29 | 0.23 | 0.48 | 0.1 | 0.3 |
| 6 | 28491950 | rs377514 | -1.3 | 1.254 | 1.569 | 1.234 | -0.043 | 0.085 | 0.085 | 0 | 0.2 | 0.29 | 0.23 | 0.48 | 0.1 | 0.3 |
| 6 | 28494327 | rs445870 | -2.476 | 1.253 | 1.606 | 1.23 | 0.025 | 0.071 | 0.071 | 0 | 0.28 | 0.59 | 0.37 | 0.51 | 0.31 | 0.35 |
| 6 | 28500235 | rs769189 | -0.607 | 1.174 | 1.486 | 1.252 | 0.03 | -0.052 | -0.052 | 0.643 | 0.16 | 0.08 | 0.21 | 0.42 | 0.09 | 0.27 |
| 6 | 28500449 | rs380879 | -1.289 | 1.175 | 1.503 | 1.246 | -0.044 | 0.088 | 0.088 | 0 | 0.2 | 0.3 | 0.23 | 0.48 | 0.1 | 0.31 |
| 6 | 28501006 | rs448450 | -1.289 | 1.175 | 1.503 | 1.246 | -0.044 | 0.088 | 0.088 | 0 | 0.2 | 0.3 | 0.23 | 0.48 | 0.1 | 0.31 |
| 6 | 28502550 | rs451774 | -2.708 | 1.219 | 1.293 | 1.222 | 0.002 | -0.029 | -0.029 | 0.162 | 0.34 | 0.74 | 0.39 | 0.5 | 0.29 | 0.34 |
| 6 | 28502794 | rs13215054 | -0.105 | 0.35 | 1.189 | 1.008 | 0.002 | -0.001 | -0.001 | 0 | 0.17 | 0.18 | 0.18 | 0.15 | 0.22 | 0.21 |
| 6 | 28504639 | rs392230 | -1.263 | 0.169 | 1.334 | 1.218 | -0.007 | 0.024 | 0.024 | 0 | 0.21 | 0.24 | 0.23 | 0.52 | 0.15 | 0.32 |
| 6 | 28512017 | rs2394102 | 0.14 | 0.173 | 1.256 | 1.178 | -0.01 | -0.001 | -0.001 | 0 | 0.16 | 0.16 | 0.15 | 0.13 | 0.1 | 0.2 |
| 6 | 28514583 | rs4711173 | -1.235 | 0.328 | 1.168 | 1.24 | -0.003 | -0.042 | -0.042 | 0 | 0.2 | 0.17 | 0.27 | 0.47 | 0.2 | 0.3 |
| 6 | 28521316 | rs393414 | -1.748 | 0.353 | 1.375 | 1.217 | 0.005 | 0.128 | 0.128 | 0 | 0.25 | 0.48 | 0.31 | 0.53 | 0.24 | 0.33 |
| 6 | 28522695 | rs6456825 | -2.537 | 0.285 | 1.207 | 1.131 | 0.004 | 0.124 | 0.124 | 0.239 | 0.5 | 0.93 | 0.54 | 0.67 | 0.44 | 0.55 |
| 6 | 28526655 | rs414745 | -1.726 | 0.284 | 1.415 | 1.16 | -0.002 | -0.002 | -0.002 | 0 | 0.32 | 0.69 | 0.31 | 0.21 | 0.28 | 0.29 |
| 6 | 28532960 | rs4711174 | 0.201 | 0.243 | 1.448 | 1.311 | -0.008 | -0.042 | -0.042 | 0 | 0.1 | 0.03 | 0.16 | 0.36 | 0.11 | 0.13 |
| 6 | 28533946 | rs418092 | -1.712 | 0.269 | 1.466 | 1.198 | -0.002 | -0.002 | -0.002 | 0 | 0.32 | 0.69 | 0.31 | 0.21 | 0.28 | 0.29 |
| 6 | 28542424 | rs450630 | -2.672 | 0.263 | 1.452 | 1.193 | 0.004 | 0.129 | 0.129 | 0.239 | 0.5 | 0.93 | 0.55 | 0.67 | 0.44 | 0.56 |
| 6 | 28543264 | rs17336532 | 0.105 | 0.221 | 1.646 | 1.147 | -0.001 | -0.001 | -0.001 | 0 | 0.16 | 0.19 | 0.17 | 0.14 | 0.17 | 0.21 |
| 6 | 28546952 | rs16901848 | -1.125 | 0.241 | 1.603 | 1.153 | -0.035 | -0.024 | -0.024 | 0.21 | 0.26 | 0.44 | 0.23 | 0.15 | 0.2 | 0.23 |
| 6 | 28565827 | rs6901724 | -0.319 | 0.141 | 1.479 | 1.079 | -0.042 | -0.06 | -0.06 | 0.116 | 0.15 | 0.32 | 0.13 | 0.04 | 0.1 | 0.14 |
| 6 | 28568528 | rs418914 | -1.999 | 0.19 | 1.412 | 1.108 | -0.003 | -0.003 | -0.003 | 0 | 0.32 | 0.68 | 0.32 | 0.21 | 0.28 | 0.28 |
| 6 | 28569949 | rs380914 | -2.937 | 0.158 | 1.255 | 1.139 | 0.009 | 0.127 | 0.127 | 0.332 | 0.51 | 0.93 | 0.56 | 0.67 | 0.44 | 0.55 |
| 6 | 28573006 | rs442439 | -2.236 | 0.159 | 1.249 | 1.129 | 0.008 | 0.034 | 0.034 | 0.753 | 0.35 | 0.89 | 0.35 | 0.22 | 0.3 | 0.3 |
| 6 | 28574415 | rs911178 | 0.446 | 0.197 | 1.315 | 1.149 | 0.049 | 0.023 | 0.023 | 0 | 0.94 | 0.76 | 0.92 | 0.94 | 0.92 | 0.95 |
| 6 | 28579471 | rs9501180 | 2.69 | 0.167 | 1.175 | 1.157 | 0.006 | 0.126 | 0.126 | 0.332 | 0.49 | 0.07 | 0.44 | 0.33 | 0.56 | 0.45 |
| 6 | 28584775 | rs34556183 | -1.538 | 0.172 | 1.312 | 1.15 | -0.008 | -0.014 | -0.014 | 0 | 0.26 | 0.41 | 0.23 | 0.15 | 0.21 | 0.2 |
| 6 | 28600492 | rs7742658 | 0.127 | 0.118 | 1.342 | 1.174 | -0.02 | -0.01 | -0.01 | 0 | 0.07 | 0.17 | 0.04 | 0.01 | 0.02 | 0 |
| 6 | 28602272 | rs16894171 | -0.814 | 0.194 | 1.019 | 1.293 | 0.005 | -0.015 | -0.015 | 0 | 0.18 | 0.14 | 0.26 | 0.46 | 0.18 | 0.31 |
| 6 | 28616327 | rs732395 | 1.54 | 0.134 | 1.178 | 1.27 | -0.038 | 0.144 | 0.144 | 0 | 0.76 | 0.5 | 0.67 | 0.49 | 0.78 | 0.66 |
| 6 | 28627469 | rs9295775 | -1.234 | 0.136 | 1.174 | 1.27 | -0.023 | 0.108 | 0.108 | 0 | 0.21 | 0.34 | 0.3 | 0.46 | 0.2 | 0.31 |
| 6 | 28630793 | rs7767008 | -1.437 | 0.285 | 1.044 | 1.122 | -0.005 | -0.017 | -0.017 | 2.629 | 0.26 | 0.43 | 0.24 | 0.15 | 0.22 | 0.22 |
| 6 | 28631731 | rs16894184 | 0.186 | 0.373 | 0.769 | 1.134 | -0.012 | -0.023 | -0.023 | 0.369 | 0.08 | 0.02 | 0.11 | 0.11 | 0.08 | 0.1 |
| 6 | 28641957 | rs3864294 | 2.367 | 0.041 | 0.042 | 0.436 | -0.143 | 0.152 | 0.152 | 0.905 | 0.29 | 0.02 | 0.18 | 0.05 | 0.4 | 0.16 |
| 6 | 28645170 | rs9393921 | 2.137 | 0.19 | 0.583 | 0.966 | 0.002 | 0.144 | 0.144 | 0.469 | 0.5 | 0.07 | 0.43 | 0.32 | 0.55 | 0.43 |
| 6 | 28650353 | rs9885928 | 2.154 | 0.333 | 0.585 | 0.969 | -0.015 | 0.14 | 0.14 | 0.434 | 0.49 | 0.07 | 0.44 | 0.32 | 0.55 | 0.43 |
| 6 | 28662521 | rs1319075 | 1.997 | 0.311 | 0.721 | 0.967 | 0.026 | 0.073 | 0.073 | 0.519 | 0.52 | 0.26 | 0.47 | 0.34 | 0.57 | 0.43 |
| 6 | 28664631 | rs4143771 | -1.916 | 0.333 | 0.783 | 1.068 | -0.003 | -0.009 | -0.009 | 0 | 0.44 | 0.58 | 0.49 | 0.62 | 0.41 | 0.53 |
| 6 | 28665109 | rs7382112 | -1.858 | 0.335 | 0.788 | 1.007 | -0.004 | -0.01 | -0.01 | 0 | 0.44 | 0.56 | 0.49 | 0.61 | 0.41 | 0.53 |
| 6 | 28668480 | rs7383248 | -1.915 | 0.144 | 0.651 | 1.05 | 0 | 0 | 0 | 0 | 0.44 | 0.6 | 0.5 | 0.61 | 0.41 | 0.53 |
| 6 | 28669315 | rs6939576 | -1.735 | -0.121 | 0.746 | 1.11 | -0.022 | -0.006 | -0.006 | 0 | 0.34 | 0.52 | 0.29 | 0.29 | 0.25 | 0.37 |
| 6 | 28671422 | rs1107160 | -0.379 | -0.231 | 0.433 | 0.929 | 0.04 | 0.003 | 0.003 | 0 | 0.13 | 0.04 | 0.05 | 0.08 | 0.05 | 0.16 |

| | | | | | | | | | | | | | | | | |
|---|---|---|---|---|---|---|---|---|---|---|---|---|---|---|---|---|
| 6 | 28671723 | rs1107159 | -0.379 | -0.208 | 0.433 | 0.953 | 0.034 | -0.002 | -0.002 | 0 | 0.13 | 0.04 | 0.06 | 0.08 | 0.06 | 0.16 |
| 6 | 28671898 | rs2893974 | -0.378 | -0.208 | 0.433 | 0.933 | 0.034 | 0 | 0 | 0 | 0.13 | 0.04 | 0.06 | 0.08 | 0.06 | 0.16 |
| 6 | 28675630 | rs6901325 | -0.393 | -0.215 | 0.429 | 0.929 | 0.035 | 0 | 0 | 0 | 0.14 | 0.04 | 0.06 | 0.08 | 0.06 | 0.16 |
| 6 | 28679945 | rs1539584 | -0.142 | -0.392 | 0.259 | 0.832 | 0.003 | -0.012 | -0.012 | 0 | 0.22 | 0.21 | 0.31 | 0.45 | 0.22 | 0.24 |
| 6 | 28688921 | rs6909960 | -0.189 | -0.389 | 0.348 | 0.786 | 0.006 | 0.002 | 0.002 | 0 | 0.14 | 0.09 | 0.06 | 0.08 | 0.06 | 0.16 |
| 6 | 28689672 | rs9468413 | 0.635 | -0.436 | 0.818 | 0.671 | 0.016 | -0.074 | -0.074 | 0 | 0.83 | 0.49 | 0.84 | 0.98 | 0.84 | 0.91 |
| 6 | 28695668 | rs7772682 | -0.145 | -0.36 | 0.727 | 0.714 | 0.006 | 0.002 | 0.002 | 0 | 0.14 | 0.09 | 0.06 | 0.08 | 0.06 | 0.16 |
| 6 | 28697086 | rs2893973 | -0.26 | -0.303 | 0.952 | 0.766 | 0 | 0.008 | 0.008 | 3.522 | 0.79 | 0.8 | 0.74 | 0.56 | 0.81 | 0.8 |
| 6 | 28697625 | rs17280797 | -0.111 | -0.393 | 0.905 | 0.729 | 0.005 | 0.003 | 0.003 | 0 | 0.14 | 0.09 | 0.05 | 0.08 | 0.05 | 0.16 |
| 6 | 28699691 | rs10498733 | -0.108 | -0.396 | 0.905 | 0.729 | 0.004 | 0.003 | 0.003 | 0 | 0.14 | 0.09 | 0.05 | 0.08 | 0.04 | 0.16 |
| 6 | 28703281 | rs2893970 | 0.061 | -0.419 | 0.966 | 0.734 | 0.012 | -0.05 | -0.05 | 0 | 0.21 | 0.16 | 0.25 | 0.44 | 0.18 | 0.2 |
| 6 | 28703432 | rs7766599 | -0.125 | -0.353 | 0.819 | 0.737 | 0.005 | 0.003 | 0.003 | 0 | 0.14 | 0.09 | 0.05 | 0.08 | 0.05 | 0.16 |
| 6 | 28706912 | rs880638 | -1.874 | -0.352 | 0.9 | 0.701 | 0.056 | 0.049 | 0.049 | 0.275 | 0.51 | 0.76 | 0.47 | 0.55 | 0.39 | 0.45 |
| 6 | 28711074 | rs11754985 | -0.319 | -0.342 | 0.881 | 0.612 | -0.01 | -0.036 | -0.036 | 0 | 0.07 | 0.18 | 0.04 | 0 | 0.03 | 0.05 |
| 6 | 28711334 | rs3905240 | -0.119 | -0.296 | 0.582 | 0.673 | 0.005 | 0.003 | 0.003 | 0 | 0.14 | 0.09 | 0.05 | 0.08 | 0.05 | 0.16 |
| 6 | 28716345 | rs1233585 | -0.215 | -0.307 | 0.604 | 0.649 | 0.006 | -0.006 | -0.006 | 0 | 0.14 | 0.18 | 0.09 | 0.08 | 0.13 | 0.16 |
| 6 | 28720982 | rs1233591 | -2.433 | -0.291 | 0.405 | 0.648 | -0.101 | 0.07 | 0.07 | 0.815 | 0.59 | 0.92 | 0.6 | 0.64 | 0.49 | 0.58 |
| 6 | 28722799 | rs17338569 | -0.619 | -0.283 | 0.465 | 0.689 | 0.002 | -0.006 | -0.006 | 0.031 | 0.17 | 0.11 | 0.15 | 0.17 | 0.13 | 0.24 |
| 6 | 28727673 | rs1233596 | -2.396 | -0.305 | 0.395 | 0.623 | -0.101 | 0.07 | 0.07 | 0.705 | 0.59 | 0.92 | 0.6 | 0.64 | 0.49 | 0.58 |
| 6 | 28732364 | rs3852213 | -0.042 | -0.31 | 0.586 | 0.694 | 0.004 | 0.001 | 0.001 | 0 | 0.13 | 0.09 | 0.06 | 0.08 | 0.06 | 0.16 |
| 6 | 28733403 | rs1233602 | -2.35 | 0.154 | 0.487 | 0.641 | -0.086 | 0.08 | 0.08 | 0.161 | 0.57 | 0.92 | 0.6 | 0.64 | 0.5 | 0.58 |
| 6 | 28749012 | rs7758798 | -0.034 | 0.163 | 0.624 | 0.66 | 0.004 | 0.001 | 0.001 | 0 | 0.13 | 0.09 | 0.06 | 0.08 | 0.06 | 0.16 |
| 6 | 28751727 | rs1233627 | -2.363 | 0.14 | 0.574 | 0.641 | -0.082 | 0.083 | 0.083 | 0.166 | 0.57 | 0.92 | 0.6 | 0.64 | 0.5 | 0.57 |
| 6 | 28752883 | rs7751451 | -0.021 | 0.158 | 0.588 | 0.639 | 0.004 | 0.001 | 0.001 | 0 | 0.13 | 0.09 | 0.06 | 0.08 | 0.06 | 0.16 |
| 6 | 28754520 | rs9257181 | -0.464 | 0.053 | 0.965 | 0.649 | 0.004 | 0.011 | 0.011 | 0.321 | 0.25 | 0.28 | 0.42 | 0.54 | 0.32 | 0.32 |
| 6 | 28754704 | rs11963531 | -0.064 | 0.065 | 1.075 | 0.58 | -0.035 | -0.014 | -0.014 | 0 | 0.08 | 0.22 | 0.04 | 0.01 | 0.02 | 0.03 |
| 6 | 28769671 | rs2765220 | 0.322 | -0.004 | 0.944 | 0.546 | 0.023 | -0.034 | -0.034 | 0 | 0.11 | 0.17 | 0.07 | 0.03 | 0.12 | 0.1 |
| 6 | 28770349 | rs2765219 | -2.558 | -0.176 | 0.801 | 0.47 | -0.09 | 0.07 | 0.07 | 0.571 | 0.58 | 0.93 | 0.6 | 0.65 | 0.5 | 0.57 |
| 6 | 28774593 | rs9404976 | -0.322 | -0.158 | 0.893 | 0.52 | 0.001 | -0.018 | -0.018 | 0 | 0.25 | 0.23 | 0.33 | 0.51 | 0.25 | 0.29 |
| 6 | 28775241 | rs2754767 | -2.514 | -0.11 | 0.772 | 0.509 | -0.09 | 0.07 | 0.07 | 0.571 | 0.58 | 0.93 | 0.6 | 0.65 | 0.5 | 0.57 |
| 6 | 28784031 | rs398795 | 0.492 | -0.146 | 0.796 | 0.511 | -0.004 | -0.005 | -0.005 | 0 | 0.1 | 0.08 | 0.05 | 0.03 | 0.04 | 0.1 |
| 6 | 28784396 | rs386628 | -2.531 | -0.142 | 0.781 | 0.796 | -0.083 | 0.017 | 0.017 | 0.612 | 0.58 | 0.92 | 0.6 | 0.6 | 0.5 | 0.57 |
| 6 | 28784741 | rs377392 | -2.531 | -0.142 | 0.781 | 0.821 | -0.083 | 0.013 | 0.013 | 0.612 | 0.58 | 0.92 | 0.6 | 0.6 | 0.5 | 0.57 |
| 6 | 28786884 | rs172330 | 0.498 | -0.137 | 0.798 | 0.797 | -0.004 | -0.005 | -0.005 | 0.318 | 0.1 | 0.08 | 0.05 | 0.03 | 0.04 | 0.1 |
| 6 | 28791289 | rs209184 | -2.55 | -0.18 | 0.746 | 0.774 | -0.085 | 0.014 | 0.014 | 0.573 | 0.58 | 0.92 | 0.6 | 0.6 | 0.5 | 0.61 |
| 6 | 28792477 | rs209181 | 0.42 | -0.122 | 0.853 | 0.482 | 0.022 | -0.032 | -0.032 | 0 | 0.1 | 0.17 | 0.07 | 0.02 | 0.11 | 0.1 |
| 6 | 28803103 | rs209177 | -1.06 | -0.179 | 0.941 | 0.507 | 0.006 | -0.008 | -0.008 | 0 | 0.35 | 0.3 | 0.37 | 0.53 | 0.29 | 0.41 |
| 6 | 28804186 | rs209176 | -2.661 | -0.2 | 0.913 | 0.483 | -0.086 | 0.067 | 0.067 | 0.64 | 0.58 | 0.92 | 0.59 | 0.65 | 0.5 | 0.62 |
| 6 | 28806318 | rs6904278 | -1.461 | -0.216 | 1.178 | 0.472 | -0.018 | -0.011 | -0.011 | 0 | 0.19 | 0.43 | 0.18 | 0.11 | 0.12 | 0.12 |
| 6 | 28810817 | rs422331 | -0.872 | -0.215 | 1.183 | 0.473 | 0.004 | 0.013 | 0.013 | 0.658 | 0.35 | 0.39 | 0.4 | 0.53 | 0.37 | 0.41 |
| 6 | 28815213 | rs526983 | -2.523 | -0.216 | 1.158 | 0.469 | -0.021 | 0.09 | 0.09 | 0.488 | 0.54 | 0.82 | 0.57 | 0.65 | 0.49 | 0.53 |
| 6 | 28824958 | rs9501112 | -0.613 | -0.24 | 1.24 | 0.466 | -0.057 | -0.033 | -0.033 | 0 | 0.11 | 0.31 | 0.06 | 0.01 | 0.02 | 0.03 |
| 6 | 28826262 | rs9257274 | -0.302 | -0.216 | 1.126 | 0.481 | 0.001 | -0.016 | -0.016 | 0 | 0.25 | 0.23 | 0.32 | 0.51 | 0.25 | 0.31 |
| 6 | 28826405 | rs176461 | 0.547 | -0.204 | 1.126 | 0.479 | -0.006 | -0.006 | -0.006 | 0 | 0.1 | 0.08 | 0.04 | 0.02 | 0.03 | 0.1 |
| 6 | 28833507 | rs9257280 | -0.775 | -0.208 | 1.061 | 0.503 | 0.007 | -0.026 | -0.026 | 2.134 | 0.29 | 0.25 | 0.34 | 0.51 | 0.27 | 0.31 |
| 6 | 28836456 | rs209164 | -2.328 | -0.174 | 0.96 | 0.48 | -0.021 | 0.09 | 0.09 | 0.447 | 0.54 | 0.82 | 0.57 | 0.65 | 0.49 | 0.54 |
| 6 | 28839879 | rs209160 | -2.751 | -0.298 | 0.762 | 0.411 | -0.16 | 0.017 | 0.017 | 0.681 | 0.6 | 0.97 | 0.6 | 0.65 | 0.51 | 0.63 |
| 6 | 28846249 | rs209153 | -2.626 | -0.262 | 0.777 | 0.43 | -0.086 | 0.068 | 0.068 | 0.592 | 0.58 | 0.92 | 0.6 | 0.65 | 0.5 | 0.62 |
| 6 | 28846462 | rs209152 | -2.328 | -0.2 | 0.806 | 0.469 | -0.021 | 0.09 | 0.09 | 0.447 | 0.54 | 0.82 | 0.57 | 0.64 | 0.49 | 0.53 |
| 6 | 28849790 | rs172326 | -1.141 | -0.193 | 0.838 | 0.51 | -0.001 | 0.001 | 0.001 | 0.298 | 0.38 | 0.4 | 0.4 | 0.53 | 0.37 | 0.41 |
| 6 | 28850382 | rs2006758 | -0.336 | -0.264 | 0.79 | 0.542 | 0.001 | -0.016 | -0.016 | 0 | 0.25 | 0.22 | 0.32 | 0.51 | 0.25 | 0.3 |
| 6 | 28851637 | rs9257319 | 0.171 | -0.262 | 0.746 | 0.603 | -0.006 | -0.069 | -0.069 | 0.028 | 0.19 | 0.1 | 0.24 | 0.45 | 0.18 | 0.19 |
| 6 | 28851908 | rs429369 | -1.107 | -0.265 | 0.829 | 0.518 | -0.004 | 0.001 | 0.001 | 0.214 | 0.38 | 0.39 | 0.38 | 0.53 | 0.3 | 0.41 |
| 6 | 28852425 | rs6930087 | -1.094 | -0.279 | 0.921 | 0.517 | 0.025 | 0.048 | 0.048 | 0 | 0.14 | 0.42 | 0.15 | 0.13 | 0.1 | 0.13 |
| 6 | 28853157 | rs7750338 | -1.107 | -0.262 | 0.798 | 0.554 | -0.004 | 0.001 | 0.001 | 0.214 | 0.38 | 0.39 | 0.38 | 0.53 | 0.3 | 0.41 |
| 6 | 28856572 | rs169679 | -2.515 | -0.194 | 0.569 | 0.456 | -0.086 | 0.071 | 0.071 | 0.574 | 0.58 | 0.92 | 0.6 | 0.67 | 0.5 | 0.62 |
| 6 | 28857269 | rs209148 | -2.515 | -0.194 | 0.569 | 0.456 | -0.086 | 0.071 | 0.071 | 0.574 | 0.58 | 0.92 | 0.6 | 0.67 | 0.5 | 0.62 |
| 6 | 28858850 | rs209146 | -1.088 | -0.268 | 0.755 | 0.436 | -0.003 | 0.001 | 0.001 | 0.184 | 0.38 | 0.39 | 0.37 | 0.53 | 0.3 | 0.4 |
| 6 | 28860499 | rs6456867 | 0.477 | -0.291 | 1 | 0.43 | 0.023 | 0.051 | 0.051 | 0 | 0.05 | 0.25 | 0.02 | 0.01 | 0.01 | 0.03 |
| 6 | 28863483 | rs209139 | 0.294 | -0.359 | 0.973 | 0.399 | 0.003 | 0.002 | 0.002 | 0.032 | 0.12 | 0.11 | 0.06 | 0.02 | 0.04 | 0.1 |
| 6 | 28867755 | rs209131 | -1.673 | -0.348 | 0.795 | 0.487 | 0.055 | 0.01 | 0.01 | 0 | 0.45 | 0.68 | 0.43 | 0.52 | 0.32 | 0.41 |
| 6 | 28867800 | rs209130 | -2.676 | -0.456 | 0.679 | 0.39 | -0.072 | 0.106 | 0.106 | 0.035 | 0.54 | 0.92 | 0.59 | 0.67 | 0.5 | 0.57 |
| 6 | 28869173 | rs209128 | -2.783 | -0.409 | 0.71 | 0.411 | -0.086 | 0.071 | 0.071 | 0.574 | 0.58 | 0.92 | 0.59 | 0.67 | 0.5 | 0.62 |
| 6 | 28870975 | rs1056032 | -0.609 | -0.408 | 0.746 | 0.415 | 0 | -0.001 | -0.001 | 0.005 | 0.07 | 0.12 | 0.12 | 0.12 | 0.09 | 0.09 |
| 6 | 28872446 | rs209126 | -2.803 | -0.416 | 0.769 | 0.408 | -0.09 | 0.069 | 0.069 | 0.577 | 0.59 | 0.92 | 0.59 | 0.67 | 0.5 | 0.62 |
| 6 | 28873060 | rs209123 | -2.787 | -0.414 | 0.766 | 0.408 | -0.077 | 0.073 | 0.073 | 0.574 | 0.58 | 0.91 | 0.59 | 0.67 | 0.5 | 0.62 |
| 6 | 28875502 | rs209122 | -2.787 | -0.414 | 0.766 | 0.408 | -0.077 | 0.073 | 0.073 | 0.574 | 0.58 | 0.91 | 0.59 | 0.67 | 0.5 | 0.62 |
| 6 | 28876509 | rs2269553 | -0.413 | -0.437 | 0.867 | 0.405 | 0.001 | -0.016 | -0.016 | 0 | 0.25 | 0.22 | 0.31 | 0.5 | 0.25 | 0.3 |
| 6 | 28881660 | rs209121 | -1.649 | -0.217 | 0.436 | 0.392 | 0.063 | 0.023 | 0.023 | 0 | 0.46 | 0.67 | 0.42 | 0.54 | 0.32 | 0.44 |

| | | | | | | | | | | | | | | | | |
|---|---|---|---|---|---|---|---|---|---|---|---|---|---|---|---|---|
| 6 | 28885974 | rs3132377 | -1.684 | -0.209 | 0.456 | 0.396 | 0.067 | 0.025 | 0.025 | 0 | 0.46 | 0.67 | 0.42 | 0.54 | 0.32 | 0.44 |
| 6 | 28890219 | rs9295782 | -0.619 | -0.195 | 0.453 | 0.396 | 0 | -0.001 | -0.001 | 0.019 | 0.07 | 0.12 | 0.12 | 0.12 | 0.09 | 0.09 |
| 6 | 28891458 | rs1061625 | -1.67 | -0.199 | 0.45 | 0.392 | 0.065 | 0.024 | 0.024 | 0 | 0.46 | 0.67 | 0.42 | 0.54 | 0.32 | 0.44 |
| 6 | 28893927 | rs2894066 | -1.754 | -0.197 | 0.466 | 0.393 | 0.032 | 0.028 | 0.028 | 0 | 0.47 | 0.71 | 0.45 | 0.54 | 0.39 | 0.45 |
| 6 | 28894311 | rs3763338 | -0.682 | -0.196 | 0.442 | 0.394 | 0 | -0.001 | -0.001 | 0 | 0.07 | 0.12 | 0.12 | 0.12 | 0.09 | 0.09 |
| 6 | 28894344 | rs1237485 | -2.71 | -0.197 | 0.442 | 0.416 | -0.088 | 0.078 | 0.078 | 0.455 | 0.58 | 0.92 | 0.59 | 0.67 | 0.5 | 0.62 |
| 6 | 28897633 | rs1233508 | -2.649 | -0.116 | 0.439 | 0.407 | -0.085 | 0.059 | 0.059 | 0.708 | 0.57 | 0.92 | 0.57 | 0.63 | 0.47 | 0.58 |
| 6 | 28901034 | rs932776 | -2.743 | -0.114 | 0.441 | 0.394 | -0.089 | 0.071 | 0.071 | 0.574 | 0.58 | 0.92 | 0.59 | 0.67 | 0.5 | 0.62 |
| 6 | 28901565 | rs2032502 | -0.501 | -0.114 | 0.442 | 0.394 | 0 | 0 | 0 | 0 | 0.07 | 0.12 | 0.12 | 0.12 | 0.09 | 0.09 |
| 6 | 28902133 | rs1004062 | -2.734 | -0.11 | 0.43 | 0.394 | -0.089 | 0.071 | 0.071 | 0.574 | 0.58 | 0.92 | 0.59 | 0.67 | 0.5 | 0.62 |
| 6 | 28904162 | rs6912843 | -1.715 | -0.112 | 0.441 | 0.402 | 0.069 | 0.026 | 0.026 | 0 | 0.47 | 0.68 | 0.42 | 0.54 | 0.32 | 0.45 |
| 6 | 28909465 | rs4713186 | -2.442 | -0.099 | 0.302 | 0.367 | -0.034 | 0.084 | 0.084 | 0.022 | 0.57 | 0.92 | 0.55 | 0.66 | 0.42 | 0.61 |
| 6 | 28909779 | rs7761745 | -1.375 | -0.119 | 0.414 | 0.363 | 0.005 | 0.002 | 0.002 | 0 | 0.12 | 0.24 | 0.14 | 0.12 | 0.1 | 0.17 |
| 6 | 28910235 | rs7762289 | -1.375 | -0.119 | 0.414 | 0.363 | 0.005 | 0.002 | 0.002 | 0 | 0.12 | 0.24 | 0.14 | 0.12 | 0.1 | 0.17 |
| 6 | 28911802 | rs2071790 | -2.41 | -0.068 | 0.311 | 0.378 | -0.086 | 0.073 | 0.073 | 0.561 | 0.58 | 0.92 | 0.58 | 0.66 | 0.5 | 0.62 |
| 6 | 28912076 | rs2071789 | 0.076 | -0.054 | 0.347 | 0.508 | -0.002 | -0.061 | -0.061 | 0 | 0.2 | 0.11 | 0.23 | 0.44 | 0.18 | 0.19 |
| 6 | 28915108 | rs763009 | -2.542 | -0.023 | 0.175 | 0.366 | -0.161 | 0.031 | 0.031 | 0.606 | 0.6 | 0.98 | 0.62 | 0.66 | 0.51 | 0.64 |
| 6 | 28916252 | rs4947339 | -2.355 | 0.051 | 0.262 | 0.367 | -0.049 | 0.072 | 0.072 | 0.474 | 0.58 | 0.92 | 0.56 | 0.66 | 0.43 | 0.62 |
| 6 | 28918936 | rs6456876 | -2.355 | 0.048 | 0.269 | 0.367 | -0.05 | 0.073 | 0.073 | 0.474 | 0.58 | 0.91 | 0.56 | 0.66 | 0.43 | 0.61 |
| 6 | 28921940 | rs4947256 | -0.496 | 0.029 | 0.253 | 0.409 | 0.001 | 0.001 | 0.001 | 0 | 0.07 | 0.12 | 0.12 | 0.12 | 0.09 | 0.09 |
| 6 | 28922719 | rs1476016 | -2.398 | 0.029 | 0.252 | 0.409 | -0.086 | 0.072 | 0.072 | 0.554 | 0.58 | 0.92 | 0.59 | 0.66 | 0.5 | 0.62 |
| 6 | 28923980 | rs9380100 | -0.584 | -0.04 | 0.302 | 0.452 | 0.005 | 0.005 | 0.005 | 0 | 0.07 | 0.13 | 0.12 | 0.12 | 0.09 | 0.08 |
| 6 | 28928167 | rs12386522 | 1.104 | -0.183 | 0.258 | 0.397 | 0.018 | 0.019 | 0.019 | 0 | 0.07 | 0.01 | 0.04 | 0.02 | 0.03 | 0.1 |
| 6 | 28936556 | rs9257425 | -1.512 | -0.177 | 0.349 | 0.438 | 0.044 | 0.003 | 0.003 | 0 | 0.43 | 0.64 | 0.39 | 0.52 | 0.31 | 0.42 |
| 6 | 28941688 | rs6933672 | -1.753 | -0.137 | 0.669 | 0.295 | -0.078 | -0.085 | -0.085 | 0.744 | 0.24 | 0.53 | 0.17 | 0.05 | 0.12 | 0.24 |
| 6 | 28949904 | rs968722 | -1.738 | 0.011 | 0.684 | 0.431 | -0.04 | -0.024 | -0.024 | 0 | 0.25 | 0.47 | 0.2 | 0.14 | 0.15 | 0.24 |
| 6 | 28952035 | rs6904975 | -2.465 | -0.242 | 0.312 | 0.386 | -0.083 | 0.036 | 0.036 | 1.003 | 0.59 | 0.94 | 0.56 | 0.65 | 0.43 | 0.59 |
| 6 | 28956416 | rs3129792 | -2.465 | -0.165 | 0.312 | 0.386 | -0.136 | 0.032 | 0.032 | 1.003 | 0.59 | 0.96 | 0.6 | 0.65 | 0.51 | 0.6 |
| 6 | 28961819 | rs16894540 | -0.385 | -0.135 | 0.419 | 0.645 | -0.032 | 0.059 | 0.059 | 0.766 | 0.24 | 0.36 | 0.29 | 0.4 | 0.19 | 0.21 |
| 6 | 28963248 | rs6456880 | -2.667 | -0.116 | 0.418 | 0.645 | -0.095 | 0.032 | 0.032 | 1.003 | 0.59 | 0.95 | 0.57 | 0.65 | 0.44 | 0.6 |
| 6 | 28964847 | rs2269555 | -2.763 | -0.362 | 0.243 | 0.493 | -0.11 | 0.026 | 0.026 | 1.827 | 0.61 | 0.95 | 0.57 | 0.66 | 0.44 | 0.61 |
| 6 | 28967038 | rs6920392 | -2.655 | -0.359 | 0.263 | 0.347 | -0.099 | 0.032 | 0.032 | 1.003 | 0.59 | 0.95 | 0.57 | 0.65 | 0.44 | 0.6 |
| 6 | 28968146 | rs6906909 | -2.655 | -0.359 | 0.263 | 0.348 | -0.099 | 0.023 | 0.023 | 1.003 | 0.59 | 0.96 | 0.57 | 0.65 | 0.44 | 0.6 |
| 6 | 28968930 | rs9257453 | 0.145 | -0.371 | 0.3 | 0.341 | 0 | 0 | 0 | 0 | 0.09 | 0.07 | 0.07 | 0.11 | 0.09 | 0.12 |
| 6 | 28969522 | rs6933976 | -2.108 | -0.359 | 0.286 | 0.348 | -0.005 | -0.005 | -0.005 | 0.257 | 0.36 | 0.58 | 0.27 | 0.25 | 0.24 | 0.39 |
| 6 | 28970176 | rs3129793 | -2.627 | -0.362 | 0.282 | 0.346 | -0.136 | 0.025 | 0.025 | 1.136 | 0.59 | 0.96 | 0.59 | 0.65 | 0.5 | 0.61 |
| 6 | 28970369 | rs6456883 | -2.636 | -0.362 | 0.282 | 0.346 | -0.099 | 0.024 | 0.024 | 1.056 | 0.59 | 0.95 | 0.57 | 0.65 | 0.43 | 0.61 |
| 6 | 28972089 | rs9257456 | -0.005 | -0.36 | 0.284 | 0.347 | 0.001 | -0.002 | -0.002 | 0.332 | 0.1 | 0.11 | 0.07 | 0.12 | 0.09 | 0.15 |
| 6 | 28973215 | rs6925178 | -2.118 | -0.362 | 0.29 | 0.346 | -0.004 | -0.004 | -0.004 | 0 | 0.36 | 0.59 | 0.28 | 0.25 | 0.24 | 0.39 |
| 6 | 28976071 | rs6923005 | -2.144 | -0.378 | 0.282 | 0.338 | -0.003 | -0.003 | -0.003 | 0.04 | 0.36 | 0.59 | 0.28 | 0.25 | 0.24 | 0.39 |
| 6 | 28977925 | rs9257461 | -2.648 | -0.377 | 0.281 | 0.338 | -0.099 | 0.024 | 0.024 | 1.212 | 0.59 | 0.95 | 0.57 | 0.65 | 0.43 | 0.61 |
| 6 | 28981244 | rs6930903 | -2.124 | -0.419 | 0.717 | 0.317 | -0.004 | -0.004 | -0.004 | 0 | 0.36 | 0.59 | 0.28 | 0.25 | 0.24 | 0.39 |
| 6 | 28982287 | rs9357075 | -1.707 | -0.419 | 0.71 | 0.317 | -0.043 | -0.027 | -0.027 | 0 | 0.25 | 0.48 | 0.2 | 0.14 | 0.16 | 0.24 |
| 6 | 28984163 | rs6916923 | 0.01 | -0.405 | 0.672 | 0.32 | 0 | 0 | 0 | 0.195 | 0.1 | 0.1 | 0.07 | 0.11 | 0.09 | 0.15 |
| 6 | 28984512 | rs9468464 | 0.109 | -0.437 | 0.652 | 0.32 | -0.006 | -0.001 | -0.001 | 0 | 0.12 | 0.14 | 0.06 | 0.03 | 0.05 | 0.12 |
| 6 | 28984755 | rs3129795 | -2.33 | -0.599 | -0.279 | -0.585 | -0.101 | 0.104 | 0.104 | 0.258 | 0.81 | 0.99 | 0.88 | 0.96 | 0.73 | 0.93 |
| 6 | 28988583 | rs6941946 | -2.481 | 0.057 | 0.566 | 0.219 | -0.085 | 0.026 | 0.026 | 2.36 | 0.59 | 0.95 | 0.56 | 0.65 | 0.42 | 0.61 |
| 6 | 28990195 | rs7768299 | -2.481 | 0.057 | 0.566 | 0.219 | -0.085 | 0.026 | 0.026 | 2.36 | 0.59 | 0.95 | 0.56 | 0.65 | 0.42 | 0.6 |
| 6 | 28990618 | rs1543796 | -2.455 | 0.067 | 0.566 | 0.219 | -0.146 | 0.028 | 0.028 | 2.251 | 0.59 | 0.96 | 0.59 | 0.65 | 0.49 | 0.61 |
| 6 | 28992184 | rs2394512 | -2.473 | 0.025 | 0.565 | 0.218 | -0.084 | 0.027 | 0.027 | 2.383 | 0.59 | 0.95 | 0.57 | 0.65 | 0.42 | 0.61 |
| 6 | 28994570 | rs7741520 | -1.924 | -0.015 | 0.641 | 0.212 | -0.006 | -0.005 | -0.005 | 0.552 | 0.36 | 0.6 | 0.28 | 0.25 | 0.23 | 0.39 |
| 6 | 28995719 | rs10484544 | -0.272 | -0.021 | 0.638 | 0.209 | -0.033 | 0.061 | 0.061 | 0.775 | 0.24 | 0.35 | 0.28 | 0.39 | 0.18 | 0.21 |
| 6 | 28999218 | rs9468469 | 0.513 | -0.04 | 0.688 | 0.202 | 0.013 | 0.04 | 0.04 | 0 | 0.06 | 0.23 | 0.02 | 0.01 | 0.01 | 0.03 |
| 6 | 28999826 | rs16894557 | 0.777 | -0.079 | 0.596 | 0.281 | 0.017 | -0.032 | -0.032 | 0 | 0.14 | 0.09 | 0.18 | 0.33 | 0.1 | 0.1 |
| 6 | 29002062 | rs7755402 | -0.305 | -0.032 | 0.552 | 0.231 | -0.033 | 0.061 | 0.061 | 0.775 | 0.24 | 0.35 | 0.28 | 0.39 | 0.18 | 0.21 |
| 6 | 29004091 | rs9468471 | -1.853 | 0.027 | 0.614 | 0.251 | -0.006 | -0.005 | -0.005 | 0 | 0.36 | 0.59 | 0.28 | 0.25 | 0.23 | 0.39 |
| 6 | 29006250 | rs9468473 | -1.853 | 0.027 | 0.609 | 0.251 | -0.006 | -0.005 | -0.005 | 0 | 0.36 | 0.6 | 0.28 | 0.25 | 0.23 | 0.39 |
| 6 | 29006844 | rs7743837 | -1.853 | 0.027 | 0.609 | 0.251 | -0.005 | -0.005 | -0.005 | 0 | 0.36 | 0.6 | 0.28 | 0.26 | 0.24 | 0.39 |
| 6 | 29009568 | rs6919044 | -1.852 | 0.019 | 0.609 | 0.251 | -0.006 | -0.005 | -0.005 | 0 | 0.36 | 0.6 | 0.28 | 0.25 | 0.23 | 0.39 |
| 6 | 29010153 | rs6924824 | -1.852 | 0.019 | 0.609 | 0.251 | -0.006 | -0.005 | -0.005 | 0.622 | 0.36 | 0.6 | 0.28 | 0.25 | 0.23 | 0.39 |
| 6 | 29010824 | rs6456889 | -1.852 | 0.019 | 0.609 | 0.251 | -0.006 | -0.005 | -0.005 | 0.032 | 0.36 | 0.6 | 0.28 | 0.25 | 0.23 | 0.39 |
| 6 | 29011927 | rs2050230 | -2.422 | 0.056 | 0.436 | 0.245 | -0.146 | 0.028 | 0.028 | 2.195 | 0.59 | 0.96 | 0.59 | 0.65 | 0.49 | 0.6 |
| 6 | 29011960 | rs9391651 | -1.452 | 0.003 | 0.622 | 0.235 | -0.043 | -0.027 | -0.027 | 0 | 0.25 | 0.48 | 0.2 | 0.14 | 0.16 | 0.24 |
| 6 | 29012116 | rs7341218 | -0.226 | 0.016 | 0.68 | 0.24 | -0.03 | 0.066 | 0.066 | 0.367 | 0.23 | 0.35 | 0.28 | 0.39 | 0.19 | 0.2 |
| 6 | 29013133 | rs9468474 | 1.564 | -0.032 | 0.624 | 0.229 | 0.013 | 0.014 | 0.014 | 0 | 0.07 | 0.01 | 0.03 | 0.02 | 0.03 | 0.11 |
| 6 | 29014026 | rs2143574 | -2.542 | -0.005 | 0.632 | 0.235 | -0.084 | 0.028 | 0.028 | 2.195 | 0.59 | 0.95 | 0.57 | 0.65 | 0.42 | 0.6 |
| 6 | 29016587 | rs6456891 | -2.541 | -0.008 | 0.641 | 0.234 | -0.084 | 0.028 | 0.028 | 2.195 | 0.59 | 0.95 | 0.57 | 0.65 | 0.42 | 0.6 |
| 6 | 29017101 | rs3130756 | -2.541 | -0.006 | 0.641 | 0.234 | -0.146 | 0.028 | 0.028 | 2.195 | 0.59 | 0.96 | 0.59 | 0.65 | 0.49 | 0.6 |
| 6 | 29021017 | rs9501294 | -1.479 | -0.007 | 0.657 | 0.226 | -0.042 | -0.026 | -0.026 | 0.002 | 0.25 | 0.49 | 0.2 | 0.14 | 0.15 | 0.24 |

| | | | | | | | | | | | | | | | | |
|---|---|---|---|---|---|---|---|---|---|---|---|---|---|---|---|---|
| 6 | 29025680 | rs7747023 | -1.573 | 0.001 | 0.66 | 0.23 | -0.04 | -0.024 | -0.024 | 0 | 0.25 | 0.48 | 0.2 | 0.14 | 0.15 | 0.24 |
| 6 | 29033053 | rs6904527 | 0.065 | 0.007 | 0.592 | 0.238 | 0 | -0.002 | -0.002 | 0 | 0.11 | 0.11 | 0.07 | 0.12 | 0.07 | 0.15 |
| 6 | 29039895 | rs6456895 | -2.495 | -0.005 | 0.539 | 0.228 | -0.084 | 0.028 | 0.028 | 2.195 | 0.59 | 0.95 | 0.57 | 0.65 | 0.42 | 0.6 |
| 6 | 29039977 | rs3131083 | -2.495 | 0.022 | 0.539 | 0.235 | -0.146 | 0.023 | 0.023 | 2.195 | 0.59 | 0.96 | 0.59 | 0.65 | 0.49 | 0.6 |
| 6 | 29042937 | rs3130758 | -2.495 | 0.022 | 0.539 | 0.229 | -0.146 | 0.028 | 0.028 | 2.195 | 0.59 | 0.96 | 0.59 | 0.65 | 0.49 | 0.6 |
| 6 | 29045632 | rs9393941 | -2.503 | 0.01 | 0.536 | 0.226 | -0.084 | 0.027 | 0.027 | 2.157 | 0.59 | 0.95 | 0.57 | 0.65 | 0.42 | 0.6 |
| 6 | 29047203 | rs9348821 | -1.567 | -0.007 | 0.638 | 0.242 | -0.041 | -0.024 | -0.024 | 0.69 | 0.25 | 0.49 | 0.2 | 0.14 | 0.15 | 0.24 |
| 6 | 29050714 | rs11758255 | -1.996 | -0.013 | 0.644 | 0.242 | -0.006 | -0.005 | -0.005 | 0 | 0.36 | 0.6 | 0.28 | 0.25 | 0.23 | 0.4 |
| 6 | 29052572 | rs6903771 | -1.29 | -0.024 | 0.639 | 0.237 | -0.03 | -0.016 | -0.016 | 0 | 0.23 | 0.49 | 0.2 | 0.14 | 0.15 | 0.21 |
| 6 | 29053998 | rs7752270 | 0.083 | -0.015 | 0.641 | 0.241 | 0 | -0.002 | -0.002 | 0.144 | 0.11 | 0.11 | 0.07 | 0.12 | 0.07 | 0.15 |
| 6 | 29055442 | rs6456901 | -2.527 | -0.015 | 0.638 | 0.241 | -0.084 | 0.028 | 0.028 | 2.195 | 0.59 | 0.95 | 0.57 | 0.65 | 0.42 | 0.6 |
| 6 | 29057971 | rs9380109 | -2.527 | -0.015 | 0.638 | 0.241 | -0.084 | 0.028 | 0.028 | 2.195 | 0.59 | 0.95 | 0.57 | 0.65 | 0.42 | 0.6 |
| 6 | 29064772 | rs2142905 | -0.269 | -0.033 | 0.635 | 0.234 | -0.029 | 0.062 | 0.062 | 0.58 | 0.23 | 0.35 | 0.29 | 0.4 | 0.19 | 0.24 |
| 6 | 29067772 | rs3129111 | 0.087 | -0.012 | 0.623 | 0.242 | 0 | -0.002 | -0.002 | 6.11 | 0.11 | 0.11 | 0.07 | 0.12 | 0.07 | 0.15 |
| 6 | 29073094 | rs3131091 | 2.537 | -0.008 | 0.57 | 0.24 | -0.146 | 0.028 | 0.028 | 2.195 | 0.41 | 0.04 | 0.41 | 0.35 | 0.51 | 0.4 |
| 6 | 29075233 | rs9461507 | 0.765 | -0.031 | 0.637 | 0.235 | 0.013 | 0.012 | 0.012 | 0 | 0.1 | 0.09 | 0.05 | 0.03 | 0.03 | 0.11 |
| 6 | 29077353 | rs3129097 | 2.554 | -0.01 | 0.608 | 0.236 | -0.084 | 0.028 | 0.028 | 2.195 | 0.41 | 0.05 | 0.43 | 0.35 | 0.58 | 0.4 |
| 6 | 29078151 | rs3116838 | 0.043 | -0.027 | 0.684 | 0.243 | -0.032 | 0.066 | 0.066 | 0.768 | 0.77 | 0.65 | 0.72 | 0.61 | 0.82 | 0.8 |
| 6 | 29079156 | rs3130763 | 2.585 | -0.017 | 0.683 | 0.302 | -0.146 | 0.041 | 0.041 | 2.195 | 0.41 | 0.04 | 0.41 | 0.34 | 0.51 | 0.4 |
| 6 | 29080344 | rs3749977 | -1.949 | -0.017 | 0.684 | 0.236 | -0.005 | -0.005 | -0.005 | 0 | 0.36 | 0.6 | 0.28 | 0.25 | 0.23 | 0.4 |
| 6 | 29080349 | rs3130764 | -0.427 | -0.043 | 0.794 | 0.221 | -0.003 | 0.013 | 0.013 | 1.056 | 0.8 | 0.77 | 0.75 | 0.61 | 0.83 | 0.8 |
| 6 | 29084232 | rs3129109 | 2.556 | -0.648 | 0.915 | 0.422 | -0.078 | 0.036 | 0.036 | 1.264 | 0.42 | 0.05 | 0.43 | 0.35 | 0.58 | 0.4 |
| 6 | 29085228 | rs3129110 | 2.556 | -0.648 | 0.915 | 0.422 | -0.078 | 0.036 | 0.036 | 1.264 | 0.42 | 0.05 | 0.43 | 0.35 | 0.58 | 0.4 |
| 6 | 29086829 | rs3130766 | 2.556 | -0.649 | 0.915 | 0.422 | -0.142 | 0.036 | 0.036 | 1.264 | 0.42 | 0.04 | 0.41 | 0.35 | 0.51 | 0.4 |
| 6 | 29096414 | rs3130774 | 2.467 | -0.676 | 1.01 | 0.416 | -0.042 | 0.062 | 0.062 | 2.073 | 0.43 | 0.09 | 0.43 | 0.35 | 0.59 | 0.39 |
| 6 | 29099579 | rs3130778 | 2.467 | -0.676 | 1.01 | 0.416 | -0.042 | 0.062 | 0.062 | 2.073 | 0.43 | 0.09 | 0.43 | 0.35 | 0.59 | 0.39 |
| 6 | 29102861 | rs3129119 | -1.53 | -0.634 | 1.128 | 0.409 | 0.016 | -0.022 | -0.022 | 0.399 | 0.25 | 0.48 | 0.23 | 0.14 | 0.22 | 0.25 |
| 6 | 29104672 | rs2223363 | 0.23 | -0.633 | 1.044 | 0.437 | -0.001 | 0.002 | 0.002 | 0 | 0.1 | 0.07 | 0.07 | 0.12 | 0.07 | 0.14 |
| 6 | 29107052 | rs9468487 | 0.488 | -0.608 | 1.122 | 0.456 | -0.012 | -0.006 | -0.006 | 0 | 0.12 | 0.14 | 0.06 | 0.03 | 0.05 | 0.13 |
| 6 | 29114616 | rs3116846 | 2.358 | -0.585 | 1.05 | 0.455 | -0.089 | 0.072 | 0.072 | 0.644 | 0.43 | 0.08 | 0.41 | 0.35 | 0.52 | 0.39 |
| 6 | 29114621 | rs3130718 | 2.513 | -0.611 | 1.036 | 0.453 | -0.143 | 0.034 | 0.034 | 1.335 | 0.42 | 0.04 | 0.4 | 0.35 | 0.51 | 0.39 |
| 6 | 29118530 | rs3130724 | -0.347 | -0.625 | 1.211 | 0.44 | 0.09 | 0.022 | 0.022 | 0 | 0.13 | 0.33 | 0.18 | 0.1 | 0.18 | 0.12 |
| 6 | 29118813 | rs3116847 | -0.037 | -0.671 | 1.267 | 0.445 | -0.032 | 0.08 | 0.08 | 0.088 | 0.78 | 0.63 | 0.72 | 0.61 | 0.83 | 0.8 |
| 6 | 29119159 | rs3129123 | 1.734 | -0.658 | 1.214 | 0.416 | -0.033 | 0.053 | 0.053 | 0.571 | 0.54 | 0.17 | 0.46 | 0.39 | 0.56 | 0.51 |
| 6 | 29125402 | rs3129131 | 1.922 | -0.662 | 1.208 | 0.414 | -0.061 | 0.077 | 0.077 | 0.844 | 0.52 | 0.11 | 0.45 | 0.38 | 0.55 | 0.51 |
| 6 | 29125747 | rs3130729 | 2.585 | -0.665 | 1.205 | 0.412 | -0.142 | 0.035 | 0.035 | 1.485 | 0.42 | 0.04 | 0.4 | 0.35 | 0.51 | 0.39 |
| 6 | 29129983 | rs2142906 | 1.926 | -0.642 | 1.221 | 0.425 | -0.061 | 0.077 | 0.077 | 0.844 | 0.52 | 0.11 | 0.45 | 0.38 | 0.55 | 0.51 |
| 6 | 29137220 | rs2394520 | 1.137 | -0.643 | 1.255 | 0.427 | -0.03 | 0.05 | 0.05 | 1.931 | 0.66 | 0.52 | 0.64 | 0.49 | 0.74 | 0.65 |
| 6 | 29138315 | rs3129151 | -0.35 | -0.638 | 1.259 | 0.401 | 0.09 | 0.022 | 0.022 | 0 | 0.13 | 0.33 | 0.18 | 0.1 | 0.18 | 0.12 |
| 6 | 29138777 | rs3129152 | -0.835 | -0.639 | 1.257 | 0.4 | 0.089 | 0.029 | 0.029 | 0 | 0.15 | 0.39 | 0.19 | 0.11 | 0.19 | 0.13 |
| 6 | 29142476 | rs2206042 | 1.928 | -0.65 | 1.265 | 0.389 | -0.06 | 0.082 | 0.082 | 0.781 | 0.51 | 0.11 | 0.45 | 0.38 | 0.55 | 0.51 |
| 6 | 29144127 | rs3116857 | 1.934 | -0.657 | 1.286 | 0.386 | -0.06 | 0.083 | 0.083 | 0.625 | 0.51 | 0.11 | 0.45 | 0.38 | 0.55 | 0.51 |
| 6 | 29148619 | rs3130745 | 1.747 | -0.642 | 1.298 | 0.394 | -0.032 | 0.057 | 0.057 | 0.568 | 0.53 | 0.17 | 0.46 | 0.39 | 0.56 | 0.51 |
| 6 | 29149561 | rs3116817 | 1.927 | -0.642 | 1.291 | 0.393 | -0.06 | 0.082 | 0.082 | 0.781 | 0.51 | 0.11 | 0.45 | 0.38 | 0.55 | 0.51 |
| 6 | 29151671 | rs3116820 | 1.746 | -0.642 | 1.298 | 0.394 | -0.032 | 0.057 | 0.057 | 0.568 | 0.53 | 0.17 | 0.46 | 0.38 | 0.56 | 0.51 |
| 6 | 29155749 | rs3129171 | -0.838 | -0.634 | 1.358 | 0.408 | 0.096 | 0.033 | 0.033 | 0 | 0.15 | 0.39 | 0.2 | 0.12 | 0.2 | 0.13 |
| 6 | 29161435 | rs3116826 | 1.767 | -0.63 | 1.331 | 0.379 | -0.032 | 0.058 | 0.058 | 0.662 | 0.53 | 0.17 | 0.46 | 0.38 | 0.56 | 0.51 |
| 6 | 29170261 | rs3129093 | -0.35 | -0.622 | 1.367 | 0.366 | 0.091 | 0.023 | 0.023 | 0 | 0.13 | 0.33 | 0.18 | 0.1 | 0.18 | 0.12 |
| 6 | 29171039 | rs760804 | -0.366 | -0.621 | 1.368 | 0.367 | 0.09 | 0.022 | 0.022 | 0 | 0.13 | 0.33 | 0.18 | 0.1 | 0.18 | 0.12 |
| 6 | 29176296 | rs736466 | 1.739 | -0.656 | 1.339 | 0.368 | -0.021 | 0.062 | 0.062 | 0.316 | 0.53 | 0.18 | 0.47 | 0.38 | 0.57 | 0.53 |
| 6 | 29177102 | rs1883329 | 1.095 | -0.656 | 1.348 | 0.369 | -0.034 | 0.046 | 0.046 | 1.425 | 0.66 | 0.52 | 0.66 | 0.49 | 0.75 | 0.65 |
| 6 | 29178514 | rs2206040 | 2.599 | -0.693 | 1.322 | 0.348 | -0.145 | 0.038 | 0.038 | 1.295 | 0.41 | 0.04 | 0.4 | 0.35 | 0.51 | 0.39 |
| 6 | 29178606 | rs2206041 | 2.599 | -0.693 | 1.322 | 0.348 | -0.145 | 0.038 | 0.038 | 1.295 | 0.41 | 0.04 | 0.4 | 0.35 | 0.51 | 0.39 |
| 6 | 29179237 | rs3130817 | -0.356 | -0.627 | 1.368 | 0.384 | 0.09 | 0.022 | 0.022 | 0 | 0.13 | 0.33 | 0.18 | 0.1 | 0.18 | 0.12 |
| 6 | 29181314 | rs3130826 | 1.105 | -0.629 | 1.368 | 0.384 | -0.034 | 0.046 | 0.046 | 1.425 | 0.66 | 0.52 | 0.66 | 0.49 | 0.75 | 0.65 |
| 6 | 29182308 | rs2894083 | 1.74 | -0.627 | 1.364 | 0.384 | -0.021 | 0.062 | 0.062 | 0.316 | 0.53 | 0.18 | 0.47 | 0.38 | 0.57 | 0.53 |
| 6 | 29183346 | rs3129096 | 1.74 | -0.627 | 1.364 | 0.384 | -0.021 | 0.062 | 0.062 | 0.316 | 0.53 | 0.18 | 0.47 | 0.38 | 0.57 | 0.53 |
| 6 | 29184660 | rs3116837 | 1.098 | -0.631 | 1.366 | 0.384 | -0.034 | 0.047 | 0.047 | 1.541 | 0.66 | 0.52 | 0.66 | 0.49 | 0.75 | 0.65 |
| 6 | 29189155 | rs3130803 | 1.69 | -0.65 | 1.137 | 0.407 | -0.032 | 0.061 | 0.061 | 0.281 | 0.53 | 0.17 | 0.46 | 0.38 | 0.56 | 0.51 |
| 6 | 29191411 | rs714470 | 1.681 | -0.653 | 1.138 | 0.408 | -0.021 | 0.061 | 0.061 | 0.261 | 0.53 | 0.18 | 0.47 | 0.38 | 0.57 | 0.53 |
| 6 | 29191548 | rs12660419 | 0.947 | -0.698 | 1.163 | 0.479 | 0.013 | -0.026 | -0.026 | 0 | 0.13 | 0.1 | 0.18 | 0.33 | 0.1 | 0.11 |
| 6 | 29193785 | rs3130812 | 1.619 | -0.639 | 1.011 | 0.463 | -0.018 | 0.061 | 0.061 | 0.261 | 0.53 | 0.18 | 0.47 | 0.38 | 0.57 | 0.53 |
| 6 | 29195011 | rs3130813 | 1.619 | -0.616 | 0.962 | 0.442 | -0.021 | 0.061 | 0.061 | 0.261 | 0.53 | 0.18 | 0.47 | 0.38 | 0.57 | 0.53 |
| 6 | 29199824 | rs2207338 | 1.127 | -0.689 | 1.036 | 0.43 | -0.002 | -0.013 | -0.013 | 0 | 0.64 | 0.3 | 0.55 | 0.49 | 0.64 | 0.66 |
| 6 | 29203911 | rs3130816 | 2.518 | -0.793 | 0.933 | 0.358 | -0.154 | 0.017 | 0.017 | 1.456 | 0.39 | 0.04 | 0.4 | 0.33 | 0.51 | 0.38 |
| 6 | 29213070 | rs3129105 | 2.517 | -0.771 | 0.933 | 0.358 | -0.158 | 0.017 | 0.017 | 1.456 | 0.39 | 0.04 | 0.4 | 0.33 | 0.51 | 0.38 |
| 6 | 29225795 | rs3117330 | 1.139 | -0.782 | 0.982 | 0.364 | -0.001 | -0.013 | -0.013 | 2.37 | 0.63 | 0.3 | 0.55 | 0.49 | 0.64 | 0.66 |
| 6 | 29227117 | rs2024679 | 2.141 | -0.727 | 1.017 | 0.391 | -0.056 | 0.057 | 0.057 | 0.302 | 0.44 | 0.16 | 0.43 | 0.34 | 0.52 | 0.38 |
| 6 | 29227643 | rs3117329 | 2.141 | -0.727 | 1.017 | 0.391 | -0.051 | 0.057 | 0.057 | 0.302 | 0.44 | 0.17 | 0.44 | 0.35 | 0.53 | 0.4 |

| | | | | | | | | | | | | | | | | |
|---|---|---|---|---|---|---|---|---|---|---|---|---|---|---|---|---|
| 6 | 29230577 | rs3117328 | 2.141 | -0.727 | 1.017 | 0.391 | -0.051 | 0.057 | 0.057 | 0.302 | 0.44 | 0.17 | 0.44 | 0.35 | 0.53 | 0.4 |
| 6 | 29230683 | rs3130827 | 1.07 | -0.723 | 1.026 | 0.396 | -0.002 | -0.013 | -0.013 | 0 | 0.64 | 0.31 | 0.55 | 0.49 | 0.64 | 0.66 |
| 6 | 29231172 | rs9257657 | -0.299 | -0.761 | 1.224 | 0.405 | -0.007 | 0.025 | 0.025 | 0.009 | 0.8 | 0.75 | 0.75 | 0.59 | 0.83 | 0.78 |
| 6 | 29235900 | rs3130829 | 2.584 | -0.702 | 1.085 | 0.245 | -0.161 | 0.017 | 0.017 | 1.494 | 0.39 | 0.04 | 0.4 | 0.33 | 0.51 | 0.38 |
| 6 | 29236645 | rs7766902 | -0.25 | -0.688 | 1.244 | 0.242 | 0.021 | 0.012 | 0.012 | 0 | 0.13 | 0.26 | 0.15 | 0.1 | 0.12 | 0.12 |
| 6 | 29236722 | rs3130830 | 2.115 | -0.67 | 1.193 | 0.193 | -0.053 | 0.057 | 0.057 | 0.302 | 0.44 | 0.17 | 0.44 | 0.35 | 0.53 | 0.4 |
| 6 | 29238015 | rs6942318 | 0.336 | -0.683 | 1.204 | 0.188 | 0.001 | 0.003 | 0.003 | 0 | 0.1 | 0.08 | 0.07 | 0.11 | 0.07 | 0.14 |
| 6 | 29239150 | rs3117327 | 2.104 | -0.619 | 0.672 | 0.187 | -0.053 | 0.057 | 0.057 | 0.302 | 0.44 | 0.17 | 0.44 | 0.35 | 0.53 | 0.4 |
| 6 | 29253756 | rs9468508 | 1.19 | -0.666 | 0.692 | 0.163 | 0.019 | 0.025 | 0.025 | 0 | 0.09 | 0.01 | 0.04 | 0.04 | 0.04 | 0.12 |
| 6 | 29256420 | rs1884123 | 1.074 | -0.664 | 0.775 | 0.193 | -0.002 | -0.009 | -0.009 | 0 | 0.63 | 0.3 | 0.52 | 0.49 | 0.63 | 0.66 |
| 6 | 29257772 | rs1033569 | 1.279 | -0.487 | 0.715 | 0.246 | 0.007 | 0.012 | 0.012 | 0 | 0.57 | 0.26 | 0.5 | 0.45 | 0.6 | 0.54 |
| 6 | 29257967 | rs1033568 | 1.279 | -0.487 | 0.715 | 0.246 | 0.007 | 0.012 | 0.012 | 0 | 0.57 | 0.26 | 0.5 | 0.45 | 0.6 | 0.54 |
| 6 | 29259623 | rs4446587 | 2.35 | -0.436 | 0.56 | 0.265 | -0.163 | 0.031 | 0.031 | 1.458 | 0.39 | 0.04 | 0.37 | 0.32 | 0.5 | 0.37 |
| 6 | 29269066 | rs7754926 | 0.106 | -0.223 | 0.823 | 0.325 | 0.011 | 0.012 | 0.012 | 0 | 0.14 | 0.12 | 0.09 | 0.12 | 0.08 | 0.15 |
| 6 | 29269143 | rs7755211 | 0.106 | -0.223 | 0.823 | 0.325 | 0.011 | 0.012 | 0.012 | 0 | 0.14 | 0.12 | 0.09 | 0.12 | 0.08 | 0.15 |
| 6 | 29270498 | rs4594993 | 0.106 | -0.223 | 0.823 | 0.325 | 0.011 | 0.012 | 0.012 | 0 | 0.14 | 0.12 | 0.09 | 0.12 | 0.08 | 0.15 |
| 6 | 29271126 | rs6930876 | 0.073 | -0.241 | 0.852 | 0.323 | -0.001 | -0.001 | -0.001 | 0 | 0.15 | 0.15 | 0.09 | 0.12 | 0.08 | 0.15 |
| 6 | 29272012 | rs3117426 | -0.58 | 0.215 | 1.068 | 0.364 | 0.173 | 0.048 | 0.048 | 0 | 0.1 | 0.29 | 0.18 | 0.08 | 0.18 | 0.11 |
| 6 | 29274356 | rs7774255 | -1.467 | 0.216 | 0.339 | 0.534 | -0.027 | 0.057 | 0.057 | 0 | 0.55 | 0.21 | 0.41 | 0.33 | 0.53 | 0.39 |
| 6 | 29274486 | rs9257694 | 1.394 | 0.234 | 0.202 | 0.572 | -0.012 | 0.066 | 0.066 | 0 | 0.46 | 0.25 | 0.43 | 0.32 | 0.53 | 0.37 |
| 6 | 29275507 | rs9257696 | -1.133 | -0.329 | 0.253 | 0.512 | 0.007 | 0.002 | 0.002 | 0.251 | 0.16 | 0.21 | 0.2 | 0.15 | 0.16 | 0.11 |
| 6 | 29284518 | rs720831 | 1.984 | -0.525 | 0.253 | 0.8 | -0.007 | -0.014 | -0.014 | 0 | 0.64 | 0.25 | 0.61 | 0.64 | 0.65 | 0.65 |
| 6 | 29295487 | rs6937882 | 0.235 | -0.476 | 0.202 | 0.792 | 0.034 | -0.035 | -0.035 | 0 | 0.26 | 0.17 | 0.25 | 0.4 | 0.17 | 0.29 |
| 6 | 29301222 | rs6930435 | 0.416 | -0.452 | 0.296 | 0.79 | 0.002 | 0.027 | 0.027 | 0 | 0.8 | 0.46 | 0.81 | 0.79 | 0.8 | 0.75 |
| 6 | 29302030 | rs6903989 | 0.564 | -0.456 | 0.302 | 0.788 | 0.006 | 0.033 | 0.033 | 0 | 0.8 | 0.47 | 0.81 | 0.79 | 0.8 | 0.75 |
| 6 | 29303364 | rs3117433 | 1.522 | -0.458 | 0.446 | 0.78 | 0.048 | -0.025 | -0.025 | 1.67 | 0.07 | 0.13 | 0.06 | 0.01 | 0.1 | 0.11 |
| 6 | 29306656 | rs9257745 | -1.077 | -0.338 | -0.213 | 0.832 | 0.006 | 0 | 0 | 0.365 | 0.16 | 0.21 | 0.19 | 0.15 | 0.16 | 0.1 |
| 6 | 29308393 | rs9348827 | -0.469 | -0.34 | -0.194 | 0.831 | 0.016 | 0.012 | 0.012 | 0.193 | 0.11 | 0.17 | 0.17 | 0.11 | 0.13 | 0.08 |
| 6 | 29310681 | rs1014258 | 0.15 | -0.328 | -0.29 | 0.838 | 0.002 | 0.027 | 0.027 | 0 | 0.8 | 0.46 | 0.81 | 0.79 | 0.8 | 0.75 |
| 6 | 29323655 | rs7761746 | 0.363 | -0.324 | -0.254 | 0.833 | -0.028 | -0.004 | -0.004 | 0 | 0.07 | 0.18 | 0.05 | 0.01 | 0.02 | 0 |
| 6 | 29323905 | rs6930033 | 0.04 | -0.112 | -0.182 | 0.963 | 0.035 | -0.035 | -0.035 | 0.257 | 0.26 | 0.17 | 0.25 | 0.4 | 0.17 | 0.29 |
| 6 | 29324599 | rs10484546 | 0.04 | -0.112 | -0.182 | 0.963 | 0.035 | -0.035 | -0.035 | 0 | 0.26 | 0.17 | 0.25 | 0.4 | 0.17 | 0.29 |
| 6 | 29327079 | rs9257775 | 0.728 | 0.045 | -0.753 | 0.521 | -0.008 | 0.081 | 0.081 | 0 | 0.48 | 0.27 | 0.43 | 0.28 | 0.52 | 0.37 |
| 6 | 29334439 | rs9257792 | -0.58 | -0.003 | -0.241 | 0.596 | -0.03 | -0.021 | -0.021 | 0 | 0.11 | 0.21 | 0.07 | 0.06 | 0.05 | 0.02 |
| 6 | 29335661 | rs9257794 | 0.25 | 0.13 | -0.494 | 1.091 | -0.002 | -0.01 | -0.01 | 0 | 0.21 | 0.17 | 0.25 | 0.27 | 0.21 | 0.25 |
| 6 | 29336054 | rs4713211 | 1.011 | -0.496 | -0.163 | 1.171 | 0.015 | -0.018 | -0.018 | 0 | 0.62 | 0.33 | 0.6 | 0.52 | 0.69 | 0.6 |
| 6 | 29336147 | rs9295804 | 2.258 | -0.706 | -0.424 | 1.095 | -0.136 | 0.061 | 0.061 | 1.067 | 0.38 | 0.07 | 0.36 | 0.25 | 0.49 | 0.35 |
| 6 | 29337500 | rs6934993 | 0.607 | -0.611 | -0.366 | 1.08 | -0.002 | -0.01 | -0.01 | 0 | 0.21 | 0.17 | 0.25 | 0.27 | 0.21 | 0.25 |
| 6 | 29339271 | rs9257799 | -0.302 | -0.496 | -0.561 | 1.071 | -0.009 | -0.003 | -0.003 | 2.142 | 0.1 | 0.08 | 0.16 | 0.14 | 0.13 | 0.1 |
| 6 | 29341007 | rs7772982 | -1.327 | -0.538 | -0.589 | 1.073 | -0.001 | -0.001 | -0.001 | 0 | 0.25 | 0.26 | 0.23 | 0.26 | 0.19 | 0.25 |
| 6 | 29341308 | rs7753474 | -1.327 | -0.538 | -0.589 | 1.073 | -0.001 | -0.001 | -0.001 | 0 | 0.25 | 0.27 | 0.23 | 0.26 | 0.19 | 0.25 |
| 6 | 29341364 | rs7773534 | -1.327 | -0.538 | -0.589 | 1.088 | -0.001 | 0 | 0 | 0 | 0.25 | 0.26 | 0.23 | 0.26 | 0.19 | 0.25 |
| 6 | 29341449 | rs10456372 | 2.107 | -0.606 | -0.695 | 0.933 | -0.135 | 0.053 | 0.053 | 1.239 | 0.38 | 0.07 | 0.36 | 0.26 | 0.49 | 0.37 |
| 6 | 29341623 | rs7774029 | 0.676 | -0.602 | -0.644 | 0.955 | 0.005 | 0.002 | 0.002 | 0 | 0.1 | 0.04 | 0.05 | 0.11 | 0.05 | 0.15 |
| 6 | 29342236 | rs9380122 | 0.348 | -0.663 | -0.455 | 1.004 | -0.015 | 0.039 | 0.039 | 0 | 0.08 | 0.13 | 0.04 | 0.22 | 0.03 | 0.11 |
| 6 | 29342825 | rs3749970 | 2.112 | -0.379 | -0.568 | 0.865 | -0.136 | 0.052 | 0.052 | 1.067 | 0.38 | 0.07 | 0.36 | 0.26 | 0.49 | 0.35 |
| 6 | 29343355 | rs9257802 | -0.681 | -0.315 | -0.373 | 0.881 | 0.001 | -0.015 | -0.015 | 4.566 | 0.2 | 0.27 | 0.13 | 0.16 | 0.16 | 0.16 |
| 6 | 29344036 | rs5003265 | 1.752 | -0.095 | -0.439 | 0.825 | -0.121 | 0.062 | 0.062 | 1.191 | 0.39 | 0.07 | 0.37 | 0.25 | 0.51 | 0.37 |
| 6 | 29344083 | rs5003266 | 1.752 | -0.095 | -0.455 | 0.825 | -0.134 | 0.057 | 0.057 | 1.191 | 0.39 | 0.07 | 0.37 | 0.26 | 0.5 | 0.37 |
| 6 | 29344395 | rs9257803 | -0.374 | -0.145 | -0.322 | 0.786 | 0.029 | -0.024 | -0.024 | 1.392 | 0.11 | 0.25 | 0.08 | 0.04 | 0.1 | 0.02 |
| 6 | 29344704 | rs4598109 | 1.825 | -0.209 | -0.369 | 0.774 | -0.135 | 0.058 | 0.058 | 1.056 | 0.38 | 0.07 | 0.37 | 0.24 | 0.5 | 0.36 |
| 6 | 29344939 | rs2144425 | 1.802 | -0.237 | -0.365 | 0.758 | -0.111 | 0.069 | 0.069 | 0.317 | 0.38 | 0.09 | 0.38 | 0.25 | 0.52 | 0.37 |
| 6 | 29346329 | rs9257805 | -0.334 | -0.206 | -0.332 | 0.773 | 0.03 | -0.024 | -0.024 | 1.724 | 0.11 | 0.25 | 0.08 | 0.04 | 0.1 | 0.02 |
| 6 | 29346363 | rs238882 | 1.735 | -0.316 | -0.357 | 0.731 | -0.11 | 0.072 | 0.072 | 0.84 | 0.39 | 0.09 | 0.38 | 0.25 | 0.52 | 0.37 |
| 6 | 29349091 | rs3117442 | 1.68 | -0.434 | 0.265 | 0.728 | -0.038 | 0.063 | 0.063 | 1.492 | 0.41 | 0.2 | 0.39 | 0.25 | 0.52 | 0.39 |
| 6 | 29351873 | rs238872 | 0.946 | -0.276 | 0.397 | 0.676 | -0.003 | -0.005 | -0.005 | 0 | 0.52 | 0.32 | 0.44 | 0.49 | 0.54 | 0.44 |
| 6 | 29352943 | rs3117444 | 0.736 | -0.224 | 0.26 | 0.702 | 0 | 0 | 0 | 0.046 | 0.55 | 0.45 | 0.45 | 0.49 | 0.55 | 0.46 |
| 6 | 29353408 | rs3094556 | 0.743 | -0.222 | 0.253 | 0.703 | 0 | 0.001 | 0.001 | 0.058 | 0.55 | 0.45 | 0.45 | 0.5 | 0.55 | 0.46 |
| 6 | 29354383 | rs3129681 | 0.663 | -0.265 | 0.232 | 0.681 | 0 | 0 | 0 | 0.09 | 0.55 | 0.45 | 0.45 | 0.49 | 0.55 | 0.46 |
| 6 | 29354799 | rs3094551 | 0.618 | -0.338 | 0.185 | 0.672 | 0.008 | 0.011 | 0.011 | 0.024 | 0.61 | 0.46 | 0.48 | 0.5 | 0.59 | 0.59 |
| 6 | 29354809 | rs3094550 | 0.618 | -0.338 | 0.185 | 0.672 | 0.008 | 0.011 | 0.011 | 0.024 | 0.61 | 0.46 | 0.48 | 0.5 | 0.59 | 0.59 |
| 6 | 29355113 | rs238869 | 0.962 | -0.27 | 0.166 | 0.855 | 0.01 | 0.01 | 0.01 | 0.134 | 0.56 | 0.38 | 0.57 | 0.64 | 0.67 | 0.6 |
| 6 | 29355148 | rs3094549 | 0.762 | -0.186 | 0.161 | 0.886 | -0.004 | 0.003 | 0.003 | 0 | 0.62 | 0.48 | 0.51 | 0.56 | 0.66 | 0.7 |
| 6 | 29355519 | rs1419638 | -0.296 | 0.547 | -0.567 | 0.57 | -0.011 | -0.008 | -0.008 | 0.532 | 0.29 | 0.24 | 0.32 | 0.3 | 0.35 | 0.29 |
| 6 | 29355918 | rs1419635 | 1.09 | 0.167 | -0.876 | 0.422 | -0.042 | -0.032 | -0.032 | 0 | 0.2 | 0.1 | 0.15 | 0.24 | 0.31 | 0.1 |
| 6 | 29356046 | rs4711185 | 0.31 | 0.221 | -0.847 | 0.428 | -0.041 | -0.025 | -0.025 | 0 | 0.27 | 0.19 | 0.31 | 0.32 | 0.41 | 0.15 |
| 6 | 29356587 | rs379157 | 1.078 | -0.306 | -1.334 | 0.155 | -0.022 | 0.009 | 0.009 | 0.782 | 0.49 | 0.13 | 0.37 | 0.41 | 0.56 | 0.42 |
| 6 | 29356734 | rs9257811 | -0.073 | -0.694 | -1.414 | -0.016 | -0.053 | -0.013 | -0.013 | 1.754 | 0.37 | 0.24 | 0.36 | 0.36 | 0.46 | 0.29 |
| 6 | 29357362 | rs9257813 | 0.329 | -0.724 | -1.433 | -0.033 | -0.064 | -0.024 | -0.024 | 0 | 0.35 | 0.19 | 0.34 | 0.36 | 0.45 | 0.29 |

| | | | | | | | | | | | | | | | | |
|---|---|---|---|---|---|---|---|---|---|---|---|---|---|---|---|---|
| 6 | 29357645 | rs9393956 | 1.491 | -1.022 | -1.634 | -0.11 | -0.01 | 0.007 | 0.007 | 0 | 0.07 | 0.03 | 0.06 | 0.07 | 0.03 | 0.12 |
| 6 | 29357996 | rs4713214 | 0.447 | -0.998 | -1.607 | -0.091 | -0.064 | -0.024 | -0.024 | 0.034 | 0.35 | 0.19 | 0.35 | 0.36 | 0.45 | 0.29 |
| 6 | 29360183 | rs9257819 | 0.447 | -0.998 | -1.607 | -0.084 | -0.064 | -0.026 | -0.026 | 0 | 0.35 | 0.19 | 0.35 | 0.37 | 0.45 | 0.29 |
| 6 | 29360677 | rs1362065 | 0.447 | -0.998 | -1.574 | -0.085 | -0.064 | -0.024 | -0.024 | 3.747 | 0.35 | 0.19 | 0.35 | 0.36 | 0.45 | 0.29 |
| 6 | 29360776 | rs1362064 | 0.029 | -1.011 | -1.588 | -0.092 | -0.053 | -0.013 | -0.013 | 0 | 0.38 | 0.24 | 0.36 | 0.36 | 0.46 | 0.29 |
| 6 | 29361707 | rs16894836 | 2.792 | -1.446 | -2.295 | -0.401 | 0.008 | -0.01 | -0.01 | 0 | 0.17 | 0.01 | 0.14 | 0.17 | 0.11 | 0.29 |
| 6 | 29361735 | rs9257823 | 0.728 | -1.225 | -2.039 | -0.25 | -0.064 | -0.024 | -0.024 | 0 | 0.35 | 0.19 | 0.35 | 0.36 | 0.45 | 0.29 |
| 6 | 29362293 | rs1362063 | 0.569 | -1.257 | -1.921 | -0.26 | -0.052 | -0.017 | -0.017 | 0 | 0.36 | 0.21 | 0.35 | 0.37 | 0.46 | 0.29 |
| 6 | 29363697 | rs9257828 | 0.176 | -1.269 | -1.733 | -0.257 | -0.025 | -0.009 | -0.009 | 0 | 0.4 | 0.31 | 0.37 | 0.38 | 0.47 | 0.29 |
| 6 | 29364147 | rs4713216 | 0.751 | -1.269 | -1.744 | -0.257 | -0.064 | -0.024 | -0.024 | 0 | 0.35 | 0.19 | 0.35 | 0.36 | 0.45 | 0.29 |
| 6 | 29364325 | rs4713217 | 0.751 | -1.232 | -1.798 | -0.256 | -0.064 | -0.024 | -0.024 | 0.771 | 0.35 | 0.19 | 0.35 | 0.36 | 0.45 | 0.29 |
| 6 | 29364615 | rs9257834 | 0.751 | -1.232 | -1.798 | -0.256 | -0.064 | -0.024 | -0.024 | 0.023 | 0.35 | 0.19 | 0.35 | 0.36 | 0.45 | 0.29 |
| 6 | 29364951 | rs2073151 | 0.751 | -1.265 | -1.773 | -0.264 | -0.064 | -0.024 | -0.024 | 0 | 0.35 | 0.19 | 0.35 | 0.36 | 0.45 | 0.29 |
| 6 | 29365241 | rs2073150 | 0.422 | -1.265 | -1.773 | -0.264 | -0.057 | -0.015 | -0.015 | 0 | 0.37 | 0.23 | 0.36 | 0.36 | 0.45 | 0.29 |
| 6 | 29365974 | rs4713218 | 0.745 | -1.244 | -1.768 | -0.252 | -0.064 | -0.024 | -0.024 | 0 | 0.35 | 0.19 | 0.35 | 0.36 | 0.45 | 0.29 |
| 6 | 29366265 | rs2158279 | 0.745 | -1.244 | -1.768 | -0.252 | -0.064 | -0.024 | -0.024 | 0.774 | 0.35 | 0.19 | 0.35 | 0.36 | 0.45 | 0.29 |
| 6 | 29366377 | rs1024470 | 0.745 | -1.244 | -1.768 | -0.252 | -0.064 | -0.024 | -0.024 | 0 | 0.35 | 0.19 | 0.35 | 0.36 | 0.45 | 0.29 |
| 6 | 29367399 | rs1028411 | 0.745 | -1.244 | -1.768 | -0.252 | -0.064 | -0.024 | -0.024 | 0 | 0.35 | 0.19 | 0.35 | 0.36 | 0.45 | 0.29 |
| 6 | 29367590 | rs4711187 | 0.426 | -1.253 | -1.768 | -0.257 | -0.057 | -0.015 | -0.015 | 0 | 0.37 | 0.23 | 0.36 | 0.36 | 0.45 | 0.29 |
| 6 | 29368021 | rs9257837 | 2.276 | -1.248 | -1.81 | -0.28 | -0.017 | 0.005 | 0.005 | 0 | 0.11 | 0.07 | 0.19 | 0.08 | 0.14 | 0.16 |
| 6 | 29368813 | rs9405124 | 2.041 | -1.247 | -1.811 | -0.276 | -0.041 | -0.033 | -0.033 | 0 | 0.2 | 0.1 | 0.15 | 0.24 | 0.31 | 0.09 |
| 6 | 29369672 | rs1419634 | 0.397 | -1.262 | -1.769 | -0.259 | -0.056 | -0.014 | -0.014 | 0 | 0.38 | 0.23 | 0.36 | 0.36 | 0.45 | 0.29 |
| 6 | 29371415 | rs1362060 | 0.44 | -1.258 | -1.409 | -0.257 | -0.057 | -0.015 | -0.015 | 0.036 | 0.37 | 0.23 | 0.36 | 0.36 | 0.45 | 0.29 |
| 6 | 29373213 | rs1419633 | 0.207 | -1.293 | -1.316 | -0.263 | -0.02 | -0.007 | -0.007 | 0.598 | 0.4 | 0.33 | 0.38 | 0.43 | 0.48 | 0.31 |
| 6 | 29373249 | rs1419632 | 0.162 | -1.295 | -1.308 | -0.264 | -0.019 | -0.006 | -0.006 | 0 | 0.4 | 0.33 | 0.38 | 0.43 | 0.48 | 0.32 |
| 6 | 29379891 | rs12192194 | 0.87 | -1.249 | -1.301 | -0.244 | 0 | 0.001 | 0.001 | 0 | 0.29 | 0.29 | 0.2 | 0.35 | 0.34 | 0.15 |
| 6 | 29381906 | rs1544403 | 2.06 | -1.317 | -1.333 | -0.299 | -0.009 | 0.006 | 0.006 | 0.054 | 0.06 | 0.01 | 0.05 | 0.07 | 0.03 | 0.12 |
| 6 | 29383320 | rs1362073 | 0.908 | -1.291 | -1.315 | -0.252 | 0 | 0.001 | 0.001 | 0 | 0.29 | 0.29 | 0.2 | 0.35 | 0.34 | 0.15 |
| 6 | 29385237 | rs3131024 | -1.636 | -1.632 | -0.912 | -0.133 | -0.004 | 0.024 | 0.024 | 0 | 0.89 | 0.85 | 0.77 | 0.77 | 0.87 | 0.9 |
| 6 | 29390440 | rs2523421 | 2.56 | -1.647 | -1.174 | -0.279 | -0.017 | 0.043 | 0.043 | 0.021 | 0.1 | 0 | 0.04 | 0 | 0.11 | 0.02 |
| 6 | 29391936 | rs1419643 | 1.922 | -1.386 | -0.685 | 0.05 | -0.046 | 0.078 | 0.078 | 0.107 | 0.31 | 0.06 | 0.25 | 0.18 | 0.33 | 0.33 |
| 6 | 29392235 | rs1419642 | 1.771 | -1.39 | -0.582 | 0.048 | -0.032 | 0.086 | 0.086 | 0.009 | 0.32 | 0.09 | 0.25 | 0.18 | 0.33 | 0.33 |
| 6 | 29394427 | rs2074470 | 0.079 | -1.287 | -0.211 | 0.081 | 0.002 | 0.011 | 0.011 | 0 | 0.68 | 0.65 | 0.51 | 0.62 | 0.69 | 0.58 |
| 6 | 29394926 | rs9257857 | 2.031 | -1.253 | -0.334 | 0.087 | -0.041 | 0.009 | 0.009 | 0 | 0.13 | 0.05 | 0.19 | 0.08 | 0.15 | 0.17 |
| 6 | 29395057 | rs16894898 | 1.718 | -1.261 | -0.255 | 0.037 | 0.01 | 0.01 | 0.01 | 0 | 0.06 | 0.05 | 0.02 | 0.04 | 0.01 | 0.01 |
| 6 | 29395747 | rs7770592 | 0.878 | -1.277 | -0.217 | 0.043 | 0 | 0.001 | 0.001 | 0 | 0.29 | 0.29 | 0.21 | 0.35 | 0.34 | 0.15 |
| 6 | 29397779 | rs9295805 | 2.239 | -1.289 | -0.196 | 0.047 | -0.041 | -0.033 | -0.033 | 0 | 0.2 | 0.1 | 0.16 | 0.24 | 0.31 | 0.09 |
| 6 | 29398000 | rs2394609 | 0.895 | -1.269 | -0.211 | 0.051 | 0 | 0 | 0 | 0 | 0.29 | 0.29 | 0.21 | 0.35 | 0.34 | 0.15 |
| 6 | 29398815 | rs3131025 | 0.019 | -1.118 | -0.171 | 0.108 | 0.003 | 0.021 | 0.021 | 0 | 0.68 | 0.61 | 0.51 | 0.62 | 0.69 | 0.58 |
| 6 | 29398859 | rs7349863 | 1.997 | -0.808 | -0.434 | 0.217 | -0.016 | -0.013 | -0.013 | 0 | 0.2 | 0.15 | 0.16 | 0.24 | 0.31 | 0.09 |
| 6 | 29402435 | rs16894908 | -0.955 | -1.275 | -0.041 | -0.156 | -0.069 | -0.063 | -0.063 | 0 | 0.14 | 0.29 | 0.08 | 0.1 | 0.04 | 0.06 |
| 6 | 29404157 | rs3094561 | -0.91 | -1.481 | -0.063 | 0.229 | -0.044 | -0.044 | -0.044 | 0 | 0.64 | 0.78 | 0.59 | 0.62 | 0.55 | 0.66 |
| 6 | 29407970 | rs2074469 | 1.569 | -1.364 | -0.181 | 0.267 | -0.048 | -0.054 | -0.054 | 0 | 0.15 | 0.08 | 0.25 | 0.25 | 0.18 | 0.19 |
| 6 | 29408476 | rs2074465 | 0.761 | -1.389 | 0.133 | 0.277 | -0.036 | -0.031 | -0.031 | 0 | 0.1 | 0.15 | 0.06 | 0.05 | 0.02 | 0.01 |
| 6 | 29410093 | rs7765791 | 0.761 | -1.382 | 0.133 | 0.281 | -0.033 | -0.028 | -0.028 | 0 | 0.1 | 0.15 | 0.07 | 0.08 | 0.03 | 0.04 |
| 6 | 29412222 | rs16894919 | -1.067 | -1.406 | 0.186 | 0.294 | -0.085 | -0.024 | -0.024 | 0.011 | 0.16 | 0.34 | 0.09 | 0.18 | 0.04 | 0.11 |
| 6 | 29414286 | rs7738722 | -0.952 | -1.443 | 0.134 | 0.236 | -0.076 | -0.067 | -0.067 | 0 | 0.15 | 0.29 | 0.08 | 0.11 | 0.04 | 0.09 |
| 6 | 29414346 | rs7738742 | 0.436 | -1.443 | 0.134 | 0.236 | -0.036 | -0.028 | -0.028 | 0 | 0.12 | 0.15 | 0.06 | 0.05 | 0.02 | 0.04 |
| 6 | 29414613 | rs7739243 | -0.917 | -1.437 | 0.108 | 0.239 | -0.076 | -0.067 | -0.067 | 0 | 0.15 | 0.29 | 0.08 | 0.11 | 0.04 | 0.09 |
| 6 | 29415636 | rs6906897 | -0.917 | -1.437 | 0.108 | 0.239 | -0.076 | -0.067 | -0.067 | 1.845 | 0.15 | 0.29 | 0.08 | 0.11 | 0.04 | 0.09 |
| 6 | 29416652 | rs6935708 | 0.917 | -1.44 | 0.109 | 0.237 | -0.044 | -0.035 | -0.035 | 0 | 0.1 | 0.16 | 0.05 | 0.05 | 0.02 | 0.04 |
| 6 | 29417817 | rs2523442 | 2.006 | -1.59 | -0.09 | 0.154 | 0.005 | 0.086 | 0.086 | 0.113 | 0.3 | 0.03 | 0.2 | 0.16 | 0.26 | 0.32 |
| 6 | 29421209 | rs7738548 | -0.54 | -1.581 | -0.018 | 0.127 | -0.065 | -0.059 | -0.059 | 0.885 | 0.13 | 0.29 | 0.08 | 0.1 | 0.04 | 0.06 |
| 6 | 29422092 | rs3094574 | -0.986 | -1.542 | 0.008 | 0.217 | -0.042 | -0.042 | -0.042 | 0.041 | 0.64 | 0.78 | 0.58 | 0.62 | 0.55 | 0.67 |
| 6 | 29425584 | rs9257863 | 2.311 | -1.531 | -0.032 | 0.237 | -0.041 | -0.055 | -0.055 | 0 | 0.12 | 0.05 | 0.21 | 0.25 | 0.15 | 0.17 |
| 6 | 29426008 | rs3117194 | -1.009 | -1.539 | 0.006 | 0.236 | -0.046 | -0.047 | -0.047 | 0 | 0.64 | 0.8 | 0.59 | 0.62 | 0.55 | 0.66 |
| 6 | 29426203 | rs6456951 | -0.552 | -1.527 | 0.076 | 0.211 | -0.065 | -0.059 | -0.059 | 0 | 0.13 | 0.29 | 0.08 | 0.1 | 0.04 | 0.06 |
| 6 | 29428831 | rs7768854 | 1.421 | -1.523 | 0.132 | 0.214 | -0.024 | -0.019 | -0.019 | 0 | 0.07 | 0.1 | 0.04 | 0.05 | 0.01 | 0.01 |
| 6 | 29430170 | rs2073148 | -1.011 | -1.576 | 0.162 | 0.229 | -0.083 | -0.022 | -0.022 | 0 | 0.16 | 0.33 | 0.09 | 0.18 | 0.04 | 0.11 |
| 6 | 29432390 | rs1345228 | 2.163 | -1.544 | 0.058 | 0.259 | -0.036 | -0.049 | -0.049 | 0 | 0.13 | 0.07 | 0.22 | 0.25 | 0.16 | 0.17 |
| 6 | 29433942 | rs2107192 | 2.107 | -1.612 | 0.009 | 0.207 | 0.013 | 0.076 | 0.076 | 0.055 | 0.3 | 0.05 | 0.21 | 0.17 | 0.26 | 0.32 |
| 6 | 29434295 | rs2107191 | -1.031 | -1.557 | 0.095 | 0.223 | -0.046 | -0.047 | -0.047 | 0 | 0.64 | 0.8 | 0.6 | 0.61 | 0.55 | 0.66 |
| 6 | 29439105 | rs11961170 | 0.225 | -1.481 | 0.184 | 0.214 | -0.02 | -0.021 | -0.021 | 0 | 0.06 | 0.19 | 0.04 | 0.06 | 0.02 | 0.05 |
| 6 | 29440257 | rs16894948 | -0.993 | -1.511 | 0.212 | 0.211 | -0.084 | -0.022 | -0.022 | 0.98 | 0.16 | 0.33 | 0.09 | 0.19 | 0.04 | 0.11 |
| 6 | 29440741 | rs12206077 | 2.781 | -1.491 | 0.038 | 0.268 | -0.1 | -0.046 | -0.046 | 0 | 0.17 | 0.02 | 0.13 | 0.12 | 0.3 | 0.08 |
| 6 | 29441169 | rs1362076 | 0.901 | -1.481 | 0.122 | 0.189 | -0.034 | 0.029 | 0.029 | 0 | 0.07 | 0.2 | 0.02 | 0.13 | 0.01 | 0.09 |
| 6 | 29441618 | rs7751705 | 0.172 | -1.475 | 0.121 | 0.184 | -0.056 | -0.045 | -0.045 | 0 | 0.11 | 0.26 | 0.06 | 0.1 | 0.03 | 0.06 |
| 6 | 29444121 | rs1233495 | 1.803 | -1.522 | 0.392 | 0.189 | 0.068 | 0.046 | 0.046 | 0 | 0.14 | 0.24 | 0.26 | 0.25 | 0.24 | 0.18 |
| 6 | 29446632 | rs3094573 | -0.997 | -1.578 | 0.326 | 0.18 | -0.046 | -0.044 | -0.044 | 0 | 0.64 | 0.8 | 0.59 | 0.63 | 0.55 | 0.67 |

| | | | | | | | | | | | | | | | | |
|---|---|---|---|---|---|---|---|---|---|---|---|---|---|---|---|---|
| 6 | 29448038 | rs7771335 | -0.897 | -1.588 | 0.254 | 0.172 | -0.088 | -0.022 | -0.022 | 0.018 | 0.16 | 0.33 | 0.08 | 0.18 | 0.03 | 0.11 |
| 6 | 29451168 | rs2107189 | -0.897 | -1.588 | 0.24 | 0.172 | -0.087 | -0.022 | -0.022 | 0 | 0.16 | 0.33 | 0.08 | 0.18 | 0.03 | 0.11 |
| 6 | 29452411 | rs17415601 | 3.609 | -1.583 | 0.214 | 0.165 | 0 | 0.023 | 0.023 | 0 | 0.07 | 0.01 | 0.03 | 0.01 | 0.04 | 0.14 |
| 6 | 29454489 | rs2066951 | -0.889 | -1.604 | 0.274 | 0.161 | -0.087 | -0.022 | -0.022 | 4.572 | 0.16 | 0.33 | 0.08 | 0.18 | 0.03 | 0.11 |
| 6 | 29455599 | rs17184114 | -0.251 | -1.588 | 0.232 | 0.169 | -0.067 | -0.01 | -0.01 | 0 | 0.13 | 0.27 | 0.06 | 0.13 | 0.03 | 0.07 |
| 6 | 29458477 | rs1233492 | 2.109 | -1.538 | 0.202 | 0.172 | 0.018 | 0.075 | 0.075 | 0.022 | 0.3 | 0.05 | 0.21 | 0.17 | 0.26 | 0.33 |
| 6 | 29461914 | rs1233490 | 2.12 | -1.545 | 0.192 | 0.169 | 0.004 | 0.062 | 0.062 | 0.1 | 0.29 | 0.03 | 0.21 | 0.17 | 0.26 | 0.32 |
| 6 | 29463892 | rs6906307 | 0.254 | -1.508 | 0.289 | 0.19 | -0.052 | -0.043 | -0.043 | 0.035 | 0.11 | 0.22 | 0.05 | 0.1 | 0.02 | 0.06 |
| 6 | 29463968 | rs6926506 | 0.254 | -1.508 | 0.289 | 0.188 | -0.052 | -0.047 | -0.047 | 0 | 0.11 | 0.22 | 0.05 | 0.07 | 0.02 | 0.04 |
| 6 | 29464365 | rs6907220 | 0.254 | -1.508 | 0.289 | 0.188 | -0.052 | -0.047 | -0.047 | 0 | 0.11 | 0.22 | 0.05 | 0.07 | 0.02 | 0.04 |
| 6 | 29466956 | rs1345227 | -1.509 | -1.471 | 0.236 | 0.246 | -0.04 | -0.05 | -0.05 | 0 | 0.71 | 0.83 | 0.65 | 0.69 | 0.59 | 0.83 |
| 6 | 29468809 | rs1233487 | 1.687 | -1.14 | 0.225 | 0.361 | 0.062 | 0.042 | 0.042 | 0 | 0.14 | 0.24 | 0.26 | 0.25 | 0.24 | 0.17 |
| 6 | 29469652 | rs1233486 | 1.942 | -1.287 | -0.126 | 0.335 | 0.024 | 0.081 | 0.081 | 0.044 | 0.31 | 0.05 | 0.21 | 0.17 | 0.26 | 0.33 |
| 6 | 29470985 | rs3131019 | -1.068 | -1.314 | 0.043 | 0.381 | -0.004 | -0.006 | -0.006 | 0 | 0.67 | 0.69 | 0.63 | 0.69 | 0.57 | 0.83 |
| 6 | 29472916 | rs757256 | 0.405 | -1.308 | 0.183 | 0.379 | -0.001 | -0.004 | -0.004 | 1.009 | 0.47 | 0.49 | 0.38 | 0.35 | 0.41 | 0.58 |
| 6 | 29474385 | rs3130860 | 0.423 | -1.286 | 0.105 | 0.372 | -0.001 | -0.004 | -0.004 | 1.033 | 0.47 | 0.49 | 0.39 | 0.34 | 0.42 | 0.58 |
| 6 | 29474662 | rs734959 | -1.17 | -1.274 | 0.086 | 0.383 | -0.004 | -0.005 | -0.005 | 0.026 | 0.68 | 0.69 | 0.63 | 0.69 | 0.57 | 0.82 |
| 6 | 29475142 | rs734960 | 2.399 | -1.234 | 0.072 | 0.404 | -0.017 | -0.025 | -0.025 | 0.346 | 0.12 | 0.08 | 0.2 | 0.25 | 0.14 | 0.19 |
| 6 | 29475467 | rs1002187 | -0.21 | -1.259 | 0.054 | 0.397 | -0.001 | 0 | 0 | 0 | 0.59 | 0.56 | 0.58 | 0.59 | 0.56 | 0.76 |
| 6 | 29475682 | rs1233482 | 1.754 | -1.283 | -0.041 | 0.342 | 0.024 | 0.081 | 0.081 | 0.044 | 0.31 | 0.05 | 0.21 | 0.17 | 0.26 | 0.33 |
| 6 | 29475902 | rs3131020 | 0.197 | -1.191 | 0.07 | 0.39 | 0.01 | 0.026 | 0.026 | 0 | 0.49 | 0.55 | 0.39 | 0.37 | 0.41 | 0.6 |
| 6 | 29476142 | rs3131021 | -0.385 | -1.139 | 0.105 | 0.422 | 0 | 0.001 | 0.001 | 0 | 0.61 | 0.6 | 0.59 | 0.62 | 0.56 | 0.78 |
| 6 | 29476325 | rs3128847 | -0.703 | -1.112 | 0.046 | 0.609 | -0.031 | 0.046 | 0.046 | 0 | 0.85 | 0.7 | 0.76 | 0.81 | 0.87 | 0.9 |
| 6 | 29476508 | rs9257877 | 0.927 | -0.805 | 0.318 | 0.399 | 0.036 | 0.053 | 0.053 | 0 | 0.07 | 0.12 | 0.18 | 0.18 | 0.1 | 0.07 |
| 6 | 29477821 | rs1233478 | -0.345 | -0.869 | -0.294 | 0.254 | 0.102 | -0.026 | -0.026 | 0.001 | 0.14 | 0.45 | 0.18 | 0.09 | 0.16 | 0.2 |
| 6 | 29478301 | rs2285791 | 1.923 | -0.936 | -0.388 | 0.272 | -0.025 | -0.04 | -0.04 | 0 | 0.13 | 0.07 | 0.2 | 0.27 | 0.15 | 0.21 |
| 6 | 29480993 | rs3094572 | -1.155 | -1.062 | -0.448 | 0.204 | -0.002 | -0.002 | -0.002 | 0 | 0.66 | 0.68 | 0.63 | 0.69 | 0.57 | 0.82 |
| 6 | 29481948 | rs1010409 | 0.295 | -1.13 | -0.462 | 0.196 | 0.001 | 0.006 | 0.006 | 0 | 0.46 | 0.55 | 0.41 | 0.29 | 0.42 | 0.55 |
| 6 | 29482128 | rs1362075 | -1.179 | -1.129 | -0.481 | 0.197 | -0.002 | -0.003 | -0.003 | 0 | 0.66 | 0.69 | 0.63 | 0.69 | 0.57 | 0.82 |
| 6 | 29482432 | rs2523433 | 1.665 | -1.148 | -0.534 | 0.168 | 0.01 | 0.077 | 0.077 | 0.089 | 0.31 | 0.04 | 0.23 | 0.17 | 0.27 | 0.33 |
| 6 | 29483911 | rs1592411 | 0.437 | -1.019 | -0.375 | 0.319 | -0.002 | -0.019 | -0.019 | 0 | 0.46 | 0.49 | 0.4 | 0.27 | 0.42 | 0.53 |
| 6 | 29483968 | rs1592410 | 0.437 | -1.019 | -0.375 | 0.319 | -0.002 | -0.019 | -0.019 | 0.1 | 0.46 | 0.49 | 0.4 | 0.27 | 0.42 | 0.53 |
| 6 | 29485289 | rs9257887 | 0.803 | -1.013 | -0.378 | 0.338 | 0 | -0.055 | -0.055 | 0 | 0.2 | 0.12 | 0.22 | 0.42 | 0.16 | 0.3 |
| 6 | 29485373 | rs1015868 | 0.436 | -1.013 | -0.392 | 0.326 | -0.002 | -0.019 | -0.019 | 0 | 0.45 | 0.49 | 0.4 | 0.27 | 0.41 | 0.53 |
| 6 | 29486345 | rs3094564 | -1.013 | -0.797 | -0.58 | 0.402 | -0.001 | -0.001 | -0.001 | 0.012 | 0.66 | 0.68 | 0.63 | 0.69 | 0.57 | 0.82 |
| 6 | 29486908 | rs9257890 | 2.07 | -0.752 | -0.773 | 0.519 | -0.04 | -0.053 | -0.053 | 0 | 0.12 | 0.03 | 0.2 | 0.25 | 0.14 | 0.19 |
| 6 | 29488249 | rs2745400 | 0.041 | -1.1 | -0.633 | 0.348 | -0.013 | 0.002 | 0.002 | 0 | 0.45 | 0.63 | 0.4 | 0.49 | 0.5 | 0.32 |
| 6 | 29489048 | rs724078 | -0.204 | -1.515 | -0.579 | 0.264 | 0.029 | 0.031 | 0.031 | 0 | 0.53 | 0.6 | 0.39 | 0.28 | 0.34 | 0.44 |
| 6 | 29494897 | rs969931 | 0.083 | -1.191 | -0.528 | 0.515 | -0.012 | 0.002 | 0.002 | 0 | 0.46 | 0.61 | 0.42 | 0.49 | 0.49 | 0.33 |
| 6 | 29495280 | rs6933331 | 1.004 | -1.368 | -0.327 | 0.61 | -0.024 | 0.004 | 0.004 | 0 | 0.09 | 0.12 | 0.04 | 0.13 | 0.01 | 0.06 |
| 6 | 29496664 | rs1233427 | 1.433 | -1.532 | -0.095 | 0.535 | -0.032 | -0.022 | -0.022 | 1.915 | 0.31 | 0.22 | 0.33 | 0.31 | 0.46 | 0.21 |
| 6 | 29496776 | rs9501676 | 1.38 | -1.533 | -0.039 | 0.533 | 0.034 | 0.057 | 0.057 | 0.169 | 0.05 | 0.27 | 0.04 | 0.01 | 0.01 | 0 |
| 6 | 29497002 | rs9501677 | -0.069 | -1.534 | -0.087 | 0.535 | -0.084 | 0.027 | 0.027 | 0 | 0.14 | 0.4 | 0.09 | 0.16 | 0.03 | 0.09 |
| 6 | 29498102 | rs1233426 | 1.4 | -1.444 | -0.21 | 0.555 | -0.032 | -0.022 | -0.022 | 0 | 0.31 | 0.22 | 0.33 | 0.31 | 0.46 | 0.21 |
| 6 | 29498423 | rs6908631 | 1.014 | -1.492 | -0.164 | 0.551 | -0.024 | 0.004 | 0.004 | 0 | 0.08 | 0.12 | 0.04 | 0.13 | 0.01 | 0.06 |
| 6 | 29499416 | rs362546 | 1.42 | -1.495 | -0.189 | 0.551 | -0.032 | -0.022 | -0.022 | 0.007 | 0.31 | 0.22 | 0.33 | 0.31 | 0.46 | 0.21 |
| 6 | 29499505 | rs909968 | 1.649 | -1.525 | -0.161 | 0.537 | -0.066 | -0.046 | -0.046 | 0 | 0.3 | 0.14 | 0.32 | 0.3 | 0.46 | 0.16 |
| 6 | 29499537 | rs909967 | -0.039 | -1.567 | -0.078 | 0.522 | -0.013 | 0.004 | 0.004 | 0 | 0.46 | 0.62 | 0.42 | 0.5 | 0.49 | 0.33 |
| 6 | 29500225 | rs886381 | 1.367 | -1.745 | -0.369 | 0.386 | 0.005 | 0.062 | 0.062 | 0.587 | 0.29 | 0.02 | 0.21 | 0.17 | 0.25 | 0.33 |
| 6 | 29500412 | rs1233422 | 1.486 | -1.637 | -0.242 | 0.448 | -0.032 | -0.022 | -0.022 | 4.57 | 0.32 | 0.23 | 0.33 | 0.33 | 0.46 | 0.24 |
| 6 | 29501422 | rs9461533 | 1.185 | -1.503 | -0.145 | 0.48 | 0.021 | 0.049 | 0.049 | 0.106 | 0.06 | 0.27 | 0.04 | 0.06 | 0.01 | 0.05 |
| 6 | 29501514 | rs1233421 | 0.38 | -1.537 | -0.203 | 0.464 | 0.012 | 0.006 | 0.006 | 0 | 0.4 | 0.51 | 0.39 | 0.31 | 0.5 | 0.17 |
| 6 | 29503323 | rs1233420 | -0.226 | -1.618 | -0.24 | 0.52 | -0.007 | 0.012 | 0.012 | 0 | 0.5 | 0.72 | 0.45 | 0.48 | 0.51 | 0.33 |
| 6 | 29507310 | rs362514 | 0.286 | -1.588 | -0.226 | 0.518 | -0.015 | -0.017 | -0.017 | 0 | 0.43 | 0.36 | 0.39 | 0.41 | 0.5 | 0.23 |
| 6 | 29507426 | rs446198 | 0.929 | -1.675 | -0.206 | 0.522 | -0.035 | -0.015 | -0.015 | 0.03 | 0.35 | 0.25 | 0.35 | 0.29 | 0.49 | 0.17 |
| 6 | 29508273 | rs10484548 | 1.516 | -1.695 | -0.156 | 0.514 | 0.034 | 0.057 | 0.057 | 0.169 | 0.05 | 0.27 | 0.04 | 0.01 | 0.01 | 0 |
| 6 | 29510250 | rs362541 | 1.516 | -1.695 | -0.156 | 0.514 | 0.034 | 0.057 | 0.057 | 0.169 | 0.05 | 0.27 | 0.04 | 0.01 | 0.01 | 0 |
| 6 | 29510325 | rs362540 | -0.317 | -1.691 | -0.202 | 0.515 | -0.004 | 0.01 | 0.01 | 0 | 0.5 | 0.72 | 0.46 | 0.44 | 0.52 | 0.29 |
| 6 | 29511216 | rs3025657 | 1.105 | -2.086 | -0.194 | 0.518 | -0.021 | 0.003 | 0.003 | 0 | 0.09 | 0.12 | 0.04 | 0.13 | 0.01 | 0.06 |
| 6 | 29511341 | rs2534795 | 1.392 | -2.155 | -0.303 | 0.466 | 0.007 | 0.059 | 0.059 | 0.552 | 0.29 | 0.02 | 0.21 | 0.18 | 0.25 | 0.33 |
| 6 | 29511850 | rs1233417 | -0.246 | -2.148 | -0.288 | 0.472 | 0.001 | 0.017 | 0.017 | 0.004 | 0.71 | 0.62 | 0.63 | 0.54 | 0.76 | 0.59 |
| 6 | 29513807 | rs407161 | -0.157 | -2.126 | -0.291 | 0.476 | -0.003 | 0.006 | 0.006 | 0 | 0.49 | 0.64 | 0.45 | 0.48 | 0.52 | 0.28 |
| 6 | 29515802 | rs3094575 | -0.464 | -2.143 | -0.183 | 0.5 | -0.001 | -0.008 | -0.008 | 0 | 0.72 | 0.71 | 0.64 | 0.55 | 0.76 | 0.59 |
| 6 | 29516180 | rs387659 | 0.32 | -2.156 | -0.153 | 0.913 | -0.015 | 0.001 | 0.001 | 0 | 0.44 | 0.37 | 0.4 | 0.37 | 0.51 | 0.25 |
| 6 | 29516242 | rs3094576 | -0.662 | -2.154 | -0.282 | 0.982 | 0.031 | 0.056 | 0.056 | 0 | 0.92 | 0.87 | 0.78 | 0.76 | 0.85 | 0.89 |
| 6 | 29517093 | rs376681 | 1.016 | -2.146 | -0.23 | 0.91 | -0.045 | 0.006 | 0.006 | 0 | 0.35 | 0.24 | 0.35 | 0.23 | 0.49 | 0.19 |
| 6 | 29518442 | rs2745411 | 0.307 | -2.225 | -0.221 | 0.859 | -0.013 | -0.003 | -0.003 | 0 | 0.45 | 0.62 | 0.43 | 0.37 | 0.49 | 0.25 |
| 6 | 29518472 | rs417374 | -0.248 | -2.215 | -0.191 | 0.867 | -0.006 | 0.012 | 0.012 | 0 | 0.5 | 0.71 | 0.46 | 0.38 | 0.51 | 0.3 |
| 6 | 29519848 | rs453658 | -1.148 | -2.181 | -0.233 | 0.87 | -0.045 | 0.006 | 0.006 | 0 | 0.65 | 0.76 | 0.65 | 0.77 | 0.51 | 0.81 |

| | | | | | | | | | | | | | | | | |
|---|---|---|---|---|---|---|---|---|---|---|---|---|---|---|---|---|
| 6 | 29522588 | rs6915177 | -0.191 | -2.217 | -0.16 | 0.845 | -0.094 | 0.019 | 0.019 | 0.314 | 0.15 | 0.48 | 0.11 | 0.15 | 0.03 | 0.12 |
| 6 | 29523872 | rs2076486 | -0.191 | -2.217 | -0.16 | 0.845 | -0.094 | 0.019 | 0.019 | 0.314 | 0.15 | 0.48 | 0.11 | 0.15 | 0.03 | 0.12 |
| 6 | 29523952 | rs2076485 | 1.443 | -2.262 | -0.226 | 0.818 | 0.007 | 0.06 | 0.06 | 0.566 | 0.29 | 0.03 | 0.21 | 0.18 | 0.25 | 0.3 |
| 6 | 29524003 | rs2076484 | -0.166 | -2.217 | -0.23 | 0.819 | -0.094 | 0.019 | 0.019 | 0.314 | 0.15 | 0.48 | 0.11 | 0.15 | 0.03 | 0.12 |
| 6 | 29524168 | rs2534790 | 1.45 | -2.223 | -0.236 | 0.814 | 0.003 | 0.058 | 0.058 | 0.559 | 0.28 | 0.03 | 0.21 | 0.18 | 0.25 | 0.3 |
| 6 | 29524401 | rs2745412 | 2.402 | -2.473 | -0.42 | 0.442 | 0.005 | 0.034 | 0.034 | 1.09 | 0.08 | 0 | 0.02 | 0 | 0.08 | 0.02 |
| 6 | 29526940 | rs362536 | 0.231 | -2.288 | -0.247 | 0.588 | -0.006 | 0.013 | 0.013 | 0 | 0.5 | 0.29 | 0.54 | 0.62 | 0.49 | 0.7 |
| 6 | 29527267 | rs64036 | 0.143 | -2.215 | -0.201 | 0.645 | -0.08 | 0.03 | 0.03 | 0 | 0.14 | 0.39 | 0.09 | 0.14 | 0.03 | 0.07 |
| 6 | 29527599 | rs2272991 | -0.151 | -2.12 | -0.39 | 0.692 | -0.094 | 0.019 | 0.019 | 0.314 | 0.15 | 0.48 | 0.11 | 0.15 | 0.03 | 0.12 |
| 6 | 29528318 | rs362513 | -1.764 | -2.024 | -0.618 | 0.698 | -0.067 | -0.014 | -0.014 | 0.055 | 0.7 | 0.86 | 0.68 | 0.77 | 0.54 | 0.82 |
| 6 | 29530413 | rs362532 | 0.619 | -1.753 | -0.339 | 0.71 | -0.049 | 0.013 | 0.013 | 0 | 0.1 | 0.25 | 0.07 | 0.14 | 0.01 | 0.12 |
| 6 | 29530455 | rs362531 | 0.646 | -1.741 | -0.434 | 0.718 | -0.042 | 0.008 | 0.008 | 0 | 0.1 | 0.2 | 0.06 | 0.14 | 0.01 | 0.12 |
| 6 | 29530628 | rs362512 | 0.105 | -1.767 | -0.69 | 0.745 | -0.009 | 0.013 | 0.013 | 0 | 0.49 | 0.29 | 0.53 | 0.51 | 0.49 | 0.65 |
| 6 | 29530850 | rs362511 | -0.335 | -1.73 | -0.581 | 0.531 | -0.014 | 0.007 | 0.007 | 0 | 0.26 | 0.38 | 0.29 | 0.29 | 0.22 | 0.21 |
| 6 | 29530868 | rs362529 | 1.338 | -1.798 | -0.509 | 0.331 | -0.011 | 0.021 | 0.021 | 0.042 | 0.2 | 0.37 | 0.18 | 0.22 | 0.15 | 0.33 |
| 6 | 29531253 | rs459970 | 1.078 | -1.621 | -0.844 | 0.446 | -0.007 | 0.015 | 0.015 | 0 | 0.06 | 0.11 | 0.05 | 0.09 | 0.09 | 0.06 |
| 6 | 29531818 | rs362527 | 1.176 | -1.593 | -1.108 | 0.185 | 0.03 | 0.046 | 0.046 | 0.248 | 0.13 | 0.05 | 0.05 | 0 | 0.05 | 0.03 |
| 6 | 29533295 | rs388234 | -0.645 | -1.919 | -1.149 | 0.289 | 0.008 | 0.005 | 0.005 | 0 | 0.48 | 0.49 | 0.49 | 0.57 | 0.4 | 0.52 |
| 6 | 29533870 | rs3094577 | -1.714 | -1.795 | -1.231 | 0.194 | 0.001 | -0.022 | -0.022 | 0.054 | 0.66 | 0.71 | 0.73 | 0.88 | 0.62 | 0.79 |
| 6 | 29534240 | rs3117072 | -1.654 | -1.769 | -1.261 | 0.2 | 0.001 | -0.033 | -0.033 | 0 | 0.65 | 0.67 | 0.73 | 0.88 | 0.62 | 0.79 |
| 6 | 29535217 | rs12525160 | 1.847 | -1.513 | -1.669 | 0.424 | -0.001 | -0.001 | -0.001 | 0 | 0.2 | 0.2 | 0.22 | 0.22 | 0.19 | 0.26 |
| 6 | 29536685 | rs1119080 | -0.857 | -1.522 | -1.676 | 0.428 | 0.008 | 0.005 | 0.005 | 0 | 0.49 | 0.5 | 0.49 | 0.58 | 0.41 | 0.52 |
| 6 | 29537975 | rs1233400 | 1.474 | -1.45 | -1.717 | 0.498 | 0.024 | 0.027 | 0.027 | 0 | 0.09 | 0.12 | 0.19 | 0.21 | 0.16 | 0.07 |
| 6 | 29539765 | rs3025652 | 1.793 | -1.364 | -1.709 | 0.507 | -0.002 | -0.001 | -0.001 | 0 | 0.2 | 0.21 | 0.22 | 0.22 | 0.19 | 0.26 |
| 6 | 29540745 | rs362510 | 0.823 | -1.427 | -1.702 | 0.494 | -0.005 | -0.003 | -0.003 | 0.861 | 0.2 | 0.16 | 0.22 | 0.22 | 0.2 | 0.09 |
| 6 | 29542947 | rs3025650 | 1.549 | -1.28 | -1.649 | 0.548 | 0.002 | 0.021 | 0.021 | 0 | 0.07 | 0.18 | 0.09 | 0.08 | 0.07 | 0.05 |
| 6 | 29543646 | rs362525 | 1.448 | -1.32 | -1.668 | 0.558 | 0.001 | -0.001 | -0.001 | 0 | 0.21 | 0.22 | 0.22 | 0.22 | 0.19 | 0.26 |
| 6 | 29544825 | rs3117073 | -2.425 | -1.146 | -2.136 | 0.171 | -0.097 | 0.057 | 0.057 | 0 | 0.83 | 0.98 | 0.87 | 0.93 | 0.72 | 0.93 |
| 6 | 29545208 | rs362522 | 1.648 | -1.244 | -2.084 | 0.376 | -0.01 | -0.022 | -0.022 | 0 | 0.17 | 0.13 | 0.28 | 0.3 | 0.25 | 0.32 |
| 6 | 29545284 | rs362523 | 0.954 | -1.275 | -2.013 | 0.408 | 0.022 | 0.072 | 0.072 | 0 | 0.12 | 0.17 | 0.2 | 0.35 | 0.16 | 0.18 |
| 6 | 29545715 | rs1233397 | 0.866 | -1.411 | -2.064 | 0.341 | 0.004 | -0.022 | -0.022 | 0 | 0.28 | 0.22 | 0.32 | 0.49 | 0.21 | 0.36 |
| 6 | 29549261 | rs1233394 | 1.5 | -1.492 | -2.047 | 0.395 | 0.019 | 0.029 | 0.029 | 0 | 0.1 | 0.13 | 0.19 | 0.24 | 0.15 | 0.08 |
| 6 | 29549443 | rs12530421 | 2.384 | -1.53 | -2.065 | 0.382 | 0.055 | 0.016 | 0.016 | 0 | 0.16 | 0.05 | 0.12 | 0.13 | 0.05 | 0.2 |
| 6 | 29550416 | rs1233392 | 1.507 | -1.516 | -2.06 | 0.389 | 0.019 | 0.029 | 0.029 | 0 | 0.1 | 0.13 | 0.19 | 0.24 | 0.15 | 0.08 |
| 6 | 29550618 | rs1233391 | 0.826 | -1.532 | -2.064 | 0.38 | 0.006 | -0.023 | -0.023 | 0 | 0.28 | 0.22 | 0.32 | 0.49 | 0.2 | 0.36 |
| 6 | 29551861 | rs1233390 | 1.044 | -1.505 | -2.049 | 0.384 | 0.021 | 0.075 | 0.075 | 0 | 0.12 | 0.17 | 0.19 | 0.35 | 0.15 | 0.16 |
| 6 | 29553727 | rs1233388 | 0.832 | -1.523 | -2.06 | 0.341 | 0.005 | -0.02 | -0.02 | 0.189 | 0.28 | 0.23 | 0.32 | 0.48 | 0.21 | 0.36 |
| 6 | 29555778 | rs2235698 | 1.104 | -1.519 | -2.108 | 0.283 | -0.003 | -0.027 | -0.027 | 0.16 | 0.22 | 0.16 | 0.3 | 0.34 | 0.26 | 0.34 |
| 6 | 29556759 | rs362521 | 2.921 | -1.382 | -2.221 | 0.45 | 0.03 | -0.001 | -0.001 | 0 | 0.11 | 0.01 | 0.1 | 0.1 | 0.04 | 0.17 |
| 6 | 29558190 | rs1233386 | 1.278 | -1.239 | -2.143 | 0.524 | 0.019 | 0.031 | 0.031 | 0 | 0.1 | 0.13 | 0.19 | 0.24 | 0.15 | 0.08 |
| 6 | 29559295 | rs1233384 | 1.278 | -1.239 | -2.143 | 0.524 | 0.019 | 0.031 | 0.031 | 0 | 0.1 | 0.13 | 0.19 | 0.24 | 0.15 | 0.08 |
| 6 | 29563067 | rs17508548 | 1.033 | -1.64 | -2.577 | 0.188 | 0.049 | -0.013 | -0.013 | 0.003 | 0.22 | 0.11 | 0.16 | 0.25 | 0.11 | 0.32 |
| 6 | 29566369 | rs3095273 | -1.981 | -1.587 | -2.693 | 0.074 | -0.098 | 0.051 | 0.051 | 0 | 0.83 | 0.98 | 0.87 | 0.93 | 0.7 | 0.93 |
| 6 | 29566842 | rs3025646 | 1.556 | -2.47 | -1.759 | 0.722 | 0.047 | 0.012 | 0.012 | 0 | 0.14 | 0.02 | 0.11 | 0.1 | 0.05 | 0.18 |
| 6 | 29567638 | rs362518 | 0.56 | -2.484 | -1.759 | 0.736 | 0.049 | -0.013 | -0.013 | 0.174 | 0.22 | 0.11 | 0.16 | 0.25 | 0.11 | 0.32 |
| 6 | 29569045 | rs362517 | 0.923 | -2.441 | -1.736 | 0.748 | 0 | 0.003 | 0.003 | 0 | 0.06 | 0.05 | 0.04 | 0.12 | 0.05 | 0.12 |
| 6 | 29569532 | rs362515 | 0.923 | -2.441 | -1.736 | 0.748 | 0 | 0.003 | 0.003 | 0 | 0.06 | 0.05 | 0.04 | 0.12 | 0.05 | 0.12 |
| 6 | 29570665 | rs3025640 | 0.387 | -2.615 | -1.819 | 0.626 | 0.045 | 0.053 | 0.053 | 0.513 | 0.15 | 0.06 | 0.06 | 0.03 | 0.05 | 0.04 |
| 6 | 29570841 | rs2267633 | 0.616 | -2.605 | -1.812 | 0.648 | 0.049 | -0.013 | -0.013 | 0 | 0.22 | 0.11 | 0.16 | 0.25 | 0.11 | 0.32 |
| 6 | 29571545 | rs2076483 | 0.616 | -2.605 | -1.812 | 0.648 | 0.049 | -0.013 | -0.013 | 0.009 | 0.22 | 0.11 | 0.16 | 0.25 | 0.11 | 0.32 |
| 6 | 29571609 | rs10946999 | 1.589 | -2.517 | -1.823 | 0.75 | -0.02 | -0.069 | -0.069 | 0 | 0.11 | 0.01 | 0.15 | 0.28 | 0.11 | 0.26 |
| 6 | 29574631 | rs740884 | -0.221 | -2.188 | -1.263 | 0.248 | 0.032 | 0.032 | 0.032 | 0.017 | 0.17 | 0.13 | 0.07 | 0.05 | 0.06 | 0.1 |
| 6 | 29575003 | rs29267 | 0.313 | -2.165 | -1.269 | 0.324 | 0.049 | -0.013 | -0.013 | 0 | 0.22 | 0.11 | 0.16 | 0.25 | 0.11 | 0.32 |
| 6 | 29575456 | rs740882 | 0.011 | -2.203 | -1.291 | 0.292 | 0.024 | 0.005 | 0.005 | 0 | 0.16 | 0.09 | 0.08 | 0.12 | 0.1 | 0.13 |
| 6 | 29576047 | rs881284 | 0.011 | -2.203 | -1.291 | 0.292 | 0.024 | 0.005 | 0.005 | 0 | 0.16 | 0.09 | 0.08 | 0.12 | 0.1 | 0.13 |
| 6 | 29576393 | rs29230 | -0.551 | -2.262 | -1.319 | 0.276 | 0.092 | 0.029 | 0.029 | 0 | 0.32 | 0.15 | 0.2 | 0.25 | 0.15 | 0.33 |
| 6 | 29576592 | rs29262 | -0.119 | -1.786 | -1.124 | 0.412 | 0.024 | 0.005 | 0.005 | 0 | 0.16 | 0.09 | 0.08 | 0.12 | 0.1 | 0.13 |
| 6 | 29577617 | rs29258 | -0.119 | -1.786 | -1.124 | 0.412 | 0.024 | 0.005 | 0.005 | 0 | 0.16 | 0.09 | 0.08 | 0.12 | 0.1 | 0.13 |
| 6 | 29578030 | rs29257 | -0.119 | -1.786 | -1.124 | 0.412 | 0.024 | 0.005 | 0.005 | 0 | 0.16 | 0.09 | 0.08 | 0.12 | 0.1 | 0.13 |
| 6 | 29579544 | rs29255 | 1.757 | -1.798 | -1.129 | 0.407 | 0.033 | 0.004 | 0.004 | 0 | 0.11 | 0.02 | 0.1 | 0.1 | 0.05 | 0.17 |
| 6 | 29580436 | rs29253 | -0.173 | -1.793 | -1.127 | 0.41 | 0.026 | 0.005 | 0.005 | 0 | 0.16 | 0.09 | 0.08 | 0.12 | 0.1 | 0.15 |
| 6 | 29580588 | rs29227 | -0.12 | -1.793 | -1.127 | 0.41 | 0.024 | 0.005 | 0.005 | 0 | 0.16 | 0.09 | 0.08 | 0.12 | 0.1 | 0.13 |
| 6 | 29581011 | rs29226 | -0.173 | -1.793 | -1.127 | 0.41 | 0.026 | 0.005 | 0.005 | 0 | 0.16 | 0.09 | 0.08 | 0.12 | 0.1 | 0.15 |
| 6 | 29581041 | rs29225 | -0.12 | -1.793 | -1.127 | 0.41 | 0.024 | 0.005 | 0.005 | 0 | 0.16 | 0.09 | 0.08 | 0.12 | 0.1 | 0.13 |
| 6 | 29587874 | rs6919973 | -0.173 | -1.793 | -1.127 | 0.41 | 0.026 | 0.005 | 0.005 | 0 | 0.16 | 0.09 | 0.08 | 0.12 | 0.1 | 0.15 |
| 6 | 29588146 | rs29246 | 0.79 | -1.794 | -1.127 | 0.409 | 0.028 | 0.042 | 0.042 | 0.398 | 0.1 | 0.04 | 0.04 | 0 | 0.05 | 0.02 |
| 6 | 29588544 | rs29223 | 0.178 | -1.712 | -1.085 | 0.454 | 0.024 | 0.005 | 0.005 | 0 | 0.16 | 0.09 | 0.08 | 0.12 | 0.1 | 0.13 |
| 6 | 29592431 | rs2267635 | 1.694 | -1.504 | -1.367 | 0.494 | 0.035 | 0.005 | 0.005 | 0 | 0.11 | 0.02 | 0.1 | 0.1 | 0.05 | 0.17 |
| 6 | 29593788 | rs715044 | 1.694 | -1.504 | -1.367 | 0.494 | 0.035 | 0.005 | 0.005 | 0 | 0.11 | 0.02 | 0.1 | 0.1 | 0.05 | 0.17 |

| | | | | | | | | | | | | | | | | |
|---|---|---|---|---|---|---|---|---|---|---|---|---|---|---|---|---|
| 6 | 29598119 | rs2021749 | -0.104 | -0.962 | -1.024 | 0.357 | 0.025 | 0.005 | 0.005 | 0 | 0.16 | 0.09 | 0.08 | 0.12 | 0.1 | 0.15 |
| 6 | 29599102 | rs29243 | 0.149 | -0.483 | -0.658 | 0.494 | -0.015 | 0.013 | 0.013 | 0 | 0.05 | 0.11 | 0.03 | 0.08 | 0.02 | 0.05 |
| 6 | 29605935 | rs444189 | 0.068 | -0.476 | -0.73 | 0.397 | -0.002 | 0.005 | 0.005 | 0 | 0.39 | 0.41 | 0.41 | 0.5 | 0.32 | 0.55 |
| 6 | 29607429 | rs29218 | 1.385 | -0.584 | -0.975 | 0.583 | -0.024 | -0.064 | -0.064 | 0 | 0.18 | 0.09 | 0.27 | 0.33 | 0.17 | 0.36 |
| 6 | 29609921 | rs3117289 | 1.712 | -0.667 | -0.869 | 0.535 | -0.031 | -0.068 | -0.068 | 0 | 0.16 | 0.04 | 0.26 | 0.33 | 0.16 | 0.36 |
| 6 | 29610989 | rs29273 | 1.109 | -1.554 | -1.103 | -0.359 | 0.013 | 0.014 | 0.014 | 0 | 0.2 | 0.14 | 0.2 | 0.05 | 0.19 | 0.14 |
| 6 | 29611431 | rs29232 | 0.008 | -1.888 | -1.129 | -0.637 | 0.068 | -0.006 | -0.006 | 0 | 0.52 | 0.36 | 0.5 | 0.51 | 0.36 | 0.69 |
| 6 | 29611885 | rs3131854 | -0.483 | -1.309 | -1.075 | -0.929 | 0.01 | 0.021 | 0.021 | 0 | 0.73 | 0.77 | 0.81 | 0.84 | 0.77 | 0.79 |
| 6 | 29613218 | rs9257928 | 1.606 | -1.302 | -1.154 | -0.955 | 0.003 | -0.005 | -0.005 | 0 | 0.19 | 0.16 | 0.2 | 0.05 | 0.18 | 0.14 |
| 6 | 29615822 | rs3129073 | 2.908 | -1.257 | -1.245 | -0.912 | -0.04 | -0.079 | -0.079 | 0 | 0.16 | 0.04 | 0.19 | 0.33 | 0.16 | 0.36 |
| 6 | 29616607 | rs439812 | 1.92 | -1.356 | -1.097 | -0.884 | 0.009 | -0.001 | -0.001 | 0 | 0.11 | 0.12 | 0.26 | 0.28 | 0.25 | 0.11 |
| 6 | 29617747 | rs29269 | 1.706 | -1.396 | -1.112 | -0.944 | 0.013 | 0.014 | 0.014 | 0 | 0.2 | 0.14 | 0.2 | 0.05 | 0.19 | 0.14 |
| 6 | 29618366 | rs29272 | 0.837 | -1.549 | -1.156 | -1.018 | -0.008 | -0.024 | -0.024 | 0 | 0.08 | 0.17 | 0.09 | 0.01 | 0.05 | 0.01 |
| 6 | 29618525 | rs29231 | 1.792 | -1.55 | -1.155 | -1.018 | 0.008 | 0.002 | 0.002 | 0 | 0.21 | 0.16 | 0.21 | 0.05 | 0.19 | 0.14 |
| 6 | 29623739 | rs29228 | 2.922 | -1.139 | -1.193 | -0.739 | -0.04 | -0.08 | -0.08 | 0 | 0.16 | 0.04 | 0.19 | 0.33 | 0.16 | 0.36 |
| 6 | 29624112 | rs29234 | 0.568 | -1.045 | -0.935 | -0.761 | -0.008 | -0.024 | -0.024 | 0.063 | 0.08 | 0.17 | 0.09 | 0.01 | 0.05 | 0.01 |
| 6 | 29624894 | rs9468571 | 0.568 | -1.045 | -0.935 | -0.761 | -0.008 | -0.024 | -0.024 | 0.746 | 0.08 | 0.17 | 0.09 | 0.01 | 0.05 | 0.01 |
| 6 | 29625001 | rs3130250 | -1.739 | -0.972 | -1.173 | -0.715 | -0.023 | 0.03 | 0.03 | 0 | 0.9 | 0.95 | 0.91 | 0.97 | 0.83 | 0.96 |
| 6 | 29628437 | rs3135050 | -1.951 | -0.945 | -1.182 | -0.695 | -0.033 | 0.032 | 0.032 | 0 | 0.89 | 0.97 | 0.9 | 0.97 | 0.82 | 0.93 |
| 6 | 29632318 | rs2256266 | 1.799 | -0.728 | -0.687 | -0.629 | 0.013 | 0.003 | 0.003 | 0 | 0.11 | 0.12 | 0.26 | 0.28 | 0.24 | 0.09 |
| 6 | 29633775 | rs6905408 | 1.583 | -1.034 | -0.758 | -0.708 | 0.011 | -0.004 | -0.004 | 0 | 0.1 | 0.07 | 0.03 | 0.12 | 0.04 | 0.09 |
| 6 | 29634417 | rs2535250 | 1.194 | -1.069 | -0.821 | -0.767 | 0.001 | -0.023 | -0.023 | 0 | 0.07 | 0.2 | 0.05 | 0 | 0.03 | 0 |
| 6 | 29634927 | rs2535248 | 1.194 | -1.069 | -0.821 | -0.767 | 0.001 | -0.023 | -0.023 | 3.858 | 0.07 | 0.2 | 0.05 | 0 | 0.03 | 0 |
| 6 | 29634968 | rs2857771 | 1.194 | -1.069 | -0.821 | -0.767 | 0.001 | -0.023 | -0.023 | 0 | 0.07 | 0.2 | 0.05 | 0 | 0.03 | 0 |
| 6 | 29635317 | rs2071652 | 1.621 | -1.065 | -0.82 | -0.764 | 0.011 | -0.004 | -0.004 | 0 | 0.1 | 0.07 | 0.03 | 0.12 | 0.04 | 0.09 |
| 6 | 29635508 | rs1318631 | 1.958 | -1.063 | -0.82 | -0.764 | 0.013 | 0.003 | 0.003 | 0 | 0.11 | 0.12 | 0.19 | 0.28 | 0.24 | 0.09 |
| 6 | 29635928 | rs2071653 | -0.477 | -1.233 | -0.836 | -0.85 | 0.005 | 0.019 | 0.019 | 0 | 0.75 | 0.8 | 0.83 | 0.84 | 0.77 | 0.79 |
| 6 | 29639776 | rs9257936 | 2.434 | -1.706 | -1.154 | -1.04 | -0.002 | 0.033 | 0.033 | 0.167 | 0.09 | 0.01 | 0.06 | 0.02 | 0.12 | 0.03 |
| 6 | 29642509 | rs9257940 | 1.633 | -1.411 | -0.998 | -0.838 | 0.011 | -0.004 | -0.004 | 1.847 | 0.1 | 0.07 | 0.03 | 0.12 | 0.04 | 0.09 |
| 6 | 29644502 | rs375984 | 2.359 | -1.321 | -1.049 | -0.769 | 0.005 | 0.02 | 0.02 | 0 | 0.19 | 0.19 | 0.3 | 0.33 | 0.19 | 0.37 |
| 6 | 29645038 | rs2535238 | 2.346 | -1.329 | -1.053 | -0.773 | 0.005 | 0.019 | 0.019 | 0 | 0.2 | 0.19 | 0.3 | 0.33 | 0.19 | 0.37 |
| 6 | 29645634 | rs387642 | 2.361 | -1.326 | -1.051 | -0.771 | 0.005 | 0.02 | 0.02 | 0 | 0.19 | 0.19 | 0.3 | 0.33 | 0.19 | 0.37 |
| 6 | 29646165 | rs396660 | 1.783 | -1.62 | -1.234 | -0.911 | 0 | 0.005 | 0.005 | 0.1 | 0.26 | 0.22 | 0.38 | 0.49 | 0.26 | 0.52 |
| 6 | 29646879 | rs445150 | 1.783 | -1.62 | -1.234 | -0.911 | 0 | 0.005 | 0.005 | 0.1 | 0.26 | 0.22 | 0.38 | 0.49 | 0.26 | 0.52 |
| 6 | 29648377 | rs2747429 | 2.495 | -1.619 | -1.225 | -0.911 | 0.005 | 0.02 | 0.02 | 0 | 0.19 | 0.19 | 0.3 | 0.33 | 0.19 | 0.37 |
| 6 | 29648506 | rs2747430 | 1.793 | -1.621 | -1.234 | -0.913 | 0 | 0.005 | 0.005 | 0.1 | 0.26 | 0.22 | 0.38 | 0.49 | 0.26 | 0.52 |
| 6 | 29649056 | rs3129054 | -0.61 | -1.903 | -1.006 | -1.093 | 0.011 | 0.01 | 0.01 | 0.041 | 0.45 | 0.43 | 0.35 | 0.23 | 0.32 | 0.33 |
| 6 | 29651685 | rs2535236 | -0.6 | -1.901 | -1.005 | -1.092 | 0.01 | 0.01 | 0.01 | 0.05 | 0.45 | 0.43 | 0.34 | 0.23 | 0.32 | 0.33 |
| 6 | 29652199 | rs3117292 | -0.911 | -0.98 | -1.015 | -1.131 | 0.042 | 0.053 | 0.053 | 0 | 0.49 | 0.58 | 0.38 | 0.23 | 0.37 | 0.34 |
| 6 | 29652253 | rs9257985 | 1.548 | -2.053 | -1.206 | -1.257 | 0.03 | 0.032 | 0.032 | 1.382 | 0.13 | 0.05 | 0.04 | 0.04 | 0.06 | 0.05 |
| 6 | 29653186 | rs2747442 | -0.73 | -1.948 | -1.063 | -1.14 | 0.011 | 0.011 | 0.011 | 0.04 | 0.45 | 0.43 | 0.34 | 0.23 | 0.32 | 0.33 |
| 6 | 29653329 | rs3117294 | -0.94 | -1.944 | -1.062 | -1.142 | 0.057 | 0.05 | 0.05 | 0 | 0.48 | 0.58 | 0.37 | 0.23 | 0.35 | 0.33 |
| 6 | 29654700 | rs3117301 | 2.567 | -1.87 | -1.094 | -1.107 | 0.007 | 0.018 | 0.018 | 0 | 0.2 | 0.19 | 0.31 | 0.34 | 0.2 | 0.37 |
| 6 | 29654943 | rs2747453 | 1.808 | -2.158 | -1.037 | -1.269 | 0.005 | -0.002 | -0.002 | 0 | 0.2 | 0.16 | 0.19 | 0.04 | 0.18 | 0.11 |
| 6 | 29655113 | rs2747454 | -0.13 | -2.128 | -1.039 | -1.223 | -0.025 | -0.036 | -0.036 | 0 | 0.38 | 0.4 | 0.27 | 0.07 | 0.25 | 0.18 |
| 6 | 29656417 | rs2747457 | 2.61 | -1.989 | -1.049 | -1.165 | 0.007 | 0.018 | 0.018 | 0 | 0.2 | 0.19 | 0.31 | 0.34 | 0.2 | 0.37 |
| 6 | 29657127 | rs2747460 | 2.721 | -1.991 | -1.05 | -1.168 | 0.006 | 0.018 | 0.018 | 0 | 0.2 | 0.19 | 0.31 | 0.34 | 0.2 | 0.37 |
| 6 | 29658864 | rs3129039 | 3.183 | -1.977 | -1.096 | -1.158 | -0.056 | -0.088 | -0.088 | 0 | 0.17 | 0.03 | 0.27 | 0.34 | 0.18 | 0.37 |
| 6 | 29662443 | rs3129029 | -1.369 | -1.909 | -1.115 | -1.17 | 0.006 | 0.025 | 0.025 | 0 | 0.78 | 0.88 | 0.87 | 0.86 | 0.8 | 0.84 |
| 6 | 29663889 | rs3131888 | 1.089 | -1.884 | -1.053 | -1.159 | -0.003 | -0.003 | -0.003 | 0.231 | 0.43 | 0.4 | 0.52 | 0.38 | 0.39 | 0.48 |
| 6 | 29664452 | rs3131886 | 1.008 | -1.875 | -1.034 | -1.147 | -0.002 | -0.002 | -0.002 | 0 | 0.45 | 0.43 | 0.56 | 0.39 | 0.41 | 0.49 |
| 6 | 29666111 | rs3131875 | 1.808 | -1.847 | -0.984 | -1.123 | 0.009 | 0.009 | 0.009 | 0.128 | 0.32 | 0.34 | 0.53 | 0.62 | 0.46 | 0.48 |
| 6 | 29667273 | rs7776082 | 0.888 | -1.991 | -1.007 | -1.184 | -0.003 | -0.003 | -0.003 | 0.679 | 0.45 | 0.43 | 0.56 | 0.39 | 0.41 | 0.49 |
| 6 | 29668537 | rs3129066 | 2.601 | -1.993 | -1.004 | -1.185 | 0.009 | 0.02 | 0.02 | 0 | 0.21 | 0.22 | 0.34 | 0.35 | 0.22 | 0.39 |
| 6 | 29668913 | rs9258102 | 1.095 | -2.009 | -1.008 | -1.197 | 0.01 | 0.002 | 0.002 | 0 | 0.24 | 0.21 | 0.22 | 0.04 | 0.19 | 0.11 |
| 6 | 29669751 | rs6911622 | 1.217 | -2.021 | -0.942 | -1.133 | -0.038 | -0.027 | -0.027 | 0 | 0.14 | 0.19 | 0.05 | 0.02 | 0.06 | 0.07 |
| 6 | 29669830 | rs2107203 | 2.539 | -1.96 | -0.962 | -1.108 | 0.009 | 0.02 | 0.02 | 0 | 0.21 | 0.22 | 0.34 | 0.35 | 0.22 | 0.39 |
| 6 | 29670261 | rs3129055 | 2.539 | -1.96 | -0.962 | -1.108 | 0.009 | 0.02 | 0.02 | 0 | 0.21 | 0.22 | 0.34 | 0.35 | 0.22 | 0.39 |
| 6 | 29671043 | rs9468589 | 0.519 | -2.012 | -0.974 | -1.066 | -0.037 | 0.001 | 0.001 | 0.082 | 0.22 | 0.33 | 0.13 | 0.18 | 0.13 | 0.21 |
| 6 | 29671991 | rs3116798 | -1.18 | -1.776 | -1.074 | -1.007 | 0.001 | 0.022 | 0.022 | 0 | 0.78 | 0.88 | 0.87 | 0.85 | 0.78 | 0.81 |
| 6 | 29672686 | rs2394660 | -0.348 | -1.896 | -1.027 | -1.023 | 0.007 | 0.001 | 0.001 | 0.008 | 0.66 | 0.69 | 0.73 | 0.69 | 0.7 | 0.65 |
| 6 | 29673483 | rs3131863 | -0.348 | -1.896 | -1.04 | -1.023 | 0.007 | 0.001 | 0.001 | 0.008 | 0.66 | 0.69 | 0.73 | 0.69 | 0.7 | 0.65 |
| 6 | 29674000 | rs11962017 | 1.233 | -2.107 | -1.007 | -1.122 | -0.041 | -0.027 | -0.027 | 0.124 | 0.14 | 0.19 | 0.05 | 0.02 | 0.05 | 0.07 |
| 6 | 29674648 | rs1476573 | -0.358 | -1.942 | -1.115 | -1.036 | 0.007 | 0.001 | 0.001 | 0.008 | 0.66 | 0.69 | 0.73 | 0.69 | 0.69 | 0.65 |
| 6 | 29674741 | rs1476572 | -0.358 | -1.942 | -1.115 | -1.036 | 0.007 | 0.001 | 0.001 | 0.008 | 0.66 | 0.69 | 0.73 | 0.69 | 0.7 | 0.65 |
| 6 | 29676823 | rs7762253 | -0.629 | -1.943 | -1.189 | -1.072 | 0.007 | 0.054 | 0.054 | 3.194 | 0.48 | 0.63 | 0.43 | 0.34 | 0.5 | 0.27 |
| 6 | 29679047 | rs1610594 | -0.412 | -2.161 | -1.1 | -1.222 | 0.006 | 0.001 | 0.001 | 0.008 | 0.66 | 0.69 | 0.73 | 0.69 | 0.7 | 0.65 |
| 6 | 29679602 | rs1611364 | -1.388 | -2.191 | -1.076 | -1.268 | 0 | 0 | 0 | 0 | 0.8 | 0.81 | 0.84 | 0.85 | 0.78 | 0.8 |
| 6 | 29682667 | rs2735063 | 1.526 | -2.057 | -0.952 | -1.212 | -0.001 | -0.012 | -0.012 | 0 | 0.19 | 0.16 | 0.15 | 0.04 | 0.18 | 0.09 |

| | | | | | | | | | | | | | | | | |
|---|---|---|---|---|---|---|---|---|---|---|---|---|---|---|---|---|
| 6 | 29688501 | rs2517910 | -1.515 | -2.187 | -0.918 | -0.972 | 0.016 | 0.002 | 0.002 | 0.009 | 0.65 | 0.67 | 0.56 | 0.73 | 0.6 | 0.54 |
| 6 | 29689127 | rs3757324 | 0.38 | -2.325 | -0.794 | -0.975 | -0.019 | 0.007 | 0.007 | 0 | 0.23 | 0.28 | 0.17 | 0.29 | 0.12 | 0.29 |
| 6 | 29689590 | rs9258170 | 0.437 | -2.304 | -0.844 | -0.963 | -0.013 | 0.005 | 0.005 | 0 | 0.23 | 0.26 | 0.16 | 0.29 | 0.12 | 0.29 |
| 6 | 29691019 | rs1362126 | 0.653 | -2.539 | -1.016 | -1.128 | -0.045 | -0.019 | -0.019 | 0 | 0.31 | 0.22 | 0.35 | 0.43 | 0.44 | 0.17 |
| 6 | 29691303 | rs2076183 | 0.444 | -2.42 | -0.89 | -1.023 | -0.013 | 0.005 | 0.005 | 0 | 0.23 | 0.26 | 0.16 | 0.29 | 0.12 | 0.29 |
| 6 | 29692334 | rs9258186 | 0.444 | -2.469 | -0.89 | -1.023 | -0.014 | 0.005 | 0.005 | 0 | 0.23 | 0.26 | 0.16 | 0.29 | 0.12 | 0.29 |
| 6 | 29692562 | rs1736926 | -1.65 | -2.436 | -0.773 | -1.123 | 0.031 | -0.041 | -0.041 | 0 | 0.87 | 0.82 | 0.88 | 0.98 | 0.8 | 0.93 |
| 6 | 29693499 | rs2235383 | 0.48 | -2.507 | -0.805 | -1.113 | -0.014 | 0.005 | 0.005 | 0 | 0.23 | 0.26 | 0.16 | 0.29 | 0.12 | 0.29 |
| 6 | 29693969 | rs1736922 | 2.741 | -2.623 | -0.871 | -1.192 | -0.005 | -0.012 | -0.012 | 0.018 | 0.22 | 0.15 | 0.32 | 0.26 | 0.2 | 0.4 |
| 6 | 29695502 | rs1628578 | 2.741 | -2.621 | -0.889 | -1.202 | -0.005 | -0.012 | -0.012 | 0 | 0.22 | 0.15 | 0.32 | 0.26 | 0.2 | 0.4 |
| 6 | 29695713 | rs2523404 | -0.031 | -2.617 | -0.921 | -1.162 | -0.006 | 0.016 | 0.016 | 2.555 | 0.42 | 0.4 | 0.4 | 0.44 | 0.48 | 0.25 |
| 6 | 29695854 | rs2735059 | -0.031 | -2.656 | -0.941 | -1.184 | -0.006 | 0.016 | 0.016 | 2.555 | 0.42 | 0.4 | 0.4 | 0.44 | 0.48 | 0.25 |
| 6 | 29696245 | rs2272874 | 0.404 | -2.515 | -1.035 | -1.141 | -0.014 | 0.005 | 0.005 | 0 | 0.23 | 0.26 | 0.16 | 0.29 | 0.12 | 0.29 |
| 6 | 29696849 | rs3736694 | 0.404 | -2.5 | -1.035 | -1.141 | -0.013 | 0.005 | 0.005 | 0 | 0.23 | 0.26 | 0.16 | 0.29 | 0.12 | 0.29 |
| 6 | 29697693 | rs2735058 | 2.072 | -2.459 | -1.353 | -1.227 | 0.011 | 0.044 | 0.044 | 2.411 | 0.11 | 0.01 | 0.05 | 0.02 | 0.12 | 0.03 |
| 6 | 29698050 | rs2743925 | 1.229 | -2.475 | -1.376 | -1.198 | 0.042 | 0.084 | 0.084 | 0.879 | 0.2 | 0.05 | 0.14 | 0.05 | 0.19 | 0.09 |
| 6 | 29699156 | rs2523402 | 0.052 | -2.388 | -1.236 | -1.155 | -0.026 | 0.002 | 0.002 | 0 | 0.36 | 0.32 | 0.37 | 0.44 | 0.45 | 0.24 |
| 6 | 29700079 | rs2523399 | -1.289 | -2.569 | -1.225 | -1.209 | -0.001 | -0.001 | -0.001 | 0 | 0.58 | 0.55 | 0.54 | 0.73 | 0.57 | 0.53 |
| 6 | 29700183 | rs1610601 | 2.836 | -2.637 | -1.201 | -1.282 | -0.004 | -0.013 | -0.013 | 0 | 0.2 | 0.12 | 0.29 | 0.25 | 0.19 | 0.38 |
| 6 | 29701564 | rs2735052 | 0.073 | -2.651 | -1.223 | -1.286 | -0.03 | 0 | 0 | 0 | 0.37 | 0.32 | 0.4 | 0.44 | 0.46 | 0.25 |
| 6 | 29701800 | rs7773358 | 0.867 | -2.658 | -1.201 | -1.274 | -0.006 | 0.003 | 0.003 | 0 | 0.21 | 0.22 | 0.15 | 0.29 | 0.12 | 0.29 |
| 6 | 29702061 | rs1610602 | 1.991 | -2.715 | -1.127 | -1.315 | 0.002 | -0.002 | -0.002 | 0.065 | 0.29 | 0.24 | 0.33 | 0.25 | 0.23 | 0.4 |
| 6 | 29702510 | rs2523395 | 0.095 | -2.713 | -1.151 | -1.315 | -0.03 | 0 | 0 | 0 | 0.37 | 0.32 | 0.4 | 0.44 | 0.46 | 0.25 |
| 6 | 29702679 | rs2394159 | 0.187 | -2.709 | -1.154 | -1.315 | -0.025 | 0.002 | 0.002 | 0 | 0.36 | 0.32 | 0.37 | 0.44 | 0.45 | 0.23 |
| 6 | 29702709 | rs2248927 | -1.233 | -2.742 | -1.148 | -1.33 | -0.001 | -0.001 | -0.001 | 0 | 0.57 | 0.55 | 0.54 | 0.73 | 0.58 | 0.53 |
| 6 | 29703262 | rs2394160 | 0.104 | -2.743 | -1.149 | -1.321 | -0.03 | 0 | 0 | 0.016 | 0.37 | 0.32 | 0.4 | 0.44 | 0.46 | 0.25 |
| 6 | 29703823 | rs9258205 | 0.882 | -2.737 | -1.129 | -1.305 | -0.006 | 0.003 | 0.003 | 0 | 0.21 | 0.22 | 0.15 | 0.29 | 0.12 | 0.29 |
| 6 | 29704237 | rs9258206 | 1.598 | -2.892 | -1.194 | -1.314 | -0.032 | 0.006 | 0.006 | 0 | 0.09 | 0.16 | 0.04 | 0.12 | 0.03 | 0.09 |
| 6 | 29704316 | rs1736915 | 2.076 | -2.927 | -1.195 | -1.343 | 0.002 | -0.002 | -0.002 | 0 | 0.29 | 0.24 | 0.33 | 0.25 | 0.23 | 0.4 |
| 6 | 29704400 | rs1736913 | 0.396 | -2.947 | -1.201 | -1.361 | 0.012 | 0.01 | 0.01 | 0 | 0.57 | 0.45 | 0.69 | 0.69 | 0.65 | 0.63 |
| 6 | 29704443 | rs9258207 | 0.642 | -2.81 | -1.192 | -1.271 | -0.006 | 0.003 | 0.003 | 0 | 0.21 | 0.22 | 0.15 | 0.29 | 0.12 | 0.29 |
| 6 | 29704545 | rs9258208 | 0.624 | -2.813 | -1.193 | -1.272 | -0.005 | 0.003 | 0.003 | 0 | 0.21 | 0.22 | 0.15 | 0.29 | 0.12 | 0.3 |
| 6 | 29705402 | rs9258209 | 1.57 | -2.791 | -1.174 | -1.266 | -0.032 | 0.006 | 0.006 | 0 | 0.09 | 0.16 | 0.04 | 0.12 | 0.03 | 0.09 |
| 6 | 29705688 | rs2735051 | 0.2 | -2.791 | -1.207 | -1.282 | -0.025 | 0.002 | 0.002 | 0 | 0.36 | 0.32 | 0.37 | 0.44 | 0.45 | 0.23 |
| 6 | 29705772 | rs1736911 | -2.442 | -2.771 | -1.08 | -1.294 | -0.004 | 0.003 | 0.003 | 0 | 0.89 | 0.94 | 0.91 | 0.98 | 0.83 | 0.94 |
| 6 | 29706180 | rs9258211 | 0.652 | -2.794 | -1.185 | -1.27 | -0.006 | 0.003 | 0.003 | 0 | 0.21 | 0.22 | 0.15 | 0.29 | 0.12 | 0.29 |
| 6 | 29706277 | rs2844831 | 0.126 | -2.829 | -1.205 | -1.299 | -0.03 | 0 | 0 | 0 | 0.37 | 0.32 | 0.4 | 0.44 | 0.46 | 0.25 |
| 6 | 29706691 | rs9258213 | 1.271 | -2.843 | -1.217 | -1.298 | -0.024 | 0.004 | 0.004 | 0 | 0.1 | 0.17 | 0.09 | 0.13 | 0.05 | 0.14 |
| 6 | 29706933 | rs932338 | -1.734 | -2.742 | -1.154 | -1.301 | 0.038 | -0.05 | -0.05 | 0 | 0.86 | 0.78 | 0.88 | 0.98 | 0.8 | 0.94 |
| 6 | 29707267 | rs1633096 | 2.12 | -2.667 | -1.184 | -1.325 | 0.003 | -0.003 | -0.003 | 0 | 0.28 | 0.24 | 0.33 | 0.25 | 0.21 | 0.4 |
| 6 | 29708028 | rs9258218 | 0.79 | -2.67 | -1.155 | -1.301 | -0.006 | 0.004 | 0.004 | 0 | 0.2 | 0.22 | 0.15 | 0.29 | 0.12 | 0.29 |
| 6 | 29708222 | rs1610603 | -1.676 | -2.55 | -1.197 | -1.295 | 0.038 | -0.05 | -0.05 | 0 | 0.86 | 0.78 | 0.88 | 0.98 | 0.8 | 0.93 |
| 6 | 29709234 | rs2743951 | -0.032 | -2.517 | -1.31 | -1.242 | -0.031 | 0 | 0 | 0 | 0.38 | 0.32 | 0.44 | 0.44 | 0.46 | 0.26 |
| 6 | 29709840 | rs2517938 | 1.328 | -2.604 | -1.372 | -1.34 | 0.043 | 0.066 | 0.066 | 0.919 | 0.21 | 0.13 | 0.16 | 0.04 | 0.19 | 0.08 |
| 6 | 29710726 | rs4713240 | 1.56 | -2.552 | -1.428 | -1.309 | 0.012 | 0.009 | 0.009 | 0.169 | 0.15 | 0.23 | 0.22 | 0.39 | 0.26 | 0.15 |
| 6 | 29711747 | rs2523386 | 1.798 | -2.62 | -1.453 | -1.335 | 0.028 | 0.07 | 0.07 | 1.762 | 0.17 | 0.04 | 0.13 | 0.04 | 0.17 | 0.08 |
| 6 | 29712704 | rs9258225 | 0.583 | -2.664 | -1.35 | -1.303 | -0.011 | 0.006 | 0.006 | 0 | 0.2 | 0.22 | 0.1 | 0.28 | 0.1 | 0.24 |
| 6 | 29712866 | rs1633091 | -1.583 | -2.71 | -1.305 | -1.383 | 0.031 | -0.053 | -0.053 | 0.036 | 0.85 | 0.78 | 0.87 | 0.98 | 0.8 | 0.94 |
| 6 | 29713036 | rs2072894 | 0.625 | -2.696 | -1.311 | -1.359 | -0.011 | 0.006 | 0.006 | 0 | 0.2 | 0.22 | 0.1 | 0.28 | 0.1 | 0.24 |
| 6 | 29713926 | rs2523385 | 1.504 | -2.729 | -1.39 | -1.392 | 0.049 | 0.084 | 0.084 | 0.796 | 0.2 | 0.09 | 0.14 | 0.04 | 0.18 | 0.08 |
| 6 | 29714413 | rs1737086 | 0.757 | -2.555 | -1.292 | -1.294 | 0.05 | 0.052 | 0.052 | 0 | 0.49 | 0.32 | 0.52 | 0.3 | 0.43 | 0.53 |
| 6 | 29715332 | rs34260811 | 1.824 | -2.63 | -1.306 | -1.352 | -0.003 | -0.004 | -0.004 | 0.93 | 0.15 | 0.19 | 0.22 | 0.39 | 0.26 | 0.16 |
| 6 | 29715762 | rs1633086 | 0.727 | -2.629 | -1.307 | -1.359 | 0.043 | 0.045 | 0.045 | 0 | 0.48 | 0.32 | 0.52 | 0.3 | 0.43 | 0.47 |
| 6 | 29717980 | rs733841 | -1.628 | -2.401 | -1.186 | -1.275 | 0.037 | -0.05 | -0.05 | 0.564 | 0.86 | 0.78 | 0.88 | 0.98 | 0.8 | 0.94 |
| 6 | 29718544 | rs1610736 | -0.468 | -2.472 | -1.243 | -1.266 | 0.068 | 0.017 | 0.017 | 0 | 0.69 | 0.55 | 0.65 | 0.59 | 0.54 | 0.73 |
| 6 | 29718790 | rs1610621 | 2.524 | -2.669 | -1.258 | -1.355 | 0 | 0.006 | 0.006 | 0.104 | 0.22 | 0.21 | 0.3 | 0.27 | 0.19 | 0.33 |
| 6 | 29719561 | rs909728 | -0.404 | -2.399 | -1.196 | -1.212 | 0.066 | 0.016 | 0.016 | 0 | 0.69 | 0.55 | 0.65 | 0.59 | 0.54 | 0.72 |
| 6 | 29719991 | rs909726 | -1.636 | -2.39 | -1.177 | -1.237 | 0.037 | -0.05 | -0.05 | 1.412 | 0.86 | 0.78 | 0.88 | 0.98 | 0.8 | 0.94 |
| 6 | 29720321 | rs909725 | -1.64 | -2.38 | -1.19 | -1.232 | 0.037 | -0.05 | -0.05 | 0 | 0.86 | 0.78 | 0.88 | 0.98 | 0.8 | 0.94 |
| 6 | 29721146 | rs4713244 | 1.776 | -2.39 | -1.184 | -1.237 | 0.001 | -0.001 | -0.001 | 0 | 0.15 | 0.19 | 0.22 | 0.38 | 0.26 | 0.16 |
| 6 | 29722515 | rs9391630 | 1.992 | -2.305 | -1.206 | -1.069 | -0.073 | -0.066 | -0.066 | 1.971 | 0.12 | 0.02 | 0.15 | 0.34 | 0.24 | 0.13 |
| 6 | 29723161 | rs9258260 | 0.923 | -2.357 | -1.196 | -1.178 | 0.011 | -0.014 | -0.014 | 0 | 0.17 | 0.13 | 0.08 | 0.27 | 0.09 | 0.23 |
| 6 | 29723674 | rs4947236 | -0.13 | -2.337 | -1.187 | -1.21 | -0.013 | -0.007 | -0.007 | 5.104 | 0.35 | 0.37 | 0.34 | 0.66 | 0.37 | 0.46 |
| 6 | 29724310 | rs1633068 | -1.641 | -2.338 | -1.186 | -1.211 | 0.037 | -0.05 | -0.05 | 0 | 0.86 | 0.78 | 0.88 | 0.98 | 0.8 | 0.94 |
| 6 | 29724745 | rs1737079 | -1.641 | -2.338 | -1.186 | -1.211 | 0.037 | -0.05 | -0.05 | 0.029 | 0.86 | 0.78 | 0.88 | 0.98 | 0.8 | 0.94 |
| 6 | 29724939 | rs1737078 | 0.652 | -2.338 | -1.187 | -1.211 | 0.025 | 0.026 | 0.026 | 0 | 0.51 | 0.41 | 0.53 | 0.32 | 0.43 | 0.48 |
| 6 | 29725321 | rs1737076 | 0.652 | -2.338 | -1.187 | -1.211 | 0.025 | 0.026 | 0.026 | 0 | 0.51 | 0.41 | 0.53 | 0.32 | 0.43 | 0.48 |
| 6 | 29727023 | rs1633057 | -1.53 | -2.214 | -1.126 | -1.163 | 0.035 | -0.052 | -0.052 | 1.252 | 0.86 | 0.78 | 0.88 | 0.98 | 0.8 | 0.93 |
| 6 | 29727092 | rs1633056 | -1.604 | -2.212 | -1.124 | -1.163 | 0.038 | -0.05 | -0.05 | 0 | 0.86 | 0.78 | 0.88 | 0.98 | 0.8 | 0.94 |

| | | | | | | | | | | | | | | | |
|---|---|---|---|---|---|---|---|---|---|---|---|---|---|---|---|
| 6 | 29730504 | rs9258275 | -0.211 | -2.333 | -1.203 | -1.208 | -0.013 | -0.007 | -0.007 | 0 | 0.36 | 0.37 | 0.35 | 0.66 | 0.37 | 0.47 |
| 6 | 29730642 | rs1633048 | -1.645 | -2.359 | -1.185 | -1.23 | 0.037 | -0.05 | -0.05 | 0.311 | 0.86 | 0.78 | 0.88 | 0.98 | 0.81 | 0.94 |
| 6 | 29730730 | rs1737069 | -0.167 | -2.353 | -1.21 | -1.195 | -0.01 | -0.002 | -0.002 | 0 | 0.67 | 0.64 | 0.8 | 0.72 | 0.71 | 0.69 |
| 6 | 29731635 | rs7759474 | 1.178 | -2.198 | -1.196 | -1.209 | 0.002 | 0.001 | 0.001 | 0.069 | 0.17 | 0.22 | 0.22 | 0.39 | 0.27 | 0.21 |
| 6 | 29731787 | rs1611285 | 1.028 | -2.6 | -1.278 | -1.413 | 0.042 | 0.077 | 0.077 | 0.959 | 0.19 | 0.09 | 0.14 | 0.04 | 0.18 | 0.07 |
| 6 | 29732244 | rs1626392 | -0.055 | -2.237 | -0.761 | -1.046 | -0.001 | -0.003 | -0.003 | 0 | 0.76 | 0.76 | 0.82 | 0.87 | 0.78 | 0.83 |
| 6 | 29732302 | rs1625636 | -0.904 | -1.966 | -0.667 | -0.996 | 0.035 | -0.05 | -0.05 | 4.292 | 0.86 | 0.77 | 0.88 | 0.98 | 0.81 | 0.94 |
| 6 | 29733223 | rs1633041 | 1.314 | -2.176 | -1.002 | -0.719 | 0.008 | 0.016 | 0.016 | 0 | 0.3 | 0.25 | 0.25 | 0.11 | 0.26 | 0.3 |
| 6 | 29733375 | rs1737058 | -2.109 | -2.043 | -1.021 | -0.642 | 0.007 | -0.003 | -0.003 | 0 | 0.92 | 0.94 | 0.91 | 0.98 | 0.83 | 0.94 |
| 6 | 29733701 | rs2735048 | 1.073 | -2.266 | -1.075 | -0.543 | -0.015 | -0.007 | -0.007 | 0 | 0.25 | 0.27 | 0.35 | 0.37 | 0.39 | 0.17 |
| 6 | 29734098 | rs2735046 | 0.146 | -2.429 | -1.082 | -0.609 | -0.008 | 0 | 0 | 0.783 | 0.34 | 0.4 | 0.44 | 0.38 | 0.41 | 0.24 |
| 6 | 29736047 | rs1737043 | -0.588 | -2.556 | -1.195 | -0.615 | -0.006 | -0.057 | -0.057 | 0 | 0.7 | 0.89 | 0.72 | 0.57 | 0.69 | 0.59 |
| 6 | 29736229 | rs1737041 | 1.475 | -2.575 | -1.027 | -0.662 | 0.008 | 0.016 | 0.016 | 0 | 0.3 | 0.25 | 0.25 | 0.11 | 0.26 | 0.3 |
| 6 | 29736942 | rs11753872 | 1.851 | -2.552 | -0.953 | -0.632 | 0.03 | 0.083 | 0.083 | 0 | 0.06 | 0.24 | 0.03 | 0.08 | 0.02 | 0.05 |
| 6 | 29737481 | rs1737031 | 0.855 | -2.478 | -0.987 | -0.661 | -0.018 | -0.021 | -0.021 | 0 | 0.36 | 0.49 | 0.28 | 0.19 | 0.28 | 0.35 |
| 6 | 29737846 | rs6932033 | 1.79 | -2.251 | -0.81 | -0.514 | 0.027 | 0.079 | 0.079 | 0 | 0.06 | 0.23 | 0.04 | 0.08 | 0.02 | 0.05 |
| 6 | 29738569 | rs12662393 | 1.517 | -2.243 | -0.945 | -0.536 | -0.002 | -0.004 | -0.004 | 0 | 0.06 | 0.09 | 0.07 | 0.01 | 0.02 | 0.02 |
| 6 | 29740034 | rs7740113 | 1.517 | -2.243 | -0.945 | -0.536 | -0.002 | -0.004 | -0.004 | 0 | 0.06 | 0.09 | 0.07 | 0.01 | 0.02 | 0.02 |
| 6 | 29741338 | rs9258302 | 1.109 | -2.537 | -1.089 | -0.477 | 0.02 | -0.024 | -0.024 | 0.001 | 0.13 | 0.07 | 0.15 | 0.23 | 0.07 | 0.24 |
| 6 | 29742108 | rs1362068 | 0.254 | -2.752 | -1.135 | -0.614 | 0.031 | 0.035 | 0.035 | 0 | 0.45 | 0.6 | 0.37 | 0.2 | 0.31 | 0.38 |
| 6 | 29742299 | rs1362070 | 0.254 | -2.752 | -1.135 | -0.614 | 0.031 | 0.035 | 0.035 | 0 | 0.45 | 0.6 | 0.37 | 0.2 | 0.31 | 0.38 |
| 6 | 29742342 | rs9500983 | 1.644 | -2.706 | -1.126 | -0.595 | -0.002 | -0.004 | -0.004 | 1.576 | 0.06 | 0.09 | 0.07 | 0.01 | 0.02 | 0.02 |
| 6 | 29742809 | rs1633036 | -2.048 | -2.671 | -1.172 | -0.644 | -0.002 | 0.001 | 0.001 | 0 | 0.9 | 0.94 | 0.91 | 0.98 | 0.83 | 0.94 |
| 6 | 29742930 | rs9258303 | 1.172 | -2.653 | -1.142 | -0.606 | 0.02 | -0.024 | -0.024 | 0 | 0.13 | 0.07 | 0.15 | 0.23 | 0.07 | 0.24 |
| 6 | 29743117 | rs9500845 | 1.626 | -2.572 | -1.108 | -0.598 | -0.002 | -0.004 | -0.004 | 0.059 | 0.06 | 0.09 | 0.07 | 0.01 | 0.02 | 0.02 |
| 6 | 29745075 | rs2517930 | 1.5 | -2.521 | -1.146 | -0.61 | -0.04 | -0.022 | -0.022 | 0 | 0.23 | 0.19 | 0.28 | 0.35 | 0.35 | 0.16 |
| 6 | 29745726 | rs1615962 | 1.557 | -2.525 | -1.015 | -0.705 | 0.008 | 0.016 | 0.016 | 0 | 0.3 | 0.25 | 0.25 | 0.11 | 0.26 | 0.3 |
| 6 | 29745794 | rs1633030 | 0.904 | -2.541 | -0.983 | -0.704 | -0.019 | -0.022 | -0.022 | 0 | 0.36 | 0.48 | 0.28 | 0.19 | 0.28 | 0.35 |
| 6 | 29746367 | rs1633028 | 0.904 | -2.498 | -1.027 | -0.647 | -0.018 | -0.021 | -0.021 | 0 | 0.36 | 0.49 | 0.28 | 0.19 | 0.28 | 0.35 |
| 6 | 29746869 | rs1633021 | 1.877 | -2.664 | -1.061 | -0.737 | 0.005 | 0.009 | 0.009 | 0 | 0.26 | 0.21 | 0.22 | 0.07 | 0.21 | 0.25 |
| 6 | 29747373 | rs2735042 | 0.867 | -2.611 | -1.046 | -0.709 | -0.018 | -0.021 | -0.021 | 0 | 0.36 | 0.49 | 0.28 | 0.19 | 0.28 | 0.35 |
| 6 | 29753729 | rs1002044 | 1.498 | -2.599 | -1.063 | -0.72 | 0.009 | 0.016 | 0.016 | 0 | 0.31 | 0.25 | 0.25 | 0.11 | 0.26 | 0.3 |
| 6 | 29754016 | rs1002046 | 1.506 | -2.576 | -1.052 | -0.836 | 0.009 | 0.016 | 0.016 | 0 | 0.31 | 0.25 | 0.25 | 0.11 | 0.26 | 0.3 |
| 6 | 29755451 | rs2743941 | -0.945 | -2.572 | -1.056 | -0.818 | 0.016 | 0.031 | 0.031 | 0 | 0.77 | 0.71 | 0.84 | 0.71 | 0.74 | 0.73 |
| 6 | 29755656 | rs7758742 | 1.847 | -2.54 | -0.887 | -0.795 | 0.03 | 0.083 | 0.083 | 0 | 0.06 | 0.24 | 0.03 | 0.08 | 0.02 | 0.05 |
| 6 | 29759066 | rs1610641 | 0.301 | -2.598 | -0.93 | -0.829 | 0.014 | 0.016 | 0.016 | 0 | 0.42 | 0.57 | 0.35 | 0.19 | 0.3 | 0.38 |
| 6 | 29759445 | rs1610644 | 1.427 | -2.613 | -0.924 | -0.837 | 0.009 | 0.016 | 0.016 | 0 | 0.31 | 0.25 | 0.25 | 0.11 | 0.26 | 0.3 |
| 6 | 29759511 | rs17185132 | 1.625 | -2.59 | -0.954 | -0.828 | -0.002 | -0.004 | -0.004 | 0 | 0.06 | 0.09 | 0.07 | 0.01 | 0.02 | 0.02 |
| 6 | 29759823 | rs1611205 | 0.308 | -2.575 | -0.962 | -0.82 | 0.014 | 0.016 | 0.016 | 0 | 0.42 | 0.57 | 0.35 | 0.2 | 0.3 | 0.38 |
| 6 | 29760677 | rs1610719 | 0.308 | -2.575 | -0.962 | -0.82 | 0.014 | 0.016 | 0.016 | 0 | 0.42 | 0.57 | 0.35 | 0.19 | 0.3 | 0.38 |
| 6 | 29761473 | rs1610723 | 0.308 | -2.575 | -0.962 | -0.82 | 0.014 | 0.016 | 0.016 | 0 | 0.42 | 0.57 | 0.35 | 0.19 | 0.3 | 0.38 |
| 6 | 29761516 | rs1610724 | 0.308 | -2.575 | -0.962 | -0.82 | 0.014 | 0.016 | 0.016 | 0 | 0.42 | 0.57 | 0.35 | 0.19 | 0.3 | 0.37 |
| 6 | 29761835 | rs1611220 | 0.138 | -2.618 | -0.978 | -0.837 | 0.031 | 0.034 | 0.034 | 0 | 0.45 | 0.6 | 0.37 | 0.2 | 0.31 | 0.38 |
| 6 | 29762693 | rs1633011 | 1.425 | -2.616 | -0.95 | -0.859 | 0.009 | 0.016 | 0.016 | 0 | 0.31 | 0.25 | 0.25 | 0.11 | 0.26 | 0.3 |
| 6 | 29764443 | rs11758087 | 1.852 | -2.594 | -0.905 | -0.84 | 0.03 | 0.083 | 0.083 | 0 | 0.06 | 0.24 | 0.03 | 0.08 | 0.03 | 0.05 |
| 6 | 29764472 | rs1633005 | 2.797 | -2.874 | -0.962 | -1 | 0 | 0.007 | 0.007 | 0 | 0.17 | 0.1 | 0.13 | 0.06 | 0.15 | 0.18 |
| 6 | 29764707 | rs11751282 | 1.905 | -2.837 | -0.921 | -0.982 | 0.028 | 0.081 | 0.081 | 0 | 0.06 | 0.23 | 0.03 | 0.08 | 0.03 | 0.05 |
| 6 | 29764759 | rs1633003 | 0.042 | -2.892 | -0.964 | -1.009 | 0.031 | 0.035 | 0.035 | 0 | 0.45 | 0.6 | 0.37 | 0.2 | 0.31 | 0.38 |
| 6 | 29765001 | rs1630969 | 1.349 | -2.905 | -1.058 | -1.039 | 0.009 | 0.017 | 0.017 | 0 | 0.3 | 0.24 | 0.24 | 0.11 | 0.25 | 0.3 |
| 6 | 29765020 | rs9468623 | 1.888 | -2.65 | -0.966 | -0.946 | -0.002 | -0.002 | -0.002 | 1.111 | 0.06 | 0.09 | 0.07 | 0.01 | 0.02 | 0.02 |
| 6 | 29765087 | rs2523414 | 1.241 | -2.581 | -0.97 | -0.939 | 0 | -0.004 | -0.004 | 0 | 0.12 | 0.11 | 0.1 | 0.01 | 0.07 | 0.07 |
| 6 | 29767743 | rs28634576 | 1.772 | -2.294 | -0.916 | -0.769 | -0.002 | -0.004 | -0.004 | 2.454 | 0.06 | 0.09 | 0.07 | 0.01 | 0.02 | 0.02 |
| 6 | 29768137 | rs11751333 | 1.757 | -2.31 | -0.744 | -0.768 | 0.035 | 0.083 | 0.083 | 0 | 0.06 | 0.24 | 0.03 | 0.08 | 0.03 | 0.05 |
| 6 | 29768344 | rs1611185 | 0.787 | -2.54 | -1.018 | -0.849 | -0.043 | -0.041 | -0.041 | 2.116 | 0.32 | 0.27 | 0.38 | 0.56 | 0.45 | 0.33 |
| 6 | 29768503 | rs2017525 | 2.213 | -2.6 | -1.132 | -0.882 | -0.012 | -0.048 | -0.048 | 0 | 0.08 | 0.01 | 0.14 | 0.21 | 0.07 | 0.15 |
| 6 | 29768917 | rs1610649 | 0.167 | -2.753 | -1.219 | -0.94 | 0.014 | 0.006 | 0.006 | 0 | 0.47 | 0.51 | 0.34 | 0.39 | 0.35 | 0.51 |
| 6 | 29769497 | rs1610711 | 1.59 | -2.734 | -1.187 | -0.949 | 0.009 | 0.016 | 0.016 | 0 | 0.31 | 0.25 | 0.25 | 0.11 | 0.26 | 0.3 |
| 6 | 29769521 | rs1736982 | -0.635 | -2.724 | -1.197 | -0.947 | -0.004 | -0.048 | -0.048 | 0 | 0.7 | 0.87 | 0.72 | 0.56 | 0.7 | 0.54 |
| 6 | 29769621 | rs1632995 | 0.299 | -2.716 | -1.305 | -0.942 | 0.031 | 0.035 | 0.035 | 0 | 0.45 | 0.6 | 0.37 | 0.2 | 0.31 | 0.38 |
| 6 | 29770322 | rs1736981 | -0.631 | -2.704 | -1.31 | -0.933 | -0.004 | -0.048 | -0.048 | 0 | 0.7 | 0.87 | 0.72 | 0.56 | 0.7 | 0.54 |
| 6 | 29771066 | rs1610657 | 0.264 | -2.685 | -1.252 | -0.922 | 0.031 | 0.035 | 0.035 | 0 | 0.45 | 0.6 | 0.37 | 0.2 | 0.31 | 0.38 |
| 6 | 29771879 | rs1611192 | 1.986 | -2.694 | -1.258 | -0.933 | 0.004 | 0.007 | 0.007 | 0 | 0.25 | 0.21 | 0.22 | 0.07 | 0.21 | 0.25 |
| 6 | 29772395 | rs1632988 | 1.641 | -2.495 | -1.159 | -0.835 | 0.006 | 0.013 | 0.013 | 0 | 0.29 | 0.24 | 0.24 | 0.11 | 0.25 | 0.3 |
| 6 | 29772549 | rs1632987 | 1.68 | -2.412 | -1.153 | -0.841 | 0.009 | 0.016 | 0.016 | 0 | 0.31 | 0.25 | 0.25 | 0.11 | 0.26 | 0.3 |
| 6 | 29773509 | rs1632983 | 0.682 | -2.425 | -1.13 | -0.841 | -0.01 | -0.011 | -0.011 | 0 | 0.39 | 0.51 | 0.3 | 0.19 | 0.29 | 0.35 |
| 6 | 29773537 | rs17178826 | 1.859 | -2.419 | -1.118 | -0.836 | 0.03 | 0.083 | 0.083 | 0 | 0.06 | 0.23 | 0.03 | 0.08 | 0.03 | 0.05 |
| 6 | 29773852 | rs9258447 | 1.069 | -2.48 | -1.152 | -0.852 | 0.012 | -0.023 | -0.023 | 0.053 | 0.13 | 0.07 | 0.15 | 0.23 | 0.09 | 0.24 |
| 6 | 29773999 | rs1736976 | 0.671 | -2.447 | -1.117 | -0.86 | -0.003 | -0.003 | -0.003 | 0.022 | 0.4 | 0.54 | 0.34 | 0.19 | 0.3 | 0.37 |
| 6 | 29775525 | rs11757182 | 1.839 | -2.361 | -1.053 | -0.81 | 0.03 | 0.083 | 0.083 | 0 | 0.06 | 0.23 | 0.03 | 0.08 | 0.02 | 0.05 |

| | | | | | | | | | | | | | | | | |
|---|---|---|---|---|---|---|---|---|---|---|---|---|---|---|---|---|
| 6 | 29775662 | rs2523409 | 0.762 | -2.371 | -1.134 | -0.834 | -0.003 | -0.001 | -0.001 | 0 | 0.3 | 0.37 | 0.38 | 0.37 | 0.39 | 0.18 |
| 6 | 29775939 | rs9468632 | 1.954 | -2.356 | -1.084 | -0.853 | -0.004 | -0.005 | -0.005 | 0 | 0.06 | 0.09 | 0.07 | 0.01 | 0.02 | 0.02 |
| 6 | 29776322 | rs1736971 | 1.675 | -2.446 | -1.13 | -0.901 | 0.009 | 0.017 | 0.017 | 0 | 0.31 | 0.25 | 0.25 | 0.11 | 0.26 | 0.3 |
| 6 | 29777911 | rs1632971 | -1.908 | -2.451 | -1.145 | -0.925 | -0.002 | 0 | 0 | 0 | 0.9 | 0.94 | 0.91 | 0.98 | 0.85 | 0.94 |
| 6 | 29778626 | rs1610663 | 1.019 | -2.503 | -1.102 | -0.887 | 0.01 | 0.016 | 0.016 | 0 | 0.37 | 0.34 | 0.32 | 0.12 | 0.28 | 0.33 |
| 6 | 29778987 | rs1610699 | 0.94 | -2.506 | -1.103 | -0.828 | 0.012 | 0.018 | 0.018 | 0 | 0.37 | 0.34 | 0.32 | 0.12 | 0.28 | 0.33 |
| 6 | 29780093 | rs1610669 | 0.36 | -2.491 | -1.132 | -0.931 | 0.032 | 0.035 | 0.035 | 0 | 0.45 | 0.6 | 0.37 | 0.2 | 0.31 | 0.38 |
| 6 | 29780332 | rs3926748 | 0.993 | -2.362 | -1.102 | -0.83 | 0.012 | -0.023 | -0.023 | 0 | 0.13 | 0.07 | 0.15 | 0.23 | 0.09 | 0.24 |
| 6 | 29780698 | rs3888577 | 1.906 | -2.202 | -1.037 | -0.853 | -0.003 | -0.004 | -0.004 | 0.08 | 0.06 | 0.09 | 0.07 | 0.01 | 0.02 | 0.02 |
| 6 | 29782331 | rs11753629 | 0.179 | -2.242 | -0.988 | -0.833 | -0.039 | 0.041 | 0.041 | 0 | 0.22 | 0.35 | 0.26 | 0.3 | 0.12 | 0.23 |
| 6 | 29782470 | rs1736959 | 1.619 | -2.339 | -1.039 | -0.894 | 0.009 | 0.016 | 0.016 | 0 | 0.31 | 0.25 | 0.25 | 0.11 | 0.26 | 0.3 |
| 6 | 29782621 | rs11753654 | 1.12 | -2.357 | -1.027 | -0.901 | -0.013 | 0.096 | 0.096 | 0 | 0.13 | 0.24 | 0.17 | 0.29 | 0.09 | 0.2 |
| 6 | 29782633 | rs1736957 | 1.047 | -2.311 | -1.053 | -0.965 | -0.001 | -0.006 | -0.006 | 0 | 0.34 | 0.35 | 0.31 | 0.13 | 0.3 | 0.36 |
| 6 | 29782868 | rs1077432 | 1.563 | -2.278 | -1.09 | -0.932 | 0.009 | 0.016 | 0.016 | 0 | 0.31 | 0.25 | 0.25 | 0.11 | 0.26 | 0.3 |
| 6 | 29785149 | rs1620173 | 1.563 | -2.278 | -1.09 | -0.939 | 0.009 | 0.016 | 0.016 | 0 | 0.31 | 0.25 | 0.25 | 0.11 | 0.26 | 0.3 |
| 6 | 29785235 | rs1619379 | 0.126 | -2.155 | -1.051 | -0.833 | 0.044 | 0.036 | 0.036 | 0 | 0.51 | 0.38 | 0.38 | 0.32 | 0.37 | 0.52 |
| 6 | 29785538 | rs2735028 | 0.525 | -2.146 | -1.023 | -0.859 | 0.028 | 0.04 | 0.04 | 0 | 0.43 | 0.38 | 0.35 | 0.12 | 0.3 | 0.36 |
| 6 | 29787072 | rs2743931 | -1.759 | -1.921 | -0.987 | -0.774 | -0.002 | 0 | 0 | 0 | 0.9 | 0.94 | 0.91 | 0.98 | 0.85 | 0.95 |
| 6 | 29787369 | rs2394177 | 1.051 | -2.013 | -0.91 | -0.71 | -0.013 | 0.096 | 0.096 | 0 | 0.13 | 0.24 | 0.17 | 0.29 | 0.09 | 0.2 |
| 6 | 29788537 | rs3873245 | 1.577 | -2.071 | -0.962 | -0.718 | -0.012 | -0.049 | -0.049 | 0 | 0.08 | 0.01 | 0.04 | 0.2 | 0.06 | 0.16 |
| 6 | 29788606 | rs1610675 | 1.163 | -2.143 | -0.951 | -0.908 | 0.018 | 0.032 | 0.032 | 0 | 0.35 | 0.26 | 0.26 | 0.11 | 0.28 | 0.34 |
| 6 | 29789190 | rs1610678 | -0.532 | -1.809 | -0.909 | -0.811 | -0.002 | 0 | 0 | 0 | 0.64 | 0.62 | 0.55 | 0.61 | 0.46 | 0.72 |
| 6 | 29789999 | rs1611149 | -0.532 | -1.809 | -0.909 | -0.811 | -0.002 | 0 | 0 | 0.273 | 0.64 | 0.62 | 0.55 | 0.61 | 0.46 | 0.72 |
| 6 | 29794317 | rs1736936 | 0.06 | -1.825 | -0.91 | -0.814 | 0.001 | 0.003 | 0.003 | 0 | 0.55 | 0.51 | 0.46 | 0.6 | 0.43 | 0.7 |
| 6 | 29798425 | rs915669 | -0.433 | -1.594 | -1.039 | -0.734 | -0.002 | 0 | 0 | 0 | 0.64 | 0.62 | 0.55 | 0.61 | 0.46 | 0.73 |
| 6 | 29799103 | rs1233330 | -1.546 | -1.418 | -0.942 | -0.617 | 0.004 | -0.017 | -0.017 | 0 | 0.89 | 0.89 | 0.9 | 0.98 | 0.84 | 0.9 |
| 6 | 29801110 | rs11753296 | 0.21 | -1.588 | -0.943 | -0.56 | -0.04 | 0.03 | 0.03 | 0 | 0.22 | 0.35 | 0.26 | 0.29 | 0.13 | 0.24 |
| 6 | 29802321 | rs7772553 | 0.21 | -1.588 | -0.943 | -0.541 | -0.04 | 0.032 | 0.032 | 0 | 0.22 | 0.35 | 0.26 | 0.29 | 0.13 | 0.24 |
| 6 | 29804165 | rs9404952 | -0.18 | -1.456 | -1.141 | -0.707 | 0.006 | -0.002 | -0.002 | 0 | 0.42 | 0.55 | 0.53 | 0.64 | 0.45 | 0.38 |
| 6 | 29804176 | rs16896052 | 1.955 | -1.596 | -1.221 | -0.664 | -0.017 | -0.043 | -0.043 | 0.062 | 0.07 | 0 | 0.04 | 0.21 | 0.06 | 0.16 |
| 6 | 29804845 | rs2394178 | -0.476 | -1.966 | -1.278 | -0.923 | -0.018 | -0.005 | -0.005 | 0 | 0.38 | 0.54 | 0.38 | 0.3 | 0.23 | 0.35 |
| 6 | 29805199 | rs2394180 | 0.5 | -1.969 | -1.237 | -0.912 | -0.04 | 0.03 | 0.03 | 0 | 0.22 | 0.35 | 0.26 | 0.29 | 0.13 | 0.24 |
| 6 | 29805809 | rs2735014 | -0.456 | -1.971 | -1.275 | -0.915 | -0.02 | -0.005 | -0.005 | 0 | 0.37 | 0.54 | 0.38 | 0.3 | 0.23 | 0.35 |
| 6 | 29806622 | rs2523779 | 1.37 | -2.017 | -1.309 | -0.954 | -0.01 | -0.023 | -0.023 | 0 | 0.15 | 0.19 | 0.12 | 0.01 | 0.11 | 0.11 |
| 6 | 29807853 | rs2735009 | -0.45 | -2.023 | -1.316 | -0.952 | -0.02 | -0.005 | -0.005 | 0 | 0.37 | 0.54 | 0.38 | 0.3 | 0.23 | 0.35 |
| 6 | 29807923 | rs2743937 | -0.45 | -2.023 | -1.316 | -0.952 | -0.02 | -0.005 | -0.005 | 0 | 0.37 | 0.54 | 0.38 | 0.3 | 0.23 | 0.35 |
| 6 | 29808464 | rs2735005 | -0.801 | -2.03 | -1.335 | -0.947 | 0.006 | -0.002 | -0.002 | 0 | 0.45 | 0.54 | 0.42 | 0.51 | 0.29 | 0.51 |
| 6 | 29808634 | rs2735003 | -0.447 | -2.031 | -1.322 | -0.954 | -0.02 | -0.005 | -0.005 | 0 | 0.37 | 0.54 | 0.38 | 0.3 | 0.23 | 0.35 |
| 6 | 29808861 | rs9258537 | -0.447 | -2.031 | -1.322 | -0.954 | -0.02 | -0.005 | -0.005 | 0.059 | 0.37 | 0.54 | 0.38 | 0.3 | 0.23 | 0.35 |
| 6 | 29809087 | rs2523777 | -0.447 | -2.031 | -1.322 | -0.954 | -0.02 | -0.005 | -0.005 | 0 | 0.37 | 0.54 | 0.38 | 0.3 | 0.23 | 0.35 |
| 6 | 29809194 | rs2523776 | 1.372 | -2.022 | -1.311 | -0.955 | -0.01 | -0.023 | -0.023 | 0.014 | 0.15 | 0.19 | 0.12 | 0.01 | 0.11 | 0.11 |
| 6 | 29809382 | rs1611133 | 2.168 | -2.029 | -1.379 | -1.005 | 0.004 | 0.015 | 0.015 | 0 | 0.22 | 0.15 | 0.15 | 0.1 | 0.2 | 0.24 |
| 6 | 29809578 | rs2523774 | -0.47 | -2.021 | -1.34 | -0.931 | -0.013 | -0.004 | -0.004 | 0 | 0.4 | 0.54 | 0.38 | 0.3 | 0.23 | 0.35 |
| 6 | 29811022 | rs2523769 | -0.712 | -1.901 | -1.297 | -0.878 | 0.006 | -0.002 | -0.002 | 0.158 | 0.45 | 0.54 | 0.42 | 0.51 | 0.29 | 0.51 |
| 6 | 29811237 | rs2844827 | -0.156 | -1.895 | -1.118 | -0.826 | -0.017 | 0.021 | 0.021 | 0 | 0.33 | 0.38 | 0.32 | 0.42 | 0.26 | 0.43 |
| 6 | 29811709 | rs16896081 | 2.199 | -1.881 | -1.137 | -0.816 | -0.017 | -0.043 | -0.043 | 0.201 | 0.07 | 0 | 0.04 | 0.2 | 0.06 | 0.16 |
| 6 | 29812030 | rs2734994 | 1.099 | -1.9 | -1.106 | -0.852 | -0.011 | -0.026 | -0.026 | 0.117 | 0.14 | 0.19 | 0.12 | 0.01 | 0.1 | 0.11 |
| 6 | 29812807 | rs2523767 | -0.165 | -1.909 | -1.106 | -0.848 | -0.017 | 0.021 | 0.021 | 0 | 0.33 | 0.37 | 0.32 | 0.42 | 0.26 | 0.44 |
| 6 | 29812974 | rs2517879 | -0.164 | -1.898 | -1.102 | -0.851 | -0.017 | 0.021 | 0.021 | 0 | 0.33 | 0.38 | 0.32 | 0.42 | 0.26 | 0.43 |
| 6 | 29813942 | rs9258557 | 1.06 | -1.806 | -1.12 | -0.857 | -0.011 | -0.026 | -0.026 | 0 | 0.14 | 0.19 | 0.12 | 0.01 | 0.1 | 0.11 |
| 6 | 29814778 | rs1611750 | 1.776 | -1.728 | -1.22 | -0.844 | 0.004 | 0.01 | 0.01 | 0 | 0.23 | 0.18 | 0.16 | 0.11 | 0.2 | 0.29 |
| 6 | 29815637 | rs3128912 | -1.685 | -1.413 | -1.217 | -0.737 | -0.002 | 0 | 0 | 0 | 0.9 | 0.94 | 0.91 | 0.98 | 0.85 | 0.95 |
| 6 | 29815713 | rs3115629 | -1.685 | -1.413 | -1.217 | -0.737 | -0.002 | 0 | 0 | 0 | 0.9 | 0.94 | 0.91 | 0.98 | 0.85 | 0.95 |
| 6 | 29816371 | rs2394185 | 2.018 | -1.35 | -1.267 | -0.626 | -0.016 | -0.046 | -0.046 | 0 | 0.07 | 0.01 | 0.14 | 0.2 | 0.07 | 0.16 |
| 6 | 29816421 | rs2394186 | 1.307 | -0.541 | -1.036 | -0.586 | 0 | 0 | 0 | 0 | 0.12 | 0.13 | 0.19 | 0.2 | 0.17 | 0.17 |
| 6 | 29817244 | rs9380146 | 2.072 | -1.54 | -1.138 | -0.472 | -0.037 | -0.043 | -0.043 | 0.069 | 0.08 | 0.01 | 0.17 | 0.3 | 0.22 | 0.11 |
| 6 | 29818021 | rs3115628 | 0.441 | -2.007 | -1.148 | -0.632 | 0.071 | 0.084 | 0.084 | 0 | 0.53 | 0.36 | 0.65 | 0.82 | 0.65 | 0.65 |
| 6 | 29818662 | rs2734985 | 1.683 | -1.975 | -1.144 | -0.666 | -0.023 | -0.017 | -0.017 | 2.178 | 0.23 | 0.08 | 0.12 | 0.23 | 0.19 | 0.33 |
| 6 | 29819077 | rs2247504 | 0.366 | -2.143 | -1.037 | -0.739 | 0.017 | 0.008 | 0.008 | 0 | 0.48 | 0.39 | 0.4 | 0.44 | 0.39 | 0.61 |
| 6 | 29819093 | rs2523759 | 1.641 | -1.992 | -1.285 | -0.703 | -0.021 | -0.018 | -0.018 | 0.006 | 0.24 | 0.08 | 0.12 | 0.23 | 0.2 | 0.33 |
| 6 | 29820047 | rs2508050 | 1.491 | -2.039 | -1.268 | -0.734 | 0.006 | 0.011 | 0.011 | 0 | 0.07 | 0.07 | 0.08 | 0.06 | 0.09 | 0.06 |
| 6 | 29820278 | rs3115627 | 0.976 | -2.085 | -1.158 | -0.751 | 0.024 | 0.011 | 0.011 | 0 | 0.24 | 0.3 | 0.33 | 0.37 | 0.36 | 0.15 |
| 6 | 29820586 | rs5013093 | 1.712 | -2.105 | -1.184 | -0.758 | -0.021 | -0.018 | -0.018 | 0 | 0.24 | 0.08 | 0.12 | 0.23 | 0.2 | 0.33 |
| 6 | 29820717 | rs2523756 | 0.199 | -2.189 | -1.127 | -0.787 | 0.001 | -0.001 | -0.001 | 0 | 0.52 | 0.47 | 0.41 | 0.46 | 0.41 | 0.62 |
| 6 | 29821293 | rs9258631 | 1.353 | -2.242 | -1.126 | -0.829 | -0.037 | -0.034 | -0.034 | 0 | 0.16 | 0.27 | 0.12 | 0.03 | 0.1 | 0.12 |
| 6 | 29821692 | rs2734981 | 1.722 | -2.166 | -1.138 | -0.788 | -0.021 | -0.017 | -0.017 | 0.03 | 0.24 | 0.08 | 0.12 | 0.23 | 0.2 | 0.33 |
| 6 | 29821896 | rs2734980 | 1.736 | -2.165 | -1.137 | -0.788 | -0.021 | -0.018 | -0.018 | 0 | 0.24 | 0.08 | 0.12 | 0.23 | 0.2 | 0.33 |
| 6 | 29821937 | rs2517862 | 3.047 | -2.178 | -1.169 | -0.811 | -0.048 | 0.037 | 0.037 | 0 | 0.12 | 0.02 | 0.05 | 0.02 | 0.13 | 0.17 |
| 6 | 29821982 | rs2517861 | 1.59 | -2.188 | -1.129 | -0.809 | 0 | 0.001 | 0.001 | 0 | 0.24 | 0.15 | 0.12 | 0.23 | 0.2 | 0.33 |

| | | | | | | | | | | | | | | | | |
|---|---|---|---|---|---|---|---|---|---|---|---|---|---|---|---|---|
| 6 | 29822061 | rs3115626 | -1.626 | -2.232 | -1.142 | -0.846 | -0.002 | 0.005 | 0.005 | 0 | 0.85 | 0.88 | 0.88 | 0.94 | 0.8 | 0.84 |
| 6 | 29822521 | rs2517860 | 1.714 | -2.248 | -1.16 | -0.852 | -0.02 | -0.017 | -0.017 | 0 | 0.24 | 0.08 | 0.12 | 0.23 | 0.2 | 0.34 |
| 6 | 29823027 | rs2428510 | 0.202 | -2.262 | -1.143 | -0.863 | 0.001 | -0.001 | -0.001 | 0 | 0.52 | 0.47 | 0.41 | 0.46 | 0.41 | 0.62 |
| 6 | 29823164 | rs2517855 | 3.11 | -2.357 | -1.149 | -0.868 | -0.048 | 0.037 | 0.037 | 0 | 0.12 | 0.02 | 0.05 | 0.02 | 0.13 | 0.17 |
| 6 | 29823319 | rs2844826 | 0.199 | -2.356 | -1.148 | -0.867 | 0.001 | -0.001 | -0.001 | 0.147 | 0.52 | 0.48 | 0.41 | 0.46 | 0.4 | 0.62 |
| 6 | 29823606 | rs3132726 | -0.766 | -2.382 | -1.152 | -0.869 | 0.002 | 0.001 | 0.001 | 0.035 | 0.75 | 0.77 | 0.74 | 0.82 | 0.77 | 0.77 |
| 6 | 29823994 | rs6919513 | 0.211 | -2.378 | -1.168 | -0.867 | 0.001 | -0.001 | -0.001 | 0.364 | 0.52 | 0.47 | 0.41 | 0.46 | 0.4 | 0.62 |
| 6 | 29825434 | rs1611674 | 0.598 | -2.398 | -1.179 | -0.876 | -0.006 | -0.01 | -0.01 | 0 | 0.44 | 0.47 | 0.37 | 0.26 | 0.34 | 0.46 |
| 6 | 29825846 | rs1611684 | 0.207 | -2.346 | -1.144 | -0.852 | 0.001 | -0.001 | -0.001 | 0.133 | 0.52 | 0.47 | 0.41 | 0.46 | 0.4 | 0.62 |
| 6 | 29827390 | rs9258738 | -0.083 | -2.325 | -1.13 | -0.838 | -0.021 | 0.009 | 0.009 | 0 | 0.4 | 0.45 | 0.36 | 0.44 | 0.27 | 0.45 |
| 6 | 29827899 | rs9258740 | 0.682 | -2.338 | -1.136 | -0.845 | -0.029 | -0.001 | -0.001 | 0 | 0.27 | 0.39 | 0.29 | 0.23 | 0.21 | 0.28 |
| 6 | 29828316 | rs17185692 | 1.441 | -2.272 | -1.109 | -0.811 | 0.013 | -0.026 | -0.026 | 0.219 | 0.12 | 0.06 | 0.07 | 0.21 | 0.07 | 0.17 |
| 6 | 29828467 | rs1611704 | 1.217 | -2.174 | -1.047 | -0.808 | -0.033 | -0.019 | -0.019 | 0.015 | 0.23 | 0.21 | 0.33 | 0.37 | 0.36 | 0.16 |
| 6 | 29828660 | rs2523822 | 1.619 | -2.158 | -1.042 | -0.798 | -0.027 | -0.02 | -0.02 | 0.06 | 0.2 | 0.19 | 0.27 | 0.35 | 0.32 | 0.15 |
| 6 | 29828880 | rs9295822 | 1.753 | -2.225 | -1.08 | -0.826 | -0.041 | -0.014 | -0.014 | 0.024 | 0.11 | 0.25 | 0.1 | 0.02 | 0.05 | 0.08 |
| 6 | 29829265 | rs3094170 | -1.828 | -2.19 | -1.106 | -0.824 | -0.002 | 0 | 0 | 0 | 0.9 | 0.93 | 0.91 | 0.98 | 0.85 | 0.95 |
| 6 | 29829566 | rs2517830 | 2.667 | -2.309 | -1.131 | -0.832 | -0.005 | 0.004 | 0.004 | 1.789 | 0.08 | 0 | 0.05 | 0.04 | 0.08 | 0.03 |
| 6 | 29829577 | rs1611717 | 0.005 | -2.455 | -0.722 | -0.722 | 0.001 | -0.001 | -0.001 | 0 | 0.52 | 0.47 | 0.41 | 0.46 | 0.4 | 0.62 |
| 6 | 29829871 | rs9468643 | 1.691 | -2.479 | -0.72 | -0.731 | -0.039 | -0.012 | -0.012 | 0.695 | 0.11 | 0.25 | 0.1 | 0.02 | 0.05 | 0.08 |
| 6 | 29830074 | rs7760172 | 1.527 | -2.408 | -0.738 | -0.704 | -0.011 | 0.001 | 0.001 | 0 | 0.09 | 0.11 | 0.07 | 0.11 | 0.03 | 0.07 |
| 6 | 29830411 | rs9258766 | -0.194 | -2.357 | -0.759 | -0.698 | -0.021 | 0.009 | 0.009 | 0 | 0.4 | 0.45 | 0.36 | 0.44 | 0.27 | 0.45 |
| 6 | 29830652 | rs9258771 | -0.198 | -2.361 | -0.754 | -0.674 | -0.023 | 0.01 | 0.01 | 0 | 0.4 | 0.45 | 0.37 | 0.44 | 0.28 | 0.45 |
| 6 | 29831008 | rs1611732 | 0.003 | -2.348 | -0.761 | -0.728 | 0.001 | -0.001 | -0.001 | 3.055 | 0.52 | 0.47 | 0.41 | 0.46 | 0.4 | 0.62 |
| 6 | 29831706 | rs1611738 | 2.588 | -1.685 | -0.643 | -0.454 | -0.048 | 0.037 | 0.037 | 0 | 0.12 | 0.02 | 0.05 | 0.02 | 0.13 | 0.17 |
| 6 | 29832332 | rs2524035 | -0.608 | -1.726 | -0.527 | -0.403 | -0.001 | -0.001 | -0.001 | 4.01 | 0.8 | 0.81 | 0.83 | 0.86 | 0.81 | 0.83 |
| 6 | 29833033 | rs2508046 | 0.257 | -1.545 | -0.561 | -0.557 | -0.045 | -0.034 | -0.034 | 0.955 | 0.19 | 0.32 | 0.18 | 0.08 | 0.14 | 0.14 |
| 6 | 29833541 | rs3094165 | -0.224 | -1.571 | -0.565 | -0.539 | -0.015 | -0.012 | -0.012 | 0 | 0.67 | 0.56 | 0.63 | 0.76 | 0.68 | 0.7 |
| 6 | 29833779 | rs2734974 | 1.441 | -1.111 | -0.613 | -0.332 | -0.011 | -0.036 | -0.036 | 6.648 | 0.23 | 0.29 | 0.15 | 0.04 | 0.18 | 0.24 |
| 6 | 29833948 | rs2734973 | 0.43 | -1.137 | -0.658 | -0.362 | 0.007 | 0.029 | 0.029 | 0 | 0.13 | 0.08 | 0.1 | 0.06 | 0.13 | 0.11 |
| 6 | 29834041 | rs2517826 | 1.43 | -1.106 | -0.602 | -0.317 | -0.011 | -0.036 | -0.036 | 6.648 | 0.23 | 0.29 | 0.15 | 0.04 | 0.18 | 0.24 |
| 6 | 29835834 | rs3094159 | -0.058 | -1.097 | -0.704 | -0.179 | -0.005 | 0.009 | 0.009 | 0 | 0.61 | 0.63 | 0.62 | 0.46 | 0.64 | 0.55 |
| 6 | 29836524 | rs9258866 | 1.556 | -1.106 | -0.738 | -0.19 | 0.003 | 0.002 | 0.002 | 0 | 0.05 | 0.06 | 0.03 | 0.05 | 0.05 | 0.1 |
| 6 | 29836577 | rs3132718 | -0.048 | -1.059 | -0.727 | -0.16 | -0.005 | 0.008 | 0.008 | 0 | 0.61 | 0.63 | 0.62 | 0.46 | 0.64 | 0.55 |
| 6 | 29837188 | rs9258883 | -0.675 | -1.096 | -0.72 | -0.139 | -0.005 | 0.01 | 0.01 | 0 | 0.73 | 0.76 | 0.8 | 0.67 | 0.75 | 0.72 |
| 6 | 29839816 | rs3094654 | -0.341 | -0.668 | -0.703 | -0.363 | -0.005 | 0.008 | 0.008 | 0 | 0.61 | 0.63 | 0.62 | 0.46 | 0.64 | 0.55 |
| 6 | 29840382 | rs9258966 | -0.341 | -0.668 | -0.703 | -0.363 | -0.005 | 0.008 | 0.008 | 0 | 0.61 | 0.63 | 0.62 | 0.46 | 0.64 | 0.55 |
| 6 | 29840648 | rs3132714 | -0.341 | -0.668 | -0.703 | -0.363 | -0.005 | 0.008 | 0.008 | 0 | 0.61 | 0.63 | 0.62 | 0.46 | 0.64 | 0.55 |
| 6 | 29849619 | rs2523809 | -0.147 | -0.304 | -0.649 | -0.611 | 0.007 | 0.029 | 0.029 | 0 | 0.13 | 0.08 | 0.1 | 0.06 | 0.13 | 0.11 |
| 6 | 29849657 | rs2523808 | 0.731 | -0.198 | -0.511 | -0.597 | -0.009 | -0.036 | -0.036 | 0 | 0.22 | 0.29 | 0.15 | 0.04 | 0.18 | 0.24 |
| 6 | 29855945 | rs2734945 | 0.047 | -0.172 | -0.576 | -0.598 | -0.002 | -0.034 | -0.034 | 0.291 | 0.31 | 0.36 | 0.26 | 0.07 | 0.29 | 0.32 |
| 6 | 29868984 | rs2517904 | -0.091 | 0.126 | -0.917 | -0.113 | -0.004 | 0.031 | 0.031 | 0 | 0.08 | 0.05 | 0.09 | 0.03 | 0.09 | 0.06 |
| 6 | 29879529 | rs2524024 | -0.45 | 0.324 | -0.803 | -0.043 | -0.002 | 0 | 0 | 0 | 0.91 | 0.93 | 0.91 | 0.98 | 0.87 | 0.94 |
| 6 | 29896421 | rs9259852 | 0.769 | 0.117 | -0.858 | 0.02 | -0.003 | 0.017 | 0.017 | 0 | 0.05 | 0.02 | 0.02 | 0.01 | 0.04 | 0.04 |
| 6 | 29913067 | rs3823342 | -0.181 | -0.375 | -0.727 | 0.042 | 0.001 | -0.001 | -0.001 | 0 | 0.54 | 0.59 | 0.6 | 0.48 | 0.52 | 0.46 |
| 6 | 29913727 | rs2571382 | -0.434 | -0.43 | -0.614 | -0.093 | 0 | -0.021 | -0.021 | 0 | 0.84 | 0.79 | 0.86 | 0.97 | 0.82 | 0.94 |
| 6 | 29914270 | rs1616549 | -0.927 | -0.393 | -0.571 | -0.136 | -0.002 | 0 | 0 | 0.124 | 0.9 | 0.93 | 0.91 | 0.98 | 0.85 | 0.95 |
| 6 | 29915301 | rs2735097 | 1.103 | -1.005 | -0.444 | -0.277 | -0.013 | 0.002 | 0.002 | 0.38 | 0.3 | 0.28 | 0.42 | 0.3 | 0.36 | 0.23 |
| 6 | 29915469 | rs7745413 | 0.43 | -1.271 | -0.4 | -0.118 | -0.012 | 0.009 | 0.009 | 0 | 0.25 | 0.32 | 0.27 | 0.37 | 0.19 | 0.34 |
| 6 | 29916618 | rs1655900 | 2.135 | -0.842 | -0.639 | -0.307 | 0.001 | -0.011 | -0.011 | 0 | 0.12 | 0.08 | 0.05 | 0.02 | 0.13 | 0.17 |
| 6 | 29917439 | rs2517715 | 1.047 | -0.799 | -0.682 | -0.183 | -0.016 | 0.002 | 0.002 | 0.344 | 0.3 | 0.27 | 0.41 | 0.3 | 0.36 | 0.23 |
| 6 | 29917591 | rs2523949 | 0.367 | -0.59 | -0.583 | -0.106 | -0.001 | -0.009 | -0.009 | 0 | 0.08 | 0.1 | 0.08 | 0.02 | 0.08 | 0.05 |
| 6 | 29924831 | rs429511 | 1.775 | -0.319 | -0.532 | -0.109 | 0.003 | -0.015 | -0.015 | 0 | 0.12 | 0.08 | 0.05 | 0.02 | 0.13 | 0.16 |
| 6 | 29925086 | rs2523942 | 0.403 | -0.458 | -0.49 | -0.206 | -0.001 | -0.009 | -0.009 | 0 | 0.08 | 0.1 | 0.08 | 0.02 | 0.08 | 0.05 |
| 6 | 29926082 | rs2517700 | 0.403 | -0.458 | -0.44 | -0.206 | -0.002 | -0.015 | -0.015 | 0 | 0.08 | 0.11 | 0.09 | 0.02 | 0.08 | 0.05 |
| 6 | 29926377 | rs9260679 | 1.099 | -0.463 | -0.439 | -0.211 | -0.002 | -0.011 | -0.011 | 0 | 0.06 | 0.07 | 0.02 | 0.01 | 0.04 | 0.04 |
| 6 | 29926597 | rs1632994 | -0.198 | -0.948 | -0.396 | -0.164 | 0 | 0.002 | 0.002 | 0 | 0.51 | 0.52 | 0.59 | 0.58 | 0.53 | 0.44 |
| 6 | 29927095 | rs2735085 | -0.011 | -1.61 | -0.575 | -0.321 | 0.003 | -0.002 | -0.002 | 0.266 | 0.26 | 0.26 | 0.22 | 0.26 | 0.19 | 0.28 |
| 6 | 29928137 | rs441808 | -0.662 | -1.575 | -0.562 | -0.332 | 0 | -0.001 | -0.001 | 0 | 0.51 | 0.53 | 0.59 | 0.58 | 0.53 | 0.44 |
| 6 | 29928649 | rs2428521 | -0.493 | -1.637 | -0.482 | -0.387 | 0.001 | 0.006 | 0.006 | 0 | 0.48 | 0.46 | 0.55 | 0.57 | 0.51 | 0.44 |
| 6 | 29929253 | rs2517689 | -0.505 | -1.649 | -0.481 | -0.386 | 0.001 | 0.006 | 0.006 | 0 | 0.48 | 0.46 | 0.55 | 0.57 | 0.51 | 0.44 |
| 6 | 29929336 | rs9260706 | -1.551 | -1.711 | -0.457 | -0.496 | 0 | -0.003 | -0.003 | 1.888 | 0.91 | 0.93 | 0.92 | 0.99 | 0.89 | 0.95 |
| 6 | 29930619 | rs7739434 | -0.191 | -2.18 | -0.297 | -0.576 | -0.027 | -0.023 | -0.023 | 0.142 | 0.75 | 0.65 | 0.78 | 0.86 | 0.83 | 0.83 |
| 6 | 29931798 | rs9260722 | 0.58 | -2.267 | -0.372 | -0.625 | 0.001 | -0.006 | -0.006 | 0 | 0.1 | 0.1 | 0.08 | 0.02 | 0.08 | 0.05 |
| 6 | 29932666 | rs9260734 | 0.027 | -2.425 | -0.307 | -0.545 | -0.023 | 0.013 | 0.013 | 0.077 | 0.26 | 0.32 | 0.17 | 0.26 | 0.16 | 0.28 |
| 6 | 29932897 | rs3903160 | 1.512 | -2.305 | -0.36 | -0.536 | -0.033 | -0.012 | -0.012 | 0 | 0.23 | 0.19 | 0.35 | 0.3 | 0.33 | 0.15 |
| 6 | 29933261 | rs6457109 | 1.268 | -2.364 | -0.636 | -0.674 | -0.031 | -0.019 | -0.019 | 0 | 0.18 | 0.25 | 0.12 | 0.06 | 0.12 | 0.04 |
| 6 | 29933439 | rs2844806 | -0.262 | -2.394 | -0.545 | -0.654 | -0.014 | 0 | 0 | 0 | 0.58 | 0.65 | 0.46 | 0.55 | 0.51 | 0.67 |
| 6 | 29934163 | rs6935053 | 1.206 | -2.246 | -0.545 | -0.629 | -0.007 | -0.005 | -0.005 | 0 | 0.23 | 0.2 | 0.33 | 0.31 | 0.32 | 0.16 |
| 6 | 29934454 | rs3893259 | -1.714 | -2.343 | -0.412 | -0.677 | 0.005 | -0.017 | -0.017 | 0.077 | 0.9 | 0.89 | 0.91 | 0.99 | 0.89 | 0.95 |

| | | | | | | | | | | | | | | | | |
|---|---|---|---|---|---|---|---|---|---|---|---|---|---|---|---|---|
| 6 | 29934697 | rs4713270 | 1.235 | -2.342 | -0.414 | -0.676 | -0.007 | -0.005 | -0.005 | 0 | 0.23 | 0.2 | 0.33 | 0.31 | 0.32 | 0.16 |
| 6 | 29934891 | rs3869062 | 2.257 | -2.486 | -0.521 | -0.673 | -0.016 | -0.041 | -0.041 | 0.368 | 0.07 | 0 | 0.04 | 0.23 | 0.06 | 0.15 |
| 6 | 29935250 | rs3893464 | 0.4 | -2.644 | -0.491 | -0.715 | 0.005 | 0.008 | 0.008 | 0.318 | 0.49 | 0.41 | 0.42 | 0.62 | 0.45 | 0.66 |
| 6 | 29935372 | rs9260759 | 0.916 | -2.642 | -0.481 | -0.717 | -0.01 | -0.008 | -0.008 | 0 | 0.15 | 0.2 | 0.13 | 0.05 | 0.09 | 0.13 |
| 6 | 29935469 | rs9260761 | -1.808 | -2.59 | -0.481 | -0.697 | 0.005 | -0.017 | -0.017 | 0 | 0.9 | 0.89 | 0.91 | 0.99 | 0.89 | 0.95 |
| 6 | 29937104 | rs12193110 | 1.022 | -2.588 | -0.499 | -0.685 | 0.002 | 0.001 | 0.001 | 0.786 | 0.26 | 0.28 | 0.37 | 0.31 | 0.34 | 0.16 |
| 6 | 29937242 | rs2517673 | 0.517 | -2.649 | -0.483 | -0.716 | -0.001 | 0.001 | 0.001 | 0 | 0.18 | 0.2 | 0.16 | 0.14 | 0.11 | 0.17 |
| 6 | 29937580 | rs4713275 | 1.719 | -2.606 | -0.487 | -0.7 | -0.05 | -0.027 | -0.027 | 0.149 | 0.2 | 0.12 | 0.32 | 0.3 | 0.31 | 0.15 |
| 6 | 29937740 | rs2256539 | 0.982 | -2.646 | -0.468 | -0.737 | 0 | 0 | 0 | 0 | 0.5 | 0.48 | 0.59 | 0.56 | 0.62 | 0.49 |
| 6 | 29937833 | rs2256543 | -1.356 | -2.618 | -0.46 | -0.743 | 0.006 | -0.019 | -0.019 | 0 | 0.63 | 0.6 | 0.66 | 0.74 | 0.59 | 0.61 |
| 6 | 29937896 | rs3202637 | -0.655 | -2.721 | -0.497 | -0.785 | 0.004 | -0.009 | -0.009 | 0 | 0.74 | 0.79 | 0.82 | 0.71 | 0.77 | 0.73 |
| 6 | 29937924 | rs1061535 | -0.989 | -2.708 | -0.487 | -0.787 | -0.002 | -0.008 | -0.008 | 0.014 | 0.81 | 0.8 | 0.86 | 0.94 | 0.83 | 0.89 |
| 6 | 29938110 | rs6916422 | 1.408 | -2.64 | -0.561 | -0.75 | -0.007 | -0.005 | -0.005 | 0.089 | 0.23 | 0.19 | 0.33 | 0.31 | 0.32 | 0.16 |
| 6 | 29938258 | rs2523971 | 2.498 | -2.745 | -0.619 | -0.885 | -0.016 | -0.001 | -0.001 | 1.76 | 0.21 | 0.09 | 0.19 | 0.25 | 0.26 | 0.33 |
| 6 | 29938777 | rs2523969 | -0.008 | -2.766 | -0.604 | -0.867 | -0.011 | -0.005 | -0.005 | 0 | 0.24 | 0.3 | 0.23 | 0.15 | 0.15 | 0.24 |
| 6 | 29938932 | rs2735079 | 1.183 | -2.825 | -0.602 | -0.852 | -0.004 | -0.001 | -0.001 | 0 | 0.47 | 0.42 | 0.55 | 0.57 | 0.6 | 0.49 |
| 6 | 29939240 | rs2523966 | -1.63 | -2.876 | -0.604 | -0.89 | 0.009 | -0.015 | -0.015 | 0 | 0.66 | 0.67 | 0.68 | 0.73 | 0.61 | 0.61 |
| 6 | 29939580 | rs2523961 | 0.497 | -2.837 | -0.599 | -0.868 | -0.007 | -0.001 | -0.001 | 0 | 0.18 | 0.22 | 0.21 | 0.14 | 0.11 | 0.2 |
| 6 | 29939852 | rs435766 | 1.174 | -2.835 | -0.599 | -0.865 | -0.004 | 0 | 0 | 0 | 0.47 | 0.42 | 0.55 | 0.56 | 0.6 | 0.49 |
| 6 | 29939952 | rs2523960 | 2.081 | -2.833 | -0.598 | -0.865 | 0.003 | -0.004 | -0.004 | 0.814 | 0.25 | 0.22 | 0.22 | 0.25 | 0.28 | 0.34 |
| 6 | 29940106 | rs2256820 | -0.158 | -2.89 | -0.621 | -0.889 | -0.007 | -0.003 | -0.003 | 0 | 0.24 | 0.29 | 0.23 | 0.15 | 0.15 | 0.24 |
| 6 | 29940260 | rs2523957 | 1.127 | -2.855 | -0.605 | -0.87 | -0.003 | -0.001 | -0.001 | 0.204 | 0.48 | 0.42 | 0.55 | 0.56 | 0.6 | 0.49 |
| 6 | 29940372 | rs2256902 | 0.59 | -2.824 | -0.663 | -0.873 | -0.01 | -0.002 | -0.002 | 0 | 0.17 | 0.22 | 0.2 | 0.13 | 0.11 | 0.19 |
| 6 | 29940679 | rs2844805 | 1.699 | -2.982 | -0.674 | -0.941 | 0.005 | -0.016 | -0.016 | 0 | 0.27 | 0.36 | 0.24 | 0.25 | 0.28 | 0.33 |
| 6 | 29940750 | rs2256919 | 1.125 | -2.964 | -0.703 | -0.912 | -0.003 | -0.001 | -0.001 | 0 | 0.48 | 0.47 | 0.55 | 0.56 | 0.6 | 0.5 |
| 6 | 29941943 | rs2523946 | -0.757 | -2.962 | -0.668 | -0.929 | 0 | -0.001 | -0.001 | 0 | 0.48 | 0.55 | 0.59 | 0.46 | 0.49 | 0.39 |
| 6 | 29942083 | rs3823355 | 1.45 | -2.995 | -0.677 | -0.928 | -0.006 | -0.005 | -0.005 | 0 | 0.23 | 0.2 | 0.33 | 0.31 | 0.32 | 0.17 |
| 6 | 29942126 | rs2844804 | -0.78 | -2.996 | -0.673 | -0.928 | -0.001 | -0.001 | -0.001 | 0 | 0.48 | 0.53 | 0.59 | 0.46 | 0.49 | 0.39 |
| 6 | 29942205 | rs3823358 | 1.449 | -2.993 | -0.674 | -0.927 | -0.006 | -0.005 | -0.005 | 0.059 | 0.23 | 0.19 | 0.33 | 0.31 | 0.32 | 0.17 |
| 6 | 29942857 | rs2232238 | -0.382 | -2.993 | -0.672 | -0.926 | -0.016 | -0.01 | -0.01 | 0 | 0.25 | 0.34 | 0.26 | 0.15 | 0.16 | 0.23 |
| 6 | 29942927 | rs2523943 | -0.139 | -3.077 | -0.708 | -0.964 | -0.024 | -0.013 | -0.013 | 0.206 | 0.2 | 0.31 | 0.23 | 0.14 | 0.12 | 0.16 |
| 6 | 29943035 | rs400488 | -0.517 | -3.065 | -0.704 | -0.957 | -0.016 | -0.01 | -0.01 | 0 | 0.25 | 0.34 | 0.26 | 0.15 | 0.16 | 0.23 |
| 6 | 29943067 | rs6904029 | 1.402 | -2.854 | -0.735 | -0.851 | -0.006 | -0.005 | -0.005 | 0.001 | 0.23 | 0.19 | 0.33 | 0.31 | 0.32 | 0.17 |
| 6 | 29943490 | rs2735076 | 0.112 | -2.969 | -0.709 | -0.895 | -0.047 | -0.035 | -0.035 | 0 | 0.67 | 0.54 | 0.66 | 0.83 | 0.78 | 0.81 |
| 6 | 29943656 | rs2394250 | 0.113 | -2.952 | -0.766 | -0.88 | -0.014 | 0.001 | 0.001 | 0 | 0.42 | 0.33 | 0.42 | 0.55 | 0.47 | 0.42 |
| 6 | 29943832 | rs2893999 | 2.321 | -2.874 | -0.797 | -0.804 | -0.016 | -0.042 | -0.042 | 0 | 0.07 | 0 | 0.03 | 0.23 | 0.06 | 0.15 |
| 6 | 29944004 | rs2245961 | 1.038 | -2.887 | -0.709 | -0.847 | 0.021 | 0.002 | 0.002 | 0 | 0.51 | 0.43 | 0.57 | 0.59 | 0.65 | 0.55 |
| 6 | 29944148 | rs3823374 | 0.67 | -2.622 | -0.797 | -0.831 | 0 | 0 | 0 | 0.323 | 0.29 | 0.28 | 0.36 | 0.31 | 0.37 | 0.2 |
| 6 | 29944158 | rs3823375 | 0.702 | -2.619 | -0.795 | -0.83 | 0.001 | 0 | 0 | 0 | 0.29 | 0.28 | 0.36 | 0.31 | 0.37 | 0.2 |
| 6 | 29945270 | rs2571375 | 0.992 | -2.449 | -0.912 | -0.669 | 0 | -0.001 | -0.001 | 0 | 0.08 | 0.05 | 0.16 | 0.04 | 0.09 | 0.09 |
| 6 | 29945620 | rs2246199 | 0.181 | -2.638 | -0.79 | -0.737 | -0.043 | 0.009 | 0.009 | 0 | 0.15 | 0.26 | 0.19 | 0.15 | 0.08 | 0.13 |
| 6 | 29948365 | rs2108691 | -0.376 | -2.252 | -0.815 | -0.756 | -0.04 | -0.003 | -0.003 | 0 | 0.2 | 0.29 | 0.22 | 0.16 | 0.12 | 0.2 |
| 6 | 29949123 | rs423808 | -0.478 | -2.294 | -0.835 | -0.772 | -0.04 | -0.004 | -0.004 | 0 | 0.21 | 0.29 | 0.22 | 0.16 | 0.12 | 0.2 |
| 6 | 29949448 | rs2571430 | -0.37 | -2.28 | -0.843 | -0.767 | -0.04 | -0.003 | -0.003 | 0 | 0.2 | 0.28 | 0.22 | 0.16 | 0.12 | 0.19 |
| 6 | 29949497 | rs2246638 | -0.726 | -2.289 | -0.845 | -0.771 | -0.04 | -0.006 | -0.006 | 0 | 0.25 | 0.34 | 0.26 | 0.2 | 0.17 | 0.25 |
| 6 | 29949821 | rs2394255 | -0.405 | -2.518 | -0.893 | -0.559 | 0.001 | 0 | 0 | 0 | 0.52 | 0.49 | 0.48 | 0.5 | 0.47 | 0.62 |
| 6 | 29950140 | rs379221 | -0.568 | -2.525 | -0.959 | -0.565 | -0.022 | -0.007 | -0.007 | 0 | 0.25 | 0.3 | 0.25 | 0.16 | 0.16 | 0.23 |
| 6 | 29950210 | rs401618 | 2.162 | -2.481 | -0.905 | -0.523 | -0.012 | -0.015 | -0.015 | 0 | 0.22 | 0.11 | 0.2 | 0.29 | 0.26 | 0.33 |
| 6 | 29950583 | rs2735068 | 2.057 | -2.486 | -0.906 | -0.527 | 0.001 | 0.009 | 0.009 | 4.99 | 0.05 | 0.04 | 0.09 | 0.01 | 0.07 | 0.07 |
| 6 | 29951106 | rs2735067 | 2.1 | -2.426 | -0.867 | -0.48 | -0.012 | -0.015 | -0.015 | 0 | 0.22 | 0.11 | 0.2 | 0.29 | 0.26 | 0.34 |
| 6 | 29952162 | rs11757750 | 0.014 | -2.272 | -0.739 | -0.394 | -0.055 | 0.011 | 0.011 | 0 | 0.15 | 0.25 | 0.14 | 0.15 | 0.05 | 0.13 |
| 6 | 29952814 | rs17186937 | 0.014 | -2.272 | -0.739 | -0.394 | -0.055 | 0.011 | 0.011 | 0 | 0.15 | 0.25 | 0.14 | 0.15 | 0.04 | 0.13 |
| 6 | 29953099 | rs11755961 | 0.014 | -2.272 | -0.739 | -0.394 | -0.055 | 0.011 | 0.011 | 0 | 0.15 | 0.25 | 0.14 | 0.15 | 0.04 | 0.13 |
| 6 | 29955199 | rs5025708 | -1.994 | -2.293 | -0.886 | -0.492 | -0.029 | 0.017 | 0.017 | 0 | 0.92 | 0.99 | 0.92 | 0.98 | 0.89 | 0.95 |
| 6 | 29956842 | rs12665186 | 0.054 | -2.392 | -0.792 | -0.454 | -0.055 | 0.011 | 0.011 | 0 | 0.15 | 0.25 | 0.14 | 0.15 | 0.04 | 0.12 |
| 6 | 29957077 | rs12662611 | 0.054 | -2.353 | -0.758 | -0.436 | -0.055 | 0.012 | 0.012 | 0 | 0.15 | 0.25 | 0.14 | 0.15 | 0.04 | 0.13 |
| 6 | 29960999 | rs9260961 | -0.177 | -2.424 | -0.739 | -0.473 | -0.052 | 0.008 | 0.008 | 0 | 0.18 | 0.3 | 0.17 | 0.19 | 0.1 | 0.16 |
| 6 | 29961367 | rs9260967 | -0.177 | -2.424 | -0.739 | -0.473 | -0.052 | 0.008 | 0.008 | 0 | 0.18 | 0.3 | 0.17 | 0.19 | 0.1 | 0.16 |
| 6 | 29963065 | rs6457121 | -0.365 | -2.617 | -0.865 | -0.626 | -0.001 | 0 | 0 | 0 | 0.54 | 0.49 | 0.48 | 0.5 | 0.47 | 0.62 |
| 6 | 29963179 | rs6910898 | -0.084 | -2.517 | -0.787 | -0.565 | -0.052 | 0.008 | 0.008 | 0 | 0.18 | 0.3 | 0.17 | 0.19 | 0.1 | 0.16 |
| 6 | 29963316 | rs9260997 | -1.646 | -2.434 | -0.741 | -0.595 | -0.001 | -0.002 | -0.002 | 0 | 0.91 | 0.93 | 0.9 | 0.98 | 0.89 | 0.95 |
| 6 | 29966850 | rs6941318 | -0.081 | -2.54 | -0.67 | -0.555 | -0.052 | 0.008 | 0.008 | 0 | 0.18 | 0.3 | 0.17 | 0.19 | 0.1 | 0.16 |
| 6 | 29968275 | rs356960 | 1.791 | -2.628 | -0.742 | -0.615 | 0.007 | 0.015 | 0.015 | 6.917 | 0.07 | 0.04 | 0.08 | 0.01 | 0.07 | 0.09 |
| 6 | 29970427 | rs11760176 | 0.849 | -2.588 | -0.706 | -0.585 | -0.04 | 0.039 | 0.039 | 0 | 0.11 | 0.22 | 0.11 | 0.13 | 0.04 | 0.12 |
| 6 | 29970589 | rs7758512 | -0.097 | -2.547 | -0.71 | -0.564 | -0.052 | 0.008 | 0.008 | 0 | 0.18 | 0.3 | 0.18 | 0.19 | 0.1 | 0.16 |
| 6 | 29973392 | rs2286405 | -0.079 | -2.542 | -0.712 | -0.54 | -0.052 | 0.008 | 0.008 | 0 | 0.18 | 0.3 | 0.17 | 0.19 | 0.1 | 0.16 |
| 6 | 29973620 | rs2286404 | 2.355 | -2.641 | -0.922 | -0.501 | -0.016 | -0.042 | -0.042 | 0 | 0.07 | 0 | 0.03 | 0.19 | 0.06 | 0.15 |
| 6 | 29973925 | rs4313034 | -0.129 | -2.647 | -0.884 | -0.564 | -0.027 | -0.012 | -0.012 | 0 | 0.76 | 0.68 | 0.76 | 0.83 | 0.84 | 0.82 |
| 6 | 29974500 | rs9261105 | -0.029 | -2.583 | -0.91 | -0.496 | -0.052 | 0.008 | 0.008 | 0 | 0.18 | 0.3 | 0.17 | 0.19 | 0.1 | 0.16 |

| | | | | | | | | | | | | | | | | |
|---|---|---|---|---|---|---|---|---|---|---|---|---|---|---|---|---|
| 6 | 29975047 | rs356968 | 2.34 | -2.663 | -1.017 | -0.57 | -0.014 | -0.017 | -0.017 | 0 | 0.23 | 0.11 | 0.2 | 0.29 | 0.26 | 0.33 |
| 6 | 29975453 | rs2240619 | -0.248 | -2.646 | -1.001 | -0.597 | -0.001 | 0 | 0 | 0.075 | 0.52 | 0.49 | 0.48 | 0.5 | 0.47 | 0.62 |
| 6 | 29976024 | rs3765604 | -0.01 | -2.519 | -0.912 | -0.513 | -0.052 | 0.008 | 0.008 | 0 | 0.18 | 0.3 | 0.17 | 0.19 | 0.1 | 0.16 |
| 6 | 29977973 | rs9261115 | -0.01 | -2.519 | -0.912 | -0.513 | -0.052 | 0.008 | 0.008 | 0 | 0.18 | 0.3 | 0.17 | 0.19 | 0.1 | 0.16 |
| 6 | 29978352 | rs4713281 | 1.745 | -2.79 | -1.069 | -0.727 | -0.052 | -0.028 | -0.028 | 0.562 | 0.22 | 0.12 | 0.31 | 0.3 | 0.31 | 0.16 |
| 6 | 29979579 | rs9261129 | 0.031 | -2.741 | -0.954 | -0.688 | -0.052 | 0.008 | 0.008 | 0 | 0.18 | 0.3 | 0.17 | 0.19 | 0.1 | 0.16 |
| 6 | 29979643 | rs11753431 | 0.356 | -2.747 | -0.943 | -0.688 | -0.055 | 0.011 | 0.011 | 0 | 0.15 | 0.25 | 0.13 | 0.15 | 0.04 | 0.13 |
| 6 | 29981750 | rs16896044 | 2.451 | -2.773 | -1.075 | -0.661 | -0.016 | -0.042 | -0.042 | 1.798 | 0.07 | 0 | 0.03 | 0.19 | 0.06 | 0.15 |
| 6 | 29982429 | rs11965585 | 0.372 | -2.79 | -1.023 | -0.657 | -0.055 | 0.011 | 0.011 | 0 | 0.15 | 0.25 | 0.13 | 0.15 | 0.04 | 0.13 |
| 6 | 29983039 | rs6457126 | -0.101 | -2.919 | -1.078 | -0.764 | -0.001 | 0 | 0 | 0 | 0.53 | 0.49 | 0.48 | 0.5 | 0.47 | 0.62 |
| 6 | 29983538 | rs356954 | 2.511 | -2.935 | -1.119 | -0.776 | -0.014 | -0.017 | -0.017 | 0 | 0.23 | 0.11 | 0.2 | 0.3 | 0.26 | 0.33 |
| 6 | 29984252 | rs9357092 | 2.053 | -2.931 | -1.165 | -0.786 | -0.038 | -0.023 | -0.023 | 0 | 0.2 | 0.14 | 0.31 | 0.29 | 0.31 | 0.16 |
| 6 | 29984865 | rs9261145 | 0.084 | -2.917 | -1.158 | -0.778 | -0.052 | 0.008 | 0.008 | 0 | 0.18 | 0.3 | 0.17 | 0.19 | 0.1 | 0.16 |
| 6 | 29985849 | rs6926792 | 0.084 | -2.917 | -1.174 | -0.778 | -0.052 | 0.008 | 0.008 | 0 | 0.18 | 0.3 | 0.17 | 0.19 | 0.1 | 0.16 |
| 6 | 29986427 | rs6931776 | 0.084 | -2.917 | -1.174 | -0.778 | -0.052 | 0.008 | 0.008 | 0 | 0.18 | 0.3 | 0.17 | 0.19 | 0.1 | 0.16 |
| 6 | 29986684 | rs165256 | -0.229 | -2.919 | -1.185 | -0.782 | -0.04 | -0.005 | -0.005 | 5.794 | 0.22 | 0.29 | 0.21 | 0.16 | 0.12 | 0.22 |
| 6 | 29987184 | rs4711206 | 2.051 | -2.83 | -1.094 | -0.756 | -0.017 | -0.01 | -0.01 | 2.517 | 0.2 | 0.18 | 0.31 | 0.29 | 0.31 | 0.16 |
| 6 | 29988642 | rs3129012 | 3.541 | -2.856 | -1.485 | -0.851 | -0.051 | 0.043 | 0.043 | 0 | 0.11 | 0.03 | 0.05 | 0.02 | 0.13 | 0.17 |
| 6 | 29991699 | rs6919617 | 0.155 | -2.938 | -1.295 | -0.768 | -0.052 | 0.008 | 0.008 | 0 | 0.18 | 0.3 | 0.17 | 0.19 | 0.1 | 0.16 |
| 6 | 29992261 | rs1150743 | 2.137 | -2.91 | -1.412 | -0.748 | -0.025 | -0.063 | -0.063 | 0 | 0.1 | 0.01 | 0.14 | 0.27 | 0.12 | 0.16 |
| 6 | 29993196 | rs166325 | 1.948 | -2.775 | -1.181 | -0.725 | 0.007 | 0.016 | 0.016 | 6.774 | 0.07 | 0.04 | 0.08 | 0.01 | 0.07 | 0.09 |
| 6 | 29996855 | rs9261174 | 0.095 | -2.79 | -1.077 | -0.693 | -0.052 | 0.008 | 0.008 | 0 | 0.18 | 0.3 | 0.17 | 0.19 | 0.1 | 0.16 |
| 6 | 29999534 | rs4448133 | 0.095 | -2.79 | -1.077 | -0.693 | -0.052 | 0.008 | 0.008 | 0 | 0.18 | 0.3 | 0.17 | 0.19 | 0.1 | 0.16 |
| 6 | 30000280 | rs9261189 | 0.095 | -2.79 | -1.077 | -0.693 | -0.052 | 0.008 | 0.008 | 0 | 0.18 | 0.3 | 0.17 | 0.19 | 0.1 | 0.16 |
| 6 | 30000687 | rs16896923 | 2.504 | -2.791 | -1.198 | -0.677 | -0.016 | -0.042 | -0.042 | 0 | 0.07 | 0 | 0.03 | 0.19 | 0.06 | 0.15 |
| 6 | 30002881 | rs166327 | 0.406 | -2.876 | -1.174 | -0.765 | -0.008 | 0.003 | 0.003 | 2.574 | 0.46 | 0.45 | 0.4 | 0.49 | 0.4 | 0.53 |
| 6 | 30003797 | rs3869067 | 0.103 | -2.788 | -0.992 | -0.705 | -0.052 | 0.008 | 0.008 | 0 | 0.18 | 0.3 | 0.17 | 0.19 | 0.1 | 0.16 |
| 6 | 30004052 | rs3869068 | 0.103 | -2.788 | -0.992 | -0.705 | -0.052 | 0.008 | 0.008 | 0 | 0.18 | 0.3 | 0.17 | 0.19 | 0.1 | 0.16 |
| 6 | 30005754 | rs4711207 | 2.055 | -2.763 | -1.128 | -0.768 | -0.045 | -0.028 | -0.028 | 0 | 0.2 | 0.12 | 0.31 | 0.29 | 0.31 | 0.16 |
| 6 | 30007493 | rs259937 | 1.988 | -2.81 | -1.185 | -0.799 | 0.007 | 0.016 | 0.016 | 6.774 | 0.07 | 0.04 | 0.08 | 0.01 | 0.07 | 0.09 |
| 6 | 30008746 | rs3910312 | 2.059 | -2.833 | -0.975 | -0.814 | -0.047 | -0.029 | -0.029 | 0 | 0.2 | 0.12 | 0.31 | 0.29 | 0.31 | 0.16 |
| 6 | 30010139 | rs9261216 | 0.142 | -2.922 | -0.918 | -0.827 | -0.052 | 0.008 | 0.008 | 0 | 0.18 | 0.3 | 0.17 | 0.19 | 0.1 | 0.16 |
| 6 | 30010492 | rs9380150 | 1.903 | -2.911 | -0.962 | -0.759 | -0.02 | -0.016 | -0.016 | 0 | 0.21 | 0.18 | 0.31 | 0.33 | 0.31 | 0.17 |
| 6 | 30011581 | rs259939 | 1.904 | -3.018 | -1.083 | -0.822 | -0.019 | -0.013 | -0.013 | 0 | 0.27 | 0.12 | 0.22 | 0.3 | 0.3 | 0.37 |
| 6 | 30011934 | rs259940 | 2.496 | -3.054 | -1.147 | -0.849 | -0.014 | -0.016 | -0.016 | 0 | 0.22 | 0.11 | 0.2 | 0.29 | 0.26 | 0.33 |
| 6 | 30013887 | rs9261224 | 0.519 | -3.154 | -1.126 | -0.866 | -0.06 | 0 | 0 | 0 | 0.16 | 0.29 | 0.11 | 0.15 | 0.08 | 0.14 |
| 6 | 30015167 | rs259942 | -0.085 | -3.156 | -0.823 | -0.905 | -0.014 | -0.007 | -0.007 | 0 | 0.2 | 0.26 | 0.18 | 0.15 | 0.15 | 0.22 |
| 6 | 30015905 | rs3134879 | 1.145 | -3.134 | -0.996 | -0.89 | -0.01 | -0.006 | -0.006 | 0 | 0.36 | 0.2 | 0.37 | 0.34 | 0.39 | 0.48 |
| 6 | 30016190 | rs3132128 | -1.892 | -3.191 | -0.972 | -0.935 | 0.002 | -0.01 | -0.01 | 0.337 | 0.91 | 0.92 | 0.89 | 0.98 | 0.89 | 0.95 |
| 6 | 30016297 | rs6457138 | 0.331 | -3.228 | -0.947 | -0.94 | -0.055 | 0.011 | 0.011 | 0 | 0.15 | 0.25 | 0.14 | 0.15 | 0.04 | 0.13 |
| 6 | 30021740 | rs28570682 | 0.751 | -3.233 | -0.947 | -0.942 | -0.065 | 0.003 | 0.003 | 0 | 0.13 | 0.24 | 0.07 | 0.12 | 0.03 | 0.11 |
| 6 | 30022153 | rs7761314 | 0.446 | -3.233 | -0.947 | -0.942 | -0.06 | 0 | 0 | 0 | 0.16 | 0.29 | 0.11 | 0.15 | 0.08 | 0.14 |
| 6 | 30022808 | rs9261258 | 0.446 | -3.222 | -0.947 | -0.942 | -0.06 | 0 | 0 | 0 | 0.16 | 0.29 | 0.11 | 0.15 | 0.08 | 0.14 |
| 6 | 30023868 | rs3869070 | -0.206 | -1.92 | -0.977 | -0.919 | 0 | 0 | 0 | 0 | 0.53 | 0.5 | 0.49 | 0.5 | 0.5 | 0.62 |
| 6 | 30024365 | rs9261261 | 0.461 | -1.92 | -0.977 | -0.919 | -0.06 | 0 | 0 | 0 | 0.16 | 0.29 | 0.11 | 0.15 | 0.08 | 0.14 |
| 6 | 30025179 | rs6911724 | 0.461 | -1.92 | -0.977 | -0.919 | -0.06 | 0 | 0 | 0.053 | 0.16 | 0.29 | 0.11 | 0.15 | 0.08 | 0.14 |
| 6 | 30025503 | rs259919 | 2.509 | -1.931 | -0.99 | -0.926 | -0.029 | -0.018 | -0.018 | 0 | 0.23 | 0.12 | 0.21 | 0.3 | 0.29 | 0.33 |
| 6 | 30025759 | rs259920 | 2.136 | -2.223 | -0.992 | -0.933 | 0.007 | 0.016 | 0.016 | 6.774 | 0.07 | 0.04 | 0.08 | 0.01 | 0.07 | 0.09 |
| 6 | 30026146 | rs16896944 | 2.031 | -2.26 | -0.981 | -0.954 | -0.005 | -0.029 | -0.029 | 0 | 0.06 | 0.19 | 0.06 | 0 | 0.04 | 0.01 |
| 6 | 30026290 | rs10484549 | 2.707 | -2.258 | -1.03 | -0.944 | -0.002 | -0.02 | -0.02 | 1.407 | 0.07 | 0.02 | 0.03 | 0.19 | 0.06 | 0.15 |
| 6 | 30027413 | rs3757327 | 0.46 | -2.187 | -0.995 | -0.916 | -0.057 | 0 | 0 | 0 | 0.16 | 0.29 | 0.11 | 0.15 | 0.08 | 0.14 |
| 6 | 30028352 | rs3757328 | 0.774 | -2.194 | -0.995 | -0.916 | -0.065 | 0.003 | 0.003 | 0 | 0.13 | 0.24 | 0.07 | 0.12 | 0.03 | 0.11 |
| 6 | 30028424 | rs3757329 | 0.46 | -2.194 | -0.995 | -0.916 | -0.06 | 0 | 0 | 0 | 0.16 | 0.29 | 0.11 | 0.15 | 0.08 | 0.14 |
| 6 | 30028606 | rs3757333 | 0.774 | -2.194 | -0.995 | -0.916 | -0.065 | 0.003 | 0.003 | 0 | 0.13 | 0.24 | 0.07 | 0.12 | 0.03 | 0.11 |
| 6 | 30028800 | rs9295829 | 2.283 | -2.205 | -0.997 | -0.923 | -0.047 | -0.029 | -0.029 | 0 | 0.2 | 0.12 | 0.31 | 0.3 | 0.31 | 0.16 |
| 6 | 30028831 | rs1150742 | 2.515 | -2.169 | -0.947 | -0.894 | -0.011 | -0.014 | -0.014 | 0 | 0.23 | 0.12 | 0.2 | 0.29 | 0.26 | 0.33 |
| 6 | 30028871 | rs7770092 | 0.457 | -2.155 | -0.935 | -0.885 | -0.06 | 0 | 0 | 1.328 | 0.16 | 0.29 | 0.11 | 0.15 | 0.08 | 0.14 |
| 6 | 30030114 | rs9261269 | -1.962 | -2.16 | -0.918 | -0.887 | -0.001 | -0.002 | -0.002 | 2.607 | 0.91 | 0.93 | 0.9 | 0.98 | 0.89 | 0.95 |
| 6 | 30031091 | rs9261277 | 0.426 | -2.081 | -0.903 | -0.833 | -0.06 | 0 | 0 | 0 | 0.16 | 0.29 | 0.11 | 0.15 | 0.08 | 0.14 |
| 6 | 30031345 | rs1150739 | 0.446 | -2.075 | -0.899 | -0.831 | -0.011 | 0 | 0 | 2.128 | 0.44 | 0.45 | 0.33 | 0.45 | 0.38 | 0.51 |
| 6 | 30032673 | rs3188482 | 0.461 | -1.659 | -0.824 | -0.769 | -0.053 | 0.016 | 0.016 | 0 | 0.14 | 0.25 | 0.13 | 0.14 | 0.04 | 0.13 |
| 6 | 30033884 | rs3807031 | 2.436 | -1.125 | -1.202 | -0.714 | 0.026 | 0.009 | 0.009 | 0 | 0.09 | 0.12 | 0.25 | 0.24 | 0.23 | 0.13 |
| 6 | 30036471 | rs2074482 | 0.064 | -1.557 | -1.091 | -0.821 | -0.052 | 0.008 | 0.008 | 0 | 0.18 | 0.3 | 0.17 | 0.19 | 0.1 | 0.16 |
| 6 | 30038352 | rs1245371 | 3.754 | -1.849 | -1.335 | -0.889 | -0.014 | -0.016 | -0.016 | 7.396 | 0.22 | 0.11 | 0.2 | 0.29 | 0.26 | 0.33 |
| 6 | 30039240 | rs2301753 | 0.229 | -1.919 | -1.142 | -0.752 | -0.052 | 0.008 | 0.008 | 0 | 0.18 | 0.3 | 0.17 | 0.19 | 0.1 | 0.16 |
| 6 | 30040084 | rs9393989 | 1.546 | -1.924 | -1.157 | -0.773 | -0.017 | -0.01 | -0.01 | 0 | 0.2 | 0.18 | 0.31 | 0.29 | 0.31 | 0.16 |
| 6 | 30040810 | rs2074480 | 0.197 | -1.862 | -1.074 | -0.682 | -0.052 | 0.008 | 0.008 | 0 | 0.18 | 0.3 | 0.17 | 0.19 | 0.1 | 0.16 |
| 6 | 30041009 | rs2074479 | 0.197 | -1.862 | -1.074 | -0.682 | -0.052 | 0.008 | 0.008 | 0 | 0.18 | 0.3 | 0.17 | 0.19 | 0.1 | 0.16 |
| 6 | 30041559 | rs9261301 | -0.106 | -1.81 | -1.068 | -0.682 | 0.005 | 0.01 | 0.01 | 0 | 0.59 | 0.48 | 0.5 | 0.5 | 0.52 | 0.62 |

| | | | | | | | | | | | | | | | | |
|---|---|---|---|---|---|---|---|---|---|---|---|---|---|---|---|---|
| 6 | 30042349 | rs9261302 | 0.038 | -2.053 | -0.742 | -0.489 | -0.052 | 0.008 | 0.008 | 0 | 0.18 | 0.3 | 0.17 | 0.19 | 0.1 | 0.16 |
| 6 | 30043955 | rs3807033 | -0.144 | -2.101 | -0.727 | -0.543 | -0.086 | -0.018 | -0.018 | 4.413 | 0.21 | 0.44 | 0.19 | 0.15 | 0.08 | 0.14 |
| 6 | 30044827 | rs3807035 | -0.632 | -2.112 | -0.729 | -0.549 | -0.087 | -0.022 | -0.022 | 0 | 0.24 | 0.49 | 0.23 | 0.19 | 0.13 | 0.16 |
| 6 | 30045199 | rs1150735 | 2.705 | -2.139 | -0.944 | -0.574 | -0.04 | -0.016 | -0.016 | 0 | 0.3 | 0.12 | 0.25 | 0.31 | 0.35 | 0.38 |
| 6 | 30046246 | rs2394734 | -0.976 | -2.25 | -0.961 | -0.618 | -0.082 | -0.034 | -0.034 | 0 | 0.31 | 0.52 | 0.26 | 0.2 | 0.17 | 0.26 |
| 6 | 30047403 | rs4711209 | 2.352 | -2.24 | -1.105 | -0.521 | -0.046 | -0.031 | -0.031 | 0 | 0.19 | 0.12 | 0.31 | 0.29 | 0.31 | 0.16 |
| 6 | 30047965 | rs7382061 | -0.49 | -2.267 | -0.981 | -0.456 | -0.019 | -0.024 | -0.024 | 0 | 0.61 | 0.7 | 0.5 | 0.49 | 0.57 | 0.64 |
| 6 | 30054757 | rs2057728 | -1.382 | -2.094 | -0.909 | -0.47 | -0.028 | -0.013 | -0.013 | 2.049 | 0.71 | 0.78 | 0.52 | 0.66 | 0.58 | 0.78 |
| 6 | 30055377 | rs6904455 | -1.491 | -2.084 | -0.897 | -0.472 | -0.058 | -0.027 | -0.027 | 0.677 | 0.72 | 0.84 | 0.53 | 0.67 | 0.58 | 0.78 |
| 6 | 30057763 | rs11965797 | 2.029 | -2.06 | -0.282 | -0.692 | -0.041 | -0.009 | -0.009 | 0 | 0.08 | 0.18 | 0.04 | 0.02 | 0.02 | 0.02 |
| 6 | 30058988 | rs9261371 | -0.62 | -1.968 | -0.686 | -0.665 | -0.088 | -0.022 | -0.022 | 0 | 0.24 | 0.5 | 0.22 | 0.19 | 0.13 | 0.18 |
| 6 | 30061202 | rs17187805 | 1.157 | -1.851 | -0.696 | -0.643 | -0.064 | -0.036 | -0.036 | 0 | 0.32 | 0.45 | 0.22 | 0.17 | 0.14 | 0.32 |
| 6 | 30063496 | rs7765810 | 0.497 | -1.796 | -0.737 | -0.575 | -0.065 | -0.04 | -0.04 | 0 | 0.39 | 0.51 | 0.26 | 0.18 | 0.18 | 0.4 |
| 6 | 30064562 | rs9261394 | -0.577 | -1.339 | -1.055 | -0.168 | -0.03 | -0.015 | -0.015 | 0.219 | 0.61 | 0.76 | 0.5 | 0.6 | 0.57 | 0.63 |
| 6 | 30065575 | rs1264702 | 1.597 | -1.333 | -0.585 | -0.263 | -0.029 | -0.031 | -0.031 | 0 | 0.28 | 0.38 | 0.19 | 0.17 | 0.17 | 0.31 |
| 6 | 30066031 | rs916570 | 0.872 | -1.385 | -0.533 | -0.305 | -0.017 | -0.021 | -0.021 | 0.005 | 0.36 | 0.57 | 0.26 | 0.17 | 0.22 | 0.32 |
| 6 | 30066358 | rs1264701 | 2.356 | -1.121 | -0.87 | -0.111 | -0.058 | -0.078 | -0.078 | 0 | 0.12 | 0.02 | 0.15 | 0.38 | 0.22 | 0.17 |
| 6 | 30067177 | rs1264698 | 1.156 | -0.962 | -0.439 | -0.247 | -0.034 | -0.032 | -0.032 | 0 | 0.3 | 0.39 | 0.2 | 0.17 | 0.19 | 0.31 |
| 6 | 30067681 | rs7763159 | -2.471 | -0.878 | -0.516 | -0.339 | -0.008 | 0.011 | 0.011 | 0 | 0.92 | 0.97 | 0.92 | 0.99 | 0.89 | 0.95 |
| 6 | 30067892 | rs1264695 | 1.11 | -1.115 | -0.462 | -0.262 | -0.034 | -0.034 | -0.034 | 0.005 | 0.31 | 0.39 | 0.21 | 0.17 | 0.19 | 0.31 |
| 6 | 30069795 | rs1108052 | 1.11 | -1.043 | -0.412 | -0.249 | -0.034 | -0.034 | -0.034 | 1.608 | 0.31 | 0.39 | 0.21 | 0.17 | 0.19 | 0.31 |
| 6 | 30070937 | rs1110446 | 1.435 | -1.13 | -0.182 | -0.255 | -0.006 | -0.01 | -0.01 | 0.338 | 0.28 | 0.29 | 0.19 | 0.16 | 0.19 | 0.31 |
| 6 | 30071110 | rs2240070 | -2.029 | -0.515 | -0.477 | -0.206 | -0.008 | -0.005 | -0.005 | 0 | 0.83 | 0.85 | 0.6 | 0.83 | 0.67 | 0.81 |
| 6 | 30071279 | rs1116222 | 1.045 | -0.965 | -0.293 | -0.307 | -0.031 | -0.055 | -0.055 | 0 | 0.32 | 0.39 | 0.23 | 0.1 | 0.22 | 0.3 |
| 6 | 30071330 | rs1116221 | 1.036 | -0.985 | -0.215 | -0.303 | -0.037 | -0.063 | -0.063 | 0 | 0.32 | 0.41 | 0.22 | 0.1 | 0.22 | 0.3 |
| 6 | 30071760 | rs11964542 | 1.946 | -0.684 | -0.344 | -0.156 | -0.039 | -0.006 | -0.006 | 0 | 0.08 | 0.19 | 0.05 | 0.02 | 0.02 | 0.02 |
| 6 | 30072511 | rs2844796 | 1.766 | -0.42 | -0.643 | -0.074 | -0.025 | -0.05 | -0.05 | 0 | 0.27 | 0.34 | 0.2 | 0.09 | 0.19 | 0.22 |
| 6 | 30072721 | rs2248289 | 1.934 | -0.398 | -0.647 | -0.062 | -0.023 | -0.048 | -0.048 | 0 | 0.26 | 0.34 | 0.2 | 0.09 | 0.19 | 0.22 |
| 6 | 30073195 | rs3132680 | -1.479 | 0.696 | -1.068 | 0.496 | -0.005 | -0.004 | -0.004 | 0 | 0.83 | 0.85 | 0.6 | 0.83 | 0.68 | 0.82 |
| 6 | 30073847 | rs2844795 | -0.528 | 0.534 | -1.157 | 0.487 | 0.019 | 0.007 | 0.007 | 0 | 0.73 | 0.69 | 0.54 | 0.64 | 0.58 | 0.73 |
| 6 | 30074017 | rs3778620 | 0.969 | 0.635 | -1.465 | 0.483 | -0.014 | -0.026 | -0.026 | 0 | 0.05 | 0.01 | 0.05 | 0.22 | 0.08 | 0.08 |
| 6 | 30074028 | rs9261407 | -1.605 | 0.568 | -1.424 | -0.288 | -0.022 | 0.043 | 0.043 | 1.72 | 0.87 | 0.94 | 0.9 | 1 | 0.82 | 0.98 |
| 6 | 30075843 | rs3132679 | -0.739 | 0.319 | -0.84 | -0.258 | -0.013 | -0.034 | -0.034 | 0 | 0.85 | 0.69 | 0.85 | 0.92 | 0.86 | 0.92 |
| 6 | 30075864 | rs2023472 | 0.196 | 0.087 | -1.212 | -0.688 | 0.005 | -0.047 | -0.047 | 0 | 0.67 | 0.63 | 0.71 | 0.87 | 0.64 | 0.87 |
| 6 | 30076426 | rs3132123 | -0.947 | 0.028 | -1.129 | -0.587 | -0.013 | -0.034 | -0.034 | 0 | 0.85 | 0.69 | 0.85 | 0.92 | 0.86 | 0.92 |
| 6 | 30076630 | rs2240068 | -0.247 | -1.047 | -1.974 | -0.905 | 0 | -0.009 | -0.009 | 0.937 | 0.43 | 0.56 | 0.58 | 0.69 | 0.49 | 0.51 |
| 6 | 30076755 | rs2074483 | 1.254 | -1.01 | -1.858 | -0.885 | 0.041 | 0.098 | 0.098 | 0 | 0.42 | 0.16 | 0.26 | 0.22 | 0.37 | 0.41 |
| 6 | 30077229 | rs2523990 | 0.095 | -0.814 | -2.307 | -0.445 | -0.045 | -0.08 | -0.08 | 0.705 | 0.41 | 0.15 | 0.56 | 0.42 | 0.52 | 0.35 |
| 6 | 30077967 | rs4959041 | 1.384 | -0.924 | -1.965 | -0.637 | 0.01 | 0.001 | 0.001 | 1.953 | 0.26 | 0.27 | 0.4 | 0.23 | 0.33 | 0.15 |
| 6 | 30078275 | rs2523989 | 1.755 | -0.764 | -2.055 | -0.568 | -0.012 | 0.004 | 0.004 | 0 | 0.07 | 0.05 | 0.12 | 0.05 | 0.12 | 0.05 |
| 6 | 30078330 | rs2239529 | 1.971 | -0.762 | -2.059 | -0.567 | -0.035 | -0.001 | -0.001 | 0.799 | 0.06 | 0.01 | 0.11 | 0.05 | 0.12 | 0.05 |
| 6 | 30079307 | rs2249099 | 1.227 | -0.96 | -2.136 | -0.649 | -0.01 | 0.017 | 0.017 | 0 | 0.12 | 0.09 | 0.13 | 0.09 | 0.13 | 0.18 |
| 6 | 30080036 | rs3734837 | 2.433 | -1.141 | -2.429 | -0.471 | -0.058 | -0.075 | -0.075 | 5.849 | 0.12 | 0.01 | 0.1 | 0.35 | 0.18 | 0.24 |
| 6 | 30080231 | rs3734838 | 1.843 | -1.3 | -2.19 | -0.589 | 0.016 | -0.027 | -0.027 | 0 | 0.09 | 0.03 | 0.06 | 0.11 | 0.08 | 0.07 |
| 6 | 30080274 | rs2517598 | 1.064 | -0.917 | -1.87 | -0.761 | -0.007 | 0.012 | 0.012 | 0 | 0.12 | 0.09 | 0.13 | 0.09 | 0.13 | 0.18 |
| 6 | 30080496 | rs2844793 | 1.661 | -0.912 | -1.869 | -0.759 | -0.007 | 0.003 | 0.003 | 0 | 0.07 | 0.06 | 0.12 | 0.05 | 0.12 | 0.05 |
| 6 | 30081108 | rs9261425 | -0.768 | -1.115 | -1.823 | -0.849 | -0.002 | -0.007 | -0.007 | 0 | 0.56 | 0.56 | 0.61 | 0.75 | 0.64 | 0.52 |
| 6 | 30081334 | rs2523985 | 1.032 | -0.61 | -1.636 | -0.686 | -0.007 | 0.012 | 0.012 | 0 | 0.12 | 0.09 | 0.13 | 0.09 | 0.13 | 0.18 |
| 6 | 30082003 | rs2523984 | 1.032 | -0.61 | -1.636 | -0.686 | -0.007 | 0.012 | 0.012 | 0 | 0.12 | 0.09 | 0.13 | 0.09 | 0.13 | 0.18 |
| 6 | 30082688 | rs2245420 | 1.032 | -0.61 | -1.636 | -0.686 | -0.007 | 0.012 | 0.012 | 0 | 0.12 | 0.09 | 0.13 | 0.09 | 0.13 | 0.18 |
| 6 | 30086340 | rs1015465 | 2.532 | -1.139 | -1.396 | -0.73 | -0.046 | -0.044 | -0.044 | 0 | 0.22 | 0.3 | 0.15 | 0.11 | 0.13 | 0.2 |
| 6 | 30087776 | rs9261434 | -1.721 | -0.788 | -1.485 | -0.674 | -0.013 | 0.014 | 0.014 | 0 | 0.93 | 0.98 | 0.92 | 0.99 | 0.89 | 0.94 |
| 6 | 30089280 | rs9261438 | 1.332 | -0.867 | -1.544 | -0.73 | 0.012 | 0.001 | 0.001 | 1.424 | 0.25 | 0.26 | 0.4 | 0.23 | 0.33 | 0.15 |
| 6 | 30089428 | rs2523974 | 2.629 | -0.768 | -1.479 | -0.572 | -0.046 | -0.044 | -0.044 | 0 | 0.22 | 0.3 | 0.15 | 0.11 | 0.13 | 0.2 |
| 6 | 30090820 | rs9261440 | 1.636 | -0.563 | -1.579 | -0.551 | -0.037 | -0.035 | -0.035 | 0.232 | 0.13 | 0.07 | 0.37 | 0.22 | 0.25 | 0.11 |
| 6 | 30092707 | rs1419675 | -1.915 | -0.807 | -1.575 | -0.713 | -0.018 | -0.038 | -0.038 | 0.283 | 0.71 | 0.68 | 0.78 | 0.88 | 0.76 | 0.74 |
| 6 | 30093744 | rs9261442 | 2.057 | -0.219 | -1.306 | -0.374 | 0.005 | -0.003 | -0.003 | 0 | 0.1 | 0.09 | 0.34 | 0.22 | 0.23 | 0.09 |
| 6 | 30094788 | rs9261445 | 2.057 | -0.219 | -1.306 | -0.374 | 0.005 | -0.003 | -0.003 | 0 | 0.1 | 0.09 | 0.34 | 0.22 | 0.23 | 0.09 |
| 6 | 30095202 | rs9261447 | 0.932 | -0.568 | -1.46 | -0.544 | -0.025 | -0.02 | -0.02 | 0 | 0.19 | 0.13 | 0.38 | 0.26 | 0.25 | 0.24 |
| 6 | 30095329 | rs9261448 | 2.122 | -0.527 | -1.401 | -0.491 | -0.016 | -0.021 | -0.021 | 0.715 | 0.1 | 0.05 | 0.34 | 0.22 | 0.23 | 0.09 |
| 6 | 30096717 | rs1419673 | 2.115 | -0.524 | -1.406 | -0.489 | 0.004 | -0.003 | -0.003 | 0 | 0.1 | 0.09 | 0.34 | 0.22 | 0.23 | 0.09 |
| 6 | 30097818 | rs9261450 | 1.275 | -0.529 | -1.408 | -0.488 | 0.003 | -0.001 | -0.001 | 2.125 | 0.14 | 0.13 | 0.38 | 0.22 | 0.25 | 0.12 |
| 6 | 30098323 | rs757263 | 1.275 | -0.529 | -1.408 | -0.488 | 0.003 | -0.001 | -0.001 | 1.12 | 0.14 | 0.13 | 0.38 | 0.22 | 0.25 | 0.12 |
| 6 | 30099948 | rs9261453 | 1.426 | -0.559 | -1.022 | -0.499 | 0.001 | -0.003 | -0.003 | 0 | 0.13 | 0.12 | 0.38 | 0.22 | 0.25 | 0.11 |
| 6 | 30099996 | rs9261455 | 1.426 | -0.559 | -1.022 | -0.499 | 0.001 | -0.003 | -0.003 | 0.172 | 0.13 | 0.12 | 0.38 | 0.22 | 0.25 | 0.11 |
| 6 | 30100221 | rs9261457 | 2.081 | -0.549 | -1.02 | -0.491 | 0.002 | -0.006 | -0.006 | 0 | 0.1 | 0.08 | 0.34 | 0.22 | 0.23 | 0.09 |
| 6 | 30101656 | rs1362104 | 0.617 | -1.314 | -1.346 | -1.065 | -0.026 | -0.038 | -0.038 | 1.002 | 0.3 | 0.21 | 0.29 | 0.61 | 0.36 | 0.42 |
| 6 | 30102184 | rs2523995 | 3.777 | -1.735 | -1.214 | -1.281 | 0.009 | 0.013 | 0.013 | 0 | 0.14 | 0.12 | 0.1 | 0.04 | 0.1 | 0.17 |
| 6 | 30103394 | rs1541268 | 1.655 | -1.619 | -1.422 | -1.226 | -0.04 | -0.035 | -0.035 | 0 | 0.14 | 0.08 | 0.28 | 0.26 | 0.24 | 0.21 |

| | | | | | | | | | | | | | | | | |
|---|---|---|---|---|---|---|---|---|---|---|---|---|---|---|---|---|
| 6 | 30103840 | rs9261466 | 1.655 | -1.619 | -1.422 | -1.226 | -0.04 | -0.035 | -0.035 | 0 | 0.14 | 0.08 | 0.28 | 0.26 | 0.24 | 0.21 |
| 6 | 30103898 | rs9261467 | 2.466 | -1.617 | -1.421 | -1.225 | 0.004 | -0.005 | -0.005 | 0.199 | 0.09 | 0.08 | 0.28 | 0.22 | 0.23 | 0.08 |
| 6 | 30103923 | rs2021723 | 3.06 | -1.835 | -1.349 | -1.333 | -0.044 | -0.044 | -0.044 | 0 | 0.22 | 0.29 | 0.15 | 0.11 | 0.13 | 0.2 |
| 6 | 30104480 | rs9261468 | 2.491 | -1.774 | -1.332 | -1.287 | 0.004 | -0.005 | -0.005 | 0.59 | 0.09 | 0.08 | 0.28 | 0.22 | 0.23 | 0.08 |
| 6 | 30104783 | rs2857458 | 0.397 | -1.959 | -1.377 | -1.402 | -0.01 | -0.004 | -0.004 | 0 | 0.26 | 0.27 | 0.36 | 0.26 | 0.29 | 0.27 |
| 6 | 30104808 | rs9261470 | 1.702 | -1.901 | -1.429 | -1.349 | -0.04 | -0.035 | -0.035 | 0 | 0.14 | 0.08 | 0.28 | 0.26 | 0.24 | 0.21 |
| 6 | 30105743 | rs2107202 | -2.312 | -1.959 | -1.459 | -1.397 | -0.017 | -0.039 | -0.039 | 0 | 0.71 | 0.69 | 0.78 | 0.88 | 0.76 | 0.74 |
| 6 | 30106202 | rs2857438 | 1.49 | -1.476 | -1.559 | -1.073 | -0.02 | -0.018 | -0.018 | 0 | 0.15 | 0.12 | 0.29 | 0.26 | 0.24 | 0.21 |
| 6 | 30106493 | rs2857440 | 1.49 | -1.476 | -1.559 | -1.073 | -0.02 | -0.018 | -0.018 | 0 | 0.15 | 0.12 | 0.29 | 0.26 | 0.24 | 0.21 |
| 6 | 30108683 | rs9261484 | 0.843 | -1.497 | -1.577 | -1.094 | -0.037 | -0.027 | -0.027 | 0.213 | 0.22 | 0.14 | 0.34 | 0.26 | 0.26 | 0.25 |
| 6 | 30109412 | rs9261488 | 2.334 | -1.306 | -1.523 | -1.03 | 0.004 | -0.005 | -0.005 | 0 | 0.09 | 0.08 | 0.28 | 0.22 | 0.23 | 0.09 |
| 6 | 30109482 | rs9261489 | 2.334 | -1.306 | -1.523 | -1.03 | 0.004 | -0.005 | -0.005 | 1.355 | 0.09 | 0.08 | 0.28 | 0.22 | 0.23 | 0.09 |
| 6 | 30109831 | rs9261490 | 1.289 | -1.375 | -1.543 | -1.064 | -0.023 | -0.019 | -0.019 | 0 | 0.17 | 0.14 | 0.33 | 0.22 | 0.26 | 0.13 |
| 6 | 30110058 | rs9261492 | 1.289 | -1.375 | -1.543 | -1.064 | -0.023 | -0.019 | -0.019 | 0 | 0.17 | 0.14 | 0.33 | 0.22 | 0.26 | 0.12 |
| 6 | 30111014 | rs9261496 | 1.131 | -1.457 | -1.574 | -1.106 | -0.026 | -0.021 | -0.021 | 2.255 | 0.18 | 0.14 | 0.33 | 0.22 | 0.26 | 0.13 |
| 6 | 30111351 | rs9261498 | 0.723 | -1.524 | -1.607 | -1.137 | -0.036 | -0.025 | -0.025 | 0 | 0.23 | 0.14 | 0.34 | 0.26 | 0.26 | 0.25 |
| 6 | 30112408 | rs9261505 | 2.23 | -1.372 | -1.535 | -1.081 | 0.027 | 0.014 | 0.014 | 0 | 0.1 | 0.12 | 0.29 | 0.22 | 0.23 | 0.09 |
| 6 | 30112623 | rs9261506 | 2.005 | -1.766 | -1.778 | -0.871 | 0.016 | -0.028 | -0.028 | 0 | 0.09 | 0.03 | 0.06 | 0.13 | 0.08 | 0.07 |
| 6 | 30112704 | rs9261508 | 1.369 | -1.586 | -1.703 | -0.834 | -0.023 | -0.019 | -0.019 | 0 | 0.17 | 0.14 | 0.33 | 0.22 | 0.25 | 0.13 |
| 6 | 30113344 | rs3815086 | 1.369 | -1.586 | -1.703 | -0.834 | -0.023 | -0.019 | -0.019 | 0.143 | 0.17 | 0.14 | 0.34 | 0.22 | 0.25 | 0.13 |
| 6 | 30113955 | rs3815082 | 1.369 | -1.586 | -1.703 | -0.834 | -0.023 | -0.019 | -0.019 | 0.432 | 0.17 | 0.14 | 0.33 | 0.22 | 0.25 | 0.13 |
| 6 | 30114074 | rs3815081 | 2.42 | -1.441 | -1.928 | -0.685 | -0.058 | -0.077 | -0.077 | 0.773 | 0.12 | 0.02 | 0.1 | 0.36 | 0.18 | 0.24 |
| 6 | 30114666 | rs4959044 | 2.343 | -1.314 | -1.718 | -0.717 | 0.028 | 0.015 | 0.015 | 0 | 0.1 | 0.12 | 0.29 | 0.22 | 0.23 | 0.08 |
| 6 | 30114955 | rs757262 | 2.343 | -1.314 | -1.718 | -0.717 | 0.028 | 0.015 | 0.015 | 0 | 0.1 | 0.12 | 0.29 | 0.22 | 0.23 | 0.08 |
| 6 | 30115063 | rs757261 | 2.436 | -1.313 | -1.724 | -0.717 | 0.004 | -0.005 | -0.005 | 0 | 0.09 | 0.08 | 0.28 | 0.22 | 0.23 | 0.08 |
| 6 | 30115484 | rs9261513 | 0.994 | -0.68 | -1.858 | -0.827 | -0.035 | -0.026 | -0.026 | 0 | 0.22 | 0.14 | 0.34 | 0.26 | 0.26 | 0.25 |
| 6 | 30115542 | rs757259 | 2.376 | -0.617 | -1.828 | -0.799 | 0.028 | 0.015 | 0.015 | 0 | 0.1 | 0.12 | 0.29 | 0.22 | 0.23 | 0.08 |
| 6 | 30115965 | rs1573299 | 2.376 | -0.617 | -1.828 | -0.799 | 0.028 | 0.015 | 0.015 | 0 | 0.1 | 0.12 | 0.29 | 0.22 | 0.23 | 0.08 |
| 6 | 30116078 | rs3132676 | -1.426 | -0.732 | -1.796 | -0.966 | 0.006 | -0.019 | -0.019 | 0 | 0.89 | 0.85 | 0.9 | 0.99 | 0.86 | 0.93 |
| 6 | 30117101 | rs9261519 | 1.431 | -0.741 | -1.797 | -0.907 | -0.023 | -0.019 | -0.019 | 0.105 | 0.17 | 0.14 | 0.34 | 0.22 | 0.25 | 0.12 |
| 6 | 30118360 | rs1557611 | 2.402 | -0.714 | -1.785 | -0.894 | 0.028 | 0.015 | 0.015 | 0 | 0.1 | 0.12 | 0.29 | 0.22 | 0.23 | 0.08 |
| 6 | 30118582 | rs1557608 | 0.506 | -0.808 | -1.793 | -0.957 | -0.013 | -0.006 | -0.006 | 0 | 0.25 | 0.25 | 0.36 | 0.26 | 0.29 | 0.24 |
| 6 | 30118637 | rs1557607 | 2.394 | -0.737 | -1.754 | -0.875 | 0.028 | 0.015 | 0.015 | 0 | 0.1 | 0.12 | 0.29 | 0.22 | 0.23 | 0.08 |
| 6 | 30118707 | rs4392759 | 2.479 | -0.737 | -1.753 | -0.875 | 0.004 | -0.005 | -0.005 | 0.058 | 0.09 | 0.08 | 0.28 | 0.22 | 0.23 | 0.08 |
| 6 | 30119275 | rs9261526 | 1.665 | -0.823 | -1.757 | -0.832 | 0.004 | -0.002 | -0.002 | 0 | 0.13 | 0.12 | 0.32 | 0.22 | 0.25 | 0.11 |
| 6 | 30120268 | rs9261531 | 1.665 | -0.823 | -1.757 | -0.832 | 0.004 | -0.002 | -0.002 | 0 | 0.13 | 0.12 | 0.32 | 0.22 | 0.25 | 0.11 |
| 6 | 30120563 | rs2257914 | 3.026 | -1.37 | -1.715 | -1.142 | -0.042 | -0.044 | -0.044 | 0 | 0.22 | 0.29 | 0.15 | 0.11 | 0.13 | 0.2 |
| 6 | 30120637 | rs9468693 | 2.138 | -1.449 | -1.721 | -1.211 | -0.035 | -0.056 | -0.056 | 0 | 0.2 | 0.24 | 0.09 | 0.01 | 0.1 | 0.07 |
| 6 | 30120912 | rs2285795 | 1.238 | -1.233 | -1.636 | -1.086 | -0.002 | -0.004 | -0.004 | 0 | 0.15 | 0.13 | 0.33 | 0.27 | 0.26 | 0.14 |
| 6 | 30121048 | rs2517655 | 2.467 | -1.11 | -1.638 | -1.037 | 0.015 | 0.002 | 0.002 | 0 | 0.09 | 0.08 | 0.28 | 0.22 | 0.23 | 0.08 |
| 6 | 30121066 | rs2508031 | 1.53 | -1.112 | -1.64 | -1.037 | -0.019 | -0.016 | -0.016 | 0 | 0.16 | 0.14 | 0.33 | 0.22 | 0.25 | 0.12 |
| 6 | 30121185 | rs2517653 | 1.523 | -1.07 | -1.633 | -1.029 | -0.019 | -0.016 | -0.016 | 0 | 0.16 | 0.12 | 0.33 | 0.23 | 0.26 | 0.14 |
| 6 | 30121460 | rs2022065 | 0.891 | -1.053 | -1.643 | -0.979 | -0.027 | -0.034 | -0.034 | 0 | 0.22 | 0.16 | 0.38 | 0.34 | 0.31 | 0.15 |
| 6 | 30122623 | rs2517645 | 0.969 | -0.811 | -1.582 | -0.992 | -0.001 | 0.001 | 0.001 | 0 | 0.13 | 0.17 | 0.16 | 0.08 | 0.13 | 0.2 |
| 6 | 30125537 | rs1967 | 2.826 | -1.144 | -1.53 | -1.164 | -0.04 | -0.046 | -0.046 | 4.137 | 0.22 | 0.29 | 0.14 | 0.1 | 0.13 | 0.25 |
| 6 | 30127323 | rs9261535 | 2.712 | -0.904 | -1.593 | -0.985 | -0.041 | -0.048 | -0.048 | 0 | 0.08 | 0.02 | 0.28 | 0.21 | 0.23 | 0.08 |
| 6 | 30128442 | rs12212092 | 2.606 | -1.519 | -1.51 | -1.233 | -0.038 | -0.066 | -0.066 | 0.061 | 0.16 | 0.23 | 0.07 | 0 | 0.1 | 0.01 |
| 6 | 30129676 | rs2523734 | 2.465 | -1.078 | -1.54 | -0.945 | -0.035 | -0.039 | -0.039 | 0 | 0.25 | 0.28 | 0.15 | 0.1 | 0.13 | 0.26 |
| 6 | 30131515 | rs2523733 | 2.791 | -0.898 | -1.428 | -0.863 | -0.037 | -0.044 | -0.044 | 0 | 0.22 | 0.27 | 0.14 | 0.1 | 0.13 | 0.25 |
| 6 | 30132035 | rs2074477 | 1.885 | -0.124 | -1.603 | -0.333 | -0.058 | -0.078 | -0.078 | 0 | 0.12 | 0.01 | 0.1 | 0.36 | 0.18 | 0.25 |
| 6 | 30132827 | rs1008403 | 1.09 | -0.274 | -1.657 | -0.406 | -0.009 | -0.061 | -0.061 | 0 | 0.21 | 0.08 | 0.14 | 0.39 | 0.2 | 0.4 |
| 6 | 30135977 | rs9368624 | 1.09 | -0.348 | -1.664 | -0.425 | -0.009 | -0.061 | -0.061 | 0 | 0.21 | 0.08 | 0.14 | 0.39 | 0.2 | 0.4 |
| 6 | 30138662 | rs1029237 | 2.395 | 0.212 | -1.486 | -0.259 | -0.041 | -0.048 | -0.048 | 0 | 0.08 | 0.02 | 0.28 | 0.21 | 0.23 | 0.09 |
| 6 | 30139699 | rs929156 | 1.391 | 0.108 | -1.41 | -0.287 | -0.014 | -0.015 | -0.015 | 0.021 | 0.13 | 0.1 | 0.31 | 0.21 | 0.25 | 0.12 |
| 6 | 30140342 | rs1063280 | -1.47 | -0.14 | -1.564 | -0.494 | -0.01 | 0.012 | 0.012 | 0 | 0.9 | 0.97 | 0.86 | 0.93 | 0.87 | 0.92 |
| 6 | 30140913 | rs2844787 | 0.17 | -0.737 | -1.194 | -0.792 | -0.041 | -0.058 | -0.058 | 0 | 0.75 | 0.61 | 0.72 | 0.85 | 0.78 | 0.81 |
| 6 | 30141042 | rs9380156 | 2.143 | -0.768 | -1.544 | -0.715 | -0.058 | -0.08 | -0.08 | 0 | 0.12 | 0.01 | 0.1 | 0.38 | 0.18 | 0.25 |
| 6 | 30142690 | rs957765 | 2.616 | -0.517 | -1.307 | -0.696 | -0.043 | -0.049 | -0.049 | 0 | 0.08 | 0.02 | 0.28 | 0.19 | 0.23 | 0.08 |
| 6 | 30144420 | rs2844786 | 0.982 | -0.62 | -0.937 | -0.774 | 0.092 | 0.055 | 0.055 | 0 | 0.19 | 0.33 | 0.4 | 0.27 | 0.31 | 0.17 |
| 6 | 30145083 | rs9261539 | -0.492 | -0.72 | -0.985 | -0.885 | 0.07 | -0.002 | -0.002 | 0 | 0.31 | 0.48 | 0.48 | 0.3 | 0.43 | 0.34 |
| 6 | 30145740 | rs7774730 | -0.471 | -0.723 | -0.986 | -0.886 | 0.07 | -0.001 | -0.001 | 0 | 0.31 | 0.48 | 0.48 | 0.3 | 0.43 | 0.34 |
| 6 | 30146019 | rs9261541 | -1.674 | -0.739 | -0.994 | -0.892 | -0.01 | 0.012 | 0.012 | 0 | 0.9 | 0.97 | 0.86 | 0.93 | 0.87 | 0.92 |
| 6 | 30146650 | rs9261543 | -0.541 | -0.576 | -0.82 | -0.808 | 0.07 | -0.002 | -0.002 | 0 | 0.31 | 0.48 | 0.48 | 0.3 | 0.43 | 0.34 |
| 6 | 30147356 | rs7763162 | -0.233 | -0.472 | -0.747 | -0.762 | 0.122 | 0.022 | 0.022 | 0 | 0.26 | 0.46 | 0.47 | 0.27 | 0.43 | 0.21 |
| 6 | 30150043 | rs2844782 | 0.994 | -1.224 | -0.742 | -0.941 | 0.085 | 0.051 | 0.051 | 0 | 0.19 | 0.32 | 0.39 | 0.27 | 0.3 | 0.17 |
| 6 | 30152367 | rs7843 | 2.576 | -1.12 | -0.898 | -0.897 | -0.043 | -0.049 | -0.049 | 0 | 0.08 | 0.02 | 0.27 | 0.19 | 0.23 | 0.09 |
| 6 | 30152789 | rs1042338 | 1.697 | -1.186 | -0.905 | -0.929 | -0.043 | -0.035 | -0.035 | 0 | 0.12 | 0.05 | 0.31 | 0.19 | 0.25 | 0.12 |
| 6 | 30154199 | rs2074473 | -0.559 | -1.351 | -0.903 | -0.994 | 0.075 | 0.008 | 0.008 | 0 | 0.3 | 0.48 | 0.48 | 0.28 | 0.43 | 0.23 |
| 6 | 30154928 | rs9261547 | -2.14 | -1.264 | -0.863 | -1.065 | -0.011 | 0.007 | 0.007 | 0 | 0.94 | 0.98 | 0.92 | 0.97 | 0.89 | 0.95 |

| | | | | | | | | | | | | | | | | |
|---|---|---|---|---|---|---|---|---|---|---|---|---|---|---|---|---|
| 6 | 30155108 | rs6457164 | -0.442 | -1.186 | -0.853 | -1.056 | 0.058 | -0.001 | -0.001 | 0 | 0.31 | 0.43 | 0.46 | 0.3 | 0.43 | 0.34 |
| 6 | 30155313 | rs6457167 | -0.443 | -1.179 | -0.857 | -1.052 | 0.067 | -0.001 | -0.001 | 0 | 0.31 | 0.45 | 0.47 | 0.3 | 0.43 | 0.34 |
| 6 | 30156496 | rs2074472 | 0.647 | -1.184 | -0.839 | -1.015 | 0.059 | 0.031 | 0.031 | 0 | 0.23 | 0.34 | 0.42 | 0.27 | 0.32 | 0.2 |
| 6 | 30157654 | rs765977 | -0.404 | -1.237 | -0.861 | -0.996 | 0.067 | -0.001 | -0.001 | 0 | 0.31 | 0.45 | 0.47 | 0.3 | 0.43 | 0.34 |
| 6 | 30159854 | rs2284164 | -0.425 | -1.232 | -0.873 | -0.994 | 0.057 | -0.001 | -0.001 | 1.477 | 0.31 | 0.45 | 0.46 | 0.3 | 0.43 | 0.34 |
| 6 | 30162809 | rs4711211 | 2.634 | -1.195 | -0.527 | -1.001 | -0.043 | -0.049 | -0.049 | 0 | 0.08 | 0.02 | 0.27 | 0.19 | 0.23 | 0.09 |
| 6 | 30163955 | rs2517614 | 0.79 | -1.516 | -0.407 | -0.974 | 0.068 | -0.039 | -0.039 | 0 | 0.14 | 0.38 | 0.16 | 0.08 | 0.18 | 0.1 |
| 6 | 30165273 | rs2523722 | 0.79 | -1.516 | -0.407 | -0.96 | 0.068 | -0.039 | -0.039 | 1.414 | 0.14 | 0.38 | 0.16 | 0.08 | 0.18 | 0.1 |
| 6 | 30166266 | rs2523721 | 0.537 | -1.527 | -0.395 | -0.959 | 0.011 | -0.068 | -0.068 | 0 | 0.19 | 0.4 | 0.16 | 0.11 | 0.18 | 0.22 |
| 6 | 30166935 | rs2072107 | 3.17 | -2.286 | -1.131 | -1.22 | 0.038 | -0.02 | -0.02 | 0.409 | 0.21 | 0.04 | 0.09 | 0.15 | 0.14 | 0.24 |
| 6 | 30167294 | rs9261558 | -1.263 | -2.244 | -0.837 | -1.295 | -0.001 | 0 | 0 | 0 | 0.88 | 0.91 | 0.84 | 0.93 | 0.85 | 0.89 |
| 6 | 30168202 | rs28360013 | 1.918 | -2.44 | -0.767 | -1.311 | -0.025 | -0.011 | -0.011 | 0 | 0.1 | 0.27 | 0.11 | 0.08 | 0.07 | 0.08 |
| 6 | 30168319 | rs2523719 | 2.234 | -2.436 | -0.769 | -1.33 | 0.003 | -0.01 | -0.01 | 0.087 | 0.08 | 0.11 | 0.05 | 0.03 | 0.11 | 0.14 |
| 6 | 30169327 | rs2517611 | 1.169 | -2.531 | -0.824 | -1.302 | 0.045 | -0.048 | -0.048 | 0 | 0.15 | 0.38 | 0.15 | 0.08 | 0.18 | 0.13 |
| 6 | 30170280 | rs2517610 | 0.712 | -2.534 | -0.826 | -1.303 | -0.001 | -0.07 | -0.07 | 0.958 | 0.2 | 0.4 | 0.16 | 0.11 | 0.18 | 0.26 |
| 6 | 30170510 | rs1117490 | 0.712 | -2.534 | -0.826 | -1.303 | -0.001 | -0.07 | -0.07 | 11.124 | 0.2 | 0.4 | 0.16 | 0.11 | 0.18 | 0.26 |
| 6 | 30171827 | rs2844776 | 0.697 | -2.536 | -0.827 | -1.304 | -0.002 | -0.07 | -0.07 | 0 | 0.2 | 0.39 | 0.16 | 0.11 | 0.18 | 0.27 |
| 6 | 30171881 | rs4476873 | -1.476 | -2.533 | -0.841 | -1.306 | -0.001 | -0.002 | -0.002 | 0 | 0.89 | 0.91 | 0.85 | 0.93 | 0.87 | 0.92 |
| 6 | 30172233 | rs2284168 | 1.96 | -2.53 | -0.839 | -1.303 | -0.025 | -0.011 | -0.011 | 0 | 0.1 | 0.27 | 0.11 | 0.08 | 0.07 | 0.08 |
| 6 | 30172385 | rs2284169 | 2.924 | -2.739 | -0.887 | -1.364 | 0.042 | -0.027 | -0.027 | 0 | 0.24 | 0.13 | 0.12 | 0.21 | 0.15 | 0.25 |
| 6 | 30172513 | rs971570 | 0.76 | -2.659 | -0.857 | -1.325 | 0.001 | -0.068 | -0.068 | 0.357 | 0.2 | 0.38 | 0.16 | 0.11 | 0.18 | 0.26 |
| 6 | 30174131 | rs2021722 | 0.784 | -2.659 | -0.857 | -1.325 | -0.001 | -0.07 | -0.07 | 0 | 0.2 | 0.4 | 0.16 | 0.11 | 0.18 | 0.26 |
| 6 | 30174633 | rs885912 | 0.784 | -2.788 | -0.969 | -1.346 | -0.001 | -0.07 | -0.07 | 0.057 | 0.2 | 0.4 | 0.16 | 0.11 | 0.18 | 0.26 |
| 6 | 30175847 | rs3132672 | -0.15 | -2.762 | -1.01 | -1.349 | 0.047 | -0.005 | -0.005 | 0 | 0.33 | 0.45 | 0.46 | 0.3 | 0.43 | 0.37 |
| 6 | 30176122 | rs10947058 | 3.214 | -3.079 | -1.079 | -1.518 | -0.023 | -0.036 | -0.036 | 0 | 0.11 | 0.12 | 0.03 | 0 | 0.07 | 0.01 |
| 6 | 30178287 | rs3132671 | 0.058 | -3.078 | -1.079 | -1.518 | 0.052 | -0.004 | -0.004 | 0 | 0.32 | 0.43 | 0.46 | 0.3 | 0.43 | 0.37 |
| 6 | 30179026 | rs3130383 | 0.013 | -3.078 | -1.079 | -1.518 | 0.049 | -0.004 | -0.004 | 0 | 0.33 | 0.44 | 0.46 | 0.3 | 0.43 | 0.37 |
| 6 | 30179422 | rs2844775 | 2.388 | -3.11 | -0.953 | -1.531 | -0.036 | -0.03 | -0.03 | 0 | 0.13 | 0.06 | 0.31 | 0.19 | 0.25 | 0.11 |
| 6 | 30180246 | rs3130386 | 0.167 | -2.988 | -0.978 | -1.466 | 0.049 | -0.004 | -0.004 | 0 | 0.33 | 0.43 | 0.46 | 0.3 | 0.43 | 0.37 |
| 6 | 30181275 | rs3094614 | 0.211 | -2.924 | -1.232 | -1.453 | 0.052 | -0.004 | -0.004 | 0 | 0.32 | 0.43 | 0.46 | 0.3 | 0.43 | 0.37 |
| 6 | 30182395 | rs1345229 | 1.856 | -3.159 | -1.489 | -1.443 | 0.01 | -0.015 | -0.015 | 0.125 | 0.11 | 0.12 | 0.09 | 0.19 | 0.09 | 0.09 |
| 6 | 30184704 | rs9295837 | 3.07 | -2.704 | -0.966 | -1.278 | -0.023 | -0.036 | -0.036 | 0.496 | 0.11 | 0.12 | 0.03 | 0 | 0.07 | 0.01 |
| 6 | 30185244 | rs9261567 | 3.494 | -2.553 | -0.862 | -1.197 | 0.033 | -0.02 | -0.02 | 0 | 0.2 | 0.05 | 0.09 | 0.15 | 0.14 | 0.21 |
| 6 | 30186863 | rs2106070 | 0.015 | -2.242 | -0.705 | -1.1 | 0.052 | -0.004 | -0.004 | 0 | 0.32 | 0.43 | 0.46 | 0.3 | 0.43 | 0.37 |
| 6 | 30190640 | rs3130391 | 0.006 | -2.238 | -0.703 | -1.098 | 0.052 | -0.004 | -0.004 | 0.152 | 0.32 | 0.43 | 0.46 | 0.3 | 0.43 | 0.37 |
| 6 | 30191587 | rs3132667 | -0.005 | -2.238 | -0.703 | -1.098 | 0.051 | -0.005 | -0.005 | 0.809 | 0.32 | 0.43 | 0.46 | 0.3 | 0.43 | 0.37 |
| 6 | 30194103 | rs2040486 | 0.006 | -2.236 | -0.702 | -1.097 | 0.052 | -0.004 | -0.004 | 0.059 | 0.32 | 0.43 | 0.46 | 0.3 | 0.43 | 0.37 |
| 6 | 30196313 | rs3094140 | 0.006 | -2.152 | -0.758 | -1.093 | 0.052 | -0.004 | -0.004 | 0 | 0.32 | 0.43 | 0.46 | 0.3 | 0.43 | 0.37 |
| 6 | 30196855 | rs3129690 | 0.006 | -2.152 | -0.758 | -1.093 | 0.052 | -0.004 | -0.004 | 0 | 0.32 | 0.43 | 0.46 | 0.3 | 0.43 | 0.37 |
| 6 | 30196980 | rs9261582 | 2.417 | -2.08 | -0.689 | -1.075 | 0.026 | 0.004 | 0.004 | 0 | 0.29 | 0.25 | 0.16 | 0.21 | 0.18 | 0.25 |
| 6 | 30199016 | rs3094138 | -1.064 | -1.142 | -0.444 | -0.905 | -0.011 | 0.01 | 0.01 | 0 | 0.9 | 0.97 | 0.87 | 0.93 | 0.87 | 0.91 |
| 6 | 30200873 | rs3094137 | -1.21 | -1.129 | -0.455 | -0.902 | 0 | -0.002 | -0.002 | 0 | 0.89 | 0.91 | 0.86 | 0.93 | 0.87 | 0.91 |
| 6 | 30201955 | rs916571 | -0.105 | -0.8 | -0.568 | -0.721 | 0.094 | 0.012 | 0.012 | 0 | 0.27 | 0.43 | 0.46 | 0.27 | 0.43 | 0.25 |
| 6 | 30202429 | rs3132674 | -0.105 | -0.8 | -0.568 | -0.721 | 0.094 | 0.012 | 0.012 | 0 | 0.27 | 0.43 | 0.46 | 0.27 | 0.43 | 0.25 |
| 6 | 30202715 | rs885915 | 0.784 | -0.614 | -0.5 | -0.748 | 0.058 | 0.029 | 0.029 | 0 | 0.23 | 0.32 | 0.41 | 0.27 | 0.32 | 0.2 |
| 6 | 30202767 | rs885914 | 0.729 | -0.586 | -0.507 | -0.754 | 0.056 | 0.027 | 0.027 | 0 | 0.23 | 0.32 | 0.41 | 0.27 | 0.32 | 0.2 |
| 6 | 30203602 | rs954037 | -0.463 | -0.801 | -0.618 | -0.837 | 0.049 | -0.004 | -0.004 | 0 | 0.33 | 0.43 | 0.46 | 0.31 | 0.43 | 0.37 |
| 6 | 30204526 | rs2844780 | 0.815 | -0.747 | -0.611 | -0.778 | 0.058 | 0.027 | 0.027 | 0 | 0.23 | 0.32 | 0.41 | 0.27 | 0.32 | 0.2 |
| 6 | 30206186 | rs10456378 | 2.858 | -1.276 | -0.886 | -0.928 | 0.023 | 0.049 | 0.049 | 0.544 | 0.1 | 0.01 | 0.02 | 0 | 0.06 | 0.01 |
| 6 | 30209493 | rs2285801 | -0.122 | -1.169 | -0.742 | -0.706 | 0.077 | -0.004 | -0.004 | 0.35 | 0.3 | 0.47 | 0.43 | 0.26 | 0.44 | 0.24 |
| 6 | 30209802 | rs1362089 | -0.122 | -1.171 | -0.791 | -0.677 | 0.077 | -0.002 | -0.002 | 0.403 | 0.3 | 0.47 | 0.43 | 0.27 | 0.44 | 0.24 |
| 6 | 30210807 | rs9261588 | -2.091 | -1.326 | -0.917 | -0.783 | -0.011 | 0.007 | 0.007 | 0 | 0.94 | 0.98 | 0.93 | 0.97 | 0.89 | 0.94 |
| 6 | 30211660 | rs9261592 | -0.572 | -1.202 | -0.825 | -0.647 | 0.049 | -0.001 | -0.001 | 5.137 | 0.33 | 0.48 | 0.53 | 0.31 | 0.46 | 0.28 |
| 6 | 30211755 | rs3094636 | -0.572 | -1.202 | -0.818 | -0.647 | 0.05 | -0.001 | -0.001 | 0 | 0.33 | 0.48 | 0.53 | 0.31 | 0.46 | 0.28 |
| 6 | 30215592 | rs3132662 | -0.309 | -0.758 | -0.901 | -0.549 | 0.024 | -0.008 | -0.008 | 0 | 0.31 | 0.4 | 0.41 | 0.28 | 0.43 | 0.34 |
| 6 | 30222256 | rs2844767 | -0.309 | -0.758 | -0.901 | -0.549 | 0.024 | -0.008 | -0.008 | 0 | 0.31 | 0.4 | 0.41 | 0.28 | 0.43 | 0.34 |
| 6 | 30224305 | rs1579219 | 1.569 | -0.926 | -1.216 | -0.501 | -0.026 | -0.097 | -0.097 | 0 | 0.16 | 0.02 | 0.32 | 0.44 | 0.17 | 0.33 |
| 6 | 30224889 | rs928824 | 2.896 | -1.005 | -1.184 | -0.668 | 0.008 | 0.047 | 0.047 | 0.743 | 0.09 | 0 | 0.03 | 0 | 0.08 | 0.01 |
| 6 | 30226811 | rs962899 | 0.029 | -1.483 | -0.873 | -0.843 | 0.004 | -0.004 | -0.004 | 0.318 | 0.31 | 0.3 | 0.45 | 0.53 | 0.25 | 0.51 |
| 6 | 30227397 | rs9391806 | 1.739 | -1.545 | -0.928 | -0.87 | -0.035 | -0.107 | -0.107 | 0 | 0.16 | 0.01 | 0.32 | 0.44 | 0.16 | 0.33 |
| 6 | 30228721 | rs2516708 | -0.82 | -1.295 | -0.95 | -0.89 | 0.001 | -0.003 | -0.003 | 0.313 | 0.38 | 0.41 | 0.52 | 0.58 | 0.37 | 0.57 |
| 6 | 30229306 | rs2428507 | -0.99 | -1.36 | -0.968 | -0.924 | 0.001 | -0.003 | -0.003 | 0.011 | 0.39 | 0.43 | 0.52 | 0.58 | 0.38 | 0.58 |
| 6 | 30229694 | rs2516705 | 0.06 | -1.317 | -0.957 | -0.893 | 0.004 | -0.004 | -0.004 | 0.582 | 0.31 | 0.3 | 0.45 | 0.53 | 0.25 | 0.51 |
| 6 | 30229922 | rs2516703 | 0.06 | -1.317 | -0.957 | -0.893 | 0.004 | -0.004 | -0.004 | 0.003 | 0.31 | 0.3 | 0.45 | 0.53 | 0.25 | 0.51 |
| 6 | 30229990 | rs2516702 | 0.06 | -1.317 | -0.957 | -0.893 | 0.004 | -0.004 | -0.004 | 0.401 | 0.31 | 0.3 | 0.45 | 0.53 | 0.25 | 0.51 |
| 6 | 30230085 | rs2523746 | 0.06 | -1.317 | -0.957 | -0.893 | 0.004 | -0.004 | -0.004 | 0 | 0.31 | 0.3 | 0.45 | 0.53 | 0.25 | 0.51 |
| 6 | 30230128 | rs2523745 | 0.06 | -1.317 | -0.957 | -0.893 | 0.004 | -0.004 | -0.004 | 0 | 0.31 | 0.3 | 0.45 | 0.53 | 0.25 | 0.51 |
| 6 | 30230661 | rs2844766 | -0.793 | -1.304 | -0.957 | -0.897 | 0.001 | -0.003 | -0.003 | 0 | 0.38 | 0.41 | 0.52 | 0.58 | 0.37 | 0.57 |
| 6 | 30231108 | rs2516699 | 0.064 | -1.306 | -0.957 | -0.899 | 0.004 | -0.004 | -0.004 | 0 | 0.31 | 0.3 | 0.45 | 0.53 | 0.25 | 0.51 |

| | | | | | | | | | | | | | | | | |
|---|---|---|---|---|---|---|---|---|---|---|---|---|---|---|---|---|
| 6 | 30231224 | rs3094073 | 1.917 | -1.29 | -0.95 | -0.891 | 0.005 | 0 | 0 | 0.462 | 0.07 | 0.11 | 0.07 | 0.05 | 0.12 | 0.06 |
| 6 | 30231273 | rs3130401 | 1.917 | -1.29 | -0.95 | -0.891 | 0.005 | 0 | 0 | 0 | 0.07 | 0.11 | 0.07 | 0.05 | 0.12 | 0.06 |
| 6 | 30231330 | rs3132659 | 1.917 | -1.29 | -0.95 | -0.891 | 0.005 | 0 | 0 | 0 | 0.07 | 0.11 | 0.07 | 0.05 | 0.12 | 0.06 |
| 6 | 30231666 | rs3132658 | 1.917 | -1.29 | -0.95 | -0.891 | 0.005 | 0 | 0 | 0 | 0.07 | 0.11 | 0.07 | 0.05 | 0.12 | 0.06 |
| 6 | 30231768 | rs3094071 | 1.917 | -1.29 | -0.95 | -0.891 | 0.005 | 0 | 0 | 0 | 0.07 | 0.11 | 0.07 | 0.05 | 0.12 | 0.06 |
| 6 | 30232009 | rs3130403 | 1.935 | -1.289 | -0.951 | -0.892 | 0.006 | 0 | 0 | 0 | 0.07 | 0.11 | 0.07 | 0.05 | 0.12 | 0.06 |
| 6 | 30232374 | rs2516698 | 1.46 | -1.357 | -0.894 | -0.919 | -0.033 | -0.035 | -0.035 | 0 | 0.1 | 0.21 | 0.1 | 0.07 | 0.06 | 0.12 |
| 6 | 30232436 | rs3094630 | 1.939 | -1.281 | -0.894 | -0.884 | 0.005 | 0 | 0 | 1.596 | 0.07 | 0.11 | 0.07 | 0.05 | 0.12 | 0.06 |
| 6 | 30232514 | rs2516697 | 0.366 | -1.286 | -0.913 | -0.871 | -0.005 | 0.006 | 0.006 | 0.487 | 0.26 | 0.3 | 0.45 | 0.5 | 0.24 | 0.4 |
| 6 | 30232785 | rs3129702 | 1.941 | -1.332 | -0.91 | -0.899 | 0.005 | 0 | 0 | 0 | 0.07 | 0.11 | 0.07 | 0.05 | 0.12 | 0.06 |
| 6 | 30232953 | rs3094629 | 2.032 | -1.334 | -0.91 | -0.898 | 0.007 | -0.012 | -0.012 | 0 | 0.07 | 0.11 | 0.05 | 0 | 0.12 | 0.05 |
| 6 | 30233558 | rs3129703 | 1.941 | -1.332 | -0.91 | -0.899 | 0.005 | 0 | 0 | 0.503 | 0.07 | 0.11 | 0.07 | 0.05 | 0.12 | 0.06 |
| 6 | 30234152 | rs3130405 | 1.941 | -1.332 | -0.897 | -0.899 | -0.006 | -0.001 | -0.001 | 0.013 | 0.07 | 0.1 | 0.06 | 0.05 | 0.12 | 0.06 |
| 6 | 30234657 | rs3129832 | 1.941 | -1.332 | -0.905 | -0.899 | 0.005 | 0 | 0 | 0 | 0.07 | 0.11 | 0.07 | 0.05 | 0.12 | 0.06 |
| 6 | 30234721 | rs3129705 | 1.941 | -1.332 | -0.905 | -0.899 | 0.005 | 0 | 0 | 0.084 | 0.07 | 0.12 | 0.07 | 0.05 | 0.12 | 0.06 |
| 6 | 30235184 | rs2844763 | 0.279 | -1.402 | -0.951 | -0.875 | -0.005 | 0.006 | 0.006 | 0.469 | 0.26 | 0.3 | 0.45 | 0.51 | 0.24 | 0.41 |
| 6 | 30237454 | rs2844761 | 0.279 | -1.402 | -0.951 | -0.875 | -0.005 | 0.006 | 0.006 | 0.469 | 0.26 | 0.3 | 0.45 | 0.51 | 0.24 | 0.41 |
| 6 | 30237859 | rs9261639 | 1.749 | -1.323 | -1.259 | -0.864 | -0.094 | -0.084 | -0.084 | 0 | 0.14 | 0.04 | 0.2 | 0.24 | 0.26 | 0.1 |
| 6 | 30238559 | rs9368628 | 1.983 | -1.402 | -1.128 | -0.801 | -0.026 | -0.097 | -0.097 | 0.74 | 0.16 | 0.02 | 0.32 | 0.44 | 0.17 | 0.33 |
| 6 | 30238884 | rs2844759 | -0.744 | -1.416 | -1.122 | -0.846 | 0.02 | -0.012 | -0.012 | 0.022 | 0.37 | 0.36 | 0.49 | 0.58 | 0.29 | 0.57 |
| 6 | 30240926 | rs2097773 | 0.351 | -1.355 | -1.088 | -0.817 | -0.005 | 0.006 | 0.006 | 0.469 | 0.26 | 0.3 | 0.45 | 0.51 | 0.24 | 0.41 |
| 6 | 30241114 | rs9393999 | 1.908 | -1.325 | -1.155 | -0.794 | -0.033 | -0.108 | -0.108 | 0 | 0.16 | 0.01 | 0.32 | 0.44 | 0.16 | 0.33 |
| 6 | 30242171 | rs28780093 | 4.031 | -1.942 | -1.462 | -1.207 | 0.015 | 0.043 | 0.043 | 0 | 0.17 | 0.03 | 0.08 | 0.07 | 0.13 | 0.2 |
| 6 | 30244579 | rs968909 | -0.597 | -1.987 | -1.386 | -1.189 | 0.02 | -0.012 | -0.012 | 2.755 | 0.37 | 0.36 | 0.49 | 0.58 | 0.29 | 0.57 |
| 6 | 30245033 | rs2516725 | 0.536 | -2.007 | -1.394 | -1.193 | -0.005 | 0.006 | 0.006 | 0.469 | 0.26 | 0.3 | 0.45 | 0.51 | 0.24 | 0.41 |
| 6 | 30246212 | rs2844754 | 2.083 | -2.061 | -0.819 | -1.224 | -0.01 | -0.02 | -0.02 | 0.942 | 0.07 | 0.21 | 0.1 | 0.06 | 0.05 | 0.08 |
| 6 | 30248424 | rs2844753 | 3.524 | -2.203 | -0.969 | -1.343 | 0.008 | 0.048 | 0.048 | 0.596 | 0.09 | 0 | 0.03 | 0 | 0.08 | 0.01 |
| 6 | 30252606 | rs2394779 | 2.299 | -2.079 | -0.938 | -1.313 | -0.035 | -0.107 | -0.107 | 0.049 | 0.15 | 0.01 | 0.32 | 0.44 | 0.16 | 0.33 |
| 6 | 30253011 | rs1264626 | -0.588 | -2.087 | -0.92 | -1.322 | 0.02 | -0.012 | -0.012 | 1.495 | 0.37 | 0.36 | 0.5 | 0.58 | 0.29 | 0.57 |
| 6 | 30253339 | rs1132547 | 1.537 | -1.972 | -0.808 | -1.292 | -0.027 | -0.049 | -0.049 | 0 | 0.1 | 0.28 | 0.13 | 0.06 | 0.08 | 0.08 |
| 6 | 30256838 | rs10484551 | 2.409 | -1.922 | -0.861 | -1.25 | 0.011 | -0.007 | -0.007 | 0 | 0.07 | 0.06 | 0.04 | 0.05 | 0.04 | 0.06 |
| 6 | 30256885 | rs17194767 | 2.409 | -1.934 | -0.824 | -1.256 | 0.011 | -0.007 | -0.007 | 0 | 0.07 | 0.06 | 0.04 | 0.05 | 0.04 | 0.06 |
| 6 | 30256936 | rs1264622 | 3.318 | -1.936 | -0.801 | -1.309 | -0.015 | 0.015 | 0.015 | 0.24 | 0.1 | 0.06 | 0.05 | 0 | 0.16 | 0.01 |
| 6 | 30257693 | rs2285799 | 2.419 | -1.938 | -0.799 | -1.281 | 0.011 | -0.007 | -0.007 | 0 | 0.07 | 0.06 | 0.04 | 0.05 | 0.04 | 0.06 |
| 6 | 30258708 | rs4713309 | 2.383 | -1.94 | -0.8 | -1.282 | 0.011 | -0.007 | -0.007 | 0 | 0.07 | 0.06 | 0.04 | 0.05 | 0.04 | 0.06 |
| 6 | 30259657 | rs1045251 | 0.586 | -2.043 | -0.816 | -1.312 | -0.005 | 0.006 | 0.006 | 0.469 | 0.26 | 0.3 | 0.45 | 0.51 | 0.24 | 0.41 |
| 6 | 30260982 | rs11752362 | 3.509 | -2.304 | -0.735 | -1.433 | 0.004 | 0.004 | 0.004 | 0 | 0.22 | 0.21 | 0.12 | 0.13 | 0.15 | 0.25 |
| 6 | 30262512 | rs1048664 | 2.197 | -2.301 | -0.727 | -1.43 | -0.01 | -0.02 | -0.02 | 0 | 0.07 | 0.21 | 0.1 | 0.06 | 0.05 | 0.08 |
| 6 | 30264307 | rs1264616 | 0.687 | -2.264 | -0.746 | -1.41 | -0.005 | 0.006 | 0.006 | 0.469 | 0.26 | 0.3 | 0.45 | 0.51 | 0.24 | 0.41 |
| 6 | 30267748 | rs3094631 | -2.56 | -2.293 | -0.771 | -1.496 | -0.011 | 0.008 | 0.008 | 0 | 0.94 | 0.97 | 0.93 | 0.96 | 0.9 | 0.95 |
| 6 | 30267886 | rs7758503 | 3.6 | -2.304 | -0.771 | -1.502 | 0.004 | 0.003 | 0.003 | 0 | 0.22 | 0.21 | 0.12 | 0.13 | 0.15 | 0.25 |
| 6 | 30270080 | rs261949 | 0.303 | -2.306 | -0.848 | -1.5 | 0.004 | -0.004 | -0.004 | 0.159 | 0.31 | 0.3 | 0.45 | 0.53 | 0.25 | 0.51 |
| 6 | 30271334 | rs261946 | 2.398 | -2.289 | -0.863 | -1.495 | -0.086 | -0.078 | -0.078 | 0 | 0.13 | 0.04 | 0.2 | 0.24 | 0.26 | 0.1 |
| 6 | 30271792 | rs4713310 | 2.122 | -2.303 | -0.855 | -1.499 | 0.014 | -0.003 | -0.003 | 0 | 0.08 | 0.07 | 0.05 | 0.05 | 0.05 | 0.06 |
| 6 | 30272417 | rs261945 | -1.762 | -2.285 | -0.728 | -1.499 | -0.045 | -0.03 | -0.03 | 0 | 0.79 | 0.88 | 0.72 | 0.72 | 0.63 | 0.85 |
| 6 | 30274461 | rs9380167 | -0.034 | -2.172 | -0.933 | -1.455 | 0.005 | -0.005 | -0.005 | 0 | 0.34 | 0.38 | 0.49 | 0.55 | 0.3 | 0.47 |
| 6 | 30275246 | rs928822 | -0.892 | -1.994 | -0.938 | -1.383 | -0.007 | -0.005 | -0.005 | 0 | 0.71 | 0.77 | 0.68 | 0.71 | 0.61 | 0.83 |
| 6 | 30278112 | rs1264608 | 3.51 | -1.714 | -1.203 | -1.261 | 0.008 | 0.047 | 0.047 | 0.531 | 0.09 | 0 | 0.03 | 0 | 0.08 | 0.01 |
| 6 | 30282332 | rs3094065 | -2.118 | -1.541 | -0.982 | -0.999 | -0.01 | 0.009 | 0.009 | 0 | 0.93 | 0.97 | 0.92 | 0.96 | 0.89 | 0.95 |
| 6 | 30284375 | rs6912771 | -0.243 | -1.793 | -0.941 | -0.996 | 0.005 | -0.005 | -0.005 | 0 | 0.34 | 0.37 | 0.49 | 0.55 | 0.3 | 0.47 |
| 6 | 30285121 | rs9357097 | -0.041 | -1.88 | -0.922 | -1.008 | -0.028 | 0.008 | 0.008 | 0 | 0.32 | 0.47 | 0.39 | 0.48 | 0.29 | 0.43 |
| 6 | 30285883 | rs6924282 | -0.401 | -1.824 | -1.552 | -0.982 | -0.005 | 0.003 | 0.003 | 0 | 0.34 | 0.4 | 0.5 | 0.55 | 0.3 | 0.47 |
| 6 | 30286729 | rs9366755 | 1.993 | -1.88 | -1.627 | -1.011 | -0.035 | -0.107 | -0.107 | 2.012 | 0.16 | 0.01 | 0.32 | 0.44 | 0.16 | 0.33 |
| 6 | 30289271 | rs6900042 | -0.431 | -1.858 | -1.609 | -1.004 | -0.004 | 0.002 | 0.002 | 0 | 0.34 | 0.4 | 0.5 | 0.55 | 0.3 | 0.47 |
| 6 | 30292083 | rs12174823 | 2.479 | -1.716 | -1.609 | -0.96 | -0.04 | -0.083 | -0.083 | 0 | 0.12 | 0.01 | 0.21 | 0.41 | 0.15 | 0.29 |
| 6 | 30292147 | rs28360024 | 1.506 | -1.743 | -1.519 | -1.014 | -0.027 | -0.049 | -0.049 | 0 | 0.1 | 0.28 | 0.13 | 0.06 | 0.08 | 0.08 |
| 6 | 30293186 | rs17194824 | 2.135 | -1.729 | -1.503 | -1.002 | -0.01 | -0.02 | -0.02 | 0 | 0.07 | 0.21 | 0.1 | 0.06 | 0.05 | 0.08 |
| 6 | 30294590 | rs1264582 | 2.613 | -1.941 | -1.846 | -1.149 | 0.036 | 0.068 | 0.068 | 0 | 0.14 | 0.01 | 0.03 | 0.03 | 0.08 | 0.11 |
| 6 | 30296071 | rs1110464 | -0.829 | -1.924 | -2.038 | -1.239 | -0.123 | -0.063 | -0.063 | 0 | 0.74 | 0.89 | 0.69 | 0.72 | 0.58 | 0.8 |
| 6 | 30297505 | rs1264581 | 2.746 | -1.723 | -2.101 | -1.207 | 0.036 | 0.068 | 0.068 | 0 | 0.14 | 0.01 | 0.03 | 0.03 | 0.08 | 0.11 |
| 6 | 30297529 | rs2057727 | 0.101 | -1.129 | -1.766 | -0.882 | -0.005 | 0.003 | 0.003 | 0.625 | 0.28 | 0.32 | 0.45 | 0.51 | 0.25 | 0.41 |
| 6 | 30301600 | rs12663184 | 3.963 | -1.414 | -1.838 | -0.93 | 0.017 | 0.043 | 0.043 | 0 | 0.17 | 0.03 | 0.08 | 0.07 | 0.13 | 0.2 |
| 6 | 30302600 | rs3778624 | 0.067 | -1.177 | -1.674 | -0.804 | -0.005 | 0.003 | 0.003 | 0.633 | 0.28 | 0.32 | 0.45 | 0.51 | 0.25 | 0.41 |
| 6 | 30304816 | rs7767653 | 1.531 | -0.925 | -1.582 | -0.711 | -0.027 | -0.049 | -0.049 | 1.771 | 0.1 | 0.28 | 0.13 | 0.06 | 0.08 | 0.08 |
| 6 | 30306306 | rs3129837 | 2.038 | -0.607 | -1.587 | -0.576 | 0.055 | -0.032 | -0.032 | 1.917 | 0.07 | 0.16 | 0.05 | 0 | 0.12 | 0.05 |
| 6 | 30306553 | rs3129838 | -2.214 | -0.111 | -2.161 | -0.248 | -0.011 | 0.007 | 0.007 | 0 | 0.94 | 0.97 | 0.93 | 0.96 | 0.9 | 0.95 |
| 6 | 30310500 | rs2240058 | 1.87 | -0.11 | -2.022 | 0.09 | -0.04 | -0.083 | -0.083 | 1.123 | 0.12 | 0.01 | 0.21 | 0.41 | 0.15 | 0.29 |
| 6 | 30311932 | rs6931763 | 2.436 | 0.1 | -1.744 | 0.077 | 0.036 | 0.026 | 0.026 | 0 | 0.22 | 0.11 | 0.12 | 0.14 | 0.15 | 0.26 |
| 6 | 30315011 | rs974961 | 1.194 | 1.217 | -1.147 | 0.861 | -0.083 | -0.076 | -0.076 | 0 | 0.13 | 0.03 | 0.2 | 0.24 | 0.26 | 0.1 |

| | | | | | | | | | | | | | | | | |
|---|---|---|---|---|---|---|---|---|---|---|---|---|---|---|---|---|
| 6 | 30316739 | rs9380174 | -0.335 | 0.215 | -0.574 | 0.567 | -0.005 | 0.006 | 0.006 | 0.469 | 0.26 | 0.3 | 0.45 | 0.5 | 0.24 | 0.41 |
| 6 | 30319930 | rs6905389 | 0.501 | 0.221 | -0.089 | 0.374 | -0.038 | -0.003 | -0.003 | 0 | 0.14 | 0.3 | 0.11 | 0.07 | 0.1 | 0.06 |
| 6 | 30321361 | rs984802 | 0.213 | 1.137 | -0.528 | 0.387 | -0.013 | -0.061 | -0.061 | 0 | 0.23 | 0.11 | 0.27 | 0.41 | 0.22 | 0.38 |
| 6 | 30325531 | rs3130379 | -2.468 | 0.945 | -0.741 | -0.29 | 0 | -0.009 | -0.009 | 0 | 0.79 | 0.83 | 0.84 | 0.95 | 0.78 | 0.81 |
| 6 | 30330341 | rs2251765 | 2.154 | 0.097 | -0.836 | -0.868 | -0.006 | 0.039 | 0.039 | 0 | 0.14 | 0.02 | 0.16 | 0.04 | 0.14 | 0.08 |
| 6 | 30331027 | rs2158285 | 1.959 | 0.047 | -0.372 | -0.899 | 0.009 | 0.018 | 0.018 | 0 | 0.15 | 0.12 | 0.18 | 0.04 | 0.15 | 0.08 |
| 6 | 30332555 | rs1012411 | 0.542 | 0.002 | -0.282 | -0.294 | 0.003 | 0.003 | 0.003 | 0 | 0.28 | 0.33 | 0.3 | 0.25 | 0.31 | 0.25 |
| 6 | 30334339 | rs2040450 | 2.292 | -0.062 | -0.336 | -0.301 | -0.001 | 0.042 | 0.042 | 0 | 0.14 | 0.03 | 0.17 | 0.04 | 0.14 | 0.08 |
| 6 | 30335554 | rs3129808 | -2.933 | -0.112 | -0.356 | -0.391 | 0 | -0.01 | -0.01 | 0 | 0.8 | 0.83 | 0.84 | 0.95 | 0.79 | 0.81 |
| 6 | 30337738 | rs2022079 | 0.328 | -0.22 | -0.441 | -0.346 | 0.026 | -0.025 | -0.025 | 0 | 0.37 | 0.31 | 0.31 | 0.49 | 0.27 | 0.43 |
| 6 | 30338138 | rs2844748 | 1.811 | -0.237 | -0.587 | -0.26 | 0.021 | 0.038 | 0.038 | 0 | 0.19 | 0.04 | 0.18 | 0.09 | 0.15 | 0.21 |
| 6 | 30339293 | rs17839997 | 1.809 | -0.233 | -0.509 | -0.245 | 0.015 | 0.121 | 0.121 | 0 | 0.07 | 0.16 | 0.06 | 0.16 | 0.06 | 0.05 |
| 6 | 30339303 | rs2523724 | 1.641 | -0.253 | -0.697 | -0.371 | 0.03 | 0.046 | 0.046 | 0 | 0.19 | 0.05 | 0.18 | 0.09 | 0.16 | 0.21 |
| 6 | 30339564 | rs3130361 | -2.976 | -0.231 | -0.681 | -0.389 | 0 | -0.007 | -0.007 | 0 | 0.79 | 0.83 | 0.84 | 0.95 | 0.79 | 0.81 |
| 6 | 30339602 | rs2844746 | -1.396 | -0.218 | -0.687 | -0.39 | 0.024 | 0.084 | 0.084 | 0 | 0.61 | 0.79 | 0.66 | 0.86 | 0.63 | 0.6 |
| 6 | 30340145 | rs3132643 | -0.661 | -0.239 | -0.673 | -0.361 | 0.02 | 0.072 | 0.072 | 0 | 0.49 | 0.71 | 0.47 | 0.69 | 0.44 | 0.51 |
| 6 | 30342190 | rs3130368 | -0.655 | -0.315 | -0.677 | -0.366 | 0.022 | 0.071 | 0.071 | 0 | 0.49 | 0.71 | 0.47 | 0.69 | 0.44 | 0.51 |
| 6 | 30343183 | rs3132635 | -0.661 | -0.289 | -0.807 | -0.408 | 0.023 | 0.072 | 0.072 | 0 | 0.49 | 0.71 | 0.47 | 0.69 | 0.44 | 0.51 |
| 6 | 30343703 | rs2844745 | -1.983 | -0.292 | -0.812 | -0.413 | -0.02 | 0.012 | 0.012 | 0 | 0.68 | 0.74 | 0.64 | 0.78 | 0.59 | 0.71 |
| 6 | 30343983 | rs3130369 | -0.703 | -0.247 | -0.708 | -0.329 | 0.022 | 0.071 | 0.071 | 0 | 0.49 | 0.71 | 0.47 | 0.69 | 0.44 | 0.51 |
| 6 | 30344166 | rs3132633 | -0.703 | -0.247 | -0.708 | -0.329 | 0.022 | 0.071 | 0.071 | 0 | 0.49 | 0.71 | 0.47 | 0.69 | 0.44 | 0.51 |
| 6 | 30344690 | rs3094062 | -0.703 | -0.247 | -0.708 | -0.329 | 0.022 | 0.071 | 0.071 | 0 | 0.49 | 0.71 | 0.47 | 0.69 | 0.44 | 0.51 |
| 6 | 30345139 | rs1419653 | -0.703 | -0.321 | -0.863 | -0.314 | 0.022 | 0.071 | 0.071 | 0 | 0.49 | 0.71 | 0.47 | 0.69 | 0.44 | 0.51 |
| 6 | 30345871 | rs2524212 | -1.181 | -0.361 | -0.969 | 0.044 | 0.039 | 0.039 | 0.039 | 0 | 0.55 | 0.75 | 0.59 | 0.69 | 0.6 | 0.41 |
| 6 | 30346093 | rs3132629 | -0.728 | -0.442 | -0.724 | 0.025 | 0.025 | 0.076 | 0.076 | 0 | 0.5 | 0.72 | 0.47 | 0.69 | 0.44 | 0.51 |
| 6 | 30346618 | rs3132628 | -0.738 | -0.398 | -0.607 | 0.016 | 0.025 | 0.076 | 0.076 | 0 | 0.5 | 0.72 | 0.47 | 0.69 | 0.44 | 0.51 |
| 6 | 30347078 | rs3132626 | -0.73 | -0.39 | -0.603 | 0.02 | 0.025 | 0.076 | 0.076 | 0 | 0.5 | 0.72 | 0.47 | 0.69 | 0.44 | 0.51 |
| 6 | 30347239 | rs970269 | -0.73 | -0.39 | -0.603 | 0.02 | 0.025 | 0.076 | 0.076 | 0 | 0.5 | 0.72 | 0.47 | 0.69 | 0.44 | 0.51 |
| 6 | 30347306 | rs970270 | 2.049 | -0.375 | -0.633 | -0.002 | -0.034 | -0.022 | -0.022 | 0 | 0.11 | 0.08 | 0.19 | 0.17 | 0.19 | 0.1 |
| 6 | 30347833 | rs9404964 | 0.342 | -0.374 | -0.496 | 0.054 | 0.006 | -0.006 | -0.006 | 0 | 0.38 | 0.4 | 0.32 | 0.49 | 0.27 | 0.44 |
| 6 | 30348042 | rs1245218 | -2.891 | -0.223 | -0.595 | 0.044 | -0.001 | -0.007 | -0.007 | 0 | 0.81 | 0.85 | 0.84 | 0.95 | 0.79 | 0.81 |
| 6 | 30348090 | rs2516683 | 2.368 | -0.635 | -0.594 | 0.094 | -0.006 | 0.038 | 0.038 | 0 | 0.14 | 0.02 | 0.16 | 0.04 | 0.14 | 0.08 |
| 6 | 30348247 | rs7744914 | 2.656 | -0.754 | -0.64 | 0.086 | 0.012 | 0.001 | 0.001 | 0.015 | 0.09 | 0.1 | 0.02 | 0.06 | 0.02 | 0.06 |
| 6 | 30349382 | rs3132621 | -0.781 | -0.725 | -0.634 | 0.089 | 0.025 | 0.076 | 0.076 | 0 | 0.5 | 0.73 | 0.47 | 0.69 | 0.44 | 0.51 |
| 6 | 30350660 | rs2524211 | 2.013 | -0.728 | -0.635 | 0.088 | 0.013 | 0.03 | 0.03 | 0 | 0.18 | 0.03 | 0.17 | 0.09 | 0.15 | 0.21 |
| 6 | 30351417 | rs28780099 | 3.629 | -0.834 | -0.619 | 0.034 | 0.014 | 0.021 | 0.021 | 0 | 0.12 | 0.03 | 0.03 | 0.02 | 0.07 | 0.14 |
| 6 | 30352527 | rs3130122 | -0.618 | -0.799 | -0.589 | 0.078 | 0.026 | 0.077 | 0.077 | 0 | 0.5 | 0.73 | 0.47 | 0.69 | 0.44 | 0.51 |
| 6 | 30352986 | rs6936943 | -0.21 | -0.703 | -0.531 | 0.176 | -0.007 | 0 | 0 | 0.013 | 0.43 | 0.51 | 0.37 | 0.49 | 0.35 | 0.44 |
| 6 | 30353069 | rs3094710 | -0.52 | -0.725 | -1.247 | -0.488 | 0.022 | 0.072 | 0.072 | 0 | 0.49 | 0.73 | 0.47 | 0.69 | 0.44 | 0.51 |
| 6 | 30357780 | rs3130140 | 0.175 | -0.66 | -1.231 | -0.441 | -0.004 | -0.003 | -0.003 | 0 | 0.4 | 0.61 | 0.44 | 0.63 | 0.42 | 0.45 |
| 6 | 30358151 | rs3094051 | -0.277 | -0.162 | -1.172 | -0.434 | 0.023 | 0.074 | 0.074 | 0 | 0.49 | 0.71 | 0.47 | 0.69 | 0.44 | 0.51 |
| 6 | 30358522 | rs3130144 | -0.277 | -0.162 | -1.176 | -0.434 | 0.023 | 0.072 | 0.072 | 0 | 0.49 | 0.71 | 0.47 | 0.69 | 0.44 | 0.51 |
| 6 | 30360068 | rs3094700 | -0.277 | -0.145 | -1.161 | -0.436 | 0.023 | 0.072 | 0.072 | 0 | 0.49 | 0.71 | 0.47 | 0.69 | 0.44 | 0.51 |
| 6 | 30361193 | rs3094698 | -0.277 | -0.145 | -1.161 | -0.436 | 0.023 | 0.072 | 0.072 | 0 | 0.49 | 0.71 | 0.47 | 0.69 | 0.44 | 0.51 |
| 6 | 30361358 | rs9391683 | 0.007 | -0.145 | -1.16 | -0.436 | 0.024 | 0.021 | 0.021 | 0 | 0.47 | 0.58 | 0.4 | 0.69 | 0.34 | 0.51 |
| 6 | 30361418 | rs3094042 | -0.209 | -0.194 | -1.586 | -0.364 | 0.023 | 0.073 | 0.073 | 0 | 0.49 | 0.71 | 0.47 | 0.69 | 0.44 | 0.51 |
| 6 | 30361480 | rs3094041 | -0.209 | -0.194 | -1.591 | -0.364 | 0.023 | 0.072 | 0.072 | 0 | 0.49 | 0.71 | 0.47 | 0.69 | 0.44 | 0.51 |
| 6 | 30361608 | rs2516685 | -1.5 | -0.222 | -1.635 | -0.401 | -0.021 | 0.013 | 0.013 | 0 | 0.68 | 0.75 | 0.64 | 0.78 | 0.59 | 0.71 |
| 6 | 30361759 | rs3094040 | -0.19 | -0.188 | -1.635 | -0.383 | 0.023 | 0.072 | 0.072 | 0 | 0.49 | 0.71 | 0.47 | 0.69 | 0.44 | 0.51 |
| 6 | 30362196 | rs3130107 | -0.19 | -0.188 | -1.635 | -0.383 | 0.023 | 0.072 | 0.072 | 0 | 0.49 | 0.71 | 0.47 | 0.69 | 0.44 | 0.51 |
| 6 | 30362751 | rs3130109 | -0.19 | -0.188 | -1.635 | -0.383 | 0.023 | 0.072 | 0.072 | 0 | 0.49 | 0.71 | 0.47 | 0.69 | 0.44 | 0.51 |
| 6 | 30362961 | rs2187978 | 2.145 | -0.198 | -1.64 | -0.387 | -0.006 | 0.038 | 0.038 | 0 | 0.14 | 0.02 | 0.16 | 0.04 | 0.14 | 0.08 |
| 6 | 30362995 | rs2516684 | 1.886 | -0.121 | -1.592 | -0.348 | 0.009 | 0.025 | 0.025 | 0 | 0.17 | 0.02 | 0.17 | 0.09 | 0.15 | 0.21 |
| 6 | 30363428 | rs3130112 | -0.345 | -0.138 | -1.348 | -0.188 | 0.023 | 0.072 | 0.072 | 0 | 0.49 | 0.71 | 0.47 | 0.69 | 0.44 | 0.51 |
| 6 | 30363945 | rs3130113 | -0.345 | -0.138 | -1.348 | -0.188 | 0.023 | 0.072 | 0.072 | 0 | 0.49 | 0.71 | 0.47 | 0.69 | 0.44 | 0.51 |
| 6 | 30364508 | rs17475879 | 2.638 | 0.278 | -0.974 | -0.022 | 0.014 | 0.021 | 0.021 | 0 | 0.12 | 0.04 | 0.03 | 0.04 | 0.07 | 0.15 |
| 6 | 30366100 | rs1264567 | 1.249 | 0.53 | -1.025 | -0.246 | -0.001 | 0.034 | 0.034 | 0 | 0.18 | 0.14 | 0.19 | 0.07 | 0.2 | 0.15 |
| 6 | 30366642 | rs3131113 | 0.352 | 0.262 | -0.739 | 0.102 | 0.005 | 0.003 | 0.003 | 0 | 0.27 | 0.24 | 0.19 | 0.4 | 0.31 | 0.35 |
| 6 | 30368012 | rs12203316 | 0.612 | 0.196 | -0.703 | 0.255 | 0.027 | 0.011 | 0.011 | 0 | 0.13 | 0.21 | 0.22 | 0.16 | 0.18 | 0.1 |
| 6 | 30368635 | rs1034323 | -0.398 | 0.141 | -0.678 | 0.214 | 0.005 | 0.011 | 0.011 | 0 | 0.39 | 0.47 | 0.39 | 0.56 | 0.41 | 0.44 |
| 6 | 30369040 | rs9261743 | 0.716 | 0.191 | -0.734 | 0.226 | -0.012 | 0.008 | 0.008 | 0 | 0.26 | 0.26 | 0.16 | 0.4 | 0.23 | 0.34 |
| 6 | 30369958 | rs9261744 | 0.716 | 0.191 | -0.734 | 0.226 | -0.012 | 0.008 | 0.008 | 0 | 0.26 | 0.26 | 0.16 | 0.4 | 0.23 | 0.34 |
| 6 | 30370588 | rs1419689 | -0.817 | -0.45 | -1.004 | -0.445 | -0.002 | 0.004 | 0.004 | 0 | 0.39 | 0.63 | 0.36 | 0.25 | 0.35 | 0.28 |
| 6 | 30372278 | rs10947072 | 0.06 | -0.45 | -1.011 | -0.447 | -0.001 | 0.001 | 0.001 | 0 | 0.24 | 0.37 | 0.25 | 0.22 | 0.21 | 0.16 |
| 6 | 30372987 | rs9261761 | 0.887 | -0.257 | -0.823 | -0.427 | 0.023 | -0.045 | -0.045 | 0 | 0.15 | 0.25 | 0.11 | 0.03 | 0.15 | 0.12 |
| 6 | 30377173 | rs1264551 | -2.282 | -0.534 | -1.076 | -0.624 | -0.004 | -0.032 | -0.032 | 0.335 | 0.77 | 0.77 | 0.82 | 0.9 | 0.79 | 0.75 |
| 6 | 30378601 | rs9261817 | 0.991 | -0.422 | -0.99 | -0.545 | 0.023 | -0.045 | -0.045 | 0.156 | 0.15 | 0.25 | 0.11 | 0.03 | 0.15 | 0.12 |
| 6 | 30379074 | rs9261821 | 0.976 | -0.419 | -0.988 | -0.543 | 0.023 | -0.046 | -0.046 | 0.018 | 0.15 | 0.25 | 0.11 | 0.02 | 0.15 | 0.12 |
| 6 | 30379965 | rs9261827 | 1.085 | -0.442 | -1.016 | -0.555 | 0.028 | -0.042 | -0.042 | 0 | 0.14 | 0.25 | 0.11 | 0.03 | 0.15 | 0.12 |

| | | | | | | | | | | | | | | | | |
|---|---|---|---|---|---|---|---|---|---|---|---|---|---|---|---|---|
| 6 | 30382647 | rs9261847 | 0.951 | -0.433 | -1.028 | -0.551 | 0.024 | -0.046 | -0.046 | 0.052 | 0.15 | 0.26 | 0.11 | 0.02 | 0.15 | 0.11 |
| 6 | 30383768 | rs4713327 | -0.167 | -0.521 | -1.103 | -0.571 | 0.022 | -0.014 | -0.014 | 0 | 0.33 | 0.54 | 0.35 | 0.2 | 0.34 | 0.21 |
| 6 | 30384322 | rs2074502 | 1.138 | -0.525 | -1.109 | -0.569 | 0.005 | 0.003 | 0.003 | 0 | 0.18 | 0.28 | 0.24 | 0.18 | 0.19 | 0.1 |
| 6 | 30385057 | rs17195089 | 1.182 | -0.521 | -1.107 | -0.567 | 0.005 | 0.003 | 0.003 | 0 | 0.18 | 0.28 | 0.24 | 0.2 | 0.2 | 0.1 |
| 6 | 30385111 | rs17188743 | 1.167 | -0.524 | -1.091 | -0.568 | 0.006 | 0.004 | 0.004 | 0 | 0.18 | 0.28 | 0.24 | 0.2 | 0.19 | 0.1 |
| 6 | 30386415 | rs1573294 | 1.12 | -0.524 | -1.103 | -0.568 | 0.024 | -0.046 | -0.046 | 0 | 0.15 | 0.26 | 0.11 | 0.03 | 0.15 | 0.11 |
| 6 | 30386996 | rs1264542 | -1.598 | -0.72 | -1.173 | -0.754 | 0.016 | -0.015 | -0.015 | 0 | 0.69 | 0.66 | 0.62 | 0.79 | 0.61 | 0.7 |
| 6 | 30389258 | rs1264538 | 0.58 | -0.752 | -1.185 | -0.773 | 0.003 | 0.016 | 0.016 | 0 | 0.42 | 0.2 | 0.47 | 0.7 | 0.44 | 0.53 |
| 6 | 30389323 | rs9261895 | 1.397 | -0.79 | -1.2 | -0.79 | 0.018 | -0.049 | -0.049 | 0.234 | 0.16 | 0.26 | 0.11 | 0.02 | 0.15 | 0.11 |
| 6 | 30391723 | rs9261919 | 1.234 | -0.767 | -1.187 | -0.776 | 0.024 | -0.046 | -0.046 | 0 | 0.15 | 0.26 | 0.11 | 0.03 | 0.15 | 0.11 |
| 6 | 30392146 | rs9261922 | 0.489 | -0.772 | -1.184 | -0.779 | -0.033 | -0.058 | -0.058 | 0 | 0.26 | 0.46 | 0.14 | 0.09 | 0.17 | 0.17 |
| 6 | 30392160 | rs9261923 | 1.233 | -0.763 | -1.18 | -0.794 | 0.024 | -0.046 | -0.046 | 0 | 0.15 | 0.26 | 0.11 | 0.03 | 0.15 | 0.11 |
| 6 | 30392406 | rs9261926 | 1.233 | -0.763 | -1.18 | -0.794 | 0.024 | -0.046 | -0.046 | 0.036 | 0.15 | 0.26 | 0.11 | 0.03 | 0.15 | 0.11 |
| 6 | 30394628 | rs9261947 | 1.248 | -0.748 | -1.114 | -0.796 | 0.018 | -0.044 | -0.044 | 0.141 | 0.15 | 0.25 | 0.1 | 0.02 | 0.13 | 0.11 |
| 6 | 30394951 | rs9261950 | 1.272 | -0.766 | -1.137 | -0.802 | 0.026 | -0.045 | -0.045 | 0 | 0.15 | 0.26 | 0.11 | 0.03 | 0.15 | 0.11 |
| 6 | 30397840 | rs9501447 | 1.225 | -0.754 | -1.13 | -0.795 | 0.024 | -0.046 | -0.046 | 0 | 0.15 | 0.27 | 0.11 | 0.02 | 0.15 | 0.11 |
| 6 | 30405956 | rs1264521 | -1.158 | -0.857 | -1.03 | -0.935 | 0.004 | -0.014 | -0.014 | 0.057 | 0.88 | 0.84 | 0.87 | 0.95 | 0.85 | 0.89 |
| 6 | 30406756 | rs1079541 | 1.276 | -0.825 | -1.029 | -0.912 | 0.024 | -0.046 | -0.046 | 0.86 | 0.15 | 0.27 | 0.11 | 0.03 | 0.15 | 0.11 |
| 6 | 30407270 | rs9461604 | 1.276 | -0.825 | -1.029 | -0.912 | 0.024 | -0.046 | -0.046 | 0 | 0.15 | 0.27 | 0.11 | 0.03 | 0.15 | 0.11 |
| 6 | 30408265 | rs9295863 | 1.276 | -0.825 | -1.029 | -0.912 | 0.024 | -0.046 | -0.046 | 0 | 0.15 | 0.27 | 0.11 | 0.03 | 0.15 | 0.11 |
| 6 | 30408787 | rs1269556 | 0.779 | -0.961 | -1.111 | -0.891 | -0.008 | -0.016 | -0.016 | 0.318 | 0.4 | 0.19 | 0.46 | 0.66 | 0.43 | 0.44 |
| 6 | 30410206 | rs1264518 | 2.683 | -0.919 | -1.284 | -1.02 | -0.007 | 0.06 | 0.06 | 0 | 0.17 | 0.07 | 0.19 | 0.09 | 0.18 | 0.15 |
| 6 | 30411089 | rs9295871 | 1.294 | -0.583 | -0.968 | -0.776 | 0.024 | -0.046 | -0.046 | 0 | 0.15 | 0.27 | 0.11 | 0.03 | 0.15 | 0.11 |
| 6 | 30411546 | rs1268444 | -1.444 | -0.568 | -0.985 | -0.768 | 0.013 | -0.014 | -0.014 | 0 | 0.68 | 0.65 | 0.62 | 0.79 | 0.61 | 0.7 |
| 6 | 30412017 | rs3094716 | 0.581 | -0.548 | -1.021 | -0.739 | 0.003 | 0 | 0 | 0 | 0.42 | 0.19 | 0.46 | 0.67 | 0.44 | 0.53 |
| 6 | 30412362 | rs1264513 | 0.581 | -0.548 | -1.021 | -0.739 | 0.003 | 0 | 0 | 0 | 0.42 | 0.19 | 0.46 | 0.67 | 0.44 | 0.53 |
| 6 | 30414011 | rs1264510 | -1.575 | -0.537 | -0.96 | -0.742 | 0.014 | -0.017 | -0.017 | 1.822 | 0.91 | 0.86 | 0.87 | 0.99 | 0.86 | 0.96 |
| 6 | 30414235 | rs9295873 | 1.259 | -0.506 | -0.956 | -0.734 | 0.024 | -0.046 | -0.046 | 0 | 0.15 | 0.27 | 0.11 | 0.02 | 0.15 | 0.11 |
| 6 | 30414848 | rs1362115 | 0.54 | -0.55 | -0.958 | -0.743 | -0.033 | -0.043 | -0.043 | 0.103 | 0.26 | 0.46 | 0.15 | 0.13 | 0.17 | 0.17 |
| 6 | 30415636 | rs2105961 | 0.599 | -0.555 | -0.975 | -0.751 | 0.003 | 0 | 0 | 0 | 0.42 | 0.19 | 0.46 | 0.67 | 0.44 | 0.53 |
| 6 | 30416553 | rs4959047 | 0.52 | -0.517 | -0.944 | -0.723 | -0.034 | -0.044 | -0.044 | 0.23 | 0.26 | 0.46 | 0.15 | 0.13 | 0.17 | 0.17 |
| 6 | 30416831 | rs4959050 | 0.509 | -0.521 | -0.946 | -0.726 | -0.034 | -0.045 | -0.045 | 0.441 | 0.26 | 0.46 | 0.15 | 0.13 | 0.17 | 0.17 |
| 6 | 30417704 | rs9468768 | 1.556 | -0.512 | -0.941 | -0.723 | 0.021 | -0.047 | -0.047 | 0 | 0.15 | 0.27 | 0.11 | 0.02 | 0.15 | 0.11 |
| 6 | 30418662 | rs4947284 | 2.294 | -0.504 | -0.946 | -0.708 | -0.032 | 0.001 | 0.001 | 0 | 0.11 | 0.19 | 0.04 | 0.1 | 0.03 | 0.06 |
| 6 | 30419831 | rs1076832 | 2.517 | -0.412 | -0.955 | -0.682 | 0.028 | 0.003 | 0.003 | 0 | 0.07 | 0.11 | 0.19 | 0.1 | 0.16 | 0.04 |
| 6 | 30421397 | rs9295886 | 1.34 | -0.523 | -0.936 | -0.681 | 0.018 | -0.049 | -0.049 | 0 | 0.16 | 0.27 | 0.11 | 0.02 | 0.15 | 0.11 |
| 6 | 30421502 | rs2524198 | -1.382 | -0.541 | -0.948 | -0.699 | 0.013 | -0.014 | -0.014 | 0 | 0.68 | 0.65 | 0.62 | 0.79 | 0.61 | 0.7 |
| 6 | 30422632 | rs2844729 | -1.382 | -0.541 | -0.948 | -0.699 | 0.013 | -0.014 | -0.014 | 0 | 0.68 | 0.65 | 0.62 | 0.79 | 0.61 | 0.7 |
| 6 | 30423117 | rs17411480 | 1.265 | -0.495 | -0.95 | -0.702 | 0.025 | -0.045 | -0.045 | 2.59 | 0.15 | 0.27 | 0.11 | 0.02 | 0.15 | 0.11 |
| 6 | 30423223 | rs35407515 | 1.526 | -0.557 | -0.936 | -0.687 | 0.021 | -0.047 | -0.047 | 0 | 0.15 | 0.27 | 0.11 | 0.02 | 0.15 | 0.11 |
| 6 | 30423358 | rs34101875 | 1.492 | -0.558 | -0.936 | -0.687 | 0.02 | -0.048 | -0.048 | 1.454 | 0.15 | 0.27 | 0.11 | 0.02 | 0.15 | 0.11 |
| 6 | 30424074 | rs33986393 | 1.525 | -0.554 | -0.935 | -0.685 | 0.021 | -0.047 | -0.047 | 0 | 0.15 | 0.27 | 0.11 | 0.02 | 0.15 | 0.11 |
| 6 | 30424623 | rs2516677 | -0.051 | -0.569 | -0.969 | -0.669 | 0.025 | 0.071 | 0.071 | 0 | 0.53 | 0.39 | 0.51 | 0.77 | 0.47 | 0.59 |
| 6 | 30425157 | rs2187975 | 0.677 | -0.593 | -0.976 | -0.667 | 0.001 | -0.001 | -0.001 | 1.595 | 0.42 | 0.19 | 0.46 | 0.66 | 0.44 | 0.53 |
| 6 | 30425337 | rs2187974 | -0.155 | -0.564 | -0.971 | -0.697 | 0.02 | 0.065 | 0.065 | 0 | 0.54 | 0.39 | 0.51 | 0.77 | 0.47 | 0.59 |
| 6 | 30426130 | rs2524180 | -1.316 | -0.459 | -0.907 | -0.656 | 0.013 | -0.014 | -0.014 | 0 | 0.68 | 0.65 | 0.62 | 0.79 | 0.61 | 0.7 |
| 6 | 30426152 | rs2524179 | 0.033 | -0.488 | -0.907 | -0.684 | 0.025 | 0.071 | 0.071 | 0 | 0.53 | 0.39 | 0.51 | 0.77 | 0.47 | 0.59 |
| 6 | 30427510 | rs9501336 | 1.406 | -0.505 | -0.696 | -0.714 | 0.021 | -0.047 | -0.047 | 0 | 0.15 | 0.27 | 0.11 | 0.02 | 0.15 | 0.11 |
| 6 | 30428351 | rs2077573 | 0.409 | -0.561 | -0.724 | -0.712 | -0.03 | -0.04 | -0.04 | 0.581 | 0.25 | 0.45 | 0.16 | 0.13 | 0.17 | 0.17 |
| 6 | 30429033 | rs11966619 | 1.347 | -0.26 | -0.762 | -0.635 | 0.021 | -0.047 | -0.047 | 0.021 | 0.15 | 0.27 | 0.11 | 0.03 | 0.15 | 0.11 |
| 6 | 30430482 | rs2516676 | -0.767 | -0.223 | -0.721 | -0.552 | 0.005 | -0.017 | -0.017 | 0 | 0.88 | 0.84 | 0.86 | 0.95 | 0.85 | 0.93 |
| 6 | 30430520 | rs2516675 | -0.632 | -0.341 | -0.815 | -0.599 | -0.033 | -0.044 | -0.044 | 0.157 | 0.63 | 0.56 | 0.76 | 0.8 | 0.74 | 0.64 |
| 6 | 30433611 | rs2524172 | -1.475 | 0.486 | -0.606 | -0.212 | 0.02 | -0.012 | -0.012 | 0 | 0.93 | 0.89 | 0.88 | 0.99 | 0.87 | 0.96 |
| 6 | 30434566 | rs7764934 | -0.231 | 0.138 | -0.831 | 0.198 | 0.004 | 0.005 | 0.005 | 0 | 0.38 | 0.38 | 0.38 | 0.57 | 0.41 | 0.5 |
| 6 | 30434900 | rs7747873 | -0.246 | 0.135 | -0.833 | 0.204 | 0.003 | 0.004 | 0.004 | 0 | 0.38 | 0.38 | 0.38 | 0.58 | 0.41 | 0.5 |
| 6 | 30434999 | rs2516670 | -1.233 | 0.213 | -0.781 | 0.12 | 0.018 | -0.018 | -0.018 | 0.482 | 0.91 | 0.86 | 0.88 | 0.99 | 0.86 | 0.96 |
| 6 | 30435288 | rs6930977 | 1.954 | 0.437 | -0.806 | 0.264 | 0.007 | -0.002 | -0.002 | 0 | 0.08 | 0.09 | 0.19 | 0.1 | 0.16 | 0.08 |
| 6 | 30435801 | rs2516669 | -2.024 | 0.052 | -0.864 | 0.01 | -0.004 | -0.035 | -0.035 | 1.113 | 0.76 | 0.76 | 0.82 | 0.91 | 0.79 | 0.78 |
| 6 | 30436681 | rs12174870 | -0.038 | 0.126 | -0.855 | 0.171 | 0.005 | 0.009 | 0.009 | 0 | 0.35 | 0.37 | 0.36 | 0.57 | 0.39 | 0.42 |
| 6 | 30438226 | rs9295895 | 0.62 | 0.101 | -0.864 | 0.216 | -0.011 | 0.019 | 0.019 | 0 | 0.27 | 0.26 | 0.16 | 0.45 | 0.22 | 0.34 |
| 6 | 30446055 | rs9295897 | -0.161 | 0.104 | -0.863 | 0.213 | 0.004 | 0.008 | 0.008 | 0 | 0.36 | 0.37 | 0.36 | 0.57 | 0.39 | 0.42 |
| 6 | 30448668 | rs6909620 | 0.507 | 0.095 | -0.853 | 0.221 | -0.011 | 0.019 | 0.019 | 0 | 0.27 | 0.26 | 0.16 | 0.45 | 0.21 | 0.33 |
| 6 | 30449046 | rs9295898 | -0.057 | -0.041 | -0.881 | 0.237 | 0.005 | 0.008 | 0.008 | 0 | 0.35 | 0.37 | 0.36 | 0.57 | 0.39 | 0.42 |
| 6 | 30449482 | rs2021719 | -0.203 | -0.036 | -0.835 | 0.239 | -0.002 | -0.003 | -0.003 | 0 | 0.35 | 0.3 | 0.35 | 0.57 | 0.39 | 0.42 |
| 6 | 30451515 | rs28780106 | 2.977 | -0.011 | -1.05 | 0.575 | -0.006 | -0.016 | -0.016 | 0.035 | 0.12 | 0.08 | 0.04 | 0.04 | 0.07 | 0.15 |
| 6 | 30451904 | rs3094694 | 0.809 | 0.34 | -1.168 | 0.957 | -0.008 | -0.008 | -0.008 | 0 | 0.11 | 0.12 | 0.22 | 0.14 | 0.17 | 0.07 |
| 6 | 30453713 | rs16897572 | 1.248 | -0.038 | -1.14 | 1.023 | 0.036 | -0.075 | -0.075 | 0 | 0.11 | 0.01 | 0.05 | 0.42 | 0.08 | 0.22 |
| 6 | 30455886 | rs762324 | 0.794 | -0.106 | -1.113 | 1.08 | 0.057 | -0.057 | -0.057 | 0.483 | 0.15 | 0.02 | 0.07 | 0.28 | 0.08 | 0.22 |
| 6 | 30455920 | rs1264459 | 0.393 | 0.63 | -0.671 | 0.759 | 0.001 | -0.023 | -0.023 | 0.524 | 0.86 | 0.8 | 0.83 | 0.95 | 0.83 | 0.92 |

| | | | | | | | | | | | | | | | | |
|---|---|---|---|---|---|---|---|---|---|---|---|---|---|---|---|---|
| 6 | 30457732 | rs1059510 | 0.459 | 0.94 | -0.641 | 0.975 | 0.03 | 0.02 | 0.02 | 0 | 0.81 | 0.74 | 0.64 | 0.71 | 0.67 | 0.84 |
| 6 | 30465172 | rs2844725 | 0.474 | 0.634 | -0.684 | 0.865 | 0.008 | 0.004 | 0.004 | 0 | 0.78 | 0.77 | 0.59 | 0.62 | 0.67 | 0.77 |
| 6 | 30465512 | rs2508024 | 0.766 | 0.567 | -0.62 | 0.874 | 0.022 | 0.017 | 0.017 | 0 | 0.76 | 0.68 | 0.56 | 0.6 | 0.65 | 0.76 |
| 6 | 30468791 | rs3131115 | 0.63 | -0.047 | -0.393 | 0.455 | -0.003 | 0.002 | 0.002 | 0 | 0.35 | 0.37 | 0.19 | 0.35 | 0.3 | 0.37 |
| 6 | 30469190 | rs2844724 | 0.112 | 0.081 | -0.32 | 0.502 | 0.008 | 0.004 | 0.004 | 0 | 0.78 | 0.77 | 0.59 | 0.62 | 0.67 | 0.77 |
| 6 | 30475717 | rs2844720 | 0.098 | 0.08 | -0.321 | 0.501 | 0.008 | 0.004 | 0.004 | 0 | 0.78 | 0.77 | 0.59 | 0.62 | 0.67 | 0.77 |
| 6 | 30476034 | rs2023609 | 0.098 | 0.08 | -0.321 | 0.501 | 0.008 | 0.004 | 0.004 | 0 | 0.78 | 0.77 | 0.59 | 0.62 | 0.67 | 0.77 |
| 6 | 30476185 | rs2023608 | 0.115 | 0.12 | -0.321 | 0.501 | 0.008 | 0.004 | 0.004 | 0 | 0.78 | 0.77 | 0.59 | 0.62 | 0.67 | 0.76 |
| 6 | 30477947 | rs975195 | 0.115 | 0.105 | -0.321 | 0.501 | 0.008 | 0.004 | 0.004 | 0 | 0.78 | 0.77 | 0.59 | 0.62 | 0.67 | 0.77 |
| 6 | 30479180 | rs2844717 | 0.098 | 0.126 | -0.323 | 0.498 | 0.008 | 0.004 | 0.004 | 0 | 0.78 | 0.77 | 0.59 | 0.62 | 0.67 | 0.76 |
| 6 | 30482993 | rs2534812 | 0.546 | 0.125 | -0.323 | 0.498 | 0.039 | 0.031 | 0.031 | 0 | 0.76 | 0.61 | 0.55 | 0.6 | 0.65 | 0.76 |
| 6 | 30486456 | rs2534804 | -0.916 | 0.079 | -0.291 | 0.499 | -0.007 | -0.008 | -0.008 | 0 | 0.85 | 0.87 | 0.64 | 0.72 | 0.7 | 0.86 |
| 6 | 30495743 | rs2534825 | 0.275 | 0.209 | -0.286 | 0.474 | 0.008 | 0.004 | 0.004 | 0 | 0.78 | 0.77 | 0.59 | 0.62 | 0.67 | 0.76 |
| 6 | 30496882 | rs2516649 | 0.277 | 0.216 | -0.287 | 0.477 | 0.008 | 0.004 | 0.004 | 0 | 0.78 | 0.77 | 0.59 | 0.62 | 0.67 | 0.76 |
| 6 | 30497490 | rs2524226 | -0.441 | 0.211 | -0.282 | 0.477 | -0.004 | -0.005 | -0.005 | 0.333 | 0.82 | 0.84 | 0.62 | 0.7 | 0.69 | 0.83 |
| 6 | 30498864 | rs2534816 | 1.639 | -0.122 | -0.659 | 0.171 | -0.042 | 0.035 | 0.035 | 0 | 0.13 | 0.01 | 0.14 | 0.05 | 0.17 | 0.16 |
| 6 | 30500536 | rs3130039 | 0.097 | -0.024 | -0.502 | 0.274 | 0.008 | 0.004 | 0.004 | 0 | 0.78 | 0.77 | 0.59 | 0.62 | 0.67 | 0.76 |
| 6 | 30503945 | rs2516644 | 0.036 | -0.024 | -0.475 | 0.3 | 0.01 | 0.006 | 0.006 | 0 | 0.79 | 0.77 | 0.59 | 0.62 | 0.67 | 0.76 |
| 6 | 30506383 | rs1468079 | 0.623 | -0.043 | -0.481 | 0.291 | 0.039 | 0.031 | 0.031 | 0 | 0.75 | 0.62 | 0.55 | 0.6 | 0.66 | 0.76 |
| 6 | 30512163 | rs1058318 | -0.575 | 0.298 | -0.527 | 0.297 | -0.003 | -0.004 | -0.004 | 1.206 | 0.82 | 0.84 | 0.62 | 0.7 | 0.69 | 0.84 |
| 6 | 30518619 | rs2844714 | 0.425 | 0.265 | -0.547 | 0.3 | 0.023 | 0.018 | 0.018 | 0 | 0.76 | 0.68 | 0.56 | 0.6 | 0.66 | 0.76 |
| 6 | 30521137 | rs2074505 | 0.353 | 0.262 | -0.493 | 0.347 | 0.031 | 0.021 | 0.021 | 0 | 0.77 | 0.66 | 0.56 | 0.62 | 0.67 | 0.76 |
| 6 | 30526845 | rs2516647 | 0.636 | 0.012 | -0.082 | 0.252 | 0.034 | 0.034 | 0.034 | 0 | 0.72 | 0.62 | 0.55 | 0.59 | 0.65 | 0.76 |
| 6 | 30530496 | rs2074503 | -0.628 | 0.04 | 0.037 | 0.269 | -0.006 | -0.006 | -0.006 | 0 | 0.82 | 0.84 | 0.62 | 0.7 | 0.69 | 0.84 |
| 6 | 30532278 | rs9262113 | 1.623 | 0.138 | -0.158 | 0.181 | -0.038 | 0.027 | 0.027 | 0 | 0.14 | 0.01 | 0.14 | 0.08 | 0.17 | 0.17 |
| 6 | 30536071 | rs3130240 | -0.384 | 0.214 | -0.033 | 0.251 | -0.019 | -0.011 | -0.011 | 0 | 0.79 | 0.81 | 0.61 | 0.69 | 0.68 | 0.83 |
| 6 | 30536133 | rs3130041 | 0.139 | 0.21 | -0.023 | 0.252 | 0.001 | 0 | 0 | 0 | 0.77 | 0.72 | 0.58 | 0.67 | 0.67 | 0.83 |
| 6 | 30537339 | rs3130242 | -0.559 | 0.211 | -0.025 | 0.243 | -0.007 | -0.006 | -0.006 | 1.811 | 0.82 | 0.84 | 0.61 | 0.7 | 0.69 | 0.84 |
| 6 | 30537853 | rs3130243 | 0.138 | 0.21 | -0.023 | 0.243 | 0.001 | 0 | 0 | 0.736 | 0.77 | 0.72 | 0.58 | 0.67 | 0.67 | 0.83 |
| 6 | 30539952 | rs2269710 | 0.337 | 0.144 | -0.114 | 0.196 | -0.006 | -0.004 | -0.004 | 0 | 0.75 | 0.72 | 0.57 | 0.67 | 0.66 | 0.82 |
| 6 | 30540890 | rs2269709 | 0.087 | 0.302 | -0.059 | 0.263 | 0.002 | 0 | 0 | 0.115 | 0.78 | 0.73 | 0.58 | 0.67 | 0.67 | 0.83 |
| 6 | 30541852 | rs3132611 | -2.654 | 0.71 | -0.088 | 0.282 | -0.031 | 0.016 | 0.016 | 0.267 | 0.95 | 1 | 0.94 | 0.99 | 0.91 | 0.98 |
| 6 | 30544941 | rs732229 | -0.513 | 0.533 | -0.056 | 0.3 | 0.002 | -0.002 | -0.002 | 0 | 0.83 | 0.84 | 0.63 | 0.7 | 0.7 | 0.86 |
| 6 | 30545022 | rs3130043 | 0.991 | 0.531 | 0.066 | 0.284 | 0.059 | 0.04 | 0.04 | 0 | 0.74 | 0.53 | 0.53 | 0.6 | 0.66 | 0.79 |
| 6 | 30551286 | rs1264440 | 0.082 | 0.557 | 0.002 | 0.294 | 0.009 | 0.004 | 0.004 | 0 | 0.79 | 0.73 | 0.59 | 0.67 | 0.68 | 0.85 |
| 6 | 30552502 | rs1264439 | -0.925 | 0.554 | 0.123 | 0.306 | 0.015 | 0.003 | 0.003 | 0 | 0.86 | 0.86 | 0.64 | 0.72 | 0.7 | 0.89 |
| 6 | 30552937 | rs3130244 | 0.835 | 0.532 | 0.168 | 0.295 | 0.064 | 0.046 | 0.046 | 0 | 0.76 | 0.56 | 0.54 | 0.6 | 0.66 | 0.79 |
| 6 | 30553871 | rs1264437 | -0.521 | 0.579 | 0.001 | 0.342 | 0.003 | -0.001 | -0.001 | 0.01 | 0.83 | 0.84 | 0.63 | 0.67 | 0.7 | 0.86 |
| 6 | 30560796 | rs13195066 | 1.562 | 0.31 | -0.497 | 0.379 | -0.056 | -0.061 | -0.061 | 0 | 0.12 | 0.02 | 0.14 | 0.13 | 0.19 | 0.07 |
| 6 | 30562021 | rs1264432 | -0.725 | 0.358 | -0.408 | 0.371 | 0.003 | -0.001 | -0.001 | 0.015 | 0.83 | 0.84 | 0.63 | 0.7 | 0.7 | 0.87 |
| 6 | 30569489 | rs7759666 | 1.719 | 0.144 | -0.675 | 0.212 | -0.031 | 0.031 | 0.031 | 0 | 0.15 | 0.01 | 0.15 | 0.06 | 0.18 | 0.2 |
| 6 | 30571471 | rs1264423 | -0.38 | 0.084 | -0.636 | 0.248 | -0.055 | -0.053 | -0.053 | 0.826 | 0.74 | 0.83 | 0.49 | 0.59 | 0.52 | 0.82 |
| 6 | 30573842 | rs9262122 | 1.803 | 0.088 | -0.652 | 0.214 | -0.034 | 0.031 | 0.031 | 0 | 0.15 | 0.01 | 0.15 | 0.06 | 0.18 | 0.2 |
| 6 | 30575603 | rs1264420 | -0.853 | 0.323 | -0.692 | 0.336 | -0.021 | -0.012 | -0.012 | 0 | 0.82 | 0.88 | 0.75 | 0.83 | 0.68 | 0.88 |
| 6 | 30578335 | rs2267637 | 1.218 | -0.415 | -0.633 | 0.026 | -0.049 | 0.007 | 0.007 | 0 | 0.29 | 0.32 | 0.12 | 0.34 | 0.11 | 0.35 |
| 6 | 30579315 | rs2252745 | -1.099 | -0.16 | -0.887 | 0.082 | 0.016 | 0.004 | 0.004 | 0 | 0.85 | 0.84 | 0.63 | 0.69 | 0.7 | 0.88 |
| 6 | 30580637 | rs2267639 | 2.021 | -0.807 | -0.948 | 0.239 | 0.069 | -0.051 | -0.051 | 0 | 0.19 | 0.03 | 0.07 | 0.26 | 0.09 | 0.25 |
| 6 | 30592414 | rs2270172 | 1.123 | -0.134 | -0.709 | 0.267 | -0.011 | -0.002 | -0.002 | 0 | 0.07 | 0.13 | 0.04 | 0.05 | 0.02 | 0.03 |
| 6 | 30599702 | rs4713340 | 1.741 | -0.105 | -0.771 | 0.228 | -0.031 | 0.04 | 0.04 | 0 | 0.15 | 0.01 | 0.15 | 0.05 | 0.18 | 0.19 |
| 6 | 30608273 | rs1079687 | 0.115 | -0.061 | -0.357 | 0.344 | -0.024 | -0.005 | -0.005 | 0.164 | 0.09 | 0.22 | 0.07 | 0.08 | 0.04 | 0.09 |
| 6 | 30609629 | rs3094104 | -0.302 | -0.078 | -0.313 | 0.343 | -0.032 | 0.004 | 0.004 | 0 | 0.1 | 0.26 | 0.1 | 0.12 | 0.05 | 0.09 |
| 6 | 30609661 | rs3132608 | 0.108 | -0.02 | -0.521 | 0.34 | -0.024 | -0.005 | -0.005 | 0 | 0.09 | 0.22 | 0.07 | 0.08 | 0.04 | 0.09 |
| 6 | 30611676 | rs1140809 | -0.477 | 0.304 | -0.892 | 0.485 | -0.052 | -0.054 | -0.054 | 0 | 0.73 | 0.82 | 0.49 | 0.55 | 0.51 | 0.81 |
| 6 | 30627216 | rs1076829 | 1.366 | 0.336 | -0.87 | -0.312 | -0.017 | 0.074 | 0.074 | 0 | 0.32 | 0.16 | 0.25 | 0.09 | 0.35 | 0.36 |
| 6 | 30628082 | rs3130000 | -2.594 | 0.546 | -0.944 | -0.326 | -0.03 | 0.016 | 0.016 | 0.402 | 0.95 | 1 | 0.94 | 0.99 | 0.92 | 0.98 |
| 6 | 30628252 | rs1076828 | 1.352 | 0.398 | -0.928 | -0.36 | -0.017 | 0.074 | 0.074 | 0 | 0.32 | 0.16 | 0.25 | 0.09 | 0.35 | 0.36 |
| 6 | 30633875 | rs3094097 | -0.764 | -0.093 | -0.138 | -0.038 | -0.087 | 0.004 | 0.004 | 0 | 0.19 | 0.4 | 0.13 | 0.19 | 0.06 | 0.16 |
| 6 | 30650318 | rs2394392 | -0.146 | 0.608 | -0.233 | 0.368 | -0.021 | 0.001 | 0.001 | 0 | 0.09 | 0.22 | 0.1 | 0.08 | 0.06 | 0.09 |
| 6 | 30651587 | rs3129996 | -1.628 | 0.655 | -0.573 | 0.225 | -0.031 | 0.035 | 0.035 | 0.012 | 0.92 | 0.99 | 0.93 | 0.99 | 0.91 | 0.98 |
| 6 | 30657015 | rs3094092 | -2.438 | 0.646 | -0.49 | 0.309 | -0.026 | 0.018 | 0.018 | 0.279 | 0.95 | 1 | 0.93 | 0.99 | 0.92 | 0.98 |
| 6 | 30658239 | rs1075496 | -0.381 | 0.158 | -0.406 | 0.128 | 0.021 | 0.051 | 0.051 | 0.042 | 0.49 | 0.6 | 0.39 | 0.26 | 0.43 | 0.51 |
| 6 | 30658385 | rs2269703 | 2.244 | 0.288 | -0.62 | 0.368 | 0.016 | 0.008 | 0.008 | 0 | 0.07 | 0.03 | 0.02 | 0.06 | 0.02 | 0.09 |
| 6 | 30670700 | rs2239886 | 1.538 | 0.13 | -0.427 | 0.006 | -0.035 | -0.006 | -0.006 | 0.969 | 0.1 | 0.16 | 0.07 | 0.09 | 0.04 | 0.09 |
| 6 | 30671312 | rs28994874 | 3.128 | 0.13 | -0.466 | 0.005 | 0.008 | 0.007 | 0.007 | 0 | 0.05 | 0.03 | 0.02 | 0.02 | 0.02 | 0.07 |
| 6 | 30673625 | rs28987085 | 3.127 | 0.09 | -0.478 | 0.003 | 0.008 | 0.007 | 0.007 | 0 | 0.05 | 0.03 | 0.02 | 0.02 | 0.02 | 0.07 |
| 6 | 30673681 | rs28994868 | 3.127 | 0.09 | -0.478 | 0.003 | 0.008 | 0.007 | 0.007 | 0 | 0.05 | 0.03 | 0.02 | 0.02 | 0.02 | 0.07 |
| 6 | 30679333 | rs6929626 | 1.53 | 0.04 | -0.421 | -0.018 | -0.038 | -0.006 | -0.006 | 0 | 0.1 | 0.16 | 0.07 | 0.09 | 0.04 | 0.09 |
| 6 | 30680916 | rs9262152 | 1.261 | -0.284 | -0.832 | -0.311 | -0.031 | 0.031 | 0.031 | 0 | 0.14 | 0.01 | 0.14 | 0.06 | 0.18 | 0.19 |
| 6 | 30681477 | rs28986464 | 3.121 | -0.422 | -0.405 | -0.012 | 0.008 | 0.007 | 0.007 | 0 | 0.05 | 0.03 | 0.02 | 0.02 | 0.02 | 0.07 |

| | | | | | | | | | | | | | | | | |
|---|---|---|---|---|---|---|---|---|---|---|---|---|---|---|---|---|
| 6 | 30681980 | rs6924270 | 1.504 | -0.43 | -0.391 | -0.012 | -0.038 | -0.006 | -0.006 | 0 | 0.1 | 0.16 | 0.07 | 0.09 | 0.04 | 0.09 |
| 6 | 30685066 | rs3132587 | -2.451 | -0.223 | -0.498 | -0.055 | -0.03 | 0.016 | 0.016 | 0.402 | 0.95 | 1 | 0.94 | 0.99 | 0.92 | 0.98 |
| 6 | 30685410 | rs9461624 | 0.964 | -0.78 | -0.282 | 0.094 | -0.014 | 0.027 | 0.027 | 0 | 0.05 | 0.13 | 0.05 | 0.06 | 0.02 | 0.02 |
| 6 | 30685420 | rs4713354 | -0.846 | -1.026 | -0.207 | -0.036 | -0.095 | -0.023 | -0.023 | 0 | 0.23 | 0.5 | 0.19 | 0.19 | 0.1 | 0.17 |
| 6 | 30687714 | rs9295910 | 0.407 | -0.96 | -0.002 | 0.102 | -0.045 | 0.017 | 0.017 | 0 | 0.15 | 0.22 | 0.08 | 0.18 | 0.05 | 0.14 |
| 6 | 30688427 | rs3132584 | -0.734 | -0.704 | 0.015 | 0.087 | 0.007 | 0.005 | 0.005 | 0 | 0.2 | 0.48 | 0.18 | 0.18 | 0.18 | 0.14 |
| 6 | 30690938 | rs25527 | 0.46 | -0.705 | 0.021 | 0.095 | -0.045 | 0.018 | 0.018 | 0 | 0.14 | 0.22 | 0.08 | 0.18 | 0.05 | 0.13 |
| 6 | 30692965 | rs8233 | -0.748 | -0.701 | 0.04 | 0.077 | 0.006 | 0.004 | 0.004 | 0 | 0.2 | 0.48 | 0.18 | 0.18 | 0.18 | 0.14 |
| 6 | 30693816 | rs3095329 | -0.72 | -0.692 | 0.017 | 0.081 | 0.007 | 0.005 | 0.005 | 0 | 0.2 | 0.48 | 0.18 | 0.18 | 0.18 | 0.14 |
| 6 | 30697447 | rs3094127 | -0.901 | -0.761 | -0.017 | 0.052 | -0.006 | -0.002 | -0.002 | 0 | 0.21 | 0.49 | 0.21 | 0.22 | 0.18 | 0.16 |
| 6 | 30697740 | rs9501029 | 0.439 | -0.677 | -0.034 | 0.148 | -0.046 | 0.015 | 0.015 | 0 | 0.15 | 0.22 | 0.08 | 0.18 | 0.05 | 0.13 |
| 6 | 30698541 | rs1064627 | -0.229 | -0.685 | -0.037 | 0.139 | 0.002 | 0.01 | 0.01 | 0 | 0.17 | 0.26 | 0.15 | 0.22 | 0.15 | 0.15 |
| 6 | 30700497 | rs9262155 | 1.114 | -1.658 | -0.247 | 0.129 | 0.043 | -0.05 | -0.05 | 0 | 0.26 | 0.17 | 0.32 | 0.46 | 0.09 | 0.32 |
| 6 | 30700558 | rs9969042 | 1.429 | -0.272 | 0.071 | 0.154 | -0.019 | -0.002 | -0.002 | 0 | 0.11 | 0.16 | 0.05 | 0.08 | 0.04 | 0.11 |
| 6 | 30700695 | rs9968884 | 1.429 | -0.272 | 0.071 | 0.154 | -0.019 | -0.002 | -0.002 | 0 | 0.11 | 0.16 | 0.05 | 0.08 | 0.04 | 0.11 |
| 6 | 30708428 | rs12111310 | 1.429 | -0.272 | 0.071 | 0.156 | -0.019 | -0.003 | -0.003 | 0 | 0.11 | 0.17 | 0.05 | 0.08 | 0.04 | 0.11 |
| 6 | 30709830 | rs10947088 | 1.429 | -0.49 | 0.07 | 0.156 | -0.019 | -0.002 | -0.002 | 0 | 0.11 | 0.17 | 0.05 | 0.08 | 0.04 | 0.11 |
| 6 | 30710135 | rs10947089 | 1.429 | -0.49 | 0.07 | 0.156 | -0.019 | -0.002 | -0.002 | 0 | 0.11 | 0.17 | 0.05 | 0.08 | 0.04 | 0.11 |
| 6 | 30711031 | rs17719 | 1.429 | -0.49 | 0.07 | 0.156 | -0.019 | -0.002 | -0.002 | 0 | 0.11 | 0.17 | 0.05 | 0.08 | 0.04 | 0.11 |
| 6 | 30714479 | rs2535319 | -0.823 | -0.11 | -0.099 | 0.055 | 0.015 | -0.016 | -0.016 | 0 | 0.32 | 0.53 | 0.28 | 0.27 | 0.37 | 0.23 |
| 6 | 30717637 | rs7763502 | 0.533 | -1.506 | -1.006 | 0.216 | 0.082 | -0.067 | -0.067 | 0 | 0.35 | 0.23 | 0.48 | 0.51 | 0.21 | 0.47 |
| 6 | 30720311 | rs28780111 | 1.229 | -1.695 | -1.151 | 0.225 | 0.065 | -0.095 | -0.095 | 0 | 0.28 | 0.05 | 0.4 | 0.34 | 0.16 | 0.46 |
| 6 | 30724430 | rs2394401 | -0.226 | -1.523 | -0.878 | -0.11 | 0.001 | 0.002 | 0.002 | 0 | 0.21 | 0.19 | 0.12 | 0.13 | 0.15 | 0.14 |
| 6 | 30726748 | rs3095341 | -1.789 | -1.291 | -0.579 | -0.256 | 0.049 | -0.003 | -0.003 | 0.052 | 0.78 | 0.75 | 0.76 | 0.77 | 0.64 | 0.87 |
| 6 | 30726939 | rs3095340 | 0.102 | -1.381 | -0.552 | -0.271 | 0.004 | -0.022 | -0.022 | 0.375 | 0.14 | 0.24 | 0.09 | 0.08 | 0.14 | 0.06 |
| 6 | 30728290 | rs3131036 | -0.871 | -1.688 | -0.542 | -0.41 | -0.012 | -0.056 | -0.056 | 0 | 0.28 | 0.42 | 0.22 | 0.14 | 0.26 | 0.17 |
| 6 | 30728360 | rs3094122 | 0.116 | -1.663 | -0.547 | -0.397 | 0.001 | -0.005 | -0.005 | 0 | 0.2 | 0.22 | 0.14 | 0.13 | 0.22 | 0.12 |
| 6 | 30729487 | rs11966040 | 0.751 | -1.685 | -0.535 | -0.391 | -0.027 | 0.006 | 0.006 | 0 | 0.07 | 0.15 | 0.04 | 0.06 | 0.03 | 0.04 |
| 6 | 30730267 | rs6926530 | 0.912 | -1.811 | -0.606 | -0.343 | -0.027 | 0.002 | 0.002 | 0 | 0.08 | 0.2 | 0.04 | 0.08 | 0.03 | 0.05 |
| 6 | 30730960 | rs3094121 | 0.999 | -1.812 | -0.802 | -0.299 | -0.031 | 0.009 | 0.009 | 0 | 0.16 | 0.08 | 0.11 | 0.09 | 0.21 | 0.15 |
| 6 | 30731330 | rs9262176 | 0.737 | -1.864 | -0.776 | -0.323 | -0.033 | -0.039 | -0.039 | 0 | 0.09 | 0.21 | 0.08 | 0.01 | 0.04 | 0.04 |
| 6 | 30731505 | rs13214831 | 1.96 | -1.9 | -1.213 | -0.251 | 0.066 | -0.094 | -0.094 | 0.723 | 0.28 | 0.05 | 0.4 | 0.34 | 0.17 | 0.47 |
| 6 | 30732054 | rs3129975 | 0.935 | -1.522 | -0.91 | -0.179 | -0.031 | 0.009 | 0.009 | 0 | 0.16 | 0.08 | 0.11 | 0.09 | 0.21 | 0.15 |
| 6 | 30734080 | rs3131038 | 0.935 | -1.522 | -0.91 | -0.179 | -0.031 | 0.009 | 0.009 | 0 | 0.16 | 0.08 | 0.11 | 0.09 | 0.21 | 0.15 |
| 6 | 30734255 | rs13204228 | 1.829 | -1.473 | -0.895 | -0.167 | -0.005 | 0.002 | 0.002 | 0.015 | 0.08 | 0.01 | 0.07 | 0.07 | 0.1 | 0.07 |
| 6 | 30735105 | rs12210947 | 0.784 | -1.307 | -0.869 | -0.118 | -0.059 | -0.045 | -0.045 | 0 | 0.1 | 0.04 | 0.08 | 0.2 | 0.16 | 0.13 |
| 6 | 30735229 | rs11963870 | 0.64 | -1.392 | -0.641 | -0.111 | -0.027 | 0.005 | 0.005 | 0 | 0.07 | 0.15 | 0.04 | 0.06 | 0.03 | 0.04 |
| 6 | 30736541 | rs11964596 | 0.573 | -1.394 | -0.642 | -0.112 | -0.028 | 0.004 | 0.004 | 0 | 0.07 | 0.15 | 0.04 | 0.06 | 0.03 | 0.04 |
| 6 | 30739846 | rs6911628 | -1.05 | -1.534 | -0.646 | -0.141 | -0.028 | -0.045 | -0.045 | 0 | 0.34 | 0.47 | 0.24 | 0.18 | 0.27 | 0.21 |
| 6 | 30743014 | rs17481190 | 1.768 | -1.661 | -0.988 | -0.028 | 0.063 | -0.073 | -0.073 | 0.076 | 0.28 | 0.05 | 0.4 | 0.31 | 0.16 | 0.44 |
| 6 | 30743241 | rs3130667 | -1.688 | -1.558 | -0.777 | -0.056 | 0.046 | -0.005 | -0.005 | 0.151 | 0.78 | 0.75 | 0.76 | 0.78 | 0.64 | 0.87 |
| 6 | 30744529 | rs11757629 | 1.354 | -2.062 | -1.301 | -0.023 | -0.016 | -0.002 | -0.002 | 0 | 0.09 | 0.18 | 0.05 | 0.08 | 0.04 | 0.06 |
| 6 | 30746633 | rs12526481 | -0.031 | -2.37 | -1.478 | -0.074 | 0.114 | 0.014 | 0.014 | 0.237 | 0.52 | 0.28 | 0.54 | 0.48 | 0.33 | 0.65 |
| 6 | 30747216 | rs10947091 | -0.055 | -2.369 | -1.477 | -0.073 | 0.112 | 0.012 | 0.012 | 0.307 | 0.52 | 0.28 | 0.54 | 0.48 | 0.33 | 0.68 |
| 6 | 30747802 | rs13220225 | 1.599 | -2.434 | -1.551 | -0.064 | 0.063 | -0.073 | -0.073 | 0 | 0.28 | 0.05 | 0.4 | 0.3 | 0.17 | 0.44 |
| 6 | 30749233 | rs9468829 | -0.524 | -2.49 | -1.414 | -0.088 | -0.032 | -0.011 | -0.011 | 0.536 | 0.24 | 0.27 | 0.14 | 0.17 | 0.15 | 0.19 |
| 6 | 30749712 | rs9468830 | -0.329 | -2.457 | -1.425 | -0.085 | 0.035 | 0.001 | 0.001 | 0.741 | 0.55 | 0.37 | 0.55 | 0.5 | 0.34 | 0.69 |
| 6 | 30750682 | rs7752959 | -0.753 | -2.519 | -1.256 | 0.052 | -0.037 | -0.007 | -0.007 | 0 | 0.27 | 0.32 | 0.15 | 0.19 | 0.17 | 0.25 |
| 6 | 30753146 | rs4713360 | -0.383 | -2.483 | -1.253 | 0.058 | 0.037 | 0.002 | 0.002 | 0.676 | 0.55 | 0.37 | 0.55 | 0.5 | 0.34 | 0.69 |
| 6 | 30754540 | rs12527415 | -0.384 | -2.475 | -1.307 | 0.062 | 0.041 | 0.002 | 0.002 | 0.685 | 0.55 | 0.36 | 0.55 | 0.5 | 0.34 | 0.69 |
| 6 | 30755893 | rs16897900 | -0.384 | -2.475 | -1.307 | 0.062 | 0.041 | 0.002 | 0.002 | 0.685 | 0.55 | 0.36 | 0.55 | 0.5 | 0.34 | 0.69 |
| 6 | 30756066 | rs13201769 | -0.384 | -2.475 | -1.307 | 0.062 | 0.041 | 0.002 | 0.002 | 0.685 | 0.55 | 0.36 | 0.55 | 0.5 | 0.34 | 0.69 |
| 6 | 30756361 | rs4713366 | -0.384 | -2.475 | -1.307 | 0.082 | 0.041 | 0.001 | 0.001 | 0.685 | 0.55 | 0.36 | 0.55 | 0.49 | 0.34 | 0.69 |
| 6 | 30756744 | rs4711229 | -0.384 | -2.475 | -1.307 | 0.071 | 0.041 | 0.002 | 0.002 | 0.685 | 0.55 | 0.36 | 0.55 | 0.5 | 0.34 | 0.69 |
| 6 | 30756822 | rs4713367 | -0.384 | -2.475 | -1.307 | 0.071 | 0.041 | 0.002 | 0.002 | 0.685 | 0.55 | 0.36 | 0.55 | 0.5 | 0.34 | 0.69 |
| 6 | 30757575 | rs10947095 | -0.384 | -2.475 | -1.307 | 0.071 | 0.041 | 0.002 | 0.002 | 0.685 | 0.55 | 0.36 | 0.55 | 0.5 | 0.34 | 0.69 |
| 6 | 30758466 | rs3131043 | -0.608 | -2.349 | -1.317 | 0.051 | 0.002 | -0.002 | -0.002 | 0 | 0.59 | 0.4 | 0.58 | 0.5 | 0.43 | 0.7 |
| 6 | 30758555 | rs17189441 | 1.562 | -2.284 | -1.494 | 0.104 | 0.063 | -0.073 | -0.073 | 0 | 0.28 | 0.05 | 0.4 | 0.29 | 0.17 | 0.44 |
| 6 | 30760698 | rs7740525 | -0.293 | -2.17 | -1.381 | 0.056 | 0.019 | 0 | 0 | 0.019 | 0.56 | 0.32 | 0.57 | 0.48 | 0.42 | 0.69 |
| 6 | 30761734 | rs3094112 | 0.054 | -1.799 | -1.229 | 0.001 | -0.004 | 0.006 | 0.006 | 0 | 0.19 | 0.18 | 0.12 | 0.11 | 0.22 | 0.18 |
| 6 | 30763893 | rs3131064 | 0.302 | -1.551 | -0.992 | 0.295 | 0.013 | -0.026 | -0.026 | 0 | 0.16 | 0.29 | 0.11 | 0.09 | 0.15 | 0.1 |
| 6 | 30764420 | rs12660883 | 0.961 | -1.419 | -1.705 | 0.512 | 0.063 | -0.071 | -0.071 | 0.013 | 0.28 | 0.06 | 0.39 | 0.28 | 0.16 | 0.44 |
| 6 | 30764907 | rs1264377 | 0.454 | -0.95 | -1.305 | 0.352 | -0.008 | 0.016 | 0.016 | 0 | 0.13 | 0.1 | 0.08 | 0.03 | 0.13 | 0.11 |
| 6 | 30765579 | rs1264376 | 1.129 | -0.731 | -1.212 | 0.458 | -0.007 | 0.004 | 0.004 | 0 | 0.1 | 0.09 | 0.06 | 0.03 | 0.12 | 0.11 |
| 6 | 30765895 | rs10947096 | 0.242 | -0.163 | -1.361 | 0.525 | -0.036 | -0.018 | -0.018 | 0 | 0.06 | 0.01 | 0.06 | 0.11 | 0.14 | 0.06 |
| 6 | 30766204 | rs3129987 | -1.166 | 0.308 | -1.171 | 0.421 | 0.055 | -0.017 | -0.017 | 0 | 0.92 | 0.83 | 0.87 | 0.95 | 0.86 | 0.96 |
| 6 | 30766211 | rs1264375 | 0.465 | -0.218 | -1.478 | 0.539 | 0.009 | -0.042 | -0.042 | 1.572 | 0.32 | 0.19 | 0.48 | 0.36 | 0.27 | 0.44 |
| 6 | 30766740 | rs6905957 | 0.828 | 0.047 | -1.416 | 0.366 | 0.059 | -0.111 | -0.111 | 0 | 0.25 | 0.08 | 0.37 | 0.41 | 0.16 | 0.44 |
| 6 | 30767538 | rs4713370 | 0.206 | 0.078 | -1.418 | 0.4 | -0.021 | -0.078 | -0.078 | 0.228 | 0.31 | 0.13 | 0.47 | 0.66 | 0.31 | 0.64 |

| | | | | | | | | | | | | | | | | |
|---|---|---|---|---|---|---|---|---|---|---|---|---|---|---|---|---|
| 6 | 30767627 | rs4248149 | 0.823 | 0.073 | -1.419 | 0.4 | 0.056 | -0.107 | -0.107 | 0 | 0.25 | 0.08 | 0.37 | 0.41 | 0.16 | 0.45 |
| 6 | 30767947 | rs4573120 | 0.209 | 0.072 | -1.416 | 0.39 | -0.021 | -0.078 | -0.078 | 0 | 0.31 | 0.13 | 0.47 | 0.66 | 0.31 | 0.64 |
| 6 | 30770000 | rs9295919 | 0.199 | 0.071 | -1.412 | 0.389 | -0.019 | -0.075 | -0.075 | 5.147 | 0.31 | 0.13 | 0.47 | 0.66 | 0.31 | 0.64 |
| 6 | 30770732 | rs13198118 | 0.905 | 0.063 | -1.402 | 0.406 | 0.056 | -0.103 | -0.103 | 0 | 0.25 | 0.09 | 0.37 | 0.41 | 0.16 | 0.44 |
| 6 | 30772378 | rs3094123 | -1.217 | 0.184 | -1.309 | 0.313 | 0.039 | -0.023 | -0.023 | 0 | 0.9 | 0.81 | 0.86 | 0.95 | 0.84 | 0.94 |
| 6 | 30773314 | rs4713376 | -0.579 | 0.246 | -1.612 | 0.373 | 0.003 | -0.004 | -0.004 | 0 | 0.06 | 0.05 | 0.1 | 0.24 | 0.15 | 0.14 |
| 6 | 30774474 | rs9378109 | 0.845 | 0.157 | -1.644 | 0.359 | 0.067 | -0.107 | -0.107 | 0.162 | 0.25 | 0.08 | 0.37 | 0.41 | 0.15 | 0.44 |
| 6 | 30776483 | rs9404974 | 0.951 | 0.141 | -1.635 | 0.369 | 0.056 | -0.103 | -0.103 | 0 | 0.25 | 0.09 | 0.38 | 0.41 | 0.17 | 0.44 |
| 6 | 30776590 | rs1264362 | 1.127 | -0.246 | -1.915 | 0.117 | -0.055 | 0.105 | 0.105 | 0 | 0.28 | 0.07 | 0.17 | 0.09 | 0.31 | 0.17 |
| 6 | 30777258 | rs9380194 | 0.493 | 0.179 | -1.737 | 0.355 | 0.066 | -0.126 | -0.126 | 0 | 0.28 | 0.09 | 0.43 | 0.49 | 0.18 | 0.52 |
| 6 | 30778203 | rs9380197 | 0.354 | 0.201 | -1.732 | 0.343 | -0.021 | -0.076 | -0.076 | 0 | 0.31 | 0.13 | 0.48 | 0.65 | 0.31 | 0.58 |
| 6 | 30778786 | rs9391696 | 0.79 | 0.2 | -1.732 | 0.343 | 0.053 | -0.123 | -0.123 | 0.641 | 0.26 | 0.09 | 0.43 | 0.49 | 0.18 | 0.52 |
| 6 | 30781923 | rs4286818 | 0.369 | 0.177 | -1.705 | 0.397 | -0.021 | -0.076 | -0.076 | 0 | 0.31 | 0.13 | 0.48 | 0.65 | 0.31 | 0.58 |
| 6 | 30782105 | rs2894046 | 0.369 | 0.126 | -1.698 | 0.412 | -0.021 | -0.076 | -0.076 | 0 | 0.31 | 0.13 | 0.48 | 0.65 | 0.31 | 0.58 |
| 6 | 30782235 | rs2394412 | 0.369 | 0.126 | -1.698 | 0.412 | -0.021 | -0.076 | -0.076 | 0 | 0.31 | 0.13 | 0.48 | 0.65 | 0.31 | 0.58 |
| 6 | 30782407 | rs2894047 | 0.369 | 0.126 | -1.698 | 0.412 | -0.021 | -0.076 | -0.076 | 0.713 | 0.31 | 0.13 | 0.48 | 0.65 | 0.31 | 0.58 |
| 6 | 30785273 | rs4713380 | 0.969 | 0.125 | -1.692 | 0.407 | 0.056 | -0.099 | -0.099 | 1.942 | 0.25 | 0.09 | 0.38 | 0.4 | 0.17 | 0.44 |
| 6 | 30785886 | rs9380198 | 0.969 | -0.008 | -1.687 | 0.398 | 0.056 | -0.099 | -0.099 | 0.802 | 0.25 | 0.09 | 0.38 | 0.41 | 0.17 | 0.44 |
| 6 | 30787175 | rs4713382 | 0.969 | -0.008 | -1.687 | 0.398 | 0.056 | -0.099 | -0.099 | 2.289 | 0.25 | 0.09 | 0.38 | 0.41 | 0.17 | 0.44 |
| 6 | 30787241 | rs4713383 | 0.969 | -0.008 | -1.687 | 0.398 | 0.056 | -0.099 | -0.099 | 2.289 | 0.25 | 0.09 | 0.38 | 0.41 | 0.17 | 0.44 |
| 6 | 30787593 | rs4713385 | 0.89 | 0.003 | -1.691 | 0.403 | 0.055 | -0.107 | -0.107 | 0.949 | 0.25 | 0.09 | 0.38 | 0.41 | 0.17 | 0.44 |
| 6 | 30788191 | rs3094111 | -0.538 | 0.03 | -1.259 | 0.412 | 0.027 | -0.072 | -0.072 | 1.917 | 0.83 | 0.64 | 0.82 | 0.94 | 0.83 | 0.85 |
| 6 | 30789526 | rs16897921 | 0.113 | 0.781 | -1.991 | 0.787 | 0.004 | -0.003 | -0.003 | 0 | 0.06 | 0.05 | 0.1 | 0.24 | 0.15 | 0.14 |
| 6 | 30790481 | rs12195964 | 0.113 | 1.018 | -2.469 | 0.752 | 0.004 | -0.003 | -0.003 | 0 | 0.06 | 0.05 | 0.1 | 0.24 | 0.15 | 0.14 |
| 6 | 30790604 | rs4713389 | 0.787 | 1 | -2.484 | 0.75 | 0.056 | -0.107 | -0.107 | 0 | 0.25 | 0.08 | 0.38 | 0.41 | 0.17 | 0.44 |
| 6 | 30790798 | rs6901761 | 0.787 | 1 | -2.484 | 0.75 | 0.056 | -0.107 | -0.107 | 0.973 | 0.25 | 0.08 | 0.38 | 0.41 | 0.17 | 0.44 |
| 6 | 30791409 | rs4947289 | 0.787 | 0.991 | -2.483 | 0.748 | 0.056 | -0.107 | -0.107 | 0 | 0.25 | 0.08 | 0.38 | 0.41 | 0.17 | 0.45 |
| 6 | 30791971 | rs4713390 | 0.135 | 0.999 | -2.483 | 0.747 | -0.023 | -0.076 | -0.076 | 0 | 0.31 | 0.13 | 0.48 | 0.65 | 0.31 | 0.58 |
| 6 | 30792235 | rs4713391 | 0.117 | 1.011 | -2.475 | 0.75 | 0.004 | -0.003 | -0.003 | 0.506 | 0.06 | 0.05 | 0.1 | 0.24 | 0.15 | 0.14 |
| 6 | 30793314 | rs7751869 | 0.684 | 0.975 | -2.482 | 0.746 | 0.055 | -0.107 | -0.107 | 0 | 0.25 | 0.08 | 0.38 | 0.41 | 0.17 | 0.44 |
| 6 | 30794405 | rs4947290 | 0.771 | 0.984 | -2.477 | 0.751 | 0.057 | -0.107 | -0.107 | 0.119 | 0.25 | 0.08 | 0.38 | 0.41 | 0.17 | 0.44 |
| 6 | 30794617 | rs915664 | 0.87 | 1.004 | -2.447 | 0.466 | -0.035 | 0.095 | 0.095 | 0 | 0.27 | 0.1 | 0.18 | 0.07 | 0.32 | 0.16 |
| 6 | 30796271 | rs9380199 | -0.315 | 1.629 | -1.611 | 1.366 | -0.022 | -0.076 | -0.076 | 0 | 0.31 | 0.13 | 0.48 | 0.65 | 0.32 | 0.58 |
| 6 | 30796738 | rs3130785 | 0.346 | 1.173 | -0.801 | 0.54 | 0.022 | -0.018 | -0.018 | 0 | 0.9 | 0.86 | 0.87 | 0.95 | 0.84 | 0.94 |
| 6 | 30797991 | rs7749924 | -0.297 | 0.328 | -1.117 | 0.92 | 0.01 | -0.009 | -0.009 | 0 | 0.17 | 0.09 | 0.18 | 0.17 | 0.13 | 0.19 |
| 6 | 30799399 | rs9501030 | -0.728 | -0.302 | -1.501 | 1.072 | 0.032 | -0.028 | -0.028 | 0 | 0.28 | 0.14 | 0.38 | 0.39 | 0.19 | 0.51 |
| 6 | 30802465 | rs1264341 | 1.743 | 0.555 | -1.777 | 0.743 | -0.02 | 0.018 | 0.018 | 0 | 0.05 | 0.02 | 0.02 | 0 | 0.08 | 0.04 |
| 6 | 30803254 | rs3130649 | 0.719 | 0.504 | -1.544 | 0.749 | 0.005 | 0.021 | 0.021 | 0 | 0.47 | 0.3 | 0.32 | 0.3 | 0.53 | 0.32 |
| 6 | 30805921 | rs3095352 | 0.703 | 0.485 | -1.512 | 0.739 | 0.006 | 0.021 | 0.021 | 0 | 0.47 | 0.31 | 0.33 | 0.3 | 0.53 | 0.32 |
| 6 | 30807058 | rs11757306 | 0.783 | 0.136 | -1.13 | 0.709 | -0.02 | -0.004 | -0.004 | 0 | 0.07 | 0.13 | 0.04 | 0.04 | 0.02 | 0.08 |
| 6 | 30809864 | rs3130787 | 0.325 | -0.323 | -1.423 | 1.09 | 0.002 | 0.006 | 0.006 | 0 | 0.6 | 0.47 | 0.39 | 0.38 | 0.57 | 0.46 |
| 6 | 30810056 | rs2517580 | 1.877 | -0.09 | -2.138 | 0.825 | -0.02 | 0.053 | 0.053 | 0 | 0.26 | 0.03 | 0.18 | 0.12 | 0.27 | 0.16 |
| 6 | 30812497 | rs2535335 | 0.969 | 0.124 | -2.16 | 0.941 | -0.007 | 0.039 | 0.039 | 0 | 0.4 | 0.16 | 0.27 | 0.25 | 0.38 | 0.25 |
| 6 | 30812519 | rs7759803 | 1.109 | 0.104 | -2.154 | 0.939 | -0.02 | -0.004 | -0.004 | 1.14 | 0.07 | 0.14 | 0.04 | 0.04 | 0.03 | 0.08 |
| 6 | 30815035 | rs2517576 | 1.027 | 0.139 | -2.15 | 0.948 | 0.024 | -0.008 | -0.008 | 0 | 0.13 | 0.17 | 0.08 | 0.05 | 0.16 | 0.09 |
| 6 | 30815951 | rs2844663 | 0.98 | 0.136 | -2.155 | 0.949 | -0.007 | 0.039 | 0.039 | 0 | 0.4 | 0.15 | 0.27 | 0.25 | 0.38 | 0.25 |
| 6 | 30816270 | rs2535331 | 1.691 | 0.119 | -2.177 | 0.922 | -0.05 | -0.016 | -0.016 | 0 | 0.31 | 0.1 | 0.21 | 0.25 | 0.36 | 0.23 |
| 6 | 30816865 | rs3132578 | -0.006 | 0.113 | -2.173 | 0.923 | 0.009 | 0.016 | 0.016 | 0 | 0.53 | 0.32 | 0.35 | 0.31 | 0.55 | 0.34 |
| 6 | 30817866 | rs3095350 | -0.035 | 0.074 | -2.192 | 1.226 | 0.008 | 0.015 | 0.015 | 0 | 0.53 | 0.32 | 0.37 | 0.51 | 0.55 | 0.35 |
| 6 | 30821187 | rs2250264 | 2.272 | -0.236 | -2.507 | 0.892 | -0.045 | 0.015 | 0.015 | 0 | 0.22 | 0.01 | 0.15 | 0.12 | 0.24 | 0.16 |
| 6 | 30822413 | rs3095345 | -1.449 | 0.028 | -1.379 | 0.564 | 0.162 | 0.045 | 0.045 | 0 | 0.94 | 0.8 | 0.85 | 0.92 | 0.84 | 0.96 |
| 6 | 30822771 | rs3130653 | 0.057 | 0.004 | -1.62 | 0.569 | 0.009 | 0.016 | 0.016 | 0 | 0.53 | 0.32 | 0.35 | 0.31 | 0.55 | 0.34 |
| 6 | 30823630 | rs9295928 | 1.517 | -0.119 | -1.659 | 0.504 | 0.004 | -0.052 | -0.052 | 0 | 0.22 | 0.06 | 0.36 | 0.37 | 0.18 | 0.46 |
| 6 | 30823710 | rs3130655 | -0.228 | -0.064 | -1.631 | 0.496 | 0.009 | 0.016 | 0.016 | 0 | 0.47 | 0.32 | 0.35 | 0.3 | 0.55 | 0.34 |
| 6 | 30823760 | rs2844660 | 2.085 | -0.193 | -1.979 | 0.143 | -0.038 | 0.021 | 0.021 | 0 | 0.07 | 0.01 | 0.03 | 0 | 0.11 | 0.03 |
| 6 | 30824532 | rs2844659 | 1.268 | -0.368 | -1.745 | 0.329 | 0.025 | -0.007 | -0.007 | 0.005 | 0.13 | 0.17 | 0.08 | 0.05 | 0.16 | 0.09 |
| 6 | 30825220 | rs7772269 | 1.206 | -0.363 | -1.746 | 0.327 | 0.007 | 0.024 | 0.024 | 0 | 0.08 | 0.15 | 0.06 | 0.05 | 0.08 | 0.05 |
| 6 | 30825287 | rs9468841 | 1.847 | -0.556 | -2.008 | 0.417 | 0.014 | -0.032 | -0.032 | 0 | 0.13 | 0.03 | 0.32 | 0.26 | 0.09 | 0.2 |
| 6 | 30826904 | rs2535327 | 2.047 | -0.688 | -1.96 | 0.403 | -0.08 | 0.008 | 0.008 | 0 | 0.28 | 0.04 | 0.19 | 0.18 | 0.34 | 0.19 |
| 6 | 30829522 | rs2844657 | 1.338 | -0.648 | -1.938 | 0.424 | 0.024 | -0.006 | -0.006 | 0.767 | 0.13 | 0.17 | 0.08 | 0.05 | 0.16 | 0.09 |
| 6 | 30830292 | rs7761138 | 1.21 | -0.652 | -1.939 | 0.422 | 0.006 | 0.026 | 0.026 | 0 | 0.08 | 0.15 | 0.06 | 0.05 | 0.08 | 0.05 |
| 6 | 30831843 | rs3130791 | 0.232 | -0.643 | -1.935 | 0.421 | 0.009 | 0.016 | 0.016 | 0 | 0.53 | 0.32 | 0.35 | 0.31 | 0.54 | 0.34 |
| 6 | 30832063 | rs2535326 | 1.197 | -0.643 | -1.936 | 0.421 | -0.007 | 0.042 | 0.042 | 0 | 0.4 | 0.15 | 0.27 | 0.25 | 0.38 | 0.25 |
| 6 | 30832357 | rs3130793 | 0.241 | -0.634 | -1.924 | 0.428 | 0.009 | 0.016 | 0.016 | 0 | 0.53 | 0.32 | 0.36 | 0.31 | 0.55 | 0.34 |
| 6 | 30832810 | rs3130794 | 0.241 | -0.634 | -1.924 | 0.428 | 0.009 | 0.016 | 0.016 | 0 | 0.53 | 0.32 | 0.36 | 0.31 | 0.55 | 0.34 |
| 6 | 30834234 | rs3132575 | 0.241 | -0.634 | -1.924 | 0.428 | 0.009 | 0.016 | 0.016 | 0 | 0.53 | 0.32 | 0.36 | 0.31 | 0.55 | 0.34 |
| 6 | 30836111 | rs3095354 | 0.241 | -0.634 | -1.924 | 0.428 | 0.009 | 0.016 | 0.016 | 0 | 0.53 | 0.32 | 0.36 | 0.31 | 0.55 | 0.34 |
| 6 | 30838497 | rs2535340 | 3.296 | -0.498 | -1.63 | 0.429 | -0.02 | 0.018 | 0.018 | 0 | 0.05 | 0.02 | 0.02 | 0 | 0.08 | 0.04 |
| 6 | 30838688 | rs2844654 | 1.791 | -0.505 | -1.617 | 0.447 | -0.05 | -0.013 | -0.013 | 0 | 0.31 | 0.1 | 0.21 | 0.25 | 0.36 | 0.23 |

| | | | | | | | | | | | | | | | | |
|---|---|---|---|---|---|---|---|---|---|---|---|---|---|---|---|---|
| 6 | 30839629 | rs3130795 | 0.276 | -0.509 | -1.574 | 0.469 | 0.009 | 0.016 | 0.016 | 0 | 0.53 | 0.32 | 0.36 | 0.31 | 0.55 | 0.34 |
| 6 | 30840688 | rs3132573 | -0.155 | -0.526 | -1.551 | 0.474 | 0.002 | 0.004 | 0.004 | 0 | 0.6 | 0.46 | 0.4 | 0.35 | 0.58 | 0.45 |
| 6 | 30840950 | rs3130796 | 0.264 | -0.503 | -1.566 | 0.482 | 0.009 | 0.016 | 0.016 | 0 | 0.53 | 0.32 | 0.36 | 0.31 | 0.55 | 0.34 |
| 6 | 30844314 | rs1264333 | 1.725 | -0.421 | -1.603 | 0.434 | -0.05 | -0.013 | -0.013 | 0.066 | 0.31 | 0.1 | 0.21 | 0.25 | 0.36 | 0.24 |
| 6 | 30848253 | rs7756521 | 0.36 | -0.508 | -1.51 | 0.498 | 0.026 | -0.037 | -0.037 | 0 | 0.31 | 0.28 | 0.43 | 0.61 | 0.22 | 0.59 |
| 6 | 30849822 | rs9295930 | 1.082 | -0.552 | -1.507 | 0.595 | 0.011 | -0.032 | -0.032 | 0 | 0.22 | 0.08 | 0.36 | 0.36 | 0.18 | 0.46 |
| 6 | 30850191 | rs9501032 | 0.334 | -0.543 | -1.469 | 0.567 | 0.024 | -0.033 | -0.033 | 0.684 | 0.31 | 0.28 | 0.43 | 0.61 | 0.22 | 0.59 |
| 6 | 30852747 | rs9468842 | 0.345 | -0.548 | -1.512 | 0.565 | 0.026 | -0.037 | -0.037 | 0 | 0.31 | 0.27 | 0.43 | 0.61 | 0.22 | 0.59 |
| 6 | 30853796 | rs3131034 | -0.96 | -0.505 | -1.459 | 0.501 | 0.079 | -0.05 | -0.05 | 0 | 0.83 | 0.59 | 0.79 | 0.92 | 0.77 | 0.93 |
| 6 | 30854090 | rs6901464 | 0.376 | -0.505 | -1.459 | 0.501 | 0.027 | -0.037 | -0.037 | 0 | 0.31 | 0.28 | 0.43 | 0.61 | 0.22 | 0.59 |
| 6 | 30855251 | rs4618569 | 0.369 | -0.512 | -1.462 | 0.498 | 0.027 | -0.037 | -0.037 | 1.966 | 0.31 | 0.28 | 0.43 | 0.61 | 0.22 | 0.59 |
| 6 | 30855907 | rs1264323 | -0.559 | -0.522 | -1.493 | 0.467 | -0.001 | -0.001 | -0.001 | 0 | 0.44 | 0.45 | 0.52 | 0.67 | 0.38 | 0.68 |
| 6 | 30857542 | rs6924600 | 0.402 | -0.522 | -1.493 | 0.467 | 0.027 | -0.037 | -0.037 | 0 | 0.31 | 0.28 | 0.43 | 0.61 | 0.22 | 0.59 |
| 6 | 30857795 | rs6457281 | 2.095 | -0.53 | -1.463 | 0.469 | -0.015 | -0.001 | -0.001 | 0.042 | 0.05 | 0.1 | 0.03 | 0.02 | 0.02 | 0.07 |
| 6 | 30857894 | rs1264322 | 3.352 | -0.529 | -1.49 | 0.465 | -0.02 | 0.018 | 0.018 | 0 | 0.05 | 0.02 | 0.02 | 0 | 0.08 | 0.04 |
| 6 | 30858254 | rs7743661 | 0.348 | -0.529 | -1.49 | 0.465 | 0.027 | -0.037 | -0.037 | 0 | 0.31 | 0.28 | 0.43 | 0.61 | 0.22 | 0.59 |
| 6 | 30861225 | rs3213644 | 1.053 | -0.528 | -1.489 | 0.465 | 0.014 | -0.061 | -0.061 | 0 | 0.23 | 0.13 | 0.39 | 0.57 | 0.19 | 0.47 |
| 6 | 30862423 | rs1264319 | 1.283 | -0.329 | -1.5 | 0.473 | 0.023 | -0.007 | -0.007 | 0 | 0.13 | 0.18 | 0.1 | 0.06 | 0.16 | 0.12 |
| 6 | 30863530 | rs1264318 | 1.283 | -0.329 | -1.5 | 0.473 | 0.023 | -0.007 | -0.007 | 0.706 | 0.13 | 0.18 | 0.1 | 0.06 | 0.16 | 0.12 |
| 6 | 30864279 | rs886422 | 3.353 | -0.329 | -1.5 | 0.473 | -0.02 | 0.018 | 0.018 | 0 | 0.05 | 0.02 | 0.02 | 0 | 0.08 | 0.04 |
| 6 | 30864829 | rs1049623 | -0.629 | -0.33 | -1.5 | 0.473 | -0.001 | -0.001 | -0.001 | 0 | 0.44 | 0.45 | 0.52 | 0.67 | 0.38 | 0.68 |
| 6 | 30865115 | rs2239517 | 0.397 | -0.329 | -1.5 | 0.473 | 0.026 | -0.037 | -0.037 | 0.115 | 0.31 | 0.28 | 0.43 | 0.61 | 0.22 | 0.55 |
| 6 | 30865725 | rs2239518 | 1.053 | -0.329 | -1.5 | 0.473 | 0.014 | -0.061 | -0.061 | 0 | 0.23 | 0.13 | 0.39 | 0.57 | 0.19 | 0.47 |
| 6 | 30867106 | rs1049628 | 0.397 | -0.329 | -1.502 | 0.465 | 0.026 | -0.037 | -0.037 | 0 | 0.31 | 0.28 | 0.43 | 0.61 | 0.22 | 0.55 |
| 6 | 30867527 | rs1049633 | 3.354 | -0.329 | -1.502 | 0.465 | -0.02 | 0.018 | 0.018 | 0 | 0.05 | 0.02 | 0.02 | 0 | 0.08 | 0.04 |
| 6 | 30867958 | rs9468843 | 1.189 | -0.35 | -1.513 | 0.468 | 0.011 | -0.036 | -0.036 | 0 | 0.22 | 0.08 | 0.36 | 0.36 | 0.18 | 0.46 |
| 6 | 30868628 | rs2894055 | 0.459 | -0.33 | -1.523 | 0.379 | 0.024 | -0.036 | -0.036 | 0 | 0.3 | 0.28 | 0.43 | 0.61 | 0.22 | 0.55 |
| 6 | 30869714 | rs9295931 | 0.42 | -0.307 | -1.434 | 0.39 | 0.026 | -0.037 | -0.037 | 0 | 0.31 | 0.28 | 0.43 | 0.61 | 0.22 | 0.55 |
| 6 | 30870763 | rs9468846 | 0.42 | -0.307 | -1.434 | 0.39 | 0.026 | -0.037 | -0.037 | 0 | 0.31 | 0.28 | 0.43 | 0.61 | 0.22 | 0.55 |
| 6 | 30871619 | rs13215409 | 0.42 | -0.307 | -1.434 | 0.39 | 0.026 | -0.037 | -0.037 | 1.777 | 0.31 | 0.28 | 0.43 | 0.61 | 0.22 | 0.55 |
| 6 | 30872203 | rs3130779 | -0.966 | -0.31 | -1.435 | 0.388 | 0.077 | -0.051 | -0.051 | 2.301 | 0.83 | 0.6 | 0.79 | 0.92 | 0.77 | 0.93 |
| 6 | 30872274 | rs2894054 | 1.292 | -0.304 | -1.435 | 0.391 | 0.008 | 0.028 | 0.028 | 0 | 0.08 | 0.15 | 0.07 | 0.05 | 0.08 | 0.04 |
| 6 | 30872982 | rs1264312 | 3.386 | -0.281 | -1.449 | 0.404 | -0.019 | 0.018 | 0.018 | 0 | 0.05 | 0.02 | 0.02 | 0 | 0.08 | 0.04 |
| 6 | 30873605 | rs1264310 | 3.386 | -0.199 | -1.449 | 0.404 | -0.019 | 0.018 | 0.018 | 0 | 0.05 | 0.02 | 0.02 | 0 | 0.08 | 0.04 |
| 6 | 30874165 | rs3909130 | 0.385 | -0.333 | -1.449 | 0.405 | 0.027 | -0.037 | -0.037 | 0.944 | 0.31 | 0.28 | 0.43 | 0.61 | 0.22 | 0.55 |
| 6 | 30874308 | rs3130780 | -1.199 | -0.383 | -1.243 | 0.374 | 0.062 | -0.033 | -0.033 | 0.075 | 0.85 | 0.67 | 0.8 | 0.92 | 0.78 | 0.93 |
| 6 | 30875145 | rs2284175 | 0.355 | -0.204 | -1.195 | 0.508 | 0.026 | -0.037 | -0.037 | 0 | 0.31 | 0.28 | 0.43 | 0.61 | 0.22 | 0.55 |
| 6 | 30875622 | rs2284176 | 0.355 | -0.204 | -1.195 | 0.508 | 0.026 | -0.037 | -0.037 | 0 | 0.31 | 0.28 | 0.43 | 0.61 | 0.22 | 0.55 |
| 6 | 30875899 | rs1264309 | 1.802 | -0.224 | -1.197 | 0.497 | -0.048 | -0.012 | -0.012 | 0 | 0.31 | 0.1 | 0.21 | 0.25 | 0.36 | 0.24 |
| 6 | 30876034 | rs2074510 | 0.317 | -0.046 | -1.147 | 0.556 | 0.026 | -0.037 | -0.037 | 0 | 0.31 | 0.28 | 0.43 | 0.61 | 0.22 | 0.55 |
| 6 | 30876152 | rs1052693 | 0.317 | -0.046 | -1.147 | 0.556 | 0.026 | -0.037 | -0.037 | 0 | 0.31 | 0.28 | 0.43 | 0.61 | 0.22 | 0.55 |
| 6 | 30877202 | rs916920 | 0.317 | -0.046 | -1.148 | 0.556 | 0.026 | -0.037 | -0.037 | 0 | 0.31 | 0.28 | 0.43 | 0.61 | 0.22 | 0.55 |
| 6 | 30878769 | rs3218815 | 0.317 | -0.043 | -1.146 | 0.558 | 0.026 | -0.037 | -0.037 | 0 | 0.31 | 0.28 | 0.43 | 0.61 | 0.22 | 0.55 |
| 6 | 30878919 | rs2074512 | 0.317 | -0.458 | -1.146 | 0.558 | 0.026 | -0.037 | -0.037 | 0 | 0.31 | 0.28 | 0.43 | 0.61 | 0.22 | 0.55 |
| 6 | 30879987 | rs1264308 | 3.279 | -0.337 | -1.128 | 0.532 | -0.019 | 0.018 | 0.018 | 0 | 0.05 | 0.02 | 0.02 | 0 | 0.08 | 0.04 |
| 6 | 30880757 | rs1264307 | 1.763 | -0.505 | -1.509 | 0.517 | -0.05 | -0.014 | -0.014 | 0.054 | 0.31 | 0.1 | 0.21 | 0.25 | 0.36 | 0.24 |
| 6 | 30882277 | rs1264305 | 1.763 | -0.505 | -1.509 | 0.517 | -0.05 | -0.014 | -0.014 | 1.935 | 0.31 | 0.1 | 0.21 | 0.25 | 0.36 | 0.24 |
| 6 | 30882415 | rs1264304 | 3.327 | -0.5 | -1.512 | 0.514 | -0.019 | 0.018 | 0.018 | 0 | 0.05 | 0.02 | 0.02 | 0 | 0.08 | 0.04 |
| 6 | 30882513 | rs1264303 | 1.751 | -0.494 | -1.506 | 0.523 | -0.05 | -0.014 | -0.014 | 0 | 0.31 | 0.1 | 0.21 | 0.25 | 0.36 | 0.24 |
| 6 | 30882634 | rs1264302 | 1.751 | -0.494 | -1.506 | 0.523 | -0.05 | -0.014 | -0.014 | 0 | 0.31 | 0.1 | 0.21 | 0.25 | 0.36 | 0.24 |
| 6 | 30882689 | rs6926224 | 1.711 | -0.537 | -1.448 | 0.521 | -0.02 | 0.019 | 0.019 | 0.151 | 0.07 | 0.12 | 0.07 | 0.05 | 0.04 | 0.03 |
| 6 | 30882781 | rs1264301 | 1.809 | -0.529 | -1.453 | 0.525 | -0.05 | -0.014 | -0.014 | 0 | 0.31 | 0.1 | 0.21 | 0.25 | 0.36 | 0.24 |
| 6 | 30884436 | rs1264298 | 1.809 | -0.529 | -1.453 | 0.525 | -0.05 | -0.014 | -0.014 | 1.642 | 0.31 | 0.1 | 0.21 | 0.25 | 0.36 | 0.24 |
| 6 | 30885639 | rs9262288 | 1.809 | -0.529 | -1.453 | 0.525 | -0.05 | -0.014 | -0.014 | 0 | 0.31 | 0.1 | 0.21 | 0.25 | 0.36 | 0.24 |
| 6 | 30885821 | rs9262290 | 1.809 | -0.529 | -1.453 | 0.525 | -0.05 | -0.014 | -0.014 | 0 | 0.31 | 0.1 | 0.21 | 0.25 | 0.36 | 0.24 |
| 6 | 30887311 | rs7756286 | 1.809 | -0.529 | -1.453 | 0.525 | -0.05 | -0.014 | -0.014 | 0.249 | 0.31 | 0.1 | 0.21 | 0.25 | 0.36 | 0.24 |
| 6 | 30887344 | rs7738138 | 1.809 | -0.529 | -1.453 | 0.525 | -0.05 | -0.014 | -0.014 | 0 | 0.31 | 0.1 | 0.21 | 0.25 | 0.36 | 0.24 |
| 6 | 30887972 | rs2285319 | 0.45 | -0.557 | -1.457 | 0.51 | 0.022 | -0.038 | -0.038 | 0 | 0.28 | 0.26 | 0.42 | 0.59 | 0.2 | 0.54 |
| 6 | 30887988 | rs2249459 | 1.868 | -0.557 | -1.457 | 0.51 | -0.05 | -0.014 | -0.014 | 0 | 0.31 | 0.1 | 0.21 | 0.25 | 0.36 | 0.24 |
| 6 | 30888161 | rs2249464 | 0.561 | -0.557 | -1.445 | 0.509 | -0.018 | 0.05 | 0.05 | 0 | 0.59 | 0.36 | 0.63 | 0.84 | 0.56 | 0.78 |
| 6 | 30888169 | rs2517468 | 1.868 | -0.556 | -1.456 | 0.51 | -0.05 | -0.014 | -0.014 | 0 | 0.31 | 0.1 | 0.21 | 0.25 | 0.36 | 0.24 |
| 6 | 30889260 | rs2517467 | 0.561 | -0.564 | -1.441 | 0.512 | -0.019 | 0.052 | 0.052 | 0 | 0.59 | 0.36 | 0.63 | 0.84 | 0.56 | 0.78 |
| 6 | 30889389 | rs2074511 | 0.959 | -0.642 | -1.472 | 0.473 | 0.017 | -0.064 | -0.064 | 0 | 0.24 | 0.13 | 0.39 | 0.57 | 0.19 | 0.48 |
| 6 | 30890431 | rs885905 | -0.137 | -0.555 | -1.425 | 0.517 | -0.002 | -0.024 | -0.024 | 0 | 0.71 | 0.46 | 0.7 | 0.87 | 0.71 | 0.88 |
| 6 | 30890483 | rs2074506 | 1.883 | -0.555 | -1.425 | 0.517 | -0.05 | -0.014 | -0.014 | 0.546 | 0.31 | 0.1 | 0.21 | 0.25 | 0.36 | 0.24 |
| 6 | 30890871 | rs753725 | 0.57 | -0.555 | -1.425 | 0.517 | -0.019 | 0.052 | 0.052 | 0 | 0.59 | 0.36 | 0.63 | 0.84 | 0.56 | 0.78 |
| 6 | 30892322 | rs2532942 | 1.883 | -0.555 | -1.425 | 0.517 | -0.05 | -0.014 | -0.014 | 0 | 0.31 | 0.1 | 0.21 | 0.25 | 0.36 | 0.24 |
| 6 | 30892377 | rs2252760 | 1.872 | -0.555 | -1.425 | 0.517 | -0.05 | -0.013 | -0.013 | 0 | 0.31 | 0.1 | 0.21 | 0.25 | 0.36 | 0.24 |
| 6 | 30892521 | rs2532941 | 0.174 | -0.565 | -1.425 | 0.512 | 0 | 0 | 0 | 0 | 0.68 | 0.38 | 0.68 | 0.86 | 0.71 | 0.82 |

| | | | | | | | | | | | | | | | | |
|---|---|---|---|---|---|---|---|---|---|---|---|---|---|---|---|---|
| 6 | 30893251 | rs2252856 | 1.883 | -0.531 | -1.417 | 0.537 | -0.05 | -0.017 | -0.017 | 0.141 | 0.31 | 0.1 | 0.21 | 0.25 | 0.36 | 0.24 |
| 6 | 30893728 | rs1043483 | -0.175 | -0.533 | -1.415 | 0.528 | -0.002 | -0.027 | -0.027 | 0 | 0.71 | 0.46 | 0.7 | 0.87 | 0.71 | 0.88 |
| 6 | 30893831 | rs2532938 | 1.883 | -0.531 | -1.42 | 0.529 | -0.05 | -0.014 | -0.014 | 0 | 0.31 | 0.1 | 0.21 | 0.25 | 0.36 | 0.24 |
| 6 | 30893941 | rs4678 | 1.461 | -0.531 | -1.42 | 0.529 | -0.003 | 0.002 | 0.002 | 0 | 0.12 | 0.1 | 0.07 | 0.03 | 0.16 | 0.1 |
| 6 | 30894408 | rs2532936 | 1.853 | -0.496 | -1.432 | 0.54 | -0.05 | -0.013 | -0.013 | 0 | 0.31 | 0.1 | 0.21 | 0.25 | 0.36 | 0.24 |
| 6 | 30895680 | rs4711247 | 0.439 | -0.49 | -1.429 | 0.544 | 0.022 | -0.038 | -0.038 | 0.644 | 0.28 | 0.26 | 0.42 | 0.59 | 0.2 | 0.54 |
| 6 | 30895990 | rs3873332 | 1.015 | -0.49 | -1.429 | 0.544 | 0.015 | -0.061 | -0.061 | 0.179 | 0.23 | 0.13 | 0.39 | 0.57 | 0.19 | 0.47 |
| 6 | 30896147 | rs3873334 | 0.46 | -0.458 | -1.43 | 0.56 | 0.022 | -0.038 | -0.038 | 0 | 0.28 | 0.26 | 0.42 | 0.59 | 0.2 | 0.54 |
| 6 | 30896190 | rs2517462 | 1.775 | -0.425 | -1.445 | 0.553 | -0.05 | -0.014 | -0.014 | 0 | 0.31 | 0.1 | 0.21 | 0.25 | 0.36 | 0.24 |
| 6 | 30897022 | rs2517459 | -1.331 | -0.358 | -1.407 | 0.551 | 0.039 | -0.067 | -0.067 | 0 | 0.79 | 0.59 | 0.78 | 0.91 | 0.76 | 0.91 |
| 6 | 30897774 | rs2532929 | 1.207 | -0.52 | -1.537 | 0.89 | -0.002 | -0.097 | -0.097 | 0 | 0.42 | 0.1 | 0.5 | 0.51 | 0.38 | 0.42 |
| 6 | 30898687 | rs2844652 | 1.624 | -0.382 | -1.468 | 0.84 | -0.05 | -0.014 | -0.014 | 0 | 0.31 | 0.1 | 0.21 | 0.25 | 0.36 | 0.24 |
| 6 | 30899163 | rs2532925 | 0.269 | -0.524 | -1.389 | 0.736 | 0.004 | 0.01 | 0.01 | 0.094 | 0.66 | 0.33 | 0.66 | 0.85 | 0.7 | 0.81 |
| 6 | 30899233 | rs4713402 | 0.847 | -0.447 | -1.423 | 0.816 | 0.015 | -0.061 | -0.061 | 0 | 0.23 | 0.13 | 0.39 | 0.57 | 0.19 | 0.47 |
| 6 | 30899524 | rs3131787 | 1.527 | -0.434 | -1.433 | 0.804 | -0.002 | 0.001 | 0.001 | 0.002 | 0.12 | 0.1 | 0.07 | 0.02 | 0.16 | 0.06 |
| 6 | 30899571 | rs2286656 | 0.914 | -0.417 | -1.449 | 0.973 | 0.011 | -0.02 | -0.02 | 0 | 0.23 | 0.12 | 0.37 | 0.36 | 0.18 | 0.46 |
| 6 | 30899746 | rs2286655 | 0.869 | -0.485 | -1.444 | 0.969 | 0.014 | -0.061 | -0.061 | 0 | 0.23 | 0.13 | 0.39 | 0.57 | 0.19 | 0.47 |
| 6 | 30900150 | rs3095151 | 1.135 | -0.536 | -1.211 | 0.97 | 0.02 | -0.024 | -0.024 | 0 | 0.13 | 0.17 | 0.08 | 0.02 | 0.16 | 0.06 |
| 6 | 30900435 | rs12525616 | 2.153 | -0.58 | -1.307 | 1.026 | 0.058 | -0.062 | -0.062 | 0 | 0.11 | 0.01 | 0.3 | 0.26 | 0.03 | 0.19 |
| 6 | 30900664 | rs2844651 | 0.611 | -0.123 | -0.973 | 0.806 | -0.018 | 0.007 | 0.007 | 0 | 0.58 | 0.33 | 0.61 | 0.66 | 0.55 | 0.76 |
| 6 | 30902181 | rs10947114 | 1.176 | -0.107 | -0.973 | 0.819 | 0.013 | -0.019 | -0.019 | 0 | 0.23 | 0.12 | 0.37 | 0.37 | 0.18 | 0.46 |
| 6 | 30903948 | rs3131784 | -1.105 | -0.014 | -1.074 | 0.79 | -0.067 | 0.022 | 0.022 | 0 | 0.61 | 0.77 | 0.7 | 0.7 | 0.54 | 0.66 |
| 6 | 30905313 | rs3132571 | -0.761 | -0.016 | -0.976 | 0.802 | 0.008 | -0.004 | -0.004 | 0.03 | 0.38 | 0.62 | 0.33 | 0.34 | 0.35 | 0.2 |
| 6 | 30907176 | rs6933400 | 1.484 | 0.346 | -0.691 | 0.866 | -0.017 | -0.076 | -0.076 | 0 | 0.17 | 0.4 | 0.16 | 0.06 | 0.15 | 0.06 |
| 6 | 30907266 | rs6933909 | 1.484 | 0.346 | -0.691 | 0.866 | -0.017 | -0.076 | -0.076 | 0 | 0.17 | 0.4 | 0.16 | 0.06 | 0.15 | 0.06 |
| 6 | 30908266 | rs4711249 | 0.985 | 0.283 | -0.876 | 0.938 | 0.035 | -0.065 | -0.065 | 0.04 | 0.28 | 0.14 | 0.39 | 0.55 | 0.19 | 0.51 |
| 6 | 30912414 | rs9501035 | 0.98 | 0.19 | -0.914 | 0.999 | 0.036 | -0.065 | -0.065 | 0.257 | 0.28 | 0.14 | 0.39 | 0.55 | 0.19 | 0.52 |
| 6 | 30912481 | rs2844702 | 1.613 | 0.312 | -0.943 | 1.015 | -0.05 | -0.014 | -0.014 | 0 | 0.31 | 0.1 | 0.21 | 0.25 | 0.36 | 0.24 |
| 6 | 30912691 | rs2844701 | -0.5 | 0.383 | -0.835 | 1.037 | -0.003 | 0.002 | 0.002 | 0 | 0.92 | 0.87 | 0.91 | 0.95 | 0.89 | 0.9 |
| 6 | 30914843 | rs3130782 | 0.86 | 0.404 | -0.834 | 1.112 | 0.007 | 0.005 | 0.005 | 0 | 0.07 | 0.09 | 0.06 | 0.07 | 0.12 | 0.04 |
| 6 | 30919701 | rs2517449 | 1.483 | 0.191 | -1.098 | 0.982 | -0.049 | -0.03 | -0.03 | 0.163 | 0.31 | 0.1 | 0.21 | 0.27 | 0.36 | 0.27 |
| 6 | 30919878 | rs11970154 | 1.431 | 0.16 | -0.922 | 0.846 | -0.016 | -0.068 | -0.068 | 0.804 | 0.17 | 0.41 | 0.16 | 0.06 | 0.15 | 0.05 |
| 6 | 30920890 | rs2240804 | 1.753 | 0.215 | -1.075 | 0.887 | -0.078 | 0.014 | 0.014 | 0 | 0.28 | 0.04 | 0.2 | 0.18 | 0.34 | 0.2 |
| 6 | 30920957 | rs2240803 | 0.909 | -0.681 | -0.778 | 1.108 | 0.019 | -0.029 | -0.029 | 0 | 0.26 | 0.08 | 0.36 | 0.37 | 0.19 | 0.5 |
| 6 | 30921364 | rs12190030 | 0.394 | -0.755 | -0.757 | 1.128 | 0.048 | -0.035 | -0.035 | 0 | 0.32 | 0.14 | 0.43 | 0.42 | 0.22 | 0.54 |
| 6 | 30921882 | rs3757340 | 0.283 | -0.106 | -0.544 | 1.353 | -0.007 | -0.004 | -0.004 | 0 | 0.44 | 0.49 | 0.52 | 0.43 | 0.33 | 0.56 |
| 6 | 30925985 | rs11753326 | -0.637 | 0.305 | -0.209 | 1.576 | -0.006 | -0.004 | -0.004 | 0 | 0.11 | 0.15 | 0.12 | 0.06 | 0.09 | 0.06 |
| 6 | 30931844 | rs3131934 | 0.162 | 1.255 | -0.232 | 1.167 | 0.045 | 0.028 | 0.028 | 0 | 0.06 | 0.15 | 0.07 | 0.07 | 0.11 | 0.13 |
| 6 | 30932068 | rs3131783 | -0.9 | 1.034 | -0.172 | 1.154 | 0.037 | 0.036 | 0.036 | 1.199 | 0.09 | 0.23 | 0.09 | 0.08 | 0.12 | 0.11 |
| 6 | 30932309 | rs2844697 | -1.061 | 1.155 | -0.736 | 0.683 | -0.018 | 0.019 | 0.019 | 0.866 | 0.27 | 0.74 | 0.29 | 0.19 | 0.33 | 0.17 |
| 6 | 30932682 | rs2532924 | -0.025 | 1.135 | -1.021 | 0.539 | -0.013 | 0.032 | 0.032 | 0.096 | 0.57 | 0.76 | 0.5 | 0.32 | 0.57 | 0.49 |
| 6 | 30933794 | rs3095089 | 0.63 | 1.145 | -0.355 | 0.286 | 0.06 | -0.038 | -0.038 | 0 | 0.07 | 0.17 | 0.11 | 0 | 0.13 | 0.01 |
| 6 | 30933864 | rs3131933 | 0.268 | 0.673 | -0.92 | -0.125 | -0.022 | 0.048 | 0.048 | 0 | 0.38 | 0.28 | 0.48 | 0.19 | 0.46 | 0.22 |
| 6 | 30933975 | rs2532919 | 0.615 | 0.47 | -0.835 | 0.012 | -0.004 | 0.012 | 0.012 | 0 | 0.12 | 0.24 | 0.27 | 0.1 | 0.15 | 0.12 |
| 6 | 30936014 | rs2844695 | -0.434 | 0.744 | -0.975 | 0.381 | 0.004 | 0.011 | 0.011 | 0 | 0.23 | 0.26 | 0.36 | 0.47 | 0.31 | 0.37 |
| 6 | 30937377 | rs3095085 | 1.872 | 0.528 | -1.193 | 0.095 | -0.009 | 0.05 | 0.05 | 0 | 0.22 | 0.13 | 0.25 | 0.04 | 0.24 | 0.1 |
| 6 | 30940945 | rs2517446 | 1.201 | 0.832 | -1.139 | 0.228 | -0.001 | 0.011 | 0.011 | 0 | 0.11 | 0.15 | 0.26 | 0.12 | 0.14 | 0.12 |
| 6 | 30942284 | rs2844688 | 0.875 | 0.846 | -1.18 | 0.17 | -0.003 | 0.006 | 0.006 | 0 | 0.12 | 0.15 | 0.26 | 0.12 | 0.14 | 0.12 |
| 6 | 30942387 | rs9262315 | 1.212 | 0.86 | -1.174 | 0.177 | -0.001 | 0.011 | 0.011 | 0 | 0.11 | 0.15 | 0.26 | 0.12 | 0.14 | 0.12 |
| 6 | 30942447 | rs2517442 | 1.212 | 0.86 | -1.174 | 0.177 | -0.001 | 0.011 | 0.011 | 0 | 0.11 | 0.15 | 0.26 | 0.12 | 0.14 | 0.12 |
| 6 | 30942621 | rs2517441 | 1.212 | 0.86 | -1.174 | 0.177 | -0.001 | 0.011 | 0.011 | 0 | 0.11 | 0.15 | 0.26 | 0.12 | 0.14 | 0.12 |
| 6 | 30942929 | rs2517439 | 1.212 | 0.86 | -1.174 | 0.177 | -0.001 | 0.011 | 0.011 | 0 | 0.11 | 0.15 | 0.26 | 0.12 | 0.14 | 0.12 |
| 6 | 30942972 | rs2844686 | 1.212 | 0.86 | -1.174 | 0.177 | -0.001 | 0.011 | 0.011 | 0 | 0.11 | 0.15 | 0.26 | 0.12 | 0.14 | 0.12 |
| 6 | 30943083 | rs2844685 | 1.212 | 0.86 | -1.174 | 0.177 | -0.001 | 0.011 | 0.011 | 0 | 0.11 | 0.15 | 0.26 | 0.12 | 0.14 | 0.12 |
| 6 | 30944232 | rs2517434 | 1.212 | 0.86 | -1.174 | 0.177 | -0.001 | 0.011 | 0.011 | 0 | 0.11 | 0.15 | 0.26 | 0.12 | 0.14 | 0.12 |
| 6 | 30946148 | rs2844682 | 1.212 | 0.86 | -1.174 | 0.177 | -0.001 | 0.011 | 0.011 | 0 | 0.11 | 0.15 | 0.26 | 0.12 | 0.14 | 0.12 |
| 6 | 30946496 | rs2844680 | -0.671 | 0.34 | -1.011 | -0.137 | -0.001 | 0 | 0 | 0 | 0.53 | 0.55 | 0.64 | 0.51 | 0.57 | 0.49 |
| 6 | 30947042 | rs9394023 | 1.401 | 0.401 | -1.038 | -0.165 | -0.049 | -0.052 | -0.052 | 0.468 | 0.1 | 0.03 | 0.07 | 0.33 | 0.15 | 0.24 |
| 6 | 30947365 | rs2844679 | 1.4 | 0.387 | -1.035 | -0.162 | -0.001 | 0.011 | 0.011 | 0 | 0.11 | 0.15 | 0.26 | 0.12 | 0.14 | 0.12 |
| 6 | 30947378 | rs2517431 | -1.324 | 0.051 | -1.08 | -0.205 | -0.003 | -0.007 | -0.007 | 0.623 | 0.7 | 0.79 | 0.72 | 0.6 | 0.72 | 0.6 |
| 6 | 30948421 | rs2517430 | 1.55 | 0.064 | -1.077 | -0.22 | -0.002 | 0.011 | 0.011 | 0 | 0.11 | 0.15 | 0.26 | 0.12 | 0.15 | 0.12 |
| 6 | 30949430 | rs2517426 | 1.573 | 0.065 | -1.077 | -0.22 | -0.001 | 0.011 | 0.011 | 0 | 0.11 | 0.15 | 0.26 | 0.12 | 0.15 | 0.12 |
| 6 | 30949670 | rs2517425 | 2.294 | 0.045 | -1.059 | -0.231 | -0.001 | 0.03 | 0.03 | 0 | 0.24 | 0.17 | 0.27 | 0.04 | 0.25 | 0.11 |
| 6 | 30950589 | rs2517422 | 1.568 | 0.095 | -1.066 | -0.205 | -0.001 | 0.011 | 0.011 | 0 | 0.11 | 0.15 | 0.26 | 0.12 | 0.15 | 0.12 |
| 6 | 30951367 | rs2530700 | 1.568 | 0.095 | -1.066 | -0.205 | -0.001 | 0.011 | 0.011 | 0 | 0.11 | 0.15 | 0.26 | 0.12 | 0.15 | 0.12 |
| 6 | 30952873 | rs2517419 | 1.568 | 0.128 | -1.031 | -0.129 | -0.001 | 0.011 | 0.011 | 0 | 0.11 | 0.15 | 0.26 | 0.12 | 0.15 | 0.12 |
| 6 | 30955359 | rs2844677 | 2.344 | 0.128 | -1.047 | -0.141 | 0.007 | 0.052 | 0.052 | 0 | 0.06 | 0.15 | 0.18 | 0.11 | 0.09 | 0.11 |
| 6 | 30955681 | rs1634731 | -2.151 | -0.102 | -1.239 | -0.135 | -0.059 | -0.008 | -0.008 | 0 | 0.82 | 0.97 | 0.87 | 0.83 | 0.82 | 0.76 |
| 6 | 30955929 | rs2517416 | 1.733 | -0.018 | -0.764 | -0.266 | -0.001 | 0.011 | 0.011 | 0 | 0.11 | 0.15 | 0.26 | 0.12 | 0.15 | 0.12 |

| | | | | | | | | | | | | | | | | |
|---|---|---|---|---|---|---|---|---|---|---|---|---|---|---|---|---|
| 6 | 30956779 | rs2517415 | 1.733 | -0.018 | -0.764 | -0.266 | -0.001 | 0.011 | 0.011 | 0 | 0.11 | 0.15 | 0.26 | 0.12 | 0.15 | 0.12 |
| 6 | 30957618 | rs886403 | 2.425 | -0.03 | -0.756 | -0.272 | -0.015 | 0.065 | 0.065 | 0 | 0.22 | 0.1 | 0.26 | 0.04 | 0.24 | 0.11 |
| 6 | 30957766 | rs3873342 | 2.022 | -0.049 | -0.764 | -0.292 | 0.001 | 0.019 | 0.019 | 0 | 0.1 | 0.21 | 0.06 | 0.08 | 0.12 | 0.06 |
| 6 | 30958494 | rs2530695 | 1.742 | -0.009 | -1.101 | -0.272 | -0.001 | 0.011 | 0.011 | 0 | 0.11 | 0.15 | 0.26 | 0.12 | 0.15 | 0.12 |
| 6 | 30960267 | rs2517411 | 1.742 | -0.009 | -1.101 | -0.272 | -0.001 | 0.011 | 0.011 | 0 | 0.11 | 0.15 | 0.26 | 0.12 | 0.15 | 0.13 |
| 6 | 30961926 | rs2844673 | 1.742 | -0.009 | -1.101 | -0.19 | -0.001 | 0.019 | 0.019 | 0 | 0.11 | 0.15 | 0.26 | 0.21 | 0.15 | 0.13 |
| 6 | 30963143 | rs9468857 | 3.809 | -0.017 | -1.074 | -0.365 | 0.001 | 0 | 0 | 0 | 0.07 | 0.05 | 0.01 | 0.01 | 0.01 | 0.02 |
| 6 | 30964393 | rs2517409 | 1.706 | 0.069 | -1.104 | -0.268 | -0.001 | 0.011 | 0.011 | 0 | 0.11 | 0.15 | 0.26 | 0.12 | 0.15 | 0.12 |
| 6 | 30966304 | rs2252926 | 1.706 | 0.069 | -1.104 | -0.268 | -0.001 | 0.011 | 0.011 | 0 | 0.11 | 0.15 | 0.26 | 0.12 | 0.15 | 0.13 |
| 6 | 30966453 | rs2517408 | 1.962 | 0.066 | -1.108 | -0.251 | -0.002 | -0.007 | -0.007 | 0 | 0.1 | 0.15 | 0.2 | 0.05 | 0.13 | 0.06 |
| 6 | 30967202 | rs2530690 | 1.698 | 0.066 | -1.106 | -0.258 | -0.001 | 0.011 | 0.011 | 0 | 0.11 | 0.15 | 0.26 | 0.12 | 0.15 | 0.13 |
| 6 | 30969898 | rs1632859 | -1.117 | 0.114 | -1.027 | -0.303 | 0.007 | -0.012 | -0.012 | 0.246 | 0.66 | 0.78 | 0.71 | 0.59 | 0.73 | 0.56 |
| 6 | 30970459 | rs2253417 | 1.475 | 0.183 | -0.995 | -0.273 | -0.003 | 0.007 | 0.007 | 0 | 0.12 | 0.15 | 0.26 | 0.12 | 0.15 | 0.12 |
| 6 | 30972202 | rs380808 | 1.734 | 0.249 | -0.964 | -0.238 | -0.001 | 0.011 | 0.011 | 0 | 0.11 | 0.15 | 0.26 | 0.12 | 0.15 | 0.12 |
| 6 | 30972390 | rs436376 | 1.734 | 0.249 | -0.964 | -0.238 | -0.001 | 0.011 | 0.011 | 0 | 0.11 | 0.15 | 0.26 | 0.12 | 0.15 | 0.13 |
| 6 | 30972589 | rs1634717 | 1.172 | 0.248 | -0.963 | -0.238 | -0.004 | 0.019 | 0.019 | 0 | 0.34 | 0.27 | 0.53 | 0.16 | 0.4 | 0.23 |
| 6 | 30972865 | rs1634718 | 2.696 | 0.282 | -0.968 | -0.188 | -0.03 | 0.057 | 0.057 | 0 | 0.19 | 0.06 | 0.21 | 0.04 | 0.23 | 0.1 |
| 6 | 30974337 | rs2844667 | 1.374 | 0.979 | -0.493 | 0.37 | -0.002 | 0.011 | 0.011 | 0 | 0.11 | 0.15 | 0.26 | 0.12 | 0.15 | 0.12 |
| 6 | 30975090 | rs2523921 | 1.395 | 0.989 | -0.489 | 0.375 | -0.001 | 0.011 | 0.011 | 0 | 0.11 | 0.15 | 0.26 | 0.12 | 0.15 | 0.12 |
| 6 | 30975141 | rs2523923 | 1.395 | 0.989 | -0.489 | 0.375 | -0.001 | 0.011 | 0.011 | 0 | 0.11 | 0.15 | 0.26 | 0.12 | 0.15 | 0.12 |
| 6 | 30976781 | rs2844666 | 1.395 | 0.989 | -0.489 | 0.375 | -0.001 | 0.011 | 0.011 | 0 | 0.11 | 0.15 | 0.26 | 0.12 | 0.15 | 0.12 |
| 6 | 30977815 | rs2523926 | 1.395 | 0.17 | -0.478 | 0.4 | -0.001 | 0.011 | 0.011 | 0 | 0.11 | 0.15 | 0.26 | 0.12 | 0.15 | 0.12 |
| 6 | 30978196 | rs2530716 | 1.395 | 0.17 | -0.478 | 0.4 | -0.001 | 0.011 | 0.011 | 0 | 0.11 | 0.15 | 0.26 | 0.12 | 0.15 | 0.12 |
| 6 | 30978746 | rs12203410 | 1.269 | 0.211 | -0.604 | 0.446 | -0.025 | -0.011 | -0.011 | 0.029 | 0.18 | 0.32 | 0.09 | 0.08 | 0.16 | 0.12 |
| 6 | 30979156 | rs1634723 | -1.073 | 0.286 | -0.829 | 0.396 | -0.004 | -0.015 | -0.015 | 0 | 0.74 | 0.81 | 0.74 | 0.59 | 0.77 | 0.62 |
| 6 | 30980603 | rs12528087 | 2.344 | 0.322 | -0.318 | 1.076 | 0.015 | -0.012 | -0.012 | 0 | 0.08 | 0.03 | 0.08 | 0.07 | 0.05 | 0.12 |
| 6 | 30980747 | rs12528082 | 2.122 | 0.323 | -0.312 | 1.073 | 0.017 | -0.009 | -0.009 | 0 | 0.08 | 0.05 | 0.09 | 0.09 | 0.06 | 0.12 |
| 6 | 30982544 | rs9394026 | -0.183 | 1.03 | 0.073 | 1.057 | 0.025 | 0.053 | 0.053 | 0 | 0.2 | 0.3 | 0.14 | 0.4 | 0.27 | 0.3 |
| 6 | 30982701 | rs4713410 | 0.398 | 0.143 | 0.126 | 0.485 | -0.007 | -0.016 | -0.016 | 0 | 0.12 | 0.19 | 0.2 | 0.05 | 0.13 | 0.07 |
| 6 | 30986264 | rs3873345 | 0.398 | 0.209 | -0.085 | 0.519 | -0.007 | -0.016 | -0.016 | 0 | 0.12 | 0.19 | 0.2 | 0.05 | 0.13 | 0.07 |
| 6 | 30986396 | rs3906267 | 0.398 | 0.209 | -0.085 | 0.519 | -0.007 | -0.016 | -0.016 | 0.19 | 0.12 | 0.19 | 0.2 | 0.05 | 0.13 | 0.07 |
| 6 | 30986527 | rs9262495 | 0.398 | 0.209 | -0.085 | 0.519 | -0.007 | -0.016 | -0.016 | 0 | 0.12 | 0.19 | 0.2 | 0.05 | 0.13 | 0.07 |
| 6 | 30987327 | rs4713413 | 0.398 | 0.209 | -0.085 | 0.519 | -0.007 | -0.016 | -0.016 | 0.26 | 0.12 | 0.19 | 0.2 | 0.05 | 0.13 | 0.07 |
| 6 | 30989770 | rs9262528 | 0.398 | 0.209 | -0.085 | 0.519 | -0.007 | -0.016 | -0.016 | 0.26 | 0.12 | 0.19 | 0.2 | 0.05 | 0.13 | 0.07 |
| 6 | 30991559 | rs6903912 | -1.136 | 0.125 | 0.086 | 0.556 | -0.019 | -0.01 | -0.01 | 0 | 0.23 | 0.44 | 0.26 | 0.13 | 0.25 | 0.13 |
| 6 | 30992346 | rs12198448 | 0.042 | 0.055 | -0.073 | 0.642 | -0.014 | 0.003 | 0.003 | 0 | 0.14 | 0.29 | 0.11 | 0.08 | 0.13 | 0.06 |
| 6 | 30993499 | rs2517554 | -0.375 | 0.187 | -0.719 | 0.752 | 0.002 | -0.015 | -0.015 | 0 | 0.4 | 0.35 | 0.49 | 0.56 | 0.56 | 0.53 |
| 6 | 30993533 | rs2523898 | -0.215 | 0.06 | -1.041 | 0.677 | 0.001 | -0.011 | -0.011 | 0 | 0.44 | 0.47 | 0.48 | 0.32 | 0.51 | 0.42 |
| 6 | 30993567 | rs4713420 | 0.175 | 0.422 | -0.895 | 1.144 | 0.022 | 0.038 | 0.038 | 0 | 0.12 | 0.13 | 0.17 | 0.36 | 0.19 | 0.24 |
| 6 | 30993958 | rs2523897 | 0.239 | -0.494 | -1.238 | 1.585 | 0.015 | -0.038 | -0.038 | 0.905 | 0.82 | 0.92 | 0.78 | 0.68 | 0.85 | 0.81 |
| 6 | 30998742 | rs6457300 | 0.95 | -0.68 | -1.286 | 0.918 | -0.014 | -0.017 | -0.017 | 0 | 0.38 | 0.38 | 0.46 | 0.33 | 0.38 | 0.24 |
| 6 | 30998879 | rs6457301 | 0.95 | -0.68 | -1.286 | 0.918 | -0.014 | -0.017 | -0.017 | 0.018 | 0.38 | 0.38 | 0.46 | 0.33 | 0.38 | 0.24 |
| 6 | 30999997 | rs10947121 | 0.432 | -1.075 | -1.332 | 1.063 | -0.006 | -0.002 | -0.002 | 0 | 0.48 | 0.41 | 0.55 | 0.53 | 0.41 | 0.42 |
| 6 | 31001421 | rs3909113 | 2.101 | -0.903 | -1.741 | 1.054 | 0.033 | -0.036 | -0.036 | 1.306 | 0.1 | 0.03 | 0.1 | 0.2 | 0.02 | 0.18 |
| 6 | 31001911 | rs11754586 | 2.146 | -1.006 | -1.878 | 0.974 | 0.062 | -0.037 | -0.037 | 0 | 0.24 | 0.02 | 0.26 | 0.19 | 0.17 | 0.25 |
| 6 | 31002013 | rs6933349 | 1.435 | -0.599 | -1.373 | 1.166 | 0.004 | -0.008 | -0.008 | 0 | 0.19 | 0.22 | 0.18 | 0.15 | 0.22 | 0.12 |
| 6 | 31002101 | rs7768644 | 2.138 | -0.782 | -1.53 | 1.083 | 0.006 | 0.004 | 0.004 | 0 | 0.1 | 0.12 | 0.09 | 0.08 | 0.07 | 0.14 |
| 6 | 31002616 | rs4248154 | 1.23 | -0.565 | -1.555 | 1.252 | 0.012 | -0.034 | -0.034 | 0 | 0.22 | 0.11 | 0.22 | 0.3 | 0.18 | 0.28 |
| 6 | 31006855 | rs2844665 | -0.04 | 0.252 | -1.488 | 1.442 | -0.002 | -0.002 | -0.002 | 0 | 0.69 | 0.71 | 0.62 | 0.47 | 0.63 | 0.65 |
| 6 | 31007081 | rs4711253 | 1.997 | 0.479 | -1.542 | 1.498 | -0.001 | 0.008 | 0.008 | 0 | 0.17 | 0.12 | 0.2 | 0.17 | 0.17 | 0.15 |
| 6 | 31007590 | rs2517552 | 0.083 | -0.124 | -1.648 | 1.261 | 0.018 | 0.026 | 0.026 | 0 | 0.75 | 0.72 | 0.63 | 0.48 | 0.64 | 0.67 |
| 6 | 31008368 | rs2517550 | -0.072 | -0.073 | -1.619 | 1.285 | -0.002 | -0.002 | -0.002 | 0 | 0.69 | 0.71 | 0.62 | 0.47 | 0.63 | 0.65 |
| 6 | 31008598 | rs2517549 | 0.926 | -0.824 | -1.962 | 0.56 | 0.027 | -0.105 | -0.105 | 0 | 0.3 | 0.54 | 0.25 | 0.09 | 0.31 | 0.2 |
| 6 | 31008818 | rs2517548 | -0.219 | -0.531 | -1.938 | 0.735 | -0.001 | -0.001 | -0.001 | 0 | 0.69 | 0.71 | 0.62 | 0.47 | 0.63 | 0.65 |
| 6 | 31009302 | rs2517545 | -0.219 | -0.531 | -1.938 | 0.735 | -0.001 | -0.001 | -0.001 | 0 | 0.69 | 0.71 | 0.62 | 0.47 | 0.63 | 0.65 |
| 6 | 31011013 | rs2844647 | -0.575 | -0.831 | -2.113 | 0.67 | -0.004 | 0.001 | 0.001 | 0 | 0.44 | 0.64 | 0.37 | 0.41 | 0.43 | 0.5 |
| 6 | 31011247 | rs2248386 | -0.575 | -0.831 | -2.113 | 0.67 | -0.004 | 0.001 | 0.001 | 0 | 0.44 | 0.64 | 0.37 | 0.41 | 0.43 | 0.5 |
| 6 | 31011662 | rs10947129 | 2.978 | -0.477 | -1.924 | 0.823 | 0.047 | 0.05 | 0.05 | 0 | 0.24 | 0.07 | 0.25 | 0.07 | 0.2 | 0.15 |
| 6 | 31011845 | rs12528584 | 2.978 | -0.477 | -1.924 | 0.823 | 0.047 | 0.05 | 0.05 | 0 | 0.24 | 0.07 | 0.25 | 0.07 | 0.2 | 0.15 |
| 6 | 31012356 | rs7756294 | 3.305 | -0.467 | -1.917 | 0.76 | 0.019 | 0.008 | 0.008 | 0 | 0.08 | 0.03 | 0.06 | 0.08 | 0.05 | 0.1 |
| 6 | 31012730 | rs2523872 | 0.053 | -0.209 | -1.753 | 0.894 | 0.019 | 0.028 | 0.028 | 0 | 0.75 | 0.71 | 0.63 | 0.48 | 0.64 | 0.67 |
| 6 | 31012996 | rs3131927 | 0.131 | -0.271 | -1.804 | 0.891 | 0.014 | 0.024 | 0.024 | 0 | 0.74 | 0.72 | 0.62 | 0.47 | 0.63 | 0.67 |
| 6 | 31014116 | rs2523870 | 0.118 | -0.282 | -1.745 | 0.784 | 0.015 | 0.023 | 0.023 | 0 | 0.74 | 0.72 | 0.61 | 0.48 | 0.64 | 0.67 |
| 6 | 31016978 | rs2251830 | -0.045 | -0.501 | -1.819 | 0.387 | 0.009 | 0.021 | 0.021 | 0 | 0.49 | 0.64 | 0.35 | 0.32 | 0.41 | 0.39 |
| 6 | 31017334 | rs2517534 | 0.919 | -0.587 | -1.865 | 0.662 | 0.01 | 0.01 | 0.01 | 0 | 0.7 | 0.68 | 0.6 | 0.51 | 0.57 | 0.53 |
| 6 | 31018407 | rs2517532 | 0.919 | -0.587 | -1.865 | 0.642 | 0.01 | 0.01 | 0.01 | 0 | 0.7 | 0.68 | 0.6 | 0.51 | 0.57 | 0.53 |
| 6 | 31018546 | rs2523864 | -0.03 | -0.591 | -1.867 | 0.64 | 0.01 | 0.021 | 0.021 | 0 | 0.49 | 0.64 | 0.35 | 0.32 | 0.41 | 0.39 |
| 6 | 31019460 | rs9295947 | 3.294 | -0.589 | -1.865 | 0.641 | 0.048 | 0.012 | 0.012 | 0 | 0.22 | 0.04 | 0.26 | 0.19 | 0.17 | 0.14 |
| 6 | 31019562 | rs2394427 | 3.294 | -0.589 | -1.865 | 0.641 | 0.048 | 0.012 | 0.012 | 0 | 0.22 | 0.04 | 0.26 | 0.19 | 0.17 | 0.14 |

| | | | | | | | | | | | | | | | | |
|---|---|---|---|---|---|---|---|---|---|---|---|---|---|---|---|---|
| 6 | 31020152 | rs2428517 | -0.31 | -0.338 | -1.654 | 0.611 | 0.01 | 0.021 | 0.021 | 0 | 0.48 | 0.64 | 0.35 | 0.32 | 0.41 | 0.39 |
| 6 | 31021547 | rs2517527 | 0.054 | 0.116 | -1.324 | 0.708 | 0.026 | 0.009 | 0.009 | 0 | 0.78 | 0.74 | 0.62 | 0.65 | 0.62 | 0.72 |
| 6 | 31023840 | rs16898614 | 2.113 | -0.599 | -1.396 | 0.425 | 0.015 | 0.012 | 0.012 | 0 | 0.1 | 0.07 | 0.06 | 0.05 | 0.05 | 0.12 |
| 6 | 31024738 | rs7356893 | 2.15 | -1.264 | -1.36 | 0.109 | 0.009 | 0.014 | 0.014 | 0 | 0.28 | 0.16 | 0.31 | 0.21 | 0.25 | 0.2 |
| 6 | 31024808 | rs9262632 | 2.27 | -1.527 | -1.411 | -0.037 | 0.034 | 0.027 | 0.027 | 0 | 0.11 | 0.03 | 0.07 | 0.05 | 0.07 | 0.12 |
| 6 | 31025051 | rs2523849 | 0.688 | -1.961 | -1.359 | 0.007 | -0.018 | -0.002 | -0.002 | 0.896 | 0.33 | 0.31 | 0.21 | 0.28 | 0.17 | 0.35 |
| 6 | 31025104 | rs2523848 | 0.688 | -1.961 | -1.359 | 0.007 | -0.018 | -0.002 | -0.002 | 0.492 | 0.33 | 0.31 | 0.21 | 0.28 | 0.17 | 0.35 |
| 6 | 31025265 | rs9262633 | 2.476 | -1.944 | -1.362 | 0.004 | 0.009 | 0.014 | 0.014 | 0 | 0.28 | 0.16 | 0.31 | 0.21 | 0.25 | 0.2 |
| 6 | 31025848 | rs9262636 | 2.477 | -1.946 | -1.363 | 0.003 | 0.009 | 0.015 | 0.015 | 0 | 0.28 | 0.16 | 0.31 | 0.21 | 0.25 | 0.2 |
| 6 | 31025989 | rs9262638 | 2.375 | -1.961 | -1.357 | -0.005 | 0.01 | 0.016 | 0.016 | 0 | 0.29 | 0.17 | 0.32 | 0.21 | 0.25 | 0.2 |
| 6 | 31026009 | rs9262639 | 2.486 | -1.951 | -1.359 | 0 | 0.009 | 0.014 | 0.014 | 0 | 0.28 | 0.16 | 0.31 | 0.21 | 0.25 | 0.2 |
| 6 | 31026434 | rs2517523 | 0.417 | -1.981 | -1.352 | -0.004 | -0.001 | 0.005 | 0.005 | 0 | 0.63 | 0.52 | 0.53 | 0.49 | 0.43 | 0.56 |
| 6 | 31027031 | rs16898629 | 3.465 | -2.013 | -1.369 | -0.01 | 0.02 | 0.021 | 0.021 | 0 | 0.07 | 0.01 | 0.04 | 0.01 | 0.04 | 0.04 |
| 6 | 31028345 | rs2517518 | 1.159 | -1.862 | -1.301 | 0.007 | -0.028 | 0.001 | 0.001 | 0 | 0.3 | 0.31 | 0.21 | 0.28 | 0.17 | 0.35 |
| 6 | 31028920 | rs2517516 | 0.444 | -1.874 | -1.307 | 0.001 | -0.001 | 0.004 | 0.004 | 0 | 0.63 | 0.52 | 0.53 | 0.49 | 0.43 | 0.56 |
| 6 | 31029173 | rs9262648 | 2.445 | -1.874 | -1.307 | 0.001 | 0.009 | 0.014 | 0.014 | 0 | 0.28 | 0.16 | 0.31 | 0.21 | 0.25 | 0.2 |
| 6 | 31029455 | rs9262651 | 2.406 | -1.856 | -1.323 | -0.015 | 0.004 | 0.016 | 0.016 | 0 | 0.28 | 0.16 | 0.31 | 0.21 | 0.27 | 0.22 |
| 6 | 31029685 | rs2517512 | 2.066 | -2.2 | -1.364 | -0.102 | -0.035 | -0.048 | -0.048 | 0 | 0.21 | 0.28 | 0.17 | 0.06 | 0.14 | 0.21 |
| 6 | 31029918 | rs2517511 | 0.419 | -2.119 | -1.345 | -0.099 | -0.001 | 0.004 | 0.004 | 0 | 0.63 | 0.52 | 0.53 | 0.49 | 0.43 | 0.56 |
| 6 | 31030122 | rs2517510 | 0.419 | -2.119 | -1.345 | -0.099 | -0.001 | 0.004 | 0.004 | 0 | 0.63 | 0.52 | 0.53 | 0.49 | 0.43 | 0.56 |
| 6 | 31030283 | rs2523841 | 1.094 | -2.114 | -1.344 | -0.096 | -0.028 | 0.001 | 0.001 | 0 | 0.3 | 0.31 | 0.21 | 0.28 | 0.17 | 0.35 |
| 6 | 31031381 | rs2256436 | 0.725 | -2.054 | -1.319 | -0.076 | -0.004 | 0.008 | 0.008 | 0 | 0.61 | 0.47 | 0.52 | 0.49 | 0.42 | 0.56 |
| 6 | 31031680 | rs2517506 | 0.742 | -2.063 | -1.321 | -0.078 | -0.019 | -0.002 | -0.002 | 0 | 0.33 | 0.31 | 0.21 | 0.28 | 0.17 | 0.35 |
| 6 | 31031904 | rs3906263 | 3.169 | -2.377 | -1.368 | -0.259 | -0.006 | 0.011 | 0.011 | 0.141 | 0.22 | 0.07 | 0.29 | 0.19 | 0.24 | 0.18 |
| 6 | 31032042 | rs2394423 | 2.415 | -2.34 | -1.381 | -0.242 | 0.009 | 0.014 | 0.014 | 0 | 0.28 | 0.16 | 0.31 | 0.21 | 0.25 | 0.2 |
| 6 | 31032370 | rs2517504 | 0.358 | -2.324 | -1.375 | -0.235 | -0.001 | 0.004 | 0.004 | 0 | 0.63 | 0.52 | 0.54 | 0.49 | 0.43 | 0.56 |
| 6 | 31032937 | rs2517502 | 0.687 | -2.324 | -1.375 | -0.235 | -0.004 | 0.008 | 0.008 | 0 | 0.61 | 0.47 | 0.52 | 0.49 | 0.42 | 0.56 |
| 6 | 31033509 | rs2517500 | 0.687 | -2.324 | -1.375 | -0.235 | -0.004 | 0.008 | 0.008 | 0 | 0.61 | 0.47 | 0.53 | 0.49 | 0.42 | 0.56 |
| 6 | 31034264 | rs9262656 | 2.416 | -2.338 | -1.384 | -0.24 | 0.009 | 0.014 | 0.014 | 0 | 0.28 | 0.16 | 0.31 | 0.21 | 0.25 | 0.2 |
| 6 | 31036783 | rs2517497 | 0.416 | -1.943 | -1.513 | -0.167 | -0.001 | 0.004 | 0.004 | 0 | 0.63 | 0.52 | 0.53 | 0.49 | 0.43 | 0.56 |
| 6 | 31038976 | rs2523891 | 0.742 | -1.938 | -1.427 | -0.173 | -0.004 | 0.008 | 0.008 | 0 | 0.61 | 0.47 | 0.52 | 0.49 | 0.42 | 0.56 |
| 6 | 31041155 | rs2523884 | 0.742 | -1.938 | -1.427 | -0.173 | -0.004 | 0.008 | 0.008 | 0 | 0.61 | 0.47 | 0.52 | 0.49 | 0.42 | 0.56 |
| 6 | 31042217 | rs2523882 | 1.204 | -1.979 | -1.473 | -0.259 | -0.016 | 0.001 | 0.001 | 0.008 | 0.55 | 0.41 | 0.48 | 0.47 | 0.34 | 0.53 |
| 6 | 31042306 | rs2517489 | 0.741 | -1.891 | -1.426 | -0.19 | -0.004 | 0.008 | 0.008 | 0 | 0.61 | 0.47 | 0.52 | 0.49 | 0.42 | 0.56 |
| 6 | 31042608 | rs2523881 | 0.741 | -1.891 | -1.426 | -0.19 | -0.004 | 0.008 | 0.008 | 0 | 0.61 | 0.47 | 0.52 | 0.49 | 0.42 | 0.56 |
| 6 | 31043232 | rs2249231 | 0.415 | -1.895 | -1.424 | -0.192 | -0.001 | 0.004 | 0.004 | 0 | 0.63 | 0.52 | 0.53 | 0.49 | 0.43 | 0.56 |
| 6 | 31044468 | rs2905759 | 0.76 | -1.897 | -1.433 | -0.175 | -0.005 | 0.011 | 0.011 | 0 | 0.61 | 0.47 | 0.52 | 0.49 | 0.42 | 0.56 |
| 6 | 31044802 | rs2517478 | 0.76 | -1.897 | -1.433 | -0.175 | -0.005 | 0.011 | 0.011 | 0 | 0.61 | 0.47 | 0.52 | 0.49 | 0.42 | 0.56 |
| 6 | 31044863 | rs2517477 | 0.76 | -1.897 | -1.433 | -0.175 | -0.005 | 0.011 | 0.011 | 0 | 0.61 | 0.47 | 0.52 | 0.49 | 0.42 | 0.56 |
| 6 | 31046528 | rs2523843 | 0.75 | -1.897 | -1.433 | -0.175 | -0.005 | 0.011 | 0.011 | 0 | 0.61 | 0.48 | 0.52 | 0.49 | 0.42 | 0.56 |
| 6 | 31047145 | rs2523836 | 0.76 | -1.897 | -1.433 | -0.175 | -0.005 | 0.011 | 0.011 | 0 | 0.61 | 0.47 | 0.52 | 0.49 | 0.42 | 0.56 |
| 6 | 31047471 | rs2523895 | 0.413 | -1.898 | -1.433 | -0.175 | -0.002 | 0.007 | 0.007 | 0 | 0.63 | 0.52 | 0.53 | 0.49 | 0.43 | 0.56 |
| 6 | 31048263 | rs2106067 | 0.769 | -1.897 | -1.432 | -0.175 | -0.005 | 0.011 | 0.011 | 0 | 0.61 | 0.47 | 0.52 | 0.49 | 0.42 | 0.56 |
| 6 | 31048387 | rs2249876 | 0.422 | -1.897 | -1.432 | -0.175 | -0.002 | 0.007 | 0.007 | 0 | 0.63 | 0.52 | 0.53 | 0.49 | 0.43 | 0.56 |
| 6 | 31049655 | rs9380215 | 2.681 | -2.315 | -1.661 | -0.247 | 0.041 | -0.056 | -0.056 | 0 | 0.15 | 0.02 | 0.13 | 0.24 | 0.08 | 0.13 |
| 6 | 31050179 | rs9263473 | 0.254 | -2.129 | -1.562 | -0.201 | -0.002 | 0.007 | 0.007 | 0 | 0.63 | 0.52 | 0.53 | 0.49 | 0.43 | 0.56 |
| 6 | 31050244 | rs9263474 | 0.254 | -2.129 | -1.562 | -0.201 | -0.002 | 0.007 | 0.007 | 0 | 0.63 | 0.52 | 0.53 | 0.49 | 0.43 | 0.56 |
| 6 | 31051388 | rs2535318 | 0.264 | -2.129 | -1.562 | -0.201 | -0.002 | 0.007 | 0.007 | 0 | 0.63 | 0.52 | 0.53 | 0.49 | 0.43 | 0.56 |
| 6 | 31052127 | rs2535315 | 0.571 | -2.122 | -1.571 | -0.197 | -0.005 | 0.011 | 0.011 | 0 | 0.61 | 0.47 | 0.52 | 0.49 | 0.42 | 0.56 |
| 6 | 31052684 | rs2535311 | 0.571 | -2.122 | -1.571 | -0.197 | -0.005 | 0.011 | 0.011 | 0 | 0.61 | 0.47 | 0.52 | 0.49 | 0.42 | 0.56 |
| 6 | 31053257 | rs2535310 | 0.48 | -1.575 | -1.578 | -0.217 | 0.009 | 0.012 | 0.012 | 0 | 0.62 | 0.48 | 0.56 | 0.49 | 0.51 | 0.56 |
| 6 | 31053867 | rs2535306 | 0.582 | -1.618 | -1.578 | -0.215 | -0.005 | 0.011 | 0.011 | 0 | 0.61 | 0.47 | 0.52 | 0.49 | 0.42 | 0.56 |
| 6 | 31054511 | rs3130955 | 1.084 | -0.951 | -1.591 | -0.027 | 0.011 | 0.009 | 0.009 | 0 | 0.27 | 0.36 | 0.27 | 0.31 | 0.35 | 0.36 |
| 6 | 31054687 | rs2535304 | -0.256 | -1.863 | -1.535 | -0.1 | -0.001 | 0.004 | 0.004 | 0 | 0.63 | 0.54 | 0.53 | 0.49 | 0.43 | 0.56 |
| 6 | 31055423 | rs2535303 | -0.268 | -1.665 | -1.534 | -0.01 | 0.012 | 0.014 | 0.014 | 0 | 0.65 | 0.55 | 0.57 | 0.49 | 0.51 | 0.57 |
| 6 | 31058178 | rs4947296 | 2.669 | -1.793 | -1.603 | -0.038 | 0.038 | -0.056 | -0.056 | 0.001 | 0.14 | 0.02 | 0.1 | 0.22 | 0.08 | 0.13 |
| 6 | 31060115 | rs2517453 | -0.283 | -1.833 | -1.574 | -0.01 | 0.012 | 0.014 | 0.014 | 0 | 0.65 | 0.55 | 0.57 | 0.49 | 0.51 | 0.57 |
| 6 | 31062133 | rs2535294 | 0.082 | -1.929 | -1.618 | 0.027 | -0.001 | -0.001 | -0.001 | 0 | 0.61 | 0.55 | 0.51 | 0.4 | 0.51 | 0.47 |
| 6 | 31062363 | rs2535293 | -0.296 | -1.903 | -1.606 | 0.029 | 0.012 | 0.014 | 0.014 | 0 | 0.65 | 0.55 | 0.57 | 0.49 | 0.51 | 0.57 |
| 6 | 31062667 | rs2517448 | -0.825 | -1.942 | -1.64 | 0.068 | 0.066 | 0.03 | 0.03 | 0 | 0.83 | 0.75 | 0.63 | 0.64 | 0.63 | 0.73 |
| 6 | 31062899 | rs2535291 | -0.274 | -1.954 | -1.689 | 0.049 | 0.012 | 0.015 | 0.015 | 0 | 0.66 | 0.55 | 0.57 | 0.49 | 0.51 | 0.57 |
| 6 | 31066419 | rs9263509 | 1.628 | -2.012 | -1.71 | 0.022 | 0.013 | -0.027 | -0.027 | 0.222 | 0.23 | 0.08 | 0.14 | 0.3 | 0.17 | 0.17 |
| 6 | 31066671 | rs2894179 | 2.55 | -2.022 | -1.715 | 0.019 | -0.011 | -0.023 | -0.023 | 0 | 0.17 | 0.06 | 0.13 | 0.24 | 0.15 | 0.14 |
| 6 | 31067009 | rs2517403 | 1.4 | -2.033 | -1.71 | 0.001 | -0.017 | -0.068 | -0.068 | 0.196 | 0.38 | 0.46 | 0.37 | 0.1 | 0.34 | 0.29 |
| 6 | 31068420 | rs2535283 | -0.289 | -2.042 | -1.714 | 0.008 | 0.012 | 0.014 | 0.014 | 0 | 0.65 | 0.55 | 0.57 | 0.49 | 0.51 | 0.55 |
| 6 | 31069899 | rs6919169 | 1.362 | -2.003 | -1.693 | -0.004 | -0.017 | -0.068 | -0.068 | 0 | 0.38 | 0.46 | 0.37 | 0.1 | 0.34 | 0.28 |
| 6 | 31071645 | rs9263600 | 2.121 | -1.957 | -1.685 | 0.051 | -0.003 | -0.006 | -0.006 | 0 | 0.1 | 0.08 | 0.1 | 0.15 | 0.09 | 0.14 |
| 6 | 31073903 | rs7381897 | -0.287 | -2.01 | -1.698 | 0.025 | 0.012 | 0.014 | 0.014 | 0 | 0.65 | 0.55 | 0.57 | 0.49 | 0.51 | 0.55 |
| 6 | 31074030 | rs6457327 | -0.896 | -2.012 | -1.702 | 0.042 | 0.071 | 0.033 | 0.033 | 0 | 0.83 | 0.74 | 0.63 | 0.64 | 0.63 | 0.73 |

| | | | | | | | | | | | | | | | | |
|---|---|---|---|---|---|---|---|---|---|---|---|---|---|---|---|---|
| 6 | 31074101 | rs2517485 | 0.912 | -1.751 | -1.709 | -0.287 | -0.014 | -0.025 | -0.025 | 0 | 0.42 | 0.46 | 0.43 | 0.19 | 0.35 | 0.38 |
| 6 | 31075161 | rs9263625 | 1.008 | -1.71 | -1.69 | -0.285 | 0.039 | -0.061 | -0.061 | 0.019 | 0.28 | 0.09 | 0.2 | 0.39 | 0.17 | 0.27 |
| 6 | 31075375 | rs1064191 | -0.488 | -1.709 | -1.69 | -0.285 | 0.012 | 0.014 | 0.014 | 0 | 0.65 | 0.55 | 0.57 | 0.49 | 0.51 | 0.55 |
| 6 | 31075481 | rs2844635 | 1.182 | -1.693 | -1.665 | -0.305 | -0.017 | -0.068 | -0.068 | 0 | 0.38 | 0.46 | 0.37 | 0.1 | 0.34 | 0.29 |
| 6 | 31077405 | rs9391709 | 2.375 | -1.696 | -1.687 | -0.263 | 0.059 | -0.049 | -0.049 | 0.153 | 0.17 | 0.01 | 0.1 | 0.24 | 0.08 | 0.13 |
| 6 | 31077938 | rs1265063 | -0.485 | -1.705 | -1.691 | -0.274 | 0.012 | 0.014 | 0.014 | 0 | 0.65 | 0.55 | 0.57 | 0.49 | 0.51 | 0.55 |
| 6 | 31078005 | rs1265062 | 1.186 | -1.718 | -1.692 | -0.295 | -0.017 | -0.068 | -0.068 | 0 | 0.38 | 0.46 | 0.37 | 0.1 | 0.34 | 0.29 |
| 6 | 31078836 | rs3734854 | 2.328 | -1.627 | -1.691 | -0.169 | 0.059 | -0.049 | -0.049 | 0 | 0.17 | 0.01 | 0.1 | 0.24 | 0.08 | 0.13 |
| 6 | 31079242 | rs2233985 | 2.306 | -1.627 | -1.692 | -0.17 | 0.06 | -0.049 | -0.049 | 0.011 | 0.17 | 0.01 | 0.1 | 0.24 | 0.08 | 0.13 |
| 6 | 31079264 | rs2233984 | 2.328 | -1.626 | -1.691 | -0.169 | 0.059 | -0.049 | -0.049 | 0 | 0.17 | 0.01 | 0.1 | 0.24 | 0.08 | 0.13 |
| 6 | 31079894 | rs2233977 | 1.285 | -1.558 | -1.657 | -0.157 | 0.029 | -0.061 | -0.061 | 0 | 0.26 | 0.09 | 0.2 | 0.39 | 0.17 | 0.26 |
| 6 | 31079994 | rs2233976 | 3.345 | -1.296 | -1.465 | -0.103 | 0.006 | 0.011 | 0.011 | 0.117 | 0.1 | 0.05 | 0.15 | 0.04 | 0.12 | 0.08 |
| 6 | 31080471 | rs1265052 | -0.57 | -1.176 | -1.456 | -0.045 | 0.022 | 0.019 | 0.019 | 0 | 0.67 | 0.55 | 0.57 | 0.49 | 0.52 | 0.62 |
| 6 | 31080586 | rs2270190 | 1.502 | -1.336 | -1.587 | 0.038 | 0.021 | 0.023 | 0.023 | 0 | 0.2 | 0.11 | 0.06 | 0.11 | 0.1 | 0.27 |
| 6 | 31080718 | rs4495304 | 2.595 | -1.203 | -1.615 | 0.124 | 0.036 | -0.059 | -0.059 | 0.146 | 0.14 | 0.01 | 0.1 | 0.24 | 0.08 | 0.13 |
| 6 | 31080859 | rs2233966 | -0.344 | -1.137 | -1.518 | 0.074 | 0.008 | 0.01 | 0.01 | 0 | 0.62 | 0.52 | 0.52 | 0.49 | 0.51 | 0.55 |
| 6 | 31080899 | rs2233965 | 1.281 | -0.718 | -2.104 | 0.274 | 0.026 | -0.052 | -0.052 | 0 | 0.24 | 0.06 | 0.19 | 0.32 | 0.16 | 0.23 |
| 6 | 31080919 | rs2233964 | 0.751 | -0.707 | -2.062 | 0.279 | 0.051 | -0.017 | -0.017 | 0 | 0.28 | 0.11 | 0.2 | 0.33 | 0.16 | 0.25 |
| 6 | 31081989 | rs3130977 | 0.865 | 0.096 | -1.515 | 0.047 | 0.008 | -0.008 | -0.008 | 0 | 0.28 | 0.27 | 0.32 | 0.2 | 0.32 | 0.29 |
| 6 | 31082081 | rs17190526 | 1.704 | 0.138 | -1.625 | 0.072 | 0.005 | -0.001 | -0.001 | 0 | 0.08 | 0.08 | 0.05 | 0.06 | 0.1 | 0.06 |
| 6 | 31082285 | rs3130553 | -0.144 | -0.479 | -1.856 | -0.011 | -0.001 | -0.001 | -0.001 | 0 | 0.59 | 0.35 | 0.51 | 0.41 | 0.49 | 0.64 |
| 6 | 31082304 | rs3130554 | 0.04 | -0.442 | -1.838 | -0.014 | 0.004 | 0.009 | 0.009 | 0.019 | 0.53 | 0.34 | 0.5 | 0.39 | 0.49 | 0.59 |
| 6 | 31082361 | rs6917517 | 1.551 | -0.718 | -1.763 | 0.052 | 0.067 | 0.04 | 0.04 | 0 | 0.26 | 0.09 | 0.21 | 0.2 | 0.18 | 0.3 |
| 6 | 31082404 | rs3130980 | 0.261 | 0.076 | -1.75 | 0.26 | -0.004 | 0.008 | 0.008 | 0.205 | 0.41 | 0.62 | 0.41 | 0.53 | 0.37 | 0.32 |
| 6 | 31082987 | rs3094220 | 0.726 | 0.051 | -1.697 | 0.261 | 0.021 | 0.004 | 0.004 | 0 | 0.17 | 0.21 | 0.15 | 0.16 | 0.19 | 0.21 |
| 6 | 31083156 | rs1042147 | -0.5 | 0.042 | -1.737 | 0.237 | 0.009 | -0.001 | -0.001 | 0 | 0.66 | 0.71 | 0.61 | 0.71 | 0.55 | 0.62 |
| 6 | 31083656 | rs3094217 | -0.5 | 0.042 | -1.737 | 0.237 | 0.009 | -0.001 | -0.001 | 0.869 | 0.66 | 0.71 | 0.61 | 0.71 | 0.55 | 0.62 |
| 6 | 31083664 | rs1042134 | -0.5 | 0.042 | -1.737 | 0.237 | 0.009 | -0.001 | -0.001 | 0 | 0.66 | 0.71 | 0.61 | 0.71 | 0.55 | 0.62 |
| 6 | 31083813 | rs3130981 | -2.768 | 0.044 | -1.738 | 0.237 | -0.018 | 0.005 | 0.005 | 0 | 0.83 | 0.91 | 0.77 | 0.87 | 0.74 | 0.84 |
| 6 | 31084288 | rs1042126 | -0.5 | 0.03 | -1.742 | 0.23 | 0.009 | -0.001 | -0.001 | 0.103 | 0.66 | 0.71 | 0.61 | 0.71 | 0.55 | 0.62 |
| 6 | 31084435 | rs1062470 | 0.35 | 0.032 | -1.789 | 0.2 | -0.008 | 0.01 | 0.01 | 0.155 | 0.4 | 0.62 | 0.4 | 0.51 | 0.35 | 0.31 |
| 6 | 31084792 | rs3130983 | -0.473 | 0.083 | -1.76 | 0.215 | 0.008 | -0.001 | -0.001 | 0 | 0.66 | 0.71 | 0.61 | 0.71 | 0.55 | 0.62 |
| 6 | 31088922 | rs2239520 | 0.048 | 0.161 | -1.71 | 0.228 | 0 | 0.049 | 0.049 | 0 | 0.41 | 0.22 | 0.3 | 0.31 | 0.38 | 0.53 |
| 6 | 31093066 | rs3095302 | -0.272 | -0.008 | -1.68 | 0.275 | 0.034 | 0.051 | 0.051 | 0.642 | 0.51 | 0.75 | 0.56 | 0.61 | 0.45 | 0.43 |
| 6 | 31093190 | rs6909321 | 2.38 | -0.4 | -1.667 | 0.504 | 0.026 | -0.004 | -0.004 | 0.409 | 0.21 | 0.2 | 0.06 | 0.15 | 0.11 | 0.18 |
| 6 | 31093356 | rs3095301 | -0.225 | 0.04 | -1.358 | 0.661 | 0.022 | 0.031 | 0.031 | 0.352 | 0.5 | 0.67 | 0.55 | 0.62 | 0.45 | 0.43 |
| 6 | 31093482 | rs3131003 | -0.376 | 0.069 | -1.456 | 0.647 | 0.031 | 0.049 | 0.049 | 0.883 | 0.52 | 0.76 | 0.57 | 0.63 | 0.45 | 0.43 |
| 6 | 31093587 | rs3815087 | 0.618 | 0.075 | -1.377 | 0.69 | 0.018 | 0.001 | 0.001 | 0.295 | 0.38 | 0.36 | 0.23 | 0.26 | 0.22 | 0.36 |
| 6 | 31093699 | rs9263699 | 0.637 | 0.082 | -1.407 | 0.693 | 0.028 | 0.001 | 0.001 | 0.256 | 0.38 | 0.34 | 0.23 | 0.26 | 0.22 | 0.36 |
| 6 | 31093776 | rs3778639 | 1.718 | 0.091 | -1.452 | 0.681 | 0.061 | 0.009 | 0.009 | 0.808 | 0.27 | 0.21 | 0.06 | 0.17 | 0.11 | 0.24 |
| 6 | 31094971 | rs10947137 | 2.222 | -0.123 | -1.589 | 0.603 | 0.017 | 0.001 | 0.001 | 0.155 | 0.15 | 0.01 | 0.1 | 0.13 | 0.09 | 0.18 |
| 6 | 31095801 | rs9263715 | 0.697 | -0.226 | -1.546 | 0.639 | 0.028 | 0.002 | 0.002 | 0.296 | 0.38 | 0.34 | 0.23 | 0.27 | 0.22 | 0.36 |
| 6 | 31096575 | rs9263719 | 1.502 | -0.248 | -1.633 | 0.701 | -0.027 | 0.01 | 0.01 | 0 | 0.17 | 0.18 | 0.18 | 0.16 | 0.13 | 0.17 |
| 6 | 31097301 | rs3130559 | 0.493 | 0.673 | -1.59 | 0.944 | 0.018 | 0.07 | 0.07 | 0.246 | 0.19 | 0.39 | 0.35 | 0.39 | 0.22 | 0.23 |
| 6 | 31097453 | rs3130560 | -2.191 | 1.13 | -1.468 | 0.474 | -0.01 | 0.005 | 0.005 | 0 | 0.82 | 0.88 | 0.76 | 0.88 | 0.72 | 0.83 |
| 6 | 31097918 | rs6929464 | 0.62 | 0.834 | -1.422 | 0.397 | 0 | 0.002 | 0.002 | 0.848 | 0.38 | 0.37 | 0.18 | 0.18 | 0.19 | 0.26 |
| 6 | 31099577 | rs4959053 | 1.984 | 2.547 | -1.368 | 0.706 | 0.013 | -0.002 | -0.002 | 0 | 0.14 | 0.01 | 0.09 | 0.13 | 0.08 | 0.18 |
| 6 | 31105310 | rs1265100 | 0.765 | 2.345 | -0.681 | 1.482 | 0.01 | 0.031 | 0.031 | 0.064 | 0.13 | 0.16 | 0.23 | 0.33 | 0.17 | 0.12 |
| 6 | 31105633 | rs2074478 | -1.094 | 2.318 | -1.117 | 1.571 | -0.011 | -0.034 | -0.034 | 0 | 0.83 | 0.9 | 0.85 | 0.77 | 0.83 | 0.82 |
| 6 | 31106268 | rs3130573 | 0.364 | 1.614 | -0.958 | 0.994 | 0.004 | -0.022 | -0.022 | 0 | 0.32 | 0.49 | 0.32 | 0.18 | 0.35 | 0.32 |
| 6 | 31106499 | rs9263726 | 0.261 | 2.548 | -0.844 | 0.888 | 0.011 | -0.003 | -0.003 | 0.125 | 0.1 | 0.16 | 0.12 | 0.06 | 0.15 | 0.05 |
| 6 | 31106516 | rs9501057 | 0.41 | 2.105 | -0.267 | 1.129 | 0.003 | 0.078 | 0.078 | 0.176 | 0.05 | 0.15 | 0.02 | 0.17 | 0.04 | 0.05 |
| 6 | 31106893 | rs1265094 | -0.342 | 0.688 | -0.448 | 0.519 | -0.002 | 0.001 | 0.001 | 0 | 0.56 | 0.56 | 0.44 | 0.35 | 0.53 | 0.46 |
| 6 | 31107087 | rs3094663 | -1.455 | 0.509 | -0.63 | 0.763 | 0.001 | 0.003 | 0.003 | 0 | 0.77 | 0.74 | 0.78 | 0.7 | 0.74 | 0.7 |
| 6 | 31107187 | rs1265093 | 1.192 | 0.807 | -0.695 | 0.703 | -0.011 | -0.006 | -0.006 | 0.44 | 0.23 | 0.19 | 0.35 | 0.35 | 0.28 | 0.25 |
| 6 | 31107258 | rs2285803 | -1.453 | 0.725 | -0.678 | 0.699 | 0.001 | 0.003 | 0.003 | 0 | 0.77 | 0.74 | 0.78 | 0.7 | 0.74 | 0.7 |
| 6 | 31107361 | rs2233945 | 0.594 | 0.619 | -0.686 | 0.602 | 0.012 | -0.004 | -0.004 | 0 | 0.1 | 0.16 | 0.13 | 0.06 | 0.17 | 0.05 |
| 6 | 31107733 | rs1966 | 1.498 | 0.599 | -0.564 | 0.595 | -0.001 | -0.005 | -0.005 | 0 | 0.22 | 0.18 | 0.33 | 0.35 | 0.21 | 0.24 |
| 6 | 31108485 | rs3130454 | -1.447 | 0.566 | -0.539 | 0.581 | 0.003 | 0.007 | 0.007 | 0 | 0.77 | 0.72 | 0.77 | 0.7 | 0.74 | 0.69 |
| 6 | 31108829 | rs9263733 | 2.189 | 0.527 | -0.504 | 0.52 | 0.001 | -0.002 | -0.002 | 0 | 0.05 | 0.06 | 0.04 | 0.06 | 0.04 | 0.03 |
| 6 | 31109693 | rs16898844 | 1.124 | 0.541 | -0.483 | 0.542 | 0.01 | 0.096 | 0.096 | 0.124 | 0.05 | 0.15 | 0.02 | 0.18 | 0.04 | 0.05 |
| 6 | 31109810 | rs1265087 | 1.391 | -0.091 | -0.797 | 0.22 | 0.004 | 0.04 | 0.04 | 0 | 0.32 | 0.23 | 0.28 | 0.07 | 0.31 | 0.24 |
| 6 | 31110786 | rs12364 | 0.396 | -0.136 | -0.781 | 0.232 | -0.001 | -0.001 | -0.001 | 0 | 0.15 | 0.16 | 0.14 | 0.1 | 0.17 | 0.14 |
| 6 | 31110969 | rs2073718 | 1.171 | -0.138 | -0.763 | 0.256 | 0.01 | 0.096 | 0.096 | 0.124 | 0.05 | 0.15 | 0.02 | 0.18 | 0.04 | 0.05 |
| 6 | 31111356 | rs9263739 | 0.49 | -0.37 | -0.85 | 0.117 | -0.001 | 0 | 0 | 0 | 0.15 | 0.16 | 0.14 | 0.1 | 0.17 | 0.14 |
| 6 | 31111400 | rs9263740 | -0.735 | -0.372 | -0.858 | 0.115 | -0.006 | 0.009 | 0.009 | 0 | 0.23 | 0.31 | 0.17 | 0.28 | 0.21 | 0.22 |
| 6 | 31112075 | rs1265080 | -0.285 | -0.356 | -0.796 | 0.118 | 0.001 | -0.002 | -0.002 | 0.072 | 0.55 | 0.53 | 0.44 | 0.35 | 0.53 | 0.45 |
| 6 | 31112354 | rs11967883 | 1.148 | -0.322 | -0.727 | 0.198 | 0.01 | 0.096 | 0.096 | 0.124 | 0.05 | 0.15 | 0.02 | 0.18 | 0.04 | 0.05 |
| 6 | 31112484 | rs130072 | 3.417 | -0.731 | -0.998 | -0.204 | 0.008 | 0.009 | 0.009 | 0.31 | 0.11 | 0.06 | 0.14 | 0.03 | 0.11 | 0.08 |

| | | | | | | | | | | | | | | | | |
|---|---|---|---|---|---|---|---|---|---|---|---|---|---|---|---|---|
| 6 | 31113052 | rs3094225 | -0.991 | -0.791 | -0.856 | -0.037 | 0.049 | 0.135 | 0.135 | 0.015 | 0.79 | 0.58 | 0.79 | 0.53 | 0.75 | 0.72 |
| 6 | 31113083 | rs1265076 | 1.336 | -0.891 | -0.967 | -0.192 | 0.004 | 0.046 | 0.046 | 0 | 0.32 | 0.22 | 0.28 | 0.07 | 0.31 | 0.24 |
| 6 | 31113214 | rs1265074 | 1.799 | -0.994 | -1.032 | -0.233 | 0 | -0.002 | -0.002 | 0 | 0.22 | 0.19 | 0.33 | 0.35 | 0.21 | 0.24 |
| 6 | 31113842 | rs3132538 | -2.838 | -1.054 | -1.012 | -0.067 | -0.033 | -0.023 | -0.023 | 0 | 0.89 | 0.95 | 0.9 | 0.83 | 0.82 | 0.83 |
| 6 | 31114182 | rs746647 | 1.358 | -1.063 | -1.036 | -0.252 | 0.004 | 0.046 | 0.046 | 0 | 0.32 | 0.22 | 0.28 | 0.07 | 0.32 | 0.24 |
| 6 | 31114515 | rs9263749 | 0.665 | -1.095 | -1.036 | -0.248 | -0.001 | 0 | 0 | 0 | 0.15 | 0.16 | 0.14 | 0.1 | 0.17 | 0.14 |
| 6 | 31114573 | rs2240064 | 0.091 | -1.155 | -1.012 | -0.227 | 0.002 | 0.002 | 0.002 | 0 | 0.62 | 0.56 | 0.64 | 0.6 | 0.57 | 0.55 |
| 6 | 31114848 | rs2240061 | 1.36 | -1.138 | -0.984 | -0.2 | 0.01 | 0.096 | 0.096 | 0.124 | 0.05 | 0.15 | 0.02 | 0.18 | 0.04 | 0.05 |
| 6 | 31115441 | rs3131012 | 0.178 | -1.162 | -1.073 | -0.292 | -0.001 | -0.001 | -0.001 | 0 | 0.59 | 0.56 | 0.63 | 0.6 | 0.56 | 0.53 |
| 6 | 31116502 | rs2027937 | 2.703 | -1.267 | -1.148 | -0.289 | 0.038 | 0.054 | 0.054 | 0 | 0.16 | 0.04 | 0.1 | 0.04 | 0.08 | 0.12 |
| 6 | 31116627 | rs3131014 | 0.325 | -1.221 | -0.948 | -0.248 | -0.01 | 0.011 | 0.011 | 0 | 0.32 | 0.29 | 0.38 | 0.51 | 0.24 | 0.28 |
| 6 | 31117075 | rs1265115 | 0.072 | -0.626 | -1.086 | -0.248 | -0.001 | 0 | 0 | 0 | 0.39 | 0.36 | 0.42 | 0.56 | 0.35 | 0.51 |
| 6 | 31117188 | rs1265114 | 1.219 | -0.532 | -1.114 | -0.265 | 0.004 | 0.046 | 0.046 | 0 | 0.32 | 0.22 | 0.28 | 0.07 | 0.31 | 0.24 |
| 6 | 31118019 | rs1265112 | 1.219 | -0.532 | -1.114 | -0.265 | 0.004 | 0.046 | 0.046 | 0 | 0.32 | 0.22 | 0.28 | 0.07 | 0.31 | 0.24 |
| 6 | 31118511 | rs130067 | 1.707 | -0.593 | -1.072 | -0.16 | 0 | -0.002 | -0.002 | 0.266 | 0.22 | 0.19 | 0.33 | 0.35 | 0.21 | 0.24 |
| 6 | 31118942 | rs2517985 | 1.267 | -0.571 | -1.074 | -0.192 | 0.004 | 0.046 | 0.046 | 0 | 0.32 | 0.22 | 0.28 | 0.07 | 0.31 | 0.24 |
| 6 | 31119422 | rs1265110 | 1.947 | -0.558 | -1.061 | -0.151 | 0.065 | 0.118 | 0.118 | 0 | 0.06 | 0.15 | 0.23 | 0.3 | 0.13 | 0.12 |
| 6 | 31119589 | rs1265109 | 0.523 | -0.765 | -0.995 | -0.165 | -0.007 | 0.012 | 0.012 | 0 | 0.34 | 0.35 | 0.4 | 0.54 | 0.3 | 0.37 |
| 6 | 31120614 | rs2240059 | -1.024 | -0.595 | -0.957 | -0.13 | 0.001 | 0.008 | 0.008 | 0 | 0.77 | 0.73 | 0.8 | 0.68 | 0.78 | 0.66 |
| 6 | 31122330 | rs130077 | 0.921 | -0.514 | -0.93 | -0.162 | 0.01 | -0.003 | -0.003 | 0 | 0.1 | 0.15 | 0.13 | 0.06 | 0.17 | 0.05 |
| 6 | 31122482 | rs130076 | 2.322 | -0.754 | -1.107 | -0.358 | -0.027 | 0.032 | 0.032 | 0 | 0.17 | 0.13 | 0.09 | 0.04 | 0.19 | 0.16 |
| 6 | 31122502 | rs130075 | 1.904 | -0.739 | -0.898 | -0.213 | -0.023 | -0.016 | -0.016 | 0 | 0.09 | 0.18 | 0.04 | 0.04 | 0.04 | 0.1 |
| 6 | 31123416 | rs9263781 | 0.929 | -0.641 | -0.919 | -0.188 | 0.01 | -0.003 | -0.003 | 0 | 0.1 | 0.15 | 0.13 | 0.06 | 0.17 | 0.05 |
| 6 | 31124849 | rs3130453 | 0.161 | -0.901 | -0.867 | -0.256 | -0.001 | 0.003 | 0.003 | 0 | 0.52 | 0.5 | 0.52 | 0.4 | 0.46 | 0.48 |
| 6 | 31125158 | rs743401 | -0.146 | -0.912 | -0.916 | -0.26 | -0.007 | -0.007 | -0.007 | 0 | 0.71 | 0.72 | 0.69 | 0.61 | 0.62 | 0.63 |
| 6 | 31125407 | rs12529697 | 2.853 | -1.026 | -0.989 | -0.331 | 0.038 | 0.059 | 0.059 | 0 | 0.16 | 0.03 | 0.1 | 0.04 | 0.08 | 0.12 |
| 6 | 31125569 | rs2239524 | -0.15 | -0.948 | -0.939 | -0.278 | -0.007 | -0.007 | -0.007 | 0.612 | 0.71 | 0.72 | 0.69 | 0.61 | 0.62 | 0.63 |
| 6 | 31125777 | rs720465 | 1.069 | -1.131 | -0.979 | -0.422 | 0.006 | 0.024 | 0.024 | 0 | 0.36 | 0.33 | 0.25 | 0.08 | 0.28 | 0.28 |
| 6 | 31125823 | rs9263785 | 0.963 | -0.962 | -0.969 | -0.328 | 0.01 | -0.003 | -0.003 | 0 | 0.1 | 0.15 | 0.13 | 0.06 | 0.17 | 0.05 |
| 6 | 31126252 | rs3130456 | -0.12 | -1.01 | -0.958 | -0.333 | 0.021 | 0.02 | 0.02 | 0 | 0.69 | 0.58 | 0.68 | 0.61 | 0.61 | 0.63 |
| 6 | 31126409 | rs17197101 | 2.683 | -0.984 | -1.22 | -0.297 | 0.015 | 0.015 | 0.015 | 0.292 | 0.14 | 0.07 | 0.14 | 0.03 | 0.12 | 0.1 |
| 6 | 31126790 | rs3095239 | -0.498 | -0.999 | -1.217 | -0.306 | -0.008 | -0.007 | -0.007 | 0 | 0.6 | 0.65 | 0.55 | 0.58 | 0.5 | 0.55 |
| 6 | 31126944 | rs3094187 | -0.222 | -1.001 | -1.218 | -0.307 | -0.008 | -0.006 | -0.006 | 0 | 0.57 | 0.65 | 0.54 | 0.58 | 0.5 | 0.53 |
| 6 | 31127602 | rs6905862 | 0.258 | -1.037 | -1.216 | -0.301 | 0 | -0.003 | -0.003 | 0 | 0.45 | 0.48 | 0.48 | 0.38 | 0.41 | 0.4 |
| 6 | 31128552 | rs3130934 | -1.583 | -1.024 | -1.22 | -0.299 | -0.003 | -0.015 | -0.015 | 1.927 | 0.7 | 0.8 | 0.68 | 0.64 | 0.67 | 0.61 |
| 6 | 31129616 | rs2073721 | -1.767 | -0.941 | -1.245 | -0.288 | -0.011 | -0.02 | -0.02 | 0.232 | 0.81 | 0.87 | 0.81 | 0.68 | 0.78 | 0.68 |
| 6 | 31129707 | rs2073724 | 3.73 | -0.943 | -1.244 | -0.298 | 0.005 | 0.005 | 0.005 | 0.326 | 0.11 | 0.07 | 0.14 | 0.03 | 0.11 | 0.08 |
| 6 | 31130019 | rs9263794 | 0.788 | -0.959 | -1.24 | -0.288 | 0.009 | -0.003 | -0.003 | 0 | 0.11 | 0.16 | 0.13 | 0.07 | 0.17 | 0.12 |
| 6 | 31130078 | rs2073723 | -1.768 | -0.944 | -1.242 | -0.287 | -0.011 | -0.02 | -0.02 | 0.103 | 0.81 | 0.87 | 0.81 | 0.68 | 0.78 | 0.68 |
| 6 | 31130593 | rs1419881 | -0.161 | -0.935 | -1.238 | -0.28 | -0.002 | -0.001 | -0.001 | 1.161 | 0.5 | 0.64 | 0.5 | 0.57 | 0.45 | 0.45 |
| 6 | 31130760 | rs1841 | 2.818 | -0.893 | -1.206 | -0.262 | 0.038 | 0.059 | 0.059 | 0 | 0.16 | 0.03 | 0.1 | 0.04 | 0.08 | 0.12 |
| 6 | 31130821 | rs1044870 | 0.934 | -0.607 | -1.012 | -0.069 | 0.01 | -0.003 | -0.003 | 0 | 0.1 | 0.15 | 0.13 | 0.06 | 0.17 | 0.05 |
| 6 | 31130865 | rs17190776 | 3.649 | -0.614 | -1.013 | -0.063 | 0.005 | 0.005 | 0.005 | 0.326 | 0.11 | 0.07 | 0.14 | 0.03 | 0.11 | 0.08 |
| 6 | 31131707 | rs9501062 | 1.38 | -0.564 | -0.983 | -0.035 | 0.008 | 0.094 | 0.094 | 0.2 | 0.05 | 0.15 | 0.02 | 0.18 | 0.04 | 0.05 |
| 6 | 31132649 | rs2394882 | -1.145 | -0.463 | -1.174 | -0.081 | 0.003 | -0.014 | -0.014 | 0 | 0.67 | 0.8 | 0.67 | 0.64 | 0.67 | 0.58 |
| 6 | 31132883 | rs9263796 | 0.903 | -0.409 | -1.146 | -0.069 | 0.01 | -0.003 | -0.003 | 0 | 0.1 | 0.15 | 0.13 | 0.06 | 0.17 | 0.05 |
| 6 | 31133943 | rs3130932 | -1.138 | -0.482 | -1.186 | -0.058 | 0.003 | -0.014 | -0.014 | 4.096 | 0.67 | 0.8 | 0.67 | 0.64 | 0.67 | 0.58 |
| 6 | 31134599 | rs9263800 | 0.888 | -0.277 | -1.123 | 0.026 | 0.01 | -0.003 | -0.003 | 0 | 0.1 | 0.15 | 0.13 | 0.06 | 0.17 | 0.05 |
| 6 | 31134888 | rs3130931 | -0.878 | -0.258 | -1.25 | 0.014 | -0.029 | -0.031 | -0.031 | 0 | 0.76 | 0.86 | 0.79 | 0.66 | 0.73 | 0.54 |
| 6 | 31136453 | rs3130501 | -1.619 | -0.531 | -1.18 | 0.07 | -0.012 | -0.02 | -0.02 | 1.054 | 0.81 | 0.87 | 0.81 | 0.68 | 0.78 | 0.68 |
| 6 | 31136666 | rs3130502 | -1.234 | -0.557 | -1.18 | 0.071 | -0.01 | -0.025 | -0.025 | 0 | 0.78 | 0.87 | 0.81 | 0.68 | 0.78 | 0.66 |
| 6 | 31136832 | rs3132523 | -1.234 | -0.557 | -1.18 | 0.071 | -0.01 | -0.025 | -0.025 | 0 | 0.78 | 0.87 | 0.81 | 0.68 | 0.78 | 0.66 |
| 6 | 31137165 | rs3130503 | -2.68 | -0.449 | -1.144 | 0.167 | -0.008 | -0.006 | -0.006 | 0.051 | 0.88 | 0.91 | 0.84 | 0.71 | 0.82 | 0.78 |
| 6 | 31139452 | rs879882 | 0.574 | -1.023 | -1.281 | -0.212 | -0.013 | 0.017 | 0.017 | 0 | 0.61 | 0.52 | 0.67 | 0.45 | 0.64 | 0.54 |
| 6 | 31139490 | rs885952 | 0.843 | -1.17 | -1.238 | -0.236 | 0.004 | -0.001 | -0.001 | 0 | 0.38 | 0.43 | 0.46 | 0.38 | 0.4 | 0.39 |
| 6 | 31139504 | rs1108746 | 2.041 | -0.932 | -1.132 | -0.113 | 0.014 | 0.014 | 0.014 | 0.05 | 0.2 | 0.13 | 0.18 | 0.07 | 0.16 | 0.18 |
| 6 | 31140047 | rs1265159 | 1.731 | -0.479 | -0.885 | 0.003 | 0.002 | -0.01 | -0.01 | 0.346 | 0.19 | 0.25 | 0.13 | 0.04 | 0.19 | 0.14 |
| 6 | 31140152 | rs885950 | 0.285 | -0.488 | -0.852 | 0.111 | -0.003 | 0.004 | 0.004 | 0 | 0.43 | 0.42 | 0.47 | 0.38 | 0.41 | 0.41 |
| 6 | 31140532 | rs885948 | -0.222 | -0.345 | -0.78 | 0.309 | -0.006 | 0 | 0 | 0 | 0.54 | 0.68 | 0.57 | 0.51 | 0.49 | 0.49 |
| 6 | 31140741 | rs1265158 | -0.361 | -0.192 | -0.876 | 0.26 | -0.002 | -0.005 | -0.005 | 0 | 0.53 | 0.48 | 0.31 | 0.28 | 0.34 | 0.46 |
| 6 | 31141127 | rs2269713 | 0.367 | -0.311 | -0.511 | 0.347 | -0.002 | 0.085 | 0.085 | 0.651 | 0.07 | 0.19 | 0.04 | 0.2 | 0.05 | 0.12 |
| 6 | 31141523 | rs887468 | 1.035 | -0.23 | -0.823 | 0.189 | 0 | 0.018 | 0.018 | 0 | 0.27 | 0.28 | 0.17 | 0.05 | 0.24 | 0.23 |
| 6 | 31141836 | rs3132517 | -2.51 | 0.012 | -0.9 | 0.365 | -0.016 | -0.011 | -0.011 | 1.117 | 0.88 | 0.93 | 0.89 | 0.82 | 0.82 | 0.82 |
| 6 | 31142245 | rs3094188 | -0.519 | -0.262 | -0.893 | 0.235 | 0.012 | 0.027 | 0.027 | 0.301 | 0.62 | 0.77 | 0.72 | 0.74 | 0.67 | 0.61 |
| 6 | 31142297 | rs1265156 | 0.149 | -0.35 | -0.829 | 0.265 | 0.007 | 0.003 | 0.003 | 0 | 0.41 | 0.33 | 0.46 | 0.39 | 0.33 | 0.38 |
| 6 | 31142903 | rs9468877 | 0.916 | -0.575 | -0.838 | 0.173 | 0.021 | 0.02 | 0.02 | 0 | 0.27 | 0.16 | 0.17 | 0.08 | 0.15 | 0.25 |
| 6 | 31143511 | rs887466 | -0.096 | -0.308 | -0.758 | 0.318 | 0.001 | 0 | 0 | 0.004 | 0.43 | 0.37 | 0.51 | 0.48 | 0.41 | 0.41 |
| 6 | 31143582 | rs3131018 | -1.164 | -0.334 | -0.833 | 0.318 | 0.003 | 0 | 0 | 0 | 0.69 | 0.67 | 0.68 | 0.72 | 0.62 | 0.75 |
| 6 | 31143694 | rs1265155 | 0.005 | -0.33 | -0.842 | 0.343 | 0.01 | 0 | 0 | 0 | 0.42 | 0.33 | 0.49 | 0.48 | 0.39 | 0.41 |

| | | | | | | | | | | | | | | | | |
|---|---|---|---|---|---|---|---|---|---|---|---|---|---|---|---|---|
| 6 | 31144429 | rs4947305 | -0.011 | -0.325 | -0.839 | 0.345 | 0.011 | 0 | 0 | 0 | 0.43 | 0.33 | 0.49 | 0.48 | 0.39 | 0.4 |
| 6 | 31145271 | rs9501066 | -0.011 | -0.325 | -0.839 | 0.345 | 0.011 | 0 | 0 | 0 | 0.43 | 0.33 | 0.49 | 0.48 | 0.39 | 0.41 |
| 6 | 31145839 | rs3868542 | -0.021 | -0.327 | -0.84 | 0.345 | 0.011 | 0 | 0 | 0 | 0.43 | 0.33 | 0.5 | 0.48 | 0.39 | 0.41 |
| 6 | 31145920 | rs887464 | 0.514 | -0.269 | -0.882 | 0.251 | -0.003 | -0.002 | -0.002 | 0 | 0.33 | 0.29 | 0.48 | 0.45 | 0.42 | 0.29 |
| 6 | 31145991 | rs3871248 | -0.085 | -0.412 | -0.827 | 0.326 | 0.011 | 0 | 0 | 0 | 0.43 | 0.33 | 0.5 | 0.48 | 0.39 | 0.41 |
| 6 | 31146403 | rs1052989 | 0.751 | -0.352 | -0.805 | 0.372 | 0.05 | 0.098 | 0.098 | 0 | 0.09 | 0.16 | 0.29 | 0.39 | 0.2 | 0.08 |
| 6 | 31146584 | rs9380220 | 1.212 | -0.348 | -0.803 | 0.374 | 0.066 | 0.123 | 0.123 | 0 | 0.08 | 0.16 | 0.29 | 0.39 | 0.2 | 0.08 |
| 6 | 31146846 | rs4713438 | 0.18 | -0.63 | -0.819 | 0.26 | -0.011 | -0.008 | -0.008 | 0 | 0.35 | 0.34 | 0.24 | 0.14 | 0.2 | 0.28 |
| 6 | 31149045 | rs9263827 | 0.924 | -0.42 | -0.813 | 0.362 | -0.013 | 0.002 | 0.002 | 0 | 0.08 | 0.18 | 0.07 | 0.06 | 0.06 | 0.04 |
| 6 | 31150435 | rs1128175 | -2.407 | -0.243 | -0.989 | 0.398 | -0.004 | -0.003 | -0.003 | 0.077 | 0.78 | 0.86 | 0.78 | 0.78 | 0.76 | 0.79 |
| 6 | 31155670 | rs9263846 | 1.618 | -0.221 | -1.024 | 0.367 | 0.034 | 0.018 | 0.018 | 0.02 | 0.23 | 0.13 | 0.22 | 0.11 | 0.2 | 0.18 |
| 6 | 31157593 | rs9295957 | 1.658 | -0.07 | -0.972 | 0.539 | 0.089 | 0.155 | 0.155 | 0 | 0.06 | 0.16 | 0.28 | 0.39 | 0.19 | 0.08 |
| 6 | 31160553 | rs9461653 | 2.468 | -0.214 | -1.003 | 0.432 | 0.01 | 0.008 | 0.008 | 0 | 0.17 | 0.09 | 0.12 | 0.04 | 0.08 | 0.13 |
| 6 | 31161229 | rs1265178 | 2.881 | -0.225 | -1.124 | 0.389 | -0.067 | 0.01 | 0.01 | 0 | 0.13 | 0.07 | 0.07 | 0.04 | 0.18 | 0.14 |
| 6 | 31162154 | rs28362332 | 2.383 | -0.096 | -0.882 | 0.604 | 0.007 | 0.003 | 0.003 | 0 | 0.16 | 0.12 | 0.18 | 0.09 | 0.15 | 0.11 |
| 6 | 31162520 | rs7762933 | 0.375 | -0.085 | -0.88 | 0.61 | 0.007 | -0.002 | -0.002 | 0 | 0.47 | 0.39 | 0.62 | 0.54 | 0.47 | 0.39 |
| 6 | 31162839 | rs4713446 | 0.367 | -0.087 | -0.881 | 0.609 | 0.007 | -0.002 | -0.002 | 2.955 | 0.47 | 0.39 | 0.62 | 0.54 | 0.47 | 0.39 |
| 6 | 31162963 | rs4713447 | 0.38 | -0.079 | -0.877 | 0.613 | 0.007 | -0.002 | -0.002 | 0 | 0.47 | 0.39 | 0.62 | 0.54 | 0.47 | 0.39 |
| 6 | 31165527 | rs28362343 | 2.3 | -0.046 | -0.873 | 0.64 | 0.007 | 0.003 | 0.003 | 0 | 0.16 | 0.12 | 0.18 | 0.09 | 0.15 | 0.11 |
| 6 | 31165566 | rs3094609 | -2.561 | 0.094 | -0.876 | 0.701 | -0.021 | -0.021 | -0.021 | 0 | 0.91 | 0.96 | 0.92 | 0.84 | 0.88 | 0.87 |
| 6 | 31166828 | rs9404983 | 1.183 | -0.116 | -0.79 | 0.703 | 0.013 | 0.035 | 0.035 | 0 | 0.14 | 0.17 | 0.33 | 0.41 | 0.23 | 0.15 |
| 6 | 31167498 | rs9295961 | 2.338 | -0.233 | -0.809 | 0.621 | 0.01 | 0.008 | 0.008 | 0 | 0.17 | 0.09 | 0.11 | 0.04 | 0.08 | 0.13 |
| 6 | 31167512 | rs9501077 | 0.321 | -0.201 | -0.859 | 0.653 | 0.007 | -0.002 | -0.002 | 0 | 0.47 | 0.39 | 0.62 | 0.54 | 0.47 | 0.39 |
| 6 | 31167927 | rs4122189 | 1.123 | 0.035 | -0.775 | 0.842 | 0.013 | 0.035 | 0.035 | 0 | 0.14 | 0.17 | 0.33 | 0.4 | 0.23 | 0.15 |
| 6 | 31168029 | rs6903896 | 0.309 | -0.029 | -0.699 | 0.833 | -0.006 | 0.006 | 0.006 | 0 | 0.05 | 0.09 | 0.03 | 0.1 | 0.04 | 0.12 |
| 6 | 31170514 | rs9263870 | 1.987 | -0.609 | -0.941 | 0.47 | 0.014 | 0.005 | 0.005 | 0 | 0.17 | 0.11 | 0.18 | 0.09 | 0.15 | 0.11 |
| 6 | 31170528 | rs9263871 | 0.668 | -0.618 | -0.911 | 0.464 | 0 | -0.003 | -0.003 | 0 | 0.32 | 0.3 | 0.52 | 0.5 | 0.39 | 0.26 |
| 6 | 31170713 | rs9263873 | 0.413 | -0.741 | -0.929 | 0.361 | 0.003 | -0.001 | -0.001 | 0 | 0.47 | 0.39 | 0.61 | 0.54 | 0.47 | 0.39 |
| 6 | 31170914 | rs9263875 | 0.311 | -0.56 | -0.922 | 0.363 | 0.001 | 0.001 | 0.001 | 0 | 0.33 | 0.3 | 0.46 | 0.45 | 0.39 | 0.29 |
| 6 | 31171257 | rs6921948 | 0.171 | -0.514 | -0.861 | 0.325 | 0 | 0.001 | 0.001 | 0.042 | 0.49 | 0.42 | 0.53 | 0.39 | 0.52 | 0.38 |
| 6 | 31173096 | rs4446605 | 2.103 | -1.149 | -0.86 | 0.327 | 0.014 | 0.005 | 0.005 | 0 | 0.17 | 0.11 | 0.18 | 0.09 | 0.15 | 0.11 |
| 6 | 31173678 | rs3094606 | -2.612 | -1.217 | -0.907 | 0.356 | -0.016 | -0.016 | -0.016 | 0.235 | 0.91 | 0.94 | 0.91 | 0.84 | 0.88 | 0.87 |
| 6 | 31174527 | rs2894181 | 0.225 | -1.236 | -0.939 | 0.347 | 0 | 0.001 | 0.001 | 0.053 | 0.49 | 0.42 | 0.53 | 0.39 | 0.52 | 0.38 |
| 6 | 31176226 | rs3132506 | 1.999 | -1.537 | -0.892 | 0.232 | 0.001 | 0 | 0 | 0.334 | 0.21 | 0.26 | 0.24 | 0.22 | 0.22 | 0.18 |
| 6 | 31176335 | rs3887152 | 0.668 | -1.545 | -0.894 | 0.228 | 0.007 | -0.005 | -0.005 | 1.591 | 0.32 | 0.31 | 0.35 | 0.29 | 0.37 | 0.27 |
| 6 | 31176921 | rs4351302 | 1.455 | -1.526 | -0.888 | 0.216 | 0.013 | 0.043 | 0.043 | 0 | 0.26 | 0.13 | 0.16 | 0.06 | 0.2 | 0.21 |
| 6 | 31176928 | rs9263916 | 2.231 | -1.527 | -0.89 | 0.221 | 0.014 | 0.005 | 0.005 | 0 | 0.17 | 0.11 | 0.18 | 0.09 | 0.15 | 0.11 |
| 6 | 31177503 | rs3132505 | 1.994 | -1.513 | -0.886 | 0.225 | 0.001 | 0 | 0 | 0 | 0.21 | 0.26 | 0.24 | 0.22 | 0.22 | 0.18 |
| 6 | 31177845 | rs35718543 | 0.848 | -1.495 | -0.847 | 0.17 | -0.022 | -0.027 | -0.027 | 0 | 0.13 | 0.19 | 0.08 | 0.04 | 0.09 | 0.12 |
| 6 | 31177915 | rs3130952 | -2.552 | -1.443 | -0.999 | 0.211 | -0.019 | -0.019 | -0.019 | 0 | 0.9 | 0.95 | 0.91 | 0.84 | 0.88 | 0.87 |
| 6 | 31178743 | rs3132504 | -2.552 | -1.443 | -0.999 | 0.216 | -0.019 | -0.018 | -0.018 | 2.576 | 0.9 | 0.95 | 0.91 | 0.84 | 0.88 | 0.87 |
| 6 | 31181025 | rs9263948 | 2.142 | -1.498 | -1.008 | 0.149 | 0.014 | 0.006 | 0.006 | 0 | 0.17 | 0.11 | 0.18 | 0.1 | 0.15 | 0.11 |
| 6 | 31182383 | rs9468898 | 0.416 | -1.412 | -1.017 | 0.184 | -0.003 | 0.001 | 0.001 | 0.069 | 0.05 | 0.08 | 0.03 | 0.09 | 0.04 | 0.12 |
| 6 | 31184020 | rs9295967 | 1.317 | -1.361 | -1.049 | 0.11 | 0.013 | 0.043 | 0.043 | 0 | 0.26 | 0.13 | 0.16 | 0.06 | 0.2 | 0.21 |
| 6 | 31184196 | rs3869109 | -0.524 | -1.336 | -0.983 | 0.176 | -0.009 | -0.008 | -0.008 | 0 | 0.62 | 0.61 | 0.61 | 0.43 | 0.62 | 0.5 |
| 6 | 31184425 | rs9501522 | 2.091 | -1.26 | -0.992 | 0.162 | 0.004 | 0.001 | 0.001 | 0 | 0.17 | 0.1 | 0.08 | 0.04 | 0.08 | 0.13 |
| 6 | 31185977 | rs3132501 | -2.429 | -1.044 | -1.016 | 0.299 | -0.024 | -0.023 | -0.023 | 0 | 0.91 | 0.95 | 0.91 | 0.84 | 0.88 | 0.87 |
| 6 | 31186039 | rs9263964 | -0.4 | -1.212 | -0.997 | 0.252 | -0.006 | -0.005 | -0.005 | 0 | 0.61 | 0.59 | 0.61 | 0.41 | 0.62 | 0.49 |
| 6 | 31186534 | rs9263969 | 1.3 | -1.39 | -1.041 | 0.145 | 0.015 | 0.029 | 0.029 | 0.002 | 0.26 | 0.14 | 0.25 | 0.11 | 0.27 | 0.19 |
| 6 | 31187075 | rs3130467 | 0.552 | -1.099 | -0.874 | 0.331 | 0 | 0 | 0 | 0 | 0.29 | 0.34 | 0.29 | 0.41 | 0.26 | 0.37 |
| 6 | 31188267 | rs9263979 | 1.152 | -1.358 | -0.915 | 0.118 | 0.011 | 0.011 | 0.011 | 0 | 0.36 | 0.25 | 0.29 | 0.16 | 0.28 | 0.25 |
| 6 | 31188654 | rs28360997 | 0.857 | -1.292 | -1.037 | 0.14 | 0.016 | 0.048 | 0.048 | 0 | 0.18 | 0.07 | 0.11 | 0.05 | 0.15 | 0.2 |
| 6 | 31190538 | rs9263985 | 1.026 | -1.301 | -1.096 | 0.125 | 0.032 | 0.072 | 0.072 | 0 | 0.4 | 0.21 | 0.35 | 0.17 | 0.39 | 0.34 |
| 6 | 31190850 | rs12662501 | 2.278 | -1.423 | -1.133 | 0.081 | 0.039 | 0.053 | 0.053 | 0.047 | 0.24 | 0.09 | 0.21 | 0.08 | 0.18 | 0.2 |
| 6 | 31192766 | rs9263993 | 0.783 | -1.214 | -1.349 | 0.347 | -0.004 | -0.013 | -0.013 | 0.071 | 0.17 | 0.12 | 0.15 | 0.2 | 0.21 | 0.15 |
| 6 | 31192796 | rs6904669 | -0.936 | -1.255 | -1.295 | 0.296 | 0.003 | -0.002 | -0.002 | 0 | 0.35 | 0.58 | 0.41 | 0.33 | 0.45 | 0.28 |
| 6 | 31193155 | rs4373355 | 0.986 | -1.337 | -1.214 | 0.248 | 0.015 | 0.025 | 0.025 | 0 | 0.42 | 0.35 | 0.36 | 0.17 | 0.39 | 0.34 |
| 6 | 31195218 | rs6928399 | 0.813 | -1.168 | -1.19 | 0.302 | 0.002 | 0.006 | 0.006 | 0 | 0.38 | 0.6 | 0.45 | 0.3 | 0.38 | 0.24 |
| 6 | 31196376 | rs3130521 | -0.045 | -1.106 | -1.283 | 0.368 | -0.006 | 0.006 | 0.006 | 0 | 0.64 | 0.76 | 0.72 | 0.72 | 0.63 | 0.56 |
| 6 | 31197407 | rs28397289 | 0.015 | -1.017 | -1.187 | 0.389 | 0.106 | 0.081 | 0.081 | 0.008 | 0.13 | 0.39 | 0.23 | 0.24 | 0.21 | 0.06 |
| 6 | 31199208 | rs3130473 | 1.105 | -1.001 | -1.285 | 0.39 | -0.02 | -0.019 | -0.019 | 0.414 | 0.25 | 0.14 | 0.23 | 0.3 | 0.24 | 0.26 |
| 6 | 31199256 | rs3130474 | -0.129 | -1.035 | -1.289 | 0.352 | 0.009 | -0.025 | -0.025 | 1.894 | 0.38 | 0.3 | 0.37 | 0.48 | 0.37 | 0.44 |
| 6 | 31199460 | rs3130528 | -2.317 | -0.997 | -1.33 | 0.357 | -0.024 | -0.023 | -0.023 | 0 | 0.91 | 0.95 | 0.91 | 0.84 | 0.88 | 0.87 |
| 6 | 31199766 | rs11967684 | 0.7 | -1.065 | -1.333 | 0.292 | 0.004 | 0.021 | 0.021 | 0 | 0.39 | 0.62 | 0.47 | 0.32 | 0.41 | 0.26 |
| 6 | 31202246 | rs2394888 | -1.612 | -0.88 | -1.301 | 0.36 | -0.057 | -0.023 | -0.023 | 0 | 0.81 | 0.91 | 0.84 | 0.82 | 0.76 | 0.79 |
| 6 | 31206206 | rs3130685 | 0.358 | -1.055 | -1.307 | 0.329 | 0.001 | 0.002 | 0.002 | 0 | 0.61 | 0.57 | 0.53 | 0.54 | 0.46 | 0.58 |
| 6 | 31206979 | rs2394895 | 1.835 | -1.01 | -1.378 | 0.392 | -0.023 | -0.011 | -0.011 | 0.011 | 0.19 | 0.13 | 0.21 | 0.22 | 0.21 | 0.18 |
| 6 | 31207692 | rs3868082 | 0.303 | -1.011 | -1.32 | 0.403 | 0 | 0 | 0 | 0.133 | 0.52 | 0.59 | 0.63 | 0.49 | 0.6 | 0.51 |
| 6 | 31207920 | rs3132499 | -2.213 | -1.127 | -1.499 | 0.478 | -0.024 | -0.024 | -0.024 | 0 | 0.91 | 0.95 | 0.9 | 0.84 | 0.88 | 0.87 |

| | | | | | | | | | | | | | | | | |
|---|---|---|---|---|---|---|---|---|---|---|---|---|---|---|---|---|
| 6 | 31208340 | rs3095250 | -0.761 | -1.15 | -1.488 | 0.44 | 0.007 | -0.001 | -0.001 | 0 | 0.52 | 0.56 | 0.53 | 0.58 | 0.56 | 0.49 |
| 6 | 31208610 | rs3132496 | -0.296 | -1.183 | -1.45 | 0.449 | 0 | 0 | 0 | 0.271 | 0.45 | 0.48 | 0.55 | 0.6 | 0.53 | 0.48 |
| 6 | 31209045 | rs3130534 | -0.287 | -1.183 | -1.449 | 0.439 | 0 | -0.001 | -0.001 | 0 | 0.44 | 0.48 | 0.51 | 0.5 | 0.53 | 0.43 |
| 6 | 31209510 | rs3130712 | -0.438 | -1.212 | -1.458 | 0.404 | 0 | 0 | 0 | 0 | 0.47 | 0.55 | 0.55 | 0.65 | 0.56 | 0.45 |
| 6 | 31210216 | rs3130688 | -0.783 | -1.192 | -1.453 | 0.402 | 0.006 | 0.006 | 0.006 | 0.12 | 0.52 | 0.56 | 0.57 | 0.68 | 0.56 | 0.53 |
| 6 | 31210265 | rs6901869 | 1.955 | -1.128 | -1.483 | 0.389 | -0.032 | 0.021 | 0.021 | 0 | 0.09 | 0.03 | 0.06 | 0.02 | 0.11 | 0.08 |
| 6 | 31211050 | rs3130695 | -1.216 | -1.228 | -1.489 | 0.364 | 0 | -0.004 | -0.004 | 0 | 0.79 | 0.8 | 0.81 | 0.77 | 0.8 | 0.74 |
| 6 | 31213413 | rs4332019 | 1.866 | -1.183 | -1.45 | 0.29 | 0.001 | -0.001 | -0.001 | 0.575 | 0.27 | 0.24 | 0.24 | 0.09 | 0.23 | 0.2 |
| 6 | 31213902 | rs3095245 | 0.968 | -1.125 | -1.391 | 0.365 | 0.003 | 0.001 | 0.001 | 0 | 0.27 | 0.3 | 0.41 | 0.43 | 0.39 | 0.25 |
| 6 | 31218327 | rs3130425 | 0.196 | -1.636 | -1.511 | 0.527 | 0 | 0.004 | 0.004 | 0 | 0.39 | 0.47 | 0.53 | 0.57 | 0.52 | 0.39 |
| 6 | 31218889 | rs4084262 | 1.706 | -1.777 | -1.545 | 0.461 | 0.006 | 0.005 | 0.005 | 0 | 0.27 | 0.22 | 0.22 | 0.09 | 0.19 | 0.2 |
| 6 | 31218984 | rs3130427 | 0.14 | -1.514 | -1.434 | 0.533 | 0 | 0.004 | 0.004 | 0 | 0.39 | 0.47 | 0.53 | 0.57 | 0.52 | 0.39 |
| 6 | 31219869 | rs9264179 | 0.008 | -1.489 | -1.401 | 0.473 | -0.001 | 0.001 | 0.001 | 0.003 | 0.4 | 0.47 | 0.5 | 0.51 | 0.54 | 0.36 |
| 6 | 31219902 | rs9264180 | -0.342 | -1.171 | -1.447 | 0.529 | 0.01 | 0.001 | 0.001 | 0.141 | 0.51 | 0.56 | 0.53 | 0.55 | 0.55 | 0.46 |
| 6 | 31221039 | rs4416711 | 0.415 | -0.892 | -1.785 | 0.329 | -0.015 | -0.004 | -0.004 | 0 | 0.12 | 0.15 | 0.09 | 0.08 | 0.08 | 0.14 |
| 6 | 31221485 | rs17191559 | 1.428 | -0.585 | -1.998 | 0.258 | 0.033 | -0.03 | -0.03 | 0 | 0.16 | 0.04 | 0.15 | 0.23 | 0.09 | 0.17 |
| 6 | 31221698 | rs1793891 | 1.048 | 0.028 | -1.729 | -0.131 | -0.067 | 0.039 | 0.039 | 0 | 0.21 | 0.09 | 0.11 | 0.06 | 0.23 | 0.22 |
| 6 | 31221914 | rs9264219 | -0.041 | 0.112 | -1.176 | 0.082 | -0.006 | -0.012 | -0.012 | 0 | 0.6 | 0.48 | 0.64 | 0.74 | 0.65 | 0.65 |
| 6 | 31228410 | rs1986997 | -0.387 | 0.611 | -1.373 | 0.255 | -0.037 | 0.003 | 0.003 | 1.068 | 0.71 | 0.79 | 0.81 | 0.77 | 0.68 | 0.71 |
| 6 | 31228972 | rs2524121 | 0.824 | 0.698 | -1.356 | 0.099 | 0.002 | -0.003 | -0.003 | 0.198 | 0.1 | 0.07 | 0.12 | 0.04 | 0.12 | 0.11 |
| 6 | 31230701 | rs2394950 | 0.635 | 0.082 | -1.282 | -0.257 | -0.015 | -0.022 | -0.022 | 0.017 | 0.08 | 0.11 | 0.06 | 0.01 | 0.05 | 0.1 |
| 6 | 31230800 | rs2245822 | 1.301 | -0.158 | -1.735 | -0.559 | -0.066 | 0.038 | 0.038 | 0 | 0.21 | 0.1 | 0.11 | 0.05 | 0.23 | 0.21 |
| 6 | 31230924 | rs7747738 | 1.506 | -0.213 | -1.437 | -0.794 | 0.003 | -0.028 | -0.028 | 0 | 0.05 | 0.11 | 0.06 | 0 | 0.05 | 0.03 |
| 6 | 31231989 | rs2853961 | 0.85 | -0.678 | -1.497 | -0.897 | 0.003 | 0.012 | 0.012 | 0 | 0.4 | 0.43 | 0.32 | 0.18 | 0.4 | 0.4 |
| 6 | 31232111 | rs3130542 | -0.973 | -0.691 | -1.555 | -0.923 | -0.006 | 0.001 | 0.001 | 0.085 | 0.81 | 0.82 | 0.84 | 0.82 | 0.81 | 0.75 |
| 6 | 31232543 | rs2844623 | 1.062 | -0.63 | -1.495 | -0.415 | 0.012 | 0.011 | 0.011 | 0 | 0.12 | 0.15 | 0.18 | 0.16 | 0.16 | 0.17 |
| 6 | 31234381 | rs9264532 | -0.092 | -0.757 | -1.649 | 0.078 | 0.006 | 0.002 | 0.002 | 0.719 | 0.59 | 0.64 | 0.65 | 0.6 | 0.68 | 0.6 |
| 6 | 31236051 | rs2524099 | -0.069 | -0.756 | -1.89 | 0.298 | -0.034 | -0.005 | -0.005 | 0 | 0.58 | 0.7 | 0.62 | 0.56 | 0.52 | 0.58 |
| 6 | 31236250 | rs9394047 | 1.73 | -1.113 | -1.937 | 0.276 | 0.081 | 0.01 | 0.01 | 0.395 | 0.23 | 0.09 | 0.17 | 0.22 | 0.09 | 0.23 |
| 6 | 31236410 | rs34131062 | 1.634 | -0.323 | -1.596 | 0.61 | 0.017 | 0.039 | 0.039 | 0 | 0.05 | 0.08 | 0.13 | 0.25 | 0.13 | 0.08 |
| 6 | 31236567 | rs1049281 | -0.573 | 0.177 | -1.578 | 0.521 | 0.01 | -0.006 | -0.006 | 0 | 0.76 | 0.73 | 0.8 | 0.81 | 0.72 | 0.75 |
| 6 | 31240431 | rs2074488 | 1.584 | 0.229 | -1.598 | 0.832 | 0.017 | 0.039 | 0.039 | 0 | 0.05 | 0.08 | 0.13 | 0.25 | 0.13 | 0.08 |
| 6 | 31240721 | rs2249742 | 0.365 | -0.104 | -1.654 | 0.569 | -0.002 | -0.002 | -0.002 | 2.955 | 0.53 | 0.52 | 0.56 | 0.52 | 0.51 | 0.54 |
| 6 | 31240988 | rs7759127 | 2.094 | -0.142 | -1.541 | 0.494 | 0.038 | 0.031 | 0.031 | 0 | 0.05 | 0.09 | 0.15 | 0.1 | 0.11 | 0.05 |
| 6 | 31241032 | rs5010528 | 2.202 | -0.43 | -1.647 | 0.355 | -0.021 | -0.033 | -0.033 | 0.065 | 0.15 | 0.24 | 0.19 | 0.06 | 0.14 | 0.1 |
| 6 | 31241370 | rs6923313 | -0.488 | -0.415 | -1.775 | 0.33 | 0.014 | 0.033 | 0.033 | 0.021 | 0.57 | 0.69 | 0.76 | 0.72 | 0.66 | 0.55 |
| 6 | 31241537 | rs2844621 | -1.221 | -0.047 | -1.667 | 0.533 | 0.026 | 0.022 | 0.022 | 0 | 0.81 | 0.78 | 0.74 | 0.73 | 0.75 | 0.81 |
| 6 | 31241639 | rs2524084 | -0.444 | -0.024 | -1.602 | 0.574 | 0.004 | 0.013 | 0.013 | 0 | 0.2 | 0.22 | 0.32 | 0.52 | 0.32 | 0.25 |
| 6 | 31243170 | rs3132486 | -0.751 | -0.202 | -1.651 | 0.445 | 0.002 | -0.003 | -0.003 | 0 | 0.43 | 0.43 | 0.53 | 0.67 | 0.49 | 0.44 |
| 6 | 31243347 | rs9461680 | 0.764 | -0.11 | -1.639 | 0.523 | 0.002 | -0.002 | -0.002 | 0 | 0.24 | 0.19 | 0.19 | 0.3 | 0.18 | 0.21 |
| 6 | 31243603 | rs2524077 | 1.506 | 0.021 | -1.521 | 0.557 | 0.029 | 0.021 | 0.021 | 0 | 0.06 | 0.11 | 0.15 | 0.12 | 0.12 | 0.06 |
| 6 | 31243846 | rs2844613 | 1.201 | 0.003 | -1.591 | 0.489 | 0.012 | -0.026 | -0.026 | 0.059 | 0.08 | 0.12 | 0.08 | 0.01 | 0.14 | 0.1 |
| 6 | 31243884 | rs3130696 | -0.571 | -0.058 | -1.594 | 0.471 | 0.007 | 0.006 | 0.006 | 0 | 0.19 | 0.25 | 0.26 | 0.27 | 0.25 | 0.2 |
| 6 | 31244021 | rs2524074 | -1.244 | 0.043 | -1.65 | 0.462 | -0.011 | 0 | 0 | 0 | 0.82 | 0.83 | 0.82 | 0.82 | 0.74 | 0.78 |
| 6 | 31244068 | rs9264731 | -2.103 | -0.033 | -1.617 | 0.556 | -0.023 | -0.023 | -0.023 | 0 | 0.9 | 0.95 | 0.9 | 0.83 | 0.88 | 0.87 |
| 6 | 31244235 | rs2524073 | 2.068 | -0.017 | -1.594 | 0.564 | 0.038 | 0.031 | 0.031 | 0 | 0.05 | 0.09 | 0.15 | 0.1 | 0.11 | 0.06 |
| 6 | 31244332 | rs9468913 | 1.5 | -0.207 | -1.647 | 0.476 | 0.01 | 0.01 | 0.01 | 0 | 0.18 | 0.11 | 0.06 | 0.05 | 0.06 | 0.13 |
| 6 | 31244520 | rs2524070 | 0.967 | 0.08 | -1.542 | 0.577 | 0.054 | 0.032 | 0.032 | 0.154 | 0.08 | 0.17 | 0.17 | 0.12 | 0.16 | 0.08 |
| 6 | 31244960 | rs6919908 | -2.102 | 0.09 | -1.557 | 0.586 | 0.001 | 0 | 0 | 0 | 0.88 | 0.87 | 0.88 | 0.83 | 0.83 | 0.86 |
| 6 | 31245061 | rs2524068 | 0.967 | 0.094 | -1.547 | 0.589 | 0.054 | 0.032 | 0.032 | 0.154 | 0.08 | 0.17 | 0.17 | 0.12 | 0.16 | 0.08 |
| 6 | 31245736 | rs6906846 | -1.228 | -0.288 | -1.689 | 0.39 | -0.004 | -0.008 | -0.008 | 0 | 0.68 | 0.63 | 0.68 | 0.74 | 0.69 | 0.67 |
| 6 | 31246703 | rs7381988 | -2.432 | -0.226 | -1.723 | 0.431 | -0.032 | -0.026 | -0.026 | 2.384 | 0.89 | 0.94 | 0.9 | 0.82 | 0.86 | 0.85 |
| 6 | 31247067 | rs7382297 | -2.808 | -0.232 | -1.72 | 0.433 | -0.023 | -0.023 | -0.023 | 0 | 0.9 | 0.95 | 0.9 | 0.83 | 0.88 | 0.87 |
| 6 | 31247135 | rs4386816 | 1.714 | -0.895 | -1.588 | 0.385 | -0.019 | -0.028 | -0.028 | 0.064 | 0.21 | 0.25 | 0.2 | 0.08 | 0.16 | 0.19 |
| 6 | 31250606 | rs9468914 | 0.211 | -0.969 | -1.718 | 0.511 | 0.006 | 0.005 | 0.005 | 0.721 | 0.48 | 0.53 | 0.46 | 0.39 | 0.39 | 0.45 |
| 6 | 31250854 | rs2844603 | -1.187 | -0.97 | -1.794 | 0.524 | -0.036 | 0.011 | 0.011 | 0.193 | 0.64 | 0.78 | 0.78 | 0.71 | 0.64 | 0.66 |
| 6 | 31251312 | rs6921855 | 0.962 | -0.98 | -1.73 | 0.594 | -0.007 | 0.007 | 0.007 | 0 | 0.09 | 0.11 | 0.14 | 0.2 | 0.08 | 0.14 |
| 6 | 31253444 | rs9461684 | 2.388 | -1.149 | -1.753 | 0.461 | -0.021 | -0.034 | -0.034 | 2.034 | 0.15 | 0.26 | 0.19 | 0.03 | 0.14 | 0.08 |
| 6 | 31254088 | rs2853933 | -1.169 | -1.045 | -1.774 | 0.495 | -0.036 | 0.011 | 0.011 | 0.193 | 0.64 | 0.78 | 0.78 | 0.71 | 0.64 | 0.66 |
| 6 | 31254263 | rs7754443 | 0.571 | -1.131 | -1.797 | 0.477 | -0.011 | -0.007 | -0.007 | 0.134 | 0.34 | 0.5 | 0.32 | 0.31 | 0.34 | 0.32 |
| 6 | 31254936 | rs9468920 | 0.594 | -1.129 | -1.8 | 0.478 | -0.011 | -0.007 | -0.007 | 0.058 | 0.34 | 0.49 | 0.32 | 0.31 | 0.34 | 0.32 |
| 6 | 31256753 | rs2524044 | 1.57 | -1.107 | -1.794 | 0.49 | -0.009 | -0.008 | -0.008 | 0 | 0.18 | 0.09 | 0.28 | 0.17 | 0.16 | 0.19 |
| 6 | 31256948 | rs9468924 | 2.331 | -1.191 | -1.8 | 0.458 | -0.028 | -0.006 | -0.006 | 0 | 0.05 | 0.08 | 0.02 | 0.07 | 0.01 | 0.02 |
| 6 | 31257622 | rs3900938 | 1.493 | -1.103 | -1.758 | 0.496 | -0.004 | 0.005 | 0.005 | 0.282 | 0.25 | 0.34 | 0.25 | 0.25 | 0.27 | 0.25 |
| 6 | 31257625 | rs2524040 | -1.241 | -1.133 | -1.802 | 0.487 | -0.036 | 0.011 | 0.011 | 0.193 | 0.64 | 0.78 | 0.78 | 0.71 | 0.64 | 0.66 |
| 6 | 31258141 | rs28367594 | 2.004 | -1.151 | -1.782 | 0.467 | 0.003 | -0.029 | -0.029 | 0 | 0.05 | 0.11 | 0.06 | 0 | 0.05 | 0.03 |
| 6 | 31258837 | rs9468925 | 0.236 | -1.087 | -1.933 | 0.596 | 0.03 | 0.022 | 0.022 | 0.124 | 0.49 | 0.6 | 0.48 | 0.38 | 0.42 | 0.44 |
| 6 | 31258846 | rs2256583 | 1.538 | -1.083 | -1.938 | 0.599 | -0.009 | -0.008 | -0.008 | 0 | 0.18 | 0.09 | 0.28 | 0.17 | 0.16 | 0.19 |
| 6 | 31259579 | rs2524163 | -1.189 | -1.085 | -1.94 | 0.601 | -0.036 | 0.011 | 0.011 | 0.193 | 0.64 | 0.78 | 0.78 | 0.71 | 0.64 | 0.66 |

| | | | | | | | | | | | | | | | | |
|---|---|---|---|---|---|---|---|---|---|---|---|---|---|---|---|---|
| 6 | 31259750 | rs2524162 | 1.146 | -1.078 | -1.934 | 0.605 | 0.005 | 0.007 | 0.007 | 0 | 0.08 | 0.07 | 0.16 | 0.17 | 0.14 | 0.1 |
| 6 | 31260218 | rs9468926 | 0.524 | -1.084 | -1.934 | 0.6 | -0.003 | -0.003 | -0.003 | 0 | 0.36 | 0.57 | 0.35 | 0.32 | 0.39 | 0.32 |
| 6 | 31260345 | rs9357122 | 0.823 | -1.097 | -1.938 | 0.598 | -0.01 | 0.009 | 0.009 | 0 | 0.1 | 0.12 | 0.15 | 0.22 | 0.09 | 0.14 |
| 6 | 31260397 | rs2524156 | -1.176 | -1.08 | -1.946 | 0.584 | -0.036 | 0.011 | 0.011 | 0.193 | 0.64 | 0.78 | 0.78 | 0.71 | 0.64 | 0.66 |
| 6 | 31261137 | rs16899178 | 2.028 | -1.083 | -1.943 | 0.582 | 0.033 | 0.023 | 0.023 | 0 | 0.13 | 0.03 | 0.13 | 0.07 | 0.05 | 0.13 |
| 6 | 31261276 | rs2243868 | -1.169 | -1.068 | -1.937 | 0.588 | -0.036 | 0.011 | 0.011 | 0.193 | 0.64 | 0.78 | 0.78 | 0.71 | 0.64 | 0.66 |
| 6 | 31262169 | rs3873379 | 0.646 | -1.119 | -1.897 | 0.594 | -0.011 | -0.007 | -0.007 | 0.274 | 0.34 | 0.49 | 0.32 | 0.31 | 0.34 | 0.32 |
| 6 | 31262438 | rs3873380 | 1.041 | -1.131 | -1.904 | 0.582 | -0.02 | -0.014 | -0.014 | 0 | 0.4 | 0.45 | 0.43 | 0.29 | 0.35 | 0.38 |
| 6 | 31262461 | rs9348863 | 0.627 | -1.027 | -1.835 | 0.633 | -0.011 | 0.008 | 0.008 | 0 | 0.1 | 0.12 | 0.15 | 0.22 | 0.09 | 0.14 |
| 6 | 31263231 | rs3873381 | 0.72 | -1.017 | -1.833 | 0.638 | -0.01 | 0.009 | 0.009 | 0 | 0.1 | 0.12 | 0.15 | 0.22 | 0.09 | 0.14 |
| 6 | 31264392 | rs9380237 | 0.002 | -0.963 | -1.94 | 0.595 | 0.101 | 0.037 | 0.037 | 0 | 0.48 | 0.31 | 0.38 | 0.4 | 0.29 | 0.48 |
| 6 | 31265262 | rs2246954 | -0.538 | -0.835 | -1.957 | 0.624 | -0.011 | 0.049 | 0.049 | 0.262 | 0.51 | 0.71 | 0.64 | 0.66 | 0.59 | 0.54 |
| 6 | 31265314 | rs2524123 | -0.086 | -0.924 | -1.989 | 0.606 | 0.015 | -0.006 | -0.006 | 0 | 0.4 | 0.33 | 0.38 | 0.43 | 0.33 | 0.4 |
| 6 | 31265490 | rs2247056 | -2.911 | -0.895 | -1.996 | 0.611 | -0.011 | -0.014 | -0.014 | 1.367 | 0.9 | 0.91 | 0.88 | 0.83 | 0.8 | 0.88 |
| 6 | 31265539 | rs3905495 | 0.428 | -0.895 | -1.95 | 0.6 | 0.002 | 0.001 | 0.001 | 0.365 | 0.48 | 0.49 | 0.45 | 0.41 | 0.39 | 0.47 |
| 6 | 31265565 | rs17192463 | 2.548 | -0.854 | -2.013 | 0.601 | 0.045 | 0.014 | 0.014 | 0 | 0.11 | 0.02 | 0.13 | 0.06 | 0.03 | 0.11 |
| 6 | 31266190 | rs2853922 | -1.08 | -0.772 | -1.981 | 0.626 | -0.033 | 0.009 | 0.009 | 0.036 | 0.65 | 0.75 | 0.77 | 0.73 | 0.64 | 0.67 |
| 6 | 31266522 | rs2524089 | -1.08 | -0.772 | -1.981 | 0.626 | -0.033 | 0.009 | 0.009 | 0.036 | 0.65 | 0.75 | 0.77 | 0.73 | 0.64 | 0.67 |
| 6 | 31267618 | rs9380238 | 1.298 | -1.063 | -2.088 | 0.534 | 0.03 | -0.019 | -0.019 | 0 | 0.22 | 0.14 | 0.17 | 0.24 | 0.16 | 0.2 |
| 6 | 31268332 | rs28367646 | 0.554 | -0.742 | -1.864 | 0.679 | -0.011 | -0.007 | -0.007 | 3.346 | 0.36 | 0.5 | 0.33 | 0.34 | 0.35 | 0.4 |
| 6 | 31268832 | rs9380240 | -0.367 | -0.709 | -1.866 | 0.687 | 0.007 | -0.003 | -0.003 | 0 | 0.57 | 0.6 | 0.59 | 0.61 | 0.46 | 0.59 |
| 6 | 31269154 | rs2524066 | -1.171 | -0.657 | -1.87 | 0.695 | -0.033 | 0.009 | 0.009 | 0.049 | 0.65 | 0.75 | 0.77 | 0.73 | 0.64 | 0.66 |
| 6 | 31269173 | rs9366778 | -0.408 | -0.659 | -1.866 | 0.699 | 0.006 | -0.004 | -0.004 | 1.564 | 0.58 | 0.61 | 0.59 | 0.61 | 0.46 | 0.59 |
| 6 | 31269308 | rs3873385 | 0.819 | -0.591 | -1.778 | 0.706 | 0.006 | 0.031 | 0.031 | 0 | 0.09 | 0.09 | 0.24 | 0.31 | 0.15 | 0.1 |
| 6 | 31269348 | rs3915971 | 0.154 | -0.588 | -1.778 | 0.712 | 0.015 | 0.037 | 0.037 | 0 | 0.15 | 0.21 | 0.31 | 0.32 | 0.24 | 0.13 |
| 6 | 31269553 | rs9468935 | 0.89 | -0.754 | -1.762 | 0.639 | -0.013 | -0.016 | -0.016 | 0 | 0.32 | 0.49 | 0.32 | 0.26 | 0.33 | 0.35 |
| 6 | 31271008 | rs13200073 | 1.347 | -0.836 | -1.79 | 0.654 | 0.037 | 0.034 | 0.034 | 0 | 0.25 | 0.16 | 0.1 | 0.11 | 0.15 | 0.21 |
| 6 | 31271157 | rs7750269 | -0.323 | -0.515 | -1.595 | 0.913 | 0.006 | -0.009 | -0.009 | 0 | 0.13 | 0.09 | 0.14 | 0.29 | 0.1 | 0.13 |
| 6 | 31271601 | rs9264868 | 1.774 | -0.357 | -1.578 | 0.924 | 0.018 | 0.014 | 0.014 | 0 | 0.05 | 0.06 | 0.14 | 0.12 | 0.11 | 0.06 |
| 6 | 31271630 | rs9264869 | 1.774 | -0.357 | -1.578 | 0.924 | 0.018 | 0.014 | 0.014 | 0 | 0.05 | 0.06 | 0.14 | 0.12 | 0.11 | 0.06 |
| 6 | 31271662 | rs9264870 | 1.774 | -0.357 | -1.578 | 0.924 | 0.018 | 0.014 | 0.014 | 0 | 0.05 | 0.06 | 0.14 | 0.12 | 0.11 | 0.06 |
| 6 | 31272157 | rs9368675 | -0.32 | -0.482 | -1.607 | 0.925 | 0.007 | -0.011 | -0.011 | 0 | 0.13 | 0.09 | 0.14 | 0.29 | 0.1 | 0.13 |
| 6 | 31272209 | rs9368676 | -0.32 | -0.482 | -1.607 | 1.114 | 0.007 | -0.011 | -0.011 | 0 | 0.13 | 0.09 | 0.14 | 0.34 | 0.1 | 0.14 |
| 6 | 31272321 | rs9368677 | 0.339 | -0.464 | -1.595 | 1.035 | 0.002 | -0.004 | -0.004 | 0 | 0.09 | 0.07 | 0.12 | 0.21 | 0.08 | 0.07 |
| 6 | 31272553 | rs9264904 | 0.834 | -0.696 | -1.485 | 0.863 | -0.004 | -0.028 | -0.028 | 0 | 0.26 | 0.37 | 0.37 | 0.15 | 0.3 | 0.18 |
| 6 | 31272774 | rs9264916 | -0.123 | -0.459 | -1.696 | 1.056 | 0.001 | 0.008 | 0.008 | 0 | 0.15 | 0.13 | 0.27 | 0.32 | 0.19 | 0.14 |
| 6 | 31273030 | rs7760988 | -0.364 | -0.506 | -1.711 | 1.049 | 0.007 | -0.01 | -0.01 | 0 | 0.14 | 0.09 | 0.14 | 0.29 | 0.1 | 0.13 |
| 6 | 31273224 | rs364415 | 1.813 | -0.44 | -1.692 | 1.044 | 0.018 | 0.014 | 0.014 | 0 | 0.05 | 0.06 | 0.14 | 0.12 | 0.11 | 0.06 |
| 6 | 31273495 | rs7761965 | 0.345 | -0.927 | -1.897 | 0.911 | 0.054 | -0.021 | -0.021 | 1.067 | 0.25 | 0.11 | 0.27 | 0.36 | 0.13 | 0.24 |
| 6 | 31273745 | rs3873386 | 0.472 | -0.582 | -1.873 | 0.872 | 0.022 | 0.024 | 0.024 | 0.037 | 0.46 | 0.6 | 0.49 | 0.32 | 0.43 | 0.47 |
| 6 | 31274380 | rs9264942 | 0.086 | -1.039 | -1.955 | 0.734 | 0.021 | -0.02 | -0.02 | 0 | 0.4 | 0.27 | 0.3 | 0.39 | 0.36 | 0.52 |
| 6 | 31274441 | rs9468942 | 1.847 | -0.725 | -1.933 | 0.74 | -0.03 | -0.009 | -0.009 | 0 | 0.06 | 0.08 | 0.02 | 0.05 | 0.01 | 0.01 |
| 6 | 31274555 | rs10484554 | 1.893 | -0.628 | -2 | 0.713 | 0.005 | 0.026 | 0.026 | 0 | 0.2 | 0.08 | 0.08 | 0.05 | 0.14 | 0.2 |
| 6 | 31274693 | rs3094691 | -0.178 | -0.542 | -1.794 | 0.816 | 0.012 | 0.019 | 0.019 | 0.161 | 0.55 | 0.65 | 0.68 | 0.41 | 0.49 | 0.52 |
| 6 | 31275174 | rs396243 | -0.333 | -0.072 | -1.622 | 1.066 | -0.006 | -0.017 | -0.017 | 0 | 0.17 | 0.11 | 0.27 | 0.35 | 0.2 | 0.15 |
| 6 | 31275246 | rs2853916 | -0.449 | -0.075 | -1.611 | 1.134 | 0 | -0.004 | -0.004 | 0 | 0.18 | 0.14 | 0.27 | 0.34 | 0.19 | 0.14 |
| 6 | 31302470 | rs6914104 | 0.329 | 0.776 | -1.837 | 0.986 | -0.018 | -0.023 | -0.023 | 0 | 0.66 | 0.76 | 0.53 | 0.51 | 0.64 | 0.67 |
| 6 | 31305731 | rs28367832 | 0.574 | 0.909 | -1.99 | 0.987 | -0.011 | -0.012 | -0.012 | 0.236 | 0.57 | 0.71 | 0.5 | 0.44 | 0.55 | 0.57 |
| 6 | 31308630 | rs9265797 | -0.593 | 0.986 | -2.021 | 1.016 | -0.008 | -0.018 | -0.018 | 0 | 0.28 | 0.23 | 0.32 | 0.47 | 0.3 | 0.29 |
| 6 | 31310022 | rs28380796 | 0.894 | 0.436 | -1.826 | 0.85 | 0.023 | 0.017 | 0.017 | 0 | 0.34 | 0.27 | 0.21 | 0.21 | 0.22 | 0.32 |
| 6 | 31313101 | rs9265882 | 0.219 | 0.451 | -1.847 | 1.047 | 0.016 | 0.009 | 0.009 | 0 | 0.44 | 0.34 | 0.37 | 0.33 | 0.36 | 0.41 |
| 6 | 31314243 | rs7453967 | -2.146 | 0.523 | -1.861 | 1.031 | -0.023 | 0.021 | 0.021 | 0.184 | 0.84 | 0.94 | 0.82 | 0.91 | 0.85 | 0.85 |
| 6 | 31314781 | rs2507997 | 1.272 | 0.55 | -1.853 | 1.072 | -0.021 | -0.016 | -0.016 | 0 | 0.17 | 0.11 | 0.19 | 0.2 | 0.22 | 0.15 |
| 6 | 31317099 | rs2923006 | 0.347 | 0.469 | -1.698 | 0.99 | 0.019 | 0.028 | 0.028 | 0 | 0.38 | 0.51 | 0.44 | 0.5 | 0.5 | 0.43 |
| 6 | 31317182 | rs4540292 | 0.829 | 0.483 | -1.701 | 0.995 | 0.055 | 0.05 | 0.05 | 1.131 | 0.29 | 0.45 | 0.41 | 0.42 | 0.42 | 0.33 |
| 6 | 31317347 | rs2156875 | 0.376 | 0.39 | -1.679 | 0.962 | 0.019 | 0.028 | 0.028 | 0 | 0.38 | 0.51 | 0.44 | 0.5 | 0.5 | 0.43 |
| 6 | 31318024 | rs2844586 | 1.428 | -0.098 | -1.905 | 0.758 | -0.044 | -0.035 | -0.035 | 0 | 0.15 | 0.19 | 0.09 | 0.09 | 0.07 | 0.13 |
| 6 | 31318144 | rs2523619 | 0.903 | 0.064 | -1.836 | 0.836 | 0.026 | 0.02 | 0.02 | 0 | 0.35 | 0.28 | 0.21 | 0.21 | 0.22 | 0.35 |
| 6 | 31318177 | rs4394275 | 1.968 | 0.48 | -1.547 | 0.984 | 0.027 | 0.028 | 0.028 | 0.355 | 0.19 | 0.34 | 0.25 | 0.23 | 0.26 | 0.21 |
| 6 | 31318303 | rs2523618 | -1.797 | 0.486 | -1.588 | 0.954 | -0.023 | 0.021 | 0.021 | 0.184 | 0.84 | 0.94 | 0.82 | 0.91 | 0.86 | 0.85 |
| 6 | 31319281 | rs9391846 | 0.459 | 0.474 | -1.375 | 1.154 | 0.002 | 0.042 | 0.042 | 0 | 0.05 | 0.09 | 0.13 | 0.2 | 0.08 | 0.04 |
| 6 | 31319355 | rs2442728 | -0.064 | 0.362 | -1.627 | 0.938 | 0.022 | 0.019 | 0.019 | 0 | 0.46 | 0.32 | 0.39 | 0.33 | 0.38 | 0.45 |
| 6 | 31320810 | rs2596503 | -0.675 | 1.258 | -1.349 | 1.128 | 0.056 | 0.018 | 0.018 | 0 | 0.93 | 0.89 | 0.91 | 0.81 | 0.83 | 0.86 |
| 6 | 31321211 | rs2596501 | -0.002 | 1.187 | -1.526 | 1.197 | 0.058 | 0.067 | 0.067 | 0.117 | 0.72 | 0.66 | 0.68 | 0.47 | 0.53 | 0.64 |
| 6 | 31321685 | rs1058026 | 1.607 | -0.128 | -1.264 | 0.544 | -0.022 | -0.008 | -0.008 | 0 | 0.34 | 0.33 | 0.29 | 0.18 | 0.22 | 0.3 |
| 6 | 31325056 | rs2596487 | 1.737 | 0.564 | -1.539 | 0.393 | -0.022 | 0.027 | 0.027 | 0 | 0.14 | 0.05 | 0.15 | 0.07 | 0.15 | 0.07 |
| 6 | 31326960 | rs2523591 | 0.338 | 0.257 | -1.407 | 0.401 | 0.001 | 0 | 0 | 2.831 | 0.51 | 0.5 | 0.53 | 0.34 | 0.47 | 0.57 |
| 6 | 31327435 | rs2523586 | 0.049 | 1.319 | -1.594 | 0.481 | -0.018 | 0.003 | 0.003 | 2.231 | 0.13 | 0.09 | 0.23 | 0.12 | 0.26 | 0.24 |
| 6 | 31330546 | rs2596548 | -1.322 | 1.467 | -1.573 | 0.631 | 0.037 | 0.002 | 0.002 | 0 | 0.92 | 0.85 | 0.92 | 0.93 | 0.84 | 0.93 |

| | | | | | | | | | | | | | | | | |
|---|---|---|---|---|---|---|---|---|---|---|---|---|---|---|---|---|
| 6 | 31330788 | rs9266329 | 0.358 | 0.382 | -1.87 | 0.595 | -0.004 | -0.002 | -0.002 | 0 | 0.37 | 0.25 | 0.36 | 0.47 | 0.23 | 0.34 |
| 6 | 31331166 | rs2596549 | -1.627 | 0.815 | -1.796 | 0.619 | 0.037 | 0.002 | 0.002 | 0 | 0.92 | 0.85 | 0.92 | 0.93 | 0.84 | 0.93 |
| 6 | 31331257 | rs2523557 | -1.289 | 0.927 | -1.916 | 0.442 | 0.02 | -0.011 | -0.011 | 0 | 0.75 | 0.67 | 0.76 | 0.84 | 0.65 | 0.82 |
| 6 | 31331829 | rs2523554 | -1.276 | 1.089 | -2.002 | 0.444 | -0.01 | -0.002 | -0.002 | 0 | 0.81 | 0.78 | 0.86 | 0.82 | 0.68 | 0.84 |
| 6 | 31332239 | rs2596551 | 2.735 | 0.995 | -2.017 | 0.391 | -0.001 | -0.01 | -0.01 | 0.002 | 0.16 | 0.18 | 0.16 | 0.04 | 0.19 | 0.09 |
| 6 | 31333562 | rs2523544 | 2.346 | 0.979 | -2.01 | 0.42 | -0.031 | -0.006 | -0.006 | 0.265 | 0.11 | 0.08 | 0.07 | 0.12 | 0.16 | 0.1 |
| 6 | 31334174 | rs2596574 | 2.523 | 1.161 | -1.989 | 0.397 | -0.001 | -0.017 | -0.017 | 0 | 0.14 | 0.17 | 0.13 | 0.03 | 0.1 | 0.09 |
| 6 | 31334945 | rs2844575 | -0.429 | 0.999 | -2.103 | 0.512 | 0.001 | 0 | 0 | 0.235 | 0.53 | 0.46 | 0.42 | 0.54 | 0.48 | 0.55 |
| 6 | 31335566 | rs9266395 | 1.181 | 0.884 | -2.083 | 0.544 | -0.009 | 0.002 | 0.002 | 0.245 | 0.31 | 0.24 | 0.22 | 0.39 | 0.22 | 0.41 |
| 6 | 31335647 | rs2844571 | 2.028 | 1.054 | -2.104 | 0.528 | -0.056 | -0.005 | -0.005 | 0.05 | 0.14 | 0.08 | 0.09 | 0.12 | 0.25 | 0.1 |
| 6 | 31335707 | rs2523537 | -1.848 | 1.035 | -2.071 | 0.538 | 0.036 | 0.002 | 0.002 | 0 | 0.92 | 0.85 | 0.92 | 0.93 | 0.84 | 0.93 |
| 6 | 31336349 | rs2523534 | -0.779 | 0.926 | -2.093 | 0.573 | -0.003 | 0 | 0 | 0.731 | 0.59 | 0.49 | 0.48 | 0.62 | 0.57 | 0.62 |
| 6 | 31336568 | rs9266409 | 1.227 | 0.913 | -2.117 | 0.564 | -0.01 | 0.002 | 0.002 | 0.245 | 0.31 | 0.24 | 0.22 | 0.39 | 0.21 | 0.41 |
| 6 | 31336644 | rs2844567 | 2.673 | 0.929 | -2.127 | 0.549 | -0.001 | -0.004 | -0.004 | 0 | 0.14 | 0.17 | 0.16 | 0.05 | 0.1 | 0.1 |
| 6 | 31336870 | rs2253907 | -0.244 | 0.964 | -2.115 | 0.579 | -0.004 | -0.004 | -0.004 | 0 | 0.45 | 0.32 | 0.32 | 0.57 | 0.47 | 0.52 |
| 6 | 31336891 | rs2253908 | 2.012 | 1.12 | -2.073 | 0.581 | -0.056 | -0.005 | -0.005 | 0 | 0.14 | 0.08 | 0.09 | 0.12 | 0.25 | 0.1 |
| 6 | 31338528 | rs9295986 | 1.146 | -0.003 | -2.32 | 0.507 | -0.01 | 0 | 0 | 0.682 | 0.38 | 0.39 | 0.47 | 0.52 | 0.25 | 0.41 |
| 6 | 31340328 | rs2523522 | -1.44 | 0.278 | -2.328 | 0.587 | -0.005 | -0.006 | -0.006 | 0.063 | 0.64 | 0.51 | 0.49 | 0.71 | 0.61 | 0.7 |
| 6 | 31340477 | rs4358666 | 1.956 | 0.294 | -2.329 | 0.589 | -0.026 | -0.013 | -0.013 | 0.758 | 0.28 | 0.34 | 0.39 | 0.2 | 0.23 | 0.22 |
| 6 | 31340559 | rs2394990 | -1.44 | 0.295 | -2.329 | 0.579 | -0.005 | -0.006 | -0.006 | 0.079 | 0.64 | 0.51 | 0.48 | 0.71 | 0.61 | 0.7 |
| 6 | 31340611 | rs7770216 | 0.96 | 0.225 | -2.357 | 0.546 | -0.006 | 0.001 | 0.001 | 0.024 | 0.32 | 0.24 | 0.24 | 0.42 | 0.22 | 0.46 |
| 6 | 31340807 | rs2263311 | 2.689 | 0.21 | -2.341 | 0.509 | 0 | -0.012 | -0.012 | 0 | 0.15 | 0.17 | 0.13 | 0.03 | 0.1 | 0.09 |
| 6 | 31341156 | rs6936035 | 1.979 | 0.218 | -2.337 | 0.521 | -0.026 | -0.007 | -0.007 | 0.159 | 0.28 | 0.35 | 0.44 | 0.22 | 0.23 | 0.23 |
| 6 | 31341312 | rs9266514 | -1.47 | 0.229 | -2.337 | 0.516 | -0.005 | -0.006 | -0.006 | 0 | 0.64 | 0.51 | 0.48 | 0.71 | 0.61 | 0.7 |
| 6 | 31341357 | rs2844556 | 2.894 | 0.263 | -2.299 | 0.477 | -0.003 | -0.016 | -0.016 | 0 | 0.14 | 0.16 | 0.13 | 0.03 | 0.1 | 0.1 |
| 6 | 31342631 | rs2254556 | -1.943 | 0.325 | -2.27 | 0.527 | 0.037 | 0.002 | 0.002 | 0 | 0.92 | 0.85 | 0.92 | 0.93 | 0.84 | 0.93 |
| 6 | 31343100 | rs2507985 | -0.936 | 0.262 | -2.303 | 0.547 | -0.005 | -0.001 | -0.001 | 0 | 0.61 | 0.51 | 0.46 | 0.66 | 0.6 | 0.63 |
| 6 | 31343632 | rs6933050 | 1.144 | 0.236 | -2.32 | 0.557 | -0.01 | 0.002 | 0.002 | 0.245 | 0.31 | 0.24 | 0.22 | 0.39 | 0.21 | 0.41 |
| 6 | 31344348 | rs3957110 | -0.896 | 0.389 | -2.277 | 0.637 | -0.005 | -0.001 | -0.001 | 1.876 | 0.61 | 0.51 | 0.46 | 0.66 | 0.6 | 0.63 |
| 6 | 31345021 | rs7382258 | -0.817 | 0.353 | -2.294 | 0.64 | -0.007 | 0.007 | 0.007 | 0.414 | 0.61 | 0.5 | 0.46 | 0.56 | 0.6 | 0.57 |
| 6 | 31345229 | rs2844545 | 2.075 | 0.409 | -2.323 | 0.535 | -0.068 | -0.029 | -0.029 | 0 | 0.16 | 0.09 | 0.1 | 0.19 | 0.28 | 0.12 |
| 6 | 31345794 | rs2507983 | -0.764 | 0.247 | -2.202 | 0.52 | -0.006 | 0.003 | 0.003 | 0.426 | 0.61 | 0.5 | 0.46 | 0.61 | 0.6 | 0.61 |
| 6 | 31346414 | rs2442738 | 0.273 | 0.059 | -2.362 | 0.319 | -0.003 | 0.006 | 0.006 | 0 | 0.81 | 0.84 | 0.88 | 0.86 | 0.84 | 0.91 |
| 6 | 31346445 | rs2523627 | -1.026 | 0.507 | -2.323 | 0.384 | 0.018 | 0.027 | 0.027 | 0 | 0.52 | 0.34 | 0.41 | 0.67 | 0.57 | 0.66 |
| 6 | 31346822 | rs9266629 | 0.714 | 0.907 | -2.294 | 0.519 | -0.034 | -0.03 | -0.03 | 0 | 0.1 | 0.06 | 0.07 | 0.19 | 0.19 | 0.07 |
| 6 | 31346937 | rs9501587 | 1.978 | 0.391 | -2.33 | 0.313 | -0.034 | -0.056 | -0.056 | 0.952 | 0.33 | 0.49 | 0.37 | 0.14 | 0.26 | 0.32 |
| 6 | 31347057 | rs9266638 | 0.883 | 0.121 | -2.529 | 0.316 | 0.102 | 0.03 | 0.03 | 0 | 0.49 | 0.7 | 0.5 | 0.47 | 0.35 | 0.5 |
| 6 | 31347144 | rs3094600 | 1.173 | 0.14 | -2.62 | 0.298 | 0.042 | 0.045 | 0.045 | 0 | 0.33 | 0.09 | 0.25 | 0.19 | 0.26 | 0.24 |
| 6 | 31347451 | rs2244020 | 0.437 | 0.126 | -2.574 | 0.304 | 0.051 | -0.024 | -0.024 | 0 | 0.57 | 0.74 | 0.62 | 0.53 | 0.41 | 0.52 |
| 6 | 31347798 | rs4713460 | -0.291 | 0.106 | -2.469 | 0.337 | 0.013 | -0.004 | -0.004 | 0 | 0.43 | 0.27 | 0.42 | 0.59 | 0.3 | 0.48 |
| 6 | 31347816 | rs4713462 | 0.266 | 0.109 | -2.467 | 0.343 | 0.003 | -0.002 | -0.002 | 0 | 0.4 | 0.27 | 0.44 | 0.6 | 0.29 | 0.48 |
| 6 | 31347877 | rs2507980 | 2.805 | 0.323 | -2.2 | 0.324 | -0.004 | -0.024 | -0.024 | 0 | 0.13 | 0.17 | 0.13 | 0.03 | 0.1 | 0.09 |
| 6 | 31348077 | rs9266669 | 2.962 | 0.474 | -2.211 | 0.397 | -0.029 | -0.009 | -0.009 | 0.085 | 0.08 | 0.05 | 0.04 | 0.07 | 0.12 | 0.07 |
| 6 | 31348580 | rs9266689 | -0.399 | 0.414 | -2.113 | 0.429 | -0.001 | 0 | 0 | 0.109 | 0.62 | 0.47 | 0.66 | 0.58 | 0.65 | 0.66 |
| 6 | 31349922 | rs9266722 | 1.539 | 0.325 | -2.081 | 0.479 | -0.038 | 0.106 | 0.106 | 0 | 0.16 | 0.23 | 0.17 | 0.4 | 0.09 | 0.2 |
| 6 | 31350704 | rs1521 | -1.954 | 0.691 | -2.058 | 0.23 | 0.009 | -0.02 | -0.02 | 0 | 0.78 | 0.69 | 0.79 | 0.9 | 0.74 | 0.84 |
| 6 | 31350802 | rs2844533 | -1.954 | 0.691 | -2.058 | 0.23 | 0.009 | -0.02 | -0.02 | 0.321 | 0.78 | 0.69 | 0.79 | 0.9 | 0.74 | 0.84 |
| 6 | 31351153 | rs3016014 | -1.508 | 1.322 | -2.059 | 0.299 | 0.045 | -0.001 | -0.001 | 0 | 0.9 | 0.85 | 0.89 | 0.93 | 0.75 | 0.93 |
| 6 | 31351764 | rs2442752 | 1.12 | 0.732 | -2.287 | 0.18 | -0.001 | -0.012 | -0.012 | 0 | 0.44 | 0.51 | 0.48 | 0.25 | 0.4 | 0.4 |
| 6 | 31352040 | rs2442749 | 1.565 | 0.68 | -2.412 | 0.221 | -0.023 | -0.01 | -0.01 | 0 | 0.2 | 0.11 | 0.22 | 0.21 | 0.23 | 0.14 |
| 6 | 31352113 | rs9266772 | 1.898 | 0.035 | -2.325 | 0.127 | -0.01 | 0.017 | 0.017 | 0 | 0.21 | 0.18 | 0.23 | 0.09 | 0.2 | 0.19 |
| 6 | 31352631 | rs7741091 | 1.296 | -0.739 | -2.35 | 0.175 | 0.033 | 0.019 | 0.019 | 0 | 0.44 | 0.38 | 0.46 | 0.3 | 0.37 | 0.42 |
| 6 | 31352880 | rs9266774 | 1.637 | -0.748 | -2.311 | 0.201 | -0.034 | 0.068 | 0.068 | 0 | 0.15 | 0.21 | 0.13 | 0.34 | 0.09 | 0.18 |
| 6 | 31352902 | rs28366099 | 2.433 | -0.67 | -2.251 | 0.105 | -0.002 | -0.015 | -0.015 | 0 | 0.15 | 0.17 | 0.13 | 0.03 | 0.1 | 0.09 |
| 6 | 31353593 | rs2844529 | 0.945 | -0.734 | -2.315 | 0.105 | -0.011 | -0.009 | -0.009 | 0 | 0.45 | 0.56 | 0.55 | 0.34 | 0.39 | 0.39 |
| 6 | 31353639 | rs2523484 | 0.516 | -0.761 | -2.336 | 0.099 | -0.011 | 0 | 0 | 0 | 0.59 | 0.56 | 0.63 | 0.4 | 0.48 | 0.53 |
| 6 | 31354104 | rs2428486 | 0.592 | -0.097 | -1.965 | 0.307 | -0.011 | -0.009 | -0.009 | 0 | 0.45 | 0.56 | 0.55 | 0.34 | 0.4 | 0.39 |
| 6 | 31354504 | rs4711268 | 1.232 | -0.104 | -1.958 | 0.307 | -0.013 | -0.004 | -0.004 | 0 | 0.3 | 0.38 | 0.42 | 0.26 | 0.3 | 0.29 |
| 6 | 31354560 | rs13437082 | 1.22 | -0.102 | -1.966 | 0.291 | -0.017 | -0.001 | -0.001 | 0 | 0.31 | 0.39 | 0.43 | 0.31 | 0.3 | 0.3 |
| 6 | 31354595 | rs2596562 | 0.591 | -0.115 | -1.973 | 0.28 | -0.012 | -0.009 | -0.009 | 0 | 0.44 | 0.56 | 0.55 | 0.34 | 0.4 | 0.39 |
| 6 | 31354819 | rs4711269 | 1.188 | -0.068 | -1.944 | 0.358 | -0.017 | -0.001 | -0.001 | 0 | 0.31 | 0.39 | 0.43 | 0.31 | 0.3 | 0.3 |
| 6 | 31355119 | rs13437088 | 1.197 | -0.063 | -1.942 | 0.36 | -0.017 | -0.001 | -0.001 | 0.169 | 0.31 | 0.39 | 0.43 | 0.31 | 0.3 | 0.3 |
| 6 | 31360255 | rs7751505 | 1.34 | -0.05 | -1.956 | 0.309 | -0.015 | -0.002 | -0.002 | 1.058 | 0.3 | 0.38 | 0.39 | 0.3 | 0.3 | 0.29 |
| 6 | 31360389 | rs2523477 | 2.462 | -0.097 | -1.984 | 0.23 | -0.005 | -0.025 | -0.025 | 0 | 0.13 | 0.17 | 0.13 | 0.03 | 0.1 | 0.09 |
| 6 | 31360433 | rs7751725 | 1.386 | -0.091 | -1.981 | 0.244 | -0.015 | -0.002 | -0.002 | 0 | 0.3 | 0.38 | 0.39 | 0.3 | 0.3 | 0.3 |
| 6 | 31361424 | rs2523476 | 2.446 | -0.076 | -1.972 | 0.233 | -0.005 | -0.025 | -0.025 | 0 | 0.13 | 0.17 | 0.13 | 0.03 | 0.1 | 0.09 |
| 6 | 31361710 | rs2523475 | 0.602 | -0.101 | -1.989 | 0.231 | -0.014 | -0.013 | -0.013 | 0 | 0.44 | 0.56 | 0.52 | 0.33 | 0.4 | 0.38 |
| 6 | 31363489 | rs2523464 | 2.449 | -0.087 | -1.977 | 0.228 | -0.005 | -0.025 | -0.025 | 1.889 | 0.13 | 0.17 | 0.13 | 0.03 | 0.1 | 0.09 |
| 6 | 31364105 | rs28366135 | 2.207 | -0.091 | -1.979 | 0.225 | -0.004 | -0.022 | -0.022 | 0.035 | 0.14 | 0.17 | 0.13 | 0.03 | 0.1 | 0.09 |

| | | | | | | | | | | | | | | | | |
|---|---|---|---|---|---|---|---|---|---|---|---|---|---|---|---|---|
| 6 | 31365308 | rs2523459 | 0.659 | -0.089 | -1.988 | 0.241 | -0.015 | -0.014 | -0.014 | 0 | 0.44 | 0.56 | 0.52 | 0.33 | 0.4 | 0.38 |
| 6 | 31365707 | rs2523457 | 1.747 | 0.299 | -2.033 | 0.359 | 0.067 | 0.043 | 0.043 | 0.648 | 0.17 | 0.23 | 0.2 | 0.32 | 0.26 | 0.19 |
| 6 | 31366595 | rs2596542 | 0.893 | 0.426 | -1.971 | 0.447 | -0.018 | -0.022 | -0.022 | 0.257 | 0.43 | 0.54 | 0.52 | 0.27 | 0.4 | 0.37 |
| 6 | 31366622 | rs2428475 | 0.719 | 0.425 | -1.972 | 0.449 | -0.014 | -0.013 | -0.013 | 0 | 0.44 | 0.56 | 0.52 | 0.33 | 0.4 | 0.38 |
| 6 | 31367365 | rs2251731 | 0.888 | 0.447 | -1.968 | 0.443 | -0.018 | -0.022 | -0.022 | 0 | 0.43 | 0.54 | 0.52 | 0.27 | 0.4 | 0.37 |
| 6 | 31367865 | rs2523454 | 2.067 | 0.433 | -1.93 | 0.427 | -0.033 | -0.033 | -0.033 | 0 | 0.12 | 0.09 | 0.18 | 0.32 | 0.24 | 0.19 |
| 6 | 31368588 | rs2844523 | 2.461 | 0.302 | -1.795 | 0.458 | -0.005 | -0.025 | -0.025 | 0 | 0.13 | 0.17 | 0.13 | 0.03 | 0.1 | 0.09 |
| 6 | 31368964 | rs2844521 | 0.73 | 0.312 | -1.8 | 0.48 | -0.017 | -0.021 | -0.021 | 2.24 | 0.43 | 0.54 | 0.52 | 0.28 | 0.4 | 0.37 |
| 6 | 31371430 | rs2844519 | -1.63 | 0.81 | -1.717 | 0.841 | 0.084 | 0.038 | 0.038 | 0 | 0.93 | 0.85 | 0.91 | 0.83 | 0.83 | 0.92 |
| 6 | 31373810 | rs2263313 | 2.322 | -0.176 | -1.688 | 0.687 | -0.005 | -0.025 | -0.025 | 0 | 0.13 | 0.17 | 0.13 | 0.03 | 0.1 | 0.09 |
| 6 | 31374262 | rs2857281 | 2.265 | -0.179 | -1.684 | 0.713 | -0.008 | -0.017 | -0.017 | 0 | 0.14 | 0.18 | 0.14 | 0.08 | 0.1 | 0.1 |
| 6 | 31377047 | rs6910087 | 1.88 | -0.355 | -1.81 | 0.631 | -0.037 | -0.011 | -0.011 | 0 | 0.22 | 0.31 | 0.29 | 0.17 | 0.16 | 0.16 |
| 6 | 31378510 | rs1063632 | 0.973 | -0.45 | -1.911 | 0.566 | 0.003 | 0.001 | 0.001 | 0.074 | 0.08 | 0.08 | 0.05 | 0.07 | 0.07 | 0.04 |
| 6 | 31379109 | rs1051794 | -0.216 | -0.843 | -2.125 | 0.287 | -0.029 | -0.005 | -0.005 | 0 | 0.31 | 0.48 | 0.4 | 0.3 | 0.31 | 0.3 |
| 6 | 31379115 | rs1131896 | 1.521 | -1.099 | -2.357 | 0.262 | 0.041 | 0.014 | 0.014 | 0.781 | 0.46 | 0.29 | 0.35 | 0.42 | 0.28 | 0.41 |
| 6 | 31379198 | rs2256028 | 1.819 | -1.091 | -2.305 | 0.139 | -0.01 | -0.008 | -0.008 | 0 | 0.37 | 0.27 | 0.21 | 0.12 | 0.19 | 0.27 |
| 6 | 31380449 | rs2256175 | -1.027 | -0.668 | -2.18 | 0.322 | -0.018 | -0.026 | -0.026 | 0 | 0.75 | 0.73 | 0.65 | 0.42 | 0.53 | 0.59 |
| 6 | 31383071 | rs1131904 | 0.096 | -0.589 | -2.059 | 0.388 | -0.021 | 0.007 | 0.007 | 0.211 | 0.27 | 0.44 | 0.35 | 0.3 | 0.3 | 0.3 |
| 6 | 31383979 | rs6932236 | 0.297 | -0.595 | -2.062 | 0.385 | -0.018 | 0.012 | 0.012 | 0.044 | 0.26 | 0.44 | 0.35 | 0.3 | 0.3 | 0.3 |
| 6 | 31384479 | rs2848713 | 2.64 | -0.991 | -2.272 | 0.21 | 0.08 | 0.005 | 0.005 | 1.317 | 0.19 | 0 | 0.06 | 0.09 | 0.08 | 0.19 |
| 6 | 31384578 | rs9468992 | 1.565 | -0.469 | -1.99 | 0.376 | -0.006 | 0.013 | 0.013 | 0 | 0.16 | 0.34 | 0.28 | 0.18 | 0.18 | 0.24 |
| 6 | 31384792 | rs9266845 | 0.223 | -0.425 | -1.989 | 0.376 | -0.017 | 0.008 | 0.008 | 0.2 | 0.26 | 0.4 | 0.34 | 0.3 | 0.3 | 0.3 |
| 6 | 31387373 | rs2596530 | -1.635 | -0.202 | -2.034 | 0.473 | 0.008 | -0.001 | -0.001 | 0.012 | 0.81 | 0.74 | 0.83 | 0.78 | 0.61 | 0.8 |
| 6 | 31387557 | rs2596531 | -1.635 | -0.202 | -2.034 | 0.473 | 0.008 | -0.001 | -0.001 | 1.719 | 0.81 | 0.74 | 0.83 | 0.78 | 0.61 | 0.8 |
| 6 | 31388595 | rs9295993 | 0.092 | -0.353 | -2 | 0.455 | -0.021 | 0.007 | 0.007 | 0.211 | 0.27 | 0.44 | 0.35 | 0.3 | 0.3 | 0.3 |
| 6 | 31389784 | rs2844511 | -1.609 | -0.109 | -1.996 | 0.512 | 0.008 | -0.001 | -0.001 | 0 | 0.81 | 0.74 | 0.83 | 0.78 | 0.61 | 0.8 |
| 6 | 31390203 | rs10223568 | 1.592 | -0.248 | -1.999 | 0.5 | -0.006 | 0.013 | 0.013 | 0 | 0.16 | 0.34 | 0.28 | 0.18 | 0.18 | 0.24 |
| 6 | 31390410 | rs2516448 | -1.659 | -0.232 | -1.998 | 0.503 | 0.008 | -0.001 | -0.001 | 0 | 0.81 | 0.74 | 0.83 | 0.78 | 0.61 | 0.8 |
| 6 | 31392118 | rs9501109 | 0.537 | -0.197 | -1.962 | 0.323 | -0.017 | -0.014 | -0.014 | 3.952 | 0.35 | 0.42 | 0.38 | 0.19 | 0.24 | 0.27 |
| 6 | 31393374 | rs4947314 | 0.985 | -0.136 | -1.882 | 0.304 | -0.002 | 0.037 | 0.037 | 0 | 0.19 | 0.4 | 0.32 | 0.24 | 0.22 | 0.26 |
| 6 | 31401374 | rs4346874 | 1.458 | -0.139 | -1.88 | 0.291 | -0.006 | 0.013 | 0.013 | 0 | 0.16 | 0.34 | 0.28 | 0.18 | 0.18 | 0.24 |
| 6 | 31405604 | rs9501595 | 1.458 | -0.139 | -1.88 | 0.291 | -0.006 | 0.013 | 0.013 | 0 | 0.16 | 0.34 | 0.28 | 0.18 | 0.18 | 0.24 |
| 6 | 31407643 | rs7772549 | -0.238 | -0.24 | -1.872 | 0.205 | -0.02 | -0.01 | -0.01 | 0 | 0.4 | 0.47 | 0.43 | 0.3 | 0.33 | 0.31 |
| 6 | 31407828 | rs9469003 | 1.493 | -0.251 | -1.829 | 0.19 | -0.006 | 0.013 | 0.013 | 0 | 0.16 | 0.34 | 0.28 | 0.18 | 0.18 | 0.24 |
| 6 | 31409024 | rs9469007 | -0.22 | -0.121 | -1.852 | 0.19 | -0.02 | -0.01 | -0.01 | 0 | 0.4 | 0.47 | 0.43 | 0.3 | 0.33 | 0.31 |
| 6 | 31409605 | rs9267084 | 2.632 | 1.059 | -1.83 | 0.386 | -0.049 | -0.017 | -0.017 | 1.386 | 0.09 | 0.03 | 0.04 | 0.03 | 0.13 | 0.09 |
| 6 | 31409857 | rs28575156 | 1.39 | 0.388 | -1.717 | 0.419 | -0.006 | 0.013 | 0.013 | 0 | 0.16 | 0.34 | 0.28 | 0.19 | 0.18 | 0.24 |
| 6 | 31410408 | rs2844510 | 0.246 | 0.582 | -1.665 | 0.497 | 0.031 | 0.002 | 0.002 | 0 | 0.06 | 0.1 | 0.11 | 0.06 | 0.13 | 0.1 |
| 6 | 31412961 | rs2596464 | -0.086 | 0.463 | -2.094 | 0.233 | 0.006 | 0.007 | 0.007 | 0.72 | 0.38 | 0.34 | 0.49 | 0.47 | 0.47 | 0.41 |
| 6 | 31416156 | rs2516464 | -0.232 | 0.137 | -1.879 | -0.107 | -0.004 | 0.054 | 0.054 | 0 | 0.47 | 0.59 | 0.52 | 0.2 | 0.5 | 0.37 |
| 6 | 31416536 | rs2516463 | 1.464 | 0.121 | -1.832 | -0.156 | 0.031 | -0.036 | -0.036 | 1.412 | 0.09 | 0.23 | 0.11 | 0.01 | 0.1 | 0.07 |
| 6 | 31416639 | rs2516462 | -0.549 | 0.12 | -1.852 | -0.135 | -0.006 | 0.051 | 0.051 | 0 | 0.49 | 0.6 | 0.53 | 0.26 | 0.53 | 0.38 |
| 6 | 31416920 | rs16899646 | 1.712 | 0.078 | -1.887 | -0.168 | -0.034 | -0.01 | -0.01 | 0 | 0.24 | 0.3 | 0.28 | 0.17 | 0.15 | 0.16 |
| 6 | 31417191 | rs3128982 | 1.803 | 0.104 | -1.891 | -0.171 | -0.071 | 0.036 | 0.036 | 0 | 0.14 | 0.05 | 0.13 | 0.02 | 0.25 | 0.14 |
| 6 | 31417396 | rs9501129 | 1.39 | 0.077 | -1.886 | -0.166 | -0.024 | -0.002 | -0.002 | 0 | 0.26 | 0.32 | 0.29 | 0.23 | 0.18 | 0.18 |
| 6 | 31417510 | rs2596460 | 1.458 | 0.097 | -1.835 | -0.131 | 0.031 | -0.036 | -0.036 | 0 | 0.09 | 0.23 | 0.12 | 0.01 | 0.1 | 0.07 |
| 6 | 31417647 | rs2596458 | 1.458 | 0.097 | -1.835 | -0.131 | 0.031 | -0.036 | -0.036 | 0 | 0.09 | 0.23 | 0.11 | 0.01 | 0.1 | 0.07 |
| 6 | 31418124 | rs2523693 | -0.519 | 0.101 | -1.853 | -0.13 | -0.006 | 0.051 | 0.051 | 0 | 0.49 | 0.61 | 0.53 | 0.26 | 0.53 | 0.38 |
| 6 | 31419042 | rs2516459 | -0.489 | 0.093 | -1.835 | -0.134 | -0.006 | 0.05 | 0.05 | 0 | 0.48 | 0.61 | 0.53 | 0.26 | 0.53 | 0.38 |
| 6 | 31419387 | rs2516458 | -0.205 | 0.093 | -1.831 | -0.142 | -0.004 | 0.052 | 0.052 | 0 | 0.46 | 0.59 | 0.52 | 0.2 | 0.5 | 0.37 |
| 6 | 31420018 | rs2516456 | -0.208 | 0.054 | -1.748 | -0.162 | -0.004 | 0.051 | 0.051 | 0 | 0.46 | 0.59 | 0.52 | 0.2 | 0.5 | 0.37 |
| 6 | 31420500 | rs3131622 | 0.634 | 0.116 | -1.731 | -0.121 | -0.003 | 0.015 | 0.015 | 0 | 0.37 | 0.36 | 0.4 | 0.19 | 0.4 | 0.29 |
| 6 | 31421297 | rs2263316 | -1.368 | 0.075 | -1.769 | -0.036 | -0.003 | 0.001 | 0.001 | 0.141 | 0.6 | 0.72 | 0.67 | 0.36 | 0.61 | 0.59 |
| 6 | 31421489 | rs6920723 | 2.83 | 0.573 | -1.426 | -0.007 | 0.046 | 0.046 | 0.046 | 0.122 | 0.09 | 0.02 | 0.02 | 0 | 0.02 | 0.01 |
| 6 | 31424757 | rs2395028 | 0.744 | 1.378 | -0.956 | 0.36 | 0.031 | -0.036 | -0.036 | 0 | 0.09 | 0.23 | 0.11 | 0.01 | 0.1 | 0.07 |
| 6 | 31425499 | rs3131621 | 1.139 | 1.657 | -1.112 | 0.361 | -0.079 | 0.001 | 0.001 | 0 | 0.16 | 0.07 | 0.14 | 0.08 | 0.28 | 0.15 |
| 6 | 31425985 | rs2596480 | 0.552 | 0.606 | -0.619 | 0.368 | 0.029 | -0.037 | -0.037 | 0 | 0.09 | 0.24 | 0.12 | 0.01 | 0.1 | 0.07 |
| 6 | 31429927 | rs3094228 | 0.3 | -0.444 | -1.36 | -0.004 | 0.036 | 0.041 | 0.041 | 0 | 0.28 | 0.18 | 0.13 | 0.18 | 0.18 | 0.26 |
| 6 | 31430799 | rs2596473 | 0.384 | -0.579 | -1.284 | -0.193 | 0.002 | 0 | 0 | 0.708 | 0.49 | 0.34 | 0.48 | 0.5 | 0.5 | 0.57 |
| 6 | 31431691 | rs2255221 | 1.777 | -0.627 | -1.039 | -0.431 | 0.006 | -0.04 | -0.04 | 0 | 0.08 | 0.17 | 0.14 | 0.02 | 0.08 | 0 |
| 6 | 31431757 | rs11752262 | 0.69 | -0.839 | -1.099 | -0.384 | -0.008 | 0.001 | 0.001 | 0 | 0.14 | 0.15 | 0.07 | 0.17 | 0.06 | 0.2 |
| 6 | 31431820 | rs2243621 | 2.346 | -0.817 | -0.875 | -0.496 | -0.015 | -0.036 | -0.036 | 0 | 0.21 | 0.31 | 0.27 | 0.07 | 0.18 | 0.07 |
| 6 | 31431874 | rs2395030 | 2.581 | -1.404 | -1.388 | -0.361 | 0.06 | -0.008 | -0.008 | 0.019 | 0.14 | 0 | 0.04 | 0.08 | 0.06 | 0.1 |
| 6 | 31432125 | rs2284178 | 0.489 | -0.435 | -0.849 | -0.213 | 0.003 | 0.001 | 0.001 | 0.589 | 0.49 | 0.34 | 0.48 | 0.5 | 0.5 | 0.57 |
| 6 | 31432787 | rs11757571 | 0.495 | -0.37 | -0.771 | -0.168 | 0.013 | 0.004 | 0.004 | 0 | 0.13 | 0.08 | 0.05 | 0.16 | 0.06 | 0.13 |
| 6 | 31433073 | rs11757667 | 0.488 | -0.373 | -0.772 | -0.17 | 0.013 | 0.004 | 0.004 | 0 | 0.13 | 0.08 | 0.05 | 0.16 | 0.06 | 0.13 |
| 6 | 31433558 | rs3094014 | 0.569 | -0.311 | -0.761 | -0.18 | 0.006 | 0.007 | 0.007 | 0 | 0.2 | 0.18 | 0.1 | 0.18 | 0.15 | 0.25 |
| 6 | 31434111 | rs3094604 | 0.754 | -0.15 | -0.864 | -0.149 | 0.01 | 0.016 | 0.016 | 0 | 0.19 | 0.12 | 0.08 | 0.16 | 0.14 | 0.18 |
| 6 | 31434366 | rs3094013 | 2.875 | 0.613 | -0.702 | 0.005 | -0.01 | 0.01 | 0.01 | 0 | 0.06 | 0.03 | 0.03 | 0 | 0.08 | 0.05 |

| | | | | | | | | | | | | | | | | |
|---|---|---|---|---|---|---|---|---|---|---|---|---|---|---|---|---|
| 6 | 31435119 | rs2516518 | 0.389 | 0.58 | -0.223 | -0.053 | 0.007 | -0.055 | -0.055 | 0 | 0.1 | 0.22 | 0.16 | 0.02 | 0.1 | 0.01 |
| 6 | 31435833 | rs2244546 | 0.254 | 0.667 | -0.67 | 0.172 | -0.012 | -0.012 | -0.012 | 0.099 | 0.06 | 0.01 | 0.18 | 0.12 | 0.08 | 0.07 |
| 6 | 31435869 | rs4713466 | 0.425 | 1.631 | -0.261 | 0.626 | 0.014 | 0.034 | 0.034 | 0 | 0.1 | 0.17 | 0.17 | 0.23 | 0.12 | 0.12 |
| 6 | 31436047 | rs2518028 | 0.201 | 1.755 | -0.506 | 0.227 | 0.029 | -0.009 | -0.009 | 0 | 0.81 | 0.75 | 0.84 | 0.88 | 0.72 | 0.82 |
| 6 | 31436312 | rs2596454 | -1.413 | 1.11 | -0.968 | -0.169 | 0.001 | 0.056 | 0.056 | 0 | 0.74 | 0.86 | 0.87 | 0.93 | 0.76 | 0.8 |
| 6 | 31437305 | rs2395031 | 0.342 | 1.387 | -0.775 | -0.13 | -0.001 | -0.014 | -0.014 | 0.24 | 0.06 | 0.1 | 0.06 | 0 | 0.08 | 0.03 |
| 6 | 31437506 | rs2894220 | 0.342 | 1.387 | -0.775 | -0.13 | -0.001 | -0.014 | -0.014 | 1.214 | 0.06 | 0.1 | 0.06 | 0 | 0.08 | 0.03 |
| 6 | 31438368 | rs2244839 | 0.881 | 0.799 | -0.841 | 0.132 | -0.036 | 0 | 0 | 0.011 | 0.35 | 0.46 | 0.28 | 0.31 | 0.27 | 0.23 |
| 6 | 31438390 | rs2596449 | -1.793 | 0.606 | -1.122 | 0.014 | 0.015 | 0.076 | 0.076 | 0 | 0.71 | 0.87 | 0.86 | 0.89 | 0.75 | 0.76 |
| 6 | 31438794 | rs7758090 | 0.616 | 0.618 | -1.109 | 0.035 | 0 | -0.004 | -0.004 | 0 | 0.08 | 0.1 | 0.06 | 0.01 | 0.09 | 0.05 |
| 6 | 31439063 | rs2844505 | -1.795 | 0.61 | -1.128 | 0.017 | 0.015 | 0.076 | 0.076 | 0 | 0.71 | 0.87 | 0.86 | 0.89 | 0.75 | 0.76 |
| 6 | 31439498 | rs9267139 | 0.268 | 0.67 | -1.091 | 0.376 | 0 | 0.018 | 0.018 | 0 | 0.09 | 0.11 | 0.22 | 0.26 | 0.1 | 0.16 |
| 6 | 31439740 | rs2523663 | -0.637 | 0.634 | -1.116 | -0.02 | 0.02 | 0.081 | 0.081 | 0 | 0.63 | 0.77 | 0.81 | 0.88 | 0.66 | 0.7 |
| 6 | 31440082 | rs1055569 | 0.157 | 0.66 | -1.101 | 0.039 | 0.005 | 0.003 | 0.003 | 0 | 0.44 | 0.58 | 0.36 | 0.34 | 0.37 | 0.29 |
| 6 | 31440195 | rs2596446 | -1.834 | 0.672 | -1.141 | 0.003 | 0.016 | 0.077 | 0.077 | 0 | 0.71 | 0.87 | 0.86 | 0.89 | 0.75 | 0.76 |
| 6 | 31440497 | rs2516440 | -0.061 | 0.549 | -0.886 | 0.164 | 0.004 | 0.002 | 0.002 | 0 | 0.44 | 0.58 | 0.36 | 0.34 | 0.38 | 0.29 |
| 6 | 31440552 | rs3828886 | -0.063 | 0.184 | -0.662 | 0.177 | -0.015 | -0.005 | -0.005 | 0 | 0.4 | 0.46 | 0.28 | 0.33 | 0.29 | 0.28 |
| 6 | 31441901 | rs12665745 | 1.568 | -0.088 | -0.613 | 0.051 | 0.032 | -0.018 | -0.018 | 0 | 0.24 | 0.16 | 0.14 | 0.29 | 0.16 | 0.22 |
| 6 | 31442137 | rs6940467 | 0.454 | 0.002 | -0.505 | -0.015 | -0.029 | -0.06 | -0.06 | 0.626 | 0.15 | 0.28 | 0.14 | 0.03 | 0.12 | 0.06 |
| 6 | 31442782 | rs3131643 | 1.697 | 0.593 | -0.642 | 0.217 | 0.016 | -0.002 | -0.002 | 0 | 0.08 | 0.12 | 0.05 | 0.06 | 0.11 | 0.04 |
| 6 | 31443323 | rs12660382 | 1.186 | 0.717 | -0.83 | 0.272 | 0.019 | -0.02 | -0.02 | 0 | 0.26 | 0.18 | 0.16 | 0.31 | 0.21 | 0.31 |
| 6 | 31445216 | rs2844502 | 0.688 | 1.024 | -0.683 | 0.329 | -0.003 | -0.006 | -0.006 | 0 | 0.3 | 0.29 | 0.36 | 0.6 | 0.3 | 0.36 |
| 6 | 31445345 | rs2596534 | -0.126 | 0.868 | -0.685 | 0.128 | -0.001 | 0.001 | 0.001 | 0 | 0.38 | 0.38 | 0.42 | 0.61 | 0.4 | 0.42 |
| 6 | 31445909 | rs2395488 | 0.834 | 0.871 | -0.683 | 0.13 | -0.003 | -0.006 | -0.006 | 0 | 0.3 | 0.29 | 0.36 | 0.6 | 0.3 | 0.36 |
| 6 | 31446466 | rs2248372 | -0.438 | 0.63 | -0.97 | -0.36 | 0.003 | 0.017 | 0.017 | 0 | 0.33 | 0.25 | 0.38 | 0.22 | 0.4 | 0.37 |
| 6 | 31446546 | rs2248373 | -1.75 | 0.63 | -0.97 | -0.361 | -0.005 | -0.067 | -0.067 | 0 | 0.63 | 0.54 | 0.75 | 0.81 | 0.7 | 0.74 |
| 6 | 31446796 | rs2248462 | 1.267 | 0.635 | -1.003 | -0.346 | -0.002 | -0.002 | -0.002 | 0.006 | 0.15 | 0.14 | 0.29 | 0.15 | 0.26 | 0.23 |
| 6 | 31446820 | rs2243429 | 1.378 | 0.635 | -1.004 | -0.345 | -0.003 | -0.006 | -0.006 | 0 | 0.3 | 0.29 | 0.36 | 0.6 | 0.3 | 0.36 |
| 6 | 31447588 | rs2516513 | 1.66 | 0.216 | -0.903 | -0.541 | 0.001 | -0.005 | -0.005 | 0 | 0.23 | 0.18 | 0.3 | 0.15 | 0.26 | 0.25 |
| 6 | 31447848 | rs2596536 | 1.464 | 0.391 | -0.836 | -0.459 | -0.003 | -0.006 | -0.006 | 0 | 0.3 | 0.29 | 0.36 | 0.6 | 0.3 | 0.36 |
| 6 | 31448154 | rs2523651 | -0.656 | 0.389 | -0.836 | -0.46 | -0.003 | -0.019 | -0.019 | 0 | 0.53 | 0.52 | 0.71 | 0.76 | 0.66 | 0.65 |
| 6 | 31448315 | rs2516424 | 1.464 | 0.39 | -0.836 | -0.459 | -0.003 | -0.006 | -0.006 | 6.328 | 0.3 | 0.29 | 0.36 | 0.6 | 0.3 | 0.36 |
| 6 | 31448533 | rs2248617 | 1.464 | 0.39 | -0.836 | -0.459 | -0.003 | -0.006 | -0.006 | 2.025 | 0.3 | 0.29 | 0.36 | 0.6 | 0.3 | 0.36 |
| 6 | 31448625 | rs2516511 | 1.331 | 0.39 | -0.836 | -0.459 | -0.002 | -0.002 | -0.002 | 0 | 0.15 | 0.14 | 0.29 | 0.15 | 0.26 | 0.23 |
| 6 | 31448653 | rs3828893 | 0.939 | 0.365 | -0.805 | -0.493 | 0.001 | 0.001 | 0.001 | 0.057 | 0.08 | 0.09 | 0.06 | 0.01 | 0.1 | 0.05 |
| 6 | 31448818 | rs3749945 | 0.939 | 0.365 | -0.805 | -0.493 | 0.001 | 0.001 | 0.001 | 0.057 | 0.08 | 0.09 | 0.06 | 0.01 | 0.1 | 0.05 |
| 6 | 31448862 | rs3749946 | 1.845 | 0.192 | -0.72 | -0.583 | -0.051 | -0.047 | -0.047 | 0 | 0.18 | 0.23 | 0.11 | 0.07 | 0.07 | 0.09 |
| 6 | 31448976 | rs3099844 | 2.273 | 0.614 | -0.709 | -0.381 | 0.052 | 0.024 | 0.024 | 0 | 0.05 | 0.14 | 0.05 | 0.07 | 0.09 | 0.04 |
| 6 | 31449022 | rs2523650 | -2.399 | 0.515 | -0.833 | -0.393 | -0.004 | 0.005 | 0.005 | 0 | 0.71 | 0.75 | 0.83 | 0.83 | 0.73 | 0.74 |
| 6 | 31449327 | rs2905722 | -2.209 | 1.486 | -0.704 | 0.024 | 0.041 | -0.004 | -0.004 | 0 | 0.94 | 0.89 | 0.94 | 0.96 | 0.88 | 0.89 |
| 6 | 31449994 | rs2516509 | 0.898 | 1.295 | -0.766 | 0.005 | -0.003 | -0.002 | -0.002 | 0 | 0.16 | 0.14 | 0.29 | 0.15 | 0.26 | 0.23 |
| 6 | 31450909 | rs2523710 | 2.432 | 1.16 | -0.998 | 0.207 | 0.025 | 0.018 | 0.018 | 0 | 0.21 | 0.08 | 0.11 | 0.13 | 0.16 | 0.09 |
| 6 | 31451213 | rs2516507 | -0.749 | 1.17 | -0.846 | 0.141 | -0.002 | -0.016 | -0.016 | 0.488 | 0.52 | 0.53 | 0.69 | 0.71 | 0.62 | 0.6 |
| 6 | 31451476 | rs2905747 | -1.285 | 1.178 | -0.912 | 0.107 | 0.018 | 0 | 0 | 0 | 0.9 | 0.89 | 0.89 | 0.91 | 0.79 | 0.9 |
| 6 | 31451567 | rs2523706 | 1.201 | 1.142 | -0.859 | 0.199 | 0.001 | -0.006 | -0.006 | 0 | 0.29 | 0.29 | 0.34 | 0.54 | 0.27 | 0.31 |
| 6 | 31451680 | rs2523705 | 1.275 | 1.142 | -0.858 | 0.198 | -0.002 | -0.002 | -0.002 | 0 | 0.16 | 0.14 | 0.29 | 0.15 | 0.26 | 0.23 |
| 6 | 31451836 | rs3094011 | 2.303 | 1.144 | -0.835 | 0.259 | 0.052 | 0.024 | 0.024 | 0 | 0.05 | 0.14 | 0.05 | 0.07 | 0.09 | 0.04 |
| 6 | 31452475 | rs2516504 | 1.267 | 1.039 | -0.873 | 0.185 | 0.001 | -0.006 | -0.006 | 0 | 0.29 | 0.29 | 0.34 | 0.54 | 0.26 | 0.31 |
| 6 | 31453113 | rs2904600 | 1.235 | 1.041 | -0.873 | 0.185 | -0.002 | -0.002 | -0.002 | 0 | 0.16 | 0.14 | 0.29 | 0.15 | 0.26 | 0.23 |
| 6 | 31453618 | rs9267212 | 0.539 | 0.837 | -0.857 | 0.261 | -0.002 | -0.006 | -0.006 | 0 | 0.08 | 0.1 | 0.07 | 0.01 | 0.1 | 0.05 |
| 6 | 31453640 | rs2516500 | 1.133 | 0.866 | -0.862 | 0.303 | 0.001 | -0.006 | -0.006 | 0 | 0.29 | 0.28 | 0.33 | 0.54 | 0.26 | 0.31 |
| 6 | 31457633 | rs9267280 | 0.499 | 0.84 | -0.857 | 0.262 | -0.002 | -0.005 | -0.005 | 2.029 | 0.08 | 0.1 | 0.06 | 0.01 | 0.1 | 0.05 |
| 6 | 31458157 | rs2507964 | 1.215 | 0.82 | -0.849 | 0.317 | 0 | 0.013 | 0.013 | 0 | 0.27 | 0.29 | 0.31 | 0.54 | 0.27 | 0.29 |
| 6 | 31459742 | rs2516415 | 1.297 | 0.842 | -0.835 | 0.328 | 0.001 | 0.014 | 0.014 | 0 | 0.27 | 0.29 | 0.32 | 0.54 | 0.27 | 0.28 |
| 6 | 31459972 | rs2904783 | -0.736 | 0.872 | -0.848 | 0.141 | 0.001 | 0.001 | 0.001 | 0 | 0.53 | 0.55 | 0.69 | 0.72 | 0.63 | 0.58 |
| 6 | 31461089 | rs3130922 | -0.748 | 0.976 | -0.85 | 0.13 | 0.001 | -0.002 | -0.002 | 0.503 | 0.53 | 0.55 | 0.7 | 0.72 | 0.68 | 0.58 |
| 6 | 31461966 | rs2395043 | -1.06 | 0.717 | -0.904 | 0.024 | 0.008 | 0.012 | 0.012 | 0 | 0.62 | 0.65 | 0.74 | 0.73 | 0.72 | 0.64 |
| 6 | 31463491 | rs2516408 | 0.799 | 0.781 | -0.872 | 0.312 | -0.013 | -0.014 | -0.014 | 0 | 0.34 | 0.38 | 0.19 | 0.27 | 0.26 | 0.24 |
| 6 | 31463963 | rs2534678 | 1.068 | 0.98 | -0.924 | 0.4 | 0.004 | 0.004 | 0.004 | 0 | 0.13 | 0.14 | 0.28 | 0.17 | 0.18 | 0.23 |
| 6 | 31464739 | rs3828903 | 0.685 | 0.766 | -0.991 | 0.285 | 0.041 | -0.001 | -0.001 | 1.519 | 0.41 | 0.32 | 0.4 | 0.46 | 0.32 | 0.4 |
| 6 | 31465726 | rs3828912 | 0.693 | 0.763 | -0.993 | 0.283 | 0.041 | -0.001 | -0.001 | 0 | 0.41 | 0.32 | 0.4 | 0.46 | 0.32 | 0.4 |
| 6 | 31465819 | rs3828914 | 0.758 | 0.57 | -1.082 | 0.186 | 0.025 | -0.009 | -0.009 | 0 | 0.38 | 0.32 | 0.38 | 0.46 | 0.31 | 0.4 |
| 6 | 31468099 | rs2534667 | 1.159 | 0.834 | -0.996 | 0.319 | 0.004 | 0.004 | 0.004 | 0 | 0.13 | 0.14 | 0.28 | 0.17 | 0.18 | 0.23 |
| 6 | 31472459 | rs2534657 | 1.159 | 0.834 | -0.996 | 0.341 | 0.004 | 0.005 | 0.005 | 0 | 0.13 | 0.14 | 0.28 | 0.18 | 0.18 | 0.23 |
| 6 | 31472720 | rs2855812 | 2.429 | 0.702 | -1.01 | 0.112 | 0.016 | 0.014 | 0.014 | 0 | 0.16 | 0.24 | 0.13 | 0.21 | 0.2 | 0.17 |
| 6 | 31473286 | rs3134899 | -1.637 | 0.678 | -1.08 | 0.098 | 0.039 | 0.002 | 0.002 | 0 | 0.89 | 0.85 | 0.86 | 0.91 | 0.77 | 0.88 |
| 6 | 31474280 | rs2242955 | 2.975 | 0.648 | -1.031 | 0.13 | -0.001 | -0.001 | -0.001 | 0 | 0.11 | 0.12 | 0.09 | 0.14 | 0.11 | 0.14 |
| 6 | 31475127 | rs3131638 | -1.52 | 1.033 | -0.994 | 0.284 | 0.034 | 0.002 | 0.002 | 0 | 0.89 | 0.85 | 0.87 | 0.91 | 0.77 | 0.88 |
| 6 | 31475413 | rs3130615 | -0.95 | 1.169 | -1.015 | 0.308 | 0.036 | 0.004 | 0.004 | 0 | 0.87 | 0.82 | 0.82 | 0.88 | 0.73 | 0.78 |

| | | | | | | | | | | | | | | | | |
|---|---|---|---|---|---|---|---|---|---|---|---|---|---|---|---|---|
| 6 | 31475486 | rs3132468 | -0.95 | 1.169 | -1.015 | 0.308 | 0.036 | 0.004 | 0.004 | 0 | 0.87 | 0.82 | 0.82 | 0.88 | 0.73 | 0.78 |
| 6 | 31476854 | rs2844498 | 0.143 | 0.801 | -1.079 | 0.262 | 0.008 | 0.008 | 0.008 | 0 | 0.51 | 0.45 | 0.63 | 0.6 | 0.46 | 0.52 |
| 6 | 31478951 | rs9267431 | 0.416 | 0.74 | -1.147 | 0.277 | 0.004 | 0.01 | 0.01 | 0 | 0.22 | 0.3 | 0.37 | 0.3 | 0.26 | 0.14 |
| 6 | 31480398 | rs2844496 | 1.673 | 1.054 | -0.971 | 0.382 | 0.008 | 0.018 | 0.018 | 0 | 0.1 | 0.12 | 0.22 | 0.14 | 0.13 | 0.12 |
| 6 | 31480906 | rs2516401 | -0.288 | 1.461 | -0.911 | 0.409 | 0.043 | 0.026 | 0.026 | 0 | 0.82 | 0.76 | 0.67 | 0.77 | 0.7 | 0.77 |
| 6 | 31480923 | rs3131633 | -1.699 | 1.481 | -0.916 | 0.455 | 0.042 | 0.005 | 0.005 | 0 | 0.92 | 0.88 | 0.89 | 0.92 | 0.84 | 0.92 |
| 6 | 31481105 | rs2516400 | 0.726 | 1.058 | -1.077 | 0.262 | -0.012 | -0.014 | -0.014 | 0 | 0.36 | 0.39 | 0.19 | 0.25 | 0.27 | 0.26 |
| 6 | 31481146 | rs2246871 | 0.215 | 1.783 | -1.161 | 0.482 | 0.046 | 0.024 | 0.024 | 0 | 0.8 | 0.74 | 0.62 | 0.74 | 0.66 | 0.67 |
| 6 | 31482203 | rs2246986 | 1.256 | 1.652 | -1.156 | 0.488 | 0.008 | 0.02 | 0.02 | 0 | 0.1 | 0.12 | 0.22 | 0.14 | 0.13 | 0.12 |
| 6 | 31483415 | rs2844494 | 0.175 | 1.768 | -1.158 | 0.481 | 0.046 | 0.025 | 0.025 | 0 | 0.8 | 0.74 | 0.62 | 0.74 | 0.66 | 0.67 |
| 6 | 31484032 | rs3094001 | -0.831 | 1.765 | -1.109 | 0.362 | 0.033 | 0.006 | 0.006 | 0 | 0.91 | 0.88 | 0.89 | 0.91 | 0.83 | 0.89 |
| 6 | 31484517 | rs2395045 | -0.901 | 1.737 | -1.112 | 0.354 | 0.033 | 0.004 | 0.004 | 0 | 0.9 | 0.85 | 0.84 | 0.88 | 0.78 | 0.79 |
| 6 | 31484683 | rs3131631 | -0.819 | 1.733 | -1.114 | 0.352 | 0.032 | 0.004 | 0.004 | 0 | 0.9 | 0.85 | 0.84 | 0.88 | 0.78 | 0.79 |
| 6 | 31485354 | rs3131630 | -0.827 | 1.691 | -1.118 | 0.298 | 0.023 | -0.001 | -0.001 | 0.953 | 0.9 | 0.85 | 0.84 | 0.9 | 0.78 | 0.82 |
| 6 | 31488145 | rs3130637 | -0.629 | 1.706 | -1.139 | 0.238 | 0.016 | -0.001 | -0.001 | 0 | 0.89 | 0.84 | 0.79 | 0.87 | 0.76 | 0.8 |
| 6 | 31488191 | rs3130636 | -0.629 | 1.706 | -1.139 | 0.238 | 0.016 | -0.001 | -0.001 | 0 | 0.89 | 0.84 | 0.79 | 0.87 | 0.76 | 0.8 |
| 6 | 31489644 | rs3132454 | 0.423 | 0.74 | -1.39 | -0.097 | 0.02 | 0.007 | 0.007 | 0 | 0.43 | 0.29 | 0.41 | 0.32 | 0.38 | 0.42 |
| 6 | 31490725 | rs3093993 | -0.884 | 1.102 | -1.439 | -0.138 | 0.016 | -0.001 | -0.001 | 0.361 | 0.89 | 0.84 | 0.79 | 0.87 | 0.76 | 0.8 |
| 6 | 31490970 | rs9469021 | 3.045 | 0.804 | -1.485 | -0.298 | 0.009 | 0.008 | 0.008 | 0 | 0.09 | 0.05 | 0.04 | 0.01 | 0.06 | 0.06 |
| 6 | 31491000 | rs3095227 | -0.94 | 1.2 | -1.366 | -0.104 | 0.016 | -0.001 | -0.001 | 0 | 0.89 | 0.84 | 0.79 | 0.87 | 0.76 | 0.8 |
| 6 | 31491131 | rs3093992 | -0.94 | 1.2 | -1.366 | -0.104 | 0.016 | -0.001 | -0.001 | 2.112 | 0.89 | 0.84 | 0.79 | 0.87 | 0.76 | 0.8 |
| 6 | 31496569 | rs2516482 | 3.091 | 0.902 | -1.5 | -0.221 | -0.006 | 0.004 | 0.004 | 0.237 | 0.13 | 0.12 | 0.07 | 0.06 | 0.14 | 0.05 |
| 6 | 31496925 | rs3093983 | -1.262 | 1.149 | -1.454 | -0.158 | 0.025 | 0 | 0 | 0 | 0.92 | 0.88 | 0.88 | 0.9 | 0.83 | 0.9 |
| 6 | 31497399 | rs3130055 | -0.361 | 0.501 | -1.762 | -0.305 | -0.006 | 0.005 | 0.005 | 0 | 0.78 | 0.83 | 0.84 | 0.76 | 0.73 | 0.72 |
| 6 | 31497457 | rs3093981 | -1.334 | 0.773 | -1.648 | -0.238 | 0.025 | 0 | 0 | 0 | 0.92 | 0.88 | 0.88 | 0.9 | 0.83 | 0.9 |
| 6 | 31497478 | rs3093980 | -1.334 | 0.773 | -1.648 | -0.238 | 0.025 | 0 | 0 | 0 | 0.92 | 0.88 | 0.88 | 0.9 | 0.83 | 0.9 |
| 6 | 31497835 | rs3115537 | -1.334 | 0.773 | -1.648 | -0.238 | 0.025 | 0 | 0 | 0 | 0.92 | 0.88 | 0.88 | 0.9 | 0.83 | 0.9 |
| 6 | 31497975 | rs3219190 | 2.798 | 0.847 | -1.816 | -0.394 | -0.04 | 0.034 | 0.034 | 0 | 0.07 | 0.01 | 0.06 | 0 | 0.13 | 0.03 |
| 6 | 31498497 | rs3093978 | -1.337 | 0.686 | -1.662 | -0.244 | 0.025 | 0 | 0 | 0 | 0.92 | 0.88 | 0.88 | 0.9 | 0.83 | 0.9 |
| 6 | 31498737 | rs2516478 | 1.53 | 0.017 | -1.89 | -0.411 | 0.014 | 0.007 | 0.007 | 0 | 0.22 | 0.14 | 0.16 | 0.17 | 0.16 | 0.24 |
| 6 | 31499354 | rs3130056 | -1.397 | 0.47 | -1.719 | -0.293 | 0.025 | 0 | 0 | 0 | 0.92 | 0.88 | 0.88 | 0.9 | 0.83 | 0.9 |
| 6 | 31501413 | rs3093948 | -1.397 | 0.494 | -1.719 | -0.293 | 0.027 | 0 | 0 | 0 | 0.92 | 0.88 | 0.88 | 0.9 | 0.82 | 0.9 |
| 6 | 31501737 | rs1055388 | -0.443 | 0.08 | -1.911 | -0.419 | -0.006 | 0.005 | 0.005 | 0 | 0.78 | 0.83 | 0.84 | 0.76 | 0.73 | 0.72 |
| 6 | 31502767 | rs3131628 | -1.381 | 0.226 | -1.838 | -0.369 | 0.023 | 0 | 0 | 0 | 0.91 | 0.88 | 0.88 | 0.9 | 0.83 | 0.9 |
| 6 | 31502861 | rs3093976 | -1.411 | 0.228 | -1.836 | -0.368 | 0.024 | 0 | 0 | 0 | 0.92 | 0.88 | 0.88 | 0.9 | 0.83 | 0.9 |
| 6 | 31505887 | rs3130058 | -1.411 | 0.376 | -1.8 | -0.193 | 0.024 | 0 | 0 | 0 | 0.92 | 0.88 | 0.88 | 0.9 | 0.83 | 0.9 |
| 6 | 31506210 | rs3093974 | 1.685 | 0.018 | -1.857 | -0.277 | 0.025 | 0.04 | 0.04 | 0 | 0.3 | 0.49 | 0.33 | 0.42 | 0.31 | 0.2 |
| 6 | 31506624 | rs1129640 | -1.514 | 0.161 | -1.783 | -0.203 | 0.024 | 0 | 0 | 0 | 0.92 | 0.88 | 0.88 | 0.9 | 0.83 | 0.9 |
| 6 | 31506648 | rs933208 | -0.664 | -0.009 | -1.868 | -0.263 | -0.007 | 0.005 | 0.005 | 0 | 0.78 | 0.83 | 0.83 | 0.76 | 0.73 | 0.72 |
| 6 | 31506691 | rs2071596 | 1.569 | -0.094 | -1.856 | -0.304 | 0.032 | 0.123 | 0.123 | 0 | 0.16 | 0.33 | 0.26 | 0.37 | 0.17 | 0.14 |
| 6 | 31506744 | rs2516393 | -1.518 | -0.044 | -1.851 | -0.284 | 0.024 | 0 | 0 | 0 | 0.92 | 0.88 | 0.88 | 0.9 | 0.83 | 0.9 |
| 6 | 31506801 | rs2523512 | 1.75 | -0.233 | -1.985 | -0.395 | 0.014 | 0.007 | 0.007 | 0 | 0.22 | 0.14 | 0.16 | 0.16 | 0.16 | 0.24 |
| 6 | 31506854 | rs2523511 | -1.53 | -0.107 | -1.926 | -0.329 | 0.024 | 0 | 0 | 0 | 0.92 | 0.88 | 0.88 | 0.9 | 0.83 | 0.9 |
| 6 | 31507447 | rs2239709 | 1.738 | -0.222 | -1.934 | -0.328 | 0.037 | 0.01 | 0.01 | 0 | 0.13 | 0.05 | 0.05 | 0.14 | 0.1 | 0.18 |
| 6 | 31508761 | rs2523508 | -1.522 | -0.04 | -1.989 | -0.335 | 0.027 | 0 | 0 | 0 | 0.92 | 0.88 | 0.88 | 0.9 | 0.83 | 0.9 |
| 6 | 31509967 | rs2523506 | 2.272 | -0.48 | -2.128 | -0.5 | 0.007 | -0.003 | -0.003 | 3.343 | 0.18 | 0.11 | 0.14 | 0.16 | 0.15 | 0.23 |
| 6 | 31510858 | rs2523504 | -0.652 | -0.209 | -2.133 | -0.588 | -0.007 | 0.006 | 0.006 | 0 | 0.78 | 0.81 | 0.83 | 0.81 | 0.73 | 0.74 |
| 6 | 31510924 | rs2844509 | 1.13 | -0.608 | -2.365 | -0.667 | -0.001 | 0.001 | 0.001 | 0 | 0.24 | 0.21 | 0.36 | 0.17 | 0.27 | 0.25 |
| 6 | 31511857 | rs2251824 | 2.32 | -0.621 | -2.332 | -0.611 | 0.007 | -0.003 | -0.003 | 2.827 | 0.18 | 0.11 | 0.13 | 0.16 | 0.14 | 0.23 |
| 6 | 31512799 | rs2071593 | 1.765 | -0.645 | -2.268 | -0.557 | 0.037 | 0.015 | 0.015 | 0 | 0.13 | 0.05 | 0.05 | 0.1 | 0.1 | 0.16 |
| 6 | 31513402 | rs2239705 | 1.98 | -0.462 | -2.21 | -0.491 | 0 | -0.002 | -0.002 | 0.023 | 0.17 | 0.13 | 0.25 | 0.14 | 0.19 | 0.14 |
| 6 | 31515340 | rs2071592 | 1.665 | -0.476 | -2.211 | -0.515 | -0.004 | 0.013 | 0.013 | 0.006 | 0.36 | 0.5 | 0.34 | 0.46 | 0.29 | 0.26 |
| 6 | 31516363 | rs3219183 | 2.421 | -0.578 | -2.224 | -0.405 | 0.043 | -0.007 | -0.007 | 0 | 0.1 | 0.02 | 0.02 | 0.09 | 0.04 | 0.16 |
| 6 | 31518354 | rs2523500 | -1.071 | -0.241 | -2.273 | -0.462 | 0.004 | 0 | 0 | 0 | 0.75 | 0.75 | 0.63 | 0.76 | 0.64 | 0.76 |
| 6 | 31520426 | rs6916921 | 2.285 | -0.309 | -2.299 | -0.421 | 0.027 | -0.024 | -0.024 | 0 | 0.12 | 0.02 | 0.18 | 0.13 | 0.09 | 0.13 |
| 6 | 31522669 | rs6929796 | 1.677 | -0.226 | -2.171 | -0.395 | 0.029 | 0.127 | 0.127 | 0 | 0.16 | 0.33 | 0.26 | 0.41 | 0.17 | 0.16 |
| 6 | 31524851 | rs2857605 | -2.049 | 0.334 | -2.422 | -0.449 | -0.031 | -0.012 | -0.012 | 0.511 | 0.85 | 0.93 | 0.89 | 0.82 | 0.81 | 0.76 |
| 6 | 31525319 | rs2239707 | -1.138 | 0.121 | -2.562 | -0.449 | -0.029 | -0.023 | -0.023 | 1.705 | 0.71 | 0.89 | 0.78 | 0.7 | 0.71 | 0.69 |
| 6 | 31525448 | rs2230365 | 1.586 | -0.352 | -2.566 | -0.541 | 0.062 | 0.002 | 0.002 | 0 | 0.24 | 0.05 | 0.12 | 0.21 | 0.14 | 0.22 |
| 6 | 31526856 | rs13192469 | 2.518 | -0.294 | -2.466 | -0.487 | 0.043 | -0.007 | -0.007 | 0 | 0.1 | 0.02 | 0.02 | 0.1 | 0.04 | 0.15 |
| 6 | 31527266 | rs9469024 | 4.369 | -0.205 | -2.359 | -0.523 | 0.022 | 0.023 | 0.023 | 0 | 0.07 | 0.02 | 0.01 | 0 | 0 | 0.05 |
| 6 | 31528951 | rs17200733 | 2.942 | 0.291 | -2.324 | -0.491 | -0.042 | 0.036 | 0.036 | 0 | 0.07 | 0.01 | 0.06 | 0 | 0.13 | 0.03 |
| 6 | 31529883 | rs2516390 | -0.307 | 0.197 | -2.105 | -0.257 | -0.009 | -0.002 | -0.002 | 0 | 0.65 | 0.73 | 0.61 | 0.66 | 0.6 | 0.61 |
| 6 | 31531215 | rs928815 | -0.287 | 0.205 | -2.107 | -0.263 | -0.008 | -0.002 | -0.002 | 0.157 | 0.65 | 0.73 | 0.61 | 0.66 | 0.6 | 0.61 |
| 6 | 31532814 | rs2857709 | -2.45 | 0.531 | -2.099 | -0.228 | -0.013 | 0.009 | 0.009 | 0.57 | 0.93 | 0.97 | 0.95 | 0.97 | 0.88 | 0.95 |
| 6 | 31533378 | rs2857602 | -0.48 | -2.052 | -2.168 | -0.654 | -0.009 | -0.002 | -0.002 | 1.236 | 0.65 | 0.73 | 0.61 | 0.66 | 0.6 | 0.61 |
| 6 | 31533606 | rs2857708 | 2.399 | -1.986 | -2.141 | -0.623 | 0.001 | -0.004 | -0.004 | 0 | 0.14 | 0.04 | 0.1 | 0.12 | 0.11 | 0.07 |
| 6 | 31536224 | rs2844484 | -0.461 | -1.953 | -2.122 | -0.625 | -0.009 | -0.002 | -0.002 | 2.219 | 0.65 | 0.73 | 0.61 | 0.67 | 0.59 | 0.61 |
| 6 | 31538244 | rs2009658 | 1.822 | -1.932 | -2.069 | -0.579 | 0.007 | -0.001 | -0.001 | 0 | 0.18 | 0.11 | 0.13 | 0.16 | 0.15 | 0.23 |

| | | | | | | | | | | | | | | | | |
|---|---|---|---|---|---|---|---|---|---|---|---|---|---|---|---|---|
| 6 | 31540313 | rs909253 | 1.724 | -2.511 | -2.137 | -0.751 | 0.001 | 0.016 | 0.016 | 0 | 0.35 | 0.52 | 0.34 | 0.47 | 0.31 | 0.26 |
| 6 | 31540556 | rs2229094 | 0.374 | -2.588 | -2.022 | -0.812 | 0.003 | 0.004 | 0.004 | 0 | 0.3 | 0.27 | 0.27 | 0.2 | 0.29 | 0.35 |
| 6 | 31540757 | rs2229092 | 3.41 | -2.657 | -2.413 | -0.788 | 0.001 | 0.029 | 0.029 | 0 | 0.09 | 0 | 0.03 | 0.01 | 0.06 | 0.03 |
| 6 | 31540784 | rs1041981 | 1.591 | -2.522 | -2.259 | -0.645 | 0.002 | 0.016 | 0.016 | 0 | 0.35 | 0.51 | 0.34 | 0.47 | 0.31 | 0.26 |
| 6 | 31542308 | rs1799964 | 1.181 | -2.464 | -2.313 | -0.618 | 0.004 | 0.002 | 0.002 | 0 | 0.23 | 0.15 | 0.22 | 0.2 | 0.21 | 0.35 |
| 6 | 31542476 | rs1800630 | 1.805 | -2.477 | -2.323 | -0.627 | 0.01 | 0.001 | 0.001 | 0.055 | 0.19 | 0.1 | 0.13 | 0.16 | 0.15 | 0.24 |
| 6 | 31542482 | rs1799724 | 1.952 | -2.334 | -2.215 | -0.541 | 0.026 | -0.025 | -0.025 | 0 | 0.12 | 0.02 | 0.18 | 0.13 | 0.09 | 0.12 |
| 6 | 31543031 | rs1800629 | 2.978 | -2.011 | -1.997 | -0.6 | -0.008 | 0.006 | 0.006 | 0.254 | 0.14 | 0.12 | 0.07 | 0.06 | 0.13 | 0.05 |
| 6 | 31544189 | rs3093662 | 1.19 | -1.462 | -1.971 | -0.139 | 0 | 0.016 | 0.016 | 0 | 0.09 | 0.08 | 0.11 | 0.03 | 0.08 | 0.11 |
| 6 | 31547611 | rs769177 | 2.657 | -1.555 | -2.01 | -0.064 | 0.034 | -0.008 | -0.008 | 0 | 0.07 | 0.01 | 0.02 | 0.08 | 0.03 | 0.09 |
| 6 | 31550452 | rs3093551 | 1.622 | -1.378 | -1.922 | -0.114 | 0.009 | 0.004 | 0.004 | 0 | 0.06 | 0.05 | 0.04 | 0 | 0.03 | 0.01 |
| 6 | 31555130 | rs2256965 | -0.55 | -1.565 | -2.208 | -0.137 | 0.001 | 0.018 | 0.018 | 0 | 0.57 | 0.65 | 0.55 | 0.73 | 0.55 | 0.72 |
| 6 | 31555392 | rs2256974 | 1.148 | -1.738 | -2.326 | -0.266 | 0.035 | 0.14 | 0.14 | 0 | 0.15 | 0.38 | 0.28 | 0.41 | 0.18 | 0.15 |
| 6 | 31556581 | rs1052248 | -0.221 | -1.552 | -3.399 | -1.278 | 0.095 | 0.099 | 0.099 | 0 | 0.44 | 0.22 | 0.23 | 0.25 | 0.27 | 0.55 |
| 6 | 31556928 | rs3179003 | 1.534 | -0.966 | -3.232 | -1.002 | 0.011 | 0.006 | 0.006 | 0 | 0.07 | 0.05 | 0.04 | 0.01 | 0.03 | 0.03 |
| 6 | 31557491 | rs11575841 | 1.871 | -0.892 | -3.166 | -0.93 | 0.007 | 0.002 | 0.002 | 0 | 0.06 | 0.05 | 0.04 | 0.01 | 0.02 | 0.02 |
| 6 | 31557791 | rs11575839 | 0.735 | -0.317 | -3.024 | -0.486 | -0.004 | 0 | 0 | 0 | 0.06 | 0.09 | 0.04 | 0.04 | 0.03 | 0.15 |
| 6 | 31563240 | rs3132452 | 1.581 | -0.085 | -2.931 | -0.312 | 0.025 | -0.049 | -0.049 | 1.582 | 0.11 | 0.03 | 0.18 | 0.18 | 0.09 | 0.13 |
| 6 | 31564167 | rs2515919 | -0.468 | 0.784 | -2.402 | -0.853 | -0.012 | 0.055 | 0.055 | 0 | 0.38 | 0.17 | 0.24 | 0.25 | 0.39 | 0.49 |
| 6 | 31564821 | rs2844480 | 0.681 | 0.393 | -2.034 | -0.325 | -0.005 | 0.009 | 0.009 | 0 | 0.2 | 0.12 | 0.15 | 0.12 | 0.2 | 0.33 |
| 6 | 31566204 | rs2509217 | 0.845 | 0.897 | -1.559 | -0.052 | 0.046 | -0.002 | -0.002 | 1.047 | 0.09 | 0.01 | 0.02 | 0.11 | 0.04 | 0.11 |
| 6 | 31567422 | rs2857596 | -0.683 | 1.877 | -1.002 | 0.115 | 0.002 | -0.002 | -0.002 | 0.413 | 0.81 | 0.79 | 0.83 | 0.91 | 0.75 | 0.89 |
| 6 | 31572481 | rs2857700 | -0.901 | 1.566 | -1.095 | 0.31 | -0.011 | 0.021 | 0.021 | 0 | 0.88 | 0.93 | 0.96 | 0.99 | 0.89 | 0.93 |
| 6 | 31572927 | rs9378200 | 0.521 | 1.47 | -1.475 | 1.028 | -0.03 | -0.048 | -0.048 | 1.088 | 0.08 | 0.01 | 0.06 | 0.3 | 0.08 | 0.24 |
| 6 | 31572956 | rs2844479 | 1.085 | 1.596 | -1.662 | 0.928 | 0.069 | 0.016 | 0.016 | 0.234 | 0.49 | 0.27 | 0.44 | 0.44 | 0.37 | 0.32 |
| 6 | 31575276 | rs9348876 | 1.007 | 1.544 | -1.684 | 0.963 | -0.022 | -0.048 | -0.048 | 0 | 0.08 | 0.02 | 0.07 | 0.3 | 0.08 | 0.24 |
| 6 | 31578772 | rs2844477 | 0.86 | -0.109 | -2.602 | 0.596 | -0.071 | -0.128 | -0.128 | 0 | 0.42 | 0.04 | 0.4 | 0.6 | 0.4 | 0.57 |
| 6 | 31583155 | rs2844475 | 0.317 | -0.498 | -2.649 | 0.22 | 0.029 | -0.01 | -0.01 | 0.015 | 0.49 | 0.16 | 0.47 | 0.62 | 0.42 | 0.58 |
| 6 | 31583312 | rs2736182 | 2.293 | -0.799 | -2.116 | 0.259 | -0.058 | -0.005 | -0.005 | 0 | 0.15 | 0.22 | 0.08 | 0.15 | 0.03 | 0.07 |
| 6 | 31583827 | rs2259571 | 1.246 | -0.732 | -2.173 | 0.278 | -0.094 | -0.153 | -0.153 | 0 | 0.4 | 0.03 | 0.39 | 0.59 | 0.4 | 0.48 |
| 6 | 31583931 | rs2269475 | 0.715 | 0.417 | -1.925 | 0.545 | 0.017 | 0.019 | 0.019 | 0 | 0.05 | 0.1 | 0.19 | 0.13 | 0.16 | 0.11 |
| 6 | 31585219 | rs2857697 | 0.555 | 0.402 | -1.93 | 0.531 | 0.027 | -0.014 | -0.014 | 0 | 0.49 | 0.14 | 0.46 | 0.62 | 0.42 | 0.58 |
| 6 | 31587938 | rs3763295 | 0.674 | 0.411 | -1.903 | 0.586 | 0.017 | 0.021 | 0.021 | 0 | 0.05 | 0.1 | 0.2 | 0.13 | 0.16 | 0.11 |
| 6 | 31589676 | rs2844472 | 1.316 | 0.322 | -1.937 | 0.549 | -0.071 | -0.129 | -0.129 | 0 | 0.42 | 0.04 | 0.4 | 0.6 | 0.4 | 0.57 |
| 6 | 31590898 | rs2736172 | 1.33 | 0.686 | -1.989 | 0.524 | -0.073 | -0.13 | -0.13 | 0.039 | 0.42 | 0.04 | 0.4 | 0.6 | 0.4 | 0.57 |
| 6 | 31593476 | rs2260000 | 1.315 | 0.963 | -2.036 | 0.459 | -0.072 | -0.129 | -0.129 | 0.024 | 0.42 | 0.04 | 0.4 | 0.6 | 0.4 | 0.57 |
| 6 | 31595487 | rs2736171 | 0.567 | 0.942 | -1.994 | 0.794 | 0.031 | -0.023 | -0.023 | 0 | 0.49 | 0.16 | 0.47 | 0.58 | 0.42 | 0.58 |
| 6 | 31595882 | rs1046080 | -1.461 | 1.191 | -0.979 | 0.462 | 0.018 | -0.038 | -0.038 | 0.027 | 0.79 | 0.67 | 0.82 | 0.97 | 0.77 | 0.89 |
| 6 | 31597708 | rs2736163 | 2.611 | 1.125 | -0.799 | 0.482 | -0.054 | -0.005 | -0.005 | 0 | 0.14 | 0.23 | 0.09 | 0.15 | 0.03 | 0.06 |
| 6 | 31597753 | rs2242660 | -0.884 | 1.219 | -0.915 | 0.425 | 0.015 | 0 | 0 | 0 | 0.4 | 0.65 | 0.43 | 0.38 | 0.37 | 0.4 |
| 6 | 31597757 | rs17207225 | 1.783 | 1.184 | -0.904 | 0.42 | -0.016 | -0.022 | -0.022 | 0 | 0.07 | 0.1 | 0.06 | 0.02 | 0.02 | 0.02 |
| 6 | 31598293 | rs2736161 | 2.998 | 1.186 | -0.882 | 0.424 | -0.025 | 0 | 0 | 0.021 | 0.12 | 0.15 | 0.06 | 0.14 | 0.03 | 0.06 |
| 6 | 31601344 | rs2272593 | -1.635 | 1.256 | -0.933 | 0.413 | 0.02 | -0.039 | -0.039 | 0.383 | 0.79 | 0.68 | 0.83 | 0.97 | 0.77 | 0.87 |
| 6 | 31601513 | rs2242659 | -0.436 | 1.24 | -0.94 | 0.426 | 0.01 | -0.004 | -0.004 | 0 | 0.37 | 0.64 | 0.43 | 0.37 | 0.37 | 0.3 |
| 6 | 31602489 | rs2242657 | 0.457 | 1.289 | -0.922 | 0.446 | 0.006 | 0.01 | 0.01 | 0 | 0.06 | 0.1 | 0.19 | 0.13 | 0.16 | 0.11 |
| 6 | 31602967 | rs1046089 | 0.615 | 1.276 | -0.916 | 0.439 | -0.002 | 0.017 | 0.017 | 0.746 | 0.3 | 0.54 | 0.36 | 0.35 | 0.34 | 0.29 |
| 6 | 31603189 | rs11538264 | 1.764 | 1.287 | -0.929 | 0.443 | -0.016 | -0.022 | -0.022 | 0.335 | 0.07 | 0.1 | 0.06 | 0.02 | 0.02 | 0.02 |
| 6 | 31603591 | rs2261033 | 0.794 | 1.331 | -0.961 | 0.486 | 0.002 | -0.015 | -0.015 | 0 | 0.49 | 0.23 | 0.6 | 0.5 | 0.51 | 0.44 |
| 6 | 31606376 | rs2077491 | 0.949 | 2.39 | -0.96 | 0.425 | 0.004 | 0.025 | 0.025 | 1.568 | 0.5 | 0.3 | 0.52 | 0.62 | 0.52 | 0.62 |
| 6 | 31609813 | rs755714 | 2.294 | 2.412 | -0.745 | 0.401 | -0.036 | -0.071 | -0.071 | 0 | 0.36 | 0.02 | 0.34 | 0.35 | 0.33 | 0.31 |
| 6 | 31610529 | rs1077393 | 0.804 | 2.407 | -0.701 | 0.416 | 0 | 0.02 | 0.02 | 1.276 | 0.51 | 0.31 | 0.53 | 0.62 | 0.52 | 0.62 |
| 6 | 31610686 | rs1052486 | 0.804 | 2.407 | -0.701 | 0.416 | 0 | 0.02 | 0.02 | 1.276 | 0.51 | 0.31 | 0.53 | 0.62 | 0.52 | 0.62 |
| 6 | 31611746 | rs7770976 | 1.474 | 2.319 | -0.667 | 0.39 | -0.015 | -0.021 | -0.021 | 0.664 | 0.08 | 0.11 | 0.07 | 0.02 | 0.02 | 0.02 |
| 6 | 31613739 | rs3130048 | 0.353 | 2.056 | -0.566 | 0.011 | 0.043 | 0.056 | 0.056 | 0 | 0.17 | 0.22 | 0.14 | 0.32 | 0.25 | 0.29 |
| 6 | 31614102 | rs2242656 | -1.376 | 1.996 | -0.482 | 0.009 | -0.021 | -0.029 | -0.029 | 0 | 0.66 | 0.59 | 0.79 | 0.82 | 0.83 | 0.85 |
| 6 | 31615167 | rs2844463 | 0.775 | 2.199 | -0.212 | -0.004 | -0.057 | -0.014 | -0.014 | 0.621 | 0.31 | 0.54 | 0.22 | 0.24 | 0.13 | 0.19 |
| 6 | 31615514 | rs10484558 | 1.808 | 2.193 | -0.207 | -0.014 | -0.022 | -0.03 | -0.03 | 0 | 0.07 | 0.13 | 0.07 | 0.02 | 0.02 | 0.02 |
| 6 | 31616366 | rs805303 | -0.246 | 2.287 | -0.238 | 0.007 | 0.01 | -0.004 | -0.004 | 0 | 0.37 | 0.64 | 0.43 | 0.37 | 0.37 | 0.3 |
| 6 | 31617957 | rs805302 | -0.246 | 2.287 | -0.238 | 0.007 | 0.01 | -0.004 | -0.004 | 0 | 0.37 | 0.64 | 0.43 | 0.37 | 0.37 | 0.3 |
| 6 | 31618121 | rs805301 | -0.246 | 2.287 | -0.238 | 0.007 | 0.01 | -0.004 | -0.004 | 0 | 0.37 | 0.64 | 0.43 | 0.37 | 0.37 | 0.3 |
| 6 | 31618567 | rs805300 | 2.834 | 2.266 | -0.22 | 0.006 | -0.054 | -0.005 | -0.005 | 0.521 | 0.14 | 0.22 | 0.09 | 0.15 | 0.03 | 0.06 |
| 6 | 31618761 | rs3130050 | -3.063 | 2.317 | -0.26 | -0.002 | -0.029 | 0.032 | 0.032 | 0 | 0.88 | 0.95 | 0.96 | 0.99 | 0.88 | 0.93 |
| 6 | 31620020 | rs813115 | -0.244 | 2.266 | -0.248 | 0.013 | 0.01 | -0.004 | -0.004 | 0 | 0.37 | 0.64 | 0.43 | 0.37 | 0.37 | 0.3 |
| 6 | 31622606 | rs805297 | 2.178 | 2.262 | -0.32 | 0.001 | -0.036 | -0.071 | -0.071 | 0.253 | 0.36 | 0.02 | 0.34 | 0.35 | 0.33 | 0.31 |
| 6 | 31623873 | rs805264 | 3.222 | 2.18 | -0.239 | -0.001 | -0.025 | 0 | 0 | 0.126 | 0.12 | 0.15 | 0.06 | 0.14 | 0.03 | 0.06 |
| 6 | 31625541 | rs707921 | 2.845 | 2.168 | -0.218 | -0.007 | -0.054 | -0.005 | -0.005 | 0 | 0.14 | 0.22 | 0.09 | 0.15 | 0.03 | 0.06 |
| 6 | 31627523 | rs3130617 | -1.787 | 2.256 | -0.205 | -0.011 | 0.02 | -0.037 | -0.037 | 1.435 | 0.79 | 0.68 | 0.83 | 0.97 | 0.77 | 0.87 |
| 6 | 31628113 | rs805263 | 3.496 | 2.145 | -0.197 | 0.019 | -0.024 | -0.003 | -0.003 | 0.173 | 0.11 | 0.14 | 0.05 | 0.12 | 0.03 | 0.05 |
| 6 | 31628733 | rs805262 | 1.904 | 2.19 | -0.22 | 0.003 | -0.109 | -0.1 | -0.1 | 0 | 0.4 | 0.07 | 0.42 | 0.59 | 0.47 | 0.48 |

| | | | | | | | | | | | | | | | | |
|---|---|---|---|---|---|---|---|---|---|---|---|---|---|---|---|---|
| 6 | 31629096 | rs707920 | -0.189 | 2.127 | -0.188 | 0.021 | 0.005 | -0.002 | -0.002 | 0.139 | 0.37 | 0.61 | 0.41 | 0.37 | 0.3 | 0.3 |
| 6 | 31629499 | rs707974 | 2.68 | 1.104 | -0.916 | -0.7 | -0.042 | 0.049 | 0.049 | 0 | 0.1 | 0.01 | 0.06 | 0.02 | 0.11 | 0.03 |
| 6 | 31629953 | rs7029 | 0.16 | 1.036 | -0.865 | -0.65 | 0.005 | -0.002 | -0.002 | 0 | 0.37 | 0.61 | 0.41 | 0.37 | 0.3 | 0.3 |
| 6 | 31630241 | rs7992 | 0.16 | 1.037 | -0.864 | -0.65 | 0.005 | -0.002 | -0.002 | 0.301 | 0.37 | 0.61 | 0.41 | 0.37 | 0.3 | 0.3 |
| 6 | 31631377 | rs805259 | 3.163 | 1.051 | -0.869 | -0.64 | -0.054 | -0.005 | -0.005 | 0.051 | 0.14 | 0.22 | 0.09 | 0.15 | 0.03 | 0.06 |
| 6 | 31631907 | rs928814 | 2.043 | 1.089 | -0.865 | -0.634 | -0.023 | -0.03 | -0.03 | 0 | 0.07 | 0.13 | 0.07 | 0.02 | 0.02 | 0.02 |
| 6 | 31632686 | rs2295665 | 1.029 | 1.142 | -0.879 | -0.602 | -0.008 | -0.005 | -0.005 | 2.144 | 0.06 | 0.08 | 0.19 | 0.13 | 0.16 | 0.11 |
| 6 | 31633165 | rs2295664 | 1.029 | 1.142 | -0.879 | -0.602 | -0.008 | -0.005 | -0.005 | 0 | 0.06 | 0.08 | 0.19 | 0.13 | 0.16 | 0.11 |
| 6 | 31633552 | rs805258 | 3.53 | 0.929 | -0.915 | -0.691 | -0.025 | 0 | 0 | 1.033 | 0.12 | 0.15 | 0.06 | 0.14 | 0.03 | 0.06 |
| 6 | 31634193 | rs805257 | -0.444 | 0.968 | -0.922 | -0.72 | 0.018 | -0.002 | -0.002 | 0 | 0.39 | 0.65 | 0.43 | 0.38 | 0.37 | 0.4 |
| 6 | 31634570 | rs9267529 | 1.74 | 0.96 | -0.892 | -0.719 | -0.021 | 0.019 | 0.019 | 0.387 | 0.09 | 0.18 | 0.06 | 0.07 | 0.07 | 0.12 |
| 6 | 31635710 | rs14365 | 1.369 | 0.976 | -0.892 | -0.72 | -0.028 | 0.005 | 0.005 | 0 | 0.28 | 0.43 | 0.35 | 0.3 | 0.22 | 0.19 |
| 6 | 31636031 | rs3117578 | -3.438 | 1.056 | -0.924 | -0.743 | -0.029 | 0.032 | 0.032 | 0 | 0.88 | 0.95 | 0.96 | 0.99 | 0.88 | 0.93 |
| 6 | 31639757 | rs805267 | 3.143 | 1.019 | -0.858 | -0.708 | -0.054 | -0.005 | -0.005 | 0.171 | 0.14 | 0.22 | 0.09 | 0.15 | 0.03 | 0.06 |
| 6 | 31639845 | rs11758242 | 1.847 | 1.006 | -0.857 | -0.718 | -0.023 | -0.03 | -0.03 | 0.022 | 0.08 | 0.13 | 0.08 | 0.02 | 0.03 | 0.02 |
| 6 | 31639979 | rs9267532 | 2.499 | 0.954 | -0.734 | -0.741 | -0.008 | 0.036 | 0.036 | 0 | 0.07 | 0.1 | 0.04 | 0.06 | 0.07 | 0.11 |
| 6 | 31640518 | rs1266076 | -0.071 | 0.987 | -0.755 | -0.738 | 0.011 | -0.004 | -0.004 | 0 | 0.37 | 0.64 | 0.43 | 0.37 | 0.37 | 0.31 |
| 6 | 31641139 | rs707919 | 1.867 | 0.993 | -0.804 | -0.746 | -0.036 | -0.071 | -0.071 | 0 | 0.36 | 0.02 | 0.34 | 0.35 | 0.33 | 0.31 |
| 6 | 31641386 | rs805272 | 3.536 | 0.686 | -0.735 | -0.742 | -0.025 | 0 | 0 | 0 | 0.12 | 0.15 | 0.05 | 0.14 | 0.03 | 0.06 |
| 6 | 31643399 | rs3115667 | -2.249 | 0.726 | -0.756 | -0.76 | 0.023 | -0.039 | -0.039 | 0 | 0.8 | 0.68 | 0.83 | 0.97 | 0.77 | 0.87 |
| 6 | 31644282 | rs805269 | 3.174 | 0.677 | -0.715 | -0.716 | -0.054 | -0.005 | -0.005 | 0 | 0.14 | 0.22 | 0.09 | 0.15 | 0.03 | 0.06 |
| 6 | 31645922 | rs9267533 | 1.357 | 0.694 | -0.719 | -0.709 | -0.021 | 0.019 | 0.019 | 0.387 | 0.09 | 0.18 | 0.06 | 0.07 | 0.07 | 0.12 |
| 6 | 31646398 | rs2280800 | 1.001 | 0.72 | -0.725 | -0.703 | -0.008 | -0.005 | -0.005 | 0 | 0.06 | 0.08 | 0.19 | 0.13 | 0.16 | 0.11 |
| 6 | 31646766 | rs9267534 | 1.357 | 0.76 | -0.722 | -0.703 | -0.017 | 0.019 | 0.019 | 0.387 | 0.09 | 0.18 | 0.06 | 0.07 | 0.07 | 0.12 |
| 6 | 31648403 | rs805290 | 0.041 | 0.722 | -0.738 | -0.717 | 0.005 | -0.002 | -0.002 | 0 | 0.37 | 0.62 | 0.41 | 0.37 | 0.3 | 0.3 |
| 6 | 31651194 | rs9267536 | 2.763 | 0.72 | -0.682 | -0.69 | -0.007 | 0.037 | 0.037 | 0 | 0.06 | 0.08 | 0.04 | 0.05 | 0.05 | 0.03 |
| 6 | 31651247 | rs2844462 | 3.18 | -0.193 | -0.913 | -0.967 | 0.014 | 0.011 | 0.011 | 0 | 0.06 | 0 | 0.15 | 0.02 | 0.05 | 0.06 |
| 6 | 31655438 | rs10573 | 1.341 | 0.028 | -0.765 | -0.959 | -0.02 | 0.019 | 0.019 | 0.381 | 0.09 | 0.16 | 0.06 | 0.07 | 0.07 | 0.11 |
| 6 | 31657413 | rs1475865 | -2.104 | 0.118 | -0.766 | -1.043 | 0.027 | -0.041 | -0.041 | 0 | 0.8 | 0.68 | 0.83 | 0.97 | 0.77 | 0.87 |
| 6 | 31657924 | rs805283 | 3.002 | -0.947 | -0.621 | -0.936 | -0.053 | -0.005 | -0.005 | 0 | 0.14 | 0.22 | 0.09 | 0.15 | 0.03 | 0.06 |
| 6 | 31659731 | rs805282 | -0.439 | -1.024 | -0.926 | -0.818 | 0.017 | 0 | 0 | 0 | 0.4 | 0.62 | 0.41 | 0.38 | 0.3 | 0.4 |
| 6 | 31661489 | rs805281 | -0.291 | -1.028 | -0.927 | -0.819 | 0.006 | -0.001 | -0.001 | 0 | 0.38 | 0.61 | 0.41 | 0.37 | 0.3 | 0.3 |
| 6 | 31662957 | rs805277 | -0.636 | -1.029 | -0.925 | -0.819 | 0.015 | 0 | 0 | 0 | 0.4 | 0.61 | 0.41 | 0.38 | 0.3 | 0.4 |
| 6 | 31665194 | rs805274 | -0.31 | -0.879 | -0.874 | -0.782 | 0.006 | -0.001 | -0.001 | 0 | 0.38 | 0.61 | 0.41 | 0.37 | 0.3 | 0.3 |
| 6 | 31665452 | rs805273 | 2.077 | -1.009 | -0.868 | -0.788 | -0.086 | -0.037 | -0.037 | 0 | 0.23 | 0.35 | 0.16 | 0.17 | 0.06 | 0.07 |
| 6 | 31669496 | rs1266071 | 2.327 | -0.837 | -1 | -0.895 | -0.042 | 0.049 | 0.049 | 0 | 0.1 | 0.01 | 0.06 | 0.02 | 0.11 | 0.03 |
| 6 | 31672268 | rs34406901 | 2.658 | -1.313 | -0.525 | -0.603 | 0.017 | 0.03 | 0.03 | 0 | 0.11 | 0.05 | 0.03 | 0.01 | 0.05 | 0.1 |
| 6 | 31673436 | rs9267546 | -0.007 | -1.012 | 0.186 | -0.349 | -0.041 | -0.002 | -0.002 | 0 | 0.13 | 0.29 | 0.12 | 0.09 | 0.09 | 0.12 |
| 6 | 31675297 | rs805295 | 2.4 | -0.758 | -0.122 | -0.24 | -0.054 | -0.006 | -0.006 | 1.224 | 0.14 | 0.23 | 0.09 | 0.15 | 0.04 | 0.06 |
| 6 | 31675765 | rs2242653 | -0.027 | 0.136 | -0.788 | 0.008 | -0.008 | -0.007 | -0.007 | 0 | 0.07 | 0.09 | 0.19 | 0.14 | 0.19 | 0.16 |
| 6 | 31678730 | rs805287 | -0.875 | -0.648 | -0.932 | 0.341 | 0.078 | 0.008 | 0.008 | 0.001 | 0.43 | 0.68 | 0.43 | 0.4 | 0.32 | 0.42 |
| 6 | 31679307 | rs805286 | -1.058 | -0.741 | -0.807 | 0.273 | 0.101 | 0.013 | 0.013 | 0.056 | 0.45 | 0.71 | 0.45 | 0.38 | 0.31 | 0.43 |
| 6 | 31679874 | rs9469040 | 2.635 | -1.136 | -0.258 | 0.228 | -0.034 | -0.002 | -0.002 | 0 | 0.13 | 0.16 | 0.08 | 0.15 | 0.03 | 0.06 |
| 6 | 31681160 | rs385306 | -0.092 | -1.384 | 0.257 | 0.118 | -0.025 | 0.008 | 0.008 | 0.336 | 0.34 | 0.54 | 0.34 | 0.37 | 0.27 | 0.34 |
| 6 | 31681289 | rs442391 | 1.204 | -1.335 | 0.176 | 0.08 | -0.023 | -0.032 | -0.032 | 0 | 0.06 | 0.13 | 0.07 | 0 | 0.02 | 0 |
| 6 | 31683018 | rs9469042 | 2.554 | -1.39 | 0.132 | 0.097 | -0.032 | -0.004 | -0.004 | 0.021 | 0.16 | 0.17 | 0.08 | 0.16 | 0.03 | 0.16 |
| 6 | 31683157 | rs3749952 | 2.668 | -1.371 | 0.139 | 0.105 | -0.034 | -0.002 | -0.002 | 0 | 0.13 | 0.16 | 0.08 | 0.14 | 0.03 | 0.06 |
| 6 | 31686497 | rs15574 | -1.308 | -1.429 | 0.218 | 0.091 | -0.029 | 0.006 | 0.006 | 0.207 | 0.26 | 0.52 | 0.34 | 0.22 | 0.25 | 0.28 |
| 6 | 31688200 | rs376510 | 1.187 | -1.555 | 0.572 | 0.107 | 0.002 | 0.062 | 0.062 | 0 | 0.08 | 0.22 | 0.06 | 0.06 | 0.07 | 0.11 |
| 6 | 31688217 | rs805294 | -1.334 | -0.641 | 0.5 | 0.082 | 0.072 | -0.009 | -0.009 | 0 | 0.43 | 0.76 | 0.45 | 0.39 | 0.36 | 0.44 |
| 6 | 31688518 | rs805293 | -1.788 | 0.666 | 0.009 | 0.63 | -0.016 | 0.042 | 0.042 | 0.153 | 0.49 | 0.22 | 0.47 | 0.37 | 0.48 | 0.42 |
| 6 | 31688799 | rs4713479 | 2.827 | 0.207 | 0.084 | 0.509 | -0.034 | -0.002 | -0.002 | 0 | 0.13 | 0.17 | 0.09 | 0.14 | 0.03 | 0.06 |
| 6 | 31690009 | rs805292 | -1.526 | 0.255 | 0.083 | 0.518 | -0.031 | 0.005 | 0.005 | 0.548 | 0.27 | 0.52 | 0.34 | 0.22 | 0.26 | 0.28 |
| 6 | 31691657 | rs453098 | 0.467 | 0.277 | 0.226 | 0.502 | 0.007 | 0.017 | 0.017 | 0.581 | 0.09 | 0.31 | 0.08 | 0.04 | 0.08 | 0.1 |
| 6 | 31692163 | rs805306 | -1.219 | 0.247 | -0.091 | 0.507 | 0.076 | -0.008 | -0.008 | 0 | 0.43 | 0.78 | 0.45 | 0.39 | 0.36 | 0.44 |
| 6 | 31697558 | rs707916 | -1.228 | 0.235 | -0.119 | 0.458 | 0.077 | -0.008 | -0.008 | 0 | 0.43 | 0.77 | 0.45 | 0.39 | 0.36 | 0.44 |
| 6 | 31698088 | rs805304 | -1.228 | 0.235 | -0.119 | 0.458 | 0.077 | -0.008 | -0.008 | 0 | 0.43 | 0.78 | 0.45 | 0.39 | 0.36 | 0.44 |
| 6 | 31706474 | rs28381339 | 3.303 | -0.388 | -0.542 | 0.107 | 0.034 | 0.028 | 0.028 | 0 | 0.09 | 0.02 | 0.01 | 0.01 | 0.02 | 0.09 |
| 6 | 31708328 | rs2075789 | 0.448 | -0.29 | -0.49 | 0.219 | -0.02 | -0.028 | -0.028 | 0 | 0.06 | 0.02 | 0.16 | 0.13 | 0.1 | 0.14 |
| 6 | 31709349 | rs1144709 | -0.788 | -0.36 | -0.502 | 0.161 | 0.091 | -0.004 | -0.004 | 0 | 0.4 | 0.71 | 0.4 | 0.37 | 0.23 | 0.34 |
| 6 | 31710020 | rs1144708 | -1.194 | -0.271 | -0.526 | 0.164 | 0.073 | -0.009 | -0.009 | 0 | 0.43 | 0.78 | 0.45 | 0.39 | 0.36 | 0.43 |
| 6 | 31710968 | rs707915 | 1.197 | -0.252 | -0.283 | 0.186 | -0.004 | 0.053 | 0.053 | 0 | 0.08 | 0.24 | 0.06 | 0.06 | 0.07 | 0.11 |
| 6 | 31711124 | rs2293861 | -1.159 | -0.255 | -0.247 | 0.174 | -0.01 | 0.001 | 0.001 | 0.125 | 0.18 | 0.29 | 0.28 | 0.17 | 0.18 | 0.17 |
| 6 | 31712181 | rs2075788 | -1.159 | -0.266 | -0.224 | 0.169 | -0.01 | 0.001 | 0.001 | 0.125 | 0.18 | 0.29 | 0.28 | 0.17 | 0.18 | 0.17 |
| 6 | 31713124 | rs3749953 | -0.159 | -0.29 | -0.235 | 0.208 | -0.009 | -0.011 | -0.011 | 0 | 0.08 | 0.08 | 0.19 | 0.14 | 0.16 | 0.15 |
| 6 | 31713454 | rs3828922 | -1.043 | -0.27 | -0.264 | 0.235 | -0.01 | -0.002 | -0.002 | 0 | 0.18 | 0.29 | 0.28 | 0.16 | 0.18 | 0.17 |
| 6 | 31717692 | rs3117572 | -1.445 | 0.259 | -0.519 | 0.478 | -0.027 | -0.033 | -0.033 | 0 | 0.94 | 0.99 | 0.92 | 0.75 | 0.84 | 0.85 |
| 6 | 31717696 | rs1150793 | 1.274 | -0.32 | -0.381 | 0.42 | 0 | 0.058 | 0.058 | 0 | 0.08 | 0.24 | 0.06 | 0.06 | 0.07 | 0.11 |
| 6 | 31718602 | rs2299851 | 0.46 | -0.1 | -0.911 | 0.451 | -0.02 | -0.023 | -0.023 | 0 | 0.06 | 0.01 | 0.16 | 0.13 | 0.1 | 0.14 |

| | | | | | | | | | | | | | | | | |
|---|---|---|---|---|---|---|---|---|---|---|---|---|---|---|---|---|
| 6 | 31722262 | rs9296001 | 3.642 | -0.29 | -0.861 | 0.294 | 0.024 | -0.005 | -0.005 | 0 | 0.11 | 0.06 | 0.05 | 0.14 | 0.03 | 0.06 |
| 6 | 31726688 | rs707939 | 2.081 | -0.043 | -1.253 | 0.207 | -0.068 | -0.062 | -0.062 | 0 | 0.36 | 0.02 | 0.35 | 0.35 | 0.37 | 0.32 |
| 6 | 31727822 | rs9461718 | 3.627 | -0.713 | -0.841 | 0.265 | 0.024 | -0.006 | -0.006 | 4.878 | 0.11 | 0.06 | 0.05 | 0.14 | 0.03 | 0.07 |
| 6 | 31729359 | rs707938 | -0.705 | -1.04 | -0.261 | 0.106 | 0.048 | 0.031 | 0.031 | 0 | 0.45 | 0.69 | 0.42 | 0.35 | 0.32 | 0.47 |
| 6 | 31731014 | rs707937 | 0.835 | -1.428 | 0.076 | 0.453 | -0.009 | 0.064 | 0.064 | 0 | 0.22 | 0.32 | 0.39 | 0.49 | 0.21 | 0.24 |
| 6 | 31731881 | rs6905572 | 0.047 | -0.942 | -0.031 | 0.325 | -0.011 | -0.013 | -0.013 | 0 | 0.07 | 0.08 | 0.19 | 0.14 | 0.15 | 0.15 |
| 6 | 31733650 | rs707936 | 1.468 | -1.111 | 0.155 | 0.269 | 0 | 0.058 | 0.058 | 0 | 0.08 | 0.24 | 0.06 | 0.06 | 0.07 | 0.11 |
| 6 | 31735631 | rs707934 | 0.491 | -1.125 | -0.093 | 0.28 | -0.02 | 0.048 | 0.048 | 0.033 | 0.11 | 0.36 | 0.1 | 0.07 | 0.07 | 0.11 |
| 6 | 31738408 | rs9461721 | 3.374 | -1.063 | -0.601 | 0.266 | 0.021 | -0.004 | -0.004 | 0 | 0.11 | 0.06 | 0.04 | 0.14 | 0.03 | 0.07 |
| 6 | 31742590 | rs707928 | -1.082 | -1.127 | -1.193 | 0.21 | 0.072 | -0.005 | -0.005 | 0 | 0.41 | 0.78 | 0.46 | 0.38 | 0.34 | 0.43 |
| 6 | 31745284 | rs2736426 | -0.294 | -0.656 | -1.415 | 0.301 | -0.081 | -0.058 | -0.058 | 0.048 | 0.65 | 0.78 | 0.61 | 0.58 | 0.55 | 0.61 |
| 6 | 31745518 | rs707927 | 3.151 | -0.538 | -0.521 | 0.293 | -0.023 | -0.003 | -0.003 | 0 | 0.12 | 0.15 | 0.07 | 0.12 | 0.03 | 0.05 |
| 6 | 31745842 | rs34585796 | 3.17 | -0.425 | -1.017 | 0.287 | 0.035 | 0.029 | 0.029 | 0 | 0.09 | 0.02 | 0.01 | 0.01 | 0.02 | 0.08 |
| 6 | 31748820 | rs707926 | -0.571 | 0.254 | -0.343 | 0.644 | 0.032 | 0.116 | 0.116 | 0 | 0.14 | 0.36 | 0.26 | 0.18 | 0.2 | 0.25 |
| 6 | 31750509 | rs11531 | 3.352 | 0.12 | -0.429 | 0.618 | 0.045 | -0.007 | -0.007 | 0 | 0.1 | 0.03 | 0.04 | 0.12 | 0.02 | 0.05 |
| 6 | 31751727 | rs707924 | -0.554 | 0.157 | -0.392 | 0.639 | 0.032 | 0.116 | 0.116 | 0 | 0.14 | 0.36 | 0.26 | 0.18 | 0.2 | 0.25 |
| 6 | 31753526 | rs11751198 | 0.247 | 0.031 | -0.447 | 0.469 | -0.026 | -0.047 | -0.047 | 0 | 0.13 | 0.19 | 0.11 | 0.03 | 0.05 | 0.02 |
| 6 | 31761063 | rs9469054 | 3.307 | 0.015 | -0.428 | 0.505 | 0.036 | -0.002 | -0.002 | 0 | 0.11 | 0.05 | 0.05 | 0.12 | 0.02 | 0.05 |
| 6 | 31764899 | rs480092 | -0.536 | 0.112 | -0.33 | 0.539 | 0.03 | 0.113 | 0.113 | 0 | 0.14 | 0.36 | 0.26 | 0.18 | 0.2 | 0.25 |
| 6 | 31773821 | rs2242668 | 0.227 | 0.121 | -0.456 | 0.562 | -0.02 | -0.003 | -0.003 | 0 | 0.15 | 0.21 | 0.14 | 0.17 | 0.08 | 0.07 |
| 6 | 31778272 | rs2227956 | -1.616 | 0.979 | -0.575 | 0.785 | -0.022 | -0.032 | -0.032 | 2.26 | 0.94 | 0.99 | 0.92 | 0.77 | 0.84 | 0.84 |
| 6 | 31778529 | rs2075799 | -0.098 | 0.724 | -0.501 | 0.778 | -0.023 | -0.001 | -0.001 | 2.306 | 0.16 | 0.23 | 0.15 | 0.18 | 0.09 | 0.08 |
| 6 | 31780235 | rs7757496 | 0.274 | 0.795 | -0.519 | 0.826 | -0.019 | -0.026 | -0.026 | 0 | 0.06 | 0.02 | 0.17 | 0.13 | 0.09 | 0.14 |
| 6 | 31780594 | rs481825 | 1.389 | 0.671 | -0.413 | 0.733 | 0.034 | 0.074 | 0.074 | 0 | 0.06 | 0.28 | 0.06 | 0.04 | 0.05 | 0.1 |
| 6 | 31786365 | rs17201192 | 3.299 | 0.579 | -0.598 | 0.639 | 0.036 | 0.037 | 0.037 | 0.903 | 0.11 | 0.05 | 0.03 | 0 | 0.02 | 0.02 |
| 6 | 31794592 | rs2763979 | -1.388 | 0.739 | -0.671 | 0.645 | 0.045 | 0.026 | 0.026 | 0.157 | 0.42 | 0.71 | 0.42 | 0.26 | 0.36 | 0.4 |
| 6 | 31795550 | rs6457452 | -0.213 | 0.986 | -0.295 | 1.133 | -0.018 | -0.019 | -0.019 | 2.265 | 0.16 | 0.21 | 0.1 | 0.06 | 0.08 | 0.12 |
| 6 | 31798287 | rs35987121 | 1.832 | 1.036 | -0.273 | 1.15 | -0.008 | -0.024 | -0.024 | 0 | 0.05 | 0.09 | 0.05 | 0.02 | 0.03 | 0.01 |
| 6 | 31804729 | rs11968400 | -0.361 | 1.347 | 0.016 | 1.172 | -0.017 | -0.019 | -0.019 | 0 | 0.16 | 0.2 | 0.11 | 0.06 | 0.08 | 0.12 |
| 6 | 31807489 | rs733539 | -0.269 | 1.577 | 0.086 | 1.351 | -0.01 | -0.009 | -0.009 | 1.558 | 0.08 | 0.07 | 0.2 | 0.13 | 0.15 | 0.15 |
| 6 | 31810085 | rs3115673 | -0.468 | 1.597 | 0.03 | 1.272 | 0.025 | 0.121 | 0.121 | 0 | 0.14 | 0.36 | 0.26 | 0.18 | 0.18 | 0.25 |
| 6 | 31813774 | rs9267577 | -1.522 | 1.552 | -0.164 | 1.216 | -0.02 | -0.031 | -0.031 | 0.917 | 0.94 | 0.99 | 0.93 | 0.77 | 0.84 | 0.84 |
| 6 | 31815723 | rs12210887 | -0.272 | 1.253 | -0.104 | 1.104 | -0.02 | -0.018 | -0.018 | 0.261 | 0.14 | 0.19 | 0.08 | 0.06 | 0.07 | 0.12 |
| 6 | 31819123 | rs9501157 | 1.181 | 1.272 | -0.091 | 1.145 | -0.012 | -0.011 | -0.011 | 0.115 | 0.07 | 0.09 | 0.05 | 0.03 | 0.03 | 0.02 |
| 6 | 31819164 | rs13191375 | 0.746 | 1.555 | -0.035 | 1.304 | -0.007 | -0.002 | -0.002 | 0 | 0.08 | 0.12 | 0.03 | 0.05 | 0.04 | 0.11 |
| 6 | 31819376 | rs574914 | -0.605 | 1.999 | 0.174 | 1.533 | 0.037 | 0.134 | 0.134 | 0 | 0.14 | 0.36 | 0.26 | 0.18 | 0.19 | 0.24 |
| 6 | 31824828 | rs9267649 | -1.262 | 2.128 | -0.576 | 1.228 | -0.015 | -0.027 | -0.027 | 0 | 0.94 | 0.99 | 0.93 | 0.77 | 0.84 | 0.84 |
| 6 | 31837250 | rs660594 | 0.854 | 2.006 | 0.07 | 1.358 | 0.169 | 0.148 | 0.148 | 0 | 0.81 | 0.68 | 0.68 | 0.6 | 0.58 | 0.6 |
| 6 | 31837277 | rs660550 | 0.854 | 2.006 | 0.07 | 1.358 | 0.169 | 0.148 | 0.148 | 0 | 0.81 | 0.68 | 0.68 | 0.6 | 0.58 | 0.6 |
| 6 | 31838441 | rs644827 | 1.097 | 1.945 | 0.038 | 1.327 | 0.154 | 0.134 | 0.134 | 0 | 0.8 | 0.68 | 0.68 | 0.6 | 0.58 | 0.55 |
| 6 | 31838490 | rs644774 | 1.097 | 1.945 | 0.038 | 1.327 | 0.154 | 0.134 | 0.134 | 0 | 0.8 | 0.68 | 0.68 | 0.6 | 0.58 | 0.55 |
| 6 | 31838713 | rs494620 | 0.489 | 0.856 | -0.606 | 0.756 | 0.002 | 0 | 0 | 0 | 0.54 | 0.43 | 0.47 | 0.42 | 0.45 | 0.39 |
| 6 | 31839331 | rs2242664 | 0.714 | 0.925 | -0.572 | 0.805 | 0.154 | 0.134 | 0.134 | 0 | 0.8 | 0.68 | 0.68 | 0.6 | 0.58 | 0.55 |
| 6 | 31840625 | rs614549 | 1.047 | 0.513 | -0.797 | 0.585 | 0.007 | 0.006 | 0.006 | 0 | 0.48 | 0.45 | 0.46 | 0.39 | 0.4 | 0.36 |
| 6 | 31843924 | rs2736428 | 2.388 | 0.083 | -1.259 | 0.429 | -0.068 | -0.07 | -0.07 | 0.538 | 0.32 | 0.03 | 0.33 | 0.34 | 0.35 | 0.31 |
| 6 | 31844684 | rs6907185 | 3.069 | -0.177 | -1.288 | 0.271 | 0.006 | 0.005 | 0.005 | 0.122 | 0.1 | 0.13 | 0.07 | 0.01 | 0.07 | 0.06 |
| 6 | 31846234 | rs9267659 | -1.598 | 1.07 | -1.35 | 0.706 | -0.021 | -0.03 | -0.03 | 0.928 | 0.93 | 0.98 | 0.93 | 0.78 | 0.81 | 0.84 |
| 6 | 31847012 | rs605203 | -1.364 | 1.262 | -0.967 | 0.872 | 0.115 | 0.039 | 0.039 | 0 | 0.93 | 0.92 | 0.9 | 0.78 | 0.72 | 0.84 |
| 6 | 31848220 | rs589428 | -1.364 | 1.248 | -0.76 | 0.873 | 0.115 | 0.041 | 0.041 | 0 | 0.93 | 0.92 | 0.9 | 0.78 | 0.72 | 0.84 |
| 6 | 31850332 | rs555007 | 0.8 | 1.173 | -0.774 | 0.863 | -0.004 | -0.006 | -0.006 | 0.083 | 0.07 | 0.07 | 0.19 | 0.11 | 0.1 | 0.09 |
| 6 | 31851234 | rs652888 | 0.647 | 1.048 | -0.654 | 0.502 | 0.167 | 0.184 | 0.184 | 0 | 0.08 | 0.24 | 0.11 | 0.25 | 0.2 | 0.22 |
| 6 | 31851354 | rs3869145 | 1.835 | 0.729 | -0.76 | 0.465 | 0.009 | -0.006 | -0.006 | 0.593 | 0.06 | 0.03 | 0.06 | 0.13 | 0.03 | 0.05 |
| 6 | 31851469 | rs2844458 | 2.875 | 0.759 | -0.852 | 0.464 | -0.063 | -0.136 | -0.136 | 0.137 | 0.33 | 0.03 | 0.47 | 0.41 | 0.35 | 0.31 |
| 6 | 31856070 | rs486416 | -1.346 | 0.856 | -0.702 | 0.431 | 0.144 | 0.058 | 0.058 | 0 | 0.93 | 0.89 | 0.9 | 0.78 | 0.72 | 0.84 |
| 6 | 31859578 | rs13194087 | 0.307 | 0.487 | -0.726 | 0.319 | -0.06 | -0.043 | -0.043 | 0 | 0.19 | 0.41 | 0.11 | 0.07 | 0.09 | 0.09 |
| 6 | 31860118 | rs9501159 | 3.759 | 0.482 | -0.771 | 0.31 | 0.042 | 0.042 | 0.042 | 1.297 | 0.1 | 0.02 | 0.02 | 0 | 0.01 | 0.02 |
| 6 | 31863433 | rs2243873 | 0.783 | 0.717 | -0.74 | 0.406 | 0.184 | 0.068 | 0.068 | 0.046 | 0.8 | 0.64 | 0.78 | 0.7 | 0.59 | 0.62 |
| 6 | 31864547 | rs7887 | 2.356 | 0.603 | -0.785 | 0.419 | -0.058 | -0.157 | -0.157 | 0.028 | 0.36 | 0.03 | 0.48 | 0.55 | 0.36 | 0.36 |
| 6 | 31866418 | rs9469069 | 1.567 | 0.637 | -0.564 | 0.4 | -0.014 | -0.013 | -0.013 | 1.414 | 0.07 | 0.17 | 0.07 | 0.03 | 0.04 | 0.02 |
| 6 | 31867253 | rs9267663 | 1.349 | 0.749 | -0.846 | 0.463 | -0.022 | 0.009 | 0.009 | 0 | 0.08 | 0.18 | 0.09 | 0.04 | 0.04 | 0.09 |
| 6 | 31869673 | rs2844455 | 2.718 | 0.647 | -1.364 | 0.514 | 0.023 | -0.068 | -0.068 | 0.429 | 0.26 | 0.07 | 0.3 | 0.31 | 0.21 | 0.22 |
| 6 | 31872383 | rs10947223 | 0.2 | 0.571 | -0.715 | 0.504 | -0.056 | -0.056 | -0.056 | 0.442 | 0.18 | 0.4 | 0.11 | 0.06 | 0.08 | 0.08 |
| 6 | 31872551 | rs2763982 | 2.328 | 0.775 | -0.973 | 0.624 | -0.057 | -0.153 | -0.153 | 0 | 0.36 | 0.03 | 0.48 | 0.55 | 0.36 | 0.36 |
| 6 | 31882024 | rs2844452 | 0.601 | 0.86 | -0.576 | 0.657 | 0.024 | 0.017 | 0.017 | 0 | 0.66 | 0.5 | 0.67 | 0.63 | 0.47 | 0.45 |
| 6 | 31883679 | rs9267673 | 0.917 | 0.944 | -0.52 | 0.716 | 0.019 | 0.037 | 0.037 | 0 | 0.1 | 0.24 | 0.1 | 0.08 | 0.1 | 0.23 |
| 6 | 31883957 | rs644045 | -2.127 | 1.101 | -0.639 | 0.715 | 0.01 | 0.001 | 0.001 | 0 | 0.85 | 0.82 | 0.85 | 0.79 | 0.71 | 0.88 |
| 6 | 31888062 | rs17201403 | 3.282 | 0.872 | -0.733 | 0.528 | 0.042 | 0.042 | 0.042 | 0.597 | 0.1 | 0.02 | 0.03 | 0 | 0.01 | 0.02 |
| 6 | 31892641 | rs9267677 | 0.813 | 0.613 | -0.545 | 0.537 | 0.016 | 0.019 | 0.019 | 0 | 0.11 | 0.24 | 0.11 | 0.06 | 0.12 | 0.2 |
| 6 | 31893944 | rs2734335 | 0.902 | 0.525 | -0.701 | 0.564 | 0.162 | 0.083 | 0.083 | 0 | 0.76 | 0.54 | 0.71 | 0.67 | 0.52 | 0.55 |

| | | | | | | | | | | | | | | | | |
|---|---|---|---|---|---|---|---|---|---|---|---|---|---|---|---|---|
| 6 | 31894626 | rs3020644 | 1.807 | 0.157 | -1.353 | 0.454 | 0.034 | -0.043 | -0.043 | 0.041 | 0.48 | 0.1 | 0.54 | 0.57 | 0.39 | 0.37 |
| 6 | 31895619 | rs9332704 | 3.05 | -0.259 | -0.94 | 0.145 | 0.043 | 0.043 | 0.043 | 0.382 | 0.1 | 0.02 | 0.02 | 0 | 0.01 | 0.02 |
| 6 | 31896112 | rs9332705 | 3.262 | 0.155 | -0.724 | 0.37 | 0.008 | 0.012 | 0.012 | 0 | 0.07 | 0.03 | 0.01 | 0.01 | 0 | 0.05 |
| 6 | 31905130 | rs1042663 | -0.394 | 0.648 | -0.568 | 0.566 | 0.001 | -0.008 | -0.008 | 0 | 0.07 | 0.18 | 0.09 | 0.05 | 0.09 | 0.15 |
| 6 | 31905328 | rs2507953 | -0.394 | 0.648 | -0.568 | 0.566 | 0.001 | -0.008 | -0.008 | 0 | 0.07 | 0.18 | 0.09 | 0.05 | 0.09 | 0.15 |
| 6 | 31907147 | rs550605 | -0.394 | 0.648 | -0.568 | 0.566 | 0.001 | -0.008 | -0.008 | 0 | 0.07 | 0.18 | 0.09 | 0.05 | 0.09 | 0.15 |
| 6 | 31907168 | rs653414 | -0.394 | 0.648 | -0.568 | 0.566 | 0.001 | -0.008 | -0.008 | 0 | 0.07 | 0.18 | 0.09 | 0.05 | 0.09 | 0.15 |
| 6 | 31908761 | rs497239 | -0.394 | 0.648 | -0.567 | 0.566 | 0.001 | -0.008 | -0.008 | 0 | 0.07 | 0.18 | 0.09 | 0.06 | 0.09 | 0.15 |
| 6 | 31910162 | rs609061 | -0.394 | 0.618 | -0.567 | 0.587 | 0 | -0.006 | -0.006 | 0.09 | 0.07 | 0.17 | 0.09 | 0.06 | 0.09 | 0.16 |
| 6 | 31910929 | rs2242572 | -0.394 | 0.618 | -0.567 | 0.573 | 0 | -0.008 | -0.008 | 0 | 0.07 | 0.17 | 0.09 | 0.06 | 0.09 | 0.16 |
| 6 | 31910938 | rs547154 | -0.394 | 0.618 | -0.567 | 0.573 | 0 | -0.008 | -0.008 | 0 | 0.07 | 0.17 | 0.09 | 0.06 | 0.09 | 0.16 |
| 6 | 31914179 | rs12614 | 0.562 | 0.048 | -0.183 | 0.163 | -0.057 | -0.078 | -0.078 | 0 | 0.23 | 0.38 | 0.14 | 0.06 | 0.13 | 0.15 |
| 6 | 31914935 | rs1048709 | -1.035 | 0.708 | -0.441 | 0.889 | -0.042 | -0.027 | -0.027 | 0 | 0.93 | 0.99 | 0.78 | 0.7 | 0.85 | 0.9 |
| 6 | 31916400 | rs537160 | 0.089 | 0.806 | -0.395 | 0.967 | 0.017 | 0.029 | 0.029 | 0 | 0.87 | 0.86 | 0.68 | 0.57 | 0.75 | 0.86 |
| 6 | 31916951 | rs541862 | 0.043 | 0.797 | -0.396 | 0.967 | 0.002 | -0.005 | -0.005 | 2.138 | 0.07 | 0.18 | 0.09 | 0.06 | 0.09 | 0.15 |
| 6 | 31917540 | rs4151657 | 2.744 | 0.684 | -0.596 | 1.171 | -0.012 | 0.031 | 0.031 | 0.065 | 0.42 | 0.05 | 0.31 | 0.31 | 0.36 | 0.29 |
| 6 | 31919578 | rs2072633 | 1.592 | 0.26 | -0.515 | 1.266 | 0.025 | 0.04 | 0.04 | 0 | 0.78 | 0.67 | 0.55 | 0.47 | 0.65 | 0.71 |
| 6 | 31919917 | rs522162 | 0.11 | 0.351 | -0.509 | 1.282 | 0.002 | -0.007 | -0.007 | 0 | 0.07 | 0.18 | 0.09 | 0.06 | 0.09 | 0.15 |
| 6 | 31919956 | rs760070 | 0.11 | 0.351 | -0.509 | 1.282 | 0.002 | -0.007 | -0.007 | 0 | 0.07 | 0.18 | 0.09 | 0.06 | 0.09 | 0.15 |
| 6 | 31920687 | rs550513 | 0.11 | 0.351 | -0.607 | 1.282 | 0.002 | -0.007 | -0.007 | 0 | 0.07 | 0.18 | 0.09 | 0.06 | 0.09 | 0.15 |
| 6 | 31921475 | rs2072632 | 2.773 | 0.08 | -0.771 | 1.134 | -0.065 | -0.031 | -0.031 | 0.147 | 0.32 | 0.03 | 0.3 | 0.31 | 0.35 | 0.28 |
| 6 | 31924327 | rs9501161 | 1.796 | 0.076 | -0.655 | 1.088 | -0.011 | -0.017 | -0.017 | 0 | 0.18 | 0.19 | 0.11 | 0.05 | 0.13 | 0.14 |
| 6 | 31924880 | rs403569 | -0.03 | 0.671 | -0.614 | 1.336 | 0.004 | -0.007 | -0.007 | 0.034 | 0.07 | 0.19 | 0.09 | 0.06 | 0.09 | 0.15 |
| 6 | 31927342 | rs440454 | -1.178 | 0.377 | -0.914 | 1.316 | 0.01 | 0.017 | 0.017 | 0 | 0.9 | 0.94 | 0.75 | 0.68 | 0.78 | 0.88 |
| 6 | 31928306 | rs438999 | 0.125 | 0.352 | -0.898 | 1.289 | 0.002 | -0.007 | -0.007 | 0 | 0.07 | 0.18 | 0.09 | 0.06 | 0.09 | 0.15 |
| 6 | 31928691 | rs2280774 | 2.516 | 0.423 | -1.067 | 1.334 | -0.08 | -0.033 | -0.033 | 0.111 | 0.28 | 0.03 | 0.28 | 0.28 | 0.35 | 0.23 |
| 6 | 31928799 | rs419788 | -0.97 | 0.849 | -0.587 | 1.552 | 0.01 | 0.017 | 0.017 | 0 | 0.9 | 0.94 | 0.75 | 0.67 | 0.78 | 0.88 |
| 6 | 31929014 | rs437179 | -0.97 | 0.849 | -0.587 | 1.552 | 0.01 | 0.017 | 0.017 | 0 | 0.9 | 0.94 | 0.75 | 0.68 | 0.78 | 0.88 |
| 6 | 31930351 | rs2734331 | 0.978 | 0.323 | -0.582 | 1.028 | 0.015 | -0.013 | -0.013 | 0.581 | 0.06 | 0.01 | 0.06 | 0.13 | 0.03 | 0.05 |
| 6 | 31930462 | rs429608 | -1.08 | 0.323 | -0.502 | 0.872 | 0 | -0.002 | -0.002 | 0 | 0.13 | 0.19 | 0.12 | 0.08 | 0.15 | 0.21 |
| 6 | 31931137 | rs401775 | -0.952 | 0.326 | -0.128 | 0.997 | 0.035 | 0.033 | 0.033 | 0.112 | 0.15 | 0.4 | 0.17 | 0.15 | 0.19 | 0.34 |
| 6 | 31932177 | rs444921 | 0.208 | 0.089 | -0.196 | 0.84 | -0.013 | -0.029 | -0.029 | 0 | 0.16 | 0.35 | 0.16 | 0.1 | 0.14 | 0.21 |
| 6 | 31933161 | rs406936 | 0.191 | 0.087 | -0.196 | 0.839 | -0.013 | -0.029 | -0.029 | 0 | 0.16 | 0.34 | 0.16 | 0.1 | 0.14 | 0.21 |
| 6 | 31933518 | rs492899 | 1.373 | -0.535 | -0.43 | 0.62 | -0.038 | -0.027 | -0.027 | 0 | 0.29 | 0.32 | 0.15 | 0.09 | 0.12 | 0.15 |
| 6 | 31934372 | rs454212 | 0.275 | -0.189 | -0.281 | 0.778 | -0.013 | -0.029 | -0.029 | 0.013 | 0.16 | 0.34 | 0.16 | 0.1 | 0.14 | 0.21 |
| 6 | 31935311 | rs453821 | 0.275 | -0.189 | -0.281 | 0.8 | -0.013 | -0.029 | -0.029 | 0 | 0.16 | 0.34 | 0.16 | 0.1 | 0.14 | 0.21 |
| 6 | 31935567 | rs3911893 | 1.349 | -0.233 | -0.343 | 0.866 | 0.002 | -0.001 | -0.001 | 0 | 0.06 | 0.06 | 0.08 | 0.12 | 0.03 | 0.05 |
| 6 | 31936057 | rs34241101 | 1.944 | -0.033 | -0.223 | 0.905 | 0.055 | 0.036 | 0.036 | 0 | 0.06 | 0.17 | 0.03 | 0.03 | 0.08 | 0.19 |
| 6 | 31936668 | rs410851 | -1.115 | 0.443 | -0.441 | 1.424 | -0.013 | -0.009 | -0.009 | 0 | 0.9 | 0.95 | 0.86 | 0.69 | 0.78 | 0.88 |
| 6 | 31936679 | rs449643 | 0.263 | 0.305 | -0.39 | 1.325 | -0.013 | -0.029 | -0.029 | 0 | 0.16 | 0.34 | 0.16 | 0.1 | 0.14 | 0.21 |
| 6 | 31938107 | rs474534 | 0.989 | 0.066 | -0.643 | 1.162 | -0.031 | -0.034 | -0.034 | 0.122 | 0.23 | 0.32 | 0.16 | 0.1 | 0.15 | 0.19 |
| 6 | 31941557 | rs387608 | 0.121 | 0.542 | -0.835 | 1.421 | -0.014 | 0 | 0 | 0.195 | 0.16 | 0.35 | 0.17 | 0.15 | 0.14 | 0.21 |
| 6 | 31946614 | rs6941112 | 1.693 | 1.559 | -1.348 | 1.662 | -0.06 | -0.028 | -0.028 | 0.877 | 0.29 | 0.03 | 0.38 | 0.28 | 0.33 | 0.22 |
| 6 | 31947460 | rs389883 | -0.464 | 1.983 | -0.473 | 1.874 | -0.013 | -0.009 | -0.009 | 0 | 0.9 | 0.95 | 0.86 | 0.69 | 0.78 | 0.88 |
| 6 | 32014456 | rs34214527 | 1.132 | 0.974 | 0.074 | 1.295 | 0.072 | 0.042 | 0.042 | 0 | 0.06 | 0.2 | 0.05 | 0.03 | 0.1 | 0.19 |
| 6 | 32014575 | rs2856448 | 0.903 | 0.925 | -0.111 | 1.43 | 0.039 | 0.002 | 0.002 | 0.603 | 0.66 | 0.45 | 0.61 | 0.64 | 0.55 | 0.74 |
| 6 | 32017521 | rs10046127 | 2.142 | 0.067 | -0.528 | 0.943 | -0.023 | -0.023 | -0.023 | 0 | 0.18 | 0.17 | 0.08 | 0.04 | 0.06 | 0.08 |
| 6 | 32019769 | rs12333245 | 1.205 | -0.344 | -0.763 | 0.868 | 0.021 | 0.024 | 0.024 | 0.482 | 0.3 | 0.22 | 0.12 | 0.11 | 0.11 | 0.28 |
| 6 | 32020844 | rs2077580 | 2.024 | -0.168 | -0.647 | 0.912 | 0.064 | 0.034 | 0.034 | 0 | 0.05 | 0.16 | 0.04 | 0.03 | 0.1 | 0.19 |
| 6 | 32022898 | rs6902493 | 2.348 | 0.239 | -0.517 | 1.318 | -0.1 | 0.017 | 0.017 | 0 | 0.28 | 0.02 | 0.37 | 0.22 | 0.34 | 0.24 |
| 6 | 32026808 | rs12198173 | 1.781 | 0.241 | -0.395 | 1.326 | 0.062 | 0.032 | 0.032 | 0 | 0.06 | 0.17 | 0.04 | 0.03 | 0.1 | 0.19 |
| 6 | 32029183 | rs2269429 | 1.101 | 0.024 | -0.535 | 1.243 | 0.02 | 0.023 | 0.023 | 0.092 | 0.3 | 0.21 | 0.11 | 0.11 | 0.11 | 0.28 |
| 6 | 32030284 | rs2239689 | 2.071 | 0.3 | -0.678 | 1.309 | -0.1 | 0.026 | 0.026 | 0 | 0.29 | 0.03 | 0.37 | 0.22 | 0.34 | 0.24 |
| 6 | 32033007 | rs7766862 | 2.071 | 0.3 | -0.678 | 1.309 | -0.1 | 0.026 | 0.026 | 0 | 0.29 | 0.03 | 0.37 | 0.22 | 0.34 | 0.24 |
| 6 | 32038700 | rs2071295 | 2.257 | 1.044 | -0.57 | 1.408 | -0.101 | 0.017 | 0.017 | 0 | 0.28 | 0.02 | 0.37 | 0.22 | 0.34 | 0.24 |
| 6 | 32040993 | rs17201553 | 3.545 | 0.805 | -0.434 | 1.348 | 0.004 | 0.009 | 0.009 | 0 | 0.07 | 0.05 | 0.01 | 0 | 0 | 0.05 |
| 6 | 32050067 | rs185819 | 0.671 | 0.743 | -0.451 | 1.386 | 0.035 | 0.001 | 0.001 | 0.516 | 0.65 | 0.45 | 0.61 | 0.64 | 0.55 | 0.74 |
| 6 | 32050544 | rs3130287 | -1.601 | 1.144 | -0.601 | 1.149 | -0.035 | -0.019 | -0.019 | 0 | 0.92 | 0.99 | 0.95 | 0.92 | 0.89 | 0.91 |
| 6 | 32052898 | rs34878747 | 2.322 | 0.839 | -0.67 | 0.971 | -0.02 | -0.019 | -0.019 | 0.223 | 0.18 | 0.17 | 0.08 | 0.04 | 0.06 | 0.08 |
| 6 | 32052983 | rs169496 | 1.234 | 0.845 | -0.677 | 0.982 | -0.008 | -0.001 | -0.001 | 0 | 0.3 | 0.26 | 0.13 | 0.11 | 0.11 | 0.28 |
| 6 | 32056580 | rs204900 | 1.234 | 0.845 | -0.677 | 0.982 | -0.008 | -0.001 | -0.001 | 0 | 0.3 | 0.26 | 0.13 | 0.11 | 0.11 | 0.28 |
| 6 | 32057627 | rs204899 | 1.421 | 0.865 | -0.674 | 0.988 | -0.015 | -0.01 | -0.01 | 0 | 0.23 | 0.21 | 0.12 | 0.1 | 0.11 | 0.23 |
| 6 | 32062687 | rs2071293 | 2.631 | 0.866 | -0.87 | 0.932 | -0.106 | 0.002 | 0.002 | 0 | 0.25 | 0.02 | 0.36 | 0.22 | 0.33 | 0.25 |
| 6 | 32064098 | rs204896 | 1.333 | 0.72 | -0.871 | 0.891 | -0.013 | -0.004 | -0.004 | 2.67 | 0.29 | 0.26 | 0.12 | 0.11 | 0.11 | 0.27 |
| 6 | 32066177 | rs17421624 | 2.522 | 1 | -0.942 | 0.992 | -0.106 | 0.003 | 0.003 | 0 | 0.26 | 0.02 | 0.36 | 0.22 | 0.33 | 0.23 |
| 6 | 32066765 | rs13199524 | 2.01 | 0.962 | -0.513 | 1.03 | 0.043 | 0.019 | 0.019 | 0 | 0.05 | 0.11 | 0.04 | 0.03 | 0.1 | 0.19 |
| 6 | 32071893 | rs3134954 | -1.51 | 0.937 | -0.68 | 1.013 | -0.036 | -0.019 | -0.019 | 0 | 0.92 | 0.99 | 0.95 | 0.92 | 0.89 | 0.91 |
| 6 | 32072940 | rs439844 | 1.276 | 0.318 | -0.661 | 0.793 | -0.008 | -0.001 | -0.001 | 0 | 0.3 | 0.26 | 0.13 | 0.11 | 0.11 | 0.28 |
| 6 | 32074804 | rs12153855 | 1.697 | 0.385 | -0.43 | 0.781 | 0.061 | 0.018 | 0.018 | 0 | 0.08 | 0.17 | 0.08 | 0.04 | 0.12 | 0.19 |

| | | | | | | | | | | | | | | | | |
|---|---|---|---|---|---|---|---|---|---|---|---|---|---|---|---|---|
| 6 | 32077380 | rs411337 | 1.326 | 0.252 | -0.345 | 0.764 | 0.045 | 0.044 | 0.044 | 0 | 0.28 | 0.18 | 0.09 | 0.11 | 0.08 | 0.28 |
| 6 | 32078373 | rs3807039 | 0.832 | 0.921 | -0.262 | 0.99 | 0.003 | 0.002 | 0.002 | 0 | 0.06 | 0.07 | 0.18 | 0.1 | 0.15 | 0.09 |
| 6 | 32079567 | rs17207986 | 1.604 | 0.836 | -0.444 | 1.043 | -0.017 | -0.02 | -0.02 | 0 | 0.06 | 0.01 | 0.08 | 0.12 | 0.04 | 0.08 |
| 6 | 32080146 | rs3130342 | -1.897 | 0.718 | -0.439 | 1.07 | -0.036 | -0.017 | -0.017 | 9.262 | 0.92 | 0.99 | 0.95 | 0.92 | 0.89 | 0.91 |
| 6 | 32080383 | rs9469084 | 1.559 | 0.77 | -0.404 | 1.012 | 0.056 | 0.051 | 0.051 | 0 | 0.28 | 0.18 | 0.09 | 0.11 | 0.08 | 0.28 |
| 6 | 32081422 | rs13196724 | 2.746 | 0.624 | -0.454 | 0.944 | 0.038 | 0.035 | 0.035 | 0.075 | 0.16 | 0.09 | 0.05 | 0.04 | 0.03 | 0.08 |
| 6 | 32083175 | rs8111 | 3.081 | 0.64 | -0.54 | 1.167 | -0.061 | 0.008 | 0.008 | 0 | 0.25 | 0.02 | 0.29 | 0.18 | 0.29 | 0.17 |
| 6 | 32083300 | rs8283 | 0.902 | 0.598 | -0.351 | 1.326 | -0.045 | 0.003 | 0.003 | 0.234 | 0.36 | 0.42 | 0.19 | 0.35 | 0.21 | 0.5 |
| 6 | 32089142 | rs204888 | 1.663 | 0.673 | -0.466 | 1.129 | 0.056 | 0.051 | 0.051 | 0 | 0.28 | 0.17 | 0.09 | 0.11 | 0.08 | 0.28 |
| 6 | 32097876 | rs169494 | 1.698 | 0.603 | -0.511 | 1.123 | 0.087 | 0.071 | 0.071 | 0 | 0.28 | 0.13 | 0.09 | 0.1 | 0.08 | 0.28 |
| 6 | 32097891 | rs204892 | 1.665 | 0.896 | -0.508 | 1.176 | 0 | -0.009 | -0.009 | 0 | 0.06 | 0.12 | 0.07 | 0.02 | 0.05 | 0.01 |
| 6 | 32097983 | rs9391734 | 1.88 | 0.948 | -0.445 | 1.2 | 0.059 | 0.013 | 0.013 | 0 | 0.08 | 0.17 | 0.08 | 0.04 | 0.12 | 0.19 |
| 6 | 32101762 | rs9267803 | 2.964 | 0.926 | -0.596 | 1.184 | -0.061 | 0.007 | 0.007 | 0 | 0.24 | 0.02 | 0.29 | 0.18 | 0.29 | 0.16 |
| 6 | 32102680 | rs13211318 | 1.891 | 0.954 | -0.543 | 1.207 | 0.059 | 0.013 | 0.013 | 0 | 0.08 | 0.17 | 0.08 | 0.04 | 0.13 | 0.19 |
| 6 | 32105001 | rs4713505 | 2.621 | 0.738 | -0.759 | 1.201 | -0.052 | 0.018 | 0.018 | 0 | 0.27 | 0.02 | 0.29 | 0.18 | 0.3 | 0.17 |
| 6 | 32105512 | rs388629 | 1.642 | 0.733 | -0.469 | 1.264 | 0 | -0.009 | -0.009 | 0 | 0.06 | 0.11 | 0.07 | 0.02 | 0.05 | 0.02 |
| 6 | 32109979 | rs204999 | -0.154 | 0.844 | -0.155 | 1.23 | 0.072 | -0.032 | -0.032 | 7.277 | 0.18 | 0.4 | 0.21 | 0.08 | 0.28 | 0.22 |
| 6 | 32112626 | rs3130279 | -1.76 | 1.02 | -0.39 | 1.304 | -0.022 | -0.012 | -0.012 | 0 | 0.92 | 0.97 | 0.95 | 0.91 | 0.89 | 0.91 |
| 6 | 32113980 | rs4713506 | 2.819 | 0.873 | -0.655 | 1.149 | -0.055 | 0.008 | 0.008 | 0 | 0.25 | 0.02 | 0.29 | 0.18 | 0.28 | 0.15 |
| 6 | 32117971 | rs3134608 | 1.772 | 0.815 | -0.369 | 1.056 | 0.12 | 0.01 | 0.01 | 0 | 0.08 | 0.21 | 0.1 | 0.04 | 0.2 | 0.2 |
| 6 | 32120715 | rs1053924 | 0.289 | 0.618 | -0.267 | 1.124 | -0.004 | -0.004 | -0.004 | 1.493 | 0.75 | 0.77 | 0.61 | 0.6 | 0.69 | 0.71 |
| 6 | 32121594 | rs9267810 | 2.697 | 0.854 | -0.516 | 1.317 | -0.004 | 0.029 | 0.029 | 0 | 0.26 | 0.01 | 0.29 | 0.15 | 0.26 | 0.16 |
| 6 | 32126002 | rs3134603 | -0.926 | 0.647 | -0.669 | 1.261 | -0.012 | -0.014 | -0.014 | 0 | 0.85 | 0.97 | 0.89 | 0.79 | 0.89 | 0.83 |
| 6 | 32132233 | rs2269424 | 2.646 | 0.564 | -0.657 | 1.182 | -0.005 | 0.028 | 0.028 | 0 | 0.26 | 0.01 | 0.29 | 0.14 | 0.26 | 0.16 |
| 6 | 32134510 | rs3096697 | 1.596 | 0.782 | -0.336 | 1.237 | 0.112 | 0.007 | 0.007 | 0 | 0.08 | 0.21 | 0.1 | 0.04 | 0.2 | 0.19 |
| 6 | 32134656 | rs3130347 | 1.596 | 0.782 | -0.344 | 1.237 | 0.116 | 0.008 | 0.008 | 0 | 0.08 | 0.18 | 0.1 | 0.04 | 0.2 | 0.2 |
| 6 | 32136547 | rs1061808 | 0.592 | 0.599 | -0.208 | 1.297 | 0.015 | 0.007 | 0.007 | 0 | 0.71 | 0.68 | 0.59 | 0.59 | 0.63 | 0.69 |
| 6 | 32136838 | rs1061807 | 2.837 | 0.351 | -0.384 | 1.238 | -0.005 | 0.028 | 0.028 | 0 | 0.26 | 0.01 | 0.29 | 0.14 | 0.26 | 0.16 |
| 6 | 32138545 | rs3130283 | -2.042 | 0.418 | -0.294 | 1.207 | -0.022 | -0.007 | -0.007 | 0.031 | 0.92 | 0.97 | 0.95 | 0.92 | 0.89 | 0.91 |
| 6 | 32139565 | rs34280838 | 3.007 | 0.208 | -0.277 | 1.077 | 0.045 | 0.04 | 0.04 | 0.757 | 0.16 | 0.08 | 0.03 | 0.04 | 0.03 | 0.07 |
| 6 | 32140487 | rs3130284 | 1.587 | 0.741 | -0.048 | 1.319 | 0.112 | 0.008 | 0.008 | 0 | 0.08 | 0.21 | 0.1 | 0.04 | 0.2 | 0.19 |
| 6 | 32141883 | rs408359 | 1.689 | 0.654 | 0.022 | 1.477 | 0.038 | 0.028 | 0.028 | 0.328 | 0.3 | 0.19 | 0.1 | 0.15 | 0.09 | 0.23 |
| 6 | 32145205 | rs3134947 | 1.298 | 1.913 | 0.644 | 1.862 | 0.112 | 0.008 | 0.008 | 0 | 0.08 | 0.21 | 0.1 | 0.04 | 0.2 | 0.2 |
| 6 | 32145707 | rs2269423 | 0.953 | 1.911 | 0.642 | 1.868 | 0.014 | 0.002 | 0.002 | 0 | 0.74 | 0.71 | 0.6 | 0.65 | 0.63 | 0.75 |
| 6 | 32146492 | rs3134945 | 1.236 | 1.875 | 0.472 | 1.888 | 0.114 | 0.009 | 0.009 | 0 | 0.08 | 0.21 | 0.1 | 0.04 | 0.2 | 0.2 |
| 6 | 32147696 | rs3130349 | 1.768 | 1.571 | 0.24 | 1.987 | 0.083 | 0.046 | 0.046 | 0 | 0.05 | 0.12 | 0.09 | 0.09 | 0.16 | 0.19 |
| 6 | 32147761 | rs3134943 | -0.805 | 1.386 | 0.141 | 1.881 | -0.014 | 0.006 | 0.006 | 0 | 0.89 | 0.95 | 0.89 | 0.86 | 0.89 | 0.9 |
| 6 | 32151934 | rs3131300 | 1.565 | 0.897 | 0.291 | 1.845 | 0.088 | 0.049 | 0.049 | 0 | 0.05 | 0.14 | 0.08 | 0.09 | 0.16 | 0.19 |
| 6 | 32152442 | rs1800625 | 1.521 | 0.902 | 0.294 | 1.847 | 0.086 | 0.048 | 0.048 | 0 | 0.06 | 0.14 | 0.08 | 0.1 | 0.16 | 0.19 |
| 6 | 32154285 | rs204995 | -0.732 | 0.699 | 0.39 | 1.96 | 0.068 | 0.092 | 0.092 | 0.073 | 0.16 | 0.44 | 0.19 | 0.22 | 0.22 | 0.28 |
| 6 | 32155581 | rs204993 | 0.042 | 0.881 | 0.057 | 2.312 | 0.081 | 0.16 | 0.16 | 0 | 0.13 | 0.32 | 0.18 | 0.39 | 0.25 | 0.33 |
| 6 | 32156908 | rs204992 | 0.479 | 0.885 | 0.063 | 2.295 | 0.128 | 0.106 | 0.106 | 0 | 0.08 | 0.3 | 0.16 | 0.13 | 0.2 | 0.22 |
| 6 | 32158319 | rs176095 | 0.11 | 0.815 | 0.093 | 2.052 | 0.106 | 0.128 | 0.128 | 0 | 0.09 | 0.31 | 0.16 | 0.19 | 0.2 | 0.24 |
| 6 | 32161324 | rs12663103 | 1.147 | 0.772 | 0.092 | 1.917 | 0.022 | -0.011 | -0.011 | 0 | 0.06 | 0.1 | 0.05 | 0.06 | 0.07 | 0.14 |
| 6 | 32161366 | rs204991 | -0.015 | 0.731 | 0.164 | 2.114 | 0.026 | 0.09 | 0.09 | 0 | 0.09 | 0.29 | 0.19 | 0.3 | 0.17 | 0.19 |
| 6 | 32161430 | rs204990 | -0.015 | 0.731 | 0.164 | 2.114 | 0.026 | 0.09 | 0.09 | 0 | 0.09 | 0.29 | 0.19 | 0.3 | 0.17 | 0.19 |
| 6 | 32165444 | rs2071278 | 0.922 | 0.723 | 0.041 | 2.071 | -0.017 | -0.02 | -0.02 | 0 | 0.07 | 0.1 | 0.13 | 0.21 | 0.12 | 0.16 |
| 6 | 32165583 | rs9267820 | 1.675 | -0.293 | -0.484 | 1.294 | 0.069 | 0.103 | 0.103 | 0 | 0.4 | 0.18 | 0.25 | 0.17 | 0.29 | 0.3 |
| 6 | 32169574 | rs9267821 | 1.695 | -0.284 | -0.495 | 1.311 | 0.068 | 0.102 | 0.102 | 0 | 0.4 | 0.18 | 0.25 | 0.17 | 0.29 | 0.3 |
| 6 | 32170433 | rs2071287 | 0.405 | -0.274 | -0.473 | 1.35 | 0.005 | 0.032 | 0.032 | 0 | 0.53 | 0.37 | 0.41 | 0.38 | 0.48 | 0.58 |
| 6 | 32171075 | rs3132935 | 0.3 | -0.43 | -0.546 | 1.139 | 0.013 | 0.003 | 0.003 | 0 | 0.13 | 0.2 | 0.16 | 0.21 | 0.19 | 0.29 |
| 6 | 32171288 | rs6907458 | 1.638 | -0.413 | -0.532 | 1.121 | 0.022 | -0.011 | -0.011 | 0 | 0.06 | 0.09 | 0.05 | 0.06 | 0.07 | 0.14 |
| 6 | 32171683 | rs2071277 | 0.369 | -0.41 | -0.55 | 1.165 | 0.004 | 0.016 | 0.016 | 0 | 0.53 | 0.38 | 0.41 | 0.43 | 0.48 | 0.6 |
| 6 | 32176211 | rs206017 | -0.224 | -0.453 | -0.572 | 1.224 | 0.009 | -0.014 | -0.014 | 0.018 | 0.62 | 0.59 | 0.56 | 0.73 | 0.59 | 0.71 |
| 6 | 32177900 | rs9267833 | 1.743 | -0.608 | -0.752 | 1.103 | 0.079 | 0.135 | 0.135 | 0 | 0.39 | 0.11 | 0.24 | 0.17 | 0.28 | 0.3 |
| 6 | 32179896 | rs2071286 | 2.609 | -0.689 | -0.965 | 1.026 | -0.022 | 0.017 | 0.017 | 0 | 0.21 | 0.02 | 0.19 | 0.14 | 0.21 | 0.19 |
| 6 | 32180241 | rs206019 | 1.172 | -0.355 | -0.604 | 1.318 | 0.018 | 0.114 | 0.114 | 0.007 | 0.09 | 0.22 | 0.15 | 0.3 | 0.11 | 0.11 |
| 6 | 32181182 | rs206016 | 1.172 | -0.394 | -0.508 | 1.287 | 0.018 | 0.114 | 0.114 | 0.007 | 0.09 | 0.22 | 0.15 | 0.3 | 0.11 | 0.11 |
| 6 | 32182759 | rs206015 | 1.118 | -0.412 | -0.49 | 1.368 | 0.017 | 0.123 | 0.123 | 0.003 | 0.09 | 0.21 | 0.16 | 0.31 | 0.11 | 0.11 |
| 6 | 32183175 | rs3131290 | 1.244 | -0.668 | -0.87 | 0.996 | 0.042 | 0.134 | 0.134 | 0 | 0.44 | 0.2 | 0.29 | 0.2 | 0.38 | 0.35 |
| 6 | 32184221 | rs3134799 | 1.018 | -0.57 | -0.691 | 1.004 | 0.017 | 0.095 | 0.095 | 0 | 0.47 | 0.29 | 0.33 | 0.22 | 0.44 | 0.45 |
| 6 | 32184345 | rs404860 | 0.477 | -0.642 | -0.661 | 1.122 | 0.002 | 0.12 | 0.12 | 0 | 0.15 | 0.29 | 0.19 | 0.43 | 0.16 | 0.2 |
| 6 | 32184705 | rs3134798 | -1.573 | -0.65 | -0.631 | 0.962 | 0.034 | 0 | 0 | 0.096 | 0.88 | 0.87 | 0.69 | 0.91 | 0.73 | 0.87 |
| 6 | 32185605 | rs2854050 | 2.466 | -0.614 | -0.605 | 1.176 | -0.012 | -0.022 | -0.022 | 0.038 | 0.06 | 0.03 | 0.07 | 0.2 | 0.05 | 0.08 |
| 6 | 32186264 | rs436388 | -0.074 | -0.806 | -0.581 | 0.833 | 0.01 | -0.009 | -0.009 | 0.546 | 0.62 | 0.58 | 0.52 | 0.64 | 0.6 | 0.65 |
| 6 | 32186348 | rs379464 | 1.334 | -1.111 | -0.516 | 0.747 | -0.024 | 0.053 | 0.053 | 0 | 0.08 | 0.2 | 0.08 | 0.11 | 0.06 | 0.12 |
| 6 | 32186872 | rs2256594 | 0.401 | -0.805 | -0.744 | 0.716 | 0.001 | 0.096 | 0.096 | 0 | 0.15 | 0.29 | 0.17 | 0.38 | 0.15 | 0.18 |
| 6 | 32187023 | rs394657 | 1.129 | -0.749 | -0.827 | 0.392 | 0.017 | 0.095 | 0.095 | 0 | 0.47 | 0.29 | 0.35 | 0.22 | 0.45 | 0.46 |
| 6 | 32187605 | rs1109771 | 0.498 | -0.863 | -0.762 | -0.152 | -0.001 | 0.002 | 0.002 | 2.788 | 0.5 | 0.52 | 0.53 | 0.71 | 0.52 | 0.61 |

| | | | | | | | | | | | | | | | | |
|---|---|---|---|---|---|---|---|---|---|---|---|---|---|---|---|---|
| 6 | 32188099 | rs430916 | 1.143 | -0.357 | -1.068 | 0.06 | 0.015 | 0.089 | 0.089 | 0 | 0.3 | 0.12 | 0.21 | 0.13 | 0.28 | 0.2 |
| 6 | 32188603 | rs520803 | 0.796 | -0.312 | -1.004 | 0.143 | 0.003 | 0.041 | 0.041 | 0 | 0.34 | 0.21 | 0.26 | 0.19 | 0.35 | 0.34 |
| 6 | 32188640 | rs520692 | 0.796 | -0.312 | -1.004 | 0.143 | 0.003 | 0.041 | 0.041 | 0 | 0.34 | 0.21 | 0.26 | 0.19 | 0.35 | 0.34 |
| 6 | 32189032 | rs415929 | 0.765 | -0.68 | -1.025 | 0.208 | 0.004 | 0.032 | 0.032 | 0 | 0.35 | 0.21 | 0.26 | 0.24 | 0.35 | 0.36 |
| 6 | 32189841 | rs715299 | 0.523 | -0.935 | -0.561 | 0.406 | -0.003 | 0 | 0 | 0 | 0.4 | 0.47 | 0.34 | 0.61 | 0.27 | 0.37 |
| 6 | 32190217 | rs915895 | 0.405 | -0.862 | -0.773 | 0.399 | 0.018 | -0.014 | -0.014 | 0 | 0.42 | 0.38 | 0.33 | 0.54 | 0.31 | 0.44 |
| 6 | 32190390 | rs915894 | 0.539 | -0.064 | -0.606 | 0.579 | 0.005 | 0.01 | 0.01 | 0 | 0.3 | 0.37 | 0.33 | 0.51 | 0.35 | 0.42 |
| 6 | 32190406 | rs443198 | -0.329 | -0.795 | -0.589 | 0.406 | 0.002 | -0.004 | -0.004 | 0.684 | 0.43 | 0.47 | 0.38 | 0.56 | 0.36 | 0.5 |
| 6 | 32190620 | rs3134931 | -0.192 | -1.537 | -0.687 | 0.518 | -0.001 | 0 | 0 | 1.138 | 0.41 | 0.48 | 0.38 | 0.6 | 0.32 | 0.46 |
| 6 | 32191457 | rs375244 | 0.09 | -3.148 | -1 | 0.667 | 0.105 | -0.031 | -0.031 | 0 | 0.43 | 0.26 | 0.3 | 0.51 | 0.23 | 0.32 |
| 6 | 32191581 | rs396960 | 0.698 | -3.322 | -0.697 | 0.662 | -0.003 | 0 | 0 | 0 | 0.33 | 0.31 | 0.3 | 0.36 | 0.27 | 0.24 |
| 6 | 32191607 | rs8192565 | 1.191 | -3.411 | -0.908 | 0.676 | 0.031 | 0.019 | 0.019 | 0 | 0.14 | 0.04 | 0.09 | 0.11 | 0.09 | 0.11 |
| 6 | 32191620 | rs3134930 | 0.967 | -3.479 | -0.926 | 0.739 | -0.005 | 0.002 | 0.002 | 0 | 0.25 | 0.28 | 0.23 | 0.27 | 0.22 | 0.27 |
| 6 | 32192436 | rs391755 | 0.663 | -3.691 | -0.896 | 0.95 | -0.026 | 0.016 | 0.016 | 0 | 0.13 | 0.18 | 0.13 | 0.26 | 0.06 | 0.25 |
| 6 | 32192560 | rs2267644 | 2.884 | -3.9 | -1.155 | 0.651 | 0.031 | 0.036 | 0.036 | 0 | 0.11 | 0.05 | 0.03 | 0.02 | 0.05 | 0.09 |
| 6 | 32192617 | rs397081 | 1.567 | -3.62 | -1.012 | 0.817 | -0.012 | 0.003 | 0.003 | 0 | 0.1 | 0.1 | 0.06 | 0.15 | 0.04 | 0.17 |
| 6 | 32194339 | rs499691 | 0.239 | -4.143 | -1.005 | 0.744 | -0.019 | -0.001 | -0.001 | 0 | 0.48 | 0.49 | 0.27 | 0.57 | 0.22 | 0.5 |
| 6 | 32194854 | rs3096691 | -0.625 | -4.102 | -1.145 | 0.685 | -0.008 | -0.003 | -0.003 | 0 | 0.61 | 0.54 | 0.45 | 0.67 | 0.43 | 0.6 |
| 6 | 32194956 | rs483574 | 1.511 | -3.811 | -1.161 | 0.651 | -0.024 | 0.019 | 0.019 | 0 | 0.14 | 0.05 | 0.18 | 0.1 | 0.21 | 0.1 |
| 6 | 32195017 | rs482759 | 1.511 | -3.811 | -1.161 | 0.651 | -0.024 | 0.019 | 0.019 | 0 | 0.14 | 0.05 | 0.18 | 0.1 | 0.21 | 0.1 |
| 6 | 32195332 | rs9296012 | 0.759 | -3.778 | -0.882 | 0.812 | -0.011 | -0.001 | -0.001 | 0 | 0.38 | 0.39 | 0.21 | 0.42 | 0.18 | 0.33 |
| 6 | 32195988 | rs365053 | 1.209 | -3.83 | -1.02 | 1.016 | 0.017 | -0.027 | -0.027 | 0 | 0.27 | 0.18 | 0.33 | 0.33 | 0.25 | 0.22 |
| 6 | 32196864 | rs9378123 | 1.649 | -4.079 | -0.9 | 0.96 | -0.012 | -0.003 | -0.003 | 0 | 0.31 | 0.32 | 0.15 | 0.23 | 0.15 | 0.22 |
| 6 | 32197463 | rs495089 | 0.522 | -4.055 | -0.824 | 0.907 | -0.009 | 0.001 | 0.001 | 0 | 0.44 | 0.44 | 0.24 | 0.36 | 0.21 | 0.38 |
| 6 | 32198049 | rs1426713 | 2.378 | -3.937 | -1.405 | 0.779 | 0.004 | 0.042 | 0.042 | 0 | 0.13 | 0.06 | 0.05 | 0.03 | 0.09 | 0.19 |
| 6 | 32198056 | rs440261 | 1.279 | -3.853 | -1.382 | 0.953 | 0.013 | 0 | 0 | 0 | 0.26 | 0.18 | 0.33 | 0.27 | 0.26 | 0.21 |
| 6 | 32198867 | rs404890 | 0.595 | -3.859 | -1.42 | 0.888 | 0.013 | 0.014 | 0.014 | 0 | 0.38 | 0.26 | 0.39 | 0.32 | 0.38 | 0.36 |
| 6 | 32198936 | rs2849015 | 0.485 | -3.897 | -1.334 | 0.98 | 0.013 | -0.006 | -0.006 | 0 | 0.39 | 0.31 | 0.39 | 0.41 | 0.38 | 0.42 |
| 6 | 32199144 | rs377763 | 0.29 | -4.1 | -1.62 | 0.898 | 0.022 | 0.013 | 0.013 | 2.693 | 0.25 | 0.21 | 0.15 | 0.18 | 0.18 | 0.3 |
| 6 | 32199352 | rs9267873 | -1.653 | -4.249 | -1.619 | 0.955 | -0.034 | 0.002 | 0.002 | 0.371 | 0.87 | 0.93 | 0.67 | 0.92 | 0.6 | 0.83 |
| 6 | 32200754 | rs6936346 | 0.32 | -4.252 | -1.616 | 0.763 | 0.05 | 0.007 | 0.007 | 3.97 | 0.4 | 0.3 | 0.43 | 0.4 | 0.31 | 0.37 |
| 6 | 32202597 | rs9501398 | 2.544 | -4.36 | -1.566 | 0.737 | 0.006 | 0.018 | 0.018 | 0 | 0.14 | 0.1 | 0.05 | 0.04 | 0.1 | 0.19 |
| 6 | 32202935 | rs9267898 | -1.38 | -4.062 | -1.661 | 0.873 | -0.02 | -0.02 | -0.02 | 1.309 | 0.86 | 0.92 | 0.66 | 0.77 | 0.59 | 0.8 |
| 6 | 32203537 | rs3130299 | 1.209 | -4.085 | -1.608 | 0.742 | 0.014 | -0.002 | -0.002 | 0 | 0.26 | 0.18 | 0.33 | 0.27 | 0.25 | 0.21 |
| 6 | 32204433 | rs693797 | 0.4 | -4.076 | -1.571 | 0.752 | 0.009 | 0.013 | 0.013 | 0 | 0.38 | 0.27 | 0.39 | 0.32 | 0.39 | 0.36 |
| 6 | 32205045 | rs549182 | 1.937 | -4.116 | -1.58 | 0.736 | -0.015 | 0.001 | 0.001 | 0 | 0.08 | 0.07 | 0.05 | 0.11 | 0.02 | 0.14 |
| 6 | 32205072 | rs549270 | 0.882 | -4.092 | -1.574 | 0.748 | -0.006 | 0.002 | 0.002 | 1.995 | 0.13 | 0.12 | 0.09 | 0.13 | 0.04 | 0.16 |
| 6 | 32205110 | rs9267911 | -0.437 | -3.65 | -1.415 | 0.841 | 0.043 | 0.013 | 0.013 | 4.339 | 0.51 | 0.39 | 0.48 | 0.45 | 0.43 | 0.51 |
| 6 | 32206304 | rs6906128 | 0.707 | -3.009 | -1.344 | 0.689 | 0.089 | 0.048 | 0.048 | 0.262 | 0.42 | 0.21 | 0.45 | 0.33 | 0.29 | 0.29 |
| 6 | 32207393 | rs416352 | 0.246 | -2.324 | -0.895 | 0.897 | 0.074 | -0.025 | -0.025 | 0 | 0.57 | 0.28 | 0.4 | 0.46 | 0.31 | 0.48 |
| 6 | 32208324 | rs424232 | 0.852 | -2.547 | -0.753 | 0.863 | 0.024 | 0.005 | 0.005 | 0.013 | 0.4 | 0.31 | 0.28 | 0.36 | 0.27 | 0.37 |
| 6 | 32209027 | rs382259 | 0.196 | -2.253 | -0.6 | 0.947 | 0.033 | -0.014 | -0.014 | 0 | 0.58 | 0.36 | 0.4 | 0.47 | 0.26 | 0.48 |
| 6 | 32210799 | rs419132 | 0.051 | -2.309 | -0.214 | 0.936 | -0.034 | 0.007 | 0.007 | 2.539 | 0.6 | 0.56 | 0.42 | 0.53 | 0.27 | 0.52 |
| 6 | 32211065 | rs377743 | 0.276 | -2.383 | -0.024 | 0.953 | 0.011 | -0.004 | -0.004 | 0.052 | 0.58 | 0.37 | 0.38 | 0.55 | 0.25 | 0.52 |
| 6 | 32211085 | rs412657 | -0.737 | -2.059 | 0.197 | 1.086 | -0.001 | 0.006 | 0.006 | 1.058 | 0.69 | 0.75 | 0.51 | 0.79 | 0.41 | 0.64 |
| 6 | 32211218 | rs9267947 | -1.31 | -1.48 | -0.505 | 1.104 | 0.015 | 0.009 | 0.009 | 0 | 0.5 | 0.86 | 0.45 | 0.62 | 0.5 | 0.57 |
| 6 | 32211317 | rs411326 | -0.292 | -0.863 | -0.173 | 1.15 | 0.022 | 0.032 | 0.032 | 0 | 0.21 | 0.35 | 0.26 | 0.35 | 0.33 | 0.31 |
| 6 | 32212233 | rs9267948 | -1.487 | -1.303 | -0.754 | 0.991 | -0.13 | -0.002 | -0.002 | 0 | 0.79 | 0.94 | 0.62 | 0.86 | 0.63 | 0.79 |
| 6 | 32212264 | rs427037 | 0.572 | -2.063 | -0.573 | 0.966 | -0.027 | 0.002 | 0.002 | 0 | 0.56 | 0.41 | 0.33 | 0.47 | 0.23 | 0.49 |
| 6 | 32212985 | rs17576984 | 2.114 | -1.832 | -0.861 | 1.015 | 0.058 | -0.026 | -0.026 | 0 | 0.2 | 0.05 | 0.11 | 0.23 | 0.09 | 0.13 |
| 6 | 32213008 | rs454875 | 0.701 | -2.225 | -0.557 | 1.116 | -0.042 | 0.012 | 0.012 | 2.233 | 0.17 | 0.22 | 0.11 | 0.23 | 0.06 | 0.22 |
| 6 | 32213210 | rs454748 | 0.282 | -0.537 | -0.509 | 1.103 | 0.111 | 0.029 | 0.029 | 0 | 0.73 | 0.49 | 0.64 | 0.65 | 0.45 | 0.63 |
| 6 | 32213638 | rs9267956 | 1.373 | -0.691 | -0.72 | 1.116 | -0.002 | 0.018 | 0.018 | 0 | 0.53 | 0.17 | 0.5 | 0.36 | 0.36 | 0.38 |
| 6 | 32215063 | rs563412 | 0.936 | -0.825 | -0.521 | 1.212 | -0.019 | -0.01 | -0.01 | 0 | 0.65 | 0.32 | 0.58 | 0.56 | 0.42 | 0.57 |
| 6 | 32215796 | rs2894239 | 0.973 | -0.834 | -0.525 | 1.234 | -0.02 | -0.022 | -0.022 | 0 | 0.64 | 0.32 | 0.56 | 0.48 | 0.41 | 0.54 |
| 6 | 32215876 | rs2395110 | 0.935 | -0.829 | -0.523 | 1.227 | -0.019 | -0.01 | -0.01 | 0 | 0.65 | 0.32 | 0.58 | 0.56 | 0.42 | 0.57 |
| 6 | 32217092 | rs6936204 | -0.537 | -0.367 | 0.001 | 1.173 | 0.176 | 0.049 | 0.049 | 0 | 0.88 | 0.73 | 0.83 | 0.82 | 0.68 | 0.78 |
| 6 | 32217846 | rs2894240 | 1.337 | -0.398 | -0.04 | 1.16 | -0.02 | -0.022 | -0.022 | 0.191 | 0.64 | 0.32 | 0.56 | 0.48 | 0.41 | 0.54 |
| 6 | 32218843 | rs3115573 | 1.337 | -0.398 | -0.04 | 1.16 | -0.02 | -0.022 | -0.022 | 0 | 0.64 | 0.32 | 0.56 | 0.48 | 0.41 | 0.54 |
| 6 | 32218989 | rs9296015 | 0.158 | -0.454 | 0.046 | 1.14 | -0.011 | 0.035 | 0.035 | 0 | 0.18 | 0.27 | 0.24 | 0.3 | 0.22 | 0.19 |
| 6 | 32219725 | rs4959089 | 0.158 | -0.454 | 0.046 | 1.14 | -0.011 | 0.035 | 0.035 | 0 | 0.18 | 0.27 | 0.24 | 0.3 | 0.22 | 0.19 |
| 6 | 32219838 | rs9357138 | 2.507 | -0.878 | -0.276 | 0.982 | 0.093 | -0.015 | -0.015 | 0 | 0.27 | 0.08 | 0.18 | 0.24 | 0.14 | 0.22 |
| 6 | 32220484 | rs3115572 | 0.014 | -0.527 | 0.012 | 1.063 | 0.147 | 0.023 | 0.023 | 0.041 | 0.82 | 0.6 | 0.8 | 0.78 | 0.63 | 0.73 |
| 6 | 32220685 | rs3130315 | 1.453 | -0.526 | 0.002 | 1.064 | -0.02 | -0.022 | -0.022 | 0 | 0.64 | 0.32 | 0.56 | 0.48 | 0.41 | 0.54 |
| 6 | 32221228 | rs3130316 | 0.464 | -0.341 | -0.023 | 0.986 | 0.133 | 0.014 | 0.014 | 0 | 0.81 | 0.59 | 0.8 | 0.78 | 0.63 | 0.73 |
| 6 | 32221782 | rs3096700 | 0.369 | -0.331 | -0.018 | 0.991 | 0.141 | 0.019 | 0.019 | 0 | 0.81 | 0.59 | 0.8 | 0.78 | 0.63 | 0.73 |
| 6 | 32230608 | rs9268055 | -0.427 | 0.106 | 0.382 | 0.993 | 0.079 | 0.043 | 0.043 | 0.715 | 0.15 | 0.35 | 0.14 | 0.24 | 0.24 | 0.26 |
| 6 | 32233547 | rs2395114 | 1.99 | 0.019 | 0.2 | 0.98 | 0.005 | 0.02 | 0.02 | 0.083 | 0.52 | 0.13 | 0.5 | 0.36 | 0.37 | 0.38 |
| 6 | 32235177 | rs3096681 | -0.353 | 0.089 | 0.316 | 0.97 | 0.079 | 0.043 | 0.043 | 0.715 | 0.15 | 0.35 | 0.14 | 0.24 | 0.24 | 0.26 |

| | | | | | | | | | | | | | | | | |
|---|---|---|---|---|---|---|---|---|---|---|---|---|---|---|---|---|
| 6 | 32235757 | rs17201917 | 1.717 | 0.073 | 0.366 | 1.013 | -0.002 | -0.002 | -0.002 | 0 | 0.08 | 0.1 | 0.16 | 0.12 | 0.18 | 0.09 |
| 6 | 32235895 | rs3132931 | 0.414 | 0.044 | 0.427 | 0.99 | 0.074 | 0.039 | 0.039 | 1.287 | 0.12 | 0.22 | 0.1 | 0.22 | 0.22 | 0.25 |
| 6 | 32236054 | rs3132933 | 1.49 | -0.027 | 0.5 | 1 | -0.017 | 0.004 | 0.004 | 0.244 | 0.11 | 0.15 | 0.07 | 0.12 | 0.06 | 0.16 |
| 6 | 32236142 | rs3115560 | -0.419 | 0.076 | 0.509 | 0.98 | 0.08 | 0.044 | 0.044 | 0.715 | 0.15 | 0.36 | 0.14 | 0.24 | 0.24 | 0.26 |
| 6 | 32238013 | rs3096673 | -0.419 | 0.076 | 0.501 | 0.98 | 0.079 | 0.043 | 0.043 | 0.715 | 0.15 | 0.35 | 0.14 | 0.24 | 0.24 | 0.26 |
| 6 | 32238219 | rs3096674 | 0.417 | 0.076 | 0.501 | 0.98 | 0.074 | 0.039 | 0.039 | 1.287 | 0.12 | 0.22 | 0.1 | 0.22 | 0.22 | 0.25 |
| 6 | 32238680 | rs3132945 | -0.419 | 0.076 | 0.501 | 0.98 | 0.079 | 0.043 | 0.043 | 0.715 | 0.15 | 0.36 | 0.14 | 0.24 | 0.24 | 0.26 |
| 6 | 32239651 | rs3115557 | 0.417 | 0.077 | 0.499 | 0.98 | 0.074 | 0.039 | 0.039 | 1.287 | 0.12 | 0.22 | 0.1 | 0.22 | 0.22 | 0.25 |
| 6 | 32244627 | rs3130340 | -0.416 | 0.076 | 0.496 | 0.98 | 0.079 | 0.043 | 0.043 | 0.715 | 0.15 | 0.35 | 0.14 | 0.24 | 0.24 | 0.26 |
| 6 | 32245370 | rs9268103 | -0.875 | 0.065 | 0.49 | 0.98 | 0.029 | -0.002 | -0.002 | 0 | 0.87 | 0.78 | 0.87 | 0.89 | 0.85 | 0.92 |
| 6 | 32245827 | rs3115553 | -0.421 | 0.067 | 0.445 | 0.985 | 0.079 | 0.043 | 0.043 | 0.715 | 0.15 | 0.35 | 0.14 | 0.24 | 0.24 | 0.26 |
| 6 | 32246156 | rs3115552 | -0.421 | 0.067 | 0.445 | 0.985 | 0.079 | 0.043 | 0.043 | 0.715 | 0.15 | 0.35 | 0.14 | 0.24 | 0.24 | 0.26 |
| 6 | 32252678 | rs9268125 | 0.383 | 0.07 | 0.443 | 0.987 | 0.073 | 0.041 | 0.041 | 1.38 | 0.12 | 0.22 | 0.1 | 0.22 | 0.22 | 0.25 |
| 6 | 32254452 | rs9268131 | 0.412 | 0.071 | 0.443 | 0.987 | 0.074 | 0.039 | 0.039 | 1.287 | 0.12 | 0.22 | 0.1 | 0.22 | 0.22 | 0.25 |
| 6 | 32255230 | rs9268135 | 0.376 | 0.071 | 0.443 | 0.987 | 0.074 | 0.039 | 0.039 | 1.287 | 0.12 | 0.22 | 0.1 | 0.22 | 0.22 | 0.25 |
| 6 | 32255269 | rs9268137 | 0.376 | 0.071 | 0.443 | 0.987 | 0.074 | 0.039 | 0.039 | 1.287 | 0.12 | 0.22 | 0.1 | 0.22 | 0.22 | 0.25 |
| 6 | 32255410 | rs7751896 | -0.451 | 0.068 | 0.436 | 0.986 | 0.079 | 0.043 | 0.043 | 0.715 | 0.15 | 0.35 | 0.14 | 0.24 | 0.24 | 0.26 |
| 6 | 32256418 | rs9268142 | 0.008 | 0.032 | 0.429 | 0.963 | 0.056 | 0.023 | 0.023 | 0.037 | 0.14 | 0.23 | 0.13 | 0.22 | 0.23 | 0.25 |
| 6 | 32257337 | rs4713518 | 2.094 | -0.071 | 0.329 | 0.978 | 0.005 | 0.02 | 0.02 | 0.083 | 0.52 | 0.13 | 0.5 | 0.36 | 0.37 | 0.38 |
| 6 | 32257566 | rs926070 | -0.248 | -0.099 | 0.415 | 0.998 | 0.036 | 0.009 | 0.009 | 0 | 0.79 | 0.68 | 0.71 | 0.77 | 0.67 | 0.82 |
| 6 | 32259527 | rs9268148 | -0.819 | -0.051 | 0.4 | 1.001 | 0.029 | -0.002 | -0.002 | 0 | 0.87 | 0.78 | 0.87 | 0.89 | 0.85 | 0.92 |
| 6 | 32260350 | rs6935269 | -0.394 | -0.027 | 0.247 | 0.976 | 0.079 | 0.043 | 0.043 | 0.715 | 0.15 | 0.35 | 0.14 | 0.24 | 0.24 | 0.26 |
| 6 | 32261507 | rs3749966 | -0.362 | 0.021 | 0.244 | 0.996 | 0.08 | 0.044 | 0.044 | 0.631 | 0.15 | 0.35 | 0.14 | 0.24 | 0.24 | 0.26 |
| 6 | 32261771 | rs560505 | 2.066 | -0.286 | 0.011 | 0.853 | -0.022 | -0.005 | -0.005 | 0 | 0.43 | 0.13 | 0.5 | 0.36 | 0.37 | 0.38 |
| 6 | 32266535 | rs565571 | 2.057 | -0.271 | 0.005 | 0.888 | -0.021 | 0 | 0 | 0 | 0.43 | 0.13 | 0.5 | 0.36 | 0.37 | 0.38 |
| 6 | 32268501 | rs537757 | -2.087 | -0.084 | -0.025 | 0.911 | 0.005 | 0.02 | 0.02 | 0.083 | 0.48 | 0.14 | 0.5 | 0.36 | 0.37 | 0.38 |
| 6 | 32268701 | rs6909427 | -0.378 | 0.055 | 0.101 | 0.935 | 0.08 | 0.044 | 0.044 | 0.631 | 0.15 | 0.35 | 0.14 | 0.24 | 0.24 | 0.26 |
| 6 | 32270500 | rs477005 | 1.922 | -0.07 | -0.176 | 0.905 | -0.021 | -0.004 | -0.004 | 0 | 0.43 | 0.13 | 0.5 | 0.36 | 0.37 | 0.38 |
| 6 | 32271807 | rs3864300 | -0.038 | 0.167 | 0.103 | 1.024 | 0.056 | 0.03 | 0.03 | 0.285 | 0.13 | 0.22 | 0.12 | 0.24 | 0.23 | 0.26 |
| 6 | 32272510 | rs9268168 | -0.038 | 0.167 | 0.103 | 1.024 | 0.056 | 0.03 | 0.03 | 0.285 | 0.13 | 0.22 | 0.12 | 0.24 | 0.23 | 0.26 |
| 6 | 32273765 | rs6457536 | -0.038 | 0.167 | 0.103 | 1.024 | 0.056 | 0.03 | 0.03 | 0.285 | 0.13 | 0.22 | 0.12 | 0.24 | 0.23 | 0.26 |
| 6 | 32275194 | rs7341328 | -0.038 | 0.167 | 0.103 | 1.024 | 0.056 | 0.03 | 0.03 | 0.285 | 0.13 | 0.22 | 0.12 | 0.24 | 0.23 | 0.26 |
| 6 | 32277211 | rs9268192 | -0.038 | 0.167 | 0.138 | 1.024 | 0.056 | 0.03 | 0.03 | 0.285 | 0.13 | 0.22 | 0.12 | 0.24 | 0.23 | 0.26 |
| 6 | 32278266 | rs502626 | 1.952 | 0.073 | 0.073 | 0.982 | 0.006 | 0.021 | 0.021 | 0.07 | 0.52 | 0.14 | 0.5 | 0.36 | 0.37 | 0.38 |
| 6 | 32278635 | rs9268199 | -2.656 | -0.068 | -0.076 | 0.909 | -0.047 | 0.004 | 0.004 | 0 | 0.85 | 0.99 | 0.87 | 0.93 | 0.85 | 0.9 |
| 6 | 32278670 | rs9268200 | -0.165 | 0.221 | 0.223 | 1.029 | 0.063 | 0.035 | 0.035 | 0.421 | 0.14 | 0.24 | 0.12 | 0.24 | 0.23 | 0.26 |
| 6 | 32278792 | rs3864302 | -0.484 | 0.171 | 0.24 | 1.057 | 0.065 | 0.068 | 0.068 | 0 | 0.16 | 0.35 | 0.14 | 0.29 | 0.24 | 0.28 |
| 6 | 32279340 | rs9268202 | 0.486 | 0.224 | 0.197 | 1.021 | 0.079 | 0.043 | 0.043 | 0.598 | 0.12 | 0.22 | 0.1 | 0.22 | 0.22 | 0.25 |
| 6 | 32279902 | rs9501173 | 1.671 | 0.256 | 0.237 | 1.024 | 0 | 0.027 | 0.027 | 0 | 0.07 | 0.1 | 0.03 | 0.2 | 0.07 | 0.11 |
| 6 | 32280182 | rs6939410 | 0.362 | 0.241 | 0.211 | 1.023 | 0.079 | 0.043 | 0.043 | 0.973 | 0.12 | 0.22 | 0.1 | 0.22 | 0.22 | 0.25 |
| 6 | 32282033 | rs547261 | 1.86 | 0.023 | 0.21 | 0.957 | 0 | 0.015 | 0.015 | 0.037 | 0.52 | 0.14 | 0.5 | 0.36 | 0.37 | 0.38 |
| 6 | 32282854 | rs6910071 | 1.68 | 0.289 | 0.227 | 0.984 | 0.047 | -0.003 | -0.003 | 0 | 0.05 | 0.04 | 0.23 | 0.05 | 0.19 | 0.07 |
| 6 | 32282979 | rs6909790 | -0.189 | 0.153 | 0.387 | 1.002 | 0.054 | 0.029 | 0.029 | 0.285 | 0.13 | 0.21 | 0.12 | 0.24 | 0.23 | 0.26 |
| 6 | 32283394 | rs493136 | 1.551 | 0.055 | 0.466 | 1.045 | -0.017 | 0.004 | 0.004 | 0.313 | 0.11 | 0.15 | 0.07 | 0.12 | 0.06 | 0.15 |
| 6 | 32283494 | rs6915455 | -0.176 | 0.131 | 0.416 | 1.057 | 0.056 | 0.03 | 0.03 | 0.285 | 0.13 | 0.22 | 0.12 | 0.24 | 0.23 | 0.26 |
| 6 | 32283844 | rs3749967 | 1.676 | 0.15 | 0.441 | 1.061 | 0 | 0.027 | 0.027 | 0 | 0.07 | 0.1 | 0.03 | 0.2 | 0.07 | 0.11 |
| 6 | 32284340 | rs9268220 | 1.13 | 0.19 | 0.419 | 1.032 | 0.101 | -0.005 | -0.005 | 1.914 | 0.05 | 0.12 | 0.07 | 0.02 | 0.15 | 0.14 |
| 6 | 32285596 | rs9296020 | 2.104 | 0.055 | 0.525 | 1.071 | -0.022 | 0.028 | 0.028 | 0 | 0.05 | 0.1 | 0.02 | 0.1 | 0.01 | 0.06 |
| 6 | 32286102 | rs531094 | 2.073 | 0.025 | 0.439 | 1.053 | 0.013 | 0.027 | 0.027 | 0.068 | 0.52 | 0.15 | 0.51 | 0.36 | 0.37 | 0.38 |
| 6 | 32286761 | rs498422 | 0.898 | -0.007 | 0.58 | 1.076 | -0.046 | 0.034 | 0.034 | 0.657 | 0.14 | 0.32 | 0.1 | 0.18 | 0.06 | 0.18 |
| 6 | 32288089 | rs6920338 | -1.045 | 0.106 | 0.551 | 1.037 | 0.028 | 0.008 | 0.008 | 0 | 0.9 | 0.85 | 0.92 | 0.89 | 0.88 | 0.92 |
| 6 | 32288190 | rs574710 | 1.884 | -0.049 | 0.458 | 1.052 | -0.006 | 0.001 | 0.001 | 0.257 | 0.53 | 0.14 | 0.52 | 0.38 | 0.38 | 0.39 |
| 6 | 32289318 | rs547077 | 1.884 | -0.206 | 0.37 | 1.068 | -0.006 | 0.001 | 0.001 | 0.257 | 0.53 | 0.14 | 0.52 | 0.38 | 0.38 | 0.4 |
| 6 | 32289594 | rs570963 | 0.1 | -0.204 | 0.433 | 1.083 | -0.047 | 0.075 | 0.075 | 0.05 | 0.21 | 0.42 | 0.13 | 0.38 | 0.14 | 0.29 |
| 6 | 32290927 | rs4279480 | -0.371 | -0.252 | 0.328 | 1.105 | 0.036 | 0.009 | 0.009 | 0 | 0.79 | 0.68 | 0.71 | 0.77 | 0.67 | 0.82 |
| 6 | 32290954 | rs485774 | 2.091 | -0.261 | 0.323 | 1.103 | 0.005 | 0.02 | 0.02 | 0.083 | 0.52 | 0.14 | 0.5 | 0.36 | 0.37 | 0.38 |
| 6 | 32291190 | rs496360 | 1.195 | -0.267 | 0.347 | 1.114 | -0.026 | 0.001 | 0.001 | 0.766 | 0.13 | 0.19 | 0.08 | 0.12 | 0.06 | 0.15 |
| 6 | 32291359 | rs4348357 | -0.345 | -0.226 | 0.052 | 1.077 | 0.036 | 0.009 | 0.009 | 0 | 0.79 | 0.68 | 0.71 | 0.77 | 0.67 | 0.82 |
| 6 | 32291542 | rs742697 | -0.345 | -0.226 | 0.052 | 1.077 | 0.036 | 0.009 | 0.009 | 0 | 0.79 | 0.68 | 0.71 | 0.77 | 0.67 | 0.82 |
| 6 | 32291955 | rs9501611 | 2.121 | -0.299 | -0.089 | 1.057 | -0.022 | 0.028 | 0.028 | 0 | 0.05 | 0.09 | 0.02 | 0.1 | 0.01 | 0.06 |
| 6 | 32292084 | rs525607 | 1.571 | -0.235 | -0.277 | 1.003 | -0.006 | 0.001 | 0.001 | 0.257 | 0.53 | 0.14 | 0.52 | 0.38 | 0.38 | 0.4 |
| 6 | 32292715 | rs552339 | 1.746 | -0.209 | -0.264 | 1.004 | 0.005 | 0.02 | 0.02 | 0.083 | 0.52 | 0.14 | 0.5 | 0.36 | 0.37 | 0.38 |
| 6 | 32293152 | rs533885 | 1.746 | -0.209 | -0.264 | 1.004 | 0.005 | 0.02 | 0.02 | 0.083 | 0.52 | 0.14 | 0.5 | 0.37 | 0.37 | 0.38 |
| 6 | 32294563 | rs3129905 | -0.468 | -0.068 | 0.064 | 1.027 | 0.036 | 0.009 | 0.009 | 0 | 0.79 | 0.68 | 0.71 | 0.77 | 0.67 | 0.83 |
| 6 | 32294566 | rs501545 | 1.801 | -0.169 | -0.039 | 1.049 | 0.005 | 0.02 | 0.02 | 0.083 | 0.52 | 0.13 | 0.5 | 0.36 | 0.37 | 0.38 |
| 6 | 32295357 | rs524578 | 1.628 | -0.189 | -0.048 | 1.049 | -0.006 | 0.001 | 0.001 | 0.257 | 0.53 | 0.13 | 0.52 | 0.38 | 0.38 | 0.4 |
| 6 | 32297337 | rs9368713 | 1.815 | -0.189 | -0.048 | 1.049 | 0.005 | 0.02 | 0.02 | 0.083 | 0.52 | 0.14 | 0.5 | 0.36 | 0.37 | 0.38 |
| 6 | 32297901 | rs3132958 | -0.813 | -0.081 | -0.05 | 1.083 | 0.039 | -0.011 | -0.011 | 0 | 0.83 | 0.7 | 0.8 | 0.87 | 0.78 | 0.9 |
| 6 | 32298372 | rs9405090 | 1.963 | -0.081 | -0.051 | 1.083 | 0.005 | 0.02 | 0.02 | 0.083 | 0.52 | 0.14 | 0.5 | 0.36 | 0.37 | 0.38 |

| | | | | | | | | | | | | | | | | |
|---|---|---|---|---|---|---|---|---|---|---|---|---|---|---|---|---|
| 6 | 32298814 | rs3129949 | 0.441 | -0.093 | 0.188 | 1.145 | 0.026 | -0.007 | -0.007 | 0 | 0.18 | 0.31 | 0.15 | 0.13 | 0.21 | 0.29 |
| 6 | 32298942 | rs3132959 | -0.83 | -0.093 | 0.188 | 1.145 | 0.039 | -0.011 | -0.011 | 0 | 0.83 | 0.7 | 0.8 | 0.87 | 0.78 | 0.9 |
| 6 | 32299592 | rs1003879 | 0.847 | -0.557 | -0.087 | 1.023 | -0.012 | -0.001 | -0.001 | 0 | 0.46 | 0.53 | 0.35 | 0.43 | 0.35 | 0.54 |
| 6 | 32299715 | rs742582 | 1.977 | -0.711 | -0.092 | 0.993 | 0.005 | 0.02 | 0.02 | 0.083 | 0.52 | 0.14 | 0.5 | 0.36 | 0.37 | 0.38 |
| 6 | 32299822 | rs1003878 | 0.536 | -0.71 | -0.091 | 0.993 | 0.026 | -0.007 | -0.007 | 0 | 0.18 | 0.31 | 0.15 | 0.13 | 0.21 | 0.29 |
| 6 | 32300152 | rs1076712 | -1.038 | -0.667 | -0.092 | 0.993 | 0.013 | 0.024 | 0.024 | 1.037 | 0.3 | 0.56 | 0.28 | 0.43 | 0.4 | 0.47 |
| 6 | 32304031 | rs6930681 | 1.976 | -0.686 | -0.111 | 0.998 | 0.005 | 0.02 | 0.02 | 0.083 | 0.52 | 0.14 | 0.5 | 0.36 | 0.37 | 0.38 |
| 6 | 32306419 | rs2273019 | -1.022 | -0.699 | -0.105 | 1.035 | 0.006 | 0.019 | 0.019 | 0.482 | 0.31 | 0.56 | 0.3 | 0.46 | 0.41 | 0.49 |
| 6 | 32307260 | rs2022533 | 1.809 | -0.785 | -0.087 | 1.026 | -0.005 | 0.001 | 0.001 | 0.249 | 0.53 | 0.14 | 0.51 | 0.38 | 0.38 | 0.39 |
| 6 | 32307382 | rs1033500 | 1.97 | -0.778 | -0.084 | 1.024 | 0.005 | 0.02 | 0.02 | 0.069 | 0.52 | 0.14 | 0.5 | 0.36 | 0.37 | 0.38 |
| 6 | 32307869 | rs1033498 | 2.031 | -0.802 | -0.095 | 1.012 | 0.007 | 0.023 | 0.023 | 0.109 | 0.51 | 0.14 | 0.5 | 0.36 | 0.37 | 0.38 |
| 6 | 32309003 | rs2143468 | -1.997 | -0.804 | -0.085 | 0.987 | -0.001 | 0.014 | 0.014 | 0.078 | 0.47 | 0.14 | 0.5 | 0.36 | 0.38 | 0.38 |
| 6 | 32309352 | rs2143465 | 1.967 | -0.742 | -0.183 | 1.002 | 0.005 | 0.02 | 0.02 | 0.076 | 0.52 | 0.14 | 0.5 | 0.36 | 0.37 | 0.38 |
| 6 | 32309911 | rs3117137 | 0.513 | -0.742 | -0.182 | 1.002 | 0.027 | -0.007 | -0.007 | 1.168 | 0.18 | 0.32 | 0.15 | 0.13 | 0.21 | 0.29 |
| 6 | 32311459 | rs3132954 | -0.493 | -0.848 | -0.495 | 0.892 | 0 | -0.001 | -0.001 | 0 | 0.41 | 0.57 | 0.37 | 0.5 | 0.44 | 0.56 |
| 6 | 32312082 | rs6904320 | -0.487 | -0.444 | -0.269 | 1.017 | 0.025 | 0.012 | 0.012 | 0 | 0.81 | 0.69 | 0.78 | 0.79 | 0.78 | 0.86 |
| 6 | 32312796 | rs4576282 | 1.932 | -0.442 | -0.285 | 1.018 | 0.005 | 0.02 | 0.02 | 0.076 | 0.52 | 0.14 | 0.5 | 0.36 | 0.37 | 0.38 |
| 6 | 32313550 | rs3117134 | -0.923 | -0.439 | -0.284 | 1.02 | 0.011 | 0.023 | 0.023 | 1.047 | 0.3 | 0.56 | 0.29 | 0.43 | 0.4 | 0.47 |
| 6 | 32313653 | rs3117133 | -0.518 | -0.439 | -0.284 | 1.02 | 0.025 | 0.012 | 0.012 | 0 | 0.81 | 0.7 | 0.78 | 0.79 | 0.78 | 0.86 |
| 6 | 32314676 | rs4959096 | 1.93 | -0.442 | -0.304 | 1.016 | 0.006 | 0.021 | 0.021 | 0.078 | 0.52 | 0.14 | 0.5 | 0.36 | 0.37 | 0.38 |
| 6 | 32315727 | rs910049 | -0.509 | -0.428 | -0.271 | 1.018 | 0.025 | 0.012 | 0.012 | 0 | 0.81 | 0.7 | 0.78 | 0.79 | 0.78 | 0.85 |
| 6 | 32317276 | rs2076540 | 1.966 | -0.495 | -0.507 | 0.957 | -0.022 | -0.005 | -0.005 | 0 | 0.43 | 0.13 | 0.49 | 0.36 | 0.37 | 0.37 |
| 6 | 32317471 | rs2076538 | 1.844 | -0.324 | -0.42 | 1.038 | 0.006 | 0.021 | 0.021 | 0.078 | 0.52 | 0.13 | 0.49 | 0.36 | 0.37 | 0.38 |
| 6 | 32317973 | rs761188 | -1.008 | -0.251 | -0.135 | 1.092 | 0.011 | 0.023 | 0.023 | 1.047 | 0.3 | 0.56 | 0.29 | 0.43 | 0.4 | 0.47 |
| 6 | 32318036 | rs761187 | 2.021 | -0.425 | -0.296 | 1.013 | -0.022 | -0.005 | -0.005 | 0 | 0.43 | 0.13 | 0.49 | 0.36 | 0.37 | 0.37 |
| 6 | 32318610 | rs3117119 | -1.039 | -0.158 | -0.099 | 1.152 | 0.025 | -0.015 | -0.015 | 0 | 0.87 | 0.78 | 0.88 | 0.9 | 0.86 | 0.92 |
| 6 | 32319364 | rs1265763 | -0.487 | -0.143 | 0.171 | 1.141 | -0.03 | 0.057 | 0.057 | 0.119 | 0.25 | 0.44 | 0.22 | 0.42 | 0.25 | 0.34 |
| 6 | 32320153 | rs3132963 | -1.053 | -0.145 | 0.172 | 1.136 | 0.024 | -0.015 | -0.015 | 0 | 0.87 | 0.78 | 0.88 | 0.9 | 0.86 | 0.92 |
| 6 | 32321004 | rs2022544 | 0.775 | -0.186 | 0.284 | 1.207 | 0.001 | 0.018 | 0.018 | 0 | 0.11 | 0.11 | 0.12 | 0.3 | 0.19 | 0.19 |
| 6 | 32321115 | rs1265762 | 1.442 | -0.188 | 0.28 | 1.216 | -0.003 | -0.031 | -0.031 | 0.174 | 0.58 | 0.26 | 0.58 | 0.46 | 0.53 | 0.56 |
| 6 | 32321597 | rs1265761 | 1.175 | -0.17 | 0.342 | 1.259 | -0.026 | 0 | 0 | 0.769 | 0.13 | 0.2 | 0.08 | 0.12 | 0.06 | 0.15 |
| 6 | 32321872 | rs1265760 | 1.449 | -0.168 | 0.34 | 1.257 | -0.003 | -0.031 | -0.031 | 0.171 | 0.58 | 0.26 | 0.58 | 0.46 | 0.53 | 0.56 |
| 6 | 32322393 | rs1265759 | 1.449 | -0.168 | 0.34 | 1.257 | -0.003 | -0.031 | -0.031 | 0.053 | 0.58 | 0.26 | 0.58 | 0.46 | 0.53 | 0.56 |
| 6 | 32323529 | rs1265758 | 1.279 | -0.168 | 0.34 | 1.257 | -0.002 | -0.01 | -0.01 | 0 | 0.6 | 0.39 | 0.6 | 0.45 | 0.54 | 0.56 |
| 6 | 32323745 | rs3129907 | -0.663 | -0.202 | 0.324 | 1.239 | 0.037 | -0.011 | -0.011 | 0 | 0.83 | 0.7 | 0.8 | 0.87 | 0.79 | 0.9 |
| 6 | 32324945 | rs6907322 | -0.284 | -0.192 | 0.329 | 1.248 | -0.015 | 0.032 | 0.032 | 0.117 | 0.23 | 0.31 | 0.2 | 0.42 | 0.25 | 0.34 |
| 6 | 32325835 | rs2894249 | -0.685 | -0.192 | 0.329 | 1.248 | 0.038 | -0.011 | -0.011 | 0 | 0.83 | 0.7 | 0.8 | 0.87 | 0.78 | 0.9 |
| 6 | 32326045 | rs2395150 | 1.271 | -0.192 | 0.329 | 1.248 | -0.002 | -0.01 | -0.01 | 1.576 | 0.6 | 0.39 | 0.6 | 0.46 | 0.54 | 0.56 |
| 6 | 32326165 | rs9268303 | -0.284 | -0.193 | 0.325 | 1.249 | -0.015 | 0.032 | 0.032 | 0.117 | 0.23 | 0.31 | 0.2 | 0.42 | 0.25 | 0.34 |
| 6 | 32327781 | rs6904636 | 1.44 | -0.189 | 0.299 | 1.248 | -0.003 | -0.031 | -0.031 | 0 | 0.58 | 0.26 | 0.58 | 0.46 | 0.53 | 0.56 |
| 6 | 32328403 | rs3129911 | 1.44 | -0.189 | 0.299 | 1.248 | -0.003 | -0.031 | -0.031 | 0.078 | 0.58 | 0.26 | 0.58 | 0.46 | 0.53 | 0.56 |
| 6 | 32330153 | rs9268326 | 2.261 | -0.2 | 0.283 | 1.242 | 0.007 | 0.022 | 0.022 | 0.073 | 0.52 | 0.14 | 0.5 | 0.36 | 0.37 | 0.38 |
| 6 | 32330306 | rs9268330 | -0.614 | -0.202 | 0.296 | 1.243 | -0.03 | 0.057 | 0.057 | 0.119 | 0.25 | 0.44 | 0.22 | 0.41 | 0.25 | 0.34 |
| 6 | 32330695 | rs6457553 | 0.892 | -0.202 | 0.298 | 1.245 | 0.002 | 0.02 | 0.02 | 0 | 0.11 | 0.11 | 0.12 | 0.3 | 0.19 | 0.19 |
| 6 | 32331998 | rs9268347 | -0.677 | -0.231 | 0.285 | 1.279 | -0.031 | 0.056 | 0.056 | 0.062 | 0.25 | 0.44 | 0.22 | 0.42 | 0.26 | 0.34 |
| 6 | 32333098 | rs3129922 | 1.434 | -0.213 | 0.293 | 1.277 | -0.003 | -0.031 | -0.031 | 0.059 | 0.58 | 0.26 | 0.58 | 0.46 | 0.53 | 0.56 |
| 6 | 32333425 | rs9268364 | -0.613 | -0.214 | 0.294 | 1.277 | -0.03 | 0.057 | 0.057 | 0.119 | 0.25 | 0.44 | 0.22 | 0.42 | 0.25 | 0.34 |
| 6 | 32333439 | rs9268365 | -0.582 | -0.218 | 0.291 | 1.275 | -0.029 | 0.059 | 0.059 | 0.133 | 0.24 | 0.44 | 0.22 | 0.42 | 0.25 | 0.34 |
| 6 | 32333955 | rs9268368 | 2.254 | -0.213 | 0.291 | 1.274 | 0.007 | 0.022 | 0.022 | 0.073 | 0.52 | 0.13 | 0.5 | 0.36 | 0.37 | 0.38 |
| 6 | 32334754 | rs7770048 | 0.897 | -0.213 | 0.293 | 1.275 | 0.002 | 0.02 | 0.02 | 0 | 0.11 | 0.11 | 0.12 | 0.3 | 0.19 | 0.19 |
| 6 | 32335204 | rs2143462 | 0.093 | -0.463 | 0.178 | 1.147 | 0.042 | 0.01 | 0.01 | 0.016 | 0.08 | 0.12 | 0.09 | 0.1 | 0.16 | 0.18 |
| 6 | 32335433 | rs2073048 | 0.937 | -0.374 | 0.218 | 1.192 | 0.002 | 0.02 | 0.02 | 0 | 0.11 | 0.11 | 0.12 | 0.3 | 0.19 | 0.19 |
| 6 | 32335516 | rs3129931 | -0.786 | -0.347 | 0.146 | 1.192 | 0.038 | -0.011 | -0.011 | 0.398 | 0.83 | 0.7 | 0.8 | 0.87 | 0.78 | 0.9 |
| 6 | 32336074 | rs2073046 | 2.181 | -0.36 | 0.129 | 1.555 | 0.007 | 0.022 | 0.022 | 0.073 | 0.52 | 0.13 | 0.5 | 0.36 | 0.37 | 0.38 |
| 6 | 32336187 | rs3129934 | -1.182 | -0.356 | 0.133 | 1.554 | 0.025 | -0.015 | -0.015 | 0 | 0.87 | 0.78 | 0.88 | 0.9 | 0.86 | 0.92 |
| 6 | 32336364 | rs3129937 | 0.391 | -0.429 | -0.002 | 1.559 | 0.064 | 0.023 | 0.023 | 0 | 0.07 | 0.12 | 0.09 | 0.1 | 0.16 | 0.18 |
| 6 | 32336586 | rs9268384 | 2.088 | -0.435 | -0.011 | 1.558 | 0.007 | 0.022 | 0.022 | 0.073 | 0.52 | 0.13 | 0.5 | 0.36 | 0.37 | 0.38 |
| 6 | 32336766 | rs3129939 | 0.372 | -0.405 | 0.024 | 1.53 | 0.064 | 0.023 | 0.023 | 0 | 0.07 | 0.12 | 0.09 | 0.1 | 0.16 | 0.18 |
| 6 | 32337630 | rs2273017 | 1.269 | -0.571 | 0.003 | 1.493 | 0.011 | 0.011 | 0.011 | 0.004 | 0.59 | 0.26 | 0.59 | 0.61 | 0.57 | 0.59 |
| 6 | 32337686 | rs3129941 | -0.731 | -0.266 | -0.021 | 1.465 | 0.038 | -0.011 | -0.011 | 0 | 0.83 | 0.7 | 0.8 | 0.87 | 0.78 | 0.9 |
| 6 | 32338283 | rs3129942 | 0.185 | -0.356 | -0.048 | 1.568 | 0.073 | 0.07 | 0.07 | 1.541 | 0.16 | 0.32 | 0.15 | 0.26 | 0.25 | 0.27 |
| 6 | 32338400 | rs3117111 | -0.788 | -0.297 | -0.019 | 1.44 | 0.038 | -0.011 | -0.011 | 1.466 | 0.83 | 0.7 | 0.8 | 0.87 | 0.78 | 0.9 |
| 6 | 32338695 | rs3129943 | -0.473 | -0.122 | -0.197 | 1.545 | 0.03 | 0.072 | 0.072 | 1.042 | 0.2 | 0.32 | 0.17 | 0.37 | 0.25 | 0.37 |
| 6 | 32338986 | rs2073044 | 1.5 | 0.266 | -1.465 | 1.107 | -0.052 | -0.032 | -0.032 | 0 | 0.27 | 0.12 | 0.29 | 0.23 | 0.23 | 0.22 |
| 6 | 32339076 | rs2050190 | 0.344 | 0.004 | -0.699 | 1.069 | 0.001 | 0.028 | 0.028 | 0 | 0.28 | 0.48 | 0.27 | 0.41 | 0.32 | 0.35 |
| 6 | 32339548 | rs2073045 | 1.171 | 0.641 | -0.832 | 0.733 | 0.035 | -0.013 | -0.013 | 2.087 | 0.23 | 0.3 | 0.32 | 0.11 | 0.35 | 0.24 |
| 6 | 32339647 | rs2050189 | 1.257 | -0.075 | -1.539 | 0.871 | 0.037 | -0.001 | -0.001 | 0.437 | 0.31 | 0.04 | 0.32 | 0.36 | 0.18 | 0.32 |
| 6 | 32339897 | rs2050188 | -0.622 | 0.224 | -0.587 | 1.233 | -0.006 | -0.002 | -0.002 | 0 | 0.53 | 0.52 | 0.65 | 0.59 | 0.67 | 0.63 |
| 6 | 32341398 | rs9391858 | 0.652 | 0.758 | -0.622 | 1.268 | 0.004 | 0 | 0 | 0.017 | 0.08 | 0.1 | 0.11 | 0.16 | 0.19 | 0.18 |

| | | | | | | | | | | | | | | | | |
|---|---|---|---|---|---|---|---|---|---|---|---|---|---|---|---|---|
| 6 | 32341473 | rs9268403 | 1.268 | 0.602 | -0.757 | 1.143 | -0.04 | 0.018 | 0.018 | 0 | 0.22 | 0.17 | 0.31 | 0.11 | 0.27 | 0.22 |
| 6 | 32341719 | rs9268404 | 1.268 | 0.602 | -0.757 | 1.143 | -0.04 | 0.018 | 0.018 | 0 | 0.22 | 0.17 | 0.31 | 0.11 | 0.27 | 0.22 |
| 6 | 32342537 | rs3129945 | 0.662 | 0.113 | -0.146 | 1.457 | -0.03 | 0.019 | 0.019 | 1.703 | 0.25 | 0.39 | 0.19 | 0.33 | 0.21 | 0.19 |
| 6 | 32344376 | rs9268425 | 1.402 | 0.249 | -0.18 | 1.416 | -0.04 | 0.018 | 0.018 | 0 | 0.22 | 0.17 | 0.32 | 0.11 | 0.27 | 0.22 |
| 6 | 32344973 | rs9268428 | 1.402 | 0.249 | -0.18 | 1.416 | -0.04 | 0.018 | 0.018 | 0 | 0.22 | 0.17 | 0.31 | 0.11 | 0.27 | 0.22 |
| 6 | 32347490 | rs8180664 | 1.226 | 0.435 | -0.12 | 1.498 | -0.041 | 0.02 | 0.02 | 0 | 0.23 | 0.17 | 0.31 | 0.11 | 0.27 | 0.22 |
| 6 | 32348145 | rs2395157 | 1.414 | 0.484 | -0.097 | 1.523 | -0.04 | 0.017 | 0.017 | 0 | 0.22 | 0.17 | 0.31 | 0.12 | 0.27 | 0.22 |
| 6 | 32349711 | rs9268454 | 1.394 | 0.477 | -0.072 | 1.52 | -0.036 | 0.016 | 0.016 | 0 | 0.22 | 0.19 | 0.32 | 0.11 | 0.27 | 0.22 |
| 6 | 32349946 | rs9268456 | 1.394 | 0.477 | -0.092 | 1.52 | -0.038 | 0.017 | 0.017 | 0 | 0.22 | 0.19 | 0.32 | 0.11 | 0.27 | 0.22 |
| 6 | 32351283 | rs9268460 | 0.066 | 0.431 | -0.089 | 1.591 | -0.007 | -0.001 | -0.001 | 0 | 0.33 | 0.33 | 0.48 | 0.27 | 0.44 | 0.42 |
| 6 | 32351901 | rs9268461 | 1.441 | 0.429 | -0.136 | 1.586 | -0.04 | 0.017 | 0.017 | 0 | 0.22 | 0.17 | 0.31 | 0.11 | 0.27 | 0.22 |
| 6 | 32354428 | rs4424066 | 0.066 | 0.409 | 0.034 | 1.581 | -0.007 | -0.001 | -0.001 | 0 | 0.33 | 0.33 | 0.48 | 0.27 | 0.44 | 0.42 |
| 6 | 32355605 | rs9268472 | 0.066 | 0.409 | 0.034 | 1.581 | -0.007 | -0.001 | -0.001 | 2.409 | 0.33 | 0.33 | 0.48 | 0.27 | 0.44 | 0.42 |
| 6 | 32357133 | rs17423649 | 0.538 | 0.418 | 0.039 | 1.573 | 0.054 | 0.018 | 0.018 | 0 | 0.06 | 0.14 | 0.15 | 0.04 | 0.16 | 0.08 |
| 6 | 32357165 | rs9268474 | 1.446 | 0.417 | 0.037 | 1.574 | -0.04 | 0.017 | 0.017 | 0 | 0.22 | 0.17 | 0.31 | 0.11 | 0.27 | 0.22 |
| 6 | 32357489 | rs17202259 | 0.537 | 0.411 | 0.042 | 1.585 | 0.054 | 0.021 | 0.021 | 0 | 0.06 | 0.14 | 0.14 | 0.04 | 0.16 | 0.08 |
| 6 | 32358270 | rs3117099 | 0.584 | 0.241 | 0.03 | 1.585 | -0.035 | 0.049 | 0.049 | 0.369 | 0.26 | 0.4 | 0.2 | 0.43 | 0.2 | 0.3 |
| 6 | 32358513 | rs3117098 | -1.646 | 0.27 | 0.021 | 1.591 | 0.031 | 0.037 | 0.037 | 1.857 | 0.63 | 0.81 | 0.71 | 0.74 | 0.71 | 0.76 |
| 6 | 32359460 | rs16870123 | 0.32 | 0.402 | 0.024 | 1.596 | 0.052 | 0.02 | 0.02 | 0 | 0.06 | 0.16 | 0.15 | 0.05 | 0.17 | 0.09 |
| 6 | 32361111 | rs3817973 | 0.057 | 0.401 | 0.144 | 1.617 | -0.007 | -0.001 | -0.001 | 0.027 | 0.33 | 0.33 | 0.48 | 0.27 | 0.44 | 0.42 |
| 6 | 32361388 | rs3817969 | 0.708 | 0.406 | 0.158 | 1.628 | 0.008 | 0.026 | 0.026 | 0 | 0.1 | 0.16 | 0.17 | 0.16 | 0.16 | 0.2 |
| 6 | 32363955 | rs2076529 | 0.067 | 0.348 | 0.121 | 1.623 | -0.008 | -0.001 | -0.001 | 0 | 0.33 | 0.33 | 0.48 | 0.27 | 0.44 | 0.42 |
| 6 | 32365580 | rs3129954 | -1.555 | 0.544 | 0.106 | 1.615 | 0.033 | 0.04 | 0.04 | 1.857 | 0.63 | 0.82 | 0.71 | 0.74 | 0.71 | 0.76 |
| 6 | 32365840 | rs3129955 | -1.555 | 0.544 | 0.106 | 1.615 | 0.033 | 0.04 | 0.04 | 1.857 | 0.63 | 0.82 | 0.71 | 0.74 | 0.71 | 0.76 |
| 6 | 32366421 | rs4248166 | 0.136 | 0.754 | 0.115 | 1.778 | 0.067 | 0.096 | 0.096 | 0 | 0.07 | 0.16 | 0.21 | 0.22 | 0.21 | 0.1 |
| 6 | 32367259 | rs2294884 | 0.049 | 0.754 | 0.116 | 1.778 | 0.06 | 0.086 | 0.086 | 0 | 0.07 | 0.14 | 0.21 | 0.22 | 0.21 | 0.13 |
| 6 | 32367515 | rs2294882 | 0.181 | 0.74 | 0.109 | 1.777 | 0.014 | 0.064 | 0.064 | 0 | 0.11 | 0.16 | 0.23 | 0.34 | 0.22 | 0.24 |
| 6 | 32367604 | rs2294881 | 0.181 | 0.74 | 0.109 | 1.777 | 0.014 | 0.064 | 0.064 | 0 | 0.11 | 0.16 | 0.23 | 0.34 | 0.22 | 0.24 |
| 6 | 32367795 | rs2294878 | 0.117 | 0.773 | 0.082 | 1.727 | 0.004 | 0 | 0 | 0 | 0.29 | 0.31 | 0.52 | 0.33 | 0.49 | 0.34 |
| 6 | 32368087 | rs3817963 | 0.73 | 0.779 | 0.061 | 1.758 | -0.041 | -0.007 | -0.007 | 2.028 | 0.27 | 0.17 | 0.33 | 0.23 | 0.28 | 0.33 |
| 6 | 32369355 | rs3763304 | 0.113 | 0.792 | 0.079 | 1.771 | 0.061 | 0.088 | 0.088 | 0 | 0.07 | 0.14 | 0.21 | 0.22 | 0.21 | 0.13 |
| 6 | 32370616 | rs2076525 | 1.238 | -0.782 | 0.146 | 1.746 | -0.041 | 0.018 | 0.018 | 0 | 0.23 | 0.17 | 0.31 | 0.11 | 0.27 | 0.22 |
| 6 | 32370684 | rs2076524 | 1.238 | -0.782 | 0.146 | 1.746 | -0.041 | 0.018 | 0.018 | 0 | 0.23 | 0.17 | 0.31 | 0.11 | 0.27 | 0.22 |
| 6 | 32370835 | rs2076523 | 0.092 | -0.776 | 0.148 | 1.759 | -0.003 | -0.003 | -0.003 | 0 | 0.31 | 0.32 | 0.47 | 0.43 | 0.37 | 0.43 |
| 6 | 32371619 | rs3793126 | 1.22 | -0.773 | 0.149 | 1.754 | -0.041 | 0.018 | 0.018 | 0 | 0.23 | 0.17 | 0.31 | 0.11 | 0.27 | 0.22 |
| 6 | 32371915 | rs3793127 | 1.774 | -0.762 | 0.143 | 1.749 | -0.048 | 0.027 | 0.027 | 0 | 0.19 | 0.09 | 0.23 | 0.08 | 0.2 | 0.2 |
| 6 | 32373232 | rs10947261 | 0.751 | -0.792 | 0.158 | 1.757 | -0.011 | 0.091 | 0.091 | 0 | 0.09 | 0.14 | 0.15 | 0.32 | 0.1 | 0.22 |
| 6 | 32373312 | rs10947262 | 0.723 | -0.762 | 0.13 | 1.753 | -0.011 | 0.093 | 0.093 | 0 | 0.09 | 0.16 | 0.16 | 0.32 | 0.1 | 0.22 |
| 6 | 32373801 | rs3806157 | 0.437 | -0.696 | 0.15 | 1.661 | 0.009 | 0.004 | 0.004 | 0 | 0.27 | 0.3 | 0.45 | 0.31 | 0.37 | 0.32 |
| 6 | 32375330 | rs9268493 | -0.112 | -0.895 | 0.037 | 1.482 | -0.001 | -0.001 | -0.001 | 0 | 0.34 | 0.36 | 0.34 | 0.36 | 0.31 | 0.4 |
| 6 | 32375352 | rs9268494 | -0.399 | -0.952 | -0.001 | 1.519 | 0 | -0.006 | -0.006 | 0 | 0.38 | 0.36 | 0.35 | 0.47 | 0.31 | 0.52 |
| 6 | 32375745 | rs6926737 | -1.103 | -1.064 | 0.064 | 1.432 | 0.001 | 0.003 | 0.003 | 0.512 | 0.57 | 0.65 | 0.59 | 0.69 | 0.57 | 0.71 |
| 6 | 32375973 | rs3763309 | 1.881 | -1.008 | -0.076 | 1.433 | -0.048 | 0.027 | 0.027 | 0 | 0.19 | 0.09 | 0.23 | 0.08 | 0.2 | 0.2 |
| 6 | 32376176 | rs3763311 | 0.212 | -1.049 | 0.004 | 1.442 | -0.005 | -0.003 | -0.003 | 0 | 0.3 | 0.28 | 0.32 | 0.33 | 0.29 | 0.35 |
| 6 | 32376348 | rs3763312 | 1.91 | -0.996 | -0.184 | 1.434 | -0.048 | 0.026 | 0.026 | 0 | 0.18 | 0.08 | 0.23 | 0.08 | 0.2 | 0.2 |
| 6 | 32376471 | rs3763313 | 1.686 | -1.014 | -0.151 | 1.445 | 0.003 | 0 | 0 | 0 | 0.18 | 0.21 | 0.21 | 0.22 | 0.25 | 0.19 |
| 6 | 32376746 | rs3763316 | 1.232 | -0.839 | -0.501 | 1.387 | -0.041 | 0.018 | 0.018 | 0 | 0.23 | 0.17 | 0.31 | 0.11 | 0.27 | 0.22 |
| 6 | 32376788 | rs3763317 | -0.83 | -0.876 | -0.392 | 1.407 | 0 | -0.002 | -0.002 | 0.733 | 0.53 | 0.55 | 0.5 | 0.66 | 0.5 | 0.69 |
| 6 | 32377061 | rs9268503 | -0.97 | -0.915 | -0.37 | 1.404 | 0 | 0.001 | 0.001 | 0.45 | 0.57 | 0.63 | 0.59 | 0.69 | 0.57 | 0.71 |
| 6 | 32377284 | rs9380293 | 1.296 | -0.93 | -0.378 | 1.442 | 0.002 | -0.004 | -0.004 | 0 | 0.09 | 0.08 | 0.03 | 0.24 | 0.02 | 0.18 |
| 6 | 32378866 | rs5007265 | -0.971 | -0.916 | -0.366 | 1.417 | 0 | 0.001 | 0.001 | 0.45 | 0.57 | 0.63 | 0.59 | 0.69 | 0.57 | 0.71 |
| 6 | 32378982 | rs5007263 | -0.971 | -0.916 | -0.366 | 1.417 | 0 | 0.001 | 0.001 | 0.45 | 0.57 | 0.63 | 0.59 | 0.69 | 0.57 | 0.71 |
| 6 | 32379101 | rs5007259 | -0.971 | -0.916 | -0.366 | 1.417 | 0 | 0.001 | 0.001 | 0.45 | 0.57 | 0.63 | 0.59 | 0.69 | 0.57 | 0.71 |
| 6 | 32379489 | rs9268516 | 1.081 | -0.891 | -0.43 | 1.405 | -0.043 | 0.022 | 0.022 | 0 | 0.23 | 0.17 | 0.32 | 0.12 | 0.28 | 0.22 |
| 6 | 32379736 | rs9405098 | 0.892 | -0.327 | -0.066 | 1.773 | 0.001 | -0.003 | -0.003 | 0 | 0.09 | 0.08 | 0.03 | 0.24 | 0.02 | 0.18 |
| 6 | 32380190 | rs6932810 | -0.478 | -0.126 | -0.134 | 1.528 | 0 | 0.001 | 0.001 | 0.45 | 0.57 | 0.63 | 0.59 | 0.69 | 0.57 | 0.71 |
| 6 | 32380208 | rs3129963 | 0.171 | -0.047 | -0.068 | 1.569 | 0.074 | 0.059 | 0.059 | 0 | 0.06 | 0.16 | 0.12 | 0.05 | 0.14 | 0.05 |
| 6 | 32380262 | rs6932542 | -0.477 | -0.108 | -0.146 | 1.546 | 0 | 0.001 | 0.001 | 0.45 | 0.57 | 0.63 | 0.59 | 0.69 | 0.57 | 0.71 |
| 6 | 32383108 | rs9268528 | 1.431 | -0.106 | -0.419 | 1.326 | 0.015 | -0.028 | -0.028 | 0.184 | 0.55 | 0.2 | 0.45 | 0.45 | 0.37 | 0.45 |
| 6 | 32384527 | rs9268541 | 2.181 | -0.433 | -0.479 | 1.411 | 0.022 | 0.005 | 0.005 | 0 | 0.17 | 0.08 | 0.09 | 0.13 | 0.06 | 0.15 |
| 6 | 32384721 | rs9268542 | 1.368 | -0.36 | -0.486 | 0.842 | 0.013 | -0.028 | -0.028 | 7.575 | 0.55 | 0.2 | 0.45 | 0.45 | 0.37 | 0.45 |
| 6 | 32387809 | rs2395163 | 2.184 | -0.39 | -0.584 | 0.793 | -0.041 | 0.059 | 0.059 | 2.054 | 0.25 | 0.09 | 0.23 | 0.09 | 0.2 | 0.21 |
| 6 | 32388144 | rs2395165 | -0.765 | 0.019 | -0.096 | 0.791 | 0.056 | -0.031 | -0.031 | 0 | 0.88 | 0.74 | 0.84 | 0.97 | 0.79 | 0.93 |
| 6 | 32388275 | rs2395166 | 0.399 | -0.097 | 0.019 | 0.894 | 0.068 | -0.008 | -0.008 | 0.272 | 0.79 | 0.58 | 0.7 | 0.82 | 0.61 | 0.7 |
| 6 | 32389305 | rs9268557 | 1.342 | -0.04 | -0.318 | 0.925 | -0.024 | -0.002 | -0.002 | 1.638 | 0.58 | 0.31 | 0.55 | 0.52 | 0.5 | 0.47 |
| 6 | 32389648 | rs3135363 | 0.492 | 0.332 | -0.313 | 1.175 | 0 | 0.016 | 0.016 | 0 | 0.18 | 0.15 | 0.26 | 0.3 | 0.24 | 0.18 |
| 6 | 32392945 | rs3135351 | -0.494 | 1.438 | 0.334 | 1.127 | 0.036 | 0.033 | 0.033 | 0 | 0.05 | 0.05 | 0.11 | 0.12 | 0.13 | 0.08 |
| 6 | 32395036 | rs3135344 | 0.336 | 1.515 | 0.33 | 0.987 | 0.055 | -0.019 | -0.019 | 0 | 0.86 | 0.72 | 0.82 | 0.94 | 0.69 | 0.81 |
| 6 | 32396475 | rs3129846 | -0.708 | 1.355 | 0.346 | 1.082 | 0.027 | 0.027 | 0.027 | 0 | 0.15 | 0.22 | 0.23 | 0.24 | 0.28 | 0.19 |

| | | | | | | | | | | | | | | | | |
|---|---|---|---|---|---|---|---|---|---|---|---|---|---|---|---|---|
| 6 | 32396615 | rs3135342 | -0.555 | 1.372 | 0.338 | 1.062 | 0.003 | -0.001 | -0.001 | 0 | 0.14 | 0.15 | 0.22 | 0.14 | 0.27 | 0.16 |
| 6 | 32397049 | rs3129848 | -0.555 | 1.372 | 0.338 | 1.051 | 0.003 | -0.001 | -0.001 | 0 | 0.14 | 0.15 | 0.22 | 0.18 | 0.27 | 0.17 |
| 6 | 32398202 | rs9268589 | 2.238 | 1.155 | 0.395 | 1.034 | -0.003 | -0.003 | -0.003 | 0 | 0.62 | 0.34 | 0.46 | 0.52 | 0.38 | 0.47 |
| 6 | 32399833 | rs9268605 | 2.238 | 1.154 | 0.403 | 1.027 | -0.003 | -0.003 | -0.003 | 1.265 | 0.62 | 0.34 | 0.46 | 0.52 | 0.38 | 0.47 |
| 6 | 32399842 | rs2395172 | -0.468 | 1.195 | 0.427 | 1.03 | 0.003 | -0.001 | -0.001 | 0 | 0.14 | 0.15 | 0.22 | 0.18 | 0.27 | 0.17 |
| 6 | 32400070 | rs9268606 | 2.285 | 1.195 | 0.427 | 1.03 | -0.003 | -0.003 | -0.003 | 0.663 | 0.62 | 0.34 | 0.46 | 0.52 | 0.38 | 0.47 |
| 6 | 32400088 | rs984778 | 1.28 | 1.195 | 0.427 | 1.03 | 0.062 | 0.005 | 0.005 | 0 | 0.76 | 0.49 | 0.68 | 0.69 | 0.66 | 0.64 |
| 6 | 32400344 | rs9501626 | -0.303 | 1.214 | 0.522 | 1.033 | 0.011 | 0.008 | 0.008 | 0 | 0.11 | 0.3 | 0.21 | 0.16 | 0.14 | 0.14 |
| 6 | 32400520 | rs3129858 | -0.748 | 1.208 | 0.513 | 1.034 | -0.002 | -0.001 | -0.001 | 0 | 0.15 | 0.15 | 0.22 | 0.18 | 0.27 | 0.17 |
| 6 | 32401079 | rs3129860 | -1.077 | 1.331 | 0.575 | 1.072 | -0.022 | -0.017 | -0.017 | 0.014 | 0.92 | 0.97 | 0.94 | 0.88 | 0.88 | 0.87 |
| 6 | 32401217 | rs3135338 | 1.449 | 1.329 | 0.579 | 1.071 | 0.062 | 0.005 | 0.005 | 0 | 0.76 | 0.49 | 0.68 | 0.69 | 0.66 | 0.64 |
| 6 | 32402464 | rs7773756 | 2.409 | 1.297 | 0.568 | 1.071 | -0.003 | -0.003 | -0.003 | 0 | 0.62 | 0.34 | 0.46 | 0.52 | 0.38 | 0.47 |
| 6 | 32402705 | rs2027856 | -0.294 | 1.299 | 0.58 | 1.071 | 0.011 | 0.008 | 0.008 | 0 | 0.11 | 0.3 | 0.21 | 0.16 | 0.14 | 0.14 |
| 6 | 32402889 | rs9268615 | 2.363 | 1.282 | 0.454 | 1.075 | -0.003 | -0.003 | -0.003 | 0 | 0.62 | 0.34 | 0.46 | 0.52 | 0.38 | 0.47 |
| 6 | 32404220 | rs3129867 | 1.16 | 1.275 | 0.471 | 1.075 | 0.072 | 0.017 | 0.017 | 0 | 0.77 | 0.51 | 0.69 | 0.71 | 0.68 | 0.64 |
| 6 | 32404377 | rs3129868 | -1.064 | 1.289 | 0.326 | 1.088 | -0.022 | -0.017 | -0.017 | 0 | 0.92 | 0.97 | 0.94 | 0.88 | 0.88 | 0.87 |
| 6 | 32404859 | rs2395173 | 1.305 | 1.254 | 0.321 | 1.079 | 0.061 | 0.005 | 0.005 | 0 | 0.75 | 0.49 | 0.68 | 0.69 | 0.66 | 0.64 |
| 6 | 32404878 | rs2395174 | -0.494 | 1.283 | 0.328 | 1.091 | 0.003 | -0.001 | -0.001 | 0 | 0.14 | 0.15 | 0.22 | 0.18 | 0.27 | 0.17 |
| 6 | 32405192 | rs3135395 | 1.313 | 1.442 | 0.485 | 1.289 | 0.062 | 0.005 | 0.005 | 0 | 0.76 | 0.49 | 0.68 | 0.69 | 0.66 | 0.64 |
| 6 | 32405865 | rs9268628 | 1.335 | 1.041 | 0.294 | 1.204 | 0.096 | -0.02 | -0.02 | 0 | 0.34 | 0.16 | 0.22 | 0.37 | 0.18 | 0.24 |
| 6 | 32406342 | rs3129871 | 1.317 | 1.202 | 0.408 | 1.247 | 0.027 | 0.007 | 0.007 | 0 | 0.74 | 0.45 | 0.65 | 0.67 | 0.63 | 0.6 |
| 6 | 32408012 | rs3129876 | -0.568 | 1.275 | 0.363 | 1.249 | 0.021 | 0.021 | 0.021 | 0 | 0.17 | 0.23 | 0.23 | 0.24 | 0.27 | 0.19 |
| 6 | 32408044 | rs9268644 | 0.519 | 1.326 | 0.349 | 1.243 | -0.009 | -0.001 | -0.001 | 0.287 | 0.81 | 0.82 | 0.73 | 0.82 | 0.6 | 0.81 |
| 6 | 32408735 | rs3129878 | -1.317 | 1.184 | 0.349 | 1.255 | 0 | 0.007 | 0.007 | 0 | 0.24 | 0.27 | 0.28 | 0.36 | 0.29 | 0.32 |
| 6 | 32408842 | rs3135393 | 0.675 | 1.512 | 0.42 | 0.923 | 0.007 | 0.007 | 0.007 | 1.741 | 0.08 | 0.05 | 0.12 | 0.13 | 0.14 | 0.09 |
| 6 | 32409484 | rs3129881 | -0.335 | 1.504 | 0.487 | 0.869 | 0.012 | -0.004 | -0.004 | 0 | 0.15 | 0.17 | 0.22 | 0.1 | 0.27 | 0.13 |
| 6 | 32409530 | rs3129882 | -0.332 | 1.434 | 0.45 | 0.84 | 0.01 | 0.006 | 0.006 | 4.592 | 0.52 | 0.57 | 0.58 | 0.37 | 0.55 | 0.47 |
| 6 | 32409996 | rs6911777 | 0.204 | 2.337 | 1.075 | 0.994 | 0.047 | 0.044 | 0.044 | 0 | 0.06 | 0.19 | 0.09 | 0.12 | 0.07 | 0.11 |
| 6 | 32410137 | rs3129883 | 0.598 | 1.803 | 0.785 | 1.13 | 0.044 | 0 | 0 | 0 | 0.85 | 0.72 | 0.8 | 0.8 | 0.73 | 0.7 |
| 6 | 32410576 | rs3129886 | 0.598 | 1.822 | 0.78 | 1.13 | 0.047 | 0 | 0 | 0 | 0.85 | 0.72 | 0.8 | 0.8 | 0.73 | 0.7 |
| 6 | 32410716 | rs9268658 | 2.133 | 1.745 | 0.791 | 1.138 | 0.063 | 0.013 | 0.013 | 0 | 0.68 | 0.5 | 0.58 | 0.63 | 0.46 | 0.55 |
| 6 | 32411035 | rs8084 | 1.691 | 1.778 | 0.796 | 1.138 | 0.079 | 0.035 | 0.035 | 0 | 0.75 | 0.6 | 0.67 | 0.68 | 0.59 | 0.63 |
| 6 | 32411307 | rs2239806 | -0.095 | 1.779 | 0.808 | 1.141 | 0.023 | 0.04 | 0.04 | 2.405 | 0.1 | 0.13 | 0.13 | 0.12 | 0.15 | 0.07 |
| 6 | 32411376 | rs2239805 | 0.281 | 1.778 | 0.802 | 1.145 | 0.008 | 0 | 0 | 0 | 0.08 | 0.05 | 0.12 | 0.06 | 0.14 | 0.05 |
| 6 | 32411523 | rs2239804 | 2.15 | 1.788 | 0.801 | 1.113 | 0.063 | 0.013 | 0.013 | 0 | 0.68 | 0.5 | 0.58 | 0.63 | 0.46 | 0.55 |
| 6 | 32411646 | rs7192 | 1.708 | 1.692 | 0.802 | 1.112 | 0.056 | 0.039 | 0.039 | 0.013 | 0.76 | 0.6 | 0.69 | 0.69 | 0.63 | 0.72 |
| 6 | 32411726 | rs3129888 | -0.121 | 1.686 | 0.822 | 1.11 | 0.08 | -0.023 | -0.023 | 0 | 0.92 | 0.8 | 0.85 | 0.96 | 0.79 | 0.97 |
| 6 | 32411833 | rs2239803 | 1.932 | 1.735 | 0.671 | 1.125 | 0.104 | 0.028 | 0.028 | 0 | 0.72 | 0.54 | 0.59 | 0.65 | 0.47 | 0.6 |
| 6 | 32411846 | rs2239802 | 0.5 | 1.722 | 0.683 | 1.139 | 0.027 | 0 | 0 | 0 | 0.86 | 0.73 | 0.82 | 0.81 | 0.78 | 0.79 |
| 6 | 32412435 | rs3177928 | 0.145 | 1.734 | 0.686 | 1.145 | -0.022 | 0.009 | 0.009 | 0 | 0.08 | 0.1 | 0.11 | 0.05 | 0.18 | 0.17 |
| 6 | 32412539 | rs7195 | 1.605 | 1.705 | 0.704 | 1.159 | 0.056 | 0.039 | 0.039 | 0.013 | 0.76 | 0.6 | 0.69 | 0.69 | 0.63 | 0.72 |
| 6 | 32412809 | rs1041885 | -0.104 | 1.706 | 0.702 | 1.159 | 0.022 | 0.038 | 0.038 | 2.405 | 0.1 | 0.13 | 0.13 | 0.12 | 0.15 | 0.07 |
| 6 | 32413094 | rs2213586 | 1.597 | 1.703 | 0.699 | 1.159 | 0.056 | 0.039 | 0.039 | 0.013 | 0.76 | 0.6 | 0.69 | 0.69 | 0.63 | 0.72 |
| 6 | 32413150 | rs2213585 | 1.597 | 1.703 | 0.699 | 1.159 | 0.056 | 0.039 | 0.039 | 0.013 | 0.76 | 0.6 | 0.69 | 0.69 | 0.63 | 0.72 |
| 6 | 32413317 | rs2395182 | 0.498 | 1.702 | 0.698 | 1.155 | 0.028 | 0.001 | 0.001 | 0 | 0.86 | 0.73 | 0.82 | 0.81 | 0.78 | 0.79 |
| 6 | 32413459 | rs2227139 | 1.579 | 1.704 | 0.699 | 1.158 | 0.056 | 0.039 | 0.039 | 0.013 | 0.76 | 0.6 | 0.69 | 0.69 | 0.63 | 0.72 |
| 6 | 32413557 | rs3763326 | 0.777 | 1.7 | 0.679 | 1.153 | -0.016 | -0.002 | -0.002 | 0.685 | 0.07 | 0.05 | 0.03 | 0.05 | 0.03 | 0.05 |
| 6 | 32414273 | rs3129890 | -1.119 | 1.641 | 0.457 | 1.082 | -0.044 | 0.018 | 0.018 | 0 | 0.26 | 0.38 | 0.26 | 0.33 | 0.25 | 0.26 |
| 6 | 32418031 | rs6937545 | 0.115 | 1.478 | 0.232 | 1.033 | 0.02 | 0.016 | 0.016 | 0 | 0.76 | 0.73 | 0.8 | 0.73 | 0.72 | 0.79 |
| 6 | 32418848 | rs28895095 | 0.058 | 1.611 | -0.133 | 0.989 | -0.021 | 0.008 | 0.008 | 0 | 0.08 | 0.1 | 0.11 | 0.05 | 0.18 | 0.16 |
| 6 | 32420179 | rs7754768 | 0.777 | 1.508 | -0.104 | 1.005 | -0.001 | 0.004 | 0.004 | 0 | 0.66 | 0.48 | 0.67 | 0.54 | 0.62 | 0.6 |
| 6 | 32423021 | rs28895171 | -0.255 | 1.958 | -0.419 | 0.91 | -0.021 | 0.008 | 0.008 | 0 | 0.08 | 0.1 | 0.11 | 0.05 | 0.18 | 0.17 |
| 6 | 32424380 | rs28895187 | -0.255 | 1.958 | -0.419 | 0.91 | -0.021 | 0.008 | 0.008 | 0 | 0.08 | 0.1 | 0.11 | 0.05 | 0.18 | 0.17 |
| 6 | 32424882 | rs7763262 | 0.365 | 1.681 | -0.054 | 1.04 | 0.02 | 0.016 | 0.016 | 0 | 0.76 | 0.73 | 0.8 | 0.73 | 0.72 | 0.79 |
| 6 | 32427748 | rs9268831 | 1.012 | 1.358 | -0.161 | 0.983 | 0.01 | 0.003 | 0.003 | 0 | 0.59 | 0.51 | 0.63 | 0.52 | 0.52 | 0.44 |
| 6 | 32427789 | rs9268832 | 0.804 | 1.246 | 0.078 | 0.92 | 0.012 | 0.036 | 0.036 | 0 | 0.7 | 0.58 | 0.69 | 0.54 | 0.63 | 0.61 |
| 6 | 32428186 | rs6923504 | -0.818 | 0.824 | 0.005 | 1.041 | 0.008 | 0.001 | 0.001 | 3.274 | 0.58 | 0.62 | 0.77 | 0.62 | 0.7 | 0.63 |
| 6 | 32428285 | rs6903608 | -0.811 | 0.822 | 0.004 | 1.048 | 0.008 | 0.001 | 0.001 | 3.438 | 0.58 | 0.62 | 0.77 | 0.61 | 0.7 | 0.63 |
| 6 | 32429643 | rs9268853 | 1.128 | 0.856 | 0.103 | 1.037 | -0.055 | -0.026 | -0.026 | 0 | 0.33 | 0.19 | 0.41 | 0.31 | 0.32 | 0.3 |
| 6 | 32430167 | rs9268862 | -0.924 | 0.823 | 0.184 | 1.017 | -0.001 | 0.018 | 0.018 | 0.066 | 0.19 | 0.39 | 0.28 | 0.26 | 0.25 | 0.26 |
| 6 | 32430752 | rs7766854 | -0.924 | 0.823 | 0.184 | 1.017 | -0.001 | 0.018 | 0.018 | 0.066 | 0.19 | 0.39 | 0.28 | 0.26 | 0.25 | 0.26 |
| 6 | 32431129 | rs7767493 | -0.924 | 0.823 | 0.184 | 1.036 | -0.001 | 0.021 | 0.021 | 0.066 | 0.19 | 0.39 | 0.28 | 0.27 | 0.25 | 0.26 |
| 6 | 32431147 | rs9268877 | -0.398 | 0.837 | 0.136 | 1.034 | 0.015 | 0.008 | 0.008 | 6.413 | 0.52 | 0.58 | 0.69 | 0.57 | 0.57 | 0.56 |
| 6 | 32431358 | rs9268880 | -0.857 | 0.835 | 0.181 | 1.029 | 0.016 | 0.003 | 0.003 | 3.472 | 0.58 | 0.63 | 0.77 | 0.61 | 0.7 | 0.63 |
| 6 | 32432406 | rs9268911 | -0.893 | 0.826 | 0.2 | 1.038 | -0.001 | 0.018 | 0.018 | 0.066 | 0.19 | 0.39 | 0.27 | 0.26 | 0.24 | 0.26 |
| 6 | 32433167 | rs2395185 | 1.131 | 0.834 | 0.196 | 1.039 | -0.057 | -0.027 | -0.027 | 0.253 | 0.33 | 0.19 | 0.41 | 0.31 | 0.32 | 0.3 |
| 6 | 32434939 | rs9268977 | -0.891 | 0.83 | 0.207 | 1.038 | -0.001 | 0.018 | 0.018 | 0.066 | 0.19 | 0.39 | 0.28 | 0.26 | 0.25 | 0.26 |
| 6 | 32435044 | rs9268979 | -0.409 | 0.831 | 0.204 | 1.039 | 0.015 | 0.008 | 0.008 | 6.413 | 0.52 | 0.58 | 0.69 | 0.57 | 0.57 | 0.56 |
| 6 | 32435123 | rs9268980 | -0.894 | 0.829 | 0.21 | 1.037 | -0.001 | 0.018 | 0.018 | 0.066 | 0.19 | 0.39 | 0.28 | 0.26 | 0.25 | 0.25 |

| | | | | | | | | | | | | | | | | |
|---|---|---|---|---|---|---|---|---|---|---|---|---|---|---|---|---|
| 6 | 32438542 | rs9368726 | 1.129 | 0.835 | 0.191 | 1.07 | -0.055 | -0.024 | -0.024 | 0.035 | 0.33 | 0.19 | 0.41 | 0.31 | 0.32 | 0.3 |
| 6 | 32438598 | rs9269043 | -0.888 | 0.831 | 0.204 | 1.061 | -0.001 | 0.018 | 0.018 | 0.066 | 0.19 | 0.39 | 0.28 | 0.26 | 0.25 | 0.26 |
| 6 | 32438648 | rs9405108 | 1.133 | 0.832 | 0.201 | 1.062 | -0.055 | -0.026 | -0.026 | 8.926 | 0.33 | 0.19 | 0.41 | 0.31 | 0.32 | 0.3 |
| 6 | 32439323 | rs2157338 | -0.89 | 0.83 | 0.206 | 1.061 | -0.001 | 0.018 | 0.018 | 0.066 | 0.19 | 0.39 | 0.28 | 0.26 | 0.25 | 0.26 |
| 6 | 32439508 | rs2187823 | -0.89 | 0.83 | 0.206 | 1.08 | -0.001 | 0.021 | 0.021 | 0.066 | 0.19 | 0.36 | 0.27 | 0.26 | 0.24 | 0.25 |
| 6 | 32439828 | rs9286790 | 1.03 | 0.994 | 0.102 | 1.119 | -0.055 | -0.026 | -0.026 | 0.966 | 0.33 | 0.2 | 0.41 | 0.31 | 0.32 | 0.3 |
| 6 | 32448763 | rs7748472 | 0.657 | 0.97 | 0.212 | 1.178 | -0.012 | 0 | 0 | 0 | 0.06 | 0.12 | 0.04 | 0.12 | 0.04 | 0.07 |
| 6 | 32448904 | rs7749057 | 0.652 | 1.076 | 0.318 | 1.201 | 0.066 | 0.031 | 0.031 | 0.871 | 0.07 | 0.14 | 0.13 | 0.04 | 0.13 | 0.05 |
| 6 | 32449411 | rs1964995 | 0.429 | 1.075 | 0.038 | 1.174 | -0.04 | -0.02 | -0.02 | 0 | 0.39 | 0.24 | 0.49 | 0.35 | 0.45 | 0.37 |
| 6 | 32501715 | rs33997707 | 0.245 | 1.083 | 0.037 | 1.132 | -0.007 | 0.012 | 0.012 | 0 | 0.1 | 0.1 | 0.12 | 0.04 | 0.15 | 0.08 |
| 6 | 32512613 | rs35571839 | 0.988 | 1.555 | 0.092 | 1.137 | 0.002 | -0.001 | -0.001 | 0 | 0.05 | 0.07 | 0.13 | 0.02 | 0.15 | 0.03 |
| 6 | 32519005 | rs28490179 | -0.449 | 1.358 | 0.27 | 1.365 | 0.001 | 0 | 0 | 0.998 | 0.81 | 0.84 | 0.88 | 0.76 | 0.85 | 0.72 |
| 6 | 32560859 | rs28366298 | 1.201 | 1.298 | 0.052 | 1.256 | -0.069 | -0.042 | -0.042 | 0 | 0.29 | 0.14 | 0.36 | 0.31 | 0.31 | 0.26 |
| 6 | 32561334 | rs35265698 | 1.262 | 1.408 | -0.32 | 1.283 | -0.075 | -0.026 | -0.026 | 1.762 | 0.14 | 0.04 | 0.21 | 0.12 | 0.16 | 0.08 |
| 6 | 32566232 | rs9270665 | 1.638 | 1.177 | -0.027 | 1.307 | -0.031 | -0.02 | -0.02 | 0 | 0.28 | 0.35 | 0.22 | 0.12 | 0.22 | 0.13 |
| 6 | 32569687 | rs28605404 | 1.298 | 1.422 | -0.142 | 1.217 | -0.001 | 0.001 | 0.001 | 0 | 0.05 | 0.06 | 0.11 | 0.02 | 0.13 | 0.03 |
| 6 | 32569691 | rs477515 | 0.829 | 1.403 | -0.104 | 1.451 | -0.032 | 0.019 | 0.019 | 3.765 | 0.3 | 0.21 | 0.34 | 0.17 | 0.32 | 0.29 |
| 6 | 32570400 | rs2516049 | 0.829 | 1.874 | -0.097 | 1.506 | -0.022 | 0.033 | 0.033 | 3.765 | 0.3 | 0.18 | 0.29 | 0.14 | 0.27 | 0.24 |
| 6 | 32572251 | rs2858870 | 1.706 | 1.518 | -0.107 | 1.398 | -0.002 | 0.014 | 0.014 | 2.472 | 0.15 | 0.11 | 0.11 | 0.04 | 0.13 | 0.18 |
| 6 | 32573562 | rs602457 | 1.387 | 1.531 | -0.151 | 1.454 | -0.013 | 0.009 | 0.009 | 0.139 | 0.13 | 0.09 | 0.21 | 0.06 | 0.16 | 0.08 |
| 6 | 32574060 | rs9270986 | -0.753 | 1.352 | -0.181 | 1.575 | 0.002 | 0.003 | 0.003 | 0.925 | 0.81 | 0.86 | 0.9 | 0.79 | 0.86 | 0.74 |
| 6 | 32575369 | rs9271055 | -0.753 | 1.352 | -0.181 | 1.575 | 0.002 | 0.003 | 0.003 | 0.925 | 0.81 | 0.86 | 0.9 | 0.79 | 0.86 | 0.73 |
| 6 | 32576478 | rs9271100 | -0.225 | 1.399 | -0.172 | 1.567 | -0.004 | 0.006 | 0.006 | 3.291 | 0.76 | 0.8 | 0.79 | 0.77 | 0.73 | 0.71 |
| 6 | 32577380 | rs660895 | 1.279 | 1.44 | -0.133 | 1.708 | -0.029 | -0.014 | -0.014 | 0 | 0.16 | 0.13 | 0.29 | 0.24 | 0.22 | 0.16 |
| 6 | 32577889 | rs9271170 | -0.265 | 1.356 | -0.13 | 1.704 | -0.004 | 0.006 | 0.006 | 3.291 | 0.76 | 0.8 | 0.79 | 0.77 | 0.73 | 0.71 |
| 6 | 32578052 | rs532098 | -0.08 | 1.338 | -0.003 | 1.801 | -0.002 | -0.001 | -0.001 | 0 | 0.41 | 0.38 | 0.49 | 0.39 | 0.47 | 0.44 |
| 6 | 32581096 | rs35759989 | 1.519 | 1.293 | 0.008 | 1.796 | -0.004 | 0.006 | 0.006 | 0 | 0.05 | 0.09 | 0.02 | 0.1 | 0.04 | 0.02 |
| 6 | 32581889 | rs4530903 | 1.367 | 1.387 | 0.001 | 1.731 | -0.001 | 0.001 | 0.001 | 0 | 0.05 | 0.06 | 0.11 | 0.02 | 0.13 | 0.03 |
| 6 | 32582075 | rs642093 | 1.632 | 1.186 | 0.062 | 1.85 | -0.029 | 0.005 | 0.005 | 0 | 0.28 | 0.34 | 0.28 | 0.26 | 0.23 | 0.14 |
| 6 | 32582796 | rs13204672 | 1.611 | 1.415 | 0.016 | 1.596 | 0 | -0.005 | -0.005 | 0 | 0.15 | 0.12 | 0.12 | 0.04 | 0.15 | 0.2 |
| 6 | 32583742 | rs9271348 | -0.988 | 1.248 | -0.071 | 1.548 | 0.01 | -0.007 | -0.007 | 0 | 0.74 | 0.76 | 0.81 | 0.79 | 0.72 | 0.54 |
| 6 | 32586854 | rs9271366 | -1.222 | 0.937 | -0.022 | 1.522 | 0.01 | 0.004 | 0.004 | 0 | 0.89 | 0.87 | 0.93 | 0.84 | 0.88 | 0.74 |
| 6 | 32589000 | rs9271488 | 0.914 | 0.991 | -0.272 | 1.647 | -0.043 | -0.026 | -0.026 | 4.339 | 0.3 | 0.19 | 0.38 | 0.32 | 0.31 | 0.29 |
| 6 | 32590722 | rs17533090 | 0.191 | 0.951 | -0.201 | 1.613 | 0.002 | -0.001 | -0.001 | 0.078 | 0.13 | 0.15 | 0.15 | 0.13 | 0.18 | 0.15 |
| 6 | 32590925 | rs3129763 | 1.712 | 0.727 | 0.083 | 1.576 | 0.001 | 0.002 | 0.002 | 0 | 0.28 | 0.26 | 0.24 | 0.12 | 0.21 | 0.13 |
| 6 | 32590953 | rs9271588 | 0.018 | 0.865 | 0.035 | 1.627 | 0.005 | -0.015 | -0.015 | 0 | 0.43 | 0.34 | 0.53 | 0.45 | 0.49 | 0.45 |
| 6 | 32590990 | rs35399661 | 0.181 | 0.953 | 0.095 | 1.592 | 0.002 | -0.001 | -0.001 | 0.312 | 0.13 | 0.15 | 0.15 | 0.13 | 0.18 | 0.15 |
| 6 | 32594328 | rs9271775 | -0.737 | 0.946 | 0.191 | 1.621 | 0.06 | 0.059 | 0.059 | 0 | 0.87 | 0.67 | 0.83 | 0.73 | 0.82 | 0.65 |
| 6 | 32599999 | rs9272105 | 0.038 | 0.905 | 0.088 | 1.617 | 0.014 | 0.003 | 0.003 | 0 | 0.51 | 0.58 | 0.53 | 0.47 | 0.55 | 0.43 |
| 6 | 32602269 | rs9272219 | 1.381 | 0.588 | -0.088 | 1.545 | -0.019 | -0.009 | -0.009 | 3.014 | 0.29 | 0.32 | 0.33 | 0.23 | 0.29 | 0.2 |
| 6 | 32602482 | rs3104369 | 0.035 | 0.621 | -0.059 | 1.53 | -0.001 | -0.01 | -0.01 | 0 | 0.74 | 0.63 | 0.78 | 0.67 | 0.74 | 0.6 |
| 6 | 32605884 | rs2187668 | 1.932 | -0.134 | -0.573 | 1.179 | 0.013 | 0.02 | 0.02 | 0 | 0.14 | 0.05 | 0.12 | 0.05 | 0.11 | 0.08 |
| 6 | 32611641 | rs9273012 | 0.847 | -0.148 | -0.506 | 1.206 | -0.019 | -0.009 | -0.009 | 0 | 0.29 | 0.3 | 0.32 | 0.22 | 0.29 | 0.16 |
| 6 | 32620283 | rs17843604 | 0.463 | 0.069 | -0.607 | 1.364 | 0 | -0.003 | -0.003 | 0 | 0.57 | 0.36 | 0.59 | 0.43 | 0.53 | 0.33 |
| 6 | 32626086 | rs7744001 | 0.642 | 0.283 | -1.051 | 1.499 | -0.015 | 0.004 | 0.004 | 0 | 0.35 | 0.43 | 0.39 | 0.36 | 0.34 | 0.42 |
| 6 | 32626249 | rs6928482 | 0.278 | 0.305 | -1.11 | 1.53 | -0.001 | 0 | 0 | 0 | 0.45 | 0.46 | 0.64 | 0.51 | 0.49 | 0.42 |
| 6 | 32626272 | rs9273363 | 1.361 | 0.31 | -1.112 | 1.518 | -0.039 | -0.044 | -0.044 | 0 | 0.28 | 0.09 | 0.33 | 0.32 | 0.26 | 0.26 |
| 6 | 32626311 | rs6906021 | -0.026 | 0.113 | -1.079 | 1.513 | -0.003 | 0.005 | 0.005 | 0.29 | 0.59 | 0.51 | 0.47 | 0.5 | 0.46 | 0.54 |
| 6 | 32627714 | rs1063355 | 0.344 | 0.175 | -0.995 | 1.444 | -0.002 | -0.001 | -0.001 | 2.226 | 0.58 | 0.51 | 0.71 | 0.54 | 0.6 | 0.49 |
| 6 | 32628428 | rs2854275 | 2.074 | 0.586 | -0.843 | 1.447 | 0.012 | 0.024 | 0.024 | 0 | 0.14 | 0.05 | 0.07 | 0.03 | 0.1 | 0.07 |
| 6 | 32635949 | rs2647025 | 0.898 | 0.735 | -0.792 | 1.6 | -0.01 | -0.01 | -0.01 | 0 | 0.28 | 0.21 | 0.45 | 0.33 | 0.31 | 0.29 |
| 6 | 32637994 | rs9274741 | -0.311 | 0.804 | -0.666 | 1.178 | 0.011 | -0.035 | -0.035 | 0 | 0.61 | 0.51 | 0.68 | 0.8 | 0.54 | 0.72 |
| 6 | 32650809 | rs17212090 | 0.766 | 0.894 | -0.709 | 1.403 | -0.002 | 0.004 | 0.004 | 9.058 | 0.26 | 0.29 | 0.27 | 0.46 | 0.25 | 0.26 |
| 6 | 32651117 | rs9275141 | -0.155 | 0.913 | -0.709 | 1.409 | 0.01 | -0.022 | -0.022 | 0 | 0.58 | 0.5 | 0.66 | 0.75 | 0.52 | 0.64 |
| 6 | 32651894 | rs2856695 | -0.139 | 0.911 | -0.709 | 1.407 | 0.01 | -0.015 | -0.015 | 0 | 0.58 | 0.5 | 0.66 | 0.66 | 0.52 | 0.6 |
| 6 | 32652359 | rs3021058 | -0.155 | 0.913 | -0.709 | 1.386 | 0.01 | -0.022 | -0.022 | 0 | 0.58 | 0.48 | 0.66 | 0.75 | 0.51 | 0.63 |
| 6 | 32653070 | rs4947342 | 0.84 | 0.902 | -0.726 | 1.405 | 0.004 | -0.007 | -0.007 | 0 | 0.32 | 0.22 | 0.41 | 0.31 | 0.26 | 0.39 |
| 6 | 32653385 | rs2856692 | 0.84 | 0.902 | -0.726 | 1.405 | 0.004 | -0.007 | -0.007 | 1.281 | 0.32 | 0.21 | 0.39 | 0.31 | 0.26 | 0.39 |
| 6 | 32657543 | rs4642516 | -0.158 | 0.91 | -0.71 | 1.31 | 0.01 | -0.023 | -0.023 | 0 | 0.58 | 0.5 | 0.67 | 0.76 | 0.52 | 0.64 |
| 6 | 32657578 | rs7774434 | -0.199 | 1.049 | -0.583 | 1.255 | -0.002 | 0.002 | 0.002 | 0 | 0.44 | 0.4 | 0.53 | 0.41 | 0.43 | 0.55 |
| 6 | 32658079 | rs7775228 | 1.397 | 0.975 | -0.6 | 1.315 | -0.006 | 0.004 | 0.004 | 0.231 | 0.2 | 0.18 | 0.21 | 0.23 | 0.16 | 0.31 |
| 6 | 32658310 | rs9469220 | -0.362 | 1.007 | -0.614 | 1.307 | 0 | 0.003 | 0.003 | 5.279 | 0.54 | 0.6 | 0.6 | 0.65 | 0.54 | 0.61 |
| 6 | 32659878 | rs9275224 | -0.259 | 0.982 | -0.57 | 1.34 | -0.001 | -0.002 | -0.002 | 0 | 0.49 | 0.58 | 0.68 | 0.65 | 0.55 | 0.44 |
| 6 | 32660375 | rs2858324 | -0.872 | 0.942 | -0.585 | 1.325 | 0.004 | -0.012 | -0.012 | 0 | 0.67 | 0.67 | 0.79 | 0.81 | 0.65 | 0.73 |
| 6 | 32660943 | rs9275245 | -0.255 | 0.963 | -0.579 | 1.331 | -0.001 | -0.002 | -0.002 | 0.434 | 0.49 | 0.58 | 0.68 | 0.65 | 0.55 | 0.44 |
| 6 | 32663564 | rs5000634 | 0.251 | 0.981 | -0.568 | 1.334 | -0.002 | -0.012 | -0.012 | 2.375 | 0.39 | 0.31 | 0.56 | 0.5 | 0.41 | 0.3 |
| 6 | 32663851 | rs6457617 | -0.223 | 0.979 | -0.562 | 1.335 | 0 | 0 | 0 | 4.023 | 0.49 | 0.54 | 0.68 | 0.56 | 0.54 | 0.39 |
| 6 | 32664163 | rs6457622 | 0.246 | 0.984 | -0.568 | 1.337 | -0.002 | -0.012 | -0.012 | 0.268 | 0.39 | 0.37 | 0.56 | 0.5 | 0.41 | 0.3 |
| 6 | 32664458 | rs2647012 | -0.907 | 0.972 | -0.572 | 1.304 | 0.005 | -0.013 | -0.013 | 0 | 0.67 | 0.68 | 0.79 | 0.81 | 0.65 | 0.73 |

| | | | | | | | | | | | | | | | | |
|---|---|---|---|---|---|---|---|---|---|---|---|---|---|---|---|---|
| 6 | 32664880 | rs2647003 | -0.907 | 0.972 | -0.572 | 1.304 | 0.005 | -0.013 | -0.013 | 0 | 0.67 | 0.67 | 0.79 | 0.81 | 0.65 | 0.73 |
| 6 | 32664960 | rs9357152 | 0.919 | 0.943 | -0.598 | 1.379 | 0.003 | -0.008 | -0.008 | 0.457 | 0.25 | 0.25 | 0.27 | 0.37 | 0.24 | 0.21 |
| 6 | 32665616 | rs2856664 | 0.407 | 1.076 | -0.559 | 1.31 | 0.021 | -0.009 | -0.009 | 0 | 0.43 | 0.35 | 0.37 | 0.53 | 0.34 | 0.51 |
| 6 | 32665728 | rs9275312 | 1.119 | 1.153 | -0.549 | 1.321 | 0.001 | 0.002 | 0.002 | 0.753 | 0.15 | 0.15 | 0.29 | 0.23 | 0.17 | 0.14 |
| 6 | 32666738 | rs2856725 | -0.848 | 1.342 | -0.526 | 1.269 | 0.005 | -0.014 | -0.014 | 0.676 | 0.67 | 0.67 | 0.79 | 0.81 | 0.65 | 0.73 |
| 6 | 32666822 | rs9275328 | 1.127 | 1.348 | -0.524 | 1.306 | 0.001 | 0.002 | 0.002 | 0.753 | 0.15 | 0.15 | 0.29 | 0.23 | 0.17 | 0.14 |
| 6 | 32666875 | rs9275330 | 1.127 | 1.348 | -0.524 | 1.315 | 0.001 | 0.001 | 0.001 | 0.753 | 0.15 | 0.15 | 0.29 | 0.23 | 0.17 | 0.14 |
| 6 | 32666968 | rs9275333 | 1.127 | 1.348 | -0.524 | 1.311 | 0.001 | 0.002 | 0.002 | 0.753 | 0.15 | 0.15 | 0.29 | 0.23 | 0.17 | 0.14 |
| 6 | 32666997 | rs35998847 | 0.874 | 1.333 | -0.504 | 1.293 | 0.004 | -0.01 | -0.01 | 0.105 | 0.09 | 0.17 | 0.12 | 0.06 | 0.14 | 0.09 |
| 6 | 32667119 | rs3135006 | -0.888 | 1.169 | -0.504 | 1.214 | 0.044 | -0.005 | -0.005 | 0 | 0.85 | 0.74 | 0.87 | 0.9 | 0.79 | 0.89 |
| 6 | 32667230 | rs2858312 | -0.588 | 1.308 | -0.814 | 1.137 | 0.025 | -0.012 | -0.012 | 0 | 0.58 | 0.36 | 0.37 | 0.52 | 0.31 | 0.42 |
| 6 | 32667364 | rs17427599 | -0.294 | 1.55 | -0.751 | 1.204 | 0.053 | 0.041 | 0.041 | 0.15 | 0.14 | 0.3 | 0.25 | 0.22 | 0.28 | 0.16 |
| 6 | 32668296 | rs9275371 | 0.064 | 1.453 | -0.78 | 1.156 | 0.007 | 0 | 0 | 0 | 0.24 | 0.32 | 0.41 | 0.28 | 0.31 | 0.22 |
| 6 | 32668526 | rs9275374 | 0.064 | 1.453 | -0.78 | 1.156 | 0.007 | 0 | 0 | 0 | 0.24 | 0.32 | 0.41 | 0.28 | 0.31 | 0.22 |
| 6 | 32669018 | rs1612904 | -1.223 | 1.384 | -0.814 | 1.116 | 0.004 | -0.018 | -0.018 | 0 | 0.7 | 0.69 | 0.81 | 0.86 | 0.69 | 0.81 |
| 6 | 32669084 | rs9275388 | 0.151 | 1.379 | -0.812 | 1.117 | 0.007 | 0 | 0 | 0 | 0.24 | 0.32 | 0.41 | 0.28 | 0.31 | 0.22 |
| 6 | 32669156 | rs9275390 | 0.151 | 1.379 | -0.812 | 1.117 | 0.007 | 0 | 0 | 0 | 0.24 | 0.32 | 0.41 | 0.28 | 0.31 | 0.22 |
| 6 | 32669439 | rs9275393 | 0.151 | 1.379 | -0.812 | 1.117 | 0.007 | 0 | 0 | 0 | 0.24 | 0.32 | 0.41 | 0.28 | 0.31 | 0.22 |
| 6 | 32669767 | rs2647050 | 0.335 | 1.382 | -0.823 | 1.157 | 0.02 | -0.01 | -0.01 | 0 | 0.43 | 0.36 | 0.38 | 0.53 | 0.35 | 0.51 |
| 6 | 32669955 | rs9275406 | 0.16 | 1.383 | -0.823 | 1.157 | 0.007 | 0 | 0 | 0 | 0.24 | 0.32 | 0.41 | 0.28 | 0.31 | 0.22 |
| 6 | 32670000 | rs2858308 | 1.723 | 1.183 | -0.881 | 1.235 | 0.014 | -0.003 | -0.003 | 0 | 0.18 | 0.1 | 0.11 | 0.16 | 0.1 | 0.29 |
| 6 | 32670037 | rs9275407 | 0.176 | 1.28 | -0.838 | 1.201 | 0.007 | 0 | 0 | 0 | 0.24 | 0.32 | 0.41 | 0.28 | 0.31 | 0.22 |
| 6 | 32670244 | rs9275418 | 0.176 | 1.28 | -0.838 | 1.201 | 0.007 | 0 | 0 | 0 | 0.24 | 0.32 | 0.41 | 0.28 | 0.31 | 0.22 |
| 6 | 32670255 | rs2856718 | 0.31 | 1.279 | -0.839 | 1.201 | 0.02 | -0.01 | -0.01 | 0.91 | 0.43 | 0.36 | 0.38 | 0.53 | 0.35 | 0.51 |
| 6 | 32670308 | rs2856717 | -1.026 | 1.279 | -0.837 | 1.2 | 0.005 | -0.011 | -0.011 | 0 | 0.67 | 0.68 | 0.79 | 0.81 | 0.66 | 0.73 |
| 6 | 32670464 | rs2858305 | -1.026 | 1.279 | -0.837 | 1.2 | 0.005 | -0.011 | -0.011 | 0 | 0.67 | 0.68 | 0.79 | 0.81 | 0.66 | 0.73 |
| 6 | 32670495 | rs35120848 | 0.353 | 1.344 | -0.799 | 1.263 | 0.016 | 0.009 | 0.009 | 0.001 | 0.11 | 0.25 | 0.23 | 0.12 | 0.19 | 0.1 |
| 6 | 32670576 | rs9275424 | 0.033 | 1.341 | -0.782 | 1.262 | 0.008 | 0 | 0 | 0 | 0.24 | 0.32 | 0.41 | 0.28 | 0.31 | 0.22 |
| 6 | 32670978 | rs9275428 | 0.033 | 1.341 | -0.798 | 1.262 | 0.007 | 0 | 0 | 0 | 0.24 | 0.32 | 0.41 | 0.28 | 0.31 | 0.22 |
| 6 | 32671103 | rs13192471 | 0.344 | 1.329 | -0.783 | 1.26 | 0.016 | 0.009 | 0.009 | 0.001 | 0.11 | 0.25 | 0.23 | 0.12 | 0.19 | 0.1 |
| 6 | 32671248 | rs1794275 | -0.439 | 1.269 | -0.833 | 1.23 | 0.009 | 0.005 | 0.005 | 0.158 | 0.16 | 0.26 | 0.24 | 0.17 | 0.22 | 0.18 |
| 6 | 32671521 | rs9275439 | 0.052 | 1.317 | -0.771 | 1.268 | 0.007 | 0 | 0 | 0 | 0.24 | 0.32 | 0.41 | 0.28 | 0.31 | 0.22 |
| 6 | 32672082 | rs9275464 | 0.169 | 1.323 | -0.793 | 1.232 | 0 | 0 | 0 | 0.398 | 0.23 | 0.29 | 0.41 | 0.19 | 0.3 | 0.18 |
| 6 | 32674994 | rs9275523 | 0.4 | 1.319 | -0.808 | 1.39 | -0.007 | -0.005 | -0.005 | 0 | 0.22 | 0.24 | 0.31 | 0.31 | 0.25 | 0.21 |
| 6 | 32677152 | rs16898264 | 0.271 | 1.258 | -0.827 | 1.363 | 0.02 | 0 | 0 | 1.408 | 0.43 | 0.36 | 0.38 | 0.44 | 0.35 | 0.51 |
| 6 | 32677397 | rs16898262 | 1.702 | 1.153 | -0.865 | 1.326 | 0.014 | 0.015 | 0.015 | 0 | 0.18 | 0.1 | 0.11 | 0.07 | 0.1 | 0.29 |
| 6 | 32677440 | rs10947332 | 0.785 | 1.69 | -0.729 | 1.401 | 0.004 | 0.008 | 0.008 | 0.022 | 0.09 | 0.17 | 0.12 | 0.15 | 0.14 | 0.09 |
| 6 | 32678999 | rs9275572 | -0.636 | 1.668 | -0.724 | 1.381 | 0.013 | -0.01 | -0.01 | 0.052 | 0.65 | 0.6 | 0.69 | 0.75 | 0.6 | 0.72 |
| 6 | 32680070 | rs9275582 | 0.535 | 1.686 | -0.774 | 1.379 | -0.018 | 0 | 0 | 0 | 0.21 | 0.21 | 0.3 | 0.22 | 0.24 | 0.17 |
| 6 | 32680970 | rs7745656 | 0.661 | 1.689 | -0.728 | 1.415 | 0.029 | 0.001 | 0.001 | 0.865 | 0.42 | 0.35 | 0.36 | 0.43 | 0.28 | 0.49 |
| 6 | 32681049 | rs2647087 | 0.661 | 1.669 | -0.728 | 1.415 | 0.03 | 0.001 | 0.001 | 0.865 | 0.42 | 0.34 | 0.36 | 0.43 | 0.28 | 0.49 |
| 6 | 32681085 | rs2858333 | 0.661 | 1.669 | -0.728 | 1.415 | 0.03 | 0.001 | 0.001 | 0.865 | 0.42 | 0.34 | 0.36 | 0.43 | 0.28 | 0.49 |
| 6 | 32681161 | rs2858332 | -0.57 | 1.672 | -0.719 | 1.422 | 0.03 | -0.009 | -0.009 | 0 | 0.63 | 0.59 | 0.66 | 0.74 | 0.53 | 0.7 |
| 6 | 32681277 | rs2858331 | 0.074 | 1.679 | -0.788 | 1.331 | 0.01 | 0.003 | 0.003 | 4.43 | 0.51 | 0.52 | 0.48 | 0.46 | 0.42 | 0.57 |
| 6 | 32681355 | rs9275595 | 0.549 | 1.706 | -0.788 | 1.315 | -0.018 | 0 | 0 | 0 | 0.21 | 0.21 | 0.3 | 0.22 | 0.24 | 0.17 |
| 6 | 32681530 | rs3957146 | 1.644 | 1.494 | -0.857 | 1.122 | -0.043 | 0.019 | 0.019 | 1.214 | 0.12 | 0.04 | 0.18 | 0.07 | 0.1 | 0.08 |
| 6 | 32681568 | rs2647089 | 0.596 | 1.333 | -0.784 | 1.122 | 0.028 | 0.001 | 0.001 | 0.865 | 0.42 | 0.35 | 0.36 | 0.43 | 0.28 | 0.49 |
| 6 | 32681631 | rs9275596 | -0.883 | 1.442 | -0.812 | 0.786 | 0.006 | -0.019 | -0.019 | 0.914 | 0.71 | 0.7 | 0.81 | 0.86 | 0.69 | 0.81 |
| 6 | 32682019 | rs3998159 | 1.256 | 1.652 | -0.858 | 0.845 | -0.021 | 0.02 | 0.02 | 1.193 | 0.12 | 0.04 | 0.18 | 0.07 | 0.1 | 0.08 |
| 6 | 32682137 | rs3957148 | 1.112 | 1.608 | -0.821 | 0.838 | -0.02 | 0.022 | 0.022 | 1.149 | 0.13 | 0.07 | 0.18 | 0.07 | 0.1 | 0.08 |
| 6 | 32682174 | rs3104404 | 0.035 | 0.411 | -0.841 | 0.869 | -0.002 | -0.001 | -0.001 | 0 | 0.34 | 0.35 | 0.28 | 0.44 | 0.19 | 0.51 |
| 6 | 32682308 | rs3104405 | -0.929 | 1.078 | -0.657 | 0.847 | -0.001 | 0.002 | 0.002 | 0 | 0.71 | 0.61 | 0.76 | 0.77 | 0.65 | 0.85 |
| 6 | 32682394 | rs3997848 | 0.045 | 0.613 | -0.784 | 0.565 | 0.006 | 0.006 | 0.006 | 0 | 0.42 | 0.28 | 0.3 | 0.2 | 0.32 | 0.46 |
| 6 | 32682664 | rs9275601 | -0.199 | 0.812 | -0.44 | 0.576 | 0.002 | 0.003 | 0.003 | 0 | 0.46 | 0.45 | 0.41 | 0.52 | 0.42 | 0.46 |
| 6 | 32682724 | rs3873444 | -1.255 | 0.666 | 0.152 | 0.287 | 0 | -0.041 | -0.041 | 0.432 | 0.12 | 0.25 | 0.11 | 0.06 | 0.12 | 0.17 |
| 6 | 32682862 | rs3892710 | 0.7 | -0.248 | 0.124 | -0.118 | -0.044 | -0.083 | -0.083 | 0.132 | 0.27 | 0.38 | 0.12 | 0.05 | 0.16 | 0.23 |
| 6 | 32683691 | rs6935940 | -0.176 | -0.511 | -0.333 | 0.057 | -0.022 | -0.011 | -0.011 | 0.244 | 0.36 | 0.57 | 0.34 | 0.29 | 0.24 | 0.29 |
| 6 | 32683898 | rs6915986 | 1.547 | -0.652 | -0.304 | 0.011 | -0.018 | -0.013 | -0.013 | 0 | 0.08 | 0.16 | 0.06 | 0.03 | 0.02 | 0.12 |
| 6 | 32684069 | rs6936707 | 1.547 | -0.652 | -0.304 | 0.011 | -0.018 | -0.013 | -0.013 | 0 | 0.08 | 0.16 | 0.06 | 0.03 | 0.02 | 0.12 |
| 6 | 32684257 | rs9275614 | 1.405 | -0.51 | -0.313 | 0.086 | 0.02 | 0.032 | 0.032 | 0.049 | 0.11 | 0.17 | 0.15 | 0.19 | 0.12 | 0.08 |
| 6 | 32684344 | rs5024431 | -0.186 | -0.467 | -0.509 | 0.084 | -0.022 | -0.01 | -0.01 | 2.937 | 0.36 | 0.57 | 0.34 | 0.28 | 0.24 | 0.29 |
| 6 | 32685685 | rs3104398 | 1.084 | -0.713 | -0.634 | 0.085 | 0.012 | 0.016 | 0.016 | 0 | 0.19 | 0.12 | 0.14 | 0.11 | 0.12 | 0.25 |
| 6 | 32685865 | rs9275653 | 0.022 | -0.199 | -0.428 | 0.036 | 0.001 | -0.005 | -0.005 | 0.122 | 0.62 | 0.51 | 0.6 | 0.76 | 0.64 | 0.63 |
| 6 | 32686084 | rs12177980 | 0.858 | -0.361 | -0.803 | -0.211 | 0.019 | -0.015 | -0.015 | 0 | 0.42 | 0.17 | 0.46 | 0.63 | 0.45 | 0.51 |
| 6 | 32687297 | rs9275682 | 1.99 | -0.363 | -0.79 | -0.164 | -0.001 | -0.006 | -0.006 | 0.084 | 0.2 | 0.34 | 0.14 | 0.14 | 0.19 | 0.12 |
| 6 | 32687358 | rs3104401 | 1.68 | -0.23 | -0.791 | -0.186 | 0.006 | 0.018 | 0.018 | 0 | 0.2 | 0.12 | 0.17 | 0.11 | 0.15 | 0.26 |
| 6 | 32687570 | rs9275686 | 2.005 | -0.23 | -0.791 | -0.186 | -0.001 | -0.006 | -0.006 | 0.495 | 0.2 | 0.34 | 0.14 | 0.14 | 0.19 | 0.12 |
| 6 | 32687973 | rs9275698 | 0.507 | -0.23 | -0.791 | -0.186 | 0 | -0.001 | -0.001 | 0 | 0.4 | 0.47 | 0.31 | 0.25 | 0.34 | 0.38 |
| 6 | 32689529 | rs9461799 | 0.886 | -0.229 | -0.792 | -0.186 | 0.019 | -0.021 | -0.021 | 0.011 | 0.42 | 0.17 | 0.46 | 0.62 | 0.45 | 0.51 |

| | | | | | | | | | | | | | | | | |
|---|---|---|---|---|---|---|---|---|---|---|---|---|---|---|---|---|
| 6 | 32690027 | rs9275793 | -1.593 | -0.227 | -0.79 | -0.185 | 0.094 | 0.004 | 0.004 | 0.035 | 0.82 | 0.63 | 0.77 | 0.87 | 0.78 | 0.89 |
| 6 | 32691745 | rs763026 | 2.003 | -0.231 | -0.794 | -0.183 | -0.001 | -0.006 | -0.006 | 0 | 0.2 | 0.34 | 0.14 | 0.14 | 0.19 | 0.12 |
| 6 | 32692227 | rs9469246 | 0.563 | -0.302 | -0.841 | -0.198 | 0.05 | 0.014 | 0.014 | 0.497 | 0.46 | 0.17 | 0.47 | 0.66 | 0.45 | 0.63 |
| 6 | 32693242 | rs2858892 | 0.851 | -0.256 | -0.828 | -0.186 | 0.016 | -0.02 | -0.02 | 0 | 0.42 | 0.17 | 0.46 | 0.62 | 0.45 | 0.51 |
| 6 | 32695022 | rs9276017 | -1.543 | -0.271 | -0.777 | -0.181 | 0.093 | 0.003 | 0.003 | 0.039 | 0.82 | 0.63 | 0.77 | 0.87 | 0.78 | 0.89 |
| 6 | 32698013 | rs2859112 | 0.867 | -0.319 | -0.843 | -0.189 | 0.019 | -0.015 | -0.015 | 0 | 0.42 | 0.17 | 0.46 | 0.63 | 0.45 | 0.51 |
| 6 | 32698457 | rs9276162 | -1.556 | -0.265 | -0.77 | -0.175 | 0.094 | 0.004 | 0.004 | 0.035 | 0.82 | 0.63 | 0.77 | 0.87 | 0.78 | 0.89 |
| 6 | 32698918 | rs9276171 | 1.304 | -0.333 | -0.586 | -0.204 | -0.012 | -0.014 | -0.014 | 0 | 0.23 | 0.54 | 0.18 | 0.18 | 0.2 | 0.14 |
| 6 | 32698960 | rs7751842 | -1.505 | -0.324 | -0.609 | -0.186 | 0.093 | 0.007 | 0.007 | 0.034 | 0.82 | 0.63 | 0.77 | 0.87 | 0.78 | 0.89 |
| 6 | 32698988 | rs7751856 | -0.07 | -0.524 | -0.743 | -0.234 | -0.025 | 0.014 | 0.014 | 0.585 | 0.59 | 0.28 | 0.63 | 0.72 | 0.57 | 0.76 |
| 6 | 32702449 | rs2859078 | 1.309 | -0.498 | -0.715 | -0.366 | -0.003 | -0.016 | -0.016 | 0 | 0.24 | 0.34 | 0.15 | 0.07 | 0.19 | 0.23 |
| 6 | 32702813 | rs9276291 | -1.482 | -0.53 | -0.726 | -0.377 | 0.087 | -0.001 | -0.001 | 0.69 | 0.82 | 0.63 | 0.77 | 0.87 | 0.78 | 0.89 |
| 6 | 32702930 | rs9276296 | -1.569 | -0.496 | -0.71 | -0.36 | 0.094 | 0.004 | 0.004 | 0.035 | 0.82 | 0.63 | 0.77 | 0.87 | 0.78 | 0.89 |
| 6 | 32703366 | rs2859071 | 0.058 | -0.521 | -0.74 | -0.368 | -0.016 | 0.057 | 0.057 | 0.001 | 0.57 | 0.29 | 0.62 | 0.8 | 0.59 | 0.66 |
| 6 | 32703845 | rs2858881 | 1.558 | -0.537 | -0.698 | -0.367 | -0.036 | 0.003 | 0.003 | 0 | 0.08 | 0.2 | 0.05 | 0.08 | 0.01 | 0.14 |
| 6 | 32704523 | rs9276310 | 2.273 | -0.514 | -0.87 | -0.317 | -0.001 | 0.003 | 0.003 | 0 | 0.16 | 0.12 | 0.16 | 0.07 | 0.15 | 0.14 |
| 6 | 32705276 | rs13199787 | 0.634 | -0.517 | -0.877 | -0.321 | 0.043 | 0.071 | 0.071 | 0.192 | 0.46 | 0.16 | 0.47 | 0.77 | 0.45 | 0.64 |
| 6 | 32706128 | rs6457644 | 0.943 | -0.518 | -0.877 | -0.321 | 0.012 | 0.028 | 0.028 | 0 | 0.41 | 0.17 | 0.46 | 0.73 | 0.45 | 0.52 |
| 6 | 32706234 | rs7773068 | -2 | -0.418 | -0.826 | -0.3 | 0.03 | -0.005 | -0.005 | 3.014 | 0.86 | 0.83 | 0.81 | 0.91 | 0.79 | 0.92 |
| 6 | 32706334 | rs7773694 | -2 | -0.418 | -0.826 | -0.3 | 0.03 | -0.005 | -0.005 | 0 | 0.86 | 0.83 | 0.81 | 0.91 | 0.79 | 0.92 |
| 6 | 32706398 | rs7773407 | 0.671 | -0.418 | -0.826 | -0.3 | 0.043 | 0.071 | 0.071 | 0.192 | 0.46 | 0.17 | 0.47 | 0.77 | 0.45 | 0.65 |
| 6 | 32706416 | rs9276351 | 1.317 | -0.369 | -0.774 | -0.269 | -0.016 | -0.073 | -0.073 | 0 | 0.24 | 0.54 | 0.18 | 0.07 | 0.2 | 0.13 |
| 6 | 32706480 | rs7755597 | -1.977 | -0.369 | -0.774 | -0.268 | 0.03 | -0.005 | -0.005 | 0.184 | 0.86 | 0.83 | 0.81 | 0.91 | 0.79 | 0.92 |
| 6 | 32707295 | rs9276370 | -0.309 | -0.339 | -0.868 | -0.208 | -0.021 | 0.034 | 0.034 | 0.513 | 0.62 | 0.29 | 0.63 | 0.84 | 0.59 | 0.79 |
| 6 | 32708890 | rs9276399 | 2.099 | -0.319 | -0.814 | -0.128 | -0.001 | 0.003 | 0.003 | 0 | 0.16 | 0.12 | 0.16 | 0.07 | 0.14 | 0.14 |
| 6 | 32709964 | rs4394270 | 0.654 | -0.35 | -0.798 | -0.172 | -0.002 | -0.042 | -0.042 | 0 | 0.37 | 0.51 | 0.32 | 0.13 | 0.34 | 0.26 |
| 6 | 32711178 | rs17500468 | 2.473 | -0.337 | -1.138 | -0.369 | -0.021 | 0.036 | 0.036 | 1.387 | 0.09 | 0.01 | 0.06 | 0.02 | 0.15 | 0.07 |
| 6 | 32711576 | rs2213568 | -0.666 | -0.289 | -0.988 | -0.179 | 0.002 | -0.001 | -0.001 | 0 | 0.64 | 0.63 | 0.71 | 0.69 | 0.61 | 0.66 |
| 6 | 32712818 | rs17500510 | 1.664 | -0.455 | -1.291 | -0.6 | 0.003 | 0.067 | 0.067 | 0.163 | 0.13 | 0.03 | 0.06 | 0.06 | 0.1 | 0.11 |
| 6 | 32713188 | rs2213565 | -2.27 | -0.369 | -1.175 | -0.519 | 0.024 | -0.006 | -0.006 | 0 | 0.85 | 0.83 | 0.81 | 0.91 | 0.79 | 0.92 |
| 6 | 32713267 | rs2239800 | 1.436 | -0.391 | -1.267 | 0.126 | -0.001 | 0.021 | 0.021 | 0.03 | 0.08 | 0.06 | 0.27 | 0.38 | 0.11 | 0.22 |
| 6 | 32713867 | rs9276435 | 2.8 | -0.301 | -1.087 | -0.286 | -0.033 | 0.037 | 0.037 | 0.012 | 0.13 | 0.04 | 0.07 | 0.03 | 0.17 | 0.1 |
| 6 | 32714143 | rs2071800 | 1.759 | -0.359 | -1.112 | -0.295 | 0.005 | 0.056 | 0.056 | 1.211 | 0.12 | 0.03 | 0.04 | 0.05 | 0.07 | 0.1 |
| 6 | 32717024 | rs4415152 | 1.673 | -0.232 | -0.706 | -0.279 | 0.103 | 0.014 | 0.014 | 0.113 | 0.06 | 0.27 | 0.15 | 0 | 0.08 | 0.01 |
| 6 | 32717224 | rs9276472 | 1.635 | -0.234 | -0.706 | -0.28 | 0.1 | 0.012 | 0.012 | 0.071 | 0.06 | 0.27 | 0.15 | 0 | 0.08 | 0.01 |
| 6 | 32719666 | rs9276502 | 1.635 | -0.234 | -0.706 | -0.28 | 0.1 | 0.012 | 0.012 | 0.071 | 0.06 | 0.27 | 0.15 | 0 | 0.08 | 0.01 |
| 6 | 32719804 | rs2213572 | -0.826 | -0.32 | -0.8 | -0.303 | 0 | -0.002 | -0.002 | 0 | 0.64 | 0.65 | 0.71 | 0.81 | 0.61 | 0.67 |
| 6 | 32722186 | rs10807113 | -0.275 | -0.329 | -0.791 | -0.317 | -0.009 | 0.049 | 0.049 | 0.173 | 0.58 | 0.37 | 0.57 | 0.81 | 0.53 | 0.66 |
| 6 | 32722754 | rs9276555 | 2.88 | -0.311 | -0.752 | -0.373 | 0 | -0.005 | -0.005 | 0.355 | 0.16 | 0.19 | 0.09 | 0.04 | 0.18 | 0.11 |
| 6 | 32723341 | rs7770024 | -0.41 | -0.928 | -0.795 | -0.171 | -0.009 | 0.049 | 0.049 | 0.173 | 0.58 | 0.37 | 0.57 | 0.81 | 0.53 | 0.66 |
| 6 | 32723572 | rs7769979 | -0.41 | -0.928 | -0.795 | -0.171 | -0.009 | 0.049 | 0.049 | 0.173 | 0.58 | 0.37 | 0.57 | 0.81 | 0.53 | 0.66 |
| 6 | 32723869 | rs7774452 | -0.417 | -0.932 | -0.797 | -0.173 | -0.009 | 0.048 | 0.048 | 0.228 | 0.58 | 0.37 | 0.57 | 0.81 | 0.53 | 0.66 |
| 6 | 32723917 | rs7756516 | -0.413 | -0.925 | -0.793 | -0.169 | -0.009 | 0.049 | 0.049 | 0.173 | 0.58 | 0.37 | 0.57 | 0.81 | 0.53 | 0.66 |
| 6 | 32724061 | rs9276558 | -0.964 | -0.924 | -0.768 | -0.176 | 0 | -0.002 | -0.002 | 0 | 0.64 | 0.65 | 0.71 | 0.81 | 0.61 | 0.67 |
| 6 | 32724189 | rs7774954 | 1.698 | -0.894 | -0.745 | -0.159 | 0.105 | 0.016 | 0.016 | 0.127 | 0.06 | 0.27 | 0.15 | 0 | 0.08 | 0.01 |
| 6 | 32725193 | rs2301271 | -1.538 | -0.828 | -1.063 | -0.091 | 0.016 | -0.01 | -0.01 | 0 | 0.68 | 0.65 | 0.72 | 0.84 | 0.62 | 0.79 |
| 6 | 32727905 | rs1023449 | -0.954 | -0.842 | -0.957 | -0.12 | -0.018 | 0.018 | 0.018 | 0.176 | 0.64 | 0.7 | 0.72 | 0.81 | 0.61 | 0.67 |
| 6 | 32728554 | rs4248169 | -0.944 | -0.793 | -0.936 | -0.074 | -0.015 | 0.013 | 0.013 | 0.135 | 0.64 | 0.68 | 0.7 | 0.8 | 0.61 | 0.66 |
| 6 | 32728787 | rs2006165 | -0.954 | -0.837 | -0.931 | -0.107 | -0.017 | 0.017 | 0.017 | 0.176 | 0.64 | 0.7 | 0.72 | 0.81 | 0.61 | 0.67 |
| 6 | 32729459 | rs2071551 | -1.65 | -0.9 | -0.975 | -0.017 | -0.009 | 0.006 | 0.006 | 1.344 | 0.69 | 0.7 | 0.73 | 0.87 | 0.62 | 0.81 |
| 6 | 32730012 | rs7453920 | -1.65 | -0.9 | -0.975 | -0.017 | -0.009 | 0.006 | 0.006 | 1.344 | 0.69 | 0.7 | 0.73 | 0.87 | 0.62 | 0.81 |
| 6 | 32730086 | rs2051549 | -1.67 | -0.894 | -0.972 | -0.014 | -0.008 | 0.006 | 0.006 | 1.354 | 0.69 | 0.7 | 0.73 | 0.87 | 0.62 | 0.81 |
| 6 | 32730940 | rs2071550 | 0.845 | -1.564 | -1.285 | 0.16 | -0.006 | -0.053 | -0.053 | 0 | 0.26 | 0.15 | 0.36 | 0.44 | 0.35 | 0.37 |
| 6 | 32731258 | rs1573649 | -0.79 | -1.633 | -1.31 | 0.133 | -0.004 | 0.009 | 0.009 | 0 | 0.59 | 0.63 | 0.63 | 0.71 | 0.54 | 0.56 |
| 6 | 32731439 | rs1573648 | -1.234 | -1.637 | -1.312 | 0.133 | 0 | -0.002 | -0.002 | 0 | 0.63 | 0.62 | 0.64 | 0.78 | 0.54 | 0.7 |
| 6 | 32731960 | rs6902723 | -0.776 | -1.627 | -1.273 | 0.143 | -0.003 | 0.006 | 0.006 | 0 | 0.59 | 0.61 | 0.63 | 0.71 | 0.54 | 0.57 |
| 6 | 32732210 | rs6903130 | -0.776 | -1.627 | -1.302 | 0.126 | -0.004 | 0.008 | 0.008 | 0 | 0.59 | 0.62 | 0.63 | 0.71 | 0.54 | 0.56 |
| 6 | 32732677 | rs9348894 | 0.889 | -1.638 | -1.312 | 0.133 | -0.006 | -0.053 | -0.053 | 1.56 | 0.26 | 0.15 | 0.36 | 0.44 | 0.35 | 0.37 |
| 6 | 32732937 | rs9276586 | -0.699 | -0.606 | -1.175 | -0.124 | 0 | -0.001 | -0.001 | 0.186 | 0.63 | 0.63 | 0.64 | 0.78 | 0.54 | 0.7 |
| 6 | 32733446 | rs5019296 | -0.707 | -0.61 | -1.176 | -0.126 | 0 | -0.001 | -0.001 | 0 | 0.64 | 0.63 | 0.64 | 0.78 | 0.54 | 0.7 |
| 6 | 32733823 | rs35439528 | 1.436 | -0.69 | -1.215 | -0.172 | 0.005 | 0.058 | 0.058 | 1.279 | 0.11 | 0.03 | 0.04 | 0.05 | 0.07 | 0.1 |
| 6 | 32733931 | rs9276595 | -0.337 | 0.08 | -0.71 | 0.391 | 0 | -0.001 | -0.001 | 2.263 | 0.63 | 0.63 | 0.64 | 0.78 | 0.54 | 0.7 |
| 6 | 32733987 | rs9276598 | -0.019 | 0.089 | -0.706 | 0.394 | -0.004 | 0.009 | 0.009 | 0 | 0.59 | 0.63 | 0.63 | 0.71 | 0.54 | 0.56 |
| 6 | 32734030 | rs7382794 | -0.019 | 0.089 | -0.706 | 0.394 | -0.004 | 0.009 | 0.009 | 0 | 0.59 | 0.63 | 0.63 | 0.71 | 0.54 | 0.56 |
| 6 | 32736936 | rs6901084 | -0.009 | 0.419 | -0.629 | 0.367 | -0.004 | -0.015 | -0.015 | 0 | 0.6 | 0.55 | 0.53 | 0.78 | 0.47 | 0.7 |
| 6 | 32737149 | rs6457658 | -0.009 | 0.419 | -0.629 | 0.367 | -0.004 | -0.015 | -0.015 | 0 | 0.6 | 0.55 | 0.53 | 0.78 | 0.47 | 0.7 |
| 6 | 32737494 | rs6457661 | -0.009 | 0.419 | -0.629 | 0.367 | -0.004 | -0.015 | -0.015 | 0 | 0.6 | 0.55 | 0.53 | 0.78 | 0.47 | 0.7 |
| 6 | 32739888 | rs1383265 | 0.021 | 0.046 | -0.545 | 1.013 | -0.01 | 0.01 | 0.01 | 0 | 0.29 | 0.35 | 0.22 | 0.4 | 0.21 | 0.37 |
| 6 | 32740411 | rs2857212 | -0.244 | 0.309 | -0.716 | 1.195 | -0.003 | -0.006 | -0.006 | 0.143 | 0.35 | 0.33 | 0.4 | 0.49 | 0.38 | 0.56 |

| | | | | | | | | | | | | | | | | |
|---|---|---|---|---|---|---|---|---|---|---|---|---|---|---|---|---|
| 6 | 32740442 | rs2857211 | -0.668 | 0.166 | -0.695 | 0.873 | 0.025 | 0.006 | 0.006 | 0 | 0.78 | 0.7 | 0.64 | 0.75 | 0.69 | 0.9 |
| 6 | 32740856 | rs1480383 | 0.487 | -0.396 | -0.903 | 1.01 | 0.027 | 0.017 | 0.017 | 0 | 0.4 | 0.35 | 0.22 | 0.29 | 0.3 | 0.25 |
| 6 | 32740895 | rs1480382 | 0.528 | -0.094 | -0.958 | 1.034 | 0.024 | 0.02 | 0.02 | 0 | 0.41 | 0.36 | 0.28 | 0.33 | 0.34 | 0.26 |
| 6 | 32741742 | rs2857210 | -0.518 | 0.431 | -0.959 | 0.985 | 0.021 | 0.005 | 0.005 | 0 | 0.74 | 0.7 | 0.65 | 0.71 | 0.71 | 0.73 |
| 6 | 32741868 | rs2621416 | 0.45 | 0.521 | -0.781 | 1.171 | -0.009 | -0.01 | -0.01 | 1.152 | 0.18 | 0.17 | 0.31 | 0.37 | 0.28 | 0.35 |
| 6 | 32743217 | rs9276615 | -0.919 | 0.809 | -0.774 | 1.138 | 0.043 | 0.015 | 0.015 | 0 | 0.77 | 0.7 | 0.66 | 0.74 | 0.71 | 0.83 |
| 6 | 32743298 | rs13195441 | 0.052 | 0.777 | -0.79 | 1.122 | 0.027 | 0.07 | 0.07 | 0.668 | 0.2 | 0.34 | 0.33 | 0.38 | 0.28 | 0.37 |
| 6 | 32744145 | rs2857205 | -0.925 | 0.777 | -0.787 | 1.122 | 0.043 | 0.015 | 0.015 | 0 | 0.77 | 0.7 | 0.66 | 0.74 | 0.71 | 0.83 |
| 6 | 32744347 | rs2857204 | 0.594 | 0.756 | -0.79 | 1.154 | -0.001 | -0.001 | -0.001 | 0 | 0.33 | 0.34 | 0.38 | 0.41 | 0.37 | 0.48 |
| 6 | 32745043 | rs9276644 | 0.769 | 0.942 | -0.791 | 1.148 | 0.021 | 0.03 | 0.03 | 0 | 0.41 | 0.36 | 0.28 | 0.29 | 0.34 | 0.26 |
| 6 | 32745324 | rs2170183 | -0.907 | 1.025 | -0.751 | 1.176 | 0.04 | 0.015 | 0.015 | 0 | 0.77 | 0.7 | 0.66 | 0.74 | 0.71 | 0.83 |
| 6 | 32746984 | rs2857197 | -0.907 | 1.025 | -0.751 | 1.176 | 0.04 | 0.015 | 0.015 | 0 | 0.77 | 0.7 | 0.66 | 0.74 | 0.71 | 0.83 |
| 6 | 32747060 | rs2857195 | -0.893 | 1.012 | -0.757 | 1.169 | 0.039 | 0.015 | 0.015 | 0 | 0.77 | 0.7 | 0.66 | 0.74 | 0.71 | 0.83 |
| 6 | 32747528 | rs2857193 | -1.039 | 1.055 | -0.759 | 1.171 | 0.001 | -0.001 | -0.001 | 0.321 | 0.64 | 0.7 | 0.62 | 0.71 | 0.63 | 0.72 |
| 6 | 32748398 | rs28986337 | 3.078 | 0.997 | -0.759 | 1.171 | -0.002 | 0.026 | 0.026 | 0.258 | 0.13 | 0.01 | 0.05 | 0.03 | 0.08 | 0.11 |
| 6 | 32748810 | rs28986338 | 3.078 | 0.996 | -0.756 | 1.17 | -0.002 | 0.026 | 0.026 | 0.258 | 0.13 | 0.01 | 0.05 | 0.03 | 0.08 | 0.11 |
| 6 | 32751349 | rs28986359 | 3.077 | 0.996 | -0.756 | 1.17 | -0.002 | 0.026 | 0.026 | 0.258 | 0.13 | 0.01 | 0.05 | 0.03 | 0.08 | 0.11 |
| 6 | 32751691 | rs28986364 | 3.077 | 0.996 | -0.756 | 1.17 | -0.002 | 0.026 | 0.026 | 0.258 | 0.13 | 0.01 | 0.05 | 0.03 | 0.08 | 0.11 |
| 6 | 32752049 | rs10484564 | 2.758 | 0.309 | -1.14 | 0.628 | 0.019 | 0.05 | 0.05 | 0 | 0.1 | 0 | 0.03 | 0 | 0.08 | 0.03 |
| 6 | 32752139 | rs719654 | 1.085 | 0.881 | -0.823 | 1.054 | -0.003 | -0.004 | -0.004 | 0.356 | 0.15 | 0.14 | 0.3 | 0.24 | 0.22 | 0.28 |
| 6 | 32752150 | rs28986373 | 3.288 | 0.847 | -0.881 | 0.916 | -0.002 | 0.026 | 0.026 | 0.258 | 0.13 | 0.01 | 0.05 | 0.03 | 0.08 | 0.11 |
| 6 | 32752685 | rs13204139 | 3.288 | 0.847 | -0.881 | 0.916 | -0.002 | 0.026 | 0.026 | 0.258 | 0.13 | 0.01 | 0.05 | 0.03 | 0.08 | 0.11 |
| 6 | 32752788 | rs2857178 | -1.133 | 0.844 | -0.882 | 0.916 | 0.041 | 0.016 | 0.016 | 0 | 0.77 | 0.7 | 0.66 | 0.74 | 0.71 | 0.83 |
| 6 | 32752840 | rs2857177 | -1.25 | 0.834 | -0.886 | 0.914 | 0.001 | -0.001 | -0.001 | 0 | 0.64 | 0.7 | 0.62 | 0.71 | 0.63 | 0.72 |
| 6 | 32753682 | rs1988564 | -1.039 | 0.844 | -0.882 | 0.919 | 0.039 | 0.015 | 0.015 | 0 | 0.77 | 0.7 | 0.66 | 0.74 | 0.71 | 0.83 |
| 6 | 32754430 | rs28986396 | 3.286 | 0.864 | -0.872 | 0.93 | -0.002 | 0.026 | 0.026 | 0.258 | 0.13 | 0.01 | 0.05 | 0.03 | 0.08 | 0.11 |
| 6 | 32757297 | rs9276711 | 0.265 | 0.854 | -0.876 | 0.926 | 0.037 | 0.042 | 0.042 | 0 | 0.44 | 0.35 | 0.28 | 0.33 | 0.34 | 0.36 |
| 6 | 32757453 | rs2621390 | -0.993 | 0.867 | -0.876 | 0.894 | 0.037 | 0.014 | 0.014 | 0 | 0.77 | 0.7 | 0.66 | 0.73 | 0.71 | 0.83 |
| 6 | 32757737 | rs2857165 | -1.22 | 0.867 | -0.876 | 0.894 | 0.001 | -0.001 | -0.001 | 0 | 0.64 | 0.69 | 0.62 | 0.71 | 0.63 | 0.72 |
| 6 | 32757938 | rs2621388 | -1.007 | 0.867 | -0.876 | 0.894 | 0.038 | 0.014 | 0.014 | 0 | 0.77 | 0.7 | 0.66 | 0.73 | 0.71 | 0.83 |
| 6 | 32758219 | rs2857164 | -1.22 | 0.867 | -0.876 | 0.894 | 0.001 | -0.001 | -0.001 | 0.927 | 0.64 | 0.69 | 0.62 | 0.71 | 0.63 | 0.72 |
| 6 | 32758394 | rs7758736 | 2.938 | 0.867 | -0.876 | 0.894 | -0.018 | 0.033 | 0.033 | 0.127 | 0.14 | 0.1 | 0.09 | 0.03 | 0.17 | 0.11 |
| 6 | 32758571 | rs1158785 | -1.23 | 0.769 | -0.925 | 0.929 | 0.001 | -0.001 | -0.001 | 0.867 | 0.64 | 0.69 | 0.62 | 0.71 | 0.63 | 0.72 |
| 6 | 32759273 | rs2621384 | -1.248 | 0.767 | -0.872 | 0.928 | -0.006 | 0.002 | 0.002 | 2.116 | 0.64 | 0.74 | 0.62 | 0.71 | 0.63 | 0.72 |
| 6 | 32759297 | rs2857161 | -1.259 | 0.771 | -0.988 | 0.93 | 0 | -0.001 | -0.001 | 0 | 0.65 | 0.7 | 0.63 | 0.71 | 0.64 | 0.72 |
| 6 | 32759448 | rs3948793 | 0.247 | 0.712 | -1.008 | 0.923 | 0.036 | 0.044 | 0.044 | 0 | 0.44 | 0.36 | 0.29 | 0.32 | 0.34 | 0.36 |
| 6 | 32761776 | rs2157081 | -0.674 | 0.674 | -0.975 | 0.871 | 0.045 | 0.012 | 0.012 | 0 | 0.75 | 0.69 | 0.61 | 0.69 | 0.68 | 0.82 |
| 6 | 32762103 | rs2157079 | -0.674 | 0.674 | -0.975 | 0.871 | 0.045 | 0.012 | 0.012 | 0 | 0.75 | 0.69 | 0.61 | 0.69 | 0.68 | 0.82 |
| 6 | 32762307 | rs2067098 | -0.77 | 0.666 | -0.979 | 0.867 | 0.043 | 0.011 | 0.011 | 0 | 0.75 | 0.69 | 0.61 | 0.69 | 0.67 | 0.82 |
| 6 | 32762391 | rs9276722 | 3.227 | 0.624 | -1.026 | 0.841 | -0.003 | 0.025 | 0.025 | 0.239 | 0.13 | 0 | 0.05 | 0.03 | 0.08 | 0.11 |
| 6 | 32763110 | rs2621377 | -0.964 | 0.641 | -0.99 | 0.875 | -0.002 | 0 | 0 | 0 | 0.63 | 0.69 | 0.56 | 0.67 | 0.59 | 0.71 |
| 6 | 32763514 | rs2857151 | -0.636 | 0.692 | -0.92 | 0.869 | 0.044 | 0.012 | 0.012 | 0 | 0.75 | 0.69 | 0.61 | 0.69 | 0.67 | 0.82 |
| 6 | 32763821 | rs2857150 | -0.636 | 0.692 | -0.92 | 0.869 | 0.044 | 0.012 | 0.012 | 0 | 0.75 | 0.69 | 0.61 | 0.69 | 0.67 | 0.82 |
| 6 | 32763888 | rs9276726 | 0.262 | 0.528 | -0.917 | 0.88 | 0.033 | 0.027 | 0.027 | 0 | 0.44 | 0.37 | 0.23 | 0.32 | 0.33 | 0.33 |
| 6 | 32764970 | rs9986640 | 3.255 | 0.036 | -0.917 | 0.648 | -0.026 | -0.006 | -0.006 | 0 | 0.08 | 0.07 | 0.02 | 0.03 | 0 | 0.02 |
| 6 | 32765182 | rs7767167 | -2.342 | 0.739 | -0.953 | 0.569 | 0.02 | -0.022 | -0.022 | 0 | 0.92 | 0.91 | 0.94 | 0.97 | 0.89 | 0.91 |
| 6 | 32765451 | rs1383261 | 0.008 | 0.431 | -1.012 | 0.612 | -0.012 | -0.009 | -0.009 | 0 | 0.46 | 0.54 | 0.25 | 0.32 | 0.34 | 0.35 |
| 6 | 32765671 | rs6932969 | 0.008 | 0.431 | -1.012 | 0.612 | -0.012 | -0.009 | -0.009 | 0 | 0.46 | 0.54 | 0.25 | 0.32 | 0.34 | 0.35 |
| 6 | 32766593 | rs7383606 | 0.008 | 0.431 | -1.012 | 0.612 | -0.012 | -0.009 | -0.009 | 8.292 | 0.46 | 0.54 | 0.25 | 0.32 | 0.34 | 0.35 |
| 6 | 32766854 | rs9276734 | 0.008 | 0.431 | -1.012 | 0.612 | -0.012 | -0.009 | -0.009 | 0 | 0.46 | 0.54 | 0.25 | 0.32 | 0.34 | 0.35 |
| 6 | 32767091 | rs2621371 | -0.721 | 0.489 | -1.038 | 0.667 | 0.044 | 0.012 | 0.012 | 0 | 0.75 | 0.68 | 0.61 | 0.69 | 0.67 | 0.82 |
| 6 | 32767496 | rs7382714 | 0.038 | 0.489 | -1.038 | 0.667 | -0.012 | -0.009 | -0.009 | 1.066 | 0.46 | 0.54 | 0.25 | 0.32 | 0.34 | 0.35 |
| 6 | 32767620 | rs4947350 | 1.376 | 0.269 | -1.14 | 0.561 | 0.008 | -0.008 | -0.008 | 0 | 0.17 | 0.33 | 0.11 | 0.12 | 0.17 | 0.06 |
| 6 | 32767673 | rs7381376 | -0.036 | 0.349 | -1.06 | 0.619 | -0.012 | -0.009 | -0.009 | 0 | 0.46 | 0.54 | 0.25 | 0.32 | 0.34 | 0.35 |
| 6 | 32768721 | rs2621367 | 1.335 | 0.343 | -1.063 | 0.615 | -0.019 | -0.051 | -0.051 | 0.123 | 0.3 | 0.14 | 0.36 | 0.37 | 0.34 | 0.47 |
| 6 | 32769926 | rs2199874 | 1.305 | 0.342 | -1.061 | 0.615 | -0.001 | -0.002 | -0.002 | 0.381 | 0.31 | 0.31 | 0.37 | 0.38 | 0.34 | 0.49 |
| 6 | 32770482 | rs6899857 | -0.039 | 0.342 | -1.061 | 0.615 | -0.012 | -0.009 | -0.009 | 0.083 | 0.46 | 0.54 | 0.25 | 0.32 | 0.34 | 0.35 |
| 6 | 32770808 | rs2621358 | 1.393 | 0.342 | -1.061 | 0.615 | -0.02 | -0.052 | -0.052 | 1.348 | 0.29 | 0.14 | 0.36 | 0.37 | 0.34 | 0.47 |
| 6 | 32772636 | rs2857143 | -0.769 | 0.343 | -1.061 | 0.615 | 0.044 | 0.012 | 0.012 | 0 | 0.75 | 0.68 | 0.61 | 0.69 | 0.67 | 0.82 |
| 6 | 32773172 | rs2199870 | -0.769 | 0.343 | -1.061 | 0.615 | 0.044 | 0.012 | 0.012 | 0 | 0.75 | 0.68 | 0.61 | 0.69 | 0.67 | 0.82 |
| 6 | 32774465 | rs6912414 | 0.094 | 0.345 | -1.059 | 0.617 | 0.033 | 0.027 | 0.027 | 0 | 0.44 | 0.37 | 0.23 | 0.31 | 0.33 | 0.33 |
| 6 | 32775809 | rs7382619 | -0.008 | 0.349 | -1.072 | 0.619 | -0.012 | -0.01 | -0.01 | 0 | 0.46 | 0.54 | 0.25 | 0.32 | 0.34 | 0.34 |
| 6 | 32775821 | rs2857135 | -0.876 | 0.35 | -1.072 | 0.619 | 0.043 | 0.011 | 0.011 | 0 | 0.75 | 0.68 | 0.61 | 0.69 | 0.67 | 0.82 |
| 6 | 32776087 | rs7382649 | -0.008 | 0.349 | -1.072 | 0.619 | -0.012 | -0.01 | -0.01 | 0 | 0.46 | 0.54 | 0.25 | 0.32 | 0.34 | 0.34 |
| 6 | 32776583 | rs2621338 | 1.388 | 0.351 | -1.074 | 0.62 | -0.02 | -0.052 | -0.052 | 0.859 | 0.29 | 0.14 | 0.36 | 0.37 | 0.34 | 0.47 |
| 6 | 32776832 | rs2857128 | -0.862 | 0.372 | -1.055 | 0.623 | 0.043 | 0.011 | 0.011 | 0 | 0.75 | 0.68 | 0.61 | 0.69 | 0.67 | 0.82 |
| 6 | 32776976 | rs2621336 | -0.862 | 0.371 | -1.056 | 0.623 | 0.043 | 0.011 | 0.011 | 0 | 0.75 | 0.68 | 0.61 | 0.69 | 0.67 | 0.82 |
| 6 | 32777140 | rs2857126 | -0.737 | 0.378 | -1.053 | 0.626 | 0.043 | 0.011 | 0.011 | 0 | 0.75 | 0.68 | 0.61 | 0.69 | 0.67 | 0.82 |
| 6 | 32777752 | rs12207915 | 2.211 | 0.502 | -1.141 | 0.885 | -0.017 | -0.027 | -0.027 | 0 | 0.05 | 0.01 | 0.07 | 0.18 | 0.07 | 0.12 |

| | | | | | | | | | | | | | | | | |
|---|---|---|---|---|---|---|---|---|---|---|---|---|---|---|---|---|
| 6 | 32777900 | rs2857118 | 1.424 | 0.498 | -1.022 | 0.741 | -0.02 | -0.052 | -0.052 | 0 | 0.29 | 0.14 | 0.36 | 0.37 | 0.34 | 0.47 |
| 6 | 32778281 | rs2857115 | 1.724 | 0.532 | -1.017 | 0.751 | -0.014 | -0.045 | -0.045 | 0 | 0.26 | 0.13 | 0.35 | 0.36 | 0.31 | 0.38 |
| 6 | 32778531 | rs7383433 | -0.756 | 0.572 | -0.812 | 0.749 | 0.043 | 0.011 | 0.011 | 0 | 0.75 | 0.68 | 0.61 | 0.69 | 0.67 | 0.82 |
| 6 | 32779996 | rs2857114 | -0.361 | 0.567 | -0.801 | 0.747 | 0.057 | 0.012 | 0.012 | 0 | 0.73 | 0.51 | 0.59 | 0.68 | 0.67 | 0.8 |
| 6 | 32780317 | rs2621332 | -0.754 | 0.58 | -0.823 | 0.752 | 0.043 | 0.011 | 0.011 | 0 | 0.75 | 0.68 | 0.61 | 0.69 | 0.67 | 0.82 |
| 6 | 32780470 | rs2621331 | -0.149 | 0.509 | -0.933 | 0.723 | 0.03 | 0 | 0 | 0 | 0.69 | 0.5 | 0.58 | 0.66 | 0.67 | 0.7 |
| 6 | 32780724 | rs11244 | 0.497 | 0.503 | -1.173 | 0.71 | 0.018 | 0.021 | 0.021 | 0 | 0.31 | 0.28 | 0.19 | 0.23 | 0.28 | 0.28 |
| 6 | 32781776 | rs2856997 | -0.044 | 0.527 | -0.922 | 0.812 | -0.002 | 0.009 | 0.009 | 0 | 0.43 | 0.51 | 0.45 | 0.49 | 0.44 | 0.62 |
| 6 | 32782149 | rs17501267 | 1.777 | 0.588 | -1.011 | 0.835 | -0.001 | 0.002 | 0.002 | 0 | 0.06 | 0.02 | 0.02 | 0.06 | 0.05 | 0.03 |
| 6 | 32782387 | rs2071475 | 1.182 | 0.584 | -1.079 | 0.894 | -0.008 | -0.011 | -0.011 | 1.578 | 0.16 | 0.13 | 0.3 | 0.24 | 0.22 | 0.29 |
| 6 | 32782582 | rs2071474 | 1.359 | 0.605 | -1.013 | 0.878 | -0.016 | -0.052 | -0.052 | 0 | 0.29 | 0.14 | 0.35 | 0.37 | 0.31 | 0.46 |
| 6 | 32782605 | rs2071473 | 0.5 | 0.595 | -0.859 | 0.869 | 0 | 0.012 | 0.012 | 0 | 0.36 | 0.4 | 0.41 | 0.43 | 0.39 | 0.57 |
| 6 | 32783365 | rs2857111 | 1.381 | 0.617 | -0.845 | 1.154 | 0.008 | 0.042 | 0.042 | 0 | 0.06 | 0.09 | 0.05 | 0.18 | 0.08 | 0.05 |
| 6 | 32783405 | rs1383258 | 1.831 | 0.607 | -0.836 | 1.148 | -0.001 | 0.009 | 0.009 | 0 | 0.07 | 0.19 | 0.05 | 0.06 | 0.05 | 0.1 |
| 6 | 32783896 | rs2621326 | 0.056 | 0.616 | -0.86 | 1.151 | 0 | -0.001 | -0.001 | 0 | 0.41 | 0.42 | 0.45 | 0.61 | 0.44 | 0.61 |
| 6 | 32784620 | rs2071472 | 1.338 | 0.635 | -0.944 | 1.17 | -0.016 | -0.052 | -0.052 | 0 | 0.29 | 0.14 | 0.35 | 0.37 | 0.31 | 0.46 |
| 6 | 32784783 | rs2071469 | 0.663 | 0.639 | -0.934 | 0.887 | -0.006 | -0.021 | -0.021 | 0 | 0.35 | 0.22 | 0.4 | 0.55 | 0.39 | 0.51 |
| 6 | 32785035 | rs2071468 | 1.381 | 0.475 | -0.797 | 0.877 | 0.008 | 0.042 | 0.042 | 0 | 0.06 | 0.09 | 0.05 | 0.19 | 0.08 | 0.05 |
| 6 | 32785275 | rs7742120 | 1.381 | 0.475 | -0.797 | 0.877 | 0.008 | 0.042 | 0.042 | 0 | 0.06 | 0.09 | 0.05 | 0.19 | 0.08 | 0.05 |
| 6 | 32785284 | rs7762120 | 1.381 | 0.475 | -0.797 | 0.877 | 0.008 | 0.042 | 0.042 | 0 | 0.06 | 0.09 | 0.05 | 0.19 | 0.08 | 0.05 |
| 6 | 32785515 | rs2857107 | 1.469 | 0.326 | -0.793 | 0.528 | -0.018 | -0.001 | -0.001 | 0 | 0.18 | 0.27 | 0.16 | 0.18 | 0.14 | 0.09 |
| 6 | 32786068 | rs17429444 | 2.217 | 0.255 | -1.097 | 0.385 | 0.041 | 0.078 | 0.078 | 0 | 0.17 | 0.01 | 0.08 | 0.02 | 0.11 | 0.11 |
| 6 | 32786882 | rs3763355 | 2.348 | 0.442 | -1.005 | 0.628 | -0.021 | -0.024 | -0.024 | 0 | 0.09 | 0 | 0.13 | 0.08 | 0.08 | 0.07 |
| 6 | 32786917 | rs3763354 | -1.795 | 0.34 | -0.754 | 0.621 | -0.004 | 0.004 | 0.004 | 0.574 | 0.81 | 0.89 | 0.81 | 0.89 | 0.8 | 0.84 |
| 6 | 32787036 | rs1894407 | 0.005 | 0.061 | -0.369 | 1.095 | -0.001 | 0.004 | 0.004 | 0 | 0.38 | 0.42 | 0.43 | 0.58 | 0.4 | 0.49 |
| 6 | 32787056 | rs1894406 | 0.395 | 0.08 | -0.352 | 1.127 | 0 | -0.027 | -0.027 | 0 | 0.35 | 0.23 | 0.39 | 0.55 | 0.36 | 0.49 |
| 6 | 32788707 | rs2621323 | 0.202 | 0.037 | -0.258 | 0.797 | -0.005 | 0.01 | 0.01 | 0 | 0.36 | 0.41 | 0.4 | 0.43 | 0.34 | 0.56 |
| 6 | 32788712 | rs2621322 | 0.839 | -0.024 | -0.307 | 0.816 | -0.003 | -0.015 | -0.015 | 0 | 0.16 | 0.14 | 0.29 | 0.34 | 0.19 | 0.33 |
| 6 | 32788878 | rs9784876 | 1.149 | -0.028 | -0.299 | 0.789 | 0.024 | -0.014 | -0.014 | 0 | 0.07 | 0.26 | 0.07 | 0.03 | 0.07 | 0.1 |
| 6 | 32789480 | rs2621321 | 0.768 | 0.377 | -0.132 | 1.008 | 0.003 | -0.012 | -0.012 | 0 | 0.29 | 0.14 | 0.34 | 0.27 | 0.27 | 0.43 |
| 6 | 32791299 | rs2857103 | 0.475 | 0.431 | -0.303 | 0.599 | -0.001 | 0.021 | 0.021 | 0 | 0.32 | 0.4 | 0.39 | 0.4 | 0.34 | 0.46 |
| 6 | 32792659 | rs13209654 | 3.036 | 0.332 | -0.408 | 0.519 | -0.003 | 0.024 | 0.024 | 0.175 | 0.12 | 0 | 0.05 | 0.03 | 0.09 | 0.11 |
| 6 | 32792973 | rs1894411 | 0.829 | 0.628 | -0.369 | 0.665 | -0.017 | 0 | 0 | 1.312 | 0.18 | 0.27 | 0.16 | 0.18 | 0.14 | 0.09 |
| 6 | 32794676 | rs2857101 | 0.706 | 1.027 | -0.824 | 0.972 | -0.004 | -0.062 | -0.062 | 0 | 0.28 | 0.12 | 0.3 | 0.36 | 0.27 | 0.44 |
| 6 | 32795032 | rs10484565 | 0.658 | 1.119 | -0.845 | 0.977 | -0.012 | -0.012 | -0.012 | 0 | 0.07 | 0.03 | 0.03 | 0.09 | 0.08 | 0.03 |
| 6 | 32795965 | rs241456 | 0.552 | 1.165 | -0.618 | 1.071 | 0.003 | -0.035 | -0.035 | 0 | 0.29 | 0.16 | 0.31 | 0.37 | 0.28 | 0.44 |
| 6 | 32796144 | rs241454 | 0.542 | 1.162 | -0.62 | 1.069 | 0.003 | -0.035 | -0.035 | 0 | 0.29 | 0.16 | 0.31 | 0.37 | 0.28 | 0.44 |
| 6 | 32796226 | rs241453 | 0.601 | 1.162 | -0.621 | 1.069 | -0.004 | -0.055 | -0.055 | 0 | 0.28 | 0.12 | 0.31 | 0.36 | 0.27 | 0.43 |
| 6 | 32796480 | rs241451 | 0.452 | 1.212 | -0.409 | 1.068 | 0 | 0.001 | 0.001 | 0 | 0.31 | 0.28 | 0.35 | 0.4 | 0.34 | 0.45 |
| 6 | 32796653 | rs241449 | 0.55 | 1.185 | -0.463 | 1.063 | 0.003 | -0.035 | -0.035 | 0 | 0.29 | 0.16 | 0.31 | 0.37 | 0.28 | 0.44 |
| 6 | 32796685 | rs241448 | 0.465 | 1.158 | -0.281 | 1.049 | 0.003 | -0.01 | -0.01 | 0 | 0.29 | 0.19 | 0.32 | 0.37 | 0.28 | 0.44 |
| 6 | 32796751 | rs241447 | 0.546 | 1.201 | -0.72 | 1.083 | 0.003 | -0.036 | -0.036 | 0 | 0.29 | 0.16 | 0.31 | 0.37 | 0.28 | 0.44 |
| 6 | 32796967 | rs241446 | 0.565 | 1.215 | -0.714 | 1.073 | 0.003 | -0.035 | -0.035 | 0.009 | 0.29 | 0.16 | 0.31 | 0.37 | 0.28 | 0.45 |
| 6 | 32797072 | rs241445 | 0.565 | 1.215 | -0.714 | 1.073 | 0.003 | -0.035 | -0.035 | 0 | 0.29 | 0.16 | 0.31 | 0.37 | 0.28 | 0.44 |
| 6 | 32797115 | rs241443 | 0.491 | 1.244 | -0.656 | 1.08 | 0 | 0.002 | 0.002 | 0 | 0.31 | 0.28 | 0.35 | 0.4 | 0.34 | 0.45 |
| 6 | 32797297 | rs241441 | 0.492 | 1.102 | -1.036 | 1.017 | 0.005 | -0.033 | -0.033 | 1.383 | 0.29 | 0.16 | 0.3 | 0.36 | 0.28 | 0.45 |
| 6 | 32797361 | rs241440 | 0.495 | 1.102 | -1.11 | 1.017 | 0.001 | -0.051 | -0.051 | 0 | 0.29 | 0.13 | 0.3 | 0.36 | 0.28 | 0.45 |
| 6 | 32797488 | rs4148874 | -0.723 | 1.418 | -0.988 | 0.689 | 0.034 | 0.012 | 0.012 | 0 | 0.84 | 0.7 | 0.73 | 0.83 | 0.75 | 0.84 |
| 6 | 32797620 | rs241438 | 0.55 | 0.962 | -1.634 | 0.717 | -0.01 | -0.046 | -0.046 | 0.419 | 0.31 | 0.23 | 0.38 | 0.52 | 0.36 | 0.44 |
| 6 | 32797684 | rs241437 | 0.196 | 0.978 | -1.617 | 0.709 | -0.014 | -0.042 | -0.042 | 3.851 | 0.35 | 0.24 | 0.4 | 0.54 | 0.39 | 0.53 |
| 6 | 32797876 | rs241436 | 0.197 | 0.761 | -1.72 | 0.604 | 0.042 | -0.013 | -0.013 | 0 | 0.49 | 0.22 | 0.46 | 0.55 | 0.49 | 0.52 |
| 6 | 32798283 | rs9380326 | 0.284 | 1.121 | -1.512 | 0.612 | -0.029 | -0.009 | -0.009 | 0 | 0.08 | 0.06 | 0.11 | 0.1 | 0.13 | 0.07 |
| 6 | 32798731 | rs1015166 | -0.074 | 1.08 | -1.881 | 0.665 | -0.049 | -0.016 | -0.016 | 0.318 | 0.26 | 0.18 | 0.2 | 0.29 | 0.32 | 0.2 |
| 6 | 32798795 | rs241433 | -0.857 | 0.847 | -1.818 | 0.644 | -0.002 | -0.003 | -0.003 | 0 | 0.56 | 0.48 | 0.54 | 0.55 | 0.59 | 0.58 |
| 6 | 32800224 | rs2228397 | 1.202 | 0.84 | -1.739 | 0.626 | 0.02 | 0 | 0 | 0 | 0.27 | 0.2 | 0.31 | 0.25 | 0.22 | 0.28 |
| 6 | 32800412 | rs1800454 | 1.665 | 1.604 | -1.378 | 0.759 | 0.004 | 0.003 | 0.003 | 0.155 | 0.1 | 0.15 | 0.14 | 0.19 | 0.15 | 0.09 |
| 6 | 32802203 | rs241432 | -0.524 | 1.557 | -1.484 | 0.635 | -0.002 | -0.003 | -0.003 | 0 | 0.56 | 0.48 | 0.54 | 0.55 | 0.58 | 0.58 |
| 6 | 32802807 | rs4148872 | 0.977 | 1.75 | -1.464 | 0.513 | -0.004 | 0 | 0 | 0 | 0.08 | 0.05 | 0.11 | 0.1 | 0.17 | 0.07 |
| 6 | 32802820 | rs241430 | -0.274 | 1.382 | -1.35 | 0.626 | -0.001 | 0.001 | 0.001 | 0.455 | 0.55 | 0.47 | 0.54 | 0.56 | 0.6 | 0.57 |
| 6 | 32803316 | rs4148871 | -1.273 | 0.902 | -1.028 | 0.53 | 0.004 | 0.02 | 0.02 | 0.483 | 0.73 | 0.87 | 0.78 | 0.88 | 0.78 | 0.74 |
| 6 | 32803840 | rs241429 | 0.329 | 1.133 | -0.862 | 0.886 | 0.001 | 0 | 0 | 0 | 0.65 | 0.35 | 0.53 | 0.55 | 0.58 | 0.62 |
| 6 | 32804070 | rs241428 | 1.776 | 0.522 | -0.671 | 0.735 | -0.006 | -0.004 | -0.004 | 0 | 0.07 | 0.06 | 0.06 | 0.03 | 0.04 | 0.02 |
| 6 | 32804217 | rs3819714 | 0.478 | 0.159 | -0.832 | 0.775 | 0 | 0 | 0 | 0 | 0.47 | 0.48 | 0.45 | 0.34 | 0.34 | 0.36 |
| 6 | 32804219 | rs3819715 | 0.478 | 0.159 | -0.832 | 0.775 | 0 | 0 | 0 | 0 | 0.47 | 0.48 | 0.45 | 0.34 | 0.34 | 0.36 |
| 6 | 32804414 | rs241427 | -1.087 | 0.526 | -1.428 | 0.624 | 0.002 | 0.001 | 0.001 | 0 | 0.66 | 0.78 | 0.64 | 0.67 | 0.67 | 0.66 |
| 6 | 32804553 | rs241426 | 0.571 | 0.069 | -1.504 | 0.34 | 0.03 | 0.047 | 0.047 | 0 | 0.45 | 0.74 | 0.58 | 0.66 | 0.59 | 0.62 |
| 6 | 32804570 | rs3819720 | 0.004 | 0.514 | -1.699 | 0.573 | -0.061 | 0.009 | 0.009 | 0 | 0.28 | 0.18 | 0.3 | 0.21 | 0.35 | 0.33 |
| 6 | 32804909 | rs241425 | -1.363 | 0.255 | -1.645 | 0.623 | -0.025 | 0.01 | 0.01 | 0 | 0.67 | 0.79 | 0.69 | 0.74 | 0.58 | 0.71 |
| 6 | 32804934 | rs241424 | -0.193 | 0.039 | -1.825 | 0.872 | 0.009 | -0.001 | -0.001 | 3.92 | 0.5 | 0.65 | 0.6 | 0.51 | 0.46 | 0.49 |

| | | | | | | | | | | | | | | | | |
|---|---|---|---|---|---|---|---|---|---|---|---|---|---|---|---|---|
| 6 | 32805049 | rs2239701 | -0.42 | -0.393 | -1.692 | 0.88 | 0.001 | 0.001 | 0.001 | 0 | 0.51 | 0.42 | 0.51 | 0.54 | 0.45 | 0.54 |
| 6 | 32805470 | rs2071465 | -0.689 | -0.385 | -1.526 | 0.663 | 0.002 | 0.043 | 0.043 | 0 | 0.52 | 0.4 | 0.32 | 0.34 | 0.43 | 0.48 |
| 6 | 32806121 | rs2071544 | 0.199 | -0.406 | -1.473 | 0.672 | -0.006 | -0.006 | -0.006 | 0 | 0.55 | 0.4 | 0.48 | 0.6 | 0.48 | 0.59 |
| 6 | 32806391 | rs4148870 | 0.444 | -0.352 | -1.385 | 0.736 | 0.005 | 0.022 | 0.022 | 0 | 0.48 | 0.37 | 0.46 | 0.5 | 0.44 | 0.51 |
| 6 | 32806461 | rs2071552 | 0.346 | -0.351 | -1.383 | 0.739 | 0 | 0.02 | 0.02 | 0 | 0.49 | 0.38 | 0.48 | 0.51 | 0.47 | 0.55 |
| 6 | 32806584 | rs4148868 | 0.364 | -0.363 | -1.385 | 0.698 | -0.001 | 0.041 | 0.041 | 0 | 0.49 | 0.37 | 0.44 | 0.44 | 0.47 | 0.52 |
| 6 | 32806786 | rs4713598 | 0.737 | -0.394 | -1.396 | 0.687 | -0.002 | 0.034 | 0.034 | 0 | 0.45 | 0.34 | 0.42 | 0.44 | 0.43 | 0.47 |
| 6 | 32808232 | rs3763349 | -0.055 | -0.413 | -1.134 | 0.434 | 0.002 | -0.002 | -0.002 | 0.111 | 0.55 | 0.58 | 0.54 | 0.52 | 0.54 | 0.55 |
| 6 | 32809848 | rs9357155 | 1.038 | -0.128 | -1.106 | 0.201 | -0.01 | -0.006 | -0.006 | 0 | 0.09 | 0.06 | 0.18 | 0.15 | 0.14 | 0.18 |
| 6 | 32810443 | rs9276810 | 0.393 | -0.165 | -1.104 | 0.172 | 0.003 | 0.039 | 0.039 | 0 | 0.48 | 0.37 | 0.44 | 0.45 | 0.46 | 0.48 |
| 6 | 32811383 | rs6924102 | 0.621 | -0.144 | -1.092 | 0.181 | -0.004 | 0.03 | 0.03 | 0 | 0.44 | 0.34 | 0.41 | 0.45 | 0.43 | 0.44 |
| 6 | 32811629 | rs2071543 | 0.877 | -0.18 | -1.372 | 0.276 | -0.007 | -0.004 | -0.004 | 0 | 0.1 | 0.1 | 0.19 | 0.17 | 0.15 | 0.19 |
| 6 | 32812528 | rs2071463 | 1.327 | -0.276 | -1.565 | 0.369 | 0.012 | -0.093 | -0.093 | 0 | 0.2 | 0.02 | 0.21 | 0.26 | 0.15 | 0.25 |
| 6 | 32812858 | rs2071541 | 0.43 | -0.537 | -1.321 | 0.308 | 0.025 | -0.005 | -0.005 | 0 | 0.18 | 0.07 | 0.1 | 0.13 | 0.13 | 0.2 |
| 6 | 32813279 | rs1057373 | 0.656 | -0.544 | -1.337 | 0.307 | 0.049 | -0.022 | -0.022 | 0 | 0.17 | 0.03 | 0.08 | 0.13 | 0.09 | 0.14 |
| 6 | 32813852 | rs13215 | -0.169 | -0.424 | -0.888 | 0.353 | -0.003 | -0.025 | -0.025 | 0 | 0.07 | 0.2 | 0.06 | 0 | 0.07 | 0.05 |
| 6 | 32814659 | rs4711312 | -0.29 | -0.44 | -0.9 | 0.355 | 0.006 | 0.003 | 0.003 | 0 | 0.23 | 0.24 | 0.15 | 0.13 | 0.16 | 0.21 |
| 6 | 32815488 | rs16871026 | 0.596 | -0.229 | -0.788 | 0.34 | -0.019 | -0.027 | -0.027 | 0 | 0.05 | 0.17 | 0.05 | 0 | 0.03 | 0 |
| 6 | 32815516 | rs17213812 | -0.362 | -0.224 | -0.787 | 0.344 | -0.006 | -0.02 | -0.02 | 0 | 0.07 | 0.21 | 0.07 | 0.01 | 0.07 | 0.07 |
| 6 | 32816663 | rs16871027 | 0.487 | -0.219 | -0.792 | 0.35 | -0.021 | -0.025 | -0.025 | 0 | 0.06 | 0.17 | 0.05 | 0 | 0.03 | 0 |
| 6 | 32816700 | rs2071482 | -0.126 | -0.276 | -0.708 | 0.325 | -0.007 | -0.003 | -0.003 | 0.067 | 0.22 | 0.29 | 0.14 | 0.18 | 0.16 | 0.26 |
| 6 | 32816958 | rs4148882 | -0.853 | -0.035 | -0.796 | 0.344 | -0.07 | -0.087 | -0.087 | 0.304 | 0.72 | 0.94 | 0.69 | 0.68 | 0.65 | 0.67 |
| 6 | 32816998 | rs12527715 | -0.278 | -0.148 | -0.759 | 0.231 | 0.002 | 0.001 | 0.001 | 0 | 0.25 | 0.29 | 0.15 | 0.21 | 0.18 | 0.28 |
| 6 | 32817774 | rs2395269 | 0.272 | -0.025 | -0.874 | 0.3 | 0.02 | -0.004 | -0.004 | 0 | 0.2 | 0.12 | 0.1 | 0.21 | 0.14 | 0.28 |
| 6 | 32818678 | rs2071538 | 1.896 | 0.153 | -1.165 | 0.31 | -0.066 | 0.041 | 0.041 | 0 | 0.17 | 0.01 | 0.14 | 0.05 | 0.23 | 0.12 |
| 6 | 32819478 | rs4148879 | 0.561 | -0.005 | -1.042 | 0.405 | 0.045 | -0.013 | -0.013 | 0 | 0.18 | 0.08 | 0.08 | 0.2 | 0.11 | 0.21 |
| 6 | 32819517 | rs2284190 | 0.899 | 0.098 | -1.028 | 0.42 | -0.007 | -0.005 | -0.005 | 0 | 0.1 | 0.09 | 0.19 | 0.17 | 0.15 | 0.19 |
| 6 | 32819865 | rs2071481 | 0.457 | -0.053 | -1.056 | 0.344 | 0.057 | -0.006 | -0.006 | 0 | 0.2 | 0.08 | 0.08 | 0.2 | 0.11 | 0.21 |
| 6 | 32822186 | rs4148878 | 2.132 | 0.567 | -0.873 | 0.427 | -0.026 | 0.025 | 0.025 | 0 | 0.06 | 0.02 | 0.02 | 0.02 | 0.06 | 0.01 |
| 6 | 32822269 | rs1351383 | 0.482 | 0.442 | -0.548 | 0.453 | 0.009 | 0.013 | 0.013 | 0 | 0.5 | 0.49 | 0.38 | 0.39 | 0.4 | 0.43 |
| 6 | 32822858 | rs4713600 | 0.397 | 0.438 | -0.514 | 0.453 | 0.025 | 0.036 | 0.036 | 0 | 0.5 | 0.46 | 0.37 | 0.39 | 0.4 | 0.41 |
| 6 | 32823257 | rs3763346 | 0.188 | 0.421 | -0.521 | 0.443 | 0 | 0 | 0 | 0 | 0.53 | 0.51 | 0.4 | 0.4 | 0.43 | 0.44 |
| 6 | 32823567 | rs991760 | 1.352 | 0.346 | -0.56 | 0.428 | 0.023 | -0.012 | -0.012 | 0.556 | 0.12 | 0.07 | 0.09 | 0.17 | 0.06 | 0.17 |
| 6 | 32824318 | rs2071534 | 0.176 | 0.355 | -0.553 | 0.424 | 0 | 0 | 0 | 0 | 0.53 | 0.51 | 0.4 | 0.4 | 0.43 | 0.44 |
| 6 | 32825090 | rs17587 | 1.716 | 0.301 | -0.514 | 0.377 | 0.007 | 0.027 | 0.027 | 0.004 | 0.31 | 0.23 | 0.22 | 0.21 | 0.26 | 0.19 |
| 6 | 32825348 | rs2071477 | 0.387 | 0.366 | -0.498 | 0.412 | 0.012 | 0.017 | 0.017 | 0 | 0.5 | 0.49 | 0.37 | 0.39 | 0.4 | 0.41 |
| 6 | 32825379 | rs2071476 | 0.178 | 0.366 | -0.498 | 0.412 | 0 | 0 | 0 | 0 | 0.53 | 0.51 | 0.4 | 0.4 | 0.43 | 0.44 |
| 6 | 32826233 | rs20547 | 1.192 | 0.303 | -0.519 | 0.632 | 0.013 | 0.008 | 0.008 | 0 | 0.06 | 0.02 | 0.11 | 0.21 | 0.03 | 0.16 |
| 6 | 32826450 | rs6930981 | 0.067 | 0.47 | -0.414 | 0.003 | 0 | 0 | 0 | 0 | 0.53 | 0.51 | 0.4 | 0.4 | 0.43 | 0.44 |
| 6 | 32829276 | rs9276820 | 0.084 | 0.531 | -0.342 | -0.053 | 0 | 0 | 0 | 0 | 0.53 | 0.51 | 0.4 | 0.39 | 0.43 | 0.44 |
| 6 | 32829714 | rs9276825 | 0.084 | 0.531 | -0.342 | -0.053 | 0 | 0 | 0 | 0 | 0.53 | 0.51 | 0.4 | 0.39 | 0.43 | 0.44 |
| 6 | 32830013 | rs7767288 | 0.379 | 0.531 | -0.342 | -0.054 | 0.009 | 0.013 | 0.013 | 0 | 0.5 | 0.49 | 0.38 | 0.39 | 0.4 | 0.43 |
| 6 | 32830034 | rs7766900 | 0.084 | 0.531 | -0.342 | -0.053 | 0 | 0 | 0 | 0 | 0.53 | 0.51 | 0.4 | 0.39 | 0.43 | 0.44 |
| 6 | 32832667 | rs3752430 | -0.562 | 0.643 | -0.142 | -0.046 | -0.008 | -0.008 | -0.008 | 0 | 0.65 | 0.69 | 0.58 | 0.67 | 0.57 | 0.64 |
| 6 | 32833646 | rs1383267 | 0.216 | 0.589 | -0.255 | -0.449 | 0.026 | 0.039 | 0.039 | 0 | 0.5 | 0.46 | 0.37 | 0.39 | 0.4 | 0.41 |
| 6 | 32834229 | rs1871667 | 0.28 | 0.589 | -0.254 | -0.449 | 0.01 | 0.015 | 0.015 | 0 | 0.5 | 0.5 | 0.38 | 0.39 | 0.4 | 0.43 |
| 6 | 32834732 | rs1383266 | 1.639 | 0.692 | -0.426 | -0.456 | -0.05 | 0.038 | 0.038 | 0 | 0.2 | 0.02 | 0.16 | 0.07 | 0.26 | 0.13 |
| 6 | 32835258 | rs10484566 | 1.009 | -0.256 | -0.016 | -0.346 | 0.024 | 0.021 | 0.021 | 0 | 0.12 | 0.07 | 0.01 | 0.11 | 0.06 | 0.14 |
| 6 | 32835615 | rs6931062 | 0.228 | -0.115 | 0.058 | -0.285 | 0.003 | 0.008 | 0.008 | 0 | 0.38 | 0.39 | 0.36 | 0.28 | 0.34 | 0.27 |
| 6 | 32837696 | rs1580345 | 0.19 | -0.115 | 0.059 | -0.285 | 0.003 | 0.006 | 0.006 | 0 | 0.38 | 0.4 | 0.36 | 0.28 | 0.34 | 0.29 |
| 6 | 32838584 | rs6906376 | 0.19 | -0.115 | 0.059 | -0.285 | 0.003 | 0.006 | 0.006 | 0 | 0.38 | 0.4 | 0.36 | 0.28 | 0.34 | 0.29 |
| 6 | 32838664 | rs6929078 | 0.19 | -0.115 | 0.059 | -0.285 | 0.003 | 0.006 | 0.006 | 0 | 0.38 | 0.4 | 0.36 | 0.28 | 0.34 | 0.29 |
| 6 | 32838949 | rs7453903 | 0.19 | -0.115 | 0.059 | -0.285 | 0.003 | 0.006 | 0.006 | 0 | 0.38 | 0.4 | 0.36 | 0.28 | 0.34 | 0.29 |
| 6 | 32840139 | rs9276845 | 0.19 | -0.115 | 0.059 | -0.285 | 0.003 | 0.006 | 0.006 | 0 | 0.38 | 0.4 | 0.36 | 0.28 | 0.34 | 0.29 |
| 6 | 32843238 | rs9276864 | 0.201 | -0.238 | -0.129 | -0.272 | 0.004 | 0.008 | 0.008 | 0 | 0.38 | 0.39 | 0.36 | 0.28 | 0.34 | 0.27 |
| 6 | 32845873 | rs7757767 | 0.175 | -0.263 | -0.01 | -0.23 | 0.003 | 0.006 | 0.006 | 0 | 0.38 | 0.4 | 0.36 | 0.28 | 0.34 | 0.29 |
| 6 | 32846570 | rs2621426 | 0.212 | -0.263 | -0.015 | -0.232 | 0.003 | 0.008 | 0.008 | 0 | 0.38 | 0.4 | 0.36 | 0.28 | 0.34 | 0.27 |
| 6 | 32848168 | rs9276899 | 0.195 | -0.265 | 0.035 | -0.233 | -0.001 | -0.003 | -0.003 | 0.038 | 0.38 | 0.44 | 0.36 | 0.28 | 0.34 | 0.27 |
| 6 | 32848566 | rs9276900 | 0.195 | -0.267 | 0.11 | -0.227 | -0.009 | -0.037 | -0.037 | 0 | 0.38 | 0.46 | 0.29 | 0.22 | 0.34 | 0.24 |
| 6 | 32849639 | rs2170185 | 0.193 | -0.258 | -0.158 | -0.23 | -0.001 | -0.003 | -0.003 | 0 | 0.38 | 0.44 | 0.36 | 0.28 | 0.34 | 0.27 |
| 6 | 32850839 | rs9276909 | 0.93 | -0.273 | -0.147 | -0.245 | 0.008 | 0.029 | 0.029 | 0.004 | 0.31 | 0.23 | 0.22 | 0.21 | 0.26 | 0.19 |
| 6 | 32850850 | rs2127675 | 0.218 | -0.238 | -0.221 | -0.218 | -0.001 | -0.003 | -0.003 | 0 | 0.38 | 0.44 | 0.36 | 0.28 | 0.34 | 0.27 |
| 6 | 32851507 | rs4947259 | 0.186 | -0.226 | -0.218 | -0.212 | -0.002 | -0.004 | -0.004 | 0.014 | 0.38 | 0.45 | 0.37 | 0.28 | 0.34 | 0.29 |
| 6 | 32853042 | rs7767277 | -0.054 | -0.623 | -0.02 | -0.252 | -0.017 | -0.003 | -0.003 | 2.569 | 0.18 | 0.24 | 0.13 | 0.18 | 0.1 | 0.18 |
| 6 | 32853219 | rs35571244 | 0.846 | -0.602 | -0.08 | -0.344 | -0.002 | 0.002 | 0.002 | 0 | 0.08 | 0.06 | 0.02 | 0.02 | 0.06 | 0.02 |
| 6 | 32854697 | rs10046257 | -0.106 | -0.607 | -0.044 | -0.305 | -0.014 | -0.004 | -0.004 | 1.589 | 0.19 | 0.24 | 0.15 | 0.16 | 0.12 | 0.16 |
| 6 | 32855081 | rs10046277 | -0.052 | -0.608 | -0.045 | -0.305 | -0.016 | -0.017 | -0.017 | 0 | 0.19 | 0.24 | 0.08 | 0.1 | 0.11 | 0.16 |
| 6 | 32855174 | rs10046115 | -0.052 | -0.608 | -0.045 | -0.305 | -0.016 | -0.017 | -0.017 | 0.268 | 0.19 | 0.24 | 0.08 | 0.1 | 0.11 | 0.16 |
| 6 | 32855275 | rs7740137 | -0.167 | -0.621 | -0.041 | -0.309 | -0.018 | -0.017 | -0.017 | 0 | 0.19 | 0.24 | 0.08 | 0.11 | 0.11 | 0.17 |

| | | | | | | | | | | | | | | | | |
|---|---|---|---|---|---|---|---|---|---|---|---|---|---|---|---|---|
| 6 | 32855624 | rs6457688 | -0.049 | -0.62 | -0.041 | -0.309 | -0.016 | -0.017 | -0.017 | 1.928 | 0.19 | 0.24 | 0.08 | 0.1 | 0.11 | 0.16 |
| 6 | 32855683 | rs6457689 | -0.049 | -0.62 | -0.041 | -0.309 | -0.016 | -0.017 | -0.017 | 0.277 | 0.19 | 0.24 | 0.08 | 0.1 | 0.11 | 0.16 |
| 6 | 32856208 | rs2018501 | -0.915 | -0.592 | 0.084 | -0.314 | -0.025 | -0.026 | -0.026 | 0 | 0.61 | 0.75 | 0.57 | 0.59 | 0.54 | 0.55 |
| 6 | 32857496 | rs9276913 | -0.923 | -0.626 | 0.081 | -0.317 | -0.024 | -0.026 | -0.026 | 1.939 | 0.61 | 0.75 | 0.57 | 0.59 | 0.54 | 0.55 |
| 6 | 32857536 | rs10214760 | 0.056 | -0.617 | 0.099 | -0.324 | -0.016 | -0.016 | -0.016 | 2.117 | 0.18 | 0.24 | 0.08 | 0.11 | 0.11 | 0.16 |
| 6 | 32857801 | rs241414 | 0.195 | -0.574 | 0.147 | -0.316 | 0.002 | -0.008 | -0.008 | 1.505 | 0.42 | 0.49 | 0.42 | 0.42 | 0.42 | 0.37 |
| 6 | 32859000 | rs241413 | 0.292 | -0.581 | 0.179 | -0.32 | -0.003 | 0.005 | 0.005 | 0 | 0.41 | 0.42 | 0.41 | 0.43 | 0.42 | 0.37 |
| 6 | 32860752 | rs9469300 | 0.463 | -0.605 | 0.157 | -0.432 | 0.001 | -0.001 | -0.001 | 0 | 0.16 | 0.2 | 0.08 | 0.1 | 0.11 | 0.15 |
| 6 | 32861920 | rs241409 | 0.973 | -0.558 | 0.104 | -0.411 | 0.004 | 0.001 | 0.001 | 0.146 | 0.07 | 0.09 | 0.16 | 0.1 | 0.11 | 0.1 |
| 6 | 32862682 | rs3749981 | 1.65 | -0.565 | 0.087 | -0.413 | -0.022 | -0.009 | -0.009 | 0 | 0.05 | 0.02 | 0.07 | 0.05 | 0.1 | 0.07 |
| 6 | 32862711 | rs241408 | -0.951 | -0.561 | 0.11 | -0.394 | -0.024 | -0.023 | -0.023 | 0 | 0.61 | 0.75 | 0.5 | 0.58 | 0.54 | 0.55 |
| 6 | 32862740 | rs241407 | 1.153 | -0.405 | -0.467 | -0.294 | 0.001 | 0 | 0 | 0.246 | 0.07 | 0.09 | 0.16 | 0.1 | 0.11 | 0.1 |
| 6 | 32864681 | rs11961569 | 0.525 | -0.56 | -0.866 | -0.342 | 0.034 | 0.035 | 0.035 | 0 | 0.15 | 0.09 | 0.03 | 0.1 | 0.09 | 0.15 |
| 6 | 32864890 | rs11969321 | 0.525 | -0.56 | -0.866 | -0.342 | 0.034 | 0.035 | 0.035 | 0 | 0.15 | 0.09 | 0.03 | 0.1 | 0.09 | 0.15 |
| 6 | 32865338 | rs17508331 | 0.525 | -0.547 | -0.919 | -0.34 | 0.032 | 0.035 | 0.035 | 0 | 0.15 | 0.09 | 0.03 | 0.1 | 0.09 | 0.15 |
| 6 | 32865798 | rs241405 | -0.728 | -0.586 | -0.911 | -0.355 | 0.002 | 0.001 | 0.001 | 0 | 0.58 | 0.6 | 0.45 | 0.58 | 0.51 | 0.55 |
| 6 | 32865911 | rs9276915 | 1.045 | -0.589 | -0.915 | -0.359 | 0.006 | 0.037 | 0.037 | 0.087 | 0.31 | 0.2 | 0.15 | 0.21 | 0.25 | 0.2 |
| 6 | 32865997 | rs241404 | 0.215 | -0.617 | -1.164 | -0.323 | 0.002 | -0.014 | -0.014 | 0.297 | 0.42 | 0.51 | 0.34 | 0.41 | 0.42 | 0.37 |
| 6 | 32867123 | rs9501233 | 0.475 | -0.618 | -1.204 | -0.323 | 0.035 | 0.037 | 0.037 | 0 | 0.15 | 0.09 | 0.03 | 0.1 | 0.09 | 0.15 |
| 6 | 32867859 | rs9461818 | 0.542 | -0.586 | -1.189 | -0.307 | 0.034 | 0.035 | 0.035 | 0 | 0.15 | 0.09 | 0.03 | 0.1 | 0.09 | 0.15 |
| 6 | 32869045 | rs241402 | 0.331 | -0.586 | -1.184 | -0.307 | -0.003 | 0.006 | 0.006 | 0 | 0.41 | 0.42 | 0.33 | 0.42 | 0.42 | 0.37 |
| 6 | 32869882 | rs10080898 | 0.533 | -0.568 | -1.224 | -0.306 | 0.034 | 0.035 | 0.035 | 0 | 0.15 | 0.09 | 0.03 | 0.1 | 0.09 | 0.15 |
| 6 | 32869915 | rs9296057 | 0.533 | -0.568 | -1.224 | -0.306 | 0.034 | 0.035 | 0.035 | 0 | 0.15 | 0.09 | 0.03 | 0.1 | 0.09 | 0.16 |
| 6 | 32870057 | rs3101942 | -1.477 | -0.557 | -1.245 | -0.34 | -0.076 | -0.111 | -0.111 | 0 | 0.69 | 0.93 | 0.61 | 0.61 | 0.6 | 0.57 |
| 6 | 32871317 | rs1591 | 0.342 | -0.358 | -1.033 | -0.053 | 0.034 | 0.035 | 0.035 | 0 | 0.15 | 0.09 | 0.03 | 0.1 | 0.09 | 0.15 |
| 6 | 32871558 | rs241399 | -0.957 | -0.345 | -1.014 | -0.048 | 0.003 | 0.002 | 0.002 | 0 | 0.58 | 0.6 | 0.45 | 0.58 | 0.51 | 0.55 |
| 6 | 32872266 | rs9469302 | 0.323 | -0.343 | -1.104 | -0.049 | 0.034 | 0.035 | 0.035 | 0 | 0.15 | 0.09 | 0.03 | 0.1 | 0.09 | 0.15 |
| 6 | 32874796 | rs9469310 | 0.323 | -0.343 | -1.104 | -0.049 | 0.034 | 0.035 | 0.035 | 0 | 0.15 | 0.09 | 0.03 | 0.1 | 0.09 | 0.15 |
| 6 | 32875323 | rs1585894 | 0.323 | -0.343 | -1.104 | -0.049 | 0.034 | 0.035 | 0.035 | 0 | 0.15 | 0.09 | 0.03 | 0.1 | 0.09 | 0.15 |
| 6 | 32875450 | rs241398 | 0.065 | -0.349 | -1.09 | -0.053 | -0.003 | 0.005 | 0.005 | 0 | 0.41 | 0.42 | 0.33 | 0.41 | 0.42 | 0.37 |
| 6 | 32876049 | rs9501234 | 0.634 | -0.323 | -1.272 | -0.079 | 0.022 | 0.026 | 0.026 | 0 | 0.13 | 0.08 | 0.03 | 0.1 | 0.08 | 0.14 |
| 6 | 32876514 | rs9500926 | 0.313 | -0.326 | -1.273 | -0.08 | 0.034 | 0.035 | 0.035 | 0 | 0.15 | 0.09 | 0.03 | 0.1 | 0.09 | 0.15 |
| 6 | 32877073 | rs1978030 | 0.313 | -0.326 | -1.273 | -0.08 | 0.034 | 0.035 | 0.035 | 0 | 0.15 | 0.09 | 0.03 | 0.1 | 0.09 | 0.15 |
| 6 | 32877173 | rs154989 | -0.952 | -0.327 | -1.257 | -0.072 | 0.003 | 0.002 | 0.002 | 0 | 0.58 | 0.6 | 0.44 | 0.58 | 0.51 | 0.55 |
| 6 | 32877266 | rs154988 | -0.952 | -0.327 | -1.257 | -0.072 | 0.003 | 0.002 | 0.002 | 0 | 0.58 | 0.6 | 0.44 | 0.57 | 0.51 | 0.55 |
| 6 | 32879233 | rs154986 | 0.529 | -0.473 | -1.633 | -0.146 | 0.022 | 0.025 | 0.025 | 0 | 0.13 | 0.07 | 0.03 | 0.1 | 0.08 | 0.14 |
| 6 | 32880419 | rs154984 | 0.176 | -0.475 | -1.632 | -0.145 | 0.034 | 0.035 | 0.035 | 0 | 0.15 | 0.09 | 0.03 | 0.1 | 0.09 | 0.15 |
| 6 | 32880445 | rs154983 | 0.496 | -0.404 | -1.611 | -0.115 | 0.022 | 0.026 | 0.026 | 0 | 0.13 | 0.08 | 0.03 | 0.1 | 0.08 | 0.14 |
| 6 | 32881502 | rs9276921 | 1.185 | -0.322 | -1.717 | -0.124 | 0.046 | 0.032 | 0.032 | 0.436 | 0.22 | 0.09 | 0.1 | 0.19 | 0.12 | 0.19 |
| 6 | 32886487 | rs35599935 | 1.346 | 0.076 | -1.5 | -0.038 | -0.026 | 0.022 | 0.022 | 0 | 0.05 | 0.01 | 0.02 | 0.02 | 0.06 | 0.01 |
| 6 | 32889971 | rs149392 | 0.151 | -0.119 | -1.324 | 0.128 | 0.024 | 0.029 | 0.029 | 0 | 0.13 | 0.07 | 0.03 | 0.1 | 0.08 | 0.14 |
| 6 | 32893102 | rs9296059 | -0.091 | -0.22 | -1.378 | -0.005 | 0.04 | 0.053 | 0.053 | 0 | 0.15 | 0.07 | 0.03 | 0.09 | 0.09 | 0.15 |
| 6 | 32895580 | rs9501239 | 0.229 | -0.063 | -1.204 | 0.147 | 0.013 | 0.012 | 0.012 | 0 | 0.06 | 0.07 | 0.02 | 0.06 | 0.05 | 0.07 |
| 6 | 32899139 | rs11756897 | 0.246 | 0.341 | -0.905 | 0.128 | -0.001 | -0.003 | -0.003 | 0.216 | 0.25 | 0.29 | 0.21 | 0.19 | 0.25 | 0.29 |
| 6 | 32899296 | rs154978 | 0.31 | 0.621 | -0.745 | 0.501 | 0.004 | 0.004 | 0.004 | 0 | 0.58 | 0.61 | 0.45 | 0.57 | 0.53 | 0.55 |
| 6 | 32899485 | rs3132131 | 0.117 | 1.391 | -1.36 | 0.973 | -0.069 | -0.099 | -0.099 | 0 | 0.77 | 0.96 | 0.65 | 0.64 | 0.68 | 0.73 |
| 6 | 32900642 | rs154973 | -0.107 | 0.616 | -0.742 | 0.273 | -0.007 | -0.027 | -0.027 | 2.901 | 0.14 | 0.19 | 0.15 | 0.04 | 0.11 | 0.08 |
| 6 | 32900651 | rs154972 | -0.957 | 1.123 | -0.991 | -0.517 | -0.003 | 0.004 | 0.004 | 0.193 | 0.39 | 0.32 | 0.32 | 0.08 | 0.38 | 0.25 |
| 6 | 32900718 | rs181997 | -0.97 | 1.165 | -0.366 | -0.525 | -0.003 | -0.032 | -0.032 | 0 | 0.39 | 0.39 | 0.33 | 0.1 | 0.34 | 0.24 |
| 6 | 32900787 | rs3129299 | -0.997 | 0.796 | 0.098 | 0.053 | -0.017 | -0.014 | -0.014 | 0 | 0.26 | 0.42 | 0.35 | 0.27 | 0.2 | 0.27 |
| 6 | 32901351 | rs1465651 | -0.719 | 0.474 | -0.143 | -0.226 | 0.005 | 0.004 | 0.004 | 0 | 0.24 | 0.21 | 0.18 | 0.08 | 0.14 | 0.21 |
| 6 | 32901934 | rs3132132 | -0.615 | 0.269 | -0.191 | -0.262 | 0.003 | 0 | 0 | 0 | 0.24 | 0.21 | 0.18 | 0.07 | 0.14 | 0.19 |
| 6 | 32902001 | rs12662077 | 0.676 | 0.186 | -0.447 | 0.346 | 0.007 | 0.034 | 0.034 | 0 | 0.05 | 0.05 | 0.02 | 0.21 | 0.06 | 0.06 |
| 6 | 32902583 | rs10751 | -0.29 | 0.129 | -0.497 | 0.183 | 0.017 | 0.056 | 0.056 | 0 | 0.22 | 0.11 | 0.17 | 0.07 | 0.14 | 0.19 |
| 6 | 32903637 | rs23544 | 0.657 | 0.413 | -0.702 | 0.002 | -0.003 | 0.014 | 0.014 | 0.027 | 0.4 | 0.25 | 0.41 | 0.39 | 0.46 | 0.41 |
| 6 | 32903724 | rs68600 | 0.545 | 0.364 | -0.614 | 0.01 | -0.001 | 0.014 | 0.014 | 0 | 0.42 | 0.28 | 0.44 | 0.4 | 0.47 | 0.41 |
| 6 | 32903900 | rs151719 | 0.165 | 0.625 | -0.494 | 0.192 | -0.014 | -0.005 | -0.005 | 0 | 0.71 | 0.55 | 0.75 | 0.77 | 0.77 | 0.87 |
| 6 | 32904601 | rs2071556 | 0.445 | 0.843 | -0.554 | 0.085 | 0.002 | 0.024 | 0.024 | 0 | 0.55 | 0.56 | 0.51 | 0.43 | 0.6 | 0.5 |
| 6 | 32904661 | rs171329 | 0.729 | 0.861 | -0.405 | 0.06 | 0 | 0.014 | 0.014 | 0 | 0.51 | 0.4 | 0.46 | 0.43 | 0.53 | 0.48 |
| 6 | 32904981 | rs1042337 | -0.102 | 0.632 | -0.598 | 0.253 | -0.019 | -0.039 | -0.039 | 0 | 0.77 | 0.81 | 0.73 | 0.72 | 0.79 | 0.84 |
| 6 | 32905746 | rs194675 | 1.185 | 0.31 | -0.792 | -0.037 | -0.008 | 0.013 | 0.013 | 0.261 | 0.4 | 0.22 | 0.36 | 0.37 | 0.44 | 0.44 |
| 6 | 32905811 | rs714289 | 1.596 | 0.338 | -0.658 | -0.013 | -0.001 | -0.005 | -0.005 | 0.53 | 0.08 | 0.09 | 0.04 | 0.01 | 0.05 | 0.04 |
| 6 | 32911127 | rs3101944 | 1.271 | 0.3 | -0.606 | 0.094 | -0.031 | 0.023 | 0.023 | 0 | 0.09 | 0.01 | 0.06 | 0.04 | 0.08 | 0.09 |
| 6 | 32914297 | rs3130595 | 1.282 | 0.365 | -0.438 | 0.061 | -0.031 | 0.022 | 0.022 | 0 | 0.08 | 0.01 | 0.05 | 0.04 | 0.08 | 0.09 |
| 6 | 32914330 | rs189865 | 1.753 | 0.198 | -0.787 | -0.205 | -0.004 | -0.004 | -0.004 | 0 | 0.25 | 0.17 | 0.28 | 0.35 | 0.34 | 0.35 |
| 6 | 32914613 | rs194682 | 1.674 | 0.124 | -0.811 | -0.242 | -0.001 | -0.001 | -0.001 | 0 | 0.26 | 0.2 | 0.28 | 0.35 | 0.34 | 0.35 |
| 6 | 32915823 | rs6899309 | 0.826 | 0.428 | -0.487 | 0.118 | -0.008 | 0.025 | 0.025 | 0.057 | 0.08 | 0.17 | 0.12 | 0.09 | 0.06 | 0.09 |
| 6 | 32922459 | rs9378127 | 0.626 | 0.329 | -0.7 | 0.115 | 0.009 | -0.066 | -0.066 | 1.575 | 0.16 | 0.05 | 0.19 | 0.24 | 0.12 | 0.25 |
| 6 | 32924061 | rs28724903 | 0.905 | 0.323 | -0.705 | 0.117 | -0.005 | -0.062 | -0.062 | 0.35 | 0.13 | 0.01 | 0.16 | 0.23 | 0.1 | 0.23 |

| | | | | | | | | | | | | | | | | |
|---|---|---|---|---|---|---|---|---|---|---|---|---|---|---|---|---|
| 6 | 32924584 | rs209474 | 1.401 | 0.225 | -0.677 | 0.001 | -0.001 | 0.002 | 0.002 | 0 | 0.39 | 0.3 | 0.32 | 0.37 | 0.4 | 0.41 |
| 6 | 32925585 | rs209475 | 2.039 | 0.202 | -0.685 | -0.018 | -0.007 | -0.006 | -0.006 | 0 | 0.29 | 0.18 | 0.28 | 0.36 | 0.34 | 0.36 |
| 6 | 32925692 | rs580962 | 0.166 | 0.037 | -0.784 | -0.054 | -0.035 | -0.072 | -0.072 | 0.281 | 0.59 | 0.79 | 0.53 | 0.5 | 0.59 | 0.53 |
| 6 | 32930795 | rs9276933 | 0.605 | 0.004 | -0.833 | -0.062 | 0.058 | -0.02 | -0.02 | 0.224 | 0.08 | 0.28 | 0.09 | 0.01 | 0.1 | 0.02 |
| 6 | 32932150 | rs206783 | 1.375 | 0.095 | -0.837 | -0.019 | 0 | 0.003 | 0.003 | 0 | 0.39 | 0.3 | 0.32 | 0.37 | 0.4 | 0.41 |
| 6 | 32935179 | rs206786 | 1.37 | 0.089 | -0.842 | -0.022 | 0 | 0.003 | 0.003 | 0 | 0.39 | 0.3 | 0.32 | 0.37 | 0.41 | 0.42 |
| 6 | 32936373 | rs3918149 | 0.801 | 0.194 | -1.171 | 0.077 | 0.009 | -0.066 | -0.066 | 0 | 0.16 | 0.05 | 0.19 | 0.24 | 0.12 | 0.25 |
| 6 | 32938049 | rs3135330 | 1.578 | 0.382 | -0.87 | 0.368 | -0.031 | 0.022 | 0.022 | 0 | 0.08 | 0.01 | 0.05 | 0.04 | 0.08 | 0.09 |
| 6 | 32941910 | rs620202 | 2.124 | 0.433 | -0.864 | 0.451 | -0.017 | 0.021 | 0.021 | 0 | 0.23 | 0.08 | 0.23 | 0.18 | 0.28 | 0.28 |
| 6 | 32943151 | rs635688 | 0.248 | 0.365 | -0.439 | 0.972 | -0.037 | -0.075 | -0.075 | 0.204 | 0.59 | 0.79 | 0.54 | 0.5 | 0.6 | 0.53 |
| 6 | 32944746 | rs11908 | 2.038 | 0.423 | -0.588 | 0.979 | -0.023 | 0.039 | 0.039 | 0 | 0.31 | 0.17 | 0.29 | 0.2 | 0.37 | 0.33 |
| 6 | 32946029 | rs34530779 | 2.167 | 0.274 | -0.59 | 0.907 | 0.027 | 0.029 | 0.029 | 2.885 | 0.07 | 0.02 | 0 | 0 | 0 | 0 |
| 6 | 32946133 | rs206781 | 1.834 | 0.74 | -0.457 | 1.117 | -0.023 | 0.04 | 0.04 | 0 | 0.31 | 0.17 | 0.29 | 0.2 | 0.37 | 0.33 |
| 6 | 32948462 | rs3918140 | 1.312 | 0.721 | -0.49 | 1.233 | 0.019 | -0.032 | -0.032 | 4.768 | 0.06 | 0 | 0.13 | 0.19 | 0.03 | 0.12 |
| 6 | 32950050 | rs565876 | 1.657 | 1.06 | -0.283 | 0.997 | -0.023 | 0.04 | 0.04 | 0 | 0.31 | 0.17 | 0.29 | 0.2 | 0.37 | 0.33 |
| 6 | 32952546 | rs206777 | 0.211 | 1.357 | -0.232 | 0.896 | -0.057 | -0.099 | -0.099 | 1.669 | 0.6 | 0.86 | 0.55 | 0.51 | 0.6 | 0.53 |
| 6 | 32953711 | rs206776 | -0.317 | 1.43 | -0.351 | 0.9 | -0.031 | -0.096 | -0.096 | 0 | 0.69 | 0.87 | 0.61 | 0.56 | 0.69 | 0.62 |
| 6 | 32954689 | rs3097646 | 0.964 | 1.488 | -0.303 | 0.882 | -0.031 | 0.022 | 0.022 | 0 | 0.08 | 0.01 | 0.05 | 0.04 | 0.08 | 0.09 |
| 6 | 32957419 | rs579615 | 0.093 | 1.027 | -0.422 | 0.846 | -0.061 | -0.1 | -0.1 | 6.112 | 0.6 | 0.87 | 0.55 | 0.51 | 0.6 | 0.53 |
| 6 | 32961104 | rs206769 | 0.268 | 0.984 | -0.717 | 0.844 | 0.028 | -0.013 | -0.013 | 0 | 0.28 | 0.1 | 0.28 | 0.26 | 0.22 | 0.32 |
| 6 | 32961361 | rs9500927 | 0.389 | 1.001 | -0.42 | 0.471 | -0.032 | 0.018 | 0.018 | 3.159 | 0.2 | 0.36 | 0.1 | 0.23 | 0.14 | 0.09 |
| 6 | 32961621 | rs172275 | 0.445 | 1.744 | -0.19 | 0.74 | 0 | -0.001 | -0.001 | 0 | 0.54 | 0.56 | 0.42 | 0.47 | 0.55 | 0.46 |
| 6 | 32962420 | rs206767 | 0.483 | 1.564 | -0.227 | 0.686 | 0 | -0.002 | -0.002 | 0 | 0.53 | 0.54 | 0.4 | 0.43 | 0.51 | 0.44 |
| 6 | 32964967 | rs9276964 | -0.663 | 2.001 | 0.074 | 0.367 | 0.045 | -0.042 | -0.042 | 0 | 0.22 | 0.43 | 0.25 | 0.13 | 0.27 | 0.21 |
| 6 | 32965085 | rs3128948 | 0.719 | 2.262 | 0.203 | 0.53 | 0.044 | -0.001 | -0.001 | 0 | 0.84 | 0.73 | 0.83 | 0.88 | 0.82 | 0.89 |
| 6 | 32965942 | rs176248 | -1.27 | 2.169 | 0.184 | 0.265 | 0.027 | -0.027 | -0.027 | 0 | 0.22 | 0.6 | 0.28 | 0.19 | 0.28 | 0.18 |
| 6 | 32968276 | rs2395300 | 0.688 | 1.279 | -0.652 | 0.126 | 0.058 | 0.041 | 0.041 | 0 | 0.45 | 0.21 | 0.44 | 0.3 | 0.32 | 0.43 |
| 6 | 32968397 | rs9276969 | 0.637 | 0.91 | -0.398 | -0.06 | -0.013 | -0.019 | -0.019 | 0.186 | 0.06 | 0.13 | 0.05 | 0.01 | 0.03 | 0.02 |
| 6 | 32968598 | rs12191230 | 1.09 | 0.066 | -1.001 | -0.451 | 0.031 | 0.014 | 0.014 | 0 | 0.32 | 0.16 | 0.18 | 0.25 | 0.23 | 0.34 |
| 6 | 32968693 | rs2395301 | 0.468 | 0.09 | -0.961 | -0.397 | 0.058 | 0.041 | 0.041 | 0 | 0.45 | 0.21 | 0.44 | 0.3 | 0.32 | 0.43 |
| 6 | 32969457 | rs172274 | -0.066 | 0.103 | -0.882 | -0.385 | 0.049 | 0.041 | 0.041 | 0 | 0.51 | 0.37 | 0.5 | 0.31 | 0.34 | 0.45 |
| 6 | 32969696 | rs12193085 | 1.672 | -0.279 | -1.17 | -0.635 | 0.05 | -0.034 | -0.034 | 0 | 0.22 | 0.06 | 0.12 | 0.25 | 0.13 | 0.27 |
| 6 | 32970450 | rs206762 | -0.911 | -0.485 | -1.161 | -0.063 | -0.057 | -0.011 | -0.011 | 0.002 | 0.72 | 0.82 | 0.78 | 0.67 | 0.56 | 0.7 |
| 6 | 32971708 | rs3128931 | -0.511 | -0.969 | -0.986 | -0.037 | -0.008 | 0.025 | 0.025 | 0.24 | 0.26 | 0.55 | 0.31 | 0.37 | 0.24 | 0.28 |
| 6 | 32972163 | rs3128933 | 0.564 | -0.884 | -1.031 | 0.011 | -0.009 | -0.013 | -0.013 | 0 | 0.06 | 0.15 | 0.06 | 0.01 | 0.03 | 0.02 |
| 6 | 32972404 | rs3128935 | -0.19 | -0.992 | -0.852 | 0.24 | -0.03 | 0.052 | 0.052 | 0 | 0.11 | 0.28 | 0.12 | 0.17 | 0.05 | 0.11 |
| 6 | 32972642 | rs1044429 | 1.709 | -1.353 | -1.357 | -0.13 | 0.047 | -0.048 | -0.048 | 0 | 0.22 | 0.04 | 0.12 | 0.25 | 0.13 | 0.27 |
| 6 | 32972690 | rs592625 | 1.137 | -1.823 | -1.548 | -0.29 | 0.096 | -0.023 | -0.023 | 0 | 0.29 | 0.09 | 0.13 | 0.27 | 0.14 | 0.28 |
| 6 | 32972887 | rs376892 | -0.71 | -1.59 | -0.907 | -0.508 | -0.016 | -0.019 | -0.019 | 0.006 | 0.4 | 0.49 | 0.31 | 0.24 | 0.29 | 0.31 |
| 6 | 32972903 | rs35776718 | 1.209 | -1.37 | -1.179 | -0.353 | -0.045 | -0.046 | -0.046 | 0.503 | 0.08 | 0 | 0.15 | 0.19 | 0.14 | 0.12 |
| 6 | 32973281 | rs416622 | 1.374 | -0.882 | -1.002 | -0.631 | 0 | 0.012 | 0.012 | 0.167 | 0.52 | 0.37 | 0.33 | 0.6 | 0.5 | 0.58 |
| 6 | 32973599 | rs9276975 | 0.87 | -1.347 | -1.131 | -0.689 | -0.015 | -0.028 | -0.028 | 0.321 | 0.13 | 0.06 | 0.18 | 0.2 | 0.16 | 0.16 |
| 6 | 32973743 | rs3129304 | -1.065 | -1.16 | -1.056 | -0.878 | 0.041 | -0.005 | -0.005 | 0.078 | 0.9 | 0.84 | 0.71 | 0.95 | 0.87 | 0.95 |
| 6 | 32973794 | rs9276976 | 0.87 | -1.263 | -1.056 | -0.906 | -0.003 | -0.012 | -0.012 | 0 | 0.16 | 0.09 | 0.18 | 0.19 | 0.16 | 0.16 |
| 6 | 32973878 | rs3129303 | -1.072 | -1.214 | -1.03 | -0.887 | 0.041 | -0.005 | -0.005 | 0 | 0.9 | 0.83 | 0.71 | 0.95 | 0.87 | 0.95 |
| 6 | 32974081 | rs417812 | 0.047 | -1.444 | -1.189 | -0.949 | -0.008 | -0.012 | -0.012 | 0 | 0.35 | 0.31 | 0.36 | 0.56 | 0.43 | 0.37 |
| 6 | 32974268 | rs408036 | -0.025 | -1.483 | -1.257 | -1.042 | -0.012 | -0.01 | -0.01 | 0.563 | 0.37 | 0.31 | 0.37 | 0.57 | 0.46 | 0.42 |
| 6 | 32974343 | rs3129302 | 1.322 | -1.387 | -1.198 | -1.041 | -0.001 | -0.01 | -0.01 | 0 | 0.05 | 0.08 | 0.07 | 0.01 | 0.05 | 0.03 |
| 6 | 32974401 | rs2581 | 0.125 | -1.367 | -1.209 | -1.028 | -0.01 | -0.013 | -0.013 | 4.183 | 0.36 | 0.31 | 0.37 | 0.55 | 0.45 | 0.38 |
| 6 | 32974551 | rs2582 | 2.765 | -1.527 | -1.496 | -1.091 | 0.013 | -0.058 | -0.058 | 2.119 | 0.2 | 0.01 | 0.11 | 0.22 | 0.12 | 0.27 |
| 6 | 32974843 | rs6913008 | 1.357 | -1.35 | -1.293 | -1.037 | -0.006 | -0.012 | -0.012 | 2.409 | 0.14 | 0.09 | 0.18 | 0.19 | 0.16 | 0.16 |
| 6 | 32974934 | rs378352 | 1.8 | -1.328 | -1.479 | -0.979 | -0.075 | -0.093 | -0.093 | 0 | 0.14 | 0.02 | 0.11 | 0.35 | 0.22 | 0.16 |
| 6 | 32975381 | rs453779 | -0.802 | -1.426 | -1.31 | -1.063 | 0.008 | 0.032 | 0.032 | 0 | 0.5 | 0.61 | 0.47 | 0.7 | 0.58 | 0.55 |
| 6 | 32976654 | rs2284191 | 1.132 | -1.274 | -1.301 | -0.966 | -0.019 | -0.031 | -0.031 | 0 | 0.08 | 0.03 | 0.05 | 0.13 | 0.11 | 0.1 |
| 6 | 32976759 | rs86567 | -0.863 | -1.308 | -1.284 | -1.02 | -0.021 | -0.02 | -0.02 | 2.284 | 0.74 | 0.75 | 0.56 | 0.6 | 0.6 | 0.8 |
| 6 | 32976927 | rs404557 | 1.732 | -1.354 | -1.091 | -1.035 | 0.05 | 0.003 | 0.003 | 0.016 | 0.06 | 0.24 | 0.07 | 0 | 0.04 | 0 |
| 6 | 32977315 | rs403414 | 0.3 | -1.396 | -1.208 | -1.073 | 0.001 | 0.014 | 0.014 | 0 | 0.21 | 0.2 | 0.16 | 0.14 | 0.21 | 0.16 |
| 6 | 32977420 | rs381218 | 1.747 | -1.338 | -1.204 | -1.021 | -0.06 | -0.082 | -0.082 | 0 | 0.16 | 0.05 | 0.12 | 0.35 | 0.25 | 0.17 |
| 6 | 32977535 | rs396090 | -0.51 | -1.437 | -1.116 | -1.037 | -0.001 | -0.003 | -0.003 | 0 | 0.53 | 0.47 | 0.4 | 0.58 | 0.52 | 0.52 |
| 6 | 32978178 | rs6911639 | 0.347 | -1.351 | -1.617 | -0.96 | -0.002 | -0.001 | -0.001 | 4.036 | 0.26 | 0.27 | 0.23 | 0.22 | 0.21 | 0.32 |
| 6 | 32978997 | rs3763342 | 1.893 | -1.473 | -1.839 | -0.966 | -0.047 | -0.011 | -0.011 | 0.685 | 0.26 | 0.33 | 0.19 | 0.22 | 0.14 | 0.3 |
| 6 | 32979609 | rs4713603 | 0.239 | -1.339 | -2.94 | -0.964 | 0.022 | 0.001 | 0.001 | 0 | 0.53 | 0.68 | 0.47 | 0.56 | 0.47 | 0.58 |
| 6 | 32979746 | rs4711313 | 0.26 | -1.325 | -2.935 | -0.956 | 0.023 | 0.002 | 0.002 | 0 | 0.52 | 0.68 | 0.47 | 0.56 | 0.47 | 0.58 |
| 6 | 32980606 | rs4486068 | 0.26 | -1.325 | -2.935 | -0.956 | 0.023 | 0.002 | 0.002 | 0 | 0.52 | 0.68 | 0.47 | 0.56 | 0.47 | 0.58 |
| 6 | 32980608 | rs3130603 | -1.936 | -1.25 | -2.981 | -0.97 | -0.038 | -0.009 | -0.009 | 0 | 0.94 | 0.98 | 0.73 | 0.98 | 0.87 | 0.98 |
| 6 | 32981555 | rs34354079 | 1.764 | -1.289 | -3.037 | -1.008 | 0.001 | 0.028 | 0.028 | 0 | 0.1 | 0.05 | 0.04 | 0.01 | 0.08 | 0.06 |
| 6 | 32981647 | rs6457699 | 0.331 | -1.291 | -3.044 | -1.008 | 0.023 | 0.002 | 0.002 | 0 | 0.52 | 0.68 | 0.47 | 0.56 | 0.47 | 0.58 |
| 6 | 32981937 | rs9276992 | 1.498 | -1.248 | -2.987 | -0.993 | 0.008 | 0.009 | 0.009 | 0 | 0.11 | 0.09 | 0.05 | 0.03 | 0.05 | 0.13 |
| 6 | 32982017 | rs34570277 | 1.722 | -1.207 | -2.943 | -0.968 | -0.002 | 0.032 | 0.032 | 0 | 0.1 | 0.05 | 0.04 | 0.01 | 0.08 | 0.06 |

| | | | | | | | | | | | | | | | | |
|---|---|---|---|---|---|---|---|---|---|---|---|---|---|---|---|---|
| 6 | 32982921 | rs2395304 | 0.189 | -1.201 | -2.964 | -0.951 | 0.018 | -0.002 | -0.002 | 0.09 | 0.53 | 0.68 | 0.47 | 0.56 | 0.47 | 0.6 |
| 6 | 32984011 | rs6935846 | 0.18 | -1.196 | -2.955 | -0.949 | 0.017 | -0.002 | -0.002 | 0.01 | 0.54 | 0.68 | 0.47 | 0.56 | 0.48 | 0.6 |
| 6 | 32984255 | rs9276994 | 0.714 | -1.122 | -2.871 | -0.917 | 0.043 | 0.005 | 0.005 | 0 | 0.45 | 0.63 | 0.41 | 0.42 | 0.36 | 0.49 |
| 6 | 32984788 | rs2116264 | 2.537 | -1.05 | -2.976 | -0.9 | 0.021 | -0.04 | -0.04 | 0.064 | 0.19 | 0.01 | 0.12 | 0.21 | 0.11 | 0.27 |
| 6 | 32985052 | rs3130604 | -1.732 | -0.342 | -2.337 | -0.812 | -0.038 | -0.009 | -0.009 | 1.469 | 0.94 | 0.98 | 0.73 | 0.98 | 0.87 | 0.98 |
| 6 | 32986573 | rs6457701 | 0.928 | -0.417 | -2.339 | -0.815 | 0.044 | 0.005 | 0.005 | 0 | 0.45 | 0.63 | 0.41 | 0.42 | 0.36 | 0.49 |
| 6 | 32987277 | rs3130607 | -1.26 | 0.257 | -1.618 | -0.614 | -0.04 | -0.008 | -0.008 | 0 | 0.94 | 0.98 | 0.73 | 0.98 | 0.87 | 0.98 |
| 6 | 32987774 | rs423639 | 0.227 | -0.393 | -1.513 | -0.668 | -0.004 | -0.019 | -0.019 | 0.058 | 0.15 | 0.19 | 0.11 | 0.03 | 0.09 | 0.13 |
| 6 | 32988049 | rs6457702 | 0.171 | -0.404 | -1.708 | -0.526 | -0.05 | -0.024 | -0.024 | 2.695 | 0.64 | 0.74 | 0.51 | 0.6 | 0.56 | 0.7 |
| 6 | 32988089 | rs17220695 | 0.862 | -0.382 | -1.406 | -0.336 | 0.009 | 0.009 | 0.009 | 0 | 0.1 | 0.08 | 0.03 | 0.03 | 0.04 | 0.13 |
| 6 | 32988175 | rs17214311 | 0.862 | -0.382 | -1.406 | -0.336 | 0.009 | 0.009 | 0.009 | 0 | 0.1 | 0.08 | 0.03 | 0.03 | 0.04 | 0.13 |
| 6 | 32988695 | rs9296068 | 0.656 | -0.583 | -1.417 | -0.349 | 0.017 | -0.016 | -0.016 | 0 | 0.4 | 0.56 | 0.4 | 0.51 | 0.35 | 0.5 |
| 6 | 32989521 | rs3130609 | 0.628 | -0.193 | -1.177 | -0.031 | 0.007 | 0.008 | 0.008 | 0 | 0.1 | 0.13 | 0.06 | 0.02 | 0.05 | 0.02 |
| 6 | 32989971 | rs9277003 | 0.936 | -0.052 | -0.927 | -0.115 | 0.007 | -0.013 | -0.013 | 0 | 0.39 | 0.27 | 0.37 | 0.5 | 0.37 | 0.5 |
| 6 | 32992424 | rs7738250 | 0.963 | -0.058 | -1.011 | -0.232 | 0.003 | 0.016 | 0.016 | 0 | 0.1 | 0.06 | 0.04 | 0.01 | 0.08 | 0.06 |
| 6 | 32993266 | rs9277015 | 1.046 | -0.152 | -0.842 | -0.213 | 0.008 | -0.013 | -0.013 | 0 | 0.39 | 0.27 | 0.37 | 0.5 | 0.37 | 0.5 |
| 6 | 32996014 | rs6933546 | 1.25 | -0.042 | -0.73 | -0.256 | 0.012 | -0.017 | -0.017 | 0 | 0.41 | 0.27 | 0.36 | 0.52 | 0.36 | 0.51 |
| 6 | 32997674 | rs6920606 | 0.4 | -0.04 | -0.763 | -0.291 | 0 | -0.001 | -0.001 | 0 | 0.56 | 0.43 | 0.44 | 0.58 | 0.47 | 0.71 |
| 6 | 32997871 | rs9277023 | 0.955 | -0.02 | -0.754 | -0.268 | 0.013 | -0.011 | -0.011 | 0.399 | 0.44 | 0.28 | 0.38 | 0.54 | 0.39 | 0.61 |
| 6 | 32998238 | rs9277027 | 1.791 | -0.235 | -0.944 | -0.052 | -0.008 | -0.013 | -0.013 | 0.289 | 0.33 | 0.03 | 0.18 | 0.31 | 0.27 | 0.4 |
| 6 | 32998564 | rs1431396 | -0.23 | 0.426 | -0.967 | -0.042 | 0.018 | 0.012 | 0.012 | 0.281 | 0.33 | 0.57 | 0.29 | 0.39 | 0.41 | 0.26 |
| 6 | 32998729 | rs3130171 | -0.246 | 0.431 | -0.964 | -0.039 | 0.018 | 0.012 | 0.012 | 0.25 | 0.33 | 0.57 | 0.29 | 0.39 | 0.41 | 0.26 |
| 6 | 33001735 | rs429073 | -0.31 | 0.496 | -0.875 | 0.046 | -0.017 | 0 | 0 | 0.89 | 0.22 | 0.55 | 0.28 | 0.13 | 0.2 | 0.14 |
| 6 | 33003065 | rs2858458 | -0.872 | 0.45 | -0.949 | -0.023 | -0.01 | -0.009 | -0.009 | 0 | 0.33 | 0.62 | 0.32 | 0.17 | 0.28 | 0.25 |
| 6 | 33003562 | rs423196 | -0.27 | 0.451 | -0.947 | -0.022 | -0.017 | 0 | 0 | 0.89 | 0.22 | 0.55 | 0.28 | 0.13 | 0.2 | 0.14 |
| 6 | 33003687 | rs448733 | 0.409 | 0.391 | -0.842 | -0.056 | -0.009 | -0.044 | -0.044 | 0 | 0.09 | 0.26 | 0.09 | 0 | 0.06 | 0.05 |
| 6 | 33004387 | rs763469 | -1.465 | 0.45 | -0.896 | -0.081 | -0.038 | -0.009 | -0.009 | 0.249 | 0.94 | 0.98 | 0.72 | 0.98 | 0.87 | 0.98 |
| 6 | 33004917 | rs414564 | 0.427 | 0.405 | -0.879 | -0.079 | -0.009 | -0.044 | -0.044 | 0.377 | 0.09 | 0.26 | 0.09 | 0 | 0.06 | 0.05 |
| 6 | 33004935 | rs444821 | -1.042 | 0.4 | -0.883 | -0.082 | -0.009 | -0.009 | -0.009 | 0.236 | 0.33 | 0.62 | 0.32 | 0.17 | 0.28 | 0.25 |
| 6 | 33005208 | rs439852 | -1.035 | 0.413 | -0.877 | -0.075 | -0.01 | -0.009 | -0.009 | 0.161 | 0.33 | 0.62 | 0.32 | 0.17 | 0.28 | 0.25 |
| 6 | 33005644 | rs406477 | -0.563 | 0.412 | -0.877 | -0.076 | -0.018 | -0.001 | -0.001 | 0 | 0.22 | 0.55 | 0.28 | 0.13 | 0.2 | 0.14 |
| 6 | 33005932 | rs9277052 | -0.196 | 0.44 | -0.874 | -0.06 | -0.013 | 0.005 | 0.005 | 0 | 0.24 | 0.36 | 0.24 | 0.17 | 0.21 | 0.2 |
| 6 | 33006869 | rs663310 | -1.02 | 0.445 | -0.885 | -0.057 | -0.01 | -0.009 | -0.009 | 0.032 | 0.33 | 0.62 | 0.32 | 0.17 | 0.28 | 0.25 |
| 6 | 33007192 | rs393906 | -1.02 | 0.478 | -1.011 | -0.031 | -0.01 | -0.009 | -0.009 | 0 | 0.33 | 0.62 | 0.32 | 0.17 | 0.28 | 0.25 |
| 6 | 33007752 | rs7774158 | 1.725 | 0.387 | -1.037 | -0.044 | 0.063 | 0.077 | 0.077 | 2.427 | 0.46 | 0.16 | 0.24 | 0.36 | 0.35 | 0.52 |
| 6 | 33008546 | rs441576 | 0.739 | 0.546 | -1.075 | 0.122 | -0.001 | 0.003 | 0.003 | 0.498 | 0.58 | 0.43 | 0.35 | 0.37 | 0.45 | 0.67 |
| 6 | 33008873 | rs9277059 | 1.269 | 0.492 | -1.348 | 0.2 | 0.034 | 0.054 | 0.054 | 0.506 | 0.38 | 0.1 | 0.22 | 0.32 | 0.31 | 0.5 |
| 6 | 33014050 | rs3130174 | 0.224 | 0.505 | -1.338 | 0.208 | 0 | 0 | 0 | 0 | 0.71 | 0.72 | 0.54 | 0.49 | 0.58 | 0.76 |
| 6 | 33016033 | rs3097659 | -1.179 | 0.629 | -1.916 | 0.268 | -0.038 | -0.009 | -0.009 | 0 | 0.94 | 0.98 | 0.72 | 0.98 | 0.87 | 0.98 |
| 6 | 33016728 | rs375912 | 0.884 | 0.676 | -1.737 | 0.309 | 0.021 | 0.054 | 0.054 | 0.495 | 0.39 | 0.1 | 0.22 | 0.32 | 0.32 | 0.51 |
| 6 | 33016887 | rs2105929 | 1.051 | 0.682 | -1.605 | 0.443 | 0.023 | 0.031 | 0.031 | 1.025 | 0.35 | 0.09 | 0.19 | 0.31 | 0.29 | 0.41 |
| 6 | 33018726 | rs450745 | 1.1 | 0.724 | -1.941 | 0.465 | -0.013 | -0.002 | -0.002 | 0 | 0.34 | 0.03 | 0.19 | 0.31 | 0.29 | 0.41 |
| 6 | 33019643 | rs440841 | 0.877 | 0.255 | -1.521 | 0.287 | -0.088 | -0.036 | -0.036 | 0 | 0.47 | 0.07 | 0.31 | 0.68 | 0.54 | 0.55 |
| 6 | 33020777 | rs3097662 | 0.234 | 0.725 | -1.167 | 0.504 | -0.013 | 0.002 | 0.002 | 0 | 0.79 | 0.85 | 0.64 | 0.81 | 0.81 | 0.76 |
| 6 | 33023946 | rs9277196 | -0.364 | 0.519 | -0.896 | 0.454 | -0.001 | 0.002 | 0.002 | 0.142 | 0.56 | 0.55 | 0.73 | 0.64 | 0.53 | 0.51 |
| 6 | 33024499 | rs367645 | -0.113 | -0.021 | -0.233 | 0.247 | 0.067 | -0.003 | -0.003 | 0.37 | 0.07 | 0.3 | 0.07 | 0 | 0.04 | 0.01 |
| 6 | 33024654 | rs3135402 | -0.544 | -0.899 | -0.753 | 0.081 | -0.027 | -0.025 | -0.025 | 0 | 0.38 | 0.5 | 0.45 | 0.25 | 0.26 | 0.35 |
| 6 | 33025700 | rs4551215 | 0.099 | -0.824 | -0.743 | 0.868 | 0.005 | 0.36 | 0.36 | 0 | 0.07 | 0.2 | 0.04 | 0.58 | 0.03 | 0.1 |
| 6 | 33026110 | rs422544 | -0.225 | -1.116 | -0.871 | 0.641 | 0.037 | 0.041 | 0.041 | 0 | 0.37 | 0.33 | 0.41 | 0.21 | 0.28 | 0.3 |
| 6 | 33030112 | rs9277334 | 0.175 | -1.32 | -1.02 | 0.526 | 0.001 | 0.001 | 0.001 | 0 | 0.22 | 0.24 | 0.13 | 0.16 | 0.13 | 0.22 |
| 6 | 33034596 | rs2301226 | 0.189 | -1.348 | -1.013 | 0.511 | 0 | 0.001 | 0.001 | 0 | 0.21 | 0.24 | 0.13 | 0.16 | 0.13 | 0.22 |
| 6 | 33035771 | rs1054025 | 0.539 | -1.308 | -0.977 | 0.624 | 0.012 | 0.382 | 0.382 | 0 | 0.06 | 0.2 | 0.04 | 0.55 | 0.03 | 0.1 |
| 6 | 33035860 | rs2301225 | 0.919 | -1.442 | -1.087 | 0.451 | -0.034 | -0.008 | -0.008 | 0 | 0.08 | 0.01 | 0.24 | 0.04 | 0.12 | 0.06 |
| 6 | 33036999 | rs1042190 | -0.272 | -2.368 | -1.772 | -0.15 | 0 | -0.002 | -0.002 | 0 | 0.27 | 0.58 | 0.28 | 0.65 | 0.19 | 0.36 |
| 6 | 33038369 | rs2301224 | -0.292 | -2.37 | -1.785 | -0.14 | -0.001 | 0 | 0 | 0 | 0.27 | 0.58 | 0.28 | 0.65 | 0.19 | 0.36 |
| 6 | 33039625 | rs9277341 | -0.79 | -2.424 | -1.822 | -0.177 | 0.096 | 0.139 | 0.139 | 0.397 | 0.47 | 0.74 | 0.4 | 0.84 | 0.32 | 0.59 |
| 6 | 33040715 | rs6914849 | -0.472 | -1.983 | -1.508 | 0.186 | -0.002 | 0.001 | 0.001 | 0 | 0.27 | 0.58 | 0.27 | 0.67 | 0.18 | 0.36 |
| 6 | 33041176 | rs1431400 | -0.522 | -1.983 | -1.508 | 0.176 | -0.001 | -0.001 | -0.001 | 0 | 0.28 | 0.58 | 0.28 | 0.68 | 0.19 | 0.37 |
| 6 | 33041186 | rs1431401 | -0.473 | -1.982 | -1.507 | 0.176 | -0.002 | 0 | 0 | 0 | 0.27 | 0.58 | 0.28 | 0.65 | 0.19 | 0.36 |
| 6 | 33042880 | rs987870 | 0.24 | -1.78 | -1.435 | 0.335 | -0.032 | -0.063 | -0.063 | 0 | 0.21 | 0.38 | 0.24 | 0.1 | 0.15 | 0.26 |
| 6 | 33044257 | rs2071353 | -0.314 | -1.788 | -1.46 | 0.37 | -0.005 | 0.007 | 0.007 | 0 | 0.25 | 0.56 | 0.27 | 0.66 | 0.18 | 0.31 |
| 6 | 33044388 | rs2071354 | 0.232 | -1.759 | -1.429 | 0.009 | -0.032 | -0.063 | -0.063 | 0 | 0.21 | 0.38 | 0.23 | 0.1 | 0.15 | 0.26 |
| 6 | 33048661 | rs1042151 | -0.213 | -1.519 | -2.039 | -0.182 | -0.043 | -0.085 | -0.085 | 0.719 | 0.24 | 0.42 | 0.21 | 0.12 | 0.21 | 0.25 |
| 6 | 33050279 | rs9277378 | -1.204 | -1.518 | -2.146 | 0.268 | 0.023 | -0.068 | -0.068 | 0 | 0.34 | 0.7 | 0.34 | 0.61 | 0.31 | 0.37 |
| 6 | 33050499 | rs9277386 | -0.985 | -1.472 | -2.102 | 0.325 | 0.002 | -0.008 | -0.008 | 0 | 0.32 | 0.59 | 0.32 | 0.62 | 0.31 | 0.37 |
| 6 | 33051139 | rs9277396 | -1.003 | -1.458 | -2.133 | 0.255 | 0.002 | -0.009 | -0.009 | 0 | 0.32 | 0.6 | 0.33 | 0.62 | 0.32 | 0.37 |
| 6 | 33053352 | rs9277464 | -0.666 | -1.515 | -2.134 | 0.214 | 0.006 | -0.075 | -0.075 | 0 | 0.27 | 0.19 | 0.29 | 0.62 | 0.27 | 0.3 |
| 6 | 33053468 | rs9277469 | -1.145 | -1.519 | -2.125 | 0.21 | 0.002 | -0.009 | -0.009 | 0.063 | 0.32 | 0.6 | 0.33 | 0.62 | 0.32 | 0.37 |
| 6 | 33053682 | rs9277471 | -1.145 | -1.519 | -2.125 | 0.209 | 0.002 | -0.009 | -0.009 | 0 | 0.32 | 0.6 | 0.33 | 0.62 | 0.32 | 0.37 |

| | | | | | | | | | | | | | | | | |
|---|---|---|---|---|---|---|---|---|---|---|---|---|---|---|---|---|
| 6 | 33053942 | rs9277489 | -0.664 | -1.519 | -2.125 | 0.209 | 0.006 | -0.075 | -0.075 | 0 | 0.27 | 0.19 | 0.29 | 0.62 | 0.27 | 0.3 |
| 6 | 33054721 | rs9277533 | -1.144 | -1.518 | -2.125 | 0.209 | 0.002 | -0.009 | -0.009 | 0 | 0.32 | 0.6 | 0.33 | 0.62 | 0.32 | 0.37 |
| 6 | 33055346 | rs9277546 | -1.213 | -1.537 | -2.134 | 0.202 | 0.003 | -0.011 | -0.011 | 0 | 0.33 | 0.61 | 0.33 | 0.62 | 0.31 | 0.36 |
| 6 | 33055538 | rs9277554 | -1.139 | -1.532 | -2.132 | 0.199 | 0.002 | -0.009 | -0.009 | 0.126 | 0.32 | 0.6 | 0.33 | 0.62 | 0.32 | 0.37 |
| 6 | 33055605 | rs9277555 | -0.634 | -1.54 | -2.139 | 0.242 | -0.01 | 0.038 | 0.038 | 0 | 0.27 | 0.5 | 0.26 | 0.53 | 0.27 | 0.28 |
| 6 | 33056069 | rs3117229 | -0.677 | -1.536 | -2.141 | 0.237 | 0.006 | -0.075 | -0.075 | 0 | 0.27 | 0.19 | 0.29 | 0.62 | 0.27 | 0.3 |
| 6 | 33056435 | rs3117228 | -1.161 | -1.537 | -2.141 | 0.237 | 0.002 | -0.009 | -0.009 | 0.095 | 0.32 | 0.6 | 0.33 | 0.62 | 0.32 | 0.37 |
| 6 | 33056694 | rs9277557 | -0.68 | -1.532 | -2.149 | 0.242 | 0.006 | -0.075 | -0.075 | 0 | 0.27 | 0.19 | 0.29 | 0.62 | 0.27 | 0.3 |
| 6 | 33056897 | rs9277565 | 0.025 | -1.533 | -2.154 | 0.249 | 0.005 | -0.105 | -0.105 | 0.49 | 0.22 | 0.1 | 0.22 | 0.53 | 0.23 | 0.21 |
| 6 | 33057176 | rs3130188 | -1.155 | -1.536 | -2.143 | 0.243 | 0.002 | -0.009 | -0.009 | 1.653 | 0.32 | 0.6 | 0.33 | 0.62 | 0.32 | 0.37 |
| 6 | 33057659 | rs3117226 | -0.655 | -1.543 | -2.143 | 0.251 | -0.012 | 0.028 | 0.028 | 0 | 0.27 | 0.52 | 0.25 | 0.52 | 0.26 | 0.27 |
| 6 | 33057711 | rs3117225 | -1.204 | -1.538 | -2.144 | 0.249 | 0.003 | -0.01 | -0.01 | 0 | 0.33 | 0.61 | 0.33 | 0.62 | 0.32 | 0.37 |
| 6 | 33057835 | rs3097652 | -1.27 | -1.572 | -2.159 | 0.232 | 0.004 | -0.012 | -0.012 | 0 | 0.33 | 0.61 | 0.33 | 0.62 | 0.32 | 0.37 |
| 6 | 33058774 | rs3128972 | -0.694 | -1.55 | -2.169 | 0.275 | -0.01 | 0.035 | 0.035 | 0 | 0.27 | 0.51 | 0.26 | 0.53 | 0.27 | 0.28 |
| 6 | 33058874 | rs2179920 | -0.362 | -1.55 | -2.169 | 0.276 | -0.024 | -0.064 | -0.064 | 0 | 0.25 | 0.46 | 0.23 | 0.14 | 0.23 | 0.25 |
| 6 | 33059262 | rs2179919 | -0.712 | -1.55 | -2.172 | 0.276 | -0.01 | 0.035 | 0.035 | 0 | 0.27 | 0.51 | 0.26 | 0.53 | 0.27 | 0.28 |
| 6 | 33059669 | rs2281390 | 0.287 | -1.719 | -2.558 | -0.075 | 0.001 | 0.019 | 0.019 | 0 | 0.18 | 0.15 | 0.31 | 0.06 | 0.17 | 0.11 |
| 6 | 33059796 | rs2281389 | 0.579 | -1.707 | -2.537 | -0.061 | 0.012 | 0.052 | 0.052 | 0 | 0.2 | 0.06 | 0.18 | 0.13 | 0.19 | 0.19 |
| 6 | 33059996 | rs3128917 | -0.504 | -1.712 | -2.538 | -0.063 | -0.01 | 0.034 | 0.034 | 0 | 0.28 | 0.51 | 0.27 | 0.53 | 0.27 | 0.28 |
| 6 | 33060428 | rs7763822 | 0.777 | -1.66 | -2.509 | -0.074 | 0 | 0.001 | 0.001 | 0 | 0.05 | 0.1 | 0.07 | 0.09 | 0.04 | 0.09 |
| 6 | 33060949 | rs3117222 | -0.481 | -1.668 | -2.485 | -0.069 | -0.01 | 0.035 | 0.035 | 0 | 0.27 | 0.51 | 0.26 | 0.54 | 0.27 | 0.28 |
| 6 | 33061098 | rs3128918 | 0.277 | -1.684 | -2.507 | -0.087 | 0.001 | 0.019 | 0.019 | 0 | 0.18 | 0.15 | 0.31 | 0.06 | 0.17 | 0.12 |
| 6 | 33061690 | rs3130190 | -0.553 | -1.452 | -2.389 | 0.023 | -0.01 | 0.035 | 0.035 | 0 | 0.27 | 0.51 | 0.26 | 0.53 | 0.27 | 0.28 |
| 6 | 33061871 | rs3130191 | -0.532 | -1.452 | -2.389 | 0.023 | -0.01 | 0.036 | 0.036 | 0 | 0.27 | 0.51 | 0.26 | 0.53 | 0.27 | 0.27 |
| 6 | 33061930 | rs3130192 | 1.702 | -1.463 | -2.429 | -0.012 | -0.029 | -0.016 | -0.016 | 0 | 0.06 | 0 | 0.24 | 0.04 | 0.11 | 0.05 |
| 6 | 33061947 | rs3117221 | 0.102 | -1.445 | -2.34 | 0.023 | 0.005 | -0.105 | -0.105 | 0.114 | 0.22 | 0.1 | 0.22 | 0.53 | 0.23 | 0.21 |
| 6 | 33063931 | rs3130198 | -0.569 | -1.428 | -2.337 | 0.039 | -0.009 | 0.029 | 0.029 | 0 | 0.27 | 0.52 | 0.27 | 0.53 | 0.27 | 0.28 |
| 6 | 33064605 | rs3117213 | -0.557 | -1.427 | -2.336 | 0.039 | -0.009 | 0.03 | 0.03 | 0 | 0.27 | 0.52 | 0.27 | 0.53 | 0.27 | 0.27 |
| 6 | 33071424 | rs3128924 | -0.455 | -1.56 | -2.791 | -0.161 | 0.008 | 0.041 | 0.041 | 0 | 0.44 | 0.47 | 0.54 | 0.8 | 0.37 | 0.41 |
| 6 | 33071777 | rs3117239 | -0.111 | -1.625 | -2.802 | -0.304 | -0.024 | -0.059 | -0.059 | 0.579 | 0.26 | 0.47 | 0.23 | 0.15 | 0.23 | 0.28 |
| 6 | 33072240 | rs2064479 | -0.385 | -1.621 | -2.804 | -0.245 | -0.015 | 0.044 | 0.044 | 0 | 0.27 | 0.48 | 0.26 | 0.54 | 0.26 | 0.31 |
| 6 | 33072266 | rs2064478 | -0.1 | -1.621 | -2.801 | -0.267 | -0.023 | -0.059 | -0.059 | 0 | 0.26 | 0.47 | 0.23 | 0.15 | 0.23 | 0.28 |
| 6 | 33073322 | rs2064476 | -0.938 | -1.633 | -2.821 | -0.255 | 0 | -0.001 | -0.001 | 0.879 | 0.33 | 0.57 | 0.33 | 0.63 | 0.32 | 0.4 |
| 6 | 33073984 | rs3117234 | -0.193 | -1.631 | -2.815 | -0.261 | -0.021 | -0.057 | -0.057 | 1.209 | 0.26 | 0.47 | 0.23 | 0.15 | 0.24 | 0.29 |
| 6 | 33074908 | rs3117231 | -0.332 | -1.633 | -2.816 | -0.244 | -0.015 | 0.039 | 0.039 | 0 | 0.27 | 0.48 | 0.25 | 0.52 | 0.26 | 0.3 |
| 6 | 33074963 | rs3130215 | -2.066 | -1.678 | -2.857 | -0.359 | -0.086 | 0.113 | 0.113 | 0.009 | 0.69 | 0.87 | 0.76 | 0.95 | 0.6 | 0.69 |
| 6 | 33075635 | rs3117230 | -0.37 | -1.41 | -2.32 | 0.113 | -0.023 | -0.059 | -0.059 | 0.055 | 0.26 | 0.47 | 0.23 | 0.15 | 0.23 | 0.28 |
| 6 | 33075666 | rs3128930 | -0.961 | -1.072 | -2.089 | 0.362 | -0.005 | -0.022 | -0.022 | 0 | 0.26 | 0.54 | 0.28 | 0.23 | 0.26 | 0.37 |
| 6 | 33077776 | rs6457713 | -0.059 | -1.011 | -1.813 | 0.595 | 0.09 | 0.403 | 0.403 | 0 | 0.07 | 0.46 | 0.08 | 0.39 | 0.09 | 0.08 |
| 6 | 33079812 | rs6937061 | 0.823 | -1.063 | -2.453 | 0.676 | 0.038 | 0.03 | 0.03 | 0 | 0.11 | 0.03 | 0.1 | 0.05 | 0.04 | 0.12 |
| 6 | 33081632 | rs11751659 | 0.307 | -0.673 | -2.228 | 0.402 | -0.01 | -0.004 | -0.004 | 6.096 | 0.09 | 0.05 | 0.27 | 0.05 | 0.13 | 0.07 |
| 6 | 33082109 | rs9277629 | 1.265 | -0.914 | -2.673 | -0.181 | 0.041 | 0.052 | 0.052 | 1.592 | 0.18 | 0.02 | 0.06 | 0.04 | 0.11 | 0.27 |
| 6 | 33083121 | rs1810472 | 0.242 | -0.865 | -2.988 | -0.33 | 0.038 | -0.001 | -0.001 | 0 | 0.4 | 0.18 | 0.32 | 0.36 | 0.29 | 0.38 |
| 6 | 33083134 | rs2395349 | 0.875 | -1.052 | -3.147 | -0.413 | -0.01 | -0.071 | -0.071 | 0.022 | 0.25 | 0.05 | 0.14 | 0.28 | 0.19 | 0.23 |
| 6 | 33084148 | rs9277656 | 0.875 | -1.038 | -3.147 | -0.413 | -0.006 | -0.071 | -0.071 | 2.324 | 0.25 | 0.05 | 0.14 | 0.28 | 0.19 | 0.23 |
| 6 | 33084671 | rs3129294 | -0.172 | -1.13 | -3.354 | -0.625 | 0.076 | 0.139 | 0.139 | 0 | 0.49 | 0.21 | 0.33 | 0.22 | 0.37 | 0.54 |
| 6 | 33085851 | rs3117039 | 0.149 | -1.127 | -3.399 | -0.605 | 0.036 | 0.042 | 0.042 | 0.09 | 0.51 | 0.14 | 0.26 | 0.4 | 0.4 | 0.57 |
| 6 | 33085910 | rs3117038 | 1.174 | -1.158 | -3.418 | -0.641 | 0.02 | 0.076 | 0.076 | 0.312 | 0.26 | 0.08 | 0.12 | 0.13 | 0.22 | 0.35 |
| 6 | 33086249 | rs3117035 | 0.115 | -1.155 | -3.421 | -0.637 | 0.035 | 0.019 | 0.019 | 0.074 | 0.51 | 0.14 | 0.26 | 0.41 | 0.41 | 0.58 |
| 6 | 33086448 | rs1883414 | 0.521 | -1.096 | -3.384 | -0.609 | 0.046 | 0.106 | 0.106 | 0 | 0.41 | 0.19 | 0.29 | 0.2 | 0.33 | 0.5 |
| 6 | 33087684 | rs9277682 | 0.839 | -1.337 | -3.372 | -0.586 | -0.01 | -0.071 | -0.071 | 0 | 0.25 | 0.05 | 0.14 | 0.28 | 0.19 | 0.23 |
| 6 | 33089623 | rs3117027 | -0.135 | -1.261 | -3.317 | -0.643 | 0.111 | 0.157 | 0.157 | 0 | 0.49 | 0.21 | 0.21 | 0.19 | 0.34 | 0.55 |
| 6 | 33091454 | rs9277723 | 0.029 | -1.262 | -3.293 | -0.643 | 0.091 | 0.01 | 0.01 | 0 | 0.43 | 0.17 | 0.29 | 0.36 | 0.27 | 0.38 |
| 6 | 33092049 | rs3117021 | -0.338 | -1.043 | -3.21 | -0.491 | -0.045 | -0.05 | -0.05 | 0 | 0.66 | 0.25 | 0.4 | 0.53 | 0.48 | 0.74 |
| 6 | 33093365 | rs6913896 | 0.259 | -1.135 | -3.24 | -0.518 | 0.069 | -0.006 | -0.006 | 0.822 | 0.4 | 0.17 | 0.29 | 0.36 | 0.27 | 0.38 |
| 6 | 33094258 | rs3129275 | 0.011 | -1.121 | -3.247 | -0.527 | -0.003 | -0.041 | -0.041 | 0 | 0.58 | 0.24 | 0.36 | 0.51 | 0.44 | 0.69 |
| 6 | 33094869 | rs3129274 | -1.52 | -0.834 | -2.643 | -0.677 | 0.163 | -0.094 | -0.094 | 0 | 0.83 | 0.62 | 0.77 | 0.98 | 0.69 | 0.93 |
| 6 | 33095516 | rs3117016 | -0.05 | -1.336 | -3.113 | -0.621 | 0.045 | -0.026 | -0.026 | 0.87 | 0.46 | 0.19 | 0.34 | 0.49 | 0.38 | 0.48 |
| 6 | 33096274 | rs3117008 | -0.233 | -0.973 | -2.916 | -0.419 | -0.045 | -0.05 | -0.05 | 0 | 0.66 | 0.25 | 0.4 | 0.53 | 0.48 | 0.74 |
| 6 | 33096766 | rs3117004 | 0.364 | -1.131 | -3.03 | -0.489 | -0.037 | -0.105 | -0.105 | 0.662 | 0.31 | 0.08 | 0.29 | 0.44 | 0.3 | 0.34 |
| 6 | 33097231 | rs3116998 | -0.309 | -1.141 | -3.035 | -0.499 | -0.045 | -0.05 | -0.05 | 0.535 | 0.66 | 0.25 | 0.4 | 0.53 | 0.48 | 0.74 |
| 6 | 33097382 | rs3130235 | 0.349 | -1.173 | -2.818 | -0.582 | -0.015 | -0.015 | -0.015 | 0 | 0.14 | 0.04 | 0.1 | 0.16 | 0.16 | 0.16 |
| 6 | 33097423 | rs3129270 | 1.307 | -1.176 | -2.822 | -0.584 | -0.018 | -0.014 | -0.014 | 0 | 0.06 | 0.03 | 0.06 | 0.13 | 0.11 | 0.11 |
| 6 | 33097561 | rs3130237 | 0.431 | -1.16 | -2.801 | -0.575 | -0.015 | -0.015 | -0.015 | 0 | 0.14 | 0.04 | 0.1 | 0.16 | 0.16 | 0.16 |
| 6 | 33097614 | rs3129269 | -0.2 | -1.077 | -2.742 | -0.526 | 0.116 | 0.131 | 0.131 | 0 | 0.49 | 0.22 | 0.21 | 0.25 | 0.34 | 0.56 |
| 6 | 33098276 | rs6901221 | 0.121 | -0.496 | -2.409 | -0.156 | -0.015 | -0.015 | -0.015 | 1.514 | 0.14 | 0.04 | 0.1 | 0.16 | 0.16 | 0.16 |
| 6 | 33098569 | rs9277768 | 0.559 | -0.561 | -2.674 | -0.1 | -0.049 | -0.096 | -0.096 | 5.708 | 0.23 | 0.06 | 0.25 | 0.41 | 0.26 | 0.29 |
| 6 | 33098596 | rs9277769 | -0.652 | -0.384 | -2.43 | -0.528 | 0.085 | -0.089 | -0.089 | 0 | 0.75 | 0.6 | 0.73 | 0.95 | 0.64 | 0.88 |
| 6 | 33098896 | rs3129267 | -1.187 | -0.281 | -2.407 | -0.563 | 0.159 | -0.097 | -0.097 | 0 | 0.82 | 0.58 | 0.77 | 0.98 | 0.69 | 0.93 |

| | | | | | | | | | | | | | | | | |
|---|---|---|---|---|---|---|---|---|---|---|---|---|---|---|---|---|
| 6 | 33098966 | rs2294478 | 0.153 | -1.051 | -2.723 | -0.794 | -0.055 | 0.132 | 0.132 | 0.104 | 0.56 | 0.26 | 0.62 | 0.78 | 0.48 | 0.61 |
| 6 | 33099210 | rs2294472 | -0.107 | -1.213 | -2.722 | -0.895 | -0.007 | 0.057 | 0.057 | 0 | 0.4 | 0.27 | 0.59 | 0.76 | 0.39 | 0.43 |
| 6 | 33100189 | rs2092305 | 0.319 | -1.221 | -2.715 | -0.908 | 0.004 | -0.005 | -0.005 | 0.029 | 0.11 | 0.25 | 0.1 | 0.13 | 0.13 | 0.17 |
| 6 | 33100732 | rs2395352 | 0.178 | -1.101 | -2.671 | -0.84 | -0.045 | 0.049 | 0.049 | 0 | 0.37 | 0.19 | 0.56 | 0.76 | 0.37 | 0.36 |
| 6 | 33101444 | rs7756467 | 0.178 | -1.101 | -2.671 | -0.84 | -0.045 | 0.049 | 0.049 | 0 | 0.37 | 0.19 | 0.56 | 0.76 | 0.37 | 0.36 |
| 6 | 33101602 | rs3129264 | 0.856 | -1.173 | -2.617 | -0.842 | -0.004 | -0.007 | -0.007 | 0.625 | 0.3 | 0.28 | 0.15 | 0.17 | 0.23 | 0.42 |
| 6 | 33103321 | rs7750000 | 0.475 | -1.041 | -2.626 | -0.63 | -0.022 | -0.072 | -0.072 | 0 | 0.2 | 0.16 | 0.37 | 0.46 | 0.23 | 0.18 |
| 6 | 33104175 | rs9366816 | 0.475 | -1.041 | -2.626 | -0.63 | -0.022 | -0.072 | -0.072 | 0 | 0.2 | 0.16 | 0.37 | 0.46 | 0.23 | 0.18 |
| 6 | 33105064 | rs13437000 | 0.87 | -0.955 | -2.556 | -0.541 | -0.01 | 0.006 | 0.006 | 0 | 0.12 | 0.15 | 0.34 | 0.43 | 0.19 | 0.13 |
| 6 | 33105469 | rs3129258 | 0.808 | -1.14 | -2.612 | -0.688 | -0.002 | -0.004 | -0.004 | 0 | 0.3 | 0.28 | 0.15 | 0.17 | 0.23 | 0.42 |
| 6 | 33107461 | rs4711319 | 1.814 | -1.12 | -2.677 | -0.66 | -0.024 | -0.078 | -0.078 | 0.071 | 0.17 | 0.03 | 0.19 | 0.29 | 0.15 | 0.19 |
| 6 | 33107955 | rs4713610 | 1.886 | -1.114 | -2.674 | -0.655 | -0.025 | -0.076 | -0.076 | 0 | 0.17 | 0.03 | 0.19 | 0.28 | 0.14 | 0.18 |
| 6 | 33108321 | rs9296083 | 0.9 | -1.009 | -2.615 | -0.601 | -0.014 | -0.003 | -0.003 | 0 | 0.12 | 0.14 | 0.34 | 0.44 | 0.19 | 0.13 |
| 6 | 33109556 | rs3129249 | 0.664 | -1.113 | -2.635 | -0.68 | -0.003 | -0.005 | -0.005 | 0.135 | 0.3 | 0.28 | 0.15 | 0.17 | 0.23 | 0.42 |
| 6 | 33110167 | rs9296086 | 0.927 | -1.077 | -2.632 | -0.651 | -0.014 | -0.003 | -0.003 | 0 | 0.12 | 0.14 | 0.34 | 0.44 | 0.19 | 0.13 |
| 6 | 33110834 | rs7750683 | -0.074 | -1.128 | -2.665 | -0.657 | 0.002 | 0.003 | 0.003 | 0 | 0.43 | 0.42 | 0.49 | 0.62 | 0.43 | 0.55 |
| 6 | 33110933 | rs7754200 | 0.841 | -1.148 | -2.639 | -0.68 | -0.003 | -0.005 | -0.005 | 0 | 0.3 | 0.28 | 0.15 | 0.17 | 0.23 | 0.42 |
| 6 | 33111347 | rs3129234 | 0.662 | -1.146 | -2.638 | -0.679 | -0.003 | -0.005 | -0.005 | 0.514 | 0.3 | 0.28 | 0.15 | 0.17 | 0.23 | 0.42 |
| 6 | 33112218 | rs3129231 | 0.662 | -1.146 | -2.638 | -0.679 | -0.003 | -0.005 | -0.005 | 0 | 0.3 | 0.28 | 0.15 | 0.17 | 0.23 | 0.42 |
| 6 | 33112601 | rs3129228 | 0.843 | -1.155 | -2.642 | -0.683 | -0.003 | -0.005 | -0.005 | 0 | 0.3 | 0.28 | 0.15 | 0.17 | 0.23 | 0.42 |
| 6 | 33113197 | rs3129223 | 0.665 | -1.154 | -2.642 | -0.683 | -0.003 | -0.005 | -0.005 | 0 | 0.3 | 0.28 | 0.15 | 0.17 | 0.23 | 0.42 |
| 6 | 33114171 | rs1003979 | -0.239 | -1.186 | -2.727 | -0.751 | -0.003 | 0.004 | 0.004 | 0 | 0.58 | 0.45 | 0.67 | 0.9 | 0.56 | 0.74 |
| 6 | 33115024 | rs3130151 | -0.042 | -1.009 | -2.619 | -0.622 | 0.001 | 0.002 | 0.002 | 0 | 0.42 | 0.42 | 0.49 | 0.62 | 0.42 | 0.55 |
| 6 | 33115762 | rs3130153 | -0.017 | -1.025 | -2.62 | -0.628 | 0.002 | -0.001 | -0.001 | 0.333 | 0.42 | 0.42 | 0.48 | 0.61 | 0.42 | 0.55 |
| 6 | 33116222 | rs2395365 | -0.017 | -1.025 | -2.62 | -0.628 | 0.002 | -0.001 | -0.001 | 1.676 | 0.42 | 0.42 | 0.48 | 0.61 | 0.41 | 0.55 |
| 6 | 33117258 | rs3129214 | 0.658 | -1.033 | -2.624 | -0.629 | -0.003 | -0.005 | -0.005 | 0 | 0.3 | 0.27 | 0.15 | 0.17 | 0.23 | 0.42 |
| 6 | 33117818 | rs721394 | 0.922 | -0.985 | -2.606 | -0.595 | -0.013 | 0 | 0 | 0 | 0.12 | 0.15 | 0.33 | 0.44 | 0.18 | 0.13 |
| 6 | 33121678 | rs726599 | -1.573 | -0.375 | -2.618 | -0.792 | 0.169 | -0.061 | -0.061 | 0 | 0.83 | 0.62 | 0.76 | 0.96 | 0.68 | 0.93 |
| 6 | 33122171 | rs756441 | 0.076 | -0.424 | -2.633 | -0.78 | 0.002 | -0.001 | -0.001 | 0 | 0.42 | 0.42 | 0.48 | 0.61 | 0.42 | 0.55 |
| 6 | 33122331 | rs756440 | 0.901 | -0.505 | -2.601 | -0.805 | -0.003 | -0.005 | -0.005 | 0 | 0.3 | 0.27 | 0.15 | 0.17 | 0.23 | 0.42 |
| 6 | 33124494 | rs3116959 | 0.901 | -0.505 | -2.601 | -0.805 | -0.003 | -0.005 | -0.005 | 0 | 0.3 | 0.27 | 0.15 | 0.17 | 0.23 | 0.42 |
| 6 | 33125462 | rs3129206 | -0.784 | -0.116 | -2.604 | -0.495 | 0.129 | -0.058 | -0.058 | 0 | 0.75 | 0.61 | 0.51 | 0.92 | 0.56 | 0.86 |
| 6 | 33125725 | rs9277920 | 0.832 | -0.404 | -2.746 | -0.603 | 0.095 | 0.036 | 0.036 | 0.01 | 0.52 | 0.21 | 0.31 | 0.38 | 0.33 | 0.58 |
| 6 | 33125858 | rs3130161 | 0.368 | -0.283 | -2.599 | -0.522 | 0.006 | -0.002 | -0.002 | 0 | 0.11 | 0.25 | 0.1 | 0.13 | 0.13 | 0.17 |
| 6 | 33127757 | rs3129203 | 0.409 | -0.301 | -2.61 | -0.532 | 0.01 | 0.001 | 0.001 | 0 | 0.1 | 0.25 | 0.1 | 0.13 | 0.13 | 0.18 |
| 6 | 33127987 | rs4713614 | -0.32 | -0.207 | -2.688 | -0.624 | -0.015 | 0.01 | 0.01 | 0.84 | 0.72 | 0.37 | 0.67 | 0.83 | 0.55 | 0.76 |
| 6 | 33128377 | rs9368757 | 0.624 | -0.177 | -2.698 | -0.495 | -0.014 | -0.002 | -0.002 | 0 | 0.12 | 0.15 | 0.33 | 0.43 | 0.18 | 0.16 |
| 6 | 33129071 | rs3116956 | -1.506 | -0.196 | -2.699 | -0.51 | 0.165 | -0.064 | -0.064 | 0 | 0.82 | 0.61 | 0.76 | 0.96 | 0.68 | 0.93 |
| 6 | 33129746 | rs9380351 | 0.67 | -0.166 | -2.7 | -0.508 | -0.013 | 0 | 0 | 0 | 0.12 | 0.15 | 0.33 | 0.44 | 0.18 | 0.16 |
| 6 | 33130430 | rs2235498 | 1.072 | -0.547 | -2.907 | -0.558 | 0.023 | -0.023 | -0.023 | 0 | 0.27 | 0.12 | 0.24 | 0.3 | 0.21 | 0.24 |
| 6 | 33131734 | rs2257126 | -1.371 | -0.185 | -2.598 | -0.571 | 0.163 | -0.065 | -0.065 | 0.144 | 0.82 | 0.59 | 0.75 | 0.96 | 0.68 | 0.93 |
| 6 | 33134392 | rs2855453 | -1.203 | -0.19 | -2.6 | -0.574 | 0.15 | -0.067 | -0.067 | 0 | 0.81 | 0.59 | 0.75 | 0.95 | 0.68 | 0.88 |
| 6 | 33136145 | rs986521 | -1.399 | 0.465 | -2.816 | -0.714 | -0.036 | 0.012 | 0.012 | 0 | 0.92 | 0.96 | 0.84 | 0.99 | 0.75 | 0.94 |
| 6 | 33136575 | rs2855448 | -0.777 | 0.403 | -2.644 | -0.764 | 0.17 | -0.075 | -0.075 | 0 | 0.82 | 0.58 | 0.75 | 0.95 | 0.65 | 0.9 |
| 6 | 33137403 | rs2855442 | -1.159 | 0.332 | -2.575 | -0.808 | -0.076 | 0.04 | 0.04 | 0.148 | 0.85 | 0.91 | 0.79 | 0.95 | 0.72 | 0.91 |
| 6 | 33137727 | rs2855441 | 0.203 | 0.072 | -2.627 | -0.94 | 0.023 | -0.031 | -0.031 | 0 | 0.75 | 0.39 | 0.69 | 0.94 | 0.64 | 0.89 |
| 6 | 33138021 | rs9368758 | 0.727 | 0.027 | -2.731 | -0.517 | 0 | 0.103 | 0.103 | 0 | 0.06 | 0.11 | 0.08 | 0.49 | 0.1 | 0.1 |
| 6 | 33138498 | rs2855440 | -0.945 | -0.016 | -2.726 | -0.563 | -0.038 | 0.021 | 0.021 | 0.459 | 0.82 | 0.84 | 0.76 | 0.95 | 0.69 | 0.91 |
| 6 | 33138677 | rs1799911 | 1.746 | -0.669 | -3.144 | -0.998 | 0.057 | 0.052 | 0.052 | 0.388 | 0.37 | 0.12 | 0.26 | 0.24 | 0.25 | 0.4 |
| 6 | 33138955 | rs2855437 | -0.773 | -0.573 | -3.13 | -0.877 | -0.035 | 0.012 | 0.012 | 0 | 0.8 | 0.82 | 0.75 | 0.88 | 0.69 | 0.86 |
| 6 | 33139328 | rs1799910 | 0.495 | -0.685 | -3.215 | -0.966 | 0.001 | 0.044 | 0.044 | 0 | 0.58 | 0.68 | 0.39 | 0.34 | 0.44 | 0.62 |
| 6 | 33139662 | rs2855434 | -1.6 | -0.567 | -3.145 | -0.933 | -0.073 | 0.036 | 0.036 | 0.246 | 0.85 | 0.91 | 0.79 | 0.96 | 0.72 | 0.93 |
| 6 | 33140009 | rs973233 | -1.616 | -0.564 | -3.147 | -0.934 | -0.075 | 0.057 | 0.057 | 0.231 | 0.85 | 0.92 | 0.8 | 0.98 | 0.72 | 0.93 |
| 6 | 33140983 | rs2855432 | -1.727 | -0.565 | -3.146 | -0.932 | -0.074 | 0.038 | 0.038 | 0.314 | 0.85 | 0.91 | 0.79 | 0.96 | 0.72 | 0.93 |
| 6 | 33141253 | rs9277932 | 0.254 | -0.659 | -3.169 | -0.98 | -0.015 | 0.022 | 0.022 | 0.363 | 0.63 | 0.61 | 0.65 | 0.79 | 0.54 | 0.64 |
| 6 | 33141280 | rs2855430 | 1.509 | -0.767 | -3.222 | -1.038 | -0.026 | 0.013 | 0.013 | 0 | 0.13 | 0.01 | 0.07 | 0.1 | 0.14 | 0.21 |
| 6 | 33141475 | rs970901 | 0.323 | -0.699 | -3.213 | -1.029 | -0.016 | 0.022 | 0.022 | 0.391 | 0.63 | 0.61 | 0.65 | 0.78 | 0.54 | 0.64 |
| 6 | 33142162 | rs2076312 | 0.132 | -0.673 | -3.201 | -1.02 | -0.011 | -0.012 | -0.012 | 0 | 0.22 | 0.3 | 0.14 | 0.18 | 0.17 | 0.29 |
| 6 | 33142792 | rs2071026 | 0.312 | -0.675 | -3.206 | -1.008 | -0.015 | 0.021 | 0.021 | 0.391 | 0.63 | 0.61 | 0.65 | 0.78 | 0.55 | 0.64 |
| 6 | 33144373 | rs2855425 | -1.751 | -0.637 | -3.196 | -1.015 | -0.071 | 0.038 | 0.038 | 0.328 | 0.85 | 0.91 | 0.79 | 0.96 | 0.72 | 0.93 |
| 6 | 33145369 | rs2076311 | 1.793 | -0.747 | -3.254 | -1.058 | -0.025 | -0.031 | -0.031 | 0 | 0.38 | 0.39 | 0.27 | 0.24 | 0.28 | 0.4 |
| 6 | 33145710 | rs2855423 | -1.721 | -0.632 | -3.315 | -0.96 | -0.075 | 0.056 | 0.056 | 0.221 | 0.85 | 0.92 | 0.79 | 0.98 | 0.73 | 0.93 |
| 6 | 33148878 | rs2744507 | 1.536 | -0.667 | -3.325 | -0.962 | -0.031 | 0.011 | 0.011 | 0 | 0.12 | 0.01 | 0.07 | 0.1 | 0.14 | 0.21 |
| 6 | 33158189 | rs2855429 | -1.736 | -0.556 | -3.28 | -0.916 | -0.074 | 0.038 | 0.038 | 0.314 | 0.85 | 0.91 | 0.79 | 0.96 | 0.72 | 0.93 |
| 6 | 33158898 | rs2269346 | 0.957 | -0.571 | -3.293 | -0.88 | -0.001 | 0.09 | 0.09 | 0 | 0.07 | 0.09 | 0.08 | 0.49 | 0.1 | 0.1 |
| 6 | 33159664 | rs9469378 | 2.246 | -0.628 | -3.325 | -0.926 | 0.019 | 0.028 | 0.028 | 0 | 0.08 | 0.03 | 0.03 | 0.02 | 0.04 | 0.07 |
| 6 | 33160425 | rs9277935 | -0.019 | -0.591 | -3.289 | -0.903 | -0.001 | 0.011 | 0.011 | 0 | 0.24 | 0.29 | 0.18 | 0.59 | 0.24 | 0.32 |
| 6 | 33161661 | rs1050673 | -1.671 | -0.475 | -3.264 | -0.856 | -0.075 | 0.039 | 0.039 | 0.398 | 0.85 | 0.91 | 0.79 | 0.96 | 0.72 | 0.93 |
| 6 | 33162215 | rs2744537 | -1.689 | -0.472 | -3.262 | -0.854 | -0.074 | 0.038 | 0.038 | 0.314 | 0.85 | 0.91 | 0.79 | 0.96 | 0.72 | 0.93 |

| | | | | | | | | | | | | | | | | |
|---|---|---|---|---|---|---|---|---|---|---|---|---|---|---|---|---|
| 6 | 33163451 | rs6531 | -1.533 | -0.473 | -3.262 | -0.855 | -0.073 | 0.042 | 0.042 | 0.197 | 0.84 | 0.89 | 0.79 | 0.95 | 0.72 | 0.91 |
| 6 | 33164735 | rs3117040 | -1.624 | -0.469 | -3.226 | -0.853 | -0.067 | 0.039 | 0.039 | 0.239 | 0.85 | 0.91 | 0.78 | 0.96 | 0.72 | 0.93 |
| 6 | 33166034 | rs2076310 | -0.336 | -0.492 | -3.207 | -0.841 | -0.004 | 0.023 | 0.023 | 0 | 0.28 | 0.39 | 0.22 | 0.6 | 0.27 | 0.34 |
| 6 | 33172932 | rs110662 | -1.61 | -0.434 | -3.209 | -0.845 | -0.073 | 0.022 | 0.022 | 0.193 | 0.85 | 0.92 | 0.79 | 0.91 | 0.73 | 0.87 |
| 6 | 33173842 | rs439205 | -0.376 | -0.576 | -3.124 | -0.75 | -0.004 | 0.02 | 0.02 | 0 | 0.28 | 0.38 | 0.22 | 0.6 | 0.27 | 0.34 |
| 6 | 33173998 | rs383711 | 0.843 | -0.565 | -3.122 | -0.725 | -0.001 | 0.09 | 0.09 | 0 | 0.07 | 0.1 | 0.08 | 0.49 | 0.1 | 0.1 |
| 6 | 33174783 | rs421446 | -0.647 | -0.629 | -3.161 | -0.775 | 0.004 | -0.004 | -0.004 | 0 | 0.36 | 0.41 | 0.25 | 0.62 | 0.31 | 0.39 |
| 6 | 33175575 | rs107822 | -0.42 | -0.525 | -3.084 | -0.691 | -0.004 | 0.022 | 0.022 | 0 | 0.28 | 0.39 | 0.22 | 0.6 | 0.27 | 0.34 |
| 6 | 33175824 | rs213210 | 0.804 | -0.481 | -3.061 | -0.642 | -0.001 | 0.09 | 0.09 | 0 | 0.07 | 0.1 | 0.08 | 0.49 | 0.1 | 0.1 |
| 6 | 33178010 | rs213208 | -0.186 | -0.492 | -3.039 | -0.806 | -0.004 | 0.023 | 0.023 | 0 | 0.28 | 0.39 | 0.22 | 0.61 | 0.27 | 0.35 |
| 6 | 33179689 | rs2854028 | 1.906 | -0.554 | -3.278 | -0.946 | 0.004 | 0.031 | 0.031 | 0.379 | 0.28 | 0.03 | 0.22 | 0.15 | 0.21 | 0.33 |
| 6 | 33182870 | rs1567464 | 2.037 | -0.25 | -3.013 | -0.826 | -0.036 | -0.014 | -0.014 | 0 | 0.07 | 0.01 | 0.24 | 0.04 | 0.12 | 0.09 |
| 6 | 33183730 | rs213213 | -0.795 | -0.058 | -2.248 | -0.937 | -0.031 | 0.039 | 0.039 | 0 | 0.77 | 0.85 | 0.77 | 0.91 | 0.69 | 0.87 |
| 6 | 33184894 | rs9277937 | 1.845 | -0.131 | -2.27 | -0.98 | 0.035 | 0.04 | 0.04 | 0.166 | 0.18 | 0.03 | 0.05 | 0.05 | 0.1 | 0.21 |
| 6 | 33185918 | rs213212 | -1.391 | 0.493 | -1.768 | -0.6 | -0.014 | 0.01 | 0.01 | 0.064 | 0.85 | 0.88 | 0.79 | 0.98 | 0.73 | 0.93 |
| 6 | 33192662 | rs386084 | -0.317 | 0.504 | -1.422 | -0.282 | -0.001 | 0.014 | 0.014 | 0 | 0.29 | 0.39 | 0.23 | 0.6 | 0.29 | 0.35 |
| 6 | 33195604 | rs213194 | -0.858 | 0.82 | -1.247 | -0.496 | -0.018 | 0.015 | 0.015 | 0.427 | 0.83 | 0.87 | 0.79 | 0.92 | 0.73 | 0.9 |
| 6 | 33202640 | rs213220 | -0.632 | 0.691 | -1.15 | -0.348 | -0.016 | 0.014 | 0.014 | 0 | 0.67 | 0.68 | 0.55 | 0.77 | 0.55 | 0.64 |
| 6 | 33203338 | rs213221 | -0.197 | 0.459 | -1.111 | -0.463 | -0.033 | 0.063 | 0.063 | 0.189 | 0.6 | 0.31 | 0.51 | 0.71 | 0.54 | 0.62 |
| 6 | 33204274 | rs9277952 | 0.813 | 0.361 | -1.217 | 0.039 | -0.026 | -0.066 | -0.066 | 0.023 | 0.1 | 0.01 | 0.1 | 0.38 | 0.12 | 0.22 |
| 6 | 33209310 | rs213226 | -0.072 | 0.546 | -0.548 | -0.939 | -0.013 | 0.022 | 0.022 | 0.164 | 0.6 | 0.35 | 0.52 | 0.7 | 0.55 | 0.62 |
| 6 | 33215904 | rs457565 | 0.619 | 0.337 | -0.75 | -1.232 | 0.017 | 0.01 | 0.01 | 0 | 0.41 | 0.25 | 0.37 | 0.58 | 0.39 | 0.4 |
| 6 | 33218180 | rs461338 | 1.586 | 0.358 | -0.813 | -1.24 | -0.003 | 0.037 | 0.037 | 0 | 0.18 | 0.04 | 0.13 | 0.09 | 0.15 | 0.21 |
| 6 | 33219142 | rs213216 | 0.363 | 0.388 | -0.696 | -1.088 | 0.008 | 0.013 | 0.013 | 0 | 0.4 | 0.21 | 0.37 | 0.61 | 0.39 | 0.4 |
| 6 | 33232055 | rs213202 | 0.374 | 0.511 | -0.514 | -1.022 | 0.011 | 0.007 | 0.007 | 0 | 0.41 | 0.33 | 0.38 | 0.58 | 0.38 | 0.4 |
| 6 | 33232521 | rs213201 | 0.293 | 0.495 | -0.481 | -1.03 | 0.019 | 0.012 | 0.012 | 0 | 0.42 | 0.25 | 0.37 | 0.58 | 0.38 | 0.4 |
| 6 | 33239905 | rs465658 | 1.536 | 0.34 | -0.238 | -1.2 | 0.009 | 0.047 | 0.047 | 0 | 0.19 | 0.05 | 0.13 | 0.09 | 0.15 | 0.21 |
| 6 | 33240699 | rs466886 | 1.536 | 0.34 | -0.238 | -1.2 | 0.009 | 0.047 | 0.047 | 0 | 0.19 | 0.05 | 0.13 | 0.09 | 0.15 | 0.21 |
| 6 | 33241076 | rs213204 | 0.639 | 0.353 | -0.238 | -1.241 | 0.015 | 0.008 | 0.008 | 0 | 0.41 | 0.24 | 0.37 | 0.58 | 0.38 | 0.4 |
| 6 | 33241135 | rs213205 | -1.056 | 0.366 | -0.003 | -1.262 | 0.016 | -0.013 | -0.013 | 0.046 | 0.67 | 0.61 | 0.53 | 0.71 | 0.55 | 0.62 |
| 6 | 33241169 | rs213207 | -1.056 | 0.366 | -0.003 | -1.262 | 0.016 | -0.013 | -0.013 | 0 | 0.67 | 0.61 | 0.53 | 0.71 | 0.55 | 0.62 |
| 6 | 33242825 | rs458434 | 1.538 | 0.334 | -0.184 | -1.188 | 0.007 | 0.047 | 0.047 | 0 | 0.19 | 0.05 | 0.13 | 0.09 | 0.15 | 0.21 |
| 6 | 33244641 | rs464367 | -0.969 | 0.543 | -0.114 | -1.201 | 0.019 | -0.016 | -0.016 | 0 | 0.67 | 0.54 | 0.49 | 0.72 | 0.52 | 0.63 |
| 6 | 33246518 | rs3445 | 1.395 | 0.387 | -0.416 | -1.172 | 0.009 | 0.049 | 0.049 | 0 | 0.2 | 0.05 | 0.13 | 0.09 | 0.16 | 0.21 |
| 6 | 33250476 | rs469064 | 0.354 | 0.76 | -0.55 | -1.353 | -0.035 | 0.04 | 0.04 | 0.096 | 0.45 | 0.2 | 0.5 | 0.41 | 0.5 | 0.58 |
| 6 | 33252115 | rs455567 | -0.747 | 0.533 | -0.394 | -1.457 | -0.027 | 0.017 | 0.017 | 0 | 0.58 | 0.76 | 0.56 | 0.42 | 0.52 | 0.61 |
| 6 | 33252336 | rs467508 | 1.724 | 0.477 | -0.464 | -1.524 | 0.003 | 0.036 | 0.036 | 0 | 0.19 | 0.03 | 0.11 | 0.09 | 0.13 | 0.22 |
| 6 | 33252561 | rs6906691 | 1.813 | 0.444 | -0.183 | -1.538 | -0.066 | -0.069 | -0.069 | 0.446 | 0.12 | 0.39 | 0.04 | 0.01 | 0.02 | 0.02 |
| 6 | 33252661 | rs6907034 | 2.265 | 0.446 | -0.216 | -1.537 | -0.037 | -0.038 | -0.038 | 0 | 0.09 | 0.19 | 0.03 | 0.01 | 0.02 | 0.02 |
| 6 | 33254665 | rs14398 | 2.009 | 0.494 | -0.428 | -1.518 | -0.013 | 0.046 | 0.046 | 0 | 0.18 | 0.01 | 0.11 | 0.08 | 0.12 | 0.21 |
| 6 | 33255103 | rs446735 | -0.799 | 0.503 | -0.427 | -1.526 | -0.027 | 0.017 | 0.017 | 0 | 0.58 | 0.76 | 0.56 | 0.42 | 0.52 | 0.61 |
| 6 | 33255369 | rs465151 | 2.017 | 0.506 | -0.43 | -1.552 | -0.013 | 0.046 | 0.046 | 0 | 0.18 | 0.01 | 0.11 | 0.07 | 0.12 | 0.21 |
| 6 | 33256984 | rs463260 | -0.739 | 0.975 | -0.319 | -1.444 | -0.027 | 0.018 | 0.018 | 0 | 0.59 | 0.76 | 0.56 | 0.42 | 0.52 | 0.61 |
| 6 | 33257186 | rs464865 | -0.713 | 0.527 | -0.283 | -1.419 | -0.027 | 0.019 | 0.019 | 0 | 0.58 | 0.76 | 0.56 | 0.42 | 0.52 | 0.61 |
| 6 | 33258443 | rs456261 | -0.126 | 0.497 | -0.266 | -1.433 | 0 | 0.003 | 0.003 | 0 | 0.54 | 0.52 | 0.53 | 0.38 | 0.51 | 0.59 |
| 6 | 33261871 | rs2239842 | 1.756 | 0.566 | -0.473 | -1.409 | -0.093 | -0.046 | -0.046 | 0 | 0.2 | 0.02 | 0.34 | 0.28 | 0.33 | 0.32 |
| 6 | 33267672 | rs1059288 | -0.618 | 0.531 | -0.359 | -1.324 | -0.024 | 0.023 | 0.023 | 0 | 0.58 | 0.76 | 0.57 | 0.41 | 0.52 | 0.64 |
| 6 | 33280309 | rs2282851 | 1.981 | 0.58 | -0.635 | -1.014 | -0.077 | -0.051 | -0.051 | 0 | 0.17 | 0.02 | 0.34 | 0.26 | 0.29 | 0.29 |
| 6 | 33280505 | rs3106191 | 1.965 | 0.577 | -0.636 | -1.016 | -0.077 | -0.051 | -0.051 | 0.011 | 0.17 | 0.02 | 0.34 | 0.26 | 0.29 | 0.29 |
| 6 | 33282002 | rs3106189 | -0.217 | 0.244 | -0.411 | -1.334 | 0.001 | 0.005 | 0.005 | 0 | 0.53 | 0.57 | 0.53 | 0.38 | 0.5 | 0.61 |
| 6 | 33282181 | rs3130099 | -0.217 | 0.244 | -0.405 | -1.334 | 0.001 | 0.007 | 0.007 | 0 | 0.53 | 0.57 | 0.53 | 0.38 | 0.5 | 0.61 |
| 6 | 33282338 | rs1061801 | 1.383 | 0.065 | -0.444 | -1.425 | 0.002 | 0.064 | 0.064 | 0 | 0.22 | 0.04 | 0.15 | 0.11 | 0.18 | 0.3 |
| 6 | 33282628 | rs1061783 | -0.368 | 0.116 | -0.435 | -1.379 | 0 | 0.001 | 0.001 | 0 | 0.54 | 0.57 | 0.54 | 0.39 | 0.51 | 0.62 |
| 6 | 33283766 | rs3130100 | -0.439 | -0.004 | -0.509 | -1.432 | 0 | -0.001 | -0.001 | 0 | 0.56 | 0.57 | 0.55 | 0.39 | 0.53 | 0.63 |
| 6 | 33287183 | rs2051625 | 2.05 | 0.034 | -0.276 | -1.376 | -0.04 | -0.042 | -0.042 | 0 | 0.1 | 0.21 | 0.03 | 0.01 | 0.02 | 0.02 |
| 6 | 33288075 | rs2239839 | 2.118 | 0.052 | -0.297 | -1.376 | -0.077 | -0.05 | -0.05 | 0.216 | 0.18 | 0.02 | 0.34 | 0.26 | 0.29 | 0.29 |
| 6 | 33288271 | rs1059231 | 2.118 | 0.052 | -0.297 | -1.376 | -0.077 | -0.05 | -0.05 | 0 | 0.18 | 0.02 | 0.34 | 0.26 | 0.29 | 0.29 |
| 6 | 33290402 | rs3130018 | 0.773 | 0.003 | -0.142 | -1.064 | -0.041 | 0.015 | 0.015 | 0.112 | 0.42 | 0.16 | 0.5 | 0.39 | 0.49 | 0.6 |
| 6 | 33290825 | rs2073525 | -0.478 | 0.394 | 0.05 | -0.85 | -0.021 | 0.009 | 0.009 | 0 | 0.58 | 0.76 | 0.57 | 0.43 | 0.51 | 0.64 |
| 6 | 33303989 | rs3130266 | 0.114 | 0.565 | -0.399 | -0.831 | 0.004 | 0.018 | 0.018 | 0 | 0.52 | 0.5 | 0.53 | 0.39 | 0.51 | 0.62 |
| 6 | 33308221 | rs3130270 | 0.114 | 0.565 | -0.399 | -0.855 | 0.004 | 0.018 | 0.018 | 0 | 0.52 | 0.5 | 0.53 | 0.38 | 0.51 | 0.62 |
| 6 | 33312308 | rs3130014 | 0.643 | 0.829 | -0.868 | -0.878 | -0.007 | -0.038 | -0.038 | 0 | 0.33 | 0.18 | 0.32 | 0.58 | 0.34 | 0.34 |
| 6 | 33319596 | rs211476 | 0.028 | 0.724 | -0.486 | -0.991 | -0.016 | -0.009 | -0.009 | 0.34 | 0.43 | 0.47 | 0.34 | 0.61 | 0.3 | 0.39 |
| 6 | 33320613 | rs211474 | -0.079 | 0.713 | -0.45 | -1.005 | -0.005 | -0.004 | -0.004 | 0 | 0.44 | 0.49 | 0.34 | 0.62 | 0.31 | 0.39 |
| 6 | 33324527 | rs3117324 | -0.081 | 0.833 | -0.21 | -1.032 | -0.001 | 0.018 | 0.018 | 0 | 0.55 | 0.4 | 0.56 | 0.39 | 0.59 | 0.61 |
| 6 | 33326250 | rs3106192 | 0.53 | 0.587 | -0.622 | -1.436 | -0.075 | -0.02 | -0.02 | 0 | 0.33 | 0.1 | 0.44 | 0.36 | 0.46 | 0.53 |
| 6 | 33326434 | rs9278019 | 2.485 | 0.429 | -0.693 | -1.52 | -0.031 | 0.056 | 0.056 | 0 | 0.11 | 0.01 | 0.09 | 0.02 | 0.11 | 0.06 |
| 6 | 33328518 | rs211455 | 0.435 | 0.414 | -0.606 | -1.527 | -0.022 | -0.014 | -0.014 | 0.881 | 0.43 | 0.47 | 0.34 | 0.62 | 0.3 | 0.38 |
| 6 | 33330131 | rs211453 | 0.223 | 0.562 | -0.566 | -1.654 | -0.003 | -0.007 | -0.007 | 0 | 0.49 | 0.47 | 0.39 | 0.68 | 0.36 | 0.56 |

| | | | | | | | | | | | | | | | | |
|---|---|---|---|---|---|---|---|---|---|---|---|---|---|---|---|---|
| 6 | 33330981 | rs211452 | 0.424 | 0.481 | -0.566 | -1.687 | 0.06 | 0.083 | 0.083 | 0.177 | 0.47 | 0.28 | 0.38 | 0.65 | 0.36 | 0.56 |
| 6 | 33332137 | rs3130275 | 1.235 | 0.613 | -0.805 | -1.648 | -0.099 | -0.036 | -0.036 | 0 | 0.23 | 0.04 | 0.37 | 0.3 | 0.39 | 0.34 |
| 6 | 33336930 | rs3130276 | 1.243 | 0.613 | -0.807 | -1.648 | -0.099 | -0.036 | -0.036 | 0 | 0.23 | 0.05 | 0.37 | 0.3 | 0.39 | 0.34 |
| 6 | 33340510 | rs9278027 | 2.326 | 0.613 | -0.82 | -1.654 | -0.031 | 0.056 | 0.056 | 0 | 0.11 | 0.01 | 0.09 | 0.02 | 0.12 | 0.06 |
| 6 | 33346186 | rs2747479 | 0.171 | 0.451 | -0.711 | -1.607 | 0.051 | 0.054 | 0.054 | 0.1 | 0.51 | 0.65 | 0.42 | 0.64 | 0.36 | 0.55 |
| 6 | 33349885 | rs2747477 | 0.19 | 0.436 | -0.694 | -1.614 | 0.014 | 0.015 | 0.015 | 0.003 | 0.49 | 0.53 | 0.4 | 0.64 | 0.36 | 0.55 |
| 6 | 33351696 | rs6942045 | 2.123 | 0.766 | -0.883 | -1.488 | -0.083 | -0.073 | -0.073 | 0.01 | 0.17 | 0.01 | 0.34 | 0.25 | 0.31 | 0.32 |
| 6 | 33358278 | rs456993 | 1.011 | 0.776 | -0.836 | -1.497 | -0.075 | -0.042 | -0.042 | 6.664 | 0.24 | 0.06 | 0.37 | 0.31 | 0.39 | 0.35 |
| 6 | 33359008 | rs465506 | 0.837 | 0.75 | -0.793 | -1.5 | -0.055 | -0.032 | -0.032 | 1.609 | 0.25 | 0.09 | 0.37 | 0.32 | 0.4 | 0.36 |
| 6 | 33359142 | rs465474 | 1.019 | 0.75 | -0.793 | -1.5 | -0.077 | -0.042 | -0.042 | 0 | 0.24 | 0.06 | 0.37 | 0.31 | 0.39 | 0.35 |
| 6 | 33360574 | rs461964 | 0.838 | 0.75 | -0.793 | -1.5 | -0.055 | -0.032 | -0.032 | 0.228 | 0.25 | 0.09 | 0.37 | 0.32 | 0.4 | 0.36 |
| 6 | 33380804 | rs2235982 | 2.35 | 0.857 | -0.717 | -1.616 | -0.031 | 0.056 | 0.056 | 0 | 0.11 | 0.01 | 0.09 | 0.02 | 0.11 | 0.06 |
| 6 | 33382241 | rs442745 | 1.076 | 0.9 | -0.69 | -1.578 | -0.077 | -0.042 | -0.042 | 0.285 | 0.24 | 0.06 | 0.37 | 0.31 | 0.39 | 0.35 |
| 6 | 33383974 | rs4231 | 2.345 | 0.534 | -0.672 | -1.564 | -0.031 | 0.056 | 0.056 | 0 | 0.11 | 0.01 | 0.09 | 0.02 | 0.11 | 0.06 |
| 6 | 33385953 | rs1705003 | 1.012 | 0.393 | -0.584 | -1.634 | -0.009 | -0.035 | -0.035 | 0 | 0.23 | 0.25 | 0.12 | 0.04 | 0.14 | 0.08 |
| 6 | 33389381 | rs211456 | 0.427 | 0.504 | -0.633 | -1.685 | 0.002 | 0.004 | 0.004 | 0 | 0.47 | 0.52 | 0.41 | 0.64 | 0.37 | 0.54 |
| 6 | 33389603 | rs7752202 | 1.807 | 0.504 | -0.646 | -1.642 | 0.006 | 0.029 | 0.029 | 0 | 0.15 | 0.07 | 0.1 | 0.03 | 0.13 | 0.08 |
| 6 | 33399778 | rs9394145 | 2.275 | 0.57 | -0.696 | -1.618 | -0.085 | -0.073 | -0.073 | 0.081 | 0.17 | 0.01 | 0.32 | 0.25 | 0.32 | 0.32 |
| 6 | 33403422 | rs453590 | 0.335 | 0.272 | -0.464 | -1.498 | 0.014 | 0.061 | 0.061 | 0.041 | 0.46 | 0.27 | 0.41 | 0.64 | 0.39 | 0.54 |
| 6 | 33408542 | rs411136 | 0.334 | 0.271 | -0.458 | -1.478 | 0.014 | 0.058 | 0.058 | 0.035 | 0.46 | 0.27 | 0.4 | 0.63 | 0.38 | 0.54 |
| 6 | 33416818 | rs1755044 | 1.633 | 0.333 | -0.615 | -1.452 | -0.033 | -0.034 | -0.034 | 0.519 | 0.09 | 0.17 | 0.02 | 0.01 | 0.02 | 0.02 |
| 6 | 33421577 | rs2247385 | 0.086 | 0.332 | -0.611 | -1.473 | 0.008 | 0.017 | 0.017 | 0.011 | 0.49 | 0.53 | 0.42 | 0.64 | 0.38 | 0.54 |
| 6 | 33422150 | rs2772385 | 1.612 | 0.203 | -0.405 | -1.172 | -0.03 | -0.031 | -0.031 | 0 | 0.09 | 0.16 | 0.02 | 0.01 | 0.02 | 0.02 |
| 6 | 33427350 | rs2772372 | -0.64 | 0.23 | -0.407 | -1.173 | -0.001 | -0.028 | -0.028 | 0 | 0.88 | 0.77 | 0.89 | 0.97 | 0.9 | 0.96 |
| 6 | 33429672 | rs2772373 | 0.314 | 0.271 | -0.485 | -1.195 | -0.008 | 0.016 | 0.016 | 0 | 0.33 | 0.23 | 0.38 | 0.32 | 0.41 | 0.37 |
| 6 | 33430950 | rs3130035 | 1.556 | 0.344 | 0.095 | -0.994 | 0.062 | -0.006 | -0.006 | 0 | 0.95 | 0.81 | 0.9 | 0.99 | 0.92 | 0.98 |
| 6 | 33431517 | rs2772374 | 1.143 | 0.386 | -0.228 | -0.932 | -0.033 | -0.034 | -0.034 | 0.221 | 0.09 | 0.17 | 0.02 | 0.01 | 0.02 | 0.02 |
| 6 | 33477013 | rs3857558 | -0.142 | 0.599 | -0.147 | -0.961 | 0.006 | -0.082 | -0.082 | 0.059 | 0.84 | 0.33 | 0.83 | 0.99 | 0.88 | 0.96 |
| 6 | 33478496 | rs9296092 | 0.459 | 0.618 | -0.604 | -1.171 | -0.042 | 0.132 | 0.132 | 0.188 | 0.64 | 0.18 | 0.66 | 0.93 | 0.67 | 0.86 |
| 6 | 33480435 | rs86715 | -0.822 | -0.002 | -0.6 | -1.286 | -0.009 | -0.034 | -0.034 | 0 | 0.27 | 0.71 | 0.16 | 0.06 | 0.13 | 0.09 |
| 6 | 33480738 | rs68191 | -0.04 | 0.039 | -0.5 | -1.263 | -0.031 | -0.07 | -0.07 | 2.214 | 0.21 | 0.41 | 0.13 | 0.03 | 0.11 | 0.09 |
| 6 | 33485236 | rs210179 | -0.809 | -0.018 | -0.935 | -0.492 | -0.042 | -0.061 | -0.061 | 0.235 | 0.58 | 0.8 | 0.47 | 0.55 | 0.43 | 0.46 |
| 6 | 33496714 | rs449242 | -0.201 | 0.009 | -0.963 | -0.673 | 0.007 | 0.111 | 0.111 | 0.897 | 0.4 | 0.75 | 0.31 | 0.09 | 0.3 | 0.18 |
| 6 | 33505594 | rs9469481 | -0.187 | 0.046 | -0.864 | -0.692 | 0.001 | -0.013 | -0.013 | 0 | 0.38 | 0.65 | 0.31 | 0.08 | 0.29 | 0.17 |
| 6 | 33511229 | rs210203 | -0.037 | 0.13 | -1.092 | -0.62 | -0.008 | 0.035 | 0.035 | 1.462 | 0.36 | 0.33 | 0.31 | 0.13 | 0.32 | 0.18 |
| 6 | 33515520 | rs210152 | -0.235 | 0.488 | -1.227 | -0.266 | 0.003 | 0.003 | 0.003 | 0 | 0.93 | 0.94 | 0.83 | 0.88 | 0.84 | 0.93 |
| 6 | 33525680 | rs210170 | 0.142 | 0.262 | -1.378 | -0.313 | 0 | -0.02 | -0.02 | 0 | 0.67 | 0.75 | 0.63 | 0.42 | 0.66 | 0.49 |
| 6 | 33533404 | rs210130 | 0.633 | 0.295 | -1.365 | -0.393 | -0.001 | -0.013 | -0.013 | 0 | 0.41 | 0.46 | 0.41 | 0.31 | 0.43 | 0.38 |
| 6 | 33534880 | rs6914422 | 1.71 | 0.297 | -1.383 | -0.419 | 0.003 | -0.002 | -0.002 | 0 | 0.27 | 0.33 | 0.3 | 0.25 | 0.32 | 0.3 |
| 6 | 33536670 | rs210132 | 0.748 | 0.36 | -1.515 | -0.337 | -0.001 | -0.004 | -0.004 | 0 | 0.41 | 0.48 | 0.41 | 0.35 | 0.43 | 0.38 |
| 6 | 33537802 | rs17627049 | 1.086 | 0.539 | -1.333 | -0.395 | -0.015 | 0.039 | 0.039 | 0 | 0.22 | 0.15 | 0.2 | 0.07 | 0.22 | 0.11 |
| 6 | 33540209 | rs210134 | 0.563 | 0.121 | -1.612 | -0.213 | -0.004 | -0.003 | -0.003 | 1.098 | 0.73 | 0.76 | 0.73 | 0.75 | 0.71 | 0.71 |
| 6 | 33541719 | rs513349 | 0.474 | 0.534 | -1.688 | -0.512 | 0.004 | 0.031 | 0.031 | 0.584 | 0.58 | 0.71 | 0.55 | 0.76 | 0.44 | 0.76 |
| 6 | 33542538 | rs210138 | 0.895 | 0.406 | -1.761 | -0.428 | 0.009 | -0.079 | -0.079 | 0.476 | 0.22 | 0.15 | 0.24 | 0.49 | 0.19 | 0.44 |
| 6 | 33546837 | rs210142 | 0.279 | 0.584 | -1.648 | -0.379 | -0.002 | -0.001 | -0.001 | 0 | 0.77 | 0.76 | 0.76 | 0.83 | 0.73 | 0.87 |
| 6 | 33548394 | rs5745568 | 1.58 | 0.243 | -1.646 | -0.377 | -0.004 | -0.02 | -0.02 | 0 | 0.24 | 0.08 | 0.27 | 0.24 | 0.22 | 0.3 |
| 6 | 33551514 | rs530878 | -0.184 | -0.008 | -1.472 | -0.382 | 0.012 | -0.009 | -0.009 | 0 | 0.35 | 0.64 | 0.29 | 0.53 | 0.23 | 0.49 |
| 6 | 33553580 | rs394199 | -0.051 | 0.614 | -1.722 | -0.434 | -0.006 | -0.045 | -0.045 | 0 | 0.44 | 0.25 | 0.46 | 0.56 | 0.49 | 0.53 |
| 6 | 33554147 | rs419261 | -0.356 | 1.1 | -1.675 | -0.473 | -0.028 | 0.039 | 0.039 | 0 | 0.73 | 0.82 | 0.72 | 0.9 | 0.61 | 0.85 |
| 6 | 33555348 | rs4559081 | 1.687 | 0.591 | -1.667 | -0.447 | -0.037 | 0.03 | 0.03 | 0 | 0.14 | 0.01 | 0.16 | 0.05 | 0.2 | 0.05 |
| 6 | 33555904 | rs420361 | -0.039 | 0.746 | -1.396 | -0.526 | 0 | -0.003 | -0.003 | 0 | 0.68 | 0.64 | 0.73 | 0.87 | 0.64 | 0.87 |
| 6 | 33557045 | rs396746 | 0.392 | 0.754 | -1.391 | -0.517 | 0.012 | -0.012 | -0.012 | 0 | 0.9 | 0.83 | 0.92 | 0.93 | 0.86 | 0.93 |
| 6 | 33557225 | rs430655 | -0.16 | 0.693 | -1.335 | -0.467 | 0 | -0.008 | -0.008 | 0 | 0.68 | 0.65 | 0.73 | 0.87 | 0.64 | 0.87 |
| 6 | 33557741 | rs375555 | 0.919 | 0.728 | -1.528 | -0.257 | -0.003 | -0.077 | -0.077 | 0 | 0.22 | 0.13 | 0.24 | 0.48 | 0.21 | 0.44 |
| 6 | 33560896 | rs210162 | 0.537 | 0.719 | -1.521 | -0.252 | 0.007 | -0.068 | -0.068 | 0.081 | 0.25 | 0.16 | 0.25 | 0.52 | 0.23 | 0.48 |
| 6 | 33562709 | rs10947410 | 0.156 | 0.768 | -1.006 | -0.158 | -0.002 | 0.002 | 0.002 | 0 | 0.47 | 0.4 | 0.47 | 0.36 | 0.43 | 0.43 |
| 6 | 33562720 | rs6929774 | 0.156 | 0.768 | -1.006 | -0.158 | -0.002 | 0.002 | 0.002 | 0 | 0.47 | 0.4 | 0.47 | 0.36 | 0.43 | 0.43 |
| 6 | 33564412 | rs6457736 | -0.608 | 0.888 | -1.069 | -0.211 | 0.034 | -0.036 | -0.036 | 0 | 0.24 | 0.4 | 0.21 | 0.12 | 0.21 | 0.13 |
| 6 | 33568687 | rs12213869 | 0.819 | 0.517 | -1.193 | 0.016 | 0.031 | -0.028 | -0.028 | 0 | 0.31 | 0.21 | 0.36 | 0.38 | 0.26 | 0.35 |
| 6 | 33574413 | rs210120 | 0.24 | 0.543 | -1.004 | -0.244 | 0.043 | 0.033 | 0.033 | 0 | 0.49 | 0.67 | 0.43 | 0.65 | 0.43 | 0.56 |
| 6 | 33575009 | rs444697 | 0.884 | 0.649 | -1.004 | -0.208 | 0.007 | -0.04 | -0.04 | 0.001 | 0.25 | 0.21 | 0.24 | 0.54 | 0.24 | 0.47 |
| 6 | 33576577 | rs169737 | 0.516 | 1.048 | -0.798 | -0.4 | 0.077 | -0.077 | -0.077 | 0 | 0.81 | 0.6 | 0.79 | 0.97 | 0.77 | 0.9 |
| 6 | 33581805 | rs9469525 | 1.168 | 0.73 | -1.234 | 0.029 | -0.055 | 0.001 | 0.001 | 0.265 | 0.21 | 0.02 | 0.26 | 0.14 | 0.24 | 0.28 |
| 6 | 33582090 | rs499384 | 0.684 | 0.692 | -1.172 | 0.095 | 0.001 | 0.013 | 0.013 | 0.388 | 0.11 | 0.17 | 0.06 | 0.13 | 0.09 | 0.06 |
| 6 | 33588147 | rs3748079 | 1.045 | 0.589 | -1.282 | 0.032 | -0.059 | -0.003 | -0.003 | 0 | 0.21 | 0.01 | 0.26 | 0.14 | 0.24 | 0.28 |
| 6 | 33601132 | rs9469532 | -0.229 | 0.577 | -0.635 | 0.319 | 0.003 | 0.018 | 0.018 | 0 | 0.37 | 0.55 | 0.38 | 0.53 | 0.34 | 0.56 |
| 6 | 33608379 | rs4713647 | -1.288 | 0.365 | -0.637 | 0.254 | -0.009 | -0.012 | -0.012 | 0 | 0.94 | 0.9 | 0.99 | 1 | 0.99 | 0.99 |
| 6 | 33610350 | rs4711333 | -1.027 | 0.333 | -0.488 | 0.214 | -0.019 | -0.026 | -0.026 | 0 | 0.94 | 0.86 | 0.98 | 1 | 0.99 | 0.99 |
| 6 | 33613344 | rs9357162 | 0.666 | 0.383 | -1.322 | 0.27 | -0.061 | -0.078 | -0.078 | 0 | 0.22 | 0.02 | 0.28 | 0.28 | 0.24 | 0.29 |

| | | | | | | | | | | | | | | | | |
|---|---|---|---|---|---|---|---|---|---|---|---|---|---|---|---|---|
| 6 | 33614127 | rs6935686 | 0.449 | -0.259 | -1.093 | 0.079 | -0.055 | 0.074 | 0.074 | 0 | 0.28 | 0.39 | 0.19 | 0.46 | 0.21 | 0.45 |
| 6 | 33617464 | rs6457738 | -0.056 | 0.36 | -1.19 | -0.179 | -0.011 | 0.011 | 0.011 | 0 | 0.63 | 0.64 | 0.56 | 0.7 | 0.65 | 0.62 |
| 6 | 33622933 | rs1570760 | 1.177 | 0.422 | -1.347 | -0.073 | -0.066 | -0.074 | -0.074 | 0 | 0.22 | 0.03 | 0.29 | 0.26 | 0.24 | 0.28 |
| 6 | 33624221 | rs4259245 | 0.638 | 0.305 | -1.415 | -0.254 | -0.026 | 0.036 | 0.036 | 0 | 0.33 | 0.18 | 0.33 | 0.24 | 0.39 | 0.16 |
| 6 | 33625696 | rs2274200 | 1.298 | 0.377 | -1.386 | -0.203 | -0.066 | -0.074 | -0.074 | 0.838 | 0.22 | 0.03 | 0.29 | 0.26 | 0.24 | 0.28 |
| 6 | 33627573 | rs2296341 | -0.888 | 0.157 | -0.843 | -0.468 | -0.048 | -0.029 | -0.029 | 0 | 0.73 | 0.52 | 0.76 | 0.8 | 0.79 | 0.84 |
| 6 | 33630129 | rs2147658 | 1.011 | 0.322 | -1.184 | -0.592 | -0.043 | 0.052 | 0.052 | 0 | 0.26 | 0.08 | 0.31 | 0.14 | 0.3 | 0.24 |
| 6 | 33631017 | rs1536041 | -0.126 | 0.144 | -0.935 | -0.652 | 0.002 | -0.001 | -0.001 | 0 | 0.82 | 0.72 | 0.86 | 0.83 | 0.84 | 0.94 |
| 6 | 33632014 | rs1536036 | 0.383 | -0.041 | -0.777 | -0.714 | 0 | 0.023 | 0.023 | 0 | 0.5 | 0.54 | 0.56 | 0.34 | 0.54 | 0.4 |
| 6 | 33632451 | rs12198325 | 0.82 | -0.063 | -0.852 | -0.749 | -0.034 | 0.11 | 0.11 | 0.016 | 0.44 | 0.21 | 0.51 | 0.19 | 0.52 | 0.33 |
| 6 | 33633527 | rs6901411 | 1.003 | -0.368 | -1.102 | -0.686 | -0.021 | -0.065 | -0.065 | 3.823 | 0.23 | 0.12 | 0.27 | 0.42 | 0.25 | 0.4 |
| 6 | 33633940 | rs1570759 | 0.285 | -0.304 | -1.069 | -0.664 | 0.001 | -0.05 | -0.05 | 1.385 | 0.32 | 0.17 | 0.3 | 0.48 | 0.29 | 0.53 |
| 6 | 33636907 | rs2077163 | 0.217 | -0.342 | -1.029 | -0.658 | 0.005 | -0.004 | -0.004 | 0.156 | 0.49 | 0.37 | 0.48 | 0.52 | 0.5 | 0.61 |
| 6 | 33638180 | rs2229634 | 0.344 | -0.254 | -1.11 | -0.606 | -0.005 | -0.043 | -0.043 | 0 | 0.3 | 0.16 | 0.28 | 0.47 | 0.28 | 0.52 |
| 6 | 33641956 | rs7769838 | -0.736 | -0.27 | -0.743 | -1.132 | 0.007 | 0.063 | 0.063 | 1.455 | 0.7 | 0.78 | 0.79 | 0.98 | 0.7 | 0.91 |
| 6 | 33642593 | rs942639 | 0.192 | -0.245 | -0.629 | -1.025 | 0.003 | -0.012 | -0.012 | 0 | 0.78 | 0.53 | 0.83 | 0.82 | 0.82 | 0.92 |
| 6 | 33643558 | rs2229637 | 0.797 | 0.094 | -0.688 | -0.94 | 0.001 | 0.02 | 0.02 | 0 | 0.24 | 0.19 | 0.25 | 0.07 | 0.28 | 0.11 |
| 6 | 33644033 | rs729424 | -1.132 | -0.084 | -0.712 | -0.977 | -0.005 | -0.024 | -0.024 | 0 | 0.32 | 0.48 | 0.22 | 0.26 | 0.28 | 0.25 |
| 6 | 33644253 | rs10947427 | 1.453 | 0.269 | -0.75 | -1.147 | 0.002 | 0.028 | 0.028 | 0.051 | 0.18 | 0.12 | 0.2 | 0.05 | 0.22 | 0.08 |
| 6 | 33647058 | rs10947428 | -0.679 | 0.419 | -0.461 | -1.211 | 0.065 | -0.057 | -0.057 | 0.041 | 0.17 | 0.49 | 0.15 | 0.04 | 0.2 | 0.05 |
| 6 | 33647594 | rs4713653 | -0.346 | 0.198 | -0.683 | -1.641 | -0.005 | -0.037 | -0.037 | 0 | 0.67 | 0.44 | 0.73 | 0.88 | 0.7 | 0.8 |
| 6 | 33648097 | rs9469554 | 2.281 | 0.033 | -0.783 | -1.733 | 0.011 | 0.047 | 0.047 | 0.055 | 0.15 | 0.06 | 0.17 | 0.05 | 0.18 | 0.07 |
| 6 | 33648997 | rs4713656 | 0.94 | 0.216 | -0.572 | -1.968 | 0.006 | -0.046 | -0.046 | 0.685 | 0.7 | 0.63 | 0.81 | 0.94 | 0.7 | 0.87 |
| 6 | 33650621 | rs2296330 | 1.678 | 0.254 | -1.304 | -2.269 | -0.033 | -0.076 | -0.076 | 0 | 0.21 | 0.04 | 0.21 | 0.67 | 0.23 | 0.41 |
| 6 | 33650743 | rs2296329 | 1.486 | 0.062 | -1.353 | -2.367 | -0.008 | 0.002 | 0.002 | 0.003 | 0.24 | 0.05 | 0.36 | 0.19 | 0.23 | 0.32 |
| 6 | 33650812 | rs11759011 | 2.326 | -0.022 | -1.199 | -2.605 | -0.047 | -0.016 | -0.016 | 0 | 0.15 | 0.01 | 0.31 | 0.1 | 0.2 | 0.18 |
| 6 | 33651129 | rs942636 | 1.554 | -0.009 | -1.113 | -2.599 | 0.003 | 0.008 | 0.008 | 0 | 0.2 | 0.17 | 0.34 | 0.1 | 0.2 | 0.19 |
| 6 | 33651277 | rs2296328 | 1.946 | 0.02 | -0.613 | -2.557 | -0.017 | -0.065 | -0.065 | 0 | 0.19 | 0.31 | 0.2 | 0.07 | 0.23 | 0.13 |
| 6 | 33653111 | rs942637 | -1.388 | 0.121 | -0.747 | -2.486 | 0.005 | -0.053 | -0.053 | 0.104 | 0.88 | 0.83 | 0.94 | 0.99 | 0.92 | 0.93 |
| 6 | 33653448 | rs749338 | 0.903 | 0.091 | -1.064 | -2.476 | -0.039 | 0.202 | 0.202 | 0.366 | 0.43 | 0.1 | 0.57 | 0.86 | 0.46 | 0.71 |
| 6 | 33659793 | rs2296742 | 0.543 | 0.31 | -0.956 | -2.484 | 0.001 | 0.216 | 0.216 | 0.341 | 0.51 | 0.22 | 0.61 | 0.9 | 0.49 | 0.81 |
| 6 | 33660143 | rs9296097 | 2.234 | 0.261 | -1.119 | -2.535 | 0.006 | 0.061 | 0.061 | 0.328 | 0.17 | 0.06 | 0.18 | 0.05 | 0.19 | 0.08 |
| 6 | 33661091 | rs3818528 | 1.361 | 0.296 | -1.013 | -2.515 | -0.036 | 0.128 | 0.128 | 0.129 | 0.63 | 0.17 | 0.75 | 0.91 | 0.66 | 0.85 |
| 6 | 33662667 | rs684119 | 0.202 | 0.354 | -0.848 | -2.533 | -0.001 | -0.011 | -0.011 | 0 | 0.14 | 0.17 | 0.09 | 0.03 | 0.15 | 0.04 |
| 6 | 33667965 | rs594223 | 0.298 | 0.292 | -0.815 | -2.569 | -0.013 | -0.066 | -0.066 | 0 | 0.78 | 0.39 | 0.88 | 0.94 | 0.87 | 0.86 |
| 6 | 33668618 | rs597386 | -0.308 | 0.338 | -0.83 | -2.478 | -0.001 | -0.005 | -0.005 | 0 | 0.17 | 0.23 | 0.12 | 0.05 | 0.21 | 0.06 |
| 6 | 33675642 | rs2281829 | 0.991 | 0.221 | -0.907 | -2.617 | -0.022 | 0.223 | 0.223 | 0.012 | 0.46 | 0.1 | 0.57 | 0.85 | 0.47 | 0.72 |
| 6 | 33675851 | rs542441 | 0.458 | 0.288 | -0.852 | -2.583 | -0.019 | 0.062 | 0.062 | 0.013 | 0.33 | 0.29 | 0.31 | 0.09 | 0.4 | 0.14 |
| 6 | 33684313 | rs649775 | -1.531 | 0.31 | -0.704 | -2.533 | 0.009 | -0.051 | -0.051 | 1.189 | 0.9 | 0.82 | 0.95 | 0.99 | 0.93 | 0.96 |
| 6 | 33689520 | rs2966 | 0.852 | 0.284 | -0.782 | -2.582 | -0.016 | 0.242 | 0.242 | 0.008 | 0.48 | 0.12 | 0.58 | 0.86 | 0.48 | 0.77 |
| 6 | 33689534 | rs2967 | 0.98 | 0.292 | -0.646 | -2.561 | -0.002 | -0.023 | -0.023 | 0 | 0.72 | 0.34 | 0.82 | 0.97 | 0.74 | 0.91 |
| 6 | 33689956 | rs559290 | -1.53 | 0.326 | -0.644 | -2.523 | 0.004 | -0.044 | -0.044 | 0 | 0.9 | 0.84 | 0.95 | 0.99 | 0.93 | 0.96 |
| 6 | 33690310 | rs630792 | 1.029 | 0.412 | -0.812 | -2.521 | -0.031 | 0.023 | 0.023 | 5.375 | 0.06 | 0 | 0.06 | 0.01 | 0.1 | 0.02 |
| 6 | 33697125 | rs6457740 | 1.013 | 0.336 | -0.737 | -2.573 | -0.001 | -0.029 | -0.029 | 0.426 | 0.72 | 0.37 | 0.81 | 0.96 | 0.74 | 0.87 |
| 6 | 33699899 | rs4713671 | 0.851 | 0.349 | -0.864 | -2.597 | 0 | 0.203 | 0.203 | 0.007 | 0.48 | 0.16 | 0.59 | 0.86 | 0.48 | 0.75 |
| 6 | 33702701 | rs12207382 | 0.084 | 0.413 | -0.413 | -2.673 | -0.003 | -0.06 | -0.06 | 0 | 0.35 | 0.59 | 0.28 | 0.09 | 0.34 | 0.16 |
| 6 | 33702869 | rs622917 | -0.581 | 0.382 | -0.571 | -2.737 | -0.022 | 0.106 | 0.106 | 0 | 0.45 | 0.72 | 0.35 | 0.12 | 0.43 | 0.23 |
| 6 | 33703089 | rs623813 | 0.226 | 0.45 | -0.43 | -2.661 | -0.019 | 0.032 | 0.032 | 0.88 | 0.13 | 0.07 | 0.1 | 0.02 | 0.16 | 0.05 |
| 6 | 33703230 | rs545787 | -1.425 | 0.497 | -0.595 | -2.66 | -0.01 | -0.005 | -0.005 | 0.068 | 0.3 | 0.72 | 0.24 | 0.07 | 0.33 | 0.16 |
| 6 | 33710038 | rs568901 | -0.721 | 0.473 | -0.319 | -2.643 | 0.049 | -0.094 | -0.094 | 0 | 0.23 | 0.47 | 0.18 | 0.07 | 0.26 | 0.12 |
| 6 | 33710284 | rs6942022 | -0.122 | 0.474 | -0.402 | -2.591 | 0.005 | -0.023 | -0.023 | 0 | 0.09 | 0.2 | 0.06 | 0.04 | 0.07 | 0.07 |
| 6 | 33717770 | rs9469583 | 0.186 | 0.25 | -0.526 | -2.804 | 0 | 0.109 | 0.109 | 1.434 | 0.54 | 0.32 | 0.65 | 0.91 | 0.56 | 0.79 |
| 6 | 33721933 | rs9461905 | 0.296 | -0.048 | -0.38 | -2.529 | 0.018 | -0.091 | -0.091 | 0.85 | 0.8 | 0.45 | 0.84 | 0.96 | 0.8 | 0.87 |
| 6 | 33722098 | rs4713676 | 0.959 | -0.215 | -0.578 | -2.611 | 0 | -0.024 | -0.024 | 0 | 0.72 | 0.32 | 0.8 | 0.93 | 0.78 | 0.81 |
| 6 | 33723290 | rs4304152 | 0.617 | -0.145 | -0.679 | -2.661 | 0.017 | 0.202 | 0.202 | 0 | 0.53 | 0.18 | 0.62 | 0.86 | 0.57 | 0.71 |
| 6 | 33724004 | rs6919321 | 0.158 | -0.168 | -0.677 | -2.674 | 0.024 | 0.098 | 0.098 | 0 | 0.58 | 0.28 | 0.65 | 0.87 | 0.62 | 0.75 |
| 6 | 33724794 | rs9380376 | 0.708 | -0.169 | -1.144 | -2.659 | -0.036 | -0.013 | -0.013 | 0.963 | 0.32 | 0.09 | 0.27 | 0.73 | 0.3 | 0.46 |
| 6 | 33730954 | rs4711345 | 0.71 | -0.347 | -0.85 | -2.792 | 0.002 | 0.01 | 0.01 | 0 | 0.4 | 0.39 | 0.53 | 0.19 | 0.43 | 0.33 |
| 6 | 33731408 | rs9394165 | 0.636 | -0.345 | -1.033 | -2.899 | -0.053 | 0.046 | 0.046 | 0 | 0.37 | 0.1 | 0.3 | 0.76 | 0.34 | 0.57 |
| 6 | 33731787 | rs943466 | 0.539 | -0.394 | -0.958 | -2.911 | 0.003 | -0.017 | -0.017 | 0.015 | 0.25 | 0.31 | 0.4 | 0.14 | 0.28 | 0.26 |
| 6 | 33732219 | rs6925337 | 1.369 | -0.594 | -0.765 | -2.907 | -0.05 | -0.068 | -0.068 | 0.199 | 0.25 | 0.32 | 0.2 | 0.08 | 0.17 | 0.15 |
| 6 | 33735292 | rs4711347 | 0.947 | -0.471 | -1.051 | -2.89 | -0.053 | 0.229 | 0.229 | 0.312 | 0.51 | 0.12 | 0.43 | 0.81 | 0.5 | 0.63 |
| 6 | 33736295 | rs4711348 | 0.487 | -0.421 | -0.938 | -2.866 | -0.035 | 0.156 | 0.156 | 0.081 | 0.54 | 0.19 | 0.45 | 0.84 | 0.51 | 0.7 |
| 6 | 33737800 | rs1075379 | 0.245 | -0.391 | -0.576 | -2.937 | 0.002 | -0.042 | -0.042 | 0.494 | 0.27 | 0.36 | 0.37 | 0.13 | 0.28 | 0.21 |
| 6 | 33738442 | rs943472 | -1.141 | -0.374 | -0.534 | -2.988 | 0.055 | -0.064 | -0.064 | 0 | 0.87 | 0.73 | 0.89 | 0.98 | 0.81 | 0.96 |
| 6 | 33739960 | rs10947436 | -1.133 | -0.371 | -0.532 | -2.987 | 0.054 | -0.065 | -0.065 | 0 | 0.87 | 0.73 | 0.89 | 0.98 | 0.81 | 0.96 |
| 6 | 33740025 | rs2296743 | 0.871 | -0.678 | -0.742 | -3.185 | -0.021 | 0.038 | 0.038 | 0 | 0.35 | 0.15 | 0.31 | 0.76 | 0.35 | 0.56 |
| 6 | 33741289 | rs6904716 | 0.16 | -0.697 | -0.77 | -3.189 | 0.005 | 0.01 | 0.01 | 0 | 0.48 | 0.49 | 0.56 | 0.2 | 0.46 | 0.33 |
| 6 | 33743997 | rs6933607 | 1.504 | -0.749 | -0.684 | -3.211 | -0.01 | 0.008 | 0.008 | 0 | 0.2 | 0.14 | 0.35 | 0.12 | 0.27 | 0.19 |

| | | | | | | | | | | | | | | | | |
|---|---|---|---|---|---|---|---|---|---|---|---|---|---|---|---|---|
| 6 | 33745071 | rs2296748 | 0.457 | -0.737 | -0.71 | -3.207 | -0.003 | 0.04 | 0.04 | 0 | 0.39 | 0.24 | 0.33 | 0.78 | 0.35 | 0.63 |
| 6 | 33750086 | rs943476 | 0.038 | -0.72 | -0.687 | -3.2 | 0.006 | -0.065 | -0.065 | 0 | 0.78 | 0.69 | 0.84 | 0.95 | 0.78 | 0.9 |
| 6 | 33753673 | rs2395402 | 0.688 | -0.708 | -0.855 | -3.167 | -0.012 | 0.124 | 0.124 | 0.309 | 0.53 | 0.25 | 0.46 | 0.83 | 0.5 | 0.69 |
| 6 | 33755890 | rs756137 | 1.117 | -0.948 | -1.11 | -3.005 | -0.066 | -0.046 | -0.046 | 4.197 | 0.28 | 0.09 | 0.29 | 0.66 | 0.33 | 0.46 |
| 6 | 33767727 | rs3828783 | 2.083 | -0.627 | -0.995 | -3 | -0.017 | 0.006 | 0.006 | 0 | 0.16 | 0.05 | 0.33 | 0.11 | 0.22 | 0.16 |
| 6 | 33768897 | rs2281820 | -0.084 | -0.695 | -0.847 | -3.039 | 0.041 | -0.063 | -0.063 | 0.029 | 0.65 | 0.49 | 0.52 | 0.86 | 0.54 | 0.76 |
| 6 | 33770370 | rs3806109 | 0.748 | -0.651 | -0.842 | -3.02 | -0.018 | 0.035 | 0.035 | 0 | 0.34 | 0.17 | 0.31 | 0.76 | 0.35 | 0.56 |
| 6 | 33771779 | rs2281818 | 0.437 | -0.59 | -0.601 | -3.02 | -0.017 | 0.039 | 0.039 | 0 | 0.06 | 0.19 | 0.01 | 0.09 | 0.01 | 0.07 |
| 6 | 33772761 | rs3763253 | 0.969 | -0.537 | -0.665 | -3.037 | -0.008 | 0.001 | 0.001 | 0.182 | 0.22 | 0.14 | 0.38 | 0.16 | 0.27 | 0.2 |
| 6 | 33773199 | rs9366829 | 0.508 | -0.322 | -0.84 | -3.245 | -0.003 | -0.016 | -0.016 | 0 | 0.53 | 0.48 | 0.43 | 0.78 | 0.46 | 0.65 |
| 6 | 33774948 | rs10947439 | 1.269 | -0.254 | -0.8 | -3.203 | -0.001 | 0.061 | 0.061 | 0 | 0.6 | 0.26 | 0.74 | 0.82 | 0.66 | 0.65 |
| 6 | 33775391 | rs7774407 | 1.229 | -0.259 | -0.777 | -3.205 | -0.001 | 0 | 0 | 0 | 0.62 | 0.33 | 0.76 | 0.84 | 0.66 | 0.72 |
| 6 | 33775641 | rs1547669 | 0.833 | -0.341 | -0.851 | -3.263 | -0.016 | 0.097 | 0.097 | 0 | 0.45 | 0.25 | 0.63 | 0.8 | 0.49 | 0.58 |
| 6 | 33777432 | rs2395399 | 0.039 | -0.336 | -0.809 | -3.214 | 0.014 | -0.054 | -0.054 | 2.385 | 0.76 | 0.72 | 0.83 | 0.95 | 0.73 | 0.85 |
| 6 | 33778964 | rs9394169 | 0.252 | -0.289 | -0.785 | -3.179 | 0.005 | 0.023 | 0.023 | 0 | 0.52 | 0.39 | 0.67 | 0.83 | 0.55 | 0.68 |
| 6 | 33782040 | rs9469591 | -0.675 | 0.264 | -0.729 | -2.928 | -0.004 | -0.025 | -0.025 | 0 | 0.18 | 0.29 | 0.12 | 0.14 | 0.12 | 0.24 |
| 6 | 33783883 | rs1535948 | -0.661 | 0.247 | -0.971 | -2.96 | 0.002 | -0.005 | -0.005 | 0 | 0.29 | 0.3 | 0.45 | 0.29 | 0.29 | 0.39 |
| 6 | 33784864 | rs3957165 | -0.802 | 0.321 | -0.931 | -2.948 | 0.031 | -0.065 | -0.065 | 0.646 | 0.84 | 0.71 | 0.9 | 0.97 | 0.81 | 0.93 |
| 6 | 33785827 | rs1535950 | 0.792 | 0.191 | -1.089 | -2.945 | -0.003 | -0.001 | -0.001 | 0 | 0.34 | 0.27 | 0.3 | 0.66 | 0.33 | 0.45 |
| 6 | 33787746 | rs13203745 | 2.553 | 0.085 | -1.11 | -2.808 | -0.005 | 0.082 | 0.082 | 0.743 | 0.19 | 0.01 | 0.12 | 0.02 | 0.19 | 0.09 |
| 6 | 33790415 | rs4713691 | 0.843 | 0.195 | -1.075 | -2.748 | 0.001 | -0.006 | -0.006 | 0 | 0.28 | 0.3 | 0.27 | 0.64 | 0.23 | 0.4 |
| 6 | 33791230 | rs10947443 | 1.546 | -0.404 | -1.676 | -2.575 | 0.051 | 0.072 | 0.072 | 0.906 | 0.27 | 0.1 | 0.16 | 0.14 | 0.2 | 0.24 |
| 6 | 33791515 | rs767896 | -0.236 | 0.094 | -1.35 | -2.466 | 0.012 | -0.018 | -0.018 | 0 | 0.28 | 0.27 | 0.47 | 0.18 | 0.35 | 0.26 |
| 6 | 33793760 | rs3998109 | 0.572 | 0.191 | -1.282 | -2.336 | 0.014 | 0.069 | 0.069 | 0 | 0.39 | 0.24 | 0.26 | 0.08 | 0.36 | 0.28 |
| 6 | 33794129 | rs9469597 | -1.038 | 0.235 | -1.313 | -2.157 | -0.007 | -0.013 | -0.013 | 0 | 0.34 | 0.51 | 0.48 | 0.31 | 0.35 | 0.34 |
| 6 | 33794258 | rs9469599 | 0.56 | 0.494 | -1.236 | -1.604 | 0.002 | 0.013 | 0.013 | 2.734 | 0.11 | 0.1 | 0.06 | 0.03 | 0.11 | 0.06 |
| 6 | 33797009 | rs9461917 | 0.206 | 0.374 | -1.222 | -1.607 | 0.002 | 0.032 | 0.032 | 0.329 | 0.57 | 0.37 | 0.61 | 0.29 | 0.63 | 0.45 |
| 6 | 33801510 | rs12055409 | 0.474 | 0.377 | -1.174 | -1.277 | -0.004 | -0.005 | -0.005 | 0.509 | 0.6 | 0.44 | 0.67 | 0.36 | 0.68 | 0.51 |
| 6 | 33804137 | rs7761859 | -0.87 | 0.416 | -1.477 | -1.134 | -0.061 | -0.111 | -0.111 | 2.375 | 0.84 | 0.99 | 0.79 | 0.41 | 0.8 | 0.59 |
| 6 | 33807091 | rs6901216 | 0.306 | 0.76 | -1.486 | -1.287 | -0.086 | 0.021 | 0.021 | 0.375 | 0.24 | 0.08 | 0.44 | 0.21 | 0.39 | 0.24 |
| 6 | 33813043 | rs6905353 | 0.489 | -0.144 | -0.941 | -1.108 | 0.001 | -0.005 | -0.005 | 0 | 0.37 | 0.38 | 0.24 | 0.16 | 0.3 | 0.27 |
| 6 | 33816933 | rs9469615 | 0.016 | 0.351 | -0.889 | -0.916 | -0.003 | 0.005 | 0.005 | 4.647 | 0.11 | 0.13 | 0.09 | 0.06 | 0.14 | 0.06 |
| 6 | 33817929 | rs4713693 | 0.058 | 0.365 | -0.808 | -0.755 | -0.004 | -0.012 | -0.012 | 0 | 0.48 | 0.55 | 0.36 | 0.26 | 0.46 | 0.31 |
| 6 | 33819582 | rs11753213 | 0.125 | 0.408 | -0.616 | -0.759 | 0.01 | 0.013 | 0.013 | 0.672 | 0.53 | 0.42 | 0.34 | 0.24 | 0.42 | 0.39 |
| 6 | 33820059 | rs6457742 | -0.579 | 1.103 | -0.476 | -0.975 | 0.006 | -0.011 | -0.011 | 0 | 0.33 | 0.32 | 0.33 | 0.22 | 0.41 | 0.3 |
| 6 | 33822865 | rs9296100 | -0.829 | 0.682 | -0.163 | -0.898 | 0.004 | 0.023 | 0.023 | 0 | 0.78 | 0.77 | 0.63 | 0.4 | 0.75 | 0.64 |
| 6 | 33823149 | rs9469618 | 0.583 | 0.96 | -0.643 | -1.463 | 0.027 | 0.161 | 0.161 | 0.03 | 0.49 | 0.26 | 0.42 | 0.22 | 0.49 | 0.35 |
| 6 | 33826590 | rs4713697 | -0.94 | 1.115 | -0.154 | -0.694 | 0.017 | 0.145 | 0.145 | 0 | 0.76 | 0.62 | 0.67 | 0.5 | 0.74 | 0.64 |
| 6 | 33828178 | rs9469619 | 1.266 | 0.976 | -0.488 | -1.296 | -0.004 | 0.033 | 0.033 | 0 | 0.29 | 0.19 | 0.25 | 0.1 | 0.34 | 0.13 |
| 6 | 33830362 | rs6934013 | 0.778 | 1.189 | -0.432 | -1.051 | 0 | 0 | 0 | 0.585 | 0.13 | 0.14 | 0.12 | 0.13 | 0.07 | 0.07 |
| 6 | 33833491 | rs4711358 | -0.504 | 1.14 | -0.497 | -1.112 | -0.02 | 0.033 | 0.033 | 0 | 0.34 | 0.24 | 0.38 | 0.33 | 0.33 | 0.36 |
| 6 | 33835210 | rs1555687 | -1.305 | 1.067 | -0.723 | -1.113 | 0.002 | 0.025 | 0.025 | 0 | 0.76 | 0.77 | 0.69 | 0.48 | 0.74 | 0.63 |
| 6 | 33837316 | rs6457744 | 0.266 | 1.326 | -1.302 | -0.88 | 0.004 | 0.02 | 0.02 | 0 | 0.39 | 0.37 | 0.29 | 0.15 | 0.39 | 0.27 |
| 6 | 33839359 | rs1324582 | 0.928 | 2.018 | -1.038 | -0.33 | -0.007 | 0.004 | 0.004 | 0.163 | 0.58 | 0.58 | 0.54 | 0.6 | 0.51 | 0.59 |
| 6 | 33840121 | rs10947461 | 0.062 | 2.655 | -1.122 | 0.007 | -0.055 | -0.004 | -0.004 | 0 | 0.12 | 0.03 | 0.2 | 0.12 | 0.18 | 0.12 |
| 6 | 33840657 | rs9348927 | 0.125 | 1.907 | -1.019 | 0.069 | -0.031 | 0.038 | 0.038 | 0 | 0.35 | 0.25 | 0.37 | 0.21 | 0.44 | 0.28 |
| 6 | 33841361 | rs9380384 | 0.15 | 1.619 | -0.707 | 0.237 | -0.001 | -0.001 | -0.001 | 0 | 0.89 | 0.87 | 0.86 | 0.86 | 0.87 | 0.91 |
| 6 | 33843759 | rs6926872 | 0.068 | 0.993 | -0.613 | 0.287 | 0.002 | 0.002 | 0.002 | 0 | 0.45 | 0.55 | 0.57 | 0.47 | 0.45 | 0.58 |
| 6 | 33844683 | rs16869645 | -0.428 | 0.824 | -0.55 | 0.206 | -0.006 | -0.011 | -0.011 | 0 | 0.1 | 0.14 | 0.08 | 0.04 | 0.07 | 0.09 |
| 6 | 33846143 | rs4713711 | -0.442 | 0.774 | -0.882 | -0.158 | -0.003 | -0.024 | -0.024 | 0 | 0.32 | 0.58 | 0.28 | 0.08 | 0.32 | 0.15 |
| 6 | 33847077 | rs10947463 | 1.184 | 0.818 | -0.975 | -0.128 | -0.029 | 0.038 | 0.038 | 0 | 0.15 | 0.01 | 0.14 | 0.03 | 0.19 | 0.05 |
| 6 | 33848109 | rs12214429 | 1.327 | 0.83 | -0.953 | -0.113 | -0.008 | 0.056 | 0.056 | 0 | 0.16 | 0.05 | 0.15 | 0.03 | 0.2 | 0.05 |
| 6 | 33851173 | rs16869652 | -0.069 | 0.954 | -0.518 | -0.031 | 0.003 | -0.019 | -0.019 | 0 | 0.1 | 0.17 | 0.1 | 0.04 | 0.09 | 0.07 |
| 6 | 33853719 | rs6935041 | 0.024 | 0.993 | -0.643 | 0.126 | -0.001 | -0.001 | -0.001 | 0 | 0.55 | 0.52 | 0.62 | 0.45 | 0.55 | 0.6 |
| 6 | 33854816 | rs9461924 | 0.468 | 1.17 | -1.244 | 0.276 | -0.007 | 0.013 | 0.013 | 0 | 0.27 | 0.1 | 0.33 | 0.25 | 0.27 | 0.47 |
| 6 | 33857525 | rs2274154 | -0.406 | 1.59 | -0.391 | 0.723 | 0.001 | 0.005 | 0.005 | 0 | 0.1 | 0.11 | 0.13 | 0.19 | 0.12 | 0.07 |
| 6 | 33858199 | rs6941621 | -0.86 | 1.492 | -0.416 | 0.642 | 0.025 | -0.016 | -0.016 | 0 | 0.25 | 0.51 | 0.27 | 0.26 | 0.27 | 0.18 |
| 6 | 33864237 | rs942505 | 0.431 | 1.461 | -1.208 | 1.003 | -0.06 | -0.074 | -0.074 | 3.365 | 0.86 | 0.99 | 0.91 | 0.79 | 0.81 | 0.8 |
| 6 | 33864354 | rs7741871 | -0.4 | 1.191 | -1.295 | 0.33 | 0.001 | 0.034 | 0.034 | 0 | 0.29 | 0.18 | 0.23 | 0.09 | 0.3 | 0.17 |
| 6 | 33864998 | rs3800341 | 0.905 | 1.094 | -0.375 | 0.679 | 0.053 | 0.074 | 0.074 | 0 | 0.83 | 0.46 | 0.78 | 0.77 | 0.78 | 0.86 |
| 6 | 33865343 | rs10947465 | -0.168 | 0.877 | -1.037 | 0.588 | -0.04 | 0.01 | 0.01 | 0 | 0.24 | 0.06 | 0.15 | 0.21 | 0.25 | 0.37 |
| 6 | 33865380 | rs4713714 | 0.609 | 0.393 | -1.078 | 0.977 | -0.012 | -0.008 | -0.008 | 0 | 0.2 | 0.18 | 0.39 | 0.31 | 0.33 | 0.33 |
| 6 | 33866059 | rs10485346 | 0.389 | -0.191 | -0.573 | 0.929 | -0.004 | 0.004 | 0.004 | 0 | 0.46 | 0.44 | 0.57 | 0.41 | 0.48 | 0.43 |
| 6 | 33868647 | rs12198623 | -0.029 | 0.067 | -0.479 | 1.002 | 0.001 | 0.005 | 0.005 | 0 | 0.05 | 0.05 | 0.07 | 0.07 | 0.05 | 0.05 |
| 6 | 33869806 | rs9296102 | -0.121 | -0.361 | -0.155 | 1.024 | 0.002 | -0.001 | -0.001 | 0.505 | 0.43 | 0.58 | 0.35 | 0.45 | 0.35 | 0.43 |
| 6 | 33872149 | rs6457748 | 1.011 | -0.819 | 0.279 | 0.981 | 0.006 | 0.053 | 0.053 | 0.134 | 0.71 | 0.52 | 0.58 | 0.62 | 0.67 | 0.78 |
| 6 | 33873699 | rs4563711 | 1.463 | -0.709 | 0.459 | 0.383 | 0.014 | -0.002 | -0.002 | 0.077 | 0.76 | 0.65 | 0.71 | 0.77 | 0.71 | 0.82 |
| 6 | 33874440 | rs17535965 | 0.55 | -1.138 | -0.049 | 0.206 | 0.054 | 0.06 | 0.06 | 0.614 | 0.29 | 0.08 | 0.17 | 0.13 | 0.21 | 0.11 |
| 6 | 33876082 | rs16869677 | 0.163 | -0.84 | 0.04 | 0.302 | 0.002 | -0.002 | -0.002 | 0 | 0.08 | 0.09 | 0.08 | 0.08 | 0.06 | 0.11 |

| | | | | | | | | | | | | | | | | |
|---|---|---|---|---|---|---|---|---|---|---|---|---|---|---|---|---|
| 6 | 33876434 | rs12523795 | 0.705 | -0.845 | 0.104 | 0.352 | -0.009 | 0.027 | 0.027 | 0 | 0.06 | 0.11 | 0.12 | 0.14 | 0.03 | 0.05 |
| 6 | 33880480 | rs12191571 | 0.273 | -0.808 | 0.004 | 0.316 | -0.006 | -0.002 | -0.002 | 0.285 | 0.08 | 0.14 | 0.07 | 0.06 | 0.04 | 0.08 |
| 6 | 33885802 | rs2477223 | 0.924 | -0.771 | -0.001 | 0.27 | 0.001 | -0.007 | -0.007 | 2.168 | 0.05 | 0.04 | 0.02 | 0 | 0.07 | 0.01 |
| 6 | 33886682 | rs12665045 | 0.894 | -1.067 | -0.239 | 0.152 | 0.036 | 0.049 | 0.049 | 1.797 | 0.26 | 0.05 | 0.15 | 0.12 | 0.19 | 0.1 |
| 6 | 33886844 | rs11969392 | 1.026 | -0.68 | 0.074 | 0.246 | 0.017 | 0.061 | 0.061 | 0.644 | 0.22 | 0.33 | 0.4 | 0.54 | 0.28 | 0.44 |
| 6 | 33887067 | rs4711367 | 0.38 | -0.655 | 0.072 | 0.254 | 0.021 | 0.043 | 0.043 | 0.16 | 0.28 | 0.39 | 0.46 | 0.61 | 0.32 | 0.52 |
| 6 | 33887368 | rs9368783 | 1.424 | -0.338 | 0.01 | 0.029 | -0.042 | 0.052 | 0.052 | 0 | 0.23 | 0.07 | 0.15 | 0.13 | 0.25 | 0.29 |
| 6 | 33888796 | rs9469633 | 0.424 | -0.261 | 0.016 | 0.1 | -0.004 | 0 | 0 | 0 | 0.07 | 0.12 | 0.06 | 0.06 | 0.04 | 0.08 |
| 6 | 33890451 | rs7748976 | 0.447 | -0.292 | 0.05 | 0.057 | 0.001 | -0.015 | -0.015 | 0 | 0.32 | 0.3 | 0.23 | 0.19 | 0.29 | 0.37 |
| 6 | 33890574 | rs11752328 | 1.207 | -0.206 | -0.247 | 0.122 | -0.004 | 0.011 | 0.011 | 0 | 0.24 | 0.17 | 0.16 | 0.13 | 0.25 | 0.29 |
| 6 | 33894694 | rs2477234 | 0.076 | -0.445 | -0.444 | 0.338 | -0.001 | -0.009 | -0.009 | 0 | 0.16 | 0.28 | 0.13 | 0.12 | 0.14 | 0.13 |
| 6 | 33896120 | rs12661060 | 0.822 | -0.557 | -1.254 | 0.626 | -0.003 | 0.038 | 0.038 | 0 | 0.15 | 0.01 | 0.08 | 0.04 | 0.12 | 0.05 |
| 6 | 33898869 | rs4711369 | -0.008 | -0.355 | -0.909 | 0.833 | 0 | 0.028 | 0.028 | 0 | 0.47 | 0.24 | 0.52 | 0.61 | 0.45 | 0.52 |
| 6 | 33902653 | rs6457749 | -1.021 | -0.98 | -0.223 | 1.301 | -0.027 | 0.011 | 0.011 | 0 | 0.35 | 0.53 | 0.49 | 0.45 | 0.22 | 0.27 |
| 6 | 33903529 | rs6939337 | -0.679 | -0.412 | 0.259 | 1.431 | -0.013 | 0.045 | 0.045 | 0 | 0.23 | 0.42 | 0.29 | 0.26 | 0.23 | 0.17 |
| 6 | 33905008 | rs7765294 | 0.018 | 0.033 | -0.33 | 1.132 | -0.025 | 0.012 | 0.012 | 0 | 0.67 | 0.8 | 0.57 | 0.65 | 0.59 | 0.42 |
| 6 | 33908138 | rs9394180 | 0.825 | -0.25 | 0.107 | 1.24 | 0.02 | 0.004 | 0.004 | 0 | 0.47 | 0.3 | 0.32 | 0.42 | 0.43 | 0.31 |
| 6 | 33908399 | rs9469640 | -0.37 | -0.317 | 0.654 | 1.328 | 0.006 | 0.077 | 0.077 | 0.44 | 0.07 | 0.3 | 0.02 | 0.08 | 0.02 | 0.05 |
| 6 | 33911492 | rs9469643 | -0.031 | -0.313 | 0.732 | 1.33 | 0.007 | 0.075 | 0.075 | 0 | 0.05 | 0.23 | 0.02 | 0.08 | 0.01 | 0.05 |
| 6 | 33915676 | rs7772005 | 0.204 | -0.478 | 0.655 | 1.181 | -0.014 | 0.004 | 0.004 | 0 | 0.54 | 0.68 | 0.38 | 0.51 | 0.5 | 0.35 |
| 6 | 33915771 | rs7771788 | 0.81 | -0.493 | 0.749 | 1.173 | 0 | -0.001 | -0.001 | 0 | 0.48 | 0.47 | 0.36 | 0.43 | 0.48 | 0.3 |
| 6 | 33917213 | rs2495958 | -0.043 | -0.389 | 0.678 | 1.177 | 0.006 | -0.002 | -0.002 | 0.06 | 0.88 | 0.93 | 0.82 | 0.91 | 0.89 | 0.94 |
| 6 | 33917711 | rs10947469 | 0.349 | -0.48 | 0.684 | 1.613 | -0.008 | -0.002 | -0.002 | 0 | 0.52 | 0.66 | 0.35 | 0.43 | 0.5 | 0.34 |
| 6 | 33918563 | rs2495962 | -0.018 | -0.501 | 0.695 | 1.478 | -0.008 | 0.016 | 0.016 | 0 | 0.39 | 0.31 | 0.42 | 0.39 | 0.4 | 0.6 |
| 6 | 33919050 | rs2495964 | 0.802 | -0.83 | 0.418 | 1.46 | 0.004 | -0.026 | -0.026 | 0.518 | 0.63 | 0.34 | 0.65 | 0.58 | 0.64 | 0.77 |
| 6 | 33920994 | rs9469648 | -0.706 | -0.519 | 0.548 | 1.286 | -0.031 | 0.028 | 0.028 | 0 | 0.09 | 0.29 | 0.06 | 0.1 | 0.01 | 0.06 |
| 6 | 33929065 | rs2495971 | 0.961 | -0.594 | 0.254 | 1.195 | -0.022 | 0.003 | 0.003 | 0 | 0.77 | 0.59 | 0.77 | 0.83 | 0.83 | 0.9 |
| 6 | 33932442 | rs2495972 | -0.191 | -0.604 | -0.176 | 1.025 | -0.006 | 0.004 | 0.004 | 0 | 0.45 | 0.65 | 0.47 | 0.5 | 0.41 | 0.61 |
| 6 | 33944014 | rs2495975 | 0.558 | -0.251 | -0.202 | 0.882 | 0 | 0 | 0 | 0 | 0.78 | 0.76 | 0.79 | 0.86 | 0.78 | 0.88 |
| 6 | 33945921 | rs4713731 | -0.395 | -0.232 | -0.578 | 1.067 | 0.003 | -0.002 | -0.002 | 0 | 0.3 | 0.27 | 0.45 | 0.29 | 0.27 | 0.18 |
| 6 | 33946574 | rs9380396 | -0.431 | -0.184 | -0.598 | 1.084 | 0.005 | 0.005 | 0.005 | 0 | 0.3 | 0.37 | 0.49 | 0.3 | 0.34 | 0.19 |
| 6 | 33948019 | rs6911089 | 0.414 | -0.428 | -0.38 | 0.982 | 0.003 | -0.018 | -0.018 | 0 | 0.17 | 0.33 | 0.12 | 0.12 | 0.16 | 0.1 |
| 6 | 33948624 | rs2499762 | -0.332 | -0.161 | -1.076 | 1.242 | -0.027 | -0.065 | -0.065 | 0 | 0.34 | 0.14 | 0.23 | 0.51 | 0.35 | 0.58 |
| 6 | 33953646 | rs2499766 | -0.402 | -0.261 | -0.572 | 0.974 | 0.008 | 0.008 | 0.008 | 0 | 0.63 | 0.52 | 0.42 | 0.42 | 0.58 | 0.62 |
| 6 | 33953734 | rs17531143 | -0.631 | -0.368 | -0.749 | 1.202 | 0 | 0.001 | 0.001 | 0 | 0.16 | 0.12 | 0.11 | 0.14 | 0.1 | 0.26 |
| 6 | 33956472 | rs4256433 | 0.242 | 0.248 | -0.838 | 1.419 | -0.011 | -0.05 | -0.05 | 1.416 | 0.71 | 0.84 | 0.78 | 0.55 | 0.71 | 0.59 |
| 6 | 33957134 | rs2495988 | -1.022 | -0.29 | 0.044 | 1.53 | -0.009 | 0.182 | 0.182 | 2.1 | 0.07 | 0.36 | 0.28 | 0.17 | 0.02 | 0.02 |
| 6 | 33960611 | rs2495990 | -0.465 | -0.439 | -0.579 | 1.066 | -0.014 | -0.004 | -0.004 | 0 | 0.36 | 0.46 | 0.46 | 0.34 | 0.34 | 0.27 |
| 6 | 33961441 | rs2495992 | 0.014 | -0.372 | -1.318 | 1.019 | 0.031 | 0.023 | 0.023 | 0 | 0.47 | 0.27 | 0.52 | 0.64 | 0.6 | 0.64 |
| 6 | 33963501 | rs2451370 | 0.103 | -0.174 | -1.728 | 1.268 | -0.066 | -0.111 | -0.111 | 0 | 0.72 | 0.96 | 0.67 | 0.58 | 0.68 | 0.8 |
| 6 | 33971226 | rs6900980 | -1.171 | -0.602 | -0.684 | 1.39 | -0.02 | 0.031 | 0.031 | 0 | 0.22 | 0.34 | 0.23 | 0.34 | 0.15 | 0.34 |
| 6 | 33975810 | rs9469670 | -1.247 | -0.713 | -0.233 | 1.486 | -0.02 | 0.055 | 0.055 | 0 | 0.2 | 0.32 | 0.22 | 0.37 | 0.14 | 0.32 |
| 6 | 33984420 | rs9461942 | -0.916 | -0.799 | 0.101 | 1.45 | 0.021 | 0.033 | 0.033 | 3.739 | 0.08 | 0.36 | 0.06 | 0.04 | 0.04 | 0.02 |
| 6 | 33988407 | rs9469681 | -0.968 | -0.904 | 0.621 | 1.393 | 0.037 | 0.044 | 0.044 | 3.476 | 0.08 | 0.42 | 0.07 | 0.04 | 0.04 | 0.02 |
| 6 | 33990447 | rs2229901 | -2.533 | -0.998 | 0.725 | 1.301 | -0.011 | 0.015 | 0.015 | 0.196 | 0.26 | 0.72 | 0.25 | 0.41 | 0.16 | 0.33 |
| 6 | 33991368 | rs7772436 | -1.618 | -0.938 | 0.813 | 1.172 | -0.025 | -0.028 | -0.028 | 4.272 | 0.12 | 0.68 | 0.09 | 0.04 | 0.06 | 0.03 |
| 6 | 33997375 | rs2499669 | -0.702 | -0.926 | -0.577 | 0.893 | -0.061 | 0.049 | 0.049 | 0 | 0.65 | 0.83 | 0.7 | 0.74 | 0.55 | 0.7 |
| 6 | 34007054 | rs2499677 | -1.325 | -0.556 | -0.195 | 1.35 | -0.001 | 0.001 | 0.001 | 1.663 | 0.25 | 0.76 | 0.25 | 0.46 | 0.21 | 0.3 |
| 6 | 34015765 | rs4711374 | -0.71 | -0.582 | -0.023 | 1.324 | 0.014 | 0.06 | 0.06 | 0 | 0.2 | 0.4 | 0.22 | 0.36 | 0.18 | 0.29 |
| 6 | 34016786 | rs2451328 | -1.204 | -0.677 | -0.595 | 1.024 | -0.002 | -0.001 | -0.001 | 2.557 | 0.69 | 0.85 | 0.59 | 0.75 | 0.69 | 0.72 |
| 6 | 34022740 | rs1873250 | -1.472 | -0.654 | -0.547 | 1.057 | 0.005 | -0.01 | -0.01 | 1.436 | 0.3 | 0.73 | 0.25 | 0.46 | 0.22 | 0.33 |
| 6 | 34032613 | rs10947475 | 1.348 | -0.592 | -1.075 | 0.57 | -0.055 | 0.127 | 0.127 | 0 | 0.35 | 0.1 | 0.26 | 0.11 | 0.4 | 0.37 |
| 6 | 34034441 | rs2499682 | -0.906 | -0.47 | -0.849 | 0.802 | -0.02 | -0.001 | -0.001 | 0.041 | 0.71 | 0.9 | 0.61 | 0.78 | 0.69 | 0.73 |
| 6 | 34034648 | rs10947476 | 1.376 | -0.675 | -0.887 | 0.642 | -0.062 | 0.107 | 0.107 | 0 | 0.28 | 0.07 | 0.21 | 0.08 | 0.35 | 0.3 |
| 6 | 34036446 | rs1906953 | -1.26 | -0.463 | 0.346 | 1.577 | -0.028 | 0.116 | 0.116 | 0 | 0.21 | 0.51 | 0.24 | 0.48 | 0.14 | 0.26 |
| 6 | 34041306 | rs2499689 | -1.602 | -0.581 | 0.286 | 1.492 | -0.04 | 0.028 | 0.028 | 0.093 | 0.1 | 0.26 | 0.04 | 0.12 | 0.04 | 0.04 |
| 6 | 34043298 | rs2499694 | -2.166 | -0.717 | 0.073 | 1.503 | 0.082 | -0.054 | -0.054 | 1.604 | 0.36 | 0.84 | 0.3 | 0.62 | 0.2 | 0.32 |
| 6 | 34044376 | rs6928460 | -0.778 | -0.495 | 0.414 | 1.326 | -0.027 | 0.029 | 0.029 | 0 | 0.06 | 0.19 | 0.03 | 0.1 | 0.01 | 0.01 |
| 6 | 34044918 | rs2499697 | 0.905 | -0.138 | 0.245 | 1.268 | -0.022 | -0.021 | -0.021 | 0 | 0.95 | 0.99 | 0.9 | 0.86 | 0.89 | 0.94 |
| 6 | 34046131 | rs1994582 | -2.103 | -0.203 | 0.708 | 1.18 | -0.005 | 0.002 | 0.002 | 0.119 | 0.11 | 0.33 | 0.07 | 0.1 | 0.07 | 0.04 |
| 6 | 34050917 | rs12211633 | 2.325 | 0.142 | 0.445 | 1.324 | 0.005 | -0.032 | -0.032 | 0 | 0.67 | 0.28 | 0.7 | 0.48 | 0.72 | 0.66 |
| 6 | 34051442 | rs9368793 | 2.048 | 0.131 | 0.624 | 1.335 | -0.003 | 0.009 | 0.009 | 0.274 | 0.71 | 0.35 | 0.72 | 0.53 | 0.77 | 0.71 |
| 6 | 34053424 | rs1873249 | 1.885 | 0.207 | 0.182 | 1.089 | -0.003 | 0.086 | 0.086 | 0 | 0.45 | 0.27 | 0.34 | 0.25 | 0.5 | 0.42 |
| 6 | 34054393 | rs2451357 | 1.224 | 0.542 | 0.284 | 1.321 | -0.003 | 0.003 | 0.003 | 0 | 0.92 | 0.93 | 0.9 | 0.8 | 0.85 | 0.9 |
| 6 | 34055679 | rs1472037 | 1.224 | 0.542 | 0.28 | 1.321 | -0.005 | 0 | 0 | 0.045 | 0.92 | 0.93 | 0.9 | 0.8 | 0.85 | 0.9 |
| 6 | 34059239 | rs937035 | -0.836 | 0.318 | 0.403 | 1.249 | 0.005 | 0.029 | 0.029 | 0 | 0.34 | 0.54 | 0.48 | 0.5 | 0.31 | 0.44 |
| 6 | 34062805 | rs874631 | 1.075 | 0.414 | 0.389 | 1.368 | -0.002 | 0.001 | 0.001 | 0 | 0.92 | 0.93 | 0.9 | 0.82 | 0.85 | 0.91 |
| 6 | 34065982 | rs12206652 | 0.024 | 0.383 | 0.436 | 1.332 | -0.004 | -0.006 | -0.006 | 0 | 0.25 | 0.14 | 0.37 | 0.24 | 0.24 | 0.27 |
| 6 | 34069062 | rs3887555 | 0.753 | 0.014 | 0.538 | 1.251 | -0.012 | -0.029 | -0.029 | 0 | 0.62 | 0.71 | 0.52 | 0.52 | 0.59 | 0.63 |

| | | | | | | | | | | | | | | | | |
|---|---|---|---|---|---|---|---|---|---|---|---|---|---|---|---|---|
| 6 | 34072215 | rs2499714 | -0.654 | -0.024 | 0.831 | 1.034 | -0.004 | 0.115 | 0.115 | 0.013 | 0.16 | 0.37 | 0.11 | 0.33 | 0.09 | 0.21 |
| 6 | 34072561 | rs7763695 | 0.777 | 0.3 | 1.021 | 1.107 | 0.002 | -0.001 | -0.001 | 0 | 0.59 | 0.44 | 0.47 | 0.61 | 0.59 | 0.65 |
| 6 | 34074463 | rs1109654 | 0.544 | -0.032 | 1.332 | 0.707 | 0.018 | 0.005 | 0.005 | 0 | 0.6 | 0.53 | 0.68 | 0.4 | 0.6 | 0.52 |
| 6 | 34083564 | rs13196449 | -1.596 | -0.696 | 1.366 | 0.603 | -0.055 | -0.034 | -0.034 | 0 | 0.13 | 0.28 | 0.06 | 0.04 | 0.03 | 0.02 |
| 6 | 34095696 | rs11961715 | -0.968 | -0.612 | 0.889 | 0.751 | -0.03 | 0.097 | 0.097 | 0 | 0.08 | 0.24 | 0.07 | 0.13 | 0.02 | 0.04 |
| 6 | 34102061 | rs2499730 | 1.642 | -0.448 | 0.091 | 0.754 | -0.07 | -0.003 | -0.003 | 0.012 | 0.42 | 0.13 | 0.58 | 0.37 | 0.56 | 0.46 |
| 6 | 34103662 | rs2451334 | 0.317 | -0.202 | -0.334 | 0.64 | -0.12 | -0.008 | -0.008 | 0 | 0.22 | 0.02 | 0.15 | 0.16 | 0.28 | 0.25 |
| 6 | 34103954 | rs914813 | -0.039 | -0.491 | 0.066 | 0.378 | -0.012 | -0.009 | -0.009 | 0.031 | 0.36 | 0.29 | 0.25 | 0.49 | 0.36 | 0.46 |
| 6 | 34106584 | rs2029457 | 0.834 | -0.064 | -0.752 | 0.562 | -0.096 | -0.097 | -0.097 | 0 | 0.44 | 0.1 | 0.62 | 0.47 | 0.57 | 0.47 |
| 6 | 34118586 | rs2451330 | -0.211 | 0.145 | -0.058 | 0.671 | -0.031 | -0.029 | -0.029 | 0.394 | 0.36 | 0.25 | 0.54 | 0.53 | 0.41 | 0.45 |
| 6 | 34125281 | rs7755106 | 0.527 | -0.018 | 0.341 | 0.454 | -0.013 | -0.046 | -0.046 | 0.061 | 0.58 | 0.25 | 0.48 | 0.49 | 0.53 | 0.53 |
| 6 | 34131376 | rs10947479 | 0.684 | 0.218 | 0.537 | 0.527 | -0.02 | -0.024 | -0.024 | 0 | 0.57 | 0.18 | 0.65 | 0.55 | 0.53 | 0.53 |
| 6 | 34131610 | rs10947482 | 0.724 | 0.092 | 0.549 | 0.555 | -0.037 | -0.081 | -0.081 | 0 | 0.56 | 0.12 | 0.45 | 0.47 | 0.52 | 0.5 |
| 6 | 34132951 | rs4713750 | 0.516 | -0.017 | 0.769 | 0.358 | -0.018 | -0.004 | -0.004 | 0.422 | 0.58 | 0.3 | 0.68 | 0.56 | 0.55 | 0.52 |
| 6 | 34133488 | rs4713751 | 0.128 | -0.101 | 1.352 | 0.313 | 0.02 | 0.011 | 0.011 | 0 | 0.66 | 0.61 | 0.75 | 0.56 | 0.57 | 0.53 |
| 6 | 34135235 | rs4713755 | 0.625 | -0.048 | 0.524 | 0.337 | -0.027 | -0.017 | -0.017 | 0 | 0.57 | 0.18 | 0.66 | 0.55 | 0.54 | 0.51 |
| 6 | 34143292 | rs1105695 | 0.328 | -0.039 | 0.58 | 0.404 | 0.007 | -0.001 | -0.001 | 0.909 | 0.57 | 0.34 | 0.63 | 0.56 | 0.45 | 0.49 |
| 6 | 34153441 | rs7744288 | -1.553 | -0.01 | 0.901 | 0.318 | -0.027 | -0.017 | -0.017 | 0 | 0.06 | 0.15 | 0.03 | 0.02 | 0.01 | 0.02 |
| 6 | 34165721 | rs7742369 | -1.87 | 0.049 | 1.191 | 0.185 | -0.022 | -0.044 | -0.044 | 0 | 0.23 | 0.61 | 0.4 | 0.13 | 0.18 | 0.13 |
| 6 | 34168464 | rs6922119 | -1.329 | 0.237 | 1.765 | 0.255 | 0.1 | 0.024 | 0.024 | 0 | 0.06 | 0.29 | 0.07 | 0.02 | 0.09 | 0.04 |
| 6 | 34194866 | rs1759645 | 1.93 | 0.051 | 1.898 | 0.179 | -0.01 | -0.035 | -0.035 | 0.007 | 0.8 | 0.4 | 0.61 | 0.87 | 0.84 | 0.9 |
| 6 | 34195011 | rs1776897 | 1.412 | 0.053 | 2.116 | 0.181 | -0.027 | -0.002 | -0.002 | 0 | 0.85 | 0.52 | 0.66 | 0.88 | 0.92 | 0.95 |
| 6 | 34197188 | rs1759628 | 1.263 | 0.08 | 2.15 | 0.199 | -0.049 | -0.001 | -0.001 | 0 | 0.87 | 0.62 | 0.69 | 0.9 | 0.95 | 0.97 |
| 6 | 34215919 | rs1150779 | 1.469 | 0.076 | 2.042 | 0.203 | -0.027 | 0.006 | 0.006 | 0.77 | 0.85 | 0.54 | 0.68 | 0.87 | 0.94 | 0.95 |
| 6 | 34218718 | rs9394200 | 1.339 | 0.052 | 2.023 | 0.207 | -0.064 | 0.013 | 0.013 | 0.903 | 0.86 | 0.53 | 0.69 | 0.87 | 0.97 | 0.97 |
| 6 | 34236429 | rs9469761 | -1.728 | 0.136 | 1.953 | 0.194 | -0.01 | -0.018 | -0.018 | 0 | 0.17 | 0.52 | 0.36 | 0.13 | 0.13 | 0.08 |
| 6 | 34238514 | rs6918981 | 1.749 | 0.012 | 1.672 | 0.128 | -0.003 | -0.033 | -0.033 | 0 | 0.78 | 0.42 | 0.6 | 0.83 | 0.8 | 0.87 |
| 6 | 34254178 | rs6908522 | -2.037 | -0.135 | 1.884 | -0.04 | -0.012 | -0.041 | -0.041 | 0 | 0.08 | 0.21 | 0.04 | 0 | 0.02 | 0.04 |
| 6 | 34255631 | rs16883896 | -1.199 | -0.113 | 1.96 | -0.216 | -0.002 | 0.009 | 0.009 | 0 | 0.1 | 0.08 | 0.09 | 0.04 | 0.13 | 0.1 |
| 6 | 34268684 | rs3798554 | -1.832 | -0.168 | 2.093 | -0.197 | 0 | 0.019 | 0.019 | 0 | 0.12 | 0.11 | 0.09 | 0.06 | 0.13 | 0.12 |
| 6 | 34270668 | rs6930978 | -2.946 | -0.7 | 2.274 | 1.234 | -0.041 | 0.043 | 0.043 | 0.119 | 0.32 | 0.53 | 0.48 | 0.46 | 0.21 | 0.35 |
| 6 | 34286424 | rs4711390 | -0.772 | -0.701 | 2.304 | 1.235 | 0.026 | -0.023 | -0.023 | 0 | 0.12 | 0.06 | 0.33 | 0.41 | 0.04 | 0.2 |
| 6 | 34302869 | rs206936 | -3.075 | -0.776 | 2.439 | 1.192 | -0.029 | 0.024 | 0.024 | 0 | 0.33 | 0.57 | 0.48 | 0.47 | 0.21 | 0.4 |
| 6 | 34310476 | rs3798557 | -1.132 | -0.826 | 2.281 | 1.273 | -0.012 | 0.014 | 0.014 | 0 | 0.1 | 0.03 | 0.09 | 0.05 | 0.13 | 0.1 |
| 6 | 34332179 | rs206937 | -0.214 | 0.394 | 1.449 | 1.118 | -0.085 | -0.058 | -0.058 | 0 | 0.87 | 0.98 | 0.85 | 0.76 | 0.75 | 0.76 |
| 6 | 34371707 | rs6935129 | 0.372 | 0.149 | 1.576 | 1.2 | 0.027 | -0.026 | -0.026 | 0 | 0.88 | 0.93 | 0.67 | 0.58 | 0.96 | 0.79 |
| 6 | 34382610 | rs10947500 | 1.094 | -0.323 | 0.559 | 0.216 | -0.048 | 0.071 | 0.071 | 0 | 0.16 | 0.01 | 0.09 | 0.02 | 0.17 | 0.1 |
| 6 | 34383208 | rs16889678 | -1.747 | -0.491 | 1.159 | 0.642 | -0.001 | 0.006 | 0.006 | 0.381 | 0.09 | 0.37 | 0.04 | 0.03 | 0.03 | 0.06 |
| 6 | 34383788 | rs16889690 | -1.14 | -0.466 | 1.158 | 0.637 | 0.019 | -0.005 | -0.005 | 0 | 0.07 | 0.29 | 0.04 | 0 | 0.02 | 0.04 |
| 6 | 34384852 | rs16889706 | -0.269 | -0.483 | 0.987 | 0.639 | 0.022 | -0.017 | -0.017 | 0 | 0.13 | 0.08 | 0.33 | 0.43 | 0.04 | 0.21 |
| 6 | 34397920 | rs3913010 | 1.772 | -0.408 | 1.379 | 0.619 | -0.021 | 0.016 | 0.016 | 0.288 | 0.69 | 0.42 | 0.52 | 0.51 | 0.8 | 0.64 |
| 6 | 34401378 | rs16889851 | -1.098 | -0.428 | 1.322 | 0.579 | 0.02 | -0.004 | -0.004 | 0.176 | 0.07 | 0.28 | 0.04 | 0 | 0.02 | 0.04 |
| 6 | 34405715 | rs9366856 | 1.57 | -0.46 | 1.067 | 0.784 | -0.003 | 0.001 | 0.001 | 0.164 | 0.69 | 0.34 | 0.51 | 0.5 | 0.8 | 0.65 |
| 6 | 34416388 | rs9394220 | 1.908 | -0.273 | 0.888 | 1.295 | 0.047 | -0.013 | -0.013 | 0 | 0.64 | 0.22 | 0.48 | 0.37 | 0.75 | 0.61 |
| 6 | 34416893 | rs4713791 | 1.392 | -0.23 | 1.536 | 1.486 | -0.035 | 0.013 | 0.013 | 0 | 0.77 | 0.55 | 0.87 | 0.73 | 0.83 | 0.82 |
| 6 | 34422354 | rs9394222 | 1.5 | -0.365 | 0.635 | 0.813 | 0.004 | 0.065 | 0.065 | 0 | 0.5 | 0.36 | 0.33 | 0.27 | 0.5 | 0.47 |
| 6 | 34433510 | rs6927652 | 1.872 | -0.564 | 0.071 | 0.745 | -0.02 | 0.145 | 0.145 | 0.212 | 0.5 | 0.08 | 0.54 | 0.23 | 0.51 | 0.43 |
| 6 | 34438564 | rs6914721 | -1.588 | -0.772 | 0.761 | 1.229 | -0.05 | 0.018 | 0.018 | 0 | 0.3 | 0.56 | 0.23 | 0.41 | 0.2 | 0.26 |
| 6 | 34439749 | rs9469791 | -0.339 | -0.375 | 0.724 | 1.535 | -0.013 | -0.011 | -0.011 | 1.116 | 0.13 | 0.1 | 0.17 | 0.3 | 0.27 | 0.24 |
| 6 | 34441731 | rs9394226 | -1.177 | -1.101 | 0.454 | 1.297 | -0.023 | 0.006 | 0.006 | 0 | 0.19 | 0.23 | 0.14 | 0.3 | 0.07 | 0.24 |
| 6 | 34443922 | rs3846866 | 2.155 | -0.857 | -0.239 | 0.527 | -0.044 | 0.135 | 0.135 | 2.807 | 0.5 | 0.1 | 0.52 | 0.26 | 0.47 | 0.39 |
| 6 | 34448315 | rs3846867 | -1.891 | -0.404 | 0.679 | 0.941 | 0.001 | 0 | 0 | 0 | 0.4 | 0.41 | 0.33 | 0.62 | 0.47 | 0.55 |
| 6 | 34457528 | rs3889226 | -1.226 | -0.504 | 0.577 | 0.985 | -0.091 | 0.135 | 0.135 | 0.014 | 0.25 | 0.46 | 0.15 | 0.54 | 0.13 | 0.36 |
| 6 | 34458533 | rs4713797 | 2.011 | -0.176 | 0.338 | 0.933 | -0.052 | 0.03 | 0.03 | 1.358 | 0.54 | 0.11 | 0.55 | 0.34 | 0.53 | 0.45 |
| 6 | 34461158 | rs7742005 | -1.436 | -0.07 | 0.506 | 1.03 | -0.002 | 0.005 | 0.005 | 0.001 | 0.4 | 0.53 | 0.31 | 0.58 | 0.39 | 0.54 |
| 6 | 34464010 | rs3904668 | 1.621 | 0.016 | -0.045 | 1.064 | -0.001 | 0.061 | 0.061 | 0.251 | 0.53 | 0.25 | 0.56 | 0.31 | 0.54 | 0.44 |
| 6 | 34472007 | rs4713805 | 2.024 | 0.451 | -0.248 | 1.004 | -0.038 | -0.005 | -0.005 | 0 | 0.48 | 0.06 | 0.51 | 0.38 | 0.46 | 0.42 |
| 6 | 34474629 | rs4713806 | 1.838 | 0.504 | -0.184 | 1.068 | 0.004 | 0.08 | 0.08 | 0.259 | 0.51 | 0.13 | 0.52 | 0.33 | 0.46 | 0.42 |
| 6 | 34478203 | rs2395567 | 1.785 | 0.477 | -0.207 | 0.87 | 0.013 | 0.042 | 0.042 | 0.107 | 0.5 | 0.13 | 0.51 | 0.37 | 0.46 | 0.42 |
| 6 | 34480637 | rs10947508 | -0.021 | 0.845 | 0.134 | 0.799 | -0.053 | 0.012 | 0.012 | 0 | 0.09 | 0.03 | 0.15 | 0.02 | 0.22 | 0.12 |
| 6 | 34481140 | rs3800471 | -1.138 | 0.505 | 0.536 | 0.931 | 0.073 | 0.085 | 0.085 | 0 | 0.05 | 0.18 | 0.11 | 0.1 | 0.09 | 0.03 |
| 6 | 34483015 | rs4713808 | 1.375 | 0.31 | -0.022 | 0.729 | -0.042 | 0.03 | 0.03 | 0.02 | 0.55 | 0.18 | 0.57 | 0.36 | 0.53 | 0.5 |
| 6 | 34501529 | rs913014 | 2.292 | -0.096 | 1.346 | 0.505 | -0.003 | -0.001 | -0.001 | 0.483 | 0.79 | 0.27 | 0.79 | 0.8 | 0.87 | 0.93 |
| 6 | 34503338 | rs11758426 | -1.428 | 0.185 | 1.164 | 0.576 | 0.035 | 0.024 | 0.024 | 0 | 0.08 | 0.19 | 0.13 | 0.09 | 0.09 | 0.04 |
| 6 | 34503439 | rs11756280 | -1.378 | 0.185 | 1.164 | 0.576 | 0.037 | 0.026 | 0.026 | 0 | 0.07 | 0.19 | 0.13 | 0.09 | 0.09 | 0.03 |
| 6 | 34505194 | rs7754038 | -1.724 | 0.178 | 1.291 | 0.573 | 0.025 | 0.015 | 0.015 | 0 | 0.09 | 0.19 | 0.13 | 0.09 | 0.09 | 0.03 |
| 6 | 34507311 | rs2233645 | -0.426 | 0.024 | 1.267 | 0.427 | -0.005 | -0.029 | -0.029 | 0 | 0.07 | 0.17 | 0.04 | 0.01 | 0.05 | 0.03 |
| 6 | 34513266 | rs3778076 | -1.423 | 0.071 | 1.294 | 0.482 | -0.004 | -0.001 | -0.001 | 0.115 | 0.08 | 0.16 | 0.06 | 0.08 | 0.05 | 0.03 |
| 6 | 34520267 | rs3798544 | -2.114 | -0.175 | 1.555 | 0.304 | -0.033 | -0.048 | -0.048 | 0 | 0.21 | 0.5 | 0.18 | 0.12 | 0.13 | 0.1 |

| | | | | | | | | | | | | | | | | |
|---|---|---|---|---|---|---|---|---|---|---|---|---|---|---|---|---|
| 6 | 34526326 | rs3756932 | -1.317 | -0.266 | 1.392 | 0.393 | -0.047 | -0.043 | -0.043 | 0.036 | 0.15 | 0.37 | 0.1 | 0.09 | 0.08 | 0.08 |
| 6 | 34527482 | rs9296121 | -1.821 | -0.141 | 1.042 | 0.693 | -0.019 | 0.012 | 0.012 | 0 | 0.27 | 0.43 | 0.19 | 0.34 | 0.22 | 0.19 |
| 6 | 34528517 | rs16890991 | 0.042 | -0.097 | 0.922 | 0.717 | -0.027 | -0.051 | -0.051 | 0 | 0.11 | 0.03 | 0.09 | 0.25 | 0.13 | 0.12 |
| 6 | 34530165 | rs4713813 | 1.985 | -0.155 | 1.127 | 0.705 | 0.004 | -0.004 | -0.004 | 0 | 0.67 | 0.33 | 0.73 | 0.58 | 0.77 | 0.79 |
| 6 | 34531300 | rs10947510 | -1.567 | 0.536 | 0.372 | 0.437 | 0.002 | -0.002 | -0.002 | 0 | 0.41 | 0.46 | 0.39 | 0.37 | 0.47 | 0.4 |
| 6 | 34540022 | rs205279 | -1.392 | -0.106 | 1.054 | 0.89 | -0.053 | 0.01 | 0.01 | 0.006 | 0.16 | 0.47 | 0.16 | 0.16 | 0.1 | 0.09 |
| 6 | 34542048 | rs6936773 | -1.803 | -0.039 | 0.419 | 0.609 | -0.025 | -0.003 | -0.003 | 0 | 0.32 | 0.48 | 0.21 | 0.35 | 0.26 | 0.25 |
| 6 | 34544384 | rs6917350 | -1.013 | 0.306 | 0.066 | 0.909 | 0 | -0.013 | -0.013 | 0 | 0.18 | 0.13 | 0.12 | 0.26 | 0.18 | 0.16 |
| 6 | 34546560 | rs2814982 | -1.265 | 0.552 | 0.723 | 0.659 | 0.031 | -0.003 | -0.003 | 0 | 0.11 | 0.31 | 0.08 | 0.08 | 0.1 | 0.08 |
| 6 | 34548296 | rs2814985 | -1.215 | 0.262 | 1.203 | 0.335 | 0.015 | -0.05 | -0.05 | 0.554 | 0.11 | 0.22 | 0.1 | 0.01 | 0.13 | 0.08 |
| 6 | 34551912 | rs205284 | -0.617 | -0.176 | 1.039 | 0.457 | -0.002 | 0 | 0 | 0 | 0.14 | 0.15 | 0.06 | 0.12 | 0.08 | 0.07 |
| 6 | 34553048 | rs205283 | -2.717 | -0.777 | 0.373 | 0.685 | 0.006 | 0.042 | 0.042 | 0 | 0.53 | 0.95 | 0.36 | 0.69 | 0.38 | 0.4 |
| 6 | 34555709 | rs1201872 | -0.414 | -0.445 | 0.561 | 0.589 | -0.001 | 0 | 0 | 0 | 0.14 | 0.16 | 0.06 | 0.12 | 0.08 | 0.08 |
| 6 | 34563164 | rs205262 | -2.297 | -0.515 | 0.99 | 0.52 | 0.012 | 0.019 | 0.019 | 0 | 0.36 | 0.73 | 0.23 | 0.14 | 0.28 | 0.19 |
| 6 | 34569814 | rs9469813 | -0.929 | -0.519 | 1.159 | 0.526 | -0.028 | -0.02 | -0.02 | 0 | 0.2 | 0.36 | 0.12 | 0.13 | 0.14 | 0.09 |
| 6 | 34579431 | rs2744974 | -2.615 | -0.586 | 1.057 | 0.87 | 0.022 | -0.003 | -0.003 | 0.292 | 0.42 | 0.76 | 0.31 | 0.43 | 0.34 | 0.28 |
| 6 | 34592731 | rs2744963 | -1.464 | -0.511 | 1.071 | 0.914 | 0.016 | 0.036 | 0.036 | 0 | 0.17 | 0.28 | 0.18 | 0.3 | 0.19 | 0.17 |
| 6 | 34607958 | rs17644393 | 0.84 | -0.578 | 1.001 | 0.908 | -0.005 | 0.031 | 0.031 | 0 | 0.08 | 0 | 0.03 | 0.01 | 0.05 | 0.04 |
| 6 | 34613630 | rs6909637 | -0.637 | -0.526 | 1.284 | 0.934 | -0.038 | -0.023 | -0.023 | 0 | 0.2 | 0.38 | 0.12 | 0.13 | 0.14 | 0.09 |
| 6 | 34617144 | rs2814992 | -2.394 | -0.541 | 1.228 | 0.932 | 0.027 | -0.005 | -0.005 | 0 | 0.42 | 0.77 | 0.31 | 0.43 | 0.34 | 0.28 |
| 6 | 34618893 | rs2814993 | -0.77 | -0.542 | 1.237 | 0.864 | 0.01 | -0.05 | -0.05 | 0.119 | 0.11 | 0.22 | 0.1 | 0.01 | 0.12 | 0.08 |
| 6 | 34633862 | rs2744949 | -2.412 | -0.531 | 1.248 | 0.924 | 0.026 | -0.016 | -0.016 | 0 | 0.42 | 0.77 | 0.31 | 0.45 | 0.34 | 0.28 |
| 6 | 34638847 | rs2815005 | -0.735 | -0.538 | 1.233 | 0.92 | 0.01 | -0.05 | -0.05 | 0.72 | 0.11 | 0.22 | 0.1 | 0.02 | 0.13 | 0.08 |
| 6 | 34643179 | rs2744943 | -2.433 | -0.538 | 1.233 | 0.92 | 0.028 | -0.016 | -0.016 | 0 | 0.42 | 0.77 | 0.31 | 0.45 | 0.34 | 0.28 |
| 6 | 34647978 | rs2814945 | -2.433 | 0.086 | 1.163 | 0.855 | 0.028 | -0.016 | -0.016 | 0 | 0.42 | 0.77 | 0.31 | 0.45 | 0.34 | 0.28 |
| 6 | 34659054 | rs2744972 | 0.753 | 0.197 | 1.094 | 0.897 | 0.01 | -0.05 | -0.05 | 2.837 | 0.89 | 0.78 | 0.9 | 0.99 | 0.87 | 0.92 |
| 6 | 34677103 | rs6932930 | 1.472 | 0.226 | 1.043 | 0.93 | 0.005 | 0.036 | 0.036 | 0 | 0.79 | 0.64 | 0.81 | 0.68 | 0.8 | 0.81 |
| 6 | 34694779 | rs4513788 | -2.124 | 0.185 | 0.626 | 0.891 | 0.029 | -0.018 | -0.018 | 0 | 0.42 | 0.77 | 0.31 | 0.46 | 0.34 | 0.29 |
| 6 | 34697767 | rs2814953 | -2.133 | 0.182 | 0.625 | 0.913 | 0.03 | -0.019 | -0.019 | 0.985 | 0.42 | 0.77 | 0.31 | 0.46 | 0.34 | 0.29 |
| 6 | 34704613 | rs2764211 | -2.125 | 0.18 | 0.624 | 0.893 | 0.029 | -0.018 | -0.018 | 0 | 0.42 | 0.77 | 0.31 | 0.46 | 0.34 | 0.29 |
| 6 | 34715341 | rs9462007 | -1.013 | 0.132 | 1.074 | 0.828 | 0.001 | -0.084 | -0.084 | 0 | 0.15 | 0.36 | 0.11 | 0.01 | 0.13 | 0.1 |
| 6 | 34725478 | rs7743724 | -0.57 | 0.109 | 1.135 | 0.861 | -0.006 | -0.003 | -0.003 | 0.393 | 0.19 | 0.21 | 0.11 | 0.13 | 0.14 | 0.09 |
| 6 | 34728253 | rs9462014 | -1.21 | 0.108 | 1.146 | 0.858 | 0.001 | -0.096 | -0.096 | 0 | 0.16 | 0.42 | 0.12 | 0.01 | 0.14 | 0.1 |
| 6 | 34747603 | rs11759151 | -2.402 | 0.122 | 1.106 | 0.881 | 0.029 | -0.018 | -0.018 | 0 | 0.42 | 0.77 | 0.31 | 0.46 | 0.34 | 0.29 |
| 6 | 34771820 | rs3798349 | 0.38 | 0.125 | 1.111 | 0.895 | -0.004 | 0.03 | 0.03 | 0 | 0.06 | 0.03 | 0.08 | 0.3 | 0.06 | 0.09 |
| 6 | 34797241 | rs9462027 | -1.882 | 0.114 | 1.136 | 0.842 | -0.003 | -0.014 | -0.014 | 0.118 | 0.36 | 0.67 | 0.23 | 0.15 | 0.27 | 0.2 |
| 6 | 34804204 | rs7761391 | -0.666 | 0.112 | 1.169 | 0.844 | -0.011 | -0.006 | -0.006 | 0.173 | 0.19 | 0.22 | 0.11 | 0.13 | 0.14 | 0.09 |
| 6 | 34823187 | rs3734266 | -0.679 | 0.314 | 1.165 | 0.845 | -0.012 | -0.012 | -0.012 | 0.242 | 0.19 | 0.25 | 0.11 | 0.13 | 0.15 | 0.1 |
| 6 | 34824107 | rs16894945 | 0.378 | 0.322 | 1.122 | 0.893 | -0.004 | 0.03 | 0.03 | 0 | 0.06 | 0.03 | 0.08 | 0.3 | 0.06 | 0.09 |
| 6 | 34824636 | rs11755393 | -2.483 | 0.317 | 1.145 | 0.881 | 0.018 | -0.018 | -0.018 | 0 | 0.42 | 0.77 | 0.31 | 0.46 | 0.35 | 0.29 |
| 6 | 34827085 | rs9469913 | -1.26 | 0.099 | 1.15 | 0.88 | 0.005 | -0.086 | -0.086 | 0 | 0.17 | 0.46 | 0.12 | 0.02 | 0.14 | 0.1 |
| 6 | 34828553 | rs6457796 | -2.098 | 0.056 | 1.044 | 0.871 | 0.022 | 0.035 | 0.035 | 0 | 0.37 | 0.74 | 0.24 | 0.16 | 0.28 | 0.2 |
| 6 | 34831369 | rs9469917 | -2.435 | 0.07 | 1.045 | 0.84 | 0.028 | -0.017 | -0.017 | 0.022 | 0.42 | 0.77 | 0.31 | 0.46 | 0.34 | 0.29 |
| 6 | 34831856 | rs13205210 | -0.617 | 0.092 | 1.06 | 0.846 | -0.009 | -0.005 | -0.005 | 0 | 0.19 | 0.21 | 0.11 | 0.13 | 0.14 | 0.09 |
| 6 | 34831866 | rs3734264 | -2.317 | 0.124 | 0.937 | 0.861 | 0.028 | -0.018 | -0.018 | 0.064 | 0.42 | 0.77 | 0.31 | 0.46 | 0.34 | 0.29 |
| 6 | 34838429 | rs3734263 | 0.031 | -0.24 | 0.937 | 1.052 | -0.01 | 0.007 | 0.007 | 0 | 0.06 | 0.02 | 0.09 | 0.3 | 0.07 | 0.09 |
| 6 | 34840398 | rs3734261 | -0.186 | -0.335 | 1.312 | 1.005 | -0.023 | -0.003 | -0.003 | 0 | 0.14 | 0.2 | 0.06 | 0.13 | 0.08 | 0.08 |
| 6 | 34840440 | rs3734260 | 0.067 | -0.253 | 1.12 | 1.045 | -0.01 | 0.01 | 0.01 | 0 | 0.06 | 0.03 | 0.09 | 0.29 | 0.08 | 0.09 |
| 6 | 34840515 | rs3822923 | 0.523 | -0.457 | 1.038 | 1.277 | 0.018 | -0.006 | -0.006 | 0 | 0.06 | 0.01 | 0.07 | 0.26 | 0.01 | 0.13 |
| 6 | 34841330 | rs3800453 | -0.782 | -0.641 | 1 | 1.169 | -0.022 | 0.019 | 0.019 | 0 | 0.12 | 0.14 | 0.1 | 0.27 | 0.04 | 0.15 |
| 6 | 34844629 | rs3734259 | -0.782 | -0.641 | 1 | 1.169 | -0.022 | 0.019 | 0.019 | 0 | 0.12 | 0.14 | 0.1 | 0.27 | 0.04 | 0.15 |
| 6 | 34845648 | rs2985 | -2.694 | -0.716 | 0.996 | 1.115 | -0.019 | 0.009 | 0.009 | 0.31 | 0.31 | 0.67 | 0.27 | 0.58 | 0.15 | 0.33 |
| 6 | 34854745 | rs9366871 | 0.238 | -0.495 | 0.964 | 1.215 | -0.008 | 0.018 | 0.018 | 0 | 0.06 | 0.02 | 0.09 | 0.3 | 0.07 | 0.09 |
| 6 | 34868790 | rs6907619 | -1.628 | -0.535 | 1.014 | 1.189 | -0.012 | 0.058 | 0.058 | 0 | 0.18 | 0.22 | 0.18 | 0.57 | 0.11 | 0.23 |
| 6 | 34886436 | rs9380470 | -1.481 | -0.559 | 1.083 | 1.169 | -0.049 | 0.031 | 0.031 | 0 | 0.16 | 0.26 | 0.12 | 0.26 | 0.05 | 0.16 |
| 6 | 34904488 | rs1737727 | -2.476 | -0.655 | 0.893 | 1.107 | 0.016 | -0.006 | -0.006 | 0 | 0.39 | 0.67 | 0.29 | 0.59 | 0.2 | 0.36 |
| 6 | 34904584 | rs847851 | -3.045 | -0.683 | 0.777 | 1.091 | 0.142 | 0.07 | 0.07 | 0.031 | 0.49 | 0.95 | 0.37 | 0.72 | 0.29 | 0.41 |
| 6 | 34906168 | rs847848 | -2.423 | -0.712 | 0.81 | 1.103 | -0.017 | 0.011 | 0.011 | 0.228 | 0.3 | 0.68 | 0.27 | 0.59 | 0.15 | 0.32 |
| 6 | 34909687 | rs9348955 | 0.076 | -0.685 | 0.68 | 1.113 | 0 | 0.04 | 0.04 | 0 | 0.07 | 0.05 | 0.09 | 0.31 | 0.08 | 0.09 |
| 6 | 34938809 | rs847861 | -2.052 | -0.727 | 0.725 | 1.088 | -0.078 | -0.022 | -0.022 | 0.447 | 0.32 | 0.57 | 0.2 | 0.27 | 0.13 | 0.26 |
| 6 | 34950274 | rs9394265 | -0.374 | -0.68 | 0.708 | 1.106 | 0.001 | -0.039 | -0.039 | 0 | 0.17 | 0.07 | 0.12 | 0.31 | 0.13 | 0.14 |
| 6 | 34960744 | rs6457799 | 1.693 | -0.706 | 0.567 | 1.054 | -0.106 | 0.022 | 0.022 | 0.039 | 0.78 | 0.45 | 0.83 | 0.73 | 0.92 | 0.78 |
| 6 | 34962060 | rs3823438 | 2.783 | -0.161 | 0 | 1.112 | -0.082 | 0.107 | 0.107 | 0.234 | 0.48 | 0.03 | 0.5 | 0.24 | 0.57 | 0.45 |
| 6 | 34966047 | rs9784889 | 1.187 | -0.527 | 0.375 | 1.235 | -0.061 | -0.006 | -0.006 | 0 | 0.72 | 0.56 | 0.82 | 0.73 | 0.89 | 0.76 |
| 6 | 34982599 | rs16896390 | -0.737 | -0.196 | -0.168 | 1.092 | 0.006 | -0.011 | -0.011 | 0 | 0.13 | 0.08 | 0.16 | 0.54 | 0.09 | 0.23 |
| 6 | 34990968 | rs1880090 | 1.169 | 0.184 | -0.122 | 0.681 | 0.024 | 0.04 | 0.04 | 0.182 | 0.69 | 0.23 | 0.76 | 0.83 | 0.75 | 0.84 |
| 6 | 35000453 | rs1996913 | -0.973 | 0.163 | -0.122 | 1.186 | 0.008 | -0.009 | -0.009 | 0 | 0.14 | 0.11 | 0.17 | 0.59 | 0.1 | 0.23 |
| 6 | 35000663 | rs13219307 | -0.481 | 0.062 | 0.652 | 0.99 | 0.01 | 0.043 | 0.043 | 0 | 0.12 | 0.28 | 0.11 | 0.14 | 0.11 | 0.05 |
| 6 | 35002779 | rs9462042 | 0.189 | -0.107 | 0.736 | 0.761 | -0.007 | -0.026 | -0.026 | 0.066 | 0.13 | 0.13 | 0.05 | 0.01 | 0.08 | 0.04 |

| | | | | | | | | | | | | | | | | |
|---|---|---|---|---|---|---|---|---|---|---|---|---|---|---|---|---|
| 6 | 35003603 | rs2820232 | 2.562 | 0.468 | 0.415 | 0.727 | -0.101 | 0.082 | 0.082 | 0.845 | 0.46 | 0.02 | 0.42 | 0.26 | 0.46 | 0.47 |
| 6 | 35004819 | rs12525532 | 2.571 | 0.4 | 0.34 | 0.295 | 0.009 | 0.052 | 0.052 | 0.057 | 0.4 | 0.16 | 0.38 | 0.27 | 0.39 | 0.43 |
| 6 | 35008247 | rs12204265 | -0.071 | 0.621 | 0.804 | 0.657 | 0.103 | 0.003 | 0.003 | 0 | 0.06 | 0.15 | 0.16 | 0.02 | 0.2 | 0.14 |
| 6 | 35009153 | rs2049316 | -0.47 | 0.462 | 0.746 | 0.661 | 0.026 | -0.065 | -0.065 | 0.883 | 0.22 | 0.11 | 0.2 | 0.57 | 0.14 | 0.25 |
| 6 | 35012460 | rs1568705 | 1.726 | 0.399 | 1.032 | 0.579 | -0.001 | -0.002 | -0.002 | 0.005 | 0.59 | 0.3 | 0.55 | 0.4 | 0.62 | 0.54 |
| 6 | 35017003 | rs2689091 | 2.489 | 0.447 | 0.572 | 0.584 | -0.066 | 0.115 | 0.115 | 0.916 | 0.48 | 0.03 | 0.45 | 0.26 | 0.47 | 0.47 |
| 6 | 35018461 | rs820098 | 0.885 | 0.475 | 0.862 | 0.594 | 0.022 | 0.037 | 0.037 | 0 | 0.72 | 0.58 | 0.62 | 0.41 | 0.66 | 0.56 |
| 6 | 35033854 | rs820077 | 1.83 | 0.364 | 1.016 | 0.573 | -0.022 | -0.033 | -0.033 | 0.359 | 0.77 | 0.46 | 0.83 | 0.86 | 0.82 | 0.91 |
| 6 | 35055629 | rs820103 | -0.946 | 0.439 | 1.285 | 0.633 | 0.005 | -0.004 | -0.004 | 0 | 0.15 | 0.31 | 0.14 | 0.12 | 0.15 | 0.06 |
| 6 | 35056443 | rs2005 | -0.336 | 0.375 | 1.443 | 1.145 | -0.002 | 0.013 | 0.013 | 0 | 0.86 | 0.88 | 0.92 | 0.68 | 0.9 | 0.86 |
| 6 | 35058531 | rs11552398 | -0.567 | 0.391 | 1.221 | 1.111 | 0.006 | 0.001 | 0.001 | 0 | 0.13 | 0.2 | 0.13 | 0.12 | 0.15 | 0.06 |
| 6 | 35058667 | rs2296194 | -0.174 | 0.117 | 1.12 | 1.109 | 0.014 | -0.013 | -0.013 | 0 | 0.07 | 0.04 | 0.09 | 0.25 | 0.02 | 0.11 |
| 6 | 35076076 | rs2820229 | 0.622 | 0.255 | 1.206 | 1.093 | 0.061 | 0.068 | 0.068 | 0 | 0.75 | 0.53 | 0.69 | 0.65 | 0.67 | 0.68 |
| 6 | 35078715 | rs9380489 | -0.282 | -0.588 | 1.21 | 1.065 | 0.014 | -0.013 | -0.013 | 0 | 0.07 | 0.04 | 0.09 | 0.25 | 0.02 | 0.11 |
| 6 | 35090097 | rs2234044 | -0.994 | -0.591 | 1.218 | 1.058 | 0.006 | -0.003 | -0.003 | 0 | 0.15 | 0.28 | 0.14 | 0.12 | 0.15 | 0.06 |
| 6 | 35100416 | rs9380492 | -0.492 | -0.557 | 1.029 | 1.08 | -0.005 | 0.005 | 0.005 | 0 | 0.08 | 0.05 | 0.09 | 0.25 | 0.02 | 0.11 |
| 6 | 35110705 | rs3756856 | -0.585 | -0.491 | 1.012 | 0.949 | 0.005 | 0 | 0 | 0 | 0.13 | 0.2 | 0.13 | 0.12 | 0.15 | 0.06 |
| 6 | 35113642 | rs9348966 | -0.221 | -0.521 | 0.778 | 1.118 | 0.012 | -0.013 | -0.013 | 2.186 | 0.07 | 0.04 | 0.1 | 0.21 | 0.02 | 0.11 |
| 6 | 35114911 | rs9394274 | 1.321 | -0.617 | 0.779 | 1.139 | 0.058 | -0.025 | -0.025 | 0 | 0.6 | 0.19 | 0.72 | 0.56 | 0.72 | 0.67 |
| 6 | 35115994 | rs6457802 | 0.092 | -0.647 | 0.921 | 1.128 | -0.018 | 0.044 | 0.044 | 0 | 0.78 | 0.6 | 0.86 | 0.67 | 0.83 | 0.73 |
| 6 | 35117399 | rs1886243 | 2.424 | -0.655 | 0.482 | 1.153 | 0.069 | 0.017 | 0.017 | 1.547 | 0.49 | 0.03 | 0.61 | 0.32 | 0.68 | 0.56 |
| 6 | 35118236 | rs9368849 | 1.769 | -0.628 | 0.503 | 1.125 | 0.06 | -0.051 | -0.051 | 0 | 0.55 | 0.09 | 0.7 | 0.55 | 0.69 | 0.65 |
| 6 | 35124480 | rs4711404 | 1.213 | -0.609 | 0.655 | 1.144 | 0.018 | -0.014 | -0.014 | 0.116 | 0.64 | 0.28 | 0.75 | 0.56 | 0.73 | 0.68 |
| 6 | 35129816 | rs9469953 | -1.614 | -0.517 | 0.854 | 1.123 | -0.068 | 0.048 | 0.048 | 0 | 0.24 | 0.49 | 0.22 | 0.33 | 0.12 | 0.17 |
| 6 | 35138917 | rs12191808 | 2.252 | -0.464 | 0.232 | 1.13 | 0.077 | 0.021 | 0.021 | 1.782 | 0.49 | 0.04 | 0.62 | 0.32 | 0.68 | 0.56 |
| 6 | 35140861 | rs942375 | 0.195 | -0.198 | 0.668 | 1.239 | -0.009 | 0.018 | 0.018 | 0 | 0.79 | 0.72 | 0.87 | 0.68 | 0.84 | 0.74 |
| 6 | 35155610 | rs3800400 | -0.225 | 0.002 | 0.542 | 1.52 | 0.014 | -0.008 | -0.008 | 0 | 0.07 | 0.04 | 0.1 | 0.24 | 0.02 | 0.1 |
| 6 | 35164004 | rs9357196 | -1.674 | -0.065 | 1.188 | 1.36 | -0.03 | 0.053 | 0.053 | 0 | 0.21 | 0.37 | 0.2 | 0.36 | 0.15 | 0.15 |
| 6 | 35172194 | rs1108709 | -2.711 | -0.239 | 1.358 | 1.273 | 0.014 | -0.006 | -0.006 | 0 | 0.31 | 0.73 | 0.27 | 0.36 | 0.19 | 0.17 |
| 6 | 35174642 | rs1929846 | 0.411 | -0.25 | 1.555 | 1.255 | 0 | -0.032 | -0.032 | 1.989 | 0.89 | 0.83 | 0.9 | 0.99 | 0.88 | 0.96 |
| 6 | 35174858 | rs6928769 | 1.559 | -0.251 | 1.554 | 1.257 | -0.03 | 0.05 | 0.05 | 0 | 0.79 | 0.63 | 0.8 | 0.67 | 0.85 | 0.86 |
| 6 | 35181236 | rs9368854 | -0.085 | -0.252 | 1.515 | 1.259 | 0.011 | 0.015 | 0.015 | 0 | 0.07 | 0.04 | 0.1 | 0.32 | 0.02 | 0.1 |
| 6 | 35193773 | rs763155 | 0.346 | -0.244 | 1.607 | 1.252 | 0 | -0.011 | -0.011 | 0 | 0.89 | 0.89 | 0.91 | 0.99 | 0.88 | 0.96 |
| 6 | 35203203 | rs13214290 | 2.691 | -0.259 | 1.737 | 1.25 | 0.002 | -0.002 | -0.002 | 0 | 0.7 | 0.29 | 0.73 | 0.66 | 0.82 | 0.83 |
| 6 | 35205881 | rs734538 | 0.362 | -0.258 | 1.751 | 1.249 | 0 | -0.015 | -0.015 | 1.581 | 0.89 | 0.89 | 0.91 | 0.99 | 0.88 | 0.96 |
| 6 | 35206553 | rs732594 | 1.428 | -0.015 | 1.287 | 0.931 | 0 | 0.006 | 0.006 | 0 | 0.46 | 0.45 | 0.54 | 0.37 | 0.47 | 0.62 |
| 6 | 35211211 | rs9469972 | 2.709 | 0.361 | 2.003 | 1.316 | 0.001 | -0.002 | -0.002 | 0.375 | 0.7 | 0.3 | 0.73 | 0.64 | 0.81 | 0.83 |
| 6 | 35214884 | rs16868695 | -1.207 | 0.365 | 2.04 | 1.311 | -0.01 | -0.04 | -0.04 | 0.053 | 0.08 | 0.29 | 0.06 | 0 | 0.03 | 0.02 |
| 6 | 35215904 | rs4711411 | 2.772 | 0.375 | 1.955 | 1.319 | 0 | -0.001 | -0.001 | 2.366 | 0.7 | 0.3 | 0.73 | 0.66 | 0.82 | 0.83 |
| 6 | 35216740 | rs9394287 | 0.492 | 0.462 | 1.897 | 1.34 | -0.044 | -0.069 | -0.069 | 0 | 0.14 | 0.01 | 0.09 | 0.24 | 0.14 | 0.11 |
| 6 | 35243257 | rs10947540 | 2.781 | 0.44 | 2.197 | 1.268 | 0.006 | -0.002 | -0.002 | 0 | 0.7 | 0.29 | 0.73 | 0.66 | 0.82 | 0.83 |
| 6 | 35254207 | rs743965 | -0.989 | -0.872 | 2.407 | 1.129 | 0.029 | 0.012 | 0.012 | 1.169 | 0.05 | 0.29 | 0.05 | 0 | 0.02 | 0 |
| 6 | 35258539 | rs763157 | 0.742 | -0.858 | 2.224 | 1.14 | -0.001 | -0.011 | -0.011 | 2.38 | 0.88 | 0.88 | 0.91 | 0.99 | 0.88 | 0.96 |
| 6 | 35268533 | rs1888824 | 2.603 | -0.885 | 2.019 | 1.186 | 0.006 | -0.002 | -0.002 | 0.142 | 0.7 | 0.29 | 0.73 | 0.66 | 0.82 | 0.83 |
| 6 | 35272274 | rs10807150 | 2.76 | -1.084 | 2.056 | 1.21 | 0.022 | -0.007 | -0.007 | 0.208 | 0.68 | 0.27 | 0.73 | 0.64 | 0.81 | 0.8 |
| 6 | 35278796 | rs10755687 | 2.651 | -1.088 | 2.054 | 1.211 | 0.007 | -0.004 | -0.004 | 0.004 | 0.7 | 0.28 | 0.74 | 0.64 | 0.82 | 0.8 |
| 6 | 35285720 | rs2395617 | 0.773 | -1.05 | 2.063 | 1.138 | -0.002 | -0.035 | -0.035 | 0 | 0.89 | 0.82 | 0.92 | 0.99 | 0.89 | 0.96 |
| 6 | 35289446 | rs707969 | 0.818 | -1.009 | 2.08 | 1.179 | 0.01 | -0.003 | -0.003 | 0 | 0.94 | 0.9 | 0.93 | 0.98 | 0.9 | 0.98 |
| 6 | 35298662 | rs6920432 | -2.65 | -1.182 | 2.221 | 1.303 | -0.006 | -0.003 | -0.003 | 0 | 0.26 | 0.73 | 0.21 | 0.36 | 0.09 | 0.18 |
| 6 | 35312254 | rs2267664 | -0.255 | -1.134 | 2.32 | 1.32 | 0.009 | 0.016 | 0.016 | 0 | 0.07 | 0.04 | 0.1 | 0.33 | 0.02 | 0.12 |
| 6 | 35322975 | rs7758272 | -1.159 | -1.008 | 2.487 | 1.173 | -0.001 | -0.033 | -0.033 | 0 | 0.08 | 0.26 | 0.03 | 0.01 | 0.04 | 0.03 |
| 6 | 35324741 | rs7746988 | -1.568 | -1.022 | 2.608 | 1.202 | -0.005 | 0.021 | 0.021 | 0 | 0.08 | 0.38 | 0.06 | 0.02 | 0.02 | 0.02 |
| 6 | 35327361 | rs9658077 | -0.178 | -1.059 | 2.662 | 1.176 | -0.018 | -0.026 | -0.026 | 0 | 0.07 | 0.15 | 0.04 | 0.01 | 0.03 | 0.03 |
| 6 | 35330030 | rs6901410 | -2.646 | -1.054 | 2.658 | 1.182 | -0.027 | -0.029 | -0.029 | 0 | 0.19 | 0.73 | 0.12 | 0.03 | 0.08 | 0.06 |
| 6 | 35330898 | rs6457815 | -1.877 | -1.045 | 2.711 | 1.174 | -0.045 | -0.069 | -0.069 | 0 | 0.13 | 0.52 | 0.1 | 0.01 | 0.06 | 0.03 |
| 6 | 35337197 | rs9658080 | -1.164 | -1.447 | 2.681 | 1.184 | -0.001 | -0.033 | -0.033 | 0 | 0.08 | 0.26 | 0.03 | 0.01 | 0.04 | 0.03 |
| 6 | 35337271 | rs9658081 | -0.011 | -1.468 | 2.77 | 1.174 | -0.016 | -0.024 | -0.024 | 4.177 | 0.07 | 0.14 | 0.03 | 0.01 | 0.03 | 0.03 |
| 6 | 35337467 | rs9296148 | -2.546 | -1.481 | 2.799 | 1.197 | -0.029 | -0.029 | -0.029 | 0 | 0.19 | 0.73 | 0.12 | 0.03 | 0.08 | 0.06 |
| 6 | 35339035 | rs7739752 | -2.533 | -1.47 | 2.879 | 1.194 | -0.029 | -0.031 | -0.031 | 0 | 0.19 | 0.73 | 0.11 | 0.03 | 0.07 | 0.06 |
| 6 | 35340300 | rs12173582 | -0.235 | -1.551 | 2.86 | 1.333 | 0.009 | 0.016 | 0.016 | 0 | 0.07 | 0.04 | 0.09 | 0.32 | 0.02 | 0.12 |
| 6 | 35341875 | rs6919334 | -2.521 | -1.52 | 2.847 | 1.276 | -0.029 | -0.031 | -0.031 | 0 | 0.19 | 0.73 | 0.11 | 0.03 | 0.07 | 0.06 |
| 6 | 35349739 | rs9470007 | -2.521 | -1.539 | 2.835 | 1.262 | -0.029 | -0.031 | -0.031 | 1.393 | 0.19 | 0.73 | 0.11 | 0.03 | 0.07 | 0.06 |
| 6 | 35350042 | rs7770619 | -0.6 | -1.538 | 2.844 | 1.262 | -0.014 | -0.034 | -0.034 | 0 | 0.08 | 0.2 | 0.05 | 0.01 | 0.05 | 0.03 |
| 6 | 35353523 | rs6922548 | -2.229 | -1.526 | 2.568 | 1.268 | -0.029 | -0.031 | -0.031 | 0 | 0.19 | 0.73 | 0.11 | 0.03 | 0.08 | 0.06 |
| 6 | 35356143 | rs3777744 | -2.787 | -1.558 | 2.409 | 1.315 | 0.038 | -0.021 | -0.021 | 0 | 0.28 | 0.79 | 0.22 | 0.36 | 0.1 | 0.18 |
| 6 | 35356640 | rs9658100 | -2.216 | -1.542 | 2.419 | 1.281 | -0.029 | -0.031 | -0.031 | 0 | 0.19 | 0.73 | 0.11 | 0.03 | 0.07 | 0.06 |
| 6 | 35357693 | rs3798343 | -1.137 | -1.518 | 2.416 | 1.34 | -0.034 | 0.091 | 0.091 | 0 | 0.1 | 0.18 | 0.13 | 0.33 | 0.03 | 0.12 |
| 6 | 35362848 | rs6457816 | -2.238 | -1.39 | 2.313 | 0.795 | -0.045 | -0.048 | -0.048 | 0 | 0.18 | 0.65 | 0.12 | 0.03 | 0.07 | 0.06 |
| 6 | 35365967 | rs9658119 | -2.536 | -1.461 | 2.268 | 0.632 | -0.066 | -0.125 | -0.125 | 0.022 | 0.24 | 0.42 | 0.11 | 0.02 | 0.11 | 0.06 |

| | | | | | | | | | | | | | | | | |
|---|---|---|---|---|---|---|---|---|---|---|---|---|---|---|---|---|
| 6 | 35367909 | rs1040436 | 1.895 | -1.472 | 2.057 | 0.853 | 0.021 | 0.017 | 0.017 | 0.243 | 0.62 | 0.34 | 0.78 | 0.73 | 0.76 | 0.83 |
| 6 | 35369491 | rs2267665 | 0.247 | -1.503 | 2.203 | 0.84 | 0.001 | -0.006 | -0.006 | 0 | 0.81 | 0.9 | 0.88 | 0.76 | 0.82 | 0.88 |
| 6 | 35369806 | rs1883322 | 1.905 | -1.506 | 2.276 | 0.837 | 0.004 | 0.004 | 0.004 | 0.218 | 0.63 | 0.39 | 0.78 | 0.73 | 0.76 | 0.83 |
| 6 | 35371624 | rs7751726 | 0.332 | -1.49 | 2.528 | 0.833 | -0.004 | -0.017 | -0.017 | 0 | 0.06 | 0.13 | 0.04 | 0.01 | 0.04 | 0.03 |
| 6 | 35372053 | rs9658127 | -0.669 | -1.455 | 2.536 | 0.832 | 0.015 | -0.002 | -0.002 | 0 | 0.06 | 0.28 | 0.03 | 0 | 0.01 | 0 |
| 6 | 35372524 | rs2267667 | 2.065 | -1.535 | 2.49 | 0.824 | 0.025 | 0.021 | 0.021 | 0.083 | 0.62 | 0.33 | 0.78 | 0.73 | 0.75 | 0.82 |
| 6 | 35374461 | rs2038068 | 1.728 | -1.524 | 2.027 | 0.832 | -0.005 | -0.003 | -0.003 | 0 | 0.65 | 0.41 | 0.79 | 0.73 | 0.77 | 0.84 |
| 6 | 35375526 | rs6906237 | -1.857 | -1.515 | 2.051 | 0.813 | -0.051 | -0.051 | -0.051 | 0 | 0.15 | 0.5 | 0.09 | 0.02 | 0.06 | 0.03 |
| 6 | 35376006 | rs9462081 | -0.557 | -1.517 | 2.075 | 0.805 | -0.024 | -0.03 | -0.03 | 2.109 | 0.09 | 0.21 | 0.06 | 0.01 | 0.03 | 0.03 |
| 6 | 35376287 | rs6911817 | -0.669 | -1.506 | 2.051 | 0.81 | 0.01 | -0.002 | -0.002 | 0 | 0.06 | 0.28 | 0.03 | 0 | 0.01 | 0 |
| 6 | 35377922 | rs2267668 | 0.182 | -1.519 | 1.952 | 0.824 | 0.004 | -0.001 | -0.001 | 0.167 | 0.8 | 0.9 | 0.88 | 0.77 | 0.82 | 0.88 |
| 6 | 35378124 | rs2267669 | 0.182 | -1.64 | 1.889 | 0.735 | 0.004 | -0.001 | -0.001 | 0 | 0.8 | 0.9 | 0.88 | 0.77 | 0.82 | 0.88 |
| 6 | 35380222 | rs9296153 | -0.487 | -1.705 | 1.851 | 0.771 | -0.022 | -0.028 | -0.028 | 0 | 0.08 | 0.2 | 0.06 | 0.01 | 0.03 | 0.03 |
| 6 | 35386041 | rs9462082 | 0.215 | -1.719 | 1.713 | 0.582 | 0.005 | -0.006 | -0.006 | 0 | 0.81 | 0.93 | 0.89 | 0.77 | 0.84 | 0.89 |
| 6 | 35387378 | rs9296154 | -0.446 | -1.707 | 1.695 | 0.592 | -0.022 | 0 | 0 | 0 | 0.08 | 0.2 | 0.06 | 0.02 | 0.03 | 0.03 |
| 6 | 35388384 | rs9658151 | -0.474 | -1.707 | 1.684 | 0.587 | -0.023 | -0.029 | -0.029 | 0 | 0.09 | 0.2 | 0.06 | 0.01 | 0.03 | 0.03 |
| 6 | 35388479 | rs2076169 | 0.805 | -1.688 | 1.371 | 0.602 | -0.023 | -0.058 | -0.058 | 0 | 0.11 | 0 | 0.06 | 0.22 | 0.1 | 0.1 |
| 6 | 35391787 | rs2076167 | 1.904 | -1.716 | 1.504 | 0.619 | 0.001 | 0.001 | 0.001 | 0.048 | 0.63 | 0.4 | 0.79 | 0.74 | 0.77 | 0.83 |
| 6 | 35393404 | rs2076166 | 0.23 | -1.687 | 1.447 | 0.612 | 0.007 | -0.006 | -0.006 | 0 | 0.81 | 0.91 | 0.89 | 0.76 | 0.84 | 0.88 |
| 6 | 35395010 | rs3734254 | 1.754 | -1.89 | 1.49 | 0.535 | -0.01 | -0.006 | -0.006 | 1.472 | 0.64 | 0.42 | 0.8 | 0.74 | 0.8 | 0.84 |
| 6 | 35395578 | rs1053046 | -1.757 | -1.905 | 1.504 | 0.534 | -0.065 | -0.065 | -0.065 | 0 | 0.16 | 0.5 | 0.09 | 0.02 | 0.05 | 0.05 |
| 6 | 35395618 | rs1053049 | 1.993 | -1.842 | 1.43 | 0.534 | 0.017 | 0.015 | 0.015 | 0.214 | 0.63 | 0.35 | 0.78 | 0.74 | 0.76 | 0.83 |
| 6 | 35396004 | rs760783 | 0.271 | -1.816 | 1.283 | 0.585 | 0.008 | -0.004 | -0.004 | 0 | 0.81 | 0.9 | 0.88 | 0.77 | 0.84 | 0.88 |
| 6 | 35397127 | rs7749160 | 0.169 | -1.822 | 1.255 | 0.59 | 0.005 | -0.007 | -0.007 | 0.077 | 0.81 | 0.91 | 0.89 | 0.77 | 0.84 | 0.88 |
| 6 | 35398357 | rs9470023 | -0.489 | -1.795 | 1.353 | 0.58 | -0.024 | -0.029 | -0.029 | 0 | 0.09 | 0.21 | 0.05 | 0.01 | 0.03 | 0.03 |
| 6 | 35402785 | rs4713858 | 0.033 | -1.79 | 1.036 | 0.613 | 0.003 | -0.005 | -0.005 | 0 | 0.82 | 0.9 | 0.89 | 0.77 | 0.84 | 0.87 |
| 6 | 35404437 | rs9462085 | 1.957 | -1.894 | 0.955 | 0.823 | 0.057 | -0.013 | -0.013 | 0 | 0.53 | 0.33 | 0.71 | 0.44 | 0.7 | 0.7 |
| 6 | 35406153 | rs4713859 | -1.576 | -1.995 | 0.871 | 1.212 | -0.024 | 0.013 | 0.013 | 0 | 0.2 | 0.25 | 0.16 | 0.33 | 0.06 | 0.17 |
| 6 | 35408959 | rs2894401 | -0.53 | -2.016 | 0.162 | 0.814 | -0.008 | 0.02 | 0.02 | 1.138 | 0.71 | 0.87 | 0.8 | 0.72 | 0.71 | 0.8 |
| 6 | 35417791 | rs10214770 | 1.096 | -2.116 | -0.019 | 0.98 | -0.011 | -0.055 | -0.055 | 0.455 | 0.44 | 0.12 | 0.61 | 0.51 | 0.59 | 0.51 |
| 6 | 35422754 | rs2395626 | 1.55 | -2.122 | -0.279 | 0.998 | -0.105 | -0.085 | -0.085 | 0.565 | 0.38 | 0.08 | 0.57 | 0.35 | 0.56 | 0.45 |
| 6 | 35423489 | rs7757405 | -0.763 | -2.231 | -0.302 | 0.987 | -0.021 | 0.02 | 0.02 | 0.247 | 0.7 | 0.89 | 0.82 | 0.71 | 0.68 | 0.76 |
| 6 | 35424188 | rs3800377 | -0.369 | -2.226 | -0.322 | 0.974 | -0.011 | -0.021 | -0.021 | 0 | 0.65 | 0.88 | 0.78 | 0.54 | 0.65 | 0.7 |
| 6 | 35425990 | rs3800376 | 0.138 | -2.186 | 0.023 | 0.826 | -0.004 | 0.084 | 0.084 | 0.584 | 0.06 | 0.28 | 0.04 | 0.12 | 0.01 | 0.18 |
| 6 | 35426175 | rs3823434 | 0.556 | -2.175 | -0.008 | 0.833 | 0.02 | -0.035 | -0.035 | 0.052 | 0.06 | 0.01 | 0.03 | 0.17 | 0.03 | 0.07 |
| 6 | 35430686 | rs9462088 | -0.599 | -2.003 | 0.202 | 0.767 | -0.001 | 0.076 | 0.076 | 0.898 | 0.09 | 0.44 | 0.05 | 0.12 | 0.04 | 0.18 |
| 6 | 35439932 | rs12525930 | 1.488 | -1.708 | -1.17 | 0.69 | -0.126 | -0.086 | -0.086 | 0.518 | 0.37 | 0.02 | 0.56 | 0.28 | 0.57 | 0.46 |
| 6 | 35442397 | rs2076173 | -1.893 | -1.951 | 0.018 | 0.785 | 0.122 | -0.023 | -0.023 | 5.431 | 0.35 | 0.87 | 0.26 | 0.37 | 0.12 | 0.32 |
| 6 | 35452346 | rs11756089 | -1.442 | -0.643 | 0.636 | 0.409 | -0.058 | -0.036 | -0.036 | 0 | 0.14 | 0.3 | 0.14 | 0.06 | 0.05 | 0.03 |
| 6 | 35457396 | rs7740758 | -1.47 | 0.339 | 0.876 | 0.277 | 0.062 | -0.067 | -0.067 | 0.035 | 0.17 | 0.45 | 0.13 | 0.07 | 0.18 | 0.13 |
| 6 | 35463202 | rs7744287 | -1.446 | 0.059 | 0.828 | 0.885 | 0.002 | 0.014 | 0.014 | 0.953 | 0.28 | 0.64 | 0.24 | 0.42 | 0.32 | 0.37 |
| 6 | 35465786 | rs1051952 | -1.975 | 0.008 | 0.387 | 1.084 | -0.048 | -0.038 | -0.038 | 0 | 0.41 | 0.93 | 0.47 | 0.6 | 0.43 | 0.5 |
| 6 | 35468659 | rs1883636 | -1.347 | 0.061 | 0.573 | 1.157 | -0.008 | -0.003 | -0.003 | 0.452 | 0.13 | 0.23 | 0.22 | 0.14 | 0.11 | 0.11 |
| 6 | 35473030 | rs7769417 | 0.638 | 0.31 | -0.272 | 0.62 | -0.003 | 0.005 | 0.005 | 0 | 0.53 | 0.31 | 0.65 | 0.44 | 0.47 | 0.36 |
| 6 | 35473252 | rs2395628 | -0.273 | 0.277 | 0.737 | 0.763 | 0.022 | 0.05 | 0.05 | 0.396 | 0.22 | 0.33 | 0.2 | 0.42 | 0.3 | 0.37 |
| 6 | 35476442 | rs2064319 | -1.015 | 0.208 | 0.848 | 0.687 | 0.001 | 0.006 | 0.006 | 0.842 | 0.3 | 0.34 | 0.35 | 0.53 | 0.38 | 0.47 |
| 6 | 35477152 | rs7752728 | -0.312 | 0.049 | 1.071 | 0.419 | -0.014 | -0.012 | -0.012 | 3.331 | 0.74 | 0.67 | 0.79 | 0.81 | 0.77 | 0.74 |
| 6 | 35480262 | rs1540910 | 0.079 | 0.061 | 1.19 | 0.426 | 0.002 | 0.026 | 0.026 | 0 | 0.71 | 0.85 | 0.75 | 0.76 | 0.72 | 0.65 |
| 6 | 35484790 | rs4267954 | 0.679 | 0.025 | 1.458 | 0.436 | -0.017 | -0.017 | -0.017 | 2.063 | 0.65 | 0.54 | 0.65 | 0.71 | 0.68 | 0.65 |
| 6 | 35488853 | rs1540909 | -1.036 | 0.06 | 1.408 | 0.412 | 0.006 | 0.018 | 0.018 | 0 | 0.79 | 0.9 | 0.89 | 0.87 | 0.81 | 0.76 |
| 6 | 35489854 | rs749522 | -0.327 | 0.051 | 1.409 | 0.413 | 0.017 | 0.03 | 0.03 | 0 | 0.75 | 0.89 | 0.88 | 0.83 | 0.79 | 0.71 |
| 6 | 35490238 | rs4713879 | -0.327 | 0.051 | 1.395 | 0.413 | 0.018 | 0.031 | 0.031 | 0 | 0.75 | 0.89 | 0.88 | 0.83 | 0.8 | 0.71 |
| 6 | 35495811 | rs4711420 | -0.321 | 0.061 | 1.468 | 0.421 | -0.024 | -0.035 | -0.035 | 0 | 0.74 | 0.6 | 0.85 | 0.85 | 0.77 | 0.75 |
| 6 | 35498098 | rs6935976 | -1.031 | 0.062 | 1.403 | 0.422 | 0.005 | 0.014 | 0.014 | 0 | 0.79 | 0.89 | 0.89 | 0.86 | 0.81 | 0.75 |
| 6 | 35503576 | rs7743152 | 0.834 | 0.259 | 0.994 | 0.763 | 0.009 | -0.009 | -0.009 | 0 | 0.58 | 0.55 | 0.6 | 0.64 | 0.47 | 0.45 |
| 6 | 35512955 | rs3807049 | -0.37 | -0.009 | 0.701 | 0.615 | -0.012 | 0.03 | 0.03 | 0 | 0.74 | 0.9 | 0.75 | 0.79 | 0.71 | 0.65 |
| 6 | 35518578 | rs9394305 | -0.715 | -0.014 | 0.711 | 0.614 | 0.017 | 0.04 | 0.04 | 0 | 0.78 | 0.9 | 0.88 | 0.88 | 0.81 | 0.74 |
| 6 | 35521961 | rs9470055 | -0.667 | -0.071 | 0.756 | 0.607 | 0.009 | 0.021 | 0.021 | 0 | 0.77 | 0.84 | 0.88 | 0.87 | 0.8 | 0.74 |
| 6 | 35525751 | rs6931945 | -1.415 | 0.431 | 0.999 | 0.639 | 0.123 | 0.065 | 0.065 | 0.008 | 0.15 | 0.34 | 0.34 | 0.22 | 0.34 | 0.28 |
| 6 | 35528570 | rs4713897 | -0.754 | 0.469 | 0.843 | 0.602 | 0.008 | 0.022 | 0.022 | 0 | 0.77 | 0.84 | 0.87 | 0.88 | 0.8 | 0.72 |
| 6 | 35529025 | rs6911621 | -0.449 | 0.513 | 0.929 | 0.658 | -0.01 | -0.016 | -0.016 | 0 | 0.1 | 0.07 | 0.24 | 0.16 | 0.32 | 0.27 |
| 6 | 35531205 | rs6924087 | 0.037 | 0.503 | 0.979 | 0.645 | 0 | -0.008 | -0.008 | 0 | 0.08 | 0.06 | 0.12 | 0.11 | 0.24 | 0.21 |
| 6 | 35542476 | rs3800373 | 0.173 | 0.492 | 1.074 | 0.569 | -0.011 | -0.03 | -0.03 | 0.044 | 0.71 | 0.56 | 0.72 | 0.76 | 0.71 | 0.67 |
| 6 | 35548236 | rs7757037 | 1.157 | 0.368 | 1.178 | 0.643 | -0.006 | 0.004 | 0.004 | 0 | 0.57 | 0.38 | 0.5 | 0.62 | 0.47 | 0.42 |
| 6 | 35567082 | rs9296158 | 0.186 | 0.332 | 1.079 | 0.542 | -0.017 | -0.024 | -0.024 | 0.398 | 0.68 | 0.53 | 0.64 | 0.71 | 0.7 | 0.66 |
| 6 | 35575487 | rs737054 | 2.227 | 0.399 | 0.56 | 0.429 | 0.008 | -0.008 | -0.008 | 7.803 | 0.3 | 0.06 | 0.34 | 0.24 | 0.26 | 0.15 |
| 6 | 35579002 | rs3777747 | 0.97 | 0.635 | 0.655 | 0.793 | -0.002 | 0.003 | 0.003 | 0 | 0.55 | 0.4 | 0.5 | 0.61 | 0.46 | 0.42 |
| 6 | 35579375 | rs6926133 | -0.73 | 0.657 | 0.426 | 0.883 | 0.015 | 0.023 | 0.023 | 0 | 0.76 | 0.85 | 0.85 | 0.81 | 0.8 | 0.73 |
| 6 | 35589070 | rs9380524 | 0.89 | 0.554 | 0.491 | 0.931 | -0.014 | -0.06 | -0.06 | 0.041 | 0.11 | 0.01 | 0.08 | 0.34 | 0.11 | 0.17 |

| | | | | | | | | | | | | | | | | |
|---|---|---|---|---|---|---|---|---|---|---|---|---|---|---|---|---|
| 6 | 35625147 | rs4713904 | -0.432 | 0.662 | 0.449 | 0.597 | -0.009 | 0.007 | 0.007 | 0 | 0.74 | 0.82 | 0.73 | 0.74 | 0.69 | 0.67 |
| 6 | 35646435 | rs9470080 | 0.045 | 0.692 | 0.383 | 0.598 | -0.003 | -0.009 | -0.009 | 0 | 0.69 | 0.54 | 0.64 | 0.69 | 0.67 | 0.67 |
| 6 | 35658327 | rs9380526 | 0.078 | 0.712 | 0.385 | 0.603 | -0.004 | -0.011 | -0.011 | 0.438 | 0.68 | 0.56 | 0.64 | 0.69 | 0.67 | 0.66 |
| 6 | 35671651 | rs10456432 | -0.413 | 1.14 | -0.177 | 0.659 | 0.053 | 0.007 | 0.007 | 0 | 0.08 | 0.17 | 0.12 | 0.05 | 0.19 | 0.21 |
| 6 | 35675696 | rs9380529 | 0.484 | 0.978 | -0.842 | 0.729 | -0.119 | -0.031 | -0.031 | 0 | 0.62 | 0.88 | 0.63 | 0.69 | 0.48 | 0.43 |
| 6 | 35683059 | rs9394314 | -0.071 | 0.65 | -0.691 | 0.467 | -0.001 | 0.022 | 0.022 | 0 | 0.72 | 0.82 | 0.73 | 0.77 | 0.68 | 0.65 |
| 6 | 35685490 | rs2766533 | -0.663 | 0.596 | -0.691 | 0.506 | -0.026 | 0.018 | 0.018 | 0 | 0.37 | 0.26 | 0.36 | 0.28 | 0.48 | 0.49 |
| 6 | 35685955 | rs12200498 | -0.538 | 1.004 | -1.278 | 0.499 | -0.05 | -0.013 | -0.013 | 0.003 | 0.09 | 0.04 | 0.1 | 0.06 | 0.18 | 0.15 |
| 6 | 35688619 | rs2817032 | -0.062 | 0.419 | -1.354 | 0.402 | 0.017 | 0.055 | 0.055 | 0 | 0.31 | 0.08 | 0.27 | 0.21 | 0.3 | 0.39 |
| 6 | 35701280 | rs2817037 | -0.155 | 0.687 | -1.231 | 1.062 | 0 | 0.006 | 0.006 | 0 | 0.79 | 0.81 | 0.85 | 0.57 | 0.8 | 0.77 |
| 6 | 35702887 | rs2817038 | 0.428 | 0.803 | -1.341 | 0.997 | -0.033 | -0.034 | -0.034 | 0 | 0.71 | 0.79 | 0.77 | 0.52 | 0.63 | 0.63 |
| 6 | 35705606 | rs2817040 | 0.012 | 0.293 | -1.288 | 0.621 | 0.031 | 0.028 | 0.028 | 0 | 0.29 | 0.09 | 0.26 | 0.22 | 0.29 | 0.38 |
| 6 | 35708634 | rs2766543 | -0.745 | 0.15 | -1.48 | 0.651 | -0.004 | 0.013 | 0.013 | 0.014 | 0.44 | 0.85 | 0.45 | 0.4 | 0.49 | 0.56 |
| 6 | 35713031 | rs2766545 | -0.698 | 0.124 | -1.529 | 0.64 | -0.007 | 0.019 | 0.019 | 0.036 | 0.45 | 0.87 | 0.45 | 0.38 | 0.49 | 0.56 |
| 6 | 35713426 | rs1075092 | -1.253 | 0.085 | -1.662 | 0.58 | -0.065 | -0.013 | -0.013 | 0 | 0.54 | 0.89 | 0.52 | 0.46 | 0.56 | 0.61 |
| 6 | 35718651 | rs2766551 | 0.151 | -0.089 | -0.683 | 0.615 | -0.008 | -0.001 | -0.001 | 0 | 0.3 | 0.32 | 0.26 | 0.27 | 0.26 | 0.4 |
| 6 | 35723417 | rs7752049 | -1.065 | 0.941 | 0.442 | 0.74 | 0.211 | 0.047 | 0.047 | 0 | 0.1 | 0.45 | 0.12 | 0.07 | 0.19 | 0.11 |
| 6 | 35727655 | rs2766559 | 0.571 | 0.296 | -0.079 | 0.634 | -0.023 | -0.005 | -0.005 | 0.593 | 0.66 | 0.74 | 0.48 | 0.44 | 0.52 | 0.66 |
| 6 | 35730950 | rs2766565 | -0.25 | 0.512 | -0.376 | 0.793 | -0.114 | -0.078 | -0.078 | 0.404 | 0.76 | 0.93 | 0.57 | 0.46 | 0.62 | 0.72 |
| 6 | 35734051 | rs2817056 | 0.103 | 0.368 | -0.035 | 0.751 | -0.025 | -0.011 | -0.011 | 0 | 0.72 | 0.75 | 0.57 | 0.42 | 0.62 | 0.71 |
| 6 | 35741722 | rs2766576 | 0.717 | 0.046 | 0.129 | 0.95 | -0.016 | -0.023 | -0.023 | 13.161 | 0.63 | 0.69 | 0.52 | 0.55 | 0.6 | 0.63 |
| 6 | 35742308 | rs10807154 | 0.592 | -0.539 | -0.193 | 0.59 | -0.002 | -0.002 | -0.002 | 0 | 0.43 | 0.43 | 0.33 | 0.36 | 0.37 | 0.37 |
| 6 | 35745655 | rs2478467 | -0.864 | -0.34 | -0.505 | 0.786 | -0.077 | -0.077 | -0.077 | 0 | 0.75 | 0.9 | 0.62 | 0.62 | 0.66 | 0.72 |
| 6 | 35748086 | rs7753322 | 1.021 | -0.073 | -0.368 | 0.562 | -0.004 | -0.003 | -0.003 | 0 | 0.34 | 0.33 | 0.25 | 0.32 | 0.3 | 0.28 |
| 6 | 35748415 | rs10947568 | 0.984 | -0.003 | -0.346 | 0.47 | -0.003 | -0.004 | -0.004 | 0 | 0.36 | 0.35 | 0.26 | 0.29 | 0.3 | 0.28 |
| 6 | 35750299 | rs2766581 | -0.091 | 0.07 | -0.399 | 0.294 | -0.004 | 0.013 | 0.013 | 0.179 | 0.26 | 0.19 | 0.25 | 0.16 | 0.32 | 0.35 |
| 6 | 35779879 | rs879667 | -0.393 | -0.437 | -0.54 | -0.272 | 0.039 | 0.125 | 0.125 | 0 | 0.44 | 0.75 | 0.38 | 0.24 | 0.38 | 0.48 |
| 6 | 35783497 | rs6457858 | 0.288 | -0.801 | 0.2 | -0.523 | -0.001 | 0.015 | 0.015 | 0 | 0.31 | 0.25 | 0.28 | 0.16 | 0.32 | 0.4 |
| 6 | 35787417 | rs879493 | -0.373 | -0.888 | 1.064 | -0.569 | -0.003 | -0.019 | -0.019 | 0 | 0.4 | 0.52 | 0.37 | 0.24 | 0.38 | 0.47 |
| 6 | 35788600 | rs7740886 | 0.219 | -0.865 | 1.264 | -0.586 | 0 | -0.014 | -0.014 | 0 | 0.31 | 0.31 | 0.28 | 0.16 | 0.32 | 0.39 |
| 6 | 35800540 | rs9470120 | 0.204 | -0.868 | 1.575 | -0.589 | 0 | -0.013 | -0.013 | 0 | 0.31 | 0.31 | 0.28 | 0.16 | 0.32 | 0.39 |
| 6 | 35803192 | rs1049649 | 0.204 | -0.886 | 1.575 | -0.589 | 0 | -0.013 | -0.013 | 0 | 0.31 | 0.31 | 0.28 | 0.16 | 0.32 | 0.4 |
| 6 | 35805133 | rs17704843 | 1.209 | -1.033 | 1.062 | -1.053 | -0.009 | 0.04 | 0.04 | 0 | 0.09 | 0 | 0.04 | 0.02 | 0.08 | 0.03 |
| 6 | 35806784 | rs10484264 | 0.632 | -1.136 | 1.188 | -1.069 | -0.006 | 0.025 | 0.025 | 0 | 0.28 | 0.19 | 0.27 | 0.16 | 0.31 | 0.39 |
| 6 | 35810666 | rs7758042 | 0.632 | -1.136 | 1.188 | -1.069 | -0.006 | 0.025 | 0.025 | 0 | 0.28 | 0.19 | 0.27 | 0.16 | 0.31 | 0.39 |
| 6 | 35836705 | rs9296168 | 0.275 | -1.139 | 1.21 | -1.061 | 0 | -0.039 | -0.039 | 0.056 | 0.32 | 0.42 | 0.29 | 0.19 | 0.32 | 0.42 |
| 6 | 35849985 | rs9470155 | 0.388 | -1.133 | 0.98 | -1.062 | 0 | -0.013 | -0.013 | 0 | 0.31 | 0.31 | 0.28 | 0.16 | 0.32 | 0.4 |
| 6 | 35855286 | rs11968721 | -0.754 | -0.29 | 0.003 | -0.267 | -0.123 | -0.112 | -0.112 | 0.813 | 0.81 | 0.98 | 0.61 | 0.71 | 0.69 | 0.83 |
| 6 | 35900277 | rs11755428 | 0.909 | -0.6 | 0.306 | -0.336 | -0.039 | 0.072 | 0.072 | 0.272 | 0.25 | 0.1 | 0.26 | 0.13 | 0.29 | 0.38 |
| 6 | 35902585 | rs12110812 | -1.193 | -0.48 | 0.619 | -0.394 | 0.026 | 0.039 | 0.039 | 1.614 | 0.07 | 0.45 | 0.05 | 0.02 | 0.02 | 0.03 |
| 6 | 35906389 | rs651158 | -0.211 | -0.513 | 0.299 | -0.433 | 0.005 | 0.009 | 0.009 | 0 | 0.42 | 0.6 | 0.39 | 0.23 | 0.38 | 0.46 |
| 6 | 35913096 | rs9368895 | -0.014 | -0.565 | 0.084 | 0.289 | 0.018 | -0.021 | -0.021 | 0 | 0.09 | 0.02 | 0.04 | 0.27 | 0.05 | 0.12 |
| 6 | 35923246 | rs2295852 | -0.649 | 0.082 | 0.091 | -0.836 | 0.001 | 0.021 | 0.021 | 0 | 0.4 | 0.7 | 0.29 | 0.28 | 0.35 | 0.29 |
| 6 | 35925798 | rs10456081 | 1.113 | 0.565 | -0.74 | -0.734 | -0.027 | -0.007 | -0.007 | 0 | 0.18 | 0.11 | 0.13 | 0.2 | 0.21 | 0.2 |
| 6 | 35939688 | rs10947580 | 1.199 | 0.583 | -0.731 | -0.725 | -0.027 | -0.008 | -0.008 | 0 | 0.18 | 0.11 | 0.13 | 0.2 | 0.21 | 0.2 |
| 6 | 35941099 | rs10484265 | 0.342 | 0.316 | -0.328 | -0.516 | -0.02 | 0.026 | 0.026 | 0 | 0.06 | 0 | 0.04 | 0 | 0.08 | 0.01 |
| 6 | 35949497 | rs12664249 | -0.661 | 0.244 | -0.019 | -0.396 | -0.015 | -0.017 | -0.017 | 1.47 | 0.1 | 0.19 | 0.07 | 0.06 | 0.05 | 0.08 |
| 6 | 35958871 | rs11965835 | -0.297 | 0.485 | -0.318 | -0.543 | -0.005 | 0.041 | 0.041 | 0 | 0.16 | 0.08 | 0.14 | 0.07 | 0.14 | 0.17 |
| 6 | 35960390 | rs17707331 | -0.223 | 0.493 | -0.3 | -0.538 | -0.014 | 0.064 | 0.064 | 0 | 0.16 | 0.03 | 0.14 | 0.05 | 0.14 | 0.12 |
| 6 | 35962978 | rs969659 | 1.446 | 0.484 | -0.245 | -0.904 | 0.002 | 0.082 | 0.082 | 0.023 | 0.48 | 0.33 | 0.36 | 0.27 | 0.47 | 0.56 |
| 6 | 35965810 | rs3800369 | 1.361 | 0.469 | -0.153 | -0.836 | 0.005 | 0.069 | 0.069 | 0 | 0.49 | 0.36 | 0.36 | 0.31 | 0.48 | 0.57 |
| 6 | 35967058 | rs9470191 | 1.515 | 0.445 | -0.077 | -0.849 | -0.02 | 0.139 | 0.139 | 0.185 | 0.47 | 0.18 | 0.35 | 0.27 | 0.47 | 0.56 |
| 6 | 35967772 | rs17713154 | 0.185 | 0.461 | -0.085 | -0.845 | -0.003 | 0.042 | 0.042 | 0 | 0.16 | 0.09 | 0.14 | 0.05 | 0.14 | 0.12 |
| 6 | 35969768 | rs625204 | 0.521 | 0.496 | 0.161 | -0.793 | 0.005 | -0.003 | -0.003 | 0 | 0.65 | 0.48 | 0.51 | 0.38 | 0.62 | 0.74 |
| 6 | 35980121 | rs743923 | -0.899 | 0.467 | 0.616 | -0.983 | -0.006 | -0.082 | -0.082 | 0.596 | 0.26 | 0.47 | 0.23 | 0.12 | 0.24 | 0.18 |
| 6 | 35980712 | rs7743315 | -0.899 | 0.467 | 0.622 | -0.983 | -0.006 | -0.085 | -0.085 | 0.48 | 0.26 | 0.46 | 0.23 | 0.12 | 0.24 | 0.18 |
| 6 | 35985023 | rs1100864 | 0.108 | 0.633 | 0.48 | -0.398 | -0.019 | -0.043 | -0.043 | 0.022 | 0.89 | 0.98 | 0.7 | 0.5 | 0.87 | 0.86 |
| 6 | 35987645 | rs851034 | 0.783 | 0.658 | 0.608 | -0.479 | 0.015 | 0.012 | 0.012 | 0 | 0.73 | 0.57 | 0.58 | 0.34 | 0.71 | 0.68 |
| 6 | 35991474 | rs10456082 | 0.207 | 0.556 | 0.681 | -0.452 | 0.002 | 0.004 | 0.004 | 0 | 0.14 | 0.18 | 0.11 | 0.15 | 0.16 | 0.13 |
| 6 | 35993906 | rs3761980 | 0.165 | 0.625 | 1.019 | -0.513 | 0.021 | 0.079 | 0.079 | 0.204 | 0.08 | 0.25 | 0.07 | 0.14 | 0.08 | 0.06 |
| 6 | 35994088 | rs3761979 | 0.51 | 0.673 | 0.913 | -0.454 | 0.005 | 0.043 | 0.043 | 0.012 | 0.07 | 0.02 | 0.3 | 0.5 | 0.08 | 0.11 |
| 6 | 36006220 | rs851023 | -0.316 | 0.821 | 0.938 | -0.565 | -0.006 | -0.002 | -0.002 | 0 | 0.91 | 0.93 | 0.93 | 1 | 0.87 | 0.96 |
| 6 | 36006998 | rs3804454 | -0.463 | 0.742 | 0.899 | -0.681 | 0.002 | -0.003 | -0.003 | 0 | 0.22 | 0.3 | 0.18 | 0.19 | 0.21 | 0.17 |
| 6 | 36009495 | rs12199654 | 1.471 | 0.71 | 0.838 | -0.713 | -0.026 | 0.041 | 0.041 | 0 | 0.08 | 0 | 0.05 | 0 | 0.08 | 0.07 |
| 6 | 36014161 | rs851019 | 1.912 | 0.686 | 1.125 | -0.476 | -0.044 | 0.129 | 0.129 | 0.453 | 0.56 | 0.16 | 0.42 | 0.25 | 0.56 | 0.62 |
| 6 | 36015094 | rs12665389 | 0.303 | 0.668 | 1.118 | -0.471 | -0.019 | 0.051 | 0.051 | 0 | 0.13 | 0.03 | 0.11 | 0.05 | 0.12 | 0.11 |
| 6 | 36035354 | rs2237093 | 0.409 | 0.693 | 0.961 | -0.435 | 0.003 | 0.045 | 0.045 | 0.012 | 0.07 | 0.02 | 0.3 | 0.5 | 0.08 | 0.11 |
| 6 | 36036909 | rs7775290 | -0.117 | 0.746 | 0.836 | -0.481 | 0.015 | 0.072 | 0.072 | 0.448 | 0.09 | 0.27 | 0.07 | 0.14 | 0.08 | 0.11 |
| 6 | 36054394 | rs1100858 | 0.168 | 0.731 | 0.899 | -0.539 | 0.018 | -0.048 | -0.048 | 0 | 0.88 | 0.76 | 0.92 | 0.97 | 0.85 | 0.93 |

| | | | | | | | | | | | | | | | | |
|---|---|---|---|---|---|---|---|---|---|---|---|---|---|---|---|---|
| 6 | 36065185 | rs851006 | 0.515 | 0.678 | 0.763 | -0.529 | -0.007 | -0.005 | -0.005 | 0 | 0.77 | 0.81 | 0.81 | 0.95 | 0.74 | 0.85 |
| 6 | 36070955 | rs2815806 | -0.167 | 0.691 | 0.37 | -0.518 | -0.005 | -0.004 | -0.004 | 0 | 0.91 | 0.93 | 0.93 | 1 | 0.87 | 0.96 |
| 6 | 36073831 | rs7760405 | -0.721 | 0.558 | -0.036 | -0.46 | 0.007 | 0.08 | 0.08 | 1.725 | 0.11 | 0.36 | 0.07 | 0.18 | 0.09 | 0.08 |
| 6 | 36075219 | rs9470219 | 1.468 | 0.688 | -0.203 | -0.406 | -0.031 | 0.108 | 0.108 | 0.515 | 0.57 | 0.18 | 0.42 | 0.26 | 0.56 | 0.65 |
| 6 | 36076280 | rs6457878 | -0.138 | 0.697 | -0.342 | -0.453 | 0.022 | 0.078 | 0.078 | 0.17 | 0.08 | 0.26 | 0.06 | 0.13 | 0.09 | 0.06 |
| 6 | 36076934 | rs3804452 | 0.113 | 0.695 | -0.396 | -0.456 | -0.017 | 0.052 | 0.052 | 0 | 0.13 | 0.02 | 0.11 | 0.05 | 0.12 | 0.11 |
| 6 | 36077961 | rs3804451 | -0.173 | 0.736 | -0.367 | -0.384 | 0.022 | 0.078 | 0.078 | 0.17 | 0.08 | 0.26 | 0.06 | 0.13 | 0.09 | 0.06 |
| 6 | 36085325 | rs7757672 | 0.714 | 0.609 | -1.629 | -1.126 | -0.062 | 0.113 | 0.113 | 0 | 0.3 | 0.02 | 0.27 | 0.08 | 0.32 | 0.38 |
| 6 | 36086976 | rs11759192 | -0.471 | 0.828 | -0.491 | -0.302 | -0.001 | -0.022 | -0.022 | 1.309 | 0.15 | 0.2 | 0.12 | 0.04 | 0.12 | 0.11 |
| 6 | 36094188 | rs6922865 | 0.758 | 0.957 | -0.256 | 0.216 | 0.014 | 0.024 | 0.024 | 1.374 | 0.73 | 0.41 | 0.55 | 0.4 | 0.68 | 0.78 |
| 6 | 36098410 | rs1059227 | -0.173 | 0.733 | -0.953 | -0.734 | -0.002 | 0.149 | 0.149 | 0 | 0.61 | 0.81 | 0.72 | 0.95 | 0.64 | 0.6 |
| 6 | 36100732 | rs2071864 | -0.729 | 0.46 | -0.673 | -0.861 | -0.013 | 0.002 | 0.002 | 0 | 0.29 | 0.6 | 0.43 | 0.69 | 0.25 | 0.32 |
| 6 | 36103921 | rs2252430 | -0.63 | 0.452 | -0.798 | -0.931 | -0.002 | 0.023 | 0.023 | 0 | 0.92 | 0.98 | 0.98 | 1 | 0.94 | 0.97 |
| 6 | 36107172 | rs2071863 | 0.272 | 0.302 | -0.329 | -0.775 | 0.016 | 0.113 | 0.113 | 0 | 0.08 | 0.12 | 0.33 | 0.58 | 0.16 | 0.19 |
| 6 | 36109901 | rs2859136 | -0.668 | 0.295 | -0.555 | -0.848 | -0.007 | 0.141 | 0.141 | 0 | 0.61 | 0.8 | 0.71 | 0.92 | 0.64 | 0.57 |
| 6 | 36112423 | rs2815807 | -0.262 | 0.153 | -0.031 | -0.923 | 0.021 | 0.086 | 0.086 | 0 | 0.2 | 0.39 | 0.39 | 0.66 | 0.23 | 0.29 |
| 6 | 36113598 | rs10447396 | 1.219 | 0.224 | -0.116 | -0.87 | -0.009 | 0.104 | 0.104 | 0.339 | 0.31 | 0.18 | 0.27 | 0.05 | 0.31 | 0.38 |
| 6 | 36126650 | rs2213661 | 0.851 | 0.194 | -0.035 | -0.817 | -0.055 | 0.075 | 0.075 | 0.168 | 0.35 | 0.12 | 0.29 | 0.23 | 0.4 | 0.26 |
| 6 | 36127469 | rs12199702 | 1.201 | 0.174 | -0.045 | -0.878 | -0.008 | 0.111 | 0.111 | 0.741 | 0.32 | 0.18 | 0.26 | 0.05 | 0.31 | 0.37 |
| 6 | 36144958 | rs9394333 | -0.205 | 0.336 | 0.101 | 0.308 | -0.026 | -0.003 | -0.003 | 0 | 0.55 | 0.76 | 0.45 | 0.4 | 0.52 | 0.61 |
| 6 | 36150336 | rs10807157 | 0.876 | 0.072 | 0.247 | 0.327 | 0.013 | -0.002 | -0.002 | 0.425 | 0.73 | 0.78 | 0.89 | 0.96 | 0.83 | 0.93 |
| 6 | 36151177 | rs6906612 | 0.876 | 0.072 | 0.226 | 0.327 | 0.01 | -0.008 | -0.008 | 0.168 | 0.73 | 0.77 | 0.89 | 0.96 | 0.83 | 0.93 |
| 6 | 36165213 | rs4713936 | 1.372 | 0.159 | 0.51 | 0.433 | 0.014 | 0.022 | 0.022 | 0.641 | 0.58 | 0.34 | 0.76 | 0.71 | 0.64 | 0.73 |
| 6 | 36177857 | rs2239798 | -0.175 | -0.144 | 0.672 | 0.716 | 0.047 | 0.065 | 0.065 | 0 | 0.14 | 0.31 | 0.12 | 0.19 | 0.2 | 0.17 |
| 6 | 36197373 | rs7754042 | -0.192 | -0.169 | 0.625 | 0.605 | -0.034 | -0.028 | -0.028 | 0 | 0.08 | 0.21 | 0.04 | 0.02 | 0.02 | 0.03 |
| 6 | 36203710 | rs9462164 | -0.682 | -0.105 | 0.55 | 0.67 | 0.005 | -0.004 | -0.004 | 0 | 0.24 | 0.32 | 0.17 | 0.21 | 0.29 | 0.18 |
| 6 | 36206082 | rs13196184 | 0.551 | -0.197 | 0.358 | 0.603 | 0.044 | 0.062 | 0.062 | 2.914 | 0.16 | 0.03 | 0.05 | 0.02 | 0.09 | 0.02 |
| 6 | 36207598 | rs9380559 | -1.541 | -0.281 | 0.415 | 0.69 | 0.024 | 0.033 | 0.033 | 1.131 | 0.42 | 0.69 | 0.49 | 0.73 | 0.37 | 0.37 |
| 6 | 36209623 | rs4713939 | -0.003 | 0.485 | -0.052 | 1.106 | 0.021 | -0.091 | -0.091 | 0.039 | 0.18 | 0.05 | 0.33 | 0.47 | 0.17 | 0.15 |
| 6 | 36216745 | rs4236049 | 0.102 | 0.511 | 0.141 | 1.105 | 0.051 | 0.07 | 0.07 | 0 | 0.85 | 0.65 | 0.87 | 0.8 | 0.79 | 0.84 |
| 6 | 36217160 | rs4713940 | 0.245 | 0.555 | 0.068 | 1.072 | 0.056 | 0.112 | 0.112 | 0 | 0.84 | 0.58 | 0.86 | 0.77 | 0.79 | 0.81 |
| 6 | 36226525 | rs12196677 | 0.331 | 0.446 | -0.165 | 0.361 | 0.028 | 0.065 | 0.065 | 2.925 | 0.17 | 0.05 | 0.09 | 0.02 | 0.13 | 0.03 |
| 6 | 36227308 | rs4713942 | -0.971 | 0.522 | -0.156 | 0.479 | -0.004 | -0.021 | -0.021 | 0 | 0.34 | 0.52 | 0.48 | 0.71 | 0.38 | 0.35 |
| 6 | 36229495 | rs7775107 | 0.004 | 0.238 | -0.405 | 0.629 | 0.02 | 0.009 | 0.009 | 0 | 0.48 | 0.42 | 0.56 | 0.59 | 0.4 | 0.52 |
| 6 | 36230135 | rs4713944 | -0.151 | 0.314 | -0.439 | 0.692 | -0.004 | 0.003 | 0.003 | 0 | 0.53 | 0.57 | 0.54 | 0.58 | 0.52 | 0.37 |
| 6 | 36234176 | rs11753002 | -0.399 | -0.073 | -0.015 | 0.567 | -0.01 | -0.018 | -0.018 | 0 | 0.15 | 0.18 | 0.12 | 0.1 | 0.09 | 0.1 |
| 6 | 36234259 | rs11759812 | -0.372 | -0.065 | 0.141 | 0.571 | -0.015 | -0.027 | -0.027 | 0 | 0.15 | 0.2 | 0.12 | 0.1 | 0.09 | 0.07 |
| 6 | 36242317 | rs9368920 | -0.019 | -0.657 | -0.212 | 0.591 | 0.004 | 0.004 | 0.004 | 0 | 0.51 | 0.59 | 0.32 | 0.28 | 0.32 | 0.5 |
| 6 | 36247103 | rs8180685 | 0.973 | -0.896 | -0.552 | 0.521 | 0.035 | 0.01 | 0.01 | 0.029 | 0.33 | 0.23 | 0.24 | 0.26 | 0.27 | 0.4 |
| 6 | 36248957 | rs12665222 | 0.377 | -0.436 | -0.637 | 1 | -0.053 | -0.036 | -0.036 | 0 | 0.38 | 0.12 | 0.58 | 0.37 | 0.58 | 0.33 |
| 6 | 36252839 | rs1004632 | -0.588 | -0.016 | -0.3 | 0.838 | 0.003 | 0.003 | 0.003 | 0.864 | 0.58 | 0.62 | 0.7 | 0.63 | 0.64 | 0.48 |
| 6 | 36254854 | rs6457895 | -0.481 | 0.265 | -0.559 | 0.836 | -0.001 | 0.003 | 0.003 | 0.944 | 0.53 | 0.59 | 0.61 | 0.58 | 0.52 | 0.45 |
| 6 | 36258667 | rs6914153 | 0.173 | 1.062 | -1.051 | 0.789 | -0.002 | -0.001 | -0.001 | 0 | 0.39 | 0.56 | 0.32 | 0.39 | 0.42 | 0.42 |
| 6 | 36258935 | rs9394342 | 1.672 | 1.084 | -1.433 | 0.656 | -0.09 | 0.052 | 0.052 | 0.27 | 0.26 | 0.02 | 0.23 | 0.14 | 0.31 | 0.21 |
| 6 | 36260858 | rs2239795 | 0.239 | 1.265 | -1.088 | 0.69 | -0.001 | -0.001 | -0.001 | 0 | 0.37 | 0.57 | 0.32 | 0.35 | 0.42 | 0.36 |
| 6 | 36262483 | rs9470247 | -0.218 | 1.165 | -0.899 | 0.633 | 0.003 | -0.016 | -0.016 | 0 | 0.93 | 0.92 | 0.96 | 0.99 | 0.92 | 0.82 |
| 6 | 36262991 | rs4713951 | 0.093 | 0.933 | -0.815 | 0.908 | 0.029 | 0.027 | 0.027 | 1.265 | 0.49 | 0.35 | 0.58 | 0.59 | 0.41 | 0.41 |
| 6 | 36269084 | rs732394 | 0.59 | 1.489 | -1.098 | 1.013 | -0.172 | -0.082 | -0.082 | 0 | 0.37 | 0.02 | 0.56 | 0.3 | 0.49 | 0.25 |
| 6 | 36269504 | rs10947599 | 0.609 | 1.314 | -0.667 | 1.017 | -0.032 | 0.007 | 0.007 | 0 | 0.34 | 0.2 | 0.52 | 0.26 | 0.41 | 0.25 |
| 6 | 36269603 | rs10947600 | 0.808 | 1.402 | -1.006 | 1.061 | -0.118 | -0.046 | -0.046 | 0 | 0.32 | 0.03 | 0.52 | 0.26 | 0.41 | 0.24 |
| 6 | 36270668 | rs4713955 | -1.502 | 2.275 | 0.038 | 1.295 | 0.007 | 0.026 | 0.026 | 0 | 0.11 | 0.14 | 0.14 | 0.32 | 0.17 | 0.19 |
| 6 | 36272040 | rs12189786 | -1.542 | 2.319 | 0.661 | 0.749 | 0.066 | -0.016 | -0.016 | 4.826 | 0.11 | 0.34 | 0.11 | 0.08 | 0.13 | 0.14 |
| 6 | 36272706 | rs933235 | -0.56 | 2.484 | 0.447 | 0.812 | 0.014 | -0.001 | -0.001 | 0 | 0.07 | 0.09 | 0.08 | 0.07 | 0.13 | 0.13 |
| 6 | 36275458 | rs4713956 | -1.444 | 1.893 | 0.347 | 0.793 | 0.073 | 0.012 | 0.012 | 0 | 0.51 | 0.74 | 0.66 | 0.61 | 0.64 | 0.57 |
| 6 | 36276810 | rs12662591 | -0.265 | 2.368 | 0.311 | 0.372 | 0.003 | 0.033 | 0.033 | 0 | 0.16 | 0.1 | 0.11 | 0.04 | 0.15 | 0.14 |
| 6 | 36277551 | rs7738417 | -0.779 | 3.019 | 0.951 | 0.829 | 0.073 | -0.066 | -0.066 | 0 | 0.21 | 0.33 | 0.2 | 0.12 | 0.32 | 0.25 |
| 6 | 36288745 | rs10456439 | -0.943 | 2.788 | 0.173 | 1.009 | -0.011 | 0.008 | 0.008 | 0 | 0.15 | 0.13 | 0.13 | 0.03 | 0.22 | 0.14 |
| 6 | 36289145 | rs763018 | -1.814 | 2.636 | 0.464 | 1.353 | 0.013 | 0.014 | 0.014 | 0 | 0.31 | 0.37 | 0.58 | 0.57 | 0.47 | 0.39 |
| 6 | 36289814 | rs17284939 | -0.462 | 2.626 | 0.241 | 1.282 | -0.036 | 0.029 | 0.029 | 0 | 0.13 | 0.08 | 0.17 | 0.11 | 0.24 | 0.16 |
| 6 | 36292007 | rs7751928 | -1.365 | 1.586 | 0.662 | 1.769 | -0.003 | 0.126 | 0.126 | 0 | 0.15 | 0.29 | 0.35 | 0.44 | 0.14 | 0.21 |
| 6 | 36295662 | rs1015059 | -1.378 | 1.59 | 0.347 | 1.985 | 0.009 | 0.111 | 0.111 | 0 | 0.1 | 0.19 | 0.34 | 0.39 | 0.15 | 0.12 |
| 6 | 36301499 | rs11965340 | -1.217 | 1.134 | 0.336 | 1.736 | -0.02 | 0.048 | 0.048 | 0.213 | 0.06 | 0.15 | 0.03 | 0.14 | 0.02 | 0.13 |
| 6 | 36304505 | rs12530054 | -0.792 | 1.749 | 0.964 | 2.436 | 0.066 | 0.163 | 0.163 | 0 | 0.08 | 0.22 | 0.35 | 0.3 | 0.15 | 0.09 |
| 6 | 36307044 | rs1034436 | -1.517 | 1.589 | 1.261 | 2.748 | 0.069 | 0.263 | 0.263 | 0 | 0.12 | 0.33 | 0.38 | 0.45 | 0.19 | 0.13 |
| 6 | 36312907 | rs9462177 | -0.126 | 2.09 | 0.32 | 2.241 | -0.1 | -0.007 | -0.007 | 0 | 0.24 | 0.06 | 0.24 | 0.17 | 0.38 | 0.29 |
| 6 | 36315242 | rs16888069 | -0.98 | 1.882 | 0.804 | 2.208 | 0.002 | 0.115 | 0.115 | 0.879 | 0.08 | 0.28 | 0.04 | 0.16 | 0.05 | 0.07 |
| 6 | 36318575 | rs2269563 | -0.908 | 1.549 | 0.337 | 1.891 | -0.025 | 0 | 0 | 0 | 0.38 | 0.31 | 0.33 | 0.36 | 0.49 | 0.47 |
| 6 | 36322975 | rs11966082 | -0.539 | 2.183 | -0.153 | 2.165 | 0.017 | 0.017 | 0.017 | 0 | 0.23 | 0.25 | 0.55 | 0.48 | 0.48 | 0.32 |
| 6 | 36324257 | rs10807168 | -0.527 | 2.175 | -0.158 | 2.162 | 0.014 | 0.014 | 0.014 | 0 | 0.23 | 0.25 | 0.55 | 0.46 | 0.48 | 0.32 |

| | | | | | | | | | | | | | | | |
|---|---|---|---|---|---|---|---|---|---|---|---|---|---|---|---|
| 6 | 36332927 | rs7765257 | -0.376 | 2.018 | -0.004 | 1.723 | 0.059 | 0.08 | 0.08 | 0 | 0.08 | 0.2 | 0.34 | 0.23 | 0.16 | 0.09 |
| 6 | 36333584 | rs6457903 | -0.301 | 1.925 | -0.575 | 1.421 | -0.042 | -0.044 | -0.044 | 0 | 0.93 | 1 | 0.87 | 0.81 | 0.86 | 0.89 |
| 6 | 36336281 | rs10456084 | 0.103 | 1.605 | -0.353 | 1.436 | -0.011 | 0.033 | 0.033 | 0 | 0.06 | 0 | 0.07 | 0 | 0.07 | 0.05 |
| 6 | 36341294 | rs3213530 | -0.629 | 1.375 | 0.036 | 1.678 | 0.023 | 0.047 | 0.047 | 0.176 | 0.1 | 0.22 | 0.34 | 0.24 | 0.15 | 0.18 |
| 6 | 36341693 | rs9470263 | 0.189 | 1.22 | -0.723 | 1.155 | -0.006 | 0.01 | 0.01 | 0 | 0.37 | 0.36 | 0.32 | 0.17 | 0.41 | 0.36 |
| 6 | 36342181 | rs6457907 | -0.193 | 0.732 | -1.439 | 0.689 | 0.048 | 0.147 | 0.147 | 0.467 | 0.57 | 0.86 | 0.73 | 0.8 | 0.64 | 0.69 |
| 6 | 36345977 | rs9470265 | -0.725 | 0.085 | -0.641 | 1.328 | -0.005 | 0.007 | 0.007 | 0 | 0.3 | 0.3 | 0.16 | 0.38 | 0.25 | 0.22 |
| 6 | 36348689 | rs1076928 | 0.757 | -0.219 | -0.768 | 0.658 | 0.024 | 0.073 | 0.073 | 0 | 0.52 | 0.22 | 0.37 | 0.43 | 0.53 | 0.44 |
| 6 | 36348800 | rs1076929 | -0.72 | -0.342 | -0.207 | 0.541 | -0.011 | 0.003 | 0.003 | 0.003 | 0.35 | 0.62 | 0.5 | 0.52 | 0.33 | 0.25 |
| 6 | 36355372 | rs2234071 | 1.018 | -0.399 | -0.187 | 0.483 | 0.019 | 0.066 | 0.066 | 0 | 0.53 | 0.25 | 0.37 | 0.42 | 0.53 | 0.44 |
| 6 | 36358289 | rs1998266 | 0.299 | -0.326 | 0.084 | 0.62 | 0.004 | 0.012 | 0.012 | 0 | 0.86 | 0.79 | 0.65 | 0.83 | 0.85 | 0.67 |
| 6 | 36362441 | rs12663882 | -0.571 | -0.574 | 0.574 | 0.492 | -0.003 | 0.002 | 0.002 | 0.011 | 0.29 | 0.28 | 0.23 | 0.38 | 0.27 | 0.19 |
| 6 | 36366123 | rs6927636 | 0.296 | -0.966 | 0.277 | 0.26 | -0.038 | 0.062 | 0.062 | 1.257 | 0.79 | 0.99 | 0.85 | 0.93 | 0.82 | 0.61 |
| 6 | 36370254 | rs7747099 | 0.694 | -0.931 | 0.838 | 0.331 | 0.001 | 0.007 | 0.007 | 0.47 | 0.77 | 0.71 | 0.59 | 0.8 | 0.78 | 0.61 |
| 6 | 36375304 | rs941816 | 0.359 | -0.933 | 0.843 | 0.336 | 0.001 | -0.013 | -0.013 | 0 | 0.79 | 0.85 | 0.63 | 0.8 | 0.81 | 0.61 |
| 6 | 36399120 | rs7741260 | 1.047 | -0.849 | 1.104 | 0.531 | 0.02 | 0.079 | 0.079 | 0 | 0.89 | 0.72 | 0.68 | 0.85 | 0.88 | 0.84 |
| 6 | 36422789 | rs6926522 | -0.701 | -1.185 | 1.243 | 0.436 | -0.03 | 0.05 | 0.05 | 0 | 0.25 | 0.4 | 0.21 | 0.38 | 0.22 | 0.19 |
| 6 | 36437412 | rs1003250 | 1.49 | -1.566 | 1.571 | 0.203 | -0.018 | 0 | 0 | 3.752 | 0.61 | 0.45 | 0.49 | 0.48 | 0.65 | 0.64 |
| 6 | 36439391 | rs4711453 | -1.61 | -1.56 | 1.542 | 0.208 | -0.009 | 0 | 0 | 0 | 0.39 | 0.58 | 0.5 | 0.52 | 0.32 | 0.36 |
| 6 | 36457018 | rs2146333 | -0.65 | -1.554 | 1.365 | 0.22 | -0.047 | 0.072 | 0.072 | 0 | 0.26 | 0.46 | 0.22 | 0.38 | 0.22 | 0.2 |
| 6 | 36457484 | rs1061632 | -0.668 | -1.564 | 1.36 | 0.215 | -0.048 | 0.07 | 0.07 | 0 | 0.26 | 0.46 | 0.22 | 0.38 | 0.22 | 0.2 |
| 6 | 36459466 | rs2187827 | -0.282 | -1.601 | 1.393 | 0.196 | 0.013 | -0.029 | -0.029 | 0 | 0.22 | 0.15 | 0.19 | 0.38 | 0.22 | 0.2 |
| 6 | 36462007 | rs4598 | -0.634 | -1.573 | 1.348 | 0.211 | -0.019 | 0.026 | 0.026 | 0 | 0.25 | 0.31 | 0.2 | 0.38 | 0.22 | 0.2 |
| 6 | 36466787 | rs4713978 | -0.483 | -1.41 | 1.281 | 0.15 | -0.002 | 0.003 | 0.003 | 0 | 0.13 | 0.13 | 0.28 | 0.14 | 0.1 | 0.17 |
| 6 | 36492770 | rs12193641 | 0.677 | -1.327 | 0.774 | 0.205 | 0.004 | 0.015 | 0.015 | 0 | 0.1 | 0.05 | 0.13 | 0.01 | 0.1 | 0.04 |
| 6 | 36508036 | rs6913714 | -0.339 | -1.173 | 0.583 | 0.303 | -0.019 | 0.026 | 0.026 | 0 | 0.25 | 0.32 | 0.21 | 0.38 | 0.22 | 0.2 |
| 6 | 36524075 | rs7741729 | -1.101 | -0.601 | 0.269 | 0.324 | 0.004 | 0.051 | 0.051 | 0 | 0.15 | 0.37 | 0.39 | 0.16 | 0.13 | 0.16 |
| 6 | 36525256 | rs9800473 | -0.971 | -0.473 | 0.158 | 0.36 | -0.008 | 0.042 | 0.042 | 0 | 0.15 | 0.33 | 0.37 | 0.19 | 0.12 | 0.17 |
| 6 | 36547395 | rs9296189 | -0.875 | -0.574 | -0.472 | 0.108 | -0.024 | -0.025 | -0.025 | 0.223 | 0.07 | 0.12 | 0.02 | 0 | 0.01 | 0.01 |
| 6 | 36557580 | rs2145048 | -0.595 | -0.833 | -0.729 | 0.419 | 0.019 | -0.057 | -0.057 | 0 | 0.26 | 0.17 | 0.2 | 0.39 | 0.28 | 0.21 |
| 6 | 36558964 | rs9394363 | -0.324 | -0.61 | -0.632 | 0.534 | 0.007 | -0.052 | -0.052 | 0.055 | 0.23 | 0.17 | 0.19 | 0.39 | 0.27 | 0.2 |
| 6 | 36570366 | rs7344 | -0.272 | -0.554 | -1.113 | 0.615 | 0.007 | -0.101 | -0.101 | 0 | 0.22 | 0.08 | 0.16 | 0.4 | 0.25 | 0.19 |
| 6 | 36581149 | rs13202984 | 0.411 | 0.152 | -1.526 | -0.607 | -0.112 | 0.118 | 0.118 | 0.063 | 0.23 | 0.01 | 0.22 | 0.02 | 0.34 | 0.15 |
| 6 | 36604198 | rs4713989 | -0.512 | 0.195 | -0.579 | 0.483 | -0.01 | -0.013 | -0.013 | 0.004 | 0.31 | 0.39 | 0.49 | 0.74 | 0.33 | 0.38 |
| 6 | 36613812 | rs6457931 | -0.35 | 0.293 | -0.969 | 0.667 | 0.009 | 0.025 | 0.025 | 0.009 | 0.47 | 0.67 | 0.51 | 0.6 | 0.41 | 0.45 |
| 6 | 36618821 | rs1321313 | 0.217 | 0.063 | -0.546 | 0.723 | -0.005 | 0.009 | 0.009 | 0 | 0.26 | 0.21 | 0.17 | 0.17 | 0.2 | 0.32 |
| 6 | 36621178 | rs4464789 | -0.439 | -0.391 | -0.542 | 0.527 | -0.044 | 0.122 | 0.122 | 0 | 0.23 | 0.41 | 0.35 | 0.43 | 0.22 | 0.29 |
| 6 | 36622900 | rs1321311 | 0.169 | -0.702 | -0.539 | 0.292 | 0.006 | -0.042 | -0.042 | 0 | 0.3 | 0.41 | 0.19 | 0.17 | 0.22 | 0.33 |
| 6 | 36627872 | rs9368950 | 1.14 | -1.262 | -1.196 | 0.057 | -0.145 | -0.02 | -0.02 | 0 | 0.3 | 0.07 | 0.32 | 0.26 | 0.44 | 0.31 |
| 6 | 36628042 | rs9462209 | -0.567 | -1.336 | -1.199 | 0.038 | -0.019 | 0.026 | 0.026 | 0.495 | 0.54 | 0.81 | 0.54 | 0.6 | 0.43 | 0.62 |
| 6 | 36629444 | rs10807170 | 0.105 | -1.249 | -1.06 | 0.159 | 0.002 | 0.005 | 0.005 | 0 | 0.53 | 0.45 | 0.67 | 0.69 | 0.65 | 0.61 |
| 6 | 36637597 | rs4714000 | -1.026 | -1.186 | -1.093 | 0.865 | 0.019 | 0.003 | 0.003 | 5.3 | 0.86 | 0.95 | 0.88 | 0.88 | 0.86 | 0.93 |
| 6 | 36638636 | rs1321309 | 0.79 | -1.178 | -0.978 | 0.898 | -0.095 | 0.034 | 0.034 | 0 | 0.38 | 0.12 | 0.37 | 0.28 | 0.49 | 0.49 |
| 6 | 36640157 | rs10947619 | 0.124 | -1.227 | -0.763 | 0.997 | -0.051 | 0.176 | 0.176 | 0 | 0.18 | 0.39 | 0.31 | 0.43 | 0.15 | 0.12 |
| 6 | 36642836 | rs10947623 | 0.14 | -1.224 | -0.761 | 0.999 | -0.05 | 0.178 | 0.178 | 0 | 0.18 | 0.39 | 0.31 | 0.43 | 0.15 | 0.12 |
| 6 | 36645696 | rs2395655 | -0.62 | -1.209 | -0.599 | 0.974 | 0.012 | -0.026 | -0.026 | 0 | 0.54 | 0.74 | 0.39 | 0.55 | 0.38 | 0.47 |
| 6 | 36646788 | rs3176320 | -0.131 | -1.462 | -0.54 | 0.677 | -0.006 | -0.032 | -0.032 | 0.406 | 0.43 | 0.52 | 0.32 | 0.27 | 0.34 | 0.35 |
| 6 | 36647525 | rs3176331 | 0.743 | -1.23 | -0.407 | 0.773 | -0.004 | -0.005 | -0.005 | 1.395 | 0.15 | 0.18 | 0.13 | 0.12 | 0.15 | 0.05 |
| 6 | 36651971 | rs1801270 | -0.324 | -1.457 | 0.09 | 1.132 | -0.053 | 0.151 | 0.151 | 0 | 0.15 | 0.31 | 0.25 | 0.49 | 0.07 | 0.15 |
| 6 | 36655445 | rs876581 | -0.397 | -1.505 | 0.27 | 1.2 | -0.045 | 0.111 | 0.111 | 0 | 0.15 | 0.24 | 0.24 | 0.47 | 0.06 | 0.13 |
| 6 | 36656256 | rs12207548 | -0.653 | -1.19 | -0.226 | 0.855 | 0.065 | 0.006 | 0.006 | 0 | 0.43 | 0.72 | 0.4 | 0.61 | 0.24 | 0.44 |
| 6 | 36658742 | rs12191972 | 0.28 | -0.695 | 0.011 | 0.594 | -0.002 | -0.038 | -0.038 | 0 | 0.26 | 0.32 | 0.13 | 0.12 | 0.17 | 0.32 |
| 6 | 36660453 | rs6457938 | -1.097 | -0.108 | -0.517 | 1.028 | -0.036 | -0.012 | -0.012 | 0 | 0.69 | 0.77 | 0.5 | 0.71 | 0.6 | 0.7 |
| 6 | 36663357 | rs6457940 | -0.148 | 0.273 | -0.225 | 1.031 | 0.008 | -0.012 | -0.012 | 0 | 0.25 | 0.33 | 0.25 | 0.24 | 0.27 | 0.24 |
| 6 | 36666327 | rs11750982 | 0.646 | 0.266 | -0.359 | 1.291 | -0.006 | -0.027 | -0.027 | 0 | 0.12 | 0.07 | 0.12 | 0.18 | 0.14 | 0.2 |
| 6 | 36667610 | rs6906101 | -0.896 | 0.07 | -0.371 | 1.259 | -0.001 | -0.003 | -0.003 | 0 | 0.39 | 0.38 | 0.37 | 0.45 | 0.41 | 0.45 |
| 6 | 36668768 | rs12528085 | -1.31 | -0.339 | -0.141 | 0.984 | -0.001 | 0.005 | 0.005 | 0 | 0.55 | 0.58 | 0.43 | 0.64 | 0.49 | 0.64 |
| 6 | 36670400 | rs1894559 | -1.279 | -0.409 | -0.071 | 0.953 | 0 | 0.003 | 0.003 | 0 | 0.54 | 0.57 | 0.43 | 0.64 | 0.49 | 0.64 |
| 6 | 36671392 | rs12199194 | 0.88 | -0.694 | -0.264 | 0.948 | -0.014 | -0.035 | -0.035 | 2.633 | 0.1 | 0 | 0.07 | 0.13 | 0.11 | 0.2 |
| 6 | 36672484 | rs9357223 | -0.857 | -0.744 | 0.514 | 0.828 | -0.02 | 0 | 0 | 0 | 0.32 | 0.64 | 0.22 | 0.34 | 0.22 | 0.34 |
| 6 | 36673510 | rs9380590 | 0.575 | -0.729 | 0.524 | 0.828 | -0.001 | 0.006 | 0.006 | 0 | 0.13 | 0.16 | 0.14 | 0.18 | 0.13 | 0.2 |
| 6 | 36678212 | rs9470378 | -0.408 | -0.765 | 0.54 | 0.754 | -0.081 | -0.033 | -0.033 | 2.071 | 0.2 | 0.61 | 0.1 | 0.16 | 0.09 | 0.14 |
| 6 | 36680592 | rs11969665 | 0.15 | -0.743 | 0.37 | 0.776 | -0.074 | -0.013 | -0.013 | 0 | 0.17 | 0.37 | 0.08 | 0.15 | 0.07 | 0.14 |
| 6 | 36685577 | rs236489 | 0.104 | -0.282 | -0.059 | 0.941 | -0.003 | 0.002 | 0.002 | 0 | 0.74 | 0.31 | 0.56 | 0.67 | 0.78 | 0.69 |
| 6 | 36689630 | rs236477 | 0.744 | -0.133 | -0.361 | 0.974 | 0.017 | -0.031 | -0.031 | 0 | 0.58 | 0.22 | 0.39 | 0.48 | 0.63 | 0.49 |
| 6 | 36695661 | rs236471 | 1.147 | 0.06 | -0.278 | 0.482 | 0.002 | -0.017 | -0.017 | 0 | 0.38 | 0.37 | 0.22 | 0.17 | 0.38 | 0.2 |
| 6 | 36696904 | rs236468 | 1.05 | 0.021 | -0.208 | 0.473 | 0.005 | -0.034 | -0.034 | 0 | 0.39 | 0.43 | 0.22 | 0.17 | 0.38 | 0.2 |
| 6 | 36700833 | rs236460 | 0.567 | -0.166 | -0.105 | 0.416 | -0.001 | -0.002 | -0.002 | 0 | 0.45 | 0.52 | 0.36 | 0.37 | 0.51 | 0.42 |
| 6 | 36700899 | rs236459 | 0.803 | -0.141 | -0.116 | 0.387 | 0.007 | -0.002 | -0.002 | 0 | 0.41 | 0.45 | 0.31 | 0.36 | 0.48 | 0.41 |

| | | | | | | | | | | | | | | | | |
|---|---|---|---|---|---|---|---|---|---|---|---|---|---|---|---|---|
| 6 | 36702284 | rs236452 | -0.451 | -0.304 | -0.097 | 0.408 | 0.002 | 0 | 0 | 0 | 0.69 | 0.72 | 0.55 | 0.69 | 0.76 | 0.65 |
| 6 | 36703095 | rs236448 | 1.51 | -0.173 | -0.101 | 0.361 | 0.02 | -0.017 | -0.017 | 0.065 | 0.29 | 0.32 | 0.2 | 0.17 | 0.37 | 0.21 |
| 6 | 36704787 | rs1894560 | 1.416 | -0.2 | 0.077 | 0.257 | 0.006 | -0.007 | -0.007 | 0 | 0.29 | 0.33 | 0.2 | 0.17 | 0.35 | 0.2 |
| 6 | 36704807 | rs1864750 | -0.53 | -0.467 | 0.23 | 0.243 | 0.013 | -0.001 | -0.001 | 0 | 0.7 | 0.78 | 0.56 | 0.69 | 0.76 | 0.65 |
| 6 | 36704915 | rs10807173 | 0.775 | -0.515 | 0.032 | 0.228 | -0.033 | -0.064 | -0.064 | 0 | 0.17 | 0.02 | 0.2 | 0.28 | 0.22 | 0.22 |
| 6 | 36705774 | rs11970008 | -0.332 | -0.567 | 0.179 | 0.202 | -0.002 | 0.004 | 0.004 | 0 | 0.24 | 0.32 | 0.27 | 0.3 | 0.27 | 0.23 |
| 6 | 36706157 | rs12196180 | 0.818 | -0.52 | 0.082 | 0.217 | 0 | 0 | 0 | 0 | 0.46 | 0.47 | 0.3 | 0.39 | 0.49 | 0.42 |
| 6 | 36708821 | rs1065261 | 0.776 | -0.508 | 0.077 | 0.222 | -0.002 | 0.001 | 0.001 | 0 | 0.45 | 0.44 | 0.29 | 0.39 | 0.49 | 0.42 |
| 6 | 36710269 | rs3752480 | 0.764 | -0.467 | -0.121 | 0.305 | -0.032 | -0.069 | -0.069 | 0.382 | 0.17 | 0.03 | 0.19 | 0.28 | 0.22 | 0.22 |
| 6 | 36711050 | rs7763347 | 0.486 | -0.383 | 0.173 | 0.109 | 0.003 | 0.008 | 0.008 | 0 | 0.48 | 0.64 | 0.3 | 0.39 | 0.49 | 0.42 |
| 6 | 36713720 | rs9470386 | 0.546 | 0.002 | 0.086 | -0.087 | 0 | -0.006 | -0.006 | 0.031 | 0.44 | 0.41 | 0.37 | 0.28 | 0.51 | 0.23 |
| 6 | 36713858 | rs9470387 | 1.802 | -0.027 | 0.038 | -0.125 | 0.002 | -0.002 | -0.002 | 0 | 0.25 | 0.27 | 0.16 | 0.18 | 0.31 | 0.19 |
| 6 | 36715921 | rs3213537 | 0.75 | -0.015 | -0.019 | -0.027 | 0.007 | 0.014 | 0.014 | 0.322 | 0.14 | 0.24 | 0.1 | 0.21 | 0.14 | 0.22 |
| 6 | 36716229 | rs7762245 | -0.216 | -0.107 | -0.002 | -0.061 | 0.006 | -0.01 | -0.01 | 0 | 0.32 | 0.4 | 0.29 | 0.31 | 0.33 | 0.26 |
| 6 | 36719728 | rs13193217 | -0.103 | -0.2 | -0.229 | -0.007 | 0.035 | -0.031 | -0.031 | 0 | 0.6 | 0.77 | 0.47 | 0.49 | 0.65 | 0.45 |
| 6 | 36720758 | rs3822969 | 1.266 | 0.101 | -0.153 | -0.093 | 0.002 | -0.008 | -0.008 | 0 | 0.1 | 0.08 | 0.08 | 0.02 | 0.07 | 0.02 |
| 6 | 36722842 | rs10456444 | 0.715 | 0.282 | 0.184 | 0.182 | -0.001 | 0.001 | 0.001 | 0 | 0.45 | 0.42 | 0.35 | 0.38 | 0.47 | 0.4 |
| 6 | 36722951 | rs236446 | -0.079 | 0.183 | 0.155 | 0.488 | -0.002 | 0 | 0 | 0 | 0.78 | 0.75 | 0.59 | 0.66 | 0.74 | 0.58 |
| 6 | 36723380 | rs11757521 | 0.348 | 0.125 | 0.278 | 0.575 | -0.001 | 0.007 | 0.007 | 0 | 0.1 | 0.12 | 0.12 | 0.17 | 0.1 | 0.15 |
| 6 | 36723442 | rs11755091 | -0.344 | 0.102 | 0.425 | 0.539 | -0.012 | 0.024 | 0.024 | 0 | 0.13 | 0.27 | 0.14 | 0.18 | 0.1 | 0.16 |
| 6 | 36723704 | rs236444 | -0.332 | -0.031 | 0.311 | 0.495 | 0.013 | -0.005 | -0.005 | 0 | 0.68 | 0.74 | 0.68 | 0.64 | 0.75 | 0.66 |
| 6 | 36725464 | rs916295 | 0.845 | -0.018 | 0.527 | 0.493 | 0.014 | 0.001 | 0.001 | 0.004 | 0.55 | 0.39 | 0.54 | 0.45 | 0.63 | 0.5 |
| 6 | 36726998 | rs236441 | -0.397 | -0.078 | 0.561 | 0.393 | -0.004 | -0.006 | -0.006 | 0 | 0.71 | 0.75 | 0.7 | 0.81 | 0.79 | 0.69 |
| 6 | 36727212 | rs236440 | -0.505 | -0.082 | 0.519 | 0.438 | 0.01 | 0.099 | 0.099 | 0.494 | 0.07 | 0.31 | 0.06 | 0.19 | 0.06 | 0.04 |
| 6 | 36729650 | rs236434 | 1.115 | -0.292 | 0.292 | 0.257 | 0.011 | -0.001 | -0.001 | 0.255 | 0.15 | 0.11 | 0.05 | 0 | 0.06 | 0.09 |
| 6 | 36730418 | rs236432 | -0.904 | -0.27 | 0.298 | 0.302 | -0.003 | 0.004 | 0.004 | 0 | 0.84 | 0.83 | 0.74 | 0.81 | 0.86 | 0.77 |
| 6 | 36733070 | rs3752482 | 0.783 | 0.437 | -0.093 | 0.165 | 0 | -0.001 | -0.001 | 0 | 0.15 | 0.18 | 0.1 | 0.19 | 0.15 | 0.09 |
| 6 | 36733132 | rs763048 | 0.599 | 0.603 | -0.137 | 0.143 | 0.017 | 0.023 | 0.023 | 0 | 0.52 | 0.37 | 0.58 | 0.62 | 0.62 | 0.59 |
| 6 | 36735242 | rs10947628 | 0.246 | 0.638 | -0.077 | -0.27 | 0.034 | -0.005 | -0.005 | 0.105 | 0.55 | 0.34 | 0.53 | 0.45 | 0.62 | 0.49 |
| 6 | 36737499 | rs916297 | 0.503 | 0.952 | -0.151 | 0.018 | -0.001 | -0.001 | -0.001 | 6.259 | 0.45 | 0.46 | 0.49 | 0.45 | 0.54 | 0.42 |
| 6 | 36744778 | rs236419 | -0.421 | 0.796 | -0.522 | 0.358 | 0 | -0.019 | -0.019 | 0 | 0.78 | 0.82 | 0.63 | 0.63 | 0.75 | 0.6 |
| 6 | 36745427 | rs9470391 | 0.767 | 0.645 | -0.666 | 0.109 | -0.013 | 0.008 | 0.008 | 0.424 | 0.25 | 0.18 | 0.29 | 0.2 | 0.29 | 0.15 |
| 6 | 36747406 | rs4714010 | 0.555 | 0.27 | -0.637 | -0.488 | -0.007 | 0.013 | 0.013 | 0 | 0.37 | 0.36 | 0.37 | 0.21 | 0.42 | 0.23 |
| 6 | 36747494 | rs236417 | 0.1 | -0.583 | -0.801 | -0.622 | 0.019 | 0.011 | 0.011 | 0 | 0.21 | 0.13 | 0.09 | 0.01 | 0.13 | 0.16 |
| 6 | 36749072 | rs16889125 | 0.133 | 0.232 | -0.474 | 0.353 | -0.001 | 0.034 | 0.034 | 0 | 0.11 | 0.13 | 0.07 | 0.22 | 0.11 | 0.2 |
| 6 | 36751245 | rs736348 | -0.494 | 0.303 | -0.623 | 0.316 | -0.001 | -0.003 | -0.003 | 0.093 | 0.18 | 0.16 | 0.15 | 0.3 | 0.22 | 0.23 |
| 6 | 36755220 | rs236411 | -0.089 | -0.236 | -0.613 | -0.337 | -0.035 | -0.034 | -0.034 | 0.833 | 0.4 | 0.43 | 0.42 | 0.28 | 0.25 | 0.32 |
| 6 | 36755402 | rs236410 | -0.018 | 1.333 | -0.555 | -0.501 | 0.002 | -0.034 | -0.034 | 0.016 | 0.76 | 0.76 | 0.67 | 0.98 | 0.63 | 0.86 |
| 6 | 36757266 | rs958423 | 0.242 | 0.541 | -0.739 | 0.346 | 0.02 | 0.008 | 0.008 | 0 | 0.5 | 0.58 | 0.48 | 0.57 | 0.5 | 0.6 |
| 6 | 36759741 | rs9366916 | 0.005 | 0.76 | -0.789 | 0.761 | -0.02 | 0.006 | 0.006 | 0 | 0.63 | 0.72 | 0.54 | 0.68 | 0.54 | 0.65 |
| 6 | 36760778 | rs236408 | -0.387 | 0.474 | -0.575 | 0.965 | -0.008 | 0.008 | 0.008 | 0 | 0.06 | 0.12 | 0.04 | 0.11 | 0.01 | 0.03 |
| 6 | 36766292 | rs6940728 | 0.152 | 1.107 | -0.628 | 0.961 | 0.003 | 0.001 | 0.001 | 0 | 0.7 | 0.72 | 0.72 | 0.66 | 0.6 | 0.57 |
| 6 | 36771587 | rs236403 | 0.103 | 0.093 | -1.035 | 0.933 | 0.043 | -0.007 | -0.007 | 0 | 0.45 | 0.75 | 0.53 | 0.52 | 0.38 | 0.41 |
| 6 | 36772000 | rs236402 | 0.186 | -0.292 | -0.093 | 0.875 | -0.008 | -0.008 | -0.008 | 0 | 0.28 | 0.34 | 0.42 | 0.27 | 0.23 | 0.31 |
| 6 | 36772543 | rs9296197 | 0.606 | -0.446 | 0.132 | 0.796 | 0.001 | 0.015 | 0.015 | 0 | 0.1 | 0.09 | 0.08 | 0.24 | 0.12 | 0.09 |
| 6 | 36775242 | rs110033 | -0.592 | -0.108 | 0.108 | 0.606 | 0 | -0.006 | -0.006 | 0 | 0.45 | 0.69 | 0.57 | 0.51 | 0.47 | 0.4 |
| 6 | 36776799 | rs236393 | 0.209 | -0.161 | 0.669 | 0.545 | -0.02 | -0.022 | -0.022 | 2.473 | 0.29 | 0.46 | 0.44 | 0.26 | 0.25 | 0.3 |
| 6 | 36779537 | rs236391 | 0.005 | -0.273 | 0.651 | 0.294 | -0.016 | -0.034 | -0.034 | 0 | 0.34 | 0.48 | 0.47 | 0.27 | 0.31 | 0.34 |
| 6 | 36780142 | rs7745099 | 0.243 | -0.15 | 0.6 | 0.392 | 0.019 | 0.032 | 0.032 | 0 | 0.16 | 0.26 | 0.13 | 0.27 | 0.23 | 0.11 |
| 6 | 36784429 | rs12664540 | 0.71 | -0.557 | -0.017 | 0.857 | 0.015 | -0.097 | -0.097 | 1.141 | 0.17 | 0.02 | 0.08 | 0.3 | 0.14 | 0.15 |
| 6 | 36786072 | rs1015330 | 0.423 | -0.473 | 0.667 | 0.471 | 0.006 | -0.004 | -0.004 | 0 | 0.13 | 0.15 | 0.35 | 0.13 | 0.17 | 0.18 |
| 6 | 36787859 | rs236381 | 0.166 | -0.509 | 0.897 | 0.697 | 0.007 | 0.028 | 0.028 | 0.006 | 0.17 | 0.24 | 0.14 | 0.33 | 0.22 | 0.11 |
| 6 | 36790689 | rs17374068 | 0.406 | -0.506 | 0.925 | 0.624 | 0.03 | -0.016 | -0.016 | 0 | 0.09 | 0.2 | 0.17 | 0.05 | 0.1 | 0.15 |
| 6 | 36791412 | rs4714014 | -0.96 | -0.838 | 1.076 | 0.687 | -0.022 | 0.003 | 0.003 | 0 | 0.36 | 0.52 | 0.35 | 0.39 | 0.3 | 0.3 |
| 6 | 36792006 | rs236374 | 0.276 | -0.456 | 0.689 | -0.102 | 0 | 0.005 | 0.005 | 0 | 0.51 | 0.45 | 0.53 | 0.63 | 0.51 | 0.64 |
| 6 | 36793335 | rs236371 | -1.467 | -0.266 | 0.704 | -0.117 | 0 | -0.038 | -0.038 | 0 | 0.82 | 0.85 | 0.86 | 1 | 0.8 | 0.93 |
| 6 | 36794065 | rs9380605 | -0.09 | -0.275 | 0.72 | -0.085 | 0.007 | -0.01 | -0.01 | 0.014 | 0.6 | 0.57 | 0.59 | 0.66 | 0.53 | 0.77 |
| 6 | 36796133 | rs9380607 | 1.079 | -0.045 | 0.516 | -0.024 | -0.083 | -0.091 | -0.091 | 0 | 0.16 | 0.01 | 0.24 | 0.32 | 0.27 | 0.15 |
| 6 | 36796614 | rs9470413 | 0.366 | -0.161 | 0.676 | -0.029 | -0.019 | -0.006 | -0.006 | 0 | 0.06 | 0.14 | 0.01 | 0.02 | 0.02 | 0.01 |
| 6 | 36797956 | rs16889173 | 1.175 | -0.552 | 0.339 | -0.195 | 0.034 | -0.053 | -0.053 | 0.12 | 0.2 | 0.02 | 0.09 | 0.22 | 0.15 | 0.16 |
| 6 | 36798081 | rs12524422 | 1.065 | -0.52 | 0.466 | -0.203 | 0.036 | -0.014 | -0.014 | 0.313 | 0.09 | 0.21 | 0.18 | 0.05 | 0.1 | 0.14 |
| 6 | 36798150 | rs236361 | -0.524 | -0.511 | -0.062 | -0.199 | -0.046 | -0.005 | -0.005 | 0 | 0.63 | 0.82 | 0.57 | 0.58 | 0.52 | 0.76 |
| 6 | 36800611 | rs236358 | 0.154 | -0.425 | -0.066 | -0.197 | -0.006 | -0.003 | -0.003 | 0 | 0.42 | 0.74 | 0.48 | 0.36 | 0.36 | 0.61 |
| 6 | 36804187 | rs460539 | 1.338 | -0.476 | -0.137 | -0.287 | -0.012 | -0.026 | -0.026 | 0 | 0.25 | 0.18 | 0.26 | 0.35 | 0.25 | 0.44 |
| 6 | 36804661 | rs11751844 | 1.133 | -1.051 | -0.423 | -0.16 | -0.071 | -0.099 | -0.099 | 2.282 | 0.15 | 0.01 | 0.22 | 0.34 | 0.24 | 0.15 |
| 6 | 36805753 | rs16889183 | 1.699 | -1.173 | -0.443 | -0.336 | 0.055 | -0.017 | -0.017 | 0 | 0.16 | 0.02 | 0.07 | 0.15 | 0.07 | 0.12 |
| 6 | 36805885 | rs9380610 | -0.689 | -1.056 | -0.257 | -0.44 | -0.001 | 0.003 | 0.003 | 0 | 0.52 | 0.55 | 0.52 | 0.56 | 0.52 | 0.47 |
| 6 | 36819210 | rs9380614 | 1.031 | -0.741 | -0.708 | -0.335 | -0.071 | -0.1 | -0.1 | 0 | 0.16 | 0.01 | 0.22 | 0.35 | 0.24 | 0.15 |
| 6 | 36819546 | rs236350 | 1.191 | -1.117 | -0.664 | -0.681 | -0.047 | -0.12 | -0.12 | 0 | 0.21 | 0.35 | 0.1 | 0 | 0.11 | 0.04 |

| | | | | | | | | | | | | | | | | |
|---|---|---|---|---|---|---|---|---|---|---|---|---|---|---|---|---|
| 6 | 36820565 | rs236349 | -1.555 | -1.249 | -0.658 | -0.666 | 0.001 | 0 | 0 | 0 | 0.74 | 0.74 | 0.67 | 0.66 | 0.65 | 0.52 |
| 6 | 36825573 | rs9462232 | 0.218 | -1.118 | -0.636 | -0.556 | 0.02 | 0.004 | 0.004 | 0 | 0.29 | 0.24 | 0.18 | 0.27 | 0.19 | 0.2 |
| 6 | 36831569 | rs13219913 | 0.177 | -0.954 | -0.494 | -0.441 | -0.018 | 0.061 | 0.061 | 0 | 0.13 | 0.23 | 0.12 | 0.26 | 0.11 | 0.12 |
| 6 | 36856739 | rs6910832 | -0.798 | -0.915 | -0.6 | -0.494 | -0.008 | 0.002 | 0.002 | 0.052 | 0.37 | 0.32 | 0.51 | 0.68 | 0.47 | 0.4 |
| 6 | 36860092 | rs12213912 | -0.608 | -0.639 | -0.189 | -0.146 | 0.002 | -0.007 | -0.007 | 0 | 0.21 | 0.18 | 0.24 | 0.31 | 0.22 | 0.25 |
| 6 | 36871732 | rs7762601 | 1.011 | -0.61 | -0.245 | -0.152 | -0.027 | 0.017 | 0.017 | 0 | 0.07 | 0.02 | 0.17 | 0.03 | 0.11 | 0.13 |
| 6 | 36879464 | rs6900068 | 0.62 | -0.477 | -0.132 | -0.228 | 0.03 | 0.037 | 0.037 | 0 | 0.49 | 0.37 | 0.41 | 0.35 | 0.4 | 0.25 |
| 6 | 36884681 | rs12202755 | -1.026 | -0.173 | 0.04 | 0.414 | 0.001 | -0.004 | -0.004 | 0 | 0.22 | 0.22 | 0.25 | 0.31 | 0.23 | 0.25 |
| 6 | 36887308 | rs2272276 | -1.143 | -0.165 | -0.06 | 0.41 | -0.006 | 0.02 | 0.02 | 0 | 0.23 | 0.31 | 0.25 | 0.32 | 0.23 | 0.25 |
| 6 | 36892221 | rs3074 | -1.167 | -0.167 | -0.232 | 0.41 | -0.021 | 0.028 | 0.028 | 0 | 0.25 | 0.54 | 0.27 | 0.32 | 0.23 | 0.25 |
| 6 | 36893083 | rs3734333 | -1.184 | -0.162 | -0.4 | 0.412 | -0.016 | 0.016 | 0.016 | 0.013 | 0.26 | 0.61 | 0.27 | 0.33 | 0.23 | 0.25 |
| 6 | 36894677 | rs1632649 | -0.991 | -0.083 | -1.262 | 0.124 | -0.144 | -0.016 | -0.016 | 0 | 0.65 | 0.94 | 0.74 | 0.71 | 0.57 | 0.6 |
| 6 | 36905281 | rs12213087 | -1.255 | 0.1 | -0.074 | 0.391 | -0.017 | -0.067 | -0.067 | 0 | 0.19 | 0.36 | 0.23 | 0.07 | 0.18 | 0.22 |
| 6 | 36907594 | rs9470440 | -0.887 | 0.08 | -0.71 | 0.787 | -0.021 | 0.025 | 0.025 | 0 | 0.15 | 0.22 | 0.08 | 0.25 | 0.11 | 0.12 |
| 6 | 36908382 | rs697749 | 0.582 | 1.55 | -1.462 | 0.944 | -0.092 | -0.1 | -0.1 | 0 | 0.79 | 0.92 | 0.67 | 0.68 | 0.71 | 0.83 |
| 6 | 36912256 | rs12199301 | -0.06 | 0.597 | -1.673 | -0.436 | 0.024 | 0.194 | 0.194 | 0 | 0.39 | 0.13 | 0.32 | 0.05 | 0.36 | 0.3 |
| 6 | 36922684 | rs1405069 | -0.793 | 0.604 | -1.713 | -0.293 | 0.007 | -0.016 | -0.016 | 0 | 0.47 | 0.75 | 0.57 | 0.48 | 0.45 | 0.4 |
| 6 | 36924102 | rs707542 | -1.247 | 0.639 | -1.667 | -0.309 | -0.048 | -0.068 | -0.068 | 0 | 0.66 | 0.84 | 0.68 | 0.49 | 0.62 | 0.48 |
| 6 | 36925774 | rs707998 | 0.231 | 0.412 | -0.756 | -0.376 | 0.001 | 0.02 | 0.02 | 0 | 0.23 | 0.25 | 0.31 | 0.38 | 0.24 | 0.21 |
| 6 | 36931162 | rs2296934 | -1.309 | 0.42 | -0.165 | -0.301 | 0.007 | 0.009 | 0.009 | 1.546 | 0.65 | 0.63 | 0.72 | 0.49 | 0.61 | 0.57 |
| 6 | 36932571 | rs1046279 | -0.702 | 0.323 | 0.012 | -0.377 | -0.015 | -0.036 | -0.036 | 0 | 0.37 | 0.25 | 0.51 | 0.45 | 0.44 | 0.4 |
| 6 | 36934244 | rs708006 | 0.356 | 0.022 | 0.426 | -1.096 | -0.035 | -0.093 | -0.093 | 0 | 0.28 | 0.59 | 0.24 | 0.03 | 0.19 | 0.19 |
| 6 | 36934762 | rs753635 | -0.293 | -0.049 | 0.95 | -1.086 | 0.009 | -0.043 | -0.043 | 0 | 0.12 | 0.19 | 0.19 | 0.05 | 0.13 | 0.17 |
| 6 | 36936697 | rs11548576 | -0.574 | -0.074 | 0.919 | -1.063 | -0.012 | -0.006 | -0.006 | 0 | 0.06 | 0.16 | 0.07 | 0.03 | 0.05 | 0.13 |
| 6 | 36936771 | rs7772210 | 0.181 | -0.129 | 0.789 | -1.174 | 0.011 | 0.018 | 0.018 | 0 | 0.31 | 0.25 | 0.32 | 0.09 | 0.28 | 0.26 |
| 6 | 36938432 | rs2293381 | 0.857 | -0.278 | 0.297 | -0.95 | 0.014 | -0.033 | -0.033 | 0.776 | 0.07 | 0 | 0.02 | 0.17 | 0.04 | 0.1 |
| 6 | 36939213 | rs3778027 | 0.105 | 0.273 | 0.613 | -0.591 | -0.024 | -0.053 | -0.053 | 0 | 0.28 | 0.15 | 0.38 | 0.44 | 0.33 | 0.23 |
| 6 | 36942746 | rs748408 | -0.106 | 0.054 | 0.919 | -0.865 | 0.016 | 0.019 | 0.019 | 0 | 0.31 | 0.22 | 0.3 | 0.11 | 0.22 | 0.36 |
| 6 | 36944479 | rs9296204 | 0.278 | -0.035 | 0.53 | -0.82 | -0.011 | -0.022 | -0.022 | 0 | 0.24 | 0.26 | 0.17 | 0.05 | 0.15 | 0.19 |
| 6 | 36944902 | rs9296205 | 0.852 | -0.086 | 0.272 | -0.788 | 0.004 | -0.004 | -0.004 | 0.115 | 0.21 | 0.17 | 0.11 | 0.03 | 0.09 | 0.17 |
| 6 | 36948246 | rs3778022 | -0.068 | 0.202 | 0.057 | -0.643 | -0.002 | 0.012 | 0.012 | 0 | 0.1 | 0.05 | 0.16 | 0.06 | 0.12 | 0.17 |
| 6 | 36948805 | rs3778021 | -0.733 | 0.669 | 0.107 | -0.777 | 0.003 | -0.003 | -0.003 | 0 | 0.4 | 0.66 | 0.42 | 0.12 | 0.37 | 0.37 |
| 6 | 36949527 | rs2293390 | -1.014 | 0.634 | -0.112 | -0.412 | 0.012 | 0.104 | 0.104 | 0 | 0.48 | 0.82 | 0.55 | 0.61 | 0.44 | 0.42 |
| 6 | 36951210 | rs726819 | 0.573 | 0.358 | -0.009 | -0.384 | 0.023 | 0.056 | 0.056 | 0 | 0.19 | 0.07 | 0.21 | 0.04 | 0.16 | 0.2 |
| 6 | 36955064 | rs1554721 | -0.715 | 0.409 | 0.446 | -0.232 | -0.005 | 0.013 | 0.013 | 0 | 0.35 | 0.42 | 0.43 | 0.44 | 0.28 | 0.29 |
| 6 | 36955524 | rs16868759 | 0.746 | 0.152 | 0.21 | -0.209 | 0.023 | 0.006 | 0.006 | 0 | 0.19 | 0.09 | 0.23 | 0.18 | 0.17 | 0.22 |
| 6 | 36955666 | rs13193265 | -0.568 | 0.165 | 0.476 | -0.202 | -0.009 | 0.035 | 0.035 | 0 | 0.34 | 0.43 | 0.37 | 0.41 | 0.27 | 0.2 |
| 6 | 36955888 | rs4714028 | -0.849 | 0.424 | 0.484 | -0.016 | 0.002 | 0.003 | 0.003 | 0.068 | 0.56 | 0.61 | 0.65 | 0.5 | 0.43 | 0.5 |
| 6 | 36957084 | rs12524778 | -0.906 | 0.43 | 0.204 | -0.153 | -0.002 | 0.043 | 0.043 | 0.06 | 0.56 | 0.69 | 0.67 | 0.64 | 0.46 | 0.52 |
| 6 | 36965705 | rs16889362 | 0.567 | 0.366 | -0.013 | -0.179 | 0.009 | 0.077 | 0.077 | 0 | 0.16 | 0.04 | 0.2 | 0.02 | 0.12 | 0.13 |
| 6 | 36966573 | rs13217627 | -0.498 | 0.755 | 0.519 | 0.203 | 0.008 | 0.029 | 0.029 | 0.158 | 0.53 | 0.59 | 0.63 | 0.61 | 0.44 | 0.42 |
| 6 | 36970213 | rs1318961 | -0.612 | 0.735 | 0.175 | 0.165 | -0.001 | 0.039 | 0.039 | 0 | 0.56 | 0.66 | 0.68 | 0.64 | 0.45 | 0.56 |
| 6 | 36970965 | rs1023035 | -1.426 | 0.967 | 0.847 | 0.817 | 0.028 | 0.177 | 0.177 | 0.657 | 0.11 | 0.43 | 0.28 | 0.24 | 0.13 | 0.35 |
| 6 | 36972433 | rs9380633 | -0.857 | 1.716 | 0.127 | 0.754 | 0.005 | 0.008 | 0.008 | 0 | 0.2 | 0.21 | 0.26 | 0.56 | 0.33 | 0.21 |
| 6 | 36972998 | rs7742213 | -0.449 | 0.93 | 0.589 | 0.213 | -0.018 | -0.011 | -0.011 | 0 | 0.14 | 0.38 | 0.13 | 0.09 | 0.1 | 0.04 |
| 6 | 36974212 | rs831513 | -0.933 | 0.655 | -0.463 | -0.252 | -0.021 | 0.088 | 0.088 | 0.417 | 0.52 | 0.83 | 0.54 | 0.65 | 0.47 | 0.36 |
| 6 | 36974872 | rs11967185 | -0.509 | 0.519 | 0.028 | -0.122 | 0.005 | 0.008 | 0.008 | 0 | 0.3 | 0.32 | 0.4 | 0.54 | 0.37 | 0.2 |
| 6 | 36976901 | rs6927538 | -0.327 | 0.797 | 0.48 | -0.303 | 0.007 | -0.013 | -0.013 | 0.382 | 0.23 | 0.43 | 0.18 | 0.17 | 0.21 | 0.12 |
| 6 | 36976946 | rs4714032 | -0.148 | 0.762 | 0.161 | 0.105 | 0.012 | 0.068 | 0.068 | 0 | 0.13 | 0.17 | 0.3 | 0.54 | 0.2 | 0.13 |
| 6 | 36978745 | rs1724088 | -0.454 | 0.865 | -0.049 | -0.058 | 0.004 | 0.012 | 0.012 | 0 | 0.26 | 0.27 | 0.4 | 0.55 | 0.3 | 0.23 |
| 6 | 36979583 | rs12202642 | 0 | 0.547 | -0.473 | -0.255 | 0.002 | 0.039 | 0.039 | 0 | 0.1 | 0 | 0.06 | 0.04 | 0.05 | 0.22 |
| 6 | 36982947 | rs831494 | 0.112 | 0.834 | -0.39 | -0.181 | -0.001 | -0.001 | -0.001 | 0.051 | 0.34 | 0.35 | 0.51 | 0.6 | 0.39 | 0.36 |
| 6 | 36983192 | rs12199978 | 0.748 | 1.075 | -0.421 | -0.714 | -0.014 | 0.073 | 0.073 | 0.016 | 0.32 | 0.15 | 0.21 | 0.09 | 0.36 | 0.26 |
| 6 | 36986917 | rs1224487 | -0.645 | 1.098 | -0.421 | -0.651 | -0.045 | 0.14 | 0.14 | 0.033 | 0.66 | 0.89 | 0.73 | 0.88 | 0.56 | 0.69 |
| 6 | 36988364 | rs831504 | -0.319 | 1.373 | 0.145 | 0.235 | -0.004 | -0.003 | -0.003 | 0 | 0.59 | 0.61 | 0.55 | 0.48 | 0.49 | 0.63 |
| 6 | 36989152 | rs831503 | 0.221 | 1.222 | 0.471 | 0.077 | 0.006 | 0.013 | 0.013 | 0 | 0.51 | 0.46 | 0.49 | 0.41 | 0.44 | 0.56 |
| 6 | 36992237 | rs831496 | 0.179 | 1.213 | 0.473 | 0.023 | 0.004 | 0.011 | 0.011 | 0 | 0.51 | 0.48 | 0.5 | 0.39 | 0.44 | 0.56 |
| 6 | 36995222 | rs2274587 | -0.319 | 1.317 | 0.53 | 0.2 | 0.111 | 0.412 | 0.412 | 0 | 0.05 | 0.2 | 0.22 | 0.5 | 0.11 | 0.11 |
| 6 | 36999682 | rs831489 | -0.767 | 1.276 | 0.12 | -0.498 | -0.035 | 0.128 | 0.128 | 0.293 | 0.66 | 0.89 | 0.76 | 0.88 | 0.59 | 0.7 |
| 6 | 37001267 | rs831486 | 0.081 | 1.463 | 0.44 | -0.11 | 0.003 | 0 | 0 | 0 | 0.53 | 0.57 | 0.53 | 0.53 | 0.45 | 0.59 |
| 6 | 37002012 | rs6457967 | 0.752 | 1.313 | 0.503 | -0.366 | -0.004 | -0.026 | -0.026 | 2.847 | 0.06 | 0.09 | 0.01 | 0 | 0.03 | 0.02 |
| 6 | 37002182 | rs831481 | -0.093 | 1.34 | 0.481 | -0.247 | -0.001 | -0.001 | -0.001 | 0 | 0.53 | 0.54 | 0.58 | 0.57 | 0.51 | 0.62 |
| 6 | 37030217 | rs9357234 | 0.79 | 1.076 | -0.085 | -1.599 | 0 | -0.005 | -0.005 | 0 | 0.35 | 0.37 | 0.51 | 0.73 | 0.36 | 0.31 |
| 6 | 37038432 | rs913499 | -0.403 | 0.958 | -0.255 | -1.864 | 0 | -0.05 | -0.05 | 0 | 0.59 | 0.58 | 0.73 | 0.88 | 0.56 | 0.77 |
| 6 | 37050771 | rs12204194 | 0.013 | 0.908 | -0.757 | -1.502 | -0.029 | 0.044 | 0.044 | 0 | 0.1 | 0 | 0.07 | 0.03 | 0.1 | 0.24 |
| 6 | 37060809 | rs9357239 | 0.134 | 0.851 | -0.349 | -1.516 | -0.001 | -0.04 | -0.04 | 0 | 0.4 | 0.51 | 0.59 | 0.87 | 0.38 | 0.38 |
| 6 | 37063337 | rs9296211 | -0.993 | 1.068 | 0.041 | -1.269 | 0.009 | -0.059 | -0.059 | 0 | 0.31 | 0.38 | 0.31 | 0.13 | 0.36 | 0.44 |
| 6 | 37064046 | rs7758789 | -1.091 | 1.226 | 0.166 | -1.205 | 0.031 | -0.083 | -0.083 | 0 | 0.27 | 0.47 | 0.3 | 0.08 | 0.35 | 0.31 |
| 6 | 37064487 | rs3889671 | -0.328 | 0.895 | -0.045 | -1.073 | -0.004 | 0.021 | 0.021 | 0 | 0.21 | 0.13 | 0.15 | 0.08 | 0.23 | 0.11 |

| | | | | | | | | | | | | | | | | |
|---|---|---|---|---|---|---|---|---|---|---|---|---|---|---|---|---|
| 6 | 37069885 | rs9380641 | -0.083 | 0.737 | 0.21 | -1.269 | 0 | -0.031 | -0.031 | 0.151 | 0.67 | 0.44 | 0.65 | 0.83 | 0.69 | 0.66 |
| 6 | 37079883 | rs9470483 | -0.229 | 0.895 | -0.15 | -1.083 | -0.003 | -0.017 | -0.017 | 0.391 | 0.08 | 0.14 | 0.03 | 0.03 | 0.04 | 0.18 |
| 6 | 37082809 | rs2395670 | -0.569 | 0.856 | -0.332 | -1.229 | -0.006 | 0.007 | 0.007 | 0 | 0.73 | 0.72 | 0.73 | 0.91 | 0.71 | 0.78 |
| 6 | 37087349 | rs4144258 | 0.205 | 0.638 | -0.684 | -0.294 | -0.003 | -0.002 | -0.002 | 0 | 0.39 | 0.36 | 0.52 | 0.7 | 0.41 | 0.45 |
| 6 | 37093486 | rs10807178 | 0.413 | 0.831 | -0.573 | 0.095 | 0 | -0.003 | -0.003 | 0 | 0.41 | 0.3 | 0.54 | 0.6 | 0.46 | 0.34 |
| 6 | 37098379 | rs9394410 | 0.19 | 0.736 | -0.505 | 0.07 | 0.019 | 0.011 | 0.011 | 0 | 0.48 | 0.31 | 0.5 | 0.66 | 0.44 | 0.55 |
| 6 | 37102822 | rs9380645 | 0.393 | 0.73 | -1.062 | -0.422 | -0.04 | 0.281 | 0.281 | 0 | 0.53 | 0.07 | 0.6 | 0.93 | 0.51 | 0.72 |
| 6 | 37103939 | rs11961047 | -0.655 | 1.351 | -0.652 | -0.23 | 0.02 | -0.037 | -0.037 | 0 | 0.09 | 0.14 | 0.11 | 0 | 0.16 | 0.05 |
| 6 | 37112243 | rs1766694 | -0.494 | 1.343 | -0.772 | -0.041 | -0.02 | 0.007 | 0.007 | 0 | 0.78 | 0.87 | 0.82 | 0.86 | 0.75 | 0.85 |
| 6 | 37116078 | rs262902 | -1.146 | 1.263 | -0.49 | 0.722 | 0.011 | 0.023 | 0.023 | 0.093 | 0.23 | 0.7 | 0.27 | 0.4 | 0.28 | 0.38 |
| 6 | 37118688 | rs13213436 | 0.644 | 0.624 | -1.23 | 0.734 | -0.003 | -0.03 | -0.03 | 0 | 0.22 | 0.01 | 0.3 | 0.2 | 0.14 | 0.17 |
| 6 | 37130282 | rs262908 | -1.084 | 1.16 | 0.136 | 0.89 | 0.067 | 0.014 | 0.014 | 0 | 0.2 | 0.46 | 0.21 | 0.24 | 0.26 | 0.33 |
| 6 | 37142422 | rs10507 | -0.968 | 0.993 | 0.067 | 0.638 | 0.027 | -0.078 | -0.078 | 0 | 0.24 | 0.39 | 0.29 | 0.08 | 0.28 | 0.31 |
| 6 | 37143657 | rs262928 | -1.055 | 1.105 | 0.225 | 0.847 | 0 | 0.001 | 0.001 | 1.829 | 0.27 | 0.73 | 0.35 | 0.41 | 0.29 | 0.42 |
| 6 | 37144404 | rs262929 | -0.185 | 0.969 | 0.106 | 0.835 | 0.003 | -0.006 | -0.006 | 0 | 0.81 | 0.81 | 0.84 | 0.9 | 0.8 | 0.9 |
| 6 | 37150857 | rs262918 | -1.294 | 1.133 | -0.128 | 0.638 | -0.003 | -0.016 | -0.016 | 0.783 | 0.28 | 0.77 | 0.35 | 0.42 | 0.28 | 0.42 |
| 6 | 37160049 | rs16889502 | -1.152 | 1.045 | 0.572 | 0.377 | 0.038 | 0.135 | 0.135 | 0 | 0.06 | 0.2 | 0.15 | 0.17 | 0.05 | 0.07 |
| 6 | 37161391 | rs900002 | 0.859 | 1.546 | 0.668 | -0.413 | 0.03 | -0.032 | -0.032 | 0 | 0.64 | 0.54 | 0.56 | 0.76 | 0.53 | 0.56 |
| 6 | 37169909 | rs2788075 | 0.792 | 1.437 | 0.778 | -0.694 | -0.007 | -0.008 | -0.008 | 0 | 0.66 | 0.71 | 0.7 | 0.91 | 0.56 | 0.64 |
| 6 | 37171209 | rs9462285 | 0.983 | 1.28 | 1.008 | -1.162 | -0.004 | -0.06 | -0.06 | 0.013 | 0.62 | 0.44 | 0.75 | 0.91 | 0.68 | 0.72 |
| 6 | 37172934 | rs2622890 | 1.361 | 1.287 | 1.054 | -1.236 | -0.017 | 0.084 | 0.084 | 0 | 0.55 | 0.32 | 0.61 | 0.85 | 0.55 | 0.63 |
| 6 | 37173085 | rs2776797 | 0.021 | 1.52 | 1.423 | -1.364 | 0.013 | -0.041 | -0.041 | 0 | 0.8 | 0.75 | 0.82 | 0.91 | 0.77 | 0.82 |
| 6 | 37173505 | rs1757009 | 1.723 | 1.475 | 1.38 | -1.367 | -0.013 | 0.06 | 0.06 | 0 | 0.55 | 0.35 | 0.61 | 0.85 | 0.55 | 0.64 |
| 6 | 37173793 | rs7739790 | -0.659 | 1.466 | 1.361 | -1.354 | -0.016 | -0.058 | -0.058 | 0.003 | 0.17 | 0.27 | 0.06 | 0.01 | 0.09 | 0.09 |
| 6 | 37174745 | rs1680005 | 0.706 | 1.565 | 1.269 | -1.311 | 0.056 | -0.065 | -0.065 | 0.09 | 0.73 | 0.59 | 0.67 | 0.86 | 0.63 | 0.73 |
| 6 | 37177783 | rs9470540 | -1.771 | 1.394 | 1.324 | -1.19 | 0.004 | -0.104 | -0.104 | 0 | 0.26 | 0.54 | 0.22 | 0.07 | 0.22 | 0.18 |
| 6 | 37180043 | rs1055461 | -0.642 | 1.448 | 1.085 | -1.147 | -0.025 | -0.001 | -0.001 | 0.014 | 0.07 | 0.03 | 0.14 | 0.06 | 0.13 | 0.09 |
| 6 | 37180237 | rs1224329 | 0.342 | 1.391 | 1.38 | -1.234 | 0.006 | -0.073 | -0.073 | 0 | 0.79 | 0.53 | 0.81 | 0.89 | 0.79 | 0.83 |
| 6 | 37180355 | rs1150789 | 1.72 | 1.408 | 1.288 | -1.248 | -0.014 | 0.102 | 0.102 | 0 | 0.57 | 0.22 | 0.61 | 0.82 | 0.58 | 0.65 |
| 6 | 37180523 | rs1150790 | 0.412 | 1.237 | 1.174 | -1.011 | 0.007 | -0.035 | -0.035 | 0 | 0.81 | 0.7 | 0.82 | 0.89 | 0.8 | 0.83 |
| 6 | 37181984 | rs9470543 | -0.863 | 1.423 | 0.826 | -0.683 | -0.005 | -0.011 | -0.011 | 0.492 | 0.05 | 0.22 | 0.03 | 0 | 0.01 | 0 |
| 6 | 37188494 | rs2776800 | 0.312 | 1.648 | -0.371 | -0.722 | -0.055 | 0.015 | 0.015 | 0.02 | 0.26 | 0.01 | 0.16 | 0.2 | 0.26 | 0.22 |
| 6 | 37198501 | rs262899 | 0.991 | 1.816 | 0.475 | -0.953 | 0.024 | -0.046 | -0.046 | 0 | 0.84 | 0.66 | 0.84 | 0.91 | 0.83 | 0.9 |
| 6 | 37205202 | rs166394 | 1.135 | 1.849 | 0.346 | -0.889 | -0.015 | -0.002 | -0.002 | 0.296 | 0.92 | 0.94 | 0.9 | 0.9 | 0.88 | 0.91 |
| 6 | 37221673 | rs12190959 | -0.598 | 2.401 | 0.35 | -0.916 | -0.019 | 0.014 | 0.014 | 0 | 0.14 | 0.09 | 0.21 | 0.06 | 0.22 | 0.13 |
| 6 | 37241643 | rs6917404 | 1.288 | 2.38 | 0.761 | -0.939 | 0.024 | 0.005 | 0.005 | 0 | 0.92 | 0.79 | 0.89 | 0.9 | 0.88 | 0.9 |
| 6 | 37241889 | rs9349033 | 0.49 | 2.456 | 0.393 | -1.103 | -0.042 | 0.08 | 0.08 | 0.154 | 0.22 | 0.01 | 0.13 | 0.06 | 0.18 | 0.14 |
| 6 | 37242941 | rs6457986 | 2.068 | 2.584 | 1.445 | -0.497 | 0.059 | 0.023 | 0.023 | 0.017 | 0.9 | 0.63 | 0.88 | 0.9 | 0.88 | 0.9 |
| 6 | 37252210 | rs3818136 | 2.326 | 2.567 | 1.347 | -0.506 | 0.061 | -0.045 | -0.045 | 0 | 0.75 | 0.57 | 0.68 | 0.85 | 0.66 | 0.77 |
| 6 | 37257787 | rs195764 | -0.901 | 2.512 | 1.167 | -0.484 | 0.01 | -0.008 | -0.008 | 0.149 | 0.14 | 0.14 | 0.21 | 0.06 | 0.22 | 0.14 |
| 6 | 37294875 | rs9394429 | 1.886 | 2.611 | 1.243 | -0.319 | 0.105 | 0.033 | 0.033 | 0 | 0.9 | 0.78 | 0.84 | 0.83 | 0.78 | 0.83 |
| 6 | 37295630 | rs195403 | -1.436 | 2.51 | 0.988 | -0.344 | 0.011 | -0.008 | -0.008 | 0.002 | 0.17 | 0.52 | 0.21 | 0.13 | 0.16 | 0.15 |
| 6 | 37299956 | rs195410 | -0.379 | 2.606 | 0.813 | -0.298 | 0.008 | -0.005 | -0.005 | 0 | 0.12 | 0.16 | 0.16 | 0.07 | 0.15 | 0.09 |
| 6 | 37300149 | rs2295244 | 1.242 | 2.35 | 0.801 | -0.273 | 0.012 | -0.003 | -0.003 | 3.197 | 0.78 | 0.73 | 0.69 | 0.83 | 0.66 | 0.78 |
| 6 | 37304386 | rs195371 | -1.543 | 2.389 | 0.006 | -0.267 | 0.023 | -0.033 | -0.033 | 0.7 | 0.23 | 0.57 | 0.3 | 0.13 | 0.27 | 0.2 |
| 6 | 37305622 | rs9380660 | 1.356 | 2.041 | -0.075 | -0.254 | 0.018 | -0.009 | -0.009 | 0 | 0.73 | 0.38 | 0.63 | 0.77 | 0.64 | 0.72 |
| 6 | 37315512 | rs195397 | -1.335 | 1.143 | 0.502 | 0.019 | 0.071 | 0.067 | 0.067 | 2.965 | 0.07 | 0.49 | 0.08 | 0.06 | 0.05 | 0.07 |
| 6 | 37324857 | rs2269058 | -0.01 | 1.147 | -1.169 | -0.077 | -0.074 | 0.033 | 0.033 | 0.093 | 0.44 | 0.11 | 0.58 | 0.65 | 0.51 | 0.63 |
| 6 | 37329800 | rs13203701 | 0.356 | 1.138 | -1.031 | -0.339 | -0.053 | 0.061 | 0.061 | 0 | 0.11 | 0 | 0.08 | 0 | 0.15 | 0.06 |
| 6 | 37330751 | rs9394431 | 0.497 | 1.484 | -1.386 | -0.359 | -0.064 | -0.066 | -0.066 | 0.027 | 0.33 | 0.04 | 0.43 | 0.65 | 0.37 | 0.49 |
| 6 | 37348920 | rs2284923 | -0.246 | 1.271 | -0.32 | -0.698 | 0.003 | 0.034 | 0.034 | 0 | 0.48 | 0.59 | 0.58 | 0.76 | 0.48 | 0.61 |
| 6 | 37349033 | rs2284922 | -0.115 | 1.345 | 0.371 | -0.795 | 0.018 | 0.037 | 0.037 | 0 | 0.44 | 0.56 | 0.55 | 0.74 | 0.36 | 0.6 |
| 6 | 37354435 | rs195429 | -0.997 | 1.117 | 0.644 | -0.793 | -0.021 | -0.009 | -0.009 | 0 | 0.17 | 0.71 | 0.16 | 0.11 | 0.08 | 0.14 |
| 6 | 37358022 | rs195432 | -0.804 | 1.183 | -0.167 | -1.348 | -0.113 | 0.13 | 0.13 | 1.123 | 0.69 | 0.98 | 0.76 | 0.91 | 0.66 | 0.76 |
| 6 | 37360557 | rs195434 | -1.08 | 0.852 | 0.041 | -0.716 | -0.021 | -0.008 | -0.008 | 0 | 0.18 | 0.7 | 0.16 | 0.11 | 0.08 | 0.14 |
| 6 | 37368049 | rs195436 | -0.323 | 1.115 | -0.412 | -0.574 | -0.009 | 0.009 | 0.009 | 0 | 0.73 | 0.83 | 0.56 | 0.25 | 0.69 | 0.48 |
| 6 | 37394873 | rs2173314 | -0.588 | 0.561 | 0.656 | -0.567 | -0.047 | 0.015 | 0.015 | 0 | 0.15 | 0.42 | 0.08 | 0.1 | 0.07 | 0.1 |
| 6 | 37399261 | rs2776901 | -0.655 | 0.445 | 1.224 | -0.693 | -0.057 | -0.105 | -0.105 | 1.624 | 0.15 | 0.52 | 0.07 | 0 | 0.05 | 0.06 |
| 6 | 37401003 | rs7751388 | 0.003 | 0.416 | 1.568 | -0.707 | -0.031 | -0.072 | -0.072 | 0 | 0.12 | 0.26 | 0.05 | 0 | 0.05 | 0.06 |
| 6 | 37415076 | rs9380672 | 0.83 | 0.996 | 0.646 | -0.713 | -0.069 | -0.015 | -0.015 | 0 | 0.12 | 0.02 | 0.29 | 0.11 | 0.18 | 0.11 |
| 6 | 37428577 | rs2395672 | 1.141 | 0.494 | 0.485 | -0.677 | -0.014 | -0.043 | -0.043 | 0 | 0.24 | 0 | 0.17 | 0.22 | 0.22 | 0.2 |
| 6 | 37431757 | rs2776906 | -0.227 | 0.741 | 1.155 | 0.387 | -0.037 | -0.062 | -0.062 | 0 | 0.81 | 0.86 | 0.68 | 0.46 | 0.7 | 0.58 |
| 6 | 37443950 | rs9462313 | 0.206 | 0.726 | 0.451 | 0.353 | -0.031 | -0.069 | -0.069 | 0 | 0.11 | 0.24 | 0.05 | 0 | 0.05 | 0.06 |
| 6 | 37450914 | rs10692 | -0.235 | 0.764 | -0.196 | 0.376 | -0.039 | -0.065 | -0.065 | 0.598 | 0.81 | 0.87 | 0.68 | 0.46 | 0.7 | 0.58 |
| 6 | 37451696 | rs904251 | 0.861 | 0.849 | -0.36 | 0.427 | -0.004 | -0.055 | -0.055 | 0 | 0.64 | 0.16 | 0.6 | 0.46 | 0.64 | 0.52 |
| 6 | 37464724 | rs16889848 | -0.687 | 1.555 | 0.027 | 0.829 | 0.014 | 0.025 | 0.025 | 0.17 | 0.09 | 0.13 | 0.07 | 0.13 | 0.1 | 0.16 |
| 6 | 37466079 | rs931649 | -0.254 | 1.665 | -0.202 | 0.95 | -0.029 | -0.008 | -0.008 | 0 | 0.89 | 0.93 | 0.74 | 0.46 | 0.76 | 0.6 |
| 6 | 37477262 | rs2797793 | 0.711 | 1.229 | -0.025 | -0.176 | 0.035 | 0.025 | 0.025 | 0 | 0.73 | 0.62 | 0.63 | 0.37 | 0.62 | 0.48 |
| 6 | 37485728 | rs1757181 | 0.625 | 1.034 | -0.501 | -0.361 | 0.012 | -0.027 | -0.027 | 0 | 0.68 | 0.5 | 0.61 | 0.19 | 0.64 | 0.51 |

| | | | | | | | | | | | | | | | | |
|---|---|---|---|---|---|---|---|---|---|---|---|---|---|---|---|---|
| 6 | 37487044 | rs1757171 | 0.533 | 1.01 | -0.593 | -0.336 | 0.009 | 0 | 0 | 0 | 0.69 | 0.54 | 0.62 | 0.28 | 0.65 | 0.55 |
| 6 | 37490531 | rs17636328 | 0.415 | 0.857 | -1.681 | -0.731 | -0.046 | 0.007 | 0.007 | 0.669 | 0.23 | 0.02 | 0.13 | 0.13 | 0.2 | 0.14 |
| 6 | 37491723 | rs6927590 | 0.547 | 1.485 | -1.531 | -0.703 | -0.007 | 0.052 | 0.052 | 0 | 0.24 | 0.12 | 0.17 | 0.09 | 0.21 | 0.17 |
| 6 | 37496165 | rs1776444 | -0.291 | 1.835 | -1.728 | -0.855 | -0.043 | -0.051 | -0.051 | 0 | 0.84 | 0.91 | 0.65 | 0.33 | 0.69 | 0.62 |
| 6 | 37496291 | rs2153945 | 0.176 | 1.788 | -1.333 | -0.91 | -0.012 | 0.028 | 0.028 | 0.627 | 0.58 | 0.61 | 0.46 | 0.24 | 0.48 | 0.45 |
| 6 | 37496967 | rs10498738 | 0.111 | 1.671 | -1.854 | -1.304 | -0.044 | 0.06 | 0.06 | 0 | 0.11 | 0.02 | 0.21 | 0.03 | 0.15 | 0.15 |
| 6 | 37498007 | rs11755065 | 0.513 | 1.355 | -1.529 | -0.976 | -0.059 | 0.038 | 0.038 | 0 | 0.23 | 0.02 | 0.17 | 0.09 | 0.21 | 0.17 |
| 6 | 37506415 | rs2797780 | 0.747 | 1.202 | -0.636 | -1.38 | -0.001 | -0.076 | -0.076 | 0 | 0.41 | 0.53 | 0.22 | 0.05 | 0.32 | 0.13 |
| 6 | 37508106 | rs17574699 | -0.182 | 1.676 | -0.821 | -1.017 | -0.019 | -0.018 | -0.018 | 0.347 | 0.05 | 0.04 | 0.07 | 0.1 | 0.11 | 0.08 |
| 6 | 37510161 | rs12211900 | 0.138 | 1.717 | -0.868 | -1.134 | -0.002 | 0.022 | 0.022 | 0 | 0.12 | 0.07 | 0.3 | 0.05 | 0.12 | 0.23 |
| 6 | 37510932 | rs2797777 | 0.281 | 2.036 | -0.592 | -0.334 | 0.02 | 0.076 | 0.076 | 0.029 | 0.87 | 0.78 | 0.82 | 0.37 | 0.79 | 0.7 |
| 6 | 37514461 | rs9296225 | -0.076 | 1.828 | -0.24 | -0.604 | -0.004 | 0.045 | 0.045 | 0 | 0.57 | 0.72 | 0.44 | 0.75 | 0.58 | 0.46 |
| 6 | 37516378 | rs16889902 | 1.052 | 1.449 | -0.108 | -0.81 | 0.007 | -0.037 | -0.037 | 0 | 0.34 | 0.28 | 0.16 | 0.03 | 0.26 | 0.1 |
| 6 | 37517485 | rs6457993 | 0.087 | 1.77 | -0.924 | -0.716 | -0.074 | 0.127 | 0.127 | 0 | 0.43 | 0.12 | 0.48 | 0.22 | 0.48 | 0.47 |
| 6 | 37518181 | rs914351 | -0.885 | 2.119 | -0.104 | 0.279 | 0.005 | 0.043 | 0.043 | 0.042 | 0.16 | 0.24 | 0.19 | 0.62 | 0.19 | 0.3 |
| 6 | 37526024 | rs6938002 | -1.113 | 1.935 | -0.037 | -0.488 | 0.026 | -0.01 | -0.01 | 0.074 | 0.3 | 0.39 | 0.38 | 0.77 | 0.42 | 0.48 |
| 6 | 37527977 | rs1891611 | -1.111 | 1.92 | -0.048 | -0.496 | -0.014 | 0.006 | 0.006 | 0 | 0.3 | 0.3 | 0.38 | 0.77 | 0.42 | 0.5 |
| 6 | 37529834 | rs9394437 | -0.918 | 2.249 | -0.204 | -0.207 | -0.038 | -0.031 | -0.031 | 0 | 0.18 | 0.15 | 0.29 | 0.74 | 0.31 | 0.38 |
| 6 | 37532103 | rs9394438 | -1.02 | 2.412 | -0.402 | -0.005 | -0.019 | -0.017 | -0.017 | 0 | 0.19 | 0.21 | 0.3 | 0.72 | 0.36 | 0.41 |
| 6 | 37535292 | rs1034138 | -0.398 | 1.24 | -0.946 | -0.712 | 0.008 | -0.039 | -0.039 | 0 | 0.74 | 0.9 | 0.6 | 0.76 | 0.79 | 0.64 |
| 6 | 37547262 | rs804846 | -0.369 | 1.181 | -0.789 | -0.905 | 0.009 | 0 | 0 | 0 | 0.72 | 0.9 | 0.59 | 0.8 | 0.77 | 0.72 |
| 6 | 37553195 | rs1738457 | 0.025 | 1.11 | -1.205 | -1.376 | -0.12 | 0.008 | 0.008 | 0.372 | 0.7 | 0.9 | 0.48 | 0.2 | 0.48 | 0.51 |
| 6 | 37553497 | rs1776438 | 0.815 | 0.851 | -1.04 | -1.737 | -0.003 | 0.046 | 0.046 | 0 | 0.26 | 0.15 | 0.27 | 0.07 | 0.22 | 0.19 |
| 6 | 37553949 | rs6904950 | 1.406 | 0.601 | -0.668 | -1.838 | -0.059 | -0.124 | -0.124 | 0.284 | 0.32 | 0.34 | 0.14 | 0.04 | 0.16 | 0.17 |
| 6 | 37554378 | rs6925759 | -0.587 | 0.621 | -0.62 | -1.826 | 0.002 | -0.001 | -0.001 | 2.177 | 0.07 | 0.16 | 0.11 | 0.08 | 0.12 | 0.12 |
| 6 | 37554427 | rs9462332 | 1.422 | 0.591 | -0.611 | -1.826 | -0.059 | -0.124 | -0.124 | 0.32 | 0.32 | 0.34 | 0.14 | 0.04 | 0.16 | 0.18 |
| 6 | 37554917 | rs1738443 | 0.099 | 0.685 | -0.666 | -1.754 | -0.01 | 0.006 | 0.006 | 0.072 | 0.67 | 0.64 | 0.52 | 0.19 | 0.52 | 0.51 |
| 6 | 37559282 | rs1776442 | 0.37 | 0.496 | -0.834 | -1.682 | -0.002 | 0.051 | 0.051 | 0 | 0.3 | 0.22 | 0.33 | 0.14 | 0.29 | 0.32 |
| 6 | 37561865 | rs2051072 | 0.791 | 0.405 | -0.774 | -1.735 | 0.034 | 0.037 | 0.037 | 0.565 | 0.37 | 0.66 | 0.21 | 0.13 | 0.21 | 0.3 |
| 6 | 37563079 | rs2894413 | 0.127 | 0.633 | -0.692 | -1.637 | -0.003 | 0.003 | 0.003 | 0 | 0.67 | 0.61 | 0.54 | 0.19 | 0.6 | 0.55 |
| 6 | 37563752 | rs9369011 | 0.799 | 0.714 | -0.435 | -1.653 | -0.001 | 0.012 | 0.012 | 0 | 0.26 | 0.24 | 0.28 | 0.06 | 0.23 | 0.2 |
| 6 | 37569632 | rs10510137 | 0.53 | 0.331 | -0.107 | -1.324 | -0.018 | -0.114 | -0.114 | 0 | 0.33 | 0.33 | 0.17 | 0.05 | 0.26 | 0.17 |
| 6 | 37571643 | rs804837 | 0.246 | 0.581 | -0.483 | -0.886 | -0.011 | -0.031 | -0.031 | 3.271 | 0.39 | 0.43 | 0.31 | 0.17 | 0.34 | 0.24 |
| 6 | 37572021 | rs804838 | -0.571 | 1.13 | -0.486 | -0.823 | 0.005 | -0.004 | -0.004 | 0 | 0.33 | 0.38 | 0.34 | 0.76 | 0.37 | 0.61 |
| 6 | 37572144 | rs7773994 | -0.126 | 0.616 | -0.786 | -0.468 | 0.003 | 0.005 | 0.005 | 0 | 0.41 | 0.56 | 0.5 | 0.41 | 0.42 | 0.47 |
| 6 | 37574213 | rs6939858 | -0.318 | 0.134 | -1.353 | -0.98 | -0.003 | 0.051 | 0.051 | 0 | 0.32 | 0.16 | 0.2 | 0.15 | 0.29 | 0.28 |
| 6 | 37574852 | rs9380691 | 0.13 | 0.294 | -1.446 | -0.928 | -0.022 | 0.028 | 0.028 | 0 | 0.23 | 0.08 | 0.14 | 0.13 | 0.23 | 0.2 |
| 6 | 37577683 | rs9357249 | 0.72 | 0.128 | -1.726 | -1.046 | -0.015 | 0.149 | 0.149 | 0 | 0.48 | 0.09 | 0.42 | 0.25 | 0.55 | 0.45 |
| 6 | 37582109 | rs7741051 | 0.081 | -0.022 | -1.32 | -1.155 | -0.018 | -0.059 | -0.059 | 0 | 0.12 | 0.2 | 0.05 | 0.01 | 0.05 | 0.08 |
| 6 | 37582686 | rs16890001 | 0.072 | -0.021 | -1.288 | -1.198 | -0.001 | -0.023 | -0.023 | 0 | 0.09 | 0.14 | 0.05 | 0 | 0.06 | 0.04 |
| 6 | 37584079 | rs9369012 | -0.214 | 0.032 | -1.381 | -1.213 | -0.001 | -0.007 | -0.007 | 0 | 0.29 | 0.29 | 0.15 | 0.15 | 0.24 | 0.26 |
| 6 | 37584451 | rs804820 | -0.588 | 0.21 | -1.894 | -1.287 | -0.009 | 0.047 | 0.047 | 0.047 | 0.87 | 0.95 | 0.92 | 1 | 0.87 | 0.89 |
| 6 | 37585221 | rs4714084 | -0.007 | 0.206 | -1.888 | -0.563 | -0.054 | -0.057 | -0.057 | 0.254 | 0.27 | 0.09 | 0.35 | 0.69 | 0.32 | 0.38 |
| 6 | 37588563 | rs2179782 | 0.923 | 0.211 | -1.569 | -0.416 | -0.069 | 0.154 | 0.154 | 0 | 0.31 | 0.04 | 0.24 | 0.07 | 0.37 | 0.26 |
| 6 | 37590185 | rs6918733 | 0.215 | 0.577 | -0.814 | 0.203 | 0.002 | -0.006 | -0.006 | 0 | 0.57 | 0.51 | 0.63 | 0.64 | 0.57 | 0.54 |
| 6 | 37596215 | rs6923013 | 0.382 | 0.072 | -0.587 | 0.058 | 0.001 | 0.001 | 0.001 | 0 | 0.57 | 0.38 | 0.4 | 0.29 | 0.59 | 0.58 |
| 6 | 37597129 | rs3734316 | -0.907 | -1.253 | 0.115 | -0.067 | -0.07 | -0.003 | -0.003 | 0 | 0.28 | 0.51 | 0.46 | 0.71 | 0.2 | 0.34 |
| 6 | 37599962 | rs6926735 | -1.319 | -1.442 | 0.459 | 0.957 | -0.005 | 0.137 | 0.137 | 0 | 0.12 | 0.22 | 0.19 | 0.44 | 0.12 | 0.31 |
| 6 | 37600620 | rs1738453 | -0.497 | -2.04 | 1.164 | 1.125 | -0.066 | -0.003 | -0.003 | 0.336 | 0.21 | 0.31 | 0.23 | 0.21 | 0.14 | 0.16 |
| 6 | 37600774 | rs9380696 | 0.292 | -2.109 | 0.8 | 0.654 | -0.006 | -0.07 | -0.07 | 0 | 0.87 | 0.71 | 0.92 | 0.99 | 0.9 | 0.93 |
| 6 | 37602341 | rs1738454 | -0.102 | -2.136 | 0.495 | 0.638 | -0.025 | 0.054 | 0.054 | 0 | 0.91 | 1 | 0.94 | 0.99 | 0.9 | 0.92 |
| 6 | 37602460 | rs1776447 | 0.642 | -2.202 | 0.737 | 0.567 | -0.038 | -0.129 | -0.129 | 0.02 | 0.77 | 0.64 | 0.9 | 0.99 | 0.86 | 0.88 |
| 6 | 37603516 | rs12191285 | 1.088 | -1.995 | -0.037 | -0.108 | -0.048 | -0.027 | -0.027 | 0 | 0.34 | 0.23 | 0.45 | 0.6 | 0.51 | 0.47 |
| 6 | 37603535 | rs804840 | 0.023 | -2.209 | -0.139 | -0.239 | -0.038 | -0.127 | -0.127 | 1.172 | 0.77 | 0.65 | 0.89 | 0.99 | 0.86 | 0.88 |
| 6 | 37603707 | rs12191311 | 1.761 | -2.233 | -0.306 | -0.264 | -0.18 | -0.026 | -0.026 | 0 | 0.28 | 0.03 | 0.3 | 0.17 | 0.41 | 0.24 |
| 6 | 37604489 | rs804841 | -0.06 | -2.077 | -0.119 | -0.157 | -0.032 | -0.117 | -0.117 | 0 | 0.78 | 0.68 | 0.9 | 0.99 | 0.86 | 0.88 |
| 6 | 37604620 | rs7748104 | 0.919 | -1.54 | -0.767 | 0.181 | -0.052 | -0.029 | -0.029 | 0 | 0.34 | 0.22 | 0.45 | 0.6 | 0.51 | 0.47 |
| 6 | 37606805 | rs10947686 | 0.912 | -1.603 | -0.737 | 0.065 | -0.052 | -0.046 | -0.046 | 1.47 | 0.34 | 0.22 | 0.45 | 0.59 | 0.51 | 0.47 |
| 6 | 37607290 | rs9394445 | 0.806 | -0.808 | -0.402 | 0.282 | -0.015 | 0.013 | 0.013 | 0 | 0.21 | 0.07 | 0.17 | 0.17 | 0.21 | 0.17 |
| 6 | 37610904 | rs4714089 | 0.139 | -0.897 | -0.36 | 0.196 | -0.004 | -0.023 | -0.023 | 0 | 0.6 | 0.32 | 0.59 | 0.48 | 0.61 | 0.72 |
| 6 | 37623566 | rs3846874 | 0.514 | -1.244 | -0.501 | -0.622 | 0.03 | 0.102 | 0.102 | 0 | 0.56 | 0.35 | 0.7 | 0.83 | 0.66 | 0.75 |
| 6 | 37625562 | rs10755699 | 0.411 | -1.207 | -0.137 | -0.683 | -0.039 | -0.118 | -0.118 | 0 | 0.81 | 0.65 | 0.89 | 1 | 0.88 | 0.93 |
| 6 | 37626142 | rs2273110 | 1.311 | -1.15 | -0.347 | -0.625 | -0.054 | -0.018 | -0.018 | 0.312 | 0.17 | 0.05 | 0.15 | 0.16 | 0.2 | 0.16 |
| 6 | 37627628 | rs6922589 | 0.024 | -0.987 | 0.116 | -0.516 | -0.023 | -0.077 | -0.077 | 0 | 0.84 | 0.75 | 0.9 | 1 | 0.88 | 0.93 |
| 6 | 37630020 | rs16890055 | -0.426 | -0.73 | -0.419 | 0.046 | -0.011 | 0.003 | 0.003 | 0 | 0.12 | 0.15 | 0.19 | 0.19 | 0.08 | 0.15 |
| 6 | 37631031 | rs1883901 | 0.268 | -0.489 | -0.656 | 0.187 | 0.01 | -0.011 | -0.011 | 0 | 0.09 | 0.07 | 0.19 | 0.19 | 0.07 | 0.15 |
| 6 | 37633024 | rs6925749 | -0.256 | 0.396 | -0.299 | -0.331 | -0.002 | -0.015 | -0.015 | 0 | 0.68 | 0.71 | 0.79 | 0.83 | 0.75 | 0.8 |
| 6 | 37633583 | rs9470657 | -0.274 | 0.108 | 0.181 | -0.246 | 0.018 | -0.011 | -0.011 | 0.918 | 0.05 | 0.2 | 0.03 | 0 | 0.04 | 0.03 |
| 6 | 37634933 | rs10485031 | 0.654 | -0.026 | -0.436 | -0.569 | -0.015 | 0.052 | 0.052 | 0 | 0.09 | 0.01 | 0.07 | 0 | 0.11 | 0.04 |

| | | | | | | | | | | | | | | | | |
|---|---|---|---|---|---|---|---|---|---|---|---|---|---|---|---|---|
| 6 | 37635559 | rs3846881 | 0.412 | -0.537 | -0.583 | -0.35 | -0.027 | -0.019 | -0.019 | 0 | 0.44 | 0.23 | 0.52 | 0.43 | 0.52 | 0.44 |
| 6 | 37636354 | rs9296229 | -0.965 | -0.757 | -0.062 | -0.148 | -0.034 | 0.006 | 0.006 | 0 | 0.3 | 0.42 | 0.23 | 0.39 | 0.22 | 0.35 |
| 6 | 37636600 | rs9296230 | 0.533 | -0.71 | -0.149 | 0.32 | 0.007 | -0.008 | -0.008 | 0 | 0.13 | 0.15 | 0.06 | 0.25 | 0.07 | 0.22 |
| 6 | 37638562 | rs7769372 | -0.378 | -0.648 | -0.321 | 0.155 | -0.034 | 0.038 | 0.038 | 0 | 0.2 | 0.34 | 0.15 | 0.38 | 0.11 | 0.31 |
| 6 | 37641030 | rs11759115 | 0.594 | -0.162 | -1.354 | 0.112 | 0.001 | -0.001 | -0.001 | 0.806 | 0.11 | 0.1 | 0.24 | 0.16 | 0.16 | 0.15 |
| 6 | 37643520 | rs6912334 | 0.002 | -0.134 | -1.105 | 0.437 | -0.033 | 0.081 | 0.081 | 0 | 0.2 | 0.63 | 0.16 | 0.45 | 0.12 | 0.34 |
| 6 | 37644269 | rs10947692 | 0.068 | 0.64 | -1.862 | 0.298 | -0.054 | -0.075 | -0.075 | 0.953 | 0.26 | 0.05 | 0.26 | 0.28 | 0.33 | 0.25 |
| 6 | 37647644 | rs876796 | -1.165 | 0.826 | -1.92 | -0.387 | -0.066 | 0.054 | 0.054 | 0 | 0.8 | 0.96 | 0.81 | 0.92 | 0.78 | 0.88 |
| 6 | 37649020 | rs17639655 | 0.56 | 0.412 | -2.571 | -1.086 | -0.016 | 0.105 | 0.105 | 0 | 0.17 | 0.01 | 0.11 | 0 | 0.15 | 0.05 |
| 6 | 37649344 | rs12211370 | 0.346 | 0.361 | -2.715 | -1.287 | -0.007 | 0.01 | 0.01 | 0 | 0.1 | 0.06 | 0.06 | 0.06 | 0.11 | 0.1 |
| 6 | 37650639 | rs11752386 | 1.034 | 0.507 | -3.078 | -1.316 | -0.028 | -0.005 | -0.005 | 0.55 | 0.11 | 0.02 | 0.16 | 0.12 | 0.16 | 0.18 |
| 6 | 37652186 | rs6900617 | 1.348 | 0.007 | -2.861 | -0.663 | -0.002 | 0.029 | 0.029 | 0 | 0.11 | 0.1 | 0.08 | 0.37 | 0.06 | 0.2 |
| 6 | 37652730 | rs7760531 | -0.405 | -0.061 | -1.869 | -0.787 | 0.003 | 0.012 | 0.012 | 0 | 0.36 | 0.6 | 0.45 | 0.46 | 0.4 | 0.48 |
| 6 | 37653403 | rs9357251 | -1.247 | 0.018 | -1.995 | -0.713 | -0.111 | 0.098 | 0.098 | 0.251 | 0.66 | 0.95 | 0.65 | 0.83 | 0.62 | 0.74 |
| 6 | 37657430 | rs9470668 | -0.292 | -0.161 | -0.522 | -0.284 | 0.005 | 0.083 | 0.083 | 0 | 0.24 | 0.44 | 0.38 | 0.4 | 0.26 | 0.37 |
| 6 | 37657542 | rs17374592 | 0.598 | -0.057 | -1.028 | -0.305 | 0.002 | 0.025 | 0.025 | 0 | 0.12 | 0.05 | 0.23 | 0.06 | 0.12 | 0.2 |
| 6 | 37659300 | rs3901254 | 0.395 | 0.077 | -0.827 | -0.127 | 0.003 | 0.069 | 0.069 | 0 | 0.07 | 0.1 | 0.12 | 0.27 | 0.11 | 0.12 |
| 6 | 37659584 | rs3812236 | -1.025 | -0.223 | -0.568 | -0.693 | 0 | -0.013 | -0.013 | 0 | 0.52 | 0.56 | 0.53 | 0.83 | 0.48 | 0.69 |
| 6 | 37660117 | rs6916856 | 0.02 | -0.097 | -1.1 | -0.463 | 0.026 | -0.041 | -0.041 | 0 | 0.35 | 0.17 | 0.3 | 0.47 | 0.31 | 0.37 |
| 6 | 37660867 | rs6458011 | -1.397 | 0.426 | -1.078 | -0.418 | -0.129 | 0.074 | 0.074 | 0 | 0.71 | 0.98 | 0.76 | 0.9 | 0.68 | 0.84 |
| 6 | 37661386 | rs12525618 | 0.045 | 0.371 | -0.259 | -0.091 | 0.003 | -0.041 | -0.041 | 0 | 0.11 | 0.2 | 0.14 | 0.03 | 0.12 | 0.17 |
| 6 | 37674339 | rs6938061 | -0.69 | 0.333 | -0.694 | 0.07 | -0.013 | -0.008 | -0.008 | 0 | 0.54 | 0.8 | 0.63 | 0.53 | 0.51 | 0.69 |
| 6 | 37678985 | rs3846885 | 0.095 | 0.763 | -0.196 | 0.167 | -0.029 | -0.007 | -0.007 | 0 | 0.1 | 0.02 | 0.19 | 0.1 | 0.13 | 0.19 |
| 6 | 37680294 | rs9470677 | -0.652 | 0.721 | 0.481 | 0.209 | 0.007 | -0.011 | -0.011 | 0 | 0.11 | 0.15 | 0.16 | 0.08 | 0.14 | 0.16 |
| 6 | 37683516 | rs9689490 | -0.427 | 0.54 | 0.57 | 0.102 | -0.008 | -0.02 | -0.02 | 0.236 | 0.16 | 0.17 | 0.16 | 0.08 | 0.14 | 0.16 |
| 6 | 37688151 | rs6458014 | -1.687 | 0.458 | 0.331 | 0.303 | -0.005 | -0.009 | -0.009 | 0.005 | 0.55 | 0.82 | 0.7 | 0.49 | 0.58 | 0.58 |
| 6 | 37691626 | rs742787 | -1.516 | 0.65 | 0.815 | 0.676 | 0.001 | -0.005 | -0.005 | 0 | 0.4 | 0.74 | 0.44 | 0.41 | 0.4 | 0.43 |
| 6 | 37696212 | rs2395681 | -0.654 | 0.77 | 1.257 | 0.656 | 0.014 | 0.143 | 0.143 | 0 | 0.09 | 0.32 | 0.12 | 0.23 | 0.08 | 0.09 |
| 6 | 37696933 | rs742790 | -0.03 | 0.771 | 1.365 | 0.615 | 0 | 0.017 | 0.017 | 0 | 0.07 | 0.11 | 0.11 | 0.14 | 0.08 | 0.06 |
| 6 | 37698603 | rs6458016 | -1.454 | 0.686 | 1.412 | 0.573 | -0.023 | -0.011 | -0.011 | 0.056 | 0.27 | 0.52 | 0.29 | 0.23 | 0.23 | 0.22 |
| 6 | 37699156 | rs6919087 | -1.244 | 0.807 | 1.088 | 0.47 | 0.001 | -0.006 | -0.006 | 0 | 0.31 | 0.28 | 0.3 | 0.25 | 0.32 | 0.36 |
| 6 | 37700947 | rs6908598 | -0.494 | 0.784 | 0.693 | 0.478 | -0.002 | -0.008 | -0.008 | 0.152 | 0.18 | 0.14 | 0.09 | 0.09 | 0.15 | 0.15 |
| 6 | 37704553 | rs6908743 | -0.679 | 0.787 | 0.076 | 0.59 | 0 | -0.034 | -0.034 | 0 | 0.32 | 0.45 | 0.32 | 0.26 | 0.32 | 0.37 |
| 6 | 37706992 | rs6905445 | -0.853 | 0.833 | -0.918 | 0.437 | 0.007 | -0.013 | -0.013 | 0 | 0.42 | 0.79 | 0.45 | 0.45 | 0.4 | 0.43 |
| 6 | 37710167 | rs13192185 | -0.922 | 1.423 | -0.661 | 0.779 | -0.012 | -0.005 | -0.005 | 0.211 | 0.38 | 0.62 | 0.59 | 0.57 | 0.53 | 0.47 |
| 6 | 37716124 | rs1998582 | 0.495 | 1.744 | -0.912 | 0.686 | -0.061 | 0.019 | 0.019 | 0 | 0.87 | 0.96 | 0.8 | 0.89 | 0.77 | 0.85 |
| 6 | 37719060 | rs4565301 | -0.937 | 0.762 | 0.035 | 1.302 | -0.049 | 0.182 | 0.182 | 0.57 | 0.13 | 0.37 | 0.17 | 0.33 | 0.04 | 0.24 |
| 6 | 37723353 | rs10947698 | 0.322 | 0.39 | -0.522 | -0.269 | -0.015 | 0.072 | 0.072 | 0.05 | 0.41 | 0.29 | 0.34 | 0.08 | 0.45 | 0.28 |
| 6 | 37726077 | rs10456457 | -0.008 | -0.44 | 0.151 | -0.725 | -0.002 | -0.053 | -0.053 | 1.682 | 0.48 | 0.49 | 0.35 | 0.09 | 0.45 | 0.31 |
| 6 | 37727508 | rs6940860 | 0.238 | -1.32 | 0.295 | 0.545 | 0.024 | 0.069 | 0.069 | 0 | 0.51 | 0.34 | 0.45 | 0.23 | 0.45 | 0.35 |
| 6 | 37727678 | rs6941057 | -1.062 | -0.776 | -0.174 | 0.142 | -0.036 | 0.031 | 0.031 | 0 | 0.82 | 0.91 | 0.77 | 0.94 | 0.68 | 0.8 |
| 6 | 37732700 | rs11756581 | 0.075 | -1.01 | 0.007 | 0.158 | -0.009 | -0.004 | -0.004 | 0.53 | 0.63 | 0.71 | 0.51 | 0.53 | 0.49 | 0.53 |
| 6 | 37744716 | rs6916872 | 0.183 | -1.193 | 0.074 | 0.172 | 0.009 | -0.016 | -0.016 | 0 | 0.22 | 0.16 | 0.21 | 0.4 | 0.17 | 0.25 |
| 6 | 37746668 | rs4714104 | -0.062 | -1.207 | -0.193 | -0.52 | 0 | -0.034 | -0.034 | 0.096 | 0.61 | 0.62 | 0.61 | 0.88 | 0.46 | 0.65 |
| 6 | 37746848 | rs6929224 | 0.353 | -1.559 | -0.955 | -0.237 | 0.035 | -0.029 | -0.029 | 0 | 0.24 | 0.17 | 0.11 | 0.28 | 0.13 | 0.2 |
| 6 | 37755453 | rs12529345 | 0.526 | -1.158 | -0.77 | -0.05 | 0.001 | -0.029 | -0.029 | 0 | 0.26 | 0.2 | 0.27 | 0.44 | 0.23 | 0.3 |
| 6 | 37758876 | rs6909008 | -0.499 | -1.227 | -0.275 | -0.291 | 0.016 | -0.029 | -0.029 | 0 | 0.1 | 0.31 | 0.1 | 0.02 | 0.1 | 0.06 |
| 6 | 37759821 | rs6910180 | 0.112 | -1.171 | -1.065 | -0.406 | 0.004 | 0.015 | 0.015 | 0 | 0.16 | 0.1 | 0.06 | 0.06 | 0.13 | 0.11 |
| 6 | 37761373 | rs9357255 | 0.312 | -1.29 | -1.544 | -0.173 | 0.031 | -0.106 | -0.106 | 0 | 0.3 | 0.12 | 0.4 | 0.56 | 0.22 | 0.4 |
| 6 | 37761564 | rs9369022 | 0.321 | -1.294 | -1.622 | -0.177 | 0.049 | -0.118 | -0.118 | 0 | 0.35 | 0.12 | 0.44 | 0.51 | 0.25 | 0.4 |
| 6 | 37767685 | rs6926280 | 0.206 | -1.248 | -1.63 | -0.361 | 0.081 | -0.043 | -0.043 | 0 | 0.48 | 0.14 | 0.5 | 0.54 | 0.36 | 0.46 |
| 6 | 37776962 | rs6910715 | -0.167 | -0.61 | -0.467 | -0.393 | 0.029 | -0.072 | -0.072 | 0.251 | 0.16 | 0.41 | 0.12 | 0.06 | 0.15 | 0.09 |
| 6 | 37786838 | rs11759475 | 0.98 | -0.522 | -0.439 | -0.426 | -0.003 | -0.001 | -0.001 | 2.848 | 0.05 | 0.05 | 0.05 | 0 | 0.09 | 0.02 |
| 6 | 37795279 | rs16890185 | -0.547 | -0.54 | -0.384 | -0.437 | 0.023 | -0.082 | -0.082 | 0 | 0.17 | 0.42 | 0.12 | 0.06 | 0.15 | 0.09 |
| 6 | 37851191 | rs12213835 | 1.647 | -0.595 | -0.366 | -0.596 | -0.048 | 0.034 | 0.034 | 0 | 0.19 | 0.11 | 0.28 | 0.09 | 0.36 | 0.21 |
| 6 | 37881445 | rs11752567 | 1.107 | -0.565 | -0.213 | -0.544 | 0.001 | -0.006 | -0.006 | 0 | 0.05 | 0.06 | 0.05 | 0 | 0.08 | 0.02 |
| 6 | 37894575 | rs259686 | 1.579 | -0.603 | -0.136 | -0.649 | -0.028 | 0.021 | 0.021 | 0 | 0.2 | 0.15 | 0.29 | 0.09 | 0.36 | 0.22 |
| 6 | 37909855 | rs2842512 | 0.285 | -0.779 | 0.012 | -0.358 | -0.02 | 0.231 | 0.231 | 0.013 | 0.86 | 0.57 | 0.92 | 0.66 | 0.93 | 0.79 |
| 6 | 37924728 | rs7776025 | 0.612 | -0.766 | 0.266 | -0.411 | -0.009 | -0.026 | -0.026 | 0 | 0.09 | 0.17 | 0.06 | 0.05 | 0.06 | 0.06 |
| 6 | 37924906 | rs7759792 | -0.308 | -0.604 | 0.249 | -0.404 | 0.009 | 0.072 | 0.072 | 0 | 0.15 | 0.24 | 0.11 | 0.34 | 0.15 | 0.21 |
| 6 | 37925672 | rs2842518 | 0.606 | -0.807 | 0.546 | -0.524 | 0.003 | -0.01 | -0.01 | 0 | 0.93 | 0.68 | 0.96 | 1 | 0.99 | 0.99 |
| 6 | 37949615 | rs2645124 | 1.262 | -0.777 | 0.325 | -0.683 | -0.001 | -0.003 | -0.003 | 0.377 | 0.25 | 0.23 | 0.33 | 0.12 | 0.43 | 0.26 |
| 6 | 37954692 | rs4711522 | 0.789 | -1.256 | 0.043 | -0.928 | 0.044 | -0.037 | -0.037 | 0.035 | 0.42 | 0.2 | 0.46 | 0.54 | 0.32 | 0.46 |
| 6 | 37963308 | rs12661480 | 0.799 | -1.307 | 0.122 | -0.78 | 0.005 | 0.08 | 0.08 | 0 | 0.06 | 0.08 | 0.06 | 0.29 | 0.09 | 0.15 |
| 6 | 37969064 | rs6933547 | -0.269 | -1.976 | -0.208 | -0.973 | 0 | -0.001 | -0.001 | 0 | 0.62 | 0.59 | 0.59 | 0.77 | 0.68 | 0.64 |
| 6 | 37978172 | rs4711523 | -0.492 | -1.939 | -0.356 | -0.876 | 0.011 | 0.023 | 0.023 | 0 | 0.58 | 0.35 | 0.53 | 0.83 | 0.39 | 0.64 |
| 6 | 37990758 | rs1472364 | 0.924 | -1.705 | -0.399 | -0.786 | -0.051 | -0.01 | -0.01 | 0 | 0.27 | 0.05 | 0.34 | 0.18 | 0.27 | 0.3 |
| 6 | 37999223 | rs4714120 | -1.08 | -1.691 | -0.244 | -0.728 | 0.042 | -0.075 | -0.075 | 0.67 | 0.66 | 0.48 | 0.6 | 0.83 | 0.5 | 0.68 |
| 6 | 38019720 | rs9296234 | 1.792 | -1.61 | -0.874 | -0.566 | 0.012 | -0.074 | -0.074 | 0 | 0.12 | 0.01 | 0.11 | 0.36 | 0.07 | 0.17 |

| | | | | | | | | | | | | | | | | |
|---|---|---|---|---|---|---|---|---|---|---|---|---|---|---|---|---|
| 6 | 38036420 | rs17589516 | 1.111 | -1.237 | -0.578 | -0.773 | 0.032 | 0.081 | 0.081 | 0 | 0.13 | 0 | 0.05 | 0 | 0.08 | 0.07 |
| 6 | 38057359 | rs12194275 | 0.58 | -0.412 | -0.023 | -0.358 | 0.037 | -0.048 | -0.048 | 0 | 0.32 | 0.39 | 0.38 | 0.17 | 0.48 | 0.31 |
| 6 | 38073822 | rs9470779 | -0.816 | -0.203 | -0.076 | 0.487 | 0.014 | -0.046 | -0.046 | 0.113 | 0.09 | 0.18 | 0.07 | 0 | 0.09 | 0.02 |
| 6 | 38121211 | rs3800364 | 0.003 | -0.08 | -0.232 | 0.617 | 0 | 0.066 | 0.066 | 0.044 | 0.06 | 0.07 | 0.06 | 0.29 | 0.09 | 0.15 |
| 6 | 38139482 | rs13198420 | -0.245 | -0.458 | -0.483 | 0.063 | 0.009 | -0.018 | -0.018 | 0 | 0.67 | 0.71 | 0.59 | 0.41 | 0.73 | 0.54 |
| 6 | 38139500 | rs13212638 | 0.001 | -0.892 | 0.255 | -0.441 | -0.002 | -0.017 | -0.017 | 0 | 0.6 | 0.4 | 0.55 | 0.14 | 0.73 | 0.42 |
| 6 | 38139945 | rs9470807 | -0.248 | -0.885 | 0.379 | -0.522 | 0.019 | 0.052 | 0.052 | 0 | 0.63 | 0.69 | 0.54 | 0.14 | 0.7 | 0.41 |
| 6 | 38140631 | rs737172 | 0.481 | -1 | -0.161 | -0.944 | 0.008 | 0.031 | 0.031 | 0 | 0.13 | 0.07 | 0.06 | 0.02 | 0.1 | 0.1 |
| 6 | 38140658 | rs909998 | 0.22 | -0.75 | 0.106 | -1.043 | 0.026 | 0.182 | 0.182 | 0 | 0.5 | 0.26 | 0.44 | 0.13 | 0.59 | 0.38 |
| 6 | 38141022 | rs735117 | 0.243 | -0.756 | 0.107 | -1.046 | 0.024 | 0.213 | 0.213 | 0 | 0.5 | 0.22 | 0.43 | 0.13 | 0.58 | 0.38 |
| 6 | 38141322 | rs9462399 | 0.889 | -0.732 | 0.176 | -1.04 | 0.024 | -0.025 | -0.025 | 0.529 | 0.07 | 0.06 | 0.06 | 0.01 | 0.11 | 0.03 |
| 6 | 38142004 | rs3800358 | 1.07 | -0.616 | -0.283 | -1.328 | -0.079 | 0.089 | 0.089 | 0 | 0.31 | 0.12 | 0.31 | 0.13 | 0.39 | 0.29 |
| 6 | 38142316 | rs11756301 | 0.96 | -0.454 | -0.249 | -1.238 | 0.026 | -0.028 | -0.028 | 0 | 0.07 | 0.08 | 0.06 | 0.01 | 0.11 | 0.03 |
| 6 | 38144318 | rs11754110 | 0.002 | -0.493 | -0.313 | -1.243 | 0.017 | 0.097 | 0.097 | 0 | 0.51 | 0.31 | 0.47 | 0.15 | 0.58 | 0.38 |
| 6 | 38145965 | rs17678747 | 0.823 | -0.512 | -0.37 | -1.227 | -0.056 | 0.082 | 0.082 | 0 | 0.33 | 0.15 | 0.35 | 0.14 | 0.4 | 0.3 |
| 6 | 38147745 | rs7742915 | -0.338 | -0.406 | -0.17 | -0.792 | -0.018 | -0.05 | -0.05 | 0 | 0.6 | 0.5 | 0.54 | 0.15 | 0.7 | 0.42 |
| 6 | 38157142 | rs16890428 | 1.35 | -0.415 | -0.822 | -0.14 | -0.002 | -0.071 | -0.071 | 1.823 | 0.11 | 0.01 | 0.07 | 0.41 | 0.09 | 0.17 |
| 6 | 38161207 | rs742537 | 0.125 | -0.214 | -0.498 | -1.045 | 0.006 | 0.018 | 0.018 | 0 | 0.51 | 0.43 | 0.48 | 0.15 | 0.57 | 0.38 |
| 6 | 38166511 | rs10456462 | -0.263 | 0.196 | -0.632 | -1.016 | -0.004 | -0.022 | -0.022 | 0 | 0.21 | 0.19 | 0.15 | 0.01 | 0.23 | 0.08 |
| 6 | 38173192 | rs4714129 | -0.485 | -0.258 | -0.975 | -1.627 | 0.002 | 0.041 | 0.041 | 0 | 0.77 | 0.85 | 0.85 | 0.98 | 0.79 | 0.8 |
| 6 | 38174660 | rs1931763 | 0.673 | -0.723 | -0.84 | -1.945 | 0.013 | 0 | 0 | 0.257 | 0.13 | 0.1 | 0.07 | 0 | 0.1 | 0.06 |
| 6 | 38175923 | rs12215707 | 1.167 | -1.036 | -0.833 | -2.311 | -0.023 | 0.069 | 0.069 | 0 | 0.37 | 0.23 | 0.34 | 0.1 | 0.41 | 0.35 |
| 6 | 38177857 | rs6904507 | -0.511 | -1.011 | -0.546 | -1.978 | -0.003 | -0.039 | -0.039 | 0 | 0.75 | 0.72 | 0.83 | 0.97 | 0.8 | 0.8 |
| 6 | 38192993 | rs12176364 | 1.434 | -1.179 | -0.956 | -0.671 | 0.006 | -0.062 | -0.062 | 0.513 | 0.12 | 0.02 | 0.08 | 0.44 | 0.1 | 0.14 |
| 6 | 38207636 | rs9296239 | 1.053 | -1.205 | -0.856 | 0.012 | 0.014 | -0.031 | -0.031 | 0 | 0.14 | 0.09 | 0.09 | 0.44 | 0.1 | 0.14 |
| 6 | 38214150 | rs17757975 | 0.596 | -1.062 | -0.906 | -0.232 | -0.004 | -0.006 | -0.006 | 0 | 0.1 | 0.07 | 0.09 | 0.01 | 0.14 | 0.03 |
| 6 | 38216694 | rs9366950 | 0.871 | -1.358 | -1.31 | -0.511 | -0.001 | -0.129 | -0.129 | 0 | 0.19 | 0.01 | 0.16 | 0.44 | 0.17 | 0.26 |
| 6 | 38221639 | rs726160 | -0.347 | -1.109 | -1.264 | -0.824 | -0.02 | 0.042 | 0.042 | 0 | 0.67 | 0.75 | 0.78 | 0.96 | 0.64 | 0.74 |
| 6 | 38222018 | rs10947723 | 0.378 | -1.245 | -1.423 | -0.832 | 0.004 | 0.001 | 0.001 | 0.618 | 0.5 | 0.24 | 0.48 | 0.54 | 0.53 | 0.5 |
| 6 | 38223386 | rs12191476 | 0.475 | -1.042 | -1.132 | -0.947 | -0.023 | 0.079 | 0.079 | 0.057 | 0.37 | 0.23 | 0.34 | 0.1 | 0.42 | 0.34 |
| 6 | 38237525 | rs17685625 | 0.117 | -1.317 | -1.334 | -1.169 | -0.02 | 0.098 | 0.098 | 0 | 0.39 | 0.22 | 0.33 | 0.06 | 0.42 | 0.3 |
| 6 | 38240085 | rs9470848 | -0.877 | -0.892 | -1.25 | -0.312 | 0.021 | -0.059 | -0.059 | 0 | 0.6 | 0.85 | 0.7 | 0.48 | 0.65 | 0.59 |
| 6 | 38247720 | rs7739762 | 0.147 | -0.609 | -0.759 | -0.699 | 0.003 | -0.001 | -0.001 | 0 | 0.12 | 0.14 | 0.2 | 0.08 | 0.16 | 0.07 |
| 6 | 38248195 | rs9470850 | -1.14 | -0.918 | -1.01 | -0.804 | 0.009 | -0.047 | -0.047 | 0.01 | 0.43 | 0.85 | 0.53 | 0.52 | 0.4 | 0.51 |
| 6 | 38257835 | rs12206324 | 0.956 | -1.862 | -0.096 | -0.567 | -0.111 | 0.1 | 0.1 | 0.054 | 0.25 | 0.12 | 0.27 | 0.04 | 0.42 | 0.17 |
| 6 | 38277910 | rs6905637 | -0.914 | -2.05 | 0.45 | -0.538 | 0.011 | -0.004 | -0.004 | 1.275 | 0.57 | 0.67 | 0.7 | 0.52 | 0.66 | 0.65 |
| 6 | 38287910 | rs4714146 | -0.317 | -1.644 | 0.161 | -0.466 | -0.003 | -0.017 | -0.017 | 1.183 | 0.06 | 0.01 | 0.16 | 0.09 | 0.05 | 0.11 |
| 6 | 38304855 | rs6938840 | -0.885 | -1.924 | 0.538 | -0.503 | -0.059 | 0.066 | 0.066 | 0 | 0.27 | 0.53 | 0.3 | 0.41 | 0.18 | 0.38 |
| 6 | 38304882 | rs228179 | -1.335 | -1.847 | 0.178 | -0.755 | -0.106 | 0.105 | 0.105 | 0.103 | 0.74 | 0.87 | 0.71 | 0.95 | 0.56 | 0.81 |
| 6 | 38305770 | rs12660215 | -0.167 | -1.928 | 0.336 | -0.771 | -0.01 | -0.073 | -0.073 | 0.807 | 0.17 | 0.25 | 0.18 | 0.01 | 0.18 | 0.17 |
| 6 | 38317803 | rs228187 | 1.399 | -1.834 | -0.115 | -0.564 | 0.01 | -0.1 | -0.1 | 0 | 0.43 | 0.05 | 0.27 | 0.46 | 0.35 | 0.35 |
| 6 | 38318721 | rs2179533 | -0.82 | -1.549 | 0.161 | -0.442 | -0.037 | 0.032 | 0.032 | 0.055 | 0.7 | 0.75 | 0.59 | 0.84 | 0.53 | 0.72 |
| 6 | 38320943 | rs1033480 | -1.18 | -1.379 | 0.075 | -0.678 | -0.107 | 0.103 | 0.103 | 0.096 | 0.74 | 0.87 | 0.72 | 0.96 | 0.57 | 0.81 |
| 6 | 38321529 | rs12055513 | -1.035 | -1.343 | 0.751 | -0.174 | -0.069 | 0.156 | 0.156 | 0 | 0.18 | 0.48 | 0.19 | 0.37 | 0.1 | 0.3 |
| 6 | 38323299 | rs4714148 | -0.65 | -1.253 | 0.593 | -0.891 | -0.029 | 0.077 | 0.077 | 0 | 0.59 | 0.75 | 0.71 | 0.95 | 0.58 | 0.78 |
| 6 | 38323745 | rs4573069 | -0.996 | -1.255 | 0.748 | -0.953 | 0 | -0.007 | -0.007 | 1.125 | 0.26 | 0.29 | 0.37 | 0.13 | 0.31 | 0.25 |
| 6 | 38332610 | rs9394492 | 0.459 | -1.278 | 0.769 | -1.046 | 0.001 | -0.044 | -0.044 | 1.095 | 0.44 | 0.41 | 0.54 | 0.94 | 0.4 | 0.66 |
| 6 | 38336649 | rs6907434 | -0.511 | -1.018 | 0.943 | -0.763 | 0.005 | -0.037 | -0.037 | 0 | 0.13 | 0.17 | 0.17 | 0.01 | 0.18 | 0.1 |
| 6 | 38351291 | rs7759078 | 0.669 | -1.18 | 1.217 | -0.368 | -0.017 | -0.028 | -0.028 | 0 | 0.35 | 0.6 | 0.35 | 0.84 | 0.26 | 0.54 |
| 6 | 38351981 | rs9394494 | 0.661 | -1.184 | 1.215 | -0.37 | -0.017 | -0.028 | -0.028 | 0 | 0.35 | 0.6 | 0.35 | 0.84 | 0.26 | 0.54 |
| 6 | 38361112 | rs4714156 | 0.772 | -1.172 | 1.221 | -0.365 | -0.016 | -0.028 | -0.028 | 0 | 0.35 | 0.6 | 0.35 | 0.84 | 0.26 | 0.54 |
| 6 | 38365873 | rs9357271 | 0.654 | -1.019 | 0.884 | -0.579 | -0.008 | -0.022 | -0.022 | 0.322 | 0.36 | 0.63 | 0.44 | 0.87 | 0.27 | 0.54 |
| 6 | 38374397 | rs6903060 | 0.682 | -0.954 | 0.092 | -0.614 | -0.011 | -0.026 | -0.026 | 0.389 | 0.36 | 0.61 | 0.44 | 0.87 | 0.27 | 0.54 |
| 6 | 38380804 | rs2814896 | 0.372 | -0.326 | 0.016 | -0.386 | 0.051 | 0.012 | 0.012 | 0.013 | 0.3 | 0.45 | 0.46 | 0.37 | 0.42 | 0.38 |
| 6 | 38395874 | rs13194990 | -0.035 | -0.903 | -0.406 | -0.969 | 0 | -0.03 | -0.03 | 0 | 0.1 | 0.12 | 0.06 | 0 | 0.11 | 0.05 |
| 6 | 38398803 | rs4591850 | -0.311 | -1.077 | -0.12 | -0.726 | -0.017 | -0.016 | -0.016 | 0 | 0.64 | 0.3 | 0.42 | 0.5 | 0.56 | 0.56 |
| 6 | 38400139 | rs4565302 | 0.259 | -0.906 | -0.139 | -0.759 | 0.001 | -0.012 | -0.012 | 0 | 0.09 | 0.1 | 0.13 | 0 | 0.14 | 0.07 |
| 6 | 38422752 | rs16890715 | -0.444 | -1.074 | 0.242 | -0.564 | -0.001 | 0.148 | 0.148 | 0 | 0.05 | 0.2 | 0.12 | 0.13 | 0.01 | 0.05 |
| 6 | 38436317 | rs6904723 | -0.582 | -1.012 | 0.513 | -1.023 | 0.014 | -0.079 | -0.079 | 0.401 | 0.67 | 0.6 | 0.55 | 0.96 | 0.5 | 0.74 |
| 6 | 38440970 | rs3923809 | 1.051 | -1.233 | 0.383 | -0.897 | 0.011 | 0.004 | 0.004 | 0 | 0.39 | 0.28 | 0.35 | 0.63 | 0.31 | 0.43 |
| 6 | 38442448 | rs4236058 | -1.756 | -0.741 | 0.113 | -0.957 | 0.027 | -0.045 | -0.045 | 0 | 0.79 | 0.76 | 0.76 | 0.97 | 0.69 | 0.85 |
| 6 | 38444040 | rs6920488 | 1.515 | -0.807 | -0.086 | -0.807 | 0.014 | -0.089 | -0.089 | 0.478 | 0.33 | 0.08 | 0.33 | 0.53 | 0.3 | 0.38 |
| 6 | 38444763 | rs9470884 | -0.44 | -0.318 | 0.083 | -0.982 | 0.001 | -0.035 | -0.035 | 0 | 0.1 | 0.15 | 0.07 | 0 | 0.11 | 0.04 |
| 6 | 38446634 | rs12200371 | -0.725 | -0.378 | 0.093 | -0.954 | -0.005 | -0.004 | -0.004 | 0 | 0.11 | 0.18 | 0.07 | 0.11 | 0.12 | 0.09 |
| 6 | 38463649 | rs17650092 | -0.434 | -0.209 | -0.072 | -0.982 | 0.008 | -0.019 | -0.019 | 0 | 0.13 | 0.16 | 0.23 | 0.01 | 0.23 | 0.12 |
| 6 | 38464626 | rs7770868 | -0.026 | -0.38 | 0.068 | -1.132 | 0 | -0.035 | -0.035 | 0.373 | 0.34 | 0.36 | 0.48 | 0.04 | 0.55 | 0.26 |
| 6 | 38467509 | rs9470891 | 1.227 | -0.09 | -0.051 | -0.937 | -0.001 | -0.015 | -0.015 | 0 | 0.21 | 0.2 | 0.25 | 0.03 | 0.33 | 0.15 |
| 6 | 38468887 | rs4316001 | 1.013 | -0.283 | 0.07 | -0.595 | 0.025 | 0.037 | 0.037 | 0 | 0.9 | 0.69 | 0.92 | 0.9 | 0.88 | 0.91 |
| 6 | 38470087 | rs4236060 | 0.889 | -0.709 | -0.883 | -0.286 | 0.003 | -0.131 | -0.131 | 0.509 | 0.35 | 0.02 | 0.21 | 0.4 | 0.29 | 0.37 |

| | | | | | | | | | | | | | | | | |
|---|---|---|---|---|---|---|---|---|---|---|---|---|---|---|---|---|
| 6 | 38472126 | rs9380755 | 1.787 | -0.335 | 0.139 | -0.656 | 0.001 | -0.072 | -0.072 | 0.036 | 0.85 | 0.73 | 0.79 | 1 | 0.88 | 0.94 |
| 6 | 38474138 | rs9369064 | 1.232 | -0.45 | 0.542 | -0.283 | 0.007 | 0.127 | 0.127 | 0 | 0.88 | 0.54 | 0.88 | 0.75 | 0.92 | 0.89 |
| 6 | 38474367 | rs11757985 | -0.703 | -0.934 | -0.304 | -1.442 | -0.005 | 0.014 | 0.014 | 0 | 0.23 | 0.15 | 0.21 | 0.01 | 0.22 | 0.16 |
| 6 | 38478267 | rs7740763 | 1.284 | -0.785 | -0.88 | -1.128 | 0.006 | -0.03 | -0.03 | 0 | 0.57 | 0.11 | 0.39 | 0.58 | 0.62 | 0.58 |
| 6 | 38486637 | rs9394508 | 0.767 | -0.874 | -0.467 | -1.313 | 0.007 | 0.011 | 0.011 | 0.062 | 0.47 | 0.26 | 0.47 | 0.49 | 0.51 | 0.5 |
| 6 | 38487026 | rs12525647 | -0.202 | -0.873 | -0.287 | -1.351 | 0.012 | 0.004 | 0.004 | 0 | 0.25 | 0.19 | 0.23 | 0.26 | 0.19 | 0.24 |
| 6 | 38505602 | rs13219887 | -0.144 | -0.891 | -0.288 | -1.397 | 0.01 | 0.006 | 0.006 | 0 | 0.24 | 0.18 | 0.22 | 0.23 | 0.19 | 0.21 |
| 6 | 38508509 | rs17542411 | -0.122 | -0.887 | -0.318 | -1.445 | -0.003 | 0.009 | 0.009 | 0 | 0.09 | 0.07 | 0.1 | 0 | 0.13 | 0.09 |
| 6 | 38515995 | rs6931131 | -0.526 | -0.852 | 0.161 | -1.38 | 0.001 | -0.002 | -0.002 | 0 | 0.45 | 0.48 | 0.5 | 0.51 | 0.46 | 0.48 |
| 6 | 38517566 | rs4254983 | 1.128 | -0.499 | 0.357 | -1.302 | -0.067 | 0.064 | 0.064 | 0 | 0.18 | 0.07 | 0.17 | 0.03 | 0.26 | 0.13 |
| 6 | 38522822 | rs2748153 | -0.55 | -0.546 | 0.427 | -1.285 | 0.003 | -0.026 | -0.026 | 0.253 | 0.06 | 0.25 | 0.03 | 0 | 0.03 | 0.02 |
| 6 | 38525240 | rs6910904 | -0.54 | -0.541 | 0.362 | -1.251 | -0.006 | 0.159 | 0.159 | 0 | 0.07 | 0.23 | 0.13 | 0.26 | 0.05 | 0.17 |
| 6 | 38530142 | rs2748173 | 0.16 | -0.338 | 0.156 | -1.44 | 0.043 | -0.09 | -0.09 | 0.059 | 0.24 | 0.36 | 0.2 | 0.03 | 0.3 | 0.15 |
| 6 | 38534608 | rs2748169 | -1.102 | -0.856 | 0.21 | -1.012 | 0.004 | 0.131 | 0.131 | 0 | 0.12 | 0.35 | 0.17 | 0.26 | 0.11 | 0.11 |
| 6 | 38543377 | rs17543178 | 0.624 | -0.674 | 0.117 | -1.097 | -0.007 | 0.004 | 0.004 | 0 | 0.08 | 0.05 | 0.12 | 0 | 0.14 | 0.07 |
| 6 | 38545942 | rs2748168 | -0.284 | -0.728 | 0.118 | -1.199 | 0.033 | 0 | 0 | 0 | 0.3 | 0.42 | 0.33 | 0.29 | 0.36 | 0.24 |
| 6 | 38560215 | rs2180106 | -0.209 | -1.041 | -0.136 | -1.451 | 0.005 | 0.018 | 0.018 | 0.061 | 0.26 | 0.15 | 0.19 | 0.01 | 0.26 | 0.16 |
| 6 | 38564948 | rs910516 | -1.489 | -1.099 | -0.259 | -1.106 | -0.003 | -0.068 | -0.068 | 0.266 | 0.61 | 0.86 | 0.64 | 0.57 | 0.63 | 0.56 |
| 6 | 38604754 | rs1781708 | -1.339 | -1.082 | -0.2 | -1.084 | 0.019 | -0.041 | -0.041 | 0.495 | 0.6 | 0.8 | 0.63 | 0.54 | 0.63 | 0.52 |
| 6 | 38614198 | rs1781740 | 0.349 | -1.05 | -0.227 | -1.419 | 0.02 | -0.049 | -0.049 | 0 | 0.26 | 0.35 | 0.28 | 0.09 | 0.3 | 0.18 |
| 6 | 38617720 | rs1739633 | -2.044 | -1.598 | -0.862 | -1.536 | -0.038 | -0.171 | -0.171 | 0 | 0.78 | 0.98 | 0.74 | 0.56 | 0.76 | 0.72 |
| 6 | 38626404 | rs7451595 | 0.785 | -1.213 | -0.003 | -0.893 | 0.014 | 0.048 | 0.048 | 0 | 0.13 | 0.02 | 0.07 | 0 | 0.09 | 0.04 |
| 6 | 38627279 | rs1781738 | -1.997 | -1.018 | 0.026 | -0.738 | -0.042 | -0.14 | -0.14 | 0.549 | 0.63 | 0.95 | 0.62 | 0.58 | 0.64 | 0.55 |
| 6 | 38641354 | rs6458065 | -1.325 | -1.056 | 0.246 | -0.959 | 0.007 | 0.024 | 0.024 | 1.382 | 0.53 | 0.55 | 0.55 | 0.32 | 0.55 | 0.44 |
| 6 | 38642371 | rs17622097 | 0.137 | -1.023 | -0.093 | -1.015 | -0.027 | 0.016 | 0.016 | 0 | 0.26 | 0.21 | 0.29 | 0.03 | 0.39 | 0.18 |
| 6 | 38643796 | rs7604 | 0.502 | -0.826 | -0.555 | -0.713 | 0.01 | 0.114 | 0.114 | 0 | 0.18 | 0.01 | 0.07 | 0 | 0.12 | 0.08 |
| 6 | 38650841 | rs4714175 | -1.477 | -0.601 | -1.004 | -0.614 | 0.012 | 0.05 | 0.05 | 2.029 | 0.54 | 0.72 | 0.55 | 0.4 | 0.55 | 0.52 |
| 6 | 38658883 | rs6932648 | -0.544 | -0.591 | -0.644 | -0.085 | -0.048 | -0.012 | -0.012 | 0 | 0.24 | 0.34 | 0.17 | 0.23 | 0.13 | 0.23 |
| 6 | 38664594 | rs10484854 | 0.867 | -0.432 | -0.751 | -0.144 | -0.101 | 0.068 | 0.068 | 0 | 0.15 | 0.02 | 0.16 | 0.03 | 0.25 | 0.09 |
| 6 | 38669184 | rs17622621 | -0.323 | -0.566 | -0.602 | -0.01 | -0.107 | 0.186 | 0.186 | 0 | 0.34 | 0.09 | 0.22 | 0.09 | 0.37 | 0.21 |
| 6 | 38670067 | rs2490026 | -1.153 | -0.578 | -0.625 | 0.015 | -0.001 | -0.001 | -0.001 | 0 | 0.48 | 0.63 | 0.39 | 0.51 | 0.46 | 0.44 |
| 6 | 38670837 | rs1049346 | -1.234 | -0.553 | -0.536 | 0.094 | 0.006 | 0.003 | 0.003 | 0.157 | 0.52 | 0.7 | 0.48 | 0.58 | 0.49 | 0.5 |
| 6 | 38679065 | rs1621788 | -1.258 | -0.392 | -0.676 | 0.018 | 0.001 | 0.002 | 0.002 | 0 | 0.54 | 0.66 | 0.49 | 0.59 | 0.52 | 0.5 |
| 6 | 38679676 | rs1616723 | -0.369 | 0.552 | -1.598 | -1.259 | 0.025 | 0.103 | 0.103 | 0.038 | 0.07 | 0.18 | 0.05 | 0.18 | 0.08 | 0.07 |
| 6 | 38680402 | rs12206647 | -0.679 | 0.698 | -1.814 | -1.685 | -0.11 | 0.202 | 0.202 | 0 | 0.39 | 0.05 | 0.31 | 0.16 | 0.44 | 0.28 |
| 6 | 38681778 | rs1699011 | -0.732 | 0.166 | -0.726 | -0.796 | -0.013 | -0.013 | -0.013 | 0.499 | 0.22 | 0.31 | 0.09 | 0.19 | 0.15 | 0.15 |
| 6 | 38689875 | rs1937781 | 0.361 | 0.725 | -1.224 | -1.375 | -0.01 | 0.063 | 0.063 | 0 | 0.71 | 0.82 | 0.79 | 0.99 | 0.72 | 0.73 |
| 6 | 38692437 | rs9296260 | 0.802 | 0.814 | -1.023 | -1.953 | 0.005 | -0.093 | -0.093 | 0.206 | 0.79 | 0.66 | 0.86 | 1 | 0.76 | 0.83 |
| 6 | 38693361 | rs7761629 | 0.414 | 0.335 | -1.18 | -2.169 | 0.005 | 0.004 | 0.004 | 0 | 0.09 | 0.06 | 0.05 | 0.06 | 0.06 | 0.04 |
| 6 | 38696668 | rs1698998 | -0.313 | 0.382 | -1.114 | -2.16 | 0.011 | -0.08 | -0.08 | 0 | 0.14 | 0.3 | 0.1 | 0 | 0.18 | 0.07 |
| 6 | 38698580 | rs1626976 | 0.601 | 1.187 | -1.578 | -2.241 | -0.015 | 0.016 | 0.016 | 0 | 0.83 | 0.85 | 0.73 | 0.95 | 0.79 | 0.86 |
| 6 | 38699037 | rs1631117 | -0.21 | 1.128 | -1.604 | -2.22 | 0.002 | 0.009 | 0.009 | 0.06 | 0.93 | 0.97 | 0.96 | 1 | 0.94 | 0.9 |
| 6 | 38702959 | rs17623162 | 0.85 | 1.574 | -1.856 | -2.483 | -0.095 | 0.101 | 0.101 | 0 | 0.17 | 0.01 | 0.16 | 0 | 0.24 | 0.14 |
| 6 | 38704505 | rs1937777 | 0.212 | 1.089 | -1.647 | -2.269 | -0.002 | 0.018 | 0.018 | 0 | 0.08 | 0.05 | 0.04 | 0.01 | 0.07 | 0.1 |
| 6 | 38704568 | rs7747935 | 0.248 | 1.065 | -1.594 | -2.23 | -0.007 | 0.006 | 0.006 | 0 | 0.09 | 0.07 | 0.23 | 0.01 | 0.15 | 0.04 |
| 6 | 38704943 | rs6935293 | 0.561 | 0.003 | -1.516 | -1.955 | 0.011 | -0.005 | -0.005 | 0 | 0.08 | 0.04 | 0.05 | 0.11 | 0.05 | 0.05 |
| 6 | 38706073 | rs756405 | -0.479 | -0.102 | -1.013 | -1.956 | -0.027 | -0.047 | -0.047 | 0 | 0.6 | 0.55 | 0.6 | 0.17 | 0.7 | 0.49 |
| 6 | 38712688 | rs6458068 | 0.643 | -0.444 | -0.882 | -1.583 | -0.051 | -0.022 | -0.022 | 0 | 0.31 | 0.43 | 0.26 | 0.73 | 0.21 | 0.37 |
| 6 | 38719441 | rs13209417 | -0.55 | 0.169 | -1.046 | -1.081 | 0.016 | 0.016 | 0.016 | 0 | 0.17 | 0.08 | 0.11 | 0.12 | 0.14 | 0.16 |
| 6 | 38722443 | rs769064 | 0.394 | 0.428 | -0.445 | -1.308 | -0.004 | 0.014 | 0.014 | 0 | 0.79 | 0.9 | 0.8 | 0.82 | 0.78 | 0.75 |
| 6 | 38723575 | rs9357280 | -0.7 | 0.526 | -0.216 | 0.111 | 0.019 | 0.317 | 0.317 | 0.346 | 0.07 | 0.23 | 0.16 | 0.43 | 0.06 | 0.2 |
| 6 | 38728142 | rs1678677 | -1.134 | 1.522 | -0.016 | -0.151 | 0.093 | 0.232 | 0.232 | 0.088 | 0.13 | 0.34 | 0.4 | 0.45 | 0.2 | 0.25 |
| 6 | 38734652 | rs6930268 | 0.798 | 0.746 | -0.006 | -1.289 | 0.004 | -0.039 | -0.039 | 0.018 | 0.32 | 0.29 | 0.18 | 0.01 | 0.28 | 0.26 |
| 6 | 38738537 | rs7749015 | 0.907 | 0.795 | -0.023 | -1.139 | 0.005 | 0.003 | 0.003 | 0 | 0.06 | 0.05 | 0.23 | 0.02 | 0.14 | 0.04 |
| 6 | 38742708 | rs1738264 | -0.494 | 0.78 | 0.012 | -1.117 | 0.043 | 0.123 | 0.123 | 0.092 | 0.12 | 0.23 | 0.39 | 0.45 | 0.2 | 0.25 |
| 6 | 38747581 | rs1678661 | -0.131 | 0.649 | 0.145 | -1.341 | 0.02 | 0.154 | 0.154 | 0 | 0.49 | 0.75 | 0.63 | 0.81 | 0.5 | 0.5 |
| 6 | 38750888 | rs3823430 | 0.467 | 0.549 | 0.097 | 0.524 | 0.01 | -0.022 | -0.022 | 0.315 | 0.15 | 0.11 | 0.16 | 0.37 | 0.11 | 0.18 |
| 6 | 38754502 | rs1678666 | 0.186 | 0.648 | -0.117 | 0.394 | 0.001 | 0.024 | 0.024 | 0 | 0.13 | 0.16 | 0.2 | 0.31 | 0.13 | 0.25 |
| 6 | 38757378 | rs7756191 | 0.685 | 0.524 | -0.13 | 0.175 | 0.02 | -0.009 | -0.009 | 0.222 | 0.42 | 0.23 | 0.31 | 0.39 | 0.4 | 0.39 |
| 6 | 38762555 | rs1738241 | -0.819 | 0.394 | -0.166 | -0.019 | -0.014 | 0.043 | 0.043 | 0 | 0.37 | 0.31 | 0.59 | 0.84 | 0.39 | 0.57 |
| 6 | 38762865 | rs4714183 | 0.611 | 0.377 | -0.204 | -0.001 | 0.028 | -0.072 | -0.072 | 0.402 | 0.18 | 0.08 | 0.17 | 0.51 | 0.11 | 0.27 |
| 6 | 38763733 | rs1738240 | -0.947 | 0.381 | -0.002 | -0.114 | 0.005 | 0.021 | 0.021 | 0.293 | 0.6 | 0.64 | 0.75 | 0.84 | 0.58 | 0.67 |
| 6 | 38767450 | rs12211963 | 0.818 | 0.447 | -0.28 | 0.015 | -0.029 | 0.058 | 0.058 | 1.139 | 0.35 | 0.25 | 0.41 | 0.13 | 0.43 | 0.31 |
| 6 | 38773293 | rs874808 | -0.853 | 0.496 | -0.274 | 0.359 | -0.011 | 0.008 | 0.008 | 0 | 0.34 | 0.32 | 0.58 | 0.7 | 0.38 | 0.47 |
| 6 | 38775336 | rs1738229 | -0.457 | 0.49 | -0.263 | 0.335 | 0.018 | 0.03 | 0.03 | 0 | 0.21 | 0.27 | 0.44 | 0.43 | 0.27 | 0.35 |
| 6 | 38782401 | rs1738220 | -0.652 | 0.515 | -0.172 | 0.118 | -0.008 | 0.013 | 0.013 | 0 | 0.36 | 0.32 | 0.59 | 0.8 | 0.37 | 0.52 |
| 6 | 38786528 | rs12194119 | 1.658 | 0.657 | -0.28 | -0.078 | -0.019 | 0.038 | 0.038 | 0 | 0.18 | 0.07 | 0.12 | 0 | 0.21 | 0.11 |
| 6 | 38799284 | rs9380784 | 0.943 | 0.485 | 0.106 | 0.32 | -0.005 | 0.023 | 0.023 | 0.015 | 0.16 | 0.14 | 0.16 | 0.34 | 0.11 | 0.17 |
| 6 | 38799955 | rs982705 | -1.199 | 0.374 | 0.472 | 0.173 | 0.015 | 0.095 | 0.095 | 2.214 | 0.59 | 0.75 | 0.75 | 0.84 | 0.59 | 0.67 |

| | | | | | | | | | | | | | | | | |
|---|---|---|---|---|---|---|---|---|---|---|---|---|---|---|---|---|
| 6 | 38800164 | rs9357283 | 1.023 | 0.39 | 0.529 | 0.29 | -0.005 | 0.023 | 0.023 | 0.015 | 0.16 | 0.14 | 0.16 | 0.34 | 0.11 | 0.17 |
| 6 | 38800209 | rs1678729 | 0.145 | 0.438 | 0.718 | 0.365 | 0.006 | 0.112 | 0.112 | 0.039 | 0.15 | 0.43 | 0.22 | 0.29 | 0.13 | 0.24 |
| 6 | 38800901 | rs9296264 | -0.378 | 0.466 | 0.633 | 0.428 | -0.002 | -0.006 | -0.006 | 0 | 0.26 | 0.23 | 0.42 | 0.55 | 0.25 | 0.32 |
| 6 | 38813081 | rs9369084 | -0.243 | 0.382 | 0.595 | 0.386 | -0.003 | -0.008 | -0.008 | 0 | 0.25 | 0.22 | 0.41 | 0.4 | 0.26 | 0.23 |
| 6 | 38816015 | rs1738188 | -0.391 | 0.38 | 0.777 | 0.462 | 0.052 | 0.319 | 0.319 | 1.19 | 0.07 | 0.45 | 0.1 | 0.28 | 0.04 | 0.08 |
| 6 | 38816574 | rs1678744 | -0.184 | 0.348 | 0.841 | 0.342 | 0.043 | 0.034 | 0.034 | 3.55 | 0.06 | 0.43 | 0.05 | 0 | 0.01 | 0 |
| 6 | 38820451 | rs9369085 | -0.727 | 0.407 | 0.614 | 0.292 | -0.015 | 0.006 | 0.006 | 0.153 | 0.34 | 0.24 | 0.59 | 0.7 | 0.38 | 0.47 |
| 6 | 38825329 | rs2061907 | -0.755 | 0.393 | 0.9 | 0.283 | 0.003 | -0.003 | -0.003 | 0 | 0.36 | 0.36 | 0.6 | 0.69 | 0.38 | 0.47 |
| 6 | 38827245 | rs1033524 | 1.988 | 0.126 | 0.709 | -0.082 | -0.019 | 0.131 | 0.131 | 4.807 | 0.27 | 0.08 | 0.15 | 0.01 | 0.27 | 0.21 |
| 6 | 38830410 | rs9380791 | -0.676 | 0.162 | 0.686 | -0.003 | 0.002 | -0.012 | -0.012 | 0 | 0.36 | 0.36 | 0.61 | 0.81 | 0.38 | 0.51 |
| 6 | 38839315 | rs9394546 | 0.802 | 0.088 | 0.323 | 0.098 | -0.025 | -0.129 | -0.129 | 0 | 0.22 | 0.02 | 0.22 | 0.35 | 0.2 | 0.34 |
| 6 | 38843970 | rs862427 | -1.282 | 0.091 | 0.415 | 0.061 | 0.002 | -0.002 | -0.002 | 0 | 0.87 | 0.9 | 0.92 | 0.84 | 0.86 | 0.88 |
| 6 | 38848517 | rs9394548 | 1.111 | 0.045 | 0.369 | 0.64 | -0.042 | -0.002 | -0.002 | 0.387 | 0.28 | 0.06 | 0.19 | 0.2 | 0.3 | 0.18 |
| 6 | 38848786 | rs4711560 | -0.62 | -0.209 | 0.517 | 0.787 | 0.009 | 0.015 | 0.015 | 0.002 | 0.3 | 0.19 | 0.37 | 0.8 | 0.24 | 0.52 |
| 6 | 38851669 | rs4714192 | 0.369 | 0.002 | -0.033 | 0.898 | 0.008 | -0.024 | -0.024 | 0 | 0.23 | 0.13 | 0.23 | 0.35 | 0.2 | 0.34 |
| 6 | 38852719 | rs9380804 | -0.336 | 0.047 | -0.037 | 0.6 | 0.005 | -0.007 | -0.007 | 0 | 0.28 | 0.2 | 0.36 | 0.66 | 0.24 | 0.43 |
| 6 | 38860159 | rs862435 | -0.631 | 0.102 | -0.375 | 0.463 | 0.007 | 0.023 | 0.023 | 0 | 0.52 | 0.71 | 0.7 | 0.62 | 0.49 | 0.62 |
| 6 | 38860375 | rs9366986 | 1.24 | -0.115 | -0.671 | 0.726 | -0.022 | -0.073 | -0.073 | 0 | 0.12 | 0.01 | 0.11 | 0.23 | 0.12 | 0.03 |
| 6 | 38860589 | rs862436 | 0.329 | -0.776 | -0.516 | 0.885 | -0.002 | 0.002 | 0.002 | 0 | 0.72 | 0.77 | 0.67 | 0.41 | 0.79 | 0.53 |
| 6 | 38868700 | rs9349103 | 1.196 | -0.282 | -1.09 | 0.681 | -0.109 | -0.024 | -0.024 | 0 | 0.28 | 0.02 | 0.19 | 0.21 | 0.33 | 0.14 |
| 6 | 38872930 | rs862438 | -1.191 | -0.068 | -0.761 | 1.047 | 0 | 0.109 | 0.109 | 0.84 | 0.64 | 0.9 | 0.8 | 0.85 | 0.65 | 0.69 |
| 6 | 38877605 | rs4714198 | 0.391 | -0.166 | -0.754 | 0.907 | -0.032 | -0.046 | -0.046 | 0.44 | 0.41 | 0.76 | 0.41 | 0.48 | 0.44 | 0.27 |
| 6 | 38883938 | rs7357080 | -0.996 | -0.144 | -0.761 | 0.807 | -0.048 | 0.121 | 0.121 | 0.23 | 0.6 | 0.85 | 0.6 | 0.82 | 0.55 | 0.65 |
| 6 | 38884441 | rs6458085 | 0.239 | -0.118 | -0.711 | 0.875 | -0.025 | 0.016 | 0.016 | 0.29 | 0.42 | 0.75 | 0.43 | 0.63 | 0.44 | 0.36 |
| 6 | 38886122 | rs9462464 | 0.375 | -0.149 | -0.667 | 0.926 | 0.004 | -0.003 | -0.003 | 0 | 0.13 | 0.09 | 0.15 | 0.15 | 0.1 | 0.25 |
| 6 | 38891068 | rs4714199 | -0.73 | 0.024 | -0.308 | 1.084 | 0.103 | 0.267 | 0.267 | 0.531 | 0.08 | 0.54 | 0.16 | 0.28 | 0.1 | 0.08 |
| 6 | 38893153 | rs9394554 | 1.311 | -0.005 | -0.598 | 0.877 | -0.109 | 0.021 | 0.021 | 1.015 | 0.28 | 0.02 | 0.19 | 0.17 | 0.33 | 0.14 |
| 6 | 38901236 | rs6458087 | -0.939 | 0.035 | -0.404 | 0.937 | -0.003 | 0.11 | 0.11 | 0.143 | 0.62 | 0.86 | 0.63 | 0.82 | 0.62 | 0.66 |
| 6 | 38902470 | rs12662355 | -0.211 | -0.068 | -0.224 | 0.689 | 0.006 | 0.042 | 0.042 | 0 | 0.52 | 0.32 | 0.45 | 0.37 | 0.52 | 0.48 |
| 6 | 38904298 | rs9394555 | 0.419 | 0.127 | -0.161 | 0.758 | -0.035 | 0.049 | 0.049 | 0.016 | 0.43 | 0.16 | 0.38 | 0.33 | 0.44 | 0.38 |
| 6 | 38907291 | rs16891295 | -1.016 | 0.753 | 0.374 | 0.94 | 0.039 | -0.015 | -0.015 | 0 | 0.06 | 0.16 | 0.06 | 0.01 | 0.07 | 0.07 |
| 6 | 38908311 | rs1332011 | 0.564 | 1.052 | -0.279 | 1.271 | -0.023 | -0.011 | -0.011 | 0.133 | 0.08 | 0.04 | 0.07 | 0.29 | 0.16 | 0.17 |
| 6 | 38908515 | rs579786 | -1.348 | 0.486 | -0.784 | 0.505 | -0.011 | 0.146 | 0.146 | 0 | 0.75 | 0.92 | 0.85 | 1 | 0.76 | 0.84 |
| 6 | 38910363 | rs6919432 | 1.01 | 0.301 | -0.457 | 0.625 | -0.002 | 0.031 | 0.031 | 0 | 0.34 | 0.25 | 0.26 | 0.24 | 0.33 | 0.18 |
| 6 | 38910375 | rs6919590 | 0.987 | 0.28 | -0.467 | 0.788 | 0.002 | -0.016 | -0.016 | 0.019 | 0.36 | 0.25 | 0.27 | 0.4 | 0.33 | 0.32 |
| 6 | 38913177 | rs522327 | -0.373 | 0.19 | 0.01 | 0.339 | -0.004 | -0.011 | -0.011 | 0 | 0.58 | 0.57 | 0.69 | 0.72 | 0.61 | 0.61 |
| 6 | 38921240 | rs10947767 | -0.801 | 0.091 | 0.37 | 0.593 | -0.004 | 0.014 | 0.014 | 0 | 0.12 | 0.16 | 0.22 | 0.16 | 0.1 | 0.12 |
| 6 | 38926785 | rs9380811 | 1.185 | -0.105 | 0.199 | 0.315 | -0.087 | 0.018 | 0.018 | 0 | 0.37 | 0.11 | 0.36 | 0.67 | 0.5 | 0.5 |
| 6 | 38934922 | rs9380814 | 2.104 | -0.019 | 0.045 | 0.458 | -0.111 | 0.022 | 0.022 | 0 | 0.27 | 0.02 | 0.22 | 0.17 | 0.33 | 0.17 |
| 6 | 38936956 | rs9462468 | 0.834 | -0.052 | 0.262 | 0.333 | -0.004 | -0.01 | -0.01 | 0.118 | 0.13 | 0.2 | 0.12 | 0.09 | 0.11 | 0.13 |
| 6 | 38940282 | rs9296272 | -0.217 | -0.017 | 0.324 | 0.36 | 0.004 | 0.004 | 0.004 | 0 | 0.47 | 0.53 | 0.45 | 0.25 | 0.39 | 0.36 |
| 6 | 38940796 | rs4714203 | -0.62 | -0.006 | 0.341 | 0.369 | 0 | -0.007 | -0.007 | 0.209 | 0.87 | 0.84 | 0.83 | 0.92 | 0.89 | 0.87 |
| 6 | 38946459 | rs9470956 | 0.703 | 0.049 | 0.151 | 0.434 | 0.023 | 0.044 | 0.044 | 0 | 0.35 | 0.3 | 0.23 | 0.09 | 0.3 | 0.25 |
| 6 | 38947930 | rs9394560 | 0.902 | 0.047 | 0.2 | 0.456 | 0.002 | -0.006 | -0.006 | 0 | 0.41 | 0.39 | 0.39 | 0.67 | 0.5 | 0.5 |
| 6 | 38948307 | rs6915654 | 0.351 | 0.02 | 0.227 | 0.583 | -0.005 | -0.007 | -0.007 | 0 | 0.76 | 0.69 | 0.61 | 0.76 | 0.79 | 0.75 |
| 6 | 38951998 | rs1537232 | 1.195 | 0.185 | -0.479 | 0.817 | -0.014 | 0.021 | 0.021 | 0 | 0.08 | 0.05 | 0.12 | 0.35 | 0.17 | 0.2 |
| 6 | 38953292 | rs1412264 | 0.626 | 0.171 | -0.484 | 0.768 | -0.016 | 0.019 | 0.019 | 0 | 0.1 | 0.06 | 0.14 | 0.51 | 0.17 | 0.33 |
| 6 | 38954145 | rs17336909 | 1.008 | 0.101 | -0.441 | 0.594 | 0.002 | -0.008 | -0.008 | 0.712 | 0.64 | 0.43 | 0.46 | 0.25 | 0.62 | 0.43 |
| 6 | 38956892 | rs9380816 | 0.876 | 0.097 | -0.509 | 0.553 | -0.011 | 0.038 | 0.038 | 0 | 0.42 | 0.3 | 0.34 | 0.25 | 0.44 | 0.3 |
| 6 | 38960422 | rs1412266 | -0.072 | 0.041 | -0.464 | 0.654 | -0.005 | 0.008 | 0.008 | 0 | 0.43 | 0.29 | 0.48 | 0.61 | 0.44 | 0.52 |
| 6 | 38963559 | rs16891370 | 0.623 | 0.074 | -0.477 | 0.677 | -0.006 | 0.037 | 0.037 | 0 | 0.1 | 0.08 | 0.14 | 0.51 | 0.17 | 0.33 |
| 6 | 38964131 | rs9380818 | -0.952 | -0.008 | -0.418 | 0.621 | -0.003 | 0.017 | 0.017 | 0 | 0.85 | 0.6 | 0.82 | 0.86 | 0.88 | 0.83 |
| 6 | 38964657 | rs17337012 | 0.575 | -0.001 | -0.65 | 0.636 | 0.033 | 0.109 | 0.109 | 0 | 0.32 | 0.19 | 0.33 | 0.1 | 0.26 | 0.19 |
| 6 | 38965333 | rs1412269 | 0.346 | -0.01 | -0.689 | 0.609 | 0.012 | 0.011 | 0.011 | 0 | 0.71 | 0.32 | 0.7 | 0.77 | 0.77 | 0.69 |
| 6 | 38976575 | rs12210777 | -0.503 | 0.127 | -0.71 | 0.731 | -0.001 | 0.011 | 0.011 | 0 | 0.08 | 0.11 | 0.21 | 0.18 | 0.08 | 0.17 |
| 6 | 38977970 | rs10484852 | 1.649 | 0.087 | -0.752 | 0.68 | 0.033 | 0.072 | 0.072 | 0 | 0.22 | 0.12 | 0.13 | 0.08 | 0.18 | 0.12 |
| 6 | 38980081 | rs10484847 | 1.719 | 0.079 | -0.755 | 0.682 | 0.031 | 0.073 | 0.073 | 0 | 0.22 | 0.12 | 0.12 | 0 | 0.18 | 0.11 |
| 6 | 38982244 | rs1929901 | 1.652 | 0.1 | -0.745 | 0.696 | 0.033 | 0.072 | 0.072 | 0 | 0.22 | 0.12 | 0.13 | 0.08 | 0.18 | 0.12 |
| 6 | 38983698 | rs4714207 | 0.317 | 0.353 | -0.209 | 1.202 | -0.011 | -0.024 | -0.024 | 0 | 0.17 | 0.22 | 0.13 | 0.08 | 0.12 | 0.15 |
| 6 | 38988026 | rs2281345 | 1.247 | 0.389 | -0.904 | 1.259 | 0.046 | -0.032 | -0.032 | 0 | 0.57 | 0.2 | 0.48 | 0.58 | 0.68 | 0.51 |
| 6 | 38989385 | rs2235869 | 1.092 | 0.444 | -0.878 | 1.194 | 0.034 | 0.061 | 0.061 | 0 | 0.59 | 0.2 | 0.5 | 0.73 | 0.68 | 0.63 |
| 6 | 38994185 | rs2273094 | 0.74 | 0.719 | -0.838 | 1.344 | -0.015 | 0.015 | 0.015 | 0 | 0.08 | 0.05 | 0.13 | 0.41 | 0.17 | 0.22 |
| 6 | 38996609 | rs9470963 | -0.636 | 0.37 | -0.966 | 0.825 | -0.001 | -0.003 | -0.003 | 0 | 0.23 | 0.26 | 0.32 | 0.11 | 0.19 | 0.2 |
| 6 | 38996875 | rs1332010 | 0.45 | 0.944 | -0.32 | 1.414 | 0.029 | 0.095 | 0.095 | 0 | 0.76 | 0.52 | 0.7 | 0.51 | 0.71 | 0.62 |
| 6 | 39007540 | rs7738586 | -1.127 | 0.741 | -0.86 | 0.707 | 0.001 | -0.011 | -0.011 | 0 | 0.06 | 0.1 | 0.19 | 0.02 | 0.07 | 0.05 |
| 6 | 39015118 | rs9296274 | -0.872 | 0.6 | -0.598 | 1.035 | -0.013 | 0.147 | 0.147 | 0 | 0.14 | 0.26 | 0.17 | 0.51 | 0.12 | 0.38 |
| 6 | 39020542 | rs2268657 | -0.182 | 0.856 | -1.193 | 0.329 | -0.012 | 0.046 | 0.046 | 0 | 0.45 | 0.32 | 0.42 | 0.66 | 0.49 | 0.53 |
| 6 | 39023510 | rs3799707 | 0.054 | 1.236 | -1.718 | -0.436 | -0.049 | 0.145 | 0.145 | 0.176 | 0.71 | 0.91 | 0.74 | 0.94 | 0.68 | 0.82 |
| 6 | 39023630 | rs10305432 | 0.202 | 1 | -2.549 | -0.119 | -0.071 | -0.1 | -0.1 | 0 | 0.19 | 0.02 | 0.19 | 0.29 | 0.2 | 0.16 |

| | | | | | | | | | | | | | | | | |
|---|---|---|---|---|---|---|---|---|---|---|---|---|---|---|---|---|
| 6 | 39024716 | rs10305439 | 0.129 | 0.028 | -2.473 | -0.299 | -0.109 | -0.113 | -0.113 | 0.931 | 0.31 | 0.09 | 0.34 | 0.54 | 0.45 | 0.47 |
| 6 | 39024756 | rs2143734 | 0.422 | -0.854 | -1.888 | 0.008 | -0.007 | 0.011 | 0.011 | 0 | 0.39 | 0.45 | 0.33 | 0.49 | 0.31 | 0.38 |
| 6 | 39026703 | rs9283907 | -1.203 | -0.96 | -1.966 | -0.144 | 0.001 | 0.072 | 0.072 | 0.201 | 0.79 | 0.93 | 0.86 | 0.89 | 0.83 | 0.87 |
| 6 | 39030401 | rs2268650 | 0.709 | -0.828 | -2.03 | 0.195 | -0.148 | -0.069 | -0.069 | 0 | 0.27 | 0.03 | 0.27 | 0.25 | 0.41 | 0.29 |
| 6 | 39032415 | rs910170 | -0.572 | -1.057 | -1.663 | 0.033 | -0.002 | 0.003 | 0.003 | 0 | 0.46 | 0.55 | 0.59 | 0.61 | 0.55 | 0.59 |
| 6 | 39032537 | rs874900 | -0.383 | -0.96 | -1.157 | 0.077 | 0.051 | 0.124 | 0.124 | 0.012 | 0.06 | 0.28 | 0.06 | 0.13 | 0.05 | 0.11 |
| 6 | 39034072 | rs6923761 | 1.081 | -0.652 | -1.681 | -0.366 | -0.124 | 0.118 | 0.118 | 0 | 0.2 | 0.01 | 0.17 | 0.01 | 0.33 | 0.13 |
| 6 | 39035782 | rs7766663 | -0.421 | -1.026 | -1.538 | -0.12 | -0.022 | -0.016 | -0.016 | 0 | 0.38 | 0.3 | 0.52 | 0.45 | 0.49 | 0.48 |
| 6 | 39036185 | rs12214482 | -0.499 | -1.112 | -1.4 | -0.105 | -0.001 | -0.077 | -0.077 | 0 | 0.18 | 0.07 | 0.35 | 0.44 | 0.18 | 0.35 |
| 6 | 39036345 | rs7341356 | -0.504 | -1.087 | -1.413 | -0.052 | -0.019 | -0.014 | -0.014 | 0 | 0.41 | 0.31 | 0.53 | 0.45 | 0.51 | 0.48 |
| 6 | 39040654 | rs2235868 | -0.493 | -1.091 | -1.408 | -0.047 | -0.018 | -0.014 | -0.014 | 0 | 0.41 | 0.31 | 0.54 | 0.44 | 0.52 | 0.48 |
| 6 | 39041502 | rs1042044 | -0.945 | -1.103 | -1.496 | -0.021 | 0.002 | 0.001 | 0.001 | 0.068 | 0.47 | 0.59 | 0.59 | 0.55 | 0.56 | 0.6 |
| 6 | 39042334 | rs932443 | 1.059 | -1.067 | -1.34 | 0.078 | 0 | -0.001 | -0.001 | 0.106 | 0.35 | 0.34 | 0.3 | 0.36 | 0.32 | 0.3 |
| 6 | 39043796 | rs12204668 | 0.575 | -0.814 | -1.249 | -0.036 | -0.015 | 0.013 | 0.013 | 0 | 0.25 | 0.2 | 0.23 | 0.11 | 0.38 | 0.26 |
| 6 | 39043992 | rs2268645 | 1.282 | -0.586 | -1.141 | 0.454 | -0.015 | -0.013 | -0.013 | 0 | 0.06 | 0.02 | 0.09 | 0.2 | 0.1 | 0.15 |
| 6 | 39048757 | rs7769547 | -0.157 | -0.668 | -1.008 | 0.031 | -0.005 | -0.002 | -0.002 | 0 | 0.53 | 0.48 | 0.61 | 0.54 | 0.59 | 0.61 |
| 6 | 39049205 | rs2300613 | 0.772 | -0.966 | -1.001 | 0.163 | 0.005 | -0.007 | -0.007 | 0 | 0.31 | 0.24 | 0.26 | 0.35 | 0.28 | 0.3 |
| 6 | 39050384 | rs2268640 | -0.439 | -0.912 | -1.448 | -0.244 | -0.15 | 0.179 | 0.179 | 0 | 0.62 | 0.94 | 0.71 | 0.87 | 0.52 | 0.59 |
| 6 | 39051439 | rs2206942 | -0.066 | -0.656 | -1.096 | 0.058 | 0.008 | 0.094 | 0.094 | 0 | 0.44 | 0.79 | 0.5 | 0.75 | 0.41 | 0.45 |
| 6 | 39054113 | rs10305512 | -0.58 | -0.785 | -0.072 | -0.038 | -0.007 | -0.019 | -0.019 | 0 | 0.08 | 0.3 | 0.06 | 0.02 | 0.04 | 0.12 |
| 6 | 39055485 | rs4714210 | 0.398 | -0.704 | -0.805 | 0.344 | 0.057 | 0.069 | 0.069 | 1.775 | 0.45 | 0.2 | 0.28 | 0.29 | 0.36 | 0.28 |
| 6 | 39055516 | rs10305522 | 1.55 | -0.995 | -1.024 | 0.021 | 0.013 | 0.118 | 0.118 | 0 | 0.18 | 0 | 0.06 | 0 | 0.09 | 0.08 |
| 6 | 39059679 | rs4254984 | 0.674 | -0.738 | -0.801 | 0.218 | 0.04 | 0.062 | 0.062 | 1.744 | 0.41 | 0.19 | 0.27 | 0.28 | 0.34 | 0.26 |
| 6 | 39072034 | rs10023 | 0.254 | -1.223 | -0.1 | 0.199 | -0.038 | -0.034 | -0.034 | 0 | 0.32 | 0.52 | 0.28 | 0.17 | 0.21 | 0.24 |
| 6 | 39072681 | rs7696 | 0.7 | -0.805 | -0.015 | 0.455 | 0.017 | 0.024 | 0.024 | 0.459 | 0.45 | 0.35 | 0.31 | 0.29 | 0.37 | 0.31 |
| 6 | 39074535 | rs2746304 | -0.206 | -0.612 | -0.362 | 0.317 | -0.031 | 0.076 | 0.076 | 0 | 0.61 | 0.75 | 0.65 | 0.87 | 0.54 | 0.63 |
| 6 | 39075885 | rs7757449 | -0.364 | -0.437 | -0.898 | 0.455 | -0.087 | 0.02 | 0.02 | 0 | 0.62 | 0.9 | 0.54 | 0.73 | 0.49 | 0.64 |
| 6 | 39087713 | rs9470968 | 0.613 | -0.535 | -0.491 | 0.561 | 0.038 | 0.06 | 0.06 | 0.368 | 0.4 | 0.23 | 0.28 | 0.26 | 0.32 | 0.25 |
| 6 | 39092967 | rs12202197 | 0.743 | -0.4 | -0.538 | 0.333 | 0.044 | 0.111 | 0.111 | 1.481 | 0.42 | 0.22 | 0.28 | 0.2 | 0.31 | 0.28 |
| 6 | 39093133 | rs12195232 | 0.743 | -0.4 | -0.538 | 0.333 | 0.044 | 0.111 | 0.111 | 1.481 | 0.42 | 0.22 | 0.28 | 0.2 | 0.31 | 0.28 |
| 6 | 39093551 | rs2162840 | 1.299 | -0.081 | -0.547 | 1.317 | -0.017 | -0.013 | -0.013 | 0 | 0.06 | 0 | 0.05 | 0.29 | 0.07 | 0.16 |
| 6 | 39094032 | rs9470970 | -0.724 | 0.392 | -0.482 | 0.823 | 0.001 | 0.002 | 0.002 | 0 | 0.06 | 0.08 | 0.08 | 0.11 | 0.1 | 0.1 |
| 6 | 39098344 | rs6905394 | 0.287 | 0.637 | -0.239 | 0.491 | 0.008 | -0.02 | -0.02 | 0 | 0.6 | 0.52 | 0.53 | 0.31 | 0.48 | 0.61 |
| 6 | 39099299 | rs11756949 | -0.99 | 0.554 | -0.243 | 0.484 | 0.019 | 0.045 | 0.045 | 0 | 0.13 | 0.33 | 0.21 | 0.13 | 0.1 | 0.29 |
| 6 | 39105903 | rs9462479 | 0.029 | 0.591 | -0.562 | 0.522 | 0.002 | 0.002 | 0.002 | 0 | 0.05 | 0.08 | 0.03 | 0.05 | 0.06 | 0.06 |
| 6 | 39106207 | rs4714216 | 0.738 | 0.605 | -0.576 | 0.471 | 0.022 | 0.074 | 0.074 | 0.375 | 0.53 | 0.27 | 0.42 | 0.3 | 0.43 | 0.48 |
| 6 | 39106945 | rs228815 | -0.784 | 0.78 | -0.66 | 0.84 | -0.076 | -0.001 | -0.001 | 0 | 0.75 | 0.93 | 0.74 | 0.77 | 0.65 | 0.87 |
| 6 | 39110432 | rs228819 | 1.18 | 0.605 | -0.368 | 1.15 | -0.027 | 0.022 | 0.022 | 0 | 0.07 | 0.01 | 0.16 | 0.42 | 0.1 | 0.19 |
| 6 | 39110904 | rs7757873 | -0.375 | 0.49 | -0.204 | 0.85 | -0.001 | 0.06 | 0.06 | 0 | 0.07 | 0.15 | 0.11 | 0.12 | 0.06 | 0.15 |
| 6 | 39113224 | rs1366326 | 0.958 | 0.239 | -0.663 | 0.428 | 0.013 | 0.009 | 0.009 | 0 | 0.22 | 0.13 | 0.17 | 0.13 | 0.14 | 0.21 |
| 6 | 39113807 | rs6903109 | 0.347 | 0.309 | -0.542 | 0.441 | 0.004 | 0.003 | 0.003 | 0 | 0.62 | 0.38 | 0.62 | 0.41 | 0.69 | 0.51 |
| 6 | 39115413 | rs7751419 | 1.083 | 0.103 | -1.123 | 0.203 | 0.015 | 0.075 | 0.075 | 0.083 | 0.3 | 0.15 | 0.24 | 0.18 | 0.27 | 0.26 |
| 6 | 39119814 | rs2815056 | 1.275 | 0.134 | -1.216 | 0.344 | -0.034 | 0.006 | 0.006 | 0 | 0.07 | 0.01 | 0.17 | 0.41 | 0.11 | 0.21 |
| 6 | 39119837 | rs7773754 | 0.848 | 0.126 | -1.124 | 0.248 | -0.003 | 0.062 | 0.062 | 0 | 0.33 | 0.22 | 0.27 | 0.23 | 0.33 | 0.31 |
| 6 | 39124448 | rs1544935 | 1.214 | 0.767 | -1.541 | 0.083 | -0.026 | 0.009 | 0.009 | 0 | 0.18 | 0.08 | 0.21 | 0.17 | 0.23 | 0.13 |
| 6 | 39128424 | rs9470986 | 0.688 | 0.851 | -1.037 | -0.045 | -0.001 | 0.004 | 0.004 | 0 | 0.27 | 0.21 | 0.21 | 0.18 | 0.26 | 0.22 |
| 6 | 39132818 | rs10947785 | 0.85 | 0.907 | -1.116 | -0.102 | -0.052 | 0.138 | 0.138 | 0 | 0.43 | 0.11 | 0.42 | 0.29 | 0.46 | 0.37 |
| 6 | 39134168 | rs9394575 | 0.818 | 0.947 | -0.912 | 0.146 | 0.008 | -0.001 | -0.001 | 0 | 0.3 | 0.12 | 0.21 | 0.23 | 0.26 | 0.24 |
| 6 | 39136651 | rs12214118 | 0.869 | 1.227 | -0.808 | 0.416 | -0.021 | 0.007 | 0.007 | 0 | 0.18 | 0.1 | 0.21 | 0.16 | 0.22 | 0.13 |
| 6 | 39141239 | rs2561416 | 0.025 | 1.351 | -0.761 | 0.755 | -0.043 | -0.057 | -0.057 | 1.839 | 0.1 | 0.04 | 0.25 | 0.43 | 0.14 | 0.32 |
| 6 | 39149784 | rs9394577 | 0.674 | 1.077 | -0.333 | -0.559 | 0.001 | -0.008 | -0.008 | 1.131 | 0.22 | 0.24 | 0.21 | 0.16 | 0.23 | 0.16 |
| 6 | 39150098 | rs1428373 | 0.39 | 0.904 | -0.254 | -0.74 | -0.014 | 0.017 | 0.017 | 0 | 0.41 | 0.24 | 0.49 | 0.67 | 0.41 | 0.54 |
| 6 | 39151166 | rs2815093 | -0.658 | 0.868 | 0.073 | -0.626 | -0.015 | -0.083 | -0.083 | 0 | 0.7 | 0.59 | 0.8 | 0.94 | 0.75 | 0.88 |
| 6 | 39152540 | rs1978609 | -0.865 | 1.091 | 0.385 | 0.11 | 0.008 | 0.107 | 0.107 | 0 | 0.12 | 0.21 | 0.2 | 0.53 | 0.14 | 0.29 |
| 6 | 39156108 | rs2815095 | -0.83 | 0.837 | 0.174 | -0.625 | -0.006 | -0.051 | -0.051 | 0 | 0.73 | 0.66 | 0.77 | 0.89 | 0.75 | 0.75 |
| 6 | 39157781 | rs6912288 | 0.263 | 0.645 | -0.414 | -0.486 | -0.006 | 0.121 | 0.121 | 0 | 0.52 | 0.24 | 0.48 | 0.26 | 0.53 | 0.28 |
| 6 | 39158339 | rs4714222 | 0.289 | 0.718 | -0.388 | -0.457 | -0.004 | 0.125 | 0.125 | 0 | 0.52 | 0.24 | 0.48 | 0.26 | 0.53 | 0.28 |
| 6 | 39164121 | rs751933 | 0.136 | 0.725 | -0.271 | -0.615 | -0.001 | 0.086 | 0.086 | 0 | 0.54 | 0.33 | 0.55 | 0.76 | 0.53 | 0.39 |
| 6 | 39165631 | rs2815100 | 0.908 | 0.712 | -0.553 | -0.783 | 0.013 | -0.036 | -0.036 | 0 | 0.31 | 0.15 | 0.26 | 0.55 | 0.25 | 0.2 |
| 6 | 39165965 | rs2561430 | 0.371 | 0.574 | -0.195 | -0.914 | -0.013 | -0.035 | -0.035 | 0 | 0.93 | 0.84 | 0.9 | 0.98 | 0.95 | 0.89 |
| 6 | 39166496 | rs12207689 | 0.805 | 0.713 | -0.334 | -0.91 | -0.011 | 0.01 | 0.01 | 0 | 0.22 | 0.14 | 0.27 | 0.2 | 0.27 | 0.16 |
| 6 | 39169414 | rs2815106 | 0.951 | 0.725 | -0.557 | -0.896 | 0.008 | -0.022 | -0.022 | 0.797 | 0.31 | 0.22 | 0.27 | 0.56 | 0.26 | 0.2 |
| 6 | 39174525 | rs10947787 | 0.799 | 0.527 | -0.678 | -0.809 | -0.025 | 0.01 | 0.01 | 0 | 0.22 | 0.08 | 0.26 | 0.2 | 0.26 | 0.16 |
| 6 | 39177574 | rs9462492 | 0.634 | 0.996 | -0.897 | -0.698 | -0.043 | -0.025 | -0.025 | 0 | 0.08 | 0 | 0.09 | 0.09 | 0.13 | 0.2 |
| 6 | 39177853 | rs2815113 | 0.7 | 1.038 | -0.847 | -0.582 | -0.007 | -0.029 | -0.029 | 0 | 0.32 | 0.23 | 0.3 | 0.52 | 0.3 | 0.22 |
| 6 | 39183470 | rs10456100 | 0.953 | 1.065 | -1.398 | -0.645 | -0.055 | 0.02 | 0.02 | 0 | 0.26 | 0.02 | 0.22 | 0.17 | 0.26 | 0.3 |
| 6 | 39185605 | rs2815114 | -0.551 | 1.118 | -0.427 | 0.043 | -0.004 | 0.009 | 0.009 | 0 | 0.44 | 0.37 | 0.39 | 0.61 | 0.43 | 0.43 |
| 6 | 39187168 | rs2815115 | 0.07 | 1.169 | -1.319 | 0.32 | -0.028 | -0.069 | -0.069 | 0 | 0.31 | 0.12 | 0.33 | 0.54 | 0.29 | 0.22 |
| 6 | 39191384 | rs2561401 | 0.982 | 1.002 | -0.592 | 0.378 | -0.001 | -0.002 | -0.002 | 0 | 0.78 | 0.22 | 0.72 | 0.75 | 0.79 | 0.71 |

| | | | | | | | | | | | | | | | | |
|---|---|---|---|---|---|---|---|---|---|---|---|---|---|---|---|---|
| 6 | 39192366 | rs2815122 | 0.29 | 0.893 | -0.335 | -0.411 | 0.006 | -0.049 | -0.049 | 0 | 0.58 | 0.53 | 0.5 | 0.87 | 0.51 | 0.58 |
| 6 | 39192962 | rs2815124 | 0.148 | 1.287 | -0.297 | -0.553 | -0.028 | 0.026 | 0.026 | 0.04 | 0.72 | 0.77 | 0.73 | 0.93 | 0.65 | 0.67 |
| 6 | 39195929 | rs6458102 | -0.992 | 0.852 | 0.448 | 0.927 | 0.02 | 0.197 | 0.197 | 0.327 | 0.14 | 0.38 | 0.16 | 0.36 | 0.12 | 0.24 |
| 6 | 39196662 | rs2241595 | -0.452 | 0.739 | 0.339 | 0.876 | 0.001 | 0.142 | 0.142 | 0 | 0.12 | 0.29 | 0.11 | 0.31 | 0.1 | 0.2 |
| 6 | 39202157 | rs9366999 | -0.49 | 0.481 | -0.113 | 0.001 | 0.001 | 0.013 | 0.013 | 0 | 0.33 | 0.21 | 0.4 | 0.72 | 0.32 | 0.33 |
| 6 | 39210118 | rs2115203 | 0.563 | 0.449 | -0.229 | -0.308 | 0.014 | 0.044 | 0.044 | 0 | 0.62 | 0.28 | 0.55 | 0.26 | 0.7 | 0.65 |
| 6 | 39219300 | rs1651106 | -0.78 | 0.363 | -0.357 | -0.369 | 0.029 | 0.093 | 0.093 | 0.27 | 0.39 | 0.85 | 0.39 | 0.74 | 0.3 | 0.34 |
| 6 | 39221586 | rs1744413 | -0.658 | 0.37 | -0.183 | -0.366 | 0.007 | 0.016 | 0.016 | 0.026 | 0.38 | 0.69 | 0.39 | 0.74 | 0.3 | 0.35 |
| 6 | 39223000 | rs1651114 | -0.785 | 0.514 | -1.034 | -0.314 | 0.017 | 0.142 | 0.142 | 0.761 | 0.43 | 0.91 | 0.53 | 0.76 | 0.32 | 0.42 |
| 6 | 39226854 | rs1179384 | -0.021 | 1.053 | -1.234 | -0.821 | -0.056 | 0.103 | 0.103 | 0 | 0.7 | 0.9 | 0.75 | 0.9 | 0.65 | 0.69 |
| 6 | 39227301 | rs9471032 | 0.701 | 0.231 | -1.503 | -1.28 | 0.003 | -0.01 | -0.01 | 0 | 0.18 | 0.17 | 0.13 | 0.04 | 0.12 | 0.1 |
| 6 | 39227792 | rs9380843 | 0.49 | 0.75 | -1.496 | -0.615 | -0.006 | 0.002 | 0.002 | 0 | 0.23 | 0.08 | 0.21 | 0.15 | 0.28 | 0.25 |
| 6 | 39228810 | rs1179380 | -0.184 | 1.204 | -1.706 | -0.686 | -0.063 | 0.039 | 0.039 | 0 | 0.81 | 0.95 | 0.85 | 0.85 | 0.79 | 0.68 |
| 6 | 39229521 | rs1182865 | -0.132 | 1.305 | -1.704 | -0.689 | -0.057 | 0.039 | 0.039 | 0 | 0.81 | 0.95 | 0.8 | 0.83 | 0.8 | 0.64 |
| 6 | 39230307 | rs1182864 | 0.278 | 1.216 | -2.321 | -0.122 | -0.083 | -0.152 | -0.152 | 0.017 | 0.27 | 0.02 | 0.42 | 0.48 | 0.36 | 0.41 |
| 6 | 39231013 | rs11961538 | 0.975 | 0.675 | -2.187 | -0.674 | -0.008 | 0.108 | 0.108 | 0.04 | 0.31 | 0.16 | 0.23 | 0 | 0.32 | 0.1 |
| 6 | 39239265 | rs2815022 | 0.262 | 0.777 | -1.678 | -0.473 | 0.005 | 0.025 | 0.025 | 0 | 0.51 | 0.31 | 0.45 | 0.36 | 0.51 | 0.38 |
| 6 | 39242892 | rs2395718 | 1.36 | 0.62 | -0.939 | -0.489 | -0.008 | 0.02 | 0.02 | 0 | 0.17 | 0.2 | 0.09 | 0.35 | 0.1 | 0.13 |
| 6 | 39243544 | rs2815040 | -0.075 | 0.788 | -0.384 | -0.571 | -0.001 | 0.002 | 0.002 | 0 | 0.63 | 0.42 | 0.74 | 0.49 | 0.7 | 0.56 |
| 6 | 39244185 | rs2815046 | -1.32 | 0.778 | -0.557 | -0.61 | -0.085 | 0.014 | 0.014 | 0 | 0.83 | 0.99 | 0.86 | 0.85 | 0.8 | 0.69 |
| 6 | 39244679 | rs12202455 | 0.096 | 0.74 | -0.438 | -0.68 | -0.008 | 0.004 | 0.004 | 0 | 0.15 | 0.09 | 0.24 | 0.12 | 0.19 | 0.14 |
| 6 | 39246856 | rs2758873 | 1.641 | 0.736 | -0.529 | -0.766 | 0 | 0.104 | 0.104 | 0.047 | 0.3 | 0.14 | 0.21 | 0 | 0.29 | 0.1 |
| 6 | 39247346 | rs734408 | 0.899 | 0.84 | -0.676 | -0.683 | -0.023 | -0.02 | -0.02 | 0.303 | 0.11 | 0.03 | 0.22 | 0.12 | 0.16 | 0.14 |
| 6 | 39249885 | rs2815048 | 0.443 | 0.838 | -0.232 | -0.682 | 0.001 | -0.004 | -0.004 | 0 | 0.44 | 0.44 | 0.45 | 0.12 | 0.45 | 0.24 |
| 6 | 39251648 | rs6940480 | 0.1 | 1.355 | 0.203 | 0.063 | 0.007 | 0.075 | 0.075 | 0 | 0.22 | 0.33 | 0.32 | 0.37 | 0.26 | 0.32 |
| 6 | 39261244 | rs1005995 | -0.638 | 1.244 | -0.049 | 0.027 | -0.002 | 0.024 | 0.024 | 0 | 0.33 | 0.42 | 0.47 | 0.47 | 0.38 | 0.42 |
| 6 | 39266636 | rs2815067 | -0.788 | 1.426 | -0.969 | -0.074 | -0.081 | 0.022 | 0.022 | 0 | 0.83 | 0.99 | 0.86 | 0.85 | 0.79 | 0.7 |
| 6 | 39273607 | rs10807205 | 0.376 | 1.63 | -0.93 | 0.366 | -0.079 | -0.124 | -0.124 | 0 | 0.19 | 0.02 | 0.33 | 0.39 | 0.25 | 0.31 |
| 6 | 39275282 | rs2758912 | -0.887 | 1.301 | -0.797 | 0.165 | -0.039 | -0.057 | -0.057 | 0 | 0.41 | 0.8 | 0.64 | 0.47 | 0.48 | 0.47 |
| 6 | 39279131 | rs1544050 | -0.101 | 1.111 | 0.084 | 0.729 | 0.061 | 0.043 | 0.043 | 0 | 0.75 | 0.61 | 0.56 | 0.6 | 0.58 | 0.74 |
| 6 | 39281456 | rs34179517 | 0.061 | 0.162 | -0.37 | 0.409 | 0.013 | 0.023 | 0.023 | 0 | 0.18 | 0.15 | 0.12 | 0.12 | 0.12 | 0.33 |
| 6 | 39282806 | rs11756091 | -0.601 | 0.312 | -0.541 | 0.655 | -0.139 | -0.106 | -0.106 | 0 | 0.68 | 0.89 | 0.46 | 0.47 | 0.49 | 0.48 |
| 6 | 39284184 | rs3734618 | -0.661 | 0.347 | -0.646 | 0.665 | -0.182 | -0.144 | -0.144 | 0 | 0.68 | 0.95 | 0.47 | 0.47 | 0.49 | 0.48 |
| 6 | 39289875 | rs1109798 | 0.398 | 0.27 | 0.483 | 0.749 | 0 | 0 | 0 | 0.866 | 0.47 | 0.46 | 0.4 | 0.37 | 0.36 | 0.21 |
| 6 | 39290200 | rs3734619 | 0.375 | 0.272 | 0.333 | 0.75 | 0.02 | 0.018 | 0.018 | 0.184 | 0.47 | 0.54 | 0.4 | 0.37 | 0.36 | 0.21 |
| 6 | 39291807 | rs1328384 | 0.091 | 0.244 | 0.234 | 0.785 | 0.01 | 0.011 | 0.011 | 0.959 | 0.5 | 0.54 | 0.49 | 0.63 | 0.38 | 0.29 |
| 6 | 39292084 | rs6458107 | 0.088 | -0.13 | 0.185 | 0.523 | -0.008 | -0.015 | -0.015 | 0 | 0.24 | 0.53 | 0.34 | 0.22 | 0.18 | 0.13 |
| 6 | 39292315 | rs7766608 | 0.449 | -0.18 | 0.188 | -0.07 | -0.008 | -0.131 | -0.131 | 0.916 | 0.21 | 0.34 | 0.26 | 0.01 | 0.17 | 0.05 |
| 6 | 39293898 | rs6458110 | -0.019 | -0.316 | -0.56 | -0.053 | -0.006 | -0.033 | -0.033 | 0 | 0.42 | 0.53 | 0.34 | 0.16 | 0.29 | 0.33 |
| 6 | 39294529 | rs16891930 | -0.025 | 0.336 | -0.915 | 0.835 | -0.018 | -0.061 | -0.061 | 2.836 | 0.22 | 0.12 | 0.22 | 0.63 | 0.21 | 0.23 |
| 6 | 39298450 | rs16891937 | 0.228 | -0.152 | -1.113 | 0.549 | 0.03 | -0.015 | -0.015 | 0 | 0.21 | 0.07 | 0.09 | 0.22 | 0.14 | 0.23 |
| 6 | 39301918 | rs728217 | -1.367 | -0.294 | -1.393 | 0.497 | -0.043 | 0.056 | 0.056 | 0 | 0.53 | 0.87 | 0.4 | 0.69 | 0.46 | 0.53 |
| 6 | 39302614 | rs9380848 | -0.006 | -0.405 | -1.115 | 0.506 | -0.02 | -0.051 | -0.051 | 0 | 0.24 | 0.13 | 0.25 | 0.48 | 0.25 | 0.28 |
| 6 | 39302832 | rs1887716 | -0.921 | -0.452 | -0.209 | 0.473 | 0.005 | 0.024 | 0.024 | 0 | 0.47 | 0.39 | 0.35 | 0.69 | 0.42 | 0.52 |
| 6 | 39303594 | rs9462531 | -0.719 | -0.451 | -0.143 | 0.472 | 0.015 | 0.042 | 0.042 | 0 | 0.45 | 0.24 | 0.33 | 0.68 | 0.37 | 0.5 |
| 6 | 39303870 | rs3823213 | 0.077 | -0.3 | -0.026 | 0.533 | -0.036 | -0.08 | -0.08 | 1.695 | 0.23 | 0.08 | 0.24 | 0.48 | 0.25 | 0.28 |
| 6 | 39304211 | rs3734621 | -0.64 | -0.296 | 0.157 | 0.532 | 0.074 | 0.038 | 0.038 | 2.526 | 0.06 | 0.46 | 0.05 | 0 | 0.04 | 0.01 |
| 6 | 39306063 | rs6901022 | -0.252 | -0.259 | 0.052 | 0.568 | 0.006 | 0.018 | 0.018 | 0 | 0.25 | 0.27 | 0.27 | 0.49 | 0.3 | 0.3 |
| 6 | 39308031 | rs16891961 | -0.941 | -0.58 | -0.042 | 0.584 | 0 | -0.007 | -0.007 | 0 | 0.29 | 0.7 | 0.25 | 0.46 | 0.27 | 0.29 |
| 6 | 39308207 | rs2281686 | 0.082 | -0.914 | 0.021 | 0.612 | 0.016 | -0.003 | -0.003 | 0.184 | 0.23 | 0.18 | 0.1 | 0.23 | 0.14 | 0.19 |
| 6 | 39308539 | rs9471076 | 0.272 | -0.921 | 0.146 | 0.622 | 0.017 | 0.003 | 0.003 | 0 | 0.22 | 0.18 | 0.09 | 0.2 | 0.12 | 0.19 |
| 6 | 39309875 | rs11753737 | -0.679 | -0.89 | 0.064 | 0.486 | -0.006 | -0.003 | -0.003 | 0 | 0.31 | 0.67 | 0.18 | 0.21 | 0.21 | 0.24 |
| 6 | 39310635 | rs9471078 | 0.623 | -0.786 | -0.286 | 0.625 | -0.004 | -0.047 | -0.047 | 0.59 | 0.09 | 0.01 | 0.05 | 0.19 | 0.09 | 0.18 |
| 6 | 39314976 | rs11755763 | -1.315 | -0.684 | -0.649 | 0.084 | -0.112 | -0.073 | -0.073 | 0.785 | 0.64 | 0.94 | 0.57 | 0.66 | 0.55 | 0.54 |
| 6 | 39319630 | rs9969033 | -0.083 | -0.989 | 0.82 | 0.19 | -0.015 | -0.047 | -0.047 | 0.191 | 0.24 | 0.61 | 0.16 | 0.13 | 0.19 | 0.14 |
| 6 | 39320088 | rs9471080 | 0.026 | -0.99 | 0.247 | 0.284 | -0.018 | -0.106 | -0.106 | 0 | 0.19 | 0.05 | 0.14 | 0.41 | 0.18 | 0.32 |
| 6 | 39324581 | rs6924090 | -1.018 | -0.948 | 0.405 | 0.278 | 0.024 | -0.034 | -0.034 | 2.341 | 0.43 | 0.82 | 0.33 | 0.52 | 0.36 | 0.46 |
| 6 | 39324923 | rs20456 | -1.391 | -0.985 | 0.207 | 0.226 | 0 | -0.044 | -0.044 | 0 | 0.5 | 0.91 | 0.36 | 0.53 | 0.42 | 0.5 |
| 6 | 39325078 | rs20455 | -1.049 | -0.899 | 0.279 | 0.287 | 0.021 | -0.05 | -0.05 | 0 | 0.44 | 0.85 | 0.33 | 0.52 | 0.36 | 0.46 |
| 6 | 39325549 | rs7765442 | 0.487 | -0.732 | 0.516 | 0.155 | -0.006 | -0.025 | -0.025 | 0 | 0.19 | 0.23 | 0.11 | 0.1 | 0.14 | 0.12 |
| 6 | 39329920 | rs7753481 | -0.937 | -0.669 | 0.259 | 0.151 | 0.016 | 0.025 | 0.025 | 0 | 0.47 | 0.72 | 0.35 | 0.48 | 0.41 | 0.49 |
| 6 | 39331508 | rs9394585 | -0.237 | -0.939 | 0.573 | -0.09 | 0.017 | 0 | 0 | 0 | 0.57 | 0.69 | 0.69 | 0.63 | 0.68 | 0.47 |
| 6 | 39333337 | rs9357303 | 1.029 | -0.8 | 0.332 | -0.108 | -0.108 | -0.042 | -0.042 | 0 | 0.32 | 0.08 | 0.49 | 0.37 | 0.45 | 0.23 |
| 6 | 39339013 | rs10046447 | 0.233 | -0.902 | 0.912 | -0.123 | 0.007 | -0.059 | -0.059 | 2.989 | 0.11 | 0.28 | 0.06 | 0.02 | 0.1 | 0.17 |
| 6 | 39339561 | rs9296294 | -0.338 | -0.974 | 0.645 | -0.194 | 0.02 | 0.026 | 0.026 | 0 | 0.53 | 0.65 | 0.63 | 0.4 | 0.66 | 0.44 |
| 6 | 39345527 | rs4714247 | 0.593 | -0.798 | 0.026 | -0.079 | -0.1 | -0.003 | -0.003 | 0 | 0.38 | 0.11 | 0.52 | 0.37 | 0.51 | 0.26 |
| 6 | 39348691 | rs6899860 | -0.395 | -0.907 | 0.287 | -0.097 | 0.019 | -0.004 | -0.004 | 0 | 0.56 | 0.68 | 0.68 | 0.61 | 0.66 | 0.5 |
| 6 | 39348801 | rs9394587 | 0.653 | -0.538 | -0.053 | 0.1 | -0.111 | -0.09 | -0.09 | 1.702 | 0.37 | 0.12 | 0.56 | 0.57 | 0.47 | 0.3 |
| 6 | 39349365 | rs716952 | -1.956 | -0.357 | 1.073 | 0.672 | 0.019 | 0.087 | 0.087 | 0.756 | 0.12 | 0.33 | 0.15 | 0.23 | 0.15 | 0.07 |

| | | | | | | | | | | | | | | | | |
|---|---|---|---|---|---|---|---|---|---|---|---|---|---|---|---|---|
| 6 | 39349530 | rs721755 | -0.769 | -0.307 | 0.276 | 0.696 | -0.063 | -0.145 | -0.145 | 0 | 0.72 | 0.99 | 0.78 | 0.69 | 0.75 | 0.65 |
| 6 | 39353110 | rs7774367 | -2.166 | -0.317 | 2.056 | 0.718 | 0.079 | 0.286 | 0.286 | 2.977 | 0.06 | 0.36 | 0.1 | 0.22 | 0.06 | 0.03 |
| 6 | 39356438 | rs12175497 | -2.693 | -0.453 | 2.265 | 0.66 | 0.043 | 0.183 | 0.183 | 1.617 | 0.11 | 0.45 | 0.1 | 0.22 | 0.08 | 0.06 |
| 6 | 39357257 | rs9380854 | -2.249 | -0.428 | 2.414 | 0.663 | 0.045 | 0.21 | 0.21 | 0.418 | 0.06 | 0.25 | 0.07 | 0.23 | 0.1 | 0.05 |
| 6 | 39359065 | rs9380857 | -2.003 | -0.434 | 2.434 | 0.67 | 0.024 | 0.2 | 0.2 | 1.28 | 0.06 | 0.23 | 0.05 | 0.22 | 0.05 | 0.03 |
| 6 | 39361854 | rs9296295 | -1.539 | -0.357 | 2.53 | 0.458 | 0.043 | -0.015 | -0.015 | 0.006 | 0.06 | 0.23 | 0.07 | 0.01 | 0.08 | 0.04 |
| 6 | 39364028 | rs1887714 | -2.068 | -0.714 | 2.449 | 0.29 | -0.025 | -0.007 | -0.007 | 0.428 | 0.23 | 0.62 | 0.18 | 0.21 | 0.17 | 0.1 |
| 6 | 39379854 | rs1328388 | 2.979 | -0.257 | 3.059 | 0.618 | 0.029 | 0.134 | 0.134 | 0 | 0.86 | 0.53 | 0.8 | 0.77 | 0.85 | 0.92 |
| 6 | 39383115 | rs1328389 | 2.483 | -0.219 | 3.015 | 0.629 | 0.031 | 0.106 | 0.106 | 0 | 0.88 | 0.7 | 0.82 | 0.77 | 0.85 | 0.92 |
| 6 | 39384111 | rs7759557 | -2.45 | -0.195 | 2.981 | 0.642 | -0.009 | 0.169 | 0.169 | 0 | 0.12 | 0.4 | 0.15 | 0.22 | 0.06 | 0.08 |
| 6 | 39387344 | rs9462543 | 2.678 | -0.087 | 3.231 | 0.701 | 0.064 | 0.192 | 0.192 | 0 | 0.88 | 0.55 | 0.83 | 0.78 | 0.9 | 0.93 |
| 6 | 39388887 | rs4714251 | -1.416 | -0.523 | 2.779 | 0.758 | 0.004 | 0.13 | 0.13 | 0 | 0.07 | 0.17 | 0.12 | 0.22 | 0.04 | 0.06 |
| 6 | 39389992 | rs9394591 | -2.184 | -0.517 | 2.055 | 0.755 | -0.004 | 0.136 | 0.136 | 0 | 0.12 | 0.58 | 0.17 | 0.23 | 0.1 | 0.08 |
| 6 | 39396537 | rs6458115 | -1.955 | -0.573 | 1.555 | 0.81 | -0.028 | 0.118 | 0.118 | 1.537 | 0.11 | 0.61 | 0.16 | 0.23 | 0.08 | 0.05 |
| 6 | 39399454 | rs10484825 | -0.72 | -0.697 | 1.19 | 0.201 | -0.004 | -0.033 | -0.033 | 0 | 0.07 | 0.16 | 0.05 | 0 | 0.06 | 0.02 |
| 6 | 39399659 | rs6458116 | -0.44 | -1.423 | 1.463 | -0.578 | -0.033 | -0.095 | -0.095 | 0.173 | 0.21 | 0.3 | 0.11 | 0.05 | 0.12 | 0.28 |
| 6 | 39400877 | rs1952362 | -1.203 | -1.496 | 1.321 | -0.136 | -0.022 | 0 | 0 | 0 | 0.29 | 0.31 | 0.2 | 0.28 | 0.16 | 0.38 |
| 6 | 39401745 | rs1927774 | -1.561 | -1.497 | 1.596 | -0.208 | -0.007 | -0.002 | -0.002 | 0.414 | 0.33 | 0.63 | 0.25 | 0.27 | 0.19 | 0.4 |
| 6 | 39407545 | rs9380863 | -1.418 | -1.497 | 1.626 | -0.223 | -0.015 | -0.004 | -0.004 | 0 | 0.32 | 0.62 | 0.24 | 0.27 | 0.19 | 0.4 |
| 6 | 39407831 | rs9380864 | -0.234 | -1.476 | 1.556 | -0.206 | -0.01 | -0.017 | -0.017 | 0 | 0.06 | 0.02 | 0.09 | 0.23 | 0.04 | 0.1 |
| 6 | 39412770 | rs9296298 | -2.141 | -1.751 | 1.293 | -0.635 | 0.092 | 0.125 | 0.125 | 0 | 0.5 | 0.89 | 0.35 | 0.29 | 0.33 | 0.43 |
| 6 | 39413634 | rs9471105 | -0.616 | -1.8 | 1.375 | -0.678 | -0.03 | -0.09 | -0.09 | 0.176 | 0.26 | 0.61 | 0.17 | 0.04 | 0.16 | 0.3 |
| 6 | 39430873 | rs9471112 | -1.131 | -1.679 | 1.59 | -0.515 | -0.039 | -0.127 | -0.127 | 0 | 0.21 | 0.45 | 0.11 | 0.02 | 0.14 | 0.03 |
| 6 | 39431229 | rs9471113 | -2.165 | -1.71 | 1.646 | -0.419 | -0.024 | -0.003 | -0.003 | 0.235 | 0.3 | 0.62 | 0.25 | 0.25 | 0.2 | 0.15 |
| 6 | 39451069 | rs9394599 | -0.466 | -1.22 | 1.135 | -0.292 | -0.001 | 0.058 | 0.058 | 0 | 0.07 | 0.14 | 0.11 | 0.24 | 0.04 | 0.11 |
| 6 | 39464712 | rs2073134 | -2.077 | -1.558 | 0.909 | -0.589 | 0.107 | 0.105 | 0.105 | 0 | 0.49 | 0.9 | 0.36 | 0.29 | 0.32 | 0.45 |
| 6 | 39464913 | rs875730 | -0.739 | -1.629 | 1.22 | -0.631 | -0.004 | -0.028 | -0.028 | 1.562 | 0.18 | 0.19 | 0.08 | 0.02 | 0.1 | 0.03 |
| 6 | 39474180 | rs9357307 | -1.733 | -1.669 | 1.593 | -0.64 | -0.025 | 0.003 | 0.003 | 0 | 0.27 | 0.32 | 0.13 | 0.25 | 0.16 | 0.14 |
| 6 | 39486517 | rs9380874 | -2.158 | -1.778 | 1.464 | -0.74 | 0.102 | 0.104 | 0.104 | 0 | 0.48 | 0.82 | 0.35 | 0.29 | 0.31 | 0.45 |
| 6 | 39498760 | rs13199465 | 0.356 | -1.733 | 1.491 | -0.7 | 0.012 | 0.069 | 0.069 | 0 | 0.12 | 0.01 | 0.15 | 0.01 | 0.09 | 0.21 |
| 6 | 39500579 | rs6901155 | -1.657 | -1.733 | 1.663 | -0.683 | -0.012 | 0 | 0 | 0 | 0.28 | 0.63 | 0.26 | 0.27 | 0.19 | 0.41 |
| 6 | 39507889 | rs2273063 | 0.184 | -1.655 | 1.551 | -0.639 | 0.004 | 0.024 | 0.024 | 0 | 0.06 | 0.03 | 0.02 | 0.23 | 0.03 | 0.1 |
| 6 | 39523281 | rs7751353 | -0.65 | -1.702 | 2.01 | -0.349 | -0.002 | -0.01 | -0.01 | 0 | 0.19 | 0.24 | 0.2 | 0.09 | 0.16 | 0.33 |
| 6 | 39526867 | rs1520 | -2.205 | -2.012 | 1.717 | -0.641 | 0.009 | 0.005 | 0.005 | 0 | 0.53 | 0.65 | 0.36 | 0.35 | 0.36 | 0.56 |
| 6 | 39530118 | rs4714258 | -0.634 | -1.977 | 1.689 | -0.725 | -0.013 | -0.049 | -0.049 | 0 | 0.22 | 0.44 | 0.23 | 0.09 | 0.16 | 0.33 |
| 6 | 39530428 | rs10484824 | -0.625 | -1.861 | 0.872 | -0.682 | 0.032 | 0.056 | 0.056 | 0 | 0.24 | 0.1 | 0.11 | 0.01 | 0.15 | 0.12 |
| 6 | 39542154 | rs6911091 | -1.935 | -1.812 | 0.854 | -0.564 | 0.02 | -0.004 | -0.004 | 0 | 0.56 | 0.86 | 0.39 | 0.35 | 0.38 | 0.55 |
| 6 | 39543708 | rs7764246 | -0.607 | -1.964 | 0.922 | -0.713 | 0.044 | 0.097 | 0.097 | 0 | 0.24 | 0.14 | 0.21 | 0.01 | 0.15 | 0.25 |
| 6 | 39543786 | rs11754132 | -0.499 | -1.958 | 0.89 | -0.73 | 0 | -0.001 | -0.001 | 0 | 0.07 | 0.05 | 0.05 | 0.1 | 0.08 | 0.17 |
| 6 | 39544831 | rs7769800 | -1.241 | -1.992 | 1.229 | -0.746 | 0.004 | 0.006 | 0.006 | 0 | 0.32 | 0.31 | 0.27 | 0.1 | 0.23 | 0.41 |
| 6 | 39552398 | rs725841 | -1.954 | -1.983 | 1.747 | -0.633 | 0.019 | 0.003 | 0.003 | 0 | 0.43 | 0.69 | 0.33 | 0.37 | 0.29 | 0.52 |
| 6 | 39554294 | rs9394616 | 0.25 | -1.948 | 1.712 | -0.587 | -0.006 | 0.086 | 0.086 | 0 | 0.06 | 0.12 | 0.03 | 0.26 | 0.03 | 0.09 |
| 6 | 39554898 | rs12175102 | -0.6 | -1.949 | 1.77 | -0.587 | -0.001 | 0.202 | 0.202 | 0 | 0.08 | 0.3 | 0.05 | 0.27 | 0.04 | 0.09 |
| 6 | 39564811 | rs9462557 | -0.52 | -1.976 | 1.771 | -0.648 | 0.031 | 0.035 | 0.035 | 0 | 0.25 | 0.2 | 0.21 | 0.01 | 0.15 | 0.25 |
| 6 | 39568421 | rs9296302 | -0.532 | -1.976 | 1.768 | -0.648 | 0.031 | 0.036 | 0.036 | 0 | 0.25 | 0.21 | 0.22 | 0.01 | 0.15 | 0.24 |
| 6 | 39574377 | rs9394619 | -0.575 | -1.869 | 1.588 | -0.539 | 0.002 | 0.209 | 0.209 | 0 | 0.08 | 0.3 | 0.04 | 0.27 | 0.04 | 0.09 |
| 6 | 39575866 | rs4363024 | -2.151 | -1.928 | 1.367 | -0.626 | -0.007 | -0.012 | -0.012 | 0.062 | 0.57 | 0.86 | 0.4 | 0.37 | 0.39 | 0.62 |
| 6 | 39583135 | rs2045577 | -0.801 | -1.92 | 1.562 | -0.555 | -0.002 | 0.1 | 0.1 | 0 | 0.13 | 0.28 | 0.06 | 0.28 | 0.08 | 0.1 |
| 6 | 39583661 | rs2466405 | -2.294 | -2.139 | 0.714 | -0.551 | -0.095 | -0.16 | -0.16 | 0 | 0.6 | 0.96 | 0.45 | 0.4 | 0.4 | 0.66 |
| 6 | 39587757 | rs1026526 | -0.61 | -2.081 | 1.181 | -0.569 | -0.022 | 0.034 | 0.034 | 0 | 0.28 | 0.36 | 0.23 | 0.37 | 0.16 | 0.34 |
| 6 | 39588118 | rs2446649 | -1.629 | -2.032 | 1.184 | -0.611 | -0.019 | -0.112 | -0.112 | 0.128 | 0.27 | 0.55 | 0.15 | 0.04 | 0.18 | 0.26 |
| 6 | 39590788 | rs2446653 | -2.189 | -2.018 | 1.092 | -0.597 | -0.022 | -0.088 | -0.088 | 0 | 0.56 | 0.91 | 0.39 | 0.41 | 0.36 | 0.61 |
| 6 | 39601188 | rs10498743 | 0.648 | -2.029 | 1.154 | -0.617 | 0.008 | 0.012 | 0.012 | 0 | 0.11 | 0.04 | 0.06 | 0 | 0.08 | 0.06 |
| 6 | 39606635 | rs724101 | 0.695 | -2.029 | 1.153 | -0.617 | 0.01 | 0.026 | 0.026 | 0 | 0.1 | 0.03 | 0.06 | 0 | 0.08 | 0.06 |
| 6 | 39607016 | rs12209915 | -0.558 | -2.061 | 1.224 | -0.623 | -0.023 | 0.036 | 0.036 | 0 | 0.28 | 0.36 | 0.23 | 0.37 | 0.16 | 0.34 |
| 6 | 39608028 | rs2466408 | -2.215 | -2.064 | 1.25 | -0.65 | -0.003 | -0.064 | -0.064 | 0.596 | 0.55 | 0.9 | 0.39 | 0.41 | 0.36 | 0.61 |
| 6 | 39608458 | rs10947825 | -1.393 | -2.071 | 1.186 | -0.653 | -0.003 | 0.004 | 0.004 | 0 | 0.38 | 0.38 | 0.29 | 0.41 | 0.26 | 0.53 |
| 6 | 39612095 | rs2466409 | -2.201 | -1.997 | 1.058 | -0.612 | -0.015 | -0.078 | -0.078 | 0.074 | 0.56 | 0.91 | 0.39 | 0.41 | 0.36 | 0.61 |
| 6 | 39637815 | rs302572 | 0.742 | -1.869 | 1.432 | -0.421 | 0.05 | -0.022 | -0.022 | 0.021 | 0.92 | 0.72 | 0.94 | 0.96 | 0.93 | 0.88 |
| 6 | 39639008 | rs2199171 | -0.664 | -1.217 | 0.987 | -0.319 | 0.001 | 0.03 | 0.03 | 0 | 0.43 | 0.35 | 0.41 | 0.21 | 0.4 | 0.44 |
| 6 | 39643607 | rs303678 | 1.135 | -1.195 | 0.995 | -0.313 | 0.015 | -0.062 | -0.062 | 0 | 0.87 | 0.72 | 0.92 | 0.96 | 0.9 | 0.82 |
| 6 | 39648685 | rs302562 | 0.604 | -1.031 | 1.072 | -0.223 | -0.001 | -0.097 | -0.097 | 2.883 | 0.82 | 0.65 | 0.88 | 0.96 | 0.85 | 0.82 |
| 6 | 39653406 | rs9367022 | -0.45 | -0.995 | 0.822 | -0.12 | 0 | 0 | 0 | 0 | 0.38 | 0.35 | 0.38 | 0.57 | 0.35 | 0.42 |
| 6 | 39658651 | rs10947832 | 0.239 | -0.942 | 0.182 | -0.143 | -0.012 | 0.027 | 0.027 | 0 | 0.29 | 0.14 | 0.35 | 0.21 | 0.32 | 0.33 |
| 6 | 39682934 | rs6933563 | 0.535 | -0.59 | -0.134 | -0.55 | 0.005 | 0.001 | 0.001 | 0 | 0.1 | 0.08 | 0.06 | 0 | 0.08 | 0.06 |
| 6 | 39699399 | rs303648 | 0.556 | -0.531 | -0.354 | -0.504 | -0.002 | 0.006 | 0.006 | 0 | 0.52 | 0.49 | 0.53 | 0.39 | 0.55 | 0.4 |
| 6 | 39700434 | rs2938609 | 0.883 | 0.078 | -1.301 | -0.052 | -0.085 | -0.01 | -0.01 | 1.096 | 0.37 | 0.07 | 0.46 | 0.37 | 0.45 | 0.31 |
| 6 | 39717518 | rs11751730 | -1.447 | 0.428 | -0.157 | 0.153 | 0.009 | -0.022 | -0.022 | 0 | 0.1 | 0.15 | 0.08 | 0.05 | 0.1 | 0.2 |
| 6 | 39723709 | rs9471152 | -1.001 | 0.313 | -0.419 | 0.459 | -0.015 | -0.02 | -0.02 | 2.102 | 0.05 | 0 | 0.03 | 0.08 | 0.04 | 0.12 |

| | | | | | | | | | | | | | | | | |
|---|---|---|---|---|---|---|---|---|---|---|---|---|---|---|---|---|
| 6 | 39725860 | rs2938608 | -0.148 | 0.305 | -1.373 | -0.406 | -0.033 | 0.121 | 0.121 | 0 | 0.27 | 0.09 | 0.32 | 0.05 | 0.28 | 0.39 |
| 6 | 39726757 | rs913576 | -0.752 | 0.275 | -1.076 | -0.421 | -0.003 | 0.005 | 0.005 | 0 | 0.06 | 0.08 | 0.02 | 0.08 | 0.03 | 0.12 |
| 6 | 39727833 | rs417096 | 0.973 | -0.488 | -0.65 | 0.13 | 0.034 | 0.078 | 0.078 | 0 | 0.56 | 0.31 | 0.53 | 0.26 | 0.66 | 0.39 |
| 6 | 39731430 | rs302603 | 0.869 | -0.505 | 0.256 | 0.137 | 0.004 | 0.461 | 0.461 | 0.122 | 0.91 | 0.64 | 0.88 | 0.39 | 0.96 | 0.89 |
| 6 | 39731701 | rs1018156 | 0.099 | -0.487 | 0.243 | 0.077 | -0.002 | 0.004 | 0.004 | 0 | 0.27 | 0.23 | 0.33 | 0.05 | 0.27 | 0.39 |
| 6 | 39731808 | rs372048 | 1.025 | -0.479 | 0.247 | 0.086 | -0.003 | 0.459 | 0.459 | 0.332 | 0.91 | 0.62 | 0.88 | 0.39 | 0.96 | 0.89 |
| 6 | 39732432 | rs1018155 | -0.398 | -0.454 | 0.189 | 0.033 | -0.001 | 0.001 | 0.001 | 0 | 0.33 | 0.31 | 0.34 | 0.13 | 0.29 | 0.51 |
| 6 | 39743239 | rs302615 | -1.207 | -0.19 | 0.479 | -0.492 | 0 | -0.02 | -0.02 | 0 | 0.26 | 0.29 | 0.18 | 0.1 | 0.21 | 0.25 |
| 6 | 39743801 | rs302613 | 0.564 | -0.232 | 0.672 | -0.41 | 0.036 | 0.511 | 0.511 | 0.998 | 0.95 | 0.73 | 0.89 | 0.39 | 0.98 | 0.93 |
| 6 | 39746879 | rs9296304 | 1.04 | -0.281 | 0.54 | -0.553 | -0.074 | 0.037 | 0.037 | 0 | 0.44 | 0.15 | 0.38 | 0.3 | 0.51 | 0.42 |
| 6 | 39753307 | rs9471165 | -0.005 | -0.29 | 0.554 | -0.589 | -0.013 | 0.015 | 0.015 | 0 | 0.15 | 0.1 | 0.15 | 0.02 | 0.16 | 0.06 |
| 6 | 39755999 | rs302637 | -0.227 | -0.439 | 0.625 | -0.382 | 0 | -0.057 | -0.057 | 0 | 0.46 | 0.49 | 0.5 | 0.06 | 0.45 | 0.46 |
| 6 | 39757080 | rs167325 | 1.787 | -0.476 | 0.47 | -0.384 | -0.054 | 0.15 | 0.15 | 0.005 | 0.83 | 0.6 | 0.86 | 0.29 | 0.92 | 0.75 |
| 6 | 39758362 | rs9394629 | 0.73 | -0.409 | 0.352 | -0.368 | -0.065 | 0.04 | 0.04 | 0 | 0.44 | 0.15 | 0.38 | 0.3 | 0.51 | 0.42 |
| 6 | 39758431 | rs9380895 | 1.224 | -0.439 | 0.33 | -0.365 | -0.071 | 0.052 | 0.052 | 0 | 0.38 | 0.15 | 0.37 | 0.23 | 0.48 | 0.29 |
| 6 | 39763506 | rs305851 | -0.522 | -0.25 | 0.477 | -0.42 | -0.001 | -0.031 | -0.031 | 0.727 | 0.49 | 0.55 | 0.51 | 0.07 | 0.45 | 0.49 |
| 6 | 39780248 | rs9471170 | -0.138 | 0.439 | -0.307 | 0.064 | -0.032 | -0.024 | -0.024 | 0.083 | 0.07 | 0 | 0.08 | 0.08 | 0.12 | 0.07 |
| 6 | 39784134 | rs1318002 | -0.338 | 0.274 | -0.019 | 0.015 | 0.001 | 0.013 | 0.013 | 0 | 0.47 | 0.54 | 0.6 | 0.62 | 0.48 | 0.55 |
| 6 | 39784365 | rs2395730 | -0.684 | 0.274 | 0.071 | 0.001 | 0.003 | 0.017 | 0.017 | 0.08 | 0.57 | 0.57 | 0.62 | 0.69 | 0.52 | 0.71 |
| 6 | 39786127 | rs6458124 | -1.038 | 0.091 | 0.67 | 0.48 | -0.009 | -0.004 | -0.004 | 0 | 0.31 | 0.65 | 0.3 | 0.72 | 0.24 | 0.27 |
| 6 | 39786864 | rs6908691 | 0.831 | -0.114 | 1.343 | 1.484 | -0.028 | 0.081 | 0.081 | 0.153 | 0.89 | 0.83 | 0.91 | 0.76 | 0.93 | 0.9 |
| 6 | 39788496 | rs9349128 | 1.107 | -0.025 | 0.465 | 1.333 | -0.001 | -0.046 | -0.046 | 0 | 0.57 | 0.06 | 0.63 | 0.37 | 0.68 | 0.52 |
| 6 | 39790940 | rs6914445 | -0.876 | 0.465 | 1.447 | 0.895 | -0.004 | 0.028 | 0.028 | 0 | 0.29 | 0.45 | 0.28 | 0.41 | 0.23 | 0.37 |
| 6 | 39794653 | rs9296308 | -0.946 | 0.264 | 1.261 | 1.041 | -0.008 | 0.017 | 0.017 | 0 | 0.29 | 0.56 | 0.29 | 0.41 | 0.22 | 0.39 |
| 6 | 39795793 | rs9471175 | -0.184 | 0.467 | 0.395 | 1.147 | -0.04 | 0.134 | 0.134 | 0 | 0.22 | 0.02 | 0.19 | 0.01 | 0.18 | 0.3 |
| 6 | 39800014 | rs6937133 | -0.577 | 0.513 | 0.485 | 1.25 | -0.002 | 0.016 | 0.016 | 0 | 0.26 | 0.3 | 0.24 | 0.42 | 0.2 | 0.4 |
| 6 | 39806807 | rs9367030 | -1.274 | 0.607 | 0.615 | 1.85 | 0.065 | 0.376 | 0.376 | 2.585 | 0.06 | 0.37 | 0.09 | 0.4 | 0.05 | 0.09 |
| 6 | 39809260 | rs13212273 | 0.368 | 0.823 | -0.779 | 0.922 | -0.029 | 0.131 | 0.131 | 0.637 | 0.52 | 0.14 | 0.54 | 0.22 | 0.52 | 0.39 |
| 6 | 39811883 | rs9462578 | -0.236 | 0.453 | -0.818 | 0.781 | 0.007 | 0.011 | 0.011 | 0 | 0.5 | 0.66 | 0.48 | 0.48 | 0.48 | 0.38 |
| 6 | 39815595 | rs9462580 | 0.293 | 0.082 | -1.223 | 0.449 | -0.116 | -0.06 | -0.06 | 0 | 0.27 | 0.08 | 0.38 | 0.41 | 0.4 | 0.32 |
| 6 | 39828208 | rs11755202 | 0.027 | -0.231 | -0.775 | 0.242 | -0.009 | -0.022 | -0.022 | 0 | 0.12 | 0.09 | 0.06 | 0 | 0.04 | 0.04 |
| 6 | 39829223 | rs2504097 | 0.183 | -0.305 | -0.301 | 0.277 | -0.05 | -0.02 | -0.02 | 0.003 | 0.66 | 0.74 | 0.59 | 0.55 | 0.59 | 0.67 |
| 6 | 39846137 | rs2504086 | 0.329 | -0.179 | 0.444 | 0.269 | -0.003 | 0 | 0 | 0 | 0.6 | 0.57 | 0.54 | 0.77 | 0.57 | 0.72 |
| 6 | 39851679 | rs2504082 | -0.704 | 0.157 | 0.519 | 0.255 | 0.013 | -0.032 | -0.032 | 0 | 0.4 | 0.53 | 0.45 | 0.25 | 0.42 | 0.3 |
| 6 | 39851818 | rs3003929 | 0.545 | 0.157 | 0.538 | 0.255 | 0.013 | -0.033 | -0.033 | 0 | 0.6 | 0.46 | 0.55 | 0.75 | 0.58 | 0.7 |
| 6 | 39854425 | rs3003942 | -0.383 | 0.109 | 0.555 | 0.203 | -0.082 | 0.033 | 0.033 | 0 | 0.73 | 0.88 | 0.62 | 0.79 | 0.63 | 0.74 |
| 6 | 39854901 | rs3008809 | 0.3 | 0.171 | 0.646 | 0.226 | 0.031 | -0.051 | -0.051 | 0 | 0.66 | 0.48 | 0.57 | 0.79 | 0.6 | 0.72 |
| 6 | 39856689 | rs3003946 | 0.916 | -0.331 | 0.638 | 0.663 | 0.004 | 0.005 | 0.005 | 0 | 0.51 | 0.46 | 0.45 | 0.57 | 0.49 | 0.57 |
| 6 | 39857326 | rs11759168 | 0.899 | -0.904 | -0.197 | 0.107 | 0.013 | 0.074 | 0.074 | 0 | 0.18 | 0.01 | 0.11 | 0.05 | 0.1 | 0.36 |
| 6 | 39859102 | rs9394632 | 0.093 | -0.469 | 0.21 | 0.493 | -0.003 | 0.014 | 0.014 | 0 | 0.08 | 0.12 | 0.09 | 0.18 | 0.06 | 0.12 |
| 6 | 39860030 | rs3008813 | 0.769 | -0.554 | 0.469 | 0.764 | 0.022 | 0.028 | 0.028 | 0 | 0.5 | 0.3 | 0.45 | 0.38 | 0.49 | 0.53 |
| 6 | 39860093 | rs3008814 | 0.441 | -0.534 | 0.475 | 0.757 | 0.002 | 0 | 0 | 0.001 | 0.54 | 0.3 | 0.46 | 0.42 | 0.51 | 0.54 |
| 6 | 39861595 | rs2504805 | 1.208 | -0.283 | 0.334 | 0.614 | -0.046 | 0.03 | 0.03 | 0 | 0.2 | 0.12 | 0.36 | 0.13 | 0.34 | 0.2 |
| 6 | 39863468 | rs7771410 | -0.389 | -0.335 | 0.655 | 0.776 | -0.004 | 0.189 | 0.189 | 0 | 0.07 | 0.23 | 0.09 | 0.26 | 0.03 | 0.06 |
| 6 | 39864730 | rs3008815 | 0.746 | -0.387 | 0.79 | 0.939 | 0.004 | -0.008 | -0.008 | 0 | 0.57 | 0.42 | 0.48 | 0.32 | 0.53 | 0.58 |
| 6 | 39865047 | rs3004067 | -0.743 | -0.322 | 0.685 | 0.922 | -0.045 | 0.049 | 0.049 | 0 | 0.74 | 0.83 | 0.61 | 0.87 | 0.64 | 0.77 |
| 6 | 39866290 | rs3004069 | 0.704 | -0.359 | 0.73 | 1.009 | 0 | 0 | 0 | 0 | 0.57 | 0.4 | 0.48 | 0.34 | 0.53 | 0.58 |
| 6 | 39870013 | rs3004071 | 0.128 | -0.258 | 0.012 | 0.671 | -0.054 | 0.083 | 0.083 | 0 | 0.64 | 0.75 | 0.57 | 0.86 | 0.56 | 0.68 |
| 6 | 39873146 | rs7758412 | 0.01 | -0.358 | 0.131 | 0.736 | -0.013 | 0.004 | 0.004 | 0 | 0.05 | 0.11 | 0.02 | 0.04 | 0.02 | 0.01 |
| 6 | 39874875 | rs11969769 | 0.147 | -0.352 | 0.086 | 0.739 | -0.013 | 0.001 | 0.001 | 0.008 | 0.05 | 0.08 | 0.02 | 0.04 | 0.02 | 0.01 |
| 6 | 39876433 | rs2273853 | 0.018 | -0.195 | -0.048 | 0.561 | -0.014 | 0.009 | 0.009 | 0.292 | 0.08 | 0.01 | 0.04 | 0.05 | 0.08 | 0.05 |
| 6 | 39879071 | rs1109103 | -0.042 | 0.096 | 0.061 | 0.935 | -0.004 | -0.007 | -0.007 | 0 | 0.67 | 0.69 | 0.6 | 0.54 | 0.66 | 0.73 |
| 6 | 39882750 | rs716856 | -1.276 | 0.248 | -0.128 | 0.726 | -0.017 | 0.013 | 0.013 | 0 | 0.79 | 0.83 | 0.65 | 0.86 | 0.71 | 0.81 |
| 6 | 39890267 | rs2984659 | 0.346 | 0.113 | 0.07 | 0.835 | 0 | -0.001 | -0.001 | 0.138 | 0.58 | 0.48 | 0.5 | 0.34 | 0.58 | 0.57 |
| 6 | 39892025 | rs9689513 | 0.035 | -0.153 | -0.4 | 0.557 | 0.002 | 0.005 | 0.005 | 0 | 0.28 | 0.42 | 0.24 | 0.32 | 0.27 | 0.2 |
| 6 | 39894536 | rs3008821 | 0.883 | -0.109 | -0.97 | -0.038 | -0.004 | 0.015 | 0.015 | 0 | 0.36 | 0.17 | 0.35 | 0.27 | 0.39 | 0.51 |
| 6 | 39894628 | rs2984440 | -0.259 | -0.208 | -0.985 | -0.039 | 0.043 | 0.155 | 0.155 | 0 | 0.5 | 0.68 | 0.48 | 0.81 | 0.45 | 0.63 |
| 6 | 39898112 | rs2984437 | 0.635 | 0.281 | -0.843 | 0.154 | -0.006 | 0.016 | 0.016 | 0 | 0.36 | 0.17 | 0.35 | 0.27 | 0.4 | 0.51 |
| 6 | 39904317 | rs2504080 | -0.287 | 0.593 | -0.974 | 0.101 | -0.011 | 0.02 | 0.02 | 0 | 0.71 | 0.77 | 0.61 | 0.82 | 0.7 | 0.72 |
| 6 | 39905615 | rs6904621 | -0.448 | 0.531 | -0.886 | 0.548 | 0.004 | -0.013 | -0.013 | 0 | 0.39 | 0.67 | 0.28 | 0.52 | 0.35 | 0.3 |
| 6 | 39907809 | rs2504081 | -0.065 | 0.919 | -1.178 | 0.438 | -0.034 | -0.023 | -0.023 | 3.174 | 0.25 | 0.13 | 0.37 | 0.34 | 0.38 | 0.2 |
| 6 | 39911241 | rs2475506 | -0.147 | 0.358 | -1.814 | -0.098 | -0.005 | 0.028 | 0.028 | 0 | 0.44 | 0.13 | 0.52 | 0.39 | 0.43 | 0.53 |
| 6 | 39915349 | rs10947842 | 0.818 | 0.285 | -1.898 | -0.161 | 0.019 | 0.134 | 0.134 | 0 | 0.25 | 0.02 | 0.22 | 0.05 | 0.15 | 0.38 |
| 6 | 39916614 | rs1475114 | 0.79 | 0.298 | -1.822 | -0.147 | -0.031 | -0.056 | -0.056 | 0.025 | 0.19 | 0.09 | 0.3 | 0.34 | 0.28 | 0.15 |
| 6 | 39917201 | rs9349130 | -0.35 | 0.066 | -0.676 | 0.402 | -0.029 | 0.051 | 0.051 | 0.319 | 0.08 | 0.25 | 0.07 | 0.13 | 0.03 | 0.03 |
| 6 | 39920347 | rs2475802 | -1.42 | 0.279 | -0.878 | 0.368 | -0.002 | 0.005 | 0.005 | 0 | 0.76 | 0.9 | 0.79 | 0.8 | 0.76 | 0.77 |
| 6 | 39925922 | rs880919 | -0.116 | 0.337 | -0.297 | 0.449 | 0.019 | 0.002 | 0.002 | 0 | 0.61 | 0.34 | 0.68 | 0.63 | 0.71 | 0.72 |
| 6 | 39927793 | rs2475505 | 0.828 | 0.358 | -0.348 | 0.451 | -0.009 | 0.024 | 0.024 | 0 | 0.44 | 0.12 | 0.52 | 0.39 | 0.42 | 0.53 |
| 6 | 39938471 | rs1923474 | -0.01 | 0.736 | -0.202 | 0.519 | 0.035 | 0.018 | 0.018 | 0 | 0.17 | 0.21 | 0.17 | 0.24 | 0.29 | 0.19 |

| | | | | | | | | | | | | | | | | |
|---|---|---|---|---|---|---|---|---|---|---|---|---|---|---|---|---|
| 6 | 39942114 | rs1923473 | -0.547 | 0.203 | -0.225 | 0.442 | -0.021 | -0.02 | -0.02 | 0.271 | 0.7 | 0.59 | 0.78 | 0.76 | 0.75 | 0.76 |
| 6 | 39945015 | rs9462589 | -0.42 | 0.475 | -0.118 | 0.21 | -0.003 | -0.006 | -0.006 | 0.047 | 0.47 | 0.61 | 0.33 | 0.46 | 0.48 | 0.39 |
| 6 | 39945406 | rs6912379 | -0.239 | 0.743 | -0.126 | 0.451 | 0.017 | -0.012 | -0.012 | 0 | 0.21 | 0.25 | 0.17 | 0.12 | 0.28 | 0.22 |
| 6 | 39948906 | rs12207285 | 0.85 | 0.387 | -0.17 | 0.655 | 0.008 | -0.013 | -0.013 | 0.356 | 0.24 | 0.2 | 0.13 | 0.29 | 0.19 | 0.17 |
| 6 | 39954177 | rs7451099 | 0.288 | 1.19 | 0.006 | 0.717 | 0.02 | -0.001 | -0.001 | 0.073 | 0.06 | 0.08 | 0.09 | 0.06 | 0.15 | 0.06 |
| 6 | 39961907 | rs2263328 | -0.052 | 0.794 | -0.256 | 0.744 | 0.003 | 0.023 | 0.023 | 0 | 0.14 | 0.17 | 0.28 | 0.34 | 0.18 | 0.13 |
| 6 | 39962868 | rs9369152 | 0.533 | 0.117 | -0.602 | 0.226 | 0.004 | 0.004 | 0.004 | 0 | 0.25 | 0.25 | 0.28 | 0.18 | 0.17 | 0.38 |
| 6 | 39970879 | rs2984434 | 1.705 | 0.16 | -0.969 | 0.207 | -0.067 | -0.12 | -0.12 | 0 | 0.32 | 0.02 | 0.19 | 0.33 | 0.34 | 0.21 |
| 6 | 39971284 | rs2092863 | -0.896 | -0.289 | -0.045 | 0.235 | -0.005 | 0.107 | 0.107 | 0 | 0.19 | 0.34 | 0.32 | 0.47 | 0.19 | 0.15 |
| 6 | 39980359 | rs1474870 | 0.899 | -0.384 | 0.198 | 0.11 | 0.025 | -0.001 | -0.001 | 0 | 0.54 | 0.35 | 0.4 | 0.46 | 0.65 | 0.45 |
| 6 | 39983695 | rs1080882 | 0.922 | -0.411 | 0.243 | 0.189 | 0.006 | 0.008 | 0.008 | 0 | 0.19 | 0.16 | 0.22 | 0.1 | 0.12 | 0.35 |
| 6 | 39986303 | rs3008833 | 0.968 | -0.397 | 0.198 | 0.18 | 0.039 | -0.001 | -0.001 | 0 | 0.54 | 0.33 | 0.39 | 0.46 | 0.65 | 0.45 |
| 6 | 39987315 | rs4714285 | -0.154 | -0.357 | 0.036 | 0.26 | -0.002 | 0.008 | 0.008 | 0 | 0.06 | 0.07 | 0.06 | 0.13 | 0.05 | 0.02 |
| 6 | 39999149 | rs1546965 | -0.316 | -0.369 | 0.083 | 0.215 | -0.017 | -0.002 | -0.002 | 0 | 0.33 | 0.44 | 0.49 | 0.41 | 0.26 | 0.49 |
| 6 | 40008465 | rs2983138 | 0.054 | -0.3 | 0.007 | 0.245 | -0.008 | -0.035 | -0.035 | 1.241 | 0.93 | 0.9 | 0.93 | 1 | 0.95 | 0.96 |
| 6 | 40008561 | rs12528780 | 0.052 | -0.292 | -0.056 | 0.332 | 0.001 | 0.037 | 0.037 | 0 | 0.12 | 0.17 | 0.26 | 0.34 | 0.14 | 0.12 |
| 6 | 40023088 | rs1923489 | -0.676 | -0.229 | 0.01 | 0.316 | -0.017 | 0.012 | 0.012 | 0 | 0.39 | 0.47 | 0.55 | 0.53 | 0.31 | 0.52 |
| 6 | 40045165 | rs4714299 | 0.734 | -0.197 | -0.086 | 0.267 | 0.019 | 0.013 | 0.013 | 0 | 0.22 | 0.15 | 0.26 | 0.16 | 0.13 | 0.38 |
| 6 | 40050171 | rs847769 | -0.333 | -0.183 | -0.057 | 0.307 | -0.003 | 0.003 | 0.003 | 0 | 0.34 | 0.38 | 0.52 | 0.5 | 0.27 | 0.51 |
| 6 | 40055046 | rs847779 | -0.447 | -0.137 | -0.043 | 0.338 | 0 | 0.013 | 0.013 | 0 | 0.15 | 0.17 | 0.28 | 0.37 | 0.18 | 0.14 |
| 6 | 40066652 | rs847786 | 0.191 | -0.103 | -0.04 | 0.373 | 0.001 | 0.024 | 0.024 | 0 | 0.11 | 0.14 | 0.26 | 0.34 | 0.14 | 0.14 |
| 6 | 40066677 | rs847785 | 0.191 | -0.103 | -0.04 | 0.373 | 0.001 | 0.024 | 0.024 | 0 | 0.11 | 0.14 | 0.26 | 0.34 | 0.14 | 0.14 |
| 6 | 40069436 | rs3008841 | 0.951 | -0.058 | -0.174 | 0.157 | 0.007 | 0.003 | 0.003 | 0 | 0.51 | 0.55 | 0.37 | 0.48 | 0.59 | 0.33 |
| 6 | 40084788 | rs707970 | 0.24 | -0.427 | 0.283 | 0.406 | -0.012 | -0.038 | -0.038 | 0.126 | 0.93 | 0.87 | 0.94 | 1 | 0.96 | 0.97 |
| 6 | 40085245 | rs847744 | -0.546 | -0.308 | -0.054 | 0.529 | -0.007 | 0.008 | 0.008 | 0 | 0.34 | 0.4 | 0.52 | 0.5 | 0.27 | 0.5 |
| 6 | 40096889 | rs10947855 | 0.329 | -0.159 | -0.287 | 0.597 | -0.013 | -0.008 | -0.008 | 0.409 | 0.22 | 0.26 | 0.27 | 0.16 | 0.13 | 0.37 |
| 6 | 40112662 | rs940357 | 0.42 | -0.151 | -0.643 | 0.815 | 0.01 | -0.003 | -0.003 | 0 | 0.22 | 0.19 | 0.27 | 0.22 | 0.13 | 0.38 |
| 6 | 40112732 | rs940355 | 0.118 | 0.296 | -0.407 | 0.732 | -0.011 | -0.009 | -0.009 | 1.28 | 0.91 | 0.92 | 0.76 | 0.66 | 0.87 | 0.87 |
| 6 | 40119546 | rs10498745 | -0.723 | -0.117 | -0.05 | 0.251 | -0.01 | -0.046 | -0.046 | 0.244 | 0.08 | 0.13 | 0.06 | 0 | 0.05 | 0.04 |
| 6 | 40119758 | rs9296322 | 0.403 | -0.208 | -0.579 | 0.194 | -0.023 | -0.022 | -0.022 | 2.942 | 0.23 | 0.07 | 0.25 | 0.25 | 0.22 | 0.4 |
| 6 | 40121489 | rs16893089 | -0.808 | -0.301 | -0.487 | 0.115 | 0.005 | 0.013 | 0.013 | 0 | 0.35 | 0.27 | 0.33 | 0.28 | 0.31 | 0.44 |
| 6 | 40124958 | rs7750979 | -0.911 | -0.434 | -0.452 | -0.051 | -0.014 | 0.066 | 0.066 | 0 | 0.55 | 0.68 | 0.5 | 0.63 | 0.52 | 0.77 |
| 6 | 40125459 | rs7751650 | -0.42 | -0.462 | -0.327 | -0.059 | 0.002 | 0.046 | 0.046 | 0.171 | 0.31 | 0.51 | 0.25 | 0.38 | 0.3 | 0.38 |
| 6 | 40125499 | rs1524084 | 0.785 | -0.598 | -0.237 | 0.02 | 0.008 | 0.001 | 0.001 | 0 | 0.24 | 0.15 | 0.25 | 0.25 | 0.22 | 0.39 |
| 6 | 40128253 | rs7767647 | -1.19 | -0.686 | -0.29 | -0.029 | -0.031 | 0.062 | 0.062 | 0 | 0.58 | 0.75 | 0.5 | 0.68 | 0.54 | 0.77 |
| 6 | 40128942 | rs1467313 | 0.245 | -0.829 | -0.367 | -0.016 | 0.003 | 0.003 | 0.003 | 0 | 0.31 | 0.25 | 0.31 | 0.26 | 0.26 | 0.43 |
| 6 | 40132914 | rs6930375 | 1.136 | -0.852 | -0.479 | -0.041 | -0.103 | 0.062 | 0.062 | 0 | 0.4 | 0.09 | 0.5 | 0.32 | 0.47 | 0.24 |
| 6 | 40145994 | rs940362 | -0.001 | -0.81 | -0.31 | -0.008 | 0.011 | 0.013 | 0.013 | 0 | 0.3 | 0.21 | 0.3 | 0.26 | 0.26 | 0.42 |
| 6 | 40166270 | rs3846892 | 0.884 | -0.775 | -0.071 | -0.083 | -0.024 | -0.053 | -0.053 | 0 | 0.9 | 0.72 | 0.93 | 0.99 | 0.96 | 0.96 |
| 6 | 40168513 | rs10755710 | 1.106 | -0.875 | -0.554 | -0.163 | -0.176 | 0.025 | 0.025 | 0.01 | 0.38 | 0.03 | 0.47 | 0.32 | 0.47 | 0.24 |
| 6 | 40172340 | rs9394662 | 0.963 | -0.844 | -0.448 | -0.147 | -0.096 | 0.093 | 0.093 | 0.095 | 0.42 | 0.09 | 0.49 | 0.32 | 0.47 | 0.24 |
| 6 | 40184628 | rs7746227 | 0.197 | -0.751 | 0.114 | 0.047 | 0.002 | 0.106 | 0.106 | 0 | 0.23 | 0.37 | 0.17 | 0.39 | 0.23 | 0.33 |
| 6 | 40188482 | rs11969625 | 0.662 | -0.809 | -0.254 | 0.028 | 0.012 | -0.023 | -0.023 | 0 | 0.64 | 0.31 | 0.73 | 0.57 | 0.68 | 0.63 |
| 6 | 40197201 | rs7749170 | 0.805 | -0.695 | -0.293 | 0.018 | 0 | 0 | 0 | 0 | 0.23 | 0.18 | 0.24 | 0.23 | 0.21 | 0.34 |
| 6 | 40198930 | rs10807230 | 0.866 | -1.005 | -0.341 | -0.042 | 0.012 | -0.016 | -0.016 | 0 | 0.63 | 0.31 | 0.73 | 0.55 | 0.68 | 0.6 |
| 6 | 40212720 | rs6940128 | 1.003 | -1.015 | -0.241 | 0.513 | 0.009 | -0.007 | -0.007 | 0.803 | 0.1 | 0.08 | 0.07 | 0.24 | 0.08 | 0.18 |
| 6 | 40215956 | rs12206560 | 0.381 | -0.763 | -0.11 | 0.205 | 0.001 | 0 | 0 | 0 | 0.28 | 0.2 | 0.32 | 0.31 | 0.28 | 0.37 |
| 6 | 40217930 | rs10947864 | 1.316 | -0.59 | -0.198 | 0.199 | -0.093 | 0.045 | 0.045 | 0 | 0.37 | 0.1 | 0.42 | 0.32 | 0.43 | 0.24 |
| 6 | 40218128 | rs6458146 | 0.008 | -0.55 | 0.204 | 0.238 | 0.001 | 0.001 | 0.001 | 0 | 0.65 | 0.67 | 0.66 | 0.69 | 0.69 | 0.55 |
| 6 | 40224268 | rs10498746 | 0.733 | -0.461 | -0.208 | 0.247 | 0.003 | 0.009 | 0.009 | 0 | 0.27 | 0.14 | 0.31 | 0.27 | 0.26 | 0.38 |
| 6 | 40225964 | rs1077329 | 1.091 | -0.422 | 0.36 | 0.641 | 0.023 | -0.051 | -0.051 | 0 | 0.63 | 0.22 | 0.73 | 0.6 | 0.68 | 0.61 |
| 6 | 40232327 | rs1325545 | -0.288 | -0.29 | 0.973 | 0.735 | -0.001 | 0.065 | 0.065 | 0 | 0.24 | 0.35 | 0.21 | 0.37 | 0.25 | 0.29 |
| 6 | 40233506 | rs1325548 | 1.08 | -0.258 | 0.53 | 0.727 | 0.01 | -0.024 | -0.024 | 5.749 | 0.65 | 0.27 | 0.73 | 0.59 | 0.69 | 0.64 |
| 6 | 40245822 | rs2147925 | 1.831 | -0.265 | 1.325 | 0.563 | -0.01 | -0.025 | -0.025 | 0 | 0.88 | 0.54 | 0.94 | 0.95 | 0.94 | 0.92 |
| 6 | 40250476 | rs1325555 | 0.993 | -0.228 | 1.011 | 0.583 | 0 | -0.001 | -0.001 | 2.256 | 0.94 | 0.95 | 0.97 | 0.96 | 0.94 | 0.93 |
| 6 | 40258180 | rs9394679 | 1.943 | -0.253 | 1.203 | 0.568 | -0.009 | -0.024 | -0.024 | 0 | 0.88 | 0.54 | 0.94 | 0.96 | 0.94 | 0.93 |
| 6 | 40260515 | rs9471290 | -0.041 | -0.043 | 0.575 | 0.704 | 0.008 | 0.014 | 0.014 | 0 | 0.33 | 0.4 | 0.35 | 0.43 | 0.34 | 0.45 |
| 6 | 40261080 | rs4714323 | -0.855 | 0.409 | 0.238 | 0.473 | -0.012 | -0.016 | -0.016 | 0 | 0.69 | 0.79 | 0.71 | 0.56 | 0.64 | 0.82 |
| 6 | 40267200 | rs2038865 | 0.785 | 0.061 | 0.923 | 0.256 | -0.019 | -0.012 | -0.012 | 0.279 | 0.94 | 0.82 | 0.97 | 0.99 | 0.99 | 0.99 |
| 6 | 40270047 | rs7762056 | 0.373 | -0.142 | 0.821 | 0.104 | 0.001 | 0.008 | 0.008 | 0 | 0.28 | 0.23 | 0.2 | 0.23 | 0.26 | 0.44 |
| 6 | 40271527 | rs1018631 | 0.684 | -0.092 | 0.802 | 0.198 | 0.008 | 0.044 | 0.044 | 0 | 0.57 | 0.24 | 0.62 | 0.68 | 0.59 | 0.48 |
| 6 | 40273178 | rs9462604 | 0.402 | -0.015 | 0.548 | 0.243 | -0.007 | 0 | 0 | 0 | 0.07 | 0.05 | 0.13 | 0.04 | 0.13 | 0.07 |
| 6 | 40273457 | rs4714329 | 0.448 | -0.024 | 0.643 | 0.244 | -0.002 | -0.014 | -0.014 | 0.283 | 0.59 | 0.51 | 0.63 | 0.68 | 0.6 | 0.47 |
| 6 | 40275632 | rs1886756 | 1.548 | 0.422 | 0.203 | 0.274 | -0.05 | 0.029 | 0.029 | 0 | 0.24 | 0.06 | 0.25 | 0.19 | 0.32 | 0.11 |
| 6 | 40282682 | rs1325550 | 1.355 | 0.279 | 1.087 | 0.947 | -0.027 | -0.03 | -0.03 | 0 | 0.92 | 0.71 | 0.96 | 0.99 | 0.99 | 0.99 |
| 6 | 40284376 | rs6904982 | 0.095 | 0.73 | 0.3 | 1.185 | -0.015 | 0.013 | 0.013 | 0 | 0.26 | 0.09 | 0.18 | 0.23 | 0.26 | 0.43 |
| 6 | 40287483 | rs9471302 | 0.015 | 0.735 | 0.331 | 1.144 | -0.006 | -0.007 | -0.007 | 0 | 0.26 | 0.13 | 0.18 | 0.26 | 0.26 | 0.44 |
| 6 | 40298297 | rs10498748 | 0.054 | 0.749 | 0.343 | 0.752 | -0.002 | 0.031 | 0.031 | 0 | 0.27 | 0.16 | 0.14 | 0.04 | 0.23 | 0.41 |
| 6 | 40303536 | rs4236066 | 1.257 | 0.711 | 0.467 | 1.06 | -0.021 | 0.247 | 0.247 | 0 | 0.5 | 0.23 | 0.46 | 0.09 | 0.52 | 0.26 |

| | | | | | | | | | | | | | | | | |
|---|---|---|---|---|---|---|---|---|---|---|---|---|---|---|---|---|
| 6 | 40306256 | rs4714334 | 1.319 | 0.711 | 0.495 | 1.174 | -0.013 | 0.196 | 0.196 | 0 | 0.5 | 0.24 | 0.56 | 0.19 | 0.54 | 0.26 |
| 6 | 40307960 | rs16893380 | 0.332 | 0.712 | 0.28 | 1.342 | -0.019 | -0.073 | -0.073 | 0 | 0.11 | 0 | 0.13 | 0.24 | 0.09 | 0.15 |
| 6 | 40308360 | rs9394683 | 1.206 | 0.761 | 0.474 | 0.168 | -0.004 | 0.111 | 0.111 | 0 | 0.51 | 0.33 | 0.56 | 0.19 | 0.54 | 0.26 |
| 6 | 40310573 | rs4711629 | 1.235 | 0.792 | 0.336 | 0.024 | -0.008 | 0.146 | 0.146 | 0 | 0.49 | 0.3 | 0.48 | 0.14 | 0.52 | 0.26 |
| 6 | 40310958 | rs9462608 | 0.844 | 0.805 | 0.324 | 0.068 | 0.018 | 0.097 | 0.097 | 0 | 0.54 | 0.32 | 0.59 | 0.17 | 0.6 | 0.34 |
| 6 | 40312346 | rs13203076 | 0.456 | 0.784 | -0.133 | 0.079 | 0.005 | 0.022 | 0.022 | 0 | 0.22 | 0.14 | 0.09 | 0.04 | 0.18 | 0.4 |
| 6 | 40312918 | rs17619142 | 0.456 | 0.784 | -0.133 | 0.079 | 0.005 | 0.022 | 0.022 | 0 | 0.22 | 0.14 | 0.09 | 0.04 | 0.18 | 0.4 |
| 6 | 40316295 | rs2477755 | 0.941 | 1.16 | 0.277 | 1.085 | -0.007 | -0.016 | -0.016 | 0 | 0.94 | 0.98 | 0.97 | 0.86 | 0.95 | 0.94 |
| 6 | 40321473 | rs11752896 | 1.486 | 1.139 | 0.258 | 0.919 | -0.042 | 0.039 | 0.039 | 0 | 0.64 | 0.21 | 0.58 | 0.3 | 0.6 | 0.66 |
| 6 | 40321778 | rs2477757 | 0.855 | 1.175 | 0.302 | 0.982 | -0.004 | -0.014 | -0.014 | 0 | 0.94 | 0.97 | 0.97 | 0.87 | 0.95 | 0.94 |
| 6 | 40324294 | rs966082 | 1.411 | 1.16 | 0.745 | 0.975 | 0.016 | 0.059 | 0.059 | 0.186 | 0.91 | 0.7 | 0.95 | 0.87 | 0.95 | 0.94 |
| 6 | 40327096 | rs1886753 | -1.028 | 1.26 | 0.685 | 1.076 | 0.038 | 0.052 | 0.052 | 0 | 0.27 | 0.48 | 0.37 | 0.56 | 0.35 | 0.27 |
| 6 | 40344459 | rs4714342 | -1.287 | 1.203 | 0.714 | 0.724 | 0.031 | -0.011 | -0.011 | 0 | 0.29 | 0.43 | 0.4 | 0.7 | 0.36 | 0.22 |
| 6 | 40344526 | rs4714343 | -0.89 | 0.757 | 0.555 | 0.964 | -0.003 | 0.033 | 0.033 | 0 | 0.23 | 0.58 | 0.27 | 0.63 | 0.26 | 0.2 |
| 6 | 40345698 | rs12664614 | 0.38 | -0.063 | -0.173 | 0.113 | 0.038 | 0.061 | 0.061 | 0 | 0.27 | 0.16 | 0.1 | 0.07 | 0.14 | 0.33 |
| 6 | 40345913 | rs930250 | -0.019 | 0.256 | 0.59 | 1.165 | -0.012 | 0.008 | 0.008 | 0 | 0.33 | 0.41 | 0.27 | 0.37 | 0.24 | 0.4 |
| 6 | 40347258 | rs9312 | 1.358 | -0.264 | 0.069 | 0.814 | 0.03 | -0.004 | -0.004 | 0 | 0.17 | 0.03 | 0.06 | 0.16 | 0.13 | 0.23 |
| 6 | 40347918 | rs2033530 | 0.283 | 0.359 | 0.213 | 0.862 | -0.008 | -0.004 | -0.004 | 0 | 0.37 | 0.22 | 0.25 | 0.37 | 0.4 | 0.37 |
| 6 | 40350935 | rs16863 | -1.119 | 1.047 | 1.212 | 1.641 | 0.007 | 0.002 | 0.002 | 0 | 0.47 | 0.44 | 0.65 | 0.45 | 0.63 | 0.32 |
| 6 | 40351841 | rs4145199 | 0.162 | 1.057 | 1.287 | 1.671 | 0.028 | 0.165 | 0.165 | 0 | 0.93 | 0.84 | 0.83 | 0.67 | 0.89 | 0.87 |
| 6 | 40351898 | rs4145198 | -0.233 | 1.166 | 1.477 | 1.668 | -0.018 | 0.022 | 0.022 | 0 | 0.24 | 0.17 | 0.44 | 0.14 | 0.31 | 0.11 |
| 6 | 40354316 | rs6903068 | 1.592 | 1.132 | 1.475 | 1.656 | 0.022 | 0.023 | 0.023 | 0 | 0.46 | 0.4 | 0.18 | 0.23 | 0.26 | 0.55 |
| 6 | 40360465 | rs3734557 | 0.741 | 1.179 | 1.376 | 1.687 | 0.004 | 0.007 | 0.007 | 0 | 0.69 | 0.67 | 0.39 | 0.53 | 0.58 | 0.76 |
| 6 | 40364325 | rs17619780 | -0.274 | 1.172 | 1.199 | 1.552 | -0.019 | 0.022 | 0.022 | 0 | 0.24 | 0.17 | 0.44 | 0.14 | 0.31 | 0.11 |
| 6 | 40365601 | rs2436728 | -0.26 | 1.191 | 1.059 | 1.62 | -0.039 | -0.039 | -0.039 | 0 | 0.26 | 0.18 | 0.36 | 0.55 | 0.41 | 0.27 |
| 6 | 40370499 | rs934465 | 1.53 | 0.88 | 1.415 | 1.412 | 0.059 | -0.009 | -0.009 | 0 | 0.48 | 0.2 | 0.31 | 0.49 | 0.37 | 0.64 |
| 6 | 40378321 | rs892371 | 1.782 | 0.871 | 1.915 | 1.564 | 0.031 | 0.235 | 0.235 | 0 | 0.93 | 0.71 | 0.82 | 0.74 | 0.96 | 0.86 |
| 6 | 40380338 | rs2033527 | 1.445 | 0.782 | 1.559 | 1.188 | 0.003 | 0.026 | 0.026 | 0 | 0.65 | 0.26 | 0.48 | 0.78 | 0.65 | 0.81 |
| 6 | 40381240 | rs9471341 | -0.971 | 0.945 | 1.755 | 1.109 | 0.001 | -0.001 | -0.001 | 2.432 | 0.22 | 0.23 | 0.34 | 0.14 | 0.19 | 0.08 |
| 6 | 40382265 | rs2504829 | 1.85 | 0.972 | 1.447 | 1.118 | 0.004 | 0.05 | 0.05 | 0.012 | 0.64 | 0.23 | 0.48 | 0.78 | 0.65 | 0.81 |
| 6 | 40382711 | rs2916260 | 1.814 | 1.229 | 1.633 | 1.318 | 0.079 | -0.025 | -0.025 | 0 | 0.87 | 0.49 | 0.83 | 0.93 | 0.83 | 0.9 |
| 6 | 40386975 | rs13205455 | -0.961 | 1.543 | 1.349 | 1.629 | -0.071 | -0.039 | -0.039 | 0.943 | 0.34 | 0.09 | 0.45 | 0.4 | 0.42 | 0.22 |
| 6 | 40388800 | rs6927927 | -1.496 | 1.356 | 1.802 | 1.438 | 0.102 | 0.035 | 0.035 | 0 | 0.1 | 0.46 | 0.15 | 0.09 | 0.12 | 0.07 |
| 6 | 40390291 | rs9394691 | -1.737 | 1.359 | 1.807 | 1.439 | 0.071 | 0.065 | 0.065 | 0.014 | 0.11 | 0.42 | 0.16 | 0.16 | 0.15 | 0.09 |
| 6 | 40390471 | rs9357326 | -1.759 | 1.352 | 1.677 | 1.451 | 0.021 | 0.138 | 0.138 | 0.214 | 0.05 | 0.3 | 0.05 | 0.14 | 0.03 | 0.07 |
| 6 | 40391564 | rs6925172 | -0.556 | 1.638 | 0.754 | 1.444 | 0.086 | -0.017 | -0.017 | 0 | 0.06 | 0.21 | 0.12 | 0.01 | 0.12 | 0.03 |
| 6 | 40392369 | rs13196792 | -0.644 | 1.629 | 0.42 | 1.609 | 0.006 | 0.002 | 0.002 | 0 | 0.25 | 0.25 | 0.33 | 0.39 | 0.4 | 0.17 |
| 6 | 40394175 | rs12173741 | -0.207 | 0.826 | -0.297 | 1.882 | -0.032 | -0.047 | -0.047 | 0 | 0.2 | 0.09 | 0.21 | 0.38 | 0.29 | 0.16 |
| 6 | 40405014 | rs12194543 | -1.553 | 0.145 | 0.244 | 1.497 | 0.004 | -0.053 | -0.053 | 0 | 0.14 | 0.05 | 0.27 | 0.28 | 0.11 | 0.13 |
| 6 | 40406359 | rs9394694 | -1.677 | 0.261 | 0.459 | 1.676 | -0.019 | 0.061 | 0.061 | 0 | 0.14 | 0.27 | 0.28 | 0.28 | 0.1 | 0.13 |
| 6 | 40406855 | rs4714346 | -1.735 | 0.309 | 0.449 | 1.701 | -0.019 | 0.054 | 0.054 | 0 | 0.16 | 0.32 | 0.3 | 0.28 | 0.11 | 0.12 |
| 6 | 40406923 | rs4714347 | -1.356 | 0.309 | 0.307 | 1.391 | -0.015 | 0.031 | 0.031 | 0 | 0.13 | 0.27 | 0.28 | 0.21 | 0.08 | 0.1 |
| 6 | 40411700 | rs919875 | -1.574 | 0.295 | -0.048 | 1.348 | -0.008 | 0.007 | 0.007 | 0 | 0.09 | 0.15 | 0.1 | 0.11 | 0.05 | 0.19 |
| 6 | 40416800 | rs1433734 | 0.719 | 0.236 | 0.202 | 1.442 | -0.007 | 0.018 | 0.018 | 0 | 0.89 | 0.77 | 0.84 | 0.9 | 0.92 | 0.88 |
| 6 | 40417196 | rs10807240 | 0.29 | -0.079 | -0.956 | 1.244 | 0.004 | -0.014 | -0.014 | 0 | 0.36 | 0.04 | 0.27 | 0.29 | 0.34 | 0.2 |
| 6 | 40420232 | rs1347257 | 0.402 | 0.169 | -0.408 | 1.593 | 0.002 | 0 | 0 | 0.459 | 0.49 | 0.51 | 0.45 | 0.52 | 0.54 | 0.61 |
| 6 | 40423322 | rs1433739 | -0.693 | 0.585 | -0.491 | 1.629 | -0.003 | -0.013 | -0.013 | 0 | 0.11 | 0.08 | 0.26 | 0.2 | 0.12 | 0.19 |
| 6 | 40445234 | rs16893656 | -1.518 | 0.177 | 1.051 | 1.895 | 0.051 | 0.2 | 0.2 | 2.132 | 0.07 | 0.42 | 0.09 | 0.13 | 0.03 | 0.02 |
| 6 | 40449290 | rs9394699 | -2.073 | 0.126 | 1.382 | 1.872 | 0.007 | 0.154 | 0.154 | 0.854 | 0.12 | 0.48 | 0.15 | 0.23 | 0.06 | 0.08 |
| 6 | 40452054 | rs4711637 | -0.643 | 0.194 | 0.996 | 1.798 | -0.014 | -0.007 | -0.007 | 0.015 | 0.06 | 0.02 | 0.1 | 0.06 | 0.06 | 0.16 |
| 6 | 40458259 | rs6925136 | -1.728 | 0.161 | 1.321 | 1.792 | -0.004 | 0.022 | 0.022 | 0.045 | 0.08 | 0.22 | 0.1 | 0.16 | 0.04 | 0.07 |
| 6 | 40458625 | rs2078460 | -0.565 | -0.377 | 1.325 | 1.449 | -0.028 | 0.022 | 0.022 | 0 | 0.28 | 0.41 | 0.23 | 0.34 | 0.22 | 0.29 |
| 6 | 40460226 | rs9357332 | -0.941 | -0.128 | 1.311 | 1.368 | 0.013 | 0.09 | 0.09 | 0 | 0.06 | 0.16 | 0.11 | 0.2 | 0.03 | 0.09 |
| 6 | 40461919 | rs9369207 | -0.414 | -0.096 | 1.116 | 1.36 | 0 | 0.01 | 0.01 | 0 | 0.05 | 0.03 | 0.08 | 0.17 | 0.03 | 0.07 |
| 6 | 40469438 | rs2436735 | -0.139 | 0.292 | 0.74 | 1.26 | -0.012 | -0.011 | -0.011 | 0.864 | 0.35 | 0.28 | 0.56 | 0.54 | 0.41 | 0.32 |
| 6 | 40474941 | rs2436737 | -0.137 | 0.299 | 1.156 | 1.246 | 0.027 | 0.003 | 0.003 | 0.018 | 0.36 | 0.46 | 0.63 | 0.62 | 0.44 | 0.4 |
| 6 | 40475808 | rs1036287 | -0.126 | 0.302 | 1.157 | 1.248 | 0.028 | 0.003 | 0.003 | 0.022 | 0.36 | 0.46 | 0.64 | 0.62 | 0.44 | 0.34 |
| 6 | 40486159 | rs2235705 | 0.207 | 0.265 | 1.024 | 1.025 | 0.007 | 0 | 0 | 0 | 0.63 | 0.59 | 0.33 | 0.36 | 0.51 | 0.53 |
| 6 | 40487063 | rs909982 | -0.281 | 0.389 | 1.036 | 1.008 | 0.021 | 0.014 | 0.014 | 0 | 0.69 | 0.62 | 0.41 | 0.4 | 0.6 | 0.67 |
| 6 | 40487789 | rs2281263 | 1.047 | -0.403 | 1.193 | 0.632 | 0.001 | 0.006 | 0.006 | 0 | 0.3 | 0.31 | 0.17 | 0.19 | 0.28 | 0.23 |
| 6 | 40489573 | rs909988 | 0.132 | 0.376 | 0.984 | 1.023 | 0.014 | 0.005 | 0.005 | 0 | 0.64 | 0.6 | 0.39 | 0.42 | 0.54 | 0.54 |
| 6 | 40491240 | rs382023 | -0.55 | 0.356 | 0.89 | 1.207 | -0.082 | -0.118 | -0.118 | 0.49 | 0.72 | 0.93 | 0.52 | 0.51 | 0.63 | 0.72 |
| 6 | 40493903 | rs403606 | -0.359 | 0.517 | 0.869 | 1.165 | -0.056 | -0.074 | -0.074 | 0 | 0.7 | 0.88 | 0.5 | 0.5 | 0.58 | 0.71 |
| 6 | 40494855 | rs7356878 | 0.441 | 0.394 | 0.922 | 1.106 | 0.003 | 0.006 | 0.006 | 0 | 0.58 | 0.5 | 0.37 | 0.47 | 0.49 | 0.56 |
| 6 | 40503796 | rs1433742 | 0.061 | 0.723 | 0.728 | 0.279 | -0.021 | 0.033 | 0.033 | 0 | 0.64 | 0.76 | 0.41 | 0.22 | 0.47 | 0.49 |
| 6 | 40512251 | rs7768343 | 0.428 | 0.798 | 0.494 | 0.234 | 0.011 | 0.02 | 0.02 | 0 | 0.58 | 0.68 | 0.29 | 0.22 | 0.39 | 0.45 |
| 6 | 40516220 | rs410066 | 0.055 | 1.426 | 0.292 | 0.237 | -0.051 | 0.013 | 0.013 | 0.396 | 0.65 | 0.83 | 0.42 | 0.31 | 0.56 | 0.61 |
| 6 | 40517949 | rs376285 | -0.92 | 1.521 | 0.292 | 0.223 | -0.009 | 0.011 | 0.011 | 0 | 0.06 | 0.05 | 0.14 | 0.03 | 0.16 | 0.15 |
| 6 | 40519443 | rs432959 | -0.906 | 1.522 | 0.293 | 0.224 | -0.009 | 0.011 | 0.011 | 0 | 0.06 | 0.05 | 0.14 | 0.03 | 0.16 | 0.14 |

| | | | | | | | | | | | | | | | | |
|---|---|---|---|---|---|---|---|---|---|---|---|---|---|---|---|---|
| 6 | 40522176 | rs7775867 | 0.588 | 1.457 | 0.264 | 0.183 | 0.041 | 0.052 | 0.052 | 0.014 | 0.56 | 0.7 | 0.28 | 0.24 | 0.38 | 0.38 |
| 6 | 40530612 | rs846509 | 0.075 | 1.778 | 0.476 | 0.362 | 0.001 | -0.001 | -0.001 | 0 | 0.67 | 0.7 | 0.47 | 0.32 | 0.6 | 0.62 |
| 6 | 40530800 | rs10947897 | 0.551 | 1.707 | 0.633 | 0.354 | 0.016 | -0.027 | -0.027 | 0 | 0.6 | 0.55 | 0.38 | 0.27 | 0.52 | 0.59 |
| 6 | 40536128 | rs2494938 | 0.545 | 1.718 | 0.64 | 0.328 | 0.016 | -0.027 | -0.027 | 0 | 0.61 | 0.56 | 0.38 | 0.27 | 0.52 | 0.59 |
| 6 | 40540482 | rs1737681 | 0.405 | 1.718 | 0.372 | 0.341 | -0.003 | 0.02 | 0.02 | 0 | 0.62 | 0.73 | 0.33 | 0.28 | 0.47 | 0.48 |
| 6 | 40542693 | rs2494942 | 0.466 | 1.593 | 0.222 | 0.314 | -0.001 | 0.006 | 0.006 | 0.011 | 0.58 | 0.63 | 0.33 | 0.28 | 0.49 | 0.47 |
| 6 | 40546850 | rs1890997 | 0.452 | 1.664 | 0.124 | 0.335 | -0.001 | 0.015 | 0.015 | 0.037 | 0.59 | 0.66 | 0.35 | 0.29 | 0.49 | 0.45 |
| 6 | 40549558 | rs2473592 | 0.719 | 1.642 | 0.114 | 0.322 | -0.004 | -0.02 | -0.02 | 0 | 0.56 | 0.52 | 0.31 | 0.24 | 0.46 | 0.44 |
| 6 | 40550438 | rs2473594 | 0.293 | 1.625 | 0.05 | 0.294 | -0.025 | 0.017 | 0.017 | 0.001 | 0.63 | 0.73 | 0.39 | 0.29 | 0.53 | 0.47 |
| 6 | 40557115 | rs2473596 | 0.712 | 2.244 | -0.011 | 0.265 | 0.002 | -0.019 | -0.019 | 0 | 0.57 | 0.52 | 0.33 | 0.27 | 0.5 | 0.45 |
| 6 | 40561603 | rs1340863 | 0.792 | 2.061 | -0.046 | 0.258 | -0.003 | -0.013 | -0.013 | 0 | 0.56 | 0.54 | 0.3 | 0.27 | 0.45 | 0.44 |
| 6 | 40565940 | rs846523 | 0.215 | 2.169 | -0.487 | 0.357 | -0.046 | -0.005 | -0.005 | 1.956 | 0.66 | 0.84 | 0.48 | 0.35 | 0.62 | 0.59 |
| 6 | 40572018 | rs12662332 | -0.29 | 1.797 | -0.431 | 0.36 | -0.026 | -0.016 | -0.016 | 0 | 0.05 | 0 | 0.05 | 0.05 | 0.08 | 0.03 |
| 6 | 40581254 | rs2494952 | 0.906 | 1.255 | -0.679 | -0.092 | -0.019 | -0.016 | -0.016 | 0 | 0.3 | 0.43 | 0.18 | 0.26 | 0.25 | 0.29 |
| 6 | 40582580 | rs2494956 | -0.064 | 1.268 | -0.907 | -0.309 | -0.045 | 0.015 | 0.015 | 0 | 0.63 | 0.75 | 0.37 | 0.33 | 0.54 | 0.47 |
| 6 | 40588223 | rs2494970 | -1.196 | 2.017 | -1.089 | -0.383 | -0.062 | 0.062 | 0.062 | 0.285 | 0.79 | 0.99 | 0.74 | 0.95 | 0.79 | 0.77 |
| 6 | 40588343 | rs1737675 | 0.719 | 1.996 | -1.072 | -0.4 | -0.01 | -0.018 | -0.018 | 0 | 0.36 | 0.53 | 0.25 | 0.3 | 0.33 | 0.31 |
| 6 | 40590597 | rs846537 | -0.193 | 2.203 | -1.235 | 0.097 | -0.026 | -0.024 | -0.024 | 0 | 0.71 | 0.87 | 0.65 | 0.69 | 0.7 | 0.68 |
| 6 | 40594841 | rs16893846 | 0.409 | 2.24 | -1.213 | 0.117 | -0.034 | -0.015 | -0.015 | 0 | 0.06 | 0.01 | 0.09 | 0.26 | 0.12 | 0.08 |
| 6 | 40595071 | rs17685085 | 1.054 | 2.091 | -1.141 | 0.072 | 0.018 | 0.015 | 0.015 | 0 | 0.26 | 0.16 | 0.12 | 0.03 | 0.21 | 0.16 |
| 6 | 40596239 | rs2473572 | 0.373 | 2.155 | -1.076 | 0.337 | -0.03 | 0.016 | 0.016 | 0 | 0.63 | 0.74 | 0.36 | 0.33 | 0.52 | 0.47 |
| 6 | 40598359 | rs2494974 | 0.028 | 2.188 | -1.404 | 0.215 | -0.048 | -0.038 | -0.038 | 0 | 0.29 | 0.12 | 0.38 | 0.33 | 0.34 | 0.33 |
| 6 | 40600283 | rs9394711 | -0.689 | 2.398 | -0.571 | 0.904 | -0.016 | 0.004 | 0.004 | 0 | 0.1 | 0.07 | 0.27 | 0.37 | 0.16 | 0.21 |
| 6 | 40600642 | rs9349160 | -0.763 | 2.393 | -0.497 | 0.901 | 0.008 | 0.054 | 0.054 | 0 | 0.1 | 0.12 | 0.28 | 0.37 | 0.16 | 0.21 |
| 6 | 40601537 | rs9367063 | -1.206 | 2.469 | -0.494 | 0.714 | -0.001 | -0.004 | -0.004 | 0.371 | 0.48 | 0.48 | 0.55 | 0.7 | 0.59 | 0.54 |
| 6 | 40603020 | rs7756249 | -0.782 | 1.544 | -0.703 | 0.195 | 0.019 | 0.081 | 0.081 | 0 | 0.1 | 0.16 | 0.28 | 0.37 | 0.16 | 0.2 |
| 6 | 40603057 | rs4301303 | 0.278 | 1.386 | -0.741 | 0.038 | 0.001 | -0.028 | -0.028 | 0.051 | 0.26 | 0.22 | 0.12 | 0.03 | 0.2 | 0.16 |
| 6 | 40605168 | rs9369215 | -0.293 | 1.358 | -0.787 | 0.299 | -0.002 | -0.004 | -0.004 | 0 | 0.41 | 0.37 | 0.45 | 0.43 | 0.45 | 0.39 |
| 6 | 40608248 | rs9369216 | -0.952 | 1.084 | -0.877 | -0.193 | 0.002 | 0.006 | 0.006 | 0.021 | 0.24 | 0.24 | 0.18 | 0.08 | 0.25 | 0.19 |
| 6 | 40616890 | rs10498750 | -0.599 | 1.904 | -0.856 | -0.156 | -0.009 | 0.007 | 0.007 | 0.021 | 0.2 | 0.13 | 0.17 | 0.1 | 0.3 | 0.15 |
| 6 | 40617312 | rs7765378 | 0.873 | 0.713 | -0.676 | -0.359 | 0 | -0.013 | -0.013 | 0.958 | 0.58 | 0.57 | 0.64 | 0.7 | 0.58 | 0.67 |
| 6 | 40618081 | rs12526543 | -0.101 | 0.464 | -0.709 | -0.314 | -0.03 | -0.035 | -0.035 | 0.004 | 0.21 | 0.29 | 0.18 | 0.09 | 0.12 | 0.16 |
| 6 | 40627425 | rs9394715 | -0.307 | 1.271 | -0.966 | -0.304 | -0.068 | 0.008 | 0.008 | 0.22 | 0.2 | 0.02 | 0.15 | 0.09 | 0.23 | 0.16 |
| 6 | 40628853 | rs4275056 | 0.692 | 0.791 | -0.337 | -0.309 | -0.001 | -0.016 | -0.016 | 0 | 0.63 | 0.54 | 0.61 | 0.73 | 0.64 | 0.7 |
| 6 | 40634681 | rs4560641 | 0.76 | 0.737 | -0.157 | -0.346 | -0.002 | -0.012 | -0.012 | 0 | 0.63 | 0.51 | 0.61 | 0.73 | 0.64 | 0.7 |
| 6 | 40635410 | rs16893884 | 0.763 | 1.013 | -0.444 | -0.262 | 0.01 | 0.008 | 0.008 | 0 | 0.47 | 0.4 | 0.51 | 0.64 | 0.45 | 0.56 |
| 6 | 40637653 | rs6901833 | -0.278 | 0.887 | -0.398 | 0.426 | 0 | 0.017 | 0.017 | 1.488 | 0.26 | 0.52 | 0.24 | 0.32 | 0.24 | 0.44 |
| 6 | 40638510 | rs9369219 | 0.275 | 1.056 | -0.433 | 0.404 | -0.002 | 0.004 | 0.004 | 0.051 | 0.21 | 0.19 | 0.17 | 0.1 | 0.24 | 0.16 |
| 6 | 40641396 | rs4714358 | -0.102 | 0.941 | -0.385 | 0.42 | -0.002 | 0.021 | 0.021 | 1.159 | 0.24 | 0.41 | 0.22 | 0.32 | 0.23 | 0.44 |
| 6 | 40644699 | rs12662449 | 0.191 | 1.02 | -0.63 | 0.496 | -0.043 | -0.015 | -0.015 | 0 | 0.22 | 0.09 | 0.19 | 0.19 | 0.26 | 0.2 |
| 6 | 40645737 | rs9394718 | 0.191 | 1.02 | -0.63 | 0.496 | -0.043 | -0.015 | -0.015 | 0 | 0.22 | 0.09 | 0.19 | 0.19 | 0.26 | 0.2 |
| 6 | 40648757 | rs9471397 | 0.099 | 0.994 | -0.584 | 0.483 | -0.015 | 0 | 0 | 0.065 | 0.22 | 0.11 | 0.19 | 0.19 | 0.26 | 0.2 |
| 6 | 40656881 | rs9357339 | -0.159 | 0.938 | -0.605 | 0.43 | -0.003 | 0.003 | 0.003 | 0.168 | 0.24 | 0.23 | 0.2 | 0.18 | 0.27 | 0.2 |
| 6 | 40657592 | rs7752740 | -0.162 | 0.97 | -0.624 | 0.561 | -0.002 | 0.001 | 0.001 | 0.84 | 0.24 | 0.24 | 0.21 | 0.21 | 0.27 | 0.2 |
| 6 | 40664388 | rs9367072 | 0.458 | 1.116 | -0.976 | 0.72 | -0.021 | -0.031 | -0.031 | 0.098 | 0.05 | 0 | 0.07 | 0.21 | 0.05 | 0.19 |
| 6 | 40665415 | rs6931798 | -0.354 | 1.004 | -0.847 | 0.615 | -0.008 | 0.013 | 0.013 | 2.025 | 0.24 | 0.44 | 0.21 | 0.29 | 0.21 | 0.45 |
| 6 | 40666309 | rs4588686 | -0.071 | 1.345 | -0.982 | 0.552 | -0.044 | 0.007 | 0.007 | 0 | 0.21 | 0.09 | 0.19 | 0.12 | 0.27 | 0.15 |
| 6 | 40680398 | rs6905110 | -0.716 | 1.495 | -0.8 | 0.765 | -0.04 | -0.004 | -0.004 | 0.017 | 0.21 | 0.14 | 0.2 | 0.15 | 0.27 | 0.14 |
| 6 | 40683106 | rs9462644 | -0.222 | 1.294 | -0.749 | 0.775 | -0.006 | 0.003 | 0.003 | 0 | 0.1 | 0.05 | 0.09 | 0.06 | 0.09 | 0.06 |
| 6 | 40688897 | rs4340999 | 0.406 | 0.632 | -0.26 | 0.906 | 0.01 | 0.002 | 0.002 | 0 | 0.53 | 0.57 | 0.54 | 0.53 | 0.61 | 0.55 |
| 6 | 40689730 | rs12529308 | 0.29 | 0.378 | -0.125 | 0.841 | 0.014 | -0.01 | -0.01 | 1.128 | 0.31 | 0.2 | 0.35 | 0.37 | 0.3 | 0.38 |
| 6 | 40689997 | rs4541736 | 0.069 | 0.384 | -0.043 | 0.838 | -0.005 | -0.005 | -0.005 | 0 | 0.86 | 0.82 | 0.9 | 0.9 | 0.91 | 0.93 |
| 6 | 40690493 | rs7768907 | 0.517 | 0.619 | -0.105 | 0.838 | -0.031 | -0.005 | -0.005 | 0 | 0.12 | 0.1 | 0.14 | 0.1 | 0.22 | 0.09 |
| 6 | 40691370 | rs7773686 | -0.386 | 0.517 | -0.076 | 0.827 | -0.03 | -0.023 | -0.023 | 1.289 | 0.43 | 0.27 | 0.48 | 0.46 | 0.52 | 0.46 |
| 6 | 40693403 | rs7756138 | 1.181 | 0.459 | -0.038 | 0.869 | 0.014 | 0.003 | 0.003 | 0.238 | 0.43 | 0.6 | 0.35 | 0.44 | 0.38 | 0.47 |
| 6 | 40693569 | rs7756632 | 1.325 | 0.453 | 0.095 | 0.866 | -0.003 | -0.001 | -0.001 | 0 | 0.42 | 0.51 | 0.33 | 0.44 | 0.38 | 0.47 |
| 6 | 40698365 | rs9369228 | 0.05 | 0.494 | 0.391 | 0.954 | 0.028 | 0.017 | 0.017 | 0.336 | 0.15 | 0.2 | 0.14 | 0.17 | 0.23 | 0.29 |
| 6 | 40699732 | rs4400213 | 1.398 | 0.409 | 0.34 | 0.919 | -0.004 | -0.007 | -0.007 | 0 | 0.39 | 0.39 | 0.31 | 0.36 | 0.36 | 0.28 |
| 6 | 40700237 | rs6458179 | 0.955 | 0.401 | 0.35 | 0.928 | 0.014 | 0.003 | 0.003 | 0.042 | 0.7 | 0.6 | 0.75 | 0.74 | 0.67 | 0.64 |
| 6 | 40700772 | rs7756298 | 0.289 | 0.414 | 0.207 | 0.948 | 0.007 | -0.021 | -0.021 | 0.133 | 0.32 | 0.21 | 0.44 | 0.38 | 0.31 | 0.37 |
| 6 | 40702482 | rs4308545 | -0.081 | 0.368 | 0.3 | 1.076 | 0.002 | 0.079 | 0.079 | 0 | 0.09 | 0.15 | 0.16 | 0.33 | 0.09 | 0.19 |
| 6 | 40703944 | rs9394723 | -0.166 | 0.405 | 0.296 | 1.086 | 0.003 | 0.082 | 0.082 | 0 | 0.1 | 0.14 | 0.23 | 0.34 | 0.09 | 0.2 |
| 6 | 40704337 | rs7742079 | -0.738 | 0.432 | 0.194 | 0.798 | 0.039 | 0.079 | 0.079 | 1.548 | 0.24 | 0.34 | 0.37 | 0.51 | 0.32 | 0.48 |
| 6 | 40704959 | rs6929268 | 0.275 | 0.235 | 0.492 | 0.65 | 0.006 | 0.01 | 0.01 | 0 | 0.62 | 0.64 | 0.6 | 0.87 | 0.68 | 0.77 |
| 6 | 40708366 | rs4388283 | 0.147 | 0.325 | 0.399 | 0.834 | -0.005 | -0.004 | -0.004 | 0 | 0.85 | 0.8 | 0.86 | 0.92 | 0.9 | 0.94 |
| 6 | 40713216 | rs13209441 | 1.063 | 0.35 | 0.275 | 0.811 | -0.015 | 0 | 0 | 0.442 | 0.59 | 0.33 | 0.55 | 0.42 | 0.57 | 0.44 |
| 6 | 40724131 | rs4714375 | 1.126 | 0.327 | 0.226 | 0.62 | 0.008 | 0.012 | 0.012 | 0 | 0.38 | 0.3 | 0.34 | 0.34 | 0.34 | 0.24 |
| 6 | 40725975 | rs9380991 | 0.421 | 0.506 | -0.118 | 0.8 | -0.045 | -0.033 | -0.033 | 0.468 | 0.15 | 0.09 | 0.13 | 0.19 | 0.23 | 0.32 |
| 6 | 40729065 | rs4714376 | -1.412 | 0.415 | -0.043 | 0.783 | 0.001 | 0.001 | 0.001 | 0 | 0.65 | 0.65 | 0.66 | 0.66 | 0.67 | 0.76 |

| | | | | | | | | | | | | | | | | |
|---|---|---|---|---|---|---|---|---|---|---|---|---|---|---|---|---|
| 6 | 40739924 | rs9394729 | -0.75 | 0.67 | -0.41 | 0.945 | -0.031 | -0.017 | -0.017 | 2.288 | 0.27 | 0.16 | 0.21 | 0.24 | 0.31 | 0.36 |
| 6 | 40753185 | rs9394732 | 0.153 | 0.757 | -0.231 | 0.914 | 0.014 | 0.001 | 0.001 | 0 | 0.84 | 0.57 | 0.83 | 0.9 | 0.88 | 0.89 |
| 6 | 40760413 | rs12213185 | 0.496 | 0.77 | -0.941 | 0.92 | 0.007 | 0.01 | 0.01 | 0 | 0.11 | 0.06 | 0.07 | 0.05 | 0.09 | 0.05 |
| 6 | 40760815 | rs7767835 | 1.01 | 0.72 | -1.038 | 0.885 | -0.019 | -0.006 | -0.006 | 2.499 | 0.66 | 0.25 | 0.68 | 0.71 | 0.65 | 0.56 |
| 6 | 40761873 | rs1929799 | 0.082 | 0.749 | -1.012 | 0.894 | -0.028 | -0.02 | -0.02 | 0 | 0.17 | 0.14 | 0.13 | 0.18 | 0.22 | 0.31 |
| 6 | 40765560 | rs6930345 | 0.786 | 0.64 | -1.187 | 0.815 | 0.003 | 0.028 | 0.028 | 0 | 0.25 | 0.19 | 0.26 | 0.07 | 0.27 | 0.2 |
| 6 | 40765597 | rs9471428 | 0.992 | 0.697 | -1.198 | 0.808 | -0.041 | -0.081 | -0.081 | 0 | 0.42 | 0.06 | 0.43 | 0.65 | 0.39 | 0.38 |
| 6 | 40766106 | rs1929770 | -0.187 | 0.689 | -1.137 | 0.698 | -0.01 | 0.019 | 0.019 | 0 | 0.42 | 0.39 | 0.38 | 0.25 | 0.48 | 0.5 |
| 6 | 40766466 | rs10807244 | -0.187 | 0.689 | -1.148 | 0.698 | -0.012 | 0.022 | 0.022 | 0 | 0.42 | 0.39 | 0.38 | 0.25 | 0.48 | 0.51 |
| 6 | 40770363 | rs1556452 | 0.935 | 0.771 | -2.089 | 0.455 | -0.027 | -0.007 | -0.007 | 0.455 | 0.35 | 0.04 | 0.31 | 0.32 | 0.33 | 0.18 |
| 6 | 40772971 | rs9369238 | -0.304 | 1.447 | -1.248 | 0.728 | -0.004 | 0 | 0 | 0 | 0.14 | 0.13 | 0.11 | 0.1 | 0.2 | 0.14 |
| 6 | 40776593 | rs9367079 | -0.297 | 0.808 | -1.305 | 0.875 | -0.01 | -0.05 | -0.05 | 0.186 | 0.31 | 0.21 | 0.37 | 0.47 | 0.33 | 0.56 |
| 6 | 40778147 | rs17315964 | -0.82 | 0.215 | -1.466 | 0.211 | 0.033 | 0.047 | 0.047 | 0.959 | 0.22 | 0.07 | 0.12 | 0.12 | 0.16 | 0.14 |
| 6 | 40781420 | rs962390 | -0.617 | 1.109 | -0.803 | 0.315 | 0.048 | 0.006 | 0.006 | 0.145 | 0.23 | 0.55 | 0.38 | 0.25 | 0.31 | 0.34 |
| 6 | 40783479 | rs4311505 | -0.3 | 1.034 | -1.444 | 0.027 | 0.021 | 0.042 | 0.042 | 5.642 | 0.27 | 0.08 | 0.15 | 0.13 | 0.19 | 0.16 |
| 6 | 40784861 | rs6939170 | 0.212 | 1.347 | -0.801 | -0.078 | 0.077 | 0.086 | 0.086 | 0 | 0.12 | 0.34 | 0.21 | 0.21 | 0.18 | 0.22 |
| 6 | 40785526 | rs869849 | -1.655 | 1.212 | -0.899 | 0.01 | 0.007 | 0.024 | 0.024 | 0.011 | 0.5 | 0.63 | 0.53 | 0.37 | 0.51 | 0.57 |
| 6 | 40785567 | rs12203707 | -0.206 | 1.185 | -0.894 | 0.009 | 0.021 | 0.032 | 0.032 | 5.223 | 0.28 | 0.08 | 0.15 | 0.14 | 0.2 | 0.24 |
| 6 | 40787459 | rs10738088 | -1.256 | 1.221 | -0.898 | -0.07 | 0 | -0.009 | -0.009 | 4.382 | 0.39 | 0.58 | 0.44 | 0.29 | 0.4 | 0.48 |
| 6 | 40795122 | rs9357341 | -0.961 | 1.159 | -0.544 | -0.141 | -0.004 | -0.069 | -0.069 | 0 | 0.31 | 0.5 | 0.32 | 0.15 | 0.28 | 0.26 |
| 6 | 40795151 | rs9367081 | -0.422 | 1.032 | -0.429 | -0.102 | 0.025 | -0.001 | -0.001 | 0 | 0.07 | 0.15 | 0.11 | 0.06 | 0.09 | 0.03 |
| 6 | 40796422 | rs6935623 | -1.115 | 1.018 | -0.318 | -0.115 | 0.018 | -0.01 | -0.01 | 0 | 0.1 | 0.19 | 0.18 | 0.08 | 0.12 | 0.11 |
| 6 | 40797236 | rs9357342 | -0.95 | 1.001 | -0.347 | -0.086 | -0.008 | -0.062 | -0.062 | 0.171 | 0.29 | 0.51 | 0.25 | 0.14 | 0.24 | 0.16 |
| 6 | 40801346 | rs12196942 | -0.004 | 1.01 | -0.422 | -0.082 | -0.019 | -0.053 | -0.053 | 0.054 | 0.21 | 0.33 | 0.15 | 0.07 | 0.15 | 0.13 |
| 6 | 40801982 | rs10947919 | -0.633 | 1.01 | -0.395 | -0.051 | -0.007 | -0.057 | -0.057 | 0 | 0.28 | 0.48 | 0.25 | 0.14 | 0.24 | 0.16 |
| 6 | 40802980 | rs1929779 | -1.371 | 1.374 | 0.12 | 0.354 | -0.017 | -0.009 | -0.009 | 0 | 0.29 | 0.18 | 0.33 | 0.3 | 0.34 | 0.42 |
| 6 | 40805707 | rs10947920 | -0.557 | 1.608 | 0.156 | 0.484 | -0.072 | -0.048 | -0.048 | 0 | 0.14 | 0.03 | 0.21 | 0.21 | 0.2 | 0.24 |
| 6 | 40814135 | rs6907181 | -1.65 | 0.71 | 0.749 | 0.345 | 0.006 | -0.028 | -0.028 | 0 | 0.09 | 0.41 | 0.19 | 0.02 | 0.07 | 0.11 |
| 6 | 40828945 | rs12196881 | -1.583 | 1.391 | 1.125 | 0.523 | 0.059 | 0.034 | 0.034 | 0 | 0.16 | 0.35 | 0.25 | 0.15 | 0.29 | 0.3 |
| 6 | 40841329 | rs1929762 | -0.868 | 1.421 | 0.921 | 0.504 | -0.017 | -0.004 | -0.004 | 0 | 0.09 | 0.04 | 0.13 | 0.07 | 0.19 | 0.19 |
| 6 | 40844002 | rs10807248 | -1.715 | 1.729 | 0.717 | 0.592 | 0 | 0 | 0 | 0 | 0.24 | 0.23 | 0.31 | 0.29 | 0.33 | 0.42 |
| 6 | 40852739 | rs4711652 | 2.178 | 1.441 | 1.057 | 0.388 | -0.01 | 0.012 | 0.012 | 0.056 | 0.61 | 0.3 | 0.53 | 0.66 | 0.59 | 0.44 |
| 6 | 40854152 | rs10807250 | -0.403 | 1.385 | 0.964 | 0.47 | -0.004 | -0.003 | -0.003 | 0 | 0.14 | 0.1 | 0.17 | 0.21 | 0.15 | 0.18 |
| 6 | 40863535 | rs10947929 | -0.715 | 1.368 | 1.08 | 0.465 | 0.013 | 0.009 | 0.009 | 0 | 0.16 | 0.24 | 0.2 | 0.21 | 0.17 | 0.24 |
| 6 | 40867771 | rs914509 | 1.871 | 1.456 | 1.179 | 0.494 | 0.027 | 0.01 | 0.01 | 0 | 0.69 | 0.43 | 0.58 | 0.66 | 0.59 | 0.55 |
| 6 | 40872904 | rs9369243 | 2.169 | 0.944 | 0.443 | 0.89 | -0.107 | 0.009 | 0.009 | 1.511 | 0.41 | 0.02 | 0.33 | 0.29 | 0.42 | 0.19 |
| 6 | 40889435 | rs16894135 | -0.089 | 1.367 | 1.115 | 0.965 | -0.011 | -0.009 | -0.009 | 0 | 0.13 | 0.09 | 0.17 | 0.2 | 0.15 | 0.19 |
| 6 | 40890792 | rs1929791 | 2.354 | 1.336 | 1.231 | 0.96 | 0.008 | 0.004 | 0.004 | 0.257 | 0.66 | 0.38 | 0.55 | 0.62 | 0.58 | 0.43 |
| 6 | 40891288 | rs7770120 | 1.964 | 0.99 | 1.393 | 0.963 | 0.012 | 0.012 | 0.012 | 0.011 | 0.73 | 0.41 | 0.66 | 0.68 | 0.74 | 0.6 |
| 6 | 40893601 | rs11752762 | -0.299 | 1.062 | 1.116 | 0.982 | 0.013 | 0.008 | 0.008 | 0 | 0.13 | 0.15 | 0.17 | 0.21 | 0.15 | 0.19 |
| 6 | 40894039 | rs914505 | 1.283 | 0.799 | 0.982 | 0.8 | -0.022 | -0.021 | -0.021 | 0 | 0.89 | 0.75 | 0.88 | 0.96 | 0.95 | 0.85 |
| 6 | 40896656 | rs2395760 | 1.633 | 0.814 | 0.946 | 0.809 | 0.003 | 0.007 | 0.007 | 0.092 | 0.87 | 0.72 | 0.86 | 0.89 | 0.89 | 0.83 |
| 6 | 40904543 | rs1929794 | -0.986 | 0.82 | 0.724 | 1.003 | 0.001 | -0.002 | -0.002 | 1.357 | 0.32 | 0.34 | 0.27 | 0.27 | 0.33 | 0.41 |
| 6 | 40907457 | rs6938916 | -1.087 | 0.491 | 0.968 | 0.972 | -0.029 | -0.008 | -0.008 | 0 | 0.12 | 0.23 | 0.11 | 0.07 | 0.07 | 0.18 |
| 6 | 40911805 | rs1929760 | 1.613 | 0.605 | 1.012 | 0.966 | 0.009 | 0.007 | 0.007 | 0 | 0.54 | 0.33 | 0.6 | 0.62 | 0.58 | 0.36 |
| 6 | 40914484 | rs11756334 | -0.849 | 0.656 | 1.097 | 0.993 | 0.002 | -0.001 | -0.001 | 0.063 | 0.32 | 0.34 | 0.27 | 0.27 | 0.33 | 0.41 |
| 6 | 40917816 | rs12197636 | 0.904 | 0.595 | 0.889 | 1.009 | -0.002 | 0 | 0 | 0 | 0.63 | 0.35 | 0.66 | 0.67 | 0.64 | 0.5 |
| 6 | 40921160 | rs9471447 | 1.339 | 0.591 | 0.414 | 0.985 | 0.008 | 0.007 | 0.007 | 0 | 0.54 | 0.31 | 0.6 | 0.62 | 0.58 | 0.36 |
| 6 | 40923410 | rs991762 | -0.48 | 0.692 | 0.396 | 1.248 | 0 | 0.012 | 0.012 | 0 | 0.2 | 0.23 | 0.25 | 0.36 | 0.21 | 0.21 |
| 6 | 40926958 | rs7746902 | 0.621 | 0.726 | 0.391 | 0.967 | -0.008 | 0.023 | 0.023 | 0 | 0.65 | 0.56 | 0.56 | 0.43 | 0.7 | 0.56 |
| 6 | 40927188 | rs7767329 | 0.068 | 0.824 | 0.282 | 0.893 | -0.001 | -0.004 | -0.004 | 0.046 | 0.16 | 0.22 | 0.15 | 0.09 | 0.16 | 0.17 |
| 6 | 40928923 | rs7739373 | 1.085 | 0.861 | -0.061 | 0.804 | -0.014 | 0.062 | 0.062 | 0 | 0.47 | 0.26 | 0.37 | 0.34 | 0.52 | 0.32 |
| 6 | 40939379 | rs11754308 | 1.24 | 0.623 | -0.524 | 0.613 | -0.079 | 0.129 | 0.129 | 0.616 | 0.43 | 0.07 | 0.38 | 0.22 | 0.51 | 0.26 |
| 6 | 40946626 | rs9381016 | 0.746 | 0.486 | -0.168 | 0.868 | 0.014 | -0.025 | -0.025 | 0 | 0.58 | 0.26 | 0.64 | 0.49 | 0.69 | 0.59 |
| 6 | 40949933 | rs9381017 | -0.061 | 0.431 | 0.377 | 1.27 | 0.006 | 0.073 | 0.073 | 0 | 0.13 | 0.22 | 0.18 | 0.33 | 0.14 | 0.23 |
| 6 | 40960742 | rs12664909 | -1.105 | 0.026 | 0.935 | 1.034 | -0.033 | 0.031 | 0.031 | 0 | 0.25 | 0.47 | 0.16 | 0.33 | 0.16 | 0.22 |
| 6 | 40963113 | rs714051 | -1.612 | 0.291 | 0.617 | 0.872 | 0.01 | -0.012 | -0.012 | 0 | 0.42 | 0.72 | 0.32 | 0.5 | 0.32 | 0.44 |
| 6 | 40964595 | rs9369250 | -1.433 | 0.316 | 0.623 | 0.888 | -0.001 | 0 | 0 | 0.017 | 0.37 | 0.63 | 0.29 | 0.49 | 0.29 | 0.36 |
| 6 | 40986623 | rs9381025 | -0.74 | 0.233 | 0.767 | 0.831 | -0.013 | 0.033 | 0.033 | 0 | 0.2 | 0.36 | 0.29 | 0.28 | 0.2 | 0.42 |
| 6 | 40989503 | rs9369255 | -0.048 | 0.276 | 0.74 | 0.855 | 0 | 0.047 | 0.047 | 0 | 0.16 | 0.24 | 0.25 | 0.27 | 0.17 | 0.33 |
| 6 | 40998141 | rs736795 | -0.508 | 0.237 | 0.573 | 0.833 | -0.003 | 0.025 | 0.025 | 0 | 0.18 | 0.24 | 0.27 | 0.28 | 0.2 | 0.41 |
| 6 | 40998167 | rs742493 | -0.024 | 0.46 | 0.564 | 0.875 | -0.033 | 0.025 | 0.025 | 0 | 0.09 | 0.03 | 0.09 | 0.04 | 0.12 | 0.09 |
| 6 | 41008747 | rs2073015 | -0.074 | 0.326 | 0.634 | 0.924 | 0.008 | 0.062 | 0.062 | 0 | 0.14 | 0.24 | 0.25 | 0.27 | 0.17 | 0.33 |
| 6 | 41016952 | rs2057000 | -1.234 | 0.228 | 0.711 | 0.7 | -0.04 | 0.006 | 0.006 | 0 | 0.25 | 0.49 | 0.28 | 0.27 | 0.19 | 0.3 |
| 6 | 41020510 | rs2235678 | -1.45 | 0.575 | 0.406 | 0.234 | -0.01 | -0.019 | -0.019 | 0 | 0.31 | 0.56 | 0.36 | 0.27 | 0.32 | 0.33 |
| 6 | 41027520 | rs2395754 | -2.209 | 0.534 | 0.482 | 0.393 | -0.026 | -0.006 | -0.006 | 0 | 0.45 | 0.83 | 0.5 | 0.41 | 0.42 | 0.55 |
| 6 | 41029342 | rs2076472 | 0.54 | 0.496 | 0.471 | 0.365 | -0.038 | -0.074 | -0.074 | 0.428 | 0.13 | 0.01 | 0.23 | 0.27 | 0.17 | 0.29 |
| 6 | 41033640 | rs9394761 | 0.538 | 0.489 | 0.476 | 0.37 | -0.038 | -0.074 | -0.074 | 0.881 | 0.13 | 0.01 | 0.23 | 0.27 | 0.17 | 0.29 |
| 6 | 41034858 | rs11280 | -1.181 | 0.493 | 0.501 | 0.372 | -0.037 | 0.009 | 0.009 | 0 | 0.25 | 0.51 | 0.28 | 0.27 | 0.19 | 0.3 |

| | | | | | | | | | | | | | | | | |
|---|---|---|---|---|---|---|---|---|---|---|---|---|---|---|---|---|
| 6 | 41042590 | rs2268188 | -1.48 | 0.507 | 0.495 | 0.406 | -0.011 | -0.019 | -0.019 | 0 | 0.31 | 0.56 | 0.36 | 0.27 | 0.32 | 0.32 |
| 6 | 41049508 | rs9381034 | 0.52 | 0.532 | 0.452 | 0.414 | -0.038 | -0.076 | -0.076 | 2.687 | 0.13 | 0.01 | 0.23 | 0.27 | 0.17 | 0.29 |
| 6 | 41058968 | rs9296354 | -1.388 | 0.493 | 0.498 | 0.307 | -0.022 | -0.128 | -0.128 | 1.964 | 0.19 | 0.52 | 0.13 | 0.01 | 0.15 | 0.11 |
| 6 | 41059169 | rs10080924 | -1.408 | 0.479 | 0.483 | 0.3 | -0.023 | -0.128 | -0.128 | 3.74 | 0.19 | 0.52 | 0.13 | 0.01 | 0.15 | 0.11 |
| 6 | 41059458 | rs12189627 | -1.323 | 0.487 | 0.486 | 0.303 | -0.019 | -0.131 | -0.131 | 0.505 | 0.19 | 0.52 | 0.13 | 0.01 | 0.15 | 0.04 |
| 6 | 41068423 | rs12189634 | -0.7 | 0.449 | 0.538 | 0.287 | 0.012 | -0.026 | -0.026 | 0 | 0.09 | 0.42 | 0.03 | 0 | 0.01 | 0.01 |
| 6 | 41083352 | rs10456499 | -2.511 | 0.632 | 0.152 | 0.347 | -0.075 | 0.077 | 0.077 | 1.094 | 0.57 | 0.96 | 0.61 | 0.76 | 0.57 | 0.74 |
| 6 | 41089681 | rs722442 | -2.009 | 0.158 | 0.768 | 0.451 | 0.024 | -0.013 | -0.013 | 0 | 0.36 | 0.75 | 0.41 | 0.38 | 0.32 | 0.44 |
| 6 | 41091307 | rs11969526 | 0.121 | 0.197 | 0.764 | 0.278 | -0.006 | 0.006 | 0.006 | 0 | 0.08 | 0.09 | 0.1 | 0 | 0.13 | 0.03 |
| 6 | 41116774 | rs12194477 | -1.054 | 0.114 | 0.982 | 0.392 | -0.02 | -0.058 | -0.058 | 0 | 0.11 | 0.41 | 0.03 | 0 | 0.02 | 0.01 |
| 6 | 41127972 | rs7748513 | -1.246 | 0.117 | 1.045 | 0.377 | -0.038 | -0.067 | -0.067 | 0.081 | 0.12 | 0.54 | 0.07 | 0.01 | 0.03 | 0.08 |
| 6 | 41133806 | rs7748777 | -2.005 | 0.126 | 0.123 | 0.408 | -0.028 | 0.005 | 0.005 | 0.677 | 0.51 | 0.93 | 0.52 | 0.41 | 0.43 | 0.56 |
| 6 | 41135850 | rs7759295 | -0.21 | 0.504 | 0.322 | 0.652 | -0.027 | 0.028 | 0.028 | 0 | 0.11 | 0.03 | 0.09 | 0.04 | 0.11 | 0.09 |
| 6 | 41154650 | rs9381040 | -0.623 | 0.374 | 0.015 | 0.715 | -0.001 | -0.01 | -0.01 | 0.174 | 0.24 | 0.2 | 0.36 | 0.35 | 0.3 | 0.37 |
| 6 | 41159153 | rs17328707 | -0.257 | 0.441 | -0.277 | 0.603 | -0.024 | 0.029 | 0.029 | 0 | 0.12 | 0.03 | 0.09 | 0.05 | 0.11 | 0.13 |
| 6 | 41159801 | rs9394766 | -0.141 | 0.078 | -0.232 | 0.792 | -0.049 | -0.05 | -0.05 | 0 | 0.21 | 0.04 | 0.32 | 0.25 | 0.31 | 0.34 |
| 6 | 41162060 | rs13216201 | -0.141 | 0.184 | -0.261 | 0.764 | -0.053 | -0.057 | -0.057 | 0 | 0.22 | 0.04 | 0.32 | 0.26 | 0.31 | 0.34 |
| 6 | 41162518 | rs3747742 | -0.419 | 0.16 | -0.059 | 0.721 | -0.001 | -0.011 | -0.011 | 0 | 0.25 | 0.21 | 0.37 | 0.36 | 0.31 | 0.37 |
| 6 | 41163957 | rs6914903 | -1.084 | -0.19 | 0.283 | 0.011 | -0.08 | -0.084 | -0.084 | 0.058 | 0.13 | 0.54 | 0.05 | 0 | 0.01 | 0.01 |
| 6 | 41164005 | rs6915083 | -1.041 | 0.046 | -0.41 | 0.339 | 0.038 | 0.013 | 0.013 | 0.041 | 0.43 | 0.75 | 0.48 | 0.58 | 0.4 | 0.47 |
| 6 | 41167114 | rs11752528 | 0.709 | 0.344 | -0.307 | -0.078 | 0 | -0.009 | -0.009 | 0 | 0.77 | 0.75 | 0.88 | 0.92 | 0.84 | 0.89 |
| 6 | 41168337 | rs4714431 | 1.129 | 0.501 | -0.314 | 0.2 | -0.006 | -0.002 | -0.002 | 0.302 | 0.39 | 0.23 | 0.35 | 0.4 | 0.38 | 0.38 |
| 6 | 41170895 | rs3807040 | -0.915 | 0.845 | 0.528 | 0.007 | 0.03 | -0.007 | -0.007 | 0 | 0.1 | 0.22 | 0.07 | 0.08 | 0.1 | 0.07 |
| 6 | 41171213 | rs9381042 | 0.677 | 0.701 | 0.94 | -0.268 | -0.018 | -0.067 | -0.067 | 0 | 0.85 | 0.7 | 0.92 | 0.95 | 0.89 | 0.81 |
| 6 | 41171314 | rs4714432 | 1.811 | 0.542 | 0.437 | -0.191 | -0.066 | -0.029 | -0.029 | 0 | 0.42 | 0.13 | 0.49 | 0.45 | 0.43 | 0.38 |
| 6 | 41173128 | rs6938076 | -1.024 | 0.525 | 0.74 | -0.168 | -0.001 | -0.001 | -0.001 | 0 | 0.44 | 0.43 | 0.44 | 0.5 | 0.48 | 0.55 |
| 6 | 41173220 | rs6938568 | -0.763 | 0.528 | 0.641 | -0.157 | -0.002 | -0.006 | -0.006 | 0 | 0.41 | 0.35 | 0.42 | 0.49 | 0.46 | 0.44 |
| 6 | 41176759 | rs11962624 | 0.103 | 0.553 | 0.228 | -0.133 | -0.071 | -0.167 | -0.167 | 0.36 | 0.27 | 0.04 | 0.3 | 0.4 | 0.34 | 0.35 |
| 6 | 41181489 | rs2253676 | 0.075 | 0.424 | 0.54 | -0.221 | -0.013 | -0.042 | -0.042 | 0 | 0.88 | 0.76 | 0.95 | 0.95 | 0.91 | 0.92 |
| 6 | 41185430 | rs1968871 | 1.244 | 0.43 | 0.848 | -0.178 | -0.001 | -0.042 | -0.042 | 0.305 | 0.78 | 0.57 | 0.86 | 0.87 | 0.81 | 0.84 |
| 6 | 41185458 | rs9367097 | 0.203 | 0.456 | 0.617 | -0.155 | -0.068 | -0.155 | -0.155 | 0 | 0.27 | 0.03 | 0.3 | 0.38 | 0.34 | 0.35 |
| 6 | 41185474 | rs9349180 | 1.904 | 0.052 | 0.943 | -0.287 | 0.026 | 0.08 | 0.08 | 5.38 | 0.69 | 0.17 | 0.8 | 0.84 | 0.77 | 0.73 |
| 6 | 41195903 | rs1872241 | 1.7 | 0.097 | 0.841 | -0.261 | -0.015 | -0.06 | -0.06 | 0 | 0.73 | 0.42 | 0.81 | 0.84 | 0.77 | 0.73 |
| 6 | 41196156 | rs2803496 | -0.03 | 0.088 | 0.877 | -0.267 | -0.01 | -0.041 | -0.041 | 0 | 0.88 | 0.77 | 0.95 | 0.95 | 0.91 | 0.92 |
| 6 | 41196605 | rs9369265 | 1.506 | 0.133 | 0.729 | -0.233 | -0.015 | -0.06 | -0.06 | 0 | 0.73 | 0.42 | 0.81 | 0.84 | 0.77 | 0.73 |
| 6 | 41196727 | rs2101269 | 1.506 | 0.133 | 0.731 | -0.233 | -0.015 | -0.061 | -0.061 | 0.308 | 0.73 | 0.43 | 0.81 | 0.84 | 0.77 | 0.73 |
| 6 | 41197857 | rs7769759 | 1.515 | 0.129 | 0.721 | -0.235 | -0.015 | -0.06 | -0.06 | 0 | 0.73 | 0.42 | 0.81 | 0.84 | 0.77 | 0.73 |
| 6 | 41200888 | rs9357349 | -0.316 | 0.287 | 0.534 | -0.23 | -0.004 | 0.007 | 0.007 | 0 | 0.33 | 0.36 | 0.33 | 0.39 | 0.33 | 0.3 |
| 6 | 41204805 | rs13200807 | -0.316 | 0.222 | 0.534 | -0.23 | -0.005 | 0.007 | 0.007 | 0 | 0.33 | 0.36 | 0.33 | 0.39 | 0.33 | 0.3 |
| 6 | 41205586 | rs1982755 | 2.051 | 0.198 | 0.431 | -0.271 | -0.122 | -0.081 | -0.081 | 0.29 | 0.4 | 0.08 | 0.48 | 0.46 | 0.44 | 0.39 |
| 6 | 41205690 | rs9296359 | 0.226 | 0.153 | 0.413 | -0.296 | -0.051 | -0.15 | -0.15 | 0 | 0.28 | 0.04 | 0.3 | 0.38 | 0.33 | 0.3 |
| 6 | 41205770 | rs1982756 | 0.237 | 0.171 | 0.569 | -0.298 | -0.018 | -0.061 | -0.061 | 0 | 0.85 | 0.69 | 0.93 | 0.95 | 0.9 | 0.86 |
| 6 | 41206065 | rs11753659 | 2.006 | 0.259 | -0.298 | -0.24 | -0.122 | -0.082 | -0.082 | 0.449 | 0.4 | 0.08 | 0.48 | 0.46 | 0.44 | 0.39 |
| 6 | 41206704 | rs2395767 | 1.118 | 0.248 | -0.226 | -0.225 | 0.062 | 0.009 | 0.009 | 0 | 0.5 | 0.24 | 0.57 | 0.56 | 0.56 | 0.44 |
| 6 | 41210570 | rs1484267 | 0.909 | 0.286 | -0.228 | -0.193 | 0.036 | 0.001 | 0.001 | 0 | 0.52 | 0.31 | 0.59 | 0.57 | 0.57 | 0.57 |
| 6 | 41215058 | rs9394778 | -0.481 | 0.386 | -0.247 | -0.061 | -0.024 | -0.06 | -0.06 | 3.213 | 0.37 | 0.16 | 0.38 | 0.48 | 0.44 | 0.34 |
| 6 | 41219627 | rs3997700 | 1.335 | 0.48 | -0.315 | -0.424 | -0.038 | 0.065 | 0.065 | 0 | 0.38 | 0.17 | 0.4 | 0.29 | 0.4 | 0.32 |
| 6 | 41220080 | rs4711666 | -0.503 | 0.552 | -0.134 | -0.464 | 0.002 | -0.003 | -0.003 | 0 | 0.11 | 0.14 | 0.11 | 0.11 | 0.13 | 0.06 |
| 6 | 41220384 | rs4714443 | -0.963 | 0.507 | -0.259 | -0.491 | -0.015 | -0.047 | -0.047 | 0 | 0.19 | 0.59 | 0.17 | 0.11 | 0.16 | 0.2 |
| 6 | 41224085 | rs2101270 | 0.382 | 0.462 | -0.612 | -0.342 | -0.001 | 0.043 | 0.043 | 0 | 0.57 | 0.8 | 0.57 | 0.41 | 0.56 | 0.52 |
| 6 | 41236513 | rs9369269 | 0.368 | 0.538 | -0.877 | -0.118 | -0.021 | -0.081 | -0.081 | 0.918 | 0.17 | 0.04 | 0.27 | 0.53 | 0.22 | 0.31 |
| 6 | 41238833 | rs11755124 | 1.522 | 0.893 | -1.104 | -0.494 | -0.016 | -0.025 | -0.025 | 0 | 0.34 | 0.11 | 0.34 | 0.35 | 0.33 | 0.31 |
| 6 | 41240171 | rs16894387 | -0.485 | 1.002 | -0.314 | -0.634 | 0.031 | -0.086 | -0.086 | 0 | 0.14 | 0.27 | 0.15 | 0 | 0.16 | 0.03 |
| 6 | 41241292 | rs2170887 | 0.46 | 0.988 | -0.797 | -0.398 | -0.043 | -0.133 | -0.133 | 0 | 0.18 | 0.02 | 0.25 | 0.48 | 0.22 | 0.31 |
| 6 | 41243740 | rs2234246 | 0.654 | 0.825 | -0.499 | -0.694 | -0.007 | 0.139 | 0.139 | 0 | 0.52 | 0.33 | 0.52 | 0.25 | 0.52 | 0.27 |
| 6 | 41243991 | rs2234243 | 0.37 | 0.862 | -0.245 | -0.692 | 0.028 | -0.03 | -0.03 | 0.035 | 0.06 | 0.11 | 0.09 | 0 | 0.08 | 0.02 |
| 6 | 41244020 | rs2234242 | 0.865 | 0.776 | -0.259 | -0.532 | 0.001 | 0.01 | 0.01 | 0 | 0.07 | 0.05 | 0.11 | 0.27 | 0.09 | 0.12 |
| 6 | 41246473 | rs4711668 | -0.693 | 0.677 | -0.209 | -0.644 | -0.037 | -0.131 | -0.131 | 1.098 | 0.73 | 0.92 | 0.76 | 0.54 | 0.7 | 0.54 |
| 6 | 41247764 | rs6940092 | 0.519 | 0.607 | -0.141 | -0.867 | 0.016 | 0.044 | 0.044 | 0 | 0.55 | 0.39 | 0.6 | 0.26 | 0.6 | 0.29 |
| 6 | 41250961 | rs6939973 | 0.156 | 0.964 | -0.116 | -0.876 | 0.008 | 0 | 0 | 0 | 0.09 | 0.13 | 0.09 | 0.1 | 0.17 | 0.04 |
| 6 | 41254741 | rs3789204 | 2.064 | 0.657 | -0.222 | -1.095 | -0.026 | 0.131 | 0.131 | 0.812 | 0.3 | 0.05 | 0.35 | 0.13 | 0.27 | 0.18 |
| 6 | 41279161 | rs2019392 | -1.405 | 0.656 | 0.059 | -0.596 | 0.001 | -0.001 | -0.001 | 0.565 | 0.33 | 0.62 | 0.31 | 0.34 | 0.27 | 0.31 |
| 6 | 41281398 | rs595024 | -1.021 | 0.938 | 0.051 | -0.796 | 0.006 | -0.019 | -0.019 | 0 | 0.14 | 0.21 | 0.12 | 0.08 | 0.15 | 0.16 |
| 6 | 41286253 | rs629832 | 0.692 | 0.972 | 0.2 | -0.775 | 0.01 | -0.003 | -0.003 | 0 | 0.9 | 0.9 | 0.89 | 0.92 | 0.86 | 0.84 |
| 6 | 41286507 | rs609119 | -0.626 | 0.723 | 0.187 | -0.982 | 0.004 | 0.017 | 0.017 | 0.058 | 0.49 | 0.44 | 0.4 | 0.79 | 0.48 | 0.65 |
| 6 | 41287482 | rs16894463 | 0.327 | 0.668 | -0.069 | -0.962 | -0.02 | -0.051 | -0.051 | 0 | 0.14 | 0.21 | 0.06 | 0 | 0.07 | 0.02 |
| 6 | 41287623 | rs624743 | 0.819 | 0.868 | -0.722 | -0.735 | -0.049 | -0.117 | -0.117 | 0.423 | 0.23 | 0.08 | 0.18 | 0.47 | 0.27 | 0.44 |
| 6 | 41287803 | rs612399 | 0.184 | 0.878 | -0.689 | -0.763 | -0.016 | -0.024 | -0.024 | 1.017 | 0.29 | 0.22 | 0.26 | 0.47 | 0.34 | 0.5 |
| 6 | 41290458 | rs9471552 | -0.98 | 0.438 | -0.494 | -0.619 | -0.021 | 0.008 | 0.008 | 0.123 | 0.32 | 0.58 | 0.28 | 0.4 | 0.25 | 0.31 |

| | | | | | | | | | | | | | | | | |
|---|---|---|---|---|---|---|---|---|---|---|---|---|---|---|---|---|
| 6 | 41290719 | rs9471554 | 1.569 | 0.556 | -0.733 | -0.72 | 0.003 | 0.097 | 0.097 | 0.022 | 0.33 | 0.2 | 0.41 | 0.13 | 0.32 | 0.19 |
| 6 | 41290769 | rs9471555 | 1.071 | 0.337 | -0.802 | -0.28 | -0.002 | 0.001 | 0.001 | 0.327 | 0.46 | 0.41 | 0.54 | 0.2 | 0.43 | 0.26 |
| 6 | 41293049 | rs6928260 | -0.084 | 0.25 | -0.703 | -0.274 | -0.043 | -0.09 | -0.09 | 0 | 0.14 | 0.32 | 0.07 | 0 | 0.07 | 0.02 |
| 6 | 41293326 | rs12198812 | -0.081 | 0.602 | -0.663 | 0.218 | -0.001 | -0.005 | -0.005 | 0.042 | 0.58 | 0.6 | 0.68 | 0.52 | 0.57 | 0.4 |
| 6 | 41296090 | rs9471566 | -0.172 | 0.585 | -0.647 | 0.209 | -0.006 | 0.053 | 0.053 | 0 | 0.12 | 0.18 | 0.14 | 0.31 | 0.13 | 0.14 |
| 6 | 41296679 | rs12525710 | 0.216 | 0.703 | -1.064 | 0.243 | -0.001 | -0.005 | -0.005 | 1.378 | 0.57 | 0.59 | 0.68 | 0.52 | 0.57 | 0.4 |
| 6 | 41300897 | rs9367104 | 0.19 | 0.335 | -1.988 | 0.393 | -0.007 | -0.054 | -0.054 | 0 | 0.54 | 0.14 | 0.68 | 0.56 | 0.63 | 0.58 |
| 6 | 41303207 | rs6927186 | -0.494 | -0.462 | -1.625 | 0.268 | -0.035 | 0.049 | 0.049 | 0 | 0.16 | 0.24 | 0.1 | 0.31 | 0.08 | 0.11 |
| 6 | 41305976 | rs10947963 | 1.124 | -0.548 | -2.398 | -0.268 | -0.022 | 0.081 | 0.081 | 0 | 0.21 | 0.01 | 0.34 | 0.05 | 0.26 | 0.17 |
| 6 | 41307594 | rs6929418 | 1.048 | -0.36 | -2.453 | -0.054 | -0.069 | 0.025 | 0.025 | 0.532 | 0.18 | 0.05 | 0.12 | 0.12 | 0.21 | 0.19 |
| 6 | 41309552 | rs2236369 | 0.29 | -1.165 | -1.706 | -0.175 | 0.015 | 0.002 | 0.002 | 0 | 0.48 | 0.18 | 0.68 | 0.58 | 0.59 | 0.58 |
| 6 | 41311367 | rs12193066 | -1.043 | -1.257 | -1.291 | -0.367 | 0.037 | -0.049 | -0.049 | 0 | 0.74 | 0.56 | 0.83 | 0.86 | 0.72 | 0.71 |
| 6 | 41311892 | rs12530182 | -0.335 | -1.31 | -1.291 | -0.405 | -0.002 | -0.005 | -0.005 | 0.057 | 0.66 | 0.33 | 0.76 | 0.57 | 0.68 | 0.62 |
| 6 | 41313373 | rs11757145 | 0.95 | -1.565 | -1.303 | -0.722 | -0.007 | -0.037 | -0.037 | 0 | 0.2 | 0.15 | 0.12 | 0 | 0.13 | 0.05 |
| 6 | 41313709 | rs9369272 | -0.693 | -0.542 | -0.814 | 0.024 | 0.047 | -0.051 | -0.051 | 0 | 0.74 | 0.55 | 0.82 | 0.86 | 0.7 | 0.72 |
| 6 | 41314114 | rs9369273 | 0.07 | -0.677 | -0.729 | 0.075 | 0.007 | -0.055 | -0.055 | 0 | 0.71 | 0.55 | 0.83 | 0.86 | 0.73 | 0.71 |
| 6 | 41318438 | rs2273962 | -0.49 | -0.427 | -0.659 | 0.384 | 0.021 | -0.031 | -0.031 | 0 | 0.77 | 0.49 | 0.87 | 0.86 | 0.77 | 0.75 |
| 6 | 41320465 | rs2068392 | -0.499 | -0.327 | -0.878 | 0.763 | -0.011 | 0.049 | 0.049 | 0.019 | 0.09 | 0.15 | 0.07 | 0.3 | 0.03 | 0.1 |
| 6 | 41330218 | rs2146705 | 0.317 | -0.346 | -0.715 | 0.544 | -0.004 | 0.017 | 0.017 | 0 | 0.68 | 0.37 | 0.78 | 0.59 | 0.74 | 0.65 |
| 6 | 41333340 | rs12660438 | -0.883 | -0.463 | -0.773 | 0.695 | -0.013 | 0.054 | 0.054 | 0 | 0.06 | 0.09 | 0.05 | 0.27 | 0.01 | 0.08 |
| 6 | 41336101 | rs6922617 | -0.492 | 0.01 | -0.914 | 0.707 | -0.02 | -0.015 | -0.015 | 0.022 | 0.06 | 0.02 | 0.07 | 0.25 | 0.07 | 0.1 |
| 6 | 41343599 | rs7768605 | -0.358 | -0.338 | -1.162 | -0.102 | 0.001 | 0.043 | 0.043 | 0 | 0.51 | 0.44 | 0.47 | 0.79 | 0.5 | 0.7 |
| 6 | 41345575 | rs7761534 | -0.405 | -0.79 | -1.185 | -0.492 | -0.035 | 0.027 | 0.027 | 0.097 | 0.19 | 0.33 | 0.11 | 0.25 | 0.08 | 0.19 |
| 6 | 41350821 | rs4714455 | 0.599 | -0.154 | -0.083 | -0.17 | 0.024 | -0.009 | -0.009 | 0 | 0.11 | 0.25 | 0.26 | 0.08 | 0.13 | 0.14 |
| 6 | 41351077 | rs584301 | -0.261 | -0.388 | -0.367 | -0.571 | -0.011 | -0.003 | -0.003 | 0 | 0.32 | 0.32 | 0.4 | 0.65 | 0.43 | 0.46 |
| 6 | 41351399 | rs1407789 | 0.19 | -0.206 | -0.605 | -0.435 | -0.069 | -0.103 | -0.103 | 0 | 0.25 | 0.08 | 0.34 | 0.54 | 0.38 | 0.44 |
| 6 | 41361584 | rs9471589 | 0.221 | -0.327 | -0.02 | -0.558 | 0.012 | 0.042 | 0.042 | 0.493 | 0.13 | 0.32 | 0.29 | 0.17 | 0.16 | 0.14 |
| 6 | 41362578 | rs668036 | -1.127 | -0.848 | -0.205 | -0.991 | 0.003 | -0.054 | -0.054 | 0 | 0.68 | 0.71 | 0.78 | 1 | 0.77 | 0.91 |
| 6 | 41364110 | rs1969767 | -1.604 | -0.826 | -0.056 | -0.915 | -0.007 | -0.069 | -0.069 | 0 | 0.76 | 0.77 | 0.83 | 1 | 0.82 | 0.96 |
| 6 | 41364255 | rs1535582 | -0.703 | -0.379 | -0.253 | -0.526 | 0.004 | 0.003 | 0.003 | 0.119 | 0.55 | 0.62 | 0.66 | 0.61 | 0.57 | 0.56 |
| 6 | 41366580 | rs728488 | 0.118 | -0.312 | -0.491 | -0.719 | 0.002 | -0.009 | -0.009 | 0 | 0.35 | 0.35 | 0.5 | 0.51 | 0.38 | 0.45 |
| 6 | 41366838 | rs728489 | 0.287 | 0.301 | -1.109 | -0.546 | 0.001 | -0.003 | -0.003 | 0 | 0.12 | 0.12 | 0.1 | 0.09 | 0.16 | 0.14 |
| 6 | 41371118 | rs17538671 | 1 | 0.401 | -1.292 | -0.333 | -0.099 | -0.077 | -0.077 | 0 | 0.18 | 0.03 | 0.16 | 0.27 | 0.22 | 0.23 |
| 6 | 41377992 | rs4714457 | -0.91 | -0.701 | -0.436 | -0.935 | 0.011 | 0.021 | 0.021 | 0.035 | 0.73 | 0.81 | 0.81 | 0.98 | 0.78 | 0.89 |
| 6 | 41382572 | rs9357360 | -0.834 | -0.475 | -0.369 | -0.623 | -0.01 | -0.018 | -0.018 | 0 | 0.69 | 0.65 | 0.8 | 0.89 | 0.78 | 0.86 |
| 6 | 41382620 | rs9394789 | 0.263 | -0.507 | -0.401 | -0.705 | -0.002 | -0.013 | -0.013 | 0 | 0.52 | 0.46 | 0.68 | 0.8 | 0.6 | 0.72 |
| 6 | 41382931 | rs1983798 | -0.965 | 0.026 | -0.038 | 0.142 | -0.006 | -0.028 | -0.028 | 0 | 0.33 | 0.51 | 0.39 | 0.27 | 0.31 | 0.28 |
| 6 | 41389328 | rs814836 | -1.031 | -0.356 | 0.098 | -0.002 | -0.004 | -0.057 | -0.057 | 0 | 0.13 | 0.22 | 0.09 | 0.01 | 0.08 | 0.04 |
| 6 | 41389484 | rs6912359 | -0.44 | -0.972 | 0.52 | 0.147 | -0.024 | 0.01 | 0.01 | 0.142 | 0.18 | 0.26 | 0.1 | 0.16 | 0.11 | 0.21 |
| 6 | 41392564 | rs628134 | -1.477 | -1.412 | 0.124 | 0.503 | 0.079 | 0.04 | 0.04 | 0 | 0.51 | 0.8 | 0.45 | 0.57 | 0.35 | 0.43 |
| 6 | 41392686 | rs7766624 | -0.277 | -1.472 | -0.013 | 0.48 | 0.003 | 0.108 | 0.108 | 0 | 0.13 | 0.22 | 0.26 | 0.4 | 0.15 | 0.15 |
| 6 | 41394212 | rs814839 | -1.403 | -1.32 | -0.118 | 0.486 | 0.07 | 0.008 | 0.008 | 0 | 0.51 | 0.88 | 0.38 | 0.56 | 0.35 | 0.43 |
| 6 | 41398931 | rs13219238 | -0.92 | -1.052 | 0.264 | 0.521 | -0.019 | 0.014 | 0.014 | 0 | 0.28 | 0.35 | 0.23 | 0.4 | 0.2 | 0.2 |
| 6 | 41404025 | rs13196762 | 0.214 | -1.031 | 0.072 | 0.072 | 0 | 0 | 0 | 0 | 0.77 | 0.79 | 0.83 | 0.83 | 0.8 | 0.74 |
| 6 | 41404666 | rs9369279 | 0.93 | -0.713 | 0.003 | -0.123 | -0.005 | -0.01 | -0.01 | 0 | 0.33 | 0.26 | 0.41 | 0.47 | 0.39 | 0.35 |
| 6 | 41406786 | rs766389 | 1.323 | -0.574 | 0.11 | -0.007 | -0.011 | -0.027 | -0.027 | 0.044 | 0.27 | 0.15 | 0.34 | 0.4 | 0.34 | 0.29 |
| 6 | 41412564 | rs11962670 | -0.146 | -0.709 | 0.521 | 0.062 | -0.002 | 0.012 | 0.012 | 0 | 0.24 | 0.17 | 0.19 | 0.18 | 0.23 | 0.31 |
| 6 | 41414550 | rs9381065 | -0.51 | -0.865 | 0.714 | 0.062 | 0.02 | 0.249 | 0.249 | 0 | 0.05 | 0.22 | 0.11 | 0.31 | 0.06 | 0.08 |
| 6 | 41415737 | rs912881 | 0.604 | -0.877 | 0.451 | -0.053 | -0.007 | 0.021 | 0.021 | 0 | 0.66 | 0.4 | 0.68 | 0.51 | 0.7 | 0.58 |
| 6 | 41417220 | rs1970923 | 0.427 | -0.775 | -0.075 | 0.027 | 0.047 | 0.009 | 0.009 | 0 | 0.49 | 0.28 | 0.6 | 0.45 | 0.62 | 0.55 |
| 6 | 41419988 | rs927770 | 0.641 | -0.686 | -0.505 | 0.022 | 0.027 | -0.004 | -0.004 | 0 | 0.47 | 0.27 | 0.6 | 0.47 | 0.6 | 0.48 |
| 6 | 41430550 | rs2477829 | 0.136 | -0.393 | -0.593 | -0.54 | -0.026 | 0.08 | 0.08 | 0 | 0.57 | 0.72 | 0.56 | 0.8 | 0.54 | 0.54 |
| 6 | 41430572 | rs9471595 | 0.413 | -0.226 | -0.568 | -0.375 | -0.019 | -0.019 | -0.019 | 0 | 0.25 | 0.37 | 0.2 | 0.18 | 0.2 | 0.19 |
| 6 | 41431352 | rs2496634 | -0.161 | 0.512 | -0.87 | 0.06 | -0.029 | 0.038 | 0.038 | 0 | 0.78 | 0.83 | 0.73 | 0.97 | 0.72 | 0.79 |
| 6 | 41433215 | rs9357362 | 0.378 | 0.637 | -0.909 | 0.638 | 0.001 | 0.013 | 0.013 | 0 | 0.27 | 0.22 | 0.34 | 0.56 | 0.31 | 0.29 |
| 6 | 41434817 | rs2496635 | -0.509 | 1.084 | -1.376 | 0.273 | -0.125 | 0.151 | 0.151 | 0 | 0.26 | 0.05 | 0.26 | 0.03 | 0.34 | 0.21 |
| 6 | 41443615 | rs728826 | 1.098 | 0.63 | 0.237 | 2.457 | 0.001 | 0.094 | 0.094 | 0 | 0.85 | 0.72 | 0.83 | 0.67 | 0.84 | 0.89 |
| 6 | 41443689 | rs2477831 | 1.057 | 0.641 | 0.247 | 2.463 | -0.001 | 0.064 | 0.064 | 0 | 0.86 | 0.78 | 0.84 | 0.67 | 0.84 | 0.89 |
| 6 | 41446349 | rs10947972 | -0.72 | 1.625 | 0.015 | 2.695 | -0.015 | -0.018 | -0.018 | 0.24 | 0.19 | 0.19 | 0.23 | 0.4 | 0.22 | 0.5 |
| 6 | 41447015 | rs2496639 | 0.669 | 1.417 | 0.053 | 2.651 | 0.007 | -0.011 | -0.011 | 0 | 0.88 | 0.8 | 0.85 | 0.91 | 0.86 | 0.92 |
| 6 | 41452996 | rs4714468 | -0.859 | 1.722 | -0.318 | 2.677 | -0.014 | 0.023 | 0.023 | 0.139 | 0.07 | 0.04 | 0.2 | 0.25 | 0.13 | 0.31 |
| 6 | 41455056 | rs2496637 | 0.28 | 1.365 | -0.662 | 1.593 | 0.008 | 0.036 | 0.036 | 0 | 0.47 | 0.4 | 0.34 | 0.37 | 0.45 | 0.27 |
| 6 | 41455297 | rs721313 | -1.239 | 1.682 | -0.449 | 1.522 | -0.004 | -0.001 | -0.001 | 0 | 0.14 | 0.09 | 0.3 | 0.23 | 0.17 | 0.34 |
| 6 | 41458286 | rs7742368 | 0.751 | 1.402 | -0.775 | 0.735 | -0.03 | 0.004 | 0.004 | 0 | 0.67 | 0.81 | 0.61 | 0.72 | 0.65 | 0.73 |
| 6 | 41460739 | rs16894733 | 0.138 | 0.855 | 0.022 | -0.094 | 0.003 | -0.104 | -0.104 | 0.024 | 0.21 | 0.4 | 0.19 | 0.01 | 0.22 | 0.19 |
| 6 | 41464901 | rs6913778 | -0.207 | 0.801 | 0.084 | 0.6 | 0.003 | 0.003 | 0.003 | 0 | 0.51 | 0.5 | 0.44 | 0.39 | 0.51 | 0.48 |
| 6 | 41467860 | rs6907558 | -0.307 | 0.473 | 0.147 | 0.553 | -0.009 | 0.009 | 0.009 | 0 | 0.55 | 0.57 | 0.48 | 0.39 | 0.52 | 0.59 |
| 6 | 41469071 | rs6938442 | -1.301 | 0.444 | 0.773 | 0.929 | 0.011 | 0.023 | 0.023 | 0 | 0.14 | 0.25 | 0.16 | 0.12 | 0.13 | 0.21 |
| 6 | 41474727 | rs2495232 | -1.739 | 0.465 | 0.824 | 1.026 | -0.003 | 0.049 | 0.049 | 0.73 | 0.18 | 0.45 | 0.23 | 0.2 | 0.15 | 0.25 |

| | | | | | | | | | | | | | | | | |
|---|---|---|---|---|---|---|---|---|---|---|---|---|---|---|---|---|
| 6 | 41475447 | rs2495233 | -1.545 | 0.184 | 0.927 | 1.067 | -0.027 | 0.066 | 0.066 | 0.608 | 0.15 | 0.41 | 0.21 | 0.18 | 0.1 | 0.26 |
| 6 | 41480900 | rs1886819 | 1.136 | 0.144 | 0.8 | 0.476 | -0.001 | -0.018 | -0.018 | 0 | 0.81 | 0.82 | 0.89 | 0.96 | 0.85 | 0.88 |
| 6 | 41481160 | rs2477838 | 1.736 | -0.053 | 0.928 | 0.466 | 0.009 | 0.019 | 0.019 | 0 | 0.62 | 0.37 | 0.78 | 0.8 | 0.73 | 0.57 |
| 6 | 41489441 | rs2477842 | 1.634 | -0.21 | 0.73 | 1.444 | 0.008 | -0.018 | -0.018 | 0 | 0.71 | 0.7 | 0.63 | 0.39 | 0.76 | 0.74 |
| 6 | 41490488 | rs2495239 | 1.438 | -0.441 | 0.99 | 1.741 | -0.035 | 0.125 | 0.125 | 0.707 | 0.86 | 0.68 | 0.75 | 0.57 | 0.91 | 0.83 |
| 6 | 41502669 | rs1886813 | 1.065 | -0.458 | 0.931 | 1.649 | -0.046 | 0.179 | 0.179 | 0.132 | 0.83 | 0.5 | 0.73 | 0.56 | 0.91 | 0.86 |
| 6 | 41504126 | rs2496652 | 1.069 | -0.293 | 0.062 | 1.324 | 0.086 | -0.027 | -0.027 | 0 | 0.58 | 0.19 | 0.57 | 0.38 | 0.75 | 0.51 |
| 6 | 41519522 | rs2148342 | -1.605 | 0.987 | 0.983 | 1.683 | 0.037 | 0.091 | 0.091 | 0.461 | 0.22 | 0.5 | 0.41 | 0.33 | 0.31 | 0.35 |
| 6 | 41520417 | rs7764874 | -1.063 | 1.263 | 1.397 | 1.882 | 0.01 | 0.033 | 0.033 | 0.75 | 0.1 | 0.19 | 0.13 | 0.16 | 0.13 | 0.18 |
| 6 | 41533579 | rs2104506 | -2.079 | 1.271 | 1.416 | 1.92 | 0.045 | 0.116 | 0.116 | 0.152 | 0.2 | 0.51 | 0.4 | 0.34 | 0.3 | 0.34 |
| 6 | 41536587 | rs4714485 | -2.19 | 1.218 | 1.36 | 1.922 | 0.035 | 0.106 | 0.106 | 0.098 | 0.22 | 0.51 | 0.41 | 0.35 | 0.3 | 0.34 |
| 6 | 41552005 | rs12215942 | 0.681 | 1.193 | 0.215 | 1.366 | -0.053 | -0.052 | -0.052 | 0 | 0.24 | 0.01 | 0.18 | 0.24 | 0.27 | 0.15 |
| 6 | 41552040 | rs912883 | -2.547 | 1.517 | 1.637 | 1.887 | 0.017 | 0.015 | 0.015 | 0.942 | 0.27 | 0.53 | 0.42 | 0.31 | 0.36 | 0.36 |
| 6 | 41552275 | rs11760107 | -1.474 | 1.573 | 1.716 | 1.806 | -0.001 | -0.021 | -0.021 | 0.817 | 0.12 | 0.14 | 0.11 | 0 | 0.17 | 0.14 |
| 6 | 41561354 | rs6920720 | -0.713 | 1.667 | 1.724 | 1.792 | -0.002 | -0.008 | -0.008 | 0.035 | 0.07 | 0.06 | 0.1 | 0 | 0.11 | 0.07 |
| 6 | 41562309 | rs932797 | 0.051 | 2.061 | 1.409 | 1.946 | -0.047 | -0.022 | -0.022 | 0 | 0.08 | 0.01 | 0.19 | 0.37 | 0.15 | 0.08 |
| 6 | 41568064 | rs3747745 | -2.115 | 0.966 | 1.997 | 1.22 | 0.033 | -0.029 | -0.029 | 0 | 0.18 | 0.47 | 0.23 | 0.12 | 0.23 | 0.26 |
| 6 | 41568906 | rs913075 | -2.13 | 0.968 | 1.991 | 1.22 | 0.03 | -0.031 | -0.031 | 0 | 0.19 | 0.48 | 0.23 | 0.12 | 0.23 | 0.26 |
| 6 | 41570350 | rs9381084 | -2.605 | 0.992 | 1.811 | 1.151 | 0.015 | 0.016 | 0.016 | 1.497 | 0.27 | 0.53 | 0.42 | 0.31 | 0.35 | 0.35 |
| 6 | 41573871 | rs9394816 | -1.855 | 0.918 | 1.894 | 1.114 | 0.016 | 0.159 | 0.159 | 0.19 | 0.1 | 0.27 | 0.27 | 0.27 | 0.11 | 0.11 |
| 6 | 41576250 | rs16894840 | -0.883 | 1.009 | 1.757 | 1.055 | 0.091 | -0.032 | -0.032 | 0.552 | 0.08 | 0.22 | 0.06 | 0 | 0.12 | 0.1 |
| 6 | 41576640 | rs9369298 | -2.458 | 0.782 | 2.004 | 0.89 | -0.002 | 0.002 | 0.002 | 0.282 | 0.26 | 0.56 | 0.4 | 0.28 | 0.29 | 0.26 |
| 6 | 41582464 | rs2765943 | -0.851 | 0.609 | 1.579 | 0.791 | 0.063 | -0.035 | -0.035 | 0 | 0.08 | 0.19 | 0.06 | 0 | 0.11 | 0.06 |
| 6 | 41582989 | rs2842639 | -0.851 | 0.609 | 1.579 | 0.791 | 0.063 | -0.035 | -0.035 | 0 | 0.08 | 0.19 | 0.06 | 0 | 0.11 | 0.06 |
| 6 | 41586580 | rs1028818 | -1.423 | 0.472 | 1.424 | 1.065 | -0.023 | 0.046 | 0.046 | 0.031 | 0.16 | 0.19 | 0.3 | 0.32 | 0.12 | 0.16 |
| 6 | 41593996 | rs6928533 | -0.468 | 0.671 | 0.77 | 0.881 | 0.009 | -0.024 | -0.024 | 0 | 0.05 | 0.08 | 0.06 | 0 | 0.1 | 0.05 |
| 6 | 41594613 | rs10223516 | -0.962 | 0.319 | 0.477 | 0.823 | 0 | -0.026 | -0.026 | 0 | 0.17 | 0.47 | 0.13 | 0.11 | 0.14 | 0.19 |
| 6 | 41603194 | rs1973920 | -1.045 | 0.684 | -0.154 | 0.943 | 0.011 | -0.037 | -0.037 | 0 | 0.12 | 0.04 | 0.29 | 0.29 | 0.13 | 0.1 |
| 6 | 41605830 | rs2495229 | -1.276 | 1.024 | 0.393 | 0.172 | 0.075 | -0.088 | -0.088 | 0 | 0.13 | 0.28 | 0.13 | 0.01 | 0.2 | 0.14 |
| 6 | 41608938 | rs2268411 | -0.664 | 0.575 | -0.429 | 0.607 | 0.011 | -0.032 | -0.032 | 0 | 0.11 | 0.03 | 0.07 | 0.18 | 0.09 | 0.13 |
| 6 | 41609220 | rs2268410 | -0.559 | 0.568 | -0.931 | -0.119 | -0.009 | 0.024 | 0.024 | 1.115 | 0.43 | 0.22 | 0.46 | 0.38 | 0.45 | 0.35 |
| 6 | 41609772 | rs16894871 | 0.061 | 0.511 | -0.976 | -0.863 | 0.007 | 0.046 | 0.046 | 1.67 | 0.18 | 0.1 | 0.09 | 0.01 | 0.12 | 0.11 |
| 6 | 41613772 | rs7763699 | 0.281 | 0.724 | -0.723 | -1.138 | -0.002 | -0.051 | -0.051 | 0 | 0.19 | 0.22 | 0.12 | 0.01 | 0.15 | 0.09 |
| 6 | 41614915 | rs3789208 | -0.074 | 0.823 | -1.231 | -0.62 | -0.004 | -0.016 | -0.016 | 0 | 0.38 | 0.22 | 0.61 | 0.56 | 0.45 | 0.3 |
| 6 | 41619050 | rs2842658 | -0.489 | 1.308 | -1.366 | -0.313 | -0.006 | -0.002 | -0.002 | 0 | 0.9 | 0.95 | 0.9 | 0.76 | 0.83 | 0.82 |
| 6 | 41619363 | rs4714501 | 0.066 | 0.948 | -1.334 | -0.307 | -0.006 | -0.018 | -0.018 | 0 | 0.32 | 0.23 | 0.57 | 0.54 | 0.41 | 0.27 |
| 6 | 41621271 | rs2230088 | -0.259 | 0.78 | -1.124 | -0.709 | -0.03 | 0.116 | 0.116 | 0 | 0.19 | 0.01 | 0.1 | 0 | 0.15 | 0.1 |
| 6 | 41623033 | rs2143678 | -1.1 | 1.272 | -0.094 | -0.41 | 0.048 | -0.065 | -0.065 | 0 | 0.11 | 0.28 | 0.1 | 0.01 | 0.15 | 0.05 |
| 6 | 41625429 | rs2746183 | 0.764 | 1.23 | 0.288 | -0.344 | 0.016 | -0.013 | -0.013 | 0 | 0.86 | 0.68 | 0.91 | 0.88 | 0.84 | 0.9 |
| 6 | 41625907 | rs12212842 | -0.336 | 1.514 | -0.13 | 0.082 | -0.015 | -0.024 | -0.024 | 0 | 0.09 | 0.03 | 0.06 | 0.11 | 0.11 | 0.2 |
| 6 | 41626515 | rs2179721 | 0.313 | 1.884 | -0.602 | -0.479 | -0.05 | -0.005 | -0.005 | 0 | 0.79 | 0.93 | 0.86 | 0.82 | 0.75 | 0.75 |
| 6 | 41626625 | rs2746182 | -0.228 | 1.665 | -0.916 | -0.974 | -0.015 | 0.059 | 0.059 | 0.43 | 0.83 | 0.95 | 0.9 | 0.99 | 0.85 | 0.91 |
| 6 | 41626748 | rs9357365 | 0.951 | 1.151 | -0.875 | -0.54 | -0.001 | 0.02 | 0.02 | 0 | 0.08 | 0.08 | 0.12 | 0.16 | 0.06 | 0.14 |
| 6 | 41627063 | rs1474761 | 0.274 | 0.587 | -0.241 | -1.389 | 0 | -0.041 | -0.041 | 0 | 0.61 | 0.38 | 0.72 | 0.99 | 0.66 | 0.87 |
| 6 | 41629726 | rs6458232 | -0.441 | -0.19 | -0.429 | -0.515 | -0.016 | 0.007 | 0.007 | 0 | 0.42 | 0.53 | 0.52 | 0.5 | 0.35 | 0.42 |
| 6 | 41633272 | rs4714503 | 0.559 | 0.082 | -0.56 | -0.107 | 0.003 | -0.005 | -0.005 | 1.55 | 0.11 | 0.08 | 0.17 | 0.12 | 0.06 | 0.17 |
| 6 | 41638759 | rs6458234 | -0.16 | 0.271 | -0.357 | -0.324 | -0.005 | -0.001 | -0.001 | 0 | 0.33 | 0.52 | 0.31 | 0.32 | 0.34 | 0.14 |
| 6 | 41643245 | rs1011100 | -0.414 | 0.879 | -1.308 | -0.063 | -0.064 | 0.012 | 0.012 | 0 | 0.71 | 0.93 | 0.74 | 0.74 | 0.65 | 0.62 |
| 6 | 41643386 | rs1011101 | 0.123 | 0.554 | -0.942 | -0.069 | -0.004 | -0.005 | -0.005 | 0 | 0.53 | 0.69 | 0.56 | 0.45 | 0.51 | 0.46 |
| 6 | 41643608 | rs9471624 | -0.413 | 0.454 | -0.527 | 0.478 | 0.002 | 0.027 | 0.027 | 0 | 0.22 | 0.26 | 0.27 | 0.4 | 0.24 | 0.27 |
| 6 | 41645564 | rs2842647 | 0.342 | 0.284 | -0.638 | 0.442 | -0.007 | 0.004 | 0.004 | 0 | 0.2 | 0.08 | 0.12 | 0.19 | 0.18 | 0.13 |
| 6 | 41648147 | rs4714505 | 1.668 | -0.13 | -0.736 | 0.541 | 0.019 | -0.008 | -0.008 | 4.801 | 0.06 | 0.02 | 0.1 | 0.1 | 0.03 | 0.01 |
| 6 | 41648698 | rs9349199 | 0.563 | 0.061 | -0.894 | 0.463 | -0.017 | -0.041 | -0.041 | 0.268 | 0.18 | 0.1 | 0.2 | 0.34 | 0.23 | 0.41 |
| 6 | 41650736 | rs2842643 | -1.487 | -0.069 | -1.089 | 0.018 | -0.024 | 0.023 | 0.023 | 0 | 0.72 | 0.86 | 0.67 | 0.78 | 0.7 | 0.76 |
| 6 | 41658566 | rs2273068 | 0.805 | -0.524 | -0.793 | 0.007 | 0.011 | 0.009 | 0.009 | 0 | 0.17 | 0.06 | 0.09 | 0.15 | 0.15 | 0.11 |
| 6 | 41658889 | rs1015149 | -0.787 | -0.49 | -0.298 | 0.117 | 0.002 | 0.004 | 0.004 | 0 | 0.52 | 0.57 | 0.45 | 0.44 | 0.47 | 0.34 |
| 6 | 41664501 | rs9462738 | -0.245 | -0.101 | -0.979 | 0.187 | 0.018 | 0.034 | 0.034 | 0 | 0.4 | 0.15 | 0.23 | 0.33 | 0.35 | 0.28 |
| 6 | 41665506 | rs12662389 | 0.039 | -0.272 | -0.309 | 0.729 | 0.017 | 0.015 | 0.015 | 0 | 0.19 | 0.07 | 0.11 | 0.18 | 0.15 | 0.12 |
| 6 | 41670633 | rs1467411 | 0.028 | -0.457 | -0.236 | 0.827 | 0.029 | -0.008 | -0.008 | 0.17 | 0.2 | 0.08 | 0.12 | 0.22 | 0.13 | 0.13 |
| 6 | 41670769 | rs4428488 | -0.79 | -0.362 | 0.467 | 0.652 | -0.025 | -0.034 | -0.034 | 0.212 | 0.29 | 0.49 | 0.28 | 0.23 | 0.27 | 0.19 |
| 6 | 41675740 | rs7756525 | -1.361 | -0.122 | -0.3 | 0.47 | -0.07 | 0.107 | 0.107 | 0.373 | 0.63 | 0.91 | 0.74 | 0.8 | 0.6 | 0.79 |
| 6 | 41678727 | rs1883815 | -0.723 | -0.399 | 0.456 | 1.4 | 0.028 | 0.001 | 0.001 | 0.197 | 0.44 | 0.68 | 0.46 | 0.57 | 0.31 | 0.54 |
| 6 | 41681787 | rs2057068 | -0.702 | -0.383 | 0.486 | 1.603 | 0.024 | -0.004 | -0.004 | 0 | 0.43 | 0.67 | 0.46 | 0.51 | 0.31 | 0.53 |
| 6 | 41684390 | rs9349200 | 0.767 | -0.023 | 0.384 | 1.598 | 0.002 | 0.018 | 0.018 | 0 | 0.08 | 0.07 | 0.05 | 0.21 | 0.14 | 0.2 |
| 6 | 41687366 | rs13202921 | -0.779 | -0.998 | 1.53 | 1.386 | -0.008 | 0.002 | 0.002 | 0 | 0.4 | 0.56 | 0.49 | 0.43 | 0.33 | 0.33 |
| 6 | 41688100 | rs9471635 | -0.114 | -1.2 | 1.094 | 1.564 | -0.004 | 0.07 | 0.07 | 0.765 | 0.07 | 0.17 | 0.14 | 0.16 | 0.07 | 0.19 |
| 6 | 41688990 | rs12525992 | 0.304 | -1.261 | 0.891 | 1.582 | 0.001 | -0.002 | -0.002 | 0 | 0.16 | 0.15 | 0.17 | 0.18 | 0.13 | 0.16 |
| 6 | 41694772 | rs1858584 | -0.648 | -0.972 | 1.094 | 1.416 | 0.003 | 0.096 | 0.096 | 1.207 | 0.07 | 0.22 | 0.14 | 0.16 | 0.07 | 0.19 |
| 6 | 41698134 | rs12664901 | -0.397 | -0.652 | 0.218 | 1.013 | 0.02 | -0.05 | -0.05 | 0 | 0.18 | 0.02 | 0.12 | 0.26 | 0.11 | 0.11 |

| | | | | | | | | | | | | | | | | |
|---|---|---|---|---|---|---|---|---|---|---|---|---|---|---|---|---|
| 6 | 41705612 | rs3804279 | -0.56 | -0.619 | 0.32 | 1 | -0.023 | 0.001 | 0.001 | 0 | 0.21 | 0.33 | 0.28 | 0.26 | 0.13 | 0.13 |
| 6 | 41715997 | rs2040017 | 1.179 | -0.506 | 0.434 | 1.189 | -0.024 | 0.054 | 0.054 | 0 | 0.86 | 0.7 | 0.65 | 0.76 | 0.91 | 0.8 |
| 6 | 41717705 | rs6458238 | -1.202 | -0.748 | 1.044 | 1.091 | -0.009 | 0.026 | 0.026 | 0.301 | 0.11 | 0.4 | 0.09 | 0.06 | 0.07 | 0.06 |
| 6 | 41717908 | rs6912200 | 1.163 | -1.287 | 0.303 | 0.855 | 0.075 | -0.015 | -0.015 | 0 | 0.57 | 0.23 | 0.39 | 0.5 | 0.71 | 0.45 |
| 6 | 41719110 | rs6941539 | 0.511 | -0.88 | 0.293 | 0.901 | 0.003 | -0.004 | -0.004 | 0 | 0.94 | 0.94 | 0.71 | 0.83 | 0.98 | 0.86 |
| 6 | 41726138 | rs1806719 | 0.575 | -0.634 | -0.129 | 0.001 | 0 | 0.055 | 0.055 | 0.115 | 0.43 | 0.26 | 0.51 | 0.23 | 0.4 | 0.29 |
| 6 | 41726339 | rs2395788 | 0.214 | -0.371 | -0.427 | 0.36 | -0.023 | -0.037 | -0.037 | 0 | 0.1 | 0.03 | 0.17 | 0.22 | 0.13 | 0.28 |
| 6 | 41729406 | rs12661968 | -0.141 | -0.48 | -0.164 | 0.301 | 0 | -0.004 | -0.004 | 0 | 0.12 | 0.09 | 0.18 | 0.21 | 0.13 | 0.29 |
| 6 | 41735468 | rs13197439 | 0.056 | -1.191 | -0.137 | -0.095 | -0.04 | -0.014 | -0.014 | 0.922 | 0.44 | 0.16 | 0.28 | 0.49 | 0.49 | 0.35 |
| 6 | 41744948 | rs3761781 | -0.012 | -1.424 | -0.172 | -0.197 | 0.003 | -0.001 | -0.001 | 0.008 | 0.14 | 0.11 | 0.17 | 0.22 | 0.14 | 0.2 |
| 6 | 41745989 | rs3804281 | 0.021 | -1.136 | 0.159 | -0.18 | -0.01 | -0.005 | -0.005 | 0 | 0.56 | 0.73 | 0.73 | 0.53 | 0.54 | 0.66 |
| 6 | 41748263 | rs28360509 | 0.483 | -0.6 | -0.333 | 0.49 | -0.091 | -0.094 | -0.094 | 0.393 | 0.17 | 0.02 | 0.13 | 0.26 | 0.25 | 0.15 |
| 6 | 41749692 | rs12523672 | 0.408 | -0.783 | -0.229 | 0.36 | -0.014 | -0.007 | -0.007 | 0 | 0.54 | 0.71 | 0.7 | 0.54 | 0.51 | 0.65 |
| 6 | 41753084 | rs9394831 | 0.604 | -0.895 | 0.029 | 0.565 | -0.096 | -0.111 | -0.111 | 0.01 | 0.16 | 0.02 | 0.13 | 0.3 | 0.25 | 0.15 |
| 6 | 41754576 | rs9381093 | -0.35 | -0.999 | 0.461 | 0.509 | -0.016 | -0.023 | -0.023 | 0.125 | 0.7 | 0.34 | 0.69 | 0.83 | 0.75 | 0.29 |
| 6 | 41758583 | rs3747751 | 0.997 | -1.212 | 0.315 | 0.28 | 0.02 | -0.001 | -0.001 | 0 | 0.17 | 0.09 | 0.05 | 0.15 | 0.1 | 0.1 |
| 6 | 41761495 | rs12202219 | 1.155 | -1.225 | 0.378 | 0.261 | -0.001 | -0.012 | -0.012 | 0 | 0.1 | 0.09 | 0.1 | 0.01 | 0.13 | 0.04 |
| 6 | 41764080 | rs6902650 | 1.049 | -1.233 | 0.312 | 0.227 | 0 | 0.002 | 0.002 | 0.228 | 0.51 | 0.61 | 0.68 | 0.54 | 0.49 | 0.58 |
| 6 | 41768510 | rs10498752 | 0.989 | -1.317 | 0.983 | 0.172 | -0.001 | -0.001 | -0.001 | 0 | 0.51 | 0.52 | 0.69 | 0.54 | 0.51 | 0.58 |
| 6 | 41773322 | rs2153878 | 0.655 | -1.356 | 1.114 | 0.136 | -0.004 | 0.003 | 0.003 | 2.397 | 0.13 | 0.13 | 0.16 | 0.22 | 0.13 | 0.18 |
| 6 | 41784094 | rs7753507 | 1.092 | -1.385 | 1.125 | 0.081 | 0 | 0 | 0 | 0.165 | 0.5 | 0.52 | 0.68 | 0.54 | 0.5 | 0.57 |
| 6 | 41787740 | rs9394833 | 0.05 | -1.397 | 1.137 | 0.15 | -0.016 | -0.017 | -0.017 | 0 | 0.69 | 0.66 | 0.83 | 0.82 | 0.76 | 0.8 |
| 6 | 41788380 | rs11753329 | 0.614 | -1.402 | 1.155 | 0.206 | 0.004 | -0.012 | -0.012 | 0 | 0.13 | 0.09 | 0.16 | 0.24 | 0.14 | 0.18 |
| 6 | 41792545 | rs12661667 | 0.656 | -1.348 | 1.236 | 0.2 | -0.034 | -0.041 | -0.041 | 0 | 0.17 | 0.07 | 0.13 | 0.3 | 0.25 | 0.15 |
| 6 | 41796970 | rs6899876 | 0.546 | -0.972 | 1.174 | 0.12 | -0.025 | -0.027 | -0.027 | 0.79 | 0.18 | 0.1 | 0.15 | 0.3 | 0.27 | 0.16 |
| 6 | 41814760 | rs2249703 | -1.158 | -1.139 | 1.331 | 0.153 | 0.002 | 0.003 | 0.003 | 0.089 | 0.51 | 0.58 | 0.33 | 0.46 | 0.51 | 0.43 |
| 6 | 41831143 | rs2254479 | -1.41 | -1.191 | 1.179 | 0.111 | -0.027 | 0.004 | 0.004 | 0 | 0.57 | 0.84 | 0.5 | 0.52 | 0.55 | 0.51 |
| 6 | 41869924 | rs1536701 | -0.165 | -1.195 | 1.025 | 0.081 | -0.016 | -0.023 | -0.023 | 0 | 0.31 | 0.34 | 0.17 | 0.16 | 0.24 | 0.2 |
| 6 | 41881073 | rs7746198 | 1.165 | -1.116 | 0.491 | 0.184 | 0.022 | -0.002 | -0.002 | 0 | 0.16 | 0.08 | 0.04 | 0.15 | 0.1 | 0.1 |
| 6 | 41882123 | rs11969866 | 0.581 | -1.086 | 0.428 | 0.2 | 0.004 | -0.011 | -0.011 | 0 | 0.13 | 0.09 | 0.17 | 0.23 | 0.14 | 0.2 |
| 6 | 41885319 | rs9357371 | 0.929 | -1.047 | 0.31 | 0.206 | -0.047 | -0.058 | -0.058 | 0.159 | 0.17 | 0.06 | 0.13 | 0.3 | 0.25 | 0.15 |
| 6 | 41890835 | rs2479723 | -2.44 | -1.1 | 0.131 | 0.03 | -0.088 | 0.006 | 0.006 | 0 | 0.71 | 0.97 | 0.68 | 0.76 | 0.7 | 0.73 |
| 6 | 41892809 | rs10947996 | -0.458 | -0.984 | 0.56 | 0.063 | -0.029 | -0.046 | -0.046 | 2.916 | 0.32 | 0.44 | 0.19 | 0.16 | 0.26 | 0.28 |
| 6 | 41895150 | rs2296916 | 1.313 | -0.821 | -0.181 | 0.307 | 0.027 | -0.041 | -0.041 | 0 | 0.16 | 0 | 0.04 | 0.15 | 0.1 | 0.1 |
| 6 | 41900406 | rs3828855 | 1.134 | -0.808 | -0.135 | 0.314 | 0.03 | -0.004 | -0.004 | 0.123 | 0.16 | 0.07 | 0.04 | 0.15 | 0.1 | 0.1 |
| 6 | 41902655 | rs3218108 | 0.623 | -0.383 | -0.124 | 0.547 | -0.063 | -0.075 | -0.075 | 2.091 | 0.17 | 0.05 | 0.13 | 0.3 | 0.25 | 0.14 |
| 6 | 41905275 | rs3218097 | 0.521 | -0.238 | -0.354 | 0.585 | -0.035 | -0.051 | -0.051 | 2.343 | 0.16 | 0.07 | 0.13 | 0.27 | 0.24 | 0.15 |
| 6 | 41910064 | rs3218086 | -0.222 | -0.425 | -0.553 | 0.507 | 0.013 | -0.041 | -0.041 | 0.067 | 0.18 | 0.1 | 0.33 | 0.39 | 0.16 | 0.4 |
| 6 | 41914378 | rs9349204 | 0.291 | -0.423 | -0.319 | 0.498 | -0.018 | -0.03 | -0.03 | 0 | 0.18 | 0.16 | 0.15 | 0.34 | 0.26 | 0.19 |
| 6 | 41915143 | rs6913232 | -0.607 | -0.47 | -0.261 | 0.234 | -0.004 | 0.024 | 0.024 | 0 | 0.52 | 0.76 | 0.5 | 0.51 | 0.55 | 0.45 |
| 6 | 41921241 | rs10947997 | 0.442 | -0.494 | -0.53 | 0.311 | -0.021 | -0.047 | -0.047 | 0 | 0.13 | 0.03 | 0.18 | 0.23 | 0.15 | 0.22 |
| 6 | 41924931 | rs16895130 | 1.955 | -0.434 | -0.465 | 0.134 | -0.037 | 0.096 | 0.096 | 0.174 | 0.3 | 0.1 | 0.16 | 0.11 | 0.31 | 0.14 |
| 6 | 41925304 | rs11968166 | -0.355 | -0.495 | -0.376 | 0.459 | 0.013 | -0.042 | -0.042 | 0.148 | 0.18 | 0.1 | 0.34 | 0.39 | 0.16 | 0.42 |
| 6 | 41927813 | rs4478405 | 0.012 | -0.216 | -0.194 | 0.338 | 0.001 | -0.007 | -0.007 | 0 | 0.08 | 0.11 | 0.06 | 0.05 | 0.11 | 0.1 |
| 6 | 41928382 | rs9349206 | 0.614 | -0.117 | -0.413 | -0.044 | -0.001 | 0.037 | 0.037 | 0 | 0.23 | 0.14 | 0.15 | 0.01 | 0.26 | 0.12 |
| 6 | 41931569 | rs12110546 | 0.233 | 0.136 | -0.467 | 0.249 | -0.014 | 0.09 | 0.09 | 0 | 0.11 | 0.18 | 0.19 | 0.32 | 0.04 | 0.2 |
| 6 | 41933554 | rs9357372 | 0.769 | 0.243 | -0.863 | -0.254 | -0.014 | 0.088 | 0.088 | 0 | 0.22 | 0.08 | 0.15 | 0.01 | 0.26 | 0.12 |
| 6 | 41934514 | rs4130023 | 1.57 | 0.397 | -0.64 | 0.149 | -0.02 | 0.118 | 0.118 | 0.091 | 0.29 | 0.04 | 0.13 | 0.11 | 0.25 | 0.13 |
| 6 | 41941980 | rs4415146 | -0.724 | 0.364 | -0.146 | 0.505 | -0.002 | 0.006 | 0.006 | 0 | 0.54 | 0.59 | 0.63 | 0.59 | 0.53 | 0.52 |
| 6 | 41977937 | rs11756584 | -0.275 | 0.376 | 0.035 | 0.328 | 0 | -0.001 | -0.001 | 0 | 0.31 | 0.34 | 0.35 | 0.22 | 0.32 | 0.29 |
| 6 | 41978274 | rs4607417 | 0.921 | 0.527 | 0.012 | 0.531 | 0 | 0 | 0 | 0.207 | 0.57 | 0.52 | 0.56 | 0.52 | 0.54 | 0.49 |
| 6 | 41978412 | rs4445045 | 0.512 | 0.675 | -0.082 | 0.383 | 0.006 | -0.014 | -0.014 | 0 | 0.08 | 0.1 | 0.06 | 0.05 | 0.11 | 0.12 |
| 6 | 41980666 | rs6921368 | -0.313 | 0.545 | 0.214 | 0.378 | 0 | -0.017 | -0.017 | 0 | 0.32 | 0.43 | 0.37 | 0.2 | 0.35 | 0.36 |
| 6 | 41987451 | rs4711698 | -1.124 | 0.588 | -0.431 | -0.104 | 0.007 | 0.023 | 0.023 | 0.741 | 0.7 | 0.77 | 0.85 | 0.93 | 0.74 | 0.85 |
| 6 | 41989989 | rs10948005 | 1.073 | 0.576 | -0.698 | 0.18 | -0.046 | -0.059 | -0.059 | 0 | 0.18 | 0.06 | 0.34 | 0.36 | 0.26 | 0.25 |
| 6 | 41991061 | rs4714552 | -1.246 | 0.448 | -0.456 | 0.047 | 0.006 | 0.043 | 0.043 | 0.934 | 0.71 | 0.81 | 0.85 | 0.93 | 0.74 | 0.84 |
| 6 | 41992983 | rs4714554 | -0.569 | 0.607 | -0.44 | 0.231 | -0.001 | -0.002 | -0.002 | 0 | 0.47 | 0.51 | 0.67 | 0.63 | 0.6 | 0.51 |
| 6 | 41993007 | rs9369324 | -0.248 | 0.427 | 0.08 | 0.601 | 0.007 | 0.009 | 0.009 | 0 | 0.3 | 0.4 | 0.33 | 0.27 | 0.34 | 0.27 |
| 6 | 41993688 | rs12214723 | 0.887 | 0.603 | -0.119 | 0.113 | -0.001 | -0.056 | -0.056 | 0 | 0.17 | 0.22 | 0.17 | 0.01 | 0.21 | 0.03 |
| 6 | 41993861 | rs7753265 | 0.235 | 0.684 | -0.771 | 0.21 | 0.016 | 0.02 | 0.02 | 0.072 | 0.56 | 0.34 | 0.54 | 0.69 | 0.64 | 0.68 |
| 6 | 41997926 | rs9357384 | 0.169 | 1.158 | -0.812 | 1.011 | -0.039 | -0.015 | -0.015 | 0 | 0.08 | 0.02 | 0.1 | 0.41 | 0.12 | 0.15 |
| 6 | 42002867 | rs13194688 | -0.2 | 1.045 | -0.762 | -0.485 | -0.029 | 0.075 | 0.075 | 0.011 | 0.38 | 0.22 | 0.32 | 0.03 | 0.49 | 0.22 |
| 6 | 42009583 | rs4333413 | 0.144 | 0.994 | -0.49 | -0.356 | -0.001 | -0.006 | -0.006 | 0 | 0.58 | 0.63 | 0.7 | 0.85 | 0.54 | 0.79 |
| 6 | 42012159 | rs4554318 | 0.386 | 1.12 | -1.028 | -0.134 | -0.013 | 0.083 | 0.083 | 0 | 0.55 | 0.72 | 0.48 | 0.25 | 0.47 | 0.35 |
| 6 | 42018883 | rs11967378 | -0.131 | 1.203 | -0.704 | 0.202 | -0.007 | 0.011 | 0.011 | 0 | 0.25 | 0.17 | 0.27 | 0.18 | 0.3 | 0.37 |
| 6 | 42025058 | rs6917299 | -0.185 | 1.2 | -0.538 | 0.465 | 0.013 | 0.004 | 0.004 | 0 | 0.34 | 0.44 | 0.38 | 0.33 | 0.42 | 0.24 |
| 6 | 42025318 | rs12208813 | 0.128 | 1.329 | -0.442 | 0.706 | -0.023 | -0.027 | -0.027 | 0 | 0.05 | 0.01 | 0.04 | 0.26 | 0.09 | 0.12 |
| 6 | 42026190 | rs4714560 | 1.181 | 1.153 | -0.558 | 0.554 | 0.041 | 0.033 | 0.033 | 0.001 | 0.38 | 0.04 | 0.23 | 0.22 | 0.22 | 0.3 |
| 6 | 42033440 | rs2492943 | 0.149 | 1.227 | -0.285 | 0.45 | 0.04 | 0.03 | 0.03 | 0 | 0.3 | 0.54 | 0.37 | 0.32 | 0.38 | 0.19 |

| | | | | | | | | | | | | | | | | |
|---|---|---|---|---|---|---|---|---|---|---|---|---|---|---|---|---|
| 6 | 42034685 | rs2493833 | 0.729 | 1.586 | 0.483 | 0.528 | 0.007 | 0.005 | 0.005 | 0 | 0.69 | 0.66 | 0.69 | 0.56 | 0.61 | 0.49 |
| 6 | 42037167 | rs12194513 | -1.156 | 1.243 | 0.16 | 0.645 | 0.008 | 0.052 | 0.052 | 0.035 | 0.07 | 0.13 | 0.07 | 0.26 | 0.09 | 0.12 |
| 6 | 42038925 | rs7762770 | 0.808 | 1.182 | 0.303 | -0.172 | -0.007 | -0.079 | -0.079 | 0.051 | 0.71 | 0.47 | 0.75 | 0.92 | 0.71 | 0.62 |
| 6 | 42039487 | rs9349215 | 1.213 | 1.384 | -0.045 | 0.026 | 0.04 | 0.038 | 0.038 | 1.225 | 0.47 | 0.24 | 0.47 | 0.63 | 0.41 | 0.48 |
| 6 | 42039572 | rs7763360 | 0.363 | 1.454 | 0.313 | 0.04 | 0.003 | 0.002 | 0.002 | 0 | 0.07 | 0.17 | 0.06 | 0.06 | 0.08 | 0.02 |
| 6 | 42039666 | rs2492927 | 0.011 | 1.888 | 0.654 | -0.137 | -0.001 | -0.004 | -0.004 | 1.973 | 0.79 | 0.83 | 0.82 | 0.91 | 0.78 | 0.69 |
| 6 | 42040026 | rs9394859 | -0.809 | 1.671 | 1.07 | 0.073 | 0.02 | 0.081 | 0.081 | 0 | 0.12 | 0.3 | 0.19 | 0.25 | 0.16 | 0.11 |
| 6 | 42041541 | rs9349216 | 0.954 | 1.329 | 1.022 | 0.384 | 0.009 | -0.025 | -0.025 | 1.191 | 0.6 | 0.55 | 0.64 | 0.67 | 0.59 | 0.49 |
| 6 | 42044945 | rs3800286 | 0.78 | 1.538 | 0.707 | 0.221 | -0.001 | -0.021 | -0.021 | 0 | 0.69 | 0.44 | 0.73 | 0.9 | 0.69 | 0.7 |
| 6 | 42046079 | rs3800288 | 0.255 | 1.274 | 1.099 | 0.176 | 0.018 | -0.037 | -0.037 | 0.635 | 0.77 | 0.7 | 0.75 | 0.9 | 0.71 | 0.7 |
| 6 | 42047994 | rs6906808 | 2.161 | 1.383 | 0.627 | 0.268 | -0.078 | -0.094 | -0.094 | 0.116 | 0.46 | 0.04 | 0.47 | 0.42 | 0.49 | 0.39 |
| 6 | 42048690 | rs7745547 | -1.663 | 1.351 | 0.705 | 0.281 | 0.009 | 0.086 | 0.086 | 0 | 0.25 | 0.6 | 0.27 | 0.48 | 0.22 | 0.31 |
| 6 | 42050823 | rs9381135 | -0.14 | 1.343 | -0.118 | 0.332 | -0.004 | -0.005 | -0.005 | 0 | 0.11 | 0.14 | 0.1 | 0.24 | 0.1 | 0.2 |
| 6 | 42052577 | rs9357387 | 0.977 | 1.332 | -0.199 | 0.311 | -0.001 | 0.003 | 0.003 | 0.138 | 0.6 | 0.4 | 0.59 | 0.65 | 0.59 | 0.59 |
| 6 | 42053778 | rs2492935 | -1.397 | 1.234 | -0.44 | 0.338 | 0 | 0.013 | 0.013 | 0 | 0.31 | 0.8 | 0.24 | 0.3 | 0.26 | 0.24 |
| 6 | 42054346 | rs2492937 | -1.111 | 1.324 | -0.044 | 0.111 | 0.004 | -0.026 | -0.026 | 0.514 | 0.2 | 0.68 | 0.13 | 0.07 | 0.16 | 0.1 |
| 6 | 42054380 | rs2492938 | -1.27 | 1.431 | -0.084 | 0.145 | 0.015 | 0.004 | 0.004 | 0 | 0.24 | 0.65 | 0.28 | 0.26 | 0.26 | 0.24 |
| 6 | 42060835 | rs9471759 | -0.183 | 0.343 | -0.707 | -1.21 | 0.009 | 0.044 | 0.044 | 1.226 | 0.19 | 0.11 | 0.1 | 0 | 0.15 | 0.02 |
| 6 | 42061733 | rs7749023 | -0.765 | -0.102 | -0.825 | -1.684 | 0.01 | 0.032 | 0.032 | 0.32 | 0.52 | 0.45 | 0.34 | 0.26 | 0.43 | 0.35 |
| 6 | 42066413 | rs9462769 | -0.304 | -0.905 | -0.939 | -1.978 | 0.022 | 0.121 | 0.121 | 0 | 0.29 | 0.13 | 0.17 | 0.07 | 0.27 | 0.13 |
| 6 | 42068171 | rs6458268 | -1.09 | -1.027 | -0.333 | -2.018 | 0.022 | 0.055 | 0.055 | 1.014 | 0.54 | 0.71 | 0.38 | 0.31 | 0.44 | 0.44 |
| 6 | 42068975 | rs4413610 | -0.805 | -1.088 | -0.036 | -1.947 | -0.049 | 0.012 | 0.012 | 0 | 0.25 | 0.42 | 0.22 | 0.3 | 0.19 | 0.37 |
| 6 | 42073958 | rs9394865 | -0.411 | -0.974 | 0.008 | -1.793 | 0.021 | 0.219 | 0.219 | 0 | 0.09 | 0.34 | 0.14 | 0.22 | 0.06 | 0.26 |
| 6 | 42074106 | rs9394866 | -1.274 | -0.972 | -0.074 | -2.09 | -0.049 | 0.04 | 0.04 | 5.237 | 0.61 | 0.87 | 0.42 | 0.31 | 0.47 | 0.47 |
| 6 | 42075461 | rs9381137 | 0.297 | -0.958 | -0.138 | -2.061 | -0.013 | 0.008 | 0.008 | 0 | 0.22 | 0.36 | 0.23 | 0.22 | 0.21 | 0.28 |
| 6 | 42077870 | rs9471765 | -0.731 | -1.029 | -0.146 | -2.119 | 0.013 | 0.02 | 0.02 | 2.039 | 0.43 | 0.64 | 0.32 | 0.24 | 0.33 | 0.34 |
| 6 | 42081642 | rs9471770 | 0.097 | -0.781 | -0.476 | -2.416 | 0.001 | -0.021 | -0.021 | 0 | 0.21 | 0.27 | 0.09 | 0.01 | 0.12 | 0.06 |
| 6 | 42084146 | rs10456507 | 0.827 | -0.797 | -0.982 | -2.316 | -0.021 | -0.054 | -0.054 | 0 | 0.14 | 0.02 | 0.12 | 0.18 | 0.13 | 0.2 |
| 6 | 42084516 | rs9462772 | -0.401 | -0.49 | -0.61 | -2.182 | -0.003 | 0.05 | 0.05 | 0 | 0.55 | 0.67 | 0.43 | 0.29 | 0.48 | 0.46 |
| 6 | 42091645 | rs9471773 | 0.162 | -0.143 | -0.22 | -1.812 | -0.001 | -0.07 | -0.07 | 0.048 | 0.19 | 0.22 | 0.15 | 0.03 | 0.18 | 0.12 |
| 6 | 42098095 | rs4714570 | 0.098 | -0.297 | -0.718 | -1.598 | 0.007 | 0.01 | 0.01 | 0 | 0.47 | 0.41 | 0.64 | 0.65 | 0.6 | 0.53 |
| 6 | 42098187 | rs9471779 | -0.395 | -0.561 | -0.83 | -1.781 | 0.021 | 0.052 | 0.052 | 0 | 0.24 | 0.16 | 0.16 | 0.12 | 0.19 | 0.2 |
| 6 | 42099371 | rs4714572 | -0.426 | 0.045 | -0.553 | -1.436 | 0.002 | -0.022 | -0.022 | 0 | 0.14 | 0.2 | 0.18 | 0.06 | 0.12 | 0.16 |
| 6 | 42104931 | rs17702178 | -0.37 | -0.217 | -0.917 | -1.139 | -0.005 | 0.037 | 0.037 | 0 | 0.09 | 0.01 | 0.05 | 0.05 | 0.09 | 0.13 |
| 6 | 42105449 | rs9471782 | -0.56 | -0.206 | -0.506 | -1.093 | -0.004 | -0.029 | -0.029 | 1.483 | 0.19 | 0.24 | 0.17 | 0.09 | 0.21 | 0.05 |
| 6 | 42107815 | rs2492942 | -0.446 | -0.51 | -0.547 | -0.855 | -0.008 | -0.026 | -0.026 | 0 | 0.28 | 0.32 | 0.23 | 0.09 | 0.16 | 0.17 |
| 6 | 42117366 | rs6942280 | -0.15 | -0.247 | -0.759 | -0.57 | 0.002 | 0.009 | 0.009 | 0 | 0.17 | 0.11 | 0.15 | 0.1 | 0.2 | 0.18 |
| 6 | 42125482 | rs4714574 | 0.418 | -1.369 | -0.211 | -1.355 | -0.02 | -0.043 | -0.043 | 0 | 0.57 | 0.54 | 0.65 | 0.86 | 0.72 | 0.67 |
| 6 | 42126399 | rs2236047 | 0.457 | -0.928 | 0.03 | -0.307 | 0.014 | 0.003 | 0.003 | 0.888 | 0.54 | 0.6 | 0.54 | 0.62 | 0.66 | 0.6 |
| 6 | 42126444 | rs2207729 | -0.305 | -1.018 | -0.206 | -0.348 | 0.009 | 0.028 | 0.028 | 0.261 | 0.81 | 0.89 | 0.86 | 0.92 | 0.81 | 0.83 |
| 6 | 42126529 | rs2236046 | 1.004 | -1.35 | -0.281 | -0.846 | -0.009 | -0.021 | -0.021 | 0 | 0.4 | 0.24 | 0.46 | 0.6 | 0.56 | 0.52 |
| 6 | 42126859 | rs12190581 | 0.496 | -0.958 | -0.292 | -0.545 | 0.006 | 0.006 | 0.006 | 0.348 | 0.08 | 0.03 | 0.04 | 0 | 0.08 | 0.05 |
| 6 | 42127222 | rs7745391 | 0.841 | -0.997 | -0.116 | -0.537 | -0.008 | 0.005 | 0.005 | 0 | 0.65 | 0.55 | 0.67 | 0.68 | 0.66 | 0.7 |
| 6 | 42128299 | rs2281466 | -0.718 | -0.349 | -0.223 | 0.021 | 0.004 | 0.045 | 0.045 | 0.037 | 0.14 | 0.26 | 0.18 | 0.22 | 0.15 | 0.12 |
| 6 | 42129244 | rs2281462 | -0.045 | 0.397 | -0.669 | -0.046 | -0.031 | -0.027 | -0.027 | 0 | 0.31 | 0.3 | 0.4 | 0.58 | 0.44 | 0.46 |
| 6 | 42130318 | rs2236041 | -1.09 | 0.073 | -0.997 | 0.619 | -0.013 | -0.135 | -0.135 | 0 | 0.19 | 0.01 | 0.29 | 0.4 | 0.19 | 0.26 |
| 6 | 42131875 | rs9349222 | 0.984 | -0.148 | -1.325 | -0.309 | 0.057 | 0.09 | 0.09 | 0 | 0.58 | 0.17 | 0.65 | 0.75 | 0.7 | 0.83 |
| 6 | 42134244 | rs16895258 | -0.117 | -0.302 | -0.017 | -0.696 | -0.035 | -0.024 | -0.024 | 0.622 | 0.13 | 0.3 | 0.17 | 0.09 | 0.05 | 0.01 |
| 6 | 42134517 | rs12201777 | 0.81 | -0.348 | 0.307 | -0.697 | -0.013 | -0.063 | -0.063 | 1.461 | 0.7 | 0.45 | 0.81 | 0.9 | 0.74 | 0.82 |
| 6 | 42138533 | rs4711710 | 1.246 | -0.472 | 0.299 | -0.804 | 0.075 | 0.105 | 0.105 | 0 | 0.67 | 0.19 | 0.71 | 0.91 | 0.79 | 0.92 |
| 6 | 42139825 | rs7762152 | 0.042 | -0.125 | 0.587 | -0.782 | 0.01 | 0.003 | 0.003 | 0 | 0.84 | 0.89 | 0.93 | 1 | 0.86 | 0.93 |
| 6 | 42141041 | rs13217310 | 0.514 | -0.017 | 0.512 | -0.633 | 0.004 | -0.001 | -0.001 | 0.442 | 0.09 | 0 | 0.14 | 0.08 | 0.06 | 0.01 |
| 6 | 42151235 | rs1997980 | 2.195 | 0.351 | 0.01 | -0.792 | 0.018 | -0.001 | -0.001 | 0 | 0.36 | 0.07 | 0.18 | 0.3 | 0.23 | 0.39 |
| 6 | 42152333 | rs3749920 | 0.373 | -0.127 | 0.853 | -1.356 | -0.012 | -0.008 | -0.008 | 0 | 0.1 | 0.07 | 0.15 | 0.09 | 0.06 | 0.01 |
| 6 | 42153609 | rs9471799 | 0.584 | -0.055 | 0.625 | -1.41 | -0.027 | 0.017 | 0.017 | 0 | 0.87 | 0.92 | 0.9 | 0.93 | 0.84 | 0.84 |
| 6 | 42157817 | rs7764278 | -1.035 | 0.054 | 0.924 | -1.483 | -0.035 | -0.085 | -0.085 | 0 | 0.28 | 0.37 | 0.23 | 0.09 | 0.2 | 0.07 |
| 6 | 42158003 | rs7750260 | -1.426 | 0.053 | 0.873 | -1.483 | 0.057 | 0.088 | 0.088 | 0 | 0.32 | 0.76 | 0.26 | 0.09 | 0.21 | 0.07 |
| 6 | 42162575 | rs1474867 | 0.612 | 0.065 | 0.898 | -1.481 | -0.023 | 0.01 | 0.01 | 0 | 0.87 | 0.9 | 0.89 | 0.91 | 0.84 | 0.84 |
| 6 | 42164401 | rs13217993 | 2.047 | -0.015 | 0.898 | -1.481 | 0.047 | 0.186 | 0.186 | 0 | 0.55 | 0.17 | 0.63 | 0.82 | 0.64 | 0.77 |
| 6 | 42167970 | rs2180174 | 2.007 | -0.009 | 0.935 | -1.478 | 0.045 | 0.15 | 0.15 | 0 | 0.56 | 0.21 | 0.64 | 0.82 | 0.64 | 0.77 |
| 6 | 42168542 | rs2180172 | 2.03 | -0.009 | 0.935 | -1.466 | 0.047 | 0.139 | 0.139 | 0 | 0.55 | 0.2 | 0.64 | 0.79 | 0.64 | 0.77 |
| 6 | 42174478 | rs4714579 | 1.4 | -0.023 | 1.113 | -1.473 | -0.007 | -0.095 | -0.095 | 0.626 | 0.69 | 0.52 | 0.8 | 0.91 | 0.7 | 0.78 |
| 6 | 42174558 | rs4714580 | 0.685 | 0.199 | 0.719 | -1.324 | -0.026 | 0.01 | 0.01 | 0 | 0.87 | 0.92 | 0.9 | 0.91 | 0.84 | 0.84 |
| 6 | 42179091 | rs9369350 | 1.43 | 0.194 | 0.719 | -1.322 | 0.046 | 0.136 | 0.136 | 0 | 0.55 | 0.21 | 0.64 | 0.79 | 0.64 | 0.77 |
| 6 | 42185564 | rs9471809 | -0.855 | 0.231 | 0.732 | -1.277 | 0.03 | 0.036 | 0.036 | 0 | 0.32 | 0.69 | 0.26 | 0.11 | 0.2 | 0.07 |
| 6 | 42189848 | rs7775141 | -0.565 | 0.248 | 0.948 | -1.269 | -0.043 | -0.078 | -0.078 | 0 | 0.28 | 0.39 | 0.23 | 0.11 | 0.2 | 0.07 |
| 6 | 42190446 | rs10452607 | -0.638 | 0.329 | 0.11 | -1.227 | 0.017 | 0.017 | 0.017 | 0 | 0.31 | 0.66 | 0.26 | 0.11 | 0.2 | 0.07 |
| 6 | 42194440 | rs9394889 | 0.925 | 0.417 | 0.086 | -1.179 | 0.042 | 0.139 | 0.139 | 0 | 0.56 | 0.22 | 0.63 | 0.81 | 0.64 | 0.77 |
| 6 | 42197916 | rs7755400 | 0.827 | 0.895 | 0.328 | -0.773 | 0.039 | 0.163 | 0.163 | 0.028 | 0.54 | 0.22 | 0.68 | 0.8 | 0.58 | 0.83 |

| | | | | | | | | | | | | | | | | |
|---|---|---|---|---|---|---|---|---|---|---|---|---|---|---|---|---|
| 6 | 42201863 | rs9367137 | -1.106 | 0.474 | 0.773 | -0.301 | 0.058 | 0.089 | 0.089 | 0.056 | 0.39 | 0.77 | 0.22 | 0.22 | 0.28 | 0.17 |
| 6 | 42203748 | rs9471815 | -0.755 | 0.91 | 1.261 | 0.033 | -0.033 | -0.07 | -0.07 | 0 | 0.25 | 0.57 | 0.16 | 0.14 | 0.18 | 0.07 |
| 6 | 42220472 | rs12194245 | -0.47 | 1.131 | 0.784 | 0.096 | -0.01 | -0.071 | -0.071 | 0 | 0.21 | 0.34 | 0.11 | 0.09 | 0.18 | 0.05 |
| 6 | 42232475 | rs2295274 | -0.518 | 1.209 | -0.222 | 0.799 | 0.011 | 0.012 | 0.012 | 0.169 | 0.19 | 0.14 | 0.11 | 0.12 | 0.17 | 0.21 |
| 6 | 42233518 | rs17704222 | -1.145 | 2.026 | 0.078 | 1.299 | -0.027 | 0.02 | 0.02 | 0 | 0.05 | 0 | 0.04 | 0 | 0.07 | 0.07 |
| 6 | 42240369 | rs713383 | -1.637 | 2.395 | 0.263 | 2.249 | 0 | 0.02 | 0.02 | 0.051 | 0.12 | 0.14 | 0.33 | 0.38 | 0.25 | 0.25 |
| 6 | 42252385 | rs9471831 | -1.535 | 2.389 | 0.696 | 2.752 | 0.003 | 0.032 | 0.032 | 0 | 0.16 | 0.22 | 0.32 | 0.35 | 0.22 | 0.23 |
| 6 | 42260911 | rs9394894 | -2.438 | 2.414 | 1.463 | 3.19 | 0.051 | 0.148 | 0.148 | 0.274 | 0.25 | 0.5 | 0.47 | 0.46 | 0.34 | 0.35 |
| 6 | 42264554 | rs4711712 | -1.952 | 2.007 | 1.347 | 2.67 | 0.073 | 0.106 | 0.106 | 0 | 0.08 | 0.26 | 0.14 | 0.11 | 0.11 | 0.12 |
| 6 | 42267031 | rs4711714 | -1.925 | 2.035 | 1.355 | 2.535 | 0.089 | 0.085 | 0.085 | 0 | 0.08 | 0.29 | 0.15 | 0.08 | 0.1 | 0.1 |
| 6 | 42271906 | rs7773883 | -2.076 | 1.903 | 1.316 | 2.474 | 0.065 | 0.057 | 0.057 | 0 | 0.09 | 0.33 | 0.16 | 0.08 | 0.11 | 0.12 |
| 6 | 42281967 | rs7761002 | -1.689 | 1.462 | 0.979 | 3.155 | -0.004 | 0.25 | 0.25 | 0 | 0.06 | 0.15 | 0.06 | 0.41 | 0.04 | 0.18 |
| 6 | 42285496 | rs4711717 | -1.627 | 1.812 | 0.733 | 2.641 | -0.007 | -0.011 | -0.011 | 0 | 0.2 | 0.13 | 0.42 | 0.33 | 0.25 | 0.33 |
| 6 | 42295408 | rs6920284 | -1.715 | 1.51 | 1.458 | 2.164 | 0.031 | -0.019 | -0.019 | 0 | 0.07 | 0.35 | 0.05 | 0 | 0.05 | 0.12 |
| 6 | 42305054 | rs654043 | -2.405 | 1.008 | 2.101 | 2.993 | -0.025 | 0.104 | 0.104 | 0 | 0.16 | 0.4 | 0.12 | 0.27 | 0.1 | 0.23 |
| 6 | 42310664 | rs497051 | 0.137 | 1.357 | 1.15 | 2.937 | -0.076 | -0.008 | -0.008 | 0 | 0.8 | 0.98 | 0.87 | 0.9 | 0.73 | 0.84 |
| 6 | 42319307 | rs260225 | -1.932 | 1.043 | 1.712 | 3.195 | 0.009 | 0.092 | 0.092 | 0 | 0.09 | 0.25 | 0.1 | 0.24 | 0.09 | 0.19 |
| 6 | 42330813 | rs11970679 | -1.2 | 1.043 | 1.684 | 2.72 | 0.014 | -0.03 | -0.03 | 0 | 0.07 | 0.22 | 0.05 | 0 | 0.06 | 0.09 |
| 6 | 42340131 | rs260239 | -1.954 | 1.006 | 1.413 | 3.209 | -0.019 | 0.067 | 0.067 | 2.735 | 0.12 | 0.72 | 0.13 | 0.23 | 0.07 | 0.14 |
| 6 | 42346356 | rs181990 | -1.566 | 1.035 | 1.234 | 2.909 | 0.034 | 0.1 | 0.1 | 1.526 | 0.1 | 0.54 | 0.1 | 0.14 | 0.07 | 0.08 |
| 6 | 42346554 | rs599088 | 0.718 | 1.119 | -0.4 | 2.425 | -0.064 | 0.006 | 0.006 | 0 | 0.77 | 0.98 | 0.86 | 0.88 | 0.7 | 0.78 |
| 6 | 42363916 | rs9471853 | 0.706 | 1.064 | -0.773 | 1.617 | -0.088 | 0.068 | 0.068 | 0.012 | 0.24 | 0.01 | 0.17 | 0.07 | 0.28 | 0.09 |
| 6 | 42366438 | rs9381170 | -0.264 | 1.087 | -0.394 | 2.124 | -0.057 | -0.075 | -0.075 | 0 | 0.12 | 0.01 | 0.3 | 0.32 | 0.2 | 0.22 |
| 6 | 42367757 | rs166339 | -1.312 | 0.342 | 0.631 | 1.914 | 0 | 0.082 | 0.082 | 0.855 | 0.11 | 0.58 | 0.1 | 0.13 | 0.06 | 0.05 |
| 6 | 42369308 | rs1038890 | 0.348 | 0.07 | -0.284 | 1.661 | 0.043 | -0.057 | -0.057 | 0.569 | 0.53 | 0.07 | 0.6 | 0.5 | 0.65 | 0.51 |
| 6 | 42369541 | rs260284 | -1.118 | -0.18 | 0.365 | 2.124 | 0.002 | 0.141 | 0.141 | 0.912 | 0.11 | 0.59 | 0.13 | 0.22 | 0.06 | 0.06 |
| 6 | 42372357 | rs1397175 | -1.361 | -0.264 | 0.641 | 2.128 | 0.008 | 0.044 | 0.044 | 0.041 | 0.11 | 0.27 | 0.09 | 0.14 | 0.11 | 0.17 |
| 6 | 42387426 | rs9349232 | 2.053 | -0.345 | 1.204 | 2.244 | 0 | 0.125 | 0.125 | 0.276 | 0.86 | 0.45 | 0.87 | 0.74 | 0.88 | 0.91 |
| 6 | 42388625 | rs365387 | -0.76 | -0.44 | 1.226 | 2.246 | 0.036 | 0.042 | 0.042 | 0.253 | 0.07 | 0.3 | 0.08 | 0.09 | 0.07 | 0.05 |
| 6 | 42399074 | rs9381174 | 1.141 | -0.78 | 0.551 | 1.871 | 0.021 | -0.089 | -0.089 | 0 | 0.64 | 0.08 | 0.68 | 0.59 | 0.73 | 0.63 |
| 6 | 42401082 | rs260255 | 0.539 | -0.771 | 0.731 | 1.988 | 0.009 | -0.002 | -0.002 | 0 | 0.81 | 0.84 | 0.8 | 0.82 | 0.89 | 0.8 |
| 6 | 42401992 | rs12201447 | -0.742 | -0.599 | 0.686 | 1.821 | 0.007 | 0.094 | 0.094 | 0.356 | 0.08 | 0.21 | 0.06 | 0.18 | 0.07 | 0.06 |
| 6 | 42407020 | rs694148 | 1.235 | -0.597 | 0.342 | 1.44 | -0.045 | 0.058 | 0.058 | 0 | 0.7 | 0.44 | 0.73 | 0.53 | 0.82 | 0.65 |
| 6 | 42413300 | rs7768349 | 0.321 | -0.194 | -0.099 | 1.075 | -0.001 | 0.044 | 0.044 | 0.505 | 0.72 | 0.67 | 0.68 | 0.5 | 0.73 | 0.62 |
| 6 | 42440358 | rs4714612 | 0.323 | 0.17 | -0.64 | 0.457 | -0.001 | 0 | 0 | 0 | 0.67 | 0.32 | 0.63 | 0.33 | 0.72 | 0.5 |
| 6 | 42450682 | rs9462806 | 0.115 | 0.642 | -1.827 | -0.475 | -0.037 | 0.207 | 0.207 | 0.658 | 0.3 | 0.02 | 0.15 | 0 | 0.29 | 0.1 |
| 6 | 42454850 | rs12190874 | -1.266 | 1.206 | -1.031 | 0.695 | -0.005 | -0.006 | -0.006 | 0 | 0.08 | 0.08 | 0.13 | 0.17 | 0.1 | 0.11 |
| 6 | 42455343 | rs7746290 | -0.922 | 1.703 | -0.773 | 0.033 | 0.018 | -0.042 | -0.042 | 0 | 0.15 | 0.2 | 0.15 | 0.05 | 0.17 | 0.07 |
| 6 | 42458006 | rs7743743 | 0.198 | 0.802 | -0.612 | 0.076 | -0.017 | 0.003 | 0.003 | 0.113 | 0.68 | 0.48 | 0.79 | 0.63 | 0.71 | 0.58 |
| 6 | 42458376 | rs6906363 | 0.159 | 0.633 | 0.219 | 0.173 | 0.002 | 0.003 | 0.003 | 0.243 | 0.72 | 0.71 | 0.81 | 0.72 | 0.71 | 0.63 |
| 6 | 42462055 | rs9369362 | 0.614 | 0.844 | -0.034 | 0.005 | 0.005 | -0.007 | -0.007 | 0 | 0.21 | 0.3 | 0.28 | 0.16 | 0.22 | 0.26 |
| 6 | 42467898 | rs2941399 | -0.205 | 0.699 | -0.415 | -0.801 | -0.003 | 0.008 | 0.008 | 0 | 0.59 | 0.69 | 0.65 | 0.63 | 0.53 | 0.46 |
| 6 | 42470474 | rs7760744 | 1.198 | 0.585 | -0.463 | -0.705 | 0.001 | -0.018 | -0.018 | 0 | 0.21 | 0.29 | 0.26 | 0.13 | 0.21 | 0.26 |
| 6 | 42471579 | rs12665070 | -1.116 | 0.581 | -0.554 | -0.729 | -0.004 | -0.004 | -0.004 | 0 | 0.75 | 0.8 | 0.81 | 0.69 | 0.71 | 0.65 |
| 6 | 42478124 | rs1322819 | -0.667 | 0.534 | -0.61 | -0.202 | 0.005 | -0.046 | -0.046 | 0.056 | 0.65 | 0.78 | 0.64 | 0.56 | 0.7 | 0.8 |
| 6 | 42478281 | rs9381185 | 0.242 | 0.444 | -0.577 | -0.258 | 0 | 0.004 | 0.004 | 0 | 0.48 | 0.51 | 0.47 | 0.4 | 0.5 | 0.6 |
| 6 | 42486688 | rs9381186 | 0.991 | 0.6 | -0.655 | -0.362 | -0.001 | -0.001 | -0.001 | 0 | 0.23 | 0.21 | 0.17 | 0.28 | 0.25 | 0.29 |
| 6 | 42495278 | rs16895812 | -0.047 | 0.37 | -0.801 | -0.592 | 0 | -0.003 | -0.003 | 0 | 0.21 | 0.2 | 0.19 | 0.15 | 0.22 | 0.2 |
| 6 | 42505116 | rs9381188 | -1.026 | 0.149 | -0.764 | -0.488 | -0.029 | -0.057 | -0.057 | 0 | 0.71 | 0.86 | 0.66 | 0.62 | 0.69 | 0.8 |
| 6 | 42506378 | rs4711724 | -0.36 | -0.012 | -1.032 | -0.804 | -0.009 | 0.01 | 0.01 | 0 | 0.55 | 0.8 | 0.58 | 0.62 | 0.57 | 0.69 |
| 6 | 42515372 | rs6458293 | 0.544 | 0.135 | -0.959 | -0.999 | 0.002 | 0.01 | 0.01 | 0 | 0.34 | 0.58 | 0.39 | 0.47 | 0.36 | 0.5 |
| 6 | 42531650 | rs6900726 | -1.6 | 0.289 | -1.053 | -0.989 | -0.015 | -0.083 | -0.083 | 1.92 | 0.75 | 0.87 | 0.68 | 0.53 | 0.76 | 0.85 |
| 6 | 42569086 | rs16895860 | 0.837 | 0.319 | -0.905 | -1.088 | -0.001 | -0.015 | -0.015 | 0.682 | 0.16 | 0.12 | 0.14 | 0.01 | 0.16 | 0.12 |
| 6 | 42575723 | rs9369373 | -1.625 | 0.303 | -0.892 | -0.923 | -0.013 | -0.088 | -0.088 | 0 | 0.74 | 0.87 | 0.68 | 0.54 | 0.75 | 0.81 |
| 6 | 42594053 | rs4711725 | 0.581 | 0.273 | -0.771 | -0.973 | 0.021 | 0.021 | 0.021 | 0 | 0.34 | 0.48 | 0.41 | 0.44 | 0.4 | 0.44 |
| 6 | 42598371 | rs9471889 | 1.161 | 0.33 | -0.69 | -0.983 | -0.006 | -0.003 | -0.003 | 0 | 0.25 | 0.2 | 0.22 | 0.3 | 0.32 | 0.3 |
| 6 | 42611872 | rs12198323 | 0.255 | 0.227 | -0.766 | -1.038 | -0.001 | -0.011 | -0.011 | 0 | 0.27 | 0.25 | 0.2 | 0.09 | 0.25 | 0.27 |
| 6 | 42612312 | rs7767888 | -1.291 | 0.216 | -0.767 | -1.026 | 0.009 | 0.051 | 0.051 | 0.073 | 0.85 | 0.95 | 0.91 | 1 | 0.88 | 0.88 |
| 6 | 42627434 | rs6917033 | 0.414 | 0.019 | -0.785 | -1.079 | 0.024 | 0.024 | 0.024 | 0 | 0.33 | 0.5 | 0.39 | 0.44 | 0.39 | 0.41 |
| 6 | 42629610 | rs11754231 | 1.406 | -0.164 | -0.673 | -1.287 | -0.023 | 0.028 | 0.028 | 0 | 0.05 | 0.01 | 0.06 | 0 | 0.1 | 0.01 |
| 6 | 42641704 | rs761537 | -0.003 | -0.257 | -0.63 | -1.289 | 0.025 | 0.073 | 0.073 | 0 | 0.58 | 0.74 | 0.72 | 0.91 | 0.63 | 0.61 |
| 6 | 42652667 | rs434505 | -0.661 | -0.154 | -0.401 | -1.198 | -0.014 | -0.039 | -0.039 | 2.403 | 0.75 | 0.61 | 0.72 | 0.85 | 0.8 | 0.72 |
| 6 | 42661404 | rs375435 | 0.481 | -0.149 | -0.453 | -1.212 | 0.012 | 0.073 | 0.073 | 0 | 0.48 | 0.34 | 0.52 | 0.76 | 0.55 | 0.45 |
| 6 | 42665490 | rs432753 | -0.595 | -0.11 | -0.664 | -1.128 | -0.011 | -0.027 | -0.027 | 0.015 | 0.75 | 0.69 | 0.72 | 0.85 | 0.8 | 0.72 |
| 6 | 42665888 | rs835 | 1.432 | -0.03 | -0.734 | -1.047 | -0.019 | -0.081 | -0.081 | 0 | 0.25 | 0.14 | 0.32 | 0.45 | 0.25 | 0.18 |
| 6 | 42666061 | rs434102 | -0.691 | 0.003 | -0.691 | -1.048 | -0.009 | -0.023 | -0.023 | 0 | 0.76 | 0.7 | 0.72 | 0.85 | 0.8 | 0.72 |
| 6 | 42666475 | rs433286 | -0.703 | 0.019 | -0.68 | -1.043 | -0.009 | -0.021 | -0.021 | 0.323 | 0.76 | 0.71 | 0.72 | 0.85 | 0.8 | 0.72 |
| 6 | 42671099 | rs9471904 | -0.602 | 0.361 | -0.678 | -0.717 | 0 | 0.005 | 0.005 | 0 | 0.49 | 0.54 | 0.39 | 0.44 | 0.54 | 0.55 |
| 6 | 42677167 | rs9471908 | -0.035 | 0.094 | -0.52 | -0.724 | -0.023 | -0.067 | -0.067 | 0 | 0.31 | 0.43 | 0.21 | 0.11 | 0.26 | 0.29 |

| | | | | | | | | | | | | | | | | |
|---|---|---|---|---|---|---|---|---|---|---|---|---|---|---|---|---|
| 6 | 42677753 | rs2268684 | -0.102 | 0.126 | -0.915 | -0.703 | -0.004 | 0 | 0 | 0 | 0.58 | 0.63 | 0.54 | 0.56 | 0.51 | 0.49 |
| 6 | 42689411 | rs6928781 | 1.03 | 0.059 | -0.781 | -0.641 | -0.008 | -0.044 | -0.044 | 0 | 0.34 | 0.28 | 0.38 | 0.47 | 0.34 | 0.22 |
| 6 | 42690514 | rs11754813 | 1.098 | 0.096 | -0.887 | -0.62 | -0.013 | -0.053 | -0.053 | 0 | 0.33 | 0.25 | 0.38 | 0.47 | 0.34 | 0.22 |
| 6 | 42693959 | rs9381214 | 0.171 | 0.544 | -0.507 | -0.03 | 0.005 | 0.006 | 0.006 | 0 | 0.65 | 0.66 | 0.59 | 0.56 | 0.59 | 0.51 |
| 6 | 42708628 | rs6929834 | -0.233 | 0.336 | -0.75 | 0.604 | -0.011 | -0.079 | -0.079 | 2.07 | 0.84 | 0.97 | 0.91 | 0.68 | 0.85 | 0.75 |
| 6 | 42723809 | rs11759402 | -0.389 | 0.468 | -0.562 | 0.128 | -0.001 | 0.031 | 0.031 | 0.059 | 0.17 | 0.07 | 0.15 | 0.1 | 0.18 | 0.21 |
| 6 | 42724357 | rs12195802 | -0.51 | 0.481 | -0.364 | 0.034 | 0.016 | -0.014 | -0.014 | 1.397 | 0.22 | 0.32 | 0.32 | 0.17 | 0.25 | 0.15 |
| 6 | 42731115 | rs6458307 | -0.943 | 0.43 | -0.235 | 0.102 | -0.006 | -0.028 | -0.028 | 0 | 0.35 | 0.52 | 0.39 | 0.26 | 0.31 | 0.3 |
| 6 | 42738749 | rs9381218 | 0.772 | 0.431 | -0.466 | 0.237 | -0.009 | -0.008 | -0.008 | 0 | 0.37 | 0.33 | 0.45 | 0.41 | 0.44 | 0.35 |
| 6 | 42751457 | rs3800291 | 0.616 | 0.52 | -0.223 | 0.491 | 0.014 | -0.035 | -0.035 | 0.583 | 0.57 | 0.25 | 0.67 | 0.5 | 0.61 | 0.49 |
| 6 | 42753404 | rs2894478 | -0.836 | 0.395 | -0.19 | 0.528 | 0.009 | -0.031 | -0.031 | 0.108 | 0.44 | 0.75 | 0.34 | 0.5 | 0.42 | 0.56 |
| 6 | 42778467 | rs9296399 | -0.253 | 0.269 | 0.13 | 0.642 | -0.011 | 0.034 | 0.034 | 0 | 0.07 | 0.2 | 0.04 | 0.13 | 0.04 | 0.06 |
| 6 | 42794329 | rs6924431 | -0.86 | 0.281 | -0.015 | 0.325 | 0.009 | -0.023 | -0.023 | 3.228 | 0.46 | 0.71 | 0.35 | 0.48 | 0.43 | 0.57 |
| 6 | 42810991 | rs7758869 | -0.151 | 0.313 | -0.088 | 0.322 | -0.011 | -0.034 | -0.034 | 0.079 | 0.07 | 0.21 | 0.04 | 0.01 | 0.04 | 0.05 |
| 6 | 42814291 | rs3846894 | -0.767 | 0.376 | -0.44 | 0.398 | 0.001 | -0.047 | -0.047 | 0 | 0.46 | 0.79 | 0.35 | 0.48 | 0.42 | 0.56 |
| 6 | 42821941 | rs6936589 | -0.928 | 0.082 | 0.498 | 0.266 | -0.007 | -0.001 | -0.001 | 0 | 0.08 | 0.3 | 0.04 | 0.06 | 0.04 | 0.06 |
| 6 | 42823529 | rs9381223 | -1.468 | -0.171 | 1.339 | 0.099 | -0.038 | -0.027 | -0.027 | 0 | 0.12 | 0.35 | 0.09 | 0.07 | 0.07 | 0.09 |
| 6 | 42824756 | rs9462840 | -0.483 | -0.488 | 0.7 | -1.059 | 0.009 | 0.068 | 0.068 | 0 | 0.31 | 0.19 | 0.21 | 0.01 | 0.29 | 0.22 |
| 6 | 42835487 | rs7764865 | -1.69 | -0.83 | 2.446 | -0.874 | -0.042 | 0.003 | 0.003 | 4.804 | 0.14 | 0.55 | 0.13 | 0.1 | 0.06 | 0.22 |
| 6 | 42836583 | rs12526596 | -0.825 | -0.84 | 2.432 | -0.849 | -0.003 | 0.055 | 0.055 | 0.199 | 0.07 | 0.2 | 0.08 | 0.1 | 0.04 | 0.19 |
| 6 | 42841231 | rs1150797 | -0.216 | -1.093 | 2.156 | -1 | 0.009 | 0.082 | 0.082 | 0 | 0.27 | 0.13 | 0.18 | 0.02 | 0.27 | 0.21 |
| 6 | 42845691 | rs9471938 | -1.298 | -1.244 | 2.444 | -1.076 | 0.009 | -0.012 | -0.012 | 0 | 0.08 | 0.45 | 0.05 | 0 | 0.02 | 0.02 |
| 6 | 42848071 | rs2073155 | -2.146 | -1.182 | 2.37 | -1.08 | -0.042 | -0.008 | -0.008 | 0.775 | 0.15 | 0.66 | 0.13 | 0.1 | 0.06 | 0.21 |
| 6 | 42851396 | rs2395939 | -0.243 | -1.201 | 2.245 | -1.09 | 0.011 | 0.085 | 0.085 | 0 | 0.28 | 0.14 | 0.18 | 0.02 | 0.27 | 0.22 |
| 6 | 42857827 | rs9471941 | -0.177 | -1.181 | 2.255 | -1.084 | 0.009 | 0.082 | 0.082 | 0 | 0.27 | 0.14 | 0.18 | 0.02 | 0.27 | 0.22 |
| 6 | 42869388 | rs7748409 | -1.233 | -1.084 | 2.403 | -0.846 | -0.002 | 0.079 | 0.079 | 0.446 | 0.08 | 0.25 | 0.08 | 0.13 | 0.04 | 0.19 |
| 6 | 42873823 | rs7741552 | -1.169 | -1.149 | 2.246 | -0.892 | 0.056 | 0.113 | 0.113 | 0.269 | 0.42 | 0.8 | 0.32 | 0.15 | 0.33 | 0.42 |
| 6 | 42876386 | rs7757363 | -0.966 | -1.16 | 2.093 | -0.784 | 0.004 | 0.003 | 0.003 | 0 | 0.39 | 0.36 | 0.28 | 0.14 | 0.32 | 0.4 |
| 6 | 42879289 | rs9381225 | 1.73 | -0.26 | 0.99 | -0.206 | -0.172 | -0.134 | -0.134 | 1.302 | 0.38 | 0.02 | 0.41 | 0.44 | 0.43 | 0.35 |
| 6 | 42890602 | rs9462851 | -0.444 | -0.538 | 1.46 | -0.559 | -0.012 | -0.067 | -0.067 | 0 | 0.35 | 0.4 | 0.21 | 0.02 | 0.29 | 0.22 |
| 6 | 42891022 | rs9471966 | -0.444 | -0.538 | 1.46 | -0.559 | -0.012 | -0.067 | -0.067 | 1.55 | 0.35 | 0.4 | 0.21 | 0.02 | 0.29 | 0.22 |
| 6 | 42897074 | rs2234185 | -1.601 | -0.294 | 0.724 | -0.285 | -0.121 | -0.115 | -0.115 | 0 | 0.59 | 0.97 | 0.48 | 0.4 | 0.49 | 0.64 |
| 6 | 42897712 | rs4711731 | -0.617 | -0.44 | 1.163 | -0.364 | 0.01 | 0.04 | 0.04 | 0 | 0.39 | 0.28 | 0.31 | 0.21 | 0.37 | 0.37 |
| 6 | 42902508 | rs3763236 | -0.955 | 0.195 | 1.45 | -0.969 | 0 | 0.004 | 0.004 | 0.348 | 0.5 | 0.52 | 0.53 | 0.27 | 0.52 | 0.57 |
| 6 | 42904948 | rs12525851 | 0.008 | 0.135 | 1.419 | -0.919 | -0.003 | 0.026 | 0.026 | 0 | 0.05 | 0.06 | 0.16 | 0.14 | 0.04 | 0.19 |
| 6 | 42906384 | rs9471969 | 0.086 | 0.182 | 1.428 | -1.044 | -0.001 | -0.043 | -0.043 | 1.242 | 0.32 | 0.33 | 0.21 | 0.01 | 0.27 | 0.2 |
| 6 | 42910666 | rs9462853 | -0.241 | 0.152 | 1.688 | -1.06 | -0.005 | -0.048 | -0.048 | 0 | 0.36 | 0.6 | 0.23 | 0.04 | 0.28 | 0.23 |
| 6 | 42915920 | rs6941212 | -0.363 | 0.023 | 1.675 | -0.832 | -0.003 | 0.003 | 0.003 | 0.004 | 0.44 | 0.4 | 0.35 | 0.1 | 0.46 | 0.38 |
| 6 | 42930309 | rs3800292 | -0.323 | 0.087 | 1.762 | -0.832 | -0.009 | -0.035 | -0.035 | 0 | 0.09 | 0.27 | 0.06 | 0.04 | 0.06 | 0.06 |
| 6 | 42932200 | rs1129187 | -0.331 | 0.088 | 1.203 | -0.851 | -0.005 | 0.022 | 0.022 | 0.006 | 0.44 | 0.37 | 0.35 | 0.1 | 0.46 | 0.37 |
| 6 | 42940673 | rs2395943 | -1.341 | 0.296 | 0.147 | -0.667 | -0.129 | -0.005 | -0.005 | 0 | 0.62 | 0.98 | 0.61 | 0.31 | 0.59 | 0.64 |
| 6 | 42954429 | rs3805949 | -0.663 | 0.158 | 1.286 | -0.24 | -0.014 | -0.036 | -0.036 | 0 | 0.09 | 0.25 | 0.06 | 0.04 | 0.05 | 0.07 |
| 6 | 42955524 | rs1969759 | -1.988 | 0.106 | 1.79 | -0.172 | -0.03 | 0.01 | 0.01 | 0.72 | 0.17 | 0.62 | 0.23 | 0.19 | 0.11 | 0.24 |
| 6 | 42956746 | rs4714646 | -0.996 | 0.235 | 1.977 | -0.068 | 0.073 | 0.223 | 0.223 | 0.074 | 0.05 | 0.3 | 0.15 | 0.15 | 0.03 | 0.17 |
| 6 | 42976232 | rs6458318 | -2.18 | 0.077 | 2.133 | -0.187 | -0.031 | 0.015 | 0.015 | 0.199 | 0.18 | 0.61 | 0.24 | 0.21 | 0.12 | 0.28 |
| 6 | 42978326 | rs2236356 | -2.4 | 0.044 | 2.163 | -0.217 | -0.012 | 0.001 | 0.001 | 0.351 | 0.19 | 0.71 | 0.24 | 0.21 | 0.12 | 0.28 |
| 6 | 42989242 | rs10080728 | -2.217 | 0.079 | 1.941 | -0.291 | -0.02 | 0.005 | 0.005 | 0.208 | 0.19 | 0.68 | 0.24 | 0.21 | 0.12 | 0.28 |
| 6 | 42992808 | rs3749904 | -2.354 | 0.079 | 1.884 | -0.291 | -0.013 | 0.002 | 0.002 | 0.301 | 0.19 | 0.71 | 0.24 | 0.21 | 0.12 | 0.28 |
| 6 | 43008298 | rs2273917 | -2.352 | 0.247 | 1.862 | -0.374 | -0.027 | 0.016 | 0.016 | 0.052 | 0.18 | 0.64 | 0.24 | 0.22 | 0.12 | 0.28 |
| 6 | 43013662 | rs2273918 | -1.625 | 0.262 | 2.258 | -0.356 | 0.025 | 0.162 | 0.162 | 0 | 0.08 | 0.3 | 0.18 | 0.18 | 0.06 | 0.21 |
| 6 | 43020188 | rs4711738 | -2.207 | 0.204 | 2.445 | -0.326 | -0.021 | 0.068 | 0.068 | 0.713 | 0.14 | 0.56 | 0.22 | 0.21 | 0.09 | 0.22 |
| 6 | 43026814 | rs2050367 | -2.489 | 0.152 | 2.365 | -0.353 | -0.031 | 0.051 | 0.051 | 0.822 | 0.15 | 0.58 | 0.22 | 0.2 | 0.09 | 0.23 |
| 6 | 43035007 | rs4714661 | -2.738 | 0.162 | 2.369 | -0.351 | -0.028 | 0.045 | 0.045 | 0.256 | 0.17 | 0.57 | 0.23 | 0.22 | 0.12 | 0.26 |
| 6 | 43040368 | rs13199873 | -1.393 | -0.206 | 0.967 | -1.268 | -0.153 | -0.062 | -0.062 | 0 | 0.69 | 0.95 | 0.57 | 0.33 | 0.58 | 0.68 |
| 6 | 43045462 | rs3805944 | -0.488 | -0.083 | 1.134 | -1.174 | -0.001 | -0.016 | -0.016 | 0.066 | 0.09 | 0.32 | 0.07 | 0.04 | 0.05 | 0.06 |
| 6 | 43049461 | rs2007950 | -1.55 | -0.13 | 0.911 | -1.237 | -0.163 | -0.092 | -0.092 | 0.207 | 0.72 | 0.98 | 0.59 | 0.33 | 0.6 | 0.69 |
| 6 | 43050824 | rs3729617 | -1.128 | -0.304 | 0.835 | -1.393 | 0.005 | 0.014 | 0.014 | 0.198 | 0.39 | 0.64 | 0.37 | 0.31 | 0.37 | 0.48 |
| 6 | 43063516 | rs4714664 | -0.186 | -0.298 | 0.953 | -1.034 | 0.039 | 0.236 | 0.236 | 0 | 0.05 | 0.21 | 0.15 | 0.3 | 0.04 | 0.18 |
| 6 | 43065329 | rs6458323 | -0.843 | -0.402 | 0.862 | -1.062 | 0 | 0 | 0 | 0.878 | 0.4 | 0.58 | 0.4 | 0.48 | 0.38 | 0.51 |
| 6 | 43065913 | rs3793017 | -0.062 | -0.402 | 0.858 | -1.083 | -0.001 | 0.003 | 0.003 | 0.108 | 0.32 | 0.32 | 0.23 | 0.16 | 0.31 | 0.3 |
| 6 | 43068370 | rs1034246 | 0.536 | -0.42 | 0.732 | -1.071 | -0.068 | 0.068 | 0.068 | 0 | 0.24 | 0.02 | 0.17 | 0.12 | 0.26 | 0.25 |
| 6 | 43069344 | rs9472007 | -1.278 | -0.648 | 1.735 | -0.667 | 0.024 | 0.198 | 0.198 | 0 | 0.08 | 0.26 | 0.16 | 0.32 | 0.07 | 0.21 |
| 6 | 43078066 | rs17516786 | -1.919 | -0.569 | 1.707 | -0.753 | -0.001 | 0.103 | 0.103 | 0 | 0.14 | 0.32 | 0.21 | 0.34 | 0.12 | 0.27 |
| 6 | 43078456 | rs4714666 | -1.448 | -0.579 | 1.708 | -0.747 | -0.011 | 0.142 | 0.142 | 0 | 0.12 | 0.31 | 0.18 | 0.34 | 0.07 | 0.23 |
| 6 | 43083774 | rs1105024 | -1.617 | -0.609 | 1.741 | -0.743 | -0.015 | 0.16 | 0.16 | 0 | 0.13 | 0.4 | 0.18 | 0.34 | 0.07 | 0.23 |
| 6 | 43089429 | rs3805931 | -0.212 | -0.297 | 1.491 | -0.607 | 0.001 | -0.009 | -0.009 | 0 | 0.39 | 0.37 | 0.23 | 0.07 | 0.32 | 0.24 |
| 6 | 43094144 | rs12662888 | -0.72 | -0.271 | 1.458 | -0.538 | 0.034 | 0.243 | 0.243 | 0 | 0.07 | 0.29 | 0.15 | 0.3 | 0.04 | 0.17 |
| 6 | 43095828 | rs6926032 | -1.051 | -0.296 | 1.357 | -0.59 | 0.019 | 0.096 | 0.096 | 0.295 | 0.49 | 0.81 | 0.43 | 0.36 | 0.4 | 0.45 |
| 6 | 43096199 | rs1537638 | -0.456 | -0.255 | 1.339 | -0.717 | -0.004 | -0.075 | -0.075 | 0.078 | 0.42 | 0.52 | 0.28 | 0.06 | 0.37 | 0.28 |

| | | | | | | | | | | | | | | | | |
|---|---|---|---|---|---|---|---|---|---|---|---|---|---|---|---|---|
| 6 | 43101670 | rs6941837 | -1.308 | -0.219 | 1.003 | -0.479 | -0.105 | -0.032 | -0.032 | 0 | 0.57 | 0.93 | 0.66 | 0.43 | 0.5 | 0.55 |
| 6 | 43109751 | rs6905948 | -0.602 | -0.158 | 1.441 | -0.532 | -0.002 | -0.033 | -0.033 | 0 | 0.47 | 0.48 | 0.42 | 0.1 | 0.39 | 0.29 |
| 6 | 43114702 | rs12204526 | -0.137 | -0.02 | 1.019 | -0.498 | -0.013 | 0.013 | 0.013 | 0 | 0.06 | 0 | 0.03 | 0.01 | 0.06 | 0.06 |
| 6 | 43115670 | rs6458325 | -1.004 | -0.054 | 1.278 | -0.466 | -0.002 | 0.219 | 0.219 | 0.46 | 0.51 | 0.81 | 0.48 | 0.1 | 0.45 | 0.32 |
| 6 | 43128852 | rs1051709 | -1.517 | -0.058 | 2.324 | 0.311 | 0.02 | 0.177 | 0.177 | 0 | 0.11 | 0.39 | 0.26 | 0.35 | 0.1 | 0.23 |
| 6 | 43129986 | rs7760834 | -1.848 | -0.07 | 2.403 | 0.305 | 0.024 | 0.208 | 0.208 | 0.009 | 0.12 | 0.5 | 0.27 | 0.35 | 0.1 | 0.23 |
| 6 | 43144618 | rs4711741 | -1.575 | -0.042 | 2.374 | 0.316 | 0.03 | 0.225 | 0.225 | 0 | 0.12 | 0.47 | 0.26 | 0.35 | 0.1 | 0.22 |
| 6 | 43146387 | rs6914964 | -1.023 | 0.02 | 2.343 | 0.076 | 0.055 | -0.01 | -0.01 | 0 | 0.07 | 0.32 | 0.11 | 0.01 | 0.08 | 0.07 |
| 6 | 43157908 | rs12193690 | -1.006 | -0.174 | 2.569 | -0.062 | -0.021 | -0.042 | -0.042 | 0.084 | 0.11 | 0.2 | 0.06 | 0.02 | 0.07 | 0.08 |
| 6 | 43168117 | rs9462875 | -2.286 | -0.138 | 2.726 | -0.07 | -0.014 | -0.098 | -0.098 | 0.006 | 0.17 | 0.49 | 0.17 | 0.02 | 0.14 | 0.14 |
| 6 | 43169887 | rs17209372 | -0.68 | 0.058 | 2.559 | 0.089 | 0.021 | -0.026 | -0.026 | 0 | 0.06 | 0.17 | 0.1 | 0.01 | 0.07 | 0.07 |
| 6 | 43184132 | rs2273709 | -2.615 | -0.005 | 2.491 | 0.281 | -0.004 | 0.01 | 0.01 | 0.212 | 0.21 | 0.73 | 0.22 | 0.42 | 0.17 | 0.29 |
| 6 | 43187766 | rs2651195 | -2.246 | 0.353 | 2.722 | 0.075 | 0.147 | 0.067 | 0.067 | 0.521 | 0.05 | 0.42 | 0.05 | 0 | 0.04 | 0.02 |
| 6 | 43188940 | rs6917902 | -2.191 | 0.15 | 2.404 | 0.079 | 0.021 | -0.075 | -0.075 | 0 | 0.14 | 0.49 | 0.17 | 0.02 | 0.13 | 0.12 |
| 6 | 43192269 | rs6458328 | -1.113 | 0.121 | 2.017 | 0.182 | 0.026 | 0.023 | 0.023 | 0 | 0.05 | 0.25 | 0.04 | 0.01 | 0.02 | 0.01 |
| 6 | 43199482 | rs10948067 | -0.402 | -0.253 | 1.059 | -0.224 | 0.012 | 0.013 | 0.013 | 0.001 | 0.43 | 0.32 | 0.36 | 0.46 | 0.38 | 0.35 |
| 6 | 43209739 | rs2104142 | -0.978 | -0.676 | 0.851 | -0.084 | 0.062 | 0.026 | 0.026 | 0.348 | 0.49 | 0.7 | 0.4 | 0.47 | 0.38 | 0.4 |
| 6 | 43210695 | rs12663537 | -0.304 | -0.666 | 0.581 | -0.062 | 0.013 | -0.001 | -0.001 | 1.951 | 0.39 | 0.31 | 0.36 | 0.43 | 0.34 | 0.31 |
| 6 | 43213477 | rs2651206 | -1.886 | -0.611 | 0.637 | 0.215 | 0.05 | -0.015 | -0.015 | 0 | 0.39 | 0.8 | 0.33 | 0.44 | 0.33 | 0.4 |
| 6 | 43221264 | rs12664472 | -0.656 | -0.578 | 0.798 | 0.278 | -0.021 | 0.028 | 0.028 | 0 | 0.24 | 0.33 | 0.19 | 0.36 | 0.22 | 0.22 |
| 6 | 43224755 | rs7764257 | -1.188 | -0.589 | 0.726 | 0.26 | -0.014 | 0.009 | 0.009 | 0 | 0.29 | 0.36 | 0.27 | 0.36 | 0.29 | 0.3 |
| 6 | 43225060 | rs7750845 | -0.727 | -0.597 | 0.745 | 0.281 | -0.024 | 0.031 | 0.031 | 0 | 0.25 | 0.36 | 0.19 | 0.37 | 0.22 | 0.22 |
| 6 | 43225284 | rs7769094 | -0.717 | -0.636 | 0.75 | 0.306 | -0.025 | 0.034 | 0.034 | 0 | 0.25 | 0.36 | 0.19 | 0.36 | 0.22 | 0.22 |
| 6 | 43225791 | rs7752120 | -0.621 | -0.625 | 0.751 | 0.279 | -0.022 | 0.03 | 0.03 | 0 | 0.25 | 0.36 | 0.19 | 0.36 | 0.22 | 0.22 |
| 6 | 43233266 | rs754421 | -0.853 | -0.459 | 0.74 | 0.271 | -0.004 | 0.007 | 0.007 | 0 | 0.3 | 0.4 | 0.29 | 0.36 | 0.32 | 0.31 |
| 6 | 43234316 | rs2756184 | -1.825 | -0.604 | 0.411 | 0.509 | 0.05 | -0.004 | -0.004 | 0.171 | 0.47 | 0.85 | 0.51 | 0.57 | 0.41 | 0.5 |
| 6 | 43238609 | rs2651187 | -1.112 | -0.067 | 0.963 | 0.728 | -0.019 | 0.114 | 0.114 | 0.022 | 0.14 | 0.42 | 0.23 | 0.2 | 0.08 | 0.15 |
| 6 | 43245374 | rs2651189 | -1.936 | -0.206 | 0.636 | 0.286 | 0.006 | -0.039 | -0.039 | 0 | 0.35 | 0.81 | 0.35 | 0.48 | 0.34 | 0.41 |
| 6 | 43245641 | rs2756172 | -1.269 | -0.208 | 0.823 | -0.26 | -0.058 | -0.012 | -0.012 | 0.637 | 0.17 | 0.43 | 0.11 | 0.14 | 0.07 | 0.18 |
| 6 | 43257234 | rs2756174 | -1.278 | 0.339 | 0.49 | -0.102 | 0.01 | 0.014 | 0.014 | 0 | 0.29 | 0.51 | 0.32 | 0.38 | 0.31 | 0.36 |
| 6 | 43269029 | rs1574430 | 1.214 | -0.613 | -0.785 | -1.047 | -0.049 | 0.008 | 0.008 | 0.997 | 0.43 | 0.03 | 0.5 | 0.63 | 0.58 | 0.52 |
| 6 | 43269179 | rs2841648 | 1.214 | -0.613 | -0.785 | -1.047 | -0.049 | 0.008 | 0.008 | 0.997 | 0.43 | 0.03 | 0.5 | 0.63 | 0.58 | 0.52 |
| 6 | 43273604 | rs2242416 | 1.214 | -0.646 | -0.564 | -1.049 | -0.008 | 0.037 | 0.037 | 0.997 | 0.43 | 0.07 | 0.4 | 0.62 | 0.58 | 0.52 |
| 6 | 43276390 | rs2254303 | 1.214 | -0.646 | -0.564 | -1.051 | -0.008 | 0.037 | 0.037 | 0.997 | 0.43 | 0.07 | 0.5 | 0.63 | 0.58 | 0.52 |
| 6 | 43279721 | rs4398731 | -0.722 | -0.717 | -0.206 | -1.142 | 0.067 | 0.054 | 0.054 | 0.243 | 0.48 | 0.72 | 0.38 | 0.37 | 0.33 | 0.4 |
| 6 | 43287893 | rs4714675 | 0.38 | -0.662 | -0.153 | -1.144 | 0.02 | -0.051 | -0.051 | 0 | 0.91 | 0.78 | 0.88 | 0.99 | 0.91 | 0.91 |
| 6 | 43293101 | rs9472031 | 0.39 | -1.104 | 0.271 | -1.409 | -0.038 | -0.058 | -0.058 | 0 | 0.21 | 0.34 | 0.19 | 0.05 | 0.12 | 0.11 |
| 6 | 43299058 | rs2279087 | -1.112 | -1.102 | 0.244 | -1.407 | 0.066 | 0.054 | 0.054 | 0.257 | 0.48 | 0.71 | 0.38 | 0.38 | 0.33 | 0.39 |
| 6 | 43304007 | rs7692 | -1.114 | -1.119 | 0.147 | -1.45 | 0.066 | 0.059 | 0.059 | 0.587 | 0.49 | 0.71 | 0.38 | 0.37 | 0.33 | 0.4 |
| 6 | 43322231 | rs12190755 | -0.452 | -1.159 | 0.252 | -1.428 | -0.021 | -0.001 | -0.001 | 10.184 | 0.27 | 0.37 | 0.18 | 0.32 | 0.21 | 0.28 |
| 6 | 43352898 | rs6919440 | 1.697 | -0.658 | 0.105 | -1.488 | -0.081 | 0.028 | 0.028 | 0.638 | 0.42 | 0.07 | 0.49 | 0.62 | 0.54 | 0.51 |
| 6 | 43352980 | rs1214759 | 1.032 | -0.644 | 0.151 | -1.461 | 0.05 | 0.067 | 0.067 | 0.045 | 0.5 | 0.28 | 0.61 | 0.63 | 0.63 | 0.59 |
| 6 | 43354522 | rs6458334 | -0.441 | -0.563 | 0.135 | -1.395 | -0.008 | 0.002 | 0.002 | 0 | 0.27 | 0.36 | 0.19 | 0.32 | 0.24 | 0.29 |
| 6 | 43365360 | rs12195350 | -0.473 | -0.62 | 0.104 | -1.415 | -0.009 | -0.001 | -0.001 | 0 | 0.27 | 0.36 | 0.18 | 0.31 | 0.24 | 0.29 |
| 6 | 43388054 | rs6907066 | -0.484 | -0.923 | 0.015 | -1.432 | -0.002 | -0.002 | -0.002 | 1.87 | 0.34 | 0.35 | 0.24 | 0.35 | 0.3 | 0.34 |
| 6 | 43396325 | rs1214752 | 1.181 | -1.069 | -0.049 | -1.453 | -0.004 | 0.002 | 0.002 | 0 | 0.42 | 0.37 | 0.46 | 0.59 | 0.48 | 0.47 |
| 6 | 43396856 | rs4714684 | 1.583 | -1.064 | -0.087 | -1.45 | -0.049 | -0.022 | -0.022 | 0.188 | 0.36 | 0.18 | 0.45 | 0.59 | 0.46 | 0.43 |
| 6 | 43399837 | rs700008 | -1.139 | -1.105 | -0.063 | -1.485 | 0.008 | 0.015 | 0.015 | 0 | 0.84 | 0.86 | 0.81 | 0.95 | 0.87 | 0.92 |
| 6 | 43401105 | rs9394952 | 1.77 | -1.087 | -0.085 | -1.497 | -0.147 | -0.108 | -0.108 | 0 | 0.34 | 0.03 | 0.44 | 0.59 | 0.46 | 0.43 |
| 6 | 43401826 | rs831314 | -1.123 | -1.052 | -0.157 | -1.482 | 0.007 | 0.015 | 0.015 | 0 | 0.84 | 0.86 | 0.81 | 0.95 | 0.87 | 0.92 |
| 6 | 43401888 | rs831313 | -1.326 | -1.086 | -0.156 | -1.499 | 0.008 | 0.034 | 0.034 | 0 | 0.85 | 0.92 | 0.82 | 0.95 | 0.87 | 0.92 |
| 6 | 43404511 | rs9349256 | 1.766 | -1.075 | -0.167 | -1.489 | -0.151 | -0.108 | -0.108 | 0 | 0.34 | 0.03 | 0.44 | 0.59 | 0.46 | 0.44 |
| 6 | 43405525 | rs831312 | -0.925 | -1.047 | 0.129 | -1.331 | 0.008 | 0.015 | 0.015 | 0 | 0.84 | 0.86 | 0.81 | 0.95 | 0.87 | 0.92 |
| 6 | 43406501 | rs2277122 | 1.016 | -1.001 | -0.006 | -1.163 | -0.003 | -0.034 | -0.034 | 0 | 0.06 | 0 | 0.07 | 0.11 | 0.05 | 0.19 |
| 6 | 43408005 | rs2185631 | 1.429 | -0.959 | 0.161 | -1.238 | -0.06 | -0.035 | -0.035 | 0 | 0.37 | 0.15 | 0.46 | 0.53 | 0.47 | 0.45 |
| 6 | 43409418 | rs2487663 | -0.272 | -0.58 | 0.455 | -1.388 | -0.03 | 0.012 | 0.012 | 0 | 0.71 | 0.51 | 0.67 | 0.65 | 0.77 | 0.64 |
| 6 | 43412865 | rs2125739 | -1.016 | -0.118 | 1.175 | -1.341 | 0.003 | -0.044 | -0.044 | 0 | 0.23 | 0.31 | 0.19 | 0.08 | 0.23 | 0.16 |
| 6 | 43413081 | rs9381242 | 0.051 | -0.152 | 0.622 | -1.321 | -0.027 | 0.037 | 0.037 | 0 | 0.14 | 0.03 | 0.06 | 0.07 | 0.14 | 0.05 |
| 6 | 43418691 | rs1214748 | -1.468 | -0.136 | 0.693 | -1.306 | 0 | -0.103 | -0.103 | 1.788 | 0.29 | 0.54 | 0.21 | 0.08 | 0.29 | 0.17 |
| 6 | 43423647 | rs3734687 | -1.43 | -0.087 | -0.324 | -1.276 | 0.002 | -0.003 | -0.003 | 0 | 0.31 | 0.7 | 0.23 | 0.08 | 0.29 | 0.17 |
| 6 | 43423779 | rs3734688 | -0.1 | 0 | -0.386 | -1.134 | -0.001 | 0.003 | 0.003 | 0 | 0.16 | 0.15 | 0.07 | 0.07 | 0.14 | 0.06 |
| 6 | 43427850 | rs1674850 | -0.732 | 0.066 | -0.769 | -1.118 | 0 | -0.007 | -0.007 | 0 | 0.3 | 0.69 | 0.23 | 0.08 | 0.29 | 0.17 |
| 6 | 43431000 | rs1622800 | -0.634 | -0.067 | -1.023 | -0.852 | 0.013 | 0.043 | 0.043 | 0 | 0.34 | 0.72 | 0.25 | 0.18 | 0.28 | 0.25 |
| 6 | 43434222 | rs1674870 | 0.007 | -0.444 | -0.966 | -0.724 | 0.006 | 0.008 | 0.008 | 0.204 | 0.76 | 0.84 | 0.81 | 0.81 | 0.82 | 0.74 |
| 6 | 43442651 | rs1214769 | 0.115 | -0.573 | -0.772 | -0.931 | 0.026 | 0.043 | 0.043 | 0.278 | 0.62 | 0.78 | 0.73 | 0.67 | 0.72 | 0.68 |
| 6 | 43442756 | rs16896543 | 0.047 | -0.885 | 0.141 | -0.957 | -0.002 | 0.008 | 0.008 | 1.497 | 0.07 | 0.26 | 0.04 | 0.04 | 0.03 | 0.08 |
| 6 | 43446795 | rs9472064 | 0.486 | -0.83 | -0.243 | -1.588 | -0.066 | -0.112 | -0.112 | 0 | 0.23 | 0.01 | 0.3 | 0.61 | 0.32 | 0.3 |
| 6 | 43462988 | rs9472068 | -0.345 | -0.744 | 0.408 | -1.612 | 0.001 | -0.016 | -0.016 | 0 | 0.1 | 0.41 | 0.07 | 0.04 | 0.06 | 0.12 |
| 6 | 43479067 | rs7743244 | -0.608 | -0.791 | 0.863 | -1.629 | -0.008 | -0.026 | -0.026 | 0 | 0.12 | 0.57 | 0.08 | 0.04 | 0.06 | 0.13 |

| | | | | | | | | | | | | | | | |
|---|---|---|---|---|---|---|---|---|---|---|---|---|---|---|---|
| 6 | 43490809 | rs7755135 | -0.906 | -0.768 | 0.958 | -1.702 | -0.005 | -0.044 | -0.044 | 0 | 0.13 | 0.58 | 0.1 | 0.03 | 0.1 | 0.14 |
| 6 | 43490947 | rs11077 | -0.306 | -0.798 | 0.866 | -1.943 | -0.002 | -0.028 | -0.028 | 0 | 0.39 | 0.64 | 0.44 | 0.06 | 0.43 | 0.39 |
| 6 | 43492578 | rs2257082 | 0.497 | -0.953 | 0.894 | -1.935 | 0.006 | -0.021 | -0.021 | 0.777 | 0.25 | 0.16 | 0.3 | 0.62 | 0.29 | 0.3 |
| 6 | 43496662 | rs1106841 | -0.391 | -0.838 | 1.047 | -1.701 | 0.001 | -0.037 | -0.037 | 0 | 0.37 | 0.64 | 0.42 | 0.06 | 0.39 | 0.38 |
| 6 | 43528441 | rs1096699 | 0.543 | -0.907 | 1.409 | -1.535 | -0.126 | 0.155 | 0.155 | 0.04 | 0.25 | 0.03 | 0.34 | 0.02 | 0.33 | 0.25 |
| 6 | 43540926 | rs699937 | 0.613 | -0.907 | 1.41 | -1.535 | -0.091 | 0.13 | 0.13 | 0.031 | 0.25 | 0.07 | 0.34 | 0.02 | 0.33 | 0.25 |
| 6 | 43553351 | rs9472084 | -0.871 | -0.659 | 1.668 | -1.236 | -0.005 | -0.023 | -0.023 | 0 | 0.12 | 0.56 | 0.08 | 0.03 | 0.06 | 0.13 |
| 6 | 43581586 | rs3734690 | 0.209 | -0.655 | 1.826 | -1.232 | -0.01 | -0.004 | -0.004 | 0 | 0.06 | 0.14 | 0.03 | 0.03 | 0.03 | 0.09 |
| 6 | 43582584 | rs6899628 | -0.388 | -0.655 | 1.901 | -1.224 | 0.025 | 0.037 | 0.037 | 0.326 | 0.08 | 0.37 | 0.05 | 0.03 | 0.04 | 0.09 |
| 6 | 43603468 | rs3757271 | 0.237 | -0.672 | 1.214 | -0.892 | -0.011 | 0.025 | 0.025 | 0 | 0.06 | 0.12 | 0.03 | 0.07 | 0.03 | 0.09 |
| 6 | 43608386 | rs7106 | 0.792 | -0.714 | 1.047 | -1.036 | -0.109 | 0.093 | 0.093 | 1.239 | 0.18 | 0.02 | 0.33 | 0.02 | 0.3 | 0.17 |
| 6 | 43611038 | rs12660473 | -0.011 | -1.15 | 1.68 | -0.412 | -0.011 | 0.025 | 0.025 | 0 | 0.06 | 0.12 | 0.03 | 0.07 | 0.03 | 0.09 |
| 6 | 43618026 | rs6915523 | -1.431 | -1.159 | 1.354 | -0.421 | -0.021 | 0.033 | 0.033 | 2.302 | 0.12 | 0.6 | 0.08 | 0.08 | 0.06 | 0.14 |
| 6 | 43631259 | rs7756754 | -1.528 | -1.608 | 1.218 | -0.378 | -0.05 | 0.047 | 0.047 | 0.311 | 0.2 | 0.65 | 0.11 | 0.32 | 0.1 | 0.31 |
| 6 | 43638636 | rs16896629 | -0.255 | -1.575 | 1.421 | -0.359 | -0.01 | 0.04 | 0.04 | 0 | 0.08 | 0.09 | 0.04 | 0.24 | 0.06 | 0.2 |
| 6 | 43640099 | rs2277125 | -0.558 | -2.028 | 1.06 | -0.605 | 0.018 | -0.009 | -0.009 | 0 | 0.22 | 0.15 | 0.11 | 0.31 | 0.12 | 0.29 |
| 6 | 43644800 | rs16896641 | 0.338 | -2.023 | 1.282 | -0.499 | -0.01 | 0.017 | 0.017 | 0 | 0.06 | 0.1 | 0.03 | 0.09 | 0.04 | 0.1 |
| 6 | 43645833 | rs3778492 | -1.814 | -2.095 | 1.406 | -0.53 | -0.026 | 0.02 | 0.02 | 1.215 | 0.21 | 0.73 | 0.11 | 0.34 | 0.1 | 0.31 |
| 6 | 43646401 | rs2295948 | -1.202 | -2.103 | 1.832 | -0.542 | -0.039 | 0.021 | 0.021 | 0 | 0.31 | 0.54 | 0.17 | 0.4 | 0.16 | 0.4 |
| 6 | 43649521 | rs1334600 | -1.583 | -1.519 | 2.017 | -0.245 | -0.056 | 0.034 | 0.034 | 3.927 | 0.13 | 0.62 | 0.07 | 0.09 | 0.04 | 0.1 |
| 6 | 43652770 | rs7775611 | -1.71 | -1.523 | 1.831 | -0.249 | -0.056 | 0.011 | 0.011 | 4.729 | 0.13 | 0.69 | 0.07 | 0.09 | 0.04 | 0.12 |
| 6 | 43659500 | rs16869037 | -0.842 | -1.532 | 1.814 | 0.026 | -0.024 | 0.052 | 0.052 | 0.122 | 0.1 | 0.21 | 0.06 | 0.25 | 0.06 | 0.2 |
| 6 | 43659625 | rs6908311 | 0.015 | -1.485 | 1.31 | 0.043 | -0.01 | 0.018 | 0.018 | 0 | 0.07 | 0.12 | 0.05 | 0.09 | 0.04 | 0.12 |
| 6 | 43660769 | rs9968825 | -1.79 | -1.48 | 1.288 | 0.04 | -0.043 | 0.041 | 0.041 | 1.678 | 0.21 | 0.68 | 0.11 | 0.33 | 0.1 | 0.32 |
| 6 | 43673583 | rs7764657 | -0.479 | -1.27 | 0.884 | 0.214 | -0.021 | 0.185 | 0.185 | 0 | 0.09 | 0.34 | 0.05 | 0.24 | 0.03 | 0.11 |
| 6 | 43679047 | rs1761769 | -0.545 | -0.985 | -0.745 | -0.243 | 0.002 | 0.048 | 0.048 | 0 | 0.49 | 0.77 | 0.49 | 0.35 | 0.46 | 0.6 |
| 6 | 43680462 | rs1761768 | 0.035 | -0.278 | -1.17 | -0.199 | -0.052 | 0.092 | 0.092 | 0.661 | 0.34 | 0.14 | 0.41 | 0.1 | 0.39 | 0.46 |
| 6 | 43682990 | rs752073 | 0.583 | -0.395 | -1.089 | 1.356 | -0.022 | -0.112 | -0.112 | 0 | 0.84 | 0.98 | 0.88 | 0.53 | 0.82 | 0.62 |
| 6 | 43683804 | rs833065 | 0.934 | -0.578 | -0.545 | 1.51 | -0.005 | 0.031 | 0.031 | 0 | 0.82 | 0.81 | 0.82 | 0.42 | 0.82 | 0.6 |
| 6 | 43686011 | rs7767224 | 0.776 | -0.615 | -0.764 | 1.002 | 0.029 | 0.044 | 0.044 | 0 | 0.52 | 0.28 | 0.55 | 0.35 | 0.6 | 0.41 |
| 6 | 43691873 | rs866730 | 0.464 | -0.622 | -0.123 | 0.854 | -0.001 | 0 | 0 | 0 | 0.63 | 0.64 | 0.64 | 0.53 | 0.69 | 0.68 |
| 6 | 43695675 | rs9472114 | 1.038 | -1.114 | -0.174 | 0.554 | 0.015 | -0.012 | -0.012 | 0 | 0.65 | 0.33 | 0.64 | 0.39 | 0.72 | 0.48 |
| 6 | 43705011 | rs735775 | -1.413 | -1.419 | 0.101 | 0.589 | -0.008 | -0.011 | -0.011 | 0 | 0.29 | 0.62 | 0.2 | 0.25 | 0.2 | 0.33 |
| 6 | 43710756 | rs945128 | 0.69 | -1.653 | 0.637 | 0.756 | -0.025 | 0.028 | 0.028 | 0.07 | 0.76 | 0.71 | 0.72 | 0.64 | 0.79 | 0.7 |
| 6 | 43711057 | rs7741669 | -0.858 | -1.685 | 1.213 | 0.759 | 0 | 0 | 0 | 1.47 | 0.16 | 0.28 | 0.16 | 0.2 | 0.14 | 0.23 |
| 6 | 43711459 | rs4714695 | 0.666 | -1.647 | 0.94 | 0.715 | -0.025 | 0.028 | 0.028 | 0.07 | 0.76 | 0.71 | 0.72 | 0.64 | 0.79 | 0.7 |
| 6 | 43711981 | rs1535507 | -0.196 | -1.606 | 0.355 | 0.812 | 0.005 | -0.041 | -0.041 | 0.671 | 0.86 | 0.94 | 0.85 | 0.7 | 0.84 | 0.77 |
| 6 | 43712979 | rs9357421 | 1.25 | -0.871 | -0.446 | 0.437 | -0.002 | 0.046 | 0.046 | 0 | 0.33 | 0.16 | 0.2 | 0.22 | 0.31 | 0.31 |
| 6 | 43714001 | rs7749629 | 0.298 | -0.705 | -0.218 | 0.506 | 0.001 | 0.001 | 0.001 | 0 | 0.33 | 0.61 | 0.45 | 0.41 | 0.39 | 0.34 |
| 6 | 43715994 | rs945131 | 0.384 | -0.399 | -0.88 | 0.077 | -0.006 | 0.09 | 0.09 | 0 | 0.51 | 0.33 | 0.32 | 0.27 | 0.49 | 0.38 |
| 6 | 43716429 | rs2396053 | 0.477 | -0.156 | -0.79 | 0.286 | -0.013 | 0.062 | 0.062 | 0 | 0.45 | 0.31 | 0.39 | 0.32 | 0.43 | 0.4 |
| 6 | 43719391 | rs9394963 | 0.88 | -0.121 | -0.907 | 0.325 | -0.001 | 0.044 | 0.044 | 0 | 0.37 | 0.27 | 0.22 | 0.26 | 0.36 | 0.32 |
| 6 | 43719993 | rs4714696 | 0.162 | 0.05 | -0.57 | 0.838 | 0.009 | 0.002 | 0.002 | 0.294 | 0.25 | 0.29 | 0.38 | 0.23 | 0.29 | 0.15 |
| 6 | 43723335 | rs833052 | 0.18 | -0.106 | -0.751 | 1.195 | 0.012 | -0.056 | -0.056 | 0 | 0.86 | 0.96 | 0.85 | 0.7 | 0.87 | 0.83 |
| 6 | 43726956 | rs866236 | 0.755 | -0.122 | -0.474 | 0.904 | -0.02 | 0.024 | 0.024 | 0.283 | 0.65 | 0.42 | 0.62 | 0.51 | 0.75 | 0.53 |
| 6 | 43732669 | rs699946 | 0.454 | 0.047 | -0.831 | 0.755 | 0.033 | -0.058 | -0.058 | 0.194 | 0.26 | 0.19 | 0.28 | 0.42 | 0.18 | 0.22 |
| 6 | 43737830 | rs1570360 | -0.488 | 0.906 | -1.191 | 0.751 | -0.05 | 0.026 | 0.026 | 0 | 0.74 | 0.96 | 0.8 | 0.82 | 0.68 | 0.74 |
| 6 | 43738977 | rs25648 | 0.302 | 1.308 | -1.161 | 0.72 | -0.014 | 0.01 | 0.01 | 0 | 0.13 | 0.07 | 0.14 | 0.1 | 0.17 | 0.14 |
| 6 | 43742527 | rs833068 | 0.73 | 1.235 | -0.292 | 0.642 | -0.014 | 0.007 | 0.007 | 0.977 | 0.42 | 0.49 | 0.38 | 0.43 | 0.31 | 0.25 |
| 6 | 43742579 | rs833069 | 0.609 | 1.337 | -0.507 | 0.669 | 0.005 | -0.003 | -0.003 | 0 | 0.43 | 0.63 | 0.39 | 0.43 | 0.31 | 0.25 |
| 6 | 43742626 | rs833070 | 0.246 | 1.594 | -0.415 | 0.425 | -0.02 | 0.057 | 0.057 | 0 | 0.6 | 0.76 | 0.65 | 0.74 | 0.51 | 0.61 |
| 6 | 43745095 | rs2146323 | -0.353 | 0.737 | -0.425 | 0.574 | -0.006 | 0.027 | 0.027 | 0 | 0.33 | 0.19 | 0.27 | 0.27 | 0.35 | 0.37 |
| 6 | 43747577 | rs3025010 | -0.456 | 0.936 | -0.291 | 0.677 | 0 | -0.007 | -0.007 | 0 | 0.34 | 0.38 | 0.29 | 0.31 | 0.37 | 0.38 |
| 6 | 43751359 | rs3025035 | -0.659 | 0.699 | 0.212 | 1.17 | -0.023 | 0.056 | 0.056 | 3.374 | 0.14 | 0.32 | 0.06 | 0.18 | 0.09 | 0.08 |
| 6 | 43752536 | rs3025039 | -0.521 | 1.251 | -0.192 | 1.148 | -0.009 | -0.015 | -0.015 | 0 | 0.12 | 0.08 | 0.23 | 0.17 | 0.12 | 0.11 |
| 6 | 43753212 | rs10434 | -0.007 | 1.396 | -0.342 | 0.557 | -0.016 | 0.023 | 0.023 | 0 | 0.66 | 0.67 | 0.74 | 0.78 | 0.54 | 0.55 |
| 6 | 43758485 | rs6900017 | -0.574 | 1.29 | 0.409 | 1.008 | -0.021 | 0.015 | 0.015 | 3.31 | 0.15 | 0.29 | 0.08 | 0.17 | 0.09 | 0.13 |
| 6 | 43764359 | rs12204488 | -1.102 | 1.536 | -0.045 | 0.839 | 0.006 | 0.01 | 0.01 | 2.86 | 0.32 | 0.68 | 0.18 | 0.27 | 0.23 | 0.27 |
| 6 | 43764551 | rs1358980 | 1.087 | 1.932 | -0.638 | 1.019 | -0.058 | -0.009 | -0.009 | 0.082 | 0.56 | 0.11 | 0.51 | 0.56 | 0.5 | 0.68 |
| 6 | 43767703 | rs10948095 | 1.637 | 1.677 | 0.047 | 1.335 | -0.038 | -0.021 | -0.021 | 0.159 | 0.66 | 0.17 | 0.58 | 0.63 | 0.6 | 0.69 |
| 6 | 43768435 | rs13211073 | -1.127 | 1.742 | 0.383 | 1.292 | -0.003 | -0.001 | -0.001 | 0 | 0.15 | 0.17 | 0.32 | 0.14 | 0.3 | 0.13 |
| 6 | 43770150 | rs943084 | -0.585 | 1.684 | 1.001 | 1.382 | -0.01 | 0.038 | 0.038 | 0 | 0.14 | 0.21 | 0.07 | 0.29 | 0.1 | 0.16 |
| 6 | 43770512 | rs2396081 | -1.812 | 1.748 | 0.902 | 1.35 | 0.028 | 0.036 | 0.036 | 0 | 0.32 | 0.56 | 0.4 | 0.44 | 0.39 | 0.29 |
| 6 | 43775838 | rs4507587 | -1.006 | 2.1 | 0.272 | 0.964 | -0.064 | -0.041 | -0.041 | 0 | 0.11 | 0.04 | 0.3 | 0.14 | 0.27 | 0.09 |
| 6 | 43781918 | rs10807291 | 1.679 | 1.203 | 0.779 | 0.999 | 0.002 | 0.028 | 0.028 | 4.082 | 0.86 | 0.36 | 0.72 | 0.87 | 0.87 | 0.91 |
| 6 | 43782573 | rs9462930 | 1.685 | 1.254 | 0.777 | 0.997 | 0.018 | 0.027 | 0.027 | 3.684 | 0.85 | 0.36 | 0.71 | 0.87 | 0.86 | 0.91 |
| 6 | 43789616 | rs6458350 | -0.135 | 1.173 | 0.539 | 0.309 | 0.008 | -0.055 | -0.055 | 0.064 | 0.32 | 0.42 | 0.2 | 0.17 | 0.3 | 0.14 |
| 6 | 43791080 | rs6458351 | 0.717 | 1.125 | -0.202 | 0.4 | -0.005 | -0.02 | -0.02 | 0.066 | 0.92 | 0.95 | 0.78 | 0.84 | 0.88 | 0.94 |
| 6 | 43793858 | rs9367176 | 0.182 | 1.202 | -0.172 | 0.274 | -0.059 | 0.113 | 0.113 | 0.262 | 0.29 | 0.08 | 0.18 | 0.13 | 0.29 | 0.17 |

| | | | | | | | | | | | | | | | | |
|---|---|---|---|---|---|---|---|---|---|---|---|---|---|---|---|---|
| 6 | 43798775 | rs9381256 | 0.098 | 0.367 | -0.481 | 0.877 | 0.013 | -0.027 | -0.027 | 0 | 0.19 | 0.14 | 0.12 | 0.42 | 0.13 | 0.4 |
| 6 | 43798902 | rs9381257 | 0.085 | -0.285 | -0.597 | 0.283 | 0.011 | 0.094 | 0.094 | 0 | 0.5 | 0.31 | 0.53 | 0.73 | 0.55 | 0.7 |
| 6 | 43801878 | rs2894536 | 0.334 | 0.048 | 0.168 | 0.673 | -0.04 | -0.006 | -0.006 | 0.053 | 0.8 | 0.54 | 0.7 | 0.86 | 0.82 | 0.87 |
| 6 | 43802563 | rs4714699 | -0.431 | 0.503 | -0.27 | 1.546 | -0.059 | -0.066 | -0.066 | 0.069 | 0.31 | 0.15 | 0.59 | 0.46 | 0.4 | 0.3 |
| 6 | 43804687 | rs9367177 | -0.733 | 0.261 | -0.196 | 2.147 | -0.026 | -0.042 | -0.042 | 0 | 0.08 | 0 | 0.04 | 0.25 | 0.08 | 0.04 |
| 6 | 43806609 | rs881858 | -0.564 | -0.034 | 0.011 | 1.805 | 0.025 | 0.029 | 0.029 | 0.021 | 0.62 | 0.34 | 0.76 | 0.77 | 0.69 | 0.76 |
| 6 | 43806734 | rs13201802 | -0.488 | 0.234 | -0.423 | 1.426 | -0.013 | -0.022 | -0.022 | 0 | 0.11 | 0.04 | 0.13 | 0.21 | 0.14 | 0.27 |
| 6 | 43806921 | rs1317840 | -0.273 | 0.087 | -0.108 | 1.383 | 0.022 | 0.048 | 0.048 | 0 | 0.54 | 0.71 | 0.67 | 0.65 | 0.56 | 0.49 |
| 6 | 43811268 | rs9369426 | -0.635 | 0.167 | -0.302 | 1.311 | -0.023 | -0.056 | -0.056 | 0 | 0.11 | 0.01 | 0.14 | 0.45 | 0.13 | 0.17 |
| 6 | 43811762 | rs9472138 | 0.873 | -0.073 | -0.154 | 0.782 | 0.041 | 0.088 | 0.088 | 0 | 0.34 | 0.17 | 0.19 | 0.14 | 0.3 | 0.18 |
| 6 | 43815364 | rs6937438 | -0.825 | -0.101 | 0.491 | 0.761 | -0.027 | -0.06 | -0.06 | 3.078 | 0.63 | 0.52 | 0.78 | 0.86 | 0.7 | 0.82 |
| 6 | 43816377 | rs2025704 | -0.726 | 0.005 | 0.003 | 0.86 | -0.103 | 0.068 | 0.068 | 0 | 0.62 | 0.87 | 0.56 | 0.76 | 0.55 | 0.66 |
| 6 | 43817837 | rs1536304 | -0.725 | 0.118 | 0.15 | 1.008 | -0.095 | 0.09 | 0.09 | 0 | 0.6 | 0.87 | 0.62 | 0.76 | 0.55 | 0.67 |
| 6 | 43822780 | rs1326142 | -0.582 | 0.19 | 0.665 | 0.955 | 0.017 | -0.037 | -0.037 | 0 | 0.09 | 0.14 | 0.06 | 0.01 | 0.12 | 0.17 |
| 6 | 43823689 | rs6933958 | 0.571 | 0.089 | 1.102 | 1.112 | 0.031 | 0.024 | 0.024 | 0 | 0.4 | 0.33 | 0.39 | 0.28 | 0.3 | 0.35 |
| 6 | 43825459 | rs9394969 | -0.094 | -0.001 | 0.946 | 1.024 | 0.012 | 0.036 | 0.036 | 0 | 0.48 | 0.29 | 0.39 | 0.61 | 0.47 | 0.44 |
| 6 | 43826627 | rs943080 | -0.848 | -0.002 | 0.84 | 1.058 | -0.065 | 0.07 | 0.07 | 0 | 0.59 | 0.83 | 0.52 | 0.73 | 0.51 | 0.67 |
| 6 | 43826681 | rs9472144 | 1.062 | -0.228 | 1.114 | 1.083 | 0.039 | 0.071 | 0.071 | 0 | 0.31 | 0.15 | 0.16 | 0.13 | 0.25 | 0.12 |
| 6 | 43828134 | rs2148251 | -1.499 | 0.304 | 2.314 | 1.989 | -0.011 | 0.076 | 0.076 | 0 | 0.1 | 0.4 | 0.11 | 0.12 | 0.04 | 0.22 |
| 6 | 43829251 | rs6458356 | -1.533 | 0.729 | 1.673 | 2.141 | 0.024 | 0.333 | 0.333 | 0.142 | 0.11 | 0.5 | 0.19 | 0.42 | 0.11 | 0.15 |
| 6 | 43832030 | rs4714705 | 1.511 | 0.755 | 1.097 | 1.193 | 0.008 | -0.048 | -0.048 | 0 | 0.91 | 0.75 | 0.95 | 0.99 | 0.93 | 0.84 |
| 6 | 43832177 | rs4714706 | 1.605 | 0.69 | 1.349 | 1.16 | 0.015 | -0.057 | -0.057 | 0 | 0.9 | 0.73 | 0.95 | 0.99 | 0.93 | 0.83 |
| 6 | 43832802 | rs11968561 | -1.234 | 0.775 | 0.978 | 1.395 | -0.001 | 0.004 | 0.004 | 0 | 0.18 | 0.17 | 0.26 | 0.21 | 0.19 | 0.23 |
| 6 | 43834027 | rs13192583 | -1.046 | 0.844 | 1.03 | 1.394 | 0.002 | -0.018 | -0.018 | 0 | 0.08 | 0.1 | 0.04 | 0.01 | 0.08 | 0.2 |
| 6 | 43834215 | rs9367179 | -1.221 | 0.668 | 0.87 | 2.02 | 0 | 0.013 | 0.013 | 0.002 | 0.18 | 0.16 | 0.16 | 0.4 | 0.19 | 0.31 |
| 6 | 43834251 | rs9369430 | -2.027 | 0.707 | 1.166 | 2.028 | -0.011 | 0.106 | 0.106 | 0 | 0.27 | 0.46 | 0.2 | 0.47 | 0.24 | 0.48 |
| 6 | 43836930 | rs11968152 | -1.556 | 0.73 | 0.795 | 1.483 | -0.007 | 0.025 | 0.025 | 0 | 0.08 | 0.27 | 0.05 | 0.06 | 0.05 | 0.18 |
| 6 | 43846080 | rs4714707 | 0.992 | 1.286 | 0.483 | 1.885 | 0.029 | 0.067 | 0.067 | 0 | 0.83 | 0.75 | 0.78 | 0.73 | 0.82 | 0.75 |
| 6 | 43855023 | rs1003167 | -0.023 | 1.169 | -0.303 | 1.257 | -0.017 | -0.016 | -0.016 | 0.203 | 0.38 | 0.25 | 0.47 | 0.49 | 0.47 | 0.36 |
| 6 | 43865321 | rs1570986 | 0.484 | 0.838 | 0.121 | 1.642 | 0.015 | 0.088 | 0.088 | 0 | 0.83 | 0.78 | 0.68 | 0.64 | 0.83 | 0.79 |
| 6 | 43868216 | rs7761865 | -0.53 | 0.165 | -0.496 | 0.801 | -0.001 | -0.016 | -0.016 | 0 | 0.32 | 0.25 | 0.39 | 0.55 | 0.29 | 0.41 |
| 6 | 43868986 | rs1539508 | -0.625 | 0.272 | -1.132 | 0.433 | 0.007 | 0.071 | 0.071 | 0 | 0.51 | 0.72 | 0.67 | 0.64 | 0.44 | 0.62 |
| 6 | 43870079 | rs1776723 | -0.554 | 0.233 | -1.778 | 0.274 | -0.11 | 0.159 | 0.159 | 0 | 0.61 | 0.93 | 0.73 | 0.83 | 0.51 | 0.8 |
| 6 | 43875449 | rs9381262 | -0.362 | -0.039 | -1.239 | 0.698 | 0.032 | 0.046 | 0.046 | 0 | 0.48 | 0.75 | 0.64 | 0.6 | 0.39 | 0.57 |
| 6 | 43882612 | rs9472153 | 0.441 | 0.181 | -1.136 | 1.01 | -0.058 | -0.051 | -0.051 | 0.034 | 0.35 | 0.19 | 0.37 | 0.49 | 0.43 | 0.57 |
| 6 | 43882924 | rs7747448 | -0.577 | 0.108 | -0.207 | 1.018 | -0.003 | 0.008 | 0.008 | 0 | 0.31 | 0.6 | 0.33 | 0.33 | 0.29 | 0.33 |
| 6 | 43886159 | rs721640 | 0.14 | -0.044 | -0.294 | 0.989 | -0.007 | 0.006 | 0.006 | 0 | 0.26 | 0.05 | 0.29 | 0.21 | 0.28 | 0.12 |
| 6 | 43887461 | rs7751666 | -1.129 | 0.006 | 0.954 | 1.176 | 0.046 | 0.206 | 0.206 | 1.101 | 0.05 | 0.28 | 0.04 | 0.13 | 0.03 | 0.14 |
| 6 | 43887937 | rs4426996 | -1.959 | -0.09 | 1.341 | 1.108 | 0.026 | 0.155 | 0.155 | 0.282 | 0.1 | 0.53 | 0.16 | 0.18 | 0.09 | 0.18 |
| 6 | 43888710 | rs4711757 | -2.085 | -0.087 | 1.35 | 1.106 | 0.023 | 0.129 | 0.129 | 0.054 | 0.13 | 0.47 | 0.15 | 0.17 | 0.12 | 0.18 |
| 6 | 43897727 | rs9472155 | -0.521 | -0.004 | 1.092 | 0.736 | -0.013 | -0.062 | -0.062 | 0 | 0.24 | 0.43 | 0.28 | 0.13 | 0.22 | 0.3 |
| 6 | 43898777 | rs833623 | -1.289 | 0.084 | 0.347 | 0.933 | -0.021 | -0.035 | -0.035 | 0 | 0.57 | 0.76 | 0.74 | 0.52 | 0.58 | 0.45 |
| 6 | 43899252 | rs7767550 | 0.185 | -0.13 | 0.557 | 0.903 | -0.001 | 0.003 | 0.003 | 0 | 0.21 | 0.18 | 0.26 | 0.13 | 0.22 | 0.19 |
| 6 | 43900443 | rs833621 | -0.381 | -0.059 | 0.508 | 0.931 | -0.001 | 0.004 | 0.004 | 0 | 0.34 | 0.35 | 0.46 | 0.39 | 0.36 | 0.26 |
| 6 | 43900707 | rs844294 | -0.973 | 0.204 | -0.321 | 1.036 | 0.003 | 0.074 | 0.074 | 0 | 0.53 | 0.72 | 0.59 | 0.35 | 0.51 | 0.38 |
| 6 | 43904294 | rs10948108 | -1.1 | 0.275 | -0.573 | 1.19 | 0.038 | 0.097 | 0.097 | 0 | 0.08 | 0.21 | 0.18 | 0.17 | 0.11 | 0.08 |
| 6 | 43904780 | rs11755845 | -0.241 | -0.323 | -1.347 | 0.665 | 0.011 | 0.028 | 0.028 | 0 | 0.25 | 0.12 | 0.28 | 0.18 | 0.25 | 0.18 |
| 6 | 43911551 | rs865585 | 0.701 | -0.253 | -1.192 | 0.921 | 0 | -0.001 | -0.001 | 0 | 0.76 | 0.82 | 0.84 | 0.78 | 0.77 | 0.57 |
| 6 | 43913988 | rs7764227 | -0.691 | -0.45 | -1.737 | 0.775 | 0.025 | -0.02 | -0.02 | 0 | 0.7 | 0.9 | 0.72 | 0.68 | 0.72 | 0.78 |
| 6 | 43917892 | rs729391 | -0.677 | -0.051 | -1.496 | 0.834 | -0.005 | -0.06 | -0.06 | 0 | 0.58 | 0.93 | 0.64 | 0.55 | 0.61 | 0.5 |
| 6 | 43918407 | rs9369434 | 0.078 | 0.433 | -1.588 | 0.543 | -0.132 | -0.044 | -0.044 | 1.354 | 0.3 | 0.07 | 0.5 | 0.31 | 0.42 | 0.37 |
| 6 | 43918897 | rs9472158 | -0.773 | 0.158 | -0.98 | 0.582 | 0.024 | -0.015 | -0.015 | 0 | 0.44 | 0.26 | 0.64 | 0.43 | 0.6 | 0.51 |
| 6 | 43919695 | rs9472159 | -0.329 | 0.253 | -1.301 | 0.19 | -0.122 | 0.026 | 0.026 | 0.497 | 0.36 | 0.08 | 0.54 | 0.3 | 0.48 | 0.4 |
| 6 | 43927050 | rs7767396 | -0.443 | 0.54 | -0.413 | 0.34 | 0.053 | -0.017 | -0.017 | 0 | 0.33 | 0.49 | 0.56 | 0.32 | 0.45 | 0.36 |
| 6 | 43933256 | rs9472173 | -0.61 | 0.044 | -0.526 | 0.147 | -0.028 | 0.009 | 0.009 | 0.271 | 0.42 | 0.76 | 0.6 | 0.4 | 0.48 | 0.47 |
| 6 | 43934782 | rs9472175 | 1.216 | -0.953 | 0.456 | -0.46 | -0.068 | -0.075 | -0.075 | 0.803 | 0.76 | 0.53 | 0.88 | 0.92 | 0.86 | 0.81 |
| 6 | 43935886 | rs9296424 | 1.236 | -1.025 | 0.606 | -0.515 | -0.031 | -0.035 | -0.035 | 1.692 | 0.76 | 0.69 | 0.88 | 0.92 | 0.86 | 0.8 |
| 6 | 43936789 | rs9654590 | -1.53 | -1.358 | 0.954 | -0.448 | -0.055 | -0.027 | -0.027 | 0 | 0.23 | 0.4 | 0.16 | 0.15 | 0.1 | 0.15 |
| 6 | 43937518 | rs7755431 | -1.984 | -1.391 | 0.372 | -0.693 | 0.005 | 0.005 | 0.005 | 0.446 | 0.29 | 0.74 | 0.32 | 0.18 | 0.18 | 0.21 |
| 6 | 43941645 | rs4714724 | -1.183 | -0.594 | -0.369 | -1.24 | -0.037 | 0.156 | 0.156 | 0 | 0.43 | 0.8 | 0.48 | 0.1 | 0.49 | 0.22 |
| 6 | 43942389 | rs4714729 | -1.223 | -0.59 | -0.408 | -1.215 | -0.041 | 0.163 | 0.163 | 0 | 0.43 | 0.82 | 0.5 | 0.13 | 0.49 | 0.24 |
| 6 | 43945688 | rs7770346 | 0.358 | 0.058 | -0.429 | -0.909 | -0.094 | 0.077 | 0.077 | 0 | 0.73 | 0.87 | 0.6 | 0.9 | 0.55 | 0.82 |
| 6 | 43949941 | rs1776704 | -0.916 | -0.404 | 0.045 | 0.135 | 0 | 0.001 | 0.001 | 0 | 0.45 | 0.44 | 0.45 | 0.51 | 0.47 | 0.45 |
| 6 | 43950851 | rs910611 | -0.801 | -0.628 | 0.132 | 0.396 | -0.005 | -0.014 | -0.014 | 0 | 0.08 | 0.06 | 0.09 | 0.13 | 0.08 | 0.14 |
| 6 | 43951656 | rs910609 | -1.794 | -0.687 | 0.084 | 0.371 | 0 | 0 | 0 | 0 | 0.22 | 0.24 | 0.28 | 0.23 | 0.21 | 0.28 |
| 6 | 43953358 | rs1631662 | -1.083 | -0.848 | -0.067 | 0.004 | -0.005 | -0.017 | -0.017 | 0 | 0.13 | 0.18 | 0.17 | 0.06 | 0.12 | 0.11 |
| 6 | 43960263 | rs7765917 | -1.165 | -1.532 | -0.273 | -0.33 | 0.042 | 0.047 | 0.047 | 4.173 | 0.33 | 0.72 | 0.19 | 0.08 | 0.16 | 0.21 |
| 6 | 43965165 | rs3734693 | -1.546 | -2.009 | -0.614 | 0.67 | 0.114 | -0.036 | -0.036 | 0.97 | 0.39 | 0.83 | 0.22 | 0.4 | 0.17 | 0.36 |
| 6 | 43968216 | rs16897113 | -0.946 | -1.897 | 0.791 | 0.711 | 0.001 | -0.024 | -0.024 | 0 | 0.08 | 0.37 | 0.05 | 0.01 | 0.03 | 0.09 |

| | | | | | | | | | | | | | | | | |
|---|---|---|---|---|---|---|---|---|---|---|---|---|---|---|---|---|
| 6 | 43968814 | rs2295333 | -0.916 | -1.802 | 0.935 | 0.929 | -0.003 | 0.22 | 0.22 | 0 | 0.1 | 0.23 | 0.1 | 0.36 | 0.07 | 0.18 |
| 6 | 43972845 | rs2273309 | -0.676 | -1.796 | 1.034 | 1.122 | 0.024 | 0.224 | 0.224 | 0 | 0.05 | 0.2 | 0.03 | 0.25 | 0.03 | 0.09 |
| 6 | 43978323 | rs6909859 | -1.757 | -2.341 | 0.607 | 1.274 | 0.13 | -0.082 | -0.082 | 0.159 | 0.4 | 0.86 | 0.29 | 0.45 | 0.19 | 0.26 |
| 6 | 43979345 | rs9367182 | -1.834 | -2.255 | 1.599 | 1.236 | -0.01 | -0.013 | -0.013 | 0 | 0.15 | 0.33 | 0.2 | 0.1 | 0.1 | 0.07 |
| 6 | 43980128 | rs9381275 | -1.568 | -2.245 | 1.584 | 1.298 | -0.021 | 0.026 | 0.026 | 0 | 0.07 | 0.15 | 0.04 | 0.12 | 0.02 | 0.09 |
| 6 | 43980879 | rs9472189 | -1.295 | -2.194 | 1.838 | 1.073 | 0.033 | 0.041 | 0.041 | 0 | 0.08 | 0.23 | 0.17 | 0.1 | 0.08 | 0.03 |
| 6 | 43988011 | rs7772862 | -1.182 | -2.162 | 1.435 | 0.997 | -0.016 | -0.029 | -0.029 | 0 | 0.06 | 0.15 | 0.03 | 0 | 0.02 | 0.03 |
| 6 | 43990199 | rs6907643 | -2.137 | -2.425 | 1.131 | 0.972 | 0.078 | 0.012 | 0.012 | 3.33 | 0.38 | 0.76 | 0.27 | 0.34 | 0.21 | 0.2 |
| 6 | 43992587 | rs7757024 | -1.517 | -2.415 | 1.727 | 0.888 | -0.025 | -0.024 | -0.024 | 0.005 | 0.23 | 0.29 | 0.1 | 0.11 | 0.12 | 0.08 |
| 6 | 43995668 | rs4711760 | -1.875 | -2.488 | 0.447 | 0.475 | 0.009 | 0.185 | 0.185 | 0.335 | 0.57 | 0.82 | 0.46 | 0.9 | 0.38 | 0.74 |
| 6 | 44000031 | rs4714732 | -0.229 | -2.688 | 0.461 | 1.315 | 0.007 | -0.062 | -0.062 | 0 | 0.18 | 0.13 | 0.17 | 0.42 | 0.15 | 0.43 |
| 6 | 44002077 | rs1922985 | 0.112 | -2.679 | 0.335 | 1.303 | -0.025 | -0.109 | -0.109 | 0.619 | 0.15 | 0.03 | 0.16 | 0.44 | 0.14 | 0.42 |
| 6 | 44002496 | rs9462955 | -0.314 | -2.712 | 0.682 | 1.241 | 0.001 | 0 | 0 | 0.038 | 0.16 | 0.16 | 0.04 | 0.04 | 0.06 | 0.06 |
| 6 | 44003982 | rs7757246 | -1.205 | -2.596 | 0.771 | 1.238 | -0.023 | 0.006 | 0.006 | 0.478 | 0.31 | 0.64 | 0.22 | 0.4 | 0.13 | 0.26 |
| 6 | 44007446 | rs910603 | -1.722 | -2.7 | 0.217 | 0.806 | 0.02 | 0.03 | 0.03 | 1.573 | 0.55 | 0.58 | 0.46 | 0.86 | 0.34 | 0.72 |
| 6 | 44008325 | rs9381282 | 0.058 | -2.271 | -0.64 | 1.205 | -0.01 | -0.123 | -0.123 | 0 | 0.2 | 0.05 | 0.23 | 0.51 | 0.19 | 0.43 |
| 6 | 44016989 | rs753675 | -1.24 | -2.376 | -0.507 | 0.766 | 0.067 | 0.133 | 0.133 | 0 | 0.51 | 0.68 | 0.45 | 0.81 | 0.33 | 0.66 |
| 6 | 44018156 | rs1884377 | 0.06 | -1.754 | -0.885 | 1.823 | -0.017 | -0.075 | -0.075 | 0.816 | 0.18 | 0.05 | 0.21 | 0.43 | 0.21 | 0.36 |
| 6 | 44020410 | rs9472201 | -0.852 | -1.903 | -0.746 | 1.92 | 0 | 0.007 | 0.007 | 0 | 0.08 | 0.05 | 0.03 | 0.15 | 0.04 | 0.12 |
| 6 | 44021286 | rs9472202 | -0.677 | -1.924 | -0.956 | 2.044 | 0.009 | -0.112 | -0.112 | 0 | 0.25 | 0.07 | 0.2 | 0.5 | 0.23 | 0.42 |
| 6 | 44024262 | rs1950506 | 0.146 | -1.617 | -1.958 | 1.038 | 0.02 | 0.024 | 0.024 | 0 | 0.47 | 0.17 | 0.5 | 0.6 | 0.42 | 0.66 |
| 6 | 44025275 | rs9462957 | 0.55 | -1.025 | -1.065 | 0.552 | -0.001 | -0.04 | -0.04 | 0 | 0.29 | 0.33 | 0.24 | 0.09 | 0.26 | 0.21 |
| 6 | 44025699 | rs4714735 | 0.106 | 0.822 | -0.893 | 1.097 | -0.024 | -0.036 | -0.036 | 0 | 0.07 | 0.03 | 0.12 | 0.12 | 0.12 | 0.08 |
| 6 | 44027280 | rs833803 | -0.594 | 1.519 | -0.698 | 0.979 | 0.03 | -0.015 | -0.015 | 0 | 0.21 | 0.33 | 0.26 | 0.16 | 0.34 | 0.13 |
| 6 | 44030011 | rs833805 | 0.332 | 1.398 | -1.053 | 1.316 | -0.037 | -0.035 | -0.035 | 0 | 0.93 | 0.99 | 0.93 | 0.9 | 0.87 | 0.95 |
| 6 | 44032331 | rs700004 | -0.158 | 1.027 | -0.952 | 1.397 | 0.005 | -0.016 | -0.016 | 0 | 0.8 | 0.84 | 0.77 | 0.64 | 0.79 | 0.57 |
| 6 | 44034691 | rs833810 | 0.365 | 0.999 | -0.406 | 1.046 | -0.002 | -0.016 | -0.016 | 0.888 | 0.43 | 0.57 | 0.45 | 0.32 | 0.39 | 0.28 |
| 6 | 44046143 | rs9357429 | -0.676 | 0.955 | -0.24 | 1.505 | 0.002 | 0.258 | 0.258 | 0.123 | 0.12 | 0.26 | 0.28 | 0.47 | 0.15 | 0.15 |
| 6 | 44051221 | rs6458368 | -0.304 | 0.824 | -1.079 | 0.876 | -0.06 | 0.116 | 0.116 | 0.58 | 0.6 | 0.82 | 0.7 | 0.8 | 0.54 | 0.45 |
| 6 | 44056585 | rs4714738 | -0.397 | 0.85 | -0.937 | 1.532 | -0.032 | -0.064 | -0.064 | 0.231 | 0.1 | 0.01 | 0.12 | 0.23 | 0.14 | 0.13 |
| 6 | 44073955 | rs9369453 | -0.245 | 0.869 | -0.861 | 1.608 | -0.02 | -0.028 | -0.028 | 3.997 | 0.06 | 0.01 | 0.1 | 0.23 | 0.1 | 0.12 |
| 6 | 44075807 | rs10948122 | 0.199 | 0.463 | -0.862 | 1.244 | -0.004 | -0.007 | -0.007 | 0.79 | 0.16 | 0.14 | 0.22 | 0.04 | 0.11 | 0.11 |
| 6 | 44076061 | rs877111 | -0.661 | 0.678 | -0.941 | 1.024 | -0.033 | 0.151 | 0.151 | 0.947 | 0.57 | 0.82 | 0.72 | 0.8 | 0.5 | 0.51 |
| 6 | 44077356 | rs1321097 | -0.661 | 0.685 | -0.941 | 1.024 | -0.036 | 0.151 | 0.151 | 0.947 | 0.57 | 0.81 | 0.72 | 0.8 | 0.5 | 0.51 |
| 6 | 44081510 | rs700006 | 0.42 | 0.656 | -0.939 | 1.025 | -0.004 | -0.007 | -0.007 | 0 | 0.16 | 0.14 | 0.22 | 0.04 | 0.11 | 0.11 |
| 6 | 44081718 | rs6883 | -0.55 | 0.862 | -0.866 | 1.174 | -0.033 | 0.146 | 0.146 | 1.038 | 0.57 | 0.81 | 0.72 | 0.79 | 0.5 | 0.51 |
| 6 | 44081987 | rs4711767 | -0.589 | 0.877 | -0.896 | 1.157 | -0.046 | 0.178 | 0.178 | 1.366 | 0.58 | 0.85 | 0.72 | 0.82 | 0.5 | 0.52 |
| 6 | 44084883 | rs10085205 | 0.753 | 0.719 | -0.683 | 1.145 | -0.006 | 0.069 | 0.069 | 3.796 | 0.29 | 0.56 | 0.37 | 0.53 | 0.27 | 0.25 |
| 6 | 44094305 | rs4714750 | 0.234 | 0.755 | -0.843 | 1.113 | -0.021 | -0.026 | -0.026 | 0 | 0.06 | 0.01 | 0.1 | 0.23 | 0.1 | 0.13 |
| 6 | 44097472 | rs7773700 | 1.081 | 0.736 | -0.632 | 1.168 | -0.003 | -0.004 | -0.004 | 0 | 0.27 | 0.42 | 0.22 | 0.31 | 0.23 | 0.17 |
| 6 | 44113916 | rs4714758 | -1.462 | 1.32 | -0.21 | 1.414 | -0.001 | -0.003 | -0.003 | 2.37 | 0.42 | 0.49 | 0.59 | 0.5 | 0.47 | 0.42 |
| 6 | 44115169 | rs4714759 | -0.149 | 1.251 | -1.001 | 0.662 | -0.018 | -0.025 | -0.025 | 0 | 0.06 | 0.01 | 0.1 | 0.23 | 0.1 | 0.12 |
| 6 | 44116993 | rs9369458 | -0.949 | 1.249 | -0.956 | 0.656 | 0.031 | -0.018 | -0.018 | 0 | 0.27 | 0.18 | 0.32 | 0.27 | 0.21 | 0.27 |
| 6 | 44117607 | rs4714761 | -0.396 | 2.304 | -0.841 | 1.295 | -0.018 | -0.008 | -0.008 | 0.136 | 0.05 | 0 | 0.08 | 0.23 | 0.08 | 0.12 |
| 6 | 44125996 | rs12210167 | -0.802 | 2.282 | -0.564 | 0.405 | 0.023 | -0.05 | -0.05 | 0.02 | 0.12 | 0.2 | 0.11 | 0 | 0.2 | 0.08 |
| 6 | 44133019 | rs9462966 | -1.031 | 2.74 | -0.969 | 1.051 | -0.036 | -0.025 | -0.025 | 0 | 0.18 | 0.14 | 0.33 | 0.31 | 0.39 | 0.27 |
| 6 | 44134864 | rs914501 | -0.593 | 1.395 | -2.449 | 0.634 | 0.04 | -0.103 | -0.103 | 1.446 | 0.58 | 0.95 | 0.59 | 0.6 | 0.71 | 0.62 |
| 6 | 44138022 | rs4711771 | -0.76 | 1.108 | -1.596 | 0.41 | 0.035 | 0.024 | 0.024 | 0 | 0.31 | 0.46 | 0.42 | 0.41 | 0.51 | 0.4 |
| 6 | 44138343 | rs4711773 | 0.197 | 1.25 | -1.653 | 0.461 | -0.128 | -0.092 | -0.092 | 0 | 0.16 | 0.01 | 0.29 | 0.29 | 0.39 | 0.25 |
| 6 | 44138947 | rs9369461 | -0.537 | 1.077 | -1.484 | 0.369 | 0.051 | 0.035 | 0.035 | 0 | 0.26 | 0.4 | 0.37 | 0.41 | 0.45 | 0.37 |
| 6 | 44140275 | rs1418487 | -0.81 | 0.486 | -2.015 | 0.263 | 0.028 | -0.109 | -0.109 | 0 | 0.62 | 0.94 | 0.6 | 0.62 | 0.73 | 0.63 |
| 6 | 44144798 | rs6925778 | 0.028 | 0.564 | -1.547 | 0.365 | 0.026 | 0.023 | 0.023 | 0 | 0.46 | 0.58 | 0.41 | 0.66 | 0.37 | 0.52 |
| 6 | 44146440 | rs9462971 | -0.093 | 0.362 | -1.862 | -0.259 | 0.007 | 0.026 | 0.026 | 0.007 | 0.25 | 0.15 | 0.33 | 0.15 | 0.2 | 0.25 |
| 6 | 44151490 | rs7761137 | -0.111 | 0.242 | -1.316 | -0.833 | -0.018 | -0.1 | -0.1 | 1.243 | 0.31 | 0.59 | 0.23 | 0.01 | 0.23 | 0.2 |
| 6 | 44156470 | rs1041524 | -0.503 | 1.235 | -0.453 | 0.145 | 0.019 | -0.019 | -0.019 | 0 | 0.32 | 0.42 | 0.44 | 0.23 | 0.43 | 0.26 |
| 6 | 44162146 | rs4714770 | -0.059 | 1.615 | -0.512 | 1.087 | -0.002 | -0.003 | -0.003 | 0 | 0.21 | 0.22 | 0.28 | 0.42 | 0.28 | 0.35 |
| 6 | 44163624 | rs17567624 | -0.372 | 1.451 | -0.102 | 1.04 | 0.004 | 0.094 | 0.094 | 0 | 0.24 | 0.44 | 0.29 | 0.41 | 0.24 | 0.37 |
| 6 | 44169237 | rs9369465 | 0.161 | 1.441 | -0.538 | 0.554 | 0.021 | 0.017 | 0.017 | 0 | 0.65 | 0.55 | 0.51 | 0.44 | 0.53 | 0.61 |
| 6 | 44178471 | rs2396243 | -0.075 | 1.678 | -0.823 | 0.758 | -0.011 | -0.017 | -0.017 | 0 | 0.86 | 0.91 | 0.67 | 0.72 | 0.82 | 0.79 |
| 6 | 44182720 | rs9357436 | 0.171 | 1.846 | -1.039 | 1.034 | -0.035 | -0.067 | -0.067 | 0 | 0.12 | 0 | 0.14 | 0.27 | 0.2 | 0.11 |
| 6 | 44184719 | rs2297393 | 0.269 | 1.676 | -0.84 | 0.192 | 0.003 | 0.013 | 0.013 | 0 | 0.52 | 0.4 | 0.3 | 0.3 | 0.47 | 0.31 |
| 6 | 44186390 | rs507964 | 0.567 | 1.989 | -0.923 | 0.864 | -0.037 | -0.077 | -0.077 | 0 | 0.73 | 0.89 | 0.54 | 0.45 | 0.63 | 0.69 |
| 6 | 44191920 | rs693955 | -0.245 | 1.969 | -0.93 | 0.863 | -0.063 | -0.083 | -0.083 | 0.452 | 0.87 | 0.99 | 0.7 | 0.74 | 0.82 | 0.78 |
| 6 | 44195992 | rs9394992 | -0.191 | 1.714 | -0.77 | 0.307 | -0.001 | -0.001 | -0.001 | 0 | 0.29 | 0.31 | 0.25 | 0.37 | 0.28 | 0.25 |
| 6 | 44196578 | rs324148 | -0.346 | 1.712 | -1.157 | 0.337 | -0.046 | -0.046 | -0.046 | 0 | 0.79 | 0.93 | 0.75 | 0.75 | 0.74 | 0.72 |
| 6 | 44200953 | rs760370 | 0.859 | 1.253 | -1.358 | -0.169 | 0.021 | 0.06 | 0.06 | 0 | 0.41 | 0.26 | 0.37 | 0.3 | 0.4 | 0.2 |
| 6 | 44205027 | rs666462 | 0.654 | 1.881 | -1.133 | 0.508 | -0.045 | 0.035 | 0.035 | 0.709 | 0.61 | 0.8 | 0.46 | 0.34 | 0.51 | 0.38 |
| 6 | 44206750 | rs9472236 | 0.569 | 2.081 | -1.616 | 1.029 | -0.031 | -0.104 | -0.104 | 5.371 | 0.15 | 0.02 | 0.23 | 0.39 | 0.18 | 0.29 |
| 6 | 44207117 | rs4714772 | 0.569 | 2.081 | -1.616 | 1.029 | -0.031 | -0.104 | -0.104 | 0 | 0.15 | 0.02 | 0.23 | 0.4 | 0.18 | 0.3 |

| | | | | | | | | | | | | | | | | |
|---|---|---|---|---|---|---|---|---|---|---|---|---|---|---|---|---|
| 6 | 44211867 | rs9381299 | -0.072 | 1.988 | -1.446 | 1.019 | 0.012 | 0.085 | 0.085 | 0 | 0.09 | 0.16 | 0.19 | 0.35 | 0.15 | 0.18 |
| 6 | 44214417 | rs3757283 | 0.451 | 2.324 | -2.002 | 1.191 | -0.058 | -0.11 | -0.11 | 0 | 0.15 | 0.01 | 0.19 | 0.39 | 0.2 | 0.29 |
| 6 | 44220076 | rs504697 | 0.232 | 2.341 | -1.916 | 1.02 | -0.007 | -0.009 | -0.009 | 0 | 0.76 | 0.74 | 0.66 | 0.75 | 0.72 | 0.73 |
| 6 | 44223010 | rs1875324 | -0.098 | 1.535 | -1.793 | 1.095 | -0.058 | -0.108 | -0.108 | 0 | 0.15 | 0.01 | 0.19 | 0.39 | 0.2 | 0.28 |
| 6 | 44226195 | rs2282151 | -0.202 | 1.436 | -1.769 | 1.065 | -0.037 | -0.103 | -0.103 | 0 | 0.19 | 0.04 | 0.26 | 0.41 | 0.22 | 0.29 |
| 6 | 44228452 | rs3799962 | 0.53 | 1.175 | -1.985 | 0.554 | -0.027 | -0.057 | -0.057 | 0 | 0.46 | 0.09 | 0.54 | 0.43 | 0.45 | 0.41 |
| 6 | 44232074 | rs730775 | 0.62 | 1.132 | -1.915 | 0.546 | -0.044 | -0.056 | -0.056 | 0 | 0.45 | 0.14 | 0.55 | 0.42 | 0.44 | 0.41 |
| 6 | 44235690 | rs513688 | 0.236 | 1.316 | -1.065 | 0.81 | -0.003 | -0.003 | -0.003 | 2.875 | 0.74 | 0.75 | 0.75 | 0.43 | 0.63 | 0.61 |
| 6 | 44246766 | rs12943 | 0.727 | 0.542 | -0.155 | 0.919 | -0.027 | 0.038 | 0.038 | 0.062 | 0.72 | 0.47 | 0.81 | 0.6 | 0.78 | 0.65 |
| 6 | 44250165 | rs516582 | 0.723 | 0.582 | -0.047 | 0.921 | -0.024 | 0.035 | 0.035 | 0.099 | 0.72 | 0.47 | 0.81 | 0.6 | 0.78 | 0.65 |
| 6 | 44253749 | rs324143 | -0.02 | 0.492 | 0.019 | 0.899 | -0.059 | -0.067 | -0.067 | 0 | 0.87 | 0.65 | 0.94 | 0.99 | 0.96 | 0.95 |
| 6 | 44253765 | rs2297336 | -0.121 | 0.521 | 0.001 | 0.865 | -0.021 | -0.072 | -0.072 | 0.159 | 0.14 | 0.02 | 0.13 | 0.4 | 0.18 | 0.3 |
| 6 | 44256175 | rs9381308 | -0.219 | 0.524 | 0.051 | 0.866 | -0.002 | -0.001 | -0.001 | 0 | 0.14 | 0.08 | 0.14 | 0.4 | 0.18 | 0.3 |
| 6 | 44266574 | rs1056093 | 0.332 | 0.913 | -0.334 | 0.522 | -0.062 | 0.053 | 0.053 | 0 | 0.1 | 0.02 | 0.12 | 0.01 | 0.16 | 0.1 |
| 6 | 44268371 | rs325008 | -0.025 | 0.751 | -0.101 | 0.525 | -0.063 | -0.098 | -0.098 | 0.009 | 0.86 | 0.6 | 0.95 | 1 | 0.96 | 0.95 |
| 6 | 44270936 | rs325001 | 1.561 | 0.612 | -0.16 | 0.525 | -0.011 | 0.006 | 0.006 | 0 | 0.55 | 0.22 | 0.61 | 0.41 | 0.55 | 0.48 |
| 6 | 44273546 | rs324137 | 1.453 | 0.602 | 0.025 | 0.52 | -0.008 | 0.002 | 0.002 | 0 | 0.57 | 0.28 | 0.61 | 0.42 | 0.55 | 0.48 |
| 6 | 44274797 | rs7740535 | -0.07 | 0.583 | 0.094 | 0.206 | -0.056 | -0.09 | -0.09 | 0.746 | 0.87 | 0.66 | 0.95 | 1 | 0.96 | 0.95 |
| 6 | 44275011 | rs324136 | -0.07 | 0.583 | 0.045 | 0.206 | -0.056 | -0.089 | -0.089 | 0 | 0.87 | 0.67 | 0.95 | 1 | 0.96 | 0.95 |
| 6 | 44279111 | rs9688508 | -0.796 | 1.099 | 0.001 | 0.59 | 0.059 | 0.002 | 0.002 | 0.033 | 0.18 | 0.42 | 0.21 | 0.2 | 0.22 | 0.17 |
| 6 | 44282316 | rs4714776 | -0.158 | 0.882 | -0.488 | 1.261 | 0.001 | 0.042 | 0.042 | 0 | 0.11 | 0.16 | 0.12 | 0.27 | 0.13 | 0.31 |
| 6 | 44285376 | rs2396242 | 0.255 | 0.713 | -1.359 | 0.782 | -0.037 | -0.084 | -0.084 | 0.011 | 0.4 | 0.14 | 0.52 | 0.51 | 0.42 | 0.41 |
| 6 | 44288847 | rs11962379 | -0.769 | 1.075 | -1.029 | 0.98 | -0.031 | -0.03 | -0.03 | 0 | 0.05 | 0.01 | 0.11 | 0.17 | 0.1 | 0.07 |
| 6 | 44290831 | rs9369472 | 0.662 | 0.686 | -0.702 | 0.215 | -0.005 | -0.053 | -0.053 | 0.07 | 0.82 | 0.77 | 0.91 | 0.99 | 0.86 | 0.91 |
| 6 | 44297772 | rs9472250 | 0.845 | 0.762 | -1.509 | 0.849 | -0.106 | -0.181 | -0.181 | 0.656 | 0.33 | 0.02 | 0.47 | 0.49 | 0.4 | 0.33 |
| 6 | 44297871 | rs9369476 | 0.294 | 0.292 | -0.522 | 0.717 | 0.002 | 0.003 | 0.003 | 0.124 | 0.45 | 0.43 | 0.31 | 0.32 | 0.36 | 0.47 |
| 6 | 44301107 | rs7769927 | 1.561 | 0.449 | 0.419 | 0.772 | 0.004 | -0.028 | -0.028 | 0 | 0.72 | 0.42 | 0.78 | 0.81 | 0.73 | 0.79 |
| 6 | 44303728 | rs1571773 | 1.926 | -0.282 | 1.123 | 1.578 | -0.017 | 0.106 | 0.106 | 2.113 | 0.84 | 0.45 | 0.7 | 0.72 | 0.9 | 0.84 |
| 6 | 44304376 | rs7769148 | 1.533 | -0.239 | 1.323 | 1.61 | 0.06 | 0.191 | 0.191 | 0.74 | 0.94 | 0.75 | 0.74 | 0.78 | 0.91 | 0.94 |
| 6 | 44305891 | rs9472251 | 2.431 | -0.259 | 1.452 | 1.572 | -0.012 | 0.116 | 0.116 | 1.639 | 0.85 | 0.37 | 0.69 | 0.72 | 0.9 | 0.85 |
| 6 | 44309346 | rs7757993 | 1.833 | -0.209 | 1.54 | 1.595 | 0.056 | 0.224 | 0.224 | 0.548 | 0.9 | 0.69 | 0.72 | 0.72 | 0.9 | 0.87 |
| 6 | 44316373 | rs6911823 | 1.301 | -0.516 | 1.478 | 1.695 | -0.002 | 0.001 | 0.001 | 0.703 | 0.73 | 0.61 | 0.83 | 0.7 | 0.77 | 0.66 |
| 6 | 44318072 | rs1411904 | 2.374 | -0.58 | 1.52 | 1.696 | 0.009 | -0.012 | -0.012 | 0 | 0.61 | 0.26 | 0.57 | 0.42 | 0.67 | 0.5 |
| 6 | 44324600 | rs4714784 | 1.09 | -0.485 | 1.465 | 1.701 | 0 | -0.003 | -0.003 | 0 | 0.76 | 0.77 | 0.83 | 0.71 | 0.78 | 0.67 |
| 6 | 44334388 | rs7769128 | 2.18 | -0.495 | 1.319 | 1.717 | -0.001 | 0.017 | 0.017 | 0.045 | 0.65 | 0.37 | 0.59 | 0.49 | 0.7 | 0.58 |
| 6 | 44340937 | rs13214191 | -1.007 | -0.68 | 0.995 | 1.585 | -0.007 | -0.071 | -0.071 | 0.028 | 0.19 | 0.08 | 0.18 | 0.44 | 0.19 | 0.43 |
| 6 | 44341980 | rs6918742 | 1.815 | -0.721 | 1.18 | 1.613 | -0.01 | 0.003 | 0.003 | 1.487 | 0.73 | 0.51 | 0.62 | 0.32 | 0.77 | 0.49 |
| 6 | 44346006 | rs9395010 | -1.058 | -0.676 | 0.774 | 1.603 | 0.004 | -0.04 | -0.04 | 0 | 0.19 | 0.13 | 0.18 | 0.44 | 0.18 | 0.43 |
| 6 | 44354641 | rs992160 | -0.51 | -1.064 | 0.936 | 1.063 | -0.023 | -0.01 | -0.01 | 0.934 | 0.06 | 0.05 | 0 | 0.04 | 0.01 | 0.02 |
| 6 | 44355399 | rs2297334 | -1.312 | -1.063 | 0.88 | 1.137 | -0.001 | 0.091 | 0.091 | 0.002 | 0.1 | 0.21 | 0.16 | 0.6 | 0.09 | 0.38 |
| 6 | 44355423 | rs2297333 | 2.61 | -1.189 | 1.092 | 1.013 | -0.045 | -0.009 | -0.009 | 0 | 0.73 | 0.46 | 0.64 | 0.26 | 0.84 | 0.53 |
| 6 | 44365851 | rs4143678 | 2.519 | -1.181 | 0.926 | 1.055 | -0.055 | -0.002 | -0.002 | 0 | 0.74 | 0.52 | 0.65 | 0.27 | 0.85 | 0.54 |
| 6 | 44371714 | rs6934058 | -1.533 | -0.911 | 0.346 | 1.213 | 0 | 0.066 | 0.066 | 0 | 0.11 | 0.22 | 0.16 | 0.61 | 0.09 | 0.38 |
| 6 | 44384833 | rs1888671 | -0.687 | -0.897 | 0.304 | 1.199 | -0.004 | 0.042 | 0.042 | 0 | 0.07 | 0.05 | 0.14 | 0.55 | 0.08 | 0.35 |
| 6 | 44385441 | rs13198841 | -0.071 | -1.252 | 0.046 | 0.388 | 0.018 | 0.055 | 0.055 | 0 | 0.17 | 0.02 | 0.08 | 0 | 0.15 | 0.05 |
| 6 | 44405253 | rs10484624 | -0.796 | -0.851 | 0.259 | 1.174 | -0.025 | -0.012 | -0.012 | 0 | 0.07 | 0.1 | 0.01 | 0.04 | 0.01 | 0.02 |
| 6 | 44421240 | rs9381325 | -0.658 | -0.21 | -0.19 | 1.601 | -0.005 | 0.065 | 0.065 | 0 | 0.07 | 0.05 | 0.15 | 0.54 | 0.06 | 0.29 |
| 6 | 44422466 | rs4509117 | -0.852 | -0.62 | 0.096 | 1.074 | -0.021 | 0.017 | 0.017 | 0 | 0.12 | 0.2 | 0.1 | 0.17 | 0.07 | 0.16 |
| 6 | 44424800 | rs543844 | -1.329 | -0.741 | -0.402 | 0.137 | 0.051 | 0.189 | 0.189 | 0.08 | 0.49 | 0.81 | 0.64 | 0.82 | 0.39 | 0.65 |
| 6 | 44428506 | rs16869089 | -0.782 | -0.721 | 0.015 | 0.265 | -0.001 | 0 | 0 | 0 | 0.28 | 0.32 | 0.42 | 0.65 | 0.23 | 0.41 |
| 6 | 44433330 | rs6925697 | -1.003 | -0.792 | -0.072 | -0.342 | 0.015 | 0.054 | 0.054 | 0.01 | 0.4 | 0.72 | 0.54 | 0.78 | 0.33 | 0.55 |
| 6 | 44440722 | rs9395021 | -0.159 | -0.563 | 0.054 | -0.404 | 0.001 | -0.002 | -0.002 | 0 | 0.42 | 0.43 | 0.47 | 0.51 | 0.41 | 0.55 |
| 6 | 44441178 | rs4714790 | -0.514 | -0.471 | 0.659 | -0.259 | -0.002 | 0.011 | 0.011 | 0 | 0.16 | 0.17 | 0.19 | 0.32 | 0.12 | 0.21 |
| 6 | 44441552 | rs911983 | 0.315 | -0.463 | 0.291 | -0.554 | 0.013 | -0.011 | -0.011 | 0 | 0.39 | 0.34 | 0.39 | 0.52 | 0.33 | 0.56 |
| 6 | 44443303 | rs7769305 | -0.27 | -0.101 | 0.024 | -0.671 | 0.017 | -0.059 | -0.059 | 0 | 0.11 | 0.19 | 0.1 | 0.01 | 0.12 | 0.02 |
| 6 | 44444703 | rs9296435 | -0.861 | -0.065 | 0.053 | -0.624 | -0.061 | 0.113 | 0.113 | 0.031 | 0.66 | 0.89 | 0.69 | 0.85 | 0.58 | 0.81 |
| 6 | 44445046 | rs9296436 | -0.861 | -0.045 | 0.118 | -0.733 | -0.061 | 0.113 | 0.113 | 0.031 | 0.66 | 0.89 | 0.69 | 0.85 | 0.58 | 0.8 |
| 6 | 44445990 | rs6909090 | -0.807 | 0.266 | 0.424 | -0.608 | 0.04 | -0.018 | -0.018 | 0 | 0.06 | 0.21 | 0.04 | 0 | 0.05 | 0.02 |
| 6 | 44446674 | rs2281690 | -0.816 | 0.274 | -0.064 | -0.561 | -0.066 | 0.097 | 0.097 | 0.256 | 0.69 | 0.89 | 0.7 | 0.88 | 0.59 | 0.82 |
| 6 | 44447004 | rs7770034 | -0.804 | 0.27 | -0.06 | -0.556 | -0.063 | 0.097 | 0.097 | 0.258 | 0.69 | 0.9 | 0.7 | 0.88 | 0.59 | 0.83 |
| 6 | 44449216 | rs7743369 | 0.601 | 0.673 | 0.521 | -0.142 | 0.087 | 0.081 | 0.081 | 0 | 0.89 | 0.71 | 0.85 | 0.78 | 0.85 | 0.95 |
| 6 | 44449320 | rs10948139 | 0.055 | 0.412 | 0.636 | -0.337 | 0.001 | 0.002 | 0.002 | 0 | 0.53 | 0.58 | 0.57 | 0.65 | 0.42 | 0.76 |
| 6 | 44450129 | rs4711783 | -0.046 | 0.319 | 0.596 | 0.079 | -0.002 | 0.05 | 0.05 | 0 | 0.07 | 0.17 | 0.09 | 0.11 | 0.06 | 0.14 |
| 6 | 44450880 | rs16871965 | -0.704 | 0.356 | 0.736 | -0.288 | 0.023 | -0.087 | -0.087 | 0.123 | 0.13 | 0.35 | 0.11 | 0.01 | 0.13 | 0.04 |
| 6 | 44451814 | rs7755190 | 1.1 | 0.054 | 0.531 | -0.309 | 0.009 | -0.022 | -0.022 | 0.441 | 0.72 | 0.53 | 0.77 | 0.78 | 0.74 | 0.91 |
| 6 | 44452011 | rs6458389 | 0.589 | 0.193 | 0.468 | -0.143 | 0.02 | 0.019 | 0.019 | 0 | 0.86 | 0.78 | 0.84 | 0.79 | 0.81 | 0.96 |
| 6 | 44452157 | rs7759209 | 0.633 | 0.313 | 0.121 | -0.069 | 0 | -0.005 | -0.005 | 0.854 | 0.59 | 0.43 | 0.63 | 0.64 | 0.6 | 0.7 |
| 6 | 44452606 | rs12664617 | 0.794 | 0.939 | -0.375 | 0.065 | 0.008 | -0.009 | -0.009 | 0 | 0.2 | 0.19 | 0.16 | 0.14 | 0.26 | 0.13 |
| 6 | 44454333 | rs911984 | 0.695 | 0.578 | 0.055 | 0.343 | 0.058 | 0.062 | 0.062 | 0.05 | 0.88 | 0.74 | 0.83 | 0.76 | 0.83 | 0.83 |

| | | | | | | | | | | | | | | | | |
|---|---|---|---|---|---|---|---|---|---|---|---|---|---|---|---|---|
| 6 | 44455300 | rs6932104 | -0.65 | -0.625 | -0.493 | -0.21 | -0.031 | -0.057 | -0.057 | 0 | 0.32 | 0.57 | 0.25 | 0.15 | 0.25 | 0.28 |
| 6 | 44467889 | rs12193814 | -0.415 | -0.829 | -1.189 | 0.391 | 0.062 | -0.02 | -0.02 | 0 | 0.23 | 0.05 | 0.07 | 0.18 | 0.1 | 0.09 |
| 6 | 44469029 | rs6458390 | -0.38 | -1.133 | -1.616 | 0.555 | 0.013 | -0.019 | -0.019 | 0.7 | 0.78 | 0.92 | 0.58 | 0.71 | 0.75 | 0.78 |
| 6 | 44469505 | rs6919522 | 0.827 | -0.203 | -1.084 | 0.346 | 0.033 | 0.021 | 0.021 | 0.2 | 0.27 | 0.39 | 0.23 | 0.36 | 0.37 | 0.34 |
| 6 | 44470006 | rs1322667 | 0.201 | -0.573 | -1.204 | 0.337 | 0.06 | -0.009 | -0.009 | 0 | 0.25 | 0.06 | 0.18 | 0.19 | 0.13 | 0.1 |
| 6 | 44472854 | rs9296440 | -0.342 | -0.543 | -0.803 | 0.086 | 0.026 | 0.026 | 0.026 | 0.04 | 0.43 | 0.62 | 0.4 | 0.29 | 0.31 | 0.35 |
| 6 | 44473875 | rs2145748 | 0.009 | 0.597 | -0.665 | 0.162 | 0.044 | -0.034 | -0.034 | 0 | 0.16 | 0.31 | 0.14 | 0.05 | 0.22 | 0.12 |
| 6 | 44476296 | rs9381333 | 0.887 | 0.214 | -1.487 | 0.128 | -0.014 | -0.086 | -0.086 | 0.026 | 0.34 | 0.05 | 0.3 | 0.42 | 0.37 | 0.42 |
| 6 | 44477192 | rs11757334 | 0.036 | 0.641 | -1.333 | 0.115 | 0.008 | -0.009 | -0.009 | 0.671 | 0.17 | 0.2 | 0.13 | 0.05 | 0.22 | 0.13 |
| 6 | 44482941 | rs2396256 | 0.995 | 0.16 | -2.015 | 0.313 | -0.04 | -0.111 | -0.111 | 0.364 | 0.31 | 0.03 | 0.32 | 0.41 | 0.35 | 0.38 |
| 6 | 44483052 | rs12202213 | 0.003 | 0.135 | -0.992 | 0.442 | 0.019 | -0.021 | -0.021 | 1.373 | 0.73 | 0.88 | 0.61 | 0.65 | 0.74 | 0.73 |
| 6 | 44489714 | rs2146347 | 0.85 | 0.562 | -0.991 | 0.533 | -0.003 | -0.048 | -0.048 | 1.04 | 0.61 | 0.2 | 0.61 | 0.53 | 0.65 | 0.6 |
| 6 | 44490470 | rs7774489 | -0.629 | 0.451 | -0.009 | 0.251 | 0.017 | -0.055 | -0.055 | 0 | 0.18 | 0.56 | 0.13 | 0.06 | 0.13 | 0.15 |
| 6 | 44493348 | rs11758473 | 1.247 | 0.338 | -0.279 | 0.553 | -0.012 | -0.044 | -0.044 | 0 | 0.59 | 0.13 | 0.49 | 0.46 | 0.65 | 0.6 |
| 6 | 44494370 | rs9381336 | 1.432 | 0.846 | -0.277 | 0.446 | -0.046 | 0.016 | 0.016 | 0 | 0.55 | 0.16 | 0.47 | 0.46 | 0.56 | 0.55 |
| 6 | 44494470 | rs9472294 | -0.925 | 0.813 | 0.499 | 0.326 | 0.058 | -0.031 | -0.031 | 0 | 0.13 | 0.24 | 0.11 | 0.04 | 0.18 | 0.07 |
| 6 | 44496728 | rs7740414 | -1.472 | 0.559 | 0.754 | 0.236 | -0.036 | -0.088 | -0.088 | 0 | 0.16 | 0.56 | 0.1 | 0.01 | 0.09 | 0.1 |
| 6 | 44496851 | rs2281692 | 1.987 | 1.402 | 0.526 | 0.257 | 0.032 | 0.054 | 0.054 | 0 | 0.51 | 0.25 | 0.44 | 0.44 | 0.49 | 0.49 |
| 6 | 44496950 | rs9462995 | -1.413 | 0.747 | 0.902 | 0.301 | -0.03 | -0.107 | -0.107 | 0 | 0.22 | 0.5 | 0.12 | 0.01 | 0.12 | 0.12 |
| 6 | 44498472 | rs9369494 | 0.161 | 0.344 | 0.511 | 0.301 | 0.007 | -0.014 | -0.014 | 1.884 | 0.73 | 0.82 | 0.71 | 0.52 | 0.77 | 0.74 |
| 6 | 44499803 | rs9349292 | 0.465 | 0.744 | 0.676 | 0.501 | 0.006 | 0.074 | 0.074 | 0.351 | 0.81 | 0.69 | 0.75 | 0.58 | 0.81 | 0.7 |
| 6 | 44502355 | rs7771912 | 1.099 | 0.585 | 0.408 | 0.009 | 0.027 | 0.006 | 0.006 | 0 | 0.49 | 0.57 | 0.52 | 0.49 | 0.62 | 0.55 |
| 6 | 44503246 | rs11753604 | 1.298 | 0.267 | 0.203 | 0.11 | 0.117 | -0.005 | -0.005 | 2.451 | 0.48 | 0.24 | 0.5 | 0.49 | 0.68 | 0.53 |
| 6 | 44503829 | rs9462996 | 0.297 | 0.098 | 0.216 | -0.515 | -0.025 | -0.067 | -0.067 | 0 | 0.19 | 0.25 | 0.09 | 0 | 0.11 | 0.08 |
| 6 | 44503967 | rs9367203 | 1.073 | 0.776 | 0.519 | 0.252 | -0.005 | 0.008 | 0.008 | 0 | 0.65 | 0.5 | 0.61 | 0.59 | 0.7 | 0.53 |
| 6 | 44509266 | rs1885663 | 1.171 | 1.644 | 0.131 | -0.082 | -0.024 | 0.021 | 0.021 | 0 | 0.88 | 0.89 | 0.83 | 0.96 | 0.78 | 0.9 |
| 6 | 44510011 | rs1998402 | -1.489 | 1.656 | 0.772 | 0.114 | 0.009 | 0.058 | 0.058 | 0 | 0.31 | 0.5 | 0.33 | 0.53 | 0.28 | 0.35 |
| 6 | 44518073 | rs9349293 | -0.396 | 1.641 | 0.719 | 0.083 | -0.005 | 0.159 | 0.159 | 0 | 0.2 | 0.31 | 0.26 | 0.49 | 0.15 | 0.29 |
| 6 | 44519096 | rs3734716 | 1.547 | 1.537 | 0.763 | 0.024 | 0.013 | 0.014 | 0.014 | 0 | 0.63 | 0.38 | 0.54 | 0.47 | 0.58 | 0.57 |
| 6 | 44520451 | rs1322650 | -0.864 | 1.241 | 0.606 | 0.322 | 0.013 | 0.064 | 0.064 | 0 | 0.26 | 0.36 | 0.3 | 0.55 | 0.25 | 0.33 |
| 6 | 44522398 | rs1322653 | -0.986 | 1.084 | 0.034 | -0.002 | 0.024 | 0.006 | 0.006 | 2.091 | 0.49 | 0.69 | 0.44 | 0.52 | 0.42 | 0.38 |
| 6 | 44523033 | rs990963 | 0.834 | 1.309 | 0.134 | -0.987 | 0.013 | -0.084 | -0.084 | 0.576 | 0.68 | 0.41 | 0.76 | 0.95 | 0.64 | 0.78 |
| 6 | 44523591 | rs6939369 | -0.862 | 1.052 | 0.074 | -0.243 | -0.009 | 0.115 | 0.115 | 0 | 0.11 | 0.27 | 0.2 | 0.45 | 0.1 | 0.21 |
| 6 | 44523643 | rs911982 | 0.697 | 1.333 | -0.59 | -0.338 | -0.037 | 0.026 | 0.026 | 0 | 0.92 | 0.99 | 0.92 | 0.98 | 0.89 | 0.9 |
| 6 | 44524429 | rs4711786 | 1.492 | 1.401 | -0.807 | -0.457 | -0.012 | -0.136 | -0.136 | 0 | 0.31 | 0.07 | 0.42 | 0.46 | 0.3 | 0.43 |
| 6 | 44525215 | rs4711788 | 1.258 | 1.085 | 0.355 | -0.617 | 0.023 | -0.039 | -0.039 | 0.068 | 0.88 | 0.8 | 0.86 | 0.97 | 0.82 | 0.88 |
| 6 | 44525464 | rs9472301 | -1.109 | 1.032 | 0.533 | -0.581 | 0.004 | -0.017 | -0.017 | 1.593 | 0.1 | 0.14 | 0.09 | 0.04 | 0.13 | 0.08 |
| 6 | 44525989 | rs2146349 | 0.882 | 0.918 | 0.234 | -0.585 | 0.006 | -0.016 | -0.016 | 0 | 0.45 | 0.17 | 0.55 | 0.47 | 0.54 | 0.53 |
| 6 | 44526440 | rs991234 | 0.378 | 0.533 | 0.719 | -0.512 | -0.015 | -0.001 | -0.001 | 0 | 0.53 | 0.54 | 0.62 | 0.5 | 0.66 | 0.65 |
| 6 | 44533311 | rs6923772 | 1.112 | 0.851 | 1.042 | -0.977 | 0.055 | -0.004 | -0.004 | 0.394 | 0.95 | 0.82 | 0.93 | 0.98 | 0.9 | 0.91 |
| 6 | 44534397 | rs7747763 | -1.647 | 0.465 | 1.76 | -1.044 | 0.025 | 0.079 | 0.079 | 0.316 | 0.08 | 0.47 | 0.05 | 0.09 | 0.01 | 0.04 |
| 6 | 44535078 | rs9367205 | -1.037 | 0.489 | 1.53 | -1.027 | -0.01 | 0.036 | 0.036 | 0 | 0.05 | 0.22 | 0.03 | 0.09 | 0.01 | 0.03 |
| 6 | 44538139 | rs927485 | 0.92 | 0.309 | 0.889 | -1.627 | 0.057 | 0.199 | 0.199 | 2.873 | 0.62 | 0.12 | 0.73 | 0.9 | 0.69 | 0.8 |
| 6 | 44540314 | rs7759601 | -1.126 | 1.164 | 1.016 | -1.865 | 0.02 | 0.186 | 0.186 | 1.275 | 0.49 | 0.74 | 0.32 | 0.05 | 0.46 | 0.2 |
| 6 | 44540982 | rs7764770 | -1.115 | 1.074 | 1.326 | -1.942 | 0.052 | -0.035 | -0.035 | 0 | 0.09 | 0.33 | 0.08 | 0 | 0.09 | 0.02 |
| 6 | 44541936 | rs326798 | 1.113 | 1.118 | 1.044 | -1.91 | -0.011 | 0.328 | 0.328 | 2.442 | 0.5 | 0.12 | 0.66 | 0.93 | 0.52 | 0.75 |
| 6 | 44542634 | rs1322664 | -0.674 | 1.092 | 1.321 | -1.892 | -0.003 | -0.087 | -0.087 | 0 | 0.42 | 0.59 | 0.26 | 0.04 | 0.37 | 0.17 |
| 6 | 44548984 | rs10807301 | -0.831 | 1.068 | 0.623 | -1.977 | -0.006 | 0.159 | 0.159 | 1.841 | 0.47 | 0.74 | 0.29 | 0.05 | 0.44 | 0.16 |
| 6 | 44549333 | rs11756037 | -0.373 | 1.066 | 0.376 | -2.024 | -0.003 | 0.081 | 0.081 | 0.558 | 0.39 | 0.25 | 0.22 | 0.05 | 0.37 | 0.14 |
| 6 | 44551487 | rs3997520 | -1.014 | 1.039 | -0.247 | -1.881 | -0.044 | 0.356 | 0.356 | 1.561 | 0.51 | 0.88 | 0.34 | 0.05 | 0.52 | 0.19 |
| 6 | 44558552 | rs326815 | 0.976 | 1.061 | -0.198 | -1.834 | -0.028 | 0.333 | 0.333 | 3.514 | 0.48 | 0.12 | 0.64 | 0.93 | 0.45 | 0.76 |
| 6 | 44568740 | rs618617 | 0.292 | 1.137 | 0.93 | -1.846 | 0.004 | -0.114 | -0.114 | 0.067 | 0.6 | 0.45 | 0.72 | 0.95 | 0.57 | 0.89 |
| 6 | 44572155 | rs1283946 | -0.744 | 0.844 | 0.673 | -1.343 | -0.002 | 0.001 | 0.001 | 0 | 0.54 | 0.61 | 0.48 | 0.43 | 0.54 | 0.58 |
| 6 | 44572789 | rs11755006 | 0.253 | 0.742 | 0.959 | -1.451 | -0.005 | 0.03 | 0.03 | 0.259 | 0.33 | 0.26 | 0.21 | 0.05 | 0.34 | 0.1 |
| 6 | 44575446 | rs472029 | 0.87 | 0.742 | 0.854 | -1.377 | -0.018 | -0.094 | -0.094 | 0 | 0.17 | 0.03 | 0.24 | 0.38 | 0.17 | 0.44 |
| 6 | 44577322 | rs12526438 | 1.441 | 0.702 | 0.902 | -1.5 | -0.006 | -0.046 | -0.046 | 0 | 0.35 | 0.14 | 0.45 | 0.56 | 0.34 | 0.34 |
| 6 | 44578015 | rs683425 | 0.404 | 0.721 | 1.08 | -1.499 | -0.047 | -0.106 | -0.106 | 0 | 0.2 | 0.03 | 0.28 | 0.38 | 0.26 | 0.44 |
| 6 | 44578324 | rs694927 | 0.398 | 0.615 | 1.318 | -1.67 | 0.001 | -0.036 | -0.036 | 0 | 0.56 | 0.39 | 0.74 | 0.94 | 0.6 | 0.8 |
| 6 | 44579789 | rs561160 | -0.76 | 0.527 | 1.055 | -1.552 | 0.008 | 0.02 | 0.02 | 0.393 | 0.38 | 0.58 | 0.39 | 0.5 | 0.43 | 0.59 |
| 6 | 44580672 | rs9472308 | -0.317 | 0.575 | 0.897 | -1.515 | 0.041 | -0.088 | -0.088 | 0.156 | 0.19 | 0.42 | 0.15 | 0.05 | 0.25 | 0.13 |
| 6 | 44582923 | rs9463003 | 0.299 | 0.622 | 0.575 | -1.487 | 0.013 | -0.034 | -0.034 | 0 | 0.16 | 0.24 | 0.12 | 0.04 | 0.23 | 0.06 |
| 6 | 44584387 | rs4714801 | 1.801 | 0.66 | -0.199 | -1.537 | -0.027 | -0.157 | -0.157 | 0.21 | 0.29 | 0.04 | 0.41 | 0.49 | 0.27 | 0.31 |
| 6 | 44586363 | rs6913259 | 0.798 | 0.61 | -0.086 | -1.523 | -0.007 | -0.012 | -0.012 | 0 | 0.53 | 0.28 | 0.58 | 0.49 | 0.54 | 0.38 |
| 6 | 44594159 | rs556621 | -1.49 | 0.575 | -0.5 | -1.204 | 0.003 | -0.118 | -0.118 | 0 | 0.68 | 0.92 | 0.58 | 0.51 | 0.71 | 0.7 |
| 6 | 44595064 | rs527322 | 0.644 | 0.422 | -0.745 | -1.372 | 0.004 | -0.099 | -0.099 | 0 | 0.22 | 0.07 | 0.32 | 0.46 | 0.2 | 0.53 |
| 6 | 44596053 | rs497177 | -1.371 | 0.436 | -0.616 | -1.02 | -0.023 | -0.122 | -0.122 | 0 | 0.73 | 0.94 | 0.65 | 0.55 | 0.75 | 0.76 |
| 6 | 44596435 | rs1767789 | -0.962 | 0.43 | -0.037 | -1.025 | -0.001 | 0.006 | 0.006 | 0 | 0.67 | 0.67 | 0.62 | 0.55 | 0.72 | 0.72 |
| 6 | 44597445 | rs4714802 | 0.207 | 0.592 | 0.09 | -1.078 | -0.004 | 0.004 | 0.004 | 0.553 | 0.58 | 0.52 | 0.55 | 0.45 | 0.57 | 0.4 |
| 6 | 44598555 | rs4711791 | 0.616 | 0.276 | -0.087 | -1.088 | -0.001 | -0.004 | -0.004 | 0 | 0.32 | 0.28 | 0.33 | 0.37 | 0.3 | 0.29 |

| | | | | | | | | | | | | | | | | |
|---|---|---|---|---|---|---|---|---|---|---|---|---|---|---|---|---|
| 6 | 44599143 | rs9395041 | 1.251 | 0.142 | 0.06 | -1.176 | 0.011 | -0.044 | -0.044 | 0 | 0.23 | 0.16 | 0.34 | 0.41 | 0.21 | 0.19 |
| 6 | 44599332 | rs11752906 | 0.433 | 0.423 | -0.551 | -1.841 | -0.102 | 0.099 | 0.099 | 0 | 0.2 | 0.05 | 0.18 | 0 | 0.36 | 0.11 |
| 6 | 44599507 | rs1384546 | -0.402 | -0.062 | -0.357 | -2.169 | -0.027 | 0.021 | 0.021 | 0.252 | 0.38 | 0.28 | 0.24 | 0.01 | 0.44 | 0.25 |
| 6 | 44600005 | rs6902467 | -0.766 | -0.334 | -0.184 | -2.239 | 0 | 0.046 | 0.046 | 0.923 | 0.69 | 0.62 | 0.61 | 0.46 | 0.66 | 0.47 |
| 6 | 44600814 | rs4714804 | 1.519 | -0.31 | -0.345 | -2.166 | 0.005 | -0.011 | -0.011 | 0 | 0.3 | 0.31 | 0.37 | 0.45 | 0.23 | 0.22 |
| 6 | 44601207 | rs4714805 | -1.618 | 0.048 | -0.491 | -1.832 | -0.012 | -0.103 | -0.103 | 7.3 | 0.79 | 0.95 | 0.72 | 0.46 | 0.75 | 0.54 |
| 6 | 44606357 | rs11970609 | -0.189 | 0.312 | -0.149 | -2.123 | -0.007 | -0.034 | -0.034 | 0.74 | 0.25 | 0.25 | 0.19 | 0 | 0.35 | 0.1 |
| 6 | 44609610 | rs9463005 | -0.718 | 0.199 | 0.095 | -1.969 | 0.034 | -0.081 | -0.081 | 0 | 0.12 | 0.4 | 0.11 | 0 | 0.09 | 0.05 |
| 6 | 44612363 | rs12200518 | 0.589 | 0.521 | -0.562 | -1.968 | -0.019 | -0.003 | -0.003 | 0 | 0.16 | 0.14 | 0.15 | 0 | 0.3 | 0.1 |
| 6 | 44620507 | rs9296444 | 1.733 | 0.074 | -0.92 | -1.487 | 0.006 | -0.092 | -0.092 | 0.622 | 0.22 | 0.1 | 0.29 | 0.47 | 0.2 | 0.19 |
| 6 | 44621993 | rs900404 | 0.867 | 0.175 | -0.88 | -1.504 | 0.002 | -0.049 | -0.049 | 0.346 | 0.28 | 0.17 | 0.37 | 0.49 | 0.3 | 0.48 |
| 6 | 44622298 | rs10948152 | 0.572 | 0.572 | -0.507 | -1.406 | -0.016 | -0.006 | -0.006 | 0 | 0.16 | 0.13 | 0.14 | 0 | 0.3 | 0.11 |
| 6 | 44627544 | rs1021129 | 0.786 | 0.679 | -0.616 | -1.243 | -0.066 | 0.035 | 0.035 | 0 | 0.13 | 0.03 | 0.13 | 0.03 | 0.27 | 0.08 |
| 6 | 44628216 | rs6458401 | -0.478 | 0.028 | -0.526 | -0.978 | 0.009 | -0.005 | -0.005 | 0 | 0.56 | 0.7 | 0.65 | 0.51 | 0.56 | 0.61 |
| 6 | 44631292 | rs4711793 | 1.274 | -0.132 | -0.161 | -1.088 | 0.001 | 0.077 | 0.077 | 0 | 0.25 | 0.35 | 0.47 | 0.47 | 0.22 | 0.19 |
| 6 | 44631345 | rs499374 | 1.153 | -0.322 | -0.39 | -1.106 | -0.04 | -0.058 | -0.058 | 0.365 | 0.35 | 0.1 | 0.31 | 0.44 | 0.41 | 0.35 |
| 6 | 44631987 | rs472383 | 0.558 | -0.452 | 0.277 | -1.411 | -0.05 | -0.045 | -0.045 | 0 | 0.88 | 0.72 | 0.96 | 0.96 | 0.97 | 0.96 |
| 6 | 44635740 | rs477011 | -0.362 | -0.139 | 0.341 | -1.509 | 0.069 | -0.094 | -0.094 | 0.243 | 0.25 | 0.52 | 0.25 | 0.06 | 0.39 | 0.34 |
| 6 | 44636390 | rs6940887 | 0.653 | -0.062 | 0.176 | -1.39 | 0.008 | -0.058 | -0.058 | 1.842 | 0.64 | 0.56 | 0.77 | 0.9 | 0.57 | 0.52 |
| 6 | 44637880 | rs582006 | 0.427 | 0.101 | 0.069 | -0.775 | -0.015 | 0.043 | 0.043 | 0 | 0.09 | 0.01 | 0.04 | 0 | 0.08 | 0.13 |
| 6 | 44642025 | rs489448 | -0.593 | -0.005 | 0.56 | -0.763 | -0.008 | -0.038 | -0.038 | 0 | 0.16 | 0.25 | 0.06 | 0.09 | 0.12 | 0.24 |
| 6 | 44645736 | rs649700 | -0.524 | -0.142 | 0.699 | -0.397 | -0.001 | 0 | 0 | 0 | 0.39 | 0.6 | 0.48 | 0.45 | 0.34 | 0.41 |
| 6 | 44645943 | rs660953 | -0.739 | -0.268 | 0.692 | -0.518 | 0.012 | -0.002 | -0.002 | 0 | 0.44 | 0.67 | 0.49 | 0.46 | 0.35 | 0.42 |
| 6 | 44647374 | rs9472324 | 0.97 | -0.272 | 0.717 | -0.432 | 0.003 | 0.054 | 0.054 | 0.02 | 0.23 | 0.34 | 0.42 | 0.37 | 0.21 | 0.17 |
| 6 | 44649193 | rs6901297 | 0.594 | -0.312 | 0.613 | -0.61 | -0.005 | 0.012 | 0.012 | 0 | 0.26 | 0.27 | 0.28 | 0.41 | 0.2 | 0.28 |
| 6 | 44651735 | rs1283926 | -0.532 | -0.104 | -0.043 | -0.704 | -0.006 | -0.072 | -0.072 | 0.101 | 0.16 | 0.27 | 0.07 | 0 | 0.13 | 0.12 |
| 6 | 44655191 | rs1283921 | -0.354 | -0.197 | -0.386 | -0.71 | 0.004 | -0.001 | -0.001 | 0 | 0.15 | 0.11 | 0.06 | 0.01 | 0.13 | 0.12 |
| 6 | 44659586 | rs4714811 | 1.217 | 0.001 | -0.545 | -0.49 | -0.051 | -0.101 | -0.101 | 0 | 0.2 | 0.01 | 0.27 | 0.32 | 0.2 | 0.17 |
| 6 | 44662629 | rs17335164 | 0.738 | 0.055 | -0.261 | -0.506 | -0.006 | 0.02 | 0.02 | 0 | 0.06 | 0.01 | 0.03 | 0 | 0.07 | 0.09 |
| 6 | 44671471 | rs227813 | 0.231 | 0.184 | -0.151 | -0.76 | -0.001 | -0.001 | -0.001 | 0 | 0.46 | 0.53 | 0.49 | 0.55 | 0.44 | 0.51 |
| 6 | 44674614 | rs9367207 | 1.079 | 0.086 | -0.343 | -0.784 | -0.008 | -0.03 | -0.03 | 0 | 0.34 | 0.2 | 0.42 | 0.56 | 0.33 | 0.39 |
| 6 | 44676420 | rs227819 | -1.465 | 0.209 | -0.407 | -0.884 | -0.012 | -0.036 | -0.036 | 0 | 0.79 | 0.9 | 0.71 | 0.7 | 0.79 | 0.83 |
| 6 | 44680615 | rs12205657 | 0.311 | -0.017 | -0.014 | -1.414 | 0.007 | -0.048 | -0.048 | 0 | 0.24 | 0.27 | 0.21 | 0.03 | 0.33 | 0.19 |
| 6 | 44681257 | rs227832 | 0.169 | -0.037 | -0.127 | -1.229 | 0.001 | 0.003 | 0.003 | 0 | 0.3 | 0.25 | 0.33 | 0.27 | 0.28 | 0.31 |
| 6 | 44681721 | rs1028608 | -0.368 | 0.042 | -0.364 | -1.253 | 0.002 | 0.006 | 0.006 | 0 | 0.35 | 0.66 | 0.3 | 0.18 | 0.33 | 0.3 |
| 6 | 44682316 | rs227835 | 0.509 | 0.051 | -0.667 | -1.24 | -0.01 | 0.01 | 0.01 | 0 | 0.28 | 0.11 | 0.32 | 0.27 | 0.28 | 0.3 |
| 6 | 44683123 | rs227837 | -1.394 | 0.101 | -0.806 | -1.039 | -0.046 | -0.075 | -0.075 | 0 | 0.64 | 0.88 | 0.63 | 0.49 | 0.63 | 0.61 |
| 6 | 44684124 | rs13206206 | 1.497 | -0.166 | -0.829 | -1.244 | -0.022 | -0.052 | -0.052 | 0 | 0.19 | 0.02 | 0.27 | 0.26 | 0.17 | 0.16 |
| 6 | 44684621 | rs227838 | -1.303 | -0.044 | -0.744 | -1.139 | -0.064 | -0.05 | -0.05 | 0 | 0.64 | 0.88 | 0.62 | 0.45 | 0.61 | 0.61 |
| 6 | 44699068 | rs227850 | -1.323 | -0.059 | -0.775 | -1.127 | -0.065 | -0.06 | -0.06 | 0 | 0.64 | 0.9 | 0.63 | 0.45 | 0.61 | 0.59 |
| 6 | 44710304 | rs9463017 | 1.024 | 0.012 | -0.875 | -1.195 | -0.028 | -0.049 | -0.049 | 0.494 | 0.22 | 0.02 | 0.29 | 0.25 | 0.23 | 0.14 |
| 6 | 44739949 | rs1342371 | 1.161 | -0.001 | -1.252 | -1.605 | -0.037 | 0.081 | 0.081 | 0 | 0.13 | 0.02 | 0.12 | 0 | 0.16 | 0.04 |
| 6 | 44740471 | rs2038557 | -0.292 | -0.066 | -0.78 | -1.565 | 0 | 0.074 | 0.074 | 0 | 0.5 | 0.28 | 0.52 | 0.3 | 0.56 | 0.46 |
| 6 | 44765660 | rs720988 | 1.365 | -0.191 | -0.889 | -1.5 | -0.017 | -0.061 | -0.061 | 0.365 | 0.2 | 0.01 | 0.3 | 0.24 | 0.21 | 0.14 |
| 6 | 44767072 | rs12190136 | -0.52 | -0.149 | -0.434 | -1.545 | 0 | 0.02 | 0.02 | 0 | 0.51 | 0.56 | 0.55 | 0.29 | 0.55 | 0.46 |
| 6 | 44777691 | rs10948172 | 0.828 | -0.3 | -0.437 | -1.664 | 0.032 | -0.053 | -0.053 | 3.688 | 0.18 | 0.3 | 0.18 | 0.04 | 0.29 | 0.15 |
| 6 | 44783674 | rs7349931 | 0.874 | -0.258 | -0.335 | -1.666 | 0.046 | -0.063 | -0.063 | 0 | 0.18 | 0.3 | 0.17 | 0.04 | 0.29 | 0.15 |
| 6 | 44791861 | rs12216059 | 1.3 | -0.376 | -0.51 | -1.817 | -0.016 | 0.059 | 0.059 | 0 | 0.12 | 0.02 | 0.09 | 0 | 0.13 | 0.03 |
| 6 | 44796596 | rs524492 | 0.027 | -0.457 | -0.281 | -1.716 | -0.013 | -0.013 | -0.013 | 0.104 | 0.35 | 0.36 | 0.39 | 0.25 | 0.31 | 0.33 |
| 6 | 44797088 | rs529125 | -1.02 | -0.46 | -0.206 | -1.736 | 0.034 | -0.055 | -0.055 | 0 | 0.82 | 0.69 | 0.82 | 0.96 | 0.71 | 0.85 |
| 6 | 44797271 | rs9369514 | 0.924 | -0.466 | -0.255 | -1.746 | 0 | 0.05 | 0.05 | 0 | 0.46 | 0.33 | 0.42 | 0.71 | 0.39 | 0.52 |
| 6 | 44798701 | rs3799967 | 0.727 | -0.469 | -0.262 | -1.724 | -0.001 | 0.063 | 0.063 | 0 | 0.47 | 0.33 | 0.43 | 0.71 | 0.4 | 0.53 |
| 6 | 44799657 | rs1402599 | -0.059 | -0.44 | -0.292 | -1.561 | -0.023 | -0.022 | -0.022 | 0 | 0.65 | 0.59 | 0.6 | 0.75 | 0.69 | 0.67 |
| 6 | 44800015 | rs542444 | 1.323 | -0.453 | -0.441 | -1.593 | -0.004 | -0.022 | -0.022 | 0.209 | 0.21 | 0.06 | 0.3 | 0.24 | 0.24 | 0.14 |
| 6 | 44802578 | rs9472377 | -0.259 | -0.5 | -0.396 | -1.558 | -0.045 | -0.075 | -0.075 | 1.845 | 0.14 | 0.29 | 0.09 | 0.01 | 0.07 | 0.19 |
| 6 | 44802718 | rs9463046 | 0.142 | -0.453 | -0.443 | -1.507 | -0.028 | -0.043 | -0.043 | 0 | 0.09 | 0.25 | 0.08 | 0 | 0.05 | 0.05 |
| 6 | 44810418 | rs3823252 | 1.329 | -0.343 | -0.671 | -1.448 | 0.013 | -0.024 | -0.024 | 0 | 0.21 | 0.06 | 0.29 | 0.25 | 0.2 | 0.13 |
| 6 | 44846008 | rs4714828 | 0.351 | -0.247 | -0.759 | -1.418 | -0.049 | 0.028 | 0.028 | 0.033 | 0.36 | 0.11 | 0.42 | 0.28 | 0.42 | 0.27 |
| 6 | 44863106 | rs7754378 | -0.428 | -0.293 | -0.57 | -1.43 | 0.001 | 0.045 | 0.045 | 0.007 | 0.48 | 0.34 | 0.52 | 0.28 | 0.54 | 0.32 |
| 6 | 44908728 | rs9369523 | 1.345 | -0.384 | -1.073 | -1.513 | -0.136 | -0.114 | -0.114 | 0 | 0.36 | 0.03 | 0.35 | 0.56 | 0.39 | 0.41 |
| 6 | 44925293 | rs12210292 | 1.467 | -0.048 | -1.018 | -1.61 | -0.06 | 0.051 | 0.051 | 0.097 | 0.13 | 0.02 | 0.12 | 0.02 | 0.24 | 0.08 |
| 6 | 44943982 | rs17423748 | 1.102 | -0.157 | -0.949 | -1.556 | -0.064 | 0.08 | 0.08 | 0 | 0.19 | 0.06 | 0.16 | 0.03 | 0.29 | 0.15 |
| 6 | 44944642 | rs9472412 | 0.422 | -0.327 | -0.727 | -1.657 | -0.024 | -0.057 | -0.057 | 0 | 0.12 | 0.23 | 0.11 | 0 | 0.11 | 0.05 |
| 6 | 44946463 | rs2396369 | 1.069 | -0.363 | -0.728 | -1.806 | -0.003 | -0.011 | -0.011 | 0.255 | 0.38 | 0.59 | 0.38 | 0.72 | 0.3 | 0.5 |
| 6 | 44947321 | rs10948192 | 0.381 | -0.371 | -0.63 | -1.603 | -0.049 | -0.01 | -0.01 | 0.154 | 0.36 | 0.11 | 0.46 | 0.33 | 0.43 | 0.28 |
| 6 | 44951990 | rs10948194 | 0.948 | -0.389 | -0.349 | -1.713 | -0.008 | 0.044 | 0.044 | 0 | 0.44 | 0.33 | 0.43 | 0.72 | 0.4 | 0.53 |
| 6 | 44955528 | rs4714830 | 0.316 | -0.358 | -0.444 | -1.674 | -0.034 | 0.031 | 0.031 | 0.005 | 0.36 | 0.15 | 0.42 | 0.28 | 0.42 | 0.27 |
| 6 | 44956622 | rs35687006 | 1.829 | -0.343 | -0.495 | -1.622 | -0.011 | -0.079 | -0.079 | 0.157 | 0.17 | 0.01 | 0.25 | 0.25 | 0.13 | 0.12 |
| 6 | 44972209 | rs1159131 | -0.539 | -0.326 | -0.428 | -1.555 | 0.001 | 0.042 | 0.042 | 0.007 | 0.48 | 0.34 | 0.52 | 0.29 | 0.54 | 0.33 |

| | | | | | | | | | | | | | | | | |
|---|---|---|---|---|---|---|---|---|---|---|---|---|---|---|---|---|
| 6 | 44982593 | rs12528232 | 1.7 | -0.359 | -0.496 | -1.427 | 0.006 | -0.058 | -0.058 | 0.092 | 0.17 | 0.02 | 0.26 | 0.25 | 0.13 | 0.12 |
| 6 | 45043618 | rs1411148 | -0.045 | -0.143 | -0.435 | -1.544 | -0.038 | 0.061 | 0.061 | 0 | 0.58 | 0.8 | 0.55 | 0.73 | 0.5 | 0.6 |
| 6 | 45053617 | rs11964325 | 0.323 | -0.102 | -0.426 | -1.552 | -0.032 | 0.035 | 0.035 | 0.113 | 0.37 | 0.16 | 0.42 | 0.27 | 0.44 | 0.26 |
| 6 | 45059674 | rs6922660 | 0.168 | -0.151 | -0.444 | -1.555 | -0.03 | 0.038 | 0.038 | 0 | 0.38 | 0.2 | 0.43 | 0.27 | 0.45 | 0.36 |
| 6 | 45075795 | rs1324532 | -0.489 | -0.144 | -0.421 | -1.545 | -0.038 | 0.034 | 0.034 | 0.021 | 0.64 | 0.84 | 0.58 | 0.73 | 0.56 | 0.73 |
| 6 | 45077975 | rs9381365 | -0.038 | -0.165 | -0.419 | -1.545 | -0.032 | 0.056 | 0.056 | 0 | 0.58 | 0.79 | 0.55 | 0.73 | 0.5 | 0.6 |
| 6 | 45083966 | rs9296451 | -0.193 | -0.343 | -0.205 | -1.66 | -0.036 | -0.103 | -0.103 | 0 | 0.14 | 0.47 | 0.13 | 0 | 0.12 | 0.07 |
| 6 | 45095163 | rs9395066 | 0.922 | -0.314 | -0.278 | -1.653 | 0.003 | 0.043 | 0.043 | 0 | 0.44 | 0.33 | 0.42 | 0.73 | 0.38 | 0.53 |
| 6 | 45102025 | rs1285007 | -1.54 | -0.318 | -0.44 | -1.61 | 0.009 | -0.029 | -0.029 | 0 | 0.82 | 0.94 | 0.73 | 0.75 | 0.86 | 0.79 |
| 6 | 45122247 | rs1285029 | -1.762 | -0.322 | -0.43 | -1.603 | 0.011 | -0.022 | -0.022 | 6.816 | 0.83 | 0.93 | 0.75 | 0.76 | 0.87 | 0.88 |
| 6 | 45159177 | rs6919813 | 1.035 | -0.262 | -0.465 | -1.587 | -0.073 | 0.063 | 0.063 | 0 | 0.19 | 0.06 | 0.17 | 0.04 | 0.31 | 0.15 |
| 6 | 45159336 | rs6919873 | 0.927 | -0.25 | -0.411 | -1.622 | -0.027 | 0.03 | 0.03 | 0 | 0.2 | 0.12 | 0.17 | 0.03 | 0.31 | 0.14 |
| 6 | 45159774 | rs16873215 | 0.33 | -0.343 | -0.219 | -1.665 | -0.024 | -0.062 | -0.062 | 0.227 | 0.12 | 0.23 | 0.11 | 0 | 0.12 | 0.07 |
| 6 | 45164355 | rs927657 | 0.955 | -0.531 | -0.248 | -1.611 | -0.022 | 0.049 | 0.049 | 0.172 | 0.07 | 0.21 | 0.07 | 0.16 | 0.01 | 0.11 |
| 6 | 45170466 | rs10948214 | 1.075 | -0.489 | -0.271 | -1.634 | -0.039 | 0.048 | 0.048 | 0 | 0.19 | 0.1 | 0.17 | 0.03 | 0.31 | 0.14 |
| 6 | 45174641 | rs7762957 | -0.069 | -0.554 | -0.135 | -1.625 | -0.031 | -0.097 | -0.097 | 0.278 | 0.14 | 0.44 | 0.13 | 0 | 0.12 | 0.07 |
| 6 | 45176540 | rs1284964 | -1.85 | -0.528 | -0.352 | -1.568 | 0.01 | -0.02 | -0.02 | 0 | 0.83 | 0.93 | 0.74 | 0.76 | 0.87 | 0.88 |
| 6 | 45177025 | rs2396425 | 0.823 | -0.509 | -0.379 | -1.592 | -0.002 | 0.053 | 0.053 | 0 | 0.44 | 0.32 | 0.42 | 0.73 | 0.39 | 0.53 |
| 6 | 45181340 | rs9463078 | 1.333 | -0.446 | -0.613 | -1.543 | -0.052 | -0.054 | -0.054 | 0 | 0.37 | 0.1 | 0.35 | 0.56 | 0.37 | 0.42 |
| 6 | 45205017 | rs6458432 | -1.493 | -0.084 | -0.344 | -1.373 | 0.012 | -0.029 | -0.029 | 0.27 | 0.83 | 0.94 | 0.74 | 0.76 | 0.87 | 0.88 |
| 6 | 45222674 | rs6458435 | -1.493 | -0.084 | -0.345 | -1.373 | 0.01 | -0.02 | -0.02 | 0 | 0.83 | 0.93 | 0.74 | 0.76 | 0.87 | 0.88 |
| 6 | 45246577 | rs12207402 | 1.533 | -0.016 | -0.485 | -1.524 | -0.02 | 0.036 | 0.036 | 0 | 0.09 | 0.02 | 0.07 | 0 | 0.1 | 0.03 |
| 6 | 45255230 | rs12529907 | 1.49 | 0.01 | -0.425 | -1.473 | 0.008 | -0.014 | -0.014 | 0 | 0.17 | 0.08 | 0.26 | 0.24 | 0.13 | 0.12 |
| 6 | 45302654 | rs7757282 | -0.003 | -0.031 | -0.45 | -1.593 | -0.036 | 0.036 | 0.036 | 0 | 0.37 | 0.18 | 0.43 | 0.27 | 0.45 | 0.35 |
| 6 | 45310525 | rs17209755 | 1.55 | -0.018 | -0.537 | -1.645 | -0.022 | 0.045 | 0.045 | 0 | 0.1 | 0.02 | 0.09 | 0 | 0.13 | 0.03 |
| 6 | 45321205 | rs17209769 | 0.41 | 0.304 | 0.13 | -1.112 | -0.042 | 0.054 | 0.054 | 0 | 0.19 | 0.09 | 0.17 | 0.03 | 0.31 | 0.14 |
| 6 | 45321546 | rs9472465 | -0.895 | 0.497 | 0.473 | -0.953 | -0.019 | -0.105 | -0.105 | 0 | 0.15 | 0.47 | 0.13 | 0 | 0.13 | 0.08 |
| 6 | 45339067 | rs9472472 | -0.339 | 0.509 | 0.362 | -0.939 | -0.014 | -0.063 | -0.063 | 2.294 | 0.12 | 0.23 | 0.11 | 0 | 0.13 | 0.07 |
| 6 | 45349497 | rs35565233 | 0.946 | 0.52 | 0.081 | -0.756 | -0.007 | -0.067 | -0.067 | 0 | 0.17 | 0.03 | 0.26 | 0.27 | 0.13 | 0.11 |
| 6 | 45349509 | rs9472477 | -0.016 | 0.47 | 0.528 | -0.598 | 0.003 | -0.027 | -0.027 | 0 | 0.07 | 0.27 | 0.05 | 0 | 0.07 | 0.03 |
| 6 | 45355609 | rs1321081 | -1.217 | 0.392 | -0.154 | -0.665 | -0.003 | 0.045 | 0.045 | 0.383 | 0.58 | 0.75 | 0.58 | 0.3 | 0.62 | 0.45 |
| 6 | 45358107 | rs6908650 | 1.339 | 0.424 | 0.016 | 0.178 | -0.013 | 0.002 | 0.002 | 0 | 0.45 | 0.31 | 0.41 | 0.4 | 0.44 | 0.47 |
| 6 | 45358323 | rs1321080 | -0.01 | 0.798 | 0.214 | -0.052 | -0.022 | 0.031 | 0.031 | 0.04 | 0.18 | 0.09 | 0.13 | 0.02 | 0.26 | 0.11 |
| 6 | 45366946 | rs17288355 | 0.007 | 0.846 | 0.215 | -0.203 | -0.025 | 0.009 | 0.009 | 0 | 0.11 | 0 | 0.07 | 0.05 | 0.08 | 0.17 |
| 6 | 45368797 | rs9296459 | -0.958 | 0.861 | 0.218 | -0.245 | -0.082 | 0.105 | 0.105 | 0.021 | 0.28 | 0.05 | 0.21 | 0.08 | 0.34 | 0.28 |
| 6 | 45372218 | rs17288376 | 0.012 | 0.859 | 0.24 | -0.207 | -0.025 | 0.009 | 0.009 | 0 | 0.11 | 0 | 0.09 | 0.05 | 0.08 | 0.18 |
| 6 | 45383078 | rs1467458 | -0.714 | 0.785 | 0.246 | 0.007 | -0.006 | -0.049 | -0.049 | 0.334 | 0.9 | 0.97 | 0.76 | 0.76 | 0.9 | 0.96 |
| 6 | 45389289 | rs7771980 | -0.059 | 0.859 | 0.232 | 0.045 | -0.019 | 0.007 | 0.007 | 0 | 0.11 | 0.01 | 0.09 | 0.06 | 0.09 | 0.18 |
| 6 | 45391943 | rs1321075 | -0.377 | 0.825 | 0.526 | -0.086 | -0.005 | 0.015 | 0.015 | 1.242 | 0.85 | 0.83 | 0.72 | 0.76 | 0.84 | 0.94 |
| 6 | 45402445 | rs17209895 | 0.13 | 0.902 | -0.029 | -0.191 | -0.06 | 0.082 | 0.082 | 0.178 | 0.18 | 0.04 | 0.13 | 0.02 | 0.26 | 0.1 |
| 6 | 45403774 | rs2677108 | 0.595 | 0.784 | 0.182 | -0.04 | -0.001 | 0 | 0 | 0 | 0.63 | 0.38 | 0.53 | 0.65 | 0.59 | 0.82 |
| 6 | 45404230 | rs1997992 | 0.841 | 0.784 | 0.463 | 0.107 | -0.003 | 0.016 | 0.016 | 0.149 | 0.53 | 0.37 | 0.44 | 0.6 | 0.51 | 0.65 |
| 6 | 45404528 | rs2819854 | 0.806 | 0.861 | 0.502 | -0.05 | -0.001 | 0.024 | 0.024 | 0.008 | 0.54 | 0.37 | 0.49 | 0.64 | 0.51 | 0.65 |
| 6 | 45407654 | rs2677109 | 1.072 | 1.188 | 1.267 | 0.038 | -0.012 | 0.005 | 0.005 | 0 | 0.72 | 0.82 | 0.67 | 0.79 | 0.66 | 0.7 |
| 6 | 45410312 | rs910586 | 0.841 | 1.195 | 1.258 | 0.047 | -0.017 | 0.026 | 0.026 | 0 | 0.74 | 0.85 | 0.74 | 0.96 | 0.66 | 0.72 |
| 6 | 45413479 | rs9472491 | -2.763 | 1.202 | 2.126 | -0.001 | 0.062 | -0.014 | -0.014 | 0.072 | 0.09 | 0.43 | 0.05 | 0 | 0.04 | 0.01 |
| 6 | 45424940 | rs2819861 | 0.903 | 1.256 | 1.858 | -0.01 | -0.052 | 0.093 | 0.093 | 0.013 | 0.74 | 0.92 | 0.74 | 0.96 | 0.65 | 0.72 |
| 6 | 45432214 | rs2790102 | -1.901 | 0.92 | 2.133 | 0.003 | 0.002 | -0.016 | -0.016 | 1.277 | 0.33 | 0.52 | 0.35 | 0.21 | 0.34 | 0.34 |
| 6 | 45444061 | rs2677110 | -1.157 | 1.195 | 1.746 | 0.135 | -0.033 | 0.069 | 0.069 | 0 | 0.26 | 0.11 | 0.27 | 0.04 | 0.33 | 0.28 |
| 6 | 45467765 | rs2396441 | -1.924 | 1.37 | 1.154 | 0.048 | -0.002 | 0 | 0 | 0.08 | 0.4 | 0.57 | 0.46 | 0.27 | 0.5 | 0.33 |
| 6 | 45468074 | rs6914610 | -1.628 | 1.204 | 1.195 | 0.339 | 0.07 | 0.109 | 0.109 | 2.163 | 0.17 | 0.41 | 0.21 | 0.22 | 0.19 | 0.07 |
| 6 | 45501552 | rs7748231 | -1.876 | 1.018 | 0.594 | -1.107 | -0.01 | 0.021 | 0.021 | 0.03 | 0.37 | 0.66 | 0.37 | 0.04 | 0.43 | 0.27 |
| 6 | 45506122 | rs12209785 | -0.613 | 1.104 | 0.445 | -1.07 | -0.046 | 0.109 | 0.109 | 0 | 0.21 | 0.04 | 0.22 | 0.04 | 0.27 | 0.25 |
| 6 | 45509824 | rs1928533 | -1.909 | 1.104 | 0.581 | -1.025 | -0.012 | 0.021 | 0.021 | 1.214 | 0.35 | 0.71 | 0.36 | 0.04 | 0.41 | 0.26 |
| 6 | 45511945 | rs13201287 | -1.31 | 1.356 | 0.895 | -0.703 | 0.015 | -0.129 | -0.129 | 0 | 0.25 | 0.45 | 0.24 | 0.04 | 0.28 | 0.24 |
| 6 | 45518202 | rs1200428 | -0.444 | 1.344 | 0.434 | -0.439 | -0.047 | 0.068 | 0.068 | 0 | 0.16 | 0.03 | 0.22 | 0.04 | 0.23 | 0.23 |
| 6 | 45520572 | rs6911631 | -1.269 | 1.168 | 0.492 | -0.416 | 0 | -0.016 | -0.016 | 0 | 0.3 | 0.56 | 0.24 | 0.2 | 0.28 | 0.15 |
| 6 | 45520719 | rs9472501 | -0.437 | 1.307 | 0.261 | -0.341 | -0.001 | 0.001 | 0.001 | 0.611 | 0.21 | 0.14 | 0.13 | 0.19 | 0.16 | 0.14 |
| 6 | 45521449 | rs2772395 | -1.409 | 1.474 | 0.151 | -0.406 | -0.028 | 0.027 | 0.027 | 0 | 0.35 | 0.25 | 0.38 | 0.27 | 0.4 | 0.33 |
| 6 | 45522139 | rs13191376 | -0.764 | 1.46 | -0.158 | -0.486 | -0.043 | 0.059 | 0.059 | 0.023 | 0.21 | 0.11 | 0.3 | 0.04 | 0.34 | 0.22 |
| 6 | 45523412 | rs485817 | -1.257 | 1.238 | 0.135 | -0.315 | 0 | -0.008 | -0.008 | 1.636 | 0.45 | 0.49 | 0.45 | 0.26 | 0.51 | 0.35 |
| 6 | 45526942 | rs1200426 | -0.644 | 1.335 | -0.308 | -0.328 | -0.059 | 0.092 | 0.092 | 0 | 0.18 | 0.05 | 0.21 | 0.02 | 0.23 | 0.18 |
| 6 | 45527132 | rs1200425 | -0.891 | 1.198 | 0.293 | 0.062 | 0 | 0 | 0 | 0.911 | 0.37 | 0.63 | 0.49 | 0.31 | 0.37 | 0.3 |
| 6 | 45530471 | rs12202314 | -0.751 | 0.923 | 0.5 | -0.2 | -0.001 | -0.005 | -0.005 | 0 | 0.36 | 0.59 | 0.44 | 0.25 | 0.36 | 0.21 |
| 6 | 45530668 | rs16873507 | -0.717 | 0.875 | 0.435 | -0.232 | -0.001 | -0.012 | -0.012 | 0 | 0.05 | 0.13 | 0.05 | 0.06 | 0.04 | 0.11 |
| 6 | 45532214 | rs11969213 | -0.141 | 0.918 | 0.35 | -0.377 | -0.012 | 0.015 | 0.015 | 0 | 0.07 | 0.03 | 0.05 | 0 | 0.11 | 0.04 |
| 6 | 45532251 | rs1928532 | -1.32 | 1.085 | 0.451 | -0.577 | -0.015 | 0.083 | 0.083 | 0.36 | 0.41 | 0.81 | 0.48 | 0.25 | 0.43 | 0.22 |
| 6 | 45534165 | rs1928529 | -1.015 | 0.966 | 1.331 | -0.52 | 0.014 | -0.017 | -0.017 | 0 | 0.27 | 0.56 | 0.41 | 0.25 | 0.31 | 0.19 |
| 6 | 45536359 | rs7758107 | -0.511 | 0.951 | 1.27 | -0.533 | 0.001 | 0.006 | 0.006 | 0 | 0.22 | 0.31 | 0.36 | 0.25 | 0.24 | 0.17 |

| | | | | | | | | | | | | | | | | |
|---|---|---|---|---|---|---|---|---|---|---|---|---|---|---|---|---|
| 6 | 45541920 | rs6913882 | -1.221 | 1.132 | 1.381 | -0.568 | 0.01 | -0.018 | -0.018 | 0 | 0.31 | 0.59 | 0.34 | 0.24 | 0.37 | 0.2 |
| 6 | 45545975 | rs7764616 | -1.979 | 1.388 | 0.564 | -0.922 | -0.034 | 0.208 | 0.208 | 0.359 | 0.51 | 0.94 | 0.48 | 0.28 | 0.56 | 0.44 |
| 6 | 45550070 | rs10485421 | -1.113 | 1.39 | 0.914 | -0.967 | 0.02 | -0.023 | -0.023 | 0 | 0.32 | 0.44 | 0.39 | 0.25 | 0.44 | 0.26 |
| 6 | 45556069 | rs4711819 | -1.853 | 1.363 | 0.091 | -0.862 | -0.06 | 0.199 | 0.199 | 0.347 | 0.52 | 0.95 | 0.47 | 0.28 | 0.54 | 0.45 |
| 6 | 45557327 | rs6941748 | -0.609 | 1.364 | 0.406 | -0.256 | -0.013 | -0.013 | -0.013 | 0 | 0.19 | 0.15 | 0.27 | 0.29 | 0.28 | 0.2 |
| 6 | 45558433 | rs997445 | -1.217 | 0.786 | 1.085 | -0.517 | 0 | -0.02 | -0.02 | 0.754 | 0.2 | 0.24 | 0.13 | 0.07 | 0.17 | 0.21 |
| 6 | 45569863 | rs9472521 | -2.049 | 0.694 | 1.764 | -0.511 | 0.023 | 0.011 | 0.011 | 0.019 | 0.12 | 0.39 | 0.17 | 0.12 | 0.11 | 0.16 |
| 6 | 45574144 | rs12526713 | -0.944 | 0.621 | 1.871 | -0.501 | -0.014 | 0.016 | 0.016 | 0 | 0.05 | 0.13 | 0.1 | 0.09 | 0.03 | 0.14 |
| 6 | 45574685 | rs6938758 | -2.096 | 0.672 | 1.697 | -0.561 | -0.031 | -0.063 | -0.063 | 0 | 0.26 | 0.66 | 0.19 | 0.12 | 0.15 | 0.35 |
| 6 | 45578904 | rs7760394 | -2.096 | 0.71 | 1.665 | -0.623 | -0.007 | -0.031 | -0.031 | 0 | 0.29 | 0.69 | 0.21 | 0.12 | 0.19 | 0.38 |
| 6 | 45579744 | rs16873553 | -2.158 | 0.686 | 2.006 | -0.636 | 0 | 0.039 | 0.039 | 1.222 | 0.1 | 0.5 | 0.13 | 0.08 | 0.04 | 0.1 |
| 6 | 45584868 | rs6924440 | -1.008 | 0.689 | 1.999 | -0.641 | 0.004 | -0.008 | -0.008 | 3.684 | 0.19 | 0.19 | 0.09 | 0.03 | 0.15 | 0.28 |
| 6 | 45594885 | rs6458455 | -2.452 | 0.75 | 1.886 | -0.634 | 0.003 | 0.002 | 0.002 | 0 | 0.34 | 0.71 | 0.28 | 0.15 | 0.26 | 0.39 |
| 6 | 45595446 | rs12663992 | -1.434 | 0.693 | 1.972 | -0.552 | 0.018 | 0.059 | 0.059 | 0.44 | 0.07 | 0.34 | 0.12 | 0.09 | 0.04 | 0.1 |
| 6 | 45607827 | rs7744486 | -2.421 | 0.58 | 1.61 | -0.76 | 0.022 | 0.063 | 0.063 | 0.302 | 0.39 | 0.76 | 0.3 | 0.13 | 0.28 | 0.4 |
| 6 | 45616443 | rs9472536 | -1.256 | 0.102 | 1.983 | -0.798 | -0.047 | -0.11 | -0.11 | 0.04 | 0.23 | 0.39 | 0.13 | 0.06 | 0.14 | 0.27 |
| 6 | 45618548 | rs9472537 | -1.855 | 0.128 | 1.947 | -0.815 | -0.015 | -0.073 | -0.073 | 0.075 | 0.3 | 0.62 | 0.19 | 0.06 | 0.2 | 0.29 |
| 6 | 45626855 | rs7769966 | -1.686 | 0.141 | 1.939 | 0.248 | 0.006 | 0.108 | 0.108 | 0 | 0.12 | 0.21 | 0.34 | 0.55 | 0.14 | 0.12 |
| 6 | 45647622 | rs9472548 | -2.368 | 0.111 | 1.908 | 0.21 | 0.062 | -0.007 | -0.007 | 0.012 | 0.35 | 0.78 | 0.44 | 0.61 | 0.2 | 0.37 |
| 6 | 45671069 | rs4714864 | -2.499 | 0.07 | 1.934 | 0.278 | 0.05 | 0.006 | 0.006 | 0.786 | 0.4 | 0.78 | 0.49 | 0.61 | 0.27 | 0.39 |
| 6 | 45671400 | rs6906753 | -1.087 | 0.093 | 2.271 | 0.219 | 0.031 | -0.037 | -0.037 | 1.38 | 0.08 | 0.27 | 0.07 | 0.01 | 0.08 | 0.02 |
| 6 | 45675351 | rs9472556 | -1.013 | 0.099 | 1.89 | 0.214 | -0.021 | -0.081 | -0.081 | 0.493 | 0.22 | 0.27 | 0.1 | 0.03 | 0.13 | 0.28 |
| 6 | 45677664 | rs321324 | 1.904 | 0.157 | 1.407 | 0.308 | 0.067 | -0.003 | -0.003 | 0 | 0.6 | 0.11 | 0.51 | 0.38 | 0.73 | 0.61 |
| 6 | 45687333 | rs2782660 | 1.919 | 0.433 | 1.406 | 0.344 | 0.042 | 0.012 | 0.012 | 0.211 | 0.59 | 0.13 | 0.47 | 0.34 | 0.69 | 0.6 |
| 6 | 45689356 | rs9369573 | -1.993 | 0.602 | 1.295 | 0.373 | 0 | 0.021 | 0.021 | 0 | 0.14 | 0.12 | 0.34 | 0.59 | 0.19 | 0.1 |
| 6 | 45690246 | rs6914396 | -1.025 | 0.564 | 1.471 | 0.425 | -0.04 | -0.123 | -0.123 | 0.952 | 0.21 | 0.38 | 0.11 | 0.02 | 0.13 | 0.23 |
| 6 | 45692933 | rs12525547 | -1.209 | 0.959 | 0.905 | 0.796 | -0.039 | -0.033 | -0.033 | 0 | 0.09 | 0.03 | 0.33 | 0.58 | 0.16 | 0.1 |
| 6 | 45697540 | rs2039541 | 1.824 | 0.995 | 0.869 | 0.358 | 0.024 | 0.018 | 0.018 | 0 | 0.71 | 0.55 | 0.41 | 0.35 | 0.65 | 0.64 |
| 6 | 45698116 | rs6930715 | 1.312 | 1.006 | 0.709 | 1.198 | 0.055 | 0.147 | 0.147 | 0 | 0.86 | 0.66 | 0.66 | 0.58 | 0.78 | 0.88 |
| 6 | 45701907 | rs161187 | -0.304 | 0.549 | -0.52 | -0.181 | 0.06 | 0.149 | 0.149 | 2.052 | 0.53 | 0.3 | 0.68 | 0.86 | 0.67 | 0.7 |
| 6 | 45701998 | rs321318 | -0.298 | 0.544 | -0.522 | -0.183 | 0.06 | 0.15 | 0.15 | 1.985 | 0.53 | 0.3 | 0.68 | 0.86 | 0.67 | 0.7 |
| 6 | 45702112 | rs3846898 | 0.709 | 0.564 | -0.652 | 0.113 | -0.032 | 0.012 | 0.012 | 0.054 | 0.39 | 0.24 | 0.3 | 0.33 | 0.44 | 0.59 |
| 6 | 45702779 | rs321320 | -0.812 | 0.015 | -0.367 | -0.338 | -0.047 | -0.152 | -0.152 | 0.136 | 0.68 | 0.54 | 0.87 | 0.98 | 0.79 | 0.82 |
| 6 | 45705079 | rs10948250 | 0.923 | 0.319 | -0.859 | -0.19 | -0.011 | 0.033 | 0.033 | 0 | 0.43 | 0.24 | 0.35 | 0.34 | 0.54 | 0.6 |
| 6 | 45710553 | rs829930 | -0.527 | 0.528 | -0.741 | 0.124 | -0.011 | -0.025 | -0.025 | 1.591 | 0.11 | 0.06 | 0.33 | 0.53 | 0.13 | 0.1 |
| 6 | 45719769 | rs829911 | 0.372 | 0.659 | -0.703 | -0.226 | -0.008 | 0.075 | 0.075 | 0.388 | 0.47 | 0.36 | 0.6 | 0.86 | 0.51 | 0.67 |
| 6 | 45722988 | rs321329 | -0.284 | 0.593 | -1.094 | -0.011 | -0.017 | -0.049 | -0.049 | 0 | 0.6 | 0.82 | 0.61 | 0.47 | 0.58 | 0.73 |
| 6 | 45737500 | rs1449648 | -0.526 | 0.277 | -0.818 | -0.06 | 0.02 | -0.008 | -0.008 | 2.647 | 0.55 | 0.67 | 0.65 | 0.51 | 0.61 | 0.71 |
| 6 | 45743286 | rs10498762 | -0.271 | 0.283 | -0.71 | -0.188 | 0.001 | -0.038 | -0.038 | 1.753 | 0.24 | 0.28 | 0.21 | 0.09 | 0.26 | 0.18 |
| 6 | 45748205 | rs9395117 | -0.454 | 0.176 | -0.651 | -0.176 | 0.01 | 0.032 | 0.032 | 0.03 | 0.66 | 0.46 | 0.53 | 0.58 | 0.64 | 0.46 |
| 6 | 45750575 | rs16873837 | -0.349 | 0.454 | -0.671 | 0.054 | -0.001 | -0.021 | -0.021 | 0 | 0.11 | 0.24 | 0.23 | 0.05 | 0.09 | 0.12 |
| 6 | 45756479 | rs1449645 | 0.246 | 0.574 | -1.438 | 0.261 | 0.016 | 0.003 | 0.003 | 0.217 | 0.45 | 0.31 | 0.34 | 0.45 | 0.39 | 0.29 |
| 6 | 45764178 | rs10807324 | -0.167 | 0.978 | -2.061 | 0.372 | -0.086 | -0.109 | -0.109 | 0 | 0.3 | 0.08 | 0.42 | 0.42 | 0.35 | 0.54 |
| 6 | 45764675 | rs16873881 | -0.017 | 0.946 | -2.129 | 0.338 | -0.069 | -0.109 | -0.109 | 0 | 0.26 | 0.04 | 0.28 | 0.41 | 0.3 | 0.45 |
| 6 | 45768791 | rs9472569 | -0.026 | 0.294 | -2.089 | -0.118 | 0.048 | 0.031 | 0.031 | 0 | 0.58 | 0.3 | 0.69 | 0.33 | 0.74 | 0.64 |
| 6 | 45772603 | rs4714873 | 0.174 | 0.725 | -2.456 | -0.217 | 0.072 | 0.134 | 0.134 | 0.005 | 0.52 | 0.17 | 0.53 | 0.29 | 0.68 | 0.52 |
| 6 | 45772874 | rs728398 | 0.919 | 0.81 | -2.55 | -0.262 | -0.024 | -0.004 | -0.004 | 0 | 0.06 | 0 | 0.19 | 0.03 | 0.09 | 0.09 |
| 6 | 45773324 | rs728397 | 0.496 | 1.161 | -2.613 | -0.424 | -0.043 | 0.028 | 0.028 | 0.073 | 0.09 | 0.02 | 0.13 | 0.05 | 0.19 | 0.02 |
| 6 | 45777347 | rs7746680 | -0.418 | 0.745 | -2.943 | -0.428 | 0.035 | -0.017 | -0.017 | 0 | 0.65 | 0.19 | 0.66 | 0.41 | 0.76 | 0.67 |
| 6 | 45777475 | rs4714874 | 0.766 | 0.733 | -3.032 | -0.46 | -0.063 | 0.101 | 0.101 | 0 | 0.42 | 0.12 | 0.4 | 0.24 | 0.49 | 0.49 |
| 6 | 45777794 | rs4714875 | 0.928 | 1.147 | -3.198 | -0.499 | -0.058 | 0.004 | 0.004 | 0 | 0.29 | 0.05 | 0.31 | 0.24 | 0.33 | 0.46 |
| 6 | 45779732 | rs9395123 | 0.585 | 0.713 | -2.874 | 0.304 | -0.027 | -0.081 | -0.081 | 0.604 | 0.12 | 0.01 | 0.11 | 0.35 | 0.12 | 0.06 |
| 6 | 45779946 | rs13209955 | 0.298 | 1.161 | -2.442 | -0.114 | -0.043 | 0.029 | 0.029 | 0.088 | 0.1 | 0.02 | 0.13 | 0.06 | 0.2 | 0.02 |
| 6 | 45779990 | rs16873913 | 0.059 | 0.779 | -2.105 | 0.15 | 0.016 | -0.008 | -0.008 | 0.064 | 0.12 | 0.04 | 0.13 | 0.12 | 0.07 | 0.15 |
| 6 | 45787468 | rs2894651 | -0.056 | 1.035 | -1.45 | 0.059 | -0.022 | 0.032 | 0.032 | 0 | 0.42 | 0.24 | 0.39 | 0.28 | 0.47 | 0.48 |
| 6 | 45791913 | rs12204525 | 0.203 | 0.806 | -0.939 | 0.057 | 0 | -0.006 | -0.006 | 0.52 | 0.54 | 0.5 | 0.59 | 0.32 | 0.58 | 0.66 |
| 6 | 45793972 | rs9395125 | 0.036 | 0.8 | -0.843 | 0.505 | 0 | 0.005 | 0.005 | 0 | 0.72 | 0.67 | 0.68 | 0.58 | 0.75 | 0.84 |
| 6 | 45797856 | rs10948259 | 0.846 | 0.718 | -0.618 | 0.576 | 0 | -0.002 | -0.002 | 0 | 0.56 | 0.49 | 0.6 | 0.33 | 0.6 | 0.66 |
| 6 | 45799585 | rs7770578 | -0.284 | 0.802 | -0.74 | 0.627 | 0.005 | 0.083 | 0.083 | 0 | 0.85 | 0.69 | 0.78 | 0.6 | 0.88 | 0.86 |
| 6 | 45800800 | rs9381400 | 0.039 | 0.543 | -0.939 | 0.388 | -0.036 | -0.068 | -0.068 | 0.04 | 0.13 | 0.01 | 0.06 | 0.25 | 0.13 | 0.16 |
| 6 | 45806448 | rs732080 | 0.63 | 0.338 | -1.331 | -0.057 | -0.013 | 0.061 | 0.061 | 2.526 | 0.16 | 0.05 | 0.12 | 0.06 | 0.19 | 0.02 |
| 6 | 45808357 | rs9349331 | -0.678 | 0.347 | -0.899 | 0.188 | 0.004 | 0.005 | 0.005 | 0.71 | 0.3 | 0.25 | 0.19 | 0.28 | 0.26 | 0.21 |
| 6 | 45814124 | rs6458470 | 0.299 | 0.586 | -0.807 | 0.321 | -0.012 | -0.011 | -0.011 | 1.633 | 0.36 | 0.31 | 0.44 | 0.48 | 0.35 | 0.51 |
| 6 | 45814499 | rs7767586 | 0.08 | 0.218 | -0.88 | -0.024 | 0.021 | 0.003 | 0.003 | 0 | 0.33 | 0.16 | 0.2 | 0.32 | 0.27 | 0.21 |
| 6 | 45819347 | rs1375702 | -0.798 | 0.261 | -0.452 | -0.161 | -0.004 | -0.017 | -0.017 | 0 | 0.74 | 0.76 | 0.86 | 0.92 | 0.76 | 0.81 |
| 6 | 45822614 | rs895985 | 0.884 | 0.312 | -0.353 | -0.135 | -0.005 | -0.005 | -0.005 | 0.157 | 0.34 | 0.35 | 0.43 | 0.48 | 0.34 | 0.51 |
| 6 | 45827480 | rs11965083 | 0.901 | 0.423 | -0.56 | -0.53 | 0 | -0.003 | -0.003 | 0.23 | 0.06 | 0.05 | 0.21 | 0.05 | 0.14 | 0.09 |
| 6 | 45830465 | rs1556653 | -0.591 | 0.257 | -0.572 | -0.89 | -0.013 | -0.025 | -0.025 | 0 | 0.76 | 0.76 | 0.9 | 0.92 | 0.83 | 0.81 |
| 6 | 45840604 | rs10948262 | -0.214 | 0.589 | -0.337 | -0.632 | -0.022 | 0.037 | 0.037 | 0.644 | 0.33 | 0.23 | 0.42 | 0.11 | 0.41 | 0.26 |

| | | | | | | | | | | | | | | | | |
|---|---|---|---|---|---|---|---|---|---|---|---|---|---|---|---|---|
| 6 | 45843864 | rs9381404 | 0.792 | 0.427 | -0.229 | -0.673 | 0 | -0.004 | -0.004 | 0 | 0.38 | 0.41 | 0.31 | 0.71 | 0.38 | 0.52 |
| 6 | 45845801 | rs928613 | -0.537 | 0.428 | -0.187 | -0.673 | 0.007 | 0.002 | 0.002 | 4.554 | 0.72 | 0.78 | 0.87 | 0.91 | 0.79 | 0.79 |
| 6 | 45847115 | rs873053 | 0.402 | 0.509 | -0.434 | -0.702 | -0.028 | 0.066 | 0.066 | 0.006 | 0.26 | 0.1 | 0.31 | 0.12 | 0.28 | 0.18 |
| 6 | 45852094 | rs9381406 | 0.649 | 0.129 | -0.348 | -0.541 | 0.001 | -0.004 | -0.004 | 0 | 0.38 | 0.41 | 0.32 | 0.71 | 0.37 | 0.53 |
| 6 | 45853256 | rs6934152 | 0.292 | 0.914 | -0.563 | -0.109 | -0.021 | 0.025 | 0.025 | 0.603 | 0.67 | 0.76 | 0.47 | 0.81 | 0.59 | 0.74 |
| 6 | 45862228 | rs6458474 | 0.106 | 0.491 | -0.256 | 0.421 | 0.001 | -0.006 | -0.006 | 0 | 0.44 | 0.61 | 0.35 | 0.55 | 0.41 | 0.45 |
| 6 | 45863826 | rs9463147 | 0.48 | 0.264 | 0.405 | 0.368 | -0.009 | -0.027 | -0.027 | 0.644 | 0.37 | 0.26 | 0.24 | 0.48 | 0.38 | 0.4 |
| 6 | 45867872 | rs3814071 | -0.438 | 0.044 | 0.845 | 0.168 | -0.002 | 0.056 | 0.056 | 0.004 | 0.07 | 0.29 | 0.08 | 0.1 | 0.05 | 0.13 |
| 6 | 45870148 | rs2297834 | -0.469 | 0.043 | 0.871 | 0.168 | 0.001 | 0.061 | 0.061 | 0.024 | 0.07 | 0.3 | 0.08 | 0.1 | 0.05 | 0.13 |
| 6 | 45879527 | rs3088356 | -0.792 | -0.146 | 0.641 | 0.03 | 0.018 | -0.024 | -0.024 | 1.982 | 0.74 | 0.86 | 0.72 | 0.68 | 0.77 | 0.75 |
| 6 | 45880311 | rs6458477 | -0.751 | -0.107 | 0.796 | 0.088 | -0.016 | 0.028 | 0.028 | 0.007 | 0.11 | 0.43 | 0.15 | 0.11 | 0.07 | 0.16 |
| 6 | 45880937 | rs6458478 | 0.282 | 0.145 | 0.463 | 0.083 | 0.02 | 0.025 | 0.025 | 0.014 | 0.49 | 0.71 | 0.38 | 0.59 | 0.44 | 0.56 |
| 6 | 45885673 | rs3777543 | -0.188 | 0.109 | 0.818 | 0.326 | -0.002 | 0 | 0 | 0.163 | 0.69 | 0.7 | 0.68 | 0.66 | 0.71 | 0.64 |
| 6 | 45886740 | rs12665031 | 0.238 | 0.229 | 0.385 | 0.773 | -0.014 | -0.044 | -0.044 | 0.003 | 0.3 | 0.17 | 0.2 | 0.47 | 0.3 | 0.37 |
| 6 | 45894429 | rs3777561 | 0.275 | 0.266 | -0.37 | 0.805 | -0.022 | -0.055 | -0.055 | 0.007 | 0.29 | 0.12 | 0.2 | 0.46 | 0.3 | 0.37 |
| 6 | 45897634 | rs3777565 | -0.044 | -0.157 | -0.489 | 0.784 | 0.012 | 0 | 0 | 0 | 0.88 | 0.94 | 0.83 | 0.87 | 0.92 | 0.73 |
| 6 | 45898054 | rs2396578 | -0.885 | -1.009 | 0.059 | 1.191 | -0.066 | 0.085 | 0.085 | 0.562 | 0.19 | 0.61 | 0.18 | 0.28 | 0.08 | 0.24 |
| 6 | 45902399 | rs3777573 | 0.758 | -1.05 | 0.166 | 0.887 | -0.008 | 0.018 | 0.018 | 0 | 0.28 | 0.18 | 0.17 | 0.23 | 0.27 | 0.31 |
| 6 | 45908191 | rs3822887 | 0.185 | -1.089 | 0.323 | 1.37 | -0.026 | 0.14 | 0.14 | 0 | 0.1 | 0.2 | 0.11 | 0.39 | 0.04 | 0.12 |
| 6 | 45910464 | rs3777580 | -0.466 | -1.328 | 0.354 | 1.155 | 0.033 | 0.05 | 0.05 | 0 | 0.47 | 0.71 | 0.34 | 0.68 | 0.35 | 0.59 |
| 6 | 45914090 | rs3777583 | -0.075 | -1.451 | 0.3 | 0.876 | -0.004 | 0.01 | 0.01 | 0 | 0.78 | 0.77 | 0.71 | 0.48 | 0.88 | 0.71 |
| 6 | 45914931 | rs11757345 | -0.122 | -1.453 | 0.061 | 0.892 | 0.025 | -0.046 | -0.046 | 0.02 | 0.8 | 0.86 | 0.76 | 0.48 | 0.89 | 0.78 |
| 6 | 45915088 | rs16874013 | 0.448 | -1.21 | -0.046 | 0.92 | -0.004 | 0.009 | 0.009 | 0 | 0.06 | 0.11 | 0.05 | 0.05 | 0.05 | 0.15 |
| 6 | 45918969 | rs3777585 | 0.971 | -1.017 | -0.556 | 0.483 | -0.047 | 0.06 | 0.06 | 0 | 0.23 | 0.03 | 0.14 | 0.15 | 0.22 | 0.23 |
| 6 | 45928400 | rs1555214 | 0.124 | -0.323 | -0.25 | 0.305 | 0.019 | 0.073 | 0.073 | 0 | 0.45 | 0.28 | 0.51 | 0.19 | 0.62 | 0.28 |
| 6 | 45928727 | rs4714896 | 0.337 | 0.334 | -0.425 | -0.287 | -0.002 | -0.038 | -0.038 | 0 | 0.28 | 0.26 | 0.37 | 0.03 | 0.35 | 0.16 |
| 6 | 45933157 | rs3777594 | 0.094 | -0.078 | -0.688 | -0.236 | 0.003 | 0.023 | 0.023 | 0 | 0.59 | 0.62 | 0.44 | 0.82 | 0.38 | 0.72 |
| 6 | 45936223 | rs3777595 | 0.509 | -0.713 | -1.744 | -0.485 | -0.038 | 0.055 | 0.055 | 0 | 0.3 | 0.02 | 0.32 | 0.22 | 0.26 | 0.35 |
| 6 | 45940528 | rs12662667 | 0.609 | -0.7 | -1.46 | 0.068 | 0.019 | -0.043 | -0.043 | 0.015 | 0.17 | 0.13 | 0.08 | 0.51 | 0.08 | 0.34 |
| 6 | 45943652 | rs12174264 | 0.469 | -0.744 | -1.209 | -0.108 | 0.011 | -0.018 | -0.018 | 0 | 0.18 | 0.16 | 0.09 | 0.51 | 0.08 | 0.35 |
| 6 | 45945946 | rs10484841 | 0.325 | 0.041 | -1.202 | -0.408 | -0.004 | -0.01 | -0.01 | 0 | 0.67 | 0.69 | 0.75 | 0.37 | 0.65 | 0.54 |
| 6 | 45947467 | rs4714899 | 0.339 | 0.281 | -1.268 | -0.682 | 0.002 | -0.017 | -0.017 | 0 | 0.29 | 0.62 | 0.36 | 0.1 | 0.32 | 0.2 |
| 6 | 45951349 | rs3777601 | 0.529 | 0.159 | -0.861 | -0.699 | 0.002 | -0.004 | -0.004 | 0 | 0.62 | 0.3 | 0.73 | 0.36 | 0.65 | 0.51 |
| 6 | 45951830 | rs6930840 | 0.685 | 0.392 | -0.128 | -0.453 | 0.019 | 0.059 | 0.059 | 0 | 0.39 | 0.19 | 0.39 | 0.27 | 0.34 | 0.33 |
| 6 | 45952118 | rs1338466 | -0.277 | 0.436 | -0.261 | -0.43 | 0 | 0 | 0 | 0.526 | 0.85 | 0.86 | 0.84 | 0.88 | 0.73 | 0.89 |
| 6 | 45957904 | rs3777606 | 0.844 | 0.208 | -0.521 | -0.505 | -0.056 | 0.011 | 0.011 | 0 | 0.37 | 0.02 | 0.37 | 0.27 | 0.33 | 0.33 |
| 6 | 45962212 | rs9472650 | 0.8 | 0.858 | -0.171 | -0.663 | -0.008 | 0.034 | 0.034 | 0 | 0.29 | 0.32 | 0.16 | 0.52 | 0.23 | 0.4 |
| 6 | 45968780 | rs1338472 | 0.873 | 0.858 | 0.201 | -0.69 | 0.02 | 0.05 | 0.05 | 0 | 0.4 | 0.26 | 0.4 | 0.27 | 0.34 | 0.35 |
| 6 | 45968860 | rs3777615 | 0.804 | 0.864 | 0.171 | -0.657 | -0.008 | 0.035 | 0.035 | 0 | 0.29 | 0.32 | 0.16 | 0.52 | 0.23 | 0.4 |
| 6 | 45973511 | rs2022419 | -0.149 | 1.044 | -0.083 | -0.613 | 0.007 | 0 | 0 | 0.254 | 0.74 | 0.75 | 0.59 | 0.79 | 0.63 | 0.83 |
| 6 | 45975798 | rs7771614 | 0.203 | 1.543 | -0.199 | -0.616 | 0.001 | 0.118 | 0.118 | 0 | 0.47 | 0.21 | 0.42 | 0.27 | 0.42 | 0.38 |
| 6 | 45981962 | rs3798269 | 0.015 | 1.238 | 0.073 | -0.407 | -0.001 | 0.027 | 0.027 | 0.121 | 0.53 | 0.32 | 0.58 | 0.4 | 0.55 | 0.48 |
| 6 | 45982976 | rs9472662 | -0.622 | 1.301 | 0.65 | -0.49 | 0.056 | -0.065 | -0.065 | 0.012 | 0.12 | 0.4 | 0.12 | 0.01 | 0.14 | 0.06 |
| 6 | 45985462 | rs6921518 | 0.593 | 1.353 | 0.791 | -0.474 | 0.001 | -0.028 | -0.028 | 0 | 0.07 | 0.1 | 0.05 | 0 | 0.08 | 0.01 |
| 6 | 45987653 | rs9395145 | 0.257 | 1.217 | 0.931 | -0.321 | -0.001 | 0.091 | 0.091 | 0.134 | 0.07 | 0.23 | 0.16 | 0.13 | 0.04 | 0.03 |
| 6 | 45991569 | rs2179994 | -1.367 | 1.233 | 0.907 | -0.332 | 0.021 | 0.005 | 0.005 | 3.948 | 0.17 | 0.49 | 0.27 | 0.13 | 0.18 | 0.09 |
| 6 | 45992333 | rs13328271 | -0.18 | 1.32 | 0.831 | -0.384 | 0.031 | -0.051 | -0.051 | 1.972 | 0.1 | 0.19 | 0.1 | 0.01 | 0.13 | 0.06 |
| 6 | 46000258 | rs1578005 | -0.445 | 1.326 | 0.82 | -0.377 | 0.077 | -0.057 | -0.057 | 0.508 | 0.11 | 0.4 | 0.11 | 0.01 | 0.14 | 0.05 |
| 6 | 46007845 | rs6458486 | -1.416 | 1.344 | 0.616 | -0.363 | 0.004 | -0.043 | -0.043 | 0 | 0.68 | 0.8 | 0.83 | 0.53 | 0.71 | 0.54 |
| 6 | 46009132 | rs9381416 | -0.356 | 1.254 | 0.125 | -0.638 | 0.002 | 0.044 | 0.044 | 0 | 0.28 | 0.4 | 0.35 | 0.39 | 0.31 | 0.23 |
| 6 | 46010683 | rs6936849 | 0.109 | 1.322 | 0.269 | -0.579 | 0.029 | -0.05 | -0.05 | 0 | 0.1 | 0.17 | 0.09 | 0.01 | 0.14 | 0.05 |
| 6 | 46011912 | rs9349342 | 0.652 | 1.217 | 0.207 | -0.506 | 0.032 | 0.037 | 0.037 | 0.235 | 0.1 | 0.25 | 0.12 | 0.19 | 0.18 | 0.07 |
| 6 | 46012706 | rs9472678 | 0.417 | 0.891 | -0.014 | -0.526 | 0.002 | -0.037 | -0.037 | 0.242 | 0.07 | 0.12 | 0.05 | 0 | 0.08 | 0.01 |
| 6 | 46013386 | rs9367231 | -0.51 | 0.791 | -0.24 | -0.319 | 0.003 | 0.003 | 0.003 | 0 | 0.58 | 0.5 | 0.74 | 0.53 | 0.57 | 0.49 |
| 6 | 46017626 | rs1209198 | -0.665 | 1.049 | -0.106 | 0.312 | -0.002 | -0.001 | -0.001 | 0.838 | 0.44 | 0.58 | 0.68 | 0.56 | 0.49 | 0.53 |
| 6 | 46019225 | rs10948274 | 0.231 | 1.108 | -0.739 | 0.269 | -0.002 | 0.01 | 0.01 | 0 | 0.18 | 0.1 | 0.22 | 0.13 | 0.18 | 0.38 |
| 6 | 46021435 | rs2022421 | 0.408 | 0.911 | -0.601 | 0.399 | -0.006 | 0.025 | 0.025 | 0.021 | 0.06 | 0.07 | 0.14 | 0.11 | 0.05 | 0.02 |
| 6 | 46022074 | rs1555215 | -0.655 | 0.857 | -1.064 | -0.091 | 0.007 | -0.025 | -0.025 | 0.826 | 0.58 | 0.84 | 0.78 | 0.56 | 0.63 | 0.6 |
| 6 | 46027383 | rs9296477 | -0.558 | 0.956 | -0.142 | -0.147 | 0.019 | -0.067 | -0.067 | 0.568 | 0.14 | 0.28 | 0.09 | 0.01 | 0.15 | 0.04 |
| 6 | 46030130 | rs1338463 | -0.133 | 0.533 | -0.621 | 0.105 | 0 | -0.001 | -0.001 | 0 | 0.13 | 0.14 | 0.14 | 0.05 | 0.09 | 0.06 |
| 6 | 46030540 | rs1204339 | 0.625 | 0.495 | -1.223 | 0.036 | 0.014 | 0.016 | 0.016 | 0.053 | 0.43 | 0.26 | 0.45 | 0.63 | 0.4 | 0.44 |
| 6 | 46031615 | rs12207564 | 0.979 | 0.675 | -1.522 | -0.015 | -0.047 | -0.03 | -0.03 | 1.39 | 0.15 | 0.01 | 0.15 | 0.13 | 0.15 | 0.34 |
| 6 | 46034367 | rs946223 | -0.68 | 0.193 | -0.634 | 0.144 | -0.002 | 0.001 | 0.001 | 0.175 | 0.58 | 0.63 | 0.61 | 0.66 | 0.49 | 0.5 |
| 6 | 46034491 | rs946225 | -0.335 | 0.368 | -1.211 | 0.598 | 0.013 | 0.006 | 0.006 | 0 | 0.43 | 0.32 | 0.45 | 0.61 | 0.4 | 0.43 |
| 6 | 46034726 | rs7738939 | -0.067 | 0.648 | -1.528 | 0.695 | -0.007 | -0.056 | -0.056 | 0.542 | 0.32 | 0.17 | 0.38 | 0.44 | 0.33 | 0.29 |
| 6 | 46043561 | rs7758638 | -0.008 | 0.604 | -1.626 | 0.203 | -0.019 | 0.076 | 0.076 | 0.353 | 0.36 | 0.14 | 0.32 | 0.16 | 0.34 | 0.37 |
| 6 | 46044237 | rs4714902 | -0.42 | 1.265 | -2.108 | -0.11 | -0.063 | 0.028 | 0.028 | 0 | 0.86 | 0.95 | 0.81 | 0.88 | 0.67 | 0.82 |
| 6 | 46045157 | rs6933669 | 0.373 | 1.234 | -1.585 | -0.136 | -0.008 | 0.044 | 0.044 | 0.171 | 0.36 | 0.21 | 0.32 | 0.16 | 0.35 | 0.37 |
| 6 | 46046517 | rs9381421 | 0.334 | 0.543 | -1.78 | -0.206 | -0.024 | 0.125 | 0.125 | 0 | 0.39 | 0.12 | 0.33 | 0.13 | 0.37 | 0.37 |

| | | | | | | | | | | | | | | | | |
|---|---|---|---|---|---|---|---|---|---|---|---|---|---|---|---|---|
| 6 | 46047140 | rs9381422 | 0.612 | -0.226 | -1.76 | 0.438 | 0.013 | -0.029 | -0.029 | 0 | 0.25 | 0.16 | 0.12 | 0.35 | 0.18 | 0.35 |
| 6 | 46047149 | rs9472684 | 0.9 | -0.169 | -1.852 | 0.287 | -0.018 | 0.114 | 0.114 | 0 | 0.38 | 0.12 | 0.33 | 0.14 | 0.36 | 0.34 |
| 6 | 46054285 | rs2185108 | 1.411 | -0.219 | -1.503 | 0.182 | 0.003 | 0.012 | 0.012 | 0.167 | 0.07 | 0.04 | 0.06 | 0 | 0.07 | 0.02 |
| 6 | 46058361 | rs7772635 | -0.982 | -0.165 | -1.532 | 0.248 | -0.066 | -0.009 | -0.009 | 0 | 0.87 | 0.96 | 0.8 | 0.85 | 0.68 | 0.79 |
| 6 | 46067273 | rs7754582 | 0.617 | -0.257 | -1.043 | 0.45 | -0.006 | -0.012 | -0.012 | 0.162 | 0.64 | 0.31 | 0.45 | 0.43 | 0.53 | 0.7 |
| 6 | 46076742 | rs9395154 | 0.479 | -0.44 | -0.758 | 0.741 | 0.016 | 0.024 | 0.024 | 0 | 0.72 | 0.39 | 0.51 | 0.59 | 0.61 | 0.72 |
| 6 | 46080356 | rs6921636 | 0.487 | -0.482 | -0.846 | 0.745 | 0.011 | 0.013 | 0.013 | 0 | 0.72 | 0.35 | 0.51 | 0.62 | 0.6 | 0.71 |
| 6 | 46083567 | rs7766019 | -0.321 | -0.527 | -0.422 | 0.738 | 0.087 | 0.137 | 0.137 | 0 | 0.82 | 0.56 | 0.58 | 0.62 | 0.67 | 0.74 |
| 6 | 46093667 | rs11759657 | 0.798 | -0.826 | -0.786 | 0.731 | -0.02 | -0.07 | -0.07 | 0 | 0.24 | 0.01 | 0.12 | 0.28 | 0.17 | 0.35 |
| 6 | 46096777 | rs3756804 | 1.013 | -0.84 | -0.79 | 0.666 | -0.003 | 0.002 | 0.002 | 1.136 | 0.1 | 0.13 | 0.06 | 0.17 | 0.07 | 0.01 |
| 6 | 46097569 | rs7754915 | -1.03 | -0.674 | -0.858 | 0.805 | -0.058 | -0.024 | -0.024 | 0 | 0.86 | 0.95 | 0.79 | 0.82 | 0.68 | 0.79 |
| 6 | 46097629 | rs7775082 | 0.431 | -0.815 | -0.841 | 0.788 | -0.014 | -0.029 | -0.029 | 0 | 0.62 | 0.22 | 0.45 | 0.45 | 0.53 | 0.7 |
| 6 | 46106676 | rs2143746 | 1.016 | -0.814 | -0.84 | 0.78 | -0.006 | 0.059 | 0.059 | 0 | 0.38 | 0.21 | 0.33 | 0.17 | 0.36 | 0.34 |
| 6 | 46107312 | rs11752816 | 0.547 | -0.786 | -0.94 | 0.803 | -0.016 | -0.057 | -0.057 | 0 | 0.24 | 0.01 | 0.12 | 0.28 | 0.17 | 0.35 |
| 6 | 46127626 | rs6930766 | 0.464 | -0.779 | -0.876 | 0.796 | -0.02 | -0.039 | -0.039 | 0.058 | 0.62 | 0.19 | 0.45 | 0.45 | 0.53 | 0.7 |
| 6 | 46127807 | rs2295018 | 1.229 | -0.853 | -0.937 | 0.738 | -0.014 | 0.061 | 0.061 | 0 | 0.36 | 0.18 | 0.33 | 0.17 | 0.36 | 0.33 |
| 6 | 46128269 | rs14781 | 0.472 | -0.736 | -0.866 | 0.776 | -0.02 | -0.039 | -0.039 | 2.105 | 0.62 | 0.19 | 0.45 | 0.45 | 0.53 | 0.7 |
| 6 | 46128794 | rs2235882 | 1.513 | -0.962 | -1 | 0.84 | -0.036 | -0.052 | -0.052 | 0.022 | 0.15 | 0 | 0.07 | 0.18 | 0.11 | 0.22 |
| 6 | 46129345 | rs2235881 | 0.594 | -0.851 | -0.696 | 0.756 | -0.02 | -0.04 | -0.04 | 0.193 | 0.62 | 0.19 | 0.45 | 0.45 | 0.53 | 0.7 |
| 6 | 46135489 | rs6926570 | 0.601 | -0.869 | -0.691 | 0.749 | -0.022 | -0.039 | -0.039 | 0 | 0.62 | 0.19 | 0.45 | 0.45 | 0.53 | 0.69 |
| 6 | 46142439 | rs5011271 | 0.259 | 0.047 | -0.11 | 0.817 | 0.085 | 0.115 | 0.115 | 0 | 0.81 | 0.6 | 0.57 | 0.62 | 0.64 | 0.74 |
| 6 | 46142552 | rs10807337 | 0.383 | 0.217 | -1.208 | 0.665 | 0.041 | -0.002 | -0.002 | 0 | 0.45 | 0.13 | 0.25 | 0.46 | 0.33 | 0.45 |
| 6 | 46145547 | rs2206964 | -0.091 | 0.877 | -0.835 | 0.529 | -0.056 | -0.008 | -0.008 | 1.858 | 0.84 | 0.92 | 0.78 | 0.83 | 0.64 | 0.79 |
| 6 | 46150596 | rs17466007 | -0.255 | -0.347 | -0.386 | 0.471 | -0.005 | 0.064 | 0.064 | 0 | 0.38 | 0.22 | 0.27 | 0.18 | 0.35 | 0.32 |
| 6 | 46151231 | rs6925476 | -0.408 | -0.274 | 0.054 | 0.42 | 0.011 | -0.004 | -0.004 | 0 | 0.18 | 0.26 | 0.18 | 0.14 | 0.24 | 0.2 |
| 6 | 46152939 | rs6927617 | -0.53 | -1.005 | 0.853 | 0.787 | -0.041 | 0.052 | 0.052 | 0 | 0.3 | 0.41 | 0.38 | 0.4 | 0.18 | 0.24 |
| 6 | 46153661 | rs4714904 | -0.695 | -0.938 | -0.002 | 0.821 | -0.002 | -0.001 | -0.001 | 0 | 0.33 | 0.62 | 0.4 | 0.4 | 0.18 | 0.25 |
| 6 | 46174522 | rs7744272 | -0.979 | -0.885 | 0.406 | 0.273 | -0.052 | -0.046 | -0.046 | 0 | 0.27 | 0.45 | 0.23 | 0.2 | 0.21 | 0.21 |
| 6 | 46188118 | rs2584072 | -1.133 | -1.091 | 0.807 | 0.177 | 0.017 | 0.016 | 0.016 | 0.202 | 0.3 | 0.72 | 0.24 | 0.2 | 0.2 | 0.21 |
| 6 | 46189087 | rs3799893 | -0.992 | -0.918 | 0.485 | 0.265 | -0.056 | -0.046 | -0.046 | 0 | 0.27 | 0.45 | 0.22 | 0.2 | 0.2 | 0.2 |
| 6 | 46190297 | rs3823240 | -0.906 | -0.826 | -0.199 | 0.312 | 0.03 | 0.023 | 0.023 | 0 | 0.24 | 0.11 | 0.2 | 0.2 | 0.2 | 0.2 |
| 6 | 46205342 | rs6458499 | 0.809 | -0.694 | -0.287 | 0.605 | 0.007 | -0.009 | -0.009 | 0 | 0.73 | 0.82 | 0.64 | 0.7 | 0.82 | 0.62 |
| 6 | 46214061 | rs1442222 | 0.721 | -0.774 | -0.334 | 0.524 | -0.006 | 0.04 | 0.04 | 0 | 0.81 | 0.91 | 0.81 | 0.98 | 0.86 | 0.82 |
| 6 | 46226244 | rs9296481 | 1.174 | -0.907 | 0.047 | 0.56 | -0.001 | 0 | 0 | 0 | 0.74 | 0.73 | 0.64 | 0.72 | 0.82 | 0.7 |
| 6 | 46231370 | rs2799362 | 1.084 | -0.838 | 0.008 | 0.571 | 0.001 | -0.01 | -0.01 | 0 | 0.76 | 0.82 | 0.65 | 0.72 | 0.83 | 0.72 |
| 6 | 46232325 | rs4714910 | 1.616 | -2.201 | -0.516 | 0.203 | 0.104 | -0.065 | -0.065 | 0 | 0.56 | 0.08 | 0.57 | 0.54 | 0.75 | 0.51 |
| 6 | 46251843 | rs9381438 | -0.791 | -1.958 | 0.451 | 0.198 | -0.026 | -0.001 | -0.001 | 0.299 | 0.15 | 0.23 | 0.09 | 0.18 | 0.09 | 0.2 |
| 6 | 46282207 | rs10807341 | 1.029 | -1.661 | 1.275 | 0.064 | -0.007 | 0.005 | 0.005 | 0.053 | 0.74 | 0.32 | 0.63 | 0.71 | 0.82 | 0.71 |
| 6 | 46283922 | rs16874575 | 0.94 | -1.664 | 1.776 | 0.063 | -0.013 | 0.026 | 0.026 | 0 | 0.74 | 0.44 | 0.63 | 0.71 | 0.83 | 0.72 |
| 6 | 46287917 | rs10948288 | 1.02 | -1.652 | 1.525 | 0.068 | -0.003 | 0.002 | 0.002 | 0.041 | 0.74 | 0.29 | 0.63 | 0.71 | 0.82 | 0.72 |
| 6 | 46293328 | rs2277121 | 1.059 | -1.689 | 1.464 | 0.051 | -0.007 | 0.002 | 0.002 | 0.129 | 0.73 | 0.32 | 0.62 | 0.72 | 0.82 | 0.72 |
| 6 | 46296710 | rs6906742 | 0.969 | -1.91 | 0.879 | -0.269 | 0.003 | 0.003 | 0.003 | 0 | 0.64 | 0.62 | 0.56 | 0.71 | 0.76 | 0.63 |
| 6 | 46304945 | rs6940825 | 0.244 | -1.312 | -0.168 | -0.298 | -0.072 | -0.024 | -0.024 | 0 | 0.37 | 0.16 | 0.52 | 0.6 | 0.49 | 0.42 |
| 6 | 46306927 | rs1345058 | 0.244 | -1.312 | -0.168 | -0.298 | -0.072 | -0.024 | -0.024 | 0 | 0.37 | 0.16 | 0.52 | 0.6 | 0.49 | 0.42 |
| 6 | 46321089 | rs1867015 | -0.29 | -1.943 | 0.222 | -0.672 | -0.002 | 0.034 | 0.034 | 0.019 | 0.49 | 0.38 | 0.55 | 0.6 | 0.51 | 0.59 |
| 6 | 46335169 | rs9357506 | -0.233 | -2.021 | 0.394 | -0.758 | -0.005 | 0.034 | 0.034 | 0.101 | 0.51 | 0.39 | 0.56 | 0.6 | 0.52 | 0.6 |
| 6 | 46375326 | rs2153615 | -0.45 | -2.136 | 0.476 | -0.8 | -0.023 | 0.035 | 0.035 | 0.117 | 0.54 | 0.8 | 0.59 | 0.61 | 0.52 | 0.6 |
| 6 | 46399303 | rs9357507 | 0.615 | -1.387 | -0.515 | -0.709 | -0.13 | -0.114 | -0.114 | 0.155 | 0.35 | 0.03 | 0.5 | 0.59 | 0.44 | 0.38 |
| 6 | 46410189 | rs6458505 | -0.608 | -1.297 | -0.543 | -0.854 | -0.149 | -0.118 | -0.118 | 0.809 | 0.65 | 0.97 | 0.5 | 0.41 | 0.54 | 0.63 |
| 6 | 46423922 | rs1918477 | -0.058 | -1.025 | 0.252 | -0.806 | -0.003 | 0.014 | 0.014 | 0.03 | 0.47 | 0.43 | 0.54 | 0.61 | 0.51 | 0.53 |
| 6 | 46434923 | rs876121 | 0.065 | -0.64 | -0.366 | -0.597 | -0.002 | 0.017 | 0.017 | 0.003 | 0.5 | 0.27 | 0.55 | 0.49 | 0.48 | 0.49 |
| 6 | 46437221 | rs4714937 | 0.192 | -1.161 | 0.134 | -0.525 | -0.003 | 0.061 | 0.061 | 0.064 | 0.82 | 0.71 | 0.77 | 0.67 | 0.82 | 0.81 |
| 6 | 46439907 | rs12191660 | 0.699 | -1.435 | -0.572 | -1.013 | 0.046 | 0.071 | 0.071 | 0.537 | 0.49 | 0.2 | 0.52 | 0.63 | 0.62 | 0.48 |
| 6 | 46440557 | rs1554787 | 0.518 | -1.646 | -0.191 | -1.101 | -0.019 | -0.011 | -0.011 | 0.728 | 0.09 | 0.12 | 0.21 | 0.07 | 0.03 | 0.18 |
| 6 | 46443928 | rs1534362 | -0.441 | -1.902 | 0.156 | -1.176 | 0 | -0.004 | -0.004 | 0.754 | 0.68 | 0.66 | 0.77 | 0.71 | 0.69 | 0.78 |
| 6 | 46451262 | rs16874679 | 0.735 | -2.199 | -0.202 | -1.276 | 0.017 | 0.007 | 0.007 | 0.122 | 0.12 | 0.11 | 0.06 | 0.17 | 0.05 | 0.09 |
| 6 | 46457836 | rs4711853 | -0.213 | -2.129 | -0.14 | -1.258 | -0.036 | -0.036 | -0.036 | 0.319 | 0.17 | 0.25 | 0.24 | 0.08 | 0.07 | 0.25 |
| 6 | 46459974 | rs9349363 | 0.251 | -1.921 | -0.256 | -1.218 | 0.016 | 0.018 | 0.018 | 1.627 | 0.53 | 0.36 | 0.69 | 0.53 | 0.61 | 0.64 |
| 6 | 46485635 | rs9357509 | 1.556 | -1.582 | -0.655 | -1.062 | -0.107 | -0.101 | -0.101 | 1.992 | 0.32 | 0.1 | 0.45 | 0.45 | 0.54 | 0.37 |
| 6 | 46489208 | rs4714942 | 0.8 | -1.463 | -0.239 | -0.878 | 0.018 | 0.009 | 0.009 | 0.412 | 0.67 | 0.27 | 0.61 | 0.74 | 0.76 | 0.68 |
| 6 | 46515565 | rs806511 | -0.989 | -1.476 | -0.246 | -0.882 | 0.025 | 0.014 | 0.014 | 0.43 | 0.34 | 0.78 | 0.39 | 0.26 | 0.24 | 0.33 |
| 6 | 46515745 | rs806510 | -0.747 | -1.588 | 0.16 | -0.932 | -0.067 | -0.054 | -0.054 | 0.49 | 0.17 | 0.55 | 0.23 | 0.08 | 0.05 | 0.21 |
| 6 | 46517163 | rs806509 | 0.253 | -1.623 | 0.458 | -0.951 | -0.02 | -0.035 | -0.035 | 0 | 0.07 | 0.21 | 0.02 | 0 | 0.02 | 0.02 |
| 6 | 46518407 | rs1033530 | 0.968 | -1.635 | 0.555 | -0.953 | -0.012 | -0.008 | -0.008 | 1.198 | 0.08 | 0.14 | 0.2 | 0.06 | 0.03 | 0.16 |
| 6 | 46518693 | rs699938 | -0.995 | -1.642 | 0.552 | -0.956 | -0.068 | -0.055 | -0.055 | 0 | 0.17 | 0.55 | 0.23 | 0.08 | 0.05 | 0.21 |
| 6 | 46519210 | rs806508 | -0.526 | -1.639 | 0.597 | -0.955 | -0.028 | -0.021 | -0.021 | 0 | 0.09 | 0.31 | 0.02 | 0.01 | 0.02 | 0.05 |
| 6 | 46519485 | rs806507 | -0.526 | -1.639 | 0.597 | -0.955 | -0.028 | -0.021 | -0.021 | 0.01 | 0.09 | 0.31 | 0.02 | 0.01 | 0.02 | 0.05 |
| 6 | 46520256 | rs806506 | -1.242 | -1.644 | 0.586 | -0.959 | 0.026 | 0.014 | 0.014 | 0.385 | 0.34 | 0.78 | 0.39 | 0.26 | 0.24 | 0.33 |
| 6 | 46520922 | rs806505 | -0.996 | -1.636 | 0.597 | -0.953 | -0.068 | -0.055 | -0.055 | 0 | 0.17 | 0.54 | 0.23 | 0.08 | 0.05 | 0.21 |

| | | | | | | | | | | | | | | | | |
|---|---|---|---|---|---|---|---|---|---|---|---|---|---|---|---|---|
| 6 | 46522957 | rs806502 | 0.247 | -1.6 | 0.593 | -0.722 | -0.016 | -0.036 | -0.036 | 0.863 | 0.07 | 0.21 | 0.02 | 0 | 0.02 | 0.02 |
| 6 | 46524402 | rs806501 | -0.912 | -1.642 | 0.35 | -0.717 | -0.068 | -0.055 | -0.055 | 0 | 0.17 | 0.55 | 0.23 | 0.08 | 0.05 | 0.21 |
| 6 | 46526438 | rs4714947 | 0.786 | -1.639 | 0.353 | -0.715 | -0.015 | -0.01 | -0.01 | 0.562 | 0.08 | 0.19 | 0.2 | 0.06 | 0.03 | 0.16 |
| 6 | 46528544 | rs806500 | -0.904 | -1.641 | 0.343 | -0.716 | -0.068 | -0.055 | -0.055 | 0.321 | 0.17 | 0.55 | 0.23 | 0.08 | 0.05 | 0.21 |
| 6 | 46529196 | rs3799890 | 0.694 | -1.646 | 0.347 | -0.719 | 0.006 | 0.01 | 0.01 | 0 | 0.17 | 0.21 | 0.16 | 0.19 | 0.19 | 0.1 |
| 6 | 46531648 | rs1738306 | -1.512 | -1.862 | 0.159 | -0.845 | 0.032 | 0.064 | 0.064 | 1.167 | 0.49 | 0.86 | 0.47 | 0.38 | 0.37 | 0.48 |
| 6 | 46532996 | rs806491 | -0.795 | -1.833 | 0.206 | -0.813 | -0.069 | -0.056 | -0.056 | 0 | 0.17 | 0.54 | 0.24 | 0.08 | 0.05 | 0.2 |
| 6 | 46533725 | rs9349367 | 0.771 | -1.819 | 0.213 | -0.806 | 0.006 | 0.01 | 0.01 | 0 | 0.17 | 0.21 | 0.15 | 0.19 | 0.19 | 0.1 |
| 6 | 46533903 | rs806490 | -0.763 | -1.82 | 0.213 | -0.806 | -0.068 | -0.055 | -0.055 | 0 | 0.17 | 0.54 | 0.24 | 0.08 | 0.05 | 0.2 |
| 6 | 46533919 | rs9381463 | 0.771 | -1.817 | 0.212 | -0.805 | 0.009 | 0.013 | 0.013 | 0 | 0.17 | 0.22 | 0.15 | 0.19 | 0.19 | 0.1 |
| 6 | 46536415 | rs806489 | -1.139 | -1.816 | 0.206 | -0.804 | 0.026 | 0.014 | 0.014 | 0.449 | 0.34 | 0.78 | 0.39 | 0.26 | 0.24 | 0.32 |
| 6 | 46540454 | rs9349368 | 0.765 | -1.82 | 0.204 | -0.807 | 0.006 | 0.01 | 0.01 | 0 | 0.17 | 0.21 | 0.16 | 0.19 | 0.19 | 0.1 |
| 6 | 46540908 | rs806488 | -0.303 | -1.813 | 0.229 | -0.804 | -0.031 | -0.025 | -0.025 | 0 | 0.09 | 0.3 | 0.02 | 0.01 | 0.02 | 0.05 |
| 6 | 46542226 | rs806487 | -0.277 | -1.794 | 0.224 | -0.708 | -0.028 | -0.021 | -0.021 | 0.161 | 0.09 | 0.31 | 0.02 | 0.01 | 0.02 | 0.05 |
| 6 | 46542804 | rs1527691 | 1.032 | -1.797 | 0.227 | -0.709 | -0.012 | -0.009 | -0.009 | 0 | 0.08 | 0.14 | 0.21 | 0.06 | 0.03 | 0.15 |
| 6 | 46542888 | rs1091947 | -0.278 | -1.788 | 0.151 | -0.705 | -0.028 | -0.021 | -0.021 | 0.263 | 0.09 | 0.31 | 0.02 | 0.01 | 0.02 | 0.05 |
| 6 | 46546846 | rs806485 | -0.752 | -1.789 | 0.135 | -0.705 | -0.071 | -0.057 | -0.057 | 0 | 0.17 | 0.5 | 0.23 | 0.08 | 0.05 | 0.2 |
| 6 | 46549542 | rs1727583 | -1.106 | -1.79 | -0.047 | -0.707 | 0.025 | 0.014 | 0.014 | 0.417 | 0.34 | 0.78 | 0.39 | 0.26 | 0.24 | 0.32 |
| 6 | 46549970 | rs1727595 | -1.126 | -1.768 | -0.041 | -0.695 | 0.026 | 0.014 | 0.014 | 0.449 | 0.34 | 0.78 | 0.39 | 0.26 | 0.24 | 0.32 |
| 6 | 46549976 | rs1738311 | -1.38 | -1.758 | -0.076 | -0.715 | 0.026 | 0.077 | 0.077 | 1.203 | 0.5 | 0.86 | 0.47 | 0.37 | 0.38 | 0.48 |
| 6 | 46551743 | rs1157157 | 1.191 | -1.729 | -0.063 | -0.701 | 0.001 | 0.017 | 0.017 | 0 | 0.16 | 0.08 | 0.08 | 0.11 | 0.14 | 0.17 |
| 6 | 46551858 | rs3799887 | 0.79 | -1.746 | -0.006 | -0.704 | 0.006 | 0.009 | 0.009 | 0 | 0.17 | 0.2 | 0.15 | 0.19 | 0.2 | 0.1 |
| 6 | 46552107 | rs3799886 | 0.882 | -1.676 | 0.003 | -0.668 | -0.015 | -0.01 | -0.01 | 0 | 0.08 | 0.19 | 0.21 | 0.06 | 0.03 | 0.15 |
| 6 | 46552642 | rs3799884 | 1.148 | -1.576 | -0.607 | -0.867 | 0 | 0.016 | 0.016 | 0 | 0.16 | 0.08 | 0.08 | 0.11 | 0.14 | 0.17 |
| 6 | 46553323 | rs9395203 | 0.289 | -1.448 | -0.165 | -0.554 | 0.003 | 0.006 | 0.006 | 0.188 | 0.19 | 0.23 | 0.15 | 0.19 | 0.2 | 0.1 |
| 6 | 46553670 | rs9367252 | 0.289 | -1.448 | -0.257 | -0.554 | 0.002 | 0.005 | 0.005 | 0.188 | 0.19 | 0.21 | 0.15 | 0.19 | 0.2 | 0.1 |
| 6 | 46555401 | rs737011 | -0.673 | -1.162 | -0.988 | -0.4 | 0.023 | 0.014 | 0.014 | 0.479 | 0.34 | 0.77 | 0.39 | 0.26 | 0.24 | 0.3 |
| 6 | 46560003 | rs2206926 | -0.088 | -1.376 | -0.906 | -0.833 | 0.054 | 0.03 | 0.03 | 0 | 0.21 | 0.12 | 0.1 | 0.21 | 0.1 | 0.26 |
| 6 | 46560416 | rs9472785 | -1.076 | -1.973 | -0.777 | -1.465 | 0.093 | 0.111 | 0.111 | 0 | 0.5 | 0.7 | 0.38 | 0.4 | 0.27 | 0.62 |
| 6 | 46560750 | rs9296497 | 0.687 | -2.053 | -0.034 | -1.584 | 0 | -0.019 | -0.019 | 1.696 | 0.06 | 0.18 | 0.03 | 0 | 0.02 | 0.03 |
| 6 | 46564127 | rs3799880 | 0.766 | -2.352 | 0.076 | -1.734 | -0.004 | 0.029 | 0.029 | 0.266 | 0.18 | 0.07 | 0.09 | 0.1 | 0.15 | 0.17 |
| 6 | 46564968 | rs3799879 | 0.952 | -2.424 | 0.528 | -1.764 | -0.013 | -0.017 | -0.017 | 0.523 | 0.08 | 0.14 | 0.21 | 0.06 | 0.03 | 0.16 |
| 6 | 46565064 | rs16874838 | 0.931 | -2.424 | 0.544 | -1.764 | -0.013 | -0.016 | -0.016 | 0 | 0.08 | 0.12 | 0.21 | 0.06 | 0.03 | 0.16 |
| 6 | 46565479 | rs3799878 | 0.781 | -2.41 | 0.478 | -1.76 | -0.004 | 0.028 | 0.028 | 0.266 | 0.18 | 0.07 | 0.09 | 0.1 | 0.15 | 0.17 |
| 6 | 46566795 | rs9472787 | 0.221 | -2.471 | 0.668 | -1.775 | -0.003 | -0.02 | -0.02 | 0 | 0.07 | 0.26 | 0.03 | 0 | 0.02 | 0.06 |
| 6 | 46567357 | rs1008091 | 0.789 | -2.464 | 0.716 | -1.776 | -0.002 | 0.028 | 0.028 | 0.266 | 0.18 | 0.07 | 0.09 | 0.1 | 0.15 | 0.17 |
| 6 | 46570073 | rs926665 | 0.497 | -2.465 | 0.732 | -1.776 | -0.007 | -0.022 | -0.022 | 0 | 0.06 | 0.22 | 0.03 | 0 | 0.02 | 0.06 |
| 6 | 46570116 | rs2206927 | 0.789 | -2.465 | 0.732 | -1.776 | -0.002 | 0.028 | 0.028 | 0.266 | 0.18 | 0.07 | 0.09 | 0.1 | 0.15 | 0.17 |
| 6 | 46570856 | rs6458512 | 0.209 | -2.437 | 0.74 | -1.766 | 0.006 | -0.02 | -0.02 | 0 | 0.07 | 0.26 | 0.04 | 0 | 0.03 | 0.06 |
| 6 | 46571539 | rs9463215 | 0.492 | -2.44 | 0.734 | -1.766 | -0.007 | -0.022 | -0.022 | 0 | 0.06 | 0.22 | 0.03 | 0 | 0.02 | 0.05 |
| 6 | 46574115 | rs3799877 | 0.565 | -2.552 | 0.65 | -1.823 | 0.001 | 0.035 | 0.035 | 1.112 | 0.19 | 0.07 | 0.09 | 0.1 | 0.15 | 0.17 |
| 6 | 46574447 | rs7754144 | 0.222 | -2.538 | 0.681 | -1.808 | -0.003 | -0.02 | -0.02 | 0 | 0.07 | 0.26 | 0.03 | 0 | 0.02 | 0.06 |
| 6 | 46574804 | rs7754772 | 0.503 | -2.538 | 0.679 | -1.808 | -0.007 | -0.022 | -0.022 | 0 | 0.06 | 0.22 | 0.03 | 0 | 0.02 | 0.05 |
| 6 | 46575970 | rs3799876 | -1.463 | -2.725 | 0.603 | -1.865 | 0.11 | 0.12 | 0.12 | 0.04 | 0.5 | 0.71 | 0.39 | 0.37 | 0.26 | 0.61 |
| 6 | 46578739 | rs3799875 | -0.23 | -2.735 | 0.626 | -1.869 | -0.02 | -0.081 | -0.081 | 0 | 0.26 | 0.35 | 0.12 | 0.1 | 0.17 | 0.23 |
| 6 | 46580034 | rs3799873 | -0.23 | -2.735 | 0.654 | -1.869 | -0.02 | -0.081 | -0.081 | 1.175 | 0.26 | 0.34 | 0.12 | 0.1 | 0.17 | 0.23 |
| 6 | 46580382 | rs3799872 | -1.574 | -2.751 | 0.611 | -1.877 | 0.099 | 0.109 | 0.109 | 0 | 0.52 | 0.71 | 0.39 | 0.37 | 0.27 | 0.62 |
| 6 | 46582156 | rs7758684 | -1.124 | -2.44 | 0.758 | -1.848 | 0.011 | 0.013 | 0.013 | 0.038 | 0.35 | 0.61 | 0.33 | 0.18 | 0.2 | 0.41 |
| 6 | 46587520 | rs9463217 | -0.329 | -2.267 | 0.635 | -1.792 | -0.016 | -0.069 | -0.069 | 0 | 0.26 | 0.31 | 0.12 | 0.1 | 0.17 | 0.23 |
| 6 | 46587650 | rs9472801 | -0.329 | -2.267 | 0.635 | -1.792 | -0.016 | -0.069 | -0.069 | 2.005 | 0.26 | 0.31 | 0.12 | 0.1 | 0.17 | 0.23 |
| 6 | 46587664 | rs9463218 | -0.329 | -2.267 | 0.635 | -1.792 | -0.016 | -0.069 | -0.069 | 0.572 | 0.26 | 0.31 | 0.12 | 0.1 | 0.17 | 0.23 |
| 6 | 46587916 | rs9472802 | -0.407 | -2.268 | 0.642 | -1.793 | -0.021 | -0.083 | -0.083 | 2.206 | 0.26 | 0.35 | 0.12 | 0.1 | 0.17 | 0.23 |
| 6 | 46587972 | rs9463219 | -0.329 | -2.268 | 0.639 | -1.793 | -0.016 | -0.069 | -0.069 | 0.53 | 0.26 | 0.31 | 0.12 | 0.1 | 0.17 | 0.23 |
| 6 | 46588027 | rs9463221 | -0.457 | -2.269 | 0.648 | -1.793 | -0.024 | -0.094 | -0.094 | 0.592 | 0.27 | 0.37 | 0.12 | 0.1 | 0.17 | 0.23 |
| 6 | 46588049 | rs9463222 | -0.375 | -2.262 | 0.544 | -1.79 | -0.017 | -0.07 | -0.07 | 0.412 | 0.26 | 0.32 | 0.12 | 0.1 | 0.17 | 0.23 |
| 6 | 46588362 | rs7450391 | -0.334 | -2.255 | 0.535 | -1.787 | -0.016 | -0.069 | -0.069 | 5.847 | 0.26 | 0.31 | 0.12 | 0.1 | 0.17 | 0.23 |
| 6 | 46588421 | rs2223754 | -0.334 | -2.255 | 0.535 | -1.787 | -0.016 | -0.069 | -0.069 | 0.087 | 0.26 | 0.31 | 0.12 | 0.1 | 0.17 | 0.23 |
| 6 | 46588968 | rs13199352 | 0.316 | -2.274 | 0.563 | -1.797 | 0.001 | 0.033 | 0.033 | 1.48 | 0.19 | 0.07 | 0.09 | 0.1 | 0.15 | 0.17 |
| 6 | 46589159 | rs9472805 | -0.331 | -2.261 | 0.553 | -1.789 | -0.016 | -0.069 | -0.069 | 0 | 0.26 | 0.31 | 0.12 | 0.1 | 0.17 | 0.23 |
| 6 | 46589283 | rs9296500 | -0.331 | -2.261 | 0.553 | -1.789 | -0.016 | -0.069 | -0.069 | 0.02 | 0.26 | 0.31 | 0.12 | 0.1 | 0.17 | 0.23 |
| 6 | 46589745 | rs7768852 | -0.331 | -2.261 | 0.553 | -1.789 | -0.016 | -0.069 | -0.069 | 0 | 0.26 | 0.31 | 0.12 | 0.1 | 0.17 | 0.23 |
| 6 | 46589808 | rs7768721 | -0.331 | -2.261 | 0.553 | -1.789 | -0.016 | -0.069 | -0.069 | 0.896 | 0.26 | 0.31 | 0.12 | 0.1 | 0.17 | 0.23 |
| 6 | 46590177 | rs7773028 | -0.331 | -2.196 | 0.553 | -1.789 | -0.014 | -0.069 | -0.069 | 0 | 0.26 | 0.31 | 0.12 | 0.1 | 0.17 | 0.23 |
| 6 | 46590208 | rs9472806 | -0.331 | -2.228 | 0.553 | -1.789 | -0.016 | -0.069 | -0.069 | 0 | 0.26 | 0.31 | 0.12 | 0.1 | 0.17 | 0.23 |
| 6 | 46590430 | rs9472807 | -0.331 | -2.228 | 0.553 | -1.789 | -0.016 | -0.069 | -0.069 | 0 | 0.26 | 0.31 | 0.12 | 0.1 | 0.17 | 0.23 |
| 6 | 46591039 | rs9296501 | -0.331 | -2.228 | 0.553 | -1.789 | -0.016 | -0.069 | -0.069 | 0 | 0.26 | 0.31 | 0.12 | 0.1 | 0.17 | 0.23 |
| 6 | 46591677 | rs9296503 | -0.331 | -2.228 | 0.553 | -1.789 | -0.016 | -0.069 | -0.069 | 0 | 0.26 | 0.31 | 0.12 | 0.1 | 0.17 | 0.23 |
| 6 | 46592189 | rs9472808 | 0.648 | -2.235 | 0.575 | -1.793 | -0.009 | -0.024 | -0.024 | 0.122 | 0.06 | 0.19 | 0.02 | 0 | 0.02 | 0.03 |
| 6 | 46592238 | rs9463224 | -0.342 | -2.234 | 0.551 | -1.793 | -0.016 | -0.069 | -0.069 | 2.273 | 0.26 | 0.31 | 0.12 | 0.1 | 0.17 | 0.23 |

| | | | | | | | | | | | | | | | | |
|---|---|---|---|---|---|---|---|---|---|---|---|---|---|---|---|---|
| 6 | 46592418 | rs9463226 | 0.346 | -2.238 | 0.551 | -1.795 | 0.001 | 0.032 | 0.032 | 1.175 | 0.19 | 0.07 | 0.09 | 0.1 | 0.15 | 0.17 |
| 6 | 46592879 | rs9463231 | -0.283 | -2.203 | 0.498 | -1.774 | -0.016 | -0.069 | -0.069 | 0 | 0.26 | 0.31 | 0.12 | 0.1 | 0.17 | 0.23 |
| 6 | 46596286 | rs3799866 | 1.986 | -1.326 | -0.319 | -2.196 | -0.101 | -0.085 | -0.085 | 0.142 | 0.23 | 0.05 | 0.37 | 0.56 | 0.37 | 0.2 |
| 6 | 46599187 | rs9472810 | 0.678 | -1.822 | 0.588 | -2.026 | -0.006 | -0.022 | -0.022 | 0 | 0.17 | 0.15 | 0.08 | 0.11 | 0.12 | 0.17 |
| 6 | 46603687 | rs6924295 | -0.864 | -2.017 | 0.149 | -2.331 | 0.038 | 0.038 | 0.038 | 0 | 0.48 | 0.58 | 0.26 | 0.35 | 0.28 | 0.52 |
| 6 | 46606328 | rs639434 | 0.395 | -1.662 | -0.114 | -2.299 | 0.011 | 0.032 | 0.032 | 0 | 0.68 | 0.76 | 0.81 | 0.77 | 0.73 | 0.62 |
| 6 | 46613083 | rs952884 | -1.305 | -2.024 | -0.476 | -2.34 | -0.014 | 0.032 | 0.032 | 0 | 0.62 | 0.74 | 0.5 | 0.4 | 0.4 | 0.77 |
| 6 | 46614439 | rs12215229 | -0.193 | -1.627 | -0.363 | -1.804 | 0.009 | 0.011 | 0.011 | 0 | 0.18 | 0.16 | 0.11 | 0.14 | 0.12 | 0.21 |
| 6 | 46620135 | rs9369628 | -0.55 | -1.528 | 0.02 | -1.556 | -0.04 | -0.008 | -0.008 | 0.388 | 0.08 | 0.16 | 0.21 | 0.07 | 0.02 | 0.15 |
| 6 | 46620252 | rs12192544 | 0.184 | -1.89 | -0.705 | -2.097 | -0.034 | 0.053 | 0.053 | 0 | 0.24 | 0.02 | 0.14 | 0.13 | 0.21 | 0.25 |
| 6 | 46623698 | rs3757241 | -0.76 | -1.716 | -0.923 | -2.197 | -0.038 | -0.018 | -0.018 | 0 | 0.89 | 0.97 | 0.86 | 0.91 | 0.76 | 0.89 |
| 6 | 46625274 | rs9381468 | 1.436 | -2.195 | -0.462 | -2.458 | 0.005 | -0.006 | -0.006 | 0 | 0.43 | 0.53 | 0.43 | 0.56 | 0.43 | 0.24 |
| 6 | 46625635 | rs10807344 | 0.613 | -2.2 | -0.482 | -2.46 | -0.034 | 0.058 | 0.058 | 0 | 0.24 | 0.02 | 0.15 | 0.13 | 0.21 | 0.26 |
| 6 | 46626353 | rs953062 | -0.737 | -1.97 | -0.22 | -2.36 | 0.014 | 0.019 | 0.019 | 0 | 0.46 | 0.38 | 0.43 | 0.35 | 0.33 | 0.65 |
| 6 | 46629505 | rs9296504 | 1.216 | -2.041 | -0.235 | -2.368 | 0.004 | -0.006 | -0.006 | 0 | 0.44 | 0.53 | 0.43 | 0.56 | 0.45 | 0.24 |
| 6 | 46630068 | rs9381469 | 1.25 | -2.031 | -0.231 | -2.362 | 0.004 | -0.007 | -0.007 | 0 | 0.44 | 0.53 | 0.43 | 0.56 | 0.45 | 0.24 |
| 6 | 46632006 | rs6928546 | -0.848 | -1.976 | -0.35 | -2.334 | -0.016 | -0.007 | -0.007 | 0.378 | 0.89 | 0.93 | 0.86 | 0.91 | 0.76 | 0.89 |
| 6 | 46644875 | rs2270450 | -0.467 | -2.064 | -0.131 | -2.375 | 0.054 | 0.084 | 0.084 | 0 | 0.39 | 0.23 | 0.39 | 0.22 | 0.28 | 0.52 |
| 6 | 46651055 | rs12175419 | 0.302 | -1.999 | -0.03 | -2.148 | 0.027 | -0.007 | -0.007 | 0.456 | 0.07 | 0.01 | 0.2 | 0.09 | 0.02 | 0.21 |
| 6 | 46653789 | rs9472819 | -0.75 | -1.702 | 0.094 | -2.047 | -0.037 | -0.018 | -0.018 | 0 | 0.9 | 0.97 | 0.86 | 0.91 | 0.76 | 0.89 |
| 6 | 46657057 | rs3799277 | -0.796 | -1.701 | 0.083 | -2.047 | -0.052 | -0.03 | -0.03 | 0 | 0.9 | 0.98 | 0.86 | 0.91 | 0.76 | 0.89 |
| 6 | 46670822 | rs1421371 | 0.308 | -1.786 | 0.387 | -2.09 | 0.026 | -0.011 | -0.011 | 0 | 0.83 | 0.67 | 0.84 | 0.91 | 0.76 | 0.86 |
| 6 | 46679303 | rs1805018 | 0.18 | -1.875 | -0.092 | -2.002 | -0.007 | -0.006 | -0.006 | 0 | 0.12 | 0.26 | 0.03 | 0.13 | 0.05 | 0.12 |
| 6 | 46682079 | rs1362931 | -0.622 | -1.817 | -0.241 | -2.043 | 0.013 | 0.002 | 0.002 | 0 | 0.89 | 0.9 | 0.85 | 0.91 | 0.76 | 0.89 |
| 6 | 46684222 | rs1805017 | -0.241 | -2.065 | -0.407 | -2.224 | 0.049 | 0.066 | 0.066 | 0 | 0.38 | 0.26 | 0.37 | 0.22 | 0.26 | 0.52 |
| 6 | 46685286 | rs6929105 | 0.565 | -1.896 | -0.299 | -2.107 | 0.036 | -0.004 | -0.004 | 2.056 | 0.76 | 0.57 | 0.81 | 0.78 | 0.7 | 0.74 |
| 6 | 46689529 | rs3799861 | -0.305 | -1.545 | -1.046 | -1.883 | 0.02 | 0.003 | 0.003 | 0 | 0.88 | 0.86 | 0.84 | 0.91 | 0.76 | 0.89 |
| 6 | 46689847 | rs9472831 | 1.133 | -1.645 | -1.548 | -1.95 | -0.028 | -0.002 | -0.002 | 0 | 0.39 | 0.3 | 0.43 | 0.56 | 0.45 | 0.22 |
| 6 | 46703513 | rs1421378 | -1.282 | -2.394 | -1.743 | -2.088 | -0.027 | 0.041 | 0.041 | 0.005 | 0.65 | 0.84 | 0.49 | 0.29 | 0.4 | 0.68 |
| 6 | 46705994 | rs1421379 | -0.157 | -2.599 | -1.733 | -2.196 | 0.046 | 0.064 | 0.064 | 0 | 0.31 | 0.11 | 0.33 | 0.18 | 0.19 | 0.49 |
| 6 | 46715067 | rs1421373 | -1.296 | -2.436 | -1.813 | -2.128 | -0.022 | 0.04 | 0.04 | 0.024 | 0.65 | 0.83 | 0.49 | 0.29 | 0.39 | 0.68 |
| 6 | 46717454 | rs7766818 | -0.027 | -1.996 | -1.505 | -2.177 | 0.003 | -0.015 | -0.015 | 0 | 0.35 | 0.73 | 0.17 | 0.11 | 0.21 | 0.19 |
| 6 | 46724131 | rs9381480 | 0.232 | -1.876 | -1.897 | -2.075 | -0.009 | 0.075 | 0.075 | 0 | 0.24 | 0.02 | 0.13 | 0.09 | 0.18 | 0.23 |
| 6 | 46724247 | rs9395211 | -0.513 | -1.121 | -1.169 | -1.301 | -0.019 | 0.054 | 0.054 | 0.061 | 0.64 | 0.83 | 0.5 | 0.23 | 0.4 | 0.66 |
| 6 | 46733103 | rs10948303 | -0.394 | -1.783 | -0.551 | -0.978 | -0.013 | -0.008 | -0.008 | 0 | 0.2 | 0.2 | 0.17 | 0.17 | 0.13 | 0.34 |
| 6 | 46735158 | rs7750324 | -0.962 | -2.705 | -0.427 | -1.458 | 0.015 | 0.016 | 0.016 | 0.074 | 0.42 | 0.62 | 0.25 | 0.2 | 0.18 | 0.4 |
| 6 | 46745999 | rs7760667 | -1.079 | -2.55 | -0.395 | -1.451 | 0.058 | 0.058 | 0.058 | 0.682 | 0.4 | 0.69 | 0.26 | 0.2 | 0.2 | 0.39 |
| 6 | 46748354 | rs9472853 | -0.69 | -2.876 | -0.224 | -1.581 | 0.031 | 0.032 | 0.032 | 2.35 | 0.35 | 0.7 | 0.21 | 0.17 | 0.12 | 0.34 |
| 6 | 46755352 | rs6905097 | -0.24 | -2.348 | 0.665 | -1.257 | 0.012 | 0.009 | 0.009 | 0.166 | 0.05 | 0.28 | 0.03 | 0 | 0 | 0 |
| 6 | 46765182 | rs1410820 | 0.009 | -1.855 | -0.041 | -1.816 | 0.004 | 0.099 | 0.099 | 0 | 0.34 | 0.11 | 0.36 | 0.2 | 0.34 | 0.35 |
| 6 | 46767645 | rs3799859 | 0.14 | -1.553 | -0.197 | -1.357 | 0.011 | 0.005 | 0.005 | 0 | 0.24 | 0.32 | 0.3 | 0.27 | 0.25 | 0.35 |
| 6 | 46767688 | rs3799858 | 0.882 | -1.464 | -0.401 | -1.356 | -0.036 | -0.013 | -0.013 | 0.043 | 0.11 | 0 | 0.13 | 0.1 | 0.1 | 0.2 |
| 6 | 46769628 | rs1421376 | -1.389 | -1.504 | -0.001 | -1.385 | -0.025 | -0.038 | -0.038 | 0.163 | 0.7 | 0.81 | 0.72 | 0.48 | 0.63 | 0.76 |
| 6 | 46786183 | rs7748263 | -0.258 | -1.557 | 0.018 | -1.422 | -0.002 | -0.024 | -0.024 | 1.396 | 0.47 | 0.52 | 0.42 | 0.21 | 0.38 | 0.41 |
| 6 | 46788064 | rs1410825 | -1.495 | -1.561 | 0.016 | -1.394 | -0.06 | -0.084 | -0.084 | 0 | 0.71 | 0.91 | 0.72 | 0.48 | 0.63 | 0.76 |
| 6 | 46793175 | rs6920863 | -0.287 | -1.488 | 0.08 | -1.649 | 0.003 | -0.013 | -0.013 | 0.528 | 0.49 | 0.53 | 0.43 | 0.22 | 0.38 | 0.41 |
| 6 | 46800009 | rs3799854 | 0.365 | -1.082 | -0.266 | -1.417 | 0.004 | 0.11 | 0.11 | 0 | 0.35 | 0.11 | 0.37 | 0.2 | 0.35 | 0.34 |
| 6 | 46810964 | rs220693 | 0.992 | -1.092 | -0.351 | -0.648 | 0.012 | 0.028 | 0.028 | 0 | 0.3 | 0.39 | 0.29 | 0.44 | 0.37 | 0.24 |
| 6 | 46817931 | rs6458524 | -0.898 | -1.24 | 0.047 | -0.293 | 0.041 | -0.025 | -0.025 | 0 | 0.41 | 0.78 | 0.36 | 0.37 | 0.25 | 0.41 |
| 6 | 46826844 | rs571247 | 0.968 | -1.431 | 0.306 | -0.462 | -0.054 | -0.113 | -0.113 | 0 | 0.83 | 0.56 | 0.91 | 0.99 | 0.92 | 0.94 |
| 6 | 46827126 | rs610856 | 0.902 | -1.415 | 0.313 | -0.454 | -0.049 | -0.109 | -0.109 | 0.413 | 0.84 | 0.56 | 0.91 | 0.99 | 0.92 | 0.94 |
| 6 | 46829193 | rs514313 | 0.392 | -1.443 | 0.472 | -0.469 | -0.023 | -0.06 | -0.06 | 0.381 | 0.9 | 0.71 | 0.94 | 1 | 0.96 | 0.98 |
| 6 | 46829816 | rs508818 | 1.154 | -1.47 | 0.074 | -0.442 | 0.059 | -0.014 | -0.014 | 0 | 0.55 | 0.17 | 0.62 | 0.63 | 0.72 | 0.58 |
| 6 | 46832813 | rs614826 | 1.186 | -1.491 | 0.049 | -0.445 | 0.066 | -0.032 | -0.032 | 0 | 0.55 | 0.12 | 0.62 | 0.63 | 0.72 | 0.57 |
| 6 | 46834685 | rs586024 | 0.683 | -1.328 | 0.125 | -0.44 | -0.049 | -0.109 | -0.109 | 1.921 | 0.84 | 0.56 | 0.91 | 0.99 | 0.92 | 0.94 |
| 6 | 46837390 | rs1928274 | -0.554 | -0.931 | -0.063 | -0.053 | -0.006 | 0.027 | 0.027 | 0 | 0.28 | 0.37 | 0.29 | 0.36 | 0.2 | 0.36 |
| 6 | 46840446 | rs220677 | -0.179 | -1.256 | -0.694 | -0.478 | 0.006 | 0.003 | 0.003 | 0 | 0.84 | 0.98 | 0.77 | 0.82 | 0.86 | 0.77 |
| 6 | 46840761 | rs1571442 | 0.828 | -1.293 | -1.044 | -0.704 | -0.02 | 0.003 | 0.003 | 0 | 0.09 | 0 | 0.08 | 0.05 | 0.09 | 0.06 |
| 6 | 46841806 | rs220673 | 0.608 | -1.959 | -0.292 | -0.693 | -0.017 | -0.008 | -0.008 | 0.26 | 0.67 | 0.62 | 0.62 | 0.7 | 0.78 | 0.56 |
| 6 | 46844831 | rs9472892 | -0.83 | -1.641 | -0.286 | -0.984 | 0.003 | -0.037 | -0.037 | 0 | 0.34 | 0.43 | 0.29 | 0.17 | 0.32 | 0.3 |
| 6 | 46847501 | rs2273265 | -0.419 | -1.614 | -0.188 | -1.009 | -0.001 | 0.003 | 0.003 | 0 | 0.26 | 0.22 | 0.25 | 0.16 | 0.27 | 0.26 |
| 6 | 46849057 | rs11760108 | 0.852 | -1.567 | -0.193 | -1.05 | 0.011 | -0.041 | -0.041 | 0.175 | 0.1 | 0.11 | 0.05 | 0 | 0.13 | 0.07 |
| 6 | 46854403 | rs1004018 | 0.047 | -1.58 | -0.17 | -1.017 | 0.003 | -0.003 | -0.003 | 0 | 0.17 | 0.09 | 0.19 | 0.17 | 0.15 | 0.2 |
| 6 | 46856100 | rs678312 | -0.796 | -1.804 | -0.175 | -1.005 | 0.009 | 0.026 | 0.026 | 0 | 0.84 | 0.94 | 0.86 | 0.84 | 0.86 | 0.78 |
| 6 | 46856263 | rs677439 | 2.236 | -1.949 | -0.293 | -0.936 | -0.124 | -0.129 | -0.129 | 0.092 | 0.29 | 0.06 | 0.31 | 0.52 | 0.46 | 0.24 |
| 6 | 46858993 | rs4714956 | 0.251 | -2.022 | 0.304 | -0.866 | -0.006 | 0.02 | 0.02 | 0 | 0.11 | 0.15 | 0.23 | 0.14 | 0.06 | 0.15 |
| 6 | 46867717 | rs220702 | -0.604 | -1.826 | -0.102 | -0.759 | 0.013 | -0.006 | -0.006 | 0 | 0.85 | 0.97 | 0.89 | 0.85 | 0.9 | 0.78 |
| 6 | 46868113 | rs85603 | 0.885 | -2.135 | 0.304 | -0.995 | -0.004 | -0.001 | -0.001 | 0 | 0.54 | 0.43 | 0.47 | 0.52 | 0.69 | 0.52 |
| 6 | 46868763 | rs11758970 | 0.605 | -1.704 | 0.338 | -0.89 | 0.058 | -0.033 | -0.033 | 0 | 0.06 | 0.11 | 0.09 | 0 | 0.17 | 0.06 |

| | | | | | | | | | | | | | | | | |
|---|---|---|---|---|---|---|---|---|---|---|---|---|---|---|---|---|
| 6 | 46871985 | rs6910089 | 0.714 | -1.735 | 0.496 | -0.899 | 0.055 | -0.032 | -0.032 | 0 | 0.06 | 0.11 | 0.08 | 0 | 0.15 | 0.05 |
| 6 | 46876243 | rs1328957 | -0.312 | -1.899 | 0.401 | -0.9 | -0.041 | -0.007 | -0.007 | 0 | 0.22 | 0.42 | 0.19 | 0.18 | 0.13 | 0.13 |
| 6 | 46883779 | rs17470473 | 1.086 | -1.598 | -0.087 | -0.676 | 0.001 | -0.002 | -0.002 | 0.277 | 0.11 | 0.07 | 0.14 | 0.18 | 0.09 | 0.08 |
| 6 | 46884288 | rs220715 | 2.005 | -1.28 | -0.364 | -0.436 | -0.14 | -0.153 | -0.153 | 0 | 0.3 | 0.06 | 0.32 | 0.46 | 0.47 | 0.21 |
| 6 | 46889577 | rs6927483 | -0.084 | -1.281 | 0.278 | -0.33 | 0.001 | 0.021 | 0.021 | 0 | 0.13 | 0.18 | 0.23 | 0.17 | 0.1 | 0.17 |
| 6 | 46893686 | rs6919588 | 0.571 | -1.532 | -0.004 | -0.621 | 0.002 | 0.002 | 0.002 | 0 | 0.55 | 0.56 | 0.67 | 0.69 | 0.73 | 0.61 |
| 6 | 46900508 | rs7775991 | 0.307 | -1.72 | 0.072 | -0.922 | -0.001 | -0.01 | -0.01 | 0 | 0.73 | 0.75 | 0.77 | 0.84 | 0.7 | 0.71 |
| 6 | 46910415 | rs2077055 | 1.419 | -1.892 | 0.185 | -0.946 | 0.014 | 0.056 | 0.056 | 0 | 0.5 | 0.35 | 0.49 | 0.63 | 0.57 | 0.34 |
| 6 | 46911131 | rs10948314 | 1.243 | -1.701 | -0.022 | -0.628 | -0.01 | -0.06 | -0.06 | 0.088 | 0.1 | 0.01 | 0.15 | 0.19 | 0.09 | 0.09 |
| 6 | 46914179 | rs1334968 | 0.39 | -1.844 | 0.081 | -0.836 | -0.002 | 0.006 | 0.006 | 0 | 0.73 | 0.8 | 0.78 | 0.86 | 0.71 | 0.66 |
| 6 | 46914760 | rs9367260 | 2.792 | -1.671 | -0.156 | -0.687 | -0.092 | -0.125 | -0.125 | 0 | 0.31 | 0.04 | 0.3 | 0.4 | 0.43 | 0.22 |
| 6 | 46918018 | rs761748 | 0.451 | -1.306 | 0.694 | -0.72 | 0.011 | 0.044 | 0.044 | 0 | 0.11 | 0.2 | 0.24 | 0.21 | 0.1 | 0.15 |
| 6 | 46920990 | rs4714964 | -0.326 | -1.323 | 0.443 | -0.771 | -0.011 | 0.009 | 0.009 | 0 | 0.81 | 0.89 | 0.82 | 0.85 | 0.75 | 0.67 |
| 6 | 46922061 | rs10456562 | 0.708 | -1.194 | 0.658 | -0.618 | -0.011 | -0.044 | -0.044 | 0 | 0.1 | 0.16 | 0.04 | 0 | 0.04 | 0.04 |
| 6 | 46924985 | rs1028496 | 0.583 | -0.952 | 0.895 | -0.415 | 0 | -0.022 | -0.022 | 4.338 | 0.67 | 0.72 | 0.75 | 0.86 | 0.67 | 0.6 |
| 6 | 46928442 | rs7756172 | -0.581 | -0.537 | 1.029 | 0.184 | 0.023 | 0.083 | 0.083 | 0.436 | 0.13 | 0.41 | 0.27 | 0.21 | 0.11 | 0.16 |
| 6 | 46941930 | rs10447399 | 1.177 | -0.132 | 0.046 | 0.235 | -0.012 | -0.065 | -0.065 | 1.061 | 0.3 | 0.14 | 0.26 | 0.41 | 0.34 | 0.32 |
| 6 | 46942554 | rs1334972 | 0.058 | 0.176 | 0.281 | 0.39 | 0.014 | 0.001 | 0.001 | 0.087 | 0.16 | 0.2 | 0.32 | 0.18 | 0.26 | 0.16 |
| 6 | 46942896 | rs7741079 | 0.316 | -0.043 | 0.116 | 0.114 | 0.003 | 0 | 0 | 0 | 0.44 | 0.41 | 0.34 | 0.41 | 0.41 | 0.37 |
| 6 | 46951665 | rs4714979 | 0.693 | -0.175 | 0.075 | 0.109 | 0.001 | -0.007 | -0.007 | 1.75 | 0.32 | 0.28 | 0.23 | 0.13 | 0.32 | 0.36 |
| 6 | 46953541 | rs1932033 | 0.667 | 0.079 | -0.176 | 0.389 | 0.01 | 0.007 | 0.007 | 0 | 0.64 | 0.54 | 0.51 | 0.64 | 0.67 | 0.72 |
| 6 | 46957625 | rs1226498 | -0.269 | -0.635 | -0.524 | 0.401 | 0.086 | 0.062 | 0.062 | 0.738 | 0.47 | 0.77 | 0.39 | 0.36 | 0.35 | 0.47 |
| 6 | 46966274 | rs1873318 | -0.171 | -0.915 | 0.426 | 0.382 | -0.045 | 0.051 | 0.051 | 0 | 0.12 | 0.23 | 0.14 | 0.18 | 0.03 | 0.09 |
| 6 | 46967338 | rs819497 | -0.241 | -0.91 | 0.923 | 0.454 | -0.008 | 0.201 | 0.201 | 0 | 0.08 | 0.32 | 0.13 | 0.18 | 0.01 | 0.07 |
| 6 | 46975396 | rs9369649 | 1.146 | -0.648 | 0.195 | 0.722 | -0.156 | -0.093 | -0.093 | 0.003 | 0.33 | 0.04 | 0.3 | 0.42 | 0.49 | 0.28 |
| 6 | 46977458 | rs937057 | -0.809 | -0.929 | 1.287 | 0.579 | -0.031 | 0.181 | 0.181 | 0 | 0.1 | 0.49 | 0.15 | 0.18 | 0.01 | 0.1 |
| 6 | 46979403 | rs1226478 | -0.514 | -0.941 | 1.505 | 0.572 | -0.015 | 0.199 | 0.199 | 0 | 0.09 | 0.41 | 0.14 | 0.18 | 0.01 | 0.1 |
| 6 | 46980115 | rs1226479 | -0.696 | -1.02 | 1.328 | 0.543 | 0.019 | 0.05 | 0.05 | 0.523 | 0.65 | 0.76 | 0.76 | 0.83 | 0.67 | 0.6 |
| 6 | 46980358 | rs3857591 | 0.014 | -0.965 | 1.087 | 0.565 | 0.067 | 0.032 | 0.032 | 0.506 | 0.55 | 0.26 | 0.6 | 0.65 | 0.66 | 0.5 |
| 6 | 46985955 | rs9463279 | 0.638 | -0.98 | 0.728 | 0.824 | 0.022 | 0.019 | 0.019 | 0.461 | 0.28 | 0.17 | 0.19 | 0.24 | 0.25 | 0.35 |
| 6 | 46989848 | rs1226491 | -0.857 | -0.538 | 1.339 | 0.926 | 0.019 | 0.242 | 0.242 | 0 | 0.05 | 0.33 | 0.14 | 0.19 | 0.01 | 0.1 |
| 6 | 47000661 | rs1226548 | -1.055 | -0.584 | 0.573 | 0.502 | -0.036 | 0.008 | 0.008 | 0.008 | 0.28 | 0.44 | 0.37 | 0.3 | 0.21 | 0.24 |
| 6 | 47007263 | rs1226512 | -1.224 | 0.244 | 0.497 | 0.077 | 0.033 | 0.062 | 0.062 | 0 | 0.31 | 0.53 | 0.54 | 0.46 | 0.38 | 0.34 |
| 6 | 47007748 | rs9463288 | 1.22 | 0.137 | -0.038 | -0.266 | 0.033 | 0.058 | 0.058 | 0 | 0.5 | 0.34 | 0.3 | 0.23 | 0.41 | 0.47 |
| 6 | 47009192 | rs819505 | -0.738 | 0.669 | 0.122 | -0.243 | 0.025 | 0.133 | 0.133 | 0 | 0.11 | 0.21 | 0.41 | 0.46 | 0.16 | 0.2 |
| 6 | 47010843 | rs819504 | -0.914 | 0.684 | -0.009 | -0.247 | -0.018 | -0.027 | -0.027 | 0.46 | 0.81 | 0.87 | 0.83 | 0.68 | 0.79 | 0.81 |
| 6 | 47028167 | rs1226529 | -0.63 | 0.677 | 0.339 | -0.121 | 0.003 | 0.13 | 0.13 | 0 | 0.14 | 0.22 | 0.37 | 0.39 | 0.13 | 0.18 |
| 6 | 47028451 | rs16875546 | 0.009 | 0.742 | 0.426 | -0.237 | 0.008 | -0.046 | -0.046 | 0 | 0.13 | 0.22 | 0.09 | 0 | 0.19 | 0.1 |
| 6 | 47028640 | rs1226531 | -0.158 | 0.231 | 0.261 | -0.326 | 0.005 | 0.006 | 0.006 | 0 | 0.68 | 0.63 | 0.74 | 0.63 | 0.71 | 0.73 |
| 6 | 47037469 | rs9463290 | 0.362 | 0.124 | 0.563 | -0.004 | -0.027 | -0.036 | -0.036 | 0 | 0.07 | 0.15 | 0.02 | 0 | 0.01 | 0.02 |
| 6 | 47059523 | rs13207730 | 0.486 | 0.157 | -0.024 | 0.193 | -0.008 | 0.172 | 0.172 | 0 | 0.88 | 0.77 | 0.65 | 0.61 | 0.92 | 0.87 |
| 6 | 47064338 | rs960718 | -0.189 | -0.032 | -0.695 | -0.92 | 0.037 | 0.074 | 0.074 | 1.478 | 0.49 | 0.69 | 0.29 | 0.29 | 0.36 | 0.28 |
| 6 | 47066554 | rs10948327 | 0.582 | 0.104 | -0.964 | -0.865 | -0.063 | 0.005 | 0.005 | 2.241 | 0.33 | 0.07 | 0.27 | 0.28 | 0.39 | 0.46 |
| 6 | 47068454 | rs6902883 | 0.612 | 0.067 | -0.761 | -0.815 | -0.006 | -0.001 | -0.001 | 1.915 | 0.1 | 0.09 | 0.07 | 0 | 0.12 | 0.1 |
| 6 | 47069291 | rs1532137 | -0.957 | -0.34 | -0.649 | -0.882 | -0.005 | 0.021 | 0.021 | 1.203 | 0.77 | 0.75 | 0.59 | 0.56 | 0.85 | 0.79 |
| 6 | 47073954 | rs1387879 | 0.283 | 0.18 | -1.053 | -1.015 | 0.006 | 0.053 | 0.053 | 1.481 | 0.38 | 0.27 | 0.27 | 0.22 | 0.39 | 0.5 |
| 6 | 47075799 | rs2396821 | -0.022 | -0.141 | -1.16 | -0.967 | 0.031 | -0.019 | -0.019 | 0.355 | 0.62 | 0.3 | 0.52 | 0.53 | 0.8 | 0.74 |
| 6 | 47082030 | rs9395232 | 0.051 | -0.635 | -1.07 | -0.971 | -0.004 | 0.02 | 0.02 | 0.257 | 0.4 | 0.37 | 0.33 | 0.3 | 0.55 | 0.31 |
| 6 | 47083336 | rs1738844 | -1.323 | -0.806 | -1.21 | -1.012 | -0.066 | -0.004 | -0.004 | 0 | 0.74 | 0.98 | 0.77 | 0.79 | 0.69 | 0.6 |
| 6 | 47086952 | rs937055 | 0.362 | -0.985 | -0.58 | -0.749 | -0.11 | 0.072 | 0.072 | 5.545 | 0.29 | 0.48 | 0.4 | 0.49 | 0.09 | 0.23 |
| 6 | 47089918 | rs12198742 | 0.433 | -0.65 | -1.193 | -0.731 | -0.03 | 0.033 | 0.033 | 0.007 | 0.11 | 0.04 | 0.07 | 0 | 0.15 | 0.09 |
| 6 | 47089974 | rs1738837 | -0.057 | -1.272 | -0.835 | -0.417 | -0.006 | -0.004 | -0.004 | 0 | 0.34 | 0.61 | 0.44 | 0.49 | 0.12 | 0.29 |
| 6 | 47095197 | rs1738820 | -0.439 | -0.366 | -0.818 | -0.181 | 0.004 | -0.01 | -0.01 | 0 | 0.81 | 0.85 | 0.86 | 0.75 | 0.79 | 0.75 |
| 6 | 47096004 | rs9473000 | -1.011 | -0.394 | -0.432 | -0.934 | -0.004 | -0.127 | -0.127 | 0 | 0.27 | 0.33 | 0.19 | 0.01 | 0.21 | 0.29 |
| 6 | 47100710 | rs1687134 | 0.62 | -0.499 | -0.252 | -0.966 | -0.025 | -0.043 | -0.043 | 0 | 0.7 | 0.56 | 0.76 | 0.75 | 0.75 | 0.7 |
| 6 | 47102261 | rs9463301 | -0.047 | -0.523 | -0.091 | -0.931 | -0.006 | -0.083 | -0.083 | 0.122 | 0.13 | 0.3 | 0.12 | 0 | 0.06 | 0.05 |
| 6 | 47104451 | rs1627116 | 0.136 | -0.631 | -0.272 | -0.943 | -0.008 | 0.025 | 0.025 | 0.052 | 0.73 | 0.83 | 0.83 | 0.75 | 0.72 | 0.57 |
| 6 | 47104856 | rs1630504 | 1.253 | -0.665 | -0.202 | -0.989 | -0.004 | -0.025 | -0.025 | 0 | 0.57 | 0.51 | 0.68 | 0.75 | 0.62 | 0.45 |
| 6 | 47105305 | rs9296528 | -0.392 | -0.667 | -0.143 | -0.923 | 0.011 | -0.094 | -0.094 | 0.638 | 0.13 | 0.34 | 0.13 | 0 | 0.09 | 0.06 |
| 6 | 47107045 | rs12214981 | -1.478 | -0.688 | -0.238 | -0.923 | -0.007 | -0.136 | -0.136 | 2.063 | 0.27 | 0.51 | 0.22 | 0.01 | 0.21 | 0.29 |
| 6 | 47109623 | rs1738829 | 1.434 | -0.758 | -0.061 | -0.943 | 0.004 | 0.111 | 0.111 | 0 | 0.55 | 0.26 | 0.64 | 0.74 | 0.57 | 0.46 |
| 6 | 47111842 | rs761520 | -1.304 | -0.829 | 0.075 | -1.036 | -0.002 | -0.119 | -0.119 | 0 | 0.25 | 0.55 | 0.23 | 0.01 | 0.22 | 0.26 |
| 6 | 47114409 | rs9296529 | 0.841 | -1.391 | -0.328 | -0.606 | 0.017 | 0.039 | 0.039 | 0 | 0.58 | 0.65 | 0.49 | 0.25 | 0.6 | 0.41 |
| 6 | 47114870 | rs1687137 | -0.27 | -1.576 | -0.283 | 0.072 | -0.018 | 0.007 | 0.007 | 0 | 0.17 | 0.23 | 0.09 | 0.15 | 0.12 | 0.17 |
| 6 | 47117215 | rs9367265 | -0.062 | -1.304 | -0.001 | 0.133 | -0.004 | -0.025 | -0.025 | 0.032 | 0.14 | 0.27 | 0.09 | 0.08 | 0.12 | 0.19 |
| 6 | 47117465 | rs9395235 | 1.305 | -1.335 | 0.106 | 0.179 | 0.006 | 0.021 | 0.021 | 0.549 | 0.52 | 0.49 | 0.35 | 0.33 | 0.51 | 0.43 |
| 6 | 47120281 | rs1040604 | 0.234 | -1.364 | -0.065 | 0.408 | 0.003 | 0 | 0 | 0 | 0.68 | 0.75 | 0.79 | 0.68 | 0.68 | 0.54 |
| 6 | 47121186 | rs10484958 | 0.146 | -1.347 | -0.194 | 0.326 | -0.01 | 0.037 | 0.037 | 0 | 0.16 | 0.05 | 0.09 | 0.08 | 0.14 | 0.25 |
| 6 | 47121275 | rs926774 | -0.439 | -1.332 | -0.073 | 0.369 | -0.004 | 0.142 | 0.142 | 0 | 0.13 | 0.26 | 0.38 | 0.43 | 0.16 | 0.14 |

| | | | | | | | | | | | | | | | | |
|---|---|---|---|---|---|---|---|---|---|---|---|---|---|---|---|---|
| 6 | 47124684 | rs7769886 | 0.356 | -1.232 | -0.019 | 0.761 | -0.007 | 0.003 | 0.003 | 0 | 0.68 | 0.64 | 0.55 | 0.67 | 0.68 | 0.63 |
| 6 | 47124987 | rs7774223 | -0.011 | -1.159 | -0.063 | 0.796 | 0.011 | -0.091 | -0.091 | 0.329 | 0.14 | 0.3 | 0.15 | 0 | 0.12 | 0.08 |
| 6 | 47126369 | rs6458544 | -1.113 | -1.097 | -0.553 | 1.104 | -0.042 | -0.102 | -0.102 | 0 | 0.85 | 0.99 | 0.71 | 0.67 | 0.81 | 0.73 |
| 6 | 47127323 | rs9473021 | 1.218 | -0.801 | -0.046 | 1.031 | -0.036 | 0.049 | 0.049 | 0 | 0.39 | 0.23 | 0.25 | 0.26 | 0.39 | 0.23 |
| 6 | 47128729 | rs9296534 | 0.405 | -0.528 | 0.365 | 0.494 | 0.001 | 0.005 | 0.005 | 0 | 0.54 | 0.51 | 0.64 | 0.69 | 0.56 | 0.44 |
| 6 | 47131174 | rs1884043 | -0.668 | 0.019 | 0.938 | 0.806 | 0.021 | 0.07 | 0.07 | 0 | 0.08 | 0.24 | 0.05 | 0.08 | 0.08 | 0.06 |
| 6 | 47131813 | rs6934866 | -1.363 | -0.134 | 0.949 | 0.103 | -0.001 | -0.06 | -0.06 | 0 | 0.32 | 0.42 | 0.27 | 0.07 | 0.33 | 0.27 |
| 6 | 47132114 | rs9463306 | -1.478 | -0.602 | 1.634 | 0.337 | 0.064 | -0.04 | -0.04 | 0.102 | 0.09 | 0.39 | 0.1 | 0 | 0.12 | 0.02 |
| 6 | 47136966 | rs6458546 | 0.588 | -1.047 | 1.172 | 0.571 | -0.009 | -0.002 | -0.002 | 0.11 | 0.58 | 0.56 | 0.52 | 0.32 | 0.72 | 0.45 |
| 6 | 47137735 | rs9349391 | 1.51 | -1.161 | 1.28 | 0.828 | 0.01 | 0.017 | 0.017 | 0 | 0.57 | 0.32 | 0.48 | 0.68 | 0.61 | 0.45 |
| 6 | 47142493 | rs9395238 | 2.111 | -1.193 | 1.216 | 0.843 | -0.036 | 0.003 | 0.003 | 0 | 0.42 | 0.16 | 0.33 | 0.31 | 0.53 | 0.33 |
| 6 | 47143129 | rs9381516 | 0.64 | -1.218 | 1.161 | 0.829 | 0.004 | -0.004 | -0.004 | 0 | 0.15 | 0.1 | 0.14 | 0.36 | 0.08 | 0.11 |
| 6 | 47151873 | rs4711871 | 0.073 | -1.158 | 1.226 | 0.339 | 0.008 | 0.006 | 0.006 | 0 | 0.75 | 0.79 | 0.82 | 0.78 | 0.81 | 0.57 |
| 6 | 47152278 | rs7748298 | -1.091 | -1.164 | 1.432 | 0.296 | 0.021 | -0.085 | -0.085 | 0 | 0.14 | 0.29 | 0.19 | 0 | 0.19 | 0.08 |
| 6 | 47153739 | rs6458548 | 0.091 | -1.112 | 1.275 | 0.393 | 0.008 | 0.006 | 0.006 | 0 | 0.75 | 0.8 | 0.82 | 0.78 | 0.81 | 0.57 |
| 6 | 47163013 | rs9473027 | -0.086 | -0.936 | 1.492 | 0.422 | 0.021 | -0.024 | -0.024 | 0 | 0.06 | 0.19 | 0.11 | 0 | 0.06 | 0.04 |
| 6 | 47163788 | rs2144022 | -1.151 | -1.215 | 1.533 | 0.482 | -0.033 | 0.014 | 0.014 | 0 | 0.4 | 0.42 | 0.51 | 0.54 | 0.25 | 0.32 |
| 6 | 47164010 | rs6918746 | 1.681 | -1.19 | 1.545 | 0.469 | 0.033 | 0.022 | 0.022 | 0 | 0.47 | 0.37 | 0.37 | 0.32 | 0.62 | 0.38 |
| 6 | 47167753 | rs6458551 | -0.788 | -1.109 | 1.472 | 0.471 | 0.001 | -0.001 | -0.001 | 0 | 0.88 | 0.83 | 0.89 | 0.86 | 0.87 | 0.72 |
| 6 | 47168040 | rs1884046 | -0.342 | -1.107 | 1.476 | 0.466 | 0.006 | -0.005 | -0.005 | 0 | 0.85 | 0.78 | 0.85 | 0.86 | 0.79 | 0.71 |
| 6 | 47168437 | rs1884048 | -0.797 | -1.107 | 1.454 | 0.469 | 0.001 | -0.001 | -0.001 | 0 | 0.88 | 0.83 | 0.89 | 0.86 | 0.87 | 0.72 |
| 6 | 47171810 | rs9473028 | 1.642 | -1.19 | 1.543 | 0.434 | 0.021 | 0.017 | 0.017 | 0 | 0.47 | 0.31 | 0.43 | 0.32 | 0.6 | 0.39 |
| 6 | 47177720 | rs7742235 | -0.252 | -1.065 | 1.401 | 0.59 | 0.032 | 0.03 | 0.03 | 0 | 0.2 | 0.12 | 0.11 | 0.08 | 0.12 | 0.21 |
| 6 | 47187915 | rs4629675 | 0.76 | -1.061 | 1.495 | 0.616 | 0.005 | -0.001 | -0.001 | 0 | 0.15 | 0.1 | 0.29 | 0.44 | 0.08 | 0.11 |
| 6 | 47192438 | rs9367268 | 0.255 | -0.79 | 1.48 | 0.224 | 0.013 | -0.021 | -0.021 | 0 | 0.81 | 0.65 | 0.78 | 0.84 | 0.81 | 0.68 |
| 6 | 47194617 | rs6919828 | -1.83 | -0.698 | 1.119 | 0.506 | 0.066 | -0.026 | -0.026 | 0.737 | 0.06 | 0.24 | 0.06 | 0 | 0.09 | 0.05 |
| 6 | 47205912 | rs2236039 | -1.611 | -1.178 | 1.098 | 0.785 | -0.007 | 0 | 0 | 0.02 | 0.44 | 0.5 | 0.54 | 0.58 | 0.32 | 0.36 |
| 6 | 47213667 | rs9463313 | -1.407 | -1.127 | 0.702 | 0.863 | 0.036 | -0.006 | -0.006 | 1.391 | 0.42 | 0.62 | 0.51 | 0.58 | 0.24 | 0.35 |
| 6 | 47214717 | rs1884310 | -0.911 | -0.404 | 1.251 | 1.336 | -0.031 | -0.061 | -0.061 | 0 | 0.11 | 0.32 | 0.12 | 0.01 | 0.04 | 0.08 |
| 6 | 47227915 | rs2281449 | -0.771 | 0.337 | -0.277 | 2.035 | 0 | -0.029 | -0.029 | 0 | 0.17 | 0.1 | 0.32 | 0.5 | 0.16 | 0.13 |
| 6 | 47229626 | rs714373 | -1.037 | 0.266 | -0.029 | 1.81 | 0 | 0.004 | 0.004 | 0.027 | 0.32 | 0.64 | 0.31 | 0.48 | 0.36 | 0.33 |
| 6 | 47230615 | rs2103868 | -0.518 | 0.47 | -0.232 | 2.216 | -0.034 | -0.042 | -0.042 | 0 | 0.07 | 0.01 | 0.04 | 0.23 | 0.11 | 0.14 |
| 6 | 47231741 | rs9473045 | -1.854 | -0.179 | 0.727 | 1.906 | -0.001 | 0.008 | 0.008 | 0 | 0.35 | 0.64 | 0.31 | 0.43 | 0.35 | 0.31 |
| 6 | 47236880 | rs2295261 | -1.502 | -0.142 | 0.823 | 1.954 | -0.034 | 0.001 | 0.001 | 0 | 0.25 | 0.17 | 0.23 | 0.25 | 0.36 | 0.26 |
| 6 | 47243798 | rs993612 | -2.327 | -0.606 | 0.223 | 2.197 | -0.032 | 0.1 | 0.1 | 0 | 0.51 | 0.9 | 0.6 | 0.7 | 0.47 | 0.46 |
| 6 | 47243999 | rs16875844 | 0.088 | -0.372 | 0.226 | 2.156 | -0.018 | -0.044 | -0.044 | 0 | 0.08 | 0.01 | 0.23 | 0.24 | 0.08 | 0.08 |
| 6 | 47253631 | rs11758366 | -0.035 | -0.163 | 0.491 | 1.696 | 0.001 | -0.001 | -0.001 | 0 | 0.13 | 0.14 | 0.13 | 0.07 | 0.21 | 0.12 |
| 6 | 47265841 | rs12525548 | -1.599 | -0.226 | 0.548 | 1.746 | 0.01 | -0.01 | -0.01 | 0 | 0.3 | 0.62 | 0.26 | 0.24 | 0.36 | 0.26 |
| 6 | 47272689 | rs2295259 | -1.069 | -0.633 | -0.146 | 2.16 | 0.013 | -0.067 | -0.067 | 0.074 | 0.15 | 0.05 | 0.33 | 0.46 | 0.11 | 0.2 |
| 6 | 47273413 | rs2295257 | -0.227 | -0.17 | -0.05 | 2.085 | -0.041 | -0.04 | -0.04 | 0 | 0.07 | 0.01 | 0.06 | 0.17 | 0.14 | 0.12 |
| 6 | 47275663 | rs2236035 | -1.158 | -0.402 | 0.067 | 1.931 | 0.016 | -0.046 | -0.046 | 0 | 0.3 | 0.15 | 0.21 | 0.59 | 0.26 | 0.38 |
| 6 | 47287769 | rs7744364 | -1.849 | -0.809 | -0.243 | 1.43 | 0.028 | 0.077 | 0.077 | 0 | 0.57 | 0.86 | 0.54 | 0.73 | 0.39 | 0.5 |
| 6 | 47309111 | rs6901823 | -1.097 | -0.363 | 0.299 | 1.625 | -0.004 | 0.009 | 0.009 | 0 | 0.32 | 0.35 | 0.25 | 0.54 | 0.3 | 0.38 |
| 6 | 47310107 | rs10456568 | -0.345 | -0.171 | -0.213 | 1.366 | -0.006 | 0.016 | 0.016 | 0 | 0.11 | 0.05 | 0.05 | 0 | 0.1 | 0.04 |
| 6 | 47310938 | rs9369674 | -1.032 | -0.292 | -0.215 | 1.799 | 0.01 | -0.039 | -0.039 | 0 | 0.28 | 0.19 | 0.21 | 0.47 | 0.25 | 0.31 |
| 6 | 47311005 | rs9296540 | -1.399 | -0.132 | 0.042 | 1.781 | -0.003 | -0.018 | -0.018 | 0.907 | 0.35 | 0.29 | 0.26 | 0.46 | 0.36 | 0.34 |
| 6 | 47311254 | rs9968823 | -1.154 | -0.708 | 0.131 | 1.721 | 0.016 | 0.015 | 0.015 | 0 | 0.25 | 0.19 | 0.12 | 0.14 | 0.15 | 0.18 |
| 6 | 47311304 | rs9296541 | -1.337 | -0.607 | 0.127 | 2.133 | 0.069 | 0.008 | 0.008 | 0 | 0.41 | 0.76 | 0.52 | 0.6 | 0.25 | 0.37 |
| 6 | 47311798 | rs6458559 | -1.779 | -0.497 | 1.073 | 2.708 | -0.049 | 0.302 | 0.302 | 0 | 0.12 | 0.33 | 0.3 | 0.52 | 0.03 | 0.19 |
| 6 | 47311898 | rs10948347 | -1.424 | -0.287 | 0.669 | 2.597 | 0.023 | -0.015 | -0.015 | 0 | 0.1 | 0.06 | 0.29 | 0.58 | 0.04 | 0.24 |
| 6 | 47314809 | rs9369675 | -1.226 | -0.31 | 1.233 | 2.632 | -0.023 | 0.131 | 0.131 | 0 | 0.08 | 0.15 | 0.31 | 0.6 | 0.03 | 0.14 |
| 6 | 47317925 | rs9395247 | -0.574 | 0.703 | 0.904 | 1.667 | 0 | 0 | 0 | 0 | 0.18 | 0.17 | 0.2 | 0.25 | 0.32 | 0.31 |
| 6 | 47323254 | rs9369677 | -1.027 | 0.603 | 0.853 | 1.647 | 0.011 | -0.003 | -0.003 | 0 | 0.09 | 0.05 | 0.32 | 0.6 | 0.06 | 0.15 |
| 6 | 47325161 | rs9381536 | -1.129 | 0.609 | 0.892 | 1.645 | 0.007 | 0.01 | 0.01 | 0 | 0.1 | 0.06 | 0.32 | 0.6 | 0.06 | 0.15 |
| 6 | 47329155 | rs9395250 | -0.723 | 0.387 | 0.692 | 1.279 | 0.037 | -0.085 | -0.085 | 0.407 | 0.18 | 0.04 | 0.35 | 0.58 | 0.09 | 0.2 |
| 6 | 47332204 | rs9381538 | -0.94 | 0.981 | 1.184 | 0.889 | 0.036 | -0.096 | -0.096 | 0 | 0.19 | 0.04 | 0.36 | 0.6 | 0.09 | 0.2 |
| 6 | 47338091 | rs6938470 | -0.825 | 1.044 | 1.841 | 0.322 | 0.201 | 0.043 | 0.043 | 1.68 | 0.06 | 0.39 | 0.08 | 0 | 0.11 | 0.01 |
| 6 | 47340783 | rs6458563 | -1.216 | 1.085 | 1.245 | 0.09 | -0.034 | 0.115 | 0.115 | 0 | 0.39 | 0.18 | 0.58 | 0.9 | 0.42 | 0.5 |
| 6 | 47345847 | rs6458564 | -1.552 | 0.747 | 1.006 | 0.17 | 0.008 | 0.073 | 0.073 | 0 | 0.48 | 0.66 | 0.66 | 0.91 | 0.53 | 0.53 |
| 6 | 47346320 | rs9369682 | -1.483 | 0.772 | 0.931 | 0.183 | 0.003 | 0.022 | 0.022 | 0 | 0.47 | 0.63 | 0.66 | 0.91 | 0.53 | 0.51 |
| 6 | 47361788 | rs6458566 | -1.22 | 0.718 | 0.827 | 0.149 | 0.007 | -0.046 | -0.046 | 0.014 | 0.43 | 0.5 | 0.63 | 0.84 | 0.5 | 0.5 |
| 6 | 47362455 | rs12661015 | -0.112 | 0.748 | 0.925 | 0.593 | -0.001 | 0.006 | 0.006 | 0 | 0.17 | 0.19 | 0.38 | 0.66 | 0.16 | 0.16 |
| 6 | 47363213 | rs4715004 | 1.937 | 0.435 | 0.767 | -0.36 | 0.007 | 0.127 | 0.127 | 0 | 0.43 | 0.2 | 0.27 | 0.12 | 0.44 | 0.4 |
| 6 | 47364957 | rs4715005 | 1.111 | 0.936 | 1.017 | -0.464 | -0.04 | -0.11 | -0.11 | 0.015 | 0.71 | 0.51 | 0.81 | 0.93 | 0.81 | 0.74 |
| 6 | 47369202 | rs4288204 | 2.191 | 0.952 | 0.959 | -0.484 | 0.008 | 0.128 | 0.128 | 0 | 0.43 | 0.2 | 0.27 | 0.12 | 0.44 | 0.4 |
| 6 | 47370334 | rs4711874 | 1.862 | 0.847 | 1.288 | -0.661 | 0.017 | 0.058 | 0.058 | 0 | 0.48 | 0.63 | 0.32 | 0.12 | 0.45 | 0.41 |
| 6 | 47373447 | rs1931826 | 2.019 | 0.744 | 1.584 | -0.625 | -0.004 | -0.02 | -0.02 | 0 | 0.91 | 0.46 | 0.95 | 1 | 0.99 | 0.97 |
| 6 | 47379843 | rs9369686 | 0.374 | 1.008 | 1.235 | -0.528 | -0.049 | -0.042 | -0.042 | 0.75 | 0.08 | 0.01 | 0.15 | 0.14 | 0.2 | 0.18 |
| 6 | 47382378 | rs2210711 | 0.552 | 0.737 | 1.646 | -0.423 | 0.007 | -0.003 | -0.003 | 0 | 0.72 | 0.69 | 0.44 | 0.26 | 0.63 | 0.66 |

| | | | | | | | | | | | | | | | | |
|---|---|---|---|---|---|---|---|---|---|---|---|---|---|---|---|---|
| 6 | 47387672 | rs9473100 | -1.738 | 0.662 | 1.752 | -0.411 | 0.048 | 0.039 | 0.039 | 4.651 | 0.05 | 0.39 | 0.02 | 0 | 0 | 0.03 |
| 6 | 47390536 | rs9296551 | 0.556 | 0.788 | 1.176 | -0.472 | 0.005 | -0.002 | -0.002 | 0 | 0.72 | 0.69 | 0.44 | 0.26 | 0.63 | 0.66 |
| 6 | 47397746 | rs2184397 | 1.56 | 0.511 | 0.695 | -0.66 | -0.004 | 0.162 | 0.162 | 0 | 0.51 | 0.25 | 0.27 | 0.16 | 0.5 | 0.41 |
| 6 | 47414282 | rs7775178 | -0.983 | 0.679 | 0.951 | -0.591 | 0.014 | -0.014 | -0.014 | 0 | 0.34 | 0.39 | 0.66 | 0.75 | 0.43 | 0.39 |
| 6 | 47445017 | rs1004173 | -0.504 | 1.006 | 0.826 | -0.549 | -0.016 | 0.004 | 0.004 | 0.013 | 0.15 | 0.15 | 0.22 | 0.12 | 0.25 | 0.21 |
| 6 | 47452270 | rs9296559 | -0.504 | 1.026 | 0.845 | -0.549 | -0.014 | 0.003 | 0.003 | 0.013 | 0.15 | 0.16 | 0.22 | 0.12 | 0.25 | 0.23 |
| 6 | 47466833 | rs9369704 | 1.119 | 1.093 | 1.375 | -0.69 | -0.026 | -0.004 | -0.004 | 0 | 0.87 | 0.74 | 0.89 | 0.9 | 0.93 | 0.88 |
| 6 | 47469136 | rs12523687 | -0.747 | 1.083 | 1.406 | -0.695 | 0.004 | 0 | 0 | 1.419 | 0.11 | 0.06 | 0.09 | 0.1 | 0.07 | 0.12 |
| 6 | 47494759 | rs9296564 | -0.725 | 1.072 | 1.594 | -0.726 | 0.03 | -0.011 | -0.011 | 0 | 0.17 | 0.3 | 0.23 | 0.12 | 0.26 | 0.23 |
| 6 | 47527763 | rs13191654 | 0.35 | 1.44 | 1.302 | -1.064 | -0.004 | -0.003 | -0.003 | 0 | 0.19 | 0.16 | 0.43 | 0.62 | 0.24 | 0.19 |
| 6 | 47547820 | rs10498774 | 0.28 | 1.441 | 1.324 | -1.064 | -0.001 | 0.003 | 0.003 | 0 | 0.19 | 0.17 | 0.43 | 0.62 | 0.24 | 0.19 |
| 6 | 47549629 | rs3818866 | 1.896 | 0.416 | 0.091 | -2.279 | -0.075 | 0.073 | 0.073 | 1.191 | 0.27 | 0.02 | 0.14 | 0.11 | 0.29 | 0.17 |
| 6 | 47578974 | rs12661844 | 0.766 | 0.726 | 0.567 | -1.781 | -0.001 | 0.003 | 0.003 | 0 | 0.19 | 0.17 | 0.43 | 0.62 | 0.24 | 0.19 |
| 6 | 47594002 | rs1043276 | 0.443 | 0.716 | 0.549 | -1.828 | -0.004 | -0.015 | -0.015 | 0.288 | 0.68 | 0.5 | 0.48 | 0.28 | 0.7 | 0.69 |
| 6 | 47595970 | rs923148 | 0.028 | 0.941 | 0.353 | -1.795 | 0.07 | -0.034 | -0.034 | 0 | 0.06 | 0.12 | 0.1 | 0 | 0.17 | 0.03 |
| 6 | 47596016 | rs923147 | 0.801 | 0.839 | 0.285 | -1.645 | -0.007 | -0.012 | -0.012 | 0 | 0.18 | 0.13 | 0.43 | 0.59 | 0.22 | 0.17 |
| 6 | 47597241 | rs2046465 | 0.619 | 0.802 | 0.355 | -1.745 | 0.005 | -0.02 | -0.02 | 0 | 0.67 | 0.58 | 0.44 | 0.28 | 0.66 | 0.69 |
| 6 | 47598125 | rs9395288 | 0.097 | 0.863 | 0.218 | -1.82 | 0.029 | -0.008 | -0.008 | 0 | 0.16 | 0.27 | 0.22 | 0.13 | 0.23 | 0.24 |
| 6 | 47599978 | rs16876120 | -0.468 | 0.72 | 0.137 | -1.677 | 0.004 | -0.003 | -0.003 | 0.181 | 0.13 | 0.05 | 0.1 | 0.13 | 0.12 | 0.15 |
| 6 | 47623323 | rs6916790 | 0.156 | 0.848 | 0.266 | -1.739 | 0.015 | -0.008 | -0.008 | 0.07 | 0.7 | 0.55 | 0.47 | 0.3 | 0.68 | 0.73 |
| 6 | 47626685 | rs6935668 | 0.176 | 0.822 | 0.26 | -1.752 | 0.013 | -0.009 | -0.009 | 3.114 | 0.7 | 0.54 | 0.47 | 0.3 | 0.68 | 0.72 |
| 6 | 47626833 | rs12201065 | 1.551 | 0.816 | -0.169 | -1.921 | 0.043 | 0.232 | 0.232 | 0.441 | 0.48 | 0.16 | 0.23 | 0.17 | 0.43 | 0.45 |
| 6 | 47627027 | rs4711895 | -0.118 | 0.847 | -0.08 | -1.846 | 0.014 | -0.014 | -0.014 | 0 | 0.2 | 0.28 | 0.24 | 0.13 | 0.25 | 0.27 |
| 6 | 47631622 | rs2396829 | 0.521 | 0.978 | -0.02 | -1.849 | 0.011 | -0.012 | -0.012 | 0.648 | 0.66 | 0.46 | 0.42 | 0.3 | 0.62 | 0.71 |
| 6 | 47646842 | rs6907125 | 0.181 | 0.92 | 0.086 | -1.786 | 0.021 | -0.009 | -0.009 | 1.001 | 0.68 | 0.47 | 0.44 | 0.31 | 0.63 | 0.74 |
| 6 | 47649573 | rs10807371 | 0.266 | 0.913 | 0.111 | -1.789 | 0.012 | -0.008 | -0.008 | 0.089 | 0.68 | 0.44 | 0.43 | 0.31 | 0.63 | 0.74 |
| 6 | 47649694 | rs9381594 | 0.147 | 0.912 | 0.105 | -1.79 | 0.022 | -0.009 | -0.009 | 3.841 | 0.69 | 0.48 | 0.44 | 0.31 | 0.63 | 0.75 |
| 6 | 47652617 | rs6900620 | 0.181 | 0.913 | 0.106 | -1.79 | 0.021 | -0.009 | -0.009 | 3.047 | 0.68 | 0.48 | 0.44 | 0.31 | 0.63 | 0.74 |
| 6 | 47653629 | rs10948375 | -0.094 | 0.872 | -0.045 | -1.858 | 0.006 | -0.011 | -0.011 | 1.684 | 0.2 | 0.29 | 0.19 | 0.16 | 0.19 | 0.27 |
| 6 | 47658627 | rs13194549 | 1.853 | 0.543 | -0.738 | -2.137 | -0.064 | 0.083 | 0.083 | 0.818 | 0.26 | 0.01 | 0.15 | 0.09 | 0.26 | 0.17 |
| 6 | 47664399 | rs1013501 | -0.245 | 0.929 | 0.318 | -1.521 | -0.012 | -0.001 | -0.001 | 0 | 0.28 | 0.22 | 0.53 | 0.69 | 0.37 | 0.25 |
| 6 | 47664649 | rs1013500 | 0.301 | 0.907 | 0.523 | -1.533 | 0.028 | -0.012 | -0.012 | 0 | 0.68 | 0.54 | 0.44 | 0.31 | 0.63 | 0.73 |
| 6 | 47669444 | rs1023082 | 0.326 | 0.927 | 0.441 | -1.523 | 0.022 | -0.011 | -0.011 | 0 | 0.68 | 0.49 | 0.44 | 0.31 | 0.63 | 0.74 |
| 6 | 47680450 | rs871654 | 0.801 | 0.921 | 0.364 | -1.261 | -0.003 | 0.003 | 0.003 | 0 | 0.19 | 0.18 | 0.47 | 0.56 | 0.25 | 0.12 |
| 6 | 47681494 | rs6916562 | 0.576 | 0.897 | 0.442 | -1.293 | 0.017 | 0.032 | 0.032 | 0.022 | 0.87 | 0.68 | 0.9 | 0.87 | 0.89 | 0.86 |
| 6 | 47682604 | rs9369738 | 0.618 | 0.893 | 0.44 | -1.295 | 0.015 | 0.03 | 0.03 | 0 | 0.87 | 0.68 | 0.9 | 0.87 | 0.89 | 0.85 |
| 6 | 47685655 | rs12198900 | 1.95 | 0.809 | -0.026 | -1.444 | 0.02 | 0.204 | 0.204 | 0.069 | 0.45 | 0.12 | 0.22 | 0.15 | 0.43 | 0.45 |
| 6 | 47686235 | rs11754756 | 0.876 | 0.854 | -0.031 | -1.334 | -0.002 | 0.004 | 0.004 | 0 | 0.19 | 0.17 | 0.47 | 0.56 | 0.25 | 0.12 |
| 6 | 47691208 | rs1360156 | 0.864 | 0.899 | -0.261 | -1.378 | -0.044 | 0.074 | 0.074 | 0 | 0.55 | 0.18 | 0.3 | 0.29 | 0.54 | 0.59 |
| 6 | 47702371 | rs9395316 | -0.073 | 0.709 | 0.044 | -1.534 | 0.009 | -0.009 | -0.009 | 0.22 | 0.68 | 0.45 | 0.4 | 0.29 | 0.64 | 0.71 |
| 6 | 47704736 | rs9381606 | -0.129 | 0.651 | 0.248 | -1.234 | 0.002 | -0.013 | -0.013 | 0 | 0.2 | 0.29 | 0.17 | 0.15 | 0.2 | 0.29 |
| 6 | 47709067 | rs12661182 | -0.326 | 0.714 | -0.335 | -0.663 | -0.001 | 0.002 | 0.002 | 0 | 0.32 | 0.3 | 0.57 | 0.72 | 0.35 | 0.31 |
| 6 | 47711600 | rs1890061 | -0.385 | 0.67 | -0.604 | -1.096 | 0.003 | -0.026 | -0.026 | 0.004 | 0.68 | 0.47 | 0.74 | 0.86 | 0.62 | 0.66 |
| 6 | 47711725 | rs9473167 | 0.308 | 0.455 | -1.071 | -1.478 | -0.001 | -0.009 | -0.009 | 0 | 0.27 | 0.32 | 0.14 | 0.13 | 0.27 | 0.3 |
| 6 | 47720824 | rs2039141 | -0.842 | 0.416 | -0.712 | -1.631 | 0.001 | -0.043 | -0.043 | 0.088 | 0.68 | 0.62 | 0.77 | 0.87 | 0.66 | 0.67 |
| 6 | 47722814 | rs7454987 | -0.092 | 0.361 | -1.162 | -0.87 | -0.035 | 0.035 | 0.035 | 0 | 0.28 | 0.14 | 0.45 | 0.16 | 0.34 | 0.14 |
| 6 | 47724996 | rs16876481 | 1.293 | 0.279 | -1.054 | -0.786 | 0.004 | 0.006 | 0.006 | 0 | 0.14 | 0.1 | 0.18 | 0.61 | 0.06 | 0.26 |
| 6 | 47732178 | rs1329521 | -0.193 | 0.53 | -0.975 | -1.512 | -0.012 | 0.05 | 0.05 | 0 | 0.41 | 0.28 | 0.63 | 0.76 | 0.43 | 0.36 |
| 6 | 47732716 | rs10485317 | 0.774 | 0.199 | -0.817 | -1.12 | -0.003 | 0.016 | 0.016 | 0 | 0.18 | 0.16 | 0.19 | 0.6 | 0.1 | 0.25 |
| 6 | 47739785 | rs2396835 | 0.188 | 0.359 | -0.987 | -1.613 | 0.035 | 0.022 | 0.022 | 0 | 0.36 | 0.25 | 0.31 | 0.76 | 0.3 | 0.51 |
| 6 | 47742527 | rs6912824 | 1.215 | 0.53 | -0.507 | -2.136 | 0.004 | -0.014 | -0.014 | 0 | 0.13 | 0.12 | 0.05 | 0 | 0.05 | 0.1 |
| 6 | 47748241 | rs1890063 | 0.303 | 0.808 | -0.259 | -2.431 | 0.045 | -0.11 | -0.11 | 0 | 0.24 | 0.52 | 0.18 | 0.06 | 0.31 | 0.2 |
| 6 | 47749728 | rs7741473 | -0.017 | 0.977 | -0.353 | -2.546 | 0 | -0.026 | -0.026 | 0.244 | 0.71 | 0.75 | 0.83 | 0.93 | 0.74 | 0.76 |
| 6 | 47754498 | rs1329543 | 0.494 | 1.028 | -0.079 | -2.473 | 0.008 | 0.042 | 0.042 | 0 | 0.5 | 0.4 | 0.68 | 0.87 | 0.5 | 0.55 |
| 6 | 47755080 | rs9296582 | 0.533 | 0.941 | -0.112 | -2.436 | 0 | -0.003 | -0.003 | 0 | 0.3 | 0.29 | 0.17 | 0.06 | 0.26 | 0.27 |
| 6 | 47757853 | rs9296583 | 0.51 | 0.846 | -0.168 | -2.38 | -0.001 | -0.027 | -0.027 | 1.442 | 0.29 | 0.38 | 0.17 | 0.06 | 0.25 | 0.27 |
| 6 | 47760306 | rs16876536 | -0.152 | 0.895 | -0.383 | -2.37 | 0.008 | -0.016 | -0.016 | 0 | 0.14 | 0.16 | 0.11 | 0.07 | 0.18 | 0.13 |
| 6 | 47765351 | rs9296584 | 0.523 | 0.908 | -0.508 | -2.354 | -0.004 | 0.184 | 0.184 | 0 | 0.52 | 0.22 | 0.68 | 0.87 | 0.5 | 0.58 |
| 6 | 47772079 | rs9381616 | 1.399 | 0.965 | -0.685 | -2.186 | -0.042 | 0.017 | 0.017 | 0 | 0.16 | 0.01 | 0.08 | 0.12 | 0.18 | 0.24 |
| 6 | 47776901 | rs10485320 | -1.774 | 0.984 | 0.284 | -1.703 | -0.004 | 0.01 | 0.01 | 0.181 | 0.28 | 0.21 | 0.18 | 0.21 | 0.31 | 0.3 |
| 6 | 47777529 | rs10485321 | -0.336 | 0.974 | 0.242 | -1.707 | -0.008 | 0.007 | 0.007 | 0 | 0.12 | 0.07 | 0.1 | 0.07 | 0.17 | 0.12 |
| 6 | 47777737 | rs10485322 | -0.233 | 0.975 | 0.239 | -1.707 | -0.009 | 0.007 | 0.007 | 0 | 0.12 | 0.06 | 0.1 | 0.07 | 0.17 | 0.12 |
| 6 | 47785831 | rs6907470 | -0.616 | 0.957 | 0.251 | -1.707 | 0.011 | -0.002 | -0.002 | 0.033 | 0.13 | 0.07 | 0.05 | 0.13 | 0.08 | 0.16 |
| 6 | 47788531 | rs7740768 | -1.879 | 0.973 | 0.213 | -1.706 | -0.002 | 0.004 | 0.004 | 0.339 | 0.29 | 0.29 | 0.18 | 0.21 | 0.31 | 0.3 |
| 6 | 47792575 | rs1328971 | -0.365 | 1.086 | -0.025 | -1.559 | -0.008 | 0.006 | 0.006 | 0 | 0.12 | 0.09 | 0.1 | 0.08 | 0.17 | 0.11 |
| 6 | 47793291 | rs9381619 | 1.485 | 0.907 | 0.084 | -1.506 | -0.003 | 0.068 | 0.068 | 0 | 0.42 | 0.21 | 0.63 | 0.73 | 0.43 | 0.45 |
| 6 | 47799673 | rs17624241 | -0.32 | 0.808 | 0.521 | -0.998 | 0 | 0 | 0 | 0 | 0.09 | 0.09 | 0.07 | 0.05 | 0.08 | 0.08 |
| 6 | 47804436 | rs16876602 | -0.304 | 0.8 | 0.522 | -0.997 | 0 | -0.001 | -0.001 | 0 | 0.09 | 0.09 | 0.07 | 0.05 | 0.08 | 0.08 |
| 6 | 47819564 | rs564351 | -0.572 | 0.94 | 0.744 | -1.034 | 0.024 | -0.028 | -0.028 | 0.447 | 0.86 | 0.73 | 0.9 | 0.94 | 0.81 | 0.89 |

| | | | | | | | | | | | | | | | | |
|---|---|---|---|---|---|---|---|---|---|---|---|---|---|---|---|---|
| 6 | 47824080 | rs2039122 | 1.512 | 0.887 | 0.208 | -1.077 | -0.022 | 0.095 | 0.095 | 0 | 0.46 | 0.22 | 0.68 | 0.71 | 0.47 | 0.49 |
| 6 | 47836871 | rs1360077 | 1.301 | 1.209 | 1.002 | -0.355 | 0.003 | -0.018 | -0.018 | 0.165 | 0.69 | 0.52 | 0.79 | 0.75 | 0.68 | 0.68 |
| 6 | 47841072 | rs1328981 | -0.308 | 1.207 | 1.063 | -0.363 | -0.002 | -0.003 | -0.003 | 0 | 0.08 | 0.09 | 0.05 | 0.06 | 0.07 | 0.06 |
| 6 | 47843173 | rs9369749 | -0.576 | 1.202 | 1.072 | -0.371 | 0.003 | -0.017 | -0.017 | 0 | 0.08 | 0.04 | 0.03 | 0.14 | 0.06 | 0.14 |
| 6 | 47845151 | rs6939078 | 0.999 | 1.136 | 0.929 | -0.326 | 0.003 | -0.018 | -0.018 | 1.385 | 0.69 | 0.52 | 0.79 | 0.74 | 0.68 | 0.68 |
| 6 | 47845813 | rs1887899 | 1.124 | 1.129 | 0.975 | -0.33 | -0.001 | -0.019 | -0.019 | 0.017 | 0.68 | 0.46 | 0.79 | 0.75 | 0.68 | 0.68 |
| 6 | 47852268 | rs1410829 | -0.351 | 1.133 | 0.93 | -0.321 | -0.002 | -0.003 | -0.003 | 0 | 0.08 | 0.09 | 0.05 | 0.06 | 0.07 | 0.06 |
| 6 | 47864221 | rs7742880 | 0.933 | 1.139 | 0.947 | -0.265 | 0.006 | -0.003 | -0.003 | 0 | 0.69 | 0.55 | 0.8 | 0.71 | 0.68 | 0.68 |
| 6 | 47867477 | rs9968847 | 0.326 | 1.141 | 0.969 | -0.481 | 0.011 | -0.01 | -0.01 | 1.107 | 0.76 | 0.65 | 0.84 | 0.8 | 0.73 | 0.75 |
| 6 | 47870083 | rs6458595 | 0.641 | 1.151 | 0.588 | -0.4 | 0.002 | -0.013 | -0.013 | 0.112 | 0.67 | 0.42 | 0.78 | 0.75 | 0.66 | 0.65 |
| 6 | 47878053 | rs10948390 | 0.811 | 1.267 | -0.141 | -0.429 | -0.013 | 0.023 | 0.023 | 0 | 0.65 | 0.29 | 0.75 | 0.76 | 0.59 | 0.63 |
| 6 | 47878091 | rs10948391 | 0.671 | 1.238 | -0.229 | -0.422 | 0.002 | -0.004 | -0.004 | 0.152 | 0.66 | 0.42 | 0.76 | 0.76 | 0.59 | 0.63 |
| 6 | 47881668 | rs9357565 | 0.796 | 1.185 | -0.259 | -0.422 | -0.005 | 0.01 | 0.01 | 0 | 0.65 | 0.34 | 0.75 | 0.76 | 0.59 | 0.63 |
| 6 | 47885007 | rs9381629 | 0.467 | 0.746 | -0.334 | -0.355 | 0.006 | -0.016 | -0.016 | 0 | 0.68 | 0.49 | 0.77 | 0.76 | 0.66 | 0.67 |
| 6 | 47885216 | rs9369756 | 0.549 | 0.833 | -0.506 | -0.311 | 0 | -0.007 | -0.007 | 1.087 | 0.67 | 0.41 | 0.77 | 0.76 | 0.66 | 0.64 |
| 6 | 47888700 | rs9381631 | -0.898 | 0.4 | -0.606 | -0.379 | -0.004 | -0.006 | -0.006 | 0 | 0.29 | 0.32 | 0.18 | 0.22 | 0.23 | 0.28 |
| 6 | 47891704 | rs12209155 | 0.439 | 0.296 | -1.436 | -0.893 | -0.03 | 0.011 | 0.011 | 0 | 0.15 | 0.01 | 0.06 | 0.09 | 0.14 | 0.17 |
| 6 | 47891731 | rs16876825 | 0.199 | 0.197 | -1.399 | -1.039 | -0.006 | 0.029 | 0.029 | 0 | 0.13 | 0.09 | 0.08 | 0.04 | 0.11 | 0.1 |
| 6 | 47897482 | rs1325694 | -0.087 | 0.094 | -1.376 | -1.091 | 0 | 0.013 | 0.013 | 0 | 0.15 | 0.12 | 0.08 | 0.04 | 0.11 | 0.11 |
| 6 | 47909938 | rs9349437 | 1.21 | -0.336 | -0.841 | -1.46 | -0.053 | -0.081 | -0.081 | 1.856 | 0.78 | 0.57 | 0.8 | 0.88 | 0.83 | 0.76 |
| 6 | 47916225 | rs1974051 | -1.501 | -0.391 | -0.765 | -1.465 | -0.04 | -0.05 | -0.05 | 0 | 0.12 | 0.33 | 0.13 | 0.07 | 0.08 | 0.15 |
| 6 | 47935234 | rs12192531 | -0.255 | -0.838 | -1.134 | -1.599 | 0.031 | 0.062 | 0.062 | 0 | 0.26 | 0.13 | 0.09 | 0.1 | 0.17 | 0.25 |
| 6 | 47941339 | rs2753188 | 0.789 | -0.861 | -1.124 | -1.559 | 0.006 | 0.008 | 0.008 | 0 | 0.67 | 0.66 | 0.77 | 0.82 | 0.72 | 0.66 |
| 6 | 47946823 | rs10214575 | 0.897 | -0.141 | -0.694 | -2.098 | 0.02 | -0.031 | -0.031 | 0 | 0.2 | 0.26 | 0.19 | 0.1 | 0.26 | 0.18 |
| 6 | 47949493 | rs1325713 | 0.511 | -0.494 | -0.964 | -2.108 | 0.016 | 0.053 | 0.053 | 0 | 0.24 | 0.07 | 0.08 | 0.1 | 0.17 | 0.23 |
| 6 | 47954482 | rs2180739 | 0.301 | -0.512 | -0.943 | -2.141 | 0.006 | 0.008 | 0.008 | 0 | 0.67 | 0.66 | 0.77 | 0.82 | 0.72 | 0.66 |
| 6 | 47963249 | rs1325707 | 0.99 | -0.473 | -0.973 | -2.113 | -0.002 | 0.036 | 0.036 | 0 | 0.2 | 0.08 | 0.08 | 0.1 | 0.15 | 0.19 |
| 6 | 47968236 | rs10456573 | 0.08 | -0.493 | -0.971 | -2.115 | 0.02 | 0.047 | 0.047 | 0 | 0.27 | 0.18 | 0.12 | 0.11 | 0.22 | 0.3 |
| 6 | 47970924 | rs9357566 | 0.681 | -0.842 | -1.15 | -2.22 | 0.001 | -0.003 | -0.003 | 0 | 0.55 | 0.55 | 0.73 | 0.77 | 0.67 | 0.57 |
| 6 | 47971082 | rs11751251 | 1.343 | -0.265 | -1.921 | -2.373 | -0.028 | 0.032 | 0.032 | 0 | 0.06 | 0.01 | 0.07 | 0 | 0.13 | 0.02 |
| 6 | 47971550 | rs12197671 | 0.976 | -0.456 | -1.865 | -2.3 | -0.007 | 0.044 | 0.044 | 0 | 0.22 | 0.07 | 0.1 | 0.1 | 0.2 | 0.21 |
| 6 | 47981365 | rs767548 | 0.862 | -0.053 | -1.662 | -1.777 | -0.046 | -0.107 | -0.107 | 0 | 0.27 | 0.03 | 0.41 | 0.62 | 0.23 | 0.33 |
| 6 | 47988311 | rs2180740 | 0.528 | -0.008 | -0.934 | -1.59 | -0.004 | -0.001 | -0.001 | 0 | 0.81 | 0.81 | 0.8 | 0.83 | 0.83 | 0.82 |
| 6 | 48005879 | rs12205869 | 0.597 | 0.278 | -1.341 | -1.719 | -0.016 | 0.093 | 0.093 | 0.854 | 0.27 | 0.07 | 0.11 | 0.1 | 0.25 | 0.23 |
| 6 | 48032148 | rs332571 | -0.902 | -0.052 | -0.744 | -1.987 | 0.027 | 0.045 | 0.045 | 0.195 | 0.43 | 0.7 | 0.33 | 0.28 | 0.37 | 0.46 |
| 6 | 48050299 | rs1556358 | -0.368 | -0.1 | -0.314 | -2.055 | -0.003 | -0.011 | -0.011 | 0 | 0.48 | 0.48 | 0.33 | 0.24 | 0.39 | 0.46 |
| 6 | 48050405 | rs6922920 | -0.175 | -0.108 | 0.108 | -1.74 | -0.02 | -0.009 | -0.009 | 0 | 0.17 | 0.25 | 0.18 | 0.13 | 0.13 | 0.19 |
| 6 | 48056902 | rs9473231 | 0.339 | 0.211 | -0.107 | -1.539 | 0.005 | -0.024 | -0.024 | 0.061 | 0.17 | 0.32 | 0.1 | 0.12 | 0.18 | 0.17 |
| 6 | 48063688 | rs9473239 | 0.38 | 0.229 | -0.228 | -1.416 | 0.006 | -0.01 | -0.01 | 0.022 | 0.17 | 0.27 | 0.09 | 0.13 | 0.18 | 0.17 |
| 6 | 48081318 | rs6458601 | -0.415 | -0.27 | -0.285 | -1.176 | -0.005 | 0 | 0 | 0 | 0.59 | 0.66 | 0.51 | 0.47 | 0.58 | 0.56 |
| 6 | 48081464 | rs9473247 | 0.207 | -0.375 | 0.142 | -0.933 | 0.048 | 0.017 | 0.017 | 0 | 0.12 | 0.35 | 0.14 | 0.13 | 0.17 | 0.17 |
| 6 | 48086158 | rs12198093 | 0.048 | -0.647 | -0.084 | -1.095 | -0.026 | 0 | 0 | 1.358 | 0.16 | 0.25 | 0.22 | 0.15 | 0.12 | 0.14 |
| 6 | 48087403 | rs12200178 | 0.015 | -0.653 | -0.086 | -1.098 | -0.027 | -0.001 | -0.001 | 0.054 | 0.16 | 0.26 | 0.22 | 0.15 | 0.12 | 0.14 |
| 6 | 48094245 | rs4715066 | -0.781 | -0.635 | -0.271 | -1.094 | -0.007 | -0.004 | -0.004 | 0 | 0.3 | 0.62 | 0.37 | 0.28 | 0.3 | 0.31 |
| 6 | 48102631 | rs13195572 | 0.257 | -0.671 | -0.242 | -1.285 | 0.036 | 0.035 | 0.035 | 0 | 0.11 | 0.25 | 0.13 | 0.16 | 0.17 | 0.2 |
| 6 | 48106015 | rs13198330 | 0.572 | -0.66 | -0.236 | -1.279 | 0.05 | 0.035 | 0.035 | 0 | 0.1 | 0.25 | 0.12 | 0.13 | 0.16 | 0.19 |
| 6 | 48116254 | rs998094 | 0.199 | -0.729 | -0.381 | -1.418 | -0.006 | 0.014 | 0.014 | 0 | 0.41 | 0.41 | 0.29 | 0.32 | 0.45 | 0.43 |
| 6 | 48151134 | rs9463378 | 0.267 | -0.746 | -0.43 | -1.492 | -0.05 | -0.042 | -0.042 | 0 | 0.2 | 0.32 | 0.2 | 0.11 | 0.1 | 0.2 |
| 6 | 48151180 | rs6926533 | 1.397 | -0.705 | -1.378 | -1.623 | -0.005 | 0.022 | 0.022 | 0 | 0.06 | 0.02 | 0.04 | 0 | 0.09 | 0.01 |
| 6 | 48156101 | rs10948402 | 1.697 | -0.843 | -1.408 | -1.601 | -0.087 | 0.09 | 0.09 | 0 | 0.26 | 0.01 | 0.15 | 0.16 | 0.27 | 0.21 |
| 6 | 48160865 | rs753849 | -1.08 | -0.981 | -1.457 | -1.716 | -0.097 | -0.045 | -0.045 | 0 | 0.67 | 0.91 | 0.52 | 0.42 | 0.55 | 0.71 |
| 6 | 48162231 | rs1150813 | 1.757 | -0.727 | -1.423 | -1.486 | -0.058 | -0.107 | -0.107 | 0 | 0.23 | 0.04 | 0.35 | 0.51 | 0.26 | 0.28 |
| 6 | 48205919 | rs11966702 | 1.289 | -0.156 | -0.908 | -1.13 | -0.086 | 0.091 | 0.091 | 0 | 0.26 | 0.01 | 0.15 | 0.16 | 0.28 | 0.21 |
| 6 | 48213344 | rs2496371 | 0.833 | -0.037 | -0.831 | -1.073 | -0.025 | -0.021 | -0.021 | 0 | 0.08 | 0.03 | 0.1 | 0.12 | 0.15 | 0.19 |
| 6 | 48214195 | rs2456913 | 0.162 | -0.399 | -0.608 | -0.865 | -0.002 | 0.018 | 0.018 | 0 | 0.46 | 0.46 | 0.32 | 0.32 | 0.45 | 0.52 |
| 6 | 48215008 | rs2496375 | -0.895 | -0.334 | -0.702 | -0.836 | -0.014 | 0.023 | 0.023 | 0 | 0.63 | 0.7 | 0.73 | 0.71 | 0.58 | 0.6 |
| 6 | 48231163 | rs2456924 | -0.3 | -0.364 | -1.144 | -0.831 | -0.008 | 0.057 | 0.057 | 0 | 0.54 | 0.64 | 0.69 | 0.68 | 0.55 | 0.49 |
| 6 | 48237736 | rs1828092 | -0.671 | 0.234 | -1.419 | -0.585 | -0.044 | 0.059 | 0.059 | 0 | 0.64 | 0.81 | 0.74 | 0.72 | 0.58 | 0.6 |
| 6 | 48247897 | rs7743104 | 0.061 | 0.53 | -0.875 | -0.051 | -0.022 | 0 | 0 | 0 | 0.07 | 0.04 | 0.09 | 0.07 | 0.14 | 0.01 |
| 6 | 48280842 | rs12208936 | -0.524 | -0.221 | -1.057 | -0.155 | 0.02 | -0.001 | -0.001 | 0 | 0.14 | 0 | 0.15 | 0.1 | 0.08 | 0.16 |
| 6 | 48281398 | rs9473289 | -0.817 | -0.452 | 0.033 | -0.142 | -0.052 | 0.003 | 0.003 | 0 | 0.15 | 0.38 | 0.17 | 0.11 | 0.1 | 0.17 |
| 6 | 48282842 | rs615396 | 0.767 | -0.465 | -0.2 | -0.271 | 0.003 | -0.021 | -0.021 | 0 | 0.93 | 0.86 | 0.93 | 0.97 | 0.93 | 0.94 |
| 6 | 48283538 | rs7742324 | 0.405 | -0.409 | -0.822 | -0.453 | -0.022 | 0.015 | 0.015 | 0.676 | 0.18 | 0.13 | 0.2 | 0.13 | 0.28 | 0.11 |
| 6 | 48287563 | rs688333 | 1.07 | -0.595 | -1.576 | -0.554 | -0.018 | -0.097 | -0.097 | 0 | 0.29 | 0.11 | 0.38 | 0.43 | 0.26 | 0.32 |
| 6 | 48287802 | rs689261 | 0.196 | -0.608 | -1.583 | -0.561 | -0.018 | 0.068 | 0.068 | 1.575 | 0.47 | 0.25 | 0.57 | 0.57 | 0.54 | 0.42 |
| 6 | 48288536 | rs584557 | 0.937 | -0.576 | -1.505 | -0.453 | -0.018 | -0.098 | -0.098 | 0 | 0.29 | 0.11 | 0.38 | 0.43 | 0.26 | 0.31 |
| 6 | 48289514 | rs1889757 | 0.336 | -0.64 | -0.986 | -0.451 | -0.038 | 0.03 | 0.03 | 0 | 0.13 | 0.34 | 0.18 | 0.11 | 0.09 | 0.16 |
| 6 | 48311449 | rs16877386 | 1.376 | -0.512 | -1.447 | -0.702 | -0.09 | -0.033 | -0.033 | 0 | 0.23 | 0.01 | 0.14 | 0.28 | 0.25 | 0.28 |
| 6 | 48319507 | rs161712 | 0.183 | -0.629 | -1.307 | -0.567 | 0.017 | 0.017 | 0.017 | 0.581 | 0.63 | 0.31 | 0.59 | 0.7 | 0.62 | 0.67 |

| | | | | | | | | | | | | | | | | |
|---|---|---|---|---|---|---|---|---|---|---|---|---|---|---|---|---|
| 6 | 48321014 | rs6921083 | 1.168 | -0.675 | -1.359 | -0.612 | -0.091 | 0.025 | 0.025 | 1.114 | 0.27 | 0.03 | 0.18 | 0.27 | 0.3 | 0.29 |
| 6 | 48321824 | rs325337 | 1.451 | -0.604 | -1.332 | -0.471 | -0.073 | -0.163 | -0.163 | 0 | 0.2 | 0.01 | 0.35 | 0.4 | 0.22 | 0.28 |
| 6 | 48329809 | rs7755638 | -0.167 | -0.666 | -0.487 | -0.498 | -0.06 | 0.018 | 0.018 | 0.185 | 0.15 | 0.45 | 0.18 | 0.12 | 0.09 | 0.17 |
| 6 | 48329893 | rs325041 | 0.718 | -0.635 | -0.557 | -0.285 | 0.017 | 0 | 0 | 0 | 0.63 | 0.31 | 0.59 | 0.66 | 0.62 | 0.67 |
| 6 | 48352572 | rs2166025 | -0.935 | -0.67 | -0.48 | -0.473 | -0.038 | -0.018 | -0.018 | 0 | 0.2 | 0.54 | 0.2 | 0.12 | 0.14 | 0.25 |
| 6 | 48367207 | rs9473350 | -0.503 | -0.237 | -0.874 | -0.335 | 0.005 | -0.041 | -0.041 | 0 | 0.07 | 0.2 | 0.04 | 0 | 0.05 | 0.09 |
| 6 | 48387129 | rs325286 | 0.299 | -0.127 | -1.539 | -0.182 | 0.015 | 0.08 | 0.08 | 0 | 0.53 | 0.23 | 0.54 | 0.7 | 0.51 | 0.55 |
| 6 | 48396334 | rs9473361 | -0.258 | 0.025 | -1.514 | -0.77 | -0.002 | 0.141 | 0.141 | 1.268 | 0.42 | 0.79 | 0.32 | 0.15 | 0.44 | 0.37 |
| 6 | 48406425 | rs11966364 | -0.829 | -0.506 | -0.86 | -0.876 | -0.031 | 0 | 0 | 0 | 0.13 | 0.23 | 0.1 | 0.08 | 0.07 | 0.14 |
| 6 | 48414233 | rs9395353 | 0.847 | -0.413 | -1.603 | -0.729 | -0.078 | -0.022 | -0.022 | 0 | 0.27 | 0.02 | 0.14 | 0.31 | 0.25 | 0.28 |
| 6 | 48423116 | rs16877484 | -0.894 | -0.63 | -0.999 | -0.847 | -0.057 | -0.01 | -0.01 | 1.077 | 0.16 | 0.37 | 0.11 | 0.08 | 0.07 | 0.15 |
| 6 | 48427979 | rs7764731 | -0.172 | -0.715 | -0.653 | -0.777 | 0 | 0.062 | 0.062 | 0 | 0.45 | 0.65 | 0.35 | 0.22 | 0.49 | 0.42 |
| 6 | 48441573 | rs16877498 | -0.24 | -0.712 | -0.499 | -0.765 | -0.032 | 0.019 | 0.019 | 0 | 0.11 | 0.29 | 0.09 | 0.08 | 0.06 | 0.13 |
| 6 | 48454655 | rs9473389 | 0.781 | -0.676 | -0.514 | -0.808 | 0.005 | -0.022 | -0.022 | 0 | 0.32 | 0.3 | 0.26 | 0.14 | 0.43 | 0.29 |
| 6 | 48480264 | rs2586827 | -0.943 | -0.675 | -0.522 | -0.808 | 0.009 | -0.032 | -0.032 | 0 | 0.68 | 0.67 | 0.74 | 0.86 | 0.57 | 0.71 |
| 6 | 48507718 | rs6908652 | 0.426 | -0.523 | -0.389 | -0.536 | 0.004 | -0.032 | -0.032 | 0.03 | 0.26 | 0.22 | 0.5 | 0.48 | 0.25 | 0.31 |
| 6 | 48513035 | rs9463425 | -1.014 | -0.633 | -0.652 | -0.923 | -0.002 | 0.009 | 0.009 | 0 | 0.54 | 0.63 | 0.69 | 0.87 | 0.44 | 0.71 |
| 6 | 48513746 | rs1838194 | -0.428 | -0.572 | -0.67 | -0.659 | -0.025 | -0.005 | -0.005 | 0 | 0.13 | 0.26 | 0.09 | 0.08 | 0.07 | 0.14 |
| 6 | 48515646 | rs9369787 | 0.383 | -0.255 | -0.877 | -0.376 | 0.003 | -0.032 | -0.032 | 0.038 | 0.26 | 0.22 | 0.5 | 0.47 | 0.25 | 0.3 |
| 6 | 48519267 | rs1361864 | -0.282 | -0.386 | -0.965 | -0.578 | -0.009 | 0.061 | 0.061 | 0 | 0.39 | 0.23 | 0.57 | 0.78 | 0.37 | 0.57 |
| 6 | 48530298 | rs12176541 | 0.806 | -0.68 | -0.794 | -0.082 | -0.002 | 0.066 | 0.066 | 0 | 0.05 | 0.01 | 0.37 | 0.47 | 0.05 | 0.21 |
| 6 | 48539324 | rs9395370 | 0.184 | -0.622 | -0.085 | -0.339 | 0.006 | 0.011 | 0.011 | 0 | 0.15 | 0.16 | 0.1 | 0.24 | 0.14 | 0.31 |
| 6 | 48562825 | rs10456593 | 0.524 | -0.68 | -0.291 | -0.37 | -0.049 | -0.067 | -0.067 | 0 | 0.12 | 0.01 | 0.09 | 0.3 | 0.12 | 0.28 |
| 6 | 48626275 | rs243132 | -1.06 | -1.235 | 0.853 | -0.444 | -0.051 | 0.289 | 0.289 | 0.012 | 0.15 | 0.48 | 0.43 | 0.48 | 0.09 | 0.25 |
| 6 | 48626970 | rs2069327 | -0.42 | -1.123 | 0.972 | -0.386 | -0.023 | 0.193 | 0.193 | 0 | 0.11 | 0.23 | 0.4 | 0.48 | 0.08 | 0.25 |
| 6 | 48649487 | rs243105 | -0.803 | -0.833 | 0.756 | -0.973 | 0 | -0.041 | -0.041 | 0 | 0.32 | 0.49 | 0.55 | 0.79 | 0.27 | 0.51 |
| 6 | 48674613 | rs10484857 | -0.135 | -1.312 | 0.61 | -1.117 | -0.01 | -0.009 | -0.009 | 0 | 0.15 | 0.25 | 0.09 | 0.09 | 0.08 | 0.16 |
| 6 | 48691652 | rs2208003 | -1.314 | -1.224 | 0.298 | -1.35 | -0.093 | 0.071 | 0.071 | 0 | 0.64 | 0.93 | 0.73 | 0.76 | 0.54 | 0.7 |
| 6 | 48696663 | rs9473468 | -0.054 | -1.317 | 1.036 | -1.399 | -0.006 | 0.034 | 0.034 | 0 | 0.44 | 0.34 | 0.62 | 0.67 | 0.43 | 0.51 |
| 6 | 48742066 | rs2495886 | 1.07 | -1.283 | 0.787 | -1.388 | -0.014 | 0.021 | 0.021 | 0 | 0.37 | 0.25 | 0.27 | 0.24 | 0.46 | 0.29 |
| 6 | 48751142 | rs1938193 | 1.109 | -1.279 | 0.713 | -1.384 | -0.031 | 0.043 | 0.043 | 0 | 0.36 | 0.2 | 0.27 | 0.24 | 0.46 | 0.3 |
| 6 | 48754886 | rs2490135 | 0.158 | -1.295 | 0.772 | -1.386 | 0.003 | 0.028 | 0.028 | 0 | 0.52 | 0.46 | 0.36 | 0.32 | 0.55 | 0.46 |
| 6 | 48764104 | rs7763556 | 0.829 | -1.199 | 0.692 | -1.358 | -0.02 | 0.041 | 0.041 | 0 | 0.38 | 0.21 | 0.3 | 0.23 | 0.48 | 0.29 |
| 6 | 48778106 | rs2997370 | -0.362 | -1.341 | 0.813 | -1.395 | 0.003 | -0.019 | -0.019 | 2.077 | 0.82 | 0.71 | 0.89 | 0.82 | 0.87 | 0.82 |
| 6 | 48784695 | rs13191385 | 1.096 | -1.333 | 0.697 | -1.4 | -0.039 | 0.068 | 0.068 | 0 | 0.38 | 0.15 | 0.29 | 0.23 | 0.48 | 0.29 |
| 6 | 48792612 | rs2495863 | 1.01 | -1.5 | 0.679 | -1.44 | 0.036 | 0.059 | 0.059 | 0.82 | 0.61 | 0.26 | 0.77 | 0.74 | 0.76 | 0.64 |
| 6 | 48807046 | rs12190429 | 0.838 | -1.517 | 0.483 | -1.453 | -0.054 | -0.021 | -0.021 | 0 | 0.13 | 0.01 | 0.08 | 0.18 | 0.13 | 0.18 |
| 6 | 48812072 | rs1938179 | 0.77 | -1.613 | 0.941 | -1.256 | 0.078 | 0.09 | 0.09 | 0 | 0.5 | 0.14 | 0.36 | 0.42 | 0.59 | 0.44 |
| 6 | 48812682 | rs7768096 | 0.83 | -1.526 | 1.025 | -1.249 | 0.074 | 0.084 | 0.084 | 0 | 0.5 | 0.17 | 0.36 | 0.42 | 0.59 | 0.44 |
| 6 | 48821707 | rs1938189 | 0.838 | -1.405 | 1.166 | -1.216 | -0.076 | -0.08 | -0.08 | 0.105 | 0.77 | 0.51 | 0.87 | 0.91 | 0.89 | 0.82 |
| 6 | 48830321 | rs1938201 | 0.964 | -1.121 | 1.006 | -0.976 | 0.02 | 0.012 | 0.012 | 0.733 | 0.94 | 0.71 | 0.95 | 1 | 0.98 | 0.97 |
| 6 | 48842605 | rs2154346 | -0.418 | -1.229 | 0.509 | -0.976 | 0 | -0.011 | -0.011 | 0 | 0.73 | 0.74 | 0.49 | 0.48 | 0.69 | 0.59 |
| 6 | 48851550 | rs10484429 | -0.252 | -1.387 | 0.584 | -1.082 | -0.007 | -0.006 | -0.006 | 0 | 0.14 | 0.19 | 0.07 | 0.09 | 0.08 | 0.16 |
| 6 | 48871002 | rs10484428 | 1.226 | -1.598 | 0.659 | -1.599 | -0.005 | 0.009 | 0.009 | 0.031 | 0.4 | 0.34 | 0.3 | 0.21 | 0.48 | 0.24 |
| 6 | 48880753 | rs6906787 | -0.28 | -1.674 | 0.598 | -1.354 | 0.005 | 0.006 | 0.006 | 2.624 | 0.17 | 0.17 | 0.08 | 0.07 | 0.09 | 0.2 |
| 6 | 48894648 | rs12211733 | -0.134 | -1.727 | 0.699 | -1.352 | 0 | -0.003 | -0.003 | 0 | 0.59 | 0.6 | 0.39 | 0.28 | 0.57 | 0.45 |
| 6 | 48914124 | rs2205869 | -0.237 | -1.726 | 0.624 | -0.945 | 0.006 | 0.006 | 0.006 | 2.829 | 0.14 | 0.06 | 0.06 | 0.08 | 0.07 | 0.12 |
| 6 | 48915976 | rs12196746 | 0.242 | -1.974 | 0.729 | -1.076 | 0.013 | 0.019 | 0.019 | 0.151 | 0.59 | 0.32 | 0.41 | 0.39 | 0.62 | 0.42 |
| 6 | 48919300 | rs760683 | -0.864 | -1.943 | 0.968 | -1.04 | 0 | 0.018 | 0.018 | 0 | 0.72 | 0.7 | 0.46 | 0.5 | 0.68 | 0.62 |
| 6 | 48920376 | rs12204534 | 1.553 | -1.971 | 0.943 | -1.077 | -0.022 | 0.059 | 0.059 | 0.278 | 0.4 | 0.27 | 0.3 | 0.21 | 0.48 | 0.24 |
| 6 | 48925998 | rs6458638 | 0.453 | -1.961 | 0.951 | -1.056 | -0.004 | -0.01 | -0.01 | 0.189 | 0.55 | 0.5 | 0.38 | 0.39 | 0.59 | 0.42 |
| 6 | 48928544 | rs16878481 | -0.49 | -1.917 | 0.627 | -1.024 | 0.002 | 0 | 0 | 1.945 | 0.17 | 0.2 | 0.08 | 0.11 | 0.08 | 0.2 |
| 6 | 48930453 | rs12213551 | 1.388 | -1.961 | 0.448 | -1.108 | -0.063 | 0.115 | 0.115 | 0.009 | 0.39 | 0.14 | 0.3 | 0.21 | 0.48 | 0.24 |
| 6 | 48964065 | rs12210871 | 1.626 | -1.955 | 0.364 | -1.104 | -0.121 | 0.118 | 0.118 | 0.663 | 0.38 | 0.05 | 0.29 | 0.21 | 0.48 | 0.24 |
| 6 | 48966452 | rs9357595 | 0.64 | -1.916 | 0.457 | -1 | -0.01 | -0.005 | -0.005 | 0 | 0.13 | 0.04 | 0.07 | 0.18 | 0.11 | 0.17 |
| 6 | 48995590 | rs4489146 | -0.77 | -1.853 | 1.002 | -0.941 | -0.001 | 0.03 | 0.03 | 0 | 0.72 | 0.62 | 0.47 | 0.5 | 0.68 | 0.62 |
| 6 | 49005412 | rs12212438 | 1.191 | -1.867 | 0.581 | -1.272 | -0.038 | 0.107 | 0.107 | 0 | 0.38 | 0.12 | 0.29 | 0.2 | 0.45 | 0.26 |
| 6 | 49019176 | rs12201977 | 1.261 | -1.892 | 0.569 | -1.284 | -0.041 | 0.105 | 0.105 | 0 | 0.37 | 0.11 | 0.28 | 0.2 | 0.45 | 0.26 |
| 6 | 49020666 | rs4591845 | -1.088 | -1.898 | 0.588 | -1.28 | -0.012 | -0.009 | -0.009 | 0.106 | 0.35 | 0.43 | 0.18 | 0.31 | 0.23 | 0.36 |
| 6 | 49020904 | rs4352681 | -1.105 | -1.893 | 0.595 | -1.277 | -0.001 | 0.03 | 0.03 | 0 | 0.72 | 0.62 | 0.47 | 0.5 | 0.68 | 0.62 |
| 6 | 49033696 | rs6917508 | -1.064 | -1.858 | 0.586 | -1.243 | -0.012 | -0.008 | -0.008 | 0 | 0.34 | 0.41 | 0.18 | 0.31 | 0.23 | 0.36 |
| 6 | 49035781 | rs6929570 | -0.894 | -1.853 | 0.695 | -1.137 | -0.036 | -0.009 | -0.009 | 0 | 0.2 | 0.41 | 0.1 | 0.13 | 0.1 | 0.19 |
| 6 | 49047862 | rs6927493 | -0.81 | -1.63 | 0.553 | -1.005 | -0.001 | 0.029 | 0.029 | 0 | 0.72 | 0.62 | 0.47 | 0.5 | 0.68 | 0.62 |
| 6 | 49081548 | rs12212993 | 0.203 | -1.605 | 0.187 | -0.859 | -0.013 | -0.025 | -0.025 | 1.096 | 0.12 | 0 | 0.05 | 0.09 | 0.08 | 0.12 |
| 6 | 49102186 | rs2496367 | -0.867 | -1.674 | -0.348 | -1.119 | -0.047 | 0.038 | 0.038 | 0 | 0.68 | 0.85 | 0.74 | 0.75 | 0.6 | 0.68 |
| 6 | 49107837 | rs9381738 | 0.459 | -1.705 | -0.279 | -0.936 | 0.005 | 0.008 | 0.008 | 0 | 0.18 | 0.11 | 0.09 | 0.17 | 0.13 | 0.19 |
| 6 | 49134412 | rs2516123 | 1.35 | -1.642 | -1.199 | -0.831 | -0.091 | -0.181 | -0.181 | 0.001 | 0.27 | 0.02 | 0.53 | 0.48 | 0.38 | 0.33 |
| 6 | 49162686 | rs2516114 | -0.624 | -1.607 | -0.988 | -0.943 | -0.051 | 0.057 | 0.057 | 0 | 0.67 | 0.84 | 0.74 | 0.76 | 0.62 | 0.66 |
| 6 | 49163193 | rs17722373 | 0.49 | -1.736 | -1.065 | -1.011 | 0.003 | 0.012 | 0.012 | 1.681 | 0.16 | 0.05 | 0.08 | 0.11 | 0.13 | 0.14 |

| | | | | | | | | | | | | | | | | |
|---|---|---|---|---|---|---|---|---|---|---|---|---|---|---|---|---|
| 6 | 49167053 | rs9381743 | 1.025 | -1.753 | -1.081 | -0.905 | -0.025 | -0.028 | -0.028 | 0 | 0.15 | 0.01 | 0.07 | 0.17 | 0.13 | 0.17 |
| 6 | 49172864 | rs990237 | 0.139 | -1.752 | -0.907 | -0.906 | 0.022 | 0.068 | 0.068 | 0 | 0.49 | 0.69 | 0.66 | 0.66 | 0.5 | 0.52 |
| 6 | 49173407 | rs2516112 | 1.201 | -1.602 | -0.617 | -0.822 | -0.005 | 0.021 | 0.021 | 0 | 0.32 | 0.37 | 0.56 | 0.49 | 0.36 | 0.34 |
| 6 | 49175210 | rs2516111 | -1.177 | -1.401 | -0.918 | -1.044 | -0.072 | 0.048 | 0.048 | 0 | 0.76 | 0.92 | 0.78 | 0.82 | 0.68 | 0.83 |
| 6 | 49190966 | rs4478408 | 1.056 | -1.634 | -0.602 | -0.73 | -0.012 | 0.054 | 0.054 | 0 | 0.33 | 0.55 | 0.58 | 0.47 | 0.36 | 0.3 |
| 6 | 49191974 | rs9473511 | 0.061 | -1.821 | -0.716 | -0.766 | 0.013 | -0.005 | -0.005 | 0 | 0.1 | 0.06 | 0.04 | 0.09 | 0.05 | 0.21 |
| 6 | 49198849 | rs1117179 | 1.105 | -1.658 | -0.637 | -0.765 | 0.002 | 0.027 | 0.027 | 0 | 0.33 | 0.4 | 0.57 | 0.47 | 0.39 | 0.3 |
| 6 | 49211262 | rs10456618 | 0.458 | -1.752 | -0.887 | -0.856 | 0.008 | 0.033 | 0.033 | 0.007 | 0.14 | 0.05 | 0.08 | 0.06 | 0.11 | 0.13 |
| 6 | 49212193 | rs492254 | -0.4 | -1.708 | -0.829 | -0.7 | -0.036 | 0.024 | 0.024 | 0 | 0.64 | 0.81 | 0.73 | 0.71 | 0.61 | 0.6 |
| 6 | 49218272 | rs11753211 | 1.017 | -1.669 | -0.891 | -0.685 | -0.084 | 0.037 | 0.037 | 0 | 0.26 | 0.08 | 0.22 | 0.2 | 0.34 | 0.19 |
| 6 | 49225193 | rs574321 | -1.231 | -1.71 | -0.631 | -0.422 | -0.011 | 0.035 | 0.035 | 0 | 0.68 | 0.55 | 0.42 | 0.55 | 0.64 | 0.71 |
| 6 | 49228005 | rs12191536 | 0.288 | -1.662 | -0.807 | -0.368 | 0.007 | 0.024 | 0.024 | 0.01 | 0.14 | 0.06 | 0.07 | 0.06 | 0.12 | 0.13 |
| 6 | 49235559 | rs10807405 | -0.573 | -1.725 | -0.796 | -0.396 | 0.005 | -0.003 | -0.003 | 0 | 0.3 | 0.1 | 0.15 | 0.26 | 0.25 | 0.31 |
| 6 | 49252022 | rs9349482 | 0.727 | -1.574 | -0.679 | -0.369 | -0.009 | -0.03 | -0.03 | 0 | 0.15 | 0.02 | 0.07 | 0.2 | 0.13 | 0.18 |
| 6 | 49264967 | rs9395475 | 0.438 | -1.604 | -0.569 | -0.339 | 0.002 | -0.011 | -0.011 | 0 | 0.17 | 0.12 | 0.09 | 0.2 | 0.14 | 0.2 |
| 6 | 49280970 | rs11752910 | 0.575 | -1.623 | -0.557 | -0.349 | -0.029 | -0.054 | -0.054 | 0.321 | 0.15 | 0.01 | 0.08 | 0.2 | 0.14 | 0.2 |
| 6 | 49287351 | rs12189573 | 0.702 | -1.37 | -0.672 | -0.458 | -0.028 | 0.015 | 0.015 | 0 | 0.32 | 0.19 | 0.24 | 0.27 | 0.36 | 0.38 |
| 6 | 49288384 | rs540785 | -0.386 | -1.353 | -0.563 | -0.449 | 0.008 | 0.003 | 0.003 | 0 | 0.51 | 0.55 | 0.34 | 0.47 | 0.51 | 0.58 |
| 6 | 49319025 | rs543805 | 0.237 | -1.349 | -0.53 | -0.445 | -0.008 | 0.004 | 0.004 | 0 | 0.19 | 0.26 | 0.1 | 0.2 | 0.14 | 0.2 |
| 6 | 49321223 | rs11970105 | 0.793 | -1.434 | -0.759 | -0.504 | -0.004 | 0.025 | 0.025 | 0 | 0.13 | 0.03 | 0.07 | 0.06 | 0.11 | 0.1 |
| 6 | 49329253 | rs3930746 | 0.736 | -1.45 | -0.755 | -0.514 | -0.015 | 0.009 | 0.009 | 0 | 0.32 | 0.25 | 0.25 | 0.26 | 0.36 | 0.38 |
| 6 | 49329899 | rs10484961 | 0.723 | -1.359 | -0.852 | -0.452 | -0.003 | 0.026 | 0.026 | 0 | 0.13 | 0.03 | 0.07 | 0.06 | 0.11 | 0.1 |
| 6 | 49337199 | rs503422 | -0.328 | -1.215 | -0.762 | -0.392 | 0.007 | 0.002 | 0.002 | 0 | 0.52 | 0.55 | 0.34 | 0.47 | 0.5 | 0.58 |
| 6 | 49352768 | rs9369892 | 0.556 | -1.065 | -1.159 | -0.72 | -0.008 | -0.014 | -0.014 | 0 | 0.15 | 0.06 | 0.08 | 0.18 | 0.14 | 0.17 |
| 6 | 49353357 | rs9473541 | 0.754 | -1.071 | -1.295 | -0.832 | -0.03 | 0.038 | 0.038 | 0 | 0.3 | 0.17 | 0.22 | 0.22 | 0.34 | 0.41 |
| 6 | 49354553 | rs6902615 | 0.603 | -0.864 | -1.374 | -0.679 | -0.005 | -0.011 | -0.011 | 0.098 | 0.15 | 0.07 | 0.08 | 0.18 | 0.14 | 0.17 |
| 6 | 49361187 | rs9381781 | 0.657 | -0.856 | -1.474 | -0.675 | -0.018 | -0.026 | -0.026 | 0.038 | 0.15 | 0.05 | 0.08 | 0.18 | 0.14 | 0.17 |
| 6 | 49367872 | rs10807409 | -0.145 | -0.89 | -1.455 | -0.695 | -0.01 | -0.003 | -0.003 | 0 | 0.27 | 0.12 | 0.15 | 0.25 | 0.25 | 0.27 |
| 6 | 49378668 | rs12528625 | 1.204 | -1.168 | -1.742 | -0.853 | -0.065 | -0.137 | -0.137 | 0 | 0.33 | 0.02 | 0.55 | 0.54 | 0.39 | 0.29 |
| 6 | 49388855 | rs4585547 | -0.017 | -1.166 | -1.612 | -0.881 | -0.007 | -0.001 | -0.001 | 0.012 | 0.27 | 0.14 | 0.16 | 0.25 | 0.25 | 0.25 |
| 6 | 49398115 | rs9381784 | 1.053 | -1.11 | -1.686 | -0.918 | -0.068 | -0.125 | -0.125 | 0 | 0.35 | 0.02 | 0.56 | 0.54 | 0.39 | 0.29 |
| 6 | 49420560 | rs6458692 | -0.095 | -1.023 | -1.567 | -0.853 | -0.005 | -0.003 | -0.003 | 0.04 | 0.28 | 0.12 | 0.15 | 0.27 | 0.25 | 0.25 |
| 6 | 49431569 | rs3806988 | 0.686 | -1.126 | -1.131 | -0.925 | 0.026 | -0.054 | -0.054 | 0.26 | 0.31 | 0.44 | 0.23 | 0.21 | 0.33 | 0.42 |
| 6 | 49439805 | rs4267943 | 0.76 | -1.122 | -1.137 | -0.922 | 0.026 | -0.05 | -0.05 | 0.051 | 0.3 | 0.42 | 0.23 | 0.21 | 0.33 | 0.42 |
| 6 | 49446509 | rs2251115 | 0.654 | -1.105 | -1.19 | -0.8 | 0.002 | -0.001 | -0.001 | 0.388 | 0.41 | 0.41 | 0.61 | 0.52 | 0.41 | 0.33 |
| 6 | 49459978 | rs2501968 | 0.367 | -1.046 | -1.183 | -0.803 | 0.008 | 0.006 | 0.006 | 0 | 0.44 | 0.41 | 0.67 | 0.58 | 0.43 | 0.38 |
| 6 | 49466686 | rs7755833 | 0.579 | -0.57 | -1.293 | -0.298 | -0.005 | 0.012 | 0.012 | 0 | 0.12 | 0.06 | 0.07 | 0.07 | 0.11 | 0.09 |
| 6 | 49479972 | rs7453050 | 0.409 | -0.542 | -1.367 | -0.39 | -0.014 | 0.013 | 0.013 | 0 | 0.7 | 0.38 | 0.43 | 0.45 | 0.75 | 0.62 |
| 6 | 49484201 | rs9381793 | 1.084 | -0.262 | -2.003 | -0.41 | -0.076 | 0.017 | 0.017 | 0 | 0.3 | 0.02 | 0.19 | 0.21 | 0.34 | 0.18 |
| 6 | 49485266 | rs9367358 | 0.489 | 0.273 | -1.626 | 0.048 | -0.045 | -0.125 | -0.125 | 1.503 | 0.16 | 0.01 | 0.45 | 0.5 | 0.2 | 0.26 |
| 6 | 49491305 | rs9473583 | 0.296 | 0.416 | -1.11 | -0.388 | 0.003 | -0.014 | -0.014 | 0 | 0.25 | 0.25 | 0.15 | 0.17 | 0.27 | 0.34 |
| 6 | 49494933 | rs9395504 | -0.661 | 0.172 | -1.133 | 0.333 | 0.002 | 0.006 | 0.006 | 0.492 | 0.4 | 0.33 | 0.62 | 0.54 | 0.38 | 0.46 |
| 6 | 49511969 | rs17805607 | -0.044 | -0.131 | -1.534 | 0.953 | -0.005 | -0.12 | -0.12 | 0 | 0.16 | 0.01 | 0.41 | 0.35 | 0.17 | 0.22 |
| 6 | 49513767 | rs13196352 | 0.765 | -0.075 | -1.786 | 0.103 | 0.002 | 0.061 | 0.061 | 0 | 0.27 | 0.11 | 0.17 | 0.11 | 0.27 | 0.11 |
| 6 | 49522865 | rs9367361 | -0.602 | -0.258 | -1.142 | 0.27 | -0.014 | 0.041 | 0.041 | 0.626 | 0.32 | 0.39 | 0.53 | 0.47 | 0.29 | 0.4 |
| 6 | 49534329 | rs9473594 | 0.133 | -0.351 | -1.486 | -0.32 | 0.011 | 0.062 | 0.062 | 0 | 0.51 | 0.34 | 0.33 | 0.28 | 0.48 | 0.35 |
| 6 | 49536392 | rs10948512 | 1.141 | -0.523 | -1.741 | -0.567 | -0.05 | 0.087 | 0.087 | 0 | 0.3 | 0.03 | 0.17 | 0.17 | 0.32 | 0.14 |
| 6 | 49537171 | rs9473595 | -0.213 | -0.372 | -1.507 | -0.494 | -0.016 | -0.044 | -0.044 | 2.009 | 0.2 | 0.3 | 0.15 | 0.11 | 0.16 | 0.21 |
| 6 | 49538807 | rs9381801 | 0.703 | -0.423 | -1.802 | -0.343 | -0.064 | -0.044 | -0.044 | 0 | 0.18 | 0.1 | 0.12 | 0.25 | 0.27 | 0.32 |
| 6 | 49541086 | rs6458702 | 0.66 | -0.555 | -1.479 | -0.391 | 0 | -0.001 | -0.001 | 0 | 0.58 | 0.41 | 0.71 | 0.64 | 0.56 | 0.47 |
| 6 | 49543614 | rs6927133 | 0.871 | -0.698 | -1.506 | -0.391 | -0.022 | 0.036 | 0.036 | 0.582 | 0.13 | 0.35 | 0.09 | 0.18 | 0.07 | 0.14 |
| 6 | 49545213 | rs12176464 | 0.586 | -0.349 | -1.25 | -0.348 | -0.061 | -0.043 | -0.043 | 0.73 | 0.18 | 0.1 | 0.12 | 0.25 | 0.27 | 0.32 |
| 6 | 49547357 | rs4566875 | 0.708 | -0.096 | -1.243 | -0.356 | 0.001 | -0.002 | -0.002 | 0 | 0.6 | 0.43 | 0.72 | 0.64 | 0.6 | 0.48 |
| 6 | 49547409 | rs1480620 | 0.411 | 0.417 | -1.484 | -0.023 | -0.046 | -0.045 | -0.045 | 0.044 | 0.17 | 0.09 | 0.11 | 0.25 | 0.24 | 0.31 |
| 6 | 49552535 | rs4711930 | 0.303 | 0.03 | -1.211 | -0.233 | -0.001 | -0.001 | -0.001 | 0.262 | 0.06 | 0.05 | 0.06 | 0.06 | 0.06 | 0.12 |
| 6 | 49559793 | rs10948514 | 0.379 | 0.401 | -1.366 | -0.116 | -0.04 | -0.026 | -0.026 | 0 | 0.14 | 0.08 | 0.15 | 0.2 | 0.25 | 0.3 |
| 6 | 49560784 | rs9395515 | 0.52 | -0.162 | -1.624 | -0.302 | -0.05 | 0.028 | 0.028 | 0 | 0.44 | 0.11 | 0.33 | 0.37 | 0.57 | 0.42 |
| 6 | 49575141 | rs9463517 | 0.236 | -0.105 | -0.689 | -0.159 | 0.024 | -0.012 | -0.012 | 1.635 | 0.08 | 0.27 | 0.08 | 0.03 | 0.05 | 0.04 |
| 6 | 49576183 | rs1985222 | 0.521 | 0.006 | -1.014 | -0.202 | -0.047 | -0.02 | -0.02 | 0 | 0.18 | 0.08 | 0.17 | 0.2 | 0.26 | 0.29 |
| 6 | 49577215 | rs3799675 | 0.521 | 0.006 | -1.014 | -0.202 | -0.047 | -0.02 | -0.02 | 0.022 | 0.18 | 0.08 | 0.17 | 0.2 | 0.26 | 0.29 |
| 6 | 49580194 | rs10485291 | 0.622 | -0.174 | -0.866 | -0.492 | -0.006 | 0.002 | 0.002 | 0 | 0.05 | 0.08 | 0.04 | 0.07 | 0.02 | 0.14 |
| 6 | 49580558 | rs1480617 | 0.485 | -0.538 | -0.611 | -0.799 | -0.006 | 0.017 | 0.017 | 0 | 0.33 | 0.33 | 0.5 | 0.52 | 0.29 | 0.35 |
| 6 | 49583871 | rs16879521 | 0.7 | -0.48 | -0.575 | -0.763 | -0.01 | 0.004 | 0.004 | 0 | 0.05 | 0.11 | 0.04 | 0.07 | 0.02 | 0.14 |
| 6 | 49586285 | rs1471540 | 0.941 | -0.689 | -0.291 | -0.583 | 0.006 | 0.013 | 0.013 | 0.244 | 0.17 | 0.2 | 0.11 | 0.24 | 0.15 | 0.18 |
| 6 | 49587227 | rs12524967 | 0.745 | -0.757 | -0.464 | -0.629 | -0.012 | 0.036 | 0.036 | 1.983 | 0.21 | 0.32 | 0.47 | 0.32 | 0.19 | 0.22 |
| 6 | 49592233 | rs7760223 | 0.273 | -0.751 | -0.368 | -0.647 | -0.043 | -0.048 | -0.048 | 0.471 | 0.24 | 0.12 | 0.22 | 0.37 | 0.31 | 0.34 |
| 6 | 49592298 | rs9463520 | 1.697 | -0.757 | -0.285 | -0.61 | -0.015 | 0.067 | 0.067 | 0.479 | 0.15 | 0.28 | 0.42 | 0.29 | 0.14 | 0.18 |
| 6 | 49599462 | rs6934867 | 0.189 | -0.678 | -0.182 | -0.769 | -0.002 | 0.029 | 0.029 | 0.521 | 0.23 | 0.34 | 0.15 | 0.3 | 0.17 | 0.31 |
| 6 | 49602760 | rs13198157 | 0.297 | -0.723 | -0.306 | -0.865 | -0.005 | 0.03 | 0.03 | 0 | 0.53 | 0.57 | 0.33 | 0.35 | 0.47 | 0.42 |

| | | | | | | | | | | | | | | | | |
|---|---|---|---|---|---|---|---|---|---|---|---|---|---|---|---|---|
| 6 | 49609134 | rs2518100 | -0.63 | -0.731 | -0.167 | -0.976 | 0.003 | -0.018 | -0.018 | 0 | 0.57 | 0.58 | 0.73 | 0.91 | 0.6 | 0.69 |
| 6 | 49616512 | rs2518103 | 0.071 | -0.722 | -0.187 | -0.947 | -0.008 | 0.013 | 0.013 | 0.098 | 0.39 | 0.34 | 0.62 | 0.67 | 0.45 | 0.51 |
| 6 | 49619240 | rs9395519 | 1.006 | -0.633 | -0.27 | -0.925 | -0.051 | -0.061 | -0.061 | 0.092 | 0.17 | 0.06 | 0.17 | 0.36 | 0.26 | 0.27 |
| 6 | 49620573 | rs13194498 | 1.045 | -0.75 | 0.042 | -1.235 | 0.008 | 0.059 | 0.059 | 0 | 0.32 | 0.23 | 0.21 | 0.09 | 0.31 | 0.25 |
| 6 | 49620882 | rs10080998 | -0.97 | -0.714 | -0.096 | -1.215 | -0.001 | 0.027 | 0.027 | 0.2 | 0.92 | 0.97 | 0.96 | 1 | 0.92 | 0.93 |
| 6 | 49630176 | rs13197952 | 0.748 | -0.698 | -0.029 | -1.179 | -0.004 | -0.039 | -0.039 | 0 | 0.36 | 0.39 | 0.23 | 0.09 | 0.31 | 0.25 |
| 6 | 49637027 | rs1570063 | 0.412 | -0.686 | 0.002 | -1.019 | -0.02 | -0.034 | -0.034 | 0 | 0.68 | 0.51 | 0.84 | 0.78 | 0.77 | 0.62 |
| 6 | 49641484 | rs17635077 | 0.938 | -0.606 | 0.081 | -0.962 | 0.001 | 0.003 | 0.003 | 1.805 | 0.16 | 0.22 | 0.07 | 0.18 | 0.13 | 0.17 |
| 6 | 49642763 | rs7746868 | 0.242 | -0.48 | -0.013 | -0.883 | -0.016 | 0.028 | 0.028 | 1.471 | 0.4 | 0.34 | 0.66 | 0.73 | 0.48 | 0.51 |
| 6 | 49652613 | rs13202956 | 0.918 | -0.599 | -0.047 | -0.95 | -0.003 | 0.045 | 0.045 | 1.052 | 0.42 | 0.26 | 0.23 | 0.22 | 0.42 | 0.28 |
| 6 | 49655649 | rs471198 | -0.714 | -0.601 | -0.084 | -0.981 | -0.004 | 0.025 | 0.025 | 0.076 | 0.92 | 0.97 | 0.96 | 1 | 0.92 | 0.93 |
| 6 | 49656252 | rs802071 | 0.092 | -0.721 | 0.179 | -0.929 | -0.02 | 0.019 | 0.019 | 0.663 | 0.08 | 0.2 | 0.05 | 0.08 | 0.02 | 0.15 |
| 6 | 49659871 | rs360559 | -0.429 | -0.67 | 0.002 | -1.05 | -0.002 | 0.085 | 0.085 | 1.016 | 0.5 | 0.69 | 0.74 | 0.78 | 0.51 | 0.66 |
| 6 | 49660418 | rs12624 | -0.446 | -0.67 | 0.002 | -1.05 | -0.002 | 0.087 | 0.087 | 1.127 | 0.5 | 0.69 | 0.74 | 0.78 | 0.51 | 0.66 |
| 6 | 49673195 | rs681283 | 0.24 | -0.501 | -0.018 | -1.283 | -0.004 | 0.049 | 0.049 | 0.679 | 0.81 | 0.91 | 0.88 | 0.94 | 0.84 | 0.74 |
| 6 | 49675544 | rs189664 | 0.938 | -0.461 | 0.134 | -1.106 | 0.005 | 0.004 | 0.004 | 0 | 0.6 | 0.5 | 0.65 | 0.5 | 0.58 | 0.42 |
| 6 | 49676107 | rs12203129 | 0.791 | -0.34 | 0.019 | -0.997 | -0.002 | 0.003 | 0.003 | 0 | 0.06 | 0.05 | 0.04 | 0.02 | 0.05 | 0.04 |
| 6 | 49694656 | rs6902472 | 1.765 | -0.496 | 0.04 | -0.939 | 0.005 | -0.009 | -0.009 | 0 | 0.18 | 0.13 | 0.41 | 0.28 | 0.17 | 0.21 |
| 6 | 49695232 | rs510770 | -0.631 | -0.503 | -0.043 | -1.002 | 0 | 0.031 | 0.031 | 1.329 | 0.86 | 0.94 | 0.91 | 0.95 | 0.89 | 0.79 |
| 6 | 49696473 | rs488132 | 1.328 | -0.619 | -0.077 | -1.208 | -0.002 | 0.007 | 0.007 | 1.903 | 0.16 | 0.24 | 0.06 | 0.19 | 0.11 | 0.12 |
| 6 | 49701439 | rs1864312 | 1.464 | -0.688 | -0.166 | -1.309 | 0.002 | 0.021 | 0.021 | 0 | 0.41 | 0.26 | 0.58 | 0.31 | 0.47 | 0.31 |
| 6 | 49701523 | rs495335 | -0.138 | -0.677 | -0.007 | -1.294 | 0.001 | 0.004 | 0.004 | 0 | 0.75 | 0.74 | 0.76 | 0.55 | 0.69 | 0.64 |
| 6 | 49703857 | rs597544 | 0.833 | -0.68 | 0 | -1.295 | -0.012 | 0.001 | 0.001 | 2.899 | 0.19 | 0.37 | 0.08 | 0.19 | 0.11 | 0.12 |
| 6 | 49704546 | rs509699 | -0.089 | -0.547 | -0.256 | -1.236 | 0.001 | 0.003 | 0.003 | 0 | 0.75 | 0.74 | 0.76 | 0.55 | 0.7 | 0.64 |
| 6 | 49710970 | rs13190702 | 1.604 | -0.566 | -0.253 | -1.237 | -0.005 | 0.016 | 0.016 | 0 | 0.25 | 0.17 | 0.18 | 0.03 | 0.3 | 0.1 |
| 6 | 49717197 | rs13210122 | 1.625 | -0.562 | -0.271 | -1.235 | -0.012 | 0.049 | 0.049 | 0 | 0.25 | 0.14 | 0.17 | 0.03 | 0.3 | 0.1 |
| 6 | 49722717 | rs6935473 | -0.504 | -0.548 | -0.235 | -1.181 | -0.001 | 0.003 | 0.003 | 0 | 0.81 | 0.8 | 0.8 | 0.57 | 0.74 | 0.68 |
| 6 | 49731027 | rs699986 | 0.53 | -0.626 | -0.156 | -1.122 | -0.022 | 0.043 | 0.043 | 0 | 0.35 | 0.46 | 0.47 | 0.46 | 0.28 | 0.33 |
| 6 | 49740169 | rs11755024 | 1.562 | -0.602 | -0.25 | -1.15 | -0.015 | 0.064 | 0.064 | 0 | 0.25 | 0.12 | 0.17 | 0.03 | 0.3 | 0.1 |
| 6 | 49744632 | rs1429599 | 2.058 | -0.649 | -0.121 | -1.144 | 0.015 | -0.041 | -0.041 | 0 | 0.18 | 0.08 | 0.4 | 0.27 | 0.17 | 0.21 |
| 6 | 49746284 | rs10948526 | 1.505 | -0.558 | -0.185 | -1.129 | -0.005 | 0.012 | 0.012 | 0 | 0.25 | 0.17 | 0.18 | 0.03 | 0.3 | 0.1 |
| 6 | 49752742 | rs696718 | 0.341 | -0.515 | -0.385 | -1.071 | -0.009 | 0.011 | 0.011 | 0 | 0.37 | 0.57 | 0.48 | 0.46 | 0.28 | 0.33 |
| 6 | 49756705 | rs11758714 | 1.543 | -0.054 | -0.34 | -0.893 | -0.071 | 0.126 | 0.126 | 0 | 0.22 | 0.02 | 0.16 | 0.03 | 0.29 | 0.1 |
| 6 | 49766526 | rs500032 | 0.726 | -0.152 | 0.128 | -0.504 | -0.003 | 0.006 | 0.006 | 2.874 | 0.17 | 0.24 | 0.06 | 0.19 | 0.11 | 0.12 |
| 6 | 49771994 | rs6905366 | 0.3 | -0.307 | -0.072 | -0.555 | 0.01 | -0.004 | -0.004 | 0.234 | 0.34 | 0.34 | 0.5 | 0.34 | 0.29 | 0.43 |
| 6 | 49778621 | rs9473663 | 0.441 | -0.213 | 0.033 | -0.588 | 0.014 | -0.012 | -0.012 | 0 | 0.37 | 0.29 | 0.54 | 0.37 | 0.34 | 0.46 |
| 6 | 49785074 | rs1429601 | 0.476 | 0.252 | 0.34 | -0.443 | 0.02 | 0.08 | 0.08 | 0.293 | 0.76 | 0.53 | 0.77 | 0.59 | 0.74 | 0.67 |
| 6 | 49786871 | rs780444 | 0.134 | 0.402 | 0.317 | -0.355 | 0 | 0.178 | 0.178 | 0 | 0.11 | 0.23 | 0.14 | 0.41 | 0.1 | 0.27 |
| 6 | 49801612 | rs10498779 | -0.172 | 0.698 | 0.084 | -0.701 | 0.022 | -0.031 | -0.031 | 0 | 0.08 | 0.16 | 0.07 | 0 | 0.13 | 0.05 |
| 6 | 49804908 | rs3799688 | 0.816 | 0.472 | 0.181 | -0.822 | 0.003 | 0.012 | 0.012 | 4.332 | 0.13 | 0.15 | 0.05 | 0.2 | 0.1 | 0.09 |
| 6 | 49805985 | rs12200099 | 0.481 | 0.523 | -0.362 | -1.471 | -0.043 | 0.067 | 0.067 | 0 | 0.31 | 0.21 | 0.24 | 0.04 | 0.42 | 0.17 |
| 6 | 49823814 | rs1970456 | 1.311 | 0.307 | -0.584 | -1.414 | 0.014 | -0.049 | -0.049 | 0.056 | 0.26 | 0.14 | 0.44 | 0.34 | 0.23 | 0.27 |
| 6 | 49823987 | rs1417491 | 0.365 | 0.262 | -0.667 | -1.476 | -0.015 | 0.057 | 0.057 | 2.418 | 0.3 | 0.38 | 0.21 | 0.57 | 0.23 | 0.39 |
| 6 | 49826120 | rs780781 | -0.273 | 0.491 | -0.868 | -1.72 | -0.013 | 0.032 | 0.032 | 0.871 | 0.88 | 0.96 | 0.92 | 0.95 | 0.89 | 0.8 |
| 6 | 49837603 | rs9473680 | -0.267 | 0.927 | -0.395 | -1.414 | -0.022 | 0.016 | 0.016 | 0 | 0.31 | 0.25 | 0.24 | 0.04 | 0.42 | 0.14 |
| 6 | 49869448 | rs7764242 | 0.012 | 0.881 | 0.136 | -0.593 | 0 | 0.062 | 0.062 | 1.509 | 0.26 | 0.34 | 0.18 | 0.51 | 0.2 | 0.35 |
| 6 | 49871812 | rs2792833 | 0.782 | 0.652 | -0.272 | -0.667 | 0.004 | -0.001 | -0.001 | 0 | 0.46 | 0.26 | 0.63 | 0.48 | 0.5 | 0.44 |
| 6 | 49872143 | rs13193597 | 0.031 | 0.308 | -0.06 | -0.447 | -0.009 | 0.052 | 0.052 | 0 | 0.1 | 0.01 | 0.07 | 0.02 | 0.1 | 0.05 |
| 6 | 49891258 | rs4715153 | -0.561 | 0.393 | 0.128 | -0.118 | -0.011 | -0.012 | -0.012 | 0.201 | 0.58 | 0.69 | 0.6 | 0.52 | 0.51 | 0.48 |
| 6 | 49896050 | rs7747376 | -0.057 | 0.355 | 0.371 | -0.333 | 0.023 | -0.007 | -0.007 | 1.409 | 0.46 | 0.36 | 0.51 | 0.51 | 0.37 | 0.4 |
| 6 | 49909756 | rs4715160 | -0.068 | 0.262 | 0.338 | -0.381 | 0.029 | -0.017 | -0.017 | 0 | 0.45 | 0.29 | 0.5 | 0.51 | 0.37 | 0.4 |
| 6 | 49915709 | rs9473701 | -0.34 | 0.249 | 0.444 | -0.398 | 0 | -0.01 | -0.01 | 0.069 | 0.08 | 0.08 | 0.07 | 0.01 | 0.13 | 0.08 |
| 6 | 49920177 | rs1337823 | -0.74 | 0.278 | 0.4 | -0.356 | 0.001 | 0.002 | 0.002 | 0.479 | 0.56 | 0.52 | 0.58 | 0.52 | 0.5 | 0.48 |
| 6 | 49935357 | rs1337820 | -0.219 | 0.18 | -0.059 | -0.163 | 0.032 | -0.025 | -0.025 | 0 | 0.44 | 0.23 | 0.49 | 0.51 | 0.37 | 0.4 |
| 6 | 49939494 | rs1324527 | -0.268 | 0.181 | -0.024 | -0.159 | 0.022 | -0.012 | -0.012 | 1.506 | 0.44 | 0.31 | 0.5 | 0.51 | 0.37 | 0.4 |
| 6 | 49942438 | rs1415732 | 0.006 | 0.097 | -0.686 | -0.17 | 0.02 | -0.082 | -0.082 | 0 | 0.35 | 0.13 | 0.45 | 0.51 | 0.32 | 0.33 |
| 6 | 49985939 | rs12196609 | 0.008 | -0.041 | 0.004 | -1.038 | -0.019 | -0.076 | -0.076 | 1.336 | 0.17 | 0.23 | 0.09 | 0 | 0.11 | 0.1 |
| 6 | 49989657 | rs10948540 | 0.464 | 0.18 | 0.185 | -1.031 | -0.016 | -0.009 | -0.009 | 0 | 0.05 | 0.02 | 0.13 | 0.06 | 0.11 | 0.06 |
| 6 | 50003645 | rs13199564 | 0.343 | -0.472 | -0.639 | -1.988 | 0.021 | 0.147 | 0.147 | 0.919 | 0.23 | 0.05 | 0.08 | 0 | 0.16 | 0.13 |
| 6 | 50011796 | rs628337 | -0.558 | -0.948 | 0.02 | -2.09 | 0.007 | -0.028 | -0.028 | 0 | 0.28 | 0.7 | 0.15 | 0.02 | 0.16 | 0.18 |
| 6 | 50020423 | rs12208674 | 0.731 | -0.903 | 0.339 | -2.08 | -0.025 | -0.096 | -0.096 | 0 | 0.16 | 0.29 | 0.1 | 0 | 0.11 | 0.1 |
| 6 | 50022132 | rs7764185 | 1.144 | -0.896 | 0.289 | -2.074 | -0.025 | -0.017 | -0.017 | 0.263 | 0.05 | 0 | 0.13 | 0.05 | 0.11 | 0.06 |
| 6 | 50029391 | rs2038763 | 0.694 | -1.062 | 0.486 | -2.063 | -0.018 | -0.074 | -0.074 | 0 | 0.16 | 0.23 | 0.09 | 0 | 0.11 | 0.1 |
| 6 | 50041852 | rs606106 | -0.608 | -1.119 | 0.479 | -2.058 | -0.018 | -0.062 | -0.062 | 0 | 0.12 | 0.47 | 0.05 | 0.01 | 0.05 | 0.07 |
| 6 | 50046852 | rs9381838 | 0.37 | -1.124 | 0.621 | -2.103 | 0.008 | -0.027 | -0.027 | 0.045 | 0.72 | 0.31 | 0.85 | 0.98 | 0.84 | 0.82 |
| 6 | 50055171 | rs932741 | 0.403 | -1.152 | 0.578 | -1.939 | 0.009 | -0.001 | -0.001 | 0 | 0.72 | 0.31 | 0.81 | 0.94 | 0.84 | 0.77 |
| 6 | 50077707 | rs2147372 | 0.456 | -0.753 | 0.732 | -1.508 | -0.032 | -0.074 | -0.074 | 0 | 0.75 | 0.43 | 0.85 | 0.95 | 0.85 | 0.8 |
| 6 | 50080330 | rs1324523 | -0.116 | 0.016 | 0.3 | -1.225 | -0.016 | -0.007 | -0.007 | 0 | 0.33 | 0.27 | 0.29 | 0.39 | 0.41 | 0.33 |
| 6 | 50081302 | rs6902413 | 0.769 | -0.21 | -0.218 | -1.363 | 0.015 | 0.007 | 0.007 | 0 | 0.55 | 0.38 | 0.49 | 0.53 | 0.6 | 0.6 |

| | | | | | | | | | | | | | | | | |
|---|---|---|---|---|---|---|---|---|---|---|---|---|---|---|---|---|
| 6 | 50082351 | rs9349513 | 0.942 | -0.029 | -0.726 | -0.88 | -0.05 | -0.099 | -0.099 | 0.486 | 0.13 | 0.03 | 0.35 | 0.51 | 0.18 | 0.22 |
| 6 | 50084574 | rs766512 | 0.274 | -0.127 | -0.359 | -0.978 | 0.006 | -0.023 | -0.023 | 1.211 | 0.72 | 0.79 | 0.57 | 0.53 | 0.73 | 0.73 |
| 6 | 50114275 | rs6921517 | 0.374 | -0.456 | -0.145 | -0.872 | 0.009 | -0.003 | -0.003 | 0 | 0.58 | 0.63 | 0.51 | 0.54 | 0.62 | 0.62 |
| 6 | 50123474 | rs10948547 | 0.778 | -0.494 | 0.162 | -0.926 | 0.01 | 0.003 | 0.003 | 0 | 0.52 | 0.47 | 0.54 | 0.56 | 0.62 | 0.62 |
| 6 | 50125321 | rs7750808 | 0.848 | -0.221 | 0.197 | -0.919 | 0.008 | -0.009 | -0.009 | 0 | 0.14 | 0.11 | 0.07 | 0 | 0.08 | 0.11 |
| 6 | 50135089 | rs1856857 | -0.652 | -0.214 | 0.225 | -0.928 | -0.014 | -0.075 | -0.075 | 0 | 0.83 | 0.75 | 0.9 | 1 | 0.89 | 0.85 |
| 6 | 50142856 | rs12210112 | 0.99 | -0.225 | 0.19 | -0.905 | 0.008 | -0.007 | -0.007 | 0 | 0.13 | 0.09 | 0.06 | 0 | 0.08 | 0.1 |
| 6 | 50159392 | rs1572887 | -0.84 | -0.232 | 0.294 | -0.909 | -0.029 | -0.076 | -0.076 | 0 | 0.85 | 0.77 | 0.92 | 1 | 0.92 | 0.9 |
| 6 | 50171209 | rs6914600 | 0.597 | -0.645 | 0.162 | -0.891 | 0.008 | 0.001 | 0.001 | 0 | 0.55 | 0.45 | 0.54 | 0.57 | 0.63 | 0.68 |
| 6 | 50176108 | rs2033993 | -0.32 | -0.419 | -0.261 | -0.783 | 0.019 | -0.016 | -0.016 | 0.193 | 0.37 | 0.25 | 0.42 | 0.43 | 0.34 | 0.29 |
| 6 | 50184055 | rs9369942 | 0.176 | -0.624 | -0.234 | -0.775 | -0.001 | -0.069 | -0.069 | 0.264 | 0.23 | 0.12 | 0.35 | 0.43 | 0.26 | 0.18 |
| 6 | 50186697 | rs6919348 | -0.435 | -0.625 | -0.201 | -0.756 | 0.005 | -0.004 | -0.004 | 0 | 0.38 | 0.33 | 0.43 | 0.44 | 0.34 | 0.29 |
| 6 | 50199587 | rs6922902 | 0.955 | -0.611 | -0.312 | -0.746 | -0.009 | -0.033 | -0.033 | 0 | 0.14 | 0.13 | 0.06 | 0.01 | 0.08 | 0.1 |
| 6 | 50209473 | rs1441594 | -0.022 | -0.283 | -0.511 | -0.527 | -0.001 | -0.054 | -0.054 | 0 | 0.23 | 0.14 | 0.36 | 0.43 | 0.27 | 0.18 |
| 6 | 50227356 | rs9381861 | -0.394 | -0.375 | -0.714 | -0.817 | 0.025 | 0.037 | 0.037 | 0 | 0.53 | 0.25 | 0.49 | 0.41 | 0.46 | 0.38 |
| 6 | 50236752 | rs1441601 | 0.255 | -1.011 | -0.581 | -1.483 | 0.025 | -0.025 | -0.025 | 0 | 0.62 | 0.39 | 0.83 | 1 | 0.78 | 0.78 |
| 6 | 50239686 | rs1348181 | 1.082 | -0.954 | -0.743 | -0.886 | -0.05 | -0.088 | -0.088 | 0 | 0.13 | 0.03 | 0.32 | 0.55 | 0.17 | 0.17 |
| 6 | 50241222 | rs9395576 | -0.897 | -1.199 | -0.416 | -1.137 | -0.06 | -0.168 | -0.168 | 0.004 | 0.77 | 0.56 | 0.86 | 1 | 0.83 | 0.88 |
| 6 | 50252315 | rs6923691 | 0.295 | -0.984 | -0.719 | -1.201 | -0.001 | -0.073 | -0.073 | 0 | 0.62 | 0.43 | 0.81 | 1 | 0.74 | 0.76 |
| 6 | 50256334 | rs10498782 | -0.319 | -0.513 | -0.454 | -0.524 | 0.028 | 0.027 | 0.027 | 0 | 0.16 | 0.08 | 0.04 | 0 | 0.09 | 0.12 |
| 6 | 50258198 | rs1361504 | -0.066 | -0.208 | -0.493 | -0.331 | -0.033 | -0.101 | -0.101 | 4.251 | 0.2 | 0.08 | 0.33 | 0.52 | 0.24 | 0.26 |
| 6 | 50261390 | rs993629 | 0.315 | 0.071 | 0.086 | 0.133 | 0 | -0.002 | -0.002 | 0 | 0.53 | 0.5 | 0.38 | 0.32 | 0.47 | 0.44 |
| 6 | 50261878 | rs9349529 | 0.502 | 0.054 | 0.479 | 0.082 | 0.001 | -0.023 | -0.023 | 0 | 0.55 | 0.49 | 0.42 | 0.24 | 0.52 | 0.48 |
| 6 | 50263685 | rs1492623 | -0.258 | 0.108 | 0.34 | 0.082 | -0.027 | -0.045 | -0.045 | 0 | 0.36 | 0.2 | 0.41 | 0.45 | 0.4 | 0.28 |
| 6 | 50267833 | rs1126287 | -0.584 | -0.029 | 0.355 | -0.088 | -0.004 | 0.002 | 0.002 | 0 | 0.45 | 0.28 | 0.47 | 0.46 | 0.47 | 0.3 |
| 6 | 50271595 | rs7749461 | -0.201 | 0.055 | 0.276 | 0.001 | -0.004 | -0.03 | -0.03 | 0 | 0.35 | 0.2 | 0.2 | 0.44 | 0.36 | 0.26 |
| 6 | 50278215 | rs9381865 | -0.13 | 0.098 | 0.496 | -0.123 | -0.014 | -0.025 | -0.025 | 0.213 | 0.43 | 0.23 | 0.33 | 0.49 | 0.46 | 0.31 |
| 6 | 50279484 | rs9357655 | -0.832 | 0.061 | 0.763 | -0.153 | 0 | 0 | 0 | 0.59 | 0.56 | 0.43 | 0.62 | 0.5 | 0.58 | 0.38 |
| 6 | 50294443 | rs9349534 | -0.366 | 0.067 | 0.868 | -0.152 | -0.002 | -0.002 | -0.002 | 3.782 | 0.47 | 0.46 | 0.55 | 0.51 | 0.51 | 0.33 |
| 6 | 50306801 | rs1492631 | -0.821 | -0.058 | 0.725 | -0.165 | 0.044 | 0.015 | 0.015 | 0.53 | 0.07 | 0.39 | 0.23 | 0.02 | 0.05 | 0.02 |
| 6 | 50318693 | rs7757451 | 1.084 | 0.013 | 0.419 | -0.138 | 0.001 | 0.043 | 0.043 | 0 | 0.38 | 0.26 | 0.23 | 0.19 | 0.35 | 0.41 |
| 6 | 50322852 | rs9349536 | -0.038 | -0.213 | 0.297 | -0.511 | -0.049 | 0.164 | 0.164 | 0 | 0.56 | 0.83 | 0.54 | 0.21 | 0.47 | 0.48 |
| 6 | 50334349 | rs6930610 | -1.146 | 0.168 | 0.823 | 0.219 | -0.028 | -0.101 | -0.101 | 0.134 | 0.17 | 0.47 | 0.3 | 0.02 | 0.12 | 0.07 |
| 6 | 50348137 | rs4144559 | 0.767 | 0.319 | 0.322 | 0.466 | 0.005 | -0.005 | -0.005 | 0 | 0.49 | 0.33 | 0.44 | 0.51 | 0.51 | 0.54 |
| 6 | 50352150 | rs6921067 | 0.126 | 0.167 | 0.772 | 0.297 | 0.002 | -0.025 | -0.025 | 0 | 0.63 | 0.68 | 0.75 | 0.53 | 0.63 | 0.57 |
| 6 | 50356035 | rs9473821 | 0.485 | 0.168 | 0.847 | 0.243 | 0.004 | -0.086 | -0.086 | 0 | 0.86 | 0.6 | 0.68 | 0.98 | 0.88 | 0.96 |
| 6 | 50357617 | rs1994785 | 1.493 | 0.281 | 0.607 | 0.33 | -0.008 | 0.068 | 0.068 | 0 | 0.42 | 0.23 | 0.28 | 0.17 | 0.42 | 0.32 |
| 6 | 50363775 | rs970877 | 0.768 | 0.315 | 0.726 | 0.461 | -0.024 | -0.107 | -0.107 | 0 | 0.83 | 0.5 | 0.67 | 0.98 | 0.88 | 0.96 |
| 6 | 50364717 | rs9367397 | 0.971 | 0.333 | 0.153 | 0.883 | -0.024 | -0.045 | -0.045 | 0 | 0.07 | 0 | 0.03 | 0.18 | 0.07 | 0.07 |
| 6 | 50378802 | rs7757302 | 1.359 | 0.343 | 0.494 | 0.432 | -0.006 | 0.043 | 0.043 | 0.043 | 0.76 | 0.41 | 0.5 | 0.67 | 0.78 | 0.73 |
| 6 | 50384635 | rs16880300 | -0.169 | 0.41 | -0.006 | 0.227 | -0.004 | -0.029 | -0.029 | 0 | 0.12 | 0.16 | 0.1 | 0 | 0.09 | 0.02 |
| 6 | 50389552 | rs9349542 | -0.43 | 0.801 | -0.275 | 0.578 | -0.052 | -0.142 | -0.142 | 1.879 | 0.33 | 0.1 | 0.4 | 0.5 | 0.34 | 0.43 |
| 6 | 50392475 | rs9688487 | -0.31 | 0.833 | -0.3 | 0.595 | -0.063 | -0.152 | -0.152 | 0 | 0.32 | 0.07 | 0.4 | 0.5 | 0.34 | 0.43 |
| 6 | 50402660 | rs7752955 | -0.143 | 0.799 | -0.467 | 0.599 | -0.077 | -0.167 | -0.167 | 0.835 | 0.31 | 0.04 | 0.39 | 0.5 | 0.34 | 0.44 |
| 6 | 50403568 | rs200253 | -0.66 | 0.832 | -0.198 | 0.674 | 0.005 | 0.019 | 0.019 | 0 | 0.39 | 0.48 | 0.47 | 0.5 | 0.41 | 0.46 |
| 6 | 50405920 | rs1492620 | -0.497 | 1.006 | -0.837 | 1.259 | -0.015 | 0.017 | 0.017 | 0 | 0.06 | 0.04 | 0.12 | 0.34 | 0.08 | 0.23 |
| 6 | 50406575 | rs1388930 | -0.401 | 0.935 | -0.981 | 0.897 | -0.042 | -0.12 | -0.12 | 1.061 | 0.27 | 0.05 | 0.4 | 0.55 | 0.28 | 0.4 |
| 6 | 50410433 | rs200256 | -0.874 | 1.124 | -0.063 | 0.315 | 0.027 | -0.057 | -0.057 | 0 | 0.1 | 0.2 | 0.13 | 0 | 0.15 | 0.02 |
| 6 | 50410682 | rs200257 | -0.871 | 1.12 | -0.063 | 0.317 | -0.001 | -0.05 | -0.05 | 0.259 | 0.1 | 0.18 | 0.1 | 0 | 0.1 | 0.02 |
| 6 | 50414325 | rs200232 | -1.521 | 0.375 | 0.33 | -0.299 | -0.006 | -0.073 | -0.073 | 0 | 0.25 | 0.71 | 0.2 | 0.01 | 0.24 | 0.13 |
| 6 | 50415008 | rs200234 | -1.063 | 0.436 | 0.74 | -0.17 | -0.005 | -0.117 | -0.117 | 0 | 0.17 | 0.57 | 0.13 | 0.01 | 0.16 | 0.11 |
| 6 | 50415303 | rs16880360 | -0.146 | 1.152 | 0.15 | 0.805 | -0.03 | -0.006 | -0.006 | 0 | 0.05 | 0 | 0.05 | 0.25 | 0.1 | 0.15 |
| 6 | 50429641 | rs200246 | -0.456 | 0.163 | 0.72 | -0.149 | -0.036 | -0.089 | -0.089 | 0 | 0.15 | 0.33 | 0.08 | 0 | 0.06 | 0.06 |
| 6 | 50439021 | rs1774627 | -1.335 | -0.492 | 0.942 | -0.587 | -0.031 | -0.104 | -0.104 | 0 | 0.26 | 0.65 | 0.17 | 0 | 0.15 | 0.08 |
| 6 | 50439456 | rs1832874 | 0.84 | -0.281 | -0.483 | -0.353 | -0.047 | -0.076 | -0.076 | 0 | 0.2 | 0.03 | 0.35 | 0.32 | 0.22 | 0.27 |
| 6 | 50444445 | rs1342233 | -1.793 | -0.134 | -1.25 | -1.382 | 0.004 | 0.134 | 0.134 | 0 | 0.75 | 0.96 | 0.84 | 1 | 0.73 | 0.86 |
| 6 | 50472927 | rs6906927 | 0.914 | -0.154 | -1.062 | -0.323 | -0.039 | -0.116 | -0.116 | 0.024 | 0.22 | 0.04 | 0.42 | 0.44 | 0.23 | 0.33 |
| 6 | 50516277 | rs2209903 | 0.404 | 0.021 | -0.381 | -0.364 | 0.017 | 0.023 | 0.023 | 0.769 | 0.66 | 0.42 | 0.73 | 0.56 | 0.65 | 0.66 |
| 6 | 50522228 | rs984329 | 0.656 | 0.255 | -0.446 | -0.338 | -0.055 | -0.062 | -0.062 | 0 | 0.14 | 0.06 | 0.16 | 0.44 | 0.24 | 0.21 |
| 6 | 50528436 | rs280344 | -0.743 | 0.173 | -0.284 | -0.414 | 0.023 | 0.03 | 0.03 | 0.375 | 0.33 | 0.58 | 0.28 | 0.44 | 0.36 | 0.34 |
| 6 | 50548918 | rs9367402 | 0.678 | 0.159 | -0.325 | -0.413 | -0.045 | -0.049 | -0.049 | 0 | 0.14 | 0.07 | 0.16 | 0.44 | 0.23 | 0.21 |
| 6 | 50551445 | rs280292 | 0.326 | -0.014 | 0.024 | -0.473 | -0.04 | -0.094 | -0.094 | 0 | 0.17 | 0.34 | 0.09 | 0 | 0.08 | 0.13 |
| 6 | 50567085 | rs280286 | -0.126 | -0.044 | 0.032 | -0.488 | -0.068 | -0.14 | -0.14 | 1.904 | 0.19 | 0.51 | 0.11 | 0.01 | 0.09 | 0.15 |
| 6 | 50571699 | rs10948566 | 0.862 | -0.507 | -0.55 | -0.949 | -0.009 | -0.104 | -0.104 | 0.018 | 0.28 | 0.05 | 0.44 | 0.43 | 0.31 | 0.3 |
| 6 | 50573840 | rs578024 | 0.444 | -0.333 | -0.367 | -0.913 | -0.065 | -0.118 | -0.118 | 0 | 0.18 | 0.43 | 0.1 | 0.01 | 0.08 | 0.14 |
| 6 | 50576468 | rs1161444 | -0.656 | -0.255 | -0.457 | -0.93 | 0.018 | 0.027 | 0.027 | 0.46 | 0.34 | 0.58 | 0.29 | 0.45 | 0.36 | 0.36 |
| 6 | 50596453 | rs1161397 | 0.081 | -0.392 | -0.311 | -1.048 | -0.055 | -0.139 | -0.139 | 3.627 | 0.19 | 0.51 | 0.12 | 0.01 | 0.12 | 0.15 |
| 6 | 50597061 | rs527336 | 0.211 | -0.514 | -0.394 | -1.038 | -0.068 | -0.12 | -0.12 | 0 | 0.19 | 0.44 | 0.1 | 0.01 | 0.08 | 0.14 |
| 6 | 50602212 | rs10948567 | 1.412 | -0.517 | -0.812 | -0.933 | -0.048 | -0.066 | -0.066 | 0 | 0.15 | 0.01 | 0.33 | 0.23 | 0.18 | 0.17 |

| | | | | | | | | | | | | | | | | |
|---|---|---|---|---|---|---|---|---|---|---|---|---|---|---|---|---|
| 6 | 50603992 | rs280315 | 0.291 | -1.197 | -0.531 | -1.286 | -0.068 | -0.12 | -0.12 | 1.142 | 0.19 | 0.45 | 0.1 | 0.01 | 0.08 | 0.14 |
| 6 | 50609474 | rs280308 | -1.93 | -1.487 | -0.829 | -1.647 | -0.001 | 0.112 | 0.112 | 0 | 0.78 | 0.98 | 0.89 | 1 | 0.8 | 0.86 |
| 6 | 50619699 | rs491286 | 0.582 | -1.223 | -0.308 | -1.178 | -0.045 | -0.086 | -0.086 | 0.213 | 0.16 | 0.31 | 0.08 | 0.01 | 0.08 | 0.14 |
| 6 | 50620961 | rs524066 | 0.153 | -1.205 | -0.473 | -1.168 | -0.075 | -0.131 | -0.131 | 0 | 0.19 | 0.48 | 0.1 | 0.01 | 0.08 | 0.14 |
| 6 | 50646175 | rs10948570 | 0.418 | -1.08 | -0.783 | -1.156 | -0.002 | -0.001 | -0.001 | 0.055 | 0.55 | 0.42 | 0.6 | 0.54 | 0.51 | 0.6 |
| 6 | 50664180 | rs12528998 | 1.438 | -0.696 | -0.827 | -0.93 | -0.027 | -0.072 | -0.072 | 0 | 0.13 | 0 | 0.29 | 0.23 | 0.1 | 0.16 |
| 6 | 50693607 | rs760899 | -0.509 | -0.809 | -0.544 | -1.136 | 0.056 | 0.048 | 0.048 | 0 | 0.33 | 0.56 | 0.35 | 0.44 | 0.46 | 0.37 |
| 6 | 50709707 | rs1569577 | 0.816 | -0.272 | -0.569 | -0.852 | -0.007 | 0.004 | 0.004 | 0 | 0.12 | 0.08 | 0.21 | 0.43 | 0.32 | 0.23 |
| 6 | 50715407 | rs9349553 | -1.08 | -0.701 | -0.641 | -1.126 | 0.001 | 0.049 | 0.049 | 3.07 | 0.69 | 0.85 | 0.77 | 0.83 | 0.7 | 0.7 |
| 6 | 50727940 | rs2235495 | 0.705 | 0.076 | -0.256 | -0.591 | 0.024 | 0.042 | 0.042 | 0 | 0.12 | 0.1 | 0.21 | 0.43 | 0.32 | 0.23 |
| 6 | 50746213 | rs2817374 | 0.247 | 0.027 | 0.168 | -0.612 | 0.163 | 0.211 | 0.211 | 0 | 0.14 | 0.37 | 0.23 | 0.41 | 0.32 | 0.23 |
| 6 | 50756951 | rs4383818 | 0.492 | -0.623 | -0.09 | -1.204 | 0.012 | 0.016 | 0.016 | 0 | 0.22 | 0.17 | 0.13 | 0 | 0.18 | 0.15 |
| 6 | 50765352 | rs2206272 | -0.625 | -0.655 | -0.029 | -1.059 | -0.002 | -0.004 | -0.004 | 0.975 | 0.42 | 0.63 | 0.4 | 0.43 | 0.54 | 0.37 |
| 6 | 50765751 | rs9381901 | -0.111 | -0.273 | -0.164 | -0.761 | -0.019 | -0.018 | -0.018 | 2.209 | 0.31 | 0.25 | 0.32 | 0.43 | 0.46 | 0.31 |
| 6 | 50782047 | rs9367415 | 0.73 | 0.196 | -0.265 | -0.492 | -0.002 | 0.01 | 0.01 | 0 | 0.13 | 0.09 | 0.27 | 0.43 | 0.34 | 0.22 |
| 6 | 50782834 | rs2143081 | -0.885 | -0.705 | -0.246 | -0.79 | -0.018 | -0.017 | -0.017 | 0 | 0.71 | 0.76 | 0.66 | 0.57 | 0.54 | 0.67 |
| 6 | 50798526 | rs2206277 | 0.769 | -1.243 | -0.568 | -0.785 | 0.016 | -0.01 | -0.01 | 0.021 | 0.22 | 0.13 | 0.34 | 0.23 | 0.18 | 0.24 |
| 6 | 50803050 | rs987237 | 0.843 | -1.24 | -0.62 | -1.298 | 0.013 | 0.022 | 0.022 | 0 | 0.21 | 0.09 | 0.34 | 0.15 | 0.18 | 0.24 |
| 6 | 50808745 | rs1569777 | 0.82 | -1.327 | -0.116 | -1.295 | -0.023 | 0.007 | 0.007 | 0.596 | 0.17 | 0.22 | 0.14 | 0.2 | 0.13 | 0.19 |
| 6 | 50812906 | rs2817419 | -0.033 | -1.238 | -0.332 | -1.481 | 0.015 | 0.017 | 0.017 | 0.784 | 0.65 | 0.69 | 0.76 | 0.8 | 0.74 | 0.72 |
| 6 | 50813332 | rs2817420 | -1.147 | -1.074 | -0.384 | -1.418 | -0.028 | 0.108 | 0.108 | 0 | 0.83 | 0.99 | 0.91 | 1 | 0.86 | 0.9 |
| 6 | 50820940 | rs2635727 | -0.128 | -1.427 | -0.33 | -1.471 | 0.025 | 0.025 | 0.025 | 1.616 | 0.64 | 0.72 | 0.77 | 0.8 | 0.75 | 0.72 |
| 6 | 50828072 | rs9473921 | 0.628 | -1.244 | -0.176 | -1.535 | 0.016 | -0.02 | -0.02 | 0.423 | 0.14 | 0.27 | 0.09 | 0.15 | 0.11 | 0.16 |
| 6 | 50829511 | rs4715209 | -0.886 | -1.416 | 0.064 | -1.65 | -0.023 | -0.101 | -0.101 | 0.576 | 0.81 | 0.68 | 0.89 | 1 | 0.87 | 0.91 |
| 6 | 50831194 | rs9296645 | 0.66 | -1.451 | 0.146 | -1.626 | 0.013 | -0.021 | -0.021 | 0 | 0.14 | 0.28 | 0.09 | 0.15 | 0.11 | 0.16 |
| 6 | 50834379 | rs9349560 | -0.619 | -1.605 | -0.26 | -1.619 | 0.011 | 0.002 | 0.002 | 0 | 0.52 | 0.58 | 0.54 | 0.53 | 0.41 | 0.59 |
| 6 | 50837708 | rs16880741 | 0.979 | -1.645 | -0.284 | -1.649 | 0.004 | -0.004 | -0.004 | 0 | 0.19 | 0.18 | 0.14 | 0.22 | 0.12 | 0.19 |
| 6 | 50845787 | rs11970011 | 1.386 | -1.606 | -0.3 | -1.65 | -0.097 | -0.085 | -0.085 | 1.319 | 0.29 | 0.1 | 0.35 | 0.48 | 0.47 | 0.32 |
| 6 | 50857995 | rs9473932 | 0.027 | -1.556 | -0.286 | -1.627 | -0.004 | -0.019 | -0.019 | 0.037 | 0.33 | 0.4 | 0.4 | 0.3 | 0.28 | 0.4 |
| 6 | 50865820 | rs943005 | 1.281 | -1.534 | -0.324 | -1.607 | 0.002 | 0.013 | 0.013 | 0 | 0.19 | 0.08 | 0.31 | 0.15 | 0.16 | 0.24 |
| 6 | 50911334 | rs9381917 | 1.006 | -1.197 | -0.597 | -1.236 | 0.016 | -0.012 | -0.012 | 0 | 0.17 | 0.13 | 0.13 | 0.23 | 0.12 | 0.19 |
| 6 | 50921271 | rs9369988 | -0.416 | -1.132 | -0.667 | -1.357 | 0.013 | 0.003 | 0.003 | 0 | 0.51 | 0.6 | 0.54 | 0.53 | 0.4 | 0.59 |
| 6 | 50930041 | rs6925748 | -0.29 | -0.193 | -0.221 | -1.487 | 0.008 | 0.027 | 0.027 | 0.403 | 0.5 | 0.44 | 0.48 | 0.19 | 0.4 | 0.48 |
| 6 | 50943006 | rs6940756 | -0.238 | -0.376 | 0.375 | -1.085 | -0.003 | -0.026 | -0.026 | 0 | 0.09 | 0.31 | 0.04 | 0.03 | 0.02 | 0.09 |
| 6 | 50946521 | rs4141973 | 1.035 | -0.003 | -0.523 | -1.113 | -0.031 | -0.054 | -0.054 | 0 | 0.12 | 0.01 | 0.27 | 0.18 | 0.12 | 0.21 |
| 6 | 50947295 | rs4082052 | -0.101 | 0.508 | -0.5 | -1.582 | -0.015 | 0.012 | 0.012 | 0 | 0.25 | 0.18 | 0.37 | 0.18 | 0.33 | 0.3 |
| 6 | 50951649 | rs9369991 | 1.327 | 0.396 | -1.099 | -2.084 | -0.046 | -0.04 | -0.04 | 0 | 0.27 | 0.12 | 0.33 | 0.63 | 0.33 | 0.33 |
| 6 | 50953315 | rs579544 | -1.336 | 0.036 | -1.438 | -2.351 | -0.001 | 0.105 | 0.105 | 0 | 0.83 | 0.99 | 0.92 | 1 | 0.89 | 0.91 |
| 6 | 50955081 | rs11759809 | 0.451 | 0.164 | -1.231 | -2.172 | -0.017 | 0.013 | 0.013 | 0 | 0.25 | 0.17 | 0.37 | 0.18 | 0.32 | 0.3 |
| 6 | 50960316 | rs4715215 | 0.469 | 0.17 | -1.225 | -2.169 | -0.018 | 0.013 | 0.013 | 0 | 0.25 | 0.16 | 0.37 | 0.18 | 0.32 | 0.3 |
| 6 | 50966474 | rs283548 | -0.562 | -0.211 | -0.919 | -2.424 | -0.01 | 0.026 | 0.026 | 0.431 | 0.43 | 0.61 | 0.5 | 0.37 | 0.48 | 0.47 |
| 6 | 50967007 | rs283547 | 0.227 | -0.314 | -0.704 | -2.231 | -0.009 | -0.006 | -0.006 | 0 | 0.17 | 0.44 | 0.13 | 0.19 | 0.16 | 0.17 |
| 6 | 50987682 | rs728831 | 1.749 | 0.128 | -1.152 | -2.391 | -0.032 | -0.049 | -0.049 | 0.189 | 0.24 | 0.12 | 0.29 | 0.63 | 0.26 | 0.32 |
| 6 | 50987768 | rs190759 | -0.251 | -0.34 | -1.357 | -2.745 | 0.061 | 0.08 | 0.08 | 0 | 0.7 | 0.87 | 0.85 | 0.99 | 0.83 | 0.85 |
| 6 | 50995076 | rs9395648 | -0.129 | -0.248 | -1.2 | -2.632 | 0.01 | -0.005 | -0.005 | 0.352 | 0.8 | 0.82 | 0.86 | 0.99 | 0.86 | 0.88 |
| 6 | 50997085 | rs9381926 | -0.603 | -0.313 | -1.18 | -2.608 | 0.011 | -0.004 | -0.004 | 0 | 0.83 | 0.87 | 0.89 | 0.99 | 0.9 | 0.9 |
| 6 | 50998135 | rs2770664 | -0.867 | -0.313 | -1.18 | -2.608 | 0.008 | -0.006 | -0.006 | 1.085 | 0.84 | 0.88 | 0.9 | 0.99 | 0.91 | 0.9 |
| 6 | 51007036 | rs4548008 | 0.38 | -0.358 | -0.558 | -2.671 | -0.045 | -0.068 | -0.068 | 0 | 0.71 | 0.54 | 0.78 | 0.85 | 0.76 | 0.83 |
| 6 | 51010906 | rs17541037 | 1.221 | 0.026 | -0.612 | -2.607 | -0.018 | 0.043 | 0.043 | 0.138 | 0.09 | 0.01 | 0.04 | 0 | 0.12 | 0.03 |
| 6 | 51014810 | rs6458752 | 0.636 | -0.18 | -0.349 | -2.36 | 0.015 | 0.03 | 0.03 | 0 | 0.12 | 0.31 | 0.12 | 0.15 | 0.14 | 0.08 |
| 6 | 51029764 | rs9395656 | 0.122 | -0.107 | -0.745 | -2.439 | 0.005 | 0.067 | 0.067 | 0 | 0.53 | 0.66 | 0.48 | 0.81 | 0.55 | 0.57 |
| 6 | 51049681 | rs1925216 | 0.749 | 0.037 | -0.436 | -2.306 | -0.004 | 0.012 | 0.012 | 0.022 | 0.41 | 0.31 | 0.37 | 0.6 | 0.4 | 0.4 |
| 6 | 51052686 | rs927716 | 0.765 | 0.087 | -0.416 | -2.316 | -0.001 | 0.002 | 0.002 | 0.013 | 0.39 | 0.22 | 0.34 | 0.6 | 0.35 | 0.37 |
| 6 | 51057610 | rs9370005 | 0.407 | 0.105 | -0.179 | -2.285 | 0.113 | 0.233 | 0.233 | 0 | 0.06 | 0.33 | 0.07 | 0.18 | 0.08 | 0.09 |
| 6 | 51058527 | rs10948596 | 0.862 | -0.073 | -0.502 | -2.473 | -0.025 | 0.018 | 0.018 | 0 | 0.26 | 0.15 | 0.41 | 0.19 | 0.34 | 0.34 |
| 6 | 51064520 | rs9381936 | -0.229 | 0.082 | -0.307 | -2.243 | 0.096 | 0.141 | 0.141 | 0.007 | 0.1 | 0.4 | 0.09 | 0.21 | 0.11 | 0.18 |
| 6 | 51066322 | rs12198852 | 1.145 | -0.224 | -1.053 | -2.456 | 0.018 | 0.082 | 0.082 | 0 | 0.13 | 0.01 | 0.05 | 0 | 0.07 | 0.05 |
| 6 | 51073648 | rs9381940 | -0.042 | 0.064 | -1.085 | -2.511 | 0.009 | 0.108 | 0.108 | 0 | 0.49 | 0.71 | 0.41 | 0.81 | 0.42 | 0.54 |
| 6 | 51084931 | rs1886570 | 0.577 | 0.121 | -1.058 | -2.469 | 0.001 | -0.007 | -0.007 | 0 | 0.34 | 0.62 | 0.36 | 0.81 | 0.34 | 0.51 |
| 6 | 51087083 | rs9381943 | 1.685 | 0.159 | -1.059 | -2.316 | -0.04 | -0.086 | -0.086 | 0 | 0.24 | 0.07 | 0.27 | 0.59 | 0.24 | 0.33 |
| 6 | 51087831 | rs2505276 | 0.34 | 0.19 | -0.809 | -2.228 | -0.002 | 0.096 | 0.096 | 0 | 0.5 | 0.69 | 0.44 | 0.81 | 0.46 | 0.57 |
| 6 | 51099256 | rs7762246 | 0.448 | -0.006 | -1.147 | -2.26 | 0.05 | 0.087 | 0.087 | 0 | 0.22 | 0.07 | 0.13 | 0 | 0.16 | 0.09 |
| 6 | 51100011 | rs2397027 | 0.402 | 0.307 | -0.996 | -1.979 | -0.038 | 0.012 | 0.012 | 0 | 0.26 | 0.13 | 0.43 | 0.19 | 0.38 | 0.35 |
| 6 | 51102772 | rs2505263 | -0.898 | -0.238 | -0.69 | -1.772 | 0.032 | 0.05 | 0.05 | 0 | 0.42 | 0.72 | 0.31 | 0.27 | 0.34 | 0.34 |
| 6 | 51113582 | rs6937978 | -0.233 | -0.418 | -0.434 | -1.883 | -0.006 | -0.035 | -0.035 | 0 | 0.1 | 0.16 | 0.1 | 0 | 0.09 | 0.04 |
| 6 | 51117477 | rs2505274 | 0.157 | -0.503 | -0.253 | -1.961 | -0.021 | -0.132 | -0.132 | 3.033 | 0.24 | 0.49 | 0.15 | 0.06 | 0.21 | 0.19 |
| 6 | 51119366 | rs1984814 | 0.084 | -0.5 | -0.264 | -1.955 | -0.02 | -0.111 | -0.111 | 0.069 | 0.24 | 0.53 | 0.16 | 0.07 | 0.21 | 0.19 |
| 6 | 51123503 | rs4236102 | 0.225 | -0.156 | -0.841 | -1.941 | 0.018 | -0.003 | -0.003 | 0 | 0.21 | 0.24 | 0.38 | 0.17 | 0.31 | 0.27 |
| 6 | 51130487 | rs2397030 | -1.815 | -0.245 | -1.083 | -1.761 | -0.084 | -0.08 | -0.08 | 0.699 | 0.63 | 0.95 | 0.68 | 0.44 | 0.62 | 0.59 |

| | | | | | | | | | | | | | | | | |
|---|---|---|---|---|---|---|---|---|---|---|---|---|---|---|---|---|
| 6 | 51136837 | rs12527716 | 0.436 | -0.257 | -0.1 | -1.312 | -0.01 | -0.069 | -0.069 | 1.221 | 0.19 | 0.26 | 0.11 | 0.03 | 0.18 | 0.08 |
| 6 | 51143101 | rs9463680 | 0.638 | -0.141 | -0.42 | -0.894 | 0.003 | -0.007 | -0.007 | 0 | 0.08 | 0.07 | 0.04 | 0.21 | 0.02 | 0.11 |
| 6 | 51147806 | rs17738108 | 0.04 | 0.46 | -0.423 | -1.087 | -0.001 | -0.001 | -0.001 | 0 | 0.14 | 0.12 | 0.37 | 0.19 | 0.23 | 0.32 |
| 6 | 51148435 | rs2465031 | -1.791 | 0.467 | -1.12 | -0.536 | -0.102 | -0.16 | -0.16 | 0 | 0.62 | 0.98 | 0.66 | 0.51 | 0.61 | 0.64 |
| 6 | 51150512 | rs6458760 | 0.777 | 0.598 | -0.216 | -1.171 | -0.014 | -0.087 | -0.087 | 0 | 0.2 | 0.35 | 0.1 | 0.05 | 0.17 | 0.09 |
| 6 | 51152988 | rs2478879 | -1.664 | 0.596 | -0.284 | -1.127 | -0.065 | 0.055 | 0.055 | 0 | 0.56 | 0.94 | 0.61 | 0.32 | 0.59 | 0.55 |
| 6 | 51153402 | rs2504677 | -0.634 | 0.465 | 0.322 | -1.081 | -0.006 | -0.095 | -0.095 | 0 | 0.78 | 0.42 | 0.86 | 0.95 | 0.81 | 0.89 |
| 6 | 51155243 | rs2709665 | -0.47 | 0.665 | 0.185 | -0.989 | 0.153 | 0.102 | 0.102 | 0.105 | 0.05 | 0.31 | 0.08 | 0.07 | 0.08 | 0.11 |
| 6 | 51156076 | rs17738155 | 1.21 | 0.567 | 0.03 | -1.046 | 0.003 | -0.006 | -0.006 | 0 | 0.09 | 0.05 | 0.03 | 0 | 0.08 | 0.01 |
| 6 | 51157533 | rs2397026 | 0.05 | 0.804 | -0.123 | -0.98 | -0.005 | 0.001 | 0.001 | 0.05 | 0.18 | 0.12 | 0.37 | 0.19 | 0.22 | 0.33 |
| 6 | 51160125 | rs2246447 | 1.065 | 0.921 | -0.061 | -0.738 | -0.054 | 0.083 | 0.083 | 0.014 | 0.45 | 0.1 | 0.41 | 0.69 | 0.45 | 0.46 |
| 6 | 51160604 | rs1342615 | -1.008 | 1.066 | 0.681 | -0.212 | 0.106 | 0.102 | 0.102 | 0.842 | 0.05 | 0.31 | 0.06 | 0.08 | 0.06 | 0.09 |
| 6 | 51171408 | rs1327265 | 0.303 | 0.303 | -0.399 | -0.479 | 0.003 | 0.026 | 0.026 | 0 | 0.66 | 0.72 | 0.74 | 0.81 | 0.64 | 0.6 |
| 6 | 51171829 | rs1327266 | 0.068 | -0.5 | 0.231 | -0.687 | -0.039 | -0.048 | -0.048 | 0.359 | 0.13 | 0.43 | 0.11 | 0.05 | 0.07 | 0.08 |
| 6 | 51174169 | rs2465055 | 0.571 | -1.087 | 0.052 | -0.609 | -0.031 | -0.011 | -0.011 | 0.179 | 0.68 | 0.39 | 0.63 | 0.75 | 0.81 | 0.78 |
| 6 | 51174618 | rs2796908 | 0.145 | -1.119 | 0.138 | -0.627 | -0.054 | -0.058 | -0.058 | 0 | 0.14 | 0.52 | 0.11 | 0.06 | 0.07 | 0.08 |
| 6 | 51174821 | rs2465052 | 0.585 | -1.1 | 0.13 | -0.613 | -0.031 | -0.011 | -0.011 | 0 | 0.68 | 0.39 | 0.63 | 0.75 | 0.81 | 0.78 |
| 6 | 51175375 | rs17813135 | 0.534 | -0.919 | -0.216 | -0.498 | -0.024 | 0.059 | 0.059 | 0 | 0.08 | 0.17 | 0.23 | 0.19 | 0.04 | 0.08 |
| 6 | 51175486 | rs13211684 | 0.005 | 0.05 | -0.729 | -0.244 | -0.009 | -0.004 | -0.004 | 0 | 0.17 | 0.17 | 0.2 | 0.17 | 0.32 | 0.38 |
| 6 | 51180392 | rs13202780 | 0.469 | 0.148 | -0.797 | -0.301 | -0.02 | 0.01 | 0.01 | 0 | 0.13 | 0.1 | 0.15 | 0.07 | 0.25 | 0.29 |
| 6 | 51183658 | rs2796909 | -1.62 | -0.063 | -0.87 | -0.399 | 0.015 | 0.018 | 0.018 | 0 | 0.89 | 0.95 | 0.95 | 1 | 0.92 | 0.95 |
| 6 | 51184712 | rs2478881 | 1.634 | -0.147 | -1.279 | 0.039 | -0.107 | -0.113 | -0.113 | 0 | 0.37 | 0.01 | 0.25 | 0.45 | 0.36 | 0.35 |
| 6 | 51201257 | rs2709670 | -0.578 | -0.016 | -0.755 | 0.424 | -0.05 | -0.037 | -0.037 | 0 | 0.27 | 0.18 | 0.23 | 0.32 | 0.42 | 0.41 |
| 6 | 51206168 | rs1536504 | -1.623 | -0.19 | -0.537 | 0.27 | 0.01 | 0.001 | 0.001 | 0 | 0.88 | 0.91 | 0.95 | 1 | 0.92 | 0.95 |
| 6 | 51217638 | rs6916171 | 1.659 | -0.334 | -0.553 | -0.201 | -0.026 | -0.076 | -0.076 | 0 | 0.27 | 0.34 | 0.36 | 0.13 | 0.22 | 0.22 |
| 6 | 51217642 | rs6915726 | 1.334 | -0.331 | -0.595 | -0.194 | -0.024 | -0.1 | -0.1 | 0 | 0.29 | 0.41 | 0.39 | 0.13 | 0.24 | 0.24 |
| 6 | 51222894 | rs2252998 | 0.814 | -0.186 | -0.253 | 0.169 | -0.015 | -0.002 | -0.002 | 0.002 | 0.55 | 0.62 | 0.6 | 0.55 | 0.47 | 0.48 |
| 6 | 51248219 | rs13205857 | 0.54 | 0.16 | -0.602 | 0.357 | -0.033 | -0.033 | -0.033 | 0.205 | 0.13 | 0.08 | 0.14 | 0.2 | 0.24 | 0.31 |
| 6 | 51248309 | rs1327283 | 1.023 | 0.005 | -0.487 | 0.255 | 0.001 | 0.002 | 0.002 | 0 | 0.53 | 0.52 | 0.57 | 0.55 | 0.45 | 0.47 |
| 6 | 51261134 | rs1536506 | 0.886 | 0.119 | -0.629 | -0.05 | -0.001 | -0.001 | -0.001 | 0 | 0.56 | 0.52 | 0.68 | 0.68 | 0.47 | 0.5 |
| 6 | 51264082 | rs2504703 | 0.4 | 0.132 | -0.914 | -0.064 | -0.005 | -0.003 | -0.003 | 0 | 0.67 | 0.65 | 0.72 | 0.68 | 0.56 | 0.51 |
| 6 | 51265608 | rs1409892 | -0.588 | 0.101 | -0.812 | 0.065 | -0.018 | -0.009 | -0.009 | 1.39 | 0.29 | 0.24 | 0.25 | 0.31 | 0.42 | 0.42 |
| 6 | 51266877 | rs2397045 | 0.906 | -0.255 | -0.958 | -0.137 | -0.029 | -0.025 | -0.025 | 2.875 | 0.42 | 0.46 | 0.53 | 0.26 | 0.29 | 0.34 |
| 6 | 51284262 | rs9395674 | 1.196 | -0.068 | -0.913 | -0.072 | -0.019 | -0.017 | -0.017 | 0.023 | 0.39 | 0.41 | 0.54 | 0.26 | 0.31 | 0.3 |
| 6 | 51294341 | rs1359782 | 0.143 | 0.342 | -1.102 | 0.088 | 0.009 | -0.007 | -0.007 | 0.547 | 0.66 | 0.6 | 0.78 | 0.76 | 0.61 | 0.62 |
| 6 | 51317602 | rs9367432 | -0.071 | 0.845 | -0.766 | 0.652 | -0.04 | -0.063 | -0.063 | 0 | 0.23 | 0.1 | 0.22 | 0.48 | 0.29 | 0.29 |
| 6 | 51321303 | rs3898645 | 0.878 | 1.197 | -0.466 | 1.194 | -0.067 | -0.096 | -0.096 | 0 | 0.14 | 0.01 | 0.13 | 0.41 | 0.18 | 0.16 |
| 6 | 51322232 | rs1337846 | -0.335 | 1.002 | -0.226 | 0.672 | -0.011 | -0.077 | -0.077 | 0 | 0.86 | 0.67 | 0.93 | 1 | 0.89 | 0.94 |
| 6 | 51339492 | rs4090367 | -0.242 | 1.095 | -0.103 | 0.199 | -0.013 | -0.084 | -0.084 | 0 | 0.86 | 0.64 | 0.91 | 1 | 0.89 | 0.94 |
| 6 | 51354717 | rs9474019 | 1.057 | 1.283 | -0.112 | 0.281 | -0.001 | -0.022 | -0.022 | 0.012 | 0.1 | 0.12 | 0.05 | 0 | 0.09 | 0.05 |
| 6 | 51357004 | rs4711976 | 0.052 | 1.432 | -0.428 | 0.497 | 0.045 | 0.054 | 0.054 | 0 | 0.79 | 0.6 | 0.77 | 0.62 | 0.71 | 0.75 |
| 6 | 51362605 | rs4379293 | 0.611 | 1.797 | -0.704 | 0.604 | -0.005 | -0.003 | -0.003 | 0 | 0.16 | 0.13 | 0.19 | 0.38 | 0.26 | 0.23 |
| 6 | 51362685 | rs7765090 | 0.474 | 1.672 | -0.594 | 0.432 | 0.014 | 0.013 | 0.013 | 0 | 0.71 | 0.6 | 0.76 | 0.62 | 0.65 | 0.72 |
| 6 | 51364171 | rs4113585 | 1.154 | 1.623 | -0.597 | 0.419 | 0.002 | -0.01 | -0.01 | 0.002 | 0.1 | 0.09 | 0.05 | 0 | 0.09 | 0.05 |
| 6 | 51365011 | rs10807424 | -0.125 | 1.674 | -0.614 | 0.414 | -0.013 | -0.012 | -0.012 | 0 | 0.26 | 0.22 | 0.25 | 0.38 | 0.37 | 0.29 |
| 6 | 51368689 | rs6458767 | 0.341 | 1.741 | -0.598 | 0.404 | 0.009 | 0.004 | 0.004 | 0 | 0.72 | 0.66 | 0.72 | 0.62 | 0.6 | 0.69 |
| 6 | 51370401 | rs6902943 | -0.804 | 1.729 | -0.649 | 0.379 | -0.002 | -0.022 | -0.022 | 0 | 0.9 | 0.86 | 0.95 | 1 | 0.91 | 0.95 |
| 6 | 51373205 | rs11962312 | 1.227 | 1.143 | -0.954 | 0.072 | 0.032 | 0.125 | 0.125 | 0.834 | 0.41 | 0.09 | 0.46 | 0.2 | 0.37 | 0.31 |
| 6 | 51374756 | rs4506043 | 0.679 | 1.548 | -0.565 | 0.584 | -0.072 | -0.065 | -0.065 | 0 | 0.14 | 0.01 | 0.13 | 0.23 | 0.2 | 0.15 |
| 6 | 51375080 | rs4333416 | -0.828 | 1.458 | -0.486 | 0.459 | 0 | -0.045 | -0.045 | 0 | 0.89 | 0.8 | 0.92 | 1 | 0.89 | 0.92 |
| 6 | 51386150 | rs9395687 | -0.365 | 1.389 | -0.546 | 0.577 | 0.014 | 0.008 | 0.008 | 0 | 0.29 | 0.36 | 0.28 | 0.38 | 0.4 | 0.31 |
| 6 | 51399953 | rs4260753 | 0.098 | 1.013 | -0.446 | 0.636 | 0.014 | 0.086 | 0.086 | 0.199 | 0.86 | 0.64 | 0.84 | 0.78 | 0.83 | 0.89 |
| 6 | 51405556 | rs12207656 | 1.006 | 1.492 | -1.083 | 0.355 | -0.074 | -0.033 | -0.033 | 0.659 | 0.12 | 0.02 | 0.13 | 0.13 | 0.22 | 0.1 |
| 6 | 51406848 | rs17752199 | 0.767 | 1.214 | -0.955 | 0.447 | 0 | 0 | 0 | 0 | 0.14 | 0.15 | 0.13 | 0.12 | 0.12 | 0.12 |
| 6 | 51410601 | rs7449794 | -0.276 | 1.207 | -0.823 | 0.491 | -0.023 | -0.021 | -0.021 | 0 | 0.26 | 0.21 | 0.28 | 0.34 | 0.39 | 0.21 |
| 6 | 51412786 | rs9381983 | -0.992 | 1.438 | -0.464 | 0.809 | 0.028 | 0.111 | 0.111 | 0 | 0.07 | 0.2 | 0.09 | 0.22 | 0.09 | 0.13 |
| 6 | 51421707 | rs2439032 | -0.861 | 1.457 | -0.51 | 0.682 | -0.005 | -0.005 | -0.005 | 0 | 0.34 | 0.36 | 0.36 | 0.45 | 0.43 | 0.33 |
| 6 | 51429680 | rs10948626 | 0.366 | 1.568 | -0.492 | 0.751 | -0.016 | 0.006 | 0.006 | 0 | 0.16 | 0.16 | 0.16 | 0.12 | 0.24 | 0.13 |
| 6 | 51436530 | rs2243758 | 0.884 | 1.214 | -0.508 | 0.376 | 0.008 | -0.008 | -0.008 | 0 | 0.63 | 0.52 | 0.66 | 0.35 | 0.51 | 0.51 |
| 6 | 51443715 | rs2439040 | 0.451 | 1.219 | -0.444 | 0.493 | 0.029 | 0.029 | 0.029 | 0 | 0.66 | 0.52 | 0.73 | 0.51 | 0.57 | 0.59 |
| 6 | 51444607 | rs10498789 | 1.478 | 1.067 | -0.543 | 0.378 | 0.013 | 0.02 | 0.02 | 0 | 0.47 | 0.35 | 0.53 | 0.23 | 0.38 | 0.39 |
| 6 | 51461440 | rs9296657 | 0.288 | 1.205 | -0.751 | 0.427 | -0.023 | -0.017 | -0.017 | 0 | 0.66 | 0.74 | 0.64 | 0.6 | 0.57 | 0.67 |
| 6 | 51465832 | rs2771009 | 0.108 | 1.21 | -0.773 | 0.44 | -0.05 | -0.035 | -0.035 | 0 | 0.68 | 0.84 | 0.65 | 0.61 | 0.57 | 0.67 |
| 6 | 51466152 | rs6458774 | -0.779 | 1.305 | -0.721 | 0.499 | 0.003 | 0.004 | 0.004 | 0 | 0.84 | 0.86 | 0.85 | 0.89 | 0.86 | 0.86 |
| 6 | 51469781 | rs1108203 | 0.208 | 1.286 | -0.728 | 0.558 | -0.025 | -0.065 | -0.065 | 1.434 | 0.17 | 0.1 | 0.1 | 0.36 | 0.18 | 0.28 |
| 6 | 51480486 | rs2784198 | -0.024 | 1.389 | -0.635 | 0.444 | 0 | -0.002 | -0.002 | 1.281 | 0.64 | 0.67 | 0.67 | 0.76 | 0.6 | 0.74 |
| 6 | 51481671 | rs2784200 | -0.814 | 1.393 | -0.592 | 0.531 | 0.024 | 0.062 | 0.062 | 0 | 0.05 | 0.2 | 0.09 | 0.14 | 0.09 | 0.1 |
| 6 | 51483961 | rs9381994 | 0.399 | 1.362 | -0.622 | 0.454 | 0.004 | -0.005 | -0.005 | 0 | 0.58 | 0.45 | 0.6 | 0.61 | 0.55 | 0.66 |
| 6 | 51484348 | rs9395699 | 0.594 | 1.604 | -0.537 | 0.5 | -0.03 | -0.02 | -0.02 | 0 | 0.1 | 0.06 | 0.09 | 0.36 | 0.17 | 0.23 |

| | | | | | | | | | | | | | | | | |
|---|---|---|---|---|---|---|---|---|---|---|---|---|---|---|---|---|
| 6 | 51491884 | rs4715227 | 0.541 | 0.622 | -0.345 | 0.324 | -0.001 | -0.001 | -0.001 | 0 | 0.55 | 0.46 | 0.59 | 0.58 | 0.54 | 0.64 |
| 6 | 51508011 | rs12192780 | -0.169 | 0.822 | -1.085 | 0.449 | -0.026 | -0.057 | -0.057 | 0 | 0.3 | 0.13 | 0.33 | 0.4 | 0.36 | 0.28 |
| 6 | 51514462 | rs488063 | -0.888 | 0.745 | -0.942 | 0.281 | -0.025 | 0.062 | 0.062 | 0 | 0.89 | 0.99 | 0.94 | 1 | 0.88 | 0.92 |
| 6 | 51514608 | rs624699 | 0.051 | 0.564 | -1.094 | 0.382 | -0.006 | 0.017 | 0.017 | 0 | 0.79 | 0.96 | 0.84 | 0.87 | 0.81 | 0.85 |
| 6 | 51518203 | rs2149580 | 0.679 | -0.772 | -0.859 | 0.081 | 0.011 | 0 | 0 | 0 | 0.18 | 0.13 | 0.3 | 0.24 | 0.09 | 0.2 |
| 6 | 51526931 | rs9381995 | 0.044 | -0.155 | -0.837 | 0.517 | -0.001 | 0.017 | 0.017 | 0 | 0.13 | 0.1 | 0.16 | 0.26 | 0.12 | 0.15 |
| 6 | 51528984 | rs6906559 | -0.178 | -0.223 | -0.655 | 0.482 | -0.008 | 0.056 | 0.056 | 0 | 0.14 | 0.16 | 0.16 | 0.26 | 0.12 | 0.15 |
| 6 | 51530765 | rs6937025 | 1.077 | -0.064 | -1.464 | -0.083 | -0.1 | -0.082 | -0.082 | 0 | 0.26 | 0.08 | 0.22 | 0.34 | 0.36 | 0.41 |
| 6 | 51537351 | rs2174861 | -0.518 | -0.815 | -1.262 | -0.372 | 0.13 | -0.065 | -0.065 | 0 | 0.44 | 0.84 | 0.55 | 0.5 | 0.28 | 0.41 |
| 6 | 51544360 | rs9395706 | -0.557 | -0.901 | -0.855 | -0.454 | -0.02 | 0.119 | 0.119 | 0 | 0.18 | 0.3 | 0.44 | 0.43 | 0.14 | 0.28 |
| 6 | 51545092 | rs1507912 | 0.758 | -1.033 | -0.763 | -0.599 | -0.074 | -0.068 | -0.068 | 0.964 | 0.26 | 0.37 | 0.12 | 0.12 | 0.13 | 0.16 |
| 6 | 51548481 | rs16881777 | 0.543 | -0.953 | -0.727 | -0.516 | -0.077 | -0.077 | -0.077 | 0 | 0.28 | 0.4 | 0.13 | 0.12 | 0.13 | 0.17 |
| 6 | 51552302 | rs7756268 | 0.686 | -0.702 | -1.23 | -0.277 | 0.001 | 0.004 | 0.004 | 0 | 0.1 | 0.06 | 0.37 | 0.3 | 0.1 | 0.16 |
| 6 | 51558289 | rs12203695 | 1.261 | -0.45 | -1.411 | -0.585 | -0.076 | 0.012 | 0.012 | 0.072 | 0.15 | 0.02 | 0.13 | 0.09 | 0.22 | 0.07 |
| 6 | 51560111 | rs582669 | 0.282 | -1.297 | -1.16 | -0.443 | 0.114 | -0.007 | -0.007 | 2.289 | 0.54 | 0.23 | 0.41 | 0.43 | 0.7 | 0.54 |
| 6 | 51568871 | rs481795 | -0.074 | -2.147 | -0.856 | -1.463 | 0.034 | 0.036 | 0.036 | 0.794 | 0.65 | 0.34 | 0.78 | 0.8 | 0.84 | 0.75 |
| 6 | 51570461 | rs2766124 | 0.747 | -2.038 | -1.148 | -1.432 | 0.135 | 0.064 | 0.064 | 3.859 | 0.45 | 0.23 | 0.64 | 0.63 | 0.69 | 0.66 |
| 6 | 51571944 | rs9474056 | 0.301 | -2.046 | -0.581 | -1.498 | -0.015 | -0.04 | -0.04 | 0 | 0.3 | 0.31 | 0.18 | 0.13 | 0.21 | 0.19 |
| 6 | 51572515 | rs9395707 | 0.541 | -1.631 | -0.764 | -1.538 | 0.02 | 0.014 | 0.014 | 0 | 0.3 | 0.38 | 0.49 | 0.61 | 0.34 | 0.59 |
| 6 | 51579595 | rs6921803 | -0.365 | -2.205 | -0.879 | -1.453 | -0.037 | 0.073 | 0.073 | 0.537 | 0.66 | 0.79 | 0.75 | 0.91 | 0.57 | 0.81 |
| 6 | 51581912 | rs9370043 | 0.848 | -2.339 | -0.742 | -1.504 | -0.046 | -0.024 | -0.024 | 6.654 | 0.3 | 0.17 | 0.5 | 0.61 | 0.39 | 0.6 |
| 6 | 51586771 | rs9349593 | -0.426 | -2.166 | -1.02 | -1.435 | -0.037 | 0.073 | 0.073 | 0.478 | 0.66 | 0.79 | 0.75 | 0.91 | 0.57 | 0.81 |
| 6 | 51601124 | rs2397058 | -0.507 | -2.151 | -1.003 | -1.459 | -0.037 | -0.002 | -0.002 | 0 | 0.69 | 0.8 | 0.7 | 0.74 | 0.6 | 0.79 |
| 6 | 51611470 | rs2397060 | 0 | -2.555 | -1.059 | -1.7 | 0.06 | 0.058 | 0.058 | 3.052 | 0.34 | 0.77 | 0.19 | 0.2 | 0.15 | 0.22 |
| 6 | 51613177 | rs765525 | 1.472 | -2.425 | -1.212 | -1.663 | -0.152 | -0.06 | -0.06 | 0.405 | 0.25 | 0.03 | 0.48 | 0.65 | 0.39 | 0.52 |
| 6 | 51617677 | rs1750961 | -1.297 | -2.399 | -1.257 | -1.763 | -0.059 | 0.093 | 0.093 | 4.314 | 0.79 | 0.92 | 0.83 | 0.98 | 0.7 | 0.85 |
| 6 | 51618170 | rs1326605 | -1.18 | -2.389 | -1.271 | -1.758 | -0.058 | 0.1 | 0.1 | 2.434 | 0.78 | 0.91 | 0.83 | 0.98 | 0.7 | 0.85 |
| 6 | 51618599 | rs6914483 | 1.275 | -1.864 | -1.085 | -1.463 | -0.15 | -0.053 | -0.053 | 0 | 0.25 | 0.04 | 0.47 | 0.66 | 0.39 | 0.52 |
| 6 | 51619444 | rs12661470 | 0.805 | -2.004 | -0.843 | -1.323 | -0.001 | -0.003 | -0.003 | 0.152 | 0.09 | 0.09 | 0.13 | 0.13 | 0.1 | 0.06 |
| 6 | 51630524 | rs9367451 | 0.506 | -1.948 | -0.815 | -1.453 | -0.025 | 0.102 | 0.102 | 0 | 0.38 | 0.17 | 0.62 | 0.78 | 0.5 | 0.59 |
| 6 | 51641817 | rs10498791 | 0.612 | -2.443 | -0.524 | -1.712 | -0.046 | -0.031 | -0.031 | 0 | 0.24 | 0.32 | 0.15 | 0.17 | 0.15 | 0.18 |
| 6 | 51642708 | rs1159871 | 0.075 | -2.459 | -0.557 | -1.673 | 0.003 | 0.014 | 0.014 | 1.025 | 0.6 | 0.37 | 0.68 | 0.83 | 0.63 | 0.72 |
| 6 | 51653864 | rs2182505 | -0.83 | -2.378 | -0.523 | -1.644 | 0 | -0.057 | -0.057 | 0 | 0.75 | 0.73 | 0.83 | 0.97 | 0.74 | 0.8 |
| 6 | 51664343 | rs1326567 | 1.274 | -2.17 | -0.717 | -1.517 | -0.154 | -0.052 | -0.052 | 0 | 0.27 | 0.04 | 0.5 | 0.66 | 0.42 | 0.53 |
| 6 | 51666631 | rs10498792 | 0.374 | -2.097 | -0.454 | -1.529 | -0.009 | 0.019 | 0.019 | 0 | 0.14 | 0.24 | 0.15 | 0.13 | 0.11 | 0.08 |
| 6 | 51669562 | rs1326579 | 0.027 | -2.307 | -0.55 | -1.609 | -0.006 | -0.008 | -0.008 | 0 | 0.6 | 0.49 | 0.65 | 0.66 | 0.64 | 0.66 |
| 6 | 51670597 | rs6458792 | -1.023 | -2.355 | -0.535 | -2.403 | -0.003 | -0.079 | -0.079 | 0 | 0.78 | 0.73 | 0.86 | 0.97 | 0.79 | 0.82 |
| 6 | 51672610 | rs12206227 | 0.567 | -2.227 | -0.619 | -2.352 | -0.01 | 0.069 | 0.069 | 0 | 0.12 | 0.01 | 0.05 | 0 | 0.11 | 0.02 |
| 6 | 51675083 | rs2397067 | -0.11 | -2.154 | -0.552 | -2.273 | -0.006 | -0.025 | -0.025 | 0 | 0.6 | 0.49 | 0.67 | 0.78 | 0.64 | 0.66 |
| 6 | 51676922 | rs6906626 | 0.713 | -2.136 | -0.399 | -2.158 | -0.063 | -0.051 | -0.051 | 0 | 0.19 | 0.32 | 0.1 | 0.09 | 0.11 | 0.08 |
| 6 | 51679918 | rs7760761 | 0.073 | -0.71 | -0.318 | -1.405 | -0.052 | 0.032 | 0.032 | 0 | 0.35 | 0.16 | 0.56 | 0.7 | 0.5 | 0.56 |
| 6 | 51683874 | rs4715247 | 0.617 | -0.863 | -0.05 | -1.598 | 0.041 | 0.037 | 0.037 | 1.404 | 0.95 | 0.71 | 0.97 | 1 | 1 | 0.99 |
| 6 | 51683985 | rs12208497 | 0.561 | -0.521 | -0.204 | -1.444 | 0.002 | 0.047 | 0.047 | 0 | 0.08 | 0.01 | 0.03 | 0 | 0.06 | 0.01 |
| 6 | 51684250 | rs10948653 | 0.594 | -0.547 | -0.186 | -1.476 | 0.009 | -0.017 | -0.017 | 0 | 0.43 | 0.43 | 0.24 | 0.1 | 0.35 | 0.28 |
| 6 | 51688362 | rs9463719 | 0.325 | 0.092 | -0.377 | -0.965 | -0.005 | -0.06 | -0.06 | 0 | 0.53 | 0.55 | 0.28 | 0.11 | 0.42 | 0.28 |
| 6 | 51705624 | rs788510 | -0.242 | 0.934 | -0.499 | -0.261 | -0.032 | 0.004 | 0.004 | 0 | 0.29 | 0.19 | 0.54 | 0.7 | 0.43 | 0.58 |
| 6 | 51757722 | rs12207819 | -0.954 | 1.446 | 0.322 | -0.355 | -0.026 | 0.031 | 0.031 | 0 | 0.12 | 0.08 | 0.1 | 0.08 | 0.17 | 0.08 |
| 6 | 51778076 | rs17669233 | -0.058 | 1.583 | 0.06 | -0.593 | -0.033 | 0.048 | 0.048 | 0 | 0.09 | 0 | 0.05 | 0 | 0.11 | 0.04 |
| 6 | 51797474 | rs12209427 | -1.378 | 1.794 | 0.563 | -0.72 | -0.039 | 0.057 | 0.057 | 0 | 0.21 | 0.12 | 0.16 | 0.07 | 0.29 | 0.13 |
| 6 | 51810499 | rs12194995 | -0.619 | 1.793 | 0.482 | -0.722 | -0.052 | 0.039 | 0.039 | 0 | 0.11 | 0.02 | 0.11 | 0.07 | 0.18 | 0.09 |
| 6 | 51812711 | rs1266909 | -1.337 | 1.591 | 0.812 | -0.63 | 0.022 | -0.033 | -0.033 | 0 | 0.25 | 0.36 | 0.2 | 0.19 | 0.3 | 0.17 |
| 6 | 51822297 | rs1266922 | -1.285 | 1.457 | 1.221 | -0.709 | 0.019 | -0.046 | -0.046 | 0.021 | 0.26 | 0.51 | 0.22 | 0.19 | 0.3 | 0.17 |
| 6 | 51826214 | rs1266924 | -0.555 | 1.473 | 0.244 | -0.054 | 0.003 | 0.009 | 0.009 | 0 | 0.23 | 0.35 | 0.19 | 0.7 | 0.27 | 0.42 |
| 6 | 51826281 | rs1321507 | 0.786 | 1.234 | -0.39 | -1.113 | -0.008 | 0.09 | 0.09 | 0 | 0.41 | 0.24 | 0.27 | 0.07 | 0.42 | 0.27 |
| 6 | 51826782 | rs927185 | 0.671 | 0.628 | -0.514 | 0.367 | -0.026 | -0.048 | -0.048 | 0.645 | 0.8 | 0.94 | 0.83 | 0.66 | 0.76 | 0.52 |
| 6 | 51828478 | rs1896967 | 0.703 | 0.037 | -0.084 | 0.523 | 0 | 0 | 0 | 0 | 0.63 | 0.41 | 0.72 | 0.6 | 0.72 | 0.55 |
| 6 | 51838263 | rs9296668 | 1.203 | -0.409 | -0.767 | 0.973 | -0.058 | -0.019 | -0.019 | 0.044 | 0.4 | 0.09 | 0.55 | 0.39 | 0.46 | 0.34 |
| 6 | 51838372 | rs2435327 | 0.458 | -0.449 | -0.472 | 0.939 | 0 | 0.001 | 0.001 | 0 | 0.62 | 0.43 | 0.65 | 0.47 | 0.59 | 0.49 |
| 6 | 51844205 | rs10948664 | 0.998 | -0.618 | -0.737 | 0.793 | -0.044 | -0.071 | -0.071 | 0 | 0.45 | 0.1 | 0.63 | 0.53 | 0.6 | 0.44 |
| 6 | 51859957 | rs12212492 | -0.842 | -0.552 | -0.218 | 0.633 | 0.003 | 0.012 | 0.012 | 0 | 0.07 | 0.08 | 0.06 | 0.14 | 0.1 | 0.11 |
| 6 | 51873614 | rs11968546 | 0.782 | -0.635 | -0.303 | 0.471 | 0.008 | 0.064 | 0.064 | 0.762 | 0.18 | 0.03 | 0.07 | 0.04 | 0.11 | 0.06 |
| 6 | 51875890 | rs2017134 | 0.721 | -0.713 | -0.082 | 0.704 | 0.021 | -0.001 | -0.001 | 0.247 | 0.49 | 0.24 | 0.65 | 0.49 | 0.6 | 0.51 |
| 6 | 51876267 | rs1896974 | 1.305 | -0.713 | -0.119 | 0.709 | -0.054 | 0.008 | 0.008 | 0 | 0.41 | 0.16 | 0.59 | 0.38 | 0.5 | 0.35 |
| 6 | 51890823 | rs9296669 | 1.283 | -0.725 | -0.147 | 0.707 | -0.086 | -0.015 | -0.015 | 0 | 0.4 | 0.11 | 0.59 | 0.38 | 0.5 | 0.35 |
| 6 | 51899824 | rs2499475 | 0.004 | -0.788 | 0.209 | 0.749 | -0.001 | 0.027 | 0.027 | 0 | 0.7 | 0.42 | 0.74 | 0.56 | 0.74 | 0.6 |
| 6 | 51901537 | rs4711994 | 0.355 | -0.801 | 0.479 | 0.627 | 0.018 | 0.039 | 0.039 | 0.084 | 0.21 | 0.16 | 0.08 | 0.07 | 0.14 | 0.1 |
| 6 | 51902528 | rs2435328 | 1.419 | -0.672 | 0.829 | 0.721 | 0.091 | 0.134 | 0.134 | 2.74 | 0.93 | 0.54 | 0.94 | 0.9 | 0.92 | 0.92 |
| 6 | 51912465 | rs9474138 | 0.042 | -0.755 | 0.658 | 0.763 | 0.004 | 0.035 | 0.035 | 0 | 0.27 | 0.31 | 0.23 | 0.42 | 0.25 | 0.4 |
| 6 | 51913240 | rs10948667 | 1.422 | -0.805 | 0.266 | 0.682 | -0.082 | -0.045 | -0.045 | 0.161 | 0.4 | 0.11 | 0.6 | 0.4 | 0.5 | 0.31 |

| | | | | | | | | | | | | | | | | |
|---|---|---|---|---|---|---|---|---|---|---|---|---|---|---|---|---|
| 6 | 51914956 | rs9370096 | 1.422 | -0.808 | 0.335 | 0.68 | -0.079 | -0.042 | -0.042 | 0 | 0.4 | 0.11 | 0.6 | 0.4 | 0.5 | 0.31 |
| 6 | 51919900 | rs10948668 | -0.016 | -0.872 | 0.358 | 0.591 | 0.012 | 0.064 | 0.064 | 0 | 0.24 | 0.12 | 0.09 | 0.08 | 0.17 | 0.17 |
| 6 | 51930437 | rs12529349 | 0.106 | -0.843 | 0.466 | 0.65 | 0.002 | 0.038 | 0.038 | 0 | 0.27 | 0.32 | 0.25 | 0.42 | 0.24 | 0.4 |
| 6 | 51934978 | rs4715276 | 0.433 | -0.834 | 0.313 | 0.646 | -0.002 | -0.058 | -0.058 | 0.8 | 0.65 | 0.22 | 0.7 | 0.47 | 0.68 | 0.49 |
| 6 | 51944847 | rs1892274 | 1.372 | -0.826 | 0.32 | 0.626 | -0.03 | 0.002 | 0.002 | 0 | 0.42 | 0.2 | 0.61 | 0.4 | 0.51 | 0.31 |
| 6 | 51947237 | rs9474143 | -0.032 | -0.801 | 0.413 | 0.665 | 0 | 0.063 | 0.063 | 0 | 0.28 | 0.39 | 0.25 | 0.43 | 0.25 | 0.4 |
| 6 | 51947257 | rs6901799 | 0.685 | -0.749 | 0.229 | 0.588 | 0.017 | 0.066 | 0.066 | 0.241 | 0.19 | 0.07 | 0.08 | 0.04 | 0.12 | 0.06 |
| 6 | 51948935 | rs1896978 | 1.133 | -0.592 | 0.467 | 0.697 | 0.032 | 0.061 | 0.061 | 0 | 0.94 | 0.82 | 0.96 | 0.87 | 0.9 | 0.85 |
| 6 | 51953097 | rs9357723 | 1.408 | -1.288 | -0.196 | 0.434 | -0.086 | 0.073 | 0.073 | 0.251 | 0.31 | 0.15 | 0.25 | 0.13 | 0.46 | 0.31 |
| 6 | 51965791 | rs6458809 | -0.495 | -1.426 | 0.157 | 0.46 | -0.016 | 0.033 | 0.033 | 0.193 | 0.21 | 0.33 | 0.42 | 0.41 | 0.18 | 0.26 |
| 6 | 51966414 | rs10948673 | 0.234 | -1.434 | 0.17 | 0.471 | 0.015 | -0.015 | -0.015 | 0 | 0.16 | 0.11 | 0.36 | 0.3 | 0.09 | 0.11 |
| 6 | 51970228 | rs7768816 | -0.46 | -1.419 | 0.168 | 0.487 | -0.016 | 0.032 | 0.032 | 0.22 | 0.22 | 0.34 | 0.42 | 0.41 | 0.18 | 0.26 |
| 6 | 51974289 | rs4715280 | 0.385 | -1.45 | -0.016 | 0.442 | -0.021 | 0.019 | 0.019 | 0.032 | 0.14 | 0.18 | 0.35 | 0.3 | 0.08 | 0.06 |
| 6 | 51986236 | rs12527406 | -0.139 | -1.582 | 0.041 | 0.233 | 0.02 | 0 | 0 | 0.036 | 0.32 | 0.24 | 0.41 | 0.35 | 0.18 | 0.14 |
| 6 | 51987453 | rs1937147 | -1.8 | -1.49 | 0.387 | -0.027 | 0.009 | -0.03 | -0.03 | 0.115 | 0.71 | 0.62 | 0.75 | 0.9 | 0.59 | 0.73 |
| 6 | 51990683 | rs1000094 | -0.146 | -1.536 | 0.471 | -0.043 | -0.011 | -0.006 | -0.006 | 0 | 0.37 | 0.28 | 0.34 | 0.56 | 0.41 | 0.59 |
| 6 | 51993491 | rs1937151 | -1.757 | -1.483 | 0.403 | -0.046 | 0.013 | -0.059 | -0.059 | 0.09 | 0.7 | 0.59 | 0.76 | 0.93 | 0.61 | 0.77 |
| 6 | 51999566 | rs9463760 | 0.692 | -1.641 | 0.238 | 0.156 | -0.002 | 0.011 | 0.011 | 0 | 0.12 | 0.09 | 0.14 | 0.34 | 0.13 | 0.23 |
| 6 | 52002310 | rs1413917 | 0.141 | -1.762 | -0.148 | 0.265 | -0.003 | 0.002 | 0.002 | 0.328 | 0.29 | 0.24 | 0.1 | 0.22 | 0.13 | 0.12 |
| 6 | 52002829 | rs9370101 | 1.299 | -1.675 | -0.237 | 0.266 | -0.065 | -0.06 | -0.06 | 0 | 0.13 | 0.01 | 0.42 | 0.23 | 0.16 | 0.18 |
| 6 | 52003764 | rs9370102 | 0.427 | -1.56 | 0.106 | -0.067 | 0.006 | 0.017 | 0.017 | 0 | 0.28 | 0.29 | 0.6 | 0.56 | 0.3 | 0.42 |
| 6 | 52004363 | rs17640030 | 1.013 | -1.62 | -0.01 | -0.127 | 0.016 | 0.029 | 0.029 | 0.344 | 0.17 | 0.09 | 0.07 | 0.03 | 0.11 | 0.07 |
| 6 | 52005005 | rs11966138 | -0.069 | -1.564 | -0.018 | -0.135 | 0.019 | 0.013 | 0.013 | 0.542 | 0.31 | 0.23 | 0.11 | 0.22 | 0.15 | 0.12 |
| 6 | 52008616 | rs6920648 | 1.506 | -1.498 | 0.041 | -0.419 | -0.001 | -0.018 | -0.018 | 0 | 0.3 | 0.36 | 0.24 | 0.08 | 0.44 | 0.29 |
| 6 | 52009710 | rs17578851 | 0.562 | -1.374 | -0.152 | -0.391 | 0.005 | 0.014 | 0.014 | 0.313 | 0.22 | 0.15 | 0.11 | 0.09 | 0.17 | 0.07 |
| 6 | 52011469 | rs10484882 | -0.131 | -1.089 | -0.311 | -0.22 | 0.005 | -0.019 | -0.019 | 0 | 0.12 | 0.2 | 0.07 | 0.07 | 0.11 | 0.2 |
| 6 | 52012873 | rs10948679 | 0.61 | -1.092 | -0.278 | -0.228 | -0.018 | 0.034 | 0.034 | 3.214 | 0.38 | 0.34 | 0.29 | 0.15 | 0.54 | 0.5 |
| 6 | 52018919 | rs1937146 | 0.592 | -1.155 | -0.205 | -0.061 | 0.001 | 0.04 | 0.04 | 0.391 | 0.26 | 0.31 | 0.56 | 0.46 | 0.25 | 0.36 |
| 6 | 52019663 | rs16882141 | 0.534 | -0.841 | -1.107 | 0.098 | -0.023 | 0.018 | 0.018 | 0 | 0.13 | 0.17 | 0.04 | 0.31 | 0.04 | 0.05 |
| 6 | 52022832 | rs7766405 | -0.284 | 0.169 | -1.473 | -0.729 | 0.039 | 0.234 | 0.234 | 0 | 0.5 | 0.15 | 0.34 | 0.16 | 0.61 | 0.39 |
| 6 | 52044509 | rs13208597 | 0.122 | 0.772 | -1.505 | -0.423 | -0.107 | 0.128 | 0.128 | 0.325 | 0.27 | 0.05 | 0.22 | 0.06 | 0.42 | 0.26 |
| 6 | 52050353 | rs4711998 | 0.346 | -0.134 | -0.798 | -0.05 | -0.022 | -0.019 | -0.019 | 0 | 0.68 | 0.43 | 0.75 | 0.23 | 0.74 | 0.55 |
| 6 | 52050493 | rs8193036 | 0.882 | -0.103 | -0.826 | 0.105 | -0.001 | 0.001 | 0.001 | 0.072 | 0.75 | 0.74 | 0.76 | 0.3 | 0.74 | 0.55 |
| 6 | 52050786 | rs3819024 | -0.312 | 0.402 | -0.788 | 0.123 | -0.004 | -0.009 | -0.009 | 0 | 0.3 | 0.25 | 0.24 | 0.51 | 0.38 | 0.36 |
| 6 | 52051274 | rs3819025 | 0.09 | 0.1 | -1.217 | -0.226 | 0.006 | -0.009 | -0.009 | 2.547 | 0.11 | 0.1 | 0.34 | 0.19 | 0.06 | 0.2 |
| 6 | 52051957 | rs10484879 | 0.49 | 0.697 | -1.269 | -0.415 | -0.085 | 0.124 | 0.124 | 0 | 0.19 | 0.01 | 0.15 | 0 | 0.25 | 0.12 |
| 6 | 52054249 | rs7747909 | 0.447 | 0.694 | -1.111 | -0.417 | -0.04 | 0.069 | 0.069 | 0 | 0.19 | 0.06 | 0.15 | 0 | 0.25 | 0.12 |
| 6 | 52056386 | rs16882180 | 0.036 | 0.316 | -0.976 | 0.15 | -0.047 | -0.107 | -0.107 | 0 | 0.29 | 0.09 | 0.22 | 0.48 | 0.36 | 0.38 |
| 6 | 52063101 | rs2154225 | -0.063 | -0.133 | -1.653 | -0.763 | 0.075 | 0.035 | 0.035 | 0.194 | 0.62 | 0.18 | 0.76 | 0.71 | 0.75 | 0.62 |
| 6 | 52063698 | rs11759291 | -0.096 | -0.171 | -1.378 | -0.783 | 0.057 | 0.028 | 0.028 | 0.072 | 0.62 | 0.22 | 0.75 | 0.71 | 0.75 | 0.62 |
| 6 | 52066624 | rs9370105 | 0.366 | -0.479 | -0.744 | -0.903 | 0.013 | 0.014 | 0.014 | 0 | 0.31 | 0.29 | 0.14 | 0.19 | 0.22 | 0.31 |
| 6 | 52075816 | rs2397079 | 0.165 | -0.496 | -0.264 | -0.91 | 0.031 | 0.016 | 0.016 | 0 | 0.63 | 0.31 | 0.76 | 0.7 | 0.75 | 0.62 |
| 6 | 52077984 | rs1999673 | 0.891 | -0.776 | -0.34 | -1.166 | 0.014 | 0.032 | 0.032 | 0.006 | 0.19 | 0.12 | 0.09 | 0.04 | 0.16 | 0.17 |
| 6 | 52085748 | rs591274 | -1.15 | -0.601 | -0.53 | -1.069 | -0.054 | 0.05 | 0.05 | 0.213 | 0.68 | 0.86 | 0.46 | 0.77 | 0.61 | 0.76 |
| 6 | 52088183 | rs2179560 | -0.927 | -0.641 | 0.032 | -0.156 | -0.038 | -0.043 | -0.043 | 0 | 0.35 | 0.49 | 0.16 | 0.2 | 0.23 | 0.38 |
| 6 | 52094056 | rs2064332 | 0.394 | -0.132 | -0.19 | 0.089 | -0.012 | -0.01 | -0.01 | 0.208 | 0.77 | 0.78 | 0.52 | 0.65 | 0.68 | 0.65 |
| 6 | 52097701 | rs9395771 | 1.303 | -0.799 | -0.138 | -0.075 | -0.003 | 0.012 | 0.012 | 0 | 0.54 | 0.34 | 0.35 | 0.54 | 0.53 | 0.55 |
| 6 | 52097787 | rs12207872 | 1.04 | -0.761 | -0.478 | -0.077 | -0.002 | 0.031 | 0.031 | 0 | 0.07 | 0.02 | 0.02 | 0 | 0.06 | 0.05 |
| 6 | 52098036 | rs1266831 | -0.552 | -0.787 | -0.469 | -0.004 | 0.019 | 0.002 | 0.002 | 0 | 0.85 | 0.78 | 0.56 | 0.82 | 0.74 | 0.77 |
| 6 | 52098368 | rs2064331 | 1.029 | -0.993 | -0.092 | -0.09 | -0.001 | -0.001 | -0.001 | 0 | 0.6 | 0.55 | 0.39 | 0.55 | 0.59 | 0.6 |
| 6 | 52101844 | rs2397084 | 0.784 | -0.896 | -0.537 | -0.302 | -0.006 | 0.06 | 0.06 | 0 | 0.1 | 0 | 0.05 | 0 | 0.08 | 0.05 |
| 6 | 52104689 | rs12201582 | -0.269 | -0.946 | -0.384 | -0.272 | 0.004 | 0.004 | 0.004 | 0 | 0.21 | 0.2 | 0.16 | 0.11 | 0.15 | 0.1 |
| 6 | 52105557 | rs12203582 | 1.829 | -0.932 | -0.791 | -0.254 | -0.089 | -0.047 | -0.047 | 0 | 0.42 | 0.06 | 0.27 | 0.35 | 0.45 | 0.42 |
| 6 | 52111808 | rs761167 | 1.505 | -1.194 | 0.292 | -0.259 | -0.075 | -0.019 | -0.019 | 0 | 0.44 | 0.08 | 0.3 | 0.38 | 0.5 | 0.5 |
| 6 | 52112966 | rs607175 | 0.972 | -1.314 | 0.445 | -0.183 | -0.001 | -0.004 | -0.004 | 0.666 | 0.53 | 0.47 | 0.36 | 0.57 | 0.52 | 0.61 |
| 6 | 52118025 | rs7741835 | -1.114 | -1.669 | 0.513 | -0.242 | -0.071 | -0.002 | -0.002 | 0 | 0.3 | 0.52 | 0.22 | 0.32 | 0.17 | 0.22 |
| 6 | 52121015 | rs669161 | 1.016 | -1.465 | 0.478 | -0.236 | 0.004 | 0.011 | 0.011 | 0 | 0.59 | 0.47 | 0.42 | 0.48 | 0.57 | 0.52 |
| 6 | 52124370 | rs1266836 | -0.242 | -1.326 | 0.331 | -0.605 | -0.014 | -0.006 | -0.006 | 0 | 0.75 | 0.77 | 0.51 | 0.69 | 0.66 | 0.72 |
| 6 | 52126705 | rs742552 | -0.036 | -1.27 | 0.327 | -0.196 | 0.048 | 0.116 | 0.116 | 0 | 0.77 | 0.54 | 0.51 | 0.49 | 0.69 | 0.63 |
| 6 | 52126781 | rs605134 | -0.531 | -1.26 | 0.215 | -0.187 | 0.034 | -0.009 | -0.009 | 0 | 0.82 | 0.72 | 0.55 | 0.83 | 0.71 | 0.78 |
| 6 | 52129484 | rs2230240 | 2.309 | -1.748 | -0.635 | -0.776 | -0.094 | 0.174 | 0.174 | 0.061 | 0.25 | 0.02 | 0.17 | 0 | 0.29 | 0.15 |
| 6 | 52130963 | rs2245502 | 0.825 | -1.556 | -0.43 | -0.508 | 0.019 | -0.004 | -0.004 | 0 | 0.15 | 0.01 | 0.13 | 0.12 | 0.09 | 0.06 |
| 6 | 52135494 | rs3789775 | 1.961 | -1.581 | -0.455 | -0.493 | -0.04 | -0.078 | -0.078 | 0.599 | 0.12 | 0 | 0.07 | 0.24 | 0.13 | 0.29 |
| 6 | 52141512 | rs3765447 | 0.111 | -1.368 | 0.359 | 0.321 | -0.01 | -0.039 | -0.039 | 0 | 0.1 | 0.01 | 0.04 | 0.13 | 0.07 | 0.07 |
| 6 | 52142092 | rs2307325 | 0.62 | -1.371 | 0.658 | 0.303 | 0.061 | 0.008 | 0.008 | 0 | 0.73 | 0.54 | 0.5 | 0.7 | 0.63 | 0.71 |
| 6 | 52144081 | rs2073036 | 1.863 | -1.294 | 0.289 | 0.293 | -0.08 | 0.105 | 0.105 | 0 | 0.47 | 0.09 | 0.3 | 0.24 | 0.52 | 0.49 |
| 6 | 52146272 | rs6934554 | 0.59 | -1.157 | 0.465 | 0.412 | -0.023 | 0.02 | 0.02 | 0 | 0.67 | 0.76 | 0.42 | 0.71 | 0.62 | 0.71 |
| 6 | 52147271 | rs7756989 | 0.989 | -1.409 | 0.623 | 0.433 | 0.004 | -0.01 | -0.01 | 0 | 0.58 | 0.45 | 0.47 | 0.69 | 0.54 | 0.65 |
| 6 | 52147309 | rs7774976 | -0.532 | -1.144 | 0.663 | 0.643 | -0.043 | 0.142 | 0.142 | 0 | 0.16 | 0.31 | 0.19 | 0.43 | 0.1 | 0.16 |

| | | | | | | | | | | | | | | | | |
|---|---|---|---|---|---|---|---|---|---|---|---|---|---|---|---|---|
| 6 | 52148465 | rs9474192 | -0.185 | -1.049 | 0.566 | 0.662 | 0.036 | 0.059 | 0.059 | 0 | 0.21 | 0.36 | 0.47 | 0.47 | 0.3 | 0.34 |
| 6 | 52166561 | rs9370111 | 0.002 | -1.149 | -0.122 | 0.608 | -0.004 | 0.002 | 0.002 | 0 | 0.13 | 0.1 | 0.16 | 0.17 | 0.1 | 0.14 |
| 6 | 52172655 | rs2073032 | 0.111 | -1.079 | 0.022 | 0.562 | 0.069 | -0.011 | -0.011 | 0.404 | 0.77 | 0.47 | 0.51 | 0.79 | 0.7 | 0.73 |
| 6 | 52177355 | rs1535010 | 0.802 | -1.148 | 0.044 | 0.587 | -0.003 | -0.009 | -0.009 | 0 | 0.59 | 0.28 | 0.43 | 0.48 | 0.59 | 0.61 |
| 6 | 52178604 | rs9395780 | -0.121 | -0.765 | -0.117 | 0.788 | 0.098 | 0.019 | 0.019 | 0 | 0.81 | 0.5 | 0.54 | 0.79 | 0.74 | 0.76 |
| 6 | 52182262 | rs11964170 | 0.079 | -0.762 | -0.065 | 0.77 | 0.042 | 0.066 | 0.066 | 0 | 0.69 | 0.42 | 0.45 | 0.45 | 0.65 | 0.58 |
| 6 | 52185051 | rs7750841 | -0.165 | -0.867 | -0.364 | 0.168 | 0.042 | -0.017 | -0.017 | 3.299 | 0.69 | 0.53 | 0.46 | 0.75 | 0.63 | 0.62 |
| 6 | 52187184 | rs6925628 | -0.43 | -0.699 | -1.222 | -0.668 | -0.007 | -0.044 | -0.044 | 0 | 0.18 | 0.2 | 0.09 | 0 | 0.14 | 0.1 |
| 6 | 52191257 | rs6458829 | 0.688 | -0.569 | -1.132 | -0.563 | -0.004 | -0.027 | -0.027 | 0 | 0.06 | 0.09 | 0.04 | 0 | 0.05 | 0.05 |
| 6 | 52193513 | rs11964472 | 0.003 | -0.425 | -0.924 | -0.138 | 0.01 | 0.001 | 0.001 | 0 | 0.17 | 0.12 | 0.08 | 0.13 | 0.14 | 0.11 |
| 6 | 52193550 | rs6458830 | 0.468 | 0.134 | -1.113 | -0.524 | 0 | 0.117 | 0.117 | 0 | 0.47 | 0.24 | 0.28 | 0.19 | 0.47 | 0.46 |
| 6 | 52198802 | rs560243 | 0.592 | 0.795 | -0.352 | -0.37 | 0.016 | -0.046 | -0.046 | 0.149 | 0.74 | 0.38 | 0.52 | 0.88 | 0.72 | 0.75 |
| 6 | 52203598 | rs9349621 | -0.325 | 0.296 | -0.412 | -0.739 | 0 | -0.017 | -0.017 | 0 | 0.35 | 0.4 | 0.19 | 0.11 | 0.27 | 0.38 |
| 6 | 52203832 | rs472589 | -0.164 | 1.03 | -1.066 | -0.129 | -0.066 | -0.071 | -0.071 | 0 | 0.89 | 0.99 | 0.86 | 0.77 | 0.78 | 0.77 |
| 6 | 52205426 | rs11759990 | -0.026 | 0.849 | -0.908 | -0.211 | 0 | 0.075 | 0.075 | 0 | 0.33 | 0.2 | 0.19 | 0.11 | 0.29 | 0.36 |
| 6 | 52205511 | rs7774765 | -0.341 | 0.254 | -0.15 | 0.987 | -0.065 | 0.187 | 0.187 | 0 | 0.16 | 0.39 | 0.34 | 0.42 | 0.06 | 0.14 |
| 6 | 52208065 | rs1359227 | 0.686 | 0.485 | -0.26 | 0.826 | -0.003 | 0.023 | 0.023 | 0 | 0.75 | 0.75 | 0.55 | 0.35 | 0.74 | 0.69 |
| 6 | 52209565 | rs514638 | 0.888 | 0.84 | -0.395 | 0.392 | -0.004 | -0.004 | -0.004 | 5.701 | 0.79 | 0.83 | 0.78 | 0.75 | 0.73 | 0.67 |
| 6 | 52211294 | rs567084 | -0.704 | 1.021 | -0.336 | -0.762 | 0.009 | -0.11 | -0.11 | 0 | 0.38 | 0.53 | 0.37 | 0.06 | 0.53 | 0.44 |
| 6 | 52212250 | rs570854 | -0.646 | 1.586 | -0.839 | -0.081 | -0.028 | -0.023 | -0.023 | 0 | 0.33 | 0.31 | 0.38 | 0.43 | 0.53 | 0.35 |
| 6 | 52213010 | rs6912417 | -1.29 | 1.031 | -0.343 | 0.244 | 0.053 | 0.038 | 0.038 | 0 | 0.53 | 0.64 | 0.55 | 0.64 | 0.73 | 0.65 |
| 6 | 52215788 | rs8180555 | -0.289 | 1.447 | -0.202 | 0.504 | 0.017 | 0.074 | 0.074 | 0 | 0.15 | 0.24 | 0.12 | 0.26 | 0.19 | 0.29 |
| 6 | 52217375 | rs7754194 | -0.387 | 2.592 | -0.534 | 0.714 | -0.045 | -0.044 | -0.044 | 3.426 | 0.8 | 0.92 | 0.75 | 0.76 | 0.73 | 0.84 |
| 6 | 52217699 | rs7772620 | -0.194 | 2.567 | -0.494 | 0.818 | -0.021 | -0.062 | -0.062 | 0 | 0.78 | 0.85 | 0.75 | 0.6 | 0.73 | 0.83 |
| 6 | 52220229 | rs7749979 | 0.956 | 2.423 | -0.639 | 0.357 | -0.015 | 0.004 | 0.004 | 0 | 0.68 | 0.5 | 0.72 | 0.38 | 0.72 | 0.56 |
| 6 | 52220793 | rs9382099 | 0.604 | 1.919 | -0.636 | 0.368 | -0.042 | 0.024 | 0.024 | 0.036 | 0.07 | 0 | 0.08 | 0.03 | 0.12 | 0.16 |
| 6 | 52220975 | rs9367480 | 0.976 | 1.891 | -0.641 | 0.358 | 0.013 | -0.005 | -0.005 | 0 | 0.64 | 0.49 | 0.65 | 0.38 | 0.58 | 0.54 |
| 6 | 52222164 | rs4712002 | -0.963 | 1.367 | -0.499 | 0.447 | 0.001 | -0.018 | -0.018 | 0.363 | 0.89 | 0.92 | 0.86 | 0.76 | 0.89 | 0.9 |
| 6 | 52225514 | rs2147206 | 1.575 | 1.337 | -0.53 | 0.373 | 0.013 | 0.012 | 0.012 | 0 | 0.47 | 0.48 | 0.45 | 0.36 | 0.27 | 0.33 |
| 6 | 52228225 | rs875142 | 1.463 | 1.298 | -0.517 | 0.358 | 0.026 | 0.022 | 0.022 | 0 | 0.49 | 0.46 | 0.46 | 0.36 | 0.26 | 0.32 |
| 6 | 52233226 | rs6458836 | -1.093 | 0.943 | -1.2 | -0.005 | 0.04 | 0.025 | 0.025 | 0.872 | 0.63 | 0.71 | 0.73 | 0.74 | 0.76 | 0.84 |
| 6 | 52236968 | rs17650296 | 0.5 | 1.654 | -1.149 | 0.071 | -0.055 | 0.054 | 0.054 | 0 | 0.09 | 0.01 | 0.11 | 0 | 0.19 | 0.06 |
| 6 | 52240958 | rs4712005 | -0.152 | 1.486 | -0.864 | 0.265 | -0.005 | -0.005 | -0.005 | 0 | 0.32 | 0.28 | 0.43 | 0.4 | 0.33 | 0.44 |
| 6 | 52242105 | rs10807443 | -1.161 | 1.837 | -0.457 | 0.087 | 0.004 | 0 | 0 | 0.011 | 0.28 | 0.26 | 0.31 | 0.27 | 0.54 | 0.37 |
| 6 | 52245415 | rs4712006 | -1.114 | 2 | -0.737 | -0.103 | -0.038 | 0.013 | 0.013 | 0.523 | 0.3 | 0.21 | 0.45 | 0.27 | 0.39 | 0.46 |
| 6 | 52253839 | rs10456660 | -0.347 | 2.091 | -0.256 | -0.345 | -0.088 | 0.043 | 0.043 | 0.389 | 0.15 | 0.04 | 0.15 | 0.07 | 0.29 | 0.23 |
| 6 | 52254149 | rs10484876 | 0.01 | 2.351 | -0.344 | -0.355 | -0.051 | 0.014 | 0.014 | 0.184 | 0.08 | 0.01 | 0.13 | 0.03 | 0.19 | 0.12 |
| 6 | 52255934 | rs11967792 | 0.025 | 1.232 | -0.399 | -1.038 | -0.015 | 0.008 | 0.008 | 0 | 0.39 | 0.5 | 0.31 | 0.53 | 0.29 | 0.36 |
| 6 | 52263744 | rs650342 | -0.784 | 1.28 | -0.229 | -1.409 | 0.073 | -0.065 | -0.065 | 0.072 | 0.15 | 0.33 | 0.16 | 0.03 | 0.23 | 0.13 |
| 6 | 52266511 | rs576138 | -1.255 | 0.99 | -0.563 | -1.435 | -0.009 | -0.053 | -0.053 | 0.005 | 0.9 | 1 | 0.88 | 0.53 | 0.93 | 0.86 |
| 6 | 52268008 | rs3748066 | 0.899 | 1.018 | -0.345 | -1.515 | 0.024 | 0.033 | 0.033 | 0 | 0.48 | 0.63 | 0.46 | 0.25 | 0.31 | 0.39 |
| 6 | 52268890 | rs17591381 | 1.229 | 0.68 | -0.733 | -1.658 | -0.031 | 0.042 | 0.042 | 0 | 0.17 | 0.01 | 0.12 | 0.07 | 0.16 | 0.16 |
| 6 | 52269547 | rs3180067 | 1.367 | 0.481 | -0.797 | -1.786 | 0.01 | -0.01 | -0.01 | 0 | 0.11 | 0.1 | 0.29 | 0.19 | 0.06 | 0.23 |
| 6 | 52269685 | rs3180068 | 0.556 | 0.466 | -0.806 | -1.796 | 0.007 | -0.019 | -0.019 | 0 | 0.16 | 0.11 | 0.33 | 0.22 | 0.13 | 0.29 |
| 6 | 52269936 | rs10484875 | 1.379 | 0.381 | -0.874 | -1.846 | -0.03 | 0.085 | 0.085 | 0 | 0.14 | 0.01 | 0.1 | 0 | 0.15 | 0.09 |
| 6 | 52270992 | rs10807445 | 0.571 | 0.597 | -0.88 | -2.004 | -0.082 | 0.035 | 0.035 | 0.005 | 0.13 | 0.01 | 0.12 | 0.05 | 0.25 | 0.14 |
| 6 | 52271204 | rs4715300 | 1.362 | 0.289 | -0.764 | -1.894 | 0.014 | -0.015 | -0.015 | 0.019 | 0.11 | 0.09 | 0.29 | 0.19 | 0.06 | 0.21 |
| 6 | 52272033 | rs474848 | -0.672 | 0.294 | -0.66 | -1.894 | 0.021 | -0.052 | -0.052 | 0.218 | 0.7 | 0.88 | 0.73 | 0.46 | 0.74 | 0.68 |
| 6 | 52272313 | rs1297 | 2.412 | 0.212 | -0.627 | -1.708 | 0.023 | 0.013 | 0.013 | 1.261 | 0.25 | 0.21 | 0.1 | 0.03 | 0.14 | 0.07 |
| 6 | 52283825 | rs16882521 | 1.135 | 0.37 | -0.623 | -1.477 | 0.022 | -0.032 | -0.032 | 0.686 | 0.11 | 0.05 | 0.28 | 0.2 | 0.06 | 0.2 |
| 6 | 52285957 | rs9382107 | 0.187 | 0.533 | -0.365 | -1.457 | 0.026 | 0.066 | 0.066 | 0 | 0.06 | 0.11 | 0.06 | 0.15 | 0.09 | 0.13 |
| 6 | 52290927 | rs719395 | 0.063 | 0.559 | -0.22 | -1.715 | -0.06 | 0.068 | 0.068 | 0 | 0.31 | 0.18 | 0.28 | 0.15 | 0.47 | 0.37 |
| 6 | 52316878 | rs2025226 | 0.619 | 0.599 | -0.105 | -1.662 | -0.016 | -0.016 | -0.016 | 0 | 0.16 | 0.11 | 0.38 | 0.19 | 0.19 | 0.23 |
| 6 | 52319251 | rs614570 | 0.461 | 0.467 | -0.011 | -1.757 | 0.006 | 0.036 | 0.036 | 0 | 0.51 | 0.66 | 0.56 | 0.38 | 0.45 | 0.41 |
| 6 | 52329510 | rs16882561 | 0.383 | 0.421 | -0.043 | -1.737 | 0.023 | -0.022 | -0.022 | 2.959 | 0.06 | 0.21 | 0.03 | 0 | 0.04 | 0.01 |
| 6 | 52336361 | rs2181274 | 1.36 | 0.289 | -0.104 | -1.832 | 0.026 | 0.075 | 0.075 | 0 | 0.4 | 0.3 | 0.47 | 0.23 | 0.32 | 0.3 |
| 6 | 52343899 | rs1266787 | 0.257 | 0.353 | -0.055 | -1.782 | 0.019 | -0.022 | -0.022 | 0.002 | 0.06 | 0.22 | 0.03 | 0 | 0.04 | 0.01 |
| 6 | 52347757 | rs764031 | 1.181 | 0.496 | -0.584 | -1.814 | -0.013 | 0.035 | 0.035 | 0 | 0.17 | 0.08 | 0.14 | 0.08 | 0.22 | 0.15 |
| 6 | 52357260 | rs7757370 | -2.407 | 0.238 | -0.701 | -1.823 | 0.033 | 0.038 | 0.038 | 1.845 | 0.76 | 0.84 | 0.91 | 0.97 | 0.87 | 0.94 |
| 6 | 52358629 | rs2397092 | 0.584 | 0.381 | -0.691 | -1.669 | -0.012 | -0.014 | -0.014 | 0 | 0.16 | 0.1 | 0.38 | 0.19 | 0.18 | 0.23 |
| 6 | 52363365 | rs9367486 | 0.096 | 0.439 | -0.682 | -1.668 | -0.028 | 0.002 | 0.002 | 0 | 0.06 | 0.01 | 0.13 | 0.02 | 0.18 | 0.14 |
| 6 | 52366762 | rs2073183 | 0.635 | 0.353 | -0.667 | -1.622 | -0.012 | -0.016 | -0.016 | 0.217 | 0.16 | 0.1 | 0.37 | 0.19 | 0.18 | 0.23 |
| 6 | 52366884 | rs9474230 | 0.778 | 0.351 | -0.704 | -1.547 | -0.033 | 0.017 | 0.017 | 0 | 0.1 | 0.08 | 0.1 | 0.09 | 0.19 | 0.17 |
| 6 | 52367117 | rs2073184 | -0.315 | 0.11 | -0.714 | -1.521 | -0.001 | 0.009 | 0.009 | 0 | 0.31 | 0.32 | 0.27 | 0.54 | 0.32 | 0.43 |
| 6 | 52371419 | rs2067002 | -1.603 | -0.193 | -0.809 | -1.197 | 0.011 | 0.021 | 0.021 | 0.495 | 0.73 | 0.79 | 0.82 | 0.81 | 0.79 | 0.83 |
| 6 | 52375593 | rs2268740 | -1.338 | -0.195 | -0.941 | -1.354 | -0.001 | 0.054 | 0.054 | 0.102 | 0.71 | 0.8 | 0.81 | 0.86 | 0.71 | 0.81 |
| 6 | 52376226 | rs12662071 | 0.824 | -0.432 | -1.239 | -1.288 | 0.005 | -0.123 | -0.123 | 1.709 | 0.15 | 0 | 0.14 | 0.54 | 0.1 | 0.22 |
| 6 | 52380641 | rs941968 | -1.432 | -0.261 | -0.987 | -1.383 | 0 | 0.042 | 0.042 | 0.029 | 0.71 | 0.8 | 0.82 | 0.87 | 0.73 | 0.82 |
| 6 | 52383067 | rs1033541 | -1.859 | -0.506 | -1.084 | -1.49 | 0.059 | 0.116 | 0.116 | 2.337 | 0.74 | 0.89 | 0.92 | 0.97 | 0.86 | 0.93 |

| | | | | | | | | | | | | | | | | |
|---|---|---|---|---|---|---|---|---|---|---|---|---|---|---|---|---|
| 6 | 52385064 | rs4712009 | -0.289 | -0.945 | -0.616 | -1.711 | -0.052 | -0.048 | -0.048 | 0 | 0.33 | 0.42 | 0.49 | 0.8 | 0.16 | 0.42 |
| 6 | 52389711 | rs1060912 | -0.188 | -0.935 | -0.613 | -1.695 | 0.023 | 0.025 | 0.025 | 0.405 | 0.41 | 0.35 | 0.44 | 0.24 | 0.72 | 0.52 |
| 6 | 52389776 | rs1060911 | 0.938 | -0.326 | -0.575 | -1.836 | -0.004 | 0 | 0 | 0 | 0.16 | 0.15 | 0.15 | 0.06 | 0.28 | 0.2 |
| 6 | 52393591 | rs3827788 | 1.034 | -0.479 | -0.376 | -1.629 | -0.003 | -0.002 | -0.002 | 0 | 0.13 | 0.14 | 0.3 | 0.21 | 0.05 | 0.2 |
| 6 | 52397047 | rs2268731 | 0.201 | -0.381 | -0.666 | -1.624 | -0.081 | 0.089 | 0.089 | 0.044 | 0.21 | 0.46 | 0.39 | 0.28 | 0.1 | 0.24 |
| 6 | 52399247 | rs4715305 | -0.448 | -0.508 | -1.03 | -1.775 | 0.002 | 0.005 | 0.005 | 0.243 | 0.39 | 0.59 | 0.53 | 0.28 | 0.38 | 0.46 |
| 6 | 52399307 | rs4715306 | -1.251 | -0.544 | -1.083 | -1.827 | -0.005 | 0.16 | 0.16 | 0.753 | 0.58 | 0.79 | 0.77 | 0.9 | 0.6 | 0.78 |
| 6 | 52400889 | rs1883930 | 0.173 | -0.001 | -0.816 | -1.36 | 0 | 0.007 | 0.007 | 0 | 0.19 | 0.19 | 0.24 | 0.62 | 0.22 | 0.32 |
| 6 | 52411122 | rs910244 | -0.509 | -0.033 | -0.825 | -1.545 | 0.009 | 0.008 | 0.008 | 0.109 | 0.42 | 0.39 | 0.4 | 0.17 | 0.69 | 0.52 |
| 6 | 52415023 | rs2268718 | 1.09 | 0.166 | -1.14 | -1.553 | -0.088 | 0.053 | 0.053 | 0 | 0.18 | 0.01 | 0.14 | 0.07 | 0.26 | 0.18 |
| 6 | 52418375 | rs9349634 | 0.724 | 0.152 | -1.094 | -1.57 | -0.016 | 0.018 | 0.018 | 0 | 0.19 | 0.14 | 0.16 | 0.07 | 0.29 | 0.23 |
| 6 | 52418956 | rs2268716 | 0.709 | 0.79 | -1.032 | -0.993 | -0.09 | 0.055 | 0.055 | 0 | 0.18 | 0.01 | 0.15 | 0.07 | 0.28 | 0.19 |
| 6 | 52424718 | rs4715307 | 0.021 | 0.396 | -0.819 | -1.069 | -0.016 | 0.01 | 0.01 | 1.799 | 0.6 | 0.48 | 0.69 | 0.43 | 0.7 | 0.67 |
| 6 | 52425189 | rs7744675 | -1.393 | 0.375 | -0.439 | -1.001 | 0 | -0.037 | -0.037 | 0 | 0.8 | 0.72 | 0.86 | 0.94 | 0.8 | 0.88 |
| 6 | 52427739 | rs9382120 | 0.502 | 0.008 | -0.5 | -1 | -0.001 | -0.001 | -0.001 | 0 | 0.42 | 0.41 | 0.54 | 0.36 | 0.42 | 0.48 |
| 6 | 52429462 | rs9349635 | -1.045 | 0.012 | -0.161 | -0.881 | 0.015 | 0.011 | 0.011 | 1.353 | 0.84 | 0.87 | 0.94 | 0.97 | 0.93 | 0.98 |
| 6 | 52432747 | rs9474249 | -0.415 | 0.766 | 0.093 | -0.148 | -0.001 | -0.017 | -0.017 | 0.471 | 0.06 | 0.24 | 0.04 | 0 | 0.02 | 0.01 |
| 6 | 52436648 | rs10484874 | -0.75 | 1.324 | -0.489 | 0.088 | 0.003 | -0.004 | -0.004 | 0 | 0.05 | 0.07 | 0.1 | 0.02 | 0.07 | 0.11 |
| 6 | 52439859 | rs1883931 | -1.034 | 1.556 | -0.531 | 0.035 | 0.006 | -0.003 | -0.003 | 0 | 0.18 | 0.2 | 0.15 | 0.14 | 0.25 | 0.2 |
| 6 | 52440970 | rs6928665 | -0.929 | 1.485 | -0.499 | 0.252 | 0.01 | 0.006 | 0.006 | 0 | 0.15 | 0.2 | 0.12 | 0.17 | 0.2 | 0.16 |
| 6 | 52443067 | rs9474251 | -0.513 | 0.621 | -0.966 | 0.238 | 0.064 | 0.012 | 0.012 | 2.686 | 0.5 | 0.67 | 0.38 | 0.55 | 0.59 | 0.72 |
| 6 | 52450447 | rs7746685 | -0.501 | -0.002 | -0.724 | -0.119 | -0.01 | 0.011 | 0.011 | 0 | 0.44 | 0.6 | 0.3 | 0.36 | 0.46 | 0.54 |
| 6 | 52452585 | rs2239620 | -0.63 | 0.082 | -0.522 | -0.084 | 0.16 | 0.072 | 0.072 | 0 | 0.8 | 0.58 | 0.82 | 0.73 | 0.64 | 0.63 |
| 6 | 52453220 | rs2239619 | -0.079 | -0.022 | -0.606 | -0.135 | 0.012 | -0.01 | -0.01 | 0 | 0.65 | 0.47 | 0.8 | 0.72 | 0.62 | 0.62 |
| 6 | 52454094 | rs13207432 | 0.652 | -0.584 | -1.125 | -0.123 | -0.02 | 0.007 | 0.007 | 0 | 0.32 | 0.16 | 0.28 | 0.35 | 0.36 | 0.34 |
| 6 | 52455546 | rs3823034 | 1.065 | -0.573 | -0.852 | 0.042 | 0.027 | -0.041 | -0.041 | 0 | 0.22 | 0.14 | 0.41 | 0.3 | 0.19 | 0.12 |
| 6 | 52469023 | rs9349637 | 1.211 | -0.593 | -0.958 | 0.07 | 0.028 | -0.072 | -0.072 | 0 | 0.21 | 0.08 | 0.4 | 0.28 | 0.18 | 0.11 |
| 6 | 52469904 | rs9463802 | -0.313 | -0.382 | -0.462 | -0.15 | 0.003 | 0.007 | 0.007 | 0 | 0.55 | 0.45 | 0.39 | 0.39 | 0.43 | 0.49 |
| 6 | 52476763 | rs11754064 | 0.482 | -0.387 | -0.807 | -0.18 | -0.015 | 0.014 | 0.014 | 0 | 0.34 | 0.16 | 0.26 | 0.31 | 0.35 | 0.36 |
| 6 | 52478364 | rs2063643 | 0.079 | 0.789 | 0.022 | -0.207 | 0.061 | -0.108 | -0.108 | 0.001 | 0.15 | 0.47 | 0.11 | 0.01 | 0.16 | 0.1 |
| 6 | 52479182 | rs9296685 | -0.667 | 0.773 | 0.281 | -0.124 | -0.001 | -0.01 | -0.01 | 0 | 0.67 | 0.76 | 0.47 | 0.41 | 0.63 | 0.58 |
| 6 | 52481565 | rs1513359 | 0.673 | 0.749 | 0.289 | -0.144 | 0.029 | 0.075 | 0.075 | 0 | 0.43 | 0.17 | 0.26 | 0.32 | 0.37 | 0.35 |
| 6 | 52487613 | rs7756118 | -0.355 | 0.788 | -0.138 | 0.216 | -0.001 | -0.009 | -0.009 | 0.259 | 0.66 | 0.77 | 0.47 | 0.39 | 0.63 | 0.58 |
| 6 | 52497284 | rs9474265 | -1.127 | 0.677 | -0.56 | 0.451 | 0.031 | -0.037 | -0.037 | 0 | 0.77 | 0.86 | 0.57 | 0.72 | 0.81 | 0.88 |
| 6 | 52498691 | rs6910985 | -0.123 | 0.727 | -0.34 | 0.284 | -0.005 | 0.002 | 0.002 | 0 | 0.62 | 0.68 | 0.45 | 0.39 | 0.58 | 0.58 |
| 6 | 52507285 | rs9474269 | 0.454 | 0.509 | -0.742 | 0.194 | -0.003 | 0.004 | 0.004 | 0 | 0.14 | 0.1 | 0.08 | 0.12 | 0.12 | 0.18 |
| 6 | 52508705 | rs11756750 | -1.111 | 0.169 | -0.781 | 0.305 | 0.026 | -0.018 | -0.018 | 0 | 0.76 | 0.84 | 0.58 | 0.74 | 0.82 | 0.9 |
| 6 | 52509965 | rs13205411 | 0.495 | 0.39 | -0.899 | 0.377 | -0.046 | 0.013 | 0.013 | 0 | 0.42 | 0.13 | 0.3 | 0.39 | 0.46 | 0.49 |
| 6 | 52510665 | rs10948711 | 0.269 | 0.384 | -0.87 | 0.318 | -0.055 | -0.036 | -0.036 | 0 | 0.45 | 0.13 | 0.37 | 0.49 | 0.49 | 0.6 |
| 6 | 52511650 | rs2180321 | -0.87 | -0.289 | -0.262 | 0.558 | -0.007 | 0.006 | 0.006 | 0.292 | 0.75 | 0.73 | 0.57 | 0.62 | 0.81 | 0.89 |
| 6 | 52513889 | rs9296687 | -0.303 | 0.29 | -0.317 | 0.298 | 0.008 | -0.004 | -0.004 | 0 | 0.58 | 0.49 | 0.43 | 0.39 | 0.51 | 0.42 |
| 6 | 52515078 | rs6918918 | -0.147 | -0.245 | -0.194 | 0.111 | -0.003 | -0.007 | -0.007 | 0 | 0.25 | 0.25 | 0.2 | 0.18 | 0.18 | 0.25 |
| 6 | 52519203 | rs723815 | -0.168 | -0.361 | -0.084 | 0.155 | -0.013 | -0.018 | -0.018 | 0 | 0.24 | 0.27 | 0.19 | 0.18 | 0.16 | 0.24 |
| 6 | 52524838 | rs7738712 | -0.065 | -0.053 | -0.449 | 0.375 | -0.004 | -0.008 | -0.008 | 0 | 0.09 | 0.07 | 0.13 | 0.17 | 0.11 | 0.2 |
| 6 | 52531837 | rs2397097 | 0.193 | -0.425 | -0.397 | 0.386 | -0.001 | 0 | 0 | 0 | 0.46 | 0.46 | 0.34 | 0.62 | 0.61 | 0.52 |
| 6 | 52532308 | rs9370138 | 0.91 | -0.239 | -0.472 | 0.348 | 0.002 | 0.001 | 0.001 | 0 | 0.16 | 0.16 | 0.15 | 0.19 | 0.22 | 0.11 |
| 6 | 52534369 | rs12211989 | 1.153 | -0.469 | -0.649 | 0.184 | -0.026 | 0.005 | 0.005 | 0 | 0.08 | 0 | 0.04 | 0.03 | 0.07 | 0.1 |
| 6 | 52534645 | rs767308 | 0.498 | -0.592 | -0.664 | 0.118 | -0.124 | -0.105 | -0.105 | 0 | 0.29 | 0.04 | 0.17 | 0.42 | 0.38 | 0.41 |
| 6 | 52534931 | rs1000810 | 0.013 | -1.063 | -0.01 | 0.001 | -0.01 | -0.008 | -0.008 | 0 | 0.56 | 0.52 | 0.47 | 0.77 | 0.71 | 0.73 |
| 6 | 52538171 | rs7746344 | -1.13 | -1.17 | 0.209 | 0.064 | -0.027 | 0.002 | 0.002 | 1.973 | 0.79 | 0.71 | 0.57 | 0.83 | 0.84 | 0.88 |
| 6 | 52539268 | rs2670142 | -0.355 | -0.844 | -0.137 | 0.259 | -0.052 | -0.036 | -0.036 | 1.342 | 0.63 | 0.45 | 0.49 | 0.83 | 0.78 | 0.83 |
| 6 | 52540497 | rs2670153 | -0.34 | -0.837 | -0.357 | 0.238 | -0.022 | -0.014 | -0.014 | 0 | 0.63 | 0.4 | 0.48 | 0.83 | 0.78 | 0.83 |
| 6 | 52557627 | rs2180323 | -0.28 | -0.126 | -0.366 | 0.66 | 0 | -0.004 | -0.004 | 0.02 | 0.66 | 0.63 | 0.72 | 0.53 | 0.7 | 0.67 |
| 6 | 52558017 | rs6902435 | 0.491 | 0.116 | -0.138 | 0.535 | 0.02 | -0.007 | -0.007 | 0 | 0.16 | 0.31 | 0.16 | 0.14 | 0.21 | 0.08 |
| 6 | 52560788 | rs6458856 | 0.671 | 0.107 | -0.005 | 0.526 | 0.003 | -0.003 | -0.003 | 2.98 | 0.16 | 0.19 | 0.14 | 0.14 | 0.21 | 0.08 |
| 6 | 52564490 | rs10484873 | 0.744 | 0.103 | -0.174 | 0.561 | -0.01 | 0 | 0 | 0 | 0.15 | 0.09 | 0.14 | 0.15 | 0.21 | 0.08 |
| 6 | 52565211 | rs1884459 | -1.035 | 0.019 | -0.275 | 0.783 | 0.011 | -0.003 | -0.003 | 0 | 0.69 | 0.68 | 0.81 | 0.7 | 0.76 | 0.81 |
| 6 | 52575354 | rs958202 | -0.373 | -0.199 | -0.661 | 0.803 | 0.014 | 0.007 | 0.007 | 0 | 0.52 | 0.57 | 0.67 | 0.42 | 0.65 | 0.62 |
| 6 | 52575508 | rs958204 | 0.247 | 0.184 | -1.269 | 0.747 | -0.015 | 0.012 | 0.012 | 0.246 | 0.35 | 0.25 | 0.5 | 0.27 | 0.44 | 0.53 |
| 6 | 52580355 | rs938223 | 0.831 | 0.913 | -2.328 | 0.168 | -0.017 | 0.017 | 0.017 | 0 | 0.11 | 0.03 | 0.06 | 0 | 0.15 | 0.1 |
| 6 | 52585532 | rs6905523 | -1.126 | 0.351 | -2.234 | 0.494 | 0.005 | -0.031 | -0.031 | 0 | 0.71 | 0.86 | 0.81 | 0.67 | 0.75 | 0.8 |
| 6 | 52586253 | rs4715309 | -0.033 | 0.422 | -2.167 | 0.541 | -0.091 | 0.07 | 0.07 | 1.203 | 0.35 | 0.06 | 0.48 | 0.2 | 0.44 | 0.53 |
| 6 | 52593824 | rs9296691 | 0.005 | 0.381 | -1.569 | 0.752 | -0.007 | 0.001 | 0.001 | 0 | 0.06 | 0.05 | 0.09 | 0.09 | 0.07 | 0.16 |
| 6 | 52594452 | rs2608654 | -0.09 | 0.182 | -1.759 | 0.664 | 0.026 | -0.009 | -0.009 | 0 | 0.61 | 0.8 | 0.73 | 0.58 | 0.71 | 0.64 |
| 6 | 52596516 | rs2748991 | 0.026 | 0.572 | -1.195 | 0.708 | -0.01 | -0.005 | -0.005 | 0 | 0.37 | 0.34 | 0.59 | 0.61 | 0.43 | 0.54 |
| 6 | 52598598 | rs7746808 | -0.115 | 0.333 | -1.607 | 0.094 | 0.004 | 0.089 | 0.089 | 0.58 | 0.45 | 0.17 | 0.69 | 0.67 | 0.58 | 0.63 |
| 6 | 52599135 | rs10807452 | 0.537 | 0.989 | -1.271 | 0.379 | -0.102 | -0.098 | -0.098 | 0 | 0.14 | 0.02 | 0.44 | 0.37 | 0.28 | 0.24 |
| 6 | 52602360 | rs9474314 | -0.869 | -0.153 | -0.101 | -0.973 | -0.027 | -0.039 | -0.039 | 0 | 0.18 | 0.32 | 0.1 | 0.09 | 0.14 | 0.1 |
| 6 | 52602843 | rs2144692 | 0.423 | -0.341 | -0.054 | -1.181 | -0.001 | -0.004 | -0.004 | 0 | 0.44 | 0.61 | 0.67 | 0.8 | 0.49 | 0.6 |

| | | | | | | | | | | | | | | | | |
|---|---|---|---|---|---|---|---|---|---|---|---|---|---|---|---|---|
| 6 | 52604054 | rs2180312 | 0.835 | -0.3 | -0.338 | -1.192 | -0.069 | 0.058 | 0.058 | 0.458 | 0.39 | 0.16 | 0.63 | 0.74 | 0.49 | 0.59 |
| 6 | 52609664 | rs2608615 | -0.417 | -0.401 | -0.322 | -1.222 | 0 | 0.054 | 0.054 | 2.649 | 0.13 | 0.45 | 0.08 | 0.15 | 0.1 | 0.09 |
| 6 | 52610584 | rs2207949 | -0.558 | -0.142 | -1.057 | -1.115 | -0.025 | 0.173 | 0.173 | 0.426 | 0.62 | 0.92 | 0.77 | 0.89 | 0.63 | 0.7 |
| 6 | 52611118 | rs2207951 | -0.692 | -0.131 | -1.361 | -1.109 | -0.085 | 0.154 | 0.154 | 1.178 | 0.64 | 0.98 | 0.77 | 0.89 | 0.63 | 0.7 |
| 6 | 52612632 | rs9349641 | 0.474 | 0.327 | -0.963 | -0.72 | -0.063 | 0.057 | 0.057 | 0.485 | 0.39 | 0.16 | 0.63 | 0.74 | 0.49 | 0.59 |
| 6 | 52617738 | rs2234951 | 0.365 | -0.059 | -0.584 | -0.606 | -0.005 | -0.032 | -0.032 | 0 | 0.07 | 0 | 0.02 | 0.13 | 0.05 | 0.16 |
| 6 | 52623807 | rs2144698 | -0.794 | -0.093 | -1.09 | -1.096 | -0.067 | 0.193 | 0.193 | 2.354 | 0.71 | 0.99 | 0.8 | 1 | 0.7 | 0.82 |
| 6 | 52625794 | rs2608629 | -0.16 | -0.085 | -1.063 | -1.067 | -0.005 | 0.15 | 0.15 | 0.03 | 0.63 | 0.91 | 0.77 | 0.86 | 0.65 | 0.66 |
| 6 | 52626269 | rs2608631 | -0.322 | -0.079 | -1.109 | -1.064 | -0.07 | 0.135 | 0.135 | 0.346 | 0.65 | 0.98 | 0.77 | 0.86 | 0.65 | 0.66 |
| 6 | 52631188 | rs2749029 | -0.322 | -0.135 | -0.836 | -1.093 | -0.032 | 0.148 | 0.148 | 0.346 | 0.65 | 0.94 | 0.77 | 0.86 | 0.65 | 0.66 |
| 6 | 52635331 | rs4715318 | -0.109 | -0.1 | -0.795 | -1.046 | -0.015 | 0.123 | 0.123 | 0.049 | 0.59 | 0.8 | 0.71 | 0.86 | 0.56 | 0.66 |
| 6 | 52657552 | rs4715326 | 0.001 | -0.295 | -0.316 | -1.05 | 0.003 | 0.067 | 0.067 | 0.051 | 0.58 | 0.72 | 0.71 | 0.86 | 0.58 | 0.66 |
| 6 | 52663813 | rs10948722 | 0.048 | -0.305 | -0.329 | -1.055 | -0.108 | 0.169 | 0.169 | 0.49 | 0.39 | 0.02 | 0.27 | 0.14 | 0.42 | 0.34 |
| 6 | 52666405 | rs10948723 | -0.019 | -0.302 | -0.315 | -1.054 | -0.033 | 0.168 | 0.168 | 0.328 | 0.39 | 0.09 | 0.28 | 0.14 | 0.43 | 0.34 |
| 6 | 52669185 | rs4715332 | 0.015 | -0.312 | -0.317 | -1.059 | 0.003 | 0.069 | 0.069 | 0.043 | 0.58 | 0.71 | 0.71 | 0.86 | 0.57 | 0.66 |
| 6 | 52697404 | rs7739421 | -0.15 | -0.587 | -0.198 | -0.9 | -0.058 | 0.058 | 0.058 | 1.457 | 0.15 | 0.48 | 0.08 | 0.18 | 0.06 | 0.09 |
| 6 | 52698340 | rs7745053 | 0.468 | -0.033 | -0.399 | -0.602 | -0.006 | -0.032 | -0.032 | 1.279 | 0.07 | 0 | 0.02 | 0.13 | 0.05 | 0.16 |
| 6 | 52703396 | rs4715352 | 0.951 | 0.418 | -0.571 | -0.43 | -0.067 | -0.047 | -0.047 | 3.557 | 0.1 | 0.01 | 0.37 | 0.12 | 0.17 | 0.19 |
| 6 | 52715510 | rs7773321 | -0.175 | -0.469 | -0.071 | -0.752 | 0.006 | -0.01 | -0.01 | 0 | 0.08 | 0.05 | 0.02 | 0.13 | 0.05 | 0.15 |
| 6 | 52724146 | rs11759734 | 0.013 | -0.405 | -0.271 | -0.763 | -0.102 | 0.227 | 0.227 | 2.499 | 0.32 | 0.02 | 0.25 | 0.01 | 0.37 | 0.19 |
| 6 | 52735825 | rs6910581 | -0.456 | -0.352 | -0.099 | -0.614 | -0.025 | 0.161 | 0.161 | 0.189 | 0.4 | 0.14 | 0.28 | 0.14 | 0.43 | 0.34 |
| 6 | 52739537 | rs11752349 | -0.393 | -0.388 | 0.58 | -0.006 | -0.007 | 0.038 | 0.038 | 0 | 0.31 | 0.2 | 0.26 | 0.16 | 0.34 | 0.25 |
| 6 | 52741615 | rs4715359 | 0.521 | -0.534 | 0.561 | 0.013 | 0.004 | 0.025 | 0.025 | 0 | 0.65 | 0.74 | 0.71 | 0.85 | 0.63 | 0.71 |
| 6 | 52742295 | rs7746971 | -1.024 | -0.711 | 1.276 | 0.434 | -0.038 | 0.02 | 0.02 | 0.401 | 0.11 | 0.29 | 0.06 | 0.14 | 0.04 | 0.1 |
| 6 | 52745061 | rs6937787 | 1.162 | -0.198 | 0.837 | 0.394 | -0.008 | -0.025 | -0.025 | 0 | 0.94 | 0.86 | 0.96 | 1 | 0.95 | 0.99 |
| 6 | 52753034 | rs668163 | -1.137 | -0.209 | 0.988 | 0.778 | -0.015 | 0.018 | 0.018 | 0.586 | 0.39 | 0.5 | 0.54 | 0.47 | 0.35 | 0.37 |
| 6 | 52757776 | rs10214506 | -1.163 | -0.369 | 1.534 | 0.659 | 0.001 | -0.026 | -0.026 | 0 | 0.06 | 0.21 | 0.03 | 0 | 0.03 | 0 |
| 6 | 52758076 | rs9296695 | -2.008 | -0.558 | 1.466 | 0.741 | -0.045 | 0.105 | 0.105 | 1.218 | 0.16 | 0.65 | 0.38 | 0.4 | 0.07 | 0.21 |
| 6 | 52759261 | rs9370165 | -0.564 | -0.586 | 1.774 | 0.807 | -0.018 | 0.145 | 0.145 | 0 | 0.07 | 0.11 | 0.32 | 0.34 | 0.03 | 0.19 |
| 6 | 52761898 | rs2295605 | -0.446 | -0.595 | 1.669 | 0.775 | -0.073 | 0.081 | 0.081 | 1.691 | 0.24 | 0.01 | 0.16 | 0.08 | 0.29 | 0.15 |
| 6 | 52765760 | rs557135 | -1.778 | -0.633 | 1.637 | 0.737 | 0.024 | -0.019 | -0.019 | 0 | 0.42 | 0.68 | 0.57 | 0.48 | 0.37 | 0.37 |
| 6 | 52766622 | rs9474339 | -1.054 | -0.605 | 1.678 | 0.741 | 0.002 | -0.025 | -0.025 | 0 | 0.06 | 0.21 | 0.03 | 0 | 0.03 | 0 |
| 6 | 52766983 | rs512795 | -2.549 | -0.615 | 1.682 | 0.749 | -0.043 | 0.078 | 0.078 | 0.756 | 0.18 | 0.67 | 0.4 | 0.4 | 0.08 | 0.22 |
| 6 | 52769856 | rs628263 | -1.297 | -0.502 | 1.66 | 0.811 | -0.002 | 0.442 | 0.442 | 0.484 | 0.09 | 0.46 | 0.36 | 0.4 | 0.03 | 0.19 |
| 6 | 52775872 | rs563464 | -1.358 | -0.471 | 1.316 | 0.834 | -0.023 | 0.361 | 0.361 | 1.322 | 0.1 | 0.39 | 0.35 | 0.4 | 0.03 | 0.18 |
| 6 | 52776851 | rs517029 | -0.707 | -0.487 | 1.277 | 0.802 | -0.002 | 0.048 | 0.048 | 1.347 | 0.27 | 0.18 | 0.18 | 0.07 | 0.29 | 0.16 |
| 6 | 52779005 | rs9382157 | -1.863 | -0.369 | 0.912 | 0.913 | -0.037 | 0.309 | 0.309 | 0.965 | 0.14 | 0.45 | 0.36 | 0.4 | 0.06 | 0.19 |
| 6 | 52779207 | rs12202200 | -1.844 | -0.368 | 0.913 | 0.914 | -0.035 | 0.312 | 0.312 | 1.012 | 0.14 | 0.45 | 0.36 | 0.4 | 0.06 | 0.19 |
| 6 | 52789828 | rs651733 | -1.516 | -0.361 | 0.886 | 0.886 | 0.007 | -0.025 | -0.025 | 0 | 0.35 | 0.23 | 0.52 | 0.42 | 0.34 | 0.33 |
| 6 | 52790972 | rs667173 | -0.536 | -0.361 | 0.855 | 0.875 | -0.011 | 0.073 | 0.073 | 2.481 | 0.25 | 0.09 | 0.18 | 0.08 | 0.28 | 0.16 |
| 6 | 52792854 | rs588565 | -0.608 | -0.316 | 0.499 | 0.898 | -0.04 | 0.101 | 0.101 | 2.807 | 0.25 | 0.03 | 0.16 | 0.08 | 0.28 | 0.16 |
| 6 | 52795489 | rs9349651 | -1.239 | -0.201 | 0.581 | 0.816 | -0.014 | 0.105 | 0.105 | 0.111 | 0.1 | 0.15 | 0.34 | 0.35 | 0.06 | 0.17 |
| 6 | 52799349 | rs482270 | -0.71 | -0.071 | 0.313 | 0.947 | 0.041 | 0.06 | 0.06 | 0.049 | 0.05 | 0.32 | 0.05 | 0.05 | 0.02 | 0.02 |
| 6 | 52802028 | rs373552 | -0.682 | 0.317 | -0.973 | 1.068 | -0.046 | 0.118 | 0.118 | 1.458 | 0.23 | 0.02 | 0.16 | 0.04 | 0.28 | 0.13 |
| 6 | 52803206 | rs451439 | -0.692 | 0.345 | -0.96 | 1.082 | -0.046 | 0.119 | 0.119 | 1.321 | 0.23 | 0.03 | 0.17 | 0.04 | 0.28 | 0.13 |
| 6 | 52803583 | rs9349653 | -0.92 | 0.533 | -0.393 | 1.864 | -0.007 | 0.11 | 0.11 | 0 | 0.06 | 0.07 | 0.31 | 0.34 | 0.03 | 0.16 |
| 6 | 52810653 | rs10456664 | -0.681 | 1.087 | -0.627 | 1.344 | -0.039 | -0.022 | -0.022 | 0 | 0.07 | 0.01 | 0.06 | 0.09 | 0.13 | 0.07 |
| 6 | 52810893 | rs4712026 | -1.046 | 1.249 | -0.597 | 1.974 | -0.002 | 0.014 | 0.014 | 0.066 | 0.12 | 0.11 | 0.35 | 0.3 | 0.13 | 0.15 |
| 6 | 52815766 | rs11962677 | -0.791 | 2.341 | -1.041 | 0.925 | -0.081 | 0.075 | 0.075 | 0 | 0.2 | 0.07 | 0.15 | 0.05 | 0.32 | 0.13 |
| 6 | 52817144 | rs568644 | -1.249 | 1.731 | -1.101 | 1.152 | 0.004 | 0.008 | 0.008 | 0.31 | 0.45 | 0.73 | 0.36 | 0.46 | 0.52 | 0.44 |
| 6 | 52821059 | rs316157 | -0.194 | 1.294 | -0.716 | 1.348 | 0.004 | -0.014 | -0.014 | 0 | 0.19 | 0.16 | 0.16 | 0.22 | 0.19 | 0.28 |
| 6 | 52822537 | rs521664 | -1.524 | 1.23 | -1.066 | 1.035 | 0.094 | -0.03 | -0.03 | 0 | 0.52 | 0.9 | 0.42 | 0.51 | 0.65 | 0.47 |
| 6 | 52825926 | rs316149 | -1.039 | 1.5 | -0.651 | 0.564 | -0.004 | 0.002 | 0.002 | 0 | 0.32 | 0.62 | 0.25 | 0.24 | 0.46 | 0.16 |
| 6 | 52829664 | rs9474363 | -1.358 | 0.874 | 0.25 | 0.817 | -0.011 | 0.084 | 0.084 | 0.047 | 0.12 | 0.46 | 0.07 | 0.2 | 0.04 | 0.08 |
| 6 | 52830861 | rs560788 | 1.034 | 1.244 | -0.231 | 0.184 | -0.001 | 0.014 | 0.014 | 0.033 | 0.55 | 0.34 | 0.46 | 0.76 | 0.53 | 0.59 |
| 6 | 52831785 | rs10484409 | -0.146 | 1.347 | -0.125 | 0.034 | -0.001 | -0.003 | -0.003 | 0 | 0.08 | 0.1 | 0.09 | 0.06 | 0.11 | 0.07 |
| 6 | 52832077 | rs17608022 | 0.304 | 1.543 | -0.275 | -0.07 | -0.047 | 0.012 | 0.012 | 0.02 | 0.09 | 0 | 0.05 | 0.04 | 0.13 | 0.06 |
| 6 | 52833065 | rs9463851 | -1.198 | 1.607 | -0.123 | -0.218 | -0.025 | -0.031 | -0.031 | 0 | 0.22 | 0.43 | 0.36 | 0.14 | 0.17 | 0.26 |
| 6 | 52833438 | rs396216 | 1.192 | 1.601 | -0.063 | -0.234 | -0.007 | 0.004 | 0.004 | 0 | 0.77 | 0.79 | 0.8 | 0.9 | 0.67 | 0.85 |
| 6 | 52834842 | rs1032419 | -1.174 | 1.471 | 0.013 | -0.197 | -0.021 | -0.024 | -0.024 | 0.121 | 0.22 | 0.38 | 0.35 | 0.14 | 0.17 | 0.26 |
| 6 | 52835572 | rs384505 | -2.642 | 1.788 | -0.125 | -0.102 | -0.095 | 0.08 | 0.08 | 0.786 | 0.47 | 0.98 | 0.58 | 0.3 | 0.54 | 0.42 |
| 6 | 52838158 | rs449690 | 2.243 | 1.718 | -0.025 | -0.152 | -0.002 | -0.001 | -0.001 | 1.712 | 0.55 | 0.4 | 0.46 | 0.75 | 0.48 | 0.6 |
| 6 | 52838196 | rs584124 | 2.005 | 1.702 | -0.039 | -0.198 | 0.007 | -0.032 | -0.032 | 0 | 0.59 | 0.44 | 0.48 | 0.75 | 0.54 | 0.64 |
| 6 | 52839185 | rs428957 | 2.408 | 1.702 | 0.018 | -0.198 | -0.006 | 0.021 | 0.021 | 0.053 | 0.55 | 0.35 | 0.45 | 0.75 | 0.48 | 0.6 |
| 6 | 52839888 | rs426169 | 2.408 | 1.704 | 0.018 | -0.198 | -0.005 | 0.014 | 0.014 | 0.053 | 0.55 | 0.36 | 0.45 | 0.75 | 0.48 | 0.6 |
| 6 | 52839958 | rs612483 | 2.005 | 1.704 | 0.018 | -0.197 | 0.008 | -0.032 | -0.032 | 0 | 0.59 | 0.44 | 0.48 | 0.75 | 0.53 | 0.64 |
| 6 | 52840235 | rs613764 | 2.005 | 1.704 | 0.018 | -0.197 | 0.008 | -0.032 | -0.032 | 0 | 0.59 | 0.44 | 0.48 | 0.75 | 0.53 | 0.64 |
| 6 | 52840254 | rs387853 | 2.407 | 1.705 | 0.016 | -0.197 | -0.006 | 0.021 | 0.021 | 0.053 | 0.55 | 0.35 | 0.45 | 0.75 | 0.48 | 0.6 |
| 6 | 52840745 | rs426013 | 2.196 | 1.702 | -0.187 | -0.203 | 0.005 | -0.017 | -0.017 | 0 | 0.62 | 0.49 | 0.49 | 0.75 | 0.55 | 0.64 |

| | | | | | | | | | | | | | | | | |
|---|---|---|---|---|---|---|---|---|---|---|---|---|---|---|---|---|
| 6 | 52841995 | rs419129 | 2.307 | 1.662 | -0.018 | -0.203 | 0.006 | -0.025 | -0.025 | 0 | 0.58 | 0.39 | 0.47 | 0.75 | 0.5 | 0.6 |
| 6 | 52842781 | rs405729 | 2.27 | 1.666 | -0.131 | -0.201 | 0.006 | -0.024 | -0.024 | 0 | 0.58 | 0.44 | 0.47 | 0.75 | 0.5 | 0.6 |
| 6 | 52843131 | rs367836 | 2.228 | 1.658 | -0.136 | -0.205 | 0.005 | -0.016 | -0.016 | 4.665 | 0.62 | 0.49 | 0.49 | 0.75 | 0.55 | 0.64 |
| 6 | 52843429 | rs3734431 | 0.136 | 1.674 | -0.129 | -0.216 | 0 | 0.002 | 0.002 | 0.152 | 0.08 | 0.07 | 0.09 | 0.06 | 0.11 | 0.07 |
| 6 | 52843477 | rs375887 | 2.325 | 1.673 | -0.128 | -0.214 | 0.005 | -0.019 | -0.019 | 1.097 | 0.58 | 0.45 | 0.47 | 0.75 | 0.5 | 0.6 |
| 6 | 52843524 | rs543613 | 0.032 | 1.551 | -0.192 | -0.29 | -0.003 | -0.004 | -0.004 | 3.682 | 0.95 | 0.95 | 0.95 | 1 | 0.9 | 0.99 |
| 6 | 52843796 | rs670960 | 2.263 | 1.585 | -0.168 | -0.264 | 0.005 | -0.017 | -0.017 | 0.018 | 0.62 | 0.49 | 0.49 | 0.75 | 0.56 | 0.64 |
| 6 | 52846158 | rs316141 | 2.318 | 1.58 | -0.17 | -0.264 | 0.007 | -0.024 | -0.024 | 0.157 | 0.62 | 0.45 | 0.49 | 0.75 | 0.56 | 0.64 |
| 6 | 52846188 | rs316140 | 2.318 | 1.58 | -0.17 | -0.264 | 0.007 | -0.024 | -0.024 | 0 | 0.62 | 0.45 | 0.49 | 0.75 | 0.56 | 0.64 |
| 6 | 52846474 | rs622902 | 2.318 | 1.58 | -0.17 | -0.264 | 0.007 | -0.024 | -0.024 | 0.259 | 0.62 | 0.45 | 0.49 | 0.75 | 0.56 | 0.64 |
| 6 | 52846680 | rs316137 | 2.33 | 1.58 | -0.171 | -0.265 | 0.004 | -0.017 | -0.017 | 0.094 | 0.58 | 0.45 | 0.47 | 0.75 | 0.5 | 0.6 |
| 6 | 52846769 | rs316136 | 2.351 | 1.58 | -0.171 | -0.265 | 0.004 | -0.019 | -0.019 | 0 | 0.58 | 0.45 | 0.47 | 0.75 | 0.5 | 0.6 |
| 6 | 52846777 | rs316135 | 2.351 | 1.58 | -0.17 | -0.265 | 0.006 | -0.024 | -0.024 | 0.096 | 0.58 | 0.44 | 0.47 | 0.75 | 0.5 | 0.6 |
| 6 | 52846902 | rs6904769 | -1.015 | 1.582 | -0.182 | -0.268 | -0.015 | 0 | 0 | 0 | 0.15 | 0.29 | 0.31 | 0.14 | 0.1 | 0.21 |
| 6 | 52847966 | rs316132 | 2.324 | 1.575 | -0.157 | -0.271 | 0.007 | -0.024 | -0.024 | 0 | 0.62 | 0.45 | 0.49 | 0.75 | 0.56 | 0.65 |
| 6 | 52848149 | rs316131 | 2.324 | 1.575 | -0.157 | -0.271 | 0.007 | -0.024 | -0.024 | 0 | 0.62 | 0.45 | 0.49 | 0.75 | 0.56 | 0.65 |
| 6 | 52848905 | rs316129 | 2.324 | 1.576 | -0.158 | -0.272 | 0.007 | -0.024 | -0.024 | 0 | 0.62 | 0.45 | 0.49 | 0.75 | 0.56 | 0.65 |
| 6 | 52849146 | rs316128 | 2.343 | 1.576 | -0.159 | -0.272 | 0.004 | -0.019 | -0.019 | 0 | 0.58 | 0.45 | 0.47 | 0.75 | 0.5 | 0.6 |
| 6 | 52851560 | rs34511626 | 0.611 | 1.588 | -0.166 | -0.278 | -0.051 | 0.01 | 0.01 | 0 | 0.09 | 0 | 0.05 | 0.05 | 0.14 | 0.06 |
| 6 | 52851979 | rs3756980 | -0.937 | 1.571 | -0.132 | -0.278 | -0.033 | 0.035 | 0.035 | 0 | 0.18 | 0.1 | 0.14 | 0.11 | 0.25 | 0.13 |
| 6 | 52856179 | rs6904771 | -0.485 | 0.788 | 0.132 | -0.005 | -0.041 | -0.047 | -0.047 | 0 | 0.08 | 0.3 | 0.02 | 0 | 0.01 | 0.01 |
| 6 | 52860407 | rs13197674 | -2.344 | 1.018 | 0.013 | 0.049 | -0.002 | -0.059 | -0.059 | 0 | 0.28 | 0.47 | 0.21 | 0.11 | 0.33 | 0.15 |
| 6 | 52865914 | rs2397136 | -1.493 | 0.873 | 0.023 | -0.034 | -0.015 | -0.061 | -0.061 | 0 | 0.19 | 0.34 | 0.12 | 0.06 | 0.17 | 0.13 |
| 6 | 52870320 | rs1493107 | -2.318 | 0.882 | -0.155 | -0.021 | -0.009 | -0.059 | -0.059 | 0 | 0.27 | 0.48 | 0.18 | 0.12 | 0.28 | 0.16 |
| 6 | 52879695 | rs316144 | -2.665 | 0.661 | -0.486 | 0.147 | -0.08 | -0.036 | -0.036 | 0 | 0.43 | 0.98 | 0.59 | 0.35 | 0.44 | 0.3 |
| 6 | 52885773 | rs6458895 | -1.462 | 0.924 | -0.201 | 0.235 | -0.013 | -0.056 | -0.056 | 0.008 | 0.19 | 0.34 | 0.11 | 0.06 | 0.17 | 0.11 |
| 6 | 52889275 | rs222461 | 1.703 | 0.918 | -0.203 | 0.226 | 0.001 | -0.074 | -0.074 | 0 | 0.76 | 0.62 | 0.85 | 0.94 | 0.74 | 0.89 |
| 6 | 52901032 | rs10484408 | -1.394 | 0.792 | -0.288 | 0.231 | -0.009 | -0.043 | -0.043 | 0.575 | 0.19 | 0.28 | 0.11 | 0.06 | 0.16 | 0.1 |
| 6 | 52905592 | rs10456665 | -0.054 | 0.684 | -0.128 | 0.558 | -0.027 | -0.062 | -0.062 | 0 | 0.1 | 0.01 | 0.36 | 0.19 | 0.12 | 0.13 |
| 6 | 52925277 | rs7774904 | -0.6 | 0.7 | 0.265 | 0.251 | -0.038 | -0.044 | -0.044 | 0 | 0.08 | 0.25 | 0.02 | 0 | 0.01 | 0.01 |
| 6 | 52932508 | rs1888274 | -0.003 | 0.598 | -0.041 | 0.327 | -0.026 | -0.051 | -0.051 | 0 | 0.09 | 0 | 0.07 | 0.23 | 0.1 | 0.17 |
| 6 | 52948145 | rs222439 | 2.008 | 0.456 | 0.386 | -0.082 | -0.023 | -0.086 | -0.086 | 0 | 0.74 | 0.45 | 0.84 | 0.88 | 0.78 | 0.89 |
| 6 | 52963071 | rs2876 | 1.932 | 0.487 | 0.196 | -0.059 | -0.016 | -0.063 | -0.063 | 0.157 | 0.75 | 0.49 | 0.85 | 0.87 | 0.78 | 0.89 |
| 6 | 52974638 | rs7743908 | -0.753 | 0.649 | 0.052 | -0.013 | 0 | -0.02 | -0.02 | 0.039 | 0.08 | 0.12 | 0.04 | 0 | 0.09 | 0.02 |
| 6 | 52975039 | rs2744457 | 0.718 | 0.733 | -0.006 | -0.027 | -0.007 | 0.001 | 0.001 | 0.011 | 0.93 | 0.92 | 0.94 | 0.93 | 0.86 | 0.96 |
| 6 | 52976317 | rs2816349 | 2.679 | 0.487 | -0.013 | -0.127 | 0 | -0.033 | -0.033 | 0 | 0.7 | 0.5 | 0.82 | 0.82 | 0.73 | 0.85 |
| 6 | 52977137 | rs2744458 | 1.067 | 0.528 | -0.023 | -0.117 | 0.003 | -0.028 | -0.028 | 0 | 0.88 | 0.86 | 0.92 | 1 | 0.82 | 0.97 |
| 6 | 52978635 | rs2816348 | 2.12 | 0.573 | -0.19 | -0.217 | 0.006 | -0.032 | -0.032 | 0 | 0.66 | 0.48 | 0.78 | 0.82 | 0.65 | 0.82 |
| 6 | 52987617 | rs6909766 | -0.771 | 0.108 | -0.298 | 0.011 | -0.01 | 0.024 | 0.024 | 0 | 0.11 | 0.03 | 0.08 | 0.06 | 0.12 | 0.07 |
| 6 | 52991621 | rs9474396 | -1.006 | -0.066 | 0.174 | 0.154 | -0.022 | 0.013 | 0.013 | 0 | 0.08 | 0.28 | 0.04 | 0.07 | 0.04 | 0.03 |
| 6 | 52998788 | rs2816346 | 1.411 | -0.053 | 0.053 | 0.227 | -0.007 | 0.014 | 0.014 | 0 | 0.71 | 0.54 | 0.79 | 0.58 | 0.73 | 0.73 |
| 6 | 52999489 | rs7745079 | -0.099 | 0.106 | -0.097 | 0.174 | 0.028 | -0.041 | -0.041 | 0 | 0.08 | 0.11 | 0.07 | 0 | 0.14 | 0.05 |
| 6 | 53002509 | rs2816341 | 1.398 | 0.069 | -0.109 | 0.193 | -0.001 | -0.005 | -0.005 | 0 | 0.74 | 0.73 | 0.81 | 0.59 | 0.73 | 0.73 |
| 6 | 53003574 | rs2816339 | 1.664 | -0.06 | 0.861 | 0.599 | 0 | 0.053 | 0.053 | 0 | 0.72 | 0.52 | 0.74 | 0.54 | 0.72 | 0.72 |
| 6 | 53004757 | rs2518580 | 1.962 | -0.676 | -0.373 | 0.073 | -0.138 | -0.151 | -0.151 | 0 | 0.43 | 0.02 | 0.28 | 0.44 | 0.47 | 0.54 |
| 6 | 53010182 | rs9367511 | -1.028 | -1.09 | 0.346 | 0.245 | -0.025 | 0.012 | 0.012 | 0 | 0.18 | 0.39 | 0.12 | 0.28 | 0.13 | 0.19 |
| 6 | 53016306 | rs6915699 | -1.33 | -1.171 | 0.108 | 0.263 | -0.093 | 0.016 | 0.016 | 0.301 | 0.19 | 0.57 | 0.12 | 0.17 | 0.07 | 0.05 |
| 6 | 53027389 | rs9395842 | 0.752 | -0.9 | -0.481 | 0.411 | -0.03 | -0.036 | -0.036 | 0 | 0.07 | 0 | 0.08 | 0.28 | 0.1 | 0.18 |
| 6 | 53031470 | rs12199809 | 0.36 | -1.098 | -0.582 | 0.222 | 0.012 | -0.008 | -0.008 | 0 | 0.11 | 0.02 | 0.34 | 0.11 | 0.1 | 0.2 |
| 6 | 53035505 | rs9395843 | 1.471 | -0.965 | -0.402 | 0.196 | 0.012 | -0.016 | -0.016 | 0 | 0.66 | 0.25 | 0.71 | 0.55 | 0.68 | 0.72 |
| 6 | 53044271 | rs9382180 | 1.325 | -1.093 | -0.455 | 0.153 | 0.005 | 0.001 | 0.001 | 0 | 0.67 | 0.28 | 0.74 | 0.62 | 0.7 | 0.67 |
| 6 | 53044379 | rs9382181 | 1.288 | -1.867 | -0.991 | -0.208 | 0.041 | -0.022 | -0.022 | 0 | 0.46 | 0.22 | 0.36 | 0.45 | 0.58 | 0.48 |
| 6 | 53045749 | rs4715383 | -0.466 | -1.967 | -0.915 | -0.358 | 0.032 | 0.035 | 0.035 | 0 | 0.41 | 0.66 | 0.51 | 0.22 | 0.29 | 0.26 |
| 6 | 53053945 | rs9382183 | -0.514 | -1.757 | -0.924 | 0.016 | -0.001 | -0.014 | -0.014 | 0.66 | 0.92 | 0.97 | 0.89 | 0.68 | 0.92 | 0.78 |
| 6 | 53058679 | rs6913875 | -0.12 | -1.853 | -0.471 | -0.268 | -0.057 | -0.052 | -0.052 | 0 | 0.31 | 0.44 | 0.24 | 0.18 | 0.2 | 0.22 |
| 6 | 53064313 | rs2493164 | -0.374 | -1.653 | -0.586 | -0.274 | 0.001 | -0.005 | -0.005 | 0 | 0.19 | 0.17 | 0.4 | 0.38 | 0.17 | 0.26 |
| 6 | 53068349 | rs12196615 | 0.38 | -1.974 | -0.459 | -0.486 | -0.041 | -0.04 | -0.04 | 1.179 | 0.28 | 0.32 | 0.2 | 0.13 | 0.17 | 0.19 |
| 6 | 53071349 | rs6917666 | 0.485 | -1.203 | -1.024 | -0.864 | 0 | -0.004 | -0.004 | 1.605 | 0.35 | 0.33 | 0.24 | 0.11 | 0.33 | 0.28 |
| 6 | 53072442 | rs2817114 | 0.637 | -1.702 | -0.579 | -0.665 | -0.009 | 0.042 | 0.042 | 0 | 0.12 | 0.19 | 0.11 | 0.33 | 0.1 | 0.23 |
| 6 | 53072548 | rs4487576 | 0.855 | -1.695 | -0.623 | -0.668 | -0.019 | 0.011 | 0.011 | 0 | 0.07 | 0 | 0.28 | 0.04 | 0.06 | 0.03 |
| 6 | 53072967 | rs9474440 | 0.086 | -1.805 | -0.71 | -0.793 | 0.001 | -0.063 | -0.063 | 2.959 | 0.18 | 0.33 | 0.11 | 0.03 | 0.16 | 0.11 |
| 6 | 53075109 | rs2816376 | -0.624 | -1.876 | -0.783 | -0.811 | -0.016 | 0.004 | 0.004 | 0 | 0.31 | 0.54 | 0.21 | 0.36 | 0.25 | 0.38 |
| 6 | 53075258 | rs10484663 | 0.69 | -1.782 | -0.886 | -0.584 | 0.002 | 0.001 | 0.001 | 0 | 0.12 | 0.09 | 0.1 | 0.33 | 0.09 | 0.26 |
| 6 | 53080337 | rs4715384 | 0.984 | -2.051 | -0.814 | -0.676 | -0.085 | -0.034 | -0.034 | 0 | 0.21 | 0.34 | 0.17 | 0.13 | 0.08 | 0.15 |
| 6 | 53080924 | rs2817093 | 0.226 | -1.534 | -0.821 | -0.558 | -0.001 | 0 | 0 | 0 | 0.24 | 0.23 | 0.18 | 0.34 | 0.23 | 0.36 |
| 6 | 53083892 | rs4715385 | 0.827 | -1.393 | -0.955 | -0.45 | -0.017 | -0.012 | -0.012 | 0.099 | 0.07 | 0 | 0.27 | 0.05 | 0.06 | 0.03 |
| 6 | 53085616 | rs2816372 | -0.373 | -1.612 | -0.799 | -0.626 | -0.015 | -0.019 | -0.019 | 0.846 | 0.39 | 0.51 | 0.23 | 0.34 | 0.34 | 0.42 |
| 6 | 53085914 | rs9474444 | 0.303 | -1.381 | -0.948 | -0.699 | -0.002 | 0.007 | 0.007 | 0 | 0.14 | 0.15 | 0.07 | 0 | 0.14 | 0.07 |
| 6 | 53087535 | rs7745600 | 0.434 | -1.371 | -0.953 | -0.694 | -0.004 | 0.014 | 0.014 | 0 | 0.13 | 0.11 | 0.06 | 0 | 0.14 | 0.06 |

| | | | | | | | | | | | | | | | | |
|---|---|---|---|---|---|---|---|---|---|---|---|---|---|---|---|---|
| 6 | 53088181 | rs2816369 | -1.256 | -0.695 | -1.064 | 0.05 | -0.024 | -0.032 | -0.032 | 0.011 | 0.95 | 1 | 0.95 | 0.88 | 0.9 | 0.88 |
| 6 | 53095364 | rs2816363 | 0.368 | -1.645 | -0.853 | -0.41 | -0.089 | -0.041 | -0.041 | 1.244 | 0.26 | 0.48 | 0.22 | 0.16 | 0.1 | 0.21 |
| 6 | 53095723 | rs2816362 | 0.214 | -1.684 | -0.83 | -0.432 | -0.037 | -0.015 | -0.015 | 0 | 0.27 | 0.61 | 0.22 | 0.16 | 0.1 | 0.21 |
| 6 | 53096835 | rs934150 | -0.42 | -1.69 | -1.094 | -0.55 | 0.081 | 0.101 | 0.101 | 0.277 | 0.41 | 0.78 | 0.29 | 0.16 | 0.24 | 0.28 |
| 6 | 53096986 | rs934149 | -1.285 | -1.363 | -1.194 | -0.201 | -0.091 | -0.104 | -0.104 | 0.059 | 0.68 | 0.88 | 0.69 | 0.52 | 0.5 | 0.63 |
| 6 | 53100374 | rs2562889 | -0.079 | -1.528 | -0.124 | -0.027 | -0.098 | -0.041 | -0.041 | 0.076 | 0.26 | 0.42 | 0.22 | 0.17 | 0.1 | 0.21 |
| 6 | 53101448 | rs2816356 | 0.334 | -1.445 | -0.141 | -0.089 | -0.074 | -0.03 | -0.03 | 0 | 0.2 | 0.33 | 0.18 | 0.13 | 0.07 | 0.13 |
| 6 | 53107202 | rs2252022 | -0.132 | -1.502 | -0.454 | -0.079 | -0.043 | -0.022 | -0.022 | 0.328 | 0.28 | 0.49 | 0.23 | 0.32 | 0.22 | 0.32 |
| 6 | 53111517 | rs9474466 | 0.281 | -1.266 | -0.383 | -0.149 | 0.03 | 0.012 | 0.012 | 0 | 0.08 | 0.23 | 0.27 | 0.04 | 0.08 | 0.02 |
| 6 | 53119557 | rs2817110 | -0.148 | -2.119 | -0.71 | -0.191 | 0.01 | 0.006 | 0.006 | 0 | 0.41 | 0.62 | 0.25 | 0.41 | 0.28 | 0.43 |
| 6 | 53122058 | rs9463893 | 0.894 | -1.18 | -1.327 | -0.842 | 0.005 | -0.043 | -0.043 | 0 | 0.16 | 0.18 | 0.13 | 0 | 0.23 | 0.06 |
| 6 | 53122141 | rs732648 | -1.184 | -1.387 | -1.46 | -0.897 | -0.155 | -0.083 | -0.083 | 0.82 | 0.64 | 0.95 | 0.64 | 0.46 | 0.59 | 0.49 |
| 6 | 53123118 | rs735860 | -1.153 | -1.453 | -1.904 | -0.752 | -0.199 | -0.184 | -0.184 | 0 | 0.72 | 0.98 | 0.66 | 0.52 | 0.56 | 0.61 |
| 6 | 53124501 | rs715441 | 2.22 | -0.269 | -2.088 | -0.206 | -0.023 | -0.022 | -0.022 | 0 | 0.05 | 0.01 | 0.08 | 0.18 | 0.16 | 0.12 |
| 6 | 53126794 | rs9474471 | 1.426 | -0.979 | -2.025 | -0.897 | 0.009 | -0.053 | -0.053 | 0.333 | 0.11 | 0.18 | 0.08 | 0 | 0.14 | 0.02 |
| 6 | 53128377 | rs17621311 | 1.598 | -0.998 | -2.1 | -0.857 | 0.013 | -0.029 | -0.029 | 0 | 0.12 | 0.01 | 0.05 | 0.09 | 0.07 | 0.15 |
| 6 | 53130285 | rs1429146 | -0.185 | -1.004 | -2 | -0.909 | -0.03 | 0.014 | 0.014 | 0.489 | 0.59 | 0.85 | 0.58 | 0.43 | 0.48 | 0.46 |
| 6 | 53138912 | rs2235722 | 2.18 | -1.436 | -2.309 | -1.243 | -0.038 | -0.064 | -0.064 | 0 | 0.19 | 0.01 | 0.15 | 0.34 | 0.21 | 0.29 |
| 6 | 53143554 | rs2281274 | -0.829 | -1.402 | -1.576 | -1.761 | -0.016 | -0.019 | -0.019 | 0 | 0.37 | 0.53 | 0.2 | 0.2 | 0.26 | 0.22 |
| 6 | 53159547 | rs974323 | 1.117 | -0.706 | -1.491 | -1.529 | -0.033 | -0.017 | -0.017 | 0 | 0.4 | 0.51 | 0.23 | 0.39 | 0.26 | 0.41 |
| 6 | 53166836 | rs761179 | 1.074 | -0.709 | -1.487 | -1.53 | 0.008 | 0.007 | 0.007 | 0 | 0.41 | 0.61 | 0.23 | 0.39 | 0.26 | 0.41 |
| 6 | 53175682 | rs9370194 | 1.074 | -0.709 | -1.51 | -1.53 | 0.01 | 0.008 | 0.008 | 0 | 0.41 | 0.61 | 0.23 | 0.39 | 0.26 | 0.41 |
| 6 | 53202035 | rs209510 | 0.106 | -0.466 | -2.092 | -1.556 | -0.125 | -0.052 | -0.052 | 0 | 0.38 | 0.06 | 0.4 | 0.57 | 0.5 | 0.53 |
| 6 | 53202606 | rs4562128 | 0.804 | -0.943 | -1.823 | -1.265 | 0.009 | 0.007 | 0.007 | 0.008 | 0.41 | 0.61 | 0.23 | 0.39 | 0.26 | 0.41 |
| 6 | 53203007 | rs209511 | -0.244 | -0.943 | -1.817 | -1.278 | 0 | -0.002 | -0.002 | 0.186 | 0.91 | 0.89 | 0.71 | 0.96 | 0.9 | 0.96 |
| 6 | 53203577 | rs209512 | 1.459 | -0.552 | -2.244 | -1.205 | -0.037 | 0.006 | 0.006 | 0 | 0.07 | 0.01 | 0.19 | 0.41 | 0.22 | 0.23 |
| 6 | 53205947 | rs209515 | -0.011 | -2.699 | -1.848 | -1.299 | 0.079 | 0.059 | 0.059 | 0 | 0.49 | 0.28 | 0.48 | 0.57 | 0.64 | 0.55 |
| 6 | 53210355 | rs209521 | 0.257 | -1.583 | -1.612 | -1.39 | -0.029 | -0.007 | -0.007 | 0 | 0.92 | 0.99 | 0.72 | 0.96 | 0.9 | 0.96 |
| 6 | 53220825 | rs10948748 | -0.072 | -1.63 | -1.246 | -1.26 | -0.018 | -0.021 | -0.021 | 0.215 | 0.14 | 0.01 | 0.08 | 0.09 | 0.12 | 0.21 |
| 6 | 53226311 | rs9463906 | -0.912 | -1.71 | -0.459 | -1.53 | -0.005 | -0.054 | -0.054 | 0.25 | 0.27 | 0.69 | 0.16 | 0.01 | 0.18 | 0.14 |
| 6 | 53238908 | rs7776264 | -1.495 | -1.213 | -0.415 | -1.305 | 0.05 | 0.139 | 0.139 | 1.634 | 0.36 | 0.81 | 0.25 | 0.08 | 0.33 | 0.24 |
| 6 | 53239608 | rs934147 | -0.795 | -1.215 | -0.077 | -1.313 | -0.032 | -0.095 | -0.095 | 0 | 0.22 | 0.62 | 0.16 | 0.03 | 0.18 | 0.18 |
| 6 | 53254111 | rs9296711 | -0.899 | -1.278 | -0.15 | -1.355 | -0.03 | -0.085 | -0.085 | 0 | 0.26 | 0.64 | 0.16 | 0.04 | 0.18 | 0.19 |
| 6 | 53255025 | rs9474515 | -1.49 | -1.203 | -0.559 | -1.37 | 0.068 | 0.159 | 0.159 | 0.571 | 0.39 | 0.79 | 0.26 | 0.06 | 0.33 | 0.25 |
| 6 | 53264485 | rs1429143 | 1.166 | -1.242 | -0.379 | -1.233 | 0.072 | 0.153 | 0.153 | 0.913 | 0.65 | 0.17 | 0.78 | 0.93 | 0.73 | 0.8 |
| 6 | 53276001 | rs6458931 | -0.819 | -1.029 | 0.227 | -1.094 | -0.063 | -0.157 | -0.157 | 0.132 | 0.21 | 0.51 | 0.11 | 0.01 | 0.13 | 0.15 |
| 6 | 53278242 | rs9367529 | 0.914 | -0.931 | 0.073 | -0.994 | -0.033 | -0.098 | -0.098 | 1.339 | 0.68 | 0.52 | 0.78 | 0.9 | 0.71 | 0.76 |
| 6 | 53284413 | rs10807460 | -0.388 | 0.007 | -0.047 | -0.214 | 0.011 | 0.023 | 0.023 | 0.509 | 0.28 | 0.31 | 0.46 | 0.66 | 0.3 | 0.33 |
| 6 | 53289583 | rs1356553 | -0.52 | -0.1 | 0.421 | -0.257 | 0.009 | 0.018 | 0.018 | 0.713 | 0.3 | 0.46 | 0.5 | 0.67 | 0.3 | 0.33 |
| 6 | 53291466 | rs6458934 | 0.261 | -0.068 | 0.006 | -0.568 | -0.027 | 0.022 | 0.022 | 0 | 0.9 | 0.99 | 0.94 | 0.97 | 0.88 | 0.91 |
| 6 | 53293179 | rs9370208 | 1.079 | -0.616 | 0.026 | -0.779 | 0.02 | 0.142 | 0.142 | 0.111 | 0.51 | 0.19 | 0.39 | 0.22 | 0.55 | 0.57 |
| 6 | 53293410 | rs4440459 | 0.251 | -0.679 | 0.167 | -0.789 | 0 | -0.004 | -0.004 | 0.04 | 0.88 | 0.88 | 0.94 | 0.97 | 0.88 | 0.91 |
| 6 | 53295203 | rs1518516 | 1.448 | 0.13 | -0.253 | -1.048 | -0.04 | 0.217 | 0.217 | 1.646 | 0.4 | 0.09 | 0.25 | 0.09 | 0.38 | 0.29 |
| 6 | 53296156 | rs9370209 | 0.884 | 0.685 | 0.481 | -0.356 | -0.008 | -0.051 | -0.051 | 0.711 | 0.86 | 0.7 | 0.91 | 0.97 | 0.88 | 0.91 |
| 6 | 53296986 | rs17827934 | -0.617 | 0.956 | 0.035 | -0.139 | -0.001 | 0.001 | 0.001 | 0 | 0.1 | 0.09 | 0.13 | 0.13 | 0.14 | 0.26 |
| 6 | 53298485 | rs1377392 | 0.811 | 1.297 | -0.255 | 0.497 | -0.026 | -0.039 | -0.039 | 0 | 0.8 | 0.84 | 0.59 | 0.38 | 0.71 | 0.68 |
| 6 | 53302539 | rs6918446 | -0.565 | 1.501 | -0.101 | 0.335 | 0.015 | 0.008 | 0.008 | 0 | 0.1 | 0.11 | 0.11 | 0.12 | 0.16 | 0.15 |
| 6 | 53317012 | rs4712029 | -0.679 | 1.612 | 0.136 | 0.767 | -0.015 | -0.015 | -0.015 | 0 | 0.17 | 0.14 | 0.39 | 0.57 | 0.28 | 0.38 |
| 6 | 53318038 | rs9474543 | -0.914 | 1.014 | 0.122 | 0.406 | -0.014 | 0.005 | 0.005 | 0.033 | 0.06 | 0.29 | 0.08 | 0.03 | 0.02 | 0.08 |
| 6 | 53330356 | rs2067833 | -0.887 | 1.434 | 0.044 | 0.585 | 0.029 | -0.011 | -0.011 | 0 | 0.35 | 0.38 | 0.52 | 0.74 | 0.41 | 0.62 |
| 6 | 53338597 | rs9395867 | -0.677 | 1.877 | -0.25 | 0.98 | 0.014 | 0.04 | 0.04 | 0.038 | 0.2 | 0.31 | 0.48 | 0.51 | 0.3 | 0.46 |
| 6 | 53349969 | rs510420 | -0.945 | 1.527 | -0.367 | 0.082 | 0.001 | -0.015 | -0.015 | 0.251 | 0.17 | 0.23 | 0.17 | 0.04 | 0.19 | 0.07 |
| 6 | 53349999 | rs10484785 | 0.341 | 1.236 | -0.411 | 0.036 | -0.001 | -0.003 | -0.003 | 0 | 0.49 | 0.46 | 0.3 | 0.27 | 0.38 | 0.3 |
| 6 | 53350525 | rs13214318 | -0.803 | 1.079 | -0.236 | 0.278 | 0.021 | -0.019 | -0.019 | 0 | 0.09 | 0.19 | 0.09 | 0.09 | 0.1 | 0.08 |
| 6 | 53351289 | rs1467408 | -0.333 | 0.18 | -0.752 | 0.416 | 0.008 | 0.029 | 0.029 | 0.214 | 0.57 | 0.85 | 0.61 | 0.64 | 0.38 | 0.7 |
| 6 | 53357018 | rs6922238 | -0.103 | 0.078 | -0.343 | 0.554 | 0.006 | -0.002 | -0.002 | 0 | 0.46 | 0.6 | 0.55 | 0.51 | 0.35 | 0.69 |
| 6 | 53360119 | rs2397146 | -0.368 | -0.354 | -0.285 | 0.502 | 0.001 | 0.01 | 0.01 | 0.664 | 0.34 | 0.71 | 0.23 | 0.24 | 0.27 | 0.48 |
| 6 | 53365374 | rs2066511 | 0.206 | -0.209 | 0.092 | 0.44 | 0.001 | -0.007 | -0.007 | 0 | 0.18 | 0.23 | 0.15 | 0.13 | 0.23 | 0.13 |
| 6 | 53366061 | rs12525474 | -0.379 | -0.593 | 0.265 | 0.3 | -0.012 | -0.01 | -0.01 | 0 | 0.18 | 0.21 | 0.12 | 0.17 | 0.15 | 0.18 |
| 6 | 53366989 | rs670548 | -1.644 | -0.719 | -0.178 | 0.452 | -0.133 | -0.078 | -0.078 | 2.437 | 0.75 | 0.97 | 0.79 | 0.71 | 0.62 | 0.85 |
| 6 | 53368309 | rs13437395 | 0.44 | -1.018 | 0.84 | 0.58 | -0.033 | 0.066 | 0.066 | 0.209 | 0.19 | 0.32 | 0.4 | 0.42 | 0.08 | 0.24 |
| 6 | 53369575 | rs3799694 | 0.313 | -1.007 | 0.784 | 0.53 | -0.045 | 0.089 | 0.089 | 0.329 | 0.2 | 0.38 | 0.41 | 0.42 | 0.08 | 0.24 |
| 6 | 53370014 | rs2277108 | 0.4 | -1.011 | 0.781 | 0.531 | -0.046 | 0.083 | 0.083 | 0.373 | 0.2 | 0.38 | 0.42 | 0.42 | 0.08 | 0.24 |
| 6 | 53370395 | rs524553 | 0.718 | -0.917 | 0.6 | 0.508 | 0 | -0.001 | -0.001 | 0.019 | 0.16 | 0.14 | 0.12 | 0.12 | 0.2 | 0.13 |
| 6 | 53373105 | rs9474576 | -0.118 | -0.947 | 0.704 | 0.519 | -0.028 | 0.017 | 0.017 | 0 | 0.26 | 0.34 | 0.42 | 0.42 | 0.1 | 0.24 |
| 6 | 53373771 | rs16883912 | 0.215 | -0.565 | 0.486 | 0.336 | 0 | -0.024 | -0.024 | 0 | 0.1 | 0.13 | 0.09 | 0.03 | 0.11 | 0.17 |
| 6 | 53376091 | rs642429 | -0.006 | -0.367 | 0.55 | 0.28 | 0.002 | -0.05 | -0.05 | 0 | 0.11 | 0.17 | 0.1 | 0.01 | 0.12 | 0.18 |
| 6 | 53377834 | rs574389 | -0.837 | -0.737 | 0.441 | 0.324 | 0.023 | 0.02 | 0.02 | 0 | 0.48 | 0.34 | 0.6 | 0.39 | 0.43 | 0.54 |
| 6 | 53378369 | rs648595 | -1.113 | -0.804 | 0.928 | 0.412 | 0.01 | 0.003 | 0.003 | 0 | 0.52 | 0.64 | 0.63 | 0.56 | 0.45 | 0.56 |

| | | | | | | | | | | | | | | | | |
|---|---|---|---|---|---|---|---|---|---|---|---|---|---|---|---|---|
| 6 | 53379405 | rs3736729 | -0.946 | -0.788 | 0.706 | 0.418 | 0.027 | 0.024 | 0.024 | 0 | 0.48 | 0.34 | 0.6 | 0.39 | 0.43 | 0.55 |
| 6 | 53380713 | rs553822 | -1.623 | -0.717 | 0.574 | 0.326 | -0.035 | -0.006 | -0.006 | 0 | 0.64 | 0.77 | 0.69 | 0.63 | 0.55 | 0.75 |
| 6 | 53383918 | rs617066 | 0.734 | -0.719 | 0.179 | 0.415 | 0 | -0.001 | -0.001 | 0.869 | 0.16 | 0.16 | 0.12 | 0.13 | 0.2 | 0.13 |
| 6 | 53385343 | rs761141 | 0.618 | -0.705 | 0.193 | 0.405 | 0 | -0.001 | -0.001 | 0.96 | 0.15 | 0.16 | 0.12 | 0.12 | 0.2 | 0.13 |
| 6 | 53385475 | rs2100375 | 0.125 | -0.718 | 0.144 | 0.39 | 0.019 | -0.036 | -0.036 | 0 | 0.31 | 0.2 | 0.47 | 0.37 | 0.26 | 0.36 |
| 6 | 53390709 | rs1555906 | -0.439 | -0.841 | 0.186 | 0.357 | 0.054 | -0.019 | -0.019 | 0.219 | 0.37 | 0.24 | 0.48 | 0.37 | 0.27 | 0.37 |
| 6 | 53393734 | rs4715407 | 0.007 | -0.897 | 0.467 | 0.452 | -0.025 | 0.003 | 0.003 | 0 | 0.23 | 0.3 | 0.38 | 0.25 | 0.08 | 0.23 |
| 6 | 53395363 | rs2268329 | 1 | -0.898 | 0.457 | 0.451 | 0.032 | -0.047 | -0.047 | 1.059 | 0.16 | 0.08 | 0.35 | 0.25 | 0.06 | 0.23 |
| 6 | 53396896 | rs3799696 | -0.661 | -0.913 | 0.554 | 0.421 | -0.009 | 0.007 | 0.007 | 0 | 0.35 | 0.39 | 0.45 | 0.51 | 0.2 | 0.43 |
| 6 | 53398015 | rs6934367 | 0.047 | -0.868 | 0.484 | 0.451 | -0.025 | 0.003 | 0.003 | 0 | 0.23 | 0.3 | 0.38 | 0.25 | 0.08 | 0.23 |
| 6 | 53398559 | rs502862 | -1.415 | -0.568 | 0.426 | 0.445 | -0.009 | 0.001 | 0.001 | 0.406 | 0.65 | 0.71 | 0.68 | 0.66 | 0.55 | 0.74 |
| 6 | 53401587 | rs2397147 | -0.183 | -0.321 | 0.247 | 0.277 | -0.01 | 0.016 | 0.016 | 1.396 | 0.27 | 0.2 | 0.22 | 0.16 | 0.35 | 0.31 |
| 6 | 53401999 | rs3799699 | -0.255 | -0.543 | 0.433 | 0.733 | 0.001 | -0.002 | -0.002 | 0 | 0.16 | 0.17 | 0.35 | 0.24 | 0.06 | 0.23 |
| 6 | 53402679 | rs606548 | -0.708 | -0.55 | 0.241 | 0.729 | -0.053 | 0.04 | 0.04 | 0 | 0.19 | 0.38 | 0.37 | 0.24 | 0.06 | 0.23 |
| 6 | 53404282 | rs2284650 | -0.163 | -0.253 | -0.961 | 0.862 | 0.03 | -0.041 | -0.041 | 0 | 0.15 | 0.08 | 0.35 | 0.24 | 0.05 | 0.23 |
| 6 | 53408789 | rs510088 | 0.334 | 0.413 | -0.881 | 0.16 | -0.003 | 0.016 | 0.016 | 0 | 0.75 | 0.86 | 0.79 | 0.92 | 0.68 | 0.74 |
| 6 | 53416053 | rs11758818 | 0.321 | 0.089 | -1.296 | 0.737 | -0.007 | -0.034 | -0.034 | 0 | 0.29 | 0.07 | 0.43 | 0.28 | 0.23 | 0.19 |
| 6 | 53420503 | rs4712036 | 0.136 | -0.172 | -0.482 | 0.703 | -0.037 | -0.011 | -0.011 | 0.59 | 0.34 | 0.51 | 0.45 | 0.28 | 0.21 | 0.19 |
| 6 | 53421910 | rs7738142 | -0.574 | 0.589 | -0.446 | 1.43 | -0.016 | -0.019 | -0.019 | 0 | 0.05 | 0.02 | 0.05 | 0.13 | 0.09 | 0.07 |
| 6 | 53427397 | rs13220340 | -0.944 | 0.246 | -0.19 | 1.696 | 0.001 | -0.001 | -0.001 | 4.164 | 0.13 | 0.12 | 0.32 | 0.28 | 0.14 | 0.22 |
| 6 | 53429272 | rs1883633 | -0.944 | 0.211 | -0.094 | 1.619 | -0.002 | 0.002 | 0.002 | 0.195 | 0.13 | 0.15 | 0.09 | 0.26 | 0.14 | 0.22 |
| 6 | 53432210 | rs655630 | -1.688 | 0.081 | -0.017 | 1.493 | 0.018 | 0.043 | 0.043 | 0.931 | 0.43 | 0.71 | 0.38 | 0.76 | 0.39 | 0.65 |
| 6 | 53443269 | rs809942 | -1.174 | -0.025 | 1.676 | 2.422 | 0.015 | 0.08 | 0.08 | 0 | 0.19 | 0.44 | 0.18 | 0.31 | 0.17 | 0.31 |
| 6 | 53445908 | rs10484783 | -0.513 | 0.023 | 1.576 | 2.28 | 0.018 | 0.042 | 0.042 | 0.06 | 0.11 | 0.21 | 0.08 | 0.15 | 0.11 | 0.11 |
| 6 | 53447254 | rs597655 | -2.194 | -0.048 | 1.938 | 2.243 | -0.03 | 0.065 | 0.065 | 0.044 | 0.29 | 0.43 | 0.24 | 0.52 | 0.24 | 0.55 |
| 6 | 53447934 | rs513041 | -2.677 | -0.056 | 1.679 | 2.245 | 0.013 | 0.052 | 0.052 | 1.019 | 0.42 | 0.77 | 0.38 | 0.76 | 0.39 | 0.65 |
| 6 | 53448040 | rs9349677 | -0.232 | -0.073 | 1.643 | 2.179 | 0.008 | -0.023 | -0.023 | 0 | 0.09 | 0.01 | 0.07 | 0.16 | 0.12 | 0.21 |
| 6 | 53448861 | rs543087 | -0.497 | -0.042 | 1.692 | 2.019 | -0.026 | 0.066 | 0.066 | 0 | 0.1 | 0.23 | 0.09 | 0.21 | 0.03 | 0.23 |
| 6 | 53449264 | rs627180 | -2.361 | -0.023 | 1.421 | 2.015 | -0.012 | 0.022 | 0.022 | 0 | 0.32 | 0.61 | 0.31 | 0.61 | 0.29 | 0.54 |
| 6 | 53450301 | rs13212644 | -0.469 | 0.387 | 0.781 | 1.738 | -0.033 | 0.037 | 0.037 | 0 | 0.07 | 0.01 | 0.06 | 0 | 0.13 | 0.06 |
| 6 | 53452060 | rs559102 | -0.734 | 0.192 | 1.027 | 1.978 | -0.02 | 0.036 | 0.036 | 0 | 0.09 | 0.16 | 0.09 | 0.23 | 0.02 | 0.23 |
| 6 | 53461965 | rs2179545 | 1.171 | 0.737 | 0.716 | 1.449 | 0.013 | -0.003 | -0.003 | 0 | 0.57 | 0.33 | 0.61 | 0.41 | 0.58 | 0.38 |
| 6 | 53465000 | rs9382219 | -0.114 | 0.73 | 0.671 | 1.62 | 0.001 | 0.009 | 0.009 | 0 | 0.07 | 0.08 | 0.12 | 0.21 | 0.05 | 0.16 |
| 6 | 53466800 | rs492381 | 0.426 | 0.64 | 1.4 | 1.697 | -0.004 | 0.001 | 0.001 | 0 | 0.86 | 0.81 | 0.86 | 0.88 | 0.84 | 0.85 |
| 6 | 53467311 | rs497085 | -1.532 | 0.54 | 1.527 | 1.39 | -0.018 | -0.018 | -0.018 | 0 | 0.18 | 0.08 | 0.11 | 0.26 | 0.21 | 0.3 |
| 6 | 53468079 | rs525248 | -2.081 | 0.534 | 1.908 | 1.369 | 0 | 0.001 | 0.001 | 0.033 | 0.26 | 0.26 | 0.23 | 0.47 | 0.27 | 0.47 |
| 6 | 53470671 | rs502156 | -0.425 | 0.546 | 1.714 | 1.366 | 0.009 | -0.021 | -0.021 | 1.006 | 0.1 | 0.05 | 0.04 | 0.2 | 0.08 | 0.23 |
| 6 | 53478869 | rs10948754 | 2.472 | 1.317 | 1.774 | 0.963 | 0.028 | 0.052 | 0.052 | 0.576 | 0.45 | 0.25 | 0.51 | 0.3 | 0.38 | 0.18 |
| 6 | 53479032 | rs6924228 | -1.02 | 1.336 | 2.058 | 0.98 | 0.011 | -0.005 | -0.005 | 0 | 0.1 | 0.12 | 0.06 | 0.07 | 0.13 | 0.08 |
| 6 | 53479069 | rs10948755 | 2.532 | 1.331 | 2.022 | 0.98 | 0.028 | 0.052 | 0.052 | 0.576 | 0.45 | 0.25 | 0.51 | 0.3 | 0.38 | 0.18 |
| 6 | 53480293 | rs11962209 | -1.216 | 1.331 | 2.159 | 0.973 | 0.021 | -0.008 | -0.008 | 0 | 0.1 | 0.14 | 0.07 | 0.07 | 0.13 | 0.08 |
| 6 | 53481176 | rs630152 | 0.458 | 1.305 | 2.212 | 0.969 | -0.008 | -0.012 | -0.012 | 1.556 | 0.84 | 0.62 | 0.85 | 0.91 | 0.85 | 0.86 |
| 6 | 53481989 | rs574202 | -1.983 | 1.287 | 2.249 | 0.948 | 0.007 | 0.003 | 0.003 | 0 | 0.41 | 0.53 | 0.35 | 0.61 | 0.47 | 0.69 |
| 6 | 53488119 | rs1409530 | 1.312 | 1.807 | 2.321 | 0.78 | 0.006 | -0.017 | -0.017 | 0 | 0.71 | 0.58 | 0.75 | 0.82 | 0.69 | 0.68 |
| 6 | 53489402 | rs11962201 | -1.55 | 1.697 | 2.387 | 0.804 | -0.013 | -0.019 | -0.019 | 0.125 | 0.21 | 0.18 | 0.14 | 0.32 | 0.28 | 0.34 |
| 6 | 53490644 | rs12664098 | -0.509 | 1.703 | 2.438 | 0.79 | 0.016 | -0.02 | -0.02 | 0.507 | 0.14 | 0.2 | 0.12 | 0.09 | 0.16 | 0.17 |
| 6 | 53490729 | rs12662781 | -0.126 | 1.708 | 2.233 | 0.793 | -0.051 | 0.019 | 0.019 | 0 | 0.12 | 0.03 | 0.1 | 0.09 | 0.16 | 0.17 |
| 6 | 53491730 | rs1536386 | 0.569 | 1.645 | 2.29 | 0.802 | -0.012 | -0.018 | -0.018 | 0 | 0.84 | 0.61 | 0.86 | 0.91 | 0.85 | 0.85 |
| 6 | 53493460 | rs6902510 | 2.161 | 1.744 | 2.39 | 0.887 | 0.022 | -0.001 | -0.001 | 0 | 0.48 | 0.26 | 0.61 | 0.5 | 0.41 | 0.33 |
| 6 | 53495061 | rs9474604 | 1.717 | 1.644 | 2.665 | 1.04 | -0.005 | 0.034 | 0.034 | 0 | 0.76 | 0.57 | 0.79 | 0.66 | 0.73 | 0.61 |
| 6 | 53497222 | rs1326543 | -0.771 | 1.713 | 2.611 | 1.002 | -0.019 | 0.012 | 0.012 | 0 | 0.22 | 0.19 | 0.15 | 0.16 | 0.24 | 0.24 |
| 6 | 53500789 | rs13192959 | -1.215 | 1.703 | 2.678 | 0.997 | 0.03 | -0.034 | -0.034 | 0.556 | 0.24 | 0.41 | 0.17 | 0.16 | 0.24 | 0.25 |
| 6 | 53504277 | rs13195847 | -0.769 | 1.707 | 2.496 | 0.999 | -0.019 | 0.012 | 0.012 | 0 | 0.22 | 0.19 | 0.16 | 0.16 | 0.24 | 0.24 |
| 6 | 53506487 | rs9367538 | 2.255 | 1.675 | 2.167 | 0.998 | -0.008 | -0.02 | -0.02 | 0 | 0.52 | 0.17 | 0.62 | 0.49 | 0.49 | 0.37 |
| 6 | 53508614 | rs6458950 | 0.06 | 1.753 | 1.834 | 1.046 | -0.011 | -0.013 | -0.013 | 0 | 0.94 | 0.95 | 0.88 | 0.91 | 0.88 | 0.89 |
| 6 | 53508991 | rs4142075 | -0.618 | 1.732 | 1.802 | 0.989 | -0.028 | -0.066 | -0.066 | 0 | 0.12 | 0.02 | 0.07 | 0.26 | 0.15 | 0.27 |
| 6 | 53510423 | rs7739121 | 1.954 | 1.758 | 2.042 | 1.047 | 0 | -0.003 | -0.003 | 0 | 0.53 | 0.3 | 0.63 | 0.5 | 0.49 | 0.38 |
| 6 | 53511188 | rs4715410 | 1.073 | 1.884 | 1.597 | 0.636 | -0.025 | 0.021 | 0.021 | 0 | 0.73 | 0.79 | 0.73 | 0.83 | 0.64 | 0.66 |
| 6 | 53511280 | rs4715411 | 1.416 | 1.788 | 1.572 | 0.643 | -0.001 | 0 | 0 | 0 | 0.67 | 0.65 | 0.68 | 0.75 | 0.58 | 0.61 |
| 6 | 53511561 | rs9370224 | 1.219 | 1.596 | 1.633 | 0.711 | 0.005 | -0.002 | -0.002 | 0 | 0.71 | 0.65 | 0.73 | 0.75 | 0.7 | 0.67 |
| 6 | 53513469 | rs9474613 | -0.659 | 1.248 | 1.547 | 0.231 | -0.004 | -0.023 | -0.023 | 0 | 0.2 | 0.27 | 0.09 | 0.1 | 0.14 | 0.16 |
| 6 | 53515130 | rs6941645 | 1.037 | 1.291 | 1.816 | 0.373 | 0.012 | 0.02 | 0.02 | 0.075 | 0.93 | 0.85 | 0.94 | 0.92 | 0.92 | 0.95 |
| 6 | 53515483 | rs6942249 | -1.419 | 1.245 | 2.019 | 0.469 | 0.007 | 0.001 | 0.001 | 1.322 | 0.31 | 0.69 | 0.21 | 0.32 | 0.2 | 0.35 |
| 6 | 53515759 | rs12697951 | -1.785 | 1.245 | 1.969 | 0.473 | 0.007 | -0.015 | -0.015 | 0 | 0.36 | 0.74 | 0.27 | 0.41 | 0.28 | 0.4 |
| 6 | 53517035 | rs3823111 | 0.362 | 1.311 | 2.245 | 0.491 | 0.012 | 0.091 | 0.091 | 0 | 0.06 | 0.12 | 0.09 | 0.22 | 0.06 | 0.16 |
| 6 | 53518631 | rs1326546 | 0.107 | 1.642 | 1.616 | 0.419 | 0.027 | -0.025 | -0.025 | 0 | 0.95 | 0.9 | 0.93 | 1 | 0.87 | 0.97 |
| 6 | 53519062 | rs10948758 | 1.65 | 1.64 | 1.71 | 0.418 | 0.08 | 0.108 | 0.108 | 0.055 | 0.81 | 0.53 | 0.77 | 0.7 | 0.74 | 0.75 |
| 6 | 53519605 | rs3799260 | 1.236 | 1.641 | 1.644 | 0.418 | 0.029 | -0.002 | -0.002 | 0 | 0.88 | 0.82 | 0.87 | 0.92 | 0.8 | 0.92 |
| 6 | 53519857 | rs12210809 | 1.559 | 1.649 | 1.648 | 0.425 | -0.045 | -0.014 | -0.014 | 0 | 0.59 | 0.17 | 0.67 | 0.6 | 0.59 | 0.58 |

| | | | | | | | | | | | | | | | | |
|---|---|---|---|---|---|---|---|---|---|---|---|---|---|---|---|---|
| 6 | 53522919 | rs1326548 | 0.175 | 1.3 | 1.749 | 0.343 | 0.048 | 0.18 | 0.18 | 0 | 0.07 | 0.2 | 0.09 | 0.21 | 0.06 | 0.16 |
| 6 | 53531927 | rs2478348 | 1.456 | 1.348 | 1.175 | 0.324 | -0.014 | 0.01 | 0.01 | 0.86 | 0.67 | 0.24 | 0.74 | 0.69 | 0.67 | 0.65 |
| 6 | 53535989 | rs9395875 | -0.072 | 1.078 | 0.456 | 0.619 | 0.073 | 0.229 | 0.229 | 0 | 0.07 | 0.3 | 0.1 | 0.22 | 0.06 | 0.16 |
| 6 | 53536029 | rs13203828 | -0.615 | 0.962 | -0.866 | -0.21 | 0.003 | 0.013 | 0.013 | 0 | 0.18 | 0.13 | 0.08 | 0.1 | 0.14 | 0.16 |
| 6 | 53546586 | rs9370225 | -0.343 | 1 | -1.305 | -0.389 | 0.006 | 0.061 | 0.061 | 0 | 0.51 | 0.59 | 0.41 | 0.72 | 0.45 | 0.68 |
| 6 | 53547719 | rs4289652 | 0.316 | 0.738 | -1.683 | -0.741 | 0.005 | 0.067 | 0.067 | 0 | 0.73 | 0.86 | 0.81 | 1 | 0.73 | 0.86 |
| 6 | 53564059 | rs12660527 | -0.49 | 0.878 | -1.941 | -0.113 | -0.047 | -0.04 | -0.04 | 0 | 0.2 | 0.02 | 0.18 | 0.17 | 0.21 | 0.27 |
| 6 | 53564752 | rs9463962 | -0.102 | 1.469 | -2.413 | 0.837 | -0.033 | -0.046 | -0.046 | 0 | 0.9 | 0.97 | 0.75 | 0.65 | 0.88 | 0.9 |
| 6 | 53566032 | rs4715416 | -0.053 | 1.19 | -2.491 | -0.428 | 0.017 | 0.11 | 0.11 | 0 | 0.47 | 0.27 | 0.37 | 0.18 | 0.48 | 0.38 |
| 6 | 53571485 | rs4128134 | 0.447 | 1.589 | -1.692 | -0.778 | 0.045 | -0.035 | -0.035 | 0.151 | 0.08 | 0.2 | 0.09 | 0.05 | 0.13 | 0.13 |
| 6 | 53574677 | rs4577786 | 1.198 | 1.503 | -1.615 | -0.925 | 0.016 | 0.083 | 0.083 | 0 | 0.23 | 0.08 | 0.12 | 0.02 | 0.2 | 0.11 |
| 6 | 53579392 | rs7758962 | 0.555 | 0.958 | -1.319 | -1.197 | -0.014 | 0.113 | 0.113 | 0 | 0.38 | 0.2 | 0.28 | 0.11 | 0.41 | 0.28 |
| 6 | 53582914 | rs9296718 | 0.565 | 0.883 | -0.992 | -1.236 | -0.008 | 0.075 | 0.075 | 0 | 0.38 | 0.24 | 0.27 | 0.11 | 0.41 | 0.29 |
| 6 | 53584244 | rs6920831 | 0.524 | 0.619 | -0.901 | -1.351 | 0.01 | -0.044 | -0.044 | 0.008 | 0.13 | 0.19 | 0.11 | 0.06 | 0.15 | 0.13 |
| 6 | 53585032 | rs4312987 | 0.275 | 0.608 | -0.554 | -0.954 | -0.013 | 0.021 | 0.021 | 0 | 0.59 | 0.66 | 0.48 | 0.32 | 0.58 | 0.56 |
| 6 | 53587038 | rs10948767 | 1.219 | 0.576 | -0.333 | -0.903 | -0.015 | -0.048 | -0.048 | 0.039 | 0.2 | 0.06 | 0.22 | 0.26 | 0.21 | 0.33 |
| 6 | 53587485 | rs7773640 | 0.239 | 0.625 | -0.231 | -0.99 | 0.004 | 0.007 | 0.007 | 0 | 0.51 | 0.41 | 0.39 | 0.17 | 0.56 | 0.42 |
| 6 | 53590075 | rs12201213 | 0.196 | 0.631 | -0.323 | -0.762 | -0.002 | 0.001 | 0.001 | 0.32 | 0.32 | 0.3 | 0.33 | 0.27 | 0.37 | 0.43 |
| 6 | 53593569 | rs4493745 | 0.93 | 0.553 | -0.274 | -0.821 | 0.007 | 0.028 | 0.028 | 0 | 0.43 | 0.37 | 0.29 | 0.15 | 0.38 | 0.33 |
| 6 | 53595083 | rs9463970 | 1.354 | 0.596 | -0.223 | -0.656 | -0.002 | 0.003 | 0.003 | 0 | 0.16 | 0.18 | 0.17 | 0.18 | 0.17 | 0.24 |
| 6 | 53595753 | rs11962552 | 0.551 | 0.656 | -0.001 | -0.65 | 0.023 | -0.036 | -0.036 | 0.025 | 0.16 | 0.27 | 0.16 | 0.09 | 0.19 | 0.16 |
| 6 | 53595950 | rs9357772 | 0.298 | 0.663 | -0.122 | -0.235 | -0.019 | -0.038 | -0.038 | 3.853 | 0.09 | 0.02 | 0.25 | 0.29 | 0.11 | 0.09 |
| 6 | 53599201 | rs10948768 | 0.611 | 0.749 | 0.027 | -0.351 | 0.025 | -0.021 | -0.021 | 0 | 0.16 | 0.27 | 0.16 | 0.13 | 0.2 | 0.19 |
| 6 | 53600766 | rs10456671 | 0.166 | 0.651 | 0.002 | -0.457 | 0.002 | 0.033 | 0.033 | 0 | 0.57 | 0.66 | 0.45 | 0.28 | 0.56 | 0.52 |
| 6 | 53607649 | rs7759946 | 0.402 | 0.554 | 0.354 | -0.599 | 0.016 | -0.068 | -0.068 | 0 | 0.14 | 0.34 | 0.13 | 0.04 | 0.14 | 0.16 |
| 6 | 53613398 | rs4516937 | -0.329 | 0.529 | 0.298 | -0.57 | -0.008 | -0.058 | -0.058 | 0 | 0.74 | 0.86 | 0.62 | 0.47 | 0.74 | 0.76 |
| 6 | 53617730 | rs12213392 | 1.014 | 0.522 | 0.357 | -0.602 | -0.01 | 0.111 | 0.111 | 0 | 0.43 | 0.26 | 0.28 | 0.08 | 0.42 | 0.28 |
| 6 | 53618520 | rs9474647 | 1.409 | 0.512 | 0.378 | -0.597 | -0.004 | 0.024 | 0.024 | 0 | 0.17 | 0.2 | 0.2 | 0.31 | 0.18 | 0.3 |
| 6 | 53619575 | rs6921605 | -0.081 | 0.522 | 0.411 | -0.576 | 0 | -0.005 | -0.005 | 0 | 0.72 | 0.75 | 0.59 | 0.33 | 0.73 | 0.68 |
| 6 | 53620063 | rs4400206 | -0.221 | 0.527 | 0.368 | -0.586 | -0.003 | -0.038 | -0.038 | 0.507 | 0.73 | 0.84 | 0.62 | 0.42 | 0.74 | 0.74 |
| 6 | 53620831 | rs10456674 | 0.771 | 0.505 | 0.358 | -0.714 | -0.012 | 0.013 | 0.013 | 0 | 0.14 | 0.12 | 0.15 | 0.06 | 0.2 | 0.17 |
| 6 | 53621099 | rs10456166 | -0.373 | 0.498 | 0.352 | -0.716 | 0.032 | -0.097 | -0.097 | 0 | 0.28 | 0.45 | 0.28 | 0.1 | 0.34 | 0.33 |
| 6 | 53621295 | rs10456167 | -0.373 | 0.498 | 0.352 | -0.716 | 0.032 | -0.097 | -0.097 | 0.386 | 0.28 | 0.45 | 0.28 | 0.1 | 0.34 | 0.33 |
| 6 | 53633422 | rs12216441 | 0.701 | 0.53 | 0.245 | -0.68 | 0 | -0.005 | -0.005 | 1.612 | 0.15 | 0.17 | 0.15 | 0.06 | 0.2 | 0.17 |
| 6 | 53651171 | rs4715423 | -0.417 | 0.457 | 0.309 | -0.709 | 0 | -0.02 | -0.02 | 0 | 0.44 | 0.53 | 0.6 | 0.89 | 0.44 | 0.56 |
| 6 | 53659112 | rs3813849 | -0.793 | 0.456 | 0.345 | -0.524 | 0.001 | -0.019 | -0.019 | 0 | 0.88 | 0.85 | 0.89 | 0.96 | 0.86 | 0.83 |
| 6 | 53663882 | rs3003498 | 1.227 | 0.737 | 0.615 | -0.3 | -0.002 | -0.002 | -0.002 | 0 | 0.17 | 0.17 | 0.22 | 0.25 | 0.24 | 0.26 |
| 6 | 53664707 | rs4342426 | -1.11 | 0.668 | 0.684 | -0.392 | 0 | -0.026 | -0.026 | 0 | 0.58 | 0.59 | 0.72 | 0.91 | 0.61 | 0.71 |
| 6 | 53668237 | rs9395878 | 0.067 | 0.637 | 0.617 | -0.267 | 0.019 | -0.079 | -0.079 | 0.316 | 0.25 | 0.11 | 0.33 | 0.5 | 0.19 | 0.23 |
| 6 | 53685273 | rs4486000 | 0.9 | 0.43 | 0.606 | -0.356 | 0.014 | 0.019 | 0.019 | 0 | 0.2 | 0.29 | 0.23 | 0.25 | 0.24 | 0.26 |
| 6 | 53689508 | rs6907027 | -0.685 | 0.411 | 0.6 | -0.418 | 0.001 | 0.006 | 0.006 | 0 | 0.4 | 0.34 | 0.5 | 0.66 | 0.37 | 0.45 |
| 6 | 53690779 | rs3003489 | -0.831 | 0.381 | 0.607 | -0.475 | 0.008 | -0.023 | -0.023 | 1.882 | 0.88 | 0.77 | 0.88 | 0.92 | 0.86 | 0.82 |
| 6 | 53693655 | rs4380749 | 1.931 | 0.378 | 0.61 | -0.487 | 0.011 | 0.061 | 0.061 | 0.002 | 0.28 | 0.15 | 0.15 | 0.01 | 0.24 | 0.11 |
| 6 | 53700093 | rs3001006 | 1.033 | 0.405 | 0.581 | -0.426 | 0.009 | 0.012 | 0.012 | 0 | 0.19 | 0.26 | 0.22 | 0.25 | 0.24 | 0.26 |
| 6 | 53702275 | rs10948782 | 0.685 | 0.403 | 0.586 | -0.433 | 0.014 | -0.028 | -0.028 | 0.145 | 0.14 | 0.2 | 0.14 | 0.06 | 0.18 | 0.16 |
| 6 | 53715917 | rs9382246 | 0.106 | 0.377 | 0.471 | -0.424 | 0.026 | -0.069 | -0.069 | 0 | 0.25 | 0.1 | 0.35 | 0.6 | 0.2 | 0.3 |
| 6 | 53724274 | rs9370236 | -1.201 | 0.414 | 0.452 | -0.479 | 0.002 | -0.018 | -0.018 | 0 | 0.59 | 0.63 | 0.72 | 0.95 | 0.61 | 0.73 |
| 6 | 53753255 | rs9370240 | -0.719 | 0.464 | 0.207 | -0.347 | 0.006 | 0.014 | 0.014 | 0 | 0.4 | 0.32 | 0.5 | 0.69 | 0.36 | 0.47 |
| 6 | 53780432 | rs1883632 | 0.32 | 0.481 | 0.117 | -0.277 | -0.047 | -0.007 | -0.007 | 0 | 0.3 | 0.18 | 0.34 | 0.31 | 0.41 | 0.43 |
| 6 | 53788361 | rs1543356 | -1.919 | 0.425 | 0.284 | -0.343 | 0.003 | -0.016 | -0.016 | 1.763 | 0.7 | 0.74 | 0.84 | 0.99 | 0.75 | 0.9 |
| 6 | 53789830 | rs9370243 | 0.728 | 0.444 | 0.15 | 0.174 | -0.021 | -0.023 | -0.023 | 0 | 0.06 | 0 | 0.22 | 0.27 | 0.08 | 0.08 |
| 6 | 53795480 | rs9474687 | 1.403 | 0.481 | 0.398 | 0.293 | -0.011 | -0.012 | -0.012 | 0 | 0.16 | 0.12 | 0.25 | 0.27 | 0.25 | 0.27 |
| 6 | 53796191 | rs9370244 | 0.007 | 0.409 | 0.273 | 0.211 | 0.014 | -0.043 | -0.043 | 0.109 | 0.26 | 0.16 | 0.34 | 0.53 | 0.22 | 0.25 |
| 6 | 53797560 | rs2745461 | -0.04 | 0.435 | 0.399 | 0.158 | 0.009 | 0.036 | 0.036 | 0 | 0.15 | 0.44 | 0.17 | 0.19 | 0.15 | 0.24 |
| 6 | 53799944 | rs9357777 | -1.062 | 0.472 | -0.115 | 0.106 | -0.001 | 0.015 | 0.015 | 0 | 0.49 | 0.33 | 0.54 | 0.56 | 0.5 | 0.54 |
| 6 | 53802888 | rs6909216 | -0.195 | 0.511 | -0.299 | -0.017 | -0.014 | 0.034 | 0.034 | 0 | 0.23 | 0.17 | 0.21 | 0.07 | 0.28 | 0.31 |
| 6 | 53818262 | rs1012624 | -0.408 | 0.426 | -0.263 | 0.213 | -0.013 | -0.019 | -0.019 | 0 | 0.32 | 0.17 | 0.27 | 0.33 | 0.34 | 0.42 |
| 6 | 53823067 | rs5006063 | 0.57 | 0.146 | -0.151 | 0.076 | 0.019 | 0.067 | 0.067 | 0 | 0.21 | 0.37 | 0.29 | 0.39 | 0.24 | 0.35 |
| 6 | 53823767 | rs761136 | -1.536 | 0.074 | -0.212 | -0.042 | -0.002 | -0.056 | -0.056 | 0.035 | 0.68 | 0.7 | 0.84 | 0.98 | 0.75 | 0.9 |
| 6 | 53826707 | rs6928440 | -0.464 | 0.034 | -0.194 | -0.033 | -0.008 | 0.021 | 0.021 | 0 | 0.46 | 0.3 | 0.54 | 0.6 | 0.5 | 0.54 |
| 6 | 53827200 | rs763814 | -1.538 | 0.04 | -0.196 | -0.035 | -0.002 | -0.055 | -0.055 | 0.119 | 0.68 | 0.71 | 0.84 | 0.98 | 0.74 | 0.89 |
| 6 | 53828983 | rs11751301 | 1.248 | 0.085 | -0.349 | 0.098 | -0.081 | -0.075 | -0.075 | 0 | 0.16 | 0.01 | 0.25 | 0.24 | 0.24 | 0.27 |
| 6 | 53829247 | rs9296727 | 0.608 | 0.185 | -0.262 | 0.007 | 0.02 | 0.071 | 0.071 | 0.016 | 0.21 | 0.4 | 0.29 | 0.39 | 0.24 | 0.35 |
| 6 | 53834379 | rs2397150 | -0.271 | 0.107 | -0.608 | 0.349 | 0.004 | -0.01 | -0.01 | 0 | 0.36 | 0.32 | 0.46 | 0.56 | 0.36 | 0.38 |
| 6 | 53836460 | rs12201094 | -0.137 | 0.443 | -0.517 | 0.127 | -0.025 | 0.008 | 0.008 | 0 | 0.12 | 0.07 | 0.09 | 0.08 | 0.14 | 0.19 |
| 6 | 53839730 | rs1325826 | 0.105 | 0.549 | -0.456 | 0.067 | -0.024 | 0.02 | 0.02 | 0 | 0.1 | 0.06 | 0.09 | 0.04 | 0.14 | 0.15 |
| 6 | 53847672 | rs9370251 | 0.342 | -0.085 | 0.257 | 0.401 | 0.012 | 0.021 | 0.021 | 0 | 0.67 | 0.56 | 0.56 | 0.56 | 0.63 | 0.44 |
| 6 | 53848551 | rs17554034 | 0.545 | -0.045 | 0.159 | 0.462 | -0.027 | 0.012 | 0.012 | 0 | 0.16 | 0.09 | 0.13 | 0.13 | 0.24 | 0.31 |
| 6 | 53851341 | rs1570916 | 0.834 | -0.16 | 0.631 | 0.411 | 0.003 | 0.006 | 0.006 | 0 | 0.66 | 0.42 | 0.55 | 0.58 | 0.63 | 0.44 |

| | | | | | | | | | | | | | | | | |
|---|---|---|---|---|---|---|---|---|---|---|---|---|---|---|---|---|
| 6 | 53851937 | rs17554130 | 0.676 | -0.091 | 0.08 | 0.431 | -0.033 | 0.011 | 0.011 | 0 | 0.16 | 0.08 | 0.11 | 0.11 | 0.23 | 0.31 |
| 6 | 53853589 | rs1041148 | 0.04 | 0.059 | 0.198 | 0.132 | -0.001 | 0.028 | 0.028 | 0 | 0.15 | 0.19 | 0.32 | 0.31 | 0.14 | 0.24 |
| 6 | 53857525 | rs12213836 | 0.886 | -0.575 | -0.395 | -0.536 | 0.04 | 0.2 | 0.2 | 0.011 | 0.51 | 0.17 | 0.36 | 0.2 | 0.58 | 0.34 |
| 6 | 53860598 | rs1409106 | 0.19 | -1.132 | -0.217 | -0.754 | 0.002 | 0.001 | 0.001 | 0 | 0.55 | 0.47 | 0.56 | 0.27 | 0.65 | 0.61 |
| 6 | 53863915 | rs927795 | 0.542 | -1.287 | -0.418 | -1.022 | 0.02 | 0.02 | 0.02 | 0.197 | 0.6 | 0.32 | 0.65 | 0.33 | 0.68 | 0.31 |
| 6 | 53864523 | rs9382269 | 0.598 | -1.499 | -0.41 | -1.032 | -0.001 | -0.007 | -0.007 | 0 | 0.28 | 0.3 | 0.26 | 0.21 | 0.3 | 0.18 |
| 6 | 53865667 | rs6930213 | 0.192 | -1.501 | -0.498 | -1.311 | -0.011 | 0.025 | 0.025 | 0 | 0.41 | 0.35 | 0.35 | 0.12 | 0.52 | 0.42 |
| 6 | 53866030 | rs1409105 | 1.079 | -1.356 | -0.853 | -1.347 | -0.121 | 0.077 | 0.077 | 0.322 | 0.34 | 0.07 | 0.42 | 0.23 | 0.4 | 0.21 |
| 6 | 53869221 | rs12206659 | 1.102 | -1.381 | -0.85 | -1.363 | -0.125 | 0.089 | 0.089 | 0.266 | 0.34 | 0.07 | 0.42 | 0.22 | 0.41 | 0.21 |
| 6 | 53877181 | rs9382274 | 0.955 | -1.562 | -0.662 | -1.312 | 0.07 | -0.065 | -0.065 | 1.611 | 0.19 | 0.05 | 0.2 | 0.66 | 0.07 | 0.22 |
| 6 | 53882340 | rs1409098 | 1.198 | -1.422 | -1.045 | -1.353 | -0.127 | 0.074 | 0.074 | 0.039 | 0.33 | 0.06 | 0.25 | 0.22 | 0.4 | 0.18 |
| 6 | 53882366 | rs2397153 | 0.198 | -1.467 | -0.718 | -1.44 | -0.021 | -0.024 | -0.024 | 0 | 0.61 | 0.47 | 0.67 | 0.33 | 0.69 | 0.47 |
| 6 | 53883843 | rs17625497 | 0.981 | -1.436 | -1.605 | -1.454 | -0.002 | -0.013 | -0.013 | 0 | 0.24 | 0.17 | 0.4 | 0.06 | 0.29 | 0.14 |
| 6 | 53884210 | rs10484652 | 1.065 | -1.532 | -0.627 | -0.449 | 0.047 | 0.069 | 0.069 | 2.075 | 0.57 | 0.27 | 0.65 | 0.27 | 0.69 | 0.31 |
| 6 | 53889581 | rs10080737 | 0.536 | -1.555 | -0.608 | -0.56 | -0.032 | 0.048 | 0.048 | 0 | 0.15 | 0.08 | 0.11 | 0.01 | 0.23 | 0.29 |
| 6 | 53889975 | rs9296733 | -0.06 | -1.462 | -0.56 | -0.507 | -0.01 | 0.033 | 0.033 | 0 | 0.2 | 0.16 | 0.12 | 0.06 | 0.23 | 0.46 |
| 6 | 53891008 | rs12194304 | 0.381 | -0.89 | -0.791 | -0.358 | 0.044 | 0.069 | 0.069 | 1.674 | 0.57 | 0.29 | 0.65 | 0.25 | 0.69 | 0.31 |
| 6 | 53893443 | rs2025660 | -0.201 | -0.517 | -1.014 | 0.243 | 0.032 | -0.038 | -0.038 | 0 | 0.36 | 0.15 | 0.32 | 0.54 | 0.28 | 0.41 |
| 6 | 53893446 | rs6937791 | 0.733 | -0.82 | -1.193 | -0.081 | 0.005 | 0.021 | 0.021 | 2.271 | 0.11 | 0.05 | 0.07 | 0.07 | 0.11 | 0.13 |
| 6 | 53902171 | rs16884633 | -0.117 | -1.192 | -1.086 | 0.623 | 0.031 | -0.014 | -0.014 | 1.53 | 0.3 | 0.26 | 0.22 | 0.37 | 0.18 | 0.27 |
| 6 | 53902717 | rs9382278 | -0.037 | -0.79 | -0.451 | 0.467 | 0.027 | -0.015 | -0.015 | 0 | 0.06 | 0.14 | 0.23 | 0.04 | 0.09 | 0.06 |
| 6 | 53909391 | rs13211796 | 0.928 | -0.788 | 0.145 | 0.367 | 0 | 0.01 | 0.01 | 0 | 0.07 | 0.08 | 0.04 | 0.1 | 0.05 | 0.09 |
| 6 | 53911656 | rs16884643 | 0.603 | -0.779 | 0.024 | 0.3 | -0.058 | -0.086 | -0.086 | 0 | 0.16 | 0.01 | 0.17 | 0.23 | 0.21 | 0.27 |
| 6 | 53913298 | rs9474715 | -0.235 | -1.179 | 1.269 | 0.26 | 0.014 | 0.059 | 0.059 | 0 | 0.1 | 0.38 | 0.06 | 0.09 | 0.04 | 0.13 |
| 6 | 53920024 | rs7739196 | 0.802 | -1.163 | 1.142 | 0.264 | -0.003 | 0.005 | 0.005 | 0 | 0.07 | 0.09 | 0.04 | 0.08 | 0.04 | 0.09 |
| 6 | 53921127 | rs9474720 | -0.633 | -1.289 | 1.042 | 0.242 | 0.003 | -0.005 | -0.005 | 0 | 0.34 | 0.64 | 0.43 | 0.43 | 0.21 | 0.33 |
| 6 | 53921981 | rs9367551 | 0.752 | -1.249 | 1.162 | 0.282 | -0.005 | 0.001 | 0.001 | 0 | 0.2 | 0.21 | 0.15 | 0.31 | 0.08 | 0.14 |
| 6 | 53926716 | rs9296737 | 0.991 | -1.247 | 0.901 | 0.273 | 0.052 | -0.075 | -0.075 | 0 | 0.18 | 0.05 | 0.14 | 0.31 | 0.08 | 0.14 |
| 6 | 53939724 | rs9349693 | 0.903 | -1.361 | 0.904 | 0.251 | 0.112 | -0.051 | -0.051 | 0.505 | 0.52 | 0.11 | 0.47 | 0.5 | 0.69 | 0.55 |
| 6 | 53945611 | rs16884668 | 0.914 | -1.394 | 1.125 | 0.219 | -0.005 | -0.017 | -0.017 | 1.686 | 0.12 | 0.16 | 0.08 | 0.06 | 0.1 | 0.13 |
| 6 | 53945699 | rs519844 | 0.009 | -1.452 | 1.457 | 0.208 | -0.053 | 0.034 | 0.034 | 0.039 | 0.7 | 0.46 | 0.78 | 0.6 | 0.88 | 0.74 |
| 6 | 53947087 | rs562559 | 0.25 | -1.35 | 1.811 | 0.137 | -0.012 | -0.085 | -0.085 | 0 | 0.8 | 0.42 | 0.68 | 0.9 | 0.81 | 0.8 |
| 6 | 53951333 | rs526144 | 1.065 | -1.284 | 1.939 | 0.234 | 0.055 | -0.016 | -0.016 | 0 | 0.91 | 0.58 | 0.76 | 0.96 | 0.91 | 0.94 |
| 6 | 53951857 | rs629761 | 0.502 | -1.33 | 2.034 | 0.282 | 0.028 | -0.02 | -0.02 | 0 | 0.64 | 0.26 | 0.57 | 0.57 | 0.8 | 0.69 |
| 6 | 53951990 | rs13199742 | 0.728 | -1.256 | 1.727 | 0.368 | -0.003 | 0.002 | 0.002 | 0 | 0.07 | 0.09 | 0.04 | 0.08 | 0.04 | 0.11 |
| 6 | 53955962 | rs598103 | 0.459 | -1.186 | 1.59 | 0.303 | 0.01 | -0.008 | -0.008 | 0 | 0.64 | 0.34 | 0.57 | 0.57 | 0.79 | 0.68 |
| 6 | 53956877 | rs557018 | 1.162 | -0.596 | 1.3 | 0.035 | 0.048 | -0.018 | -0.018 | 0 | 0.91 | 0.64 | 0.77 | 0.96 | 0.91 | 0.93 |
| 6 | 53960803 | rs1407239 | 0.454 | -0.588 | 0.411 | 0.226 | 0.072 | 0.082 | 0.082 | 0.833 | 0.49 | 0.2 | 0.47 | 0.68 | 0.64 | 0.63 |
| 6 | 53961513 | rs684940 | -0.297 | -0.531 | 1.426 | 0.244 | -0.016 | -0.022 | -0.022 | 0 | 0.39 | 0.48 | 0.47 | 0.26 | 0.34 | 0.26 |
| 6 | 53971401 | rs1028804 | 0.291 | -0.565 | 1.745 | 0.234 | 0.038 | 0.042 | 0.042 | 0 | 0.57 | 0.26 | 0.51 | 0.73 | 0.66 | 0.72 |
| 6 | 53975913 | rs499594 | -0.017 | -0.785 | 2.117 | 0.117 | 0.011 | -0.042 | -0.042 | 1.495 | 0.2 | 0.28 | 0.35 | 0.08 | 0.27 | 0.14 |
| 6 | 53978817 | rs486135 | -0.548 | -0.799 | 2.188 | 0.124 | -0.013 | -0.018 | -0.018 | 0 | 0.4 | 0.52 | 0.49 | 0.27 | 0.34 | 0.26 |
| 6 | 53985339 | rs1280946 | 0.512 | -0.843 | 2.254 | 0.103 | 0.035 | 0.039 | 0.039 | 0 | 0.56 | 0.29 | 0.5 | 0.73 | 0.65 | 0.72 |
| 6 | 53989526 | rs4712056 | 0.336 | -0.841 | 2.308 | 0.103 | -0.01 | -0.013 | -0.013 | 0.095 | 0.58 | 0.47 | 0.51 | 0.73 | 0.65 | 0.74 |
| 6 | 53995504 | rs816366 | 0.346 | -0.865 | 2.433 | 0.076 | 0.024 | -0.093 | -0.093 | 0 | 0.77 | 0.53 | 0.65 | 0.92 | 0.73 | 0.84 |
| 6 | 53996073 | rs816368 | -0.208 | -0.863 | 2.409 | 0.076 | 0.022 | -0.078 | -0.078 | 1.555 | 0.22 | 0.37 | 0.35 | 0.08 | 0.27 | 0.15 |
| 6 | 53996303 | rs816370 | -0.528 | -0.808 | 2.304 | 0.039 | 0.055 | -0.039 | -0.039 | 0 | 0.09 | 0.26 | 0.27 | 0.02 | 0.13 | 0.07 |
| 6 | 53999025 | rs816379 | -0.724 | -0.789 | 2.106 | 0.165 | 0.034 | 0.032 | 0.032 | 0 | 0.44 | 0.71 | 0.5 | 0.3 | 0.35 | 0.28 |
| 6 | 54002231 | rs2754779 | -0.733 | -0.734 | 2.088 | 0.146 | 0.012 | -0.073 | -0.073 | 0 | 0.24 | 0.58 | 0.35 | 0.08 | 0.27 | 0.16 |
| 6 | 54017126 | rs1407233 | -0.516 | 0.017 | 1.86 | -0.117 | 0.008 | -0.061 | -0.061 | 0 | 0.23 | 0.62 | 0.35 | 0.08 | 0.27 | 0.14 |
| 6 | 54019288 | rs12193888 | -0.516 | 0.017 | 1.86 | -0.117 | 0.008 | -0.061 | -0.061 | 4.26 | 0.23 | 0.62 | 0.35 | 0.08 | 0.27 | 0.14 |
| 6 | 54019320 | rs4380743 | -0.378 | -0.005 | 1.398 | 0.14 | -0.067 | -0.004 | -0.004 | 0.005 | 0.78 | 0.96 | 0.81 | 0.86 | 0.72 | 0.61 |
| 6 | 54025155 | rs10484648 | 1.353 | 0.038 | 1.562 | 0.298 | 0.082 | -0.052 | -0.052 | 0 | 0.19 | 0.06 | 0.14 | 0.21 | 0.07 | 0.12 |
| 6 | 54033762 | rs2145761 | 1.352 | 0.025 | 1.799 | 0.212 | 0.027 | -0.008 | -0.008 | 0 | 0.2 | 0.15 | 0.14 | 0.19 | 0.08 | 0.12 |
| 6 | 54039127 | rs816387 | 0.361 | 0.051 | 1.846 | 0.188 | 0.048 | 0.054 | 0.054 | 0 | 0.56 | 0.25 | 0.51 | 0.73 | 0.65 | 0.71 |
| 6 | 54046264 | rs2754798 | -0.608 | 0.523 | 2.418 | 0.719 | -0.079 | 0.003 | 0.003 | 0 | 0.77 | 0.61 | 0.84 | 0.81 | 0.92 | 0.88 |
| 6 | 54050219 | rs2792635 | -0.608 | 0.524 | 2.409 | 0.719 | -0.079 | 0.003 | 0.003 | 0 | 0.77 | 0.61 | 0.84 | 0.81 | 0.92 | 0.88 |
| 6 | 54061818 | rs1340665 | 0.86 | 0.552 | 2.344 | 0.733 | 0.045 | -0.017 | -0.017 | 0.451 | 0.2 | 0.1 | 0.14 | 0.18 | 0.08 | 0.12 |
| 6 | 54071032 | rs2792644 | -0.554 | 0.652 | 2.417 | 0.659 | 0.008 | -0.018 | -0.018 | 1.046 | 0.09 | 0.11 | 0.07 | 0.01 | 0.12 | 0.06 |
| 6 | 54071553 | rs9464034 | 0.986 | 0.593 | 2.271 | 0.693 | 0.076 | -0.073 | -0.073 | 0 | 0.19 | 0.01 | 0.14 | 0.19 | 0.08 | 0.12 |
| 6 | 54074482 | rs2754813 | -0.764 | 0.649 | 2.441 | 0.679 | 0.05 | -0.046 | -0.046 | 0.011 | 0.09 | 0.22 | 0.08 | 0.01 | 0.12 | 0.06 |
| 6 | 54076392 | rs11755131 | -0.124 | 0.614 | 2.198 | 0.648 | -0.073 | -0.018 | -0.018 | 0 | 0.21 | 0.04 | 0.35 | 0.15 | 0.3 | 0.38 |
| 6 | 54077679 | rs1417389 | 0.574 | 0.056 | 2.851 | 0.91 | 0.011 | 0.004 | 0.004 | 0.017 | 0.68 | 0.33 | 0.75 | 0.79 | 0.8 | 0.79 |
| 6 | 54082465 | rs16869492 | 0.405 | 0.058 | 2.697 | 0.927 | -0.058 | 0.007 | 0.007 | 0 | 0.22 | 0.31 | 0.16 | 0.18 | 0.08 | 0.12 |
| 6 | 54086108 | rs3957360 | 1.198 | 0.091 | 2.65 | 0.914 | 0.06 | -0.034 | -0.034 | 1.64 | 0.89 | 0.68 | 0.91 | 0.98 | 0.88 | 0.91 |
| 6 | 54094859 | rs816385 | 0.639 | 0.069 | 2.503 | 0.917 | 0.012 | 0.005 | 0.005 | 0.036 | 0.67 | 0.31 | 0.74 | 0.8 | 0.8 | 0.79 |
| 6 | 54095524 | rs2275769 | 0.577 | 0.063 | 2.437 | 0.921 | -0.046 | 0.007 | 0.007 | 0 | 0.21 | 0.23 | 0.15 | 0.18 | 0.08 | 0.12 |
| 6 | 54108523 | rs6458988 | -0.74 | 0.157 | 1.897 | 0.961 | 0.023 | 0.01 | 0.01 | 0.14 | 0.33 | 0.74 | 0.26 | 0.2 | 0.2 | 0.21 |
| 6 | 54111406 | rs9296740 | -0.772 | 0.172 | 1.903 | 0.992 | 0.025 | 0.009 | 0.009 | 0.056 | 0.34 | 0.74 | 0.26 | 0.22 | 0.2 | 0.2 |

| | | | | | | | | | | | | | | | | |
|---|---|---|---|---|---|---|---|---|---|---|---|---|---|---|---|---|
| 6 | 54125839 | rs9357795 | -1.585 | 1.632 | 2.671 | 1.687 | 0.015 | -0.026 | -0.026 | 0.036 | 0.1 | 0.19 | 0.08 | 0.02 | 0.11 | 0.09 |
| 6 | 54129982 | rs9296744 | -2.164 | 1.604 | 2.999 | 1.672 | 0.032 | -0.05 | -0.05 | 0 | 0.13 | 0.57 | 0.11 | 0.02 | 0.11 | 0.09 |
| 6 | 54135951 | rs9296747 | -2.099 | 1.831 | 2.877 | 2.052 | 0.046 | -0.035 | -0.035 | 0.153 | 0.13 | 0.5 | 0.1 | 0.03 | 0.11 | 0.09 |
| 6 | 54150287 | rs7774316 | -1.938 | 1.843 | 2.738 | 1.994 | 0.047 | -0.071 | -0.071 | 0.899 | 0.12 | 0.38 | 0.08 | 0.01 | 0.11 | 0.07 |
| 6 | 54164397 | rs7754752 | 0.938 | 2.287 | 1.877 | 2.19 | -0.063 | -0.042 | -0.042 | 0 | 0.85 | 0.98 | 0.68 | 0.82 | 0.81 | 0.68 |
| 6 | 54172465 | rs682088 | -0.993 | 2.246 | 1.847 | 2.306 | -0.068 | -0.013 | -0.013 | 0 | 0.15 | 0.01 | 0.32 | 0.13 | 0.19 | 0.33 |
| 6 | 54172879 | rs1340660 | -1.469 | 2.287 | 2.257 | 2.332 | 0.013 | -0.04 | -0.04 | 0 | 0.08 | 0.17 | 0.05 | 0.01 | 0.09 | 0.06 |
| 6 | 54174127 | rs1508638 | 0.924 | 2.044 | 2.183 | 2.388 | -0.011 | 0.008 | 0.008 | 0.001 | 0.84 | 0.9 | 0.67 | 0.87 | 0.81 | 0.68 |
| 6 | 54181882 | rs1995768 | 1.575 | 2.281 | 2.173 | 2.732 | 0.04 | 0.16 | 0.16 | 0.023 | 0.81 | 0.62 | 0.63 | 0.47 | 0.78 | 0.61 |
| 6 | 54185864 | rs1508642 | 1.516 | 2.059 | 1.253 | 1.998 | -0.023 | 0.001 | 0.001 | 0 | 0.77 | 0.81 | 0.61 | 0.79 | 0.7 | 0.61 |
| 6 | 54186147 | rs1058768 | 1.399 | 2.172 | 1.266 | 1.745 | -0.023 | 0.018 | 0.018 | 0 | 0.77 | 0.82 | 0.62 | 0.85 | 0.7 | 0.63 |
| 6 | 54210141 | rs9382326 | -1.784 | 2.204 | 2.037 | 1.69 | 0.052 | 0.02 | 0.02 | 0 | 0.09 | 0.24 | 0.14 | 0.09 | 0.19 | 0.17 |
| 6 | 54215710 | rs1508643 | 1.404 | 2.565 | 2.391 | 2.004 | 0.073 | 0.12 | 0.12 | 0.541 | 0.8 | 0.53 | 0.5 | 0.65 | 0.77 | 0.69 |
| 6 | 54220243 | rs17828521 | -1.178 | 2.586 | 2.309 | 2.32 | -0.006 | 0.142 | 0.142 | 0 | 0.05 | 0.09 | 0.03 | 0.31 | 0.04 | 0.05 |
| 6 | 54227991 | rs10948814 | -1.56 | 2.583 | 2.322 | 2.258 | 0.051 | 0.021 | 0.021 | 0 | 0.07 | 0.15 | 0.13 | 0.09 | 0.19 | 0.17 |
| 6 | 54234404 | rs9395949 | -1.897 | 2.623 | 2.317 | 2.274 | 0.063 | 0.03 | 0.03 | 0 | 0.08 | 0.22 | 0.13 | 0.09 | 0.19 | 0.17 |
| 6 | 54239250 | rs7758586 | -1.802 | 2.541 | 2.293 | 2.276 | -0.004 | 0.24 | 0.24 | 0 | 0.07 | 0.19 | 0.05 | 0.37 | 0.05 | 0.07 |
| 6 | 54254580 | rs3736353 | -1.439 | 2.439 | 2.253 | 2.258 | -0.006 | 0.107 | 0.107 | 0 | 0.06 | 0.1 | 0.03 | 0.31 | 0.05 | 0.05 |
| 6 | 54256028 | rs16885285 | -1.403 | 2.463 | 2.336 | 2.266 | 0.032 | -0.01 | -0.01 | 0 | 0.07 | 0.15 | 0.11 | 0.03 | 0.18 | 0.15 |
| 6 | 54256680 | rs9464069 | -2.483 | 2.453 | 2.269 | 2.256 | 0.056 | 0.125 | 0.125 | 0 | 0.13 | 0.35 | 0.15 | 0.35 | 0.23 | 0.2 |
| 6 | 54279637 | rs12192520 | -1.559 | 2.237 | 1.972 | 1.22 | 0.023 | -0.016 | -0.016 | 0.088 | 0.08 | 0.09 | 0.1 | 0.04 | 0.16 | 0.14 |
| 6 | 54290333 | rs9357804 | 1.273 | 1.927 | 1.396 | 1.553 | 0.036 | 0.108 | 0.108 | 0 | 0.85 | 0.75 | 0.56 | 0.47 | 0.79 | 0.74 |
| 6 | 54291407 | rs794268 | 0.884 | 1.993 | 0.647 | 1.587 | -0.08 | -0.097 | -0.097 | 0 | 0.87 | 0.98 | 0.59 | 0.47 | 0.8 | 0.78 |
| 6 | 54297305 | rs243815 | 0.142 | 1.464 | 0.551 | 0.844 | -0.027 | 0.06 | 0.06 | 0 | 0.88 | 0.98 | 0.95 | 1 | 0.88 | 0.94 |
| 6 | 54297588 | rs515020 | -2.1 | 1.199 | 1.665 | 1.133 | 0.058 | 0.061 | 0.061 | 4.484 | 0.14 | 0.53 | 0.16 | 0.17 | 0.12 | 0.1 |
| 6 | 54309630 | rs7757433 | -1.684 | 1.28 | 2.068 | 1.169 | 0.051 | 0.12 | 0.12 | 2.038 | 0.07 | 0.27 | 0.12 | 0.15 | 0.06 | 0.07 |
| 6 | 54310853 | rs243799 | -1.288 | 0.442 | 1.413 | 0.57 | 0.011 | 0.178 | 0.178 | 4.557 | 0.5 | 0.86 | 0.68 | 0.76 | 0.46 | 0.54 |
| 6 | 54318908 | rs243781 | -0.66 | 0.532 | 1.52 | 0.54 | 0.059 | -0.005 | -0.005 | 0.427 | 0.05 | 0.24 | 0.04 | 0.01 | 0.06 | 0.02 |
| 6 | 54350176 | rs567638 | -0.853 | 0.525 | 1.675 | 0.593 | 0.054 | 0.08 | 0.08 | 0.044 | 0.05 | 0.17 | 0.05 | 0.11 | 0.06 | 0.06 |
| 6 | 54360695 | rs1820408 | -0.852 | 0.518 | 1.665 | 0.59 | 0.051 | 0.077 | 0.077 | 0.044 | 0.05 | 0.18 | 0.06 | 0.11 | 0.06 | 0.06 |
| 6 | 54361735 | rs9395969 | -0.905 | 0.44 | 1.866 | 0.689 | -0.002 | -0.002 | -0.002 | 0 | 0.44 | 0.45 | 0.55 | 0.65 | 0.4 | 0.49 |
| 6 | 54376524 | rs181710 | -1.355 | 0.447 | 1.883 | 0.689 | 0.018 | 0.048 | 0.048 | 2.726 | 0.49 | 0.71 | 0.6 | 0.66 | 0.45 | 0.51 |
| 6 | 54378323 | rs243765 | -1.347 | 0.444 | 1.809 | 0.689 | 0.008 | 0.145 | 0.145 | 4.234 | 0.51 | 0.86 | 0.68 | 0.76 | 0.46 | 0.54 |
| 6 | 54397512 | rs243790 | -0.576 | 0.505 | 1.764 | 0.687 | 0.062 | 0.004 | 0.004 | 0.369 | 0.05 | 0.26 | 0.04 | 0.01 | 0.05 | 0.01 |
| 6 | 54403797 | rs712536 | -0.535 | 0.439 | 1.366 | 0.745 | 0.031 | -0.041 | -0.041 | 0 | 0.38 | 0.22 | 0.5 | 0.54 | 0.34 | 0.43 |
| 6 | 54403821 | rs712537 | 0.028 | 0.189 | 1.203 | 0.63 | -0.021 | -0.161 | -0.161 | 0.999 | 0.28 | 0.05 | 0.44 | 0.52 | 0.26 | 0.23 |
| 6 | 54404993 | rs569968 | -0.819 | 0.19 | 1.216 | 0.643 | 0.009 | -0.005 | -0.005 | 0.581 | 0.38 | 0.31 | 0.52 | 0.59 | 0.34 | 0.45 |
| 6 | 54411913 | rs511631 | 0.553 | 0.216 | 0.699 | 0.621 | 0.03 | -0.088 | -0.088 | 0.632 | 0.63 | 0.83 | 0.5 | 0.46 | 0.66 | 0.57 |
| 6 | 54419747 | rs9367581 | -0.368 | 0.303 | 0.66 | 0.688 | -0.05 | -0.15 | -0.15 | 2.831 | 0.74 | 0.95 | 0.56 | 0.5 | 0.69 | 0.79 |
| 6 | 54424937 | rs6459009 | -0.023 | 0.238 | 0.77 | 0.612 | -0.025 | -0.094 | -0.094 | 0 | 0.69 | 0.89 | 0.48 | 0.47 | 0.62 | 0.66 |
| 6 | 54428460 | rs9464097 | 1.506 | 0.089 | 1.186 | 0.554 | 0.007 | 0.122 | 0.122 | 3.446 | 0.48 | 0.21 | 0.31 | 0.23 | 0.51 | 0.47 |
| 6 | 54433329 | rs9395980 | -1.095 | 0.125 | 1.609 | 0.605 | 0.022 | 0.027 | 0.027 | 1.077 | 0.12 | 0.3 | 0.07 | 0.14 | 0.1 | 0.09 |
| 6 | 54438426 | rs6459012 | -2.066 | 0.133 | 1.508 | 0.605 | -0.026 | 0.022 | 0.022 | 3.613 | 0.2 | 0.64 | 0.17 | 0.24 | 0.11 | 0.19 |
| 6 | 54441359 | rs1853335 | -0.011 | 0.298 | 1.202 | 0.58 | -0.006 | -0.023 | -0.023 | 0.072 | 0.69 | 0.81 | 0.46 | 0.41 | 0.61 | 0.64 |
| 6 | 54442960 | rs9357815 | -1.303 | 0.266 | 1.335 | 0.58 | 0.016 | 0.028 | 0.028 | 1.04 | 0.12 | 0.3 | 0.07 | 0.14 | 0.09 | 0.09 |
| 6 | 54451626 | rs10484980 | -0.772 | 0.091 | 1.426 | 0.436 | -0.03 | -0.031 | -0.031 | 0.154 | 0.07 | 0.18 | 0.02 | 0 | 0.01 | 0.07 |
| 6 | 54452797 | rs10948828 | -2.311 | -0.117 | 2.073 | 0.492 | -0.047 | -0.009 | -0.009 | 0 | 0.16 | 0.61 | 0.15 | 0.14 | 0.1 | 0.25 |
| 6 | 54458991 | rs4377789 | -0.495 | -0.171 | 2.16 | 0.402 | 0.001 | -0.011 | -0.011 | 0.367 | 0.06 | 0.07 | 0.08 | 0.03 | 0.09 | 0.15 |
| 6 | 54460573 | rs12201939 | -0.887 | -0.143 | 2.218 | 0.308 | -0.001 | -0.034 | -0.034 | 0.053 | 0.08 | 0.17 | 0.03 | 0.01 | 0.06 | 0.05 |
| 6 | 54460646 | rs10484984 | 1.103 | -0.138 | 1.94 | 0.342 | 0.04 | 0.087 | 0.087 | 1.078 | 0.54 | 0.31 | 0.35 | 0.22 | 0.58 | 0.48 |
| 6 | 54467633 | rs9474919 | -2.436 | -0.144 | 2.238 | 0.389 | -0.052 | -0.087 | -0.087 | 0.063 | 0.18 | 0.53 | 0.14 | 0.07 | 0.11 | 0.28 |
| 6 | 54472452 | rs7771536 | -1.533 | -0.114 | 2.295 | 0.408 | -0.013 | -0.05 | -0.05 | 0 | 0.11 | 0.34 | 0.11 | 0.03 | 0.08 | 0.19 |
| 6 | 54479334 | rs9370322 | -1.45 | -0.083 | 2.273 | 0.613 | -0.002 | 0.015 | 0.015 | 0 | 0.11 | 0.17 | 0.17 | 0.16 | 0.11 | 0.21 |
| 6 | 54482002 | rs12203655 | -2.774 | -0.066 | 2.411 | 0.611 | -0.022 | 0.06 | 0.06 | 0.85 | 0.18 | 0.54 | 0.22 | 0.23 | 0.14 | 0.24 |
| 6 | 54483722 | rs9474935 | -1.344 | -0.046 | 2.248 | 0.521 | 0.001 | -0.003 | -0.003 | 0 | 0.11 | 0.12 | 0.1 | 0.03 | 0.11 | 0.19 |
| 6 | 54485245 | rs7746006 | 1.074 | -0.253 | 2.105 | 0.382 | 0.003 | 0.005 | 0.005 | 0.874 | 0.58 | 0.42 | 0.37 | 0.28 | 0.61 | 0.58 |
| 6 | 54488143 | rs1830662 | -1.133 | -0.233 | 2.093 | 0.428 | 0.003 | 0.023 | 0.023 | 0 | 0.05 | 0.11 | 0.09 | 0.11 | 0.06 | 0.03 |
| 6 | 54489220 | rs12213260 | -1.221 | -0.221 | 2.19 | 0.343 | -0.005 | -0.015 | -0.015 | 1.162 | 0.11 | 0.24 | 0.06 | 0.07 | 0.06 | 0.14 |
| 6 | 54491621 | rs6901995 | -2.226 | -0.221 | 2.184 | 0.343 | 0 | 0.018 | 0.018 | 0.576 | 0.16 | 0.34 | 0.15 | 0.19 | 0.11 | 0.16 |
| 6 | 54495166 | rs6936272 | -1.106 | -0.219 | 2.306 | 0.346 | 0.004 | -0.036 | -0.036 | 0 | 0.1 | 0.3 | 0.09 | 0.03 | 0.08 | 0.19 |
| 6 | 54505740 | rs511441 | 0.601 | -0.21 | 2.263 | 0.354 | -0.018 | 0.013 | 0.013 | 0.461 | 0.64 | 0.52 | 0.47 | 0.39 | 0.67 | 0.61 |
| 6 | 54509300 | rs7753256 | -1.551 | -0.209 | 2.225 | 0.332 | 0.033 | 0.044 | 0.044 | 1.788 | 0.46 | 0.72 | 0.59 | 0.68 | 0.39 | 0.52 |
| 6 | 54531949 | rs953896 | -0.562 | -0.291 | 2.139 | 0.25 | -0.008 | 0.02 | 0.02 | 0 | 0.32 | 0.36 | 0.51 | 0.62 | 0.33 | 0.38 |
| 6 | 54540252 | rs13206543 | 1.628 | -0.283 | 2.125 | 0.248 | 0.027 | 0.147 | 0.147 | 1.46 | 0.52 | 0.21 | 0.33 | 0.21 | 0.58 | 0.44 |
| 6 | 54542866 | rs9474952 | 1.819 | -0.261 | 2.073 | 0.281 | 0.003 | 0.03 | 0.03 | 1.268 | 0.87 | 0.68 | 0.86 | 0.82 | 0.91 | 0.83 |
| 6 | 54545032 | rs12528703 | 0.042 | -0.227 | 1.577 | 0.357 | 0.005 | -0.059 | -0.059 | 0 | 0.25 | 0.15 | 0.45 | 0.57 | 0.25 | 0.26 |
| 6 | 54554502 | rs4715468 | 0.087 | -0.228 | 1.633 | 0.356 | 0.004 | -0.033 | -0.033 | 1.075 | 0.24 | 0.15 | 0.43 | 0.57 | 0.25 | 0.24 |
| 6 | 54557514 | rs1395632 | -1.35 | -0.247 | 1.61 | 0.31 | 0.015 | 0.021 | 0.021 | 1.036 | 0.43 | 0.64 | 0.57 | 0.68 | 0.38 | 0.5 |
| 6 | 54565844 | rs12208090 | -1.3 | -0.259 | 1.674 | 0.348 | 0.006 | 0 | 0 | 0.184 | 0.43 | 0.61 | 0.56 | 0.61 | 0.38 | 0.5 |

| | | | | | | | | | | | | | | | | |
|---|---|---|---|---|---|---|---|---|---|---|---|---|---|---|---|---|
| 6 | 54575349 | rs925057 | 1.808 | -0.305 | 1.624 | 0.28 | 0.011 | 0.161 | 0.161 | 3.505 | 0.48 | 0.16 | 0.31 | 0.21 | 0.55 | 0.44 |
| 6 | 54576743 | rs6927382 | -1.78 | -0.301 | 1.771 | 0.269 | -0.016 | -0.068 | -0.068 | 0 | 0.16 | 0.45 | 0.09 | 0.07 | 0.09 | 0.13 |
| 6 | 54577218 | rs2048113 | -0.414 | -0.279 | 1.74 | 0.313 | 0.003 | -0.048 | -0.048 | 0.5 | 0.31 | 0.18 | 0.49 | 0.54 | 0.32 | 0.37 |
| 6 | 54580930 | rs7750348 | -1.254 | -0.197 | 1.749 | 0.085 | -0.016 | -0.06 | -0.06 | 0 | 0.14 | 0.27 | 0.09 | 0.03 | 0.12 | 0.11 |
| 6 | 54583380 | rs12525001 | 0.308 | -0.215 | 1.564 | 0.098 | 0 | -0.074 | -0.074 | 0.018 | 0.23 | 0.1 | 0.42 | 0.5 | 0.25 | 0.23 |
| 6 | 54585379 | rs12198783 | -1.893 | -0.232 | 1.561 | 0.017 | -0.011 | -0.09 | -0.09 | 0 | 0.19 | 0.44 | 0.13 | 0.07 | 0.16 | 0.16 |
| 6 | 54597298 | rs1039693 | 1.259 | -0.256 | 1.417 | 0.017 | 0.018 | 0.091 | 0.091 | 2.329 | 0.53 | 0.3 | 0.35 | 0.25 | 0.55 | 0.54 |
| 6 | 54604517 | rs10948847 | -0.245 | -0.266 | 1.399 | 0.056 | -0.004 | -0.035 | -0.035 | 0 | 0.74 | 0.87 | 0.51 | 0.39 | 0.72 | 0.71 |
| 6 | 54604829 | rs12661998 | 0.683 | -0.211 | 1.237 | 0 | 0.021 | -0.055 | -0.055 | 0 | 0.18 | 0.08 | 0.37 | 0.5 | 0.15 | 0.19 |
| 6 | 54606385 | rs7755817 | 0.079 | -0.212 | 1.236 | -0.009 | -0.004 | -0.035 | -0.035 | 0.069 | 0.74 | 0.87 | 0.51 | 0.39 | 0.72 | 0.71 |
| 6 | 54615504 | rs13220672 | -1.891 | -0.196 | 1.41 | -0.028 | -0.016 | -0.095 | -0.095 | 0 | 0.2 | 0.45 | 0.14 | 0.07 | 0.16 | 0.17 |
| 6 | 54616000 | rs1354830 | 2.11 | -0.227 | 1.394 | -0.022 | 0.015 | -0.009 | -0.009 | 0 | 0.72 | 0.49 | 0.72 | 0.75 | 0.71 | 0.73 |
| 6 | 54619663 | rs3846913 | 0.003 | -0.234 | 1.268 | -0.004 | -0.001 | 0.008 | 0.008 | 3.219 | 0.23 | 0.24 | 0.42 | 0.55 | 0.19 | 0.25 |
| 6 | 54621115 | rs9474991 | 2.108 | -0.148 | 0.846 | -0.074 | -0.017 | 0.109 | 0.109 | 1.591 | 0.45 | 0.22 | 0.28 | 0.21 | 0.49 | 0.42 |
| 6 | 54622285 | rs4715476 | 1.208 | 0.307 | 0.706 | 0.085 | -0.007 | -0.003 | -0.003 | 0 | 0.85 | 0.88 | 0.82 | 0.84 | 0.8 | 0.78 |
| 6 | 54623836 | rs17448241 | -0.448 | 0.132 | 0.772 | -0.106 | 0 | 0 | 0 | 0 | 0.07 | 0.06 | 0.06 | 0.04 | 0.07 | 0.12 |
| 6 | 54626318 | rs17545576 | -1.263 | -0.005 | 0.689 | -0.227 | -0.001 | 0.008 | 0.008 | 0 | 0.1 | 0.07 | 0.06 | 0.05 | 0.09 | 0.08 |
| 6 | 54629892 | rs2174145 | 1.944 | -0.436 | 0.857 | -0.638 | -0.059 | 0.16 | 0.16 | 1.578 | 0.43 | 0.13 | 0.26 | 0.14 | 0.47 | 0.39 |
| 6 | 54631181 | rs2397167 | -0.325 | -0.563 | 0.773 | -0.628 | -0.051 | -0.06 | -0.06 | 0 | 0.75 | 0.93 | 0.51 | 0.3 | 0.69 | 0.7 |
| 6 | 54632120 | rs2063116 | -1.058 | -0.689 | 1.49 | -0.702 | -0.009 | -0.037 | -0.037 | 0 | 0.13 | 0.18 | 0.09 | 0.04 | 0.1 | 0.22 |
| 6 | 54632536 | rs2135760 | 0.989 | -0.704 | 1.782 | -0.744 | -0.006 | -0.021 | -0.021 | 0.299 | 0.57 | 0.44 | 0.39 | 0.26 | 0.59 | 0.48 |
| 6 | 54633035 | rs16869530 | -1.385 | -0.646 | 1.69 | -0.614 | -0.005 | -0.014 | -0.014 | 3.684 | 0.14 | 0.15 | 0.08 | 0.07 | 0.09 | 0.12 |
| 6 | 54633907 | rs973020 | -0.306 | -0.197 | 1.393 | 0.085 | -0.062 | -0.093 | -0.093 | 0 | 0.74 | 0.89 | 0.45 | 0.35 | 0.64 | 0.68 |
| 6 | 54636482 | rs9474993 | -1.905 | -0.274 | 1.806 | 0.022 | -0.018 | -0.079 | -0.079 | 0 | 0.14 | 0.3 | 0.07 | 0 | 0.06 | 0.1 |
| 6 | 54648353 | rs6916633 | -0.319 | -0.32 | 1.531 | 0.034 | -0.062 | -0.092 | -0.092 | 0.207 | 0.74 | 0.93 | 0.45 | 0.35 | 0.64 | 0.65 |
| 6 | 54656673 | rs10948852 | 1.089 | -0.447 | 2.067 | -0.087 | 0 | 0.018 | 0.018 | 0 | 0.55 | 0.33 | 0.29 | 0.19 | 0.48 | 0.48 |
| 6 | 54669411 | rs12661022 | -0.434 | -0.438 | 1.974 | -0.056 | -0.007 | -0.019 | -0.019 | 0 | 0.07 | 0.03 | 0.06 | 0.11 | 0.1 | 0.05 |
| 6 | 54671090 | rs1503151 | 0.572 | -0.471 | 2.207 | -0.09 | 0.005 | -0.009 | -0.009 | 0 | 0.63 | 0.55 | 0.35 | 0.3 | 0.55 | 0.45 |
| 6 | 54690036 | rs9464141 | -1.966 | -0.256 | 2.1 | 0.628 | 0.006 | 0.084 | 0.084 | 0 | 0.13 | 0.37 | 0.11 | 0.26 | 0.14 | 0.14 |
| 6 | 54690468 | rs6934538 | 0.604 | -0.427 | 1.731 | 0.608 | -0.007 | -0.035 | -0.035 | 0.989 | 0.6 | 0.75 | 0.39 | 0.47 | 0.56 | 0.54 |
| 6 | 54695124 | rs11964644 | -1.424 | -0.358 | 1.658 | 0.715 | -0.018 | 0.083 | 0.083 | 0 | 0.13 | 0.32 | 0.07 | 0.26 | 0.09 | 0.13 |
| 6 | 54701685 | rs6926482 | 0.078 | -0.132 | 1.078 | 0.388 | -0.052 | -0.019 | -0.019 | 0.454 | 0.7 | 0.9 | 0.44 | 0.26 | 0.61 | 0.56 |
| 6 | 54704616 | rs1546321 | -1.795 | -0.266 | 1.595 | 0.279 | -0.018 | -0.054 | -0.054 | 0.003 | 0.21 | 0.4 | 0.15 | 0.09 | 0.18 | 0.16 |
| 6 | 54705511 | rs2270989 | 1.543 | 0.761 | 1.205 | -0.071 | -0.007 | -0.011 | -0.011 | 0 | 0.44 | 0.49 | 0.22 | 0.17 | 0.27 | 0.39 |
| 6 | 54706689 | rs6938005 | -2.062 | 0.072 | 1.371 | 0.085 | -0.012 | -0.058 | -0.058 | 0 | 0.24 | 0.4 | 0.17 | 0.09 | 0.21 | 0.26 |
| 6 | 54709692 | rs4712075 | 1.349 | 0.13 | 1.382 | 0.112 | 0.001 | 0.014 | 0.014 | 3.201 | 0.5 | 0.5 | 0.29 | 0.17 | 0.44 | 0.4 |
| 6 | 54720443 | rs12191386 | 1.831 | 0.078 | 0.897 | 0.092 | -0.063 | 0.14 | 0.14 | 4.239 | 0.44 | 0.11 | 0.25 | 0.17 | 0.43 | 0.38 |
| 6 | 54721617 | rs6911198 | 0.959 | 0.106 | 1.095 | 0.2 | 0 | -0.003 | -0.003 | 0 | 0.54 | 0.41 | 0.32 | 0.21 | 0.5 | 0.48 |
| 6 | 54722013 | rs9382387 | -1.323 | 0.174 | 1.219 | 0.012 | -0.017 | -0.078 | -0.078 | 0 | 0.17 | 0.52 | 0.19 | 0.05 | 0.15 | 0.17 |
| 6 | 54724405 | rs1503139 | -0.171 | 0.03 | 0.112 | 0.534 | -0.086 | -0.168 | -0.168 | 0 | 0.71 | 0.95 | 0.53 | 0.46 | 0.66 | 0.6 |
| 6 | 54726327 | rs12192659 | 1.763 | -0.261 | -0.106 | -0.554 | -0.169 | 0.137 | 0.137 | 4.154 | 0.39 | 0.04 | 0.25 | 0.1 | 0.45 | 0.33 |
| 6 | 54727075 | rs9370332 | 0.406 | -0.465 | -0.169 | -0.678 | -0.049 | -0.037 | -0.037 | 0 | 0.22 | 0.07 | 0.51 | 0.81 | 0.32 | 0.4 |
| 6 | 54727815 | rs16886072 | 0.359 | -0.512 | 0.339 | -0.768 | -0.003 | -0.02 | -0.02 | 0.348 | 0.18 | 0.22 | 0.07 | 0.01 | 0.06 | 0.17 |
| 6 | 54729949 | rs973205 | 1.967 | -0.508 | 0.841 | -0.78 | -0.017 | 0.084 | 0.084 | 3.077 | 0.42 | 0.25 | 0.31 | 0.15 | 0.51 | 0.37 |
| 6 | 54730344 | rs1155750 | 0.399 | -0.487 | 1.307 | -0.774 | -0.048 | -0.16 | -0.16 | 0 | 0.76 | 0.52 | 0.87 | 0.99 | 0.89 | 0.8 |
| 6 | 54730383 | rs1155749 | -0.605 | -0.486 | 1.282 | -0.769 | 0.004 | 0.02 | 0.02 | 0 | 0.33 | 0.26 | 0.56 | 0.84 | 0.37 | 0.43 |
| 6 | 54734123 | rs10948866 | 1.575 | -0.157 | 1.092 | -0.566 | -0.003 | 0.02 | 0.02 | 2.289 | 0.43 | 0.34 | 0.32 | 0.15 | 0.51 | 0.37 |
| 6 | 54735122 | rs1472679 | 1.561 | -0.145 | 1.098 | -0.559 | -0.003 | 0.021 | 0.021 | 2.444 | 0.43 | 0.34 | 0.32 | 0.15 | 0.51 | 0.37 |
| 6 | 54737431 | rs2221335 | 0.229 | -0.08 | 0.605 | -0.542 | 0.045 | -0.024 | -0.024 | 0 | 0.95 | 0.89 | 0.95 | 1 | 0.94 | 0.97 |
| 6 | 54744946 | rs12208341 | 1.726 | -0.119 | 0.143 | -0.553 | -0.092 | 0.169 | 0.169 | 4.33 | 0.41 | 0.09 | 0.28 | 0.12 | 0.51 | 0.34 |
| 6 | 54748051 | rs9357820 | 0.213 | -0.142 | 0.437 | -0.451 | 0.002 | -0.015 | -0.015 | 0 | 0.95 | 0.95 | 0.96 | 1 | 0.95 | 0.97 |
| 6 | 54753626 | rs7453866 | -0.778 | -0.249 | 0.828 | -0.577 | 0.016 | 0.034 | 0.034 | 0.847 | 0.44 | 0.43 | 0.6 | 0.81 | 0.42 | 0.58 |
| 6 | 54759226 | rs148109 | -1.512 | -0.229 | 0.976 | -0.602 | -0.038 | 0.154 | 0.154 | 1.197 | 0.56 | 0.89 | 0.7 | 0.84 | 0.49 | 0.65 |
| 6 | 54759929 | rs4141552 | -0.014 | -0.46 | 1.38 | -0.711 | -0.002 | 0.003 | 0.003 | 0 | 0.26 | 0.26 | 0.54 | 0.81 | 0.34 | 0.43 |
| 6 | 54764307 | rs239846 | -0.597 | -0.482 | 1.658 | -0.765 | -0.024 | -0.091 | -0.091 | 0 | 0.27 | 0.37 | 0.14 | 0.03 | 0.15 | 0.22 |
| 6 | 54764320 | rs4715491 | 0.917 | -0.399 | 1.502 | -0.198 | -0.046 | -0.064 | -0.064 | 0 | 0.17 | 0.02 | 0.21 | 0.26 | 0.23 | 0.27 |
| 6 | 54766731 | rs239839 | -0.547 | -0.384 | 1.549 | -1.056 | -0.026 | -0.095 | -0.095 | 0.06 | 0.27 | 0.37 | 0.14 | 0.03 | 0.15 | 0.22 |
| 6 | 54771019 | rs4712082 | -1.004 | -0.385 | 1.7 | -1.179 | -0.001 | -0.013 | -0.013 | 0.185 | 0.32 | 0.33 | 0.6 | 0.86 | 0.42 | 0.45 |
| 6 | 54771938 | rs239828 | -0.927 | -0.459 | 1.585 | -1.11 | 0.006 | 0.005 | 0.005 | 0.095 | 0.31 | 0.3 | 0.17 | 0.03 | 0.24 | 0.26 |
| 6 | 54773639 | rs7743188 | 0.157 | -0.522 | 1.595 | -1.071 | -0.002 | -0.02 | -0.02 | 0 | 0.22 | 0.27 | 0.09 | 0.03 | 0.09 | 0.18 |
| 6 | 54784652 | rs9367597 | 0.222 | -0.444 | 1.227 | -0.991 | -0.031 | -0.022 | -0.022 | 0 | 0.22 | 0.12 | 0.52 | 0.78 | 0.33 | 0.38 |
| 6 | 54802509 | rs239795 | 0.154 | -0.452 | 1.345 | -0.995 | 0.028 | -0.03 | -0.03 | 0.322 | 0.06 | 0.1 | 0.05 | 0 | 0.06 | 0.04 |
| 6 | 54805688 | rs239798 | -0.919 | -0.557 | 0.986 | -1.104 | 0.017 | -0.001 | -0.001 | 0.022 | 0.31 | 0.71 | 0.17 | 0.03 | 0.15 | 0.22 |
| 6 | 54806489 | rs9475077 | -0.919 | -0.557 | 0.986 | -1.104 | 0.017 | -0.001 | -0.001 | 4.646 | 0.31 | 0.71 | 0.17 | 0.03 | 0.15 | 0.22 |
| 6 | 54811210 | rs6902422 | -0.748 | -0.28 | 0.955 | -0.836 | -0.003 | 0.039 | 0.039 | 0 | 0.29 | 0.19 | 0.55 | 0.84 | 0.36 | 0.44 |
| 6 | 54823431 | rs960834 | -0.324 | -0.291 | 0.947 | -0.841 | -0.016 | 0.03 | 0.03 | 0 | 0.26 | 0.15 | 0.55 | 0.83 | 0.35 | 0.39 |
| 6 | 54841672 | rs1503135 | -0.967 | -0.265 | 1.016 | -0.853 | -0.007 | -0.034 | -0.034 | 0 | 0.32 | 0.42 | 0.58 | 0.84 | 0.36 | 0.44 |
| 6 | 54861647 | rs3125272 | -1.039 | -0.292 | 0.734 | -0.773 | 0 | 0 | 0 | 0.714 | 0.11 | 0.1 | 0.08 | 0.06 | 0.09 | 0.1 |
| 6 | 54863080 | rs7753268 | 0.168 | -0.305 | 0.396 | -1.331 | 0.032 | 0.078 | 0.078 | 0 | 0.27 | 0.17 | 0.13 | 0.05 | 0.19 | 0.27 |

| | | | | | | | | | | | | | | | | |
|---|---|---|---|---|---|---|---|---|---|---|---|---|---|---|---|---|
| 6 | 54865634 | rs2047809 | -1.209 | -0.027 | 0.637 | -1.261 | 0 | -0.026 | -0.026 | 2.932 | 0.57 | 0.52 | 0.71 | 0.81 | 0.58 | 0.66 |
| 6 | 54874576 | rs7766969 | 1.851 | -0.092 | 0.586 | -1.254 | -0.005 | 0.005 | 0.005 | 0 | 0.36 | 0.29 | 0.26 | 0.13 | 0.39 | 0.28 |
| 6 | 54885290 | rs9396029 | -0.21 | -0.114 | 0.493 | -1.145 | -0.037 | 0.007 | 0.007 | 1.963 | 0.08 | 0.17 | 0.03 | 0.06 | 0.03 | 0.06 |
| 6 | 54947384 | rs7760858 | -0.933 | -0.035 | 0.18 | -1.207 | -0.002 | 0.071 | 0.071 | 0.061 | 0.51 | 0.42 | 0.68 | 0.81 | 0.53 | 0.64 |
| 6 | 54962103 | rs9349752 | 0.901 | -0.096 | 0.04 | -1.239 | -0.049 | -0.067 | -0.067 | 1.652 | 0.18 | 0.04 | 0.49 | 0.77 | 0.25 | 0.32 |
| 6 | 54968520 | rs7741358 | -0.145 | 0.084 | 0.109 | -1.234 | -0.005 | -0.04 | -0.04 | 0 | 0.28 | 0.36 | 0.15 | 0.06 | 0.21 | 0.29 |
| 6 | 54971369 | rs4236122 | 1.878 | 0.13 | -0.072 | -1.2 | -0.051 | 0.106 | 0.106 | 0.061 | 0.34 | 0.15 | 0.25 | 0.11 | 0.37 | 0.28 |
| 6 | 55006253 | rs7766546 | 0.549 | 0.139 | -0.235 | -1.142 | 0.001 | -0.024 | -0.024 | 0 | 0.63 | 0.51 | 0.39 | 0.17 | 0.58 | 0.57 |
| 6 | 55006797 | rs7767652 | -0.395 | -0.323 | -0.171 | -1.048 | -0.002 | -0.034 | -0.034 | 0 | 0.28 | 0.33 | 0.15 | 0.04 | 0.21 | 0.3 |
| 6 | 55011730 | rs12209485 | 1.014 | 0.364 | -0.948 | -0.593 | -0.082 | 0.162 | 0.162 | 0.735 | 0.45 | 0.12 | 0.25 | 0.17 | 0.42 | 0.33 |
| 6 | 55033223 | rs13200042 | 0.338 | 0.206 | -0.759 | -0.755 | -0.001 | 0.054 | 0.054 | 5.792 | 0.19 | 0.08 | 0.15 | 0.03 | 0.22 | 0.15 |
| 6 | 55037484 | rs9382460 | 0.571 | -0.315 | -0.79 | -0.223 | 0.004 | -0.006 | -0.006 | 0 | 0.19 | 0.21 | 0.17 | 0.31 | 0.14 | 0.27 |
| 6 | 55040696 | rs10456181 | 1.458 | -0.195 | -0.558 | -0.3 | -0.047 | 0.113 | 0.113 | 1.053 | 0.4 | 0.18 | 0.23 | 0.16 | 0.39 | 0.3 |
| 6 | 55041811 | rs7738215 | 0.626 | -0.359 | -0.437 | -0.411 | 0.04 | 0.027 | 0.027 | 2.136 | 0.57 | 0.65 | 0.64 | 0.65 | 0.6 | 0.57 |
| 6 | 55044386 | rs6927478 | 1.192 | -0.211 | -0.472 | -0.343 | -0.013 | 0.075 | 0.075 | 0 | 0.42 | 0.3 | 0.46 | 0.23 | 0.41 | 0.31 |
| 6 | 55045025 | rs9370399 | -0.856 | -0.276 | -0.401 | -0.371 | -0.006 | -0.01 | -0.01 | 0.687 | 0.39 | 0.59 | 0.37 | 0.46 | 0.46 | 0.42 |
| 6 | 55050507 | rs7775745 | 1.004 | -0.159 | 0.699 | 0.136 | 0.007 | 0.015 | 0.015 | 1.535 | 0.53 | 0.47 | 0.57 | 0.26 | 0.57 | 0.43 |
| 6 | 55055198 | rs4712099 | 0.33 | -0.247 | 0.265 | 0.218 | 0.035 | -0.051 | -0.051 | 0 | 0.2 | 0.09 | 0.18 | 0.31 | 0.14 | 0.29 |
| 6 | 55065989 | rs3122148 | -1.055 | 0.668 | 1.229 | 0.986 | 0.017 | 0.032 | 0.032 | 1.299 | 0.21 | 0.45 | 0.35 | 0.24 | 0.22 | 0.22 |
| 6 | 55067627 | rs6937878 | -0.561 | 0.632 | 1.242 | 1.009 | 0.009 | 0.054 | 0.054 | 0.86 | 0.18 | 0.38 | 0.33 | 0.24 | 0.18 | 0.19 |
| 6 | 55071006 | rs3122149 | -0.993 | 0.644 | 1.12 | 1.075 | 0.015 | 0.059 | 0.059 | 0.86 | 0.21 | 0.47 | 0.35 | 0.27 | 0.22 | 0.22 |
| 6 | 55072273 | rs3134704 | 0.01 | 0.649 | 0.612 | 1.173 | -0.002 | -0.039 | -0.039 | 0 | 0.79 | 0.9 | 0.78 | 0.61 | 0.77 | 0.82 |
| 6 | 55079046 | rs9396060 | 0.199 | 0.464 | 0.685 | 0.963 | -0.034 | -0.063 | -0.063 | 0 | 0.12 | 0.03 | 0.29 | 0.27 | 0.15 | 0.15 |
| 6 | 55082807 | rs3122156 | -0.906 | 0.276 | 1.257 | 1.043 | -0.031 | -0.029 | -0.029 | 0 | 0.33 | 0.58 | 0.39 | 0.24 | 0.28 | 0.23 |
| 6 | 55084697 | rs3122160 | -0.814 | 0.22 | 1.303 | 1.043 | -0.042 | -0.039 | -0.039 | 0.059 | 0.32 | 0.54 | 0.17 | 0.19 | 0.27 | 0.22 |
| 6 | 55087410 | rs4530845 | -0.103 | 0.333 | 1.043 | 1.082 | 0 | 0.004 | 0.004 | 0 | 0.26 | 0.05 | 0.14 | 0.17 | 0.24 | 0.2 |
| 6 | 55092069 | rs9357849 | -1.298 | 0.216 | 1.374 | 0.911 | -0.017 | -0.028 | -0.028 | 0.041 | 0.39 | 0.51 | 0.21 | 0.18 | 0.31 | 0.24 |
| 6 | 55093831 | rs7771240 | -1.012 | 0.244 | 1.537 | 0.919 | -0.02 | -0.041 | -0.041 | 0 | 0.37 | 0.46 | 0.2 | 0.18 | 0.31 | 0.24 |
| 6 | 55097163 | rs9357851 | -0.024 | 0.233 | 1.527 | 0.912 | -0.012 | -0.001 | -0.001 | 0 | 0.26 | 0.05 | 0.14 | 0.18 | 0.25 | 0.2 |
| 6 | 55103892 | rs3134711 | -1.107 | 0.219 | 1.711 | 1.246 | -0.018 | -0.019 | -0.019 | 0 | 0.38 | 0.51 | 0.22 | 0.23 | 0.31 | 0.25 |
| 6 | 55105341 | rs16869572 | -0.743 | 0.217 | 2.047 | 1.199 | -0.01 | -0.045 | -0.045 | 0.06 | 0.08 | 0.24 | 0.04 | 0 | 0.03 | 0.03 |
| 6 | 55107267 | rs9382471 | -0.419 | 0.214 | 2.077 | 1.202 | 0.008 | 0.018 | 0.018 | 0 | 0.29 | 0.2 | 0.16 | 0.18 | 0.28 | 0.22 |
| 6 | 55109189 | rs9475214 | -0.97 | 0.255 | 2.028 | 1.234 | -0.02 | -0.041 | -0.041 | 0 | 0.37 | 0.46 | 0.2 | 0.18 | 0.31 | 0.24 |
| 6 | 55113411 | rs3122169 | 0.749 | 0.717 | 1.77 | 1.933 | -0.024 | -0.026 | -0.026 | 0 | 0.9 | 0.96 | 0.57 | 0.51 | 0.83 | 0.82 |
| 6 | 55121954 | rs2811239 | 1.231 | 0.719 | 1.717 | 1.999 | -0.003 | 0.013 | 0.013 | 0 | 0.87 | 0.91 | 0.76 | 0.57 | 0.84 | 0.82 |
| 6 | 55133586 | rs2653344 | 0.671 | 0.909 | 1.723 | 1.727 | 0.011 | -0.003 | -0.003 | 0 | 0.92 | 0.89 | 0.88 | 0.94 | 0.82 | 0.9 |
| 6 | 55134247 | rs2134294 | 1.342 | 0.784 | 2.109 | 1.868 | 0.014 | 0.029 | 0.029 | 0.108 | 0.53 | 0.34 | 0.43 | 0.34 | 0.53 | 0.59 |
| 6 | 55142337 | rs2653349 | 0.614 | 0.635 | 2.088 | 1.753 | 0.008 | -0.003 | -0.003 | 0 | 0.92 | 0.89 | 0.88 | 0.94 | 0.82 | 0.87 |
| 6 | 55144102 | rs2653350 | 0.952 | 0.831 | 2.365 | 2.058 | 0.002 | 0.003 | 0.003 | 0 | 0.6 | 0.53 | 0.65 | 0.56 | 0.53 | 0.64 |
| 6 | 55153427 | rs1542896 | 1.477 | 0.845 | 2.284 | 2.051 | 0.002 | 0.107 | 0.107 | 0.152 | 0.51 | 0.28 | 0.32 | 0.2 | 0.5 | 0.57 |
| 6 | 55155704 | rs2811231 | 0.843 | 0.821 | 2.352 | 2.1 | 0 | 0.01 | 0.01 | 0 | 0.61 | 0.51 | 0.65 | 0.47 | 0.57 | 0.64 |
| 6 | 55162885 | rs2653363 | 0.6 | 0.856 | 2.205 | 2.089 | -0.001 | -0.002 | -0.002 | 0 | 0.63 | 0.64 | 0.67 | 0.57 | 0.56 | 0.65 |
| 6 | 55164486 | rs9396079 | -0.179 | 0.784 | 1.858 | 2.212 | 0.008 | -0.052 | -0.052 | 0 | 0.3 | 0.21 | 0.48 | 0.57 | 0.28 | 0.25 |
| 6 | 55167015 | rs9367632 | -0.506 | 0.899 | 1.835 | 2.237 | 0 | 0.049 | 0.049 | 0.054 | 0.08 | 0.11 | 0.31 | 0.28 | 0.07 | 0.07 |
| 6 | 55171454 | rs7774110 | 1.395 | 0.888 | 0.991 | 2.02 | 0.014 | 0.104 | 0.104 | 0.31 | 0.51 | 0.28 | 0.33 | 0.2 | 0.51 | 0.55 |
| 6 | 55171842 | rs9370410 | -0.312 | 0.892 | 0.556 | 1.615 | 0.004 | -0.002 | -0.002 | 2.475 | 0.29 | 0.14 | 0.2 | 0.28 | 0.28 | 0.22 |
| 6 | 55189085 | rs10948900 | 0.978 | 1.08 | 0.632 | 1.407 | -0.001 | 0.081 | 0.081 | 0.49 | 0.54 | 0.32 | 0.35 | 0.24 | 0.51 | 0.56 |
| 6 | 55189482 | rs10948901 | 1.234 | 1.072 | 0.575 | 1.366 | -0.004 | 0.128 | 0.128 | 0.057 | 0.5 | 0.27 | 0.32 | 0.22 | 0.49 | 0.55 |
| 6 | 55190291 | rs6924570 | -0.699 | 1.35 | 1.09 | 1.689 | 0.029 | 0.06 | 0.06 | 0 | 0.08 | 0.15 | 0.11 | 0.17 | 0.15 | 0.15 |
| 6 | 55191223 | rs958626 | -1.245 | 1.972 | 1.45 | 1.169 | 0.121 | -0.056 | -0.056 | 0 | 0.1 | 0.27 | 0.09 | 0 | 0.14 | 0.07 |
| 6 | 55191559 | rs12662228 | -1.647 | 1.749 | 1.547 | 1.487 | 0.015 | 0.117 | 0.117 | 1.074 | 0.09 | 0.25 | 0.07 | 0.18 | 0.07 | 0.14 |
| 6 | 55194896 | rs1007968 | -0.143 | 2.111 | 0.531 | 1.349 | -0.171 | -0.097 | -0.097 | 0 | 0.29 | 0.04 | 0.36 | 0.37 | 0.5 | 0.23 |
| 6 | 55196587 | rs12199003 | 0.733 | 1.764 | 1.144 | 1.525 | 0.002 | 0.004 | 0.004 | 0 | 0.58 | 0.43 | 0.26 | 0.31 | 0.37 | 0.66 |
| 6 | 55204017 | rs3963 | -0.626 | 1.766 | 1.233 | 1.534 | -0.001 | 0 | 0 | 0 | 0.38 | 0.28 | 0.42 | 0.37 | 0.6 | 0.3 |
| 6 | 55216005 | rs870806 | -0.704 | 1.629 | 0.55 | 1.254 | -0.018 | -0.031 | -0.031 | 0 | 0.4 | 0.26 | 0.41 | 0.54 | 0.58 | 0.43 |
| 6 | 55217131 | rs6917737 | -0.29 | 1.782 | 0.322 | 1.091 | -0.063 | -0.017 | -0.017 | 0 | 0.32 | 0.18 | 0.36 | 0.37 | 0.52 | 0.26 |
| 6 | 55225058 | rs6933713 | -1.275 | 1.035 | 0.489 | 0.709 | 0.003 | -0.048 | -0.048 | 0 | 0.13 | 0.27 | 0.12 | 0.01 | 0.1 | 0.15 |
| 6 | 55226013 | rs6939922 | -0.98 | 1 | 0.485 | 0.693 | -0.001 | -0.012 | -0.012 | 0 | 0.11 | 0.16 | 0.08 | 0.01 | 0.07 | 0.15 |
| 6 | 55227593 | rs9475263 | -0.889 | 1.015 | 0.492 | 0.701 | -0.001 | -0.013 | -0.013 | 0 | 0.1 | 0.16 | 0.08 | 0.01 | 0.07 | 0.15 |
| 6 | 55243149 | rs973052 | 0.575 | 1.381 | 0.166 | 0.756 | -0.024 | 0.011 | 0.011 | 1.11 | 0.58 | 0.66 | 0.58 | 0.61 | 0.5 | 0.64 |
| 6 | 55255791 | rs12663965 | -0.999 | 1.129 | 0.791 | 0.794 | 0.056 | 0.099 | 0.099 | 0.364 | 0.1 | 0.33 | 0.09 | 0.15 | 0.08 | 0.19 |
| 6 | 55255828 | rs3942865 | 1.357 | 0.998 | -0.04 | 0.702 | -0.01 | 0.204 | 0.204 | 0.064 | 0.43 | 0.13 | 0.2 | 0.1 | 0.38 | 0.38 |
| 6 | 55257309 | rs9370416 | 0.376 | 0.886 | -0.12 | 1.009 | -0.056 | -0.086 | -0.086 | 0 | 0.28 | 0.03 | 0.27 | 0.37 | 0.34 | 0.21 |
| 6 | 55257935 | rs1032772 | -1.448 | 0.816 | 0.317 | 0.76 | -0.009 | -0.001 | -0.001 | 0 | 0.15 | 0.49 | 0.12 | 0.16 | 0.13 | 0.2 |
| 6 | 55259784 | rs9382489 | 0.374 | 0.821 | 0.131 | 1.042 | -0.052 | -0.022 | -0.022 | 0 | 0.28 | 0.02 | 0.24 | 0.25 | 0.33 | 0.21 |
| 6 | 55260356 | rs16886684 | -1.169 | 0.714 | 0.361 | 1.016 | -0.001 | 0.034 | 0.034 | 2.085 | 0.12 | 0.48 | 0.11 | 0.16 | 0.08 | 0.18 |
| 6 | 55263532 | rs4715541 | -0.551 | 0.075 | -0.397 | 0.276 | 0.025 | 0.031 | 0.031 | 0 | 0.59 | 0.55 | 0.81 | 0.8 | 0.73 | 0.66 |
| 6 | 55263789 | rs9396088 | -0.71 | 0.071 | -0.376 | 0.274 | 0.038 | 0.042 | 0.042 | 0 | 0.62 | 0.62 | 0.84 | 0.79 | 0.76 | 0.67 |
| 6 | 55265397 | rs1502200 | -0.554 | 0.077 | -0.402 | 0.279 | 0.017 | 0.02 | 0.02 | 0 | 0.59 | 0.54 | 0.81 | 0.8 | 0.73 | 0.66 |

| | | | | | | | | | | | | | | | | |
|---|---|---|---|---|---|---|---|---|---|---|---|---|---|---|---|---|
| 6 | 55266625 | rs9370418 | -0.554 | 0.077 | -0.408 | 0.279 | 0.025 | 0.031 | 0.031 | 0 | 0.59 | 0.55 | 0.81 | 0.8 | 0.73 | 0.66 |
| 6 | 55270134 | rs4715544 | 0.886 | 0.194 | -0.473 | 0.32 | -0.034 | -0.006 | -0.006 | 0 | 0.28 | 0.09 | 0.24 | 0.25 | 0.33 | 0.2 |
| 6 | 55272264 | rs1553329 | 0.678 | 0.17 | -0.374 | 0.464 | 0.06 | 0.048 | 0.048 | 0 | 0.38 | 0.3 | 0.15 | 0.23 | 0.24 | 0.34 |
| 6 | 55272581 | rs12203074 | 0.858 | 0.283 | -0.455 | 0.546 | -0.126 | -0.133 | -0.133 | 0.273 | 0.2 | 0.02 | 0.47 | 0.38 | 0.32 | 0.24 |
| 6 | 55274069 | rs10807492 | -1.068 | -0.056 | -0.441 | 0.419 | 0.015 | 0.013 | 0.013 | 0 | 0.67 | 0.65 | 0.85 | 0.8 | 0.84 | 0.74 |
| 6 | 55277167 | rs1979875 | -0.993 | -0.059 | -0.468 | 0.294 | 0.024 | 0.027 | 0.027 | 0 | 0.67 | 0.69 | 0.85 | 0.83 | 0.83 | 0.76 |
| 6 | 55282429 | rs1393325 | -0.615 | -0.021 | -0.526 | 0.28 | -0.015 | 0.013 | 0.013 | 0.403 | 0.41 | 0.67 | 0.62 | 0.6 | 0.51 | 0.54 |
| 6 | 55284772 | rs7746284 | 0.215 | -0.172 | -0.474 | 0.082 | 0.029 | 0.041 | 0.041 | 0 | 0.4 | 0.39 | 0.19 | 0.2 | 0.29 | 0.33 |
| 6 | 55285977 | rs17830227 | 0.25 | 0.019 | 0.093 | 0.248 | 0.025 | 0.122 | 0.122 | 0.237 | 0.06 | 0.15 | 0.05 | 0.16 | 0.05 | 0.18 |
| 6 | 55290936 | rs10498805 | -0.554 | 0.07 | -0.497 | 0.093 | -0.002 | 0.006 | 0.006 | 0 | 0.11 | 0.11 | 0.06 | 0.03 | 0.13 | 0.1 |
| 6 | 55294820 | rs7751456 | -0.093 | -0.032 | -0.731 | -0.202 | -0.02 | 0.091 | 0.091 | 1.025 | 0.25 | 0.12 | 0.19 | 0.03 | 0.31 | 0.17 |
| 6 | 55297418 | rs7767401 | 0.362 | -0.021 | -0.599 | 0.19 | 0.057 | 0.144 | 0.144 | 0 | 0.5 | 0.23 | 0.35 | 0.23 | 0.44 | 0.4 |
| 6 | 55298959 | rs1546298 | 1.332 | 0.129 | -0.785 | 0.554 | -0.015 | -0.053 | -0.053 | 0 | 0.13 | 0.02 | 0.1 | 0.24 | 0.14 | 0.18 |
| 6 | 55299220 | rs1546299 | -0.268 | 0.227 | -0.172 | 0.591 | 0.004 | 0.001 | 0.001 | 0.186 | 0.71 | 0.64 | 0.55 | 0.64 | 0.66 | 0.73 |
| 6 | 55301723 | rs1472641 | -0.429 | -0.056 | 0.153 | 0.702 | -0.004 | 0.008 | 0.008 | 0.112 | 0.4 | 0.48 | 0.49 | 0.48 | 0.39 | 0.42 |
| 6 | 55307050 | rs9370423 | 0.511 | -0.103 | 0.172 | 0.732 | 0.009 | 0.041 | 0.041 | 0 | 0.19 | 0.45 | 0.16 | 0.21 | 0.18 | 0.18 |
| 6 | 55307591 | rs9357862 | 1.05 | -0.114 | 0.036 | 0.727 | -0.011 | -0.05 | -0.05 | 0.046 | 0.15 | 0.02 | 0.11 | 0.2 | 0.15 | 0.18 |
| 6 | 55313379 | rs9464247 | -0.756 | -0.099 | 0.312 | 0.621 | -0.001 | 0.001 | 0.001 | 0.103 | 0.42 | 0.56 | 0.52 | 0.48 | 0.43 | 0.44 |
| 6 | 55317936 | rs4072373 | -0.271 | -0.103 | 0.29 | 0.703 | -0.011 | 0.07 | 0.07 | 0 | 0.26 | 0.44 | 0.31 | 0.41 | 0.26 | 0.31 |
| 6 | 55319485 | rs4256438 | 0.7 | -0.287 | 0.286 | 0.51 | 0.001 | 0.005 | 0.005 | 0 | 0.5 | 0.47 | 0.46 | 0.51 | 0.5 | 0.51 |
| 6 | 55320311 | rs4339457 | 0.931 | -0.222 | -0.125 | 0.263 | -0.037 | -0.005 | -0.005 | 0.954 | 0.14 | 0.01 | 0.18 | 0.07 | 0.16 | 0.18 |
| 6 | 55323430 | rs4286782 | -0.514 | -0.719 | -0.281 | 0.102 | -0.033 | -0.001 | -0.001 | 2.73 | 0.61 | 0.85 | 0.48 | 0.63 | 0.46 | 0.5 |
| 6 | 55331763 | rs9637974 | 1.33 | -0.964 | -0.575 | 0.124 | -0.005 | -0.098 | -0.098 | 0.352 | 0.21 | 0.01 | 0.19 | 0.26 | 0.17 | 0.2 |
| 6 | 55334381 | rs9382495 | 0.482 | -0.955 | -0.259 | 0.041 | 0.018 | -0.051 | -0.051 | 8.704 | 0.3 | 0.23 | 0.31 | 0.45 | 0.27 | 0.28 |
| 6 | 55334405 | rs7749819 | 1.03 | -0.9 | 0.099 | 0.003 | -0.072 | -0.061 | -0.061 | 0 | 0.25 | 0.47 | 0.13 | 0.17 | 0.12 | 0.2 |
| 6 | 55337062 | rs9475303 | 1.037 | -0.77 | -0.124 | 0.07 | -0.002 | -0.003 | -0.003 | 0 | 0.26 | 0.28 | 0.12 | 0.17 | 0.17 | 0.22 |
| 6 | 55344782 | rs4712105 | -0.909 | -0.331 | -0.205 | 0.257 | -0.056 | -0.004 | -0.004 | 0 | 0.76 | 0.86 | 0.66 | 0.7 | 0.61 | 0.69 |
| 6 | 55356846 | rs9464252 | -0.421 | -0.592 | -0.134 | 0.287 | -0.009 | -0.01 | -0.01 | 0 | 0.37 | 0.65 | 0.49 | 0.5 | 0.52 | 0.4 |
| 6 | 55364966 | rs6917354 | -0.262 | -0.649 | 0.494 | 0.319 | -0.002 | 0.121 | 0.121 | 0 | 0.07 | 0.27 | 0.1 | 0.19 | 0.07 | 0.08 |
| 6 | 55368815 | rs6936870 | -0.721 | -0.613 | 0.495 | 0.31 | 0.009 | 0.082 | 0.082 | 0 | 0.09 | 0.27 | 0.12 | 0.19 | 0.11 | 0.09 |
| 6 | 55377664 | rs4537142 | -0.821 | -1.014 | 0.13 | 0.142 | 0.076 | 0.021 | 0.021 | 0.154 | 0.52 | 0.67 | 0.69 | 0.58 | 0.69 | 0.59 |
| 6 | 55378314 | rs3800027 | -0.821 | -1.014 | 0.13 | 0.142 | 0.076 | 0.021 | 0.021 | 0.154 | 0.52 | 0.67 | 0.69 | 0.58 | 0.69 | 0.6 |
| 6 | 55383610 | rs12525326 | 0.863 | -0.911 | -0.118 | 0.127 | -0.003 | -0.001 | -0.001 | 0 | 0.25 | 0.19 | 0.35 | 0.31 | 0.41 | 0.32 |
| 6 | 55386477 | rs9357865 | 1.291 | -1.242 | -0.4 | 0.067 | 0.01 | -0.081 | -0.081 | 0.01 | 0.23 | 0.01 | 0.13 | 0.25 | 0.16 | 0.2 |
| 6 | 55386699 | rs9475338 | -0.424 | -0.463 | 0.22 | 0.227 | -0.001 | 0 | 0 | 0 | 0.46 | 0.45 | 0.65 | 0.38 | 0.61 | 0.51 |
| 6 | 55393286 | rs4320370 | 0.084 | -0.361 | -0.293 | 0.316 | -0.093 | -0.086 | -0.086 | 1.521 | 0.31 | 0.12 | 0.44 | 0.53 | 0.49 | 0.34 |
| 6 | 55399744 | rs9357866 | 0.493 | -0.216 | -0.236 | 0.368 | -0.001 | 0.002 | 0.002 | 0.004 | 0.06 | 0.03 | 0.1 | 0.23 | 0.09 | 0.09 |
| 6 | 55412503 | rs12201934 | 0.868 | -0.265 | -0.183 | 0.293 | -0.005 | 0.007 | 0.007 | 0 | 0.17 | 0.12 | 0.27 | 0.07 | 0.18 | 0.19 |
| 6 | 55416822 | rs4588676 | -0.613 | -0.55 | -0.235 | 0.279 | 0.057 | 0.026 | 0.026 | 0 | 0.52 | 0.41 | 0.73 | 0.63 | 0.69 | 0.65 |
| 6 | 55422618 | rs3925003 | -0.738 | 0.498 | -0.199 | 0.278 | 0 | -0.001 | -0.001 | 0 | 0.52 | 0.52 | 0.67 | 0.63 | 0.58 | 0.63 |
| 6 | 55428780 | rs4398739 | -0.703 | 0.754 | 0.11 | 0.46 | 0.012 | 0.089 | 0.089 | 0 | 0.09 | 0.2 | 0.12 | 0.23 | 0.11 | 0.09 |
| 6 | 55435041 | rs4715554 | 0.029 | 0.779 | 0.023 | -0.093 | 0.009 | -0.022 | -0.022 | 0 | 0.8 | 0.77 | 0.72 | 0.93 | 0.78 | 0.78 |
| 6 | 55443352 | rs4236134 | -0.329 | 0.627 | -0.196 | -0.382 | -0.001 | 0.005 | 0.005 | 0 | 0.55 | 0.61 | 0.39 | 0.63 | 0.49 | 0.65 |
| 6 | 55448610 | rs4715555 | -0.707 | 0.837 | -0.041 | -0.308 | -0.025 | 0.018 | 0.018 | 0 | 0.77 | 0.85 | 0.63 | 0.91 | 0.63 | 0.8 |
| 6 | 55456080 | rs4132990 | -0.556 | 0.482 | 0.112 | 0.087 | 0 | 0 | 0 | 0 | 0.46 | 0.58 | 0.32 | 0.63 | 0.35 | 0.62 |
| 6 | 55456654 | rs7771624 | -0.013 | 0.436 | -0.307 | 0.268 | 0.005 | -0.028 | -0.028 | 0.954 | 0.25 | 0.2 | 0.21 | 0.5 | 0.25 | 0.41 |
| 6 | 55466372 | rs9367646 | -0.624 | 0.759 | -0.083 | 0.047 | 0.002 | 0.001 | 0.001 | 0 | 0.45 | 0.48 | 0.31 | 0.63 | 0.35 | 0.61 |
| 6 | 55470865 | rs12332954 | -0.452 | 1.581 | 0.244 | 0.149 | -0.014 | 0.04 | 0.04 | 0.073 | 0.14 | 0.06 | 0.11 | 0.04 | 0.19 | 0.13 |
| 6 | 55480911 | rs4509122 | 0.595 | 1.41 | 0.672 | 0.298 | -0.005 | -0.001 | -0.001 | 0.884 | 0.8 | 0.75 | 0.77 | 0.77 | 0.79 | 0.71 |
| 6 | 55502257 | rs11965285 | -1.173 | 1.936 | 0.8 | 0.501 | 0.058 | 0.078 | 0.078 | 0 | 0.25 | 0.47 | 0.31 | 0.39 | 0.32 | 0.29 |
| 6 | 55525031 | rs9396128 | 1.667 | 2.079 | 0.466 | 0.484 | -0.016 | 0.01 | 0.01 | 0.115 | 0.55 | 0.27 | 0.46 | 0.38 | 0.47 | 0.42 |
| 6 | 55532767 | rs4267958 | -1.433 | 2.012 | 0.546 | 1.101 | 0.039 | 0.054 | 0.054 | 0 | 0.23 | 0.33 | 0.29 | 0.39 | 0.31 | 0.29 |
| 6 | 55539010 | rs7772313 | -0.588 | 1.807 | 0.197 | 1.372 | 0.004 | 0.002 | 0.002 | 0 | 0.09 | 0.07 | 0.17 | 0.21 | 0.16 | 0.07 |
| 6 | 55545768 | rs4712108 | -0.369 | 1.231 | 0.192 | 0.751 | 0 | 0.038 | 0.038 | 0 | 0.09 | 0.11 | 0.09 | 0.19 | 0.08 | 0.18 |
| 6 | 55559361 | rs2208606 | -0.37 | 1.244 | 0.195 | 0.702 | 0.001 | 0.035 | 0.035 | 0 | 0.09 | 0.11 | 0.1 | 0.19 | 0.09 | 0.18 |
| 6 | 55562028 | rs6927049 | -1.207 | 0.79 | 0.404 | 0.268 | -0.001 | 0 | 0 | 1.072 | 0.27 | 0.21 | 0.17 | 0.26 | 0.19 | 0.33 |
| 6 | 55563203 | rs2498529 | -1.372 | 1.463 | 0.619 | -0.15 | 0.011 | -0.02 | -0.02 | 0 | 0.33 | 0.48 | 0.35 | 0.32 | 0.37 | 0.28 |
| 6 | 55570742 | rs2475252 | -0.169 | 1.391 | 0.919 | 0.162 | -0.02 | -0.014 | -0.014 | 0 | 0.1 | 0.15 | 0.12 | 0.08 | 0.06 | 0.12 |
| 6 | 55571611 | rs9475383 | -0.487 | 1.732 | 1.117 | 0.124 | 0.102 | 0.09 | 0.09 | 0.057 | 0.07 | 0.19 | 0.14 | 0.23 | 0.18 | 0.07 |
| 6 | 55579345 | rs9396140 | 0.379 | 1.87 | 0.897 | 0.08 | -0.018 | -0.014 | -0.014 | 0 | 0.06 | 0.01 | 0.09 | 0.08 | 0.1 | 0.22 |
| 6 | 55579429 | rs12197364 | 1.868 | 2.279 | 1.366 | -0.007 | -0.004 | -0.018 | -0.018 | 1.096 | 0.57 | 0.27 | 0.48 | 0.43 | 0.47 | 0.38 |
| 6 | 55590730 | rs9357875 | -0.025 | 2.307 | 0.913 | 0.03 | -0.039 | -0.01 | -0.01 | 0 | 0.07 | 0.02 | 0.13 | 0.08 | 0.15 | 0.23 |
| 6 | 55612431 | rs1154854 | 1.204 | 2.272 | 1.014 | 0.003 | 0.006 | 0.004 | 0.004 | 0 | 0.7 | 0.65 | 0.64 | 0.67 | 0.67 | 0.5 |
| 6 | 55613365 | rs6908699 | -0.203 | 2.273 | 0.749 | 0.039 | -0.013 | -0.029 | -0.029 | 1.033 | 0.84 | 0.96 | 0.89 | 0.83 | 0.86 | 0.8 |
| 6 | 55620017 | rs11537721 | -0.948 | 2.153 | 0.715 | 0.002 | 0.001 | -0.006 | -0.006 | 0 | 0.17 | 0.19 | 0.25 | 0.16 | 0.21 | 0.34 |
| 6 | 55620251 | rs3756990 | -0.617 | 2.145 | 0.419 | 0.002 | 0.009 | 0.006 | 0.006 | 0 | 0.16 | 0.1 | 0.21 | 0.15 | 0.16 | 0.33 |
| 6 | 55626118 | rs2397190 | 1.652 | 2.17 | 0.458 | 0.014 | -0.003 | -0.01 | -0.01 | 0 | 0.57 | 0.36 | 0.49 | 0.44 | 0.47 | 0.39 |
| 6 | 55628344 | rs228140 | -1 | 1.766 | -0.263 | -0.083 | -0.003 | 0.007 | 0.007 | 0 | 0.17 | 0.15 | 0.24 | 0.15 | 0.21 | 0.34 |
| 6 | 55631762 | rs228144 | 0.284 | 1.672 | -0.663 | -0.181 | -0.009 | -0.015 | -0.015 | 16.963 | 0.81 | 0.96 | 0.87 | 0.84 | 0.84 | 0.77 |

| | | | | | | | | | | | | | | | | |
|---|---|---|---|---|---|---|---|---|---|---|---|---|---|---|---|---|
| 6 | 55636255 | rs228146 | 0.813 | 1.628 | -0.426 | -0.227 | 0.018 | 0.026 | 0.026 | 0.054 | 0.47 | 0.22 | 0.5 | 0.55 | 0.46 | 0.53 |
| 6 | 55639619 | rs1020422 | 1.027 | 1.138 | -0.703 | -0.116 | 0.043 | -0.027 | -0.027 | 0 | 0.41 | 0.19 | 0.39 | 0.49 | 0.33 | 0.3 |
| 6 | 55652914 | rs9349794 | -1.135 | 0.142 | 0.305 | -0.315 | -0.01 | 0.012 | 0.012 | 0 | 0.19 | 0.31 | 0.27 | 0.24 | 0.16 | 0.19 |
| 6 | 55656259 | rs3798836 | -1.589 | -0.344 | -0.155 | -0.837 | 0.029 | 0.033 | 0.033 | 1.626 | 0.45 | 0.9 | 0.46 | 0.4 | 0.39 | 0.34 |
| 6 | 55657453 | rs16887173 | -0.672 | -0.311 | -0.027 | -0.802 | -0.007 | 0.015 | 0.015 | 0 | 0.18 | 0.26 | 0.26 | 0.24 | 0.15 | 0.2 |
| 6 | 55675663 | rs7775292 | -0.146 | -0.395 | 0.765 | -0.888 | 0.004 | 0.025 | 0.025 | 0 | 0.22 | 0.16 | 0.16 | 0.15 | 0.23 | 0.15 |
| 6 | 55684300 | rs12174118 | -0.212 | -0.399 | 0.86 | -0.823 | 0 | 0.033 | 0.033 | 0 | 0.1 | 0.22 | 0.12 | 0.14 | 0.09 | 0.12 |
| 6 | 55687414 | rs10948955 | -1.419 | -0.427 | 1.416 | -0.855 | -0.006 | -0.006 | -0.006 | 0 | 0.43 | 0.47 | 0.45 | 0.41 | 0.41 | 0.36 |
| 6 | 55688361 | rs9475417 | -1.088 | -0.369 | 1.309 | -0.824 | -0.031 | 0.015 | 0.015 | 0.439 | 0.22 | 0.44 | 0.27 | 0.25 | 0.16 | 0.21 |
| 6 | 55689305 | rs3823039 | -0.324 | -0.316 | 1.147 | -0.818 | 0.001 | 0.034 | 0.034 | 0 | 0.23 | 0.15 | 0.18 | 0.16 | 0.26 | 0.18 |
| 6 | 55711211 | rs1470527 | -1.111 | -0.201 | 0.8 | -0.921 | 0.014 | 0.029 | 0.029 | 0 | 0.46 | 0.63 | 0.42 | 0.3 | 0.41 | 0.38 |
| 6 | 55714563 | rs7750450 | -0.919 | -0.169 | 0.8 | -1.22 | -0.008 | -0.062 | -0.062 | 0 | 0.42 | 0.52 | 0.35 | 0.16 | 0.41 | 0.27 |
| 6 | 55715715 | rs12215796 | -1.223 | -0.26 | 0.493 | -1.392 | 0 | -0.02 | -0.02 | 0 | 0.5 | 0.52 | 0.38 | 0.2 | 0.48 | 0.33 |
| 6 | 55716160 | rs3823038 | -0.968 | -0.241 | 0.233 | -1.295 | 0.007 | 0.003 | 0.003 | 0 | 0.43 | 0.62 | 0.36 | 0.15 | 0.41 | 0.27 |
| 6 | 55730212 | rs2021803 | -1.086 | -0.3 | -0.165 | -1.326 | 0.038 | 0.175 | 0.175 | 0.424 | 0.52 | 0.77 | 0.53 | 0.19 | 0.46 | 0.37 |
| 6 | 55734220 | rs9475436 | 0.179 | -0.585 | -0.231 | -1.615 | -0.018 | 0.049 | 0.049 | 0.018 | 0.19 | 0.08 | 0.29 | 0.06 | 0.24 | 0.22 |
| 6 | 55738271 | rs3798819 | -0.746 | -0.831 | 0.508 | -1.686 | -0.038 | -0.05 | -0.05 | 0 | 0.19 | 0.31 | 0.18 | 0.13 | 0.15 | 0.08 |
| 6 | 55739553 | rs3734444 | -1.409 | -0.669 | 0.368 | -1.425 | 0.027 | 0.086 | 0.086 | 0.421 | 0.5 | 0.63 | 0.53 | 0.19 | 0.46 | 0.34 |
| 6 | 55742926 | rs16887333 | -0.175 | -0.694 | 0.294 | -1.083 | 0 | 0 | 0 | 0 | 0.07 | 0.05 | 0.11 | 0.08 | 0.07 | 0.06 |
| 6 | 55744188 | rs9367669 | -0.879 | -0.676 | 0.357 | -1.078 | -0.008 | 0.005 | 0.005 | 0.296 | 0.42 | 0.33 | 0.58 | 0.45 | 0.48 | 0.63 |
| 6 | 55751496 | rs1447131 | -1.248 | -0.877 | 0.94 | -1.846 | 0.001 | -0.016 | -0.016 | 0 | 0.47 | 0.51 | 0.52 | 0.16 | 0.48 | 0.34 |
| 6 | 55751639 | rs10948958 | 0.351 | -0.984 | 1.204 | -1.841 | -0.027 | -0.058 | -0.058 | 1.116 | 0.11 | 0.23 | 0.04 | 0 | 0.06 | 0.03 |
| 6 | 55754378 | rs12195643 | -1.176 | -1.022 | 1.252 | -1.909 | -0.051 | -0.075 | -0.075 | 0 | 0.22 | 0.39 | 0.16 | 0.09 | 0.13 | 0.14 |
| 6 | 55754594 | rs12202825 | 0.27 | -1.038 | 1.365 | -1.917 | -0.029 | -0.063 | -0.063 | 0 | 0.11 | 0.27 | 0.05 | 0 | 0.06 | 0.03 |
| 6 | 55755929 | rs1374547 | -0.757 | -1.041 | 1.353 | -1.919 | 0.001 | 0.001 | 0.001 | 0 | 0.11 | 0.13 | 0.11 | 0.09 | 0.07 | 0.11 |
| 6 | 55757789 | rs12201469 | 0.584 | -1.024 | 1.297 | -1.91 | -0.021 | -0.047 | -0.047 | 0.476 | 0.1 | 0.17 | 0.04 | 0 | 0.06 | 0.03 |
| 6 | 55760688 | rs7770068 | -2.1 | -1.206 | 1.247 | -2.008 | -0.07 | 0.118 | 0.118 | 0.697 | 0.61 | 0.87 | 0.62 | 0.22 | 0.55 | 0.46 |
| 6 | 55766417 | rs9396162 | -1.092 | -1.06 | 1.454 | -2.035 | -0.044 | -0.062 | -0.062 | 0 | 0.21 | 0.36 | 0.16 | 0.1 | 0.13 | 0.14 |
| 6 | 55767358 | rs7758580 | -1.726 | -0.968 | 1.076 | -1.978 | -0.062 | 0.206 | 0.206 | 1.071 | 0.56 | 0.85 | 0.58 | 0.16 | 0.54 | 0.33 |
| 6 | 55770946 | rs1868032 | -0.535 | -0.793 | 1.519 | -1.758 | 0.011 | -0.044 | -0.044 | 0.006 | 0.3 | 0.39 | 0.39 | 0.13 | 0.36 | 0.24 |
| 6 | 55776585 | rs6902446 | 0.259 | -0.838 | 1.685 | -1.792 | -0.027 | -0.061 | -0.061 | 1.176 | 0.11 | 0.24 | 0.05 | 0 | 0.06 | 0.03 |
| 6 | 55778771 | rs7752119 | -1.465 | -0.844 | 1.575 | -1.798 | 0 | 0.108 | 0.108 | 0.028 | 0.47 | 0.74 | 0.54 | 0.17 | 0.48 | 0.35 |
| 6 | 55785780 | rs895642 | -1.282 | -0.823 | 1.527 | -1.769 | 0.001 | -0.032 | -0.032 | 0.049 | 0.42 | 0.59 | 0.54 | 0.17 | 0.48 | 0.33 |
| 6 | 55789704 | rs6906682 | -0.796 | -0.658 | 1.336 | -1.613 | -0.012 | -0.038 | -0.038 | 1.403 | 0.12 | 0.22 | 0.15 | 0.05 | 0.12 | 0.05 |
| 6 | 55796744 | rs17757400 | -0.252 | -0.658 | 1.213 | -1.459 | 0 | -0.002 | -0.002 | 0 | 0.07 | 0.07 | 0.14 | 0.14 | 0.07 | 0.05 |
| 6 | 55802644 | rs17683688 | 0.048 | -0.606 | 1.236 | -1.734 | -0.001 | -0.001 | -0.001 | 0.39 | 0.06 | 0.07 | 0.1 | 0.05 | 0.07 | 0.05 |
| 6 | 55805282 | rs12213017 | -1.256 | -0.733 | 1.38 | -1.825 | -0.025 | -0.054 | -0.054 | 0 | 0.24 | 0.31 | 0.18 | 0.09 | 0.18 | 0.11 |
| 6 | 55816727 | rs4574618 | -1.676 | -0.776 | 1.677 | -1.926 | -0.002 | 0.061 | 0.061 | 0 | 0.49 | 0.66 | 0.56 | 0.17 | 0.53 | 0.35 |
| 6 | 55816822 | rs4599631 | -1.813 | -0.814 | 1.644 | -1.946 | -0.046 | 0.241 | 0.241 | 0.144 | 0.52 | 0.86 | 0.58 | 0.17 | 0.54 | 0.35 |
| 6 | 55819362 | rs12208960 | 0.099 | -1.084 | 1.647 | -1.971 | 0.024 | 0.033 | 0.033 | 1.901 | 0.12 | 0.07 | 0.03 | 0.02 | 0.06 | 0.06 |
| 6 | 55824452 | rs6459119 | -0.097 | -1.031 | 1.814 | -1.994 | 0.017 | -0.044 | -0.044 | 0 | 0.75 | 0.64 | 0.61 | 0.92 | 0.64 | 0.79 |
| 6 | 55825851 | rs9475487 | -1.553 | -1.129 | 1.951 | -2.02 | 0.012 | 0 | 0 | 0 | 0.25 | 0.79 | 0.12 | 0.03 | 0.11 | 0.09 |
| 6 | 55828727 | rs2745745 | 0.771 | -0.92 | 1.98 | -1.974 | -0.001 | -0.021 | -0.021 | 0 | 0.63 | 0.57 | 0.57 | 0.89 | 0.58 | 0.73 |
| 6 | 55832161 | rs12190342 | 0.107 | -0.919 | 1.85 | -1.974 | 0.024 | 0.032 | 0.032 | 1.901 | 0.12 | 0.07 | 0.03 | 0.03 | 0.06 | 0.06 |
| 6 | 55836050 | rs2745739 | 0.823 | -0.881 | 1.907 | -1.947 | 0.004 | -0.067 | -0.067 | 0.712 | 0.62 | 0.44 | 0.71 | 0.9 | 0.58 | 0.7 |
| 6 | 55846527 | rs9357881 | 0.98 | -0.905 | 1.583 | -1.92 | -0.006 | 0.137 | 0.137 | 0.317 | 0.58 | 0.21 | 0.55 | 0.89 | 0.58 | 0.7 |
| 6 | 55875132 | rs4712110 | 0.038 | -0.82 | 1.421 | -1.833 | -0.009 | 0.007 | 0.007 | 0.02 | 0.75 | 0.78 | 0.63 | 0.91 | 0.65 | 0.8 |
| 6 | 55875266 | rs7357042 | -0.117 | -0.736 | 1.195 | -1.787 | -0.022 | -0.038 | -0.038 | 0 | 0.13 | 0.22 | 0.05 | 0.03 | 0.06 | 0.08 |
| 6 | 55885348 | rs9349800 | -0.451 | -0.565 | 0.57 | -1.815 | -0.027 | 0.061 | 0.061 | 0.078 | 0.78 | 0.95 | 0.7 | 0.91 | 0.75 | 0.82 |
| 6 | 55886109 | rs2206679 | 1.303 | -0.61 | 0.636 | -1.698 | 0.032 | 0.117 | 0.117 | 0.052 | 0.52 | 0.24 | 0.49 | 0.82 | 0.56 | 0.66 |
| 6 | 55891553 | rs3950156 | -0.225 | -0.377 | -0.008 | -1.387 | -0.019 | -0.039 | -0.039 | 0 | 0.13 | 0.21 | 0.04 | 0.03 | 0.07 | 0.08 |
| 6 | 55893797 | rs2223662 | 0.201 | 0.225 | -0.522 | -1.019 | -0.07 | 0.073 | 0.073 | 0.022 | 0.77 | 0.95 | 0.64 | 0.92 | 0.65 | 0.81 |
| 6 | 55913842 | rs2841001 | 0.573 | -0.225 | -0.325 | -1.061 | -0.004 | 0.013 | 0.013 | 0.105 | 0.68 | 0.68 | 0.6 | 0.82 | 0.67 | 0.7 |
| 6 | 55913867 | rs2841002 | 0.927 | -1.168 | -1.012 | -1.336 | -0.024 | 0.1 | 0.1 | 1.569 | 0.46 | 0.08 | 0.46 | 0.66 | 0.55 | 0.55 |
| 6 | 55914639 | rs4715574 | -0.59 | -1.604 | -1.338 | -1.626 | -0.051 | 0.092 | 0.092 | 0.774 | 0.74 | 0.96 | 0.65 | 0.91 | 0.71 | 0.74 |
| 6 | 55915726 | rs9296824 | -0.442 | -1.654 | -0.811 | -1.669 | -0.007 | 0.043 | 0.043 | 1.128 | 0.73 | 0.86 | 0.63 | 0.91 | 0.71 | 0.71 |
| 6 | 55916127 | rs9382578 | 0.306 | -2.052 | -0.45 | -1.862 | 0.004 | 0.034 | 0.034 | 0 | 0.56 | 0.63 | 0.47 | 0.83 | 0.54 | 0.58 |
| 6 | 55921936 | rs3823468 | 0.4 | -2.078 | -0.195 | -1.872 | 0.002 | 0.011 | 0.011 | 0 | 0.55 | 0.61 | 0.47 | 0.82 | 0.54 | 0.58 |
| 6 | 55928019 | rs13218367 | 1.613 | -2.272 | -0.363 | -1.338 | -0.146 | -0.134 | -0.134 | 0.687 | 0.33 | 0.04 | 0.23 | 0.53 | 0.41 | 0.29 |
| 6 | 55929347 | rs2235735 | 0.182 | -2.28 | -0.359 | -1.354 | 0.002 | 0.011 | 0.011 | 0 | 0.55 | 0.61 | 0.47 | 0.82 | 0.54 | 0.58 |
| 6 | 55932511 | rs9396171 | 1.574 | -2.252 | -0.426 | -1.329 | -0.166 | -0.153 | -0.153 | 0 | 0.33 | 0.03 | 0.23 | 0.53 | 0.41 | 0.29 |
| 6 | 55933942 | rs12189567 | 0.609 | -2.175 | -0.187 | -1.711 | -0.004 | -0.005 | -0.005 | 0 | 0.17 | 0.15 | 0.15 | 0.09 | 0.17 | 0.13 |
| 6 | 55937271 | rs9370479 | 1.249 | -2.224 | 0.088 | -1.73 | -0.033 | 0.063 | 0.063 | 0.48 | 0.39 | 0.19 | 0.4 | 0.74 | 0.5 | 0.5 |
| 6 | 55940735 | rs9475556 | 0.567 | -1.973 | -0.308 | -1.601 | 0.006 | 0.017 | 0.017 | 0 | 0.52 | 0.57 | 0.46 | 0.78 | 0.53 | 0.55 |
| 6 | 55948620 | rs4599643 | -0.51 | -1.952 | -0.299 | -1.59 | 0.002 | 0.022 | 0.022 | 2.583 | 0.69 | 0.73 | 0.61 | 0.87 | 0.7 | 0.7 |
| 6 | 55965671 | rs4573075 | -0.563 | -1.867 | -0.391 | -1.546 | 0 | 0.072 | 0.072 | 4.085 | 0.7 | 0.86 | 0.62 | 0.87 | 0.7 | 0.7 |
| 6 | 55988519 | rs1925165 | 1.13 | -1.519 | -0.297 | -1.254 | -0.003 | 0.009 | 0.009 | 0.099 | 0.41 | 0.33 | 0.41 | 0.74 | 0.5 | 0.5 |
| 6 | 55993245 | rs1925166 | 0.712 | -0.612 | -1.019 | -0.837 | 0.017 | 0.062 | 0.062 | 0 | 0.52 | 0.63 | 0.47 | 0.78 | 0.53 | 0.58 |
| 6 | 56025219 | rs761105 | 0.633 | -0.619 | -1.056 | -0.827 | 0.022 | 0.074 | 0.074 | 0 | 0.53 | 0.64 | 0.46 | 0.78 | 0.54 | 0.65 |

| | | | | | | | | | | | | | | | | |
|---|---|---|---|---|---|---|---|---|---|---|---|---|---|---|---|---|
| 6 | 56044578 | rs2038149 | 0.372 | -0.631 | -1.111 | -0.859 | 0.004 | -0.01 | -0.01 | 0 | 0.57 | 0.62 | 0.57 | 0.86 | 0.63 | 0.76 |
| 6 | 56064197 | rs1474698 | 0.769 | -0.706 | -1.134 | -0.829 | 0.006 | 0.023 | 0.023 | 0 | 0.52 | 0.59 | 0.44 | 0.76 | 0.52 | 0.63 |
| 6 | 56067109 | rs1925189 | 0.327 | -0.716 | -1.13 | -0.844 | 0.017 | 0.083 | 0.083 | 0 | 0.57 | 0.71 | 0.56 | 0.87 | 0.61 | 0.76 |
| 6 | 56068448 | rs4715591 | -0.183 | -0.862 | -1.22 | -0.96 | -0.021 | 0.098 | 0.098 | 3.487 | 0.71 | 0.91 | 0.61 | 0.86 | 0.69 | 0.78 |
| 6 | 56087914 | rs7771018 | 0.508 | -0.748 | -1.121 | -0.824 | -0.019 | -0.05 | -0.05 | 0 | 0.13 | 0.2 | 0.05 | 0 | 0.07 | 0.02 |
| 6 | 56102709 | rs1925152 | 0.033 | -0.439 | -1.038 | -0.536 | 0 | 0.07 | 0.07 | 0 | 0.57 | 0.71 | 0.46 | 0.8 | 0.53 | 0.66 |
| 6 | 56102780 | rs1925153 | 0.365 | 0.009 | -1.001 | 0.146 | -0.003 | 0 | 0 | 0 | 0.44 | 0.61 | 0.28 | 0.42 | 0.43 | 0.33 |
| 6 | 56104522 | rs1925154 | 0.203 | 0.773 | -1.186 | -0.06 | -0.016 | 0.005 | 0.005 | 0 | 0.66 | 0.68 | 0.6 | 0.68 | 0.53 | 0.61 |
| 6 | 56105707 | rs6903416 | -0.439 | 1.456 | -1.526 | -0.764 | -0.047 | 0.075 | 0.075 | 0 | 0.19 | 0.08 | 0.11 | 0.01 | 0.21 | 0.17 |
| 6 | 56108509 | rs7757985 | -0.548 | 0.846 | -1.31 | -1.424 | 0.006 | 0.033 | 0.033 | 0.587 | 0.45 | 0.71 | 0.29 | 0.09 | 0.38 | 0.27 |
| 6 | 56108853 | rs1925157 | 0.375 | 1.641 | -1.655 | -0.988 | 0.01 | 0.003 | 0.003 | 0 | 0.9 | 0.88 | 0.91 | 0.84 | 0.83 | 0.88 |
| 6 | 56112781 | rs6925652 | -0.248 | 1.283 | -1.911 | -1.42 | -0.019 | 0.162 | 0.162 | 0.235 | 0.36 | 0.86 | 0.21 | 0.01 | 0.32 | 0.19 |
| 6 | 56112790 | rs3807013 | 1.384 | 0.748 | -2.333 | -1.275 | 0.028 | -0.07 | -0.07 | 0 | 0.17 | 0.04 | 0.42 | 0.63 | 0.11 | 0.38 |
| 6 | 56112952 | rs9396184 | 1.294 | 0.989 | -2.424 | -1.491 | -0.027 | 0.062 | 0.062 | 0 | 0.26 | 0.02 | 0.18 | 0.13 | 0.29 | 0.24 |
| 6 | 56115894 | rs12530020 | 0.402 | 0.869 | -2.292 | -1.271 | 0.035 | -0.036 | -0.036 | 0 | 0.28 | 0.14 | 0.5 | 0.67 | 0.2 | 0.46 |
| 6 | 56120294 | rs724678 | 1.346 | 0.798 | -1.952 | -1.408 | 0.005 | 0.016 | 0.016 | 0 | 0.27 | 0.18 | 0.2 | 0.18 | 0.3 | 0.22 |
| 6 | 56120721 | rs724677 | -0.933 | 0.825 | -2.132 | -1.596 | -0.011 | 0.031 | 0.031 | 0.038 | 0.84 | 0.98 | 0.92 | 0.91 | 0.86 | 0.88 |
| 6 | 56121567 | rs9296835 | 0.11 | 0.836 | -2.003 | -1.599 | 0.023 | -0.055 | -0.055 | 0 | 0.28 | 0.62 | 0.21 | 0.07 | 0.34 | 0.18 |
| 6 | 56125726 | rs10223442 | 0.847 | 1.101 | -1.956 | -1.656 | -0.057 | 0.093 | 0.093 | 0 | 0.2 | 0.06 | 0.12 | 0.01 | 0.23 | 0.15 |
| 6 | 56126066 | rs9370495 | 0.519 | 0.675 | -1.961 | -1.649 | 0.031 | -0.02 | -0.02 | 0 | 0.3 | 0.19 | 0.51 | 0.67 | 0.22 | 0.49 |
| 6 | 56127267 | rs1925164 | 0.558 | 0.681 | -1.962 | -1.659 | -0.004 | 0.065 | 0.065 | 0 | 0.56 | 0.33 | 0.71 | 0.85 | 0.52 | 0.7 |
| 6 | 56127621 | rs1925163 | -0.599 | 0.762 | -1.718 | -1.371 | 0.015 | 0.073 | 0.073 | 0.033 | 0.57 | 0.81 | 0.72 | 0.74 | 0.56 | 0.66 |
| 6 | 56130746 | rs6904902 | 0.036 | 0.767 | -1.042 | -0.959 | 0.092 | 0.081 | 0.081 | 4.502 | 0.07 | 0.5 | 0.07 | 0.06 | 0.06 | 0.01 |
| 6 | 56150940 | rs9357902 | 0.061 | 0.513 | -0.712 | -0.993 | 0.039 | 0.014 | 0.014 | 0 | 0.61 | 0.78 | 0.44 | 0.3 | 0.73 | 0.47 |
| 6 | 56152943 | rs9464371 | -0.272 | 1.1 | -0.755 | -1.268 | -0.005 | -0.006 | -0.006 | 0.48 | 0.25 | 0.27 | 0.14 | 0.06 | 0.32 | 0.22 |
| 6 | 56155058 | rs6459143 | -0.213 | 1.241 | -0.708 | -1.261 | 0.102 | -0.043 | -0.043 | 0 | 0.18 | 0.55 | 0.13 | 0.09 | 0.27 | 0.17 |
| 6 | 56156125 | rs4715607 | 0.877 | 0.942 | -0.941 | -1.255 | -0.004 | 0.118 | 0.118 | 0.694 | 0.53 | 0.19 | 0.71 | 0.78 | 0.49 | 0.62 |
| 6 | 56176759 | rs9349816 | 0.217 | 0.609 | -0.678 | -0.859 | 0.073 | 0.132 | 0.132 | 0 | 0.55 | 0.22 | 0.4 | 0.22 | 0.69 | 0.47 |
| 6 | 56180739 | rs4636028 | 0.003 | 0.94 | 0.507 | -0.217 | 0.04 | 0.04 | 0.04 | 0 | 0.88 | 0.62 | 0.89 | 0.9 | 0.88 | 0.98 |
| 6 | 56182197 | rs9475664 | -0.525 | 1.361 | -0.325 | -0.547 | -0.002 | -0.002 | -0.002 | 0.02 | 0.26 | 0.23 | 0.22 | 0.09 | 0.46 | 0.25 |
| 6 | 56182465 | rs9475665 | -0.267 | 1.585 | -0.368 | -0.596 | 0.015 | -0.012 | -0.012 | 0 | 0.22 | 0.23 | 0.19 | 0.09 | 0.37 | 0.16 |
| 6 | 56185286 | rs9382618 | 1.368 | 0.64 | -1.387 | -0.49 | 0.021 | 0.136 | 0.136 | 0.012 | 0.39 | 0.04 | 0.21 | 0.15 | 0.32 | 0.33 |
| 6 | 56188650 | rs12662498 | 0.374 | 0.456 | -0.693 | 0.454 | 0.066 | -0.141 | -0.141 | 0 | 0.19 | 0.03 | 0.43 | 0.57 | 0.06 | 0.34 |
| 6 | 56202104 | rs7739763 | -0.162 | 1.383 | 0.337 | 0.306 | 0.053 | 0.057 | 0.057 | 0 | 0.11 | 0.39 | 0.08 | 0.1 | 0.11 | 0.02 |
| 6 | 56202243 | rs7739952 | -0.613 | 1.396 | 0.318 | 0.285 | 0.042 | 0.005 | 0.005 | 0 | 0.15 | 0.39 | 0.11 | 0.1 | 0.18 | 0.12 |
| 6 | 56203738 | rs9370523 | -0.648 | 1.281 | -0.349 | 1.178 | -0.003 | -0.004 | -0.004 | 0 | 0.16 | 0.11 | 0.23 | 0.45 | 0.17 | 0.33 |
| 6 | 56203830 | rs6931260 | -0.343 | 1.581 | -0.459 | -0.14 | 0 | 0.061 | 0.061 | 0.341 | 0.48 | 0.72 | 0.7 | 0.79 | 0.47 | 0.57 |
| 6 | 56203840 | rs6910739 | 0.532 | 1.756 | -0.701 | -0.723 | 0.024 | -0.024 | -0.024 | 0.163 | 0.8 | 0.74 | 0.87 | 0.91 | 0.72 | 0.84 |
| 6 | 56203904 | rs6935231 | 0.42 | 1.981 | -0.658 | -0.791 | 0.028 | -0.056 | -0.056 | 2.874 | 0.8 | 0.73 | 0.88 | 0.94 | 0.71 | 0.86 |
| 6 | 56203969 | rs6935271 | 0.591 | 1.798 | -0.584 | -0.811 | 0.006 | -0.029 | -0.029 | 0 | 0.81 | 0.78 | 0.88 | 0.93 | 0.74 | 0.82 |
| 6 | 56204017 | rs9382621 | 0.794 | 1.555 | -0.946 | -0.752 | 0.026 | -0.091 | -0.091 | 0 | 0.31 | 0.05 | 0.51 | 0.71 | 0.2 | 0.4 |
| 6 | 56204751 | rs17224373 | 1.286 | 1.669 | -0.764 | -0.621 | -0.014 | 0.096 | 0.096 | 1.068 | 0.3 | 0.04 | 0.21 | 0.1 | 0.26 | 0.35 |
| 6 | 56204780 | rs2038144 | 0.527 | 2.376 | 0.297 | -0.409 | 0.04 | -0.009 | -0.009 | 0 | 0.73 | 0.62 | 0.81 | 0.82 | 0.62 | 0.82 |
| 6 | 56206134 | rs10456732 | -0.513 | 2.348 | 0.728 | -0.653 | 0.087 | -0.019 | -0.019 | 0 | 0.12 | 0.29 | 0.15 | 0.06 | 0.22 | 0.11 |
| 6 | 56207222 | rs11962322 | -0.569 | 1.939 | 0.776 | -0.743 | 0.024 | -0.054 | -0.054 | 0 | 0.1 | 0.3 | 0.09 | 0.01 | 0.11 | 0.05 |
| 6 | 56208751 | rs9475676 | -1.353 | 2.158 | 0.546 | -0.86 | 0.025 | -0.052 | -0.052 | 0.258 | 0.24 | 0.66 | 0.24 | 0.09 | 0.36 | 0.11 |
| 6 | 56209919 | rs6908736 | 0.119 | 2.124 | 0.479 | -0.858 | 0.023 | -0.024 | -0.024 | 0 | 0.12 | 0.16 | 0.14 | 0.07 | 0.22 | 0.04 |
| 6 | 56210315 | rs17224911 | 0.094 | 2.101 | 0.476 | -0.889 | 0.021 | -0.028 | -0.028 | 0 | 0.12 | 0.15 | 0.13 | 0.06 | 0.22 | 0.04 |
| 6 | 56212433 | rs1543350 | 0.667 | 2.113 | 0.353 | -1.155 | 0.017 | -0.054 | -0.054 | 0 | 0.73 | 0.71 | 0.8 | 0.93 | 0.66 | 0.83 |
| 6 | 56212578 | rs2206533 | 0.889 | 1.735 | 0.203 | -1.301 | -0.002 | 0.008 | 0.008 | 0.74 | 0.37 | 0.32 | 0.3 | 0.19 | 0.38 | 0.44 |
| 6 | 56215930 | rs4715617 | 0.473 | 1.561 | 0.121 | -1.024 | 0.005 | 0.002 | 0.002 | 0.185 | 0.35 | 0.39 | 0.51 | 0.73 | 0.27 | 0.41 |
| 6 | 56217207 | rs2397220 | -0.224 | 1.553 | -0.231 | -0.952 | -0.002 | -0.013 | -0.013 | 0 | 0.47 | 0.55 | 0.64 | 0.79 | 0.49 | 0.45 |
| 6 | 56228086 | rs9475682 | -0.364 | 1.762 | 0.126 | -0.894 | 0.069 | -0.081 | -0.081 | 0 | 0.16 | 0.41 | 0.16 | 0.06 | 0.25 | 0.06 |
| 6 | 56231402 | rs9475687 | 0.003 | 1.78 | 0.262 | -0.888 | 0.057 | -0.051 | -0.051 | 0.131 | 0.13 | 0.25 | 0.14 | 0.06 | 0.21 | 0.04 |
| 6 | 56231723 | rs9370525 | -0.02 | 1.822 | 0.109 | -1.254 | -0.007 | -0.02 | -0.02 | 0 | 0.44 | 0.55 | 0.58 | 0.78 | 0.4 | 0.52 |
| 6 | 56234843 | rs9464383 | 0.151 | 1.944 | -0.24 | -1.268 | 0.003 | -0.022 | -0.022 | 0 | 0.44 | 0.46 | 0.38 | 0.21 | 0.51 | 0.36 |
| 6 | 56235411 | rs2294697 | 0.994 | 1.567 | -0.708 | -0.991 | -0.016 | 0.036 | 0.036 | 0.807 | 0.28 | 0.06 | 0.22 | 0.15 | 0.26 | 0.31 |
| 6 | 56237709 | rs7769615 | -0.555 | 1.44 | -0.115 | -0.747 | 0.028 | -0.049 | -0.049 | 0 | 0.09 | 0.26 | 0.08 | 0.01 | 0.11 | 0.05 |
| 6 | 56248348 | rs926530 | 0.086 | 1.35 | -0.952 | -0.465 | 0 | -0.041 | -0.041 | 0 | 0.32 | 0.13 | 0.51 | 0.6 | 0.29 | 0.46 |
| 6 | 56251191 | rs2397221 | -1.658 | 1.375 | -1.419 | -1.085 | 0.027 | 0.015 | 0.015 | 1.125 | 0.7 | 0.94 | 0.8 | 0.74 | 0.77 | 0.72 |
| 6 | 56253790 | rs2143349 | -0.217 | 1.751 | -1.259 | -1.24 | -0.019 | 0.076 | 0.076 | 0 | 0.47 | 0.21 | 0.59 | 0.62 | 0.47 | 0.61 |
| 6 | 56265544 | rs629506 | 0.431 | 1.879 | -1.328 | -1.263 | -0.009 | -0.026 | -0.026 | 0 | 0.33 | 0.14 | 0.53 | 0.61 | 0.35 | 0.47 |
| 6 | 56268377 | rs9475690 | -0.674 | 2.275 | -0.463 | -1.408 | -0.004 | -0.003 | -0.003 | 6.404 | 0.28 | 0.26 | 0.19 | 0.09 | 0.33 | 0.18 |
| 6 | 56268471 | rs676454 | 0.056 | 2.285 | -0.512 | -1.41 | 0.002 | -0.012 | -0.012 | 0.879 | 0.06 | 0.09 | 0.05 | 0 | 0.1 | 0.03 |
| 6 | 56271981 | rs10456189 | 0.364 | 2.268 | -0.282 | -1.43 | 0.02 | -0.036 | -0.036 | 0 | 0.14 | 0.2 | 0.12 | 0.06 | 0.17 | 0.04 |
| 6 | 56272686 | rs616516 | -1.498 | 2.2 | -0.29 | -1.502 | -0.049 | 0.12 | 0.12 | 0.759 | 0.39 | 0.79 | 0.29 | 0.13 | 0.44 | 0.23 |
| 6 | 56281321 | rs4715624 | 1.06 | 2.169 | -0.523 | -1.435 | -0.01 | -0.083 | -0.083 | 0 | 0.27 | 0.06 | 0.48 | 0.61 | 0.25 | 0.44 |
| 6 | 56283271 | rs586833 | -1.062 | 2.659 | -0.125 | -1.416 | -0.021 | 0.014 | 0.014 | 0 | 0.24 | 0.24 | 0.17 | 0.05 | 0.35 | 0.21 |
| 6 | 56285938 | rs662277 | -0.428 | 2.544 | -0.208 | -1.255 | -0.011 | 0.012 | 0.012 | 0 | 0.07 | 0.05 | 0.08 | 0.02 | 0.18 | 0.06 |

| | | | | | | | | | | | | | | | | |
|---|---|---|---|---|---|---|---|---|---|---|---|---|---|---|---|---|
| 6 | 56289743 | rs621387 | -1.221 | 2.482 | 0.215 | -1.356 | -0.001 | -0.021 | -0.021 | 0 | 0.37 | 0.41 | 0.26 | 0.1 | 0.49 | 0.24 |
| 6 | 56290289 | rs608383 | -0.882 | 2.59 | 0.216 | -1.316 | -0.011 | 0.003 | 0.003 | 0 | 0.3 | 0.27 | 0.21 | 0.09 | 0.41 | 0.23 |
| 6 | 56290873 | rs595298 | -0.345 | 2.637 | 0.824 | -0.761 | 0.03 | 0.027 | 0.027 | 0 | 0.32 | 0.35 | 0.56 | 0.55 | 0.44 | 0.47 |
| 6 | 56290948 | rs765289 | 0.932 | 2.047 | 0.77 | -1.011 | 0.044 | 0.134 | 0.134 | 0 | 0.49 | 0.18 | 0.33 | 0.31 | 0.43 | 0.35 |
| 6 | 56291176 | rs594008 | -0.631 | 1.831 | 1.225 | -0.418 | 0 | -0.003 | -0.003 | 0 | 0.45 | 0.49 | 0.58 | 0.59 | 0.49 | 0.57 |
| 6 | 56295657 | rs606894 | -1.073 | 1.551 | 1.263 | -0.339 | 0.001 | 0.004 | 0.004 | 0 | 0.62 | 0.72 | 0.72 | 0.74 | 0.68 | 0.59 |
| 6 | 56297568 | rs6911502 | 1.169 | 1.549 | 0.804 | -0.345 | 0.028 | 0.064 | 0.064 | 0 | 0.35 | 0.07 | 0.24 | 0.26 | 0.25 | 0.32 |
| 6 | 56304291 | rs10484787 | -1.761 | 1.569 | 1.334 | -0.247 | 0.083 | 0.005 | 0.005 | 0 | 0.05 | 0.21 | 0.06 | 0.01 | 0.09 | 0.1 |
| 6 | 56311121 | rs2397225 | -2.343 | 1.443 | 1.622 | -0.259 | 0.071 | 0.027 | 0.027 | 2.463 | 0.08 | 0.52 | 0.08 | 0.03 | 0.07 | 0.09 |
| 6 | 56312929 | rs7750332 | -0.581 | 1.721 | 0.801 | 0.001 | -0.034 | -0.069 | -0.069 | 3.726 | 0.65 | 0.85 | 0.46 | 0.51 | 0.59 | 0.63 |
| 6 | 56326765 | rs6459161 | 0.628 | 1.078 | 1.46 | 0.155 | -0.024 | -0.037 | -0.037 | 0 | 0.4 | 0.56 | 0.3 | 0.25 | 0.28 | 0.35 |
| 6 | 56327849 | rs11758339 | 0.969 | 1.106 | 1.179 | 0.176 | 0.039 | 0.068 | 0.068 | 0 | 0.35 | 0.08 | 0.25 | 0.24 | 0.25 | 0.33 |
| 6 | 56336406 | rs2894837 | 0.291 | 1.2 | 1.352 | 0.241 | 0.022 | 0.05 | 0.05 | 0 | 0.43 | 0.7 | 0.34 | 0.28 | 0.34 | 0.42 |
| 6 | 56342806 | rs9396214 | 0.827 | 0.73 | 1.086 | 0.172 | 0.033 | 0.053 | 0.053 | 0 | 0.36 | 0.22 | 0.27 | 0.24 | 0.27 | 0.34 |
| 6 | 56345337 | rs11753813 | 0.596 | 0.74 | 1.193 | 0.171 | 0.005 | 0.008 | 0.008 | 0 | 0.38 | 0.36 | 0.3 | 0.23 | 0.3 | 0.35 |
| 6 | 56352451 | rs1359070 | 0.968 | 0.76 | 0.774 | 0.192 | 0.03 | 0.067 | 0.067 | 0 | 0.35 | 0.08 | 0.26 | 0.25 | 0.27 | 0.34 |
| 6 | 56387256 | rs7741295 | -1.11 | 0.693 | 0.762 | 0.217 | 0.03 | 0.074 | 0.074 | 0 | 0.66 | 0.93 | 0.76 | 0.79 | 0.75 | 0.67 |
| 6 | 56401999 | rs4472350 | -0.691 | 0.717 | 1.356 | 0.384 | -0.021 | -0.009 | -0.009 | 0 | 0.62 | 0.52 | 0.71 | 0.64 | 0.73 | 0.64 |
| 6 | 56413273 | rs4712132 | -1.019 | 0.797 | 1.325 | 0.373 | 0.008 | 0.011 | 0.011 | 0 | 0.65 | 0.72 | 0.75 | 0.79 | 0.75 | 0.67 |
| 6 | 56420538 | rs11756977 | 0.688 | 0.742 | 1.176 | 0.379 | -0.03 | -0.012 | -0.012 | 0 | 0.37 | 0.39 | 0.28 | 0.35 | 0.27 | 0.36 |
| 6 | 56424232 | rs9349828 | 0.724 | 0.608 | 1.043 | 0.163 | -0.008 | -0.036 | -0.036 | 0 | 0.37 | 0.37 | 0.27 | 0.24 | 0.32 | 0.4 |
| 6 | 56424421 | rs4715634 | -0.483 | 0.307 | 1.29 | 1.064 | -0.019 | 0.108 | 0.108 | 0 | 0.12 | 0.27 | 0.18 | 0.32 | 0.11 | 0.16 |
| 6 | 56426931 | rs9367689 | 0.956 | 0.586 | 1.27 | 0.287 | 0 | 0.004 | 0.004 | 0.676 | 0.39 | 0.35 | 0.4 | 0.26 | 0.39 | 0.29 |
| 6 | 56436335 | rs9475724 | 0.718 | 0.377 | 1.326 | 0.258 | 0.001 | 0.009 | 0.009 | 0.382 | 0.39 | 0.6 | 0.39 | 0.29 | 0.38 | 0.28 |
| 6 | 56437465 | rs1014309 | 1.24 | 0.379 | 1.131 | 0.232 | -0.066 | 0.016 | 0.016 | 0.492 | 0.36 | 0.1 | 0.37 | 0.28 | 0.39 | 0.29 |
| 6 | 56445051 | rs9349832 | 0.696 | 0.451 | 1.163 | 0.097 | 0.001 | 0.011 | 0.011 | 0.645 | 0.42 | 0.58 | 0.4 | 0.25 | 0.4 | 0.29 |
| 6 | 56463410 | rs4712138 | 0.77 | 0.618 | 1.176 | 0.027 | 0 | 0.005 | 0.005 | 0.389 | 0.39 | 0.35 | 0.38 | 0.25 | 0.38 | 0.28 |
| 6 | 56470690 | rs13209548 | 0.893 | 0.597 | 1.156 | 0.01 | 0 | -0.025 | -0.025 | 0 | 0.38 | 0.5 | 0.38 | 0.24 | 0.38 | 0.23 |
| 6 | 56471402 | rs9382658 | 0.893 | 0.597 | 1.156 | 0.01 | 0 | -0.025 | -0.025 | 0 | 0.38 | 0.5 | 0.38 | 0.24 | 0.38 | 0.23 |
| 6 | 56474519 | rs2064592 | -0.169 | 0.941 | 0.76 | 0.009 | 0.007 | -0.064 | -0.064 | 0 | 0.62 | 0.86 | 0.55 | 0.53 | 0.62 | 0.61 |
| 6 | 56476023 | rs9370544 | 1.617 | 0.891 | 0.697 | -0.139 | -0.038 | 0.044 | 0.044 | 0.887 | 0.35 | 0.12 | 0.35 | 0.25 | 0.37 | 0.28 |
| 6 | 56476262 | rs7747060 | 1.036 | 0.868 | 1.069 | -0.153 | 0 | -0.023 | -0.023 | 0 | 0.4 | 0.5 | 0.38 | 0.27 | 0.4 | 0.3 |
| 6 | 56482704 | rs9396228 | 1.384 | 0.571 | 0.358 | -0.268 | 0 | 0.086 | 0.086 | 0.009 | 0.4 | 0.14 | 0.36 | 0.25 | 0.37 | 0.28 |
| 6 | 56507135 | rs1024195 | -0.055 | 0.713 | 0.623 | -0.375 | -0.081 | -0.179 | -0.179 | 0 | 0.66 | 0.98 | 0.57 | 0.55 | 0.61 | 0.65 |
| 6 | 56507489 | rs1024196 | -1.833 | 1.136 | 1.39 | -0.742 | 0.095 | 0.073 | 0.073 | 3.464 | 0.05 | 0.43 | 0.04 | 0.02 | 0.03 | 0.02 |
| 6 | 56548359 | rs4073960 | -0.753 | 1.165 | 1.382 | -0.957 | -0.037 | -0.129 | -0.129 | 0.016 | 0.25 | 0.53 | 0.17 | 0.02 | 0.21 | 0.05 |
| 6 | 56566149 | rs9918490 | 0.116 | 1.344 | 1.446 | -1.006 | 0.007 | -0.006 | -0.006 | 0 | 0.09 | 0.2 | 0.06 | 0.06 | 0.08 | 0.13 |
| 6 | 56569104 | rs16888128 | -0.738 | 1.248 | 1.622 | -0.881 | -0.023 | -0.094 | -0.094 | 0 | 0.24 | 0.32 | 0.15 | 0.02 | 0.22 | 0.05 |
| 6 | 56598426 | rs13198938 | -0.146 | 1.166 | 1.171 | -0.898 | -0.009 | -0.062 | -0.062 | 0 | 0.21 | 0.22 | 0.14 | 0 | 0.19 | 0.04 |
| 6 | 56616083 | rs12204927 | 0.382 | 1.534 | 0.788 | -0.94 | -0.076 | 0.056 | 0.056 | 0 | 0.13 | 0.01 | 0.09 | 0.03 | 0.21 | 0.2 |
| 6 | 56631976 | rs6904535 | -0.918 | 0.877 | 1.961 | -0.421 | -0.035 | -0.162 | -0.162 | 0 | 0.24 | 0.49 | 0.16 | 0.02 | 0.21 | 0.07 |
| 6 | 56666045 | rs13218913 | -0.007 | 1.082 | 2.108 | -0.301 | 0.004 | -0.071 | -0.071 | 0 | 0.15 | 0.21 | 0.13 | 0 | 0.2 | 0.03 |
| 6 | 56670263 | rs4587171 | -1.072 | 1.052 | 1.977 | -0.193 | -0.015 | -0.016 | -0.016 | 0 | 0.21 | 0.55 | 0.19 | 0.22 | 0.21 | 0.08 |
| 6 | 56684719 | rs10498813 | -0.024 | 1.177 | 1.676 | -0.305 | 0.001 | -0.052 | -0.052 | 0 | 0.15 | 0.16 | 0.13 | 0 | 0.19 | 0.03 |
| 6 | 56702253 | rs17752106 | 0.029 | 1.119 | 1.673 | -0.3 | 0.001 | -0.059 | -0.059 | 0.011 | 0.15 | 0.18 | 0.13 | 0 | 0.19 | 0.03 |
| 6 | 56702976 | rs17684878 | -0.011 | 1.127 | 1.701 | -0.296 | 0.002 | -0.074 | -0.074 | 0 | 0.16 | 0.21 | 0.14 | 0 | 0.19 | 0.03 |
| 6 | 56711276 | rs7761729 | -0.323 | 1.126 | 1.697 | -0.297 | 0 | -0.109 | -0.109 | 0 | 0.17 | 0.35 | 0.15 | 0 | 0.19 | 0.03 |
| 6 | 56722235 | rs1903332 | -0.708 | 1.11 | 1.65 | -0.3 | -0.012 | -0.137 | -0.137 | 2.142 | 0.2 | 0.49 | 0.15 | 0.02 | 0.19 | 0.03 |
| 6 | 56726819 | rs1870639 | 0.187 | 1.035 | 0.827 | -0.11 | -0.061 | -0.163 | -0.163 | 0 | 0.19 | 0.01 | 0.37 | 0.47 | 0.2 | 0.31 |
| 6 | 56732097 | rs16888189 | -0.885 | 1.043 | 1.742 | -0.031 | -0.009 | -0.006 | -0.006 | 0 | 0.19 | 0.34 | 0.14 | 0.23 | 0.19 | 0.09 |
| 6 | 56740504 | rs13191084 | -0.958 | 1.038 | 1.722 | -0.141 | -0.017 | -0.022 | -0.022 | 0.271 | 0.25 | 0.61 | 0.2 | 0.2 | 0.21 | 0.08 |
| 6 | 56756695 | rs6910217 | 0.177 | 1.041 | 1.441 | 0.05 | 0 | 0 | 0 | 0 | 0.19 | 0.18 | 0.33 | 0.46 | 0.16 | 0.36 |
| 6 | 56769980 | rs9296855 | -0.985 | 0.769 | 1.69 | 0.133 | -0.004 | -0.02 | -0.02 | 0 | 0.25 | 0.38 | 0.2 | 0.21 | 0.28 | 0.09 |
| 6 | 56779570 | rs9464407 | -1.068 | 0.766 | 1.628 | 0.132 | -0.005 | -0.027 | -0.027 | 2.215 | 0.26 | 0.56 | 0.21 | 0.21 | 0.28 | 0.09 |
| 6 | 56789311 | rs1599281 | -0.68 | 0.83 | 0.64 | 0.124 | 0.002 | -0.007 | -0.007 | 1.075 | 0.22 | 0.23 | 0.13 | 0.04 | 0.21 | 0.03 |
| 6 | 56791495 | rs17752435 | -0.454 | 0.873 | 0.436 | 0.146 | -0.011 | 0.075 | 0.075 | 0 | 0.19 | 0.09 | 0.11 | 0.04 | 0.21 | 0.03 |
| 6 | 56794154 | rs13209289 | 0.166 | 0.82 | 0.321 | 0.102 | -0.005 | -0.01 | -0.01 | 0 | 0.06 | 0 | 0.05 | 0.05 | 0.07 | 0.03 |
| 6 | 56799262 | rs2599697 | 1.412 | 0.952 | 0.112 | -0.013 | -0.07 | -0.033 | -0.033 | 0 | 0.34 | 0.02 | 0.32 | 0.25 | 0.33 | 0.27 |
| 6 | 56801774 | rs1451705 | -1.302 | 0.98 | 1.654 | 0.194 | -0.025 | 0.006 | 0.006 | 0 | 0.22 | 0.11 | 0.18 | 0.21 | 0.27 | 0.09 |
| 6 | 56811124 | rs2817573 | 0.79 | 0.789 | 1.532 | 0.246 | -0.009 | 0.081 | 0.081 | 0 | 0.83 | 0.93 | 0.9 | 1 | 0.83 | 0.97 |
| 6 | 56818146 | rs9382669 | -1.551 | 0.769 | 1.832 | 0.273 | 0.001 | 0.003 | 0.003 | 0 | 0.21 | 0.25 | 0.18 | 0.19 | 0.24 | 0.09 |
| 6 | 56820248 | rs13193465 | -0.875 | 0.777 | 1.743 | 0.279 | -0.009 | 0.081 | 0.081 | 0 | 0.17 | 0.06 | 0.1 | 0 | 0.18 | 0.03 |
| 6 | 56836145 | rs9475760 | -1.327 | 0.782 | 1.767 | 0.286 | 0.001 | 0.016 | 0.016 | 0 | 0.21 | 0.18 | 0.18 | 0.19 | 0.24 | 0.09 |
| 6 | 56843294 | rs11751461 | -0.304 | 0.776 | 1.594 | 0.269 | -0.004 | -0.011 | -0.011 | 0 | 0.05 | 0 | 0.07 | 0.04 | 0.07 | 0.02 |
| 6 | 56857665 | rs1562908 | 1.617 | 0.914 | 2.168 | 0.195 | -0.011 | -0.028 | -0.028 | 0 | 0.77 | 0.53 | 0.8 | 0.77 | 0.75 | 0.91 |
| 6 | 56877460 | rs720885 | -1.427 | 1.025 | 1.656 | 0.189 | 0 | -0.022 | -0.022 | 0.663 | 0.2 | 0.3 | 0.19 | 0.19 | 0.24 | 0.09 |
| 6 | 56900340 | rs4715646 | 1.412 | 1.17 | 2.101 | -0.182 | 0.027 | -0.062 | -0.062 | 0 | 0.8 | 0.51 | 0.8 | 0.87 | 0.75 | 0.89 |
| 6 | 56903482 | rs7753927 | 1.425 | 1.166 | 2.136 | -0.184 | 0.025 | -0.061 | -0.061 | 0.016 | 0.79 | 0.5 | 0.8 | 0.87 | 0.75 | 0.89 |
| 6 | 56910884 | rs1157713 | -1.998 | 0.897 | 2.189 | -0.153 | -0.003 | -0.066 | -0.066 | 0 | 0.22 | 0.47 | 0.2 | 0.13 | 0.24 | 0.11 |

| | | | | | | | | | | | | | | | | |
|---|---|---|---|---|---|---|---|---|---|---|---|---|---|---|---|---|
| 6 | 56920804 | rs12190575 | -0.992 | 0.896 | 2.187 | -0.173 | -0.005 | -0.032 | -0.032 | 0 | 0.16 | 0.15 | 0.09 | 0.04 | 0.15 | 0.04 |
| 6 | 56943388 | rs12190434 | -1.785 | 0.892 | 2.297 | -0.175 | 0.003 | -0.035 | -0.035 | 0 | 0.2 | 0.33 | 0.18 | 0.13 | 0.24 | 0.11 |
| 6 | 56954518 | rs6459178 | -1.037 | 0.885 | 2.195 | -0.178 | -0.005 | -0.031 | -0.031 | 0 | 0.16 | 0.15 | 0.09 | 0.04 | 0.15 | 0.04 |
| 6 | 56996688 | rs10484410 | -0.885 | 0.889 | 2.186 | -0.176 | -0.01 | 0.072 | 0.072 | 0 | 0.16 | 0.02 | 0.08 | 0.04 | 0.15 | 0.04 |
| 6 | 57012930 | rs3734738 | -1.749 | 0.891 | 2.204 | -0.176 | 0.001 | -0.046 | -0.046 | 0 | 0.21 | 0.37 | 0.19 | 0.13 | 0.24 | 0.11 |
| 6 | 57034009 | rs7756421 | -0.988 | 0.88 | 2.196 | -0.181 | -0.005 | -0.032 | -0.032 | 0 | 0.16 | 0.15 | 0.09 | 0.04 | 0.15 | 0.04 |
| 6 | 57042709 | rs3778515 | -2.066 | 0.915 | 2.155 | -0.178 | -0.004 | -0.075 | -0.075 | 0 | 0.23 | 0.5 | 0.2 | 0.13 | 0.24 | 0.11 |
| 6 | 57057183 | rs1547226 | -0.946 | 0.648 | 1.692 | -0.168 | -0.031 | 0.07 | 0.07 | 0 | 0.15 | 0.02 | 0.1 | 0.04 | 0.18 | 0.04 |
| 6 | 57063946 | rs12195875 | -1.572 | 0.346 | 2.054 | -0.167 | -0.005 | -0.02 | -0.02 | 0.67 | 0.2 | 0.28 | 0.14 | 0.1 | 0.2 | 0.06 |
| 6 | 57068940 | rs3800019 | -1.976 | 0.412 | 1.993 | -0.107 | -0.029 | -0.074 | -0.074 | 0 | 0.23 | 0.49 | 0.2 | 0.13 | 0.21 | 0.1 |
| 6 | 57069606 | rs3800018 | -0.805 | 0.539 | 1.634 | -0.043 | -0.017 | 0.071 | 0.071 | 0 | 0.15 | 0.02 | 0.09 | 0.04 | 0.16 | 0.03 |
| 6 | 57076552 | rs16888392 | -0.883 | 0.554 | 1.734 | -0.036 | -0.006 | -0.04 | -0.04 | 0.545 | 0.16 | 0.17 | 0.1 | 0.04 | 0.16 | 0.04 |
| 6 | 57077135 | rs12210597 | -1.244 | 0.556 | 1.6 | -0.02 | -0.013 | -0.051 | -0.051 | 0.081 | 0.21 | 0.39 | 0.19 | 0.13 | 0.21 | 0.1 |
| 6 | 57089942 | rs9475792 | -1.872 | 0.598 | 1.483 | 0.183 | 0.033 | 0.015 | 0.015 | 6.051 | 0.05 | 0.31 | 0.03 | 0 | 0.02 | 0.01 |
| 6 | 57098973 | rs409025 | 1.008 | 1.018 | 1.331 | 0.037 | 0.001 | 0.009 | 0.009 | 3.318 | 0.65 | 0.33 | 0.72 | 0.74 | 0.67 | 0.78 |
| 6 | 57112128 | rs7760349 | -0.984 | 1.254 | 1.263 | 0.026 | 0 | -0.018 | -0.018 | 0 | 0.46 | 0.51 | 0.61 | 0.79 | 0.44 | 0.46 |
| 6 | 57121719 | rs6927231 | -1.861 | 1.506 | 2.205 | -0.107 | 0.129 | -0.026 | -0.026 | 0 | 0.11 | 0.49 | 0.09 | 0.03 | 0.15 | 0.09 |
| 6 | 57124881 | rs1013147 | -0.75 | 1.384 | 2.145 | -0.076 | 0.003 | 0.005 | 0.005 | 0.122 | 0.41 | 0.33 | 0.59 | 0.78 | 0.35 | 0.39 |
| 6 | 57126898 | rs6459182 | 0.603 | 1.755 | 2.09 | 0.034 | 0.007 | 0.01 | 0.01 | 0 | 0.59 | 0.82 | 0.48 | 0.4 | 0.6 | 0.67 |
| 6 | 57129687 | rs10949013 | -1.209 | 1.582 | 2.368 | 0.1 | 0.022 | 0.109 | 0.109 | 0.021 | 0.07 | 0.18 | 0.15 | 0.21 | 0.12 | 0.08 |
| 6 | 57130574 | rs6906645 | -1.429 | 1.577 | 2.485 | 0.097 | 0.052 | 0.171 | 0.171 | 0.097 | 0.07 | 0.28 | 0.16 | 0.21 | 0.12 | 0.08 |
| 6 | 57133333 | rs984684 | -1.595 | 1.504 | 2.548 | 0.022 | 0.047 | 0.178 | 0.178 | 1.411 | 0.08 | 0.44 | 0.14 | 0.18 | 0.07 | 0.07 |
| 6 | 57144935 | rs7747096 | -1.434 | 1.537 | 2.114 | -0.026 | 0.053 | 0.182 | 0.182 | 1.879 | 0.49 | 0.78 | 0.66 | 0.82 | 0.43 | 0.4 |
| 6 | 57160572 | rs9367716 | -0.871 | 1.426 | 1.864 | 0.003 | 0.012 | 0.002 | 0.002 | 0.027 | 0.39 | 0.32 | 0.49 | 0.6 | 0.3 | 0.32 |
| 6 | 57161200 | rs9367717 | -1.45 | 1.34 | 0.993 | -0.319 | -0.121 | 0.169 | 0.169 | 3.371 | 0.56 | 0.94 | 0.69 | 0.81 | 0.51 | 0.51 |
| 6 | 57161932 | rs2397232 | -1.384 | 1.399 | 1.141 | -0.268 | -0.049 | 0.166 | 0.166 | 2.585 | 0.55 | 0.88 | 0.7 | 0.81 | 0.53 | 0.51 |
| 6 | 57163631 | rs6459185 | -0.897 | 1.588 | 1.75 | 1.16 | -0.016 | 0.049 | 0.049 | 1.302 | 0.26 | 0.52 | 0.24 | 0.33 | 0.24 | 0.25 |
| 6 | 57165629 | rs1860652 | 1.268 | 1.964 | 1.208 | 0.698 | -0.004 | -0.016 | -0.016 | 0.049 | 0.87 | 0.86 | 0.93 | 0.99 | 0.81 | 0.83 |
| 6 | 57165948 | rs12176102 | 1.797 | 1.852 | 1.297 | 0.845 | 0.004 | -0.002 | -0.002 | 0 | 0.48 | 0.18 | 0.4 | 0.52 | 0.48 | 0.55 |
| 6 | 57169715 | rs9382699 | -0.55 | 1.79 | 1.2 | 0.931 | 0.015 | -0.09 | -0.09 | 0 | 0.31 | 0.13 | 0.36 | 0.42 | 0.25 | 0.27 |
| 6 | 57173448 | rs7775338 | -1.559 | 1.855 | 1.612 | 0.795 | 0.027 | -0.037 | -0.037 | 0.145 | 0.15 | 0.26 | 0.19 | 0.06 | 0.2 | 0.18 |
| 6 | 57182068 | rs1888266 | 1.339 | 1.958 | 1.568 | 0.763 | 0.022 | -0.049 | -0.049 | 0 | 0.87 | 0.8 | 0.92 | 0.99 | 0.82 | 0.84 |
| 6 | 57182297 | rs1019210 | 0.841 | 2.202 | 1.464 | 0.882 | -0.019 | -0.074 | -0.074 | 0 | 0.7 | 0.82 | 0.63 | 0.57 | 0.68 | 0.74 |
| 6 | 57183734 | rs1013894 | -1.086 | 2.079 | 1.625 | 0.813 | -0.002 | 0.006 | 0.006 | 0.793 | 0.33 | 0.36 | 0.49 | 0.47 | 0.34 | 0.27 |
| 6 | 57184879 | rs1618210 | 0.83 | 2.059 | 1.618 | 0.801 | 0 | -0.001 | -0.001 | 0.062 | 0.68 | 0.68 | 0.51 | 0.53 | 0.66 | 0.73 |
| 6 | 57186294 | rs1755833 | 0.675 | 2.083 | 1.597 | 0.818 | -0.001 | -0.041 | -0.041 | 0 | 0.69 | 0.78 | 0.52 | 0.53 | 0.66 | 0.73 |
| 6 | 57192943 | rs6911353 | -0.717 | 2.01 | 1.331 | 0.878 | 0.032 | -0.052 | -0.052 | 0 | 0.1 | 0.22 | 0.06 | 0.01 | 0.11 | 0.15 |
| 6 | 57205920 | rs6459193 | 1.091 | 1.8 | 0.597 | 1.047 | 0 | 0.001 | 0.001 | 0.975 | 0.65 | 0.64 | 0.48 | 0.53 | 0.59 | 0.73 |
| 6 | 57229283 | rs4715668 | 1.476 | 2.132 | -0.117 | 0.88 | 0 | 0.016 | 0.016 | 1.382 | 0.52 | 0.18 | 0.4 | 0.5 | 0.47 | 0.57 |
| 6 | 57229990 | rs4712149 | 0.762 | 2.04 | -0.375 | 1.024 | -0.023 | -0.081 | -0.081 | 0 | 0.72 | 0.85 | 0.63 | 0.58 | 0.67 | 0.75 |
| 6 | 57230422 | rs4715669 | 0.567 | 2.116 | -0.501 | 1.302 | -0.048 | -0.052 | -0.052 | 2.067 | 0.89 | 0.98 | 0.74 | 0.66 | 0.81 | 0.79 |
| 6 | 57329013 | rs6914118 | -0.917 | 1.052 | -1.45 | 0.558 | -0.016 | -0.079 | -0.079 | 0 | 0.6 | 0.85 | 0.53 | 0.45 | 0.61 | 0.67 |
| 6 | 57329886 | rs9476005 | -0.379 | 0.954 | -1.305 | 0.299 | 0.004 | -0.02 | -0.02 | 0 | 0.37 | 0.52 | 0.35 | 0.15 | 0.45 | 0.43 |
| 6 | 57336340 | rs1570072 | -1.295 | 0.939 | -1.398 | 0.337 | -0.02 | -0.081 | -0.081 | 0.373 | 0.61 | 0.85 | 0.53 | 0.45 | 0.61 | 0.67 |
| 6 | 57359377 | rs2068287 | -1.158 | 1.175 | -1.265 | 0.505 | 0.001 | -0.062 | -0.062 | 3.93 | 0.57 | 0.83 | 0.51 | 0.45 | 0.59 | 0.67 |
| 6 | 57367388 | rs7748620 | 0.812 | 1.333 | -1.279 | 0.513 | -0.036 | 0.023 | 0.023 | 0.773 | 0.12 | 0 | 0.14 | 0.07 | 0.18 | 0.28 |
| 6 | 57381003 | rs9476059 | -0.044 | 1.288 | -1.064 | 0.62 | -0.003 | 0.007 | 0.007 | 0.028 | 0.22 | 0.26 | 0.18 | 0.3 | 0.16 | 0.24 |
| 6 | 57444439 | rs9476140 | -0.39 | 1.512 | -0.781 | 0.405 | 0.125 | 0.005 | 0.005 | 0 | 0.15 | 0.44 | 0.16 | 0.07 | 0.21 | 0.12 |
| 6 | 57535593 | rs9476225 | -0.076 | 1.349 | -0.965 | 0.459 | -0.004 | 0.007 | 0.007 | 0.004 | 0.23 | 0.29 | 0.18 | 0.29 | 0.16 | 0.24 |
| 6 | 57540722 | rs2498205 | 0.666 | 1.433 | -1.049 | 0.308 | -0.033 | 0.03 | 0.03 | 0.495 | 0.13 | 0.01 | 0.14 | 0.08 | 0.18 | 0.28 |
| 6 | 57561493 | rs7759819 | -0.367 | 1.536 | -0.828 | 0.362 | 0.102 | -0.003 | -0.003 | 1.323 | 0.15 | 0.48 | 0.17 | 0.07 | 0.22 | 0.12 |
| 6 | 57591840 | rs7759896 | -1.089 | 1.431 | -1.054 | 0.395 | -0.029 | -0.04 | -0.04 | 0 | 0.37 | 0.77 | 0.35 | 0.37 | 0.36 | 0.36 |
| 6 | 57642076 | rs12207241 | 0.634 | 1.307 | -0.944 | 0.36 | 0.009 | 0.005 | 0.005 | 0 | 0.58 | 0.58 | 0.65 | 0.61 | 0.63 | 0.46 |
| 6 | 57645576 | rs9396406 | 0.563 | 1.328 | -1.059 | 0.368 | 0.014 | 0.005 | 0.005 | 0 | 0.59 | 0.66 | 0.66 | 0.61 | 0.63 | 0.45 |
| 6 | 57665572 | rs11965101 | -1.39 | 1.387 | -1.013 | 0.383 | 0.037 | -0.007 | -0.007 | 0 | 0.24 | 0.53 | 0.32 | 0.19 | 0.29 | 0.2 |
| 6 | 57666427 | rs4440467 | 1.346 | 1.281 | -1.193 | 0.246 | -0.008 | 0.011 | 0.011 | 0.317 | 0.6 | 0.42 | 0.59 | 0.74 | 0.51 | 0.52 |
| 6 | 57670538 | rs10949183 | -0.926 | 1.19 | -1.272 | 0.269 | -0.025 | 0.03 | 0.03 | 0.374 | 0.88 | 0.98 | 0.87 | 0.92 | 0.82 | 0.72 |
| 6 | 57676198 | rs4506048 | 0.232 | 1.192 | -1.235 | 0.291 | 0 | -0.018 | -0.018 | 3.156 | 0.07 | 0.02 | 0.04 | 0.1 | 0.08 | 0.06 |
| 6 | 57687035 | rs3903046 | 0.229 | 1.192 | -1.235 | 0.291 | 0 | -0.018 | -0.018 | 0 | 0.07 | 0.02 | 0.04 | 0.1 | 0.08 | 0.06 |
| 6 | 57710238 | rs7758052 | -0.703 | 1.212 | -1.187 | 0.254 | 0.004 | 0.045 | 0.045 | 0.001 | 0.26 | 0.14 | 0.2 | 0.18 | 0.29 | 0.37 |
| 6 | 62902837 | rs482102 | -0.595 | 0.675 | -0.064 | -0.223 | -0.037 | -0.028 | -0.028 | 1.68 | 0.21 | 0.53 | 0.15 | 0.15 | 0.13 | 0.35 |
| 6 | 62922848 | rs530011 | -0.214 | 0.677 | -0.135 | -0.222 | -0.011 | -0.006 | -0.006 | 0 | 0.18 | 0.32 | 0.14 | 0.15 | 0.13 | 0.22 |
| 6 | 62979298 | rs2345326 | 2.068 | 0.716 | -0.399 | -0.344 | -0.044 | -0.055 | -0.055 | 0.804 | 0.39 | 0.04 | 0.27 | 0.42 | 0.41 | 0.26 |
| 6 | 62980430 | rs1007419 | 1.337 | 0.671 | -0.12 | -0.311 | 0.039 | -0.011 | -0.011 | 0 | 0.51 | 0.19 | 0.45 | 0.54 | 0.66 | 0.45 |
| 6 | 62985248 | rs9454825 | 0.167 | 0.57 | -0.298 | -0.34 | -0.03 | -0.008 | -0.008 | 0 | 0.13 | 0.25 | 0.06 | 0.1 | 0.05 | 0.17 |
| 6 | 62996879 | rs9446154 | -0.54 | 0.545 | -0.276 | -0.388 | -0.028 | -0.021 | -0.021 | 0 | 0.21 | 0.37 | 0.15 | 0.17 | 0.13 | 0.26 |
| 6 | 63022153 | rs523967 | -0.315 | 0.429 | -0.386 | -0.393 | -0.069 | -0.104 | -0.104 | 1.212 | 0.76 | 0.96 | 0.62 | 0.71 | 0.77 | 0.8 |
| 6 | 63044570 | rs678811 | -1.224 | 0.14 | -0.198 | -0.43 | -0.041 | -0.04 | -0.04 | 0 | 0.3 | 0.49 | 0.21 | 0.19 | 0.2 | 0.29 |
| 6 | 63049900 | rs6904711 | -0.59 | 0.153 | -0.693 | -0.425 | -0.02 | -0.015 | -0.015 | 0.321 | 0.21 | 0.3 | 0.14 | 0.17 | 0.13 | 0.26 |

| | | | | | | | | | | | | | | | | |
|---|---|---|---|---|---|---|---|---|---|---|---|---|---|---|---|---|
| 6 | 63061935 | rs12527308 | -0.406 | 0.042 | -0.801 | -0.495 | 0.008 | 0.007 | 0.007 | 0 | 0.22 | 0.19 | 0.14 | 0.15 | 0.13 | 0.23 |
| 6 | 63068054 | rs17463584 | 0.771 | -0.005 | -0.927 | -0.512 | 0 | 0.007 | 0.007 | 0.272 | 0.08 | 0.01 | 0.06 | 0.04 | 0.09 | 0.03 |
| 6 | 63069063 | rs2814651 | -0.158 | -0.049 | -0.873 | -0.469 | 0.029 | 0.026 | 0.026 | 0 | 0.21 | 0.07 | 0.13 | 0.15 | 0.13 | 0.23 |
| 6 | 63075694 | rs2345331 | -0.2 | -0.046 | -0.866 | -0.468 | 0.029 | 0.024 | 0.024 | 0 | 0.21 | 0.1 | 0.13 | 0.15 | 0.13 | 0.23 |
| 6 | 63079272 | rs2842795 | 0.868 | -0.016 | -0.918 | -0.558 | 0.015 | -0.008 | -0.008 | 0.349 | 0.58 | 0.61 | 0.51 | 0.55 | 0.73 | 0.57 |
| 6 | 63082734 | rs2842798 | -0.446 | -0.006 | -0.913 | -0.317 | 0.042 | 0.039 | 0.039 | 0 | 0.23 | 0.07 | 0.13 | 0.15 | 0.15 | 0.23 |
| 6 | 63085545 | rs9346389 | 1.819 | 0.394 | -0.844 | -0.347 | 0.002 | -0.014 | -0.014 | 0 | 0.33 | 0.17 | 0.26 | 0.42 | 0.35 | 0.25 |
| 6 | 63099756 | rs2814693 | -0.44 | 0.261 | -1.687 | -0.324 | 0.037 | 0.038 | 0.038 | 0 | 0.24 | 0.13 | 0.14 | 0.15 | 0.15 | 0.23 |
| 6 | 63102429 | rs4710686 | 1.966 | 0.288 | -1.813 | -0.277 | -0.076 | -0.103 | -0.103 | 0.158 | 0.32 | 0.03 | 0.25 | 0.42 | 0.35 | 0.25 |
| 6 | 63137637 | rs17465706 | 0.053 | 0.562 | -1.68 | -0.226 | -0.064 | -0.025 | -0.025 | 0.577 | 0.13 | 0.01 | 0.17 | 0.12 | 0.27 | 0.19 |
| 6 | 63152724 | rs646854 | 0.433 | -0.531 | -1.506 | -0.208 | 0.039 | 0.05 | 0.05 | 0 | 0.77 | 0.94 | 0.87 | 0.87 | 0.85 | 0.78 |
| 6 | 63173293 | rs676473 | -0.206 | -0.601 | -1.366 | -0.044 | 0.025 | -0.014 | -0.014 | 0.068 | 0.88 | 0.99 | 0.93 | 0.9 | 0.94 | 0.87 |
| 6 | 63173804 | rs678838 | 0.261 | -0.526 | -1.525 | 0.013 | 0.029 | -0.034 | -0.034 | 0.121 | 0.55 | 0.89 | 0.49 | 0.59 | 0.65 | 0.54 |
| 6 | 63174039 | rs678576 | 0.583 | -0.53 | -1.291 | -0.107 | 0.023 | 0.013 | 0.013 | 0 | 0.49 | 0.65 | 0.44 | 0.57 | 0.6 | 0.48 |
| 6 | 63174910 | rs9351812 | 1.666 | -0.375 | -1.22 | -0.044 | -0.06 | -0.064 | -0.064 | 0 | 0.31 | 0.12 | 0.26 | 0.44 | 0.38 | 0.3 |
| 6 | 63199731 | rs6909922 | -0.189 | -0.357 | -1.16 | -0.047 | 0.022 | -0.027 | -0.027 | 0 | 0.06 | 0.24 | 0.05 | 0.02 | 0.05 | 0.05 |
| 6 | 63209941 | rs2345981 | 1.646 | -0.355 | -1.361 | -0.025 | -0.062 | -0.066 | -0.066 | 0 | 0.31 | 0.12 | 0.26 | 0.44 | 0.38 | 0.3 |
| 6 | 63228933 | rs2065692 | 0.104 | -0.26 | -1.28 | 0.237 | 0.005 | 0.011 | 0.011 | 0 | 0.14 | 0.24 | 0.18 | 0.24 | 0.16 | 0.09 |
| 6 | 63231367 | rs7739318 | 0.871 | -0.26 | -1.244 | 0.248 | 0.002 | 0.02 | 0.02 | 0 | 0.1 | 0.17 | 0.15 | 0.23 | 0.13 | 0.05 |
| 6 | 63239542 | rs1334743 | 0.752 | -0.4 | -1.381 | 0.152 | -0.016 | 0.005 | 0.005 | 0 | 0.41 | 0.63 | 0.31 | 0.35 | 0.49 | 0.44 |
| 6 | 63250796 | rs1029158 | 1.087 | -0.635 | -1.605 | 0.098 | -0.045 | -0.101 | -0.101 | 0 | 0.22 | 0.02 | 0.36 | 0.28 | 0.19 | 0.28 |
| 6 | 63257354 | rs9446610 | 0.031 | -0.547 | -1.481 | 0.061 | 0.021 | 0.01 | 0.01 | 0.164 | 0.17 | 0.51 | 0.17 | 0.13 | 0.21 | 0.18 |
| 6 | 63270569 | rs1334737 | 0.501 | -0.639 | -1.533 | 0.05 | -0.012 | 0.023 | 0.023 | 0 | 0.45 | 0.72 | 0.34 | 0.36 | 0.52 | 0.5 |
| 6 | 63282423 | rs9446676 | 0.38 | -0.703 | -0.943 | 0.046 | 0.014 | 0.075 | 0.075 | 0.196 | 0.13 | 0.46 | 0.17 | 0.22 | 0.13 | 0.05 |
| 6 | 63286366 | rs9341378 | 0.827 | -0.73 | -1.01 | 0.025 | 0.003 | -0.003 | -0.003 | 0 | 0.42 | 0.42 | 0.32 | 0.36 | 0.52 | 0.48 |
| 6 | 63291027 | rs1572245 | 0.075 | -0.789 | -1.073 | 0.034 | 0.028 | -0.036 | -0.036 | 0 | 0.55 | 0.89 | 0.49 | 0.59 | 0.65 | 0.55 |
| 6 | 63306749 | rs1413783 | 1.079 | -0.723 | -1.102 | 0.066 | -0.053 | -0.109 | -0.109 | 0 | 0.22 | 0.01 | 0.36 | 0.28 | 0.19 | 0.27 |
| 6 | 63307884 | rs17469009 | 0.12 | -0.598 | -0.738 | 0.142 | 0.036 | 0.02 | 0.02 | 0 | 0.14 | 0.2 | 0.16 | 0.13 | 0.22 | 0.19 |
| 6 | 63319873 | rs9350474 | 1.43 | -0.564 | -0.875 | 0.157 | -0.026 | 0.017 | 0.017 | 0 | 0.28 | 0.21 | 0.16 | 0.23 | 0.31 | 0.3 |
| 6 | 63322114 | rs10943071 | 1.735 | -0.534 | -0.927 | 0.165 | -0.021 | 0.008 | 0.008 | 0 | 0.25 | 0.16 | 0.14 | 0.21 | 0.28 | 0.26 |
| 6 | 63327975 | rs10498821 | 0.871 | -0.308 | -1.065 | 0.272 | -0.027 | -0.027 | -0.027 | 0.443 | 0.07 | 0 | 0.09 | 0.09 | 0.09 | 0.14 |
| 6 | 63347849 | rs2782377 | 0.008 | -0.455 | -0.915 | 0.177 | 0.056 | 0.016 | 0.016 | 1.546 | 0.06 | 0.36 | 0.05 | 0.02 | 0.04 | 0.04 |
| 6 | 63358826 | rs10943087 | -0.111 | -0.65 | -1.218 | 0.242 | 0.035 | -0.026 | -0.026 | 0 | 0.46 | 0.12 | 0.5 | 0.42 | 0.35 | 0.46 |
| 6 | 63375122 | rs12204331 | -0.136 | -0.651 | -1.267 | 0.221 | 0.035 | -0.034 | -0.034 | 0 | 0.46 | 0.11 | 0.5 | 0.42 | 0.35 | 0.46 |
| 6 | 63376194 | rs4415141 | 1.825 | -0.581 | -1.099 | 0.23 | 0.003 | -0.004 | -0.004 | 0.482 | 0.26 | 0.25 | 0.16 | 0.2 | 0.28 | 0.28 |
| 6 | 63380661 | rs9343048 | 1.156 | -0.707 | -1.019 | 0.21 | -0.001 | -0.001 | -0.001 | 0.099 | 0.37 | 0.33 | 0.31 | 0.34 | 0.47 | 0.41 |
| 6 | 63388385 | rs12176521 | 0.377 | -0.44 | -0.649 | 0.381 | 0.055 | 0.015 | 0.015 | 2.456 | 0.93 | 0.64 | 0.95 | 0.98 | 0.96 | 0.96 |
| 6 | 63403021 | rs4431412 | 0.554 | -0.326 | -0.969 | 0.485 | -0.001 | 0.02 | 0.02 | 0 | 0.1 | 0.16 | 0.14 | 0.22 | 0.12 | 0.04 |
| 6 | 63410641 | rs2752989 | -0.726 | -0.394 | -1.396 | 0.449 | -0.073 | -0.113 | -0.113 | 0 | 0.78 | 0.98 | 0.63 | 0.7 | 0.79 | 0.71 |
| 6 | 63420430 | rs2752965 | -0.099 | -0.47 | -0.852 | 0.387 | 0.049 | 0.006 | 0.006 | 1.916 | 0.06 | 0.33 | 0.04 | 0.02 | 0.03 | 0.05 |
| 6 | 63455110 | rs9341457 | 0.535 | -0.439 | -0.883 | 0.542 | -0.001 | 0.039 | 0.039 | 0 | 0.1 | 0.22 | 0.14 | 0.22 | 0.11 | 0.04 |
| 6 | 63457733 | rs2474878 | -0.892 | -0.801 | -1.342 | 0.212 | 0.014 | 0.029 | 0.029 | 0 | 0.45 | 0.27 | 0.61 | 0.65 | 0.42 | 0.54 |
| 6 | 63458592 | rs2474879 | -0.404 | -0.862 | -1.373 | 0.229 | 0 | -0.079 | -0.079 | 0 | 0.35 | 0.08 | 0.46 | 0.43 | 0.31 | 0.49 |
| 6 | 63458776 | rs2208329 | -0.346 | -1.093 | -1.32 | 0.221 | 0.004 | -0.005 | -0.005 | 0 | 0.22 | 0.19 | 0.2 | 0.32 | 0.18 | 0.17 |
| 6 | 63460527 | rs9343126 | 0.75 | -0.891 | -1.2 | 0.302 | 0.009 | 0.024 | 0.024 | 0 | 0.1 | 0.18 | 0.15 | 0.22 | 0.13 | 0.04 |
| 6 | 63496191 | rs12207271 | 1.778 | -0.997 | -1.215 | 0.106 | 0.003 | -0.006 | -0.006 | 0 | 0.27 | 0.29 | 0.18 | 0.2 | 0.3 | 0.28 |
| 6 | 63514296 | rs1884934 | 0.397 | -1.152 | -1.235 | 0.062 | 0.041 | -0.008 | -0.008 | 0 | 0.53 | 0.86 | 0.49 | 0.57 | 0.62 | 0.49 |
| 6 | 63518796 | rs1155906 | 0.395 | -1.152 | -1.235 | 0.062 | 0.041 | -0.008 | -0.008 | 0 | 0.53 | 0.86 | 0.49 | 0.57 | 0.62 | 0.49 |
| 6 | 63520742 | rs7746043 | 0.395 | -1.152 | -1.235 | 0.062 | 0.041 | -0.008 | -0.008 | 0 | 0.53 | 0.86 | 0.49 | 0.57 | 0.62 | 0.49 |
| 6 | 63529431 | rs9343253 | 0.313 | -1.154 | -1.241 | 0.057 | 0.003 | 0.007 | 0.007 | 0 | 0.22 | 0.24 | 0.28 | 0.36 | 0.32 | 0.18 |
| 6 | 63534312 | rs504893 | -0.861 | -1.432 | -1.469 | -0.024 | -0.073 | -0.109 | -0.109 | 0.132 | 0.77 | 0.98 | 0.63 | 0.71 | 0.77 | 0.67 |
| 6 | 63543538 | rs4577773 | 0.56 | -1.153 | -1.235 | -0.059 | 0.003 | 0.001 | 0.001 | 0 | 0.49 | 0.54 | 0.46 | 0.55 | 0.62 | 0.45 |
| 6 | 63550561 | rs7741713 | 0.323 | -1.109 | -1.441 | -0.039 | 0.039 | 0.023 | 0.023 | 0 | 0.53 | 0.86 | 0.53 | 0.62 | 0.62 | 0.49 |
| 6 | 63562515 | rs12190145 | -0.048 | -1.119 | -1.33 | -0.101 | 0.016 | -0.005 | -0.005 | 0.683 | 0.34 | 0.15 | 0.37 | 0.31 | 0.24 | 0.46 |
| 6 | 63568603 | rs12524104 | -0.177 | -0.81 | -1.143 | 0.113 | -0.016 | 0.038 | 0.038 | 0 | 0.45 | 0.16 | 0.54 | 0.56 | 0.43 | 0.62 |
| 6 | 63569602 | rs6930829 | -1.158 | -0.731 | -0.915 | 0.118 | -0.001 | -0.001 | -0.001 | 0 | 0.67 | 0.69 | 0.74 | 0.78 | 0.6 | 0.72 |
| 6 | 63573800 | rs12201062 | 0.384 | -0.45 | -0.8 | 0.517 | -0.011 | -0.026 | -0.026 | 0 | 0.21 | 0.13 | 0.33 | 0.3 | 0.2 | 0.44 |
| 6 | 63585140 | rs1577822 | -0.174 | -0.324 | -1.405 | 0.348 | -0.022 | 0.016 | 0.016 | 0 | 0.41 | 0.17 | 0.53 | 0.56 | 0.43 | 0.62 |
| 6 | 63605221 | rs4391244 | -0.226 | -0.242 | -1.399 | 0.432 | -0.029 | 0.015 | 0.015 | 0 | 0.42 | 0.2 | 0.53 | 0.55 | 0.44 | 0.63 |
| 6 | 63610969 | rs10943288 | 0.437 | -0.222 | -1.525 | 0.628 | -0.032 | -0.049 | -0.049 | 0.605 | 0.2 | 0.09 | 0.33 | 0.3 | 0.2 | 0.44 |
| 6 | 63615339 | rs4605861 | 0.603 | -0.24 | -1.563 | 0.587 | -0.018 | -0.059 | -0.059 | 0 | 0.22 | 0.07 | 0.2 | 0.25 | 0.24 | 0.2 |
| 6 | 63617477 | rs6931620 | -1.089 | -0.525 | -1.248 | 0.467 | 0 | -0.011 | -0.011 | 0 | 0.59 | 0.62 | 0.72 | 0.73 | 0.59 | 0.71 |
| 6 | 63630759 | rs6934809 | -0.231 | -0.493 | -1.491 | 0.466 | -0.05 | 0.005 | 0.005 | 0 | 0.42 | 0.18 | 0.54 | 0.52 | 0.44 | 0.63 |
| 6 | 63634264 | rs4501410 | 0.748 | -0.418 | -1.454 | 0.511 | -0.025 | -0.071 | -0.071 | 0.001 | 0.22 | 0.04 | 0.2 | 0.25 | 0.24 | 0.2 |
| 6 | 63649452 | rs12200280 | -0.26 | -0.499 | -1.48 | 0.454 | -0.043 | 0.012 | 0.012 | 0 | 0.43 | 0.19 | 0.54 | 0.52 | 0.45 | 0.64 |
| 6 | 63652377 | rs4285303 | 0.661 | -0.458 | -1.47 | 0.373 | -0.03 | -0.043 | -0.043 | 0.031 | 0.21 | 0.1 | 0.33 | 0.27 | 0.2 | 0.43 |
| 6 | 63658063 | rs6901550 | 0.289 | -0.484 | -1.31 | 0.36 | -0.03 | -0.01 | -0.01 | 0.101 | 0.17 | 0.41 | 0.19 | 0.22 | 0.15 | 0.08 |
| 6 | 63665146 | rs4710390 | 0.367 | -0.102 | -1.125 | 0.375 | 0.005 | -0.063 | -0.063 | 0 | 0.36 | 0.15 | 0.37 | 0.47 | 0.39 | 0.28 |
| 6 | 63668515 | rs6931035 | -0.027 | 0.116 | -0.71 | 0.041 | 0.002 | -0.033 | -0.033 | 0 | 0.6 | 0.58 | 0.73 | 0.75 | 0.59 | 0.71 |

| | | | | | | | | | | | | | | | | |
|---|---|---|---|---|---|---|---|---|---|---|---|---|---|---|---|---|
| 6 | 63684573 | rs4504456 | -0.04 | 0.133 | -0.687 | 0.097 | 0.002 | -0.042 | -0.042 | 0 | 0.61 | 0.53 | 0.73 | 0.75 | 0.62 | 0.71 |
| 6 | 63689527 | rs9352357 | 0.512 | 0.01 | -0.812 | -0.008 | -0.106 | 0.04 | 0.04 | 0 | 0.29 | 0.03 | 0.18 | 0.18 | 0.32 | 0.24 |
| 6 | 63692564 | rs4485997 | -0.181 | 0.229 | -0.689 | 0.13 | 0.002 | -0.036 | -0.036 | 0 | 0.61 | 0.57 | 0.73 | 0.75 | 0.62 | 0.71 |
| 6 | 63702477 | rs4289625 | -0.121 | 0.162 | -0.632 | 0.199 | -0.002 | -0.028 | -0.028 | 3.22 | 0.61 | 0.51 | 0.75 | 0.75 | 0.63 | 0.72 |
| 6 | 63710146 | rs6453938 | -0.14 | 0.2 | -0.677 | 0.218 | -0.001 | -0.019 | -0.019 | 0.01 | 0.61 | 0.5 | 0.74 | 0.75 | 0.63 | 0.72 |
| 6 | 63733495 | rs9447939 | -0.192 | 0.211 | -0.658 | 0.224 | -0.001 | -0.035 | -0.035 | 0 | 0.62 | 0.58 | 0.75 | 0.75 | 0.63 | 0.72 |
| 6 | 63768896 | rs9350707 | 0.254 | 0.212 | -0.767 | 0.095 | -0.003 | -0.069 | -0.069 | 0 | 0.33 | 0.42 | 0.18 | 0.17 | 0.31 | 0.24 |
| 6 | 63771225 | rs9294075 | 1.018 | 0.026 | -0.711 | 0.099 | -0.004 | 0 | 0 | 0.428 | 0.94 | 0.91 | 0.95 | 0.92 | 0.96 | 0.94 |
| 6 | 63774385 | rs1341185 | 0.92 | 0.023 | -0.713 | 0.097 | -0.004 | 0 | 0 | 0 | 0.95 | 0.92 | 0.95 | 0.94 | 0.96 | 0.96 |
| 6 | 63803188 | rs789942 | 0.155 | 0.251 | -0.951 | 0.449 | -0.012 | -0.015 | -0.015 | 0 | 0.73 | 0.82 | 0.62 | 0.67 | 0.73 | 0.54 |
| 6 | 63806114 | rs12214367 | 0.716 | 0.66 | -1.377 | 1.042 | -0.056 | -0.05 | -0.05 | 0 | 0.09 | 0.01 | 0.24 | 0.16 | 0.13 | 0.25 |
| 6 | 63811569 | rs1841228 | 0.906 | 0.433 | -1.36 | 0.825 | -0.006 | -0.066 | -0.066 | 0 | 0.34 | 0.41 | 0.19 | 0.16 | 0.31 | 0.25 |
| 6 | 63836640 | rs1417689 | 0.688 | -0.08 | -1.672 | 0.623 | -0.064 | -0.113 | -0.113 | 0.054 | 0.24 | 0.02 | 0.26 | 0.3 | 0.31 | 0.22 |
| 6 | 63862923 | rs9352587 | -1.016 | -0.127 | -1.307 | 0.295 | -0.002 | -0.039 | -0.039 | 0 | 0.66 | 0.55 | 0.76 | 0.77 | 0.67 | 0.73 |
| 6 | 63863434 | rs9359325 | -0.921 | -0.119 | -1.283 | 0.299 | -0.004 | -0.027 | -0.027 | 0 | 0.64 | 0.51 | 0.76 | 0.77 | 0.67 | 0.72 |
| 6 | 63865624 | rs9343759 | -0.944 | -0.114 | -1.28 | 0.302 | -0.004 | -0.027 | -0.027 | 0 | 0.64 | 0.5 | 0.76 | 0.77 | 0.67 | 0.72 |
| 6 | 63871745 | rs3843509 | -0.955 | -0.13 | -1.288 | 0.293 | -0.004 | -0.027 | -0.027 | 0 | 0.65 | 0.5 | 0.76 | 0.77 | 0.67 | 0.72 |
| 6 | 63883378 | rs1948846 | -0.877 | 0.055 | -1.324 | 0.254 | -0.005 | -0.028 | -0.028 | 0 | 0.63 | 0.5 | 0.74 | 0.78 | 0.66 | 0.72 |
| 6 | 63886188 | rs9359331 | 0.717 | 0.088 | -1.268 | 0.584 | 0 | -0.005 | -0.005 | 0.045 | 0.47 | 0.46 | 0.49 | 0.27 | 0.44 | 0.45 |
| 6 | 63888126 | rs1585367 | 0.18 | -0.026 | -1.369 | 0.59 | -0.001 | 0 | 0 | 0.054 | 0.67 | 0.67 | 0.64 | 0.35 | 0.66 | 0.7 |
| 6 | 63893492 | rs9294122 | 0.635 | 0.2 | -1.585 | 0.981 | -0.06 | -0.112 | -0.112 | 0 | 0.19 | 0.02 | 0.28 | 0.36 | 0.28 | 0.21 |
| 6 | 63899051 | rs7754440 | 0.762 | 0.075 | -0.956 | 0.828 | 0 | 0 | 0 | 0 | 0.53 | 0.5 | 0.52 | 0.46 | 0.47 | 0.53 |
| 6 | 63900708 | rs7739721 | 0.763 | 0.056 | -0.914 | 0.829 | -0.022 | -0.078 | -0.078 | 0 | 0.22 | 0.06 | 0.28 | 0.36 | 0.26 | 0.21 |
| 6 | 63902525 | rs9359340 | 0.898 | 0.052 | -0.932 | 0.825 | 0.007 | 0.01 | 0.01 | 0 | 0.52 | 0.45 | 0.51 | 0.44 | 0.47 | 0.49 |
| 6 | 63903384 | rs2047899 | -0.189 | 0.039 | -0.907 | 0.822 | -0.004 | -0.011 | -0.011 | 0 | 0.22 | 0.08 | 0.18 | 0.18 | 0.26 | 0.25 |
| 6 | 63908900 | rs9361439 | 0.532 | 0.148 | -0.843 | 0.741 | 0 | 0 | 0 | 0 | 0.53 | 0.5 | 0.52 | 0.47 | 0.47 | 0.53 |
| 6 | 63910966 | rs12213121 | 1.139 | 0.262 | -0.892 | 0.566 | 0 | 0.025 | 0.025 | 0 | 0.41 | 0.37 | 0.23 | 0.3 | 0.35 | 0.27 |
| 6 | 63921853 | rs9350793 | 0.137 | 0.24 | -0.9 | 0.601 | -0.008 | 0.004 | 0.004 | 0 | 0.6 | 0.5 | 0.59 | 0.47 | 0.59 | 0.56 |
| 6 | 63922282 | rs9361465 | 0.486 | 0.352 | -0.962 | 0.561 | -0.049 | -0.047 | -0.047 | 0 | 0.1 | 0.01 | 0.28 | 0.15 | 0.12 | 0.22 |
| 6 | 63933079 | rs1870488 | -0.628 | 0.199 | -0.661 | 0.689 | 0 | 0.067 | 0.067 | 0 | 0.76 | 0.6 | 0.7 | 0.65 | 0.73 | 0.79 |
| 6 | 63936511 | rs9343864 | 0.188 | 0.122 | -0.81 | 0.571 | -0.035 | -0.021 | -0.021 | 0.023 | 0.13 | 0.04 | 0.29 | 0.15 | 0.16 | 0.22 |
| 6 | 63948305 | rs16894193 | -0.033 | 0.115 | -0.843 | 0.472 | 0.001 | 0.001 | 0.001 | 0 | 0.23 | 0.09 | 0.18 | 0.18 | 0.26 | 0.26 |
| 6 | 63951293 | rs6454102 | 1.5 | 0.032 | -0.769 | 0.45 | 0 | 0.005 | 0.005 | 0 | 0.38 | 0.4 | 0.23 | 0.31 | 0.32 | 0.27 |
| 6 | 63954459 | rs6915091 | 0.197 | 0.173 | -0.765 | 0.559 | -0.005 | 0.017 | 0.017 | 0 | 0.2 | 0.06 | 0.16 | 0.11 | 0.24 | 0.22 |
| 6 | 63958597 | rs9294140 | -0.437 | 0.077 | -0.6 | 0.594 | 0.005 | 0.021 | 0.021 | 0 | 0.72 | 0.64 | 0.65 | 0.65 | 0.64 | 0.78 |
| 6 | 63965416 | rs9352719 | 0.136 | 0.193 | -0.351 | 0.609 | -0.004 | 0.002 | 0.002 | 0 | 0.56 | 0.53 | 0.53 | 0.5 | 0.5 | 0.58 |
| 6 | 63967017 | rs6922743 | -0.101 | 0.326 | -0.175 | 0.415 | -0.001 | -0.002 | -0.002 | 0.176 | 0.55 | 0.48 | 0.55 | 0.55 | 0.49 | 0.6 |
| 6 | 63971954 | rs10943643 | 0.589 | 0.271 | -0.239 | 0.379 | 0.013 | 0.02 | 0.02 | 0 | 0.45 | 0.39 | 0.28 | 0.38 | 0.37 | 0.37 |
| 6 | 63981311 | rs9448732 | -0.125 | 0.334 | -0.668 | 0.428 | -0.008 | -0.016 | -0.016 | 1.63 | 0.57 | 0.69 | 0.6 | 0.55 | 0.49 | 0.6 |
| 6 | 63986735 | rs1320102 | -0.016 | 0.199 | -0.648 | 0.328 | 0 | -0.008 | -0.008 | 0 | 0.45 | 0.36 | 0.39 | 0.45 | 0.5 | 0.38 |
| 6 | 63988362 | rs1711932 | -0.659 | 0.212 | -0.698 | 0.235 | -0.009 | -0.017 | -0.017 | 0.119 | 0.58 | 0.48 | 0.73 | 0.67 | 0.66 | 0.7 |
| 6 | 63989264 | rs10943652 | 0.728 | 0.365 | -0.834 | 0.332 | -0.042 | -0.044 | -0.044 | 0 | 0.09 | 0.02 | 0.3 | 0.17 | 0.12 | 0.22 |
| 6 | 63994473 | rs6454130 | -0.675 | 0.182 | -0.805 | 0.219 | -0.005 | -0.01 | -0.01 | 0 | 0.58 | 0.46 | 0.72 | 0.68 | 0.66 | 0.7 |
| 6 | 64003459 | rs9343928 | 0.539 | 0.426 | -0.674 | 0.437 | -0.043 | -0.04 | -0.04 | 0.232 | 0.08 | 0.02 | 0.3 | 0.15 | 0.13 | 0.22 |
| 6 | 64005904 | rs9361556 | 0.541 | 0.415 | -0.679 | 0.463 | -0.043 | -0.041 | -0.041 | 0 | 0.08 | 0.02 | 0.3 | 0.16 | 0.13 | 0.22 |
| 6 | 64014812 | rs9352774 | 0.954 | 0.507 | -0.682 | 0.455 | -0.038 | -0.038 | -0.038 | 1.926 | 0.06 | 0.01 | 0.29 | 0.16 | 0.1 | 0.17 |
| 6 | 64024024 | rs2459569 | -0.865 | 0.347 | -0.492 | 0.387 | 0.012 | 0.013 | 0.013 | 0.988 | 0.17 | 0.28 | 0.35 | 0.21 | 0.18 | 0.3 |
| 6 | 64033048 | rs824372 | -0.288 | 0.137 | -0.518 | 0.451 | -0.015 | 0.023 | 0.023 | 3.653 | 0.1 | 0.19 | 0.05 | 0.18 | 0.06 | 0.11 |
| 6 | 64058565 | rs1741797 | -1.16 | 0.095 | -0.979 | 0.189 | -0.099 | 0.004 | 0.004 | 1.198 | 0.72 | 0.96 | 0.77 | 0.82 | 0.68 | 0.77 |
| 6 | 64078347 | rs12207361 | 0.65 | 0.071 | -0.823 | 0.281 | -0.044 | 0.033 | 0.033 | 0 | 0.23 | 0.02 | 0.12 | 0.12 | 0.28 | 0.26 |
| 6 | 64081621 | rs1779757 | -0.645 | 0.068 | 0.137 | 0.688 | -0.005 | 0.041 | 0.041 | 1.552 | 0.1 | 0.42 | 0.06 | 0.11 | 0.03 | 0.1 |
| 6 | 64084805 | rs2677024 | -0.651 | 0.085 | 0.154 | 0.701 | -0.005 | 0.044 | 0.044 | 1.693 | 0.1 | 0.38 | 0.06 | 0.11 | 0.03 | 0.08 |
| 6 | 64087275 | rs9344015 | 0.658 | 0.305 | -0.301 | 1.003 | -0.018 | -0.043 | -0.043 | 0.274 | 0.08 | 0.01 | 0.13 | 0.27 | 0.08 | 0.12 |
| 6 | 64094892 | rs1525361 | -1.336 | 0.405 | 0.379 | 0.359 | -0.03 | -0.052 | -0.052 | 1.525 | 0.15 | 0.51 | 0.1 | 0.08 | 0.07 | 0.15 |
| 6 | 64099010 | rs1525357 | 0.077 | 0.388 | 0.511 | 0.354 | -0.015 | -0.003 | -0.003 | 0 | 0.08 | 0.25 | 0.06 | 0.07 | 0.03 | 0.07 |
| 6 | 64101818 | rs1723527 | -1.471 | 0.447 | 0.52 | 0.35 | -0.022 | -0.053 | -0.053 | 0 | 0.15 | 0.52 | 0.1 | 0.08 | 0.08 | 0.15 |
| 6 | 64121251 | rs1010057 | -0.559 | 0.44 | 0.416 | 0.466 | -0.018 | -0.005 | -0.005 | 0 | 0.1 | 0.34 | 0.07 | 0.08 | 0.03 | 0.1 |
| 6 | 64136886 | rs1723535 | -0.304 | 0.736 | 0.322 | 0.423 | -0.018 | -0.005 | -0.005 | 0 | 0.6 | 0.66 | 0.5 | 0.61 | 0.59 | 0.6 |
| 6 | 64139997 | rs4445042 | 0.169 | 1.003 | 0.559 | 0.531 | 0.004 | 0.004 | 0.004 | 0 | 0.5 | 0.26 | 0.42 | 0.54 | 0.57 | 0.5 |
| 6 | 64159064 | rs9361779 | 0.235 | 0.86 | 0.703 | 0.546 | -0.012 | -0.012 | -0.012 | 0 | 0.51 | 0.19 | 0.42 | 0.53 | 0.57 | 0.49 |
| 6 | 64169534 | rs9449312 | -0.222 | 0.83 | 1.136 | 0.523 | 0.001 | 0.001 | 0.001 | 0 | 0.55 | 0.47 | 0.45 | 0.53 | 0.58 | 0.53 |
| 6 | 64186032 | rs9353016 | 0.275 | 0.848 | 1.106 | 0.484 | -0.008 | 0.014 | 0.014 | 0 | 0.5 | 0.19 | 0.42 | 0.55 | 0.58 | 0.49 |
| 6 | 64240516 | rs2622274 | -0.017 | 0.874 | 1.246 | 0.494 | 0.001 | -0.001 | -0.001 | 0.097 | 0.46 | 0.52 | 0.56 | 0.45 | 0.42 | 0.52 |
| 6 | 64273105 | rs1744156 | -0.464 | 1.003 | 1.312 | 0.645 | 0.029 | 0.102 | 0.102 | 0 | 0.07 | 0.26 | 0.11 | 0.15 | 0.06 | 0.12 |
| 6 | 64292153 | rs1062193 | -0.869 | 0.948 | 1.3 | 0.596 | 0.015 | 0.074 | 0.074 | 0 | 0.08 | 0.25 | 0.11 | 0.15 | 0.06 | 0.17 |
| 6 | 64295122 | rs13213141 | 0.579 | 1.074 | 1.354 | 0.657 | -0.012 | 0.045 | 0.045 | 0 | 0.35 | 0.2 | 0.43 | 0.26 | 0.31 | 0.35 |
| 6 | 64300240 | rs6929720 | -0.124 | 0.922 | 1.209 | 0.569 | 0.02 | 0.084 | 0.084 | 0 | 0.06 | 0.18 | 0.1 | 0.15 | 0.06 | 0.12 |
| 6 | 64302604 | rs1210839 | -0.408 | 0.922 | 1.191 | 0.569 | 0.033 | 0.113 | 0.113 | 0 | 0.07 | 0.27 | 0.11 | 0.15 | 0.06 | 0.12 |
| 6 | 64332103 | rs12523702 | 0.258 | 0.955 | 1.223 | 0.553 | 0.002 | -0.016 | -0.016 | 0 | 0.4 | 0.52 | 0.47 | 0.3 | 0.36 | 0.39 |

| | | | | | | | | | | | | | | | | |
|---|---|---|---|---|---|---|---|---|---|---|---|---|---|---|---|---|
| 6 | 64353020 | rs1681939 | -0.763 | 1.094 | 1.276 | 0.627 | 0.013 | 0.023 | 0.023 | 0 | 0.1 | 0.22 | 0.15 | 0.17 | 0.1 | 0.2 |
| 6 | 64355159 | rs7767473 | -0.383 | 1.062 | 1.491 | 0.484 | 0 | 0.01 | 0.01 | 0 | 0.49 | 0.71 | 0.59 | 0.46 | 0.41 | 0.53 |
| 6 | 64355587 | rs2225595 | -1.098 | 1.06 | 1.061 | 0.331 | -0.069 | 0.057 | 0.057 | 0 | 0.79 | 0.98 | 0.81 | 0.88 | 0.71 | 0.81 |
| 6 | 64368449 | rs4710256 | 0.729 | 1.042 | 1.244 | 0.343 | -0.001 | -0.001 | -0.001 | 0.552 | 0.39 | 0.37 | 0.43 | 0.29 | 0.32 | 0.34 |
| 6 | 64386060 | rs10485358 | -0.156 | 1.091 | 1.162 | 0.325 | -0.013 | 0.001 | 0.001 | 0 | 0.51 | 0.81 | 0.62 | 0.45 | 0.44 | 0.57 |
| 6 | 64427095 | rs1057530 | -0.138 | 1.085 | 1.184 | 0.298 | -0.011 | 0.007 | 0.007 | 0 | 0.51 | 0.76 | 0.61 | 0.46 | 0.44 | 0.57 |
| 6 | 64468738 | rs2151317 | -1.07 | 1.038 | 0.854 | 0.265 | -0.068 | 0.062 | 0.062 | 0 | 0.78 | 0.98 | 0.81 | 0.88 | 0.71 | 0.78 |
| 6 | 64470069 | rs13216525 | 0.618 | 0.978 | 1.496 | 0.498 | 0.004 | 0.013 | 0.013 | 0 | 0.35 | 0.17 | 0.39 | 0.3 | 0.28 | 0.3 |
| 6 | 64487247 | rs319924 | -0.579 | 1.035 | 1.236 | 0.598 | 0.012 | 0.011 | 0.011 | 0 | 0.5 | 0.79 | 0.59 | 0.47 | 0.39 | 0.53 |
| 6 | 64505213 | rs756274 | 0.055 | 1.092 | 1.338 | 0.606 | -0.004 | -0.014 | -0.014 | 0.552 | 0.42 | 0.51 | 0.46 | 0.3 | 0.35 | 0.35 |
| 6 | 64521068 | rs7762059 | -0.147 | 0.817 | 1.175 | 0.72 | 0.004 | 0.008 | 0.008 | 0 | 0.44 | 0.6 | 0.47 | 0.31 | 0.32 | 0.46 |
| 6 | 64547607 | rs12110344 | -0.229 | 0.791 | 1.125 | 0.771 | 0.064 | 0.126 | 0.126 | 0 | 0.06 | 0.2 | 0.13 | 0.15 | 0.07 | 0.07 |
| 6 | 64579224 | rs321498 | 0.176 | 0.701 | 0.748 | 0.426 | -0.004 | -0.01 | -0.01 | 0 | 0.56 | 0.42 | 0.53 | 0.69 | 0.67 | 0.53 |
| 6 | 64589465 | rs1482463 | -0.208 | 0.765 | 0.458 | 0.46 | 0.001 | 0.006 | 0.006 | 0 | 0.43 | 0.45 | 0.46 | 0.31 | 0.33 | 0.46 |
| 6 | 64610885 | rs12207884 | 0.024 | 0.307 | 0.277 | 0.667 | 0.004 | -0.004 | -0.004 | 0.246 | 0.32 | 0.28 | 0.4 | 0.34 | 0.26 | 0.26 |
| 6 | 64616024 | rs2200105 | -0.7 | 0.628 | 0.654 | 0.787 | 0.012 | 0.006 | 0.006 | 0 | 0.1 | 0.15 | 0.08 | 0.1 | 0.11 | 0.2 |
| 6 | 64618911 | rs12529599 | 0.032 | 0.327 | 0.507 | 0.895 | 0.004 | -0.005 | -0.005 | 0 | 0.34 | 0.3 | 0.4 | 0.37 | 0.26 | 0.32 |
| 6 | 64619150 | rs6912246 | -0.848 | 0.127 | 0.927 | 0.757 | 0.032 | -0.001 | -0.001 | 0 | 0.48 | 0.74 | 0.5 | 0.47 | 0.38 | 0.52 |
| 6 | 64625328 | rs452975 | -0.212 | 0.075 | 0.882 | 0.937 | 0.002 | -0.004 | -0.004 | 0.467 | 0.67 | 0.7 | 0.6 | 0.63 | 0.74 | 0.73 |
| 6 | 64646239 | rs1482458 | -0.807 | 0.007 | 1.002 | 0.654 | 0.03 | -0.008 | -0.008 | 0 | 0.46 | 0.73 | 0.5 | 0.42 | 0.38 | 0.47 |
| 6 | 64648161 | rs9353368 | -0.17 | 0.047 | 1.05 | 0.734 | 0.001 | 0.001 | 0.001 | 0.56 | 0.34 | 0.33 | 0.43 | 0.27 | 0.3 | 0.23 |
| 6 | 64662711 | rs4710454 | -0.103 | 0.074 | 0.94 | 0.748 | 0.007 | 0.003 | 0.003 | 0.956 | 0.34 | 0.27 | 0.42 | 0.27 | 0.3 | 0.24 |
| 6 | 64680963 | rs6936802 | -0.279 | 0.175 | 0.845 | 0.628 | 0.005 | 0.007 | 0.007 | 3.465 | 0.37 | 0.33 | 0.24 | 0.27 | 0.33 | 0.31 |
| 6 | 64693014 | rs6934646 | -0.226 | 0.203 | 1.069 | 0.744 | -0.003 | 0.001 | 0.001 | 3.867 | 0.37 | 0.37 | 0.26 | 0.33 | 0.34 | 0.38 |
| 6 | 64695023 | rs12529059 | 0.926 | 0.418 | 0.494 | 0.411 | -0.063 | 0.098 | 0.098 | 0.024 | 0.25 | 0.01 | 0.21 | 0.1 | 0.24 | 0.28 |
| 6 | 64697821 | rs1482455 | -0.597 | 0.333 | 1.413 | 0.88 | -0.009 | 0.006 | 0.006 | 2.308 | 0.39 | 0.5 | 0.27 | 0.37 | 0.34 | 0.39 |
| 6 | 64712730 | rs1038317 | -0.232 | 0.393 | 1.141 | 0.888 | -0.003 | 0.042 | 0.042 | 2.787 | 0.35 | 0.11 | 0.24 | 0.22 | 0.33 | 0.3 |
| 6 | 64712869 | rs1038316 | -0.631 | 0.389 | 1.311 | 0.897 | -0.007 | -0.002 | -0.002 | 0 | 0.4 | 0.53 | 0.27 | 0.33 | 0.34 | 0.35 |
| 6 | 64721056 | rs9342097 | -0.393 | 0.447 | 0.783 | 0.696 | 0.009 | 0.009 | 0.009 | 2.532 | 0.38 | 0.26 | 0.25 | 0.3 | 0.34 | 0.39 |
| 6 | 64728485 | rs9353437 | -0.156 | 0.516 | 0.414 | 0.647 | 0.014 | 0.038 | 0.038 | 4.039 | 0.34 | 0.1 | 0.22 | 0.22 | 0.31 | 0.32 |
| 6 | 64731257 | rs9344659 | -0.195 | 0.428 | 0.637 | 0.706 | 0.006 | 0.011 | 0.011 | 2.605 | 0.36 | 0.3 | 0.24 | 0.26 | 0.34 | 0.28 |
| 6 | 64758787 | rs7752934 | -0.657 | 0.38 | 0.927 | 0.828 | -0.018 | -0.004 | -0.004 | 0 | 0.35 | 0.48 | 0.22 | 0.29 | 0.27 | 0.29 |
| 6 | 64762195 | rs9351126 | 0.044 | 0.441 | 1.012 | 0.792 | 0.014 | 0.001 | 0.001 | 0.146 | 0.27 | 0.12 | 0.18 | 0.22 | 0.2 | 0.23 |
| 6 | 64769816 | rs3013156 | -1.909 | 0.476 | 1.164 | 0.461 | 0.039 | 0.076 | 0.076 | 1.958 | 0.06 | 0.35 | 0.03 | 0.07 | 0.04 | 0.04 |
| 6 | 64771342 | rs3003684 | -1.726 | 0.478 | 1.195 | 0.462 | 0.03 | 0.064 | 0.064 | 1.187 | 0.06 | 0.31 | 0.03 | 0.07 | 0.04 | 0.04 |
| 6 | 64789100 | rs3013148 | -0.784 | 0.359 | 1.587 | 1.197 | 0.024 | 0.006 | 0.006 | 1.814 | 0.37 | 0.75 | 0.37 | 0.56 | 0.3 | 0.33 |
| 6 | 64791408 | rs9342117 | 0.257 | 0.243 | 1.439 | 1.158 | -0.067 | -0.169 | -0.169 | 0 | 0.24 | 0.01 | 0.27 | 0.41 | 0.24 | 0.27 |
| 6 | 64850803 | rs4573063 | 0.893 | 0.59 | 0.812 | 0.405 | -0.088 | 0.226 | 0.226 | 0.296 | 0.38 | 0.03 | 0.24 | 0.1 | 0.39 | 0.42 |
| 6 | 64851149 | rs4401640 | -0.202 | 0.273 | 0.973 | 0.679 | -0.013 | 0.021 | 0.021 | 1.917 | 0.35 | 0.49 | 0.33 | 0.51 | 0.31 | 0.3 |
| 6 | 64885858 | rs13205132 | 0.19 | 0.295 | 1.028 | 0.684 | -0.004 | -0.088 | -0.088 | 9.194 | 0.3 | 0.17 | 0.29 | 0.42 | 0.29 | 0.25 |
| 6 | 64896909 | rs4326220 | 0.374 | 0.286 | 1.019 | 0.676 | -0.018 | -0.109 | -0.109 | 0.182 | 0.27 | 0.1 | 0.29 | 0.42 | 0.28 | 0.24 |
| 6 | 64907665 | rs6907750 | 0.209 | 0.246 | 1.183 | 0.674 | 0.007 | -0.003 | -0.003 | 0 | 0.64 | 0.32 | 0.63 | 0.48 | 0.69 | 0.66 |
| 6 | 64920494 | rs11757626 | -0.129 | 0.268 | 1.172 | 0.688 | 0.009 | -0.026 | -0.026 | 0 | 0.32 | 0.27 | 0.34 | 0.49 | 0.29 | 0.28 |
| 6 | 64927816 | rs11753432 | -0.395 | 0.257 | 1.167 | 0.651 | -0.005 | 0.003 | 0.003 | 6.251 | 0.35 | 0.43 | 0.36 | 0.52 | 0.31 | 0.33 |
| 6 | 64942181 | rs6916780 | 0.262 | 0.269 | 1.149 | 0.657 | 0.004 | -0.002 | -0.002 | 0 | 0.64 | 0.33 | 0.62 | 0.48 | 0.69 | 0.66 |
| 6 | 64946169 | rs9451166 | -0.145 | 0.484 | 0.818 | 0.452 | 0.011 | 0.002 | 0.002 | 0 | 0.69 | 0.46 | 0.55 | 0.62 | 0.62 | 0.59 |
| 6 | 64977136 | rs12190681 | -0.269 | 0.624 | 0.183 | 0.654 | -0.001 | -0.001 | -0.001 | 0.068 | 0.7 | 0.7 | 0.67 | 0.42 | 0.72 | 0.73 |
| 6 | 64983718 | rs10806410 | -0.205 | 0.663 | 0.341 | 0.414 | -0.007 | 0.015 | 0.015 | 1.645 | 0.68 | 0.55 | 0.65 | 0.5 | 0.71 | 0.73 |
| 6 | 64984651 | rs9353681 | 0.539 | 1.026 | 0.145 | 0.288 | -0.074 | -0.083 | -0.083 | 0 | 0.23 | 0.01 | 0.38 | 0.33 | 0.27 | 0.21 |
| 6 | 65000938 | rs6454771 | -0.724 | 0.928 | 0.26 | 0.438 | -0.029 | -0.131 | -0.131 | 0 | 0.73 | 0.93 | 0.7 | 0.58 | 0.72 | 0.75 |
| 6 | 65007530 | rs6917544 | 1.104 | 0.974 | 0.193 | 0.348 | -0.047 | 0.195 | 0.195 | 0.107 | 0.35 | 0.03 | 0.2 | 0.11 | 0.32 | 0.38 |
| 6 | 65012333 | rs6454782 | 0.037 | 0.992 | 0.942 | 1.225 | -0.001 | 0 | 0 | 2.033 | 0.7 | 0.69 | 0.67 | 0.5 | 0.72 | 0.73 |
| 6 | 65027989 | rs7770439 | 0.298 | 0.985 | 0.945 | 1.221 | -0.005 | 0.009 | 0.009 | 3.478 | 0.68 | 0.6 | 0.66 | 0.5 | 0.71 | 0.73 |
| 6 | 65028091 | rs7771804 | 0.014 | 1.019 | 0.966 | 1.235 | -0.001 | -0.041 | -0.041 | 0 | 0.71 | 0.82 | 0.67 | 0.5 | 0.72 | 0.73 |
| 6 | 65031761 | rs10944471 | 0.014 | 1.019 | 0.966 | 1.235 | -0.001 | -0.041 | -0.041 | 0 | 0.71 | 0.82 | 0.67 | 0.5 | 0.72 | 0.73 |
| 6 | 65031922 | rs6927405 | 0.689 | 1.02 | 1.01 | 1.233 | 0.001 | -0.001 | -0.001 | 0.089 | 0.63 | 0.35 | 0.63 | 0.47 | 0.69 | 0.66 |
| 6 | 65056811 | rs593493 | -0.183 | 1.032 | 0.988 | 1.241 | -0.03 | -0.125 | -0.125 | 0 | 0.73 | 0.92 | 0.7 | 0.57 | 0.72 | 0.75 |
| 6 | 65065008 | rs1922945 | -0.02 | 0.99 | 0.962 | 1.063 | -0.001 | -0.041 | -0.041 | 0.116 | 0.71 | 0.82 | 0.67 | 0.5 | 0.72 | 0.73 |
| 6 | 65067404 | rs10944484 | 0.336 | 1.067 | 0.881 | 0.992 | -0.059 | -0.077 | -0.077 | 0 | 0.11 | 0.02 | 0.33 | 0.32 | 0.19 | 0.08 |
| 6 | 65074128 | rs673676 | 0.377 | 0.538 | 1.59 | 1.419 | -0.001 | 0 | 0 | 2.033 | 0.7 | 0.69 | 0.67 | 0.5 | 0.72 | 0.73 |
| 6 | 65075236 | rs689092 | 0.377 | 0.538 | 1.59 | 1.419 | -0.001 | 0 | 0 | 2.033 | 0.7 | 0.69 | 0.67 | 0.5 | 0.72 | 0.73 |
| 6 | 65081343 | rs683901 | 0.352 | 0.538 | 1.591 | 1.419 | -0.001 | -0.041 | -0.041 | 2.551 | 0.71 | 0.82 | 0.67 | 0.5 | 0.72 | 0.73 |
| 6 | 65084617 | rs4518475 | 0.608 | 0.537 | 1.598 | 1.418 | -0.005 | 0.009 | 0.009 | 3.762 | 0.68 | 0.6 | 0.66 | 0.5 | 0.71 | 0.73 |
| 6 | 65085180 | rs607416 | 0.352 | 0.538 | 1.481 | 1.419 | -0.001 | -0.041 | -0.041 | 0 | 0.71 | 0.82 | 0.67 | 0.5 | 0.72 | 0.73 |
| 6 | 65085510 | rs609098 | 0.864 | 0.538 | 1.489 | 1.419 | -0.012 | 0.015 | 0.015 | 3.425 | 0.66 | 0.45 | 0.64 | 0.5 | 0.71 | 0.72 |
| 6 | 65092688 | rs640941 | 0.375 | 0.556 | 1.355 | 1.193 | -0.001 | 0.003 | 0.003 | 1.668 | 0.7 | 0.7 | 0.69 | 0.54 | 0.72 | 0.73 |
| 6 | 65125567 | rs16895190 | -0.529 | 0.631 | 1.06 | 1.192 | 0.015 | -0.064 | -0.064 | 0 | 0.28 | 0.18 | 0.28 | 0.46 | 0.25 | 0.27 |
| 6 | 65143008 | rs9444828 | -1.094 | 0.52 | 0.93 | 1.204 | 0.002 | -0.004 | -0.004 | 0.926 | 0.37 | 0.66 | 0.34 | 0.49 | 0.31 | 0.33 |
| 6 | 65171419 | rs9351261 | -0.934 | 0.425 | 0.241 | 1.326 | -0.002 | -0.036 | -0.036 | 0 | 0.35 | 0.24 | 0.34 | 0.44 | 0.38 | 0.35 |

| | | | | | | | | | | | | | | | | |
|---|---|---|---|---|---|---|---|---|---|---|---|---|---|---|---|---|
| 6 | 65181907 | rs9362879 | -0.684 | 0.262 | 0.311 | 1.351 | 0.006 | -0.047 | -0.047 | 0.464 | 0.31 | 0.22 | 0.33 | 0.49 | 0.33 | 0.31 |
| 6 | 65187445 | rs4142181 | -0.684 | 0.262 | 0.311 | 1.351 | 0.006 | -0.047 | -0.047 | 0.464 | 0.31 | 0.22 | 0.33 | 0.49 | 0.33 | 0.31 |
| 6 | 65199887 | rs13193668 | -0.604 | 0.266 | 0.309 | 1.369 | 0.007 | -0.052 | -0.052 | 0 | 0.3 | 0.22 | 0.32 | 0.46 | 0.32 | 0.26 |
| 6 | 65202546 | rs973428 | -1.049 | 0.197 | 0.806 | 1.3 | -0.014 | 0.034 | 0.034 | 1.269 | 0.34 | 0.5 | 0.36 | 0.49 | 0.34 | 0.31 |
| 6 | 65243245 | rs2993117 | -1.745 | 0.441 | 1.022 | 1.275 | 0.033 | 0.176 | 0.176 | 3.432 | 0.08 | 0.51 | 0.07 | 0.13 | 0.03 | 0.1 |
| 6 | 65245009 | rs2050623 | -0.811 | 0.482 | 0.415 | 1.28 | -0.009 | 0.018 | 0.018 | 0.02 | 0.24 | 0.39 | 0.27 | 0.28 | 0.21 | 0.3 |
| 6 | 65263122 | rs12205418 | -0.344 | 0.932 | 0.252 | 1.212 | -0.002 | -0.007 | -0.007 | 0 | 0.57 | 0.53 | 0.61 | 0.4 | 0.54 | 0.65 |
| 6 | 65278124 | rs17403480 | 0.125 | 1.52 | 0.338 | 1.252 | -0.019 | 0.017 | 0.017 | 0 | 0.06 | 0.03 | 0.05 | 0.02 | 0.13 | 0.1 |
| 6 | 65289746 | rs6928743 | -0.772 | 1.773 | 0.498 | 1.583 | 0.014 | 0.013 | 0.013 | 5.925 | 0.64 | 0.75 | 0.77 | 0.65 | 0.65 | 0.78 |
| 6 | 65294438 | rs12196601 | 0.684 | 1.593 | 0.365 | 1.203 | -0.021 | -0.048 | -0.048 | 0 | 0.26 | 0.09 | 0.39 | 0.31 | 0.23 | 0.29 |
| 6 | 65316377 | rs1415534 | -0.348 | 1.529 | 1.119 | 1.458 | 0 | -0.008 | -0.008 | 0 | 0.58 | 0.55 | 0.64 | 0.39 | 0.54 | 0.65 |
| 6 | 65317295 | rs953034 | -0.452 | 1.452 | 0.87 | 1.454 | -0.015 | -0.046 | -0.046 | 1.618 | 0.61 | 0.83 | 0.66 | 0.48 | 0.55 | 0.66 |
| 6 | 65318362 | rs9445030 | -0.16 | 1.571 | 0.999 | 1.303 | -0.006 | 0.016 | 0.016 | 3.293 | 0.54 | 0.46 | 0.6 | 0.37 | 0.49 | 0.66 |
| 6 | 65328120 | rs9445043 | 0.047 | 1.39 | 0.88 | 1.607 | -0.016 | -0.058 | -0.058 | 0 | 0.34 | 0.18 | 0.31 | 0.5 | 0.39 | 0.32 |
| 6 | 65334528 | rs6912039 | -0.436 | 1.341 | 1.011 | 1.628 | -0.017 | 0.024 | 0.024 | 5.684 | 0.68 | 0.62 | 0.79 | 0.56 | 0.72 | 0.81 |
| 6 | 65337075 | rs9353980 | -0.02 | 1.375 | 1.013 | 1.624 | -0.001 | 0.001 | 0.001 | 3.612 | 0.63 | 0.61 | 0.68 | 0.41 | 0.6 | 0.67 |
| 6 | 65354108 | rs2184404 | 0.556 | 1.598 | 0.711 | 1.453 | 0.001 | 0.025 | 0.025 | 1.261 | 0.52 | 0.42 | 0.61 | 0.36 | 0.45 | 0.48 |
| 6 | 65365420 | rs4710483 | 0.202 | 0.989 | 0.842 | 1.811 | -0.004 | -0.003 | -0.003 | 1.748 | 0.62 | 0.5 | 0.67 | 0.39 | 0.6 | 0.65 |
| 6 | 65376653 | rs16895644 | 0.262 | 0.563 | -0.098 | 1.737 | 0.007 | -0.081 | -0.081 | 0.74 | 0.28 | 0.05 | 0.18 | 0.34 | 0.26 | 0.18 |
| 6 | 65392942 | rs2055173 | 0.986 | 0.462 | 0.77 | 2.548 | -0.013 | 0.104 | 0.104 | 0.369 | 0.86 | 0.69 | 0.82 | 0.71 | 0.84 | 0.8 |
| 6 | 65423976 | rs9452511 | 1.891 | 0.373 | 0.978 | 2.599 | -0.005 | 0.269 | 0.269 | 0.566 | 0.92 | 0.7 | 0.9 | 0.69 | 0.93 | 0.92 |
| 6 | 65424107 | rs9363129 | 1.784 | 0.376 | 0.972 | 2.6 | -0.005 | 0.256 | 0.256 | 0.106 | 0.93 | 0.74 | 0.9 | 0.69 | 0.93 | 0.9 |
| 6 | 65435946 | rs7757694 | 0.371 | 0.293 | -0.524 | 1.854 | -0.09 | 0.005 | 0.005 | 0 | 0.29 | 0.01 | 0.42 | 0.2 | 0.3 | 0.18 |
| 6 | 65453785 | rs6934929 | 0.56 | 0.182 | -0.036 | 2.336 | -0.001 | -0.029 | -0.029 | 0 | 0.9 | 0.94 | 0.85 | 0.75 | 0.86 | 0.87 |
| 6 | 65460308 | rs1390267 | 0.87 | 0.109 | 0.832 | 2.314 | 0.017 | 0.135 | 0.135 | 0 | 0.89 | 0.62 | 0.83 | 0.75 | 0.86 | 0.87 |
| 6 | 65484442 | rs9452831 | -1.554 | -0.235 | 0.786 | 2.327 | -0.026 | 0.159 | 0.159 | 0 | 0.2 | 0.52 | 0.18 | 0.42 | 0.21 | 0.17 |
| 6 | 65485995 | rs1495542 | -1.563 | -0.24 | 0.783 | 2.324 | -0.026 | 0.158 | 0.158 | 0 | 0.2 | 0.52 | 0.18 | 0.42 | 0.21 | 0.17 |
| 6 | 65488026 | rs4710495 | 1.346 | -0.116 | -0.578 | 1.591 | -0.056 | -0.017 | -0.017 | 0 | 0.49 | 0.05 | 0.63 | 0.42 | 0.54 | 0.45 |
| 6 | 65496953 | rs6911837 | 0.666 | -0.284 | 0.078 | 1.684 | -0.009 | -0.025 | -0.025 | 0 | 0.6 | 0.43 | 0.75 | 0.77 | 0.71 | 0.59 |
| 6 | 65499982 | rs6942336 | 1.472 | -0.241 | -0.672 | 1.722 | -0.055 | -0.016 | -0.016 | 0 | 0.49 | 0.05 | 0.63 | 0.42 | 0.54 | 0.44 |
| 6 | 65503314 | rs16895780 | 0.124 | -0.228 | -0.646 | 1.743 | -0.006 | -0.062 | -0.062 | 0.743 | 0.1 | 0 | 0.06 | 0.2 | 0.09 | 0.06 |
| 6 | 65504664 | rs12663566 | 0.213 | -0.034 | -0.539 | 1.936 | -0.008 | -0.022 | -0.022 | 0.014 | 0.07 | 0.02 | 0.07 | 0.27 | 0.12 | 0.11 |
| 6 | 65506077 | rs4710283 | 0.964 | 0.048 | -0.354 | 1.438 | 0.026 | -0.014 | -0.014 | 0 | 0.53 | 0.22 | 0.67 | 0.49 | 0.59 | 0.49 |
| 6 | 65509089 | rs12215142 | 0.633 | 0.178 | -0.621 | 0.853 | -0.059 | 0.036 | 0.036 | 0 | 0.09 | 0.01 | 0.07 | 0.02 | 0.14 | 0.23 |
| 6 | 65518623 | rs9453021 | 0.992 | 0.206 | -0.615 | 1.167 | -0.005 | 0.05 | 0.05 | 0 | 0.5 | 0.17 | 0.65 | 0.43 | 0.51 | 0.44 |
| 6 | 65520297 | rs4710286 | 1.048 | 0.206 | -0.544 | 1.166 | -0.001 | 0.048 | 0.048 | 0 | 0.5 | 0.2 | 0.65 | 0.43 | 0.51 | 0.44 |
| 6 | 65525346 | rs9354152 | 1.07 | 0.216 | -0.617 | 1.172 | -0.002 | 0.052 | 0.052 | 0.021 | 0.5 | 0.18 | 0.65 | 0.43 | 0.51 | 0.44 |
| 6 | 65531993 | rs13198898 | 0.084 | -0.069 | -0.469 | 0.777 | -0.007 | -0.034 | -0.034 | 0.297 | 0.09 | 0.12 | 0.03 | 0 | 0.04 | 0.08 |
| 6 | 65558078 | rs9453089 | 1.174 | -0.149 | -0.558 | 0.574 | 0.006 | 0.012 | 0.012 | 0 | 0.58 | 0.44 | 0.61 | 0.22 | 0.59 | 0.53 |
| 6 | 65562837 | rs2146755 | 0.655 | -0.208 | -0.464 | 1.406 | -0.018 | 0.009 | 0.009 | 0 | 0.68 | 0.58 | 0.78 | 0.66 | 0.73 | 0.72 |
| 6 | 65577402 | rs9345560 | 0.552 | -0.246 | -0.584 | 1.826 | -0.001 | 0.027 | 0.027 | 0 | 0.07 | 0.05 | 0.1 | 0.31 | 0.1 | 0.15 |
| 6 | 65578285 | rs9354178 | -0.029 | -0.269 | -0.618 | 2.017 | 0 | 0.155 | 0.155 | 0 | 0.09 | 0.14 | 0.17 | 0.49 | 0.14 | 0.19 |
| 6 | 65584763 | rs16895902 | 0.329 | -0.279 | -0.341 | 1.706 | -0.01 | 0.04 | 0.04 | 0 | 0.05 | 0.08 | 0.04 | 0.15 | 0.02 | 0.02 |
| 6 | 65586419 | rs2146757 | 0.891 | -0.34 | -0.261 | 1.647 | -0.02 | -0.017 | -0.017 | 0 | 0.69 | 0.56 | 0.78 | 0.71 | 0.73 | 0.76 |
| 6 | 65588875 | rs9354184 | 1.514 | -0.067 | -0.979 | 1.172 | -0.021 | 0.129 | 0.129 | 0 | 0.49 | 0.15 | 0.66 | 0.69 | 0.55 | 0.46 |
| 6 | 65589054 | rs13217880 | 0.034 | 0.04 | -0.717 | 1.156 | -0.007 | -0.035 | -0.035 | 0 | 0.09 | 0.13 | 0.04 | 0 | 0.04 | 0.08 |
| 6 | 65591554 | rs2350284 | 0.113 | 0.292 | -1.143 | 1.583 | 0 | -0.018 | -0.018 | 0 | 0.82 | 0.8 | 0.85 | 0.87 | 0.79 | 0.85 |
| 6 | 65618894 | rs10944766 | 1.08 | 0.485 | -2.125 | 0.888 | -0.036 | 0.108 | 0.108 | 0.011 | 0.39 | 0.02 | 0.41 | 0.24 | 0.37 | 0.3 |
| 6 | 65628676 | rs2350001 | 0.721 | 0.966 | -1.75 | 0.736 | 0.009 | 0.004 | 0.004 | 0 | 0.69 | 0.66 | 0.62 | 0.42 | 0.6 | 0.61 |
| 6 | 65635522 | rs10944769 | 0.917 | 0.656 | -1.756 | 0.737 | 0.037 | -0.01 | -0.01 | 0.168 | 0.47 | 0.31 | 0.53 | 0.56 | 0.42 | 0.38 |
| 6 | 65635578 | rs10944770 | 0.731 | 0.677 | -1.755 | 0.742 | 0.046 | -0.004 | -0.004 | 0.879 | 0.49 | 0.31 | 0.53 | 0.56 | 0.42 | 0.39 |
| 6 | 65649774 | rs9351470 | 0.842 | 0.553 | -1.848 | 1.26 | -0.043 | -0.121 | -0.121 | 0.355 | 0.83 | 0.97 | 0.79 | 0.61 | 0.79 | 0.71 |
| 6 | 65650698 | rs12528109 | 1.089 | 0.539 | -1.918 | 1.068 | -0.116 | 0.032 | 0.032 | 0 | 0.49 | 0.05 | 0.55 | 0.43 | 0.51 | 0.36 |
| 6 | 65661067 | rs2144715 | 0.639 | 0.297 | -1.221 | 1.267 | 0.021 | -0.007 | -0.007 | 0.236 | 0.83 | 0.89 | 0.89 | 0.81 | 0.88 | 0.78 |
| 6 | 65662621 | rs7767572 | -0.539 | 0.547 | -1.231 | 1.246 | -0.013 | -0.015 | -0.015 | 0.647 | 0.16 | 0.11 | 0.16 | 0.22 | 0.24 | 0.17 |
| 6 | 65669109 | rs7745549 | 0.22 | 0.332 | -0.481 | 1.328 | -0.003 | -0.158 | -0.158 | 0.061 | 0.24 | 0.46 | 0.15 | 0 | 0.18 | 0.21 |
| 6 | 65676556 | rs2064701 | -0.186 | 0.544 | -1.166 | 1.545 | 0.002 | 0.012 | 0.012 | 0 | 0.11 | 0.13 | 0.11 | 0.23 | 0.17 | 0.15 |
| 6 | 65687620 | rs9351476 | -1.298 | 0.447 | -1.303 | 1.618 | -0.062 | -0.016 | -0.016 | 0 | 0.49 | 0.9 | 0.41 | 0.39 | 0.51 | 0.51 |
| 6 | 65692485 | rs16896133 | 0.337 | 0.4 | -1.223 | 1.657 | -0.014 | -0.026 | -0.026 | 0 | 0.1 | 0.14 | 0.05 | 0.01 | 0.06 | 0.09 |
| 6 | 65698357 | rs9363305 | -0.667 | 0.441 | -1.21 | 1.643 | 0.02 | 0.066 | 0.066 | 0.162 | 0.39 | 0.73 | 0.3 | 0.17 | 0.34 | 0.37 |
| 6 | 65703968 | rs9345581 | 1.524 | 0.453 | -1.301 | 1.629 | -0.121 | -0.041 | -0.041 | 0 | 0.45 | 0.04 | 0.53 | 0.47 | 0.43 | 0.38 |
| 6 | 65726738 | rs9345588 | 0.686 | 0.358 | -0.957 | 1.637 | 0 | 0 | 0 | 0 | 0.61 | 0.45 | 0.67 | 0.63 | 0.61 | 0.55 |
| 6 | 65734358 | rs7738129 | -0.131 | 0.414 | -1.066 | 1.659 | 0.002 | -0.02 | -0.02 | 0 | 0.28 | 0.57 | 0.26 | 0.16 | 0.29 | 0.28 |
| 6 | 65740172 | rs9453156 | -0.094 | 0.481 | -1.2 | 1.72 | 0.023 | 0.06 | 0.06 | 0 | 0.12 | 0.26 | 0.12 | 0.22 | 0.16 | 0.14 |
| 6 | 65743428 | rs13220430 | 0.155 | 0.613 | -1.428 | 1.751 | -0.007 | -0.15 | -0.15 | 0 | 0.26 | 0.47 | 0.16 | 0.01 | 0.2 | 0.22 |
| 6 | 65743530 | rs1121536 | 0.425 | 0.857 | -2.359 | 2.417 | -0.135 | -0.04 | -0.04 | 0.369 | 0.46 | 0.04 | 0.53 | 0.47 | 0.47 | 0.4 |
| 6 | 65759576 | rs2350516 | -1.84 | 0.826 | -0.826 | 2.775 | -0.001 | 0.083 | 0.083 | 0 | 0.06 | 0.08 | 0.14 | 0.3 | 0.11 | 0.11 |
| 6 | 65763048 | rs16896178 | -1.293 | 0.886 | -1.036 | 1.647 | -0.014 | -0.032 | -0.032 | 0.011 | 0.06 | 0.01 | 0.24 | 0.2 | 0.09 | 0.15 |
| 6 | 65765963 | rs9363329 | -1.138 | 1.297 | -1.342 | 1.106 | -0.037 | -0.018 | -0.018 | 0 | 0.07 | 0 | 0.07 | 0.08 | 0.09 | 0.08 |

| | | | | | | | | | | | | | | | | |
|---|---|---|---|---|---|---|---|---|---|---|---|---|---|---|---|---|
| 6 | 65767393 | rs10455568 | -1.464 | 1.029 | -1.586 | 0.819 | -0.031 | -0.006 | -0.006 | 0.013 | 0.11 | 0 | 0.08 | 0.06 | 0.12 | 0.09 |
| 6 | 65775758 | rs12207169 | -1.068 | 0.845 | -1.626 | 0.778 | -0.019 | -0.02 | -0.02 | 0 | 0.07 | 0 | 0.06 | 0.06 | 0.08 | 0.08 |
| 6 | 65781220 | rs12207150 | -1.257 | 1.473 | -1.946 | -0.028 | -0.035 | 0.071 | 0.071 | 0 | 0.28 | 0.15 | 0.19 | 0.1 | 0.32 | 0.35 |
| 6 | 65781547 | rs17410730 | -1.207 | 1.452 | -1.946 | -0.039 | -0.054 | 0.082 | 0.082 | 0 | 0.27 | 0.08 | 0.18 | 0.1 | 0.32 | 0.35 |
| 6 | 65801996 | rs1929724 | -1.43 | 0.968 | -0.781 | -0.599 | 0.007 | -0.097 | -0.097 | 0 | 0.35 | 0.47 | 0.19 | 0.07 | 0.36 | 0.37 |
| 6 | 65813903 | rs17637910 | -1.066 | 0.943 | -0.872 | -0.609 | -0.084 | 0.125 | 0.125 | 0.266 | 0.28 | 0.04 | 0.14 | 0.07 | 0.32 | 0.35 |
| 6 | 65816312 | rs9453182 | -1.39 | 0.903 | -0.54 | -0.633 | -0.031 | 0.112 | 0.112 | 0 | 0.32 | 0.16 | 0.15 | 0.07 | 0.33 | 0.35 |
| 6 | 65816812 | rs6915644 | -0.753 | 0.475 | -0.263 | -1.319 | 0.002 | -0.055 | -0.055 | 0 | 0.43 | 0.59 | 0.37 | 0.15 | 0.43 | 0.39 |
| 6 | 65828853 | rs9453190 | -0.158 | 0.434 | 0.008 | -1.305 | -0.023 | -0.08 | -0.08 | 0 | 0.23 | 0.47 | 0.23 | 0.1 | 0.17 | 0.1 |
| 6 | 65829390 | rs1041522 | -1.413 | 0.463 | -0.032 | -1.247 | 0 | 0.034 | 0.034 | 0 | 0.54 | 0.69 | 0.4 | 0.18 | 0.52 | 0.46 |
| 6 | 65831684 | rs9363336 | 1.607 | 0.335 | -0.081 | -0.591 | -0.042 | -0.088 | -0.088 | 0.592 | 0.28 | 0.08 | 0.23 | 0.39 | 0.32 | 0.24 |
| 6 | 65834263 | rs1000692 | 1.6 | 0.343 | -0.249 | -0.61 | -0.082 | -0.141 | -0.141 | 0.005 | 0.28 | 0.01 | 0.23 | 0.39 | 0.32 | 0.24 |
| 6 | 65839253 | rs9647608 | -0.788 | 0.443 | 0.517 | -1 | -0.005 | -0.03 | -0.03 | 0 | 0.11 | 0.19 | 0.04 | 0.03 | 0.09 | 0.07 |
| 6 | 65842061 | rs6455027 | -0.919 | 0.695 | 0.566 | -1.165 | -0.007 | -0.051 | -0.051 | 0 | 0.36 | 0.52 | 0.29 | 0.14 | 0.29 | 0.2 |
| 6 | 65857285 | rs1929723 | -0.917 | 0.721 | 0.816 | -1.168 | 0.008 | -0.047 | -0.047 | 0.242 | 0.12 | 0.25 | 0.06 | 0.03 | 0.12 | 0.08 |
| 6 | 65860539 | rs11962884 | -0.118 | 0.695 | 0.901 | -1.191 | -0.005 | -0.033 | -0.033 | 0 | 0.25 | 0.35 | 0.25 | 0.12 | 0.2 | 0.14 |
| 6 | 65871821 | rs12198932 | -1.066 | 0.698 | 0.875 | -1.194 | -0.003 | -0.042 | -0.042 | 0 | 0.36 | 0.6 | 0.31 | 0.14 | 0.32 | 0.22 |
| 6 | 65878609 | rs13210192 | 0.496 | 0.849 | 0.861 | -1.293 | -0.029 | 0.052 | 0.052 | 1.514 | 0.2 | 0.07 | 0.12 | 0.04 | 0.25 | 0.26 |
| 6 | 65882186 | rs1145943 | -1.118 | 0.325 | 1.134 | -0.958 | 0.014 | -0.052 | -0.052 | 0 | 0.12 | 0.29 | 0.06 | 0.03 | 0.12 | 0.08 |
| 6 | 65903328 | rs9453222 | -0.334 | 0.109 | 0.834 | -1.09 | -0.007 | 0.002 | 0.002 | 0 | 0.12 | 0.11 | 0.21 | 0.11 | 0.16 | 0.07 |
| 6 | 65910220 | rs475994 | -1.27 | 0.076 | 1.038 | -1.066 | -0.002 | -0.066 | -0.066 | 0 | 0.35 | 0.54 | 0.31 | 0.16 | 0.33 | 0.21 |
| 6 | 65924421 | rs2093342 | -0.525 | 0.116 | 1.074 | -1.077 | -0.001 | 0.003 | 0.003 | 0 | 0.24 | 0.26 | 0.26 | 0.11 | 0.22 | 0.13 |
| 6 | 65933316 | rs11966927 | -0.385 | 0.082 | 1.061 | -1.226 | -0.023 | 0.061 | 0.061 | 0 | 0.23 | 0.08 | 0.19 | 0.11 | 0.22 | 0.13 |
| 6 | 65947059 | rs12209387 | 0.226 | 0.118 | 0.929 | -1.187 | -0.023 | 0.004 | 0.004 | 0 | 0.07 | 0 | 0.02 | 0.04 | 0.08 | 0.07 |
| 6 | 65972053 | rs697305 | -1.419 | 0.108 | 0.811 | -1.221 | -0.001 | -0.03 | -0.03 | 0.089 | 0.38 | 0.62 | 0.27 | 0.15 | 0.34 | 0.22 |
| 6 | 65986503 | rs7750139 | 0.856 | 0.215 | 0.78 | -1.396 | 0 | 0.002 | 0.002 | 0 | 0.64 | 0.32 | 0.71 | 0.91 | 0.67 | 0.79 |
| 6 | 65996855 | rs9354216 | -0.7 | 0.363 | 0.446 | -1.564 | -0.013 | 0.032 | 0.032 | 0 | 0.19 | 0.06 | 0.2 | 0.14 | 0.21 | 0.28 |
| 6 | 66001749 | rs10498832 | -0.441 | 0.158 | 1.152 | -1.457 | 0 | -0.042 | -0.042 | 0 | 0.08 | 0.16 | 0.02 | 0 | 0.05 | 0.07 |
| 6 | 66006148 | rs9342465 | 1.48 | 0.163 | 0.761 | -1.479 | -0.006 | 0.155 | 0.155 | 0.104 | 0.56 | 0.2 | 0.71 | 0.89 | 0.59 | 0.71 |
| 6 | 66021416 | rs6937429 | 1.198 | 0.3 | 0.769 | -1.331 | -0.001 | 0.011 | 0.011 | 0.155 | 0.6 | 0.27 | 0.72 | 0.89 | 0.61 | 0.71 |
| 6 | 66030650 | rs7754286 | -1.052 | 0.352 | 0.699 | -1.332 | 0.001 | -0.097 | -0.097 | 0 | 0.31 | 0.44 | 0.23 | 0.11 | 0.31 | 0.2 |
| 6 | 66032226 | rs17412215 | -1.495 | 0.297 | 0.528 | -1.22 | 0 | -0.055 | -0.055 | 0.075 | 0.39 | 0.6 | 0.28 | 0.11 | 0.39 | 0.29 |
| 6 | 66042231 | rs12207746 | -0.949 | 0.263 | 0.515 | -0.4 | -0.005 | -0.047 | -0.047 | 0.057 | 0.11 | 0.25 | 0.05 | 0.04 | 0.08 | 0.08 |
| 6 | 66050797 | rs9445507 | -0.722 | 0.336 | -0.076 | 0.205 | 0.001 | 0 | 0 | 0 | 0.26 | 0.16 | 0.23 | 0.26 | 0.22 | 0.34 |
| 6 | 66055189 | rs9354231 | 0.916 | 0.664 | -0.938 | 0.264 | -0.102 | -0.017 | -0.017 | 0.204 | 0.22 | 0.01 | 0.26 | 0.18 | 0.3 | 0.16 |
| 6 | 66059086 | rs13202000 | -0.041 | 0.632 | -0.948 | 0.201 | 0.001 | 0 | 0 | 0.408 | 0.27 | 0.17 | 0.23 | 0.26 | 0.22 | 0.34 |
| 6 | 66064126 | rs1393685 | 1.003 | -0.014 | -0.397 | 0.495 | 0.003 | -0.003 | -0.003 | 0 | 0.39 | 0.31 | 0.36 | 0.38 | 0.42 | 0.36 |
| 6 | 66067950 | rs1502957 | -0.314 | 0.067 | 0.049 | 0.597 | -0.013 | 0.091 | 0.091 | 0.742 | 0.89 | 0.66 | 0.85 | 0.78 | 0.94 | 0.86 |
| 6 | 66070703 | rs12195773 | 1.209 | 0.062 | 0.061 | 0.593 | -0.097 | 0.028 | 0.028 | 0.979 | 0.21 | 0.01 | 0.25 | 0.15 | 0.3 | 0.16 |
| 6 | 66071980 | rs4580839 | -0.357 | 0.063 | 0.051 | 0.593 | -0.003 | -0.001 | -0.001 | 0.708 | 0.68 | 0.65 | 0.61 | 0.63 | 0.65 | 0.7 |
| 6 | 66075988 | rs9342468 | 1.198 | 0.059 | -0.016 | 0.601 | -0.102 | 0.025 | 0.025 | 0.979 | 0.21 | 0.01 | 0.18 | 0.15 | 0.3 | 0.16 |
| 6 | 66077188 | rs9345609 | -0.098 | 0.03 | 0.307 | 0.614 | -0.016 | 0.074 | 0.074 | 0.252 | 0.91 | 0.77 | 0.85 | 0.78 | 0.95 | 0.86 |
| 6 | 66083394 | rs998829 | 0.451 | -0.199 | 0.065 | 0.502 | -0.004 | -0.008 | -0.008 | 0 | 0.21 | 0.12 | 0.22 | 0.26 | 0.21 | 0.32 |
| 6 | 66083780 | rs9345612 | -0.85 | -0.313 | 0.298 | 0.657 | -0.003 | 0 | 0 | 0 | 0.54 | 0.47 | 0.51 | 0.45 | 0.54 | 0.52 |
| 6 | 66091351 | rs13196323 | 0.133 | -0.264 | -0.081 | 0.696 | -0.016 | -0.026 | -0.026 | 0 | 0.43 | 0.18 | 0.47 | 0.45 | 0.5 | 0.48 |
| 6 | 66106319 | rs9453294 | -0.666 | -0.254 | -0.025 | 0.697 | 0.015 | -0.001 | -0.001 | 0 | 0.52 | 0.35 | 0.5 | 0.46 | 0.54 | 0.51 |
| 6 | 66106499 | rs9363355 | 0.354 | -0.159 | -0.07 | 0.614 | -0.017 | 0.036 | 0.036 | 0.081 | 0.31 | 0.24 | 0.28 | 0.16 | 0.33 | 0.19 |
| 6 | 66115654 | rs11968993 | 0.091 | -0.189 | 0.016 | 0.649 | -0.008 | -0.024 | -0.024 | 0 | 0.08 | 0.12 | 0.03 | 0 | 0.04 | 0.03 |
| 6 | 66125723 | rs441770 | 0.465 | -0.141 | 0.213 | 0.878 | -0.014 | 0.044 | 0.044 | 1.37 | 0.09 | 0.21 | 0.13 | 0.18 | 0.05 | 0.11 |
| 6 | 66126554 | rs9453298 | -0.59 | -0.127 | -0.264 | 0.837 | 0.02 | 0.021 | 0.021 | 0 | 0.52 | 0.35 | 0.5 | 0.37 | 0.54 | 0.55 |
| 6 | 66133281 | rs10944796 | 0.395 | -0.119 | -0.373 | 0.795 | -0.025 | 0.043 | 0.043 | 0.121 | 0.3 | 0.17 | 0.28 | 0.16 | 0.32 | 0.23 |
| 6 | 66137260 | rs1502991 | -0.425 | -0.107 | -0.126 | 0.874 | 0.013 | 0.013 | 0.013 | 0 | 0.52 | 0.4 | 0.5 | 0.37 | 0.54 | 0.55 |
| 6 | 66139220 | rs9342476 | 0.131 | -0.028 | 0.004 | 1.102 | 0.023 | 0.015 | 0.015 | 0.451 | 0.34 | 0.46 | 0.49 | 0.59 | 0.38 | 0.42 |
| 6 | 66141023 | rs17254301 | 0.037 | -0.009 | -0.733 | 0.863 | -0.048 | -0.014 | -0.014 | 0 | 0.17 | 0.01 | 0.21 | 0.08 | 0.18 | 0.18 |
| 6 | 66143990 | rs9445545 | -0.302 | 0.12 | -0.912 | 1.216 | -0.018 | -0.045 | -0.045 | 0 | 0.17 | 0.08 | 0.18 | 0.34 | 0.18 | 0.23 |
| 6 | 66144068 | rs9345625 | 1.401 | -0.107 | -1.119 | 0.956 | -0.094 | -0.036 | -0.036 | 0 | 0.23 | 0.01 | 0.25 | 0.21 | 0.26 | 0.2 |
| 6 | 66164344 | rs12206730 | 0.755 | -0.405 | -0.86 | 0.872 | -0.016 | -0.031 | -0.031 | 0 | 0.3 | 0.12 | 0.27 | 0.28 | 0.27 | 0.26 |
| 6 | 66177078 | rs10944808 | -0.114 | -0.747 | -0.343 | 0.656 | 0.004 | 0.005 | 0.005 | 0 | 0.49 | 0.42 | 0.45 | 0.32 | 0.41 | 0.44 |
| 6 | 66187107 | rs10484939 | 0.736 | -0.751 | -0.278 | 0.637 | -0.006 | 0.019 | 0.019 | 0.54 | 0.16 | 0 | 0.17 | 0.04 | 0.13 | 0.18 |
| 6 | 66188166 | rs1988374 | 1.549 | -0.513 | -0.062 | 0.805 | -0.086 | -0.067 | -0.067 | 0 | 0.24 | 0.01 | 0.25 | 0.24 | 0.26 | 0.2 |
| 6 | 66204945 | rs12193967 | 1.643 | -0.316 | -0.085 | 0.781 | -0.074 | -0.071 | -0.071 | 0 | 0.23 | 0.01 | 0.24 | 0.24 | 0.24 | 0.2 |
| 6 | 66207218 | rs10944813 | 0.662 | -0.269 | 0.124 | 0.81 | -0.004 | -0.001 | -0.001 | 0 | 0.33 | 0.4 | 0.28 | 0.28 | 0.27 | 0.24 |
| 6 | 66214813 | rs4710525 | -0.002 | 0.024 | -0.225 | 0.999 | -0.031 | -0.038 | -0.038 | 0.188 | 0.15 | 0.09 | 0.2 | 0.36 | 0.21 | 0.23 |
| 6 | 66215549 | rs7449561 | 0.858 | -0.273 | 0.393 | 0.673 | -0.004 | -0.006 | -0.006 | 0 | 0.3 | 0.34 | 0.28 | 0.26 | 0.25 | 0.26 |
| 6 | 66232692 | rs16896746 | -0.084 | 0.054 | 0.329 | 0.722 | 0.023 | 0.036 | 0.036 | 0 | 0.09 | 0.24 | 0.15 | 0.12 | 0.09 | 0.09 |
| 6 | 66235307 | rs9453319 | -1.028 | 0.412 | 0.543 | 0.814 | 0.022 | 0.036 | 0.036 | 0.152 | 0.27 | 0.63 | 0.37 | 0.51 | 0.29 | 0.34 |
| 6 | 66237309 | rs1034137 | -1.672 | 0.283 | 0.091 | 0.341 | -0.046 | 0.085 | 0.085 | 0 | 0.84 | 0.96 | 0.84 | 1 | 0.73 | 0.93 |
| 6 | 66241604 | rs7776123 | -1.013 | 0.469 | 0.424 | 0.636 | 0.02 | 0.027 | 0.027 | 0.204 | 0.28 | 0.64 | 0.39 | 0.51 | 0.33 | 0.34 |
| 6 | 66245983 | rs7767963 | 0.34 | 0.279 | 0.732 | 0.692 | 0.015 | 0.056 | 0.056 | 0 | 0.08 | 0.24 | 0.14 | 0.11 | 0.06 | 0.08 |

| | | | | | | | | | | | | | | | | |
|---|---|---|---|---|---|---|---|---|---|---|---|---|---|---|---|---|
| 6 | 66258778 | rs1331197 | -0.571 | 0.028 | 0.023 | 0.503 | -0.032 | 0.072 | 0.072 | 0 | 0.75 | 0.92 | 0.8 | 0.84 | 0.67 | 0.78 |
| 6 | 66259508 | rs6932966 | 0.103 | -0.203 | 0.078 | 0.481 | -0.003 | -0.006 | -0.006 | 0 | 0.55 | 0.54 | 0.57 | 0.72 | 0.55 | 0.56 |
| 6 | 66267533 | rs6926034 | 0.099 | -0.207 | 0.085 | 0.48 | -0.005 | -0.008 | -0.008 | 0 | 0.55 | 0.54 | 0.57 | 0.72 | 0.55 | 0.56 |
| 6 | 66269399 | rs1331205 | 0.093 | -0.034 | 0.084 | 0.626 | 0.01 | 0.011 | 0.011 | 0 | 0.17 | 0.18 | 0.18 | 0.36 | 0.24 | 0.24 |
| 6 | 66278575 | rs13203580 | 0.821 | -0.306 | -0.266 | 0.555 | 0.027 | -0.021 | -0.021 | 0 | 0.17 | 0.01 | 0.17 | 0.08 | 0.12 | 0.19 |
| 6 | 66284186 | rs9360139 | 2.275 | -0.447 | -0.472 | 0.482 | -0.065 | -0.062 | -0.062 | 0 | 0.26 | 0.01 | 0.23 | 0.25 | 0.26 | 0.22 |
| 6 | 66284808 | rs1590967 | -1.001 | -0.224 | -0.057 | 0.521 | 0.001 | -0.001 | -0.001 | 0.119 | 0.44 | 0.56 | 0.49 | 0.54 | 0.41 | 0.5 |
| 6 | 66285958 | rs9351511 | -0.137 | -0.226 | -0.027 | 0.55 | 0.014 | 0.102 | 0.102 | 0.782 | 0.58 | 0.89 | 0.63 | 0.76 | 0.55 | 0.59 |
| 6 | 66287365 | rs354349 | 1.52 | -0.171 | 0.219 | 0.703 | -0.007 | 0.003 | 0.003 | 2.13 | 0.32 | 0.34 | 0.31 | 0.31 | 0.27 | 0.28 |
| 6 | 66298313 | rs6902867 | 0.398 | 0.138 | 0.071 | 0.642 | -0.005 | -0.004 | -0.004 | 0 | 0.18 | 0.21 | 0.17 | 0.09 | 0.15 | 0.2 |
| 6 | 66304163 | rs354364 | -0.305 | 0.591 | 0.143 | 0.899 | 0.086 | 0.148 | 0.148 | 0 | 0.17 | 0.4 | 0.23 | 0.41 | 0.21 | 0.26 |
| 6 | 66307143 | rs2183675 | 0.181 | -0.088 | -0.294 | 0.684 | 0.012 | 0.029 | 0.029 | 0 | 0.51 | 0.59 | 0.5 | 0.66 | 0.49 | 0.46 |
| 6 | 66308325 | rs354342 | -0.795 | -0.63 | 0.038 | 1.352 | -0.02 | 0.018 | 0.018 | 0.044 | 0.34 | 0.45 | 0.39 | 0.59 | 0.22 | 0.47 |
| 6 | 66314880 | rs354401 | 0.7 | -0.253 | -0.258 | 1.313 | 0.01 | 0.024 | 0.024 | 0.041 | 0.84 | 0.79 | 0.82 | 0.63 | 0.8 | 0.78 |
| 6 | 66321191 | rs13211922 | 0.388 | -0.049 | -0.497 | 1.209 | -0.036 | -0.004 | -0.004 | 0 | 0.09 | 0.01 | 0.05 | 0.1 | 0.1 | 0.14 |
| 6 | 66333105 | rs6900513 | 1.875 | -0.293 | -0.752 | 1.158 | -0.157 | 0.018 | 0.018 | 1.965 | 0.39 | 0.02 | 0.38 | 0.22 | 0.51 | 0.24 |
| 6 | 66333672 | rs902287 | 0.644 | -0.509 | -0.421 | 1.243 | 0.009 | 0.079 | 0.079 | 0.773 | 0.78 | 0.62 | 0.68 | 0.46 | 0.78 | 0.63 |
| 6 | 66342056 | rs13195796 | -0.85 | -0.267 | -0.294 | 1.065 | 0.015 | 0.019 | 0.019 | 0 | 0.4 | 0.61 | 0.31 | 0.25 | 0.28 | 0.39 |
| 6 | 66344855 | rs1873291 | -1.351 | -0.266 | -0.274 | 1.096 | -0.005 | 0.071 | 0.071 | 0 | 0.49 | 0.68 | 0.42 | 0.28 | 0.45 | 0.48 |
| 6 | 66346347 | rs1586658 | 0.949 | -0.25 | -0.147 | 1.386 | 0.01 | 0.005 | 0.005 | 0 | 0.57 | 0.31 | 0.53 | 0.45 | 0.62 | 0.53 |
| 6 | 66362817 | rs958706 | 0.917 | -0.312 | -0.032 | 1.489 | 0.011 | 0.037 | 0.037 | 0 | 0.61 | 0.27 | 0.6 | 0.79 | 0.65 | 0.64 |
| 6 | 66366047 | rs9918344 | -0.004 | -0.542 | -0.235 | 1.498 | 0.01 | 0 | 0 | 0 | 0.19 | 0.2 | 0.19 | 0.25 | 0.13 | 0.32 |
| 6 | 66366331 | rs1602235 | -0.71 | -0.718 | 0.241 | 1.379 | -0.004 | -0.001 | -0.001 | 0 | 0.79 | 0.8 | 0.79 | 0.85 | 0.83 | 0.79 |
| 6 | 66369304 | rs9345666 | 2.025 | -0.719 | 0.189 | 1.367 | -0.152 | -0.144 | -0.144 | 0 | 0.41 | 0.05 | 0.42 | 0.54 | 0.52 | 0.32 |
| 6 | 66370779 | rs2171902 | -0.634 | -0.728 | 0.666 | 1.326 | -0.007 | -0.002 | -0.002 | 0 | 0.78 | 0.77 | 0.77 | 0.85 | 0.82 | 0.78 |
| 6 | 66374179 | rs6455050 | -0.768 | -0.7 | 0.725 | 1.347 | 0.034 | -0.032 | -0.032 | 0.567 | 0.17 | 0.43 | 0.17 | 0.06 | 0.17 | 0.14 |
| 6 | 66377946 | rs923536 | -1.561 | -0.69 | 0.683 | 1.369 | 0.003 | -0.002 | -0.002 | 0 | 0.4 | 0.68 | 0.43 | 0.66 | 0.33 | 0.55 |
| 6 | 66395381 | rs9294646 | -1.549 | -0.687 | 0.676 | 1.371 | 0.001 | -0.001 | -0.001 | 0.432 | 0.4 | 0.65 | 0.43 | 0.66 | 0.33 | 0.55 |
| 6 | 66397288 | rs16897066 | 0.89 | -0.719 | 1.088 | 1.323 | -0.003 | -0.009 | -0.009 | 0 | 0.05 | 0.18 | 0.02 | 0 | 0.01 | 0.03 |
| 6 | 66403767 | rs34483511 | 0.176 | -0.724 | 0.948 | 1.759 | 0.014 | -0.035 | -0.035 | 4.316 | 0.19 | 0.14 | 0.2 | 0.42 | 0.13 | 0.36 |
| 6 | 66405466 | rs7450869 | -0.799 | -0.725 | 0.998 | 1.487 | 0.032 | -0.031 | -0.031 | 0.018 | 0.17 | 0.41 | 0.17 | 0.06 | 0.17 | 0.14 |
| 6 | 66408758 | rs1873294 | 1.306 | -0.743 | 0.791 | 1.43 | 0.014 | 0.01 | 0.01 | 0.185 | 0.56 | 0.23 | 0.52 | 0.44 | 0.62 | 0.55 |
| 6 | 66410718 | rs9351517 | -1.312 | -0.721 | 0.917 | 1.458 | 0.017 | 0.128 | 0.128 | 0.883 | 0.22 | 0.48 | 0.25 | 0.41 | 0.21 | 0.23 |
| 6 | 66414828 | rs12526729 | -0.002 | -0.761 | 0.816 | 1.416 | 0 | 0 | 0 | 0 | 0.2 | 0.22 | 0.18 | 0.25 | 0.12 | 0.32 |
| 6 | 66418352 | rs1490129 | -1.437 | -0.722 | 1.029 | 1.439 | 0.008 | 0.103 | 0.103 | 0.509 | 0.23 | 0.49 | 0.23 | 0.41 | 0.21 | 0.24 |
| 6 | 66423317 | rs2636131 | -0.445 | -0.768 | 0.574 | 1.393 | -0.009 | -0.004 | -0.004 | 2.195 | 0.78 | 0.75 | 0.77 | 0.85 | 0.82 | 0.79 |
| 6 | 66424345 | rs9342497 | -1.453 | -0.772 | 0.574 | 1.404 | 0.002 | 0.078 | 0.078 | 1.447 | 0.23 | 0.56 | 0.25 | 0.41 | 0.21 | 0.23 |
| 6 | 66424507 | rs2636133 | -0.378 | -0.708 | 0.546 | 1.121 | -0.013 | -0.005 | -0.005 | 0 | 0.78 | 0.74 | 0.77 | 0.85 | 0.82 | 0.78 |
| 6 | 66429923 | rs2492640 | 2.158 | -0.589 | 0.286 | 1.087 | -0.171 | 0.049 | 0.049 | 0.896 | 0.36 | 0.02 | 0.35 | 0.19 | 0.49 | 0.22 |
| 6 | 66434217 | rs1032310 | 0.301 | -0.804 | 0.493 | 1.07 | -0.03 | 0.009 | 0.009 | 0 | 0.73 | 0.5 | 0.71 | 0.81 | 0.82 | 0.75 |
| 6 | 66438702 | rs9354281 | -0.943 | -0.791 | 0.387 | 1.069 | 0 | -0.006 | -0.006 | 1.58 | 0.37 | 0.45 | 0.36 | 0.62 | 0.33 | 0.53 |
| 6 | 66442496 | rs2814142 | 0.282 | -0.682 | 0.862 | 1.02 | -0.039 | 0.031 | 0.031 | 0 | 0.75 | 0.57 | 0.72 | 0.81 | 0.82 | 0.75 |
| 6 | 66443150 | rs2814140 | 0.548 | -0.72 | 0.654 | 1.028 | -0.021 | 0.005 | 0.005 | 0 | 0.73 | 0.46 | 0.71 | 0.81 | 0.82 | 0.75 |
| 6 | 66449211 | rs7742881 | -0.294 | -0.719 | 0.878 | 1.061 | -0.019 | 0.053 | 0.053 | 0.208 | 0.08 | 0.29 | 0.06 | 0.05 | 0.01 | 0.06 |
| 6 | 66456442 | rs12526256 | 0.199 | -0.559 | 0.758 | 0.924 | -0.002 | -0.003 | -0.003 | 1.649 | 0.11 | 0.12 | 0.09 | 0.03 | 0.13 | 0.11 |
| 6 | 66478480 | rs9453400 | -1.34 | -0.7 | 0.43 | 1.105 | 0.001 | 0.022 | 0.022 | 0.25 | 0.45 | 0.72 | 0.49 | 0.56 | 0.41 | 0.47 |
| 6 | 66491056 | rs9453403 | -0.646 | -0.722 | 0.577 | 1.21 | -0.026 | 0.216 | 0.216 | 0 | 0.11 | 0.26 | 0.13 | 0.38 | 0.04 | 0.16 |
| 6 | 66491725 | rs13191280 | -1.073 | -0.697 | 0.576 | 1.177 | -0.001 | 0.099 | 0.099 | 0 | 0.21 | 0.35 | 0.22 | 0.41 | 0.17 | 0.26 |
| 6 | 66496110 | rs2251342 | -0.664 | -0.699 | 0.419 | 1.152 | 0 | -0.001 | -0.001 | 0 | 0.78 | 0.78 | 0.74 | 0.84 | 0.76 | 0.79 |
| 6 | 66502214 | rs2802036 | 1.179 | -0.731 | 0.239 | 1.075 | 0 | 0.023 | 0.023 | 0.668 | 0.54 | 0.26 | 0.51 | 0.43 | 0.58 | 0.53 |
| 6 | 66503818 | rs2802038 | 1.23 | -0.751 | 0.075 | 1.082 | -0.012 | 0.009 | 0.009 | 1.445 | 0.54 | 0.15 | 0.51 | 0.44 | 0.59 | 0.57 |
| 6 | 66506140 | rs10498834 | 0.264 | -0.941 | -0.164 | 1.122 | 0.021 | -0.016 | -0.016 | 0.002 | 0.21 | 0.15 | 0.28 | 0.3 | 0.14 | 0.36 |
| 6 | 66510926 | rs1938105 | 0.714 | -0.836 | -0.189 | 1.19 | -0.007 | 0.005 | 0.005 | 0 | 0.1 | 0.07 | 0.1 | 0.1 | 0.13 | 0.11 |
| 6 | 66511710 | rs2814112 | 2.306 | -0.869 | -0.288 | 1.143 | -0.153 | 0.086 | 0.086 | 1.095 | 0.33 | 0.01 | 0.24 | 0.14 | 0.45 | 0.21 |
| 6 | 66516749 | rs9363448 | -0.698 | -0.974 | 0.102 | 1.236 | -0.064 | 0.008 | 0.008 | 0.47 | 0.3 | 0.5 | 0.35 | 0.34 | 0.15 | 0.38 |
| 6 | 66519756 | rs2245704 | -0.727 | -0.723 | -0.092 | 1.328 | 0.004 | 0 | 0 | 0 | 0.78 | 0.75 | 0.75 | 0.84 | 0.77 | 0.79 |
| 6 | 66524243 | rs2802047 | 0.501 | -0.635 | -0.556 | 1.111 | 0.005 | 0.002 | 0.002 | 0 | 0.53 | 0.62 | 0.42 | 0.55 | 0.55 | 0.4 |
| 6 | 66527803 | rs13216907 | -0.003 | -0.544 | -0.466 | 1.12 | 0.006 | -0.007 | -0.007 | 0.379 | 0.25 | 0.12 | 0.33 | 0.28 | 0.21 | 0.39 |
| 6 | 66528177 | rs4299807 | 0.219 | -0.489 | -0.166 | 1.073 | 0.005 | -0.016 | -0.016 | 0 | 0.12 | 0.26 | 0.11 | 0.11 | 0.13 | 0.11 |
| 6 | 66528870 | rs10498836 | 0.628 | -0.496 | -0.254 | 1.069 | -0.013 | 0.01 | 0.01 | 0 | 0.1 | 0.05 | 0.09 | 0.11 | 0.13 | 0.11 |
| 6 | 66529457 | rs9445592 | -1.436 | -0.674 | -0.302 | 1.097 | -0.02 | 0.05 | 0.05 | 0 | 0.25 | 0.61 | 0.24 | 0.42 | 0.18 | 0.22 |
| 6 | 66541181 | rs11969957 | -0.014 | -1.13 | -0.59 | 0.62 | 0.004 | 0.005 | 0.005 | 0 | 0.13 | 0.11 | 0.09 | 0.11 | 0.1 | 0.12 |
| 6 | 66541198 | rs2000382 | -0.791 | -1.248 | -0.724 | 0.634 | 0.023 | 0.015 | 0.015 | 0 | 0.36 | 0.19 | 0.4 | 0.38 | 0.31 | 0.5 |
| 6 | 66542099 | rs10498837 | 1.194 | -1.349 | -0.83 | 0.801 | -0.047 | -0.084 | -0.084 | 0 | 0.21 | 0.09 | 0.27 | 0.42 | 0.28 | 0.25 |
| 6 | 66542891 | rs9354305 | -1.403 | -1.328 | -0.734 | 0.813 | 0.001 | 0.026 | 0.026 | 0 | 0.58 | 0.43 | 0.69 | 0.79 | 0.59 | 0.78 |
| 6 | 66544648 | rs12529685 | 0.549 | -1.374 | -0.382 | 0.843 | 0.005 | 0.006 | 0.006 | 0 | 0.12 | 0.08 | 0.08 | 0.11 | 0.1 | 0.11 |
| 6 | 66545656 | rs1938060 | 0.303 | -1.328 | -0.386 | 0.833 | -0.008 | -0.025 | -0.025 | 0 | 0.24 | 0.07 | 0.32 | 0.27 | 0.21 | 0.39 |
| 6 | 66546219 | rs2802056 | 2.171 | -1.214 | -0.188 | 0.662 | -0.131 | 0.159 | 0.159 | 0.125 | 0.29 | 0.01 | 0.18 | 0.07 | 0.37 | 0.17 |
| 6 | 66549080 | rs9363457 | -1.177 | -1.06 | -0.227 | 0.736 | 0 | 0.026 | 0.026 | 0 | 0.58 | 0.43 | 0.69 | 0.79 | 0.59 | 0.78 |

| | | | | | | | | | | | | | | | | |
|---|---|---|---|---|---|---|---|---|---|---|---|---|---|---|---|---|
| 6 | 66593133 | rs1499704 | -1.156 | -1.17 | -0.425 | 0.622 | 0.01 | -0.037 | -0.037 | 0 | 0.59 | 0.55 | 0.68 | 0.75 | 0.51 | 0.69 |
| 6 | 66594482 | rs4464770 | 0.49 | -1.246 | -0.533 | 0.967 | -0.045 | -0.079 | -0.079 | 0.237 | 0.28 | 0.14 | 0.32 | 0.43 | 0.34 | 0.29 |
| 6 | 66609594 | rs9354336 | 0.235 | -1.257 | -0.61 | 0.993 | 0.025 | -0.049 | -0.049 | 0 | 0.22 | 0.15 | 0.3 | 0.32 | 0.14 | 0.37 |
| 6 | 66610017 | rs9453479 | -0.077 | -1.134 | -0.472 | 1.041 | 0.013 | 0.044 | 0.044 | 0.019 | 0.32 | 0.38 | 0.36 | 0.43 | 0.36 | 0.3 |
| 6 | 66622346 | rs13193633 | 0.744 | -1.028 | -0.817 | 0.931 | -0.031 | 0.01 | 0.01 | 0 | 0.09 | 0.02 | 0.15 | 0.1 | 0.09 | 0.17 |
| 6 | 66624682 | rs9453509 | 1.1 | -0.94 | -1.075 | 0.648 | 0.014 | -0.005 | -0.005 | 0 | 0.14 | 0.28 | 0.1 | 0.07 | 0.12 | 0.05 |
| 6 | 66628392 | rs4424062 | -0.121 | -0.923 | -1.605 | 0.591 | 0.003 | 0.01 | 0.01 | 0.592 | 0.6 | 0.5 | 0.55 | 0.5 | 0.54 | 0.57 |
| 6 | 66633603 | rs12214948 | 2.393 | -1.13 | -1.807 | 0.477 | -0.099 | 0.1 | 0.1 | 0 | 0.23 | 0.01 | 0.14 | 0.08 | 0.3 | 0.14 |
| 6 | 66636204 | rs6939967 | -0.266 | -0.987 | -1.698 | 0.386 | -0.014 | 0.012 | 0.012 | 0.134 | 0.35 | 0.42 | 0.4 | 0.42 | 0.24 | 0.43 |
| 6 | 66646546 | rs767069 | 1.176 | -1.022 | -1.797 | 0.716 | -0.05 | -0.066 | -0.066 | 7.721 | 0.21 | 0.08 | 0.3 | 0.34 | 0.32 | 0.25 |
| 6 | 66652098 | rs9294676 | 0.589 | -1.318 | -1.278 | 0.941 | 0.046 | -0.052 | -0.052 | 0 | 0.11 | 0.04 | 0.14 | 0.26 | 0.03 | 0.22 |
| 6 | 66655301 | rs13191573 | 0.282 | -0.459 | -0.837 | 0.326 | -0.032 | 0.014 | 0.014 | 0 | 0.09 | 0.02 | 0.16 | 0.1 | 0.09 | 0.15 |
| 6 | 66656097 | rs1903406 | 0.145 | -0.597 | -0.503 | 0.453 | -0.038 | -0.01 | -0.01 | 0 | 0.81 | 0.65 | 0.86 | 0.84 | 0.87 | 0.82 |
| 6 | 66660054 | rs9453525 | -0.044 | -0.088 | -1.062 | 0.331 | -0.029 | 0.018 | 0.018 | 0 | 0.11 | 0.03 | 0.08 | 0.11 | 0.14 | 0.19 |
| 6 | 66664477 | rs9453530 | -0.469 | -0.672 | -0.32 | 0.167 | -0.011 | -0.018 | -0.018 | 0.012 | 0.26 | 0.36 | 0.3 | 0.24 | 0.26 | 0.26 |
| 6 | 66667351 | rs9363500 | 1.045 | -0.302 | -0.759 | 0.448 | -0.04 | -0.036 | -0.036 | 0 | 0.17 | 0.06 | 0.28 | 0.29 | 0.28 | 0.23 |
| 6 | 66668793 | rs981469 | -1.346 | -0.927 | -0.648 | 0.056 | -0.025 | -0.04 | -0.04 | 0 | 0.91 | 0.88 | 0.96 | 1 | 0.97 | 0.93 |
| 6 | 66669464 | rs2061578 | -0.643 | -1.27 | -0.745 | 0.02 | 0 | 0 | 0 | 0.002 | 0.66 | 0.65 | 0.76 | 0.71 | 0.81 | 0.65 |
| 6 | 66672155 | rs1037883 | -1.611 | -1.244 | -0.851 | 0.023 | -0.025 | -0.04 | -0.04 | 0 | 0.9 | 0.88 | 0.96 | 0.99 | 0.97 | 0.93 |
| 6 | 66672165 | rs1037882 | 1.168 | -1.312 | -0.848 | 0.002 | 0.019 | -0.048 | -0.048 | 0 | 0.24 | 0.13 | 0.2 | 0.28 | 0.16 | 0.28 |
| 6 | 66674314 | rs9345734 | 1.845 | -1.157 | -0.777 | 0.039 | -0.028 | -0.025 | -0.025 | 1.804 | 0.16 | 0.08 | 0.28 | 0.29 | 0.28 | 0.23 |
| 6 | 66676508 | rs9345736 | -0.196 | -1.273 | -0.656 | -0.064 | -0.038 | -0.055 | -0.055 | 0 | 0.64 | 0.49 | 0.66 | 0.76 | 0.71 | 0.66 |
| 6 | 66687508 | rs10944868 | 2.254 | -1.333 | -1.171 | 0.01 | -0.038 | 0.03 | 0.03 | 0 | 0.19 | 0.08 | 0.17 | 0.1 | 0.26 | 0.14 |
| 6 | 66688490 | rs1484576 | -1.627 | -1.371 | -1.044 | 0.149 | -0.025 | -0.04 | -0.04 | 0 | 0.9 | 0.88 | 0.96 | 0.99 | 0.97 | 0.93 |
| 6 | 66689252 | rs9345745 | 1.383 | -1.348 | -1.551 | 0.211 | 0.027 | -0.034 | -0.034 | 2.05 | 0.13 | 0.09 | 0.14 | 0.29 | 0.05 | 0.23 |
| 6 | 66690190 | rs12529678 | 1.636 | -1.139 | -1.333 | 0.107 | 0.012 | 0.002 | 0.002 | 0 | 0.11 | 0.04 | 0.06 | 0.11 | 0.11 | 0.1 |
| 6 | 66694905 | rs10498842 | 1.143 | -0.911 | -1.249 | 0.19 | -0.028 | -0.002 | -0.002 | 0 | 0.12 | 0.01 | 0.19 | 0.13 | 0.14 | 0.17 |
| 6 | 66697003 | rs7766730 | -0.038 | -0.686 | -0.681 | 0.443 | -0.006 | 0.003 | 0.003 | 2.459 | 0.47 | 0.39 | 0.38 | 0.47 | 0.43 | 0.42 |
| 6 | 66701759 | rs6455065 | 0.471 | -0.686 | -0.925 | 0.85 | -0.006 | -0.061 | -0.061 | 0 | 0.22 | 0.08 | 0.22 | 0.37 | 0.17 | 0.26 |
| 6 | 66712662 | rs9453563 | 0.773 | -0.529 | -1.124 | 0.656 | -0.003 | -0.022 | -0.022 | 0 | 0.21 | 0.1 | 0.25 | 0.25 | 0.22 | 0.25 |
| 6 | 66713643 | rs10498843 | 1.301 | -0.642 | -1.391 | 0.436 | -0.018 | 0.003 | 0.003 | 0 | 0.11 | 0.01 | 0.09 | 0.07 | 0.14 | 0.1 |
| 6 | 66719597 | rs6899720 | 0.077 | -0.938 | -0.964 | 0.078 | 0.014 | 0.029 | 0.029 | 3.455 | 0.54 | 0.37 | 0.61 | 0.72 | 0.59 | 0.64 |
| 6 | 66722969 | rs6455069 | 1.92 | -0.835 | -1.192 | 0.093 | -0.035 | 0.054 | 0.054 | 3.817 | 0.1 | 0.02 | 0.06 | 0 | 0.11 | 0.04 |
| 6 | 66739530 | rs10455596 | 0.545 | -0.563 | -1.148 | 0.183 | -0.078 | -0.008 | -0.008 | 0 | 0.38 | 0.11 | 0.41 | 0.33 | 0.49 | 0.41 |
| 6 | 66742205 | rs13200955 | 0.363 | -1.112 | -0.666 | 0.116 | -0.006 | 0.066 | 0.066 | 1.543 | 0.33 | 0.26 | 0.34 | 0.19 | 0.37 | 0.32 |
| 6 | 66754285 | rs6913344 | 0.502 | -0.967 | -0.85 | 0.173 | -0.027 | -0.037 | -0.037 | 0.174 | 0.33 | 0.22 | 0.34 | 0.47 | 0.35 | 0.39 |
| 6 | 66757302 | rs10498844 | 1.124 | -0.785 | -1.046 | 0.267 | -0.009 | -0.001 | -0.001 | 0 | 0.18 | 0.08 | 0.15 | 0.14 | 0.2 | 0.17 |
| 6 | 66760443 | rs12202343 | 0.441 | -0.814 | -0.944 | 0.252 | -0.066 | 0.054 | 0.054 | 2.532 | 0.37 | 0.14 | 0.32 | 0.22 | 0.47 | 0.33 |
| 6 | 66763069 | rs1385225 | -1.666 | -0.871 | -0.548 | 0.451 | -0.011 | 0.001 | 0.001 | 0.373 | 0.9 | 0.89 | 0.87 | 0.88 | 0.95 | 0.89 |
| 6 | 66764927 | rs17683993 | 0.862 | -0.573 | -0.558 | 0.445 | -0.006 | 0.001 | 0.001 | 4.432 | 0.19 | 0.11 | 0.15 | 0.14 | 0.2 | 0.17 |
| 6 | 66781190 | rs9354365 | 0.964 | -0.722 | -0.382 | 0.781 | -0.001 | -0.003 | -0.003 | 0 | 0.12 | 0.14 | 0.18 | 0.3 | 0.09 | 0.23 |
| 6 | 66796668 | rs9453585 | -0.639 | -0.829 | -0.707 | 0.474 | -0.023 | -0.01 | -0.01 | 0 | 0.68 | 0.77 | 0.65 | 0.65 | 0.65 | 0.61 |
| 6 | 66814819 | rs10080512 | -0.21 | -0.973 | -0.606 | 0.51 | 0.031 | 0.033 | 0.033 | 2 | 0.5 | 0.65 | 0.51 | 0.42 | 0.46 | 0.44 |
| 6 | 66817912 | rs9363524 | 0.991 | -0.856 | -0.489 | 0.838 | -0.004 | 0.008 | 0.008 | 0 | 0.12 | 0.2 | 0.16 | 0.18 | 0.09 | 0.14 |
| 6 | 66826432 | rs9453597 | 0.718 | -0.961 | -0.532 | -0.22 | 0 | 0.023 | 0.023 | 1.608 | 0.22 | 0.17 | 0.16 | 0.02 | 0.23 | 0.15 |
| 6 | 66828794 | rs3900131 | 0.539 | -1.062 | -0.721 | -0.184 | 0.003 | 0.024 | 0.024 | 0 | 0.23 | 0.15 | 0.29 | 0.19 | 0.27 | 0.28 |
| 6 | 66830696 | rs10214532 | -0.271 | -1.073 | -0.113 | 0.012 | 0.003 | 0.011 | 0.011 | 2.379 | 0.49 | 0.61 | 0.54 | 0.58 | 0.48 | 0.56 |
| 6 | 66840711 | rs9354379 | -0.274 | -1.005 | -0.027 | 0.016 | 0.003 | 0.011 | 0.011 | 2.287 | 0.49 | 0.61 | 0.54 | 0.58 | 0.48 | 0.56 |
| 6 | 66843649 | rs9354382 | -0.309 | -0.984 | -0.065 | 0.032 | 0.006 | 0.019 | 0.019 | 2.445 | 0.5 | 0.64 | 0.54 | 0.58 | 0.48 | 0.56 |
| 6 | 66846732 | rs2797690 | 0.176 | -1.019 | -0.152 | 0.109 | -0.002 | -0.007 | -0.007 | 0.06 | 0.32 | 0.27 | 0.36 | 0.38 | 0.35 | 0.34 |
| 6 | 66852482 | rs1103371 | 0.818 | -0.795 | -0.526 | 0.172 | -0.001 | -0.003 | -0.003 | 0 | 0.13 | 0.1 | 0.22 | 0.17 | 0.16 | 0.17 |
| 6 | 66890454 | rs6928834 | 0.412 | -0.815 | -0.464 | 0.111 | -0.005 | 0.004 | 0.004 | 0 | 0.27 | 0.34 | 0.31 | 0.35 | 0.24 | 0.38 |
| 6 | 66891318 | rs7751095 | -0.529 | -0.84 | -0.557 | -0.021 | -0.033 | 0.033 | 0.033 | 0.702 | 0.67 | 0.83 | 0.61 | 0.76 | 0.57 | 0.65 |
| 6 | 66894095 | rs1998226 | 1.502 | -0.828 | -0.962 | 0.055 | 0.006 | -0.001 | -0.001 | 0.107 | 0.1 | 0.08 | 0.12 | 0.12 | 0.05 | 0.09 |
| 6 | 66897470 | rs208436 | -0.776 | -0.808 | -1 | 0.023 | -0.029 | 0.053 | 0.053 | 0 | 0.7 | 0.9 | 0.73 | 0.82 | 0.69 | 0.74 |
| 6 | 66905763 | rs12111378 | 0.656 | -0.753 | -0.898 | 0.107 | 0.006 | 0.009 | 0.009 | 0 | 0.29 | 0.33 | 0.19 | 0.29 | 0.28 | 0.19 |
| 6 | 66916257 | rs6919921 | -0.242 | -0.686 | -0.966 | 0.099 | -0.004 | 0.019 | 0.019 | 0 | 0.58 | 0.73 | 0.51 | 0.64 | 0.55 | 0.58 |
| 6 | 66919338 | rs208476 | 0.449 | -0.709 | -0.967 | 0.097 | -0.009 | 0.001 | 0.001 | 0 | 0.24 | 0.25 | 0.33 | 0.3 | 0.2 | 0.3 |
| 6 | 66922606 | rs9360188 | 0.388 | -0.717 | -0.944 | 0.084 | -0.004 | 0 | 0 | 0 | 0.29 | 0.36 | 0.31 | 0.35 | 0.26 | 0.41 |
| 6 | 66925793 | rs2106555 | -0.146 | -0.575 | -1.245 | -0.307 | -0.004 | 0.019 | 0.019 | 0 | 0.58 | 0.74 | 0.51 | 0.64 | 0.55 | 0.58 |
| 6 | 66927774 | rs208490 | 0.395 | -0.562 | -1.19 | -0.26 | -0.01 | 0.002 | 0.002 | 0 | 0.24 | 0.24 | 0.33 | 0.29 | 0.2 | 0.3 |
| 6 | 66931208 | rs6903678 | -0.195 | 0.174 | -1.099 | -0.093 | -0.006 | 0.005 | 0.005 | 0.051 | 0.68 | 0.71 | 0.62 | 0.77 | 0.58 | 0.67 |
| 6 | 66931399 | rs208497 | 0.854 | 0.006 | -1.064 | -0.054 | 0.001 | -0.003 | -0.003 | 0 | 0.13 | 0.11 | 0.21 | 0.16 | 0.15 | 0.2 |
| 6 | 66940263 | rs208507 | 0.981 | 0.012 | -1.092 | -0.051 | -0.003 | -0.013 | -0.013 | 0 | 0.13 | 0.06 | 0.21 | 0.16 | 0.15 | 0.2 |
| 6 | 66942478 | rs208510 | 0.591 | 0.015 | -1.13 | -0.049 | -0.005 | 0.003 | 0.003 | 0 | 0.15 | 0.19 | 0.22 | 0.17 | 0.15 | 0.2 |
| 6 | 66945749 | rs208515 | -1.332 | 0.062 | -1.212 | -0.074 | -0.089 | 0.028 | 0.028 | 0 | 0.84 | 0.99 | 0.85 | 0.93 | 0.75 | 0.88 |
| 6 | 66948909 | rs6455090 | -0.251 | 0.048 | -1.169 | -0.074 | 0 | -0.001 | -0.001 | 0 | 0.68 | 0.69 | 0.62 | 0.77 | 0.6 | 0.67 |
| 6 | 66952385 | rs1029400 | -0.261 | 0.066 | -1.16 | -0.081 | 0 | -0.001 | -0.001 | 0.835 | 0.68 | 0.69 | 0.62 | 0.77 | 0.6 | 0.69 |
| 6 | 66952828 | rs208520 | -1.35 | 0.137 | -1.142 | -0.033 | -0.026 | 0.025 | 0.025 | 0 | 0.85 | 0.9 | 0.85 | 0.93 | 0.77 | 0.88 |

| | | | | | | | | | | | | | | | | |
|---|---|---|---|---|---|---|---|---|---|---|---|---|---|---|---|---|
| 6 | 66953453 | rs4710584 | -0.416 | 0.144 | -1.258 | -0.001 | 0.002 | -0.003 | -0.003 | 0 | 0.7 | 0.69 | 0.63 | 0.77 | 0.62 | 0.69 |
| 6 | 66957869 | rs9453647 | 0.194 | 0.038 | -1.443 | 0.081 | 0.005 | 0.006 | 0.006 | 0 | 0.28 | 0.2 | 0.3 | 0.28 | 0.26 | 0.42 |
| 6 | 66981393 | rs13209666 | 0.708 | 0.013 | -1.588 | -0.019 | 0.003 | 0.074 | 0.074 | 0 | 0.14 | 0.01 | 0.11 | 0.03 | 0.15 | 0.19 |
| 6 | 66987179 | rs9453670 | 0.196 | 0.074 | -1.499 | 0.049 | 0.003 | 0.005 | 0.005 | 0 | 0.28 | 0.23 | 0.3 | 0.27 | 0.26 | 0.4 |
| 6 | 66995395 | rs2078901 | 0.437 | 0.395 | -1.962 | 0.105 | -0.003 | 0.004 | 0.004 | 0.338 | 0.42 | 0.48 | 0.33 | 0.52 | 0.36 | 0.29 |
| 6 | 66999917 | rs12206488 | 0.768 | 1.12 | -2.439 | -0.03 | -0.072 | 0.013 | 0.013 | 0 | 0.14 | 0.01 | 0.13 | 0.07 | 0.21 | 0.11 |
| 6 | 67002076 | rs7449962 | -0.701 | 1.132 | -2.27 | 0.333 | -0.005 | -0.044 | -0.044 | 0 | 0.78 | 0.89 | 0.67 | 0.61 | 0.75 | 0.76 |
| 6 | 67004518 | rs13217132 | 0.259 | 1.409 | -2.032 | -0.053 | 0.024 | -0.012 | -0.012 | 0 | 0.22 | 0.08 | 0.17 | 0.21 | 0.15 | 0.22 |
| 6 | 67005173 | rs11760090 | -0.218 | 2.281 | -1.117 | 0.246 | -0.019 | 0.04 | 0.04 | 0 | 0.09 | 0.01 | 0.09 | 0.03 | 0.15 | 0.14 |
| 6 | 67008286 | rs2106553 | -0.595 | 2.354 | -0.407 | 0.648 | 0.036 | -0.1 | -0.1 | 0 | 0.14 | 0.35 | 0.09 | 0 | 0.12 | 0.05 |
| 6 | 67017543 | rs9345809 | -0.635 | 2.418 | -1.368 | 0.959 | -0.048 | -0.045 | -0.045 | 0 | 0.07 | 0.01 | 0.12 | 0.16 | 0.18 | 0.08 |
| 6 | 67039740 | rs1398803 | 0.511 | 2.2 | -1.659 | 0.163 | -0.069 | 0.082 | 0.082 | 0 | 0.72 | 0.89 | 0.71 | 0.93 | 0.65 | 0.8 |
| 6 | 67044431 | rs1708552 | 0.754 | 1.335 | -1.236 | 0.667 | -0.011 | 0.009 | 0.009 | 1.363 | 0.56 | 0.69 | 0.59 | 0.62 | 0.55 | 0.76 |
| 6 | 67049406 | rs1708558 | -0.55 | -0.052 | -0.84 | 0.435 | -0.032 | 0.018 | 0.018 | 0 | 0.27 | 0.33 | 0.16 | 0.41 | 0.13 | 0.17 |
| 6 | 67054884 | rs851860 | -0.715 | 0.52 | -1.154 | 0.099 | -0.03 | 0.047 | 0.047 | 0.04 | 0.82 | 0.95 | 0.79 | 0.95 | 0.82 | 0.89 |
| 6 | 67054961 | rs851859 | -0.04 | 0.493 | -1.017 | -0.204 | 0.038 | 0.049 | 0.049 | 0 | 0.5 | 0.59 | 0.49 | 0.76 | 0.38 | 0.64 |
| 6 | 67060189 | rs3846814 | 1.249 | 0.846 | -0.878 | -0.468 | -0.003 | -0.003 | -0.003 | 0 | 0.07 | 0.06 | 0.12 | 0.17 | 0.1 | 0.1 |
| 6 | 67060771 | rs17702764 | 1.43 | 0.739 | -0.557 | -0.618 | 0.009 | -0.024 | -0.024 | 0 | 0.07 | 0.09 | 0.05 | 0 | 0.09 | 0.03 |
| 6 | 67063913 | rs851880 | 0.802 | 0.344 | -0.505 | -0.73 | -0.01 | 0.003 | 0.003 | 0 | 0.21 | 0.24 | 0.14 | 0.32 | 0.12 | 0.11 |
| 6 | 67070206 | rs9363581 | 1.086 | 0.362 | -0.526 | -0.72 | -0.015 | 0.044 | 0.044 | 0 | 0.07 | 0.17 | 0.03 | 0.1 | 0.02 | 0.05 |
| 6 | 67072560 | rs12192666 | 0.514 | 0.275 | -0.681 | -0.786 | -0.088 | 0.086 | 0.086 | 0 | 0.16 | 0.01 | 0.13 | 0.01 | 0.25 | 0.12 |
| 6 | 67079357 | rs2882826 | 1.744 | 0.083 | -0.785 | -0.868 | -0.024 | -0.12 | -0.12 | 0 | 0.21 | 0.02 | 0.31 | 0.35 | 0.22 | 0.47 |
| 6 | 67081110 | rs1101054 | 0.813 | 0.269 | -0.579 | -1.072 | 0.018 | 0.013 | 0.013 | 0 | 0.43 | 0.39 | 0.45 | 0.67 | 0.34 | 0.59 |
| 6 | 67088132 | rs6900214 | -0.801 | 0.136 | -0.709 | -1.135 | -0.006 | 0.015 | 0.015 | 0 | 0.72 | 0.79 | 0.66 | 0.79 | 0.71 | 0.79 |
| 6 | 67092438 | rs9363588 | -0.005 | 0.409 | -0.38 | -0.927 | 0.017 | -0.006 | -0.006 | 0 | 0.13 | 0.29 | 0.08 | 0.1 | 0.1 | 0.08 |
| 6 | 67095924 | rs3846816 | -0.994 | 0.539 | -0.41 | -0.957 | 0.004 | -0.012 | -0.012 | 0 | 0.29 | 0.3 | 0.2 | 0.11 | 0.36 | 0.21 |
| 6 | 67098218 | rs851854 | -0.894 | 0.507 | -0.203 | -0.814 | -0.004 | -0.043 | -0.043 | 0.105 | 0.19 | 0.41 | 0.11 | 0.1 | 0.14 | 0.16 |
| 6 | 67098337 | rs9363590 | 0.047 | 0.462 | -0.405 | -0.731 | 0.006 | -0.003 | -0.003 | 0 | 0.09 | 0.06 | 0.03 | 0.1 | 0.05 | 0.08 |
| 6 | 67102644 | rs3914329 | 0.216 | 0.811 | -0.398 | -0.582 | -0.008 | 0.001 | 0.001 | 0 | 0.13 | 0.11 | 0.16 | 0.15 | 0.17 | 0.14 |
| 6 | 67106679 | rs9445701 | -1.463 | 0.692 | -0.49 | -0.73 | -0.009 | 0.011 | 0.011 | 0.544 | 0.33 | 0.67 | 0.33 | 0.28 | 0.36 | 0.33 |
| 6 | 67110160 | rs7754676 | 0.329 | 0.873 | -0.409 | -0.649 | 0.011 | -0.002 | -0.002 | 0.505 | 0.14 | 0.14 | 0.16 | 0.14 | 0.18 | 0.14 |
| 6 | 67115389 | rs17703383 | 0.588 | 0.88 | -0.443 | -0.652 | -0.016 | 0.021 | 0.021 | 0 | 0.05 | 0.03 | 0.06 | 0.01 | 0.14 | 0.03 |
| 6 | 67123130 | rs9283828 | -0.007 | 0.863 | -0.104 | -0.661 | 0.087 | -0.031 | -0.031 | 0 | 0.07 | 0.18 | 0.07 | 0.01 | 0.14 | 0.03 |
| 6 | 67123755 | rs1464072 | -0.969 | 0.855 | -0.028 | -0.589 | 0.062 | -0.01 | -0.01 | 0 | 0.25 | 0.39 | 0.28 | 0.28 | 0.32 | 0.2 |
| 6 | 67123794 | rs1567793 | -0.85 | 0.596 | -0.296 | -0.112 | 0.028 | 0.012 | 0.012 | 0.008 | 0.32 | 0.39 | 0.43 | 0.45 | 0.39 | 0.36 |
| 6 | 67128840 | rs10484416 | 0.449 | 0.59 | -0.322 | -0.027 | 0.067 | -0.023 | -0.023 | 0 | 0.06 | 0.13 | 0.06 | 0.01 | 0.14 | 0.03 |
| 6 | 67139880 | rs6455106 | -1.026 | 0.581 | -0.684 | -0.103 | -0.007 | 0.001 | 0.001 | 0.569 | 0.42 | 0.66 | 0.49 | 0.45 | 0.47 | 0.5 |
| 6 | 67143902 | rs2812152 | -0.261 | 0.882 | -1.014 | 0.272 | -0.005 | -0.006 | -0.006 | 0 | 0.23 | 0.22 | 0.38 | 0.44 | 0.27 | 0.33 |
| 6 | 67149979 | rs4710604 | -0.446 | 0.553 | -1.123 | 0.954 | -0.048 | -0.111 | -0.111 | 0 | 0.79 | 0.98 | 0.8 | 0.69 | 0.75 | 0.78 |
| 6 | 67154418 | rs9342552 | 0.433 | 0.476 | -0.727 | 0.218 | -0.04 | -0.017 | -0.017 | 0 | 0.4 | 0.19 | 0.43 | 0.4 | 0.48 | 0.41 |
| 6 | 67157506 | rs6924193 | 0.217 | 0.221 | 0.518 | -0.699 | 0.025 | -0.117 | -0.117 | 0 | 0.77 | 0.5 | 0.83 | 0.96 | 0.78 | 0.81 |
| 6 | 67167633 | rs2670379 | -0.09 | 0.233 | 0.536 | -0.728 | -0.017 | 0.032 | 0.032 | 0 | 0.83 | 0.92 | 0.87 | 0.96 | 0.8 | 0.83 |
| 6 | 67179287 | rs9345835 | -0.405 | 0.194 | 0.609 | -0.732 | 0.001 | 0.043 | 0.043 | 0.008 | 0.88 | 1 | 0.93 | 0.98 | 0.93 | 0.86 |
| 6 | 67185712 | rs9345837 | 1.469 | -0.068 | 0.733 | -1.034 | 0.004 | 0.056 | 0.056 | 0.345 | 0.69 | 0.22 | 0.78 | 0.96 | 0.73 | 0.74 |
| 6 | 67187759 | rs6936132 | 0.621 | -0.184 | 1.123 | -0.984 | -0.01 | -0.036 | -0.036 | 3.282 | 0.07 | 0.13 | 0.04 | 0 | 0.05 | 0.07 |
| 6 | 67188306 | rs2812171 | 0.664 | -0.211 | 1.495 | -1.011 | -0.028 | -0.066 | -0.066 | 0.626 | 0.77 | 0.66 | 0.87 | 0.97 | 0.88 | 0.78 |
| 6 | 67189717 | rs4710610 | -0.124 | -0.15 | 1.049 | -0.959 | 0.012 | 0.013 | 0.013 | 0 | 0.85 | 0.87 | 0.92 | 0.97 | 0.94 | 0.86 |
| 6 | 67194108 | rs10484419 | -0.436 | -0.288 | 0.936 | -0.908 | -0.019 | -0.011 | -0.011 | 0.023 | 0.15 | 0.24 | 0.05 | 0.1 | 0.09 | 0.08 |
| 6 | 67194835 | rs10484420 | -0.673 | -0.231 | 0.936 | -0.91 | 0.013 | -0.014 | -0.014 | 0 | 0.17 | 0.25 | 0.1 | 0.11 | 0.21 | 0.11 |
| 6 | 67196809 | rs6914260 | -0.694 | -0.234 | 0.97 | -0.923 | 0.02 | -0.02 | -0.02 | 0 | 0.17 | 0.27 | 0.1 | 0.11 | 0.21 | 0.11 |
| 6 | 67197596 | rs1398800 | 1.209 | -0.472 | 0.599 | -0.948 | 0.035 | 0.121 | 0.121 | 0 | 0.58 | 0.21 | 0.75 | 0.85 | 0.66 | 0.67 |
| 6 | 67206225 | rs2812184 | -0.726 | -0.48 | 0.715 | -0.993 | 0.007 | -0.049 | -0.049 | 0 | 0.27 | 0.36 | 0.14 | 0.12 | 0.27 | 0.2 |
| 6 | 67224061 | rs12212253 | 1.377 | -0.673 | 0.209 | -0.83 | 0.021 | 0.112 | 0.112 | 0 | 0.57 | 0.12 | 0.66 | 0.8 | 0.65 | 0.67 |
| 6 | 67228452 | rs2812143 | -0.312 | -0.735 | 0.323 | -0.886 | 0.002 | -0.004 | -0.004 | 0 | 0.92 | 0.95 | 0.85 | 0.92 | 0.96 | 0.9 |
| 6 | 67231094 | rs2670371 | -1.259 | -0.686 | 0.385 | -0.882 | 0.001 | 0.096 | 0.096 | 0 | 0.34 | 0.79 | 0.19 | 0.11 | 0.31 | 0.23 |
| 6 | 67238014 | rs12660876 | 0.259 | -0.6 | 0.203 | -0.905 | 0.023 | -0.016 | -0.016 | 0 | 0.08 | 0.07 | 0.15 | 0.08 | 0.04 | 0.1 |
| 6 | 67240297 | rs1224521 | -0.71 | -0.247 | 0.525 | -1.156 | -0.018 | -0.026 | -0.026 | 0 | 0.24 | 0.3 | 0.1 | 0.11 | 0.14 | 0.14 |
| 6 | 67245184 | rs1355799 | -0.032 | -0.323 | 0.4 | -0.953 | -0.012 | -0.009 | -0.009 | 0.55 | 0.18 | 0.19 | 0.06 | 0.1 | 0.08 | 0.11 |
| 6 | 67249532 | rs1224526 | -0.246 | 0.398 | 0.226 | -1.335 | 0.012 | -0.032 | -0.032 | 0 | 0.91 | 0.86 | 0.89 | 1 | 0.87 | 0.97 |
| 6 | 67250008 | rs1224524 | 0.162 | 0.056 | 0.147 | -1.139 | -0.001 | 0 | 0 | 0 | 0.18 | 0.16 | 0.06 | 0.1 | 0.08 | 0.11 |
| 6 | 67262816 | rs1028802 | 0.006 | 0.205 | -0.177 | -1.726 | -0.001 | -0.019 | -0.019 | 2.978 | 0.34 | 0.33 | 0.18 | 0.09 | 0.3 | 0.26 |
| 6 | 67265164 | rs7771415 | -0.37 | -0.43 | -0.154 | -2.029 | 0.014 | 0.013 | 0.013 | 1.923 | 0.47 | 0.6 | 0.21 | 0.13 | 0.34 | 0.41 |
| 6 | 67266198 | rs6899803 | 0.313 | -0.403 | -0.103 | -2.017 | -0.004 | -0.031 | -0.031 | 0 | 0.34 | 0.35 | 0.18 | 0.09 | 0.3 | 0.28 |
| 6 | 67266253 | rs9354442 | 0.924 | -0.208 | -0.273 | -1.495 | 0.009 | 0.028 | 0.028 | 0 | 0.18 | 0.21 | 0.27 | 0.38 | 0.24 | 0.2 |
| 6 | 67269926 | rs16898533 | 0.856 | -0.256 | -0.197 | -1.54 | -0.015 | -0.024 | -0.024 | 0.001 | 0.08 | 0.15 | 0.02 | 0.02 | 0.04 | 0.11 |
| 6 | 67270731 | rs1925793 | 0.576 | 0.079 | -0.299 | -1.653 | 0.001 | -0.025 | -0.025 | 1.341 | 0.37 | 0.37 | 0.21 | 0.09 | 0.41 | 0.3 |
| 6 | 67272216 | rs3920611 | 0.037 | 0.358 | -0.426 | -1.47 | 0.005 | 0.058 | 0.058 | 1.688 | 0.5 | 0.65 | 0.25 | 0.13 | 0.46 | 0.44 |
| 6 | 67272582 | rs9363611 | 0.435 | 0.106 | -0.118 | -1.132 | 0.025 | 0.052 | 0.052 | 0 | 0.19 | 0.29 | 0.28 | 0.38 | 0.27 | 0.21 |
| 6 | 67276790 | rs2148014 | 0.897 | 0.396 | -0.451 | -1.357 | -0.047 | -0.082 | -0.082 | 0 | 0.26 | 0.11 | 0.39 | 0.47 | 0.33 | 0.42 |

| | | | | | | | | | | | | | | | | |
|---|---|---|---|---|---|---|---|---|---|---|---|---|---|---|---|---|
| 6 | 67277460 | rs2352501 | 0.181 | 0.291 | -0.025 | -1.141 | 0.039 | 0.038 | 0.038 | 0 | 0.71 | 0.56 | 0.57 | 0.56 | 0.62 | 0.74 |
| 6 | 67278814 | rs10806542 | 0.508 | 0.146 | -0.136 | -1.034 | 0.005 | 0.02 | 0.02 | 0 | 0.19 | 0.22 | 0.33 | 0.44 | 0.28 | 0.21 |
| 6 | 67284077 | rs4142334 | 0.462 | 0.179 | -0.095 | -1.02 | 0.003 | 0.019 | 0.019 | 0 | 0.19 | 0.2 | 0.32 | 0.4 | 0.26 | 0.21 |
| 6 | 67285555 | rs13194883 | 1.125 | -0.018 | -0.421 | -1.296 | 0.006 | 0 | 0 | 0 | 0.22 | 0.21 | 0.09 | 0.04 | 0.12 | 0.17 |
| 6 | 67293317 | rs9453836 | 0.664 | 0.746 | -0.756 | -0.751 | -0.021 | -0.038 | -0.038 | 6.512 | 0.17 | 0.1 | 0.31 | 0.4 | 0.26 | 0.21 |
| 6 | 67305307 | rs1555821 | 0.771 | 0.468 | -0.995 | -0.835 | -0.064 | -0.108 | -0.108 | 0 | 0.32 | 0.1 | 0.43 | 0.49 | 0.41 | 0.48 |
| 6 | 67308320 | rs1925804 | 0.135 | 0.199 | -0.734 | -1.071 | 0.022 | 0.007 | 0.007 | 0 | 0.64 | 0.59 | 0.56 | 0.57 | 0.58 | 0.72 |
| 6 | 67309864 | rs2504159 | -0.24 | 0.831 | -0.822 | -0.773 | -0.011 | -0.009 | -0.009 | 0 | 0.6 | 0.68 | 0.46 | 0.51 | 0.47 | 0.5 |
| 6 | 67322556 | rs388232 | -0.108 | 1.729 | -0.324 | -0.04 | 0.043 | -0.045 | -0.045 | 0 | 0.92 | 0.85 | 0.9 | 1 | 0.88 | 0.98 |
| 6 | 67325681 | rs1455090 | -0.339 | 1.25 | -0.899 | 0.39 | -0.014 | -0.017 | -0.017 | 0 | 0.22 | 0.15 | 0.34 | 0.38 | 0.29 | 0.24 |
| 6 | 67330025 | rs9345858 | 0.163 | 0.637 | -1.025 | -0.052 | 0.008 | 0.06 | 0.06 | 0.082 | 0.49 | 0.3 | 0.68 | 0.62 | 0.48 | 0.55 |
| 6 | 67335750 | rs995522 | 0.748 | 0.335 | -0.866 | -0.359 | -0.005 | -0.02 | -0.02 | 0 | 0.06 | 0.05 | 0.02 | 0 | 0.03 | 0.05 |
| 6 | 67335843 | rs183895 | -0.526 | 0.312 | -0.779 | -0.43 | 0.029 | 0.059 | 0.059 | 0.043 | 0.48 | 0.69 | 0.28 | 0.36 | 0.46 | 0.43 |
| 6 | 67339041 | rs1563300 | -0.051 | 0.034 | -1.247 | -0.743 | -0.009 | 0.036 | 0.036 | 0 | 0.42 | 0.64 | 0.51 | 0.75 | 0.59 | 0.56 |
| 6 | 67348139 | rs1563301 | -0.175 | -0.167 | -0.58 | -0.796 | -0.032 | -0.042 | -0.042 | 0 | 0.77 | 0.69 | 0.92 | 0.9 | 0.9 | 0.82 |
| 6 | 67350269 | rs1869677 | 0.941 | -0.078 | -0.796 | -0.805 | -0.008 | 0.073 | 0.073 | 0 | 0.37 | 0.21 | 0.41 | 0.15 | 0.32 | 0.27 |
| 6 | 67360369 | rs1349981 | 1.064 | 0.223 | -0.661 | -0.546 | -0.008 | -0.002 | -0.002 | 0 | 0.16 | 0.12 | 0.31 | 0.49 | 0.23 | 0.3 |
| 6 | 67368150 | rs17527707 | 0.801 | 0.146 | -0.823 | -1.185 | -0.058 | -0.055 | -0.055 | 0 | 0.17 | 0.06 | 0.16 | 0.26 | 0.28 | 0.23 |
| 6 | 67372800 | rs12527276 | 1.221 | -0.005 | -0.99 | -1.557 | 0.001 | -0.033 | -0.033 | 0.577 | 0.13 | 0.14 | 0.08 | 0 | 0.13 | 0.08 |
| 6 | 67382664 | rs7738998 | -0.333 | -0.238 | -1.093 | -1.651 | 0.031 | 0.134 | 0.134 | 0.476 | 0.5 | 0.69 | 0.54 | 0.76 | 0.62 | 0.62 |
| 6 | 67384568 | rs9453873 | 1.236 | -0.253 | -0.592 | -1.675 | -0.001 | -0.028 | -0.028 | 0 | 0.06 | 0.12 | 0.03 | 0.01 | 0.07 | 0.03 |
| 6 | 67385829 | rs1869678 | 1.282 | -0.492 | -0.761 | -1.868 | -0.003 | 0.018 | 0.018 | 0.555 | 0.21 | 0.07 | 0.33 | 0.14 | 0.17 | 0.17 |
| 6 | 67387453 | rs9363624 | 1.608 | -0.346 | -0.484 | -1.591 | -0.014 | -0.016 | -0.016 | 0 | 0.15 | 0.12 | 0.31 | 0.49 | 0.22 | 0.3 |
| 6 | 67391355 | rs9294716 | 0.595 | -0.463 | -0.455 | -1.695 | -0.003 | 0.029 | 0.029 | 0 | 0.41 | 0.34 | 0.44 | 0.15 | 0.39 | 0.29 |
| 6 | 67392200 | rs2168248 | -1.118 | -0.447 | -0.155 | -1.616 | -0.008 | -0.004 | -0.004 | 2.797 | 0.84 | 0.84 | 0.95 | 0.89 | 0.94 | 0.88 |
| 6 | 67395750 | rs9453879 | 0.45 | -0.485 | -0.172 | -1.682 | -0.003 | -0.033 | -0.033 | 0 | 0.12 | 0.13 | 0.04 | 0.01 | 0.1 | 0.08 |
| 6 | 67398716 | rs2352603 | 0.591 | -0.471 | -0.179 | -1.678 | 0.007 | 0.04 | 0.04 | 0 | 0.22 | 0.26 | 0.34 | 0.5 | 0.3 | 0.33 |
| 6 | 67402631 | rs4493734 | -0.048 | -0.545 | -0.179 | -1.705 | -0.003 | 0.001 | 0.001 | 0 | 0.62 | 0.56 | 0.61 | 0.4 | 0.68 | 0.52 |
| 6 | 67403398 | rs1455097 | -1.148 | -0.553 | -0.185 | -1.705 | 0.037 | 0.028 | 0.028 | 3.131 | 0.85 | 0.89 | 0.96 | 0.89 | 0.94 | 0.88 |
| 6 | 67403591 | rs1455096 | 0.399 | -0.522 | 0.093 | -1.903 | -0.026 | -0.012 | -0.012 | 0.038 | 0.23 | 0.13 | 0.18 | 0.26 | 0.36 | 0.27 |
| 6 | 67403879 | rs7747972 | 0.23 | -0.622 | -0.013 | -1.646 | 0.004 | 0 | 0 | 0 | 0.57 | 0.56 | 0.6 | 0.4 | 0.6 | 0.49 |
| 6 | 67410283 | rs9294717 | -0.337 | -0.693 | -0.084 | -1.567 | 0.008 | 0.017 | 0.017 | 0 | 0.48 | 0.43 | 0.54 | 0.78 | 0.65 | 0.63 |
| 6 | 67410666 | rs9345882 | 1.104 | -0.715 | -0.244 | -1.513 | -0.005 | 0.002 | 0.002 | 0 | 0.17 | 0.19 | 0.28 | 0.46 | 0.24 | 0.29 |
| 6 | 67416255 | rs1378301 | 0.064 | -0.746 | -0.223 | -1.544 | -0.009 | 0.042 | 0.042 | 0 | 0.4 | 0.33 | 0.52 | 0.78 | 0.61 | 0.57 |
| 6 | 67419244 | rs6455132 | -0.908 | -0.906 | -0.274 | -1.62 | 0.015 | -0.002 | -0.002 | 0 | 0.9 | 0.9 | 0.92 | 0.91 | 0.96 | 0.93 |
| 6 | 67419300 | rs6942076 | 1.328 | -0.208 | -0.288 | -1.504 | -0.009 | -0.03 | -0.03 | 1.562 | 0.16 | 0.08 | 0.13 | 0.3 | 0.22 | 0.21 |
| 6 | 67428031 | rs1349979 | 0.434 | -0.198 | -0.298 | -1.506 | -0.055 | -0.035 | -0.035 | 0 | 0.23 | 0.09 | 0.23 | 0.32 | 0.37 | 0.28 |
| 6 | 67429337 | rs9342576 | -0.587 | -0.228 | -0.28 | -1.528 | 0.019 | 0.045 | 0.045 | 0 | 0.48 | 0.41 | 0.54 | 0.78 | 0.65 | 0.63 |
| 6 | 67433274 | rs1455082 | 0.338 | -0.015 | -0.462 | -1.397 | -0.028 | -0.017 | -0.017 | 0.056 | 0.23 | 0.14 | 0.24 | 0.32 | 0.37 | 0.29 |
| 6 | 67436761 | rs12197251 | 0.996 | 0.009 | -0.596 | -1.387 | -0.043 | 0.033 | 0.033 | 0 | 0.07 | 0.01 | 0.11 | 0.02 | 0.15 | 0.07 |
| 6 | 67441714 | rs4286759 | 1.199 | -0.135 | -0.512 | -1.311 | -0.008 | -0.027 | -0.027 | 0.121 | 0.16 | 0.08 | 0.13 | 0.3 | 0.22 | 0.22 |
| 6 | 67446248 | rs7746440 | 0.729 | -0.139 | -0.356 | -1.089 | -0.001 | 0.016 | 0.016 | 0 | 0.17 | 0.2 | 0.28 | 0.46 | 0.24 | 0.29 |
| 6 | 67448232 | rs6916073 | 0.9 | -0.083 | -0.73 | -1.06 | -0.029 | -0.053 | -0.053 | 0 | 0.16 | 0.13 | 0.28 | 0.46 | 0.24 | 0.29 |
| 6 | 67452650 | rs966984 | 0.114 | -0.145 | -0.473 | -1.092 | -0.011 | -0.018 | -0.018 | 0.139 | 0.3 | 0.28 | 0.35 | 0.49 | 0.4 | 0.41 |
| 6 | 67458231 | rs10447347 | 0.781 | -0.156 | -0.677 | -1.111 | -0.038 | 0.041 | 0.041 | 0 | 0.08 | 0.01 | 0.06 | 0.01 | 0.13 | 0.07 |
| 6 | 67467355 | rs4601103 | 0.353 | -0.48 | -0.588 | -1.46 | -0.003 | -0.009 | -0.009 | 1.303 | 0.29 | 0.31 | 0.35 | 0.13 | 0.22 | 0.21 |
| 6 | 67468362 | rs9351595 | 0.951 | -0.157 | -0.813 | -1.27 | -0.066 | -0.133 | -0.133 | 0 | 0.21 | 0.07 | 0.29 | 0.48 | 0.27 | 0.35 |
| 6 | 67473138 | rs11754594 | 0.512 | -0.252 | -0.67 | -1.389 | -0.019 | 0.005 | 0.005 | 0 | 0.28 | 0.12 | 0.2 | 0.29 | 0.34 | 0.31 |
| 6 | 67478540 | rs4458660 | 0.893 | -0.385 | -1.065 | -1.581 | -0.046 | 0.05 | 0.05 | 0 | 0.09 | 0 | 0.07 | 0 | 0.13 | 0.01 |
| 6 | 67480516 | rs13212321 | 0.463 | -0.388 | -0.807 | -1.359 | -0.03 | 0.002 | 0.002 | 0 | 0.28 | 0.09 | 0.19 | 0.29 | 0.34 | 0.31 |
| 6 | 67488602 | rs12201073 | 0.873 | -0.737 | -0.71 | -1.203 | -0.075 | -0.023 | -0.023 | 3.705 | 0.25 | 0.07 | 0.19 | 0.29 | 0.36 | 0.3 |
| 6 | 67488728 | rs12201163 | 0.115 | -0.89 | -0.684 | -1.285 | 0.034 | 0.109 | 0.109 | 0 | 0.47 | 0.18 | 0.48 | 0.77 | 0.64 | 0.65 |
| 6 | 67492398 | rs9360232 | 0.023 | -0.796 | -0.255 | -1.118 | -0.001 | -0.002 | -0.002 | 0.034 | 0.36 | 0.31 | 0.36 | 0.48 | 0.38 | 0.44 |
| 6 | 67501501 | rs4546468 | -0.828 | -0.916 | -0.123 | -1.168 | -0.004 | -0.001 | -0.001 | 0 | 0.69 | 0.69 | 0.59 | 0.77 | 0.77 | 0.75 |
| 6 | 67505713 | rs9354478 | 0.374 | -0.104 | -0.098 | -0.828 | -0.041 | -0.078 | -0.078 | 0.556 | 0.22 | 0.13 | 0.31 | 0.49 | 0.29 | 0.43 |
| 6 | 67519114 | rs12209528 | 0.833 | -0.175 | -0.134 | -0.941 | -0.048 | -0.008 | -0.008 | 0 | 0.25 | 0.1 | 0.18 | 0.28 | 0.37 | 0.25 |
| 6 | 67529319 | rs7748379 | -0.759 | -0.277 | -0.294 | -0.93 | -0.035 | -0.02 | -0.02 | 1.284 | 0.63 | 0.5 | 0.57 | 0.76 | 0.76 | 0.78 |
| 6 | 67541144 | rs4523069 | 0.158 | 0.317 | -0.779 | -0.658 | -0.005 | 0.009 | 0.009 | 0 | 0.4 | 0.32 | 0.26 | 0.28 | 0.48 | 0.35 |
| 6 | 67546906 | rs4395708 | 0.704 | 0.482 | -1.16 | -0.514 | -0.056 | -0.1 | -0.1 | 2.544 | 0.19 | 0.06 | 0.29 | 0.49 | 0.27 | 0.4 |
| 6 | 67548872 | rs13218643 | 0.463 | -0.267 | -0.538 | -0.899 | 0.001 | 0.016 | 0.016 | 0 | 0.13 | 0.04 | 0.06 | 0 | 0.09 | 0.1 |
| 6 | 67556311 | rs9354481 | 0.364 | -0.243 | -0.168 | -0.168 | -0.005 | -0.003 | -0.003 | 0.02 | 0.61 | 0.55 | 0.54 | 0.74 | 0.72 | 0.74 |
| 6 | 67557245 | rs11967591 | -0.906 | 0.321 | 0.375 | -0.305 | 0.028 | -0.078 | -0.078 | 0 | 0.12 | 0.38 | 0.08 | 0 | 0.1 | 0.03 |
| 6 | 67557621 | rs4554308 | -0.688 | 1.116 | -0.257 | -0.133 | -0.006 | 0 | 0 | 0 | 0.15 | 0.14 | 0.24 | 0.21 | 0.24 | 0.18 |
| 6 | 67557893 | rs4644033 | 1.166 | 0.707 | 0.151 | -0.526 | 0.002 | -0.019 | -0.019 | 0 | 0.69 | 0.35 | 0.68 | 0.82 | 0.66 | 0.66 |
| 6 | 67558004 | rs11757875 | 0.641 | -0.716 | -1.032 | -1.254 | 0.075 | 0.051 | 0.051 | 0 | 0.23 | 0.02 | 0.06 | 0.14 | 0.12 | 0.16 |
| 6 | 67559927 | rs9345900 | 0.004 | 0.089 | -1.155 | -1.578 | -0.009 | 0.038 | 0.038 | 0.088 | 0.39 | 0.29 | 0.58 | 0.79 | 0.52 | 0.52 |
| 6 | 67561944 | rs4544889 | -0.664 | -0.414 | -1.002 | -1.857 | 0.059 | 0.141 | 0.141 | 0.252 | 0.57 | 0.83 | 0.67 | 0.79 | 0.65 | 0.61 |
| 6 | 67563022 | rs9354486 | 0.003 | -0.425 | -0.852 | -1.869 | 0 | 0.007 | 0.007 | 0.036 | 0.43 | 0.39 | 0.6 | 0.78 | 0.52 | 0.52 |
| 6 | 67569562 | rs6911607 | -0.17 | -0.3 | -0.721 | -1.818 | -0.019 | 0.044 | 0.044 | 0.085 | 0.42 | 0.73 | 0.42 | 0.63 | 0.43 | 0.45 |

| | | | | | | | | | | | | | | | | |
|---|---|---|---|---|---|---|---|---|---|---|---|---|---|---|---|---|
| 6 | 67575005 | rs10455623 | 0.82 | -0.171 | -0.87 | -1.641 | -0.068 | -0.018 | -0.018 | 0.304 | 0.13 | 0.02 | 0.23 | 0.16 | 0.22 | 0.17 |
| 6 | 67575965 | rs9351606 | 0.412 | -0.123 | -0.678 | -1.596 | 0 | 0 | 0 | 0.195 | 0.29 | 0.27 | 0.35 | 0.63 | 0.3 | 0.36 |
| 6 | 67576472 | rs9445785 | -0.21 | 0.077 | -0.432 | -1.578 | 0.007 | -0.024 | -0.024 | 0 | 0.41 | 0.54 | 0.56 | 0.73 | 0.48 | 0.55 |
| 6 | 67576843 | rs7763651 | -0.035 | 0.342 | -0.406 | -0.843 | 0.01 | 0.05 | 0.05 | 0 | 0.77 | 0.66 | 0.69 | 0.52 | 0.76 | 0.67 |
| 6 | 67577304 | rs13196633 | 0.476 | 0.371 | -0.367 | 0.175 | 0 | 0.047 | 0.047 | 0 | 0.16 | 0.24 | 0.24 | 0.36 | 0.15 | 0.31 |
| 6 | 67577530 | rs9363649 | -0.789 | 0.316 | -0.282 | 0.112 | -0.004 | -0.028 | -0.028 | 0.007 | 0.95 | 0.86 | 0.97 | 1 | 0.96 | 0.98 |
| 6 | 67578852 | rs9453907 | 0.552 | 0.241 | -0.257 | 0.195 | 0.002 | 0.046 | 0.046 | 0 | 0.14 | 0.17 | 0.28 | 0.38 | 0.15 | 0.29 |
| 6 | 67582625 | rs9345915 | 1.332 | -0.348 | -0.717 | -0.304 | 0.01 | -0.004 | -0.004 | 0 | 0.17 | 0.08 | 0.08 | 0.22 | 0.14 | 0.16 |
| 6 | 67588332 | rs12207263 | 1.838 | -0.498 | -1.126 | -0.659 | -0.044 | -0.083 | -0.083 | 0 | 0.16 | 0.01 | 0.24 | 0.2 | 0.19 | 0.13 |
| 6 | 67588454 | rs9363656 | 0.336 | -0.184 | -0.62 | -0.843 | 0 | -0.002 | -0.002 | 0 | 0.43 | 0.36 | 0.38 | 0.42 | 0.44 | 0.36 |
| 6 | 67588475 | rs4544890 | -1.083 | -0.3 | -0.768 | -1.043 | 0.008 | -0.002 | -0.002 | 0 | 0.83 | 0.82 | 0.72 | 0.8 | 0.71 | 0.85 |
| 6 | 67589752 | rs9445788 | 0.79 | -0.308 | -0.73 | -1.047 | 0.036 | -0.044 | -0.044 | 1.089 | 0.08 | 0.25 | 0.06 | 0 | 0.1 | 0.06 |
| 6 | 67590453 | rs4631258 | -1.05 | -0.286 | -0.909 | -0.983 | 0.008 | -0.002 | -0.002 | 0.054 | 0.83 | 0.82 | 0.72 | 0.8 | 0.71 | 0.85 |
| 6 | 67621368 | rs6919455 | -0.069 | -0.434 | -0.945 | -1.008 | -0.007 | 0.001 | 0.001 | 0 | 0.4 | 0.4 | 0.34 | 0.38 | 0.28 | 0.5 |
| 6 | 67624339 | rs7753849 | 0.047 | -0.472 | -1.336 | -0.996 | 0 | 0.022 | 0.022 | 1.155 | 0.58 | 0.69 | 0.62 | 0.77 | 0.59 | 0.65 |
| 6 | 67625027 | rs7762782 | 1.861 | -0.903 | -1.23 | -0.871 | -0.043 | -0.083 | -0.083 | 0 | 0.16 | 0.01 | 0.24 | 0.2 | 0.19 | 0.14 |
| 6 | 67625265 | rs6455175 | -0.968 | -0.969 | -0.986 | -0.773 | -0.024 | 0.004 | 0.004 | 0 | 0.66 | 0.76 | 0.65 | 0.58 | 0.57 | 0.69 |
| 6 | 67626861 | rs6912062 | 0.154 | -0.96 | -0.942 | -0.976 | 0.016 | 0.005 | 0.005 | 0.196 | 0.26 | 0.15 | 0.1 | 0.02 | 0.13 | 0.2 |
| 6 | 67628407 | rs9445791 | -0.149 | -0.692 | -0.756 | -0.619 | 0.005 | 0.005 | 0.005 | 0 | 0.39 | 0.6 | 0.47 | 0.43 | 0.44 | 0.48 |
| 6 | 67635684 | rs4257844 | -0.14 | -1.617 | -0.718 | -0.786 | -0.025 | 0.027 | 0.027 | 0 | 0.76 | 0.68 | 0.63 | 0.41 | 0.88 | 0.59 |
| 6 | 67641300 | rs6912709 | 0.265 | -1.405 | -1.04 | -1.146 | 0.013 | 0.033 | 0.033 | 0.012 | 0.56 | 0.36 | 0.66 | 0.89 | 0.61 | 0.69 |
| 6 | 67642950 | rs7739520 | -0.059 | -1.589 | -1.046 | -1.165 | -0.031 | -0.085 | -0.085 | 0 | 0.23 | 0.36 | 0.14 | 0.02 | 0.15 | 0.1 |
| 6 | 67646913 | rs4481415 | 1.085 | -0.964 | -1.881 | -0.938 | -0.163 | -0.045 | -0.045 | 0 | 0.3 | 0.05 | 0.25 | 0.26 | 0.43 | 0.26 |
| 6 | 67648681 | rs4489139 | -0.11 | -1.644 | -1.827 | -1.212 | 0.046 | 0.001 | 0.001 | 0 | 0.58 | 0.79 | 0.49 | 0.4 | 0.67 | 0.52 |
| 6 | 67650186 | rs4573065 | 1.683 | -1.48 | -1.754 | -1.235 | -0.009 | -0.012 | -0.012 | 0 | 0.12 | 0.02 | 0.15 | 0.12 | 0.14 | 0.14 |
| 6 | 67651277 | rs4524594 | -0.929 | -1.762 | -1.841 | -1.146 | 0.013 | 0.047 | 0.047 | 0 | 0.74 | 0.83 | 0.84 | 0.88 | 0.78 | 0.78 |
| 6 | 67652222 | rs13208784 | 1.593 | -1.095 | -1.199 | -0.866 | -0.034 | -0.019 | -0.019 | 0.147 | 0.2 | 0.14 | 0.21 | 0.27 | 0.37 | 0.2 |
| 6 | 67653117 | rs12529449 | 0.679 | -1.369 | -0.715 | -1.02 | 0.037 | -0.003 | -0.003 | 0 | 0.3 | 0.57 | 0.28 | 0.27 | 0.45 | 0.24 |
| 6 | 67654492 | rs4314479 | -0.159 | -1.824 | -0.557 | -1.199 | -0.035 | 0.035 | 0.035 | 0 | 0.28 | 0.37 | 0.45 | 0.54 | 0.17 | 0.39 |
| 6 | 67655625 | rs4298326 | -0.806 | -1.808 | -0.773 | -1.517 | 0.017 | 0.055 | 0.055 | 0 | 0.69 | 0.78 | 0.8 | 0.89 | 0.73 | 0.71 |
| 6 | 67656128 | rs9360252 | 1.765 | -1.479 | -0.908 | -1.368 | -0.033 | -0.02 | -0.02 | 0.001 | 0.12 | 0.01 | 0.13 | 0.12 | 0.16 | 0.11 |
| 6 | 67660858 | rs6930795 | -0.901 | -1.499 | -0.491 | -1.299 | 0 | -0.062 | -0.062 | 0 | 0.88 | 0.73 | 0.9 | 1 | 0.88 | 0.98 |
| 6 | 67664566 | rs9345935 | 0.739 | -1.748 | -0.502 | -1.442 | 0.014 | 0.041 | 0.041 | 0 | 0.4 | 0.25 | 0.32 | 0.23 | 0.4 | 0.43 |
| 6 | 67666442 | rs4422596 | -1.096 | -1.597 | -0.416 | -1.205 | -0.003 | -0.001 | -0.001 | 0 | 0.89 | 0.91 | 0.92 | 1 | 0.88 | 0.98 |
| 6 | 67672896 | rs4373347 | 1.631 | -1.791 | -0.44 | -1.208 | -0.025 | -0.01 | -0.01 | 0 | 0.18 | 0.12 | 0.18 | 0.24 | 0.34 | 0.2 |
| 6 | 67674622 | rs6908019 | -0.923 | -1.741 | -0.577 | -1.393 | -0.001 | -0.042 | -0.042 | 0.269 | 0.89 | 0.83 | 0.92 | 1 | 0.88 | 0.98 |
| 6 | 67680571 | rs9354514 | 0.173 | -1.824 | -0.881 | -1.218 | -0.04 | 0.052 | 0.052 | 0 | 0.37 | 0.19 | 0.55 | 0.77 | 0.42 | 0.5 |
| 6 | 67681572 | rs9345943 | 0.165 | -1.824 | -0.881 | -1.218 | -0.04 | 0.052 | 0.052 | 0 | 0.37 | 0.19 | 0.55 | 0.77 | 0.42 | 0.5 |
| 6 | 67688283 | rs4416661 | 1.206 | -2.106 | -0.588 | -0.679 | -0.021 | 0.005 | 0.005 | 0 | 0.42 | 0.25 | 0.31 | 0.42 | 0.49 | 0.4 |
| 6 | 67689857 | rs9453947 | -0.933 | -2.482 | -0.014 | -0.194 | -0.008 | 0.005 | 0.005 | 4.142 | 0.08 | 0.54 | 0.03 | 0.02 | 0.01 | 0.05 |
| 6 | 67700383 | rs4610540 | -0.444 | -2.144 | -0.678 | -0.038 | 0.001 | -0.002 | -0.002 | 0 | 0.9 | 0.93 | 0.88 | 0.88 | 0.89 | 0.87 |
| 6 | 67705734 | rs9453960 | -0.101 | -2.012 | -0.646 | 0.1 | 0.058 | -0.089 | -0.089 | 0.212 | 0.2 | 0.06 | 0.34 | 0.43 | 0.09 | 0.28 |
| 6 | 67711014 | rs9354520 | -0.146 | -2.008 | -0.664 | 0.103 | 0.061 | -0.094 | -0.094 | 0 | 0.2 | 0.05 | 0.34 | 0.43 | 0.09 | 0.28 |
| 6 | 67720985 | rs9445803 | -0.389 | -1.466 | 0.186 | -0.269 | -0.021 | -0.014 | -0.014 | 0.05 | 0.08 | 0.28 | 0.03 | 0.03 | 0.02 | 0.03 |
| 6 | 67724162 | rs9445804 | -0.015 | -1.42 | -0.264 | 0.169 | 0.003 | -0.096 | -0.096 | 0 | 0.16 | 0.05 | 0.36 | 0.44 | 0.12 | 0.3 |
| 6 | 67726176 | rs10944944 | 1.258 | -1.331 | -0.386 | -0.44 | -0.043 | 0.04 | 0.04 | 0 | 0.43 | 0.25 | 0.28 | 0.39 | 0.45 | 0.39 |
| 6 | 67730693 | rs10944945 | 0.967 | -1.348 | -0.354 | -0.543 | 0.033 | 0.076 | 0.076 | 0 | 0.51 | 0.32 | 0.36 | 0.39 | 0.55 | 0.4 |
| 6 | 67736539 | rs2504074 | 0.88 | -1.208 | -0.369 | -0.463 | -0.013 | 0.027 | 0.027 | 0 | 0.08 | 0.04 | 0.07 | 0 | 0.1 | 0.02 |
| 6 | 67738061 | rs4235984 | 0.763 | -1.481 | -0.168 | -0.586 | 0.017 | 0.081 | 0.081 | 0.038 | 0.59 | 0.29 | 0.66 | 0.83 | 0.6 | 0.67 |
| 6 | 67742720 | rs11963560 | -0.182 | -1.541 | 0.104 | -0.625 | -0.029 | -0.017 | -0.017 | 0 | 0.09 | 0.35 | 0.03 | 0.03 | 0.02 | 0.03 |
| 6 | 67744470 | rs12205099 | 1.586 | -1.503 | -0.034 | -0.596 | -0.036 | 0.039 | 0.039 | 0 | 0.43 | 0.24 | 0.38 | 0.4 | 0.45 | 0.39 |
| 6 | 67753435 | rs7765154 | 0.852 | -1.332 | 0.272 | -0.739 | 0.01 | 0.039 | 0.039 | 0 | 0.6 | 0.37 | 0.67 | 0.83 | 0.6 | 0.67 |
| 6 | 67757689 | rs2503955 | -0.108 | -1.075 | 0.168 | -0.673 | -0.005 | -0.025 | -0.025 | 0 | 0.19 | 0.15 | 0.17 | 0 | 0.28 | 0.15 |
| 6 | 67770505 | rs9354531 | 0.855 | -1.11 | 0.188 | -0.645 | 0.002 | -0.048 | -0.048 | 0 | 0.68 | 0.55 | 0.69 | 0.84 | 0.65 | 0.68 |
| 6 | 67771375 | rs2502289 | -0.704 | -1.112 | 0.147 | -0.639 | -0.007 | -0.042 | -0.042 | 0.178 | 0.91 | 1 | 0.88 | 0.86 | 0.9 | 0.85 |
| 6 | 67775437 | rs2503947 | -0.606 | -1.035 | 0.253 | -0.676 | 0.024 | -0.13 | -0.13 | 0 | 0.23 | 0.46 | 0.19 | 0.03 | 0.26 | 0.17 |
| 6 | 67783032 | rs2503925 | 0.602 | -1.045 | -0.391 | -0.569 | -0.006 | 0.019 | 0.019 | 0 | 0.08 | 0.04 | 0.08 | 0 | 0.11 | 0.02 |
| 6 | 67789537 | rs2444664 | -0.645 | -1.095 | -0.463 | -0.508 | -0.011 | -0.042 | -0.042 | 0 | 0.91 | 1 | 0.88 | 0.86 | 0.9 | 0.84 |
| 6 | 67792729 | rs2502294 | -0.387 | -1.275 | -0.859 | -0.593 | 0.001 | -0.022 | -0.022 | 0 | 0.4 | 0.22 | 0.43 | 0.55 | 0.39 | 0.52 |
| 6 | 67793569 | rs2444659 | -0.801 | -1.303 | -0.56 | -0.655 | -0.001 | 0.01 | 0.01 | 0 | 0.52 | 0.49 | 0.53 | 0.59 | 0.54 | 0.65 |
| 6 | 67801176 | rs9363696 | 0.722 | -1.385 | -0.825 | -0.502 | -0.007 | -0.056 | -0.056 | 0 | 0.16 | 0.08 | 0.26 | 0.38 | 0.16 | 0.25 |
| 6 | 67801858 | rs1349813 | 1.235 | -1.281 | -0.722 | -0.562 | -0.015 | 0.024 | 0.024 | 0 | 0.11 | 0.04 | 0.08 | 0.04 | 0.15 | 0.14 |
| 6 | 67803212 | rs9351629 | -0.335 | -1.328 | -0.877 | -0.594 | -0.011 | 0.015 | 0.015 | 0.105 | 0.43 | 0.22 | 0.43 | 0.59 | 0.44 | 0.64 |
| 6 | 67804416 | rs9294736 | 0.214 | -1.257 | -0.883 | -0.499 | 0.004 | -0.067 | -0.067 | 4.699 | 0.32 | 0.16 | 0.35 | 0.55 | 0.29 | 0.5 |
| 6 | 67822272 | rs10944969 | 1.491 | -1.147 | -0.571 | -0.463 | -0.04 | 0.013 | 0.013 | 0.047 | 0.29 | 0.15 | 0.31 | 0.27 | 0.34 | 0.17 |
| 6 | 67840862 | rs10944973 | 0.704 | -1.164 | -0.665 | -0.471 | -0.003 | 0.011 | 0.011 | 0.662 | 0.39 | 0.6 | 0.35 | 0.27 | 0.36 | 0.19 |
| 6 | 67848752 | rs12662125 | 0.043 | -1.13 | -1.073 | -0.212 | -0.024 | -0.086 | -0.086 | 0 | 0.23 | 0.09 | 0.33 | 0.39 | 0.26 | 0.29 |
| 6 | 67856537 | rs4639317 | 1.434 | -1.222 | -1.149 | -0.363 | -0.099 | 0.009 | 0.009 | 0.558 | 0.31 | 0.09 | 0.32 | 0.28 | 0.36 | 0.27 |
| 6 | 67856976 | rs9360266 | -0.404 | -1.676 | -0.795 | 0.085 | -0.003 | -0.04 | -0.04 | 0 | 0.37 | 0.18 | 0.42 | 0.57 | 0.38 | 0.44 |

| | | | | | | | | | | | | | | | | |
|---|---|---|---|---|---|---|---|---|---|---|---|---|---|---|---|---|
| 6 | 67858985 | rs1453882 | -0.812 | -1.403 | -0.375 | 0.249 | -0.001 | 0.003 | 0.003 | 0.02 | 0.33 | 0.26 | 0.47 | 0.6 | 0.35 | 0.45 |
| 6 | 67866601 | rs674659 | -0.355 | -1.54 | 0.185 | -0.038 | -0.038 | -0.013 | -0.013 | 0 | 0.17 | 0.38 | 0.16 | 0.13 | 0.11 | 0.15 |
| 6 | 67876797 | rs9363705 | 0.512 | -1.609 | -0.029 | 0.077 | 0.01 | -0.093 | -0.093 | 0.373 | 0.14 | 0.01 | 0.23 | 0.39 | 0.11 | 0.29 |
| 6 | 67877233 | rs7740616 | 0.552 | -1.554 | 0.241 | 0.007 | -0.048 | -0.028 | -0.028 | 0 | 0.82 | 0.51 | 0.83 | 0.88 | 0.88 | 0.84 |
| 6 | 67877245 | rs11968408 | 0.307 | -1.299 | 0.497 | 0.096 | -0.001 | -0.017 | -0.017 | 0 | 0.06 | 0.21 | 0.02 | 0 | 0.02 | 0 |
| 6 | 67885038 | rs9354552 | -0.112 | -1.11 | 0.023 | 0.269 | -0.014 | -0.08 | -0.08 | 0.82 | 0.22 | 0.1 | 0.33 | 0.4 | 0.25 | 0.31 |
| 6 | 67885212 | rs16899152 | 0.279 | -1.295 | 0.036 | 0.103 | -0.029 | -0.032 | -0.032 | 0 | 0.24 | 0.34 | 0.13 | 0.17 | 0.17 | 0.23 |
| 6 | 67887156 | rs16899159 | -0.763 | -1.21 | -0.239 | 0.142 | 0 | -0.063 | -0.063 | 1.557 | 0.33 | 0.17 | 0.46 | 0.53 | 0.35 | 0.48 |
| 6 | 67893198 | rs9351642 | -0.854 | -1.26 | -0.238 | 0.16 | 0.003 | -0.014 | -0.014 | 0 | 0.34 | 0.29 | 0.47 | 0.53 | 0.35 | 0.48 |
| 6 | 67895080 | rs9360275 | -0.083 | -0.968 | -0.318 | 0.239 | -0.014 | -0.08 | -0.08 | 0 | 0.22 | 0.1 | 0.33 | 0.4 | 0.25 | 0.34 |
| 6 | 67897583 | rs4609013 | -0.615 | -0.468 | -0.115 | -0.358 | -0.007 | 0.024 | 0.024 | 0.302 | 0.57 | 0.63 | 0.6 | 0.7 | 0.51 | 0.68 |
| 6 | 67904039 | rs9294750 | -0.51 | -0.368 | -0.429 | -0.334 | 0 | -0.013 | -0.013 | 0.023 | 0.56 | 0.51 | 0.59 | 0.71 | 0.5 | 0.68 |
| 6 | 67905710 | rs17541229 | -0.724 | 0.055 | -0.599 | -0.247 | -0.001 | 0.005 | 0.005 | 0 | 0.09 | 0.07 | 0.07 | 0.05 | 0.08 | 0.02 |
| 6 | 67907952 | rs12197620 | -0.63 | 0.117 | -0.355 | -0.429 | 0.006 | -0.014 | -0.014 | 0.821 | 0.07 | 0.12 | 0.11 | 0.04 | 0.09 | 0.1 |
| 6 | 67921402 | rs9354570 | 1.303 | 0.184 | 0.194 | -0.568 | 0.032 | -0.025 | -0.025 | 0 | 0.91 | 0.79 | 0.86 | 0.96 | 0.89 | 0.91 |
| 6 | 67931943 | rs7766725 | 0.376 | 0.974 | -0.465 | -0.885 | -0.007 | 0.003 | 0.003 | 0 | 0.69 | 0.69 | 0.5 | 0.71 | 0.51 | 0.65 |
| 6 | 67948758 | rs9342631 | 0.513 | 0.781 | -0.423 | -1.27 | 0.008 | -0.08 | -0.08 | 2.743 | 0.86 | 0.72 | 0.88 | 1 | 0.88 | 0.94 |
| 6 | 67950785 | rs12205444 | 1.028 | 0.158 | -0.894 | -1.644 | 0.027 | -0.035 | -0.035 | 0 | 0.32 | 0.13 | 0.22 | 0.6 | 0.23 | 0.38 |
| 6 | 67951164 | rs12202269 | 0.908 | 0.144 | -0.833 | -1.651 | 0.026 | -0.021 | -0.021 | 0 | 0.33 | 0.19 | 0.23 | 0.6 | 0.23 | 0.38 |
| 6 | 67952979 | rs9345994 | -0.212 | 0.148 | -0.827 | -1.654 | 0.003 | 0.003 | 0.003 | 0.205 | 0.54 | 0.57 | 0.66 | 0.4 | 0.65 | 0.57 |
| 6 | 67955735 | rs9360281 | 0.284 | 0.112 | -0.93 | -1.728 | 0.018 | 0.08 | 0.08 | 0.129 | 0.57 | 0.3 | 0.6 | 0.85 | 0.62 | 0.63 |
| 6 | 67959832 | rs9363732 | 1.793 | -0.229 | -1.328 | -1.954 | 0.02 | -0.141 | -0.141 | 0 | 0.2 | 0.01 | 0.17 | 0.55 | 0.14 | 0.33 |
| 6 | 67964764 | rs10498851 | 0.661 | -0.199 | -1.333 | -1.998 | -0.086 | 0.013 | 0.013 | 0 | 0.29 | 0.16 | 0.47 | 0.28 | 0.48 | 0.33 |
| 6 | 67965720 | rs1903058 | 0.183 | -0.248 | -1.291 | -2.004 | 0 | -0.005 | -0.005 | 0 | 0.57 | 0.51 | 0.41 | 0.72 | 0.41 | 0.61 |
| 6 | 67967125 | rs10498853 | 1.744 | -0.265 | -1.308 | -1.941 | 0.02 | -0.142 | -0.142 | 0 | 0.2 | 0.01 | 0.18 | 0.55 | 0.14 | 0.33 |
| 6 | 67968844 | rs6903890 | 1.166 | -0.294 | -1.109 | -1.962 | -0.038 | -0.037 | -0.037 | 0 | 0.25 | 0.38 | 0.19 | 0.13 | 0.17 | 0.24 |
| 6 | 67973067 | rs13218852 | 1.149 | -0.195 | -1.26 | -1.997 | -0.071 | 0.004 | 0.004 | 0 | 0.24 | 0.12 | 0.37 | 0.25 | 0.39 | 0.25 |
| 6 | 67977326 | rs4358597 | 0.772 | -0.303 | -1.225 | -1.98 | -0.014 | -0.013 | -0.013 | 0.633 | 0.45 | 0.45 | 0.37 | 0.67 | 0.31 | 0.56 |
| 6 | 67984800 | rs9346003 | 1.092 | -0.234 | -1.191 | -1.908 | -0.05 | -0.048 | -0.048 | 1.207 | 0.25 | 0.41 | 0.19 | 0.12 | 0.17 | 0.24 |
| 6 | 67986138 | rs961195 | 0.247 | 0.126 | -1.624 | -1.841 | -0.079 | 0.046 | 0.046 | 0 | 0.36 | 0.17 | 0.4 | 0.3 | 0.48 | 0.29 |
| 6 | 67996590 | rs12210059 | -0.026 | 0.021 | -1.604 | -1.838 | -0.01 | 0.058 | 0.058 | 0 | 0.4 | 0.21 | 0.52 | 0.32 | 0.57 | 0.38 |
| 6 | 68014487 | rs12525981 | 0.196 | 0.13 | -0.994 | -2.142 | -0.005 | 0.03 | 0.03 | 0 | 0.13 | 0.06 | 0.04 | 0.04 | 0.09 | 0.05 |
| 6 | 68024278 | rs12199119 | 1.29 | 0.565 | -1.378 | -2.427 | -0.004 | 0.048 | 0.048 | 0 | 0.35 | 0.03 | 0.45 | 0.8 | 0.34 | 0.47 |
| 6 | 68025246 | rs12207946 | 0.741 | 0.579 | -1.366 | -2.543 | 0.007 | 0.16 | 0.16 | 0 | 0.45 | 0.1 | 0.53 | 0.8 | 0.44 | 0.51 |
| 6 | 68026073 | rs13220457 | 0.073 | 0.687 | -1.193 | -2.743 | -0.025 | 0.018 | 0.018 | 0 | 0.18 | 0.16 | 0.14 | 0.05 | 0.27 | 0.16 |
| 6 | 68029041 | rs9363741 | 0.583 | 0.501 | -0.845 | -2.91 | -0.01 | 0.044 | 0.044 | 0 | 0.27 | 0.14 | 0.19 | 0.13 | 0.25 | 0.27 |
| 6 | 68030203 | rs12207358 | 1.193 | 0.518 | -0.866 | -2.884 | 0.007 | 0.16 | 0.16 | 0 | 0.45 | 0.1 | 0.53 | 0.8 | 0.44 | 0.51 |
| 6 | 68033877 | rs13201396 | 0.707 | 0.693 | -0.646 | -2.71 | 0.013 | -0.01 | -0.01 | 0 | 0.11 | 0.15 | 0.11 | 0.05 | 0.2 | 0.12 |
| 6 | 68050150 | rs6903899 | 1.286 | 0.576 | -0.096 | -2.741 | -0.035 | 0.04 | 0.04 | 0 | 0.66 | 0.32 | 0.7 | 0.89 | 0.61 | 0.75 |
| 6 | 68057505 | rs7743494 | -0.656 | 0.564 | -0.097 | -2.626 | 0.002 | 0.009 | 0.009 | 0 | 0.2 | 0.15 | 0.11 | 0 | 0.16 | 0.08 |
| 6 | 68060896 | rs4272197 | 1.46 | 0.604 | 0.06 | -2.602 | 0.019 | -0.052 | -0.052 | 0 | 0.63 | 0.54 | 0.61 | 0.84 | 0.55 | 0.71 |
| 6 | 68062743 | rs12211418 | 1.862 | 0.453 | -0.151 | -2.663 | -0.031 | 0.04 | 0.04 | 0 | 0.37 | 0.03 | 0.37 | 0.76 | 0.36 | 0.48 |
| 6 | 68064253 | rs13200842 | 0.853 | 0.684 | 0.111 | -2.532 | 0.008 | -0.003 | -0.003 | 1.022 | 0.1 | 0.14 | 0.11 | 0.04 | 0.19 | 0.12 |
| 6 | 68072138 | rs7765600 | 0.65 | 0.629 | 0.084 | -2.555 | 0.047 | -0.023 | -0.023 | 5.282 | 0.77 | 0.67 | 0.7 | 0.84 | 0.69 | 0.78 |
| 6 | 68076933 | rs7740795 | 0.101 | -0.474 | 0.356 | -2.141 | -0.002 | -0.029 | -0.029 | 2.984 | 0.11 | 0.12 | 0.08 | 0 | 0.1 | 0.05 |
| 6 | 68084705 | rs9454120 | 0.627 | -0.357 | 0.363 | -2.038 | 0.062 | 0.121 | 0.121 | 0.082 | 0.92 | 0.75 | 0.81 | 0.85 | 0.9 | 0.9 |
| 6 | 68089887 | rs4431416 | 1.355 | -0.495 | 0.228 | -2.228 | -0.004 | 0.201 | 0.201 | 0.285 | 0.48 | 0.14 | 0.49 | 0.81 | 0.56 | 0.59 |
| 6 | 68103139 | rs1374556 | 0.629 | -0.336 | 0.323 | -2.029 | 0.062 | 0.121 | 0.121 | 0.082 | 0.92 | 0.75 | 0.81 | 0.85 | 0.9 | 0.9 |
| 6 | 68111529 | rs12192510 | -1.197 | -0.712 | 0.45 | -1.919 | 0.052 | 0.185 | 0.185 | 0.608 | 0.06 | 0.29 | 0.1 | 0.15 | 0.03 | 0.09 |
| 6 | 68118383 | rs7759951 | -1.246 | -0.804 | 0.165 | -2.095 | 0.065 | 0.253 | 0.253 | 0 | 0.48 | 0.78 | 0.34 | 0.08 | 0.35 | 0.32 |
| 6 | 68125255 | rs9363753 | -1.708 | -0.783 | 0.534 | -1.804 | -0.016 | -0.015 | -0.015 | 0.047 | 0.29 | 0.54 | 0.21 | 0.21 | 0.2 | 0.18 |
| 6 | 68140783 | rs9351662 | -1.577 | -0.607 | 0.677 | -1.802 | 0.008 | 0.146 | 0.146 | 1.212 | 0.49 | 0.9 | 0.43 | 0.22 | 0.41 | 0.41 |
| 6 | 68156538 | rs9454149 | 0 | -0.649 | 0.705 | -1.866 | -0.015 | -0.087 | -0.087 | 0.055 | 0.24 | 0.32 | 0.23 | 0.04 | 0.23 | 0.24 |
| 6 | 68157010 | rs7739784 | -1.623 | -0.656 | 0.705 | -1.876 | 0.022 | 0.138 | 0.138 | 1.191 | 0.48 | 0.86 | 0.43 | 0.22 | 0.41 | 0.4 |
| 6 | 68161428 | rs9294768 | -1.4 | -0.624 | 0.796 | -2.104 | 0.007 | 0.038 | 0.038 | 0 | 0.44 | 0.59 | 0.41 | 0.16 | 0.41 | 0.38 |
| 6 | 68166974 | rs6914736 | -0.01 | -0.519 | 0.323 | -1.993 | -0.015 | -0.091 | -0.091 | 0.261 | 0.24 | 0.33 | 0.24 | 0.04 | 0.23 | 0.24 |
| 6 | 68171159 | rs11751851 | -0.01 | -0.519 | 0.323 | -1.994 | -0.015 | -0.087 | -0.087 | 1.057 | 0.24 | 0.33 | 0.24 | 0.04 | 0.23 | 0.24 |
| 6 | 68200464 | rs9454157 | -0.046 | -0.83 | 0.465 | -1.885 | -0.012 | -0.083 | -0.083 | 0.55 | 0.25 | 0.33 | 0.23 | 0.04 | 0.23 | 0.24 |
| 6 | 68201044 | rs12194129 | -0.484 | -0.668 | -0.463 | -1.766 | -0.004 | 0.001 | 0.001 | 0 | 0.05 | 0.03 | 0.09 | 0.12 | 0.07 | 0.08 |
| 6 | 68211808 | rs7756876 | -1.301 | -0.798 | -0.487 | -1.889 | 0.062 | 0.146 | 0.146 | 0.832 | 0.46 | 0.85 | 0.42 | 0.19 | 0.38 | 0.39 |
| 6 | 68216688 | rs9445881 | -1.176 | -0.801 | -0.487 | -1.887 | 0.06 | 0.155 | 0.155 | 0.896 | 0.44 | 0.81 | 0.34 | 0.07 | 0.34 | 0.32 |
| 6 | 68221990 | rs4710656 | -1.544 | -0.787 | -0.788 | -1.861 | 0.026 | 0.205 | 0.205 | 0.468 | 0.49 | 0.89 | 0.44 | 0.19 | 0.41 | 0.4 |
| 6 | 68228212 | rs13213906 | 0.022 | -0.701 | -0.72 | -1.834 | -0.001 | 0.091 | 0.091 | 0 | 0.23 | 0.1 | 0.22 | 0.04 | 0.23 | 0.24 |
| 6 | 68232042 | rs9363764 | -1.125 | -0.765 | -0.943 | -1.562 | 0.05 | 0.128 | 0.128 | 0.428 | 0.53 | 0.87 | 0.42 | 0.24 | 0.38 | 0.42 |
| 6 | 68237791 | rs2028958 | -1.096 | -0.276 | -0.782 | -1.491 | -0.018 | 0.192 | 0.192 | 0.543 | 0.49 | 0.92 | 0.44 | 0.22 | 0.4 | 0.4 |
| 6 | 68250118 | rs7775598 | -0.341 | -0.316 | 0.279 | -1.558 | 0.019 | -0.082 | -0.082 | 0 | 0.17 | 0.21 | 0.14 | 0 | 0.2 | 0.1 |
| 6 | 68269956 | rs1418903 | 1.275 | -0.618 | 0.829 | -1.729 | 0.008 | 0.191 | 0.191 | 0.234 | 0.52 | 0.14 | 0.57 | 0.79 | 0.61 | 0.59 |
| 6 | 68276861 | rs856817 | 0.559 | -0.508 | 0.975 | -1.68 | 0.028 | -0.034 | -0.034 | 0 | 0.79 | 0.68 | 0.79 | 0.89 | 0.77 | 0.84 |
| 6 | 68277173 | rs1343032 | -0.298 | -0.571 | 0.622 | -1.45 | 0.03 | 0.074 | 0.074 | 0 | 0.22 | 0.1 | 0.19 | 0.08 | 0.17 | 0.23 |

| | | | | | | | | | | | | | | | | |
|---|---|---|---|---|---|---|---|---|---|---|---|---|---|---|---|---|
| 6 | 68282178 | rs17782592 | -0.321 | -0.509 | 0.561 | -1.478 | 0.02 | 0.039 | 0.039 | 0 | 0.23 | 0.12 | 0.19 | 0.08 | 0.16 | 0.23 |
| 6 | 68283502 | rs17709890 | -0.308 | -0.467 | 0.561 | -1.407 | 0.02 | 0.037 | 0.037 | 0 | 0.23 | 0.12 | 0.18 | 0.09 | 0.17 | 0.23 |
| 6 | 68284080 | rs856831 | 1.018 | -0.335 | 0.734 | -1.55 | -0.003 | -0.026 | -0.026 | 0 | 0.58 | 0.56 | 0.6 | 0.79 | 0.61 | 0.6 |
| 6 | 68285094 | rs13220356 | -0.305 | -0.416 | 0.869 | -1.602 | -0.016 | -0.039 | -0.039 | 0 | 0.23 | 0.26 | 0.2 | 0.09 | 0.16 | 0.25 |
| 6 | 68298216 | rs2144100 | -0.358 | -0.413 | 0.87 | -1.528 | -0.007 | 0.022 | 0.022 | 0 | 0.88 | 0.94 | 0.89 | 1 | 0.85 | 0.91 |
| 6 | 68301978 | rs2818075 | 0.352 | -0.413 | 0.876 | -1.527 | -0.003 | 0.002 | 0.002 | 0 | 0.84 | 0.84 | 0.82 | 0.89 | 0.81 | 0.85 |
| 6 | 68302141 | rs2747148 | -0.36 | -0.413 | 0.846 | -1.531 | -0.007 | 0.022 | 0.022 | 0 | 0.88 | 0.94 | 0.89 | 1 | 0.85 | 0.91 |
| 6 | 68317796 | rs6911763 | -0.481 | -0.225 | 0.875 | -1.508 | -0.006 | -0.035 | -0.035 | 0.133 | 0.26 | 0.26 | 0.21 | 0.09 | 0.19 | 0.26 |
| 6 | 68322659 | rs2747160 | 1.11 | -0.239 | 0.654 | -1.484 | -0.004 | -0.031 | -0.031 | 0 | 0.57 | 0.52 | 0.6 | 0.8 | 0.62 | 0.58 |
| 6 | 68351062 | rs2747173 | -0.199 | -0.136 | 0.155 | -1.14 | -0.039 | 0.067 | 0.067 | 0 | 0.89 | 0.99 | 0.88 | 1 | 0.82 | 0.91 |
| 6 | 68364866 | rs9342657 | -0.429 | -0.195 | 0.113 | -1.15 | -0.065 | -0.017 | -0.017 | 10.027 | 0.15 | 0.01 | 0.2 | 0.12 | 0.23 | 0.15 |
| 6 | 68368610 | rs11964693 | -0.117 | -0.812 | 0.517 | -1.233 | -0.023 | -0.019 | -0.019 | 1.06 | 0.19 | 0.16 | 0.17 | 0.06 | 0.09 | 0.19 |
| 6 | 68372070 | rs2818105 | 1.091 | -1.156 | 1.105 | -1.125 | -0.059 | -0.007 | -0.007 | 0.919 | 0.71 | 0.47 | 0.72 | 0.77 | 0.85 | 0.72 |
| 6 | 68379334 | rs12524310 | -0.258 | -1.2 | 1.006 | -1.326 | -0.031 | -0.022 | -0.022 | 0.016 | 0.25 | 0.25 | 0.2 | 0.08 | 0.12 | 0.2 |
| 6 | 68380125 | rs2816903 | 0.901 | -1.219 | 1.174 | -1.482 | -0.046 | -0.029 | -0.029 | 0 | 0.67 | 0.48 | 0.7 | 0.8 | 0.83 | 0.72 |
| 6 | 68384198 | rs12200729 | -0.635 | -1.171 | 1.063 | -1.456 | 0.004 | 0.05 | 0.05 | 0 | 0.08 | 0.2 | 0.1 | 0.14 | 0.05 | 0.08 |
| 6 | 68388841 | rs4498332 | -0.097 | -1.143 | 0.84 | -1.466 | 0.025 | 0.024 | 0.024 | 0 | 0.23 | 0.11 | 0.19 | 0.06 | 0.12 | 0.2 |
| 6 | 68397567 | rs9346057 | 0.091 | -1.194 | 1.144 | -1.463 | -0.059 | -0.07 | -0.07 | 0 | 0.76 | 0.69 | 0.79 | 0.94 | 0.88 | 0.8 |
| 6 | 68400534 | rs13209120 | 1.624 | -0.636 | -0.006 | -1.372 | -0.083 | -0.029 | -0.029 | 1.246 | 0.25 | 0.02 | 0.23 | 0.24 | 0.3 | 0.21 |
| 6 | 68423329 | rs12214799 | 0.319 | -0.193 | 0.379 | -1.398 | -0.02 | -0.035 | -0.035 | 0 | 0.06 | 0 | 0.09 | 0.15 | 0.05 | 0.08 |
| 6 | 68435871 | rs9454257 | 0.133 | -0.358 | 0.776 | -1.51 | -0.076 | -0.097 | -0.097 | 4.842 | 0.23 | 0.4 | 0.22 | 0.06 | 0.12 | 0.19 |
| 6 | 68438093 | rs9363799 | 0.366 | -0.316 | 0.74 | -1.467 | -0.02 | -0.035 | -0.035 | 0 | 0.06 | 0 | 0.09 | 0.14 | 0.05 | 0.08 |
| 6 | 68445577 | rs13219398 | -0.254 | 0.317 | 0.674 | -1.846 | -0.03 | 0.023 | 0.023 | 0.435 | 0.25 | 0.14 | 0.19 | 0 | 0.35 | 0.18 |
| 6 | 68447698 | rs9454264 | -1.317 | 0.295 | 0.886 | -1.673 | 0.003 | 0.006 | 0.006 | 0 | 0.5 | 0.5 | 0.47 | 0.2 | 0.46 | 0.42 |
| 6 | 68452823 | rs7766667 | -0.348 | 0.732 | 0.814 | -1.524 | -0.01 | 0.005 | 0.005 | 0 | 0.08 | 0.05 | 0.06 | 0 | 0.14 | 0.06 |
| 6 | 68453011 | rs12193750 | -0.563 | 0.568 | 0.824 | -1.449 | 0.004 | 0.036 | 0.036 | 0 | 0.08 | 0.2 | 0.1 | 0.13 | 0.05 | 0.08 |
| 6 | 68459626 | rs12200788 | 0.31 | 0.606 | 0.6 | -1.429 | -0.02 | -0.034 | -0.034 | 0 | 0.06 | 0 | 0.09 | 0.13 | 0.05 | 0.08 |
| 6 | 68472310 | rs12203810 | 0.31 | 0.606 | 0.6 | -1.429 | -0.02 | -0.034 | -0.034 | 0 | 0.06 | 0 | 0.09 | 0.13 | 0.05 | 0.08 |
| 6 | 68491969 | rs17450573 | -0.252 | 0.139 | 0.557 | -1.709 | -0.033 | 0.028 | 0.028 | 0.018 | 0.21 | 0.12 | 0.17 | 0 | 0.29 | 0.15 |
| 6 | 68492448 | rs4131870 | -0.608 | 0.044 | 0.525 | -1.757 | -0.027 | 0.028 | 0.028 | 1.777 | 0.23 | 0.18 | 0.17 | 0 | 0.29 | 0.15 |
| 6 | 68494531 | rs12195466 | 0.381 | 0.064 | 0.475 | -1.703 | -0.02 | -0.026 | -0.026 | 0 | 0.06 | 0 | 0.17 | 0.12 | 0.05 | 0.05 |
| 6 | 68503143 | rs6920238 | -0.366 | -0.173 | 0.583 | -1.646 | -0.021 | 0.004 | 0.004 | 0.11 | 0.21 | 0.15 | 0.17 | 0 | 0.29 | 0.15 |
| 6 | 68526511 | rs1913477 | -1.483 | -0.389 | 0.639 | -1.562 | 0.011 | 0.059 | 0.059 | 0 | 0.51 | 0.63 | 0.53 | 0.19 | 0.44 | 0.39 |
| 6 | 68530199 | rs12524210 | 0.456 | -0.267 | 0.559 | -1.574 | -0.006 | 0.008 | 0.008 | 0.576 | 0.16 | 0.09 | 0.12 | 0 | 0.18 | 0.09 |
| 6 | 68532965 | rs7760466 | -0.039 | -0.305 | 0.585 | -1.569 | -0.093 | -0.09 | -0.09 | 0.297 | 0.25 | 0.49 | 0.21 | 0.07 | 0.12 | 0.2 |
| 6 | 68533069 | rs13205069 | 0.363 | -0.225 | 0.335 | -1.551 | -0.027 | 0.026 | 0.026 | 0 | 0.05 | 0.02 | 0.05 | 0 | 0.11 | 0.06 |
| 6 | 68533625 | rs6925414 | 0.1 | -0.236 | 0.461 | -1.534 | -0.042 | -0.04 | -0.04 | 0.1 | 0.23 | 0.31 | 0.21 | 0.07 | 0.12 | 0.2 |
| 6 | 68538382 | rs1399230 | -0.943 | -0.039 | 0.016 | -1.45 | 0.013 | 0.014 | 0.014 | 0 | 0.45 | 0.58 | 0.48 | 0.18 | 0.32 | 0.32 |
| 6 | 68538877 | rs9363809 | -1.396 | -0.143 | -0.481 | -1.251 | -0.01 | 0.065 | 0.065 | 0 | 0.54 | 0.75 | 0.56 | 0.26 | 0.46 | 0.39 |
| 6 | 68542695 | rs12663045 | -0.272 | -0.441 | -0.605 | -0.234 | -0.018 | -0.038 | -0.038 | 0 | 0.07 | 0.01 | 0.16 | 0.19 | 0.05 | 0.05 |
| 6 | 68545861 | rs16899546 | -0.147 | -0.605 | -0.258 | -0.611 | 0.002 | 0.002 | 0.002 | 0 | 0.18 | 0.14 | 0.2 | 0.07 | 0.12 | 0.2 |
| 6 | 68547726 | rs9360306 | -0.763 | -0.385 | 0.119 | -0.354 | -0.033 | -0.01 | -0.01 | 0.361 | 0.16 | 0.33 | 0.17 | 0.09 | 0.09 | 0.11 |
| 6 | 68548907 | rs2203381 | -0.823 | 0.239 | -0.292 | -0.8 | -0.01 | 0.011 | 0.011 | 0 | 0.16 | 0.11 | 0.08 | 0.02 | 0.17 | 0.13 |
| 6 | 68549473 | rs1399224 | 0.432 | 0.583 | -1.329 | -0.518 | -0.091 | -0.089 | -0.089 | 0 | 0.6 | 0.88 | 0.61 | 0.44 | 0.49 | 0.7 |
| 6 | 68554386 | rs1399228 | 0.466 | 0.141 | -0.657 | -0.617 | -0.003 | -0.002 | -0.002 | 0.745 | 0.29 | 0.27 | 0.35 | 0.21 | 0.27 | 0.34 |
| 6 | 68574726 | rs9346081 | 0.835 | -0.011 | -1.212 | -0.812 | -0.035 | -0.053 | -0.053 | 0 | 0.23 | 0.03 | 0.32 | 0.21 | 0.24 | 0.33 |
| 6 | 68586809 | rs9354637 | -0.277 | -0.239 | -1.208 | -0.921 | 0.011 | -0.074 | -0.074 | 0 | 0.39 | 0.85 | 0.52 | 0.39 | 0.33 | 0.36 |
| 6 | 68607744 | rs10945038 | 0.719 | -0.189 | -1.297 | -1.427 | 0.023 | 0.138 | 0.138 | 0 | 0.3 | 0.12 | 0.15 | 0.05 | 0.24 | 0.42 |
| 6 | 68609591 | rs6933411 | -1.144 | -0.042 | -0.999 | -1.334 | -0.025 | -0.062 | -0.062 | 0 | 0.74 | 0.88 | 0.7 | 0.44 | 0.64 | 0.83 |
| 6 | 68609707 | rs6912774 | -0.196 | -0.076 | -0.61 | -1.328 | 0.005 | 0.023 | 0.023 | 0 | 0.43 | 0.67 | 0.33 | 0.22 | 0.37 | 0.49 |
| 6 | 68631478 | rs9454382 | 0.363 | -0.092 | 0.434 | -1.346 | 0.007 | 0.071 | 0.071 | 0 | 0.35 | 0.22 | 0.18 | 0.06 | 0.31 | 0.46 |
| 6 | 68633976 | rs12198681 | -0.111 | -0.03 | 0.174 | -1.296 | 0.003 | 0.015 | 0.015 | 0 | 0.43 | 0.66 | 0.33 | 0.23 | 0.37 | 0.49 |
| 6 | 68649501 | rs6940958 | -0.201 | -0.032 | 0.141 | -1.325 | 0.006 | 0.033 | 0.033 | 0 | 0.44 | 0.67 | 0.34 | 0.23 | 0.37 | 0.49 |
| 6 | 68653644 | rs7748806 | 0.296 | -0.06 | 0.161 | -1.329 | -0.004 | -0.056 | -0.056 | 0 | 0.37 | 0.62 | 0.19 | 0.05 | 0.24 | 0.42 |
| 6 | 68663856 | rs9346107 | -0.858 | -0.044 | 0.174 | -1.326 | -0.017 | -0.045 | -0.045 | 0 | 0.71 | 0.82 | 0.68 | 0.44 | 0.61 | 0.82 |
| 6 | 68665889 | rs7749767 | -0.072 | 0.055 | 0.321 | -1.326 | 0.004 | -0.026 | -0.026 | 0 | 0.39 | 0.64 | 0.21 | 0.06 | 0.31 | 0.46 |
| 6 | 68673028 | rs1877775 | -0.307 | 0.069 | 0.272 | -1.317 | 0.007 | 0.034 | 0.034 | 0 | 0.44 | 0.67 | 0.34 | 0.23 | 0.37 | 0.49 |
| 6 | 68686425 | rs9454404 | 0.57 | -0.054 | 0.389 | -1.288 | -0.017 | -0.08 | -0.08 | 1.408 | 0.32 | 0.39 | 0.16 | 0.05 | 0.22 | 0.41 |
| 6 | 68687348 | rs1877779 | -1.454 | 0.246 | 0.376 | -1.139 | 0.041 | 0.047 | 0.047 | 0 | 0.14 | 0.31 | 0.19 | 0.17 | 0.17 | 0.07 |
| 6 | 68691634 | rs11758293 | -0.796 | 0.253 | -0.034 | -1.328 | -0.005 | 0.002 | 0.002 | 0 | 0.11 | 0.09 | 0.1 | 0.04 | 0.2 | 0.1 |
| 6 | 68694445 | rs9363842 | 0.827 | 0.001 | -0.215 | -1.397 | -0.017 | -0.021 | -0.021 | 0.03 | 0.25 | 0.1 | 0.43 | 0.23 | 0.26 | 0.32 |
| 6 | 68697160 | rs12193625 | -0.301 | -0.256 | 0.198 | -1.541 | 0.001 | -0.016 | -0.016 | 0 | 0.43 | 0.4 | 0.25 | 0.08 | 0.39 | 0.49 |
| 6 | 68699843 | rs7775008 | -0.648 | -0.266 | -0.296 | -1.543 | 0.027 | 0.251 | 0.251 | 0 | 0.48 | 0.84 | 0.28 | 0.08 | 0.4 | 0.5 |
| 6 | 68705526 | rs1355931 | -0.707 | -0.262 | -0.358 | -1.556 | 0.023 | 0.301 | 0.301 | 0 | 0.48 | 0.88 | 0.29 | 0.08 | 0.4 | 0.49 |
| 6 | 68707131 | rs4707250 | -0.757 | -0.461 | -0.206 | -1.381 | 0.03 | 0.044 | 0.044 | 0 | 0.69 | 0.81 | 0.86 | 0.96 | 0.81 | 0.6 |
| 6 | 68717033 | rs6930772 | -0.088 | -0.321 | -0.456 | -1.368 | 0.045 | 0.088 | 0.088 | 0 | 0.6 | 0.25 | 0.77 | 0.95 | 0.74 | 0.56 |
| 6 | 68718726 | rs9360312 | 1.146 | 0.069 | -0.884 | -1.113 | -0.018 | -0.111 | -0.111 | 3.047 | 0.16 | 0.01 | 0.2 | 0.68 | 0.15 | 0.12 |
| 6 | 68722068 | rs9354655 | -0.426 | -0.014 | -0.435 | -1.523 | -0.012 | -0.117 | -0.117 | 0 | 0.7 | 0.41 | 0.81 | 0.96 | 0.75 | 0.62 |
| 6 | 68722543 | rs13193425 | 0.077 | 0.058 | -0.744 | -1.309 | -0.041 | -0.137 | -0.137 | 0.643 | 0.39 | 0.08 | 0.31 | 0.6 | 0.38 | 0.3 |

| | | | | | | | | | | | | | | | | |
|---|---|---|---|---|---|---|---|---|---|---|---|---|---|---|---|---|
| 6 | 68727910 | rs12153928 | 1.257 | 0.237 | -0.883 | -1.416 | -0.02 | -0.034 | -0.034 | 0 | 0.17 | 0.06 | 0.38 | 0.22 | 0.17 | 0.21 |
| 6 | 68728797 | rs1020689 | 0.07 | 0.003 | -0.328 | -1.552 | -0.048 | -0.104 | -0.104 | 0 | 0.83 | 0.6 | 0.94 | 0.99 | 0.93 | 0.92 |
| 6 | 68729130 | rs4706314 | 0.115 | 0.103 | -0.812 | -1.565 | 0.012 | 0.089 | 0.089 | 0 | 0.49 | 0.28 | 0.69 | 0.82 | 0.54 | 0.47 |
| 6 | 68731635 | rs2340741 | 0.006 | 0.359 | -0.707 | -1.395 | -0.003 | -0.005 | -0.005 | 0 | 0.35 | 0.31 | 0.44 | 0.23 | 0.27 | 0.32 |
| 6 | 68738618 | rs10945048 | -0.031 | 0.379 | -0.133 | -1.248 | 0 | -0.014 | -0.014 | 0 | 0.63 | 0.34 | 0.73 | 0.83 | 0.6 | 0.55 |
| 6 | 68739602 | rs9351689 | -0.043 | 0.387 | -0.139 | -1.25 | 0.002 | -0.029 | -0.029 | 0 | 0.63 | 0.38 | 0.73 | 0.83 | 0.6 | 0.55 |
| 6 | 68747569 | rs1378455 | -0.335 | 0.395 | -0.123 | -1.444 | -0.026 | -0.148 | -0.148 | 0 | 0.69 | 0.42 | 0.78 | 0.96 | 0.72 | 0.65 |
| 6 | 68753540 | rs17563853 | 0.385 | 0.533 | -0.21 | -1.638 | -0.002 | -0.015 | -0.015 | 0.496 | 0.09 | 0.08 | 0.17 | 0 | 0.12 | 0.06 |
| 6 | 68764690 | rs9454422 | 0.827 | 0.516 | -0.643 | -1.266 | -0.005 | 0.043 | 0.043 | 0.156 | 0.2 | 0.28 | 0.22 | 0.25 | 0.13 | 0.24 |
| 6 | 68765982 | rs7768996 | 0.252 | 0.455 | -0.71 | -1.337 | 0.005 | 0.038 | 0.038 | 0 | 0.54 | 0.32 | 0.57 | 0.83 | 0.53 | 0.55 |
| 6 | 68774543 | rs12198964 | -0.398 | 0.532 | -0.076 | -1.192 | -0.003 | -0.057 | -0.057 | 0 | 0.15 | 0.19 | 0.18 | 0 | 0.14 | 0.1 |
| 6 | 68776313 | rs9354665 | -0.45 | 0.517 | -0.027 | -1.103 | -0.019 | 0.007 | 0.007 | 0.765 | 0.75 | 0.42 | 0.73 | 0.8 | 0.83 | 0.73 |
| 6 | 68783499 | rs6900918 | 0.648 | 0.503 | -0.006 | -1.023 | -0.012 | 0.017 | 0.017 | 0 | 0.22 | 0.28 | 0.22 | 0.21 | 0.14 | 0.29 |
| 6 | 68799400 | rs3936209 | -0.475 | 0.803 | -0.6 | -0.948 | 0.024 | 0.047 | 0.047 | 0 | 0.53 | 0.38 | 0.61 | 0.73 | 0.64 | 0.44 |
| 6 | 68800238 | rs4707839 | 0.052 | 0.882 | -0.98 | -0.673 | -0.04 | 0.05 | 0.05 | 0.656 | 0.4 | 0.16 | 0.39 | 0.73 | 0.46 | 0.37 |
| 6 | 68803176 | rs9354670 | 0.594 | 1.017 | -1.094 | -0.302 | -0.037 | -0.112 | -0.112 | 0 | 0.33 | 0.09 | 0.34 | 0.52 | 0.37 | 0.3 |
| 6 | 68804067 | rs13219267 | 0.33 | 0.993 | -0.994 | -0.191 | -0.029 | -0.073 | -0.073 | 0.257 | 0.38 | 0.12 | 0.37 | 0.52 | 0.41 | 0.33 |
| 6 | 68804218 | rs9354671 | 0.271 | 0.858 | -0.953 | -0.658 | -0.031 | 0.049 | 0.049 | 0.581 | 0.4 | 0.18 | 0.39 | 0.73 | 0.46 | 0.37 |
| 6 | 68810538 | rs897405 | 0.103 | 0.86 | -0.718 | -0.332 | 0.016 | 0.017 | 0.017 | 0.142 | 0.45 | 0.31 | 0.32 | 0.34 | 0.37 | 0.58 |
| 6 | 68813247 | rs10945056 | -0.636 | 1.183 | -0.327 | -0.344 | 0.027 | 0.034 | 0.034 | 0 | 0.07 | 0.24 | 0.11 | 0.12 | 0.1 | 0.04 |
| 6 | 68815483 | rs701716 | 0.291 | 1.303 | -1.103 | -1.258 | 0.004 | 0.194 | 0.194 | 0.381 | 0.49 | 0.22 | 0.46 | 0.83 | 0.41 | 0.58 |
| 6 | 68823251 | rs9454429 | 1.01 | 1.152 | -1.147 | -0.696 | 0.002 | -0.019 | -0.019 | 0 | 0.16 | 0.1 | 0.15 | 0.26 | 0.12 | 0.18 |
| 6 | 68827634 | rs10945059 | 0.122 | 1.087 | -0.904 | -0.997 | 0.001 | 0 | 0 | 0 | 0.31 | 0.69 | 0.25 | 0.2 | 0.29 | 0.28 |
| 6 | 68828279 | rs13216403 | 0.128 | 1.088 | -0.904 | -0.997 | 0.001 | 0 | 0 | 0 | 0.31 | 0.68 | 0.25 | 0.2 | 0.29 | 0.28 |
| 6 | 68833136 | rs777632 | 1.165 | 1.288 | -1.294 | -1.079 | -0.067 | -0.132 | -0.132 | 0 | 0.28 | 0.02 | 0.29 | 0.56 | 0.27 | 0.36 |
| 6 | 68836838 | rs9454430 | 0.619 | 1.292 | -0.81 | -1.422 | -0.02 | 0.017 | 0.017 | 0 | 0.21 | 0.16 | 0.29 | 0 | 0.3 | 0.17 |
| 6 | 68838350 | rs7744782 | 0.213 | 0.644 | -0.063 | -1.045 | -0.006 | -0.043 | -0.043 | 0.856 | 0.23 | 0.23 | 0.14 | 0.08 | 0.21 | 0.25 |
| 6 | 68838783 | rs11963711 | -0.04 | 0.669 | -0.126 | -1.033 | -0.033 | -0.127 | -0.127 | 0 | 0.26 | 0.51 | 0.16 | 0.08 | 0.22 | 0.25 |
| 6 | 68854620 | rs1083339 | -0.069 | 1.238 | -0.367 | -0.331 | -0.004 | -0.01 | -0.01 | 0 | 0.4 | 0.38 | 0.32 | 0.5 | 0.5 | 0.47 |
| 6 | 68860309 | rs10455651 | -0.156 | 1.69 | -0.44 | -0.368 | -0.011 | 0.001 | 0.001 | 0 | 0.13 | 0.11 | 0.26 | 0.1 | 0.21 | 0.14 |
| 6 | 68860756 | rs701713 | -0.638 | 1.608 | -0.424 | -0.405 | 0.02 | 0.008 | 0.008 | 0 | 0.5 | 0.44 | 0.51 | 0.5 | 0.64 | 0.57 |
| 6 | 68865718 | rs777648 | -0.668 | 1.279 | -0.161 | -0.573 | -0.022 | -0.053 | -0.053 | 2.899 | 0.87 | 0.72 | 0.94 | 0.96 | 0.95 | 0.91 |
| 6 | 68867750 | rs4357093 | 1.219 | 1.257 | -0.151 | -0.587 | 0.011 | -0.038 | -0.038 | 0.213 | 0.35 | 0.16 | 0.42 | 0.46 | 0.29 | 0.32 |
| 6 | 68868332 | rs777649 | 0.584 | 1.342 | -0.389 | -0.564 | 0.001 | -0.01 | -0.01 | 0 | 0.37 | 0.33 | 0.25 | 0.4 | 0.44 | 0.43 |
| 6 | 68873924 | rs12211024 | 0.741 | 1.305 | -0.686 | -0.182 | 0.006 | -0.008 | -0.008 | 0 | 0.36 | 0.28 | 0.43 | 0.46 | 0.29 | 0.32 |
| 6 | 68879790 | rs1869714 | 0.641 | 1.403 | -0.808 | -0.188 | 0.01 | 0.006 | 0.006 | 0.302 | 0.66 | 0.56 | 0.61 | 0.63 | 0.52 | 0.48 |
| 6 | 68882437 | rs12207132 | 0.235 | 1.601 | -1.247 | -0.188 | -0.019 | 0.015 | 0.015 | 0.009 | 0.1 | 0.05 | 0.19 | 0 | 0.15 | 0.1 |
| 6 | 68882785 | rs9363862 | 1.127 | 0.752 | -1.92 | -0.323 | 0.044 | -0.154 | -0.154 | 0 | 0.29 | 0.02 | 0.28 | 0.47 | 0.18 | 0.29 |
| 6 | 68883862 | rs921816 | 0.225 | 1.119 | -1.428 | 0.038 | -0.026 | -0.076 | -0.076 | 0 | 0.18 | 0.08 | 0.19 | 0.33 | 0.25 | 0.37 |
| 6 | 68884304 | rs945823 | -0.427 | 1.287 | -0.794 | -0.77 | -0.001 | -0.006 | -0.006 | 0.38 | 0.73 | 0.72 | 0.81 | 0.86 | 0.74 | 0.86 |
| 6 | 68887832 | rs778008 | -0.565 | 1.554 | -0.563 | -0.824 | 0.052 | 0 | 0 | 0 | 0.08 | 0.23 | 0.11 | 0.03 | 0.15 | 0.08 |
| 6 | 68890028 | rs701662 | -0.162 | 1.561 | -0.555 | -0.807 | -0.003 | 0 | 0 | 0 | 0.07 | 0.07 | 0.09 | 0.03 | 0.15 | 0.08 |
| 6 | 68894710 | rs701665 | -1.193 | 2.341 | -0.956 | -1.087 | 0.043 | -0.02 | -0.02 | 3.68 | 0.19 | 0.25 | 0.34 | 0.07 | 0.38 | 0.21 |
| 6 | 68896299 | rs1455441 | 0.42 | 2.473 | -1.163 | -1.051 | -0.033 | 0.03 | 0.03 | 0 | 0.06 | 0 | 0.17 | 0 | 0.14 | 0.07 |
| 6 | 68897315 | rs9354683 | 0.637 | 1.706 | -1.465 | -1.194 | 0.021 | -0.037 | -0.037 | 0 | 0.32 | 0.22 | 0.3 | 0.52 | 0.26 | 0.28 |
| 6 | 68897853 | rs1823908 | 0.671 | 1.803 | -1.121 | -0.898 | -0.033 | 0.03 | 0.03 | 0 | 0.06 | 0 | 0.17 | 0 | 0.14 | 0.07 |
| 6 | 68915890 | rs7454318 | 0.51 | 1.099 | -1.97 | -1.052 | 0.038 | 0.007 | 0.007 | 0 | 0.49 | 0.23 | 0.42 | 0.52 | 0.46 | 0.32 |
| 6 | 68921177 | rs9346129 | 0.684 | 1.682 | -1.349 | -1.062 | -0.026 | 0.018 | 0.018 | 0 | 0.68 | 0.26 | 0.58 | 0.79 | 0.58 | 0.72 |
| 6 | 68930628 | rs9360322 | 0.592 | 1.256 | -1.888 | -1.361 | -0.008 | -0.109 | -0.109 | 1.376 | 0.25 | 0.02 | 0.25 | 0.69 | 0.23 | 0.3 |
| 6 | 68930738 | rs9445980 | 0.247 | 1.708 | -1.754 | -1.734 | -0.029 | 0.024 | 0.024 | 0 | 0.75 | 0.22 | 0.77 | 0.88 | 0.65 | 0.81 |
| 6 | 68936060 | rs1993329 | 0.372 | 1.607 | -1.218 | -1.94 | 0.071 | -0.076 | -0.076 | 0 | 0.78 | 0.43 | 0.81 | 0.93 | 0.67 | 0.85 |
| 6 | 68936391 | rs1993330 | -0.138 | 1.262 | -1.293 | -1.987 | 0.005 | -0.047 | -0.047 | 3.261 | 0.63 | 0.42 | 0.73 | 0.93 | 0.54 | 0.81 |
| 6 | 68937082 | rs12174660 | 0.56 | 1.21 | -1.509 | -1.434 | 0.013 | -0.009 | -0.009 | 0 | 0.25 | 0.1 | 0.4 | 0.18 | 0.19 | 0.46 |
| 6 | 68944408 | rs2603687 | -0.566 | 1.482 | -0.925 | -1.186 | 0.004 | 0.029 | 0.029 | 0.234 | 0.19 | 0.11 | 0.16 | 0.14 | 0.2 | 0.19 |
| 6 | 68945419 | rs2603688 | -0.791 | 1.843 | -0.453 | -1.588 | 0 | 0.002 | 0.002 | 0 | 0.57 | 0.55 | 0.53 | 0.77 | 0.58 | 0.57 |
| 6 | 68950223 | rs10455654 | 0.138 | 1.777 | -0.52 | -1.607 | 0 | 0.004 | 0.004 | 0 | 0.14 | 0.11 | 0.11 | 0.09 | 0.13 | 0.1 |
| 6 | 68950298 | rs2786577 | -0.702 | 1.814 | -0.441 | -1.474 | -0.001 | -0.018 | -0.018 | 0 | 0.57 | 0.49 | 0.53 | 0.77 | 0.58 | 0.57 |
| 6 | 68985256 | rs9454457 | 0.815 | 2.045 | -0.981 | -1.49 | 0.032 | 0.047 | 0.047 | 0 | 0.39 | 0.15 | 0.42 | 0.2 | 0.32 | 0.4 |
| 6 | 69029483 | rs1388052 | 0.661 | 2.001 | -1.064 | -1.643 | -0.076 | 0.156 | 0.156 | 0.102 | 0.63 | 0.2 | 0.69 | 0.83 | 0.52 | 0.64 |
| 6 | 69032817 | rs2341484 | 0.152 | 1.992 | -0.863 | -1.72 | 0.063 | -0.079 | -0.079 | 0.032 | 0.82 | 0.55 | 0.85 | 0.88 | 0.76 | 0.87 |
| 6 | 69041360 | rs11751599 | 0.327 | 1.989 | -0.856 | -1.712 | 0.033 | -0.101 | -0.101 | 0 | 0.19 | 0.35 | 0.16 | 0.05 | 0.24 | 0.23 |
| 6 | 69041861 | rs4612125 | 0.375 | 1.397 | -0.7 | -0.899 | -0.033 | -0.138 | -0.138 | 2.616 | 0.23 | 0.03 | 0.26 | 0.6 | 0.19 | 0.25 |
| 6 | 69052097 | rs1907146 | 0.497 | 1.175 | -0.581 | -0.852 | 0.004 | -0.011 | -0.011 | 2.214 | 0.6 | 0.53 | 0.59 | 0.28 | 0.57 | 0.62 |
| 6 | 69074135 | rs12203903 | -0.701 | 0.92 | -0.787 | -0.557 | -0.001 | -0.012 | -0.012 | 0 | 0.13 | 0.14 | 0.11 | 0.1 | 0.14 | 0.09 |
| 6 | 69077820 | rs9342701 | -0.088 | 0.732 | -1.08 | -0.537 | 0.017 | 0.004 | 0.004 | 0 | 0.43 | 0.27 | 0.44 | 0.65 | 0.42 | 0.28 |
| 6 | 69077987 | rs2029516 | -0.517 | 1.084 | -0.989 | -0.899 | 0 | 0.005 | 0.005 | 0 | 0.55 | 0.38 | 0.54 | 0.66 | 0.6 | 0.33 |
| 6 | 69082145 | rs1907149 | 0.653 | 0.901 | -1.665 | -1.008 | -0.122 | -0.073 | -0.073 | 0.637 | 0.27 | 0.07 | 0.43 | 0.29 | 0.43 | 0.58 |
| 6 | 69084796 | rs9354724 | 0.103 | 0.456 | -1.194 | -1.558 | 0.032 | 0.054 | 0.054 | 0 | 0.5 | 0.28 | 0.43 | 0.65 | 0.44 | 0.22 |
| 6 | 69085833 | rs9342703 | 0.499 | 0.126 | -1.359 | -1.723 | 0.034 | -0.002 | -0.002 | 0 | 0.39 | 0.17 | 0.35 | 0.65 | 0.33 | 0.19 |

| | | | | | | | | | | | | | | | | |
|---|---|---|---|---|---|---|---|---|---|---|---|---|---|---|---|---|
| 6 | 69108473 | rs9342706 | 0.694 | -0.391 | -1.58 | -1.619 | -0.03 | -0.024 | -0.024 | 0 | 0.43 | 0.09 | 0.54 | 0.36 | 0.58 | 0.67 |
| 6 | 69109884 | rs12196984 | 0.272 | -0.891 | -1.333 | -1.409 | 0.116 | 0.023 | 0.023 | 4.069 | 0.52 | 0.19 | 0.63 | 0.37 | 0.7 | 0.71 |
| 6 | 69121834 | rs6937202 | 0.237 | -0.858 | -1.335 | -1.393 | 0.105 | 0.021 | 0.021 | 4.262 | 0.53 | 0.21 | 0.63 | 0.36 | 0.7 | 0.71 |
| 6 | 69124811 | rs9360341 | 0.31 | -0.695 | -1.152 | -0.999 | 0.02 | -0.031 | -0.031 | 0 | 0.27 | 0.22 | 0.23 | 0.57 | 0.19 | 0.11 |
| 6 | 69132263 | rs4132691 | 0.054 | 0.176 | -1.293 | -1.352 | -0.037 | 0.049 | 0.049 | 0 | 0.11 | 0.01 | 0.07 | 0.03 | 0.11 | 0.11 |
| 6 | 69135009 | rs17576708 | -0.297 | 0.26 | -1.384 | -1.462 | -0.008 | 0.009 | 0.009 | 0 | 0.07 | 0.04 | 0.07 | 0.04 | 0.1 | 0.09 |
| 6 | 69142008 | rs9363918 | 0.683 | -0.13 | -1.555 | -2.342 | -0.001 | 0.101 | 0.101 | 0.329 | 0.43 | 0.22 | 0.55 | 0.85 | 0.38 | 0.54 |
| 6 | 69149444 | rs11756671 | 1.033 | 0.093 | -1.825 | -2.115 | -0.071 | 0.008 | 0.008 | 0.142 | 0.27 | 0.13 | 0.33 | 0.22 | 0.33 | 0.39 |
| 6 | 69151721 | rs9346180 | 1.378 | 0.023 | -1.682 | -1.92 | 0.01 | -0.062 | -0.062 | 0.004 | 0.16 | 0.05 | 0.28 | 0.69 | 0.13 | 0.24 |
| 6 | 69152472 | rs7771529 | -0.1 | -0.092 | -1.42 | -2.01 | -0.019 | -0.086 | -0.086 | 0 | 0.24 | 0.3 | 0.17 | 0.04 | 0.22 | 0.16 |
| 6 | 69156120 | rs9354741 | -0.023 | 0.11 | -1.235 | -1.87 | 0.005 | 0.006 | 0.006 | 0 | 0.4 | 0.35 | 0.45 | 0.73 | 0.35 | 0.4 |
| 6 | 69175865 | rs7749220 | 0.116 | 0.376 | -1.052 | -1.829 | 0.004 | -0.082 | -0.082 | 0 | 0.61 | 0.44 | 0.7 | 0.9 | 0.6 | 0.66 |
| 6 | 69185047 | rs13207908 | 0.985 | 0.32 | -1.547 | -1.87 | -0.083 | 0.045 | 0.045 | 0 | 0.32 | 0.07 | 0.34 | 0.24 | 0.38 | 0.37 |
| 6 | 69185660 | rs4708157 | 0.854 | 0.469 | -1.606 | -1.884 | -0.032 | -0.001 | -0.001 | 0 | 0.07 | 0.01 | 0.1 | 0.06 | 0.09 | 0.12 |
| 6 | 69189605 | rs9454527 | 0.615 | 0.132 | -0.946 | -2.113 | 0.001 | 0.095 | 0.095 | 1.311 | 0.47 | 0.29 | 0.41 | 0.24 | 0.48 | 0.4 |
| 6 | 69193580 | rs352517 | -0.294 | 0.132 | -0.62 | -1.904 | 0.003 | -0.014 | -0.014 | 0 | 0.28 | 0.38 | 0.31 | 0.17 | 0.26 | 0.31 |
| 6 | 69197250 | rs9342711 | 0.171 | -0.124 | -0.662 | -1.736 | -0.02 | 0.006 | 0.006 | 0 | 0.42 | 0.46 | 0.34 | 0.55 | 0.32 | 0.32 |
| 6 | 69198837 | rs352523 | -1.044 | -0.067 | -0.346 | -1.629 | -0.018 | 0.182 | 0.182 | 0 | 0.8 | 0.56 | 0.78 | 0.5 | 0.83 | 0.75 |
| 6 | 69199797 | rs9363933 | 1.261 | -0.115 | -0.535 | -1.599 | -0.012 | 0.049 | 0.049 | 0 | 0.18 | 0.21 | 0.19 | 0.5 | 0.12 | 0.24 |
| 6 | 69210493 | rs10806600 | 1.059 | 0.207 | -0.833 | -1.761 | -0.178 | -0.168 | -0.168 | 0.226 | 0.32 | 0.03 | 0.48 | 0.44 | 0.43 | 0.49 |
| 6 | 69235100 | rs9342717 | 1.008 | -0.031 | -0.429 | -1.884 | -0.063 | 0.168 | 0.168 | 0 | 0.2 | 0.5 | 0.21 | 0.5 | 0.11 | 0.23 |
| 6 | 69237307 | rs4481397 | -1.535 | -0.135 | -0.53 | -2.019 | -0.006 | 0.038 | 0.038 | 0.101 | 0.86 | 0.93 | 0.92 | 0.95 | 0.85 | 0.93 |
| 6 | 69240370 | rs12208189 | 0.374 | -0.251 | -0.479 | -2.034 | 0.011 | -0.053 | -0.053 | 0 | 0.09 | 0.25 | 0.07 | 0 | 0.09 | 0.03 |
| 6 | 69241954 | rs965056 | 1.924 | -0.029 | -0.57 | -2.008 | -0.099 | -0.086 | -0.086 | 0.004 | 0.23 | 0.01 | 0.29 | 0.33 | 0.28 | 0.32 |
| 6 | 69249384 | rs13194034 | 1.242 | -0.078 | 0.098 | -1.417 | 0 | 0.005 | 0.005 | 0.129 | 0.1 | 0.06 | 0.07 | 0 | 0.11 | 0.05 |
| 6 | 69250345 | rs9354778 | 1.126 | -0.147 | 0.132 | -1.36 | -0.009 | 0.043 | 0.043 | 0 | 0.17 | 0.19 | 0.19 | 0.5 | 0.11 | 0.23 |
| 6 | 69256487 | rs12194811 | 0.576 | -0.135 | -0.007 | -1.358 | 0.007 | -0.033 | -0.033 | 0 | 0.07 | 0.12 | 0.06 | 0 | 0.09 | 0.03 |
| 6 | 69260172 | rs6929667 | 0.542 | -0.162 | 0.009 | -1.3 | -0.063 | 0.162 | 0.162 | 0 | 0.21 | 0.5 | 0.21 | 0.5 | 0.11 | 0.23 |
| 6 | 69273529 | rs16900061 | 0.502 | 0.454 | -0.419 | -1.001 | -0.04 | -0.041 | -0.041 | 0 | 0.09 | 0.03 | 0.19 | 0.12 | 0.15 | 0.17 |
| 6 | 69282291 | rs13209526 | -0.74 | -0.311 | 0.106 | -1.039 | -0.022 | 0.02 | 0.02 | 0 | 0.36 | 0.5 | 0.33 | 0.45 | 0.3 | 0.3 |
| 6 | 69285887 | rs9354787 | -0.664 | -0.195 | -0.803 | -0.978 | 0.009 | -0.016 | -0.016 | 0 | 0.33 | 0.27 | 0.32 | 0.45 | 0.3 | 0.3 |
| 6 | 69287242 | rs7754996 | 0.868 | -0.216 | -0.82 | -0.995 | 0.014 | -0.016 | -0.016 | 0 | 0.16 | 0.1 | 0.18 | 0.45 | 0.11 | 0.23 |
| 6 | 69289549 | rs4416643 | -0.286 | 0.094 | -1.22 | -1.037 | 0 | -0.035 | -0.035 | 0 | 0.17 | 0.17 | 0.14 | 0 | 0.19 | 0.08 |
| 6 | 69291365 | rs7748176 | 0.407 | 0.436 | -1.226 | -0.757 | 0.033 | -0.046 | -0.046 | 0 | 0.91 | 0.84 | 0.92 | 1 | 0.86 | 0.97 |
| 6 | 69291570 | rs9346223 | 0.595 | 0.332 | -1.791 | -0.484 | 0.011 | 0.033 | 0.033 | 0.21 | 0.12 | 0.03 | 0.06 | 0.09 | 0.1 | 0.06 |
| 6 | 69294401 | rs9346224 | -0.038 | 0.783 | -1.491 | -0.208 | 0.029 | -0.004 | -0.004 | 0 | 0.49 | 0.3 | 0.57 | 0.49 | 0.54 | 0.63 |
| 6 | 69324546 | rs9351732 | -0.382 | 1.792 | -0.229 | -0.913 | 0.013 | 0.003 | 0.003 | 0 | 0.11 | 0.16 | 0.13 | 0.14 | 0.14 | 0.31 |
| 6 | 69328787 | rs3121773 | -0.665 | 0.874 | 0.025 | -0.576 | 0.003 | -0.024 | -0.024 | 0 | 0.35 | 0.48 | 0.44 | 0.2 | 0.37 | 0.24 |
| 6 | 69334292 | rs2585619 | -0.572 | 0.815 | -0.143 | -0.61 | -0.009 | 0.008 | 0.008 | 0.036 | 0.3 | 0.24 | 0.41 | 0.19 | 0.34 | 0.18 |
| 6 | 69337256 | rs2802705 | -0.607 | 0.809 | -0.144 | -0.613 | -0.006 | 0.011 | 0.011 | 0 | 0.31 | 0.27 | 0.41 | 0.2 | 0.34 | 0.18 |
| 6 | 69345483 | rs2256337 | -0.452 | 0.719 | -0.188 | -0.668 | -0.012 | 0.014 | 0.014 | 0 | 0.31 | 0.23 | 0.41 | 0.19 | 0.34 | 0.19 |
| 6 | 69357668 | rs9360356 | 0.964 | 0.38 | -0.428 | -0.905 | -0.004 | -0.03 | -0.03 | 0.07 | 0.15 | 0.07 | 0.15 | 0.63 | 0.16 | 0.19 |
| 6 | 69358141 | rs2585626 | -1.055 | 0.254 | -0.13 | -1.123 | 0.012 | 0.018 | 0.018 | 0.035 | 0.47 | 0.8 | 0.67 | 0.37 | 0.54 | 0.58 |
| 6 | 69364038 | rs2802691 | -1.119 | 0.143 | 0.012 | -1.173 | 0.023 | 0.019 | 0.019 | 0 | 0.51 | 0.68 | 0.68 | 0.37 | 0.56 | 0.58 |
| 6 | 69372017 | rs2253759 | -0.614 | 0.149 | 0.12 | -1.201 | 0.001 | 0.005 | 0.005 | 0 | 0.46 | 0.41 | 0.64 | 0.36 | 0.53 | 0.51 |
| 6 | 69384471 | rs1033520 | 0.733 | 0.184 | 0.055 | -0.968 | -0.001 | 0.055 | 0.055 | 0.174 | 0.1 | 0.07 | 0.14 | 0.61 | 0.14 | 0.18 |
| 6 | 69393510 | rs4607402 | 0.213 | 0.057 | 0.254 | -0.78 | 0.028 | -0.068 | -0.068 | 1.287 | 0.14 | 0.01 | 0.15 | 0.17 | 0.06 | 0.06 |
| 6 | 69404850 | rs2746128 | -0.04 | 0.25 | 0.567 | -0.719 | 0.095 | 0.076 | 0.076 | 0 | 0.87 | 0.61 | 0.74 | 0.83 | 0.81 | 0.68 |
| 6 | 69405498 | rs9342729 | 1.528 | 0.211 | 0.428 | -0.83 | 0.019 | -0.04 | -0.04 | 0 | 0.07 | 0.01 | 0.11 | 0.13 | 0.03 | 0.01 |
| 6 | 69417357 | rs9346234 | -0.429 | 0.436 | 0.615 | -0.864 | 0.001 | 0.002 | 0.002 | 0 | 0.41 | 0.61 | 0.44 | 0.21 | 0.37 | 0.26 |
| 6 | 69425811 | rs2064389 | -0.392 | 0.483 | 0.742 | -0.782 | 0.004 | -0.001 | -0.001 | 0 | 0.91 | 0.89 | 0.84 | 0.86 | 0.81 | 0.68 |
| 6 | 69439107 | rs9363963 | -0.484 | 0.483 | 0.739 | -0.782 | 0.026 | 0.005 | 0.005 | 0 | 0.9 | 0.84 | 0.84 | 0.86 | 0.81 | 0.68 |
| 6 | 69439577 | rs10945138 | 0.887 | 0.436 | 0.872 | -0.833 | 0.018 | 0.048 | 0.048 | 0 | 0.5 | 0.25 | 0.42 | 0.66 | 0.47 | 0.48 |
| 6 | 69440758 | rs7768591 | 0.737 | 0.464 | 0.915 | -0.819 | 0 | 0.001 | 0.001 | 0 | 0.53 | 0.47 | 0.43 | 0.66 | 0.47 | 0.48 |
| 6 | 69451081 | rs2024562 | 0.202 | 0.596 | 0.787 | -0.837 | 0.077 | 0.007 | 0.007 | 0 | 0.87 | 0.68 | 0.82 | 0.86 | 0.79 | 0.67 |
| 6 | 69459942 | rs9346237 | 0.179 | 0.578 | 0.542 | -0.889 | -0.042 | 0.041 | 0.041 | 0 | 0.32 | 0.07 | 0.37 | 0.17 | 0.33 | 0.18 |
| 6 | 69460974 | rs6908159 | -0.225 | 0.674 | 0.765 | -0.886 | -0.007 | -0.026 | -0.026 | 0 | 0.38 | 0.56 | 0.4 | 0.2 | 0.34 | 0.18 |
| 6 | 69464386 | rs10484791 | 0.796 | 0.617 | 0.442 | -0.841 | 0.019 | 0.047 | 0.047 | 0 | 0.51 | 0.25 | 0.42 | 0.66 | 0.46 | 0.48 |
| 6 | 69468150 | rs2073135 | 1.136 | -0.043 | -0.023 | -1.398 | 0.023 | -0.013 | -0.013 | 0 | 0.33 | 0.18 | 0.3 | 0.66 | 0.26 | 0.37 |
| 6 | 69474494 | rs9346239 | 0.412 | 0.073 | 0.2 | -1.49 | 0.007 | -0.028 | -0.028 | 0.073 | 0.68 | 0.39 | 0.68 | 0.85 | 0.59 | 0.54 |
| 6 | 69480408 | rs1523932 | 0.584 | -0.016 | 0.147 | -1.502 | -0.022 | 0.023 | 0.023 | 0 | 0.32 | 0.12 | 0.36 | 0.19 | 0.34 | 0.17 |
| 6 | 69483708 | rs1523947 | 0.39 | 0.326 | 0.815 | -1.212 | 0.05 | -0.01 | -0.01 | 0 | 0.85 | 0.72 | 0.8 | 0.85 | 0.75 | 0.61 |
| 6 | 69504861 | rs1033521 | 0.417 | 0.318 | 0.79 | -1.257 | -0.009 | -0.104 | -0.104 | 0 | 0.17 | 0.32 | 0.12 | 0 | 0.15 | 0.07 |
| 6 | 69508067 | rs2223724 | 0.589 | 0.59 | 0.357 | -0.77 | 0.039 | -0.009 | -0.009 | 0.113 | 0.86 | 0.75 | 0.82 | 0.86 | 0.78 | 0.67 |
| 6 | 69520768 | rs9294811 | 0.694 | 0.566 | 0.197 | -0.663 | 0.037 | -0.011 | -0.011 | 0 | 0.85 | 0.66 | 0.8 | 0.86 | 0.76 | 0.6 |
| 6 | 69520926 | rs9346246 | 0.779 | 0.576 | 0.105 | -0.681 | 0.019 | -0.016 | -0.016 | 0 | 0.81 | 0.67 | 0.78 | 0.86 | 0.73 | 0.6 |
| 6 | 69521978 | rs582962 | 0.528 | 0.283 | -0.289 | -0.293 | -0.007 | 0.008 | 0.008 | 0.412 | 0.63 | 0.45 | 0.6 | 0.58 | 0.7 | 0.51 |
| 6 | 69525365 | rs9360362 | -0.859 | 1.457 | -0.824 | 0.241 | -0.093 | -0.095 | -0.095 | 0.317 | 0.15 | 0.03 | 0.33 | 0.68 | 0.27 | 0.34 |
| 6 | 69527865 | rs492444 | -0.457 | 1.825 | -1.072 | -0.058 | -0.026 | 0.058 | 0.058 | 0.802 | 0.65 | 0.77 | 0.6 | 0.85 | 0.59 | 0.49 |

| | | | | | | | | | | | | | | | | |
|---|---|---|---|---|---|---|---|---|---|---|---|---|---|---|---|---|
| 6 | 69531957 | rs2022212 | -0.384 | 1.59 | -0.938 | 0.54 | -0.019 | -0.03 | -0.03 | 0 | 0.1 | 0.03 | 0.29 | 0.64 | 0.13 | 0.19 |
| 6 | 69540950 | rs571299 | -0.732 | 1.889 | -0.629 | 0.611 | 0.004 | 0.039 | 0.039 | 0 | 0.13 | 0.15 | 0.32 | 0.58 | 0.21 | 0.23 |
| 6 | 69541114 | rs514518 | -0.137 | 1.506 | -0.492 | 0.067 | -0.001 | -0.022 | -0.022 | 0 | 0.41 | 0.76 | 0.44 | 0.58 | 0.39 | 0.39 |
| 6 | 69548012 | rs544495 | -0.495 | 1.714 | -0.474 | 0.329 | 0.001 | 0.043 | 0.043 | 0 | 0.92 | 0.89 | 0.76 | 0.71 | 0.92 | 0.74 |
| 6 | 69549897 | rs493768 | -0.537 | 1.74 | -0.21 | 0.36 | 0.004 | 0.037 | 0.037 | 0 | 0.14 | 0.16 | 0.32 | 0.58 | 0.2 | 0.22 |
| 6 | 69560086 | rs13207774 | 0.731 | 1.424 | -0.706 | -0.288 | -0.003 | 0.028 | 0.028 | 2.893 | 0.18 | 0.06 | 0.12 | 0.09 | 0.19 | 0.12 |
| 6 | 69560985 | rs500125 | 0.236 | 1.529 | -0.591 | -0.338 | -0.007 | -0.008 | -0.008 | 0 | 0.42 | 0.31 | 0.47 | 0.59 | 0.53 | 0.5 |
| 6 | 69572657 | rs10223798 | 0.322 | 1.544 | -0.26 | -0.502 | 0.058 | 0.12 | 0.12 | 2.826 | 0.49 | 0.63 | 0.28 | 0.12 | 0.39 | 0.24 |
| 6 | 69581991 | rs551216 | -0.54 | 1.634 | -0.37 | -0.455 | -0.021 | -0.021 | -0.021 | 0.619 | 0.21 | 0.19 | 0.38 | 0.59 | 0.35 | 0.34 |
| 6 | 69595671 | rs12199028 | 2.087 | 1.618 | -0.296 | -0.472 | -0.004 | 0.109 | 0.109 | 0 | 0.2 | 0.02 | 0.09 | 0 | 0.18 | 0.16 |
| 6 | 69596283 | rs4706710 | 0.606 | 1.624 | -0.285 | -0.471 | -0.018 | -0.018 | -0.018 | 0 | 0.78 | 0.8 | 0.62 | 0.41 | 0.65 | 0.66 |
| 6 | 69596592 | rs4706061 | 0.131 | 1.669 | -0.267 | -0.394 | 0.003 | 0.034 | 0.034 | 0 | 0.86 | 0.85 | 0.68 | 0.42 | 0.8 | 0.78 |
| 6 | 69598003 | rs12209015 | 2.078 | 1.679 | -0.263 | -0.389 | -0.005 | 0.108 | 0.108 | 0 | 0.2 | 0.02 | 0.09 | 0 | 0.18 | 0.16 |
| 6 | 69598297 | rs1403926 | 0.193 | 1.753 | -0.011 | -0.081 | 0.004 | 0.038 | 0.038 | 0 | 0.87 | 0.85 | 0.68 | 0.42 | 0.8 | 0.78 |
| 6 | 69610281 | rs2342761 | 0.387 | 1.127 | 0.28 | -0.09 | 0 | -0.003 | -0.003 | 0 | 0.4 | 0.57 | 0.42 | 0.3 | 0.41 | 0.54 |
| 6 | 69615168 | rs10485436 | 0.88 | 0.909 | 0.023 | -0.046 | 0.004 | 0.004 | 0.004 | 8.285 | 0.13 | 0.05 | 0.11 | 0.09 | 0.14 | 0.07 |
| 6 | 69615615 | rs10485435 | 0.868 | 1.032 | 0.362 | 0.202 | -0.016 | -0.005 | -0.005 | 0 | 0.31 | 0.39 | 0.34 | 0.29 | 0.27 | 0.42 |
| 6 | 69619007 | rs1283951 | -0.397 | 1.231 | 0.288 | 1.482 | -0.028 | 0.017 | 0.017 | 0 | 0.95 | 0.99 | 0.85 | 0.64 | 0.91 | 0.88 |
| 6 | 69629187 | rs7747549 | 0.645 | 1.167 | 0.616 | 1.068 | 0.014 | 0.19 | 0.19 | 0 | 0.88 | 0.74 | 0.71 | 0.43 | 0.86 | 0.79 |
| 6 | 69632227 | rs10945147 | 1.237 | 1.029 | 0.764 | 0.995 | 0 | 0.216 | 0.216 | 0 | 0.84 | 0.56 | 0.69 | 0.43 | 0.86 | 0.79 |
| 6 | 69647244 | rs2342763 | 1.308 | 1.016 | 0.758 | 0.988 | -0.003 | 0.209 | 0.209 | 0 | 0.84 | 0.55 | 0.69 | 0.43 | 0.86 | 0.79 |
| 6 | 69652994 | rs7748808 | 0.191 | 1.382 | 0.782 | 0.732 | 0.072 | -0.036 | -0.036 | 0 | 0.09 | 0.21 | 0.08 | 0.01 | 0.15 | 0.12 |
| 6 | 69654650 | rs7759377 | 0.386 | 1.408 | 0.668 | 0.746 | 0.056 | -0.031 | -0.031 | 1.774 | 0.08 | 0.15 | 0.07 | 0.01 | 0.15 | 0.12 |
| 6 | 69660841 | rs1932616 | -0.756 | 1.135 | 0.815 | 1.065 | -0.014 | 0.192 | 0.192 | 0.016 | 0.13 | 0.28 | 0.29 | 0.56 | 0.1 | 0.2 |
| 6 | 69663880 | rs1889879 | -0.897 | 1.179 | 0.588 | 0.059 | 0.001 | -0.06 | -0.06 | 1.122 | 0.67 | 0.63 | 0.78 | 0.92 | 0.65 | 0.79 |
| 6 | 69665063 | rs12207529 | 0.506 | 1.179 | 0.464 | 0.604 | 0 | -0.002 | -0.002 | 0.224 | 0.09 | 0.07 | 0.17 | 0.28 | 0.09 | 0.28 |
| 6 | 69666520 | rs7757138 | 0.904 | 0.747 | 0.267 | 0.333 | -0.001 | -0.031 | -0.031 | 0 | 0.33 | 0.33 | 0.19 | 0.05 | 0.31 | 0.2 |
| 6 | 69666684 | rs1932618 | -1.306 | 0.813 | 0.171 | 0.182 | -0.002 | 0.002 | 0.002 | 2.678 | 0.89 | 0.91 | 0.9 | 0.92 | 0.86 | 0.94 |
| 6 | 69667040 | rs1932619 | 0.64 | 0.82 | 0.228 | 0.132 | -0.016 | -0.019 | -0.019 | 0 | 0.7 | 0.73 | 0.65 | 0.63 | 0.61 | 0.55 |
| 6 | 69667472 | rs3823061 | -1.41 | 0.831 | 0.14 | -0.285 | 0.001 | 0.013 | 0.013 | 3.231 | 0.9 | 0.95 | 0.91 | 0.97 | 0.87 | 0.95 |
| 6 | 69669063 | rs9454659 | 0.814 | 0.72 | 0.151 | -0.265 | -0.006 | -0.065 | -0.065 | 0 | 0.34 | 0.4 | 0.19 | 0.05 | 0.31 | 0.2 |
| 6 | 69672039 | rs3798971 | 0.769 | 0.701 | 0.074 | 0.288 | 0.019 | -0.057 | -0.057 | 0.573 | 0.35 | 0.1 | 0.35 | 0.55 | 0.32 | 0.35 |
| 6 | 69675838 | rs3798974 | 0.612 | 0.708 | 0.162 | 0.306 | 0.015 | -0.026 | -0.026 | 0 | 0.37 | 0.27 | 0.37 | 0.61 | 0.33 | 0.36 |
| 6 | 69676937 | rs1556754 | -0.069 | 0.629 | 0.179 | 0.211 | -0.005 | 0.039 | 0.039 | 0 | 0.81 | 0.73 | 0.63 | 0.67 | 0.8 | 0.67 |
| 6 | 69677322 | rs1006997 | -0.721 | 0.551 | 0.158 | 0.184 | -0.003 | -0.018 | -0.018 | 0 | 0.88 | 0.92 | 0.7 | 0.71 | 0.85 | 0.7 |
| 6 | 69677917 | rs1953613 | 0.753 | 0.602 | 0.076 | 0.423 | -0.012 | -0.003 | -0.003 | 0 | 0.7 | 0.74 | 0.4 | 0.33 | 0.62 | 0.45 |
| 6 | 69678117 | rs3798977 | 0.771 | 0.786 | 0.273 | -0.152 | 0 | -0.051 | -0.051 | 0 | 0.35 | 0.42 | 0.19 | 0.12 | 0.32 | 0.21 |
| 6 | 69681617 | rs3823064 | -0.395 | 0.751 | -0.151 | -0.141 | 0.021 | 0.046 | 0.046 | 0 | 0.52 | 0.7 | 0.45 | 0.7 | 0.41 | 0.41 |
| 6 | 69684850 | rs2275209 | -0.209 | 0.843 | 0.192 | -0.192 | 0.105 | 0.019 | 0.019 | 0 | 0.12 | 0.4 | 0.1 | 0.07 | 0.15 | 0.14 |
| 6 | 69687698 | rs1959407 | 1.233 | 0.8 | -0.061 | -0.344 | 0.002 | 0.041 | 0.041 | 0 | 0.31 | 0.19 | 0.14 | 0.04 | 0.27 | 0.17 |
| 6 | 69688098 | rs3798983 | 0.596 | 0.792 | -0.194 | -0.216 | 0.013 | -0.052 | -0.052 | 2.908 | 0.13 | 0.03 | 0.25 | 0.58 | 0.1 | 0.19 |
| 6 | 69688165 | rs3798984 | 1.829 | 0.889 | -0.201 | -0.243 | -0.034 | 0.015 | 0.015 | 0 | 0.06 | 0.01 | 0.05 | 0.01 | 0.13 | 0.11 |
| 6 | 69690567 | rs12665384 | 0.586 | 0.712 | -0.173 | -0.203 | 0.013 | -0.052 | -0.052 | 0.129 | 0.13 | 0.03 | 0.25 | 0.58 | 0.1 | 0.19 |
| 6 | 69698538 | rs3798992 | -0.661 | 0.652 | -0.078 | -0.42 | 0.009 | 0.017 | 0.017 | 0 | 0.57 | 0.66 | 0.5 | 0.7 | 0.54 | 0.51 |
| 6 | 69702416 | rs9454667 | 0.665 | 0.645 | -0.148 | -0.467 | 0.001 | 0.009 | 0.009 | 0 | 0.33 | 0.28 | 0.16 | 0.05 | 0.29 | 0.18 |
| 6 | 69706344 | rs9354812 | 0.473 | 0.461 | 0.176 | -0.181 | -0.019 | 0.113 | 0.113 | 0 | 0.82 | 0.52 | 0.66 | 0.74 | 0.85 | 0.68 |
| 6 | 69708219 | rs3798997 | 0.601 | 0.458 | -0.049 | -0.15 | 0.013 | -0.043 | -0.043 | 0 | 0.12 | 0.02 | 0.24 | 0.6 | 0.1 | 0.19 |
| 6 | 69708918 | rs6908180 | 0.487 | 0.45 | 0.01 | -0.348 | -0.009 | -0.028 | -0.028 | 1.666 | 0.1 | 0.23 | 0.09 | 0.03 | 0.09 | 0.03 |
| 6 | 69710575 | rs1999787 | -0.515 | 0.447 | 0.254 | -0.298 | 0 | 0.017 | 0.017 | 0 | 0.87 | 0.89 | 0.7 | 0.73 | 0.85 | 0.7 |
| 6 | 69713057 | rs13191563 | 0.072 | 0.343 | -0.411 | -1.082 | 0 | 0.013 | 0.013 | 0 | 0.25 | 0.24 | 0.15 | 0.03 | 0.24 | 0.13 |
| 6 | 69714947 | rs3798999 | 0.435 | 0.552 | -0.354 | -1.005 | 0.001 | 0.001 | 0.001 | 0 | 0.6 | 0.43 | 0.49 | 0.64 | 0.56 | 0.47 |
| 6 | 69715521 | rs3799003 | -0.003 | 0.588 | -0.424 | -1.161 | 0.001 | 0.034 | 0.034 | 0 | 0.75 | 0.89 | 0.44 | 0.09 | 0.74 | 0.49 |
| 6 | 69717102 | rs1336638 | -0.382 | 0.67 | -0.493 | -1.163 | -0.044 | -0.04 | -0.04 | 0 | 0.81 | 0.94 | 0.64 | 0.75 | 0.73 | 0.59 |
| 6 | 69718158 | rs12213405 | 0.887 | 0.64 | -0.354 | -1.452 | 0.004 | 0.016 | 0.016 | 0 | 0.47 | 0.41 | 0.25 | 0.04 | 0.46 | 0.28 |
| 6 | 69723045 | rs3799007 | 1.499 | 0.565 | -0.543 | -1.257 | -0.006 | -0.007 | -0.007 | 0.535 | 0.06 | 0.1 | 0.06 | 0.03 | 0.06 | 0.03 |
| 6 | 69728452 | rs10485430 | 1.492 | 0.531 | -0.47 | -1.359 | 0.004 | 0.015 | 0.015 | 0 | 0.17 | 0.12 | 0.1 | 0.06 | 0.15 | 0.07 |
| 6 | 69729678 | rs1889877 | 0.822 | 0.396 | -0.145 | -1.508 | 0 | -0.074 | -0.074 | 0 | 0.48 | 0.49 | 0.26 | 0.04 | 0.46 | 0.28 |
| 6 | 69745723 | rs2225803 | 0.549 | 0.451 | -0.534 | -1.533 | 0 | 0.004 | 0.004 | 0 | 0.21 | 0.22 | 0.13 | 0.04 | 0.2 | 0.13 |
| 6 | 69766509 | rs634371 | -0.677 | 0.554 | -0.627 | -1.43 | 0.005 | 0.171 | 0.171 | 0.261 | 0.48 | 0.22 | 0.71 | 0.94 | 0.54 | 0.7 |
| 6 | 69767587 | rs9446089 | -0.363 | 0.591 | -0.612 | -1.316 | 0.004 | 0.031 | 0.031 | 0 | 0.74 | 0.86 | 0.44 | 0.09 | 0.73 | 0.49 |
| 6 | 69772650 | rs3734465 | 0.991 | 0.55 | -0.536 | -1.284 | -0.012 | -0.027 | -0.027 | 0 | 0.08 | 0.15 | 0.06 | 0.03 | 0.06 | 0.03 |
| 6 | 69794037 | rs3823070 | 0.95 | 0.545 | -0.826 | -1.052 | 0.013 | -0.082 | -0.082 | 0 | 0.13 | 0.01 | 0.1 | 0.63 | 0.08 | 0.23 |
| 6 | 69794079 | rs11964764 | 1.309 | 0.824 | -0.9 | -1.81 | -0.006 | 0.03 | 0.03 | 0 | 0.18 | 0.07 | 0.12 | 0.04 | 0.19 | 0.09 |
| 6 | 69802200 | rs17481523 | 1.264 | 0.824 | -1.075 | -2.23 | -0.016 | 0.016 | 0.016 | 0 | 0.12 | 0.07 | 0.06 | 0 | 0.14 | 0.09 |
| 6 | 69802502 | rs10455681 | 0.899 | 0.714 | -1.198 | -2.233 | -0.005 | -0.043 | -0.043 | 0.053 | 0.18 | 0.02 | 0.29 | 0.83 | 0.15 | 0.49 |
| 6 | 69803462 | rs6913677 | -0.006 | 0.718 | -1.067 | -2.242 | 0.006 | 0.095 | 0.095 | 0 | 0.36 | 0.13 | 0.41 | 0.88 | 0.34 | 0.57 |
| 6 | 69804537 | rs2128290 | 0.276 | 0.486 | -0.727 | -2.373 | 0.015 | -0.146 | -0.146 | 0 | 0.73 | 0.5 | 0.69 | 0.97 | 0.7 | 0.86 |
| 6 | 69808955 | rs9454693 | 2.047 | 0.538 | -0.702 | -2.323 | -0.006 | -0.088 | -0.088 | 1.034 | 0.29 | 0.3 | 0.14 | 0 | 0.26 | 0.15 |
| 6 | 69813713 | rs2046251 | -0.362 | 0.689 | -0.682 | -2.155 | 0.019 | -0.097 | -0.097 | 0 | 0.8 | 0.65 | 0.75 | 1 | 0.76 | 0.88 |

| | | | | | | | | | | | | | | | | |
|---|---|---|---|---|---|---|---|---|---|---|---|---|---|---|---|---|
| 6 | 69816451 | rs10485427 | 0.701 | 0.775 | -0.862 | -2.2 | -0.034 | 0.057 | 0.057 | 0 | 0.23 | 0.09 | 0.15 | 0.04 | 0.27 | 0.2 |
| 6 | 69816863 | rs964664 | 0.294 | 0.596 | -0.575 | -2.315 | 0.009 | -0.137 | -0.137 | 0 | 0.72 | 0.42 | 0.69 | 0.96 | 0.7 | 0.86 |
| 6 | 69818593 | rs9363983 | 0.798 | 0.595 | -0.693 | -2.155 | -0.01 | -0.026 | -0.026 | 0 | 0.19 | 0.05 | 0.31 | 0.83 | 0.19 | 0.49 |
| 6 | 69826507 | rs3799039 | 0.655 | 0.547 | -0.695 | -2.279 | 0.032 | 0.084 | 0.084 | 0 | 0.49 | 0.33 | 0.54 | 0.92 | 0.43 | 0.66 |
| 6 | 69840109 | rs9360376 | -0.186 | 0.564 | -0.615 | -2.791 | 0.009 | -0.085 | -0.085 | 0 | 0.67 | 0.43 | 0.78 | 0.96 | 0.65 | 0.81 |
| 6 | 69844768 | rs1482326 | 0.537 | 0.566 | -0.669 | -2.824 | -0.09 | 0.006 | 0.006 | 0.832 | 0.26 | 0.05 | 0.36 | 0.87 | 0.3 | 0.6 |
| 6 | 69847967 | rs67807 | 0.887 | 0.563 | -0.502 | -2.786 | -0.002 | -0.024 | -0.024 | 1.019 | 0.14 | 0.13 | 0.07 | 0 | 0.18 | 0.11 |
| 6 | 69848733 | rs1336633 | -0.11 | 0.609 | -0.717 | -2.746 | 0.008 | -0.034 | -0.034 | 0.54 | 0.67 | 0.42 | 0.69 | 0.87 | 0.65 | 0.8 |
| 6 | 69849203 | rs11759629 | 1.606 | 0.593 | -0.684 | -2.954 | -0.002 | 0.008 | 0.008 | 0 | 0.14 | 0.09 | 0.11 | 0.04 | 0.17 | 0.06 |
| 6 | 69850406 | rs9454705 | 1.804 | 0.422 | -0.539 | -2.878 | 0.015 | -0.017 | -0.017 | 0 | 0.31 | 0.22 | 0.14 | 0 | 0.24 | 0.15 |
| 6 | 69851816 | rs1912988 | -0.057 | 0.588 | -0.088 | -2.173 | 0.034 | -0.139 | -0.139 | 0 | 0.81 | 0.52 | 0.89 | 1 | 0.82 | 0.87 |
| 6 | 69877758 | rs314215 | -0.495 | 0.617 | -0.431 | -2.275 | -0.001 | -0.053 | -0.053 | 0 | 0.29 | 0.3 | 0.32 | 0.01 | 0.33 | 0.19 |
| 6 | 69898036 | rs12528123 | 1.772 | 0.612 | -0.623 | -2.447 | -0.003 | 0.01 | 0.01 | 0 | 0.12 | 0.07 | 0.09 | 0.03 | 0.13 | 0.04 |
| 6 | 69900102 | rs779484 | -0.69 | 0.664 | -0.983 | -2.515 | -0.096 | 0.021 | 0.021 | 2.044 | 0.74 | 0.97 | 0.66 | 0.12 | 0.7 | 0.37 |
| 6 | 69900836 | rs9354826 | 1.123 | 0.556 | -0.803 | -1.862 | -0.003 | -0.084 | -0.084 | 0 | 0.14 | 0.02 | 0.11 | 0.58 | 0.12 | 0.27 |
| 6 | 69912542 | rs1751710 | -0.851 | 0.52 | -0.174 | -2.522 | 0 | -0.065 | -0.065 | 0 | 0.31 | 0.57 | 0.44 | 0.09 | 0.32 | 0.2 |
| 6 | 69924282 | rs779473 | 0.607 | 0.738 | -0.158 | -2.557 | 0 | -0.027 | -0.027 | 0 | 0.13 | 0.14 | 0.09 | 0 | 0.16 | 0.1 |
| 6 | 69928844 | rs779471 | 0.137 | 0.722 | -0.104 | -2.559 | 0.028 | -0.018 | -0.018 | 0 | 0.16 | 0.43 | 0.21 | 0.08 | 0.16 | 0.1 |
| 6 | 69931419 | rs779467 | 1.111 | 0.853 | -0.461 | -2.552 | -0.001 | -0.019 | -0.019 | 0 | 0.1 | 0.11 | 0.05 | 0 | 0.1 | 0.07 |
| 6 | 69944232 | rs1512229 | 0.08 | 0.79 | -0.491 | -2.458 | -0.001 | -0.028 | -0.028 | 0 | 0.15 | 0.15 | 0.23 | 0 | 0.16 | 0.06 |
| 6 | 69983212 | rs11965487 | 0.959 | 0.916 | -0.293 | -2.383 | -0.003 | -0.012 | -0.012 | 0 | 0.1 | 0.11 | 0.05 | 0 | 0.1 | 0.07 |
| 6 | 69989413 | rs9454725 | 0.789 | 0.604 | -0.16 | -2.793 | -0.011 | -0.079 | -0.079 | 0 | 0.34 | 0.35 | 0.18 | 0.03 | 0.28 | 0.14 |
| 6 | 70003389 | rs9454727 | 1.39 | 0.106 | -0.438 | -3.122 | -0.002 | 0.126 | 0.126 | 0 | 0.27 | 0.07 | 0.15 | 0.02 | 0.27 | 0.18 |
| 6 | 70003609 | rs1321980 | 0.067 | 0.346 | 0.029 | -3.068 | -0.003 | -0.001 | -0.001 | 0 | 0.12 | 0.16 | 0.27 | 0.07 | 0.11 | 0.05 |
| 6 | 70003822 | rs16900620 | -0.08 | 0.36 | -0.007 | -3.109 | 0.002 | -0.083 | -0.083 | 0 | 0.14 | 0.26 | 0.06 | 0 | 0.1 | 0.07 |
| 6 | 70010225 | rs13191298 | -0.518 | 0.529 | 0.013 | -3.108 | 0.006 | -0.013 | -0.013 | 0 | 0.15 | 0.3 | 0.27 | 0.07 | 0.13 | 0.05 |
| 6 | 70013594 | rs4602695 | -1.009 | 0.648 | -0.627 | -3.165 | 0.002 | -0.033 | -0.033 | 0.079 | 0.33 | 0.37 | 0.39 | 0.13 | 0.33 | 0.14 |
| 6 | 70013768 | rs4348277 | -1.035 | 0.624 | -0.614 | -3.15 | 0.004 | -0.027 | -0.027 | 0 | 0.36 | 0.37 | 0.43 | 0.13 | 0.38 | 0.15 |
| 6 | 70018250 | rs13195278 | 0.562 | 0.521 | -0.395 | -2.606 | -0.01 | 0.009 | 0.009 | 0 | 0.09 | 0.06 | 0.04 | 0 | 0.1 | 0.07 |
| 6 | 70030617 | rs7769609 | 0.804 | 0.526 | -0.36 | -2.568 | 0.003 | -0.004 | -0.004 | 0 | 0.07 | 0.09 | 0.15 | 0.03 | 0.07 | 0.04 |
| 6 | 70038147 | rs1283468 | -0.916 | 0.443 | -0.297 | -2.62 | 0.007 | -0.081 | -0.081 | 0 | 0.2 | 0.35 | 0.2 | 0.03 | 0.16 | 0.11 |
| 6 | 70042045 | rs17748715 | -0.311 | 0.494 | -0.377 | -2.594 | 0.001 | -0.013 | -0.013 | 0 | 0.16 | 0.17 | 0.19 | 0.03 | 0.17 | 0.11 |
| 6 | 70045307 | rs1297415 | 0.628 | 0.533 | -0.807 | -2.567 | -0.007 | 0.006 | 0.006 | 0 | 0.09 | 0.08 | 0.04 | 0 | 0.1 | 0.07 |
| 6 | 70049915 | rs9454732 | -0.483 | 0.356 | -1.057 | -2.698 | 0.01 | 0.087 | 0.087 | 0.306 | 0.4 | 0.76 | 0.45 | 0.15 | 0.36 | 0.21 |
| 6 | 70071173 | rs913543 | 1.065 | -0.06 | -1.298 | -2.837 | -0.034 | 0.08 | 0.08 | 0 | 0.38 | 0.17 | 0.46 | 0.83 | 0.45 | 0.61 |
| 6 | 70075331 | rs7745125 | 0.246 | 0.193 | -1.129 | -2.648 | 0.011 | 0.075 | 0.075 | 0.309 | 0.59 | 0.24 | 0.55 | 0.79 | 0.64 | 0.78 |
| 6 | 70076733 | rs1410700 | -0.125 | 0.168 | -1.041 | -2.631 | 0.015 | -0.082 | -0.082 | 0.004 | 0.81 | 0.64 | 0.8 | 0.97 | 0.83 | 0.89 |
| 6 | 70082863 | rs3799095 | 0.957 | 0.299 | -2.232 | -2.558 | -0.03 | 0.071 | 0.071 | 0 | 0.39 | 0.17 | 0.46 | 0.77 | 0.45 | 0.6 |
| 6 | 70088968 | rs9351751 | 1.069 | 0.148 | -1.34 | -2.014 | -0.004 | 0.031 | 0.031 | 0 | 0.17 | 0.1 | 0.13 | 0.12 | 0.16 | 0.08 |
| 6 | 70095690 | rs1952435 | 0.597 | 0.242 | -1.413 | -2.036 | 0.011 | 0.084 | 0.084 | 0.995 | 0.6 | 0.24 | 0.66 | 0.85 | 0.66 | 0.81 |
| 6 | 70102584 | rs12525747 | 0.645 | 0.197 | -1.386 | -1.232 | -0.014 | 0.031 | 0.031 | 0 | 0.17 | 0.04 | 0.12 | 0.12 | 0.16 | 0.08 |
| 6 | 70104023 | rs9346273 | 0.687 | 0.209 | -1.381 | -1.254 | -0.015 | 0.034 | 0.034 | 0 | 0.17 | 0.03 | 0.12 | 0.12 | 0.16 | 0.08 |
| 6 | 70108848 | rs1328729 | 0.658 | 0.156 | -1.756 | -1.702 | -0.057 | 0.085 | 0.085 | 0 | 0.18 | 0.01 | 0.09 | 0.02 | 0.19 | 0.19 |
| 6 | 70112998 | rs9454742 | -0.302 | -0.072 | -1.373 | -1.534 | -0.007 | -0.003 | -0.003 | 0.168 | 0.43 | 0.58 | 0.52 | 0.58 | 0.52 | 0.49 |
| 6 | 70124911 | rs9454746 | -0.085 | -0.334 | -1.536 | -1.697 | -0.026 | -0.029 | -0.029 | 0 | 0.83 | 0.64 | 0.85 | 0.92 | 0.9 | 0.87 |
| 6 | 70130357 | rs1328717 | -0.725 | -0.274 | -1.475 | -1.682 | 0.008 | 0.006 | 0.006 | 0 | 0.9 | 0.95 | 0.97 | 1 | 0.95 | 0.93 |
| 6 | 70136445 | rs7762967 | -0.361 | -0.218 | -1.332 | -1.584 | 0.023 | 0.078 | 0.078 | 0 | 0.07 | 0.26 | 0.12 | 0.08 | 0.04 | 0.07 |
| 6 | 70137886 | rs10945154 | 0.95 | -0.267 | -1.492 | -1.687 | -0.092 | -0.029 | -0.029 | 0 | 0.41 | 0.06 | 0.33 | 0.34 | 0.38 | 0.39 |
| 6 | 70138140 | rs16867531 | 0.531 | -0.287 | -0.961 | -1.842 | 0.015 | -0.114 | -0.114 | 5.724 | 0.21 | 0.39 | 0.15 | 0.02 | 0.23 | 0.2 |
| 6 | 70139179 | rs2026018 | -0.988 | -0.257 | -0.968 | -1.667 | 0.016 | 0.033 | 0.033 | 0 | 0.87 | 0.96 | 0.97 | 1 | 0.93 | 0.92 |
| 6 | 70165296 | rs2770310 | 0.002 | -0.235 | -0.936 | -1.648 | -0.023 | -0.034 | -0.034 | 0 | 0.8 | 0.69 | 0.85 | 0.92 | 0.89 | 0.85 |
| 6 | 70166530 | rs9360386 | 1.625 | -0.094 | -1.266 | -1.748 | -0.08 | -0.095 | -0.095 | 0 | 0.2 | 0.02 | 0.22 | 0.28 | 0.2 | 0.32 |
| 6 | 70168558 | rs7755341 | 0.051 | -0.054 | -0.853 | -1.688 | 0.005 | -0.037 | -0.037 | 0 | 0.25 | 0.59 | 0.27 | 0.08 | 0.29 | 0.22 |
| 6 | 70169182 | rs17694107 | 2.076 | -0.192 | -1.008 | -1.768 | -0.021 | 0.044 | 0.044 | 0 | 0.15 | 0.01 | 0.09 | 0.06 | 0.15 | 0.05 |
| 6 | 70176916 | rs2094596 | -0.298 | 0.129 | -0.241 | -1.099 | 0.018 | 0.032 | 0.032 | 0 | 0.84 | 0.88 | 0.96 | 0.97 | 0.92 | 0.87 |
| 6 | 70189837 | rs516295 | -0.064 | 0.221 | 0.018 | -1.009 | 0.03 | -0.126 | -0.126 | 0 | 0.78 | 0.56 | 0.84 | 0.98 | 0.74 | 0.81 |
| 6 | 70221457 | rs2146066 | 1.42 | 0.123 | -0.483 | -0.905 | -0.06 | -0.028 | -0.028 | 0.161 | 0.41 | 0.06 | 0.32 | 0.35 | 0.36 | 0.39 |
| 6 | 70233708 | rs9342757 | 1.019 | 0.194 | 0.339 | -0.632 | 0.006 | 0.013 | 0.013 | 0 | 0.5 | 0.47 | 0.48 | 0.41 | 0.45 | 0.49 |
| 6 | 70233881 | rs1293044 | -0.707 | 0.262 | 0.219 | -0.693 | -0.064 | 0.055 | 0.055 | 0.036 | 0.86 | 0.99 | 0.9 | 0.98 | 0.8 | 0.82 |
| 6 | 70246201 | rs10046309 | -1.136 | 0.203 | 0.341 | -0.596 | 0.046 | 0.143 | 0.143 | 0 | 0.07 | 0.41 | 0.13 | 0.09 | 0.04 | 0.07 |
| 6 | 70260055 | rs1631595 | -0.642 | 0.069 | 0.673 | -0.941 | 0 | 0.012 | 0.012 | 0 | 0.34 | 0.39 | 0.41 | 0.57 | 0.36 | 0.36 |
| 6 | 70261307 | rs1743789 | -0.737 | 0.072 | 1.029 | -0.97 | -0.008 | 0.011 | 0.011 | 0.056 | 0.85 | 0.92 | 0.89 | 0.98 | 0.8 | 0.82 |
| 6 | 70262569 | rs6455327 | 1.284 | 0.124 | 0.89 | -0.939 | 0.005 | 0.023 | 0.023 | 0 | 0.48 | 0.33 | 0.48 | 0.41 | 0.44 | 0.45 |
| 6 | 70269974 | rs980223 | -0.32 | 0.191 | 0.693 | -0.906 | 0.001 | 0.015 | 0.015 | 0 | 0.31 | 0.37 | 0.41 | 0.57 | 0.35 | 0.33 |
| 6 | 70280617 | rs6914524 | 1.006 | 0.169 | 0.433 | -0.855 | 0.005 | 0.03 | 0.03 | 0 | 0.51 | 0.34 | 0.48 | 0.41 | 0.45 | 0.49 |
| 6 | 70305899 | rs1817255 | -0.803 | 0.457 | 0.369 | -0.612 | 0.007 | -0.001 | -0.001 | 0 | 0.37 | 0.58 | 0.5 | 0.63 | 0.49 | 0.48 |
| 6 | 70315758 | rs10455225 | -0.417 | 0.545 | 0.566 | -0.503 | 0.011 | -0.056 | -0.056 | 0.102 | 0.21 | 0.41 | 0.14 | 0.07 | 0.21 | 0.21 |
| 6 | 70324397 | rs12204227 | -0.417 | 0.545 | 0.566 | -0.503 | 0.011 | -0.056 | -0.056 | 6.184 | 0.21 | 0.41 | 0.14 | 0.07 | 0.21 | 0.21 |
| 6 | 70324562 | rs13207307 | 0.175 | 0.549 | 0.608 | -0.518 | 0.004 | -0.027 | -0.027 | 0 | 0.16 | 0.16 | 0.11 | 0.02 | 0.2 | 0.18 |

| | | | | | | | | | | | | | | | | |
|---|---|---|---|---|---|---|---|---|---|---|---|---|---|---|---|---|
| 6 | 70329779 | rs6455330 | -0.049 | 0.598 | 0.215 | -0.495 | 0.024 | -0.079 | -0.079 | 0.175 | 0.18 | 0.36 | 0.13 | 0.03 | 0.2 | 0.18 |
| 6 | 70345513 | rs1563605 | -0.105 | 0.618 | 0.016 | -0.505 | -0.004 | -0.01 | -0.01 | 0 | 0.69 | 0.72 | 0.6 | 0.41 | 0.65 | 0.67 |
| 6 | 70349202 | rs13194186 | -0.602 | 0.654 | 0.479 | -0.62 | 0.025 | -0.091 | -0.091 | 0 | 0.18 | 0.41 | 0.13 | 0.03 | 0.2 | 0.18 |
| 6 | 70362734 | rs959519 | 0.139 | 0.614 | -1.512 | -0.541 | -0.013 | 0.056 | 0.056 | 0 | 0.66 | 0.91 | 0.75 | 0.77 | 0.72 | 0.58 |
| 6 | 70363194 | rs9351766 | 0.434 | 1.225 | -1.278 | -0.793 | 0.02 | -0.018 | -0.018 | 0.095 | 0.82 | 0.32 | 0.74 | 0.9 | 0.77 | 0.86 |
| 6 | 70364668 | rs9354871 | 0.436 | 1.354 | -1.279 | -0.701 | 0.021 | 0.004 | 0.004 | 0.812 | 0.84 | 0.29 | 0.75 | 0.85 | 0.8 | 0.88 |
| 6 | 70364817 | rs9354872 | 0.741 | 1.099 | -1.889 | -0.644 | -0.008 | -0.127 | -0.127 | 0.603 | 0.16 | 0.03 | 0.27 | 0.56 | 0.13 | 0.26 |
| 6 | 70365156 | rs7752231 | 0.107 | 1.246 | -1.062 | -0.706 | 0.037 | -0.01 | -0.01 | 0 | 0.11 | 0.46 | 0.09 | 0.07 | 0.08 | 0.03 |
| 6 | 70367148 | rs1114785 | -0.253 | 1.267 | -0.984 | -0.701 | 0.004 | 0 | 0 | 0 | 0.94 | 0.93 | 0.83 | 0.91 | 0.84 | 0.88 |
| 6 | 70367724 | rs9454860 | 0.76 | 1.512 | -1.622 | -1.35 | -0.015 | 0.057 | 0.057 | 0.031 | 0.37 | 0.11 | 0.29 | 0.25 | 0.33 | 0.39 |
| 6 | 70369159 | rs6918453 | 0.153 | 1.442 | -1.643 | -1.381 | -0.021 | 0.084 | 0.084 | 0.436 | 0.25 | 0.48 | 0.35 | 0.63 | 0.21 | 0.29 |
| 6 | 70369712 | rs6924031 | 0.638 | 1.335 | -1.384 | -1.417 | 0.049 | 0.057 | 0.057 | 2.673 | 0.05 | 0.4 | 0.04 | 0.01 | 0 | 0 |
| 6 | 70370323 | rs1585187 | 0.792 | 1.38 | -1.435 | -1.415 | -0.023 | 0.057 | 0.057 | 0.031 | 0.37 | 0.1 | 0.29 | 0.25 | 0.33 | 0.39 |
| 6 | 70371977 | rs12197211 | 1.527 | 1.363 | -1.803 | -1.798 | -0.012 | 0.104 | 0.104 | 0 | 0.2 | 0.03 | 0.08 | 0 | 0.16 | 0.12 |
| 6 | 70372422 | rs2061312 | 0.283 | 1.311 | -1.764 | -1.679 | -0.027 | 0.107 | 0.107 | 0.458 | 0.25 | 0.48 | 0.35 | 0.57 | 0.2 | 0.29 |
| 6 | 70373470 | rs6930785 | 0.087 | 1.251 | -1.768 | -1.741 | -0.003 | 0.004 | 0.004 | 0 | 0.54 | 0.57 | 0.53 | 0.61 | 0.51 | 0.48 |
| 6 | 70373515 | rs17765037 | 1.22 | 1.252 | -1.885 | -1.731 | -0.014 | 0.103 | 0.103 | 0 | 0.29 | 0.1 | 0.17 | 0.04 | 0.31 | 0.2 |
| 6 | 70375796 | rs2345237 | 0.126 | 1.232 | -1.737 | -1.663 | -0.002 | 0.029 | 0.029 | 0 | 0.52 | 0.55 | 0.42 | 0.36 | 0.48 | 0.48 |
| 6 | 70377693 | rs9354874 | -0.727 | 1.339 | -1.495 | -1.554 | -0.003 | 0.004 | 0.004 | 0 | 0.72 | 0.71 | 0.74 | 0.91 | 0.67 | 0.76 |
| 6 | 70380290 | rs6912049 | 0.255 | 1.44 | -1.569 | -1.463 | 0.08 | 0.118 | 0.118 | 3.084 | 0.06 | 0.4 | 0.04 | 0.05 | 0.02 | 0.07 |
| 6 | 70385709 | rs9185 | -0.004 | 1.387 | -1.682 | -1.405 | -0.009 | 0.027 | 0.027 | 0 | 0.59 | 0.66 | 0.64 | 0.74 | 0.54 | 0.65 |
| 6 | 70389764 | rs12111306 | 0.62 | 1.362 | -1.588 | -1.375 | 0.085 | 0.122 | 0.122 | 1.476 | 0.05 | 0.34 | 0.04 | 0.05 | 0.02 | 0.07 |
| 6 | 70391640 | rs4706273 | 1.033 | 1.408 | -1.87 | -1.21 | -0.013 | -0.041 | -0.041 | 0 | 0.43 | 0.12 | 0.43 | 0.56 | 0.43 | 0.47 |
| 6 | 70398402 | rs17503505 | 2.279 | 1.402 | -2.121 | -1.469 | 0.009 | 0.073 | 0.073 | 0 | 0.17 | 0 | 0.09 | 0.02 | 0.12 | 0.05 |
| 6 | 70442866 | rs1031631 | 0.737 | 2.301 | -1.345 | -0.819 | 0.001 | 0.028 | 0.028 | 0 | 0.52 | 0.63 | 0.51 | 0.61 | 0.47 | 0.54 |
| 6 | 70463509 | rs9354887 | 1.176 | 2.246 | -1.337 | -0.818 | -0.002 | -0.029 | -0.029 | 0 | 0.45 | 0.14 | 0.44 | 0.56 | 0.43 | 0.47 |
| 6 | 70466875 | rs13197020 | 1.304 | 2.737 | -1.563 | -0.63 | -0.017 | 0.021 | 0.021 | 0 | 0.05 | 0.02 | 0.05 | 0 | 0.07 | 0.02 |
| 6 | 70483082 | rs7749324 | 0.789 | 2.601 | -1.488 | -0.531 | 0.011 | 0.03 | 0.03 | 0 | 0.52 | 0.63 | 0.48 | 0.61 | 0.44 | 0.54 |
| 6 | 70493832 | rs17583400 | 0.617 | 2.628 | -1.606 | -0.321 | -0.032 | 0.008 | 0.008 | 0.254 | 0.05 | 0.01 | 0.08 | 0.02 | 0.14 | 0.07 |
| 6 | 70494213 | rs9454892 | 0.487 | 2.048 | -1.369 | -0.146 | -0.009 | -0.012 | -0.012 | 1.516 | 0.08 | 0.03 | 0.15 | 0.13 | 0.08 | 0.11 |
| 6 | 70499868 | rs1457498 | 0.584 | 1.99 | -1.354 | -0.179 | -0.002 | 0.002 | 0.002 | 0 | 0.07 | 0.04 | 0.15 | 0.04 | 0.08 | 0.1 |
| 6 | 70500518 | rs2305840 | 0.714 | 1.867 | -1.274 | 0.046 | -0.003 | -0.044 | -0.044 | 2.67 | 0.07 | 0 | 0.05 | 0.17 | 0.06 | 0.08 |
| 6 | 70509357 | rs1563610 | -0.134 | 1.859 | -1.225 | -0.052 | -0.023 | -0.038 | -0.038 | 0 | 0.13 | 0.08 | 0.22 | 0.26 | 0.22 | 0.21 |
| 6 | 70514750 | rs9364063 | 0.269 | 1.336 | -0.96 | -0.091 | 0 | 0 | 0 | 0 | 0.48 | 0.52 | 0.48 | 0.57 | 0.43 | 0.47 |
| 6 | 70520830 | rs13207318 | -0.515 | 1.843 | -0.749 | -0.07 | 0.006 | -0.021 | -0.021 | 0 | 0.08 | 0.12 | 0.11 | 0 | 0.14 | 0.02 |
| 6 | 70525533 | rs7743815 | -0.432 | 1.534 | -0.791 | 0.093 | 0.001 | -0.002 | -0.002 | 0 | 0.15 | 0.13 | 0.17 | 0.19 | 0.11 | 0.16 |
| 6 | 70530581 | rs9454900 | -0.508 | 1.686 | -0.718 | -0.091 | 0.003 | 0.006 | 0.006 | 0 | 0.18 | 0.1 | 0.09 | 0.09 | 0.16 | 0.18 |
| 6 | 70535032 | rs2487397 | 1.475 | 2.403 | -0.703 | 0.11 | -0.007 | 0.004 | 0.004 | 0.122 | 0.56 | 0.2 | 0.54 | 0.66 | 0.45 | 0.54 |
| 6 | 70535799 | rs7773772 | -1.163 | 2.319 | 0.206 | 0.168 | 0.001 | -0.015 | -0.015 | 0 | 0.32 | 0.59 | 0.23 | 0.2 | 0.31 | 0.22 |
| 6 | 70538315 | rs13197316 | -1.057 | 2.454 | 0.11 | 0.362 | -0.013 | -0.011 | -0.011 | 0 | 0.06 | 0.04 | 0.2 | 0.15 | 0.14 | 0.15 |
| 6 | 70541265 | rs1340969 | 0.262 | 2.373 | 0.978 | 0.499 | -0.001 | -0.015 | -0.015 | 0.837 | 0.74 | 0.69 | 0.85 | 0.85 | 0.72 | 0.75 |
| 6 | 70547652 | rs2487410 | 1.235 | 2.325 | 1.677 | 0.636 | 0.019 | 0.026 | 0.026 | 0.282 | 0.66 | 0.46 | 0.56 | 0.49 | 0.58 | 0.56 |
| 6 | 70550975 | rs9351778 | 0.726 | 2.266 | 1.905 | 0.581 | -0.006 | 0.01 | 0.01 | 0.087 | 0.71 | 0.39 | 0.76 | 0.63 | 0.71 | 0.71 |
| 6 | 70551688 | rs2502586 | 0.307 | 2.27 | 1.874 | 0.59 | -0.002 | 0.001 | 0.001 | 0 | 0.74 | 0.69 | 0.78 | 0.73 | 0.72 | 0.72 |
| 6 | 70562435 | rs13202110 | -2.222 | 2.161 | 1.869 | 0.495 | 0.126 | 0.139 | 0.139 | 0 | 0.13 | 0.38 | 0.26 | 0.25 | 0.19 | 0.23 |
| 6 | 70562615 | rs951494 | 1.626 | 2.117 | 1.865 | 0.472 | -0.04 | -0.036 | -0.036 | 0.006 | 0.6 | 0.23 | 0.51 | 0.49 | 0.52 | 0.5 |
| 6 | 70563502 | rs1361517 | 0.319 | 2.103 | 1.453 | 0.394 | -0.003 | -0.001 | -0.001 | 0 | 0.73 | 0.61 | 0.77 | 0.73 | 0.72 | 0.73 |
| 6 | 70566269 | rs2502570 | 0.365 | 1.929 | 1.235 | 0.329 | -0.002 | -0.001 | -0.001 | 0.237 | 0.72 | 0.69 | 0.77 | 0.73 | 0.71 | 0.7 |
| 6 | 70575948 | rs1340960 | 0.335 | 1.93 | 1.21 | 0.356 | -0.002 | -0.001 | -0.001 | 0 | 0.72 | 0.7 | 0.77 | 0.73 | 0.71 | 0.7 |
| 6 | 70579104 | rs1572898 | -1.719 | 1.601 | 0.887 | 0.647 | 0.051 | 0.081 | 0.081 | 0 | 0.1 | 0.18 | 0.23 | 0.25 | 0.16 | 0.24 |
| 6 | 70586332 | rs10498870 | -0.674 | 1.654 | 0.483 | 0.158 | 0.034 | 0.017 | 0.017 | 0 | 0.21 | 0.3 | 0.3 | 0.25 | 0.29 | 0.32 |
| 6 | 70586816 | rs1934897 | -0.103 | 0.923 | -0.115 | -0.674 | -0.001 | 0.026 | 0.026 | 0 | 0.49 | 0.59 | 0.43 | 0.34 | 0.47 | 0.45 |
| 6 | 70593803 | rs13200171 | -0.686 | 0.683 | -0.497 | -0.897 | 0.019 | 0.031 | 0.031 | 0 | 0.09 | 0.11 | 0.23 | 0.25 | 0.16 | 0.25 |
| 6 | 70594414 | rs2502562 | 1.256 | -0.04 | -0.526 | -1.225 | 0.001 | -0.032 | -0.032 | 0 | 0.11 | 0.14 | 0.07 | 0 | 0.13 | 0.07 |
| 6 | 70597502 | rs7757078 | 1.071 | -0.287 | -0.572 | -1.022 | 0.046 | 0.046 | 0.046 | 2.159 | 0.27 | 0.11 | 0.12 | 0 | 0.17 | 0.12 |
| 6 | 70597721 | rs10755538 | 1.329 | -0.169 | -0.75 | -0.95 | 0.024 | 0.103 | 0.103 | 2.482 | 0.2 | 0.03 | 0.12 | 0 | 0.17 | 0.12 |
| 6 | 70605160 | rs2096059 | -0.025 | 0.004 | -0.206 | -0.585 | -0.016 | -0.095 | -0.095 | 3.384 | 0.18 | 0.35 | 0.11 | 0 | 0.14 | 0.12 |
| 6 | 70612132 | rs9454912 | -0.87 | 0.157 | -0.091 | -0.192 | -0.003 | 0.035 | 0.035 | 0.02 | 0.26 | 0.5 | 0.23 | 0.29 | 0.2 | 0.33 |
| 6 | 70612818 | rs9446169 | -0.782 | 0.164 | -0.244 | -0.161 | 0.003 | 0.037 | 0.037 | 0 | 0.07 | 0.08 | 0.13 | 0.2 | 0.06 | 0.2 |
| 6 | 70622425 | rs17689448 | 0.503 | 0.133 | -0.206 | -0.192 | -0.002 | -0.082 | -0.082 | 0 | 0.15 | 0.34 | 0.1 | 0 | 0.13 | 0.1 |
| 6 | 70625166 | rs16868360 | -0.208 | 0.097 | -0.15 | -0.177 | -0.023 | -0.057 | -0.057 | 0.088 | 0.2 | 0.33 | 0.11 | 0.09 | 0.14 | 0.13 |
| 6 | 70641461 | rs3805959 | -1.046 | 0.096 | -0.15 | -0.166 | -0.007 | 0.021 | 0.021 | 0.066 | 0.27 | 0.59 | 0.22 | 0.29 | 0.2 | 0.31 |
| 6 | 70643018 | rs2273428 | 0.383 | 0.38 | -0.833 | -0.174 | -0.134 | -0.102 | -0.102 | 0 | 0.23 | 0.01 | 0.25 | 0.31 | 0.38 | 0.3 |
| 6 | 70655786 | rs9454921 | -0.447 | 0.538 | -0.32 | -0.683 | -0.018 | -0.069 | -0.069 | 0 | 0.25 | 0.33 | 0.13 | 0.04 | 0.2 | 0.14 |
| 6 | 70667927 | rs3805971 | -0.526 | 0.634 | -0.288 | 0.106 | -0.017 | -0.025 | -0.025 | 0 | 0.05 | 0 | 0.1 | 0.08 | 0.04 | 0.09 |
| 6 | 70681382 | rs17766479 | -1.112 | 0.601 | -0.281 | 0.022 | 0.001 | 0.033 | 0.033 | 0 | 0.06 | 0.06 | 0.12 | 0.2 | 0.06 | 0.18 |
| 6 | 70700121 | rs1934904 | -0.086 | 0.465 | -0.377 | -0.26 | 0.001 | 0.005 | 0.005 | 1.76 | 0.57 | 0.39 | 0.64 | 0.72 | 0.61 | 0.71 |
| 6 | 70728672 | rs978291 | -1.499 | 0.84 | -0.592 | 0.592 | 0 | 0.003 | 0.003 | 0 | 0.14 | 0.15 | 0.16 | 0.2 | 0.13 | 0.15 |
| 6 | 70739788 | rs3793038 | -1.12 | 0.971 | -0.634 | 0.849 | -0.003 | 0.011 | 0.011 | 0 | 0.07 | 0.06 | 0.14 | 0.19 | 0.1 | 0.12 |

| | | | | | | | | | | | | | | | | |
|---|---|---|---|---|---|---|---|---|---|---|---|---|---|---|---|---|
| 6 | 70740033 | rs3828766 | -1.264 | 0.662 | -0.404 | 0.984 | 0.001 | 0.047 | 0.047 | 0 | 0.12 | 0.19 | 0.16 | 0.19 | 0.1 | 0.14 |
| 6 | 70740151 | rs16868442 | -0.902 | 0.678 | -0.43 | 0.901 | -0.001 | 0.075 | 0.075 | 0 | 0.11 | 0.18 | 0.16 | 0.23 | 0.09 | 0.14 |
| 6 | 70740306 | rs1156404 | -0.361 | 1.099 | -1.005 | -0.056 | -0.057 | 0.09 | 0.09 | 0 | 0.38 | 0.15 | 0.56 | 0.83 | 0.45 | 0.67 |
| 6 | 70740773 | rs9342782 | -0.337 | 0.934 | -0.641 | -0.102 | -0.005 | 0.008 | 0.008 | 0.02 | 0.41 | 0.39 | 0.58 | 0.8 | 0.52 | 0.72 |
| 6 | 70743251 | rs3828767 | 0.13 | 1.024 | -1.072 | 0.281 | -0.007 | -0.02 | -0.02 | 0.274 | 0.58 | 0.72 | 0.46 | 0.45 | 0.54 | 0.49 |
| 6 | 70746249 | rs10945192 | -0.297 | 1.043 | -1.293 | -0.995 | -0.049 | 0.082 | 0.082 | 0 | 0.36 | 0.19 | 0.24 | 0.01 | 0.52 | 0.31 |
| 6 | 70748686 | rs6939166 | -0.362 | 1.316 | -0.618 | -1.119 | 0.055 | -0.048 | -0.048 | 3.187 | 0.09 | 0.26 | 0.09 | 0 | 0.14 | 0.06 |
| 6 | 70754134 | rs6937838 | -0.803 | 1.086 | -0.815 | -1.054 | 0.036 | 0.093 | 0.093 | 0 | 0.53 | 0.73 | 0.37 | 0.27 | 0.61 | 0.45 |
| 6 | 70755533 | rs6921039 | -0.129 | 1.056 | -0.98 | -1.13 | -0.126 | 0.218 | 0.218 | 0.231 | 0.37 | 0.06 | 0.27 | 0.05 | 0.51 | 0.31 |
| 6 | 70761121 | rs7747377 | -1.227 | 0.047 | -1.38 | -0.969 | -0.025 | -0.024 | -0.024 | 0.014 | 0.64 | 0.9 | 0.5 | 0.35 | 0.63 | 0.54 |
| 6 | 70766504 | rs3805994 | -0.356 | 0.185 | -1.333 | -1.049 | 0.02 | 0.137 | 0.137 | 0 | 0.49 | 0.73 | 0.33 | 0.24 | 0.56 | 0.43 |
| 6 | 70767756 | rs3806000 | -0.652 | 0.16 | -1.337 | -1.052 | 0.044 | 0.127 | 0.127 | 0 | 0.54 | 0.8 | 0.39 | 0.27 | 0.62 | 0.45 |
| 6 | 70776980 | rs11753925 | 0.957 | 0.176 | -1.307 | -1.042 | -0.11 | 0.173 | 0.173 | 0 | 0.27 | 0.01 | 0.14 | 0.01 | 0.34 | 0.23 |
| 6 | 70778297 | rs7349861 | 0.977 | 0.168 | -1.308 | -1.047 | -0.105 | 0.173 | 0.173 | 0 | 0.27 | 0.01 | 0.14 | 0.01 | 0.34 | 0.23 |
| 6 | 70778481 | rs16868507 | 1.904 | -0.016 | -1.296 | -1.011 | -0.006 | -0.001 | -0.001 | 0.724 | 0.06 | 0.06 | 0.08 | 0.03 | 0.01 | 0.05 |
| 6 | 70781701 | rs12206771 | 0.433 | 0.045 | -1.301 | -1.047 | -0.122 | 0.216 | 0.216 | 0.012 | 0.34 | 0.06 | 0.22 | 0.01 | 0.46 | 0.29 |
| 6 | 70784746 | rs3793048 | 0.361 | -0.58 | -1.563 | -1.075 | -0.03 | 0.018 | 0.018 | 0 | 0.19 | 0.67 | 0.31 | 0.3 | 0.11 | 0.32 |
| 6 | 70785382 | rs17711463 | 0.864 | -0.251 | -1.679 | -1.194 | -0.1 | 0.184 | 0.184 | 0 | 0.31 | 0.05 | 0.21 | 0.01 | 0.42 | 0.24 |
| 6 | 70794142 | rs12194223 | 0.489 | -0.301 | -1.651 | -1.181 | -0.073 | 0.136 | 0.136 | 0 | 0.35 | 0.14 | 0.26 | 0.05 | 0.47 | 0.26 |
| 6 | 70800617 | rs4707627 | 1.106 | -0.228 | -1.584 | -1.063 | 0.006 | 0.008 | 0.008 | 2.272 | 0.09 | 0.07 | 0.1 | 0.03 | 0.05 | 0.1 |
| 6 | 70801387 | rs6903986 | 1.305 | -0.163 | -1.854 | -1.392 | -0.003 | -0.013 | -0.013 | 0 | 0.12 | 0.08 | 0.12 | 0 | 0.18 | 0.05 |
| 6 | 70802259 | rs3806021 | -0.585 | -0.427 | -1.562 | -1.202 | 0.002 | 0.009 | 0.009 | 0.638 | 0.66 | 0.67 | 0.68 | 0.42 | 0.64 | 0.73 |
| 6 | 70802956 | rs2208608 | -0.381 | -0.47 | -1.387 | -1.329 | -0.017 | -0.001 | -0.001 | 0 | 0.38 | 0.49 | 0.5 | 0.37 | 0.3 | 0.54 |
| 6 | 70803453 | rs3806022 | 1.134 | -0.497 | -1.385 | -1.326 | -0.027 | 0.069 | 0.069 | 0 | 0.29 | 0.18 | 0.19 | 0.05 | 0.35 | 0.19 |
| 6 | 70804987 | rs9454968 | -0.613 | -0.58 | -1.294 | -1.319 | -0.006 | 0 | 0 | 0 | 0.42 | 0.55 | 0.54 | 0.41 | 0.35 | 0.56 |
| 6 | 70806443 | rs9446195 | 0.061 | -0.441 | -1.606 | -1.346 | 0.009 | 0.022 | 0.022 | 0 | 0.21 | 0.14 | 0.21 | 0.08 | 0.19 | 0.21 |
| 6 | 70808020 | rs12190708 | 1.069 | -0.297 | -1.561 | -1.064 | -0.037 | 0.107 | 0.107 | 0 | 0.25 | 0.1 | 0.14 | 0.01 | 0.29 | 0.18 |
| 6 | 70812123 | rs7772672 | -0.779 | -0.769 | -1.413 | 0.379 | 0.02 | -0.056 | -0.056 | 0.186 | 0.9 | 0.99 | 0.83 | 0.55 | 0.96 | 0.86 |
| 6 | 70813146 | rs10945197 | -0.04 | -0.631 | -1.225 | -0.437 | 0.001 | 0.031 | 0.031 | 0 | 0.52 | 0.34 | 0.46 | 0.43 | 0.49 | 0.62 |
| 6 | 70815691 | rs3806028 | 0.996 | -0.643 | -1.343 | -0.529 | -0.004 | 0.001 | 0.001 | 0 | 0.15 | 0.17 | 0.27 | 0.24 | 0.1 | 0.32 |
| 6 | 70816378 | rs17712300 | 1.352 | -0.78 | -1.431 | -0.751 | -0.008 | 0.063 | 0.063 | 1.746 | 0.17 | 0.01 | 0.03 | 0.04 | 0.11 | 0.1 |
| 6 | 70816928 | rs3793055 | -0.128 | -0.782 | -1.246 | -0.775 | 0.029 | 0.034 | 0.034 | 0.314 | 0.35 | 0.3 | 0.32 | 0.28 | 0.22 | 0.43 |
| 6 | 70817620 | rs3793056 | 1.427 | -0.622 | -1.195 | -0.676 | -0.038 | -0.018 | -0.018 | 0.105 | 0.16 | 0.05 | 0.14 | 0.15 | 0.26 | 0.19 |
| 6 | 70819800 | rs3793057 | 0.237 | -0.471 | 0.091 | -0.958 | -0.006 | 0.006 | 0.006 | 0 | 0.16 | 0.13 | 0.16 | 0.05 | 0.24 | 0.12 |
| 6 | 70822009 | rs3806030 | 1.095 | -0.553 | 0.127 | -0.966 | -0.008 | -0.009 | -0.009 | 1.125 | 0.13 | 0.17 | 0.26 | 0.1 | 0.09 | 0.24 |
| 6 | 70824403 | rs7751501 | -0.401 | -0.577 | 0.231 | -1.006 | 0.007 | 0.007 | 0.007 | 0 | 0.34 | 0.35 | 0.32 | 0.28 | 0.21 | 0.43 |
| 6 | 70827134 | rs3806032 | -1.277 | -0.611 | 0.203 | -0.969 | -0.002 | 0.085 | 0.085 | 0 | 0.53 | 0.77 | 0.5 | 0.34 | 0.45 | 0.55 |
| 6 | 70828336 | rs3806033 | -0.987 | -0.523 | 0.091 | -0.924 | 0.003 | 0.017 | 0.017 | 0 | 0.5 | 0.48 | 0.48 | 0.34 | 0.45 | 0.55 |
| 6 | 70836769 | rs10945199 | 0.985 | -0.38 | -0.63 | -0.877 | 0.006 | 0.07 | 0.07 | 0 | 0.17 | 0.07 | 0.04 | 0.03 | 0.11 | 0.11 |
| 6 | 70841325 | rs3806037 | -0.818 | -0.091 | -0.403 | -0.693 | 0.005 | 0.036 | 0.036 | 0 | 0.51 | 0.61 | 0.49 | 0.32 | 0.45 | 0.55 |
| 6 | 70842620 | rs6938971 | -0.35 | -0.528 | -0.312 | -0.364 | -0.026 | -0.032 | -0.032 | 0 | 0.34 | 0.47 | 0.33 | 0.27 | 0.2 | 0.43 |
| 6 | 70847385 | rs9446206 | -0.734 | -0.386 | -0.299 | 0.024 | -0.018 | 0.019 | 0.019 | 0 | 0.12 | 0.24 | 0.07 | 0.17 | 0.09 | 0.16 |
| 6 | 70858863 | rs2025283 | -1.278 | -0.277 | -1.05 | 0.027 | -0.045 | -0.122 | -0.122 | 2.793 | 0.73 | 0.94 | 0.64 | 0.46 | 0.73 | 0.76 |
| 6 | 70860725 | rs3806040 | 1.203 | -0.699 | -0.557 | -0.453 | -0.014 | -0.014 | -0.014 | 0.008 | 0.16 | 0.31 | 0.29 | 0.1 | 0.1 | 0.19 |
| 6 | 70860739 | rs3806041 | -1.052 | -0.648 | -0.876 | -0.787 | -0.005 | 0.068 | 0.068 | 0 | 0.54 | 0.78 | 0.49 | 0.31 | 0.46 | 0.52 |
| 6 | 70861487 | rs12204438 | 0.845 | -0.379 | -0.839 | -0.631 | -0.067 | 0.123 | 0.123 | 0 | 0.24 | 0.02 | 0.12 | 0.03 | 0.26 | 0.17 |
| 6 | 70861994 | rs6928404 | -0.779 | -0.425 | -0.744 | -0.621 | -0.03 | 0.026 | 0.026 | 0 | 0.6 | 0.72 | 0.51 | 0.31 | 0.47 | 0.53 |
| 6 | 70866609 | rs2229799 | -0.783 | -0.424 | -0.648 | -0.621 | -0.023 | 0.021 | 0.021 | 0 | 0.6 | 0.69 | 0.51 | 0.31 | 0.47 | 0.53 |
| 6 | 70869087 | rs3806043 | 0.548 | -0.173 | -0.5 | -0.45 | -0.008 | 0.028 | 0.028 | 0 | 0.06 | 0.16 | 0.05 | 0.04 | 0.02 | 0.01 |
| 6 | 70873331 | rs2296010 | -0.137 | -0.501 | -0.745 | -0.836 | 0.007 | 0.056 | 0.056 | 0 | 0.45 | 0.34 | 0.33 | 0.25 | 0.41 | 0.44 |
| 6 | 70874168 | rs802172 | -1.827 | -0.494 | -0.969 | -0.412 | -0.039 | -0.077 | -0.077 | 0 | 0.79 | 0.94 | 0.58 | 0.58 | 0.75 | 0.78 |
| 6 | 70875234 | rs802173 | -1.264 | -0.551 | -0.871 | -0.463 | -0.029 | -0.053 | -0.053 | 0 | 0.75 | 0.91 | 0.55 | 0.57 | 0.7 | 0.72 |
| 6 | 70887942 | rs12192237 | 1.126 | -0.507 | -1.118 | -0.701 | -0.06 | 0.072 | 0.072 | 0 | 0.19 | 0.01 | 0.11 | 0.03 | 0.24 | 0.15 |
| 6 | 70891740 | rs10945200 | -0.185 | -0.361 | -0.717 | -0.423 | 0.005 | 0.044 | 0.044 | 0 | 0.49 | 0.39 | 0.36 | 0.26 | 0.46 | 0.51 |
| 6 | 70894894 | rs2296006 | 1.256 | -0.361 | -0.607 | -0.285 | -0.012 | 0 | 0 | 0.008 | 0.23 | 0.15 | 0.14 | 0.19 | 0.27 | 0.25 |
| 6 | 70903546 | rs1200562 | -0.382 | -0.419 | -0.45 | -0.247 | 0.009 | 0.092 | 0.092 | 0.105 | 0.1 | 0.4 | 0.1 | 0.14 | 0.07 | 0.08 |
| 6 | 70910711 | rs550152 | 0.372 | -0.462 | -0.414 | -0.297 | -0.01 | 0.012 | 0.012 | 0 | 0.31 | 0.6 | 0.23 | 0.32 | 0.29 | 0.26 |
| 6 | 70911776 | rs610473 | -0.065 | -0.481 | -0.448 | -0.321 | -0.006 | 0.005 | 0.005 | 0 | 0.36 | 0.61 | 0.25 | 0.44 | 0.35 | 0.38 |
| 6 | 70912335 | rs613109 | -0.139 | -0.506 | -0.347 | -0.305 | 0.029 | 0.19 | 0.19 | 1.402 | 0.07 | 0.42 | 0.09 | 0.13 | 0.02 | 0.02 |
| 6 | 70912835 | rs10498871 | 1.025 | -0.483 | -0.526 | -0.318 | -0.001 | -0.013 | -0.013 | 0.018 | 0.25 | 0.17 | 0.14 | 0.3 | 0.28 | 0.31 |
| 6 | 70916998 | rs1736 | 0.379 | -0.509 | -0.609 | -0.429 | 0.001 | 0.023 | 0.023 | 0 | 0.42 | 0.34 | 0.33 | 0.13 | 0.39 | 0.41 |
| 6 | 70919443 | rs552628 | 0.699 | -0.453 | -0.59 | -0.39 | 0.01 | 0.031 | 0.031 | 0 | 0.28 | 0.35 | 0.46 | 0.43 | 0.3 | 0.26 |
| 6 | 70919597 | rs529805 | 0.874 | -0.457 | -0.574 | -0.393 | 0.004 | 0.014 | 0.014 | 0 | 0.26 | 0.29 | 0.46 | 0.43 | 0.3 | 0.26 |
| 6 | 70919860 | rs9346371 | 1.614 | -0.478 | -0.585 | -0.396 | -0.036 | -0.082 | -0.082 | 0 | 0.22 | 0.06 | 0.42 | 0.42 | 0.25 | 0.2 |
| 6 | 70920612 | rs624243 | -1.618 | -0.497 | -0.694 | -0.39 | -0.064 | 0.05 | 0.05 | 0 | 0.8 | 0.99 | 0.87 | 0.93 | 0.74 | 0.79 |
| 6 | 70921389 | rs997953 | 0.659 | -0.385 | -0.667 | -0.322 | 0.011 | 0.034 | 0.034 | 0 | 0.27 | 0.35 | 0.46 | 0.43 | 0.3 | 0.26 |
| 6 | 70923469 | rs6933155 | 0.421 | -0.284 | -0.77 | -0.536 | 0.001 | 0.042 | 0.042 | 0 | 0.42 | 0.3 | 0.33 | 0.13 | 0.39 | 0.41 |
| 6 | 70926334 | rs1064250 | 0.255 | -0.167 | -0.872 | -0.468 | -0.001 | -0.006 | -0.006 | 0 | 0.46 | 0.46 | 0.39 | 0.27 | 0.41 | 0.43 |
| 6 | 70932535 | rs1210977 | -1.241 | 0.042 | -1.237 | -0.364 | -0.065 | 0.055 | 0.055 | 0 | 0.8 | 0.99 | 0.87 | 0.94 | 0.74 | 0.78 |

| | | | | | | | | | | | | | | | | |
|---|---|---|---|---|---|---|---|---|---|---|---|---|---|---|---|---|
| 6 | 70940652 | rs10498872 | 0.461 | 0.812 | -0.932 | -0.521 | 0.013 | -0.042 | -0.042 | 0 | 0.17 | 0.19 | 0.13 | 0.01 | 0.25 | 0.11 |
| 6 | 70940822 | rs544179 | -0.801 | 0.352 | -0.84 | -0.564 | -0.001 | 0.002 | 0.002 | 0 | 0.44 | 0.59 | 0.32 | 0.29 | 0.52 | 0.34 |
| 6 | 70943041 | rs12154117 | 0.262 | 0.413 | -1.256 | -0.37 | -0.044 | 0.092 | 0.092 | 0 | 0.22 | 0.07 | 0.14 | 0.04 | 0.3 | 0.22 |
| 6 | 70944465 | rs12191701 | -0.243 | 0.25 | -1.201 | -0.391 | -0.087 | 0.17 | 0.17 | 0 | 0.36 | 0.1 | 0.23 | 0.04 | 0.5 | 0.27 |
| 6 | 70957772 | rs701693 | -0.653 | 0.257 | -1.042 | -0.251 | 0.009 | -0.02 | -0.02 | 0 | 0.42 | 0.56 | 0.31 | 0.3 | 0.49 | 0.33 |
| 6 | 70958346 | rs3793071 | 1.775 | 0.031 | -1.355 | -0.014 | -0.024 | -0.085 | -0.085 | 0.166 | 0.12 | 0.01 | 0.35 | 0.42 | 0.11 | 0.17 |
| 6 | 70961833 | rs1135056 | -0.455 | -0.049 | -0.892 | -0.653 | -0.004 | 0.003 | 0.003 | 0 | 0.37 | 0.36 | 0.25 | 0.29 | 0.43 | 0.26 |
| 6 | 70962044 | rs519068 | -0.771 | -0.09 | -0.973 | -0.693 | 0.009 | -0.024 | -0.024 | 0.01 | 0.42 | 0.53 | 0.31 | 0.3 | 0.49 | 0.32 |
| 6 | 70963124 | rs9346373 | 1.156 | -0.465 | -0.974 | -0.889 | -0.043 | -0.044 | -0.044 | 0.052 | 0.21 | 0.29 | 0.2 | 0.1 | 0.1 | 0.24 |
| 6 | 70963232 | rs701689 | -0.513 | -0.192 | -0.879 | -0.943 | 0.016 | -0.024 | -0.024 | 1.389 | 0.38 | 0.51 | 0.28 | 0.29 | 0.43 | 0.26 |
| 6 | 70963769 | rs7755873 | 1.013 | -0.039 | -0.933 | -1.014 | -0.019 | 0.009 | 0.009 | 0 | 0.14 | 0.08 | 0.1 | 0 | 0.2 | 0.05 |
| 6 | 70968276 | rs555663 | -0.44 | -0.039 | -0.915 | -0.964 | 0.014 | -0.023 | -0.023 | 0 | 0.35 | 0.42 | 0.27 | 0.18 | 0.41 | 0.21 |
| 6 | 70970743 | rs3806089 | -0.014 | 0.041 | -0.983 | -1.151 | 0 | 0 | 0 | 0 | 0.07 | 0.1 | 0.04 | 0.13 | 0.09 | 0.11 |
| 6 | 70972814 | rs12194091 | 1.01 | 0.022 | -0.86 | -1.177 | -0.002 | -0.023 | -0.023 | 3.21 | 0.16 | 0.14 | 0.11 | 0 | 0.19 | 0.05 |
| 6 | 70975512 | rs558485 | -1.398 | -0.18 | -1.033 | -1.262 | -0.03 | -0.026 | -0.026 | 0 | 0.63 | 0.86 | 0.5 | 0.39 | 0.59 | 0.55 |
| 6 | 70975782 | rs604896 | -0.153 | -0.266 | -0.78 | -1.027 | 0.022 | 0.257 | 0.257 | 0 | 0.05 | 0.28 | 0.07 | 0.26 | 0.03 | 0.06 |
| 6 | 70976538 | rs883708 | 0.456 | -0.172 | -0.971 | -1.016 | 0.014 | -0.026 | -0.026 | 0 | 0.18 | 0.22 | 0.11 | 0.12 | 0.21 | 0.1 |
| 6 | 70979193 | rs550675 | -0.555 | -0.312 | -1.207 | -0.879 | 0.007 | -0.02 | -0.02 | 0 | 0.42 | 0.56 | 0.3 | 0.29 | 0.48 | 0.32 |
| 6 | 70980388 | rs518558 | -0.491 | -0.091 | -1.402 | -0.459 | 0.005 | 0.027 | 0.027 | 0.209 | 0.5 | 0.85 | 0.51 | 0.41 | 0.43 | 0.5 |
| 6 | 70984436 | rs592121 | -0.381 | -0.266 | -1.004 | -0.488 | 0.02 | 0.08 | 0.08 | 0 | 0.48 | 0.72 | 0.43 | 0.31 | 0.36 | 0.49 |
| 6 | 70987200 | rs495558 | -0.584 | -0.29 | -1.062 | -0.493 | -0.004 | -0.013 | -0.013 | 0 | 0.51 | 0.88 | 0.48 | 0.45 | 0.38 | 0.5 |
| 6 | 70987242 | rs3793073 | 1.056 | -0.408 | -1.191 | -0.969 | 0.009 | -0.064 | -0.064 | 0 | 0.15 | 0.01 | 0.17 | 0.19 | 0.07 | 0.25 |
| 6 | 70987976 | rs3806095 | 0.451 | -0.292 | -0.67 | -0.942 | 0 | -0.014 | -0.014 | 0 | 0.35 | 0.34 | 0.32 | 0.19 | 0.29 | 0.39 |
| 6 | 70988686 | rs616642 | -0.241 | -0.102 | -0.822 | -0.81 | 0.006 | 0.021 | 0.021 | 0 | 0.08 | 0.28 | 0.1 | 0.12 | 0.07 | 0.08 |
| 6 | 70989075 | rs13199337 | 0.193 | 0.143 | -1.476 | -0.937 | -0.004 | 0.012 | 0.012 | 0 | 0.14 | 0.09 | 0.08 | 0.03 | 0.18 | 0.14 |
| 6 | 70989703 | rs3806099 | -0.466 | 0.14 | -1.275 | -0.846 | 0.025 | 0.108 | 0.108 | 0 | 0.49 | 0.78 | 0.43 | 0.31 | 0.36 | 0.49 |
| 6 | 70991630 | rs677152 | -0.138 | -0.114 | -0.932 | -1.622 | 0.003 | -0.068 | -0.068 | 0 | 0.53 | 0.54 | 0.66 | 0.96 | 0.4 | 0.56 |
| 6 | 70993935 | rs701682 | 0.923 | 0.084 | -0.932 | -1.609 | 0.02 | -0.044 | -0.044 | 0.993 | 0.09 | 0.22 | 0.08 | 0.01 | 0.11 | 0.1 |
| 6 | 70996590 | rs495823 | -0.586 | 0 | -1.027 | -1.667 | -0.009 | -0.034 | -0.034 | 0 | 0.63 | 0.71 | 0.73 | 0.97 | 0.51 | 0.66 |
| 6 | 70998563 | rs1321061 | 0.892 | -0.229 | -1.135 | -1.152 | -0.031 | 0.053 | 0.053 | 0.128 | 0.22 | 0.35 | 0.24 | 0.45 | 0.09 | 0.28 |
| 6 | 71008012 | rs9455034 | 0.486 | -0.175 | -1.228 | -1.179 | -0.017 | 0.042 | 0.042 | 0 | 0.24 | 0.37 | 0.27 | 0.45 | 0.15 | 0.29 |
| 6 | 71008339 | rs564031 | -0.952 | 0.58 | -0.874 | -0.812 | -0.002 | -0.01 | -0.01 | 0 | 0.61 | 0.63 | 0.61 | 0.51 | 0.6 | 0.52 |
| 6 | 71008745 | rs9354912 | 0.691 | 0.225 | -1.304 | -0.387 | -0.007 | -0.154 | -0.154 | 0.714 | 0.22 | 0.02 | 0.3 | 0.46 | 0.19 | 0.21 |
| 6 | 71010101 | rs12210123 | 0.212 | 0.443 | -0.831 | -0.514 | 0.015 | -0.103 | -0.103 | 0 | 0.2 | 0.27 | 0.12 | 0 | 0.18 | 0.14 |
| 6 | 71011596 | rs9294863 | -0.37 | 0.423 | -0.76 | 0.08 | 0.005 | -0.004 | -0.004 | 0.053 | 0.07 | 0.07 | 0.06 | 0.04 | 0.06 | 0.06 |
| 6 | 71011831 | rs2242589 | -0.122 | 0.661 | -0.692 | 0.017 | -0.003 | 0.002 | 0.002 | 0 | 0.52 | 0.39 | 0.52 | 0.52 | 0.49 | 0.42 |
| 6 | 71012575 | rs2242588 | -1.141 | 0.628 | -0.03 | -0.123 | 0.034 | 0.044 | 0.044 | 0 | 0.17 | 0.35 | 0.14 | 0.21 | 0.18 | 0.25 |
| 6 | 71013640 | rs6903040 | -0.408 | -0.03 | -0.314 | -0.72 | -0.044 | -0.068 | -0.068 | 7.265 | 0.3 | 0.43 | 0.23 | 0.12 | 0.17 | 0.25 |
| 6 | 71013722 | rs4706422 | 0.209 | 0.365 | -1.233 | -0.467 | -0.035 | -0.149 | -0.149 | 0 | 0.24 | 0.02 | 0.33 | 0.59 | 0.25 | 0.21 |
| 6 | 71020330 | rs498373 | -0.105 | -0.187 | -0.27 | -0.795 | -0.026 | -0.063 | -0.063 | 0 | 0.32 | 0.41 | 0.24 | 0.12 | 0.21 | 0.33 |
| 6 | 71024741 | rs652462 | -0.317 | -0.105 | -1.027 | -0.933 | 0.027 | 0.067 | 0.067 | 0 | 0.4 | 0.7 | 0.31 | 0.13 | 0.3 | 0.4 |
| 6 | 71025669 | rs509417 | -0.018 | 1.296 | -0.808 | -0.569 | -0.017 | 0.004 | 0.004 | 0 | 0.72 | 0.82 | 0.7 | 0.78 | 0.6 | 0.63 |
| 6 | 71026324 | rs6920980 | -0.774 | 0.964 | -0.819 | -0.349 | -0.001 | -0.04 | -0.04 | 0 | 0.09 | 0.15 | 0.06 | 0 | 0.09 | 0.06 |
| 6 | 71035791 | rs682946 | 0.341 | 1.464 | -1.37 | -0.058 | -0.055 | -0.03 | -0.03 | 0 | 0.79 | 0.86 | 0.72 | 0.75 | 0.66 | 0.63 |
| 6 | 71037050 | rs13216351 | -0.339 | 1.075 | -1.459 | -0.272 | -0.063 | 0.034 | 0.034 | 0 | 0.4 | 0.21 | 0.42 | 0.3 | 0.47 | 0.42 |
| 6 | 71047198 | rs1739672 | -0.497 | 0.681 | -1.173 | -0.42 | -0.02 | -0.03 | -0.03 | 0.175 | 0.14 | 0.56 | 0.1 | 0.06 | 0.1 | 0.13 |
| 6 | 71048944 | rs1711725 | -0.904 | 0.723 | -1.275 | -0.412 | 0.015 | 0.032 | 0.032 | 0.362 | 0.54 | 0.77 | 0.52 | 0.36 | 0.57 | 0.56 |
| 6 | 71051702 | rs1739663 | -0.984 | 0.481 | -0.326 | -0.138 | -0.018 | -0.027 | -0.027 | 0 | 0.14 | 0.59 | 0.1 | 0.06 | 0.1 | 0.13 |
| 6 | 71060566 | rs9455066 | -0.125 | 0.498 | -0.384 | -0.146 | 0.003 | 0.004 | 0.004 | 0 | 0.49 | 0.43 | 0.48 | 0.19 | 0.43 | 0.39 |
| 6 | 71062163 | rs1321057 | -0.131 | 0.486 | -0.355 | -0.149 | 0.005 | 0.011 | 0.011 | 0 | 0.49 | 0.42 | 0.48 | 0.19 | 0.43 | 0.39 |
| 6 | 71064295 | rs7772055 | 0.457 | 0.781 | -0.075 | 0.636 | -0.004 | -0.01 | -0.01 | 0.748 | 0.41 | 0.34 | 0.43 | 0.55 | 0.45 | 0.35 |
| 6 | 71065942 | rs9455070 | -0.465 | 0.699 | -0.015 | 0.653 | 0.004 | 0.012 | 0.012 | 0 | 0.1 | 0.14 | 0.08 | 0.2 | 0.12 | 0.25 |
| 6 | 71067423 | rs1418611 | 0.602 | 0.606 | 0.346 | 0.401 | 0.014 | 0.022 | 0.022 | 0 | 0.41 | 0.26 | 0.22 | 0.01 | 0.39 | 0.33 |
| 6 | 71075085 | rs36000024 | -0.067 | 0.685 | 0.206 | 0.519 | -0.009 | -0.007 | -0.007 | 0 | 0.09 | 0.08 | 0.07 | 0.21 | 0.12 | 0.24 |
| 6 | 71076673 | rs9455077 | 0.471 | 0.197 | 0.117 | 0.15 | -0.001 | 0.083 | 0.083 | 0 | 0.37 | 0.21 | 0.22 | 0.01 | 0.38 | 0.28 |
| 6 | 71076785 | rs6916429 | 1.05 | 0.078 | 0.679 | 0.511 | -0.043 | 0.196 | 0.196 | 0 | 0.86 | 0.56 | 0.72 | 0.73 | 0.94 | 0.88 |
| 6 | 71079888 | rs9342788 | 0.664 | 0.207 | 0.03 | 0.565 | -0.043 | -0.053 | -0.053 | 0.01 | 0.39 | 0.1 | 0.35 | 0.5 | 0.43 | 0.35 |
| 6 | 71085959 | rs10945224 | -0.167 | 0.407 | 0.008 | 0.388 | -0.019 | -0.015 | -0.015 | 0 | 0.07 | 0.04 | 0.12 | 0.22 | 0.14 | 0.26 |
| 6 | 71087091 | rs3909058 | -0.375 | 0.228 | 0.014 | -0.357 | -0.004 | 0.01 | 0.01 | 0 | 0.61 | 0.67 | 0.75 | 0.93 | 0.59 | 0.76 |
| 6 | 71096450 | rs9346388 | 0.031 | 0.473 | -0.611 | 0.1 | -0.089 | -0.153 | -0.153 | 1.374 | 0.18 | 0.01 | 0.18 | 0.48 | 0.29 | 0.26 |
| 6 | 71097590 | rs4707802 | 0.733 | -0.012 | -0.497 | -0.526 | 0.039 | 0.058 | 0.058 | 2.716 | 0.73 | 0.15 | 0.74 | 0.91 | 0.83 | 0.83 |
| 6 | 71109059 | rs16869156 | -0.361 | 0.092 | 0.288 | -0.451 | -0.016 | -0.031 | -0.031 | 0 | 0.21 | 0.29 | 0.22 | 0.09 | 0.17 | 0.17 |
| 6 | 71180505 | rs13218183 | 1.635 | 0.072 | -0.338 | -0.924 | -0.054 | 0.091 | 0.091 | 0 | 0.18 | 0.03 | 0.13 | 0.02 | 0.2 | 0.08 |
| 6 | 71189074 | rs17634863 | 1.903 | -0.214 | -0.806 | -1.318 | 0.001 | 0.129 | 0.129 | 0 | 0.26 | 0.01 | 0.07 | 0.06 | 0.18 | 0.13 |
| 6 | 71193800 | rs17713105 | 0.751 | -0.008 | -0.502 | -1.142 | -0.005 | 0.014 | 0.014 | 0 | 0.13 | 0.07 | 0.1 | 0 | 0.15 | 0.14 |
| 6 | 71214998 | rs2262681 | -0.361 | 0.162 | -0.548 | -0.892 | -0.036 | -0.034 | -0.034 | 2.226 | 0.91 | 0.96 | 0.65 | 0.66 | 0.85 | 0.74 |
| 6 | 71219464 | rs6907049 | 0.486 | 0.073 | -0.237 | -0.942 | -0.021 | -0.023 | -0.023 | 0 | 0.05 | 0.22 | 0.02 | 0 | 0.01 | 0 |
| 6 | 71236216 | rs7740873 | -0.768 | 0.024 | -0.021 | -0.943 | -0.046 | -0.088 | -0.088 | 0 | 0.24 | 0.52 | 0.24 | 0.09 | 0.17 | 0.17 |
| 6 | 71238105 | rs2747701 | 1.19 | 0.018 | -0.168 | -0.934 | -0.1 | -0.112 | -0.112 | 1.937 | 0.45 | 0.02 | 0.25 | 0.54 | 0.48 | 0.48 |

| | | | | | | | | | | | | | | | | |
|---|---|---|---|---|---|---|---|---|---|---|---|---|---|---|---|---|
| 6 | 71238377 | rs2842063 | -1.384 | -0.455 | 0.049 | -1.663 | 0.063 | 0.119 | 0.119 | 3.658 | 0.28 | 0.9 | 0.27 | 0.09 | 0.18 | 0.17 |
| 6 | 71251421 | rs9294876 | -1.146 | -0.268 | 0.19 | -1.567 | -0.05 | -0.092 | -0.092 | 0.336 | 0.25 | 0.52 | 0.23 | 0.09 | 0.18 | 0.17 |
| 6 | 71252760 | rs7756774 | -0.253 | -0.212 | 0.192 | -1.527 | -0.018 | -0.049 | -0.049 | 0.739 | 0.18 | 0.3 | 0.21 | 0.09 | 0.16 | 0.17 |
| 6 | 71269459 | rs16869371 | 0.08 | -0.141 | 0.113 | -1.512 | -0.002 | -0.007 | -0.007 | 0 | 0.17 | 0.17 | 0.21 | 0.09 | 0.16 | 0.16 |
| 6 | 71269632 | rs16869373 | 0.015 | -0.16 | 0.107 | -1.511 | -0.004 | -0.011 | -0.011 | 0 | 0.17 | 0.19 | 0.21 | 0.09 | 0.16 | 0.17 |
| 6 | 71273253 | rs6455371 | -0.619 | -0.111 | 0.027 | -1.533 | 0.003 | -0.004 | -0.004 | 0.143 | 0.43 | 0.59 | 0.37 | 0.11 | 0.38 | 0.26 |
| 6 | 71288659 | rs12203089 | 0.282 | -0.15 | -0.215 | -1.459 | -0.005 | 0.046 | 0.046 | 0 | 0.14 | 0.05 | 0.1 | 0.01 | 0.16 | 0.14 |
| 6 | 71289189 | rs1048886 | -0.911 | -0.265 | 0.008 | -1.466 | -0.029 | -0.045 | -0.045 | 8.205 | 0.24 | 0.32 | 0.22 | 0.08 | 0.18 | 0.16 |
| 6 | 71296182 | rs2691498 | -0.937 | 0.094 | -0.129 | -1.347 | 0.005 | -0.034 | -0.034 | 0 | 0.43 | 0.53 | 0.54 | 0.82 | 0.55 | 0.68 |
| 6 | 71297763 | rs10945231 | 0.226 | 0.539 | -0.308 | -1.135 | -0.008 | 0.043 | 0.043 | 0 | 0.13 | 0.05 | 0.1 | 0.01 | 0.16 | 0.14 |
| 6 | 71299649 | rs9446271 | -0.807 | 0.472 | -0.504 | -1.176 | 0 | -0.012 | -0.012 | 0 | 0.48 | 0.56 | 0.4 | 0.13 | 0.43 | 0.26 |
| 6 | 71307347 | rs12198920 | -0.655 | 0.152 | -0.277 | -0.964 | -0.004 | -0.034 | -0.034 | 0 | 0.44 | 0.55 | 0.37 | 0.13 | 0.38 | 0.25 |
| 6 | 71317960 | rs9455190 | 0.097 | 0.666 | -0.106 | -0.653 | 0.051 | -0.044 | -0.044 | 0.244 | 0.08 | 0.16 | 0.08 | 0 | 0.14 | 0.01 |
| 6 | 71318071 | rs2691511 | -1.06 | 0.397 | -0.303 | -0.973 | -0.006 | -0.106 | -0.106 | 0 | 0.3 | 0.39 | 0.19 | 0.02 | 0.33 | 0.17 |
| 6 | 71325285 | rs2691473 | -0.29 | 0.879 | -0.511 | -0.363 | 0.002 | 0.018 | 0.018 | 0 | 0.36 | 0.34 | 0.31 | 0.16 | 0.38 | 0.35 |
| 6 | 71329265 | rs1608696 | -0.002 | 0.663 | -0.479 | -0.232 | 0.006 | 0.026 | 0.026 | 0 | 0.3 | 0.28 | 0.27 | 0.12 | 0.32 | 0.2 |
| 6 | 71330911 | rs2691480 | -0.045 | 0.848 | -0.508 | 0.703 | 0.001 | -0.014 | -0.014 | 0.026 | 0.29 | 0.28 | 0.33 | 0.32 | 0.31 | 0.21 |
| 6 | 71331467 | rs2691482 | -0.915 | 0.664 | -0.543 | 0.827 | -0.008 | -0.006 | -0.006 | 0.325 | 0.49 | 0.71 | 0.73 | 0.48 | 0.52 | 0.43 |
| 6 | 71340647 | rs2494976 | -0.192 | 0.806 | -0.84 | 0.888 | -0.041 | 0.052 | 0.052 | 0 | 0.33 | 0.16 | 0.34 | 0.14 | 0.45 | 0.19 |
| 6 | 71340880 | rs9446284 | 0.297 | 0.605 | -0.199 | 1.16 | 0.007 | -0.021 | -0.021 | 0.122 | 0.07 | 0.08 | 0.08 | 0 | 0.13 | 0.01 |
| 6 | 71343073 | rs9455200 | -0.659 | 0.527 | -0.21 | 1.219 | -0.011 | 0.009 | 0.009 | 0.247 | 0.15 | 0.09 | 0.36 | 0.11 | 0.18 | 0.21 |
| 6 | 71358164 | rs781994 | -0.019 | 0.52 | -0.652 | 0.983 | -0.052 | 0.021 | 0.021 | 0 | 0.24 | 0.08 | 0.17 | 0.23 | 0.34 | 0.32 |
| 6 | 71360155 | rs16869446 | -0.181 | -0.001 | -0.587 | 1.712 | -0.006 | 0.077 | 0.077 | 0 | 0.09 | 0.1 | 0.13 | 0.34 | 0.06 | 0.06 |
| 6 | 71367566 | rs782002 | -0.028 | -0.88 | -0.994 | 0.665 | 0.023 | 0.037 | 0.037 | 0 | 0.51 | 0.36 | 0.33 | 0.37 | 0.46 | 0.57 |
| 6 | 71382765 | rs4706458 | 0.92 | -0.856 | -1.271 | -0.037 | -0.088 | 0.054 | 0.054 | 0 | 0.23 | 0.03 | 0.25 | 0.1 | 0.33 | 0.31 |
| 6 | 71383021 | rs13196394 | 0.715 | -0.688 | -1.556 | -0.281 | -0.023 | -0.03 | -0.03 | 0 | 0.15 | 0.02 | 0.2 | 0.16 | 0.15 | 0.18 |
| 6 | 71392411 | rs16880774 | 1.167 | -1.152 | -1.604 | -0.52 | 0.026 | 0.021 | 0.021 | 0 | 0.14 | 0.05 | 0.04 | 0.11 | 0.07 | 0.13 |
| 6 | 71398202 | rs781967 | 0.353 | -0.868 | -1.088 | -0.399 | 0.012 | 0.04 | 0.04 | 0 | 0.48 | 0.38 | 0.34 | 0.23 | 0.47 | 0.54 |
| 6 | 71399151 | rs1084382 | -1.3 | -1.044 | -1.232 | -0.557 | -0.103 | 0.13 | 0.13 | 0.059 | 0.8 | 0.98 | 0.8 | 1 | 0.72 | 0.9 |
| 6 | 71433076 | rs9354939 | -0.511 | -1.044 | -1.155 | -0.532 | -0.112 | 0.102 | 0.102 | 0.112 | 0.65 | 0.91 | 0.6 | 0.84 | 0.57 | 0.72 |
| 6 | 71457053 | rs10498874 | 0.551 | -1.018 | -1.128 | -0.527 | 0.013 | 0.039 | 0.039 | 0 | 0.48 | 0.38 | 0.34 | 0.23 | 0.47 | 0.54 |
| 6 | 71463508 | rs9360426 | 0.507 | -1.004 | -1.094 | -0.398 | 0.022 | 0.407 | 0.407 | 0 | 0.09 | 0.33 | 0.13 | 0.53 | 0.05 | 0.05 |
| 6 | 71472879 | rs9354940 | -0.558 | -1.015 | -1.127 | -0.426 | -0.114 | 0.093 | 0.093 | 0.164 | 0.65 | 0.91 | 0.59 | 0.83 | 0.57 | 0.71 |
| 6 | 71498986 | rs6927575 | 0.113 | -0.993 | -1.099 | -0.598 | -0.002 | 0.002 | 0.002 | 0.052 | 0.56 | 0.57 | 0.46 | 0.29 | 0.52 | 0.66 |
| 6 | 71507158 | rs9364106 | -0.048 | -0.945 | -1.027 | -0.482 | -0.009 | 0.017 | 0.017 | 0.029 | 0.24 | 0.35 | 0.33 | 0.7 | 0.2 | 0.24 |
| 6 | 71531711 | rs4706464 | 0.33 | -1.066 | -0.958 | -0.505 | 0.001 | 0.002 | 0.002 | 0 | 0.5 | 0.48 | 0.36 | 0.23 | 0.5 | 0.55 |
| 6 | 71546702 | rs2273566 | -0.248 | -1.121 | -0.847 | -0.064 | -0.012 | 0.251 | 0.251 | 0 | 0.13 | 0.33 | 0.19 | 0.55 | 0.07 | 0.17 |
| 6 | 71558805 | rs12190869 | 0.541 | -1.091 | -1.741 | -0.277 | 0.058 | 0.05 | 0.05 | 0 | 0.24 | 0.09 | 0.07 | 0.13 | 0.14 | 0.23 |
| 6 | 71568841 | rs3734372 | 0.041 | -0.202 | -1.856 | 0.289 | -0.034 | -0.074 | -0.074 | 0 | 0.18 | 0.02 | 0.33 | 0.29 | 0.18 | 0.32 |
| 6 | 71569090 | rs576516 | -0.619 | -0.11 | -1.668 | 0.062 | -0.001 | 0.036 | 0.036 | 0 | 0.45 | 0.35 | 0.58 | 0.3 | 0.5 | 0.43 |
| 6 | 71569129 | rs3734369 | 0.989 | -0.385 | -1.94 | 0.282 | -0.001 | 0.124 | 0.124 | 0 | 0.06 | 0.04 | 0.07 | 0.48 | 0.05 | 0.05 |
| 6 | 71570641 | rs9680 | 0.377 | -0.314 | -1.939 | -0.297 | -0.022 | -0.047 | -0.047 | 0.346 | 0.16 | 0.04 | 0.24 | 0.23 | 0.16 | 0.2 |
| 6 | 71573109 | rs2038477 | -0.19 | -0.291 | -1.716 | -0.117 | -0.025 | 0.062 | 0.062 | 0 | 0.42 | 0.28 | 0.47 | 0.26 | 0.45 | 0.4 |
| 6 | 71583804 | rs2504748 | -0.209 | -0.291 | -1.743 | -0.044 | -0.039 | 0.054 | 0.054 | 0 | 0.4 | 0.24 | 0.46 | 0.28 | 0.43 | 0.4 |
| 6 | 71588806 | rs2460699 | -0.115 | -0.212 | -2.041 | -0.065 | -0.078 | 0.033 | 0.033 | 0 | 0.38 | 0.15 | 0.53 | 0.3 | 0.43 | 0.41 |
| 6 | 71594785 | rs659925 | -0.501 | -1.263 | -2.082 | -0.149 | 0.115 | -0.03 | -0.03 | 0 | 0.39 | 0.81 | 0.2 | 0.57 | 0.17 | 0.37 |
| 6 | 71595571 | rs13212642 | 0.5 | -1.137 | -2.386 | -0.361 | 0.011 | -0.037 | -0.037 | 0 | 0.21 | 0.11 | 0.34 | 0.32 | 0.2 | 0.2 |
| 6 | 71598545 | rs1619936 | -0.087 | -1.638 | -2.485 | -0.608 | 0.076 | -0.051 | -0.051 | 0 | 0.31 | 0.79 | 0.17 | 0.57 | 0.15 | 0.36 |
| 6 | 71598571 | rs9446305 | 0.484 | -1.389 | -2.34 | -0.495 | 0.014 | -0.037 | -0.037 | 0.167 | 0.22 | 0.15 | 0.34 | 0.32 | 0.2 | 0.18 |
| 6 | 71599788 | rs9784769 | 0.5 | -1.306 | -2.286 | -0.637 | -0.008 | -0.024 | -0.024 | 0 | 0.08 | 0.08 | 0.01 | 0 | 0.03 | 0.07 |
| 6 | 71599826 | rs10447360 | 0.959 | -0.506 | -2.696 | -0.867 | -0.123 | 0.146 | 0.146 | 0.813 | 0.22 | 0.01 | 0.16 | 0 | 0.3 | 0.09 |
| 6 | 71599887 | rs651984 | -0.034 | -0.757 | -1.539 | 0.075 | -0.026 | 0.254 | 0.254 | 0.083 | 0.12 | 0.3 | 0.06 | 0.4 | 0.05 | 0.08 |
| 6 | 71600335 | rs639448 | -0.439 | -0.588 | -2.095 | -0.63 | 0.03 | 0.276 | 0.276 | 0.169 | 0.56 | 0.94 | 0.53 | 0.89 | 0.35 | 0.51 |
| 6 | 71600923 | rs490887 | -0.152 | -0.06 | -1.428 | -0.115 | 0.04 | -0.01 | -0.01 | 0 | 0.86 | 0.77 | 0.8 | 0.88 | 0.78 | 0.76 |
| 6 | 71602420 | rs9294881 | -0.13 | -0.162 | -1.882 | 0.404 | -0.009 | -0.078 | -0.078 | 0 | 0.3 | 0.11 | 0.42 | 0.45 | 0.3 | 0.35 |
| 6 | 71606987 | rs495433 | -0.491 | -0.8 | -0.908 | 0.64 | -0.044 | 0.036 | 0.036 | 0.084 | 0.25 | 0.57 | 0.14 | 0.39 | 0.11 | 0.23 |
| 6 | 71609049 | rs7750238 | -0.569 | -0.137 | -1.089 | 0.491 | 0.002 | 0.002 | 0.002 | 0 | 0.1 | 0.07 | 0.06 | 0.11 | 0.09 | 0.08 |
| 6 | 71609384 | rs521116 | 0.224 | -0.057 | -0.921 | 0.61 | 0.005 | 0 | 0 | 0 | 0.62 | 0.6 | 0.41 | 0.41 | 0.52 | 0.5 |
| 6 | 71609707 | rs4707862 | -1.05 | -0.558 | -1.325 | 0.434 | -0.007 | -0.004 | -0.004 | 0 | 0.14 | 0.02 | 0.14 | 0.11 | 0.12 | 0.1 |
| 6 | 71610796 | rs7765174 | -1.116 | -1.022 | -0.478 | 0.375 | -0.029 | -0.022 | -0.022 | 0 | 0.18 | 0.32 | 0.14 | 0.12 | 0.11 | 0.16 |
| 6 | 71613747 | rs2474918 | -1.171 | -1.367 | -0.598 | 0.239 | 0.007 | 0.007 | 0.007 | 0.014 | 0.46 | 0.6 | 0.58 | 0.69 | 0.36 | 0.38 |
| 6 | 71618168 | rs2255286 | -1.215 | -1.466 | -0.699 | 0.039 | 0.006 | 0.008 | 0.008 | 0.061 | 0.47 | 0.61 | 0.6 | 0.72 | 0.39 | 0.4 |
| 6 | 71631823 | rs6907179 | 1.716 | -1.069 | -0.65 | -0.523 | -0.011 | -0.018 | -0.018 | 1.491 | 0.24 | 0.18 | 0.14 | 0 | 0.28 | 0.14 |
| 6 | 71650172 | rs9294882 | -0.996 | -1.121 | -0.607 | -0.503 | 0.013 | 0.019 | 0.019 | 0 | 0.49 | 0.61 | 0.61 | 0.72 | 0.4 | 0.4 |
| 6 | 71660111 | rs6922893 | -1.497 | -1.056 | -0.716 | -0.041 | -0.004 | -0.006 | -0.006 | 0 | 0.73 | 0.8 | 0.75 | 0.64 | 0.68 | 0.55 |
| 6 | 71660233 | rs9446311 | -0.399 | -1.353 | -0.608 | -0.022 | 0.009 | -0.045 | -0.045 | 0.373 | 0.36 | 0.21 | 0.47 | 0.51 | 0.36 | 0.33 |
| 6 | 71660752 | rs6900853 | 0.849 | -1.326 | -0.587 | -0.035 | 0.002 | -0.043 | -0.043 | 1.066 | 0.34 | 0.58 | 0.28 | 0.14 | 0.32 | 0.22 |
| 6 | 71666146 | rs1320315 | -1.487 | -1.208 | -0.543 | 0.039 | -0.003 | -0.001 | -0.001 | 0 | 0.73 | 0.8 | 0.75 | 0.72 | 0.68 | 0.54 |
| 6 | 71676933 | rs1111994 | 0.866 | -1.345 | -0.603 | 0.217 | -0.005 | -0.033 | -0.033 | 0.099 | 0.16 | 0.08 | 0.22 | 0.36 | 0.16 | 0.16 |

| | | | | | | | | | | | | | | | | |
|---|---|---|---|---|---|---|---|---|---|---|---|---|---|---|---|---|
| 6 | 71680129 | rs6928051 | -0.711 | -1.354 | -0.253 | -0.013 | -0.045 | 0.015 | 0.015 | 0 | 0.23 | 0.45 | 0.29 | 0.25 | 0.13 | 0.18 |
| 6 | 71683625 | rs6930956 | -0.723 | -1.337 | -0.27 | -0.049 | -0.043 | 0.014 | 0.014 | 0 | 0.23 | 0.44 | 0.29 | 0.25 | 0.13 | 0.18 |
| 6 | 71683895 | rs6455394 | -0.139 | -1.286 | -0.389 | -0.098 | 0.006 | 0.027 | 0.027 | 0 | 0.48 | 0.68 | 0.43 | 0.33 | 0.41 | 0.32 |
| 6 | 71698630 | rs9354950 | -0.571 | -1.168 | -0.154 | -0.5 | 0.004 | -0.119 | -0.119 | 0 | 0.64 | 0.52 | 0.76 | 0.96 | 0.62 | 0.78 |
| 6 | 71704432 | rs9283838 | -1.078 | -1.26 | -0.172 | -0.532 | 0.008 | -0.068 | -0.068 | 0 | 0.69 | 0.62 | 0.85 | 0.93 | 0.71 | 0.86 |
| 6 | 71716067 | rs4706474 | -0.362 | -1.27 | -0.213 | -0.546 | 0 | 0.007 | 0.007 | 0.198 | 0.61 | 0.45 | 0.76 | 0.89 | 0.63 | 0.78 |
| 6 | 71718143 | rs11757574 | 1.4 | -1.027 | -0.415 | -0.552 | -0.002 | -0.044 | -0.044 | 0 | 0.25 | 0.22 | 0.14 | 0 | 0.28 | 0.13 |
| 6 | 71722821 | rs6910674 | 1.015 | -1.321 | -0.737 | -0.183 | -0.05 | -0.034 | -0.034 | 0 | 0.24 | 0.05 | 0.25 | 0.28 | 0.3 | 0.46 |
| 6 | 71723119 | rs9351798 | -0.093 | -1.662 | -0.457 | -0.304 | 0.011 | 0.099 | 0.099 | 0 | 0.49 | 0.29 | 0.59 | 0.82 | 0.48 | 0.69 |
| 6 | 71723390 | rs9346415 | 0.62 | -1.585 | -0.302 | -0.211 | 0.001 | -0.001 | -0.001 | 0 | 0.22 | 0.17 | 0.24 | 0.42 | 0.16 | 0.18 |
| 6 | 71726234 | rs2682188 | -0.237 | -1.578 | -0.056 | -0.231 | -0.052 | 0.076 | 0.076 | 0 | 0.29 | 0.4 | 0.36 | 0.55 | 0.18 | 0.22 |
| 6 | 71729002 | rs1079987 | 1.339 | -1.573 | -0.021 | -0.366 | -0.015 | -0.012 | -0.012 | 0 | 0.3 | 0.23 | 0.17 | 0 | 0.34 | 0.15 |
| 6 | 71731535 | rs2682194 | -0.06 | -1.777 | 0.488 | 0.389 | -0.044 | 0.105 | 0.105 | 0 | 0.26 | 0.49 | 0.32 | 0.51 | 0.16 | 0.19 |
| 6 | 71732366 | rs2748324 | 0.192 | -1.585 | -0.152 | 0.206 | -0.001 | 0.017 | 0.017 | 0 | 0.87 | 0.9 | 0.88 | 0.97 | 0.85 | 0.89 |
| 6 | 71733002 | rs9446323 | -0.819 | -1.604 | 0.066 | 0.655 | -0.044 | 0.031 | 0.031 | 0 | 0.29 | 0.46 | 0.43 | 0.64 | 0.18 | 0.23 |
| 6 | 71733748 | rs2092836 | -0.348 | -1.221 | -0.04 | 0.6 | -0.026 | 0.08 | 0.08 | 0 | 0.24 | 0.34 | 0.32 | 0.51 | 0.16 | 0.19 |
| 6 | 71741216 | rs6455400 | -0.547 | -0.748 | -0.364 | -0.292 | 0.013 | -0.037 | -0.037 | 0 | 0.65 | 0.61 | 0.66 | 0.86 | 0.57 | 0.79 |
| 6 | 71742118 | rs9360434 | -0.239 | -1.118 | -0.303 | -0.077 | 0.009 | -0.016 | -0.016 | 0 | 0.58 | 0.43 | 0.61 | 0.72 | 0.54 | 0.66 |
| 6 | 71743738 | rs1983678 | 0.821 | -1.621 | -0.528 | -0.219 | 0.001 | 0.008 | 0.008 | 0 | 0.31 | 0.35 | 0.4 | 0.38 | 0.35 | 0.41 |
| 6 | 71743943 | rs16869719 | 0.949 | -1.636 | -0.492 | -0.222 | -0.012 | 0.031 | 0.031 | 0 | 0.07 | 0.16 | 0.05 | 0.15 | 0.03 | 0.12 |
| 6 | 71744351 | rs1983676 | 0.133 | -1.615 | -0.51 | -0.264 | -0.005 | 0.008 | 0.008 | 0 | 0.38 | 0.51 | 0.45 | 0.53 | 0.37 | 0.53 |
| 6 | 71746446 | rs12055622 | -0.189 | -1.722 | -0.403 | -0.272 | 0.021 | -0.026 | -0.026 | 0 | 0.29 | 0.18 | 0.31 | 0.47 | 0.2 | 0.27 |
| 6 | 71750120 | rs1342761 | -0.742 | -1.722 | -0.136 | -0.295 | -0.006 | -0.053 | -0.053 | 0 | 0.61 | 0.64 | 0.73 | 1 | 0.53 | 0.79 |
| 6 | 71751997 | rs13208889 | 0.058 | -1.58 | -0.708 | -0.354 | 0.033 | -0.063 | -0.063 | 0 | 0.32 | 0.07 | 0.33 | 0.47 | 0.26 | 0.28 |
| 6 | 71752093 | rs9351800 | -0.768 | -1.553 | -0.693 | -0.306 | 0 | -0.011 | -0.011 | 0 | 0.54 | 0.42 | 0.68 | 0.86 | 0.51 | 0.67 |
| 6 | 71752504 | rs13208627 | 0.616 | -0.969 | -0.359 | 0.121 | 0.003 | 0.054 | 0.054 | 0.984 | 0.2 | 0.26 | 0.34 | 0.38 | 0.25 | 0.39 |
| 6 | 71754687 | rs7772103 | -0.474 | -1.161 | -0.547 | -0.608 | -0.023 | -0.043 | -0.043 | 0.285 | 0.23 | 0.33 | 0.1 | 0.11 | 0.13 | 0.15 |
| 6 | 71756160 | rs9354960 | 0.452 | -1.374 | -1.25 | -0.628 | 0.001 | -0.046 | -0.046 | 0 | 0.24 | 0.19 | 0.37 | 0.43 | 0.29 | 0.47 |
| 6 | 71761066 | rs1891523 | -0.636 | -1.479 | -1.086 | -1.119 | 0.037 | -0.053 | -0.053 | 0 | 0.35 | 0.18 | 0.41 | 0.46 | 0.26 | 0.3 |
| 6 | 71763859 | rs1342757 | 0.202 | -1.56 | -1.126 | -1.151 | 0.004 | -0.005 | -0.005 | 0.17 | 0.2 | 0.15 | 0.37 | 0.46 | 0.18 | 0.22 |
| 6 | 71772828 | rs2144402 | 0.598 | -1.501 | -0.506 | -1.403 | -0.001 | 0.001 | 0.001 | 0.051 | 0.43 | 0.5 | 0.42 | 0.46 | 0.43 | 0.6 |
| 6 | 71780591 | rs1033609 | -0.219 | -1.811 | -0.261 | -1.697 | 0.005 | -0.072 | -0.072 | 0 | 0.59 | 0.55 | 0.77 | 0.92 | 0.6 | 0.83 |
| 6 | 71781342 | rs6911809 | -0.539 | -1.721 | -0.563 | -1.653 | -0.028 | 0.012 | 0.012 | 0 | 0.66 | 0.78 | 0.8 | 1 | 0.6 | 0.83 |
| 6 | 71785901 | rs12195809 | 0.635 | -1.747 | -0.656 | -1.701 | -0.008 | 0.078 | 0.078 | 0 | 0.13 | 0.02 | 0.04 | 0 | 0.09 | 0.07 |
| 6 | 71788265 | rs1418677 | -1.861 | -1.779 | -0.669 | -1.722 | -0.122 | 0.135 | 0.135 | 0.246 | 0.8 | 0.98 | 0.84 | 1 | 0.69 | 0.91 |
| 6 | 71788468 | rs1739807 | -0.425 | -1.755 | -0.646 | -1.707 | -0.025 | 0.033 | 0.033 | 0 | 0.64 | 0.75 | 0.78 | 0.92 | 0.57 | 0.81 |
| 6 | 71793334 | rs718356 | 0.81 | -1.431 | -0.495 | -1.157 | -0.034 | -0.076 | -0.076 | 0 | 0.3 | 0.13 | 0.36 | 0.45 | 0.36 | 0.55 |
| 6 | 71812751 | rs9446337 | -0.322 | -1.15 | -0.138 | -1.241 | 0.006 | -0.015 | -0.015 | 0 | 0.18 | 0.17 | 0.06 | 0 | 0.13 | 0.11 |
| 6 | 71814944 | rs9364129 | -0.352 | -1.233 | -0.027 | -1.022 | -0.012 | 0.009 | 0.009 | 0.056 | 0.53 | 0.29 | 0.47 | 0.58 | 0.53 | 0.7 |
| 6 | 71815111 | rs7774503 | -1.204 | -1.295 | 1.036 | -1.62 | -0.04 | -0.081 | -0.081 | 0 | 0.25 | 0.47 | 0.09 | 0.09 | 0.14 | 0.1 |
| 6 | 71815318 | rs995863 | -0.309 | -0.523 | 0.979 | -1.591 | 0.005 | -0.053 | -0.053 | 0 | 0.73 | 0.66 | 0.7 | 0.91 | 0.71 | 0.89 |
| 6 | 71819535 | rs9446338 | -0.976 | -1.007 | 1.802 | -0.627 | -0.032 | -0.089 | -0.089 | 0.842 | 0.16 | 0.3 | 0.04 | 0 | 0.07 | 0.02 |
| 6 | 71820001 | rs992998 | -2.157 | -1.034 | 1.726 | -0.558 | -0.023 | -0.01 | -0.01 | 0.003 | 0.25 | 0.62 | 0.21 | 0.19 | 0.15 | 0.1 |
| 6 | 71832412 | rs1775549 | -1.962 | -0.918 | 1.705 | -0.765 | -0.016 | -0.084 | -0.084 | 0.092 | 0.24 | 0.64 | 0.1 | 0 | 0.13 | 0.08 |
| 6 | 71833162 | rs9455336 | -1.952 | -0.889 | 1.721 | -0.75 | -0.017 | -0.087 | -0.087 | 0 | 0.23 | 0.66 | 0.1 | 0 | 0.13 | 0.09 |
| 6 | 71835518 | rs1857224 | -1.978 | -0.84 | 1.669 | -0.796 | -0.01 | -0.078 | -0.078 | 0 | 0.26 | 0.66 | 0.12 | 0 | 0.18 | 0.1 |
| 6 | 71860000 | rs10945289 | 1.501 | -0.856 | 1.94 | -0.72 | -0.02 | -0.091 | -0.091 | 0 | 0.83 | 0.39 | 0.93 | 1 | 0.91 | 0.96 |
| 6 | 71861955 | rs12206074 | -0.454 | -0.844 | 2.037 | -0.711 | -0.004 | -0.037 | -0.037 | 0 | 0.11 | 0.11 | 0.03 | 0 | 0.07 | 0.02 |
| 6 | 71867477 | rs1418684 | 0.617 | -0.836 | 1.987 | -0.707 | -0.009 | -0.06 | -0.06 | 0.164 | 0.88 | 0.81 | 0.97 | 1 | 0.93 | 0.98 |
| 6 | 71868834 | rs1935796 | 1.229 | -0.842 | 2.138 | -0.716 | -0.004 | -0.082 | -0.082 | 0 | 0.86 | 0.68 | 0.95 | 1 | 0.9 | 0.96 |
| 6 | 71883338 | rs7775859 | 0.633 | -0.793 | 1.282 | -0.688 | -0.007 | -0.045 | -0.045 | 0 | 0.88 | 0.82 | 0.97 | 1 | 0.93 | 0.98 |
| 6 | 71891609 | rs9364137 | 1.926 | -0.917 | 0.908 | -0.603 | 0.03 | 0.04 | 0.04 | 0 | 0.65 | 0.23 | 0.75 | 0.79 | 0.76 | 0.81 |
| 6 | 71902656 | rs1342747 | 0.759 | -0.93 | 0.93 | -0.781 | -0.01 | -0.044 | -0.044 | 0 | 0.91 | 0.8 | 0.96 | 0.99 | 0.93 | 0.94 |
| 6 | 71912141 | rs12200732 | -0.209 | -1.05 | 0.453 | -0.718 | 0.01 | -0.05 | -0.05 | 0 | 0.1 | 0.01 | 0.1 | 0.17 | 0.06 | 0.16 |
| 6 | 71916420 | rs4585520 | 2.12 | -1.069 | 0.593 | -0.908 | 0.042 | 0.084 | 0.084 | 0 | 0.59 | 0.2 | 0.75 | 0.82 | 0.71 | 0.71 |
| 6 | 71917597 | rs12661758 | 1.586 | -0.934 | 1.076 | -0.935 | -0.022 | -0.103 | -0.103 | 0 | 0.79 | 0.37 | 0.91 | 0.99 | 0.88 | 0.94 |
| 6 | 71919099 | rs9360442 | 0.876 | -0.919 | 1.201 | -0.924 | 0.028 | -0.081 | -0.081 | 0.049 | 0.87 | 0.48 | 0.93 | 0.99 | 0.92 | 0.97 |
| 6 | 71922715 | rs9346446 | 1.834 | -0.887 | 0.89 | -0.905 | 0.009 | -0.003 | -0.003 | 0 | 0.69 | 0.24 | 0.85 | 0.99 | 0.77 | 0.86 |
| 6 | 71929868 | rs9342822 | 1.94 | -0.838 | 0.885 | -0.874 | 0.005 | -0.019 | -0.019 | 2.059 | 0.7 | 0.27 | 0.86 | 0.99 | 0.77 | 0.86 |
| 6 | 71933272 | rs9294888 | 1.611 | -0.994 | 1.004 | -0.606 | -0.024 | -0.012 | -0.012 | 0 | 0.73 | 0.38 | 0.79 | 0.8 | 0.83 | 0.8 |
| 6 | 71936292 | rs12193343 | -0.237 | -0.959 | 1.503 | -0.59 | 0.036 | -0.04 | -0.04 | 0.011 | 0.08 | 0.28 | 0.04 | 0 | 0.07 | 0.02 |
| 6 | 71937060 | rs12665552 | 1.689 | -0.935 | 1.316 | -0.566 | 0.022 | 0.021 | 0.021 | 0.29 | 0.64 | 0.3 | 0.76 | 0.8 | 0.79 | 0.75 |
| 6 | 71937998 | rs4707902 | 1.803 | -0.923 | 1.106 | -0.562 | 0.04 | 0.048 | 0.048 | 0.564 | 0.63 | 0.2 | 0.74 | 0.79 | 0.76 | 0.75 |
| 6 | 71959728 | rs7758727 | -1.418 | -0.553 | 1.366 | -0.152 | 0.056 | 0.131 | 0.131 | 1.891 | 0.07 | 0.33 | 0.06 | 0.13 | 0.06 | 0.09 |
| 6 | 71963483 | rs9455394 | -1.739 | -0.61 | 1.397 | -0.235 | 0.002 | 0.011 | 0.011 | 0.64 | 0.16 | 0.4 | 0.09 | 0.14 | 0.12 | 0.14 |
| 6 | 71968427 | rs9446361 | -2.004 | -0.715 | 1.48 | -0.138 | -0.011 | -0.001 | -0.001 | 0 | 0.17 | 0.75 | 0.14 | 0.16 | 0.1 | 0.14 |
| 6 | 71979329 | rs6914937 | -1.268 | -0.831 | 0.718 | -0.312 | 0.042 | 0.085 | 0.085 | 0.287 | 0.5 | 0.83 | 0.7 | 0.63 | 0.44 | 0.42 |
| 6 | 71993701 | rs7748559 | 0.488 | -0.944 | 0.907 | 0.507 | -0.013 | -0.056 | -0.056 | 0 | 0.2 | 0.1 | 0.4 | 0.35 | 0.25 | 0.17 |
| 6 | 71997278 | rs2208845 | -0.133 | -1.02 | 1.307 | 0.455 | 0 | 0.104 | 0.104 | 0 | 0.14 | 0.37 | 0.22 | 0.27 | 0.1 | 0.13 |

| | | | | | | | | | | | | | | | | |
|---|---|---|---|---|---|---|---|---|---|---|---|---|---|---|---|---|
| 6 | 71999404 | rs1885674 | 0.425 | -0.976 | 1.01 | 0.456 | -0.001 | -0.011 | -0.011 | 0 | 0.2 | 0.17 | 0.52 | 0.57 | 0.22 | 0.18 |
| 6 | 72005794 | rs9354987 | -1.784 | -1.001 | 0.885 | 0.369 | 0.048 | 0.07 | 0.07 | 0.322 | 0.49 | 0.83 | 0.69 | 0.63 | 0.42 | 0.42 |
| 6 | 72008997 | rs7766995 | -1.947 | -0.608 | 0.034 | -0.099 | -0.121 | -0.068 | -0.068 | 0.355 | 0.68 | 0.98 | 0.81 | 0.68 | 0.67 | 0.57 |
| 6 | 72011086 | rs16880821 | -0.632 | -0.541 | 0.829 | 0.348 | 0.009 | 0.011 | 0.011 | 0 | 0.14 | 0.12 | 0.05 | 0.01 | 0.09 | 0.12 |
| 6 | 72029515 | rs4707910 | 0.321 | -0.511 | 0.818 | 0.523 | 0 | 0.031 | 0.031 | 0 | 0.2 | 0.27 | 0.41 | 0.37 | 0.22 | 0.17 |
| 6 | 72030119 | rs13215762 | -0.505 | -0.441 | 0.661 | -0.09 | 0.008 | 0.066 | 0.066 | 0 | 0.13 | 0.07 | 0.04 | 0 | 0.09 | 0.06 |
| 6 | 72030526 | rs9354993 | -0.174 | -0.408 | 0.996 | 0.002 | -0.018 | 0.067 | 0.067 | 0 | 0.13 | 0.23 | 0.19 | 0.26 | 0.08 | 0.12 |
| 6 | 72034337 | rs7753757 | -0.892 | -0.404 | 1.053 | -0.04 | -0.04 | 0.019 | 0.019 | 0 | 0.22 | 0.36 | 0.23 | 0.27 | 0.13 | 0.23 |
| 6 | 72037276 | rs2295956 | 0.314 | -0.38 | 0.913 | 0.006 | 0.001 | -0.009 | -0.009 | 0 | 0.19 | 0.2 | 0.27 | 0.08 | 0.23 | 0.12 |
| 6 | 72039761 | rs9364162 | -1.195 | -0.339 | 0.819 | 0.02 | -0.005 | -0.001 | -0.001 | 0 | 0.26 | 0.29 | 0.26 | 0.27 | 0.19 | 0.25 |
| 6 | 72040511 | rs1322760 | -1.788 | -0.421 | 0.61 | 0.013 | 0.016 | 0.081 | 0.081 | 0.599 | 0.5 | 0.9 | 0.56 | 0.35 | 0.42 | 0.37 |
| 6 | 72043897 | rs828630 | -0.252 | -0.409 | 0.667 | 0.02 | 0.003 | -0.055 | -0.055 | 0 | 0.25 | 0.39 | 0.3 | 0.08 | 0.27 | 0.13 |
| 6 | 72048250 | rs866142 | -1.295 | -0.202 | -0.135 | 0.178 | 0.035 | 0.06 | 0.06 | 0.418 | 0.48 | 0.72 | 0.54 | 0.34 | 0.4 | 0.36 |
| 6 | 72056910 | rs828655 | -0.148 | -0.251 | -0.18 | 0.242 | 0.007 | 0.033 | 0.033 | 0 | 0.23 | 0.14 | 0.25 | 0.1 | 0.23 | 0.13 |
| 6 | 72059524 | rs6903710 | -0.679 | -0.501 | 0.667 | 0.586 | -0.019 | 0.13 | 0.13 | 0 | 0.16 | 0.45 | 0.25 | 0.3 | 0.1 | 0.12 |
| 6 | 72061460 | rs978766 | 1.541 | -0.038 | 0.215 | 0.43 | -0.015 | -0.027 | -0.027 | 0 | 0.46 | 0.2 | 0.38 | 0.48 | 0.46 | 0.58 |
| 6 | 72072140 | rs1407593 | 1.052 | 0.032 | 0.602 | 0.56 | -0.011 | 0.049 | 0.049 | 0 | 0.78 | 0.54 | 0.71 | 0.7 | 0.83 | 0.85 |
| 6 | 72073567 | rs1192048 | 1.799 | -0.017 | 0.485 | 0.552 | -0.006 | 0.001 | 0.001 | 0.724 | 0.71 | 0.34 | 0.6 | 0.68 | 0.81 | 0.76 |
| 6 | 72081794 | rs828644 | 0.807 | 0.09 | 0.884 | 0.407 | 0.016 | -0.072 | -0.072 | 0 | 0.86 | 0.72 | 0.89 | 0.98 | 0.88 | 0.88 |
| 6 | 72087785 | rs9455449 | 0.075 | 0.084 | 0.919 | 0.408 | 0.019 | -0.018 | -0.018 | 0 | 0.06 | 0.2 | 0.04 | 0 | 0.04 | 0.02 |
| 6 | 72090956 | rs1192041 | 1.476 | -0.048 | 1.519 | 0.295 | 0.001 | -0.121 | -0.121 | 1.608 | 0.83 | 0.59 | 0.9 | 1 | 0.87 | 0.92 |
| 6 | 72093537 | rs1322764 | 1.094 | -0.116 | 0.577 | 0.145 | 0.015 | 0.064 | 0.064 | 0 | 0.38 | 0.22 | 0.33 | 0.23 | 0.38 | 0.47 |
| 6 | 72095766 | rs16881192 | 1.754 | -0.047 | 0.286 | 0.441 | -0.048 | -0.061 | -0.061 | 0 | 0.16 | 0.04 | 0.2 | 0.28 | 0.24 | 0.17 |
| 6 | 72099296 | rs10455727 | 1.432 | -0.055 | 0.496 | 0.444 | -0.003 | -0.039 | -0.039 | 1.794 | 0.5 | 0.13 | 0.5 | 0.51 | 0.6 | 0.62 |
| 6 | 72100307 | rs9364164 | 0.454 | -0.387 | 0.677 | 0.642 | -0.006 | 0.032 | 0.032 | 0 | 0.1 | 0.13 | 0.21 | 0.28 | 0.05 | 0.09 |
| 6 | 72100877 | rs1727650 | -0.344 | -0.417 | 0.75 | 0.598 | 0 | -0.004 | -0.004 | 1.416 | 0.32 | 0.31 | 0.38 | 0.5 | 0.28 | 0.24 |
| 6 | 72105557 | rs1527714 | -0.977 | -0.443 | 0.819 | 0.289 | -0.017 | -0.01 | -0.01 | 0 | 0.8 | 0.85 | 0.83 | 0.8 | 0.75 | 0.87 |
| 6 | 72108896 | rs9351811 | -0.259 | -0.483 | 0.624 | 0.399 | -0.006 | 0.032 | 0.032 | 0.467 | 0.14 | 0.15 | 0.23 | 0.42 | 0.07 | 0.16 |
| 6 | 72112591 | rs13202946 | 1.886 | -0.469 | 0.112 | 0.292 | -0.056 | 0.18 | 0.18 | 0 | 0.32 | 0.02 | 0.32 | 0.1 | 0.34 | 0.41 |
| 6 | 72113712 | rs2222722 | -1.889 | -0.47 | 0.653 | 0.564 | -0.031 | 0.036 | 0.036 | 1.475 | 0.29 | 0.54 | 0.33 | 0.44 | 0.17 | 0.3 |
| 6 | 72118362 | rs852933 | 0.433 | -0.389 | 0.676 | 0.462 | 0.01 | 0.003 | 0.003 | 0 | 0.22 | 0.3 | 0.19 | 0.2 | 0.24 | 0.14 |
| 6 | 72124994 | rs765466 | -0.038 | -0.563 | 0.556 | 0.657 | 0 | -0.031 | -0.031 | 0 | 0.11 | 0.11 | 0.06 | 0 | 0.1 | 0.05 |
| 6 | 72131926 | rs6903167 | -1.767 | -0.499 | 0.142 | 0.709 | -0.009 | 0.008 | 0.008 | 1.141 | 0.29 | 0.63 | 0.31 | 0.43 | 0.17 | 0.27 |
| 6 | 72136315 | rs9346470 | -0.157 | -0.143 | -0.166 | 1.025 | 0 | 0.055 | 0.055 | 0.046 | 0.08 | 0.08 | 0.19 | 0.42 | 0.07 | 0.15 |
| 6 | 72137905 | rs9346471 | -0.917 | -0.224 | 0.262 | 1.049 | -0.008 | 0.175 | 0.175 | 0.025 | 0.14 | 0.28 | 0.22 | 0.52 | 0.11 | 0.18 |
| 6 | 72143093 | rs852953 | -0.542 | -0.163 | -0.283 | 0.457 | -0.001 | -0.011 | -0.011 | 0 | 0.36 | 0.62 | 0.32 | 0.13 | 0.33 | 0.19 |
| 6 | 72154332 | rs9360453 | 1.526 | -0.239 | -0.752 | 0.155 | -0.01 | 0.191 | 0.191 | 0.158 | 0.35 | 0.03 | 0.31 | 0.11 | 0.32 | 0.45 |
| 6 | 72157229 | rs852937 | 0.583 | -0.209 | -0.289 | 0.191 | 0.005 | -0.053 | -0.053 | 0 | 0.62 | 0.42 | 0.55 | 0.16 | 0.59 | 0.57 |
| 6 | 72157939 | rs9446392 | 1.008 | -0.003 | -0.259 | 0.22 | -0.001 | -0.003 | -0.003 | 0.112 | 0.06 | 0.07 | 0.04 | 0.01 | 0.06 | 0.03 |
| 6 | 72164390 | rs852944 | -1.309 | -0.105 | -0.202 | 0.483 | -0.021 | -0.032 | -0.032 | 0 | 0.47 | 0.81 | 0.54 | 0.48 | 0.43 | 0.36 |
| 6 | 72179118 | rs701698 | -0.907 | -0.216 | -0.189 | 0.193 | -0.01 | 0.072 | 0.072 | 0.19 | 0.42 | 0.77 | 0.36 | 0.23 | 0.37 | 0.25 |
| 6 | 72186990 | rs852961 | -0.286 | -0.205 | -0.111 | 0.184 | 0.022 | -0.031 | -0.031 | 3.04 | 0.3 | 0.48 | 0.27 | 0.19 | 0.31 | 0.2 |
| 6 | 72191860 | rs504983 | 0.47 | -0.194 | -0.254 | 0.153 | 0.049 | -0.061 | -0.061 | 0.019 | 0.23 | 0.38 | 0.25 | 0.08 | 0.27 | 0.16 |
| 6 | 72193707 | rs9351814 | 0.841 | -0.269 | -0.616 | 0.389 | 0.042 | -0.029 | -0.029 | 1.115 | 0.43 | 0.24 | 0.5 | 0.54 | 0.38 | 0.56 |
| 6 | 72195981 | rs493265 | 0.045 | -0.205 | -0.577 | 0.376 | 0.02 | -0.066 | -0.066 | 0 | 0.32 | 0.49 | 0.36 | 0.11 | 0.32 | 0.2 |
| 6 | 72197919 | rs543827 | -1.241 | -0.072 | -0.651 | 0.494 | -0.05 | -0.057 | -0.057 | 0 | 0.82 | 0.92 | 0.82 | 0.74 | 0.74 | 0.79 |
| 6 | 72202349 | rs473757 | -0.606 | -0.091 | -0.627 | 0.431 | -0.004 | -0.008 | -0.008 | 0.083 | 0.75 | 0.74 | 0.79 | 0.63 | 0.7 | 0.76 |
| 6 | 72203005 | rs9351817 | 0.62 | -0.12 | -1.101 | 0.471 | 0.042 | -0.035 | -0.035 | 0 | 0.42 | 0.23 | 0.43 | 0.53 | 0.38 | 0.56 |
| 6 | 72205044 | rs558092 | -0.83 | 0.446 | -1.45 | 0.508 | -0.066 | -0.026 | -0.026 | 0 | 0.8 | 0.96 | 0.83 | 0.84 | 0.72 | 0.79 |
| 6 | 72205132 | rs12195791 | 1.434 | -0.276 | -1.964 | 0.247 | -0.043 | -0.041 | -0.041 | 0 | 0.22 | 0.02 | 0.23 | 0.24 | 0.27 | 0.42 |
| 6 | 72205797 | rs489755 | -0.961 | 0.067 | -1.596 | 0.455 | -0.065 | -0.057 | -0.057 | 2.957 | 0.82 | 0.93 | 0.83 | 0.79 | 0.74 | 0.81 |
| 6 | 72210423 | rs547136 | 1.294 | -0.009 | -1.407 | 0.199 | -0.004 | 0.013 | 0.013 | 0 | 0.06 | 0.03 | 0.04 | 0 | 0.07 | 0.03 |
| 6 | 72213306 | rs182106 | -0.077 | -0.227 | -1.019 | 0.066 | -0.001 | -0.002 | -0.002 | 0 | 0.58 | 0.58 | 0.45 | 0.59 | 0.49 | 0.65 |
| 6 | 72214719 | rs199622 | 0.869 | -0.434 | -0.983 | 0.081 | 0.023 | -0.025 | -0.025 | 0.015 | 0.4 | 0.26 | 0.34 | 0.51 | 0.38 | 0.6 |
| 6 | 72223787 | rs199644 | -0.504 | -0.423 | -1.189 | 0.017 | -0.012 | -0.015 | -0.015 | 0.069 | 0.46 | 0.82 | 0.52 | 0.47 | 0.42 | 0.35 |
| 6 | 72226003 | rs199635 | -1.517 | -0.384 | -1.537 | 0.044 | -0.051 | -0.062 | -0.062 | 0 | 0.85 | 0.97 | 0.83 | 0.75 | 0.78 | 0.81 |
| 6 | 72232928 | rs742713 | 0.837 | -0.414 | -1.521 | 0.018 | 0.017 | 0.053 | 0.053 | 0.28 | 0.38 | 0.07 | 0.3 | 0.28 | 0.36 | 0.48 |
| 6 | 72235703 | rs2129327 | 1.416 | -0.261 | -1.657 | 0.094 | -0.05 | -0.021 | -0.021 | 1.193 | 0.29 | 0.03 | 0.26 | 0.25 | 0.33 | 0.44 |
| 6 | 72237028 | rs185634 | -1.268 | 0.172 | -1.218 | 0.191 | -0.039 | -0.047 | -0.047 | 0.055 | 0.85 | 0.95 | 0.83 | 0.75 | 0.78 | 0.83 |
| 6 | 72247218 | rs4707917 | 0.516 | 0.367 | -1.133 | -0.065 | -0.062 | 0.03 | 0.03 | 0 | 0.18 | 0.09 | 0.27 | 0.08 | 0.26 | 0.15 |
| 6 | 72250141 | rs199634 | -0.635 | -0.464 | -0.586 | -0.162 | -0.031 | -0.015 | -0.015 | 0.075 | 0.2 | 0.65 | 0.17 | 0.13 | 0.09 | 0.05 |
| 6 | 72252391 | rs199633 | -0.635 | -0.464 | -0.586 | -0.162 | -0.031 | -0.015 | -0.015 | 0 | 0.2 | 0.65 | 0.17 | 0.13 | 0.09 | 0.05 |
| 6 | 72254897 | rs9342842 | 0.902 | -0.615 | -1.174 | -0.289 | 0.023 | -0.032 | -0.032 | 0 | 0.36 | 0.21 | 0.33 | 0.52 | 0.33 | 0.55 |
| 6 | 72277032 | rs11967789 | 0.277 | -0.644 | -0.535 | -0.271 | -0.003 | 0.09 | 0.09 | 0.234 | 0.08 | 0.29 | 0.12 | 0.13 | 0.03 | 0.02 |
| 6 | 72279021 | rs9358960 | -0.044 | -0.647 | -0.506 | -0.273 | 0 | 0.096 | 0.096 | 0.279 | 0.92 | 0.67 | 0.87 | 0.87 | 0.97 | 0.98 |
| 6 | 72279876 | rs12207816 | 0.573 | -0.695 | -1.099 | -0.403 | 0 | 0.045 | 0.045 | 2.218 | 0.12 | 0.01 | 0.07 | 0.03 | 0.1 | 0.1 |
| 6 | 72281932 | rs4707922 | 0.781 | -0.681 | -1.16 | -0.411 | -0.125 | -0.062 | -0.062 | 0 | 0.33 | 0.09 | 0.42 | 0.33 | 0.49 | 0.3 |
| 6 | 72286838 | rs6914250 | 1.189 | -0.69 | -0.68 | -0.083 | 0.094 | -0.09 | -0.09 | 0 | 0.57 | 0.1 | 0.66 | 0.55 | 0.74 | 0.69 |
| 6 | 72289651 | rs6927647 | 0.398 | -0.341 | 0.077 | 0.141 | -0.016 | 0.151 | 0.151 | 0.377 | 0.12 | 0.33 | 0.1 | 0.29 | 0.07 | 0.16 |

| | | | | | | | | | | | | | | | | |
|---|---|---|---|---|---|---|---|---|---|---|---|---|---|---|---|---|
| 6 | 72291534 | rs9446418 | -0.758 | -0.099 | 0.096 | -0.195 | -0.005 | 0.012 | 0.012 | 1.591 | 0.23 | 0.48 | 0.21 | 0.24 | 0.15 | 0.1 |
| 6 | 72293273 | rs1032166 | 0.073 | -0.406 | 0.349 | -0.047 | -0.033 | 0.07 | 0.07 | 0.032 | 0.79 | 0.56 | 0.84 | 0.69 | 0.88 | 0.76 |
| 6 | 72300534 | rs12193019 | 0.708 | -0.717 | -0.239 | -0.593 | 0.009 | 0.007 | 0.007 | 0 | 0.29 | 0.26 | 0.29 | 0.26 | 0.28 | 0.39 |
| 6 | 72303339 | rs9351832 | 0.758 | -0.748 | -0.004 | -0.571 | 0.007 | 0.131 | 0.131 | 0 | 0.13 | 0.31 | 0.11 | 0.28 | 0.09 | 0.16 |
| 6 | 72303559 | rs1486605 | -0.554 | -0.884 | -0.088 | -0.634 | 0.001 | 0.001 | 0.001 | 5.544 | 0.88 | 0.85 | 0.83 | 0.93 | 0.97 | 0.92 |
| 6 | 72319782 | rs982254 | 0.238 | -0.686 | -0.203 | -0.95 | 0.013 | 0.03 | 0.03 | 0.695 | 0.46 | 0.56 | 0.46 | 0.67 | 0.55 | 0.46 |
| 6 | 72321902 | rs720742 | -0.085 | -0.698 | -0.419 | -0.98 | 0.07 | -0.001 | -0.001 | 0.022 | 0.51 | 0.74 | 0.58 | 0.61 | 0.39 | 0.64 |
| 6 | 72323850 | rs982564 | 0.655 | -0.761 | -0.492 | -0.96 | 0.034 | 0.013 | 0.013 | 1.297 | 0.57 | 0.28 | 0.54 | 0.57 | 0.69 | 0.69 |
| 6 | 72323926 | rs982565 | 0.846 | -0.715 | -0.524 | -0.934 | -0.082 | -0.073 | -0.073 | 0 | 0.35 | 0.17 | 0.35 | 0.4 | 0.46 | 0.29 |
| 6 | 72329609 | rs6453558 | 0.491 | -0.777 | -0.33 | -0.961 | -0.022 | -0.05 | -0.05 | 2.896 | 0.7 | 0.55 | 0.65 | 0.83 | 0.78 | 0.85 |
| 6 | 72337017 | rs10942957 | 1.803 | -0.77 | -0.492 | -0.975 | 0.008 | 0.026 | 0.026 | 0 | 0.21 | 0.05 | 0.18 | 0.17 | 0.23 | 0.4 |
| 6 | 72351393 | rs717519 | 0.049 | -0.81 | -0.417 | -0.949 | 0.049 | -0.007 | -0.007 | 0 | 0.48 | 0.25 | 0.42 | 0.4 | 0.6 | 0.35 |
| 6 | 72363324 | rs9351841 | 1.634 | -0.8 | -0.382 | -0.931 | 0.011 | 0.026 | 0.026 | 0 | 0.22 | 0.1 | 0.19 | 0.17 | 0.24 | 0.4 |
| 6 | 72365557 | rs272213 | -1.219 | -0.824 | -0.29 | -0.928 | 0.011 | 0.1 | 0.1 | 0 | 0.87 | 0.72 | 0.89 | 0.74 | 0.91 | 0.84 |
| 6 | 72366400 | rs1407200 | 1.093 | -0.846 | -0.467 | -0.97 | -0.105 | -0.1 | -0.1 | 0 | 0.32 | 0.12 | 0.34 | 0.4 | 0.45 | 0.29 |
| 6 | 72367279 | rs9293848 | -0.668 | -0.789 | -0.377 | -0.942 | -0.013 | 0.005 | 0.005 | 0 | 0.64 | 0.5 | 0.7 | 0.57 | 0.67 | 0.44 |
| 6 | 72372229 | rs2220290 | 1.801 | -0.718 | -0.934 | -0.934 | -0.032 | 0.001 | 0.001 | 0.008 | 0.21 | 0.02 | 0.18 | 0.17 | 0.24 | 0.39 |
| 6 | 72378469 | rs272229 | -1.414 | -0.517 | -0.706 | -0.776 | 0 | 0.016 | 0.016 | 0 | 0.88 | 0.91 | 0.88 | 0.73 | 0.89 | 0.85 |
| 6 | 72384595 | rs6918162 | 0.602 | -0.596 | -0.612 | -0.821 | -0.007 | -0.004 | -0.004 | 0 | 0.37 | 0.35 | 0.35 | 0.34 | 0.43 | 0.3 |
| 6 | 72385311 | rs4706488 | -0.083 | -0.608 | -0.578 | -0.855 | 0.017 | 0.036 | 0.036 | 0.188 | 0.49 | 0.4 | 0.53 | 0.36 | 0.46 | 0.54 |
| 6 | 72406319 | rs4707930 | 0.296 | -0.323 | -0.877 | -0.779 | -0.024 | 0.026 | 0.026 | 2.471 | 0.67 | 0.48 | 0.59 | 0.53 | 0.8 | 0.74 |
| 6 | 72408820 | rs4415130 | 0.44 | -0.224 | -1.332 | -0.758 | -0.031 | -0.014 | -0.014 | 0 | 0.38 | 0.29 | 0.38 | 0.35 | 0.5 | 0.31 |
| 6 | 72413312 | rs2754048 | -1.155 | 0.009 | -1.548 | -0.497 | 0 | 0.01 | 0.01 | 0 | 0.88 | 0.91 | 0.89 | 0.71 | 0.89 | 0.84 |
| 6 | 72417543 | rs1003260 | 1.564 | 0.159 | -1.691 | -0.461 | -0.055 | -0.068 | -0.068 | 0 | 0.11 | 0.02 | 0.19 | 0.34 | 0.22 | 0.16 |
| 6 | 72427371 | rs1204328 | 0.753 | -0.458 | -1.396 | -0.344 | -0.024 | 0.005 | 0.005 | 0.652 | 0.09 | 0.23 | 0.12 | 0.09 | 0.04 | 0.02 |
| 6 | 72430922 | rs9358971 | 1.464 | -0.405 | -2.706 | -0.419 | -0.02 | -0.024 | -0.024 | 0 | 0.19 | 0.02 | 0.15 | 0.18 | 0.2 | 0.41 |
| 6 | 72437006 | rs9293851 | 0.878 | -0.243 | -2.576 | -0.247 | -0.012 | -0.012 | -0.012 | 0 | 0.3 | 0.25 | 0.31 | 0.34 | 0.38 | 0.27 |
| 6 | 72437128 | rs829470 | 0.602 | 0.399 | -3.263 | -0.065 | -0.033 | -0.053 | -0.053 | 0 | 0.43 | 0.75 | 0.45 | 0.43 | 0.42 | 0.29 |
| 6 | 72437718 | rs829472 | 0.059 | 0.697 | -2.29 | 0.496 | -0.021 | -0.027 | -0.027 | 0.013 | 0.81 | 0.9 | 0.82 | 0.71 | 0.76 | 0.82 |
| 6 | 72440951 | rs829473 | 0.155 | 0.836 | -2.061 | 0.538 | -0.016 | -0.031 | -0.031 | 0.963 | 0.69 | 0.85 | 0.63 | 0.61 | 0.68 | 0.74 |
| 6 | 72450580 | rs1999083 | -0.711 | 0.54 | -1.447 | 0.987 | 0.007 | -0.046 | -0.046 | 0.029 | 0.18 | 0.09 | 0.11 | 0.25 | 0.11 | 0.21 |
| 6 | 72451423 | rs1328693 | 0.215 | 0.206 | -1.47 | 0.074 | 0.018 | 0.043 | 0.043 | 0 | 0.61 | 0.69 | 0.8 | 0.83 | 0.7 | 0.75 |
| 6 | 72453228 | rs11757535 | 0.158 | 0.729 | -1.548 | 0.46 | -0.012 | -0.022 | -0.022 | 0.348 | 0.11 | 0.05 | 0.14 | 0.34 | 0.16 | 0.16 |
| 6 | 72453286 | rs1328706 | 0.413 | 0.668 | -1.709 | -0.358 | -0.009 | 0.011 | 0.011 | 0 | 0.46 | 0.26 | 0.44 | 0.42 | 0.43 | 0.54 |
| 6 | 72453383 | rs1328705 | 0.862 | 0.6 | -2.177 | -0.729 | -0.102 | -0.102 | -0.102 | 0.305 | 0.28 | 0.05 | 0.36 | 0.38 | 0.31 | 0.44 |
| 6 | 72453473 | rs1328704 | -0.213 | 0.231 | -2.204 | -0.932 | -0.006 | -0.001 | -0.001 | 0 | 0.57 | 0.28 | 0.49 | 0.43 | 0.51 | 0.65 |
| 6 | 72453533 | rs1328703 | -0.368 | 0.345 | -2.157 | -1.018 | -0.001 | -0.004 | -0.004 | 0 | 0.59 | 0.32 | 0.5 | 0.43 | 0.53 | 0.66 |
| 6 | 72454176 | rs1932327 | -0.182 | 0.291 | -2.164 | -0.994 | -0.007 | -0.001 | -0.001 | 1.258 | 0.57 | 0.28 | 0.49 | 0.43 | 0.51 | 0.65 |
| 6 | 72455336 | rs17722725 | 1.545 | 0.445 | -2.039 | -1.024 | -0.01 | -0.011 | -0.011 | 0 | 0.05 | 0.04 | 0.18 | 0.09 | 0.09 | 0.1 |
| 6 | 72456393 | rs6935403 | 0.073 | -0.04 | -1.914 | -1.178 | 0.005 | 0.009 | 0.009 | 0 | 0.34 | 0.61 | 0.44 | 0.45 | 0.39 | 0.31 |
| 6 | 72457298 | rs9446462 | 0.246 | -0.146 | -1.418 | -1.005 | -0.005 | -0.018 | -0.018 | 0.013 | 0.18 | 0.16 | 0.08 | 0.05 | 0.12 | 0.1 |
| 6 | 72469672 | rs872209 | 0.07 | -0.055 | -0.834 | -0.943 | 0.003 | -0.075 | -0.075 | 3.66 | 0.86 | 0.66 | 0.93 | 0.97 | 0.88 | 0.84 |
| 6 | 72471656 | rs9360492 | 0.577 | -0.112 | -0.683 | -1.12 | 0.008 | -0.047 | -0.047 | 0 | 0.35 | 0.1 | 0.22 | 0.4 | 0.26 | 0.28 |
| 6 | 72473612 | rs4707941 | 0.609 | -0.148 | -0.488 | -1.139 | 0.008 | -0.044 | -0.044 | 0.305 | 0.34 | 0.1 | 0.23 | 0.4 | 0.26 | 0.28 |
| 6 | 72476816 | rs17722853 | 1.171 | 0.013 | -0.572 | -1.144 | -0.005 | 0.015 | 0.015 | 0.031 | 0.1 | 0.04 | 0.05 | 0.03 | 0.1 | 0.15 |
| 6 | 72477412 | rs17779253 | 0.35 | 0.102 | -0.036 | -1.17 | -0.031 | 0.028 | 0.028 | 0 | 0.33 | 0.23 | 0.39 | 0.24 | 0.45 | 0.19 |
| 6 | 72479263 | rs2881955 | 0.483 | -0.017 | -0.012 | -1.365 | -0.001 | -0.049 | -0.049 | 0.558 | 0.31 | 0.28 | 0.17 | 0.09 | 0.25 | 0.26 |
| 6 | 72481492 | rs10498879 | 1.673 | 0.121 | -0.237 | -1.109 | 0 | -0.089 | -0.089 | 0 | 0.16 | 0.04 | 0.12 | 0.34 | 0.12 | 0.18 |
| 6 | 72482458 | rs10455248 | 1.27 | 0.288 | -0.239 | -1.278 | -0.028 | 0.038 | 0.038 | 0 | 0.2 | 0.11 | 0.3 | 0.1 | 0.29 | 0.13 |
| 6 | 72488446 | rs2105067 | 0.806 | -0.118 | 0.287 | -1.216 | -0.021 | -0.117 | -0.117 | 0 | 0.25 | 0.4 | 0.13 | 0.06 | 0.15 | 0.1 |
| 6 | 72489033 | rs9442714 | 0.121 | -0.122 | 0.268 | -1.266 | -0.011 | -0.108 | -0.108 | 0 | 0.35 | 0.43 | 0.18 | 0.09 | 0.26 | 0.24 |
| 6 | 72489836 | rs9351853 | 0.812 | -0.122 | 0.241 | -1.266 | -0.014 | -0.087 | -0.087 | 2.096 | 0.24 | 0.32 | 0.13 | 0.06 | 0.15 | 0.1 |
| 6 | 72494265 | rs9446473 | -0.237 | -0.078 | 0.393 | -1.156 | 0.002 | 0.013 | 0.013 | 0 | 0.49 | 0.56 | 0.47 | 0.4 | 0.42 | 0.6 |
| 6 | 72504270 | rs9360497 | -0.176 | -0.081 | 0.398 | -1.155 | 0 | 0.004 | 0.004 | 0 | 0.49 | 0.52 | 0.46 | 0.41 | 0.42 | 0.6 |
| 6 | 72514270 | rs1410664 | 1.12 | -0.038 | 0.059 | -1.161 | -0.055 | -0.101 | -0.101 | 2.24 | 0.14 | 0.01 | 0.28 | 0.33 | 0.17 | 0.34 |
| 6 | 72515280 | rs10737976 | -0.564 | -0.123 | 0.194 | -1.2 | -0.007 | 0.013 | 0.013 | 0 | 0.65 | 0.65 | 0.59 | 0.74 | 0.54 | 0.76 |
| 6 | 72518644 | rs16881855 | 0.202 | -0.085 | 0.29 | -1.116 | 0.001 | 0.005 | 0.005 | 0 | 0.25 | 0.22 | 0.34 | 0.35 | 0.27 | 0.48 |
| 6 | 72525221 | rs9358982 | 1.007 | -0.13 | 0.232 | -1.02 | -0.002 | -0.064 | -0.064 | 0 | 0.22 | 0.27 | 0.12 | 0.06 | 0.16 | 0.1 |
| 6 | 72535853 | rs4235866 | 0.078 | -0.204 | -0.154 | -1 | 0.006 | 0.042 | 0.042 | 0 | 0.49 | 0.21 | 0.67 | 0.59 | 0.61 | 0.58 |
| 6 | 72556396 | rs12665263 | 0.42 | -0.247 | -0.061 | -1.03 | -0.006 | -0.002 | -0.002 | 0 | 0.39 | 0.5 | 0.26 | 0.4 | 0.28 | 0.28 |
| 6 | 72560565 | rs9293853 | 0.131 | -0.221 | -0.198 | -1.029 | -0.011 | 0.037 | 0.037 | 0 | 0.49 | 0.17 | 0.67 | 0.59 | 0.61 | 0.58 |
| 6 | 72560664 | rs17723887 | 1.472 | -0.234 | -0.204 | -1.037 | -0.024 | 0.045 | 0.045 | 0.054 | 0.1 | 0.01 | 0.05 | 0.01 | 0.1 | 0.14 |
| 6 | 72561091 | rs7766423 | 0.045 | -0.206 | -0.16 | -1.011 | 0.004 | 0.047 | 0.047 | 0 | 0.49 | 0.22 | 0.68 | 0.59 | 0.61 | 0.58 |
| 6 | 72567430 | rs9358983 | -0.436 | -0.168 | -0.008 | -1.001 | -0.02 | -0.112 | -0.112 | 3.625 | 0.7 | 0.5 | 0.84 | 0.96 | 0.79 | 0.7 |
| 6 | 72572521 | rs9351873 | 0.785 | -0.144 | 0.015 | -0.776 | -0.023 | -0.047 | -0.047 | 0 | 0.11 | 0.02 | 0.09 | 0.16 | 0.17 | 0.11 |
| 6 | 72580090 | rs4706494 | 0.662 | 0.076 | 0.284 | -0.446 | 0 | 0.041 | 0.041 | 0 | 0.22 | 0.34 | 0.17 | 0.38 | 0.18 | 0.11 |
| 6 | 72582893 | rs13210478 | 0.946 | 0.099 | 0.04 | -0.358 | -0.013 | -0.058 | -0.058 | 0 | 0.17 | 0.18 | 0.09 | 0.02 | 0.1 | 0.17 |
| 6 | 72590002 | rs4707954 | -0.278 | 0.512 | -0.834 | 0.067 | 0.016 | 0.011 | 0.011 | 0 | 0.54 | 0.41 | 0.56 | 0.65 | 0.63 | 0.34 |
| 6 | 72595779 | rs9358985 | 0.16 | 0.845 | -0.663 | 0.481 | -0.002 | -0.002 | -0.002 | 0.699 | 0.2 | 0.2 | 0.16 | 0.36 | 0.2 | 0.12 |

| | | | | | | | | | | | | | | | | |
|---|---|---|---|---|---|---|---|---|---|---|---|---|---|---|---|---|
| 6 | 72602564 | rs1328710 | 0.1 | 0.417 | -0.521 | 0.419 | 0.001 | 0.023 | 0.023 | 0 | 0.52 | 0.37 | 0.56 | 0.42 | 0.61 | 0.63 |
| 6 | 72603664 | rs2040055 | -0.034 | 0.317 | -0.472 | 0.626 | 0.006 | -0.011 | -0.011 | 0 | 0.09 | 0.08 | 0.17 | 0.19 | 0.06 | 0.07 |
| 6 | 72604896 | rs1753505 | -0.253 | 0.695 | 0.324 | 0.151 | 0.054 | 0.067 | 0.067 | 0 | 0.15 | 0.38 | 0.22 | 0.22 | 0.2 | 0.25 |
| 6 | 72607594 | rs7760432 | -0.52 | 0.856 | -0.517 | -0.378 | -0.044 | -0.033 | -0.033 | 0 | 0.35 | 0.23 | 0.36 | 0.6 | 0.46 | 0.34 |
| 6 | 72615220 | rs1192250 | -0.601 | 0.503 | -0.821 | -0.445 | 0.019 | 0.041 | 0.041 | 0.021 | 0.6 | 0.72 | 0.62 | 0.62 | 0.67 | 0.68 |
| 6 | 72617655 | rs1343227 | 0.755 | 0.436 | -0.699 | -0.46 | 0.017 | 0.056 | 0.056 | 0 | 0.34 | 0.15 | 0.3 | 0.18 | 0.33 | 0.37 |
| 6 | 72622271 | rs10484972 | 1.233 | 0.428 | -0.805 | -0.793 | -0.029 | 0.051 | 0.051 | 0 | 0.1 | 0.01 | 0.06 | 0.01 | 0.14 | 0.15 |
| 6 | 72624376 | rs2185876 | -1.873 | 0.353 | -0.856 | -0.864 | -0.011 | 0.121 | 0.121 | 0.247 | 0.78 | 0.96 | 0.88 | 0.99 | 0.85 | 0.82 |
| 6 | 72624622 | rs1936017 | 0.488 | 0.404 | -0.526 | -0.611 | -0.003 | 0.01 | 0.01 | 0 | 0.18 | 0.23 | 0.24 | 0.34 | 0.18 | 0.13 |
| 6 | 72627481 | rs12215297 | 0.418 | 0.534 | -0.463 | -0.575 | 0.005 | 0 | 0 | 0 | 0.18 | 0.14 | 0.13 | 0.16 | 0.13 | 0.29 |
| 6 | 72631747 | rs6931045 | -0.362 | 0.858 | -0.297 | -0.489 | 0.046 | 0.091 | 0.091 | 0 | 0.25 | 0.55 | 0.34 | 0.47 | 0.34 | 0.32 |
| 6 | 72636355 | rs17489346 | 0.599 | 0.596 | -0.765 | -0.679 | -0.032 | 0.062 | 0.062 | 0 | 0.11 | 0.01 | 0.07 | 0.01 | 0.15 | 0.17 |
| 6 | 72646391 | rs9341357 | -1.032 | 0.527 | -0.676 | -0.983 | 0 | 0.057 | 0.057 | 0 | 0.6 | 0.8 | 0.76 | 0.99 | 0.63 | 0.74 |
| 6 | 72651482 | rs6924133 | -0.017 | 0.572 | 0.08 | -0.051 | 0.025 | 0.163 | 0.163 | 0 | 0.12 | 0.36 | 0.22 | 0.39 | 0.13 | 0.11 |
| 6 | 72654432 | rs1936019 | -0.46 | 0.811 | -0.521 | 0.171 | -0.003 | -0.036 | -0.036 | 0 | 0.24 | 0.18 | 0.36 | 0.44 | 0.26 | 0.24 |
| 6 | 72662508 | rs10214880 | 0.53 | 0.717 | 0.049 | 0.496 | 0.001 | 0.034 | 0.034 | 0 | 0.08 | 0.06 | 0.07 | 0.25 | 0.1 | 0.09 |
| 6 | 72663571 | rs761177 | 0.73 | 0.913 | -0.186 | 0.43 | -0.013 | 0.026 | 0.026 | 0 | 0.1 | 0.02 | 0.07 | 0 | 0.1 | 0.06 |
| 6 | 72666352 | rs1936022 | 0.612 | 0.888 | -0.392 | -0.109 | -0.002 | -0.031 | -0.031 | 8.888 | 0.36 | 0.31 | 0.43 | 0.52 | 0.42 | 0.39 |
| 6 | 72666847 | rs13206801 | 1.302 | 1.132 | -0.654 | 0.114 | -0.031 | -0.038 | -0.038 | 0 | 0.07 | 0.01 | 0.15 | 0.29 | 0.12 | 0.09 |
| 6 | 72671565 | rs1482567 | -0.453 | 1.053 | -0.303 | -0.37 | -0.002 | -0.012 | -0.012 | 0.719 | 0.58 | 0.64 | 0.73 | 0.73 | 0.64 | 0.54 |
| 6 | 72674436 | rs9350450 | 0.229 | 1.027 | -0.416 | -0.403 | -0.006 | -0.011 | -0.011 | 0 | 0.45 | 0.44 | 0.47 | 0.54 | 0.51 | 0.43 |
| 6 | 72677981 | rs2348944 | -1.875 | 0.97 | -0.425 | -0.646 | 0 | -0.09 | -0.09 | 0.1 | 0.82 | 0.78 | 0.89 | 1 | 0.88 | 0.83 |
| 6 | 72686090 | rs12663081 | 0.696 | 0.923 | -0.657 | -0.484 | -0.026 | -0.057 | -0.057 | 0.063 | 0.1 | 0.02 | 0.21 | 0.16 | 0.1 | 0.07 |
| 6 | 72686211 | rs2348945 | -0.876 | 0.868 | -0.508 | -0.661 | -0.013 | -0.123 | -0.123 | 0 | 0.66 | 0.67 | 0.8 | 0.98 | 0.73 | 0.62 |
| 6 | 72686255 | rs7739862 | 0.489 | 0.987 | -0.586 | -0.704 | 0.001 | -0.019 | -0.019 | 0 | 0.46 | 0.58 | 0.52 | 0.83 | 0.53 | 0.48 |
| 6 | 72689212 | rs1564608 | -1.706 | 0.889 | -0.607 | -0.623 | -0.004 | -0.071 | -0.071 | 0 | 0.82 | 0.86 | 0.9 | 1 | 0.88 | 0.83 |
| 6 | 72692125 | rs7743295 | -1.706 | 0.889 | -0.669 | -0.623 | -0.003 | -0.086 | -0.086 | 0 | 0.82 | 0.8 | 0.89 | 1 | 0.88 | 0.83 |
| 6 | 72692282 | rs6453569 | -1.768 | 0.93 | -1.828 | -0.601 | -0.034 | 0.11 | 0.11 | 0 | 0.83 | 0.99 | 0.91 | 1 | 0.88 | 0.83 |
| 6 | 72693502 | rs9442725 | 0.539 | 1.08 | -1.066 | 0.301 | 0.02 | 0.112 | 0.112 | 0 | 0.09 | 0.15 | 0.09 | 0.28 | 0.11 | 0.09 |
| 6 | 72699513 | rs1384390 | 0.691 | 0.694 | -1.432 | -0.357 | -0.005 | 0.089 | 0.089 | 0 | 0.16 | 0.03 | 0.08 | 0.01 | 0.15 | 0.2 |
| 6 | 72706488 | rs2057022 | 0.657 | 0.655 | -1.413 | -0.348 | -0.008 | 0.079 | 0.079 | 0 | 0.14 | 0.02 | 0.08 | 0.01 | 0.14 | 0.2 |
| 6 | 72727772 | rs519470 | -0.576 | 0.665 | -1.762 | -0.437 | -0.014 | 0.012 | 0.012 | 0 | 0.43 | 0.21 | 0.48 | 0.33 | 0.49 | 0.52 |
| 6 | 72729528 | rs557365 | 0.914 | 0.697 | -1.845 | -0.444 | -0.057 | -0.133 | -0.133 | 0 | 0.21 | 0.03 | 0.35 | 0.32 | 0.25 | 0.3 |
| 6 | 72734047 | rs7761468 | 0.9 | 0.681 | -1.704 | -0.463 | 0.009 | 0.028 | 0.028 | 0 | 0.14 | 0.1 | 0.09 | 0.01 | 0.14 | 0.19 |
| 6 | 72749978 | rs481951 | -1.042 | 0.719 | -1.753 | -0.389 | -0.057 | -0.133 | -0.133 | 1.879 | 0.79 | 0.97 | 0.65 | 0.69 | 0.75 | 0.71 |
| 6 | 72758831 | rs9446555 | 0.156 | 0.764 | -1.6 | -0.517 | 0.004 | 0.017 | 0.017 | 0 | 0.21 | 0.14 | 0.12 | 0.01 | 0.22 | 0.21 |
| 6 | 72763339 | rs6919752 | 0.705 | 0.786 | -1.584 | -0.484 | 0.002 | -0.028 | -0.028 | 0 | 0.07 | 0.1 | 0.04 | 0 | 0.08 | 0.03 |
| 6 | 72769390 | rs17569784 | 0.932 | 0.768 | -1.636 | -0.501 | -0.001 | 0.074 | 0.074 | 0 | 0.14 | 0.04 | 0.08 | 0.01 | 0.14 | 0.19 |
| 6 | 72771098 | rs1012112 | 0.725 | 0.896 | -1.543 | -0.426 | 0 | -0.015 | -0.015 | 0 | 0.07 | 0.05 | 0.04 | 0 | 0.08 | 0.03 |
| 6 | 72801288 | rs1471640 | 0.002 | 0.63 | -1.947 | -0.52 | -0.021 | -0.072 | -0.072 | 0 | 0.38 | 0.07 | 0.44 | 0.32 | 0.4 | 0.5 |
| 6 | 72805811 | rs2496543 | 0.835 | 0.683 | -1.569 | -0.878 | -0.007 | 0.01 | 0.01 | 0 | 0.44 | 0.61 | 0.44 | 0.68 | 0.43 | 0.41 |
| 6 | 72806978 | rs1528505 | -0.247 | 0.66 | -1.754 | -0.9 | 0.01 | 0.03 | 0.03 | 0.01 | 0.47 | 0.2 | 0.49 | 0.32 | 0.49 | 0.54 |
| 6 | 72809986 | rs9293862 | -0.861 | 0.668 | -1.724 | -0.88 | -0.013 | 0.016 | 0.016 | 0 | 0.55 | 0.3 | 0.56 | 0.32 | 0.57 | 0.58 |
| 6 | 72812882 | rs1482574 | 0.11 | 0.824 | -1.58 | -0.616 | -0.03 | -0.058 | -0.058 | 0 | 0.29 | 0.1 | 0.4 | 0.31 | 0.34 | 0.34 |
| 6 | 72822523 | rs9341360 | -0.5 | 0.795 | -1.568 | -0.64 | -0.018 | -0.006 | -0.006 | 0 | 0.38 | 0.26 | 0.47 | 0.31 | 0.43 | 0.39 |
| 6 | 72825419 | rs9351889 | 0.092 | 0.775 | -1.786 | -0.583 | -0.051 | -0.047 | -0.047 | 0.097 | 0.29 | 0.05 | 0.39 | 0.27 | 0.33 | 0.33 |
| 6 | 72845455 | rs17781511 | 0.235 | 0.959 | -1.83 | -0.799 | -0.083 | 0.032 | 0.032 | 0 | 0.28 | 0.06 | 0.28 | 0.17 | 0.35 | 0.29 |
| 6 | 72851032 | rs2496551 | -0.319 | 0.917 | -1.429 | -0.857 | -0.005 | -0.038 | -0.038 | 0 | 0.92 | 0.88 | 0.97 | 1 | 0.93 | 0.97 |
| 6 | 72861276 | rs1969530 | 0.452 | 0.923 | -1.464 | -0.883 | -0.033 | 0.078 | 0.078 | 0 | 0.61 | 0.78 | 0.67 | 0.83 | 0.57 | 0.66 |
| 6 | 72875512 | rs1852705 | 0.947 | 0.596 | -1.439 | -1.246 | -0.002 | 0.053 | 0.053 | 0 | 0.44 | 0.66 | 0.59 | 0.82 | 0.42 | 0.47 |
| 6 | 72875873 | rs2349095 | 1.236 | 0.678 | -1.513 | -1.283 | -0.057 | -0.014 | -0.014 | 0 | 0.21 | 0.04 | 0.22 | 0.17 | 0.26 | 0.25 |
| 6 | 72882414 | rs2496495 | -0.461 | 0.668 | -1.495 | -1.285 | -0.034 | 0.04 | 0.04 | 0 | 0.7 | 0.84 | 0.71 | 0.83 | 0.65 | 0.7 |
| 6 | 72886012 | rs9442750 | 0.393 | 0.639 | -1.493 | -1.309 | 0.016 | 0.022 | 0.022 | 0 | 0.26 | 0.15 | 0.12 | 0.01 | 0.23 | 0.23 |
| 6 | 72888264 | rs9446589 | 0.674 | 0.613 | -1.527 | -1.311 | -0.003 | -0.027 | -0.027 | 0 | 0.09 | 0.1 | 0.03 | 0 | 0.07 | 0.03 |
| 6 | 72889472 | rs2249021 | -0.582 | 0.63 | -1.668 | -1.306 | -0.032 | 0.042 | 0.042 | 0 | 0.7 | 0.83 | 0.71 | 0.83 | 0.65 | 0.7 |
| 6 | 72890762 | rs11759297 | 1.085 | 0.304 | -2.059 | -1.696 | 0.029 | 0.127 | 0.127 | 0 | 0.23 | 0.04 | 0.13 | 0.01 | 0.18 | 0.21 |
| 6 | 72902964 | rs1830338 | 1.805 | 0.725 | -1.99 | -1.618 | -0.049 | -0.05 | -0.05 | 0.767 | 0.15 | 0.03 | 0.2 | 0.17 | 0.22 | 0.24 |
| 6 | 72903035 | rs17782317 | 1.231 | 0.689 | -1.972 | -1.601 | -0.003 | 0.146 | 0.146 | 0 | 0.25 | 0.04 | 0.11 | 0.01 | 0.24 | 0.22 |
| 6 | 72907197 | rs2249751 | 1.478 | 0.623 | -1.909 | -2.153 | 0.008 | 0.016 | 0.016 | 0 | 0.18 | 0.07 | 0.09 | 0 | 0.11 | 0.15 |
| 6 | 72908238 | rs1819062 | -0.855 | 0.707 | -1.966 | -2 | -0.019 | 0.027 | 0.027 | 0 | 0.73 | 0.83 | 0.71 | 0.83 | 0.68 | 0.7 |
| 6 | 72908366 | rs1852702 | -0.227 | 0.827 | -2.007 | -2.031 | -0.014 | 0.142 | 0.142 | 0 | 0.45 | 0.79 | 0.59 | 0.82 | 0.42 | 0.47 |
| 6 | 72911454 | rs1819061 | -0.739 | 0.945 | -1.826 | -1.8 | -0.02 | 0.028 | 0.028 | 0 | 0.73 | 0.83 | 0.71 | 0.83 | 0.68 | 0.7 |
| 6 | 72918620 | rs9351900 | 0.998 | 1.106 | -1.731 | -0.672 | -0.019 | -0.027 | -0.027 | 0.11 | 0.09 | 0.04 | 0.19 | 0.34 | 0.15 | 0.13 |
| 6 | 72927964 | rs2496540 | -0.437 | 0.719 | -1.311 | -1.159 | 0.006 | 0.027 | 0.027 | 0 | 0.42 | 0.63 | 0.57 | 0.81 | 0.37 | 0.46 |
| 6 | 72934518 | rs2807535 | -1.036 | 0.751 | -1.543 | -1.451 | -0.027 | 0.171 | 0.171 | 0 | 0.64 | 0.87 | 0.8 | 0.99 | 0.62 | 0.72 |
| 6 | 72941467 | rs1971561 | 1.764 | 0.694 | -1.659 | -1.566 | -0.057 | 0.16 | 0.16 | 0 | 0.24 | 0.03 | 0.13 | 0.01 | 0.29 | 0.22 |
| 6 | 72949319 | rs2697446 | -1 | 0.693 | -1.589 | -1.499 | -0.026 | 0.171 | 0.171 | 0 | 0.64 | 0.87 | 0.8 | 0.99 | 0.62 | 0.72 |
| 6 | 72974063 | rs12211702 | 1.061 | 0.589 | -1.59 | -1.266 | 0.007 | -0.02 | -0.02 | 0 | 0.1 | 0.05 | 0.15 | 0.16 | 0.11 | 0.14 |
| 6 | 72993334 | rs6905900 | -0.982 | 0.566 | -1.341 | -1.41 | -0.009 | 0.073 | 0.073 | 0 | 0.63 | 0.72 | 0.78 | 0.99 | 0.61 | 0.72 |

| | | | | | | | | | | | | | | | | |
|---|---|---|---|---|---|---|---|---|---|---|---|---|---|---|---|---|
| 6 | 72994975 | rs10455087 | 0.772 | 0.659 | -1.204 | -1.03 | 0.005 | -0.009 | -0.009 | 0.285 | 0.11 | 0.09 | 0.19 | 0.19 | 0.12 | 0.15 |
| 6 | 72998372 | rs988096 | -0.691 | 0.751 | -0.713 | -0.53 | 0.008 | 0.017 | 0.017 | 0 | 0.72 | 0.72 | 0.65 | 0.55 | 0.62 | 0.72 |
| 6 | 73029875 | rs9442769 | 0.865 | 0.838 | -1.194 | -1.033 | -0.007 | 0.052 | 0.052 | 0 | 0.34 | 0.23 | 0.27 | 0.17 | 0.39 | 0.37 |
| 6 | 73051884 | rs12202907 | 1.075 | 0.966 | -1.441 | -1.118 | -0.054 | 0.136 | 0.136 | 0 | 0.33 | 0.04 | 0.25 | 0.16 | 0.37 | 0.4 |
| 6 | 73063011 | rs1970373 | -2.063 | 1.044 | -1.578 | -0.946 | -0.072 | -0.061 | -0.061 | 13.148 | 0.86 | 0.98 | 0.74 | 0.82 | 0.76 | 0.77 |
| 6 | 73070497 | rs9446627 | 0.025 | 0.981 | -0.885 | -1.407 | -0.011 | 0.117 | 0.117 | 0 | 0.46 | 0.29 | 0.34 | 0.16 | 0.51 | 0.46 |
| 6 | 73086145 | rs12201232 | 0.028 | 0.987 | -1.117 | -1.393 | -0.062 | 0.02 | 0.02 | 1.412 | 0.18 | 0.01 | 0.21 | 0.14 | 0.22 | 0.18 |
| 6 | 73101361 | rs2807521 | 1.322 | 0.902 | -0.932 | -1.454 | 0.008 | 0.021 | 0.021 | 0 | 0.28 | 0.22 | 0.11 | 0.01 | 0.24 | 0.3 |
| 6 | 73109948 | rs7738055 | -0.112 | -0.039 | -0.443 | -1.447 | -0.002 | 0.011 | 0.011 | 0 | 0.49 | 0.46 | 0.35 | 0.16 | 0.5 | 0.49 |
| 6 | 73111454 | rs2815736 | -0.29 | -0.182 | -0.25 | -1.534 | 0.003 | 0.048 | 0.048 | 0 | 0.52 | 0.63 | 0.3 | 0.16 | 0.53 | 0.5 |
| 6 | 73111492 | rs9446638 | 0.724 | 0.014 | -0.618 | -1.288 | -0.001 | -0.001 | -0.001 | 0 | 0.09 | 0.08 | 0.05 | 0.15 | 0.13 | 0.14 |
| 6 | 73112380 | rs1055270 | 0.71 | 0.029 | -0.603 | -1.307 | 0.007 | 0.003 | 0.003 | 0 | 0.09 | 0.11 | 0.06 | 0.15 | 0.13 | 0.14 |
| 6 | 73115584 | rs11754392 | 1.266 | -0.118 | -0.906 | -1.356 | -0.058 | 0.177 | 0.177 | 0 | 0.26 | 0.02 | 0.11 | 0.01 | 0.26 | 0.3 |
| 6 | 73118850 | rs9446644 | -0.361 | -0.275 | -0.291 | -0.865 | -0.001 | -0.005 | -0.005 | 0.056 | 0.62 | 0.64 | 0.37 | 0.36 | 0.63 | 0.54 |
| 6 | 73120271 | rs6908357 | -0.229 | -0.095 | -0.103 | -0.969 | 0.001 | -0.071 | -0.071 | 2.16 | 0.11 | 0.36 | 0.08 | 0 | 0.1 | 0.02 |
| 6 | 73120320 | rs9442780 | 0.18 | 0.447 | -0.709 | -0.869 | -0.007 | -0.003 | -0.003 | 0 | 0.11 | 0.07 | 0.09 | 0 | 0.13 | 0.08 |
| 6 | 73120671 | rs763762 | -0.116 | 0.071 | -0.566 | -0.263 | -0.002 | -0.005 | -0.005 | 0 | 0.8 | 0.84 | 0.7 | 0.74 | 0.75 | 0.84 |
| 6 | 73125530 | rs2492970 | 0.583 | -0.936 | -0.26 | -0.447 | -0.025 | 0.097 | 0.097 | 0 | 0.87 | 0.54 | 0.89 | 0.84 | 0.94 | 0.92 |
| 6 | 73130696 | rs16882378 | 0.385 | -1.315 | -0.315 | -0.178 | 0 | 0.01 | 0.01 | 0 | 0.15 | 0.13 | 0.32 | 0.33 | 0.17 | 0.13 |
| 6 | 73131012 | rs9446649 | 1.335 | -1.422 | -0.672 | -1.345 | 0.029 | 0.117 | 0.117 | 0 | 0.33 | 0.13 | 0.1 | 0.01 | 0.24 | 0.27 |
| 6 | 73135441 | rs7740241 | 0.798 | -1.315 | -0.703 | -1.252 | 0.017 | 0.122 | 0.122 | 0 | 0.45 | 0.22 | 0.3 | 0.18 | 0.41 | 0.4 |
| 6 | 73143697 | rs2789588 | 1.526 | -1.209 | -0.79 | -1.269 | -0.035 | -0.049 | -0.049 | 0 | 0.2 | 0.14 | 0.22 | 0.39 | 0.31 | 0.33 |
| 6 | 73146011 | rs2486267 | -0.087 | -1.115 | -0.578 | -1.1 | -0.062 | -0.011 | -0.011 | 0 | 0.79 | 0.54 | 0.83 | 0.83 | 0.89 | 0.91 |
| 6 | 73147684 | rs2492964 | 0.633 | -1.077 | -0.586 | -1.084 | -0.025 | -0.043 | -0.043 | 0 | 0.72 | 0.49 | 0.77 | 0.83 | 0.8 | 0.84 |
| 6 | 73153314 | rs12190935 | 0.774 | -0.497 | -1.413 | -0.774 | -0.012 | -0.01 | -0.01 | 0 | 0.09 | 0.05 | 0.19 | 0.16 | 0.13 | 0.09 |
| 6 | 73154699 | rs9446655 | -0.164 | -0.589 | -1.481 | -0.913 | 0.003 | 0.018 | 0.018 | 0 | 0.22 | 0.36 | 0.39 | 0.26 | 0.26 | 0.17 |
| 6 | 73155197 | rs6915301 | 0.381 | -0.633 | -1.405 | -0.818 | -0.015 | 0.036 | 0.036 | 0 | 0.17 | 0.31 | 0.33 | 0.26 | 0.17 | 0.13 |
| 6 | 73156951 | rs12211844 | 0.997 | -0.408 | -1.556 | -0.669 | -0.042 | -0.032 | -0.032 | 0.018 | 0.08 | 0.01 | 0.19 | 0.15 | 0.12 | 0.08 |
| 6 | 73158346 | rs1339229 | -0.067 | -0.418 | -1.576 | -1.021 | 0.002 | -0.011 | -0.011 | 0.073 | 0.55 | 0.48 | 0.49 | 0.28 | 0.5 | 0.48 |
| 6 | 73160811 | rs1361310 | -0.209 | -0.315 | -1.522 | -1.034 | 0.017 | -0.021 | -0.021 | 0 | 0.61 | 0.48 | 0.51 | 0.28 | 0.52 | 0.49 |
| 6 | 73167827 | rs2781567 | 0.223 | -0.111 | -1.03 | -1.732 | 0.013 | 0.116 | 0.116 | 0 | 0.15 | 0.35 | 0.25 | 0.31 | 0.12 | 0.12 |
| 6 | 73168479 | rs12525276 | 0.364 | -0.177 | -1.507 | -1.86 | -0.03 | 0.158 | 0.158 | 0 | 0.32 | 0.09 | 0.15 | 0.01 | 0.29 | 0.29 |
| 6 | 73169177 | rs985564 | 0.236 | 0.203 | -1.034 | -1.168 | 0.001 | 0.022 | 0.022 | 0 | 0.48 | 0.54 | 0.41 | 0.32 | 0.42 | 0.42 |
| 6 | 73171594 | rs1572208 | 0.356 | 0.432 | -0.916 | -0.818 | -0.001 | -0.004 | -0.004 | 0 | 0.5 | 0.45 | 0.41 | 0.34 | 0.4 | 0.39 |
| 6 | 73171881 | rs1572209 | -0.05 | 0.799 | -1.477 | -0.402 | -0.069 | -0.009 | -0.009 | 0 | 0.7 | 0.86 | 0.57 | 0.35 | 0.58 | 0.47 |
| 6 | 73172623 | rs2781562 | -0.452 | 0.009 | -0.877 | -0.97 | -0.033 | -0.078 | -0.078 | 0 | 0.36 | 0.45 | 0.2 | 0.16 | 0.22 | 0.2 |
| 6 | 73177863 | rs2781564 | -0.757 | 0.624 | -1.641 | -0.347 | -0.034 | -0.05 | -0.05 | 0.345 | 0.53 | 0.92 | 0.57 | 0.57 | 0.59 | 0.56 |
| 6 | 73188091 | rs2815717 | -0.554 | 0.717 | -0.674 | -0.779 | -0.034 | 0.074 | 0.074 | 0 | 0.26 | 0.45 | 0.27 | 0.39 | 0.22 | 0.32 |
| 6 | 73190006 | rs7760670 | -0.014 | 1.304 | -0.131 | -1.013 | -0.029 | 0.052 | 0.052 | 0 | 0.21 | 0.09 | 0.16 | 0.01 | 0.29 | 0.19 |
| 6 | 73204913 | rs9360565 | -1.361 | 0.986 | -0.299 | -1.242 | 0.005 | -0.035 | -0.035 | 0.205 | 0.41 | 0.54 | 0.35 | 0.19 | 0.48 | 0.3 |
| 6 | 73217364 | rs9350463 | -1.222 | 0.993 | -0.235 | -1.27 | -0.001 | 0 | 0 | 0 | 0.2 | 0.15 | 0.16 | 0.17 | 0.18 | 0.12 |
| 6 | 73228407 | rs9446692 | 0.211 | 1.187 | -0.086 | -1.446 | 0.032 | -0.062 | -0.062 | 0.037 | 0.12 | 0.25 | 0.1 | 0.01 | 0.17 | 0.1 |
| 6 | 73234147 | rs1328854 | -0.056 | 1.084 | 0.352 | -1.431 | 0.01 | -0.03 | -0.03 | 0 | 0.06 | 0.18 | 0.06 | 0.01 | 0.05 | 0.04 |
| 6 | 73234224 | rs1328853 | -1.328 | 1.043 | 0.433 | -1.48 | 0.025 | -0.067 | -0.067 | 0 | 0.11 | 0.43 | 0.11 | 0.01 | 0.12 | 0.09 |
| 6 | 73234827 | rs9442812 | -0.852 | 1.027 | 0.208 | -1.541 | 0.005 | -0.012 | -0.012 | 0 | 0.18 | 0.1 | 0.06 | 0.01 | 0.12 | 0.08 |
| 6 | 73237751 | rs16882514 | 0.306 | 1.135 | 0.338 | -1.32 | -0.001 | -0.013 | -0.013 | 0 | 0.05 | 0.05 | 0.06 | 0.01 | 0.05 | 0.04 |
| 6 | 73258102 | rs9293890 | -2.412 | 1.976 | 0.048 | -1.234 | 0.004 | -0.026 | -0.026 | 0 | 0.52 | 0.97 | 0.59 | 0.62 | 0.62 | 0.61 |
| 6 | 73267169 | rs1202116 | 1.76 | 2.314 | 0.457 | -1.426 | 0.015 | -0.118 | -0.118 | 1.035 | 0.64 | 0.44 | 0.73 | 0.97 | 0.58 | 0.73 |
| 6 | 73270104 | rs10943033 | -0.92 | 2.279 | 0.303 | -1.279 | -0.006 | -0.004 | -0.004 | 0 | 0.1 | 0.08 | 0.06 | 0.01 | 0.13 | 0.07 |
| 6 | 73289040 | rs12205054 | -0.542 | 2.335 | 0.189 | -1.261 | -0.048 | 0.043 | 0.043 | 0 | 0.08 | 0.01 | 0.06 | 0.01 | 0.14 | 0.08 |
| 6 | 73303078 | rs997542 | -1.473 | 2.357 | 0.322 | -1.299 | 0.025 | -0.071 | -0.071 | 0 | 0.17 | 0.24 | 0.13 | 0.02 | 0.18 | 0.12 |
| 6 | 73306528 | rs1202100 | 1.829 | 2.339 | 0.286 | -1.327 | 0.008 | -0.056 | -0.056 | 2.609 | 0.79 | 0.77 | 0.83 | 0.98 | 0.73 | 0.83 |
| 6 | 73312081 | rs1202109 | 1.923 | 2.332 | 0.504 | -1.456 | 0.001 | -0.07 | -0.07 | 1.039 | 0.64 | 0.38 | 0.73 | 0.97 | 0.59 | 0.73 |
| 6 | 73315247 | rs1147572 | 0.69 | 2.322 | 0.539 | -0.492 | -0.02 | -0.064 | -0.064 | 0 | 0.88 | 0.97 | 0.74 | 0.51 | 0.78 | 0.71 |
| 6 | 73316732 | rs7749187 | -1.315 | 1.677 | 0.896 | -1.36 | 0.051 | 0.037 | 0.037 | 0.884 | 0.05 | 0.39 | 0.03 | 0 | 0.01 | 0 |
| 6 | 73318299 | rs12189795 | 0.368 | 1.857 | 0.237 | -1.309 | -0.051 | 0.055 | 0.055 | 0 | 0.1 | 0.01 | 0.07 | 0.01 | 0.13 | 0.1 |
| 6 | 73322054 | rs10943040 | -1.512 | 1.84 | 0.505 | -1.235 | 0.032 | -0.129 | -0.129 | 0 | 0.27 | 0.51 | 0.19 | 0.05 | 0.28 | 0.21 |
| 6 | 73325585 | rs12524568 | -1.255 | 1.915 | -0.005 | -0.855 | -0.031 | 0.015 | 0.015 | 0 | 0.08 | 0.04 | 0.06 | 0.03 | 0.15 | 0.07 |
| 6 | 73336288 | rs17692121 | 1.022 | 1.074 | 0.082 | -1.171 | -0.022 | 0.026 | 0.026 | 0 | 0.47 | 0.21 | 0.32 | 0.45 | 0.5 | 0.33 |
| 6 | 73343796 | rs6913237 | 1.02 | 1.074 | 0.078 | -1.154 | -0.021 | 0.023 | 0.023 | 0 | 0.47 | 0.21 | 0.32 | 0.45 | 0.5 | 0.33 |
| 6 | 73351870 | rs12200468 | 0.131 | 0.629 | 0.118 | -1.072 | -0.026 | 0.055 | 0.055 | 0 | 0.15 | 0.05 | 0.07 | 0.01 | 0.14 | 0.14 |
| 6 | 73360633 | rs9442831 | -0.204 | 0.289 | 0.24 | -1.057 | -0.067 | 0.162 | 0.162 | 0 | 0.2 | 0.45 | 0.43 | 0.53 | 0.12 | 0.43 |
| 6 | 73365274 | rs4706509 | 0.556 | 0.505 | 0.045 | -1.179 | 0 | -0.001 | -0.001 | 0 | 0.1 | 0.06 | 0.11 | 0.22 | 0.16 | 0.06 |
| 6 | 73370429 | rs9360584 | -0.126 | 0.267 | 0.113 | -1.146 | -0.061 | 0.146 | 0.146 | 0 | 0.2 | 0.43 | 0.43 | 0.54 | 0.12 | 0.4 |
| 6 | 73373640 | rs10943050 | -0.57 | 0.323 | -0.237 | -1.32 | 0.002 | 0.088 | 0.088 | 0 | 0.25 | 0.11 | 0.11 | 0.01 | 0.2 | 0.19 |
| 6 | 73373916 | rs10943051 | 0.598 | 0.313 | -0.25 | -1.331 | -0.024 | 0.066 | 0.066 | 0 | 0.15 | 0.05 | 0.07 | 0.01 | 0.13 | 0.17 |
| 6 | 73376868 | rs12194741 | -0.686 | 0.332 | -0.238 | -1.306 | 0.008 | 0.101 | 0.101 | 0 | 0.27 | 0.11 | 0.11 | 0.01 | 0.2 | 0.21 |
| 6 | 73377551 | rs12201380 | -0.658 | 0.333 | -0.238 | -1.334 | 0.008 | 0.098 | 0.098 | 0 | 0.27 | 0.11 | 0.11 | 0.01 | 0.2 | 0.21 |
| 6 | 73388290 | rs768790 | -0.147 | 0.354 | -0.217 | -1.277 | 0.011 | 0.02 | 0.02 | 0 | 0.11 | 0.07 | 0.06 | 0.02 | 0.07 | 0.05 |

| | | | | | | | | | | | | | | | | |
|---|---|---|---|---|---|---|---|---|---|---|---|---|---|---|---|---|
| 6 | 73396263 | rs9442834 | -0.751 | 0.345 | -0.037 | -1.267 | -0.022 | 0.02 | 0.02 | 0 | 0.35 | 0.47 | 0.5 | 0.54 | 0.25 | 0.56 |
| 6 | 73400219 | rs9360588 | -1.56 | 0.299 | -0.134 | -1.311 | 0.091 | 0.035 | 0.035 | 0 | 0.49 | 0.88 | 0.58 | 0.56 | 0.33 | 0.61 |
| 6 | 73405265 | rs17729195 | 0.462 | 0.387 | -0.11 | -1.308 | -0.018 | 0.063 | 0.063 | 0 | 0.17 | 0.07 | 0.08 | 0.01 | 0.13 | 0.19 |
| 6 | 73412888 | rs6453597 | -0.236 | 0.334 | 0.161 | -1.271 | -0.019 | -0.066 | -0.066 | 0 | 0.11 | 0.29 | 0.06 | 0 | 0.07 | 0.01 |
| 6 | 73419514 | rs6904964 | -1.253 | 0.555 | -0.345 | -1.286 | 0.029 | -0.003 | -0.003 | 2.551 | 0.25 | 0.5 | 0.19 | 0.23 | 0.23 | 0.1 |
| 6 | 73430216 | rs6906977 | -1.34 | -0.284 | -0.779 | -1.057 | 0.091 | 0.048 | 0.048 | 0 | 0.51 | 0.91 | 0.59 | 0.58 | 0.33 | 0.6 |
| 6 | 73437355 | rs12205374 | 0.352 | -0.779 | -0.733 | -1.041 | 0.002 | 0.058 | 0.058 | 0 | 0.16 | 0.07 | 0.07 | 0.01 | 0.11 | 0.17 |
| 6 | 73438218 | rs9442837 | -0.83 | -0.836 | -0.665 | -1.032 | 0.047 | -0.02 | -0.02 | 3.164 | 0.33 | 0.76 | 0.51 | 0.57 | 0.19 | 0.42 |
| 6 | 73441990 | rs12211580 | 0.214 | -0.581 | -0.698 | -0.925 | 0.007 | 0.025 | 0.025 | 0 | 0.17 | 0.11 | 0.07 | 0.01 | 0.11 | 0.17 |
| 6 | 73442239 | rs9351938 | -0.71 | -0.471 | -0.011 | -0.294 | -0.011 | 0.008 | 0.008 | 0 | 0.27 | 0.33 | 0.46 | 0.55 | 0.18 | 0.41 |
| 6 | 73448086 | rs4706511 | -0.731 | -0.805 | -0.872 | -0.521 | 0.094 | 0 | 0 | 0 | 0.43 | 0.77 | 0.54 | 0.54 | 0.27 | 0.45 |
| 6 | 73449742 | rs4235869 | -0.539 | -0.008 | -0.687 | -0.904 | -0.025 | 0.014 | 0.014 | 0.111 | 0.26 | 0.12 | 0.5 | 0.76 | 0.33 | 0.37 |
| 6 | 73459390 | rs2840794 | 1.018 | -0.683 | 0.41 | -1.128 | -0.029 | -0.099 | -0.099 | 2.468 | 0.76 | 0.64 | 0.88 | 0.95 | 0.85 | 0.78 |
| 6 | 73459762 | rs9446746 | -0.482 | -0.741 | 0.508 | -1.166 | 0.001 | 0.018 | 0.018 | 0 | 0.17 | 0.08 | 0.07 | 0.02 | 0.14 | 0.16 |
| 6 | 73463894 | rs7765507 | -0.803 | -0.826 | 0.628 | -1.198 | 0.046 | 0.043 | 0.043 | 1.139 | 0.05 | 0.44 | 0.04 | 0 | 0 | 0 |
| 6 | 73470432 | rs9293911 | -1.191 | -0.784 | 0.139 | -1.187 | -0.014 | -0.071 | -0.071 | 0.223 | 0.24 | 0.68 | 0.15 | 0.02 | 0.14 | 0.16 |
| 6 | 73471683 | rs2982751 | 1.278 | -0.723 | -0.197 | -1.153 | 0.026 | 0.032 | 0.032 | 0.142 | 0.71 | 0.19 | 0.84 | 0.95 | 0.84 | 0.79 |
| 6 | 73475153 | rs7758723 | -1.243 | -0.81 | 0.398 | -0.954 | -0.071 | -0.079 | -0.079 | 1.206 | 0.19 | 0.57 | 0.09 | 0.07 | 0.08 | 0.11 |
| 6 | 73478625 | rs7775641 | 0.992 | -0.074 | -0.049 | -0.167 | -0.007 | -0.019 | -0.019 | 0 | 0.64 | 0.62 | 0.64 | 0.56 | 0.58 | 0.67 |
| 6 | 73480021 | rs12213639 | 1.317 | -0.468 | -0.982 | -0.625 | -0.027 | 0.021 | 0.021 | 0 | 0.23 | 0.1 | 0.17 | 0.21 | 0.25 | 0.39 |
| 6 | 73490084 | rs16882872 | 0.101 | -0.3 | -0.779 | -0.364 | 0.003 | 0.003 | 0.003 | 0 | 0.15 | 0.18 | 0.04 | 0.07 | 0.08 | 0.13 |
| 6 | 73495941 | rs9360599 | -0.527 | -0.069 | -0.964 | -0.203 | -0.092 | -0.111 | -0.111 | 0.279 | 0.81 | 0.97 | 0.72 | 0.63 | 0.66 | 0.8 |
| 6 | 73506104 | rs9341388 | 0.402 | -0.047 | -1.08 | -0.014 | -0.023 | -0.02 | -0.02 | 0 | 0.06 | 0.01 | 0.22 | 0.28 | 0.09 | 0.07 |
| 6 | 73514264 | rs6453606 | -0.085 | -0.028 | -1.164 | -0.017 | -0.115 | -0.108 | -0.108 | 2.372 | 0.7 | 0.94 | 0.68 | 0.61 | 0.58 | 0.72 |
| 6 | 73524224 | rs4583938 | 0.646 | -0.002 | -0.109 | -0.397 | 0.002 | 0.004 | 0.004 | 0 | 0.56 | 0.45 | 0.6 | 0.53 | 0.47 | 0.66 |
| 6 | 73525349 | rs9351947 | 0.945 | -0.456 | -0.459 | -0.636 | 0.029 | -0.001 | -0.001 | 0 | 0.32 | 0.17 | 0.4 | 0.33 | 0.27 | 0.27 |
| 6 | 73527070 | rs9446761 | 0.268 | -0.343 | -0.108 | -0.673 | 0.061 | -0.024 | -0.024 | 0 | 0.1 | 0.51 | 0.09 | 0.01 | 0.08 | 0.01 |
| 6 | 73530851 | rs10498884 | 0.528 | -0.417 | -0.72 | -0.662 | 0.019 | 0.025 | 0.025 | 0 | 0.12 | 0.05 | 0.03 | 0.02 | 0.06 | 0.08 |
| 6 | 73531612 | rs17742423 | 1.42 | -0.218 | -0.839 | -0.545 | -0.024 | 0.021 | 0.021 | 0 | 0.22 | 0.06 | 0.16 | 0.19 | 0.21 | 0.38 |
| 6 | 73531644 | rs10943068 | 0.675 | -0.327 | -0.531 | -0.356 | 0.022 | -0.001 | -0.001 | 0 | 0.32 | 0.22 | 0.41 | 0.33 | 0.27 | 0.26 |
| 6 | 73533851 | rs6928101 | 0.312 | -0.142 | -0.345 | -0.151 | 0 | 0 | 0 | 0 | 0.53 | 0.46 | 0.55 | 0.51 | 0.49 | 0.65 |
| 6 | 73535037 | rs2840803 | 0.945 | -0.757 | -0.569 | 0.014 | -0.011 | 0.007 | 0.007 | 0.155 | 0.66 | 0.42 | 0.57 | 0.6 | 0.79 | 0.78 |
| 6 | 73535759 | rs1935515 | 0.887 | -0.549 | -0.093 | -0.125 | -0.016 | -0.006 | -0.006 | 0 | 0.66 | 0.5 | 0.59 | 0.69 | 0.74 | 0.78 |
| 6 | 73535931 | rs9442858 | 0.877 | -0.327 | -0.564 | -0.618 | 0.004 | 0.04 | 0.04 | 0.444 | 0.47 | 0.65 | 0.29 | 0.29 | 0.44 | 0.54 |
| 6 | 73536997 | rs3005767 | 0.975 | 0.175 | -0.64 | -0.011 | -0.081 | -0.036 | -0.036 | 0 | 0.21 | 0.02 | 0.3 | 0.26 | 0.3 | 0.25 |
| 6 | 73544500 | rs6915355 | 1.422 | 0.088 | -0.681 | -0.162 | 0.024 | 0 | 0 | 0.179 | 0.32 | 0.18 | 0.15 | 0.28 | 0.21 | 0.47 |
| 6 | 73544735 | rs4708000 | -0.033 | 0.886 | -0.081 | 0.323 | 0.018 | 0.034 | 0.034 | 0.256 | 0.31 | 0.42 | 0.44 | 0.45 | 0.37 | 0.21 |
| 6 | 73546750 | rs6927484 | 0.368 | 0.906 | -0.089 | -0.304 | 0.024 | -0.098 | -0.098 | 0.023 | 0.24 | 0.46 | 0.16 | 0.03 | 0.26 | 0.13 |
| 6 | 73561338 | rs9342975 | -0.024 | 1.107 | -0.178 | 0.579 | 0.003 | -0.002 | -0.002 | 0 | 0.8 | 0.8 | 0.69 | 0.77 | 0.71 | 0.74 |
| 6 | 73571223 | rs2840796 | 0.989 | 0.951 | -0.002 | 0.784 | -0.008 | 0.037 | 0.037 | 0 | 0.72 | 0.46 | 0.59 | 0.47 | 0.76 | 0.84 |
| 6 | 73572196 | rs1316976 | 1.57 | 0.895 | -0.053 | 0.275 | -0.003 | 0.104 | 0.104 | 0 | 0.46 | 0.32 | 0.31 | 0.21 | 0.5 | 0.56 |
| 6 | 73573279 | rs9351948 | 1.55 | 0.88 | 0.06 | 0.27 | -0.006 | 0.087 | 0.087 | 0 | 0.47 | 0.38 | 0.31 | 0.21 | 0.5 | 0.56 |
| 6 | 73575403 | rs2840795 | 0.342 | 0.841 | 0.198 | 0.762 | 0.004 | 0 | 0 | 0 | 0.25 | 0.2 | 0.28 | 0.25 | 0.26 | 0.28 |
| 6 | 73575709 | rs9342979 | 1.502 | 0.708 | 0.03 | 0.523 | -0.022 | 0.105 | 0.105 | 0 | 0.43 | 0.26 | 0.31 | 0.23 | 0.49 | 0.54 |
| 6 | 73578415 | rs4706515 | 1.407 | 1.183 | 0.059 | 0.57 | -0.015 | 0.136 | 0.136 | 0 | 0.44 | 0.24 | 0.32 | 0.21 | 0.43 | 0.51 |
| 6 | 73592597 | rs6930157 | 0.139 | 1.18 | 0.187 | 0.944 | 0 | 0.001 | 0.001 | 0 | 0.26 | 0.19 | 0.33 | 0.25 | 0.32 | 0.22 |
| 6 | 73596433 | rs9283796 | -0.732 | 1.082 | 0.393 | 0.931 | 0.011 | 0.05 | 0.05 | 0 | 0.25 | 0.33 | 0.3 | 0.47 | 0.23 | 0.16 |
| 6 | 73606783 | rs7775087 | -1.808 | 1.062 | 0.127 | 0.766 | -0.008 | 0.117 | 0.117 | 0 | 0.57 | 0.74 | 0.69 | 0.79 | 0.58 | 0.5 |
| 6 | 73616236 | rs9360616 | 0.337 | 1.034 | 0.043 | 0.836 | -0.035 | -0.033 | -0.033 | 0 | 0.24 | 0.06 | 0.33 | 0.27 | 0.33 | 0.23 |
| 6 | 73622234 | rs10498886 | 0.562 | 1.092 | -0.016 | 0.892 | -0.023 | -0.053 | -0.053 | 0 | 0.1 | 0.01 | 0.15 | 0.31 | 0.1 | 0.06 |
| 6 | 73626639 | rs951762 | -1.848 | 0.949 | 0.174 | 0.493 | -0.007 | 0.118 | 0.118 | 0 | 0.57 | 0.74 | 0.58 | 0.77 | 0.59 | 0.51 |
| 6 | 73626650 | rs9342986 | 0.201 | 1.053 | 0.317 | 0.5 | 0.006 | -0.001 | -0.001 | 0 | 0.26 | 0.18 | 0.23 | 0.25 | 0.32 | 0.23 |
| 6 | 73627045 | rs9446790 | -0.884 | 1.007 | 0.515 | 0.505 | 0.024 | 0.053 | 0.053 | 0 | 0.26 | 0.37 | 0.33 | 0.46 | 0.27 | 0.17 |
| 6 | 73635005 | rs1316784 | -0.722 | 0.659 | 0.346 | 0.846 | -0.025 | -0.008 | -0.008 | 0 | 0.32 | 0.43 | 0.37 | 0.32 | 0.32 | 0.34 |
| 6 | 73641100 | rs2882328 | -0.136 | 0.872 | -0.39 | 0.925 | 0.005 | -0.007 | -0.007 | 0 | 0.25 | 0.15 | 0.34 | 0.26 | 0.31 | 0.24 |
| 6 | 73642777 | rs6924280 | -0.125 | 0.877 | -0.377 | 0.941 | 0.005 | -0.006 | -0.006 | 0 | 0.25 | 0.17 | 0.34 | 0.27 | 0.31 | 0.24 |
| 6 | 73649330 | rs1317537 | 0.278 | 0.834 | -0.654 | 0.694 | -0.01 | 0.016 | 0.016 | 0 | 0.59 | 0.66 | 0.56 | 0.66 | 0.49 | 0.52 |
| 6 | 73650662 | rs2153830 | 0.829 | 0.291 | -0.053 | 2 | -0.012 | 0.009 | 0.009 | 1.845 | 0.64 | 0.09 | 0.78 | 0.71 | 0.74 | 0.69 |
| 6 | 73652005 | rs11758798 | -0.704 | -0.327 | 0.714 | 2.222 | 0.012 | -0.007 | -0.007 | 0 | 0.36 | 0.47 | 0.23 | 0.3 | 0.33 | 0.3 |
| 6 | 73652443 | rs6453613 | -1.116 | -1.543 | 1.709 | 3.36 | 0.003 | 0.003 | 0.003 | 0 | 0.37 | 0.67 | 0.33 | 0.34 | 0.29 | 0.28 |
| 6 | 73653891 | rs12202465 | -1.028 | -1.315 | 1.951 | 3.41 | 0 | 0.013 | 0.013 | 0 | 0.09 | 0.09 | 0.11 | 0.23 | 0.09 | 0.03 |
| 6 | 73660234 | rs885247 | -1.252 | -1.308 | 2.083 | 3.692 | 0.002 | 0.015 | 0.015 | 0 | 0.07 | 0.08 | 0.15 | 0.27 | 0.15 | 0.19 |
| 6 | 73660344 | rs1891395 | 0.842 | -1.321 | 2.246 | 3.817 | 0.019 | 0.116 | 0.116 | 0 | 0.92 | 0.78 | 0.79 | 0.78 | 0.93 | 0.87 |
| 6 | 73663745 | rs13206617 | -1.288 | -1.329 | 2.251 | 3.791 | -0.003 | -0.004 | -0.004 | 0.254 | 0.12 | 0.11 | 0.17 | 0.27 | 0.18 | 0.2 |
| 6 | 73664251 | rs1539336 | 0.869 | -1.289 | 2.155 | 3.609 | 0.023 | 0.126 | 0.126 | 0 | 0.92 | 0.75 | 0.79 | 0.78 | 0.93 | 0.87 |
| 6 | 73672548 | rs12203617 | -0.758 | -1.271 | 2.047 | 3.357 | 0.005 | 0.037 | 0.037 | 0 | 0.07 | 0.11 | 0.11 | 0.22 | 0.08 | 0.03 |
| 6 | 73681783 | rs9360623 | 0.79 | -1.277 | 2.139 | 3.244 | 0.031 | 0.117 | 0.117 | 0 | 0.92 | 0.77 | 0.8 | 0.81 | 0.93 | 0.87 |
| 6 | 73687752 | rs7747839 | -1.637 | -1.545 | 2.357 | 3.23 | -0.003 | 0.068 | 0.068 | 0.01 | 0.16 | 0.47 | 0.24 | 0.21 | 0.09 | 0.14 |
| 6 | 73697858 | rs9446807 | -0.228 | -1.506 | 2.117 | 2.997 | 0.01 | -0.028 | -0.028 | 0.515 | 0.08 | 0.26 | 0.03 | 0.02 | 0.04 | 0.01 |

| | | | | | | | | | | | | | | | | |
|---|---|---|---|---|---|---|---|---|---|---|---|---|---|---|---|---|
| 6 | 73717568 | rs7747368 | -1.961 | -1.498 | 1.942 | 3.235 | -0.003 | 0.037 | 0.037 | 0 | 0.24 | 0.65 | 0.36 | 0.44 | 0.18 | 0.21 |
| 6 | 73719982 | rs10943075 | -1.337 | -1.36 | 1.6 | 3.372 | 0.006 | 0.016 | 0.016 | 0 | 0.09 | 0.1 | 0.15 | 0.28 | 0.15 | 0.21 |
| 6 | 73727741 | rs10943076 | 1.828 | -1.54 | 1.384 | 3.289 | -0.08 | -0.028 | -0.028 | 19.185 | 0.45 | 0.02 | 0.38 | 0.26 | 0.56 | 0.51 |
| 6 | 73730617 | rs4706522 | 1.413 | -0.585 | 2.883 | 4.336 | -0.01 | -0.002 | -0.002 | 0 | 0.8 | 0.46 | 0.84 | 0.8 | 0.87 | 0.93 |
| 6 | 73733188 | rs11963228 | 2.141 | -0.589 | 2.203 | 4.182 | 0.016 | -0.038 | -0.038 | 0 | 0.69 | 0.18 | 0.62 | 0.56 | 0.81 | 0.77 |
| 6 | 73743509 | rs6453619 | -2.323 | -0.499 | 2.872 | 4.165 | -0.01 | 0.209 | 0.209 | 0 | 0.18 | 0.38 | 0.36 | 0.57 | 0.18 | 0.31 |
| 6 | 73746516 | rs9446827 | -2.968 | -0.779 | 2.91 | 3.485 | -0.04 | 0.082 | 0.082 | 0.708 | 0.21 | 0.6 | 0.3 | 0.48 | 0.09 | 0.16 |
| 6 | 73749861 | rs9351963 | -2.112 | -0.454 | 3.297 | 3.691 | -0.012 | 0.293 | 0.293 | 0 | 0.13 | 0.46 | 0.25 | 0.34 | 0.05 | 0.12 |
| 6 | 73749874 | rs16883236 | -2.156 | -0.248 | 3.081 | 2.296 | -0.008 | 0.008 | 0.008 | 0.056 | 0.06 | 0.08 | 0.05 | 0.1 | 0.03 | 0.02 |
| 6 | 73789370 | rs9446833 | -2.361 | -0.135 | 2.987 | 2.112 | -0.028 | -0.03 | -0.03 | 0.121 | 0.13 | 0.66 | 0.07 | 0.06 | 0.05 | 0.08 |
| 6 | 73794270 | rs12111515 | -1.704 | -0.128 | 3.069 | 2.105 | 0.025 | -0.022 | -0.022 | 0 | 0.1 | 0.44 | 0.05 | 0.02 | 0.05 | 0.07 |
| 6 | 73798609 | rs1539346 | 2.382 | -0.209 | 2.235 | 2.146 | -0.013 | -0.003 | -0.003 | 0 | 0.83 | 0.28 | 0.86 | 0.81 | 0.89 | 0.89 |
| 6 | 73811074 | rs6933440 | 2.763 | -0.115 | 2.257 | 2.262 | -0.038 | 0.069 | 0.069 | 6.521 | 0.88 | 0.28 | 0.72 | 0.75 | 0.93 | 0.94 |
| 6 | 73822453 | rs9446844 | 2.705 | -0.255 | 3.101 | 2.411 | -0.042 | 0.09 | 0.09 | 6.072 | 0.88 | 0.32 | 0.72 | 0.75 | 0.93 | 0.94 |
| 6 | 73845875 | rs2796051 | -2.176 | -0.195 | 2.509 | 2.469 | 0.015 | 0.071 | 0.071 | 0.208 | 0.07 | 0.37 | 0.06 | 0.05 | 0.04 | 0.04 |
| 6 | 73851242 | rs2797089 | -1.6 | -0.074 | 2.066 | 2.324 | -0.014 | 0.012 | 0.012 | 0.387 | 0.05 | 0.12 | 0.03 | 0.05 | 0.04 | 0.01 |
| 6 | 73853568 | rs6911751 | -2.251 | -0.101 | 1.891 | 2.429 | -0.01 | 0.327 | 0.327 | 3.397 | 0.1 | 0.43 | 0.26 | 0.25 | 0.06 | 0.09 |
| 6 | 73855954 | rs7764373 | -1.872 | -0.155 | 1.323 | 2.079 | -0.024 | 0.129 | 0.129 | 7.597 | 0.07 | 0.69 | 0.24 | 0.13 | 0.01 | 0.04 |
| 6 | 73861530 | rs947746 | 0.637 | 1.023 | -0.047 | 1.164 | -0.09 | 0.075 | 0.075 | 1.953 | 0.84 | 0.99 | 0.9 | 0.98 | 0.77 | 0.77 |
| 6 | 73863598 | rs9343012 | 0.945 | 0.486 | -0.793 | 1.072 | -0.013 | 0.07 | 0.07 | 2.216 | 0.43 | 0.13 | 0.38 | 0.31 | 0.41 | 0.39 |
| 6 | 73865196 | rs6453644 | 0.189 | 0.291 | -0.553 | 1.23 | -0.04 | 0.058 | 0.058 | 0.563 | 0.75 | 0.9 | 0.85 | 0.89 | 0.66 | 0.6 |
| 6 | 73865758 | rs7765352 | 1.281 | 0.299 | -0.16 | 1.275 | 0.009 | 0.064 | 0.064 | 2.103 | 0.44 | 0.19 | 0.38 | 0.34 | 0.39 | 0.39 |
| 6 | 73868831 | rs6909792 | -0.105 | 0.199 | -0.084 | 1.614 | -0.005 | -0.01 | -0.01 | 0 | 0.1 | 0.11 | 0.05 | 0.09 | 0.09 | 0.04 |
| 6 | 73871994 | rs952880 | 1.029 | 0.098 | -0.186 | 1.467 | -0.054 | 0.086 | 0.086 | 1.194 | 0.58 | 0.78 | 0.76 | 0.8 | 0.5 | 0.54 |
| 6 | 73872276 | rs6939575 | 0.398 | 0.084 | 0.207 | 1.464 | -0.029 | 0.057 | 0.057 | 0.617 | 0.7 | 0.81 | 0.81 | 0.9 | 0.6 | 0.58 |
| 6 | 73875255 | rs878523 | -0.054 | -0.225 | 1.282 | 2.289 | -0.031 | 0.076 | 0.076 | 0 | 0.1 | 0.16 | 0.18 | 0.3 | 0.07 | 0.13 |
| 6 | 73880130 | rs3799278 | -0.278 | -0.178 | 1.187 | 2.23 | -0.007 | -0.011 | -0.011 | 0 | 0.09 | 0.1 | 0.07 | 0.1 | 0.1 | 0.04 |
| 6 | 73883238 | rs3823118 | -1.129 | -0.871 | 1.144 | 2.52 | -0.04 | 0.047 | 0.047 | 0 | 0.22 | 0.66 | 0.41 | 0.5 | 0.1 | 0.16 |
| 6 | 73883761 | rs1970549 | -2.137 | -0.828 | 1.044 | 2.463 | 0.05 | 0.04 | 0.04 | 0 | 0.39 | 0.91 | 0.55 | 0.69 | 0.3 | 0.24 |
| 6 | 73886305 | rs7756501 | -1.985 | -0.902 | 1.587 | 2.407 | 0.034 | 0.029 | 0.029 | 0 | 0.39 | 0.72 | 0.54 | 0.68 | 0.3 | 0.26 |
| 6 | 73888451 | rs3799280 | -1.792 | -0.906 | 1.777 | 2.441 | -0.01 | -0.001 | -0.001 | 0.022 | 0.37 | 0.57 | 0.51 | 0.59 | 0.27 | 0.23 |
| 6 | 73893197 | rs3778268 | -1.787 | -0.777 | 1.725 | 2.441 | -0.002 | 0.012 | 0.012 | 0.265 | 0.25 | 0.71 | 0.22 | 0.29 | 0.22 | 0.12 |
| 6 | 73893677 | rs3757105 | -0.388 | -0.962 | 1.75 | 2.374 | -0.014 | 0.006 | 0.006 | 0 | 0.15 | 0.13 | 0.18 | 0.29 | 0.09 | 0.13 |
| 6 | 73895541 | rs16883476 | 1.157 | -0.91 | 1.097 | 1.698 | 0.041 | 0.065 | 0.065 | 8.291 | 0.36 | 0.14 | 0.24 | 0.23 | 0.27 | 0.15 |
| 6 | 73899356 | rs9343016 | -0.498 | -0.568 | 1.233 | 1.559 | -0.023 | -0.003 | -0.003 | 0 | 0.61 | 0.72 | 0.43 | 0.41 | 0.5 | 0.27 |
| 6 | 73900091 | rs9341413 | 1.211 | -0.662 | 1.05 | 1.691 | 0.043 | 0.107 | 0.107 | 10.235 | 0.36 | 0.14 | 0.18 | 0.1 | 0.27 | 0.12 |
| 6 | 73907138 | rs10455270 | -1.3 | -0.572 | 1.358 | 1.799 | 0.006 | 0.036 | 0.036 | 0.179 | 0.24 | 0.41 | 0.23 | 0.31 | 0.22 | 0.15 |
| 6 | 73914319 | rs6929988 | -0.772 | -0.552 | 1.072 | 1.672 | -0.177 | -0.112 | -0.112 | 1.974 | 0.65 | 0.97 | 0.66 | 0.58 | 0.5 | 0.28 |
| 6 | 73916577 | rs10943080 | -0.323 | -0.533 | 1.334 | 1.67 | 0.003 | -0.009 | -0.009 | 0 | 0.58 | 0.47 | 0.38 | 0.36 | 0.47 | 0.22 |
| 6 | 73922654 | rs9688431 | -0.592 | -0.42 | 0.987 | 1.08 | -0.003 | 0.05 | 0.05 | 0 | 0.08 | 0.2 | 0.11 | 0.2 | 0.06 | 0.04 |
| 6 | 73930950 | rs12190643 | -1.166 | -0.409 | 0.926 | 0.977 | 0 | -0.001 | -0.001 | 0 | 0.22 | 0.28 | 0.2 | 0.26 | 0.21 | 0.12 |
| 6 | 73931120 | rs12173399 | -0.258 | -0.352 | 0.943 | 1.032 | 0.008 | -0.002 | -0.002 | 0 | 0.59 | 0.53 | 0.42 | 0.41 | 0.49 | 0.27 |
| 6 | 73933401 | rs2282448 | -1.1 | -0.209 | 0.791 | 1.164 | -0.001 | 0.01 | 0.01 | 0 | 0.08 | 0.14 | 0.11 | 0.2 | 0.06 | 0.04 |
| 6 | 73948140 | rs2350753 | -1.414 | -0.246 | 0.994 | 1.198 | -0.005 | 0.091 | 0.091 | 0 | 0.11 | 0.26 | 0.14 | 0.24 | 0.07 | 0.05 |
| 6 | 73948431 | rs6911421 | 0.914 | -0.321 | 0.875 | 0.743 | 0.027 | 0.051 | 0.051 | 7.581 | 0.36 | 0.2 | 0.21 | 0.14 | 0.3 | 0.15 |
| 6 | 73951025 | rs1891494 | -2.622 | -0.049 | 1.187 | 0.867 | -0.003 | 0.001 | 0.001 | 0 | 0.32 | 0.72 | 0.29 | 0.31 | 0.22 | 0.22 |
| 6 | 73952103 | rs6453660 | -2.617 | 0.017 | 1.352 | 0.737 | -0.012 | -0.006 | -0.006 | 0.051 | 0.29 | 0.7 | 0.27 | 0.25 | 0.2 | 0.17 |
| 6 | 73958442 | rs2124006 | 2.546 | 0.417 | 1.31 | 0.676 | 0 | 0 | 0 | 0 | 0.67 | 0.27 | 0.68 | 0.72 | 0.73 | 0.78 |
| 6 | 73965968 | rs2603472 | -1.779 | 0.127 | 1.701 | 0.903 | 0.005 | 0.013 | 0.013 | 0 | 0.15 | 0.29 | 0.18 | 0.16 | 0.14 | 0.11 |
| 6 | 73969722 | rs10943084 | -1.086 | 0.146 | 1.727 | 0.888 | 0.013 | -0.039 | -0.039 | 0.072 | 0.08 | 0.23 | 0.08 | 0.01 | 0.08 | 0.02 |
| 6 | 73975404 | rs4708042 | -3.174 | 0.079 | 1.944 | 0.825 | 0 | 0.041 | 0.041 | 0 | 0.24 | 0.52 | 0.26 | 0.29 | 0.21 | 0.2 |
| 6 | 73984720 | rs10943088 | -1.086 | 0.136 | 1.973 | 1.107 | 0.002 | -0.002 | -0.002 | 0 | 0.07 | 0.07 | 0.08 | 0.06 | 0.09 | 0.08 |
| 6 | 74000604 | rs947753 | -0.653 | 0.143 | 1.898 | 1.126 | -0.011 | -0.021 | -0.021 | 0 | 0.06 | 0.06 | 0.1 | 0.14 | 0.06 | 0.09 |
| 6 | 74013156 | rs2124003 | -1.446 | 0.106 | 1.697 | 1.123 | 0 | 0.009 | 0.009 | 0.048 | 0.09 | 0.13 | 0.11 | 0.19 | 0.08 | 0.14 |
| 6 | 74014468 | rs555779 | 3.125 | 0.059 | 1.859 | 1.097 | -0.01 | 0.014 | 0.014 | 0.284 | 0.78 | 0.24 | 0.75 | 0.71 | 0.84 | 0.81 |
| 6 | 74015765 | rs7769905 | -2.233 | 0.342 | 1.011 | 1.278 | 0.005 | 0.134 | 0.134 | 0.575 | 0.11 | 0.32 | 0.15 | 0.27 | 0.09 | 0.16 |
| 6 | 74016113 | rs7753960 | -1.501 | 0.362 | 0.766 | 1.287 | 0 | 0.009 | 0.009 | 0.048 | 0.09 | 0.13 | 0.11 | 0.19 | 0.08 | 0.14 |
| 6 | 74019376 | rs16883573 | -1.485 | 0.337 | 0.77 | 0.949 | 0.009 | 0.038 | 0.038 | 0 | 0.11 | 0.3 | 0.12 | 0.14 | 0.1 | 0.1 |
| 6 | 74041165 | rs554727 | -0.154 | 0.936 | 0.127 | 1.029 | 0.04 | 0.03 | 0.03 | 3.727 | 0.32 | 0.42 | 0.49 | 0.41 | 0.4 | 0.61 |
| 6 | 74041424 | rs9360664 | -0.164 | 0.735 | -0.171 | 0.747 | -0.043 | 0.049 | 0.049 | 0 | 0.2 | 0.03 | 0.1 | 0.15 | 0.19 | 0.19 |
| 6 | 74047002 | rs11753467 | -0.037 | 0.758 | -0.106 | 0.968 | -0.004 | -0.027 | -0.027 | 0 | 0.14 | 0.03 | 0.14 | 0.21 | 0.11 | 0.15 |
| 6 | 74058006 | rs526434 | 1.185 | 0.633 | 0.05 | 0.771 | -0.001 | 0.015 | 0.015 | 0 | 0.48 | 0.37 | 0.41 | 0.54 | 0.49 | 0.56 |
| 6 | 74062007 | rs6906976 | 0.102 | 0.36 | 0.105 | 0.568 | -0.02 | -0.018 | -0.018 | 0 | 0.19 | 0.25 | 0.16 | 0.11 | 0.13 | 0.1 |
| 6 | 74064396 | rs17755984 | -0.048 | 0.537 | -0.013 | 0.509 | -0.027 | 0.058 | 0.058 | 0 | 0.13 | 0.05 | 0.07 | 0.03 | 0.11 | 0.16 |
| 6 | 74073885 | rs2295761 | 0.277 | 0.573 | 0.019 | 0.531 | -0.027 | 0.045 | 0.045 | 0 | 0.11 | 0.04 | 0.07 | 0.01 | 0.11 | 0.12 |
| 6 | 74075199 | rs9360669 | 0.269 | 0.415 | 0.114 | 0.505 | 0.01 | 0.01 | 0.01 | 0 | 0.15 | 0.13 | 0.12 | 0.05 | 0.1 | 0.08 |
| 6 | 74092923 | rs17812683 | -1.101 | 0.587 | -0.099 | 0.866 | -0.054 | 0.062 | 0.062 | 0 | 0.25 | 0.08 | 0.2 | 0.13 | 0.3 | 0.21 |
| 6 | 74098703 | rs488114 | -0.529 | 1.058 | -0.203 | 0.751 | -0.046 | 0.032 | 0.032 | 0 | 0.09 | 0.01 | 0.06 | 0.03 | 0.14 | 0.09 |
| 6 | 74104299 | rs6917933 | -0.015 | 0.538 | 0.09 | 1.001 | -0.034 | 0.049 | 0.049 | 0.814 | 0.09 | 0.01 | 0.09 | 0 | 0.14 | 0.06 |

| | | | | | | | | | | | | | | | | |
|---|---|---|---|---|---|---|---|---|---|---|---|---|---|---|---|---|
| 6 | 74123363 | rs9442907 | -0.644 | 0.521 | 0.116 | 1.049 | -0.007 | 0.013 | 0.013 | 0 | 0.14 | 0.08 | 0.12 | 0.1 | 0.18 | 0.09 |
| 6 | 74125875 | rs311686 | 1.113 | 0.365 | 0.653 | 1.135 | 0.017 | 0.026 | 0.026 | 0 | 0.52 | 0.31 | 0.45 | 0.54 | 0.57 | 0.63 |
| 6 | 74125888 | rs311685 | 1.113 | 0.365 | 0.653 | 1.137 | 0.017 | 0.028 | 0.028 | 0 | 0.52 | 0.31 | 0.45 | 0.55 | 0.57 | 0.63 |
| 6 | 74126084 | rs311684 | 0.986 | 0.36 | 0.757 | 1.128 | 0.004 | 0.018 | 0.018 | 0 | 0.54 | 0.41 | 0.62 | 0.71 | 0.57 | 0.66 |
| 6 | 74126941 | rs311683 | 1.089 | 0.412 | 0.409 | 0.966 | 0.018 | 0.027 | 0.027 | 0 | 0.52 | 0.31 | 0.45 | 0.54 | 0.57 | 0.63 |
| 6 | 74134199 | rs311680 | 0.268 | 0.117 | 0.427 | 0.931 | -0.007 | -0.01 | -0.01 | 1.079 | 0.86 | 0.84 | 0.87 | 0.93 | 0.9 | 0.89 |
| 6 | 74136427 | rs311677 | 0.22 | 0.15 | 0.487 | 0.849 | -0.017 | -0.035 | -0.035 | 0 | 0.86 | 0.8 | 0.87 | 0.95 | 0.91 | 0.93 |
| 6 | 74166098 | rs9293930 | 0.479 | 0.106 | 1.013 | 1.022 | 0.004 | 0.004 | 0.004 | 0.065 | 0.83 | 0.88 | 0.86 | 0.95 | 0.86 | 0.92 |
| 6 | 74167406 | rs9341423 | 1.524 | -0.017 | 0.985 | 1.183 | -0.02 | -0.006 | -0.006 | 0 | 0.61 | 0.44 | 0.59 | 0.57 | 0.66 | 0.66 |
| 6 | 74170860 | rs9341424 | -0.122 | -0.221 | 1.277 | 1.148 | -0.01 | -0.02 | -0.02 | 0.319 | 0.86 | 0.86 | 0.89 | 0.95 | 0.93 | 0.93 |
| 6 | 74179395 | rs7453831 | -1.183 | -0.381 | 1.255 | 1.112 | -0.003 | 0.003 | 0.003 | 0 | 0.18 | 0.27 | 0.18 | 0.21 | 0.18 | 0.17 |
| 6 | 74179788 | rs12529857 | -1.125 | -0.384 | 1.265 | 1.111 | -0.001 | 0.001 | 0.001 | 0 | 0.18 | 0.23 | 0.17 | 0.21 | 0.18 | 0.17 |
| 6 | 74191545 | rs535902 | -0.362 | -0.392 | 1.273 | 1.102 | 0.005 | 0.002 | 0.002 | 0 | 0.86 | 0.89 | 0.89 | 0.95 | 0.93 | 0.93 |
| 6 | 74192422 | rs12526844 | -0.876 | -0.424 | 1.237 | 1.106 | -0.039 | -0.05 | -0.05 | 0 | 0.14 | 0.01 | 0.15 | 0.19 | 0.18 | 0.17 |
| 6 | 74194928 | rs7768980 | -0.789 | -0.438 | 1.301 | 1.173 | -0.022 | 0.033 | 0.033 | 0 | 0.08 | 0.14 | 0.12 | 0.19 | 0.03 | 0.04 |
| 6 | 74200726 | rs577372 | 1.731 | -0.496 | 1.342 | 1.536 | -0.002 | -0.001 | -0.001 | 0 | 0.58 | 0.38 | 0.4 | 0.46 | 0.7 | 0.69 |
| 6 | 74217278 | rs674708 | -0.346 | -0.494 | 1.301 | 1.469 | -0.007 | -0.016 | -0.016 | 0.52 | 0.86 | 0.87 | 0.9 | 0.95 | 0.93 | 0.94 |
| 6 | 74226353 | rs11755449 | -0.465 | -0.398 | 1.23 | 1.51 | -0.043 | -0.049 | -0.049 | 0 | 0.12 | 0.01 | 0.16 | 0.2 | 0.17 | 0.17 |
| 6 | 74228006 | rs1874229 | 1.491 | -0.467 | 1.302 | 1.584 | 0.007 | -0.003 | -0.003 | 0 | 0.64 | 0.65 | 0.62 | 0.53 | 0.72 | 0.74 |
| 6 | 74228049 | rs1874230 | 1.783 | -0.489 | 1.454 | 1.844 | 0.01 | -0.003 | -0.003 | 0 | 0.58 | 0.33 | 0.39 | 0.42 | 0.69 | 0.68 |
| 6 | 74229405 | rs4708060 | -0.479 | -0.486 | 1.367 | 1.835 | 0.006 | 0.003 | 0.003 | 0 | 0.86 | 0.89 | 0.9 | 0.95 | 0.93 | 0.94 |
| 6 | 74229792 | rs2073465 | -1.166 | -0.474 | 1.346 | 1.844 | -0.016 | -0.025 | -0.025 | 0 | 0.18 | 0.07 | 0.18 | 0.23 | 0.2 | 0.18 |
| 6 | 74238511 | rs9352004 | 1.717 | -0.507 | 1.575 | 1.849 | 0.007 | -0.003 | -0.003 | 0 | 0.58 | 0.36 | 0.39 | 0.42 | 0.69 | 0.68 |
| 6 | 74238608 | rs9352005 | 1.153 | -0.495 | 1.449 | 1.847 | 0.018 | 0.02 | 0.02 | 0.095 | 0.68 | 0.76 | 0.72 | 0.73 | 0.74 | 0.76 |
| 6 | 74246392 | rs9343053 | 2.444 | 0.165 | 1.244 | 1.473 | -0.025 | -0.036 | -0.036 | 0 | 0.4 | 0.11 | 0.23 | 0.41 | 0.37 | 0.54 |
| 6 | 74270574 | rs642722 | -1.441 | 0.099 | 1.205 | 1.358 | -0.036 | 0.001 | 0.001 | 0.044 | 0.39 | 0.51 | 0.39 | 0.45 | 0.28 | 0.27 |
| 6 | 74275405 | rs4708064 | -0.55 | 0.148 | 1.204 | 1.346 | 0.061 | -0.017 | -0.017 | 0 | 0.21 | 0.36 | 0.37 | 0.12 | 0.35 | 0.2 |
| 6 | 74279910 | rs606263 | 0.525 | 0.108 | 1.124 | 1.411 | -0.003 | -0.006 | -0.006 | 0 | 0.14 | 0.15 | 0.09 | 0.07 | 0.07 | 0.08 |
| 6 | 74280026 | rs9442924 | -1.559 | 0.188 | 1.016 | 1.5 | -0.015 | 0.029 | 0.029 | 0 | 0.24 | 0.34 | 0.29 | 0.38 | 0.22 | 0.2 |
| 6 | 74290291 | rs583443 | -1.053 | 0.606 | 0.871 | 0.907 | 0.001 | -0.011 | -0.011 | 0 | 0.38 | 0.62 | 0.48 | 0.17 | 0.42 | 0.3 |
| 6 | 74305276 | rs12528608 | -0.285 | 0.628 | 0.735 | 1.466 | -0.022 | -0.042 | -0.042 | 0 | 0.12 | 0.02 | 0.21 | 0.26 | 0.2 | 0.12 |
| 6 | 74309401 | rs364734 | -0.04 | 0.439 | 0.944 | 1.111 | 0.017 | 0.008 | 0.008 | 0 | 0.19 | 0.15 | 0.16 | 0.18 | 0.07 | 0.08 |
| 6 | 74315347 | rs2351120 | -0.391 | 0.703 | 1.045 | 1.078 | 0.088 | -0.083 | -0.083 | 0 | 0.17 | 0.33 | 0.17 | 0.02 | 0.31 | 0.15 |
| 6 | 74329524 | rs12214877 | -0.232 | 0.684 | 1.115 | 1.105 | 0.042 | -0.056 | -0.056 | 0 | 0.15 | 0.21 | 0.13 | 0.02 | 0.25 | 0.15 |
| 6 | 74340159 | rs4032316 | -1.301 | 0.456 | 1.263 | 1.106 | 0.002 | 0.008 | 0.008 | 0 | 0.4 | 0.62 | 0.52 | 0.27 | 0.37 | 0.29 |
| 6 | 74341221 | rs685491 | -0.358 | 0.411 | 1.228 | 1.122 | -0.056 | -0.022 | -0.022 | 0 | 0.22 | 0.38 | 0.17 | 0.18 | 0.08 | 0.08 |
| 6 | 74341589 | rs687210 | -0.358 | 0.411 | 1.228 | 1.122 | -0.056 | -0.022 | -0.022 | 0.032 | 0.22 | 0.38 | 0.17 | 0.18 | 0.08 | 0.08 |
| 6 | 74351976 | rs10455281 | -0.137 | 0.489 | 1.23 | 1.104 | 0.042 | -0.056 | -0.056 | 0.642 | 0.15 | 0.21 | 0.13 | 0.02 | 0.25 | 0.15 |
| 6 | 74354175 | rs472294 | -0.009 | 0.451 | 1.259 | 1.099 | 0.005 | 0.004 | 0.004 | 0 | 0.19 | 0.16 | 0.16 | 0.18 | 0.07 | 0.08 |
| 6 | 74360511 | rs13212809 | -0.129 | 0.484 | 1.066 | 1.114 | 0.042 | -0.056 | -0.056 | 0.229 | 0.15 | 0.21 | 0.13 | 0.02 | 0.25 | 0.15 |
| 6 | 74360695 | rs1703467 | 1.285 | 0.442 | 0.869 | 1.188 | 0.02 | 0.054 | 0.054 | 0 | 0.57 | 0.21 | 0.46 | 0.68 | 0.63 | 0.71 |
| 6 | 74371910 | rs4708072 | 0.259 | 0.301 | 0.513 | 1.312 | -0.047 | -0.015 | -0.015 | 0 | 0.79 | 0.66 | 0.83 | 0.8 | 0.92 | 0.92 |
| 6 | 74377465 | rs3846752 | -0.987 | 0.862 | 0.269 | 1.723 | 0.001 | 0 | 0 | 0.223 | 0.49 | 0.54 | 0.56 | 0.5 | 0.57 | 0.37 |
| 6 | 74378808 | rs9446975 | -0.233 | 1.154 | 0.094 | 1.664 | -0.006 | -0.016 | -0.016 | 0.267 | 0.37 | 0.41 | 0.26 | 0.26 | 0.32 | 0.26 |
| 6 | 74381776 | rs10498893 | -0.269 | 1.179 | -0.024 | 1.551 | -0.009 | -0.014 | -0.014 | 0 | 0.37 | 0.46 | 0.29 | 0.32 | 0.32 | 0.27 |
| 6 | 74381926 | rs9446976 | -0.873 | 0.951 | -0.675 | 1.301 | -0.007 | 0.009 | 0.009 | 0 | 0.55 | 0.8 | 0.59 | 0.58 | 0.59 | 0.39 |
| 6 | 74382213 | rs2351127 | -1.073 | 0.996 | -0.351 | 1.296 | -0.01 | -0.02 | -0.02 | 0.131 | 0.06 | 0.01 | 0.21 | 0.07 | 0.06 | 0.08 |
| 6 | 74422722 | rs4708076 | 1.353 | 1.253 | 1.277 | 2.768 | -0.021 | 0.187 | 0.187 | 0 | 0.87 | 0.67 | 0.79 | 0.56 | 0.91 | 0.67 |
| 6 | 74424069 | rs9293937 | 0.318 | 1.089 | 0.43 | 1.624 | 0 | 0.003 | 0.003 | 0 | 0.34 | 0.26 | 0.35 | 0.26 | 0.31 | 0.16 |
| 6 | 74430495 | rs9446996 | 0.483 | 1.182 | 0.717 | 1.489 | 0.006 | 0.017 | 0.017 | 0 | 0.32 | 0.17 | 0.31 | 0.26 | 0.29 | 0.16 |
| 6 | 74430673 | rs7747167 | 0.961 | 0.502 | 0.192 | 0.882 | -0.091 | 0.201 | 0.201 | 0.495 | 0.38 | 0.05 | 0.29 | 0.1 | 0.45 | 0.4 |
| 6 | 74432636 | rs9442951 | 0.909 | 0.277 | 0.228 | 0.874 | -0.026 | 0.079 | 0.079 | 0.254 | 0.41 | 0.25 | 0.39 | 0.2 | 0.53 | 0.48 |
| 6 | 74433212 | rs4706552 | 1.223 | 0.158 | 0.237 | 1.018 | -0.035 | 0.058 | 0.058 | 0.493 | 0.74 | 0.33 | 0.72 | 0.5 | 0.86 | 0.64 |
| 6 | 74438648 | rs6935547 | -0.023 | 0.162 | 0.155 | 0.875 | 0.006 | 0.002 | 0.002 | 0.159 | 0.1 | 0.08 | 0.06 | 0.06 | 0.04 | 0.02 |
| 6 | 74449960 | rs12203043 | 0.918 | 0.167 | 0.336 | 0.903 | -0.018 | -0.009 | -0.009 | 0.833 | 0.32 | 0.16 | 0.33 | 0.35 | 0.31 | 0.17 |
| 6 | 74450775 | rs2351311 | -0.475 | 0.169 | 0.776 | 0.875 | 0.024 | 0.006 | 0.006 | 0 | 0.5 | 0.73 | 0.51 | 0.52 | 0.5 | 0.28 |
| 6 | 74458737 | rs9447004 | -0.803 | 0.022 | 0.604 | 0.95 | 0.015 | -0.003 | -0.003 | 0.172 | 0.52 | 0.66 | 0.51 | 0.54 | 0.49 | 0.36 |
| 6 | 74468860 | rs7774119 | 1.076 | -0.022 | 0.746 | 0.908 | -0.004 | -0.002 | -0.002 | 0 | 0.3 | 0.22 | 0.33 | 0.31 | 0.29 | 0.16 |
| 6 | 74477478 | rs9359043 | 0.434 | -0.037 | 0.468 | 0.985 | 0.025 | 0.004 | 0.004 | 0 | 0.06 | 0 | 0.16 | 0.31 | 0.01 | 0.21 |
| 6 | 74489384 | rs9343078 | -1.072 | -0.037 | 0.524 | 0.941 | -0.001 | 0.084 | 0.084 | 0.279 | 0.58 | 0.73 | 0.68 | 0.84 | 0.51 | 0.55 |
| 6 | 74489482 | rs9447017 | -0.08 | 0.093 | 0.731 | 0.876 | -0.006 | 0.001 | 0.001 | 0.04 | 0.43 | 0.47 | 0.41 | 0.42 | 0.34 | 0.22 |
| 6 | 74493432 | rs10455097 | -0.59 | 0.091 | 0.744 | 0.879 | 0.019 | -0.005 | -0.005 | 0.018 | 0.49 | 0.66 | 0.51 | 0.53 | 0.5 | 0.34 |
| 6 | 74497009 | rs2351528 | 0.034 | -0.04 | 0.587 | 0.903 | -0.005 | 0.004 | 0.004 | 0 | 0.41 | 0.47 | 0.4 | 0.41 | 0.34 | 0.22 |
| 6 | 74497152 | rs5023688 | 0.071 | -0.049 | 0.582 | 0.899 | -0.004 | 0.004 | 0.004 | 0 | 0.41 | 0.46 | 0.4 | 0.41 | 0.34 | 0.22 |
| 6 | 74517352 | rs4708090 | -1.652 | 0.123 | 0.789 | 1.383 | -0.001 | 0.05 | 0.05 | 0 | 0.2 | 0.3 | 0.28 | 0.39 | 0.19 | 0.36 |
| 6 | 74521947 | rs2917862 | 0.886 | -0.317 | 0.09 | 0.629 | 0.026 | -0.009 | -0.009 | 0.955 | 0.51 | 0.2 | 0.47 | 0.46 | 0.49 | 0.51 |
| 6 | 74525643 | rs10943133 | 0.751 | -0.386 | 0.143 | 0.588 | -0.032 | -0.036 | -0.036 | 0.3 | 0.26 | 0.14 | 0.35 | 0.37 | 0.29 | 0.22 |
| 6 | 74529922 | rs3012571 | 0.935 | -0.369 | 0.31 | 0.571 | 0.019 | -0.006 | -0.006 | 0.042 | 0.52 | 0.3 | 0.48 | 0.46 | 0.49 | 0.51 |
| 6 | 74531720 | rs3012572 | 0.173 | -0.378 | 0.44 | 0.658 | 0 | -0.001 | -0.001 | 0 | 0.6 | 0.52 | 0.58 | 0.6 | 0.64 | 0.62 |

| | | | | | | | | | | | | | | | | |
|---|---|---|---|---|---|---|---|---|---|---|---|---|---|---|---|---|
| 6 | 74533192 | rs2917887 | 0.416 | -0.381 | 0.408 | 0.666 | -0.044 | -0.053 | -0.053 | 1.189 | 0.86 | 0.66 | 0.93 | 0.95 | 0.93 | 0.84 |
| 6 | 74534122 | rs6907830 | -0.069 | -0.49 | 0.127 | 0.621 | 0 | -0.036 | -0.036 | 0 | 0.09 | 0.01 | 0.18 | 0.4 | 0.06 | 0.19 |
| 6 | 74536510 | rs3012519 | 0.679 | -0.451 | 0.221 | 0.69 | -0.07 | -0.017 | -0.017 | 0 | 0.83 | 0.51 | 0.88 | 0.9 | 0.93 | 0.83 |
| 6 | 74538823 | rs4708093 | 0.773 | -0.259 | -0.175 | 0.757 | -0.071 | -0.047 | -0.047 | 0.923 | 0.24 | 0.08 | 0.31 | 0.32 | 0.29 | 0.2 |
| 6 | 74541452 | rs1992141 | 0.095 | -0.276 | -0.06 | 0.768 | -0.029 | -0.026 | -0.026 | 1.353 | 0.31 | 0.25 | 0.46 | 0.48 | 0.45 | 0.32 |
| 6 | 74542387 | rs1437579 | 0.524 | -0.481 | 0.066 | 0.649 | -0.07 | -0.089 | -0.089 | 0.557 | 0.83 | 0.52 | 0.92 | 0.94 | 0.93 | 0.89 |
| 6 | 74544376 | rs7753766 | 0.601 | -0.349 | -0.274 | 0.739 | -0.012 | -0.013 | -0.013 | 0 | 0.26 | 0.22 | 0.37 | 0.36 | 0.3 | 0.22 |
| 6 | 74564596 | rs2196622 | -0.139 | -0.474 | -0.547 | 0.474 | -0.018 | -0.021 | -0.021 | 4.823 | 0.16 | 0.22 | 0.07 | 0.08 | 0.09 | 0.14 |
| 6 | 74570251 | rs3012553 | 1.152 | -0.359 | -1.354 | 0.55 | -0.036 | -0.105 | -0.105 | 0 | 0.44 | 0.05 | 0.42 | 0.44 | 0.44 | 0.5 |
| 6 | 74571797 | rs3012557 | 1.761 | -0.299 | -1.339 | 0.367 | -0.053 | 0.22 | 0.22 | 0 | 0.35 | 0.04 | 0.25 | 0.05 | 0.38 | 0.27 |
| 6 | 74581042 | rs2882704 | -0.064 | 0.169 | -0.807 | 0.865 | -0.077 | -0.048 | -0.048 | 0.827 | 0.31 | 0.08 | 0.34 | 0.35 | 0.33 | 0.25 |
| 6 | 74586805 | rs920234 | -1.12 | -0.052 | -0.486 | 0.957 | -0.001 | -0.001 | -0.001 | 5.058 | 0.53 | 0.51 | 0.57 | 0.58 | 0.56 | 0.48 |
| 6 | 74587661 | rs2917891 | 1.311 | -0.045 | -0.56 | 1.046 | 0.017 | 0.003 | 0.003 | 0 | 0.4 | 0.36 | 0.33 | 0.29 | 0.37 | 0.4 |
| 6 | 74589109 | rs9359045 | -0.183 | 0.925 | -0.983 | 0.839 | -0.067 | 0.027 | 0.027 | 0.048 | 0.11 | 0.02 | 0.16 | 0.08 | 0.27 | 0.13 |
| 6 | 74589713 | rs2917888 | 0.794 | 0.412 | -0.737 | 0.933 | 0.051 | 0.003 | 0.003 | 0.72 | 0.51 | 0.36 | 0.41 | 0.43 | 0.39 | 0.51 |
| 6 | 74605806 | rs3005511 | 0.363 | 0.561 | -0.815 | 0.969 | -0.005 | -0.022 | -0.022 | 0 | 0.61 | 0.33 | 0.57 | 0.51 | 0.66 | 0.64 |
| 6 | 74606039 | rs4708101 | -0.357 | 0.647 | -0.932 | 0.856 | -0.034 | 0.024 | 0.024 | 0 | 0.13 | 0.07 | 0.16 | 0.08 | 0.27 | 0.15 |
| 6 | 74606438 | rs3005512 | 1.645 | 0.219 | -1.129 | 0.847 | 0.023 | -0.007 | -0.007 | 0 | 0.38 | 0.18 | 0.38 | 0.33 | 0.37 | 0.38 |
| 6 | 74612516 | rs2917870 | 0.792 | 0.208 | -1.13 | 0.844 | 0.088 | -0.003 | -0.003 | 3.284 | 0.49 | 0.26 | 0.41 | 0.43 | 0.39 | 0.49 |
| 6 | 74625056 | rs4235876 | -0.29 | 0.255 | -1.068 | 0.701 | -0.034 | 0.024 | 0.024 | 0 | 0.13 | 0.07 | 0.16 | 0.08 | 0.27 | 0.15 |
| 6 | 74637913 | rs2217987 | -0.251 | 0.048 | -1.036 | 0.727 | 0.008 | 0 | 0 | 0 | 0.11 | 0.07 | 0.03 | 0.1 | 0.03 | 0.11 |
| 6 | 74639586 | rs12530086 | -0.251 | 0.048 | -1.033 | 0.727 | 0.004 | 0 | 0 | 0 | 0.11 | 0.07 | 0.03 | 0.1 | 0.03 | 0.11 |
| 6 | 74648481 | rs6453705 | 0.12 | 0.259 | -1.037 | 0.946 | 0 | -0.006 | -0.006 | 0.504 | 0.7 | 0.38 | 0.75 | 0.8 | 0.75 | 0.68 |
| 6 | 74662900 | rs3012521 | -0.623 | 0.304 | -0.498 | 0.832 | 0.056 | -0.014 | -0.014 | 0 | 0.9 | 0.59 | 0.93 | 0.96 | 0.93 | 0.91 |
| 6 | 74666559 | rs4551140 | -0.002 | 0.199 | -0.528 | 0.843 | -0.046 | 0.001 | 0.001 | 0.109 | 0.08 | 0.19 | 0.02 | 0.04 | 0.01 | 0.09 |
| 6 | 74673166 | rs12212272 | -0.131 | 0.36 | -0.58 | 0.834 | -0.033 | 0.007 | 0.007 | 0 | 0.12 | 0.07 | 0.16 | 0.12 | 0.26 | 0.15 |
| 6 | 74674332 | rs12194422 | -1.007 | 0.345 | -0.572 | 0.83 | 0.01 | -0.008 | -0.008 | 0 | 0.21 | 0.26 | 0.18 | 0.16 | 0.27 | 0.24 |
| 6 | 74675686 | rs3003177 | -1.617 | 0.184 | -0.677 | 0.731 | 0.024 | 0.113 | 0.113 | 1.97 | 0.5 | 0.78 | 0.42 | 0.32 | 0.52 | 0.56 |
| 6 | 74675731 | rs3003178 | -0.46 | 0.215 | -0.646 | 0.743 | 0.002 | -0.053 | -0.053 | 0 | 0.3 | 0.5 | 0.24 | 0.16 | 0.24 | 0.32 |
| 6 | 74677240 | rs3003180 | -1.57 | 0.19 | -0.671 | 1.002 | 0.027 | 0.068 | 0.068 | 2.699 | 0.49 | 0.68 | 0.42 | 0.35 | 0.52 | 0.49 |
| 6 | 74678421 | rs6916197 | -0.716 | -0.041 | -0.74 | 1.072 | -0.031 | 0.073 | 0.073 | 0.041 | 0.17 | 0.25 | 0.21 | 0.39 | 0.1 | 0.13 |
| 6 | 74680038 | rs3003187 | -1.66 | 0.07 | -0.734 | 1.118 | 0.029 | 0.033 | 0.033 | 0 | 0.48 | 0.72 | 0.4 | 0.57 | 0.34 | 0.39 |
| 6 | 74680740 | rs3003190 | -0.327 | 0.068 | -0.727 | 1.116 | 0.017 | -0.054 | -0.054 | 0.024 | 0.28 | 0.47 | 0.19 | 0.18 | 0.23 | 0.27 |
| 6 | 74682273 | rs4421160 | -0.807 | 0.059 | -0.747 | 1.123 | -0.02 | 0.042 | 0.042 | 0.089 | 0.2 | 0.25 | 0.22 | 0.39 | 0.11 | 0.13 |
| 6 | 74693189 | rs4235882 | -2.136 | 0.444 | -0.767 | 1.094 | -0.001 | 0.053 | 0.053 | 0.002 | 0.59 | 0.76 | 0.55 | 0.69 | 0.54 | 0.57 |
| 6 | 74693901 | rs7775279 | -1.377 | 0.461 | -0.74 | 1.107 | 0 | -0.011 | -0.011 | 0 | 0.3 | 0.27 | 0.32 | 0.49 | 0.31 | 0.25 |
| 6 | 74694400 | rs9447089 | 0.744 | 0.463 | -0.54 | 1.082 | -0.013 | 0.022 | 0.022 | 0.133 | 0.18 | 0.08 | 0.12 | 0.14 | 0.16 | 0.18 |
| 6 | 74694463 | rs12195936 | 0.438 | 0.302 | -0.548 | 1.16 | -0.05 | -0.013 | -0.013 | 0 | 0.1 | 0.02 | 0.1 | 0.1 | 0.19 | 0.12 |
| 6 | 74699350 | rs7764706 | -2.136 | 0.293 | -0.54 | 1.134 | -0.013 | 0.048 | 0.048 | 0 | 0.6 | 0.83 | 0.56 | 0.69 | 0.55 | 0.61 |
| 6 | 74706111 | rs9442995 | -0.226 | 0.165 | -0.43 | 1.147 | 0.018 | -0.058 | -0.058 | 0 | 0.27 | 0.45 | 0.18 | 0.18 | 0.23 | 0.27 |
| 6 | 74725759 | rs4352650 | 0.277 | 0.232 | -0.696 | 1.091 | 0.005 | -0.004 | -0.004 | 0 | 0.06 | 0.14 | 0.08 | 0.05 | 0.08 | 0.04 |
| 6 | 74727651 | rs9447104 | -1.94 | 0.267 | -0.71 | 1.313 | -0.005 | 0.034 | 0.034 | 0.015 | 0.59 | 0.78 | 0.55 | 0.66 | 0.54 | 0.58 |
| 6 | 74728216 | rs9360723 | 0.179 | 0.215 | -0.813 | 1.578 | 0.002 | 0.013 | 0.013 | 5.122 | 0.08 | 0.08 | 0.15 | 0.28 | 0.05 | 0.03 |
| 6 | 74728719 | rs4706565 | -0.098 | 0.295 | -0.78 | 1.588 | -0.011 | -0.044 | -0.044 | 0 | 0.16 | 0.09 | 0.19 | 0.28 | 0.14 | 0.08 |
| 6 | 74734641 | rs7763484 | -0.136 | 0.358 | -0.504 | 1.652 | -0.035 | 0.012 | 0.012 | 0 | 0.1 | 0.14 | 0.05 | 0.12 | 0.03 | 0.09 |
| 6 | 74739868 | rs9343101 | -0.525 | 0.515 | -0.173 | 1.566 | -0.002 | -0.021 | -0.021 | 0 | 0.36 | 0.29 | 0.36 | 0.54 | 0.34 | 0.35 |
| 6 | 74746153 | rs6453717 | -0.912 | 0.468 | -0.1 | 1.498 | -0.024 | -0.008 | -0.008 | 0.297 | 0.23 | 0.56 | 0.16 | 0.15 | 0.13 | 0.2 |
| 6 | 74753516 | rs13200097 | -0.252 | 0.431 | -0.103 | 1.471 | 0.002 | -0.002 | -0.002 | 0 | 0.36 | 0.3 | 0.29 | 0.41 | 0.27 | 0.25 |
| 6 | 74754551 | rs9447121 | 0.18 | 0.389 | -0.136 | 1.429 | 0.065 | -0.027 | -0.027 | 1.119 | 0.13 | 0.39 | 0.11 | 0.04 | 0.1 | 0.13 |
| 6 | 74767624 | rs9360732 | 0.038 | 0.41 | -0.468 | 1.412 | 0.022 | -0.039 | -0.039 | 0 | 0.31 | 0.16 | 0.26 | 0.41 | 0.23 | 0.23 |
| 6 | 74768985 | rs13215297 | -0.269 | 0.47 | -0.592 | 1.34 | 0.011 | -0.01 | -0.01 | 0 | 0.12 | 0.08 | 0.11 | 0.14 | 0.1 | 0.16 |
| 6 | 74771648 | rs6915086 | -1.855 | 0.625 | -0.613 | 1.382 | -0.032 | 0.041 | 0.041 | 0.738 | 0.64 | 0.82 | 0.69 | 0.73 | 0.57 | 0.6 |
| 6 | 74774958 | rs4708114 | -2.454 | 0.647 | -0.787 | 1.35 | -0.089 | -0.025 | -0.025 | 0 | 0.7 | 0.98 | 0.77 | 0.75 | 0.65 | 0.62 |
| 6 | 74781080 | rs4708116 | -2.181 | 0.311 | -0.585 | 1.088 | -0.086 | -0.078 | -0.078 | 1.648 | 0.67 | 0.94 | 0.69 | 0.61 | 0.59 | 0.49 |
| 6 | 74781737 | rs7751176 | -1.39 | 0.225 | -0.379 | 1.123 | -0.024 | 0.004 | 0.004 | 0.334 | 0.59 | 0.76 | 0.57 | 0.39 | 0.45 | 0.35 |
| 6 | 74786250 | rs9352063 | 0.218 | 0.133 | -0.186 | 1.026 | 0.021 | -0.008 | -0.008 | 0 | 0.3 | 0.21 | 0.37 | 0.35 | 0.2 | 0.2 |
| 6 | 74795428 | rs4706570 | -1.723 | 0.13 | -0.122 | 1.005 | -0.034 | -0.013 | -0.013 | 8.523 | 0.61 | 0.75 | 0.6 | 0.59 | 0.5 | 0.48 |
| 6 | 74795933 | rs7746248 | -0.697 | 0.13 | -0.083 | 1.031 | -0.016 | -0.026 | -0.026 | 1.685 | 0.23 | 0.45 | 0.14 | 0.15 | 0.17 | 0.2 |
| 6 | 74800529 | rs6921555 | -0.326 | 0.169 | -0.227 | 1.037 | 0.006 | 0 | 0 | 0 | 0.37 | 0.36 | 0.46 | 0.37 | 0.29 | 0.22 |
| 6 | 74818731 | rs2251494 | 2.078 | 0.326 | -0.523 | 1.186 | -0.085 | -0.077 | -0.077 | 0.043 | 0.33 | 0.06 | 0.31 | 0.39 | 0.41 | 0.53 |
| 6 | 74828682 | rs2501180 | 1.799 | 0.323 | -0.495 | 1.156 | -0.049 | -0.08 | -0.08 | 0 | 0.35 | 0.12 | 0.34 | 0.52 | 0.41 | 0.57 |
| 6 | 74849280 | rs12202674 | -0.233 | 0.467 | -0.421 | 0.929 | -0.007 | -0.004 | -0.004 | 0.02 | 0.06 | 0.05 | 0.08 | 0.09 | 0.14 | 0.07 |
| 6 | 74851053 | rs10943177 | -0.233 | 0.467 | -0.421 | 0.929 | -0.007 | -0.004 | -0.004 | 0 | 0.06 | 0.05 | 0.08 | 0.09 | 0.14 | 0.07 |
| 6 | 74851591 | rs12216455 | -0.233 | 0.467 | -0.421 | 0.929 | -0.007 | -0.004 | -0.004 | 0 | 0.06 | 0.05 | 0.08 | 0.09 | 0.14 | 0.07 |
| 6 | 74851650 | rs12189582 | -0.233 | 0.467 | -0.421 | 0.929 | -0.007 | -0.004 | -0.004 | 0 | 0.06 | 0.05 | 0.08 | 0.09 | 0.14 | 0.07 |
| 6 | 74853727 | rs9447166 | -1.207 | 0.362 | -0.444 | 0.902 | -0.011 | 0.037 | 0.037 | 0.441 | 0.35 | 0.73 | 0.32 | 0.16 | 0.39 | 0.23 |
| 6 | 74855431 | rs1320559 | -0.143 | -0.195 | -0.436 | 1.029 | 0.024 | -0.031 | -0.031 | 0 | 0.69 | 0.78 | 0.62 | 0.55 | 0.8 | 0.76 |
| 6 | 74858096 | rs4708129 | 0.217 | -0.144 | -0.42 | 0.982 | 0.018 | -0.036 | -0.036 | 0.822 | 0.66 | 0.77 | 0.58 | 0.49 | 0.73 | 0.7 |
| 6 | 74858456 | rs7775575 | -0.244 | -0.212 | -0.499 | 0.961 | 0.017 | 0.037 | 0.037 | 0.474 | 0.73 | 0.83 | 0.75 | 0.82 | 0.77 | 0.79 |

| | | | | | | | | | | | | | | | | |
|---|---|---|---|---|---|---|---|---|---|---|---|---|---|---|---|---|
| 6 | 74863350 | rs10455295 | 2.091 | -0.141 | -0.561 | 1.013 | -0.126 | -0.087 | -0.087 | 0 | 0.35 | 0.05 | 0.34 | 0.41 | 0.46 | 0.55 |
| 6 | 74870820 | rs6453739 | -0.106 | -0.081 | -0.051 | 0.767 | 0.001 | -0.086 | -0.086 | 0.088 | 0.2 | 0.31 | 0.11 | 0.02 | 0.17 | 0.13 |
| 6 | 74876690 | rs12523779 | -0.23 | 0.07 | 0.022 | 0.976 | 0.029 | 0.03 | 0.03 | 0 | 0.07 | 0.28 | 0.12 | 0.05 | 0.09 | 0.03 |
| 6 | 74877900 | rs4708135 | 0.2 | 0.059 | -0.028 | 1.104 | 0 | 0.047 | 0.047 | 0.433 | 0.07 | 0.05 | 0.17 | 0.35 | 0.04 | 0.09 |
| 6 | 74879308 | rs9360745 | 0.181 | 0.022 | -0.039 | 1.042 | 0 | 0.036 | 0.036 | 1.198 | 0.07 | 0.05 | 0.17 | 0.34 | 0.04 | 0.09 |
| 6 | 74881307 | rs9293954 | -0.065 | -0.056 | -0.075 | 1.018 | 0.003 | 0.02 | 0.02 | 0.873 | 0.09 | 0.06 | 0.17 | 0.36 | 0.04 | 0.15 |
| 6 | 74882723 | rs9293955 | 0.255 | -0.1 | -0.119 | 0.834 | -0.012 | -0.101 | -0.101 | 0 | 0.57 | 0.55 | 0.79 | 0.94 | 0.65 | 0.73 |
| 6 | 74891239 | rs1934016 | 1.827 | -0.075 | -0.017 | 1.125 | -0.144 | -0.119 | -0.119 | 0 | 0.36 | 0.04 | 0.45 | 0.45 | 0.47 | 0.51 |
| 6 | 74891934 | rs1316495 | -0.211 | -0.082 | 0.065 | 1.113 | 0.029 | 0.026 | 0.026 | 0 | 0.07 | 0.28 | 0.12 | 0.05 | 0.09 | 0.03 |
| 6 | 74894737 | rs10455296 | 0.608 | -0.092 | 0.062 | 1.119 | 0.012 | 0.01 | 0.01 | 0.299 | 0.5 | 0.5 | 0.63 | 0.58 | 0.63 | 0.59 |
| 6 | 74920235 | rs2184234 | 1.116 | -0.043 | -0.134 | 1.09 | -0.024 | -0.032 | -0.032 | 0 | 0.43 | 0.27 | 0.51 | 0.53 | 0.54 | 0.57 |
| 6 | 74944004 | rs2788978 | 1.789 | -0.035 | -0.245 | 1.16 | -0.18 | -0.109 | -0.109 | 0 | 0.35 | 0.02 | 0.42 | 0.41 | 0.48 | 0.46 |
| 6 | 74961837 | rs1319624 | 1.268 | -0.073 | -0.167 | 1.126 | -0.037 | -0.043 | -0.043 | 0.094 | 0.42 | 0.23 | 0.5 | 0.53 | 0.54 | 0.57 |
| 6 | 74963264 | rs1767861 | 1.232 | -0.074 | -0.162 | 1.125 | -0.029 | -0.034 | -0.034 | 0.378 | 0.42 | 0.26 | 0.5 | 0.53 | 0.54 | 0.57 |
| 6 | 74970787 | rs2635117 | 1.768 | -0.039 | -0.249 | 1.209 | -0.15 | -0.134 | -0.134 | 0 | 0.35 | 0.04 | 0.45 | 0.47 | 0.47 | 0.51 |
| 6 | 74976178 | rs9341459 | -1.034 | 0.134 | 0.596 | 1.62 | 0.018 | 0.308 | 0.308 | 0 | 0.09 | 0.32 | 0.27 | 0.39 | 0.09 | 0.17 |
| 6 | 74982935 | rs4708140 | -1.752 | 0.13 | 0.605 | 1.596 | 0.018 | 0.301 | 0.301 | 0.001 | 0.14 | 0.43 | 0.32 | 0.47 | 0.16 | 0.23 |
| 6 | 74997392 | rs9443027 | -0.252 | -0.123 | 0.981 | 1.311 | -0.002 | -0.038 | -0.038 | 0 | 0.26 | 0.28 | 0.17 | 0.07 | 0.21 | 0.18 |
| 6 | 74998457 | rs9360769 | -1.336 | 0.232 | 1.238 | 1.942 | 0.006 | 0.236 | 0.236 | 0 | 0.09 | 0.3 | 0.27 | 0.39 | 0.1 | 0.18 |
| 6 | 75001722 | rs9343128 | -1.492 | 0.231 | 1.286 | 1.942 | 0.029 | 0.355 | 0.355 | 0 | 0.1 | 0.39 | 0.27 | 0.39 | 0.1 | 0.18 |
| 6 | 75018941 | rs6914400 | -0.742 | 0.279 | 1.189 | 1.922 | 0.004 | 0.007 | 0.007 | 0 | 0.05 | 0.12 | 0.05 | 0.07 | 0.07 | 0.05 |
| 6 | 75035349 | rs1552568 | -1.06 | 0.373 | 0.397 | 1.993 | 0.002 | 0.214 | 0.214 | 0 | 0.07 | 0.15 | 0.22 | 0.44 | 0.08 | 0.18 |
| 6 | 75038067 | rs12523889 | -0.357 | 0.426 | 0.311 | 1.444 | 0.014 | 0.005 | 0.005 | 0 | 0.06 | 0.19 | 0.1 | 0.05 | 0.09 | 0.06 |
| 6 | 75056242 | rs524762 | 1.287 | 0.22 | -0.315 | 1.873 | 0.004 | -0.043 | -0.043 | 0.083 | 0.84 | 0.86 | 0.71 | 0.52 | 0.84 | 0.73 |
| 6 | 75056542 | rs599121 | 1.417 | -0.116 | -1.032 | 1.512 | -0.081 | -0.02 | -0.02 | 0 | 0.43 | 0.06 | 0.45 | 0.37 | 0.57 | 0.52 |
| 6 | 75058189 | rs6936857 | -0.392 | -0.047 | -0.94 | 0.988 | 0.04 | 0.087 | 0.087 | 2.123 | 0.41 | 0.73 | 0.25 | 0.15 | 0.27 | 0.22 |
| 6 | 75061324 | rs278351 | 0.324 | 0.467 | -0.528 | 1.292 | -0.003 | -0.002 | -0.002 | 0 | 0.9 | 0.89 | 0.91 | 0.94 | 0.91 | 0.92 |
| 6 | 75073124 | rs526770 | 1.593 | 0.512 | -0.466 | 1.711 | -0.005 | 0.035 | 0.035 | 0 | 0.81 | 0.74 | 0.85 | 0.61 | 0.82 | 0.73 |
| 6 | 75077150 | rs2635108 | 0.926 | 0.487 | -0.452 | 1.735 | -0.012 | 0.008 | 0.008 | 0 | 0.88 | 0.83 | 0.96 | 0.73 | 0.96 | 0.81 |
| 6 | 75080425 | rs9447193 | -0.057 | 0.299 | -0.325 | 1.47 | 0.004 | -0.009 | -0.009 | 0.963 | 0.19 | 0.56 | 0.16 | 0.13 | 0.15 | 0.07 |
| 6 | 75082067 | rs2756339 | 1.47 | 0.289 | -0.295 | 1.476 | 0.002 | -0.002 | -0.002 | 0.017 | 0.8 | 0.78 | 0.86 | 0.61 | 0.82 | 0.73 |
| 6 | 75093523 | rs278343 | 1.767 | 0.113 | -0.541 | 1.164 | -0.008 | 0.014 | 0.014 | 0 | 0.66 | 0.55 | 0.72 | 0.57 | 0.68 | 0.62 |
| 6 | 75094359 | rs278341 | 1.227 | 0.155 | -0.635 | 1.294 | -0.005 | 0.044 | 0.044 | 0 | 0.81 | 0.76 | 0.85 | 0.61 | 0.82 | 0.76 |
| 6 | 75102073 | rs686017 | 1.199 | 0.146 | -0.681 | 1.285 | 0.002 | -0.006 | -0.006 | 0 | 0.82 | 0.82 | 0.86 | 0.61 | 0.82 | 0.76 |
| 6 | 75112715 | rs795857 | 1.844 | 0.403 | -0.886 | 1.193 | -0.131 | -0.053 | -0.053 | 0.529 | 0.46 | 0.03 | 0.57 | 0.44 | 0.54 | 0.59 |
| 6 | 75119049 | rs1358880 | 1.687 | 0.387 | 0.042 | 1.553 | 0 | 0.005 | 0.005 | 0 | 0.81 | 0.77 | 0.85 | 0.61 | 0.82 | 0.76 |
| 6 | 75125541 | rs7774793 | 1.153 | 0.377 | 0.072 | 1.519 | -0.015 | -0.005 | -0.005 | 0 | 0.86 | 0.77 | 0.88 | 0.9 | 0.87 | 0.86 |
| 6 | 75140143 | rs716314 | 2.011 | 0.601 | -0.06 | 1.572 | -0.002 | 0.015 | 0.015 | 0 | 0.78 | 0.72 | 0.83 | 0.58 | 0.77 | 0.77 |
| 6 | 75140978 | rs9447201 | 1.966 | 0.613 | -0.102 | 1.579 | -0.001 | 0.009 | 0.009 | 0 | 0.78 | 0.75 | 0.83 | 0.58 | 0.77 | 0.77 |
| 6 | 75142356 | rs4706582 | 0.796 | 0.572 | -0.126 | 1.565 | -0.021 | -0.013 | -0.013 | 0 | 0.9 | 0.96 | 0.89 | 0.88 | 0.85 | 0.9 |
| 6 | 75144576 | rs9443037 | 2.079 | 0.587 | -0.12 | 1.55 | 0 | 0.007 | 0.007 | 0 | 0.78 | 0.75 | 0.83 | 0.58 | 0.77 | 0.77 |
| 6 | 75175449 | rs9343171 | -1.762 | 0.744 | 0.133 | 1.536 | 0.003 | 0.064 | 0.064 | 0 | 0.17 | 0.26 | 0.16 | 0.43 | 0.22 | 0.21 |
| 6 | 75184522 | rs539255 | -0.502 | 1.168 | 0.504 | 1.09 | 0.043 | 0.001 | 0.001 | 0 | 0.16 | 0.47 | 0.13 | 0.09 | 0.18 | 0.04 |
| 6 | 75184612 | rs9918354 | -0.165 | 1.199 | 0.066 | 0.793 | -0.002 | -0.004 | -0.004 | 0 | 0.15 | 0.13 | 0.16 | 0.07 | 0.19 | 0.09 |
| 6 | 75217944 | rs6902731 | -1.169 | 1.056 | 0.021 | 1.687 | 0.012 | 0.204 | 0.204 | 0.345 | 0.07 | 0.18 | 0.16 | 0.43 | 0.07 | 0.12 |
| 6 | 75229656 | rs9293973 | 1.526 | 0.584 | 0.039 | 1.733 | -0.02 | 0.179 | 0.179 | 0 | 0.86 | 0.72 | 0.78 | 0.54 | 0.88 | 0.8 |
| 6 | 75236060 | rs1931691 | 2.145 | 0.778 | -0.127 | 1.508 | -0.002 | 0.082 | 0.082 | 0 | 0.79 | 0.68 | 0.73 | 0.54 | 0.79 | 0.78 |
| 6 | 75238840 | rs6923011 | 1.392 | 0.747 | -0.102 | 1.553 | -0.018 | 0.184 | 0.184 | 0 | 0.87 | 0.73 | 0.79 | 0.54 | 0.88 | 0.8 |
| 6 | 75264996 | rs1577217 | -0.809 | 0.744 | -0.105 | 1.553 | 0.017 | -0.023 | -0.023 | 0 | 0.11 | 0.07 | 0.13 | 0.17 | 0.09 | 0.15 |
| 6 | 75291866 | rs2992983 | -1.071 | 0.759 | -0.172 | 1.181 | -0.021 | 0.268 | 0.268 | 0 | 0.1 | 0.26 | 0.19 | 0.45 | 0.07 | 0.13 |
| 6 | 75310669 | rs2785687 | -2.254 | 0.189 | -1.474 | 0.211 | -0.036 | 0.119 | 0.119 | 0.543 | 0.79 | 0.99 | 0.89 | 0.98 | 0.81 | 0.88 |
| 6 | 75316346 | rs12111118 | -0.032 | -0.034 | -0.975 | 0.651 | -0.018 | 0.021 | 0.021 | 0 | 0.07 | 0.11 | 0.1 | 0.15 | 0.03 | 0.08 |
| 6 | 75319716 | rs1167367 | -2.18 | -0.152 | -1.718 | 0.42 | -0.092 | 0.004 | 0.004 | 0 | 0.51 | 0.96 | 0.53 | 0.6 | 0.54 | 0.31 |
| 6 | 75320927 | rs528961 | -2.128 | -0.112 | -1.395 | 0.287 | -0.018 | 0.029 | 0.029 | 0 | 0.47 | 0.79 | 0.51 | 0.58 | 0.54 | 0.3 |
| 6 | 75324453 | rs1336764 | 0.288 | 0.093 | -1.298 | 0.538 | -0.068 | 0.057 | 0.057 | 0 | 0.12 | 0.01 | 0.15 | 0.04 | 0.18 | 0.06 |
| 6 | 75331064 | rs1167363 | -1.061 | 0.073 | -1.265 | 0.565 | -0.007 | -0.032 | -0.032 | 0.02 | 0.23 | 0.17 | 0.33 | 0.5 | 0.27 | 0.24 |
| 6 | 75340459 | rs2485140 | -1.002 | 0.114 | -1.248 | 0.545 | -0.008 | -0.017 | -0.017 | 0.112 | 0.23 | 0.17 | 0.29 | 0.37 | 0.26 | 0.21 |
| 6 | 75341929 | rs1336763 | -0.577 | 0.369 | -0.989 | 0.554 | 0.02 | 0.005 | 0.005 | 0 | 0.17 | 0.25 | 0.2 | 0.17 | 0.2 | 0.14 |
| 6 | 75346425 | rs6932497 | -1.484 | -0.382 | -0.833 | 0.384 | -0.025 | -0.075 | -0.075 | 0.896 | 0.22 | 0.6 | 0.17 | 0.05 | 0.16 | 0.06 |
| 6 | 75347593 | rs12215324 | 2.157 | -0.502 | -1.279 | 0.226 | -0.132 | -0.113 | -0.113 | 0.033 | 0.36 | 0.03 | 0.45 | 0.47 | 0.42 | 0.5 |
| 6 | 75349672 | rs2502496 | -1.92 | -0.371 | -0.004 | 0.065 | 0.034 | 0.019 | 0.019 | 0 | 0.21 | 0.36 | 0.23 | 0.26 | 0.26 | 0.2 |
| 6 | 75351286 | rs2485075 | -2.729 | -0.367 | -0.013 | 0.03 | -0.014 | 0.014 | 0.014 | 0 | 0.36 | 0.87 | 0.34 | 0.3 | 0.35 | 0.22 |
| 6 | 75351505 | rs6910395 | -0.736 | -0.387 | 0.054 | 0.064 | 0.014 | -0.029 | -0.029 | 1.097 | 0.11 | 0.45 | 0.07 | 0.02 | 0.06 | 0.01 |
| 6 | 75352934 | rs10498900 | -1.173 | -0.324 | -0.046 | -0.018 | -0.032 | -0.045 | -0.045 | 0 | 0.15 | 0.5 | 0.12 | 0.05 | 0.09 | 0.02 |
| 6 | 75353931 | rs2485077 | -0.743 | -0.171 | -0.007 | 0.057 | 0.032 | -0.009 | -0.009 | 0 | 0.12 | 0.21 | 0.12 | 0.12 | 0.18 | 0.11 |
| 6 | 75367272 | rs2502473 | -1.026 | -0.238 | 0 | 0.066 | 0.061 | -0.024 | -0.024 | 0.436 | 0.14 | 0.28 | 0.13 | 0.13 | 0.21 | 0.12 |
| 6 | 75405874 | rs2485116 | -1.416 | -0.254 | 0.004 | 0.157 | 0.016 | -0.012 | -0.012 | 0 | 0.2 | 0.29 | 0.18 | 0.16 | 0.24 | 0.14 |
| 6 | 75409434 | rs2502521 | -2.582 | -0.34 | -0.063 | 0.09 | -0.007 | 0 | 0 | 0 | 0.37 | 0.87 | 0.34 | 0.31 | 0.35 | 0.22 |
| 6 | 75415209 | rs1932833 | -0.5 | -0.215 | 0.13 | -0.037 | 0.021 | 0.046 | 0.046 | 6.301 | 0.07 | 0.5 | 0.05 | 0.02 | 0.02 | 0 |

| | | | | | | | | | | | | | | | | |
|---|---|---|---|---|---|---|---|---|---|---|---|---|---|---|---|---|
| 6 | 75427451 | rs9360837 | 0.667 | 0.083 | 0.026 | 0.151 | 0.011 | 0.005 | 0.005 | 0 | 0.86 | 0.83 | 0.87 | 0.83 | 0.79 | 0.88 |
| 6 | 75438427 | rs9343216 | 1.859 | -0.34 | 0.104 | 0.177 | 0.006 | -0.007 | -0.007 | 0 | 0.73 | 0.67 | 0.77 | 0.8 | 0.7 | 0.82 |
| 6 | 75440634 | rs9443057 | 1.449 | -0.154 | 0.088 | 0.308 | -0.004 | 0.007 | 0.007 | 0 | 0.8 | 0.77 | 0.78 | 0.79 | 0.83 | 0.85 |
| 6 | 75447088 | rs7752646 | -1.114 | -0.139 | 0.247 | 0.291 | 0.028 | -0.001 | -0.001 | 0 | 0.09 | 0.14 | 0.06 | 0.1 | 0.14 | 0.07 |
| 6 | 75451924 | rs2502463 | -1.054 | -0.363 | 0.249 | 0.293 | -0.002 | -0.005 | -0.005 | 0 | 0.17 | 0.22 | 0.17 | 0.09 | 0.16 | 0.11 |
| 6 | 75452066 | rs4112823 | -0.11 | -0.508 | 0.139 | 0.097 | -0.051 | -0.118 | -0.118 | 0 | 0.2 | 0.48 | 0.08 | 0.02 | 0.08 | 0.12 |
| 6 | 75454012 | rs945817 | -0.549 | -0.363 | 0.272 | 0.25 | -0.036 | -0.052 | -0.052 | 0 | 0.28 | 0.48 | 0.19 | 0.14 | 0.17 | 0.21 |
| 6 | 75455385 | rs2502465 | -1.287 | -0.333 | 0.233 | 0.271 | 0.005 | 0.016 | 0.016 | 0 | 0.21 | 0.37 | 0.18 | 0.19 | 0.21 | 0.13 |
| 6 | 75457103 | rs885463 | -1.856 | -0.326 | 0.019 | 0.273 | 0.038 | 0.047 | 0.047 | 0 | 0.49 | 0.86 | 0.37 | 0.33 | 0.37 | 0.34 |
| 6 | 75457183 | rs2485453 | -0.226 | 0.249 | 0.252 | 0.203 | 0.008 | -0.001 | -0.001 | 0.044 | 0.1 | 0.15 | 0.12 | 0.06 | 0.12 | 0.09 |
| 6 | 75459754 | rs16885535 | -1.017 | -0.189 | 0.035 | 0.263 | -0.016 | 0.018 | 0.018 | 0 | 0.1 | 0.19 | 0.06 | 0.13 | 0.08 | 0.04 |
| 6 | 75461596 | rs2248047 | -0.98 | -0.148 | -0.455 | 0.055 | 0.027 | 0.063 | 0.063 | 0.034 | 0.36 | 0.74 | 0.2 | 0.16 | 0.27 | 0.19 |
| 6 | 75461657 | rs2248046 | -0.982 | -0.394 | -0.44 | 0.035 | 0.041 | 0.058 | 0.058 | 0.05 | 0.37 | 0.73 | 0.19 | 0.16 | 0.21 | 0.2 |
| 6 | 75461769 | rs2502467 | -1 | -0.44 | -0.065 | 0.141 | -0.015 | -0.017 | -0.017 | 0 | 0.36 | 0.61 | 0.22 | 0.22 | 0.22 | 0.29 |
| 6 | 75462448 | rs7756912 | -1.025 | 0.262 | 0.395 | 1.26 | 0.014 | 0.052 | 0.052 | 0 | 0.16 | 0.24 | 0.26 | 0.33 | 0.16 | 0.05 |
| 6 | 75477971 | rs2883136 | -1.973 | 0.032 | 0.877 | 1.194 | -0.003 | -0.018 | -0.018 | 0.078 | 0.39 | 0.57 | 0.44 | 0.74 | 0.33 | 0.25 |
| 6 | 75484531 | rs4706586 | -1.094 | -0.023 | 1.106 | 1.499 | -0.003 | -0.034 | -0.034 | 0 | 0.21 | 0.16 | 0.27 | 0.46 | 0.19 | 0.06 |
| 6 | 75485472 | rs10943218 | -1.107 | -0.028 | 1.366 | 1.498 | 0 | 0.027 | 0.027 | 0 | 0.21 | 0.24 | 0.27 | 0.46 | 0.19 | 0.06 |
| 6 | 75490041 | rs756258 | -1.135 | -0.12 | 1.767 | 1.274 | -0.005 | 0.021 | 0.021 | 0 | 0.18 | 0.31 | 0.17 | 0.28 | 0.14 | 0.19 |
| 6 | 75507330 | rs2353325 | -1.002 | -0.046 | 1.834 | 1.308 | -0.017 | 0.043 | 0.043 | 0 | 0.16 | 0.33 | 0.15 | 0.27 | 0.11 | 0.16 |
| 6 | 75516703 | rs12526043 | -0.909 | -0.24 | 1.906 | 1.344 | -0.028 | 0.019 | 0.019 | 0 | 0.07 | 0.19 | 0.03 | 0.11 | 0.02 | 0.07 |
| 6 | 75517725 | rs4084884 | -2.303 | -0.231 | 1.915 | 1.425 | -0.001 | -0.003 | -0.003 | 0 | 0.38 | 0.63 | 0.43 | 0.71 | 0.33 | 0.23 |
| 6 | 75531422 | rs9343235 | -1.139 | -0.211 | 1.812 | 1.352 | -0.011 | 0.204 | 0.204 | 1.263 | 0.14 | 0.38 | 0.22 | 0.43 | 0.1 | 0.04 |
| 6 | 75532577 | rs6918679 | -1.121 | -0.209 | 1.813 | 1.353 | -0.01 | 0.206 | 0.206 | 1.112 | 0.14 | 0.38 | 0.22 | 0.43 | 0.1 | 0.04 |
| 6 | 75533420 | rs1890091 | -0.93 | -0.272 | 1.902 | 1.286 | -0.03 | 0.017 | 0.017 | 0 | 0.08 | 0.2 | 0.03 | 0.11 | 0.02 | 0.07 |
| 6 | 75533452 | rs1890090 | -2.305 | -0.231 | 1.65 | 1.206 | -0.001 | -0.004 | -0.004 | 0.08 | 0.38 | 0.61 | 0.41 | 0.7 | 0.33 | 0.23 |
| 6 | 75536823 | rs16879413 | -2.256 | -0.214 | 1.368 | 1.349 | -0.005 | -0.018 | -0.018 | 0 | 0.33 | 0.31 | 0.34 | 0.49 | 0.3 | 0.17 |
| 6 | 75536940 | rs17786760 | -0.554 | -0.156 | 1.201 | 1.184 | -0.062 | 0.062 | 0.062 | 0 | 0.13 | 0.03 | 0.11 | 0.02 | 0.19 | 0.06 |
| 6 | 75542900 | rs4397194 | -1.312 | -0.224 | 1.677 | 1.759 | -0.018 | 0.186 | 0.186 | 1.11 | 0.15 | 0.33 | 0.33 | 0.57 | 0.12 | 0.08 |
| 6 | 75543827 | rs7748489 | -1.302 | -0.225 | 1.685 | 1.758 | -0.017 | 0.18 | 0.18 | 1.109 | 0.15 | 0.32 | 0.33 | 0.57 | 0.12 | 0.08 |
| 6 | 75544130 | rs7752681 | -2.021 | -0.199 | 1.945 | 1.761 | -0.032 | 0.105 | 0.105 | 0.459 | 0.21 | 0.41 | 0.35 | 0.68 | 0.14 | 0.16 |
| 6 | 75549552 | rs2883138 | -1.997 | -0.199 | 1.945 | 1.756 | -0.027 | 0.126 | 0.126 | 0.497 | 0.2 | 0.39 | 0.3 | 0.45 | 0.14 | 0.12 |
| 6 | 75551717 | rs9343246 | -2.041 | -0.198 | 1.945 | 1.734 | -0.032 | 0.106 | 0.106 | 0.614 | 0.21 | 0.41 | 0.35 | 0.67 | 0.14 | 0.16 |
| 6 | 75553805 | rs9443079 | -2.091 | -0.209 | 1.94 | 1.739 | -0.033 | 0.105 | 0.105 | 0.631 | 0.21 | 0.41 | 0.35 | 0.67 | 0.14 | 0.16 |
| 6 | 75557870 | rs9447314 | -1.648 | -0.255 | 1.966 | 1.726 | 0.02 | 0.244 | 0.244 | 0 | 0.06 | 0.29 | 0.09 | 0.32 | 0.03 | 0.11 |
| 6 | 75558759 | rs9293982 | -2.413 | -0.328 | 2.225 | 1.485 | -0.005 | -0.01 | -0.01 | 0 | 0.37 | 0.8 | 0.48 | 0.7 | 0.32 | 0.23 |
| 6 | 75559326 | rs6932300 | -1.389 | -0.299 | 2.157 | 1.465 | 0.017 | 0.074 | 0.074 | 0 | 0.06 | 0.26 | 0.04 | 0.11 | 0.03 | 0.08 |
| 6 | 75567726 | rs9447322 | -2.472 | -0.301 | 2.428 | 1.698 | 0.006 | 0.178 | 0.178 | 8.561 | 0.07 | 0.59 | 0.08 | 0.2 | 0.01 | 0.05 |
| 6 | 75585492 | rs16885806 | -0.004 | -0.312 | 2.037 | 1.628 | -0.028 | -0.059 | -0.059 | 0.015 | 0.14 | 0.01 | 0.18 | 0.19 | 0.14 | 0.07 |
| 6 | 75585795 | rs7757844 | -1.794 | -0.124 | 2.418 | 1.762 | 0.031 | 0.027 | 0.027 | 7.127 | 0.06 | 0.38 | 0.03 | 0 | 0 | 0 |
| 6 | 75587451 | rs9443096 | -1.829 | -0.122 | 2.372 | 1.799 | 0.007 | 0.171 | 0.171 | 6.301 | 0.06 | 0.33 | 0.06 | 0.2 | 0 | 0.05 |
| 6 | 75600804 | rs7743423 | -1.619 | 0.044 | 1.059 | 1.154 | -0.08 | -0.115 | -0.115 | 1.035 | 0.71 | 0.98 | 0.67 | 0.72 | 0.68 | 0.51 |
| 6 | 75610722 | rs9447347 | -1.708 | -0.533 | 1.941 | 1.084 | -0.046 | 0.043 | 0.043 | 1.301 | 0.16 | 0.68 | 0.19 | 0.33 | 0.07 | 0.15 |
| 6 | 75620466 | rs9447358 | -2.022 | -0.349 | 2.205 | 0.828 | 0.025 | 0.204 | 0.204 | 8.578 | 0.07 | 0.53 | 0.08 | 0.2 | 0.01 | 0.05 |
| 6 | 75680571 | rs6932000 | -1.706 | -0.659 | 2.243 | 0.863 | 0.028 | 0.223 | 0.223 | 5.842 | 0.06 | 0.54 | 0.07 | 0.21 | 0.01 | 0.05 |
| 6 | 75700158 | rs4495246 | -2.14 | -0.688 | 2.064 | 0.848 | 0.011 | 0.195 | 0.195 | 4.605 | 0.07 | 0.53 | 0.07 | 0.21 | 0.01 | 0.06 |
| 6 | 75701667 | rs10943230 | 1.104 | -0.823 | 1.714 | 0.599 | 0.06 | -0.039 | -0.039 | 0 | 0.49 | 0.1 | 0.54 | 0.47 | 0.65 | 0.66 |
| 6 | 75708289 | rs9447400 | -1.91 | -0.141 | 2.734 | 0.944 | 0.032 | 0.089 | 0.089 | 6.909 | 0.06 | 0.56 | 0.04 | 0.08 | 0.01 | 0.05 |
| 6 | 75737968 | rs4291062 | 1.92 | -0.733 | 2.985 | 1.334 | -0.029 | -0.012 | -0.012 | 0 | 0.82 | 0.38 | 0.79 | 0.82 | 0.89 | 0.87 |
| 6 | 75741836 | rs9360872 | 1.988 | -0.956 | 2.793 | 1.408 | 0.01 | -0.017 | -0.017 | 0 | 0.78 | 0.19 | 0.74 | 0.66 | 0.85 | 0.88 |
| 6 | 75750133 | rs6453809 | 1.391 | -0.713 | 3.07 | 1.361 | -0.018 | -0.018 | -0.018 | 0 | 0.85 | 0.41 | 0.83 | 0.89 | 0.9 | 0.92 |
| 6 | 75752445 | rs4569931 | -1.301 | -0.705 | 2.85 | 1.369 | -0.01 | -0.007 | -0.007 | 0.512 | 0.14 | 0.3 | 0.15 | 0.11 | 0.1 | 0.08 |
| 6 | 75775381 | rs9343264 | 1.949 | -0.732 | 2.301 | 1.343 | 0.008 | -0.012 | -0.012 | 0 | 0.78 | 0.23 | 0.76 | 0.66 | 0.84 | 0.86 |
| 6 | 75779343 | rs6453812 | -1.354 | -0.437 | 2.922 | 1.581 | 0.163 | 0.31 | 0.31 | 0.573 | 0.05 | 0.42 | 0.08 | 0.21 | 0.05 | 0.05 |
| 6 | 75789526 | rs554152 | 0.892 | -0.242 | 2.006 | 1.236 | 0.029 | 0.017 | 0.017 | 0.086 | 0.88 | 0.78 | 0.85 | 0.85 | 0.86 | 0.88 |
| 6 | 75797302 | rs970547 | 0.948 | -1.115 | 1.775 | 1.366 | 0.006 | 0 | 0 | 0 | 0.65 | 0.68 | 0.77 | 0.68 | 0.77 | 0.68 |
| 6 | 75808390 | rs860569 | 0.707 | -0.541 | 2.012 | 1.621 | 0 | 0 | 0 | 0 | 0.89 | 0.9 | 0.92 | 0.86 | 0.9 | 0.89 |
| 6 | 75811880 | rs556674 | 1.783 | -0.647 | 2.384 | 1.949 | -0.003 | 0.099 | 0.099 | 0 | 0.82 | 0.54 | 0.84 | 0.67 | 0.88 | 0.83 |
| 6 | 75815840 | rs7758152 | -1.346 | -0.63 | 2.294 | 1.943 | 0.03 | 0.18 | 0.18 | 0 | 0.06 | 0.28 | 0.07 | 0.19 | 0.02 | 0.06 |
| 6 | 75825181 | rs2300795 | -1.728 | -0.636 | 2.259 | 2.014 | 0.027 | 0.254 | 0.254 | 0.045 | 0.07 | 0.35 | 0.08 | 0.23 | 0.02 | 0.06 |
| 6 | 75829373 | rs636885 | 0.86 | -0.572 | 1.469 | 1.89 | 0.001 | -0.001 | -0.001 | 0.005 | 0.88 | 0.87 | 0.91 | 0.86 | 0.89 | 0.88 |
| 6 | 75834248 | rs51630 | 1.239 | -0.825 | 1.17 | 1.851 | 0.015 | 0.031 | 0.031 | 0 | 0.66 | 0.84 | 0.73 | 0.7 | 0.72 | 0.8 |
| 6 | 75835305 | rs6453815 | 1.413 | -0.692 | 1.452 | 1.945 | -0.011 | 0.048 | 0.048 | 0 | 0.78 | 0.45 | 0.83 | 0.64 | 0.84 | 0.82 |
| 6 | 75838596 | rs11966644 | -0.733 | -0.752 | 0.992 | 1.668 | -0.001 | 0 | 0 | 0 | 0.23 | 0.22 | 0.26 | 0.22 | 0.17 | 0.08 |
| 6 | 75840120 | rs1412709 | 1.093 | -0.325 | 1.115 | 1.983 | -0.006 | 0.022 | 0.022 | 0 | 0.83 | 0.69 | 0.82 | 0.76 | 0.87 | 0.88 |
| 6 | 75843360 | rs240727 | 0.99 | -0.329 | 0.388 | 1.614 | 0.008 | 0.006 | 0.006 | 0 | 0.85 | 0.88 | 0.9 | 0.85 | 0.89 | 0.88 |
| 6 | 75843845 | rs1332778 | 1.048 | -0.283 | 0.439 | 1.747 | -0.018 | 0.09 | 0.09 | 0.094 | 0.74 | 0.52 | 0.71 | 0.4 | 0.79 | 0.77 |
| 6 | 75844600 | rs1332020 | -0.373 | 0.472 | 0.14 | 1.554 | 0.011 | 0.022 | 0.022 | 0 | 0.31 | 0.45 | 0.49 | 0.53 | 0.42 | 0.43 |
| 6 | 75844924 | rs2239643 | 0.781 | 0.023 | 0.536 | 1.759 | -0.001 | 0.14 | 0.14 | 0 | 0.81 | 0.67 | 0.79 | 0.61 | 0.8 | 0.68 |

| | | | | | | | | | | | | | | | | |
|---|---|---|---|---|---|---|---|---|---|---|---|---|---|---|---|---|
| 6 | 75845113 | rs240731 | 0.231 | -0.432 | 0.241 | 0.141 | -0.001 | -0.023 | -0.023 | 0.03 | 0.48 | 0.53 | 0.74 | 0.83 | 0.63 | 0.6 |
| 6 | 75845184 | rs240732 | 0.893 | -0.68 | 0.59 | 0.668 | -0.036 | -0.013 | -0.013 | 0 | 0.9 | 0.78 | 0.96 | 0.95 | 0.97 | 0.91 |
| 6 | 75847036 | rs240734 | -0.558 | -0.408 | 0.408 | 0.929 | -0.003 | 0.001 | 0.001 | 0.638 | 0.34 | 0.58 | 0.33 | 0.31 | 0.29 | 0.32 |
| 6 | 75848181 | rs240736 | -0.256 | -0.146 | 0.297 | 1.11 | 0.007 | -0.041 | -0.041 | 0 | 0.28 | 0.35 | 0.29 | 0.15 | 0.29 | 0.22 |
| 6 | 75852488 | rs240724 | -0.21 | -0.119 | -0.136 | 0.69 | 0.002 | -0.005 | -0.005 | 0 | 0.28 | 0.27 | 0.29 | 0.27 | 0.29 | 0.25 |
| 6 | 75876767 | rs4706592 | 0.57 | -0.289 | -1.18 | 0.665 | -0.075 | -0.191 | -0.191 | 0 | 0.28 | 0.01 | 0.47 | 0.46 | 0.33 | 0.36 |
| 6 | 75878425 | rs9447453 | -0.246 | -0.585 | -0.639 | 0.198 | 0.004 | 0.014 | 0.014 | 0.435 | 0.14 | 0.1 | 0.08 | 0 | 0.11 | 0.12 |
| 6 | 75879135 | rs240721 | -0.936 | -0.689 | -0.244 | 0.734 | -0.047 | 0.053 | 0.053 | 0.203 | 0.13 | 0.29 | 0.1 | 0.23 | 0.04 | 0.14 |
| 6 | 75880623 | rs3777505 | -0.785 | -0.623 | -0.44 | 0.768 | -0.01 | 0.01 | 0.01 | 0.02 | 0.12 | 0.2 | 0.1 | 0.23 | 0.04 | 0.13 |
| 6 | 75889208 | rs12215389 | -1.09 | -0.934 | -0.516 | 0.505 | 0.005 | -0.006 | -0.006 | 0.071 | 0.2 | 0.25 | 0.13 | 0.24 | 0.07 | 0.22 |
| 6 | 75915356 | rs240366 | 0.992 | -1.041 | 0.109 | 0.489 | -0.035 | -0.016 | -0.016 | 1.938 | 0.81 | 0.76 | 0.87 | 0.89 | 0.93 | 0.8 |
| 6 | 75917093 | rs240367 | 0.21 | -1.093 | 0.146 | 0.551 | -0.024 | 0.031 | 0.031 | 0 | 0.9 | 0.82 | 0.96 | 0.86 | 0.96 | 0.88 |
| 6 | 75919484 | rs240369 | -0.069 | -1.092 | 0.124 | 0.541 | 0.004 | -0.002 | -0.002 | 0.198 | 0.91 | 0.93 | 0.98 | 0.99 | 0.96 | 0.88 |
| 6 | 75924559 | rs240370 | 1.1 | -1.115 | 0.377 | 0.514 | -0.053 | 0.022 | 0.022 | 0.35 | 0.79 | 0.69 | 0.85 | 0.75 | 0.93 | 0.77 |
| 6 | 75932139 | rs13198422 | -0.039 | -0.186 | -0.228 | -0.062 | -0.074 | 0.056 | 0.056 | 0 | 0.13 | 0.02 | 0.11 | 0.05 | 0.25 | 0.08 |
| 6 | 75945548 | rs436898 | 0.726 | -0.562 | -0.038 | 0.065 | -0.034 | -0.014 | -0.014 | 0 | 0.83 | 0.76 | 0.88 | 0.88 | 0.93 | 0.81 |
| 6 | 75946422 | rs431116 | 0.785 | -0.562 | -0.058 | 0.065 | -0.041 | -0.018 | -0.018 | 0.97 | 0.83 | 0.74 | 0.88 | 0.88 | 0.93 | 0.81 |
| 6 | 75953705 | rs9360898 | 0.732 | -0.31 | 0.012 | 0.06 | -0.004 | 0.004 | 0.004 | 1.529 | 0.22 | 0.16 | 0.23 | 0.24 | 0.2 | 0.21 |
| 6 | 75963751 | rs654428 | -0.08 | -0.489 | 0.222 | 0.139 | -0.021 | -0.036 | -0.036 | 0.277 | 0.9 | 0.84 | 0.97 | 0.98 | 0.96 | 0.89 |
| 6 | 75963951 | rs381141 | -0.08 | -0.459 | 0.222 | 0.139 | -0.019 | -0.036 | -0.036 | 2.504 | 0.9 | 0.84 | 0.97 | 0.98 | 0.96 | 0.89 |
| 6 | 75964976 | rs638590 | -0.08 | -0.483 | 0.222 | 0.139 | -0.021 | -0.036 | -0.036 | 0.516 | 0.9 | 0.84 | 0.97 | 0.98 | 0.96 | 0.89 |
| 6 | 75968720 | rs147523 | -0.057 | -0.484 | 0.264 | 0.138 | -0.027 | -0.045 | -0.045 | 0 | 0.9 | 0.82 | 0.97 | 0.98 | 0.96 | 0.89 |
| 6 | 75973264 | rs240390 | 0.465 | -0.617 | 0.381 | 0.208 | -0.043 | 0 | 0 | 0.356 | 0.86 | 0.71 | 0.94 | 0.88 | 0.95 | 0.87 |
| 6 | 75978209 | rs173274 | 0.746 | -0.699 | 0.908 | 0.2 | -0.079 | 0.029 | 0.029 | 1.223 | 0.8 | 0.58 | 0.84 | 0.77 | 0.92 | 0.79 |
| 6 | 75980587 | rs240399 | 0.855 | -0.807 | 0.789 | 0.171 | -0.092 | 0.053 | 0.053 | 1.01 | 0.79 | 0.55 | 0.84 | 0.75 | 0.92 | 0.79 |
| 6 | 75992476 | rs7741596 | -1.008 | -0.432 | 0.504 | 0.018 | -0.016 | -0.003 | -0.003 | 0.036 | 0.13 | 0.27 | 0.09 | 0.1 | 0.08 | 0.02 |
| 6 | 75994151 | rs240374 | -1.06 | -0.493 | 0.835 | 0.021 | -0.018 | 0.01 | 0.01 | 0.535 | 0.13 | 0.35 | 0.09 | 0.11 | 0.08 | 0.03 |
| 6 | 76014987 | rs2748254 | -0.174 | -0.561 | 0.805 | 0.13 | 0.001 | -0.001 | -0.001 | 0.657 | 0.84 | 0.84 | 0.85 | 0.84 | 0.89 | 0.76 |
| 6 | 76016421 | rs13220419 | 1.462 | -0.326 | 0.117 | -0.206 | -0.009 | -0.043 | -0.043 | 0 | 0.26 | 0.15 | 0.36 | 0.42 | 0.32 | 0.31 |
| 6 | 76019007 | rs10806035 | -0.534 | -0.281 | 0.233 | -0.077 | -0.001 | 0 | 0 | 0 | 0.62 | 0.6 | 0.56 | 0.46 | 0.6 | 0.67 |
| 6 | 76022082 | rs9352202 | -0.529 | -0.276 | 0.235 | -0.075 | -0.001 | 0 | 0 | 0 | 0.62 | 0.6 | 0.56 | 0.46 | 0.6 | 0.67 |
| 6 | 76029760 | rs4275021 | -0.669 | -0.285 | 0.193 | 0.002 | -0.001 | -0.011 | -0.011 | 0.277 | 0.63 | 0.68 | 0.57 | 0.47 | 0.6 | 0.68 |
| 6 | 76038894 | rs16886475 | 0.321 | -0.224 | 0.02 | -0.187 | 0.006 | 0.005 | 0.005 | 0 | 0.21 | 0.18 | 0.18 | 0.17 | 0.16 | 0.24 |
| 6 | 76053386 | rs6929270 | -0.324 | -0.036 | 0.293 | -0.051 | -0.006 | 0.012 | 0.012 | 0.022 | 0.62 | 0.5 | 0.55 | 0.48 | 0.6 | 0.68 |
| 6 | 76067117 | rs7451911 | -1.219 | -0.057 | 0.118 | -0.076 | -0.001 | 0.001 | 0.001 | 0 | 0.72 | 0.7 | 0.61 | 0.48 | 0.68 | 0.69 |
| 6 | 76074442 | rs2808188 | -1.617 | -0.058 | -0.271 | -0.045 | -0.037 | -0.09 | -0.09 | 0 | 0.75 | 0.91 | 0.64 | 0.59 | 0.68 | 0.71 |
| 6 | 76077948 | rs13211961 | 0.602 | -0.004 | -0.3 | -0.182 | 0 | -0.003 | -0.003 | 0 | 0.08 | 0.05 | 0.04 | 0 | 0.07 | 0.01 |
| 6 | 76096230 | rs9293999 | 1.396 | 0.132 | -0.366 | -0.307 | -0.022 | -0.017 | -0.017 | 0 | 0.21 | 0.09 | 0.26 | 0.23 | 0.23 | 0.19 |
| 6 | 76125691 | rs2808203 | -0.58 | 0.165 | -0.188 | 0.024 | -0.045 | -0.094 | -0.094 | 0.249 | 0.66 | 0.87 | 0.58 | 0.48 | 0.61 | 0.68 |
| 6 | 76127504 | rs9359122 | -0.23 | 0.201 | 0.113 | 0.045 | -0.006 | 0.01 | 0.01 | 0.161 | 0.61 | 0.49 | 0.55 | 0.48 | 0.61 | 0.68 |
| 6 | 76130173 | rs2808206 | -0.749 | 0.604 | -0.432 | 0.645 | -0.003 | -0.093 | -0.093 | 2.209 | 0.69 | 0.84 | 0.52 | 0.49 | 0.65 | 0.64 |
| 6 | 76136268 | rs2951928 | 0.436 | 0.531 | -0.195 | 0.341 | 0.016 | -0.015 | -0.015 | 0 | 0.38 | 0.48 | 0.31 | 0.26 | 0.46 | 0.35 |
| 6 | 76160101 | rs2703714 | -0.901 | 0.517 | -0.434 | 0.571 | -0.032 | -0.065 | -0.065 | 0.369 | 0.7 | 0.92 | 0.55 | 0.62 | 0.65 | 0.69 |
| 6 | 76169472 | rs2998365 | -0.783 | 0.537 | -0.537 | 0.47 | -0.01 | -0.078 | -0.078 | 0.048 | 0.7 | 0.82 | 0.51 | 0.44 | 0.65 | 0.67 |
| 6 | 76171513 | rs2998367 | -1.074 | 0.529 | -0.604 | 0.51 | -0.043 | -0.097 | -0.097 | 0.047 | 0.72 | 0.92 | 0.54 | 0.58 | 0.65 | 0.72 |
| 6 | 76174682 | rs11754006 | 0.522 | 0.363 | -0.304 | 0.277 | -0.04 | 0.081 | 0.081 | 0.117 | 0.4 | 0.21 | 0.28 | 0.15 | 0.5 | 0.4 |
| 6 | 76174857 | rs2951916 | -0.076 | 0.271 | 0.121 | 0.3 | 0.002 | 0.058 | 0.058 | 0 | 0.48 | 0.6 | 0.32 | 0.22 | 0.5 | 0.51 |
| 6 | 76186658 | rs9360910 | -0.024 | 0.245 | 0.127 | 0.213 | -0.024 | 0.002 | 0.002 | 0 | 0.25 | 0.39 | 0.21 | 0.27 | 0.15 | 0.21 |
| 6 | 76194007 | rs2312929 | 1.191 | 0.617 | -0.294 | 0.068 | -0.1 | 0.161 | 0.161 | 0.3 | 0.31 | 0.03 | 0.21 | 0.05 | 0.36 | 0.27 |
| 6 | 76194895 | rs1932523 | -1.074 | 0.702 | -0.026 | 0.176 | 0.001 | -0.011 | -0.011 | 0.244 | 0.69 | 0.72 | 0.51 | 0.45 | 0.65 | 0.67 |
| 6 | 76195635 | rs12176377 | 1.021 | 0.721 | -0.252 | 0.186 | -0.029 | -0.113 | -0.113 | 0.062 | 0.29 | 0.11 | 0.48 | 0.55 | 0.35 | 0.33 |
| 6 | 76200514 | rs9359124 | 0.309 | 0.314 | -0.172 | 0.064 | 0.011 | -0.014 | -0.014 | 1.147 | 0.2 | 0.11 | 0.1 | 0.24 | 0.13 | 0.22 |
| 6 | 76202656 | rs9343292 | 0.341 | 0.397 | -0.553 | 0.153 | -0.028 | -0.022 | -0.022 | 0 | 0.09 | 0.07 | 0.07 | 0.11 | 0.13 | 0.13 |
| 6 | 76218758 | rs10943254 | 1.673 | 0.995 | -0.793 | 0.165 | -0.062 | -0.101 | -0.101 | 0 | 0.27 | 0.03 | 0.51 | 0.41 | 0.32 | 0.29 |
| 6 | 76222009 | rs10806042 | 0.223 | 1.001 | -0.645 | 0.189 | -0.064 | -0.057 | -0.057 | 0.09 | 0.42 | 0.18 | 0.63 | 0.52 | 0.51 | 0.5 |
| 6 | 76228952 | rs11752842 | 1.223 | 1.069 | -0.684 | 0.132 | -0.1 | 0.156 | 0.156 | 0.227 | 0.31 | 0.01 | 0.19 | 0.05 | 0.33 | 0.25 |
| 6 | 76236061 | rs9343298 | -0.033 | 1.054 | -0.318 | 0.204 | -0.002 | 0.01 | 0.01 | 0.013 | 0.53 | 0.48 | 0.33 | 0.29 | 0.48 | 0.48 |
| 6 | 76247512 | rs7756065 | 0.389 | 1.128 | -0.153 | 0.26 | 0.061 | 0.244 | 0.244 | 0.432 | 0.87 | 0.63 | 0.89 | 0.7 | 0.86 | 0.85 |
| 6 | 76265642 | rs9360921 | 0.243 | 1.154 | -0.38 | 0.162 | -0.006 | 0.006 | 0.006 | 0 | 0.1 | 0.11 | 0.07 | 0.04 | 0.12 | 0.13 |
| 6 | 76271141 | rs1414596 | 0.995 | 1.205 | -0.503 | 0.079 | -0.025 | 0.119 | 0.119 | 0.023 | 0.32 | 0.13 | 0.21 | 0.05 | 0.33 | 0.24 |
| 6 | 76272584 | rs6911868 | -1.081 | 1.172 | -0.464 | 0.116 | 0.006 | 0.037 | 0.037 | 0 | 0.67 | 0.67 | 0.44 | 0.46 | 0.61 | 0.67 |
| 6 | 76320846 | rs646967 | -0.823 | 1.146 | -0.404 | 0.14 | -0.006 | -0.011 | -0.011 | 0 | 0.34 | 0.48 | 0.2 | 0.28 | 0.26 | 0.4 |
| 6 | 76357426 | rs9352231 | 0.228 | 1.17 | -0.451 | 0.07 | -0.017 | 0.034 | 0.034 | 0 | 0.1 | 0.06 | 0.07 | 0.04 | 0.12 | 0.12 |
| 6 | 76367201 | rs766244 | 1.035 | 1.242 | -0.455 | 0.016 | -0.014 | 0.085 | 0.085 | 0.018 | 0.32 | 0.18 | 0.21 | 0.05 | 0.33 | 0.24 |
| 6 | 76375321 | rs2647415 | 0.928 | 1.24 | -0.596 | 0.123 | 0.007 | 0.037 | 0.037 | 0 | 0.33 | 0.34 | 0.57 | 0.54 | 0.4 | 0.33 |
| 6 | 76376552 | rs17414687 | 1.169 | 1.231 | -0.671 | 0.041 | -0.028 | 0.117 | 0.117 | 0.082 | 0.31 | 0.1 | 0.2 | 0.05 | 0.33 | 0.24 |
| 6 | 76390077 | rs2647399 | -1.457 | 1.228 | -0.673 | 0.045 | -0.049 | -0.051 | -0.051 | 0 | 0.69 | 0.93 | 0.45 | 0.58 | 0.62 | 0.68 |
| 6 | 76408189 | rs7770824 | 0.015 | 1.226 | -0.641 | -0.126 | -0.016 | 0 | 0 | 0 | 0.24 | 0.35 | 0.13 | 0.24 | 0.14 | 0.28 |
| 6 | 76413709 | rs17792478 | 1.101 | 1.292 | -0.698 | -0.1 | -0.028 | 0.117 | 0.117 | 0.076 | 0.31 | 0.1 | 0.2 | 0.05 | 0.33 | 0.24 |

| | | | | | | | | | | | | | | | | |
|---|---|---|---|---|---|---|---|---|---|---|---|---|---|---|---|---|
| 6 | 76416316 | rs17792496 | 1.793 | 1.318 | -0.848 | -0.254 | -0.012 | 0.039 | 0.039 | 0 | 0.08 | 0.01 | 0.05 | 0 | 0.1 | 0.01 |
| 6 | 76416655 | rs3798461 | -0.851 | 1.311 | -0.448 | 0.172 | -0.006 | -0.011 | -0.011 | 0 | 0.34 | 0.48 | 0.2 | 0.28 | 0.26 | 0.4 |
| 6 | 76425594 | rs7385 | 0.694 | 1.464 | -0.468 | 0.184 | -0.005 | -0.01 | -0.01 | 0.687 | 0.66 | 0.53 | 0.8 | 0.72 | 0.74 | 0.6 |
| 6 | 76447204 | rs2842556 | 0.391 | 1.611 | -1.207 | 0.364 | 0.022 | -0.043 | -0.043 | 2.329 | 0.72 | 0.23 | 0.82 | 0.49 | 0.83 | 0.71 |
| 6 | 76450023 | rs2647405 | 0.347 | 1.621 | -1.202 | 0.369 | 0.021 | -0.043 | -0.043 | 0 | 0.73 | 0.23 | 0.82 | 0.49 | 0.83 | 0.71 |
| 6 | 76458378 | rs2842549 | 1.106 | 1.839 | -1.175 | 0.466 | -0.023 | -0.049 | -0.049 | 0.504 | 0.3 | 0.06 | 0.52 | 0.4 | 0.33 | 0.38 |
| 6 | 76462711 | rs3798458 | 1.26 | 1.261 | -1.148 | 0.452 | -0.036 | -0.05 | -0.05 | 0 | 0.09 | 0 | 0.07 | 0.13 | 0.13 | 0.02 |
| 6 | 76469115 | rs276695 | -1.003 | 1.313 | -0.905 | 0.558 | -0.044 | 0.19 | 0.19 | 0.225 | 0.65 | 0.99 | 0.82 | 0.95 | 0.7 | 0.75 |
| 6 | 76489774 | rs6906615 | -0.251 | 1.283 | -0.448 | 0.813 | -0.005 | 0.083 | 0.083 | 0 | 0.21 | 0.6 | 0.14 | 0.38 | 0.14 | 0.25 |
| 6 | 76517896 | rs721264 | -0.49 | 1.276 | -0.469 | 0.863 | -0.002 | 0.009 | 0.009 | 0 | 0.22 | 0.72 | 0.17 | 0.51 | 0.16 | 0.27 |
| 6 | 76532719 | rs2273859 | 0.875 | 1.426 | -0.563 | 0.763 | -0.002 | -0.019 | -0.019 | 0.39 | 0.34 | 0.27 | 0.57 | 0.54 | 0.4 | 0.41 |
| 6 | 76536080 | rs3798447 | 1.041 | 1.533 | -0.564 | 0.691 | -0.014 | -0.007 | -0.007 | 0 | 0.32 | 0.16 | 0.55 | 0.4 | 0.38 | 0.39 |
| 6 | 76536333 | rs1280049 | -0.312 | 1.638 | -0.914 | 0.724 | -0.02 | 0.125 | 0.125 | 0 | 0.53 | 0.79 | 0.69 | 0.77 | 0.52 | 0.63 |
| 6 | 76550730 | rs12203461 | 0.176 | 1.997 | -0.668 | 0.908 | -0.017 | 0.018 | 0.018 | 0 | 0.11 | 0.11 | 0.1 | 0.04 | 0.17 | 0.09 |
| 6 | 76576527 | rs6453845 | 0.488 | 1.385 | -0.089 | 1.383 | 0.01 | 0.116 | 0.116 | 0.235 | 0.82 | 0.4 | 0.85 | 0.63 | 0.86 | 0.77 |
| 6 | 76601992 | rs2295938 | -0.129 | 1.207 | -0.431 | 0.402 | 0.022 | -0.016 | -0.016 | 0.074 | 0.89 | 0.64 | 0.94 | 0.93 | 0.93 | 0.87 |
| 6 | 76607654 | rs720863 | 0.024 | 1.246 | -1.401 | 0.597 | -0.011 | -0.005 | -0.005 | 0.255 | 0.32 | 0.16 | 0.54 | 0.39 | 0.38 | 0.39 |
| 6 | 76625079 | rs699186 | 0.024 | 1.246 | -1.401 | 0.58 | -0.011 | -0.003 | -0.003 | 0 | 0.32 | 0.16 | 0.54 | 0.39 | 0.38 | 0.39 |
| 6 | 76633385 | rs3778005 | 0.008 | 1.309 | -1.631 | 0.213 | -0.082 | 0.078 | 0.078 | 0 | 0.21 | 0.02 | 0.23 | 0.05 | 0.26 | 0.12 |
| 6 | 76635743 | rs9343330 | -0.518 | 0.879 | -2.159 | 0.292 | -0.043 | 0.084 | 0.084 | 0 | 0.52 | 0.17 | 0.38 | 0.29 | 0.48 | 0.35 |
| 6 | 76640781 | rs3734313 | 0.464 | 0.895 | -2.122 | 0.756 | -0.013 | -0.019 | -0.019 | 0 | 0.17 | 0.13 | 0.2 | 0.25 | 0.18 | 0.22 |
| 6 | 76640803 | rs10943299 | 0.987 | 0.71 | -2.36 | 0.495 | 0.027 | 0.081 | 0.081 | 0 | 0.23 | 0.09 | 0.08 | 0 | 0.14 | 0.14 |
| 6 | 76643642 | rs11754389 | 0.955 | 0.897 | -2.094 | 1.194 | -0.075 | 0.071 | 0.071 | 0.07 | 0.21 | 0.02 | 0.22 | 0.06 | 0.25 | 0.12 |
| 6 | 76645803 | rs9352247 | -0.081 | 0.832 | -1.352 | 1.184 | -0.055 | -0.058 | -0.058 | 0 | 0.29 | 0.09 | 0.3 | 0.28 | 0.33 | 0.22 |
| 6 | 76647200 | rs6453850 | 0.526 | 0.554 | -0.082 | 1.776 | -0.001 | 0.002 | 0.002 | 0 | 0.28 | 0.26 | 0.36 | 0.32 | 0.34 | 0.42 |
| 6 | 76650059 | rs2313752 | -0.56 | 0.684 | -0.598 | 1.592 | -0.002 | 0.005 | 0.005 | 0 | 0.69 | 0.38 | 0.59 | 0.54 | 0.65 | 0.57 |
| 6 | 76652443 | rs2313754 | -0.454 | 0.701 | -0.388 | 1.899 | -0.001 | 0.004 | 0.004 | 0 | 0.08 | 0.05 | 0.08 | 0.23 | 0.08 | 0.1 |
| 6 | 76662813 | rs4708201 | -0.831 | 0.619 | -0.083 | 2.201 | 0.003 | 0.152 | 0.152 | 0 | 0.07 | 0.16 | 0.13 | 0.34 | 0.07 | 0.13 |
| 6 | 76672441 | rs6924084 | -0.839 | 0.368 | -0.044 | 1.658 | -0.006 | 0.015 | 0.015 | 0 | 0.34 | 0.4 | 0.45 | 0.52 | 0.34 | 0.37 |
| 6 | 76676893 | rs9352252 | -0.943 | 0.259 | 0.027 | 1.927 | -0.002 | 0.018 | 0.018 | 0.031 | 0.1 | 0.13 | 0.17 | 0.21 | 0.09 | 0.22 |
| 6 | 76680290 | rs6933014 | -0.648 | -0.007 | 0.795 | 1.797 | -0.014 | 0.013 | 0.013 | 0 | 0.36 | 0.42 | 0.52 | 0.58 | 0.33 | 0.38 |
| 6 | 76681611 | rs7451737 | -1.167 | -0.069 | 0.577 | 1.626 | 0.023 | 0.025 | 0.025 | 0 | 0.48 | 0.6 | 0.62 | 0.66 | 0.4 | 0.48 |
| 6 | 76683635 | rs12527817 | -1.098 | 0.004 | 0.604 | 1.638 | -0.003 | -0.008 | -0.008 | 0 | 0.47 | 0.47 | 0.62 | 0.7 | 0.42 | 0.5 |
| 6 | 76683832 | rs7768845 | 0.213 | 0.073 | 0.599 | 1.308 | 0 | 0.001 | 0.001 | 0 | 0.09 | 0.08 | 0.09 | 0.06 | 0.07 | 0.1 |
| 6 | 76687433 | rs2794286 | 0.437 | 0.205 | 0.625 | 0.69 | 0 | -0.04 | -0.04 | 0 | 0.18 | 0.18 | 0.11 | 0 | 0.17 | 0.12 |
| 6 | 76698468 | rs12197455 | -0.582 | -0.295 | 0.738 | 1.221 | 0.009 | -0.011 | -0.011 | 0 | 0.43 | 0.35 | 0.56 | 0.54 | 0.39 | 0.43 |
| 6 | 76710341 | rs17802413 | 0.434 | -0.572 | 0.465 | 1.087 | -0.02 | -0.012 | -0.012 | 0 | 0.09 | 0 | 0.08 | 0.06 | 0.06 | 0.1 |
| 6 | 76714364 | rs7769507 | -0.097 | -0.804 | 0.743 | 1.041 | 0.013 | -0.013 | -0.013 | 1 | 0.35 | 0.28 | 0.42 | 0.48 | 0.33 | 0.32 |
| 6 | 76715299 | rs17802616 | 0.192 | -0.448 | 0.485 | 0.835 | -0.028 | 0.005 | 0.005 | 0 | 0.1 | 0.02 | 0.09 | 0.07 | 0.1 | 0.11 |
| 6 | 76715926 | rs1926098 | 0.919 | -0.662 | 0.908 | 0.741 | 0.019 | 0.021 | 0.021 | 0 | 0.49 | 0.34 | 0.44 | 0.36 | 0.56 | 0.56 |
| 6 | 76720204 | rs9447587 | 0.476 | -1.181 | 1.059 | 0.75 | 0.01 | -0.057 | -0.057 | 0.014 | 0.13 | 0.17 | 0.1 | 0 | 0.15 | 0.11 |
| 6 | 76723011 | rs724941 | 0.474 | -1.216 | 1.319 | 1.215 | 0.004 | -0.004 | -0.004 | 0.011 | 0.59 | 0.35 | 0.52 | 0.43 | 0.66 | 0.67 |
| 6 | 76730285 | rs4708205 | -0.538 | -1.617 | 1.179 | 1.355 | -0.009 | 0.022 | 0.022 | 0 | 0.08 | 0.13 | 0.08 | 0.16 | 0.07 | 0.16 |
| 6 | 76736996 | rs2148261 | 0.801 | -1.68 | 1.204 | 1.252 | -0.014 | -0.002 | -0.002 | 0 | 0.58 | 0.48 | 0.56 | 0.37 | 0.64 | 0.63 |
| 6 | 76741709 | rs1326172 | 0.64 | -1.81 | 1.077 | 1.27 | 0.01 | -0.009 | -0.009 | 0.018 | 0.59 | 0.64 | 0.6 | 0.46 | 0.64 | 0.64 |
| 6 | 76744298 | rs2274348 | 0.233 | -1.786 | 0.479 | 1.166 | 0.021 | -0.031 | -0.031 | 2.774 | 0.24 | 0.04 | 0.23 | 0.31 | 0.23 | 0.13 |
| 6 | 76755221 | rs943086 | 0.859 | -1.712 | 0.745 | 1.105 | 0.05 | 0.002 | 0.002 | 0 | 0.55 | 0.26 | 0.56 | 0.39 | 0.64 | 0.64 |
| 6 | 76763390 | rs3798588 | 0.385 | -1.373 | 0.688 | 0.897 | -0.026 | -0.02 | -0.02 | 0 | 0.06 | 0 | 0.09 | 0.07 | 0.07 | 0.07 |
| 6 | 76767224 | rs1933483 | 1.296 | -1.498 | 1.416 | 0.944 | 0.02 | 0.074 | 0.074 | 1.236 | 0.89 | 0.48 | 0.84 | 0.85 | 0.93 | 0.92 |
| 6 | 76793338 | rs2094199 | -0.113 | -1.428 | 1.168 | 0.926 | 0.009 | 0.004 | 0.004 | 0 | 0.07 | 0.14 | 0.09 | 0.07 | 0.07 | 0.07 |
| 6 | 76796339 | rs1581836 | -0.571 | -1.892 | 0.947 | 1.163 | 0.026 | 0.012 | 0.012 | 0 | 0.33 | 0.22 | 0.28 | 0.26 | 0.3 | 0.27 |
| 6 | 76830541 | rs6922651 | -0.948 | -1.763 | 0.94 | 0.741 | 0.014 | 0.017 | 0.017 | 0 | 0.38 | 0.66 | 0.42 | 0.27 | 0.3 | 0.28 |
| 6 | 76833742 | rs1890311 | 0.239 | -1.4 | 1.152 | 1.11 | -0.037 | 0.037 | 0.037 | 0 | 0.8 | 0.69 | 0.78 | 0.67 | 0.89 | 0.84 |
| 6 | 76839113 | rs9359158 | -0.479 | -1.691 | 0.782 | 1.052 | -0.034 | 0.025 | 0.025 | 0.171 | 0.23 | 0.05 | 0.16 | 0.15 | 0.28 | 0.21 |
| 6 | 76857008 | rs2029151 | 0.772 | -1.708 | 1.383 | 1.175 | -0.05 | 0.08 | 0.08 | 0 | 0.73 | 0.42 | 0.65 | 0.52 | 0.82 | 0.76 |
| 6 | 76863568 | rs6453856 | -0.034 | -1.696 | 1.469 | 1.199 | -0.023 | 0.024 | 0.024 | 0 | 0.8 | 0.71 | 0.77 | 0.67 | 0.87 | 0.82 |
| 6 | 76876104 | rs6900948 | -0.533 | -1.675 | 0.964 | 1.129 | -0.009 | -0.034 | -0.034 | 1.461 | 0.24 | 0.35 | 0.15 | 0.15 | 0.25 | 0.2 |
| 6 | 76894472 | rs9343358 | 1.629 | -1.722 | 0.911 | 1.14 | 0.039 | 0.017 | 0.017 | 0.276 | 0.49 | 0.06 | 0.5 | 0.37 | 0.57 | 0.55 |
| 6 | 76897563 | rs9341552 | 0.771 | -1.643 | 1.041 | 1.2 | -0.05 | 0.084 | 0.084 | 0 | 0.73 | 0.42 | 0.65 | 0.52 | 0.82 | 0.75 |
| 6 | 76908536 | rs1486039 | 0.344 | -1.612 | 1.123 | 1.055 | -0.028 | 0.056 | 0.056 | 0 | 0.77 | 0.62 | 0.71 | 0.6 | 0.82 | 0.76 |
| 6 | 76923089 | rs2314203 | 0.649 | -1.607 | 1.09 | 1.058 | -0.055 | 0.072 | 0.072 | 0 | 0.74 | 0.49 | 0.7 | 0.6 | 0.82 | 0.76 |
| 6 | 76925211 | rs1976108 | 0.671 | -1.606 | 1.172 | 1.058 | -0.046 | 0.054 | 0.054 | 0 | 0.74 | 0.45 | 0.7 | 0.6 | 0.82 | 0.76 |
| 6 | 76926454 | rs1825651 | 1.071 | -1.812 | 0.515 | 1.036 | 0.038 | -0.015 | -0.015 | 0.976 | 0.55 | 0.36 | 0.6 | 0.5 | 0.62 | 0.63 |
| 6 | 76927353 | rs9343377 | 0.263 | -1.308 | 0.983 | 1.344 | -0.055 | 0.089 | 0.089 | 0 | 0.8 | 0.57 | 0.77 | 0.68 | 0.87 | 0.82 |
| 6 | 76934173 | rs4562099 | 1.385 | -1.331 | 0.622 | 1.307 | 0.016 | -0.124 | -0.124 | 0.193 | 0.58 | 0.07 | 0.6 | 0.47 | 0.61 | 0.61 |
| 6 | 76934563 | rs13193039 | 1.385 | -1.331 | 0.622 | 1.307 | 0.016 | -0.124 | -0.124 | 0 | 0.58 | 0.07 | 0.6 | 0.47 | 0.61 | 0.61 |
| 6 | 76937256 | rs16887322 | -0.667 | -1.322 | 0.814 | 1.312 | 0.004 | -0.014 | -0.014 | 0 | 0.21 | 0.3 | 0.17 | 0.14 | 0.26 | 0.2 |
| 6 | 76938442 | rs9361005 | 0.348 | -1.331 | 0.845 | 1.334 | -0.046 | 0.072 | 0.072 | 0 | 0.79 | 0.44 | 0.77 | 0.61 | 0.87 | 0.81 |
| 6 | 76947598 | rs12205161 | -0.701 | -1.296 | 0.827 | 1.25 | -0.003 | 0.006 | 0.006 | 0 | 0.2 | 0.19 | 0.16 | 0.14 | 0.26 | 0.2 |

| | | | | | | | | | | | | | | | | |
|---|---|---|---|---|---|---|---|---|---|---|---|---|---|---|---|---|
| 6 | 76948675 | rs6912202 | 1.596 | -1.39 | 0.806 | 1.258 | 0.049 | -0.055 | -0.055 | 0 | 0.56 | 0.08 | 0.55 | 0.42 | 0.61 | 0.54 |
| 6 | 76960326 | rs1564966 | 1.594 | -1.395 | 0.82 | 1.256 | 0.039 | -0.071 | -0.071 | 0.931 | 0.56 | 0.08 | 0.55 | 0.42 | 0.61 | 0.54 |
| 6 | 76969607 | rs985022 | 0.284 | -1.397 | 0.996 | 1.274 | -0.054 | 0.103 | 0.103 | 0 | 0.79 | 0.56 | 0.73 | 0.63 | 0.87 | 0.77 |
| 6 | 76999636 | rs967231 | 0.124 | -1.395 | 1.045 | 1.277 | -0.032 | 0.057 | 0.057 | 0 | 0.81 | 0.68 | 0.79 | 0.69 | 0.87 | 0.82 |
| 6 | 77000409 | rs6918782 | 1.33 | -1.377 | 0.518 | 1.28 | 0.04 | -0.05 | -0.05 | 0 | 0.58 | 0.2 | 0.56 | 0.51 | 0.62 | 0.57 |
| 6 | 77020848 | rs7773032 | 1.172 | -1.375 | 0.547 | 1.284 | 0.015 | -0.015 | -0.015 | 0 | 0.6 | 0.29 | 0.61 | 0.54 | 0.62 | 0.6 |
| 6 | 77023797 | rs9341562 | 1.239 | -1.375 | 0.541 | 1.284 | 0.035 | -0.043 | -0.043 | 0 | 0.59 | 0.2 | 0.59 | 0.54 | 0.62 | 0.6 |
| 6 | 77059064 | rs2451592 | 0.048 | -1.322 | 0.508 | 1.306 | -0.048 | 0.069 | 0.069 | 0 | 0.8 | 0.57 | 0.78 | 0.69 | 0.87 | 0.81 |
| 6 | 77059861 | rs2451593 | 1.451 | -1.279 | 0.025 | 1.317 | 0.04 | -0.101 | -0.101 | 0 | 0.56 | 0.08 | 0.55 | 0.46 | 0.62 | 0.55 |
| 6 | 77088201 | rs1490200 | 1.486 | -1.308 | 0.006 | 1.301 | 0.048 | -0.053 | -0.053 | 0.792 | 0.54 | 0.06 | 0.54 | 0.41 | 0.62 | 0.55 |
| 6 | 77097008 | rs2505024 | 1.443 | -1.242 | 0.034 | 1.322 | 0.043 | -0.047 | -0.047 | 0 | 0.56 | 0.1 | 0.55 | 0.41 | 0.62 | 0.55 |
| 6 | 77128063 | rs9689844 | -0.688 | -1.272 | 0.107 | 1.291 | -0.009 | 0.018 | 0.018 | 0 | 0.2 | 0.15 | 0.15 | 0.11 | 0.25 | 0.18 |
| 6 | 77133082 | rs9343441 | -1.592 | -1.258 | 0.064 | 1.282 | 0.046 | -0.04 | -0.04 | 0 | 0.44 | 0.87 | 0.44 | 0.59 | 0.38 | 0.45 |
| 6 | 77137597 | rs9341575 | -1.284 | -1.285 | 0.072 | 1.221 | 0.038 | -0.055 | -0.055 | 0 | 0.37 | 0.83 | 0.41 | 0.55 | 0.3 | 0.39 |
| 6 | 77142000 | rs951993 | -1.153 | -1.268 | 0.081 | 1.229 | 0.031 | -0.055 | -0.055 | 0.018 | 0.36 | 0.79 | 0.4 | 0.45 | 0.3 | 0.38 |
| 6 | 77157446 | rs9443242 | -1.418 | -1.312 | 0.122 | 1.229 | 0.026 | -0.015 | -0.015 | 0 | 0.41 | 0.77 | 0.36 | 0.43 | 0.38 | 0.37 |
| 6 | 77159214 | rs9352339 | -1.395 | -1.317 | 0.131 | 1.224 | -0.024 | 0.013 | 0.013 | 0 | 0.11 | 0.18 | 0.12 | 0.19 | 0.09 | 0.15 |
| 6 | 77159321 | rs12660098 | -1.207 | -1.304 | 0.133 | 1.185 | 0.01 | 0.006 | 0.006 | 0 | 0.1 | 0.07 | 0.1 | 0.09 | 0.09 | 0.13 |
| 6 | 77159742 | rs7759390 | -1.111 | -1.272 | 0.154 | 1.157 | 0.006 | -0.01 | -0.01 | 0 | 0.34 | 0.72 | 0.32 | 0.39 | 0.3 | 0.31 |
| 6 | 77173321 | rs7756845 | -1.037 | -1.272 | 0.143 | 1.157 | -0.007 | 0.011 | 0.011 | 0 | 0.33 | 0.62 | 0.32 | 0.39 | 0.3 | 0.3 |
| 6 | 77179910 | rs1387845 | -1.343 | -1.26 | 0.038 | 1.163 | 0.028 | -0.032 | -0.032 | 0 | 0.38 | 0.78 | 0.37 | 0.45 | 0.3 | 0.37 |
| 6 | 77181896 | rs1907045 | -1.381 | -1.26 | 0.005 | 1.163 | 0.044 | -0.064 | -0.064 | 0 | 0.38 | 0.85 | 0.38 | 0.45 | 0.3 | 0.37 |
| 6 | 77208816 | rs7756385 | -0.542 | -1.241 | -0.049 | 1.215 | -0.063 | 0.089 | 0.089 | 0 | 0.22 | 0.47 | 0.24 | 0.37 | 0.13 | 0.26 |
| 6 | 77226819 | rs4513750 | -1.532 | -1.077 | -0.05 | 1.239 | 0.06 | -0.01 | -0.01 | 0 | 0.47 | 0.89 | 0.42 | 0.62 | 0.41 | 0.46 |
| 6 | 77242311 | rs10046327 | -0.438 | -0.916 | -0.037 | 1.335 | -0.044 | 0.03 | 0.03 | 0 | 0.26 | 0.48 | 0.25 | 0.38 | 0.24 | 0.28 |
| 6 | 77242394 | rs10046429 | -1.19 | -0.918 | -0.117 | 1.213 | 0.014 | -0.013 | -0.013 | 0.707 | 0.41 | 0.69 | 0.36 | 0.47 | 0.4 | 0.38 |
| 6 | 77242538 | rs9447723 | -0.419 | -0.907 | -0.139 | 1.157 | -0.038 | 0.044 | 0.044 | 0 | 0.26 | 0.42 | 0.24 | 0.4 | 0.24 | 0.28 |
| 6 | 77243034 | rs4329074 | -0.595 | -0.525 | -0.93 | 0.944 | 0.021 | -0.015 | -0.015 | 0 | 0.42 | 0.74 | 0.37 | 0.57 | 0.4 | 0.4 |
| 6 | 77259118 | rs6453912 | -0.794 | -0.507 | -1.633 | 0.894 | 0.06 | -0.02 | -0.02 | 0 | 0.47 | 0.93 | 0.43 | 0.63 | 0.41 | 0.46 |
| 6 | 77266188 | rs4235898 | -0.176 | 0.167 | -2.307 | 0.152 | -0.006 | 0.136 | 0.136 | 0.115 | 0.59 | 0.89 | 0.69 | 0.79 | 0.61 | 0.71 |
| 6 | 77268990 | rs7760456 | -0.629 | 0.975 | -1.731 | 0.557 | -0.015 | -0.017 | -0.017 | 0 | 0.13 | 0.07 | 0.09 | 0.16 | 0.17 | 0.17 |
| 6 | 77280443 | rs4441926 | -0.303 | 1.114 | -1.349 | -0.237 | -0.006 | 0.015 | 0.015 | 0 | 0.59 | 0.6 | 0.67 | 0.87 | 0.54 | 0.7 |
| 6 | 77282477 | rs9352350 | -0.322 | 0.926 | -1.305 | 0.343 | -0.018 | 0.025 | 0.025 | 0.006 | 0.33 | 0.44 | 0.26 | 0.33 | 0.3 | 0.28 |
| 6 | 77284830 | rs9359208 | 0.579 | 0.9 | -1.427 | 0.331 | -0.034 | -0.116 | -0.116 | 14.318 | 0.25 | 0.07 | 0.4 | 0.53 | 0.23 | 0.43 |
| 6 | 77284946 | rs9343467 | -0.57 | 0.9 | -0.711 | 0.207 | 0.003 | 0.011 | 0.011 | 0 | 0.52 | 0.41 | 0.6 | 0.82 | 0.42 | 0.61 |
| 6 | 77288517 | rs9343469 | -0.217 | 0.697 | -0.713 | 0.216 | -0.005 | 0.014 | 0.014 | 0.115 | 0.41 | 0.6 | 0.58 | 0.74 | 0.42 | 0.57 |
| 6 | 77291161 | rs4706623 | -1.009 | 0.904 | -0.699 | -0.518 | -0.001 | 0.028 | 0.028 | 0 | 0.64 | 0.72 | 0.74 | 0.93 | 0.67 | 0.77 |
| 6 | 77291948 | rs4580829 | -1.271 | 0.775 | -1.053 | -0.805 | -0.06 | 0.155 | 0.155 | 0 | 0.69 | 0.97 | 0.76 | 0.93 | 0.7 | 0.81 |
| 6 | 77299771 | rs4128345 | -1.181 | 0.81 | -1.05 | -1.009 | -0.027 | 0.184 | 0.184 | 0.661 | 0.67 | 0.94 | 0.81 | 1 | 0.68 | 0.8 |
| 6 | 77299933 | rs12110507 | -0.192 | 1.137 | 0.209 | -0.034 | 0.023 | 0.052 | 0.052 | 0.085 | 0.12 | 0.22 | 0.16 | 0.2 | 0.15 | 0.06 |
| 6 | 77315788 | rs1457972 | 0.25 | 0.925 | 0.453 | -0.257 | -0.045 | -0.08 | -0.08 | 0.014 | 0.74 | 0.5 | 0.85 | 0.89 | 0.85 | 0.81 |
| 6 | 77315831 | rs2500200 | 0.711 | 0.987 | 0.403 | -0.233 | -0.013 | -0.029 | -0.029 | 0 | 0.71 | 0.39 | 0.81 | 0.84 | 0.78 | 0.72 |
| 6 | 77320030 | rs2256364 | 1.363 | 1.091 | 0.666 | -0.201 | -0.002 | 0.006 | 0.006 | 0.074 | 0.58 | 0.15 | 0.63 | 0.63 | 0.63 | 0.65 |
| 6 | 77322488 | rs2500202 | 1.52 | 1.275 | 0.687 | -0.057 | 0.06 | 0.08 | 0.08 | 0 | 0.77 | 0.46 | 0.65 | 0.7 | 0.65 | 0.74 |
| 6 | 77324522 | rs12199162 | 0.035 | 1.262 | 0.456 | 0.09 | 0.014 | 0.071 | 0.071 | 0.101 | 0.12 | 0.25 | 0.16 | 0.21 | 0.13 | 0.05 |
| 6 | 77326802 | rs1563663 | 1.153 | 1.075 | 0.359 | -0.394 | -0.004 | 0.038 | 0.038 | 0.066 | 0.6 | 0.16 | 0.58 | 0.74 | 0.62 | 0.6 |
| 6 | 77327833 | rs1379675 | 1.139 | 1.129 | 1.316 | -0.132 | 0.026 | 0.004 | 0.004 | 1.328 | 0.86 | 0.4 | 0.84 | 0.89 | 0.86 | 0.9 |
| 6 | 77341104 | rs9361100 | -0.298 | 1.312 | 1.222 | -0.039 | 0.076 | 0.049 | 0.049 | 0 | 0.11 | 0.37 | 0.14 | 0.12 | 0.14 | 0.1 |
| 6 | 77350529 | rs6453926 | -1.366 | 1.554 | 1.196 | -0.084 | 0.064 | 0.038 | 0.038 | 0 | 0.2 | 0.48 | 0.33 | 0.19 | 0.27 | 0.26 |
| 6 | 77356681 | rs1457930 | -0.421 | 1.215 | 0.339 | -1.091 | 0.001 | -0.069 | -0.069 | 0 | 0.6 | 0.57 | 0.67 | 0.94 | 0.59 | 0.6 |
| 6 | 77356936 | rs1870262 | 0.853 | 1.944 | 0.393 | -0.142 | -0.043 | -0.044 | -0.044 | 0 | 0.07 | 0.01 | 0.2 | 0.16 | 0.15 | 0.07 |
| 6 | 77357356 | rs16887867 | -0.212 | 1.84 | 0.823 | -0.336 | 0.012 | -0.032 | -0.032 | 0 | 0.11 | 0.27 | 0.09 | 0.04 | 0.1 | 0.06 |
| 6 | 77357678 | rs2504437 | -0.868 | 1.895 | 0.874 | -0.628 | -0.014 | -0.062 | -0.062 | 0.156 | 0.88 | 0.77 | 0.94 | 1 | 0.88 | 0.9 |
| 6 | 77358648 | rs2446092 | -0.122 | 1.893 | 0.974 | -0.659 | 0.034 | -0.097 | -0.097 | 0 | 0.82 | 0.58 | 0.78 | 0.98 | 0.76 | 0.8 |
| 6 | 77364722 | rs1457940 | -0.459 | 1.798 | 0.443 | -0.68 | 0 | -0.006 | -0.006 | 0 | 0.77 | 0.76 | 0.85 | 0.89 | 0.73 | 0.78 |
| 6 | 77369341 | rs3004002 | 1.432 | 1.676 | 0.406 | -0.64 | 0.018 | 0.072 | 0.072 | 0 | 0.52 | 0.22 | 0.39 | 0.67 | 0.37 | 0.56 |
| 6 | 77375719 | rs2061367 | -0.981 | 1.473 | 0.555 | -0.706 | 0.018 | -0.036 | -0.036 | 0 | 0.28 | 0.53 | 0.34 | 0.16 | 0.36 | 0.28 |
| 6 | 77377283 | rs2504430 | 0.156 | 1.499 | 0.905 | -0.556 | 0.01 | -0.012 | -0.012 | 0.688 | 0.81 | 0.52 | 0.86 | 0.85 | 0.83 | 0.86 |
| 6 | 77379689 | rs2504432 | 1.238 | 1.705 | 0.691 | -0.46 | -0.006 | 0.04 | 0.04 | 0 | 0.56 | 0.23 | 0.42 | 0.67 | 0.41 | 0.54 |
| 6 | 77382995 | rs7750507 | 0.185 | 1.618 | 0.82 | -0.49 | 0.02 | 0.022 | 0.022 | 0 | 0.07 | 0.25 | 0.03 | 0.04 | 0.03 | 0.02 |
| 6 | 77383344 | rs7751153 | 0.333 | 1.699 | 0.611 | -0.515 | -0.004 | -0.022 | -0.022 | 0 | 0.7 | 0.4 | 0.8 | 0.85 | 0.71 | 0.76 |
| 6 | 77386915 | rs17695396 | 0.946 | 1.814 | 0.202 | -0.434 | -0.011 | -0.01 | -0.01 | 0 | 0.09 | 0.05 | 0.24 | 0.16 | 0.18 | 0.12 |
| 6 | 77387073 | rs1902852 | 0.781 | 1.795 | 0.229 | -0.17 | -0.014 | 0.019 | 0.019 | 0.322 | 0.65 | 0.29 | 0.61 | 0.75 | 0.59 | 0.65 |
| 6 | 77391594 | rs9361112 | -0.894 | 1.858 | 0.388 | -0.096 | 0.096 | 0.087 | 0.087 | 0 | 0.22 | 0.46 | 0.47 | 0.3 | 0.32 | 0.25 |
| 6 | 77391826 | rs12529543 | -0.056 | 1.844 | 0.625 | -0.101 | 0.076 | 0.08 | 0.08 | 0.193 | 0.09 | 0.32 | 0.22 | 0.1 | 0.12 | 0.11 |
| 6 | 77393108 | rs9343476 | 0.06 | 1.721 | 0.493 | -0.056 | 0.018 | 0.014 | 0.014 | 0 | 0.13 | 0.14 | 0.26 | 0.2 | 0.2 | 0.14 |
| 6 | 77394093 | rs1457963 | -1.132 | 1.455 | 0.107 | 0.298 | -0.014 | 0.033 | 0.033 | 0.067 | 0.93 | 0.83 | 0.85 | 0.89 | 0.97 | 0.99 |
| 6 | 77394271 | rs1457964 | 1.243 | 1.522 | -0.069 | 0.289 | -0.05 | -0.046 | -0.046 | 0.003 | 0.08 | 0.01 | 0.22 | 0.15 | 0.17 | 0.11 |
| 6 | 77394934 | rs9352373 | 1.244 | 0.662 | -0.136 | -0.022 | 0.052 | 0.014 | 0.014 | 0 | 0.44 | 0.22 | 0.19 | 0.45 | 0.32 | 0.44 |

| | | | | | | | | | | | | | | | | |
|---|---|---|---|---|---|---|---|---|---|---|---|---|---|---|---|---|
| 6 | 77397112 | rs9343478 | -0.796 | 0.931 | -0.258 | 0.27 | 0.025 | -0.006 | -0.006 | 0.241 | 0.38 | 0.48 | 0.56 | 0.32 | 0.51 | 0.44 |
| 6 | 77399578 | rs9361114 | -0.311 | 1.17 | -0.247 | 0.45 | 0.053 | 0.035 | 0.035 | 0 | 0.26 | 0.38 | 0.47 | 0.31 | 0.36 | 0.34 |
| 6 | 77400852 | rs9343481 | 0.105 | 0.36 | -1.477 | 0.264 | 0.038 | -0.067 | -0.067 | 2.466 | 0.32 | 0.05 | 0.13 | 0.44 | 0.26 | 0.39 |
| 6 | 77404440 | rs6934358 | -0.609 | 0.684 | -0.73 | -0.052 | -0.002 | -0.039 | -0.039 | 0 | 0.66 | 0.6 | 0.79 | 0.85 | 0.67 | 0.8 |
| 6 | 77406068 | rs9361115 | 0.699 | 1.642 | -1.085 | 0.378 | -0.061 | -0.057 | -0.057 | 0 | 0.09 | 0.01 | 0.22 | 0.22 | 0.23 | 0.12 |
| 6 | 77406237 | rs17763139 | 0.845 | 1.176 | -0.831 | -0.332 | 0 | 0 | 0 | 0 | 0.12 | 0.09 | 0.1 | 0.12 | 0.16 | 0.08 |
| 6 | 77408514 | rs1457956 | -0.607 | 0.973 | -0.751 | -0.124 | -0.006 | -0.014 | -0.014 | 1.323 | 0.63 | 0.59 | 0.66 | 0.77 | 0.67 | 0.77 |
| 6 | 77410315 | rs987672 | -0.285 | 0.606 | -0.572 | 0.02 | -0.018 | -0.013 | -0.013 | 0.054 | 0.7 | 0.8 | 0.67 | 0.31 | 0.61 | 0.49 |
| 6 | 77412066 | rs1457955 | -0.579 | 0.376 | -0.027 | 0.075 | -0.009 | -0.065 | -0.065 | 0 | 0.19 | 0.36 | 0.25 | 0.06 | 0.14 | 0.15 |
| 6 | 77419782 | rs10498907 | -0.194 | 0.883 | -0.519 | 1.05 | -0.017 | -0.005 | -0.005 | 0 | 0.07 | 0.07 | 0.13 | 0.06 | 0.12 | 0.06 |
| 6 | 77422630 | rs1016555 | -0.02 | 0.679 | -0.37 | 0.999 | -0.006 | 0.013 | 0.013 | 0 | 0.3 | 0.38 | 0.39 | 0.63 | 0.28 | 0.53 |
| 6 | 77422644 | rs1016554 | -0.346 | 0.385 | 0.239 | 0.847 | -0.006 | 0.005 | 0.005 | 0 | 0.33 | 0.66 | 0.41 | 0.63 | 0.28 | 0.53 |
| 6 | 77422929 | rs9341592 | 0.077 | 0.371 | 0.173 | 0.884 | -0.007 | 0.014 | 0.014 | 0 | 0.28 | 0.37 | 0.17 | 0.42 | 0.27 | 0.39 |
| 6 | 77424462 | rs1457974 | -0.641 | 0.372 | -0.348 | 0.177 | 0.001 | -0.001 | -0.001 | 1.053 | 0.66 | 0.79 | 0.7 | 0.68 | 0.65 | 0.8 |
| 6 | 77432018 | rs1457962 | -1.351 | 0.363 | -0.506 | -0.033 | -0.003 | -0.03 | -0.03 | 0.078 | 0.78 | 0.84 | 0.9 | 0.99 | 0.86 | 0.92 |
| 6 | 77434121 | rs9294021 | 0.816 | 0.436 | -0.543 | 0.041 | -0.064 | -0.077 | -0.077 | 0 | 0.11 | 0.01 | 0.2 | 0.31 | 0.21 | 0.11 |
| 6 | 77436710 | rs9352377 | -0.017 | 0.346 | -0.402 | -0.045 | -0.007 | 0.017 | 0.017 | 0 | 0.41 | 0.37 | 0.52 | 0.77 | 0.49 | 0.64 |
| 6 | 77461188 | rs11757291 | 0.502 | 0.317 | -0.398 | -0.129 | 0.006 | -0.019 | -0.019 | 0 | 0.36 | 0.46 | 0.38 | 0.19 | 0.37 | 0.28 |
| 6 | 77462539 | rs9447791 | 0.888 | 0.238 | -0.318 | -0.259 | -0.001 | -0.004 | -0.004 | 0 | 0.31 | 0.31 | 0.25 | 0.16 | 0.26 | 0.25 |
| 6 | 77465384 | rs1598867 | -1.127 | 0.393 | 0.012 | 0.1 | -0.003 | -0.029 | -0.029 | 0 | 0.78 | 0.84 | 0.9 | 0.99 | 0.86 | 0.91 |
| 6 | 77472335 | rs2061649 | -0.863 | 0.397 | 0.04 | 0.099 | -0.049 | -0.119 | -0.119 | 0.83 | 0.76 | 0.63 | 0.88 | 0.99 | 0.86 | 0.91 |
| 6 | 77476663 | rs987008 | -0.863 | 0.397 | 0.052 | 0.099 | -0.05 | -0.121 | -0.121 | 0 | 0.76 | 0.63 | 0.88 | 0.99 | 0.86 | 0.91 |
| 6 | 77476992 | rs9343494 | 0.815 | 0.402 | -0.482 | 0.221 | -0.03 | -0.092 | -0.092 | 1.542 | 0.22 | 0.03 | 0.13 | 0.43 | 0.26 | 0.38 |
| 6 | 77478249 | rs1457946 | 0.173 | 0.517 | -0.236 | 0.24 | 0.007 | 0.001 | 0.001 | 0.383 | 0.61 | 0.37 | 0.65 | 0.66 | 0.65 | 0.79 |
| 6 | 77481958 | rs1466410 | 0.048 | 1.055 | -0.098 | 0.623 | -0.023 | 0.064 | 0.064 | 0 | 0.43 | 0.69 | 0.49 | 0.21 | 0.41 | 0.42 |
| 6 | 77482314 | rs1381674 | 0.473 | 1.06 | -0.381 | 0.714 | -0.049 | -0.07 | -0.07 | 3.399 | 0.11 | 0.01 | 0.2 | 0.33 | 0.19 | 0.13 |
| 6 | 77483034 | rs4351216 | -0.563 | 0.852 | -0.224 | 0.567 | 0.046 | -0.014 | -0.014 | 0.501 | 0.54 | 0.77 | 0.7 | 0.51 | 0.61 | 0.55 |
| 6 | 77484430 | rs9343496 | -0.577 | 0.858 | -0.276 | 0.38 | 0.047 | 0.044 | 0.044 | 0 | 0.54 | 0.78 | 0.77 | 0.64 | 0.61 | 0.56 |
| 6 | 77485310 | rs1350406 | -0.569 | 0.857 | -0.276 | 0.38 | 0.048 | 0.045 | 0.045 | 0 | 0.54 | 0.78 | 0.77 | 0.64 | 0.61 | 0.56 |
| 6 | 77490717 | rs985454 | 0.387 | 1.048 | -0.099 | 0.027 | 0.002 | -0.03 | -0.03 | 0 | 0.32 | 0.31 | 0.19 | 0.01 | 0.24 | 0.25 |
| 6 | 77502291 | rs7756324 | -0.767 | 1.973 | 0.895 | 1.485 | 0.155 | 0.046 | 0.046 | 0.329 | 0.12 | 0.49 | 0.36 | 0.17 | 0.19 | 0.19 |
| 6 | 77524070 | rs1462472 | -0.386 | 1.967 | 1.122 | 1.499 | 0.081 | 0.026 | 0.026 | 0.019 | 0.1 | 0.26 | 0.34 | 0.17 | 0.17 | 0.19 |
| 6 | 77524451 | rs1350941 | -0.252 | 1.833 | 0.964 | 1.446 | -0.034 | -0.004 | -0.004 | 0.358 | 0.06 | 0.01 | 0.13 | 0.27 | 0.14 | 0.14 |
| 6 | 77534953 | rs9341603 | -0.938 | 1.762 | 1.213 | 1.75 | 0.062 | 0.181 | 0.181 | 0 | 0.07 | 0.26 | 0.15 | 0.27 | 0.14 | 0.14 |
| 6 | 77553864 | rs12190467 | 0.755 | 0.669 | 0.629 | 0.384 | 0.049 | 0.106 | 0.106 | 0.252 | 0.2 | 0.01 | 0.07 | 0.01 | 0.11 | 0.09 |
| 6 | 77574184 | rs9294030 | -0.545 | 1.305 | 1.223 | 0.917 | 0.1 | 0.057 | 0.057 | 0.647 | 0.12 | 0.39 | 0.3 | 0.17 | 0.17 | 0.19 |
| 6 | 77577055 | rs9447838 | -0.808 | 1.321 | 1.028 | 0.92 | 0.135 | 0.079 | 0.079 | 1.134 | 0.13 | 0.53 | 0.31 | 0.17 | 0.17 | 0.2 |
| 6 | 77587889 | rs2050427 | 0.031 | 0.85 | 0.087 | 0.926 | 0.006 | -0.014 | -0.014 | 0 | 0.25 | 0.15 | 0.22 | 0.29 | 0.28 | 0.23 |
| 6 | 77599333 | rs7767310 | 0.477 | 1.776 | 0.354 | 1.237 | -0.006 | 0.002 | 0.002 | 0 | 0.64 | 0.66 | 0.59 | 0.32 | 0.48 | 0.52 |
| 6 | 77605748 | rs12208442 | -0.068 | 1.756 | 0.421 | 1.24 | -0.065 | -0.055 | -0.055 | 0 | 0.1 | 0.01 | 0.16 | 0.4 | 0.24 | 0.25 |
| 6 | 77606341 | rs9443309 | -0.062 | 1.543 | 1.14 | 1.145 | -0.022 | 0.004 | 0.004 | 0.114 | 0.28 | 0.52 | 0.25 | 0.28 | 0.28 | 0.23 |
| 6 | 77608445 | rs10455323 | -0.066 | 1.216 | 0.386 | 0.379 | -0.019 | -0.012 | -0.012 | 0 | 0.36 | 0.28 | 0.5 | 0.59 | 0.45 | 0.56 |
| 6 | 77624386 | rs9443323 | 0.184 | 1.127 | 0.455 | 0.686 | -0.023 | 0.045 | 0.045 | 0.076 | 0.29 | 0.52 | 0.32 | 0.4 | 0.28 | 0.24 |
| 6 | 77663444 | rs6453953 | 0.803 | 1.091 | 0.337 | 0.825 | -0.001 | 0 | 0 | 0 | 0.21 | 0.2 | 0.15 | 0.26 | 0.25 | 0.22 |
| 6 | 77682370 | rs323260 | 0.563 | 1.168 | 0.277 | 0.774 | 0.009 | -0.026 | -0.026 | 0 | 0.62 | 0.45 | 0.53 | 0.2 | 0.48 | 0.5 |
| 6 | 77687606 | rs323264 | 1.866 | 0.542 | -0.212 | 0.313 | 0.018 | 0.069 | 0.069 | 0.177 | 0.15 | 0.01 | 0.07 | 0 | 0.11 | 0.06 |
| 6 | 77698853 | rs323239 | -0.132 | 0.602 | 0.016 | 0.514 | -0.011 | 0.005 | 0.005 | 0.139 | 0.71 | 0.34 | 0.73 | 0.67 | 0.71 | 0.76 |
| 6 | 77700022 | rs10943400 | 0.997 | 0.732 | -0.285 | 0.558 | 0.003 | -0.009 | -0.009 | 0.032 | 0.21 | 0.16 | 0.15 | 0.26 | 0.25 | 0.21 |
| 6 | 77711970 | rs9350687 | 0.359 | 1.021 | -0.455 | 0.75 | -0.067 | -0.059 | -0.059 | 1.832 | 0.1 | 0.01 | 0.16 | 0.38 | 0.25 | 0.25 |
| 6 | 77718243 | rs9443334 | 0.488 | 0.825 | -0.664 | 0.418 | 0.017 | 0.086 | 0.086 | 0 | 0.25 | 0.07 | 0.2 | 0.07 | 0.21 | 0.13 |
| 6 | 77730422 | rs784090 | 0.013 | 0.768 | -0.484 | 0.317 | 0.004 | 0.006 | 0.006 | 0 | 0.52 | 0.54 | 0.65 | 0.67 | 0.55 | 0.65 |
| 6 | 77740109 | rs1322100 | 1.119 | 0.86 | -0.629 | 0.41 | 0.002 | -0.016 | -0.016 | 0.974 | 0.2 | 0.12 | 0.14 | 0.25 | 0.24 | 0.22 |
| 6 | 77766235 | rs4403243 | 0.252 | 0.934 | -0.308 | 0.404 | 0.001 | -0.026 | -0.026 | 0 | 0.27 | 0.39 | 0.35 | 0.19 | 0.19 | 0.34 |
| 6 | 77773428 | rs13196188 | 0.653 | 0.904 | -0.827 | 0.481 | 0 | 0 | 0 | 0 | 0.09 | 0.09 | 0.04 | 0.18 | 0.06 | 0.2 |
| 6 | 77780168 | rs1072272 | 0.345 | 0.979 | -0.713 | 0.413 | 0.001 | 0.004 | 0.004 | 0 | 0.25 | 0.3 | 0.4 | 0.28 | 0.19 | 0.36 |
| 6 | 77798214 | rs10943413 | -0.161 | 1.394 | -0.627 | 0.648 | 0.159 | 0.262 | 0.262 | 0 | 0.12 | 0.41 | 0.23 | 0.3 | 0.24 | 0.23 |
| 6 | 77803487 | rs16888579 | 0.918 | 1.207 | -0.879 | 0.494 | 0.018 | -0.014 | -0.014 | 0 | 0.19 | 0.08 | 0.17 | 0.19 | 0.16 | 0.11 |
| 6 | 77805227 | rs2206819 | 0.232 | 0.999 | -1.228 | 0.19 | 0.005 | -0.023 | -0.023 | 0 | 0.15 | 0.07 | 0.08 | 0.22 | 0.14 | 0.22 |
| 6 | 77807794 | rs10943414 | 0.467 | 1.007 | -1.222 | 0.154 | 0.001 | 0 | 0 | 0 | 0.27 | 0.31 | 0.36 | 0.28 | 0.2 | 0.37 |
| 6 | 77814697 | rs978308 | -0.48 | 1.006 | -1.268 | 0.14 | -0.001 | 0 | 0 | 0 | 0.41 | 0.45 | 0.44 | 0.5 | 0.34 | 0.59 |
| 6 | 77820495 | rs1548240 | -1.375 | 0.799 | -1.547 | -0.111 | -0.003 | 0.15 | 0.15 | 1.201 | 0.73 | 0.93 | 0.83 | 1 | 0.74 | 0.93 |
| 6 | 77825472 | rs9443353 | -0.329 | 0.736 | -0.978 | 0.663 | 0.006 | -0.012 | -0.012 | 0 | 0.14 | 0.1 | 0.08 | 0.22 | 0.13 | 0.22 |
| 6 | 77852276 | rs4708318 | -0.381 | 1.059 | -0.754 | 0.471 | 0.009 | 0.014 | 0.014 | 3.263 | 0.51 | 0.27 | 0.33 | 0.47 | 0.39 | 0.59 |
| 6 | 77861942 | rs9447983 | -0.308 | 1.079 | -0.734 | 0.583 | 0.012 | 0.021 | 0.021 | 2.606 | 0.48 | 0.26 | 0.31 | 0.4 | 0.37 | 0.52 |
| 6 | 77877447 | rs11752111 | -0.002 | 1.362 | -1.125 | 0.542 | 0 | -0.019 | -0.019 | 0 | 0.27 | 0.19 | 0.21 | 0.37 | 0.23 | 0.31 |
| 6 | 77884287 | rs7747803 | -0.649 | 1.624 | -1.142 | 0.588 | 0.003 | 0.029 | 0.029 | 2.868 | 0.5 | 0.24 | 0.37 | 0.41 | 0.45 | 0.54 |
| 6 | 77890344 | rs9343581 | -0.493 | 1.332 | -1.184 | 0.582 | -0.008 | 0.003 | 0.003 | 0.613 | 0.08 | 0.03 | 0.17 | 0.11 | 0.11 | 0.14 |
| 6 | 77903893 | rs4075571 | 0.277 | 0.943 | -0.553 | 0.522 | -0.007 | -0.016 | -0.016 | 0 | 0.65 | 0.36 | 0.65 | 0.57 | 0.67 | 0.6 |
| 6 | 77914024 | rs12206214 | 1.096 | 0.549 | -0.739 | 0.253 | -0.027 | -0.044 | -0.044 | 0 | 0.39 | 0.21 | 0.47 | 0.49 | 0.5 | 0.33 |

| | | | | | | | | | | | | | | | | |
|---|---|---|---|---|---|---|---|---|---|---|---|---|---|---|---|---|
| 6 | 77918905 | rs12202018 | 1.769 | 0.695 | -0.817 | 0.01 | -0.054 | 0.195 | 0.195 | 0.354 | 0.32 | 0.02 | 0.18 | 0.07 | 0.33 | 0.18 |
| 6 | 77921902 | rs4265000 | 1.371 | 0.765 | -0.77 | 0.122 | 0.024 | 0.091 | 0.091 | 0.266 | 0.36 | 0.19 | 0.24 | 0.16 | 0.34 | 0.19 |
| 6 | 77926495 | rs13196496 | 0.625 | 0.587 | -0.808 | 0.083 | -0.027 | 0.116 | 0.116 | 0 | 0.21 | 0.05 | 0.11 | 0.01 | 0.17 | 0.19 |
| 6 | 77933661 | rs4463226 | -0.194 | 0.687 | -0.193 | 0.584 | 0.016 | 0.026 | 0.026 | 0.499 | 0.71 | 0.53 | 0.73 | 0.67 | 0.71 | 0.66 |
| 6 | 77936784 | rs9448005 | -1.241 | 0.379 | 0.037 | 0.42 | -0.042 | 0.016 | 0.016 | 0.113 | 0.13 | 0.34 | 0.12 | 0.13 | 0.06 | 0.07 |
| 6 | 77941912 | rs6919231 | -0.615 | 0.861 | 0.1 | 0.5 | 0.023 | 0.007 | 0.007 | 0 | 0.31 | 0.42 | 0.3 | 0.33 | 0.37 | 0.39 |
| 6 | 77942237 | rs6940394 | -0.592 | 1.059 | 0.077 | 0.434 | -0.012 | -0.103 | -0.103 | 4.632 | 0.28 | 0.41 | 0.18 | 0.07 | 0.23 | 0.3 |
| 6 | 77944195 | rs9448010 | 0.351 | 0.871 | 0.117 | 0.417 | -0.011 | -0.004 | -0.004 | 0 | 0.06 | 0.08 | 0.1 | 0.03 | 0.01 | 0.04 |
| 6 | 77956116 | rs6906798 | -0.566 | 0.713 | 0.333 | 0.349 | 0.002 | 0.014 | 0.014 | 0 | 0.5 | 0.44 | 0.44 | 0.38 | 0.5 | 0.56 |
| 6 | 77956816 | rs1418591 | -0.04 | 0.876 | 0.452 | 0.493 | -0.013 | -0.024 | -0.024 | 0 | 0.26 | 0.37 | 0.14 | 0.2 | 0.19 | 0.3 |
| 6 | 77969725 | rs9294068 | -0.32 | 0.797 | 0.333 | 0.364 | 0.023 | 0.049 | 0.049 | 0 | 0.29 | 0.37 | 0.27 | 0.46 | 0.33 | 0.4 |
| 6 | 77969910 | rs7762813 | -1.729 | 0.691 | 0.037 | 0.129 | 0.011 | 0.124 | 0.124 | 0.201 | 0.69 | 0.91 | 0.82 | 0.93 | 0.71 | 0.81 |
| 6 | 77991734 | rs6942008 | -1.477 | 1.375 | 0.824 | 0.639 | 0.089 | 0.178 | 0.178 | 0 | 0.13 | 0.34 | 0.38 | 0.35 | 0.21 | 0.16 |
| 6 | 77997341 | rs1342626 | 0.501 | 1.074 | 0.813 | 0.423 | -0.014 | -0.016 | -0.016 | 0 | 0.25 | 0.2 | 0.29 | 0.37 | 0.33 | 0.37 |
| 6 | 78011282 | rs9784885 | 0.48 | 1.113 | 0.869 | 0.44 | -0.006 | -0.007 | -0.007 | 0.327 | 0.25 | 0.21 | 0.3 | 0.36 | 0.32 | 0.36 |
| 6 | 78016134 | rs11750960 | 0.95 | 0.941 | 0.794 | 0.303 | 0.004 | 0.012 | 0.012 | 0 | 0.56 | 0.24 | 0.47 | 0.43 | 0.61 | 0.56 |
| 6 | 78020351 | rs17263843 | 1.919 | 0.878 | 0.823 | 0.453 | -0.031 | 0.168 | 0.168 | 1.06 | 0.31 | 0.02 | 0.18 | 0.06 | 0.29 | 0.19 |
| 6 | 78028507 | rs9448062 | -0.849 | 0.861 | 1.395 | 0.649 | 0.024 | 0.034 | 0.034 | 0 | 0.08 | 0.26 | 0.16 | 0.09 | 0.09 | 0.05 |
| 6 | 78031111 | rs9352465 | 0.103 | 0.824 | 1.501 | 0.644 | -0.061 | -0.043 | -0.043 | 0.053 | 0.68 | 0.54 | 0.84 | 0.78 | 0.82 | 0.7 |
| 6 | 78033996 | rs9343598 | 0.398 | 0.841 | 1.573 | 0.679 | -0.053 | -0.025 | -0.025 | 0 | 0.67 | 0.52 | 0.75 | 0.73 | 0.82 | 0.71 |
| 6 | 78035270 | rs9443399 | 0.071 | 0.742 | 1.693 | 0.627 | -0.033 | -0.016 | -0.016 | 0.003 | 0.27 | 0.3 | 0.13 | 0.22 | 0.16 | 0.28 |
| 6 | 78036137 | rs9443400 | -0.706 | 0.777 | 1.523 | 0.615 | 0.016 | 0.024 | 0.024 | 0 | 0.08 | 0.21 | 0.16 | 0.09 | 0.09 | 0.05 |
| 6 | 78045886 | rs13204564 | 0.747 | 0.695 | 1.413 | 0.534 | 0.013 | -0.005 | -0.005 | 0.012 | 0.21 | 0.11 | 0.11 | 0.01 | 0.13 | 0.17 |
| 6 | 78060981 | rs6902991 | 1.222 | 0.816 | 1.399 | 0.633 | 0.011 | 0.015 | 0.015 | 0 | 0.56 | 0.23 | 0.48 | 0.43 | 0.62 | 0.56 |
| 6 | 78066685 | rs1539580 | 1.124 | 0.874 | 1.423 | 0.67 | 0.007 | 0.009 | 0.009 | 0 | 0.57 | 0.28 | 0.48 | 0.43 | 0.62 | 0.56 |
| 6 | 78080049 | rs13208046 | 0.378 | 0.971 | 1.343 | 0.625 | 0.033 | 0.044 | 0.044 | 0 | 0.23 | 0.11 | 0.18 | 0.07 | 0.12 | 0.16 |
| 6 | 78081840 | rs10943451 | 0.732 | 1.063 | 1.453 | 0.738 | 0 | 0.002 | 0.002 | 0 | 0.64 | 0.41 | 0.52 | 0.63 | 0.65 | 0.7 |
| 6 | 78082075 | rs9443403 | -1.015 | 1.067 | 1.519 | 0.771 | 0.094 | 0.164 | 0.164 | 0 | 0.1 | 0.26 | 0.28 | 0.31 | 0.23 | 0.14 |
| 6 | 78082678 | rs1342633 | 0.707 | 1.064 | 1.414 | 0.746 | 0 | 0.001 | 0.001 | 0 | 0.65 | 0.4 | 0.52 | 0.62 | 0.64 | 0.7 |
| 6 | 78089035 | rs7770590 | -0.211 | 1.102 | 1.077 | 0.826 | 0.014 | 0.017 | 0.017 | 0 | 0.07 | 0.13 | 0.15 | 0.09 | 0.09 | 0.04 |
| 6 | 78106701 | rs4708333 | 0.583 | 1.046 | 0.773 | 0.714 | -0.07 | -0.061 | -0.061 | 0.039 | 0.24 | 0.07 | 0.3 | 0.33 | 0.3 | 0.37 |
| 6 | 78107089 | rs7772582 | 1.151 | 0.894 | 1.211 | 0.871 | -0.01 | 0.008 | 0.008 | 0 | 0.6 | 0.27 | 0.49 | 0.39 | 0.61 | 0.58 |
| 6 | 78108747 | rs1777807 | -0.236 | 0.405 | 1.282 | 0.269 | 0.031 | 0.252 | 0.252 | 0.974 | 0.06 | 0.33 | 0.03 | 0.21 | 0.02 | 0.11 |
| 6 | 78112044 | rs1777802 | -0.349 | 0.57 | 0.373 | 0.02 | 0.029 | 0.04 | 0.04 | 0 | 0.77 | 0.89 | 0.82 | 0.92 | 0.87 | 0.84 |
| 6 | 78122482 | rs2223832 | -1.228 | 0.767 | 0.951 | 0.37 | 0.008 | 0.141 | 0.141 | 0 | 0.16 | 0.42 | 0.26 | 0.36 | 0.15 | 0.17 |
| 6 | 78125662 | rs1777766 | -0.674 | 0.612 | 0.729 | 0.062 | 0.025 | 0.019 | 0.019 | 0 | 0.79 | 0.85 | 0.9 | 0.99 | 0.88 | 0.85 |
| 6 | 78127675 | rs12110491 | -0.081 | 0.619 | 0.451 | 0.086 | 0.002 | -0.001 | -0.001 | 0 | 0.07 | 0.08 | 0.14 | 0.07 | 0.12 | 0.04 |
| 6 | 78129122 | rs9359267 | 1.099 | 1.261 | 0.135 | 0.023 | -0.004 | 0.001 | 0.001 | 0 | 0.53 | 0.36 | 0.49 | 0.54 | 0.55 | 0.47 |
| 6 | 78134695 | rs12209650 | 0.671 | 1.594 | -0.217 | 0.286 | -0.056 | -0.025 | -0.025 | 0 | 0.09 | 0.01 | 0.16 | 0.09 | 0.19 | 0.22 |
| 6 | 78137057 | rs11757592 | -1.189 | 1.575 | -0.067 | 0.179 | 0.001 | 0.082 | 0.082 | 0 | 0.16 | 0.33 | 0.25 | 0.36 | 0.14 | 0.17 |
| 6 | 78139832 | rs11753559 | -0.471 | 2.051 | -0.289 | 0.409 | 0.039 | 0.015 | 0.015 | 0 | 0.08 | 0.2 | 0.14 | 0.07 | 0.12 | 0.04 |
| 6 | 78142473 | rs9343614 | 0.103 | 1.929 | -1.139 | 0.147 | -0.007 | -0.043 | -0.043 | 0.37 | 0.2 | 0.12 | 0.29 | 0.45 | 0.24 | 0.3 |
| 6 | 78144371 | rs17272780 | 0.298 | 2.209 | -1.153 | 0.242 | -0.056 | -0.025 | -0.025 | 0 | 0.09 | 0.01 | 0.15 | 0.09 | 0.19 | 0.22 |
| 6 | 78149229 | rs6453981 | 0.073 | 1.905 | -1.166 | 0.251 | -0.066 | -0.026 | -0.026 | 0.028 | 0.1 | 0.01 | 0.13 | 0.09 | 0.22 | 0.19 |
| 6 | 78160933 | rs1343485 | 0.837 | 2.001 | -1.207 | 0.201 | -0.061 | 0.154 | 0.154 | 0.918 | 0.32 | 0.04 | 0.27 | 0.12 | 0.36 | 0.17 |
| 6 | 78162536 | rs9352481 | 0.021 | 0.844 | -0.77 | 0.301 | 0.004 | 0.001 | 0.001 | 0 | 0.52 | 0.38 | 0.56 | 0.54 | 0.58 | 0.51 |
| 6 | 78166120 | rs9359271 | -0.729 | 0.81 | -0.405 | 0.59 | -0.004 | 0.046 | 0.046 | 0.125 | 0.3 | 0.54 | 0.36 | 0.48 | 0.26 | 0.47 |
| 6 | 78172992 | rs6298 | -0.216 | 0.983 | -0.387 | 0.668 | -0.005 | -0.009 | -0.009 | 0 | 0.26 | 0.24 | 0.4 | 0.51 | 0.26 | 0.33 |
| 6 | 78176009 | rs10806098 | -0.115 | 1.009 | -0.403 | 0.657 | -0.014 | -0.027 | -0.027 | 0 | 0.25 | 0.18 | 0.4 | 0.51 | 0.27 | 0.32 |
| 6 | 78184375 | rs1936158 | -1.087 | 0.674 | -1.079 | 0.149 | -0.072 | 0.089 | 0.089 | 0.804 | 0.43 | 0.9 | 0.54 | 0.73 | 0.37 | 0.46 |
| 6 | 78185001 | rs6908090 | 0.761 | 0.787 | -1.114 | -0.272 | -0.001 | 0 | 0 | 0 | 0.1 | 0.09 | 0.06 | 0.06 | 0.13 | 0.07 |
| 6 | 78190633 | rs4543330 | 1.817 | 0.522 | -1.719 | -0.815 | 0.025 | 0.122 | 0.122 | 0 | 0.22 | 0.05 | 0.08 | 0.03 | 0.13 | 0.18 |
| 6 | 78191262 | rs1343491 | 1.292 | 0.513 | -1.728 | -0.761 | -0.019 | -0.028 | -0.028 | 0 | 0.07 | 0.01 | 0.05 | 0.15 | 0.08 | 0.08 |
| 6 | 78191811 | rs1213352 | -1.531 | 0.791 | -1.794 | -0.92 | 0.009 | 0.122 | 0.122 | 0.211 | 0.78 | 0.97 | 0.78 | 0.92 | 0.76 | 0.76 |
| 6 | 78198521 | rs2798950 | -0.526 | 0.818 | -1.024 | -0.889 | 0.013 | -0.021 | -0.021 | 0.245 | 0.48 | 0.56 | 0.67 | 0.89 | 0.62 | 0.56 |
| 6 | 78204530 | rs1213356 | -0.515 | 0.471 | -0.959 | -0.969 | 0.014 | 0 | 0 | 0.078 | 0.48 | 0.57 | 0.62 | 0.86 | 0.62 | 0.55 |
| 6 | 78209820 | rs17209149 | 1.224 | 0.554 | -1.206 | -0.907 | -0.033 | -0.008 | -0.008 | 0.322 | 0.1 | 0.01 | 0.05 | 0.06 | 0.13 | 0.05 |
| 6 | 78211113 | rs4235903 | -1.629 | 0.31 | -1.28 | -1.037 | 0.003 | 0.113 | 0.113 | 0.249 | 0.79 | 0.97 | 0.79 | 0.92 | 0.77 | 0.76 |
| 6 | 78211608 | rs2798952 | -1.068 | 0.489 | -0.831 | -0.83 | 0.075 | 0.151 | 0.151 | 0.404 | 0.54 | 0.73 | 0.64 | 0.86 | 0.64 | 0.58 |
| 6 | 78215569 | rs2798953 | -0.635 | 0.608 | -0.74 | -0.766 | 0.016 | 0.005 | 0.005 | 0.094 | 0.49 | 0.53 | 0.61 | 0.86 | 0.63 | 0.55 |
| 6 | 78226917 | rs1213378 | 0.267 | 0.831 | -0.64 | -0.6 | 0.061 | -0.001 | -0.001 | 0 | 0.32 | 0.48 | 0.5 | 0.65 | 0.42 | 0.43 |
| 6 | 78236296 | rs10943465 | 0.82 | 1.087 | -0.364 | -0.391 | -0.004 | -0.007 | -0.007 | 0 | 0.26 | 0.22 | 0.39 | 0.49 | 0.27 | 0.34 |
| 6 | 78241057 | rs1419162 | -1.133 | 0.554 | -0.733 | -0.766 | 0.075 | 0.179 | 0.179 | 0.577 | 0.55 | 0.81 | 0.65 | 0.86 | 0.64 | 0.58 |
| 6 | 78246469 | rs10943467 | 0.359 | 0.699 | -0.841 | -0.688 | 0.059 | -0.001 | -0.001 | 0 | 0.32 | 0.48 | 0.5 | 0.65 | 0.43 | 0.43 |
| 6 | 78247104 | rs4706652 | -0.554 | 0.312 | -0.726 | -0.725 | 0.013 | -0.002 | -0.002 | 0.044 | 0.49 | 0.52 | 0.61 | 0.86 | 0.63 | 0.56 |
| 6 | 78251854 | rs2609063 | -0.545 | 0.329 | -0.71 | -0.716 | 0.012 | -0.004 | -0.004 | 0 | 0.48 | 0.52 | 0.61 | 0.86 | 0.63 | 0.55 |
| 6 | 78255395 | rs898038 | 1.233 | 0.218 | -0.678 | -0.541 | -0.016 | -0.026 | -0.026 | 0 | 0.07 | 0.02 | 0.05 | 0.15 | 0.08 | 0.07 |
| 6 | 78258171 | rs2798976 | -0.521 | 0.337 | -0.661 | -0.754 | 0.01 | -0.01 | -0.01 | 0.044 | 0.49 | 0.52 | 0.62 | 0.88 | 0.62 | 0.56 |
| 6 | 78259489 | rs2446287 | -1.569 | 0.384 | -1.194 | -0.625 | -0.016 | 0.091 | 0.091 | 0 | 0.75 | 0.95 | 0.85 | 0.94 | 0.82 | 0.8 |

| | | | | | | | | | | | | | | | | |
|---|---|---|---|---|---|---|---|---|---|---|---|---|---|---|---|---|
| 6 | 78261702 | rs1778296 | -0.918 | 0.159 | -1.318 | -0.705 | 0.011 | 0.082 | 0.082 | 0.28 | 0.61 | 0.93 | 0.81 | 0.79 | 0.74 | 0.72 |
| 6 | 78262337 | rs1824282 | -0.225 | 0.372 | -1.017 | -0.619 | 0.027 | 0.011 | 0.011 | 0.19 | 0.45 | 0.74 | 0.58 | 0.62 | 0.42 | 0.59 |
| 6 | 78264479 | rs1458405 | -0.225 | 0.372 | -1.017 | -0.619 | 0.027 | 0.011 | 0.011 | 0.19 | 0.45 | 0.74 | 0.58 | 0.62 | 0.42 | 0.58 |
| 6 | 78265478 | rs1458406 | 0.485 | 0.104 | -0.547 | -0.383 | 0 | 0.055 | 0.055 | 0 | 0.27 | 0.5 | 0.37 | 0.57 | 0.23 | 0.36 |
| 6 | 78270696 | rs12523809 | 0.841 | 0.104 | -0.488 | -0.323 | -0.001 | 0.001 | 0.001 | 0 | 0.24 | 0.19 | 0.32 | 0.5 | 0.22 | 0.34 |
| 6 | 78291914 | rs2504264 | -1.888 | -0.164 | -0.78 | -0.605 | -0.012 | 0.101 | 0.101 | 0.042 | 0.75 | 0.97 | 0.86 | 0.95 | 0.83 | 0.81 |
| 6 | 78294880 | rs1902970 | -0.851 | 0.238 | -0.831 | -0.705 | -0.015 | 0.221 | 0.221 | 0.288 | 0.5 | 0.81 | 0.56 | 0.82 | 0.42 | 0.52 |
| 6 | 78297902 | rs12527883 | 0.809 | 0.222 | -0.451 | -0.659 | 0.001 | 0.034 | 0.034 | 0 | 0.25 | 0.27 | 0.35 | 0.57 | 0.22 | 0.36 |
| 6 | 78298165 | rs2252216 | -1.009 | 0.304 | -0.271 | -0.752 | -0.022 | 0.197 | 0.197 | 0.013 | 0.52 | 0.73 | 0.57 | 0.89 | 0.45 | 0.53 |
| 6 | 78303603 | rs2085590 | -1.024 | 0.301 | -0.272 | -0.754 | -0.023 | 0.195 | 0.195 | 0.008 | 0.52 | 0.73 | 0.57 | 0.89 | 0.45 | 0.53 |
| 6 | 78315289 | rs7764654 | 0.073 | 0.264 | -0.072 | -0.767 | 0.012 | 0.005 | 0.005 | 0 | 0.32 | 0.45 | 0.47 | 0.67 | 0.33 | 0.44 |
| 6 | 78316625 | rs2446248 | 0.186 | 0.377 | 0.004 | -0.674 | -0.013 | 0.008 | 0.008 | 0 | 0.19 | 0.27 | 0.1 | 0.22 | 0.13 | 0.09 |
| 6 | 78319616 | rs12199701 | 0.753 | 0.423 | -0.147 | -0.719 | -0.009 | 0.003 | 0.003 | 0 | 0.23 | 0.24 | 0.28 | 0.07 | 0.36 | 0.29 |
| 6 | 78326093 | rs1598929 | 0.838 | 0.451 | -0.098 | -0.449 | 0.01 | -0.002 | -0.002 | 0 | 0.15 | 0.08 | 0.06 | 0.15 | 0.09 | 0.07 |
| 6 | 78330693 | rs1304885 | 0.811 | 0.293 | -0.095 | -0.38 | -0.001 | 0.002 | 0.002 | 0 | 0.23 | 0.19 | 0.35 | 0.57 | 0.21 | 0.35 |
| 6 | 78332893 | rs2504298 | 0.262 | 0.242 | -0.242 | -0.297 | 0.01 | -0.014 | -0.014 | 0 | 0.18 | 0.12 | 0.08 | 0.22 | 0.13 | 0.09 |
| 6 | 78336649 | rs1160571 | -0.28 | -0.034 | -0.013 | -0.33 | -0.01 | 0 | 0 | 0 | 0.38 | 0.48 | 0.38 | 0.62 | 0.28 | 0.42 |
| 6 | 78336806 | rs2504295 | 1.018 | -0.002 | -0.197 | 0.544 | -0.012 | 0.073 | 0.073 | 0 | 0.12 | 0.2 | 0.14 | 0.29 | 0.07 | 0.21 |
| 6 | 78351916 | rs898040 | 0.507 | -0.017 | -0.558 | 0.237 | -0.024 | 0.037 | 0.037 | 0 | 0.27 | 0.18 | 0.3 | 0.2 | 0.39 | 0.34 |
| 6 | 78352070 | rs898039 | 0.053 | 0.053 | -0.502 | 0.239 | -0.014 | 0.008 | 0.008 | 0 | 0.32 | 0.33 | 0.42 | 0.29 | 0.49 | 0.41 |
| 6 | 78353362 | rs2504284 | -1.739 | -0.252 | -0.437 | -0.021 | 0.005 | 0.019 | 0.019 | 0 | 0.7 | 0.81 | 0.81 | 0.91 | 0.77 | 0.83 |
| 6 | 78354757 | rs12206237 | 1.168 | -0.275 | -0.496 | -0.059 | -0.06 | 0.091 | 0.091 | 0.161 | 0.21 | 0.06 | 0.28 | 0.05 | 0.38 | 0.3 |
| 6 | 78354813 | rs2504283 | -0.548 | -0.156 | -0.301 | 0.086 | 0.004 | -0.018 | -0.018 | 0 | 0.39 | 0.44 | 0.47 | 0.71 | 0.38 | 0.47 |
| 6 | 78355600 | rs7752338 | -0.246 | -0.307 | -0.569 | 0.024 | 0 | 0.014 | 0.014 | 0 | 0.38 | 0.42 | 0.39 | 0.26 | 0.45 | 0.39 |
| 6 | 78356011 | rs12662855 | 1.019 | -0.037 | -0.314 | 0.294 | -0.01 | 0.086 | 0.086 | 0 | 0.13 | 0.2 | 0.12 | 0.27 | 0.08 | 0.22 |
| 6 | 78359807 | rs9352507 | 0.413 | 0.175 | -0.562 | -0.136 | -0.014 | -0.002 | -0.002 | 0 | 0.09 | 0.13 | 0.04 | 0.06 | 0.04 | 0.03 |
| 6 | 78381291 | rs2504270 | -1.146 | 0.252 | -1.096 | -0.484 | -0.027 | -0.016 | -0.016 | 0 | 0.71 | 0.89 | 0.79 | 0.76 | 0.73 | 0.76 |
| 6 | 78381834 | rs9343638 | -0.579 | 0.424 | -0.306 | -0.622 | 0 | -0.066 | -0.066 | 2.564 | 0.32 | 0.42 | 0.33 | 0.14 | 0.39 | 0.28 |
| 6 | 78392084 | rs12055464 | -0.264 | 0.703 | -0.51 | -0.313 | 0.003 | 0.048 | 0.048 | 0 | 0.2 | 0.09 | 0.11 | 0.07 | 0.18 | 0.07 |
| 6 | 78393124 | rs483602 | -0.561 | 0.973 | 0.477 | 0.13 | 0.04 | 0.227 | 0.227 | 2.32 | 0.16 | 0.54 | 0.3 | 0.39 | 0.14 | 0.2 |
| 6 | 78400093 | rs11969443 | 0.281 | 0.954 | 0.822 | 0.282 | 0.019 | 0.073 | 0.073 | 0 | 0.09 | 0.15 | 0.23 | 0.31 | 0.13 | 0.15 |
| 6 | 78401868 | rs12203229 | -0.363 | 0.819 | 0.359 | -0.259 | -0.099 | -0.092 | -0.092 | 0.013 | 0.29 | 0.11 | 0.41 | 0.5 | 0.48 | 0.45 |
| 6 | 78412532 | rs10943480 | -2.243 | 0.567 | 0.996 | -0.312 | -0.007 | -0.013 | -0.013 | 0.255 | 0.64 | 0.67 | 0.78 | 0.87 | 0.8 | 0.69 |
| 6 | 78420432 | rs7761742 | -1.103 | 0.578 | 0.993 | -0.339 | 0.06 | -0.017 | -0.017 | 0 | 0.49 | 0.22 | 0.52 | 0.56 | 0.66 | 0.53 |
| 6 | 78422191 | rs7752164 | -2.226 | 0.539 | 0.974 | -0.358 | -0.008 | -0.014 | -0.014 | 0 | 0.65 | 0.68 | 0.79 | 0.87 | 0.8 | 0.69 |
| 6 | 78439540 | rs16889371 | 0.545 | 0.815 | 0.675 | -0.207 | -0.013 | -0.028 | -0.028 | 0 | 0.15 | 0.02 | 0.08 | 0.18 | 0.11 | 0.18 |
| 6 | 78439841 | rs6939080 | -0.114 | 1.122 | 0.841 | -0.187 | 0.013 | 0.057 | 0.057 | 0 | 0.23 | 0.35 | 0.28 | 0.44 | 0.28 | 0.14 |
| 6 | 78446870 | rs4708362 | -1.322 | 1.144 | 0.834 | -0.211 | -0.033 | 0.069 | 0.069 | 1.047 | 0.39 | 0.78 | 0.55 | 0.75 | 0.42 | 0.3 |
| 6 | 78451324 | rs1157630 | -0.032 | 1.155 | 0.934 | -0.216 | 0.035 | 0.117 | 0.117 | 1.568 | 0.14 | 0.31 | 0.25 | 0.31 | 0.14 | 0.16 |
| 6 | 78456439 | rs10943488 | -0.085 | 1.156 | 0.883 | -0.216 | 0.047 | 0.147 | 0.147 | 1.584 | 0.14 | 0.34 | 0.25 | 0.31 | 0.14 | 0.16 |
| 6 | 78470331 | rs1483047 | -0.247 | 1.124 | 0.93 | -0.238 | 0.025 | 0.114 | 0.114 | 0 | 0.25 | 0.48 | 0.29 | 0.44 | 0.28 | 0.14 |
| 6 | 78476694 | rs7748475 | -2.736 | 1.066 | 0.285 | -0.354 | 0.07 | 0.199 | 0.199 | 0.266 | 0.67 | 0.97 | 0.84 | 0.95 | 0.8 | 0.72 |
| 6 | 78482257 | rs13210329 | -0.466 | 1.234 | 0.395 | -0.183 | -0.047 | 0.059 | 0.059 | 0 | 0.27 | 0.1 | 0.29 | 0.21 | 0.37 | 0.41 |
| 6 | 78494764 | rs17777705 | 0.609 | 1.429 | 0.483 | -0.054 | -0.055 | 0.059 | 0.059 | 0 | 0.12 | 0.02 | 0.19 | 0.02 | 0.26 | 0.23 |
| 6 | 78524718 | rs9294087 | -0.21 | 1.317 | 0.999 | 0.228 | 0.051 | 0.155 | 0.155 | 1.024 | 0.14 | 0.36 | 0.25 | 0.31 | 0.14 | 0.16 |
| 6 | 78526794 | rs9443479 | -0.418 | 1.312 | 1.008 | 0.225 | 0.065 | 0.195 | 0.195 | 1.454 | 0.15 | 0.43 | 0.25 | 0.31 | 0.14 | 0.16 |
| 6 | 78531689 | rs9359294 | -0.21 | 1.165 | 0.94 | 0.155 | 0.01 | 0.044 | 0.044 | 0 | 0.23 | 0.3 | 0.28 | 0.44 | 0.28 | 0.14 |
| 6 | 78543069 | rs9294089 | -0.666 | 0.531 | 1.113 | 0.138 | -0.034 | -0.103 | -0.103 | 0 | 0.26 | 0.43 | 0.12 | 0.07 | 0.18 | 0.07 |
| 6 | 78547147 | rs9361296 | -0.883 | 0.691 | 1.311 | 0.254 | 0.026 | 0.045 | 0.045 | 0.32 | 0.29 | 0.54 | 0.35 | 0.38 | 0.31 | 0.21 |
| 6 | 78548866 | rs12197987 | -0.322 | 0.572 | 1.052 | 0.079 | -0.048 | 0.058 | 0.058 | 0 | 0.27 | 0.1 | 0.29 | 0.2 | 0.37 | 0.41 |
| 6 | 78549160 | rs12211471 | -0.813 | 0.526 | 1.107 | 0.061 | -0.026 | -0.026 | -0.026 | 0.136 | 0.35 | 0.34 | 0.53 | 0.51 | 0.51 | 0.55 |
| 6 | 78558734 | rs4056845 | -0.374 | 0.451 | 1.009 | -0.017 | -0.045 | 0.059 | 0.059 | 0 | 0.27 | 0.11 | 0.29 | 0.2 | 0.38 | 0.41 |
| 6 | 78562076 | rs1877531 | -0.917 | 0.414 | 1.007 | -0.039 | 0 | 0.002 | 0.002 | 1.834 | 0.33 | 0.62 | 0.35 | 0.38 | 0.32 | 0.21 |
| 6 | 78568478 | rs12200754 | -0.47 | 0.41 | 0.984 | -0.158 | -0.045 | 0.058 | 0.058 | 0 | 0.27 | 0.11 | 0.29 | 0.21 | 0.37 | 0.41 |
| 6 | 78605490 | rs12208156 | -0.646 | 0.423 | 0.984 | -0.151 | -0.026 | 0.055 | 0.055 | 0 | 0.29 | 0.17 | 0.29 | 0.2 | 0.38 | 0.41 |
| 6 | 78620417 | rs12189558 | 0.495 | 0.854 | 0.606 | -0.023 | -0.049 | 0.05 | 0.05 | 0 | 0.09 | 0.01 | 0.17 | 0.02 | 0.23 | 0.22 |
| 6 | 78633116 | rs9448258 | -0.342 | 0.355 | 1.156 | 0.104 | -0.019 | -0.032 | -0.032 | 0 | 0.08 | 0.33 | 0.04 | 0 | 0.01 | 0.02 |
| 6 | 78644509 | rs9359304 | 0.275 | 0.394 | 0.723 | 0.239 | 0.001 | 0.048 | 0.048 | 0.667 | 0.17 | 0.04 | 0.08 | 0.06 | 0.15 | 0.05 |
| 6 | 78657290 | rs9361318 | -0.792 | 0.552 | 0.989 | 0.27 | 0.006 | 0.02 | 0.02 | 0.523 | 0.3 | 0.57 | 0.35 | 0.4 | 0.29 | 0.2 |
| 6 | 78663663 | rs9448264 | -1.652 | 0.553 | 0.839 | 0.058 | 0.027 | 0.065 | 0.065 | 0 | 0.57 | 0.72 | 0.75 | 0.88 | 0.68 | 0.63 |
| 6 | 78684899 | rs9361324 | -0.721 | 0.392 | 1.144 | 0.133 | 0.014 | 0.035 | 0.035 | 0.654 | 0.29 | 0.42 | 0.33 | 0.38 | 0.29 | 0.21 |
| 6 | 78686454 | rs9352548 | -2.241 | 0.396 | 0.834 | -0.021 | -0.007 | 0.224 | 0.224 | 0.893 | 0.64 | 0.95 | 0.81 | 0.96 | 0.7 | 0.68 |
| 6 | 78708695 | rs6932080 | -0.675 | 0.626 | 0.9 | 0.26 | -0.003 | 0.003 | 0.003 | 0 | 0.3 | 0.3 | 0.3 | 0.2 | 0.39 | 0.39 |
| 6 | 78718367 | rs4463222 | -0.368 | 0.708 | 0.405 | 0.337 | -0.104 | -0.154 | -0.154 | 0 | 0.27 | 0.07 | 0.39 | 0.51 | 0.4 | 0.43 |
| 6 | 78722248 | rs4532411 | -0.76 | 0.725 | 0.946 | 0.273 | -0.012 | -0.025 | -0.025 | 0 | 0.37 | 0.29 | 0.52 | 0.53 | 0.57 | 0.56 |
| 6 | 78723311 | rs9343714 | -1.829 | 0.545 | 1.134 | -0.036 | 0.048 | 0.081 | 0.081 | 0.138 | 0.59 | 0.75 | 0.78 | 0.96 | 0.71 | 0.69 |
| 6 | 78724973 | rs9448269 | -0.427 | 0.747 | 1.463 | 0.139 | -0.02 | -0.034 | -0.034 | 0 | 0.19 | 0.34 | 0.15 | 0.12 | 0.14 | 0.12 |
| 6 | 78735368 | rs1888778 | -0.287 | 0.893 | 1.249 | 0.07 | -0.037 | -0.028 | -0.028 | 4.31 | 0.25 | 0.19 | 0.43 | 0.39 | 0.43 | 0.44 |
| 6 | 78751523 | rs1590999 | -0.926 | 0.624 | 1.186 | 0.077 | -0.003 | 0.001 | 0.001 | 0 | 0.41 | 0.38 | 0.66 | 0.88 | 0.56 | 0.62 |

| | | | | | | | | | | | | | | | | |
|---|---|---|---|---|---|---|---|---|---|---|---|---|---|---|---|---|
| 6 | 78753675 | rs2223968 | -0.97 | 0.623 | 1.187 | 0.116 | -0.006 | -0.009 | -0.009 | 0.768 | 0.4 | 0.33 | 0.51 | 0.56 | 0.56 | 0.55 |
| 6 | 78753960 | rs9361343 | 1.035 | 0.605 | 1.2 | -0.03 | 0.001 | 0.001 | 0.001 | 0 | 0.06 | 0.09 | 0.03 | 0.07 | 0.07 | 0.03 |
| 6 | 78764662 | rs9294096 | -1.244 | 0.633 | 1.057 | -0.044 | 0.003 | -0.059 | -0.059 | 0 | 0.47 | 0.47 | 0.69 | 0.96 | 0.62 | 0.65 |
| 6 | 78797479 | rs9361356 | -0.338 | 0.495 | 0.698 | 0.338 | -0.001 | -0.004 | -0.004 | 0.258 | 0.14 | 0.12 | 0.08 | 0.07 | 0.13 | 0.04 |
| 6 | 78801824 | rs9343725 | 0.673 | 0.595 | 0.747 | 0.12 | -0.001 | -0.001 | -0.001 | 1.907 | 0.06 | 0.04 | 0.02 | 0.11 | 0.07 | 0.04 |
| 6 | 78811146 | rs6938098 | -1.056 | 0.459 | 1.159 | 0.013 | -0.012 | 0.008 | 0.008 | 0 | 0.14 | 0.27 | 0.08 | 0.12 | 0.12 | 0.04 |
| 6 | 78811713 | rs10080897 | -1.217 | 0.436 | 1.279 | 0.001 | -0.016 | 0.008 | 0.008 | 0 | 0.15 | 0.37 | 0.09 | 0.12 | 0.12 | 0.05 |
| 6 | 78813219 | rs1011709 | -0.636 | 0.508 | 1.269 | 0.042 | -0.016 | -0.026 | -0.026 | 0.003 | 0.35 | 0.3 | 0.51 | 0.51 | 0.55 | 0.52 |
| 6 | 78817347 | rs11752049 | 0.86 | 0.672 | 1.307 | 0.053 | 0.049 | 0.113 | 0.113 | 0 | 0.07 | 0.12 | 0.17 | 0.28 | 0.14 | 0.15 |
| 6 | 78821232 | rs9294097 | -1.718 | 0.633 | 1.319 | -0.013 | 0.065 | 0.04 | 0.04 | 0 | 0.5 | 0.65 | 0.6 | 0.63 | 0.67 | 0.56 |
| 6 | 78825444 | rs1331804 | -1.051 | 0.642 | 1.309 | 0 | -0.015 | 0.009 | 0.009 | 0 | 0.15 | 0.37 | 0.09 | 0.12 | 0.12 | 0.04 |
| 6 | 78834409 | rs4706686 | -1.922 | 0.587 | 1.175 | -0.067 | 0.05 | 0.013 | 0.013 | 0.493 | 0.57 | 0.74 | 0.61 | 0.63 | 0.67 | 0.58 |
| 6 | 78844153 | rs9361364 | -0.51 | 0.735 | 0.848 | -0.123 | -0.06 | 0.079 | 0.079 | 0 | 0.83 | 0.94 | 0.74 | 0.95 | 0.69 | 0.77 |
| 6 | 78848097 | rs6914238 | -0.747 | -0.057 | 1.1 | 0.103 | -0.001 | 0.015 | 0.015 | 0 | 0.18 | 0.22 | 0.11 | 0.27 | 0.16 | 0.14 |
| 6 | 78863599 | rs6454046 | -0.416 | 0 | 1.095 | 0.107 | -0.046 | -0.025 | -0.025 | 1.877 | 0.74 | 0.82 | 0.68 | 0.65 | 0.56 | 0.58 |
| 6 | 78866052 | rs9448304 | 0.418 | 0.074 | 1.103 | 0.278 | 0.003 | 0.008 | 0.008 | 0 | 0.08 | 0.1 | 0.06 | 0.15 | 0.09 | 0.09 |
| 6 | 78866182 | rs9343742 | -0.229 | 0.058 | 1.165 | 0.281 | -0.001 | 0.006 | 0.006 | 0 | 0.11 | 0.16 | 0.08 | 0.11 | 0.11 | 0.05 |
| 6 | 78872236 | rs9448308 | 0.223 | 0.089 | 1.019 | 0.311 | 0 | 0 | 0 | 0 | 0.1 | 0.09 | 0.08 | 0.16 | 0.1 | 0.1 |
| 6 | 78876920 | rs10080585 | 0.231 | 0.12 | 0.99 | 0.312 | -0.001 | -0.001 | -0.001 | 0 | 0.1 | 0.1 | 0.09 | 0.15 | 0.11 | 0.11 |
| 6 | 78912889 | rs9443542 | 0.99 | -0.181 | 0.93 | 0.102 | 0.017 | 0.157 | 0.157 | 1.947 | 0.48 | 0.24 | 0.32 | 0.2 | 0.42 | 0.44 |
| 6 | 78923047 | rs7748524 | 0.123 | -0.125 | 0.809 | 0.146 | -0.002 | -0.001 | -0.001 | 0 | 0.31 | 0.28 | 0.33 | 0.35 | 0.45 | 0.43 |
| 6 | 78927086 | rs10806128 | 0.241 | -0.137 | 0.807 | 0.145 | -0.038 | -0.019 | -0.019 | 0 | 0.3 | 0.21 | 0.33 | 0.35 | 0.44 | 0.43 |
| 6 | 78934100 | rs2050689 | 0.177 | -0.152 | 0.685 | 0.082 | -0.001 | -0.002 | -0.002 | 0.58 | 0.11 | 0.1 | 0.09 | 0.16 | 0.11 | 0.12 |
| 6 | 78942566 | rs10455344 | -0.516 | -0.127 | 0.77 | 0.085 | -0.001 | 0.003 | 0.003 | 0 | 0.42 | 0.46 | 0.42 | 0.51 | 0.56 | 0.55 |
| 6 | 78945906 | rs12234129 | 0.586 | 0.179 | 0.502 | 0.342 | 0 | 0.013 | 0.013 | 0 | 0.07 | 0.06 | 0.03 | 0.11 | 0.07 | 0.04 |
| 6 | 78946797 | rs6901674 | -0.529 | 0.24 | 0.515 | 0.466 | 0.029 | 0.437 | 0.437 | 0.055 | 0.06 | 0.3 | 0.22 | 0.33 | 0.02 | 0.07 |
| 6 | 78953246 | rs6911371 | 1.003 | 0.38 | -0.09 | 0.07 | 0.024 | 0.129 | 0.129 | 0.884 | 0.43 | 0.2 | 0.26 | 0.16 | 0.37 | 0.36 |
| 6 | 78955138 | rs7741242 | -1.035 | 0.555 | 0.553 | 1.214 | 0.002 | 0.268 | 0.268 | 0 | 0.16 | 0.41 | 0.35 | 0.51 | 0.14 | 0.19 |
| 6 | 78958666 | rs818228 | -0.409 | 0.709 | -0.176 | 0.687 | -0.021 | -0.037 | -0.037 | 0 | 0.09 | 0.02 | 0.05 | 0.12 | 0.12 | 0.08 |
| 6 | 78959888 | rs818237 | 0.281 | 0.571 | -0.269 | -0.552 | 0.01 | 0.035 | 0.035 | 0.692 | 0.63 | 0.31 | 0.72 | 0.96 | 0.64 | 0.78 |
| 6 | 78962319 | rs699171 | -0.438 | 0.472 | 0.443 | -0.776 | 0 | -0.047 | -0.047 | 0 | 0.79 | 0.78 | 0.81 | 0.96 | 0.73 | 0.81 |
| 6 | 79039234 | rs818316 | 0.871 | 0.237 | 0.852 | -0.245 | 0.014 | 0.053 | 0.053 | 0.499 | 0.61 | 0.31 | 0.56 | 0.77 | 0.62 | 0.75 |
| 6 | 79042157 | rs9443553 | -0.446 | 0.285 | 1.179 | -0.169 | -0.012 | 0.017 | 0.017 | 0 | 0.14 | 0.2 | 0.21 | 0.21 | 0.11 | 0.06 |
| 6 | 79042990 | rs818314 | 0.847 | 0.255 | 1.225 | -0.186 | 0.015 | 0.062 | 0.062 | 0.624 | 0.62 | 0.27 | 0.56 | 0.75 | 0.62 | 0.75 |
| 6 | 79047537 | rs818276 | 0.863 | 0.353 | 1.018 | -0.161 | 0.009 | 0.069 | 0.069 | 0.628 | 0.62 | 0.27 | 0.61 | 0.75 | 0.61 | 0.75 |
| 6 | 79051952 | rs818269 | -0.109 | 0.389 | 1.187 | -0.105 | 0.013 | -0.003 | -0.003 | 0 | 0.77 | 0.72 | 0.67 | 0.75 | 0.67 | 0.77 |
| 6 | 79052420 | rs235461 | -1.033 | 0.523 | 0.426 | -0.939 | -0.043 | 0.068 | 0.068 | 0 | 0.39 | 0.3 | 0.5 | 0.75 | 0.51 | 0.48 |
| 6 | 79055410 | rs235456 | -0.49 | 1.111 | 0.418 | -0.969 | -0.054 | 0.02 | 0.02 | 0.007 | 0.28 | 0.23 | 0.48 | 0.75 | 0.48 | 0.45 |
| 6 | 79080410 | rs2493224 | -1.058 | 1.101 | 0.726 | -0.913 | -0.006 | 0.035 | 0.035 | 0 | 0.41 | 0.37 | 0.52 | 0.75 | 0.55 | 0.48 |
| 6 | 79090822 | rs12213776 | 0.728 | 1.054 | 0.608 | -0.933 | -0.023 | 0.056 | 0.056 | 0.187 | 0.2 | 0.11 | 0.19 | 0.07 | 0.27 | 0.19 |
| 6 | 79091268 | rs16889911 | 0.682 | 1.326 | 0.52 | 0.037 | -0.016 | -0.027 | -0.027 | 0.659 | 0.06 | 0.01 | 0.18 | 0.24 | 0.06 | 0.13 |
| 6 | 79095072 | rs2493237 | -1.193 | 1.134 | 0.614 | -0.377 | -0.005 | 0.03 | 0.03 | 0 | 0.41 | 0.37 | 0.52 | 0.75 | 0.55 | 0.48 |
| 6 | 79115925 | rs236226 | 0.178 | 1.277 | 0.532 | 0.053 | -0.002 | 0.002 | 0.002 | 0 | 0.1 | 0.09 | 0.06 | 0.08 | 0.13 | 0.07 |
| 6 | 79129963 | rs7451761 | 0.175 | 1.103 | 0.596 | -0.274 | -0.008 | 0.005 | 0.005 | 0 | 0.22 | 0.17 | 0.21 | 0.08 | 0.3 | 0.23 |
| 6 | 79134092 | rs12202927 | -0.534 | 0.837 | 0.932 | -0.479 | -0.001 | 0 | 0 | 0.083 | 0.3 | 0.26 | 0.23 | 0.08 | 0.32 | 0.24 |
| 6 | 79138641 | rs10455346 | 0.498 | 0.866 | 0.859 | -0.458 | -0.033 | 0.059 | 0.059 | 0 | 0.2 | 0.1 | 0.19 | 0.08 | 0.29 | 0.21 |
| 6 | 79155534 | rs4706705 | -0.771 | 0.788 | 1.079 | -0.24 | -0.005 | -0.046 | -0.046 | 0 | 0.37 | 0.48 | 0.39 | 0.26 | 0.39 | 0.28 |
| 6 | 79157921 | rs12214772 | -0.205 | 0.904 | 1.288 | -0.168 | 0.011 | -0.012 | -0.012 | 0.323 | 0.27 | 0.32 | 0.35 | 0.26 | 0.37 | 0.26 |
| 6 | 79158574 | rs9361406 | -0.276 | 0.899 | 1.345 | -0.171 | 0.02 | -0.021 | -0.021 | 0 | 0.27 | 0.4 | 0.36 | 0.26 | 0.37 | 0.26 |
| 6 | 79163463 | rs11754966 | -0.07 | 0.866 | 1.457 | -0.218 | 0.029 | -0.128 | -0.128 | 0 | 0.25 | 0.47 | 0.24 | 0.07 | 0.34 | 0.23 |
| 6 | 79165394 | rs2103720 | -0.104 | 0.865 | 1.269 | -0.217 | 0 | -0.029 | -0.029 | 0 | 0.26 | 0.27 | 0.23 | 0.07 | 0.35 | 0.24 |
| 6 | 79167366 | rs742715 | -0.051 | 1.402 | 0.885 | 0.368 | 0.01 | 0.038 | 0.038 | 0 | 0.14 | 0.17 | 0.25 | 0.45 | 0.29 | 0.24 |
| 6 | 79171938 | rs10943540 | -0.084 | 0.912 | 1.208 | 0.002 | 0.027 | -0.018 | -0.018 | 2.047 | 0.26 | 0.47 | 0.34 | 0.26 | 0.35 | 0.26 |
| 6 | 79172613 | rs6915719 | -0.77 | 0.777 | 0.424 | -0.122 | 0.02 | 0.11 | 0.11 | 0.617 | 0.42 | 0.77 | 0.4 | 0.26 | 0.41 | 0.3 |
| 6 | 79173398 | rs7758061 | -1.124 | 0.525 | 0.494 | 0.13 | -0.003 | 0.007 | 0.007 | 0 | 0.27 | 0.6 | 0.23 | 0.3 | 0.23 | 0.33 |
| 6 | 79174120 | rs9294112 | -0.785 | 0.063 | -0.541 | -0.945 | 0.063 | 0.112 | 0.112 | 0 | 0.5 | 0.68 | 0.59 | 0.68 | 0.6 | 0.7 |
| 6 | 79174637 | rs6927933 | -0.688 | 0.156 | -0.634 | -1.109 | 0.007 | 0.111 | 0.111 | 0 | 0.49 | 0.65 | 0.5 | 0.7 | 0.5 | 0.7 |
| 6 | 79175007 | rs7768031 | 0.654 | 0.353 | -0.041 | -0.994 | -0.012 | -0.034 | -0.034 | 0.325 | 0.07 | 0.1 | 0.02 | 0 | 0.03 | 0.04 |
| 6 | 79175661 | rs2068816 | -0.113 | 0.077 | 0.072 | -1.126 | -0.036 | -0.083 | -0.083 | 1.317 | 0.15 | 0.31 | 0.09 | 0.01 | 0.06 | 0.18 |
| 6 | 79176271 | rs6939797 | 0.583 | 0.143 | 0.126 | -1.039 | -0.017 | -0.041 | -0.041 | 0.433 | 0.07 | 0.19 | 0.03 | 0 | 0.03 | 0.03 |
| 6 | 79184742 | rs16890052 | -0.302 | 0.248 | 0.021 | -1.113 | 0.007 | -0.094 | -0.094 | 0.46 | 0.18 | 0.33 | 0.19 | 0.01 | 0.17 | 0.18 |
| 6 | 79185106 | rs600188 | 0.811 | -0.352 | -0.534 | -1.081 | 0.015 | 0.078 | 0.078 | 0 | 0.32 | 0.17 | 0.26 | 0.15 | 0.34 | 0.24 |
| 6 | 79185214 | rs12199335 | 0.692 | -0.363 | -0.497 | -1.086 | 0.016 | 0.049 | 0.049 | 0 | 0.33 | 0.22 | 0.26 | 0.15 | 0.34 | 0.24 |
| 6 | 79188906 | rs649612 | 0.275 | -0.25 | -0.365 | -1.008 | 0.007 | 0.009 | 0.009 | 0 | 0.4 | 0.39 | 0.3 | 0.16 | 0.37 | 0.28 |
| 6 | 79190209 | rs1474755 | 0.642 | -0.241 | 0.101 | -0.293 | 0.001 | 0.047 | 0.047 | 0 | 0.17 | 0.24 | 0.18 | 0.41 | 0.16 | 0.28 |
| 6 | 79193292 | rs6454062 | 0.164 | -0.09 | 0.043 | -1.059 | -0.006 | -0.062 | -0.062 | 0 | 0.33 | 0.42 | 0.38 | 0.15 | 0.27 | 0.22 |
| 6 | 79196788 | rs7759740 | -0.653 | -0.108 | 0.01 | -0.869 | -0.005 | -0.004 | -0.004 | 0 | 0.38 | 0.59 | 0.47 | 0.62 | 0.31 | 0.52 |
| 6 | 79206005 | rs11961883 | 1.543 | -0.101 | -0.034 | -0.868 | 0.003 | 0.011 | 0.011 | 0 | 0.18 | 0.14 | 0.21 | 0.14 | 0.14 | 0.07 |
| 6 | 79206340 | rs9361408 | 0.243 | -0.106 | 0.191 | -0.879 | -0.01 | 0.009 | 0.009 | 0.272 | 0.29 | 0.34 | 0.34 | 0.62 | 0.22 | 0.5 |

| | | | | | | | | | | | | | | | | |
|---|---|---|---|---|---|---|---|---|---|---|---|---|---|---|---|---|
| 6 | 79212957 | rs4632856 | 0.362 | 0.021 | 0.153 | -0.85 | 0.026 | -0.042 | -0.042 | 0.555 | 0.08 | 0.21 | 0.13 | 0 | 0.1 | 0.01 |
| 6 | 79226369 | rs1321596 | -1.013 | -0.096 | 0.071 | -1.002 | -0.013 | 0.035 | 0.035 | 0 | 0.57 | 0.69 | 0.68 | 0.76 | 0.47 | 0.6 |
| 6 | 79227466 | rs2057298 | 0.545 | -0.125 | 0.172 | -1.001 | -0.002 | 0.006 | 0.006 | 0 | 0.24 | 0.27 | 0.3 | 0.61 | 0.2 | 0.37 |
| 6 | 79227797 | rs1358817 | 1.606 | -0.016 | 0.184 | -1.036 | 0.007 | 0.02 | 0.02 | 0 | 0.18 | 0.11 | 0.21 | 0.14 | 0.14 | 0.07 |
| 6 | 79231334 | rs941684 | 0.29 | -0.055 | -0.074 | -1.18 | -0.003 | -0.002 | -0.002 | 0 | 0.32 | 0.37 | 0.36 | 0.68 | 0.27 | 0.46 |
| 6 | 79233635 | rs9294116 | -0.981 | 0.282 | -0.38 | -1.027 | -0.021 | 0.048 | 0.048 | 0 | 0.64 | 0.74 | 0.74 | 0.83 | 0.54 | 0.67 |
| 6 | 79235696 | rs10943546 | -0.065 | 0.546 | -0.189 | -0.527 | 0.012 | -0.032 | -0.032 | 0 | 0.07 | 0.12 | 0.12 | 0 | 0.1 | 0.02 |
| 6 | 79241525 | rs7744328 | 0.088 | 0.316 | -0.209 | -0.695 | -0.002 | -0.064 | -0.064 | 0.559 | 0.17 | 0.24 | 0.11 | 0.01 | 0.13 | 0.14 |
| 6 | 79242701 | rs1395452 | 1.035 | 0.187 | -0.696 | -0.219 | -0.01 | -0.062 | -0.062 | 0.078 | 0.16 | 0.08 | 0.17 | 0.39 | 0.17 | 0.27 |
| 6 | 79248588 | rs686492 | 0.589 | 0.6 | -0.47 | -1.051 | -0.009 | 0.023 | 0.023 | 0 | 0.32 | 0.22 | 0.39 | 0.24 | 0.37 | 0.2 |
| 6 | 79251664 | rs586228 | 0.645 | 0.584 | -0.485 | -1.06 | -0.011 | 0.025 | 0.025 | 0 | 0.32 | 0.24 | 0.4 | 0.24 | 0.37 | 0.21 |
| 6 | 79259290 | rs9443592 | 0.721 | 0.49 | -0.567 | -1.109 | 0.01 | 0.014 | 0.014 | 0.112 | 0.15 | 0.12 | 0.1 | 0.08 | 0.1 | 0.23 |
| 6 | 79268815 | rs625065 | 0.182 | 0.415 | -0.16 | -1.193 | -0.002 | -0.048 | -0.048 | 0 | 0.37 | 0.43 | 0.41 | 0.25 | 0.37 | 0.2 |
| 6 | 79268831 | rs625051 | -0.781 | 0.512 | -0.192 | -1.122 | 0.003 | 0.035 | 0.035 | 0.053 | 0.52 | 0.55 | 0.5 | 0.32 | 0.47 | 0.43 |
| 6 | 79272439 | rs1395451 | 0.857 | 0.532 | -0.203 | -0.843 | 0 | 0.003 | 0.003 | 0.014 | 0.08 | 0.02 | 0.05 | 0.07 | 0.07 | 0.1 |
| 6 | 79293967 | rs11751885 | 0.236 | 0.632 | 0.104 | -0.959 | -0.012 | 0.014 | 0.014 | 0 | 0.08 | 0.04 | 0.05 | 0 | 0.1 | 0.01 |
| 6 | 79298664 | rs236867 | -1.064 | 0.617 | 0.028 | -1.16 | 0 | 0.018 | 0.018 | 0 | 0.58 | 0.59 | 0.56 | 0.32 | 0.57 | 0.45 |
| 6 | 79301526 | rs7776020 | 1.756 | 0.667 | -0.265 | -1.362 | -0.017 | 0.005 | 0.005 | 0 | 0.16 | 0.01 | 0.2 | 0.16 | 0.13 | 0.07 |
| 6 | 79304684 | rs236875 | -0.109 | 1.047 | -0.071 | -1.241 | -0.011 | 0.033 | 0.033 | 0.659 | 0.43 | 0.32 | 0.48 | 0.33 | 0.5 | 0.3 |
| 6 | 79305484 | rs236877 | -0.879 | 0.85 | -0.193 | -0.989 | -0.001 | 0.016 | 0.016 | 0 | 0.58 | 0.59 | 0.6 | 0.33 | 0.57 | 0.44 |
| 6 | 79306231 | rs11756326 | 0.429 | 0.753 | -0.033 | -0.636 | -0.007 | 0 | 0 | 0 | 0.06 | 0.04 | 0.05 | 0 | 0.1 | 0.01 |
| 6 | 79315357 | rs1570075 | -0.475 | -0.209 | -0.165 | -0.27 | -0.04 | 0.014 | 0.014 | 0 | 0.63 | 0.52 | 0.6 | 0.66 | 0.76 | 0.66 |
| 6 | 79316177 | rs12200556 | 0.217 | -0.476 | -0.396 | -0.309 | -0.028 | -0.004 | -0.004 | 0.525 | 0.14 | 0.06 | 0.12 | 0.15 | 0.23 | 0.16 |
| 6 | 79316230 | rs10806133 | -0.291 | -0.141 | -0.488 | -0.637 | 0.001 | 0.006 | 0.006 | 0 | 0.49 | 0.38 | 0.45 | 0.65 | 0.47 | 0.5 |
| 6 | 79318288 | rs1995650 | -0.683 | 0.118 | -0.402 | -0.528 | -0.001 | 0.001 | 0.001 | 0 | 0.59 | 0.44 | 0.53 | 0.65 | 0.58 | 0.64 |
| 6 | 79322411 | rs236889 | 0.475 | -0.074 | -0.235 | -0.57 | -0.018 | -0.058 | -0.058 | 0 | 0.1 | 0.21 | 0.1 | 0 | 0.05 | 0.03 |
| 6 | 79326617 | rs16890234 | 0.812 | -0.123 | -0.809 | -0.412 | 0 | 0 | 0 | 0 | 0.13 | 0.11 | 0.13 | 0.36 | 0.14 | 0.26 |
| 6 | 79337035 | rs12200526 | -0.111 | -0.12 | -1.125 | -0.527 | -0.095 | -0.141 | -0.141 | 0 | 0.26 | 0.05 | 0.28 | 0.52 | 0.37 | 0.43 |
| 6 | 79341891 | rs236855 | -1.623 | -0.119 | -0.894 | -0.718 | -0.036 | -0.071 | -0.071 | 2.766 | 0.73 | 0.83 | 0.63 | 0.65 | 0.63 | 0.69 |
| 6 | 79345146 | rs512778 | -0.114 | -0.407 | -0.609 | -0.772 | 0.018 | 0.022 | 0.022 | 0 | 0.52 | 0.62 | 0.68 | 0.51 | 0.62 | 0.52 |
| 6 | 79369333 | rs12210702 | 1.725 | -0.239 | -0.975 | -0.701 | -0.051 | -0.102 | -0.102 | 0 | 0.23 | 0.11 | 0.32 | 0.35 | 0.32 | 0.31 |
| 6 | 79379460 | rs1954659 | 0.017 | -0.499 | -0.518 | -0.606 | -0.006 | -0.004 | -0.004 | 1.391 | 0.46 | 0.49 | 0.39 | 0.49 | 0.32 | 0.42 |
| 6 | 79379599 | rs11756825 | 1.417 | -0.147 | -0.769 | -0.688 | -0.036 | 0.047 | 0.047 | 0 | 0.09 | 0.02 | 0.08 | 0.01 | 0.11 | 0.14 |
| 6 | 79396968 | rs9352611 | 1.801 | -0.191 | -0.118 | -0.419 | -0.04 | -0.057 | -0.057 | 0 | 0.32 | 0.12 | 0.34 | 0.3 | 0.36 | 0.45 |
| 6 | 79401413 | rs639370 | 0.159 | -0.317 | 0.286 | -0.517 | -0.009 | -0.001 | -0.001 | 0.781 | 0.56 | 0.51 | 0.61 | 0.46 | 0.69 | 0.6 |
| 6 | 79403793 | rs605868 | -0.426 | -0.488 | -0.051 | -0.598 | 0.027 | 0.073 | 0.073 | 0 | 0.7 | 0.86 | 0.74 | 0.82 | 0.81 | 0.85 |
| 6 | 79410392 | rs237115 | 1.082 | -0.465 | 0.166 | -0.558 | -0.007 | -0.024 | -0.024 | 0 | 0.07 | 0.1 | 0.1 | 0 | 0.05 | 0.05 |
| 6 | 79419656 | rs1782783 | -0.477 | -0.79 | -0.065 | -0.621 | 0.005 | 0.007 | 0.007 | 0 | 0.91 | 0.92 | 0.84 | 0.94 | 0.95 | 0.92 |
| 6 | 79420799 | rs17642139 | 0.53 | -0.915 | -0.156 | -0.747 | 0.011 | 0.045 | 0.045 | 0 | 0.18 | 0.07 | 0.09 | 0.02 | 0.15 | 0.22 |
| 6 | 79422789 | rs9341740 | 1.167 | -0.696 | -0.106 | -0.633 | 0.026 | 0 | 0 | 0 | 0.48 | 0.25 | 0.46 | 0.43 | 0.55 | 0.53 |
| 6 | 79428024 | rs1180819 | -0.059 | -0.407 | -0.174 | -0.902 | -0.001 | 0 | 0 | 0 | 0.18 | 0.16 | 0.2 | 0.15 | 0.24 | 0.12 |
| 6 | 79429138 | rs1543482 | 0.723 | -0.413 | -0.27 | -1.191 | 0.003 | 0.016 | 0.016 | 0 | 0.13 | 0.05 | 0.08 | 0.12 | 0.11 | 0.06 |
| 6 | 79432926 | rs1180823 | 0.809 | -0.298 | -0.264 | -0.968 | -0.018 | -0.02 | -0.02 | 0 | 0.72 | 0.72 | 0.64 | 0.39 | 0.65 | 0.77 |
| 6 | 79455282 | rs9343798 | 1.52 | -0.408 | -0.46 | -0.989 | -0.009 | 0.02 | 0.02 | 0 | 0.38 | 0.2 | 0.42 | 0.26 | 0.38 | 0.48 |
| 6 | 79457222 | rs7760429 | 0.056 | -0.388 | -0.533 | -0.927 | -0.018 | -0.03 | -0.03 | 1.348 | 0.13 | 0.07 | 0.14 | 0.55 | 0.2 | 0.14 |
| 6 | 79461603 | rs9350774 | 1.016 | -0.405 | -0.44 | -0.873 | 0.005 | 0.064 | 0.064 | 0 | 0.46 | 0.23 | 0.45 | 0.26 | 0.43 | 0.49 |
| 6 | 79479882 | rs9448560 | -0.029 | -0.333 | -0.009 | -1.824 | -0.011 | -0.053 | -0.053 | 0 | 0.14 | 0.17 | 0.13 | 0 | 0.1 | 0.05 |
| 6 | 79483130 | rs12663824 | 0.145 | -0.185 | -0.026 | -1.821 | -0.004 | -0.004 | -0.004 | 0 | 0.11 | 0.12 | 0.08 | 0.13 | 0.17 | 0.08 |
| 6 | 79483474 | rs1507152 | -1.278 | -0.36 | 0.395 | -1.975 | -0.009 | -0.026 | -0.026 | 0 | 0.35 | 0.47 | 0.37 | 0.2 | 0.32 | 0.24 |
| 6 | 79484386 | rs1507153 | 2.179 | -0.414 | 0.147 | -2.158 | -0.062 | -0.018 | -0.018 | 0 | 0.32 | 0.09 | 0.39 | 0.67 | 0.36 | 0.52 |
| 6 | 79489676 | rs9361440 | -0.415 | -0.691 | 0.67 | -2.316 | -0.004 | -0.009 | -0.009 | 0 | 0.81 | 0.82 | 0.9 | 0.87 | 0.89 | 0.92 |
| 6 | 79506885 | rs6454084 | -0.027 | -0.731 | 0.717 | -2.351 | -0.012 | -0.024 | -0.024 | 1.01 | 0.15 | 0.23 | 0.27 | 0.05 | 0.1 | 0.11 |
| 6 | 79519842 | rs906320 | -1.024 | -0.743 | 0.75 | -2.351 | 0.003 | 0.004 | 0.004 | 0 | 0.85 | 0.89 | 0.91 | 0.87 | 0.89 | 0.92 |
| 6 | 79532610 | rs955765 | 0.331 | -0.639 | 0.731 | -2.387 | 0.007 | 0.007 | 0.007 | 0 | 0.11 | 0.17 | 0.08 | 0.13 | 0.17 | 0.1 |
| 6 | 79536282 | rs6902294 | -0.635 | -0.722 | 0.649 | -2.404 | -0.006 | -0.021 | -0.021 | 0.012 | 0.23 | 0.26 | 0.29 | 0.05 | 0.15 | 0.12 |
| 6 | 79554272 | rs17741785 | 1.138 | -0.588 | 0.666 | -2.439 | -0.006 | 0.037 | 0.037 | 0.014 | 0.1 | 0.04 | 0.06 | 0.01 | 0.11 | 0.14 |
| 6 | 79562655 | rs12190108 | 0.53 | -0.553 | 0.589 | -2.441 | -0.043 | -0.047 | -0.047 | 0 | 0.1 | 0.01 | 0.07 | 0.13 | 0.17 | 0.08 |
| 6 | 79572678 | rs9352668 | -1.57 | -0.616 | 0.894 | -2.545 | 0 | -0.008 | -0.008 | 0.059 | 0.4 | 0.42 | 0.41 | 0.19 | 0.41 | 0.21 |
| 6 | 79574875 | rs6923327 | 0.096 | -0.617 | 0.929 | -2.553 | -0.011 | -0.03 | -0.03 | 0.449 | 0.08 | 0.12 | 0.11 | 0 | 0.04 | 0.03 |
| 6 | 79591805 | rs2174743 | 1.728 | -0.699 | 1.003 | -2.618 | -0.006 | 0.024 | 0.024 | 0.81 | 0.44 | 0.31 | 0.47 | 0.69 | 0.48 | 0.67 |
| 6 | 79595097 | rs6908105 | -1.224 | -0.697 | 1.014 | -2.615 | -0.005 | 0.015 | 0.015 | 0.091 | 0.34 | 0.29 | 0.31 | 0.19 | 0.37 | 0.18 |
| 6 | 79621214 | rs9361468 | -1.173 | -0.676 | 0.726 | -2.6 | 0.009 | 0.022 | 0.022 | 0 | 0.85 | 0.9 | 0.9 | 0.88 | 0.89 | 0.9 |
| 6 | 79654655 | rs9688399 | 1.04 | -0.686 | 0.794 | -2.612 | 0.006 | -0.069 | -0.069 | 0.668 | 0.11 | 0.2 | 0.07 | 0.01 | 0.11 | 0.14 |
| 6 | 79655477 | rs10943605 | 1.749 | -0.689 | 0.715 | -2.616 | -0.008 | 0.026 | 0.026 | 1.058 | 0.44 | 0.3 | 0.46 | 0.68 | 0.48 | 0.67 |
| 6 | 79657042 | rs9361473 | -0.172 | -0.683 | 0.718 | -2.63 | -0.009 | -0.012 | -0.012 | 0 | 0.2 | 0.12 | 0.23 | 0.18 | 0.23 | 0.15 |
| 6 | 79657391 | rs1984195 | 1.718 | -0.781 | 0.748 | -2.598 | -0.008 | 0.026 | 0.026 | 1.058 | 0.44 | 0.3 | 0.46 | 0.68 | 0.48 | 0.67 |
| 6 | 79674929 | rs10943608 | 2.481 | -0.757 | 0.687 | -2.581 | -0.046 | 0.007 | 0.007 | 0.125 | 0.32 | 0.12 | 0.38 | 0.68 | 0.37 | 0.53 |
| 6 | 79675701 | rs9350797 | -0.354 | -0.752 | 0.696 | -2.51 | -0.023 | -0.001 | -0.001 | 0.019 | 0.23 | 0.17 | 0.28 | 0.19 | 0.32 | 0.16 |
| 6 | 79691175 | rs10943611 | 0.929 | -0.745 | 0.66 | -2.468 | 0.008 | 0.027 | 0.027 | 0 | 0.14 | 0.07 | 0.1 | 0.12 | 0.11 | 0.1 |

| | | | | | | | | | | | | | | | | |
|---|---|---|---|---|---|---|---|---|---|---|---|---|---|---|---|---|
| 6 | 79700038 | rs1415862 | -0.118 | -0.643 | 0.653 | -2.403 | -0.002 | -0.036 | -0.036 | 0 | 0.93 | 0.83 | 0.88 | 0.99 | 0.95 | 0.95 |
| 6 | 79745107 | rs10943613 | 1.181 | -0.705 | 0.531 | -2.418 | 0 | 0.02 | 0.02 | 0.263 | 0.52 | 0.36 | 0.49 | 0.69 | 0.53 | 0.69 |
| 6 | 79754360 | rs7739298 | -0.443 | -0.408 | 0.122 | -2.149 | -0.006 | -0.027 | -0.027 | 0 | 0.95 | 0.87 | 0.89 | 1 | 0.96 | 0.95 |
| 6 | 79822314 | rs1415310 | 1.973 | -0.314 | -0.086 | -2.212 | -0.072 | 0.002 | 0.002 | 0 | 0.33 | 0.09 | 0.4 | 0.68 | 0.39 | 0.53 |
| 6 | 79823371 | rs12208017 | 0.138 | -0.297 | -0.154 | -2.242 | -0.06 | -0.053 | -0.053 | 0 | 0.14 | 0.01 | 0.09 | 0.14 | 0.23 | 0.11 |
| 6 | 79823541 | rs10943616 | 1.803 | -0.857 | 0.048 | -2.11 | -0.07 | 0.001 | 0.001 | 0 | 0.33 | 0.08 | 0.4 | 0.68 | 0.37 | 0.53 |
| 6 | 79826148 | rs9361491 | -1.572 | -0.616 | 0.498 | -1.489 | -0.033 | 0.018 | 0.018 | 0.009 | 0.58 | 0.78 | 0.48 | 0.38 | 0.56 | 0.42 |
| 6 | 79831660 | rs7747540 | 0.646 | -0.655 | 0.718 | -1.51 | -0.001 | -0.043 | -0.043 | 0 | 0.11 | 0.11 | 0.06 | 0.01 | 0.11 | 0.14 |
| 6 | 79835010 | rs6454096 | -0.989 | -0.579 | 0.512 | -1.472 | -0.001 | 0.016 | 0.016 | 0 | 0.43 | 0.55 | 0.39 | 0.2 | 0.47 | 0.31 |
| 6 | 79855431 | rs6904674 | -1.564 | -0.533 | 0.233 | -1.442 | -0.035 | 0.041 | 0.041 | 0.152 | 0.58 | 0.78 | 0.48 | 0.3 | 0.56 | 0.4 |
| 6 | 79856630 | rs10484946 | 0.024 | 0.087 | -0.259 | -0.577 | 0.006 | -0.002 | -0.002 | 0 | 0.14 | 0.1 | 0.08 | 0.19 | 0.09 | 0.09 |
| 6 | 79859877 | rs9352701 | -0.64 | 0.564 | 0.148 | -0.566 | -0.032 | -0.015 | -0.015 | 0 | 0.18 | 0.09 | 0.27 | 0.19 | 0.3 | 0.15 |
| 6 | 79861169 | rs9294130 | -0.942 | 0.803 | -0.095 | -0.367 | 0.011 | -0.022 | -0.022 | 1.139 | 0.3 | 0.49 | 0.33 | 0.19 | 0.35 | 0.17 |
| 6 | 79872281 | rs1335533 | 0.281 | 0.659 | -0.354 | -0.956 | -0.002 | -0.037 | -0.037 | 0 | 0.56 | 0.52 | 0.77 | 0.87 | 0.59 | 0.57 |
| 6 | 79875143 | rs2874649 | -0.028 | 0.344 | -0.587 | -1.441 | -0.08 | 0.04 | 0.04 | 0 | 0.27 | 0.05 | 0.16 | 0.13 | 0.33 | 0.31 |
| 6 | 79876976 | rs12206378 | -0.338 | 0.287 | -0.147 | -1.47 | 0.003 | -0.01 | -0.01 | 0 | 0.3 | 0.32 | 0.18 | 0.13 | 0.34 | 0.31 |
| 6 | 79878217 | rs6454097 | 0.399 | 0.222 | -0.318 | -1.388 | -0.01 | -0.045 | -0.045 | 0 | 0.88 | 0.84 | 0.96 | 1 | 0.92 | 0.89 |
| 6 | 79880555 | rs4276463 | 0.686 | 0.312 | -0.339 | -1.458 | 0.003 | -0.011 | -0.011 | 0 | 0.87 | 0.92 | 0.95 | 1 | 0.9 | 0.89 |
| 6 | 79881053 | rs12214432 | -0.107 | 0.412 | -0.553 | -1.31 | -0.059 | -0.014 | -0.014 | 0 | 0.24 | 0.06 | 0.15 | 0.17 | 0.3 | 0.29 |
| 6 | 79886077 | rs1577792 | 0.653 | -0.045 | 0.034 | -1.817 | 0.001 | 0.052 | 0.052 | 0 | 0.55 | 0.35 | 0.76 | 0.84 | 0.61 | 0.65 |
| 6 | 79887872 | rs6454101 | -0.127 | -0.12 | 0.737 | -1.988 | -0.006 | 0.025 | 0.025 | 0 | 0.35 | 0.24 | 0.19 | 0.16 | 0.34 | 0.34 |
| 6 | 79892520 | rs16890450 | -0.883 | -0.121 | 1.056 | -1.981 | 0.012 | -0.039 | -0.039 | 0 | 0.08 | 0.27 | 0.04 | 0 | 0.05 | 0.01 |
| 6 | 79902254 | rs7750289 | -0.829 | -0.135 | 1.131 | -2.031 | 0 | -0.032 | -0.032 | 0 | 0.11 | 0.15 | 0.06 | 0 | 0.11 | 0.1 |
| 6 | 79910280 | rs9343883 | 1.063 | -0.195 | 1.37 | -1.826 | -0.009 | 0.11 | 0.11 | 0 | 0.49 | 0.29 | 0.71 | 0.8 | 0.55 | 0.55 |
| 6 | 79910318 | rs9352704 | -0.073 | -0.233 | 1.649 | -1.848 | -0.004 | 0.014 | 0.014 | 0 | 0.36 | 0.26 | 0.2 | 0.2 | 0.34 | 0.34 |
| 6 | 79911290 | rs9448657 | -0.9 | -0.201 | 1.508 | -1.828 | 0 | -0.046 | -0.046 | 0 | 0.12 | 0.18 | 0.06 | 0.01 | 0.11 | 0.11 |
| 6 | 79912189 | rs2236543 | 1.17 | -0.319 | 0.385 | -2.174 | -0.062 | -0.134 | -0.134 | 1.918 | 0.22 | 0.02 | 0.44 | 0.7 | 0.25 | 0.35 |
| 6 | 79914733 | rs10943623 | 0.416 | -0.375 | 0.486 | -2.211 | -0.012 | 0.103 | 0.103 | 0 | 0.48 | 0.28 | 0.71 | 0.8 | 0.55 | 0.52 |
| 6 | 79922256 | rs13210296 | 0.348 | -0.463 | 0.552 | -2.005 | 0.002 | 0.014 | 0.014 | 0 | 0.07 | 0.04 | 0.04 | 0.01 | 0.06 | 0.1 |
| 6 | 79934569 | rs13209338 | 0.103 | -0.489 | 0.829 | -2.017 | -0.005 | -0.032 | -0.032 | 0 | 0.07 | 0.1 | 0.04 | 0.01 | 0.06 | 0.1 |
| 6 | 79942484 | rs1335535 | 0.533 | -0.539 | 1.132 | -2.082 | 0.014 | -0.036 | -0.036 | 0 | 0.93 | 0.76 | 0.96 | 1 | 0.95 | 0.99 |
| 6 | 79942929 | rs9294133 | 1.177 | -0.565 | 1.144 | -2.097 | -0.002 | -0.099 | -0.099 | 0 | 0.85 | 0.63 | 0.92 | 1 | 0.89 | 0.88 |
| 6 | 79944994 | rs1060419 | 0.575 | -0.712 | 0.78 | -2.631 | 0.006 | -0.021 | -0.021 | 0 | 0.39 | 0.41 | 0.3 | 0.13 | 0.44 | 0.32 |
| 6 | 79948276 | rs12214161 | 0.635 | -0.813 | 0.646 | -2.657 | -0.017 | 0.076 | 0.076 | 0 | 0.41 | 0.16 | 0.6 | 0.79 | 0.44 | 0.5 |
| 6 | 79959909 | rs2322217 | 0.671 | -0.615 | 0.799 | -2.643 | -0.006 | 0.01 | 0.01 | 0 | 0.38 | 0.32 | 0.29 | 0.13 | 0.43 | 0.31 |
| 6 | 79979700 | rs4499887 | 0.789 | -0.664 | 0.56 | -2.668 | -0.079 | 0.095 | 0.095 | 0 | 0.36 | 0.1 | 0.28 | 0.13 | 0.43 | 0.31 |
| 6 | 79979927 | rs1414283 | -0.252 | -0.504 | 0.574 | -2.564 | 0.012 | -0.03 | -0.03 | 0.02 | 0.94 | 0.85 | 0.97 | 1 | 0.95 | 0.99 |
| 6 | 79985624 | rs12208128 | 1.097 | -0.687 | 0.329 | -2.592 | -0.016 | 0.025 | 0.025 | 0 | 0.22 | 0.13 | 0.27 | 0.09 | 0.31 | 0.15 |
| 6 | 79990109 | rs6913297 | 0.466 | -0.233 | 0.109 | -2.344 | 0.008 | -0.044 | -0.044 | 0 | 0.92 | 0.73 | 0.95 | 1 | 0.95 | 0.98 |
| 6 | 79991629 | rs7757062 | 0.491 | -0.233 | 0.102 | -2.344 | 0.01 | -0.044 | -0.044 | 0 | 0.92 | 0.72 | 0.95 | 1 | 0.95 | 0.98 |
| 6 | 80002450 | rs9341767 | -0.176 | -0.324 | -0.241 | -2.488 | -0.003 | -0.053 | -0.053 | 0.506 | 0.67 | 0.57 | 0.69 | 0.83 | 0.69 | 0.7 |
| 6 | 80002510 | rs9341768 | 0.935 | -0.126 | -1.009 | -2.545 | -0.098 | 0.053 | 0.053 | 0 | 0.25 | 0.03 | 0.22 | 0.09 | 0.29 | 0.22 |
| 6 | 80010581 | rs10943629 | 0.162 | -0.032 | -0.963 | -2.823 | 0 | 0.002 | 0.002 | 0 | 0.43 | 0.32 | 0.33 | 0.17 | 0.43 | 0.42 |
| 6 | 80012425 | rs4706767 | 1.271 | -0.197 | -0.977 | -2.822 | 0.004 | -0.059 | -0.059 | 0 | 0.19 | 0.1 | 0.36 | 0.7 | 0.19 | 0.37 |
| 6 | 80016681 | rs4706768 | -0.006 | -0.161 | -1.306 | -2.855 | 0.025 | 0.047 | 0.047 | 0 | 0.44 | 0.77 | 0.49 | 0.74 | 0.4 | 0.49 |
| 6 | 80027915 | rs4706090 | -0.558 | -0.064 | -1.965 | -2.825 | -0.096 | 0.069 | 0.069 | 0 | 0.62 | 0.93 | 0.58 | 0.81 | 0.54 | 0.66 |
| 6 | 80035972 | rs12200596 | 0.444 | -0.079 | -1.796 | -2.857 | -0.013 | -0.071 | -0.071 | 0 | 0.24 | 0.67 | 0.12 | 0.04 | 0.21 | 0.11 |
| 6 | 80036131 | rs4706772 | -1.129 | 0.345 | -1.88 | -2.862 | -0.056 | -0.012 | -0.012 | 0 | 0.87 | 0.98 | 0.8 | 0.9 | 0.83 | 0.89 |
| 6 | 80046198 | rs2184425 | -0.759 | -0.275 | -1.554 | -2.639 | -0.094 | 0.082 | 0.082 | 0 | 0.61 | 0.92 | 0.58 | 0.81 | 0.53 | 0.65 |
| 6 | 80046945 | rs12198122 | 0.535 | -0.332 | -1.328 | -2.634 | -0.005 | -0.108 | -0.108 | 0 | 0.22 | 0.5 | 0.11 | 0.04 | 0.21 | 0.11 |
| 6 | 80047861 | rs10484944 | 0.296 | -0.143 | -1.67 | -2.396 | -0.012 | -0.02 | -0.02 | 0 | 0.18 | 0.17 | 0.1 | 0.08 | 0.14 | 0.18 |
| 6 | 80054021 | rs4706776 | 0.315 | 0.309 | -1.723 | -1.967 | 0.009 | -0.028 | -0.028 | 0 | 0.22 | 0.12 | 0.37 | 0.7 | 0.2 | 0.4 |
| 6 | 80063980 | rs7740673 | -0.067 | 0.658 | -1.818 | -1.932 | 0.019 | 0.088 | 0.088 | 0 | 0.46 | 0.82 | 0.5 | 0.74 | 0.42 | 0.51 |
| 6 | 80070120 | rs12191986 | -0.348 | 1.413 | -1.381 | -0.775 | 0.009 | 0.012 | 0.012 | 0.509 | 0.05 | 0.1 | 0.07 | 0.09 | 0.1 | 0.04 |
| 6 | 80079559 | rs10943641 | 0.061 | 1.13 | -2.006 | -1.247 | -0.035 | 0.153 | 0.153 | 0.002 | 0.26 | 0.01 | 0.14 | 0.09 | 0.21 | 0.26 |
| 6 | 80082927 | rs725750 | 0.318 | 2.244 | -1.596 | -0.777 | -0.036 | 0.013 | 0.013 | 0 | 0.9 | 0.95 | 0.9 | 0.96 | 0.82 | 0.9 |
| 6 | 80085887 | rs4055090 | 0.397 | 1.337 | -1.292 | -0.935 | 0.035 | 0.016 | 0.016 | 0.581 | 0.45 | 0.33 | 0.52 | 0.7 | 0.3 | 0.53 |
| 6 | 80086885 | rs3857448 | 0.195 | 1.297 | -0.8 | -1.018 | 0.007 | 0 | 0 | 0 | 0.18 | 0.21 | 0.14 | 0.14 | 0.25 | 0.13 |
| 6 | 80094171 | rs17799269 | 0.683 | 1.003 | -0.517 | -1.291 | -0.014 | 0.03 | 0.03 | 0 | 0.17 | 0.08 | 0.19 | 0.08 | 0.19 | 0.21 |
| 6 | 80095914 | rs16890620 | 0.819 | 0.9 | -0.431 | -1.341 | 0.004 | 0.024 | 0.024 | 0 | 0.11 | 0.05 | 0.03 | 0.04 | 0.09 | 0.13 |
| 6 | 80102138 | rs3843504 | 0.83 | 0.802 | -0.627 | -1.346 | -0.011 | -0.036 | -0.036 | 0 | 0.65 | 0.54 | 0.75 | 0.79 | 0.7 | 0.62 |
| 6 | 80107679 | rs17743190 | 0.596 | 1.2 | -0.728 | -1.038 | -0.006 | 0.032 | 0.032 | 0.088 | 0.13 | 0.04 | 0.09 | 0.1 | 0.1 | 0.12 |
| 6 | 80111104 | rs9294143 | 1.049 | 1.296 | -0.573 | -1.063 | 0.004 | -0.001 | -0.001 | 0 | 0.45 | 0.37 | 0.5 | 0.58 | 0.41 | 0.34 |
| 6 | 80113449 | rs196669 | -0.73 | 1.267 | -0.34 | -1.031 | -0.019 | -0.061 | -0.061 | 0 | 0.3 | 0.53 | 0.19 | 0.15 | 0.24 | 0.24 |
| 6 | 80113595 | rs196670 | -0.364 | 1.239 | -0.131 | -1.057 | -0.014 | -0.076 | -0.076 | 0.021 | 0.19 | 0.39 | 0.16 | 0.1 | 0.16 | 0.13 |
| 6 | 80116301 | rs3846737 | -0.232 | 1.258 | -0.186 | -0.95 | 0.008 | 0.02 | 0.02 | 0.219 | 0.25 | 0.17 | 0.12 | 0.16 | 0.19 | 0.23 |
| 6 | 80126245 | rs3843505 | 0.796 | 1.416 | -0.025 | -0.683 | 0.005 | -0.03 | -0.03 | 0.656 | 0.75 | 0.47 | 0.73 | 0.8 | 0.74 | 0.74 |
| 6 | 80129933 | rs196686 | 0.711 | 1.408 | 0.054 | -0.674 | -0.005 | -0.019 | -0.019 | 0 | 0.77 | 0.57 | 0.77 | 0.8 | 0.78 | 0.76 |
| 6 | 80130929 | rs196687 | 1.087 | 1.617 | 0.328 | -0.776 | 0.018 | -0.029 | -0.029 | 0 | 0.72 | 0.48 | 0.64 | 0.79 | 0.68 | 0.69 |

| | | | | | | | | | | | | | | | | |
|---|---|---|---|---|---|---|---|---|---|---|---|---|---|---|---|---|
| 6 | 80131140 | rs16890716 | 0.093 | 1.278 | 0.277 | -0.595 | 0 | -0.021 | -0.021 | 0.83 | 0.15 | 0.2 | 0.13 | 0.11 | 0.13 | 0.13 |
| 6 | 80131636 | rs196689 | 0.651 | 1.282 | 0.071 | -0.528 | 0 | -0.011 | -0.011 | 0 | 0.17 | 0.14 | 0.13 | 0.2 | 0.17 | 0.22 |
| 6 | 80132292 | rs16890720 | -0.306 | 1.173 | 0.377 | -0.177 | -0.013 | -0.055 | -0.055 | 0 | 0.18 | 0.32 | 0.13 | 0.1 | 0.15 | 0.12 |
| 6 | 80133500 | rs196691 | 0.599 | 1.137 | 0.16 | -0.065 | 0.015 | 0.002 | 0.002 | 0 | 0.77 | 0.57 | 0.66 | 0.78 | 0.74 | 0.74 |
| 6 | 80136327 | rs4706786 | 0.486 | 0.746 | -1.092 | 0.089 | -0.013 | -0.129 | -0.129 | 0 | 0.16 | 0.01 | 0.27 | 0.58 | 0.16 | 0.21 |
| 6 | 80136466 | rs3846739 | 0.017 | 0.514 | -0.296 | -0.677 | 0.01 | 0.001 | 0.001 | 0 | 0.85 | 0.7 | 0.81 | 0.88 | 0.85 | 0.78 |
| 6 | 80136577 | rs3846740 | 0.531 | 0.413 | -0.41 | -0.967 | -0.011 | -0.049 | -0.049 | 0 | 0.69 | 0.57 | 0.72 | 0.81 | 0.7 | 0.66 |
| 6 | 80137259 | rs9361533 | -0.316 | 0.168 | -0.363 | -0.604 | -0.008 | -0.008 | -0.008 | 0 | 0.37 | 0.34 | 0.49 | 0.67 | 0.33 | 0.42 |
| 6 | 80138778 | rs196705 | -0.915 | -0.15 | -0.584 | -0.936 | 0.008 | -0.039 | -0.039 | 0.314 | 0.89 | 0.95 | 0.86 | 0.83 | 0.93 | 0.84 |
| 6 | 80139673 | rs9443669 | 0.738 | 0.031 | -0.27 | -1.87 | 0.001 | -0.112 | -0.112 | 0 | 0.36 | 0.39 | 0.21 | 0.06 | 0.39 | 0.2 |
| 6 | 80141077 | rs12213051 | 0.123 | -0.495 | -0.343 | -2.19 | 0.016 | 0.044 | 0.044 | 0 | 0.55 | 0.63 | 0.51 | 0.72 | 0.45 | 0.44 |
| 6 | 80141139 | rs11754404 | 0.515 | -0.009 | -0.437 | -2.357 | -0.031 | 0.06 | 0.06 | 2.094 | 0.28 | 0.18 | 0.28 | 0.1 | 0.36 | 0.38 |
| 6 | 80143500 | rs10498920 | 1.384 | -0.165 | -0.198 | -2.366 | -0.002 | -0.006 | -0.006 | 0 | 0.32 | 0.25 | 0.18 | 0.02 | 0.35 | 0.19 |
| 6 | 80147187 | rs196701 | -1.507 | -0.167 | -0.225 | -2.322 | 0.01 | -0.042 | -0.042 | 0.085 | 0.88 | 0.95 | 0.86 | 0.83 | 0.93 | 0.83 |
| 6 | 80147522 | rs9448710 | 1.205 | -0.131 | 0.099 | -2.431 | -0.001 | -0.03 | -0.03 | 0 | 0.33 | 0.29 | 0.19 | 0.06 | 0.36 | 0.19 |
| 6 | 80149183 | rs196698 | -0.573 | -0.393 | 0.135 | -2.401 | -0.002 | -0.01 | -0.01 | 0.482 | 0.7 | 0.75 | 0.56 | 0.17 | 0.85 | 0.59 |
| 6 | 80155789 | rs3846742 | 0.766 | -0.243 | -0.236 | -2.41 | -0.018 | 0.087 | 0.087 | 1.71 | 0.26 | 0.12 | 0.16 | 0.1 | 0.28 | 0.36 |
| 6 | 80159327 | rs2803185 | 0.019 | -0.301 | -0.056 | -2.298 | 0.023 | 0.035 | 0.035 | 0.646 | 0.56 | 0.65 | 0.53 | 0.25 | 0.63 | 0.4 |
| 6 | 80160375 | rs2803191 | 0.783 | -0.244 | -0.055 | -2.333 | -0.002 | -0.05 | -0.05 | 0 | 0.41 | 0.58 | 0.29 | 0.07 | 0.5 | 0.2 |
| 6 | 80162372 | rs10484941 | 1.038 | 0.019 | -0.17 | -1.587 | -0.021 | -0.039 | -0.039 | 0 | 0.13 | 0.06 | 0.23 | 0.17 | 0.13 | 0.2 |
| 6 | 80166444 | rs9343914 | 0.649 | -0.055 | -0.248 | -1.567 | 0.036 | -0.041 | -0.041 | 0.196 | 0.19 | 0.1 | 0.3 | 0.66 | 0.1 | 0.25 |
| 6 | 80170708 | rs2437701 | -0.683 | 0.849 | 0.324 | -1.083 | 0.063 | 0.035 | 0.035 | 0.356 | 0.24 | 0.49 | 0.36 | 0.23 | 0.26 | 0.24 |
| 6 | 80176353 | rs17727505 | -0.041 | 1.083 | 0.079 | -1.109 | 0.043 | -0.026 | -0.026 | 0 | 0.05 | 0.11 | 0.09 | 0.01 | 0.12 | 0.03 |
| 6 | 80176497 | rs1485376 | 0.488 | 0.41 | 0.155 | -1.701 | 0.001 | -0.007 | -0.007 | 0 | 0.55 | 0.45 | 0.45 | 0.3 | 0.56 | 0.41 |
| 6 | 80177371 | rs1485373 | -0.635 | 0.141 | 0.041 | -1.612 | -0.054 | 0.031 | 0.031 | 0.444 | 0.81 | 0.94 | 0.83 | 0.85 | 0.73 | 0.68 |
| 6 | 80178048 | rs2803181 | -0.635 | 0.141 | 0.042 | -1.632 | -0.057 | 0.034 | 0.034 | 0.444 | 0.81 | 0.95 | 0.83 | 0.85 | 0.73 | 0.68 |
| 6 | 80178342 | rs1485371 | 1.171 | -0.063 | -0.147 | -1.845 | -0.001 | 0.003 | 0.003 | 0.265 | 0.49 | 0.45 | 0.43 | 0.26 | 0.51 | 0.37 |
| 6 | 80178365 | rs4706789 | 0.66 | -0.236 | 0.075 | -1.858 | -0.007 | 0.161 | 0.161 | 0 | 0.07 | 0.13 | 0.25 | 0.53 | 0.05 | 0.19 |
| 6 | 80178392 | rs2803182 | -0.395 | -0.31 | -0.091 | -1.892 | -0.013 | 0.105 | 0.105 | 1.301 | 0.7 | 0.93 | 0.81 | 0.85 | 0.72 | 0.64 |
| 6 | 80178863 | rs2803183 | -0.836 | -0.026 | 0.225 | -1.515 | 0.05 | 0.188 | 0.188 | 0.134 | 0.14 | 0.39 | 0.35 | 0.55 | 0.17 | 0.22 |
| 6 | 80187982 | rs16890791 | -0.184 | -0.21 | 0.455 | -1.801 | -0.003 | -0.025 | -0.025 | 0.016 | 0.33 | 0.51 | 0.34 | 0.23 | 0.25 | 0.28 |
| 6 | 80191532 | rs9294147 | 0.106 | -0.285 | 0.243 | -2.028 | 0.005 | -0.007 | -0.007 | 0 | 0.25 | 0.17 | 0.27 | 0.25 | 0.21 | 0.27 |
| 6 | 80192120 | rs2655668 | -0.789 | -0.3 | 0.013 | -2.001 | 0 | -0.002 | -0.002 | 0 | 0.43 | 0.57 | 0.38 | 0.27 | 0.33 | 0.34 |
| 6 | 80195421 | rs16890805 | 1.195 | 0.064 | 0.227 | -1.721 | 0.031 | 0.008 | 0.008 | 0 | 0.06 | 0 | 0.02 | 0.04 | 0.03 | 0.04 |
| 6 | 80196848 | rs1875845 | -0.652 | 0.049 | 0.276 | -1.729 | 0.007 | -0.006 | -0.006 | 0 | 0.3 | 0.55 | 0.34 | 0.27 | 0.27 | 0.27 |
| 6 | 80216840 | rs9352743 | 0.534 | -0.136 | 0.013 | -1.595 | 0.084 | -0.108 | -0.108 | 0 | 0.21 | 0.04 | 0.28 | 0.57 | 0.09 | 0.25 |
| 6 | 80228541 | rs2655655 | -0.753 | 0.22 | -0.137 | -1.86 | -0.036 | 0.044 | 0.044 | 0 | 0.86 | 0.95 | 0.88 | 0.97 | 0.79 | 0.84 |
| 6 | 80228673 | rs2803195 | -0.753 | 0.22 | -0.137 | -1.86 | -0.036 | 0.044 | 0.044 | 0 | 0.86 | 0.95 | 0.88 | 0.97 | 0.79 | 0.84 |
| 6 | 80233948 | rs2655703 | -0.038 | 0.317 | 0.133 | -1.899 | 0.054 | -0.044 | -0.044 | 0.003 | 0.08 | 0.24 | 0.09 | 0.01 | 0.11 | 0.02 |
| 6 | 80237507 | rs1827628 | 1.454 | 0.394 | 0.076 | -2.012 | 0.012 | 0.08 | 0.08 | 0 | 0.4 | 0.26 | 0.26 | 0.17 | 0.43 | 0.35 |
| 6 | 80240951 | rs6927389 | 1.456 | 0.606 | -0.035 | -1.814 | 0.015 | -0.024 | -0.024 | 0.066 | 0.08 | 0.11 | 0.08 | 0.01 | 0.15 | 0.04 |
| 6 | 80242512 | rs9352745 | -0.366 | 0.143 | 0.048 | -1.436 | 0.01 | 0.018 | 0.018 | 0 | 0.37 | 0.32 | 0.53 | 0.79 | 0.26 | 0.48 |
| 6 | 80252423 | rs11753205 | 1.152 | 0.469 | -0.48 | -1.203 | -0.038 | -0.045 | -0.045 | 0 | 0.15 | 0.07 | 0.24 | 0.22 | 0.19 | 0.23 |
| 6 | 80258802 | rs9359383 | 1.072 | 0.411 | -0.491 | -1.231 | 0.047 | -0.084 | -0.084 | 2.155 | 0.15 | 0.04 | 0.27 | 0.54 | 0.05 | 0.21 |
| 6 | 80269343 | rs7743617 | 0.812 | 0.865 | -0.13 | -0.989 | -0.024 | -0.023 | -0.023 | 0 | 0.15 | 0.1 | 0.24 | 0.18 | 0.19 | 0.23 |
| 6 | 80278470 | rs6454124 | -0.07 | 0.755 | 0.144 | -1.036 | 0.06 | -0.033 | -0.033 | 0.345 | 0.06 | 0.16 | 0.09 | 0.01 | 0.11 | 0.02 |
| 6 | 80286973 | rs1354452 | 0.964 | 0.395 | -0.257 | -0.987 | 0.024 | -0.033 | -0.033 | 0 | 0.16 | 0.09 | 0.27 | 0.55 | 0.06 | 0.2 |
| 6 | 80301114 | rs2814380 | 0.407 | 0.491 | -0.229 | -1.501 | 0.013 | 0.059 | 0.059 | 0.069 | 0.55 | 0.35 | 0.51 | 0.77 | 0.45 | 0.57 |
| 6 | 80307921 | rs6939569 | 0.121 | 0.613 | -0.376 | -1.488 | -0.013 | 0.019 | 0.019 | 0 | 0.1 | 0.04 | 0.1 | 0.05 | 0.14 | 0.05 |
| 6 | 80311361 | rs1394289 | 1.288 | 0.888 | -0.302 | -1.572 | -0.02 | -0.008 | -0.008 | 0.797 | 0.15 | 0.11 | 0.24 | 0.15 | 0.19 | 0.24 |
| 6 | 80313297 | rs12208969 | 1.104 | 0.686 | -0.344 | -1.612 | 0.014 | 0.071 | 0.071 | 0.112 | 0.39 | 0.25 | 0.24 | 0.21 | 0.38 | 0.38 |
| 6 | 80322737 | rs2655699 | 1.253 | 0.592 | -0.38 | -1.522 | 0.013 | -0.015 | -0.015 | 0 | 0.17 | 0.11 | 0.27 | 0.56 | 0.06 | 0.19 |
| 6 | 80322860 | rs12523953 | 1.091 | 1.264 | -0.264 | -1.646 | -0.006 | -0.003 | -0.003 | 0 | 0.15 | 0.15 | 0.24 | 0.16 | 0.19 | 0.24 |
| 6 | 80323834 | rs2812683 | 0.747 | 1.334 | 0.012 | -1.671 | -0.004 | -0.028 | -0.028 | 0.021 | 0.56 | 0.51 | 0.52 | 0.76 | 0.45 | 0.57 |
| 6 | 80325234 | rs2812681 | -0.524 | 0.84 | 0.054 | -1.495 | 0.018 | -0.083 | -0.083 | 0.36 | 0.82 | 0.71 | 0.86 | 0.97 | 0.77 | 0.87 |
| 6 | 80325857 | rs2245410 | 0.19 | 0.82 | 0.008 | -1.245 | 0.007 | -0.024 | -0.024 | 0.202 | 0.6 | 0.53 | 0.6 | 0.78 | 0.56 | 0.59 |
| 6 | 80330183 | rs2655696 | 0.334 | 0.634 | 0.137 | -0.839 | -0.013 | -0.032 | -0.032 | 0 | 0.68 | 0.61 | 0.48 | 0.22 | 0.76 | 0.57 |
| 6 | 80339088 | rs2655684 | 0.59 | 1.422 | 0.412 | -1.067 | 0.035 | -0.068 | -0.068 | 0 | 0.78 | 0.7 | 0.74 | 0.94 | 0.69 | 0.9 |
| 6 | 80340234 | rs997226 | -0.183 | 1.576 | 0.27 | -0.698 | 0.012 | 0.021 | 0.021 | 0 | 0.2 | 0.28 | 0.31 | 0.32 | 0.25 | 0.27 |
| 6 | 80343642 | rs2134856 | 0.684 | 1.307 | 0.271 | -0.692 | 0.061 | 0.009 | 0.009 | 0 | 0.7 | 0.5 | 0.54 | 0.66 | 0.54 | 0.69 |
| 6 | 80344948 | rs10455361 | 1.838 | 1.103 | -0.191 | -0.659 | 0.031 | 0.156 | 0.156 | 0 | 0.41 | 0.17 | 0.23 | 0.2 | 0.37 | 0.45 |
| 6 | 80350534 | rs2655685 | 1.336 | 0.966 | -0.05 | -0.513 | 0.032 | -0.04 | -0.04 | 0 | 0.15 | 0.1 | 0.19 | 0.41 | 0.06 | 0.18 |
| 6 | 80353483 | rs2812719 | 1.435 | 1.034 | -0.123 | -0.705 | 0.054 | 0.048 | 0.048 | 0.251 | 0.54 | 0.29 | 0.4 | 0.61 | 0.38 | 0.62 |
| 6 | 80357321 | rs12527526 | 0.786 | 0.609 | -0.126 | -0.641 | -0.037 | 0.029 | 0.029 | 0 | 0.09 | 0 | 0.11 | 0.04 | 0.1 | 0.06 |
| 6 | 80364814 | rs10455362 | 0.616 | 0.894 | -0.049 | -0.784 | 0.047 | 0.101 | 0.101 | 0 | 0.57 | 0.32 | 0.44 | 0.24 | 0.64 | 0.57 |
| 6 | 80366145 | rs2746313 | 0.541 | 1.033 | -0.212 | -0.693 | 0.019 | -0.106 | -0.106 | 0 | 0.24 | 0.12 | 0.3 | 0.45 | 0.17 | 0.24 |
| 6 | 80366532 | rs2812713 | -0.615 | 1.381 | 0.195 | -0.763 | 0.03 | -0.069 | -0.069 | 0.126 | 0.87 | 0.8 | 0.86 | 0.98 | 0.79 | 0.95 |
| 6 | 80374293 | rs2746314 | -0.634 | 1.396 | 0.201 | -0.732 | 0.031 | -0.064 | -0.064 | 0 | 0.87 | 0.8 | 0.86 | 0.98 | 0.79 | 0.95 |
| 6 | 80377842 | rs2812710 | 0.375 | 1.346 | 0.086 | -0.775 | 0.021 | -0.114 | -0.114 | 0.083 | 0.75 | 0.62 | 0.74 | 0.96 | 0.68 | 0.9 |

| | | | | | | | | | | | | | | | | |
|---|---|---|---|---|---|---|---|---|---|---|---|---|---|---|---|---|
| 6 | 80381609 | rs17584185 | 0.496 | 1.436 | -0.509 | -0.606 | -0.019 | 0.017 | 0.017 | 0 | 0.08 | 0.07 | 0.09 | 0 | 0.16 | 0.06 |
| 6 | 80389433 | rs9359387 | 0.245 | 1.215 | -0.501 | -0.37 | 0.001 | 0.002 | 0.002 | 0 | 0.1 | 0.08 | 0.21 | 0.32 | 0.09 | 0.2 |
| 6 | 80400034 | rs10943658 | 1.094 | 0.997 | -0.713 | -0.742 | -0.073 | 0.198 | 0.198 | 0 | 0.46 | 0.12 | 0.33 | 0.21 | 0.47 | 0.55 |
| 6 | 80408882 | rs2235880 | -1.003 | 0.794 | -0.5 | -0.856 | 0.001 | -0.025 | -0.025 | 3.658 | 0.85 | 0.84 | 0.87 | 0.98 | 0.77 | 0.94 |
| 6 | 80410876 | rs2255090 | 0.986 | 0.349 | -0.758 | -0.459 | 0.06 | -0.078 | -0.078 | 0 | 0.19 | 0.11 | 0.2 | 0.4 | 0.06 | 0.25 |
| 6 | 80410956 | rs2295015 | 1.164 | 0.342 | -0.88 | -0.329 | 0 | 0.014 | 0.014 | 0 | 0.07 | 0.08 | 0.11 | 0.22 | 0.06 | 0.07 |
| 6 | 80411396 | rs1062793 | 1.604 | 0.464 | -1.238 | -0.539 | -0.012 | 0.103 | 0.103 | 0 | 0.36 | 0.1 | 0.22 | 0.21 | 0.35 | 0.45 |
| 6 | 80411511 | rs1503 | 1.558 | 0.505 | -1.235 | -0.54 | -0.007 | 0.093 | 0.093 | 0 | 0.36 | 0.11 | 0.22 | 0.22 | 0.35 | 0.47 |
| 6 | 80411572 | rs1507 | 0.889 | 0.782 | -0.91 | -0.545 | 0.078 | -0.111 | -0.111 | 0.01 | 0.18 | 0.06 | 0.2 | 0.4 | 0.06 | 0.25 |
| 6 | 80412925 | rs1044204 | 0.665 | 0.905 | -0.827 | -0.54 | -0.013 | 0.013 | 0.013 | 0 | 0.07 | 0.05 | 0.09 | 0 | 0.16 | 0.05 |
| 6 | 80414202 | rs6899814 | 0.589 | 0.18 | -0.354 | -0.274 | -0.001 | -0.001 | -0.001 | 0 | 0.54 | 0.48 | 0.43 | 0.43 | 0.57 | 0.59 |
| 6 | 80415428 | rs2746319 | 0.085 | -0.195 | -0.58 | 0.514 | 0.08 | -0.112 | -0.112 | 0 | 0.18 | 0.05 | 0.2 | 0.39 | 0.06 | 0.25 |
| 6 | 80419573 | rs2812677 | 0.67 | 0.503 | -0.351 | 0.675 | -0.014 | -0.031 | -0.031 | 0 | 0.84 | 0.93 | 0.58 | 0.51 | 0.79 | 0.72 |
| 6 | 80421665 | rs2812720 | -0.163 | 0.597 | -0.427 | 0.616 | -0.062 | 0.008 | 0.008 | 0.125 | 0.9 | 0.99 | 0.71 | 0.9 | 0.84 | 0.9 |
| 6 | 80432965 | rs7745524 | -0.471 | 0.331 | -0.166 | 0.043 | 0.009 | 0.003 | 0.003 | 0 | 0.48 | 0.5 | 0.43 | 0.56 | 0.4 | 0.63 |
| 6 | 80436297 | rs9343931 | 0.044 | 0.178 | -0.137 | -0.041 | 0.032 | -0.012 | -0.012 | 0 | 0.41 | 0.35 | 0.3 | 0.51 | 0.33 | 0.59 |
| 6 | 80438871 | rs9350824 | -1.067 | 0.17 | -0.034 | -0.185 | -0.056 | -0.009 | -0.009 | 0.503 | 0.59 | 0.79 | 0.63 | 0.61 | 0.52 | 0.7 |
| 6 | 80441475 | rs9443691 | 0.106 | 0.218 | 0.378 | -0.19 | 0.054 | -0.024 | -0.024 | 0 | 0.11 | 0.19 | 0.21 | 0.09 | 0.15 | 0.1 |
| 6 | 80441913 | rs9352760 | -0.275 | 0.217 | 0.399 | -0.195 | 0.061 | -0.033 | -0.033 | 0.314 | 0.13 | 0.2 | 0.31 | 0.1 | 0.19 | 0.11 |
| 6 | 80442333 | rs2207369 | -1.422 | 0.132 | 0.002 | -0.225 | -0.092 | -0.026 | -0.026 | 0 | 0.63 | 0.88 | 0.66 | 0.64 | 0.55 | 0.73 |
| 6 | 80443859 | rs13220519 | 0.89 | 0.02 | 0.047 | -0.03 | 0.031 | 0.033 | 0.033 | 0 | 0.19 | 0.13 | 0.06 | 0 | 0.09 | 0.12 |
| 6 | 80449232 | rs9294155 | -1.674 | 0.45 | 0.056 | 0.433 | -0.127 | -0.091 | -0.091 | 1.063 | 0.68 | 0.95 | 0.68 | 0.65 | 0.61 | 0.76 |
| 6 | 80454831 | rs2092745 | -0.288 | 0.421 | 0.291 | 0.477 | 0.062 | 0.01 | 0.01 | 0 | 0.49 | 0.69 | 0.45 | 0.55 | 0.42 | 0.63 |
| 6 | 80458862 | rs6914970 | -1.348 | 0.417 | 0.546 | 0.019 | 0.131 | 0.026 | 0.026 | 0 | 0.07 | 0.33 | 0.15 | 0.01 | 0.09 | 0.03 |
| 6 | 80466124 | rs6925255 | -1.148 | 0.428 | 0.23 | 0.296 | -0.098 | -0.099 | -0.099 | 0 | 0.63 | 0.89 | 0.59 | 0.57 | 0.56 | 0.73 |
| 6 | 80472811 | rs2038293 | -0.167 | 0.462 | 0.328 | 0.372 | 0.026 | 0 | 0 | 0 | 0.51 | 0.36 | 0.41 | 0.53 | 0.45 | 0.68 |
| 6 | 80483731 | rs9341791 | -0.282 | 0.235 | 0 | 0.457 | 0.007 | -0.002 | -0.002 | 7.306 | 0.52 | 0.46 | 0.42 | 0.53 | 0.45 | 0.68 |
| 6 | 80486611 | rs714702 | -0.698 | 0.408 | -0.987 | 0.425 | -0.07 | -0.072 | -0.072 | 0 | 0.59 | 0.85 | 0.57 | 0.58 | 0.53 | 0.72 |
| 6 | 80489544 | rs9448821 | -0.256 | 0.714 | 0.385 | 1.142 | -0.041 | 0.043 | 0.043 | 0.386 | 0.12 | 0.1 | 0.05 | 0.31 | 0.04 | 0.19 |
| 6 | 80492072 | rs9294160 | -1.396 | 0.722 | 0.481 | 1.061 | -0.058 | 0.086 | 0.086 | 0.271 | 0.18 | 0.31 | 0.19 | 0.36 | 0.1 | 0.23 |
| 6 | 80494345 | rs9294163 | -0.637 | 0.572 | 0.233 | 1.182 | -0.053 | 0.093 | 0.093 | 0.135 | 0.13 | 0.2 | 0.08 | 0.35 | 0.06 | 0.21 |
| 6 | 80495689 | rs9341793 | -1.039 | 0.672 | -0.186 | 1.215 | -0.013 | 0.013 | 0.013 | 0.817 | 0.19 | 0.16 | 0.08 | 0.35 | 0.11 | 0.22 |
| 6 | 80501763 | rs12525074 | -0.221 | 1.279 | -0.535 | 0.829 | -0.049 | 0.044 | 0.044 | 0 | 0.08 | 0 | 0.11 | 0 | 0.13 | 0.09 |
| 6 | 80511814 | rs12525600 | -0.453 | 1.051 | -0.554 | 1.454 | -0.029 | -0.066 | -0.066 | 0.131 | 0.12 | 0.03 | 0.26 | 0.32 | 0.16 | 0.24 |
| 6 | 80512765 | rs9448824 | -1.57 | 0.906 | 0.026 | 1.232 | 0 | 0.002 | 0.002 | 0 | 0.38 | 0.63 | 0.49 | 0.52 | 0.41 | 0.64 |
| 6 | 80516121 | rs9341795 | -0.607 | 0.901 | 0.398 | 1.146 | 0.01 | -0.034 | -0.034 | 0 | 0.22 | 0.38 | 0.2 | 0.16 | 0.25 | 0.39 |
| 6 | 80520724 | rs6903123 | -1.365 | 0.871 | 0.476 | 1.161 | 0.003 | 0.004 | 0.004 | 0 | 0.36 | 0.46 | 0.47 | 0.48 | 0.41 | 0.63 |
| 6 | 80522040 | rs9361566 | -0.311 | 0.74 | 0.177 | 1.398 | 0.006 | -0.007 | -0.007 | 0 | 0.12 | 0.08 | 0.17 | 0.38 | 0.11 | 0.23 |
| 6 | 80522080 | rs9448826 | -0.817 | 0.667 | 0.736 | 0.754 | 0.014 | -0.07 | -0.07 | 0.016 | 0.3 | 0.51 | 0.21 | 0.16 | 0.3 | 0.39 |
| 6 | 80522780 | rs12194503 | 1.212 | 0.474 | 0.033 | 0.697 | -0.001 | 0.029 | 0.029 | 0 | 0.39 | 0.18 | 0.5 | 0.34 | 0.4 | 0.3 |
| 6 | 80526386 | rs4601089 | -0.973 | 0.72 | 0.717 | 0.917 | -0.003 | 0.104 | 0.104 | 0.126 | 0.21 | 0.55 | 0.27 | 0.42 | 0.19 | 0.26 |
| 6 | 80527359 | rs10943664 | 0.13 | 0.922 | 0.772 | 0.817 | 0.014 | -0.028 | -0.028 | 0 | 0.07 | 0.08 | 0.03 | 0 | 0.1 | 0.04 |
| 6 | 80529597 | rs9343937 | 0.322 | 0.825 | 0.61 | 0.725 | -0.04 | 0.019 | 0.019 | 1.56 | 0.64 | 0.78 | 0.61 | 0.71 | 0.58 | 0.64 |
| 6 | 80530332 | rs11759038 | 0.577 | 0.795 | 0.741 | 0.88 | -0.005 | 0.103 | 0.103 | 0 | 0.09 | 0.1 | 0.19 | 0.41 | 0.07 | 0.21 |
| 6 | 80533187 | rs1321847 | 0.326 | 0.836 | 0.863 | 0.956 | -0.001 | 0.02 | 0.02 | 0 | 0.92 | 0.87 | 0.93 | 0.84 | 0.9 | 0.96 |
| 6 | 80533953 | rs9343939 | 0.622 | 0.793 | 0.772 | 0.936 | -0.003 | 0.074 | 0.074 | 0.498 | 0.1 | 0.1 | 0.19 | 0.42 | 0.08 | 0.24 |
| 6 | 80535882 | rs13207543 | 0.68 | 0.904 | 0.21 | 0.859 | -0.046 | 0.009 | 0.009 | 0.072 | 0.17 | 0.01 | 0.19 | 0.06 | 0.15 | 0.17 |
| 6 | 80539454 | rs9352773 | -0.586 | 1.186 | 0.331 | 0.756 | 0.031 | 0.158 | 0.158 | 0.168 | 0.21 | 0.39 | 0.26 | 0.42 | 0.21 | 0.29 |
| 6 | 80540223 | rs1590244 | 0.234 | 1.015 | -0.144 | 0.722 | -0.001 | 0.014 | 0.014 | 0 | 0.93 | 0.89 | 0.93 | 0.84 | 0.9 | 0.96 |
| 6 | 80546242 | rs9341796 | 0.219 | 1.035 | -0.402 | 0.213 | -0.012 | 0.087 | 0.087 | 3.108 | 0.78 | 0.5 | 0.7 | 0.41 | 0.79 | 0.69 |
| 6 | 80550286 | rs2145572 | -0.284 | 1.442 | -0.801 | 0.049 | -0.016 | 0.037 | 0.037 | 0 | 0.24 | 0.12 | 0.22 | 0.05 | 0.25 | 0.21 |
| 6 | 80555040 | rs6931690 | -0.361 | 1.357 | -0.579 | 0.683 | -0.005 | 0.003 | 0.003 | 0.291 | 0.11 | 0.14 | 0.14 | 0.07 | 0.16 | 0.12 |
| 6 | 80558480 | rs12205684 | -0.641 | 1.311 | -1.087 | 1.076 | -0.057 | -0.076 | -0.076 | 0.729 | 0.21 | 0.06 | 0.33 | 0.28 | 0.31 | 0.17 |
| 6 | 80564519 | rs9341799 | 0.7 | 0.883 | -1.798 | 0.251 | -0.013 | 0.007 | 0.007 | 0 | 0.42 | 0.2 | 0.46 | 0.58 | 0.37 | 0.34 |
| 6 | 80564836 | rs346291 | -0.016 | 0.98 | -1.893 | 0.292 | 0.009 | 0.066 | 0.066 | 0 | 0.38 | 0.74 | 0.26 | 0.2 | 0.36 | 0.5 |
| 6 | 80564945 | rs1010938 | 0.492 | 0.966 | -2.24 | 0.508 | 0.003 | 0.011 | 0.011 | 0 | 0.14 | 0.12 | 0.18 | 0.41 | 0.12 | 0.14 |
| 6 | 80566234 | rs9343946 | 0.921 | 0.797 | -1.831 | 0.169 | -0.005 | 0 | 0 | 0.067 | 0.07 | 0.11 | 0.04 | 0.12 | 0.04 | 0.29 |
| 6 | 80569885 | rs103194 | -0.6 | 0.802 | -1.909 | 0.16 | -0.04 | -0.026 | -0.026 | 0.451 | 0.8 | 0.95 | 0.72 | 0.78 | 0.74 | 0.84 |
| 6 | 80572483 | rs9448842 | 1.057 | 0.222 | -1.465 | 0.157 | 0.009 | 0.062 | 0.062 | 0.009 | 0.37 | 0.2 | 0.31 | 0.19 | 0.32 | 0.22 |
| 6 | 80572548 | rs1321844 | 0.737 | 0.188 | -1.014 | 0.355 | -0.008 | 0.052 | 0.052 | 0 | 0.15 | 0.17 | 0.19 | 0.39 | 0.12 | 0.14 |
| 6 | 80576353 | rs346305 | -0.456 | 0.245 | -0.306 | 0.11 | 0.002 | -0.005 | -0.005 | 3.03 | 0.75 | 0.66 | 0.68 | 0.78 | 0.69 | 0.81 |
| 6 | 80576744 | rs346306 | -0.615 | 0.324 | -0.291 | 0.172 | 0.001 | 0.003 | 0.003 | 0 | 0.38 | 0.49 | 0.37 | 0.59 | 0.37 | 0.58 |
| 6 | 80576812 | rs346307 | 0.922 | 0.435 | -0.224 | -0.014 | -0.012 | 0.005 | 0.005 | 0 | 0.14 | 0.09 | 0.14 | 0.07 | 0.21 | 0.14 |
| 6 | 80578610 | rs1742485 | 0.026 | 0.407 | -0.077 | -0.002 | 0.02 | -0.005 | -0.005 | 0 | 0.21 | 0.26 | 0.19 | 0.2 | 0.25 | 0.45 |
| 6 | 80582193 | rs346315 | 0.627 | 0.261 | -0.452 | 0.279 | 0.02 | 0.036 | 0.036 | 0 | 0.48 | 0.21 | 0.4 | 0.31 | 0.54 | 0.33 |
| 6 | 80590664 | rs2874814 | 0.606 | 0.56 | -0.124 | 0.183 | 0.007 | -0.023 | -0.023 | 0.018 | 0.63 | 0.51 | 0.63 | 0.7 | 0.62 | 0.48 |
| 6 | 80591625 | rs343688 | -0.043 | 0.494 | 0.243 | 0.125 | -0.006 | -0.007 | -0.007 | 1.41 | 0.84 | 0.83 | 0.93 | 0.85 | 0.88 | 0.67 |
| 6 | 80591852 | rs6911924 | 0.317 | 0.538 | 0.37 | 0.148 | 0.004 | -0.024 | -0.024 | 1.751 | 0.7 | 0.65 | 0.78 | 0.79 | 0.69 | 0.57 |
| 6 | 80597728 | rs1512117 | 1.211 | 0.563 | 0.219 | 0.273 | -0.004 | -0.002 | -0.002 | 0 | 0.08 | 0.06 | 0.15 | 0.08 | 0.07 | 0.09 |

| | | | | | | | | | | | | | | | | |
|---|---|---|---|---|---|---|---|---|---|---|---|---|---|---|---|---|
| 6 | 80598166 | rs1512116 | 0.754 | 0.21 | 0.455 | -0.193 | 0.004 | 0.034 | 0.034 | 0 | 0.49 | 0.47 | 0.56 | 0.67 | 0.45 | 0.43 |
| 6 | 80604428 | rs16891209 | 1.246 | 0.076 | 0.445 | 0.143 | -0.002 | 0.044 | 0.044 | 1.669 | 0.09 | 0.09 | 0.16 | 0.36 | 0.08 | 0.1 |
| 6 | 80604524 | rs343681 | 0.043 | 0.036 | 0.776 | -0.102 | -0.021 | -0.075 | -0.075 | 3.734 | 0.7 | 0.57 | 0.73 | 0.89 | 0.7 | 0.86 |
| 6 | 80606507 | rs10943676 | 1.575 | -0.195 | 0.356 | -0.333 | -0.012 | 0.011 | 0.011 | 0 | 0.31 | 0.2 | 0.24 | 0.24 | 0.3 | 0.23 |
| 6 | 80606537 | rs163981 | -0.306 | -0.489 | 0.443 | -0.229 | 0.013 | -0.001 | -0.001 | 0 | 0.37 | 0.45 | 0.48 | 0.64 | 0.39 | 0.58 |
| 6 | 80609178 | rs9361573 | 0.931 | -0.236 | 0.271 | -0.801 | -0.01 | -0.015 | -0.015 | 0.095 | 0.15 | 0.06 | 0.17 | 0.21 | 0.12 | 0.39 |
| 6 | 80610839 | rs2164166 | 0.4 | -0.248 | 0.462 | -0.816 | 0.008 | -0.02 | -0.02 | 0 | 0.62 | 0.49 | 0.62 | 0.71 | 0.6 | 0.46 |
| 6 | 80611616 | rs1887571 | 1.234 | 0.195 | -0.085 | -0.872 | 0.041 | -0.025 | -0.025 | 0 | 0.36 | 0.17 | 0.34 | 0.6 | 0.3 | 0.3 |
| 6 | 80614816 | rs9448846 | -0.355 | 0.522 | 0.489 | -0.884 | 0.021 | -0.012 | -0.012 | 0 | 0.33 | 0.5 | 0.35 | 0.3 | 0.34 | 0.52 |
| 6 | 80618523 | rs10806180 | 1.284 | 0.604 | -0.196 | -0.914 | -0.066 | -0.014 | -0.014 | 0 | 0.11 | 0.01 | 0.14 | 0.07 | 0.2 | 0.08 |
| 6 | 80624656 | rs2991 | 0.754 | 0.498 | 0.215 | -0.949 | -0.004 | 0.002 | 0.002 | 1.62 | 0.17 | 0.2 | 0.19 | 0.23 | 0.12 | 0.41 |
| 6 | 80625658 | rs16891260 | 0.703 | 0.495 | 0.357 | -0.951 | -0.006 | 0.006 | 0.006 | 1.347 | 0.17 | 0.25 | 0.2 | 0.23 | 0.12 | 0.41 |
| 6 | 80626375 | rs3812153 | 0.703 | 0.509 | 0.357 | -0.951 | -0.008 | 0.006 | 0.006 | 1.347 | 0.17 | 0.25 | 0.2 | 0.23 | 0.12 | 0.41 |
| 6 | 80631931 | rs1007416 | 0.703 | 0.488 | 0.357 | -0.951 | -0.006 | 0.006 | 0.006 | 1.347 | 0.17 | 0.25 | 0.2 | 0.23 | 0.12 | 0.41 |
| 6 | 80639321 | rs9448860 | 0.76 | 0.275 | 0.219 | -0.858 | -0.05 | 0.057 | 0.057 | 0 | 0.21 | 0.06 | 0.16 | 0.1 | 0.27 | 0.16 |
| 6 | 80647285 | rs9448863 | 0.768 | 0.074 | 0.332 | -0.721 | -0.004 | 0.002 | 0.002 | 1.853 | 0.17 | 0.2 | 0.19 | 0.23 | 0.12 | 0.41 |
| 6 | 80652229 | rs239520 | 0.347 | 0.06 | -0.31 | -0.728 | -0.017 | 0.003 | 0.003 | 1.763 | 0.19 | 0.55 | 0.22 | 0.23 | 0.12 | 0.42 |
| 6 | 80654633 | rs239516 | -1.042 | -0.117 | -0.469 | -0.728 | 0.039 | -0.022 | -0.022 | 0 | 0.59 | 0.89 | 0.56 | 0.41 | 0.63 | 0.71 |
| 6 | 80658793 | rs1512120 | 0.366 | -0.842 | -1.171 | -1.809 | 0.04 | 0.041 | 0.041 | 0 | 0.63 | 0.3 | 0.75 | 0.71 | 0.77 | 0.52 |
| 6 | 80661020 | rs151584 | -1.234 | -0.769 | -1.156 | -1.855 | 0.016 | 0.01 | 0.01 | 0.263 | 0.9 | 0.95 | 0.98 | 1 | 0.93 | 0.99 |
| 6 | 80664928 | rs6915142 | 0.156 | -0.65 | -1.007 | -1.757 | -0.016 | -0.028 | -0.028 | 0 | 0.66 | 0.43 | 0.77 | 0.77 | 0.77 | 0.54 |
| 6 | 80670767 | rs12207890 | 1.873 | -0.431 | -1.779 | -1.739 | -0.049 | 0.008 | 0.008 | 0.877 | 0.17 | 0.02 | 0.08 | 0.12 | 0.2 | 0.03 |
| 6 | 80674294 | rs9343953 | 1.437 | -0.102 | -1.229 | -1.92 | -0.055 | -0.057 | -0.057 | 0.077 | 0.34 | 0.07 | 0.49 | 0.59 | 0.35 | 0.25 |
| 6 | 80681241 | rs4541708 | 0.787 | -0.009 | -0.993 | -1.838 | 0.005 | 0 | 0 | 0 | 0.18 | 0.1 | 0.16 | 0.24 | 0.13 | 0.44 |
| 6 | 80683573 | rs149293 | -0.368 | 0.359 | -0.732 | -1.522 | -0.007 | -0.007 | -0.007 | 0.143 | 0.67 | 0.67 | 0.78 | 0.71 | 0.77 | 0.53 |
| 6 | 80683626 | rs10455125 | 0.745 | 0.837 | -2.042 | -1.783 | -0.049 | 0.109 | 0.109 | 0 | 0.18 | 0.01 | 0.12 | 0.01 | 0.18 | 0.14 |
| 6 | 80688583 | rs9448870 | 0.293 | 0.779 | -1.327 | -1.1 | -0.03 | 0.011 | 0.011 | 0 | 0.12 | 0.06 | 0.14 | 0.09 | 0.23 | 0.14 |
| 6 | 80688614 | rs3857457 | 0.206 | 0.254 | -1.177 | -0.793 | -0.01 | 0.066 | 0.066 | 0 | 0.54 | 0.36 | 0.62 | 0.68 | 0.54 | 0.4 |
| 6 | 80692007 | rs17239632 | -0.806 | -0.677 | -1.183 | -0.716 | 0.045 | 0.055 | 0.055 | 0 | 0.24 | 0.09 | 0.15 | 0.09 | 0.15 | 0.23 |
| 6 | 80715838 | rs240226 | -0.452 | -0.563 | -0.028 | -0.862 | -0.015 | 0.012 | 0.012 | 0 | 0.62 | 0.71 | 0.78 | 0.71 | 0.56 | 0.63 |
| 6 | 80724666 | rs12528948 | 0.02 | -0.517 | -0.27 | -1.079 | -0.013 | 0.077 | 0.077 | 0 | 0.14 | 0.01 | 0.09 | 0.01 | 0.14 | 0.15 |
| 6 | 80730966 | rs239576 | -0.801 | -0.352 | -0.149 | -1.168 | -0.033 | 0.045 | 0.045 | 0 | 0.64 | 0.8 | 0.8 | 0.78 | 0.59 | 0.69 |
| 6 | 80732242 | rs17175251 | -0.049 | -0.464 | 0 | -1.245 | 0.003 | 0.082 | 0.082 | 0 | 0.15 | 0.03 | 0.11 | 0.01 | 0.15 | 0.16 |
| 6 | 80750290 | rs608962 | -0.47 | -0.468 | 0.048 | -1.253 | 0.004 | -0.005 | -0.005 | 0.007 | 0.61 | 0.63 | 0.78 | 0.71 | 0.55 | 0.63 |
| 6 | 80751942 | rs17254634 | -0.092 | -0.524 | -0.073 | -1.268 | 0.004 | 0.085 | 0.085 | 0 | 0.16 | 0.03 | 0.1 | 0.01 | 0.15 | 0.16 |
| 6 | 80754346 | rs16891457 | 0.128 | -0.554 | 0.063 | -1.223 | -0.007 | -0.005 | -0.005 | 2.768 | 0.11 | 0.14 | 0.08 | 0.08 | 0.06 | 0.11 |
| 6 | 80765111 | rs9294169 | 0.581 | -0.429 | -0.021 | -1.17 | -0.019 | 0.014 | 0.014 | 0 | 0.36 | 0.26 | 0.21 | 0.29 | 0.41 | 0.37 |
| 6 | 80775330 | rs3812152 | 0.556 | -0.472 | 0.152 | -1.214 | -0.017 | 0.013 | 0.013 | 0 | 0.36 | 0.25 | 0.21 | 0.3 | 0.41 | 0.38 |
| 6 | 80777467 | rs3828761 | 0.096 | -0.583 | 0.142 | -1.216 | 0.01 | 0.075 | 0.075 | 0 | 0.45 | 0.24 | 0.3 | 0.29 | 0.5 | 0.43 |
| 6 | 80781605 | rs1410521 | 1.212 | -0.641 | 0.068 | -1.244 | -0.004 | -0.05 | -0.05 | 0 | 0.34 | 0.1 | 0.55 | 0.61 | 0.34 | 0.35 |
| 6 | 80784426 | rs10943688 | 1.123 | -0.647 | 0.301 | -1.247 | 0.013 | -0.004 | -0.004 | 0 | 0.35 | 0.27 | 0.56 | 0.61 | 0.34 | 0.35 |
| 6 | 80793667 | rs7772677 | 0.209 | -0.787 | -0.068 | -1.48 | -0.053 | 0.075 | 0.075 | 0 | 0.37 | 0.18 | 0.26 | 0.22 | 0.44 | 0.19 |
| 6 | 80799439 | rs2322626 | 0.776 | -0.87 | 0.083 | -1.526 | 0.029 | 0.097 | 0.097 | 0 | 0.45 | 0.22 | 0.6 | 0.77 | 0.41 | 0.65 |
| 6 | 80801693 | rs2505951 | -0.315 | -0.741 | -0.212 | -1.345 | -0.097 | 0.07 | 0.07 | 0 | 0.64 | 0.86 | 0.74 | 0.79 | 0.58 | 0.8 |
| 6 | 80803557 | rs9448883 | 0.142 | -0.755 | 0.128 | -1.145 | 0.008 | 0.018 | 0.018 | 0 | 0.11 | 0.07 | 0.11 | 0.04 | 0.08 | 0.1 |
| 6 | 80804273 | rs17807445 | -0.159 | -0.71 | 0.363 | -1.033 | 0.007 | 0.018 | 0.018 | 0 | 0.1 | 0.06 | 0.08 | 0.01 | 0.12 | 0.13 |
| 6 | 80804686 | rs10943692 | -1.06 | -0.853 | 0.865 | -0.855 | -0.051 | 0.018 | 0.018 | 0.051 | 0.16 | 0.52 | 0.06 | 0.17 | 0.06 | 0.33 |
| 6 | 80805796 | rs10452561 | -1.404 | -0.746 | 0.975 | -0.714 | -0.045 | -0.01 | -0.01 | 0 | 0.21 | 0.4 | 0.09 | 0.16 | 0.13 | 0.42 |
| 6 | 80807454 | rs2256531 | -0.224 | -0.382 | 0.389 | -0.365 | -0.065 | -0.025 | -0.025 | 0 | 0.71 | 0.87 | 0.76 | 0.72 | 0.64 | 0.82 |
| 6 | 80810013 | rs7453746 | 1.211 | -0.402 | 0.398 | -0.34 | -0.09 | -0.072 | -0.072 | 0.124 | 0.49 | 0.07 | 0.64 | 0.56 | 0.51 | 0.41 |
| 6 | 80814802 | rs2490232 | 0.669 | -0.446 | 0.715 | -0.348 | -0.019 | -0.022 | -0.022 | 2.194 | 0.6 | 0.75 | 0.73 | 0.58 | 0.58 | 0.49 |
| 6 | 80839315 | rs4706113 | 0.646 | -0.489 | 0.901 | -0.351 | -0.03 | 0.033 | 0.033 | 0 | 0.59 | 0.24 | 0.65 | 0.7 | 0.51 | 0.71 |
| 6 | 80851191 | rs2505940 | -0.163 | -0.492 | 1.121 | -0.361 | -0.015 | -0.001 | -0.001 | 0 | 0.69 | 0.69 | 0.71 | 0.71 | 0.6 | 0.81 |
| 6 | 80854470 | rs10080237 | 1.274 | -0.498 | 0.91 | -0.339 | -0.037 | -0.054 | -0.054 | 0 | 0.49 | 0.06 | 0.62 | 0.56 | 0.46 | 0.42 |
| 6 | 80880138 | rs6931421 | 1.764 | -0.689 | 0.656 | 0.036 | -0.034 | -0.084 | -0.084 | 0 | 0.36 | 0.03 | 0.42 | 0.38 | 0.32 | 0.29 |
| 6 | 80915986 | rs3805914 | -0.409 | -0.069 | 1.562 | -0.022 | -0.017 | 0.067 | 0.067 | 0 | 0.1 | 0.27 | 0.05 | 0.14 | 0.05 | 0.29 |
| 6 | 80919632 | rs7757610 | 0.048 | 0.039 | 1.443 | -0.049 | -0.003 | -0.023 | -0.023 | 1.501 | 0.08 | 0.11 | 0.04 | 0.01 | 0.08 | 0.1 |
| 6 | 80939228 | rs688867 | 0.243 | 0.077 | 1.44 | -0.067 | -0.058 | -0.027 | -0.027 | 1.532 | 0.25 | 0.15 | 0.27 | 0.3 | 0.36 | 0.19 |
| 6 | 80940719 | rs671614 | 0.179 | 0.006 | 1.476 | -0.121 | -0.01 | -0.065 | -0.065 | 0 | 0.88 | 0.74 | 0.94 | 0.99 | 0.9 | 0.9 |
| 6 | 80956146 | rs1359239 | 0.942 | 0.009 | 1.558 | -0.111 | 0.009 | -0.012 | -0.012 | 0 | 0.6 | 0.48 | 0.66 | 0.69 | 0.51 | 0.71 |
| 6 | 80956281 | rs636499 | 0.018 | -0.025 | 1.134 | -0.083 | 0.006 | 0.011 | 0.011 | 0 | 0.89 | 0.92 | 0.95 | 0.99 | 0.9 | 0.89 |
| 6 | 80962957 | rs3812123 | -0.401 | 0.009 | 1.223 | -0.034 | -0.012 | 0.017 | 0.017 | 0 | 0.09 | 0.12 | 0.03 | 0.13 | 0.06 | 0.29 |
| 6 | 80975741 | rs604732 | 0.098 | -0.006 | 1.159 | -0.023 | 0.002 | 0.003 | 0.003 | 0 | 0.88 | 0.9 | 0.95 | 0.93 | 0.9 | 0.89 |
| 6 | 80978928 | rs613588 | 0.28 | 0.108 | 1.139 | -0.02 | -0.081 | -0.017 | -0.017 | 0 | 0.25 | 0.14 | 0.26 | 0.24 | 0.36 | 0.18 |
| 6 | 80980808 | rs638056 | -0.805 | 0.081 | 1.443 | -0.02 | 0.012 | 0.009 | 0.009 | 0.226 | 0.37 | 0.56 | 0.31 | 0.44 | 0.44 | 0.48 |
| 6 | 80985539 | rs3805896 | -0.569 | 0.044 | 1.58 | -0.007 | -0.004 | 0.045 | 0.045 | 0 | 0.11 | 0.19 | 0.04 | 0.19 | 0.08 | 0.3 |
| 6 | 81026899 | rs806843 | 1.668 | -0.03 | 1.614 | -0.06 | -0.029 | -0.004 | -0.004 | 0 | 0.77 | 0.55 | 0.89 | 0.79 | 0.81 | 0.6 |
| 6 | 81030941 | rs806850 | -0.315 | 0.143 | 1.568 | 0.103 | 0.005 | 0.007 | 0.007 | 0 | 0.94 | 0.98 | 0.98 | 0.99 | 0.95 | 0.93 |
| 6 | 81034730 | rs806847 | 1.646 | 0.111 | 1.753 | 0.111 | -0.022 | 0.021 | 0.021 | 0 | 0.82 | 0.67 | 0.93 | 0.8 | 0.87 | 0.63 |

| | | | | | | | | | | | | | | | | |
|---|---|---|---|---|---|---|---|---|---|---|---|---|---|---|---|---|
| 6 | 81034777 | rs806846 | -0.275 | 0.099 | 1.724 | 0.097 | -0.007 | -0.018 | -0.018 | 0 | 0.94 | 0.92 | 0.98 | 0.99 | 0.95 | 0.93 |
| 6 | 81038412 | rs1341279 | -0.176 | 0.58 | 1.8 | 0.082 | 0.018 | -0.011 | -0.011 | 0 | 0.26 | 0.38 | 0.18 | 0.21 | 0.31 | 0.12 |
| 6 | 81040302 | rs1040596 | 1.864 | 0.573 | 1.49 | 0.08 | -0.008 | 0.025 | 0.025 | 0 | 0.8 | 0.66 | 0.83 | 0.74 | 0.82 | 0.57 |
| 6 | 81050936 | rs947223 | 0.046 | 0.621 | 1.574 | 0.091 | -0.032 | 0.006 | 0.006 | 0 | 0.24 | 0.19 | 0.17 | 0.21 | 0.3 | 0.12 |
| 6 | 81053642 | rs926733 | 1.923 | 0.641 | 1.303 | 0.174 | -0.008 | 0.024 | 0.024 | 0.008 | 0.8 | 0.66 | 0.83 | 0.74 | 0.82 | 0.57 |
| 6 | 81054515 | rs4502885 | 1.939 | 0.64 | 1.302 | 0.173 | -0.009 | 0.023 | 0.023 | 0 | 0.79 | 0.66 | 0.83 | 0.74 | 0.82 | 0.57 |
| 6 | 81054965 | rs3805848 | 1.939 | 0.64 | 1.302 | 0.173 | -0.009 | 0.023 | 0.023 | 0 | 0.79 | 0.66 | 0.83 | 0.74 | 0.82 | 0.57 |
| 6 | 81055299 | rs7758761 | 1.939 | 0.64 | 1.302 | 0.173 | -0.009 | 0.023 | 0.023 | 0 | 0.79 | 0.66 | 0.83 | 0.74 | 0.82 | 0.57 |
| 6 | 81055331 | rs7740932 | 1.939 | 0.64 | 1.302 | 0.173 | -0.009 | 0.023 | 0.023 | 0 | 0.79 | 0.66 | 0.83 | 0.74 | 0.82 | 0.57 |
| 6 | 81055396 | rs7759304 | 1.939 | 0.64 | 1.302 | 0.173 | -0.009 | 0.023 | 0.023 | 0 | 0.79 | 0.66 | 0.83 | 0.74 | 0.82 | 0.57 |
| 6 | 81055413 | rs7740958 | 1.939 | 0.64 | 1.302 | 0.173 | -0.009 | 0.023 | 0.023 | 0 | 0.79 | 0.66 | 0.83 | 0.74 | 0.82 | 0.57 |
| 6 | 81055792 | rs15062 | 1.939 | 0.64 | 1.302 | 0.173 | -0.009 | 0.023 | 0.023 | 0 | 0.79 | 0.66 | 0.83 | 0.74 | 0.82 | 0.57 |
| 6 | 81055803 | rs10864 | 1.939 | 0.64 | 1.302 | 0.173 | -0.009 | 0.023 | 0.023 | 0 | 0.79 | 0.66 | 0.83 | 0.74 | 0.82 | 0.57 |
| 6 | 81056360 | rs723587 | -0.929 | 0.76 | 1.426 | 0.251 | 0.002 | 0.076 | 0.076 | 0.096 | 0.12 | 0.26 | 0.13 | 0.22 | 0.09 | 0.26 |
| 6 | 81057918 | rs9352825 | -1.7 | 0.765 | 1.316 | 0.239 | 0.013 | 0.052 | 0.052 | 0.39 | 0.17 | 0.35 | 0.16 | 0.25 | 0.15 | 0.41 |
| 6 | 81059188 | rs9350851 | 1.223 | 0.377 | 0.284 | 0.146 | -0.1 | -0.086 | -0.086 | 0.007 | 0.51 | 0.04 | 0.63 | 0.53 | 0.51 | 0.45 |
| 6 | 81070372 | rs6906638 | 0.882 | 0.272 | 0.778 | 0.325 | 0.018 | 0.006 | 0.006 | 0.038 | 0.48 | 0.38 | 0.62 | 0.57 | 0.44 | 0.45 |
| 6 | 81072063 | rs7754957 | -0.147 | -0.008 | 0.823 | 0.296 | -0.055 | 0.089 | 0.089 | 0 | 0.38 | 0.18 | 0.29 | 0.19 | 0.53 | 0.22 |
| 6 | 81075751 | rs9448941 | 0.754 | 0.054 | 0.779 | 0.407 | 0.054 | 0.06 | 0.06 | 0.128 | 0.49 | 0.21 | 0.64 | 0.67 | 0.41 | 0.5 |
| 6 | 81075912 | rs9448942 | 0.815 | 0.061 | 0.763 | 0.411 | 0.05 | 0.056 | 0.056 | 0.509 | 0.48 | 0.18 | 0.64 | 0.67 | 0.41 | 0.5 |
| 6 | 81076339 | rs1572941 | 1.342 | -0.073 | 0.502 | 0.422 | -0.002 | -0.142 | -0.142 | 0 | 0.39 | 0.04 | 0.52 | 0.55 | 0.34 | 0.36 |
| 6 | 81122793 | rs9352835 | 0.174 | 0.67 | 1.096 | 0.49 | -0.007 | -0.007 | -0.007 | 0 | 0.05 | 0.03 | 0.12 | 0.12 | 0.12 | 0.08 |
| 6 | 81150078 | rs9352838 | 0.718 | 0.43 | 1.707 | 0.381 | -0.006 | 0.002 | 0.002 | 0 | 0.91 | 0.76 | 0.93 | 0.94 | 0.92 | 0.79 |
| 6 | 81153486 | rs560670 | 0.087 | 0.359 | 1.075 | 0.344 | -0.14 | -0.004 | -0.004 | 0 | 0.35 | 0.04 | 0.28 | 0.26 | 0.52 | 0.27 |
| 6 | 81159931 | rs9344003 | 0.257 | 0.393 | 1.099 | 0.364 | -0.008 | -0.007 | -0.007 | 0 | 0.05 | 0.04 | 0.12 | 0.12 | 0.12 | 0.08 |
| 6 | 81166178 | rs13199780 | 1.413 | 0.06 | 1.158 | 0.32 | 0.069 | -0.071 | -0.071 | 0 | 0.43 | 0.07 | 0.47 | 0.57 | 0.28 | 0.44 |
| 6 | 81171274 | rs574098 | 2.238 | 0.282 | 1.853 | 0.107 | -0.03 | -0.025 | -0.025 | 0 | 0.85 | 0.24 | 0.88 | 0.94 | 0.92 | 0.78 |
| 6 | 81177227 | rs515347 | 0.996 | 0.448 | 1.464 | 0.191 | -0.008 | -0.001 | -0.001 | 0 | 0.91 | 0.77 | 0.93 | 0.94 | 0.93 | 0.79 |
| 6 | 81198826 | rs1865917 | -0.891 | 0.46 | 1.402 | 0.202 | -0.008 | -0.001 | -0.001 | 0 | 0.09 | 0.22 | 0.07 | 0.06 | 0.07 | 0.21 |
| 6 | 81208541 | rs1436868 | -1.137 | 0.48 | 1.553 | 0.214 | 0.011 | 0.021 | 0.021 | 0 | 0.1 | 0.37 | 0.08 | 0.06 | 0.07 | 0.2 |
| 6 | 81215558 | rs632422 | -0.155 | 0.834 | 0.778 | 0.308 | -0.158 | -0.062 | -0.062 | 0 | 0.68 | 0.95 | 0.74 | 0.69 | 0.52 | 0.79 |
| 6 | 81216957 | rs998759 | 0.001 | 0.945 | 0.751 | 0.359 | -0.025 | -0.009 | -0.009 | 0 | 0.07 | 0.02 | 0.13 | 0.12 | 0.14 | 0.1 |
| 6 | 81218067 | rs11751803 | 1.5 | 0.77 | 0.672 | 0.287 | 0.067 | -0.134 | -0.134 | 0.087 | 0.45 | 0.04 | 0.5 | 0.52 | 0.3 | 0.49 |
| 6 | 81228527 | rs1436876 | -1.554 | 1.123 | 1.811 | 0.506 | 0.076 | 0.057 | 0.057 | 0 | 0.12 | 0.41 | 0.17 | 0.18 | 0.16 | 0.12 |
| 6 | 81237839 | rs16891947 | -1.241 | 1.076 | 1.889 | 0.503 | 0.06 | 0.091 | 0.091 | 0.134 | 0.05 | 0.38 | 0.04 | 0.04 | 0.01 | 0.03 |
| 6 | 81253907 | rs10498923 | -0.077 | 1.156 | 1.472 | 0.511 | -0.042 | -0.032 | -0.032 | 0 | 0.07 | 0.01 | 0.13 | 0.14 | 0.15 | 0.1 |
| 6 | 81262941 | rs2322840 | 0.439 | 0.912 | 1.495 | 0.374 | 0.021 | -0.009 | -0.009 | 0 | 0.62 | 0.44 | 0.64 | 0.67 | 0.53 | 0.71 |
| 6 | 81266567 | rs953500 | 0.389 | 0.514 | 1.434 | 0.346 | 0.027 | -0.023 | -0.023 | 0 | 0.59 | 0.75 | 0.55 | 0.53 | 0.38 | 0.61 |
| 6 | 81274336 | rs10943720 | 0.646 | 0.506 | 1.469 | 0.353 | 0.016 | -0.007 | -0.007 | 0 | 0.57 | 0.61 | 0.53 | 0.49 | 0.38 | 0.59 |
| 6 | 81276112 | rs9449074 | 0.394 | 0.51 | 0.932 | 0.34 | -0.021 | -0.1 | -0.1 | 0 | 0.61 | 0.94 | 0.56 | 0.53 | 0.38 | 0.61 |
| 6 | 81293837 | rs2503725 | -1.437 | 0.434 | 1.289 | 0.302 | 0.07 | 0.008 | 0.008 | 0.42 | 0.5 | 0.74 | 0.49 | 0.52 | 0.63 | 0.42 |
| 6 | 81297890 | rs722715 | -0.434 | 0.422 | 0.782 | 0.299 | 0.001 | -0.078 | -0.078 | 0 | 0.4 | 0.09 | 0.44 | 0.48 | 0.62 | 0.39 |
| 6 | 81298688 | rs1454446 | 1.456 | 0.45 | 0.752 | 0.334 | 0.082 | -0.109 | -0.109 | 0 | 0.45 | 0.04 | 0.45 | 0.47 | 0.31 | 0.46 |
| 6 | 81309680 | rs6926124 | 1.559 | 0.507 | 1.179 | 0.394 | 0.084 | -0.072 | -0.072 | 0 | 0.45 | 0.09 | 0.5 | 0.46 | 0.34 | 0.44 |
| 6 | 81313076 | rs9449095 | -0.761 | 0.847 | 1.628 | 0.386 | 0.029 | -0.022 | -0.022 | 0.213 | 0.1 | 0.3 | 0.08 | 0.05 | 0.09 | 0.08 |
| 6 | 81315235 | rs2444827 | -0.478 | 0.811 | 1.838 | 0.341 | -0.037 | -0.035 | -0.035 | 0 | 0.37 | 0.22 | 0.37 | 0.46 | 0.53 | 0.34 |
| 6 | 81315597 | rs9443804 | 0.728 | 0.814 | 1.768 | 0.344 | 0.003 | -0.003 | -0.003 | 1.232 | 0.56 | 0.39 | 0.59 | 0.53 | 0.43 | 0.53 |
| 6 | 81322363 | rs12216014 | 1.271 | 0.848 | 1.509 | 0.4 | 0.075 | 0.054 | 0.054 | 1.289 | 0.5 | 0.12 | 0.6 | 0.61 | 0.39 | 0.51 |
| 6 | 81341175 | rs2444830 | -0.298 | 1.253 | 1.748 | 0.44 | -0.032 | -0.007 | -0.007 | 0.041 | 0.27 | 0.13 | 0.24 | 0.31 | 0.35 | 0.24 |
| 6 | 81342885 | rs2496689 | -0.732 | 1.243 | 1.823 | 0.475 | -0.006 | -0.002 | -0.002 | 0 | 0.29 | 0.24 | 0.26 | 0.32 | 0.37 | 0.37 |
| 6 | 81344004 | rs1377979 | -1.032 | 1.264 | 1.917 | 0.451 | 0 | 0 | 0 | 0 | 0.33 | 0.3 | 0.3 | 0.32 | 0.45 | 0.42 |
| 6 | 81346033 | rs1902066 | 1.59 | 1.375 | 1.862 | 0.572 | -0.009 | -0.027 | -0.027 | 0 | 0.59 | 0.27 | 0.63 | 0.5 | 0.54 | 0.49 |
| 6 | 81346504 | rs2496690 | 1.681 | 1.528 | 1.849 | 0.592 | -0.01 | -0.014 | -0.014 | 0 | 0.66 | 0.28 | 0.71 | 0.57 | 0.59 | 0.6 |
| 6 | 81349756 | rs2503749 | 2.196 | 1.499 | 1.829 | 0.576 | -0.016 | -0.034 | -0.034 | 0 | 0.52 | 0.14 | 0.56 | 0.46 | 0.43 | 0.44 |
| 6 | 81352515 | rs9294177 | -0.817 | 1.497 | 1.892 | 0.558 | -0.001 | 0.003 | 0.003 | 0 | 0.14 | 0.12 | 0.15 | 0.11 | 0.15 | 0.16 |
| 6 | 81352611 | rs2444818 | 2.389 | 1.155 | 1.605 | 0.78 | 0.004 | 0.067 | 0.067 | 0.425 | 0.44 | 0.11 | 0.46 | 0.3 | 0.38 | 0.27 |
| 6 | 81364570 | rs2444825 | 0.786 | 1.178 | 2.036 | 0.812 | 0.03 | 0.101 | 0.101 | 0 | 0.77 | 0.63 | 0.77 | 0.56 | 0.72 | 0.64 |
| 6 | 81365907 | rs2503761 | 1.029 | 1.176 | 1.995 | 0.809 | 0.018 | 0.074 | 0.074 | 0 | 0.74 | 0.5 | 0.74 | 0.53 | 0.71 | 0.63 |
| 6 | 81366484 | rs16892086 | -0.011 | 0.51 | 2.074 | 0.982 | -0.007 | -0.026 | -0.026 | 0 | 0.05 | 0.14 | 0.05 | 0 | 0.06 | 0.01 |
| 6 | 81380831 | rs2503718 | 1.011 | 0.518 | 2.25 | 1.02 | 0.018 | 0.03 | 0.03 | 0 | 0.94 | 0.76 | 0.95 | 0.95 | 0.95 | 0.84 |
| 6 | 81385896 | rs1454456 | 1.27 | 0.608 | 2.04 | 1.023 | 0.008 | -0.013 | -0.013 | 0 | 0.82 | 0.78 | 0.8 | 0.88 | 0.75 | 0.79 |
| 6 | 81387450 | rs9344029 | -0.841 | 0.641 | 1.767 | 1.005 | -0.056 | 0.035 | 0.035 | 0.241 | 0.15 | 0.04 | 0.17 | 0.08 | 0.23 | 0.2 |
| 6 | 81392005 | rs9361640 | -1.351 | 0.698 | 1.974 | 1.139 | -0.002 | -0.002 | -0.002 | 0 | 0.26 | 0.23 | 0.31 | 0.49 | 0.3 | 0.35 |
| 6 | 81400749 | rs9344031 | 1.119 | 0.57 | 2.013 | 0.924 | -0.004 | -0.03 | -0.03 | 0.213 | 0.89 | 0.54 | 0.91 | 0.98 | 0.9 | 0.85 |
| 6 | 81406662 | rs1584897 | -0.756 | 0.625 | 1.812 | 0.953 | -0.001 | 0.004 | 0.004 | 0 | 0.13 | 0.1 | 0.15 | 0.08 | 0.16 | 0.14 |
| 6 | 81423391 | rs1545557 | -1.386 | 0.498 | 1.991 | 1.016 | -0.04 | 0.012 | 0.012 | 0 | 0.19 | 0.47 | 0.15 | 0.18 | 0.14 | 0.31 |
| 6 | 81427214 | rs9791244 | -0.07 | 0.592 | 1.863 | 1.174 | 0.001 | 0.034 | 0.034 | 0 | 0.13 | 0.13 | 0.16 | 0.42 | 0.15 | 0.21 |
| 6 | 81428336 | rs2123993 | -2.475 | 0.641 | 1.953 | 1.15 | -0.009 | 0.103 | 0.103 | 0.017 | 0.5 | 0.81 | 0.52 | 0.68 | 0.53 | 0.7 |
| 6 | 81428748 | rs985737 | -1.862 | 0.608 | 1.979 | 1.167 | -0.022 | -0.07 | -0.07 | 0 | 0.24 | 0.57 | 0.26 | 0.1 | 0.26 | 0.29 |

| | | | | | | | | | | | | | | | | |
|---|---|---|---|---|---|---|---|---|---|---|---|---|---|---|---|---|
| 6 | 81429073 | rs7754084 | -0.1 | 0.682 | 1.572 | 1.251 | 0.001 | 0.034 | 0.034 | 0 | 0.13 | 0.13 | 0.16 | 0.42 | 0.15 | 0.21 |
| 6 | 81432886 | rs12663591 | 0.434 | 0.726 | 1.449 | 1.161 | -0.007 | -0.048 | -0.048 | 0.413 | 0.08 | 0 | 0.06 | 0.18 | 0.05 | 0.16 |
| 6 | 81435634 | rs9361645 | -1.884 | 0.683 | 1.825 | 1.273 | 0.003 | 0.007 | 0.007 | 0 | 0.34 | 0.61 | 0.32 | 0.38 | 0.4 | 0.46 |
| 6 | 81435835 | rs2039768 | -0.61 | 0.674 | 2.125 | 1.259 | 0 | -0.006 | -0.006 | 1.777 | 0.13 | 0.13 | 0.18 | 0.08 | 0.16 | 0.13 |
| 6 | 81439973 | rs1856524 | -1.964 | 0.673 | 2.07 | 1.27 | 0.003 | 0.008 | 0.008 | 0 | 0.34 | 0.61 | 0.32 | 0.38 | 0.4 | 0.46 |
| 6 | 81446168 | rs9350868 | -2.094 | 0.678 | 2.086 | 1.256 | -0.001 | -0.001 | -0.001 | 0 | 0.42 | 0.61 | 0.38 | 0.56 | 0.45 | 0.62 |
| 6 | 81447367 | rs1339377 | -2.326 | 0.687 | 2.093 | 1.241 | -0.007 | 0.005 | 0.005 | 0 | 0.47 | 0.69 | 0.42 | 0.56 | 0.53 | 0.67 |
| 6 | 81450225 | rs9352859 | -1.78 | 0.705 | 1.987 | 1.251 | 0.011 | 0.038 | 0.038 | 0 | 0.33 | 0.44 | 0.23 | 0.49 | 0.37 | 0.54 |
| 6 | 81452534 | rs9341825 | -0.478 | 0.704 | 1.848 | 1.213 | -0.014 | 0.017 | 0.017 | 0 | 0.13 | 0.05 | 0.16 | 0.08 | 0.16 | 0.13 |
| 6 | 81460295 | rs11752339 | -0.365 | 0.663 | 1.923 | 1.309 | -0.007 | -0.012 | -0.012 | 0 | 0.13 | 0.09 | 0.09 | 0.18 | 0.13 | 0.21 |
| 6 | 81463045 | rs13197404 | -0.365 | 0.663 | 1.923 | 1.309 | -0.007 | -0.012 | -0.012 | 0 | 0.13 | 0.09 | 0.09 | 0.18 | 0.13 | 0.21 |
| 6 | 81466614 | rs9359428 | -1.018 | 0.666 | 2.049 | 1.33 | 0.016 | 0.055 | 0.055 | 0 | 0.19 | 0.35 | 0.14 | 0.31 | 0.25 | 0.33 |
| 6 | 81478607 | rs10485285 | -1.888 | 0.669 | 1.72 | 1.322 | 0.007 | 0.025 | 0.025 | 0 | 0.33 | 0.44 | 0.23 | 0.49 | 0.37 | 0.54 |
| 6 | 81508411 | rs9344046 | -0.576 | 0.092 | 1.825 | 1.329 | 0.009 | 0.027 | 0.027 | 0 | 0.15 | 0.21 | 0.11 | 0.3 | 0.21 | 0.18 |
| 6 | 81517302 | rs9341828 | -0.556 | 0.097 | 1.793 | 1.329 | 0.006 | 0.022 | 0.022 | 0 | 0.15 | 0.18 | 0.11 | 0.29 | 0.21 | 0.18 |
| 6 | 81522847 | rs9361660 | -0.125 | 0.099 | 1.711 | 1.33 | -0.015 | -0.024 | -0.024 | 0 | 0.11 | 0.08 | 0.06 | 0.29 | 0.15 | 0.18 |
| 6 | 81526092 | rs1556902 | -1.251 | -0.009 | 1.893 | 1.366 | -0.024 | 0.046 | 0.046 | 0.194 | 0.07 | 0.15 | 0.13 | 0.13 | 0.04 | 0.17 |
| 6 | 81529339 | rs6935889 | -1.219 | 0 | 1.906 | 1.371 | -0.023 | 0.049 | 0.049 | 0.654 | 0.07 | 0.15 | 0.13 | 0.13 | 0.04 | 0.17 |
| 6 | 81536262 | rs1017741 | -2.571 | -0.114 | 1.696 | 1.385 | -0.01 | 0.166 | 0.166 | 0.965 | 0.49 | 0.86 | 0.54 | 0.75 | 0.48 | 0.7 |
| 6 | 81536714 | rs13193950 | -0.777 | -0.131 | 1.796 | 1.434 | 0.008 | -0.007 | -0.007 | 0 | 0.14 | 0.1 | 0.05 | 0.2 | 0.1 | 0.35 |
| 6 | 81543810 | rs9283799 | -1.862 | -0.13 | 2.016 | 1.457 | -0.001 | 0.022 | 0.022 | 0 | 0.3 | 0.33 | 0.16 | 0.49 | 0.31 | 0.53 |
| 6 | 81564185 | rs7769005 | -1.69 | -0.128 | 2.026 | 1.44 | 0.007 | -0.007 | -0.007 | 0 | 0.27 | 0.21 | 0.21 | 0.29 | 0.25 | 0.48 |
| 6 | 81578293 | rs481777 | -1.18 | -0.152 | 2.015 | 1.402 | -0.003 | -0.008 | -0.008 | 0.057 | 0.2 | 0.18 | 0.13 | 0.3 | 0.26 | 0.34 |
| 6 | 81585206 | rs4142360 | -0.358 | -0.186 | 1.987 | 1.389 | -0.021 | -0.007 | -0.007 | 0.358 | 0.12 | 0.05 | 0.17 | 0.09 | 0.15 | 0.13 |
| 6 | 81589369 | rs9361676 | -1.515 | -0.1 | 2.148 | 1.543 | 0 | 0.02 | 0.02 | 0 | 0.22 | 0.31 | 0.14 | 0.3 | 0.27 | 0.34 |
| 6 | 81593880 | rs9361677 | -1.645 | -0.102 | 2.169 | 1.542 | 0.004 | 0.059 | 0.059 | 0 | 0.23 | 0.45 | 0.15 | 0.3 | 0.27 | 0.34 |
| 6 | 81597546 | rs9449170 | -1.303 | -0.047 | 2.151 | 1.596 | -0.041 | 0.141 | 0.141 | 0.661 | 0.15 | 0.43 | 0.15 | 0.3 | 0.05 | 0.21 |
| 6 | 81607383 | rs12527737 | -1.402 | -0.043 | 2.17 | 1.598 | -0.051 | 0.125 | 0.125 | 1.73 | 0.16 | 0.5 | 0.15 | 0.3 | 0.05 | 0.21 |
| 6 | 81615591 | rs3846743 | -1.206 | -0.05 | 2.142 | 1.595 | -0.042 | 0.141 | 0.141 | 0.722 | 0.15 | 0.43 | 0.15 | 0.3 | 0.05 | 0.21 |
| 6 | 81617682 | rs562466 | -2.155 | 0.008 | 1.949 | 1.61 | -0.008 | 0.036 | 0.036 | 1.364 | 0.3 | 0.45 | 0.25 | 0.59 | 0.26 | 0.39 |
| 6 | 81619103 | rs12525920 | -1.108 | -0.046 | 1.981 | 1.635 | -0.032 | 0.074 | 0.074 | 0.085 | 0.14 | 0.29 | 0.15 | 0.3 | 0.05 | 0.22 |
| 6 | 81648445 | rs534800 | -0.7 | 0.097 | 1.723 | 1.701 | 0.009 | -0.037 | -0.037 | 0 | 0.14 | 0.08 | 0.07 | 0.28 | 0.12 | 0.15 |
| 6 | 81650550 | rs1017742 | -1.135 | -0.001 | 1.733 | 1.638 | 0.017 | 0.057 | 0.057 | 0 | 0.09 | 0.12 | 0.16 | 0.3 | 0.14 | 0.21 |
| 6 | 81656521 | rs9361688 | -1.773 | -0.059 | 1.797 | 1.574 | 0.002 | 0.021 | 0.021 | 0 | 0.25 | 0.32 | 0.24 | 0.58 | 0.26 | 0.35 |
| 6 | 81658974 | rs492269 | -1.773 | -0.059 | 1.797 | 1.574 | 0.002 | 0.021 | 0.021 | 0 | 0.25 | 0.32 | 0.24 | 0.59 | 0.26 | 0.35 |
| 6 | 81662008 | rs1350686 | -1.366 | 0.043 | 1.675 | 1.574 | -0.004 | -0.005 | -0.005 | 0 | 0.22 | 0.24 | 0.14 | 0.48 | 0.26 | 0.33 |
| 6 | 81667079 | rs9350889 | -1.984 | -0.562 | 1.779 | 1.515 | 0 | -0.001 | -0.001 | 0.018 | 0.34 | 0.36 | 0.29 | 0.55 | 0.36 | 0.45 |
| 6 | 81667151 | rs9344072 | -1.984 | -0.562 | 1.779 | 1.515 | 0 | -0.001 | -0.001 | 0 | 0.34 | 0.36 | 0.29 | 0.55 | 0.36 | 0.45 |
| 6 | 81667881 | rs7770169 | -1.984 | -0.562 | 1.779 | 1.515 | 0 | -0.001 | -0.001 | 0 | 0.34 | 0.36 | 0.29 | 0.55 | 0.36 | 0.45 |
| 6 | 81668203 | rs7770677 | -1.984 | -0.562 | 1.779 | 1.515 | 0 | -0.001 | -0.001 | 0 | 0.34 | 0.36 | 0.29 | 0.55 | 0.36 | 0.45 |
| 6 | 81670380 | rs7771176 | -0.441 | -0.589 | 1.777 | 1.511 | -0.001 | -0.001 | -0.001 | 0 | 0.14 | 0.12 | 0.17 | 0.09 | 0.13 | 0.13 |
| 6 | 81675870 | rs9352903 | 0.357 | -0.584 | 1.514 | 1.627 | 0.022 | -0.028 | -0.028 | 0 | 0.08 | 0.02 | 0.01 | 0.28 | 0.03 | 0.14 |
| 6 | 81678658 | rs9352906 | -1.031 | -0.59 | 1.509 | 1.626 | -0.031 | -0.083 | -0.083 | 0 | 0.17 | 0.07 | 0.1 | 0.47 | 0.23 | 0.32 |
| 6 | 81682916 | rs11969591 | -0.872 | -0.843 | 1.495 | 1.52 | -0.003 | 0.008 | 0.008 | 1.06 | 0.18 | 0.17 | 0.19 | 0.4 | 0.09 | 0.34 |
| 6 | 81685863 | rs1418866 | -0.844 | -0.768 | 1.226 | 1.384 | -0.006 | -0.005 | -0.005 | 0 | 0.12 | 0.15 | 0.1 | 0.19 | 0.2 | 0.18 |
| 6 | 81687677 | rs1075102 | -2.615 | -0.839 | 1.338 | 1.362 | 0.042 | 0.148 | 0.148 | 1.543 | 0.49 | 0.85 | 0.52 | 0.72 | 0.43 | 0.66 |
| 6 | 81689287 | rs9352909 | -1.73 | -0.223 | 1.33 | 0.994 | 0 | 0 | 0 | 0 | 0.26 | 0.34 | 0.28 | 0.28 | 0.33 | 0.29 |
| 6 | 81698459 | rs9449207 | -1.529 | -0.203 | 1.091 | 1.004 | -0.04 | -0.02 | -0.02 | 0.727 | 0.24 | 0.14 | 0.26 | 0.28 | 0.33 | 0.29 |
| 6 | 81706529 | rs2874936 | -2.206 | -0.249 | 1.367 | 1.019 | -0.007 | 0.008 | 0.008 | 0.019 | 0.37 | 0.51 | 0.41 | 0.39 | 0.38 | 0.49 |
| 6 | 81708701 | rs9449212 | -1.708 | -0.26 | 1.584 | 1.022 | -0.054 | 0 | 0 | 1.433 | 0.26 | 0.5 | 0.32 | 0.21 | 0.19 | 0.31 |
| 6 | 81710053 | rs7751304 | -0.961 | -0.242 | 1.523 | 1.033 | 0.007 | 0.004 | 0.004 | 0 | 0.12 | 0.16 | 0.1 | 0.19 | 0.2 | 0.18 |
| 6 | 81713515 | rs6910151 | -0.865 | -0.284 | 1.527 | 1.024 | -0.044 | 0.018 | 0.018 | 0.9 | 0.12 | 0.23 | 0.14 | 0.12 | 0.06 | 0.2 |
| 6 | 81720652 | rs9449218 | -0.44 | -0.283 | 1.519 | 1.025 | -0.022 | 0.01 | 0.01 | 0.778 | 0.1 | 0.12 | 0.14 | 0.12 | 0.06 | 0.2 |
| 6 | 81736311 | rs9449223 | -0.778 | -0.267 | 1.379 | 1.033 | -0.05 | 0.024 | 0.024 | 1.428 | 0.12 | 0.28 | 0.15 | 0.12 | 0.06 | 0.2 |
| 6 | 81740994 | rs11757828 | -0.954 | -0.202 | 0.845 | 1.053 | 0.008 | 0.004 | 0.004 | 0 | 0.12 | 0.16 | 0.1 | 0.19 | 0.2 | 0.17 |
| 6 | 81745735 | rs2120524 | -1.835 | -0.08 | 0.853 | 1.004 | 0.003 | -0.004 | -0.004 | 0 | 0.38 | 0.66 | 0.42 | 0.4 | 0.39 | 0.49 |
| 6 | 81769538 | rs6918133 | -0.813 | -0.009 | 0.341 | 0.725 | -0.002 | -0.007 | -0.007 | 0.846 | 0.13 | 0.2 | 0.17 | 0.09 | 0.12 | 0.12 |
| 6 | 81777045 | rs9361722 | -0.86 | -0.392 | -0.126 | 0.524 | 0.012 | 0.018 | 0.018 | 0 | 0.19 | 0.15 | 0.19 | 0.09 | 0.14 | 0.15 |
| 6 | 81778014 | rs7774156 | -0.64 | -0.984 | 0.24 | 1.365 | -0.026 | 0.153 | 0.153 | 0 | 0.1 | 0.22 | 0.06 | 0.35 | 0.04 | 0.16 |
| 6 | 81782500 | rs310385 | 0.359 | -1.193 | 0.599 | 1.249 | 0.006 | 0.032 | 0.032 | 0 | 0.89 | 0.81 | 0.82 | 0.78 | 0.88 | 0.87 |
| 6 | 81784903 | rs9352922 | 0.258 | -1.209 | 0.486 | 1.482 | -0.012 | 0.15 | 0.15 | 0 | 0.07 | 0.2 | 0.05 | 0.25 | 0.03 | 0.14 |
| 6 | 81786684 | rs9449234 | -0.283 | -1.191 | 0.347 | 1.408 | 0.012 | 0.018 | 0.018 | 0 | 0.19 | 0.14 | 0.19 | 0.09 | 0.14 | 0.15 |
| 6 | 81792413 | rs310420 | 2.284 | -0.95 | 0.302 | 1.034 | -0.016 | 0.081 | 0.081 | 0.072 | 0.45 | 0.23 | 0.48 | 0.29 | 0.54 | 0.36 |
| 6 | 81797984 | rs310410 | -0.498 | -0.956 | 0.34 | 1.001 | -0.003 | 0.003 | 0.003 | 0 | 0.94 | 0.98 | 0.94 | 1 | 0.92 | 0.99 |
| 6 | 81798980 | rs7765033 | 0.055 | -0.961 | 0.286 | 1.036 | 0.009 | 0.013 | 0.013 | 0 | 0.19 | 0.15 | 0.2 | 0.09 | 0.14 | 0.15 |
| 6 | 81800362 | rs310405 | 2.12 | -0.945 | 0.268 | 1.038 | -0.018 | 0.081 | 0.081 | 0.085 | 0.45 | 0.23 | 0.48 | 0.29 | 0.54 | 0.37 |
| 6 | 81818875 | rs1418862 | -1.803 | -1.042 | 0.315 | 1.042 | 0.051 | 0.038 | 0.038 | 0.329 | 0.44 | 0.7 | 0.45 | 0.66 | 0.29 | 0.44 |
| 6 | 81820941 | rs9350904 | -1.615 | -1.121 | 0.291 | 1.006 | 0.04 | 0.027 | 0.027 | 0.209 | 0.42 | 0.68 | 0.44 | 0.66 | 0.29 | 0.44 |
| 6 | 81828364 | rs9344109 | -1.831 | -1.015 | 0.302 | 1.023 | 0.052 | 0.038 | 0.038 | 0.258 | 0.44 | 0.69 | 0.44 | 0.66 | 0.29 | 0.44 |

| | | | | | | | | | | | | | | | | |
|---|---|---|---|---|---|---|---|---|---|---|---|---|---|---|---|---|
| 6 | 81831574 | rs7748211 | -1.819 | -1.049 | 0.293 | 1.013 | 0.053 | 0.038 | 0.038 | 0.33 | 0.44 | 0.69 | 0.44 | 0.66 | 0.29 | 0.44 |
| 6 | 81837758 | rs12660400 | -0.209 | -0.864 | 0.346 | 1.095 | 0.01 | 0.05 | 0.05 | 0 | 0.11 | 0.21 | 0.25 | 0.24 | 0.12 | 0.13 |
| 6 | 81842393 | rs9359456 | -1.005 | -0.852 | 0.217 | 1.11 | -0.011 | 0.001 | 0.001 | 0 | 0.31 | 0.36 | 0.37 | 0.33 | 0.25 | 0.28 |
| 6 | 81843363 | rs6908031 | -1.499 | -0.955 | -0.853 | 0.44 | -0.128 | -0.066 | -0.066 | 0 | 0.8 | 0.98 | 0.7 | 0.78 | 0.67 | 0.78 |
| 6 | 81850755 | rs9359458 | 0.338 | -1.046 | -0.205 | 0.9 | 0.004 | 0.005 | 0.005 | 0 | 0.15 | 0.09 | 0.1 | 0.09 | 0.13 | 0.15 |
| 6 | 81856665 | rs6935142 | -0.321 | -1.163 | -0.073 | 0.949 | 0 | 0.054 | 0.054 | 0.44 | 0.13 | 0.29 | 0.26 | 0.24 | 0.12 | 0.15 |
| 6 | 81865348 | rs7769553 | -1.408 | -1.216 | -0.019 | 0.899 | 0.014 | -0.015 | -0.015 | 0 | 0.41 | 0.31 | 0.24 | 0.47 | 0.35 | 0.51 |
| 6 | 81872212 | rs6924063 | -1.518 | -1.119 | -0.008 | 1 | -0.002 | 0.031 | 0.031 | 0 | 0.35 | 0.53 | 0.4 | 0.53 | 0.34 | 0.49 |
| 6 | 81878755 | rs7773268 | -1.575 | -1.116 | -0.053 | 0.935 | 0.005 | -0.003 | -0.003 | 0 | 0.44 | 0.45 | 0.25 | 0.47 | 0.35 | 0.52 |
| 6 | 81889500 | rs7738750 | 0.348 | -0.992 | -0.11 | 0.997 | 0 | 0.001 | 0.001 | 0.018 | 0.17 | 0.19 | 0.1 | 0.1 | 0.13 | 0.15 |
| 6 | 81900828 | rs4706882 | -1.59 | -0.728 | -0.406 | 1.059 | -0.007 | 0.052 | 0.052 | 0.165 | 0.56 | 0.71 | 0.59 | 0.74 | 0.47 | 0.69 |
| 6 | 81904308 | rs9344125 | -0.896 | -0.579 | -0.676 | 1.295 | 0 | 0.001 | 0.001 | 0 | 0.24 | 0.23 | 0.15 | 0.36 | 0.22 | 0.36 |
| 6 | 81907559 | rs9344126 | -1.403 | -0.155 | -1.305 | 0.665 | -0.038 | 0.063 | 0.063 | 0.607 | 0.58 | 0.79 | 0.64 | 0.73 | 0.52 | 0.68 |
| 6 | 81909920 | rs13197911 | -0.937 | 0.358 | -0.966 | 1.032 | -0.021 | -0.008 | -0.008 | 0 | 0.14 | 0.06 | 0.18 | 0.15 | 0.18 | 0.24 |
| 6 | 81934605 | rs7762518 | -0.839 | 0.253 | -0.709 | 1.396 | -0.002 | -0.003 | -0.003 | 0 | 0.29 | 0.28 | 0.25 | 0.36 | 0.35 | 0.27 |
| 6 | 81960107 | rs9341865 | -1.409 | -0.429 | -0.773 | 1.376 | -0.025 | 0.032 | 0.032 | 0.222 | 0.6 | 0.74 | 0.6 | 0.67 | 0.57 | 0.75 |
| 6 | 81985033 | rs1334636 | -0.629 | -0.856 | -0.505 | 1.844 | 0 | -0.002 | -0.002 | 4.213 | 0.36 | 0.38 | 0.36 | 0.55 | 0.37 | 0.46 |
| 6 | 82006729 | rs16892963 | -0.762 | -0.954 | -0.453 | 1.783 | 0.007 | 0.156 | 0.156 | 0.01 | 0.09 | 0.23 | 0.21 | 0.31 | 0.09 | 0.21 |
| 6 | 82007386 | rs9361785 | -0.762 | -0.954 | -0.453 | 1.783 | 0.007 | 0.156 | 0.156 | 0.01 | 0.09 | 0.23 | 0.21 | 0.31 | 0.09 | 0.21 |
| 6 | 82010784 | rs12199170 | -0.203 | -0.982 | -0.479 | 1.789 | -0.011 | -0.001 | -0.001 | 1.36 | 0.26 | 0.1 | 0.15 | 0.24 | 0.28 | 0.24 |
| 6 | 82012892 | rs7741560 | -0.881 | -0.966 | -0.491 | 1.713 | 0 | -0.005 | -0.005 | 0.019 | 0.35 | 0.34 | 0.36 | 0.55 | 0.37 | 0.46 |
| 6 | 82015654 | rs9344146 | -0.005 | -0.822 | -0.433 | 1.521 | 0.001 | 0.005 | 0.005 | 0 | 0.05 | 0.06 | 0.14 | 0.08 | 0.07 | 0.1 |
| 6 | 82017878 | rs10485394 | -0.093 | -0.902 | -0.491 | 1.645 | -0.009 | -0.012 | -0.012 | 0 | 0.24 | 0.16 | 0.15 | 0.29 | 0.29 | 0.26 |
| 6 | 82018209 | rs1343236 | 1.076 | -0.588 | 0.157 | 1.039 | 0.043 | 0.033 | 0.033 | 1.321 | 0.89 | 0.57 | 0.78 | 0.93 | 0.91 | 0.87 |
| 6 | 82019857 | rs1562068 | 0.719 | -1.371 | -0.202 | 0.985 | -0.001 | -0.001 | -0.001 | 0 | 0.66 | 0.61 | 0.73 | 0.64 | 0.73 | 0.65 |
| 6 | 82019874 | rs9359479 | 0.28 | -1.25 | -0.421 | 0.899 | -0.058 | 0.017 | 0.017 | 0 | 0.25 | 0.05 | 0.15 | 0.2 | 0.29 | 0.25 |
| 6 | 82022657 | rs12055792 | 0.286 | -1.314 | -0.373 | 0.876 | -0.023 | 0.02 | 0.02 | 0 | 0.26 | 0.13 | 0.16 | 0.22 | 0.29 | 0.25 |
| 6 | 82027161 | rs1343238 | 0.062 | -1.354 | -0.313 | 0.837 | -0.013 | 0.027 | 0.027 | 0 | 0.27 | 0.17 | 0.16 | 0.21 | 0.29 | 0.25 |
| 6 | 82034903 | rs9361791 | -0.037 | -1.367 | -0.308 | 0.83 | -0.004 | 0.012 | 0.012 | 0 | 0.28 | 0.25 | 0.16 | 0.21 | 0.29 | 0.24 |
| 6 | 82036764 | rs1413720 | -0.453 | -1.459 | -0.026 | 0.817 | -0.018 | 0.014 | 0.014 | 0 | 0.28 | 0.31 | 0.23 | 0.36 | 0.21 | 0.31 |
| 6 | 82064465 | rs962983 | 0.799 | -1.523 | -0.035 | 0.764 | -0.039 | 0.006 | 0.006 | 0.472 | 0.68 | 0.48 | 0.75 | 0.69 | 0.76 | 0.65 |
| 6 | 82075973 | rs6454218 | 0.728 | -1.726 | -0.089 | 0.645 | -0.041 | -0.004 | -0.004 | 0 | 0.64 | 0.48 | 0.72 | 0.69 | 0.75 | 0.65 |
| 6 | 82078440 | rs1419027 | 1.065 | -1.536 | -0.215 | 0.625 | 0 | 0 | 0 | 0.06 | 0.6 | 0.38 | 0.71 | 0.69 | 0.67 | 0.64 |
| 6 | 82087297 | rs996136 | 1.552 | -1.537 | -0.277 | 0.65 | -0.011 | -0.006 | -0.006 | 0 | 0.37 | 0.3 | 0.57 | 0.45 | 0.46 | 0.41 |
| 6 | 82092004 | rs2166042 | -0.123 | -1.762 | -0.179 | 0.705 | -0.016 | 0.012 | 0.012 | 0.042 | 0.2 | 0.31 | 0.19 | 0.25 | 0.16 | 0.28 |
| 6 | 82093262 | rs2120535 | -1.314 | -1.736 | -0.27 | 0.696 | 0.001 | 0 | 0 | 0.815 | 0.57 | 0.61 | 0.4 | 0.54 | 0.49 | 0.55 |
| 6 | 82099191 | rs2323252 | 1.695 | -1.737 | -0.273 | 0.675 | -0.009 | -0.005 | -0.005 | 0.526 | 0.38 | 0.3 | 0.57 | 0.45 | 0.46 | 0.41 |
| 6 | 82099375 | rs991112 | -0.464 | -1.835 | -0.305 | 0.719 | -0.016 | -0.041 | -0.041 | 0.055 | 0.41 | 0.08 | 0.26 | 0.38 | 0.41 | 0.31 |
| 6 | 82109125 | rs11756740 | 0.839 | -1.634 | -0.296 | 0.696 | -0.031 | -0.082 | -0.082 | 0 | 0.19 | 0.02 | 0.12 | 0.26 | 0.19 | 0.21 |
| 6 | 82115320 | rs7751405 | 1.827 | -1.609 | -0.174 | 0.9 | 0.002 | -0.001 | -0.001 | 0 | 0.32 | 0.28 | 0.46 | 0.36 | 0.43 | 0.29 |
| 6 | 82115934 | rs1891700 | -0.442 | -1.631 | -0.265 | 0.679 | -0.034 | 0.034 | 0.034 | 0.688 | 0.21 | 0.43 | 0.2 | 0.27 | 0.17 | 0.28 |
| 6 | 82120647 | rs9359489 | -0.199 | -1.525 | -0.419 | 0.671 | 0.015 | 0.042 | 0.042 | 0 | 0.39 | 0.16 | 0.29 | 0.28 | 0.35 | 0.3 |
| 6 | 82126750 | rs16893126 | -0.662 | -1.464 | -0.136 | 0.751 | -0.053 | 0.022 | 0.022 | 2.987 | 0.25 | 0.46 | 0.2 | 0.27 | 0.17 | 0.29 |
| 6 | 82128917 | rs7741650 | -1.355 | -1.544 | -0.25 | 0.745 | 0.002 | 0.001 | 0.001 | 0.399 | 0.62 | 0.62 | 0.45 | 0.59 | 0.49 | 0.57 |
| 6 | 82130505 | rs1159703 | -0.119 | -1.639 | -0.19 | 0.756 | -0.03 | 0.016 | 0.016 | 0 | 0.39 | 0.07 | 0.31 | 0.32 | 0.4 | 0.31 |
| 6 | 82140311 | rs1088022 | -0.87 | -1.589 | -0.008 | 0.863 | -0.01 | 0.014 | 0.014 | 0.048 | 0.57 | 0.3 | 0.4 | 0.41 | 0.52 | 0.43 |
| 6 | 82149482 | rs2120533 | 1.882 | -1.554 | -0.023 | 0.847 | 0.02 | 0.013 | 0.013 | 4.276 | 0.31 | 0.35 | 0.49 | 0.41 | 0.4 | 0.39 |
| 6 | 82159696 | rs475930 | -0.914 | -1.48 | -0.161 | 0.821 | -0.009 | 0.015 | 0.015 | 0.049 | 0.57 | 0.3 | 0.4 | 0.41 | 0.52 | 0.43 |
| 6 | 82169305 | rs796350 | 0.141 | -1.048 | 0.361 | 0.983 | -0.022 | 0.021 | 0.021 | 0 | 0.94 | 0.98 | 0.91 | 1 | 0.88 | 0.92 |
| 6 | 82186156 | rs6454221 | 0.021 | -1.803 | 0.492 | 0.536 | -0.021 | -0.002 | -0.002 | 0 | 0.93 | 0.91 | 0.96 | 0.93 | 0.95 | 0.95 |
| 6 | 82193109 | rs17530068 | 0.038 | -1.908 | -0.086 | 0.565 | -0.033 | -0.057 | -0.057 | 0 | 0.18 | 0.04 | 0.24 | 0.25 | 0.23 | 0.18 |
| 6 | 82195125 | rs12213217 | 1.537 | -2.062 | 0.242 | 0.559 | 0.003 | -0.001 | -0.001 | 0.742 | 0.52 | 0.41 | 0.5 | 0.5 | 0.6 | 0.62 |
| 6 | 82203213 | rs6905459 | 0.188 | -2.036 | 0.319 | 0.547 | -0.018 | -0.02 | -0.02 | 0 | 0.05 | 0.24 | 0.02 | 0 | 0 | 0 |
| 6 | 82221302 | rs9449340 | 1.565 | -2.078 | 0.301 | 0.55 | 0.004 | -0.001 | -0.001 | 0.003 | 0.52 | 0.4 | 0.49 | 0.5 | 0.6 | 0.62 |
| 6 | 82222692 | rs6937455 | -1.132 | -1.919 | 0.416 | 0.633 | 0.004 | 0.001 | 0.001 | 0.743 | 0.09 | 0.32 | 0.05 | 0.05 | 0.04 | 0.04 |
| 6 | 82228814 | rs9361832 | -1.384 | -1.948 | 0.44 | 0.637 | -0.01 | 0.003 | 0.003 | 4.003 | 0.41 | 0.51 | 0.48 | 0.43 | 0.36 | 0.35 |
| 6 | 82238322 | rs9344194 | -0.159 | -1.922 | 0.439 | 0.573 | -0.027 | -0.062 | -0.062 | 2.35 | 0.2 | 0.07 | 0.26 | 0.3 | 0.26 | 0.16 |
| 6 | 82244147 | rs1983728 | -0.554 | -1.933 | 0.466 | 0.566 | -0.002 | -0.001 | -0.001 | 0.847 | 0.94 | 0.96 | 0.97 | 0.92 | 0.96 | 0.94 |
| 6 | 82253194 | rs16893209 | -0.073 | -1.824 | 0.441 | 0.553 | -0.001 | -0.006 | -0.006 | 0.483 | 0.06 | 0.04 | 0.03 | 0.01 | 0.04 | 0.04 |
| 6 | 82268753 | rs4448140 | -1.291 | -1.824 | 0.244 | 0.562 | 0.037 | -0.029 | -0.029 | 0 | 0.41 | 0.19 | 0.46 | 0.45 | 0.36 | 0.32 |
| 6 | 82271163 | rs9294233 | -1.408 | -1.815 | 0.248 | 0.568 | 0.04 | -0.014 | -0.014 | 0 | 0.42 | 0.27 | 0.47 | 0.45 | 0.36 | 0.32 |
| 6 | 82286331 | rs1917065 | 1.568 | -1.815 | 0.207 | 0.574 | 0.009 | -0.004 | -0.004 | 0.213 | 0.53 | 0.38 | 0.49 | 0.49 | 0.59 | 0.64 |
| 6 | 82304235 | rs13437553 | 0.226 | -1.461 | 0.283 | 0.761 | 0.002 | 0.002 | 0.002 | 0 | 0.16 | 0.18 | 0.25 | 0.22 | 0.24 | 0.15 |
| 6 | 82321632 | rs6906639 | 0.385 | -1.397 | 0.427 | 0.673 | 0.063 | 0.073 | 0.073 | 0.671 | 0.93 | 0.73 | 0.95 | 0.94 | 0.94 | 0.95 |
| 6 | 82321973 | rs16893263 | 0.471 | -1.444 | 0.438 | 0.637 | 0.008 | 0.011 | 0.011 | 0 | 0.12 | 0.1 | 0.18 | 0.09 | 0.07 | 0.12 |
| 6 | 82325529 | rs9344208 | 0.135 | -1.463 | 0.405 | 0.688 | 0.004 | 0.009 | 0.009 | 0 | 0.17 | 0.2 | 0.24 | 0.24 | 0.23 | 0.15 |
| 6 | 82328546 | rs16893279 | 0.514 | -1.508 | 0.354 | 0.664 | 0.017 | 0 | 0 | 0 | 0.07 | 0.02 | 0.04 | 0.08 | 0.04 | 0.05 |
| 6 | 82333380 | rs2181364 | 0.544 | -1.519 | 0.357 | 0.657 | 0.012 | 0.017 | 0.017 | 0 | 0.11 | 0.08 | 0.18 | 0.09 | 0.07 | 0.12 |
| 6 | 82335012 | rs10943824 | -1.127 | -1.419 | 0.372 | 0.696 | -0.005 | 0.015 | 0.015 | 0.521 | 0.33 | 0.4 | 0.44 | 0.43 | 0.32 | 0.29 |

| | | | | | | | | | | | | | | | | |
|---|---|---|---|---|---|---|---|---|---|---|---|---|---|---|---|---|
| 6 | 82346997 | rs9344210 | -0.818 | -1.419 | 0.369 | 0.699 | 0 | 0 | 0 | 0 | 0.29 | 0.31 | 0.42 | 0.37 | 0.31 | 0.27 |
| 6 | 82355896 | rs12214288 | -0.112 | -1.034 | 0.144 | 0.646 | -0.004 | -0.024 | -0.024 | 0 | 0.2 | 0.13 | 0.25 | 0.3 | 0.24 | 0.17 |
| 6 | 82357304 | rs16893308 | -0.179 | -0.457 | 0.239 | 0.737 | 0.022 | 0.018 | 0.018 | 0.337 | 0.19 | 0.13 | 0.21 | 0.17 | 0.11 | 0.17 |
| 6 | 82379575 | rs1341230 | 0.314 | 0.592 | -0.427 | 1.142 | -0.01 | -0.005 | -0.005 | 0 | 0.37 | 0.3 | 0.38 | 0.39 | 0.52 | 0.43 |
| 6 | 82394320 | rs1156970 | -0.925 | 0.374 | -0.075 | 1.852 | -0.007 | 0.158 | 0.158 | 0 | 0.08 | 0.11 | 0.17 | 0.38 | 0.05 | 0.28 |
| 6 | 82399911 | rs10943829 | 0.57 | 0.784 | -0.729 | 1.456 | -0.039 | 0.02 | 0.02 | 0 | 0.41 | 0.2 | 0.39 | 0.28 | 0.49 | 0.38 |
| 6 | 82399967 | rs940510 | 0.393 | 1.203 | 0.068 | 1.214 | 0.086 | -0.013 | -0.013 | 0.445 | 0.94 | 0.79 | 0.94 | 0.98 | 0.9 | 0.97 |
| 6 | 82403301 | rs9350935 | -0.138 | 0.721 | -0.453 | 0.875 | -0.001 | -0.071 | -0.071 | 0 | 0.17 | 0.05 | 0.22 | 0.37 | 0.11 | 0.29 |
| 6 | 82406730 | rs194876 | 0.018 | 0.382 | 0.119 | 0.175 | -0.004 | -0.004 | -0.004 | 0 | 0.23 | 0.23 | 0.28 | 0.21 | 0.22 | 0.26 |
| 6 | 82409797 | rs1144175 | -0.775 | 0.59 | 0.297 | 0.16 | 0.015 | -0.038 | -0.038 | 0.155 | 0.18 | 0.4 | 0.12 | 0.14 | 0.19 | 0.09 |
| 6 | 82413747 | rs194881 | -0.432 | 0.486 | -0.217 | 0.169 | -0.035 | -0.01 | -0.01 | 0 | 0.12 | 0.02 | 0.18 | 0.12 | 0.15 | 0.22 |
| 6 | 82422740 | rs1276895 | -0.548 | 0.203 | 0.936 | 0.155 | 0.018 | 0.009 | 0.009 | 0 | 0.16 | 0.49 | 0.18 | 0.18 | 0.1 | 0.13 |
| 6 | 82426477 | rs194905 | -1.528 | 0.578 | 0.775 | 0.176 | 0.009 | 0.001 | 0.001 | 0 | 0.28 | 0.52 | 0.38 | 0.33 | 0.26 | 0.39 |
| 6 | 82427049 | rs1145865 | -0.488 | 0.571 | 0.776 | 0.178 | 0.017 | 0.008 | 0.008 | 0 | 0.16 | 0.49 | 0.18 | 0.18 | 0.1 | 0.13 |
| 6 | 82431631 | rs194910 | -0.242 | 0.584 | 0.752 | 0.179 | -0.037 | -0.028 | -0.028 | 0 | 0.12 | 0.02 | 0.2 | 0.15 | 0.17 | 0.26 |
| 6 | 82432826 | rs194914 | -2.104 | 0.395 | 0.961 | 0.015 | -0.018 | 0.037 | 0.037 | 0 | 0.43 | 0.73 | 0.47 | 0.43 | 0.41 | 0.52 |
| 6 | 82439009 | rs1014901 | -0.142 | 0.46 | 1.043 | 0.019 | 0.033 | -0.044 | -0.044 | 0.109 | 0.13 | 0.39 | 0.17 | 0.08 | 0.09 | 0.1 |
| 6 | 82441337 | rs194933 | -2.024 | -0.02 | 0.342 | 0.125 | -0.072 | -0.039 | -0.039 | 0.022 | 0.56 | 0.88 | 0.63 | 0.55 | 0.51 | 0.57 |
| 6 | 82441832 | rs194935 | -2.002 | -0.053 | 0.403 | 0.028 | -0.034 | 0.087 | 0.087 | 0 | 0.46 | 0.85 | 0.55 | 0.41 | 0.41 | 0.51 |
| 6 | 82443499 | rs194943 | -0.848 | -0.116 | 1.007 | -0.024 | 0.023 | -0.022 | -0.022 | 0 | 0.18 | 0.36 | 0.34 | 0.16 | 0.19 | 0.29 |
| 6 | 82443708 | rs10484871 | -0.422 | -0.04 | 1.048 | 0.027 | -0.022 | -0.01 | -0.01 | 0 | 0.17 | 0.27 | 0.06 | 0.15 | 0.08 | 0.1 |
| 6 | 82445375 | rs1033825 | -0.093 | -0.04 | 0.894 | 0.026 | -0.012 | -0.014 | -0.014 | 0 | 0.15 | 0.2 | 0.05 | 0.12 | 0.08 | 0.09 |
| 6 | 82447223 | rs13199056 | -2.042 | -0.03 | 0.965 | 0.011 | -0.001 | -0.002 | -0.002 | 0 | 0.33 | 0.69 | 0.33 | 0.27 | 0.25 | 0.26 |
| 6 | 82448870 | rs7741215 | -0.156 | -0.003 | 0.853 | -0.001 | -0.014 | 0.016 | 0.016 | 0 | 0.09 | 0.05 | 0.13 | 0.08 | 0.12 | 0.06 |
| 6 | 82452566 | rs16893407 | -0.221 | -0.078 | 0.872 | 0.046 | -0.012 | -0.014 | -0.014 | 0 | 0.15 | 0.2 | 0.06 | 0.12 | 0.08 | 0.09 |
| 6 | 82458639 | rs7742628 | -1.025 | 0.087 | 0.763 | -0.097 | -0.026 | -0.044 | -0.044 | 0 | 0.2 | 0.41 | 0.08 | 0.14 | 0.1 | 0.16 |
| 6 | 82463376 | rs915125 | -0.976 | 0.577 | 0.655 | 0.165 | -0.022 | -0.002 | -0.002 | 0 | 0.3 | 0.44 | 0.34 | 0.3 | 0.29 | 0.38 |
| 6 | 82464216 | rs9361858 | -1.575 | 0.628 | 0.223 | 0.249 | 0.002 | 0.045 | 0.045 | 0 | 0.45 | 0.77 | 0.6 | 0.38 | 0.45 | 0.48 |
| 6 | 82484634 | rs9353027 | -1.541 | 0.582 | 1.07 | 1.225 | -0.021 | 0.099 | 0.099 | 0.025 | 0.17 | 0.39 | 0.3 | 0.34 | 0.17 | 0.21 |
| 6 | 82496434 | rs194615 | -1.351 | 0.49 | 1.308 | 1.192 | -0.011 | 0.08 | 0.08 | 0 | 0.15 | 0.61 | 0.23 | 0.3 | 0.16 | 0.17 |
| 6 | 82496625 | rs16893491 | -0.218 | 0.424 | 1.059 | 1.143 | 0.001 | 0.014 | 0.014 | 0 | 0.1 | 0.14 | 0.06 | 0.18 | 0.08 | 0.09 |
| 6 | 82497480 | rs191786 | -2.527 | 0.473 | 0.247 | 0.832 | 0.004 | -0.088 | -0.088 | 0 | 0.42 | 0.87 | 0.49 | 0.52 | 0.38 | 0.37 |
| 6 | 82503792 | rs16893522 | -0.201 | 0.51 | 0.37 | 0.771 | 0.002 | 0.011 | 0.011 | 0 | 0.13 | 0.09 | 0.12 | 0.07 | 0.12 | 0.12 |
| 6 | 82505705 | rs1291465 | -0.11 | 1.43 | 0.265 | 0.663 | 0.051 | -0.031 | -0.031 | 0.04 | 0.14 | 0.28 | 0.27 | 0.09 | 0.24 | 0.25 |
| 6 | 82507123 | rs11961442 | -0.203 | 1.347 | 0.164 | 0.714 | -0.001 | 0.017 | 0.017 | 0 | 0.11 | 0.15 | 0.15 | 0.19 | 0.08 | 0.11 |
| 6 | 82515315 | rs16893526 | 0.116 | 1.368 | 0.174 | 0.654 | 0.001 | 0.006 | 0.006 | 0 | 0.1 | 0.09 | 0.05 | 0.16 | 0.08 | 0.09 |
| 6 | 82521379 | rs9353030 | -1.086 | 1.388 | 0.14 | 0.713 | -0.001 | 0.005 | 0.005 | 0 | 0.16 | 0.15 | 0.28 | 0.33 | 0.14 | 0.21 |
| 6 | 82521957 | rs194635 | -2.586 | 1.369 | -0.001 | 0.691 | -0.006 | -0.054 | -0.054 | 0.406 | 0.54 | 0.86 | 0.73 | 0.57 | 0.61 | 0.62 |
| 6 | 82527317 | rs1931625 | -0.052 | 1.41 | 0.289 | 0.653 | -0.006 | -0.021 | -0.021 | 0 | 0.09 | 0.26 | 0.06 | 0.05 | 0.1 | 0.03 |
| 6 | 82531367 | rs194606 | -0.649 | 1.504 | 0.039 | 0.617 | 0.044 | -0.056 | -0.056 | 0.125 | 0.18 | 0.57 | 0.28 | 0.1 | 0.25 | 0.26 |
| 6 | 82535868 | rs1981090 | -1.235 | 1.351 | -0.345 | 0.712 | 0.008 | -0.018 | -0.018 | 1.123 | 0.26 | 0.17 | 0.38 | 0.31 | 0.28 | 0.26 |
| 6 | 82544288 | rs12194273 | 2.064 | 1.358 | -0.533 | 0.659 | -0.011 | -0.067 | -0.067 | 0.782 | 0.45 | 0.12 | 0.27 | 0.45 | 0.37 | 0.37 |
| 6 | 82546753 | rs9449373 | -0.146 | 1.434 | -0.507 | 0.567 | -0.013 | 0.039 | 0.039 | 0 | 0.14 | 0.04 | 0.11 | 0.06 | 0.16 | 0.11 |
| 6 | 82555016 | rs3812145 | 0.097 | 1.567 | -0.009 | 0.522 | 0.022 | -0.021 | -0.021 | 0 | 0.86 | 0.83 | 0.74 | 0.92 | 0.73 | 0.73 |
| 6 | 82563225 | rs9361863 | -0.222 | 1.381 | 0.066 | 0.676 | -0.002 | 0.013 | 0.013 | 0 | 0.14 | 0.1 | 0.2 | 0.07 | 0.15 | 0.11 |
| 6 | 82569146 | rs12214571 | 2.252 | 1.312 | -0.216 | 0.863 | -0.068 | -0.056 | -0.056 | 3.314 | 0.4 | 0.02 | 0.23 | 0.32 | 0.36 | 0.33 |
| 6 | 82571236 | rs9443939 | 0.603 | 1.263 | 0.175 | 0.581 | 0.014 | -0.01 | -0.01 | 0 | 0.69 | 0.53 | 0.52 | 0.73 | 0.62 | 0.6 |
| 6 | 82575450 | rs6940643 | 2.05 | 1.315 | 0.064 | 0.616 | -0.016 | -0.078 | -0.078 | 0.387 | 0.44 | 0.12 | 0.28 | 0.46 | 0.39 | 0.37 |
| 6 | 82575594 | rs6940839 | 1.161 | 1.318 | 0.109 | 0.608 | 0.003 | -0.006 | -0.006 | 0.436 | 0.58 | 0.57 | 0.36 | 0.66 | 0.47 | 0.49 |
| 6 | 82578196 | rs1931643 | 0.41 | 1.356 | -0.534 | 0.634 | 0.011 | 0 | 0 | 0 | 0.72 | 0.68 | 0.54 | 0.73 | 0.62 | 0.6 |
| 6 | 82585857 | rs12203649 | 1.873 | 1.574 | -1.746 | 0.676 | -0.026 | -0.085 | -0.085 | 0 | 0.44 | 0.11 | 0.27 | 0.46 | 0.39 | 0.37 |
| 6 | 82587813 | rs9361865 | 1.294 | 1.471 | -1.392 | 0.852 | -0.045 | -0.042 | -0.042 | 0 | 0.57 | 0.15 | 0.44 | 0.53 | 0.53 | 0.48 |
| 6 | 82588292 | rs1931646 | -0.556 | 1.039 | -0.954 | 0.851 | 0 | 0.001 | 0.001 | 0 | 0.1 | 0.09 | 0.06 | 0.17 | 0.08 | 0.11 |
| 6 | 82590868 | rs9344227 | 0.875 | 1.345 | -1.607 | 0.679 | -0.036 | 0.083 | 0.083 | 0 | 0.5 | 0.06 | 0.39 | 0.61 | 0.47 | 0.55 |
| 6 | 82599523 | rs7450340 | -1.191 | 1.631 | -0.527 | 1.113 | -0.002 | -0.001 | -0.001 | 0.019 | 0.46 | 0.59 | 0.51 | 0.4 | 0.56 | 0.51 |
| 6 | 82611058 | rs9361875 | -0.859 | 1.737 | -0.327 | 0.992 | -0.029 | 0.015 | 0.015 | 0 | 0.33 | 0.22 | 0.36 | 0.28 | 0.46 | 0.43 |
| 6 | 82614505 | rs9344229 | -1.615 | 2.297 | -0.203 | 1.515 | 0.049 | 0.124 | 0.124 | 0.878 | 0.24 | 0.48 | 0.34 | 0.42 | 0.3 | 0.43 |
| 6 | 82616083 | rs2875087 | -1.646 | 1.943 | 0.215 | 1.478 | 0.016 | -0.001 | -0.001 | 0 | 0.46 | 0.78 | 0.54 | 0.52 | 0.57 | 0.52 |
| 6 | 82616216 | rs9344230 | -1.344 | 1.88 | 0.256 | 1.688 | -0.006 | -0.006 | -0.006 | 0 | 0.22 | 0.25 | 0.35 | 0.25 | 0.3 | 0.34 |
| 6 | 82622131 | rs2050304 | -1.301 | 1.342 | 0.033 | 1.429 | 0.003 | 0.002 | 0.002 | 0 | 0.27 | 0.32 | 0.36 | 0.24 | 0.31 | 0.36 |
| 6 | 82635257 | rs6909813 | -1.509 | 1.257 | 0.284 | 1.716 | 0.025 | 0.037 | 0.037 | 0 | 0.55 | 0.72 | 0.72 | 0.61 | 0.61 | 0.63 |
| 6 | 82647162 | rs9294240 | 0.114 | 1.498 | 0.3 | 1.535 | -0.031 | 0.086 | 0.086 | 0 | 0.26 | 0.1 | 0.2 | 0.07 | 0.34 | 0.17 |
| 6 | 82651663 | rs11963665 | -0.837 | 1.369 | 0.213 | 1.645 | -0.013 | 0.018 | 0.018 | 0 | 0.21 | 0.11 | 0.19 | 0.11 | 0.21 | 0.26 |
| 6 | 82657079 | rs9341905 | -0.688 | 1.228 | 0.599 | 1.967 | 0.023 | 0.023 | 0.023 | 0.013 | 0.33 | 0.48 | 0.5 | 0.37 | 0.43 | 0.38 |
| 6 | 82660374 | rs2050990 | -0.774 | 1.272 | 0.403 | 1.829 | 0.005 | 0.018 | 0.018 | 0 | 0.35 | 0.62 | 0.53 | 0.48 | 0.43 | 0.39 |
| 6 | 82672457 | rs1361685 | -1.615 | 0.866 | 0.549 | 1.941 | -0.012 | 0.064 | 0.064 | 1.022 | 0.22 | 0.63 | 0.38 | 0.37 | 0.15 | 0.41 |
| 6 | 82672570 | rs1361683 | -1.642 | 0.959 | 0.693 | 2.159 | 0.009 | 0.059 | 0.059 | 0 | 0.28 | 0.43 | 0.39 | 0.42 | 0.25 | 0.41 |
| 6 | 82685299 | rs10806234 | 0.005 | 1.176 | 0.74 | 2.226 | 0.022 | 0.028 | 0.028 | 0.058 | 0.68 | 0.51 | 0.56 | 0.56 | 0.58 | 0.58 |
| 6 | 82692160 | rs4706916 | 1.426 | 1.06 | 0.54 | 2.262 | 0.016 | 0.02 | 0.02 | 0.12 | 0.44 | 0.32 | 0.29 | 0.39 | 0.37 | 0.38 |

| | | | | | | | | | | | | | | | | |
|---|---|---|---|---|---|---|---|---|---|---|---|---|---|---|---|---|
| 6 | 82699370 | rs4054489 | -0.947 | 1.127 | 0.214 | 1.965 | -0.06 | -0.001 | -0.001 | 0.308 | 0.2 | 0.01 | 0.25 | 0.11 | 0.19 | 0.19 |
| 6 | 82723807 | rs4540297 | -0.198 | 1.043 | 0.62 | 1.668 | 0.003 | -0.027 | -0.027 | 0 | 0.71 | 0.73 | 0.72 | 0.87 | 0.63 | 0.75 |
| 6 | 82725168 | rs6902034 | -1.659 | 0.965 | 0.489 | 2.264 | 0.002 | 0.051 | 0.051 | 0 | 0.27 | 0.6 | 0.44 | 0.45 | 0.28 | 0.31 |
| 6 | 82725586 | rs7741179 | 0.662 | 1.11 | 0.402 | 1.968 | -0.01 | -0.01 | -0.01 | 0 | 0.54 | 0.27 | 0.32 | 0.52 | 0.47 | 0.4 |
| 6 | 82729032 | rs1418306 | -0.974 | 0.858 | 0.449 | 2.285 | 0.017 | 0.18 | 0.18 | 0.841 | 0.15 | 0.36 | 0.38 | 0.37 | 0.14 | 0.29 |
| 6 | 82730936 | rs1361694 | 1.037 | 0.87 | 0.454 | 2.282 | 0.036 | 0.01 | 0.01 | 0 | 0.46 | 0.32 | 0.26 | 0.47 | 0.32 | 0.36 |
| 6 | 82731411 | rs1573033 | -0.115 | 0.91 | 0.412 | 2.251 | -0.012 | 0.015 | 0.015 | 0.011 | 0.61 | 0.71 | 0.75 | 0.86 | 0.48 | 0.7 |
| 6 | 82735843 | rs9353048 | 1.014 | 0.862 | -0.212 | 2.28 | 0.062 | 0.006 | 0.006 | 0 | 0.45 | 0.21 | 0.25 | 0.47 | 0.32 | 0.36 |
| 6 | 82739863 | rs945855 | 0.691 | 1.055 | -0.405 | 1.748 | 0.002 | 0.002 | 0.002 | 0 | 0.5 | 0.54 | 0.4 | 0.74 | 0.37 | 0.55 |
| 6 | 82750075 | rs2152844 | -0.557 | 0.91 | -0.904 | 1.323 | 0.002 | -0.008 | -0.008 | 0 | 0.09 | 0.04 | 0.23 | 0.1 | 0.09 | 0.11 |
| 6 | 82753898 | rs9359512 | 0.156 | 0.786 | -1.206 | 1.084 | 0.004 | 0.011 | 0.011 | 0 | 0.31 | 0.21 | 0.17 | 0.35 | 0.25 | 0.26 |
| 6 | 82755854 | rs1342194 | -0.086 | 1.271 | -1.709 | 0.842 | -0.084 | 0.07 | 0.07 | 0.406 | 0.34 | 0.05 | 0.39 | 0.17 | 0.41 | 0.3 |
| 6 | 82758463 | rs4706928 | -0.121 | 1.647 | -1.699 | 1.495 | -0.014 | -0.014 | -0.014 | 0.019 | 0.11 | 0.08 | 0.2 | 0.25 | 0.2 | 0.24 |
| 6 | 82767410 | rs884656 | -0.218 | 1.727 | -1.101 | 2.014 | 0.008 | 0.008 | 0.008 | 0 | 0.11 | 0.16 | 0.21 | 0.25 | 0.2 | 0.24 |
| 6 | 82769430 | rs1361682 | -0.879 | 0.877 | -0.331 | 1.67 | 0.051 | 0.018 | 0.018 | 0.325 | 0.55 | 0.77 | 0.69 | 0.65 | 0.68 | 0.64 |
| 6 | 82776291 | rs9449392 | -0.236 | 1.097 | -0.67 | 1.351 | -0.011 | 0.083 | 0.083 | 0.088 | 0.43 | 0.22 | 0.59 | 0.28 | 0.56 | 0.33 |
| 6 | 82778405 | rs9344244 | 0.506 | 1.212 | -0.254 | 1.537 | 0.001 | 0.015 | 0.015 | 0 | 0.73 | 0.44 | 0.63 | 0.62 | 0.74 | 0.62 |
| 6 | 82778502 | rs9361886 | 0.488 | 1.363 | -0.067 | 1.702 | -0.001 | -0.01 | -0.01 | 0 | 0.56 | 0.36 | 0.54 | 0.48 | 0.49 | 0.48 |
| 6 | 82778549 | rs12526513 | -0.439 | 1.369 | -0.323 | 1.962 | -0.034 | -0.028 | -0.028 | 0 | 0.39 | 0.22 | 0.59 | 0.54 | 0.55 | 0.38 |
| 6 | 82780595 | rs11758730 | -0.946 | 1.075 | 0.041 | 1.917 | -0.005 | 0.009 | 0.009 | 0 | 0.16 | 0.19 | 0.3 | 0.24 | 0.13 | 0.18 |
| 6 | 82782672 | rs7454792 | -0.931 | 0.827 | 0.04 | 1.544 | 0.003 | 0.009 | 0.009 | 0 | 0.57 | 0.62 | 0.72 | 0.63 | 0.6 | 0.66 |
| 6 | 82787023 | rs10943852 | -0.491 | 0.798 | 0.112 | 1.555 | 0.005 | 0.002 | 0.002 | 0 | 0.14 | 0.13 | 0.2 | 0.1 | 0.11 | 0.15 |
| 6 | 82794111 | rs1984257 | -0.347 | 0.79 | 0.213 | 1.552 | 0.004 | 0.002 | 0.002 | 0 | 0.14 | 0.08 | 0.19 | 0.1 | 0.12 | 0.14 |
| 6 | 82794239 | rs2039669 | -1.306 | 0.729 | 0.357 | 1.656 | 0.004 | -0.004 | -0.004 | 0 | 0.42 | 0.65 | 0.6 | 0.55 | 0.38 | 0.51 |
| 6 | 82794594 | rs1538138 | -1.702 | 0.711 | 0.323 | 1.604 | 0.015 | 0.008 | 0.008 | 0.156 | 0.66 | 0.74 | 0.85 | 0.73 | 0.76 | 0.71 |
| 6 | 82796643 | rs9449397 | 0.586 | 1.188 | 0.227 | 0.865 | 0.008 | 0 | 0 | 0 | 0.08 | 0.12 | 0.13 | 0.1 | 0.16 | 0.05 |
| 6 | 82797161 | rs16893863 | 0.689 | 1.135 | 0.206 | 0.447 | 0.009 | -0.009 | -0.009 | 0 | 0.16 | 0.15 | 0.17 | 0.21 | 0.11 | 0.27 |
| 6 | 82799872 | rs7775215 | 0.848 | 0.691 | 0.014 | 0.348 | -0.059 | 0.031 | 0.031 | 0.022 | 0.25 | 0.09 | 0.25 | 0.15 | 0.38 | 0.21 |
| 6 | 82804125 | rs9443948 | -0.354 | 0.665 | 0.213 | 0.341 | 0.003 | -0.005 | -0.005 | 0 | 0.25 | 0.36 | 0.28 | 0.21 | 0.25 | 0.33 |
| 6 | 82805292 | rs1342196 | -0.357 | 0.662 | 0.258 | 0.339 | 0.003 | -0.004 | -0.004 | 0.882 | 0.25 | 0.35 | 0.28 | 0.21 | 0.25 | 0.34 |
| 6 | 82807521 | rs9449409 | -0.597 | 0.412 | 0.206 | 0.384 | 0.01 | 0.007 | 0.007 | 0.609 | 0.51 | 0.5 | 0.54 | 0.36 | 0.65 | 0.55 |
| 6 | 82820392 | rs1933023 | 0.119 | 0.48 | 0.125 | 0.193 | -0.015 | 0.028 | 0.028 | 0.083 | 0.35 | 0.3 | 0.37 | 0.15 | 0.54 | 0.29 |
| 6 | 82822343 | rs9294244 | -1.199 | 0.222 | 0.193 | 0.091 | -0.002 | -0.001 | -0.001 | 0 | 0.61 | 0.63 | 0.61 | 0.36 | 0.73 | 0.61 |
| 6 | 82824453 | rs9443956 | -0.491 | 0.259 | 0.339 | -0.011 | 0.009 | 0.013 | 0.013 | 0.016 | 0.45 | 0.42 | 0.43 | 0.18 | 0.63 | 0.34 |
| 6 | 82825655 | rs10485090 | 0.614 | 0.238 | 0.245 | 0.006 | 0.003 | 0.023 | 0.023 | 0 | 0.16 | 0.1 | 0.19 | 0.06 | 0.15 | 0.14 |
| 6 | 82826548 | rs1361094 | -0.852 | 0.195 | 0.253 | 0.024 | -0.014 | -0.007 | -0.007 | 0 | 0.55 | 0.55 | 0.58 | 0.36 | 0.71 | 0.6 |
| 6 | 82830438 | rs6941911 | -0.042 | 0.796 | 0.321 | 0.25 | 0.02 | -0.078 | -0.078 | 0 | 0.16 | 0.3 | 0.14 | 0 | 0.17 | 0.07 |
| 6 | 82831311 | rs13208556 | 0.612 | 0.897 | -0.196 | 0.369 | -0.059 | 0.034 | 0.034 | 0.049 | 0.25 | 0.09 | 0.26 | 0.18 | 0.4 | 0.22 |
| 6 | 82844294 | rs13211186 | 0.744 | 0.97 | -0.379 | 0.315 | -0.093 | 0.02 | 0.02 | 0 | 0.23 | 0.04 | 0.26 | 0.18 | 0.39 | 0.21 |
| 6 | 82852544 | rs9341911 | 0.211 | -0.191 | -0.185 | 0.685 | 0.031 | 0.024 | 0.024 | 0.461 | 0.51 | 0.37 | 0.45 | 0.64 | 0.35 | 0.44 |
| 6 | 82855424 | rs9361892 | -0.42 | -1.121 | -0.148 | 0.979 | -0.05 | 0.048 | 0.048 | 0 | 0.28 | 0.39 | 0.31 | 0.44 | 0.15 | 0.22 |
| 6 | 82856409 | rs16893934 | 0.447 | -0.767 | -0.313 | 0.85 | 0 | -0.002 | -0.002 | 0 | 0.06 | 0 | 0.11 | 0.06 | 0.04 | 0.07 |
| 6 | 82857374 | rs9449437 | 0.306 | -0.43 | -0.417 | 0.472 | -0.008 | 0.012 | 0.012 | 0.013 | 0.17 | 0.15 | 0.19 | 0.07 | 0.2 | 0.11 |
| 6 | 82857479 | rs1890118 | 1.303 | -0.517 | -0.512 | 0.516 | -0.009 | -0.01 | -0.01 | 0 | 0.23 | 0.21 | 0.17 | 0.3 | 0.3 | 0.28 |
| 6 | 82870823 | rs6904953 | 0.016 | -0.856 | -0.28 | -0.386 | 0.022 | 0.046 | 0.046 | 0 | 0.49 | 0.4 | 0.41 | 0.77 | 0.41 | 0.51 |
| 6 | 82883836 | rs197246 | 0.797 | -0.395 | 0.887 | -0.635 | 0.019 | -0.075 | -0.075 | 0.402 | 0.24 | 0.24 | 0.25 | 0.03 | 0.33 | 0.21 |
| 6 | 82888064 | rs6908373 | 0.457 | -0.389 | 0.815 | -0.716 | 0.029 | -0.002 | -0.002 | 0 | 0.1 | 0.18 | 0.18 | 0.07 | 0.19 | 0.1 |
| 6 | 82893577 | rs2132288 | 0.9 | -0.548 | 0.994 | -0.796 | 0.031 | 0.086 | 0.086 | 0.051 | 0.52 | 0.3 | 0.41 | 0.76 | 0.42 | 0.52 |
| 6 | 82900811 | rs9449444 | -0.446 | -0.355 | 0.749 | -0.678 | 0.022 | 0.015 | 0.015 | 0 | 0.16 | 0.25 | 0.31 | 0.2 | 0.23 | 0.26 |
| 6 | 82926778 | rs7743511 | -0.474 | -0.354 | 0.749 | -0.678 | 0.021 | 0.014 | 0.014 | 0 | 0.16 | 0.29 | 0.31 | 0.2 | 0.23 | 0.26 |
| 6 | 82938817 | rs655345 | -0.999 | -0.272 | 0.468 | -0.76 | -0.009 | -0.01 | -0.01 | 0 | 0.76 | 0.83 | 0.75 | 0.97 | 0.67 | 0.79 |
| 6 | 82958246 | rs9294254 | 0.789 | -0.398 | 0.323 | -0.412 | -0.01 | 0.034 | 0.034 | 0 | 0.45 | 0.4 | 0.51 | 0.67 | 0.51 | 0.53 |
| 6 | 82973801 | rs521805 | 0.262 | -0.914 | 0.541 | -0.2 | -0.059 | 0.08 | 0.08 | 0 | 0.34 | 0.12 | 0.43 | 0.14 | 0.54 | 0.3 |
| 6 | 82974089 | rs12664788 | 0.232 | -1.15 | 0.879 | -0.175 | -0.047 | 0.011 | 0.011 | 0 | 0.18 | 0.3 | 0.1 | 0.21 | 0.08 | 0.09 |
| 6 | 82980983 | rs540814 | -0.711 | -0.502 | 1.121 | 0.204 | 0.005 | -0.01 | -0.01 | 0.308 | 0.62 | 0.67 | 0.7 | 0.5 | 0.67 | 0.56 |
| 6 | 83002810 | rs9344267 | -0.223 | -0.328 | 1.174 | 0.306 | 0.027 | -0.007 | -0.007 | 0 | 0.59 | 0.32 | 0.67 | 0.43 | 0.69 | 0.54 |
| 6 | 83010246 | rs511002 | -0.785 | -0.283 | 1.032 | 0.333 | -0.004 | 0.005 | 0.005 | 0.542 | 0.66 | 0.62 | 0.73 | 0.57 | 0.71 | 0.7 |
| 6 | 83010602 | rs507500 | -0.246 | -0.304 | 1.209 | 0.351 | -0.019 | 0.002 | 0.002 | 0.033 | 0.61 | 0.47 | 0.59 | 0.42 | 0.69 | 0.55 |
| 6 | 83010705 | rs517187 | -0.246 | -0.304 | 1.209 | 0.351 | -0.019 | 0.002 | 0.002 | 0.033 | 0.61 | 0.47 | 0.59 | 0.42 | 0.69 | 0.55 |
| 6 | 83030053 | rs526833 | -1.118 | -0.286 | 1.171 | 0.345 | 0.01 | -0.031 | -0.031 | 0 | 0.7 | 0.75 | 0.75 | 0.58 | 0.72 | 0.72 |
| 6 | 83037925 | rs16894082 | -0.16 | -0.456 | 1.263 | 0.233 | -0.007 | 0.026 | 0.026 | 0 | 0.05 | 0.08 | 0.13 | 0.16 | 0.03 | 0.14 |
| 6 | 83065888 | rs770898 | -0.324 | -0.273 | 1.121 | 0.114 | 0.01 | 0.007 | 0.007 | 0.003 | 0.59 | 0.61 | 0.57 | 0.4 | 0.64 | 0.54 |
| 6 | 83073446 | rs4590314 | -1.171 | -0.23 | 0.984 | 0.153 | 0.004 | -0.032 | -0.032 | 0.108 | 0.68 | 0.74 | 0.72 | 0.58 | 0.69 | 0.71 |
| 6 | 83074365 | rs770911 | -0.169 | -0.303 | 1.039 | 0.126 | -0.002 | -0.002 | -0.002 | 0 | 0.57 | 0.43 | 0.55 | 0.4 | 0.64 | 0.56 |
| 6 | 83075914 | rs700494 | -1.13 | -0.22 | 0.472 | 0.105 | 0.002 | -0.023 | -0.023 | 0.204 | 0.68 | 0.73 | 0.72 | 0.58 | 0.69 | 0.71 |
| 6 | 83096577 | rs9449462 | 0.712 | -0.118 | 0.544 | -0.83 | 0 | 0 | 0 | 0 | 0.39 | 0.39 | 0.33 | 0.61 | 0.34 | 0.49 |
| 6 | 83105313 | rs714133 | 0.4 | 0.086 | 0.44 | -0.734 | -0.007 | -0.006 | -0.006 | 0 | 0.44 | 0.59 | 0.36 | 0.61 | 0.35 | 0.51 |
| 6 | 83126449 | rs10943875 | 1.239 | 0.909 | 0.346 | -0.633 | -0.063 | 0.101 | 0.101 | 0 | 0.54 | 0.12 | 0.39 | 0.7 | 0.46 | 0.63 |
| 6 | 83132239 | rs3011884 | 0.957 | 0.891 | 1.074 | -0.668 | 0.046 | -0.014 | -0.014 | 0.09 | 0.67 | 0.52 | 0.45 | 0.7 | 0.51 | 0.69 |

| | | | | | | | | | | | | | | | | |
|---|---|---|---|---|---|---|---|---|---|---|---|---|---|---|---|---|
| 6 | 83156490 | rs2983882 | 1.253 | 1.008 | 1.323 | -0.762 | 0.025 | -0.014 | -0.014 | 0 | 0.67 | 0.45 | 0.57 | 0.72 | 0.51 | 0.69 |
| 6 | 83170897 | rs9341923 | -0.014 | 1.578 | 1.206 | -0.745 | -0.044 | -0.008 | -0.008 | 0 | 0.08 | 0.01 | 0.15 | 0.06 | 0.15 | 0.08 |
| 6 | 83191207 | rs2983865 | 1.005 | 1.709 | 1.565 | -0.796 | 0.058 | -0.002 | -0.002 | 0.588 | 0.77 | 0.66 | 0.69 | 0.78 | 0.64 | 0.81 |
| 6 | 83197587 | rs1853144 | 0.169 | 1.563 | 0.953 | -0.762 | -0.066 | 0.004 | 0.004 | 0 | 0.87 | 0.99 | 0.87 | 0.93 | 0.8 | 0.93 |
| 6 | 83204488 | rs9344283 | -1.054 | 1.351 | 1.239 | -0.627 | 0.001 | -0.035 | -0.035 | 0 | 0.1 | 0.21 | 0.11 | 0.06 | 0.12 | 0.12 |
| 6 | 83223691 | rs6899828 | -0.399 | 1.521 | 0.996 | -0.567 | -0.01 | 0.013 | 0.013 | 0 | 0.07 | 0.02 | 0.08 | 0.06 | 0.12 | 0.11 |
| 6 | 83228260 | rs4546539 | -0.989 | 1.746 | 1.274 | -0.644 | -0.023 | -0.002 | -0.002 | 2.021 | 0.23 | 0.16 | 0.4 | 0.22 | 0.36 | 0.18 |
| 6 | 83230032 | rs4706167 | -2.003 | 1.415 | 1.508 | -0.744 | 0.006 | -0.028 | -0.028 | 0 | 0.39 | 0.57 | 0.48 | 0.16 | 0.5 | 0.32 |
| 6 | 83231888 | rs4351310 | -2.008 | 1.389 | 1.622 | -0.74 | 0.002 | -0.006 | -0.006 | 0.054 | 0.41 | 0.57 | 0.54 | 0.3 | 0.52 | 0.34 |
| 6 | 83252450 | rs2117966 | -1.572 | 1.472 | 1.566 | -0.716 | 0.035 | -0.017 | -0.017 | 3.09 | 0.33 | 0.44 | 0.5 | 0.28 | 0.47 | 0.31 |
| 6 | 83260390 | rs7747067 | -1.612 | 1.498 | 1.413 | -0.711 | 0.042 | -0.023 | -0.023 | 0 | 0.35 | 0.51 | 0.52 | 0.29 | 0.49 | 0.31 |
| 6 | 83276413 | rs2917488 | -2.34 | 1.363 | 1.499 | -0.599 | -0.005 | 0.01 | 0.01 | 0.669 | 0.34 | 0.7 | 0.47 | 0.26 | 0.39 | 0.33 |
| 6 | 83282876 | rs1836837 | -2.335 | 1.121 | 1.525 | -0.486 | -0.007 | 0.023 | 0.023 | 1.225 | 0.36 | 0.83 | 0.48 | 0.3 | 0.38 | 0.37 |
| 6 | 83284573 | rs9449501 | -1.035 | 1.155 | 1.707 | -0.431 | -0.016 | 0.039 | 0.039 | 0 | 0.1 | 0.32 | 0.05 | 0.1 | 0.05 | 0.08 |
| 6 | 83293061 | rs367890 | -1.526 | 1.166 | 1.332 | -0.482 | -0.003 | 0.001 | 0.001 | 0.136 | 0.21 | 0.17 | 0.39 | 0.18 | 0.32 | 0.28 |
| 6 | 83300948 | rs285990 | -1.595 | 1.135 | 1.35 | -0.459 | 0.001 | 0.002 | 0.002 | 0 | 0.13 | 0.14 | 0.25 | 0.17 | 0.19 | 0.22 |
| 6 | 83309943 | rs285984 | -0.955 | 0.859 | 1.36 | -0.458 | -0.011 | 0.054 | 0.054 | 0 | 0.06 | 0.12 | 0.16 | 0.14 | 0.04 | 0.1 |
| 6 | 83311136 | rs285982 | -2.397 | 0.829 | 1.552 | -0.489 | 0.011 | 0.017 | 0.017 | 1.137 | 0.36 | 0.73 | 0.45 | 0.29 | 0.34 | 0.35 |
| 6 | 83316216 | rs285979 | -1.649 | 0.88 | 1.16 | -0.356 | -0.002 | 0.002 | 0.002 | 0 | 0.13 | 0.13 | 0.22 | 0.17 | 0.17 | 0.21 |
| 6 | 83337469 | rs7356857 | -1.56 | 0.645 | 1.492 | -0.659 | -0.012 | -0.05 | -0.05 | 0 | 0.24 | 0.42 | 0.2 | 0.12 | 0.18 | 0.16 |
| 6 | 83355550 | rs17771989 | -0.124 | 1.491 | 0.566 | -0.733 | -0.036 | -0.028 | -0.028 | 0 | 0.06 | 0 | 0.1 | 0.07 | 0.15 | 0.08 |
| 6 | 83356887 | rs6454284 | -1.675 | 1.3 | 0.781 | -0.855 | 0.014 | 0.028 | 0.028 | 1.782 | 0.27 | 0.81 | 0.23 | 0.12 | 0.18 | 0.16 |
| 6 | 83366282 | rs1369869 | -2.275 | 1.264 | 0.586 | -0.989 | -0.042 | 0.141 | 0.141 | 0.092 | 0.4 | 0.87 | 0.38 | 0.17 | 0.43 | 0.34 |
| 6 | 83370452 | rs376096 | 0.54 | 1.252 | 0.592 | -1.071 | -0.002 | 0.007 | 0.007 | 0.893 | 0.86 | 0.93 | 0.86 | 0.99 | 0.88 | 0.92 |
| 6 | 83372225 | rs2989879 | 2.452 | 1.115 | 0.49 | -1.113 | -0.075 | 0.145 | 0.145 | 0.35 | 0.56 | 0.07 | 0.58 | 0.79 | 0.54 | 0.59 |
| 6 | 83378048 | rs10455408 | -0.723 | 1.113 | 0.953 | -1.119 | -0.006 | -0.019 | -0.019 | 0 | 0.05 | 0.17 | 0.04 | 0.02 | 0.04 | 0.06 |
| 6 | 83380203 | rs992014 | 2.031 | 1.086 | 1.41 | -1.118 | 0.006 | -0.046 | -0.046 | 1.132 | 0.65 | 0.59 | 0.66 | 0.88 | 0.59 | 0.66 |
| 6 | 83385150 | rs1369867 | 1.827 | 1.104 | 1.37 | -1.103 | -0.016 | 0.031 | 0.031 | 0 | 0.71 | 0.84 | 0.7 | 0.9 | 0.63 | 0.73 |
| 6 | 83385527 | rs9344292 | -0.435 | 1.115 | 1.361 | -1.081 | -0.003 | 0.006 | 0.006 | 0.001 | 0.05 | 0.02 | 0.05 | 0.02 | 0.09 | 0.09 |
| 6 | 83394432 | rs10498932 | -1.204 | 1.112 | 1.442 | -1.062 | 0.026 | 0.08 | 0.08 | 0.27 | 0.09 | 0.51 | 0.08 | 0.1 | 0.05 | 0.08 |
| 6 | 83397096 | rs287822 | 2.09 | 1.132 | 1.329 | -1.089 | -0.005 | 0.019 | 0.019 | 0.674 | 0.66 | 0.32 | 0.67 | 0.82 | 0.68 | 0.74 |
| 6 | 83401393 | rs9361958 | -1.51 | 1.197 | 1.354 | -1.065 | 0.003 | -0.043 | -0.043 | 0 | 0.11 | 0.28 | 0.09 | 0.04 | 0.14 | 0.16 |
| 6 | 83403744 | rs7760846 | -0.141 | 1.173 | 1.927 | -1.078 | 0.049 | 0.1 | 0.1 | 0 | 0.05 | 0.23 | 0.05 | 0.1 | 0.05 | 0.08 |
| 6 | 83405526 | rs3846777 | -1.501 | 1.179 | 1.667 | -1.081 | 0.004 | -0.043 | -0.043 | 0 | 0.11 | 0.29 | 0.09 | 0.04 | 0.14 | 0.16 |
| 6 | 83407023 | rs2492419 | 2.203 | 1.182 | 1.814 | -1.103 | 0.001 | -0.021 | -0.021 | 0.512 | 0.64 | 0.42 | 0.64 | 0.87 | 0.58 | 0.67 |
| 6 | 83410478 | rs9361960 | -0.727 | 1.18 | 1.826 | -1.089 | 0.046 | 0.094 | 0.094 | 0 | 0.07 | 0.37 | 0.06 | 0.1 | 0.05 | 0.08 |
| 6 | 83414014 | rs11752723 | -0.158 | 1.242 | 1.667 | -1.092 | -0.002 | -0.004 | -0.004 | 0 | 0.06 | 0.04 | 0.1 | 0.07 | 0.15 | 0.08 |
| 6 | 83416600 | rs287810 | 1.984 | 1.18 | 1.641 | -1.079 | -0.005 | 0.014 | 0.014 | 0.508 | 0.67 | 0.35 | 0.68 | 0.81 | 0.67 | 0.75 |
| 6 | 83423397 | rs2253003 | 2.271 | 1.212 | 0.724 | -1.045 | -0.114 | 0.116 | 0.116 | 0.175 | 0.56 | 0.05 | 0.58 | 0.78 | 0.54 | 0.57 |
| 6 | 83434226 | rs1757826 | 1.605 | 1.284 | 1.084 | -1.168 | -0.002 | -0.086 | -0.086 | 0 | 0.7 | 0.47 | 0.69 | 0.93 | 0.71 | 0.75 |
| 6 | 83435717 | rs9350960 | 2.294 | 1.591 | 0.65 | -1.349 | -0.075 | 0.189 | 0.189 | 0.331 | 0.52 | 0.08 | 0.44 | 0.77 | 0.47 | 0.58 |
| 6 | 83435849 | rs9341941 | 2.294 | 1.591 | 0.65 | -1.349 | -0.075 | 0.189 | 0.189 | 0.331 | 0.52 | 0.08 | 0.44 | 0.77 | 0.47 | 0.58 |
| 6 | 83440063 | rs700497 | -0.17 | 1.603 | 0.589 | -1.425 | -0.069 | 0.091 | 0.091 | 0.143 | 0.15 | 0.02 | 0.19 | 0.01 | 0.22 | 0.11 |
| 6 | 83440417 | rs7769325 | 1.306 | 1.604 | 0.813 | -1.35 | 0.004 | -0.09 | -0.09 | 0 | 0.68 | 0.5 | 0.55 | 0.89 | 0.63 | 0.76 |
| 6 | 83443347 | rs6909889 | -1.746 | 1.393 | 1.022 | -1.113 | 0.019 | -0.014 | -0.014 | 0 | 0.16 | 0.43 | 0.12 | 0.12 | 0.17 | 0.17 |
| 6 | 83445290 | rs9444011 | -0.732 | 1.512 | 1.209 | -0.919 | 0.049 | 0.092 | 0.092 | 0 | 0.07 | 0.34 | 0.05 | 0.09 | 0.05 | 0.08 |
| 6 | 83446800 | rs9359536 | 1.599 | 1.942 | 0.81 | -1.208 | -0.031 | 0.079 | 0.079 | 0 | 0.64 | 0.17 | 0.52 | 0.8 | 0.56 | 0.66 |
| 6 | 83447417 | rs6933883 | 1.981 | 1.376 | 0.993 | -1.006 | 0.002 | 0.073 | 0.073 | 0.282 | 0.52 | 0.26 | 0.44 | 0.71 | 0.46 | 0.6 |
| 6 | 83447512 | rs287833 | 2.229 | 1.729 | 1.675 | -0.702 | 0.066 | 0.118 | 0.118 | 0.702 | 0.9 | 0.45 | 0.9 | 0.85 | 0.85 | 0.89 |
| 6 | 83447830 | rs7773826 | -1.769 | 1.45 | 1.725 | -0.66 | -0.008 | 0 | 0 | 0 | 0.16 | 0.3 | 0.23 | 0.15 | 0.15 | 0.17 |
| 6 | 83453859 | rs287849 | -1.271 | 1.968 | 1.781 | -0.758 | -0.019 | 0.014 | 0.014 | 0 | 0.29 | 0.19 | 0.34 | 0.15 | 0.43 | 0.25 |
| 6 | 83454017 | rs287848 | -2.294 | 1.21 | 1.636 | -1.09 | 0.024 | 0.039 | 0.039 | 0 | 0.51 | 0.7 | 0.58 | 0.31 | 0.62 | 0.48 |
| 6 | 83461302 | rs2458538 | -1.331 | 1.78 | 1.926 | -0.739 | -0.016 | 0.013 | 0.013 | 0 | 0.29 | 0.23 | 0.34 | 0.16 | 0.43 | 0.26 |
| 6 | 83473150 | rs1331785 | 1.869 | 1.542 | 2.293 | -0.737 | 0.012 | 0.057 | 0.057 | 0.653 | 0.86 | 0.46 | 0.77 | 0.83 | 0.86 | 0.82 |
| 6 | 83473573 | rs1040046 | -1.158 | 1.618 | 1.933 | -0.731 | 0.01 | 0.016 | 0.016 | 0 | 0.11 | 0.25 | 0.21 | 0.13 | 0.14 | 0.13 |
| 6 | 83479174 | rs10455410 | -0.838 | 1.694 | 1.763 | -0.679 | -0.001 | -0.001 | -0.001 | 0.761 | 0.1 | 0.12 | 0.21 | 0.13 | 0.15 | 0.14 |
| 6 | 83481903 | rs2323770 | 2.428 | 1.779 | 1.636 | -0.558 | 0.018 | 0.047 | 0.047 | 0.016 | 0.52 | 0.14 | 0.4 | 0.63 | 0.42 | 0.58 |
| 6 | 83482938 | rs17181739 | -0.485 | 1.964 | 1.711 | -0.619 | -0.055 | 0.072 | 0.072 | 0 | 0.22 | 0.06 | 0.26 | 0.07 | 0.27 | 0.11 |
| 6 | 83484534 | rs9344309 | -0.777 | 1.948 | 2.053 | -0.621 | 0.068 | 0.06 | 0.06 | 0 | 0.09 | 0.25 | 0.09 | 0.13 | 0.17 | 0.12 |
| 6 | 83485920 | rs4391233 | 2.725 | 1.946 | 2.005 | -0.623 | 0.017 | 0.046 | 0.046 | 0.018 | 0.53 | 0.14 | 0.4 | 0.63 | 0.42 | 0.58 |
| 6 | 83487064 | rs6931603 | 1.636 | 1.947 | 2.023 | -0.617 | 0.017 | -0.026 | -0.026 | 0 | 0.64 | 0.53 | 0.5 | 0.76 | 0.59 | 0.7 |
| 6 | 83491868 | rs6909405 | 1.742 | 1.89 | 1.985 | -0.53 | -0.004 | 0.022 | 0.022 | 0.076 | 0.84 | 0.55 | 0.76 | 0.83 | 0.85 | 0.81 |
| 6 | 83498310 | rs2203283 | -0.696 | 1.921 | 1.624 | -0.519 | 0.02 | 0.016 | 0.016 | 0 | 0.08 | 0.09 | 0.08 | 0.13 | 0.17 | 0.12 |
| 6 | 83536070 | rs9359543 | -1.021 | 2.06 | 1.511 | -0.599 | 0.015 | 0.007 | 0.007 | 0 | 0.1 | 0.09 | 0.09 | 0.13 | 0.19 | 0.14 |
| 6 | 83539461 | rs2057026 | 2.439 | 2.035 | 1.548 | -0.219 | -0.001 | -0.003 | -0.003 | 0 | 0.53 | 0.17 | 0.4 | 0.62 | 0.42 | 0.57 |
| 6 | 83541333 | rs209411 | -1.527 | 2.046 | 1.827 | -0.194 | 0.004 | 0.023 | 0.023 | 0.224 | 0.14 | 0.39 | 0.23 | 0.12 | 0.14 | 0.13 |
| 6 | 83542389 | rs985802 | -0.937 | 2.082 | 1.835 | -0.136 | 0.03 | 0.064 | 0.064 | 0 | 0.09 | 0.15 | 0.09 | 0.19 | 0.16 | 0.18 |
| 6 | 83545752 | rs2783168 | -1.185 | 2.015 | 1.279 | -0.184 | 0.008 | 0.023 | 0.023 | 0 | 0.13 | 0.33 | 0.22 | 0.12 | 0.12 | 0.13 |
| 6 | 83546026 | rs12197652 | 2.242 | 1.969 | 0.637 | -0.207 | -0.026 | -0.027 | -0.027 | 0 | 0.52 | 0.07 | 0.4 | 0.61 | 0.44 | 0.57 |

| | | | | | | | | | | | | | | | | |
|---|---|---|---|---|---|---|---|---|---|---|---|---|---|---|---|---|
| 6 | 83550056 | rs1932217 | -0.341 | 1.986 | 0.736 | -0.723 | -0.022 | 0.023 | 0.023 | 0.073 | 0.13 | 0.09 | 0.11 | 0.07 | 0.21 | 0.16 |
| 6 | 83551482 | rs7760612 | -1.006 | 2.056 | 0.749 | -0.821 | 0.119 | -0.001 | -0.001 | 0.286 | 0.17 | 0.43 | 0.15 | 0.14 | 0.27 | 0.19 |
| 6 | 83551624 | rs9294264 | -0.35 | 1.963 | 0.675 | -0.811 | 0.001 | -0.001 | -0.001 | 0 | 0.09 | 0.11 | 0.19 | 0.08 | 0.07 | 0.07 |
| 6 | 83570155 | rs553094 | -2.502 | 1.996 | 0.4 | -0.799 | 0.038 | 0.025 | 0.025 | 1.031 | 0.56 | 0.93 | 0.63 | 0.32 | 0.67 | 0.5 |
| 6 | 83571914 | rs9344322 | -0.635 | 2.223 | 0.633 | -0.781 | -0.068 | -0.014 | -0.014 | 0 | 0.14 | 0.03 | 0.11 | 0.13 | 0.26 | 0.17 |
| 6 | 83573836 | rs975001 | -0.794 | 2.087 | 0.681 | -0.75 | 0.093 | 0.011 | 0.011 | 0 | 0.16 | 0.41 | 0.13 | 0.14 | 0.23 | 0.18 |
| 6 | 83574391 | rs1040544 | -1.626 | 1.775 | 0.02 | -0.685 | -0.012 | 0.068 | 0.068 | 0 | 0.43 | 0.72 | 0.42 | 0.23 | 0.55 | 0.41 |
| 6 | 83576815 | rs209420 | -1.988 | 1.785 | -0.208 | -0.515 | -0.066 | 0.031 | 0.031 | 0.447 | 0.46 | 0.94 | 0.66 | 0.31 | 0.55 | 0.43 |
| 6 | 83580759 | rs209401 | -2.001 | 1.778 | -0.21 | -0.741 | -0.06 | 0.09 | 0.09 | 0.51 | 0.47 | 0.94 | 0.66 | 0.28 | 0.54 | 0.43 |
| 6 | 83582258 | rs9688767 | -0.823 | 1.788 | 0.638 | -0.632 | 0.004 | 0.022 | 0.022 | 0 | 0.13 | 0.24 | 0.4 | 0.12 | 0.14 | 0.12 |
| 6 | 83584304 | rs9361981 | -0.318 | 1.937 | 0.763 | -0.881 | 0.049 | -0.015 | -0.015 | 0.485 | 0.1 | 0.16 | 0.09 | 0.09 | 0.17 | 0.11 |
| 6 | 83588696 | rs4317380 | -0.788 | 1.183 | 0.443 | -0.78 | 0.001 | -0.003 | -0.003 | 0 | 0.12 | 0.16 | 0.39 | 0.11 | 0.13 | 0.12 |
| 6 | 83590891 | rs13206195 | -0.289 | 1.176 | 0.107 | -0.73 | 0.001 | -0.013 | -0.013 | 2.821 | 0.1 | 0.13 | 0.1 | 0.01 | 0.14 | 0.11 |
| 6 | 83594304 | rs10484711 | -0.271 | 1.187 | -0.132 | -0.724 | -0.024 | 0.035 | 0.035 | 0 | 0.1 | 0.05 | 0.1 | 0.01 | 0.14 | 0.11 |
| 6 | 83595589 | rs159045 | -2.064 | 0.9 | -0.626 | -0.982 | -0.035 | 0.103 | 0.103 | 0.74 | 0.39 | 0.95 | 0.59 | 0.18 | 0.35 | 0.37 |
| 6 | 83598857 | rs1328892 | -1.817 | 0.428 | 0.797 | -0.724 | -0.028 | -0.027 | -0.027 | 0 | 0.26 | 0.57 | 0.49 | 0.19 | 0.22 | 0.29 |
| 6 | 83602583 | rs4706975 | -1.88 | 0.425 | 0.803 | -0.726 | -0.03 | -0.029 | -0.029 | 0 | 0.26 | 0.58 | 0.49 | 0.19 | 0.22 | 0.29 |
| 6 | 83608890 | rs1981129 | -1.104 | 0.367 | 0.998 | -0.69 | -0.001 | 0.005 | 0.005 | 0 | 0.14 | 0.12 | 0.32 | 0.13 | 0.17 | 0.15 |
| 6 | 83609239 | rs9294265 | -1.382 | 0.454 | 0.957 | -0.709 | -0.007 | 0.018 | 0.018 | 0.041 | 0.13 | 0.46 | 0.18 | 0.11 | 0.06 | 0.19 |
| 6 | 83610839 | rs159054 | -2.821 | 0.43 | 0.784 | -0.721 | 0.005 | 0.006 | 0.006 | 0.99 | 0.3 | 0.94 | 0.52 | 0.24 | 0.23 | 0.34 |
| 6 | 83612460 | rs7738961 | -0.974 | 0.473 | 0.953 | -0.725 | -0.014 | 0.022 | 0.022 | 0 | 0.1 | 0.26 | 0.15 | 0.11 | 0.05 | 0.16 |
| 6 | 83613259 | rs7744090 | -1.671 | 0.492 | 0.926 | -0.729 | 0.001 | -0.003 | -0.003 | 0 | 0.19 | 0.28 | 0.46 | 0.2 | 0.19 | 0.3 |
| 6 | 83613680 | rs9449559 | -1.598 | 0.49 | 0.93 | -0.727 | -0.004 | -0.002 | -0.002 | 0 | 0.19 | 0.29 | 0.44 | 0.2 | 0.18 | 0.27 |
| 6 | 83615910 | rs9444019 | -1.976 | 0.612 | 1.011 | -0.842 | -0.005 | -0.039 | -0.039 | 0.046 | 0.2 | 0.66 | 0.15 | 0.05 | 0.16 | 0.17 |
| 6 | 83616522 | rs2324167 | -1.914 | 0.546 | 1.017 | -0.836 | -0.011 | -0.039 | -0.039 | 1.592 | 0.19 | 0.66 | 0.14 | 0.05 | 0.15 | 0.14 |
| 6 | 83630591 | rs293498 | 1.451 | 0.463 | 1.079 | -0.731 | -0.004 | -0.042 | -0.042 | 0 | 0.85 | 0.72 | 0.81 | 0.93 | 0.87 | 0.92 |
| 6 | 83666480 | rs12201727 | -1.17 | 0.341 | 1.027 | -0.727 | -0.001 | -0.001 | -0.001 | 0.444 | 0.14 | 0.13 | 0.13 | 0.13 | 0.14 | 0.15 |
| 6 | 83667045 | rs7739323 | -1.387 | 0.349 | 1.084 | -0.748 | -0.002 | -0.003 | -0.003 | 0 | 0.16 | 0.18 | 0.14 | 0.13 | 0.16 | 0.17 |
| 6 | 83679599 | rs6922073 | -1.364 | 0.347 | 1.171 | -0.762 | -0.004 | -0.007 | -0.007 | 0 | 0.15 | 0.21 | 0.14 | 0.12 | 0.16 | 0.16 |
| 6 | 83698770 | rs2149537 | 2.348 | 0.32 | 1.301 | -0.806 | -0.014 | -0.047 | -0.047 | 0 | 0.7 | 0.52 | 0.75 | 0.81 | 0.72 | 0.76 |
| 6 | 83704533 | rs11758364 | -0.974 | 0.397 | 1.281 | -0.741 | 0.001 | -0.037 | -0.037 | 0 | 0.12 | 0.17 | 0.09 | 0.03 | 0.14 | 0.11 |
| 6 | 83710287 | rs12212188 | -1.551 | 0.539 | 1.124 | -0.633 | 0.004 | 0.007 | 0.007 | 0 | 0.15 | 0.21 | 0.19 | 0.17 | 0.19 | 0.12 |
| 6 | 83718503 | rs9359554 | 2.213 | -0.063 | 0.895 | -1.167 | -0.013 | -0.036 | -0.036 | 0 | 0.66 | 0.47 | 0.71 | 0.78 | 0.69 | 0.69 |
| 6 | 83733785 | rs4235816 | 2.137 | -0.261 | 0.836 | -1.223 | -0.015 | -0.035 | -0.035 | 0 | 0.64 | 0.47 | 0.7 | 0.78 | 0.68 | 0.68 |
| 6 | 83746059 | rs12209871 | -0.681 | -0.019 | 0.775 | -1.17 | -0.004 | 0.02 | 0.02 | 0 | 0.12 | 0.05 | 0.09 | 0.03 | 0.15 | 0.1 |
| 6 | 83748563 | rs6934537 | 0.798 | -0.043 | 0.861 | -1.18 | -0.001 | -0.008 | -0.008 | 0.346 | 0.86 | 0.88 | 0.9 | 0.95 | 0.88 | 0.93 |
| 6 | 83758364 | rs4706978 | 1.592 | -0.063 | 0.725 | -1.169 | 0 | 0.011 | 0.011 | 3.422 | 0.74 | 0.81 | 0.8 | 0.9 | 0.73 | 0.8 |
| 6 | 83769908 | rs1538356 | 1.849 | 0.091 | 0.87 | -0.739 | 0.009 | 0.011 | 0.011 | 0 | 0.75 | 0.45 | 0.69 | 0.75 | 0.72 | 0.77 |
| 6 | 83782301 | rs4515348 | 1.304 | 0.371 | 1.05 | -0.61 | 0.02 | 0.002 | 0.002 | 0 | 0.82 | 0.78 | 0.8 | 0.81 | 0.72 | 0.84 |
| 6 | 83784497 | rs7766560 | 1.694 | 0.357 | 1.167 | -0.616 | 0.046 | 0.002 | 0.002 | 0.108 | 0.78 | 0.52 | 0.74 | 0.79 | 0.7 | 0.82 |
| 6 | 83817591 | rs6906567 | 1.907 | 0.34 | 1.157 | -0.61 | 0.017 | 0.021 | 0.021 | 0.509 | 0.77 | 0.4 | 0.72 | 0.71 | 0.7 | 0.77 |
| 6 | 83828254 | rs2148011 | 1.239 | 0.338 | 1.203 | -0.609 | 0.008 | -0.001 | -0.001 | 1.364 | 0.82 | 0.82 | 0.8 | 0.82 | 0.72 | 0.86 |
| 6 | 83838673 | rs4706980 | -0.22 | 0.359 | 1.077 | -0.61 | -0.038 | -0.035 | -0.035 | 0 | 0.09 | 0 | 0.09 | 0.15 | 0.14 | 0.08 |
| 6 | 83859627 | rs13208108 | 0.559 | 0.189 | 0.375 | -1.251 | -0.068 | 0.109 | 0.109 | 0 | 0.23 | 0.01 | 0.24 | 0.06 | 0.25 | 0.1 |
| 6 | 83873386 | rs12190346 | 0.705 | 0.647 | 0.376 | -1.27 | -0.047 | 0.056 | 0.056 | 0 | 0.11 | 0.01 | 0.09 | 0 | 0.14 | 0.07 |
| 6 | 83877496 | rs495944 | -0.757 | 0.383 | 1.031 | -0.63 | 0.018 | 0.002 | 0.002 | 0 | 0.19 | 0.23 | 0.2 | 0.19 | 0.28 | 0.16 |
| 6 | 83880167 | rs473267 | -0.988 | 0.43 | 0.484 | -0.606 | 0.015 | 0.023 | 0.023 | 0.432 | 0.23 | 0.59 | 0.28 | 0.29 | 0.3 | 0.22 |
| 6 | 83881740 | rs542948 | -1.007 | 0.433 | 0.437 | -0.604 | 0.013 | 0.022 | 0.022 | 0.444 | 0.23 | 0.6 | 0.28 | 0.29 | 0.3 | 0.23 |
| 6 | 83894382 | rs628629 | -0.737 | 0.518 | -0.651 | -0.45 | 0.002 | 0.075 | 0.075 | 0 | 0.51 | 0.88 | 0.57 | 0.37 | 0.58 | 0.38 |
| 6 | 83908738 | rs6902437 | -1.049 | 1.134 | 0.241 | 0.479 | 0.03 | 0.039 | 0.039 | 0 | 0.17 | 0.29 | 0.25 | 0.28 | 0.28 | 0.2 |
| 6 | 83911334 | rs6910678 | -1.64 | 0.488 | 0.537 | 0.858 | -0.016 | 0.089 | 0.089 | 0 | 0.11 | 0.32 | 0.13 | 0.26 | 0.1 | 0.1 |
| 6 | 83915414 | rs9444043 | -0.79 | -0.066 | -0.023 | 0.213 | 0.052 | 0.042 | 0.042 | 1.524 | 0.47 | 0.79 | 0.48 | 0.41 | 0.4 | 0.34 |
| 6 | 83916069 | rs13219666 | 0.215 | -0.312 | -0.084 | -0.014 | -0.022 | 0.144 | 0.144 | 4.971 | 0.29 | 0.08 | 0.27 | 0.05 | 0.28 | 0.09 |
| 6 | 83925555 | rs1180242 | -1.633 | -0.047 | -0.402 | -0.103 | -0.135 | -0.111 | -0.111 | 0.43 | 0.59 | 0.98 | 0.59 | 0.45 | 0.53 | 0.43 |
| 6 | 83932070 | rs1180241 | 0.183 | -0.189 | 1.095 | -0.079 | -0.012 | 0.075 | 0.075 | 4.431 | 0.31 | 0.17 | 0.28 | 0.06 | 0.31 | 0.13 |
| 6 | 83932731 | rs9449593 | -1.666 | -0.208 | 1.26 | -0.052 | -0.029 | 0.111 | 0.111 | 0 | 0.14 | 0.44 | 0.18 | 0.29 | 0.1 | 0.17 |
| 6 | 83943928 | rs1180237 | -1.047 | -0.177 | 1.261 | -0.055 | 0.001 | 0.004 | 0.004 | 2.409 | 0.45 | 0.6 | 0.47 | 0.35 | 0.42 | 0.29 |
| 6 | 83954264 | rs983087 | 0.037 | -0.214 | 1.17 | -0.05 | -0.008 | 0.052 | 0.052 | 4.256 | 0.32 | 0.16 | 0.33 | 0.2 | 0.33 | 0.16 |
| 6 | 83956724 | rs1180204 | -0.931 | -0.221 | 1.374 | -0.051 | -0.001 | -0.004 | -0.004 | 0 | 0.43 | 0.49 | 0.45 | 0.34 | 0.42 | 0.29 |
| 6 | 83960339 | rs2179847 | -1.043 | -0.199 | 1.374 | -0.041 | -0.019 | 0.006 | 0.006 | 0 | 0.08 | 0.22 | 0.07 | 0.11 | 0.05 | 0.08 |
| 6 | 83960862 | rs1180195 | -0.959 | -0.215 | 1.597 | -0.003 | 0.017 | 0.024 | 0.024 | 4.481 | 0.44 | 0.68 | 0.44 | 0.38 | 0.38 | 0.28 |
| 6 | 83964709 | rs1180192 | 0.083 | -0.427 | 1.159 | -0.069 | 0.012 | 0.034 | 0.034 | 6.023 | 0.29 | 0.17 | 0.3 | 0.19 | 0.27 | 0.13 |
| 6 | 83967454 | rs1180187 | -1.285 | -0.44 | 0.881 | -0.041 | 0.044 | 0.02 | 0.02 | 3.875 | 0.47 | 0.95 | 0.48 | 0.4 | 0.36 | 0.32 |
| 6 | 83972313 | rs1954537 | -1.299 | -0.118 | 1.182 | 0.119 | -0.008 | 0.01 | 0.01 | 0 | 0.1 | 0.19 | 0.11 | 0.16 | 0.09 | 0.13 |
| 6 | 83976802 | rs767144 | -0.154 | -0.193 | 0.98 | 0.138 | 0.005 | 0.048 | 0.048 | 5.725 | 0.3 | 0.08 | 0.31 | 0.21 | 0.27 | 0.15 |
| 6 | 83995864 | rs13192303 | -0.538 | -0.086 | 0.922 | 0.11 | -0.033 | 0.003 | 0.003 | 0.056 | 0.07 | 0.01 | 0.1 | 0.06 | 0.1 | 0.06 |
| 6 | 84005867 | rs1143796 | -0.791 | -0.175 | 1.603 | 0.096 | 0.005 | 0.018 | 0.018 | 3.16 | 0.4 | 0.34 | 0.43 | 0.27 | 0.37 | 0.21 |
| 6 | 84017921 | rs1144184 | -1.205 | -0.219 | 1.77 | 0.18 | 0.038 | 0.03 | 0.03 | 2.797 | 0.46 | 0.77 | 0.55 | 0.42 | 0.39 | 0.31 |
| 6 | 84020324 | rs1170348 | -1.073 | -0.507 | 2.079 | 0.147 | -0.01 | -0.014 | -0.014 | 0.298 | 0.43 | 0.55 | 0.52 | 0.36 | 0.36 | 0.26 |

| | | | | | | | | | | | | | | | | |
|---|---|---|---|---|---|---|---|---|---|---|---|---|---|---|---|---|
| 6 | 84024620 | rs1144180 | 0.05 | -0.354 | 2.377 | 0.828 | -0.004 | 0.036 | 0.036 | 6.202 | 0.28 | 0.12 | 0.34 | 0.18 | 0.31 | 0.12 |
| 6 | 84030582 | rs750385 | 0.749 | -0.804 | 1.957 | 0.27 | -0.021 | 0.063 | 0.063 | 4.435 | 0.14 | 0.03 | 0.1 | 0 | 0.17 | 0.03 |
| 6 | 84038677 | rs1170485 | -1.116 | -0.628 | 2.263 | 0.396 | 0.008 | 0.017 | 0.017 | 3.915 | 0.39 | 0.34 | 0.49 | 0.26 | 0.33 | 0.2 |
| 6 | 84051050 | rs1535588 | -1.116 | -0.628 | 2.263 | 0.396 | 0.008 | 0.017 | 0.017 | 3.915 | 0.39 | 0.34 | 0.49 | 0.26 | 0.33 | 0.2 |
| 6 | 84064190 | rs6917851 | -0.776 | -0.629 | 2.145 | 0.402 | 0.005 | 0.151 | 0.151 | 4.969 | 0.36 | 0.1 | 0.37 | 0.12 | 0.32 | 0.14 |
| 6 | 84087659 | rs13205856 | -0.32 | -0.587 | 1.878 | 0.361 | -0.025 | 0.001 | 0.001 | 0.065 | 0.07 | 0 | 0.11 | 0.06 | 0.08 | 0.05 |
| 6 | 84102550 | rs12191369 | -0.717 | -0.589 | 1.894 | 0.371 | 0.006 | 0.152 | 0.152 | 4.888 | 0.36 | 0.1 | 0.37 | 0.11 | 0.32 | 0.13 |
| 6 | 84119361 | rs3798890 | -0.724 | -0.535 | 1.934 | 0.407 | 0.007 | 0.138 | 0.138 | 4.583 | 0.36 | 0.1 | 0.37 | 0.12 | 0.32 | 0.14 |
| 6 | 84138485 | rs6941094 | -0.937 | -0.544 | 2.106 | 0.443 | -0.008 | -0.019 | -0.019 | 0.393 | 0.4 | 0.44 | 0.5 | 0.26 | 0.33 | 0.2 |
| 6 | 84162042 | rs12190945 | -0.889 | -0.565 | 2.025 | 0.483 | 0.026 | 0.05 | 0.05 | 4.697 | 0.37 | 0.25 | 0.48 | 0.27 | 0.31 | 0.2 |
| 6 | 84169965 | rs2000375 | -1.443 | -0.705 | 2.198 | 0.487 | 0.016 | 0.009 | 0.009 | 1.28 | 0.46 | 0.58 | 0.54 | 0.41 | 0.34 | 0.3 |
| 6 | 84174401 | rs12203860 | -1.021 | -0.709 | 2.309 | 0.466 | -0.013 | -0.04 | -0.04 | 0 | 0.41 | 0.43 | 0.28 | 0.15 | 0.31 | 0.18 |
| 6 | 84175552 | rs6454330 | -0.683 | -0.714 | 2.216 | 0.463 | 0.014 | 0.039 | 0.039 | 6.745 | 0.36 | 0.22 | 0.26 | 0.15 | 0.3 | 0.17 |
| 6 | 84176481 | rs13201274 | -0.032 | -0.717 | 2.057 | 0.436 | 0.023 | 0.111 | 0.111 | 5.7 | 0.28 | 0.07 | 0.14 | 0.06 | 0.22 | 0.08 |
| 6 | 84182057 | rs9444076 | -2.193 | -0.712 | 2.118 | 0.532 | -0.029 | 0.076 | 0.076 | 0 | 0.18 | 0.53 | 0.24 | 0.31 | 0.12 | 0.18 |
| 6 | 84184472 | rs4706178 | -1.644 | -0.831 | 1.995 | 0.44 | 0.061 | 0.05 | 0.05 | 2.45 | 0.48 | 0.67 | 0.4 | 0.39 | 0.34 | 0.31 |
| 6 | 84197019 | rs9449628 | -2.658 | -0.673 | 2.084 | 0.71 | -0.023 | 0.051 | 0.051 | 0 | 0.19 | 0.67 | 0.29 | 0.35 | 0.12 | 0.23 |
| 6 | 84198622 | rs7747101 | -1.921 | -0.589 | 2.035 | 0.576 | -0.005 | -0.01 | -0.01 | 0 | 0.16 | 0.5 | 0.14 | 0.15 | 0.11 | 0.12 |
| 6 | 84204107 | rs11752474 | -0.902 | -0.329 | 2.039 | 0.535 | 0.042 | -0.029 | -0.029 | 0 | 0.1 | 0.2 | 0.12 | 0.05 | 0.14 | 0.07 |
| 6 | 84211936 | rs10943944 | -1.973 | -0.34 | 1.934 | 0.559 | 0.049 | 0.031 | 0.031 | 0 | 0.17 | 0.49 | 0.19 | 0.19 | 0.17 | 0.16 |
| 6 | 84217168 | rs7753918 | -1.029 | -0.386 | 1.739 | 0.478 | 0.061 | -0.056 | -0.056 | 0.034 | 0.11 | 0.29 | 0.1 | 0.03 | 0.14 | 0.07 |
| 6 | 84217876 | rs607259 | 1.366 | -0.67 | 1.213 | 0.401 | 0.05 | 0.048 | 0.048 | 2.945 | 0.5 | 0.2 | 0.54 | 0.58 | 0.61 | 0.67 |
| 6 | 84219312 | rs6928830 | -1.946 | -0.409 | 1.471 | 0.583 | 0.018 | 0.072 | 0.072 | 0 | 0.18 | 0.42 | 0.28 | 0.27 | 0.15 | 0.2 |
| 6 | 84220165 | rs637974 | 0.27 | -0.835 | 0.931 | 0.069 | 0.041 | 0.061 | 0.061 | 2.172 | 0.63 | 0.78 | 0.84 | 0.85 | 0.76 | 0.86 |
| 6 | 84220405 | rs12204318 | -0.737 | -0.374 | 1.122 | 1.347 | 0.055 | 0.09 | 0.09 | 0 | 0.09 | 0.23 | 0.1 | 0.29 | 0.13 | 0.09 |
| 6 | 84220770 | rs9341961 | -1.084 | -0.386 | 1.147 | 1.39 | 0.05 | 0.143 | 0.143 | 0 | 0.11 | 0.3 | 0.17 | 0.34 | 0.13 | 0.13 |
| 6 | 84223563 | rs592911 | 0.664 | -0.911 | 0.484 | 0.433 | 0.001 | 0.005 | 0.005 | 0.602 | 0.47 | 0.51 | 0.69 | 0.57 | 0.56 | 0.59 |
| 6 | 84224999 | rs579599 | 0.034 | -1.201 | 0.914 | 0.284 | 0.013 | 0.015 | 0.015 | 2.593 | 0.62 | 0.72 | 0.81 | 0.83 | 0.73 | 0.86 |
| 6 | 84227646 | rs1171114 | 0.779 | -1.067 | 0.845 | 0.209 | -0.001 | -0.003 | -0.003 | 0.077 | 0.48 | 0.6 | 0.69 | 0.62 | 0.57 | 0.61 |
| 6 | 84229114 | rs568129 | 0.058 | -1.071 | 0.832 | 0.174 | 0.026 | 0.044 | 0.044 | 2.199 | 0.61 | 0.77 | 0.8 | 0.85 | 0.71 | 0.84 |
| 6 | 84234144 | rs3812141 | -0.077 | -1.408 | 1.587 | 0.24 | 0.009 | 0.032 | 0.032 | 0 | 0.14 | 0.26 | 0.11 | 0.22 | 0.13 | 0.25 |
| 6 | 84240606 | rs645252 | -0.178 | -1.339 | 1.657 | 0.873 | 0.026 | -0.001 | -0.001 | 0.438 | 0.6 | 0.76 | 0.63 | 0.62 | 0.68 | 0.79 |
| 6 | 84250330 | rs7741310 | 0.856 | -1.294 | 1.678 | 0.892 | -0.002 | -0.007 | -0.007 | 0 | 0.46 | 0.58 | 0.69 | 0.6 | 0.57 | 0.61 |
| 6 | 84264333 | rs1158059 | 1.29 | -1.481 | 1.163 | 0.668 | -0.099 | -0.047 | -0.047 | 1.734 | 0.33 | 0.14 | 0.42 | 0.39 | 0.52 | 0.54 |
| 6 | 84268666 | rs3798878 | 0.035 | -1.585 | 0.76 | 0.517 | 0.004 | 0.003 | 0.003 | 0 | 0.23 | 0.17 | 0.14 | 0.24 | 0.17 | 0.26 |
| 6 | 84282761 | rs2023238 | -0.39 | -1.49 | 0.658 | 0.538 | 0.042 | 0.003 | 0.003 | 1.935 | 0.58 | 0.8 | 0.6 | 0.62 | 0.68 | 0.79 |
| 6 | 84286477 | rs11756746 | 0.369 | -0.998 | 0.499 | 0.543 | -0.018 | 0.009 | 0.009 | 0.41 | 0.23 | 0.13 | 0.15 | 0.23 | 0.23 | 0.26 |
| 6 | 84313180 | rs11754643 | 0.534 | -0.991 | 0.355 | 0.465 | -0.08 | -0.005 | -0.005 | 0 | 0.22 | 0.01 | 0.14 | 0.2 | 0.23 | 0.27 |
| 6 | 84314423 | rs1546977 | 1.272 | -1.101 | 0.889 | 0.459 | 0.018 | 0.022 | 0.022 | 0.238 | 0.3 | 0.39 | 0.42 | 0.42 | 0.44 | 0.52 |
| 6 | 84322084 | rs9362001 | -0.861 | -1.326 | 1.03 | 0.405 | -0.014 | -0.014 | -0.014 | 0.039 | 0.62 | 0.64 | 0.67 | 0.77 | 0.75 | 0.87 |
| 6 | 84325184 | rs12193281 | -0.153 | -1.3 | 1.045 | 0.417 | -0.03 | -0.022 | -0.022 | 0 | 0.28 | 0.16 | 0.24 | 0.34 | 0.31 | 0.36 |
| 6 | 84335881 | rs217344 | -0.898 | -1.326 | 1.076 | 0.395 | 0.064 | 0.066 | 0.066 | 4.013 | 0.62 | 0.88 | 0.69 | 0.76 | 0.76 | 0.87 |
| 6 | 84347714 | rs217328 | 1.19 | -1.144 | 1.087 | 0.322 | -0.004 | -0.004 | -0.004 | 0 | 0.31 | 0.33 | 0.42 | 0.41 | 0.44 | 0.52 |
| 6 | 84348873 | rs3798865 | 0.404 | -1.197 | 0.836 | 0.319 | -0.053 | -0.103 | -0.103 | 0 | 0.23 | 0.03 | 0.3 | 0.39 | 0.24 | 0.29 |
| 6 | 84354101 | rs217323 | 1.053 | -1.093 | 0.952 | 0.155 | -0.06 | -0.069 | -0.069 | 0 | 0.3 | 0.16 | 0.4 | 0.42 | 0.44 | 0.52 |
| 6 | 84365478 | rs6940118 | 0.735 | -0.996 | 1.004 | 0.207 | 0 | 0.004 | 0.004 | 0 | 0.05 | 0.07 | 0.04 | 0.09 | 0.06 | 0.06 |
| 6 | 84369869 | rs217308 | 1.152 | -0.986 | 0.986 | 0.207 | -0.04 | -0.045 | -0.045 | 0 | 0.28 | 0.22 | 0.41 | 0.43 | 0.44 | 0.52 |
| 6 | 84393942 | rs217291 | 1.104 | -0.892 | 0.172 | 0.257 | -0.025 | -0.028 | -0.028 | 0 | 0.29 | 0.24 | 0.41 | 0.43 | 0.44 | 0.52 |
| 6 | 84399772 | rs7749272 | 0.381 | -0.638 | 0.364 | 0.894 | 0 | 0.006 | 0.006 | 0 | 0.05 | 0.08 | 0.04 | 0.09 | 0.06 | 0.06 |
| 6 | 84405335 | rs12201719 | -0.7 | -0.696 | 0.26 | 0.791 | -0.026 | -0.069 | -0.069 | 0 | 0.28 | 0.16 | 0.39 | 0.52 | 0.3 | 0.36 |
| 6 | 84410228 | rs813882 | -0.64 | -0.746 | 0.292 | 1.053 | 0.001 | 0.004 | 0.004 | 0 | 0.26 | 0.28 | 0.37 | 0.57 | 0.27 | 0.4 |
| 6 | 84412693 | rs522115 | 0.101 | -1.287 | 0.602 | 1.025 | -0.002 | 0.043 | 0.043 | 0 | 0.07 | 0.12 | 0.05 | 0.13 | 0.06 | 0.15 |
| 6 | 84419793 | rs624076 | -1.471 | -1.895 | 0.632 | 0.371 | 0.039 | 0.013 | 0.013 | 0 | 0.34 | 0.72 | 0.21 | 0.27 | 0.18 | 0.29 |
| 6 | 84423854 | rs484056 | -0.045 | -2.07 | 0.664 | 0.339 | -0.002 | 0.004 | 0.004 | 0 | 0.14 | 0.15 | 0.1 | 0.19 | 0.12 | 0.2 |
| 6 | 84428280 | rs586661 | -1.563 | -2.014 | 0.579 | 0.361 | 0.018 | 0.006 | 0.006 | 0 | 0.34 | 0.67 | 0.2 | 0.27 | 0.17 | 0.29 |
| 6 | 84437425 | rs660956 | -1.35 | -1.912 | 0.232 | 0.41 | -0.06 | -0.011 | -0.011 | 0 | 0.31 | 0.49 | 0.18 | 0.27 | 0.17 | 0.29 |
| 6 | 84469645 | rs623263 | -0.046 | -1.725 | -0.121 | 0.443 | -0.012 | 0.053 | 0.053 | 0 | 0.15 | 0.28 | 0.12 | 0.19 | 0.13 | 0.2 |
| 6 | 84479999 | rs1853802 | 1.394 | -2.022 | -0.93 | 0.118 | 0.119 | -0.085 | -0.085 | 1.757 | 0.61 | 0.09 | 0.57 | 0.48 | 0.78 | 0.67 |
| 6 | 84490881 | rs626674 | 0.006 | -2.046 | -1.11 | 0.125 | -0.011 | 0.073 | 0.073 | 0 | 0.19 | 0.27 | 0.33 | 0.47 | 0.16 | 0.25 |
| 6 | 84493078 | rs676282 | -0.689 | -2.097 | -1.093 | 0.078 | -0.02 | 0.056 | 0.056 | 0 | 0.23 | 0.36 | 0.36 | 0.48 | 0.17 | 0.28 |
| 6 | 84501600 | rs592188 | -0.3 | -2 | -1.218 | 0.139 | 0.022 | 0.031 | 0.031 | 0 | 0.9 | 0.96 | 0.97 | 0.96 | 0.96 | 0.95 |
| 6 | 84504749 | rs671367 | 0.1 | -1.84 | -1.279 | 0.215 | 0.009 | -0.054 | -0.054 | 0 | 0.18 | 0.14 | 0.32 | 0.47 | 0.17 | 0.24 |
| 6 | 84520520 | rs1570843 | 0.409 | -1.381 | -1.397 | 0.422 | 0.079 | -0.162 | -0.162 | 1.682 | 0.63 | 0.05 | 0.59 | 0.49 | 0.79 | 0.7 |
| 6 | 84525784 | rs13211072 | 0.204 | -0.297 | -1.417 | 0.162 | -0.121 | 0.057 | 0.057 | 0 | 0.32 | 0.02 | 0.22 | 0.17 | 0.38 | 0.26 |
| 6 | 84533341 | rs9359570 | -1.203 | -0.953 | 0.31 | 0.902 | -0.033 | 0.221 | 0.221 | 0.001 | 0.13 | 0.57 | 0.27 | 0.26 | 0.08 | 0.06 |
| 6 | 84534357 | rs4707016 | 1.425 | -0.964 | -0.09 | 0.526 | -0.151 | 0.001 | 0.001 | 2.919 | 0.33 | 0.03 | 0.26 | 0.24 | 0.4 | 0.36 |
| 6 | 84539678 | rs927737 | -0.077 | -1.21 | 0.821 | 0.474 | -0.01 | 0.022 | 0.022 | 0 | 0.2 | 0.11 | 0.29 | 0.15 | 0.22 | 0.2 |
| 6 | 84553478 | rs9353141 | -1.195 | -1.048 | 1.164 | 0.685 | -0.036 | 0.207 | 0.207 | 0.04 | 0.13 | 0.58 | 0.27 | 0.26 | 0.08 | 0.06 |
| 6 | 84560151 | rs4707017 | -0.877 | -1.048 | 1.163 | 0.686 | 0.003 | 0.377 | 0.377 | 1.074 | 0.09 | 0.52 | 0.25 | 0.26 | 0.04 | 0.05 |

| | | | | | | | | | | | | | | | | |
|---|---|---|---|---|---|---|---|---|---|---|---|---|---|---|---|---|
| 6 | 84568924 | rs2296412 | -0.919 | -1.051 | 1.161 | 0.685 | -0.001 | 0.37 | 0.37 | 1.356 | 0.09 | 0.52 | 0.22 | 0.26 | 0.04 | 0.05 |
| 6 | 84594547 | rs9353149 | -1.668 | -1.047 | 1.3 | 0.264 | -0.043 | 0.152 | 0.152 | 0.124 | 0.14 | 0.66 | 0.25 | 0.24 | 0.08 | 0.05 |
| 6 | 84594649 | rs9341967 | 0.002 | -1.199 | 1.204 | 0.183 | -0.006 | 0.049 | 0.049 | 0 | 0.74 | 0.92 | 0.69 | 0.8 | 0.75 | 0.71 |
| 6 | 84603226 | rs9449718 | -2.153 | -0.977 | 2.6 | 0.998 | -0.034 | 0.239 | 0.239 | 0.096 | 0.13 | 0.55 | 0.24 | 0.25 | 0.08 | 0.05 |
| 6 | 84603422 | rs3757003 | -2.655 | -1.01 | 2.363 | 0.986 | 0.005 | -0.008 | -0.008 | 0 | 0.16 | 0.84 | 0.26 | 0.25 | 0.08 | 0.05 |
| 6 | 84635498 | rs9341969 | -1.704 | -0.961 | 2.242 | 0.973 | -0.028 | 0.274 | 0.274 | 2.881 | 0.1 | 0.58 | 0.23 | 0.24 | 0.04 | 0.07 |
| 6 | 84641238 | rs9344377 | -2.205 | -0.957 | 1.729 | 0.973 | -0.041 | 0.055 | 0.055 | 0.901 | 0.16 | 0.71 | 0.26 | 0.24 | 0.08 | 0.08 |
| 6 | 84660034 | rs3778183 | -1.107 | -0.578 | 0.587 | 1.183 | -0.015 | 0.093 | 0.093 | 0 | 0.08 | 0.14 | 0.2 | 0.24 | 0.04 | 0.13 |
| 6 | 84675796 | rs7774773 | -1.277 | -0.513 | 0.965 | 0.177 | -0.028 | -0.064 | -0.064 | 0 | 0.11 | 0.6 | 0.07 | 0 | 0.05 | 0.01 |
| 6 | 84693623 | rs215973 | -1.179 | -0.577 | 0.078 | 0.045 | -0.032 | 0.049 | 0.049 | 1.538 | 0.5 | 0.85 | 0.45 | 0.57 | 0.48 | 0.37 |
| 6 | 84695944 | rs9359577 | -2.046 | -0.781 | 0.627 | 0.574 | -0.02 | 0.022 | 0.022 | 1.484 | 0.17 | 0.79 | 0.21 | 0.29 | 0.08 | 0.08 |
| 6 | 84729122 | rs1753990 | -0.787 | -0.132 | 1.39 | -0.675 | -0.016 | 0.009 | 0.009 | 1.216 | 0.17 | 0.12 | 0.13 | 0.16 | 0.27 | 0.21 |
| 6 | 84736254 | rs4707031 | 1.501 | -0.205 | 1.284 | -0.751 | -0.03 | 0.051 | 0.051 | 1.028 | 0.75 | 0.14 | 0.8 | 0.84 | 0.72 | 0.7 |
| 6 | 84738434 | rs9341971 | 1.294 | -0.472 | 1.558 | -0.92 | -0.003 | -0.077 | -0.077 | 0 | 0.71 | 0.47 | 0.84 | 0.85 | 0.71 | 0.62 |
| 6 | 84740396 | rs2480181 | -0.756 | -0.497 | 1.272 | -0.994 | -0.031 | -0.039 | -0.039 | 0 | 0.09 | 0.21 | 0.04 | 0.01 | 0.03 | 0.09 |
| 6 | 84759647 | rs9362026 | 0.066 | -0.467 | 0.788 | -1.043 | -0.079 | 0.037 | 0.037 | 0.036 | 0.84 | 0.99 | 0.77 | 0.9 | 0.76 | 0.69 |
| 6 | 84762822 | rs2497108 | -1.681 | -1.029 | 1.889 | -0.906 | -0.024 | -0.027 | -0.027 | 0 | 0.37 | 0.56 | 0.21 | 0.17 | 0.23 | 0.32 |
| 6 | 84765609 | rs9449760 | -1.763 | -1.04 | 1.939 | -0.911 | 0.007 | 0.007 | 0.007 | 0 | 0.38 | 0.62 | 0.22 | 0.16 | 0.26 | 0.32 |
| 6 | 84767124 | rs17729906 | -0.732 | -1.125 | 2.01 | -0.949 | -0.011 | 0.014 | 0.014 | 0 | 0.23 | 0.06 | 0.15 | 0.19 | 0.21 | 0.21 |
| 6 | 84767152 | rs2497113 | -1.74 | -1.057 | 1.973 | -0.937 | 0.003 | 0.001 | 0.001 | 0 | 0.38 | 0.6 | 0.22 | 0.19 | 0.26 | 0.32 |
| 6 | 84769478 | rs2497115 | -1.831 | -1.068 | 1.75 | -0.991 | 0.07 | 0.098 | 0.098 | 0.019 | 0.41 | 0.79 | 0.23 | 0.18 | 0.27 | 0.33 |
| 6 | 84769939 | rs2480191 | -1.777 | -0.767 | 1.92 | -0.838 | 0.007 | 0.007 | 0.007 | 0 | 0.38 | 0.66 | 0.22 | 0.17 | 0.27 | 0.33 |
| 6 | 84776051 | rs1556111 | -0.827 | -0.773 | 1.779 | -0.889 | 0.004 | 0.007 | 0.007 | 0 | 0.2 | 0.17 | 0.13 | 0.19 | 0.17 | 0.12 |
| 6 | 84790544 | rs2497139 | -1.058 | -0.686 | 1.931 | -0.862 | -0.014 | -0.039 | -0.039 | 0 | 0.1 | 0.2 | 0.11 | 0 | 0.06 | 0.08 |
| 6 | 84796363 | rs2480196 | -1.525 | -0.708 | 2.285 | -0.877 | -0.031 | -0.074 | -0.074 | 0 | 0.13 | 0.48 | 0.13 | 0.01 | 0.07 | 0.09 |
| 6 | 84796475 | rs6928551 | -0.597 | -0.809 | 2.253 | -0.891 | 0 | 0.001 | 0.001 | 0 | 0.16 | 0.13 | 0.12 | 0.16 | 0.16 | 0.13 |
| 6 | 84799059 | rs9449776 | -1.876 | -0.859 | 2.359 | -0.917 | -0.055 | -0.095 | -0.095 | 1.501 | 0.16 | 0.58 | 0.15 | 0.01 | 0.07 | 0.09 |
| 6 | 84799185 | rs2875382 | -2.114 | -0.86 | 2.358 | -0.917 | 0.004 | 0.003 | 0.003 | 1.168 | 0.32 | 0.71 | 0.27 | 0.17 | 0.23 | 0.22 |
| 6 | 84799673 | rs2497129 | -1.885 | -0.857 | 2.37 | -0.903 | -0.055 | -0.095 | -0.095 | 0 | 0.16 | 0.58 | 0.15 | 0.01 | 0.07 | 0.09 |
| 6 | 84800063 | rs2497128 | -1.885 | -0.857 | 2.37 | -0.903 | -0.055 | -0.095 | -0.095 | 0 | 0.16 | 0.58 | 0.15 | 0.01 | 0.07 | 0.09 |
| 6 | 84800166 | rs1045649 | -1.885 | -0.857 | 2.37 | -0.903 | -0.055 | -0.095 | -0.095 | 0.059 | 0.16 | 0.58 | 0.15 | 0.01 | 0.07 | 0.09 |
| 6 | 84800378 | rs3219 | -1.513 | -0.857 | 2.39 | -0.903 | -0.033 | -0.08 | -0.08 | 0 | 0.13 | 0.51 | 0.14 | 0.01 | 0.07 | 0.09 |
| 6 | 84800492 | rs8348 | -1.788 | -0.851 | 2.044 | -0.9 | -0.056 | -0.096 | -0.096 | 0 | 0.16 | 0.58 | 0.15 | 0.01 | 0.07 | 0.09 |
| 6 | 84802783 | rs2497123 | -2.097 | -0.775 | 1.889 | -0.972 | 0.011 | 0.015 | 0.015 | 0.706 | 0.35 | 0.71 | 0.29 | 0.21 | 0.26 | 0.24 |
| 6 | 84829622 | rs7741955 | -1 | -0.739 | 2.01 | -0.588 | -0.021 | -0.051 | -0.051 | 0.236 | 0.1 | 0.25 | 0.11 | 0.01 | 0.05 | 0.08 |
| 6 | 84851397 | rs1886254 | -0.469 | -0.64 | 1.529 | -0.577 | -0.035 | -0.038 | -0.038 | 0.452 | 0.13 | 0.02 | 0.12 | 0.19 | 0.15 | 0.11 |
| 6 | 84853623 | rs9449802 | -1.768 | -0.7 | 1.745 | -0.525 | -0.026 | -0.043 | -0.043 | 2.011 | 0.16 | 0.74 | 0.16 | 0.01 | 0.07 | 0.09 |
| 6 | 84908967 | rs9283803 | -1.648 | -0.595 | 0.628 | -0.415 | -0.03 | -0.039 | -0.039 | 0.339 | 0.15 | 0.74 | 0.16 | 0.02 | 0.06 | 0.08 |
| 6 | 84911336 | rs2274092 | -1.661 | -0.153 | 0.312 | -0.479 | 0.018 | 0.103 | 0.103 | 1.373 | 0.41 | 0.84 | 0.34 | 0.23 | 0.34 | 0.4 |
| 6 | 84926060 | rs9449841 | -1.473 | -0.235 | 1.052 | 0.018 | -0.033 | -0.055 | -0.055 | 0.3 | 0.12 | 0.55 | 0.14 | 0.02 | 0.04 | 0.07 |
| 6 | 84935635 | rs4707035 | 1.349 | 0.352 | 0.25 | 0.221 | -0.013 | 0.013 | 0.013 | 0.278 | 0.76 | 0.74 | 0.81 | 0.84 | 0.74 | 0.81 |
| 6 | 84935694 | rs9449850 | -1.762 | -0.089 | 0.997 | 0.098 | -0.021 | -0.037 | -0.037 | 0.163 | 0.25 | 0.63 | 0.19 | 0.11 | 0.17 | 0.17 |
| 6 | 84943627 | rs6922633 | -2.431 | 0.343 | 1.545 | 1.175 | 0.008 | 0.025 | 0.025 | 0.021 | 0.29 | 0.47 | 0.4 | 0.56 | 0.34 | 0.26 |
| 6 | 84951849 | rs735641 | 1.344 | 0.365 | 1.425 | 1.217 | -0.004 | 0.036 | 0.036 | 0 | 0.9 | 0.9 | 0.71 | 0.53 | 0.78 | 0.89 |
| 6 | 84959435 | rs7739877 | 2.176 | 0.44 | 1.619 | 1.202 | 0.002 | 0.004 | 0.004 | 0 | 0.74 | 0.74 | 0.6 | 0.44 | 0.66 | 0.74 |
| 6 | 84965282 | rs1324089 | 1.84 | 0.454 | 1.572 | 1.69 | 0 | -0.005 | -0.005 | 0.051 | 0.78 | 0.79 | 0.78 | 0.6 | 0.75 | 0.78 |
| 6 | 84971363 | rs6910017 | 2.374 | 0.457 | 1.535 | 1.687 | 0.01 | 0.024 | 0.024 | 0.058 | 0.72 | 0.49 | 0.58 | 0.44 | 0.66 | 0.74 |
| 6 | 84972805 | rs1408274 | -0.719 | 0.635 | 1.104 | 1.574 | 0.007 | 0.128 | 0.128 | 0 | 0.06 | 0.1 | 0.11 | 0.31 | 0.13 | 0.07 |
| 6 | 84982208 | rs12174666 | -1.787 | 0.391 | 0.903 | 1.345 | 0.002 | 0.04 | 0.04 | 0 | 0.23 | 0.27 | 0.23 | 0.39 | 0.27 | 0.15 |
| 6 | 84985239 | rs13214117 | -2.054 | 0.32 | 0.914 | 1.252 | 0.004 | 0.029 | 0.029 | 0.839 | 0.31 | 0.53 | 0.41 | 0.45 | 0.35 | 0.19 |
| 6 | 84997747 | rs2324570 | -2.007 | 0.431 | 0.65 | 1.272 | 0.012 | 0.035 | 0.035 | 0.647 | 0.31 | 0.51 | 0.42 | 0.52 | 0.36 | 0.21 |
| 6 | 85012209 | rs13200720 | -1.009 | 0.015 | 0.521 | 0.707 | -0.024 | 0.006 | 0.006 | 0 | 0.14 | 0.03 | 0.13 | 0.09 | 0.13 | 0.08 |
| 6 | 85014975 | rs7767211 | -0.951 | 0.353 | -0.398 | 0.886 | -0.018 | 0.013 | 0.013 | 0 | 0.13 | 0.02 | 0.12 | 0.08 | 0.11 | 0.07 |
| 6 | 85034419 | rs9449866 | 0.359 | 0.274 | -0.53 | 0.605 | -0.002 | 0.001 | 0.001 | 0 | 0.44 | 0.39 | 0.6 | 0.49 | 0.51 | 0.59 |
| 6 | 85034682 | rs2223793 | -0.901 | -0.159 | -0.108 | 0.367 | 0.008 | 0.038 | 0.038 | 0.38 | 0.11 | 0.52 | 0.13 | 0.11 | 0.09 | 0.12 |
| 6 | 85034839 | rs12190678 | -0.884 | -0.384 | -0.12 | 0.239 | -0.002 | -0.002 | -0.002 | 0.009 | 0.15 | 0.18 | 0.13 | 0.12 | 0.13 | 0.13 |
| 6 | 85036285 | rs2206984 | 1.217 | -1.168 | -0.098 | 0.106 | -0.004 | -0.001 | -0.001 | 0.173 | 0.85 | 0.85 | 0.9 | 0.84 | 0.9 | 0.93 |
| 6 | 85038293 | rs9444158 | -0.702 | -1.132 | -0.223 | -0.187 | -0.003 | 0.002 | 0.002 | 0 | 0.15 | 0.14 | 0.27 | 0.12 | 0.21 | 0.17 |
| 6 | 85038395 | rs9350994 | 0.206 | -0.976 | -0.282 | -0.222 | 0.001 | 0.015 | 0.015 | 0 | 0.51 | 0.42 | 0.62 | 0.6 | 0.49 | 0.5 |
| 6 | 85039069 | rs6907261 | 0.777 | -1.153 | 0.282 | -0.266 | -0.028 | -0.005 | -0.005 | 0 | 0.85 | 0.74 | 0.9 | 0.88 | 0.89 | 0.95 |
| 6 | 85043476 | rs1535060 | -0.675 | -1.273 | 0.053 | -0.299 | 0.006 | 0.01 | 0.01 | 0 | 0.18 | 0.29 | 0.31 | 0.22 | 0.21 | 0.28 |
| 6 | 85043919 | rs9359585 | 1.325 | -1.531 | 0.001 | -0.512 | 0.023 | -0.047 | -0.047 | 0 | 0.3 | 0.21 | 0.32 | 0.49 | 0.22 | 0.17 |
| 6 | 85052572 | rs13191052 | -0.569 | -1.306 | 0.065 | -0.466 | 0.036 | 0.07 | 0.07 | 0 | 0.07 | 0.28 | 0.15 | 0.11 | 0.08 | 0.07 |
| 6 | 85054553 | rs12663831 | -0.383 | -1.312 | 0.161 | -0.469 | 0.04 | 0.075 | 0.075 | 0 | 0.06 | 0.27 | 0.14 | 0.11 | 0.07 | 0.07 |
| 6 | 85054710 | rs1883897 | 0.562 | -1.488 | 0.119 | -0.943 | 0.046 | 0.041 | 0.041 | 0.006 | 0.49 | 0.25 | 0.52 | 0.61 | 0.4 | 0.47 |
| 6 | 85055653 | rs12208987 | 1.583 | -1.228 | -0.017 | -1.376 | -0.085 | -0.023 | -0.023 | 0 | 0.2 | 0.04 | 0.16 | 0.2 | 0.32 | 0.36 |
| 6 | 85056975 | rs10455426 | 1.536 | -1.252 | 0.199 | -1.385 | -0.032 | -0.002 | -0.002 | 0.208 | 0.21 | 0.1 | 0.17 | 0.2 | 0.32 | 0.36 |
| 6 | 85059005 | rs7775532 | 0.085 | -1.315 | 0.485 | -1.465 | 0.002 | -0.004 | -0.004 | 0.037 | 0.61 | 0.66 | 0.71 | 0.72 | 0.53 | 0.58 |
| 6 | 85059296 | rs7758189 | 1.063 | -1.267 | 0.518 | -1.481 | 0.009 | -0.033 | -0.033 | 0 | 0.06 | 0.12 | 0.05 | 0 | 0.05 | 0.03 |

| | | | | | | | | | | | | | | | | |
|---|---|---|---|---|---|---|---|---|---|---|---|---|---|---|---|---|
| 6 | 85062826 | rs6928921 | 0.783 | -1.3 | 0.534 | -1.499 | -0.023 | -0.076 | -0.076 | 0.229 | 0.76 | 0.63 | 0.82 | 0.91 | 0.79 | 0.91 |
| 6 | 85065248 | rs6903593 | 0.192 | -1.541 | 0.396 | -1.721 | -0.03 | 0.035 | 0.035 | 0.311 | 0.41 | 0.18 | 0.37 | 0.32 | 0.48 | 0.66 |
| 6 | 85066581 | rs761409 | 0.065 | -1.534 | 0.431 | -1.708 | -0.022 | -0.034 | -0.034 | 0 | 0.83 | 0.75 | 0.87 | 0.91 | 0.84 | 0.94 |
| 6 | 85066620 | rs6454364 | 1.966 | -1.478 | 0.236 | -1.67 | -0.006 | 0.002 | 0.002 | 0 | 0.36 | 0.45 | 0.46 | 0.6 | 0.31 | 0.25 |
| 6 | 85071341 | rs7745556 | 1.638 | -1.428 | 0.174 | -1.664 | -0.032 | -0.002 | -0.002 | 0.221 | 0.21 | 0.1 | 0.17 | 0.2 | 0.32 | 0.36 |
| 6 | 85080622 | rs6901000 | 0.486 | -1.467 | 0.132 | -1.497 | -0.002 | 0.035 | 0.035 | 0 | 0.19 | 0.06 | 0.19 | 0.12 | 0.17 | 0.3 |
| 6 | 85082479 | rs2206989 | 2.3 | -1.472 | 0.023 | -1.449 | 0.027 | -0.044 | -0.044 | 0 | 0.29 | 0.17 | 0.39 | 0.58 | 0.24 | 0.18 |
| 6 | 85084256 | rs7760461 | -0.592 | -1.454 | 0.142 | -1.48 | 0 | -0.003 | -0.003 | 2.381 | 0.65 | 0.68 | 0.74 | 0.8 | 0.59 | 0.61 |
| 6 | 85085336 | rs11756826 | -0.131 | -1.287 | -0.331 | -1.046 | -0.006 | 0.001 | 0.001 | 0 | 0.1 | 0.15 | 0.09 | 0.1 | 0.12 | 0.06 |
| 6 | 85089473 | rs1080140 | 2.256 | -1.367 | -0.861 | -1.087 | 0.027 | -0.049 | -0.049 | 0.895 | 0.25 | 0.17 | 0.37 | 0.57 | 0.19 | 0.17 |
| 6 | 85089511 | rs10455146 | -0.496 | -0.305 | -0.523 | -0.962 | 0.001 | -0.004 | -0.004 | 0 | 0.12 | 0.13 | 0.11 | 0.1 | 0.16 | 0.06 |
| 6 | 85091805 | rs2324578 | 0.649 | -0.279 | -0.515 | -0.94 | -0.001 | -0.004 | -0.004 | 0 | 0.82 | 0.84 | 0.87 | 0.9 | 0.8 | 0.93 |
| 6 | 85101180 | rs6454367 | 1.056 | -0.351 | -0.897 | -1.044 | 0.006 | 0.023 | 0.023 | 0.11 | 0.7 | 0.88 | 0.7 | 0.74 | 0.72 | 0.63 |
| 6 | 85103530 | rs11759310 | 0.33 | -0.175 | -0.883 | -0.986 | -0.041 | -0.001 | -0.001 | 0.324 | 0.15 | 0.03 | 0.18 | 0.12 | 0.16 | 0.28 |
| 6 | 85105960 | rs9359587 | 1.564 | -0.751 | -0.6 | -1.244 | 0.003 | -0.004 | -0.004 | 0.505 | 0.3 | 0.3 | 0.37 | 0.59 | 0.22 | 0.16 |
| 6 | 85107756 | rs9362046 | 0.026 | -0.293 | -0.245 | -1.474 | 0 | -0.004 | -0.004 | 0 | 0.53 | 0.49 | 0.62 | 0.71 | 0.45 | 0.47 |
| 6 | 85107882 | rs10943979 | 0.965 | -0.281 | 0.067 | -1.605 | 0.002 | -0.008 | -0.008 | 0 | 0.31 | 0.26 | 0.23 | 0.15 | 0.37 | 0.4 |
| 6 | 85109553 | rs9449891 | 0.093 | -0.271 | 0.136 | -1.552 | 0.008 | -0.019 | -0.019 | 0 | 0.06 | 0.17 | 0.06 | 0.01 | 0.07 | 0.08 |
| 6 | 85109688 | rs2206985 | 0.827 | -0.305 | 0.04 | -1.632 | 0.008 | 0.017 | 0.017 | 0 | 0.5 | 0.47 | 0.6 | 0.71 | 0.4 | 0.46 |
| 6 | 85110264 | rs6899777 | 0.112 | 0.191 | 0.238 | -1.336 | -0.001 | 0.002 | 0.002 | 0.346 | 0.87 | 0.9 | 0.89 | 0.88 | 0.85 | 0.95 |
| 6 | 85110283 | rs9341975 | 0.82 | 0.21 | 0.303 | -1.314 | 0 | -0.008 | -0.008 | 0 | 0.83 | 0.75 | 0.83 | 0.87 | 0.78 | 0.86 |
| 6 | 85111127 | rs2023558 | -1.207 | 0.206 | 0.579 | -1.481 | 0.014 | -0.065 | -0.065 | 1.015 | 0.21 | 0.39 | 0.29 | 0.08 | 0.21 | 0.3 |
| 6 | 85111931 | rs12206828 | -1.142 | 0.446 | 1.041 | -1.39 | 0.01 | -0.018 | -0.018 | 1.903 | 0.19 | 0.47 | 0.19 | 0.16 | 0.22 | 0.23 |
| 6 | 85112992 | rs13437417 | -0.443 | 0.531 | 0.834 | -1.34 | 0.009 | 0.006 | 0.006 | 0 | 0.15 | 0.3 | 0.17 | 0.16 | 0.2 | 0.22 |
| 6 | 85113293 | rs9449894 | -0.687 | 0.53 | 0.836 | -1.338 | 0.012 | -0.003 | -0.003 | 0 | 0.17 | 0.3 | 0.18 | 0.16 | 0.22 | 0.17 |
| 6 | 85114269 | rs12197418 | -0.206 | 0.519 | 0.87 | -1.324 | 0.001 | 0.013 | 0.013 | 0 | 0.13 | 0.24 | 0.13 | 0.16 | 0.17 | 0.15 |
| 6 | 85116484 | rs9362047 | 0.627 | 1.084 | 0.197 | -1.889 | 0 | -0.01 | -0.01 | 3.465 | 0.58 | 0.43 | 0.68 | 0.78 | 0.54 | 0.5 |
| 6 | 85118284 | rs7763862 | -1.042 | 1.347 | 0.33 | -1.777 | -0.001 | -0.002 | -0.002 | 0 | 0.33 | 0.34 | 0.4 | 0.24 | 0.37 | 0.44 |
| 6 | 85119504 | rs9444174 | -0.108 | 1.703 | 0.233 | -1.762 | 0.051 | 0.007 | 0.007 | 0.529 | 0.09 | 0.32 | 0.06 | 0.05 | 0.11 | 0.22 |
| 6 | 85119727 | rs9362048 | 0.185 | 1.435 | -0.434 | -1.689 | -0.04 | 0.037 | 0.037 | 0.224 | 0.17 | 0.02 | 0.23 | 0.09 | 0.17 | 0.23 |
| 6 | 85121532 | rs4706203 | 0.928 | 1.506 | -0.307 | -1.553 | 0.029 | 0.066 | 0.066 | 0.731 | 0.44 | 0.18 | 0.44 | 0.7 | 0.37 | 0.27 |
| 6 | 85122601 | rs17578634 | -0.452 | 1.223 | -0.01 | -1.575 | 0.005 | -0.014 | -0.014 | 0 | 0.3 | 0.32 | 0.37 | 0.18 | 0.38 | 0.59 |
| 6 | 85125094 | rs10943983 | 0.122 | 0.837 | 0.06 | -1.396 | 0 | -0.013 | -0.013 | 0 | 0.07 | 0.07 | 0.05 | 0 | 0.05 | 0.04 |
| 6 | 85127929 | rs16874544 | -0.601 | 0.961 | 0.432 | -1.441 | 0.102 | -0.016 | -0.016 | 0.048 | 0.09 | 0.3 | 0.08 | 0.03 | 0.14 | 0.21 |
| 6 | 85130409 | rs6937168 | -0.498 | 1.078 | 0.051 | -1.648 | -0.046 | 0.061 | 0.061 | 0.019 | 0.74 | 0.86 | 0.81 | 0.91 | 0.67 | 0.86 |
| 6 | 85135586 | rs713170 | 0.428 | 0.68 | -0.02 | -1.505 | 0 | -0.016 | -0.016 | 0.659 | 0.58 | 0.44 | 0.69 | 0.88 | 0.47 | 0.62 |
| 6 | 85140399 | rs3734451 | 0.133 | 0.612 | -0.214 | -1.205 | -0.001 | -0.006 | -0.006 | 0 | 0.28 | 0.25 | 0.21 | 0.13 | 0.26 | 0.35 |
| 6 | 85141439 | rs6906061 | -0.945 | 0.707 | -0.313 | -1.309 | 0.033 | 0.019 | 0.019 | 0.018 | 0.6 | 0.76 | 0.55 | 0.33 | 0.65 | 0.76 |
| 6 | 85147411 | rs9294298 | -0.439 | 0.985 | -0.064 | -1.3 | -0.004 | 0.009 | 0.009 | 0 | 0.37 | 0.67 | 0.42 | 0.24 | 0.41 | 0.61 |
| 6 | 85163946 | rs12215178 | 0.783 | 0.895 | -0.052 | -1.264 | 0 | 0 | 0 | 0.162 | 0.22 | 0.19 | 0.3 | 0.22 | 0.23 | 0.33 |
| 6 | 85165330 | rs9449923 | -0.723 | 0.871 | -0.348 | -1.392 | 0.055 | 0.052 | 0.052 | 0.444 | 0.53 | 0.78 | 0.55 | 0.33 | 0.63 | 0.72 |
| 6 | 85171535 | rs4707051 | 2.123 | 0.545 | -0.752 | -1.11 | 0.004 | -0.176 | -0.176 | 0 | 0.25 | 0.01 | 0.28 | 0.56 | 0.19 | 0.2 |
| 6 | 85173502 | rs4618489 | -0.953 | 1.266 | -0.251 | -1.199 | -0.015 | 0.02 | 0.02 | 0.538 | 0.79 | 0.87 | 0.86 | 0.91 | 0.75 | 0.86 |
| 6 | 85178268 | rs9362054 | 0.071 | 1.423 | -0.358 | -1.116 | -0.002 | 0.005 | 0.005 | 0 | 0.54 | 0.47 | 0.53 | 0.69 | 0.51 | 0.54 |
| 6 | 85181754 | rs9353186 | 0.662 | 1.479 | -0.957 | -1.061 | 0.06 | 0.057 | 0.057 | 0.474 | 0.47 | 0.17 | 0.44 | 0.67 | 0.36 | 0.29 |
| 6 | 85194010 | rs11756328 | -0.246 | 2.085 | -0.529 | -0.392 | 0.042 | -0.01 | -0.01 | 0.122 | 0.06 | 0.15 | 0.07 | 0.02 | 0.14 | 0.22 |
| 6 | 85200684 | rs10214636 | -0.394 | 2.085 | -0.524 | -0.393 | 0.053 | -0.012 | -0.012 | 0 | 0.06 | 0.18 | 0.07 | 0.02 | 0.14 | 0.22 |
| 6 | 85202005 | rs10498941 | -0.164 | 1.037 | -0.725 | -0.242 | -0.001 | -0.001 | -0.001 | 0 | 0.22 | 0.21 | 0.31 | 0.22 | 0.23 | 0.31 |
| 6 | 85205713 | rs10498942 | -0.86 | 1.225 | -0.832 | -0.037 | -0.007 | 0.017 | 0.017 | 0 | 0.12 | 0.07 | 0.09 | 0 | 0.14 | 0.05 |
| 6 | 85211340 | rs7741380 | -0.662 | 1.404 | -1.052 | 0.738 | -0.047 | -0.082 | -0.082 | 0.461 | 0.28 | 0.15 | 0.22 | 0.45 | 0.33 | 0.49 |
| 6 | 85211681 | rs13193467 | -0.531 | 1.393 | -1.379 | 0.323 | -0.016 | 0.035 | 0.035 | 0 | 0.14 | 0.06 | 0.16 | 0.04 | 0.17 | 0.09 |
| 6 | 85216471 | rs6933537 | 1.518 | 0.848 | -1.346 | 0.227 | 0.05 | 0.043 | 0.043 | 0 | 0.36 | 0.17 | 0.22 | 0.28 | 0.26 | 0.31 |
| 6 | 85225179 | rs9344403 | -0.318 | 0.839 | 0.084 | 0.265 | 0.008 | 0.113 | 0.113 | 0 | 0.08 | 0.19 | 0.27 | 0.23 | 0.06 | 0.11 |
| 6 | 85225727 | rs9449951 | -0.64 | 0.836 | 0.481 | 0.05 | 0.008 | -0.065 | -0.065 | 0 | 0.26 | 0.38 | 0.21 | 0.04 | 0.33 | 0.1 |
| 6 | 85228896 | rs6936385 | -0.685 | 0.446 | 0.673 | 0.483 | -0.012 | 0.005 | 0.005 | 0 | 0.16 | 0.23 | 0.11 | 0.2 | 0.12 | 0.18 |
| 6 | 85232898 | rs4510639 | 0.603 | 0.724 | 0.25 | 0.537 | -0.11 | -0.104 | -0.104 | 0 | 0.21 | 0.02 | 0.18 | 0.3 | 0.3 | 0.36 |
| 6 | 85237857 | rs9344405 | 1.156 | -0.002 | 0.535 | 0.46 | -0.002 | -0.011 | -0.011 | 0.113 | 0.66 | 0.33 | 0.72 | 0.8 | 0.64 | 0.77 |
| 6 | 85246086 | rs12526313 | 1.476 | 0.062 | 0.181 | 0.477 | 0.007 | -0.013 | -0.013 | 0 | 0.57 | 0.11 | 0.45 | 0.58 | 0.58 | 0.66 |
| 6 | 85246447 | rs13214308 | 0.211 | 0.108 | 0.385 | 0.333 | 0.005 | -0.044 | -0.044 | 0.368 | 0.16 | 0.21 | 0.14 | 0 | 0.24 | 0.05 |
| 6 | 85266769 | rs9294306 | 0.748 | 0.145 | 0.138 | 0.445 | -0.01 | -0.036 | -0.036 | 0 | 0.67 | 0.39 | 0.77 | 0.84 | 0.7 | 0.79 |
| 6 | 85266965 | rs17792917 | -0.018 | 1.018 | -0.294 | 0.92 | -0.048 | -0.068 | -0.068 | 0 | 0.19 | 0.05 | 0.16 | 0.29 | 0.23 | 0.34 |
| 6 | 85286434 | rs16874800 | -0.009 | 0.971 | -0.3 | 0.918 | -0.041 | -0.088 | -0.088 | 2.194 | 0.2 | 0.06 | 0.24 | 0.37 | 0.24 | 0.39 |
| 6 | 85298904 | rs13215124 | -0.354 | 0.869 | 0.75 | 0.482 | -0.002 | -0.019 | -0.019 | 0 | 0.18 | 0.15 | 0.13 | 0.01 | 0.24 | 0.11 |
| 6 | 85328621 | rs10498943 | 1.758 | 0.979 | 0.575 | 0.12 | -0.029 | 0.264 | 0.264 | 0.956 | 0.41 | 0.07 | 0.25 | 0.09 | 0.37 | 0.35 |
| 6 | 85342772 | rs4707082 | 1.323 | 0.918 | 1.066 | 0.226 | 0.111 | 0.248 | 0.248 | 2.38 | 0.93 | 0.66 | 0.73 | 0.77 | 0.9 | 0.85 |
| 6 | 85343652 | rs7770869 | 1.429 | 0.971 | 1.355 | 0.227 | -0.001 | 0.213 | 0.213 | 0.019 | 0.48 | 0.21 | 0.32 | 0.11 | 0.47 | 0.39 |
| 6 | 85347257 | rs13217066 | -0.731 | 0.968 | 1.63 | 0.235 | -0.001 | 0.005 | 0.005 | 0 | 0.4 | 0.34 | 0.36 | 0.66 | 0.4 | 0.5 |
| 6 | 85354880 | rs4349787 | -1.341 | 0.889 | 1.622 | 0.062 | 0.008 | 0.003 | 0.003 | 0 | 0.51 | 0.61 | 0.71 | 0.88 | 0.55 | 0.63 |
| 6 | 85367282 | rs1034027 | -0.477 | 1.007 | 1.783 | 0.49 | 0.001 | -0.057 | -0.057 | 0 | 0.79 | 0.89 | 0.8 | 0.62 | 0.84 | 0.64 |

| | | | | | | | | | | | | | | | | |
|---|---|---|---|---|---|---|---|---|---|---|---|---|---|---|---|---|
| 6 | 85370513 | rs7772438 | -0.572 | 1.111 | 2.135 | 0.313 | -0.009 | -0.062 | -0.062 | 0 | 0.12 | 0.23 | 0.06 | 0.01 | 0.11 | 0.12 |
| 6 | 85371011 | rs9449999 | 1.242 | 1.081 | 2.114 | 0.295 | -0.001 | 0.001 | 0.001 | 0 | 0.6 | 0.4 | 0.41 | 0.38 | 0.62 | 0.41 |
| 6 | 85371398 | rs7759428 | 0.941 | 1.061 | 2.077 | 0.301 | -0.015 | 0.019 | 0.019 | 0 | 0.65 | 0.54 | 0.72 | 0.61 | 0.73 | 0.52 |
| 6 | 85376381 | rs9450002 | 0.979 | 1.078 | 1.995 | 0.171 | 0.006 | -0.076 | -0.076 | 0 | 0.58 | 0.47 | 0.33 | 0.13 | 0.53 | 0.47 |
| 6 | 85379474 | rs13211475 | 0.009 | 0.974 | 1.959 | 0.228 | 0.023 | 0.086 | 0.086 | 0 | 0.13 | 0.33 | 0.14 | 0.26 | 0.17 | 0.04 |
| 6 | 85381928 | rs6454393 | -0.935 | 0.97 | 1.912 | 0.073 | -0.012 | -0.086 | -0.086 | 0 | 0.14 | 0.34 | 0.07 | 0.01 | 0.11 | 0.12 |
| 6 | 85384335 | rs1327756 | 0.654 | 1.008 | 1.71 | 0.179 | -0.002 | -0.009 | -0.009 | 0.386 | 0.65 | 0.55 | 0.65 | 0.36 | 0.67 | 0.61 |
| 6 | 85386138 | rs1887505 | 0.421 | 0.854 | 1.439 | 0.093 | 0.003 | 0.012 | 0.012 | 0 | 0.65 | 0.8 | 0.67 | 0.36 | 0.68 | 0.59 |
| 6 | 85394092 | rs10498945 | 2.05 | 0.968 | 1.252 | -0.089 | -0.017 | 0.234 | 0.234 | 0.374 | 0.38 | 0.07 | 0.2 | 0.09 | 0.34 | 0.32 |
| 6 | 85396512 | rs2025866 | 1.822 | 1.014 | 1.421 | -0.091 | -0.042 | 0.076 | 0.076 | 0.058 | 0.42 | 0.16 | 0.51 | 0.33 | 0.45 | 0.43 |
| 6 | 85397988 | rs10806281 | 2.078 | 0.958 | 1.347 | -0.137 | -0.023 | 0.24 | 0.24 | 0.489 | 0.38 | 0.06 | 0.19 | 0.09 | 0.34 | 0.32 |
| 6 | 85404156 | rs13212866 | 0.528 | 0.932 | 1.393 | 0.056 | -0.01 | -0.019 | -0.019 | 0 | 0.11 | 0.04 | 0.11 | 0.25 | 0.16 | 0.03 |
| 6 | 85407811 | rs9450007 | -1.342 | 0.892 | 1.691 | -0.001 | -0.02 | -0.089 | -0.089 | 0.032 | 0.18 | 0.66 | 0.15 | 0.03 | 0.15 | 0.06 |
| 6 | 85417046 | rs13215600 | 0.53 | 0.899 | 1.668 | 0.007 | -0.011 | -0.022 | -0.022 | 0 | 0.11 | 0.04 | 0.11 | 0.25 | 0.16 | 0.03 |
| 6 | 85418565 | rs2182799 | 0.516 | 0.895 | 1.686 | 0.004 | -0.006 | 0.011 | 0.011 | 0 | 0.66 | 0.88 | 0.68 | 0.36 | 0.68 | 0.58 |
| 6 | 85429204 | rs9362096 | 1.659 | 0.907 | 1.673 | 0.011 | 0.001 | 0.091 | 0.091 | 0 | 0.47 | 0.2 | 0.53 | 0.33 | 0.53 | 0.51 |
| 6 | 85434491 | rs9362097 | 1.659 | 0.907 | 1.673 | 0.011 | 0.001 | 0.091 | 0.091 | 0 | 0.47 | 0.2 | 0.53 | 0.33 | 0.53 | 0.51 |
| 6 | 85435275 | rs9294310 | 0.182 | 0.927 | 1.821 | 0.025 | -0.005 | -0.03 | -0.03 | 0 | 0.06 | 0.15 | 0.03 | 0 | 0.04 | 0.02 |
| 6 | 85444089 | rs1738363 | -0.61 | 0.962 | 1.483 | 0.056 | -0.01 | -0.019 | -0.019 | 0.306 | 0.89 | 0.96 | 0.89 | 0.75 | 0.84 | 0.97 |
| 6 | 85453782 | rs2015519 | 1.036 | 0.863 | 1.476 | 0.022 | -0.002 | -0.004 | -0.004 | 0 | 0.56 | 0.43 | 0.62 | 0.36 | 0.64 | 0.55 |
| 6 | 85461800 | rs6936473 | -2.146 | 0.833 | 1.49 | 0.034 | 0.001 | 0.001 | 0.001 | 1.032 | 0.24 | 0.78 | 0.47 | 0.26 | 0.28 | 0.19 |
| 6 | 85462640 | rs860844 | -0.345 | 0.88 | 1.487 | 0.195 | 0.004 | 0.035 | 0.035 | 0 | 0.88 | 0.83 | 0.87 | 0.73 | 0.85 | 0.97 |
| 6 | 85466286 | rs215939 | 0.639 | 0.881 | 1.526 | 0.145 | 0.054 | 0.166 | 0.166 | 0 | 0.82 | 0.64 | 0.54 | 0.51 | 0.74 | 0.85 |
| 6 | 85471464 | rs6942044 | 0.109 | 0.849 | 1.253 | 0.127 | 0.001 | -0.008 | -0.008 | 0.006 | 0.08 | 0.1 | 0.07 | 0.04 | 0.11 | 0.04 |
| 6 | 85487246 | rs215934 | 0.392 | 0.743 | 1.032 | 0.048 | -0.013 | -0.111 | -0.111 | 0 | 0.19 | 0.03 | 0.18 | 0.38 | 0.14 | 0.39 |
| 6 | 85489234 | rs141127 | 0.085 | 0.849 | 1.422 | 0.242 | 0.001 | -0.059 | -0.059 | 0.988 | 0.86 | 0.77 | 0.88 | 1 | 0.86 | 0.98 |
| 6 | 85489520 | rs16875134 | 0.524 | 0.901 | 1.045 | 0.208 | 0.013 | 0 | 0 | 0 | 0.05 | 0.1 | 0.06 | 0.04 | 0.08 | 0.03 |
| 6 | 85516100 | rs16875178 | 0.524 | 0.901 | 1.045 | 0.208 | 0.013 | 0 | 0 | 0 | 0.05 | 0.1 | 0.06 | 0.04 | 0.08 | 0.04 |
| 6 | 85516598 | rs215921 | -0.483 | 0.877 | 1.008 | 0.187 | -0.008 | 0.017 | 0.017 | 0 | 0.89 | 0.97 | 0.91 | 1 | 0.86 | 0.98 |
| 6 | 85518398 | rs9450039 | -0.902 | 0.768 | 1.033 | 0.139 | -0.005 | -0.042 | -0.042 | 0 | 0.13 | 0.17 | 0.06 | 0 | 0.14 | 0.1 |
| 6 | 85520268 | rs9450041 | -0.184 | 0.756 | 0.878 | 0.159 | -0.003 | -0.018 | -0.018 | 0.113 | 0.06 | 0.1 | 0.03 | 0 | 0.06 | 0.02 |
| 6 | 85521035 | rs215922 | 2.007 | 0.752 | 0.706 | 0.201 | -0.09 | -0.089 | -0.089 | 0.938 | 0.59 | 0.08 | 0.38 | 0.48 | 0.51 | 0.72 |
| 6 | 85523135 | rs858738 | -0.202 | 0.659 | 0.986 | 0.495 | 0.003 | 0.003 | 0.003 | 0 | 0.08 | 0.1 | 0.13 | 0.08 | 0.1 | 0.02 |
| 6 | 85555632 | rs12526113 | 1.542 | 0.382 | 0.025 | 0.017 | -0.076 | -0.004 | -0.004 | 0 | 0.54 | 0.1 | 0.3 | 0.42 | 0.47 | 0.57 |
| 6 | 85556282 | rs6454399 | -0.032 | 0.88 | -0.026 | -0.28 | -0.027 | -0.023 | -0.023 | 0 | 0.93 | 1 | 0.87 | 0.92 | 0.9 | 0.98 |
| 6 | 85563069 | rs2224211 | 0.111 | 0.992 | -0.18 | -0.274 | 0.007 | 0.006 | 0.006 | 0.276 | 0.12 | 0.11 | 0.31 | 0.23 | 0.24 | 0.24 |
| 6 | 85572537 | rs7741631 | 0.422 | 0.458 | -0.23 | -0.651 | -0.002 | -0.033 | -0.033 | 0.023 | 0.11 | 0.14 | 0.08 | 0 | 0.09 | 0.01 |
| 6 | 85572756 | rs9450065 | 0.092 | 0.443 | -0.13 | -0.649 | 0.013 | -0.036 | -0.036 | 0.193 | 0.07 | 0.15 | 0.05 | 0 | 0.09 | 0.14 |
| 6 | 85575780 | rs2024846 | 0.904 | 0.413 | -0.153 | -0.611 | 0.002 | -0.039 | -0.039 | 0 | 0.74 | 0.7 | 0.74 | 0.91 | 0.72 | 0.82 |
| 6 | 85580439 | rs13199506 | -0.048 | 0.215 | -0.51 | -0.546 | 0.001 | -0.019 | -0.019 | 2.034 | 0.13 | 0.12 | 0.09 | 0 | 0.09 | 0.03 |
| 6 | 85582396 | rs17692425 | 1.136 | 0.169 | -1.12 | -0.79 | -0.035 | 0.147 | 0.147 | 0 | 0.36 | 0.08 | 0.22 | 0.14 | 0.34 | 0.21 |
| 6 | 85588632 | rs1884478 | -0.738 | -0.261 | -0.53 | -0.339 | -0.001 | -0.001 | -0.001 | 0.157 | 0.47 | 0.45 | 0.5 | 0.5 | 0.49 | 0.67 |
| 6 | 85588678 | rs9450069 | -0.067 | -0.427 | -0.406 | -0.455 | 0.002 | -0.006 | -0.006 | 0 | 0.14 | 0.12 | 0.21 | 0.37 | 0.12 | 0.12 |
| 6 | 85591387 | rs11758273 | 0.817 | -0.365 | -0.749 | -0.173 | -0.039 | -0.066 | -0.066 | 0 | 0.1 | 0.01 | 0.15 | 0.26 | 0.13 | 0.16 |
| 6 | 85593031 | rs763725 | -2.275 | -0.406 | -0.6 | -0.157 | -0.08 | 0.155 | 0.155 | 0 | 0.65 | 0.92 | 0.73 | 0.87 | 0.62 | 0.78 |
| 6 | 85594528 | rs9344437 | 0.727 | -0.424 | -0.528 | -0.37 | -0.028 | -0.027 | -0.027 | 0 | 0.11 | 0.05 | 0.24 | 0.22 | 0.2 | 0.31 |
| 6 | 85602929 | rs9362110 | -1.412 | -0.441 | -0.281 | -0.392 | 0.027 | 0.041 | 0.041 | 0 | 0.29 | 0.53 | 0.48 | 0.59 | 0.32 | 0.42 |
| 6 | 85604347 | rs9450078 | -1.412 | -0.441 | -0.281 | -0.392 | 0.027 | 0.041 | 0.041 | 0 | 0.29 | 0.53 | 0.48 | 0.59 | 0.32 | 0.42 |
| 6 | 85625729 | rs9444253 | -0.13 | -0.519 | -0.233 | -0.204 | -0.002 | 0.022 | 0.022 | 0 | 0.15 | 0.16 | 0.21 | 0.37 | 0.12 | 0.12 |
| 6 | 85631896 | rs4620088 | -0.904 | -0.537 | 0.211 | -0.325 | -0.028 | -0.01 | -0.01 | 0 | 0.37 | 0.5 | 0.3 | 0.3 | 0.3 | 0.33 |
| 6 | 85641642 | rs10944052 | -0.082 | -0.346 | -0.237 | -0.2 | -0.015 | -0.047 | -0.047 | 0 | 0.21 | 0.1 | 0.22 | 0.3 | 0.24 | 0.31 |
| 6 | 85642512 | rs9444258 | 0.543 | -0.353 | -0.371 | -0.292 | -0.006 | 0.029 | 0.029 | 0 | 0.1 | 0.06 | 0.05 | 0 | 0.11 | 0.05 |
| 6 | 85643917 | rs1322880 | -1.802 | -0.481 | -0.085 | -0.226 | 0.009 | 0.015 | 0.015 | 0 | 0.41 | 0.68 | 0.46 | 0.67 | 0.37 | 0.47 |
| 6 | 85652904 | rs16875357 | -0.107 | -0.284 | -0.595 | -0.152 | -0.024 | -0.063 | -0.063 | 0 | 0.21 | 0.07 | 0.22 | 0.3 | 0.24 | 0.31 |
| 6 | 85662325 | rs9351022 | -1.054 | -0.501 | -0.442 | -0.453 | 0 | 0.039 | 0.039 | 0 | 0.39 | 0.27 | 0.36 | 0.23 | 0.39 | 0.56 |
| 6 | 85665162 | rs9450098 | 0.291 | -0.587 | -0.68 | -0.499 | -0.012 | 0.024 | 0.024 | 0 | 0.1 | 0.07 | 0.05 | 0 | 0.11 | 0.05 |
| 6 | 85674483 | rs4707103 | -2.411 | -0.558 | -0.911 | -0.793 | -0.052 | 0.167 | 0.167 | 0.007 | 0.65 | 0.86 | 0.76 | 0.89 | 0.64 | 0.8 |
| 6 | 85679883 | rs9362119 | -1.114 | -0.651 | -0.73 | -0.793 | 0.005 | 0.067 | 0.067 | 0.856 | 0.41 | 0.72 | 0.4 | 0.22 | 0.39 | 0.5 |
| 6 | 85680000 | rs11964358 | 0.614 | -0.537 | -0.744 | -0.539 | -0.014 | 0.023 | 0.023 | 0 | 0.1 | 0.06 | 0.05 | 0.02 | 0.12 | 0.05 |
| 6 | 85680478 | rs11754342 | -0.55 | -0.751 | -0.826 | -0.649 | 0.003 | -0.041 | -0.041 | 0 | 0.26 | 0.08 | 0.36 | 0.66 | 0.26 | 0.3 |
| 6 | 85683405 | rs962054 | -0.407 | -0.601 | -0.735 | -0.549 | -0.006 | -0.045 | -0.045 | 0.418 | 0.24 | 0.08 | 0.36 | 0.66 | 0.25 | 0.3 |
| 6 | 85684033 | rs2208880 | -0.908 | -0.795 | -0.656 | -0.744 | 0.006 | -0.012 | -0.012 | 0 | 0.33 | 0.61 | 0.38 | 0.22 | 0.36 | 0.34 |
| 6 | 85686384 | rs1359013 | 0.152 | -1.554 | -0.78 | -0.344 | -0.032 | -0.077 | -0.077 | 0 | 0.19 | 0.05 | 0.22 | 0.3 | 0.25 | 0.22 |
| 6 | 85688541 | rs9359617 | 0.671 | -1.348 | -0.264 | -0.381 | 0 | 0.024 | 0.024 | 0 | 0.17 | 0.26 | 0.26 | 0.23 | 0.17 | 0.28 |
| 6 | 85689003 | rs10485066 | 0.171 | -1.336 | -0.452 | -0.29 | -0.011 | 0.031 | 0.031 | 0 | 0.15 | 0.04 | 0.12 | 0.09 | 0.16 | 0.12 |
| 6 | 85690306 | rs1322868 | -2.643 | -1.509 | -0.535 | -0.51 | -0.087 | 0.167 | 0.167 | 0.016 | 0.64 | 0.85 | 0.73 | 0.88 | 0.57 | 0.8 |
| 6 | 85690595 | rs2324764 | -1.148 | -1.516 | -0.442 | -0.502 | 0.019 | 0.031 | 0.031 | 0.674 | 0.39 | 0.7 | 0.42 | 0.23 | 0.31 | 0.5 |
| 6 | 85691102 | rs2324765 | -1.06 | -1.255 | -0.365 | -0.221 | -0.004 | -0.049 | -0.049 | 0.537 | 0.25 | 0.09 | 0.31 | 0.65 | 0.26 | 0.3 |
| 6 | 85697194 | rs10806286 | 0.199 | -1.125 | -0.144 | -0.053 | -0.004 | 0.077 | 0.077 | 2.361 | 0.1 | 0.23 | 0.12 | 0.21 | 0.1 | 0.17 |

| | | | | | | | | | | | | | | | | |
|---|---|---|---|---|---|---|---|---|---|---|---|---|---|---|---|---|
| 6 | 85705312 | rs4707110 | -0.996 | -1.277 | -0.151 | -0.201 | 0.002 | 0.085 | 0.085 | 0.051 | 0.24 | 0.47 | 0.24 | 0.37 | 0.23 | 0.22 |
| 6 | 85706347 | rs16875429 | 0.848 | -1.295 | -0.219 | -0.163 | -0.016 | -0.04 | -0.04 | 0 | 0.07 | 0 | 0.11 | 0.22 | 0.09 | 0.16 |
| 6 | 85706748 | rs9359618 | 0.091 | -1.394 | -0.405 | -0.3 | -0.051 | -0.154 | -0.154 | 0 | 0.22 | 0.02 | 0.39 | 0.43 | 0.26 | 0.44 |
| 6 | 85707337 | rs9444276 | 0.18 | -1.396 | -0.159 | -0.308 | -0.04 | -0.071 | -0.071 | 0 | 0.11 | 0.3 | 0.05 | 0 | 0.04 | 0.04 |
| 6 | 85708695 | rs9444279 | -0.479 | -1.499 | -0.226 | -0.373 | -0.1 | -0.151 | -0.151 | 0.636 | 0.19 | 0.41 | 0.07 | 0 | 0.07 | 0.08 |
| 6 | 85710457 | rs9444281 | 0.305 | -1.446 | -0.44 | -0.351 | 0.001 | 0.029 | 0.029 | 0.016 | 0.11 | 0.14 | 0.19 | 0.38 | 0.19 | 0.17 |
| 6 | 85716701 | rs2208876 | -0.613 | -1.652 | -0.838 | -0.453 | -0.051 | -0.085 | -0.085 | 0 | 0.21 | 0.62 | 0.09 | 0 | 0.07 | 0.07 |
| 6 | 85718178 | rs9344451 | 0.698 | -1.074 | -0.938 | -0.042 | -0.039 | -0.066 | -0.066 | 0 | 0.14 | 0.02 | 0.29 | 0.21 | 0.17 | 0.28 |
| 6 | 85720325 | rs1322863 | -0.198 | -1.051 | -0.69 | -0.196 | 0.008 | 0.01 | 0.01 | 0 | 0.12 | 0.04 | 0.07 | 0.08 | 0.07 | 0.1 |
| 6 | 85727796 | rs7773154 | -1.543 | -1.029 | -0.334 | 0.185 | -0.019 | 0.021 | 0.021 | 0 | 0.3 | 0.54 | 0.3 | 0.35 | 0.26 | 0.43 |
| 6 | 85732319 | rs16875549 | 0.148 | -1.129 | -0.324 | 0.2 | 0.011 | -0.006 | -0.006 | 0.017 | 0.11 | 0.06 | 0.07 | 0.09 | 0.05 | 0.12 |
| 6 | 85732409 | rs4707117 | -2.032 | -1.227 | -0.441 | 0.336 | 0.029 | 0.029 | 0.029 | 0 | 0.43 | 0.83 | 0.44 | 0.65 | 0.37 | 0.48 |
| 6 | 85732548 | rs6936512 | -0.872 | -1.238 | -0.385 | 0.347 | -0.031 | 0.058 | 0.058 | 0.654 | 0.23 | 0.47 | 0.22 | 0.29 | 0.15 | 0.28 |
| 6 | 85737235 | rs7739166 | 0.186 | -1.188 | -0.679 | 0.374 | 0 | 0.014 | 0.014 | 0 | 0.11 | 0.09 | 0.12 | 0.3 | 0.11 | 0.05 |
| 6 | 85739871 | rs10806287 | -1.803 | -1.124 | -0.82 | 0.195 | 0.03 | 0.03 | 0.03 | 0 | 0.43 | 0.83 | 0.44 | 0.65 | 0.37 | 0.48 |
| 6 | 85744722 | rs10498949 | 0.899 | -0.967 | -0.803 | 0.241 | -0.005 | 0.008 | 0.008 | 0 | 0.07 | 0.04 | 0.08 | 0.06 | 0.1 | 0.15 |
| 6 | 85746881 | rs1885632 | -1.851 | -1.16 | -1.041 | 0.153 | 0.029 | 0.116 | 0.116 | 0 | 0.5 | 0.9 | 0.46 | 0.71 | 0.4 | 0.51 |
| 6 | 85750834 | rs1923597 | -0.759 | -1.028 | -0.488 | 0.315 | -0.003 | 0.039 | 0.039 | 0 | 0.19 | 0.48 | 0.22 | 0.25 | 0.2 | 0.3 |
| 6 | 85760635 | rs7760395 | -0.845 | -1.509 | -0.568 | 0.238 | -0.015 | 0.047 | 0.047 | 0 | 0.27 | 0.34 | 0.24 | 0.39 | 0.18 | 0.19 |
| 6 | 85761559 | rs725227 | 0.228 | -1.443 | -0.513 | 0.329 | -0.001 | 0.007 | 0.007 | 0 | 0.12 | 0.09 | 0.12 | 0.3 | 0.12 | 0.05 |
| 6 | 85766507 | rs6938017 | -0.295 | -1.26 | -0.244 | 0.5 | 0.012 | 0.103 | 0.103 | 0.781 | 0.12 | 0.42 | 0.14 | 0.2 | 0.1 | 0.15 |
| 6 | 85770652 | rs6930127 | -0.295 | -1.26 | -0.226 | 0.492 | 0.013 | 0.1 | 0.1 | 0.781 | 0.12 | 0.42 | 0.14 | 0.19 | 0.1 | 0.15 |
| 6 | 85772144 | rs11965782 | 0.864 | -1.142 | -0.509 | 0.577 | -0.011 | -0.029 | -0.029 | 1.849 | 0.06 | 0.01 | 0.1 | 0.2 | 0.09 | 0.15 |
| 6 | 85774866 | rs9344458 | 0.754 | -1.156 | -0.84 | 0.08 | -0.037 | -0.083 | -0.083 | 0 | 0.14 | 0.02 | 0.29 | 0.24 | 0.18 | 0.29 |
| 6 | 85777822 | rs7765284 | -0.367 | -1.011 | -0.325 | 0.077 | -0.023 | -0.008 | -0.008 | 0 | 0.14 | 0.25 | 0.11 | 0.09 | 0.07 | 0.14 |
| 6 | 85782095 | rs10498951 | -0.997 | -1.161 | -0.357 | 0.009 | -0.012 | -0.021 | -0.021 | 0.676 | 0.28 | 0.35 | 0.18 | 0.18 | 0.19 | 0.23 |
| 6 | 85787115 | rs1923589 | -0.638 | -1.257 | -0.315 | -0.248 | 0 | -0.005 | -0.005 | 0 | 0.83 | 0.85 | 0.87 | 0.94 | 0.84 | 0.81 |
| 6 | 85790460 | rs1325973 | -1.957 | -1.472 | -0.395 | -0.059 | -0.016 | 0.067 | 0.067 | 0 | 0.57 | 0.83 | 0.5 | 0.7 | 0.44 | 0.58 |
| 6 | 85793173 | rs6931497 | -0.904 | -1.205 | -0.122 | -0.092 | -0.034 | 0.085 | 0.085 | 0 | 0.29 | 0.47 | 0.33 | 0.53 | 0.25 | 0.36 |
| 6 | 85797051 | rs1409191 | 0.218 | -1.04 | -0.367 | -0.354 | 0.001 | 0.011 | 0.011 | 0 | 0.21 | 0.14 | 0.17 | 0.06 | 0.24 | 0.26 |
| 6 | 85797118 | rs7750062 | -0.659 | -1.074 | -0.204 | -0.167 | -0.005 | -0.032 | -0.032 | 0 | 0.29 | 0.37 | 0.28 | 0.19 | 0.31 | 0.4 |
| 6 | 85800575 | rs10485301 | -0.629 | -1.042 | -0.297 | -0.187 | -0.009 | 0.044 | 0.044 | 0 | 0.31 | 0.43 | 0.35 | 0.51 | 0.31 | 0.42 |
| 6 | 85802566 | rs6454417 | -1.107 | -1.226 | -0.461 | -0.516 | -0.004 | 0.004 | 0.004 | 0 | 0.45 | 0.5 | 0.41 | 0.51 | 0.44 | 0.47 |
| 6 | 85810298 | rs12527709 | 0.235 | -1.113 | -0.835 | -0.48 | -0.002 | 0.016 | 0.016 | 0 | 0.14 | 0.07 | 0.06 | 0 | 0.13 | 0.05 |
| 6 | 85818790 | rs9450163 | -1.077 | -1.019 | -0.715 | -0.257 | 0.024 | -0.03 | -0.03 | 0 | 0.49 | 0.78 | 0.45 | 0.51 | 0.45 | 0.47 |
| 6 | 85818874 | rs16875673 | 0.251 | -1.304 | -0.118 | -0.045 | 0.001 | 0.028 | 0.028 | 1.224 | 0.08 | 0.2 | 0.12 | 0.13 | 0.08 | 0.14 |
| 6 | 85821243 | rs10485303 | -0.942 | -1.517 | -0.902 | -0.192 | 0.022 | 0.02 | 0.02 | 0 | 0.23 | 0.12 | 0.19 | 0.21 | 0.17 | 0.33 |
| 6 | 85821694 | rs867744 | -0.99 | -1.676 | -0.589 | -0.286 | -0.012 | -0.013 | -0.013 | 0 | 0.19 | 0.54 | 0.22 | 0.17 | 0.15 | 0.29 |
| 6 | 85823394 | rs1536277 | -0.081 | -2.074 | -1.016 | -0.24 | 0.041 | -0.02 | -0.02 | 2.111 | 0.32 | 0.16 | 0.21 | 0.38 | 0.22 | 0.27 |
| 6 | 85825144 | rs10485305 | -0.362 | -1.8 | -1.124 | -0.149 | -0.005 | -0.002 | -0.002 | 0 | 0.15 | 0.17 | 0.16 | 0.14 | 0.12 | 0.14 |
| 6 | 85827524 | rs1359602 | -1.256 | -1.668 | -1.566 | -0.162 | 0.001 | -0.046 | -0.046 | 0.022 | 0.59 | 0.85 | 0.45 | 0.58 | 0.38 | 0.59 |
| 6 | 85831810 | rs941956 | 0.238 | -0.964 | -1.055 | 0.135 | -0.008 | -0.026 | -0.026 | 0.131 | 0.08 | 0.02 | 0.14 | 0.14 | 0.1 | 0.11 |
| 6 | 85833802 | rs10944063 | -0.535 | -1.221 | -1.107 | -0.076 | -0.005 | -0.002 | -0.002 | 0 | 0.15 | 0.08 | 0.21 | 0.15 | 0.19 | 0.1 |
| 6 | 85834204 | rs1998393 | -1.039 | -1.873 | -0.923 | -0.167 | 0.005 | -0.004 | -0.004 | 0.495 | 0.37 | 0.63 | 0.4 | 0.54 | 0.3 | 0.34 |
| 6 | 85840403 | rs595890 | -0.673 | -1.915 | -0.902 | -1.132 | 0.004 | -0.04 | -0.04 | 0 | 0.45 | 0.54 | 0.63 | 0.92 | 0.38 | 0.53 |
| 6 | 85842716 | rs526225 | -1.035 | -0.801 | -0.998 | -1.511 | 0.017 | -0.117 | -0.117 | 0 | 0.73 | 0.58 | 0.72 | 0.95 | 0.64 | 0.87 |
| 6 | 85845189 | rs562034 | 1.074 | -0.985 | -1.117 | -0.814 | -0.003 | -0.032 | -0.032 | 0.175 | 0.14 | 0.06 | 0.13 | 0.34 | 0.12 | 0.14 |
| 6 | 85849736 | rs11759993 | 1.083 | -1.072 | -1.198 | -0.735 | -0.015 | -0.037 | -0.037 | 2.628 | 0.09 | 0.02 | 0.19 | 0.19 | 0.1 | 0.15 |
| 6 | 85850774 | rs1407160 | 0.034 | -1.164 | -0.948 | -0.935 | -0.021 | 0.041 | 0.041 | 0.235 | 0.29 | 0.39 | 0.52 | 0.55 | 0.26 | 0.39 |
| 6 | 85854434 | rs563365 | 1.105 | -0.935 | -0.858 | -1.046 | 0.001 | -0.025 | -0.025 | 0 | 0.15 | 0.1 | 0.15 | 0.35 | 0.13 | 0.15 |
| 6 | 85867042 | rs1321737 | -0.432 | -0.942 | -0.76 | -1.152 | 0.006 | -0.006 | -0.006 | 0.56 | 0.59 | 0.49 | 0.63 | 0.61 | 0.52 | 0.73 |
| 6 | 85870792 | rs1332371 | 1.453 | -0.759 | -1.116 | -1.236 | -0.038 | 0.07 | 0.07 | 0.113 | 0.2 | 0.08 | 0.09 | 0.04 | 0.22 | 0.27 |
| 6 | 85870984 | rs7768362 | -0.048 | -0.916 | -1.038 | -1.282 | -0.032 | 0.015 | 0.015 | 0.155 | 0.32 | 0.49 | 0.54 | 0.61 | 0.27 | 0.43 |
| 6 | 85872082 | rs1407163 | 1.285 | -0.852 | -1.058 | -1.201 | -0.005 | -0.009 | -0.009 | 0 | 0.22 | 0.18 | 0.15 | 0.35 | 0.22 | 0.29 |
| 6 | 85876291 | rs515583 | -1.003 | -0.64 | -1.195 | -1.338 | -0.008 | 0.028 | 0.028 | 0 | 0.68 | 0.77 | 0.78 | 0.96 | 0.62 | 0.83 |
| 6 | 85880616 | rs1570933 | -0.162 | -0.645 | -1.151 | -1.289 | -0.001 | 0.086 | 0.086 | 0 | 0.51 | 0.65 | 0.68 | 0.94 | 0.48 | 0.7 |
| 6 | 85881189 | rs1321742 | 0.059 | -0.665 | -1.1 | -1.199 | -0.022 | 0.031 | 0.031 | 0 | 0.28 | 0.43 | 0.53 | 0.61 | 0.27 | 0.42 |
| 6 | 85881718 | rs581170 | 1.126 | -0.626 | -1.111 | -1.336 | -0.005 | 0.012 | 0.012 | 0 | 0.14 | 0.08 | 0.09 | 0 | 0.13 | 0.12 |
| 6 | 85890143 | rs1407167 | -0.204 | -0.657 | -1.14 | -1.37 | 0 | -0.002 | -0.002 | 0 | 0.51 | 0.58 | 0.67 | 0.94 | 0.48 | 0.7 |
| 6 | 85892529 | rs10498939 | 0.885 | -0.665 | -1.088 | -0.854 | 0 | 0.01 | 0.01 | 0.154 | 0.1 | 0.09 | 0.2 | 0.32 | 0.09 | 0.13 |
| 6 | 85894338 | rs9362141 | 0.756 | -0.71 | -1.129 | -1.073 | -0.007 | 0.053 | 0.053 | 0 | 0.34 | 0.24 | 0.24 | 0.06 | 0.39 | 0.17 |
| 6 | 85901263 | rs1321749 | -0.637 | -0.464 | -1.106 | -0.953 | -0.037 | 0.074 | 0.074 | 0.444 | 0.63 | 0.78 | 0.8 | 0.96 | 0.61 | 0.59 |
| 6 | 85901618 | rs9294329 | 0.688 | -0.472 | -0.882 | -0.331 | -0.014 | 0.121 | 0.121 | 0 | 0.17 | 0.35 | 0.33 | 0.29 | 0.17 | 0.26 |
| 6 | 85906208 | rs12213732 | 0.724 | -0.17 | -1.189 | -0.234 | -0.005 | -0.006 | -0.006 | 0 | 0.24 | 0.25 | 0.16 | 0.34 | 0.23 | 0.3 |
| 6 | 85941871 | rs16875886 | 0.718 | 0.11 | -1.457 | -0.161 | -0.004 | -0.047 | -0.047 | 0 | 0.09 | 0.02 | 0.19 | 0.33 | 0.09 | 0.12 |
| 6 | 85948591 | rs9351054 | 0.684 | -0.198 | -1.663 | -0.663 | -0.02 | -0.027 | -0.027 | 0 | 0.35 | 0.2 | 0.47 | 0.31 | 0.43 | 0.38 |
| 6 | 85951214 | rs1547702 | -0.954 | -0.409 | -1.56 | -1.084 | -0.023 | 0.053 | 0.053 | 0.337 | 0.65 | 0.81 | 0.83 | 0.95 | 0.66 | 0.61 |
| 6 | 85953339 | rs9444314 | 1.3 | -0.265 | -1.265 | -0.891 | 0.074 | 0.022 | 0.022 | 0.096 | 0.08 | 0.35 | 0.07 | 0.02 | 0.09 | 0.05 |
| 6 | 85954619 | rs9450204 | 0.815 | -0.272 | -1.266 | -0.892 | 0.054 | -0.009 | -0.009 | 0 | 0.09 | 0.36 | 0.09 | 0.03 | 0.1 | 0.06 |

| | | | | | | | | | | | | | | | | |
|---|---|---|---|---|---|---|---|---|---|---|---|---|---|---|---|---|
| 6 | 85962470 | rs9294331 | 1.574 | -0.376 | -1.554 | -0.969 | -0.053 | 0.085 | 0.085 | 0 | 0.22 | 0.08 | 0.19 | 0.05 | 0.34 | 0.16 |
| 6 | 85963188 | rs944669 | 0.037 | -0.448 | -1.499 | -0.873 | 0 | -0.001 | -0.001 | 0 | 0.44 | 0.56 | 0.55 | 0.34 | 0.52 | 0.44 |
| 6 | 85972492 | rs1037163 | -0.842 | -0.461 | -1.582 | -0.742 | -0.004 | 0.002 | 0.002 | 0.493 | 0.64 | 0.68 | 0.77 | 0.66 | 0.66 | 0.6 |
| 6 | 85979455 | rs9362176 | 1.248 | -0.472 | -1.831 | -0.784 | 0.022 | -0.083 | -0.083 | 0.085 | 0.13 | 0.02 | 0.27 | 0.26 | 0.08 | 0.22 |
| 6 | 85982579 | rs1885016 | 1.477 | -0.164 | -1.772 | -1.617 | -0.027 | 0.058 | 0.058 | 0 | 0.22 | 0.12 | 0.1 | 0.05 | 0.21 | 0.28 |
| 6 | 85985075 | rs7749042 | 1.063 | 0.34 | -1.562 | -1.385 | -0.029 | 0.041 | 0.041 | 0 | 0.23 | 0.17 | 0.21 | 0.05 | 0.35 | 0.16 |
| 6 | 85987004 | rs519667 | -1.148 | 0.27 | -1.481 | -1.304 | -0.001 | 0.011 | 0.011 | 0 | 0.87 | 0.95 | 0.93 | 1 | 0.87 | 0.89 |
| 6 | 85987239 | rs521621 | 0.071 | 0.022 | -1.581 | -1.301 | -0.002 | 0.037 | 0.037 | 0 | 0.46 | 0.37 | 0.46 | 0.37 | 0.55 | 0.36 |
| 6 | 85991153 | rs313233 | -0.785 | 0.069 | -1.671 | -1.251 | 0.012 | -0.016 | -0.016 | 0 | 0.66 | 0.82 | 0.57 | 0.42 | 0.72 | 0.46 |
| 6 | 85991961 | rs313232 | 0.314 | -0.017 | -1.521 | -1.29 | -0.008 | 0.055 | 0.055 | 0.799 | 0.11 | 0.2 | 0.2 | 0.28 | 0.11 | 0.12 |
| 6 | 85992024 | rs9359645 | 0.678 | 0.247 | -1.657 | -1.557 | -0.012 | 0.053 | 0.053 | 0 | 0.33 | 0.25 | 0.22 | 0.07 | 0.33 | 0.26 |
| 6 | 85992684 | rs3903689 | 1.276 | 0.222 | -1.637 | -1.556 | 0.074 | 0.053 | 0.053 | 0 | 0.08 | 0.37 | 0.07 | 0.07 | 0.09 | 0.02 |
| 6 | 85995036 | rs313230 | -0.467 | 0.28 | -2.048 | -1.571 | -0.007 | 0.06 | 0.06 | 0 | 0.52 | 0.8 | 0.49 | 0.42 | 0.53 | 0.4 |
| 6 | 85995936 | rs10498952 | 0.861 | 0.462 | -2.207 | -1.498 | 0.001 | 0.001 | 0.001 | 1.181 | 0.09 | 0.07 | 0.18 | 0.28 | 0.09 | 0.11 |
| 6 | 85996401 | rs313229 | 0.69 | 0.59 | -2.13 | -1.419 | 0.005 | 0.014 | 0.014 | 0 | 0.42 | 0.61 | 0.5 | 0.65 | 0.37 | 0.56 |
| 6 | 85997097 | rs12199725 | 1.425 | 0.952 | -2.142 | -1.485 | -0.048 | 0.039 | 0.039 | 0.158 | 0.11 | 0.04 | 0.05 | 0.07 | 0.16 | 0.28 |
| 6 | 85997241 | rs2016 | 0.687 | 0.871 | -1.916 | -1.295 | -0.014 | 0.049 | 0.049 | 0 | 0.19 | 0.1 | 0.15 | 0.07 | 0.21 | 0.16 |
| 6 | 85997900 | rs532570 | 0.687 | 0.871 | -1.916 | -1.295 | -0.014 | 0.049 | 0.049 | 0 | 0.19 | 0.1 | 0.15 | 0.07 | 0.21 | 0.16 |
| 6 | 85998177 | rs3857479 | 0.387 | 0.77 | -1.376 | -0.707 | -0.009 | 0.019 | 0.019 | 0 | 0.14 | 0.08 | 0.07 | 0 | 0.12 | 0.1 |
| 6 | 86004204 | rs567513 | 0.109 | 1.108 | -1.129 | 0.192 | -0.027 | -0.001 | -0.001 | 0 | 0.33 | 0.19 | 0.29 | 0.34 | 0.37 | 0.47 |
| 6 | 86004627 | rs499915 | 0.092 | 1.095 | -0.943 | 0.185 | -0.018 | 0.001 | 0.001 | 0 | 0.33 | 0.23 | 0.3 | 0.34 | 0.37 | 0.47 |
| 6 | 86008097 | rs7759585 | 0.905 | 0.884 | -0.789 | 0.39 | 0.018 | 0.013 | 0.013 | 0.7 | 0.36 | 0.3 | 0.38 | 0.22 | 0.21 | 0.27 |
| 6 | 86008956 | rs313210 | -0.084 | 1.072 | -0.761 | 0.268 | 0.037 | 0.093 | 0.093 | 0 | 0.78 | 0.6 | 0.71 | 0.56 | 0.69 | 0.82 |
| 6 | 86011079 | rs9362180 | 0.251 | 1.033 | -0.879 | 0.238 | -0.114 | 0.02 | 0.02 | 0.009 | 0.41 | 0.08 | 0.31 | 0.35 | 0.48 | 0.55 |
| 6 | 86025172 | rs1358939 | 1.053 | 0.97 | -0.746 | 0.212 | 0.046 | 0.03 | 0.03 | 0.466 | 0.35 | 0.24 | 0.38 | 0.22 | 0.21 | 0.27 |
| 6 | 86028681 | rs11752804 | 0.604 | 1.152 | -0.771 | -0.145 | -0.025 | 0.045 | 0.045 | 0 | 0.09 | 0.01 | 0.03 | 0 | 0.11 | 0.07 |
| 6 | 86029090 | rs9450234 | 0.65 | 1.268 | -0.713 | 0.01 | 0.037 | 0.038 | 0.038 | 0 | 0.1 | 0.35 | 0.08 | 0.09 | 0.11 | 0.04 |
| 6 | 86029970 | rs1830630 | 0.274 | 1.601 | -1.33 | 0.154 | -0.081 | -0.038 | -0.038 | 0.311 | 0.32 | 0.07 | 0.28 | 0.35 | 0.37 | 0.48 |
| 6 | 86032269 | rs313178 | 0.327 | 1.546 | -0.529 | 0.173 | -0.03 | 0.056 | 0.056 | 0 | 0.89 | 0.98 | 0.96 | 1 | 0.87 | 0.92 |
| 6 | 86042637 | rs188750 | 0.576 | 1.575 | -0.741 | 1.279 | -0.026 | -0.067 | -0.067 | 0 | 0.84 | 0.94 | 0.64 | 0.56 | 0.78 | 0.79 |
| 6 | 86042869 | rs9444325 | 0.318 | 1.215 | -0.502 | 0.697 | 0.001 | 0.005 | 0.005 | 0 | 0.44 | 0.43 | 0.49 | 0.26 | 0.42 | 0.3 |
| 6 | 86049040 | rs4707190 | 0.096 | 1.377 | -0.195 | 0.604 | 0 | -0.001 | -0.001 | 3.731 | 0.41 | 0.43 | 0.34 | 0.32 | 0.44 | 0.38 |
| 6 | 86054042 | rs9359652 | 0.21 | 1.442 | 0.009 | 0.501 | -0.019 | -0.003 | -0.003 | 0 | 0.36 | 0.23 | 0.27 | 0.33 | 0.36 | 0.37 |
| 6 | 86056783 | rs313243 | 0.814 | 0.944 | -0.307 | -0.087 | 0.003 | -0.019 | -0.019 | 0 | 0.16 | 0.13 | 0.08 | 0.04 | 0.11 | 0.22 |
| 6 | 86057368 | rs313242 | -0.555 | 0.946 | -0.377 | 0.128 | -0.013 | 0.002 | 0.002 | 0.219 | 0.93 | 0.97 | 0.83 | 0.57 | 0.89 | 0.85 |
| 6 | 86059902 | rs9450247 | 0.595 | 0.831 | 0.058 | -0.358 | 0.061 | -0.032 | -0.032 | 0 | 0.07 | 0.22 | 0.06 | 0 | 0.14 | 0.05 |
| 6 | 86060171 | rs9450248 | 0.217 | 0.79 | 0.023 | -0.33 | -0.002 | -0.023 | -0.023 | 0 | 0.39 | 0.44 | 0.46 | 0.2 | 0.43 | 0.26 |
| 6 | 86067547 | rs505923 | -0.35 | 0.861 | -0.049 | -0.28 | -0.011 | 0.034 | 0.034 | 0 | 0.56 | 0.64 | 0.55 | 0.24 | 0.54 | 0.48 |
| 6 | 86071757 | rs9362197 | 0.63 | 0.796 | -0.37 | 0.083 | 0.012 | 0.001 | 0.001 | 0 | 0.32 | 0.15 | 0.4 | 0.25 | 0.29 | 0.21 |
| 6 | 86073707 | rs586364 | -0.701 | 1.109 | -0.448 | 0.156 | -0.01 | -0.025 | -0.025 | 0 | 0.74 | 0.76 | 0.71 | 0.51 | 0.7 | 0.65 |
| 6 | 86078534 | rs6454463 | -0.448 | 0.845 | -0.136 | 0.255 | 0.008 | 0.015 | 0.015 | 0 | 0.27 | 0.34 | 0.24 | 0.3 | 0.3 | 0.23 |
| 6 | 86091724 | rs4707202 | 0.522 | 0.688 | -0.118 | 0.179 | -0.001 | -0.001 | -0.001 | 0 | 0.25 | 0.27 | 0.43 | 0.21 | 0.25 | 0.2 |
| 6 | 86091803 | rs6914591 | 0.769 | 0.92 | -0.9 | 0.1 | -0.078 | 0.069 | 0.069 | 0 | 0.3 | 0.1 | 0.26 | 0.11 | 0.38 | 0.26 |
| 6 | 86092879 | rs545969 | -1.126 | 0.683 | -0.705 | 0.445 | -0.002 | 0.003 | 0.003 | 0 | 0.85 | 0.82 | 0.85 | 0.76 | 0.89 | 0.81 |
| 6 | 86094187 | rs10498955 | 0.951 | 0.46 | -1.139 | 0.415 | -0.023 | -0.09 | -0.09 | 1.598 | 0.13 | 0.01 | 0.07 | 0.37 | 0.14 | 0.19 |
| 6 | 86100832 | rs577388 | -0.193 | 0.437 | -1.04 | 0.379 | 0.006 | -0.038 | -0.038 | 0 | 0.72 | 0.79 | 0.78 | 0.37 | 0.75 | 0.61 |
| 6 | 86104456 | rs11753050 | 0.861 | 0.462 | -1.138 | 0.314 | -0.078 | 0.068 | 0.068 | 0 | 0.3 | 0.1 | 0.26 | 0.11 | 0.38 | 0.26 |
| 6 | 86106092 | rs16876172 | 0.83 | 0.465 | -1 | 0.509 | -0.025 | -0.093 | -0.093 | 0 | 0.13 | 0.01 | 0.07 | 0.39 | 0.14 | 0.2 |
| 6 | 86113775 | rs184332 | -0.755 | 0.215 | -0.181 | 0.321 | 0 | -0.003 | -0.003 | 0 | 0.61 | 0.64 | 0.44 | 0.55 | 0.64 | 0.61 |
| 6 | 86115244 | rs554113 | 0.299 | 0.029 | 0.254 | 0.257 | -0.036 | -0.059 | -0.059 | 0 | 0.15 | 0.39 | 0.09 | 0.06 | 0.1 | 0.14 |
| 6 | 86118728 | rs9362210 | 0.565 | 0.95 | -0.351 | 0.662 | -0.072 | -0.101 | -0.101 | 0.15 | 0.17 | 0.02 | 0.12 | 0.37 | 0.24 | 0.22 |
| 6 | 86132363 | rs16876230 | 1.166 | 0.098 | 0.012 | 0.683 | -0.033 | 0.018 | 0.018 | 0 | 0.12 | 0.21 | 0.11 | 0.22 | 0.04 | 0.18 |
| 6 | 86136433 | rs9362215 | 0.763 | -0.319 | -0.466 | 0.239 | 0.004 | 0.057 | 0.057 | 0 | 0.16 | 0.01 | 0.08 | 0.05 | 0.14 | 0.17 |
| 6 | 86137888 | rs9450273 | 1.138 | 0.012 | -0.227 | 0.4 | 0.007 | 0.031 | 0.031 | 0 | 0.49 | 0.25 | 0.61 | 0.32 | 0.52 | 0.47 |
| 6 | 86144875 | rs6931295 | -0.915 | 0.137 | -0.318 | 0.462 | -0.072 | -0.077 | -0.077 | 0 | 0.82 | 0.97 | 0.84 | 0.6 | 0.7 | 0.83 |
| 6 | 86161269 | rs3734442 | 1.449 | -0.215 | 0.032 | 0.419 | -0.054 | 0.047 | 0.047 | 0 | 0.12 | 0.3 | 0.11 | 0.23 | 0.03 | 0.18 |
| 6 | 86168265 | rs6942065 | 0.182 | -0.045 | 0.37 | 0.259 | -0.018 | -0.058 | -0.058 | 0.033 | 0.18 | 0.29 | 0.09 | 0.05 | 0.14 | 0.16 |
| 6 | 86178324 | rs4707206 | 1.266 | -0.257 | 0.195 | 0.448 | -0.032 | 0.012 | 0.012 | 0 | 0.12 | 0.18 | 0.11 | 0.24 | 0.04 | 0.17 |
| 6 | 86180732 | rs10944128 | 0.88 | -0.152 | 0.122 | 0.382 | 0.018 | 0.006 | 0.006 | 0.442 | 0.54 | 0.27 | 0.65 | 0.32 | 0.57 | 0.51 |
| 6 | 86181931 | rs4593336 | 0.36 | -0.105 | 0.301 | 0.45 | -0.001 | 0.033 | 0.033 | 0.102 | 0.67 | 0.54 | 0.71 | 0.37 | 0.66 | 0.64 |
| 6 | 86203963 | rs6913634 | 0.752 | 0.006 | 0.207 | 0.424 | -0.022 | -0.001 | -0.001 | 0 | 0.08 | 0.26 | 0.05 | 0.05 | 0.05 | 0.1 |
| 6 | 86205323 | rs6922 | -0.469 | -0.149 | -0.206 | 0.341 | -0.128 | -0.112 | -0.112 | 0.219 | 0.77 | 0.99 | 0.83 | 0.6 | 0.67 | 0.79 |
| 6 | 86208525 | rs9450286 | 0.444 | -0.145 | 0.106 | 0.246 | -0.019 | -0.038 | -0.038 | 1.477 | 0.13 | 0.27 | 0.06 | 0.05 | 0.09 | 0.13 |
| 6 | 86214507 | rs6900270 | 0.382 | -0.286 | 0.27 | 0.081 | -0.007 | 0.018 | 0.018 | 0.936 | 0.08 | 0.38 | 0.06 | 0.05 | 0.05 | 0.1 |
| 6 | 86221771 | rs6911143 | -0.189 | -0.436 | -1.047 | -0.158 | 0.01 | -0.033 | -0.033 | 0 | 0.52 | 0.95 | 0.43 | 0.41 | 0.39 | 0.44 |
| 6 | 86228109 | rs6909767 | -0.29 | -0.211 | -0.433 | -0.348 | 0 | 0.021 | 0.021 | 0 | 0.63 | 0.54 | 0.7 | 0.37 | 0.64 | 0.62 |
| 6 | 86232543 | rs9353319 | 1.163 | -0.198 | -0.415 | -0.321 | -0.007 | 0.017 | 0.017 | 0.336 | 0.07 | 0.3 | 0.05 | 0.05 | 0.05 | 0.1 |
| 6 | 86234835 | rs17440806 | 1.076 | -0.133 | -0.835 | -0.468 | -0.017 | 0.014 | 0.014 | 0 | 0.23 | 0.04 | 0.19 | 0.11 | 0.23 | 0.19 |
| 6 | 86240951 | rs7452605 | -1.515 | -0.157 | -0.786 | -0.389 | -0.089 | -0.077 | -0.077 | 1.522 | 0.78 | 0.96 | 0.83 | 0.6 | 0.67 | 0.79 |

| | | | | | | | | | | | | | | | | |
|---|---|---|---|---|---|---|---|---|---|---|---|---|---|---|---|---|
| 6 | 86269039 | rs4626396 | 1.67 | -0.174 | -0.895 | -0.411 | -0.015 | -0.028 | -0.028 | 0 | 0.15 | 0.02 | 0.27 | 0.14 | 0.12 | 0.21 |
| 6 | 86279737 | rs4296866 | 0.17 | -0.029 | -0.558 | -0.286 | -0.007 | 0.07 | 0.07 | 0.795 | 0.5 | 0.27 | 0.56 | 0.24 | 0.48 | 0.46 |
| 6 | 86295858 | rs2842608 | 1.249 | -0.95 | -0.484 | -0.059 | -0.044 | -0.038 | -0.038 | 0 | 0.22 | 0.08 | 0.17 | 0.4 | 0.33 | 0.21 |
| 6 | 86297719 | rs12190637 | 1.942 | -1.115 | -0.515 | -0.088 | 0.006 | -0.013 | -0.013 | 0.451 | 0.15 | 0.03 | 0.13 | 0.11 | 0.12 | 0.16 |
| 6 | 86300851 | rs3900740 | 1.461 | -1.031 | -0.353 | -0.093 | -0.025 | 0.007 | 0.007 | 0 | 0.12 | 0.17 | 0.11 | 0.23 | 0.03 | 0.18 |
| 6 | 86340237 | rs13202606 | 1.878 | -1.04 | -0.384 | -0.127 | 0.037 | 0.035 | 0.035 | 0 | 0.1 | 0.04 | 0.03 | 0.02 | 0.03 | 0.17 |
| 6 | 86342679 | rs12212560 | 0.297 | -1.024 | -0.418 | -0.124 | -0.001 | -0.007 | -0.007 | 0.576 | 0.36 | 0.32 | 0.29 | 0.63 | 0.37 | 0.38 |
| 6 | 86400900 | rs9444361 | -1.447 | -0.932 | -0.652 | -0.203 | -0.097 | -0.069 | -0.069 | 0 | 0.78 | 0.96 | 0.84 | 0.62 | 0.67 | 0.8 |
| 6 | 86410651 | rs7752502 | -0.369 | -0.98 | -0.564 | -0.263 | -0.001 | 0.026 | 0.026 | 0 | 0.63 | 0.52 | 0.71 | 0.4 | 0.63 | 0.62 |
| 6 | 86415236 | rs2475787 | -0.346 | -1.052 | -1.088 | -0.235 | -0.001 | 0 | 0 | 0.047 | 0.62 | 0.63 | 0.7 | 0.4 | 0.6 | 0.61 |
| 6 | 86475814 | rs9450325 | 0.049 | -0.324 | -0.893 | 0.102 | 0.001 | -0.006 | -0.006 | 0.227 | 0.61 | 0.56 | 0.65 | 0.32 | 0.59 | 0.65 |
| 6 | 86500316 | rs1414201 | 0.316 | 0.03 | -0.633 | 0.5 | 0.013 | 0.049 | 0.049 | 0 | 0.22 | 0.33 | 0.23 | 0.58 | 0.3 | 0.18 |
| 6 | 86521582 | rs17442622 | 1.342 | 0.101 | -0.823 | 0.433 | -0.056 | -0.042 | -0.042 | 0 | 0.1 | 0.02 | 0.12 | 0.11 | 0.13 | 0.09 |
| 6 | 86576854 | rs2121594 | 0.738 | -0.009 | -0.652 | 0.558 | -0.01 | -0.009 | -0.009 | 0 | 0.18 | 0.12 | 0.14 | 0.36 | 0.29 | 0.16 |
| 6 | 86578833 | rs9342047 | 0.849 | -0.257 | -0.552 | 0.447 | 0.004 | 0.026 | 0.026 | 0.308 | 0.08 | 0.34 | 0.05 | 0.05 | 0.05 | 0.1 |
| 6 | 86589152 | rs1447157 | 0.876 | -0.257 | -0.559 | 0.447 | 0.001 | 0.022 | 0.022 | 0.273 | 0.07 | 0.32 | 0.05 | 0.05 | 0.05 | 0.1 |
| 6 | 86589290 | rs13206945 | 1.694 | -0.475 | -1.293 | 0.325 | 0.026 | -0.013 | -0.013 | 0 | 0.21 | 0.01 | 0.3 | 0.16 | 0.13 | 0.36 |
| 6 | 86592226 | rs10455449 | 0.729 | 0.052 | -0.624 | 0.475 | -0.097 | -0.089 | -0.089 | 0.621 | 0.17 | 0.01 | 0.13 | 0.35 | 0.29 | 0.16 |
| 6 | 86592362 | rs6454505 | -0.725 | 0.105 | -0.5 | 0.5 | -0.003 | -0.028 | -0.028 | 0 | 0.3 | 0.78 | 0.28 | 0.62 | 0.35 | 0.26 |
| 6 | 86656653 | rs12200330 | 0.499 | 0.477 | -0.583 | 0.661 | -0.019 | -0.017 | -0.017 | 0 | 0.18 | 0.1 | 0.14 | 0.35 | 0.29 | 0.16 |
| 6 | 86681475 | rs17443200 | 0.478 | 0.545 | -0.847 | 0.481 | -0.039 | 0.081 | 0.081 | 0.074 | 0.14 | 0.03 | 0.1 | 0 | 0.15 | 0.08 |
| 6 | 86683779 | rs806606 | -0.045 | 0.422 | -0.162 | 1.271 | 0 | 0.01 | 0.01 | 0 | 0.77 | 0.94 | 0.81 | 0.9 | 0.78 | 0.79 |
| 6 | 86726221 | rs12199516 | -0.09 | 0.967 | -0.198 | 0.959 | 0.006 | 0.005 | 0.005 | 0 | 0.19 | 0.18 | 0.14 | 0.35 | 0.29 | 0.16 |
| 6 | 86767686 | rs6923394 | -0.655 | -0.025 | -0.103 | 0.637 | 0 | 0.01 | 0.01 | 0 | 0.23 | 0.06 | 0.19 | 0.1 | 0.22 | 0.21 |
| 6 | 86783258 | rs9450383 | -0.573 | 0.206 | 0.819 | 0.345 | 0.008 | 0.029 | 0.029 | 0.309 | 0.07 | 0.34 | 0.05 | 0.05 | 0.05 | 0.1 |
| 6 | 86797650 | rs220430 | 0.321 | 0.556 | 0.215 | 0.471 | -0.022 | 0.032 | 0.032 | 0.826 | 0.06 | 0.01 | 0.08 | 0 | 0.08 | 0.11 |
| 6 | 86855532 | rs61875 | 1.278 | 0.434 | 1.126 | 0.707 | 0.022 | -0.025 | -0.025 | 0 | 0.9 | 0.72 | 0.91 | 0.94 | 0.88 | 0.92 |
| 6 | 86857389 | rs220428 | 1.356 | 0.434 | 0.666 | 0.709 | 0.003 | 0.035 | 0.035 | 1.366 | 0.85 | 0.3 | 0.79 | 0.78 | 0.86 | 0.88 |
| 6 | 86865514 | rs6931546 | 0.294 | -0.142 | -0.21 | 0.293 | 0.006 | -0.001 | -0.001 | 0 | 0.3 | 0.24 | 0.22 | 0.31 | 0.19 | 0.34 |
| 6 | 86870117 | rs11756369 | 0.493 | -0.207 | -0.796 | -0.23 | 0.021 | 0.085 | 0.085 | 0 | 0.25 | 0.1 | 0.27 | 0.06 | 0.24 | 0.14 |
| 6 | 86870628 | rs220409 | -0.111 | 0.033 | -0.301 | 0.128 | -0.002 | 0.005 | 0.005 | 0.125 | 0.42 | 0.63 | 0.46 | 0.65 | 0.51 | 0.55 |
| 6 | 86874123 | rs220402 | 0.037 | 0.661 | -0.188 | -0.038 | 0.05 | 0.062 | 0.062 | 1.603 | 0.28 | 0.48 | 0.39 | 0.58 | 0.39 | 0.4 |
| 6 | 86874355 | rs2143772 | -0.199 | 0.894 | -0.008 | 0.214 | 0.009 | 0.12 | 0.12 | 1.471 | 0.1 | 0.37 | 0.14 | 0.22 | 0.09 | 0.16 |
| 6 | 86885509 | rs220394 | -0.173 | 0.863 | -0.137 | 0.068 | -0.005 | 0.02 | 0.02 | 0 | 0.8 | 0.87 | 0.89 | 0.94 | 0.81 | 0.83 |
| 6 | 86886644 | rs1555171 | 1.032 | 0.824 | -0.079 | 0.133 | 0.033 | 0.038 | 0.038 | 0 | 0.51 | 0.31 | 0.49 | 0.35 | 0.42 | 0.44 |
| 6 | 86887829 | rs6454524 | 0.977 | 0.844 | -0.132 | 0.143 | 0.013 | 0.015 | 0.015 | 0 | 0.52 | 0.37 | 0.5 | 0.35 | 0.42 | 0.44 |
| 6 | 86893950 | rs1577237 | -0.948 | 0.888 | -0.061 | 0.239 | -0.065 | -0.074 | -0.074 | 0 | 0.83 | 0.96 | 0.75 | 0.64 | 0.7 | 0.77 |
| 6 | 86895945 | rs6926588 | 0.633 | 0.88 | -0.048 | 0.223 | 0.043 | 0.024 | 0.024 | 0.246 | 0.67 | 0.47 | 0.56 | 0.41 | 0.55 | 0.59 |
| 6 | 86896768 | rs9359682 | -0.896 | 0.877 | 0.011 | 0.302 | -0.046 | -0.052 | -0.052 | 2.329 | 0.83 | 0.93 | 0.75 | 0.64 | 0.7 | 0.77 |
| 6 | 86898957 | rs3949571 | -0.371 | 0.805 | 0.012 | 0.3 | 0.015 | 0.158 | 0.158 | 3.894 | 0.1 | 0.45 | 0.14 | 0.22 | 0.09 | 0.17 |
| 6 | 86913307 | rs9359686 | -0.868 | 0.96 | -0.438 | 0.332 | -0.065 | -0.074 | -0.074 | 0 | 0.83 | 0.96 | 0.75 | 0.64 | 0.7 | 0.78 |
| 6 | 86950469 | rs9359692 | -0.83 | 0.744 | -0.232 | 0.195 | -0.002 | -0.01 | -0.01 | 0.376 | 0.3 | 0.44 | 0.24 | 0.29 | 0.28 | 0.33 |
| 6 | 86953342 | rs12199696 | -0.385 | 0.719 | -0.666 | 0.134 | -0.04 | -0.026 | -0.026 | 0 | 0.38 | 0.14 | 0.37 | 0.42 | 0.49 | 0.4 |
| 6 | 86956248 | rs9344588 | 0.298 | 0.133 | -0.877 | 0.077 | 0.022 | -0.037 | -0.037 | 0 | 0.24 | 0.16 | 0.24 | 0.43 | 0.17 | 0.43 |
| 6 | 86956777 | rs4707263 | 0.258 | -0.098 | -1.195 | -0.353 | -0.012 | -0.066 | -0.066 | 0 | 0.43 | 0.11 | 0.4 | 0.51 | 0.51 | 0.43 |
| 6 | 86957149 | rs4707264 | 0.502 | 0.91 | -1.206 | -0.409 | 0.016 | 0.003 | 0.003 | 0 | 0.05 | 0.05 | 0.09 | 0.09 | 0.13 | 0.02 |
| 6 | 86957952 | rs2988685 | -0.36 | 0.794 | -0.971 | -0.497 | 0.043 | 0.004 | 0.004 | 0 | 0.09 | 0.15 | 0.13 | 0.09 | 0.19 | 0.04 |
| 6 | 86959447 | rs2988682 | -0.666 | 0.68 | -0.557 | -0.564 | -0.026 | -0.098 | -0.098 | 0.861 | 0.71 | 0.54 | 0.69 | 0.94 | 0.74 | 0.89 |
| 6 | 86963127 | rs3000069 | 0.337 | 0.753 | -1.075 | -0.363 | -0.077 | -0.057 | -0.057 | 0 | 0.38 | 0.06 | 0.31 | 0.41 | 0.38 | 0.42 |
| 6 | 86966457 | rs2798530 | 1.141 | 0.947 | -1.035 | -0.275 | -0.047 | -0.072 | -0.072 | 0 | 0.18 | 0.04 | 0.25 | 0.34 | 0.24 | 0.24 |
| 6 | 86968616 | rs2816545 | 0.403 | 0.674 | -0.942 | -0.215 | -0.016 | -0.032 | -0.032 | 0 | 0.37 | 0.18 | 0.38 | 0.44 | 0.41 | 0.53 |
| 6 | 86968841 | rs2324898 | -0.022 | 0.656 | -0.813 | -0.226 | 0.008 | -0.001 | -0.001 | 0.307 | 0.39 | 0.65 | 0.5 | 0.39 | 0.36 | 0.27 |
| 6 | 86976150 | rs4707265 | -0.597 | 0.641 | -0.301 | -0.323 | -0.02 | 0.109 | 0.109 | 0.348 | 0.2 | 0.5 | 0.25 | 0.34 | 0.17 | 0.21 |
| 6 | 86983160 | rs2798535 | 0.719 | 0.603 | -0.332 | -0.351 | -0.001 | 0.008 | 0.008 | 1.673 | 0.78 | 0.26 | 0.68 | 0.57 | 0.74 | 0.79 |
| 6 | 86987698 | rs2211493 | -0.315 | 0.4 | -0.19 | -0.394 | -0.04 | 0.153 | 0.153 | 0 | 0.18 | 0.44 | 0.23 | 0.36 | 0.14 | 0.19 |
| 6 | 87000772 | rs1573190 | -0.092 | 0.343 | 0.39 | -0.42 | -0.011 | 0.183 | 0.183 | 0 | 0.13 | 0.33 | 0.21 | 0.36 | 0.11 | 0.18 |
| 6 | 87001570 | rs17541873 | 1.043 | 0.273 | 0.018 | -0.669 | 0.008 | 0.079 | 0.079 | 0 | 0.21 | 0.06 | 0.06 | 0.06 | 0.14 | 0.19 |
| 6 | 87006947 | rs9444400 | 1.583 | 0.353 | 0.082 | -0.594 | -0.013 | -0.029 | -0.029 | 0 | 0.05 | 0.1 | 0.02 | 0 | 0.03 | 0.01 |
| 6 | 87008721 | rs2816567 | -0.476 | 0.356 | -0.139 | -0.737 | 0.018 | 0.131 | 0.131 | 0 | 0.58 | 0.77 | 0.66 | 0.89 | 0.64 | 0.74 |
| 6 | 87018295 | rs2816579 | 1.271 | 0.427 | -0.408 | -0.649 | -0.021 | -0.023 | -0.023 | 0 | 0.21 | 0.11 | 0.32 | 0.43 | 0.36 | 0.29 |
| 6 | 87028147 | rs7772098 | -0.27 | -0.184 | -0.167 | -0.554 | -0.014 | 0.072 | 0.072 | 0 | 0.13 | 0.24 | 0.21 | 0.36 | 0.11 | 0.18 |
| 6 | 87041092 | rs13213204 | 1.411 | -0.124 | -0.55 | -0.875 | -0.003 | 0.009 | 0.009 | 0 | 0.16 | 0.02 | 0.08 | 0.1 | 0.11 | 0.25 |
| 6 | 87041910 | rs7751415 | 0.689 | -0.042 | -0.264 | -0.907 | -0.002 | 0.04 | 0.04 | 2.921 | 0.2 | 0.14 | 0.28 | 0.04 | 0.21 | 0.07 |
| 6 | 87047727 | rs7741866 | 0.299 | -0.007 | -0.247 | -0.882 | 0.004 | 0.009 | 0.009 | 3.139 | 0.25 | 0.21 | 0.29 | 0.04 | 0.23 | 0.07 |
| 6 | 87062176 | rs13211518 | -0.064 | 0.218 | -0.094 | -0.127 | 0 | -0.001 | -0.001 | 0 | 0.08 | 0.1 | 0.12 | 0.16 | 0.08 | 0.09 |
| 6 | 87070493 | rs7741755 | -0.398 | -0.207 | -0.435 | -0.8 | 0 | -0.011 | -0.011 | 0 | 0.29 | 0.29 | 0.26 | 0.08 | 0.19 | 0.17 |
| 6 | 87078175 | rs9353392 | 0.921 | 0.417 | -0.905 | -0.336 | -0.054 | -0.063 | -0.063 | 2.725 | 0.22 | 0.07 | 0.27 | 0.59 | 0.34 | 0.23 |
| 6 | 87090015 | rs1361848 | 0.03 | 0.555 | -0.786 | -0.33 | -0.002 | -0.012 | -0.012 | 0 | 0.29 | 0.32 | 0.29 | 0.08 | 0.18 | 0.2 |
| 6 | 87094486 | rs12193553 | 1.356 | 0.417 | -0.662 | -0.614 | 0.005 | -0.014 | -0.014 | 0 | 0.24 | 0.2 | 0.1 | 0.05 | 0.16 | 0.18 |

| | | | | | | | | | | | | | | | | |
|---|---|---|---|---|---|---|---|---|---|---|---|---|---|---|---|---|
| 6 | 87097985 | rs9353399 | 1.395 | 0.732 | -0.953 | -0.351 | -0.025 | -0.102 | -0.102 | 0 | 0.2 | 0.04 | 0.22 | 0.51 | 0.25 | 0.22 |
| 6 | 87100832 | rs6900662 | 1.265 | 0.594 | -1.011 | -0.5 | -0.01 | 0.053 | 0.053 | 0.039 | 0.09 | 0.01 | 0.05 | 0 | 0.08 | 0.05 |
| 6 | 87101329 | rs9342070 | 0.156 | 0.706 | -0.535 | -0.336 | -0.012 | 0.093 | 0.093 | 0 | 0.44 | 0.24 | 0.54 | 0.79 | 0.56 | 0.61 |
| 6 | 87113172 | rs6919366 | 0.117 | 0.578 | -0.411 | -0.416 | 0.02 | 0.041 | 0.041 | 0 | 0.45 | 0.34 | 0.6 | 0.86 | 0.66 | 0.62 |
| 6 | 87115977 | rs4707276 | 0.971 | 0.902 | -0.573 | -0.126 | -0.038 | -0.041 | -0.041 | 0 | 0.22 | 0.11 | 0.28 | 0.6 | 0.34 | 0.23 |
| 6 | 87119490 | rs4053720 | 1.304 | 0.978 | -0.551 | 0.315 | -0.023 | -0.095 | -0.095 | 1.484 | 0.2 | 0.04 | 0.2 | 0.41 | 0.24 | 0.24 |
| 6 | 87122261 | rs9450448 | 1.071 | 0.323 | -0.521 | -0.214 | -0.047 | -0.079 | -0.079 | 0.326 | 0.22 | 0.1 | 0.3 | 0.52 | 0.33 | 0.25 |
| 6 | 87124225 | rs16876987 | 0.738 | 0.104 | -0.397 | -0.346 | -0.012 | 0.005 | 0.005 | 0 | 0.12 | 0.19 | 0.18 | 0.1 | 0.06 | 0.05 |
| 6 | 87124569 | rs3857481 | 0.647 | 0.18 | -0.839 | -0.386 | -0.008 | -0.004 | -0.004 | 0 | 0.23 | 0.2 | 0.32 | 0.28 | 0.31 | 0.37 |
| 6 | 87125624 | rs9353406 | 0.408 | -0.445 | -0.713 | -0.567 | 0.017 | 0.037 | 0.037 | 0 | 0.45 | 0.37 | 0.62 | 0.82 | 0.64 | 0.62 |
| 6 | 87126861 | rs6928430 | 0.889 | -0.375 | -1.402 | -0.638 | 0.042 | 0.078 | 0.078 | 0.665 | 0.25 | 0.12 | 0.11 | 0.06 | 0.17 | 0.19 |
| 6 | 87130477 | rs9450450 | -0.055 | -0.148 | -1.404 | -0.526 | 0.06 | 0.156 | 0.156 | 0 | 0.39 | 0.17 | 0.17 | 0.09 | 0.29 | 0.35 |
| 6 | 87132262 | rs6927637 | 0.436 | 0.635 | -1.093 | 0.18 | -0.003 | 0.032 | 0.032 | 0 | 0.11 | 0.05 | 0.06 | 0 | 0.1 | 0.05 |
| 6 | 87140364 | rs1508245 | 0.476 | 1.272 | -1.112 | 0.686 | -0.07 | -0.09 | -0.09 | 0.033 | 0.25 | 0.08 | 0.33 | 0.51 | 0.38 | 0.27 |
| 6 | 87140674 | rs1508246 | -0.173 | 1.169 | -0.183 | 0.643 | 0.036 | 0.047 | 0.047 | 0 | 0.79 | 0.63 | 0.69 | 0.71 | 0.74 | 0.7 |
| 6 | 87152611 | rs1508233 | -0.25 | 0.31 | -0.003 | 0.555 | -0.017 | 0.063 | 0.063 | 0 | 0.13 | 0.22 | 0.19 | 0.33 | 0.08 | 0.17 |
| 6 | 87153030 | rs1858038 | -0.177 | 0.542 | -0.179 | 0.148 | 0.017 | -0.043 | -0.043 | 0 | 0.37 | 0.52 | 0.37 | 0.12 | 0.38 | 0.28 |
| 6 | 87165643 | rs9784836 | -0.252 | 0.161 | -0.461 | -0.084 | -0.026 | 0 | 0 | 0.231 | 0.67 | 0.57 | 0.71 | 0.59 | 0.75 | 0.63 |
| 6 | 87173621 | rs6899410 | 0.037 | 0.28 | -0.353 | -0.205 | 0.016 | -0.04 | -0.04 | 0 | 0.37 | 0.5 | 0.36 | 0.12 | 0.38 | 0.28 |
| 6 | 87174156 | rs9450458 | 1.127 | 0.212 | -0.391 | -0.176 | -0.058 | -0.144 | -0.144 | 1.083 | 0.31 | 0.04 | 0.35 | 0.5 | 0.37 | 0.35 |
| 6 | 87178252 | rs994732 | 0.122 | 0.428 | -0.304 | -0.23 | 0.042 | -0.032 | -0.032 | 0 | 0.24 | 0.46 | 0.3 | 0.09 | 0.27 | 0.14 |
| 6 | 87183511 | rs9351099 | 0.433 | 0.307 | -0.281 | -0.389 | -0.016 | 0.061 | 0.061 | 0 | 0.12 | 0.22 | 0.18 | 0.3 | 0.08 | 0.15 |
| 6 | 87208546 | rs2102421 | -0.395 | 0.332 | -0.485 | -0.406 | -0.024 | -0.003 | -0.003 | 2.162 | 0.68 | 0.49 | 0.71 | 0.62 | 0.75 | 0.65 |
| 6 | 87212587 | rs1022023 | 0.024 | 0.509 | -0.539 | 0.092 | 0.021 | 0.016 | 0.016 | 0 | 0.24 | 0.39 | 0.29 | 0.22 | 0.28 | 0.14 |
| 6 | 87213096 | rs9344620 | 0.368 | 0.404 | -0.473 | 0.052 | -0.035 | -0.01 | -0.01 | 1.151 | 0.31 | 0.44 | 0.29 | 0.34 | 0.24 | 0.36 |
| 6 | 87219320 | rs9342077 | 1.523 | 0.321 | -0.66 | -0.009 | 0.011 | 0.061 | 0.061 | 0.322 | 0.18 | 0.07 | 0.09 | 0.07 | 0.17 | 0.2 |
| 6 | 87220627 | rs1948991 | -0.409 | 0.559 | -0.714 | 0.047 | 0.004 | -0.002 | -0.002 | 0 | 0.37 | 0.44 | 0.36 | 0.26 | 0.39 | 0.29 |
| 6 | 87220776 | rs1112672 | 1.059 | 0.732 | -0.805 | -0.028 | -0.027 | -0.042 | -0.042 | 0.547 | 0.1 | 0.06 | 0.15 | 0.22 | 0.12 | 0.05 |
| 6 | 87229518 | rs16877346 | 0.711 | 0.73 | -0.663 | -0.066 | 0.007 | -0.048 | -0.048 | 0 | 0.14 | 0.19 | 0.13 | 0 | 0.16 | 0.1 |
| 6 | 87243828 | rs1338715 | -0.113 | 0.685 | -0.689 | -0.086 | 0.032 | 0.038 | 0.038 | 0.879 | 0.55 | 0.34 | 0.63 | 0.63 | 0.64 | 0.48 |
| 6 | 87245600 | rs1567266 | 1.06 | 0.918 | -0.494 | 0.12 | -0.001 | -0.013 | -0.013 | 0 | 0.13 | 0.13 | 0.08 | 0.03 | 0.11 | 0.14 |
| 6 | 87251194 | rs952571 | 1.099 | 0.866 | -0.541 | 0.144 | -0.03 | -0.047 | -0.047 | 0 | 0.3 | 0.08 | 0.34 | 0.41 | 0.36 | 0.34 |
| 6 | 87265335 | rs4428472 | -0.156 | 0.709 | -0.505 | 0.036 | 0 | 0 | 0 | 1.345 | 0.57 | 0.42 | 0.55 | 0.44 | 0.63 | 0.58 |
| 6 | 87294410 | rs12527613 | 0.227 | 0.772 | -0.818 | 0.357 | -0.015 | -0.001 | -0.001 | 0.846 | 0.43 | 0.23 | 0.42 | 0.44 | 0.47 | 0.48 |
| 6 | 87298528 | rs12212922 | 0.725 | 0.74 | -1.101 | 0.369 | -0.067 | -0.121 | -0.121 | 0 | 0.28 | 0.05 | 0.37 | 0.49 | 0.35 | 0.3 |
| 6 | 87304060 | rs10944251 | 1.003 | -0.4 | -0.875 | 0.037 | 0.001 | 0 | 0 | 0.956 | 0.17 | 0.13 | 0.18 | 0.14 | 0.15 | 0.24 |
| 6 | 87306370 | rs9450485 | -1.147 | -0.087 | -1.19 | -0.069 | -0.07 | -0.06 | -0.06 | 0.858 | 0.7 | 0.88 | 0.53 | 0.54 | 0.54 | 0.6 |
| 6 | 87314539 | rs4707306 | -0.139 | -0.58 | -0.879 | -0.479 | 0.012 | 0.013 | 0.013 | 0 | 0.58 | 0.37 | 0.69 | 0.77 | 0.69 | 0.6 |
| 6 | 87317857 | rs11755770 | 0.772 | -0.652 | -0.87 | -0.52 | -0.001 | -0.015 | -0.015 | 0 | 0.15 | 0.18 | 0.09 | 0.06 | 0.12 | 0.14 |
| 6 | 87318870 | rs6906784 | 1.231 | -0.616 | -0.852 | -0.385 | 0.005 | 0.036 | 0.036 | 0.893 | 0.17 | 0.05 | 0.18 | 0.14 | 0.15 | 0.24 |
| 6 | 87319151 | rs10944254 | -0.438 | -0.216 | -0.618 | -0.092 | 0.001 | -0.004 | -0.004 | 0 | 0.58 | 0.39 | 0.7 | 0.77 | 0.71 | 0.6 |
| 6 | 87333241 | rs9294358 | 0.28 | 0.008 | -0.213 | 0.444 | -0.009 | 0.035 | 0.035 | 0 | 0.72 | 0.59 | 0.76 | 0.69 | 0.77 | 0.8 |
| 6 | 87338453 | rs1538407 | 0.227 | -0.052 | -0.185 | 0.351 | 0 | 0.093 | 0.093 | 0 | 0.17 | 0.28 | 0.17 | 0.3 | 0.15 | 0.16 |
| 6 | 87338626 | rs1538408 | -0.067 | -0.073 | -0.336 | 0.391 | 0.001 | -0.066 | -0.066 | 0 | 0.14 | 0.07 | 0.15 | 0.24 | 0.11 | 0.06 |
| 6 | 87339591 | rs7758873 | 0.336 | 0.036 | 0.023 | 0.299 | -0.001 | -0.03 | -0.03 | 1.118 | 0.14 | 0.21 | 0.08 | 0.06 | 0.12 | 0.14 |
| 6 | 87345576 | rs16877559 | 0.372 | -0.077 | -0.189 | 0.236 | -0.003 | -0.03 | -0.03 | 0 | 0.15 | 0.21 | 0.09 | 0.06 | 0.12 | 0.13 |
| 6 | 87349163 | rs2211074 | 0.824 | -0.23 | -0.819 | 0.156 | -0.128 | -0.09 | -0.09 | 0 | 0.3 | 0.05 | 0.47 | 0.42 | 0.48 | 0.4 |
| 6 | 87351573 | rs1954290 | -0.136 | -0.3 | -0.625 | 0.198 | 0.015 | -0.002 | -0.002 | 0.778 | 0.46 | 0.35 | 0.57 | 0.5 | 0.6 | 0.54 |
| 6 | 87353474 | rs1935606 | 0.522 | -0.207 | -0.607 | 0.18 | -0.001 | -0.008 | -0.008 | 0.033 | 0.16 | 0.21 | 0.09 | 0.06 | 0.12 | 0.13 |
| 6 | 87358929 | rs16877616 | 0.54 | -0.181 | -0.745 | 0.205 | -0.003 | 0.036 | 0.036 | 0 | 0.14 | 0.1 | 0.08 | 0.06 | 0.12 | 0.13 |
| 6 | 87395567 | rs1325183 | 0.613 | -0.276 | -0.948 | 0.472 | -0.134 | -0.053 | -0.053 | 0 | 0.31 | 0.05 | 0.53 | 0.64 | 0.49 | 0.45 |
| 6 | 87408274 | rs592805 | -0.779 | -0.262 | -0.929 | 0.444 | -0.014 | 0.042 | 0.042 | 0 | 0.86 | 0.92 | 0.93 | 0.94 | 0.88 | 0.87 |
| 6 | 87412973 | rs1066335 | -0.787 | -0.266 | -0.861 | 0.441 | 0 | 0.003 | 0.003 | 0 | 0.85 | 0.82 | 0.92 | 0.94 | 0.88 | 0.87 |
| 6 | 87424388 | rs499254 | -0.062 | -0.253 | -0.869 | 0.639 | 0.008 | -0.016 | -0.016 | 0 | 0.57 | 0.72 | 0.46 | 0.51 | 0.41 | 0.44 |
| 6 | 87430465 | rs1325182 | 0.583 | -0.262 | -1.133 | 0.664 | -0.13 | -0.047 | -0.047 | 0 | 0.31 | 0.06 | 0.53 | 0.64 | 0.49 | 0.45 |
| 6 | 87432577 | rs1292672 | 1.345 | -0.788 | -1.241 | 0.344 | 0.01 | -0.002 | -0.002 | 0 | 0.17 | 0.15 | 0.18 | 0.2 | 0.15 | 0.24 |
| 6 | 87439714 | rs1325181 | 0.748 | -0.546 | -1.431 | 0.448 | -0.134 | -0.05 | -0.05 | 0 | 0.31 | 0.05 | 0.53 | 0.64 | 0.49 | 0.45 |
| 6 | 87449564 | rs654118 | -0.27 | -0.559 | -1.432 | 0.45 | -0.009 | -0.05 | -0.05 | 0 | 0.58 | 0.85 | 0.47 | 0.51 | 0.41 | 0.44 |
| 6 | 87456331 | rs13202251 | 1.011 | -0.546 | -1.451 | 0.454 | -0.125 | -0.095 | -0.095 | 0 | 0.28 | 0.05 | 0.46 | 0.43 | 0.47 | 0.42 |
| 6 | 87464604 | rs476014 | 0.009 | -0.561 | -1.372 | 0.472 | 0.026 | 0.061 | 0.061 | 0.704 | 0.53 | 0.69 | 0.38 | 0.28 | 0.39 | 0.41 |
| 6 | 87466492 | rs493187 | -0.182 | -0.41 | -1.291 | 0.199 | 0.01 | -0.014 | -0.014 | 0 | 0.57 | 0.75 | 0.46 | 0.51 | 0.41 | 0.44 |
| 6 | 87474883 | rs9342091 | 1.609 | -0.141 | -1.688 | 0.393 | -0.016 | -0.016 | -0.016 | 0 | 0.06 | 0 | 0.21 | 0.29 | 0.05 | 0.09 |
| 6 | 87478447 | rs9362361 | 1.586 | -0.479 | -1.783 | 0.068 | -0.019 | 0.071 | 0.071 | 0.43 | 0.15 | 0.01 | 0.09 | 0.05 | 0.13 | 0.18 |
| 6 | 87479890 | rs503887 | 0.097 | -0.197 | -1.487 | 0.326 | 0.011 | -0.007 | -0.007 | 0 | 0.56 | 0.7 | 0.45 | 0.51 | 0.41 | 0.44 |
| 6 | 87483694 | rs494991 | -0.547 | -0.215 | -1.583 | 0.449 | -0.027 | 0.041 | 0.041 | 0 | 0.86 | 0.95 | 0.92 | 0.93 | 0.88 | 0.87 |
| 6 | 87486840 | rs9353427 | 0.143 | -0.11 | -1.638 | 0.468 | 0.006 | 0.034 | 0.034 | 0 | 0.47 | 0.31 | 0.57 | 0.69 | 0.48 | 0.59 |
| 6 | 87491157 | rs12198149 | 0.736 | -0.392 | -1.92 | 0.673 | -0.069 | -0.072 | -0.072 | 2.283 | 0.27 | 0.08 | 0.37 | 0.45 | 0.36 | 0.38 |
| 6 | 87492241 | rs584256 | -1.183 | -0.369 | -2.064 | 0.616 | -0.03 | 0.032 | 0.032 | 0 | 0.87 | 0.95 | 0.91 | 0.94 | 0.87 | 0.86 |
| 6 | 87494690 | rs9351108 | 0.69 | -0.317 | -1.823 | 0.752 | -0.069 | -0.072 | -0.072 | 0.189 | 0.27 | 0.08 | 0.36 | 0.45 | 0.36 | 0.38 |

| | | | | | | | | | | | | | | | | |
|---|---|---|---|---|---|---|---|---|---|---|---|---|---|---|---|---|
| 6 | 87500445 | rs6454568 | -0.397 | -0.506 | -1.67 | 0.773 | 0.008 | 0.032 | 0.032 | 0 | 0.5 | 0.37 | 0.59 | 0.72 | 0.5 | 0.66 |
| 6 | 87500866 | rs6911565 | 0.156 | -0.854 | -1.066 | 0.808 | -0.017 | 0.005 | 0.005 | 0.075 | 0.2 | 0.21 | 0.21 | 0.24 | 0.12 | 0.2 |
| 6 | 87509113 | rs9344653 | 0.227 | -0.756 | -1.239 | 0.803 | 0.057 | 0.002 | 0.002 | 0.388 | 0.22 | 0.04 | 0.13 | 0.14 | 0.13 | 0.21 |
| 6 | 87532216 | rs6918275 | 0.682 | 0.498 | -1.045 | 1.081 | -0.145 | -0.05 | -0.05 | 0 | 0.25 | 0.02 | 0.19 | 0.26 | 0.37 | 0.34 |
| 6 | 87533862 | rs1358781 | -0.173 | 0.313 | -0.544 | 1.175 | -0.03 | 0.061 | 0.061 | 0 | 0.87 | 0.97 | 0.92 | 0.97 | 0.88 | 0.87 |
| 6 | 87540023 | rs7751208 | 1.401 | 0.289 | 0.15 | 1.48 | 0.003 | 0.034 | 0.034 | 1.619 | 0.76 | 0.42 | 0.6 | 0.66 | 0.74 | 0.78 |
| 6 | 87541825 | rs941660 | 0.834 | 0.433 | -0.197 | 1.22 | 0.002 | 0 | 0 | 0.022 | 0.81 | 0.8 | 0.81 | 0.76 | 0.76 | 0.81 |
| 6 | 87541925 | rs941661 | 0.796 | 0.442 | -0.237 | 1.224 | -0.002 | -0.001 | -0.001 | 0 | 0.81 | 0.83 | 0.81 | 0.76 | 0.76 | 0.81 |
| 6 | 87545780 | rs2145049 | -0.416 | 0.521 | -0.206 | 1.316 | -0.004 | 0.023 | 0.023 | 0 | 0.06 | 0.05 | 0.11 | 0.21 | 0.09 | 0.05 |
| 6 | 87557277 | rs9444441 | -0.315 | 0.413 | 0.071 | 0.685 | -0.008 | -0.058 | -0.058 | 0.145 | 0.14 | 0.25 | 0.07 | 0.03 | 0.08 | 0.15 |
| 6 | 87562096 | rs10944276 | 0.886 | 0.407 | -0.21 | 0.574 | -0.078 | -0.005 | -0.005 | 0 | 0.33 | 0.08 | 0.3 | 0.31 | 0.39 | 0.36 |
| 6 | 87564247 | rs9450536 | -0.161 | 0.439 | -0.158 | 0.536 | -0.01 | -0.059 | -0.059 | 0 | 0.15 | 0.25 | 0.07 | 0.03 | 0.08 | 0.15 |
| 6 | 87565559 | rs16877863 | 0.151 | 0.47 | -0.124 | 0.529 | 0.001 | -0.037 | -0.037 | 0 | 0.14 | 0.18 | 0.08 | 0.04 | 0.11 | 0.14 |
| 6 | 87566319 | rs9450537 | -0.741 | 0.408 | 0.003 | 0.685 | -0.001 | -0.001 | -0.001 | 0 | 0.57 | 0.61 | 0.7 | 0.57 | 0.62 | 0.58 |
| 6 | 87569383 | rs12525091 | -0.651 | 0.452 | 0.556 | 0.769 | -0.004 | 0.004 | 0.004 | 0 | 0.06 | 0.06 | 0.12 | 0.15 | 0.1 | 0.05 |
| 6 | 87599076 | rs1535228 | -0.723 | 0.422 | 0.509 | 0.806 | 0 | -0.076 | -0.076 | 0 | 0.16 | 0.31 | 0.1 | 0.04 | 0.12 | 0.16 |
| 6 | 87615970 | rs9450576 | -1.845 | 0.415 | 0.559 | 0.916 | 0.008 | 0.037 | 0.037 | 0.457 | 0.24 | 0.48 | 0.4 | 0.27 | 0.24 | 0.22 |
| 6 | 87621929 | rs745172 | -1.944 | 0.397 | 0.551 | 0.879 | 0.002 | 0.027 | 0.027 | 1.195 | 0.25 | 0.55 | 0.41 | 0.27 | 0.24 | 0.22 |
| 6 | 87626449 | rs16877989 | -0.861 | 0.501 | 0.167 | 0.731 | -0.002 | 0.006 | 0.006 | 0 | 0.06 | 0.08 | 0.12 | 0.15 | 0.1 | 0.05 |
| 6 | 87626774 | rs16877991 | -0.562 | 0.465 | 0.108 | 0.744 | 0.001 | -0.064 | -0.064 | 0.459 | 0.16 | 0.27 | 0.09 | 0.04 | 0.12 | 0.16 |
| 6 | 87628559 | rs16877997 | -0.386 | 0.471 | 0.106 | 0.746 | 0.001 | -0.038 | -0.038 | 0 | 0.14 | 0.18 | 0.08 | 0.04 | 0.11 | 0.14 |
| 6 | 87630007 | rs11966339 | -0.349 | 0.504 | -0.009 | 0.768 | -0.026 | -0.025 | -0.025 | 0.326 | 0.05 | 0.01 | 0.12 | 0.15 | 0.1 | 0.05 |
| 6 | 87638339 | rs9450594 | -0.739 | 0.406 | 0.103 | 0.678 | -0.003 | -0.078 | -0.078 | 0 | 0.17 | 0.32 | 0.1 | 0.04 | 0.12 | 0.16 |
| 6 | 87646919 | rs11964560 | -0.795 | 0.636 | -0.282 | 1.138 | 0.001 | 0.021 | 0.021 | 0.049 | 0.05 | 0.08 | 0.08 | 0.15 | 0.09 | 0.05 |
| 6 | 87648471 | rs828361 | -0.85 | -0.016 | -0.259 | 0.664 | 0 | -0.002 | -0.002 | 0 | 0.22 | 0.23 | 0.3 | 0.11 | 0.16 | 0.14 |
| 6 | 87651742 | rs10944288 | -0.425 | 0.271 | -0.281 | 0.867 | 0.002 | 0.023 | 0.023 | 0.422 | 0.66 | 0.59 | 0.73 | 0.55 | 0.66 | 0.64 |
| 6 | 87664609 | rs1886333 | -1.014 | 0.225 | -0.039 | 0.911 | -0.019 | 0.02 | 0.02 | 0 | 0.24 | 0.39 | 0.22 | 0.27 | 0.18 | 0.16 |
| 6 | 87666493 | rs16878041 | -0.835 | 0.314 | 0.033 | 1.021 | -0.011 | 0.026 | 0.026 | 0 | 0.22 | 0.37 | 0.22 | 0.28 | 0.17 | 0.15 |
| 6 | 87676707 | rs942472 | -0.693 | 0.206 | -0.036 | 1.17 | 0.002 | 0.024 | 0.024 | 3.322 | 0.71 | 0.65 | 0.77 | 0.6 | 0.7 | 0.74 |
| 6 | 87682125 | rs6922679 | -0.68 | 0.407 | 0.014 | 1.21 | -0.013 | 0.017 | 0.017 | 0 | 0.18 | 0.25 | 0.17 | 0.29 | 0.12 | 0.13 |
| 6 | 87692655 | rs722763 | -0.486 | 0.137 | 0.066 | 1.255 | -0.023 | 0.068 | 0.068 | 0 | 0.15 | 0.32 | 0.17 | 0.27 | 0.1 | 0.11 |
| 6 | 87709146 | rs6936751 | -0.599 | -0.027 | -0.026 | 1.251 | -0.061 | 0.078 | 0.078 | 0 | 0.15 | 0.33 | 0.14 | 0.28 | 0.05 | 0.1 |
| 6 | 87722604 | rs9344662 | -1.336 | 0.042 | -0.071 | 1.256 | -0.028 | 0.048 | 0.048 | 0 | 0.2 | 0.36 | 0.18 | 0.31 | 0.13 | 0.16 |
| 6 | 87729868 | rs11961178 | -0.547 | 0.149 | -0.182 | 1.096 | -0.057 | 0.072 | 0.072 | 0 | 0.15 | 0.3 | 0.14 | 0.28 | 0.05 | 0.1 |
| 6 | 87730855 | rs16878151 | -0.423 | 0.15 | -0.156 | 1.097 | -0.042 | 0.048 | 0.048 | 0 | 0.14 | 0.23 | 0.13 | 0.28 | 0.05 | 0.1 |
| 6 | 87732983 | rs12195969 | -0.676 | -0.046 | -0.475 | 0.917 | -0.014 | 0.002 | 0.002 | 0 | 0.29 | 0.34 | 0.24 | 0.31 | 0.23 | 0.34 |
| 6 | 87740064 | rs16878170 | -0.551 | 0.073 | -0.429 | 0.807 | 0.004 | -0.021 | -0.021 | 2.537 | 0.06 | 0.14 | 0.06 | 0 | 0.08 | 0.06 |
| 6 | 87749815 | rs7739124 | 0.777 | -0.016 | -0.436 | 0.874 | -0.009 | 0.001 | 0.001 | 0.008 | 0.67 | 0.58 | 0.72 | 0.67 | 0.7 | 0.65 |
| 6 | 87757319 | rs7752881 | -0.209 | 0.187 | -0.474 | 0.81 | -0.025 | 0.024 | 0.024 | 0 | 0.06 | 0.04 | 0.06 | 0 | 0.12 | 0.08 |
| 6 | 87763956 | rs7761538 | 0.819 | 0.195 | -0.337 | 0.987 | 0 | 0.004 | 0.004 | 0 | 0.67 | 0.61 | 0.72 | 0.61 | 0.63 | 0.68 |
| 6 | 87772911 | rs11964150 | 0.584 | -0.048 | -0.912 | 0.894 | 0.016 | -0.027 | -0.027 | 0.004 | 0.38 | 0.28 | 0.55 | 0.52 | 0.31 | 0.38 |
| 6 | 87774205 | rs9342101 | -0.367 | 0.125 | -1.056 | 0.508 | 0 | -0.008 | -0.008 | 0 | 0.78 | 0.89 | 0.87 | 0.76 | 0.76 | 0.7 |
| 6 | 87780737 | rs9359727 | 0.767 | -0.225 | -0.922 | 0.573 | -0.005 | -0.055 | -0.055 | 10.59 | 0.33 | 0.15 | 0.48 | 0.44 | 0.31 | 0.39 |
| 6 | 87788479 | rs9342103 | 0.525 | -0.214 | -0.846 | 0.558 | 0.008 | -0.03 | -0.03 | 5.221 | 0.35 | 0.22 | 0.49 | 0.44 | 0.31 | 0.39 |
| 6 | 87789164 | rs17449571 | -0.044 | -0.07 | -1.156 | 0.333 | -0.036 | 0.027 | 0.027 | 0 | 0.08 | 0.03 | 0.07 | 0.02 | 0.14 | 0.06 |
| 6 | 87794580 | rs9359730 | 1.026 | -0.247 | -1.215 | 0.392 | -0.025 | -0.106 | -0.106 | 0 | 0.28 | 0.07 | 0.46 | 0.46 | 0.28 | 0.37 |
| 6 | 87795310 | rs1055409 | -0.388 | -0.15 | -0.086 | 0.041 | 0.037 | -0.032 | -0.032 | 0 | 0.16 | 0.37 | 0.14 | 0.1 | 0.23 | 0.14 |
| 6 | 87805592 | rs2031365 | -1.474 | -0.557 | -0.245 | -0.087 | 0.018 | 0.026 | 0.026 | 0.95 | 0.37 | 0.74 | 0.28 | 0.27 | 0.29 | 0.3 |
| 6 | 87809402 | rs981087 | 0.129 | -0.173 | 0.046 | 0.069 | 0.008 | -0.002 | -0.002 | 0 | 0.43 | 0.57 | 0.6 | 0.61 | 0.5 | 0.5 |
| 6 | 87813843 | rs6909630 | -0.514 | 0.282 | 0.202 | -0.148 | 0.013 | -0.007 | -0.007 | 0 | 0.16 | 0.23 | 0.14 | 0.17 | 0.22 | 0.14 |
| 6 | 87814241 | rs6910649 | -0.53 | 0.26 | 0.198 | -0.14 | 0.01 | -0.007 | -0.007 | 0.039 | 0.16 | 0.23 | 0.14 | 0.17 | 0.22 | 0.14 |
| 6 | 87816905 | rs4574603 | -0.074 | -0.345 | 0.421 | -0.352 | -0.001 | -0.002 | -0.002 | 0.013 | 0.54 | 0.53 | 0.65 | 0.6 | 0.47 | 0.62 |
| 6 | 87821501 | rs2031522 | 0.737 | -0.41 | 0.76 | -0.356 | 0.01 | -0.017 | -0.017 | 0 | 0.43 | 0.25 | 0.58 | 0.47 | 0.37 | 0.51 |
| 6 | 87834068 | rs6454581 | -0.672 | -0.584 | 0.779 | -0.543 | -0.108 | -0.048 | -0.048 | 0.628 | 0.63 | 0.88 | 0.71 | 0.6 | 0.51 | 0.65 |
| 6 | 87843365 | rs9353455 | -0.028 | -0.597 | 0.939 | -0.546 | -0.001 | 0.002 | 0.002 | 0 | 0.54 | 0.56 | 0.66 | 0.6 | 0.47 | 0.62 |
| 6 | 87856003 | rs6907805 | -0.078 | -0.607 | 1.186 | -0.551 | -0.006 | 0.008 | 0.008 | 0.004 | 0.56 | 0.63 | 0.67 | 0.6 | 0.47 | 0.62 |
| 6 | 87870819 | rs1886751 | -0.665 | -0.604 | 1.083 | -0.597 | -0.099 | -0.041 | -0.041 | 0.015 | 0.63 | 0.87 | 0.71 | 0.61 | 0.51 | 0.66 |
| 6 | 87872702 | rs9444479 | -0.683 | -0.6 | 1.084 | -0.596 | -0.099 | -0.042 | -0.042 | 0.208 | 0.63 | 0.87 | 0.71 | 0.6 | 0.51 | 0.66 |
| 6 | 87874389 | rs9362399 | 0.107 | -0.637 | 1.261 | -0.605 | -0.008 | 0.003 | 0.003 | 0 | 0.11 | 0.2 | 0.06 | 0.13 | 0.09 | 0.11 |
| 6 | 87885952 | rs10085214 | -0.191 | -0.832 | 1.285 | -0.624 | -0.006 | -0.001 | -0.001 | 0.77 | 0.14 | 0.11 | 0.16 | 0.11 | 0.11 | 0.11 |
| 6 | 87886186 | rs7739034 | -0.819 | -0.802 | 1.209 | -0.617 | -0.105 | -0.046 | -0.046 | 0 | 0.63 | 0.89 | 0.71 | 0.6 | 0.51 | 0.66 |
| 6 | 87887245 | rs4706289 | -0.819 | -0.802 | 1.209 | -0.617 | -0.099 | -0.042 | -0.042 | 0.286 | 0.63 | 0.87 | 0.71 | 0.6 | 0.51 | 0.66 |
| 6 | 87891960 | rs6904310 | -0.819 | -0.802 | 1.209 | -0.617 | -0.099 | -0.042 | -0.042 | 2.785 | 0.63 | 0.87 | 0.71 | 0.6 | 0.51 | 0.66 |
| 6 | 87920555 | rs4707349 | 1.709 | -0.868 | 1.355 | -0.673 | 0.002 | -0.028 | -0.028 | 0 | 0.31 | 0.15 | 0.43 | 0.37 | 0.27 | 0.4 |
| 6 | 87922320 | rs4099965 | 1.161 | -0.894 | 1.181 | -0.7 | -0.007 | 0.013 | 0.013 | 1.25 | 0.06 | 0.02 | 0.06 | 0.02 | 0.08 | 0.02 |
| 6 | 87930169 | rs9450636 | 0.561 | -0.867 | 1.124 | -0.763 | 0.013 | -0.012 | -0.012 | 0.543 | 0.44 | 0.26 | 0.58 | 0.46 | 0.37 | 0.51 |
| 6 | 87937025 | rs7774603 | -0.278 | -0.875 | 1.177 | -0.766 | -0.004 | 0.005 | 0.005 | 0 | 0.56 | 0.62 | 0.67 | 0.6 | 0.47 | 0.63 |
| 6 | 87965528 | rs6908834 | 0.564 | -0.875 | 1.173 | -0.765 | 0.013 | -0.012 | -0.012 | 0 | 0.44 | 0.26 | 0.58 | 0.46 | 0.37 | 0.51 |
| 6 | 87967636 | rs6941356 | 0.532 | -0.88 | 1.132 | -0.767 | -0.009 | -0.027 | -0.027 | 0 | 0.08 | 0.03 | 0.05 | 0.14 | 0.1 | 0.11 |

| | | | | | | | | | | | | | | | | |
|---|---|---|---|---|---|---|---|---|---|---|---|---|---|---|---|---|
| 6 | 87968565 | rs9362415 | -0.275 | -0.866 | 1.142 | -0.761 | -0.004 | 0.005 | 0.005 | 0 | 0.56 | 0.62 | 0.67 | 0.6 | 0.47 | 0.63 |
| 6 | 87969341 | rs16878809 | -0.743 | -0.871 | 1.207 | -0.77 | 0.007 | -0.034 | -0.034 | 0 | 0.07 | 0.22 | 0.04 | 0 | 0.04 | 0.03 |
| 6 | 87969737 | rs3734187 | -0.272 | -0.871 | 1.186 | -0.77 | -0.004 | 0.005 | 0.005 | 0 | 0.56 | 0.62 | 0.67 | 0.6 | 0.47 | 0.63 |
| 6 | 87976073 | rs4707356 | -0.923 | -0.871 | 1.167 | -0.768 | -0.115 | -0.06 | -0.06 | 0.006 | 0.63 | 0.9 | 0.71 | 0.6 | 0.51 | 0.66 |
| 6 | 87977775 | rs9450645 | -0.186 | -0.873 | 1.171 | -0.756 | 0.01 | -0.002 | -0.002 | 0.056 | 0.53 | 0.55 | 0.62 | 0.46 | 0.41 | 0.54 |
| 6 | 87991236 | rs6916994 | -0.21 | -0.864 | 1.082 | -0.773 | -0.002 | 0.003 | 0.003 | 0 | 0.55 | 0.61 | 0.66 | 0.6 | 0.47 | 0.63 |
| 6 | 87993090 | rs2240461 | 1.873 | -0.816 | 1.106 | -0.791 | -0.03 | -0.074 | -0.074 | 0 | 0.29 | 0.07 | 0.41 | 0.37 | 0.26 | 0.4 |
| 6 | 87993761 | rs3812130 | -0.175 | -0.824 | 1.188 | -0.726 | -0.002 | 0.002 | 0.002 | 0 | 0.55 | 0.59 | 0.66 | 0.6 | 0.47 | 0.63 |
| 6 | 87994015 | rs6934603 | -0.175 | -0.824 | 1.188 | -0.726 | -0.002 | 0.002 | 0.002 | 0 | 0.55 | 0.59 | 0.66 | 0.6 | 0.47 | 0.63 |
| 6 | 87994572 | rs4707358 | -0.86 | -0.806 | 0.928 | -0.715 | -0.105 | -0.052 | -0.052 | 0 | 0.63 | 0.89 | 0.71 | 0.6 | 0.51 | 0.66 |
| 6 | 88008168 | rs10498960 | -0.986 | -0.649 | 0.919 | -0.646 | -0.004 | -0.043 | -0.043 | 0 | 0.08 | 0.21 | 0.04 | 0 | 0.05 | 0.03 |
| 6 | 88018313 | rs242263 | 0.041 | -0.741 | 0.864 | -0.704 | 0.013 | 0.002 | 0.002 | 0 | 0.48 | 0.41 | 0.37 | 0.54 | 0.58 | 0.47 |
| 6 | 88021118 | rs1145712 | -0.192 | -0.78 | 0.849 | -0.692 | 0.027 | 0.003 | 0.003 | 0 | 0.51 | 0.41 | 0.39 | 0.54 | 0.63 | 0.49 |
| 6 | 88029197 | rs16878999 | -0.528 | -0.562 | 0.774 | -0.554 | 0 | -0.024 | -0.024 | 0 | 0.06 | 0.21 | 0.02 | 0 | 0.02 | 0.01 |
| 6 | 88030587 | rs2875533 | 1.782 | -0.62 | 0.234 | -0.609 | -0.037 | -0.077 | -0.077 | 0 | 0.25 | 0.06 | 0.41 | 0.36 | 0.24 | 0.4 |
| 6 | 88039779 | rs9359746 | 0.004 | -0.586 | 0.4 | -0.57 | 0 | -0.001 | -0.001 | 0 | 0.1 | 0.13 | 0.08 | 0.13 | 0.12 | 0.13 |
| 6 | 88045963 | rs242275 | -0.63 | -0.667 | 0.478 | -0.628 | 0.007 | -0.012 | -0.012 | 0 | 0.58 | 0.73 | 0.41 | 0.54 | 0.63 | 0.54 |
| 6 | 88046800 | rs1048893 | 0.117 | -0.568 | 0.32 | -0.549 | -0.001 | -0.004 | -0.004 | 0 | 0.09 | 0.1 | 0.07 | 0.13 | 0.12 | 0.13 |
| 6 | 88051052 | rs397516 | -0.647 | -0.726 | 0.274 | -0.743 | 0.001 | -0.018 | -0.018 | 0 | 0.61 | 0.74 | 0.41 | 0.57 | 0.63 | 0.49 |
| 6 | 88057130 | rs381798 | -0.17 | -0.204 | 0.622 | -0.292 | 0.007 | -0.002 | -0.002 | 0.038 | 0.53 | 0.66 | 0.4 | 0.54 | 0.61 | 0.52 |
| 6 | 88058675 | rs9450659 | 1.132 | -0.246 | 0.64 | -0.244 | 0 | -0.008 | -0.008 | 0 | 0.31 | 0.18 | 0.43 | 0.33 | 0.27 | 0.42 |
| 6 | 88061884 | rs242268 | -0.003 | -0.179 | 0.415 | -0.212 | 0.012 | 0 | 0 | 0 | 0.52 | 0.47 | 0.38 | 0.54 | 0.61 | 0.49 |
| 6 | 88067157 | rs450712 | 0.168 | -0.023 | 0.156 | -0.019 | -0.005 | -0.001 | -0.001 | 0 | 0.85 | 0.88 | 0.83 | 0.87 | 0.88 | 0.94 |
| 6 | 88075122 | rs9353482 | -0.206 | 0.232 | 0.166 | 0.116 | -0.002 | -0.003 | -0.003 | 0 | 0.1 | 0.11 | 0.07 | 0.14 | 0.11 | 0.13 |
| 6 | 88078615 | rs6931037 | -0.361 | 0.2 | 0.199 | 0.083 | -0.001 | -0.001 | -0.001 | 0 | 0.11 | 0.12 | 0.07 | 0.13 | 0.12 | 0.13 |
| 6 | 88089160 | rs1884320 | 0.982 | -0.017 | 0.567 | 0.023 | -0.01 | 0.001 | 0.001 | 0.036 | 0.75 | 0.65 | 0.76 | 0.73 | 0.78 | 0.81 |
| 6 | 88097764 | rs4707364 | 1.369 | -0.033 | 0.772 | -0.143 | 0.001 | 0.008 | 0.008 | 0 | 0.28 | 0.29 | 0.44 | 0.33 | 0.26 | 0.35 |
| 6 | 88109304 | rs1051550 | 1.196 | 0.269 | 0.923 | -0.145 | -0.002 | -0.005 | -0.005 | 0.066 | 0.7 | 0.61 | 0.73 | 0.76 | 0.7 | 0.79 |
| 6 | 88126423 | rs16879281 | 0.089 | 0.296 | 0.879 | -0.137 | -0.005 | -0.001 | -0.001 | 0.118 | 0.09 | 0.09 | 0.07 | 0.1 | 0.12 | 0.12 |
| 6 | 88127101 | rs16879289 | 0.778 | 0.26 | 0.903 | -0.162 | 0.001 | -0.004 | -0.004 | 0 | 0.09 | 0.07 | 0.12 | 0.12 | 0.09 | 0.05 |
| 6 | 88128267 | rs6454618 | -1.621 | 0.197 | 1.07 | -0.182 | -0.01 | -0.024 | -0.024 | 0 | 0.32 | 0.55 | 0.29 | 0.24 | 0.3 | 0.21 |
| 6 | 88129314 | rs6900397 | -1.735 | 0.174 | 1.066 | -0.194 | -0.01 | -0.02 | -0.02 | 0 | 0.33 | 0.61 | 0.29 | 0.24 | 0.3 | 0.21 |
| 6 | 88135708 | rs728582 | 1.479 | 0.227 | 0.849 | -0.064 | -0.004 | -0.014 | -0.014 | 0 | 0.28 | 0.15 | 0.43 | 0.3 | 0.26 | 0.41 |
| 6 | 88142979 | rs6940716 | -0.963 | 0.229 | 0.89 | -0.067 | -0.001 | -0.001 | -0.001 | 0.09 | 0.24 | 0.27 | 0.25 | 0.23 | 0.26 | 0.2 |
| 6 | 88145338 | rs9450681 | 1.423 | 0.234 | 0.807 | -0.041 | -0.001 | -0.008 | -0.008 | 1.289 | 0.28 | 0.21 | 0.44 | 0.3 | 0.26 | 0.4 |
| 6 | 88152535 | rs9294379 | -0.44 | 0.388 | 0.705 | 0.143 | 0 | 0.001 | 0.001 | 0 | 0.59 | 0.58 | 0.7 | 0.53 | 0.55 | 0.61 |
| 6 | 88166279 | rs2179845 | -0.319 | 0.515 | 0.099 | 0.31 | 0 | -0.001 | -0.001 | 0 | 0.15 | 0.16 | 0.17 | 0.13 | 0.14 | 0.07 |
| 6 | 88171627 | rs10944312 | 0.439 | 0.89 | -0.314 | 0.304 | -0.051 | 0.007 | 0.007 | 0.247 | 0.14 | 0.01 | 0.08 | 0.08 | 0.22 | 0.09 |
| 6 | 88182439 | rs2273129 | -0.487 | 0.888 | 0.124 | 0.468 | -0.019 | -0.012 | -0.012 | 0 | 0.58 | 0.76 | 0.7 | 0.52 | 0.53 | 0.59 |
| 6 | 88184912 | rs12524205 | 0.969 | 0.778 | 0.62 | 0.389 | 0.001 | 0.002 | 0.002 | 0 | 0.32 | 0.24 | 0.43 | 0.3 | 0.3 | 0.37 |
| 6 | 88185777 | rs2268996 | 0.448 | 0.76 | 1.092 | 0.342 | -0.002 | -0.005 | -0.005 | 0.013 | 0.37 | 0.42 | 0.49 | 0.31 | 0.35 | 0.39 |
| 6 | 88188253 | rs2300900 | 1.064 | 0.754 | 1.027 | 0.344 | 0 | 0.001 | 0.001 | 0 | 0.31 | 0.22 | 0.43 | 0.3 | 0.3 | 0.37 |
| 6 | 88202841 | rs9450716 | 0.433 | 0.753 | 1.256 | 0.353 | -0.004 | -0.014 | -0.014 | 0 | 0.38 | 0.5 | 0.5 | 0.31 | 0.35 | 0.39 |
| 6 | 88221892 | rs1051148 | -0.84 | 0.738 | 1.213 | 0.444 | -0.034 | -0.008 | -0.008 | 0 | 0.59 | 0.78 | 0.7 | 0.56 | 0.53 | 0.59 |
| 6 | 88224164 | rs8802 | 0.188 | 0.703 | 1.408 | 0.466 | -0.011 | 0.013 | 0.013 | 4.166 | 0.12 | 0.16 | 0.14 | 0.15 | 0.1 | 0.1 |
| 6 | 88239266 | rs17850652 | -0.134 | 0.708 | 1.371 | 0.556 | 0.003 | 0.002 | 0.002 | 0 | 0.09 | 0.12 | 0.06 | 0.11 | 0.09 | 0.1 |
| 6 | 88240047 | rs2295085 | -1.48 | 0.694 | 1.534 | 0.543 | -0.012 | 0.016 | 0.016 | 1.277 | 0.26 | 0.44 | 0.27 | 0.26 | 0.24 | 0.22 |
| 6 | 88250589 | rs12663400 | 0.299 | 0.756 | 1.523 | 0.672 | 0 | -0.001 | -0.001 | 0 | 0.41 | 0.45 | 0.5 | 0.4 | 0.39 | 0.47 |
| 6 | 88250620 | rs12660483 | 0.051 | 0.76 | 1.478 | 0.671 | -0.008 | -0.007 | -0.007 | 0.183 | 0.08 | 0.04 | 0.05 | 0.11 | 0.09 | 0.1 |
| 6 | 88256092 | rs7757636 | -0.985 | 0.717 | 1.544 | 0.648 | -0.081 | -0.044 | -0.044 | 0 | 0.6 | 0.89 | 0.71 | 0.56 | 0.54 | 0.59 |
| 6 | 88273969 | rs6900057 | 0.491 | 0.759 | 1.621 | 0.635 | 0.005 | -0.02 | -0.02 | 0 | 0.05 | 0.14 | 0.05 | 0.01 | 0.05 | 0.02 |
| 6 | 88291081 | rs9450749 | 0.955 | 0.735 | 1.507 | 0.66 | -0.001 | -0.001 | -0.001 | 0 | 0.33 | 0.34 | 0.44 | 0.3 | 0.3 | 0.37 |
| 6 | 88299310 | rs9450751 | 1.118 | 0.733 | 1.535 | 0.647 | 0 | 0.001 | 0.001 | 0 | 0.32 | 0.29 | 0.43 | 0.3 | 0.3 | 0.37 |
| 6 | 88311975 | rs4707380 | 0.36 | 0.734 | 1.494 | 0.649 | 0 | 0 | 0 | 0 | 0.41 | 0.42 | 0.49 | 0.4 | 0.39 | 0.48 |
| 6 | 88325270 | rs7769144 | 0.179 | 0.751 | 1.467 | 0.639 | -0.006 | 0.005 | 0.005 | 5.179 | 0.12 | 0.13 | 0.14 | 0.15 | 0.1 | 0.1 |
| 6 | 88329942 | rs13213285 | 0.534 | 0.762 | 1.506 | 0.64 | 0.004 | -0.019 | -0.019 | 0 | 0.05 | 0.11 | 0.05 | 0.01 | 0.05 | 0.02 |
| 6 | 88331299 | rs7772561 | -0.854 | 0.875 | 1.563 | 0.653 | -0.01 | 0 | 0 | 0.301 | 0.58 | 0.74 | 0.7 | 0.56 | 0.54 | 0.6 |
| 6 | 88343318 | rs9444530 | -0.44 | 0.887 | 1.512 | 0.662 | -0.006 | 0.011 | 0.011 | 0 | 0.53 | 0.62 | 0.65 | 0.55 | 0.49 | 0.58 |
| 6 | 88345048 | rs9450765 | 0.319 | 0.934 | 1.275 | 0.634 | 0.001 | 0 | 0 | 0 | 0.41 | 0.43 | 0.5 | 0.41 | 0.39 | 0.48 |
| 6 | 88346369 | rs16879646 | -0.094 | 1 | 0.93 | 0.631 | -0.003 | -0.003 | -0.003 | 0 | 0.08 | 0.07 | 0.06 | 0.11 | 0.09 | 0.1 |
| 6 | 88362943 | rs2307393 | 0.42 | 1.274 | 1.07 | 0.83 | 0.003 | -0.016 | -0.016 | 0 | 0.05 | 0.1 | 0.05 | 0.01 | 0.05 | 0.02 |
| 6 | 88366708 | rs2307373 | -0.796 | 1.364 | 0.733 | 1.346 | -0.033 | -0.059 | -0.059 | 0 | 0.32 | 0.16 | 0.4 | 0.48 | 0.43 | 0.28 |
| 6 | 88370957 | rs715207 | -0.01 | 1.539 | 1.725 | 1 | -0.009 | 0.008 | 0.008 | 4.025 | 0.12 | 0.12 | 0.13 | 0.13 | 0.1 | 0.1 |
| 6 | 88382859 | rs2754245 | -0.896 | 1.44 | 1.623 | 0.961 | -0.031 | -0.007 | -0.007 | 0 | 0.56 | 0.77 | 0.69 | 0.55 | 0.54 | 0.57 |
| 6 | 88391734 | rs2787917 | -1.148 | 1.415 | 1.908 | 0.924 | -0.004 | 0.032 | 0.032 | 3.717 | 0.2 | 0.55 | 0.22 | 0.24 | 0.19 | 0.24 |
| 6 | 88393705 | rs2250795 | -1.255 | 1.494 | 1.901 | 0.899 | 0.011 | 0.021 | 0.021 | 3.704 | 0.21 | 0.56 | 0.31 | 0.24 | 0.23 | 0.25 |
| 6 | 88397273 | rs2787919 | -1.162 | 1.086 | 1.94 | 0.96 | -0.004 | 0.032 | 0.032 | 3.717 | 0.2 | 0.55 | 0.22 | 0.24 | 0.19 | 0.24 |
| 6 | 88397755 | rs2250276 | -0.361 | 1.155 | 1.804 | 0.901 | 0.053 | 0.05 | 0.05 | 0 | 0.05 | 0.13 | 0.05 | 0.11 | 0.09 | 0.1 |
| 6 | 88398718 | rs4707385 | 0.89 | 1.167 | 1.674 | 0.895 | -0.009 | -0.003 | -0.003 | 0.056 | 0.31 | 0.2 | 0.43 | 0.3 | 0.3 | 0.37 |

| | | | | | | | | | | | | | | | | |
|---|---|---|---|---|---|---|---|---|---|---|---|---|---|---|---|---|
| 6 | 88411768 | rs9474 | 0.813 | 1.251 | 1.137 | 0.927 | -0.001 | 0 | 0 | 0 | 0.31 | 0.29 | 0.44 | 0.29 | 0.3 | 0.38 |
| 6 | 88412786 | rs2787927 | -0.81 | 1.106 | 0.831 | 0.956 | -0.032 | 0.015 | 0.015 | 0.727 | 0.55 | 0.85 | 0.7 | 0.6 | 0.52 | 0.62 |
| 6 | 88420902 | rs2787933 | -2.22 | 1.284 | 1.954 | 0.935 | -0.004 | 0.009 | 0.009 | 0.39 | 0.27 | 0.57 | 0.26 | 0.31 | 0.25 | 0.25 |
| 6 | 88427097 | rs12190021 | -0.224 | 1.267 | 1.792 | 0.844 | -0.002 | -0.01 | -0.01 | 0.208 | 0.05 | 0.05 | 0.04 | 0.01 | 0.04 | 0.01 |
| 6 | 88432049 | rs2787937 | -1.8 | 1.339 | 2.15 | 0.925 | 0.13 | 0.061 | 0.061 | 0 | 0.09 | 0.39 | 0.08 | 0.11 | 0.13 | 0.13 |
| 6 | 88441663 | rs6902334 | 1.42 | 1.306 | 1.703 | 0.957 | 0.053 | 0.018 | 0.018 | 0 | 0.93 | 0.8 | 0.93 | 0.89 | 0.88 | 0.87 |
| 6 | 88453491 | rs2325077 | -1.11 | 1.359 | 1.587 | 0.859 | 0 | -0.001 | -0.001 | 0 | 0.31 | 0.28 | 0.29 | 0.25 | 0.34 | 0.24 |
| 6 | 88455481 | rs9342120 | 1.568 | 1.326 | 1.409 | 0.829 | -0.06 | 0.012 | 0.012 | 0.138 | 0.58 | 0.16 | 0.61 | 0.64 | 0.53 | 0.63 |
| 6 | 88462545 | rs9344717 | 1.171 | 1.129 | 1.894 | 0.801 | 0.082 | 0.037 | 0.037 | 0 | 0.95 | 0.69 | 0.95 | 0.98 | 0.94 | 0.95 |
| 6 | 88466990 | rs4707403 | 1.44 | 0.857 | 1.209 | 0.815 | -0.028 | 0.033 | 0.033 | 1.498 | 0.62 | 0.16 | 0.64 | 0.73 | 0.6 | 0.71 |
| 6 | 88482324 | rs7744213 | -1.662 | 0.553 | 1.767 | 0.732 | 0.007 | -0.037 | -0.037 | 0 | 0.1 | 0.34 | 0.08 | 0.02 | 0.05 | 0.06 |
| 6 | 88486333 | rs6454648 | 1.824 | 0.501 | 1.504 | 1.174 | -0.01 | 0.071 | 0.071 | 0.164 | 0.84 | 0.46 | 0.77 | 0.79 | 0.86 | 0.87 |
| 6 | 88492526 | rs10944332 | -1.323 | 0.492 | 1.008 | 1.244 | 0 | -0.011 | -0.011 | 1.986 | 0.13 | 0.08 | 0.14 | 0.22 | 0.14 | 0.11 |
| 6 | 88495214 | rs7747510 | -1.455 | 0.207 | 1.722 | 0.992 | 0.022 | 0.012 | 0.012 | 1.998 | 0.07 | 0.41 | 0.05 | 0 | 0.01 | 0.05 |
| 6 | 88497286 | rs9344726 | 1.596 | 0.536 | 0.398 | 1.063 | -0.057 | -0.074 | -0.074 | 1.801 | 0.57 | 0.12 | 0.53 | 0.54 | 0.47 | 0.67 |
| 6 | 88498604 | rs2245604 | 1.589 | -0.088 | 0.891 | 0.781 | -0.045 | -0.002 | -0.002 | 0 | 0.76 | 0.65 | 0.83 | 0.8 | 0.85 | 0.87 |
| 6 | 88499661 | rs9294390 | 0.759 | 0.218 | 0.643 | 0.838 | -0.021 | -0.021 | -0.021 | 0 | 0.64 | 0.72 | 0.7 | 0.55 | 0.51 | 0.71 |
| 6 | 88501091 | rs9344728 | 0.987 | 0.323 | 0.131 | 0.819 | -0.03 | -0.017 | -0.017 | 0.554 | 0.57 | 0.26 | 0.67 | 0.57 | 0.53 | 0.69 |
| 6 | 88503309 | rs2787885 | -0.174 | 0.14 | -0.681 | -0.034 | -0.092 | 0.15 | 0.15 | 3.16 | 0.67 | 0.95 | 0.69 | 0.92 | 0.5 | 0.7 |
| 6 | 88503850 | rs2787887 | 0.393 | -0.35 | -0.461 | -0.29 | 0.048 | 0.134 | 0.134 | 2.318 | 0.49 | 0.76 | 0.68 | 0.91 | 0.41 | 0.61 |
| 6 | 88505654 | rs13202853 | -0.084 | -0.495 | -0.762 | 0.569 | -0.017 | 0.132 | 0.132 | 8.947 | 0.08 | 0.21 | 0.18 | 0.28 | 0.04 | 0.07 |
| 6 | 88505746 | rs2787889 | 0.496 | 0.065 | -0.713 | -0.329 | -0.021 | -0.16 | -0.16 | 0 | 0.75 | 0.63 | 0.86 | 0.99 | 0.76 | 0.85 |
| 6 | 88516052 | rs722903 | 0.444 | 0.277 | -0.872 | -0.009 | 0.013 | -0.008 | -0.008 | 0.269 | 0.3 | 0.43 | 0.25 | 0.23 | 0.38 | 0.31 |
| 6 | 88520263 | rs6922280 | 1.27 | 0.41 | -0.026 | -0.659 | -0.009 | -0.131 | -0.131 | 0.926 | 0.7 | 0.56 | 0.82 | 0.98 | 0.73 | 0.88 |
| 6 | 88523217 | rs1535986 | 0.377 | 0.422 | -0.31 | -0.682 | -0.012 | 0.012 | 0.012 | 0.072 | 0.89 | 0.92 | 0.93 | 0.99 | 0.87 | 0.97 |
| 6 | 88526850 | rs2787897 | -1.094 | -0.012 | -0.032 | -0.116 | -0.041 | -0.127 | -0.127 | 0 | 0.19 | 0.29 | 0.1 | 0.01 | 0.13 | 0.09 |
| 6 | 88541569 | rs2787901 | 0.08 | 0.646 | -0.862 | 0.489 | -0.014 | -0.019 | -0.019 | 0.785 | 0.56 | 0.74 | 0.45 | 0.45 | 0.45 | 0.53 |
| 6 | 88542614 | rs6926419 | -0.426 | 0.848 | -0.9 | 0.261 | -0.011 | 0.029 | 0.029 | 0 | 0.14 | 0.07 | 0.14 | 0.04 | 0.14 | 0.1 |
| 6 | 88549795 | rs718220 | -1.268 | 0.713 | 0.242 | 0.389 | 0.007 | -0.004 | -0.004 | 0.06 | 0.24 | 0.47 | 0.27 | 0.23 | 0.22 | 0.22 |
| 6 | 88556885 | rs1324689 | -1.459 | 0.34 | 1.223 | 0.623 | 0.029 | 0.208 | 0.208 | 1.143 | 0.08 | 0.43 | 0.14 | 0.23 | 0.07 | 0.12 |
| 6 | 88560672 | rs7764222 | -1.011 | 1.209 | 1.139 | 1.014 | 0.028 | 0.078 | 0.078 | 0 | 0.27 | 0.55 | 0.44 | 0.55 | 0.35 | 0.31 |
| 6 | 88571309 | rs6454657 | 1.099 | 1.038 | 0.944 | 0.285 | -0.001 | -0.005 | -0.005 | 0 | 0.49 | 0.56 | 0.35 | 0.24 | 0.46 | 0.41 |
| 6 | 88573884 | rs7753922 | 0.536 | 1.014 | 0.947 | 0.41 | -0.01 | 0.014 | 0.014 | 0 | 0.68 | 0.68 | 0.65 | 0.57 | 0.75 | 0.62 |
| 6 | 88577115 | rs9342127 | 0.224 | 0.619 | 0.778 | 0.201 | -0.026 | 0.092 | 0.092 | 0 | 0.81 | 0.71 | 0.79 | 0.6 | 0.88 | 0.74 |
| 6 | 88579560 | rs4129674 | 0.601 | 0.639 | 0.745 | 0.092 | 0.005 | -0.013 | -0.013 | 0 | 0.64 | 0.59 | 0.49 | 0.28 | 0.59 | 0.54 |
| 6 | 88585913 | rs9444562 | 1.281 | 0.843 | 0.763 | -0.01 | 0.009 | -0.039 | -0.039 | 0.448 | 0.49 | 0.51 | 0.26 | 0.01 | 0.4 | 0.33 |
| 6 | 88593579 | rs9359760 | 0.124 | 0.868 | 0.721 | 0.165 | -0.001 | 0.002 | 0.002 | 0.119 | 0.16 | 0.23 | 0.23 | 0.26 | 0.18 | 0.2 |
| 6 | 88605278 | rs4706301 | 0.762 | 0.928 | 0.594 | 0.141 | -0.059 | -0.055 | -0.055 | 0.681 | 0.16 | 0.05 | 0.3 | 0.31 | 0.31 | 0.21 |
| 6 | 88609666 | rs9362459 | -0.673 | 0.841 | 0.859 | 0.212 | 0.005 | 0.007 | 0.007 | 0 | 0.36 | 0.42 | 0.55 | 0.53 | 0.49 | 0.46 |
| 6 | 88612769 | rs6925312 | 0.626 | 0.852 | 0.739 | 0.039 | -0.014 | 0.021 | 0.021 | 0 | 0.65 | 0.74 | 0.49 | 0.24 | 0.58 | 0.53 |
| 6 | 88619412 | rs9344740 | 1.243 | 0.454 | 0.654 | 0.123 | 0.067 | 0.041 | 0.041 | 2.079 | 0.56 | 0.16 | 0.54 | 0.35 | 0.69 | 0.4 |
| 6 | 88620460 | rs12528768 | -0.907 | 0.367 | 1.117 | 0.07 | -0.004 | 0.003 | 0.003 | 0.45 | 0.28 | 0.35 | 0.38 | 0.65 | 0.24 | 0.47 |
| 6 | 88620665 | rs12528790 | -1.144 | 0.382 | 1.175 | 0.077 | -0.018 | 0.004 | 0.004 | 1.892 | 0.32 | 0.46 | 0.39 | 0.65 | 0.24 | 0.53 |
| 6 | 88624437 | rs9450829 | -0.183 | 0.386 | 0.911 | -0.181 | -0.005 | -0.036 | -0.036 | 0 | 0.91 | 0.88 | 0.97 | 1 | 0.96 | 0.96 |
| 6 | 88626861 | rs4076053 | -0.019 | -0.248 | -0.187 | -0.83 | 0.01 | 0.133 | 0.133 | 1.723 | 0.49 | 0.32 | 0.68 | 0.93 | 0.59 | 0.66 |
| 6 | 88628864 | rs9450833 | 0.35 | -0.286 | -0.93 | -0.918 | -0.035 | -0.138 | -0.138 | 1.472 | 0.7 | 0.53 | 0.83 | 0.97 | 0.85 | 0.74 |
| 6 | 88629773 | rs6910293 | 1.167 | -0.447 | -1.007 | -1 | 0.004 | -0.043 | -0.043 | 0 | 0.25 | 0.25 | 0.18 | 0.04 | 0.28 | 0.18 |
| 6 | 88630632 | rs9344744 | -0.191 | -0.439 | -1.273 | -0.965 | -0.027 | -0.127 | -0.127 | 0.732 | 0.74 | 0.57 | 0.85 | 0.97 | 0.86 | 0.83 |
| 6 | 88633009 | rs4428477 | -0.175 | -0.431 | -1.266 | -0.96 | -0.03 | -0.126 | -0.126 | 1.301 | 0.75 | 0.57 | 0.85 | 0.97 | 0.87 | 0.83 |
| 6 | 88635440 | rs9344745 | 0.652 | -0.191 | -2.446 | -0.532 | -0.073 | -0.088 | -0.088 | 0.09 | 0.37 | 0.17 | 0.47 | 0.51 | 0.49 | 0.38 |
| 6 | 88636038 | rs12195946 | 1.075 | -0.015 | -2.421 | -0.827 | -0.016 | 0.049 | 0.049 | 0.163 | 0.23 | 0.09 | 0.16 | 0.04 | 0.27 | 0.18 |
| 6 | 88641724 | rs13204059 | -0.407 | 0.407 | -2.486 | -0.994 | 0.017 | 0.07 | 0.07 | 0.003 | 0.3 | 0.2 | 0.21 | 0.03 | 0.24 | 0.29 |
| 6 | 88643072 | rs4235835 | -0.269 | 0.404 | -2.289 | -0.482 | 0.023 | 0.029 | 0.029 | 0 | 0.58 | 0.23 | 0.6 | 0.35 | 0.62 | 0.53 |
| 6 | 88652580 | rs2509458 | 0.138 | 0.907 | -2.101 | -0.506 | 0.006 | 0.029 | 0.029 | 0 | 0.18 | 0.07 | 0.08 | 0 | 0.1 | 0.08 |
| 6 | 88654437 | rs9353513 | -0.338 | 1.863 | -2.057 | 0.464 | -0.141 | -0.148 | -0.148 | 0.178 | 0.23 | 0.06 | 0.38 | 0.49 | 0.39 | 0.29 |
| 6 | 88655566 | rs2282521 | -0.168 | 1.346 | -1.995 | 0.513 | -0.037 | -0.12 | -0.12 | 1.477 | 0.18 | 0.05 | 0.31 | 0.38 | 0.2 | 0.19 |
| 6 | 88655801 | rs2282519 | -0.639 | 1.179 | -1.402 | -0.052 | -0.011 | 0.002 | 0.002 | 3.674 | 0.42 | 0.67 | 0.49 | 0.55 | 0.48 | 0.47 |
| 6 | 88657347 | rs10944335 | 0.356 | 0.848 | -1.21 | -0.828 | -0.034 | 0.078 | 0.078 | 0 | 0.35 | 0.26 | 0.26 | 0.06 | 0.41 | 0.27 |
| 6 | 88658834 | rs6908365 | -0.454 | 0.276 | -0.635 | -0.495 | -0.004 | -0.016 | -0.016 | 0 | 0.4 | 0.44 | 0.52 | 0.76 | 0.38 | 0.57 |
| 6 | 88660007 | rs1940893 | -0.014 | 0.288 | -1.038 | -0.499 | -0.006 | 0.012 | 0.012 | 0 | 0.3 | 0.13 | 0.44 | 0.71 | 0.3 | 0.49 |
| 6 | 88660328 | rs2499503 | -0.402 | 0.266 | -0.594 | -0.734 | 0.006 | -0.022 | -0.022 | 0.078 | 0.94 | 0.89 | 0.96 | 0.97 | 0.94 | 0.98 |
| 6 | 88661106 | rs7749166 | -0.338 | 0.146 | -1.012 | -0.948 | -0.012 | 0.137 | 0.137 | 0 | 0.37 | 0.13 | 0.57 | 0.79 | 0.34 | 0.52 |
| 6 | 88661802 | rs2509453 | 0.628 | -0.448 | -0.378 | -0.545 | -0.024 | 0.001 | 0.001 | 0 | 0.71 | 0.6 | 0.82 | 0.61 | 0.83 | 0.62 |
| 6 | 88663601 | rs4707428 | 1.373 | 0.057 | -0.956 | -0.834 | -0.08 | 0.152 | 0.152 | 0 | 0.32 | 0.1 | 0.37 | 0.05 | 0.42 | 0.18 |
| 6 | 88663964 | rs1940890 | 0.769 | -0.124 | -0.664 | -0.541 | -0.03 | 0.098 | 0.098 | 0 | 0.46 | 0.24 | 0.47 | 0.31 | 0.5 | 0.43 |
| 6 | 88671201 | rs16879998 | -0.126 | -0.199 | -0.105 | -0.503 | -0.003 | -0.024 | -0.024 | 0.628 | 0.13 | 0.07 | 0.11 | 0.27 | 0.09 | 0.24 |
| 6 | 88671941 | rs632385 | 0.118 | -0.377 | -0.012 | -0.624 | -0.027 | -0.033 | -0.033 | 0.004 | 0.75 | 0.43 | 0.85 | 0.83 | 0.83 | 0.84 |
| 6 | 88675020 | rs677120 | 1.441 | -0.244 | -0.086 | -0.663 | -0.013 | 0.048 | 0.048 | 0 | 0.39 | 0.3 | 0.4 | 0.05 | 0.45 | 0.27 |
| 6 | 88676599 | rs1321501 | 0.434 | -0.123 | 0.001 | -0.311 | -0.03 | -0.099 | -0.099 | 0.256 | 0.23 | 0.07 | 0.33 | 0.51 | 0.28 | 0.32 |

| | | | | | | | | | | | | | | | | |
|---|---|---|---|---|---|---|---|---|---|---|---|---|---|---|---|---|
| 6 | 88678075 | rs543049 | 0.877 | -0.119 | 0.009 | -0.409 | 0.02 | 0.067 | 0.067 | 0 | 0.52 | 0.33 | 0.51 | 0.33 | 0.55 | 0.46 |
| 6 | 88678534 | rs648114 | 0.382 | -0.064 | -0.139 | -0.204 | -0.005 | 0.028 | 0.028 | 0 | 0.85 | 0.96 | 0.9 | 0.9 | 0.87 | 0.88 |
| 6 | 88679631 | rs489468 | 0.996 | -0.131 | 0.302 | 0.119 | -0.01 | 0.03 | 0.03 | 0 | 0.7 | 0.44 | 0.59 | 0.45 | 0.71 | 0.64 |
| 6 | 88680458 | rs2466413 | -0.655 | 0.117 | 0.714 | -0.813 | 0.011 | -0.049 | -0.049 | 0 | 0.09 | 0.22 | 0.04 | 0.01 | 0.08 | 0.04 |
| 6 | 88681691 | rs2474884 | -1.412 | 0.058 | 1.035 | -0.928 | 0.012 | -0.041 | -0.041 | 0.049 | 0.16 | 0.48 | 0.12 | 0.1 | 0.15 | 0.15 |
| 6 | 88686618 | rs269442 | 0.435 | 0.3 | 0.252 | -0.566 | -0.018 | -0.092 | -0.092 | 0 | 0.14 | 0.03 | 0.33 | 0.4 | 0.17 | 0.24 |
| 6 | 88688679 | rs11970470 | -0.541 | 0.385 | 1.039 | -1.137 | 0.032 | -0.093 | -0.093 | 0 | 0.14 | 0.32 | 0.13 | 0.01 | 0.14 | 0.07 |
| 6 | 88691312 | rs7760757 | -1.705 | 0.252 | 1.023 | -1.333 | 0.013 | -0.091 | -0.091 | 0.513 | 0.26 | 0.42 | 0.22 | 0.09 | 0.25 | 0.28 |
| 6 | 88695068 | rs11758530 | -0.296 | 0.231 | 0.897 | -1.315 | 0.01 | -0.059 | -0.059 | 0 | 0.13 | 0.19 | 0.12 | 0.01 | 0.14 | 0.07 |
| 6 | 88699915 | rs9450849 | -1.24 | 0.225 | 0.882 | -1.345 | 0.006 | -0.114 | -0.114 | 0.578 | 0.21 | 0.35 | 0.14 | 0.01 | 0.18 | 0.1 |
| 6 | 88702545 | rs580486 | 1.621 | -0.203 | 1.006 | -1.368 | -0.001 | -0.043 | -0.043 | 0 | 0.69 | 0.39 | 0.7 | 0.85 | 0.71 | 0.67 |
| 6 | 88705241 | rs17202238 | -0.211 | -0.191 | 0.488 | -1.346 | -0.063 | 0.081 | 0.081 | 0 | 0.14 | 0.03 | 0.14 | 0.02 | 0.19 | 0.08 |
| 6 | 88715087 | rs589644 | 0.501 | -0.227 | 0.617 | -1.221 | -0.005 | -0.045 | -0.045 | 0 | 0.89 | 0.99 | 0.89 | 0.87 | 0.9 | 0.77 |
| 6 | 88717766 | rs366676 | 0.422 | -0.337 | 0.312 | -2.003 | -0.003 | 0.017 | 0.017 | 0.003 | 0.41 | 0.34 | 0.57 | 0.81 | 0.45 | 0.51 |
| 6 | 88729447 | rs269432 | 0.337 | -0.42 | 0.142 | -1.66 | -0.012 | -0.054 | -0.054 | 0 | 0.91 | 0.99 | 0.89 | 0.83 | 0.9 | 0.77 |
| 6 | 88731608 | rs269431 | 0.764 | -0.432 | 0.198 | -1.666 | -0.014 | 0.054 | 0.054 | 0.187 | 0.88 | 0.74 | 0.87 | 0.83 | 0.9 | 0.76 |
| 6 | 88743121 | rs621935 | -0.228 | -0.655 | 0.07 | -1.881 | 0.013 | 0.082 | 0.082 | 0 | 0.6 | 0.73 | 0.74 | 0.82 | 0.66 | 0.59 |
| 6 | 88743501 | rs12205892 | 0.175 | -0.639 | 0.065 | -1.876 | 0.002 | 0.006 | 0.006 | 0.032 | 0.39 | 0.39 | 0.26 | 0.18 | 0.35 | 0.4 |
| 6 | 88745769 | rs269424 | 0.79 | -0.757 | -0.086 | -1.768 | -0.077 | -0.071 | -0.071 | 0 | 0.36 | 0.1 | 0.5 | 0.48 | 0.45 | 0.34 |
| 6 | 88777778 | rs806441 | 0.637 | -0.818 | 0.346 | -2.353 | -0.016 | -0.053 | -0.053 | 0 | 0.9 | 0.99 | 0.89 | 0.84 | 0.9 | 0.76 |
| 6 | 88780388 | rs1090709 | 0.229 | -0.883 | 0.47 | -2.32 | 0.008 | 0.034 | 0.034 | 0 | 0.59 | 0.62 | 0.73 | 0.83 | 0.65 | 0.59 |
| 6 | 88793304 | rs9450874 | -0.005 | -0.215 | 0.269 | -2.485 | -0.021 | 0.105 | 0.105 | 0.041 | 0.33 | 0.12 | 0.17 | 0.01 | 0.3 | 0.17 |
| 6 | 88797199 | rs10944342 | -0.386 | -0.23 | 0.541 | -2.493 | -0.005 | -0.154 | -0.154 | 0.154 | 0.38 | 0.46 | 0.2 | 0.01 | 0.3 | 0.17 |
| 6 | 88797697 | rs9450877 | -0.102 | -0.094 | -0.192 | -2.423 | -0.006 | 0.072 | 0.072 | 0 | 0.35 | 0.21 | 0.18 | 0.01 | 0.3 | 0.17 |
| 6 | 88803599 | rs806454 | 1.427 | -0.165 | 0.414 | -1.281 | -0.002 | -0.063 | -0.063 | 2.709 | 0.78 | 0.72 | 0.86 | 1 | 0.78 | 0.87 |
| 6 | 88817786 | rs6908755 | -1.091 | -0.067 | 0.185 | -0.859 | -0.028 | -0.087 | -0.087 | 0 | 0.19 | 0.35 | 0.15 | 0.04 | 0.16 | 0.18 |
| 6 | 88818965 | rs11968144 | -1.095 | -0.065 | 0.126 | -0.858 | -0.029 | -0.089 | -0.089 | 0 | 0.19 | 0.35 | 0.15 | 0.04 | 0.16 | 0.18 |
| 6 | 88821736 | rs1408260 | 1.273 | 0.86 | -0.931 | -0.316 | -0.012 | 0.012 | 0.012 | 0 | 0.94 | 0.96 | 0.95 | 0.99 | 0.91 | 0.94 |
| 6 | 88825342 | rs4706308 | 0.41 | 0.892 | -1.612 | -0.351 | -0.055 | 0.029 | 0.029 | 1.046 | 0.67 | 0.89 | 0.56 | 0.23 | 0.65 | 0.61 |
| 6 | 88826061 | rs9353523 | -0.619 | 0.531 | -1.658 | -0.484 | 0.016 | 0.026 | 0.026 | 0.196 | 0.09 | 0.02 | 0.03 | 0.05 | 0.04 | 0.06 |
| 6 | 88826436 | rs9359765 | -1.058 | 0.502 | -1.958 | -1.264 | 0.007 | 0.022 | 0.022 | 0.411 | 0.27 | 0.77 | 0.24 | 0.11 | 0.29 | 0.18 |
| 6 | 88829036 | rs3857490 | -1.171 | 0.834 | -2.559 | -1.908 | -0.037 | 0.164 | 0.164 | 0 | 0.44 | 0.8 | 0.38 | 0.14 | 0.56 | 0.44 |
| 6 | 88830721 | rs9450891 | -0.688 | 1.102 | -1.512 | -2.111 | 0.037 | -0.115 | -0.115 | 0 | 0.28 | 0.43 | 0.26 | 0.08 | 0.44 | 0.36 |
| 6 | 88836054 | rs6938321 | -0.716 | 1.038 | -1.359 | -2.081 | 0 | 0.019 | 0.019 | 0 | 0.15 | 0.13 | 0.11 | 0.07 | 0.12 | 0.07 |
| 6 | 88843377 | rs9353525 | 0.775 | 0.378 | -2.048 | -1.61 | 0.057 | -0.104 | -0.104 | 0.709 | 0.26 | 0.07 | 0.29 | 0.53 | 0.12 | 0.23 |
| 6 | 88843390 | rs10485171 | -0.297 | 0.345 | -2.058 | -1.683 | 0.035 | -0.108 | -0.108 | 4.657 | 0.29 | 0.45 | 0.27 | 0.09 | 0.44 | 0.36 |
| 6 | 88845949 | rs806365 | -0.665 | 0.316 | -2.024 | -1.681 | -0.023 | 0.008 | 0.008 | 3.327 | 0.57 | 0.68 | 0.58 | 0.61 | 0.56 | 0.6 |
| 6 | 88847589 | rs806366 | -0.542 | 0.224 | -1.809 | -1.706 | -0.013 | 0.02 | 0.02 | 1.315 | 0.54 | 0.79 | 0.56 | 0.58 | 0.49 | 0.47 |
| 6 | 88850100 | rs806368 | 0.717 | 0.293 | -1.849 | -1.664 | 0.016 | -0.111 | -0.111 | 12.804 | 0.32 | 0.08 | 0.36 | 0.49 | 0.22 | 0.3 |
| 6 | 88851751 | rs4707436 | 0.035 | 0.794 | -1.491 | -1.366 | 0.092 | -0.04 | -0.04 | 0.101 | 0.15 | 0.42 | 0.18 | 0.08 | 0.26 | 0.18 |
| 6 | 88853635 | rs1049353 | 0.197 | 0.95 | -2.045 | -1.224 | -0.047 | 0.011 | 0.011 | 0.363 | 0.1 | 0.03 | 0.15 | 0.08 | 0.26 | 0.17 |
| 6 | 88856178 | rs806369 | -0.715 | 0.495 | -1.975 | -1.167 | -0.036 | -0.057 | -0.057 | 0 | 0.71 | 0.88 | 0.68 | 0.58 | 0.69 | 0.7 |
| 6 | 88857320 | rs806374 | -0.001 | 0.433 | -0.984 | -0.85 | 0.001 | -0.001 | -0.001 | 0 | 0.42 | 0.42 | 0.4 | 0.51 | 0.33 | 0.33 |
| 6 | 88858648 | rs806376 | -0.59 | 0.619 | -0.554 | -0.609 | 0.009 | 0.062 | 0.062 | 0 | 0.49 | 0.69 | 0.35 | 0.25 | 0.46 | 0.39 |
| 6 | 88859551 | rs806378 | 0.353 | 0.518 | -0.812 | -0.976 | -0.044 | 0.042 | 0.042 | 0.396 | 0.27 | 0.06 | 0.18 | 0.18 | 0.27 | 0.21 |
| 6 | 88861698 | rs6928813 | -0.152 | 0.48 | -0.058 | -0.982 | 0.008 | -0.079 | -0.079 | 0 | 0.16 | 0.37 | 0.14 | 0.02 | 0.16 | 0.13 |
| 6 | 88864653 | rs806380 | -0.131 | 0.448 | -0.291 | -1.072 | -0.044 | 0.08 | 0.08 | 0 | 0.32 | 0.08 | 0.21 | 0.19 | 0.32 | 0.26 |
| 6 | 88865901 | rs806381 | -0.532 | 0.301 | -0.028 | -0.765 | 0.009 | 0.016 | 0.016 | 0.008 | 0.38 | 0.3 | 0.32 | 0.31 | 0.32 | 0.3 |
| 6 | 88866376 | rs7752758 | 0.155 | 0.359 | 0.214 | -0.807 | 0.013 | -0.043 | -0.043 | 0.333 | 0.1 | 0.24 | 0.09 | 0.01 | 0.11 | 0.09 |
| 6 | 88867488 | rs12528858 | 0.579 | 0.736 | 0.032 | -0.766 | -0.023 | -0.003 | -0.003 | 0 | 0.07 | 0.01 | 0.03 | 0.04 | 0.07 | 0.02 |
| 6 | 88867925 | rs12205430 | 0.609 | 0.745 | -0.001 | -0.784 | -0.036 | 0.001 | 0.001 | 0.5 | 0.22 | 0.01 | 0.13 | 0.18 | 0.19 | 0.19 |
| 6 | 88871049 | rs6454673 | -0.52 | 0.874 | 0.139 | -0.725 | 0.001 | 0.009 | 0.009 | 0.616 | 0.32 | 0.33 | 0.2 | 0.22 | 0.29 | 0.25 |
| 6 | 88872930 | rs6454674 | -0.785 | 0.89 | 0.145 | -0.657 | 0.004 | 0.008 | 0.008 | 0.072 | 0.36 | 0.35 | 0.31 | 0.29 | 0.31 | 0.3 |
| 6 | 88877952 | rs2180619 | 0.834 | 1.31 | 1.116 | -0.332 | -0.007 | 0.045 | 0.045 | 0 | 0.57 | 0.33 | 0.5 | 0.75 | 0.56 | 0.57 |
| 6 | 88879928 | rs9344757 | 0.368 | 1.422 | 1.07 | -0.324 | -0.002 | -0.009 | -0.009 | 1.773 | 0.66 | 0.41 | 0.55 | 0.75 | 0.68 | 0.61 |
| 6 | 88880701 | rs9353528 | 0.986 | 1.434 | 1.042 | -0.335 | -0.017 | 0.064 | 0.064 | 0 | 0.55 | 0.25 | 0.49 | 0.71 | 0.55 | 0.55 |
| 6 | 88882652 | rs10485170 | -0.399 | 1.051 | 1.361 | -0.019 | 0.055 | -0.023 | -0.023 | 0 | 0.07 | 0.29 | 0.08 | 0 | 0.09 | 0.07 |
| 6 | 88883522 | rs9450907 | -0.701 | 1.054 | 0.929 | -0.017 | 0.077 | -0.019 | -0.019 | 0.773 | 0.09 | 0.46 | 0.1 | 0 | 0.09 | 0.07 |
| 6 | 88884821 | rs1406977 | -0.08 | 0.943 | -0.002 | 0.313 | -0.001 | 0.003 | 0.003 | 0 | 0.75 | 0.88 | 0.65 | 0.75 | 0.77 | 0.68 |
| 6 | 88887215 | rs6906184 | -1.182 | 1.064 | 1.007 | 0.37 | 0.162 | 0.112 | 0.112 | 1.501 | 0.06 | 0.45 | 0.09 | 0.05 | 0.13 | 0.03 |
| 6 | 88890620 | rs7770704 | -0.975 | 0.87 | 1.131 | 0.648 | 0.06 | 0.12 | 0.12 | 0.159 | 0.06 | 0.28 | 0.08 | 0.1 | 0.12 | 0.04 |
| 6 | 88891040 | rs7757075 | 0.343 | 0.784 | 0.475 | -0.483 | -0.008 | 0.036 | 0.036 | 0 | 0.6 | 0.31 | 0.7 | 0.89 | 0.54 | 0.67 |
| 6 | 88893001 | rs9351138 | 0.964 | 0.614 | -0.572 | -0.06 | -0.09 | -0.134 | -0.134 | 0 | 0.25 | 0.02 | 0.26 | 0.43 | 0.26 | 0.17 |
| 6 | 88894422 | rs1535257 | -0.867 | 0.043 | -0.029 | -0.022 | -0.014 | 0.02 | 0.02 | 2.539 | 0.2 | 0.62 | 0.24 | 0.19 | 0.2 | 0.24 |
| 6 | 88898611 | rs9294397 | -1.063 | -0.032 | -0.444 | -0.215 | -0.042 | 0.152 | 0.152 | 0 | 0.72 | 0.94 | 0.82 | 0.99 | 0.72 | 0.81 |
| 6 | 88900035 | rs12209554 | 0.305 | -0.088 | -0.29 | -0.083 | 0.005 | 0.137 | 0.137 | 0 | 0.51 | 0.22 | 0.58 | 0.81 | 0.52 | 0.58 |
| 6 | 88900739 | rs4498322 | 0.527 | -0.118 | -0.328 | -0.083 | 0.013 | -0.001 | -0.001 | 0 | 0.11 | 0.07 | 0.13 | 0.08 | 0.03 | 0.08 |
| 6 | 88902564 | rs12213790 | 0.85 | -0.116 | -0.769 | -0.115 | -0.139 | -0.131 | -0.131 | 0 | 0.33 | 0.03 | 0.32 | 0.51 | 0.4 | 0.36 |
| 6 | 88903855 | rs4707441 | 0.6 | -0.132 | -0.486 | -0.002 | -0.002 | 0.01 | 0.01 | 0.02 | 0.16 | 0.18 | 0.26 | 0.3 | 0.12 | 0.24 |

| | | | | | | | | | | | | | | | | |
|---|---|---|---|---|---|---|---|---|---|---|---|---|---|---|---|---|
| 6 | 88903993 | rs9353530 | -0.371 | -0.173 | -0.028 | -0.093 | -0.027 | 0.079 | 0.079 | 0.943 | 0.17 | 0.5 | 0.23 | 0.17 | 0.15 | 0.1 |
| 6 | 88904043 | rs12660255 | -0.532 | 0.315 | -0.299 | -0.045 | 0.006 | 0.079 | 0.079 | 0 | 0.59 | 0.71 | 0.67 | 0.91 | 0.64 | 0.63 |
| 6 | 88904498 | rs4707442 | -0.021 | 0.265 | -0.319 | 0.655 | -0.002 | 0.011 | 0.011 | 0.012 | 0.16 | 0.18 | 0.26 | 0.31 | 0.12 | 0.24 |
| 6 | 88906439 | rs11967553 | 0.76 | 0.578 | -0.569 | 0.656 | -0.021 | -0.021 | -0.021 | 0 | 0.86 | 0.91 | 0.86 | 0.86 | 0.75 | 0.8 |
| 6 | 88906784 | rs13201863 | 0.262 | 0.58 | -0.261 | 0.405 | 0.005 | 0.012 | 0.012 | 0 | 0.31 | 0.31 | 0.32 | 0.48 | 0.4 | 0.25 |
| 6 | 88907388 | rs9450916 | -0.05 | 0.307 | -0.088 | 0.119 | 0.003 | -0.005 | -0.005 | 0.488 | 0.62 | 0.58 | 0.61 | 0.36 | 0.53 | 0.62 |
| 6 | 88907598 | rs9450917 | 0.225 | -0.18 | 0.11 | -0.16 | -0.001 | -0.011 | -0.011 | 0.601 | 0.55 | 0.51 | 0.59 | 0.36 | 0.53 | 0.61 |
| 6 | 88907735 | rs9450918 | 0.158 | -0.358 | 0.213 | -0.453 | -0.019 | -0.097 | -0.097 | 0.24 | 0.15 | 0.33 | 0.07 | 0 | 0.07 | 0.09 |
| 6 | 88914554 | rs7766691 | 0.292 | -0.42 | 0.078 | -0.423 | 0.001 | 0.017 | 0.017 | 0.011 | 0.48 | 0.41 | 0.55 | 0.36 | 0.47 | 0.56 |
| 6 | 88914904 | rs7752742 | 0.347 | -0.441 | 0.112 | -0.434 | 0.001 | 0.028 | 0.028 | 0.015 | 0.48 | 0.37 | 0.55 | 0.36 | 0.47 | 0.56 |
| 6 | 88921181 | rs7774945 | 0.191 | -0.531 | 0.481 | -0.241 | -0.004 | -0.037 | -0.037 | 0 | 0.12 | 0.14 | 0.06 | 0 | 0.07 | 0.07 |
| 6 | 88922785 | rs1555340 | -0.08 | -0.55 | 0.119 | 0.036 | -0.01 | -0.1 | -0.1 | 0 | 0.28 | 0.09 | 0.24 | 0.53 | 0.23 | 0.2 |
| 6 | 88931764 | rs6927294 | -0.093 | -0.577 | 0.067 | -0.196 | -0.013 | -0.078 | -0.078 | 0 | 0.12 | 0.28 | 0.06 | 0 | 0.07 | 0.09 |
| 6 | 88942948 | rs2325102 | -0.526 | -0.404 | -0.31 | 0.042 | -0.009 | 0.002 | 0.002 | 0.117 | 0.43 | 0.54 | 0.31 | 0.53 | 0.3 | 0.31 |
| 6 | 88947667 | rs9362473 | 0.33 | -0.061 | -0.554 | 1.086 | -0.024 | -0.053 | -0.053 | 0 | 0.1 | 0.01 | 0.15 | 0.33 | 0.11 | 0.16 |
| 6 | 88950505 | rs7769940 | -1.136 | 0.105 | -0.13 | 0.673 | -0.006 | -0.004 | -0.004 | 0 | 0.2 | 0.68 | 0.23 | 0.2 | 0.2 | 0.26 |
| 6 | 88955689 | rs3929734 | -0.586 | 0.147 | 0.447 | 0.621 | 0.016 | 0.02 | 0.02 | 0.546 | 0.13 | 0.46 | 0.18 | 0.14 | 0.12 | 0.21 |
| 6 | 88970587 | rs1408701 | -0.855 | 0.077 | 0.354 | 0.586 | -0.013 | -0.01 | -0.01 | 0.33 | 0.15 | 0.57 | 0.2 | 0.14 | 0.13 | 0.21 |
| 6 | 88973751 | rs1408702 | -1.088 | 0.271 | -0.509 | 0.547 | 0.004 | -0.026 | -0.026 | 0 | 0.41 | 0.81 | 0.44 | 0.56 | 0.39 | 0.49 |
| 6 | 88978312 | rs10944353 | 0.82 | 0.146 | -0.775 | -0.189 | 0.015 | 0.037 | 0.037 | 0.39 | 0.17 | 0.08 | 0.09 | 0 | 0.1 | 0.07 |
| 6 | 88984253 | rs10944354 | 0.544 | 0.122 | -0.774 | -0.067 | -0.008 | 0.066 | 0.066 | 0.016 | 0.21 | 0.08 | 0.15 | 0.08 | 0.2 | 0.26 |
| 6 | 88987617 | rs9444597 | -0.27 | 0.315 | -0.713 | -0.285 | 0.004 | 0.077 | 0.077 | 0 | 0.48 | 0.68 | 0.52 | 0.32 | 0.44 | 0.53 |
| 6 | 88989087 | rs9362478 | 0.806 | 0.286 | -0.373 | -0.309 | 0.008 | -0.009 | -0.009 | 0 | 0.23 | 0.38 | 0.34 | 0.24 | 0.23 | 0.25 |
| 6 | 88997402 | rs12524684 | 0.94 | 0.327 | -0.864 | -0.285 | 0.001 | 0.015 | 0.015 | 0 | 0.22 | 0.14 | 0.16 | 0.08 | 0.2 | 0.26 |
| 6 | 89009765 | rs6931165 | -0.485 | 0.386 | -1.157 | 0.048 | -0.067 | 0.025 | 0.025 | 0 | 0.54 | 0.89 | 0.64 | 0.42 | 0.52 | 0.61 |
| 6 | 89027593 | rs6932835 | 0.91 | 0.432 | -1.412 | 0.105 | -0.071 | -0.145 | -0.145 | 0 | 0.27 | 0.03 | 0.25 | 0.44 | 0.32 | 0.26 |
| 6 | 89033459 | rs9362481 | -0.134 | 0.312 | -0.924 | -0.392 | 0.002 | 0.042 | 0.042 | 0 | 0.62 | 0.7 | 0.69 | 0.84 | 0.64 | 0.58 |
| 6 | 89033722 | rs9344779 | -0.864 | 0.329 | -1.12 | -0.288 | 0 | 0.007 | 0.007 | 0 | 0.91 | 0.96 | 0.95 | 1 | 0.92 | 0.93 |
| 6 | 89045060 | rs6934116 | 0.4 | 0.398 | -1.056 | -0.115 | -0.007 | 0.001 | 0.001 | 0 | 0.38 | 0.13 | 0.33 | 0.6 | 0.4 | 0.33 |
| 6 | 89051075 | rs6938352 | 0.099 | 0.549 | -1.05 | -0.106 | -0.002 | -0.002 | -0.002 | 0 | 0.45 | 0.31 | 0.36 | 0.39 | 0.43 | 0.43 |
| 6 | 89053621 | rs992426 | -0.871 | 0.538 | -1.384 | -0.469 | -0.035 | 0.11 | 0.11 | 0 | 0.71 | 0.9 | 0.81 | 0.93 | 0.71 | 0.71 |
| 6 | 89053706 | rs1324960 | 1.365 | 0.527 | -1.495 | -0.196 | -0.045 | -0.091 | -0.091 | 0.224 | 0.21 | 0.03 | 0.26 | 0.32 | 0.27 | 0.22 |
| 6 | 89071574 | rs9353544 | 1.053 | 0.584 | -1.35 | -0.204 | -0.022 | -0.019 | -0.019 | 0 | 0.22 | 0.07 | 0.3 | 0.28 | 0.2 | 0.26 |
| 6 | 89073338 | rs4707460 | -0.513 | 0.559 | -0.998 | -0.26 | 0.018 | -0.042 | -0.042 | 0 | 0.92 | 0.83 | 0.95 | 1 | 0.93 | 0.94 |
| 6 | 89074336 | rs990060 | -0.052 | 0.597 | -1.289 | -0.218 | -0.014 | -0.002 | -0.002 | 0 | 0.43 | 0.23 | 0.5 | 0.57 | 0.42 | 0.46 |
| 6 | 89075687 | rs10485166 | 0.82 | 0.801 | -1.513 | -0.084 | -0.006 | -0.064 | -0.064 | 0 | 0.2 | 0.11 | 0.18 | 0.29 | 0.19 | 0.19 |
| 6 | 89077257 | rs9353546 | 1.117 | 0.971 | -1.318 | -0.774 | -0.052 | -0.056 | -0.056 | 0.09 | 0.17 | 0.01 | 0.28 | 0.27 | 0.17 | 0.25 |
| 6 | 89078189 | rs12529670 | 0.349 | 0.821 | -0.495 | -1.002 | -0.013 | 0.07 | 0.07 | 0 | 0.24 | 0.1 | 0.15 | 0.07 | 0.22 | 0.26 |
| 6 | 89078417 | rs989625 | 1.038 | 0.753 | 0.506 | -0.215 | 0.006 | -0.067 | -0.067 | 0 | 0.83 | 0.47 | 0.9 | 0.95 | 0.86 | 0.92 |
| 6 | 89084901 | rs9362494 | -0.433 | 0.619 | 0.139 | 0.171 | 0.004 | 0.005 | 0.005 | 0.103 | 0.09 | 0.32 | 0.05 | 0.05 | 0.07 | 0.03 |
| 6 | 89086057 | rs12215322 | -0.621 | 0.747 | -0.604 | -0.343 | -0.006 | 0.032 | 0.032 | 0 | 0.18 | 0.07 | 0.1 | 0.03 | 0.17 | 0.14 |
| 6 | 89086152 | rs723787 | -0.863 | 1.065 | -0.538 | 0.193 | -0.007 | -0.009 | -0.009 | 0.069 | 0.08 | 0.04 | 0.06 | 0.15 | 0.11 | 0.23 |
| 6 | 89086860 | rs16880525 | 0.096 | 0.128 | -0.487 | 0.012 | -0.007 | 0.001 | 0.001 | 1.086 | 0.22 | 0.26 | 0.16 | 0.25 | 0.15 | 0.23 |
| 6 | 89094542 | rs11962482 | -0.396 | -0.113 | -0.124 | 0.026 | 0.034 | 0.12 | 0.12 | 0 | 0.05 | 0.15 | 0.07 | 0.22 | 0.09 | 0.16 |
| 6 | 89094735 | rs4707461 | 0.139 | -0.127 | -0.849 | -0.324 | -0.063 | -0.02 | -0.02 | 0 | 0.59 | 0.15 | 0.69 | 0.6 | 0.52 | 0.53 |
| 6 | 89096817 | rs9344795 | 1.431 | -0.305 | -1.052 | -0.113 | -0.004 | -0.067 | -0.067 | 0 | 0.11 | 0 | 0.31 | 0.17 | 0.11 | 0.17 |
| 6 | 89098112 | rs6936494 | 0.138 | -0.151 | -0.919 | -0.225 | 0.034 | 0.12 | 0.12 | 0 | 0.05 | 0.15 | 0.07 | 0.21 | 0.09 | 0.16 |
| 6 | 89108830 | rs6927539 | -0.846 | -0.051 | -0.551 | -0.42 | -0.002 | -0.029 | -0.029 | 0.247 | 0.88 | 0.83 | 0.93 | 0.98 | 0.89 | 0.93 |
| 6 | 89111410 | rs10455163 | 0.254 | -0.015 | -0.858 | -0.371 | -0.062 | -0.02 | -0.02 | 0 | 0.59 | 0.15 | 0.69 | 0.6 | 0.52 | 0.53 |
| 6 | 89112817 | rs10485165 | 1.65 | -0.161 | -1.139 | -0.554 | -0.037 | -0.101 | -0.101 | 4.031 | 0.2 | 0.01 | 0.23 | 0.34 | 0.16 | 0.22 |
| 6 | 89140027 | rs9353553 | 0.968 | -0.249 | -0.958 | -0.571 | 0.019 | -0.024 | -0.024 | 0 | 0.22 | 0.13 | 0.34 | 0.25 | 0.16 | 0.23 |
| 6 | 89141561 | rs16880562 | 0.369 | -0.216 | -1.236 | -0.639 | -0.003 | 0.006 | 0.006 | 0 | 0.25 | 0.2 | 0.19 | 0.22 | 0.28 | 0.24 |
| 6 | 89150753 | rs2325118 | -1.123 | -0.041 | -1.031 | -0.593 | -0.001 | 0.008 | 0.008 | 0 | 0.89 | 0.92 | 0.93 | 0.98 | 0.89 | 0.92 |
| 6 | 89152479 | rs936495 | -1.05 | -0.052 | -1.02 | -0.599 | 0 | 0.007 | 0.007 | 0 | 0.89 | 0.92 | 0.93 | 0.98 | 0.89 | 0.92 |
| 6 | 89155606 | rs749420 | 0.209 | 0.029 | -0.922 | -0.465 | -0.018 | -0.021 | -0.021 | 0.01 | 0.43 | 0.29 | 0.43 | 0.56 | 0.48 | 0.48 |
| 6 | 89157027 | rs1870832 | -0.255 | 0.343 | -0.889 | -0.268 | -0.003 | 0.004 | 0.004 | 0.296 | 0.66 | 0.67 | 0.79 | 0.81 | 0.64 | 0.71 |
| 6 | 89162999 | rs7751158 | 0.236 | 0.381 | -0.88 | -0.199 | 0.004 | -0.005 | -0.005 | 0.468 | 0.45 | 0.57 | 0.54 | 0.47 | 0.43 | 0.47 |
| 6 | 89169157 | rs7763511 | 0.405 | 0.367 | -0.804 | 0.291 | -0.018 | -0.052 | -0.052 | 0 | 0.21 | 0.08 | 0.24 | 0.34 | 0.21 | 0.24 |
| 6 | 89173758 | rs6454694 | 0.408 | -0.266 | -0.264 | -0.504 | -0.013 | -0.043 | -0.043 | 3.499 | 0.83 | 0.77 | 0.89 | 0.98 | 0.86 | 0.88 |
| 6 | 89173954 | rs10944384 | -0.142 | 0.128 | -0.631 | -0.219 | -0.061 | -0.049 | -0.049 | 0 | 0.34 | 0.16 | 0.4 | 0.6 | 0.43 | 0.56 |
| 6 | 89176668 | rs6939339 | -0.174 | -0.028 | -0.754 | -0.342 | -0.021 | -0.014 | -0.014 | 0.335 | 0.36 | 0.69 | 0.45 | 0.6 | 0.44 | 0.57 |
| 6 | 89179019 | rs11757134 | 0.299 | 0.345 | -1.012 | -0.199 | 0.001 | -0.003 | -0.003 | 0 | 0.07 | 0.11 | 0.03 | 0.04 | 0.08 | 0.07 |
| 6 | 89181414 | rs9450988 | -0.526 | 0.222 | -1.054 | -0.255 | -0.039 | -0.02 | -0.02 | 0 | 0.4 | 0.75 | 0.44 | 0.6 | 0.44 | 0.58 |
| 6 | 89181634 | rs11756329 | -0.325 | 0.23 | -1.122 | -0.24 | 0 | 0.002 | 0.002 | 0.39 | 0.11 | 0.12 | 0.03 | 0.04 | 0.08 | 0.07 |
| 6 | 89183007 | rs12195030 | -0.101 | 0.14 | -1.455 | -0.483 | -0.015 | 0.139 | 0.139 | 0 | 0.45 | 0.25 | 0.26 | 0.21 | 0.42 | 0.3 |
| 6 | 89183666 | rs16880601 | -0.239 | -0.028 | -1.335 | 0.186 | 0.01 | -0.023 | -0.023 | 0 | 0.16 | 0.13 | 0.21 | 0.2 | 0.13 | 0.17 |
| 6 | 89186177 | rs1408734 | 0.095 | 0.007 | -1.329 | 0.254 | -0.001 | -0.054 | -0.054 | 0 | 0.12 | 0.05 | 0.19 | 0.2 | 0.11 | 0.19 |
| 6 | 89192819 | rs9450990 | -0.748 | -0.177 | -1.428 | -0.369 | -0.001 | -0.007 | -0.007 | 2.035 | 0.18 | 0.7 | 0.11 | 0.09 | 0.17 | 0.11 |
| 6 | 89193082 | rs10944386 | -0.672 | -0.18 | -1.405 | -0.37 | -0.001 | -0.01 | -0.01 | 0.056 | 0.18 | 0.68 | 0.11 | 0.08 | 0.17 | 0.1 |

| | | | | | | | | | | | | | | | | |
|---|---|---|---|---|---|---|---|---|---|---|---|---|---|---|---|---|
| 6 | 89195922 | rs6650988 | -0.595 | -0.916 | -2.224 | -0.211 | 0.025 | -0.074 | -0.074 | 0 | 0.82 | 0.94 | 0.85 | 0.69 | 0.88 | 0.72 |
| 6 | 89196308 | rs6650989 | 0.675 | -0.946 | -2.179 | -0.231 | 0.034 | -0.013 | -0.013 | 0 | 0.63 | 0.21 | 0.73 | 0.61 | 0.71 | 0.63 |
| 6 | 89197310 | rs2031449 | 0.675 | -0.946 | -2.182 | -0.231 | 0.034 | -0.015 | -0.015 | 0 | 0.63 | 0.21 | 0.73 | 0.61 | 0.71 | 0.63 |
| 6 | 89200648 | rs9344819 | 0.936 | -0.979 | -1.636 | -0.24 | -0.017 | -0.074 | -0.074 | 0 | 0.12 | 0.01 | 0.18 | 0.2 | 0.11 | 0.19 |
| 6 | 89200843 | rs9344820 | 0.249 | -0.699 | -2.466 | -0.916 | -0.049 | 0.006 | 0.006 | 0.393 | 0.36 | 0.09 | 0.32 | 0.26 | 0.32 | 0.32 |
| 6 | 89201960 | rs9359790 | 0.365 | -0.66 | -2.401 | -0.912 | 0.012 | -0.054 | -0.054 | 0 | 0.16 | 0.53 | 0.09 | 0.08 | 0.17 | 0.1 |
| 6 | 89208252 | rs4707477 | 0.441 | -0.563 | -2.564 | -0.822 | -0.059 | -0.003 | -0.003 | 0.095 | 0.46 | 0.84 | 0.52 | 0.43 | 0.56 | 0.41 |
| 6 | 89211957 | rs1325008 | -0.851 | -0.493 | -2.512 | -0.789 | 0.03 | -0.049 | -0.049 | 0.077 | 0.81 | 0.91 | 0.84 | 0.69 | 0.88 | 0.72 |
| 6 | 89216444 | rs17576557 | 1.593 | -0.35 | -2.662 | -1.002 | -0.031 | 0.083 | 0.083 | 1.061 | 0.15 | 0.02 | 0.1 | 0.03 | 0.15 | 0.04 |
| 6 | 89219177 | rs12193211 | 1.365 | -0.301 | -2.583 | -0.925 | -0.018 | 0.022 | 0.022 | 0.143 | 0.08 | 0.02 | 0.04 | 0.04 | 0.06 | 0.09 |
| 6 | 89223096 | rs9353572 | 0.48 | -0.442 | -1.947 | -1.028 | 0.02 | -0.012 | -0.012 | 0 | 0.1 | 0.15 | 0.07 | 0.08 | 0.16 | 0.1 |
| 6 | 89226654 | rs9344826 | 0.297 | -0.635 | -1.759 | -1.058 | 0.002 | 0.021 | 0.021 | 0.054 | 0.55 | 0.65 | 0.67 | 0.58 | 0.57 | 0.58 |
| 6 | 89227341 | rs10498962 | 1.598 | -0.663 | -1.898 | -1.035 | -0.094 | -0.079 | -0.079 | 1.323 | 0.29 | 0.05 | 0.4 | 0.33 | 0.39 | 0.3 |
| 6 | 89238395 | rs17577049 | 1.462 | -0.603 | -1.814 | -1.013 | -0.02 | 0.023 | 0.023 | 0.087 | 0.09 | 0.02 | 0.04 | 0.04 | 0.06 | 0.09 |
| 6 | 89239967 | rs17492709 | 0.824 | -0.651 | -1.835 | -1.031 | -0.016 | 0.056 | 0.056 | 0 | 0.48 | 0.17 | 0.56 | 0.64 | 0.51 | 0.57 |
| 6 | 89241026 | rs9353578 | -0.024 | -0.577 | -1.639 | -0.903 | -0.012 | 0.006 | 0.006 | 3.048 | 0.27 | 0.62 | 0.27 | 0.25 | 0.18 | 0.29 |
| 6 | 89246061 | rs17577546 | 1.283 | 0.036 | -1.815 | -0.997 | -0.059 | -0.077 | -0.077 | 0.017 | 0.27 | 0.08 | 0.35 | 0.33 | 0.32 | 0.28 |
| 6 | 89250297 | rs7763299 | 0.745 | 0.097 | -0.714 | -1.088 | -0.001 | -0.032 | -0.032 | 0 | 0.16 | 0.12 | 0.13 | 0.03 | 0.21 | 0.06 |
| 6 | 89252152 | rs964888 | 0.637 | 0.056 | -1.004 | -0.61 | -0.005 | -0.005 | -0.005 | 0 | 0.45 | 0.64 | 0.47 | 0.49 | 0.52 | 0.44 |
| 6 | 89253131 | rs7741087 | 0.717 | 0.125 | -1.211 | -0.3 | 0.016 | -0.048 | -0.048 | 0 | 0.17 | 0.09 | 0.22 | 0.3 | 0.12 | 0.24 |
| 6 | 89256093 | rs2757721 | 0.187 | 0.494 | -1.19 | -0.62 | -0.056 | 0.036 | 0.036 | 1.757 | 0.87 | 0.98 | 0.83 | 0.92 | 0.82 | 0.78 |
| 6 | 89256465 | rs2610763 | 0.24 | 0.487 | -1.052 | -0.623 | -0.019 | 0.025 | 0.025 | 1.848 | 0.87 | 0.94 | 0.82 | 0.9 | 0.82 | 0.77 |
| 6 | 89259374 | rs2279083 | -0.269 | 0.101 | -0.538 | -0.53 | 0.004 | -0.04 | -0.04 | 0 | 0.27 | 0.37 | 0.2 | 0.11 | 0.27 | 0.17 |
| 6 | 89260886 | rs4320332 | -0.19 | -0.024 | -1.097 | -0.466 | 0.027 | 0.021 | 0.021 | 0.047 | 0.58 | 0.27 | 0.64 | 0.68 | 0.7 | 0.73 |
| 6 | 89261177 | rs4706323 | 1.305 | -0.087 | -1.188 | -0.616 | -0.01 | -0.01 | -0.01 | 0 | 0.24 | 0.09 | 0.24 | 0.21 | 0.21 | 0.16 |
| 6 | 89262470 | rs2875553 | 0.504 | -0.058 | -1.336 | -0.8 | -0.07 | -0.04 | -0.04 | 0 | 0.28 | 0.15 | 0.38 | 0.41 | 0.48 | 0.41 |
| 6 | 89264146 | rs9362529 | 0.634 | -0.154 | -0.968 | -0.796 | 0.008 | -0.007 | -0.007 | 0.196 | 0.42 | 0.33 | 0.44 | 0.48 | 0.32 | 0.46 |
| 6 | 89264558 | rs4707489 | -0.043 | -0.166 | -1.033 | -0.784 | -0.028 | -0.076 | -0.076 | 0.961 | 0.7 | 0.49 | 0.82 | 0.89 | 0.8 | 0.86 |
| 6 | 89265214 | rs13191303 | 0.988 | 0.497 | -1.506 | -0.82 | -0.104 | -0.083 | -0.083 | 0 | 0.22 | 0.04 | 0.19 | 0.31 | 0.32 | 0.2 |
| 6 | 89268509 | rs6918869 | 1.303 | 0.305 | -1.267 | -0.929 | 0.001 | -0.006 | -0.006 | 0 | 0.06 | 0.04 | 0.03 | 0 | 0.06 | 0.02 |
| 6 | 89268715 | rs1923415 | 0.59 | 0.289 | -0.91 | -0.973 | -0.019 | -0.069 | -0.069 | 0 | 0.18 | 0.29 | 0.11 | 0.03 | 0.14 | 0.06 |
| 6 | 89268935 | rs2757728 | -0.376 | 0.476 | -1.405 | -0.884 | -0.022 | -0.009 | -0.009 | 0 | 0.7 | 0.72 | 0.61 | 0.62 | 0.51 | 0.6 |
| 6 | 89270624 | rs2757732 | -0.461 | 0.591 | -0.782 | -1.349 | 0.002 | -0.052 | -0.052 | 0.505 | 0.82 | 0.75 | 0.83 | 0.97 | 0.77 | 0.92 |
| 6 | 89271778 | rs9294404 | -0.175 | 0.593 | -1.518 | -1.128 | 0.025 | 0.077 | 0.077 | 0 | 0.65 | 0.21 | 0.64 | 0.87 | 0.66 | 0.73 |
| 6 | 89272006 | rs10498963 | -0.164 | 0.812 | -0.998 | -0.461 | -0.033 | -0.029 | -0.029 | 0 | 0.28 | 0.1 | 0.28 | 0.31 | 0.25 | 0.26 |
| 6 | 89293745 | rs12527253 | -0.029 | 1.359 | -0.954 | -1.036 | -0.092 | 0.057 | 0.057 | 0 | 0.21 | 0.02 | 0.17 | 0.08 | 0.31 | 0.41 |
| 6 | 89293829 | rs2756380 | -0.249 | 0.379 | -0.668 | -1.349 | 0.022 | 0.064 | 0.064 | 0 | 0.66 | 0.83 | 0.45 | 0.16 | 0.67 | 0.54 |
| 6 | 89295741 | rs9444632 | -1.23 | 0.804 | 0.963 | -0.85 | 0.037 | -0.025 | -0.025 | 0.703 | 0.1 | 0.3 | 0.1 | 0.04 | 0.1 | 0.03 |
| 6 | 89296870 | rs9294406 | -1.286 | 0.813 | 1.054 | -0.767 | 0.034 | 0.001 | 0.001 | 0.207 | 0.09 | 0.2 | 0.13 | 0.07 | 0.11 | 0.03 |
| 6 | 89299046 | rs744270 | -1.83 | 0.376 | 1.11 | -0.722 | 0.039 | 0.034 | 0.034 | 3.852 | 0.1 | 0.49 | 0.12 | 0.06 | 0.07 | 0.03 |
| 6 | 89304760 | rs16879934 | -0.884 | 0.528 | 0.257 | -0.586 | -0.006 | 0.007 | 0.007 | 0 | 0.05 | 0.13 | 0.08 | 0.06 | 0.05 | 0.06 |
| 6 | 89310475 | rs10085363 | -0.302 | 0.364 | 0.003 | -0.869 | 0.001 | 0.001 | 0.001 | 0 | 0.16 | 0.14 | 0.09 | 0.09 | 0.14 | 0.06 |
| 6 | 89317670 | rs6940660 | -0.533 | 0.21 | -0.682 | -1.51 | 0.007 | 0.047 | 0.047 | 0 | 0.63 | 0.83 | 0.61 | 0.3 | 0.67 | 0.53 |
| 6 | 89318640 | rs2610768 | -0.685 | -0.236 | 0.283 | -1.654 | -0.038 | -0.039 | -0.039 | 0.042 | 0.24 | 0.37 | 0.2 | 0.17 | 0.17 | 0.13 |
| 6 | 89320589 | rs2756367 | -0.601 | -0.245 | 0.284 | -1.656 | -0.033 | -0.032 | -0.032 | 0 | 0.23 | 0.35 | 0.2 | 0.17 | 0.16 | 0.12 |
| 6 | 89329939 | rs16880966 | 0.103 | -0.092 | -0.103 | -1.583 | -0.054 | -0.007 | -0.007 | 0 | 0.1 | 0.01 | 0.09 | 0.07 | 0.18 | 0.13 |
| 6 | 89337530 | rs3798787 | 0.148 | -0.301 | 0.048 | -1.737 | 0.003 | 0.032 | 0.032 | 0 | 0.16 | 0.1 | 0.09 | 0.08 | 0.13 | 0.1 |
| 6 | 89339534 | rs2756374 | -0.329 | 0.047 | -0.139 | -1.859 | -0.055 | 0.083 | 0.083 | 0.356 | 0.6 | 0.89 | 0.48 | 0.27 | 0.54 | 0.47 |
| 6 | 89349438 | rs3778172 | -0.511 | -0.014 | 0.105 | -1.824 | 0.004 | 0.001 | 0.001 | 0 | 0.06 | 0.1 | 0.06 | 0.08 | 0.07 | 0.03 |
| 6 | 89421027 | rs9353592 | -0.255 | -0.714 | -0.255 | -2.345 | 0.047 | 0.047 | 0.047 | 0.089 | 0.3 | 0.72 | 0.11 | 0.08 | 0.12 | 0.19 |
| 6 | 89432362 | rs6901395 | -0.461 | -0.352 | -0.483 | -2.136 | -0.051 | 0.103 | 0.103 | 0.049 | 0.57 | 0.8 | 0.41 | 0.26 | 0.46 | 0.43 |
| 6 | 89460542 | rs7747355 | 0.228 | -0.044 | -0.207 | -2.224 | 0.03 | 0.056 | 0.056 | 0 | 0.25 | 0.12 | 0.08 | 0.08 | 0.14 | 0.21 |
| 6 | 89468856 | rs9351179 | 0.311 | -0.1 | -0.018 | -2.254 | 0.016 | 0.022 | 0.022 | 0 | 0.25 | 0.15 | 0.09 | 0.08 | 0.14 | 0.21 |
| 6 | 89511628 | rs10455494 | -0.587 | 0.048 | -0.015 | -2.181 | 0.007 | -0.002 | -0.002 | 0 | 0.07 | 0.12 | 0.09 | 0.08 | 0.09 | 0.04 |
| 6 | 89522122 | rs12200224 | -0.462 | 0.044 | -0.054 | -2.183 | 0.003 | -0.001 | -0.001 | 0 | 0.07 | 0.1 | 0.09 | 0.08 | 0.1 | 0.05 |
| 6 | 89620255 | rs9451101 | -1.103 | 0.022 | -0.179 | -2.294 | -0.053 | -0.009 | -0.009 | 0.005 | 0.65 | 0.89 | 0.5 | 0.36 | 0.57 | 0.48 |
| 6 | 89633046 | rs10944413 | 0.98 | -0.039 | -0.407 | -2.412 | -0.101 | 0.014 | 0.014 | 0.318 | 0.26 | 0.05 | 0.29 | 0.19 | 0.3 | 0.22 |
| 6 | 89633891 | rs16881113 | -1 | -0.387 | 0.355 | -1.981 | 0.002 | -0.006 | -0.006 | 0.502 | 0.08 | 0.17 | 0.07 | 0.08 | 0.09 | 0.04 |
| 6 | 89642071 | rs9359829 | -0.712 | -0.424 | 0.127 | -1.87 | 0.008 | 0.007 | 0.007 | 0.276 | 0.35 | 0.67 | 0.16 | 0.17 | 0.21 | 0.28 |
| 6 | 89673861 | rs10223700 | -0.554 | -0.129 | 0.264 | -1.356 | 0.002 | -0.035 | -0.035 | 1.187 | 0.12 | 0.23 | 0.1 | 0.06 | 0.16 | 0.22 |
| 6 | 89676206 | rs11751805 | -0.059 | -0.057 | -0.047 | -1.921 | 0.009 | 0.026 | 0.026 | 0 | 0.36 | 0.25 | 0.31 | 0.22 | 0.3 | 0.3 |
| 6 | 89682207 | rs2207554 | 1.1 | -0.273 | -0.074 | -2.048 | 0.004 | 0.049 | 0.049 | 0 | 0.47 | 0.39 | 0.55 | 0.72 | 0.5 | 0.4 |
| 6 | 89683536 | rs6454727 | 0.491 | -0.237 | -0.079 | -2.033 | -0.01 | 0.055 | 0.055 | 0 | 0.85 | 0.94 | 0.88 | 0.94 | 0.82 | 0.71 |
| 6 | 89695108 | rs2207553 | 1.196 | -0.301 | -0.14 | -1.957 | 0.001 | 0.036 | 0.036 | 0 | 0.46 | 0.42 | 0.55 | 0.71 | 0.5 | 0.4 |
| 6 | 89698024 | rs9451118 | 0.325 | -0.316 | -0.109 | -1.954 | 0.014 | 0.03 | 0.03 | 0 | 0.58 | 0.43 | 0.63 | 0.78 | 0.65 | 0.62 |
| 6 | 89698052 | rs2325168 | 0.294 | -0.311 | -0.421 | -1.955 | 0.014 | 0.058 | 0.058 | 0 | 0.33 | 0.13 | 0.28 | 0.21 | 0.28 | 0.29 |
| 6 | 89709564 | rs12208298 | -0.259 | -0.191 | -0.397 | -1.914 | -0.015 | 0.066 | 0.066 | 0 | 0.14 | 0.06 | 0.09 | 0.05 | 0.16 | 0.22 |
| 6 | 89728922 | rs17277859 | 0.879 | -0.594 | -0.59 | -2.073 | 0.02 | 0.027 | 0.027 | 0 | 0.19 | 0.11 | 0.06 | 0.04 | 0.11 | 0.19 |
| 6 | 89730740 | rs2748178 | 0.69 | -0.581 | -0.328 | -1.792 | 0.02 | 0.031 | 0.031 | 0 | 0.16 | 0.26 | 0.28 | 0.18 | 0.19 | 0.11 |

| | | | | | | | | | | | | | | | | |
|---|---|---|---|---|---|---|---|---|---|---|---|---|---|---|---|---|
| 6 | 89732101 | rs12201676 | 0.064 | -0.556 | -0.82 | -1.789 | 0.031 | 0.089 | 0.089 | 0.017 | 0.3 | 0.14 | 0.14 | 0.09 | 0.25 | 0.39 |
| 6 | 89732364 | rs16881241 | 2.006 | -0.399 | -1.176 | -1.569 | -0.159 | -0.124 | -0.124 | 0 | 0.37 | 0.03 | 0.45 | 0.59 | 0.45 | 0.34 |
| 6 | 89737500 | rs12190865 | -0.201 | -0.601 | -0.956 | -1.626 | 0.058 | 0.152 | 0.152 | 0.14 | 0.35 | 0.11 | 0.16 | 0.1 | 0.28 | 0.42 |
| 6 | 89737774 | rs6903926 | -0.194 | -0.574 | -0.546 | -1.511 | 0.024 | 0.102 | 0.102 | 0.533 | 0.08 | 0.35 | 0.06 | 0.13 | 0.06 | 0.1 |
| 6 | 89739221 | rs9359837 | -0.605 | -0.591 | -0.593 | -1.559 | 0.008 | 0.006 | 0.006 | 0 | 0.41 | 0.46 | 0.22 | 0.16 | 0.33 | 0.51 |
| 6 | 89740625 | rs205231 | -0.623 | -0.426 | -0.531 | -1.475 | 0.005 | -0.005 | -0.005 | 0.086 | 0.94 | 0.91 | 0.94 | 0.98 | 0.94 | 0.95 |
| 6 | 89741014 | rs205228 | -0.835 | -0.642 | -0.538 | -1.102 | -0.002 | -0.004 | -0.004 | 0 | 0.52 | 0.5 | 0.46 | 0.34 | 0.47 | 0.53 |
| 6 | 89743077 | rs205226 | -1.501 | -0.664 | -0.672 | -1.1 | 0.027 | 0.061 | 0.061 | 0 | 0.51 | 0.85 | 0.48 | 0.38 | 0.46 | 0.59 |
| 6 | 89743455 | rs12205662 | 0.602 | -0.635 | -0.864 | -1.128 | 0.005 | 0.01 | 0.01 | 0 | 0.15 | 0.1 | 0.1 | 0.07 | 0.17 | 0.25 |
| 6 | 89743488 | rs205223 | -0.222 | -0.671 | -0.799 | -1.12 | -0.007 | -0.01 | -0.01 | 0 | 0.28 | 0.39 | 0.34 | 0.25 | 0.26 | 0.28 |
| 6 | 89745218 | rs110337 | -0.874 | -0.321 | -0.907 | -1.116 | -0.009 | -0.019 | -0.019 | 0.12 | 0.44 | 0.48 | 0.44 | 0.32 | 0.43 | 0.54 |
| 6 | 89749189 | rs205216 | 0.364 | 0.362 | -1.159 | -1.129 | -0.07 | -0.01 | -0.01 | 0 | 0.17 | 0.03 | 0.22 | 0.12 | 0.21 | 0.12 |
| 6 | 89750761 | rs9344901 | 0.727 | 0.219 | -1.137 | -0.859 | -0.048 | -0.045 | -0.045 | 0 | 0.15 | 0.03 | 0.18 | 0.17 | 0.16 | 0.13 |
| 6 | 89751272 | rs172221 | -0.025 | 0.506 | -1.485 | -1.414 | -0.023 | -0.026 | -0.026 | 0.715 | 0.68 | 0.9 | 0.54 | 0.36 | 0.71 | 0.63 |
| 6 | 89754750 | rs855575 | 0.947 | 0.649 | -0.44 | -1.433 | 0.02 | 0.017 | 0.017 | 0 | 0.14 | 0.3 | 0.18 | 0.13 | 0.15 | 0.11 |
| 6 | 89757972 | rs9362606 | 0.005 | 0.33 | -0.645 | -1.395 | -0.015 | -0.057 | -0.057 | 0 | 0.22 | 0.09 | 0.4 | 0.67 | 0.21 | 0.34 |
| 6 | 89759640 | rs17291400 | 2.184 | 0.569 | -0.559 | -1.712 | 0 | 0.042 | 0.042 | 0 | 0.21 | 0.09 | 0.12 | 0 | 0.21 | 0.1 |
| 6 | 89763711 | rs6921997 | 0.538 | 0.146 | 0.169 | -1.628 | 0 | -0.012 | -0.012 | 3.163 | 0.44 | 0.48 | 0.55 | 0.69 | 0.43 | 0.43 |
| 6 | 89769624 | rs9294419 | -0.364 | 0.21 | 0.228 | -1.678 | -0.014 | -0.057 | -0.057 | 0.005 | 0.32 | 0.57 | 0.21 | 0.17 | 0.3 | 0.39 |
| 6 | 89770216 | rs6932023 | 0.03 | 0.126 | 0.289 | -1.727 | 0 | -0.051 | -0.051 | 0 | 0.27 | 0.36 | 0.18 | 0.11 | 0.3 | 0.32 |
| 6 | 89772615 | rs10447337 | 0.02 | 0.164 | 0.083 | -1.711 | 0.001 | -0.027 | -0.027 | 0.009 | 0.27 | 0.34 | 0.19 | 0.11 | 0.33 | 0.32 |
| 6 | 89773599 | rs6907604 | -0.851 | 0.174 | -0.332 | -1.725 | -0.025 | 0.024 | 0.024 | 0 | 0.75 | 0.9 | 0.76 | 0.84 | 0.74 | 0.83 |
| 6 | 89778440 | rs9451140 | -0.634 | 0.173 | -0.1 | -1.728 | 0.002 | 0.002 | 0.002 | 0 | 0.34 | 0.72 | 0.23 | 0.17 | 0.33 | 0.4 |
| 6 | 89781378 | rs10080656 | -0.644 | 0.142 | -0.1 | -1.729 | 0.002 | 0.003 | 0.003 | 0 | 0.34 | 0.72 | 0.23 | 0.17 | 0.32 | 0.39 |
| 6 | 89782384 | rs1573857 | 1.045 | 0.117 | -0.261 | -1.747 | -0.02 | 0.026 | 0.026 | 0 | 0.41 | 0.16 | 0.52 | 0.68 | 0.42 | 0.43 |
| 6 | 89799737 | rs10455501 | 0.073 | 0.186 | -0.358 | -1.846 | -0.003 | 0.028 | 0.028 | 0 | 0.28 | 0.23 | 0.17 | 0.12 | 0.31 | 0.34 |
| 6 | 89801278 | rs7745708 | -0.998 | 0.235 | -0.557 | -1.844 | -0.051 | -0.002 | -0.002 | 0 | 0.77 | 0.97 | 0.77 | 0.85 | 0.78 | 0.85 |
| 6 | 89804646 | rs1929267 | 0.784 | 0.266 | -0.166 | -1.696 | -0.004 | 0.012 | 0.012 | 0 | 0.41 | 0.29 | 0.52 | 0.67 | 0.41 | 0.43 |
| 6 | 89805888 | rs1054973 | -0.004 | 0.4 | -0.345 | -1.873 | 0.015 | 0.091 | 0.091 | 0 | 0.16 | 0.28 | 0.4 | 0.67 | 0.17 | 0.31 |
| 6 | 89806689 | rs9362615 | 0.678 | 0.392 | -0.33 | -1.858 | -0.005 | 0.011 | 0.011 | 0 | 0.4 | 0.29 | 0.53 | 0.67 | 0.41 | 0.43 |
| 6 | 89820477 | rs7773548 | -0.554 | 0.375 | -0.218 | -1.884 | -0.001 | 0.016 | 0.016 | 0.019 | 0.7 | 0.74 | 0.72 | 0.83 | 0.7 | 0.78 |
| 6 | 89832079 | rs387177 | -0.086 | 0.53 | -0.631 | -1.865 | 0.013 | 0.116 | 0.116 | 0 | 0.16 | 0.28 | 0.32 | 0.62 | 0.17 | 0.3 |
| 6 | 89834020 | rs1796723 | -0.039 | 0.554 | -0.62 | -1.873 | 0.015 | 0.12 | 0.12 | 0 | 0.16 | 0.27 | 0.32 | 0.62 | 0.17 | 0.3 |
| 6 | 89854436 | rs12215916 | 0.102 | 0.539 | -0.677 | -1.972 | -0.068 | 0.142 | 0.142 | 0 | 0.27 | 0.01 | 0.17 | 0.06 | 0.29 | 0.28 |
| 6 | 89862725 | rs6937095 | 0.171 | 0.622 | -0.466 | -1.915 | -0.005 | 0.064 | 0.064 | 0 | 0.26 | 0.15 | 0.16 | 0.13 | 0.29 | 0.31 |
| 6 | 89864679 | rs4053639 | 0.048 | 0.502 | -0.372 | -1.812 | -0.007 | -0.004 | -0.004 | 0.03 | 0.33 | 0.57 | 0.39 | 0.26 | 0.3 | 0.27 |
| 6 | 89879168 | rs7775554 | -0.853 | 1.275 | 0.376 | -1.143 | 0 | -0.006 | -0.006 | 0 | 0.42 | 0.5 | 0.51 | 0.79 | 0.47 | 0.67 |
| 6 | 89885206 | rs13202074 | -0.443 | 1.253 | 0.454 | -1.05 | 0.03 | 0.079 | 0.079 | 0 | 0.22 | 0.34 | 0.4 | 0.62 | 0.3 | 0.34 |
| 6 | 89885237 | rs416115 | -0.443 | 1.253 | 0.454 | -1.05 | 0.03 | 0.079 | 0.079 | 0 | 0.22 | 0.34 | 0.4 | 0.62 | 0.3 | 0.34 |
| 6 | 89888443 | rs3734201 | -0.64 | 1.144 | 0.424 | -1.586 | 0.001 | 0.022 | 0.022 | 0 | 0.44 | 0.43 | 0.45 | 0.27 | 0.43 | 0.45 |
| 6 | 89888744 | rs1796743 | -0.576 | 1.008 | 0.451 | -1.629 | 0.021 | 0.048 | 0.048 | 0 | 0.27 | 0.5 | 0.43 | 0.63 | 0.34 | 0.36 |
| 6 | 89890375 | rs407221 | -1.897 | 1.02 | 0.604 | -1.618 | -0.031 | 0.148 | 0.148 | 0 | 0.67 | 0.96 | 0.85 | 0.89 | 0.73 | 0.79 |
| 6 | 89894119 | rs9362624 | 0.68 | 1.238 | 0.855 | -1.46 | -0.012 | -0.019 | -0.019 | 0 | 0.17 | 0.09 | 0.28 | 0.2 | 0.18 | 0.14 |
| 6 | 89894968 | rs407206 | 1.778 | 1.269 | 0.827 | -1.501 | -0.03 | 0.181 | 0.181 | 0 | 0.37 | 0.07 | 0.19 | 0.11 | 0.33 | 0.22 |
| 6 | 89895207 | rs364936 | 1.3 | 1.562 | 1.106 | -1.302 | -0.009 | 0.062 | 0.062 | 0 | 0.54 | 0.21 | 0.46 | 0.31 | 0.51 | 0.36 |
| 6 | 89900599 | rs453503 | -0.079 | 1.686 | 1.26 | -1.219 | -0.006 | 0.112 | 0.112 | 0 | 0.17 | 0.28 | 0.34 | 0.62 | 0.18 | 0.28 |
| 6 | 89905239 | rs6930407 | -0.606 | 1.753 | 1.339 | -1.924 | 0.058 | -0.035 | -0.035 | 0 | 0.08 | 0.25 | 0.07 | 0.01 | 0.13 | 0.07 |
| 6 | 89907015 | rs439912 | -1.193 | 1.815 | 1.19 | -2.017 | 0 | -0.033 | -0.033 | 0.008 | 0.52 | 0.55 | 0.75 | 0.94 | 0.56 | 0.7 |
| 6 | 89907873 | rs422751 | 0.359 | 1.631 | 1.365 | -1.775 | -0.017 | -0.074 | -0.074 | 0 | 0.79 | 0.64 | 0.88 | 0.95 | 0.84 | 0.74 |
| 6 | 89913520 | rs2150817 | -1.171 | 1.664 | 0.827 | -1.622 | 0.004 | -0.017 | -0.017 | 4.279 | 0.58 | 0.67 | 0.8 | 0.93 | 0.67 | 0.69 |
| 6 | 89914187 | rs2183797 | -0.908 | 1.908 | 0.541 | -1.408 | 0.007 | -0.004 | -0.004 | 0.73 | 0.33 | 0.45 | 0.49 | 0.72 | 0.42 | 0.4 |
| 6 | 89914763 | rs11756847 | 0.553 | 1.916 | 0.043 | -1.148 | -0.063 | -0.042 | -0.042 | 0 | 0.19 | 0.03 | 0.29 | 0.21 | 0.24 | 0.28 |
| 6 | 89916641 | rs7758893 | 0 | 1.726 | 0.342 | -0.907 | -0.006 | -0.022 | -0.022 | 0 | 0.85 | 0.92 | 0.65 | 0.37 | 0.8 | 0.77 |
| 6 | 89916843 | rs9444675 | 0.818 | 1.687 | 0.758 | -1.286 | 0.017 | -0.026 | -0.026 | 0 | 0.49 | 0.48 | 0.24 | 0.03 | 0.32 | 0.3 |
| 6 | 89918342 | rs13215029 | -0.864 | 1.997 | 1.039 | -0.931 | 0.192 | 0.067 | 0.067 | 0 | 0.07 | 0.35 | 0.08 | 0.04 | 0.15 | 0.05 |
| 6 | 89918386 | rs13215160 | -0.393 | 1.913 | 1.041 | -0.921 | 0.192 | 0.108 | 0.108 | 0 | 0.05 | 0.35 | 0.06 | 0.04 | 0.1 | 0.04 |
| 6 | 89918562 | rs4707529 | -0.184 | 1.444 | 0.268 | -0.816 | -0.004 | -0.018 | -0.018 | 0 | 0.86 | 0.93 | 0.67 | 0.37 | 0.82 | 0.78 |
| 6 | 89921623 | rs9353653 | 0.61 | 1.405 | 0.349 | -1.09 | 0.034 | 0.011 | 0.011 | 0 | 0.68 | 0.5 | 0.53 | 0.35 | 0.6 | 0.67 |
| 6 | 89930208 | rs914479 | -0.952 | 1.456 | 0.143 | -0.729 | 0.056 | -0.094 | -0.094 | 0.039 | 0.28 | 0.54 | 0.22 | 0.09 | 0.38 | 0.25 |
| 6 | 89932204 | rs282114 | 0.413 | 1.379 | -0.811 | -0.582 | -0.083 | -0.074 | -0.074 | 1.426 | 0.23 | 0.04 | 0.31 | 0.29 | 0.26 | 0.3 |
| 6 | 89950960 | rs282136 | 0.636 | 1.214 | -0.477 | -0.703 | 0.033 | 0.05 | 0.05 | 0.395 | 0.51 | 0.24 | 0.52 | 0.64 | 0.45 | 0.5 |
| 6 | 89951335 | rs9451186 | -0.72 | 1.341 | -0.343 | -0.796 | 0.001 | -0.02 | -0.02 | 0 | 0.25 | 0.72 | 0.18 | 0.08 | 0.26 | 0.21 |
| 6 | 89953068 | rs9444678 | -0.773 | 1.431 | -0.232 | -0.728 | 0.008 | -0.016 | -0.016 | 0 | 0.08 | 0.16 | 0.08 | 0.02 | 0.12 | 0.08 |
| 6 | 89954461 | rs282133 | -0.642 | 1.467 | -0.363 | -0.795 | -0.034 | -0.034 | -0.034 | 0 | 0.75 | 0.91 | 0.71 | 0.71 | 0.73 | 0.67 |
| 6 | 89954672 | rs17740802 | -0.187 | 1.548 | -0.38 | -0.828 | 0.007 | -0.008 | -0.008 | 0.002 | 0.06 | 0.13 | 0.06 | 0.02 | 0.1 | 0.08 |
| 6 | 89959665 | rs282125 | 0.11 | 1.118 | -0.064 | -1.324 | -0.013 | 0.051 | 0.051 | 0 | 0.57 | 0.72 | 0.34 | 0.09 | 0.54 | 0.4 |
| 6 | 89960200 | rs282124 | 1.72 | 0.854 | -0.05 | -1.409 | 0.081 | 0.079 | 0.079 | 0.3 | 0.3 | 0.11 | 0.11 | 0 | 0.18 | 0.16 |
| 6 | 89960222 | rs282123 | 0.492 | 1.032 | -0.159 | -1.346 | 0.023 | 0.021 | 0.021 | 0 | 0.5 | 0.58 | 0.26 | 0.03 | 0.43 | 0.3 |
| 6 | 89963796 | rs16881632 | -0.03 | 1.819 | 0.059 | -0.501 | -0.019 | -0.071 | -0.071 | 0 | 0.16 | 0.05 | 0.35 | 0.63 | 0.18 | 0.26 |

| | | | | | | | | | | | | | | | | |
|---|---|---|---|---|---|---|---|---|---|---|---|---|---|---|---|---|
| 6 | 89964478 | rs3734197 | -0.631 | 1.889 | 0.546 | -0.395 | 0.005 | -0.008 | -0.008 | 0.23 | 0.06 | 0.11 | 0.06 | 0.02 | 0.1 | 0.12 |
| 6 | 89967498 | rs282129 | -1.011 | 1.622 | 0.905 | -0.17 | 0.096 | -0.012 | -0.012 | 0 | 0.19 | 0.46 | 0.17 | 0.14 | 0.25 | 0.15 |
| 6 | 89968712 | rs12205041 | -0.601 | 1.889 | 0.294 | -0.084 | 0.075 | -0.002 | -0.002 | 0 | 0.16 | 0.27 | 0.14 | 0.12 | 0.25 | 0.18 |
| 6 | 89974063 | rs723041 | 1.039 | 1.23 | -0.152 | 0.925 | -0.001 | 0.016 | 0.016 | 0 | 0.85 | 0.83 | 0.72 | 0.72 | 0.87 | 0.84 |
| 6 | 89974066 | rs958854 | -0.123 | 0.504 | -0.625 | 0.915 | 0.017 | -0.027 | -0.027 | 1.028 | 0.16 | 0.11 | 0.15 | 0.36 | 0.15 | 0.24 |
| 6 | 89974626 | rs12206367 | -0.197 | 1.123 | -0.492 | 0.578 | 0.015 | -0.011 | -0.011 | 0 | 0.05 | 0.06 | 0.05 | 0 | 0.1 | 0.05 |
| 6 | 89976372 | rs9451192 | -0.427 | 1.208 | -0.942 | 0.145 | 0.002 | -0.068 | -0.068 | 0.982 | 0.32 | 0.21 | 0.38 | 0.6 | 0.34 | 0.47 |
| 6 | 89977906 | rs4707535 | -1.164 | 0.924 | -1.275 | -0.247 | -0.012 | -0.046 | -0.046 | 0.087 | 0.73 | 0.9 | 0.68 | 0.72 | 0.78 | 0.87 |
| 6 | 89980083 | rs282121 | -0.919 | 0.3 | -0.087 | -0.411 | -0.023 | 0.036 | 0.036 | 0 | 0.57 | 0.82 | 0.56 | 0.73 | 0.58 | 0.69 |
| 6 | 89981413 | rs282117 | -0.883 | 0.292 | -0.091 | -0.415 | -0.022 | 0.038 | 0.038 | 0 | 0.57 | 0.82 | 0.55 | 0.73 | 0.58 | 0.69 |
| 6 | 89985343 | rs9444682 | 0.233 | 0.222 | 0.25 | -0.268 | 0.003 | -0.01 | -0.01 | 0 | 0.22 | 0.24 | 0.2 | 0.42 | 0.24 | 0.29 |
| 6 | 89985379 | rs9451194 | -0.187 | 0.266 | 0.324 | -0.342 | -0.002 | -0.008 | -0.008 | 0 | 0.38 | 0.38 | 0.44 | 0.66 | 0.43 | 0.55 |
| 6 | 89985498 | rs9353660 | -0.847 | 0.116 | 0.471 | -0.383 | -0.001 | -0.008 | -0.008 | 1.998 | 0.53 | 0.58 | 0.52 | 0.73 | 0.58 | 0.68 |
| 6 | 89992200 | rs3777522 | -0.032 | 0.301 | 0.673 | -0.119 | -0.006 | -0.055 | -0.055 | 0.615 | 0.18 | 0.45 | 0.11 | 0.07 | 0.15 | 0.13 |
| 6 | 89995299 | rs12196758 | -0.947 | 0.422 | 0.598 | -0.162 | 0.071 | -0.013 | -0.013 | 0 | 0.09 | 0.34 | 0.07 | 0.01 | 0.09 | 0.08 |
| 6 | 89996405 | rs3798256 | -0.676 | 0.329 | 0.19 | -0.208 | 0 | -0.006 | -0.006 | 0 | 0.47 | 0.6 | 0.51 | 0.72 | 0.53 | 0.69 |
| 6 | 89999296 | rs964626 | -1.035 | 0.044 | 0.138 | 0.141 | -0.004 | 0.006 | 0.006 | 0 | 0.26 | 0.26 | 0.16 | 0.47 | 0.19 | 0.35 |
| 6 | 90000554 | rs9451196 | -0.274 | 0.157 | 0.17 | 0.213 | -0.021 | 0.061 | 0.061 | 0 | 0.18 | 0.25 | 0.18 | 0.5 | 0.11 | 0.28 |
| 6 | 90001465 | rs7762767 | -0.345 | 0.804 | 0.182 | -0.405 | 0.006 | -0.018 | -0.018 | 0 | 0.06 | 0.13 | 0.04 | 0.01 | 0.07 | 0.09 |
| 6 | 90005925 | rs2148174 | 0.348 | 1.204 | 0.212 | -0.879 | 0.016 | -0.07 | -0.07 | 0 | 0.76 | 0.72 | 0.82 | 0.97 | 0.71 | 0.75 |
| 6 | 90006421 | rs9344921 | 0.686 | 1.132 | -0.231 | -0.877 | -0.035 | 0.053 | 0.053 | 0 | 0.71 | 0.81 | 0.8 | 0.96 | 0.63 | 0.67 |
| 6 | 90008700 | rs7762714 | 0.261 | 0.681 | 0.311 | -0.858 | 0.013 | -0.088 | -0.088 | 1.021 | 0.86 | 0.72 | 0.94 | 1 | 0.91 | 0.93 |
| 6 | 90008744 | rs7742664 | -0.279 | 0.996 | 0.168 | -0.253 | 0.012 | 0.043 | 0.043 | 0 | 0.24 | 0.3 | 0.24 | 0.38 | 0.26 | 0.3 |
| 6 | 90009311 | rs1570932 | 0.569 | 0.83 | -0.589 | -0.081 | -0.006 | -0.001 | -0.001 | 0 | 0.62 | 0.43 | 0.7 | 0.62 | 0.64 | 0.61 |
| 6 | 90018361 | rs6454752 | -0.878 | 0.635 | -0.45 | -0.289 | -0.001 | -0.008 | -0.008 | 0.506 | 0.23 | 0.27 | 0.23 | 0.13 | 0.23 | 0.28 |
| 6 | 90019967 | rs9294432 | -1.183 | 0.463 | -1.564 | -0.405 | -0.138 | -0.005 | -0.005 | 0 | 0.61 | 0.92 | 0.52 | 0.74 | 0.46 | 0.83 |
| 6 | 90022921 | rs7764923 | -0.61 | -0.19 | -1.102 | -0.282 | 0.075 | 0.09 | 0.09 | 0 | 0.42 | 0.69 | 0.3 | 0.19 | 0.23 | 0.39 |
| 6 | 90025112 | rs2236204 | 0.05 | -0.589 | -1.36 | -0.505 | -0.002 | 0 | 0 | 0 | 0.23 | 0.04 | 0.21 | 0.19 | 0.19 | 0.28 |
| 6 | 90026797 | rs9362636 | -0.114 | -0.672 | -0.195 | -0.19 | -0.091 | -0.037 | -0.037 | 0.01 | 0.19 | 0.53 | 0.09 | 0.12 | 0.04 | 0.12 |
| 6 | 90033736 | rs12195070 | -0.236 | -0.661 | -0.263 | -0.201 | 0.037 | 0.036 | 0.036 | 0.243 | 0.28 | 0.12 | 0.22 | 0.19 | 0.18 | 0.29 |
| 6 | 90036511 | rs13215418 | 0.41 | -0.396 | -0.427 | -0.149 | -0.021 | -0.079 | -0.079 | 0 | 0.13 | 0.04 | 0.17 | 0.43 | 0.14 | 0.37 |
| 6 | 90039670 | rs10502 | -1.359 | -0.354 | -0.601 | -0.27 | 0.017 | 0.059 | 0.059 | 0.625 | 0.55 | 0.89 | 0.49 | 0.74 | 0.36 | 0.81 |
| 6 | 90043137 | rs1998576 | 0.263 | 0.006 | -0.514 | -0.408 | -0.001 | -0.006 | -0.006 | 1.097 | 0.19 | 0.21 | 0.14 | 0.09 | 0.2 | 0.25 |
| 6 | 90046404 | rs12189673 | 1.546 | 0.376 | -1.654 | -0.786 | -0.095 | -0.082 | -0.082 | 0 | 0.23 | 0.03 | 0.36 | 0.25 | 0.31 | 0.13 |
| 6 | 90051915 | rs16881711 | -0.225 | 0.322 | -1.308 | -0.187 | -0.033 | -0.066 | -0.066 | 0 | 0.22 | 0.11 | 0.22 | 0.43 | 0.24 | 0.44 |
| 6 | 90052604 | rs7760851 | -0.896 | 0.285 | -0.941 | -0.414 | -0.036 | 0.051 | 0.051 | 0 | 0.58 | 0.77 | 0.53 | 0.72 | 0.51 | 0.79 |
| 6 | 90068458 | rs9362644 | -0.967 | 0.62 | -1.023 | -0.043 | -0.056 | 0.007 | 0.007 | 0 | 0.49 | 0.9 | 0.4 | 0.34 | 0.44 | 0.45 |
| 6 | 90071201 | rs10944445 | -1.449 | 1.02 | -1.216 | -0.015 | -0.009 | -0.016 | -0.016 | 0 | 0.59 | 0.92 | 0.57 | 0.67 | 0.59 | 0.7 |
| 6 | 90075359 | rs1062108 | -1.433 | 1.04 | -1.217 | -0.008 | -0.012 | -0.011 | -0.011 | 0 | 0.59 | 0.92 | 0.57 | 0.67 | 0.58 | 0.7 |
| 6 | 90076384 | rs1555402 | -0.12 | 1.04 | -0.505 | -0.256 | -0.019 | -0.085 | -0.085 | 0.688 | 0.3 | 0.44 | 0.17 | 0.07 | 0.24 | 0.28 |
| 6 | 90076993 | rs2274087 | 0.827 | 1.14 | -0.506 | -0.218 | -0.049 | -0.056 | -0.056 | 0.266 | 0.1 | 0.02 | 0.16 | 0.32 | 0.15 | 0.24 |
| 6 | 90078402 | rs16881774 | -0.018 | 1.251 | 0.003 | -0.123 | 0.044 | -0.087 | -0.087 | 0 | 0.15 | 0.22 | 0.1 | 0 | 0.2 | 0.13 |
| 6 | 90089677 | rs9451211 | -0.259 | 1.274 | 0.063 | -0.001 | 0 | -0.006 | -0.006 | 0 | 0.41 | 0.47 | 0.34 | 0.4 | 0.39 | 0.54 |
| 6 | 90100643 | rs2875567 | 0.105 | 1.135 | 0.271 | 0.403 | 0.033 | -0.066 | -0.066 | 6.897 | 0.11 | 0.19 | 0.05 | 0 | 0.11 | 0.09 |
| 6 | 90113275 | rs6922408 | -1.152 | 0.744 | -0.075 | 0.993 | 0.057 | 0.259 | 0.259 | 0.293 | 0.05 | 0.21 | 0.16 | 0.27 | 0.07 | 0.02 |
| 6 | 90119043 | rs1885070 | -0.812 | -0.08 | -0.675 | 0.057 | 0.032 | 0.008 | 0.008 | 0 | 0.34 | 0.74 | 0.3 | 0.38 | 0.27 | 0.44 |
| 6 | 90120991 | rs1358852 | -0.817 | -0.076 | -0.599 | 0.05 | -0.007 | -0.004 | -0.004 | 0 | 0.35 | 0.58 | 0.3 | 0.38 | 0.27 | 0.44 |
| 6 | 90123995 | rs854919 | -0.058 | -0.095 | -1.349 | -0.67 | 0.03 | 0.14 | 0.14 | 0 | 0.69 | 0.89 | 0.81 | 0.94 | 0.77 | 0.72 |
| 6 | 90127390 | rs854917 | 0.37 | -0.165 | -1.425 | -0.715 | 0.033 | 0.151 | 0.151 | 0 | 0.64 | 0.88 | 0.78 | 0.88 | 0.71 | 0.6 |
| 6 | 90131156 | rs10455510 | -0.521 | 0.556 | -0.636 | -0.103 | 0.014 | 0.009 | 0.009 | 1.624 | 0.24 | 0.32 | 0.22 | 0.28 | 0.3 | 0.24 |
| 6 | 90134102 | rs2093237 | -0.502 | 0.565 | -0.557 | -0.076 | 0.009 | 0.012 | 0.012 | 2.285 | 0.24 | 0.35 | 0.22 | 0.29 | 0.29 | 0.24 |
| 6 | 90140564 | rs854915 | 0.402 | 0.589 | -0.465 | -0.011 | 0.015 | 0.019 | 0.019 | 0 | 0.16 | 0.2 | 0.17 | 0.21 | 0.24 | 0.11 |
| 6 | 90142093 | rs854916 | 0.46 | 0.593 | -0.547 | -0.009 | -0.002 | -0.002 | -0.002 | 0 | 0.16 | 0.13 | 0.17 | 0.21 | 0.24 | 0.11 |
| 6 | 90149275 | rs12201016 | 0.554 | 0.281 | -1.063 | -0.513 | -0.023 | 0.071 | 0.071 | 0 | 0.15 | 0.01 | 0.08 | 0.03 | 0.14 | 0.22 |
| 6 | 90162180 | rs9342199 | -0.045 | 0.622 | -0.287 | 0.512 | 0.032 | 0.271 | 0.271 | 0.474 | 0.06 | 0.23 | 0.28 | 0.4 | 0.08 | 0.03 |
| 6 | 90166419 | rs854864 | 0.81 | 1.24 | -0.514 | 0.376 | -0.074 | -0.06 | -0.06 | 0.041 | 0.16 | 0.04 | 0.18 | 0.2 | 0.27 | 0.11 |
| 6 | 90167977 | rs9451226 | 0.663 | 0.862 | -0.477 | -0.513 | -0.006 | 0.001 | 0.001 | 0.399 | 0.29 | 0.29 | 0.26 | 0.06 | 0.3 | 0.25 |
| 6 | 90170989 | rs854865 | -0.034 | 0.431 | -0.761 | -0.725 | -0.053 | -0.065 | -0.065 | 0 | 0.39 | 0.13 | 0.36 | 0.49 | 0.47 | 0.49 |
| 6 | 90179545 | rs1209055 | -0.427 | 0.191 | -0.475 | -0.608 | 0.018 | 0.194 | 0.194 | 1.732 | 0.13 | 0.33 | 0.32 | 0.45 | 0.13 | 0.18 |
| 6 | 90197586 | rs7745260 | -2.164 | 0.047 | -0.842 | -0.829 | 0.038 | 0.12 | 0.12 | 0 | 0.81 | 0.99 | 0.95 | 1 | 0.91 | 0.92 |
| 6 | 90200219 | rs6914963 | 0.942 | 0.223 | -0.665 | -0.687 | -0.001 | -0.006 | -0.006 | 0 | 0.06 | 0.05 | 0.03 | 0.03 | 0.06 | 0.14 |
| 6 | 90207300 | rs2224727 | 0.163 | 0.317 | -0.637 | -0.792 | -0.014 | -0.011 | -0.011 | 0 | 0.37 | 0.67 | 0.57 | 0.46 | 0.42 | 0.28 |
| 6 | 90212737 | rs7769333 | 0.959 | 0.275 | -0.694 | -0.8 | -0.001 | 0.008 | 0.008 | 0.357 | 0.06 | 0.04 | 0.04 | 0.03 | 0.06 | 0.13 |
| 6 | 90218106 | rs942115 | 0.335 | 1.175 | -0.648 | -0.911 | -0.004 | 0.015 | 0.015 | 2.371 | 0.33 | 0.59 | 0.53 | 0.46 | 0.31 | 0.18 |
| 6 | 90228498 | rs10944450 | 0.833 | 0.702 | -0.818 | -0.841 | 0.014 | -0.038 | -0.038 | 0 | 0.21 | 0.08 | 0.16 | 0.28 | 0.18 | 0.38 |
| 6 | 90240886 | rs13209503 | 1.695 | 0.738 | -0.888 | -0.963 | -0.041 | -0.049 | -0.049 | 0 | 0.16 | 0.07 | 0.16 | 0.22 | 0.22 | 0.11 |
| 6 | 90244235 | rs2026651 | 0.765 | 0.744 | -0.667 | -1.018 | 0.012 | -0.056 | -0.056 | 0.048 | 0.29 | 0.35 | 0.29 | 0.06 | 0.34 | 0.25 |
| 6 | 90261621 | rs11965790 | 0.686 | 0.422 | -1.132 | -0.771 | 0.022 | -0.037 | -0.037 | 1.477 | 0.22 | 0.07 | 0.16 | 0.24 | 0.18 | 0.38 |
| 6 | 90261899 | rs1887585 | 1.094 | 0.36 | -1.073 | -0.722 | 0.013 | 0.043 | 0.043 | 0 | 0.36 | 0.12 | 0.21 | 0.24 | 0.32 | 0.2 |

| | | | | | | | | | | | | | | | | |
|---|---|---|---|---|---|---|---|---|---|---|---|---|---|---|---|---|
| 6 | 90285588 | rs7745254 | 0.19 | 0.282 | -0.704 | -0.909 | 0.038 | 0.045 | 0.045 | 0 | 0.49 | 0.34 | 0.27 | 0.27 | 0.34 | 0.61 |
| 6 | 90294088 | rs9451249 | -0.298 | 0.6 | -0.285 | -0.71 | 0.002 | 0.057 | 0.057 | 0 | 0.19 | 0.37 | 0.28 | 0.27 | 0.16 | 0.26 |
| 6 | 90296588 | rs9362662 | -0.353 | 0.582 | -0.311 | -1.063 | 0.012 | 0.005 | 0.005 | 0.366 | 0.26 | 0.32 | 0.49 | 0.61 | 0.51 | 0.29 |
| 6 | 90298371 | rs9344946 | -0.709 | 0.116 | -0.304 | -1.175 | 0.003 | -0.038 | -0.038 | 0 | 0.82 | 0.7 | 0.81 | 0.88 | 0.83 | 0.66 |
| 6 | 90300010 | rs6912680 | 0.322 | 0.041 | -0.415 | -1.212 | -0.004 | 0 | 0 | 0 | 0.68 | 0.38 | 0.66 | 0.64 | 0.78 | 0.45 |
| 6 | 90310966 | rs6454764 | -0.356 | 0.005 | -0.151 | -1.134 | -0.07 | 0.144 | 0.144 | 0 | 0.85 | 0.57 | 0.78 | 0.74 | 0.95 | 0.78 |
| 6 | 90315789 | rs3748085 | -0.723 | 0.099 | -0.312 | -1.219 | 0.004 | -0.053 | -0.053 | 0 | 0.82 | 0.68 | 0.87 | 0.91 | 0.83 | 0.67 |
| 6 | 90316178 | rs6454765 | -0.216 | -0.204 | -0.6 | -1.703 | 0.038 | 0.043 | 0.043 | 0 | 0.46 | 0.31 | 0.61 | 0.64 | 0.64 | 0.34 |
| 6 | 90321146 | rs9451253 | 0.024 | -0.323 | -0.731 | -1.875 | -0.046 | -0.01 | -0.01 | 0 | 0.35 | 0.39 | 0.33 | 0.29 | 0.18 | 0.28 |
| 6 | 90321196 | rs1179901 | 0.959 | -0.234 | -0.714 | -1.846 | 0.015 | -0.052 | -0.052 | 0 | 0.17 | 0.32 | 0.13 | 0.09 | 0.18 | 0.32 |
| 6 | 90321409 | rs4707554 | -1.141 | -0.485 | -0.914 | -2.051 | 0.014 | 0.032 | 0.032 | 0 | 0.55 | 0.67 | 0.79 | 0.91 | 0.66 | 0.56 |
| 6 | 90321545 | rs1179902 | 1.436 | -0.43 | -0.865 | -1.966 | 0.001 | -0.011 | -0.011 | 0 | 0.13 | 0.13 | 0.11 | 0.05 | 0.15 | 0.18 |
| 6 | 90321955 | rs2073272 | 0.558 | 0.152 | -0.904 | -1.971 | -0.013 | -0.019 | -0.019 | 0 | 0.2 | 0.17 | 0.4 | 0.6 | 0.31 | 0.15 |
| 6 | 90322507 | rs4706346 | -2.249 | -0.25 | -0.863 | -2.076 | 0.022 | 0.03 | 0.03 | 0 | 0.76 | 0.86 | 0.91 | 1 | 0.84 | 0.83 |
| 6 | 90322573 | rs1179904 | 0.295 | 0.312 | -0.16 | -1.431 | 0.002 | -0.057 | -0.057 | 0 | 0.23 | 0.29 | 0.15 | 0.08 | 0.21 | 0.34 |
| 6 | 90322922 | rs1179905 | -0.023 | 0.226 | 0.121 | -1.304 | -0.012 | -0.11 | -0.11 | 0.435 | 0.26 | 0.44 | 0.17 | 0.09 | 0.21 | 0.37 |
| 6 | 90326360 | rs2273238 | 0.27 | 0.211 | -0.145 | -0.995 | -0.005 | 0.249 | 0.249 | 0 | 0.09 | 0.35 | 0.19 | 0.26 | 0.04 | 0.23 |
| 6 | 90337633 | rs13214731 | 0.405 | 0.45 | -0.711 | -1.307 | -0.052 | 0.106 | 0.106 | 0 | 0.19 | 0.04 | 0.19 | 0.03 | 0.23 | 0.07 |
| 6 | 90340276 | rs3210511 | -0.969 | 0.508 | -0.422 | -1.355 | -0.001 | -0.024 | -0.024 | 0 | 0.44 | 0.57 | 0.7 | 0.91 | 0.47 | 0.39 |
| 6 | 90341607 | rs9351210 | -0.491 | 0.509 | -0.461 | -0.985 | -0.011 | 0.006 | 0.006 | 0 | 0.31 | 0.42 | 0.57 | 0.69 | 0.3 | 0.3 |
| 6 | 90347511 | rs34012596 | 0.533 | 0.631 | -0.691 | -1.189 | -0.017 | 0.02 | 0.02 | 0 | 0.14 | 0.12 | 0.11 | 0.04 | 0.21 | 0.33 |
| 6 | 90361532 | rs761727 | -2.686 | 0.483 | -0.71 | -1.327 | 0.022 | 0.018 | 0.018 | 0 | 0.74 | 0.87 | 0.92 | 1 | 0.84 | 0.9 |
| 6 | 90371986 | rs3818132 | -0.754 | 0.518 | -0.784 | -1.397 | 0.001 | -0.011 | -0.011 | 0 | 0.45 | 0.59 | 0.7 | 0.91 | 0.47 | 0.38 |
| 6 | 90374283 | rs16882046 | 1.739 | 0.505 | -0.811 | -1.409 | 0.024 | 0.029 | 0.029 | 2.458 | 0.09 | 0.01 | 0.04 | 0.03 | 0.03 | 0.14 |
| 6 | 90378808 | rs761731 | -1.379 | 0.549 | -0.769 | -1.416 | 0.002 | 0.019 | 0.019 | 0 | 0.59 | 0.68 | 0.79 | 0.96 | 0.63 | 0.56 |
| 6 | 90382150 | rs715203 | 1.57 | 0.598 | -0.76 | -1.399 | 0.025 | -0.007 | -0.007 | 0 | 0.06 | 0.1 | 0.07 | 0.01 | 0.18 | 0.19 |
| 6 | 90390443 | rs9353689 | -0.765 | 0.483 | -0.759 | -1.402 | 0.001 | 0.004 | 0.004 | 0 | 0.45 | 0.61 | 0.7 | 0.91 | 0.47 | 0.39 |
| 6 | 90398903 | rs6914142 | 1.23 | 0.509 | -0.766 | -1.333 | -0.011 | 0.027 | 0.027 | 0 | 0.13 | 0.06 | 0.09 | 0.05 | 0.16 | 0.18 |
| 6 | 90402482 | rs9294445 | -0.845 | 0.488 | -0.763 | -1.4 | 0.001 | -0.012 | -0.012 | 0 | 0.44 | 0.6 | 0.7 | 0.91 | 0.46 | 0.38 |
| 6 | 90408589 | rs3736984 | 0.471 | 0.436 | -0.794 | -1.423 | 0.004 | 0.064 | 0.064 | 0 | 0.22 | 0.08 | 0.12 | 0.08 | 0.19 | 0.32 |
| 6 | 90408742 | rs12530146 | 0.779 | 0.337 | -0.642 | -1.226 | -0.007 | 0.012 | 0.012 | 0 | 0.11 | 0.1 | 0.1 | 0.21 | 0.06 | 0.2 |
| 6 | 90418262 | rs2026634 | -2.765 | 0.464 | -0.98 | -1.438 | 0.013 | -0.012 | -0.012 | 0 | 0.73 | 0.8 | 0.9 | 1 | 0.84 | 0.9 |
| 6 | 90425648 | rs4707561 | -1.817 | 0.468 | -0.982 | -1.442 | 0.015 | 0.04 | 0.04 | 0 | 0.6 | 0.74 | 0.82 | 0.95 | 0.68 | 0.72 |
| 6 | 90437699 | rs7767196 | -2.729 | 0.448 | -0.965 | -1.405 | 0.01 | -0.019 | -0.019 | 4.691 | 0.73 | 0.79 | 0.9 | 1 | 0.84 | 0.9 |
| 6 | 90442119 | rs17294425 | 0.599 | 0.486 | -0.975 | -1.408 | -0.02 | 0.027 | 0.027 | 0 | 0.16 | 0.11 | 0.12 | 0.04 | 0.22 | 0.33 |
| 6 | 90448092 | rs4140446 | -2.691 | 0.411 | -0.939 | -1.36 | 0.012 | -0.016 | -0.016 | 0 | 0.73 | 0.8 | 0.9 | 1 | 0.84 | 0.9 |
| 6 | 90450135 | rs2180140 | -0.32 | 0.441 | -0.917 | -1.225 | -0.013 | 0.011 | 0.011 | 0 | 0.31 | 0.42 | 0.56 | 0.69 | 0.28 | 0.29 |
| 6 | 90455084 | rs9359861 | -2.811 | 0.46 | -1.54 | -1.238 | 0.006 | 0.182 | 0.182 | 0 | 0.74 | 0.98 | 0.93 | 1 | 0.84 | 0.9 |
| 6 | 90461123 | rs4707563 | 0.626 | 0.8 | -1.308 | -1.008 | 0.001 | 0.014 | 0.014 | 0 | 0.13 | 0.18 | 0.14 | 0.22 | 0.18 | 0.09 |
| 6 | 90465626 | rs6942026 | 0.287 | 0.829 | -1.328 | -1.078 | -0.001 | 0.007 | 0.007 | 0 | 0.64 | 0.72 | 0.73 | 0.7 | 0.63 | 0.59 |
| 6 | 90473260 | rs4707565 | -0.126 | 0.876 | -1.301 | -1.071 | 0.019 | 0.008 | 0.008 | 0.222 | 0.2 | 0.3 | 0.22 | 0.22 | 0.37 | 0.29 |
| 6 | 90476276 | rs6910277 | 0.77 | 0.841 | -1.3 | -1.05 | -0.002 | 0.016 | 0.016 | 0.038 | 0.57 | 0.62 | 0.65 | 0.69 | 0.44 | 0.39 |
| 6 | 90476452 | rs9362681 | 1.312 | 0.759 | -1.314 | -1.053 | 0.026 | 0.01 | 0.01 | 0 | 0.42 | 0.59 | 0.48 | 0.67 | 0.25 | 0.34 |
| 6 | 90482397 | rs12110451 | 0.488 | 1.047 | -1.275 | -0.702 | 0.001 | 0.013 | 0.013 | 0 | 0.13 | 0.18 | 0.14 | 0.22 | 0.18 | 0.09 |
| 6 | 90497589 | rs4707569 | 0.658 | 1.055 | -1.336 | -0.697 | -0.005 | -0.006 | -0.006 | 0 | 0.13 | 0.14 | 0.14 | 0.22 | 0.18 | 0.09 |
| 6 | 90499502 | rs1036853 | 1.068 | 1.028 | -1.411 | -0.753 | -0.05 | 0.046 | 0.046 | 0 | 0.13 | 0.01 | 0.09 | 0.05 | 0.16 | 0.18 |
| 6 | 90520848 | rs7744538 | 0.137 | 1.043 | -0.862 | -0.614 | 0.004 | 0.051 | 0.051 | 0.007 | 0.14 | 0.27 | 0.14 | 0.22 | 0.17 | 0.09 |
| 6 | 90529746 | rs12207432 | 2.129 | 0.847 | -1.13 | -0.895 | 0.039 | 0.128 | 0.128 | 0 | 0.27 | 0.08 | 0.09 | 0 | 0.17 | 0.11 |
| 6 | 90540008 | rs11967579 | 1.113 | 1.093 | -1.052 | -0.648 | -0.05 | 0.038 | 0.038 | 0 | 0.1 | 0.01 | 0.09 | 0.04 | 0.15 | 0.14 |
| 6 | 90544486 | rs12661409 | 0.626 | 1.119 | -1.213 | -0.616 | 0.001 | -0.003 | -0.003 | 0.198 | 0.12 | 0.12 | 0.1 | 0.21 | 0.12 | 0.08 |
| 6 | 90565242 | rs369328 | -1.081 | 1.063 | -1.149 | -0.645 | -0.007 | 0.028 | 0.028 | 0 | 0.43 | 0.35 | 0.33 | 0.3 | 0.54 | 0.59 |
| 6 | 90584424 | rs9344968 | -0.112 | 1.085 | -1.102 | -0.829 | -0.013 | -0.008 | -0.008 | 0.021 | 0.22 | 0.22 | 0.18 | 0.22 | 0.31 | 0.31 |
| 6 | 90586725 | rs453752 | 0.739 | 1.079 | -1.037 | -0.867 | -0.002 | -0.013 | -0.013 | 1.418 | 0.1 | 0.1 | 0.06 | 0.04 | 0.08 | 0.13 |
| 6 | 90592223 | rs441577 | 0.745 | 1.084 | -1.035 | -0.863 | -0.002 | -0.013 | -0.013 | 0.795 | 0.1 | 0.1 | 0.06 | 0.04 | 0.08 | 0.13 |
| 6 | 90597428 | rs11755610 | 0.071 | 1.38 | -1.152 | -0.925 | -0.065 | 0.07 | 0.07 | 0.75 | 0.2 | 0.1 | 0.17 | 0.04 | 0.34 | 0.38 |
| 6 | 90602268 | rs17764308 | 0.71 | 1.253 | -1.069 | -0.818 | -0.027 | 0.032 | 0.032 | 0 | 0.11 | 0.05 | 0.12 | 0.03 | 0.19 | 0.16 |
| 6 | 90603505 | rs292241 | -0.181 | 1.178 | -1.001 | -0.92 | -0.006 | 0.007 | 0.007 | 0 | 0.22 | 0.17 | 0.16 | 0.07 | 0.25 | 0.29 |
| 6 | 90606331 | rs292240 | -1.259 | 1.151 | -1.275 | -1.196 | -0.033 | 0.2 | 0.2 | 0 | 0.58 | 0.85 | 0.78 | 0.96 | 0.59 | 0.71 |
| 6 | 90609718 | rs4706352 | -0.622 | 1.446 | -0.848 | -0.476 | -0.002 | 0.004 | 0.004 | 0 | 0.39 | 0.67 | 0.6 | 0.74 | 0.43 | 0.38 |
| 6 | 90615280 | rs9342215 | -0.199 | 1.596 | -1.369 | -0.021 | -0.026 | -0.037 | -0.037 | 0 | 0.21 | 0.12 | 0.18 | 0.25 | 0.26 | 0.23 |
| 6 | 90622187 | rs367274 | -0.802 | 1.491 | -1.131 | 0.007 | -0.014 | 0.03 | 0.03 | 0 | 0.42 | 0.34 | 0.36 | 0.32 | 0.47 | 0.49 |
| 6 | 90622723 | rs366655 | -0.246 | 1.525 | -1.016 | -0.166 | 0.005 | 0.046 | 0.046 | 0.237 | 0.23 | 0.14 | 0.22 | 0.1 | 0.21 | 0.19 |
| 6 | 90627452 | rs176765 | -0.341 | 0.661 | -1.052 | 0.034 | -0.006 | 0.012 | 0.012 | 0.174 | 0.26 | 0.23 | 0.34 | 0.14 | 0.32 | 0.22 |
| 6 | 90629110 | rs9351229 | -0.75 | 0.453 | -1.422 | 0.514 | -0.021 | 0.038 | 0.038 | 0 | 0.43 | 0.24 | 0.58 | 0.67 | 0.5 | 0.36 |
| 6 | 90631132 | rs2304895 | -0.649 | 0.364 | -0.534 | 1.498 | -0.002 | 0.01 | 0.01 | 0 | 0.14 | 0.18 | 0.12 | 0.21 | 0.13 | 0.11 |
| 6 | 90631504 | rs2304894 | -0.622 | 0.515 | -0.576 | 1.481 | -0.026 | 0.085 | 0.085 | 0 | 0.08 | 0.19 | 0.04 | 0.2 | 0.03 | 0.15 |
| 6 | 90634552 | rs4707581 | -0.756 | 0.852 | -1.31 | 1.27 | -0.036 | -0.12 | -0.12 | 0 | 0.14 | 0.01 | 0.18 | 0.45 | 0.15 | 0.26 |
| 6 | 90634912 | rs17513180 | -0.467 | 1.218 | -0.855 | 0.462 | 0.005 | -0.017 | -0.017 | 0 | 0.07 | 0.08 | 0.09 | 0.03 | 0.11 | 0.02 |
| 6 | 90638553 | rs1065272 | -0.079 | 0.553 | -0.961 | 0.095 | 0 | -0.004 | -0.004 | 1.566 | 0.53 | 0.51 | 0.65 | 0.73 | 0.45 | 0.38 |

| | | | | | | | | | | | | | | | | |
|---|---|---|---|---|---|---|---|---|---|---|---|---|---|---|---|---|
| 6 | 90638963 | rs1055403 | -0.119 | 0.898 | -1.217 | 0.221 | 0.006 | -0.006 | -0.006 | 0.035 | 0.19 | 0.07 | 0.17 | 0.23 | 0.17 | 0.28 |
| 6 | 90640491 | rs2289577 | -0.434 | 1.11 | -0.69 | 0.633 | 0.032 | 0.018 | 0.018 | 0 | 0.07 | 0.17 | 0.08 | 0.09 | 0.1 | 0.03 |
| 6 | 90641522 | rs10455512 | -0.248 | 0.692 | -1.54 | 0.99 | -0.035 | -0.151 | -0.151 | 0 | 0.19 | 0.01 | 0.27 | 0.4 | 0.2 | 0.24 |
| 6 | 90641851 | rs17765013 | 0.18 | 0.954 | -1.352 | 0.671 | -0.031 | 0.02 | 0.02 | 0 | 0.05 | 0.01 | 0.08 | 0.03 | 0.1 | 0.04 |
| 6 | 90642358 | rs3798789 | 0.264 | 0.636 | -1.398 | 0.211 | 0.004 | 0.008 | 0.008 | 0 | 0.17 | 0.15 | 0.07 | 0.07 | 0.11 | 0.09 |
| 6 | 90642900 | rs292258 | 0.557 | 0.335 | -1.102 | 0.517 | 0.004 | 0.006 | 0.006 | 0.802 | 0.61 | 0.51 | 0.54 | 0.41 | 0.61 | 0.54 |
| 6 | 90642998 | rs292257 | 0.651 | 0.552 | -1.471 | 0.528 | 0.027 | 0.001 | 0.001 | 0.622 | 0.55 | 0.28 | 0.42 | 0.53 | 0.43 | 0.37 |
| 6 | 90643281 | rs292255 | 0.133 | 0.517 | -1.152 | 0.54 | 0.009 | 0.019 | 0.019 | 0.031 | 0.69 | 0.43 | 0.51 | 0.53 | 0.66 | 0.62 |
| 6 | 90643949 | rs17708487 | 0.714 | 0.745 | -0.979 | 0.457 | 0.017 | -0.032 | -0.032 | 0.643 | 0.14 | 0.14 | 0.09 | 0 | 0.23 | 0.27 |
| 6 | 90644783 | rs9359876 | 0.781 | 0.244 | -1.13 | 0.593 | 0.001 | -0.006 | -0.006 | 7.978 | 0.19 | 0.2 | 0.17 | 0.2 | 0.17 | 0.12 |
| 6 | 90646226 | rs292251 | 0.435 | 0.19 | -0.792 | 0.692 | -0.008 | -0.008 | -0.008 | 0 | 0.59 | 0.48 | 0.53 | 0.29 | 0.67 | 0.56 |
| 6 | 90649238 | rs4706353 | -0.103 | 0.101 | -0.443 | 0.742 | 0 | -0.001 | -0.001 | 0.034 | 0.72 | 0.71 | 0.77 | 0.73 | 0.73 | 0.72 |
| 6 | 90651086 | rs3734662 | 0.789 | 0.085 | -0.416 | 0.491 | 0.001 | 0.027 | 0.027 | 0 | 0.48 | 0.32 | 0.44 | 0.09 | 0.52 | 0.44 |
| 6 | 90652950 | rs9362704 | 0.348 | 0.311 | -0.479 | 0.58 | -0.003 | -0.004 | -0.004 | 0.544 | 0.59 | 0.45 | 0.39 | 0.47 | 0.54 | 0.53 |
| 6 | 90654861 | rs3966851 | 1.469 | 0.36 | -0.492 | 0.669 | 0.006 | -0.015 | -0.015 | 0 | 0.09 | 0.06 | 0.07 | 0.25 | 0.06 | 0.07 |
| 6 | 90655622 | rs9351230 | 0.176 | 0.406 | -0.433 | 0.333 | 0.038 | -0.06 | -0.06 | 0 | 0.66 | 0.63 | 0.62 | 0.86 | 0.61 | 0.65 |
| 6 | 90659596 | rs210059 | -0.091 | 1.021 | -0.242 | 0.79 | 0 | 0 | 0 | 1.606 | 0.22 | 0.21 | 0.15 | 0.25 | 0.18 | 0.1 |
| 6 | 90660319 | rs9451298 | -0.055 | 1.099 | 0.014 | 1.066 | 0.003 | 0.016 | 0.016 | 0 | 0.25 | 0.43 | 0.19 | 0.24 | 0.28 | 0.35 |
| 6 | 90663436 | rs17708991 | 0.951 | 1.194 | -0.423 | 0.823 | -0.034 | -0.015 | -0.015 | 0.585 | 0.08 | 0.01 | 0.21 | 0.07 | 0.1 | 0.07 |
| 6 | 90664582 | rs12213272 | -0.576 | 0.96 | 0.185 | 1.007 | -0.001 | -0.005 | -0.005 | 0 | 0.26 | 0.31 | 0.27 | 0.4 | 0.26 | 0.32 |
| 6 | 90665718 | rs12208074 | -0.531 | 0.969 | 0.184 | 0.996 | -0.001 | -0.008 | -0.008 | 0.054 | 0.26 | 0.31 | 0.26 | 0.38 | 0.26 | 0.3 |
| 6 | 90665969 | rs10806414 | -0.138 | 1.008 | 0.079 | 0.729 | -0.007 | -0.004 | -0.004 | 0.028 | 0.2 | 0.22 | 0.19 | 0.22 | 0.2 | 0.25 |
| 6 | 90669325 | rs9362708 | 0.923 | 1.018 | 0.1 | 0.732 | -0.001 | 0.002 | 0.002 | 0 | 0.13 | 0.15 | 0.1 | 0.18 | 0.15 | 0.17 |
| 6 | 90671114 | rs176713 | -0.036 | 1.029 | 0.19 | 0.357 | 0.024 | -0.034 | -0.034 | 2.021 | 0.09 | 0.19 | 0.09 | 0.04 | 0.11 | 0.04 |
| 6 | 90673538 | rs12213197 | -0.443 | 0.885 | 0.079 | 0.422 | 0.002 | 0.013 | 0.013 | 0 | 0.34 | 0.45 | 0.29 | 0.41 | 0.36 | 0.43 |
| 6 | 90674451 | rs16882302 | 1.428 | 0.956 | -0.09 | 0.554 | -0.001 | 0.003 | 0.003 | 0 | 0.06 | 0.04 | 0.07 | 0.21 | 0.06 | 0.04 |
| 6 | 90674918 | rs12525504 | -0.022 | 0.927 | 0.237 | -0.355 | -0.001 | -0.027 | -0.027 | 0.28 | 0.1 | 0.12 | 0.07 | 0.01 | 0.09 | 0.13 |
| 6 | 90677126 | rs6911941 | -0.139 | 0.817 | 0.206 | -0.87 | 0.001 | -0.04 | -0.04 | 0 | 0.53 | 0.46 | 0.31 | 0.08 | 0.47 | 0.36 |
| 6 | 90677176 | rs6454789 | 0.734 | 0.949 | 0.229 | -0.161 | -0.023 | -0.004 | -0.004 | 0 | 0.62 | 0.37 | 0.42 | 0.56 | 0.56 | 0.5 |
| 6 | 90678323 | rs6941452 | -0.202 | 1.129 | 0.319 | -0.953 | 0.051 | -0.081 | -0.081 | 0 | 0.74 | 0.54 | 0.81 | 0.84 | 0.69 | 0.71 |
| 6 | 90681466 | rs6917758 | -0.123 | 0.451 | 0.406 | -1.02 | 0.02 | 0.11 | 0.11 | 0 | 0.28 | 0.1 | 0.18 | 0.03 | 0.27 | 0.27 |
| 6 | 90683044 | rs11969265 | -1.102 | 0.589 | 0.762 | -0.741 | 0 | 0.004 | 0.004 | 1.401 | 0.5 | 0.5 | 0.53 | 0.37 | 0.52 | 0.48 |
| 6 | 90683587 | rs6910129 | -0.382 | 0.983 | 0.849 | -0.945 | 0.033 | 0.073 | 0.073 | 1.268 | 0.05 | 0.16 | 0.24 | 0.17 | 0.07 | 0.01 |
| 6 | 90684355 | rs6936055 | -0.485 | 0.972 | 0.573 | -1.016 | 0.013 | 0.084 | 0.084 | 0.396 | 0.2 | 0.48 | 0.36 | 0.31 | 0.2 | 0.16 |
| 6 | 90687073 | rs6454792 | -1.019 | 1.071 | 0.258 | -1.136 | 0 | 0.034 | 0.034 | 3.704 | 0.5 | 0.6 | 0.56 | 0.31 | 0.53 | 0.4 |
| 6 | 90695246 | rs10806415 | -1.411 | 0.885 | 0.521 | -0.645 | 0 | 0.001 | 0.001 | 2.124 | 0.29 | 0.61 | 0.44 | 0.31 | 0.31 | 0.3 |
| 6 | 90695766 | rs6908310 | -0.77 | 0.47 | 0.124 | -0.599 | -0.002 | -0.023 | -0.023 | 0 | 0.08 | 0.13 | 0.06 | 0 | 0.08 | 0.12 |
| 6 | 90698125 | rs7761142 | -0.752 | 0.364 | 0.442 | -0.378 | -0.029 | 0.048 | 0.048 | 0 | 0.19 | 0.52 | 0.35 | 0.27 | 0.18 | 0.18 |
| 6 | 90698820 | rs10806416 | -0.504 | 0.661 | 0.165 | -1.438 | 0 | -0.017 | -0.017 | 0.529 | 0.37 | 0.28 | 0.23 | 0.06 | 0.35 | 0.31 |
| 6 | 90700549 | rs6926007 | 0.489 | 0.801 | 0.36 | -1.455 | -0.007 | -0.019 | -0.019 | 0 | 0.66 | 0.46 | 0.67 | 0.73 | 0.71 | 0.75 |
| 6 | 90701109 | rs3857494 | -0.149 | 0.946 | 0.603 | -1.579 | 0.016 | 0.045 | 0.045 | 0 | 0.76 | 0.86 | 0.88 | 1 | 0.8 | 0.82 |
| 6 | 90707030 | rs4707585 | 0.677 | 0.884 | 0.667 | -1.536 | -0.005 | -0.013 | -0.013 | 0.079 | 0.67 | 0.45 | 0.69 | 0.73 | 0.71 | 0.75 |
| 6 | 90707282 | rs4707586 | 1.391 | 1.386 | 0.11 | -2.091 | -0.065 | 0.084 | 0.084 | 0.939 | 0.43 | 0.05 | 0.33 | 0.67 | 0.42 | 0.6 |
| 6 | 90713018 | rs6454796 | -0.63 | 1.276 | 0.504 | -2.013 | 0.041 | 0.025 | 0.025 | 1.17 | 0.33 | 0.68 | 0.32 | 0.31 | 0.21 | 0.24 |
| 6 | 90714054 | rs9444725 | 0.088 | 1.43 | 0.848 | -1.962 | -0.035 | -0.048 | -0.048 | 0 | 0.25 | 0.45 | 0.26 | 0.2 | 0.19 | 0.21 |
| 6 | 90717525 | rs9451313 | 0.077 | 1.41 | 1.121 | -1.993 | 0.001 | -0.001 | -0.001 | 0 | 0.27 | 0.28 | 0.13 | 0.09 | 0.13 | 0.18 |
| 6 | 90718877 | rs13206749 | 1.123 | 1.297 | 0.421 | -2.154 | -0.062 | -0.025 | -0.025 | 1.061 | 0.16 | 0.01 | 0.15 | 0.12 | 0.18 | 0.07 |
| 6 | 90719164 | rs3857496 | -0.512 | 1.21 | 1.199 | -2.062 | 0.096 | -0.065 | -0.065 | 0 | 0.24 | 0.44 | 0.44 | 0.21 | 0.36 | 0.18 |
| 6 | 90724855 | rs9451316 | -1.205 | 0.581 | 1.404 | -1.618 | 0.015 | 0.033 | 0.033 | 0 | 0.44 | 0.65 | 0.47 | 0.28 | 0.33 | 0.27 |
| 6 | 90730438 | rs9444729 | -0.844 | 1.752 | 1.575 | -0.898 | 0.003 | -0.078 | -0.078 | 0.66 | 0.14 | 0.35 | 0.22 | 0.01 | 0.12 | 0.06 |
| 6 | 90734082 | rs12663434 | -1.787 | 1.739 | 1.467 | -0.922 | 0.01 | -0.056 | -0.056 | 0.193 | 0.23 | 0.56 | 0.39 | 0.19 | 0.23 | 0.11 |
| 6 | 90734908 | rs4388268 | -1.685 | 1.81 | 0.947 | -0.886 | 0.022 | -0.071 | -0.071 | 0.22 | 0.23 | 0.47 | 0.4 | 0.19 | 0.23 | 0.11 |
| 6 | 90735222 | rs4053616 | 1.51 | 1.8 | 0.915 | -0.866 | 0.053 | -0.081 | -0.081 | 0.238 | 0.75 | 0.53 | 0.57 | 0.81 | 0.69 | 0.86 |
| 6 | 90735491 | rs16882364 | -1.285 | 1.197 | 0.465 | -0.269 | 0.026 | 0.018 | 0.018 | 0 | 0.09 | 0.16 | 0.16 | 0.18 | 0.11 | 0.05 |
| 6 | 90738735 | rs6931936 | -0.781 | 0.945 | 0.255 | -0.541 | -0.006 | -0.051 | -0.051 | 0 | 0.3 | 0.46 | 0.42 | 0.25 | 0.25 | 0.13 |
| 6 | 90739172 | rs3857498 | -0.465 | 1.15 | 0.476 | 0.185 | 0.022 | 0.024 | 0.024 | 0 | 0.25 | 0.37 | 0.22 | 0.32 | 0.28 | 0.3 |
| 6 | 90739827 | rs9362711 | -0.963 | 0.755 | 0.531 | 0.375 | 0.005 | 0.052 | 0.052 | 0 | 0.17 | 0.28 | 0.25 | 0.64 | 0.17 | 0.4 |
| 6 | 90756453 | rs9342219 | -1.575 | 0.552 | 0.265 | -0.065 | -0.009 | -0.021 | -0.021 | 0.099 | 0.25 | 0.24 | 0.29 | 0.65 | 0.3 | 0.51 |
| 6 | 90772650 | rs4053606 | 0.197 | -0.154 | -0.397 | -0.508 | 0.047 | 0.005 | 0.005 | 9.133 | 0.15 | 0.01 | 0.11 | 0.09 | 0.06 | 0.06 |
| 6 | 90776990 | rs6932747 | 0.17 | 0.566 | -0.176 | -0.428 | -0.01 | 0.076 | 0.076 | 0 | 0.86 | 0.97 | 0.79 | 0.99 | 0.85 | 0.89 |
| 6 | 90777594 | rs13199026 | -0.647 | 0.617 | 0.362 | 0.205 | -0.03 | 0.045 | 0.045 | 0.188 | 0.09 | 0.02 | 0.06 | 0 | 0.14 | 0.11 |
| 6 | 90795343 | rs4707594 | -1.482 | 0.73 | 0.282 | 0.279 | -0.007 | -0.013 | -0.013 | 0 | 0.22 | 0.17 | 0.28 | 0.65 | 0.27 | 0.44 |
| 6 | 90800438 | rs17767336 | -0.872 | 0.786 | -0.632 | -0.641 | -0.036 | 0.049 | 0.049 | 1.58 | 0.1 | 0.03 | 0.11 | 0 | 0.18 | 0.15 |
| 6 | 90814199 | rs6454802 | -0.045 | 0.407 | -0.869 | -1.161 | -0.026 | 0.101 | 0.101 | 0.492 | 0.34 | 0.18 | 0.33 | 0.01 | 0.4 | 0.3 |
| 6 | 90816437 | rs207260 | -0.182 | 0.314 | -0.805 | -0.881 | 0.002 | -0.003 | -0.003 | 0 | 0.07 | 0.07 | 0.11 | 0.08 | 0.05 | 0.08 |
| 6 | 90820581 | rs207264 | -0.361 | -0.25 | 0.043 | -1.095 | -0.001 | -0.02 | -0.02 | 0 | 0.51 | 0.52 | 0.55 | 0.13 | 0.48 | 0.46 |
| 6 | 90828169 | rs207269 | -0.562 | -0.559 | 0.209 | -1.052 | -0.001 | 0 | 0 | 0 | 0.6 | 0.64 | 0.62 | 0.22 | 0.52 | 0.55 |
| 6 | 90849635 | rs2501726 | -1.143 | -0.762 | 1.124 | -0.705 | -0.031 | -0.021 | -0.021 | 3.523 | 0.12 | 0.38 | 0.08 | 0.08 | 0.05 | 0.1 |
| 6 | 90859267 | rs2208039 | -0.553 | -0.642 | 0.569 | -0.592 | -0.02 | 0.006 | 0.006 | 0.164 | 0.06 | 0.13 | 0.07 | 0.07 | 0.02 | 0.02 |

| | | | | | | | | | | | | | | | | |
|---|---|---|---|---|---|---|---|---|---|---|---|---|---|---|---|---|
| 6 | 90865557 | rs7748973 | -0.479 | -0.658 | 0.395 | -0.601 | -0.013 | 0 | 0 | 0 | 0.06 | 0.08 | 0.06 | 0.08 | 0.02 | 0.06 |
| 6 | 90885698 | rs2501718 | -1.113 | -0.832 | 0.927 | -0.69 | 0.003 | 0.013 | 0.013 | 0 | 0.09 | 0.31 | 0.11 | 0.07 | 0.06 | 0.13 |
| 6 | 90887951 | rs9344992 | 0.218 | -0.94 | -0.322 | 0.223 | 0.007 | -0.042 | -0.042 | 1.11 | 0.13 | 0.05 | 0.14 | 0.4 | 0.1 | 0.17 |
| 6 | 90899561 | rs2501720 | -1.562 | -1.019 | -0.81 | -0.42 | 0.07 | -0.045 | -0.045 | 0 | 0.44 | 0.89 | 0.41 | 0.56 | 0.27 | 0.5 |
| 6 | 90909157 | rs17712288 | 1.012 | -0.621 | -1.226 | -1.082 | -0.021 | 0.198 | 0.198 | 0 | 0.43 | 0.14 | 0.42 | 0.15 | 0.45 | 0.39 |
| 6 | 90912594 | rs12209546 | 0.535 | -0.659 | -1.248 | -1.096 | -0.055 | -0.031 | -0.031 | 0 | 0.55 | 0.13 | 0.56 | 0.56 | 0.55 | 0.57 |
| 6 | 90916345 | rs3734660 | 1.092 | -0.647 | -1.245 | -1.098 | -0.035 | 0.202 | 0.202 | 0.086 | 0.42 | 0.11 | 0.42 | 0.15 | 0.45 | 0.39 |
| 6 | 90921740 | rs370409 | -1.379 | -0.685 | -1.356 | -1.062 | -0.101 | -0.111 | -0.111 | 0.108 | 0.83 | 0.99 | 0.8 | 0.63 | 0.69 | 0.79 |
| 6 | 90926612 | rs10806425 | 1.344 | -0.687 | -1.203 | -1.144 | -0.003 | 0.143 | 0.143 | 0 | 0.41 | 0.18 | 0.38 | 0.09 | 0.43 | 0.36 |
| 6 | 90927728 | rs206916 | -0.538 | -0.654 | -1.249 | -1.097 | -0.058 | -0.09 | -0.09 | 0 | 0.75 | 0.87 | 0.7 | 0.56 | 0.65 | 0.71 |
| 6 | 90929301 | rs9344996 | 0.902 | -0.574 | -1.544 | -0.777 | -0.002 | -0.08 | -0.08 | 0 | 0.14 | 0.02 | 0.14 | 0.38 | 0.11 | 0.18 |
| 6 | 90929696 | rs12205059 | 0.535 | -0.598 | -1.469 | -0.81 | -0.012 | 0.02 | 0.02 | 0.05 | 0.11 | 0.04 | 0.07 | 0 | 0.09 | 0.15 |
| 6 | 90936894 | rs206913 | -0.496 | -0.564 | -1.541 | -0.776 | -0.095 | -0.113 | -0.113 | 0.077 | 0.75 | 0.95 | 0.72 | 0.55 | 0.63 | 0.66 |
| 6 | 90941240 | rs11757155 | 1.876 | -0.37 | -1.493 | -0.828 | -0.035 | 0.16 | 0.16 | 0.149 | 0.33 | 0.06 | 0.29 | 0.04 | 0.36 | 0.23 |
| 6 | 90946479 | rs4707609 | 1.625 | -0.373 | -1.463 | -0.786 | -0.016 | 0.129 | 0.129 | 0.571 | 0.36 | 0.09 | 0.34 | 0.11 | 0.38 | 0.26 |
| 6 | 90957463 | rs3757247 | 0.987 | -0.359 | -1.374 | -0.774 | 0.004 | -0.015 | -0.015 | 0 | 0.5 | 0.18 | 0.49 | 0.49 | 0.49 | 0.43 |
| 6 | 90973159 | rs1847472 | 1.571 | -0.025 | -1.415 | -0.718 | -0.024 | 0.144 | 0.144 | 0 | 0.33 | 0.09 | 0.29 | 0.04 | 0.37 | 0.24 |
| 6 | 90975999 | rs661713 | -0.133 | -0.02 | -1.251 | -0.391 | -0.094 | -0.137 | -0.137 | 0 | 0.74 | 0.94 | 0.67 | 0.49 | 0.62 | 0.64 |
| 6 | 90981653 | rs9111 | -1.044 | 0.031 | -1.274 | -0.402 | -0.1 | -0.109 | -0.109 | 0.025 | 0.83 | 0.99 | 0.81 | 0.63 | 0.69 | 0.79 |
| 6 | 90987872 | rs2174281 | 0.95 | -0.035 | -1.085 | -0.685 | -0.011 | 0.096 | 0.096 | 0.215 | 0.45 | 0.22 | 0.37 | 0.04 | 0.46 | 0.39 |
| 6 | 91002494 | rs597325 | 0.052 | 1.079 | -1.21 | 0.122 | -0.107 | -0.161 | -0.161 | 0 | 0.74 | 0.96 | 0.68 | 0.48 | 0.62 | 0.65 |
| 6 | 91019654 | rs12195431 | -0.91 | 1.029 | -0.624 | -1.09 | 0.002 | 0.005 | 0.005 | 0 | 0.05 | 0.06 | 0.11 | 0.09 | 0.08 | 0.02 |
| 6 | 91020169 | rs285644 | 0.686 | 1.06 | -0.443 | -1.295 | 0.015 | -0.037 | -0.037 | 0 | 0.91 | 0.88 | 0.92 | 1 | 0.89 | 0.89 |
| 6 | 91028797 | rs285640 | -0.193 | 0.547 | -1.42 | -1.843 | -0.084 | 0.009 | 0.009 | 0 | 0.28 | 0.04 | 0.39 | 0.79 | 0.33 | 0.29 |
| 6 | 91037023 | rs1847473 | 0.532 | 0.381 | -0.945 | -2.711 | -0.007 | -0.033 | -0.033 | 0 | 0.33 | 0.36 | 0.34 | 0.15 | 0.29 | 0.2 |
| 6 | 91039808 | rs285630 | -0.714 | 0.406 | -0.038 | -2.967 | 0 | -0.061 | -0.061 | 0 | 0.12 | 0.36 | 0.09 | 0.01 | 0.08 | 0.1 |
| 6 | 91046297 | rs4143444 | 0.131 | 0.834 | -0.165 | -3.027 | 0.026 | -0.041 | -0.041 | 0.05 | 0.91 | 0.84 | 0.93 | 1 | 0.88 | 0.89 |
| 6 | 91050550 | rs285621 | 1.148 | 0.406 | -0.023 | -3.312 | 0.003 | -0.023 | -0.023 | 0.091 | 0.76 | 0.7 | 0.81 | 0.87 | 0.73 | 0.73 |
| 6 | 91054195 | rs285619 | 0.899 | 0.591 | -0.197 | -3.505 | -0.002 | 0 | 0 | 0 | 0.73 | 0.76 | 0.7 | 0.88 | 0.68 | 0.71 |
| 6 | 91056784 | rs2064960 | 1.264 | 0.803 | -0.468 | -3.618 | -0.035 | 0.006 | 0.006 | 0 | 0.19 | 0.07 | 0.4 | 0.84 | 0.27 | 0.31 |
| 6 | 91061153 | rs9444758 | 1.119 | -0.272 | -0.412 | -3.924 | -0.001 | -0.049 | -0.049 | 0 | 0.21 | 0.22 | 0.13 | 0 | 0.18 | 0.11 |
| 6 | 91063151 | rs9451389 | -0.03 | -0.505 | -0.379 | -4.403 | 0.007 | 0.001 | 0.001 | 0 | 0.33 | 0.28 | 0.15 | 0.11 | 0.24 | 0.34 |
| 6 | 91064554 | rs717893 | 0.672 | -0.045 | -0.403 | -4.299 | -0.018 | 0.057 | 0.057 | 0 | 0.35 | 0.19 | 0.5 | 0.87 | 0.38 | 0.43 |
| 6 | 91064774 | rs285651 | -0.489 | 0.185 | -0.43 | -4.192 | 0.034 | -0.045 | -0.045 | 0 | 0.89 | 0.8 | 0.81 | 0.99 | 0.82 | 0.87 |
| 6 | 91067623 | rs6899522 | 1.207 | 0.081 | -0.995 | -4.252 | -0.001 | -0.037 | -0.037 | 0 | 0.17 | 0.18 | 0.11 | 0 | 0.15 | 0.1 |
| 6 | 91069230 | rs11754391 | 0.568 | 0.153 | -1.334 | -4.008 | 0.018 | 0.118 | 0.118 | 0 | 0.24 | 0.07 | 0.11 | 0.01 | 0.21 | 0.22 |
| 6 | 91082136 | rs4707614 | -0.082 | 0.268 | -1.275 | -3.998 | -0.019 | 0.126 | 0.126 | 0.202 | 0.65 | 0.83 | 0.69 | 0.99 | 0.6 | 0.66 |
| 6 | 91084452 | rs1570168 | 0.292 | 0.007 | -0.819 | -3.721 | -0.015 | -0.083 | -0.083 | 0 | 0.28 | 0.52 | 0.18 | 0.1 | 0.23 | 0.17 |
| 6 | 91086462 | rs790595 | -0.269 | 0.373 | -0.987 | -3.522 | 0.004 | 0.045 | 0.045 | 0 | 0.54 | 0.66 | 0.32 | 0.11 | 0.47 | 0.41 |
| 6 | 91091844 | rs790585 | -0.286 | 0.59 | -0.753 | -2.719 | -0.001 | -0.01 | -0.01 | 0 | 0.38 | 0.4 | 0.38 | 0.16 | 0.34 | 0.28 |
| 6 | 91092352 | rs790584 | 0.057 | 0.662 | -0.694 | -2.701 | -0.001 | -0.087 | -0.087 | 0 | 0.54 | 0.42 | 0.64 | 0.98 | 0.57 | 0.63 |
| 6 | 91094257 | rs790583 | -0.057 | 0.592 | -0.743 | -0.937 | 0.004 | 0.007 | 0.007 | 0 | 0.52 | 0.6 | 0.72 | 0.76 | 0.6 | 0.71 |
| 6 | 91098430 | rs790576 | 0.173 | 0.831 | -0.962 | -1.04 | -0.002 | 0.031 | 0.031 | 0.383 | 0.62 | 0.34 | 0.69 | 0.74 | 0.61 | 0.73 |
| 6 | 91105896 | rs790605 | -0.008 | 0.676 | -0.88 | -1.547 | 0.002 | 0.052 | 0.052 | 0 | 0.53 | 0.7 | 0.64 | 0.68 | 0.52 | 0.6 |
| 6 | 91106000 | rs9353722 | 0.236 | 0.534 | -0.885 | -1.908 | -0.049 | 0.015 | 0.015 | 0 | 0.39 | 0.17 | 0.57 | 0.67 | 0.46 | 0.46 |
| 6 | 91108519 | rs9444762 | -0.967 | 0.469 | -0.788 | -2.131 | -0.064 | 0.093 | 0.093 | 0.408 | 0.82 | 0.97 | 0.87 | 0.95 | 0.78 | 0.8 |
| 6 | 91112237 | rs12208250 | 0.661 | 0.657 | -0.762 | -2.04 | -0.03 | -0.018 | -0.018 | 0.028 | 0.34 | 0.12 | 0.51 | 0.67 | 0.36 | 0.44 |
| 6 | 91114488 | rs1451071 | 0.828 | 0.482 | -0.845 | -1.979 | -0.028 | -0.076 | -0.076 | 0 | 0.27 | 0.02 | 0.21 | 0.26 | 0.26 | 0.2 |
| 6 | 91117088 | rs12190699 | -0.185 | 0.622 | -0.286 | -1.559 | 0.007 | 0.001 | 0.001 | 0.015 | 0.09 | 0.19 | 0.08 | 0.12 | 0.11 | 0.12 |
| 6 | 91121017 | rs12195320 | 0.134 | 0.646 | -0.416 | -1.543 | -0.004 | -0.002 | -0.002 | 0 | 0.08 | 0.09 | 0.06 | 0.12 | 0.11 | 0.12 |
| 6 | 91122641 | rs9451413 | -0.43 | 0.48 | 0.068 | -1.671 | 0.072 | 0.045 | 0.045 | 0.016 | 0.06 | 0.38 | 0.04 | 0 | 0.02 | 0.04 |
| 6 | 91122971 | rs9353726 | -0.235 | 0.438 | -0.526 | -2.037 | -0.025 | 0.071 | 0.071 | 0 | 0.46 | 0.26 | 0.62 | 0.68 | 0.52 | 0.49 |
| 6 | 91123042 | rs9353727 | -0.273 | 0.857 | -0.729 | -2.158 | -0.019 | 0.042 | 0.042 | 0 | 0.42 | 0.25 | 0.43 | 0.68 | 0.44 | 0.45 |
| 6 | 91123268 | rs7745444 | 0.067 | 0.598 | -0.848 | -2.167 | -0.013 | 0.014 | 0.014 | 0 | 0.14 | 0.09 | 0.13 | 0 | 0.17 | 0.07 |
| 6 | 91123494 | rs7745771 | -0.749 | 0.675 | -0.761 | -2.134 | -0.007 | 0.035 | 0.035 | 0.337 | 0.49 | 0.8 | 0.62 | 0.56 | 0.48 | 0.46 |
| 6 | 91129879 | rs10944485 | 0.651 | 0.536 | -0.725 | -0.923 | -0.058 | -0.084 | -0.084 | 0.412 | 0.13 | 0.01 | 0.28 | 0.24 | 0.16 | 0.29 |
| 6 | 91131746 | rs1221552 | -0.042 | 0.696 | -0.531 | -1.125 | -0.001 | -0.03 | -0.03 | 0 | 0.07 | 0.13 | 0.03 | 0.01 | 0.03 | 0.07 |
| 6 | 91133398 | rs9362733 | 0.946 | 0.824 | -0.528 | -1.284 | 0.001 | 0.008 | 0.008 | 0 | 0.46 | 0.45 | 0.42 | 0.33 | 0.47 | 0.33 |
| 6 | 91135896 | rs1451070 | 0.39 | 0.75 | -0.42 | -1.139 | -0.022 | -0.108 | -0.108 | 0.088 | 0.27 | 0.08 | 0.39 | 0.55 | 0.26 | 0.34 |
| 6 | 91137151 | rs1668657 | -0.874 | 0.775 | -0.447 | -1.17 | 0.001 | 0.003 | 0.003 | 0 | 0.45 | 0.54 | 0.51 | 0.67 | 0.41 | 0.58 |
| 6 | 91138902 | rs1668702 | -0.597 | 0.774 | -0.444 | -1.174 | -0.006 | 0 | 0 | 0.064 | 0.41 | 0.52 | 0.47 | 0.56 | 0.33 | 0.46 |
| 6 | 91141142 | rs9451422 | -0.148 | 0.729 | -0.696 | -1.081 | -0.035 | -0.03 | -0.03 | 0 | 0.32 | 0.13 | 0.44 | 0.66 | 0.34 | 0.46 |
| 6 | 91158691 | rs7773219 | -0.028 | 0.614 | -0.628 | -0.959 | 0.006 | -0.023 | -0.023 | 0.023 | 0.29 | 0.23 | 0.4 | 0.55 | 0.26 | 0.33 |
| 6 | 91165896 | rs958656 | -0.883 | 0.601 | -0.413 | -0.964 | -0.032 | -0.072 | -0.072 | 2.077 | 0.14 | 0.62 | 0.08 | 0.01 | 0.07 | 0.12 |
| 6 | 91168261 | rs791039 | 0.168 | 0.638 | -0.162 | -0.928 | 0.009 | -0.018 | -0.018 | 0 | 0.07 | 0.28 | 0.04 | 0 | 0.02 | 0.07 |
| 6 | 91168953 | rs1145721 | -0.639 | 0.71 | -1.162 | -0.997 | 0.018 | -0.017 | -0.017 | 0 | 0.39 | 0.67 | 0.49 | 0.56 | 0.35 | 0.45 |
| 6 | 91169306 | rs1144152 | -0.963 | 0.865 | -1.307 | -1.008 | -0.011 | 0.004 | 0.004 | 0.268 | 0.92 | 0.94 | 0.93 | 1 | 0.88 | 0.92 |
| 6 | 91176901 | rs1353227 | 1.154 | 0.649 | -1.629 | -1.228 | 0.013 | 0.002 | 0.002 | 0 | 0.33 | 0.19 | 0.22 | 0.27 | 0.28 | 0.23 |
| 6 | 91180389 | rs9353731 | 1.328 | 0.249 | -1.656 | -1.109 | 0.014 | -0.07 | -0.07 | 0 | 0.1 | 0.01 | 0.06 | 0.3 | 0.08 | 0.04 |

| | | | | | | | | | | | | | | | | |
|---|---|---|---|---|---|---|---|---|---|---|---|---|---|---|---|---|
| 6 | 91184183 | rs1144157 | 0.313 | 0.467 | -1.326 | -1.554 | -0.005 | 0.003 | 0.003 | 0.869 | 0.66 | 0.34 | 0.69 | 0.93 | 0.67 | 0.72 |
| 6 | 91185262 | rs791057 | 0.678 | 0.488 | -1.364 | -1.547 | -0.021 | 0.117 | 0.117 | 1.741 | 0.58 | 0.27 | 0.63 | 0.82 | 0.56 | 0.56 |
| 6 | 91186769 | rs1221561 | -0.333 | 0.603 | -1.388 | -1.59 | -0.028 | 0.051 | 0.051 | 1.663 | 0.74 | 0.84 | 0.75 | 0.83 | 0.72 | 0.69 |
| 6 | 91187510 | rs1145724 | -0.756 | 0.477 | -1.362 | -1.664 | -0.009 | -0.069 | -0.069 | 1.282 | 0.18 | 0.66 | 0.13 | 0.01 | 0.16 | 0.13 |
| 6 | 91187609 | rs1231690 | -0.484 | 0.627 | -1.326 | -1.571 | -0.024 | 0.075 | 0.075 | 1.863 | 0.77 | 0.88 | 0.78 | 0.94 | 0.76 | 0.77 |
| 6 | 91188927 | rs6911698 | -0.203 | 0.352 | -0.748 | -1.138 | 0.049 | -0.05 | -0.05 | 0.014 | 0.1 | 0.51 | 0.09 | 0.01 | 0.09 | 0.05 |
| 6 | 91192230 | rs4707623 | 1.593 | 0.23 | -1.516 | -1.034 | 0.018 | -0.071 | -0.071 | 0.038 | 0.21 | 0.05 | 0.18 | 0.29 | 0.17 | 0.19 |
| 6 | 91192557 | rs791059 | -0.111 | 0.276 | -1.187 | -1.084 | 0.017 | 0.003 | 0.003 | 0.006 | 0.44 | 0.6 | 0.35 | 0.43 | 0.4 | 0.44 |
| 6 | 91194149 | rs1221220 | -0.37 | 0.524 | -1.542 | -0.591 | -0.049 | -0.084 | -0.084 | 0 | 0.67 | 0.86 | 0.5 | 0.54 | 0.62 | 0.64 |
| 6 | 91200145 | rs157686 | -0.95 | 0.486 | -1.544 | -0.686 | -0.035 | -0.098 | -0.098 | 1.801 | 0.77 | 0.9 | 0.52 | 0.54 | 0.7 | 0.69 |
| 6 | 91204215 | rs806272 | -0.189 | 0.335 | -0.946 | -0.216 | -0.023 | 0.071 | 0.071 | 0.154 | 0.58 | 0.72 | 0.27 | 0.22 | 0.44 | 0.47 |
| 6 | 91205726 | rs2325302 | 1.113 | 0.387 | -0.834 | -0.288 | -0.015 | 0.046 | 0.046 | 0 | 0.23 | 0.15 | 0.11 | 0.04 | 0.21 | 0.14 |
| 6 | 91207091 | rs806275 | -0.276 | 0.297 | -0.77 | -0.36 | 0.014 | -0.088 | -0.088 | 0 | 0.13 | 0.5 | 0.09 | 0.01 | 0.09 | 0.1 |
| 6 | 91207351 | rs806276 | -0.125 | 0.626 | -0.792 | -0.043 | -0.01 | 0.022 | 0.022 | 0.008 | 0.39 | 0.68 | 0.34 | 0.26 | 0.37 | 0.29 |
| 6 | 91208083 | rs9451435 | 0.859 | 0.536 | -0.558 | -0.738 | 0.139 | 0.052 | 0.052 | 0.246 | 0.05 | 0.31 | 0.05 | 0 | 0.05 | 0.01 |
| 6 | 91208376 | rs284509 | -0.474 | 0.338 | -1.464 | -0.548 | -0.031 | 0.029 | 0.029 | 0 | 0.63 | 0.77 | 0.32 | 0.32 | 0.51 | 0.57 |
| 6 | 91208693 | rs284513 | 0.024 | 0.043 | -1.314 | 0.328 | 0.035 | 0.009 | 0.009 | 0 | 0.28 | 0.13 | 0.11 | 0.28 | 0.21 | 0.33 |
| 6 | 91210407 | rs205664 | 0.055 | 0.364 | -0.881 | 0.531 | -0.001 | 0.001 | 0.001 | 0 | 0.4 | 0.39 | 0.51 | 0.51 | 0.42 | 0.38 |
| 6 | 91210594 | rs205662 | -0.331 | 0.123 | -0.598 | 0.213 | -0.101 | -0.095 | -0.095 | 0 | 0.59 | 0.93 | 0.45 | 0.45 | 0.43 | 0.52 |
| 6 | 91210640 | rs205661 | -0.957 | -0.472 | -0.437 | 0.183 | -0.059 | 0.048 | 0.048 | 0 | 0.18 | 0.42 | 0.18 | 0.22 | 0.1 | 0.24 |
| 6 | 91212086 | rs9362744 | 0.506 | -0.425 | -1.184 | -0.269 | -0.005 | 0.061 | 0.061 | 0 | 0.16 | 0.01 | 0.05 | 0.09 | 0.12 | 0.14 |
| 6 | 91218473 | rs157695 | -0.48 | -0.568 | -0.716 | -0.331 | 0.019 | 0.038 | 0.038 | 3.739 | 0.49 | 0.59 | 0.41 | 0.34 | 0.38 | 0.51 |
| 6 | 91219007 | rs157694 | -1.406 | -0.272 | -1.032 | -0.542 | 0.001 | 0.005 | 0.005 | 2.09 | 0.77 | 0.78 | 0.58 | 0.5 | 0.71 | 0.71 |
| 6 | 91220555 | rs711264 | 0.761 | -0.196 | -1.087 | -0.564 | -0.001 | 0.041 | 0.041 | 0.16 | 0.26 | 0.18 | 0.16 | 0.11 | 0.31 | 0.16 |
| 6 | 91221810 | rs806287 | 0.849 | -0.221 | -1.093 | -0.577 | -0.023 | 0.08 | 0.08 | 0.171 | 0.25 | 0.08 | 0.15 | 0.11 | 0.31 | 0.16 |
| 6 | 91225135 | rs157705 | 0.451 | -0.407 | -0.912 | -0.817 | -0.01 | -0.02 | -0.02 | 0.635 | 0.4 | 0.49 | 0.32 | 0.33 | 0.31 | 0.43 |
| 6 | 91260867 | rs157688 | -0.709 | -0.658 | -0.696 | -0.75 | 0.004 | 0.009 | 0.009 | 0.417 | 0.73 | 0.7 | 0.56 | 0.45 | 0.69 | 0.65 |
| 6 | 91278668 | rs1145736 | 1.414 | -0.76 | -0.707 | -0.849 | -0.008 | 0.04 | 0.04 | 0.13 | 0.23 | 0.13 | 0.15 | 0.11 | 0.3 | 0.14 |
| 6 | 91280109 | rs1143794 | 1.473 | -0.748 | -0.776 | -0.847 | -0.035 | 0.07 | 0.07 | 0.155 | 0.23 | 0.05 | 0.15 | 0.11 | 0.3 | 0.14 |
| 6 | 91284990 | rs157432 | 0.374 | -0.785 | -0.634 | -0.826 | -0.006 | -0.012 | -0.012 | 0 | 0.4 | 0.45 | 0.31 | 0.33 | 0.31 | 0.43 |
| 6 | 91301364 | rs282065 | -0.917 | -0.745 | -0.724 | -0.791 | 0 | 0.004 | 0.004 | 1.027 | 0.74 | 0.73 | 0.59 | 0.5 | 0.73 | 0.7 |
| 6 | 91305608 | rs6454815 | 0.101 | -0.672 | -0.885 | -1.273 | -0.004 | 0.027 | 0.027 | 0 | 0.38 | 0.29 | 0.3 | 0.14 | 0.46 | 0.29 |
| 6 | 91311839 | rs7747353 | -0.188 | -0.798 | -0.725 | -0.826 | -0.001 | -0.002 | -0.002 | 0.812 | 0.67 | 0.35 | 0.54 | 0.82 | 0.71 | 0.68 |
| 6 | 91314637 | rs1999090 | 0.485 | -0.812 | -1.065 | -0.906 | 0.026 | -0.003 | -0.003 | 0 | 0.36 | 0.14 | 0.31 | 0.7 | 0.28 | 0.45 |
| 6 | 91317922 | rs1389762 | -0.859 | -0.895 | -1.068 | -0.95 | -0.035 | 0.004 | 0.004 | 1.209 | 0.74 | 0.91 | 0.58 | 0.83 | 0.71 | 0.69 |
| 6 | 91318722 | rs401147 | -0.59 | -0.98 | -0.803 | -0.753 | 0.046 | 0.091 | 0.091 | 1.347 | 0.49 | 0.81 | 0.41 | 0.73 | 0.37 | 0.53 |
| 6 | 91319984 | rs6454818 | 0.365 | -0.965 | -0.751 | -0.649 | 0.032 | 0.001 | 0.001 | 0.24 | 0.36 | 0.15 | 0.31 | 0.7 | 0.27 | 0.45 |
| 6 | 91327303 | rs427647 | -0.313 | -0.97 | -0.11 | -0.95 | 0.071 | 0.072 | 0.072 | 1.353 | 0.05 | 0.45 | 0.03 | 0.01 | 0 | 0.01 |
| 6 | 91344010 | rs17590916 | 0.302 | -0.839 | -0.18 | -1.084 | 0.008 | 0.028 | 0.028 | 0 | 0.39 | 0.32 | 0.25 | 0.13 | 0.4 | 0.21 |
| 6 | 91344491 | rs1556091 | -0.737 | -0.78 | -0.241 | -1.06 | 0.057 | -0.057 | -0.057 | 0.352 | 0.75 | 0.46 | 0.59 | 0.88 | 0.69 | 0.7 |
| 6 | 91345657 | rs9362761 | 0.098 | -0.926 | -0.468 | -0.732 | 0.02 | 0.057 | 0.057 | 0.476 | 0.36 | 0.16 | 0.34 | 0.75 | 0.3 | 0.49 |
| 6 | 91352508 | rs2750451 | -1.204 | -0.694 | -0.985 | -0.052 | -0.056 | 0.008 | 0.008 | 0.93 | 0.84 | 0.96 | 0.67 | 0.89 | 0.8 | 0.77 |
| 6 | 91356629 | rs16883304 | 0.34 | -0.808 | -0.767 | 0.587 | 0.022 | -0.081 | -0.081 | 2.693 | 0.14 | 0 | 0.05 | 0.27 | 0.08 | 0.14 |
| 6 | 91360144 | rs1923087 | 0.654 | -0.99 | -0.782 | -0.014 | 0.065 | 0.042 | 0.042 | 0.789 | 0.71 | 0.87 | 0.67 | 0.21 | 0.77 | 0.51 |
| 6 | 91362112 | rs1126353 | 0.107 | -1.026 | 0.105 | 0.066 | -0.032 | 0.001 | 0.001 | 0.123 | 0.78 | 0.74 | 0.77 | 0.22 | 0.87 | 0.59 |
| 6 | 91363390 | rs932571 | -0.128 | -1.003 | 0.022 | 0.078 | 0.045 | -0.023 | -0.023 | 0.799 | 0.8 | 0.91 | 0.79 | 0.22 | 0.87 | 0.59 |
| 6 | 91365419 | rs9451456 | 1.025 | -0.989 | 0.264 | -0.326 | -0.004 | -0.04 | -0.04 | 0.373 | 0.21 | 0.35 | 0.35 | 0.08 | 0.18 | 0.17 |
| 6 | 91366039 | rs2750440 | 0.466 | -1.041 | 0.233 | -0.372 | 0.023 | 0.03 | 0.03 | 0 | 0.52 | 0.57 | 0.37 | 0.13 | 0.64 | 0.36 |
| 6 | 91368633 | rs2616023 | 0.585 | -1.051 | 0.209 | -0.362 | 0.043 | 0.08 | 0.08 | 0 | 0.51 | 0.37 | 0.35 | 0.13 | 0.63 | 0.36 |
| 6 | 91374106 | rs1474964 | 0.211 | -0.927 | 0.041 | -0.335 | 0.028 | 0.031 | 0.031 | 2.269 | 0.71 | 0.91 | 0.7 | 0.2 | 0.76 | 0.49 |
| 6 | 91376033 | rs1923075 | 0.215 | -0.927 | 0.041 | -0.335 | 0.029 | 0.032 | 0.032 | 2.287 | 0.71 | 0.91 | 0.7 | 0.2 | 0.76 | 0.49 |
| 6 | 91376392 | rs1884839 | 0.805 | -0.957 | 0.024 | -0.379 | 0.03 | 0.152 | 0.152 | 0 | 0.48 | 0.3 | 0.33 | 0.12 | 0.59 | 0.32 |
| 6 | 91376578 | rs1923076 | 0.825 | -0.599 | 0.049 | -0.162 | -0.003 | -0.049 | -0.049 | 0 | 0.21 | 0.38 | 0.35 | 0.08 | 0.17 | 0.17 |
| 6 | 91385628 | rs2750473 | -0.493 | -0.561 | -0.201 | 0.291 | 0.002 | -0.004 | -0.004 | 0 | 0.79 | 0.83 | 0.86 | 0.72 | 0.82 | 0.73 |
| 6 | 91386419 | rs2236329 | 0.027 | -0.238 | 0.144 | 0.969 | -0.002 | 0.077 | 0.077 | 0.238 | 0.11 | 0.18 | 0.21 | 0.32 | 0.1 | 0.2 |
| 6 | 91388868 | rs2750423 | 0.732 | 0.031 | -0.077 | 0.263 | 0.004 | 0.01 | 0.01 | 0.052 | 0.31 | 0.3 | 0.19 | 0.11 | 0.33 | 0.21 |
| 6 | 91395059 | rs4141907 | -1.282 | 0.061 | 0.009 | 1.192 | -0.009 | 0.026 | 0.026 | 0 | 0.18 | 0.25 | 0.28 | 0.37 | 0.16 | 0.25 |
| 6 | 91396592 | rs9444778 | -0.178 | 0.1 | -0.027 | 1.214 | 0 | 0.095 | 0.095 | 0.211 | 0.1 | 0.18 | 0.23 | 0.34 | 0.1 | 0.2 |
| 6 | 91398349 | rs9345044 | -1.713 | 0.197 | -0.27 | 1.14 | 0.006 | 0.019 | 0.019 | 0 | 0.35 | 0.41 | 0.41 | 0.54 | 0.37 | 0.38 |
| 6 | 91399141 | rs13217057 | -0.836 | 1.17 | -0.841 | 1.051 | -0.044 | -0.026 | -0.026 | 1.738 | 0.14 | 0.05 | 0.12 | 0.16 | 0.21 | 0.13 |
| 6 | 91401234 | rs6916490 | -0.859 | 0.366 | -0.449 | 0.91 | 0.001 | 0 | 0 | 0 | 0.39 | 0.41 | 0.45 | 0.62 | 0.4 | 0.45 |
| 6 | 91404252 | rs1547015 | -0.941 | -0.055 | 0.16 | 1.158 | 0.001 | 0.002 | 0.002 | 0 | 0.43 | 0.44 | 0.58 | 0.43 | 0.52 | 0.55 |
| 6 | 91406809 | rs9362770 | -0.599 | 0.074 | 0.217 | 1.271 | -0.088 | 0.049 | 0.049 | 0.03 | 0.28 | 0.1 | 0.44 | 0.2 | 0.39 | 0.37 |
| 6 | 91412651 | rs1145743 | -1.911 | -0.737 | 0.51 | 1.094 | 0.003 | 0.062 | 0.062 | 1.681 | 0.6 | 0.74 | 0.77 | 0.76 | 0.58 | 0.87 |
| 6 | 91414247 | rs1145744 | -0.684 | -0.876 | 0.621 | 1.04 | 0.016 | -0.038 | -0.038 | 0 | 0.26 | 0.2 | 0.3 | 0.49 | 0.19 | 0.49 |
| 6 | 91417393 | rs2325312 | -0.631 | -0.876 | 1.298 | 1.032 | 0.002 | -0.002 | -0.002 | 0 | 0.32 | 0.33 | 0.45 | 0.27 | 0.39 | 0.36 |
| 6 | 91424218 | rs1145754 | -0.733 | -0.837 | 1.427 | 1.26 | 0.025 | -0.086 | -0.086 | 0.268 | 0.25 | 0.14 | 0.29 | 0.49 | 0.19 | 0.5 |
| 6 | 91424763 | rs9353758 | -0.384 | -0.684 | 1.656 | 1.277 | 0.035 | 0.404 | 0.404 | 1.254 | 0.05 | 0.3 | 0.05 | 0.23 | 0.01 | 0.03 |
| 6 | 91428616 | rs6931375 | -0.493 | -0.683 | 1.726 | 1.221 | 0.026 | 0.364 | 0.364 | 2.495 | 0.06 | 0.3 | 0.05 | 0.22 | 0.01 | 0.03 |

| | | | | | | | | | | | | | | | | |
|---|---|---|---|---|---|---|---|---|---|---|---|---|---|---|---|---|
| 6 | 91435173 | rs1231500 | -2.046 | -1.11 | 1.338 | 0.793 | -0.002 | 0.006 | 0.006 | 0.077 | 0.65 | 0.67 | 0.78 | 0.77 | 0.6 | 0.89 |
| 6 | 91454709 | rs1231483 | -0.475 | -1.126 | 1.109 | 0.717 | 0.022 | -0.078 | -0.078 | 3.266 | 0.24 | 0.11 | 0.29 | 0.53 | 0.19 | 0.45 |
| 6 | 91455178 | rs1231482 | -0.475 | -1.126 | 1.109 | 0.717 | 0.022 | -0.078 | -0.078 | 0.362 | 0.24 | 0.11 | 0.29 | 0.53 | 0.19 | 0.45 |
| 6 | 91455484 | rs1231481 | -2.441 | -1.083 | 1.143 | 0.469 | -0.013 | 0.028 | 0.028 | 0.076 | 0.66 | 0.77 | 0.81 | 1 | 0.61 | 0.92 |
| 6 | 91463437 | rs9362780 | -0.507 | -0.812 | 1.181 | 0.836 | -0.051 | 0.049 | 0.049 | 0 | 0.29 | 0.09 | 0.47 | 0.2 | 0.34 | 0.39 |
| 6 | 91467387 | rs9353762 | -0.566 | -0.816 | 1.353 | 0.87 | -0.021 | 0.024 | 0.024 | 0 | 0.3 | 0.16 | 0.47 | 0.27 | 0.34 | 0.39 |
| 6 | 91470459 | rs9451477 | -1.557 | -0.967 | 1.542 | 0.834 | -0.003 | 0.05 | 0.05 | 1.64 | 0.54 | 0.32 | 0.76 | 0.92 | 0.54 | 0.85 |
| 6 | 91470905 | rs4706375 | -1.036 | -1.063 | 1.905 | 0.782 | -0.002 | 0.002 | 0.002 | 0 | 0.38 | 0.42 | 0.51 | 0.28 | 0.39 | 0.42 |
| 6 | 91472289 | rs2204742 | -0.524 | -0.997 | 1.452 | 0.854 | 0.017 | -0.071 | -0.071 | 0 | 0.23 | 0.1 | 0.25 | 0.5 | 0.19 | 0.42 |
| 6 | 91474545 | rs2141484 | -1.579 | -0.991 | 1.424 | 0.954 | -0.004 | 0.046 | 0.046 | 1.406 | 0.54 | 0.32 | 0.73 | 0.77 | 0.53 | 0.81 |
| 6 | 91475449 | rs1358602 | -0.633 | -1.061 | 1.448 | 0.601 | -0.001 | 0.001 | 0.001 | 0 | 0.31 | 0.29 | 0.49 | 0.27 | 0.35 | 0.4 |
| 6 | 91480768 | rs7341365 | -1.678 | -0.895 | 1.761 | 1.143 | -0.029 | -0.03 | -0.03 | 0 | 0.35 | 0.48 | 0.38 | 0.73 | 0.25 | 0.51 |
| 6 | 91487213 | rs2840191 | -1.391 | -0.805 | 1.18 | 1.157 | 0.002 | -0.001 | -0.001 | 0.063 | 0.32 | 0.31 | 0.34 | 0.64 | 0.24 | 0.49 |
| 6 | 91490710 | rs1145756 | -1.764 | -0.728 | 0.76 | 1.43 | 0.029 | -0.055 | -0.055 | 0 | 0.38 | 0.77 | 0.38 | 0.56 | 0.32 | 0.52 |
| 6 | 91490914 | rs12526019 | 0.091 | -0.77 | 0.824 | 1.724 | 0.001 | 0.038 | 0.038 | 0 | 0.07 | 0.04 | 0.15 | 0.3 | 0.04 | 0.18 |
| 6 | 91494166 | rs6904240 | 1.671 | -0.675 | 0.703 | 1.295 | 0.031 | -0.039 | -0.039 | 0.914 | 0.61 | 0.23 | 0.58 | 0.41 | 0.68 | 0.47 |
| 6 | 91500926 | rs16883515 | -1.018 | -0.982 | 1 | 0.771 | -0.001 | -0.02 | -0.02 | 0 | 0.2 | 0.21 | 0.14 | 0.09 | 0.22 | 0.21 |
| 6 | 91502205 | rs17523287 | -0.022 | -0.723 | 0.683 | 0.817 | -0.021 | 0.026 | 0.026 | 0.021 | 0.12 | 0.05 | 0.07 | 0 | 0.18 | 0.1 |
| 6 | 91505115 | rs12525150 | 0.188 | -0.583 | 0.381 | 0.962 | 0.004 | -0.042 | -0.042 | 0.144 | 0.16 | 0.07 | 0.33 | 0.34 | 0.17 | 0.27 |
| 6 | 91505386 | rs9451486 | 0.298 | -0.639 | -0.06 | 0.978 | -0.002 | -0.02 | -0.02 | 0 | 0.14 | 0.07 | 0.19 | 0.18 | 0.13 | 0.19 |
| 6 | 91505411 | rs10485409 | -1.173 | -0.662 | 0.224 | 0.992 | -0.01 | 0.024 | 0.024 | 0.075 | 0.37 | 0.46 | 0.47 | 0.49 | 0.38 | 0.51 |
| 6 | 91505957 | rs10485408 | 0.14 | -0.428 | -0.503 | 1.109 | -0.01 | -0.092 | -0.092 | 0 | 0.15 | 0.02 | 0.32 | 0.33 | 0.17 | 0.26 |
| 6 | 91508722 | rs9451487 | 0.267 | -0.128 | -0.24 | 1.43 | 0.024 | -0.023 | -0.023 | 0.596 | 0.56 | 0.26 | 0.48 | 0.47 | 0.57 | 0.44 |
| 6 | 91510922 | rs6899740 | 0.858 | 0.114 | -0.357 | 1.141 | -0.082 | 0.084 | 0.084 | 1.232 | 0.42 | 0.16 | 0.29 | 0.3 | 0.45 | 0.26 |
| 6 | 91517782 | rs9294467 | 0.863 | 0.022 | -1.208 | 1.195 | -0.082 | 0.072 | 0.072 | 1.363 | 0.42 | 0.17 | 0.27 | 0.3 | 0.45 | 0.24 |
| 6 | 91518694 | rs4707649 | -0.891 | -0.143 | -0.221 | 1.609 | -0.02 | 0.044 | 0.044 | 0.035 | 0.23 | 0.34 | 0.26 | 0.36 | 0.16 | 0.34 |
| 6 | 91529722 | rs9353764 | -0.865 | 0.012 | -0.816 | 1.673 | 0.007 | -0.03 | -0.03 | 0 | 0.17 | 0.12 | 0.27 | 0.3 | 0.16 | 0.11 |
| 6 | 91531693 | rs9351245 | -0.834 | 0.189 | -1.412 | 1.601 | 0.003 | 0.015 | 0.015 | 0 | 0.42 | 0.24 | 0.49 | 0.6 | 0.41 | 0.49 |
| 6 | 91532613 | rs1542650 | -0.581 | 0.232 | -1.374 | 0.783 | -0.002 | 0 | 0 | 0 | 0.74 | 0.76 | 0.79 | 0.89 | 0.72 | 0.79 |
| 6 | 91534091 | rs12529724 | 1.073 | 0.193 | -1.448 | 0.872 | -0.015 | -0.002 | -0.002 | 0 | 0.22 | 0.13 | 0.16 | 0.23 | 0.24 | 0.36 |
| 6 | 91543216 | rs10485405 | 0.321 | -0.315 | -1.118 | 0.725 | -0.016 | 0.034 | 0.034 | 0 | 0.34 | 0.11 | 0.31 | 0.26 | 0.36 | 0.32 |
| 6 | 91544513 | rs1528983 | 0.617 | -0.3 | -1.489 | 0.797 | -0.048 | 0.009 | 0.009 | 0.112 | 0.38 | 0.14 | 0.33 | 0.31 | 0.43 | 0.51 |
| 6 | 91547290 | rs721025 | -1.094 | -0.173 | -1.175 | 0.741 | 0.007 | -0.031 | -0.031 | 0 | 0.79 | 0.74 | 0.86 | 0.94 | 0.76 | 0.87 |
| 6 | 91550774 | rs7452644 | -0.251 | -0.251 | -0.406 | 0.77 | 0.008 | -0.036 | -0.036 | 0.058 | 0.72 | 0.41 | 0.72 | 0.83 | 0.7 | 0.84 |
| 6 | 91552785 | rs11965033 | -0.018 | -0.373 | -0.678 | 0.653 | 0.002 | -0.039 | -0.039 | 0.202 | 0.07 | 0.01 | 0.24 | 0.25 | 0.07 | 0.16 |
| 6 | 91560367 | rs12192521 | 1.03 | -0.459 | -0.577 | 0.493 | 0.003 | 0.083 | 0.083 | 0.542 | 0.19 | 0.05 | 0.12 | 0 | 0.18 | 0.09 |
| 6 | 91561402 | rs1145761 | 0.129 | -0.389 | -0.402 | 0.514 | 0 | -0.009 | -0.009 | 2.138 | 0.29 | 0.67 | 0.28 | 0.16 | 0.29 | 0.16 |
| 6 | 91565841 | rs1385411 | -0.55 | -0.413 | -0.198 | 0.623 | -0.025 | 0.16 | 0.16 | 0 | 0.09 | 0.25 | 0.26 | 0.35 | 0.05 | 0.08 |
| 6 | 91566322 | rs1220315 | -0.556 | -0.387 | -0.209 | 0.642 | -0.018 | 0.034 | 0.034 | 0.75 | 0.54 | 0.72 | 0.54 | 0.41 | 0.51 | 0.31 |
| 6 | 91567851 | rs6913163 | 1.119 | -0.389 | -0.287 | 0.548 | -0.024 | -0.001 | -0.001 | 6.699 | 0.35 | 0.14 | 0.31 | 0.29 | 0.38 | 0.44 |
| 6 | 91580571 | rs1220319 | 0.899 | -0.529 | 0.639 | 0.619 | 0.016 | -0.048 | -0.048 | 0 | 0.21 | 0.32 | 0.13 | 0.06 | 0.24 | 0.14 |
| 6 | 91581760 | rs1145775 | 0.798 | -0.56 | 0.866 | 0.602 | 0.029 | -0.076 | -0.076 | 1.065 | 0.22 | 0.39 | 0.14 | 0.06 | 0.24 | 0.13 |
| 6 | 91584932 | rs698162 | -0.704 | -0.563 | 0.783 | 0.592 | -0.018 | 0.082 | 0.082 | 1.493 | 0.54 | 0.73 | 0.46 | 0.31 | 0.51 | 0.31 |
| 6 | 91587146 | rs1775875 | 0.4 | 0.102 | 0.848 | 1.077 | -0.014 | 0.016 | 0.016 | 0 | 0.2 | 0.13 | 0.18 | 0.14 | 0.24 | 0.15 |
| 6 | 91596456 | rs1342840 | -1.614 | 0.186 | 1.111 | 1.428 | -0.001 | 0.009 | 0.009 | 0 | 0.38 | 0.58 | 0.42 | 0.47 | 0.38 | 0.36 |
| 6 | 91601691 | rs7750922 | -0.956 | 0.397 | 0.88 | 1.239 | -0.012 | 0.014 | 0.014 | 0 | 0.17 | 0.28 | 0.21 | 0.23 | 0.16 | 0.21 |
| 6 | 91603644 | rs12211262 | -0.082 | 0.776 | 1.019 | 1.264 | -0.003 | 0.005 | 0.005 | 0 | 0.1 | 0.07 | 0.07 | 0 | 0.11 | 0.1 |
| 6 | 91607588 | rs1394245 | -0.653 | 0.75 | 1.039 | 1.318 | -0.001 | 0 | 0 | 0 | 0.09 | 0.11 | 0.18 | 0.1 | 0.06 | 0.07 |
| 6 | 91611628 | rs711274 | -0.46 | 0.795 | 1.161 | 1.531 | 0.061 | 0.189 | 0.189 | 0.329 | 0.15 | 0.38 | 0.23 | 0.44 | 0.19 | 0.2 |
| 6 | 91611925 | rs12191179 | -0.848 | 0.763 | 0.74 | 1.152 | -0.004 | -0.054 | -0.054 | 0.623 | 0.2 | 0.27 | 0.2 | 0.03 | 0.18 | 0.15 |
| 6 | 91612079 | rs9451525 | -0.405 | 1.071 | 0.296 | 1.791 | 0.009 | 0.069 | 0.069 | 0.239 | 0.08 | 0.14 | 0.13 | 0.21 | 0.09 | 0.18 |
| 6 | 91620319 | rs7739445 | -0.07 | 1.268 | -0.155 | 1.208 | 0.034 | -0.013 | -0.013 | 0 | 0.09 | 0.17 | 0.07 | 0.05 | 0.14 | 0.09 |
| 6 | 91620431 | rs1145784 | 0.742 | 1.015 | -0.741 | 2.019 | -0.008 | -0.026 | -0.026 | 0 | 0.91 | 0.95 | 0.82 | 0.57 | 0.91 | 0.86 |
| 6 | 91622357 | rs12200299 | 1.348 | 1.329 | -1.17 | 1.041 | 0.074 | 0.13 | 0.13 | 0.009 | 0.47 | 0.14 | 0.27 | 0.2 | 0.38 | 0.45 |
| 6 | 91622657 | rs1145787 | -0.104 | 1.843 | -1.207 | 0.849 | -0.079 | 0.111 | 0.111 | 0 | 0.8 | 0.96 | 0.87 | 1 | 0.72 | 0.88 |
| 6 | 91634701 | rs1145805 | -0.644 | 1.313 | -0.008 | 1.908 | 0.017 | 0.206 | 0.206 | 0.178 | 0.12 | 0.31 | 0.33 | 0.46 | 0.14 | 0.25 |
| 6 | 91636139 | rs2325337 | -0.87 | 0.903 | 0.051 | 1.473 | 0.037 | 0.097 | 0.097 | 0.279 | 0.2 | 0.45 | 0.26 | 0.33 | 0.2 | 0.18 |
| 6 | 91638813 | rs1016075 | 0.495 | 0.513 | -0.559 | 0.894 | 0.007 | 0.013 | 0.013 | 0.006 | 0.58 | 0.39 | 0.67 | 0.64 | 0.61 | 0.59 |
| 6 | 91639322 | rs1145807 | -0.225 | 0.322 | -0.494 | 0.509 | 0 | 0.027 | 0.027 | 0 | 0.54 | 0.67 | 0.5 | 0.61 | 0.54 | 0.7 |
| 6 | 91642307 | rs711280 | -0.542 | 0.003 | -0.32 | 0.813 | -0.004 | 0.016 | 0.016 | 0 | 0.09 | 0.1 | 0.09 | 0.19 | 0.07 | 0.11 |
| 6 | 91644580 | rs1220321 | -0.021 | -0.121 | -0.569 | 0.731 | 0.008 | -0.018 | -0.018 | 0 | 0.41 | 0.28 | 0.43 | 0.4 | 0.36 | 0.4 |
| 6 | 91644742 | rs1220322 | -0.021 | -0.121 | -0.405 | 0.746 | 0.007 | -0.015 | -0.015 | 0 | 0.41 | 0.31 | 0.43 | 0.4 | 0.36 | 0.41 |
| 6 | 91645630 | rs17533654 | 0.592 | -0.077 | -0.464 | 0.742 | -0.004 | 0.015 | 0.015 | 0 | 0.31 | 0.18 | 0.34 | 0.21 | 0.28 | 0.29 |
| 6 | 91651283 | rs711285 | -0.855 | 0.118 | -0.355 | 0.826 | -0.002 | -0.007 | -0.007 | 0.13 | 0.18 | 0.13 | 0.15 | 0.22 | 0.18 | 0.33 |
| 6 | 91655112 | rs1238132 | -0.411 | -0.47 | 0.54 | 0.661 | 0.01 | 0.011 | 0.011 | 0 | 0.59 | 0.54 | 0.51 | 0.43 | 0.49 | 0.62 |
| 6 | 91656112 | rs1220445 | -0.131 | -0.511 | 0.321 | 0.626 | -0.013 | -0.01 | -0.01 | 0 | 0.54 | 0.33 | 0.49 | 0.43 | 0.48 | 0.59 |
| 6 | 91659604 | rs1145813 | -0.424 | -0.504 | 0.608 | 0.415 | -0.005 | 0.002 | 0.002 | 0 | 0.64 | 0.67 | 0.62 | 0.62 | 0.56 | 0.7 |
| 6 | 91660540 | rs1145814 | 0.203 | -0.581 | 0.64 | 0.428 | -0.001 | 0.002 | 0.002 | 0 | 0.56 | 0.57 | 0.59 | 0.62 | 0.5 | 0.64 |
| 6 | 91660833 | rs9451543 | 0.094 | -0.315 | -0.176 | 0.463 | 0 | -0.016 | -0.016 | 0 | 0.09 | 0.08 | 0.05 | 0 | 0.06 | 0.07 |

| | | | | | | | | | | | | | | | | |
|---|---|---|---|---|---|---|---|---|---|---|---|---|---|---|---|---|
| 6 | 91669341 | rs7775358 | -0.417 | -0.336 | -0.095 | 0.549 | -0.006 | 0.002 | 0.002 | 0 | 0.66 | 0.67 | 0.62 | 0.63 | 0.57 | 0.71 |
| 6 | 91670249 | rs2008577 | -0.722 | -0.383 | -0.125 | 0.464 | -0.013 | -0.007 | -0.007 | 0.086 | 0.69 | 0.72 | 0.62 | 0.61 | 0.58 | 0.71 |
| 6 | 91675072 | rs10806440 | -0.279 | -0.18 | -0.229 | 0.291 | -0.001 | -0.003 | -0.003 | 0 | 0.61 | 0.59 | 0.58 | 0.45 | 0.55 | 0.63 |
| 6 | 91683972 | rs6914621 | -0.11 | 0.228 | -0.596 | 0.614 | 0.001 | -0.001 | -0.001 | 0 | 0.5 | 0.4 | 0.53 | 0.46 | 0.49 | 0.54 |
| 6 | 91695600 | rs1485042 | 0.002 | 0.627 | -0.875 | 0.186 | -0.031 | 0.052 | 0.052 | 0 | 0.21 | 0.07 | 0.15 | 0.08 | 0.25 | 0.21 |
| 6 | 91695668 | rs1144168 | -0.776 | -0.109 | 0.124 | -0.25 | -0.009 | -0.048 | -0.048 | 0 | 0.32 | 0.61 | 0.2 | 0.1 | 0.27 | 0.29 |
| 6 | 91697611 | rs9345064 | 0.101 | 0.372 | -0.233 | 0.059 | 0 | -0.001 | -0.001 | 0 | 0.17 | 0.17 | 0.15 | 0.22 | 0.27 | 0.21 |
| 6 | 91698265 | rs1144171 | -1.291 | -0.237 | -0.479 | 0.183 | 0.002 | -0.055 | -0.055 | 0.403 | 0.58 | 0.85 | 0.38 | 0.5 | 0.59 | 0.56 |
| 6 | 91703486 | rs10485402 | -0.142 | -0.589 | 0.859 | 0.068 | -0.022 | -0.031 | -0.031 | 0.134 | 0.08 | 0.22 | 0.02 | 0.02 | 0.02 | 0.07 |
| 6 | 91704358 | rs9351248 | -0.132 | -0.403 | 0.49 | 0.18 | 0.011 | -0.005 | -0.005 | 1.92 | 0.1 | 0.07 | 0.03 | 0.18 | 0.05 | 0.07 |
| 6 | 91714276 | rs9345068 | -0.273 | -0.755 | 0.53 | 0.31 | 0.009 | -0.012 | -0.012 | 0 | 0.1 | 0.07 | 0.02 | 0.23 | 0.01 | 0.1 |
| 6 | 91717296 | rs1836137 | 0.644 | -0.75 | 0.648 | 0.31 | -0.066 | 0.066 | 0.066 | 0 | 0.88 | 0.67 | 0.96 | 0.75 | 0.99 | 0.88 |
| 6 | 91722948 | rs9353776 | -0.906 | -0.856 | 0.13 | 0.28 | -0.077 | 0.022 | 0.022 | 0 | 0.22 | 0.59 | 0.08 | 0.28 | 0.05 | 0.11 |
| 6 | 91723068 | rs16883886 | 0.11 | -0.213 | -0.775 | 0.305 | -0.041 | -0.015 | -0.015 | 0 | 0.13 | 0.09 | 0.15 | 0.15 | 0.26 | 0.22 |
| 6 | 91724888 | rs11756310 | 0.623 | -0.193 | -1.214 | -0.211 | -0.092 | 0.133 | 0.133 | 0 | 0.28 | 0.07 | 0.27 | 0.06 | 0.41 | 0.31 |
| 6 | 91725260 | rs9345070 | -0.558 | -1.348 | -0.894 | 0.392 | 0.037 | -0.134 | -0.134 | 0 | 0.66 | 0.95 | 0.54 | 0.48 | 0.72 | 0.64 |
| 6 | 91728217 | rs12211003 | -0.029 | -0.972 | -0.616 | 0.537 | -0.072 | -0.184 | -0.184 | 0 | 0.24 | 0.01 | 0.44 | 0.43 | 0.29 | 0.3 |
| 6 | 91751775 | rs16883933 | -1.213 | -1.237 | 0.845 | 1.062 | -0.016 | 0.232 | 0.232 | 2.289 | 0.06 | 0.21 | 0.07 | 0.4 | 0.04 | 0.07 |
| 6 | 91753027 | rs6936812 | -1.861 | -1.178 | 1.462 | 0.951 | 0.014 | 0.268 | 0.268 | 2.696 | 0.12 | 0.39 | 0.12 | 0.4 | 0.09 | 0.16 |
| 6 | 91756461 | rs9362810 | -1.43 | -1.259 | 1.098 | 0.861 | 0.006 | 0.098 | 0.098 | 6.067 | 0.09 | 0.45 | 0.09 | 0.11 | 0.04 | 0.05 |
| 6 | 91772347 | rs11963862 | -2.13 | -1.402 | -0.058 | 0.651 | 0.047 | -0.072 | -0.072 | 0 | 0.22 | 0.84 | 0.21 | 0.45 | 0.11 | 0.2 |
| 6 | 91778114 | rs9359916 | -1.843 | -1.563 | 0.144 | 0.702 | -0.009 | 0.002 | 0.002 | 1.575 | 0.2 | 0.69 | 0.17 | 0.44 | 0.07 | 0.14 |
| 6 | 91780338 | rs7766974 | -1.773 | -1.758 | -0.058 | -0.413 | 0.036 | 0.041 | 0.041 | 0 | 0.3 | 0.71 | 0.18 | 0.09 | 0.14 | 0.3 |
| 6 | 91781177 | rs16883970 | -1.22 | -1.83 | 0.349 | -0.337 | 0.011 | 0.008 | 0.008 | 0 | 0.2 | 0.16 | 0.12 | 0.11 | 0.1 | 0.26 |
| 6 | 91783441 | rs16883972 | -0.693 | -1.792 | 0.345 | -0.299 | 0.004 | 0 | 0 | 0.191 | 0.14 | 0.11 | 0.09 | 0.11 | 0.07 | 0.11 |
| 6 | 91786112 | rs9451570 | -0.904 | -1.657 | 0.832 | -0.325 | 0.017 | -0.005 | -0.005 | 0 | 0.06 | 0.25 | 0.03 | 0 | 0.02 | 0.02 |
| 6 | 91790493 | rs7754397 | -1.721 | -1.762 | 0.465 | -0.428 | 0.029 | 0.026 | 0.026 | 0 | 0.28 | 0.73 | 0.14 | 0.05 | 0.11 | 0.25 |
| 6 | 91793109 | rs10944520 | -0.49 | -2.424 | -0.025 | 0.124 | 0.033 | -0.018 | -0.018 | 0.18 | 0.08 | 0.04 | 0.09 | 0.59 | 0.04 | 0.12 |
| 6 | 91797486 | rs9451576 | -0.736 | -2.579 | 0.379 | -0.017 | 0 | -0.022 | -0.022 | 0 | 0.18 | 0.18 | 0.07 | 0 | 0.05 | 0.09 |
| 6 | 91798500 | rs1534777 | -1.998 | -2.854 | -0.336 | -0.811 | 0.082 | 0.176 | 0.176 | 0 | 0.54 | 0.89 | 0.33 | 0.74 | 0.28 | 0.46 |
| 6 | 91801885 | rs2178373 | -0.751 | -2.429 | 0.412 | -0.525 | 0.027 | 0.043 | 0.043 | 0.278 | 0.11 | 0.55 | 0.12 | 0.11 | 0.09 | 0.19 |
| 6 | 91802492 | rs9345089 | -0.805 | -2.45 | 0.402 | -0.529 | 0.033 | -0.038 | -0.038 | 0 | 0.25 | 0.16 | 0.14 | 0.59 | 0.13 | 0.18 |
| 6 | 91805713 | rs9362825 | -1.897 | -2.425 | 0.303 | -0.528 | 0.048 | 0.065 | 0.065 | 0.207 | 0.38 | 0.78 | 0.27 | 0.74 | 0.22 | 0.38 |
| 6 | 91816034 | rs6924365 | -0.834 | -1.892 | 0.796 | -0.207 | -0.031 | 0.087 | 0.087 | 0.02 | 0.2 | 0.36 | 0.13 | 0.58 | 0.08 | 0.1 |
| 6 | 91837498 | rs6902540 | -1.471 | -1.758 | 0.028 | -0.215 | -0.012 | -0.003 | -0.003 | 0 | 0.27 | 0.39 | 0.21 | 0.68 | 0.17 | 0.29 |
| 6 | 91840179 | rs9359925 | -0.315 | -1.616 | -0.402 | -0.122 | 0.011 | -0.048 | -0.048 | 0 | 0.16 | 0.08 | 0.11 | 0.57 | 0.08 | 0.09 |
| 6 | 91841370 | rs4598041 | 0.774 | -0.207 | -0.86 | -1.16 | -0.09 | 0.108 | 0.108 | 0 | 0.27 | 0.08 | 0.37 | 0.11 | 0.44 | 0.25 |
| 6 | 91841769 | rs12661308 | -0.883 | -0.027 | -0.135 | -0.299 | 0.037 | 0.037 | 0.037 | 0 | 0.09 | 0.25 | 0.1 | 0.1 | 0.09 | 0.2 |
| 6 | 91842612 | rs10806442 | -0.497 | 0.013 | -0.961 | -0.524 | -0.035 | 0.066 | 0.066 | 2.112 | 0.11 | 0.01 | 0.05 | 0 | 0.09 | 0.04 |
| 6 | 91844387 | rs12207238 | 0.082 | 0.162 | -1.478 | -1.024 | -0.01 | 0.125 | 0.125 | 0 | 0.44 | 0.17 | 0.38 | 0.17 | 0.53 | 0.32 |
| 6 | 91844509 | rs9345098 | -0.045 | -0.014 | -0.928 | -0.806 | -0.005 | 0.02 | 0.02 | 0 | 0.45 | 0.38 | 0.42 | 0.32 | 0.5 | 0.41 |
| 6 | 91845134 | rs9451582 | 0.138 | -0.29 | -1.3 | -0.924 | 0.029 | 0.165 | 0.165 | 0 | 0.49 | 0.18 | 0.4 | 0.19 | 0.56 | 0.34 |
| 6 | 91845407 | rs4424056 | 0.16 | -0.085 | -0.528 | -1.207 | -0.003 | -0.071 | -0.071 | 0 | 0.76 | 0.67 | 0.85 | 0.96 | 0.75 | 0.88 |
| 6 | 91845703 | rs4495251 | 0.576 | -0.102 | -0.421 | -1.238 | -0.001 | -0.081 | -0.081 | 0 | 0.72 | 0.62 | 0.84 | 0.96 | 0.72 | 0.86 |
| 6 | 91846329 | rs9353802 | 0.566 | 0.771 | -0.708 | -1.482 | -0.022 | -0.023 | -0.023 | 0 | 0.18 | 0.15 | 0.29 | 0.34 | 0.3 | 0.29 |
| 6 | 91848874 | rs4469264 | -0.683 | 0.508 | 0.145 | -1.589 | 0.072 | -0.023 | -0.023 | 0 | 0.87 | 0.55 | 0.9 | 0.94 | 0.83 | 0.92 |
| 6 | 91859454 | rs10806443 | 0.145 | 0.52 | 0.065 | -1.627 | 0.011 | -0.046 | -0.046 | 0 | 0.16 | 0.26 | 0.1 | 0.04 | 0.21 | 0.1 |
| 6 | 91860131 | rs9359928 | 0.361 | 0.551 | -0.286 | -1.965 | -0.015 | 0.103 | 0.103 | 0 | 0.44 | 0.31 | 0.71 | 0.92 | 0.49 | 0.72 |
| 6 | 91862425 | rs4612131 | -0.903 | 0.785 | -0.218 | -1.602 | 0.043 | 0.038 | 0.038 | 0 | 0.84 | 0.69 | 0.71 | 0.72 | 0.74 | 0.73 |
| 6 | 91865681 | rs4707657 | -0.419 | 0.059 | -0.676 | -1.92 | 0.021 | 0.085 | 0.085 | 0 | 0.48 | 0.27 | 0.56 | 0.67 | 0.49 | 0.57 |
| 6 | 91868414 | rs9353805 | 0.116 | -0.041 | -0.94 | -0.87 | -0.016 | -0.034 | -0.034 | 0.021 | 0.22 | 0.13 | 0.32 | 0.38 | 0.32 | 0.39 |
| 6 | 91872869 | rs12191736 | 0.367 | 0.173 | -0.691 | -0.739 | -0.007 | 0.033 | 0.033 | 0 | 0.2 | 0.1 | 0.1 | 0.03 | 0.22 | 0.12 |
| 6 | 91873522 | rs9451593 | 0.208 | -0.188 | -0.244 | -0.151 | 0.017 | 0.031 | 0.031 | 0 | 0.67 | 0.51 | 0.48 | 0.43 | 0.6 | 0.64 |
| 6 | 91874505 | rs9345104 | -0.275 | 0.311 | -0.002 | -0.369 | 0.078 | 0.027 | 0.027 | 0 | 0.82 | 0.57 | 0.68 | 0.71 | 0.7 | 0.76 |
| 6 | 91882335 | rs6942007 | -0.191 | -0.057 | 0.022 | -0.527 | 0.034 | -0.027 | -0.027 | 0 | 0.94 | 0.84 | 0.95 | 1 | 0.88 | 0.93 |
| 6 | 91887566 | rs4513756 | 0.573 | -0.596 | -0.233 | -0.414 | 0.022 | -0.034 | -0.034 | 0 | 0.15 | 0.1 | 0.19 | 0.28 | 0.1 | 0.13 |
| 6 | 91894594 | rs9362844 | 0.675 | -0.265 | 0.134 | -0.555 | 0.056 | 0.013 | 0.013 | 0 | 0.17 | 0.26 | 0.31 | 0.28 | 0.33 | 0.24 |
| 6 | 91905655 | rs4574605 | -0.026 | -0.545 | -0.194 | -0.863 | 0.022 | -0.015 | -0.015 | 0 | 0.27 | 0.13 | 0.45 | 0.65 | 0.25 | 0.46 |
| 6 | 91909129 | rs10498972 | 0.884 | -0.472 | -0.153 | -0.851 | 0.045 | 0.009 | 0.009 | 0 | 0.17 | 0.23 | 0.31 | 0.27 | 0.32 | 0.24 |
| 6 | 91909164 | rs2820350 | 0.868 | -0.574 | -0.212 | -0.884 | 0.026 | -0.034 | -0.034 | 0.015 | 0.06 | 0.16 | 0.04 | 0.01 | 0.11 | 0.08 |
| 6 | 91914423 | rs4707660 | 0.881 | -0.579 | -0.521 | -0.733 | -0.037 | -0.061 | -0.061 | 0.045 | 0.1 | 0.01 | 0.24 | 0.38 | 0.14 | 0.3 |
| 6 | 91939989 | rs6928795 | 1.027 | -0.763 | -0.273 | -1.131 | 0.025 | 0.009 | 0.009 | 0 | 0.16 | 0.19 | 0.42 | 0.31 | 0.32 | 0.24 |
| 6 | 91942743 | rs7758604 | 0.283 | -0.916 | -0.383 | -1.076 | 0.027 | -0.018 | -0.018 | 6.255 | 0.26 | 0.13 | 0.3 | 0.65 | 0.2 | 0.44 |
| 6 | 91944524 | rs4235845 | 0.023 | -0.98 | -0.049 | -1.109 | -0.01 | 0.001 | 0.001 | 0 | 0.28 | 0.29 | 0.31 | 0.65 | 0.2 | 0.44 |
| 6 | 91948582 | rs9294483 | -0.302 | -1.605 | -0.232 | -1.671 | 0.003 | 0.054 | 0.054 | 0 | 0.57 | 0.67 | 0.77 | 0.96 | 0.56 | 0.74 |
| 6 | 91957965 | rs9345117 | 0.215 | -1.641 | -0.267 | -1.596 | -0.02 | -0.008 | -0.008 | 0 | 0.4 | 0.43 | 0.34 | 0.65 | 0.25 | 0.49 |
| 6 | 91962247 | rs9362860 | 0.528 | -1.235 | -0.546 | -1.375 | 0.022 | -0.053 | -0.053 | 0 | 0.24 | 0.08 | 0.3 | 0.64 | 0.19 | 0.43 |
| 6 | 91970752 | rs4626401 | 1.451 | -0.954 | -0.441 | -1.232 | -0.01 | -0.012 | -0.012 | 0 | 0.16 | 0.12 | 0.42 | 0.31 | 0.31 | 0.24 |
| 6 | 91978768 | rs10498973 | 1.256 | -1.035 | -0.589 | -0.581 | -0.023 | -0.054 | -0.054 | 3.289 | 0.09 | 0.01 | 0.21 | 0.31 | 0.08 | 0.3 |

| | | | | | | | | | | | | | | | | |
|---|---|---|---|---|---|---|---|---|---|---|---|---|---|---|---|---|
| 6 | 91981841 | rs10755495 | -0.059 | -1.365 | -0.761 | -0.943 | 0.054 | 0.064 | 0.064 | 0 | 0.52 | 0.72 | 0.6 | 0.35 | 0.65 | 0.47 |
| 6 | 91994620 | rs10944534 | 0.996 | -0.82 | -0.731 | -0.23 | 0.002 | -0.003 | -0.003 | 0.325 | 0.1 | 0.07 | 0.22 | 0.31 | 0.08 | 0.3 |
| 6 | 92009172 | rs2069205 | -0.587 | -0.814 | -1.039 | -0.528 | -0.055 | 0.013 | 0.013 | 0 | 0.69 | 0.85 | 0.51 | 0.69 | 0.58 | 0.7 |
| 6 | 92009946 | rs4707671 | -0.306 | -1.903 | -0.745 | -0.288 | -0.018 | -0.001 | -0.001 | 0 | 0.34 | 0.6 | 0.3 | 0.31 | 0.17 | 0.39 |
| 6 | 92010992 | rs12174540 | -0.204 | -1.619 | -1.179 | -0.165 | -0.045 | 0.065 | 0.065 | 0 | 0.2 | 0.39 | 0.26 | 0.33 | 0.09 | 0.37 |
| 6 | 92011208 | rs9345126 | 0.668 | -0.996 | -1.721 | 0.189 | 0.001 | 0.019 | 0.019 | 0 | 0.1 | 0.11 | 0.23 | 0.31 | 0.06 | 0.32 |
| 6 | 92012025 | rs2811659 | -0.177 | 0.037 | -1.894 | 0.392 | 0.004 | 0.006 | 0.006 | 0 | 0.45 | 0.29 | 0.27 | 0.49 | 0.51 | 0.37 |
| 6 | 92012043 | rs2748279 | -0.296 | 1.244 | -2.11 | 0.452 | 0.013 | 0.036 | 0.036 | 0.15 | 0.22 | 0.04 | 0.11 | 0.15 | 0.19 | 0.12 |
| 6 | 92014180 | rs2748276 | 0.386 | 2.252 | -1.252 | 0.897 | -0.126 | -0.062 | -0.062 | 0 | 0.84 | 0.99 | 0.59 | 0.85 | 0.67 | 0.82 |
| 6 | 92019086 | rs2224063 | 0.306 | 0.898 | -0.835 | 0.896 | -0.044 | -0.02 | -0.02 | 0 | 0.67 | 0.82 | 0.48 | 0.55 | 0.6 | 0.68 |
| 6 | 92026031 | rs13211543 | -0.276 | -0.05 | -0.481 | 0.536 | -0.018 | -0.032 | -0.032 | 0 | 0.1 | 0.1 | 0.03 | 0 | 0.03 | 0.14 |
| 6 | 92041904 | rs2144363 | -0.377 | -0.425 | 0.92 | 1.186 | -0.026 | 0.181 | 0.181 | 0 | 0.08 | 0.18 | 0.08 | 0.28 | 0.03 | 0.15 |
| 6 | 92045797 | rs9342274 | -1.005 | -0.311 | 0.947 | 0.959 | 0.016 | 0.06 | 0.06 | 0 | 0.53 | 0.73 | 0.62 | 0.73 | 0.41 | 0.61 |
| 6 | 92054323 | rs7772106 | -0.408 | -0.071 | 1.197 | 1.011 | -0.002 | 0.008 | 0.008 | 0 | 0.44 | 0.39 | 0.55 | 0.45 | 0.42 | 0.45 |
| 6 | 92075429 | rs2017790 | -0.844 | 0.102 | 0.887 | 0.526 | -0.009 | 0.059 | 0.059 | 0 | 0.53 | 0.74 | 0.63 | 0.71 | 0.45 | 0.58 |
| 6 | 92086748 | rs7738757 | -0.827 | 0.099 | 0.911 | 0.525 | -0.005 | 0.046 | 0.046 | 0 | 0.53 | 0.71 | 0.63 | 0.71 | 0.45 | 0.58 |
| 6 | 92096750 | rs9294489 | -0.796 | 0.151 | 0.793 | 0.525 | -0.003 | 0.045 | 0.045 | 0 | 0.53 | 0.68 | 0.64 | 0.74 | 0.45 | 0.61 |
| 6 | 92097137 | rs9294491 | -0.315 | -0.007 | 1.185 | 1.064 | -0.001 | -0.008 | -0.008 | 0.123 | 0.43 | 0.32 | 0.57 | 0.57 | 0.42 | 0.49 |
| 6 | 92119129 | rs6937560 | -1.03 | 0.298 | 1.36 | 1.266 | 0.002 | 0.012 | 0.012 | 1.272 | 0.34 | 0.65 | 0.55 | 0.55 | 0.35 | 0.48 |
| 6 | 92132799 | rs9359938 | -0.561 | 0.303 | 1.647 | 1.128 | 0.003 | 0.001 | 0.001 | 1.24 | 0.27 | 0.28 | 0.51 | 0.33 | 0.33 | 0.37 |
| 6 | 92136903 | rs6938848 | -0.634 | 0.298 | 1.658 | 1.126 | 0.002 | 0.001 | 0.001 | 1.623 | 0.27 | 0.29 | 0.51 | 0.33 | 0.33 | 0.37 |
| 6 | 92146299 | rs2748285 | -1.196 | 0.182 | 1.956 | 0.847 | 0.001 | 0.003 | 0.003 | 0 | 0.44 | 0.48 | 0.61 | 0.56 | 0.46 | 0.43 |
| 6 | 92148779 | rs2818290 | -1.299 | 0.174 | 2.038 | 0.841 | -0.004 | -0.007 | -0.007 | 0 | 0.47 | 0.7 | 0.63 | 0.56 | 0.47 | 0.42 |
| 6 | 92149334 | rs2818291 | -0.266 | 0.181 | 1.853 | 0.893 | -0.018 | -0.003 | -0.003 | 0 | 0.19 | 0.25 | 0.12 | 0.23 | 0.15 | 0.06 |
| 6 | 92164986 | rs9362875 | -0.934 | 0.219 | 1.527 | 0.801 | -0.001 | -0.003 | -0.003 | 0.17 | 0.33 | 0.33 | 0.57 | 0.56 | 0.38 | 0.35 |
| 6 | 92170010 | rs9342279 | 0.161 | 0.391 | 0.92 | 0.804 | -0.053 | -0.066 | -0.066 | 0 | 0.14 | 0.05 | 0.44 | 0.33 | 0.23 | 0.29 |
| 6 | 92184987 | rs13208509 | -0.238 | 0.122 | 1.157 | 0.326 | -0.001 | 0.001 | 0.001 | 0 | 0.1 | 0.05 | 0.04 | 0 | 0.09 | 0.08 |
| 6 | 92190410 | rs4130902 | -1.336 | -0.336 | 0.949 | 0.077 | 0.05 | 0.089 | 0.089 | 0 | 0.39 | 0.8 | 0.22 | 0.22 | 0.26 | 0.22 |
| 6 | 92190651 | rs6924108 | -1.357 | -0.315 | 1.325 | -0.067 | -0.016 | 0.011 | 0.011 | 0 | 0.19 | 0.44 | 0.09 | 0.22 | 0.16 | 0.17 |
| 6 | 92194948 | rs9351269 | -1.286 | 0.192 | 0.847 | -0.365 | 0.002 | 0.008 | 0.008 | 0 | 0.59 | 0.61 | 0.69 | 0.91 | 0.63 | 0.62 |
| 6 | 92200553 | rs9362878 | -0.975 | 0.111 | 0.911 | -0.383 | 0.003 | 0.004 | 0.004 | 0 | 0.4 | 0.63 | 0.22 | 0.24 | 0.3 | 0.23 |
| 6 | 92208127 | rs9353843 | -1.409 | 0.08 | 0.611 | -0.41 | -0.012 | 0.056 | 0.056 | 0 | 0.56 | 0.63 | 0.67 | 0.91 | 0.55 | 0.62 |
| 6 | 92208884 | rs16884411 | -0.006 | 0.23 | 0.044 | 0.386 | -0.02 | -0.053 | -0.053 | 0 | 0.12 | 0.04 | 0.06 | 0.32 | 0.15 | 0.21 |
| 6 | 92208995 | rs9359945 | 0.558 | 0.126 | 0.343 | 0.592 | -0.004 | 0.002 | 0.002 | 0.311 | 0.12 | 0.11 | 0.43 | 0.34 | 0.2 | 0.23 |
| 6 | 92210391 | rs4385282 | 0.188 | -0.034 | 0.327 | 0.271 | 0.02 | 0.043 | 0.043 | 2.641 | 0.18 | 0.1 | 0.1 | 0.01 | 0.12 | 0.07 |
| 6 | 92210672 | rs9362881 | -0.802 | 0.214 | 0.76 | 0.692 | 0.006 | 0.058 | 0.058 | 0.427 | 0.23 | 0.33 | 0.51 | 0.59 | 0.28 | 0.33 |
| 6 | 92214481 | rs4422587 | -0.684 | 0.194 | 0.637 | 0.98 | 0.003 | 0.055 | 0.055 | 0 | 0.07 | 0.1 | 0.04 | 0.26 | 0.06 | 0.09 |
| 6 | 92221558 | rs4707675 | 1.854 | 0.138 | 0.029 | 0.231 | -0.049 | 0.101 | 0.101 | 0 | 0.36 | 0.17 | 0.32 | 0.09 | 0.41 | 0.34 |
| 6 | 92224243 | rs16884430 | -0.571 | 0.161 | 0.371 | 0.457 | 0.013 | 0.072 | 0.072 | 0 | 0.08 | 0.21 | 0.05 | 0.19 | 0.06 | 0.1 |
| 6 | 92224496 | rs12529119 | 0.277 | 0.33 | 0.348 | 0.426 | 0.001 | -0.007 | -0.007 | 0.157 | 0.15 | 0.11 | 0.06 | 0.19 | 0.15 | 0.11 |
| 6 | 92226306 | rs16884433 | -0.374 | 0.935 | 0.217 | 0.354 | 0.016 | 0.077 | 0.077 | 0 | 0.08 | 0.21 | 0.05 | 0.19 | 0.06 | 0.1 |
| 6 | 92229590 | rs6913021 | 1.049 | 0.852 | 0.115 | 0.361 | 0 | 0.047 | 0.047 | 0 | 0.78 | 0.46 | 0.88 | 0.61 | 0.83 | 0.77 |
| 6 | 92233404 | rs6904895 | -0.652 | 0.853 | -0.166 | 0.286 | -0.01 | -0.092 | -0.092 | 0 | 0.4 | 0.12 | 0.55 | 0.51 | 0.42 | 0.43 |
| 6 | 92233748 | rs17555108 | 0.427 | 0.876 | -0.166 | 0.321 | -0.026 | -0.038 | -0.038 | 0 | 0.14 | 0.01 | 0.06 | 0.19 | 0.16 | 0.11 |
| 6 | 92240790 | rs4296871 | 0.491 | 0.91 | 0.232 | 0.327 | 0.007 | 0.035 | 0.035 | 0.092 | 0.86 | 0.72 | 0.93 | 0.79 | 0.88 | 0.87 |
| 6 | 92247872 | rs12173279 | -0.045 | 0.92 | -0.225 | 0.338 | -0.004 | -0.061 | -0.061 | 0 | 0.26 | 0.09 | 0.49 | 0.32 | 0.27 | 0.33 |
| 6 | 92250788 | rs9359951 | 0.584 | 0.878 | -0.327 | 0.294 | -0.028 | -0.099 | -0.099 | 0 | 0.15 | 0.01 | 0.42 | 0.31 | 0.19 | 0.24 |
| 6 | 92254620 | rs9345148 | 1.83 | 0.854 | -0.221 | 0.215 | -0.035 | 0.069 | 0.069 | 0 | 0.45 | 0.25 | 0.36 | 0.27 | 0.46 | 0.45 |
| 6 | 92256169 | rs9294508 | 1.438 | 0.851 | -0.096 | 0.189 | 0.003 | -0.008 | -0.008 | 0 | 0.62 | 0.37 | 0.81 | 0.41 | 0.68 | 0.66 |
| 6 | 92264022 | rs4573056 | -0.155 | 0.832 | -0.272 | 0.169 | -0.002 | -0.063 | -0.063 | 0 | 0.26 | 0.09 | 0.48 | 0.32 | 0.27 | 0.33 |
| 6 | 92270124 | rs9345152 | -0.54 | 0.758 | -0.19 | 0.19 | -0.004 | -0.054 | -0.054 | 2.632 | 0.24 | 0.08 | 0.1 | 0.37 | 0.21 | 0.22 |
| 6 | 92288357 | rs11760152 | 0.883 | 0.743 | -0.201 | 0.111 | -0.001 | 0.009 | 0.009 | 0.363 | 0.1 | 0.07 | 0.06 | 0 | 0.08 | 0.01 |
| 6 | 92289144 | rs9353857 | 1.248 | 0.717 | -0.085 | 0.191 | 0.006 | -0.017 | -0.017 | 0 | 0.62 | 0.34 | 0.8 | 0.42 | 0.67 | 0.67 |
| 6 | 92298780 | rs1556623 | 2.276 | 1.226 | -0.432 | -0.125 | -0.05 | 0.146 | 0.146 | 0 | 0.36 | 0.13 | 0.32 | 0.07 | 0.39 | 0.33 |
| 6 | 92313979 | rs9353859 | -0.947 | 1.175 | -0.074 | -0.066 | 0 | -0.001 | -0.001 | 0 | 0.32 | 0.3 | 0.5 | 0.72 | 0.35 | 0.49 |
| 6 | 92314646 | rs4613794 | -1.349 | 0.857 | -0.593 | -0.732 | 0.002 | 0.017 | 0.017 | 0 | 0.6 | 0.62 | 0.64 | 0.91 | 0.6 | 0.6 |
| 6 | 92318594 | rs9359954 | -1.575 | 0.345 | -0.509 | -0.607 | -0.028 | 0.14 | 0.14 | 0.355 | 0.55 | 0.73 | 0.63 | 0.92 | 0.51 | 0.6 |
| 6 | 92321000 | rs4112305 | -1.471 | 0.507 | -0.411 | -0.522 | -0.004 | 0.074 | 0.074 | 0.262 | 0.52 | 0.62 | 0.6 | 0.92 | 0.5 | 0.59 |
| 6 | 92330487 | rs369910 | 0.41 | 0.874 | -0.483 | -0.497 | 0.005 | -0.046 | -0.046 | 0 | 0.15 | 0.25 | 0.1 | 0.01 | 0.16 | 0.08 |
| 6 | 92330602 | rs10485101 | 0.518 | 0.861 | -0.54 | 0.029 | -0.01 | -0.02 | -0.02 | 0 | 0.09 | 0.06 | 0.05 | 0.23 | 0.09 | 0.13 |
| 6 | 92340802 | rs1201867 | -1.659 | 0.72 | -0.502 | -0.617 | -0.008 | 0.11 | 0.11 | 0 | 0.65 | 0.82 | 0.69 | 0.89 | 0.67 | 0.67 |
| 6 | 92348148 | rs984739 | -0.532 | 0.872 | 0.23 | 0.167 | -0.003 | 0.039 | 0.039 | 0.074 | 0.27 | 0.35 | 0.46 | 0.55 | 0.29 | 0.37 |
| 6 | 92349020 | rs1931821 | -1.155 | 0.913 | 0.223 | -0.143 | -0.005 | -0.031 | -0.031 | 0 | 0.33 | 0.63 | 0.19 | 0.2 | 0.29 | 0.2 |
| 6 | 92349598 | rs1334633 | -0.85 | 0.666 | 0.689 | -0.093 | -0.011 | 0.011 | 0.011 | 0 | 0.17 | 0.42 | 0.11 | 0.18 | 0.12 | 0.14 |
| 6 | 92357468 | rs915167 | -1.666 | 0.746 | 0.388 | -0.288 | -0.002 | 0.013 | 0.013 | 0 | 0.38 | 0.75 | 0.24 | 0.37 | 0.35 | 0.33 |
| 6 | 92365086 | rs16884621 | -0.295 | 0.742 | 0.743 | -0.323 | 0.014 | 0.008 | 0.008 | 0.033 | 0.13 | 0.2 | 0.11 | 0.17 | 0.14 | 0.23 |
| 6 | 92367643 | rs7453029 | 0.022 | 0.747 | 0.321 | -0.346 | -0.024 | -0.003 | -0.003 | 1.131 | 0.18 | 0.09 | 0.1 | 0.19 | 0.21 | 0.13 |
| 6 | 92368944 | rs4707684 | 0.84 | 0.998 | 0.148 | -0.054 | 0.001 | 0.09 | 0.09 | 0.126 | 0.09 | 0.08 | 0.31 | 0.47 | 0.14 | 0.15 |
| 6 | 92377118 | rs1160726 | -1.645 | 0.805 | -0.313 | -0.14 | 0.001 | 0.017 | 0.017 | 0 | 0.39 | 0.74 | 0.34 | 0.36 | 0.36 | 0.36 |

| | | | | | | | | | | | | | | | | |
|---|---|---|---|---|---|---|---|---|---|---|---|---|---|---|---|---|
| 6 | 92381402 | rs7774651 | -1.15 | 0.917 | -0.331 | -0.148 | -0.007 | 0.013 | 0.013 | 0 | 0.35 | 0.27 | 0.44 | 0.72 | 0.4 | 0.39 |
| 6 | 92382537 | rs9345169 | 0.998 | 0.972 | -0.403 | 0.036 | -0.01 | 0.004 | 0.004 | 0.247 | 0.08 | 0.01 | 0.29 | 0.45 | 0.1 | 0.11 |
| 6 | 92388426 | rs4707685 | -1.977 | 1.572 | 0.7 | 0.197 | 0.016 | 0.21 | 0.21 | 0.015 | 0.51 | 0.79 | 0.66 | 0.89 | 0.54 | 0.58 |
| 6 | 92390455 | rs9362902 | -0.84 | 1.926 | 0.938 | 1 | 0.006 | 0.018 | 0.018 | 2.325 | 0.22 | 0.22 | 0.39 | 0.6 | 0.27 | 0.33 |
| 6 | 92393787 | rs565665 | -1.712 | 0.966 | 1.322 | 1.136 | 0.005 | 0.1 | 0.1 | 3.756 | 0.13 | 0.47 | 0.16 | 0.19 | 0.08 | 0.14 |
| 6 | 92393990 | rs529607 | -2.059 | 1.06 | 1.154 | 1.126 | 0.003 | 0.069 | 0.069 | 0.109 | 0.26 | 0.5 | 0.24 | 0.4 | 0.23 | 0.23 |
| 6 | 92395675 | rs570258 | -1.654 | 1.171 | 1.246 | 1.07 | 0.014 | 0.037 | 0.037 | 0.065 | 0.13 | 0.31 | 0.08 | 0.19 | 0.13 | 0.16 |
| 6 | 92401171 | rs1761783 | -1.499 | 1.21 | 1.203 | 1.164 | 0.029 | 0.087 | 0.087 | 0 | 0.12 | 0.28 | 0.15 | 0.23 | 0.12 | 0.16 |
| 6 | 92412540 | rs574892 | -2.036 | 1.277 | 0.791 | 1.227 | 0.028 | 0.091 | 0.091 | 0 | 0.26 | 0.48 | 0.33 | 0.43 | 0.27 | 0.22 |
| 6 | 92424784 | rs13206653 | -0.529 | 1.247 | -0.179 | 1.336 | -0.01 | -0.005 | -0.005 | 0.646 | 0.06 | 0.01 | 0.1 | 0.05 | 0.06 | 0.07 |
| 6 | 92424803 | rs9345171 | -0.61 | 1.183 | -0.186 | 1.244 | 0.001 | -0.013 | -0.013 | 0 | 0.18 | 0.11 | 0.37 | 0.52 | 0.19 | 0.29 |
| 6 | 92426566 | rs9451764 | -0.577 | 1.251 | -0.151 | 1.347 | -0.033 | -0.038 | -0.038 | 0 | 0.12 | 0.03 | 0.07 | 0.2 | 0.16 | 0.07 |
| 6 | 92446395 | rs2064943 | -1.863 | 1.308 | 0.441 | 1.422 | 0.008 | 0.026 | 0.026 | 0 | 0.29 | 0.61 | 0.35 | 0.43 | 0.3 | 0.27 |
| 6 | 92462541 | rs9451793 | -0.787 | 1.305 | 0.381 | 1.54 | -0.051 | -0.053 | -0.053 | 0.213 | 0.13 | 0.01 | 0.16 | 0.2 | 0.19 | 0.11 |
| 6 | 92492833 | rs12660904 | -0.116 | 1.095 | 0.351 | 1.916 | 0.008 | -0.026 | -0.026 | 0 | 0.08 | 0.02 | 0.31 | 0.4 | 0.1 | 0.11 |
| 6 | 92493227 | rs1979796 | -1.98 | 1.172 | 0.628 | 1.714 | 0.017 | 0.048 | 0.048 | 0 | 0.3 | 0.53 | 0.36 | 0.43 | 0.33 | 0.31 |
| 6 | 92518321 | rs2501636 | -2.116 | 1.179 | 0.776 | 1.716 | 0.024 | 0.063 | 0.063 | 0 | 0.29 | 0.51 | 0.27 | 0.43 | 0.33 | 0.31 |
| 6 | 92525977 | rs904308 | -2.039 | 1.146 | 0.873 | 1.775 | 0.011 | 0.122 | 0.122 | 0.297 | 0.23 | 0.52 | 0.22 | 0.46 | 0.22 | 0.3 |
| 6 | 92526806 | rs976832 | -1.249 | 0.848 | 1.223 | 1.201 | 0.012 | 0.011 | 0.011 | 0.015 | 0.13 | 0.2 | 0.09 | 0.12 | 0.17 | 0.17 |
| 6 | 92552875 | rs2501657 | 1.436 | 0.948 | 1.73 | 0.998 | 0.06 | -0.084 | -0.084 | 0 | 0.86 | 0.59 | 0.87 | 0.97 | 0.8 | 0.86 |
| 6 | 92562268 | rs2101980 | -1.313 | 1.203 | 1.714 | 1.011 | 0.006 | 0.003 | 0.003 | 0.178 | 0.18 | 0.2 | 0.13 | 0.21 | 0.22 | 0.27 |
| 6 | 92569711 | rs2454242 | -0.184 | 1.22 | 1.422 | 1.003 | -0.003 | -0.006 | -0.006 | 0 | 0.09 | 0.1 | 0.06 | 0 | 0.13 | 0.05 |
| 6 | 92572932 | rs9342302 | 2.286 | 1.185 | 1.606 | 0.954 | 0.004 | -0.01 | -0.01 | 0.176 | 0.66 | 0.36 | 0.75 | 0.76 | 0.57 | 0.57 |
| 6 | 92577468 | rs9451847 | -1.047 | 1.238 | 1.69 | 0.981 | 0.038 | -0.016 | -0.016 | 0 | 0.05 | 0.18 | 0.06 | 0.03 | 0.07 | 0.09 |
| 6 | 92582660 | rs6920127 | 1.999 | 1.156 | 1.587 | 0.966 | 0.018 | -0.028 | -0.028 | 0 | 0.67 | 0.45 | 0.75 | 0.76 | 0.58 | 0.58 |
| 6 | 92590109 | rs4318847 | 2.336 | 1.119 | 1.433 | 1.288 | -0.007 | -0.005 | -0.005 | 0.03 | 0.58 | 0.38 | 0.68 | 0.48 | 0.45 | 0.46 |
| 6 | 92595284 | rs9345186 | -1.167 | 1.21 | 1.304 | 1.382 | 0.001 | 0.027 | 0.027 | 0 | 0.13 | 0.17 | 0.11 | 0.31 | 0.17 | 0.2 |
| 6 | 92599727 | rs9353872 | -1.465 | 1.213 | 1.355 | 1.378 | 0.008 | 0.053 | 0.053 | 0 | 0.14 | 0.25 | 0.12 | 0.31 | 0.18 | 0.22 |
| 6 | 92604703 | rs7742254 | 0.827 | 0.97 | 1.355 | 1.188 | 0.002 | -0.001 | -0.001 | 0.438 | 0.8 | 0.81 | 0.87 | 0.8 | 0.75 | 0.72 |
| 6 | 92606067 | rs9451855 | 1.439 | 0.843 | 1.273 | 1.128 | 0.013 | -0.031 | -0.031 | 0.299 | 0.69 | 0.62 | 0.8 | 0.8 | 0.62 | 0.67 |
| 6 | 92611525 | rs1498258 | -1.272 | 0.822 | 0.912 | 1.058 | -0.002 | -0.002 | -0.002 | 0 | 0.17 | 0.17 | 0.13 | 0.31 | 0.2 | 0.22 |
| 6 | 92615457 | rs1391502 | -1.123 | 0.719 | 0.717 | 0.357 | 0.02 | -0.03 | -0.03 | 0.517 | 0.24 | 0.1 | 0.49 | 0.71 | 0.23 | 0.27 |
| 6 | 92616418 | rs9451856 | -1.971 | 0.712 | 0.863 | 0.314 | 0 | 0.001 | 0.001 | 0 | 0.39 | 0.45 | 0.59 | 0.77 | 0.4 | 0.44 |
| 6 | 92620641 | rs991880 | -1.028 | 0.487 | 0.679 | 0.398 | -0.004 | -0.062 | -0.062 | 0 | 0.24 | 0.05 | 0.49 | 0.71 | 0.24 | 0.27 |
| 6 | 92627946 | rs1498260 | -1.167 | 0.351 | 0.626 | 0.355 | 0.032 | -0.015 | -0.015 | 0.45 | 0.27 | 0.14 | 0.5 | 0.69 | 0.24 | 0.27 |
| 6 | 92641939 | rs1995145 | 2.299 | 0.488 | 0.657 | 0.306 | -0.001 | -0.026 | -0.026 | 3.74 | 0.41 | 0.38 | 0.3 | 0.09 | 0.35 | 0.34 |
| 6 | 92645170 | rs9351294 | -1.435 | 0.142 | 0.39 | 0.486 | 0.023 | 0.024 | 0.024 | 0 | 0.36 | 0.32 | 0.55 | 0.72 | 0.33 | 0.38 |
| 6 | 92647085 | rs2030055 | 1.806 | 0.362 | 0.428 | 0.026 | -0.024 | 0.223 | 0.223 | 1.577 | 0.48 | 0.21 | 0.44 | 0.11 | 0.44 | 0.45 |
| 6 | 92650235 | rs1498252 | 0.395 | 0.594 | 0.777 | 0.695 | -0.004 | -0.004 | -0.004 | 0 | 0.79 | 0.82 | 0.86 | 0.68 | 0.69 | 0.79 |
| 6 | 92669376 | rs12526373 | -1.263 | 0.421 | 1.056 | 1.021 | -0.023 | 0.121 | 0.121 | 1.423 | 0.11 | 0.5 | 0.16 | 0.15 | 0.05 | 0.11 |
| 6 | 92670118 | rs4432695 | -0.266 | 0.454 | 1.241 | 1.038 | 0.041 | 0.213 | 0.213 | 1.251 | 0.06 | 0.27 | 0.15 | 0.15 | 0.03 | 0.09 |
| 6 | 92684706 | rs6930755 | -0.172 | 0.469 | 0.632 | 0.838 | -0.002 | -0.008 | -0.008 | 0 | 0.1 | 0.13 | 0.09 | 0.01 | 0.12 | 0.12 |
| 6 | 92686777 | rs1493558 | 1.998 | 0.472 | 0.579 | 0.781 | -0.013 | 0.192 | 0.192 | 0.053 | 0.51 | 0.24 | 0.44 | 0.12 | 0.44 | 0.48 |
| 6 | 92687036 | rs13194228 | -1.937 | 0.461 | 0.632 | 1.125 | -0.012 | 0.046 | 0.046 | 0.091 | 0.27 | 0.62 | 0.4 | 0.48 | 0.25 | 0.31 |
| 6 | 92687857 | rs2201302 | 1.876 | 0.416 | 0.541 | 1.084 | -0.017 | 0.183 | 0.183 | 0.02 | 0.52 | 0.24 | 0.45 | 0.12 | 0.46 | 0.48 |
| 6 | 92689878 | rs1493560 | -0.602 | 0.272 | 1.112 | 1.666 | 0.003 | 0.01 | 0.01 | 0 | 0.09 | 0.08 | 0.17 | 0.34 | 0.09 | 0.1 |
| 6 | 92712552 | rs12664176 | -0.602 | 0.272 | 1.11 | 1.666 | 0.002 | 0.019 | 0.019 | 0 | 0.09 | 0.09 | 0.18 | 0.34 | 0.09 | 0.1 |
| 6 | 92712609 | rs12661313 | -0.602 | 0.272 | 1.11 | 1.666 | 0.002 | 0.019 | 0.019 | 0 | 0.09 | 0.09 | 0.18 | 0.34 | 0.09 | 0.1 |
| 6 | 92715572 | rs13216475 | -1.083 | 0.414 | 0.919 | 1.567 | -0.001 | -0.017 | -0.017 | 0.229 | 0.14 | 0.14 | 0.1 | 0.01 | 0.16 | 0.13 |
| 6 | 92722564 | rs11756114 | -0.481 | 0.515 | 0.632 | 1.605 | -0.012 | 0.016 | 0.016 | 0.03 | 0.11 | 0.09 | 0.1 | 0.02 | 0.15 | 0.12 |
| 6 | 92725708 | rs2452987 | 1.031 | 0.542 | -0.234 | 1.624 | 0.032 | -0.008 | -0.008 | 0 | 0.57 | 0.18 | 0.4 | 0.43 | 0.6 | 0.57 |
| 6 | 92730285 | rs6942324 | -0.927 | 1.008 | -0.649 | 1.181 | 0.001 | -0.003 | -0.003 | 0 | 0.36 | 0.39 | 0.45 | 0.4 | 0.37 | 0.34 |
| 6 | 92749256 | rs10944560 | -0.918 | 2.341 | -1.105 | 1.257 | -0.104 | -0.064 | -0.064 | 1 | 0.15 | 0.02 | 0.15 | 0.19 | 0.26 | 0.16 |
| 6 | 92750174 | rs2249818 | 1.12 | 1.779 | -0.895 | 1.257 | 0.007 | 0.003 | 0.003 | 0 | 0.5 | 0.36 | 0.6 | 0.47 | 0.47 | 0.53 |
| 6 | 92750444 | rs12213860 | -0.382 | 1.649 | -0.535 | 1.295 | 0.006 | -0.006 | -0.006 | 0 | 0.2 | 0.16 | 0.09 | 0.23 | 0.17 | 0.19 |
| 6 | 92750490 | rs2201306 | -1.071 | 1.746 | -0.044 | 1.203 | -0.01 | 0.019 | 0.019 | 0.183 | 0.26 | 0.38 | 0.23 | 0.4 | 0.26 | 0.29 |
| 6 | 92751145 | rs4707716 | -0.156 | 1.934 | 0.08 | 1.372 | 0.019 | 0.023 | 0.023 | 0 | 0.07 | 0.22 | 0.15 | 0.13 | 0.09 | 0.1 |
| 6 | 92752666 | rs2452974 | 1.881 | 1.747 | 0.026 | 1.329 | -0.004 | -0.002 | -0.002 | 0 | 0.55 | 0.34 | 0.61 | 0.43 | 0.5 | 0.53 |
| 6 | 92755089 | rs2452975 | 1.296 | 1.748 | 0.123 | 1.324 | -0.023 | -0.031 | -0.031 | 0.064 | 0.71 | 0.77 | 0.8 | 0.56 | 0.63 | 0.68 |
| 6 | 92755341 | rs10944561 | -0.335 | 1.741 | 0.136 | 1.332 | -0.001 | 0.009 | 0.009 | 0 | 0.14 | 0.15 | 0.07 | 0.23 | 0.13 | 0.16 |
| 6 | 92756602 | rs2503662 | 0.174 | 1.859 | 0.227 | 1.12 | 0.008 | 0 | 0 | 0 | 0.87 | 0.87 | 0.87 | 0.84 | 0.78 | 0.86 |
| 6 | 92760362 | rs2503661 | 0.861 | 1.839 | 0.234 | 1.022 | -0.018 | -0.003 | -0.003 | 0 | 0.82 | 0.87 | 0.75 | 0.76 | 0.69 | 0.76 |
| 6 | 92760762 | rs9353880 | -0.24 | 1.793 | 0.766 | 1.11 | -0.003 | -0.004 | -0.004 | 1.354 | 0.1 | 0.16 | 0.05 | 0.09 | 0.1 | 0.13 |
| 6 | 92766359 | rs2503654 | 1.983 | 1.679 | 0.918 | 1.083 | -0.005 | 0.002 | 0.002 | 0 | 0.67 | 0.33 | 0.66 | 0.67 | 0.58 | 0.64 |
| 6 | 92773778 | rs2452961 | 0.123 | 1.735 | 0.917 | 1.078 | -0.002 | 0 | 0 | 0 | 0.87 | 0.9 | 0.88 | 0.88 | 0.78 | 0.86 |
| 6 | 92777420 | rs6454889 | -1.264 | 1.65 | 0.903 | 0.924 | -0.021 | -0.116 | -0.116 | 0 | 0.17 | 0.4 | 0.17 | 0 | 0.15 | 0.13 |
| 6 | 92786021 | rs2503647 | 1.236 | 1.701 | 0.841 | 1.008 | 0.02 | 0 | 0 | 0 | 0.76 | 0.72 | 0.82 | 0.72 | 0.68 | 0.72 |
| 6 | 92817698 | rs11961411 | -0.373 | 1.537 | 0.829 | 0.92 | -0.002 | -0.038 | -0.038 | 0 | 0.09 | 0.14 | 0.04 | 0.02 | 0.08 | 0.09 |
| 6 | 92819644 | rs2452944 | 1.763 | 1.615 | 1.065 | 0.889 | -0.017 | 0.036 | 0.036 | 2.481 | 0.6 | 0.31 | 0.62 | 0.69 | 0.53 | 0.62 |

| | | | | | | | | | | | | | | | | |
|---|---|---|---|---|---|---|---|---|---|---|---|---|---|---|---|---|
| 6 | 92819879 | rs2503639 | 0.14 | 1.661 | 1.116 | 0.926 | -0.005 | -0.004 | -0.004 | 0 | 0.85 | 0.87 | 0.84 | 0.76 | 0.76 | 0.84 |
| 6 | 92820925 | rs13217394 | 2.083 | 1.524 | 0.607 | 0.97 | 0.065 | -0.025 | -0.025 | 0 | 0.41 | 0.17 | 0.32 | 0.38 | 0.27 | 0.39 |
| 6 | 92822870 | rs2452940 | 1.194 | 1.639 | 0.814 | 0.642 | 0.008 | -0.003 | -0.003 | 0 | 0.7 | 0.38 | 0.75 | 0.68 | 0.64 | 0.72 |
| 6 | 92823967 | rs10944562 | -0.246 | 1.655 | 0.953 | 0.649 | 0.063 | 0.033 | 0.033 | 0.021 | 0.84 | 0.71 | 0.82 | 0.76 | 0.76 | 0.84 |
| 6 | 92824700 | rs9342313 | -0.704 | 1.377 | 0.726 | 0.469 | -0.001 | -0.031 | -0.031 | 0 | 0.13 | 0.31 | 0.07 | 0.09 | 0.11 | 0.12 |
| 6 | 92827954 | rs9451925 | -0.523 | 1.432 | 0.679 | 0.46 | 0.004 | -0.085 | -0.085 | 0 | 0.14 | 0.29 | 0.16 | 0 | 0.14 | 0.13 |
| 6 | 92829488 | rs4446534 | 1.473 | 1.791 | 0.551 | 0.541 | 0.062 | 0.055 | 0.055 | 0.35 | 0.48 | 0.34 | 0.52 | 0.67 | 0.34 | 0.54 |
| 6 | 92830228 | rs4334953 | -0.485 | 1.994 | 0.461 | 0.502 | -0.073 | -0.066 | -0.066 | 0 | 0.86 | 0.95 | 0.84 | 0.77 | 0.76 | 0.85 |
| 6 | 92836208 | rs4073532 | 1.74 | 1.88 | 0.532 | 0.463 | 0.096 | 0.068 | 0.068 | 0.435 | 0.46 | 0.23 | 0.49 | 0.67 | 0.32 | 0.52 |
| 6 | 92838614 | rs7759851 | 0.602 | 2.066 | 0.695 | 0.496 | -0.026 | -0.003 | -0.003 | 0 | 0.72 | 0.77 | 0.67 | 0.71 | 0.58 | 0.71 |
| 6 | 92843692 | rs13192548 | -0.132 | 1.983 | 0.781 | 0.13 | 0.01 | -0.074 | -0.074 | 0 | 0.23 | 0.29 | 0.15 | 0.05 | 0.26 | 0.19 |
| 6 | 92844213 | rs9345201 | -0.116 | 2.042 | 0.723 | 0.136 | 0.008 | -0.03 | -0.03 | 0 | 0.13 | 0.18 | 0.17 | 0.05 | 0.16 | 0.13 |
| 6 | 92849774 | rs7773056 | -0.662 | 2.048 | 0.628 | 0.552 | -0.079 | -0.095 | -0.095 | 0 | 0.85 | 0.95 | 0.69 | 0.65 | 0.75 | 0.84 |
| 6 | 92853369 | rs7745179 | 0.127 | 1.788 | 0.866 | 1.076 | -0.001 | 0.013 | 0.013 | 0 | 0.84 | 0.79 | 0.81 | 0.72 | 0.82 | 0.76 |
| 6 | 92859874 | rs13202708 | 1.841 | 1.795 | 0.56 | 0.935 | 0.076 | 0.103 | 0.103 | 0.585 | 0.39 | 0.15 | 0.31 | 0.28 | 0.26 | 0.38 |
| 6 | 92867219 | rs4502906 | -0.998 | 2.116 | 0.223 | 1.104 | -0.008 | -0.003 | -0.003 | 0 | 0.38 | 0.32 | 0.44 | 0.38 | 0.5 | 0.34 |
| 6 | 92889890 | rs12212424 | -0.089 | 2.232 | -0.786 | 1.059 | -0.054 | -0.083 | -0.083 | 0 | 0.19 | 0.1 | 0.32 | 0.41 | 0.28 | 0.15 |
| 6 | 92890665 | rs1373376 | 0.116 | 2.291 | -0.971 | 1.09 | -0.101 | -0.135 | -0.135 | 0.214 | 0.18 | 0.01 | 0.31 | 0.41 | 0.28 | 0.15 |
| 6 | 92895021 | rs10498978 | 0.015 | 1.988 | -0.64 | 0.188 | -0.061 | 0.07 | 0.07 | 0 | 0.14 | 0.03 | 0.13 | 0.01 | 0.2 | 0.18 |
| 6 | 92897624 | rs9359974 | 0.955 | 1.34 | 0.029 | 0.397 | -0.002 | -0.083 | -0.083 | 0.018 | 0.85 | 0.71 | 0.92 | 0.97 | 0.9 | 0.92 |
| 6 | 92904799 | rs10944570 | -0.476 | 1.483 | -0.611 | 0.676 | -0.029 | -0.021 | -0.021 | 0 | 0.08 | 0.02 | 0.05 | 0.14 | 0.09 | 0.13 |
| 6 | 92912828 | rs9359977 | 1.076 | 1.695 | -0.906 | 0.62 | -0.028 | 0.058 | 0.058 | 0.033 | 0.57 | 0.26 | 0.71 | 0.68 | 0.56 | 0.51 |
| 6 | 92913083 | rs9345216 | 0.19 | 1.812 | -0.496 | 0.432 | -0.018 | 0.024 | 0.024 | 0 | 0.73 | 0.8 | 0.83 | 0.94 | 0.71 | 0.76 |
| 6 | 92918288 | rs1444133 | 0.221 | 1.353 | -0.96 | 0.538 | -0.003 | -0.006 | -0.006 | 0.454 | 0.2 | 0.16 | 0.17 | 0.29 | 0.24 | 0.18 |
| 6 | 92919416 | rs2120546 | -0.113 | 0.676 | -0.544 | 0.755 | 0.012 | -0.019 | -0.019 | 0 | 0.65 | 0.54 | 0.66 | 0.7 | 0.67 | 0.77 |
| 6 | 92922779 | rs12210356 | 0.473 | 0.833 | -0.742 | 1.06 | -0.024 | -0.038 | -0.038 | 0 | 0.19 | 0.07 | 0.16 | 0.29 | 0.24 | 0.18 |
| 6 | 92922807 | rs1444131 | -1.195 | 0.497 | -0.97 | 0.675 | -0.026 | 0.054 | 0.054 | 0 | 0.43 | 0.72 | 0.53 | 0.74 | 0.49 | 0.56 |
| 6 | 92929307 | rs7775910 | -0.633 | 0.503 | -0.431 | 0.726 | 0.008 | 0.014 | 0.014 | 0.55 | 0.17 | 0.48 | 0.1 | 0.18 | 0.15 | 0.28 |
| 6 | 92938341 | rs9294529 | -0.953 | 0.668 | -0.968 | 0.487 | 0.002 | 0.005 | 0.005 | 0.072 | 0.34 | 0.64 | 0.17 | 0.21 | 0.28 | 0.35 |
| 6 | 92941472 | rs9451963 | -0.953 | 0.668 | -0.968 | 0.487 | 0.002 | 0.005 | 0.005 | 0.072 | 0.34 | 0.64 | 0.17 | 0.21 | 0.28 | 0.35 |
| 6 | 92942447 | rs2325434 | -1.111 | 0.405 | -1.251 | 0.381 | 0.008 | 0.05 | 0.05 | 0.45 | 0.61 | 0.75 | 0.6 | 0.79 | 0.64 | 0.65 |
| 6 | 92954254 | rs10944578 | 0.838 | 0.864 | -1.048 | 0.736 | -0.053 | -0.063 | -0.063 | 0.113 | 0.11 | 0.01 | 0.09 | 0.18 | 0.17 | 0.08 |
| 6 | 92958861 | rs11963376 | -0.408 | 0.389 | -0.476 | 0.396 | -0.011 | -0.023 | -0.023 | 0 | 0.23 | 0.51 | 0.14 | 0.13 | 0.18 | 0.2 |
| 6 | 92960855 | rs13205488 | 0.584 | 0.467 | -0.399 | 0.328 | -0.015 | 0.03 | 0.03 | 1.168 | 0.35 | 0.5 | 0.41 | 0.5 | 0.32 | 0.44 |
| 6 | 92964208 | rs9362961 | 0.321 | 0.047 | -0.565 | 0.088 | -0.002 | -0.03 | -0.03 | 0 | 0.6 | 0.48 | 0.58 | 0.74 | 0.61 | 0.65 |
| 6 | 92965297 | rs1822589 | 1.411 | 0.098 | -0.878 | 0.13 | -0.108 | 0.024 | 0.024 | 3.106 | 0.26 | 0.01 | 0.18 | 0.13 | 0.31 | 0.19 |
| 6 | 92966484 | rs7747017 | 0.829 | -0.179 | -0.707 | 0.261 | -0.012 | 0.017 | 0.017 | 0 | 0.31 | 0.41 | 0.35 | 0.4 | 0.26 | 0.38 |
| 6 | 92966839 | rs6937033 | 0.365 | -0.344 | -0.491 | 0.108 | -0.002 | 0.004 | 0.004 | 0 | 0.22 | 0.2 | 0.19 | 0.1 | 0.25 | 0.28 |
| 6 | 92967813 | rs7753136 | 0.938 | -0.375 | -0.469 | 0.114 | -0.024 | 0.034 | 0.034 | 0.233 | 0.3 | 0.44 | 0.35 | 0.4 | 0.24 | 0.33 |
| 6 | 92972984 | rs13204161 | -0.126 | -0.287 | -0.435 | 0.167 | 0.004 | 0.002 | 0.002 | 0.693 | 0.51 | 0.6 | 0.53 | 0.47 | 0.49 | 0.61 |
| 6 | 92973406 | rs9353901 | 1.234 | -0.247 | -0.604 | 0.214 | -0.058 | -0.009 | -0.009 | 0 | 0.3 | 0.09 | 0.23 | 0.3 | 0.37 | 0.32 |
| 6 | 92976132 | rs6937171 | 1.13 | -0.255 | -0.609 | -0.181 | -0.058 | -0.092 | -0.092 | 0.15 | 0.31 | 0.09 | 0.39 | 0.52 | 0.37 | 0.33 |
| 6 | 92985361 | rs6930450 | 0.049 | -0.111 | -0.241 | -0.11 | 0.008 | 0.001 | 0.001 | 0.718 | 0.48 | 0.34 | 0.51 | 0.47 | 0.49 | 0.6 |
| 6 | 92986363 | rs10944582 | 0.165 | -0.138 | -0.347 | -0.123 | -0.013 | -0.026 | -0.026 | 0.56 | 0.47 | 0.2 | 0.51 | 0.47 | 0.49 | 0.6 |
| 6 | 92988226 | rs12663329 | 1.851 | -0.365 | -0.463 | -0.16 | 0.022 | -0.036 | -0.036 | 0 | 0.21 | 0.15 | 0.26 | 0.37 | 0.12 | 0.26 |
| 6 | 92996311 | rs9353905 | -0.828 | -0.37 | -0.191 | -0.456 | 0.005 | -0.132 | -0.132 | 0 | 0.83 | 0.56 | 0.91 | 0.99 | 0.86 | 0.94 |
| 6 | 92998916 | rs1025933 | -0.644 | -0.376 | -0.242 | -0.303 | -0.004 | 0.042 | 0.042 | 0 | 0.81 | 0.5 | 0.75 | 0.78 | 0.85 | 0.93 |
| 6 | 92998990 | rs1963461 | 0.036 | -0.259 | -0.871 | -0.32 | -0.024 | -0.017 | -0.017 | 0.006 | 0.45 | 0.18 | 0.5 | 0.46 | 0.47 | 0.59 |
| 6 | 93005995 | rs1015455 | -0.935 | 0.175 | -0.971 | -0.078 | 0.003 | 0.044 | 0.044 | 0.013 | 0.83 | 0.63 | 0.75 | 0.78 | 0.86 | 0.93 |
| 6 | 93013237 | rs12195418 | 0.406 | 0.514 | -1.598 | -0.276 | -0.063 | 0.088 | 0.088 | 0.071 | 0.19 | 0.02 | 0.17 | 0.06 | 0.22 | 0.26 |
| 6 | 93014024 | rs9362967 | 0.944 | 0.417 | -1.068 | 0.188 | -0.077 | -0.051 | -0.051 | 0.972 | 0.27 | 0.06 | 0.24 | 0.32 | 0.38 | 0.3 |
| 6 | 93050990 | rs2067050 | 0.706 | -0.411 | -0.321 | -0.275 | -0.022 | -0.073 | -0.073 | 0 | 0.17 | 0.2 | 0.07 | 0.01 | 0.12 | 0.05 |
| 6 | 93071015 | rs6927251 | -0.436 | -0.549 | -0.738 | -0.311 | 0.006 | 0.024 | 0.024 | 1.078 | 0.48 | 0.36 | 0.44 | 0.38 | 0.46 | 0.41 |
| 6 | 93073425 | rs609578 | -1.723 | -0.616 | -0.901 | -0.181 | -0.097 | -0.038 | -0.038 | 0 | 0.68 | 0.92 | 0.73 | 0.68 | 0.62 | 0.67 |
| 6 | 93077500 | rs634940 | 0.917 | -0.507 | -0.957 | -0.288 | -0.02 | 0.008 | 0.008 | 0 | 0.16 | 0.1 | 0.14 | 0.01 | 0.24 | 0.14 |
| 6 | 93081335 | rs582993 | 0.257 | -0.421 | -0.731 | -0.12 | -0.003 | -0.004 | -0.004 | 0.128 | 0.33 | 0.29 | 0.37 | 0.37 | 0.34 | 0.36 |
| 6 | 93101639 | rs650849 | -1.501 | -0.274 | -0.966 | -0.079 | -0.063 | -0.072 | -0.072 | 0 | 0.7 | 0.89 | 0.76 | 0.64 | 0.66 | 0.65 |
| 6 | 93105260 | rs600466 | 0.38 | -0.659 | -0.92 | -0.477 | 0.016 | -0.004 | -0.004 | 0.521 | 0.23 | 0.14 | 0.1 | 0.01 | 0.17 | 0.07 |
| 6 | 93105629 | rs602548 | 0.38 | -0.659 | -0.92 | -0.477 | 0.016 | -0.004 | -0.004 | 0 | 0.23 | 0.14 | 0.1 | 0.01 | 0.17 | 0.07 |
| 6 | 93106944 | rs12193330 | 0.918 | -0.667 | -0.671 | -0.396 | -0.018 | -0.006 | -0.006 | 0 | 0.27 | 0.16 | 0.34 | 0.3 | 0.3 | 0.32 |
| 6 | 93115912 | rs2757023 | 0.46 | -0.674 | -0.839 | -0.348 | 0 | -0.005 | -0.005 | 0.056 | 0.12 | 0.1 | 0.12 | 0.09 | 0.18 | 0.24 |
| 6 | 93119314 | rs2555757 | -0.6 | -0.722 | -0.922 | -0.327 | 0 | 0.013 | 0.013 | 0.368 | 0.46 | 0.75 | 0.66 | 0.63 | 0.5 | 0.58 |
| 6 | 93125582 | rs2730853 | -0.012 | -0.605 | -1.03 | -0.194 | -0.023 | 0.059 | 0.059 | 0 | 0.16 | 0.04 | 0.16 | 0.09 | 0.24 | 0.24 |
| 6 | 93126584 | rs13205064 | 1.561 | -0.582 | -1.056 | -0.08 | -0.01 | -0.047 | -0.047 | 0.375 | 0.08 | 0.01 | 0.2 | 0.28 | 0.04 | 0.17 |
| 6 | 93127953 | rs663809 | -0.559 | -0.56 | -0.679 | -0.278 | 0.022 | 0.018 | 0.018 | 0.465 | 0.37 | 0.27 | 0.4 | 0.34 | 0.36 | 0.32 |
| 6 | 93144865 | rs9345231 | 1.023 | -0.081 | -0.055 | 0.169 | -0.001 | 0.001 | 0.001 | 1.548 | 0.43 | 0.34 | 0.27 | 0.36 | 0.38 | 0.42 |
| 6 | 93147787 | rs2818142 | 0.191 | -0.123 | 0.162 | 0.01 | 0.001 | -0.004 | -0.004 | 0 | 0.59 | 0.55 | 0.52 | 0.67 | 0.51 | 0.47 |
| 6 | 93150800 | rs2730846 | 0.164 | -0.296 | 0.23 | 0.085 | 0.005 | -0.01 | -0.01 | 0 | 0.15 | 0.14 | 0.25 | 0.31 | 0.13 | 0.04 |
| 6 | 93157828 | rs879174 | 1.313 | -0.536 | 0.346 | -0.165 | -0.001 | -0.001 | -0.001 | 0.137 | 0.6 | 0.6 | 0.4 | 0.34 | 0.63 | 0.56 |

| | | | | | | | | | | | | | | | | |
|---|---|---|---|---|---|---|---|---|---|---|---|---|---|---|---|---|
| 6 | 93163594 | rs1427138 | 0.417 | 0.136 | 0.209 | -0.234 | 0.002 | -0.018 | -0.018 | 0 | 0.14 | 0.12 | 0.08 | 0.03 | 0.12 | 0.03 |
| 6 | 93166766 | rs12524154 | 1.951 | 0.254 | -0.016 | -0.226 | -0.009 | -0.025 | -0.025 | 0 | 0.36 | 0.1 | 0.21 | 0.33 | 0.33 | 0.43 |
| 6 | 93172063 | rs2555779 | 0.038 | -0.028 | 0.199 | 0.162 | 0.011 | 0.003 | 0.003 | 0.333 | 0.92 | 0.96 | 0.96 | 0.94 | 0.93 | 0.9 |
| 6 | 93178019 | rs7740559 | 1.617 | 0.013 | 0.394 | 0.18 | 0.005 | 0.014 | 0.014 | 0 | 0.56 | 0.32 | 0.35 | 0.34 | 0.57 | 0.55 |
| 6 | 93178604 | rs1427144 | 1.105 | 0.087 | 0.06 | 0.251 | -0.004 | 0.008 | 0.008 | 0 | 0.76 | 0.74 | 0.65 | 0.65 | 0.76 | 0.6 |
| 6 | 93179624 | rs10944594 | 0.467 | 0.479 | -0.371 | 0.569 | -0.014 | -0.036 | -0.036 | 0 | 0.07 | 0.01 | 0.2 | 0.28 | 0.05 | 0.16 |
| 6 | 93183255 | rs1427142 | 0.276 | 0.628 | 0.003 | 0.248 | 0.011 | -0.005 | -0.005 | 0 | 0.7 | 0.49 | 0.63 | 0.68 | 0.67 | 0.66 |
| 6 | 93183933 | rs2555773 | 1.379 | 0.357 | -0.315 | 0.06 | 0.003 | -0.009 | -0.009 | 0 | 0.66 | 0.34 | 0.47 | 0.43 | 0.69 | 0.58 |
| 6 | 93185340 | rs2730856 | 0.299 | 0.875 | -0.438 | 0.124 | -0.005 | -0.007 | -0.007 | 0 | 0.11 | 0.09 | 0.08 | 0.03 | 0.12 | 0.04 |
| 6 | 93185955 | rs2555772 | 1.14 | 0.67 | -0.142 | 0.449 | 0.004 | 0.018 | 0.018 | 0 | 0.86 | 0.85 | 0.73 | 0.68 | 0.81 | 0.68 |
| 6 | 93199724 | rs6935299 | 0.437 | 0.141 | -0.184 | 0.36 | -0.014 | 0.093 | 0.093 | 0 | 0.12 | 0.24 | 0.19 | 0.29 | 0.07 | 0.12 |
| 6 | 93212002 | rs958847 | 0.1 | 0.465 | -0.359 | 0.449 | -0.003 | -0.018 | -0.018 | 0.594 | 0.33 | 0.31 | 0.39 | 0.52 | 0.35 | 0.3 |
| 6 | 93243960 | rs784065 | 0.225 | 0.52 | -0.547 | 0.305 | -0.001 | -0.04 | -0.04 | 0 | 0.37 | 0.27 | 0.39 | 0.52 | 0.35 | 0.36 |
| 6 | 93255714 | rs783856 | -0.38 | 1.111 | -0.48 | 0.343 | 0.001 | 0.004 | 0.004 | 0 | 0.19 | 0.18 | 0.37 | 0.31 | 0.27 | 0.25 |
| 6 | 93267755 | rs1999063 | 0.992 | 0.989 | -0.897 | -0.166 | -0.008 | 0.083 | 0.083 | 0.555 | 0.3 | 0.08 | 0.19 | 0.14 | 0.28 | 0.3 |
| 6 | 93274584 | rs9362985 | 1.476 | 1.034 | -1.251 | -0.203 | -0.079 | -0.043 | -0.043 | 0.406 | 0.25 | 0.03 | 0.17 | 0.22 | 0.29 | 0.23 |
| 6 | 93277987 | rs12216384 | 0.617 | 0.8 | -0.81 | -0.104 | -0.032 | 0.06 | 0.06 | 0 | 0.37 | 0.16 | 0.24 | 0.26 | 0.38 | 0.35 |
| 6 | 93296511 | rs783845 | -0.225 | 0.718 | -0.094 | 0.158 | 0.026 | 0.076 | 0.076 | 0 | 0.2 | 0.34 | 0.39 | 0.34 | 0.26 | 0.25 |
| 6 | 93341099 | rs783616 | 0.195 | 0.899 | -0.51 | 0.117 | -0.042 | -0.045 | -0.045 | 1.008 | 0.11 | 0.03 | 0.36 | 0.34 | 0.22 | 0.21 |
| 6 | 93348842 | rs10498982 | 0.633 | 0.772 | -0.586 | -0.03 | -0.054 | 0.052 | 0.052 | 0 | 0.37 | 0.13 | 0.24 | 0.26 | 0.39 | 0.35 |
| 6 | 93349763 | rs1841258 | -1.049 | 0.69 | -0.312 | -0.056 | -0.015 | 0.14 | 0.14 | 0 | 0.89 | 0.67 | 0.81 | 0.74 | 0.94 | 0.9 |
| 6 | 93351530 | rs6926880 | 0.525 | 0.895 | -0.294 | -0.123 | -0.054 | 0.052 | 0.052 | 0 | 0.37 | 0.13 | 0.24 | 0.26 | 0.39 | 0.35 |
| 6 | 93368951 | rs13203066 | -0.474 | 1.279 | -0.199 | 0.116 | -0.009 | -0.004 | -0.004 | 0 | 0.13 | 0.12 | 0.31 | 0.35 | 0.24 | 0.23 |
| 6 | 93369655 | rs6918777 | 0.105 | 1.024 | -0.096 | -0.48 | -0.001 | -0.002 | -0.002 | 0 | 0.44 | 0.44 | 0.66 | 0.69 | 0.45 | 0.44 |
| 6 | 93372070 | rs12175910 | -0.381 | 1.185 | -0.476 | -0.411 | 0.007 | 0.055 | 0.055 | 1.281 | 0.14 | 0.17 | 0.32 | 0.39 | 0.18 | 0.17 |
| 6 | 93378241 | rs9345265 | -0.029 | 1.169 | -0.768 | -0.507 | 0 | -0.019 | -0.019 | 0 | 0.39 | 0.3 | 0.48 | 0.5 | 0.39 | 0.4 |
| 6 | 93378969 | rs1459810 | -0.146 | 1.405 | -0.897 | -0.238 | 0.007 | 0.057 | 0.057 | 1.462 | 0.12 | 0.17 | 0.32 | 0.39 | 0.16 | 0.15 |
| 6 | 93396190 | rs12528064 | 0.973 | 1.042 | -1.347 | -0.418 | 0.014 | 0.032 | 0.032 | 0.255 | 0.24 | 0.15 | 0.16 | 0.09 | 0.22 | 0.26 |
| 6 | 93402547 | rs9345268 | 0.636 | 1.963 | -1.453 | -0.067 | -0.007 | 0.005 | 0.005 | 0 | 0.27 | 0.16 | 0.19 | 0.22 | 0.29 | 0.35 |
| 6 | 93403758 | rs17654546 | 0.927 | 2.024 | -1.498 | -0.085 | -0.026 | 0.031 | 0.031 | 0 | 0.06 | 0.01 | 0.06 | 0 | 0.1 | 0.09 |
| 6 | 93408386 | rs2045591 | -0.522 | 1.983 | -1.219 | 0.172 | -0.074 | -0.097 | -0.097 | 0 | 0.13 | 0.02 | 0.34 | 0.43 | 0.21 | 0.16 |
| 6 | 93409510 | rs9345269 | -0.093 | 1.201 | -0.581 | -0.583 | -0.002 | 0.008 | 0.008 | 0 | 0.29 | 0.2 | 0.19 | 0.22 | 0.31 | 0.35 |
| 6 | 93429797 | rs1824423 | -0.207 | 0.883 | -0.456 | -1.682 | -0.003 | -0.061 | -0.061 | 0 | 0.34 | 0.33 | 0.21 | 0.05 | 0.34 | 0.31 |
| 6 | 93443329 | rs6908465 | 0.833 | 0.694 | -0.711 | -2.412 | 0.028 | 0.058 | 0.058 | 0 | 0.31 | 0.18 | 0.1 | 0.03 | 0.21 | 0.31 |
| 6 | 93452526 | rs12199478 | 0.11 | 0.277 | -0.645 | -2.191 | -0.004 | -0.058 | -0.058 | 0 | 0.34 | 0.32 | 0.21 | 0.05 | 0.34 | 0.31 |
| 6 | 93467158 | rs12211463 | 0.561 | 0.635 | -1.134 | -2.32 | -0.005 | 0.054 | 0.054 | 0 | 0.21 | 0.1 | 0.23 | 0.08 | 0.2 | 0.2 |
| 6 | 93473885 | rs6454911 | 0.253 | 0.38 | -0.929 | -2.063 | 0.017 | 0.104 | 0.104 | 0.251 | 0.27 | 0.1 | 0.15 | 0.04 | 0.28 | 0.32 |
| 6 | 93487547 | rs2674381 | -0.264 | 0.628 | -0.673 | -1.467 | 0.003 | 0.022 | 0.022 | 0 | 0.15 | 0.1 | 0.1 | 0.04 | 0.14 | 0.07 |
| 6 | 93488364 | rs16870237 | -0.53 | 0.514 | -0.219 | -1.331 | -0.006 | -0.009 | -0.009 | 0 | 0.17 | 0.23 | 0.1 | 0.12 | 0.15 | 0.1 |
| 6 | 93491572 | rs12190936 | 0.516 | 0.539 | -0.839 | -2.035 | -0.035 | 0.093 | 0.093 | 0.607 | 0.24 | 0.07 | 0.4 | 0.08 | 0.27 | 0.27 |
| 6 | 93494267 | rs9452114 | 0.933 | 0.334 | -0.253 | -2.265 | -0.015 | -0.074 | -0.074 | 1.323 | 0.2 | 0.28 | 0.1 | 0.02 | 0.2 | 0.17 |
| 6 | 93494372 | rs7454934 | 0.312 | 1.062 | -0.346 | -2.028 | 0.035 | 0.103 | 0.103 | 0 | 0.07 | 0.11 | 0.24 | 0.71 | 0.13 | 0.14 |
| 6 | 93512325 | rs9345274 | 0.104 | 1.036 | -0.311 | -2.094 | 0.041 | 0.105 | 0.105 | 0.067 | 0.07 | 0.14 | 0.24 | 0.73 | 0.14 | 0.14 |
| 6 | 93518927 | rs6899693 | -0.324 | 0.295 | -0.406 | -2.47 | 0.006 | 0.004 | 0.004 | 1.118 | 0.4 | 0.61 | 0.22 | 0.14 | 0.35 | 0.29 |
| 6 | 93520257 | rs9294536 | -0.267 | 0.313 | -0.407 | -2.451 | 0.005 | 0.004 | 0.004 | 0.737 | 0.39 | 0.6 | 0.22 | 0.14 | 0.35 | 0.29 |
| 6 | 93524382 | rs13205868 | 0.016 | 0.279 | -0.101 | -2.26 | -0.033 | -0.058 | -0.058 | 0 | 0.27 | 0.53 | 0.14 | 0.11 | 0.24 | 0.21 |
| 6 | 93530757 | rs9445025 | 0.016 | 0.279 | -0.078 | -2.26 | -0.033 | -0.058 | -0.058 | 0.601 | 0.27 | 0.53 | 0.14 | 0.11 | 0.24 | 0.21 |
| 6 | 93532022 | rs9445026 | 0.809 | 0.234 | 0.09 | -2.289 | -0.015 | -0.095 | -0.095 | 1.406 | 0.19 | 0.34 | 0.11 | 0.02 | 0.19 | 0.16 |
| 6 | 93532328 | rs16870296 | 0.022 | 0.445 | -0.187 | -2.05 | -0.008 | 0.016 | 0.016 | 0.574 | 0.07 | 0.14 | 0.04 | 0.09 | 0.06 | 0.05 |
| 6 | 93543673 | rs9452143 | 0.425 | 0.417 | -0.558 | -2.539 | 0.002 | -0.005 | -0.005 | 0 | 0.3 | 0.26 | 0.15 | 0.06 | 0.24 | 0.24 |
| 6 | 93545700 | rs6919358 | 0.788 | 0.422 | -0.867 | -2.95 | -0.004 | 0.117 | 0.117 | 0 | 0.33 | 0.14 | 0.22 | 0.04 | 0.32 | 0.26 |
| 6 | 93553425 | rs951556 | -0.414 | 0.472 | -0.273 | -2.983 | 0.003 | 0.013 | 0.013 | 1.445 | 0.49 | 0.51 | 0.73 | 0.85 | 0.54 | 0.6 |
| 6 | 93555944 | rs6909411 | 0.79 | 0.458 | -0.276 | -3.008 | 0.003 | -0.005 | -0.005 | 0 | 0.3 | 0.26 | 0.15 | 0.06 | 0.24 | 0.26 |
| 6 | 93561006 | rs9359995 | -1.28 | 0.399 | -0.196 | -2.825 | 0 | -0.022 | -0.022 | 0 | 0.8 | 0.77 | 0.89 | 0.98 | 0.8 | 0.85 |
| 6 | 93561150 | rs1040155 | -0.396 | 0.398 | -0.388 | -3.031 | -0.013 | -0.042 | -0.042 | 0 | 0.69 | 0.55 | 0.83 | 0.87 | 0.71 | 0.82 |
| 6 | 93564260 | rs9294541 | -1.075 | 0.492 | -0.376 | -2.97 | -0.002 | -0.007 | -0.007 | 0 | 0.78 | 0.72 | 0.86 | 0.9 | 0.76 | 0.83 |
| 6 | 93566103 | rs9294543 | 0.222 | 0.51 | -0.324 | -3.048 | 0.013 | -0.033 | -0.033 | 0 | 0.35 | 0.37 | 0.61 | 0.82 | 0.39 | 0.56 |
| 6 | 93566127 | rs9294544 | -0.79 | 0.522 | -0.702 | -2.677 | -0.003 | -0.013 | -0.013 | 0 | 0.77 | 0.69 | 0.86 | 0.9 | 0.76 | 0.83 |
| 6 | 93571654 | rs9351335 | 0.063 | 0.671 | -1.024 | -2.719 | -0.05 | 0.239 | 0.239 | 0 | 0.44 | 0.12 | 0.48 | 0.89 | 0.5 | 0.45 |
| 6 | 93575149 | rs4707762 | 0.076 | 0.667 | -0.988 | -2.699 | -0.056 | 0.159 | 0.159 | 0 | 0.44 | 0.15 | 0.47 | 0.81 | 0.48 | 0.45 |
| 6 | 93576022 | rs1810303 | 1.224 | 0.454 | -1.134 | -2.676 | -0.034 | 0.088 | 0.088 | 0.05 | 0.19 | 0.07 | 0.09 | 0.01 | 0.19 | 0.23 |
| 6 | 93577492 | rs9445034 | 0.21 | 0.457 | -1.114 | -2.637 | -0.039 | 0.232 | 0.232 | 0 | 0.45 | 0.15 | 0.48 | 0.89 | 0.5 | 0.45 |
| 6 | 93582968 | rs1572626 | 0.33 | 0.302 | -0.706 | -2.714 | -0.021 | -0.043 | -0.043 | 0 | 0.1 | 0.43 | 0.04 | 0 | 0.02 | 0 |
| 6 | 93584284 | rs9363010 | 0.606 | 0.52 | -0.833 | -2.766 | -0.013 | 0 | 0 | 0 | 0.07 | 0.02 | 0.25 | 0.73 | 0.13 | 0.23 |
| 6 | 93587834 | rs1591740 | -1.282 | 0.342 | -0.789 | -2.824 | 0.003 | 0 | 0 | 0 | 0.76 | 0.74 | 0.85 | 0.95 | 0.74 | 0.75 |
| 6 | 93588687 | rs9351340 | 0.2 | 0.345 | -0.743 | -2.768 | -0.02 | 0.186 | 0.186 | 0 | 0.45 | 0.18 | 0.48 | 0.89 | 0.5 | 0.45 |
| 6 | 93596223 | rs9351341 | 0.306 | 0.345 | -0.749 | -2.763 | -0.067 | 0.159 | 0.159 | 0 | 0.44 | 0.12 | 0.46 | 0.81 | 0.48 | 0.45 |
| 6 | 93596338 | rs2506908 | -0.602 | 0.319 | -0.721 | -2.781 | -0.011 | 0.101 | 0.101 | 0 | 0.57 | 0.68 | 0.74 | 0.86 | 0.53 | 0.52 |
| 6 | 93596433 | rs4707766 | 0.268 | 0.347 | -0.769 | -2.762 | -0.035 | 0.135 | 0.135 | 0 | 0.44 | 0.18 | 0.47 | 0.81 | 0.48 | 0.45 |

| | | | | | | | | | | | | | | | | |
|---|---|---|---|---|---|---|---|---|---|---|---|---|---|---|---|---|
| 6 | 93601661 | rs9353964 | 0.305 | 0.348 | -0.822 | -2.762 | -0.067 | 0.159 | 0.159 | 0 | 0.44 | 0.12 | 0.46 | 0.81 | 0.48 | 0.45 |
| 6 | 93606144 | rs2506929 | -1.196 | 0.355 | -0.777 | -2.765 | 0 | -0.011 | -0.011 | 0 | 0.75 | 0.69 | 0.63 | 0.92 | 0.74 | 0.74 |
| 6 | 93615450 | rs2506933 | -0.883 | 0.365 | -0.505 | -2.446 | -0.04 | 0.006 | 0.006 | 3.859 | 0.14 | 0.61 | 0.3 | 0.12 | 0.06 | 0.07 |
| 6 | 93616295 | rs2225922 | -0.571 | 0.409 | -0.539 | -2.476 | -0.001 | 0.034 | 0.034 | 0 | 0.54 | 0.55 | 0.51 | 0.81 | 0.5 | 0.45 |
| 6 | 93628360 | rs9363019 | 0.311 | 0.621 | -1.068 | -2.423 | -0.012 | 0.001 | 0.001 | 0 | 0.07 | 0.02 | 0.25 | 0.73 | 0.13 | 0.22 |
| 6 | 93629457 | rs2185056 | -0.978 | 0.642 | -1.149 | -2.576 | -0.011 | 0.13 | 0.13 | 0 | 0.58 | 0.72 | 0.76 | 0.93 | 0.54 | 0.52 |
| 6 | 93632765 | rs9452157 | 0.491 | 0.58 | -1.161 | -2.577 | -0.055 | 0.221 | 0.221 | 0.119 | 0.37 | 0.08 | 0.22 | 0.08 | 0.36 | 0.22 |
| 6 | 93636781 | rs2506950 | -0.652 | 0.606 | -0.936 | -2.227 | -0.006 | 0.131 | 0.131 | 0 | 0.58 | 0.72 | 0.76 | 0.93 | 0.55 | 0.52 |
| 6 | 93643091 | rs7742301 | -0.008 | 0.337 | -0.148 | -2.125 | 0.008 | 0.016 | 0.016 | 0 | 0.39 | 0.29 | 0.23 | 0.15 | 0.33 | 0.22 |
| 6 | 93646297 | rs1338266 | -0.692 | 0.523 | -0.506 | -2.573 | -0.037 | 0.078 | 0.078 | 0 | 0.64 | 0.79 | 0.74 | 0.95 | 0.58 | 0.65 |
| 6 | 93646679 | rs6923057 | 0.703 | 0.373 | -0.369 | -1.59 | 0.016 | 0.043 | 0.043 | 0.137 | 0.13 | 0.04 | 0.26 | 0.05 | 0.11 | 0.18 |
| 6 | 93674438 | rs17528777 | 0.214 | 0.596 | -0.222 | -1.369 | -0.017 | 0.016 | 0.016 | 0 | 0.07 | 0.04 | 0.13 | 0.02 | 0.14 | 0.06 |
| 6 | 93674832 | rs17437983 | 0.062 | 0.274 | -0.608 | -1.812 | 0.039 | 0.184 | 0.184 | 0.025 | 0.4 | 0.12 | 0.33 | 0.16 | 0.33 | 0.27 |
| 6 | 93678513 | rs554089 | 0.674 | 0.149 | -0.472 | -1.864 | -0.036 | 0.095 | 0.095 | 0 | 0.09 | 0.16 | 0.19 | 0.71 | 0.03 | 0.2 |
| 6 | 93684282 | rs17738153 | 0.491 | -0.297 | -0.625 | -1.885 | 0.017 | 0.134 | 0.134 | 0 | 0.26 | 0.06 | 0.13 | 0.04 | 0.21 | 0.12 |
| 6 | 93686244 | rs655096 | -1.114 | -0.085 | -0.535 | -1.568 | -0.003 | -0.037 | -0.037 | 1.236 | 0.62 | 0.64 | 0.7 | 0.93 | 0.53 | 0.58 |
| 6 | 93689951 | rs17529284 | 0.764 | 0.319 | -1.231 | -0.891 | 0.011 | 0.054 | 0.054 | 0.429 | 0.17 | 0.02 | 0.2 | 0.07 | 0.12 | 0.18 |
| 6 | 93690281 | rs10485369 | 0.434 | 0.335 | -1.198 | -0.852 | 0.005 | 0.026 | 0.026 | 0.197 | 0.21 | 0.04 | 0.24 | 0.11 | 0.16 | 0.24 |
| 6 | 93696324 | rs11751768 | -0.687 | 1.111 | -0.769 | -0.459 | -0.021 | 0.058 | 0.058 | 0.047 | 0.27 | 0.13 | 0.26 | 0.1 | 0.33 | 0.14 |
| 6 | 93697732 | rs2485800 | -1.089 | 0.972 | -0.385 | -0.062 | 0.001 | 0 | 0 | 0 | 0.1 | 0.11 | 0.14 | 0.13 | 0.1 | 0.08 |
| 6 | 93702141 | rs677242 | 0.284 | 0.755 | -0.371 | 0.162 | 0.012 | -0.011 | -0.011 | 0 | 0.41 | 0.3 | 0.38 | 0.51 | 0.39 | 0.41 |
| 6 | 93713805 | rs652520 | -0.578 | 0.653 | -0.235 | 0.072 | -0.034 | 0.002 | 0.002 | 0 | 0.57 | 0.85 | 0.55 | 0.66 | 0.49 | 0.57 |
| 6 | 93716579 | rs10455181 | 0.474 | 0.627 | -0.407 | -0.217 | -0.003 | 0.021 | 0.021 | 0 | 0.05 | 0.03 | 0.05 | 0 | 0.07 | 0.03 |
| 6 | 93718740 | rs9918469 | 0.304 | 0.74 | -0.181 | 0.018 | -0.001 | -0.034 | -0.034 | 0.415 | 0.08 | 0.12 | 0.04 | 0 | 0.06 | 0.05 |
| 6 | 93720269 | rs2787751 | 0.55 | 0.728 | -0.704 | 0.126 | 0.016 | -0.03 | -0.03 | 0 | 0.41 | 0.24 | 0.47 | 0.61 | 0.39 | 0.44 |
| 6 | 93722247 | rs1969811 | -1.352 | 0.673 | 0.147 | 0.055 | 0.016 | 0.007 | 0.007 | 0 | 0.16 | 0.58 | 0.17 | 0.15 | 0.1 | 0.16 |
| 6 | 93726713 | rs1324136 | -1.195 | 0.496 | -0.682 | -0.265 | 0.011 | 0.028 | 0.028 | 0 | 0.14 | 0.07 | 0.07 | 0 | 0.11 | 0.08 |
| 6 | 93728200 | rs9452184 | -0.475 | 0.673 | -1.522 | -1.258 | -0.03 | 0.224 | 0.224 | 0.738 | 0.52 | 0.87 | 0.35 | 0.14 | 0.52 | 0.33 |
| 6 | 93735074 | rs1590392 | -0.809 | 0.412 | -1.367 | -1.449 | 0.008 | 0.018 | 0.018 | 0 | 0.36 | 0.67 | 0.32 | 0.29 | 0.33 | 0.32 |
| 6 | 93735832 | rs2476880 | -0.763 | 0.328 | -0.767 | -1.51 | -0.015 | -0.048 | -0.048 | 0 | 0.3 | 0.49 | 0.28 | 0.23 | 0.28 | 0.3 |
| 6 | 93749161 | rs9294552 | -0.362 | 0.371 | -0.732 | -1.48 | 0 | 0.007 | 0.007 | 0 | 0.06 | 0.13 | 0.04 | 0.06 | 0.05 | 0.02 |
| 6 | 93749165 | rs1408296 | 0.208 | 0.27 | -0.859 | -1.618 | -0.006 | 0.045 | 0.045 | 0 | 0.08 | 0.02 | 0.07 | 0.01 | 0.09 | 0.1 |
| 6 | 93750416 | rs9452212 | -1.017 | 0.203 | -0.202 | -1.656 | -0.02 | -0.045 | -0.045 | 0.115 | 0.16 | 0.3 | 0.12 | 0.07 | 0.14 | 0.12 |
| 6 | 93752014 | rs7760335 | -0.773 | 0.201 | -0.107 | -1.634 | -0.023 | -0.043 | -0.043 | 0 | 0.13 | 0.32 | 0.11 | 0.06 | 0.11 | 0.11 |
| 6 | 93753233 | rs9452215 | -0.756 | 0.201 | -0.097 | -1.636 | -0.022 | -0.077 | -0.077 | 0.615 | 0.13 | 0.29 | 0.11 | 0.01 | 0.11 | 0.11 |
| 6 | 93755285 | rs9445061 | -0.612 | 0.156 | 0.023 | -1.642 | 0.003 | -0.015 | -0.015 | 2.438 | 0.22 | 0.26 | 0.2 | 0.23 | 0.24 | 0.21 |
| 6 | 93755554 | rs9445062 | -1.485 | 0.175 | -0.065 | -1.628 | 0.006 | 0.014 | 0.014 | 0 | 0.36 | 0.66 | 0.32 | 0.29 | 0.35 | 0.32 |
| 6 | 93759334 | rs6924566 | 0.576 | 0.163 | -0.152 | -1.618 | 0 | -0.002 | -0.002 | 0.192 | 0.07 | 0.09 | 0.03 | 0.03 | 0.05 | 0.03 |
| 6 | 93760118 | rs12216260 | -0.95 | 0.138 | -0.082 | -1.643 | -0.011 | -0.031 | -0.031 | 0.109 | 0.18 | 0.29 | 0.12 | 0.09 | 0.15 | 0.13 |
| 6 | 93765656 | rs576798 | 0.722 | 0.066 | -0.566 | -1.938 | 0 | -0.009 | -0.009 | 0 | 0.15 | 0.17 | 0.29 | 0.09 | 0.12 | 0.18 |
| 6 | 93781036 | rs2057541 | 0.819 | 0.066 | -0.519 | -2.04 | -0.011 | -0.044 | -0.044 | 0 | 0.66 | 0.4 | 0.57 | 0.8 | 0.72 | 0.7 |
| 6 | 93787786 | rs9342349 | -1.03 | -0.059 | -0.591 | -2.057 | 0.005 | -0.006 | -0.006 | 0 | 0.88 | 0.83 | 0.72 | 0.91 | 0.89 | 0.82 |
| 6 | 93790248 | rs1324115 | 0.849 | 0.015 | -0.55 | -2.032 | -0.016 | -0.065 | -0.065 | 0 | 0.67 | 0.44 | 0.57 | 0.8 | 0.72 | 0.7 |
| 6 | 93802038 | rs9353982 | 1.876 | 0.451 | -0.904 | -2.045 | -0.092 | -0.124 | -0.124 | 2.187 | 0.26 | 0.03 | 0.23 | 0.57 | 0.28 | 0.38 |
| 6 | 93806820 | rs9452240 | 0.34 | 0.453 | -0.888 | -2.047 | 0 | -0.015 | -0.015 | 0.054 | 0.4 | 0.38 | 0.34 | 0.23 | 0.45 | 0.32 |
| 6 | 93808577 | rs1359236 | -0.538 | 0.278 | -0.373 | -1.694 | 0 | -0.009 | -0.009 | 0.014 | 0.86 | 0.83 | 0.71 | 0.9 | 0.89 | 0.82 |
| 6 | 93812283 | rs1408289 | 0.112 | 0.277 | 0.054 | -1.328 | -0.003 | -0.02 | -0.02 | 0 | 0.09 | 0.16 | 0.05 | 0.04 | 0.07 | 0.03 |
| 6 | 93819929 | rs9445074 | -0.189 | 0.4 | -0.122 | -1.241 | 0.019 | 0.002 | 0.002 | 0 | 0.16 | 0.3 | 0.18 | 0.2 | 0.19 | 0.18 |
| 6 | 93821257 | rs9294559 | -0.852 | 0.33 | -0.154 | -0.937 | -0.019 | -0.064 | -0.064 | 0.059 | 0.19 | 0.42 | 0.14 | 0.11 | 0.16 | 0.12 |
| 6 | 93825256 | rs2209258 | -0.326 | 0.554 | -0.765 | -0.208 | 0.001 | -0.014 | -0.014 | 0 | 0.27 | 0.29 | 0.19 | 0.56 | 0.25 | 0.21 |
| 6 | 93826168 | rs768382 | -0.699 | 0.187 | -1.163 | -0.013 | 0.014 | -0.025 | -0.025 | 0 | 0.77 | 0.89 | 0.63 | 0.71 | 0.82 | 0.57 |
| 6 | 93833213 | rs10944649 | 0.498 | 0.507 | -1.463 | -0.34 | -0.016 | 0.018 | 0.018 | 4.265 | 0.22 | 0.12 | 0.28 | 0.16 | 0.21 | 0.15 |
| 6 | 93833436 | rs6938042 | 0.943 | 0.479 | -1.84 | -0.715 | -0.017 | -0.06 | -0.06 | 0.507 | 0.28 | 0.14 | 0.18 | 0.45 | 0.32 | 0.27 |
| 6 | 93834075 | rs16870805 | 1.483 | 0.399 | -1.304 | -0.926 | -0.006 | -0.023 | -0.023 | 0.011 | 0.11 | 0.13 | 0.06 | 0.04 | 0.08 | 0.08 |
| 6 | 93836577 | rs1575540 | 1.836 | 0.502 | -1.456 | -1.113 | -0.001 | 0.006 | 0.006 | 0 | 0.06 | 0.03 | 0.04 | 0 | 0.06 | 0.02 |
| 6 | 93837991 | rs1924474 | -0.588 | 0.436 | -1.506 | -1.077 | 0.003 | 0.007 | 0.007 | 3.018 | 0.38 | 0.64 | 0.35 | 0.26 | 0.3 | 0.23 |
| 6 | 93838942 | rs11961866 | 0.696 | 0.422 | -1.444 | -1.084 | 0.005 | -0.036 | -0.036 | 0.109 | 0.25 | 0.47 | 0.29 | 0.16 | 0.22 | 0.15 |
| 6 | 93842533 | rs1930934 | 0.11 | 0.376 | -1.502 | -0.979 | 0.003 | 0.007 | 0.007 | 3.018 | 0.62 | 0.36 | 0.65 | 0.74 | 0.7 | 0.77 |
| 6 | 93845633 | rs2147222 | 0.782 | 0.481 | -1.568 | -0.915 | -0.013 | 0.011 | 0.011 | 1.848 | 0.34 | 0.2 | 0.47 | 0.29 | 0.38 | 0.5 |
| 6 | 93847197 | rs571484 | 0.891 | 0.357 | -1.829 | -1.116 | -0.026 | 0.04 | 0.04 | 0.001 | 0.44 | 0.17 | 0.42 | 0.65 | 0.45 | 0.56 |
| 6 | 93848400 | rs538751 | -1.004 | 0.259 | -1.717 | -1.058 | -0.002 | 0.017 | 0.017 | 0.729 | 0.74 | 0.83 | 0.71 | 0.81 | 0.76 | 0.77 |
| 6 | 93849608 | rs7772178 | 1.469 | 0.521 | -1.736 | -1.109 | 0.001 | -0.022 | -0.022 | 0 | 0.12 | 0.14 | 0.07 | 0.04 | 0.12 | 0.08 |
| 6 | 93851245 | rs532830 | -1.583 | 0.512 | -1.771 | -1.1 | -0.016 | -0.036 | -0.036 | 0 | 0.86 | 0.96 | 0.78 | 0.84 | 0.88 | 0.85 |
| 6 | 93851328 | rs533741 | 0.612 | 0.524 | -1.748 | -1.131 | -0.012 | 0.034 | 0.034 | 0 | 0.55 | 0.63 | 0.47 | 0.65 | 0.48 | 0.57 |
| 6 | 93851462 | rs12215808 | 0.082 | 0.511 | -1.697 | -1.043 | -0.017 | 0.017 | 0.017 | 0.173 | 0.19 | 0.16 | 0.24 | 0.1 | 0.29 | 0.2 |
| 6 | 93854735 | rs9342351 | 1.927 | 0.22 | -1.57 | -0.944 | 0.011 | -0.01 | -0.01 | 0 | 0.16 | 0.05 | 0.22 | 0.17 | 0.11 | 0.26 |
| 6 | 93854804 | rs6935980 | 0.247 | 0.335 | -1.373 | -1.056 | -0.001 | 0 | 0 | 0 | 0.42 | 0.52 | 0.41 | 0.59 | 0.41 | 0.49 |
| 6 | 93856156 | rs9353991 | 1.168 | 0.448 | -1.265 | -1.089 | 0 | -0.023 | -0.023 | 0.101 | 0.3 | 0.4 | 0.32 | 0.26 | 0.27 | 0.36 |
| 6 | 93858168 | rs1324098 | 0.807 | 0.437 | -1.383 | -1.341 | -0.001 | 0.038 | 0.038 | 0.098 | 0.22 | 0.16 | 0.23 | 0.1 | 0.25 | 0.15 |

| | | | | | | | | | | | | | | | | |
|---|---|---|---|---|---|---|---|---|---|---|---|---|---|---|---|---|
| 6 | 93859226 | rs9445080 | 2.01 | 0.401 | -1.402 | -1.377 | 0.004 | -0.004 | -0.004 | 0 | 0.11 | 0.03 | 0.15 | 0.09 | 0.1 | 0.1 |
| 6 | 93859659 | rs998776 | 0.508 | 0.29 | -1.305 | -1.315 | -0.009 | 0.045 | 0.045 | 0.18 | 0.23 | 0.17 | 0.24 | 0.1 | 0.28 | 0.19 |
| 6 | 93867544 | rs7774264 | 1.238 | 0.271 | -1.28 | -1.293 | -0.002 | 0.115 | 0.115 | 0 | 0.14 | 0.23 | 0.09 | 0.38 | 0.14 | 0.14 |
| 6 | 93869122 | rs9353992 | -1.873 | 0.533 | -1.778 | -1.52 | -0.034 | -0.05 | -0.05 | 0 | 0.86 | 0.98 | 0.77 | 0.81 | 0.85 | 0.83 |
| 6 | 93875254 | rs9452269 | 1.477 | 0.633 | -1.77 | -1.579 | -0.007 | 0.008 | 0.008 | 0 | 0.14 | 0.03 | 0.18 | 0.09 | 0.18 | 0.1 |
| 6 | 93876819 | rs1541123 | -0.647 | 0.798 | -1.779 | -1.483 | -0.007 | 0.016 | 0.016 | 3.947 | 0.67 | 0.82 | 0.65 | 0.75 | 0.69 | 0.69 |
| 6 | 93880914 | rs16870964 | 1.724 | 0.82 | -1.534 | -1.199 | 0.002 | -0.019 | -0.019 | 0 | 0.06 | 0.12 | 0.05 | 0.01 | 0.07 | 0.05 |
| 6 | 93880964 | rs16870967 | 1.217 | 0.776 | -1.637 | -0.983 | 0 | -0.011 | -0.011 | 0 | 0.13 | 0.13 | 0.1 | 0.04 | 0.16 | 0.09 |
| 6 | 93881538 | rs4288174 | 0.633 | 0.449 | -1.781 | -0.517 | 0.014 | -0.042 | -0.042 | 2.101 | 0.29 | 0.19 | 0.31 | 0.55 | 0.25 | 0.41 |
| 6 | 93882387 | rs6915708 | -0.007 | 0.824 | -1.326 | -0.311 | -0.001 | 0.01 | 0.01 | 0 | 0.18 | 0.13 | 0.11 | 0.07 | 0.16 | 0.14 |
| 6 | 93886912 | rs914924 | -0.457 | 0.954 | -1.567 | 0.29 | -0.065 | -0.03 | -0.03 | 0 | 0.83 | 0.99 | 0.76 | 0.8 | 0.82 | 0.78 |
| 6 | 93890176 | rs11966984 | 0.554 | 0.269 | -1.522 | -0.088 | 0.012 | 0.015 | 0.015 | 0 | 0.18 | 0.16 | 0.09 | 0.05 | 0.13 | 0.07 |
| 6 | 93892299 | rs6921600 | 0.232 | 0.219 | -1.133 | -0.194 | -0.021 | -0.044 | -0.044 | 0.097 | 0.37 | 0.53 | 0.19 | 0.16 | 0.28 | 0.17 |
| 6 | 93898633 | rs7772899 | -0.324 | 0.185 | -1.631 | -0.009 | 0.002 | 0.043 | 0.043 | 0 | 0.53 | 0.9 | 0.43 | 0.35 | 0.41 | 0.45 |
| 6 | 93898896 | rs13205265 | 0.363 | 0.131 | -1.078 | -0.023 | -0.023 | -0.074 | -0.074 | 10.694 | 0.36 | 0.49 | 0.19 | 0.08 | 0.28 | 0.17 |
| 6 | 93901016 | rs1324103 | 0.176 | 0.395 | -1.573 | 0.413 | -0.057 | -0.098 | -0.098 | 0 | 0.32 | 0.11 | 0.41 | 0.45 | 0.42 | 0.33 |
| 6 | 93906373 | rs6931820 | 0.718 | 0.621 | -1.777 | 0.347 | -0.049 | -0.073 | -0.073 | 0 | 0.16 | 0.06 | 0.33 | 0.42 | 0.28 | 0.23 |
| 6 | 93906936 | rs4391235 | -0.862 | 0.116 | -1.111 | 0.157 | -0.004 | 0.021 | 0.021 | 0 | 0.54 | 0.56 | 0.52 | 0.65 | 0.55 | 0.4 |
| 6 | 93909751 | rs7745734 | 0.526 | 0.546 | -0.946 | 0.974 | -0.031 | -0.038 | -0.038 | 0 | 0.11 | 0.05 | 0.19 | 0.28 | 0.18 | 0.12 |
| 6 | 93909926 | rs1575536 | -0.222 | -0.009 | -0.517 | 0.563 | -0.038 | 0.01 | 0.01 | 0 | 0.21 | 0.38 | 0.12 | 0.17 | 0.15 | 0.16 |
| 6 | 93913423 | rs7776099 | 0.376 | 0.022 | -0.588 | 0.559 | -0.019 | 0.062 | 0.062 | 0 | 0.22 | 0.45 | 0.12 | 0.27 | 0.17 | 0.15 |
| 6 | 93915413 | rs635899 | 0.4 | 0.073 | -1.072 | -0.233 | -0.028 | 0.049 | 0.049 | 0 | 0.14 | 0.01 | 0.09 | 0.04 | 0.16 | 0.1 |
| 6 | 93917260 | rs12210498 | 0.568 | 0.125 | -1.397 | -0.473 | -0.018 | -0.054 | -0.054 | 4.571 | 0.25 | 0.15 | 0.36 | 0.53 | 0.28 | 0.33 |
| 6 | 93918878 | rs364774 | -1.022 | 0.089 | -1.354 | -0.677 | -0.012 | 0.001 | 0.001 | 0 | 0.76 | 0.83 | 0.7 | 0.77 | 0.71 | 0.83 |
| 6 | 93918959 | rs388461 | 1.512 | -0.25 | -0.971 | -0.823 | -0.027 | -0.087 | -0.087 | 0.219 | 0.18 | 0.25 | 0.08 | 0 | 0.12 | 0.02 |
| 6 | 93919659 | rs1486761 | -0.061 | 0.088 | -0.979 | -0.618 | -0.001 | 0 | 0 | 0 | 0.42 | 0.45 | 0.52 | 0.66 | 0.44 | 0.7 |
| 6 | 93920448 | rs7747056 | 0.835 | 0.083 | -0.87 | -0.913 | -0.001 | -0.026 | -0.026 | 0.59 | 0.17 | 0.32 | 0.16 | 0.13 | 0.16 | 0.37 |
| 6 | 93921556 | rs631175 | -0.224 | 0.174 | -1.206 | -1.046 | -0.002 | 0.051 | 0.051 | 2.224 | 0.48 | 0.3 | 0.54 | 0.71 | 0.49 | 0.77 |
| 6 | 93922926 | rs782706 | 0.375 | 0.27 | -1.189 | -1.307 | -0.042 | 0.009 | 0.009 | 0.208 | 0.45 | 0.14 | 0.44 | 0.58 | 0.46 | 0.57 |
| 6 | 93928345 | rs782714 | 0.335 | 0.262 | -1.161 | -1.311 | -0.015 | 0.021 | 0.021 | 0.124 | 0.46 | 0.21 | 0.45 | 0.58 | 0.46 | 0.57 |
| 6 | 93931092 | rs12189899 | 0.737 | 0.269 | -1.369 | -1.494 | -0.025 | 0.077 | 0.077 | 1.085 | 0.24 | 0.04 | 0.14 | 0.12 | 0.21 | 0.22 |
| 6 | 93931365 | rs17543146 | 2.14 | 0.317 | -1.333 | -1.521 | -0.002 | 0.009 | 0.009 | 0 | 0.11 | 0.08 | 0.06 | 0.01 | 0.12 | 0.07 |
| 6 | 93941861 | rs1486757 | 0.683 | 0.799 | -1.579 | -1.204 | 0.01 | 0.029 | 0.029 | 1.485 | 0.22 | 0.09 | 0.19 | 0.15 | 0.21 | 0.45 |
| 6 | 93942185 | rs1486756 | -0.834 | 0.878 | -1.484 | -1.127 | -0.001 | 0 | 0 | 2.06 | 0.7 | 0.32 | 0.59 | 0.7 | 0.68 | 0.79 |
| 6 | 93943117 | rs1572102 | 1.036 | 1.151 | -1.72 | -0.88 | -0.058 | -0.125 | -0.125 | 1.667 | 0.23 | 0.05 | 0.25 | 0.42 | 0.26 | 0.13 |
| 6 | 93947044 | rs9354001 | 0.585 | 1.308 | -0.964 | -0.556 | 0.007 | -0.003 | -0.003 | 0 | 0.13 | 0.08 | 0.07 | 0.14 | 0.13 | 0.34 |
| 6 | 93947307 | rs9354002 | 0.693 | 1.459 | -0.747 | -0.323 | -0.009 | 0.03 | 0.03 | 1.983 | 0.23 | 0.09 | 0.21 | 0.15 | 0.24 | 0.46 |
| 6 | 93947347 | rs17446388 | 0.563 | 1.601 | -0.629 | -0.258 | -0.005 | 0.016 | 0.016 | 0.04 | 0.12 | 0.06 | 0.09 | 0.05 | 0.16 | 0.06 |
| 6 | 93956769 | rs450238 | -1.084 | 1.626 | -0.393 | -0.192 | 0.005 | -0.003 | -0.003 | 0 | 0.78 | 0.69 | 0.7 | 0.83 | 0.72 | 0.86 |
| 6 | 93959361 | rs3823231 | 0.306 | 1.712 | -0.961 | -0.191 | -0.022 | 0.015 | 0.015 | 0.078 | 0.57 | 0.28 | 0.47 | 0.65 | 0.5 | 0.73 |
| 6 | 93961571 | rs1386274 | -0.835 | 1.611 | -0.589 | 0.096 | 0.003 | -0.013 | -0.013 | 2.128 | 0.74 | 0.41 | 0.69 | 0.82 | 0.72 | 0.86 |
| 6 | 93973569 | rs345725 | 0.516 | 1.613 | -1.768 | 0.8 | 0.005 | 0.04 | 0.04 | 0 | 0.06 | 0.11 | 0.13 | 0.13 | 0.07 | 0.07 |
| 6 | 93997137 | rs12663435 | 0.065 | 0.93 | -1.715 | 0.625 | 0.01 | -0.071 | -0.071 | 0 | 0.35 | 0.18 | 0.27 | 0.49 | 0.26 | 0.27 |
| 6 | 94004485 | rs346462 | -0.201 | 1.329 | -1.356 | 0.505 | -0.007 | -0.002 | -0.002 | 0 | 0.3 | 0.21 | 0.33 | 0.31 | 0.31 | 0.5 |
| 6 | 94005099 | rs17448051 | 0.307 | 1.667 | -1.579 | 0.273 | -0.03 | 0.012 | 0.012 | 0.058 | 0.07 | 0.01 | 0.08 | 0.03 | 0.14 | 0.04 |
| 6 | 94008535 | rs9363058 | -0.185 | 1.465 | -2.019 | -0.106 | -0.049 | 0.08 | 0.08 | 0 | 0.3 | 0.06 | 0.24 | 0.17 | 0.3 | 0.3 |
| 6 | 94023695 | rs164290 | -0.284 | 0.853 | -0.713 | 0.288 | -0.013 | 0.024 | 0.024 | 0 | 0.33 | 0.56 | 0.25 | 0.48 | 0.3 | 0.29 |
| 6 | 94025584 | rs164294 | -0.9 | 0.906 | -1.264 | -0.048 | -0.031 | -0.031 | -0.031 | 0 | 0.76 | 0.9 | 0.64 | 0.73 | 0.71 | 0.75 |
| 6 | 94028965 | rs164298 | 0.301 | 0.787 | -0.947 | 0.523 | -0.02 | -0.003 | -0.003 | 0 | 0.33 | 0.47 | 0.24 | 0.31 | 0.3 | 0.29 |
| 6 | 94029815 | rs164301 | -0.513 | 0.617 | -1.138 | 0.307 | -0.003 | 0.031 | 0.031 | 0 | 0.54 | 0.74 | 0.56 | 0.65 | 0.56 | 0.51 |
| 6 | 94031097 | rs1492964 | 1.309 | 0.583 | -1.128 | 0.3 | -0.051 | -0.069 | -0.069 | 0.063 | 0.16 | 0.03 | 0.3 | 0.34 | 0.22 | 0.21 |
| 6 | 94031651 | rs164302 | 0.408 | 0.589 | -1.046 | 0.339 | -0.012 | -0.003 | -0.003 | 0 | 0.33 | 0.56 | 0.25 | 0.31 | 0.3 | 0.29 |
| 6 | 94034201 | rs7754087 | 0.715 | 0.571 | -0.985 | 0.35 | 0.015 | -0.019 | -0.019 | 0.888 | 0.28 | 0.17 | 0.2 | 0.31 | 0.25 | 0.28 |
| 6 | 94037400 | rs164556 | -0.758 | 0.409 | -1.105 | 0.375 | -0.008 | -0.002 | -0.002 | 0 | 0.67 | 0.77 | 0.61 | 0.69 | 0.66 | 0.71 |
| 6 | 94041109 | rs16871170 | 0.819 | 0.153 | -1.116 | 0.286 | -0.016 | -0.05 | -0.05 | 0 | 0.09 | 0.01 | 0.03 | 0.16 | 0.06 | 0.09 |
| 6 | 94049334 | rs13200186 | -0.06 | 0.344 | -0.889 | 0.258 | 0.003 | 0.016 | 0.016 | 0.079 | 0.61 | 0.37 | 0.68 | 0.74 | 0.63 | 0.66 |
| 6 | 94051719 | rs13200697 | 0.508 | 0.412 | -1.092 | 0.229 | -0.027 | 0.082 | 0.082 | 0 | 0.37 | 0.22 | 0.32 | 0.16 | 0.42 | 0.34 |
| 6 | 94051765 | rs7751214 | 1.034 | 0.5 | -1.143 | 0.624 | -0.002 | -0.034 | -0.034 | 0 | 0.21 | 0.12 | 0.28 | 0.32 | 0.21 | 0.24 |
| 6 | 94051912 | rs13203837 | 1.067 | 0.958 | -1.033 | 0.233 | -0.068 | 0.064 | 0.064 | 0.323 | 0.2 | 0.04 | 0.25 | 0.07 | 0.27 | 0.14 |
| 6 | 94057477 | rs164533 | 0.093 | 0.815 | -0.564 | 0.406 | 0.01 | 0.072 | 0.072 | 0.356 | 0.11 | 0.31 | 0.13 | 0.21 | 0.08 | 0.09 |
| 6 | 94060369 | rs164538 | 0.488 | 0.771 | -1.454 | 0.195 | -0.01 | 0.011 | 0.011 | 0 | 0.34 | 0.59 | 0.41 | 0.56 | 0.26 | 0.35 |
| 6 | 94061405 | rs164539 | 0.104 | 0.774 | -1.503 | 0.123 | 0.034 | -0.002 | -0.002 | 0.658 | 0.45 | 0.68 | 0.44 | 0.6 | 0.33 | 0.43 |
| 6 | 94062054 | rs17688085 | 1.568 | 0.822 | -1.51 | 0.105 | -0.001 | 0.031 | 0.031 | 0 | 0.09 | 0.02 | 0.02 | 0.03 | 0.07 | 0.07 |
| 6 | 94064406 | rs164540 | -0.254 | 1.133 | -1.775 | 0.297 | -0.076 | 0.145 | 0.145 | 0.834 | 0.6 | 0.89 | 0.65 | 0.83 | 0.55 | 0.65 |
| 6 | 94065683 | rs2450423 | 0.553 | 1.113 | -1.739 | 0.318 | 0.031 | -0.002 | -0.002 | 0.309 | 0.44 | 0.68 | 0.44 | 0.6 | 0.33 | 0.43 |
| 6 | 94067981 | rs164544 | 0.078 | 1.05 | -2.203 | -0.258 | -0.077 | 0.147 | 0.147 | 0.835 | 0.6 | 0.89 | 0.65 | 0.83 | 0.55 | 0.65 |
| 6 | 94071608 | rs164548 | -0.008 | 0.771 | -1.487 | 0.205 | 0.01 | 0.063 | 0.063 | 0 | 0.78 | 0.89 | 0.9 | 0.92 | 0.82 | 0.8 |
| 6 | 94076418 | rs4707796 | -0.08 | 0.582 | -1.124 | 0.844 | 0.007 | -0.023 | -0.023 | 0.005 | 0.12 | 0.09 | 0.13 | 0.31 | 0.1 | 0.16 |
| 6 | 94082008 | rs2006999 | 1.035 | 0.771 | -1.632 | 0.405 | -0.036 | 0.104 | 0.104 | 0 | 0.25 | 0.05 | 0.12 | 0.07 | 0.22 | 0.16 |

| | | | | | | | | | | | | | | | | |
|---|---|---|---|---|---|---|---|---|---|---|---|---|---|---|---|---|
| 6 | 94083123 | rs351328 | -0.965 | 0.642 | -1.634 | 0.225 | 0.002 | -0.002 | -0.002 | 0 | 0.92 | 0.94 | 0.94 | 1 | 0.92 | 0.98 |
| 6 | 94089337 | rs16880204 | 1.363 | 0.789 | -1.779 | 0.23 | -0.03 | 0.089 | 0.089 | 0 | 0.23 | 0.02 | 0.09 | 0.07 | 0.17 | 0.15 |
| 6 | 94103626 | rs2052636 | 0.227 | 0.596 | -1.57 | 0.327 | 0.001 | 0.007 | 0.007 | 0 | 0.13 | 0.14 | 0.14 | 0.31 | 0.1 | 0.18 |
| 6 | 94104527 | rs549965 | -0.352 | 0.978 | -1.507 | 0.503 | -0.016 | -0.006 | -0.006 | 0 | 0.57 | 0.81 | 0.49 | 0.59 | 0.52 | 0.48 |
| 6 | 94108873 | rs564158 | -0.116 | 1.264 | -0.678 | 0.744 | -0.029 | 0.037 | 0.037 | 0 | 0.24 | 0.56 | 0.22 | 0.31 | 0.19 | 0.19 |
| 6 | 94112802 | rs625822 | 0.198 | 2.023 | -0.921 | 0.647 | -0.087 | 0.065 | 0.065 | 0 | 0.79 | 0.98 | 0.77 | 0.9 | 0.73 | 0.8 |
| 6 | 94117546 | rs3799792 | -0.337 | 1.773 | -0.066 | 1.355 | 0.025 | 0.051 | 0.051 | 0 | 0.07 | 0.13 | 0.16 | 0.19 | 0.11 | 0.12 |
| 6 | 94130024 | rs351331 | -0.444 | 1.578 | -0.127 | 1.58 | -0.055 | -0.117 | -0.117 | 2.896 | 0.26 | 0.09 | 0.5 | 0.42 | 0.36 | 0.38 |
| 6 | 94133927 | rs519420 | -0.418 | 1.571 | 0.947 | 1.558 | 0.004 | 0.017 | 0.017 | 0 | 0.25 | 0.27 | 0.51 | 0.43 | 0.36 | 0.38 |
| 6 | 94143528 | rs481992 | 0 | 1.913 | 1.088 | 1.295 | -0.044 | 0.016 | 0.016 | 0 | 0.89 | 0.98 | 0.88 | 0.91 | 0.79 | 0.8 |
| 6 | 94162298 | rs591356 | -0.652 | 1.284 | 1.41 | 1.283 | 0.021 | 0.022 | 0.022 | 0.009 | 0.31 | 0.46 | 0.56 | 0.51 | 0.42 | 0.45 |
| 6 | 94206244 | rs9452333 | 0.762 | 1.372 | 1.435 | 1.37 | -0.003 | -0.007 | -0.007 | 0.411 | 0.63 | 0.3 | 0.43 | 0.6 | 0.57 | 0.64 |
| 6 | 94216545 | rs967879 | -1.349 | 1.851 | 1.996 | 1.394 | 0.085 | 0.043 | 0.043 | 0.417 | 0.1 | 0.59 | 0.19 | 0.12 | 0.12 | 0.1 |
| 6 | 94240619 | rs9452355 | -1.184 | 1.884 | 2.246 | 1.426 | 0.124 | 0.084 | 0.084 | 0.027 | 0.09 | 0.51 | 0.18 | 0.12 | 0.12 | 0.1 |
| 6 | 94246104 | rs7746862 | -0.098 | 1.924 | 2.131 | 1.481 | -0.021 | -0.034 | -0.034 | 0 | 0.78 | 0.87 | 0.61 | 0.74 | 0.68 | 0.75 |
| 6 | 94269760 | rs12196821 | -1.133 | 1.909 | 2.062 | 1.466 | 0.13 | 0.089 | 0.089 | 0.02 | 0.09 | 0.46 | 0.18 | 0.12 | 0.12 | 0.1 |
| 6 | 94274392 | rs6940694 | -0.84 | 1.908 | 2.077 | 1.45 | 0.168 | 0.062 | 0.062 | 0.722 | 0.05 | 0.39 | 0.15 | 0 | 0.06 | 0.02 |
| 6 | 94287959 | rs2200875 | 0.226 | 1.969 | 1.583 | 1.591 | -0.005 | -0.037 | -0.037 | 0.002 | 0.73 | 0.8 | 0.52 | 0.59 | 0.67 | 0.65 |
| 6 | 94306412 | rs1906965 | 0.171 | 1.696 | 0.223 | 1.403 | -0.004 | -0.014 | -0.014 | 1.335 | 0.75 | 0.77 | 0.56 | 0.59 | 0.67 | 0.67 |
| 6 | 94316889 | rs9351362 | -0.365 | 1.953 | -0.251 | 1.317 | -0.039 | -0.043 | -0.043 | 0 | 0.17 | 0.06 | 0.19 | 0.25 | 0.25 | 0.34 |
| 6 | 94327480 | rs9363099 | 0.425 | 1.224 | 0.111 | 1.161 | -0.005 | -0.024 | -0.024 | 0 | 0.74 | 0.76 | 0.56 | 0.59 | 0.67 | 0.67 |
| 6 | 94333920 | rs11966981 | -0.843 | 0.588 | 0.458 | 0.273 | -0.03 | -0.121 | -0.121 | 0.223 | 0.22 | 0.33 | 0.19 | 0.01 | 0.16 | 0.07 |
| 6 | 94348038 | rs1906980 | -0.344 | 1.198 | -0.024 | 0.377 | -0.075 | -0.011 | -0.011 | 0.059 | 0.27 | 0.15 | 0.3 | 0.31 | 0.47 | 0.34 |
| 6 | 94369637 | rs7757023 | -0.792 | 0.958 | 0.558 | 0.912 | 0.025 | 0.052 | 0.052 | 0 | 0.34 | 0.5 | 0.38 | 0.48 | 0.54 | 0.41 |
| 6 | 94403778 | rs1577595 | -1.674 | 0.658 | 0.664 | 1.377 | 0.067 | 0.069 | 0.069 | 0 | 0.58 | 0.96 | 0.72 | 0.72 | 0.72 | 0.56 |
| 6 | 94408714 | rs6933279 | 0.021 | 0.651 | 0.819 | 1.43 | -0.024 | -0.103 | -0.103 | 0.119 | 0.21 | 0.31 | 0.19 | 0.01 | 0.17 | 0.07 |
| 6 | 94419927 | rs6454977 | -1.638 | 0.674 | 0.428 | 1.468 | 0.068 | 0.069 | 0.069 | 0 | 0.58 | 0.96 | 0.72 | 0.72 | 0.72 | 0.55 |
| 6 | 94437933 | rs6931842 | -0.269 | 0.987 | 0.568 | 1.575 | -0.076 | -0.043 | -0.043 | 0 | 0.28 | 0.14 | 0.32 | 0.39 | 0.47 | 0.36 |
| 6 | 94460497 | rs1414614 | -0.837 | 1.11 | 0.534 | 1.129 | 0.05 | 0.075 | 0.075 | 0.013 | 0.28 | 0.47 | 0.34 | 0.47 | 0.3 | 0.45 |
| 6 | 94470429 | rs1360909 | -1.828 | -0.176 | -0.643 | 0.487 | 0.059 | 0.153 | 0.153 | 0 | 0.73 | 0.97 | 0.89 | 0.96 | 0.8 | 0.84 |
| 6 | 94481162 | rs9345394 | -0.341 | -0.136 | -0.43 | 0.649 | -0.025 | -0.037 | -0.037 | 0 | 0.27 | 0.19 | 0.33 | 0.48 | 0.3 | 0.46 |
| 6 | 94487858 | rs9354042 | -1.87 | 0.001 | -0.332 | 0.513 | 0.057 | 0.121 | 0.121 | 0 | 0.74 | 0.92 | 0.88 | 0.97 | 0.8 | 0.84 |
| 6 | 94490153 | rs6939009 | -0.497 | -0.39 | 0.107 | 0.382 | 0.008 | -0.009 | -0.009 | 0 | 0.32 | 0.57 | 0.31 | 0.31 | 0.31 | 0.4 |
| 6 | 94491539 | rs1414615 | 0.21 | -0.085 | -0.209 | 0.562 | -0.092 | -0.111 | -0.111 | 0 | 0.36 | 0.11 | 0.49 | 0.48 | 0.48 | 0.37 |
| 6 | 94500169 | rs1537834 | 1.101 | -0.539 | -0.062 | 0.342 | 0.105 | -0.012 | -0.012 | 0 | 0.42 | 0.16 | 0.42 | 0.41 | 0.32 | 0.27 |
| 6 | 94567726 | rs1937359 | -0.179 | 0.038 | 0.293 | 0.216 | 0.002 | -0.033 | -0.033 | 0 | 0.4 | 0.19 | 0.45 | 0.57 | 0.53 | 0.68 |
| 6 | 94568431 | rs7765999 | -0.459 | 0.013 | 0.327 | 0.231 | 0.015 | -0.002 | -0.002 | 1.068 | 0.43 | 0.34 | 0.49 | 0.57 | 0.57 | 0.69 |
| 6 | 94624568 | rs11753100 | -0.227 | 0.035 | 0.13 | 0.005 | 0.006 | 0.012 | 0.012 | 0.908 | 0.4 | 0.2 | 0.46 | 0.62 | 0.55 | 0.69 |
| 6 | 94635123 | rs1219031 | -0.306 | 0.043 | 0.199 | -0.023 | 0.011 | 0.022 | 0.022 | 1.004 | 0.41 | 0.29 | 0.47 | 0.63 | 0.55 | 0.7 |
| 6 | 94640147 | rs9354063 | -0.297 | -0.231 | 0.411 | -0.09 | -0.009 | 0.043 | 0.043 | 0 | 0.09 | 0.12 | 0.12 | 0.33 | 0.05 | 0.07 |
| 6 | 94646136 | rs12525634 | -0.232 | -0.289 | 0.435 | -0.134 | 0.013 | 0.02 | 0.02 | 0.87 | 0.41 | 0.25 | 0.48 | 0.63 | 0.55 | 0.71 |
| 6 | 94652023 | rs2633615 | -1.729 | -0.713 | -0.394 | -0.454 | 0.022 | 0.181 | 0.181 | 0 | 0.7 | 0.97 | 0.84 | 0.97 | 0.75 | 0.81 |
| 6 | 94660447 | rs7449883 | -0.167 | 0.318 | 0.561 | -0.293 | -0.03 | -0.015 | -0.015 | 3.284 | 0.93 | 0.9 | 0.99 | 0.98 | 0.99 | 0.99 |
| 6 | 94673014 | rs4706440 | 0.989 | 0.498 | 0.776 | -0.27 | 0.027 | 0.039 | 0.039 | 1.537 | 0.51 | 0.39 | 0.5 | 0.32 | 0.44 | 0.29 |
| 6 | 94677177 | rs1488309 | 1.014 | 0.5 | 0.6 | -0.276 | 0.067 | 0.098 | 0.098 | 2.698 | 0.49 | 0.11 | 0.48 | 0.35 | 0.44 | 0.29 |
| 6 | 94708712 | rs9452527 | -0.988 | 1.093 | 0.68 | -0.332 | -0.019 | 0.015 | 0.015 | 0.276 | 0.41 | 0.24 | 0.46 | 0.63 | 0.49 | 0.6 |
| 6 | 94715324 | rs1488307 | 0.646 | 1.529 | 0.821 | -0.37 | -0.004 | 0 | 0 | 0 | 0.73 | 0.75 | 0.81 | 0.9 | 0.65 | 0.76 |
| 6 | 94720600 | rs9351398 | 0.181 | 1.435 | 1.591 | -0.358 | -0.004 | 0.042 | 0.042 | 0 | 0.1 | 0.13 | 0.11 | 0.3 | 0.08 | 0.18 |
| 6 | 94726087 | rs1352380 | 1.792 | 1.275 | 1.594 | -0.439 | -0.104 | -0.038 | -0.038 | 0 | 0.57 | 0.08 | 0.67 | 0.56 | 0.56 | 0.47 |
| 6 | 94743025 | rs679881 | 0.204 | 1.396 | 1.598 | -0.395 | -0.135 | 0.062 | 0.062 | 0 | 0.78 | 0.97 | 0.83 | 0.91 | 0.66 | 0.77 |
| 6 | 94748716 | rs9345428 | 0.205 | 1.263 | 1.799 | -0.309 | -0.006 | 0.044 | 0.044 | 0 | 0.11 | 0.15 | 0.11 | 0.3 | 0.08 | 0.18 |
| 6 | 94756060 | rs12663518 | 0.301 | 1.538 | 1.468 | -0.35 | -0.054 | -0.044 | -0.044 | 0 | 0.1 | 0.02 | 0.29 | 0.53 | 0.18 | 0.25 |
| 6 | 94757719 | rs16871829 | 0.251 | 1.362 | 1.811 | -0.533 | 0.004 | 0.036 | 0.036 | 0 | 0.06 | 0.32 | 0.02 | 0.05 | 0.02 | 0.11 |
| 6 | 94761761 | rs667243 | 0.422 | 1.559 | 1.443 | -0.586 | -0.046 | 0.04 | 0.04 | 0 | 0.77 | 0.87 | 0.83 | 0.92 | 0.67 | 0.77 |
| 6 | 94771185 | rs16871880 | 0.501 | 1.437 | 1.585 | -0.559 | -0.027 | -0.004 | -0.004 | 0 | 0.06 | 0.12 | 0.01 | 0.05 | 0.02 | 0.12 |
| 6 | 94773306 | rs1338341 | 0.508 | 1.533 | 1.566 | -0.641 | 0.036 | -0.039 | -0.039 | 0.013 | 0.77 | 0.75 | 0.82 | 0.92 | 0.67 | 0.78 |
| 6 | 94805705 | rs647683 | 1.276 | 1.515 | 1.107 | -0.424 | -0.006 | -0.003 | -0.003 | 0 | 0.66 | 0.68 | 0.68 | 0.6 | 0.59 | 0.58 |
| 6 | 94811175 | rs574793 | 1.837 | 1.487 | 1.015 | -0.593 | 0.045 | 0.093 | 0.093 | 0.274 | 0.53 | 0.69 | 0.42 | 0.03 | 0.41 | 0.24 |
| 6 | 94824086 | rs640252 | 1.322 | 1.887 | 0.915 | -0.351 | 0.031 | 0.012 | 0.012 | 0 | 0.68 | 0.43 | 0.67 | 0.61 | 0.6 | 0.61 |
| 6 | 94837882 | rs665947 | -0.065 | 2.349 | 0.61 | -0.537 | -0.041 | 0.046 | 0.046 | 0 | 0.84 | 0.93 | 0.88 | 0.95 | 0.79 | 0.79 |
| 6 | 94839093 | rs2535669 | 1.103 | 1.952 | 0.848 | -0.463 | 0.053 | 0.01 | 0.01 | 0 | 0.7 | 0.47 | 0.7 | 0.67 | 0.62 | 0.63 |
| 6 | 94842774 | rs537092 | -0.723 | 1.853 | 0.419 | -0.333 | 0.001 | 0.039 | 0.039 | 0 | 0.14 | 0.19 | 0.27 | 0.63 | 0.16 | 0.32 |
| 6 | 94845639 | rs515814 | 0.307 | 1.758 | 0.49 | -0.308 | -0.026 | 0.012 | 0.012 | 0 | 0.06 | 0.15 | 0.01 | 0.07 | 0.01 | 0.1 |
| 6 | 94856048 | rs525413 | 0.151 | 1.898 | 0.519 | -0.348 | 0.01 | -0.013 | -0.013 | 0 | 0.95 | 0.88 | 0.94 | 0.98 | 0.89 | 0.96 |
| 6 | 94861241 | rs491003 | -0.045 | 1.115 | 0.131 | -0.811 | -0.003 | -0.011 | -0.011 | 0 | 0.28 | 0.31 | 0.17 | 0.11 | 0.23 | 0.24 |
| 6 | 94861991 | rs16872039 | 0.493 | 1.3 | -0.454 | -0.639 | -0.037 | -0.016 | -0.016 | 0 | 0.08 | 0.02 | 0.26 | 0.56 | 0.15 | 0.23 |
| 6 | 94885246 | rs9345447 | 0.68 | 0.938 | -0.475 | -0.564 | 0.003 | 0.076 | 0.076 | 0 | 0.21 | 0.09 | 0.16 | 0.04 | 0.22 | 0.14 |
| 6 | 94887219 | rs1750596 | 0.06 | 0.928 | -0.303 | -0.706 | 0.027 | 0.008 | 0.008 | 0 | 0.74 | 0.64 | 0.71 | 0.67 | 0.66 | 0.61 |
| 6 | 94888278 | rs1626647 | -0.435 | 0.423 | -0.692 | -0.642 | 0.025 | 0.052 | 0.052 | 0 | 0.27 | 0.2 | 0.16 | 0.09 | 0.22 | 0.23 |

| | | | | | | | | | | | | | | | | |
|---|---|---|---|---|---|---|---|---|---|---|---|---|---|---|---|---|
| 6 | 94891562 | rs1029882 | 0.033 | 0.595 | -0.569 | -0.374 | -0.024 | -0.005 | -0.005 | 0.029 | 0.06 | 0.11 | 0.01 | 0.04 | 0.01 | 0.09 |
| 6 | 94919013 | rs7750925 | -0.24 | 1.159 | -0.592 | -0.344 | 0.006 | 0.006 | 0.006 | 0 | 0.27 | 0.18 | 0.17 | 0.04 | 0.27 | 0.16 |
| 6 | 94920611 | rs1468639 | -0.558 | 1.711 | -0.764 | 0.658 | -0.053 | -0.056 | -0.056 | 0.055 | 0.09 | 0.01 | 0.25 | 0.67 | 0.16 | 0.24 |
| 6 | 94921408 | rs9363161 | -0.795 | 1.743 | -0.859 | 1.42 | -0.059 | -0.073 | -0.073 | 0 | 0.11 | 0.01 | 0.13 | 0.31 | 0.18 | 0.19 |
| 6 | 94922837 | rs452930 | 1.07 | 1.574 | -1.418 | 0.668 | -0.026 | -0.057 | -0.057 | 10.08 | 0.27 | 0.11 | 0.49 | 0.39 | 0.32 | 0.24 |
| 6 | 94925228 | rs416145 | 0.533 | 1.271 | -0.786 | 0.143 | -0.005 | -0.01 | -0.01 | 0 | 0.3 | 0.34 | 0.15 | 0.23 | 0.22 | 0.33 |
| 6 | 94925987 | rs423246 | -0.939 | 1.145 | -0.558 | -0.063 | 0 | 0 | 0 | 0 | 0.61 | 0.41 | 0.4 | 0.57 | 0.59 | 0.65 |
| 6 | 94935308 | rs7742539 | 1.03 | 0.967 | -0.974 | -0.287 | -0.03 | -0.003 | -0.003 | 0.508 | 0.19 | 0.02 | 0.12 | 0.16 | 0.15 | 0.23 |
| 6 | 94937733 | rs2021314 | -0.027 | 1.07 | -0.906 | -0.203 | -0.044 | -0.045 | -0.045 | 0 | 0.31 | 0.1 | 0.25 | 0.34 | 0.37 | 0.31 |
| 6 | 94938658 | rs433793 | -1.407 | 1.016 | -0.299 | -0.235 | -0.007 | -0.015 | -0.015 | 0 | 0.64 | 0.71 | 0.41 | 0.57 | 0.59 | 0.64 |
| 6 | 94939515 | rs6454990 | 0.887 | 0.678 | -0.055 | -0.21 | 0.008 | 0.011 | 0.011 | 0 | 0.21 | 0.15 | 0.13 | 0.04 | 0.19 | 0.13 |
| 6 | 94940269 | rs9360055 | 0.174 | 0.913 | -0.056 | -0.032 | -0.058 | -0.073 | -0.073 | 0 | 0.11 | 0.01 | 0.13 | 0.3 | 0.17 | 0.19 |
| 6 | 94958322 | rs220944 | -0.843 | 0.679 | 0.219 | -0.092 | 0.038 | -0.016 | -0.016 | 0.359 | 0.11 | 0.47 | 0.12 | 0.04 | 0.09 | 0.12 |
| 6 | 94963758 | rs9351412 | 2.082 | 0.904 | -0.091 | -0.13 | -0.012 | -0.053 | -0.053 | 0 | 0.26 | 0.12 | 0.43 | 0.39 | 0.28 | 0.23 |
| 6 | 94978674 | rs1408937 | 0.6 | 0.792 | -0.118 | -0.147 | 0.01 | 0.094 | 0.094 | 0.062 | 0.55 | 0.21 | 0.68 | 0.72 | 0.65 | 0.53 |
| 6 | 94979161 | rs9363169 | 1.1 | 0.845 | -0.075 | -0.155 | 0.007 | -0.003 | -0.003 | 0.424 | 0.45 | 0.2 | 0.55 | 0.43 | 0.48 | 0.35 |
| 6 | 94980584 | rs220971 | -0.325 | 1.173 | 0.45 | 0.163 | 0.024 | -0.039 | -0.039 | 0.279 | 0.07 | 0.2 | 0.1 | 0 | 0.09 | 0.09 |
| 6 | 94990974 | rs6935280 | 1.02 | 1.134 | 0.255 | 0.171 | -0.025 | -0.003 | -0.003 | 0 | 0.55 | 0.22 | 0.58 | 0.43 | 0.54 | 0.43 |
| 6 | 94991591 | rs7770720 | 1.028 | 1.1 | 0.148 | 0.074 | -0.019 | 0.157 | 0.157 | 6.098 | 0.27 | 0.02 | 0.13 | 0.05 | 0.26 | 0.2 |
| 6 | 94994034 | rs979568 | 0.056 | 1.107 | 0.428 | 0.15 | 0 | -0.041 | -0.041 | 2.116 | 0.69 | 0.57 | 0.73 | 0.8 | 0.72 | 0.64 |
| 6 | 94999281 | rs380842 | -0.711 | 1.114 | 0.86 | 0.157 | -0.004 | 0.006 | 0.006 | 0 | 0.77 | 0.82 | 0.82 | 0.77 | 0.8 | 0.77 |
| 6 | 95002441 | rs9345459 | 0.338 | 1.229 | 0.623 | 0.319 | -0.056 | -0.058 | -0.058 | 0.017 | 0.09 | 0.01 | 0.13 | 0.29 | 0.18 | 0.18 |
| 6 | 95002750 | rs7763538 | 0.904 | 1.168 | 0.736 | 0.305 | -0.015 | -0.011 | -0.011 | 0 | 0.56 | 0.28 | 0.58 | 0.46 | 0.54 | 0.44 |
| 6 | 95003444 | rs12529767 | 1.822 | 1.221 | 0.644 | 0.357 | -0.003 | -0.041 | -0.041 | 0 | 0.26 | 0.12 | 0.44 | 0.39 | 0.28 | 0.23 |
| 6 | 95004561 | rs434310 | -0.748 | 1.433 | 0.7 | 0.564 | -0.002 | 0 | 0 | 0 | 0.16 | 0.15 | 0.22 | 0.29 | 0.26 | 0.26 |
| 6 | 95005684 | rs371345 | -0.32 | 1.309 | 0.872 | 0.262 | -0.002 | -0.014 | -0.014 | 0.261 | 0.74 | 0.56 | 0.81 | 0.76 | 0.8 | 0.72 |
| 6 | 95014649 | rs422906 | -0.455 | 1.355 | 0.912 | 0.292 | 0 | -0.004 | -0.004 | 0 | 0.76 | 0.73 | 0.82 | 0.79 | 0.79 | 0.74 |
| 6 | 95014933 | rs366115 | -1.294 | 1.455 | 1.303 | 0.419 | 0.049 | -0.014 | -0.014 | 0 | 0.1 | 0.41 | 0.11 | 0.03 | 0.1 | 0.11 |
| 6 | 95024737 | rs7769230 | 0.5 | 1.361 | 0.774 | 0.591 | -0.012 | 0.008 | 0.008 | 0.059 | 0.39 | 0.22 | 0.59 | 0.61 | 0.45 | 0.41 |
| 6 | 95026172 | rs9452620 | -0.448 | 1.304 | 0.109 | 0.363 | -0.004 | -0.015 | -0.015 | 0 | 0.05 | 0.06 | 0.02 | 0 | 0.04 | 0.04 |
| 6 | 95028476 | rs1325478 | 0.491 | 1.379 | -0.079 | 0.273 | -0.021 | 0.146 | 0.146 | 4.407 | 0.33 | 0.1 | 0.16 | 0.1 | 0.3 | 0.28 |
| 6 | 95034121 | rs416366 | -0.323 | 1.531 | 0.289 | 0.421 | 0.001 | -0.012 | -0.012 | 0 | 0.77 | 0.68 | 0.82 | 0.79 | 0.8 | 0.76 |
| 6 | 95035288 | rs1325479 | 0.483 | 1.526 | 0.365 | 0.408 | 0.008 | 0.061 | 0.061 | 3.394 | 0.34 | 0.24 | 0.17 | 0.14 | 0.31 | 0.28 |
| 6 | 95036897 | rs385368 | 0.439 | 1.529 | 0.197 | 0.398 | 0.003 | -0.016 | -0.016 | 0 | 0.43 | 0.44 | 0.65 | 0.65 | 0.49 | 0.48 |
| 6 | 95042308 | rs7770285 | 0.449 | 1.559 | -0.225 | 0.443 | -0.003 | 0.129 | 0.129 | 3.524 | 0.33 | 0.14 | 0.17 | 0.1 | 0.31 | 0.27 |
| 6 | 95043459 | rs9342406 | -0.141 | 1.603 | -0.001 | 1.17 | -0.045 | -0.053 | -0.053 | 0 | 0.1 | 0.03 | 0.13 | 0.33 | 0.16 | 0.16 |
| 6 | 95049615 | rs459393 | -0.833 | 1.609 | 0.166 | 1.1 | 0.01 | 0.045 | 0.045 | 0 | 0.14 | 0.15 | 0.19 | 0.34 | 0.22 | 0.24 |
| 6 | 95056999 | rs462497 | 0.331 | 1.167 | 0.377 | 0.566 | -0.002 | 0 | 0 | 0 | 0.39 | 0.53 | 0.56 | 0.41 | 0.38 | 0.38 |
| 6 | 95060183 | rs885717 | 0.55 | 1.114 | 0.343 | 0.61 | 0.002 | -0.002 | -0.002 | 0.011 | 0.36 | 0.61 | 0.51 | 0.41 | 0.33 | 0.3 |
| 6 | 95060430 | rs4571535 | 0.961 | 1.211 | 0.089 | 0.554 | -0.002 | -0.001 | -0.001 | 0.013 | 0.31 | 0.29 | 0.48 | 0.31 | 0.32 | 0.26 |
| 6 | 95061816 | rs467349 | 0.426 | 2.232 | -0.084 | 1.322 | -0.035 | -0.034 | -0.034 | 0 | 0.69 | 0.77 | 0.71 | 0.42 | 0.64 | 0.62 |
| 6 | 95064614 | rs13204646 | -1.004 | 2.4 | -0.087 | 1.929 | -0.03 | -0.044 | -0.044 | 2.215 | 0.11 | 0.04 | 0.13 | 0.38 | 0.18 | 0.21 |
| 6 | 95065361 | rs461442 | 1.554 | 2.736 | -0.761 | 1.682 | -0.037 | 0.167 | 0.167 | 0 | 0.51 | 0.06 | 0.58 | 0.19 | 0.46 | 0.57 |
| 6 | 95065664 | rs12204226 | -1.343 | 2.643 | 0.138 | 1.951 | 0.021 | 0.111 | 0.111 | 0 | 0.16 | 0.31 | 0.17 | 0.4 | 0.21 | 0.22 |
| 6 | 95071919 | rs447142 | 1.569 | 2.762 | -0.285 | 1.788 | -0.055 | 0.13 | 0.13 | 0 | 0.51 | 0.04 | 0.57 | 0.23 | 0.46 | 0.53 |
| 6 | 95078646 | rs449578 | 0.892 | 3.093 | 0.106 | 2.264 | -0.022 | -0.091 | -0.091 | 0 | 0.67 | 0.15 | 0.73 | 0.51 | 0.69 | 0.73 |
| 6 | 95080580 | rs9354114 | -0.876 | 3.185 | 0.047 | 2.29 | -0.027 | -0.046 | -0.046 | 0 | 0.16 | 0.07 | 0.15 | 0.33 | 0.23 | 0.22 |
| 6 | 95081321 | rs404945 | -0.009 | 3.037 | 0.74 | 2.199 | 0.009 | 0.003 | 0.003 | 0 | 0.78 | 0.7 | 0.82 | 0.75 | 0.75 | 0.8 |
| 6 | 95088393 | rs10080541 | -0.937 | 2.829 | 0.918 | 2.169 | 0.022 | 0.182 | 0.182 | 0 | 0.06 | 0.19 | 0.06 | 0.21 | 0.05 | 0.04 |
| 6 | 95089937 | rs2076704 | -0.821 | 2.834 | 0.825 | 2.149 | 0.012 | 0.142 | 0.142 | 0 | 0.06 | 0.16 | 0.06 | 0.2 | 0.05 | 0.04 |
| 6 | 95090000 | rs384105 | 1.612 | 2.875 | 0.267 | 1.607 | -0.065 | 0.195 | 0.195 | 0 | 0.51 | 0.06 | 0.57 | 0.18 | 0.46 | 0.51 |
| 6 | 95090661 | rs390661 | 0.425 | 2.684 | 1.283 | 1.84 | 0.019 | 0.018 | 0.018 | 0 | 0.78 | 0.63 | 0.8 | 0.7 | 0.76 | 0.83 |
| 6 | 95092853 | rs2325585 | -1.459 | 2.523 | 1.197 | 1.571 | -0.001 | -0.006 | -0.006 | 0 | 0.43 | 0.46 | 0.49 | 0.79 | 0.55 | 0.48 |
| 6 | 95094319 | rs9445241 | -0.903 | 2.767 | 1.755 | 1.802 | 0.048 | 0.109 | 0.109 | 0.05 | 0.11 | 0.41 | 0.1 | 0.23 | 0.13 | 0.12 |
| 6 | 95096690 | rs12196652 | -0.987 | 2.903 | 1.85 | 1.784 | 0.041 | 0.037 | 0.037 | 0 | 0.14 | 0.32 | 0.12 | 0.23 | 0.19 | 0.12 |
| 6 | 95098682 | rs9342409 | -0.785 | 3.077 | 1.536 | 1.815 | -0.016 | -0.029 | -0.029 | 0.017 | 0.13 | 0.06 | 0.2 | 0.54 | 0.16 | 0.29 |
| 6 | 95101041 | rs11753129 | 0.081 | 3.142 | 1.59 | 2.028 | -0.052 | 0.071 | 0.071 | 0 | 0.16 | 0.03 | 0.16 | 0.02 | 0.2 | 0.06 |
| 6 | 95104559 | rs9363197 | -0.694 | 3.091 | 1.884 | 2.086 | -0.002 | 0.033 | 0.033 | 0 | 0.11 | 0.08 | 0.19 | 0.54 | 0.12 | 0.29 |
| 6 | 95106958 | rs9445244 | -1.851 | 3.365 | 2.08 | 1.994 | -0.01 | 0.006 | 0.006 | 0 | 0.35 | 0.67 | 0.33 | 0.29 | 0.46 | 0.2 |
| 6 | 95112651 | rs1033581 | -1.13 | 3.285 | 2.074 | 2.064 | 0.07 | 0.038 | 0.038 | 0 | 0.15 | 0.37 | 0.13 | 0.23 | 0.22 | 0.13 |
| 6 | 95145322 | rs9452723 | -0.648 | 2.854 | 2.058 | 2.045 | 0.061 | -0.057 | -0.057 | 0.034 | 0.22 | 0.46 | 0.32 | 0.13 | 0.27 | 0.14 |
| 6 | 95147902 | rs9354118 | -0.817 | 2.237 | 0.975 | 2.153 | 0.003 | 0.126 | 0.126 | 0 | 0.11 | 0.18 | 0.19 | 0.54 | 0.12 | 0.29 |
| 6 | 95172841 | rs6454998 | 0.919 | 1.815 | 0.526 | 1.398 | -0.007 | 0.002 | 0.002 | 0 | 0.57 | 0.37 | 0.49 | 0.32 | 0.5 | 0.49 |
| 6 | 95176359 | rs1322422 | -0.831 | 1.791 | 0.575 | 1.282 | 0.002 | 0.001 | 0.001 | 0 | 0.35 | 0.37 | 0.48 | 0.5 | 0.41 | 0.45 |
| 6 | 95176640 | rs7454136 | -0.907 | 1.273 | 0.566 | 0.991 | -0.001 | -0.009 | -0.009 | 0 | 0.34 | 0.46 | 0.4 | 0.4 | 0.32 | 0.37 |
| 6 | 95184421 | rs9363211 | -0.317 | 0.208 | -0.247 | 0.213 | 0.028 | 0.039 | 0.039 | 4.762 | 0.22 | 0.08 | 0.25 | 0.08 | 0.15 | 0.16 |
| 6 | 95188014 | rs9445284 | -1.008 | 0.292 | 0.163 | -0.112 | -0.004 | -0.018 | -0.018 | 0.028 | 0.3 | 0.27 | 0.32 | 0.08 | 0.2 | 0.19 |
| 6 | 95188273 | rs9445285 | -1.013 | 0 | 0.304 | -0.086 | -0.017 | -0.033 | -0.033 | 0 | 0.29 | 0.3 | 0.32 | 0.1 | 0.2 | 0.21 |
| 6 | 95197882 | rs9445291 | -0.761 | 0.228 | 0.328 | -0.305 | 0.013 | -0.026 | -0.026 | 0 | 0.12 | 0.22 | 0.09 | 0.04 | 0.17 | 0.03 |

| | | | | | | | | | | | | | | | |
|---|---|---|---|---|---|---|---|---|---|---|---|---|---|---|---|
| 6 | 95202197 | rs1022787 | 1.653 | -0.394 | -0.519 | 0.116 | -0.024 | -0.021 | -0.021 | 0 | 0.46 | 0.07 | 0.71 | 0.35 | 0.57 | 0.41 |
| 6 | 95211042 | rs9452812 | 0.116 | -0.305 | -0.4 | 0.132 | 0.017 | -0.029 | -0.029 | 0 | 0.06 | 0.17 | 0.06 | 0 | 0.08 | 0.01 |
| 6 | 95211341 | rs222547 | -0.279 | -0.711 | 0.307 | 0.164 | -0.017 | 0.166 | 0.166 | 0 | 0.14 | 0.28 | 0.09 | 0.53 | 0.11 | 0.38 |
| 6 | 95215623 | rs222556 | -0.102 | -0.733 | 0.348 | 0.166 | -0.02 | 0.2 | 0.2 | 0 | 0.12 | 0.26 | 0.06 | 0.53 | 0.07 | 0.38 |
| 6 | 95219795 | rs9363216 | 0.11 | -0.793 | 0.047 | 0.001 | 0.045 | 0.053 | 0.053 | 1.733 | 0.25 | 0.09 | 0.1 | 0.08 | 0.15 | 0.17 |
| 6 | 95234099 | rs222534 | -1.425 | -0.847 | -0.022 | 0.057 | 0.122 | -0.045 | -0.045 | 0.43 | 0.48 | 0.91 | 0.23 | 0.58 | 0.3 | 0.57 |
| 6 | 95244829 | rs222519 | -1.495 | -0.865 | 0.53 | -0.011 | 0.022 | 0.014 | 0.014 | 0.233 | 0.5 | 0.59 | 0.23 | 0.63 | 0.39 | 0.59 |
| 6 | 95247573 | rs9363217 | 0.356 | -0.919 | 0.414 | -0.027 | 0.036 | 0.065 | 0.065 | 1.695 | 0.23 | 0.06 | 0.09 | 0.06 | 0.15 | 0.17 |
| 6 | 95252573 | rs16873343 | 0.523 | -0.888 | 0.379 | -0.008 | -0.03 | -0.009 | -0.009 | 6.068 | 0.05 | 0.01 | 0.03 | 0.04 | 0.09 | 0.02 |
| 6 | 95255735 | rs9360071 | 0.644 | -1.144 | 0.207 | -0.187 | 0.002 | 0.01 | 0.01 | 0.741 | 0.13 | 0 | 0.07 | 0.05 | 0.11 | 0.12 |
| 6 | 95256763 | rs2145067 | 0.794 | -0.933 | -0.508 | -0.682 | -0.017 | -0.045 | -0.045 | 0 | 0.14 | 0.04 | 0.41 | 0.31 | 0.18 | 0.17 |
| 6 | 95257918 | rs2208173 | -0.851 | -1.047 | -0.341 | -0.642 | 0.02 | -0.003 | -0.003 | 0 | 0.45 | 0.61 | 0.21 | 0.59 | 0.3 | 0.57 |
| 6 | 95262849 | rs2819966 | -0.522 | -1.021 | -0.451 | -0.62 | -0.017 | 0.112 | 0.112 | 0 | 0.18 | 0.34 | 0.11 | 0.52 | 0.14 | 0.39 |
| 6 | 95269855 | rs2750033 | -1.25 | -0.901 | -1.129 | -0.564 | 0.015 | -0.017 | -0.017 | 0 | 0.53 | 0.93 | 0.26 | 0.63 | 0.39 | 0.59 |
| 6 | 95278429 | rs2750031 | -0.095 | -1.05 | -0.902 | -0.491 | 0.075 | -0.052 | -0.052 | 0 | 0.36 | 0.18 | 0.15 | 0.61 | 0.22 | 0.57 |
| 6 | 95285511 | rs2224317 | -1.752 | -0.9 | -0.258 | -0.619 | -0.049 | 0.094 | 0.094 | 0.351 | 0.66 | 0.87 | 0.65 | 0.96 | 0.56 | 0.75 |
| 6 | 95296005 | rs2078604 | -0.516 | -0.847 | -0.472 | -0.491 | 0.047 | 0.206 | 0.206 | 0 | 0.48 | 0.22 | 0.55 | 0.91 | 0.4 | 0.72 |
| 6 | 95309419 | rs2501985 | 0.663 | -0.915 | -0.613 | -0.271 | 0.029 | 0.059 | 0.059 | 0.914 | 0.24 | 0.03 | 0.09 | 0.09 | 0.17 | 0.19 |
| 6 | 95320818 | rs993200 | -1.421 | -0.518 | -0.336 | -0.184 | -0.14 | 0.189 | 0.189 | 1.299 | 0.67 | 0.96 | 0.66 | 0.96 | 0.57 | 0.74 |
| 6 | 95332279 | rs7738219 | 0.801 | -0.656 | 0.51 | 1.344 | 0.071 | -0.065 | -0.065 | 2.092 | 0.56 | 0.08 | 0.63 | 0.51 | 0.69 | 0.55 |
| 6 | 95335012 | rs4299796 | 0.228 | -0.813 | 1.069 | 0.84 | 0.006 | 0.001 | 0.001 | 1.485 | 0.65 | 0.65 | 0.88 | 0.64 | 0.79 | 0.59 |
| 6 | 95336020 | rs9354130 | 0.218 | -0.781 | 0.887 | 0.863 | 0.005 | 0.001 | 0.001 | 1.508 | 0.65 | 0.65 | 0.88 | 0.64 | 0.79 | 0.59 |
| 6 | 95342961 | rs6455001 | 0.118 | -0.767 | 0.831 | 0.87 | -0.008 | -0.002 | -0.002 | 0 | 0.65 | 0.63 | 0.87 | 0.64 | 0.79 | 0.59 |
| 6 | 95345767 | rs6924653 | 0.118 | -0.767 | 0.831 | 0.87 | -0.008 | -0.002 | -0.002 | 0 | 0.65 | 0.63 | 0.87 | 0.64 | 0.79 | 0.59 |
| 6 | 95350702 | rs13207128 | -0.847 | -0.11 | 1.068 | 0.738 | -0.004 | 0.006 | 0.006 | 0 | 0.05 | 0.06 | 0.05 | 0 | 0.07 | 0.02 |
| 6 | 95351548 | rs6928079 | -1.649 | -0.009 | 1.402 | 0.82 | 0.045 | 0.046 | 0.046 | 0.131 | 0.08 | 0.46 | 0.05 | 0.06 | 0.05 | 0.04 |
| 6 | 95355112 | rs10944752 | 0.594 | -0.486 | 0.471 | 0.897 | 0.087 | -0.024 | -0.024 | 0.333 | 0.53 | 0.1 | 0.56 | 0.51 | 0.66 | 0.51 |
| 6 | 95362698 | rs9452884 | -0.269 | -0.58 | 0.974 | 0.805 | -0.019 | -0.028 | -0.028 | 0.199 | 0.06 | 0.16 | 0.02 | 0 | 0.02 | 0.02 |
| 6 | 95364959 | rs2502026 | -0.044 | -0.786 | 1.265 | 0.82 | 0.004 | -0.002 | -0.002 | 0 | 0.33 | 0.34 | 0.17 | 0.11 | 0.24 | 0.23 |
| 6 | 95367184 | rs4552698 | 0.819 | -0.426 | 0.944 | 0.783 | -0.127 | -0.125 | -0.125 | 0.639 | 0.23 | 0.02 | 0.31 | 0.31 | 0.32 | 0.2 |
| 6 | 95367435 | rs2502037 | 1.598 | -1.071 | 2.05 | 0.974 | -0.011 | 0.172 | 0.172 | 0.611 | 0.85 | 0.46 | 0.72 | 0.62 | 0.9 | 0.74 |
| 6 | 95368911 | rs9445337 | -1.551 | -1.023 | 1.93 | 0.972 | 0.007 | -0.057 | -0.057 | 0 | 0.1 | 0.28 | 0.07 | 0 | 0.07 | 0.03 |
| 6 | 95372264 | rs2502042 | 0.192 | -1.015 | 1.127 | 0.974 | 0.018 | 0.044 | 0.044 | 0.941 | 0.27 | 0.15 | 0.15 | 0.11 | 0.22 | 0.21 |
| 6 | 95373184 | rs2493968 | -0.2 | -0.652 | 0.73 | 1.112 | -0.042 | -0.053 | -0.053 | 0 | 0.34 | 0.52 | 0.18 | 0.18 | 0.24 | 0.23 |
| 6 | 95380245 | rs2502045 | 0.51 | -0.462 | -0.392 | 1.221 | -0.014 | 0.065 | 0.065 | 0 | 0.92 | 0.86 | 0.79 | 0.69 | 0.96 | 0.77 |
| 6 | 95390186 | rs10498989 | 0.725 | -0.518 | -0.434 | 1.229 | 0.091 | -0.032 | -0.032 | 0.533 | 0.53 | 0.1 | 0.55 | 0.51 | 0.66 | 0.51 |
| 6 | 95403650 | rs12210972 | 0.34 | -0.76 | -0.186 | 0.538 | 0.032 | 0.035 | 0.035 | 0.29 | 0.63 | 0.27 | 0.81 | 0.82 | 0.76 | 0.76 |
| 6 | 95421874 | rs6931706 | -1.48 | -0.071 | 0.583 | 1.247 | -0.008 | 0.057 | 0.057 | 0.905 | 0.12 | 0.34 | 0.23 | 0.23 | 0.1 | 0.25 |
| 6 | 95435352 | rs9363234 | -0.882 | -0.057 | 0.341 | 1.379 | -0.009 | -0.039 | -0.039 | 0 | 0.07 | 0.02 | 0.09 | 0.21 | 0.06 | 0.21 |
| 6 | 95456914 | rs6918852 | -1.348 | -0.405 | 1.118 | 1.118 | 0.002 | -0.013 | -0.013 | 0.368 | 0.05 | 0.22 | 0.02 | 0 | 0.02 | 0.01 |
| 6 | 95458317 | rs794672 | -2.648 | -1.652 | 0.948 | 2.103 | 0.103 | -0.108 | -0.108 | 0 | 0.28 | 0.89 | 0.17 | 0.57 | 0.1 | 0.34 |
| 6 | 95464758 | rs11969409 | -1.686 | -1.65 | 1.999 | 2.039 | 0.038 | 0.017 | 0.017 | 1.602 | 0.07 | 0.43 | 0.03 | 0 | 0.02 | 0.01 |
| 6 | 95502730 | rs9363237 | -2.584 | -1.704 | 2.448 | 3.093 | -0.049 | 0.047 | 0.047 | 3.545 | 0.14 | 0.75 | 0.06 | 0.28 | 0.03 | 0.07 |
| 6 | 95504340 | rs794669 | -3.127 | -1.659 | 2.366 | 3.094 | 0.099 | -0.099 | -0.099 | 0 | 0.27 | 0.89 | 0.17 | 0.57 | 0.1 | 0.34 |
| 6 | 95524427 | rs9452967 | -1.301 | -1.7 | 2.988 | 3.081 | 0.015 | -0.001 | -0.001 | 0 | 0.05 | 0.3 | 0.03 | 0 | 0.02 | 0.01 |
| 6 | 95540600 | rs197883 | -3.024 | -1.736 | 3.426 | 3.252 | -0.038 | 0.004 | 0.004 | 0.47 | 0.2 | 0.73 | 0.14 | 0.28 | 0.06 | 0.27 |
| 6 | 95544333 | rs9632477 | -1.581 | -1.731 | 3.706 | 3.227 | 0.027 | 0.054 | 0.054 | 0.503 | 0.06 | 0.33 | 0.03 | 0.05 | 0.02 | 0.02 |
| 6 | 95546028 | rs9445398 | -2.539 | -1.731 | 3.577 | 3.227 | -0.061 | -0.045 | -0.045 | 0.064 | 0.13 | 0.67 | 0.05 | 0.05 | 0.03 | 0.05 |
| 6 | 95552683 | rs9360079 | -1.387 | -1.345 | 3.059 | 3.035 | -0.026 | -0.054 | -0.054 | 0 | 0.1 | 0.02 | 0.13 | 0.22 | 0.12 | 0.25 |
| 6 | 95556864 | rs197870 | -2.641 | -1.391 | 3.491 | 2.954 | -0.053 | 0.028 | 0.028 | 2.941 | 0.19 | 0.74 | 0.07 | 0.35 | 0.03 | 0.12 |
| 6 | 95572362 | rs197892 | -1.519 | -1.178 | 3.729 | 3.209 | -0.025 | 0.262 | 0.262 | 0 | 0.08 | 0.28 | 0.09 | 0.43 | 0.03 | 0.21 |
| 6 | 95597741 | rs12195016 | -1.766 | -1.193 | 3.649 | 3.044 | 0.008 | -0.022 | -0.022 | 0 | 0.06 | 0.14 | 0.06 | 0 | 0.07 | 0.05 |
| 6 | 95610355 | rs9354149 | -1.947 | -0.928 | 3.794 | 3.392 | -0.009 | 0.469 | 0.469 | 0.035 | 0.1 | 0.46 | 0.1 | 0.48 | 0.03 | 0.23 |
| 6 | 95633301 | rs9453067 | -0.671 | -0.979 | 3.788 | 3.503 | -0.033 | 0.149 | 0.149 | 0 | 0.07 | 0.17 | 0.02 | 0.25 | 0.01 | 0.08 |
| 6 | 95643248 | rs6905617 | -2.465 | -1.219 | 3.609 | 3.417 | -0.007 | 0.137 | 0.137 | 0.151 | 0.15 | 0.37 | 0.13 | 0.46 | 0.11 | 0.28 |
| 6 | 95660707 | rs9363255 | -0.474 | -1.215 | 3.139 | 3.424 | 0.004 | 0.076 | 0.076 | 0 | 0.06 | 0.06 | 0.07 | 0.46 | 0.03 | 0.22 |
| 6 | 95662219 | rs12663902 | -0.144 | -1.218 | 3.074 | 3.443 | -0.017 | 0.053 | 0.053 | 0 | 0.06 | 0.06 | 0.01 | 0.24 | 0.01 | 0.08 |
| 6 | 95668509 | rs1028640 | -3.03 | -1.245 | 3.245 | 3.406 | -0.017 | 0.011 | 0.011 | 4.149 | 0.24 | 0.66 | 0.16 | 0.71 | 0.11 | 0.37 |
| 6 | 95676106 | rs12664570 | -0.224 | -1.091 | 2.681 | 3.048 | -0.014 | 0.053 | 0.053 | 0 | 0.06 | 0.06 | 0.01 | 0.25 | 0.01 | 0.08 |
| 6 | 95836160 | rs2252451 | -0.096 | -0.919 | 1.618 | 1.313 | -0.003 | 0.078 | 0.078 | 0 | 0.71 | 0.85 | 0.82 | 0.95 | 0.69 | 0.74 |
| 6 | 95841116 | rs2380218 | -0.658 | -0.446 | 1.669 | 1.009 | 0.014 | -0.017 | -0.017 | 0 | 0.29 | 0.45 | 0.43 | 0.23 | 0.32 | 0.2 |
| 6 | 95857781 | rs9490554 | -0.682 | -0.031 | 1.704 | 0.845 | -0.022 | -0.07 | -0.07 | 0.102 | 0.2 | 0.41 | 0.09 | 0.1 | 0.16 | 0.18 |
| 6 | 95871752 | rs7746362 | -0.223 | -0.039 | 1.695 | 0.84 | -0.033 | -0.07 | -0.07 | 0 | 0.17 | 0.37 | 0.09 | 0.07 | 0.12 | 0.1 |
| 6 | 95872321 | rs12216275 | -0.627 | 0.426 | 1.657 | 0.825 | 0.004 | -0.001 | -0.001 | 2.137 | 0.15 | 0.15 | 0.36 | 0.19 | 0.26 | 0.19 |
| 6 | 95874120 | rs2613542 | 0.923 | 0.35 | 1.449 | 0.763 | 0.008 | 0.007 | 0.007 | 0 | 0.38 | 0.32 | 0.37 | 0.69 | 0.31 | 0.45 |
| 6 | 95874258 | rs2613543 | 1.127 | 0.307 | 1.054 | 0.741 | 0.021 | 0.02 | 0.02 | 0 | 0.37 | 0.16 | 0.36 | 0.69 | 0.31 | 0.45 |
| 6 | 95880011 | rs2716081 | -0.805 | 0.34 | 0.904 | 0.51 | -0.009 | 0.054 | 0.054 | 0 | 0.69 | 0.82 | 0.81 | 0.92 | 0.65 | 0.66 |
| 6 | 95891910 | rs9399724 | 0.299 | 0.967 | 0.966 | 0.091 | 0.017 | -0.06 | -0.06 | 0 | 0.61 | 0.48 | 0.5 | 0.86 | 0.55 | 0.73 |
| 6 | 95894399 | rs2613565 | 0.412 | 1.357 | 1.008 | -0.024 | 0.039 | -0.09 | -0.09 | 0.383 | 0.26 | 0.41 | 0.41 | 0.08 | 0.34 | 0.21 |

| | | | | | | | | | | | | | | | | |
|---|---|---|---|---|---|---|---|---|---|---|---|---|---|---|---|---|
| 6 | 95937046 | rs877096 | 1.271 | 1.385 | 0.902 | 0.596 | 0.025 | 0.065 | 0.065 | 0 | 0.19 | 0.04 | 0.26 | 0.08 | 0.14 | 0.16 |
| 6 | 95951267 | rs12204802 | 1.702 | 1.815 | -0.5 | -0.659 | -0.014 | 0.07 | 0.07 | 0 | 0.12 | 0.01 | 0.11 | 0 | 0.12 | 0.04 |
| 6 | 95952089 | rs7449934 | 1.53 | 2.027 | -0.548 | -0.376 | -0.038 | -0.014 | -0.014 | 2.221 | 0.29 | 0.03 | 0.27 | 0.28 | 0.27 | 0.32 |
| 6 | 95956614 | rs2380214 | 0.091 | 2.364 | 0.005 | -0.073 | 0.001 | 0.009 | 0.009 | 0 | 0.88 | 0.93 | 0.92 | 0.94 | 0.9 | 0.94 |
| 6 | 95963764 | rs7749777 | 0.472 | 2.252 | 0.042 | 0.351 | -0.001 | -0.001 | -0.001 | 0 | 0.57 | 0.41 | 0.45 | 0.49 | 0.55 | 0.62 |
| 6 | 95968991 | rs729278 | 0.169 | 2.311 | 0.019 | 0.34 | -0.008 | 0.007 | 0.007 | 0.045 | 0.62 | 0.6 | 0.5 | 0.89 | 0.54 | 0.74 |
| 6 | 95974657 | rs13196543 | -0.058 | 2.194 | -0.124 | 0.359 | -0.004 | 0.013 | 0.013 | 0 | 0.15 | 0.1 | 0.11 | 0.06 | 0.13 | 0.06 |
| 6 | 95980623 | rs9400397 | -1.39 | 2.056 | -0.139 | 0.224 | 0.009 | 0.046 | 0.046 | 0 | 0.78 | 0.89 | 0.64 | 0.92 | 0.79 | 0.83 |
| 6 | 95988605 | rs4448101 | -1.005 | 2.189 | -0.11 | 0.221 | -0.011 | 0.005 | 0.005 | 0.01 | 0.74 | 0.74 | 0.58 | 0.92 | 0.64 | 0.79 |
| 6 | 96000122 | rs9400554 | 0.474 | 1.549 | 0.069 | 0.555 | -0.001 | 0.001 | 0.001 | 0.084 | 0.53 | 0.46 | 0.43 | 0.53 | 0.47 | 0.57 |
| 6 | 96007343 | rs10782175 | -0.449 | 1.438 | -0.18 | 0.234 | 0 | 0.065 | 0.065 | 0.013 | 0.67 | 0.86 | 0.57 | 0.86 | 0.68 | 0.76 |
| 6 | 96008082 | rs7748108 | -1.241 | 1.48 | -0.19 | 0.183 | -0.002 | 0.048 | 0.048 | 0 | 0.76 | 0.9 | 0.64 | 0.92 | 0.76 | 0.81 |
| 6 | 96014686 | rs6932267 | -0.395 | 1.604 | 0.132 | 0.189 | -0.007 | -0.005 | -0.005 | 0 | 0.68 | 0.69 | 0.56 | 0.87 | 0.57 | 0.76 |
| 6 | 96025446 | rs9320526 | 0.752 | 1.533 | 0.289 | 0.546 | 0.006 | 0.015 | 0.015 | 0 | 0.52 | 0.35 | 0.36 | 0.43 | 0.48 | 0.58 |
| 6 | 96034050 | rs13205436 | 0.603 | 1.557 | 0.02 | 0.594 | -0.018 | -0.015 | -0.015 | 0 | 0.08 | 0 | 0.06 | 0.06 | 0.08 | 0.04 |
| 6 | 96035175 | rs9374586 | -0.674 | 1.556 | 0.186 | 0.6 | -0.007 | 0 | 0 | 0 | 0.7 | 0.72 | 0.59 | 0.87 | 0.62 | 0.76 |
| 6 | 96035410 | rs12661585 | 0.085 | 1.535 | 0.186 | 0.611 | -0.015 | 0.252 | 0.252 | 0 | 0.07 | 0.23 | 0.13 | 0.37 | 0.02 | 0.13 |
| 6 | 96036952 | rs12525680 | 0.453 | 1.626 | 0.077 | 0.395 | 0.05 | -0.023 | -0.023 | 0.025 | 0.09 | 0.18 | 0.07 | 0.05 | 0.18 | 0.06 |
| 6 | 96042338 | rs4498363 | -1.594 | 1.534 | -0.028 | 0.4 | 0.006 | 0.078 | 0.078 | 0 | 0.79 | 0.94 | 0.67 | 0.91 | 0.81 | 0.83 |
| 6 | 96063401 | rs9387603 | -0.499 | 1.068 | 0.028 | 0.506 | 0.001 | -0.023 | -0.023 | 0 | 0.67 | 0.65 | 0.56 | 0.87 | 0.57 | 0.75 |
| 6 | 96067121 | rs12333032 | 0.335 | 1.048 | 0.028 | 0.506 | 0.001 | -0.023 | -0.023 | 0 | 0.33 | 0.35 | 0.44 | 0.13 | 0.43 | 0.24 |
| 6 | 96070254 | rs4383824 | 0.394 | 0.767 | -0.182 | 1.18 | -0.01 | 0.264 | 0.264 | 0.002 | 0.93 | 0.76 | 0.93 | 0.64 | 0.98 | 0.87 |
| 6 | 96078320 | rs9482027 | 0.661 | 0.475 | -0.39 | 0.808 | 0.013 | 0.031 | 0.031 | 0 | 0.18 | 0.07 | 0.11 | 0.1 | 0.17 | 0.15 |
| 6 | 96080915 | rs4327706 | 0.513 | 0.669 | -0.138 | 1.304 | -0.01 | 0.235 | 0.235 | 0 | 0.93 | 0.75 | 0.87 | 0.66 | 0.98 | 0.87 |
| 6 | 96083087 | rs4388292 | 1.214 | 0.649 | -0.185 | 1.135 | -0.009 | 0.101 | 0.101 | 0 | 0.82 | 0.67 | 0.77 | 0.6 | 0.86 | 0.82 |
| 6 | 96085046 | rs4074107 | 0.567 | 0.481 | -0.254 | 1.207 | -0.011 | 0.077 | 0.077 | 0 | 0.85 | 0.75 | 0.81 | 0.6 | 0.91 | 0.83 |
| 6 | 96086583 | rs9490131 | 0.577 | 0.702 | -0.383 | 0.648 | 0 | -0.005 | -0.005 | 0 | 0.39 | 0.56 | 0.3 | 0.38 | 0.38 | 0.44 |
| 6 | 96089739 | rs4438965 | 0.626 | 0.9 | -0.787 | 0.277 | 0.027 | 0.132 | 0.132 | 0 | 0.39 | 0.07 | 0.45 | 0.18 | 0.34 | 0.33 |
| 6 | 96090419 | rs12525999 | 0.481 | 1.135 | -0.834 | 0.279 | -0.006 | 0.008 | 0.008 | 0 | 0.06 | 0.03 | 0.05 | 0 | 0.15 | 0.05 |
| 6 | 96091123 | rs4620131 | -0.448 | 0.949 | -0.881 | 0.594 | -0.079 | 0.024 | 0.024 | 0 | 0.59 | 0.88 | 0.43 | 0.71 | 0.5 | 0.6 |
| 6 | 96100775 | rs9375155 | 1.02 | 1.504 | -0.5 | 0.728 | -0.002 | -0.004 | -0.004 | 0 | 0.91 | 0.93 | 0.87 | 0.88 | 0.83 | 0.93 |
| 6 | 96111372 | rs6907357 | 0.533 | 1.39 | -0.376 | 0.861 | 0.039 | 0.018 | 0.018 | 0 | 0.46 | 0.33 | 0.52 | 0.39 | 0.39 | 0.44 |
| 6 | 96124293 | rs4839952 | -0.387 | 1.493 | 0.009 | 0.792 | 0.023 | 0.026 | 0.026 | 0.151 | 0.62 | 0.55 | 0.63 | 0.48 | 0.52 | 0.52 |
| 6 | 96149449 | rs13215970 | -0.157 | 1.52 | 0.068 | 0.933 | -0.005 | -0.017 | -0.017 | 0.153 | 0.58 | 0.35 | 0.61 | 0.43 | 0.59 | 0.53 |
| 6 | 96159101 | rs4382270 | 0.903 | 1.485 | 0.167 | 0.912 | 0.003 | 0.009 | 0.009 | 0 | 0.29 | 0.33 | 0.25 | 0.4 | 0.29 | 0.39 |
| 6 | 96164110 | rs9482958 | 0.928 | 1.491 | 0.158 | 0.92 | 0.006 | 0.016 | 0.016 | 0 | 0.29 | 0.36 | 0.25 | 0.4 | 0.29 | 0.39 |
| 6 | 96193945 | rs4839992 | -0.326 | 1.408 | 0.148 | 0.989 | -0.002 | -0.001 | -0.001 | 0.301 | 0.86 | 0.88 | 0.86 | 0.77 | 0.85 | 0.83 |
| 6 | 96196122 | rs13208734 | 0.38 | 1.509 | 0.363 | 1.259 | 0.011 | -0.015 | -0.015 | 0 | 0.1 | 0 | 0.07 | 0.07 | 0.06 | 0.07 |
| 6 | 96212413 | rs9321415 | -0.263 | 1.635 | 0.569 | 1.278 | 0.001 | -0.001 | -0.001 | 0.085 | 0.4 | 0.37 | 0.35 | 0.42 | 0.47 | 0.38 |
| 6 | 96218908 | rs9493940 | 0.386 | 1.329 | 1.016 | 1.496 | 0.004 | 0.007 | 0.007 | 0 | 0.26 | 0.4 | 0.24 | 0.36 | 0.28 | 0.27 |
| 6 | 96228904 | rs13203167 | 0.155 | 1.149 | 1.017 | 1.454 | -0.033 | -0.005 | -0.005 | 2.004 | 0.58 | 0.23 | 0.56 | 0.38 | 0.52 | 0.59 |
| 6 | 96233216 | rs12530302 | 0.198 | 1.082 | 1.135 | 1.539 | -0.015 | -0.007 | -0.007 | 0 | 0.6 | 0.25 | 0.6 | 0.38 | 0.57 | 0.6 |
| 6 | 96245540 | rs13205985 | 0.029 | 1.095 | 1.106 | 1.527 | 0.012 | 0.008 | 0.008 | 0 | 0.62 | 0.53 | 0.61 | 0.38 | 0.57 | 0.6 |
| 6 | 96246431 | rs9494677 | 0.042 | 1.101 | 0.77 | 1.49 | 0.001 | 0 | 0 | 0 | 0.36 | 0.38 | 0.32 | 0.45 | 0.4 | 0.38 |
| 6 | 96259105 | rs9399238 | 0.014 | 1.159 | -0.616 | 1.245 | -0.016 | -0.015 | -0.015 | 0 | 0.34 | 0.23 | 0.3 | 0.48 | 0.4 | 0.39 |
| 6 | 96276408 | rs6936274 | -0.08 | 0.673 | -0.203 | 0.887 | -0.011 | -0.004 | -0.004 | 0 | 0.43 | 0.43 | 0.37 | 0.35 | 0.35 | 0.4 |
| 6 | 96283525 | rs17055551 | 0.215 | 0.445 | -0.351 | 0.734 | -0.003 | 0.002 | 0.002 | 0 | 0.31 | 0.42 | 0.31 | 0.29 | 0.3 | 0.35 |
| 6 | 96283669 | rs6570428 | 0.445 | 0.796 | -0.98 | 0.231 | -0.02 | 0.047 | 0.047 | 0.663 | 0.39 | 0.2 | 0.48 | 0.37 | 0.42 | 0.45 |
| 6 | 96289600 | rs9376622 | 0.028 | 0.615 | -0.939 | -0.124 | 0.008 | 0.04 | 0.04 | 0 | 0.23 | 0.07 | 0.12 | 0.12 | 0.18 | 0.16 |
| 6 | 96289803 | rs4434479 | 0.517 | 0.439 | -0.907 | -0.396 | 0.005 | 0.041 | 0.041 | 0 | 0.29 | 0.15 | 0.22 | 0.18 | 0.28 | 0.37 |
| 6 | 96291232 | rs1544136 | 0.418 | 0.429 | -0.932 | -0.373 | 0.008 | 0.04 | 0.04 | 0 | 0.23 | 0.06 | 0.12 | 0.12 | 0.18 | 0.16 |
| 6 | 96291278 | rs1544137 | 1.103 | 0.85 | -0.415 | -0.756 | 0.001 | 0.008 | 0.008 | 0.008 | 0.26 | 0.29 | 0.32 | 0.66 | 0.29 | 0.24 |
| 6 | 96291800 | rs10499007 | -0.02 | 0.778 | -0.772 | -1.183 | -0.094 | 0.114 | 0.114 | 0.72 | 0.24 | 0.07 | 0.37 | 0.03 | 0.38 | 0.25 |
| 6 | 96293482 | rs10499006 | 0.053 | 0.593 | -0.256 | -1 | 0.024 | 0.08 | 0.08 | 0 | 0.49 | 0.34 | 0.52 | 0.18 | 0.57 | 0.52 |
| 6 | 96293530 | rs17718892 | 1.081 | 0.632 | -0.598 | -1.012 | -0.013 | 0.069 | 0.069 | 0 | 0.33 | 0.11 | 0.23 | 0.18 | 0.33 | 0.38 |
| 6 | 96294640 | rs1336959 | 0.525 | 1.011 | -0.741 | -0.851 | -0.062 | 0.063 | 0.063 | 0 | 0.14 | 0.04 | 0.28 | 0 | 0.25 | 0.15 |
| 6 | 96296327 | rs2785917 | 0.521 | 0.752 | -0.556 | -0.782 | 0.013 | 0.043 | 0.043 | 0 | 0.23 | 0.12 | 0.12 | 0.09 | 0.17 | 0.16 |
| 6 | 96301487 | rs7754801 | 1.256 | 0.877 | -0.41 | -0.492 | -0.006 | 0.009 | 0.009 | 0 | 0.21 | 0.14 | 0.23 | 0.23 | 0.29 | 0.18 |
| 6 | 96302858 | rs2785905 | 0.5 | 0.803 | -0.507 | -0.897 | 0.013 | 0.037 | 0.037 | 0 | 0.23 | 0.12 | 0.12 | 0.11 | 0.17 | 0.16 |
| 6 | 96308231 | rs916498 | 0.42 | 1.024 | -0.381 | -0.702 | -0.003 | -0.022 | -0.022 | 0.393 | 0.14 | 0.12 | 0.23 | 0 | 0.17 | 0.14 |
| 6 | 96309043 | rs2799665 | -0.971 | 1.022 | -0.333 | -0.655 | 0.003 | 0.003 | 0.003 | 0 | 0.77 | 0.75 | 0.64 | 0.32 | 0.68 | 0.75 |
| 6 | 96314776 | rs17719229 | -0.048 | 0.857 | -0.222 | -0.383 | -0.006 | -0.014 | -0.014 | 0 | 0.14 | 0.11 | 0.23 | 0 | 0.17 | 0.14 |
| 6 | 96316201 | rs1537982 | -0.249 | 0.094 | -0.327 | -0.245 | 0.014 | 0.039 | 0.039 | 0 | 0.23 | 0.12 | 0.12 | 0.11 | 0.17 | 0.16 |
| 6 | 96317644 | rs2785895 | -1.058 | -0.135 | 0.387 | -0.388 | -0.007 | -0.074 | -0.074 | 0 | 0.27 | 0.49 | 0.26 | 0.13 | 0.21 | 0.31 |
| 6 | 96323966 | rs1857484 | 0.956 | -0.291 | 0.799 | 0.038 | -0.001 | 0 | 0 | 0.015 | 0.59 | 0.59 | 0.48 | 0.45 | 0.63 | 0.61 |
| 6 | 96324391 | rs17055573 | 0.173 | -0.36 | 0.833 | 0.032 | 0.033 | 0.043 | 0.043 | 0 | 0.15 | 0.07 | 0.07 | 0.03 | 0.07 | 0.08 |
| 6 | 96324667 | rs17719564 | 0.204 | -0.307 | 0.843 | 0.018 | -0.002 | -0.017 | -0.017 | 0 | 0.14 | 0.08 | 0.17 | 0 | 0.12 | 0.13 |
| 6 | 96326035 | rs2785918 | 0.273 | -0.303 | 0.921 | 0.067 | 0.025 | 0.125 | 0.125 | 0 | 0.88 | 0.76 | 0.72 | 0.49 | 0.82 | 0.81 |
| 6 | 96332211 | rs9390350 | 1.46 | -0.429 | 0.939 | 0.002 | 0.01 | -0.003 | -0.003 | 0 | 0.65 | 0.24 | 0.59 | 0.41 | 0.74 | 0.65 |

| | | | | | | | | | | | | | | | | |
|---|---|---|---|---|---|---|---|---|---|---|---|---|---|---|---|---|
| 6 | 96332506 | rs2785910 | 0.829 | -0.247 | 0.964 | 0.118 | 0.016 | 0.034 | 0.034 | 0 | 0.73 | 0.65 | 0.55 | 0.49 | 0.69 | 0.69 |
| 6 | 96350191 | rs1336951 | -0.01 | -0.112 | 1.233 | 0.449 | -0.007 | 0.048 | 0.048 | 0 | 0.05 | 0.23 | 0.21 | 0.1 | 0.02 | 0.09 |
| 6 | 96353696 | rs2785919 | 0.847 | -0.384 | 1.332 | 0.326 | -0.001 | 0.091 | 0.091 | 0 | 0.8 | 0.54 | 0.67 | 0.45 | 0.81 | 0.74 |
| 6 | 96357307 | rs1015924 | 1.35 | -0.44 | 1.255 | 0.239 | 0.008 | 0.023 | 0.023 | 0 | 0.67 | 0.46 | 0.51 | 0.45 | 0.69 | 0.6 |
| 6 | 96360452 | rs13215966 | 0.237 | -0.417 | 1.036 | -0.094 | -0.003 | -0.019 | -0.019 | 0.079 | 0.13 | 0.08 | 0.17 | 0.01 | 0.13 | 0.15 |
| 6 | 96362604 | rs12190516 | 1.67 | -0.515 | 0.328 | -0.151 | 0.032 | 0.057 | 0.057 | 0 | 0.49 | 0.19 | 0.41 | 0.42 | 0.61 | 0.51 |
| 6 | 96363607 | rs601581 | -0.124 | -0.356 | -0.002 | 0.362 | 0.039 | 0.051 | 0.051 | 0 | 0.85 | 0.95 | 0.92 | 0.97 | 0.93 | 0.92 |
| 6 | 96370232 | rs7765008 | -0.68 | 0.032 | 0.172 | 0.698 | 0.006 | 0.069 | 0.069 | 0 | 0.14 | 0.22 | 0.28 | 0.55 | 0.17 | 0.24 |
| 6 | 96373191 | rs654659 | 0.557 | 0.073 | -0.228 | -0.059 | 0.025 | 0.112 | 0.112 | 0.047 | 0.52 | 0.33 | 0.65 | 0.23 | 0.56 | 0.51 |
| 6 | 96374858 | rs2785926 | 0.42 | -0.461 | -0.146 | 0.478 | -0.02 | 0.003 | 0.003 | 0 | 0.74 | 0.33 | 0.61 | 0.6 | 0.9 | 0.74 |
| 6 | 96379508 | rs526939 | -0.282 | -0.554 | 0.454 | 0.478 | 0.007 | 0.007 | 0.007 | 0 | 0.83 | 0.88 | 0.91 | 0.87 | 0.93 | 0.86 |
| 6 | 96380002 | rs12204999 | 1.684 | -0.304 | 0.024 | 0.092 | -0.082 | 0.182 | 0.182 | 0 | 0.36 | 0.06 | 0.27 | 0.12 | 0.42 | 0.19 |
| 6 | 96384515 | rs6902603 | 0.217 | -0.271 | 0.233 | 0.264 | -0.014 | -0.023 | -0.023 | 1.236 | 0.26 | 0.21 | 0.18 | 0.48 | 0.36 | 0.39 |
| 6 | 96400830 | rs2143437 | -0.884 | -0.313 | 0.215 | 0.167 | -0.008 | 0.022 | 0.022 | 0 | 0.44 | 0.4 | 0.56 | 0.59 | 0.5 | 0.63 |
| 6 | 96407271 | rs584453 | 0.341 | -0.276 | 0.143 | 0.138 | -0.006 | -0.008 | -0.008 | 0 | 0.13 | 0.09 | 0.16 | 0 | 0.12 | 0.17 |
| 6 | 96409424 | rs10484741 | 0.189 | -0.398 | 0.278 | 0.206 | -0.022 | -0.031 | -0.031 | 0 | 0.27 | 0.21 | 0.19 | 0.49 | 0.36 | 0.38 |
| 6 | 96410114 | rs659115 | -0.662 | -0.463 | 0.089 | 0.12 | 0.054 | 0.011 | 0.011 | 0 | 0.86 | 0.99 | 0.94 | 0.88 | 0.94 | 0.86 |
| 6 | 96418766 | rs9386276 | -0.531 | -0.292 | 0.347 | 0.271 | -0.002 | 0.015 | 0.015 | 0 | 0.34 | 0.73 | 0.3 | 0.65 | 0.37 | 0.4 |
| 6 | 96426952 | rs590973 | -0.11 | 0.175 | 0.347 | 0.47 | 0.011 | 0.004 | 0.004 | 0.013 | 0.12 | 0.17 | 0.24 | 0.17 | 0.13 | 0.15 |
| 6 | 96427122 | rs655865 | -0.985 | 0.1 | 0.352 | 0.45 | -0.003 | -0.002 | -0.002 | 0 | 0.31 | 0.32 | 0.54 | 0.39 | 0.26 | 0.43 |
| 6 | 96427525 | rs501817 | -0.514 | 0.365 | 0.059 | 0.598 | -0.059 | -0.107 | -0.107 | 0 | 0.75 | 0.89 | 0.84 | 0.55 | 0.66 | 0.62 |
| 6 | 96438320 | rs1108409 | 0.265 | 0.35 | 0.006 | 0.598 | 0.011 | 0.036 | 0.036 | 0 | 0.18 | 0.05 | 0.12 | 0.13 | 0.12 | 0.17 |
| 6 | 96442356 | rs13211985 | 0.481 | 0.362 | -0.016 | 0.52 | 0.001 | 0.024 | 0.024 | 0 | 0.17 | 0.01 | 0.12 | 0.13 | 0.12 | 0.17 |
| 6 | 96444545 | rs612612 | -0.048 | 0.525 | 0.136 | 0.649 | -0.005 | 0.002 | 0.002 | 0 | 0.71 | 0.71 | 0.54 | 0.29 | 0.58 | 0.5 |
| 6 | 96459814 | rs1325076 | 0.994 | 0.282 | 0.162 | 0.507 | 0.005 | 0.013 | 0.013 | 0 | 0.45 | 0.6 | 0.36 | 0.32 | 0.4 | 0.21 |
| 6 | 96460364 | rs6911579 | 1.097 | 0.304 | 0.186 | 0.504 | -0.001 | -0.021 | -0.021 | 1.166 | 0.44 | 0.56 | 0.3 | 0.16 | 0.4 | 0.2 |
| 6 | 96460471 | rs9404127 | 1.026 | 0.314 | 0.142 | 0.524 | 0.004 | 0.013 | 0.013 | 0 | 0.44 | 0.6 | 0.36 | 0.33 | 0.4 | 0.21 |
| 6 | 96462168 | rs1546898 | 0.272 | 0.136 | 0.518 | 0.635 | 0.004 | 0.035 | 0.035 | 0 | 0.18 | 0.01 | 0.11 | 0.13 | 0.12 | 0.18 |
| 6 | 96475105 | rs1325067 | 1.031 | 0.397 | -0.017 | 0.425 | -0.013 | 0.103 | 0.103 | 0 | 0.42 | 0.26 | 0.3 | 0.17 | 0.44 | 0.22 |
| 6 | 96477293 | rs9377313 | -0.614 | 0.517 | 0.321 | 1.044 | 0.006 | 0.047 | 0.047 | 0 | 0.12 | 0.23 | 0.36 | 0.26 | 0.07 | 0.25 |
| 6 | 96479399 | rs4365940 | -0.265 | 0.527 | 0.084 | 1.007 | -0.002 | -0.001 | -0.001 | 0 | 0.27 | 0.35 | 0.21 | 0.28 | 0.26 | 0.38 |
| 6 | 96479757 | rs6924617 | -1.004 | 0.554 | -0.335 | 1.03 | 0.024 | 0.141 | 0.141 | 0.169 | 0.45 | 0.88 | 0.46 | 0.75 | 0.36 | 0.54 |
| 6 | 96481452 | rs9322615 | -0.681 | 1.195 | 0.022 | 0.474 | 0.007 | -0.007 | -0.007 | 0 | 0.38 | 0.43 | 0.41 | 0.33 | 0.42 | 0.51 |
| 6 | 96486869 | rs7747031 | -0.428 | 0.997 | 0.172 | 0.643 | -0.017 | 0.014 | 0.014 | 2.958 | 0.12 | 0.31 | 0.06 | 0.08 | 0.06 | 0.07 |
| 6 | 96488054 | rs9485570 | -0.224 | 1.403 | -0.426 | 0.516 | 0 | 0 | 0 | 0 | 0.27 | 0.3 | 0.2 | 0.25 | 0.25 | 0.37 |
| 6 | 96489816 | rs12195970 | -0.899 | 1.721 | -0.675 | 0.097 | -0.042 | 0.047 | 0.047 | 0 | 0.13 | 0.05 | 0.12 | 0 | 0.21 | 0.1 |
| 6 | 96491897 | rs11156165 | 0.065 | 1.798 | -0.303 | 0.101 | 0.02 | -0.025 | -0.025 | 0 | 0.07 | 0.08 | 0.07 | 0 | 0.13 | 0.04 |
| 6 | 96493233 | rs2248641 | -0.608 | 1.567 | -0.512 | -0.24 | 0.004 | 0.025 | 0.025 | 0 | 0.61 | 0.73 | 0.54 | 0.7 | 0.6 | 0.7 |
| 6 | 96503490 | rs9386302 | 0.175 | 1.538 | -0.338 | -0.047 | -0.006 | 0.007 | 0.007 | 0.312 | 0.38 | 0.41 | 0.33 | 0.55 | 0.31 | 0.54 |
| 6 | 96505219 | rs10499003 | 1.077 | 1.638 | -0.388 | -0.107 | 0 | -0.004 | -0.004 | 0.411 | 0.07 | 0.04 | 0.07 | 0 | 0.13 | 0.04 |
| 6 | 96505920 | rs1325072 | -0.603 | 1.457 | -0.381 | -0.052 | -0.002 | -0.005 | -0.005 | 0 | 0.56 | 0.5 | 0.51 | 0.65 | 0.54 | 0.68 |
| 6 | 96513935 | rs4634465 | 1.145 | 1.364 | -0.907 | -0.137 | -0.051 | 0.033 | 0.033 | 0 | 0.33 | 0.07 | 0.43 | 0.21 | 0.36 | 0.18 |
| 6 | 96521701 | rs12214839 | -0.357 | 1.561 | -0.211 | 0.314 | 0.06 | 0.018 | 0.018 | 0 | 0.18 | 0.31 | 0.35 | 0.19 | 0.26 | 0.1 |
| 6 | 96544414 | rs12200614 | -0.045 | 1.588 | -0.245 | 0.267 | 0.027 | -0.021 | -0.021 | 0.702 | 0.12 | 0.16 | 0.09 | 0.09 | 0.18 | 0.17 |
| 6 | 96547654 | rs12175153 | 0.911 | 1.738 | -0.221 | 0.325 | 0.001 | 0.011 | 0.011 | 0 | 0.07 | 0.07 | 0.23 | 0.18 | 0.1 | 0.05 |
| 6 | 96551023 | rs17056117 | 1.393 | 1.156 | -0.52 | 0.138 | 0.038 | -0.052 | -0.052 | 0 | 0.2 | 0.08 | 0.13 | 0.36 | 0.17 | 0.39 |
| 6 | 96554218 | rs17056121 | 1.876 | 1.024 | -0.574 | -0.171 | -0.042 | 0.033 | 0.033 | 0 | 0.31 | 0.07 | 0.33 | 0.21 | 0.36 | 0.18 |
| 6 | 96556603 | rs7768089 | -0.426 | 0.94 | -0.209 | -0.203 | 0.009 | 0.034 | 0.034 | 0 | 0.49 | 0.38 | 0.53 | 0.65 | 0.41 | 0.64 |
| 6 | 96566217 | rs9377383 | 0.936 | 0.782 | 0.093 | -0.318 | 0 | 0.001 | 0.001 | 0.097 | 0.9 | 0.63 | 0.96 | 0.92 | 0.94 | 0.87 |
| 6 | 96569905 | rs7772067 | 0.758 | 0.833 | -0.07 | -0.311 | 0.022 | -0.026 | -0.026 | 0.067 | 0.8 | 0.46 | 0.86 | 0.86 | 0.77 | 0.81 |
| 6 | 96571137 | rs9485739 | -0.216 | 0.807 | -0.204 | -0.179 | 0.005 | 0.012 | 0.012 | 0.638 | 0.41 | 0.36 | 0.44 | 0.73 | 0.37 | 0.68 |
| 6 | 96572628 | rs6571057 | 0.371 | 0.678 | -0.212 | -0.32 | 0 | -0.002 | -0.002 | 0.19 | 0.31 | 0.32 | 0.4 | 0.64 | 0.31 | 0.45 |
| 6 | 96574671 | rs9404263 | 0.347 | 0.681 | -0.209 | -0.317 | 0 | -0.002 | -0.002 | 0.482 | 0.32 | 0.32 | 0.4 | 0.64 | 0.33 | 0.45 |
| 6 | 96577879 | rs9390903 | 0.557 | 0.811 | -0.162 | -0.468 | -0.004 | -0.067 | -0.067 | 0.483 | 0.76 | 0.43 | 0.86 | 0.9 | 0.81 | 0.72 |
| 6 | 96578287 | rs4472357 | 1.448 | 0.671 | -0.354 | -0.485 | -0.06 | 0.116 | 0.116 | 0 | 0.41 | 0.08 | 0.43 | 0.2 | 0.44 | 0.26 |
| 6 | 96578439 | rs4555922 | -0.037 | 0.516 | 0.142 | -0.358 | -0.048 | -0.08 | -0.08 | 0 | 0.88 | 0.63 | 0.96 | 0.99 | 0.97 | 0.9 |
| 6 | 96586116 | rs12214860 | -0.093 | 0.77 | -0.298 | -0.341 | -0.023 | -0.07 | -0.07 | 0.531 | 0.24 | 0.05 | 0.23 | 0.31 | 0.21 | 0.28 |
| 6 | 96593695 | rs6900374 | 0.757 | 0.993 | -0.071 | -0.443 | -0.005 | 0.001 | 0.001 | 0.094 | 0.63 | 0.37 | 0.66 | 0.69 | 0.62 | 0.65 |
| 6 | 96595020 | rs4132260 | 0.407 | 0.985 | -0.046 | -0.425 | 0.002 | -0.002 | -0.002 | 0.003 | 0.67 | 0.39 | 0.67 | 0.78 | 0.63 | 0.78 |
| 6 | 96599728 | rs4385319 | -0.079 | 1.022 | 0.112 | -0.568 | 0.064 | -0.072 | -0.072 | 0 | 0.88 | 0.46 | 0.89 | 0.99 | 0.86 | 0.93 |
| 6 | 96606573 | rs6571086 | 0.878 | 1.042 | 0.241 | -0.558 | 0.077 | -0.04 | -0.04 | 0.312 | 0.08 | 0.19 | 0.09 | 0 | 0.14 | 0.04 |
| 6 | 96608866 | rs4599653 | 1.315 | 0.73 | -0.123 | -0.536 | 0.034 | -0.048 | -0.048 | 0.022 | 0.23 | 0.09 | 0.13 | 0.37 | 0.18 | 0.37 |
| 6 | 96622433 | rs17725092 | 1.814 | 0.928 | -0.177 | -0.611 | -0.032 | 0.032 | 0.032 | 0 | 0.06 | 0 | 0.04 | 0 | 0.07 | 0.02 |
| 6 | 96625462 | rs12661236 | 1.01 | 0.813 | -0.212 | -0.669 | 0.022 | -0.025 | -0.025 | 0 | 0.56 | 0.19 | 0.41 | 0.6 | 0.49 | 0.72 |
| 6 | 96628270 | rs9377473 | 0.646 | 0.798 | -0.151 | -0.553 | 0.009 | 0.002 | 0.002 | 0.121 | 0.12 | 0.19 | 0.27 | 0.15 | 0.15 | 0.06 |
| 6 | 96635942 | rs9391015 | 0.458 | 0.787 | -0.035 | -0.559 | 0.02 | 0.006 | 0.006 | 0.024 | 0.12 | 0.24 | 0.27 | 0.15 | 0.15 | 0.06 |
| 6 | 96658670 | rs17056459 | 1.479 | 0.751 | -0.189 | -0.554 | 0.034 | -0.074 | -0.074 | 2.112 | 0.23 | 0.09 | 0.15 | 0.46 | 0.2 | 0.38 |
| 6 | 96660070 | rs17056460 | 0.395 | 1.071 | 0.035 | -0.354 | 0.021 | 0.006 | 0.006 | 0.044 | 0.12 | 0.24 | 0.27 | 0.15 | 0.15 | 0.06 |
| 6 | 96666863 | rs9373728 | 0.41 | 1.054 | -0.101 | -0.388 | 0.012 | 0.004 | 0.004 | 0.106 | 0.13 | 0.26 | 0.24 | 0.15 | 0.13 | 0.07 |

| | | | | | | | | | | | | | | | | |
|---|---|---|---|---|---|---|---|---|---|---|---|---|---|---|---|---|
| 6 | 96669605 | rs10498999 | 1.63 | 1.002 | -0.18 | -0.42 | 0.093 | 0.003 | 0.003 | 0 | 0.51 | 0.13 | 0.39 | 0.49 | 0.43 | 0.58 |
| 6 | 96672687 | rs4590278 | -0.102 | 1.24 | -0.353 | -0.375 | -0.019 | -0.008 | -0.008 | 0.026 | 0.27 | 0.2 | 0.29 | 0.24 | 0.36 | 0.17 |
| 6 | 96674755 | rs7768506 | -1.183 | 1.249 | -0.448 | -0.368 | -0.002 | 0.004 | 0.004 | 0 | 0.41 | 0.49 | 0.56 | 0.39 | 0.51 | 0.25 |
| 6 | 96681831 | rs9322777 | 0.437 | 1.363 | -1.002 | -0.214 | 0.002 | -0.001 | -0.001 | 0 | 0.13 | 0.16 | 0.27 | 0.15 | 0.15 | 0.09 |
| 6 | 96689890 | rs9404497 | 0.621 | 1.365 | -1.283 | -0.148 | -0.063 | -0.086 | -0.086 | 0 | 0.18 | 0.04 | 0.21 | 0.24 | 0.22 | 0.13 |
| 6 | 96693765 | rs17056489 | 1.099 | 1.108 | -1.281 | -0.285 | 0.031 | 0.005 | 0.005 | 0 | 0.55 | 0.15 | 0.4 | 0.51 | 0.48 | 0.71 |
| 6 | 96695140 | rs11752646 | 1.415 | 0.958 | -1.295 | -0.652 | 0.034 | 0.154 | 0.154 | 0 | 0.29 | 0.05 | 0.23 | 0.04 | 0.24 | 0.2 |
| 6 | 96700328 | rs9322788 | -1.121 | 1.255 | -1.226 | -0.469 | -0.038 | -0.018 | -0.018 | 0 | 0.34 | 0.8 | 0.37 | 0.32 | 0.41 | 0.23 |
| 6 | 96703681 | rs11754959 | 1.423 | 1.029 | -1.245 | -0.383 | 0.008 | 0.155 | 0.155 | 0.055 | 0.27 | 0.04 | 0.22 | 0.03 | 0.25 | 0.21 |
| 6 | 96711264 | rs6938549 | 0.298 | 1.235 | -0.615 | -0.02 | -0.008 | -0.009 | -0.009 | 0 | 0.18 | 0.13 | 0.23 | 0.22 | 0.23 | 0.13 |
| 6 | 96721908 | rs6903469 | -1.491 | 1.291 | -0.592 | 0.01 | -0.031 | -0.047 | -0.047 | 0 | 0.28 | 0.82 | 0.34 | 0.42 | 0.35 | 0.3 |
| 6 | 96723784 | rs9399876 | -0.169 | 1.488 | -0.646 | -0.045 | -0.04 | -0.034 | -0.034 | 0 | 0.18 | 0.11 | 0.31 | 0.24 | 0.3 | 0.21 |
| 6 | 96739393 | rs2263190 | -1.231 | 0.772 | -1.009 | -0.55 | 0.031 | 0.171 | 0.171 | 0.038 | 0.71 | 0.97 | 0.83 | 0.96 | 0.75 | 0.77 |
| 6 | 96746620 | rs7754354 | -0.106 | 0.693 | -0.557 | -0.175 | -0.017 | -0.022 | -0.022 | 1.198 | 0.17 | 0.13 | 0.29 | 0.31 | 0.24 | 0.18 |
| 6 | 96746717 | rs17056546 | -0.215 | 0.706 | -0.256 | 0.027 | 0.028 | 0.04 | 0.04 | 0 | 0.16 | 0.26 | 0.29 | 0.26 | 0.19 | 0.14 |
| 6 | 96751756 | rs2387154 | -0.758 | 0.636 | -0.425 | -0.006 | 0.005 | 0.013 | 0.013 | 0 | 0.24 | 0.68 | 0.32 | 0.31 | 0.24 | 0.19 |
| 6 | 96756254 | rs9391219 | 0.049 | 0.622 | -0.366 | 0.022 | 0.018 | 0.032 | 0.032 | 0 | 0.13 | 0.21 | 0.26 | 0.25 | 0.17 | 0.12 |
| 6 | 96762799 | rs4839824 | 0.25 | 0.623 | -0.445 | 0.04 | -0.03 | -0.028 | -0.028 | 0 | 0.12 | 0.08 | 0.25 | 0.17 | 0.16 | 0.09 |
| 6 | 96764612 | rs7763613 | -1.626 | 0.581 | -0.467 | 0.082 | -0.073 | -0.029 | -0.029 | 0.386 | 0.38 | 0.88 | 0.61 | 0.57 | 0.43 | 0.31 |
| 6 | 96796050 | rs1983551 | -1.252 | 0.075 | -0.541 | -0.116 | 0.056 | 0.121 | 0.121 | 0 | 0.77 | 0.97 | 0.86 | 0.97 | 0.84 | 0.81 |
| 6 | 96796630 | rs9322829 | -0.032 | 0.713 | 0.012 | 0.435 | -0.008 | -0.008 | -0.008 | 0.166 | 0.13 | 0.11 | 0.25 | 0.17 | 0.16 | 0.09 |
| 6 | 96825758 | rs2472918 | -0.017 | 0.711 | 0.003 | 0.313 | 0.008 | 0.008 | 0.008 | 0 | 0.12 | 0.15 | 0.26 | 0.17 | 0.16 | 0.09 |
| 6 | 96830805 | rs9386441 | 0.299 | 0.963 | -0.319 | 0.362 | -0.036 | -0.057 | -0.057 | 0 | 0.12 | 0.04 | 0.17 | 0.25 | 0.15 | 0.09 |
| 6 | 96832495 | rs2472921 | -1.451 | 0.868 | -0.013 | 0.303 | -0.005 | -0.006 | -0.006 | 0 | 0.31 | 0.72 | 0.47 | 0.48 | 0.34 | 0.23 |
| 6 | 96834665 | rs9386443 | -0.121 | 0.867 | 0.041 | 0.305 | -0.021 | -0.036 | -0.036 | 0 | 0.14 | 0.1 | 0.18 | 0.31 | 0.18 | 0.14 |
| 6 | 96835536 | rs2472923 | -0.962 | 0.682 | 0.328 | 0.243 | 0.022 | 0.023 | 0.023 | 0.03 | 0.17 | 0.61 | 0.29 | 0.17 | 0.16 | 0.09 |
| 6 | 96838735 | rs767618 | -0.143 | 0.943 | -0.048 | 0.342 | 0.017 | 0.016 | 0.016 | 0 | 0.13 | 0.17 | 0.26 | 0.17 | 0.16 | 0.09 |
| 6 | 96846264 | rs2499792 | -1.165 | 0.432 | -0.229 | -0.072 | 0.01 | 0.031 | 0.031 | 0 | 0.81 | 0.9 | 0.85 | 0.97 | 0.81 | 0.84 |
| 6 | 96846975 | rs9384578 | -0.198 | 0.629 | -0.408 | 0.326 | -0.047 | -0.064 | -0.064 | 1.788 | 0.16 | 0.06 | 0.3 | 0.34 | 0.22 | 0.16 |
| 6 | 96847181 | rs6939383 | 0.816 | 0.373 | 0.028 | 0.307 | 0 | 0.022 | 0.022 | 0 | 0.51 | 0.38 | 0.4 | 0.36 | 0.51 | 0.54 |
| 6 | 96849679 | rs976357 | -1.532 | 0.246 | 0.152 | -0.141 | 0.011 | 0.032 | 0.032 | 0 | 0.81 | 0.9 | 0.85 | 0.97 | 0.81 | 0.84 |
| 6 | 96850379 | rs9399923 | -0.6 | 0.263 | 0.199 | -0.137 | -0.008 | -0.02 | -0.02 | 1.654 | 0.72 | 0.67 | 0.82 | 0.78 | 0.76 | 0.75 |
| 6 | 96853579 | rs6924957 | -0.697 | 0.253 | 0.231 | -0.202 | -0.006 | -0.019 | -0.019 | 0 | 0.7 | 0.67 | 0.81 | 0.78 | 0.75 | 0.74 |
| 6 | 96858902 | rs760620 | -0.039 | 0.268 | 0.355 | -0.151 | -0.018 | 0.002 | 0.002 | 2.409 | 0.11 | 0.25 | 0.04 | 0.14 | 0.05 | 0.14 |
| 6 | 96861476 | rs6568392 | -0.546 | 0.216 | 0.434 | -0.23 | -0.039 | -0.114 | -0.114 | 0 | 0.72 | 0.49 | 0.77 | 0.92 | 0.76 | 0.79 |
| 6 | 96894328 | rs4839835 | 0.925 | 0.317 | 0.427 | -0.244 | 0.029 | 0.049 | 0.049 | 0.104 | 0.95 | 0.77 | 0.96 | 0.96 | 0.94 | 0.94 |
| 6 | 96938227 | rs6934392 | 0.66 | 0.125 | 0.42 | 0.081 | 0 | 0.006 | 0.006 | 0 | 0.49 | 0.4 | 0.45 | 0.45 | 0.51 | 0.52 |
| 6 | 96969694 | rs13210597 | 1.247 | -0.156 | 0.433 | -0.339 | 0.025 | 0.064 | 0.064 | 0 | 0.23 | 0.1 | 0.18 | 0.04 | 0.22 | 0.16 |
| 6 | 96999725 | rs1127175 | 0.056 | -0.082 | 0.462 | -0.296 | 0.013 | 0.02 | 0.02 | 0 | 0.42 | 0.33 | 0.29 | 0.31 | 0.37 | 0.33 |
| 6 | 97003725 | rs9384654 | 0.777 | -0.084 | 0.36 | -0.336 | -0.005 | -0.039 | -0.039 | 0 | 0.15 | 0.07 | 0.07 | 0.27 | 0.13 | 0.16 |
| 6 | 97012746 | rs11153058 | 1.234 | -0.089 | 0.351 | -0.35 | 0.025 | 0.061 | 0.061 | 0 | 0.23 | 0.1 | 0.18 | 0.04 | 0.21 | 0.17 |
| 6 | 97052754 | rs2064947 | 1.217 | -0.087 | 0.472 | -0.349 | 0.024 | 0.057 | 0.057 | 0 | 0.23 | 0.11 | 0.18 | 0.04 | 0.21 | 0.16 |
| 6 | 97058553 | rs2273621 | -0.252 | -0.086 | 0.607 | -0.331 | -0.004 | -0.003 | -0.003 | 0 | 0.45 | 0.55 | 0.3 | 0.35 | 0.35 | 0.36 |
| 6 | 97058574 | rs2252816 | 0.499 | -0.16 | -0.084 | -0.677 | -0.142 | -0.153 | -0.153 | 0 | 0.34 | 0.04 | 0.5 | 0.56 | 0.39 | 0.35 |
| 6 | 97061940 | rs6923212 | 1.459 | -0.327 | 0.085 | -0.542 | 0.025 | 0.065 | 0.065 | 0 | 0.23 | 0.1 | 0.18 | 0.04 | 0.21 | 0.16 |
| 6 | 97063555 | rs9398148 | 0.167 | -0.383 | 0.19 | -0.561 | -0.005 | -0.003 | -0.003 | 0 | 0.45 | 0.5 | 0.3 | 0.35 | 0.35 | 0.35 |
| 6 | 97063724 | rs9398152 | 0.959 | -0.257 | -0.257 | -0.43 | 0.005 | -0.021 | -0.021 | 0 | 0.16 | 0.08 | 0.07 | 0.27 | 0.11 | 0.15 |
| 6 | 97069838 | rs211168 | -1.185 | -0.069 | -0.54 | -0.528 | -0.09 | 0.026 | 0.026 | 0.589 | 0.82 | 0.99 | 0.83 | 0.89 | 0.72 | 0.69 |
| 6 | 97070100 | rs211170 | -1.522 | 0.042 | 0.354 | 0.28 | -0.059 | 0.192 | 0.192 | 0 | 0.59 | 0.89 | 0.64 | 0.86 | 0.52 | 0.53 |
| 6 | 97073320 | rs211175 | -0.203 | 0.003 | 0.521 | 0.32 | 0 | 0 | 0 | 0 | 0.35 | 0.36 | 0.52 | 0.54 | 0.37 | 0.31 |
| 6 | 97075910 | rs927271 | -0.411 | 0.291 | -0.161 | 0.109 | 0.004 | -0.001 | -0.001 | 0 | 0.41 | 0.73 | 0.54 | 0.58 | 0.37 | 0.37 |
| 6 | 97080198 | rs12200560 | 1.079 | 0.49 | -0.641 | -0.095 | -0.059 | 0.19 | 0.19 | 0 | 0.41 | 0.11 | 0.36 | 0.14 | 0.48 | 0.47 |
| 6 | 97083485 | rs12197804 | 1.077 | 1.176 | -0.191 | 0.514 | 0.007 | 0.206 | 0.206 | 0 | 0.47 | 0.18 | 0.35 | 0.14 | 0.47 | 0.44 |
| 6 | 97084774 | rs9374010 | 0.125 | 1.073 | -0.522 | 0.79 | 0 | -0.004 | -0.004 | 0 | 0.17 | 0.13 | 0.06 | 0.24 | 0.14 | 0.16 |
| 6 | 97085238 | rs4839850 | -0.095 | 1.343 | -0.279 | 1.636 | -0.061 | -0.117 | -0.117 | 2.415 | 0.16 | 0.01 | 0.25 | 0.38 | 0.21 | 0.24 |
| 6 | 97093037 | rs1034240 | 0.335 | 1.09 | 0.342 | 0.847 | 0.012 | 0.012 | 0.012 | 0 | 0.64 | 0.45 | 0.41 | 0.38 | 0.6 | 0.6 |
| 6 | 97094092 | rs1413524 | 1.145 | 1.149 | 0.319 | 0.717 | 0.019 | -0.01 | -0.01 | 0 | 0.93 | 0.83 | 0.92 | 0.97 | 0.92 | 0.91 |
| 6 | 97094444 | rs1334328 | 0.561 | 1.48 | 0.085 | 0.492 | 0.017 | -0.018 | -0.018 | 0.284 | 0.62 | 0.44 | 0.62 | 0.72 | 0.53 | 0.44 |
| 6 | 97096065 | rs9400212 | -0.214 | 1.332 | -0.383 | 0.353 | -0.004 | -0.023 | -0.023 | 0 | 0.37 | 0.21 | 0.24 | 0.41 | 0.36 | 0.37 |
| 6 | 97099220 | rs9398164 | 0.204 | 1.25 | 0.363 | -0.006 | 0.027 | -0.058 | -0.058 | 0.229 | 0.9 | 0.66 | 0.89 | 1 | 0.9 | 0.95 |
| 6 | 97102615 | rs1413522 | 0.043 | 1.689 | 0.253 | -0.138 | 0.004 | 0.018 | 0.018 | 0 | 0.34 | 0.34 | 0.32 | 0.44 | 0.33 | 0.38 |
| 6 | 97115136 | rs17798014 | 0.018 | 1.488 | -0.326 | -0.613 | 0 | 0.026 | 0.026 | 0 | 0.17 | 0.16 | 0.14 | 0.07 | 0.17 | 0.14 |
| 6 | 97115959 | rs7755528 | -0.256 | 1.557 | 0.101 | -0.721 | -0.002 | -0.003 | -0.003 | 2.285 | 0.57 | 0.46 | 0.58 | 0.57 | 0.57 | 0.57 |
| 6 | 97119260 | rs4839692 | 1.134 | 1.541 | -0.117 | -0.898 | 0.001 | -0.004 | -0.004 | 0 | 0.21 | 0.15 | 0.12 | 0.37 | 0.16 | 0.18 |
| 6 | 97120254 | rs996059 | -0.136 | 1.499 | -0.126 | -0.822 | 0.005 | -0.054 | -0.054 | 0 | 0.9 | 0.78 | 0.9 | 1 | 0.9 | 0.95 |
| 6 | 97120756 | rs17741645 | 0.469 | 1.498 | -0.441 | -0.862 | -0.003 | 0.008 | 0.008 | 0 | 0.1 | 0.1 | 0.08 | 0.07 | 0.11 | 0.1 |
| 6 | 97120870 | rs9400228 | 1.318 | 1.192 | -0.598 | -0.997 | -0.034 | -0.132 | -0.132 | 0.992 | 0.31 | 0.03 | 0.39 | 0.49 | 0.31 | 0.36 |
| 6 | 97125641 | rs9486890 | -0.354 | 1.222 | -0.09 | -1.21 | 0.001 | -0.025 | -0.025 | 0 | 0.12 | 0.17 | 0.12 | 0.04 | 0.12 | 0.08 |
| 6 | 97126729 | rs4839861 | 1.443 | 1.171 | -0.107 | -1.595 | 0.003 | -0.003 | -0.003 | 0 | 0.66 | 0.35 | 0.72 | 0.96 | 0.68 | 0.7 |

| | | | | | | | | | | | | | | | | |
|---|---|---|---|---|---|---|---|---|---|---|---|---|---|---|---|---|
| 6 | 97127920 | rs9386728 | 1.459 | 1.186 | -0.11 | -1.585 | 0.003 | 0.003 | 0.003 | 0 | 0.66 | 0.34 | 0.63 | 0.93 | 0.68 | 0.69 |
| 6 | 97130909 | rs13218880 | -0.087 | 1.138 | -0.308 | -1.293 | 0 | -0.003 | -0.003 | 0.843 | 0.11 | 0.1 | 0.12 | 0.04 | 0.12 | 0.08 |
| 6 | 97135177 | rs933042 | 0.111 | 0.808 | -1.119 | -1.398 | -0.026 | 0.02 | 0.02 | 0 | 0.4 | 0.23 | 0.43 | 0.38 | 0.43 | 0.42 |
| 6 | 97136981 | rs7746875 | -0.427 | 0.679 | -0.741 | -1.574 | 0.002 | -0.095 | -0.095 | 0 | 0.17 | 0.36 | 0.2 | 0.03 | 0.14 | 0.26 |
| 6 | 97145600 | rs9386745 | 1.497 | 0.302 | -1.373 | -1.01 | 0.017 | -0.123 | -0.123 | 1.159 | 0.27 | 0.03 | 0.17 | 0.34 | 0.27 | 0.23 |
| 6 | 97159223 | rs538052 | 0.413 | 0.515 | -1.147 | -0.976 | 0.032 | 0.001 | 0.001 | 0 | 0.44 | 0.65 | 0.29 | 0.38 | 0.38 | 0.36 |
| 6 | 97160140 | rs13207096 | -0.307 | 1.329 | -1.392 | -0.778 | -0.101 | 0.052 | 0.052 | 0.013 | 0.21 | 0.04 | 0.18 | 0.07 | 0.27 | 0.25 |
| 6 | 97162969 | rs381952 | 0.307 | 1.444 | -0.981 | -0.849 | 0.022 | -0.001 | -0.001 | 0 | 0.76 | 0.74 | 0.65 | 0.75 | 0.69 | 0.63 |
| 6 | 97163187 | rs1334327 | 0.243 | 1.426 | -0.718 | -1.008 | 0.04 | -0.027 | -0.027 | 0 | 0.07 | 0.25 | 0.08 | 0.01 | 0.09 | 0.05 |
| 6 | 97164468 | rs9374051 | 1.903 | 0.928 | -1.288 | -1.199 | 0.004 | -0.142 | -0.142 | 0.591 | 0.25 | 0.04 | 0.19 | 0.38 | 0.27 | 0.22 |
| 6 | 97164505 | rs394490 | 0.831 | 1.033 | -1.2 | -1.262 | -0.03 | 0.037 | 0.037 | 0 | 0.09 | 0.03 | 0.08 | 0 | 0.12 | 0.11 |
| 6 | 97166640 | rs9480890 | -0.147 | 0.868 | -1.095 | -1.223 | -0.001 | -0.065 | -0.065 | 0.208 | 0.14 | 0.31 | 0.11 | 0.01 | 0.12 | 0.09 |
| 6 | 97171823 | rs11756135 | 1.267 | 0.802 | -1.131 | -1.173 | 0.002 | -0.004 | -0.004 | 0.062 | 0.07 | 0.07 | 0.03 | 0 | 0.03 | 0.04 |
| 6 | 97174304 | rs17056960 | 0.992 | 0.784 | -1.032 | -1.169 | -0.035 | 0.011 | 0.011 | 0 | 0.28 | 0.37 | 0.27 | 0.35 | 0.16 | 0.22 |
| 6 | 97186347 | rs13217314 | 2.057 | 0.62 | -1.316 | -1.227 | 0 | -0.121 | -0.121 | 0.16 | 0.23 | 0.05 | 0.19 | 0.39 | 0.26 | 0.2 |
| 6 | 97191923 | rs9386770 | 0.68 | 0.674 | -1.016 | -1.284 | -0.022 | -0.002 | -0.002 | 0.131 | 0.34 | 0.54 | 0.34 | 0.37 | 0.25 | 0.26 |
| 6 | 97192188 | rs12661366 | 1.075 | 1.189 | -1.044 | -1.235 | -0.041 | -0.001 | -0.001 | 0 | 0.07 | 0.01 | 0.04 | 0.04 | 0.11 | 0.1 |
| 6 | 97194665 | rs12201147 | 0.097 | 0.45 | -0.837 | -1.174 | -0.005 | 0.013 | 0.013 | 0.352 | 0.16 | 0.12 | 0.11 | 0 | 0.16 | 0.14 |
| 6 | 97208361 | rs2206573 | 0.604 | 0.475 | -0.853 | -1.193 | -0.001 | 0.011 | 0.011 | 0 | 0.62 | 0.68 | 0.51 | 0.75 | 0.47 | 0.49 |
| 6 | 97209235 | rs4839868 | 0.202 | 0.634 | -0.831 | -1.174 | 0.005 | -0.002 | -0.002 | 0 | 0.69 | 0.72 | 0.58 | 0.75 | 0.58 | 0.6 |
| 6 | 97213142 | rs6937313 | 0.828 | 0.222 | -0.831 | -1.14 | -0.009 | 0.001 | 0.001 | 0 | 0.29 | 0.3 | 0.28 | 0.36 | 0.17 | 0.21 |
| 6 | 97215370 | rs9374085 | 1.79 | 0.309 | -1.129 | -0.527 | -0.019 | 0.005 | 0.005 | 0 | 0.06 | 0.03 | 0.12 | 0.25 | 0.14 | 0.08 |
| 6 | 97222294 | rs12528520 | 1 | 0.305 | -0.91 | -0.617 | 0.001 | -0.001 | -0.001 | 0.018 | 0.06 | 0.05 | 0.08 | 0.03 | 0.09 | 0.03 |
| 6 | 97229612 | rs6941115 | 1.013 | 0.017 | -0.792 | -0.708 | -0.038 | 0.007 | 0.007 | 0 | 0.3 | 0.6 | 0.28 | 0.38 | 0.16 | 0.24 |
| 6 | 97230516 | rs213858 | 0.858 | 0.221 | -0.939 | -0.596 | -0.022 | -0.059 | -0.059 | 0.091 | 0.33 | 0.12 | 0.28 | 0.37 | 0.39 | 0.32 |
| 6 | 97234008 | rs213860 | 0.122 | 0.127 | -0.821 | -0.517 | -0.016 | -0.007 | -0.007 | 0 | 0.4 | 0.31 | 0.35 | 0.4 | 0.48 | 0.39 |
| 6 | 97234433 | rs9487145 | -0.105 | -0.044 | -0.551 | -0.725 | -0.015 | -0.044 | -0.044 | 0 | 0.14 | 0.37 | 0.06 | 0.06 | 0.09 | 0.11 |
| 6 | 97235589 | rs850435 | -1.054 | 0.212 | -0.713 | -0.785 | 0.002 | -0.001 | -0.001 | 0 | 0.78 | 0.79 | 0.69 | 0.8 | 0.73 | 0.65 |
| 6 | 97238122 | rs437346 | 0.808 | 0.294 | -0.984 | -0.7 | -0.01 | -0.037 | -0.037 | 0 | 0.35 | 0.15 | 0.29 | 0.38 | 0.41 | 0.35 |
| 6 | 97239429 | rs11153198 | 2.242 | -0.11 | -1.263 | -0.774 | -0.031 | -0.093 | -0.093 | 0.163 | 0.17 | 0.01 | 0.25 | 0.34 | 0.15 | 0.15 |
| 6 | 97240665 | rs1928967 | 1.746 | 0.102 | -1.116 | -0.75 | -0.01 | -0.11 | -0.11 | 0.497 | 0.23 | 0.06 | 0.19 | 0.38 | 0.27 | 0.24 |
| 6 | 97281052 | rs1328130 | -1.018 | 0.295 | -0.908 | -0.847 | -0.029 | 0.052 | 0.052 | 0 | 0.89 | 0.97 | 0.92 | 1 | 0.87 | 0.9 |
| 6 | 97283783 | rs6568629 | 0.828 | 0.232 | -0.68 | -0.985 | 0.013 | -0.007 | -0.007 | 0 | 0.25 | 0.29 | 0.33 | 0.2 | 0.29 | 0.34 |
| 6 | 97314345 | rs9374164 | 1.446 | -0.027 | -0.747 | -1.064 | 0 | -0.001 | -0.001 | 0 | 0.26 | 0.23 | 0.34 | 0.36 | 0.22 | 0.22 |
| 6 | 97316278 | rs11967227 | 1.261 | 0.006 | -0.843 | -1.05 | -0.007 | 0.005 | 0.005 | 0 | 0.27 | 0.3 | 0.34 | 0.36 | 0.22 | 0.22 |
| 6 | 97318206 | rs12528971 | -0.693 | -0.033 | -0.917 | -0.951 | -0.001 | -0.001 | -0.001 | 0 | 0.62 | 0.63 | 0.7 | 0.61 | 0.56 | 0.62 |
| 6 | 97320164 | rs7749140 | 0.363 | 0.35 | -0.617 | -1.189 | 0.009 | 0.001 | 0.001 | 0.121 | 0.25 | 0.3 | 0.33 | 0.25 | 0.3 | 0.35 |
| 6 | 97332805 | rs6568649 | 0.3 | 0.506 | -1.258 | -1.149 | -0.03 | 0.107 | 0.107 | 0.23 | 0.81 | 0.96 | 0.89 | 1 | 0.82 | 0.85 |
| 6 | 97338341 | rs11759571 | 0.836 | 0.504 | -1.26 | -1.151 | 0.005 | 0.039 | 0.039 | 0 | 0.08 | 0.01 | 0.03 | 0 | 0.04 | 0.06 |
| 6 | 97339078 | rs6684 | 0.875 | 0.595 | -0.961 | -0.994 | 0.013 | 0.054 | 0.054 | 0 | 0.51 | 0.37 | 0.54 | 0.76 | 0.49 | 0.47 |
| 6 | 97339088 | rs11402 | 0.074 | 0.96 | -1.061 | -0.591 | -0.028 | 0.046 | 0.046 | 0 | 0.89 | 0.97 | 0.92 | 1 | 0.87 | 0.9 |
| 6 | 97360419 | rs4839882 | 0.245 | 0.613 | -0.785 | -0.12 | -0.005 | 0.014 | 0.014 | 0 | 0.44 | 0.36 | 0.45 | 0.35 | 0.51 | 0.23 |
| 6 | 97365239 | rs850577 | -1.048 | 0.692 | -0.921 | -0.289 | -0.009 | -0.062 | -0.062 | 0 | 0.7 | 0.83 | 0.69 | 0.53 | 0.72 | 0.83 |
| 6 | 97370194 | rs850583 | -0.499 | 0.208 | -1.047 | -0.38 | 0.031 | 0.08 | 0.08 | 0 | 0.5 | 0.7 | 0.55 | 0.27 | 0.57 | 0.6 |
| 6 | 97371612 | rs850586 | 0.835 | -0.069 | -1.111 | -0.529 | 0.006 | 0.003 | 0.003 | 0 | 0.09 | 0.07 | 0.04 | 0.07 | 0.04 | 0.14 |
| 6 | 97377572 | rs1359931 | -0.408 | 0.054 | -0.958 | -0.778 | -0.002 | 0.019 | 0.019 | 0 | 0.25 | 0.22 | 0.19 | 0.07 | 0.27 | 0.25 |
| 6 | 97381858 | rs982124 | 1.161 | 0.705 | -0.663 | -0.678 | -0.008 | 0.022 | 0.022 | 0 | 0.09 | 0.02 | 0.08 | 0.04 | 0.09 | 0.05 |
| 6 | 97387349 | rs7770996 | 0.349 | 0.721 | -0.659 | -0.727 | -0.006 | -0.012 | -0.012 | 0 | 0.14 | 0.15 | 0.13 | 0 | 0.21 | 0.11 |
| 6 | 97387465 | rs12662765 | -0.662 | 0.343 | 0.146 | -0.08 | 0.032 | -0.001 | -0.001 | 0 | 0.26 | 0.43 | 0.37 | 0.2 | 0.31 | 0.35 |
| 6 | 97388179 | rs7761614 | 0.246 | 0.351 | 0.172 | -0.073 | 0.052 | 0.037 | 0.037 | 0.016 | 0.16 | 0.33 | 0.28 | 0.17 | 0.21 | 0.3 |
| 6 | 97397823 | rs4140566 | 0.302 | -0.042 | -0.232 | -0.004 | 0.032 | -0.037 | -0.037 | 0.374 | 0.19 | 0.03 | 0.13 | 0.26 | 0.14 | 0.22 |
| 6 | 97405915 | rs7762923 | 0.354 | 0.011 | -0.069 | -0.022 | -0.003 | -0.019 | -0.019 | 0.524 | 0.11 | 0.14 | 0.09 | 0.03 | 0.1 | 0.06 |
| 6 | 97409378 | rs12525889 | 0.755 | 0.038 | -0.078 | -0.008 | -0.001 | 0.006 | 0.006 | 0 | 0.09 | 0.05 | 0.09 | 0.03 | 0.09 | 0.06 |
| 6 | 97419598 | rs974417 | 0.077 | 0.164 | 0.18 | 0.126 | 0.068 | 0.08 | 0.08 | 0 | 0.14 | 0.37 | 0.25 | 0.18 | 0.17 | 0.3 |
| 6 | 97422930 | rs2179537 | -0.288 | 0.042 | 0.265 | 0.076 | -0.001 | 0.002 | 0.002 | 0 | 0.42 | 0.42 | 0.47 | 0.47 | 0.4 | 0.57 |
| 6 | 97423683 | rs11756272 | -0.424 | 0.12 | 0.065 | 0.117 | -0.002 | 0.002 | 0.002 | 0 | 0.44 | 0.51 | 0.47 | 0.47 | 0.4 | 0.57 |
| 6 | 97430103 | rs12523684 | -0.495 | 0.148 | -0.276 | 0.131 | 0.009 | -0.006 | -0.006 | 0 | 0.45 | 0.61 | 0.48 | 0.46 | 0.4 | 0.57 |
| 6 | 97438289 | rs1933459 | 0.162 | -0.327 | 0.363 | 0.157 | -0.022 | -0.051 | -0.051 | 1.406 | 0.27 | 0.39 | 0.17 | 0.11 | 0.22 | 0.21 |
| 6 | 97441555 | rs1206112 | -1.035 | -0.4 | 0.554 | 0.454 | -0.01 | 0.036 | 0.036 | 0.435 | 0.29 | 0.59 | 0.49 | 0.51 | 0.28 | 0.42 |
| 6 | 97449042 | rs9320375 | -0.576 | -0.427 | 0.617 | 0.492 | -0.028 | 0.174 | 0.174 | 0 | 0.18 | 0.4 | 0.4 | 0.51 | 0.15 | 0.34 |
| 6 | 97459945 | rs7766446 | -0.257 | -0.316 | 0.709 | 0.645 | 0.072 | 0.332 | 0.332 | 0 | 0.05 | 0.36 | 0.28 | 0.18 | 0.03 | 0.09 |
| 6 | 97461663 | rs12055565 | 0.041 | -0.37 | 0.388 | 0.638 | 0.011 | -0.052 | -0.052 | 0 | 0.12 | 0.04 | 0.12 | 0.33 | 0.12 | 0.26 |
| 6 | 97463090 | rs9387043 | 0.298 | -0.373 | 0.457 | 0.698 | 0.014 | 0.011 | 0.011 | 0 | 0.1 | 0.05 | 0.06 | 0.07 | 0.08 | 0.15 |
| 6 | 97466656 | rs1206097 | 0.263 | -0.312 | 0.543 | 0.63 | -0.004 | -0.007 | -0.007 | 0 | 0.08 | 0.05 | 0.08 | 0 | 0.13 | 0.08 |
| 6 | 97469390 | rs1206100 | -1.159 | -0.305 | 0.171 | 0.503 | -0.13 | -0.07 | -0.07 | 4.907 | 0.6 | 0.94 | 0.69 | 0.57 | 0.54 | 0.71 |
| 6 | 97476678 | rs13211055 | 0.41 | -0.382 | 0.761 | 0.852 | 0.008 | 0.002 | 0.002 | 0 | 0.29 | 0.27 | 0.23 | 0.29 | 0.25 | 0.29 |
| 6 | 97489770 | rs1766483 | -0.981 | 0.268 | 1.515 | 1.423 | 0.034 | -0.002 | -0.002 | 0.151 | 0.12 | 0.3 | 0.11 | 0.07 | 0.15 | 0.11 |
| 6 | 97498501 | rs4839708 | -1.502 | 0.19 | 1.659 | 1.242 | 0.021 | 0.089 | 0.089 | 1.28 | 0.15 | 0.49 | 0.36 | 0.19 | 0.16 | 0.16 |
| 6 | 97501179 | rs1766467 | -0.433 | 0.26 | 1.504 | 1.237 | 0.008 | -0.044 | -0.044 | 0.25 | 0.09 | 0.15 | 0.08 | 0 | 0.13 | 0.08 |

| | | | | | | | | | | | | | | | | |
|---|---|---|---|---|---|---|---|---|---|---|---|---|---|---|---|---|
| 6 | 97501957 | rs1737659 | -0.692 | 0.258 | 1.555 | 1.253 | 0.019 | 0.001 | 0.001 | 0 | 0.1 | 0.21 | 0.1 | 0.08 | 0.14 | 0.1 |
| 6 | 97508112 | rs1206117 | -1.327 | 0.247 | 1.595 | 1.269 | 0.049 | 0.13 | 0.13 | 0 | 0.14 | 0.48 | 0.36 | 0.19 | 0.16 | 0.16 |
| 6 | 97518003 | rs1933463 | 1.117 | 0.036 | 1.124 | 0.844 | -0.011 | -0.052 | -0.052 | 0 | 0.8 | 0.7 | 0.86 | 0.93 | 0.79 | 0.78 |
| 6 | 97518987 | rs9374329 | -0.199 | 0.051 | 1.102 | 0.876 | -0.025 | -0.011 | -0.011 | 0 | 0.11 | 0.2 | 0.06 | 0.07 | 0.07 | 0.15 |
| 6 | 97519369 | rs11153367 | 0.415 | 0.438 | 0.914 | 0.669 | -0.002 | -0.004 | -0.004 | 1.887 | 0.46 | 0.44 | 0.36 | 0.36 | 0.39 | 0.47 |
| 6 | 97528325 | rs963402 | 0.147 | 0.702 | 0.835 | 0.633 | 0.009 | -0.031 | -0.031 | 0 | 0.91 | 0.91 | 0.91 | 1 | 0.85 | 0.94 |
| 6 | 97534341 | rs9320410 | -0.013 | 0.789 | 0.829 | 0.651 | 0.006 | -0.023 | -0.023 | 0.013 | 0.92 | 0.93 | 0.91 | 1 | 0.85 | 0.94 |
| 6 | 97537000 | rs879384 | 0.464 | 0.674 | 0.222 | 0.918 | -0.052 | -0.063 | -0.063 | 0 | 0.4 | 0.13 | 0.27 | 0.47 | 0.43 | 0.38 |
| 6 | 97537182 | rs1997933 | 0.28 | 0.736 | 0.279 | 0.594 | 0.018 | 0.015 | 0.015 | 0 | 0.45 | 0.37 | 0.37 | 0.42 | 0.39 | 0.51 |
| 6 | 97541008 | rs1206131 | -0.673 | 1.038 | 0.209 | 0.432 | 0.019 | -0.042 | -0.042 | 0 | 0.09 | 0.19 | 0.1 | 0 | 0.15 | 0.07 |
| 6 | 97544329 | rs946120 | 0.074 | 0.847 | -0.114 | 0.665 | 0.008 | 0.009 | 0.009 | 0 | 0.53 | 0.45 | 0.44 | 0.44 | 0.55 | 0.58 |
| 6 | 97546036 | rs1206153 | -0.036 | 0.945 | 0.133 | 0.638 | -0.001 | 0 | 0 | 0 | 0.55 | 0.58 | 0.39 | 0.45 | 0.53 | 0.58 |
| 6 | 97549206 | rs1206155 | -0.418 | 0.861 | -0.184 | 0.714 | 0.001 | -0.002 | -0.002 | 0 | 0.74 | 0.88 | 0.66 | 0.75 | 0.72 | 0.8 |
| 6 | 97551806 | rs1206157 | -0.174 | 0.718 | 0.018 | 0.572 | -0.003 | -0.002 | -0.002 | 1.252 | 0.69 | 0.69 | 0.63 | 0.74 | 0.7 | 0.83 |
| 6 | 97565173 | rs1206147 | 0.376 | 0.661 | -0.342 | 0.536 | -0.023 | 0.18 | 0.18 | 0 | 0.53 | 0.84 | 0.41 | 0.25 | 0.5 | 0.57 |
| 6 | 97566938 | rs1206149 | 1.114 | 0.47 | 0.045 | 0.273 | -0.015 | 0.048 | 0.048 | 0 | 0.43 | 0.27 | 0.36 | 0.25 | 0.47 | 0.52 |
| 6 | 97576742 | rs1766530 | -0.631 | 0.55 | -0.448 | 0.432 | -0.011 | 0.017 | 0.017 | 0 | 0.71 | 0.9 | 0.63 | 0.76 | 0.67 | 0.81 |
| 6 | 97588710 | rs11966316 | -0.379 | 0.287 | -0.285 | 0.507 | -0.007 | 0.012 | 0.012 | 0 | 0.07 | 0.28 | 0.12 | 0.06 | 0.03 | 0.08 |
| 6 | 97591680 | rs1206178 | 0.387 | 0.324 | -0.374 | 0.442 | -0.007 | 0.155 | 0.155 | 0 | 0.51 | 0.72 | 0.41 | 0.25 | 0.49 | 0.59 |
| 6 | 97599612 | rs3822908 | 0.154 | 0.291 | -0.389 | 0.462 | 0 | -0.053 | -0.053 | 0 | 0.19 | 0.13 | 0.22 | 0.52 | 0.17 | 0.25 |
| 6 | 97608206 | rs1206141 | 0.292 | 0.454 | -0.447 | -0.402 | -0.008 | 0.155 | 0.155 | 0 | 0.51 | 0.72 | 0.4 | 0.24 | 0.49 | 0.59 |
| 6 | 97613299 | rs10484830 | -0.076 | 0.487 | -0.386 | -0.393 | 0.009 | -0.013 | -0.013 | 0 | 0.07 | 0.29 | 0.03 | 0 | 0.03 | 0.05 |
| 6 | 97616717 | rs1206125 | 0.272 | 0.495 | -0.382 | -0.389 | -0.006 | 0.153 | 0.153 | 0 | 0.51 | 0.72 | 0.41 | 0.25 | 0.5 | 0.59 |
| 6 | 97626374 | rs1737145 | 0.638 | 0.548 | -0.532 | -0.353 | 0.002 | -0.027 | -0.027 | 0 | 0.44 | 0.42 | 0.37 | 0.25 | 0.47 | 0.54 |
| 6 | 97644935 | rs12207304 | 0.937 | 0.539 | -0.732 | -0.49 | 0.004 | -0.045 | -0.045 | 2.157 | 0.41 | 0.42 | 0.35 | 0.24 | 0.41 | 0.5 |
| 6 | 97645628 | rs1474620 | -1.5 | 0.174 | -0.188 | -0.108 | 0 | -0.035 | -0.035 | 0 | 0.41 | 0.6 | 0.52 | 0.76 | 0.41 | 0.54 |
| 6 | 97677118 | rs9481410 | 1.372 | -0.829 | 0.572 | 0.593 | -0.035 | -0.064 | -0.064 | 0 | 0.69 | 0.47 | 0.55 | 0.24 | 0.77 | 0.65 |
| 6 | 97722634 | rs9372393 | -1.628 | -0.79 | 0.425 | 0.614 | -0.031 | -0.068 | -0.068 | 0.014 | 0.33 | 0.55 | 0.46 | 0.77 | 0.24 | 0.42 |
| 6 | 97740651 | rs17057619 | -0.581 | -0.708 | 0.279 | 0.592 | -0.007 | -0.025 | -0.025 | 0 | 0.07 | 0.23 | 0.02 | 0 | 0.04 | 0.05 |
| 6 | 97746730 | rs9400737 | -1.09 | -0.326 | -0.348 | 1.04 | 0.006 | -0.007 | -0.007 | 0 | 0.25 | 0.22 | 0.44 | 0.63 | 0.21 | 0.37 |
| 6 | 97749365 | rs9387227 | -1.09 | -0.336 | -0.348 | 1.04 | 0.005 | -0.007 | -0.007 | 0 | 0.25 | 0.22 | 0.44 | 0.63 | 0.21 | 0.37 |
| 6 | 97751624 | rs2388040 | -0.99 | -0.303 | -0.192 | 0.906 | 0 | -0.002 | -0.002 | 0 | 0.2 | 0.22 | 0.41 | 0.67 | 0.18 | 0.37 |
| 6 | 97751948 | rs2388041 | -0.97 | -0.397 | -0.246 | 0.04 | -0.008 | -0.044 | -0.044 | 7.03 | 0.34 | 0.49 | 0.55 | 0.82 | 0.31 | 0.39 |
| 6 | 97764445 | rs10499010 | -0.559 | -0.198 | -0.821 | 0.26 | -0.044 | 0.054 | 0.054 | 1.694 | 0.28 | 0.06 | 0.21 | 0.17 | 0.27 | 0.29 |
| 6 | 97771510 | rs9398398 | -1.044 | -0.495 | -0.629 | 0.591 | -0.017 | 0.04 | 0.04 | 0.022 | 0.11 | 0.18 | 0.09 | 0.25 | 0.09 | 0.3 |
| 6 | 97779994 | rs6901184 | -1.155 | -0.636 | -0.501 | 0.544 | 0.016 | -0.003 | -0.003 | 0 | 0.4 | 0.3 | 0.31 | 0.42 | 0.36 | 0.6 |
| 6 | 97846571 | rs7771567 | -1.332 | -0.645 | -0.448 | 0.565 | 0.017 | -0.003 | -0.003 | 0.056 | 0.4 | 0.31 | 0.3 | 0.43 | 0.34 | 0.56 |
| 6 | 97887474 | rs17057750 | -0.8 | -0.667 | -0.455 | 0.578 | -0.003 | -0.002 | -0.002 | 3.99 | 0.08 | 0.06 | 0.07 | 0.08 | 0.04 | 0.23 |
| 6 | 97888120 | rs200825 | -1.381 | -0.636 | -0.356 | 0.525 | 0.012 | -0.002 | -0.002 | 0 | 0.4 | 0.32 | 0.31 | 0.43 | 0.36 | 0.6 |
| 6 | 97921380 | rs4145406 | -1.076 | -0.534 | -0.478 | 0.685 | 0.023 | -0.038 | -0.038 | 1.631 | 0.09 | 0.04 | 0.07 | 0.25 | 0.07 | 0.28 |
| 6 | 97964889 | rs9489047 | -0.313 | -0.746 | 0.182 | 0.762 | -0.024 | 0.224 | 0.224 | 0.041 | 0.1 | 0.26 | 0.35 | 0.48 | 0.07 | 0.08 |
| 6 | 97967913 | rs7775139 | 1.257 | -0.87 | 0.091 | 0.485 | -0.005 | -0.043 | -0.043 | 0.326 | 0.58 | 0.5 | 0.38 | 0.12 | 0.61 | 0.39 |
| 6 | 98014794 | rs13213903 | -0.093 | -0.747 | -0.412 | 0.542 | -0.078 | 0.01 | 0.01 | 1.231 | 0.21 | 0.03 | 0.18 | 0.17 | 0.24 | 0.29 |
| 6 | 98015352 | rs9398455 | -0.811 | -0.689 | -0.21 | 0.59 | 0.021 | -0.025 | -0.025 | 0.112 | 0.08 | 0.03 | 0.08 | 0.23 | 0.08 | 0.23 |
| 6 | 98033236 | rs7356837 | -0.884 | -0.991 | 0.251 | 0.511 | -0.007 | 0.005 | 0.005 | 0 | 0.37 | 0.45 | 0.51 | 0.58 | 0.33 | 0.39 |
| 6 | 98044632 | rs4391266 | 0.35 | -1.682 | 0.305 | 0.369 | -0.013 | -0.049 | -0.049 | 0 | 0.12 | 0.03 | 0.08 | 0.18 | 0.13 | 0.12 |
| 6 | 98066596 | rs9374686 | 1.423 | -1.924 | 0.867 | 0.162 | 0.065 | 0.096 | 0.096 | 0 | 0.52 | 0.33 | 0.4 | 0.08 | 0.65 | 0.44 |
| 6 | 98068593 | rs3924094 | 0.572 | -1.868 | 1.017 | 0.265 | -0.057 | 0.014 | 0.014 | 0.543 | 0.69 | 0.47 | 0.55 | 0.3 | 0.82 | 0.69 |
| 6 | 98094026 | rs6930802 | 0.922 | -1.871 | 1.078 | 0.26 | -0.046 | -0.004 | -0.004 | 0.016 | 0.66 | 0.46 | 0.49 | 0.27 | 0.79 | 0.55 |
| 6 | 98104948 | rs4397233 | 1.249 | -1.6 | 0.981 | 0.439 | 0.11 | -0.04 | -0.04 | 1.916 | 0.53 | 0.12 | 0.7 | 0.55 | 0.67 | 0.46 |
| 6 | 98125710 | rs9387583 | -0.09 | -1.51 | 1.192 | 0.424 | -0.005 | 0.007 | 0.007 | 0 | 0.18 | 0.17 | 0.14 | 0.21 | 0.17 | 0.3 |
| 6 | 98161552 | rs6938329 | -0.288 | -1.178 | 0.932 | 0.73 | -0.026 | 0.055 | 0.055 | 3.067 | 0.19 | 0.41 | 0.13 | 0.27 | 0.12 | 0.21 |
| 6 | 98169324 | rs9387620 | -0.132 | -1.324 | 0.629 | 0.711 | -0.031 | -0.006 | -0.006 | 0.538 | 0.26 | 0.36 | 0.17 | 0.2 | 0.2 | 0.33 |
| 6 | 98171790 | rs9320672 | -0.603 | -0.328 | -0.98 | 0.625 | 0.069 | -0.055 | -0.055 | 4.58 | 0.48 | 0.87 | 0.34 | 0.47 | 0.39 | 0.61 |
| 6 | 98183112 | rs9401123 | -0.837 | -0.628 | -0.365 | 1.179 | 0.039 | -0.035 | -0.035 | 0 | 0.34 | 0.77 | 0.35 | 0.43 | 0.23 | 0.37 |
| 6 | 98200131 | rs12527293 | -0.534 | -0.673 | 0.416 | 1.548 | -0.025 | 0.102 | 0.102 | 0.457 | 0.06 | 0.18 | 0.14 | 0.15 | 0 | 0.07 |
| 6 | 98202741 | rs636234 | 0.072 | -0.396 | 0.214 | 0.589 | -0.013 | 0.034 | 0.034 | 0 | 0.63 | 0.76 | 0.72 | 0.79 | 0.6 | 0.49 |
| 6 | 98204209 | rs9387644 | 0.414 | -0.487 | 0.098 | 0.76 | -0.002 | 0.006 | 0.006 | 0.328 | 0.12 | 0.14 | 0.09 | 0.22 | 0.11 | 0.04 |
| 6 | 98206805 | rs600613 | 0.525 | -0.51 | 0.229 | 0.743 | 0 | 0.002 | 0.002 | 0 | 0.5 | 0.46 | 0.64 | 0.57 | 0.5 | 0.45 |
| 6 | 98209349 | rs1338551 | -0.065 | -0.199 | -0.197 | 0.472 | -0.013 | 0.034 | 0.034 | 0 | 0.63 | 0.76 | 0.72 | 0.79 | 0.6 | 0.49 |
| 6 | 98212409 | rs6569077 | 0.46 | -0.074 | -0.046 | 0.666 | 0.01 | 0.02 | 0.02 | 0 | 0.37 | 0.44 | 0.46 | 0.55 | 0.44 | 0.32 |
| 6 | 98214395 | rs9489706 | -0.196 | -0.517 | 0.246 | 0.556 | 0.003 | -0.002 | -0.002 | 0 | 0.22 | 0.19 | 0.09 | 0.22 | 0.16 | 0.27 |
| 6 | 98240442 | rs595074 | -0.096 | -0.609 | 0.138 | 0.232 | -0.002 | 0 | 0 | 0 | 0.7 | 0.75 | 0.57 | 0.69 | 0.72 | 0.6 |
| 6 | 98242041 | rs9489770 | 0.231 | -0.908 | 0.837 | 0.228 | 0.005 | -0.01 | -0.01 | 0 | 0.21 | 0.16 | 0.08 | 0.05 | 0.16 | 0.22 |
| 6 | 98251106 | rs17058043 | -0.014 | -0.79 | 1.045 | 0.301 | -0.008 | 0.003 | 0.003 | 0 | 0.27 | 0.39 | 0.41 | 0.31 | 0.26 | 0.39 |
| 6 | 98263613 | rs4839713 | 0.665 | -0.476 | 0.612 | 0.565 | 0.02 | -0.012 | -0.012 | 0 | 0.47 | 0.19 | 0.5 | 0.47 | 0.58 | 0.31 |
| 6 | 98274551 | rs9401196 | 0.635 | -0.458 | 0.778 | 0.56 | 0.025 | -0.003 | -0.003 | 0.172 | 0.48 | 0.27 | 0.5 | 0.47 | 0.58 | 0.3 |
| 6 | 98318226 | rs6922758 | -0.225 | -0.149 | 0.815 | 0.418 | -0.004 | 0.021 | 0.021 | 0 | 0.85 | 0.77 | 0.79 | 0.76 | 0.88 | 0.85 |
| 6 | 98328774 | rs9401295 | 0.079 | -0.228 | 1.022 | 0.272 | 0.003 | 0.006 | 0.006 | 0 | 0.56 | 0.61 | 0.52 | 0.65 | 0.61 | 0.43 |

| | | | | | | | | | | | | | | | | |
|---|---|---|---|---|---|---|---|---|---|---|---|---|---|---|---|---|
| 6 | 98331498 | rs9320760 | 1.186 | -0.184 | 0.914 | 0.322 | -0.007 | -0.026 | -0.026 | 0 | 0.11 | 0.04 | 0.19 | 0.17 | 0.11 | 0.1 |
| 6 | 98332567 | rs10484867 | 0.988 | -0.177 | 0.924 | 0.311 | -0.015 | -0.006 | -0.006 | 0 | 0.41 | 0.35 | 0.31 | 0.41 | 0.49 | 0.27 |
| 6 | 98337903 | rs6928545 | 0.075 | -0.173 | 0.424 | 0.292 | 0.001 | 0 | 0 | 0.185 | 0.34 | 0.31 | 0.38 | 0.34 | 0.31 | 0.42 |
| 6 | 98342812 | rs9320768 | -0.667 | 0.325 | 0.042 | 0.758 | -0.001 | 0.023 | 0.023 | 0.187 | 0.88 | 0.79 | 0.8 | 0.83 | 0.89 | 0.9 |
| 6 | 98344031 | rs9372625 | 1.002 | 0.796 | -0.773 | 0.674 | -0.134 | -0.136 | -0.136 | 3.511 | 0.32 | 0.05 | 0.25 | 0.4 | 0.41 | 0.23 |
| 6 | 98361795 | rs6934784 | -0.565 | 0.737 | -0.09 | 0.518 | -0.003 | 0.008 | 0.008 | 0 | 0.54 | 0.57 | 0.54 | 0.4 | 0.5 | 0.68 |
| 6 | 98369230 | rs4339469 | -1.056 | 0.769 | -0.048 | 0.615 | -0.01 | -0.014 | -0.014 | 0 | 0.66 | 0.7 | 0.74 | 0.57 | 0.61 | 0.78 |
| 6 | 98379027 | rs6940021 | -0.483 | 0.756 | -0.014 | 0.546 | 0.004 | 0.022 | 0.022 | 0 | 0.54 | 0.63 | 0.48 | 0.4 | 0.47 | 0.65 |
| 6 | 98395421 | rs9372650 | -0.802 | 0.749 | 0.317 | 0.463 | 0.012 | 0.008 | 0.008 | 0 | 0.62 | 0.49 | 0.64 | 0.57 | 0.55 | 0.73 |
| 6 | 98419104 | rs9387954 | 0.92 | 0.753 | 0.283 | 0.497 | -0.001 | -0.003 | -0.003 | 1.17 | 0.12 | 0.11 | 0.2 | 0.18 | 0.11 | 0.12 |
| 6 | 98435125 | rs10872224 | -1.168 | 0.719 | 0.451 | 0.469 | -0.035 | -0.039 | -0.039 | 0 | 0.65 | 0.79 | 0.68 | 0.58 | 0.59 | 0.78 |
| 6 | 98468284 | rs4131042 | 1.035 | 0.714 | 0.39 | 0.493 | -0.009 | -0.029 | -0.029 | 0 | 0.11 | 0.04 | 0.2 | 0.18 | 0.11 | 0.13 |
| 6 | 98470233 | rs17813462 | 1.141 | 0.081 | -0.108 | 0.084 | 0.02 | 0.026 | 0.026 | 0 | 0.16 | 0.03 | 0.26 | 0.12 | 0.12 | 0.14 |
| 6 | 98481755 | rs7760876 | 1.33 | 0.145 | 0.152 | 0.089 | -0.034 | -0.026 | -0.026 | 0.072 | 0.37 | 0.3 | 0.34 | 0.42 | 0.44 | 0.25 |
| 6 | 98489654 | rs4839936 | -0.153 | -0.069 | 0.363 | -0.073 | 0.025 | 0.01 | 0.01 | 0.019 | 0.33 | 0.12 | 0.35 | 0.32 | 0.32 | 0.38 |
| 6 | 98495952 | rs2505059 | -0.815 | 0.267 | 0.339 | 0.237 | 0.007 | 0.021 | 0.021 | 0.014 | 0.53 | 0.59 | 0.47 | 0.34 | 0.47 | 0.62 |
| 6 | 98517272 | rs12214442 | -0.468 | 0.261 | 0.362 | 0.235 | -0.001 | -0.002 | -0.002 | 0.871 | 0.5 | 0.44 | 0.45 | 0.34 | 0.43 | 0.59 |
| 6 | 98528404 | rs4458695 | 1.348 | 0.278 | 0.184 | 0.232 | -0.058 | -0.06 | -0.06 | 0 | 0.33 | 0.17 | 0.31 | 0.42 | 0.4 | 0.22 |
| 6 | 98547488 | rs9490542 | -0.36 | 0.534 | 0.323 | 0.495 | -0.008 | -0.021 | -0.021 | 0.019 | 0.43 | 0.42 | 0.31 | 0.25 | 0.4 | 0.47 |
| 6 | 98559692 | rs13213457 | 0.087 | 0.836 | 0.511 | 0.35 | -0.001 | 0.012 | 0.012 | 0 | 0.16 | 0.26 | 0.22 | 0.23 | 0.15 | 0.17 |
| 6 | 98560826 | rs1906251 | 0.278 | 0.888 | 0.574 | 0.33 | 0 | 0 | 0 | 1.355 | 0.58 | 0.46 | 0.57 | 0.65 | 0.66 | 0.42 |
| 6 | 98562720 | rs9375195 | 0.988 | 0.939 | 0.485 | 0.401 | -0.027 | -0.024 | -0.024 | 0 | 0.45 | 0.21 | 0.37 | 0.49 | 0.54 | 0.3 |
| 6 | 98597689 | rs1487452 | -0.552 | 0.829 | 0.545 | 0.374 | 0.008 | 0.011 | 0.011 | 1.359 | 0.5 | 0.53 | 0.6 | 0.37 | 0.55 | 0.64 |
| 6 | 98597809 | rs2505065 | 0.819 | 0.88 | 0.458 | 0.445 | 0.007 | 0.017 | 0.017 | 0 | 0.12 | 0.21 | 0.2 | 0.18 | 0.12 | 0.11 |
| 6 | 98611300 | rs1487439 | 1.203 | 0.842 | 0.389 | 0.428 | 0.033 | -0.037 | -0.037 | 0.086 | 0.34 | 0.18 | 0.18 | 0.41 | 0.31 | 0.21 |
| 6 | 98622703 | rs9490788 | 0.321 | 0.887 | 0.417 | 0.442 | 0.001 | 0 | 0 | 0 | 0.14 | 0.16 | 0.21 | 0.16 | 0.15 | 0.14 |
| 6 | 98626561 | rs9372752 | 0.436 | 0.873 | 0.466 | 0.414 | 0.019 | 0.033 | 0.033 | 1.441 | 0.5 | 0.44 | 0.4 | 0.65 | 0.46 | 0.37 |
| 6 | 98652595 | rs1376499 | 0.593 | 0.914 | 0.327 | 0.417 | 0.018 | 0.03 | 0.03 | 1.634 | 0.47 | 0.41 | 0.39 | 0.65 | 0.44 | 0.35 |
| 6 | 98677699 | rs10223664 | 0.877 | 0.976 | 0.598 | 0.474 | 0.01 | 0.013 | 0.013 | 0 | 0.12 | 0.15 | 0.2 | 0.16 | 0.12 | 0.11 |
| 6 | 98678145 | rs10499016 | 0.733 | 0.846 | 0.22 | 0.166 | 0.001 | 0.031 | 0.031 | 0.458 | 0.23 | 0.03 | 0.33 | 0.14 | 0.16 | 0.14 |
| 6 | 98701016 | rs1349485 | -0.368 | 0.65 | 0.832 | 0.381 | 0 | 0 | 0 | 2.201 | 0.54 | 0.43 | 0.61 | 0.38 | 0.58 | 0.65 |
| 6 | 98725758 | rs12206944 | 0.071 | 0.618 | 0.822 | 0.357 | 0.019 | 0.045 | 0.045 | 1.786 | 0.48 | 0.32 | 0.58 | 0.33 | 0.54 | 0.54 |
| 6 | 98735611 | rs1376505 | -0.961 | 0.918 | 0.264 | 0.37 | 0.03 | -0.056 | -0.056 | 0 | 0.68 | 0.86 | 0.83 | 0.59 | 0.71 | 0.79 |
| 6 | 98739867 | rs9375351 | 0.875 | 0.907 | 0.264 | 0.364 | 0.026 | -0.047 | -0.047 | 0 | 0.32 | 0.18 | 0.17 | 0.41 | 0.29 | 0.21 |
| 6 | 98742599 | rs1451336 | -0.155 | 0.925 | 0.252 | 0.358 | 0.004 | 0 | 0 | 2.243 | 0.56 | 0.6 | 0.62 | 0.44 | 0.59 | 0.67 |
| 6 | 98749329 | rs7754741 | 0.822 | 0.808 | 0.087 | 0.215 | 0.025 | -0.04 | -0.04 | 0 | 0.32 | 0.19 | 0.17 | 0.41 | 0.29 | 0.21 |
| 6 | 98760918 | rs6940828 | -0.154 | 0.755 | 0.228 | 0.278 | -0.001 | 0 | 0 | 0 | 0.53 | 0.52 | 0.58 | 0.49 | 0.57 | 0.66 |
| 6 | 98767943 | rs6941784 | 0.114 | 0.894 | 0.21 | 0.248 | 0 | -0.004 | -0.004 | 0 | 0.46 | 0.53 | 0.44 | 0.41 | 0.49 | 0.59 |
| 6 | 98770554 | rs6569407 | -0.711 | 0.947 | 0.021 | 0.277 | -0.007 | -0.044 | -0.044 | 1.27 | 0.6 | 0.87 | 0.65 | 0.53 | 0.61 | 0.7 |
| 6 | 98781853 | rs17058404 | 0.251 | 1.046 | 0.322 | -0.406 | 0.003 | -0.049 | -0.049 | 0 | 0.09 | 0.19 | 0.07 | 0 | 0.07 | 0 |
| 6 | 98789325 | rs12529268 | -0.481 | 0.983 | 0.253 | -0.393 | 0.011 | 0.011 | 0.011 | 0 | 0.13 | 0.24 | 0.19 | 0.12 | 0.11 | 0.06 |
| 6 | 98795332 | rs17058417 | 0.267 | 0.982 | 0.09 | -0.467 | -0.003 | -0.037 | -0.037 | 0 | 0.09 | 0.14 | 0.05 | 0 | 0.06 | 0 |
| 6 | 98798280 | rs13437130 | -0.785 | 1.009 | -0.068 | 0.006 | 0.017 | 0.059 | 0.059 | 0 | 0.06 | 0.16 | 0.16 | 0.12 | 0.06 | 0.07 |
| 6 | 98819800 | rs980899 | -0.272 | 0.828 | -0.488 | -0.514 | -0.004 | 0.013 | 0.013 | 0 | 0.7 | 0.77 | 0.74 | 0.73 | 0.69 | 0.8 |
| 6 | 98824077 | rs1507774 | -0.196 | 0.861 | -0.474 | -0.496 | -0.007 | -0.015 | -0.015 | 0 | 0.69 | 0.56 | 0.8 | 0.72 | 0.73 | 0.78 |
| 6 | 98829949 | rs7767825 | -0.026 | 1.204 | -0.945 | -0.116 | -0.038 | 0.003 | 0.003 | 0 | 0.11 | 0.01 | 0.09 | 0.06 | 0.13 | 0.12 |
| 6 | 98892120 | rs1494777 | -0.172 | 0.87 | -0.945 | 0.034 | -0.026 | 0.069 | 0.069 | 3.079 | 0.42 | 0.08 | 0.26 | 0.29 | 0.38 | 0.26 |
| 6 | 98893182 | rs3123339 | -0.177 | 0.868 | -0.999 | 0.037 | -0.046 | 0.065 | 0.065 | 3.059 | 0.42 | 0.07 | 0.26 | 0.29 | 0.39 | 0.26 |
| 6 | 98895069 | rs3125575 | -0.308 | 0.827 | -0.336 | 0.026 | -0.004 | -0.014 | -0.014 | 0 | 0.43 | 0.46 | 0.27 | 0.28 | 0.39 | 0.29 |
| 6 | 98904572 | rs17761570 | 0.458 | 0.828 | -0.288 | -0.089 | -0.007 | 0.095 | 0.095 | 0 | 0.45 | 0.27 | 0.3 | 0.22 | 0.44 | 0.39 |
| 6 | 98905933 | rs9388482 | 0.573 | 0.116 | 0.251 | 0.511 | 0.015 | -0.016 | -0.016 | 0 | 0.64 | 0.31 | 0.49 | 0.58 | 0.73 | 0.55 |
| 6 | 98931278 | rs210400 | -0.78 | 0.374 | 0.129 | 0.642 | -0.011 | 0.028 | 0.028 | 0.094 | 0.29 | 0.27 | 0.39 | 0.42 | 0.21 | 0.41 |
| 6 | 98931716 | rs169750 | -0.703 | 0.439 | -0.08 | 0.676 | 0 | 0.001 | 0.001 | 0.14 | 0.28 | 0.21 | 0.39 | 0.41 | 0.21 | 0.41 |
| 6 | 98935310 | rs9398813 | -0.907 | 1.002 | -0.152 | 0.285 | 0.001 | -0.001 | -0.001 | 0 | 0.05 | 0.09 | 0.11 | 0.05 | 0.04 | 0.08 |
| 6 | 98938023 | rs17058578 | -1.181 | 1.251 | -0.299 | 0.308 | 0.008 | -0.003 | -0.003 | 0.245 | 0.06 | 0.12 | 0.1 | 0.07 | 0.1 | 0.04 |
| 6 | 98938237 | rs9401901 | -0.029 | 1.307 | -1.31 | -1.253 | -0.046 | 0.186 | 0.186 | 0 | 0.59 | 0.82 | 0.67 | 0.99 | 0.57 | 0.83 |
| 6 | 98938249 | rs9388508 | 0.008 | 0.831 | -1.217 | -0.599 | 0.001 | -0.003 | -0.003 | 0 | 0.18 | 0.19 | 0.14 | 0.26 | 0.17 | 0.29 |
| 6 | 98938834 | rs2227121 | -0.092 | 0.789 | -1.084 | -0.639 | 0.003 | 0.032 | 0.032 | 0.467 | 0.49 | 0.41 | 0.6 | 0.66 | 0.49 | 0.39 |
| 6 | 98949550 | rs158773 | 0.08 | 1.03 | -0.72 | -0.61 | 0.005 | -0.013 | -0.013 | 0 | 0.63 | 0.49 | 0.78 | 0.67 | 0.61 | 0.54 |
| 6 | 98953633 | rs158774 | -0.463 | 1.015 | -0.624 | -0.724 | -0.003 | 0.003 | 0.003 | 0 | 0.27 | 0.25 | 0.18 | 0.21 | 0.3 | 0.26 |
| 6 | 98959605 | rs158777 | 0.444 | 0.748 | -0.609 | -0.566 | 0.009 | 0.042 | 0.042 | 0 | 0.5 | 0.37 | 0.59 | 0.65 | 0.48 | 0.49 |
| 6 | 98962591 | rs150396 | -1.181 | 0.226 | 0.329 | 0.069 | 0.011 | 0.035 | 0.035 | 0 | 0.28 | 0.53 | 0.29 | 0.38 | 0.27 | 0.35 |
| 6 | 98971740 | rs1481449 | -0.372 | 0.158 | 0.779 | 0.276 | -0.024 | 0.103 | 0.103 | 0.218 | 0.08 | 0.25 | 0.15 | 0.21 | 0.02 | 0.02 |
| 6 | 98975810 | rs211215 | -1.683 | 0.423 | 0.951 | 0.673 | 0.022 | 0.048 | 0.048 | 0 | 0.29 | 0.55 | 0.31 | 0.4 | 0.3 | 0.35 |
| 6 | 98978496 | rs11961327 | -0.444 | 0.333 | 0.977 | 0.666 | -0.025 | 0.098 | 0.098 | 0.251 | 0.08 | 0.25 | 0.15 | 0.21 | 0.02 | 0.02 |
| 6 | 98980147 | rs211222 | -0.433 | -0.115 | 0.586 | 0.397 | 0.029 | 0.031 | 0.031 | 0 | 0.75 | 0.86 | 0.89 | 0.82 | 0.8 | 0.79 |
| 6 | 98991464 | rs17058667 | -1.069 | 0.024 | 0.668 | 0.468 | 0.01 | -0.034 | -0.034 | 0.084 | 0.27 | 0.46 | 0.16 | 0.19 | 0.27 | 0.31 |
| 6 | 98992369 | rs183316 | -1.175 | 0.028 | 0.534 | 0.438 | 0.007 | -0.061 | -0.061 | 0 | 0.28 | 0.54 | 0.16 | 0.17 | 0.27 | 0.34 |
| 6 | 99001499 | rs211227 | -0.714 | 0.15 | 0.625 | 0.379 | -0.031 | 0.116 | 0.116 | 0.062 | 0.17 | 0.47 | 0.2 | 0.29 | 0.1 | 0.19 |

| | | | | | | | | | | | | | | | | |
|---|---|---|---|---|---|---|---|---|---|---|---|---|---|---|---|---|
| 6 | 99004191 | rs9372864 | 1.345 | 0.077 | 0.726 | 0.138 | -0.004 | 0.003 | 0.003 | 0 | 0.45 | 0.27 | 0.6 | 0.42 | 0.53 | 0.44 |
| 6 | 99013740 | rs2046174 | 1.021 | 0.03 | 0.718 | -0.334 | -0.012 | -0.027 | -0.027 | 0 | 0.81 | 0.57 | 0.9 | 0.89 | 0.85 | 0.79 |
| 6 | 99017732 | rs17829357 | -0.798 | 0.266 | 0.54 | -0.76 | 0.002 | 0.009 | 0.009 | 0.167 | 0.14 | 0.08 | 0.06 | 0.08 | 0.1 | 0.21 |
| 6 | 99019431 | rs9491794 | -0.433 | 0.345 | 0.875 | -0.831 | -0.008 | -0.012 | -0.012 | 0 | 0.32 | 0.42 | 0.16 | 0.25 | 0.24 | 0.26 |
| 6 | 99032723 | rs1481440 | 0.138 | 0.313 | 0.86 | -0.885 | -0.021 | -0.016 | -0.016 | 0.396 | 0.69 | 0.33 | 0.86 | 0.74 | 0.79 | 0.77 |
| 6 | 99033489 | rs11752460 | -0.717 | 0.436 | 0.793 | -0.927 | -0.006 | 0.026 | 0.026 | 0 | 0.16 | 0.08 | 0.09 | 0.08 | 0.13 | 0.24 |
| 6 | 99034313 | rs1481438 | 1.347 | 0.447 | 0.792 | -0.928 | 0.018 | 0.052 | 0.052 | 0 | 0.53 | 0.25 | 0.77 | 0.66 | 0.66 | 0.53 |
| 6 | 99045749 | rs6929790 | -0.324 | 0.344 | 0.859 | -1.041 | -0.02 | -0.019 | -0.019 | 0 | 0.3 | 0.46 | 0.13 | 0.26 | 0.21 | 0.23 |
| 6 | 99050825 | rs17830067 | -0.93 | 0.366 | 0.367 | -1.16 | -0.01 | 0.041 | 0.041 | 0 | 0.17 | 0.08 | 0.1 | 0.08 | 0.13 | 0.23 |
| 6 | 99058170 | rs969540 | 0.983 | 0.371 | 0.388 | -1.157 | 0 | -0.026 | -0.026 | 1.116 | 0.83 | 0.72 | 0.9 | 0.92 | 0.86 | 0.76 |
| 6 | 99076746 | rs17058737 | -0.953 | 0.385 | 0.136 | -1.151 | -0.003 | 0.019 | 0.019 | 0 | 0.17 | 0.15 | 0.1 | 0.08 | 0.14 | 0.24 |
| 6 | 99080658 | rs11757142 | -1.033 | 0.385 | 0.06 | -1.15 | -0.002 | 0.016 | 0.016 | 0 | 0.17 | 0.15 | 0.1 | 0.08 | 0.13 | 0.24 |
| 6 | 99092539 | rs17058761 | -0.958 | 0.442 | -0.121 | -1.063 | -0.01 | 0.03 | 0.03 | 0 | 0.14 | 0.09 | 0.08 | 0.07 | 0.12 | 0.24 |
| 6 | 99124184 | rs11752997 | 0.266 | 0.243 | -0.444 | -1.107 | -0.005 | -0.052 | -0.052 | 0 | 0.17 | 0 | 0.07 | 0.19 | 0.13 | 0.16 |
| 6 | 99126082 | rs4472356 | 0.827 | 0.851 | -0.335 | -1.17 | -0.014 | 0.101 | 0.101 | 0.069 | 0.56 | 0.15 | 0.65 | 0.72 | 0.64 | 0.45 |
| 6 | 99138875 | rs12204275 | 0.826 | 0.878 | -0.44 | -1.389 | -0.068 | 0.095 | 0.095 | 0.705 | 0.55 | 0.04 | 0.65 | 0.73 | 0.63 | 0.45 |
| 6 | 99139350 | rs6939572 | 0.713 | 1.015 | -0.043 | -1.284 | 0.004 | 0.083 | 0.083 | 0 | 0.57 | 0.24 | 0.67 | 0.73 | 0.64 | 0.44 |
| 6 | 99140492 | rs6924761 | 1.357 | 1.58 | 0.69 | -0.965 | 0.184 | 0.061 | 0.061 | 0 | 0.95 | 0.76 | 0.86 | 0.95 | 0.89 | 0.82 |
| 6 | 99170039 | rs9375604 | 1.434 | 0.71 | 0.644 | -0.956 | 0.023 | 0.009 | 0.009 | 0 | 0.91 | 0.72 | 0.96 | 0.93 | 0.93 | 0.74 |
| 6 | 99173425 | rs4529305 | -0.745 | 0.726 | 0.18 | -1.143 | -0.021 | -0.013 | -0.013 | 0.193 | 0.07 | 0 | 0.04 | 0.07 | 0.08 | 0.19 |
| 6 | 99175625 | rs9388676 | 0.716 | 1.041 | 0.692 | -1.252 | 0.088 | -0.042 | -0.042 | 0 | 0.9 | 0.67 | 0.9 | 1 | 0.87 | 0.92 |
| 6 | 99195548 | rs11154467 | 0.946 | 0.911 | 0.038 | -2.15 | -0.001 | -0.039 | -0.039 | 0 | 0.27 | 0.23 | 0.25 | 0.02 | 0.29 | 0.08 |
| 6 | 99203222 | rs4406241 | 0.866 | 1.098 | 0.098 | -1.756 | 0.076 | -0.004 | -0.004 | 0 | 0.87 | 0.56 | 0.9 | 0.91 | 0.86 | 0.74 |
| 6 | 99214884 | rs7762570 | 0.214 | 0.812 | -0.435 | -1.692 | -0.005 | 0.004 | 0.004 | 0.15 | 0.6 | 0.64 | 0.39 | 0.27 | 0.5 | 0.48 |
| 6 | 99218958 | rs4839976 | -0.238 | 0.653 | -0.638 | -1.903 | 0.001 | 0.048 | 0.048 | 0 | 0.52 | 0.37 | 0.64 | 0.93 | 0.52 | 0.59 |
| 6 | 99219522 | rs4398748 | 0.636 | 0.251 | -0.341 | -1.593 | -0.017 | -0.071 | -0.071 | 0 | 0.86 | 0.74 | 0.94 | 0.98 | 0.92 | 0.85 |
| 6 | 99221669 | rs9385493 | -0.57 | 0.339 | -0.884 | -2.247 | -0.003 | -0.004 | -0.004 | 0 | 0.65 | 0.38 | 0.67 | 0.94 | 0.61 | 0.84 |
| 6 | 99226729 | rs10499023 | -0.276 | 0.546 | -1.538 | -2.663 | -0.041 | 0.109 | 0.109 | 0 | 0.24 | 0.09 | 0.14 | 0.02 | 0.25 | 0.22 |
| 6 | 99229574 | rs4839977 | 0.344 | 0.203 | -1.549 | -2.385 | 0.007 | 0.007 | 0.007 | 0 | 0.28 | 0.18 | 0.46 | 0.76 | 0.23 | 0.43 |
| 6 | 99230880 | rs4555920 | -0.683 | 0.134 | -1.329 | -2.135 | 0.003 | 0.088 | 0.088 | 0 | 0.55 | 0.28 | 0.62 | 0.78 | 0.54 | 0.66 |
| 6 | 99235909 | rs9375642 | -0.593 | -0.355 | -1.69 | -0.85 | 0.05 | -0.019 | -0.019 | 0 | 0.35 | 0.17 | 0.24 | 0.36 | 0.25 | 0.35 |
| 6 | 99236649 | rs9402154 | -0.281 | -0.383 | -1.189 | -1.565 | 0.006 | -0.003 | -0.003 | 0.201 | 0.37 | 0.39 | 0.53 | 0.63 | 0.44 | 0.55 |
| 6 | 99237315 | rs9388719 | -1.895 | -0.372 | -1.183 | -1.652 | 0.028 | -0.17 | -0.17 | 0.408 | 0.72 | 0.56 | 0.77 | 0.99 | 0.69 | 0.9 |
| 6 | 99237479 | rs12174549 | 0.104 | -0.67 | -1.42 | -1.835 | 0.014 | 0.096 | 0.096 | 0.224 | 0.24 | 0.09 | 0.11 | 0.04 | 0.21 | 0.22 |
| 6 | 99242592 | rs7762652 | -2.069 | -0.648 | -1.181 | -1.822 | 0.019 | -0.13 | -0.13 | 1.83 | 0.72 | 0.6 | 0.77 | 0.99 | 0.69 | 0.9 |
| 6 | 99250916 | rs9385517 | 0.107 | -0.628 | -1.288 | -1.826 | 0.007 | 0.098 | 0.098 | 0.204 | 0.24 | 0.1 | 0.11 | 0.04 | 0.21 | 0.22 |
| 6 | 99251754 | rs4839986 | -1.216 | -0.655 | -1.163 | -1.636 | -0.001 | -0.022 | -0.022 | 0 | 0.62 | 0.51 | 0.65 | 0.68 | 0.65 | 0.75 |
| 6 | 99258445 | rs6569636 | -2.082 | -0.607 | -1.062 | -1.723 | 0.02 | -0.129 | -0.129 | 0 | 0.73 | 0.62 | 0.78 | 0.99 | 0.69 | 0.9 |
| 6 | 99270438 | rs9375688 | 0.138 | -0.794 | -0.987 | -0.948 | 0.005 | 0.003 | 0.003 | 0.028 | 0.28 | 0.26 | 0.48 | 0.36 | 0.38 | 0.36 |
| 6 | 99270533 | rs9375689 | -2.012 | -0.622 | -0.585 | -0.988 | 0.008 | -0.084 | -0.084 | 0 | 0.73 | 0.67 | 0.78 | 0.99 | 0.69 | 0.9 |
| 6 | 99275290 | rs2444935 | -0.346 | -0.692 | -0.834 | -0.858 | -0.003 | 0.031 | 0.031 | 0.096 | 0.85 | 0.89 | 0.91 | 0.95 | 0.85 | 0.83 |
| 6 | 99277867 | rs1869641 | 1.25 | -0.595 | -0.856 | -0.701 | 0 | -0.007 | -0.007 | 0.674 | 0.38 | 0.36 | 0.31 | 0.15 | 0.41 | 0.26 |
| 6 | 99279449 | rs1883306 | 0.859 | -0.595 | -0.9 | -0.291 | 0.01 | -0.01 | -0.01 | 0 | 0.13 | 0.11 | 0.13 | 0.38 | 0.04 | 0.15 |
| 6 | 99284532 | rs3823036 | 0.156 | -0.431 | -0.877 | -0.812 | -0.007 | -0.008 | -0.008 | 0 | 0.29 | 0.2 | 0.5 | 0.73 | 0.3 | 0.43 |
| 6 | 99290334 | rs195853 | -0.358 | -0.384 | -0.864 | -1.201 | -0.001 | 0.023 | 0.023 | 0.016 | 0.85 | 0.9 | 0.92 | 0.95 | 0.85 | 0.84 |
| 6 | 99290592 | rs195852 | 1.253 | -0.429 | -0.501 | -1.173 | 0 | -0.058 | -0.058 | 0.233 | 0.33 | 0.31 | 0.26 | 0.02 | 0.38 | 0.17 |
| 6 | 99294322 | rs195851 | 0.703 | -0.34 | -0.436 | -0.977 | 0 | -0.084 | -0.084 | 0 | 0.44 | 0.44 | 0.3 | 0.02 | 0.44 | 0.24 |
| 6 | 99309057 | rs174447 | -0.176 | -0.271 | 0.071 | 0.332 | -0.02 | 0.102 | 0.102 | 0 | 0.11 | 0.28 | 0.2 | 0.2 | 0.11 | 0.16 |
| 6 | 99320715 | rs9388789 | 0.378 | -0.3 | 0.038 | 1.133 | -0.004 | 0.123 | 0.123 | 0 | 0.11 | 0.13 | 0.13 | 0.49 | 0.07 | 0.23 |
| 6 | 99323424 | rs11537982 | 0.666 | -0.484 | -0.155 | 0.731 | 0.006 | 0.011 | 0.011 | 0 | 0.08 | 0.05 | 0.04 | 0 | 0.06 | 0.04 |
| 6 | 99326490 | rs9375728 | -0.339 | -0.478 | -0.158 | 0.739 | -0.003 | 0.031 | 0.031 | 0.118 | 0.19 | 0.18 | 0.16 | 0.5 | 0.13 | 0.27 |
| 6 | 99328730 | rs195831 | -0.395 | -0.212 | -0.555 | -0.038 | -0.04 | 0.012 | 0.012 | 0.668 | 0.72 | 0.87 | 0.63 | 0.73 | 0.63 | 0.64 |
| 6 | 99336279 | rs10484609 | 0.114 | -0.202 | -0.496 | -0.03 | -0.005 | 0.041 | 0.041 | 0.118 | 0.19 | 0.21 | 0.16 | 0.5 | 0.13 | 0.26 |
| 6 | 99346145 | rs7739884 | 1.557 | -0.286 | -0.393 | -0.188 | -0.001 | -0.029 | -0.029 | 0 | 0.41 | 0.38 | 0.27 | 0.03 | 0.39 | 0.21 |
| 6 | 99351047 | rs195824 | -0.796 | -0.325 | -0.37 | -0.095 | -0.036 | -0.02 | -0.02 | 0.107 | 0.31 | 0.49 | 0.36 | 0.7 | 0.24 | 0.42 |
| 6 | 99432747 | rs196959 | -0.302 | -0.173 | -0.777 | 0.448 | -0.046 | 0.003 | 0.003 | 0.087 | 0.73 | 0.88 | 0.63 | 0.72 | 0.63 | 0.63 |
| 6 | 99433614 | rs196960 | -0.392 | -0.45 | -0.557 | 0.449 | -0.025 | 0.074 | 0.074 | 0 | 0.22 | 0.33 | 0.32 | 0.38 | 0.18 | 0.3 |
| 6 | 99437566 | rs2747734 | 1.023 | -0.268 | -0.591 | 0.277 | 0 | 0.002 | 0.002 | 1.284 | 0.5 | 0.48 | 0.31 | 0.33 | 0.43 | 0.31 |
| 6 | 99440369 | rs2572109 | 1.223 | -0.329 | -0.259 | 0.21 | 0.003 | 0.076 | 0.076 | 0 | 0.1 | 0.15 | 0.05 | 0.3 | 0.06 | 0.13 |
| 6 | 99441083 | rs12200990 | 1.502 | -0.265 | -0.357 | 0.045 | 0.001 | 0.021 | 0.021 | 0.194 | 0.39 | 0.33 | 0.26 | 0.03 | 0.37 | 0.19 |
| 6 | 99450081 | rs2180046 | -0.674 | 0.173 | -0.56 | 0.054 | -0.016 | 0.022 | 0.022 | 0.125 | 0.86 | 0.91 | 0.9 | 0.98 | 0.83 | 0.87 |
| 6 | 99462075 | rs2747739 | 0.253 | 0.301 | -0.367 | 0.196 | 0.026 | 0.071 | 0.071 | 0 | 0.22 | 0.29 | 0.31 | 0.55 | 0.28 | 0.32 |
| 6 | 99462324 | rs9402354 | 1.371 | 0.338 | -0.447 | 0.27 | -0.027 | -0.03 | -0.03 | 0 | 0.11 | 0.06 | 0.27 | 0.26 | 0.21 | 0.19 |
| 6 | 99463771 | rs9375844 | -0.14 | -0.102 | -0.663 | -0.033 | -0.006 | 0.009 | 0.009 | 0 | 0.27 | 0.15 | 0.31 | 0.27 | 0.29 | 0.3 |
| 6 | 99464542 | rs2207446 | 0.18 | -0.238 | -0.487 | 0.096 | -0.001 | 0.005 | 0.005 | 0.184 | 0.74 | 0.66 | 0.7 | 0.77 | 0.72 | 0.7 |
| 6 | 99467661 | rs12173555 | 1.477 | -0.235 | -0.581 | 0.098 | -0.033 | -0.062 | -0.062 | 0 | 0.1 | 0.01 | 0.04 | 0.29 | 0.07 | 0.14 |
| 6 | 99468942 | rs9388950 | -0.858 | -0.183 | -0.531 | 0.109 | 0.002 | -0.006 | -0.006 | 0 | 0.39 | 0.39 | 0.53 | 0.51 | 0.42 | 0.46 |
| 6 | 99473410 | rs2092772 | 2.029 | -0.212 | -0.406 | 0.048 | 0.003 | 0.046 | 0.046 | 0.073 | 0.36 | 0.24 | 0.33 | 0.17 | 0.34 | 0.27 |
| 6 | 99478174 | rs12201236 | 2.765 | -0.274 | -0.672 | -0.399 | -0.01 | 0.053 | 0.053 | 0 | 0.28 | 0.17 | 0.22 | 0.02 | 0.31 | 0.12 |

| | | | | | | | | | | | | | | | | |
|---|---|---|---|---|---|---|---|---|---|---|---|---|---|---|---|---|
| 6 | 99478304 | rs4839737 | 0.631 | -0.217 | -0.659 | -0.108 | -0.024 | 0.07 | 0.07 | 0 | 0.13 | 0.22 | 0.21 | 0.25 | 0.12 | 0.17 |
| 6 | 99479842 | rs2747748 | 0.079 | 0.089 | -0.689 | -0.352 | -0.01 | 0.105 | 0.105 | 0 | 0.73 | 0.9 | 0.86 | 0.98 | 0.76 | 0.79 |
| 6 | 99480633 | rs7756447 | 0.691 | 0.059 | -0.651 | -0.274 | -0.043 | 0.061 | 0.061 | 0 | 0.61 | 0.77 | 0.59 | 0.71 | 0.54 | 0.59 |
| 6 | 99485266 | rs11756151 | 0.459 | -0.056 | -0.646 | -0.263 | 0.028 | 0.029 | 0.029 | 0.052 | 0.14 | 0.07 | 0.05 | 0 | 0.07 | 0.08 |
| 6 | 99498599 | rs12173832 | 1.193 | 0.255 | -0.671 | -0.022 | -0.036 | -0.012 | -0.012 | 0 | 0.11 | 0.03 | 0.15 | 0.15 | 0.21 | 0.18 |
| 6 | 99500963 | rs2144241 | -1.662 | 0.032 | -0.66 | -0.461 | -0.035 | 0.085 | 0.085 | 0 | 0.59 | 0.72 | 0.68 | 0.96 | 0.52 | 0.72 |
| 6 | 99501709 | rs6916751 | -0.028 | -0.107 | -0.501 | -0.214 | 0.009 | 0.025 | 0.025 | 0 | 0.28 | 0.3 | 0.24 | 0.46 | 0.3 | 0.39 |
| 6 | 99504377 | rs9493282 | -0.402 | -0.106 | -0.501 | 0.081 | -0.008 | 0 | 0 | 0.254 | 0.29 | 0.29 | 0.24 | 0.27 | 0.2 | 0.22 |
| 6 | 99515134 | rs12660289 | 0.304 | 0.309 | -0.25 | 0.227 | 0.031 | 0.06 | 0.06 | 0 | 0.21 | 0.32 | 0.22 | 0.37 | 0.25 | 0.35 |
| 6 | 99521031 | rs2207445 | -0.183 | 0.27 | -0.362 | -0.075 | 0 | -0.007 | -0.007 | 0.454 | 0.75 | 0.79 | 0.68 | 0.62 | 0.76 | 0.77 |
| 6 | 99524062 | rs7764372 | -0.328 | 0.365 | -0.284 | -0.376 | 0.023 | 0.015 | 0.015 | 0 | 0.28 | 0.44 | 0.26 | 0.36 | 0.3 | 0.42 |
| 6 | 99524281 | rs9375894 | 0.526 | -0.027 | -0.058 | -0.149 | -0.012 | 0.118 | 0.118 | 0 | 0.13 | 0.32 | 0.08 | 0.29 | 0.06 | 0.18 |
| 6 | 99531172 | rs12212298 | 1.722 | -0.135 | -0.409 | -0.467 | -0.012 | 0.026 | 0.026 | 0 | 0.3 | 0.21 | 0.27 | 0.02 | 0.36 | 0.13 |
| 6 | 99531774 | rs9385615 | 0.827 | -0.342 | 0.109 | 0.359 | -0.008 | 0.085 | 0.085 | 0.136 | 0.11 | 0.23 | 0.07 | 0.27 | 0.05 | 0.17 |
| 6 | 99538132 | rs2572098 | 0.5 | -0.108 | -0.41 | 0.073 | -0.012 | 0.026 | 0.026 | 0.036 | 0.07 | 0.02 | 0.16 | 0.02 | 0.1 | 0.07 |
| 6 | 99539153 | rs9402444 | 0.545 | -0.442 | -0.15 | 0.463 | -0.01 | 0.059 | 0.059 | 0.01 | 0.15 | 0.15 | 0.16 | 0.54 | 0.08 | 0.29 |
| 6 | 99550464 | rs6902772 | -0.399 | 0.307 | -0.473 | 0.124 | -0.03 | -0.009 | -0.009 | 0 | 0.34 | 0.25 | 0.35 | 0.63 | 0.42 | 0.52 |
| 6 | 99551169 | rs6940049 | -1.137 | -0.061 | -0.291 | -0.314 | 0.003 | 0.004 | 0.004 | 0 | 0.53 | 0.49 | 0.53 | 0.66 | 0.57 | 0.69 |
| 6 | 99551765 | rs3860236 | 0.266 | -0.072 | -0.357 | -0.354 | -0.003 | -0.029 | -0.029 | 0.334 | 0.18 | 0.24 | 0.18 | 0.04 | 0.16 | 0.16 |
| 6 | 99551970 | rs10457592 | 2.261 | -0.395 | -0.231 | -0.672 | 0.025 | 0.023 | 0.023 | 0.801 | 0.27 | 0.13 | 0.15 | 0.01 | 0.22 | 0.14 |
| 6 | 99552868 | rs12208335 | 0.239 | -0.195 | -0.303 | -0.432 | -0.006 | -0.051 | -0.051 | 0 | 0.19 | 0.28 | 0.19 | 0.03 | 0.16 | 0.16 |
| 6 | 99557664 | rs6938641 | -0.233 | -0.013 | -0.196 | -0.176 | 0 | 0.033 | 0.033 | 0.056 | 0.81 | 0.74 | 0.7 | 0.67 | 0.79 | 0.83 |
| 6 | 99559773 | rs1081025 | 0.775 | -0.127 | -0.484 | -0.023 | 0.019 | -0.016 | -0.016 | 0 | 0.17 | 0.11 | 0.13 | 0.32 | 0.06 | 0.2 |
| 6 | 99565215 | rs12526079 | 1.117 | 0.16 | -0.578 | -0.258 | -0.115 | -0.001 | -0.001 | 0 | 0.17 | 0.03 | 0.2 | 0.09 | 0.36 | 0.23 |
| 6 | 99569709 | rs7769941 | -0.945 | -0.079 | -0.545 | -0.269 | -0.019 | 0.02 | 0.02 | 0 | 0.93 | 0.98 | 0.84 | 0.98 | 0.89 | 0.93 |
| 6 | 99574363 | rs2388839 | 0.68 | -0.412 | -0.52 | -0.537 | 0.012 | 0.015 | 0.015 | 0 | 0.13 | 0.09 | 0.09 | 0.03 | 0.1 | 0.11 |
| 6 | 99575874 | rs4454147 | 1.206 | -0.43 | -0.4 | -0.525 | -0.003 | 0.082 | 0.082 | 0 | 0.44 | 0.26 | 0.33 | 0.1 | 0.54 | 0.32 |
| 6 | 99583773 | rs2029964 | 0.162 | -0.353 | -0.334 | -0.408 | -0.078 | -0.017 | -0.017 | 0 | 0.34 | 0.14 | 0.32 | 0.39 | 0.41 | 0.42 |
| 6 | 99584658 | rs728758 | 2.275 | -0.336 | -0.341 | -0.559 | 0.004 | -0.038 | -0.038 | 0.029 | 0.26 | 0.19 | 0.13 | 0.01 | 0.19 | 0.13 |
| 6 | 99592404 | rs11154718 | 0.789 | -0.518 | -0.064 | -0.101 | -0.001 | 0.002 | 0.002 | 0.034 | 0.61 | 0.42 | 0.46 | 0.61 | 0.6 | 0.64 |
| 6 | 99593258 | rs12190591 | 0.621 | 0.364 | -0.381 | -0.188 | 0.02 | -0.029 | -0.029 | 0 | 0.06 | 0.09 | 0.12 | 0 | 0.09 | 0.06 |
| 6 | 99600083 | rs9321394 | -0.226 | 0.08 | -0.471 | -0.005 | -0.007 | 0.005 | 0.005 | 0 | 0.33 | 0.39 | 0.45 | 0.64 | 0.27 | 0.39 |
| 6 | 99607499 | rs11754157 | 0.167 | 1.179 | -0.837 | -0.331 | -0.089 | 0.002 | 0.002 | 0.668 | 0.16 | 0.02 | 0.18 | 0.09 | 0.26 | 0.19 |
| 6 | 99610074 | rs4839999 | 0.45 | 0.633 | -0.567 | -0.376 | 0.006 | 0.008 | 0.008 | 0 | 0.17 | 0.14 | 0.27 | 0.11 | 0.13 | 0.25 |
| 6 | 99613136 | rs12206927 | 0.311 | 0.722 | -0.124 | -1.075 | -0.001 | -0.025 | -0.025 | 0 | 0.47 | 0.46 | 0.33 | 0.09 | 0.55 | 0.3 |
| 6 | 99629252 | rs12110525 | 0.517 | 0.757 | -0.011 | -1.152 | -0.017 | 0.014 | 0.014 | 1.038 | 0.17 | 0.13 | 0.17 | 0.09 | 0.25 | 0.17 |
| 6 | 99630038 | rs9373057 | 0.39 | 0.641 | 0.15 | -0.769 | -0.001 | 0.004 | 0.004 | 3.703 | 0.24 | 0.23 | 0.4 | 0.5 | 0.25 | 0.22 |
| 6 | 99633011 | rs9399068 | -1.565 | 0.527 | 0.527 | -0.877 | -0.003 | -0.091 | -0.091 | 0 | 0.72 | 0.69 | 0.84 | 1 | 0.71 | 0.88 |
| 6 | 99637123 | rs9375977 | 1.538 | 0.51 | 0.543 | -0.886 | -0.01 | 0.095 | 0.095 | 0 | 0.27 | 0.1 | 0.15 | 0 | 0.29 | 0.12 |
| 6 | 99640610 | rs7745052 | 0.458 | 0.495 | 0.42 | -0.869 | 0.023 | -0.025 | -0.025 | 0.123 | 0.28 | 0.17 | 0.27 | 0.41 | 0.2 | 0.48 |
| 6 | 99641248 | rs1496971 | 1.419 | 0.528 | 0.465 | -0.927 | 0.001 | 0.006 | 0.006 | 0 | 0.28 | 0.2 | 0.15 | 0 | 0.29 | 0.12 |
| 6 | 99659802 | rs7767885 | -0.391 | 0.539 | -0.103 | -0.895 | 0.001 | 0.043 | 0.043 | 0.303 | 0.52 | 0.36 | 0.66 | 0.9 | 0.45 | 0.69 |
| 6 | 99662681 | rs7769752 | 0.819 | 0.751 | 0.186 | -0.411 | -0.002 | -0.056 | -0.056 | 0.004 | 0.32 | 0.43 | 0.27 | 0.09 | 0.33 | 0.15 |
| 6 | 99662743 | rs17059246 | 0.202 | 0.841 | -0.521 | -0.346 | 0.004 | -0.021 | -0.021 | 0.267 | 0.46 | 0.25 | 0.44 | 0.5 | 0.44 | 0.64 |
| 6 | 99664294 | rs9389116 | -0.19 | 1.055 | -0.708 | 0.673 | -0.015 | -0.045 | -0.045 | 0 | 0.21 | 0.13 | 0.3 | 0.41 | 0.23 | 0.2 |
| 6 | 99669676 | rs9373072 | 0.172 | 1.033 | -1.158 | 0.334 | -0.009 | -0.005 | -0.005 | 0 | 0.43 | 0.19 | 0.36 | 0.44 | 0.37 | 0.44 |
| 6 | 99672014 | rs2132683 | -0.223 | 1.121 | -0.887 | 0.663 | 0.03 | 0.133 | 0.133 | 0 | 0.24 | 0.46 | 0.47 | 0.47 | 0.32 | 0.32 |
| 6 | 99673152 | rs9375997 | -0.123 | 0.799 | -1.97 | 0.374 | -0.001 | 0.022 | 0.022 | 1.353 | 0.44 | 0.06 | 0.29 | 0.41 | 0.32 | 0.44 |
| 6 | 99673466 | rs9402556 | -0.04 | 0.874 | -1.947 | 0.569 | -0.041 | -0.08 | -0.08 | 1.723 | 0.17 | 0.01 | 0.08 | 0.3 | 0.14 | 0.15 |
| 6 | 99675908 | rs12528619 | -0.545 | 1.962 | -1.67 | 0.379 | 0.008 | -0.001 | -0.001 | 0 | 0.1 | 0.06 | 0.42 | 0.09 | 0.21 | 0.12 |
| 6 | 99678424 | rs1566116 | -0.609 | 1.406 | -1.476 | 0.451 | -0.001 | -0.003 | -0.003 | 0 | 0.47 | 0.51 | 0.61 | 0.61 | 0.53 | 0.44 |
| 6 | 99679578 | rs4840017 | -0.476 | 1.095 | -1.361 | 0.389 | -0.04 | 0.057 | 0.057 | 0 | 0.36 | 0.26 | 0.55 | 0.2 | 0.45 | 0.29 |
| 6 | 99679902 | rs9402564 | 0.553 | 1.052 | -1.484 | 0.241 | 0.002 | 0.001 | 0.001 | 0 | 0.1 | 0.11 | 0.05 | 0.12 | 0.07 | 0.07 |
| 6 | 99683802 | rs2029965 | -0.925 | 0.937 | -1.205 | 0.164 | 0.01 | 0 | 0 | 0 | 0.4 | 0.44 | 0.57 | 0.6 | 0.47 | 0.4 |
| 6 | 99690090 | rs9376014 | 0.013 | 1.044 | -1.218 | 0.402 | -0.011 | -0.029 | -0.029 | 0 | 0.2 | 0.13 | 0.13 | 0.49 | 0.24 | 0.26 |
| 6 | 99690449 | rs12193060 | -0.175 | 0.774 | -1.131 | 0.293 | -0.013 | 0.082 | 0.082 | 0 | 0.23 | 0.13 | 0.12 | 0 | 0.22 | 0.13 |
| 6 | 99694494 | rs1874538 | 0.068 | 0.926 | -1.041 | 0.49 | -0.004 | -0.015 | -0.015 | 0.565 | 0.19 | 0.13 | 0.11 | 0.49 | 0.19 | 0.25 |
| 6 | 99717329 | rs9483707 | -0.092 | 0.949 | -0.783 | 0.003 | -0.009 | -0.04 | -0.04 | 0 | 0.7 | 0.79 | 0.71 | 0.47 | 0.64 | 0.64 |
| 6 | 99721049 | rs1045728 | 0.061 | 0.39 | -0.947 | -0.408 | -0.007 | 0.087 | 0.087 | 0 | 0.54 | 0.66 | 0.49 | 0.17 | 0.55 | 0.35 |
| 6 | 99724713 | rs910034 | -1.036 | 0.279 | -1.019 | -0.451 | -0.009 | -0.02 | -0.02 | 1.259 | 0.76 | 0.82 | 0.75 | 0.47 | 0.69 | 0.66 |
| 6 | 99727653 | rs12660321 | 1.064 | 0.276 | -1.03 | -0.456 | 0.005 | -0.027 | -0.027 | 0 | 0.16 | 0.12 | 0.22 | 0.29 | 0.08 | 0.28 |
| 6 | 99728442 | rs1496979 | 0.881 | 0.019 | -1.021 | -0.389 | 0.015 | -0.058 | -0.058 | 0 | 0.34 | 0.4 | 0.23 | 0.12 | 0.43 | 0.16 |
| 6 | 99729901 | rs6933093 | -0.049 | 0.003 | -1.034 | -0.399 | -0.006 | 0.074 | 0.074 | 0 | 0.54 | 0.67 | 0.49 | 0.22 | 0.56 | 0.36 |
| 6 | 99733939 | rs9373105 | 0.716 | 0.084 | -0.884 | -0.172 | -0.011 | -0.022 | -0.022 | 0 | 0.2 | 0.26 | 0.25 | 0.1 | 0.13 | 0.2 |
| 6 | 99734185 | rs1496980 | 0.854 | 0.694 | -0.965 | 0.285 | 0.018 | -0.072 | -0.072 | 0 | 0.21 | 0.08 | 0.11 | 0.49 | 0.15 | 0.31 |
| 6 | 99740492 | rs12207550 | 1.363 | 0.735 | -0.9 | 0.253 | 0.014 | 0.021 | 0.021 | 0 | 0.07 | 0.03 | 0.04 | 0 | 0.05 | 0.02 |
| 6 | 99744107 | rs221582 | 0.133 | 0.683 | -0.77 | 0.272 | -0.003 | -0.017 | -0.017 | 1.107 | 0.69 | 0.71 | 0.7 | 0.51 | 0.64 | 0.64 |
| 6 | 99747633 | rs221578 | -0.16 | 1.139 | -0.49 | 0.475 | 0.015 | -0.034 | -0.034 | 0 | 0.9 | 0.83 | 0.82 | 1 | 0.79 | 0.95 |
| 6 | 99753499 | rs221530 | 0.56 | 1.113 | -0.449 | 0.899 | 0.009 | 0.022 | 0.022 | 0 | 0.86 | 0.8 | 0.79 | 0.63 | 0.77 | 0.83 |

| | | | | | | | | | | | | | | | | |
|---|---|---|---|---|---|---|---|---|---|---|---|---|---|---|---|---|
| 6 | 99763266 | rs6913076 | 1.08 | 1.268 | 0.364 | 1.022 | 0.188 | 0.347 | 0.347 | 0 | 0.93 | 0.67 | 0.83 | 0.57 | 0.81 | 0.84 |
| 6 | 99768894 | rs182613 | -0.418 | 0.955 | -0.318 | 0.4 | 0.01 | 0.09 | 0.09 | 0 | 0.55 | 0.75 | 0.61 | 0.81 | 0.37 | 0.7 |
| 6 | 99772374 | rs13206094 | 0.805 | 1.014 | -0.684 | 0.13 | -0.016 | -0.034 | -0.034 | 0 | 0.24 | 0.01 | 0.12 | 0.14 | 0.25 | 0.19 |
| 6 | 99773417 | rs4840031 | 1.16 | 0.954 | -0.743 | 0.37 | -0.023 | -0.034 | -0.034 | 0 | 0.08 | 0 | 0.19 | 0.12 | 0.08 | 0.11 |
| 6 | 99776986 | rs6922449 | 0.143 | 0.919 | -0.528 | 0.438 | -0.014 | 0.011 | 0.011 | 0 | 0.45 | 0.29 | 0.29 | 0.61 | 0.5 | 0.4 |
| 6 | 99781218 | rs11757364 | 0.366 | 0.917 | -0.544 | 0.419 | -0.01 | 0.074 | 0.074 | 0 | 0.43 | 0.24 | 0.26 | 0.19 | 0.47 | 0.27 |
| 6 | 99782879 | rs17059400 | 0.681 | 0.535 | -0.415 | 0.154 | -0.012 | 0.012 | 0.012 | 0 | 0.19 | 0.24 | 0.3 | 0.26 | 0.08 | 0.29 |
| 6 | 99793615 | rs11963108 | 0.234 | 0.714 | -0.211 | 0.301 | -0.004 | 0.031 | 0.031 | 0 | 0.35 | 0.55 | 0.22 | 0.54 | 0.32 | 0.33 |
| 6 | 99801698 | rs9402701 | 0.909 | 0.636 | 0.147 | 0.411 | 0.039 | 0.269 | 0.269 | 0 | 0.15 | 0.39 | 0.12 | 0.52 | 0.2 | 0.16 |
| 6 | 99803269 | rs12198238 | 0.1 | 0.766 | -0.07 | 0.212 | 0.037 | 0.033 | 0.033 | 0 | 0.51 | 0.33 | 0.56 | 0.37 | 0.35 | 0.59 |
| 6 | 99804157 | rs13194648 | 1.346 | 0.812 | -0.192 | -0.04 | -0.026 | -0.023 | -0.023 | 0.133 | 0.12 | 0.03 | 0.07 | 0.09 | 0.17 | 0.06 |
| 6 | 99806408 | rs17059457 | 0.868 | 0.736 | -0.239 | -0.105 | -0.021 | -0.026 | -0.026 | 0 | 0.09 | 0 | 0.14 | 0.11 | 0.08 | 0.15 |
| 6 | 99819379 | rs6925344 | -0.653 | 0.635 | 0.181 | 0.089 | 0.007 | 0.049 | 0.049 | 0 | 0.85 | 0.71 | 0.91 | 0.86 | 0.83 | 0.94 |
| 6 | 99826265 | rs12193590 | 0.884 | 0.372 | -0.148 | -0.196 | 0.022 | 0.16 | 0.16 | 0.411 | 0.38 | 0.12 | 0.32 | 0.13 | 0.3 | 0.27 |
| 6 | 99827314 | rs9402716 | -0.86 | 0.648 | -0.043 | -0.2 | 0.132 | 0.073 | 0.073 | 0 | 0.84 | 0.6 | 0.77 | 0.86 | 0.67 | 0.91 |
| 6 | 99828941 | rs4840038 | 0.777 | 0.507 | -0.423 | -0.267 | 0.02 | 0.168 | 0.168 | 0.478 | 0.38 | 0.09 | 0.32 | 0.13 | 0.3 | 0.27 |
| 6 | 99842056 | rs9376137 | 0.954 | 1.01 | -0.192 | 0.351 | -0.011 | -0.059 | -0.059 | 0 | 0.23 | 0.14 | 0.32 | 0.5 | 0.22 | 0.36 |
| 6 | 99842138 | rs9376138 | 0.033 | 0.9 | -0.086 | 0.242 | 0.006 | 0.026 | 0.026 | 0 | 0.37 | 0.27 | 0.39 | 0.79 | 0.32 | 0.6 |
| 6 | 99847260 | rs4144165 | -0.162 | 0.975 | 0.425 | 0.301 | 0.011 | -0.03 | -0.03 | 0 | 0.4 | 0.48 | 0.54 | 0.8 | 0.48 | 0.63 |
| 6 | 99857773 | rs9321534 | 0.21 | 0.87 | 0.647 | 0.114 | 0.007 | -0.027 | -0.027 | 0 | 0.58 | 0.46 | 0.46 | 0.2 | 0.52 | 0.37 |
| 6 | 99871010 | rs12198321 | -0.167 | 0.885 | 0.348 | 0.123 | 0.011 | -0.031 | -0.031 | 0 | 0.41 | 0.48 | 0.54 | 0.8 | 0.48 | 0.63 |
| 6 | 99877219 | rs6923983 | -0.367 | 0.914 | -0.027 | 0.052 | -0.003 | 0.119 | 0.119 | 0 | 0.48 | 0.31 | 0.47 | 0.88 | 0.49 | 0.66 |
| 6 | 99882563 | rs17785525 | 1.602 | 0.792 | -0.003 | -0.207 | -0.044 | -0.024 | -0.024 | 0 | 0.11 | 0 | 0.07 | 0.08 | 0.18 | 0.06 |
| 6 | 99883137 | rs6570064 | -0.304 | 0.865 | 0.181 | -0.259 | -0.001 | 0.08 | 0.08 | 0 | 0.48 | 0.33 | 0.47 | 0.88 | 0.49 | 0.67 |
| 6 | 99883694 | rs9402791 | 0.974 | 0.736 | -0.43 | -0.048 | -0.016 | -0.076 | -0.076 | 0 | 0.22 | 0.11 | 0.32 | 0.5 | 0.22 | 0.37 |
| 6 | 99889915 | rs4839748 | 0.32 | 0.528 | -0.512 | -0.104 | 0.01 | 0.17 | 0.17 | 0.281 | 0.41 | 0.16 | 0.36 | 0.11 | 0.34 | 0.29 |
| 6 | 99893527 | rs9494471 | 0.324 | 0.523 | 0.32 | 0.55 | 0.04 | 0.296 | 0.296 | 0.329 | 0.08 | 0.29 | 0.08 | 0.35 | 0.09 | 0.16 |
| 6 | 99893938 | rs4504482 | 0.628 | 0.749 | -0.047 | -0.104 | 0.003 | -0.038 | -0.038 | 0.01 | 0.37 | 0.2 | 0.37 | 0.49 | 0.33 | 0.42 |
| 6 | 99905558 | rs6916603 | 0.43 | 0.566 | 0.222 | -0.341 | 0.008 | 0.017 | 0.017 | 0 | 0.39 | 0.32 | 0.35 | 0.09 | 0.32 | 0.27 |
| 6 | 99916182 | rs10457650 | -0.033 | 0.56 | 0.237 | -0.158 | 0.014 | 0.046 | 0.046 | 0 | 0.48 | 0.33 | 0.54 | 0.23 | 0.57 | 0.37 |
| 6 | 99921822 | rs7745012 | 0.045 | 0.553 | 0.207 | -0.179 | 0.121 | -0.028 | -0.028 | 0 | 0.91 | 0.74 | 0.83 | 0.99 | 0.81 | 0.95 |
| 6 | 99933680 | rs9483935 | 0.485 | 0.428 | 0.148 | -0.112 | -0.024 | 0.081 | 0.081 | 0 | 0.14 | 0.29 | 0.08 | 0.28 | 0.09 | 0.22 |
| 6 | 99934446 | rs6918880 | 0.791 | 0.424 | 0.128 | -0.247 | 0.003 | -0.04 | -0.04 | 0 | 0.37 | 0.22 | 0.37 | 0.49 | 0.33 | 0.42 |
| 6 | 99939045 | rs12717185 | -0.164 | 0.665 | 0.22 | -0.083 | -0.016 | 0.089 | 0.089 | 0 | 0.2 | 0.3 | 0.12 | 0.41 | 0.16 | 0.28 |
| 6 | 99964012 | rs2209157 | 0.32 | 0.938 | 0.37 | -0.433 | 0.128 | -0.045 | -0.045 | 0 | 0.9 | 0.66 | 0.83 | 0.99 | 0.81 | 0.95 |
| 6 | 99971429 | rs9402863 | 1.056 | 0.801 | 0.025 | -0.357 | -0.007 | -0.014 | -0.014 | 0 | 0.71 | 0.29 | 0.6 | 0.63 | 0.61 | 0.66 |
| 6 | 99973893 | rs10223892 | -0.079 | 0.576 | 0.045 | -0.48 | -0.014 | -0.006 | -0.006 | 0 | 0.24 | 0.31 | 0.12 | 0.14 | 0.19 | 0.21 |
| 6 | 99974948 | rs17224695 | 0.687 | 0.635 | -0.938 | -0.997 | -0.047 | 0.057 | 0.057 | 0.211 | 0.21 | 0.06 | 0.2 | 0.09 | 0.24 | 0.14 |
| 6 | 99980252 | rs2057517 | 0.579 | 0.128 | -0.564 | -0.875 | -0.012 | 0.035 | 0.035 | 0 | 0.13 | 0.21 | 0.18 | 0.24 | 0.12 | 0.2 |
| 6 | 99986195 | rs7754710 | 0.079 | 0.636 | -0.32 | -1.102 | -0.005 | -0.001 | -0.001 | 0 | 0.71 | 0.74 | 0.66 | 0.9 | 0.61 | 0.82 |
| 6 | 99990856 | rs1054227 | 0.778 | 0.624 | -0.346 | -1.108 | -0.026 | 0.061 | 0.061 | 0.202 | 0.21 | 0.05 | 0.19 | 0.09 | 0.19 | 0.14 |
| 6 | 99993268 | rs2296154 | 0.791 | 0.628 | -0.344 | -1.106 | -0.026 | 0.06 | 0.06 | 0.112 | 0.21 | 0.05 | 0.19 | 0.09 | 0.19 | 0.14 |
| 6 | 99995320 | rs13205324 | 0.526 | 0.576 | -0.187 | -1.048 | -0.021 | 0.035 | 0.035 | 0 | 0.27 | 0.34 | 0.22 | 0.37 | 0.2 | 0.29 |
| 6 | 100005775 | rs543967 | 0.746 | 0.63 | -0.174 | -1.008 | 0.001 | -0.004 | -0.004 | 0 | 0.44 | 0.39 | 0.44 | 0.53 | 0.41 | 0.52 |
| 6 | 100022125 | rs1590359 | 0.202 | 0.398 | 0.049 | -0.857 | -0.021 | 0.032 | 0.032 | 0 | 0.27 | 0.34 | 0.22 | 0.37 | 0.2 | 0.29 |
| 6 | 100031037 | rs514769 | 0.319 | 0.443 | -0.679 | -0.701 | -0.015 | -0.013 | -0.013 | 0 | 0.39 | 0.34 | 0.44 | 0.61 | 0.44 | 0.53 |
| 6 | 100046845 | rs1552855 | 0.481 | 0.497 | -1.474 | -0.691 | -0.094 | -0.119 | -0.119 | 0 | 0.37 | 0.07 | 0.42 | 0.57 | 0.43 | 0.52 |
| 6 | 100046933 | rs9484083 | 0.103 | 0.208 | -0.755 | -1.16 | 0.004 | -0.035 | -0.035 | 0 | 0.25 | 0.19 | 0.18 | 0.01 | 0.16 | 0.12 |
| 6 | 100053626 | rs7741279 | 0.287 | 0.739 | -0.94 | -0.809 | 0.006 | 0.027 | 0.027 | 1.015 | 0.16 | 0.09 | 0.15 | 0.02 | 0.14 | 0.13 |
| 6 | 100060761 | rs330843 | 1.282 | 1.126 | -1.192 | -1.199 | -0.031 | 0.048 | 0.048 | 0 | 0.17 | 0.07 | 0.11 | 0 | 0.25 | 0.07 |
| 6 | 100062766 | rs3734346 | 0.557 | 1.153 | -1.102 | -1.046 | 0 | -0.012 | -0.012 | 0 | 0.05 | 0.04 | 0.04 | 0.01 | 0.06 | 0.05 |
| 6 | 100081441 | rs6927488 | 0.404 | 1.388 | -0.872 | -1.156 | -0.008 | -0.05 | -0.05 | 0 | 0.15 | 0.17 | 0.13 | 0.02 | 0.1 | 0.12 |
| 6 | 100082983 | rs594231 | 1.341 | 1.545 | -1.136 | -1.32 | -0.021 | 0.032 | 0.032 | 0 | 0.18 | 0.08 | 0.11 | 0 | 0.24 | 0.06 |
| 6 | 100088902 | rs9402954 | 0.158 | 1.129 | -1.076 | -0.37 | 0.008 | -0.019 | -0.019 | 0.263 | 0.11 | 0.05 | 0.22 | 0.46 | 0.08 | 0.35 |
| 6 | 100092390 | rs1339203 | 0.887 | 0.582 | -0.973 | -1.115 | 0.015 | 0.035 | 0.035 | 0 | 0.33 | 0.23 | 0.17 | 0.11 | 0.24 | 0.22 |
| 6 | 100099703 | rs546567 | 0.592 | 0.379 | -1.303 | -1.022 | -0.007 | 0.03 | 0.03 | 0.272 | 0.31 | 0.21 | 0.23 | 0.02 | 0.35 | 0.18 |
| 6 | 100108500 | rs13193313 | 0.106 | 0.205 | -1.121 | 0.214 | 0.025 | -0.035 | -0.035 | 0.372 | 0.18 | 0.07 | 0.06 | 0.15 | 0.11 | 0.12 |
| 6 | 100116092 | rs9321659 | -1.062 | 0.339 | -0.102 | 0.32 | 0.004 | 0.085 | 0.085 | 0 | 0.15 | 0.59 | 0.14 | 0.25 | 0.1 | 0.17 |
| 6 | 100129570 | rs9373217 | 0.205 | 0.957 | -0.282 | 0.462 | -0.029 | 0.114 | 0.114 | 0 | 0.11 | 0.17 | 0.18 | 0.42 | 0.05 | 0.35 |
| 6 | 100129959 | rs503649 | 1.231 | 1.244 | -0.921 | -0.32 | -0.008 | 0.007 | 0.007 | 0 | 0.19 | 0.09 | 0.13 | 0.01 | 0.24 | 0.07 |
| 6 | 100130991 | rs472977 | 1.288 | 1.06 | -0.936 | -0.337 | -0.002 | 0.003 | 0.003 | 0 | 0.18 | 0.09 | 0.12 | 0.01 | 0.23 | 0.07 |
| 6 | 100131219 | rs9495145 | -0.201 | 1.13 | 0.133 | -0.107 | 0.025 | -0.018 | -0.018 | 0.443 | 0.08 | 0.32 | 0.05 | 0.01 | 0.04 | 0.07 |
| 6 | 100132920 | rs9376355 | 0.656 | 1.089 | -0.036 | -0.071 | 0.023 | -0.054 | -0.054 | 0 | 0.1 | 0.02 | 0.17 | 0.42 | 0.05 | 0.34 |
| 6 | 100161035 | rs12661094 | 0.335 | 0.847 | -0.459 | -0.582 | -0.003 | -0.005 | -0.005 | 0 | 0.57 | 0.33 | 0.46 | 0.56 | 0.49 | 0.61 |
| 6 | 100172277 | rs12211649 | 0.514 | 0.855 | -0.287 | -0.449 | 0.008 | 0.059 | 0.059 | 0 | 0.07 | 0.18 | 0.09 | 0.2 | 0.06 | 0.09 |
| 6 | 100173173 | rs4839755 | 1.072 | 0.634 | -0.374 | -0.476 | 0.01 | -0.018 | -0.018 | 1.035 | 0.26 | 0.12 | 0.24 | 0.22 | 0.22 | 0.21 |
| 6 | 100173832 | rs9389645 | 0.937 | 0.873 | -0.058 | -0.177 | 0.167 | 0.053 | 0.053 | 0 | 0.83 | 0.49 | 0.71 | 0.78 | 0.71 | 0.81 |
| 6 | 100174464 | rs6916754 | 0.408 | 0.711 | -0.325 | 0.01 | -0.026 | 0.101 | 0.101 | 0 | 0.11 | 0.17 | 0.18 | 0.42 | 0.05 | 0.34 |
| 6 | 100187748 | rs12527523 | -0.068 | 1.26 | -0.465 | 0.128 | 0.082 | 0.053 | 0.053 | 0 | 0.09 | 0.19 | 0.24 | 0.24 | 0.25 | 0.12 |

| | | | | | | | | | | | | | | | | |
|---|---|---|---|---|---|---|---|---|---|---|---|---|---|---|---|---|
| 6 | 100192215 | rs12525414 | -0.082 | 1.258 | -0.466 | 0.127 | 0.076 | 0.049 | 0.049 | 0 | 0.09 | 0.19 | 0.24 | 0.24 | 0.25 | 0.12 |
| 6 | 100195578 | rs17826560 | 0.881 | 0.888 | -0.902 | -0.098 | -0.032 | -0.02 | -0.02 | 0.276 | 0.26 | 0.02 | 0.26 | 0.17 | 0.24 | 0.21 |
| 6 | 100197285 | rs7742890 | -0.229 | -0.154 | -1.054 | -0.545 | 0.06 | -0.009 | -0.009 | 0 | 0.59 | 0.34 | 0.65 | 0.55 | 0.71 | 0.45 |
| 6 | 100197712 | rs7356874 | 0.107 | -0.3 | -0.536 | -0.668 | 0.021 | -0.026 | -0.026 | 0 | 0.08 | 0.3 | 0.05 | 0.01 | 0.05 | 0.06 |
| 6 | 100206681 | rs10485226 | 1.348 | -0.287 | -1.16 | -0.687 | -0.05 | 0.036 | 0.036 | 0 | 0.24 | 0.02 | 0.15 | 0.14 | 0.21 | 0.13 |
| 6 | 100207838 | rs4388294 | -0.108 | -0.37 | -1.03 | -0.626 | 0.08 | -0.015 | -0.015 | 1.159 | 0.59 | 0.3 | 0.65 | 0.56 | 0.71 | 0.45 |
| 6 | 100208624 | rs17059765 | 0.654 | -0.297 | -0.781 | -0.262 | 0.034 | -0.039 | -0.039 | 0 | 0.09 | 0.02 | 0.17 | 0.38 | 0.04 | 0.35 |
| 6 | 100213515 | rs4431442 | -0.671 | -0.27 | -0.547 | -0.545 | 0.034 | -0.004 | -0.004 | 0.05 | 0.45 | 0.68 | 0.41 | 0.56 | 0.33 | 0.55 |
| 6 | 100228256 | rs9403091 | -1.037 | -0.35 | -0.226 | -0.639 | -0.016 | 0.064 | 0.064 | 0 | 0.26 | 0.51 | 0.29 | 0.57 | 0.18 | 0.44 |
| 6 | 100231618 | rs9376435 | -1.338 | 0.197 | -0.465 | -0.78 | -0.009 | 0.128 | 0.128 | 0.308 | 0.54 | 0.84 | 0.58 | 0.76 | 0.54 | 0.64 |
| 6 | 100233278 | rs9385869 | -0.098 | -0.271 | -0.322 | -0.864 | 0.006 | -0.01 | -0.01 | 0 | 0.26 | 0.23 | 0.29 | 0.38 | 0.18 | 0.37 |
| 6 | 100233870 | rs9399284 | -1.446 | -0.363 | -0.569 | -0.958 | -0.013 | 0.009 | 0.009 | 0 | 0.39 | 0.71 | 0.5 | 0.61 | 0.43 | 0.5 |
| 6 | 100236473 | rs12663112 | -0.047 | -0.405 | -0.574 | -0.892 | 0.019 | -0.043 | -0.043 | 0 | 0.22 | 0.16 | 0.11 | 0.33 | 0.17 | 0.25 |
| 6 | 100236866 | rs4596491 | 0.929 | -0.389 | -0.59 | -0.88 | 0.006 | -0.022 | -0.022 | 0 | 0.14 | 0.02 | 0.07 | 0.15 | 0.11 | 0.14 |
| 6 | 100237464 | rs12198721 | -0.942 | -0.249 | -0.691 | -1.248 | 0.021 | 0.038 | 0.038 | 0 | 0.5 | 0.36 | 0.26 | 0.34 | 0.43 | 0.38 |
| 6 | 100237880 | rs6936455 | -2.349 | -0.478 | -1.133 | -1.519 | 0 | 0.006 | 0.006 | 0 | 0.76 | 0.88 | 0.74 | 0.78 | 0.73 | 0.75 |
| 6 | 100240267 | rs9403103 | -2.421 | -0.482 | -1.227 | -1.522 | -0.014 | 0.002 | 0.002 | 0 | 0.77 | 0.94 | 0.74 | 0.78 | 0.73 | 0.75 |
| 6 | 100242158 | rs6929006 | -0.932 | -0.208 | -0.724 | -1.161 | -0.048 | -0.012 | -0.012 | 0 | 0.41 | 0.78 | 0.48 | 0.43 | 0.47 | 0.43 |
| 6 | 100242366 | rs6929428 | -2.111 | -0.253 | -0.843 | -1.451 | -0.018 | 0.08 | 0.08 | 0.01 | 0.63 | 0.93 | 0.59 | 0.77 | 0.64 | 0.67 |
| 6 | 100243033 | rs17236659 | -0.086 | -0.925 | -0.574 | -1.467 | 0.009 | 0.038 | 0.038 | 0 | 0.24 | 0.14 | 0.09 | 0.15 | 0.21 | 0.19 |
| 6 | 100247048 | rs6931227 | 0.047 | -0.739 | -0.267 | -1.897 | 0.007 | 0.027 | 0.027 | 0 | 0.19 | 0.09 | 0.13 | 0.01 | 0.16 | 0.1 |
| 6 | 100249106 | rs7763983 | -0.81 | -0.767 | -0.105 | -1.931 | -0.009 | -0.114 | -0.114 | 0 | 0.32 | 0.58 | 0.18 | 0.01 | 0.26 | 0.13 |
| 6 | 100250740 | rs17059858 | 0.198 | -0.397 | -0.322 | -1.689 | 0 | -0.02 | -0.02 | 0 | 0.17 | 0.11 | 0.12 | 0 | 0.17 | 0.09 |
| 6 | 100255471 | rs9373250 | -1.092 | -0.401 | -0.324 | -1.74 | -0.024 | -0.049 | -0.049 | 0 | 0.63 | 0.52 | 0.61 | 0.78 | 0.71 | 0.7 |
| 6 | 100257066 | rs12216503 | 0.876 | 0.185 | -0.814 | -1.213 | -0.012 | 0.06 | 0.06 | 0.128 | 0.12 | 0.02 | 0.02 | 0 | 0.1 | 0.05 |
| 6 | 100257668 | rs4495279 | 0.06 | 0.72 | -0.427 | -0.546 | 0.12 | 0.245 | 0.245 | 0 | 0.1 | 0.25 | 0.3 | 0.44 | 0.2 | 0.28 |
| 6 | 100265121 | rs6934621 | -0.465 | -0.115 | -1.083 | -0.689 | 0.023 | -0.033 | -0.033 | 1.613 | 0.25 | 0.21 | 0.2 | 0.34 | 0.22 | 0.27 |
| 6 | 100266123 | rs7760502 | -0.945 | -0.032 | -1.238 | -0.714 | -0.02 | -0.039 | -0.039 | 0.027 | 0.3 | 0.2 | 0.26 | 0.34 | 0.35 | 0.31 |
| 6 | 100268849 | rs17059881 | 0.279 | -0.052 | -1.315 | -0.652 | 0.003 | -0.014 | -0.014 | 0 | 0.17 | 0.02 | 0.09 | 0.16 | 0.15 | 0.18 |
| 6 | 100277840 | rs13204333 | -1.012 | 0.651 | -0.739 | -0.789 | 0 | 0 | 0 | 0 | 0.15 | 0.19 | 0.19 | 0.17 | 0.2 | 0.14 |
| 6 | 100279956 | rs9403141 | 0.169 | -0.121 | -0.615 | -0.994 | 0.015 | 0.002 | 0.002 | 0 | 0.37 | 0.34 | 0.26 | 0.67 | 0.27 | 0.5 |
| 6 | 100280032 | rs9385894 | 0.063 | 0.356 | -0.456 | -0.76 | 0.011 | 0.01 | 0.01 | 0 | 0.49 | 0.63 | 0.33 | 0.81 | 0.34 | 0.69 |
| 6 | 100281817 | rs7453413 | 0.344 | 1.411 | 0.156 | -0.045 | -0.002 | -0.026 | -0.026 | 0 | 0.6 | 0.62 | 0.48 | 0.83 | 0.45 | 0.74 |
| 6 | 100282075 | rs3922542 | 0.414 | 1.438 | -0.187 | -0.08 | -0.002 | -0.005 | -0.005 | 0 | 0.59 | 0.52 | 0.48 | 0.71 | 0.45 | 0.6 |
| 6 | 100286127 | rs9321763 | 0.302 | 1.32 | -0.477 | -0.052 | 0.012 | -0.027 | -0.027 | 0 | 0.59 | 0.55 | 0.52 | 0.82 | 0.53 | 0.8 |
| 6 | 100288317 | rs4370372 | -0.61 | 1.388 | -0.763 | -0.249 | -0.012 | 0.017 | 0.017 | 0 | 0.29 | 0.2 | 0.2 | 0.18 | 0.33 | 0.27 |
| 6 | 100292091 | rs9495653 | 0.752 | 0.727 | -0.467 | -0.68 | -0.009 | -0.04 | -0.04 | 0 | 0.88 | 0.81 | 0.95 | 1 | 0.91 | 0.9 |
| 6 | 100293531 | rs9495657 | -0.299 | 0.312 | -0.28 | -1.962 | -0.013 | -0.166 | -0.166 | 0.92 | 0.34 | 0.52 | 0.19 | 0 | 0.29 | 0.18 |
| 6 | 100296024 | rs4990843 | -0.49 | 0.292 | -0.324 | -1.85 | -0.007 | -0.073 | -0.073 | 0 | 0.4 | 0.45 | 0.25 | 0.12 | 0.36 | 0.33 |
| 6 | 100297481 | rs9403163 | 0.478 | 0.786 | -0.652 | -1.742 | -0.021 | -0.01 | -0.01 | 0 | 0.18 | 0.15 | 0.24 | 0.22 | 0.3 | 0.24 |
| 6 | 100317146 | rs12197810 | 0.652 | 0.376 | -0.807 | -2.612 | -0.015 | 0.089 | 0.089 | 0.091 | 0.27 | 0.11 | 0.29 | 0.04 | 0.26 | 0.13 |
| 6 | 100319031 | rs9403180 | 0.379 | 0.207 | -0.35 | -2.493 | -0.006 | -0.033 | -0.033 | 0 | 0.42 | 0.42 | 0.3 | 0.2 | 0.36 | 0.46 |
| 6 | 100321734 | rs2397664 | 0.512 | 0.153 | -0.497 | -2.386 | 0 | 0 | 0 | 0 | 0.07 | 0.06 | 0.12 | 0.19 | 0.07 | 0.28 |
| 6 | 100324464 | rs9403186 | 0.434 | 0.201 | -0.439 | -2.37 | -0.013 | 0.018 | 0.018 | 1.354 | 0.3 | 0.24 | 0.39 | 0.76 | 0.37 | 0.42 |
| 6 | 100326214 | rs9495745 | 0.721 | 0.79 | -0.417 | -2.339 | 0.014 | 0.082 | 0.082 | 0.497 | 0.24 | 0.08 | 0.25 | 0.04 | 0.19 | 0.11 |
| 6 | 100328198 | rs4840097 | -0.389 | 0.798 | 0.014 | -2.274 | -0.001 | -0.099 | -0.099 | 0 | 0.64 | 0.41 | 0.79 | 1 | 0.66 | 0.8 |
| 6 | 100329446 | rs9389810 | 0.675 | 0.834 | -0.158 | -2.21 | -0.037 | 0.082 | 0.082 | 0 | 0.29 | 0.04 | 0.42 | 0.15 | 0.32 | 0.31 |
| 6 | 100345238 | rs2397678 | -0.704 | 0.824 | -0.015 | -2.244 | 0.006 | -0.085 | -0.085 | 0.515 | 0.69 | 0.45 | 0.75 | 0.92 | 0.68 | 0.72 |
| 6 | 100345403 | rs6902801 | -0.24 | 0.712 | 0.278 | -2.204 | -0.004 | -0.043 | -0.043 | 0.071 | 0.41 | 0.48 | 0.39 | 0.84 | 0.36 | 0.48 |
| 6 | 100353268 | rs7751620 | -0.621 | 0.613 | 0.386 | -2.196 | -0.025 | -0.081 | -0.081 | 4.498 | 0.7 | 0.44 | 0.82 | 0.91 | 0.77 | 0.86 |
| 6 | 100354935 | rs9403208 | 0.047 | 0.904 | 0.093 | -2.195 | -0.027 | 0.038 | 0.038 | 1.468 | 0.37 | 0.25 | 0.37 | 0.77 | 0.44 | 0.48 |
| 6 | 100366635 | rs4560657 | 1.444 | 0.795 | 0.314 | -2.207 | -0.004 | -0.072 | -0.072 | 0 | 0.22 | 0.24 | 0.1 | 0 | 0.19 | 0.08 |
| 6 | 100369284 | rs13206575 | 0.739 | 0.785 | 0.401 | -2.218 | -0.012 | -0.102 | -0.102 | 0.325 | 0.27 | 0.31 | 0.17 | 0 | 0.22 | 0.1 |
| 6 | 100391872 | rs4559096 | -0.676 | 0.813 | 0.391 | -2.232 | -0.001 | -0.004 | -0.004 | 0 | 0.66 | 0.66 | 0.58 | 0.78 | 0.66 | 0.6 |
| 6 | 100392532 | rs12215494 | 0.89 | 0.822 | 0.045 | -2.223 | -0.032 | 0.114 | 0.114 | 0 | 0.31 | 0.07 | 0.4 | 0.11 | 0.34 | 0.36 |
| 6 | 100393761 | rs13195863 | 0.5 | 0.679 | -0.013 | -2.06 | -0.019 | -0.128 | -0.128 | 0 | 0.28 | 0.38 | 0.17 | 0 | 0.21 | 0.1 |
| 6 | 100397526 | rs10499026 | 1.234 | 0.809 | -0.396 | -1.941 | -0.004 | -0.076 | -0.076 | 0 | 0.22 | 0.24 | 0.1 | 0 | 0.19 | 0.08 |
| 6 | 100403514 | rs4840106 | -0.093 | 0.989 | -0.747 | -1.271 | -0.014 | 0.038 | 0.038 | 1.646 | 0.32 | 0.12 | 0.33 | 0.78 | 0.3 | 0.41 |
| 6 | 100427015 | rs7739904 | -0.709 | 1.24 | -0.349 | -0.312 | -0.002 | 0 | 0 | 0 | 0.35 | 0.35 | 0.34 | 0.72 | 0.4 | 0.4 |
| 6 | 100428029 | rs9496070 | -0.312 | 1.262 | -0.692 | -0.323 | -0.008 | -0.005 | -0.005 | 0 | 0.43 | 0.59 | 0.51 | 0.6 | 0.54 | 0.36 |
| 6 | 100428313 | rs13212643 | 0.668 | 0.948 | -0.688 | -0.329 | 0.03 | 0.091 | 0.091 | 0.174 | 0.5 | 0.12 | 0.39 | 0.28 | 0.39 | 0.56 |
| 6 | 100430062 | rs12203515 | 0.732 | 1.119 | -0.356 | -0.444 | 0.056 | -0.117 | -0.117 | 0 | 0.17 | 0.34 | 0.22 | 0.01 | 0.2 | 0.1 |
| 6 | 100430803 | rs9496085 | 0.011 | 1.06 | -0.191 | 0.048 | -0.035 | -0.057 | -0.057 | 0.378 | 0.27 | 0.18 | 0.31 | 0.59 | 0.4 | 0.32 |
| 6 | 100431321 | rs9389934 | 0.286 | 1.073 | -0.273 | -0.027 | 0.008 | 0 | 0 | 0 | 0.66 | 0.52 | 0.6 | 0.29 | 0.58 | 0.64 |
| 6 | 100439009 | rs2001456 | 0.236 | 0.827 | -0.842 | -0.287 | 0.028 | -0.028 | -0.028 | 0 | 0.3 | 0.09 | 0.3 | 0.29 | 0.22 | 0.46 |
| 6 | 100442268 | rs6925272 | 0.147 | 0.917 | -0.238 | -0.641 | 0 | -0.056 | -0.056 | 0 | 0.39 | 0.48 | 0.33 | 0.12 | 0.37 | 0.21 |
| 6 | 100448788 | rs3763374 | 0.075 | 1.221 | -0.62 | 0.543 | -0.029 | -0.008 | -0.008 | 0 | 0.07 | 0 | 0.1 | 0.1 | 0.1 | 0.04 |
| 6 | 100449301 | rs9385975 | 1.209 | 0.733 | 0.473 | 1.408 | -0.011 | 0.218 | 0.218 | 1.134 | 0.84 | 0.52 | 0.88 | 0.56 | 0.87 | 0.78 |
| 6 | 100457411 | rs9389951 | 0.368 | 0.984 | -0.104 | 0.958 | -0.011 | 0.018 | 0.018 | 0.299 | 0.39 | 0.33 | 0.45 | 0.24 | 0.48 | 0.24 |

| | | | | | | | | | | | | | | | | |
|---|---|---|---|---|---|---|---|---|---|---|---|---|---|---|---|---|
| 6 | 100480906 | rs4240585 | 0.949 | 0.535 | -0.541 | 0.526 | 0.023 | -0.028 | -0.028 | 0 | 0.33 | 0.12 | 0.37 | 0.33 | 0.31 | 0.42 |
| 6 | 100482016 | rs6925134 | 0.4 | 0.935 | -0.628 | 0.45 | -0.006 | -0.005 | -0.005 | 0.79 | 0.1 | 0.11 | 0.16 | 0.12 | 0.13 | 0.04 |
| 6 | 100492824 | rs12202476 | 0.353 | 0.824 | -0.114 | 0.368 | 0.007 | 0.007 | 0.007 | 0 | 0.17 | 0.14 | 0.1 | 0.08 | 0.13 | 0.1 |
| 6 | 100494186 | rs3890820 | 0.602 | 0.638 | 0.132 | 0.326 | -0.047 | 0.033 | 0.033 | 0 | 0.72 | 0.62 | 0.84 | 0.56 | 0.82 | 0.6 |
| 6 | 100510685 | rs6903499 | 0.376 | 0.948 | 0.477 | 0.624 | -0.002 | 0.087 | 0.087 | 0 | 0.91 | 0.89 | 0.94 | 0.65 | 0.92 | 0.75 |
| 6 | 100514432 | rs11155243 | 0.614 | 0.927 | 0.544 | 0.565 | -0.014 | 0.029 | 0.029 | 0 | 0.76 | 0.68 | 0.85 | 0.56 | 0.81 | 0.67 |
| 6 | 100521649 | rs6938191 | 1.085 | 1.111 | 0.375 | 0.274 | -0.008 | 0.003 | 0.003 | 0 | 0.29 | 0.22 | 0.38 | 0.24 | 0.35 | 0.2 |
| 6 | 100523912 | rs4144262 | 0.451 | 0.893 | 0.229 | 0.295 | 0.004 | 0.002 | 0.002 | 0 | 0.16 | 0.13 | 0.09 | 0.08 | 0.11 | 0.08 |
| 6 | 100534262 | rs2474265 | 0.016 | 0.985 | 0.28 | 0.679 | 0 | 0.016 | 0.016 | 0 | 0.07 | 0.05 | 0.06 | 0.28 | 0.08 | 0.25 |
| 6 | 100539396 | rs6910833 | -0.997 | 1.215 | 0.387 | 0.489 | 0.004 | -0.004 | -0.004 | 0.837 | 0.67 | 0.69 | 0.59 | 0.76 | 0.63 | 0.79 |
| 6 | 100548788 | rs17060130 | 0.176 | 1.267 | 0.19 | 0.309 | 0.006 | -0.038 | -0.038 | 0 | 0.1 | 0.17 | 0.07 | 0 | 0.13 | 0.02 |
| 6 | 100560563 | rs9386011 | 1.208 | 1.167 | -0.077 | 0.392 | 0.017 | -0.052 | -0.052 | 0.087 | 0.35 | 0.12 | 0.3 | 0.4 | 0.32 | 0.45 |
| 6 | 100562360 | rs2474258 | -0.071 | 1.181 | 0.112 | 0.768 | -0.008 | 0.01 | 0.01 | 0 | 0.08 | 0.06 | 0.15 | 0.29 | 0.13 | 0.26 |
| 6 | 100574090 | rs9969059 | 0.242 | 0.749 | 0.229 | 0.418 | -0.003 | -0.011 | -0.011 | 0.12 | 0.14 | 0.15 | 0.07 | 0.08 | 0.09 | 0.07 |
| 6 | 100575272 | rs932511 | -0.291 | 0.829 | 0.015 | 0.412 | -0.006 | -0.007 | -0.007 | 0 | 0.6 | 0.66 | 0.44 | 0.48 | 0.5 | 0.54 |
| 6 | 100583430 | rs4839772 | 0.083 | 1.145 | -0.743 | 0.785 | 0.02 | -0.002 | -0.002 | 0 | 0.44 | 0.23 | 0.35 | 0.4 | 0.4 | 0.46 |
| 6 | 100592593 | rs9390065 | 0.076 | 1.702 | -0.432 | 1.314 | 0.021 | 0.083 | 0.083 | 0.7 | 0.72 | 0.47 | 0.58 | 0.52 | 0.71 | 0.69 |
| 6 | 100592868 | rs2474254 | -1.12 | 1.911 | -1.267 | 1.06 | -0.031 | -0.007 | -0.007 | 0 | 0.24 | 0.05 | 0.18 | 0.26 | 0.19 | 0.3 |
| 6 | 100593022 | rs11155282 | 0.506 | 1.84 | -1.237 | 0.807 | 0.019 | 0.049 | 0.049 | 2.267 | 0.44 | 0.23 | 0.38 | 0.32 | 0.49 | 0.38 |
| 6 | 100602931 | rs2503095 | -0.31 | 1.862 | -0.888 | 1.445 | 0.018 | 0.036 | 0.036 | 0 | 0.09 | 0.18 | 0.29 | 0.2 | 0.17 | 0.19 |
| 6 | 100603029 | rs761714 | -0.243 | 1.865 | -0.936 | 1.31 | -0.006 | -0.037 | -0.037 | 0.527 | 0.18 | 0.09 | 0.18 | 0.31 | 0.21 | 0.16 |
| 6 | 100606568 | rs2503097 | -0.872 | 1.926 | -0.622 | 0.707 | 0.013 | 0.019 | 0.019 | 0 | 0.11 | 0.18 | 0.29 | 0.2 | 0.2 | 0.23 |
| 6 | 100612700 | rs17789452 | -0.306 | 2.007 | -0.601 | 0.583 | -0.001 | -0.008 | -0.008 | 0.207 | 0.29 | 0.26 | 0.22 | 0.11 | 0.29 | 0.29 |
| 6 | 100626148 | rs761711 | -0.856 | 2.084 | -0.641 | 0.713 | -0.005 | 0.007 | 0.007 | 0 | 0.42 | 0.34 | 0.52 | 0.32 | 0.49 | 0.51 |
| 6 | 100627435 | rs13205707 | -0.62 | 1.423 | -0.891 | 0.057 | -0.011 | 0.039 | 0.039 | 0 | 0.33 | 0.17 | 0.23 | 0.12 | 0.32 | 0.33 |
| 6 | 100631103 | rs7741147 | 0.689 | 1.319 | -0.601 | 0.855 | 0.001 | 0.003 | 0.003 | 0 | 0.69 | 0.34 | 0.7 | 0.56 | 0.66 | 0.58 |
| 6 | 100633338 | rs12201659 | 0.776 | 0.939 | -1.256 | 0.349 | -0.029 | 0.103 | 0.103 | 0.272 | 0.41 | 0.15 | 0.32 | 0.2 | 0.42 | 0.23 |
| 6 | 100634009 | rs6899958 | 0.045 | 0.814 | -0.387 | 0.137 | 0.038 | -0.077 | -0.077 | 0.241 | 0.87 | 0.57 | 0.92 | 0.98 | 0.89 | 0.86 |
| 6 | 100636619 | rs7754315 | 0.275 | 0.903 | -0.583 | 0.848 | 0.01 | 0.006 | 0.006 | 0 | 0.52 | 0.35 | 0.41 | 0.46 | 0.56 | 0.43 |
| 6 | 100642867 | rs6570574 | 0.84 | 1.033 | -0.511 | 1.306 | -0.009 | -0.01 | -0.01 | 0 | 0.86 | 0.89 | 0.84 | 0.72 | 0.76 | 0.9 |
| 6 | 100650387 | rs9496769 | 0.047 | 0.039 | -0.769 | 1.086 | 0.068 | 0.034 | 0.034 | 0.083 | 0.43 | 0.29 | 0.36 | 0.28 | 0.28 | 0.35 |
| 6 | 100661336 | rs9484823 | 0.628 | -0.517 | -1.07 | 1.109 | 0.042 | 0.029 | 0.029 | 0.224 | 0.3 | 0.21 | 0.27 | 0.15 | 0.18 | 0.17 |
| 6 | 100668277 | rs9390146 | -0.06 | -0.488 | -1.026 | 1.147 | 0 | -0.021 | -0.021 | 0 | 0.12 | 0.03 | 0.09 | 0.13 | 0.1 | 0.17 |
| 6 | 100674018 | rs6928245 | 1.287 | -0.628 | -1.041 | 0.999 | -0.011 | -0.025 | -0.025 | 0.033 | 0.72 | 0.79 | 0.73 | 0.59 | 0.64 | 0.77 |
| 6 | 100674799 | rs7749433 | 0.063 | -0.537 | -1.005 | 1.218 | -0.016 | 0.003 | 0.003 | 0 | 0.84 | 0.91 | 0.87 | 0.82 | 0.75 | 0.9 |
| 6 | 100676593 | rs730223 | 1.039 | -0.654 | -0.796 | 1.207 | -0.001 | 0.002 | 0.002 | 0.6 | 0.44 | 0.55 | 0.47 | 0.44 | 0.48 | 0.6 |
| 6 | 100680565 | rs2073265 | -1.049 | -0.744 | -0.527 | 1.529 | 0.034 | 0.044 | 0.044 | 0.002 | 0.49 | 0.35 | 0.47 | 0.61 | 0.45 | 0.4 |
| 6 | 100693008 | rs4140483 | 0.355 | -0.832 | -0.626 | 1.541 | 0.052 | 0.031 | 0.031 | 0 | 0.29 | 0.16 | 0.26 | 0.14 | 0.16 | 0.18 |
| 6 | 100698564 | rs9390162 | -0.036 | -0.851 | -0.364 | 1.597 | 0.01 | -0.009 | -0.009 | 0 | 0.34 | 0.31 | 0.38 | 0.35 | 0.21 | 0.24 |
| 6 | 100706809 | rs12175103 | -0.849 | -0.797 | -0.284 | 1.53 | 0.045 | 0.048 | 0.048 | 0 | 0.47 | 0.29 | 0.46 | 0.61 | 0.43 | 0.4 |
| 6 | 100707145 | rs6908458 | -0.842 | -0.69 | -0.335 | 1.584 | -0.011 | -0.022 | -0.022 | 0.287 | 0.18 | 0.13 | 0.21 | 0.44 | 0.28 | 0.16 |
| 6 | 100722593 | rs7750764 | 0.273 | -0.64 | -0.369 | 1.648 | -0.011 | -0.017 | -0.017 | 0 | 0.09 | 0.04 | 0.06 | 0.21 | 0.12 | 0.06 |
| 6 | 100734930 | rs9399482 | -0.844 | -0.815 | -0.004 | 1.534 | 0.001 | 0.001 | 0.001 | 0 | 0.51 | 0.53 | 0.49 | 0.61 | 0.45 | 0.4 |
| 6 | 100744281 | rs10499030 | -1.479 | -0.333 | 0.141 | 1.724 | 0.013 | 0.083 | 0.083 | 0 | 0.22 | 0.34 | 0.24 | 0.47 | 0.3 | 0.22 |
| 6 | 100752002 | rs9390223 | -1.087 | -0.158 | -0.432 | 1.843 | -0.001 | 0.009 | 0.009 | 0 | 0.19 | 0.14 | 0.22 | 0.47 | 0.28 | 0.15 |
| 6 | 100758142 | rs11752981 | -0.505 | -0.086 | -0.779 | 1.516 | -0.033 | -0.036 | -0.036 | 0 | 0.15 | 0.05 | 0.1 | 0.25 | 0.23 | 0.1 |
| 6 | 100760042 | rs9376867 | -1.103 | 0.036 | -1.115 | 1.598 | -0.057 | -0.081 | -0.081 | 0 | 0.22 | 0.05 | 0.22 | 0.39 | 0.3 | 0.17 |
| 6 | 100760170 | rs13196561 | -0.445 | 0.102 | -1.198 | 1.488 | -0.055 | -0.052 | -0.052 | 0 | 0.15 | 0.02 | 0.09 | 0.24 | 0.22 | 0.1 |
| 6 | 100760409 | rs3957418 | 0.409 | 0.038 | -1.017 | 0.914 | 0.008 | -0.018 | -0.018 | 0 | 0.29 | 0.32 | 0.21 | 0.14 | 0.38 | 0.41 |
| 6 | 100763213 | rs924974 | 0.643 | -0.822 | -1.148 | 1.421 | -0.008 | -0.007 | -0.007 | 5.051 | 0.61 | 0.54 | 0.71 | 0.64 | 0.65 | 0.78 |
| 6 | 100769678 | rs10046239 | 0.57 | -0.89 | -1.251 | 1.441 | 0.001 | 0.036 | 0.036 | 0 | 0.92 | 0.86 | 0.85 | 0.81 | 0.91 | 0.91 |
| 6 | 100773433 | rs1506075 | 0.528 | -1.057 | -1.443 | 1.841 | -0.016 | -0.068 | -0.068 | 0 | 0.11 | 0.01 | 0.26 | 0.19 | 0.13 | 0.14 |
| 6 | 100779276 | rs2841300 | 0.78 | -1.014 | -1.196 | 1.313 | -0.002 | -0.001 | -0.001 | 0 | 0.61 | 0.59 | 0.71 | 0.64 | 0.65 | 0.78 |
| 6 | 100779692 | rs9390276 | 0.049 | -0.939 | -1.239 | 1.358 | -0.017 | -0.029 | -0.029 | 0 | 0.07 | 0.01 | 0.14 | 0.2 | 0.09 | 0.07 |
| 6 | 100785588 | rs2841296 | 0.034 | -1.061 | -1.247 | 1.268 | -0.025 | 0 | 0 | 0.003 | 0.77 | 0.69 | 0.82 | 0.81 | 0.88 | 0.84 |
| 6 | 100791267 | rs1395118 | -0.089 | -1.019 | -1.399 | 1.23 | -0.006 | 0.005 | 0.005 | 0 | 0.83 | 0.86 | 0.89 | 0.83 | 0.78 | 0.91 |
| 6 | 100793016 | rs7749859 | 0.859 | -1.107 | -1.463 | 1.132 | -0.027 | -0.046 | -0.046 | 0 | 0.14 | 0.16 | 0.03 | 0 | 0.03 | 0.07 |
| 6 | 100793443 | rs2841292 | 0.837 | -0.804 | -2.317 | 1.376 | -0.001 | 0 | 0 | 0.022 | 0.74 | 0.76 | 0.74 | 0.6 | 0.69 | 0.81 |
| 6 | 100797890 | rs1876155 | -0.9 | -0.874 | -2.665 | 1.446 | -0.04 | -0.049 | -0.049 | 4.37 | 0.09 | 0.01 | 0.18 | 0.17 | 0.16 | 0.13 |
| 6 | 100798472 | rs1506079 | -0.202 | -0.728 | -2.794 | 1.516 | -0.005 | -0.111 | -0.111 | 0 | 0.14 | 0.01 | 0.33 | 0.48 | 0.11 | 0.16 |
| 6 | 100800122 | rs2841287 | -1.097 | -0.873 | -2.853 | 1.399 | 0 | 0.007 | 0.007 | 0.014 | 0.37 | 0.05 | 0.54 | 0.64 | 0.31 | 0.32 |
| 6 | 100802068 | rs9403683 | -0.504 | -1.018 | -2.454 | 1.088 | 0.038 | -0.031 | -0.031 | 0 | 0.23 | 0.03 | 0.17 | 0.16 | 0.13 | 0.16 |
| 6 | 100810976 | rs9403695 | -0.984 | -0.346 | -2.11 | 1.437 | -0.022 | -0.04 | -0.04 | 0 | 0.08 | 0.01 | 0.16 | 0.15 | 0.1 | 0.09 |
| 6 | 100811392 | rs17060420 | -0.777 | -0.297 | -1.996 | 1.537 | -0.016 | -0.014 | -0.014 | 0.089 | 0.06 | 0.01 | 0.13 | 0.17 | 0.09 | 0.06 |
| 6 | 100811533 | rs17792983 | -0.634 | -0.303 | -2.244 | 0.895 | -0.018 | 0.02 | 0.02 | 0.367 | 0.22 | 0.08 | 0.21 | 0.2 | 0.28 | 0.15 |
| 6 | 100813188 | rs1506084 | -1.975 | -0.45 | -1.743 | 1.367 | -0.005 | 0.015 | 0.015 | 0.007 | 0.31 | 0.24 | 0.44 | 0.74 | 0.35 | 0.33 |
| 6 | 100816614 | rs13213077 | -1.707 | -0.499 | -1.76 | 1.628 | -0.001 | -0.087 | -0.087 | 0 | 0.24 | 0.12 | 0.33 | 0.57 | 0.26 | 0.25 |
| 6 | 100818915 | rs9376940 | -1.767 | -0.52 | -1.691 | 1.617 | 0.003 | -0.059 | -0.059 | 0 | 0.24 | 0.15 | 0.33 | 0.57 | 0.26 | 0.25 |
| 6 | 100827450 | rs9403718 | -1.812 | -0.525 | -1.695 | 1.662 | -0.017 | -0.061 | -0.061 | 0 | 0.2 | 0.09 | 0.31 | 0.57 | 0.24 | 0.26 |

| | | | | | | | | | | | | | | | | |
|---|---|---|---|---|---|---|---|---|---|---|---|---|---|---|---|---|
| 6 | 100827834 | rs17060460 | -1.837 | -0.523 | -1.702 | 1.663 | -0.034 | -0.087 | -0.087 | 0 | 0.19 | 0.06 | 0.3 | 0.57 | 0.23 | 0.25 |
| 6 | 100829135 | rs453530 | 0.392 | -0.829 | -1.85 | 1.457 | 0.067 | 0.085 | 0.085 | 2.073 | 0.14 | 0 | 0.01 | 0 | 0.02 | 0.05 |
| 6 | 100831997 | rs9403732 | -1.319 | 0.603 | -1.073 | 2.341 | -0.027 | -0.103 | -0.103 | 0 | 0.14 | 0 | 0.21 | 0.4 | 0.16 | 0.18 |
| 6 | 100836784 | rs13201004 | -0.582 | 0.65 | -1.037 | 2.344 | -0.001 | -0.006 | -0.006 | 0.722 | 0.09 | 0.08 | 0.12 | 0.17 | 0.1 | 0.07 |
| 6 | 100857869 | rs7747189 | -2.003 | 0.641 | -0.974 | 2.303 | 0 | -0.034 | -0.034 | 0 | 0.23 | 0.17 | 0.34 | 0.57 | 0.25 | 0.26 |
| 6 | 100859159 | rs718268 | -2.205 | 0.646 | -1.004 | 2.305 | -0.002 | 0.019 | 0.019 | 0 | 0.25 | 0.26 | 0.35 | 0.57 | 0.25 | 0.26 |
| 6 | 100859947 | rs17060530 | -2.638 | 0.682 | -1 | 2.306 | 0 | -0.003 | -0.003 | 0 | 0.31 | 0.33 | 0.44 | 0.74 | 0.34 | 0.33 |
| 6 | 100862051 | rs17060534 | -1.459 | 1.051 | -1.39 | 2.538 | -0.002 | 0.024 | 0.024 | 0 | 0.06 | 0.06 | 0.1 | 0.18 | 0.08 | 0.07 |
| 6 | 100868779 | rs3734354 | -1.092 | 0.971 | -1.421 | 1.946 | -0.037 | -0.062 | -0.062 | 0.054 | 0.1 | 0 | 0.21 | 0.41 | 0.15 | 0.19 |
| 6 | 100872787 | rs714378 | -1.463 | 0.888 | -1.315 | 1.895 | -0.006 | -0.023 | -0.023 | 0 | 0.16 | 0.13 | 0.28 | 0.57 | 0.18 | 0.23 |
| 6 | 100875516 | rs3778033 | -1.272 | 0.949 | -1.342 | 1.783 | -0.017 | -0.051 | -0.051 | 0 | 0.17 | 0.07 | 0.25 | 0.41 | 0.21 | 0.22 |
| 6 | 100878267 | rs3798498 | 0.164 | 0.835 | -1.42 | 1.413 | 0.006 | 0.001 | 0.001 | 0 | 0.55 | 0.33 | 0.45 | 0.38 | 0.6 | 0.68 |
| 6 | 100879651 | rs241816 | 0.084 | 0.576 | -0.849 | 1.842 | -0.001 | 0.004 | 0.004 | 0 | 0.61 | 0.43 | 0.55 | 0.55 | 0.65 | 0.78 |
| 6 | 100880696 | rs1847912 | -0.389 | 0.44 | -1.034 | 1.536 | 0.023 | 0.082 | 0.082 | 0 | 0.24 | 0.58 | 0.4 | 0.44 | 0.25 | 0.17 |
| 6 | 100888625 | rs3798492 | -0.167 | 0.272 | -0.891 | 0.945 | -0.021 | -0.029 | -0.029 | 0 | 0.29 | 0.13 | 0.38 | 0.39 | 0.29 | 0.19 |
| 6 | 100894587 | rs1857859 | -0.239 | 0.242 | -0.678 | 0.93 | -0.013 | -0.02 | -0.02 | 0.583 | 0.29 | 0.18 | 0.38 | 0.39 | 0.29 | 0.19 |
| 6 | 100895344 | rs397662 | -0.499 | 0.173 | -0.674 | 0.822 | -0.002 | 0.083 | 0.083 | 0 | 0.82 | 0.97 | 0.94 | 0.98 | 0.86 | 0.94 |
| 6 | 100910798 | rs241819 | 0.66 | 0.393 | -0.393 | 1.3 | -0.032 | -0.014 | -0.014 | 0 | 0.53 | 0.85 | 0.53 | 0.5 | 0.54 | 0.74 |
| 6 | 100932409 | rs1972281 | -0.225 | 0.383 | -0.123 | 1.474 | -0.011 | -0.036 | -0.036 | 0 | 0.81 | 0.97 | 0.89 | 0.88 | 0.81 | 0.93 |
| 6 | 100936361 | rs6570772 | -0.234 | 0.406 | -0.13 | 1.466 | -0.006 | -0.037 | -0.037 | 0 | 0.82 | 0.97 | 0.89 | 0.88 | 0.82 | 0.93 |
| 6 | 100936628 | rs10457800 | 1.166 | 0.329 | 0.131 | 1.695 | -0.007 | 0.058 | 0.058 | 0 | 0.46 | 0.33 | 0.48 | 0.33 | 0.51 | 0.67 |
| 6 | 100942138 | rs7764271 | -0.79 | 0.355 | 0.114 | 1.933 | 0.048 | 0.361 | 0.361 | 0.76 | 0.05 | 0.32 | 0.1 | 0.32 | 0.01 | 0.08 |
| 6 | 100949822 | rs7751526 | 0.494 | 0.302 | -0.02 | 1.64 | -0.047 | 0.104 | 0.104 | 0.019 | 0.55 | 0.87 | 0.65 | 0.83 | 0.54 | 0.82 |
| 6 | 100953148 | rs240764 | 0.737 | 0.449 | 0.204 | 1.404 | -0.002 | -0.003 | -0.003 | 0 | 0.48 | 0.52 | 0.5 | 0.71 | 0.47 | 0.76 |
| 6 | 100954036 | rs240766 | -0.027 | 0.408 | 0.054 | 1.192 | -0.093 | 0.033 | 0.033 | 1.392 | 0.6 | 0.93 | 0.59 | 0.79 | 0.54 | 0.8 |
| 6 | 100955752 | rs12374612 | 1.137 | 0.347 | 0.21 | 1.341 | 0.014 | 0.086 | 0.086 | 0 | 0.45 | 0.29 | 0.38 | 0.27 | 0.47 | 0.63 |
| 6 | 100957344 | rs240768 | -0.869 | 0.325 | 0.62 | 1.489 | -0.013 | 0.064 | 0.064 | 0.368 | 0.08 | 0.23 | 0.11 | 0.16 | 0.05 | 0.09 |
| 6 | 100961198 | rs401864 | -1.48 | 0.321 | 0.77 | 1.495 | -0.018 | 0.067 | 0.067 | 0 | 0.11 | 0.28 | 0.12 | 0.2 | 0.07 | 0.09 |
| 6 | 100966006 | rs3213542 | -0.804 | 0.33 | 0.757 | 1.496 | -0.015 | 0.044 | 0.044 | 0.349 | 0.08 | 0.19 | 0.11 | 0.16 | 0.05 | 0.09 |
| 6 | 101027440 | rs6931919 | 0.879 | 0.33 | 0.735 | 1.496 | -0.015 | 0.045 | 0.045 | 0.345 | 0.92 | 0.82 | 0.89 | 0.84 | 0.95 | 0.91 |
| 6 | 101039215 | rs17245314 | 0.45 | 0.52 | 0.473 | 1.263 | -0.098 | 0.116 | 0.116 | 0.24 | 0.27 | 0.02 | 0.2 | 0.07 | 0.28 | 0.14 |
| 6 | 101072307 | rs240150 | -1.154 | 0.491 | 0.655 | 1.34 | -0.002 | 0.084 | 0.084 | 0 | 0.54 | 0.7 | 0.5 | 0.73 | 0.51 | 0.36 |
| 6 | 101091434 | rs1392969 | -1.712 | 0.31 | 0.77 | 1.522 | 0.001 | 0.388 | 0.388 | 0 | 0.07 | 0.27 | 0.15 | 0.51 | 0.03 | 0.15 |
| 6 | 101094554 | rs239239 | -1.269 | 0.342 | 0.763 | 1.439 | 0.004 | 0.021 | 0.021 | 0 | 0.55 | 0.6 | 0.62 | 0.73 | 0.54 | 0.37 |
| 6 | 101109175 | rs17306522 | 1.395 | -0.245 | 0.533 | 0.69 | 0.022 | 0.039 | 0.039 | 0 | 0.19 | 0.12 | 0.18 | 0.09 | 0.14 | 0.38 |
| 6 | 101147095 | rs12201157 | 1.098 | -0.01 | 0.552 | 0.433 | -0.003 | -0.017 | -0.017 | 0 | 0.09 | 0.02 | 0.07 | 0.16 | 0.09 | 0.13 |
| 6 | 101149540 | rs9386233 | 0.468 | 0.083 | 0.632 | 0.419 | -0.016 | 0.016 | 0.016 | 0.837 | 0.45 | 0.28 | 0.5 | 0.52 | 0.47 | 0.28 |
| 6 | 101157916 | rs6904959 | -0.007 | 0.025 | 0.546 | 0.421 | 0.006 | 0.031 | 0.031 | 0 | 0.47 | 0.41 | 0.39 | 0.31 | 0.48 | 0.61 |
| 6 | 101166809 | rs7748644 | -0.164 | 0.053 | 0.513 | 0.398 | -0.002 | -0.002 | -0.002 | 0 | 0.49 | 0.48 | 0.45 | 0.48 | 0.48 | 0.7 |
| 6 | 101174245 | rs12525128 | 0.883 | 0.039 | 0.112 | 0.382 | -0.123 | 0.026 | 0.026 | 1.341 | 0.4 | 0.06 | 0.41 | 0.22 | 0.46 | 0.2 |
| 6 | 101191397 | rs1039033 | -0.375 | 0.071 | 0.418 | 0.361 | -0.003 | 0.018 | 0.018 | 0.896 | 0.53 | 0.56 | 0.44 | 0.27 | 0.53 | 0.63 |
| 6 | 101196080 | rs12212435 | -0.242 | 0.116 | 0.483 | 0.464 | -0.001 | -0.001 | -0.001 | 1.132 | 0.49 | 0.5 | 0.45 | 0.44 | 0.48 | 0.7 |
| 6 | 101234788 | rs12664423 | 0.583 | -0.104 | 0.269 | 0.334 | 0.02 | 0.051 | 0.051 | 0 | 0.3 | 0.13 | 0.26 | 0.25 | 0.24 | 0.5 |
| 6 | 101247503 | rs3811078 | -0.45 | 0.099 | 0.362 | 0.49 | 0 | 0.042 | 0.042 | 0.193 | 0.52 | 0.66 | 0.53 | 0.74 | 0.49 | 0.78 |
| 6 | 101263516 | rs7740783 | 0.16 | 0.245 | 0.454 | 0.636 | -0.007 | -0.012 | -0.012 | 0 | 0.47 | 0.23 | 0.52 | 0.43 | 0.51 | 0.31 |
| 6 | 101275792 | rs705588 | -0.715 | 0.274 | 0.855 | 0.897 | 0.027 | 0.144 | 0.144 | 0.352 | 0.21 | 0.55 | 0.31 | 0.48 | 0.26 | 0.24 |
| 6 | 101295873 | rs9399696 | -0.411 | 0.268 | 0.999 | 0.972 | -0.001 | 0.16 | 0.16 | 0 | 0.08 | 0.24 | 0.13 | 0.31 | 0.05 | 0.09 |
| 6 | 101296389 | rs9390698 | 1.159 | 0.102 | 0.395 | 0.556 | -0.12 | 0.006 | 0.006 | 1.245 | 0.39 | 0.06 | 0.42 | 0.23 | 0.45 | 0.17 |
| 6 | 101304411 | rs9386259 | 0.484 | 0.331 | 0.197 | 0.27 | 0.001 | 0.049 | 0.049 | 0 | 0.2 | 0.26 | 0.22 | 0.47 | 0.21 | 0.14 |
| 6 | 101304829 | rs705599 | -1.161 | 0.319 | 0.372 | 0.042 | -0.003 | -0.005 | -0.005 | 0 | 0.66 | 0.68 | 0.74 | 0.75 | 0.73 | 0.44 |
| 6 | 101312424 | rs998254 | 1.685 | 0.349 | 0.223 | 0.091 | -0.06 | 0.127 | 0.127 | 0.617 | 0.27 | 0.06 | 0.32 | 0.08 | 0.29 | 0.13 |
| 6 | 101329173 | rs12530388 | 0.226 | -0.012 | 0.853 | 0.238 | 0.001 | 0.018 | 0.018 | 0.181 | 0.52 | 0.44 | 0.61 | 0.75 | 0.51 | 0.34 |
| 6 | 101330188 | rs4840159 | 0.503 | -0.017 | 0.833 | 0.254 | 0.005 | 0.045 | 0.045 | 0.022 | 0.51 | 0.42 | 0.61 | 0.71 | 0.5 | 0.35 |
| 6 | 101344511 | rs612117 | 0.635 | 0.097 | 0.916 | 0.202 | 0.059 | -0.057 | -0.057 | 0.818 | 0.1 | 0.28 | 0.1 | 0 | 0.17 | 0.06 |
| 6 | 101351390 | rs2398247 | 1.417 | -0.026 | 0.798 | 0.327 | -0.112 | -0.005 | -0.005 | 0 | 0.38 | 0.06 | 0.4 | 0.24 | 0.44 | 0.18 |
| 6 | 101351852 | rs625745 | 0.306 | -0.042 | 0.897 | 0.297 | 0.038 | -0.065 | -0.065 | 0 | 0.13 | 0.24 | 0.12 | 0 | 0.22 | 0.08 |
| 6 | 101351896 | rs626120 | 0.221 | -0.058 | 0.923 | 0.363 | 0.026 | -0.009 | -0.009 | 0 | 0.55 | 0.37 | 0.61 | 0.44 | 0.69 | 0.35 |
| 6 | 101354510 | rs4840161 | 1.401 | -0.053 | 0.865 | 0.365 | -0.127 | -0.015 | -0.015 | 0.294 | 0.4 | 0.06 | 0.42 | 0.28 | 0.47 | 0.18 |
| 6 | 101363967 | rs705619 | -0.203 | -0.015 | 1.055 | 0.338 | 0.045 | 0.013 | 0.013 | 0.66 | 0.16 | 0.38 | 0.19 | 0.16 | 0.22 | 0.16 |
| 6 | 101371187 | rs10485300 | -1.558 | -0.023 | 1.181 | 0.342 | 0.021 | 0.169 | 0.169 | 0 | 0.06 | 0.24 | 0.09 | 0.2 | 0.03 | 0.09 |
| 6 | 101371644 | rs661142 | -0.505 | -0.096 | 1.365 | 0.328 | -0.023 | -0.033 | -0.033 | 0.001 | 0.64 | 0.6 | 0.73 | 0.76 | 0.73 | 0.44 |
| 6 | 101371999 | rs673119 | -0.554 | -0.07 | 1.313 | 0.335 | 0.042 | 0.013 | 0.013 | 0.105 | 0.18 | 0.38 | 0.21 | 0.2 | 0.25 | 0.17 |
| 6 | 101380707 | rs7739574 | -0.555 | -0.153 | 1.402 | 0.363 | 0.018 | 0.025 | 0.025 | 0 | 0.66 | 0.75 | 0.74 | 0.76 | 0.74 | 0.44 |
| 6 | 101392348 | rs595422 | -0.049 | -0.125 | 1.343 | 0.373 | 0.001 | -0.001 | -0.001 | 0 | 0.61 | 0.63 | 0.64 | 0.53 | 0.71 | 0.35 |
| 6 | 101406380 | rs6925884 | -1.614 | -0.107 | 1.238 | 0.441 | 0 | 0.148 | 0.148 | 0 | 0.12 | 0.33 | 0.14 | 0.34 | 0.09 | 0.1 |
| 6 | 101409649 | rs817229 | 0.387 | -0.147 | 0.851 | 0.36 | 0.095 | 0.056 | 0.056 | 0 | 0.53 | 0.21 | 0.54 | 0.28 | 0.69 | 0.27 |
| 6 | 101426759 | rs4240589 | 1.38 | -0.09 | 0.743 | 0.342 | -0.058 | 0.021 | 0.021 | 0.134 | 0.38 | 0.16 | 0.41 | 0.24 | 0.43 | 0.18 |
| 6 | 101449041 | rs2244658 | -0.531 | -0.072 | 0.978 | 0.36 | 0.018 | 0.023 | 0.023 | 0 | 0.66 | 0.75 | 0.73 | 0.74 | 0.73 | 0.44 |
| 6 | 101463320 | rs9404070 | 1.102 | -0.004 | 0.764 | 0.452 | -0.057 | -0.053 | -0.053 | 3.825 | 0.44 | 0.14 | 0.49 | 0.4 | 0.46 | 0.27 |

| | | | | | | | | | | | | | | | | |
|---|---|---|---|---|---|---|---|---|---|---|---|---|---|---|---|---|
| 6 | 101468591 | rs1321204 | -1.832 | 0.028 | 1.18 | 0.565 | -0.017 | 0.217 | 0.217 | 0.175 | 0.12 | 0.33 | 0.17 | 0.46 | 0.06 | 0.17 |
| 6 | 101471710 | rs1575100 | 1.062 | 0.045 | 0.86 | 0.523 | -0.014 | -0.009 | -0.009 | 0 | 0.46 | 0.31 | 0.49 | 0.5 | 0.48 | 0.21 |
| 6 | 101481014 | rs1765075 | 0.641 | 0.012 | 0.668 | 0.448 | 0.006 | -0.025 | -0.025 | 0 | 0.11 | 0.08 | 0.11 | 0 | 0.2 | 0.09 |
| 6 | 101482379 | rs13211508 | 1.436 | -0.342 | 0.286 | 0.325 | 0.004 | 0.029 | 0.029 | 0 | 0.16 | 0.01 | 0.17 | 0.08 | 0.12 | 0.41 |
| 6 | 101487483 | rs12528419 | 1.425 | -0.248 | 0.449 | 0.397 | -0.071 | -0.047 | -0.047 | 0.409 | 0.43 | 0.12 | 0.45 | 0.46 | 0.45 | 0.2 |
| 6 | 101488697 | rs10499032 | -0.632 | -0.335 | 0.724 | 0.334 | -0.006 | -0.031 | -0.031 | 0 | 0.06 | 0.15 | 0.05 | 0 | 0.04 | 0.03 |
| 6 | 101490341 | rs9390717 | 1.512 | -0.223 | 0.608 | 0.404 | -0.097 | -0.071 | -0.071 | 0 | 0.42 | 0.09 | 0.45 | 0.46 | 0.45 | 0.2 |
| 6 | 101508921 | rs9498467 | -1.382 | -0.108 | 1.164 | 0.688 | 0.013 | 0.255 | 0.255 | 0 | 0.08 | 0.3 | 0.1 | 0.35 | 0.05 | 0.04 |
| 6 | 101540875 | rs9390728 | 0.918 | -0.175 | 1.294 | 0.626 | 0.029 | 0.139 | 0.139 | 0.283 | 0.78 | 0.57 | 0.72 | 0.53 | 0.75 | 0.75 |
| 6 | 101556289 | rs9404082 | 0.387 | -0.338 | 0.955 | 0.523 | 0.021 | -0.034 | -0.034 | 0 | 0.28 | 0.14 | 0.25 | 0.32 | 0.23 | 0.51 |
| 6 | 101569185 | rs9399711 | -1.348 | -0.021 | 1.188 | 0.792 | 0.004 | 0.248 | 0.248 | 0 | 0.08 | 0.26 | 0.1 | 0.35 | 0.04 | 0.04 |
| 6 | 101573555 | rs2770333 | -0.839 | 0.045 | 1.238 | 0.736 | 0.042 | 0.081 | 0.081 | 0 | 0.19 | 0.34 | 0.21 | 0.34 | 0.25 | 0.13 |
| 6 | 101576535 | rs943910 | 0.355 | 0.091 | 0.972 | 0.659 | 0.009 | -0.032 | -0.032 | 0 | 0.12 | 0.11 | 0.11 | 0 | 0.2 | 0.09 |
| 6 | 101583773 | rs1928070 | -0.55 | 0.053 | 1.188 | 0.732 | 0.047 | 0.088 | 0.088 | 0 | 0.18 | 0.33 | 0.2 | 0.35 | 0.25 | 0.13 |
| 6 | 101584291 | rs2813651 | -1.308 | -0.402 | 1.007 | 0.539 | -0.006 | 0.049 | 0.049 | 0 | 0.32 | 0.5 | 0.33 | 0.52 | 0.33 | 0.25 |
| 6 | 101594795 | rs9377263 | -0.171 | 0.184 | 1.047 | 0.978 | 0 | -0.004 | -0.004 | 0 | 0.16 | 0.17 | 0.12 | 0.24 | 0.15 | 0.15 |
| 6 | 101598363 | rs9390733 | 0.985 | 0.207 | 0.747 | 0.997 | -0.027 | -0.056 | -0.056 | 0 | 0.1 | 0 | 0.07 | 0.24 | 0.1 | 0.11 |
| 6 | 101608851 | rs1149271 | -0.173 | 0.339 | 0.973 | 0.779 | 0.014 | 0.007 | 0.007 | 0.131 | 0.87 | 0.89 | 0.83 | 0.84 | 0.8 | 0.84 |
| 6 | 101615409 | rs1775644 | -0.173 | 0.335 | 0.969 | 0.826 | 0.014 | 0.007 | 0.007 | 0.131 | 0.87 | 0.89 | 0.83 | 0.83 | 0.8 | 0.84 |
| 6 | 101650474 | rs1408771 | 1.73 | 0.305 | 0.987 | 0.75 | 0 | 0.002 | 0.002 | 0.194 | 0.73 | 0.32 | 0.68 | 0.48 | 0.7 | 0.76 |
| 6 | 101658891 | rs3849200 | -1.66 | 0.017 | 1.063 | 0.897 | -0.032 | 0.239 | 0.239 | 0.098 | 0.15 | 0.57 | 0.21 | 0.52 | 0.1 | 0.15 |
| 6 | 101659388 | rs2764272 | -0.048 | -0.036 | 0.502 | 0.758 | 0.032 | -0.016 | -0.016 | 0 | 0.33 | 0.18 | 0.27 | 0.34 | 0.27 | 0.59 |
| 6 | 101688127 | rs7754461 | -0.781 | 0.767 | 0.897 | 1.138 | 0.005 | 0.137 | 0.137 | 0 | 0.08 | 0.16 | 0.11 | 0.3 | 0.05 | 0.07 |
| 6 | 101726103 | rs643101 | -0.146 | 1.243 | 0.601 | 0.539 | 0.004 | -0.021 | -0.021 | 0.323 | 0.13 | 0.11 | 0.12 | 0 | 0.22 | 0.09 |
| 6 | 101727949 | rs2252114 | -0.799 | 1.191 | 0.2 | 0.704 | -0.035 | -0.069 | -0.069 | 0 | 0.89 | 1 | 0.93 | 0.76 | 0.89 | 0.89 |
| 6 | 101731443 | rs283210 | -1.594 | 1.041 | 0.665 | 0.727 | 0 | 0.004 | 0.004 | 1.374 | 0.27 | 0.66 | 0.33 | 0.51 | 0.34 | 0.24 |
| 6 | 101734063 | rs775258 | 0.6 | 0.642 | 0.637 | 0.504 | -0.024 | -0.023 | -0.023 | 0.065 | 0.16 | 0.24 | 0.17 | 0.09 | 0.13 | 0.4 |
| 6 | 101792444 | rs689383 | -0.435 | 1.168 | 0.844 | 0.773 | 0.011 | 0.075 | 0.075 | 0.204 | 0.06 | 0.24 | 0.1 | 0.17 | 0.03 | 0.08 |
| 6 | 101800051 | rs2629983 | -1.794 | 1.35 | 0.945 | 0.673 | 0.049 | -0.019 | -0.019 | 0.494 | 0.23 | 0.43 | 0.25 | 0.22 | 0.31 | 0.24 |
| 6 | 101801581 | rs661064 | 0.222 | 1.385 | 0.954 | 0.668 | 0.052 | -0.039 | -0.039 | 0 | 0.09 | 0.13 | 0.1 | 0 | 0.21 | 0.07 |
| 6 | 101805148 | rs2852494 | 0.319 | 1.668 | 0.627 | 0.771 | -0.021 | 0.008 | 0.008 | 0 | 0.94 | 0.99 | 0.88 | 0.6 | 0.92 | 0.88 |
| 6 | 101807507 | rs1485833 | -2.096 | 1.394 | 0.949 | 0.893 | 0.018 | 0.051 | 0.051 | 0.351 | 0.26 | 0.61 | 0.33 | 0.59 | 0.32 | 0.3 |
| 6 | 101815243 | rs1385764 | -0.942 | 1.514 | 0.638 | -0.415 | 0.092 | 0.002 | 0.002 | 0.434 | 0.15 | 0.32 | 0.21 | 0.17 | 0.24 | 0.16 |
| 6 | 101820085 | rs17812498 | -0.247 | 1.559 | 0.589 | -0.386 | 0.004 | 0 | 0 | 0 | 0.07 | 0.03 | 0.05 | 0.05 | 0.06 | 0.09 |
| 6 | 101828259 | rs4103516 | -1.486 | 1.53 | 0.607 | -0.392 | 0.052 | -0.017 | -0.017 | 1.145 | 0.22 | 0.43 | 0.25 | 0.22 | 0.31 | 0.25 |
| 6 | 101839863 | rs6930752 | -0.164 | 1.693 | -0.011 | -0.96 | 0.015 | 0.041 | 0.041 | 0 | 0.44 | 0.64 | 0.4 | 0.86 | 0.35 | 0.71 |
| 6 | 101846343 | rs1415946 | -1.219 | 2.227 | 0.792 | 0.06 | 0.13 | -0.038 | -0.038 | 0.344 | 0.14 | 0.4 | 0.15 | 0.05 | 0.26 | 0.08 |
| 6 | 101854633 | rs2579931 | -1.094 | 1.648 | 0.002 | 0.29 | -0.001 | 0 | 0 | 0 | 0.08 | 0.06 | 0.06 | 0.09 | 0.1 | 0.13 |
| 6 | 101855808 | rs7755912 | -1.299 | 1.688 | 0.233 | 0.383 | 0.004 | 0.015 | 0.015 | 0.011 | 0.23 | 0.39 | 0.21 | 0.24 | 0.24 | 0.2 |
| 6 | 101856136 | rs1338162 | -1.892 | 1.666 | 0.185 | 0.373 | 0.018 | 0.005 | 0.005 | 1.319 | 0.31 | 0.45 | 0.27 | 0.34 | 0.34 | 0.34 |
| 6 | 101859505 | rs12195965 | -1.422 | 1.271 | 0.05 | 0.419 | 0.003 | 0.007 | 0.007 | 0 | 0.23 | 0.35 | 0.21 | 0.24 | 0.24 | 0.2 |
| 6 | 101860248 | rs2225912 | -0.747 | 1.567 | -0.389 | 0.011 | 0 | -0.015 | -0.015 | 0 | 0.08 | 0.08 | 0.05 | 0 | 0.11 | 0.11 |
| 6 | 101866368 | rs1856309 | 0.329 | 1.018 | -0.525 | -0.778 | -0.004 | -0.076 | -0.076 | 0 | 0.63 | 0.59 | 0.74 | 0.9 | 0.68 | 0.67 |
| 6 | 101868479 | rs2852517 | -0.418 | 0.803 | -0.691 | -0.868 | 0.007 | -0.002 | -0.002 | 2.069 | 0.08 | 0.01 | 0.02 | 0.07 | 0.04 | 0.17 |
| 6 | 101871923 | rs12194114 | 0.954 | 0.901 | -1.212 | -0.286 | 0.001 | 0.007 | 0.007 | 0 | 0.54 | 0.58 | 0.64 | 0.6 | 0.58 | 0.61 |
| 6 | 101874585 | rs1890276 | -0.529 | 0.982 | -0.934 | 0.356 | -0.012 | -0.013 | -0.013 | 0.534 | 0.09 | 0.12 | 0.02 | 0.06 | 0.03 | 0.16 |
| 6 | 101875714 | rs2852525 | 0.017 | 0.612 | -1.497 | -0.486 | -0.012 | 0.117 | 0.117 | 0 | 0.56 | 0.77 | 0.61 | 0.93 | 0.55 | 0.69 |
| 6 | 101880376 | rs1465169 | 0.719 | 0.622 | -1.558 | -0.495 | 0.003 | -0.034 | -0.034 | 0.425 | 0.44 | 0.55 | 0.57 | 0.87 | 0.51 | 0.55 |
| 6 | 101883637 | rs2787565 | 0.106 | 0.622 | -1.523 | -0.494 | 0.001 | 0.024 | 0.024 | 0 | 0.55 | 0.65 | 0.62 | 0.93 | 0.57 | 0.69 |
| 6 | 101889424 | rs997279 | 0.064 | 0.6 | -1.624 | -0.289 | -0.001 | -0.018 | -0.018 | 0 | 0.37 | 0.32 | 0.35 | 0.07 | 0.38 | 0.28 |
| 6 | 101889443 | rs2518238 | -0.536 | 0.573 | -1.643 | -0.292 | -0.005 | 0.02 | 0.02 | 0 | 0.94 | 1 | 0.96 | 0.99 | 0.95 | 0.98 |
| 6 | 101890395 | rs2788274 | -0.087 | 0.517 | -1.63 | -0.326 | -0.003 | 0.04 | 0.04 | 0 | 0.38 | 0.25 | 0.35 | 0.07 | 0.38 | 0.28 |
| 6 | 101891573 | rs17061967 | -0.135 | 0.391 | -1.622 | -0.439 | 0 | -0.015 | -0.015 | 0 | 0.1 | 0.15 | 0.11 | 0.03 | 0.09 | 0.11 |
| 6 | 101897534 | rs1856310 | -0.455 | 0.308 | -1.676 | -0.248 | 0.004 | 0.006 | 0.006 | 0.117 | 0.42 | 0.33 | 0.44 | 0.4 | 0.38 | 0.33 |
| 6 | 101900114 | rs2518160 | 0.646 | 0.327 | -1.784 | -0.321 | -0.045 | 0.127 | 0.127 | 0 | 0.27 | 0.06 | 0.24 | 0.03 | 0.3 | 0.17 |
| 6 | 101901973 | rs1338167 | 0.799 | 0.035 | -1.58 | -0.305 | 0.005 | 0.028 | 0.028 | 0 | 0.49 | 0.53 | 0.52 | 0.58 | 0.54 | 0.54 |
| 6 | 101903439 | rs10485268 | 0.441 | 0.132 | -1.665 | -0.228 | 0 | 0 | 0 | 0 | 0.06 | 0.07 | 0.1 | 0.07 | 0.07 | 0.1 |
| 6 | 101906652 | rs2518224 | 0.418 | -0.06 | -2.077 | -0.675 | -0.019 | 0.024 | 0.024 | 0 | 0.09 | 0.03 | 0.13 | 0 | 0.12 | 0.05 |
| 6 | 101913238 | rs2788278 | 0.872 | 0.363 | -2.459 | -0.338 | -0.046 | -0.051 | -0.051 | 0 | 0.12 | 0.01 | 0.14 | 0.12 | 0.16 | 0.1 |
| 6 | 101916765 | rs2788279 | -0.18 | 0.278 | -2.264 | -0.222 | 0.01 | 0.023 | 0.023 | 0.076 | 0.36 | 0.1 | 0.32 | 0.37 | 0.31 | 0.3 |
| 6 | 101927338 | rs2749065 | -0.248 | 0.541 | -2.176 | -0.298 | 0.002 | -0.007 | -0.007 | 0 | 0.13 | 0.15 | 0.14 | 0.08 | 0.13 | 0.13 |
| 6 | 101929203 | rs990386 | -0.999 | 0.553 | -2.085 | -0.309 | 0.018 | 0.056 | 0.056 | 0 | 0.49 | 0.31 | 0.46 | 0.43 | 0.44 | 0.44 |
| 6 | 101929286 | rs2252682 | -0.091 | 0.471 | -1.925 | -0.19 | -0.005 | 0.033 | 0.033 | 0.208 | 0.31 | 0.06 | 0.26 | 0.29 | 0.27 | 0.19 |
| 6 | 101933087 | rs989467 | -0.598 | 0.458 | -1.749 | -0.184 | 0.005 | 0.016 | 0.016 | 0.692 | 0.4 | 0.37 | 0.36 | 0.32 | 0.35 | 0.25 |
| 6 | 101933529 | rs989466 | -0.936 | 0.766 | -1.765 | 0.084 | 0.007 | 0.012 | 0.012 | 0.647 | 0.48 | 0.43 | 0.44 | 0.46 | 0.43 | 0.43 |
| 6 | 101939743 | rs2399711 | 0.491 | 0.673 | -1.667 | -0.241 | 0.005 | 0.026 | 0.026 | 0 | 0.2 | 0.09 | 0.19 | 0.01 | 0.17 | 0.13 |
| 6 | 101941240 | rs2749060 | -1.223 | 0.698 | -1.306 | -0.137 | -0.038 | -0.006 | -0.006 | 0 | 0.63 | 0.8 | 0.62 | 0.6 | 0.53 | 0.8 |
| 6 | 101941460 | rs2579942 | -1.231 | 0.699 | -1.306 | -0.137 | -0.038 | -0.006 | -0.006 | 0 | 0.64 | 0.8 | 0.62 | 0.6 | 0.53 | 0.8 |
| 6 | 101941632 | rs9399721 | -0.236 | 0.567 | -1.049 | -0.034 | -0.006 | 0.003 | 0.003 | 0 | 0.43 | 0.48 | 0.39 | 0.5 | 0.33 | 0.59 |

| | | | | | | | | | | | | | | | | |
|---|---|---|---|---|---|---|---|---|---|---|---|---|---|---|---|---|
| 6 | 101948938 | rs6930522 | -0.331 | 0.439 | -1.031 | 0.053 | 0.011 | 0.076 | 0.076 | 0 | 0.4 | 0.28 | 0.34 | 0.19 | 0.34 | 0.37 |
| 6 | 101953003 | rs4840191 | 0.792 | 0.441 | -1.042 | 0.036 | -0.004 | 0.031 | 0.031 | 0 | 0.28 | 0.24 | 0.29 | 0.04 | 0.26 | 0.2 |
| 6 | 101953217 | rs2144706 | -1.745 | 0.478 | -1.049 | 0.085 | -0.039 | -0.002 | -0.002 | 0 | 0.63 | 0.8 | 0.61 | 0.6 | 0.52 | 0.8 |
| 6 | 101953555 | rs3808 | 0.092 | 0.428 | -0.473 | 0.774 | -0.017 | 0.117 | 0.117 | 0 | 0.22 | 0.38 | 0.19 | 0.49 | 0.16 | 0.46 |
| 6 | 101957291 | rs7747072 | 0.66 | 1.112 | 0.184 | 1.274 | 0.035 | 0.259 | 0.259 | 0 | 0.11 | 0.38 | 0.14 | 0.35 | 0.08 | 0.31 |
| 6 | 101959098 | rs12215960 | 0.047 | 1.355 | -0.138 | 1.182 | 0.002 | 0.03 | 0.03 | 0 | 0.09 | 0.07 | 0.08 | 0.31 | 0.13 | 0.08 |
| 6 | 101962940 | rs2749056 | 0.161 | 1.128 | -0.186 | 0.658 | -0.002 | -0.003 | -0.003 | 0 | 0.11 | 0.11 | 0.13 | 0.06 | 0.11 | 0.14 |
| 6 | 101964843 | rs7760831 | -0.02 | 1.282 | -0.031 | 0.718 | 0.017 | -0.011 | -0.011 | 3.288 | 0.08 | 0.2 | 0.1 | 0.03 | 0.09 | 0.07 |
| 6 | 101964914 | rs9390754 | -0.121 | 1.053 | 0.189 | 1.2 | 0.028 | 0.242 | 0.242 | 0 | 0.14 | 0.44 | 0.16 | 0.35 | 0.12 | 0.35 |
| 6 | 101966253 | rs2247210 | -1.068 | 1.099 | -0.18 | 0.763 | -0.005 | -0.002 | -0.002 | 0 | 0.44 | 0.42 | 0.52 | 0.44 | 0.46 | 0.61 |
| 6 | 101966333 | rs10485270 | 0.314 | 1.273 | -0.737 | 0.972 | -0.041 | -0.054 | -0.054 | 0 | 0.09 | 0.01 | 0.08 | 0.31 | 0.12 | 0.12 |
| 6 | 101966553 | rs2247218 | -0.239 | 1.411 | -0.717 | 1.289 | -0.025 | 0.056 | 0.056 | 0 | 0.31 | 0.09 | 0.34 | 0.15 | 0.33 | 0.27 |
| 6 | 101967710 | rs2518245 | 0.174 | 1.398 | -0.792 | 1.215 | -0.004 | 0.027 | 0.027 | 0 | 0.15 | 0.03 | 0.14 | 0.06 | 0.15 | 0.14 |
| 6 | 101976291 | rs2518251 | -0.669 | 1.395 | -0.562 | 1.235 | 0 | 0.002 | 0.002 | 0 | 0.28 | 0.31 | 0.32 | 0.35 | 0.28 | 0.47 |
| 6 | 101984119 | rs2749081 | 1.437 | 1.274 | -0.364 | 1.165 | 0.005 | -0.006 | -0.006 | 1.962 | 0.6 | 0.53 | 0.59 | 0.35 | 0.59 | 0.42 |
| 6 | 101987410 | rs2255306 | 0.341 | 1.307 | -0.318 | 1.218 | -0.055 | -0.104 | -0.104 | 0 | 0.77 | 0.97 | 0.78 | 0.64 | 0.74 | 0.77 |
| 6 | 101988513 | rs9390757 | -0.527 | 1.302 | -0.333 | 1.159 | 0.004 | 0.008 | 0.008 | 0 | 0.31 | 0.43 | 0.37 | 0.38 | 0.31 | 0.49 |
| 6 | 101988652 | rs2518216 | 2.054 | 1.3 | -0.35 | 1.127 | -0.005 | -0.017 | -0.017 | 5.184 | 0.45 | 0.49 | 0.4 | 0.26 | 0.42 | 0.28 |
| 6 | 101988727 | rs17760780 | 0.829 | 1.293 | -0.233 | 1.411 | -0.036 | 0.008 | 0.008 | 0 | 0.16 | 0.04 | 0.19 | 0.09 | 0.17 | 0.14 |
| 6 | 101989945 | rs1040711 | 1.932 | 1.314 | -0.087 | 1.38 | 0 | -0.002 | -0.002 | 0.579 | 0.49 | 0.46 | 0.41 | 0.26 | 0.45 | 0.3 |
| 6 | 101990259 | rs2023169 | 1.901 | 1.327 | -0.259 | 1.38 | 0.006 | 0.024 | 0.024 | 0 | 0.49 | 0.53 | 0.41 | 0.26 | 0.45 | 0.3 |
| 6 | 101991728 | rs2518220 | 0.533 | 1.229 | -0.276 | 1.538 | -0.013 | -0.05 | -0.05 | 0 | 0.81 | 0.93 | 0.79 | 0.64 | 0.76 | 0.79 |
| 6 | 102002092 | rs2852571 | 1.516 | 1.235 | -0.18 | 1.604 | 0.002 | 0 | 0 | 0 | 0.65 | 0.59 | 0.61 | 0.35 | 0.62 | 0.44 |
| 6 | 102002356 | rs2245817 | -0.596 | 1.37 | -0.331 | 1.696 | 0 | 0.008 | 0.008 | 0 | 0.89 | 0.94 | 0.87 | 0.94 | 0.88 | 0.9 |
| 6 | 102002591 | rs2245825 | 0.351 | 1.16 | -0.292 | 1.541 | -0.008 | -0.04 | -0.04 | 0 | 0.8 | 0.89 | 0.79 | 0.64 | 0.76 | 0.79 |
| 6 | 102005291 | rs9404119 | -0.186 | 0.349 | 0.01 | 1.425 | 0.013 | 0.046 | 0.046 | 0 | 0.13 | 0.24 | 0.17 | 0.29 | 0.14 | 0.35 |
| 6 | 102005370 | rs9373614 | -0.186 | 0.375 | 0.01 | 1.425 | 0.014 | 0.046 | 0.046 | 0 | 0.13 | 0.24 | 0.18 | 0.29 | 0.14 | 0.35 |
| 6 | 102008823 | rs17828670 | 0.215 | 0.012 | -0.252 | 0.828 | 0 | 0 | 0 | 0 | 0.13 | 0.04 | 0.16 | 0.07 | 0.12 | 0.08 |
| 6 | 102021152 | rs627428 | -1.709 | -0.224 | 0.448 | 0.786 | 0 | -0.004 | -0.004 | 0.548 | 0.13 | 0.54 | 0.07 | 0.09 | 0.08 | 0.18 |
| 6 | 102026214 | rs493632 | -1.493 | -0.162 | 0.774 | 0.652 | 0.048 | 0.012 | 0.012 | 0 | 0.06 | 0.22 | 0.04 | 0.05 | 0.06 | 0.12 |
| 6 | 102026932 | rs487083 | -1.493 | -0.162 | 0.766 | 0.652 | 0.046 | 0.011 | 0.011 | 0 | 0.06 | 0.22 | 0.04 | 0.05 | 0.06 | 0.12 |
| 6 | 102033541 | rs2021678 | -0.731 | -0.299 | 0.153 | 0.339 | -0.003 | -0.045 | -0.045 | 0 | 0.32 | 0.36 | 0.3 | 0.15 | 0.29 | 0.26 |
| 6 | 102037354 | rs6908225 | -0.998 | -0.337 | -0.026 | 0.814 | -0.003 | 0.003 | 0.003 | 0 | 0.36 | 0.34 | 0.39 | 0.33 | 0.33 | 0.26 |
| 6 | 102037684 | rs6936552 | -1.004 | -0.759 | 0.12 | 0.992 | -0.011 | 0.128 | 0.128 | 0 | 0.15 | 0.32 | 0.16 | 0.24 | 0.11 | 0.18 |
| 6 | 102042953 | rs1415484 | -1.269 | -0.79 | 0.158 | 0.946 | -0.021 | 0.06 | 0.06 | 0 | 0.26 | 0.35 | 0.33 | 0.33 | 0.22 | 0.24 |
| 6 | 102045546 | rs479851 | 0.534 | -0.778 | 0.253 | 0.906 | 0.002 | 0.001 | 0.001 | 0 | 0.12 | 0.12 | 0.19 | 0.09 | 0.11 | 0.07 |
| 6 | 102053814 | rs2224202 | -1.42 | -0.754 | 0.34 | 0.912 | 0.006 | 0.092 | 0.092 | 0 | 0.2 | 0.39 | 0.19 | 0.29 | 0.18 | 0.3 |
| 6 | 102054465 | rs611348 | -1.801 | -1.168 | 0.163 | 0.703 | -0.057 | 0.111 | 0.111 | 0 | 0.62 | 0.84 | 0.63 | 0.81 | 0.56 | 0.75 |
| 6 | 102054910 | rs609531 | -1.701 | -1.206 | 0.871 | 1.19 | -0.005 | 0.017 | 0.017 | 0 | 0.32 | 0.56 | 0.38 | 0.37 | 0.29 | 0.38 |
| 6 | 102058583 | rs2210968 | -1.333 | -1.569 | 0.837 | 1.384 | -0.012 | 0.056 | 0.056 | 0 | 0.27 | 0.37 | 0.35 | 0.36 | 0.25 | 0.32 |
| 6 | 102058891 | rs513216 | -0.792 | -1.244 | 1.132 | 1.833 | 0.019 | -0.004 | -0.004 | 0.692 | 0.38 | 0.66 | 0.33 | 0.58 | 0.3 | 0.49 |
| 6 | 102065160 | rs9390764 | -1.155 | -0.751 | 1.793 | 2.268 | 0.001 | 0.13 | 0.13 | 0 | 0.15 | 0.29 | 0.16 | 0.28 | 0.14 | 0.26 |
| 6 | 102066487 | rs512051 | -1.419 | -0.769 | 1.782 | 2.27 | 0 | 0.185 | 0.185 | 0 | 0.17 | 0.44 | 0.17 | 0.29 | 0.15 | 0.31 |
| 6 | 102077056 | rs1856137 | -0.929 | -0.321 | 1.07 | 1.466 | 0.044 | 0.017 | 0.017 | 0.24 | 0.09 | 0.26 | 0.2 | 0.09 | 0.13 | 0.12 |
| 6 | 102111184 | rs9485526 | 1.124 | -1.213 | 0.712 | 1.845 | 0.024 | -0.038 | -0.038 | 0 | 0.66 | 0.78 | 0.63 | 0.48 | 0.76 | 0.61 |
| 6 | 102113164 | rs1415483 | -0.738 | -1.453 | 0.622 | 1.942 | 0.072 | 0.018 | 0.018 | 0 | 0.21 | 0.05 | 0.09 | 0.19 | 0.07 | 0.24 |
| 6 | 102114780 | rs9485528 | 1.411 | -1.466 | 0.653 | 2.071 | 0.021 | 0.012 | 0.012 | 0 | 0.71 | 0.79 | 0.75 | 0.74 | 0.84 | 0.69 |
| 6 | 102116069 | rs7739084 | -0.944 | -1.453 | 0.646 | 2.069 | 0.052 | 0.027 | 0.027 | 0 | 0.22 | 0.08 | 0.1 | 0.19 | 0.09 | 0.24 |
| 6 | 102119829 | rs9485530 | 2.077 | -1.713 | 1.826 | 1.977 | 0.026 | -0.019 | -0.019 | 0 | 0.6 | 0.3 | 0.6 | 0.5 | 0.77 | 0.61 |
| 6 | 102120244 | rs9399725 | -0.417 | -1.745 | 1.783 | 1.983 | 0.013 | 0.001 | 0.001 | 0 | 0.07 | 0.13 | 0.15 | 0.07 | 0.1 | 0.05 |
| 6 | 102135266 | rs2518298 | 1.531 | -1.771 | 2.562 | 1.966 | 0.036 | 0.01 | 0.01 | 1.441 | 0.89 | 0.51 | 0.82 | 0.93 | 0.9 | 0.95 |
| 6 | 102138254 | rs2065938 | 1.949 | -1.868 | 2.553 | 1.993 | 0.02 | -0.016 | -0.016 | 0 | 0.62 | 0.35 | 0.63 | 0.51 | 0.79 | 0.62 |
| 6 | 102145638 | rs9498669 | 2.212 | -1.892 | 2.541 | 2.146 | -0.013 | 0.052 | 0.052 | 1.155 | 0.83 | 0.38 | 0.72 | 0.78 | 0.86 | 0.91 |
| 6 | 102151300 | rs1337412 | 2.037 | -1.878 | 2.572 | 2.139 | 0.006 | -0.006 | -0.006 | 0 | 0.61 | 0.37 | 0.62 | 0.5 | 0.79 | 0.62 |
| 6 | 102157669 | rs1856134 | -0.446 | -1.82 | 2.506 | 2.108 | 0.008 | 0.002 | 0.002 | 0 | 0.07 | 0.14 | 0.16 | 0.07 | 0.1 | 0.05 |
| 6 | 102161798 | rs17062378 | -1.09 | -2.025 | 2.787 | 2.234 | 0.043 | -0.058 | -0.058 | 0 | 0.22 | 0.01 | 0.11 | 0.27 | 0.09 | 0.27 |
| 6 | 102165329 | rs1222955 | -1.754 | -2.171 | 3.21 | 2.294 | 0 | -0.005 | -0.005 | 0 | 0.28 | 0.24 | 0.21 | 0.42 | 0.11 | 0.3 |
| 6 | 102166559 | rs1222956 | -2.137 | -1.622 | 3.21 | 2.457 | 0.028 | 0.198 | 0.198 | 1.113 | 0.08 | 0.4 | 0.13 | 0.16 | 0.04 | 0.04 |
| 6 | 102168372 | rs7770500 | -1.39 | -1.838 | 2.787 | 2.443 | -0.05 | 0.01 | 0.01 | 0 | 0.22 | 0.3 | 0.15 | 0.29 | 0.06 | 0.29 |
| 6 | 102169607 | rs1222964 | -2.057 | -1.975 | 2.546 | 2.445 | -0.002 | -0.007 | -0.007 | 0 | 0.3 | 0.72 | 0.27 | 0.43 | 0.09 | 0.32 |
| 6 | 102170855 | rs942666 | -1.925 | -1.918 | 2.455 | 2.475 | -0.065 | 0.046 | 0.046 | 1.098 | 0.29 | 0.61 | 0.27 | 0.45 | 0.09 | 0.33 |
| 6 | 102178732 | rs9322604 | -1.854 | -1.607 | 2.81 | 2.986 | -0.083 | 0.13 | 0.13 | 1.313 | 0.25 | 0.45 | 0.28 | 0.47 | 0.14 | 0.35 |
| 6 | 102183265 | rs1925227 | -1.298 | -1.543 | 2.404 | 2.535 | -0.01 | -0.001 | -0.001 | 1.374 | 0.17 | 0.15 | 0.14 | 0.18 | 0.07 | 0.23 |
| 6 | 102198018 | rs9498687 | -1.436 | -1.551 | 2.428 | 2.424 | -0.066 | -0.006 | -0.006 | 2.625 | 0.19 | 0.3 | 0.16 | 0.16 | 0.08 | 0.25 |
| 6 | 102198738 | rs9386290 | -1.898 | -1.018 | 2.653 | 3.152 | -0.033 | 0.284 | 0.284 | 0.643 | 0.11 | 0.4 | 0.17 | 0.51 | 0.07 | 0.25 |
| 6 | 102250872 | rs9498712 | -1.323 | -1.001 | 2.959 | 2.324 | 0.014 | -0.005 | -0.005 | 0 | 0.07 | 0.27 | 0.09 | 0.03 | 0.08 | 0.04 |
| 6 | 102259114 | rs2852573 | -1.81 | -0.915 | 2.75 | 3.064 | -0.007 | 0.115 | 0.115 | 0 | 0.07 | 0.09 | 0.19 | 0.61 | 0.06 | 0.31 |
| 6 | 102261917 | rs12333317 | -1.166 | -0.971 | 2.756 | 2.751 | -0.005 | -0.008 | -0.008 | 0 | 0.07 | 0.11 | 0.06 | 0.03 | 0.06 | 0.04 |
| 6 | 102291342 | rs1547384 | -1.396 | -0.157 | 2.647 | 2.768 | 0.032 | 0.05 | 0.05 | 0 | 0.05 | 0.08 | 0.22 | 0.14 | 0.13 | 0.1 |

| | | | | | | | | | | | | | | | | |
|---|---|---|---|---|---|---|---|---|---|---|---|---|---|---|---|---|
| 6 | 102293249 | rs2791829 | 1.138 | -0.305 | 2.712 | 2.339 | -0.007 | -0.01 | -0.01 | 0.426 | 0.94 | 0.83 | 0.94 | 0.97 | 0.95 | 0.97 |
| 6 | 102310726 | rs2000937 | 1.098 | -0.197 | 2.637 | 2.201 | -0.001 | 0.006 | 0.006 | 0 | 0.91 | 0.93 | 0.91 | 0.97 | 0.89 | 0.97 |
| 6 | 102318339 | rs2153445 | 1.636 | -0.175 | 2.541 | 2.23 | 0.049 | -0.039 | -0.039 | 0 | 0.83 | 0.45 | 0.82 | 0.92 | 0.78 | 0.88 |
| 6 | 102323696 | rs4120910 | -1.87 | -0.078 | 1.951 | 3.002 | -0.028 | 0.238 | 0.238 | 0 | 0.08 | 0.24 | 0.25 | 0.52 | 0.03 | 0.19 |
| 6 | 102343078 | rs9485557 | -1.501 | -0.041 | 1.325 | 1.824 | -0.004 | 0.026 | 0.026 | 0.219 | 0.08 | 0.16 | 0.19 | 0.08 | 0.05 | 0.04 |
| 6 | 102343504 | rs17784776 | 0.45 | -0.126 | -0.06 | 0.495 | -0.078 | 0.18 | 0.18 | 0 | 0.31 | 0.02 | 0.25 | 0.06 | 0.37 | 0.23 |
| 6 | 102350747 | rs1475922 | 1.094 | 0.175 | 0.784 | 0.506 | 0.041 | 0.027 | 0.027 | 0.287 | 0.93 | 0.75 | 0.94 | 0.95 | 0.94 | 0.92 |
| 6 | 102354341 | rs17062661 | 0.389 | 0.008 | 0.186 | 1.12 | -0.062 | -0.104 | -0.104 | 0.229 | 0.38 | 0.04 | 0.4 | 0.6 | 0.42 | 0.49 |
| 6 | 102355883 | rs7772343 | 1.215 | -0.106 | 0.726 | 1.005 | 0.013 | 0.021 | 0.021 | 0 | 0.94 | 0.82 | 0.93 | 0.95 | 0.94 | 0.95 |
| 6 | 102358855 | rs1475919 | 0.231 | 0.014 | 0.339 | 0.336 | -0.001 | -0.027 | -0.027 | 0.94 | 0.79 | 0.73 | 0.88 | 0.95 | 0.84 | 0.92 |
| 6 | 102388612 | rs1340270 | 0.926 | 0.071 | 0.256 | 0.383 | -0.073 | -0.105 | -0.105 | 1.288 | 0.4 | 0.04 | 0.37 | 0.59 | 0.45 | 0.53 |
| 6 | 102399279 | rs1413970 | 0.324 | 0.324 | 0.692 | 0.16 | 0.002 | -0.045 | -0.045 | 1.163 | 0.83 | 0.71 | 0.89 | 0.95 | 0.86 | 0.93 |
| 6 | 102407708 | rs988376 | 0.509 | 0.233 | 1.049 | 0.392 | 0 | -0.006 | -0.006 | 0.014 | 0.47 | 0.68 | 0.42 | 0.52 | 0.47 | 0.57 |
| 6 | 102408680 | rs1335035 | 0.687 | 0.215 | 1.352 | 0.406 | 0 | -0.001 | -0.001 | 0 | 0.45 | 0.55 | 0.42 | 0.51 | 0.47 | 0.56 |
| 6 | 102411802 | rs1223045 | 0.649 | 0.234 | 1.203 | 0.551 | 0.003 | 0.005 | 0.005 | 0 | 0.46 | 0.6 | 0.41 | 0.45 | 0.45 | 0.53 |
| 6 | 102423363 | rs1223052 | 0.511 | -0.053 | 0.67 | 0.169 | -0.005 | 0.094 | 0.094 | 0.019 | 0.32 | 0.12 | 0.21 | 0 | 0.35 | 0.15 |
| 6 | 102445951 | rs7745267 | 0.855 | -0.048 | 0.738 | 0.308 | -0.012 | -0.008 | -0.008 | 2.728 | 0.42 | 0.22 | 0.37 | 0.44 | 0.43 | 0.51 |
| 6 | 102455878 | rs10485277 | 0.103 | 0.012 | 0.89 | 0.5 | 0.002 | -0.004 | -0.004 | 0.083 | 0.05 | 0.07 | 0.17 | 0.09 | 0.03 | 0.04 |
| 6 | 102457961 | rs13208276 | 0.288 | -0.029 | 0.664 | 0.192 | -0.002 | -0.03 | -0.03 | 0 | 0.28 | 0.14 | 0.29 | 0.36 | 0.32 | 0.27 |
| 6 | 102480523 | rs1335026 | -0.075 | 0.383 | 1.077 | 0.27 | -0.003 | 0.005 | 0.005 | 0 | 0.06 | 0.1 | 0.18 | 0.08 | 0.04 | 0.04 |
| 6 | 102482848 | rs995640 | 0.275 | 0.135 | 1.455 | 0.267 | -0.007 | 0.027 | 0.027 | 0 | 0.27 | 0.48 | 0.29 | 0.33 | 0.32 | 0.29 |
| 6 | 102492342 | rs2852611 | -0.432 | -0.173 | 1.434 | 0.003 | 0 | 0 | 0 | 0.022 | 0.42 | 0.44 | 0.35 | 0.32 | 0.44 | 0.29 |
| 6 | 102501605 | rs954764 | 0.005 | -0.104 | 1.001 | -0.243 | -0.007 | 0.081 | 0.081 | 1.003 | 0.15 | 0.02 | 0.11 | 0 | 0.11 | 0.02 |
| 6 | 102501774 | rs954765 | -0.38 | -0.068 | 1.273 | -0.112 | -0.004 | 0.008 | 0.008 | 0 | 0.42 | 0.24 | 0.41 | 0.36 | 0.44 | 0.25 |
| 6 | 102505857 | rs2252510 | -0.528 | -0.148 | 1.351 | -0.153 | 0.001 | 0.008 | 0.008 | 0 | 0.43 | 0.35 | 0.42 | 0.36 | 0.44 | 0.25 |
| 6 | 102512506 | rs2253370 | -0.471 | -0.11 | 1.328 | -0.158 | 0.001 | 0.008 | 0.008 | 0 | 0.43 | 0.32 | 0.42 | 0.36 | 0.44 | 0.26 |
| 6 | 102513865 | rs1034253 | 0.571 | -0.131 | 1.282 | -0.155 | -0.001 | -0.001 | -0.001 | 1.04 | 0.27 | 0.2 | 0.3 | 0.36 | 0.33 | 0.22 |
| 6 | 102514974 | rs1034254 | -0.358 | -0.112 | 1.256 | -0.139 | -0.002 | 0.007 | 0.007 | 0 | 0.42 | 0.28 | 0.41 | 0.36 | 0.44 | 0.24 |
| 6 | 102526478 | rs2782900 | 0.811 | -0.188 | 0.87 | -0.362 | -0.013 | 0.027 | 0.027 | 0 | 0.24 | 0.13 | 0.23 | 0.08 | 0.33 | 0.12 |
| 6 | 102530252 | rs2782901 | -1.545 | -0.14 | 0.038 | -0.464 | -0.106 | 0.184 | 0.184 | 0.034 | 0.65 | 0.99 | 0.75 | 0.92 | 0.6 | 0.75 |
| 6 | 102594313 | rs1232239 | -0.367 | 0.273 | 0.994 | 0.072 | -0.001 | 0.007 | 0.007 | 0 | 0.46 | 0.53 | 0.47 | 0.35 | 0.47 | 0.29 |
| 6 | 102595588 | rs1232241 | -0.249 | 0.287 | 0.839 | 0.103 | -0.001 | -0.002 | -0.002 | 0 | 0.45 | 0.53 | 0.46 | 0.35 | 0.46 | 0.28 |
| 6 | 102601177 | rs3849205 | -0.374 | 0.626 | 0.629 | 0.301 | 0.014 | -0.003 | -0.003 | 0 | 0.53 | 0.6 | 0.62 | 0.46 | 0.61 | 0.46 |
| 6 | 102603013 | rs3995732 | 0.107 | 0.951 | 0.817 | 0.278 | 0.016 | 0.132 | 0.132 | 0 | 0.44 | 0.23 | 0.3 | 0.1 | 0.44 | 0.33 |
| 6 | 102609098 | rs9399737 | -0.048 | 1.074 | 0.848 | 0.326 | 0.009 | 0.028 | 0.028 | 0 | 0.48 | 0.37 | 0.32 | 0.1 | 0.45 | 0.33 |
| 6 | 102614062 | rs9373629 | -0.704 | 1.013 | 0.607 | 0.328 | -0.052 | -0.08 | -0.08 | 0 | 0.58 | 0.9 | 0.62 | 0.46 | 0.61 | 0.49 |
| 6 | 102616731 | rs7749939 | -1.768 | 1.032 | 1.704 | 1.229 | 0.11 | 0.239 | 0.239 | 4.928 | 0.08 | 0.36 | 0.25 | 0.27 | 0.15 | 0.12 |
| 6 | 102626292 | rs9485630 | -1.528 | 1.07 | 1.806 | 1.249 | 0.125 | 0.258 | 0.258 | 2.551 | 0.07 | 0.34 | 0.25 | 0.27 | 0.15 | 0.12 |
| 6 | 102629815 | rs2895436 | -1.51 | 1.171 | 0.903 | 1.139 | 0.135 | 0.336 | 0.336 | 2.31 | 0.1 | 0.46 | 0.31 | 0.36 | 0.17 | 0.16 |
| 6 | 102635841 | rs7750307 | -0.519 | 0.418 | 0.278 | -0.313 | -0.03 | -0.136 | -0.136 | 0 | 0.34 | 0.43 | 0.2 | 0.05 | 0.27 | 0.27 |
| 6 | 102639153 | rs6909610 | -0.484 | 0.542 | -0.043 | -0.123 | -0.007 | -0.01 | -0.01 | 2.425 | 0.52 | 0.73 | 0.56 | 0.51 | 0.52 | 0.63 |
| 6 | 102642021 | rs6937225 | 0.632 | 0.279 | -0.543 | -0.081 | 0.001 | -0.036 | -0.036 | 0 | 0.16 | 0.03 | 0.17 | 0.22 | 0.18 | 0.23 |
| 6 | 102653273 | rs6929339 | -1.196 | 0.723 | 0.093 | 0.713 | -0.096 | -0.085 | -0.085 | 0 | 0.55 | 0.94 | 0.57 | 0.52 | 0.52 | 0.64 |
| 6 | 102660014 | rs12206631 | -0.066 | 0.597 | 0.726 | 0.667 | -0.005 | -0.02 | -0.02 | 0 | 0.3 | 0.37 | 0.34 | 0.16 | 0.27 | 0.28 |
| 6 | 102669512 | rs1232226 | 0.961 | 0.739 | 0.36 | 0.25 | -0.018 | -0.114 | -0.114 | 0 | 0.24 | 0.04 | 0.25 | 0.48 | 0.22 | 0.29 |
| 6 | 102670220 | rs11156175 | 0.341 | 0.744 | 0.672 | 0.206 | 0 | 0.006 | 0.006 | 0 | 0.52 | 0.48 | 0.52 | 0.17 | 0.53 | 0.36 |
| 6 | 102674605 | rs10872745 | 1.032 | 0.719 | 0.514 | 0.177 | -0.054 | 0.066 | 0.066 | 0 | 0.32 | 0.1 | 0.39 | 0.12 | 0.36 | 0.2 |
| 6 | 102680964 | rs9386305 | -0.514 | 0.925 | 1.2 | 0.898 | 0.003 | 0.051 | 0.051 | 0.008 | 0.24 | 0.53 | 0.24 | 0.36 | 0.26 | 0.36 |
| 6 | 102682418 | rs9986468 | -0.986 | 1.023 | 1.472 | 0.976 | 0.116 | 0.127 | 0.127 | 2.078 | 0.06 | 0.37 | 0.05 | 0.12 | 0.07 | 0.12 |
| 6 | 102701920 | rs1232212 | -0.15 | 0.972 | 1.34 | 0.898 | 0.01 | 0.023 | 0.023 | 0.055 | 0.32 | 0.59 | 0.31 | 0.61 | 0.28 | 0.43 |
| 6 | 102706463 | rs9498995 | 0.726 | 0.864 | 1.331 | 0.831 | 0.001 | 0.041 | 0.041 | 0.768 | 0.52 | 0.35 | 0.52 | 0.16 | 0.55 | 0.36 |
| 6 | 102713169 | rs9373633 | -0.005 | 0.874 | 1.325 | 0.806 | 0.013 | 0.027 | 0.027 | 0.004 | 0.3 | 0.42 | 0.31 | 0.6 | 0.29 | 0.4 |
| 6 | 102727741 | rs2226908 | -0.905 | 0.864 | 1.218 | 0.9 | -0.001 | 0.029 | 0.029 | 0.349 | 0.48 | 0.65 | 0.5 | 0.79 | 0.47 | 0.65 |
| 6 | 102728121 | rs12173739 | 0.632 | 0.955 | 1.004 | 1.481 | -0.035 | 0.032 | 0.032 | 0.613 | 0.24 | 0.02 | 0.18 | 0.16 | 0.21 | 0.31 |
| 6 | 102731735 | rs3911831 | 0.425 | 0.966 | 1.074 | 1.313 | 0.001 | -0.004 | -0.004 | 0 | 0.26 | 0.22 | 0.19 | 0.25 | 0.21 | 0.31 |
| 6 | 102733010 | rs2994284 | 0.118 | 0.966 | 1.031 | 1.313 | -0.004 | 0.001 | 0.001 | 0 | 0.28 | 0.33 | 0.2 | 0.25 | 0.21 | 0.31 |
| 6 | 102733537 | rs2153395 | -0.137 | 0.998 | 1.05 | 1.396 | 0.007 | 0.022 | 0.022 | 0.087 | 0.31 | 0.43 | 0.29 | 0.57 | 0.28 | 0.41 |
| 6 | 102738413 | rs1361417 | -0.016 | 1.101 | 0.848 | 1.493 | -0.005 | -0.006 | -0.006 | 5.317 | 0.81 | 0.8 | 0.7 | 0.45 | 0.74 | 0.65 |
| 6 | 102739061 | rs2994291 | 0.017 | 1.095 | 0.846 | 1.49 | -0.005 | -0.008 | -0.008 | 0 | 0.81 | 0.79 | 0.7 | 0.45 | 0.74 | 0.65 |
| 6 | 102739084 | rs2994292 | -0.432 | 1.046 | 0.856 | 1.353 | 0 | -0.001 | -0.001 | 1.569 | 0.83 | 0.81 | 0.78 | 0.77 | 0.81 | 0.73 |
| 6 | 102748329 | rs9322644 | 0.288 | 1.095 | 0.791 | 0.968 | 0 | 0.004 | 0.004 | 0.163 | 0.59 | 0.63 | 0.46 | 0.42 | 0.59 | 0.47 |
| 6 | 102766711 | rs9322648 | 0.324 | 1.021 | 0.529 | 1.034 | -0.049 | -0.095 | -0.095 | 0.424 | 0.25 | 0.06 | 0.31 | 0.49 | 0.28 | 0.33 |
| 6 | 102779335 | rs954551 | 0.032 | 1.1 | 0.586 | 1.005 | -0.03 | -0.055 | -0.055 | 0 | 0.8 | 0.88 | 0.53 | 0.31 | 0.73 | 0.68 |
| 6 | 102800381 | rs4334996 | -0.869 | 0.826 | 0.872 | 0.709 | -0.001 | -0.011 | -0.011 | 1.48 | 0.48 | 0.59 | 0.67 | 0.89 | 0.49 | 0.6 |
| 6 | 102814820 | rs2399799 | 0.408 | 0.862 | 0.652 | 0.92 | 0.005 | -0.068 | -0.068 | 0 | 0.18 | 0.11 | 0.35 | 0.38 | 0.2 | 0.25 |
| 6 | 102819959 | rs17493048 | 0.533 | 0.938 | 0.648 | 1.004 | -0.009 | 0.001 | 0.001 | 0.104 | 0.24 | 0.08 | 0.18 | 0.18 | 0.22 | 0.26 |
| 6 | 102833855 | rs9322655 | 0.144 | 0.943 | 0.774 | 0.978 | -0.004 | -0.006 | -0.006 | 0.021 | 0.27 | 0.24 | 0.32 | 0.51 | 0.29 | 0.35 |
| 6 | 102835601 | rs1538381 | 0.055 | 0.94 | 0.789 | 0.977 | 0.011 | 0.035 | 0.035 | 0 | 0.28 | 0.3 | 0.32 | 0.51 | 0.29 | 0.35 |
| 6 | 102840130 | rs9404204 | 0.387 | 0.85 | 0.468 | 0.96 | 0.012 | -0.047 | -0.047 | 2.134 | 0.18 | 0.08 | 0.33 | 0.39 | 0.2 | 0.24 |

| | | | | | | | | | | | | | | | | |
|---|---|---|---|---|---|---|---|---|---|---|---|---|---|---|---|---|
| 6 | 102847110 | rs17063233 | 0.629 | 0.88 | 0.343 | 0.979 | -0.042 | -0.027 | -0.027 | 0 | 0.23 | 0.03 | 0.18 | 0.19 | 0.22 | 0.26 |
| 6 | 102849336 | rs1338657 | -0.542 | 0.841 | 0.84 | 0.945 | 0.003 | 0.004 | 0.004 | 0.028 | 0.43 | 0.34 | 0.51 | 0.58 | 0.42 | 0.5 |
| 6 | 102856592 | rs9499070 | 0.425 | 0.947 | 0.726 | 0.985 | 0.001 | -0.003 | -0.003 | 0.05 | 0.25 | 0.23 | 0.18 | 0.19 | 0.22 | 0.25 |
| 6 | 102860663 | rs7773696 | 0.299 | 0.876 | 0.733 | 1.017 | 0.015 | -0.052 | -0.052 | 0 | 0.18 | 0.12 | 0.33 | 0.4 | 0.2 | 0.24 |
| 6 | 102865390 | rs9377361 | -0.091 | 0.826 | 0.742 | 0.295 | 0.001 | 0.022 | 0.022 | 0 | 0.22 | 0.35 | 0.47 | 0.71 | 0.27 | 0.28 |
| 6 | 102884588 | rs6915057 | -0.502 | 0.816 | 0.432 | -0.328 | -0.002 | 0.015 | 0.015 | 0 | 0.47 | 0.6 | 0.66 | 0.9 | 0.49 | 0.54 |
| 6 | 102896943 | rs4475333 | -0.526 | 0.524 | -0.026 | -0.891 | 0.001 | -0.053 | -0.053 | 0.235 | 0.52 | 0.53 | 0.71 | 0.92 | 0.58 | 0.62 |
| 6 | 102901092 | rs17501289 | 0.686 | 0.403 | -0.206 | -1.045 | 0.005 | 0.029 | 0.029 | 1.746 | 0.06 | 0 | 0.03 | 0 | 0.05 | 0.04 |
| 6 | 102903773 | rs6928093 | 0.408 | 0.689 | 0.369 | -0.462 | 0 | 0 | 0 | 0 | 0.26 | 0.16 | 0.19 | 0.28 | 0.29 | 0.25 |
| 6 | 102904579 | rs4445064 | -0.851 | 0.85 | 0.502 | -0.25 | 0.001 | -0.006 | -0.006 | 0 | 0.49 | 0.43 | 0.68 | 0.8 | 0.51 | 0.59 |
| 6 | 102922642 | rs601907 | 0.848 | 0.836 | 0.527 | -0.262 | -0.001 | -0.076 | -0.076 | 0 | 0.48 | 0.49 | 0.29 | 0.08 | 0.42 | 0.38 |
| 6 | 102931736 | rs643863 | -0.431 | 0.759 | 0.599 | -0.213 | -0.001 | -0.03 | -0.03 | 0 | 0.8 | 0.88 | 0.52 | 0.5 | 0.79 | 0.71 |
| 6 | 102953010 | rs9499109 | -0.231 | 0.75 | 0.637 | -0.213 | 0.005 | 0.011 | 0.011 | 0 | 0.32 | 0.42 | 0.23 | 0.44 | 0.38 | 0.33 |
| 6 | 102963131 | rs17063343 | 0.169 | 0.744 | 0.57 | -0.214 | -0.03 | -0.041 | -0.041 | 0 | 0.28 | 0.17 | 0.22 | 0.43 | 0.37 | 0.29 |
| 6 | 102966628 | rs9390844 | 0.845 | 0.737 | 0.51 | -0.243 | 0.004 | -0.037 | -0.037 | 0.006 | 0.49 | 0.53 | 0.29 | 0.06 | 0.41 | 0.38 |
| 6 | 102969306 | rs1157234 | 0.466 | 0.83 | -0.05 | -0.188 | -0.015 | -0.025 | -0.025 | 0 | 0.26 | 0.08 | 0.19 | 0.29 | 0.29 | 0.26 |
| 6 | 102970894 | rs1934007 | -0.413 | 0.861 | 0.055 | 0.066 | -0.044 | -0.138 | -0.138 | 0 | 0.81 | 0.98 | 0.52 | 0.5 | 0.79 | 0.71 |
| 6 | 103005548 | rs13211916 | 1.188 | 0.229 | -0.304 | -0.708 | 0.047 | 0.164 | 0.164 | 2.907 | 0.25 | 0.02 | 0.1 | 0 | 0.14 | 0.07 |
| 6 | 103007392 | rs1933995 | -0.128 | 0.408 | 0.411 | -0.425 | 0 | 0.081 | 0.081 | 0.644 | 0.55 | 0.74 | 0.34 | 0.22 | 0.52 | 0.45 |
| 6 | 103016670 | rs9322665 | 0.703 | 0.244 | 0.8 | -0.548 | -0.009 | 0.16 | 0.16 | 1.613 | 0.45 | 0.22 | 0.28 | 0.07 | 0.42 | 0.33 |
| 6 | 103018974 | rs7752919 | 0.022 | 0.316 | 0.526 | -0.415 | -0.002 | 0.115 | 0.115 | 1.099 | 0.55 | 0.79 | 0.34 | 0.23 | 0.52 | 0.43 |
| 6 | 103028147 | rs7758751 | 0.456 | 0.871 | 0.866 | 0.443 | -0.001 | 0.034 | 0.034 | 0.235 | 0.53 | 0.38 | 0.38 | 0.29 | 0.46 | 0.43 |
| 6 | 103030420 | rs6907131 | 0.237 | 1.101 | 1.116 | 0.837 | 0 | -0.002 | -0.002 | 2.514 | 0.27 | 0.24 | 0.2 | 0.29 | 0.28 | 0.26 |
| 6 | 103041607 | rs9485706 | 0.131 | 1.089 | 1.439 | 0.831 | -0.002 | 0.006 | 0.006 | 0 | 0.27 | 0.34 | 0.21 | 0.29 | 0.28 | 0.26 |
| 6 | 103052113 | rs7771843 | 0.252 | 1.09 | 1.462 | 0.831 | 0 | -0.006 | -0.006 | 0.044 | 0.27 | 0.19 | 0.2 | 0.29 | 0.28 | 0.26 |
| 6 | 103067505 | rs12206844 | 0.193 | 1.087 | 1.437 | 0.831 | -0.001 | 0.003 | 0.003 | 0 | 0.27 | 0.28 | 0.2 | 0.29 | 0.28 | 0.26 |
| 6 | 103067777 | rs7454288 | -0.305 | 1.162 | 1.438 | 1.025 | -0.005 | 0.039 | 0.039 | 0.019 | 0.19 | 0.33 | 0.28 | 0.27 | 0.18 | 0.24 |
| 6 | 103071746 | rs9322668 | 0.442 | 1.465 | 1.052 | 0.823 | -0.033 | -0.062 | -0.062 | 0 | 0.1 | 0.02 | 0.14 | 0.27 | 0.12 | 0.13 |
| 6 | 103084016 | rs6571039 | -0.88 | 1.589 | 1.693 | 1.194 | 0.002 | 0.003 | 0.003 | 0 | 0.46 | 0.43 | 0.47 | 0.57 | 0.45 | 0.52 |
| 6 | 103095006 | rs11156202 | 1.123 | 1.564 | 1.798 | 1.174 | 0.004 | 0.084 | 0.084 | 0.428 | 0.51 | 0.29 | 0.44 | 0.3 | 0.46 | 0.4 |
| 6 | 103111984 | rs9654652 | -0.174 | 1.349 | 1.871 | 1.122 | -0.002 | -0.002 | -0.002 | 0 | 0.28 | 0.4 | 0.23 | 0.29 | 0.28 | 0.28 |
| 6 | 103119441 | rs9399760 | -0.551 | 1.369 | 1.735 | 1.132 | -0.009 | 0.06 | 0.06 | 0 | 0.19 | 0.37 | 0.27 | 0.27 | 0.18 | 0.23 |
| 6 | 103124343 | rs7769811 | -0.227 | 1.369 | 1.47 | 1.007 | -0.001 | -0.001 | -0.001 | 0 | 0.28 | 0.3 | 0.22 | 0.29 | 0.28 | 0.28 |
| 6 | 103135634 | rs10457108 | -0.196 | 1.411 | 0.972 | 1.031 | 0.001 | 0.002 | 0.002 | 0 | 0.28 | 0.2 | 0.21 | 0.29 | 0.28 | 0.28 |
| 6 | 103146074 | rs9377381 | -0.845 | 1.378 | 1.15 | 0.847 | 0.002 | 0.007 | 0.007 | 0.005 | 0.47 | 0.57 | 0.49 | 0.62 | 0.44 | 0.55 |
| 6 | 103157617 | rs1523414 | -1.03 | 1.477 | 1.381 | 0.776 | 0.027 | 0.066 | 0.066 | 0.305 | 0.47 | 0.65 | 0.56 | 0.71 | 0.58 | 0.65 |
| 6 | 103159056 | rs13215862 | -0.695 | 1.492 | 1.857 | 0.812 | -0.022 | 0.017 | 0.017 | 0.028 | 0.42 | 0.26 | 0.5 | 0.63 | 0.49 | 0.5 |
| 6 | 103169903 | rs6902640 | -1.11 | 1.479 | 1.871 | 0.822 | 0.029 | 0.077 | 0.077 | 0.266 | 0.48 | 0.71 | 0.57 | 0.71 | 0.58 | 0.65 |
| 6 | 103175290 | rs13195313 | -0.719 | 1.478 | 1.942 | 0.832 | -0.018 | 0.015 | 0.015 | 0.046 | 0.43 | 0.27 | 0.51 | 0.64 | 0.49 | 0.51 |
| 6 | 103179548 | rs1560455 | -0.924 | 1.48 | 2.067 | 0.814 | 0.006 | -0.012 | -0.012 | 0.147 | 0.44 | 0.43 | 0.52 | 0.66 | 0.5 | 0.54 |
| 6 | 103188348 | rs17505047 | 0.195 | 1.562 | 1.578 | 1.09 | -0.095 | -0.06 | -0.06 | 0 | 0.24 | 0.03 | 0.24 | 0.33 | 0.33 | 0.27 |
| 6 | 103198856 | rs1431213 | -1.118 | 1.515 | 1.724 | 1.047 | 0.011 | 0.022 | 0.022 | 0.527 | 0.47 | 0.56 | 0.55 | 0.69 | 0.58 | 0.62 |
| 6 | 103210765 | rs951197 | -1.272 | 1.472 | 1.543 | 1.101 | 0.016 | 0.082 | 0.082 | 0.5 | 0.47 | 0.7 | 0.55 | 0.72 | 0.55 | 0.65 |
| 6 | 103225448 | rs12216167 | -0.035 | 1.307 | 0.044 | 1.164 | -0.095 | -0.06 | -0.06 | 0 | 0.24 | 0.03 | 0.23 | 0.33 | 0.33 | 0.27 |
| 6 | 103229346 | rs1431210 | -0.555 | 1.139 | 0.481 | 0.742 | -0.013 | 0.015 | 0.015 | 0.027 | 0.42 | 0.29 | 0.48 | 0.64 | 0.46 | 0.5 |
| 6 | 103242059 | rs9377388 | -1.745 | 0.952 | 1.325 | 0.682 | 0.072 | 0.122 | 0.122 | 2.47 | 0.06 | 0.42 | 0.11 | 0.1 | 0.03 | 0.13 |
| 6 | 103261515 | rs9390890 | -2.127 | 0.834 | 1.416 | 0.622 | 0.036 | 0.083 | 0.083 | 4.232 | 0.07 | 0.51 | 0.12 | 0.1 | 0.03 | 0.14 |
| 6 | 103287274 | rs9390898 | -0.833 | 1.621 | 1.039 | 0.268 | -0.01 | 0.005 | 0.005 | 0.291 | 0.22 | 0.23 | 0.29 | 0.17 | 0.32 | 0.36 |
| 6 | 103288401 | rs9377400 | -0.675 | 1.618 | 1.044 | 0.265 | -0.032 | 0.022 | 0.022 | 0.319 | 0.2 | 0.14 | 0.2 | 0.11 | 0.3 | 0.28 |
| 6 | 103292745 | rs1367645 | -0.348 | 1.533 | 1.268 | -0.099 | 0.007 | -0.045 | -0.045 | 0.001 | 0.08 | 0.2 | 0.05 | 0 | 0.08 | 0.02 |
| 6 | 103294020 | rs6571064 | -0.348 | 1.533 | 1.283 | -0.099 | 0.005 | -0.045 | -0.045 | 0.687 | 0.08 | 0.19 | 0.05 | 0 | 0.08 | 0.02 |
| 6 | 103299213 | rs12196138 | -1.249 | 1.571 | 0.764 | -0.295 | -0.009 | 0.005 | 0.005 | 0 | 0.3 | 0.37 | 0.35 | 0.19 | 0.43 | 0.41 |
| 6 | 103303052 | rs7766397 | -0.974 | 1.803 | 0.842 | -0.399 | 0.007 | -0.01 | -0.01 | 0 | 0.24 | 0.3 | 0.25 | 0.13 | 0.34 | 0.27 |
| 6 | 103303838 | rs9485742 | 0.393 | 1.845 | 0.659 | -0.381 | -0.007 | 0.014 | 0.014 | 0 | 0.07 | 0.03 | 0.04 | 0.02 | 0.08 | 0.03 |
| 6 | 103314799 | rs9399770 | -0.976 | 1.234 | 0.828 | 0.149 | 0.006 | 0.003 | 0.003 | 0 | 0.31 | 0.3 | 0.54 | 0.66 | 0.38 | 0.6 |
| 6 | 103317153 | rs9499255 | 0.521 | 0.456 | 0.176 | -0.897 | 0.002 | -0.03 | -0.03 | 0 | 0.32 | 0.38 | 0.19 | 0.06 | 0.33 | 0.22 |
| 6 | 103325998 | rs9377407 | 0.09 | 1.185 | -0.082 | -0.775 | -0.085 | 0.023 | 0.023 | 0.001 | 0.16 | 0.03 | 0.2 | 0.11 | 0.26 | 0.24 |
| 6 | 103327607 | rs1431218 | -0.548 | 0.773 | -0.031 | -0.9 | -0.005 | -0.009 | -0.009 | 0.058 | 0.64 | 0.7 | 0.53 | 0.43 | 0.63 | 0.55 |
| 6 | 103338050 | rs1359053 | -0.065 | 0.758 | 0.473 | -1.01 | -0.001 | 0.001 | 0.001 | 0 | 0.54 | 0.49 | 0.45 | 0.4 | 0.52 | 0.44 |
| 6 | 103341545 | rs9404275 | 0.298 | 0.603 | 0.498 | -1.084 | -0.034 | 0.109 | 0.109 | 0.848 | 0.49 | 0.12 | 0.43 | 0.38 | 0.51 | 0.43 |
| 6 | 103346215 | rs17063732 | 0.796 | 0.621 | 0.556 | -1.102 | -0.005 | -0.002 | -0.002 | 0 | 0.08 | 0.07 | 0.06 | 0 | 0.11 | 0.02 |
| 6 | 103352202 | rs13213644 | 0.166 | 0.537 | 0.659 | -0.92 | -0.036 | 0.107 | 0.107 | 0.739 | 0.49 | 0.12 | 0.43 | 0.38 | 0.51 | 0.43 |
| 6 | 103352728 | rs6914436 | 0.689 | 0.742 | 0.496 | -0.999 | 0.05 | 0.092 | 0.092 | 0.208 | 0.39 | 0.08 | 0.27 | 0.26 | 0.3 | 0.17 |
| 6 | 103357748 | rs9377412 | -0.368 | 0.875 | 0.666 | -0.901 | -0.016 | 0.007 | 0.007 | 0 | 0.57 | 0.66 | 0.49 | 0.44 | 0.56 | 0.55 |
| 6 | 103375422 | rs7775194 | -0.115 | 0.896 | 0.947 | -0.881 | -0.002 | 0.005 | 0.005 | 0 | 0.55 | 0.4 | 0.46 | 0.42 | 0.55 | 0.53 |
| 6 | 103375639 | rs1323203 | 1.528 | 0.452 | 0.539 | -1.237 | -0.018 | 0.11 | 0.11 | 0 | 0.17 | 0.01 | 0.1 | 0.01 | 0.16 | 0.05 |
| 6 | 103376636 | rs1323205 | -0.335 | 0.955 | 1.263 | -0.619 | -0.042 | -0.007 | -0.007 | 0 | 0.59 | 0.77 | 0.48 | 0.44 | 0.56 | 0.54 |
| 6 | 103380970 | rs6571070 | -0.129 | 1.107 | 0.907 | -0.53 | -0.008 | 0.006 | 0.006 | 0 | 0.57 | 0.59 | 0.47 | 0.44 | 0.55 | 0.53 |
| 6 | 103403853 | rs6913900 | -0.028 | 1.139 | 0.671 | -0.514 | 0 | -0.002 | -0.002 | 0 | 0.56 | 0.49 | 0.47 | 0.44 | 0.55 | 0.53 |

| | | | | | | | | | | | | | | | |
|---|---|---|---|---|---|---|---|---|---|---|---|---|---|---|---|
| 6 | 103438452 | rs9373671 | 0.402 | 1.511 | 0.513 | -0.154 | -0.034 | 0 | 0 | 0 | 0.11 | 0.04 | 0.14 | 0.12 | 0.2 | 0.27 |
| 6 | 103467054 | rs13212776 | -0.002 | 1.133 | 0.505 | -0.143 | 0.009 | 0.016 | 0.016 | 0 | 0.66 | 0.54 | 0.5 | 0.46 | 0.61 | 0.57 |
| 6 | 103467436 | rs9322695 | -0.126 | 1.213 | 0.341 | -0.384 | 0.022 | -0.03 | -0.03 | 0.009 | 0.2 | 0.23 | 0.18 | 0.11 | 0.25 | 0.26 |
| 6 | 103468941 | rs9404299 | 0.166 | 1.316 | 0.134 | 0.372 | 0.005 | -0.002 | -0.002 | 0 | 0.25 | 0.62 | 0.23 | 0.21 | 0.25 | 0.33 |
| 6 | 103475623 | rs6900581 | 0.297 | 1.302 | -0.42 | 0.372 | -0.035 | 0.037 | 0.037 | 0.708 | 0.19 | 0.08 | 0.15 | 0.12 | 0.22 | 0.32 |
| 6 | 103481466 | rs6900275 | -0.431 | 1.314 | -0.445 | 0.643 | 0 | 0.048 | 0.048 | 0 | 0.86 | 0.84 | 0.67 | 0.64 | 0.82 | 0.82 |
| 6 | 103483952 | rs9404303 | 0.183 | 1.051 | -0.691 | 0.425 | 0.009 | 0.003 | 0.003 | 0 | 0.25 | 0.33 | 0.24 | 0.24 | 0.29 | 0.3 |
| 6 | 103484412 | rs9390939 | 0.293 | 0.443 | -1.256 | 0.346 | 0.043 | 0.074 | 0.074 | 0.235 | 0.39 | 0.16 | 0.26 | 0.27 | 0.31 | 0.2 |
| 6 | 103485282 | rs9485766 | -0.925 | 0.817 | -0.387 | 0.96 | 0.015 | 0.102 | 0.102 | 0 | 0.22 | 0.51 | 0.23 | 0.38 | 0.23 | 0.23 |
| 6 | 103485557 | rs9404304 | 0.204 | 0.424 | -1.37 | 0.828 | -0.022 | -0.131 | -0.131 | 0.107 | 0.21 | 0.02 | 0.32 | 0.41 | 0.2 | 0.18 |
| 6 | 103496197 | rs2399886 | -0.124 | 0.334 | -1.355 | -0.22 | -0.032 | 0.111 | 0.111 | 0 | 0.49 | 0.14 | 0.53 | 0.65 | 0.51 | 0.57 |
| 6 | 103519750 | rs9377433 | -0.876 | 0.276 | -0.643 | -0.56 | -0.005 | 0.063 | 0.063 | 0.062 | 0.65 | 0.85 | 0.7 | 0.77 | 0.65 | 0.76 |
| 6 | 103521720 | rs9647631 | -0.778 | 0.267 | -0.646 | -0.563 | -0.023 | 0.024 | 0.024 | 0.017 | 0.63 | 0.83 | 0.67 | 0.67 | 0.59 | 0.7 |
| 6 | 103528930 | rs12207731 | 0.617 | 0.517 | -0.789 | -0.699 | -0.091 | 0.005 | 0.005 | 0.108 | 0.41 | 0.04 | 0.47 | 0.65 | 0.41 | 0.54 |
| 6 | 103553252 | rs2205680 | -0.428 | 0.781 | -0.256 | -0.371 | -0.002 | -0.009 | -0.009 | 0 | 0.92 | 0.94 | 0.95 | 1 | 0.91 | 0.97 |
| 6 | 103555683 | rs7745925 | 0.177 | 0.744 | 0.101 | -0.36 | 0.018 | -0.035 | -0.035 | 0 | 0.07 | 0.12 | 0.11 | 0 | 0.08 | 0.05 |
| 6 | 103563630 | rs7740615 | 0.497 | 0.715 | 0.004 | -0.512 | 0.002 | -0.032 | -0.032 | 1.122 | 0.86 | 0.87 | 0.84 | 1 | 0.83 | 0.91 |
| 6 | 103573865 | rs12206675 | 0.89 | 0.465 | -0.1 | -0.884 | -0.012 | 0.131 | 0.131 | 0 | 0.33 | 0.14 | 0.27 | 0.08 | 0.33 | 0.22 |
| 6 | 103575112 | rs9404329 | 0.708 | 0.373 | -0.223 | -0.712 | -0.072 | -0.018 | -0.018 | 0 | 0.44 | 0.04 | 0.48 | 0.63 | 0.42 | 0.54 |
| 6 | 103576712 | rs6571081 | -0.135 | 0.401 | 0.174 | -0.715 | 0.018 | 0.041 | 0.041 | 0 | 0.89 | 0.82 | 0.89 | 0.74 | 0.83 | 0.86 |
| 6 | 103577654 | rs2179079 | 0.157 | 0.317 | 0.697 | -0.801 | -0.001 | -0.056 | -0.056 | 0.369 | 0.42 | 0.51 | 0.39 | 0.11 | 0.41 | 0.27 |
| 6 | 103580795 | rs7775708 | 0.779 | 0.392 | 0.526 | -0.769 | 0.001 | 0.068 | 0.068 | 0 | 0.34 | 0.22 | 0.27 | 0.08 | 0.34 | 0.22 |
| 6 | 103583945 | rs6915549 | -0.997 | 0.58 | 0.819 | -0.42 | 0.094 | 0.021 | 0.021 | 0.696 | 0.12 | 0.48 | 0.17 | 0.08 | 0.14 | 0.17 |
| 6 | 103600715 | rs505347 | -0.306 | 0.431 | 0.56 | -0.582 | 0.001 | 0.022 | 0.022 | 0.735 | 0.56 | 0.6 | 0.65 | 0.75 | 0.57 | 0.73 |
| 6 | 103601517 | rs4946850 | 0.576 | 0.373 | 0.004 | -0.259 | -0.032 | -0.07 | -0.07 | 0 | 0.43 | 0.09 | 0.48 | 0.57 | 0.41 | 0.47 |
| 6 | 103604578 | rs6915705 | -0.416 | 0.733 | 0.695 | -0.118 | 0.025 | -0.044 | -0.044 | 0 | 0.08 | 0.23 | 0.11 | 0 | 0.08 | 0.1 |
| 6 | 103609631 | rs479238 | 0.214 | 0.452 | 0.129 | -0.093 | 0.013 | -0.031 | -0.031 | 0 | 0.3 | 0.17 | 0.36 | 0.62 | 0.25 | 0.32 |
| 6 | 103612659 | rs522447 | 0.282 | 0.779 | 0.427 | 0.044 | 0.006 | 0.027 | 0.027 | 0 | 0.9 | 0.81 | 0.91 | 0.8 | 0.9 | 0.96 |
| 6 | 103620601 | rs578419 | 0.738 | 0.731 | 0.82 | 0.043 | 0.028 | 0.068 | 0.068 | 0 | 0.87 | 0.7 | 0.89 | 0.82 | 0.85 | 0.77 |
| 6 | 103622001 | rs522737 | 0.089 | 0.641 | 0.203 | -0.3 | -0.001 | -0.003 | -0.003 | 0 | 0.19 | 0.18 | 0.17 | 0.22 | 0.21 | 0.29 |
| 6 | 103623456 | rs2294216 | 0.462 | 0.727 | 0.224 | -0.302 | -0.012 | -0.011 | -0.011 | 2.295 | 0.73 | 0.78 | 0.75 | 0.68 | 0.66 | 0.63 |
| 6 | 103624403 | rs17623275 | 0.808 | 0.484 | 0.013 | -0.68 | -0.05 | 0.196 | 0.196 | 0 | 0.39 | 0.16 | 0.31 | 0.04 | 0.42 | 0.25 |
| 6 | 103624808 | rs494028 | -0.241 | 0.713 | 0.018 | -0.368 | -0.025 | -0.01 | -0.01 | 0 | 0.91 | 0.95 | 0.92 | 0.9 | 0.86 | 0.92 |
| 6 | 103625805 | rs6912940 | 0.443 | 0.713 | 0.042 | -0.375 | -0.016 | -0.011 | -0.011 | 0 | 0.73 | 0.78 | 0.75 | 0.68 | 0.65 | 0.63 |
| 6 | 103634233 | rs472570 | 0.57 | 0.72 | -0.296 | -0.273 | -0.053 | -0.049 | -0.049 | 0 | 0.14 | 0.01 | 0.13 | 0.11 | 0.17 | 0.31 |
| 6 | 103635120 | rs12213111 | 0.642 | 0.705 | 0.052 | -0.17 | -0.002 | -0.015 | -0.015 | 0 | 0.3 | 0.24 | 0.46 | 0.55 | 0.29 | 0.27 |
| 6 | 103652391 | rs501241 | 0.43 | 0.622 | 0.924 | -0.147 | 0.015 | 0.066 | 0.066 | 0 | 0.88 | 0.6 | 0.89 | 0.81 | 0.87 | 0.92 |
| 6 | 103674148 | rs4636062 | 0.419 | 0.615 | 0.632 | -0.535 | -0.001 | 0.139 | 0.139 | 1.141 | 0.47 | 0.19 | 0.62 | 0.76 | 0.52 | 0.67 |
| 6 | 103684742 | rs6919280 | 0.505 | 0.508 | 0.947 | -0.592 | 0.008 | 0.105 | 0.105 | 0.156 | 0.37 | 0.19 | 0.26 | 0.04 | 0.35 | 0.21 |
| 6 | 103716267 | rs721383 | -0.286 | 0.521 | 0.844 | -0.568 | 0.014 | 0.083 | 0.083 | 0.343 | 0.51 | 0.68 | 0.37 | 0.24 | 0.48 | 0.29 |
| 6 | 103720661 | rs7746268 | -0.27 | 0.512 | 0.83 | -0.572 | 0.017 | 0.099 | 0.099 | 0.438 | 0.5 | 0.71 | 0.37 | 0.24 | 0.48 | 0.29 |
| 6 | 103726563 | rs2208332 | -0.609 | 0.641 | 0.793 | -0.499 | -0.04 | 0.064 | 0.064 | 1.518 | 0.56 | 0.84 | 0.42 | 0.34 | 0.54 | 0.42 |
| 6 | 103728158 | rs1008965 | 0.365 | 0.651 | 0.79 | -0.495 | -0.005 | 0.051 | 0.051 | 0.723 | 0.47 | 0.28 | 0.6 | 0.66 | 0.47 | 0.62 |
| 6 | 103767356 | rs4945830 | 0.384 | 0.651 | 0.787 | -0.513 | -0.006 | 0.051 | 0.051 | 0.914 | 0.47 | 0.29 | 0.6 | 0.67 | 0.46 | 0.62 |
| 6 | 103774790 | rs12207319 | 0.495 | 0.749 | 0.692 | -0.681 | 0.026 | 0.137 | 0.137 | 0.157 | 0.35 | 0.12 | 0.24 | 0.04 | 0.3 | 0.18 |
| 6 | 103775306 | rs7768059 | -0.759 | 1.2 | 0.895 | -0.335 | -0.021 | 0.034 | 0.034 | 0 | 0.94 | 0.97 | 0.95 | 1 | 0.91 | 0.97 |
| 6 | 103784799 | rs9322714 | 0.011 | 1.056 | 1.177 | -0.497 | -0.001 | -0.017 | -0.017 | 0 | 0.6 | 0.6 | 0.62 | 0.87 | 0.56 | 0.67 |
| 6 | 103788127 | rs9404359 | -0.695 | 1.072 | 1.231 | -0.469 | 0.016 | -0.036 | -0.036 | 0 | 0.93 | 0.83 | 0.95 | 1 | 0.91 | 0.97 |
| 6 | 103793130 | rs11156266 | 0.366 | 1.013 | 1.023 | -0.527 | -0.008 | 0.043 | 0.043 | 0 | 0.29 | 0.15 | 0.25 | 0.13 | 0.32 | 0.28 |
| 6 | 103797506 | rs12202509 | 0.719 | 1.049 | 1.03 | -0.531 | 0.005 | 0.034 | 0.034 | 0 | 0.26 | 0.14 | 0.17 | 0.02 | 0.26 | 0.19 |
| 6 | 103797987 | rs7759361 | -0.029 | 1.025 | 1.212 | -0.373 | -0.003 | 0.002 | 0.002 | 0 | 0.35 | 0.59 | 0.3 | 0.39 | 0.36 | 0.33 |
| 6 | 103799046 | rs994988 | 0.671 | 1.029 | 1.214 | -0.4 | -0.009 | 0.03 | 0.03 | 0 | 0.53 | 0.15 | 0.56 | 0.61 | 0.52 | 0.61 |
| 6 | 103799673 | rs9404361 | -0.416 | 1.021 | 1.178 | -0.382 | 0.038 | 0.355 | 0.355 | 2.797 | 0.07 | 0.44 | 0.05 | 0.26 | 0.04 | 0.05 |
| 6 | 103801640 | rs1155126 | 0.471 | 1.334 | 0.405 | -0.831 | 0.022 | -0.061 | -0.061 | 0 | 0.87 | 0.77 | 0.85 | 0.99 | 0.83 | 0.91 |
| 6 | 103816260 | rs7743013 | 0.194 | 1.232 | 0.129 | -0.718 | -0.005 | 0.036 | 0.036 | 0 | 0.29 | 0.16 | 0.25 | 0.14 | 0.32 | 0.27 |
| 6 | 103822130 | rs12524438 | -0.054 | 1.24 | 0.089 | -0.701 | -0.012 | 0.037 | 0.037 | 0 | 0.35 | 0.24 | 0.29 | 0.14 | 0.41 | 0.3 |
| 6 | 103827338 | rs12196235 | 0.35 | 1.036 | -0.378 | -0.652 | -0.003 | 0.125 | 0.125 | 0 | 0.27 | 0.08 | 0.16 | 0.03 | 0.27 | 0.19 |
| 6 | 103840702 | rs17624545 | -0.713 | 1.568 | -0.145 | -0.403 | 0.003 | 0 | 0 | 0 | 0.13 | 0.12 | 0.2 | 0.11 | 0.19 | 0.17 |
| 6 | 103842567 | rs9377478 | -0.698 | 1.699 | -0.18 | -0.019 | 0.029 | 0.014 | 0.014 | 0.221 | 0.62 | 0.79 | 0.76 | 0.65 | 0.65 | 0.5 |
| 6 | 103843861 | rs9499413 | -1.036 | 2.136 | -0.563 | 0.216 | 0.006 | -0.009 | -0.009 | 0 | 0.23 | 0.25 | 0.26 | 0.11 | 0.3 | 0.2 |
| 6 | 103846166 | rs7771431 | -0.078 | 1.737 | -0.502 | 0.317 | 0.001 | 0.012 | 0.012 | 0 | 0.11 | 0.06 | 0.12 | 0 | 0.09 | 0.06 |
| 6 | 103863554 | rs17624721 | -0.273 | 1.771 | -0.579 | 0.312 | 0.008 | -0.035 | -0.035 | 0 | 0.12 | 0.15 | 0.14 | 0 | 0.14 | 0.07 |
| 6 | 103865241 | rs12197896 | 0.063 | 1.768 | -0.578 | 0.312 | -0.002 | -0.037 | -0.037 | 0 | 0.1 | 0.14 | 0.12 | 0 | 0.09 | 0.07 |
| 6 | 103866950 | rs12201549 | 0.542 | 1.827 | -0.937 | 0.311 | -0.18 | -0.098 | -0.098 | 2.222 | 0.36 | 0.03 | 0.39 | 0.32 | 0.41 | 0.3 |
| 6 | 103873242 | rs9391003 | -1.329 | 1.889 | 0.414 | 1.02 | 0.042 | 0.257 | 0.257 | 5.956 | 0.08 | 0.43 | 0.12 | 0.31 | 0.08 | 0.07 |
| 6 | 103882949 | rs2895553 | -1.705 | 1.886 | 0.415 | 1.017 | 0 | 0.08 | 0.08 | 3.188 | 0.18 | 0.5 | 0.25 | 0.31 | 0.17 | 0.14 |
| 6 | 103886531 | rs2223814 | -1.908 | 1.889 | 0.415 | 1.017 | 0.044 | 0.065 | 0.065 | 3.534 | 0.19 | 0.5 | 0.27 | 0.31 | 0.22 | 0.14 |
| 6 | 103889026 | rs4945866 | -1.339 | 1.558 | 0.388 | 1.038 | 0.042 | 0.257 | 0.257 | 5.956 | 0.08 | 0.43 | 0.13 | 0.31 | 0.08 | 0.07 |
| 6 | 103895135 | rs2207016 | -0.227 | 1.497 | 0.204 | 0.997 | 0 | -0.01 | -0.01 | 2.134 | 0.1 | 0.08 | 0.12 | 0 | 0.09 | 0.07 |

| | | | | | | | | | | | | | | | | |
|---|---|---|---|---|---|---|---|---|---|---|---|---|---|---|---|---|
| 6 | 103907058 | rs6941032 | -1.58 | 1.489 | 0.168 | 1.159 | 0.011 | 0.066 | 0.066 | 1.292 | 0.23 | 0.5 | 0.33 | 0.35 | 0.21 | 0.18 |
| 6 | 103913327 | rs4551211 | -1.864 | 1.572 | 0.191 | 1.148 | 0.025 | 0.05 | 0.05 | 1.564 | 0.24 | 0.54 | 0.36 | 0.35 | 0.24 | 0.19 |
| 6 | 103916711 | rs6910997 | -1.313 | 1.611 | -0.115 | 1.177 | -0.005 | -0.062 | -0.062 | 0.011 | 0.19 | 0.38 | 0.22 | 0.06 | 0.14 | 0.12 |
| 6 | 103917357 | rs9499444 | -1.345 | 1.606 | -0.189 | 1.151 | 0.004 | -0.033 | -0.033 | 0.486 | 0.19 | 0.29 | 0.24 | 0.06 | 0.17 | 0.12 |
| 6 | 103924015 | rs9499449 | -0.826 | 1.401 | -0.246 | 1.232 | -0.029 | 0.017 | 0.017 | 0.431 | 0.07 | 0.27 | 0.1 | 0.06 | 0.01 | 0.04 |
| 6 | 103924200 | rs9391023 | -1.35 | 1.437 | -0.301 | 1.223 | 0.005 | 0.058 | 0.058 | 1.099 | 0.24 | 0.51 | 0.34 | 0.38 | 0.22 | 0.18 |
| 6 | 103945290 | rs9373700 | -0.898 | 1.483 | -0.307 | 1.237 | 0.021 | 0.02 | 0.02 | 1.01 | 0.2 | 0.3 | 0.26 | 0.32 | 0.24 | 0.14 |
| 6 | 103945668 | rs2506721 | -0.462 | 1.744 | -1.252 | 1.278 | -0.049 | 0.008 | 0.008 | 0.232 | 0.67 | 0.95 | 0.83 | 0.76 | 0.68 | 0.79 |
| 6 | 103948927 | rs9386361 | -1.345 | 1.994 | 0.049 | 1.787 | 0.022 | 0.033 | 0.033 | 0.831 | 0.19 | 0.32 | 0.24 | 0.32 | 0.21 | 0.14 |
| 6 | 103982636 | rs9404414 | -1.335 | 1.593 | -0.679 | 1.95 | -0.045 | -0.051 | -0.051 | 0 | 0.08 | 0.02 | 0.09 | 0.3 | 0.15 | 0.04 |
| 6 | 103990472 | rs9499550 | -0.112 | 1.241 | -0.486 | 1.388 | -0.001 | 0.002 | 0.002 | 0 | 0.55 | 0.42 | 0.64 | 0.65 | 0.59 | 0.72 |
| 6 | 103996305 | rs2506754 | -1.012 | 1.297 | 0.087 | 1.414 | 0.073 | 0.19 | 0.19 | 0.435 | 0.08 | 0.39 | 0.07 | 0.17 | 0.07 | 0.06 |
| 6 | 103996442 | rs2506753 | -0.57 | 1.058 | 0.335 | 1.261 | -0.005 | 0.077 | 0.077 | 0 | 0.64 | 0.82 | 0.78 | 0.85 | 0.66 | 0.79 |
| 6 | 103998972 | rs7356965 | 0.062 | 1.254 | -0.148 | 1.575 | -0.001 | -0.001 | -0.001 | 0.657 | 0.33 | 0.34 | 0.49 | 0.37 | 0.36 | 0.53 |
| 6 | 104002330 | rs2503298 | -0.064 | 1.321 | 0.433 | 1.707 | 0.013 | 0.028 | 0.028 | 0 | 0.36 | 0.5 | 0.54 | 0.45 | 0.41 | 0.59 |
| 6 | 104003082 | rs2506739 | -0.826 | 1.074 | 0.224 | 1.333 | -0.024 | -0.027 | -0.027 | 0 | 0.61 | 0.55 | 0.82 | 0.77 | 0.76 | 0.7 |
| 6 | 104004347 | rs4133417 | -0.525 | 0.682 | 1.044 | 1.294 | 0.004 | 0.015 | 0.015 | 0 | 0.36 | 0.43 | 0.39 | 0.51 | 0.36 | 0.31 |
| 6 | 104015974 | rs10457116 | -0.559 | 0.806 | 1.333 | 1.371 | 0 | 0.001 | 0.001 | 0 | 0.34 | 0.37 | 0.39 | 0.48 | 0.36 | 0.3 |
| 6 | 104016730 | rs6923962 | -1.358 | 0.811 | 1.335 | 1.311 | -0.006 | -0.037 | -0.037 | 0 | 0.61 | 0.6 | 0.79 | 0.88 | 0.69 | 0.82 |
| 6 | 104016969 | rs7452689 | 0.694 | 0.92 | 1.074 | 1.415 | 0 | 0.015 | 0.015 | 0.353 | 0.11 | 0.11 | 0.3 | 0.31 | 0.15 | 0.34 |
| 6 | 104023932 | rs4130441 | 0.421 | 0.9 | 1.108 | 1.503 | -0.034 | -0.03 | -0.03 | 0 | 0.24 | 0.07 | 0.26 | 0.27 | 0.28 | 0.22 |
| 6 | 104031495 | rs9391068 | 0.066 | 0.929 | 1.17 | 1.514 | 0 | 0.003 | 0.003 | 0 | 0.28 | 0.3 | 0.28 | 0.4 | 0.29 | 0.23 |
| 6 | 104032936 | rs9499572 | -0.827 | 0.914 | 1.138 | 1.347 | 0.061 | 0.154 | 0.154 | 0.69 | 0.49 | 0.22 | 0.69 | 0.77 | 0.65 | 0.66 |
| 6 | 104045013 | rs9377531 | 0.015 | 0.878 | 1.156 | 0.953 | -0.004 | 0.002 | 0.002 | 0 | 0.28 | 0.28 | 0.22 | 0.29 | 0.32 | 0.26 |
| 6 | 104055272 | rs9404438 | 0.165 | 0.867 | 1.136 | 0.937 | 0 | -0.005 | -0.005 | 0 | 0.27 | 0.29 | 0.19 | 0.2 | 0.26 | 0.21 |
| 6 | 104058040 | rs4945902 | -1.219 | 0.603 | 1.063 | 0.906 | -0.005 | -0.006 | -0.006 | 1.011 | 0.57 | 0.49 | 0.63 | 0.68 | 0.64 | 0.77 |
| 6 | 104060926 | rs9391079 | -1.09 | 0.606 | 1.052 | 0.908 | 0.001 | 0.001 | 0.001 | 0 | 0.54 | 0.47 | 0.62 | 0.67 | 0.63 | 0.64 |
| 6 | 104065330 | rs13206744 | -0.09 | 0.614 | 1.008 | 0.925 | -0.005 | -0.036 | -0.036 | 0 | 0.26 | 0.15 | 0.4 | 0.39 | 0.31 | 0.39 |
| 6 | 104071986 | rs9499616 | -0.337 | 0.611 | 1.028 | 0.916 | -0.015 | -0.033 | -0.033 | 0 | 0.28 | 0.19 | 0.44 | 0.47 | 0.36 | 0.44 |
| 6 | 104072159 | rs4327729 | -1.06 | 0.591 | 1.037 | 0.892 | 0.001 | 0.001 | 0.001 | 0 | 0.54 | 0.47 | 0.62 | 0.67 | 0.62 | 0.64 |
| 6 | 104072481 | rs2501185 | -1.342 | 0.412 | 1.144 | 0.759 | -0.04 | -0.064 | -0.064 | 0 | 0.12 | 0.34 | 0.1 | 0.01 | 0.04 | 0.16 |
| 6 | 104075466 | rs12212731 | 0.633 | 0.454 | 0.975 | 0.746 | -0.054 | 0.124 | 0.124 | 0.243 | 0.23 | 0.05 | 0.17 | 0.1 | 0.25 | 0.2 |
| 6 | 104076572 | rs9399841 | -1.09 | 0.54 | 1.02 | 0.736 | 0 | 0.001 | 0.001 | 0.013 | 0.54 | 0.49 | 0.57 | 0.6 | 0.51 | 0.71 |
| 6 | 104077204 | rs9391087 | -1.135 | 0.556 | 1.017 | 0.733 | 0.003 | 0.006 | 0.006 | 0 | 0.57 | 0.49 | 0.59 | 0.6 | 0.59 | 0.72 |
| 6 | 104091611 | rs9499625 | -0.479 | 0.571 | 0.895 | 0.808 | -0.015 | -0.033 | -0.033 | 1.284 | 0.28 | 0.19 | 0.44 | 0.47 | 0.36 | 0.44 |
| 6 | 104097573 | rs9499627 | -0.048 | 0.466 | 0.807 | 0.781 | 0.008 | -0.033 | -0.033 | 0 | 0.24 | 0.15 | 0.37 | 0.38 | 0.24 | 0.38 |
| 6 | 104100800 | rs12210804 | 0.051 | 0.464 | 0.819 | 0.659 | -0.008 | -0.025 | -0.025 | 0.755 | 0.06 | 0.11 | 0.08 | 0 | 0.02 | 0.03 |
| 6 | 104102329 | rs13213991 | 0.991 | 0.474 | 0.636 | 0.727 | -0.004 | -0.014 | -0.014 | 0 | 0.11 | 0.01 | 0.07 | 0.07 | 0.09 | 0.05 |
| 6 | 104111556 | rs9499632 | -1.473 | 0.527 | 1.319 | 0.612 | 0.001 | -0.089 | -0.089 | 0 | 0.13 | 0.45 | 0.12 | 0 | 0.08 | 0.04 |
| 6 | 104112259 | rs9373721 | 0.53 | 0.592 | 1.018 | 0.648 | -0.026 | 0.04 | 0.04 | 0.008 | 0.24 | 0.09 | 0.18 | 0.22 | 0.28 | 0.21 |
| 6 | 104123344 | rs13197871 | 0.345 | 0.576 | 1.011 | 0.731 | -0.018 | -0.061 | -0.061 | 0 | 0.25 | 0.09 | 0.39 | 0.37 | 0.29 | 0.4 |
| 6 | 104123688 | rs9399846 | 0.474 | 0.65 | 1.072 | 0.72 | -0.026 | 0.041 | 0.041 | 0 | 0.24 | 0.09 | 0.18 | 0.22 | 0.28 | 0.21 |
| 6 | 104136200 | rs9386386 | -0.996 | 0.633 | 1.242 | 0.731 | 0.006 | 0.045 | 0.045 | 0.843 | 0.42 | 0.73 | 0.34 | 0.3 | 0.38 | 0.42 |
| 6 | 104151513 | rs4634493 | -0.879 | 0.909 | 1.123 | 0.848 | 0.039 | 0.032 | 0.032 | 0.638 | 0.09 | 0.29 | 0.12 | 0.07 | 0.08 | 0.07 |
| 6 | 104155910 | rs9391107 | -0.054 | 1.077 | 0.94 | 0.968 | -0.004 | 0.029 | 0.029 | 0 | 0.27 | 0.19 | 0.19 | 0.22 | 0.3 | 0.32 |
| 6 | 104163538 | rs9377569 | 0.474 | 1.104 | 0.726 | 1.03 | -0.068 | 0.007 | 0.007 | 0 | 0.22 | 0.05 | 0.18 | 0.25 | 0.28 | 0.19 |
| 6 | 104180114 | rs9377572 | -0.985 | 1.184 | 0.848 | 0.979 | -0.001 | -0.001 | -0.001 | 0 | 0.6 | 0.58 | 0.61 | 0.66 | 0.64 | 0.8 |
| 6 | 104181365 | rs6929257 | -0.97 | 1.2 | 0.838 | 0.993 | -0.001 | -0.001 | -0.001 | 0 | 0.59 | 0.55 | 0.57 | 0.59 | 0.59 | 0.74 |
| 6 | 104192796 | rs9377576 | 0.25 | 1.142 | 0.619 | 0.796 | -0.012 | 0.05 | 0.05 | 0 | 0.25 | 0.08 | 0.17 | 0.26 | 0.26 | 0.33 |
| 6 | 104194389 | rs12662867 | -0.014 | 1.281 | 0.698 | 0.865 | 0.004 | 0 | 0 | 0.794 | 0.33 | 0.25 | 0.38 | 0.3 | 0.31 | 0.42 |
| 6 | 104195078 | rs11756082 | 0.034 | 1.293 | 0.664 | 0.871 | 0.005 | 0 | 0 | 0.092 | 0.32 | 0.23 | 0.39 | 0.3 | 0.31 | 0.42 |
| 6 | 104198180 | rs9391114 | 0.256 | 1.309 | 0.729 | 0.909 | -0.002 | -0.018 | -0.018 | 0 | 0.26 | 0.3 | 0.18 | 0.26 | 0.27 | 0.32 |
| 6 | 104198552 | rs9404464 | -0.863 | 0.941 | -0.008 | 0.75 | 0 | -0.002 | -0.002 | 0 | 0.59 | 0.56 | 0.59 | 0.68 | 0.6 | 0.8 |
| 6 | 104199348 | rs9499661 | -0.663 | 1.563 | -0.104 | 0.706 | 0.027 | -0.034 | -0.034 | 0 | 0.06 | 0.17 | 0.08 | 0 | 0.08 | 0.03 |
| 6 | 104203193 | rs12208114 | -0.649 | 1.283 | -0.652 | 1.028 | -0.028 | -0.014 | -0.014 | 0 | 0.26 | 0.17 | 0.2 | 0.37 | 0.32 | 0.38 |
| 6 | 104214242 | rs9373729 | -0.727 | 1.287 | -0.485 | 0.854 | 0.001 | -0.002 | -0.002 | 0 | 0.4 | 0.5 | 0.4 | 0.65 | 0.38 | 0.53 |
| 6 | 104220230 | rs4351290 | -1.227 | 1.008 | 0.245 | 1.068 | -0.013 | 0.096 | 0.096 | 0.324 | 0.22 | 0.64 | 0.31 | 0.46 | 0.17 | 0.21 |
| 6 | 104222812 | rs7760658 | 0.562 | 1.792 | -0.39 | 0.604 | -0.056 | 0.012 | 0.012 | 0 | 0.74 | 0.88 | 0.67 | 0.72 | 0.61 | 0.57 |
| 6 | 104223877 | rs4610597 | -0.726 | 1.515 | 0.642 | 0.491 | 0.005 | -0.04 | -0.04 | 0 | 0.08 | 0.24 | 0.04 | 0 | 0.04 | 0.01 |
| 6 | 104224562 | rs4479968 | -0.256 | 1.498 | 0.75 | 0.731 | -0.011 | 0.215 | 0.215 | 0 | 0.13 | 0.38 | 0.19 | 0.34 | 0.12 | 0.15 |
| 6 | 104230129 | rs4291121 | 0.285 | 2.349 | 0.636 | 0.517 | -0.041 | 0.007 | 0.007 | 0 | 0.83 | 0.93 | 0.73 | 0.79 | 0.7 | 0.63 |
| 6 | 104230815 | rs12111484 | 1.075 | 2.35 | 0.779 | 0.625 | -0.026 | -0.009 | -0.009 | 0.048 | 0.75 | 0.82 | 0.64 | 0.56 | 0.6 | 0.55 |
| 6 | 104232520 | rs9499667 | -1.858 | 1.986 | 1.086 | 0.406 | -0.023 | 0.043 | 0.043 | 0 | 0.3 | 0.55 | 0.34 | 0.43 | 0.25 | 0.26 |
| 6 | 104234792 | rs9404475 | 1.674 | 2.434 | 0.908 | 0.367 | -0.002 | -0.007 | -0.007 | 1.212 | 0.66 | 0.35 | 0.49 | 0.57 | 0.62 | 0.69 |
| 6 | 104236919 | rs11156348 | -1.092 | 2.047 | 0.884 | 0.677 | -0.009 | 0.003 | 0.003 | 0 | 0.1 | 0.09 | 0.17 | 0.36 | 0.11 | 0.12 |
| 6 | 104240245 | rs4623274 | 0.565 | 2.048 | 0.896 | 0.669 | -0.016 | -0.011 | -0.011 | 0 | 0.81 | 0.85 | 0.64 | 0.67 | 0.68 | 0.58 |
| 6 | 104245438 | rs4632930 | 0.009 | 2.043 | 0.776 | 0.701 | -0.015 | -0.022 | -0.022 | 0 | 0.06 | 0.03 | 0.07 | 0.23 | 0.07 | 0.07 |
| 6 | 104245631 | rs13213026 | 1.305 | 1.994 | 1.098 | 0.634 | 0.045 | 0.034 | 0.034 | 0 | 0.72 | 0.62 | 0.56 | 0.44 | 0.6 | 0.53 |
| 6 | 104263238 | rs6904256 | -0.739 | 1.837 | 0.986 | 0.562 | -0.014 | -0.018 | -0.018 | 0 | 0.12 | 0.21 | 0.13 | 0.05 | 0.08 | 0.16 |

| | | | | | | | | | | | | | | | | |
|---|---|---|---|---|---|---|---|---|---|---|---|---|---|---|---|---|
| 6 | 104276001 | rs9499672 | 0.689 | 2.104 | 0.961 | 0.559 | 0.032 | -0.022 | -0.022 | 3.128 | 0.78 | 0.74 | 0.71 | 0.8 | 0.71 | 0.65 |
| 6 | 104279864 | rs9485899 | 1.183 | 1.937 | 0.919 | 0.604 | 0.034 | 0.003 | 0.003 | 0 | 0.71 | 0.67 | 0.62 | 0.57 | 0.61 | 0.57 |
| 6 | 104298866 | rs5011511 | -1.14 | 1.649 | 0.821 | 0.705 | 0.001 | -0.005 | -0.005 | 0 | 0.14 | 0.13 | 0.2 | 0.17 | 0.1 | 0.19 |
| 6 | 104302477 | rs4639362 | -0.801 | 1.488 | 0.794 | 0.65 | 0.006 | 0.011 | 0.011 | 0.001 | 0.12 | 0.07 | 0.13 | 0.05 | 0.09 | 0.14 |
| 6 | 104305047 | rs11156349 | -1.126 | 1.199 | 1.075 | 0.568 | -0.017 | 0.074 | 0.074 | 0 | 0.22 | 0.44 | 0.28 | 0.62 | 0.2 | 0.24 |
| 6 | 104307953 | rs9386391 | -1.558 | 1.921 | 0.948 | 0.628 | 0.011 | -0.01 | -0.01 | 0.003 | 0.29 | 0.63 | 0.33 | 0.31 | 0.31 | 0.5 |
| 6 | 104308569 | rs6571142 | 0.125 | 2.1 | 0.616 | 0.363 | -0.069 | 0.078 | 0.078 | 0 | 0.85 | 0.98 | 0.85 | 1 | 0.76 | 0.89 |
| 6 | 104315701 | rs3922556 | -1.054 | 2.122 | 1.024 | 0.562 | 0.016 | -0.08 | -0.08 | 5.131 | 0.18 | 0.43 | 0.2 | 0.07 | 0.18 | 0.28 |
| 6 | 104317458 | rs12190404 | 0.321 | 1.999 | 0.819 | 0.868 | -0.007 | 0.021 | 0.021 | 0 | 0.67 | 0.8 | 0.75 | 0.69 | 0.66 | 0.72 |
| 6 | 104321907 | rs7774633 | 1.835 | 1.89 | 0.489 | 0.962 | -0.026 | 0.171 | 0.171 | 0 | 0.45 | 0.16 | 0.31 | 0.24 | 0.43 | 0.37 |
| 6 | 104325755 | rs7761721 | 1.803 | 1.901 | 0.505 | 0.968 | -0.009 | 0.155 | 0.155 | 0 | 0.45 | 0.23 | 0.31 | 0.24 | 0.43 | 0.37 |
| 6 | 104347952 | rs11970747 | -0.848 | 1.891 | 1.202 | 1.792 | 0.031 | 0.265 | 0.265 | 6.893 | 0.07 | 0.43 | 0.12 | 0.17 | 0.02 | 0.04 |
| 6 | 104351361 | rs4489183 | -1.021 | 1.929 | 1.133 | 1.84 | 0.015 | 0.129 | 0.129 | 0 | 0.06 | 0.08 | 0.26 | 0.46 | 0.13 | 0.19 |
| 6 | 104353431 | rs7753976 | -0.745 | 1.829 | 1.225 | 1.707 | 0.023 | 0.055 | 0.055 | 6.358 | 0.07 | 0.39 | 0.03 | 0.04 | 0.01 | 0.03 |
| 6 | 104359363 | rs4463284 | -1.183 | 2.126 | 0.605 | 0.882 | -0.089 | -0.023 | -0.023 | 0.184 | 0.17 | 0.08 | 0.35 | 0.22 | 0.28 | 0.4 |
| 6 | 104363975 | rs11965133 | -0.278 | 2.123 | 0.633 | 0.876 | -0.035 | 0.019 | 0.019 | 0 | 0.12 | 0.06 | 0.23 | 0.06 | 0.18 | 0.23 |
| 6 | 104367936 | rs12526433 | -0.045 | 1.396 | 0.638 | 0.758 | -0.019 | 0.059 | 0.059 | 0 | 0.23 | 0.09 | 0.16 | 0.1 | 0.25 | 0.19 |
| 6 | 104377656 | rs4270819 | 1.013 | 1.411 | 0.945 | 0.896 | 0.025 | 0.196 | 0.196 | 5.905 | 0.92 | 0.56 | 0.93 | 0.86 | 0.98 | 0.94 |
| 6 | 104384978 | rs9377585 | 1.055 | 1.444 | 0.904 | 0.914 | 0.022 | 0.188 | 0.188 | 5.687 | 0.92 | 0.54 | 0.93 | 0.86 | 0.98 | 0.95 |
| 6 | 104403825 | rs4144638 | 0.425 | 1.751 | 0.491 | 0.942 | 0.041 | -0.004 | -0.004 | 0 | 0.78 | 0.66 | 0.63 | 0.74 | 0.72 | 0.57 |
| 6 | 104407049 | rs4495295 | 1.267 | 1.75 | -0.582 | 0.922 | 0.03 | 0.014 | 0.014 | 0.3 | 0.52 | 0.32 | 0.43 | 0.53 | 0.45 | 0.36 |
| 6 | 104407723 | rs6921682 | -0.15 | 1.7 | -0.903 | 0.298 | -0.004 | 0.047 | 0.047 | 0 | 0.24 | 0.15 | 0.16 | 0.1 | 0.25 | 0.19 |
| 6 | 104423624 | rs4357170 | 1.022 | 1.835 | -0.274 | 0.852 | 0.052 | -0.01 | -0.01 | 0 | 0.76 | 0.48 | 0.63 | 0.73 | 0.72 | 0.57 |
| 6 | 104445367 | rs9499708 | 0.084 | 1.375 | -1.542 | 1.03 | -0.15 | -0.092 | -0.092 | 6.34 | 0.69 | 0.97 | 0.73 | 0.69 | 0.63 | 0.53 |
| 6 | 104459663 | rs2205749 | 0.417 | 0.657 | -1.15 | 1.384 | -0.005 | 0.004 | 0.004 | 0 | 0.5 | 0.89 | 0.55 | 0.48 | 0.46 | 0.42 |
| 6 | 104462555 | rs2205748 | -0.188 | 0.238 | -0.636 | 1.537 | 0.006 | 0.008 | 0.008 | 0 | 0.54 | 0.76 | 0.59 | 0.64 | 0.45 | 0.35 |
| 6 | 104466287 | rs2205746 | -0.163 | 0.191 | -0.245 | 1.383 | 0 | 0.017 | 0.017 | 0 | 0.53 | 0.64 | 0.63 | 0.73 | 0.46 | 0.39 |
| 6 | 104469545 | rs6915190 | -0.206 | 0.259 | 0.166 | 1.29 | -0.008 | 0.046 | 0.046 | 0 | 0.54 | 0.7 | 0.63 | 0.73 | 0.47 | 0.39 |
| 6 | 104475450 | rs13210163 | 0.693 | -0.056 | -0.144 | 1.171 | 0.026 | 0.064 | 0.064 | 0 | 0.32 | 0.06 | 0.25 | 0.24 | 0.23 | 0.17 |
| 6 | 104478130 | rs2056928 | 0.747 | -0.126 | -0.276 | 1.137 | -0.006 | 0.038 | 0.038 | 0 | 0.31 | 0.03 | 0.24 | 0.24 | 0.23 | 0.17 |
| 6 | 104483487 | rs9399862 | -0.092 | -0.069 | -0.053 | 1.326 | 0.013 | 0.041 | 0.041 | 0 | 0.19 | 0.41 | 0.31 | 0.22 | 0.21 | 0.16 |
| 6 | 104484800 | rs9499726 | -0.092 | -0.069 | -0.196 | 1.326 | 0.02 | 0.054 | 0.054 | 0 | 0.19 | 0.44 | 0.31 | 0.22 | 0.21 | 0.17 |
| 6 | 104493098 | rs6919443 | -0.146 | 0.146 | -0.426 | 1.241 | 0.004 | 0.005 | 0.005 | 0 | 0.52 | 0.6 | 0.57 | 0.46 | 0.44 | 0.32 |
| 6 | 104496518 | rs1736577 | -0.607 | 0.586 | -1.302 | 0.929 | -0.128 | -0.004 | -0.004 | 0 | 0.72 | 0.97 | 0.73 | 0.84 | 0.61 | 0.54 |
| 6 | 104497838 | rs859985 | -0.128 | 0.404 | -0.858 | 1.15 | -0.018 | -0.004 | -0.004 | 0 | 0.58 | 0.7 | 0.63 | 0.54 | 0.5 | 0.42 |
| 6 | 104499121 | rs9404502 | 0.569 | 0.061 | -1.319 | 0.946 | 0.004 | 0.021 | 0.021 | 0 | 0.22 | 0.1 | 0.18 | 0.15 | 0.16 | 0.16 |
| 6 | 104500232 | rs859982 | 0.525 | -0.019 | -1.701 | 0.758 | -0.075 | -0.044 | -0.044 | 0 | 0.25 | 0.03 | 0.39 | 0.35 | 0.3 | 0.19 |
| 6 | 104501198 | rs859980 | -0.59 | -0.085 | -0.996 | 1.086 | -0.002 | 0.012 | 0.012 | 0 | 0.57 | 0.42 | 0.67 | 0.8 | 0.52 | 0.46 |
| 6 | 104503032 | rs6906370 | -0.131 | -0.133 | -0.915 | 1.488 | -0.018 | 0.092 | 0.092 | 0.046 | 0.1 | 0.28 | 0.05 | 0.2 | 0.04 | 0.08 |
| 6 | 104512664 | rs859969 | -0.515 | -0.007 | -1.212 | 1.27 | 0.007 | -0.003 | -0.003 | 0 | 0.56 | 0.26 | 0.66 | 0.62 | 0.55 | 0.44 |
| 6 | 104512842 | rs859967 | 0.058 | -0.006 | -1.355 | 1.119 | -0.092 | -0.055 | -0.055 | 0 | 0.45 | 0.07 | 0.62 | 0.54 | 0.47 | 0.35 |
| 6 | 104513200 | rs11156355 | 0.461 | 0.007 | -1.337 | 0.843 | -0.08 | -0.013 | -0.013 | 0 | 0.26 | 0.03 | 0.35 | 0.31 | 0.31 | 0.19 |
| 6 | 104515333 | rs9499734 | -0.201 | 0.063 | -0.574 | 0.814 | -0.005 | -0.034 | -0.034 | 0.7 | 0.11 | 0.15 | 0.03 | 0 | 0.06 | 0.05 |
| 6 | 104517073 | rs9391148 | -0.232 | 0.375 | -0.518 | 1.372 | -0.023 | 0 | 0 | 0 | 0.56 | 0.75 | 0.65 | 0.56 | 0.46 | 0.37 |
| 6 | 104518126 | rs9377599 | 0.164 | 0.271 | -0.139 | 1.469 | 0 | 0 | 0 | 0 | 0.46 | 0.43 | 0.62 | 0.4 | 0.42 | 0.29 |
| 6 | 104518954 | rs12110847 | -0.846 | 0.416 | 0.283 | 1.364 | 0.064 | 0.138 | 0.138 | 1.529 | 0.07 | 0.32 | 0.04 | 0.09 | 0.05 | 0.03 |
| 6 | 104519786 | rs9386402 | -0.327 | 0.46 | -0.322 | 1.17 | -0.024 | 0.054 | 0.054 | 0 | 0.57 | 0.74 | 0.67 | 0.66 | 0.47 | 0.37 |
| 6 | 104520128 | rs11156357 | -1.364 | 0.289 | 0.374 | 1.597 | -0.003 | 0.172 | 0.172 | 0.717 | 0.16 | 0.56 | 0.09 | 0.26 | 0.13 | 0.09 |
| 6 | 104521604 | rs9499736 | -0.48 | 0.263 | 0.66 | 1.118 | -0.003 | -0.024 | -0.024 | 0.508 | 0.1 | 0.18 | 0.05 | 0.05 | 0.1 | 0.08 |
| 6 | 104523342 | rs12203539 | -0.391 | 0.084 | 0.387 | 0.424 | -0.002 | -0.009 | -0.009 | 0 | 0.67 | 0.87 | 0.68 | 0.29 | 0.68 | 0.54 |
| 6 | 104524060 | rs2399978 | 0.547 | 0.295 | 0.09 | 0.007 | 0.017 | 0.094 | 0.094 | 0.265 | 0.51 | 0.31 | 0.46 | 0.16 | 0.55 | 0.48 |
| 6 | 104539777 | rs9322793 | 0.547 | 0.295 | 0.09 | 0.007 | 0.017 | 0.094 | 0.094 | 0.265 | 0.51 | 0.31 | 0.46 | 0.16 | 0.55 | 0.48 |
| 6 | 104545642 | rs4945574 | 1.117 | 0.251 | -0.2 | 0.422 | -0.038 | -0.06 | -0.06 | 2.064 | 0.09 | 0.01 | 0.19 | 0.26 | 0.12 | 0.26 |
| 6 | 104554681 | rs9391161 | 1.058 | 0.166 | 0.08 | -0.488 | -0.062 | 0.169 | 0.169 | 0.015 | 0.41 | 0.13 | 0.41 | 0.12 | 0.45 | 0.4 |
| 6 | 104559148 | rs9386407 | -0.09 | 0.207 | 0.097 | -0.439 | 0.004 | 0.049 | 0.049 | 0.123 | 0.55 | 0.64 | 0.49 | 0.16 | 0.56 | 0.49 |
| 6 | 104560332 | rs9485936 | -0.09 | 0.207 | 0.097 | -0.439 | 0.004 | 0.049 | 0.049 | 0.123 | 0.55 | 0.64 | 0.49 | 0.17 | 0.56 | 0.49 |
| 6 | 104583607 | rs13203780 | -0.039 | 0.23 | 0.257 | -0.317 | 0.004 | -0.03 | -0.03 | 0.272 | 0.11 | 0.32 | 0.05 | 0.05 | 0.1 | 0.08 |
| 6 | 104587040 | rs11156367 | 0.965 | 0.136 | 0.038 | -0.539 | -0.078 | 0.155 | 0.155 | 0.046 | 0.39 | 0.12 | 0.41 | 0.12 | 0.44 | 0.37 |
| 6 | 104592769 | rs4946271 | 0.438 | 0.15 | -0.038 | -0.477 | 0.002 | -0.034 | -0.034 | 0 | 0.44 | 0.56 | 0.54 | 0.16 | 0.45 | 0.37 |
| 6 | 104604684 | rs9499779 | -0.986 | 0.162 | -0.737 | -0.878 | -0.041 | 0.107 | 0.107 | 0 | 0.43 | 0.81 | 0.57 | 0.17 | 0.52 | 0.29 |
| 6 | 104608299 | rs1365683 | 1.425 | 0.199 | -0.824 | -0.795 | -0.028 | -0.032 | -0.032 | 0 | 0.1 | 0.01 | 0.08 | 0.08 | 0.11 | 0.04 |
| 6 | 104625659 | rs6915167 | 0.173 | 0.122 | 0.086 | -0.854 | 0.004 | 0.017 | 0.017 | 0.32 | 0.12 | 0.4 | 0.14 | 0.09 | 0.1 | 0.09 |
| 6 | 104628972 | rs2744161 | -0.615 | 0.118 | 0.003 | -0.928 | -0.026 | 0.058 | 0.058 | 0 | 0.34 | 0.76 | 0.45 | 0.07 | 0.42 | 0.2 |
| 6 | 104652538 | rs9485946 | 0.581 | 0.139 | -0.069 | -0.834 | -0.006 | 0.002 | 0.002 | 0 | 0.1 | 0.2 | 0.13 | 0.09 | 0.1 | 0.09 |
| 6 | 104653067 | rs2744204 | -1.019 | 0.364 | -0.191 | -0.899 | -0.041 | 0.11 | 0.11 | 0 | 0.42 | 0.81 | 0.56 | 0.16 | 0.51 | 0.29 |
| 6 | 104654028 | rs1158956 | -1.869 | 0.041 | -0.594 | -0.977 | -0.014 | 0.028 | 0.028 | 0 | 0.79 | 0.98 | 0.89 | 0.89 | 0.85 | 0.8 |
| 6 | 104676937 | rs2820247 | 0.194 | 0.51 | 0.619 | -0.829 | 0.026 | -0.078 | -0.078 | 0.015 | 0.19 | 0.29 | 0.33 | 0.01 | 0.29 | 0.17 |
| 6 | 104681487 | rs2820263 | -1.081 | 0.464 | 0.495 | -0.77 | 0.002 | -0.008 | -0.008 | 0 | 0.4 | 0.61 | 0.54 | 0.18 | 0.51 | 0.3 |
| 6 | 104683494 | rs2744190 | -1.526 | 0.462 | -0.177 | -0.803 | -0.036 | 0.128 | 0.128 | 0.002 | 0.43 | 0.88 | 0.56 | 0.18 | 0.5 | 0.29 |

| | | | | | | | | | | | | | | | | |
|---|---|---|---|---|---|---|---|---|---|---|---|---|---|---|---|---|
| 6 | 104684669 | rs11156374 | 0.308 | 0.681 | 0.092 | -0.765 | 0.009 | -0.039 | -0.039 | 0 | 0.15 | 0.17 | 0.32 | 0.01 | 0.27 | 0.16 |
| 6 | 104690621 | rs201200 | 0.189 | 0.393 | 0.022 | -0.269 | -0.009 | -0.044 | -0.044 | 0 | 0.07 | 0 | 0.05 | 0.22 | 0.06 | 0.13 |
| 6 | 104697453 | rs2168261 | 0.077 | 0.35 | 0.558 | -0.583 | -0.014 | -0.002 | -0.002 | 0 | 0.1 | 0.22 | 0.13 | 0.09 | 0.1 | 0.1 |
| 6 | 104698658 | rs2744184 | 0.622 | 0.389 | -0.231 | -0.586 | -0.003 | -0.001 | -0.001 | 0.503 | 0.11 | 0.04 | 0.08 | 0.05 | 0.12 | 0.03 |
| 6 | 104703974 | rs2744183 | -1.631 | 0.373 | -0.266 | -0.733 | -0.024 | 0.118 | 0.118 | 0.142 | 0.4 | 0.84 | 0.53 | 0.15 | 0.49 | 0.27 |
| 6 | 104706579 | rs6571174 | -0.385 | 0.364 | 0.735 | -0.079 | 0.017 | -0.057 | -0.057 | 0.789 | 0.15 | 0.21 | 0.29 | 0.01 | 0.28 | 0.14 |
| 6 | 104730184 | rs9499822 | -1.092 | -0.229 | 1.148 | 0.552 | -0.033 | -0.004 | -0.004 | 0 | 0.13 | 0.59 | 0.15 | 0.08 | 0.09 | 0.09 |
| 6 | 104740181 | rs201202 | -0.41 | -0.296 | 0.327 | 0.576 | 0.019 | 0.048 | 0.048 | 0 | 0.32 | 0.14 | 0.28 | 0.23 | 0.31 | 0.26 |
| 6 | 104742765 | rs6942087 | -0.442 | -0.149 | 0.961 | 0.805 | 0.016 | 0.016 | 0.016 | 0 | 0.12 | 0.18 | 0.15 | 0.13 | 0.18 | 0.26 |
| 6 | 104745414 | rs9391179 | 1.281 | -0.224 | 1.39 | 0.7 | 0.017 | 0.038 | 0.038 | 0 | 0.49 | 0.35 | 0.54 | 0.64 | 0.51 | 0.46 |
| 6 | 104750318 | rs2604500 | -0.456 | -0.507 | 1.334 | 0.686 | 0.042 | 0.067 | 0.067 | 0 | 0.34 | 0.16 | 0.29 | 0.23 | 0.3 | 0.31 |
| 6 | 104756256 | rs2198337 | -0.141 | -0.843 | 1.585 | 0.712 | -0.034 | 0 | 0 | 0 | 0.27 | 0.36 | 0.23 | 0.28 | 0.15 | 0.22 |
| 6 | 104756449 | rs2218984 | -1.023 | -0.787 | 1.532 | 0.733 | -0.025 | 0 | 0 | 0 | 0.41 | 0.53 | 0.34 | 0.4 | 0.33 | 0.41 |
| 6 | 104760556 | rs9386412 | 0.206 | -0.796 | 1.337 | 0.734 | 0.067 | -0.023 | -0.023 | 0 | 0.25 | 0.13 | 0.21 | 0.28 | 0.15 | 0.22 |
| 6 | 104761978 | rs736429 | -1.074 | -0.776 | 1.486 | 0.731 | -0.001 | 0 | 0 | 0.969 | 0.42 | 0.58 | 0.34 | 0.4 | 0.33 | 0.41 |
| 6 | 104765523 | rs1341131 | -0.034 | -0.683 | 1.135 | 0.629 | -0.018 | 0.02 | 0.02 | 0 | 0.1 | 0.03 | 0.17 | 0.01 | 0.16 | 0.09 |
| 6 | 104765922 | rs404900 | 0.116 | -0.762 | 1.234 | 0.769 | 0.003 | -0.002 | -0.002 | 0 | 0.71 | 0.69 | 0.78 | 0.7 | 0.84 | 0.77 |
| 6 | 104778803 | rs9404535 | 0.257 | -0.743 | 0.88 | 0.781 | 0.074 | -0.034 | -0.034 | 0 | 0.25 | 0.12 | 0.21 | 0.29 | 0.15 | 0.22 |
| 6 | 104783184 | rs9377625 | 1.602 | -0.562 | 1.009 | 0.826 | -0.032 | -0.038 | -0.038 | 0 | 0.43 | 0.18 | 0.44 | 0.56 | 0.49 | 0.48 |
| 6 | 104787528 | rs2054105 | 2.11 | -0.487 | 0.634 | 1.457 | -0.115 | 0.025 | 0.025 | 0 | 0.35 | 0.04 | 0.37 | 0.31 | 0.39 | 0.39 |
| 6 | 104792647 | rs270695 | 0.645 | -1.461 | 1.136 | 1.088 | 0.025 | -0.014 | -0.014 | 2.122 | 0.63 | 0.7 | 0.69 | 0.59 | 0.68 | 0.58 |
| 6 | 104792894 | rs12215393 | 1.747 | -1.462 | 1.214 | 1.079 | 0.002 | -0.006 | -0.006 | 0 | 0.45 | 0.32 | 0.45 | 0.56 | 0.51 | 0.46 |
| 6 | 104796488 | rs270690 | 0.495 | -1.47 | 1.286 | 1.076 | 0.043 | -0.043 | -0.043 | 0.039 | 0.63 | 0.81 | 0.71 | 0.59 | 0.68 | 0.58 |
| 6 | 104801107 | rs156149 | -1.183 | -1.594 | 1.095 | 0.981 | -0.003 | -0.107 | -0.107 | 0.095 | 0.23 | 0.53 | 0.26 | 0.03 | 0.21 | 0.15 |
| 6 | 104803190 | rs1417667 | 0.481 | -1.663 | 1.005 | 1.124 | 0.046 | -0.04 | -0.04 | 0.01 | 0.18 | 0.13 | 0.18 | 0.29 | 0.11 | 0.17 |
| 6 | 104817535 | rs443578 | 1.516 | -1.139 | 1.283 | 0.719 | 0.004 | -0.028 | -0.028 | 0 | 0.68 | 0.46 | 0.65 | 0.85 | 0.65 | 0.67 |
| 6 | 104817775 | rs17459015 | -0.422 | -1.125 | 1.076 | 1.325 | -0.054 | -0.03 | -0.03 | 0 | 0.11 | 0.01 | 0.1 | 0.12 | 0.18 | 0.2 |
| 6 | 104829526 | rs156135 | 1.597 | -1.44 | 1.33 | 1.444 | -0.015 | 0.001 | 0.001 | 0 | 0.65 | 0.41 | 0.79 | 0.67 | 0.75 | 0.68 |
| 6 | 104831503 | rs4946463 | -1.44 | -1.387 | 1.177 | 1.477 | 0.021 | -0.069 | -0.069 | 0.699 | 0.3 | 0.1 | 0.2 | 0.33 | 0.29 | 0.33 |
| 6 | 104853996 | rs9499858 | -1.024 | -1.282 | 1.621 | 1.563 | -0.017 | -0.021 | -0.021 | 0 | 0.22 | 0.38 | 0.09 | 0.14 | 0.19 | 0.21 |
| 6 | 104854997 | rs565729 | 1.171 | -1.538 | 1.577 | 1.833 | 0.004 | -0.002 | -0.002 | 0.251 | 0.59 | 0.39 | 0.71 | 0.61 | 0.66 | 0.62 |
| 6 | 104856170 | rs7740010 | 0.435 | -1.526 | 1.55 | 1.791 | 0.019 | -0.067 | -0.067 | 0.136 | 0.17 | 0.04 | 0.17 | 0.26 | 0.15 | 0.17 |
| 6 | 104856640 | rs267333 | 1.351 | -1.335 | 1.682 | 1.786 | 0.056 | 0.059 | 0.059 | 0 | 0.52 | 0.29 | 0.61 | 0.4 | 0.59 | 0.55 |
| 6 | 104872729 | rs6918717 | -0.279 | -1.253 | 1.083 | 1.867 | -0.006 | 0.037 | 0.037 | 0 | 0.07 | 0.09 | 0.11 | 0.19 | 0.07 | 0.07 |
| 6 | 104875654 | rs267336 | 1.339 | -1.383 | 0.861 | 1.738 | 0.099 | -0.004 | -0.004 | 0 | 0.51 | 0.14 | 0.67 | 0.6 | 0.64 | 0.57 |
| 6 | 104896294 | rs6928812 | -1.111 | -1.229 | 1.152 | 1.673 | -0.01 | 0.008 | 0.008 | 0 | 0.14 | 0.21 | 0.13 | 0.2 | 0.1 | 0.11 |
| 6 | 104896918 | rs3851208 | -1.68 | -1.274 | 1.473 | 1.672 | -0.034 | 0.02 | 0.02 | 0 | 0.18 | 0.32 | 0.16 | 0.21 | 0.11 | 0.13 |
| 6 | 104910848 | rs2084907 | -1.059 | -1.156 | 1.457 | 1.588 | -0.01 | 0.007 | 0.007 | 0 | 0.13 | 0.17 | 0.13 | 0.19 | 0.09 | 0.11 |
| 6 | 104913044 | rs408949 | 0.469 | -1.313 | 1.771 | 1.656 | 0.027 | 0.042 | 0.042 | 0 | 0.86 | 0.66 | 0.9 | 0.82 | 0.83 | 0.8 |
| 6 | 104917557 | rs156192 | 1.004 | -1.445 | 1.788 | 1.737 | -0.008 | 0.005 | 0.005 | 0 | 0.61 | 0.43 | 0.82 | 0.59 | 0.67 | 0.61 |
| 6 | 104919013 | rs1901708 | -0.39 | -1.319 | 1.716 | 1.749 | 0.024 | 0.036 | 0.036 | 0 | 0.14 | 0.28 | 0.09 | 0.18 | 0.17 | 0.2 |
| 6 | 104921314 | rs13207723 | -0.276 | -1.312 | 1.638 | 1.764 | 0.006 | 0.008 | 0.008 | 0 | 0.13 | 0.18 | 0.09 | 0.17 | 0.17 | 0.2 |
| 6 | 104921322 | rs12197133 | -0.85 | -1.275 | 1.055 | 1.818 | -0.013 | 0.003 | 0.003 | 0 | 0.12 | 0.18 | 0.11 | 0.14 | 0.09 | 0.11 |
| 6 | 104924823 | rs94802 | 1.736 | -1.203 | 0.662 | 1.72 | 0.038 | 0.071 | 0.071 | 0.072 | 0.46 | 0.11 | 0.66 | 0.4 | 0.55 | 0.49 |
| 6 | 104926159 | rs995291 | -0.093 | -1.228 | 0.688 | 1.749 | 0.027 | -0.01 | -0.01 | 0 | 0.25 | 0.16 | 0.1 | 0.24 | 0.17 | 0.19 |
| 6 | 104930625 | rs271870 | 1.708 | -0.288 | 1.401 | 1.719 | -0.013 | 0.047 | 0.047 | 0.12 | 0.72 | 0.38 | 0.76 | 0.63 | 0.72 | 0.69 |
| 6 | 104943423 | rs1341129 | -1.181 | -0.364 | 0.099 | 1.557 | 0.001 | 0.001 | 0.001 | 0 | 0.1 | 0.11 | 0.12 | 0.17 | 0.08 | 0.08 |
| 6 | 104948139 | rs271854 | 1.095 | 0.158 | 0.652 | 1.311 | 0.019 | 0.018 | 0.018 | 1.227 | 0.83 | 0.45 | 0.81 | 0.79 | 0.86 | 0.84 |
| 6 | 104952471 | rs6918769 | -0.163 | -0.752 | -0.192 | 2.215 | 0.054 | -0.099 | -0.099 | 0 | 0.16 | 0 | 0.05 | 0.25 | 0.1 | 0.08 |
| 6 | 104957450 | rs6906183 | -1.619 | -1.512 | 0.653 | 1.844 | 0.005 | -0.002 | -0.002 | 0.094 | 0.43 | 0.57 | 0.31 | 0.49 | 0.31 | 0.33 |
| 6 | 104966905 | rs450996 | 1.502 | -1.526 | 0.472 | 1.956 | 0.098 | -0.01 | -0.01 | 0 | 0.56 | 0.22 | 0.68 | 0.49 | 0.71 | 0.66 |
| 6 | 104972955 | rs9386420 | 0.408 | -1.559 | 0.535 | 1.954 | 0.018 | 0.014 | 0.014 | 1.451 | 0.84 | 0.42 | 0.83 | 0.83 | 0.86 | 0.83 |
| 6 | 104974611 | rs2185204 | 1.433 | -1.495 | 0.311 | 1.874 | 0.097 | -0.011 | -0.011 | 0 | 0.56 | 0.22 | 0.68 | 0.49 | 0.71 | 0.66 |
| 6 | 104978218 | rs7747489 | -1.06 | -2.149 | 0.517 | 0.978 | 0.002 | 0.002 | 0.002 | 0 | 0.27 | 0.26 | 0.29 | 0.2 | 0.2 | 0.23 |
| 6 | 104980478 | rs4142560 | 1.94 | -1.142 | -0.243 | 0.441 | -0.144 | -0.132 | -0.132 | 2.989 | 0.34 | 0.03 | 0.35 | 0.59 | 0.39 | 0.51 |
| 6 | 104980550 | rs1890512 | -0.202 | -1.324 | 0.349 | 0.039 | 0.011 | -0.052 | -0.052 | 0 | 0.16 | 0.21 | 0.2 | 0 | 0.25 | 0.07 |
| 6 | 104983139 | rs12204422 | -1.137 | -1.597 | 0.921 | 0.049 | 0.001 | -0.048 | -0.048 | 0.175 | 0.09 | 0.3 | 0.1 | 0 | 0.07 | 0.05 |
| 6 | 104990836 | rs9404560 | -1.051 | -1.579 | 0.933 | 0.072 | 0.038 | 0.033 | 0.033 | 0.029 | 0.44 | 0.31 | 0.35 | 0.31 | 0.33 | 0.4 |
| 6 | 104993712 | rs17521744 | 0.075 | -1.319 | 0.932 | 0.049 | 0.01 | 0.008 | 0.008 | 0.629 | 0.12 | 0.15 | 0.15 | 0.16 | 0.13 | 0.16 |
| 6 | 105033213 | rs4946608 | -0.98 | -1.467 | 1.323 | 0.038 | 0.008 | 0.008 | 0.008 | 0 | 0.37 | 0.3 | 0.34 | 0.16 | 0.26 | 0.27 |
| 6 | 105054947 | rs1541166 | -1.593 | -1.437 | 1.456 | 0.075 | 0.046 | 0.05 | 0.05 | 0 | 0.5 | 0.62 | 0.49 | 0.33 | 0.4 | 0.41 |
| 6 | 105060069 | rs1933500 | -0.944 | -1.439 | 1.179 | 0.048 | -0.01 | -0.039 | -0.039 | 0 | 0.08 | 0.12 | 0.08 | 0 | 0.07 | 0.05 |
| 6 | 105062381 | rs9391217 | -1.531 | -1.223 | 0.502 | -0.368 | -0.001 | -0.002 | -0.002 | 0 | 0.67 | 0.67 | 0.66 | 0.45 | 0.66 | 0.45 |
| 6 | 105066456 | rs7761328 | -1.611 | -1.493 | 0.809 | -0.318 | 0.055 | 0.068 | 0.068 | 0 | 0.5 | 0.69 | 0.5 | 0.34 | 0.4 | 0.41 |
| 6 | 105069088 | rs4946623 | -1.385 | -1.472 | 0.927 | -0.388 | -0.018 | -0.02 | -0.02 | 1.085 | 0.4 | 0.54 | 0.37 | 0.17 | 0.28 | 0.28 |
| 6 | 105073676 | rs1416041 | -0.555 | -1.123 | 0.473 | -0.256 | 0.05 | 0.011 | 0.011 | 0 | 0.22 | 0.08 | 0.23 | 0.15 | 0.13 | 0.18 |
| 6 | 105078118 | rs2185061 | -0.482 | -1.182 | 0.558 | -0.375 | -0.026 | -0.08 | -0.08 | 0.845 | 0.16 | 0.22 | 0.11 | 0 | 0.13 | 0.1 |
| 6 | 105079446 | rs6914213 | -1.79 | -0.902 | -0.308 | -0.052 | 0.026 | 0.011 | 0.011 | 0 | 0.48 | 0.85 | 0.52 | 0.42 | 0.41 | 0.4 |
| 6 | 105086125 | rs6940468 | -0.65 | -0.495 | 0.226 | 0.08 | 0.109 | 0.079 | 0.079 | 0.013 | 0.06 | 0.15 | 0.15 | 0.14 | 0.15 | 0.1 |

| | | | | | | | | | | | | | | | | |
|---|---|---|---|---|---|---|---|---|---|---|---|---|---|---|---|---|
| 6 | 105091850 | rs9377661 | -1.133 | -0.579 | 0.187 | 0.089 | 0.071 | 0.043 | 0.043 | 0 | 0.08 | 0.14 | 0.16 | 0.17 | 0.17 | 0.16 |
| 6 | 105101208 | rs4946632 | -0.4 | -0.866 | -0.3 | -0.028 | 0.021 | -0.018 | -0.018 | 0 | 0.14 | 0.07 | 0.19 | 0.13 | 0.1 | 0.16 |
| 6 | 105105089 | rs1074289 | -1.818 | -1.074 | -0.276 | -0.177 | 0.058 | 0.022 | 0.022 | 0 | 0.5 | 0.77 | 0.49 | 0.43 | 0.42 | 0.47 |
| 6 | 105131101 | rs6937026 | -0.527 | -0.977 | 0.545 | -0.271 | -0.032 | -0.094 | -0.094 | 0 | 0.19 | 0.29 | 0.12 | 0 | 0.13 | 0.16 |
| 6 | 105131373 | rs4946640 | -1.999 | -0.282 | -0.111 | -0.403 | 0.096 | 0.132 | 0.132 | 0 | 0.49 | 0.68 | 0.36 | 0.18 | 0.34 | 0.35 |
| 6 | 105133622 | rs6910034 | -2.319 | -0.224 | -0.366 | -0.638 | -0.031 | 0.081 | 0.081 | 0 | 0.56 | 0.84 | 0.54 | 0.31 | 0.48 | 0.46 |
| 6 | 105136569 | rs13194505 | -0.777 | -0.458 | -0.01 | -0.78 | 0.034 | 0.129 | 0.129 | 0.581 | 0.24 | 0.04 | 0.19 | 0.02 | 0.18 | 0.14 |
| 6 | 105153614 | rs9499927 | -0.424 | -0.171 | 0.666 | -0.595 | -0.006 | -0.048 | -0.048 | 0.137 | 0.25 | 0.3 | 0.14 | 0 | 0.15 | 0.13 |
| 6 | 105154521 | rs9404573 | -0.557 | 0.218 | 1.092 | -0.47 | 0.082 | 0.063 | 0.063 | 0 | 0.15 | 0.3 | 0.2 | 0.27 | 0.2 | 0.17 |
| 6 | 105155782 | rs7772567 | -2.173 | 0.194 | 0.854 | -0.309 | -0.049 | 0.001 | 0.001 | 0.036 | 0.53 | 0.89 | 0.52 | 0.43 | 0.47 | 0.43 |
| 6 | 105164613 | rs9499934 | -0.447 | -0.098 | 1.15 | -0.365 | -0.058 | -0.134 | -0.134 | 0 | 0.26 | 0.42 | 0.15 | 0 | 0.15 | 0.14 |
| 6 | 105171129 | rs7757634 | -0.309 | 0.167 | 1.066 | -0.166 | 0.002 | -0.011 | -0.011 | 0 | 0.1 | 0.07 | 0.18 | 0.15 | 0.12 | 0.14 |
| 6 | 105173534 | rs7755772 | -0.246 | 0.066 | 1.255 | -0.255 | 0.028 | 0.02 | 0.02 | 0 | 0.24 | 0.19 | 0.13 | 0 | 0.15 | 0.14 |
| 6 | 105180893 | rs17065282 | -0.352 | 0.072 | 1.042 | -0.238 | 0.003 | -0.011 | -0.011 | 0 | 0.1 | 0.07 | 0.18 | 0.16 | 0.12 | 0.14 |
| 6 | 105191814 | rs6942231 | -1.581 | 0.247 | 1.15 | -0.232 | -0.012 | 0.067 | 0.067 | 0 | 0.41 | 0.77 | 0.35 | 0.26 | 0.37 | 0.29 |
| 6 | 105211360 | rs6907016 | -2.186 | 0.257 | 1.328 | -0.254 | -0.024 | 0.035 | 0.035 | 0.178 | 0.51 | 0.85 | 0.53 | 0.41 | 0.48 | 0.43 |
| 6 | 105216405 | rs6922441 | -2.211 | 0.261 | 1.318 | -0.237 | -0.023 | 0.031 | 0.031 | 0.13 | 0.51 | 0.85 | 0.53 | 0.42 | 0.48 | 0.43 |
| 6 | 105230729 | rs7769270 | -2.171 | 0.253 | 1.313 | -0.293 | -0.02 | 0.042 | 0.042 | 0.277 | 0.51 | 0.85 | 0.53 | 0.41 | 0.48 | 0.42 |
| 6 | 105232233 | rs9322817 | -1.469 | 0.26 | 1.349 | -0.29 | -0.004 | 0.015 | 0.015 | 0 | 0.4 | 0.64 | 0.34 | 0.26 | 0.37 | 0.28 |
| 6 | 105235902 | rs4409213 | -0.214 | 0.248 | 1.486 | -0.313 | -0.013 | -0.062 | -0.062 | 0.475 | 0.24 | 0.25 | 0.13 | 0 | 0.15 | 0.13 |
| 6 | 105235975 | rs1378720 | -1.71 | 0.259 | 1.227 | -0.302 | 0.058 | 0.062 | 0.062 | 0.266 | 0.26 | 0.46 | 0.39 | 0.41 | 0.34 | 0.29 |
| 6 | 105239352 | rs17065321 | 0.376 | 0.299 | 0.406 | -0.271 | 0.001 | 0.023 | 0.023 | 5.87 | 0.07 | 0.04 | 0.04 | 0.03 | 0.07 | 0.09 |
| 6 | 105254000 | rs9486015 | -2.113 | 0.305 | 0.406 | -0.268 | -0.017 | 0.045 | 0.045 | 0.33 | 0.5 | 0.84 | 0.52 | 0.41 | 0.48 | 0.42 |
| 6 | 105266118 | rs4079063 | -2.113 | 0.305 | 0.406 | -0.268 | -0.017 | 0.049 | 0.049 | 0.33 | 0.5 | 0.84 | 0.52 | 0.41 | 0.48 | 0.42 |
| 6 | 105270890 | rs2486144 | 0.356 | 0.373 | 0.184 | -0.267 | 0.002 | -0.011 | -0.011 | 0.863 | 0.9 | 0.92 | 0.82 | 0.85 | 0.88 | 0.86 |
| 6 | 105288125 | rs6927608 | -0.227 | 0.373 | 0.184 | -0.267 | 0.002 | -0.011 | -0.011 | 0 | 0.1 | 0.07 | 0.18 | 0.15 | 0.12 | 0.14 |
| 6 | 105302222 | rs13215256 | -0.283 | 0.374 | 0.211 | -0.31 | 0.045 | 0.062 | 0.062 | 0 | 0.25 | 0.15 | 0.14 | 0 | 0.15 | 0.14 |
| 6 | 105311616 | rs6911031 | -0.264 | 0.677 | 0.13 | -0.041 | 0.009 | -0.001 | -0.001 | 1.821 | 0.12 | 0.12 | 0.18 | 0.19 | 0.21 | 0.15 |
| 6 | 105313205 | rs11156426 | -0.446 | 0.425 | -0.152 | 0.055 | 0.002 | -0.011 | -0.011 | 0 | 0.1 | 0.07 | 0.18 | 0.16 | 0.12 | 0.14 |
| 6 | 105315369 | rs12206525 | -0.462 | 0.424 | -0.152 | 0.046 | 0.003 | -0.011 | -0.011 | 0.861 | 0.1 | 0.07 | 0.18 | 0.16 | 0.12 | 0.14 |
| 6 | 105332814 | rs9404586 | 1.357 | 0.461 | -0.158 | -0.019 | -0.023 | 0.03 | 0.03 | 0.235 | 0.49 | 0.16 | 0.47 | 0.58 | 0.52 | 0.57 |
| 6 | 105347297 | rs9373782 | -0.826 | 0.202 | 0.804 | -0.265 | 0.013 | 0.016 | 0.016 | 0 | 0.17 | 0.3 | 0.17 | 0.22 | 0.16 | 0.14 |
| 6 | 105348544 | rs2153127 | 1.91 | 0.16 | 0.521 | -0.435 | -0.069 | 0.011 | 0.011 | 0.171 | 0.45 | 0.12 | 0.43 | 0.59 | 0.47 | 0.55 |
| 6 | 105348705 | rs6923490 | -0.885 | 0.256 | 1.013 | -0.463 | 0.02 | -0.003 | -0.003 | 0 | 0.2 | 0.25 | 0.21 | 0.23 | 0.24 | 0.17 |
| 6 | 105352035 | rs4945713 | -1.576 | 0.246 | 1.55 | -0.459 | -0.004 | -0.006 | -0.006 | 0 | 0.45 | 0.52 | 0.49 | 0.38 | 0.4 | 0.33 |
| 6 | 105378954 | rs7759938 | 1.585 | 0.139 | 1.679 | -0.574 | -0.001 | -0.004 | -0.004 | 0.682 | 0.64 | 0.41 | 0.72 | 0.69 | 0.65 | 0.7 |
| 6 | 105384492 | rs11156431 | -0.371 | 0.063 | 1.569 | -0.7 | -0.001 | -0.029 | -0.029 | 0.025 | 0.14 | 0.16 | 0.06 | 0 | 0.12 | 0.1 |
| 6 | 105401013 | rs9404590 | 0.175 | 0.022 | 1.415 | -0.646 | 0 | 0.001 | 0.001 | 0 | 0.12 | 0.12 | 0.16 | 0.27 | 0.18 | 0.11 |
| 6 | 105402083 | rs314279 | -0.09 | 0.058 | 1.039 | -0.656 | -0.015 | 0.02 | 0.02 | 1.506 | 0.93 | 0.96 | 0.94 | 0.97 | 0.88 | 0.91 |
| 6 | 105404198 | rs12194974 | 1.371 | 0.425 | 0.69 | -0.889 | -0.023 | 0.033 | 0.033 | 0 | 0.1 | 0.03 | 0.06 | 0 | 0.12 | 0.03 |
| 6 | 105406274 | rs17065417 | -0.007 | -0.226 | 1.559 | -0.078 | -0.003 | -0.033 | -0.033 | 0.423 | 0.11 | 0.13 | 0.04 | 0 | 0.1 | 0.08 |
| 6 | 105407662 | rs314277 | 0.691 | -0.214 | 1.436 | -0.064 | -0.057 | -0.126 | -0.126 | 0.03 | 0.77 | 0.54 | 0.88 | 0.96 | 0.84 | 0.85 |
| 6 | 105436053 | rs314286 | 0.776 | -0.203 | 1.435 | 0.16 | -0.001 | 0 | 0 | 0 | 0.66 | 0.65 | 0.7 | 0.69 | 0.65 | 0.72 |
| 6 | 105444058 | rs369065 | 0.769 | -0.174 | 0.914 | 0.42 | 0.002 | 0.011 | 0.011 | 0 | 0.65 | 0.48 | 0.68 | 0.62 | 0.64 | 0.72 |
| 6 | 105462004 | rs314272 | 1.525 | -0.118 | 0.461 | -0.065 | -0.007 | 0.122 | 0.122 | 0.001 | 0.51 | 0.13 | 0.64 | 0.69 | 0.55 | 0.61 |
| 6 | 105478927 | rs2057391 | -1.564 | -0.364 | 1.576 | 0.245 | -0.023 | -0.014 | -0.014 | 0 | 0.2 | 0.47 | 0.22 | 0.14 | 0.18 | 0.14 |
| 6 | 105489760 | rs7750532 | -1.609 | -0.325 | 1.718 | 0.367 | -0.016 | -0.002 | -0.002 | 0 | 0.18 | 0.62 | 0.23 | 0.14 | 0.15 | 0.09 |
| 6 | 105493055 | rs160595 | 1.675 | -0.541 | 1.897 | 0.529 | -0.058 | -0.008 | -0.008 | 0 | 0.78 | 0.56 | 0.77 | 0.83 | 0.85 | 0.86 |
| 6 | 105506242 | rs221626 | 1.416 | -0.495 | 1.6 | 0.532 | -0.006 | -0.001 | -0.001 | 0.241 | 0.8 | 0.78 | 0.78 | 0.83 | 0.85 | 0.87 |
| 6 | 105520630 | rs9404597 | -0.861 | 0.141 | 1.069 | 0.843 | -0.054 | -0.095 | -0.095 | 0.889 | 0.14 | 0 | 0.21 | 0.38 | 0.16 | 0.16 |
| 6 | 105538504 | rs221645 | 1.763 | -0.779 | 1.796 | -0.397 | -0.083 | -0.059 | -0.059 | 14.909 | 0.81 | 0.57 | 0.91 | 0.93 | 0.9 | 0.87 |
| 6 | 105541159 | rs221649 | 1.755 | -0.752 | 1.815 | -0.378 | -0.076 | -0.054 | -0.054 | 3.195 | 0.81 | 0.58 | 0.91 | 0.92 | 0.9 | 0.87 |
| 6 | 105544119 | rs17065477 | -1.519 | -0.906 | 1.94 | -0.548 | -0.016 | -0.019 | -0.019 | 0 | 0.13 | 0.22 | 0.05 | 0.05 | 0.07 | 0.09 |
| 6 | 105545057 | rs17774009 | -1.062 | -1.064 | 1.912 | -0.586 | 0.005 | 0.018 | 0.018 | 0.024 | 0.1 | 0.05 | 0.03 | 0.03 | 0.06 | 0.09 |
| 6 | 105546452 | rs221652 | 2.704 | -1.326 | 2.872 | -0.693 | 0.016 | 0.017 | 0.017 | 2.065 | 0.69 | 0.24 | 0.87 | 0.89 | 0.85 | 0.81 |
| 6 | 105548850 | rs221657 | 0.822 | -1.307 | 2.691 | -0.644 | 0.022 | -0.014 | -0.014 | 0 | 0.91 | 1 | 0.97 | 0.93 | 0.96 | 0.91 |
| 6 | 105552470 | rs221661 | 1.135 | -1.442 | 2.896 | -0.819 | -0.039 | 0.001 | 0.001 | 0 | 0.88 | 0.74 | 0.96 | 0.91 | 0.96 | 0.91 |
| 6 | 105553236 | rs6911494 | -2.182 | -1.482 | 3.218 | -0.848 | 0.055 | -0.007 | -0.007 | 1.091 | 0.08 | 0.52 | 0.06 | 0 | 0.05 | 0.03 |
| 6 | 105562120 | rs2001119 | 2.583 | -1.504 | 3.241 | -0.861 | 0.003 | 0.003 | 0.003 | 2.177 | 0.69 | 0.27 | 0.87 | 0.88 | 0.84 | 0.8 |
| 6 | 105565429 | rs964730 | 0.971 | -1.504 | 3.277 | -0.861 | -0.025 | -0.01 | -0.01 | 0 | 0.89 | 0.81 | 0.96 | 0.94 | 0.96 | 0.91 |
| 6 | 105566414 | rs9500032 | 2.541 | -1.495 | 3.253 | -0.857 | 0.003 | 0.002 | 0.002 | 2.124 | 0.69 | 0.27 | 0.87 | 0.88 | 0.84 | 0.8 |
| 6 | 105568685 | rs9404601 | 2.104 | -1.496 | 3.263 | -0.856 | -0.056 | -0.041 | -0.041 | 0 | 0.75 | 0.42 | 0.9 | 0.88 | 0.89 | 0.81 |
| 6 | 105573104 | rs17524448 | -0.079 | -1.149 | 3.007 | -1.124 | -0.059 | 0.077 | 0.077 | 0.003 | 0.14 | 0.01 | 0.13 | 0 | 0.24 | 0.03 |
| 6 | 105585511 | rs9404605 | 2.473 | -1.766 | 3.763 | -0.388 | -0.043 | -0.044 | -0.044 | 0 | 0.74 | 0.42 | 0.9 | 0.89 | 0.86 | 0.8 |
| 6 | 105587627 | rs1890228 | 1.583 | -2.066 | 3.813 | -0.349 | -0.055 | -0.026 | -0.026 | 0 | 0.87 | 0.62 | 0.95 | 0.94 | 0.96 | 0.9 |
| 6 | 105588638 | rs1338019 | 1.583 | -2.066 | 3.813 | -0.349 | -0.055 | -0.026 | -0.026 | 0.387 | 0.87 | 0.62 | 0.95 | 0.94 | 0.96 | 0.9 |
| 6 | 105596610 | rs6905202 | -1.296 | -2.065 | 3.81 | -0.349 | -0.004 | -0.013 | -0.013 | 0 | 0.12 | 0.12 | 0.05 | 0.03 | 0.07 | 0.09 |
| 6 | 105598184 | rs1190278 | 2.884 | -2.067 | 3.816 | -0.35 | 0.003 | 0.002 | 0.002 | 2.214 | 0.69 | 0.26 | 0.87 | 0.89 | 0.84 | 0.8 |
| 6 | 105599671 | rs4626463 | -1.439 | -2.065 | 3.806 | -0.349 | -0.011 | -0.014 | -0.014 | 0.029 | 0.12 | 0.2 | 0.05 | 0.05 | 0.07 | 0.09 |

| | | | | | | | | | | | | | | | | |
|---|---|---|---|---|---|---|---|---|---|---|---|---|---|---|---|---|
| 6 | 105605865 | rs1190281 | 1.366 | -2.064 | 3.777 | -0.348 | -0.022 | -0.01 | -0.01 | 0 | 0.88 | 0.82 | 0.96 | 0.94 | 0.95 | 0.91 |
| 6 | 105606018 | rs768781 | 2.023 | -2.06 | 3.571 | -0.358 | -0.049 | -0.023 | -0.023 | 0 | 0.84 | 0.48 | 0.92 | 0.92 | 0.91 | 0.89 |
| 6 | 105610946 | rs1190286 | 2.965 | -2.061 | 3.379 | -0.359 | 0.073 | 0.083 | 0.083 | 3.242 | 0.69 | 0.11 | 0.86 | 0.89 | 0.84 | 0.8 |
| 6 | 105612220 | rs11962089 | -2.059 | -2.017 | 3.101 | -0.339 | -0.003 | -0.068 | -0.068 | 0 | 0.16 | 0.35 | 0.08 | 0.03 | 0.11 | 0.1 |
| 6 | 105615994 | rs1190295 | 2.803 | -2.022 | 3.047 | -0.339 | -0.001 | -0.002 | -0.002 | 0 | 0.7 | 0.29 | 0.87 | 0.89 | 0.84 | 0.8 |
| 6 | 105618017 | rs923405 | 2.445 | -1.917 | 2.483 | -0.282 | -0.039 | -0.029 | -0.029 | 0.573 | 0.75 | 0.63 | 0.91 | 0.89 | 0.89 | 0.82 |
| 6 | 105624415 | rs1190270 | 1.823 | -1.661 | 1.851 | 0.021 | -0.048 | -0.023 | -0.023 | 0 | 0.84 | 0.5 | 0.92 | 0.92 | 0.91 | 0.9 |
| 6 | 105635404 | rs7745815 | -1.139 | -1.401 | 0.955 | -0.014 | 0.009 | 0.006 | 0.006 | 0 | 0.08 | 0.05 | 0.02 | 0.05 | 0.04 | 0.02 |
| 6 | 105637762 | rs1190307 | 0.998 | -1.575 | 0.945 | 0.263 | -0.005 | 0.011 | 0.011 | 0 | 0.93 | 0.91 | 0.85 | 0.85 | 0.96 | 0.91 |
| 6 | 105643136 | rs1341268 | 2.382 | -1.965 | 0.764 | 0.481 | 0.091 | -0.079 | -0.079 | 0 | 0.7 | 0.08 | 0.73 | 0.61 | 0.86 | 0.73 |
| 6 | 105647791 | rs1765924 | 1.414 | -1.6 | 1.38 | 0.484 | -0.016 | 0.055 | 0.055 | 0 | 0.91 | 0.77 | 0.83 | 0.82 | 0.95 | 0.87 |
| 6 | 105656760 | rs1149297 | 1.828 | -1.307 | 1.672 | 0.079 | -0.013 | -0.01 | -0.01 | 0 | 0.83 | 0.65 | 0.9 | 0.87 | 0.85 | 0.85 |
| 6 | 105660039 | rs7771542 | -1.706 | -1.332 | 1.225 | 0.062 | -0.008 | -0.003 | -0.003 | 0 | 0.11 | 0.2 | 0.04 | 0.08 | 0.07 | 0.07 |
| 6 | 105661450 | rs6911417 | -1.914 | -1.333 | 1.228 | 0.072 | -0.008 | -0.003 | -0.003 | 2.036 | 0.14 | 0.21 | 0.05 | 0.08 | 0.08 | 0.07 |
| 6 | 105670495 | rs3995549 | -1.362 | -1.263 | 1.158 | 0.073 | 0.003 | 0.053 | 0.053 | 0 | 0.11 | 0.3 | 0.08 | 0.15 | 0.09 | 0.11 |
| 6 | 105675144 | rs1190023 | -2.124 | -1.16 | 0.056 | 0.019 | 0.004 | -0.026 | -0.026 | 0 | 0.33 | 0.94 | 0.24 | 0.33 | 0.29 | 0.33 |
| 6 | 105682496 | rs947213 | -1.373 | -0.943 | 0.607 | -0.103 | 0.012 | -0.031 | -0.031 | 0 | 0.06 | 0.13 | 0.03 | 0 | 0.06 | 0.03 |
| 6 | 105691778 | rs1190027 | -1.773 | -1.783 | -0.283 | -0.834 | 0.025 | 0.073 | 0.073 | 0 | 0.49 | 0.88 | 0.31 | 0.38 | 0.41 | 0.4 |
| 6 | 105699964 | rs1149301 | -0.947 | -1.726 | -0.126 | -0.81 | 0.011 | -0.032 | -0.032 | 0 | 0.06 | 0.13 | 0.04 | 0 | 0.06 | 0.04 |
| 6 | 105701475 | rs17065667 | -0.763 | -2.059 | 0.051 | -0.962 | 0.025 | 0.008 | 0.008 | 4.99 | 0.23 | 0.13 | 0.12 | 0.24 | 0.16 | 0.24 |
| 6 | 105704168 | rs1189924 | -2.354 | -2.047 | -0.046 | -0.957 | 0.009 | 0.066 | 0.066 | 0.581 | 0.5 | 0.94 | 0.32 | 0.38 | 0.41 | 0.4 |
| 6 | 105704658 | rs17065694 | -0.364 | -2.301 | 0.461 | -0.991 | -0.003 | 0.004 | 0.004 | 0 | 0.07 | 0.03 | 0.09 | 0.07 | 0.08 | 0.03 |
| 6 | 105720483 | rs17065714 | -0.068 | -2.354 | 0.859 | -1.054 | 0.002 | -0.007 | -0.007 | 0 | 0.12 | 0.07 | 0.05 | 0 | 0.08 | 0.09 |
| 6 | 105724549 | rs1190052 | -1.675 | -2.547 | 1.604 | -1.177 | -0.03 | -0.065 | -0.065 | 0 | 0.2 | 0.47 | 0.14 | 0.1 | 0.14 | 0.17 |
| 6 | 105726036 | rs1051484 | -2.01 | -2.813 | 1.228 | -1.138 | 0.013 | 0.009 | 0.009 | 1.14 | 0.29 | 0.73 | 0.21 | 0.2 | 0.17 | 0.25 |
| 6 | 105732207 | rs1358878 | -1.56 | -2.846 | 1.48 | -1.107 | -0.038 | -0.028 | -0.028 | 0 | 0.25 | 0.62 | 0.19 | 0.14 | 0.14 | 0.21 |
| 6 | 105735164 | rs3799999 | -1.479 | -2.813 | 1.017 | -1.1 | -0.025 | -0.039 | -0.039 | 0.012 | 0.25 | 0.63 | 0.18 | 0.1 | 0.14 | 0.18 |
| 6 | 105741030 | rs2793399 | -1.199 | -2.992 | 0.795 | -1.167 | -0.007 | -0.005 | -0.005 | 0 | 0.25 | 0.3 | 0.13 | 0.15 | 0.16 | 0.22 |
| 6 | 105742642 | rs1190050 | -1.98 | -2.856 | 0.26 | -0.978 | 0.142 | 0.135 | 0.135 | 1.001 | 0.51 | 0.88 | 0.3 | 0.27 | 0.3 | 0.43 |
| 6 | 105745188 | rs1190053 | -1.074 | -2.048 | 0.641 | -0.689 | -0.005 | -0.005 | -0.005 | 0 | 0.1 | 0.13 | 0.06 | 0.07 | 0.08 | 0.12 |
| 6 | 105756525 | rs1149316 | -1.345 | -2.469 | 0.894 | -0.886 | -0.049 | 0.009 | 0.009 | 7.63 | 0.21 | 0.7 | 0.12 | 0.25 | 0.07 | 0.18 |
| 6 | 105757070 | rs1617445 | -2.271 | -3.085 | 0.024 | -1.512 | 0.068 | 0.042 | 0.042 | 0.387 | 0.53 | 0.96 | 0.31 | 0.33 | 0.29 | 0.43 |
| 6 | 105760925 | rs1149319 | -1.428 | -2.961 | 1.103 | -1.531 | -0.022 | -0.024 | -0.024 | 0 | 0.06 | 0.27 | 0.01 | 0 | 0 | 0.05 |
| 6 | 105761860 | rs1201442 | -1.401 | -2.963 | 1.162 | -1.532 | -0.026 | -0.028 | -0.028 | 0 | 0.05 | 0.2 | 0.01 | 0 | 0 | 0.05 |
| 6 | 105768250 | rs1149321 | -2.345 | -3.222 | 1.17 | -1.685 | 0.125 | 0.097 | 0.097 | 0.769 | 0.42 | 0.85 | 0.22 | 0.25 | 0.21 | 0.4 |
| 6 | 105771520 | rs2253957 | -1.332 | -3.263 | 1.354 | -1.696 | -0.021 | -0.024 | -0.024 | 0 | 0.21 | 0.74 | 0.08 | 0.07 | 0.07 | 0.2 |
| 6 | 105775152 | rs12189723 | -1.332 | -3.258 | 1.368 | -1.659 | 0.011 | 0.008 | 0.008 | 0 | 0.21 | 0.16 | 0.1 | 0.02 | 0.14 | 0.2 |
| 6 | 105776792 | rs6902415 | -0.963 | -3.273 | 1.416 | -1.665 | -0.05 | -0.055 | -0.055 | 0.395 | 0.18 | 0.62 | 0.07 | 0.07 | 0.07 | 0.15 |
| 6 | 105776866 | rs12192054 | -1.294 | -3.245 | 1.361 | -1.676 | 0.019 | 0.039 | 0.039 | 0 | 0.21 | 0.11 | 0.1 | 0.02 | 0.14 | 0.2 |
| 6 | 105780539 | rs7759457 | -2.314 | -3.19 | 1.201 | -1.632 | 0.134 | 0.106 | 0.106 | 1.485 | 0.43 | 0.87 | 0.22 | 0.25 | 0.21 | 0.42 |
| 6 | 105781538 | rs4532474 | -1.38 | -3.094 | 1.966 | -1.708 | -0.019 | -0.007 | -0.007 | 0.002 | 0.25 | 0.29 | 0.13 | 0.2 | 0.15 | 0.22 |
| 6 | 105783150 | rs4512258 | -2.019 | -3.036 | 2.23 | -1.711 | -0.032 | -0.033 | -0.033 | 0 | 0.37 | 0.53 | 0.19 | 0.2 | 0.21 | 0.29 |
| 6 | 105792819 | rs6936682 | -1.148 | -2.748 | 1.78 | -1.818 | -0.018 | -0.068 | -0.068 | 0 | 0.12 | 0.24 | 0.06 | 0 | 0.06 | 0.07 |
| 6 | 105809919 | rs17065857 | -0.684 | -2.751 | 1.619 | -1.889 | -0.012 | -0.055 | -0.055 | 0.035 | 0.15 | 0.17 | 0.05 | 0 | 0.08 | 0.12 |
| 6 | 105811285 | rs12191229 | -1.866 | -3.033 | 1.967 | -2.145 | -0.028 | -0.07 | -0.07 | 0.005 | 0.23 | 0.46 | 0.13 | 0.12 | 0.17 | 0.22 |
| 6 | 105811308 | rs4374856 | -2.428 | -3.086 | 1.508 | -2.164 | 0.094 | 0.236 | 0.236 | 2.176 | 0.43 | 0.9 | 0.21 | 0.12 | 0.24 | 0.34 |
| 6 | 105812640 | rs1078726 | -1.726 | -2.899 | 2.019 | -2.155 | -0.033 | -0.111 | -0.111 | 0.177 | 0.26 | 0.32 | 0.12 | 0 | 0.16 | 0.26 |
| 6 | 105815882 | rs9322836 | -2.361 | -2.983 | 1.859 | -2.071 | 0.057 | 0.117 | 0.117 | 5.309 | 0.33 | 0.82 | 0.16 | 0.09 | 0.21 | 0.29 |
| 6 | 105817378 | rs717324 | -1.516 | -2.877 | 2.221 | -2.048 | -0.018 | -0.167 | -0.167 | 2.81 | 0.22 | 0.49 | 0.09 | 0 | 0.14 | 0.14 |
| 6 | 105821449 | rs9486069 | -2.209 | -3.172 | 0.95 | -2.631 | 0.07 | 0.202 | 0.202 | 0.511 | 0.43 | 0.82 | 0.2 | 0 | 0.29 | 0.23 |
| 6 | 105821512 | rs9500087 | -2.152 | -3.341 | 1.117 | -2.672 | -0.009 | -0.031 | -0.031 | 0 | 0.31 | 0.62 | 0.15 | 0.09 | 0.21 | 0.29 |
| 6 | 105831571 | rs6900098 | -0.454 | -3.34 | 1.074 | -2.65 | 0.003 | -0.026 | -0.026 | 0 | 0.17 | 0.15 | 0.08 | 0 | 0.12 | 0.06 |
| 6 | 105841041 | rs7741545 | -2.709 | -3.398 | 0.981 | -2.586 | 0.128 | 0.241 | 0.241 | 0 | 0.49 | 0.83 | 0.23 | 0.15 | 0.33 | 0.38 |
| 6 | 105841952 | rs11758609 | -1.746 | -3.4 | 1.13 | -2.666 | -0.007 | -0.146 | -0.146 | 0 | 0.21 | 0.52 | 0.09 | 0.01 | 0.15 | 0.13 |
| 6 | 105845414 | rs9320114 | -1.349 | -3.116 | 0.427 | -2.629 | 0.013 | 0.005 | 0.005 | 0 | 0.29 | 0.24 | 0.14 | 0.06 | 0.21 | 0.23 |
| 6 | 105845906 | rs6910880 | -2.028 | -2.733 | 0.124 | -2.41 | 0.095 | 0.259 | 0.259 | 0 | 0.5 | 0.88 | 0.26 | 0.19 | 0.34 | 0.44 |
| 6 | 105854300 | rs6927596 | -1.625 | -2.316 | 0.672 | -2.585 | 0 | -0.076 | -0.076 | 0 | 0.45 | 0.55 | 0.23 | 0.03 | 0.36 | 0.43 |
| 6 | 105860107 | rs6920614 | -1.637 | -1.626 | 0.52 | -2.159 | 0.021 | 0.04 | 0.04 | 0 | 0.4 | 0.75 | 0.24 | 0.27 | 0.35 | 0.4 |
| 6 | 105875614 | rs6909720 | -2.04 | -1.955 | 1.7 | -2.619 | -0.001 | -0.114 | -0.114 | 0.482 | 0.18 | 0.57 | 0.11 | 0.03 | 0.17 | 0.17 |
| 6 | 105879268 | rs17065917 | -1.884 | -1.969 | 2.017 | -2.627 | 0.005 | -0.104 | -0.104 | 0.03 | 0.17 | 0.41 | 0.1 | 0.03 | 0.17 | 0.17 |
| 6 | 105888195 | rs17065925 | -1.696 | -1.973 | 2.181 | -2.629 | 0.002 | -0.075 | -0.075 | 1.595 | 0.16 | 0.29 | 0.09 | 0.03 | 0.17 | 0.16 |
| 6 | 105889825 | rs9486097 | -2.396 | -2.001 | 2.487 | -2.643 | -0.01 | -0.071 | -0.071 | 0.456 | 0.19 | 0.69 | 0.11 | 0.03 | 0.17 | 0.16 |
| 6 | 105898952 | rs10080910 | -2.141 | -1.475 | 2.278 | -2.561 | -0.007 | -0.06 | -0.06 | 0.437 | 0.19 | 0.25 | 0.09 | 0.03 | 0.18 | 0.17 |
| 6 | 105902893 | rs1325419 | -2.789 | -1.656 | 1.837 | -2.678 | -0.003 | -0.058 | -0.058 | 0 | 0.34 | 0.63 | 0.19 | 0.03 | 0.27 | 0.41 |
| 6 | 105906485 | rs9486103 | -2.778 | -1.767 | 1.886 | -2.657 | -0.008 | -0.072 | -0.072 | 0.308 | 0.34 | 0.62 | 0.18 | 0.03 | 0.27 | 0.36 |
| 6 | 105907955 | rs12196063 | -0.909 | -1.584 | 1.42 | -2.567 | 0.029 | 0.07 | 0.07 | 0 | 0.16 | 0.05 | 0.09 | 0 | 0.11 | 0.23 |
| 6 | 105914910 | rs9486111 | -2.916 | -1.58 | 1.459 | -2.569 | 0.052 | 0.152 | 0.152 | 0.77 | 0.35 | 0.85 | 0.2 | 0.03 | 0.26 | 0.36 |
| 6 | 105923060 | rs9486112 | -1.888 | -1.503 | 1.478 | -2.537 | 0 | -0.042 | -0.042 | 0 | 0.25 | 0.19 | 0.11 | 0 | 0.21 | 0.28 |
| 6 | 105926104 | rs2159209 | -1.728 | -1.222 | 1.769 | -2.24 | -0.004 | -0.072 | -0.072 | 0 | 0.14 | 0.69 | 0.1 | 0.03 | 0.08 | 0.1 |

| | | | | | | | | | | | | | | | | |
|---|---|---|---|---|---|---|---|---|---|---|---|---|---|---|---|---|
| 6 | 105933744 | rs7748192 | -1.281 | -0.937 | 1.98 | -1.902 | -0.07 | -0.119 | -0.119 | 0 | 0.2 | 0.37 | 0.09 | 0 | 0.08 | 0.21 |
| 6 | 105948763 | rs9486125 | -1.834 | -0.579 | 0.627 | -1.72 | -0.001 | -0.032 | -0.032 | 0 | 0.2 | 0.65 | 0.12 | 0.05 | 0.18 | 0.16 |
| 6 | 105951839 | rs1269287 | -0.926 | -0.327 | 0.281 | -1.284 | 0.016 | 0.033 | 0.033 | 0 | 0.26 | 0.21 | 0.15 | 0.06 | 0.22 | 0.32 |
| 6 | 105953724 | rs9486131 | -1.508 | 0.183 | 1.398 | -0.308 | 0.042 | 0.026 | 0.026 | 0.044 | 0.07 | 0.22 | 0.06 | 0.06 | 0.08 | 0.04 |
| 6 | 105960712 | rs7752055 | -2.535 | -0.123 | 0.981 | -0.598 | 0.024 | 0.054 | 0.054 | 4.157 | 0.19 | 0.84 | 0.15 | 0.06 | 0.11 | 0.28 |
| 6 | 105974727 | rs9399916 | 0.842 | 0.17 | 0.805 | 0.34 | 0.017 | -0.072 | -0.072 | 0 | 0.48 | 0.07 | 0.35 | 0.45 | 0.38 | 0.38 |
| 6 | 105978666 | rs9480578 | -1.723 | 0.193 | 1.615 | 0.336 | -0.004 | -0.09 | -0.09 | 2.892 | 0.14 | 0.56 | 0.12 | 0 | 0.1 | 0.25 |
| 6 | 105983239 | rs6919641 | -2.111 | 0.429 | 2.148 | 1.095 | -0.012 | 0.062 | 0.062 | 3.322 | 0.1 | 0.65 | 0.26 | 0.14 | 0.03 | 0.14 |
| 6 | 105987536 | rs9480580 | -0.661 | 0.681 | 1.827 | 1.207 | 0.015 | -0.026 | -0.026 | 0.234 | 0.05 | 0.16 | 0.07 | 0 | 0.03 | 0.14 |
| 6 | 105988260 | rs9373790 | 0.128 | 0.79 | 1.358 | 1.43 | -0.007 | -0.008 | -0.008 | 0 | 0.62 | 0.67 | 0.42 | 0.49 | 0.47 | 0.63 |
| 6 | 105991031 | rs4946672 | 0.219 | 0.914 | 0.757 | 0.769 | 0.011 | -0.008 | -0.008 | 0 | 0.61 | 0.48 | 0.57 | 0.6 | 0.47 | 0.62 |
| 6 | 105991917 | rs9372090 | 0.144 | 0.822 | 0.577 | 0.733 | 0.023 | -0.012 | -0.012 | 0 | 0.62 | 0.54 | 0.59 | 0.65 | 0.5 | 0.64 |
| 6 | 105997617 | rs6934122 | -0.868 | 0.79 | 0.71 | 1.646 | 0.039 | 0.322 | 0.322 | 0 | 0.06 | 0.16 | 0.11 | 0.49 | 0.06 | 0.12 |
| 6 | 106000830 | rs4946673 | -0.31 | 0.444 | 0.146 | 0.548 | 0.02 | 0.268 | 0.268 | 0.738 | 0.49 | 0.85 | 0.52 | 0.91 | 0.4 | 0.58 |
| 6 | 106006230 | rs9386455 | 0.584 | 0.194 | 0.446 | 0.661 | 0.024 | -0.004 | -0.004 | 0 | 0.34 | 0.15 | 0.34 | 0.26 | 0.26 | 0.34 |
| 6 | 106006902 | rs6914292 | -0.717 | 0.062 | 0.209 | 0.438 | -0.136 | 0.286 | 0.286 | 2.956 | 0.56 | 0.98 | 0.55 | 0.91 | 0.43 | 0.58 |
| 6 | 106007726 | rs9320121 | -1.567 | 0.056 | 1.312 | 0.519 | 0.024 | 0.23 | 0.23 | 3.733 | 0.16 | 0.48 | 0.17 | 0.65 | 0.12 | 0.23 |
| 6 | 106009443 | rs9398033 | -0.238 | 0.103 | 1.6 | 0.503 | 0.028 | 0.167 | 0.167 | 0.265 | 0.47 | 0.78 | 0.5 | 0.84 | 0.36 | 0.5 |
| 6 | 106011498 | rs17215314 | 0.292 | 0.061 | 1.495 | 0.573 | 0.009 | 0.04 | 0.04 | 0.144 | 0.4 | 0.29 | 0.45 | 0.76 | 0.33 | 0.47 |
| 6 | 106013013 | rs12214850 | -1.617 | 0.344 | 1.769 | 0.873 | 0.053 | 0.429 | 0.429 | 1.625 | 0.11 | 0.43 | 0.14 | 0.58 | 0.1 | 0.17 |
| 6 | 106014983 | rs9791195 | -0.467 | 0.433 | 1.346 | 0.888 | 0.023 | 0.294 | 0.294 | 0 | 0.06 | 0.14 | 0.12 | 0.5 | 0.06 | 0.11 |
| 6 | 106017976 | rs10457128 | -0.359 | 0.221 | 0.874 | -0.164 | 0.02 | 0.134 | 0.134 | 0 | 0.57 | 0.79 | 0.54 | 0.19 | 0.64 | 0.55 |
| 6 | 106035017 | rs7752220 | -0.816 | 0.281 | 1.392 | 0.034 | 0.011 | -0.029 | -0.029 | 1.346 | 0.08 | 0.31 | 0.04 | 0 | 0.04 | 0.12 |
| 6 | 106036533 | rs9386458 | 0.148 | 0.229 | 0.253 | -0.205 | -0.046 | 0.06 | 0.06 | 1.894 | 0.41 | 0.21 | 0.44 | 0.25 | 0.53 | 0.47 |
| 6 | 106040613 | rs2013092 | 0.063 | 0.227 | 0.258 | -0.193 | -0.032 | 0.066 | 0.066 | 0.952 | 0.42 | 0.25 | 0.44 | 0.23 | 0.52 | 0.46 |
| 6 | 106042885 | rs714530 | 0.468 | 0.233 | 0.159 | -0.176 | -0.139 | 0.092 | 0.092 | 1.454 | 0.35 | 0.04 | 0.37 | 0.16 | 0.46 | 0.39 |
| 6 | 106042948 | rs763131 | -0.931 | 0.306 | 0.247 | -0.207 | 0.009 | -0.015 | -0.015 | 1.224 | 0.15 | 0.63 | 0.1 | 0.09 | 0.12 | 0.19 |
| 6 | 106051024 | rs13207480 | 0.429 | 0.541 | -0.661 | -0.766 | -0.067 | 0.032 | 0.032 | 0.984 | 0.24 | 0.07 | 0.34 | 0.14 | 0.38 | 0.45 |
| 6 | 106051738 | rs6933379 | -0.55 | 0.298 | -0.626 | -0.423 | 0.053 | 0.148 | 0.148 | 0 | 0.53 | 0.75 | 0.5 | 0.16 | 0.61 | 0.58 |
| 6 | 106059644 | rs2027873 | 0.212 | 0.662 | -0.893 | 0.01 | -0.013 | -0.019 | -0.019 | 0.755 | 0.62 | 0.7 | 0.79 | 0.58 | 0.56 | 0.65 |
| 6 | 106060602 | rs1041914 | 0.178 | 0.642 | -0.706 | 0.262 | -0.016 | -0.036 | -0.036 | 0.338 | 0.63 | 0.71 | 0.79 | 0.52 | 0.56 | 0.64 |
| 6 | 106065730 | rs9372096 | 0.561 | 0.359 | -0.616 | -0.426 | -0.013 | 0.023 | 0.023 | 0.003 | 0.38 | 0.49 | 0.62 | 0.51 | 0.31 | 0.48 |
| 6 | 106073530 | rs17805224 | 0.247 | 0.385 | -0.623 | -0.54 | -0.001 | -0.035 | -0.035 | 0.121 | 0.25 | 0.26 | 0.16 | 0.06 | 0.23 | 0.16 |
| 6 | 106075247 | rs11152903 | 0.837 | 0.212 | -1.11 | -0.431 | -0.071 | 0.043 | 0.043 | 0.269 | 0.28 | 0.02 | 0.5 | 0.15 | 0.27 | 0.27 |
| 6 | 106080487 | rs4946676 | -1.063 | 0.329 | -1.328 | 0.045 | -0.017 | -0.051 | -0.051 | 0 | 0.88 | 0.96 | 0.92 | 0.74 | 0.86 | 0.89 |
| 6 | 106089212 | rs12663565 | -0.101 | 0.147 | -0.492 | -0.069 | 0.003 | 0.257 | 0.257 | 0.211 | 0.06 | 0.25 | 0.1 | 0.29 | 0.03 | 0.11 |
| 6 | 106091250 | rs1011659 | -0.214 | -0.149 | -0.549 | -0.574 | 0.001 | -0.001 | -0.001 | 0 | 0.38 | 0.61 | 0.28 | 0.42 | 0.33 | 0.39 |
| 6 | 106093733 | rs6931024 | -0.74 | 0.062 | -0.938 | -0.721 | -0.006 | -0.063 | -0.063 | 0 | 0.74 | 0.86 | 0.85 | 0.57 | 0.75 | 0.71 |
| 6 | 106094057 | rs9386463 | 1.048 | 0.114 | -1.341 | -1.189 | -0.058 | 0.184 | 0.184 | 0.078 | 0.45 | 0.12 | 0.63 | 0.21 | 0.52 | 0.44 |
| 6 | 106098475 | rs12206056 | 1.861 | 0.154 | -1.328 | -1.125 | -0.067 | 0.015 | 0.015 | 0 | 0.23 | 0.02 | 0.49 | 0.15 | 0.27 | 0.26 |
| 6 | 106101469 | rs6919834 | -0.037 | -0.001 | -0.821 | -1.032 | -0.04 | 0.094 | 0.094 | 0.055 | 0.2 | 0.56 | 0.18 | 0.35 | 0.15 | 0.25 |
| 6 | 106103769 | rs7771901 | -0.652 | 0.063 | -0.825 | -1.036 | -0.021 | 0.024 | 0.024 | 0.348 | 0.28 | 0.57 | 0.23 | 0.35 | 0.24 | 0.21 |
| 6 | 106109980 | rs11754094 | 1.447 | 0.271 | -0.673 | -0.677 | -0.081 | 0.028 | 0.028 | 0 | 0.25 | 0.04 | 0.5 | 0.15 | 0.3 | 0.27 |
| 6 | 106114885 | rs6568387 | -0.473 | 0.251 | -0.345 | -0.599 | 0.011 | 0.022 | 0.022 | 0 | 0.48 | 0.66 | 0.35 | 0.42 | 0.45 | 0.45 |
| 6 | 106115795 | rs7751764 | 0.398 | 0.411 | -0.393 | -0.858 | -0.001 | -0.048 | -0.048 | 0 | 0.34 | 0.31 | 0.2 | 0.06 | 0.35 | 0.25 |
| 6 | 106117118 | rs6568389 | 0.368 | 0.435 | -0.436 | -0.846 | -0.001 | -0.072 | -0.072 | 0 | 0.34 | 0.36 | 0.21 | 0.06 | 0.35 | 0.25 |
| 6 | 106123662 | rs1270380 | -0.488 | 0.501 | -0.715 | -0.401 | -0.01 | -0.016 | -0.016 | 0 | 0.62 | 0.71 | 0.43 | 0.54 | 0.6 | 0.62 |
| 6 | 106124324 | rs845858 | -0.674 | 0.559 | -0.875 | -0.412 | 0.006 | 0.034 | 0.034 | 0 | 0.83 | 0.77 | 0.9 | 0.71 | 0.81 | 0.85 |
| 6 | 106125974 | rs845860 | -0.887 | 0.555 | -0.911 | -0.387 | -0.004 | 0.053 | 0.053 | 0 | 0.84 | 0.75 | 0.91 | 0.71 | 0.85 | 0.86 |
| 6 | 106126253 | rs845861 | -0.16 | 0.496 | -0.734 | -0.518 | -0.013 | 0.089 | 0.089 | 2.782 | 0.28 | 0.48 | 0.23 | 0.46 | 0.23 | 0.3 |
| 6 | 106129782 | rs11152909 | 1.701 | 0.53 | -0.768 | -0.565 | -0.048 | 0.012 | 0.012 | 0.003 | 0.21 | 0.04 | 0.46 | 0.14 | 0.21 | 0.23 |
| 6 | 106132323 | rs845864 | 0.27 | 0.211 | 0.022 | -0.249 | -0.004 | 0.013 | 0.013 | 0 | 0.17 | 0.44 | 0.11 | 0.17 | 0.13 | 0.12 |
| 6 | 106140281 | rs1029321 | -0.466 | 0.304 | -0.202 | -0.388 | -0.009 | 0.076 | 0.076 | 1.439 | 0.3 | 0.49 | 0.25 | 0.51 | 0.26 | 0.33 |
| 6 | 106145993 | rs845854 | -0.348 | 0.391 | -0.238 | -0.804 | -0.007 | -0.085 | -0.085 | 0.742 | 0.27 | 0.39 | 0.18 | 0.04 | 0.24 | 0.29 |
| 6 | 106147299 | rs7749157 | 0.904 | 0.421 | -0.566 | -1.077 | -0.002 | 0.046 | 0.046 | 0 | 0.3 | 0.18 | 0.19 | 0.07 | 0.33 | 0.27 |
| 6 | 106149846 | rs2202768 | 1.053 | 0.486 | -0.78 | -0.753 | -0.095 | 0.052 | 0.052 | 0 | 0.27 | 0.03 | 0.49 | 0.14 | 0.29 | 0.28 |
| 6 | 106159033 | rs4946680 | 0.081 | 0.79 | -1.144 | -0.558 | -0.006 | 0.016 | 0.016 | 7.236 | 0.66 | 0.69 | 0.75 | 0.89 | 0.61 | 0.56 |
| 6 | 106160279 | rs9372100 | 0.863 | 0.809 | -1.426 | 0.071 | -0.066 | -0.14 | -0.14 | 0 | 0.33 | 0.05 | 0.53 | 0.45 | 0.33 | 0.38 |
| 6 | 106183999 | rs9480594 | -0.253 | 0.71 | -0.064 | 0.833 | 0.035 | 0.139 | 0.139 | 0 | 0.07 | 0.22 | 0.04 | 0.16 | 0.07 | 0.03 |
| 6 | 106189232 | rs7747094 | -0.339 | 0.685 | -0.072 | 0.694 | 0.024 | 0.157 | 0.157 | 0.819 | 0.16 | 0.46 | 0.18 | 0.33 | 0.17 | 0.2 |
| 6 | 106198018 | rs10782150 | 0.934 | 0.692 | -0.508 | 0.72 | -0.001 | -0.002 | -0.002 | 0 | 0.59 | 0.37 | 0.69 | 0.59 | 0.56 | 0.61 |
| 6 | 106205297 | rs1948999 | 1.134 | 0.7 | -0.545 | 0.855 | 0 | -0.001 | -0.001 | 0.745 | 0.58 | 0.32 | 0.69 | 0.45 | 0.54 | 0.49 |
| 6 | 106208888 | rs2136058 | -0.645 | 0.79 | -0.713 | 1.202 | 0.019 | -0.058 | -0.058 | 0 | 0.87 | 0.98 | 0.95 | 0.8 | 0.91 | 0.83 |
| 6 | 106212994 | rs1876551 | 1.502 | 0.746 | -1.192 | 0.936 | -0.002 | -0.003 | -0.003 | 0 | 0.38 | 0.37 | 0.23 | 0.3 | 0.27 | 0.19 |
| 6 | 106213051 | rs1876550 | 0.23 | 0.753 | -1.149 | 0.94 | -0.036 | -0.039 | -0.039 | 0.077 | 0.57 | 0.84 | 0.42 | 0.5 | 0.54 | 0.34 |
| 6 | 106220531 | rs1876548 | -0.936 | 0.73 | -1.261 | 0.972 | 0.017 | -0.026 | -0.026 | 0.301 | 0.87 | 0.94 | 0.95 | 0.79 | 0.92 | 0.83 |
| 6 | 106228704 | rs11152918 | 0.896 | 0.744 | -1.318 | 0.924 | -0.101 | 0.021 | 0.021 | 0.132 | 0.26 | 0.03 | 0.51 | 0.17 | 0.35 | 0.29 |
| 6 | 106229598 | rs7758554 | 0.305 | 0.839 | -1.093 | 0.935 | -0.006 | 0.082 | 0.082 | 0 | 0.79 | 0.71 | 0.87 | 0.66 | 0.78 | 0.78 |
| 6 | 106229781 | rs9486186 | 0.344 | 0.871 | -1.378 | 0.911 | 0.06 | 0.098 | 0.098 | 0 | 0.08 | 0.22 | 0.08 | 0.14 | 0.13 | 0.05 |

| | | | | | | | | | | | | | | | | |
|---|---|---|---|---|---|---|---|---|---|---|---|---|---|---|---|---|
| 6 | 106237505 | rs2202766 | 0.945 | 0.859 | -1.409 | 0.852 | -0.002 | -0.033 | -0.033 | 0.021 | 0.11 | 0.25 | 0.11 | 0.06 | 0.15 | 0.1 |
| 6 | 106239385 | rs2063201 | 1.015 | 0.907 | -1.44 | 0.878 | -0.008 | 0.045 | 0.045 | 0 | 0.68 | 0.48 | 0.75 | 0.6 | 0.63 | 0.69 |
| 6 | 106244548 | rs17066312 | 1.41 | 0.788 | -1.392 | 0.791 | -0.01 | -0.02 | -0.02 | 0 | 0.37 | 0.4 | 0.22 | 0.28 | 0.27 | 0.2 |
| 6 | 106244980 | rs1508371 | -0.348 | 0.758 | -1.341 | 0.861 | 0.011 | 0.023 | 0.023 | 0.018 | 0.18 | 0.47 | 0.18 | 0.22 | 0.24 | 0.12 |
| 6 | 106246658 | rs9320132 | -0.325 | 0.789 | -1.453 | 0.826 | -0.09 | -0.059 | -0.059 | 0 | 0.6 | 0.94 | 0.43 | 0.67 | 0.57 | 0.54 |
| 6 | 106247394 | rs1876560 | -0.536 | 0.735 | -1.473 | 0.814 | 0.003 | -0.01 | -0.01 | 0.193 | 0.42 | 0.52 | 0.68 | 0.37 | 0.56 | 0.46 |
| 6 | 106250151 | rs9373811 | 0.205 | 1.31 | -1.039 | 0.796 | 0.005 | -0.032 | -0.032 | 0.403 | 0.13 | 0.26 | 0.13 | 0.06 | 0.17 | 0.13 |
| 6 | 106250400 | rs9384583 | -0.74 | 1.271 | -1.116 | 0.837 | 0.032 | 0.006 | 0.006 | 0.001 | 0.22 | 0.5 | 0.21 | 0.23 | 0.28 | 0.17 |
| 6 | 106252343 | rs1876558 | -0.789 | 1.271 | -1.087 | 0.938 | -0.044 | -0.068 | -0.068 | 0.711 | 0.62 | 0.95 | 0.46 | 0.66 | 0.6 | 0.57 |
| 6 | 106252945 | rs1355024 | -0.504 | 1.273 | -1.053 | 0.925 | 0.041 | 0.028 | 0.028 | 0 | 0.19 | 0.48 | 0.2 | 0.23 | 0.26 | 0.16 |
| 6 | 106252999 | rs1355023 | -0.795 | 1.315 | -1.107 | 0.931 | 0.04 | 0.083 | 0.083 | 0 | 0.24 | 0.55 | 0.23 | 0.41 | 0.31 | 0.38 |
| 6 | 106253136 | rs1876555 | -0.757 | 1.331 | -1.282 | 0.934 | -0.045 | -0.061 | -0.061 | 0.316 | 0.62 | 0.95 | 0.46 | 0.68 | 0.6 | 0.57 |
| 6 | 106257080 | rs6937783 | 0.318 | 0.833 | -0.7 | -0.029 | -0.01 | -0.032 | -0.032 | 0 | 0.39 | 0.45 | 0.23 | 0.26 | 0.29 | 0.2 |
| 6 | 106261769 | rs17066362 | -0.299 | 1.132 | -0.383 | 0.432 | 0.005 | -0.032 | -0.032 | 1.019 | 0.11 | 0.25 | 0.11 | 0.06 | 0.13 | 0.11 |
| 6 | 106263076 | rs7738222 | 0.659 | 1.095 | -0.528 | 0.843 | 0.002 | 0.014 | 0.014 | 0 | 0.65 | 0.57 | 0.46 | 0.53 | 0.65 | 0.77 |
| 6 | 106264634 | rs1007664 | 1.185 | 0.477 | -0.99 | 0.779 | -0.038 | 0.09 | 0.09 | 0.107 | 0.46 | 0.05 | 0.33 | 0.24 | 0.42 | 0.36 |
| 6 | 106264730 | rs7748267 | -0.545 | 0.267 | -0.132 | 0.459 | -0.023 | -0.056 | -0.056 | 0 | 0.1 | 0.23 | 0.06 | 0 | 0.11 | 0.04 |
| 6 | 106266373 | rs2895618 | -1.415 | 0.655 | 0.232 | 0.833 | 0.001 | -0.003 | -0.003 | 0 | 0.44 | 0.5 | 0.52 | 0.47 | 0.51 | 0.57 |
| 6 | 106270505 | rs9386476 | -1.532 | 0.703 | 0.248 | 0.845 | 0.001 | -0.003 | -0.003 | 0 | 0.44 | 0.5 | 0.52 | 0.46 | 0.51 | 0.57 |
| 6 | 106282402 | rs7766016 | -1.848 | 1.208 | 0.672 | 1.442 | -0.009 | 0.052 | 0.052 | 0 | 0.29 | 0.45 | 0.41 | 0.51 | 0.31 | 0.33 |
| 6 | 106282540 | rs1567268 | -0.335 | 0.924 | 0.839 | 1.213 | 0.006 | 0 | 0 | 0.649 | 0.18 | 0.22 | 0.19 | 0.26 | 0.28 | 0.29 |
| 6 | 106282631 | rs9384586 | -0.928 | -0.463 | 1.208 | 1.003 | 0.016 | -0.018 | -0.018 | 0.012 | 0.13 | 0.07 | 0.23 | 0.2 | 0.07 | 0.21 |
| 6 | 106289336 | rs9486199 | -0.848 | -0.111 | 0.258 | 0.786 | 0.002 | -0.037 | -0.037 | 0 | 0.09 | 0.15 | 0.08 | 0 | 0.15 | 0.04 |
| 6 | 106296457 | rs6917664 | -1.122 | -0.329 | 0.443 | 1.511 | -0.029 | 0.071 | 0.071 | 2.314 | 0.2 | 0.28 | 0.28 | 0.32 | 0.11 | 0.17 |
| 6 | 106298461 | rs7765357 | -0.673 | -0.196 | 0.477 | 1.209 | -0.001 | 0 | 0 | 0.811 | 0.31 | 0.66 | 0.2 | 0.37 | 0.23 | 0.25 |
| 6 | 106303274 | rs11966261 | -0.695 | -0.626 | 0.249 | 1.379 | 0.087 | -0.109 | -0.109 | 0 | 0.33 | 0.89 | 0.23 | 0.42 | 0.23 | 0.31 |
| 6 | 106316554 | rs12199117 | -0.767 | 1.174 | -0.591 | 0.485 | -0.078 | 0.113 | 0.113 | 0 | 0.27 | 0.12 | 0.22 | 0 | 0.35 | 0.14 |
| 6 | 106317196 | rs9372110 | -1.345 | 1.032 | -0.19 | 2.021 | 0.005 | 0.122 | 0.122 | 0.626 | 0.05 | 0.08 | 0.25 | 0.39 | 0.07 | 0.1 |
| 6 | 106319854 | rs9486210 | -1.183 | 0.862 | 0.05 | 1.187 | -0.004 | -0.057 | -0.057 | 0.215 | 0.1 | 0.24 | 0.08 | 0 | 0.09 | 0.05 |
| 6 | 106320260 | rs9399953 | -1.366 | 1.507 | -0.282 | 1.623 | 0.01 | 0.016 | 0.016 | 0.023 | 0.25 | 0.33 | 0.55 | 0.56 | 0.29 | 0.45 |
| 6 | 106321833 | rs9486212 | -1.253 | 1.48 | -0.181 | 2.142 | -0.01 | -0.011 | -0.011 | 0 | 0.08 | 0.05 | 0.13 | 0.2 | 0.09 | 0.17 |
| 6 | 106321844 | rs9480610 | -1.489 | 1.173 | -0.396 | 1.887 | 0.002 | 0.024 | 0.024 | 0 | 0.24 | 0.32 | 0.48 | 0.46 | 0.2 | 0.43 |
| 6 | 106322750 | rs13437093 | -1.636 | 1.199 | -0.458 | 1.878 | 0.001 | 0.008 | 0.008 | 0 | 0.32 | 0.39 | 0.54 | 0.47 | 0.29 | 0.45 |
| 6 | 106325399 | rs7749121 | -0.838 | 0.52 | -0.549 | 1.146 | 0.014 | 0 | 0 | 0.82 | 0.39 | 0.65 | 0.56 | 0.63 | 0.3 | 0.46 |
| 6 | 106326030 | rs6917182 | -1.359 | 1.133 | -0.019 | 0.973 | 0.042 | 0.014 | 0.014 | 0 | 0.06 | 0.2 | 0.09 | 0.05 | 0.11 | 0.08 |
| 6 | 106328676 | rs6934413 | -0.641 | 1.269 | -0.711 | -0.178 | -0.023 | 0.073 | 0.073 | 0 | 0.29 | 0.18 | 0.17 | 0.03 | 0.33 | 0.2 |
| 6 | 106329055 | rs9386485 | -0.9 | 0.648 | -0.718 | 0.215 | 0.003 | 0.004 | 0.004 | 0 | 0.52 | 0.61 | 0.37 | 0.45 | 0.49 | 0.44 |
| 6 | 106329966 | rs4568493 | -0.172 | 0.302 | -0.312 | 0.177 | -0.011 | 0.049 | 0.049 | 0 | 0.29 | 0.19 | 0.17 | 0.1 | 0.34 | 0.24 |
| 6 | 106330503 | rs4072698 | 0.061 | 0.225 | 0.696 | 0.647 | 0.063 | 0.183 | 0.183 | 0.289 | 0.05 | 0.26 | 0.07 | 0.13 | 0.04 | 0.03 |
| 6 | 106331317 | rs4345426 | 0.65 | 0.146 | 0.612 | 0.189 | 0.013 | -0.055 | -0.055 | 1.023 | 0.9 | 0.59 | 0.92 | 1 | 0.92 | 0.98 |
| 6 | 106332553 | rs4075042 | 0.117 | 0.219 | 0.225 | 0.033 | -0.003 | 0.031 | 0.031 | 0 | 0.17 | 0.04 | 0.14 | 0.09 | 0.16 | 0.21 |
| 6 | 106337211 | rs7452412 | -0.321 | 0.204 | 0.784 | 0.059 | -0.004 | 0.005 | 0.005 | 0 | 0.21 | 0.21 | 0.18 | 0.22 | 0.19 | 0.24 |
| 6 | 106339446 | rs4530899 | 1.848 | 0.317 | -0.077 | 0.463 | -0.059 | -0.107 | -0.107 | 1.41 | 0.23 | 0.02 | 0.37 | 0.31 | 0.24 | 0.34 |
| 6 | 106345057 | rs7772593 | 0.239 | 0.272 | 0.558 | 0.434 | 0.016 | -0.043 | -0.043 | 1.654 | 0.09 | 0.29 | 0.08 | 0 | 0.08 | 0.02 |
| 6 | 106345283 | rs11152931 | 0.852 | 0.169 | 0.388 | 0.516 | -0.026 | -0.021 | -0.021 | 0 | 0.42 | 0.13 | 0.53 | 0.44 | 0.42 | 0.53 |
| 6 | 106346382 | rs7740356 | -0.231 | 0.252 | 0.589 | 0.583 | 0.021 | 0.036 | 0.036 | 0.363 | 0.11 | 0.39 | 0.11 | 0.08 | 0.1 | 0.04 |
| 6 | 106348246 | rs4946705 | 0.116 | 0.225 | 0.307 | 0.76 | -0.004 | 0.004 | 0.004 | 0 | 0.53 | 0.58 | 0.63 | 0.6 | 0.5 | 0.57 |
| 6 | 106350616 | rs9398051 | 0.861 | -0.028 | -0.027 | 0.801 | -0.07 | -0.011 | -0.011 | 0 | 0.23 | 0.07 | 0.16 | 0.2 | 0.3 | 0.19 |
| 6 | 106350725 | rs9386491 | 0.861 | -0.026 | -0.027 | 0.802 | -0.07 | -0.011 | -0.011 | 0 | 0.23 | 0.06 | 0.16 | 0.2 | 0.3 | 0.19 |
| 6 | 106353761 | rs10457138 | -0.617 | -0.086 | 0.458 | 0.901 | -0.009 | -0.003 | -0.003 | 0 | 0.31 | 0.59 | 0.28 | 0.36 | 0.29 | 0.23 |
| 6 | 106359032 | rs4945734 | 0.635 | -0.237 | 0.332 | 1.12 | 0.025 | -0.011 | -0.011 | 0 | 0.57 | 0.35 | 0.69 | 0.6 | 0.64 | 0.75 |
| 6 | 106367446 | rs12202386 | -0.38 | -0.287 | 0.474 | 1.435 | 0.037 | 0.048 | 0.048 | 0 | 0.37 | 0.13 | 0.21 | 0.22 | 0.32 | 0.22 |
| 6 | 106368709 | rs9386492 | 1.614 | -0.408 | 0.179 | 1.261 | -0.1 | -0.033 | -0.033 | 0 | 0.36 | 0.07 | 0.52 | 0.44 | 0.4 | 0.55 |
| 6 | 106369045 | rs6913299 | -0.752 | -0.121 | 0.793 | 1.476 | 0.043 | 0.044 | 0.044 | 0.466 | 0.08 | 0.35 | 0.07 | 0.05 | 0.05 | 0.07 |
| 6 | 106371088 | rs11152936 | 0.387 | -0.125 | 0.237 | 1.486 | 0.004 | -0.016 | -0.016 | 2.418 | 0.74 | 0.36 | 0.78 | 0.8 | 0.72 | 0.78 |
| 6 | 106373715 | rs7766801 | -0.087 | -0.256 | 0.11 | 1.449 | 0.021 | 0.018 | 0.018 | 0 | 0.29 | 0.12 | 0.15 | 0.2 | 0.22 | 0.2 |
| 6 | 106374015 | rs4946713 | -0.72 | -0.272 | 0.13 | 1.503 | 0.008 | 0.007 | 0.007 | 0 | 0.46 | 0.32 | 0.34 | 0.51 | 0.45 | 0.37 |
| 6 | 106374868 | rs7772171 | -0.252 | -0.282 | 0.148 | 1.521 | 0.002 | -0.004 | -0.004 | 0 | 0.3 | 0.28 | 0.21 | 0.36 | 0.23 | 0.22 |
| 6 | 106378009 | rs6913325 | -0.664 | -0.3 | 0.056 | 1.102 | 0.001 | 0.044 | 0.044 | 0 | 0.45 | 0.19 | 0.28 | 0.35 | 0.45 | 0.36 |
| 6 | 106379169 | rs11152938 | 1.193 | -0.218 | 0.079 | 1.153 | -0.037 | -0.023 | -0.023 | 0 | 0.46 | 0.14 | 0.61 | 0.56 | 0.51 | 0.56 |
| 6 | 106379699 | rs7760686 | 0.071 | -0.125 | 0.307 | 0.816 | -0.009 | -0.017 | -0.017 | 0 | 0.11 | 0.15 | 0.06 | 0.04 | 0.07 | 0.07 |
| 6 | 106385525 | rs6924145 | 0.529 | 0.021 | 0.259 | 0.846 | -0.003 | -0.003 | -0.003 | 0 | 0.67 | 0.4 | 0.77 | 0.73 | 0.71 | 0.72 |
| 6 | 106396686 | rs9486233 | -0.262 | 0.314 | 0.572 | 0.839 | -0.006 | -0.005 | -0.005 | 0 | 0.2 | 0.34 | 0.13 | 0.24 | 0.19 | 0.24 |
| 6 | 106397748 | rs6568420 | 0.683 | 0.881 | 0.412 | 1.064 | -0.004 | -0.004 | -0.004 | 0 | 0.53 | 0.33 | 0.65 | 0.51 | 0.51 | 0.57 |
| 6 | 106397895 | rs6926851 | -0.098 | 1.021 | 0.438 | 1.176 | 0.004 | 0.001 | 0.001 | 0 | 0.19 | 0.24 | 0.1 | 0.24 | 0.18 | 0.2 |
| 6 | 106399648 | rs6938669 | 0.562 | 0.88 | 0.11 | 1.034 | 0.027 | 0.112 | 0.112 | 0 | 0.76 | 0.43 | 0.78 | 0.57 | 0.74 | 0.77 |
| 6 | 106403729 | rs9320141 | -0.018 | 0.894 | -0.139 | 0.525 | -0.001 | 0.014 | 0.014 | 0 | 0.7 | 0.73 | 0.79 | 0.84 | 0.7 | 0.72 |
| 6 | 106406999 | rs9373821 | 0.579 | 0.881 | -0.329 | 0.875 | 0.008 | 0.005 | 0.005 | 0 | 0.42 | 0.45 | 0.62 | 0.61 | 0.45 | 0.48 |
| 6 | 106407386 | rs9386496 | -0.062 | 1.341 | -0.53 | 0.54 | -0.017 | 0.027 | 0.027 | 0.153 | 0.72 | 0.79 | 0.77 | 0.9 | 0.63 | 0.76 |

| | | | | | | | | | | | | | | | | |
|---|---|---|---|---|---|---|---|---|---|---|---|---|---|---|---|---|
| 6 | 106411339 | rs9373823 | -0.969 | 1.032 | -0.219 | 1.655 | -0.01 | 0.349 | 0.349 | 0 | 0.05 | 0.21 | 0.05 | 0.44 | 0.02 | 0.08 |
| 6 | 106414990 | rs9480625 | -1.678 | 1.492 | -0.443 | 1.342 | 0.009 | 0.062 | 0.062 | 0 | 0.19 | 0.3 | 0.18 | 0.5 | 0.19 | 0.22 |
| 6 | 106418389 | rs9480628 | -1.297 | 1.754 | -0.567 | 0.624 | 0.005 | 0.014 | 0.014 | 0 | 0.17 | 0.27 | 0.16 | 0.22 | 0.17 | 0.12 |
| 6 | 106425101 | rs7746706 | -2.123 | 1.996 | -0.655 | 0.535 | -0.01 | -0.006 | -0.006 | 0 | 0.24 | 0.3 | 0.24 | 0.26 | 0.28 | 0.22 |
| 6 | 106427506 | rs4945738 | -1.674 | 1.643 | -0.692 | 0.567 | -0.03 | -0.032 | -0.032 | 0 | 0.2 | 0.18 | 0.21 | 0.25 | 0.23 | 0.2 |
| 6 | 106434279 | rs9320142 | 1.301 | 1.475 | -0.427 | 0.335 | -0.001 | 0.002 | 0.002 | 0 | 0.84 | 0.82 | 0.87 | 0.84 | 0.86 | 0.89 |
| 6 | 106438420 | rs4318909 | 1.446 | 1.674 | -0.096 | 0.458 | 0.036 | 0.048 | 0.048 | 0 | 0.87 | 0.73 | 0.83 | 0.78 | 0.8 | 0.85 |
| 6 | 106456807 | rs6913174 | 0.997 | 1.714 | -0.388 | 0.3 | 0.004 | 0.003 | 0.003 | 0 | 0.88 | 0.85 | 0.83 | 0.8 | 0.81 | 0.88 |
| 6 | 106461835 | rs4275097 | 0.37 | 1.615 | -1.321 | -0.605 | -0.032 | 0.142 | 0.142 | 0.037 | 0.37 | 0.15 | 0.22 | 0.03 | 0.37 | 0.25 |
| 6 | 106468486 | rs9480634 | 0.006 | 1.175 | -1.236 | -0.553 | -0.037 | 0.134 | 0.134 | 2.556 | 0.56 | 0.79 | 0.7 | 0.82 | 0.57 | 0.77 |
| 6 | 106481498 | rs11152954 | 0.118 | 1.194 | -1.229 | -0.506 | -0.013 | -0.019 | -0.019 | 0 | 0.9 | 0.92 | 0.87 | 0.82 | 0.88 | 0.94 |
| 6 | 106482901 | rs6909087 | -0.005 | 1.204 | -1.267 | -0.5 | -0.039 | -0.048 | -0.048 | 0 | 0.9 | 0.97 | 0.88 | 0.82 | 0.88 | 0.94 |
| 6 | 106487292 | rs7769470 | 0.543 | 1.197 | -1.281 | -0.562 | -0.01 | 0.106 | 0.106 | 0.094 | 0.34 | 0.11 | 0.18 | 0 | 0.31 | 0.18 |
| 6 | 106504727 | rs4576290 | 0.415 | 1.245 | -0.809 | 0.212 | -0.006 | -0.008 | -0.008 | 0 | 0.9 | 0.9 | 0.87 | 0.82 | 0.88 | 0.94 |
| 6 | 106510995 | rs4555967 | 0.104 | 1.06 | -0.486 | -0.584 | -0.001 | -0.033 | -0.033 | 0.333 | 0.44 | 0.45 | 0.68 | 0.83 | 0.45 | 0.76 |
| 6 | 106511888 | rs6903235 | -1.116 | 1.138 | -0.534 | -0.926 | 0.004 | 0.07 | 0.07 | 0.204 | 0.71 | 0.89 | 0.87 | 1 | 0.73 | 0.9 |
| 6 | 106511932 | rs6925611 | -0.859 | 1.078 | 0.37 | 0.095 | 0.001 | 0.003 | 0.003 | 0.169 | 0.16 | 0.31 | 0.13 | 0.16 | 0.17 | 0.11 |
| 6 | 106514039 | rs9398059 | 0.32 | 0.988 | 0.233 | 0.29 | 0.006 | -0.009 | -0.009 | 0 | 0.05 | 0.02 | 0.04 | 0.1 | 0.04 | 0.06 |
| 6 | 106519665 | rs9398061 | -0.67 | 0.966 | 0.264 | 0.215 | -0.004 | 0.005 | 0.005 | 0 | 0.15 | 0.25 | 0.11 | 0.16 | 0.11 | 0.08 |
| 6 | 106521653 | rs9386510 | -0.917 | 0.901 | 0.306 | 0.419 | 0.007 | -0.003 | -0.003 | 0.453 | 0.25 | 0.38 | 0.18 | 0.2 | 0.23 | 0.12 |
| 6 | 106524830 | rs9320148 | -0.809 | 0.88 | 0.384 | 0.419 | 0.002 | -0.001 | -0.001 | 1.749 | 0.12 | 0.33 | 0.09 | 0.1 | 0.11 | 0.09 |
| 6 | 106525156 | rs4946721 | -1.174 | 0.933 | -0.389 | 0.463 | -0.001 | 0.004 | 0.004 | 0.152 | 0.37 | 0.7 | 0.27 | 0.3 | 0.35 | 0.21 |
| 6 | 106536253 | rs2185379 | -0.362 | 0.607 | -0.277 | 0.651 | -0.008 | 0.001 | 0.001 | 0 | 0.06 | 0.07 | 0.05 | 0.08 | 0.03 | 0.07 |
| 6 | 106537750 | rs6923419 | -0.825 | 0.956 | 0.418 | 1.357 | 0.086 | 0.201 | 0.201 | 0.463 | 0.09 | 0.4 | 0.1 | 0.2 | 0.11 | 0.07 |
| 6 | 106540276 | rs4946725 | -0.172 | 1.048 | -0.187 | 1.21 | -0.033 | -0.043 | -0.043 | 0 | 0.1 | 0.01 | 0.1 | 0.18 | 0.12 | 0.07 |
| 6 | 106544560 | rs12175062 | -0.236 | 1.121 | -0.157 | 1.218 | -0.017 | -0.038 | -0.038 | 0 | 0.11 | 0.04 | 0.1 | 0.23 | 0.13 | 0.07 |
| 6 | 106548730 | rs535780 | -1.168 | 1.018 | -0.383 | 0.925 | 0 | 0 | 0 | 0 | 0.38 | 0.35 | 0.3 | 0.34 | 0.4 | 0.42 |
| 6 | 106549398 | rs486522 | 0.001 | 0.857 | -0.431 | 0.358 | -0.007 | 0.008 | 0.008 | 0 | 0.17 | 0.09 | 0.15 | 0.02 | 0.2 | 0.22 |
| 6 | 106552031 | rs573869 | -0.641 | 0.842 | -0.476 | 0.465 | -0.001 | 0.002 | 0.002 | 0 | 0.26 | 0.22 | 0.19 | 0.11 | 0.27 | 0.35 |
| 6 | 106563554 | rs12526490 | 0.206 | -0.116 | -1.088 | 0.572 | 0.023 | -0.006 | -0.006 | 0 | 0.26 | 0.11 | 0.37 | 0.24 | 0.24 | 0.3 |
| 6 | 106564434 | rs535490 | 0.388 | -0.311 | -0.796 | -0.17 | -0.002 | -0.025 | -0.025 | 0 | 0.37 | 0.38 | 0.25 | 0.08 | 0.28 | 0.29 |
| 6 | 106568864 | rs693612 | -0.064 | -0.066 | -0.304 | 0.769 | 0.011 | 0.003 | 0.003 | 0 | 0.48 | 0.59 | 0.35 | 0.46 | 0.39 | 0.38 |
| 6 | 106570056 | rs526531 | -0.8 | 0.334 | -0.296 | 0.867 | -0.062 | -0.044 | -0.044 | 0 | 0.82 | 0.95 | 0.81 | 0.72 | 0.7 | 0.81 |
| 6 | 106574892 | rs9480642 | 0.656 | -0.418 | -0.435 | 0.742 | -0.042 | -0.02 | -0.02 | 0 | 0.25 | 0.51 | 0.15 | 0.21 | 0.18 | 0.13 |
| 6 | 106576986 | rs7761791 | -0.291 | -0.484 | -0.452 | 0.69 | 0.023 | -0.017 | -0.017 | 0 | 0.42 | 0.71 | 0.31 | 0.49 | 0.35 | 0.31 |
| 6 | 106583418 | rs1008945 | -0.866 | -0.66 | -0.611 | 0.297 | 0.042 | -0.003 | -0.003 | 0 | 0.45 | 0.35 | 0.34 | 0.56 | 0.37 | 0.48 |
| 6 | 106583964 | rs9320151 | 0.48 | -0.747 | -0.73 | 0.391 | 0.007 | -0.087 | -0.087 | 0.115 | 0.19 | 0.03 | 0.17 | 0.36 | 0.16 | 0.33 |
| 6 | 106585095 | rs9373835 | 1.262 | -0.716 | -0.59 | 0.242 | -0.002 | 0.004 | 0.004 | 0 | 0.39 | 0.42 | 0.51 | 0.45 | 0.4 | 0.47 |
| 6 | 106588806 | rs6568431 | -0.084 | -0.711 | -0.565 | 0.237 | 0.031 | 0 | 0 | 0 | 0.65 | 0.59 | 0.67 | 0.65 | 0.61 | 0.62 |
| 6 | 106589496 | rs6901662 | 1.462 | -0.795 | -0.598 | 0.182 | 0.003 | -0.033 | -0.033 | 0.295 | 0.37 | 0.18 | 0.46 | 0.43 | 0.39 | 0.45 |
| 6 | 106590363 | rs4946728 | -0.818 | -0.83 | -0.821 | 0.081 | 0.095 | -0.153 | -0.153 | 0 | 0.79 | 0.56 | 0.81 | 0.99 | 0.75 | 0.9 |
| 6 | 106591318 | rs17586975 | 0.638 | -0.923 | -0.945 | -0.032 | -0.035 | -0.044 | -0.044 | 0.077 | 0.11 | 0.01 | 0.15 | 0.2 | 0.12 | 0.07 |
| 6 | 106591448 | rs17625558 | -0.411 | -0.933 | -0.946 | -0.026 | -0.026 | -0.131 | -0.131 | 0 | 0.3 | 0.04 | 0.31 | 0.56 | 0.29 | 0.41 |
| 6 | 106591781 | rs6929396 | 0.775 | -0.918 | -0.899 | -0.048 | 0.02 | 0.008 | 0.008 | 0 | 0.15 | 0.13 | 0.05 | 0.01 | 0.09 | 0.1 |
| 6 | 106594224 | rs742109 | 0.164 | -0.852 | -0.958 | -0.023 | 0.025 | -0.042 | -0.042 | 0 | 0.62 | 0.54 | 0.66 | 0.78 | 0.58 | 0.86 |
| 6 | 106597639 | rs1012894 | -0.227 | -0.726 | -0.331 | 0.261 | 0.003 | -0.001 | -0.001 | 0 | 0.64 | 0.66 | 0.69 | 0.66 | 0.61 | 0.59 |
| 6 | 106598023 | rs1040411 | 0.281 | 0.016 | -0.736 | 0.753 | -0.006 | 0.023 | 0.023 | 1.447 | 0.46 | 0.3 | 0.6 | 0.63 | 0.51 | 0.49 |
| 6 | 106599710 | rs9386512 | -1.106 | 0.086 | -0.323 | 1.871 | -0.001 | 0.148 | 0.148 | 0 | 0.05 | 0.05 | 0.09 | 0.47 | 0.07 | 0.23 |
| 6 | 106601165 | rs9398069 | -1.252 | -0.436 | -0.225 | 0.887 | 0.027 | -0.002 | -0.002 | 0.327 | 0.47 | 0.63 | 0.41 | 0.51 | 0.39 | 0.53 |
| 6 | 106601936 | rs7772706 | -0.052 | -0.587 | -0.458 | 0.606 | -0.025 | -0.024 | -0.024 | 0 | 0.12 | 0.01 | 0.13 | 0.15 | 0.11 | 0.03 |
| 6 | 106604136 | rs12110990 | 0.118 | -0.735 | -0.024 | 0.649 | -0.011 | 0.046 | 0.046 | 0 | 0.06 | 0.23 | 0.03 | 0.11 | 0.03 | 0.06 |
| 6 | 106613091 | rs4946730 | -1.505 | -0.945 | 0.024 | 0.477 | -0.033 | -0.008 | -0.008 | 0 | 0.32 | 0.48 | 0.24 | 0.35 | 0.24 | 0.2 |
| 6 | 106616379 | rs9373836 | -0.154 | -0.85 | -0.144 | 0.485 | 0.001 | 0.003 | 0.003 | 0.118 | 0.16 | 0.1 | 0.18 | 0.14 | 0.16 | 0.34 |
| 6 | 106618785 | rs9486298 | -0.249 | -0.733 | 0.002 | 0.455 | -0.019 | -0.018 | -0.018 | 0 | 0.13 | 0.23 | 0.09 | 0.08 | 0.11 | 0.12 |
| 6 | 106623391 | rs9398070 | -1.421 | -0.488 | -0.333 | 0.577 | 0.015 | 0.005 | 0.005 | 0 | 0.48 | 0.58 | 0.42 | 0.5 | 0.39 | 0.53 |
| 6 | 106627910 | rs9398071 | 0.687 | 0.386 | -0.558 | -0.073 | 0.003 | 0.045 | 0.045 | 1.246 | 0.74 | 0.83 | 0.66 | 0.92 | 0.79 | 0.78 |
| 6 | 106631780 | rs1322178 | -0.362 | 1.793 | -0.368 | -0.108 | 0.034 | -0.008 | -0.008 | 0 | 0.06 | 0.06 | 0.1 | 0.03 | 0.19 | 0.19 |
| 6 | 106631860 | rs10484575 | -0.831 | 1.458 | -0.038 | -0.201 | -0.011 | -0.033 | -0.033 | 0 | 0.13 | 0.19 | 0.11 | 0.05 | 0.08 | 0.16 |
| 6 | 106642208 | rs9486302 | 0.202 | 1.219 | 0.626 | -0.368 | -0.013 | -0.035 | -0.035 | 0.012 | 0.16 | 0.27 | 0.13 | 0.07 | 0.13 | 0.15 |
| 6 | 106665523 | rs2299860 | 0.084 | 1.114 | 0.512 | 0.508 | -0.001 | -0.001 | -0.001 | 0 | 0.12 | 0.09 | 0.31 | 0.13 | 0.07 | 0.03 |
| 6 | 106697392 | rs12526431 | 0.319 | 1.106 | 0.429 | 0.504 | -0.012 | -0.026 | -0.026 | 0 | 0.11 | 0.01 | 0.29 | 0.13 | 0.07 | 0.03 |
| 6 | 106734040 | rs633724 | -2.259 | 1.34 | 0.577 | 0.557 | 0.027 | 0.032 | 0.032 | 0 | 0.34 | 0.54 | 0.54 | 0.44 | 0.35 | 0.4 |
| 6 | 106734408 | rs484621 | -2.4 | 1.301 | 0.536 | 0.525 | 0.006 | -0.011 | -0.011 | 0 | 0.49 | 0.8 | 0.67 | 0.51 | 0.48 | 0.55 |
| 6 | 106735484 | rs2787538 | -0.527 | 1.337 | 0.567 | 0.419 | -0.01 | -0.034 | -0.034 | 0 | 0.13 | 0.2 | 0.12 | 0.05 | 0.09 | 0.17 |
| 6 | 106740458 | rs490010 | -2.233 | 1.402 | 0.329 | 0.386 | -0.002 | 0.005 | 0.005 | 0.758 | 0.5 | 0.84 | 0.67 | 0.55 | 0.48 | 0.55 |
| 6 | 106749308 | rs12212740 | 1.277 | 1.09 | -0.142 | -0.017 | -0.023 | 0.046 | 0.046 | 0 | 0.14 | 0.01 | 0.08 | 0.08 | 0.13 | 0.05 |
| 6 | 106764866 | rs573775 | -1.401 | 1.064 | 0.177 | 0.088 | 0.06 | 0.084 | 0.084 | 0 | 0.23 | 0.53 | 0.25 | 0.35 | 0.27 | 0.37 |
| 6 | 106774030 | rs510432 | -1.72 | 0.982 | -0.078 | -0.068 | -0.023 | -0.006 | -0.006 | 0.374 | 0.45 | 0.81 | 0.64 | 0.56 | 0.43 | 0.55 |
| 6 | 106776081 | rs688810 | -0.77 | 0.81 | 0.061 | 0.024 | -0.019 | 0.07 | 0.07 | 0 | 0.26 | 0.48 | 0.23 | 0.38 | 0.21 | 0.3 |

| | | | | | | | | | | | | | | | | |
|---|---|---|---|---|---|---|---|---|---|---|---|---|---|---|---|---|
| 6 | 106788486 | rs9384603 | 0.059 | 0.987 | 0.145 | 0.1 | 0.051 | -0.017 | -0.017 | 0.537 | 0.06 | 0.1 | 0.1 | 0.03 | 0.14 | 0.23 |
| 6 | 106792518 | rs2787513 | -0.572 | 0.876 | 0.046 | -0.074 | -0.001 | 0 | 0 | 0 | 0.41 | 0.58 | 0.35 | 0.38 | 0.45 | 0.42 |
| 6 | 106795331 | rs6924580 | 1.651 | 0.735 | -0.096 | -0.087 | -0.02 | -0.03 | -0.03 | 0 | 0.39 | 0.21 | 0.28 | 0.45 | 0.44 | 0.39 |
| 6 | 106799104 | rs683456 | -0.167 | 0.747 | -0.035 | -0.049 | -0.005 | 0.002 | 0.002 | 0 | 0.35 | 0.44 | 0.24 | 0.35 | 0.31 | 0.19 |
| 6 | 106800346 | rs9384604 | -0.157 | 0.79 | -0.064 | -0.089 | 0 | -0.001 | -0.001 | 0.284 | 0.36 | 0.44 | 0.29 | 0.35 | 0.4 | 0.24 |
| 6 | 106800661 | rs11152977 | 1.305 | 0.796 | -0.309 | 0.008 | -0.002 | -0.027 | -0.027 | 0 | 0.38 | 0.24 | 0.24 | 0.45 | 0.35 | 0.35 |
| 6 | 106803325 | rs503518 | -0.429 | 0.716 | 0.067 | -0.293 | 0.032 | -0.019 | -0.019 | 0 | 0.07 | 0.13 | 0.1 | 0.03 | 0.14 | 0.23 |
| 6 | 106815036 | rs3862833 | 0.905 | 0.391 | 0.082 | -0.275 | 0.01 | 0.033 | 0.033 | 0 | 0.62 | 0.74 | 0.64 | 0.74 | 0.64 | 0.6 |
| 6 | 106817806 | rs4946744 | 1.029 | 0.327 | 0.18 | -0.307 | 0.009 | 0.024 | 0.024 | 0 | 0.61 | 0.69 | 0.64 | 0.74 | 0.64 | 0.6 |
| 6 | 106817850 | rs4946745 | 0.34 | 0.256 | 0.105 | -0.373 | -0.01 | 0.017 | 0.017 | 0.728 | 0.75 | 0.79 | 0.73 | 0.89 | 0.69 | 0.69 |
| 6 | 106821165 | rs17251215 | -0.058 | 0.349 | -0.032 | -0.401 | -0.018 | 0.042 | 0.042 | 0.129 | 0.2 | 0.09 | 0.24 | 0.08 | 0.27 | 0.22 |
| 6 | 106821393 | rs17066910 | 0.365 | 0.373 | 0.161 | -0.331 | 0 | 0 | 0 | 2.593 | 0.07 | 0.1 | 0.09 | 0.15 | 0.03 | 0.02 |
| 6 | 106821428 | rs2787520 | 0.747 | 0.442 | 0.726 | -0.358 | 0.017 | 0.04 | 0.04 | 0 | 0.58 | 0.28 | 0.6 | 0.74 | 0.63 | 0.6 |
| 6 | 106823539 | rs3851212 | -0.284 | 0.674 | 0.412 | 0.027 | 0.048 | 0.009 | 0.009 | 0.084 | 0.08 | 0.37 | 0.07 | 0.09 | 0.06 | 0.02 |
| 6 | 106824214 | rs7755014 | -0.711 | 0.519 | -0.586 | 0.067 | -0.037 | 0.03 | 0.03 | 0.141 | 0.42 | 0.16 | 0.44 | 0.35 | 0.49 | 0.47 |
| 6 | 106829172 | rs803520 | 1.104 | 0.559 | -0.319 | 0.412 | -0.034 | -0.016 | -0.016 | 0.032 | 0.42 | 0.21 | 0.33 | 0.47 | 0.44 | 0.56 |
| 6 | 106829537 | rs6568433 | -0.824 | 0.709 | 0.102 | 0.387 | 0.025 | -0.001 | -0.001 | 0.064 | 0.51 | 0.57 | 0.57 | 0.48 | 0.42 | 0.34 |
| 6 | 106831666 | rs7740539 | -0.608 | 0.494 | 0.357 | 0.392 | 0.04 | 0.027 | 0.027 | 0 | 0.35 | 0.17 | 0.31 | 0.26 | 0.29 | 0.31 |
| 6 | 106831766 | rs7759046 | -0.583 | 0.488 | 0.408 | 0.389 | 0.043 | 0.023 | 0.023 | 0 | 0.35 | 0.1 | 0.3 | 0.26 | 0.29 | 0.31 |
| 6 | 106831875 | rs803532 | 1.469 | 0.61 | 0.578 | 0.433 | -0.021 | -0.007 | -0.007 | 0 | 0.42 | 0.29 | 0.34 | 0.47 | 0.44 | 0.56 |
| 6 | 106833696 | rs9399991 | -0.08 | 0.397 | 0.336 | 0.031 | -0.014 | 0.011 | 0.011 | 0 | 0.11 | 0.16 | 0.14 | 0.14 | 0.09 | 0.06 |
| 6 | 106836299 | rs2749083 | 0.533 | 0.544 | 0.618 | -0.153 | 0.009 | -0.012 | -0.012 | 0 | 0.91 | 0.83 | 0.92 | 0.94 | 0.87 | 0.89 |
| 6 | 106836645 | rs17317556 | 0.856 | 0.52 | 1.035 | -0.149 | 0.007 | 0.027 | 0.027 | 0 | 0.13 | 0.2 | 0.26 | 0.22 | 0.13 | 0.02 |
| 6 | 106838862 | rs6922072 | -0.191 | 0.498 | 0.807 | -0.352 | 0.011 | 0.024 | 0.024 | 0 | 0.26 | 0.18 | 0.21 | 0.07 | 0.22 | 0.26 |
| 6 | 106839721 | rs803523 | 0.985 | 0.625 | 0.657 | -0.191 | 0.005 | 0.094 | 0.094 | 0 | 0.5 | 0.26 | 0.44 | 0.66 | 0.52 | 0.6 |
| 6 | 106840296 | rs2353594 | -0.94 | 0.681 | 0.818 | -0.061 | 0 | 0.041 | 0.041 | 0 | 0.51 | 0.66 | 0.61 | 0.4 | 0.47 | 0.38 |
| 6 | 106841948 | rs803522 | -0.548 | 0.649 | 0.993 | 0.036 | 0.007 | 0.009 | 0.009 | 0 | 0.84 | 0.95 | 0.91 | 0.84 | 0.85 | 0.92 |
| 6 | 106852349 | rs910688 | 0.383 | 0.663 | 2.261 | 0.159 | -0.002 | -0.006 | -0.006 | 0 | 0.65 | 0.66 | 0.7 | 0.75 | 0.65 | 0.77 |
| 6 | 106860712 | rs987377 | -0.047 | 0.645 | 2.166 | 0.157 | -0.003 | 0.037 | 0.037 | 1.2 | 0.68 | 0.91 | 0.73 | 0.74 | 0.66 | 0.79 |
| 6 | 106862821 | rs11152984 | -0.452 | 0.748 | 2.126 | 0.149 | 0.003 | -0.008 | -0.008 | 0 | 0.1 | 0.14 | 0.08 | 0.08 | 0.12 | 0.14 |
| 6 | 106863710 | rs2353592 | 0.679 | 0.796 | 2.03 | 0.243 | -0.001 | 0.007 | 0.007 | 1.136 | 0.58 | 0.45 | 0.56 | 0.6 | 0.55 | 0.73 |
| 6 | 106865294 | rs9384609 | 0.621 | 0.778 | 2.038 | 0.247 | 0 | -0.012 | -0.012 | 0 | 0.58 | 0.52 | 0.56 | 0.6 | 0.55 | 0.73 |
| 6 | 106869488 | rs1340611 | -0.288 | 0.9 | 1.536 | -0.173 | 0.041 | 0.028 | 0.028 | 0 | 0.07 | 0.16 | 0.13 | 0.15 | 0.1 | 0.05 |
| 6 | 106877696 | rs4946752 | -0.232 | 0.853 | 1.429 | -0.197 | 0.004 | 0.001 | 0.001 | 0 | 0.07 | 0.14 | 0.07 | 0.08 | 0.09 | 0.13 |
| 6 | 106879750 | rs1884462 | 0.619 | 0.941 | 1.098 | -0.327 | -0.004 | -0.046 | -0.046 | 0 | 0.55 | 0.55 | 0.65 | 0.82 | 0.57 | 0.66 |
| 6 | 106879930 | rs1884461 | 1.066 | 0.988 | 0.483 | -0.231 | 0.001 | 0.099 | 0.099 | 1.013 | 0.46 | 0.27 | 0.49 | 0.68 | 0.43 | 0.62 |
| 6 | 106881308 | rs7767036 | 0.389 | 0.687 | 0.297 | -0.247 | 0.001 | 0.009 | 0.009 | 0.704 | 0.65 | 0.75 | 0.74 | 0.9 | 0.67 | 0.79 |
| 6 | 106885785 | rs927016 | 0.305 | 0.649 | 0.302 | -0.222 | 0 | -0.028 | -0.028 | 0 | 0.59 | 0.58 | 0.69 | 0.81 | 0.53 | 0.55 |
| 6 | 106887924 | rs9400005 | -0.423 | 0.591 | 0.191 | 0.171 | -0.003 | -0.006 | -0.006 | 0.345 | 0.22 | 0.19 | 0.23 | 0.26 | 0.19 | 0.14 |
| 6 | 106890953 | rs7753319 | 1.07 | 0.472 | 0.452 | 1.058 | -0.012 | -0.015 | -0.015 | 0 | 0.38 | 0.28 | 0.52 | 0.53 | 0.41 | 0.42 |
| 6 | 106893041 | rs6908605 | -0.397 | 0.545 | 0.78 | 0.731 | 0.011 | 0.055 | 0.055 | 0.474 | 0.39 | 0.29 | 0.26 | 0.11 | 0.35 | 0.24 |
| 6 | 106893687 | rs9320165 | 0.103 | 0.452 | 0.533 | 0.723 | 0.002 | 0.023 | 0.023 | 0 | 0.15 | 0.09 | 0.08 | 0.08 | 0.12 | 0.12 |
| 6 | 106894342 | rs9480671 | -0.363 | 0.432 | 0.524 | 0.713 | 0.013 | 0.039 | 0.039 | 0 | 0.18 | 0.1 | 0.08 | 0.08 | 0.12 | 0.13 |
| 6 | 106895837 | rs9373866 | 0.669 | 0.582 | 0.767 | 0.743 | -0.017 | 0.022 | 0.022 | 0 | 0.76 | 0.81 | 0.82 | 0.85 | 0.71 | 0.72 |
| 6 | 106896650 | rs4530901 | 0.164 | 0.589 | 0.769 | 0.724 | 0.033 | -0.019 | -0.019 | 0 | 0.9 | 0.84 | 0.89 | 0.93 | 0.82 | 0.85 |
| 6 | 106898421 | rs2236073 | 1.461 | 0.508 | 0.967 | 0.74 | -0.012 | 0.014 | 0.014 | 0.097 | 0.57 | 0.24 | 0.53 | 0.37 | 0.57 | 0.5 |
| 6 | 106899226 | rs1417352 | 0.787 | 0.585 | 0.536 | 0.815 | 0.012 | 0.011 | 0.011 | 0 | 0.77 | 0.72 | 0.72 | 0.65 | 0.7 | 0.63 |
| 6 | 106900429 | rs11751463 | -0.035 | 1.055 | 0.191 | 0.955 | 0.001 | 0.024 | 0.024 | 0 | 0.15 | 0.1 | 0.08 | 0.08 | 0.12 | 0.13 |
| 6 | 106900902 | rs9386545 | 1.199 | 1.328 | 0.01 | 1.099 | -0.018 | -0.043 | -0.043 | 0 | 0.33 | 0.09 | 0.35 | 0.36 | 0.34 | 0.38 |
| 6 | 106901586 | rs9486351 | -0.669 | 1.075 | 0.074 | 1.133 | -0.029 | -0.007 | -0.007 | 0.363 | 0.58 | 0.81 | 0.54 | 0.57 | 0.49 | 0.47 |
| 6 | 106907560 | rs6919213 | 0.791 | 1.08 | 0.287 | 1.046 | -0.001 | -0.001 | -0.001 | 0.003 | 0.35 | 0.36 | 0.37 | 0.28 | 0.34 | 0.35 |
| 6 | 106909194 | rs6908055 | -0.657 | 1.861 | 0.383 | 1.115 | -0.05 | 0.03 | 0.03 | 0 | 0.15 | 0.01 | 0.07 | 0.15 | 0.16 | 0.28 |
| 6 | 106914005 | rs6905896 | 1.331 | 2.065 | 0.738 | 1.38 | 0.016 | 0.003 | 0.003 | 0 | 0.7 | 0.66 | 0.6 | 0.59 | 0.58 | 0.56 |
| 6 | 106914582 | rs1340616 | -0.773 | 2.108 | 0.755 | 1.384 | 0.01 | 0.014 | 0.014 | 0 | 0.21 | 0.22 | 0.3 | 0.38 | 0.23 | 0.27 |
| 6 | 106915933 | rs1340618 | 0.795 | 1.944 | 0.873 | 1.241 | 0.046 | -0.02 | -0.02 | 0 | 0.9 | 0.79 | 0.89 | 0.97 | 0.81 | 0.83 |
| 6 | 106921200 | rs9400010 | -0.962 | 1.117 | 0.74 | 1.474 | 0.01 | 0.013 | 0.013 | 0 | 0.2 | 0.21 | 0.29 | 0.39 | 0.23 | 0.27 |
| 6 | 106921538 | rs9386551 | -1.041 | 1.138 | 0.572 | 1.484 | 0.025 | 0.031 | 0.031 | 0 | 0.21 | 0.33 | 0.3 | 0.39 | 0.23 | 0.27 |
| 6 | 106925710 | rs6925883 | -1.458 | 0.665 | 0.86 | 1.916 | 0.034 | 0.256 | 0.256 | 0.04 | 0.13 | 0.44 | 0.29 | 0.37 | 0.12 | 0.12 |
| 6 | 106936937 | rs9372132 | -1.496 | 0.512 | 0.825 | 1.843 | -0.001 | 0.013 | 0.013 | 0 | 0.09 | 0.12 | 0.25 | 0.28 | 0.12 | 0.08 |
| 6 | 106937904 | rs7751527 | -1.538 | -0.65 | 0.566 | 1.029 | -0.016 | -0.009 | -0.009 | 0 | 0.28 | 0.59 | 0.32 | 0.25 | 0.18 | 0.34 |
| 6 | 106940336 | rs9480675 | -0.874 | -0.731 | 0.857 | 1.314 | 0.002 | 0.05 | 0.05 | 0 | 0.1 | 0.32 | 0.13 | 0.1 | 0.07 | 0.04 |
| 6 | 106941334 | rs12209167 | -1.835 | -1.107 | 0.99 | 1.066 | -0.002 | -0.008 | -0.008 | 0 | 0.29 | 0.47 | 0.43 | 0.27 | 0.28 | 0.19 |
| 6 | 106942339 | rs12211147 | -0.562 | -1.073 | 1.147 | 1.023 | 0.006 | -0.004 | -0.004 | 0 | 0.12 | 0.11 | 0.09 | 0.07 | 0.16 | 0.13 |
| 6 | 106943311 | rs1181557 | -0.243 | -1.281 | -0.126 | 0.912 | 0.042 | 0.008 | 0.008 | 0 | 0.61 | 0.96 | 0.72 | 0.69 | 0.67 | 0.57 |
| 6 | 106944820 | rs7760555 | -0.752 | -1.255 | -0.125 | 0.956 | 0.024 | -0.032 | -0.032 | 0 | 0.67 | 0.97 | 0.77 | 0.69 | 0.72 | 0.58 |
| 6 | 106944901 | rs9480677 | 0.566 | -0.405 | -0.095 | 0.564 | 0.021 | -0.048 | -0.048 | 0.733 | 0.17 | 0.3 | 0.16 | 0.06 | 0.23 | 0.05 |
| 6 | 106945125 | rs9480678 | -0.555 | -0.396 | -0.293 | 0.582 | 0.057 | -0.015 | -0.015 | 0 | 0.56 | 0.86 | 0.62 | 0.59 | 0.61 | 0.54 |
| 6 | 106945563 | rs11152998 | 0.234 | -0.105 | 0.039 | 0.784 | 0.005 | 0.003 | 0.003 | 0.011 | 0.13 | 0.11 | 0.12 | 0.19 | 0.16 | 0.14 |
| 6 | 106946212 | rs17067117 | 1.107 | -0.095 | 0.014 | 0.795 | 0.002 | 0.025 | 0.025 | 0 | 0.07 | 0.1 | 0.11 | 0.1 | 0.07 | 0.04 |

| | | | | | | | | | | | | | | | | |
|---|---|---|---|---|---|---|---|---|---|---|---|---|---|---|---|---|
| 6 | 106948651 | rs1340625 | -0.035 | -0.048 | -0.214 | 0.607 | 0.008 | 0.005 | 0.005 | 0 | 0.14 | 0.15 | 0.12 | 0.19 | 0.16 | 0.13 |
| 6 | 106950833 | rs6928575 | 1.272 | -0.165 | -0.462 | 0.797 | -0.051 | 0.053 | 0.053 | 0 | 0.22 | 0.05 | 0.32 | 0.11 | 0.22 | 0.32 |
| 6 | 106954144 | rs783412 | -0.58 | -0.522 | 0.028 | 1.023 | -0.009 | 0.001 | 0.001 | 2.042 | 0.66 | 0.68 | 0.65 | 0.65 | 0.78 | 0.71 |
| 6 | 106955602 | rs7752730 | 1.071 | -0.144 | -0.148 | 0.432 | -0.035 | 0.11 | 0.11 | 0.149 | 0.3 | 0.1 | 0.25 | 0.05 | 0.31 | 0.39 |
| 6 | 106955838 | rs783409 | 0.403 | -0.21 | 0.506 | 0.617 | -0.005 | 0.027 | 0.027 | 0 | 0.1 | 0.34 | 0.15 | 0.11 | 0.07 | 0.05 |
| 6 | 106956635 | rs783407 | 0.684 | -0.193 | 0.471 | 0.618 | -0.003 | 0.02 | 0.02 | 0 | 0.08 | 0.24 | 0.14 | 0.11 | 0.07 | 0.05 |
| 6 | 106957120 | rs1417356 | 0.022 | -0.69 | 0.38 | 0.641 | -0.048 | 0.029 | 0.029 | 0.09 | 0.39 | 0.2 | 0.37 | 0.31 | 0.46 | 0.45 |
| 6 | 106957474 | rs1417357 | -0.057 | -0.703 | 0.381 | 0.582 | -0.047 | 0.007 | 0.007 | 0.01 | 0.39 | 0.22 | 0.46 | 0.57 | 0.47 | 0.49 |
| 6 | 106957506 | rs783403 | -1.063 | -0.887 | 0.677 | 0.697 | -0.033 | 0.048 | 0.048 | 0.377 | 0.84 | 0.74 | 0.9 | 0.73 | 0.89 | 0.95 |
| 6 | 106958744 | rs783423 | 0.478 | -0.736 | 0.573 | 0.751 | -0.016 | 0.011 | 0.011 | 0 | 0.1 | 0.29 | 0.15 | 0.12 | 0.07 | 0.06 |
| 6 | 106959424 | rs4946758 | 0.937 | -0.59 | 0.045 | 0.67 | 0.015 | -0.05 | -0.05 | 0.505 | 0.22 | 0.09 | 0.3 | 0.51 | 0.17 | 0.31 |
| 6 | 106959961 | rs4946759 | 0.278 | -0.586 | 0.396 | 0.733 | -0.042 | 0.017 | 0.017 | 0.662 | 0.72 | 0.43 | 0.72 | 0.61 | 0.78 | 0.89 |
| 6 | 106961655 | rs1340624 | 0.709 | -0.428 | 0.255 | 0.697 | 0.01 | -0.052 | -0.052 | 0.771 | 0.25 | 0.14 | 0.23 | 0.49 | 0.22 | 0.34 |
| 6 | 106962186 | rs13218830 | 1.987 | -0.309 | 0.287 | 0.255 | -0.01 | 0.037 | 0.037 | 0 | 0.27 | 0.12 | 0.19 | 0.05 | 0.25 | 0.35 |
| 6 | 106964724 | rs6938452 | 0.326 | -0.081 | 0.47 | 0.328 | 0.01 | 0.013 | 0.013 | 0.504 | 0.5 | 0.41 | 0.36 | 0.17 | 0.55 | 0.53 |
| 6 | 106967185 | rs1159148 | 2.028 | 0.472 | 0.451 | 0.315 | 0.001 | -0.035 | -0.035 | 1.076 | 0.26 | 0.2 | 0.19 | 0.05 | 0.24 | 0.35 |
| 6 | 106967778 | rs3747787 | 0.32 | 0.634 | 0.2 | 0.476 | -0.011 | 0.016 | 0.016 | 0 | 0.2 | 0.18 | 0.14 | 0.12 | 0.25 | 0.17 |
| 6 | 106973667 | rs9320172 | 0.8 | 0.48 | 0.086 | 0.677 | -0.019 | -0.007 | -0.007 | 0 | 0.24 | 0.1 | 0.18 | 0.27 | 0.27 | 0.31 |
| 6 | 106975536 | rs11153000 | 0.868 | 0.449 | 0.095 | 0.557 | -0.011 | -0.069 | -0.069 | 0 | 0.23 | 0.1 | 0.27 | 0.46 | 0.24 | 0.32 |
| 6 | 106976289 | rs783390 | -1.284 | 0.486 | 0.42 | 0.314 | 0.001 | -0.03 | -0.03 | 0.577 | 0.6 | 0.68 | 0.72 | 0.88 | 0.68 | 0.59 |
| 6 | 106980324 | rs13212772 | 0.983 | 0.268 | 0.396 | 0.556 | 0.017 | 0.002 | 0.002 | 0 | 0.53 | 0.28 | 0.47 | 0.51 | 0.52 | 0.71 |
| 6 | 106981188 | rs6929649 | 0.424 | 0.162 | 0.608 | 0.64 | -0.022 | 0.016 | 0.016 | 0 | 0.7 | 0.54 | 0.59 | 0.62 | 0.73 | 0.86 |
| 6 | 106982488 | rs1462147 | 0.049 | 0.153 | 0.711 | 0.594 | -0.002 | -0.014 | -0.014 | 0 | 0.33 | 0.21 | 0.4 | 0.59 | 0.35 | 0.39 |
| 6 | 106982560 | rs1111830 | -0.689 | 0.121 | 0.752 | 0.338 | 0.009 | 0.033 | 0.033 | 0 | 0.82 | 0.92 | 0.88 | 0.93 | 0.85 | 0.97 |
| 6 | 106982672 | rs898894 | 0.899 | 0.133 | 0.849 | 0.396 | -0.002 | -0.002 | -0.002 | 1.086 | 0.46 | 0.43 | 0.32 | 0.15 | 0.48 | 0.53 |
| 6 | 106983180 | rs936497 | 0.942 | 0.174 | 0.749 | 0.519 | -0.003 | 0 | 0 | 0 | 0.09 | 0.09 | 0.13 | 0.12 | 0.1 | 0.07 |
| 6 | 106984739 | rs11153007 | 0.468 | 0.121 | 0.768 | 0.398 | -0.004 | 0.005 | 0.005 | 0 | 0.16 | 0.13 | 0.12 | 0.1 | 0.19 | 0.18 |
| 6 | 106987370 | rs783396 | -1.482 | -0.198 | 0.8 | 0.292 | 0.011 | 0.013 | 0.013 | 0 | 0.87 | 0.94 | 0.92 | 0.93 | 0.9 | 0.97 |
| 6 | 106988018 | rs783394 | -1.076 | -0.147 | 0.762 | 0.241 | 0.001 | -0.014 | -0.014 | 1.74 | 0.55 | 0.59 | 0.71 | 0.88 | 0.62 | 0.58 |
| 6 | 106989212 | rs783392 | -0.663 | -0.14 | 0.761 | 0.248 | -0.001 | -0.003 | -0.003 | 0 | 0.51 | 0.56 | 0.67 | 0.88 | 0.55 | 0.57 |
| 6 | 106992103 | rs2027561 | 0.327 | 0.01 | 0.561 | 0.646 | -0.017 | -0.015 | -0.015 | 0 | 0.31 | 0.17 | 0.33 | 0.42 | 0.41 | 0.34 |
| 6 | 106993436 | rs1770720 | -0.206 | -1.493 | 1.025 | 1.101 | -0.013 | 0.033 | 0.033 | 0 | 0.81 | 0.64 | 0.78 | 0.76 | 0.84 | 0.93 |
| 6 | 106994326 | rs2219667 | 1.28 | -1.544 | 1.037 | 0.7 | 0.014 | 0.007 | 0.007 | 0.004 | 0.6 | 0.3 | 0.62 | 0.64 | 0.69 | 0.76 |
| 6 | 106995863 | rs1616375 | -0.268 | -1.166 | 1.254 | 0.621 | -0.01 | -0.066 | -0.066 | 0.505 | 0.82 | 0.67 | 0.87 | 0.93 | 0.84 | 0.95 |
| 6 | 106998973 | rs4946762 | -0.12 | -1.359 | 1.342 | 0.63 | -0.072 | -0.107 | -0.107 | 0 | 0.81 | 0.42 | 0.88 | 0.92 | 0.9 | 0.97 |
| 6 | 107002149 | rs1676016 | -0.051 | -1.299 | 1.286 | 0.779 | -0.033 | -0.024 | -0.024 | 0.318 | 0.79 | 0.62 | 0.88 | 0.83 | 0.85 | 0.87 |
| 6 | 107004132 | rs2054366 | 0.514 | -1.313 | 1.395 | 0.772 | -0.055 | -0.042 | -0.042 | 3.645 | 0.77 | 0.5 | 0.86 | 0.83 | 0.85 | 0.87 |
| 6 | 107004452 | rs2615206 | -0.132 | -1.306 | 1.39 | 0.776 | -0.017 | -0.012 | -0.012 | 0 | 0.8 | 0.72 | 0.89 | 0.83 | 0.85 | 0.87 |
| 6 | 107014734 | rs9486403 | -0.147 | -1.303 | 1.471 | 0.778 | -0.027 | -0.018 | -0.018 | 0 | 0.2 | 0.33 | 0.11 | 0.17 | 0.15 | 0.13 |
| 6 | 107016343 | rs3747789 | 1.051 | -1.327 | 1.464 | 0.765 | 0.021 | 0.05 | 0.05 | 0 | 0.13 | 0.01 | 0.07 | 0.07 | 0.1 | 0.03 |
| 6 | 107016731 | rs3747790 | 0.092 | -1.243 | 1.449 | 1.159 | -0.003 | 0.004 | 0.004 | 0 | 0.2 | 0.22 | 0.23 | 0.35 | 0.18 | 0.15 |
| 6 | 107018446 | rs11758586 | 0.104 | -1.244 | 1.449 | 1.155 | -0.005 | -0.003 | -0.003 | 0 | 0.2 | 0.21 | 0.1 | 0.17 | 0.15 | 0.13 |
| 6 | 107019287 | rs9320182 | 0.072 | -1.244 | 1.45 | 1.169 | -0.023 | -0.014 | -0.014 | 0 | 0.2 | 0.3 | 0.11 | 0.17 | 0.15 | 0.13 |
| 6 | 107019635 | rs9320183 | 0.104 | -1.244 | 1.441 | 1.154 | -0.005 | -0.003 | -0.003 | 0 | 0.2 | 0.21 | 0.1 | 0.17 | 0.15 | 0.13 |
| 6 | 107022395 | rs11968064 | 0.905 | -1.221 | 1.423 | 1.088 | -0.023 | 0.018 | 0.018 | 0 | 0.72 | 0.4 | 0.7 | 0.65 | 0.75 | 0.83 |
| 6 | 107027481 | rs4946768 | 0.905 | -1.22 | 1.205 | 1.429 | -0.034 | 0.086 | 0.086 | 0 | 0.72 | 0.44 | 0.61 | 0.49 | 0.75 | 0.81 |
| 6 | 107043751 | rs4945759 | 0.41 | -1.218 | 1.169 | 1.422 | -0.047 | 0.05 | 0.05 | 0 | 0.76 | 0.54 | 0.77 | 0.67 | 0.84 | 0.84 |
| 6 | 107056401 | rs6938201 | 1.147 | -1.019 | 1.096 | 1.482 | 0.012 | 0.194 | 0.194 | 0 | 0.92 | 0.75 | 0.72 | 0.65 | 0.91 | 0.94 |
| 6 | 107063162 | rs898896 | 0.053 | -1.088 | 0.733 | 1.047 | 0 | -0.001 | -0.001 | 0.119 | 0.71 | 0.67 | 0.59 | 0.35 | 0.69 | 0.67 |
| 6 | 107063540 | rs898898 | 1.049 | -1.928 | 0.85 | 1.171 | -0.014 | 0.019 | 0.019 | 0 | 0.66 | 0.38 | 0.61 | 0.54 | 0.7 | 0.76 |
| 6 | 107069273 | rs1870867 | -0.726 | -2.135 | 0.76 | 1.028 | -0.031 | 0.016 | 0.016 | 0.028 | 0.3 | 0.6 | 0.47 | 0.63 | 0.25 | 0.24 |
| 6 | 107073939 | rs7773795 | 0.279 | -1.793 | 0.201 | 0.729 | 0 | 0.001 | 0.001 | 0 | 0.07 | 0.09 | 0.13 | 0.11 | 0.08 | 0.08 |
| 6 | 107081022 | rs3804340 | -0.262 | -1.902 | -0.142 | 0.468 | 0.014 | 0.009 | 0.009 | 0 | 0.86 | 0.9 | 0.91 | 0.99 | 0.92 | 0.91 |
| 6 | 107084941 | rs1419054 | 0.255 | -1.869 | 0.162 | 0.867 | -0.019 | 0.033 | 0.033 | 0 | 0.19 | 0.22 | 0.23 | 0.35 | 0.17 | 0.15 |
| 6 | 107086815 | rs12530128 | 0.231 | -2.246 | -0.296 | 0.242 | -0.003 | 0.111 | 0.111 | 1.63 | 0.18 | 0.01 | 0.11 | 0.01 | 0.16 | 0.18 |
| 6 | 107087870 | rs3804344 | 1.054 | -2.085 | -0.076 | 0.405 | -0.023 | -0.042 | -0.042 | 0 | 0.14 | 0.19 | 0.08 | 0.07 | 0.09 | 0.02 |
| 6 | 107089289 | rs7750365 | 0.909 | -2.167 | 0.161 | 0.023 | 0.001 | -0.08 | -0.08 | 0.717 | 0.44 | 0.46 | 0.36 | 0.13 | 0.47 | 0.3 |
| 6 | 107090369 | rs9400038 | 0.503 | -2.228 | 0.128 | 0.291 | -0.001 | -0.001 | -0.001 | 0 | 0.51 | 0.5 | 0.61 | 0.42 | 0.58 | 0.4 |
| 6 | 107091220 | rs9400041 | 0.55 | -2.227 | 0.129 | 0.217 | 0 | -0.018 | -0.018 | 0 | 0.51 | 0.5 | 0.48 | 0.23 | 0.55 | 0.38 |
| 6 | 107098989 | rs11752776 | 1.537 | -1.776 | 0.238 | 0.547 | -0.009 | -0.016 | -0.016 | 0 | 0.3 | 0.27 | 0.28 | 0.06 | 0.38 | 0.28 |
| 6 | 107099191 | rs12333016 | 0.64 | -1.774 | 0.22 | 0.576 | -0.009 | -0.013 | -0.013 | 0 | 0.06 | 0.01 | 0.12 | 0.1 | 0.08 | 0.08 |
| 6 | 107103157 | rs6910496 | -0.407 | -1.829 | 0.413 | 0.996 | -0.004 | 0.089 | 0.089 | 0 | 0.07 | 0.14 | 0.12 | 0.27 | 0.06 | 0.11 |
| 6 | 107104055 | rs9400043 | 0.526 | -1.608 | 0.396 | 0.633 | -0.006 | -0.007 | -0.007 | 0 | 0.07 | 0.03 | 0.12 | 0.1 | 0.08 | 0.08 |
| 6 | 107113715 | rs2015205 | 0.773 | -1.647 | 0.406 | 0.645 | -0.011 | 0.023 | 0.023 | 0 | 0.37 | 0.31 | 0.4 | 0.16 | 0.46 | 0.36 |
| 6 | 107114361 | rs1026619 | 1.313 | -1.62 | 0.384 | 0.628 | -0.021 | 0.007 | 0.007 | 0 | 0.3 | 0.25 | 0.28 | 0.06 | 0.38 | 0.28 |
| 6 | 107115394 | rs3101493 | 1.455 | -1.612 | 0.39 | 0.456 | -0.022 | 0.009 | 0.009 | 0 | 0.3 | 0.25 | 0.28 | 0.06 | 0.38 | 0.28 |
| 6 | 107123879 | rs3121445 | 1.238 | -1.75 | 0.375 | 0.446 | -0.019 | 0.003 | 0.003 | 0.217 | 0.3 | 0.25 | 0.28 | 0.05 | 0.39 | 0.26 |
| 6 | 107126673 | rs9400046 | -0.008 | -1.97 | 0.378 | 0.719 | 0.022 | 0.032 | 0.032 | 0 | 0.08 | 0.19 | 0.16 | 0.18 | 0.09 | 0.08 |
| 6 | 107129085 | rs3121433 | 0.515 | -2.455 | 0.232 | 1.017 | 0.037 | 0 | 0 | 0.011 | 0.45 | 0.64 | 0.61 | 0.48 | 0.6 | 0.4 |

| | | | | | | | | | | | | | | | | |
|---|---|---|---|---|---|---|---|---|---|---|---|---|---|---|---|---|
| 6 | 107133719 | rs3127660 | 0.289 | -2.604 | 0.562 | 1.351 | -0.049 | 0.077 | 0.077 | 0 | 0.78 | 0.62 | 0.64 | 0.51 | 0.88 | 0.77 |
| 6 | 107135667 | rs9386579 | -0.091 | -2.974 | 0.505 | 0.988 | -0.023 | -0.011 | -0.011 | 0 | 0.45 | 0.47 | 0.33 | 0.63 | 0.32 | 0.44 |
| 6 | 107136274 | rs9398102 | -0.316 | -2.996 | 0.476 | 0.773 | -0.005 | -0.019 | -0.019 | 0 | 0.48 | 0.5 | 0.51 | 0.81 | 0.33 | 0.47 |
| 6 | 107137735 | rs2354552 | -0.1 | -2.573 | 0.026 | 0.762 | -0.006 | 0.062 | 0.062 | 0 | 0.8 | 0.89 | 0.83 | 0.98 | 0.78 | 0.78 |
| 6 | 107139657 | rs3104037 | 0.737 | -3.519 | 0.12 | 1.163 | 0.038 | 0.003 | 0.003 | 0 | 0.52 | 0.54 | 0.54 | 0.48 | 0.64 | 0.57 |
| 6 | 107139868 | rs12525297 | 0.077 | -3.626 | -0.12 | 1.253 | -0.023 | -0.01 | -0.01 | 0 | 0.24 | 0.08 | 0.15 | 0.18 | 0.24 | 0.29 |
| 6 | 107140356 | rs17067644 | 0.317 | -3.658 | 0.116 | 1.31 | -0.007 | 0.009 | 0.009 | 0 | 0.15 | 0.13 | 0.1 | 0.21 | 0.13 | 0.25 |
| 6 | 107141246 | rs6940205 | -0.254 | -3.808 | 0.348 | 1.195 | -0.031 | 0.037 | 0.037 | 0 | 0.3 | 0.32 | 0.3 | 0.41 | 0.24 | 0.31 |
| 6 | 107141376 | rs9320187 | 1.536 | -3.837 | 0.132 | 0.916 | 0.007 | -0.009 | -0.009 | 0 | 0.13 | 0.08 | 0.05 | 0 | 0.09 | 0.03 |
| 6 | 107141927 | rs3127661 | 0.567 | -3.449 | 0.362 | 1.174 | -0.013 | -0.013 | -0.013 | 0 | 0.59 | 0.4 | 0.58 | 0.65 | 0.7 | 0.67 |
| 6 | 107142323 | rs2354549 | 0.693 | -3.444 | 0.364 | 1.229 | -0.008 | 0.002 | 0.002 | 0 | 0.58 | 0.37 | 0.43 | 0.44 | 0.67 | 0.65 |
| 6 | 107144796 | rs2254924 | -0.572 | -3.446 | 0.361 | 1.25 | -0.019 | -0.008 | -0.008 | 0.074 | 0.37 | 0.54 | 0.38 | 0.38 | 0.32 | 0.31 |
| 6 | 107151522 | rs910528 | 1.668 | -3.098 | 0.389 | 1.401 | -0.054 | -0.046 | -0.046 | 0 | 0.3 | 0.18 | 0.29 | 0.37 | 0.46 | 0.36 |
| 6 | 107154445 | rs2798361 | 0.182 | -3.124 | 0.466 | 1.207 | -0.033 | -0.018 | -0.018 | 0 | 0.59 | 0.57 | 0.66 | 0.66 | 0.7 | 0.65 |
| 6 | 107160488 | rs2748293 | 0.217 | -3.114 | 0.083 | 1.17 | 0.044 | 0.035 | 0.035 | 0.895 | 0.61 | 0.79 | 0.8 | 0.71 | 0.73 | 0.67 |
| 6 | 107161532 | rs2642465 | -0.424 | -3.076 | 0.138 | 1.169 | -0.017 | -0.011 | -0.011 | 0.38 | 0.71 | 0.75 | 0.84 | 0.77 | 0.78 | 0.71 |
| 6 | 107162032 | rs2748289 | -0.092 | -2.726 | 0.303 | 1.157 | -0.006 | -0.006 | -0.006 | 0 | 0.13 | 0.14 | 0.06 | 0.06 | 0.06 | 0.05 |
| 6 | 107162648 | rs2642467 | -0.837 | -2.576 | -0.091 | 0.619 | 0.017 | 0.01 | 0.01 | 0 | 0.4 | 0.36 | 0.42 | 0.36 | 0.3 | 0.32 |
| 6 | 107174515 | rs6915445 | -0.694 | -2.652 | -0.133 | 0.714 | -0.034 | 0.008 | 0.008 | 0 | 0.35 | 0.48 | 0.37 | 0.38 | 0.24 | 0.28 |
| 6 | 107178504 | rs10872009 | -0.271 | -2.041 | -0.488 | 0.684 | 0.074 | 0.025 | 0.025 | 0 | 0.46 | 0.7 | 0.55 | 0.38 | 0.35 | 0.33 |
| 6 | 107179509 | rs12196331 | -0.433 | -1.443 | -0.092 | 0.642 | -0.001 | 0.042 | 0.042 | 1.811 | 0.08 | 0.24 | 0.07 | 0.11 | 0.06 | 0.04 |
| 6 | 107182045 | rs2354223 | -0.265 | -1.326 | -1.274 | 0.563 | 0.018 | -0.08 | -0.08 | 0 | 0.52 | 0.97 | 0.64 | 0.53 | 0.59 | 0.56 |
| 6 | 107187910 | rs9486454 | -0.086 | -1.326 | -0.459 | 0.319 | 0.002 | -0.049 | -0.049 | 0.172 | 0.09 | 0.15 | 0.06 | 0 | 0.07 | 0.07 |
| 6 | 107189647 | rs1022630 | 0.698 | -1.177 | -0.32 | 0.334 | -0.034 | -0.024 | -0.024 | 0 | 0.24 | 0.19 | 0.26 | 0.31 | 0.38 | 0.32 |
| 6 | 107190424 | rs9480703 | -1.009 | -1.348 | 0.568 | 0.352 | 0.038 | 0.133 | 0.133 | 0.538 | 0.09 | 0.54 | 0.26 | 0.16 | 0.08 | 0.11 |
| 6 | 107192985 | rs3117391 | -0.757 | -1.743 | -0.224 | 0.222 | -0.016 | -0.046 | -0.046 | 0 | 0.83 | 0.99 | 0.91 | 0.77 | 0.85 | 0.76 |
| 6 | 107193215 | rs9480704 | -0.397 | -1.209 | 0.676 | 0.277 | 0.052 | 0.097 | 0.097 | 0 | 0.06 | 0.29 | 0.18 | 0.11 | 0.06 | 0.07 |
| 6 | 107193720 | rs9480705 | -1.214 | -1.384 | 0.028 | 0.143 | -0.019 | -0.017 | -0.017 | 0.899 | 0.22 | 0.67 | 0.33 | 0.15 | 0.18 | 0.26 |
| 6 | 107196789 | rs11968442 | -0.46 | -0.964 | -0.385 | 0.137 | 0.005 | 0.008 | 0.008 | 0 | 0.14 | 0.09 | 0.08 | 0.05 | 0.12 | 0.19 |
| 6 | 107200239 | rs3117389 | 0.002 | -0.836 | -0.294 | 0.304 | -0.021 | -0.017 | -0.017 | 0 | 0.36 | 0.24 | 0.35 | 0.37 | 0.5 | 0.41 |
| 6 | 107210671 | rs6568450 | -0.142 | -0.905 | -0.228 | 0.197 | 0 | -0.003 | -0.003 | 0 | 0.42 | 0.34 | 0.52 | 0.41 | 0.55 | 0.46 |
| 6 | 107213273 | rs7772155 | 0.454 | -1.3 | -0.053 | -0.161 | 0.006 | 0.007 | 0.007 | 0 | 0.49 | 0.45 | 0.58 | 0.62 | 0.54 | 0.41 |
| 6 | 107216553 | rs3117418 | -0.33 | -1.895 | 0.517 | -0.615 | -0.014 | -0.094 | -0.094 | 0 | 0.31 | 0.52 | 0.21 | 0.05 | 0.27 | 0.22 |
| 6 | 107216674 | rs11757353 | 0.585 | -2.053 | -0.128 | -0.57 | -0.002 | -0.016 | -0.016 | 1.744 | 0.1 | 0.01 | 0.06 | 0.08 | 0.09 | 0.14 |
| 6 | 107220091 | rs3117416 | -0.474 | -1.704 | 0.506 | -0.631 | -0.011 | -0.107 | -0.107 | 1.275 | 0.3 | 0.41 | 0.2 | 0.05 | 0.27 | 0.22 |
| 6 | 107226021 | rs6920626 | -0.104 | -2.109 | -0.186 | -0.41 | 0.018 | -0.008 | -0.008 | 0 | 0.36 | 0.61 | 0.2 | 0.39 | 0.21 | 0.34 |
| 6 | 107231570 | rs1884270 | 0.508 | -2.102 | -0.278 | -0.352 | 0.018 | -0.016 | -0.016 | 0 | 0.27 | 0.22 | 0.13 | 0.3 | 0.15 | 0.31 |
| 6 | 107236494 | rs7775079 | -0.085 | -1.079 | 0.222 | -0.696 | 0.012 | 0.068 | 0.068 | 0.017 | 0.21 | 0.08 | 0.13 | 0.04 | 0.19 | 0.2 |
| 6 | 107237052 | rs6902983 | -0.856 | -1.038 | 0.194 | -0.683 | -0.005 | -0.039 | -0.039 | 0 | 0.31 | 0.6 | 0.2 | 0.12 | 0.25 | 0.24 |
| 6 | 107237748 | rs3117414 | -0.551 | -0.936 | -0.342 | -0.467 | -0.003 | -0.028 | -0.028 | 0.839 | 0.31 | 0.34 | 0.26 | 0.11 | 0.22 | 0.25 |
| 6 | 107245957 | rs3127623 | 0.061 | -0.279 | -0.634 | 0.195 | -0.021 | -0.032 | -0.032 | 0 | 0.35 | 0.18 | 0.47 | 0.45 | 0.36 | 0.39 |
| 6 | 107248110 | rs3127646 | -0.104 | -0.305 | 0.229 | 0.3 | 0 | 0.001 | 0.001 | 0 | 0.37 | 0.39 | 0.54 | 0.51 | 0.37 | 0.45 |
| 6 | 107249301 | rs17067756 | 0.043 | -0.873 | 0.314 | -0.148 | -0.018 | -0.042 | -0.042 | 0.459 | 0.11 | 0.15 | 0.04 | 0 | 0.08 | 0.01 |
| 6 | 107249554 | rs910532 | -0.097 | -0.83 | 0.722 | -0.044 | 0.014 | 0.002 | 0.002 | 1.051 | 0.46 | 0.3 | 0.51 | 0.51 | 0.43 | 0.45 |
| 6 | 107252236 | rs11751550 | -0.373 | -0.56 | 1.102 | -0.222 | 0.037 | 0.01 | 0.01 | 0 | 0.07 | 0.28 | 0.05 | 0.06 | 0.07 | 0.08 |
| 6 | 107252668 | rs910529 | -0.373 | -0.563 | 1.104 | -0.224 | 0.037 | 0.01 | 0.01 | 0 | 0.07 | 0.28 | 0.05 | 0.06 | 0.07 | 0.08 |
| 6 | 107256695 | rs1358725 | -0.761 | -0.989 | 0.679 | -0.031 | 0.001 | -0.001 | -0.001 | 0.21 | 0.67 | 0.64 | 0.67 | 0.7 | 0.58 | 0.58 |
| 6 | 107266807 | rs4946795 | -0.642 | -1.226 | 0.649 | -0.032 | -0.014 | -0.014 | -0.014 | 0.812 | 0.18 | 0.22 | 0.16 | 0.11 | 0.15 | 0.09 |
| 6 | 107272331 | rs3117397 | -0.483 | -1.006 | 0.581 | 0.103 | -0.062 | -0.058 | -0.058 | 0 | 0.63 | 0.82 | 0.66 | 0.61 | 0.55 | 0.53 |
| 6 | 107291157 | rs7767043 | -0.479 | -1.169 | 1.789 | 0.308 | -0.053 | 0.007 | 0.007 | 1.827 | 0.13 | 0.38 | 0.14 | 0.11 | 0.07 | 0.08 |
| 6 | 107292036 | rs2354157 | -0.503 | -1.141 | 1.606 | 0.322 | -0.054 | 0.006 | 0.006 | 1.697 | 0.13 | 0.38 | 0.14 | 0.11 | 0.07 | 0.08 |
| 6 | 107294875 | rs9486475 | -0.133 | -1.081 | 1.347 | 0.41 | -0.03 | 0.008 | 0.008 | 2.813 | 0.12 | 0.19 | 0.14 | 0.12 | 0.07 | 0.07 |
| 6 | 107295198 | rs272586 | -0.567 | -1.394 | 1.052 | 0.33 | 0.016 | 0.028 | 0.028 | 0 | 0.49 | 0.65 | 0.65 | 0.67 | 0.46 | 0.4 |
| 6 | 107298061 | rs272583 | -0.371 | -1.801 | 1.136 | 0.353 | -0.007 | 0.025 | 0.025 | 0 | 0.34 | 0.56 | 0.43 | 0.47 | 0.3 | 0.35 |
| 6 | 107299417 | rs272580 | 0.511 | -1.067 | 0.747 | 0.514 | -0.017 | -0.049 | -0.049 | 0 | 0.22 | 0.14 | 0.36 | 0.4 | 0.26 | 0.29 |
| 6 | 107300725 | rs272578 | -0.28 | -1.42 | 0.691 | 0.196 | -0.004 | -0.024 | -0.024 | 0 | 0.56 | 0.52 | 0.65 | 0.82 | 0.46 | 0.68 |
| 6 | 107304447 | rs1714631 | -0.578 | -1.31 | 0.337 | -0.052 | -0.003 | -0.008 | -0.008 | 0.108 | 0.62 | 0.63 | 0.69 | 0.9 | 0.52 | 0.7 |
| 6 | 107307003 | rs908819 | 0.219 | -1.004 | -0.022 | 0.708 | 0.005 | -0.022 | -0.022 | 0 | 0.22 | 0.19 | 0.34 | 0.39 | 0.21 | 0.31 |
| 6 | 107307786 | rs9486478 | -0.314 | -0.88 | -0.12 | 0.351 | -0.006 | 0 | 0 | 0 | 0.39 | 0.52 | 0.52 | 0.59 | 0.35 | 0.38 |
| 6 | 107310131 | rs12197404 | -0.242 | -1.117 | -0.487 | -0.076 | 0.01 | 0.012 | 0.012 | 0.09 | 0.19 | 0.16 | 0.09 | 0.06 | 0.11 | 0.11 |
| 6 | 107310381 | rs9486484 | -0.482 | -1.349 | -0.389 | -0.111 | -0.044 | -0.037 | -0.037 | 0.271 | 0.34 | 0.46 | 0.16 | 0.11 | 0.18 | 0.21 |
| 6 | 107313977 | rs1900601 | -0.203 | -0.267 | -1.149 | -0.148 | -0.02 | 0.067 | 0.067 | 0.539 | 0.25 | 0.07 | 0.11 | 0.17 | 0.25 | 0.25 |
| 6 | 107328319 | rs7746539 | -0.826 | -0.453 | -0.278 | 0.864 | 0.003 | 0.151 | 0.151 | 0 | 0.08 | 0.16 | 0.06 | 0.34 | 0.07 | 0.19 |
| 6 | 107329095 | rs3099605 | -0.092 | 0.29 | -0.623 | 0.89 | -0.044 | 0.006 | 0.006 | 1.602 | 0.33 | 0.17 | 0.35 | 0.23 | 0.35 | 0.36 |
| 6 | 107329406 | rs1665914 | -1.199 | -0.113 | -0.15 | 0.815 | 0 | -0.003 | -0.003 | 0 | 0.34 | 0.64 | 0.46 | 0.69 | 0.38 | 0.45 |
| 6 | 107329420 | rs1665913 | 0.29 | -0.144 | 0.293 | 0.758 | 0.022 | 0.041 | 0.041 | 0 | 0.17 | 0.25 | 0.37 | 0.35 | 0.28 | 0.24 |
| 6 | 107348589 | rs2783031 | 0.24 | -0.14 | 0.309 | 0.807 | 0.018 | 0.037 | 0.037 | 0 | 0.17 | 0.24 | 0.35 | 0.35 | 0.28 | 0.24 |
| 6 | 107351798 | rs9373908 | -1.011 | -0.594 | 0.558 | 0.413 | 0.006 | 0.173 | 0.173 | 1.237 | 0.1 | 0.23 | 0.15 | 0.35 | 0.07 | 0.2 |
| 6 | 107370460 | rs9400081 | -1.038 | -0.645 | 0.426 | 0.401 | -0.008 | 0.051 | 0.051 | 0.581 | 0.1 | 0.15 | 0.14 | 0.34 | 0.07 | 0.19 |

| | | | | | | | | | | | | | | | | |
|---|---|---|---|---|---|---|---|---|---|---|---|---|---|---|---|---|
| 6 | 107374147 | rs2484325 | 1.103 | -0.617 | 0.534 | 0.371 | -0.052 | 0.097 | 0.097 | 0.905 | 0.82 | 0.5 | 0.91 | 0.66 | 0.92 | 0.81 |
| 6 | 107380733 | rs6568461 | 0.079 | -0.244 | 0.877 | -0.527 | -0.002 | -0.121 | -0.121 | 0.5 | 0.23 | 0.39 | 0.14 | 0.05 | 0.16 | 0.13 |
| 6 | 107380837 | rs6920629 | 0.76 | -0.037 | 0.491 | -0.627 | -0.011 | 0.206 | 0.206 | 0 | 0.5 | 0.19 | 0.42 | 0.14 | 0.48 | 0.36 |
| 6 | 107381245 | rs3763212 | 0.446 | -0.008 | 0.364 | -0.229 | -0.018 | -0.068 | -0.068 | 0.578 | 0.21 | 0.1 | 0.36 | 0.46 | 0.3 | 0.3 |
| 6 | 107381609 | rs3763215 | -0.023 | 0.06 | 0.61 | -0.239 | 0.001 | 0.022 | 0.022 | 0 | 0.73 | 0.44 | 0.79 | 0.67 | 0.78 | 0.71 |
| 6 | 107383411 | rs12199744 | 1.2 | -0.133 | -0.117 | -0.71 | -0.092 | 0.103 | 0.103 | 0 | 0.23 | 0.01 | 0.27 | 0.06 | 0.28 | 0.17 |
| 6 | 107386233 | rs9486513 | 0.193 | -0.606 | 0.816 | -0.079 | -0.004 | -0.037 | -0.037 | 0 | 0.2 | 0.25 | 0.14 | 0.07 | 0.17 | 0.14 |
| 6 | 107386793 | rs1060369 | -0.222 | -0.4 | 0.094 | 0.223 | -0.001 | 0.012 | 0.012 | 0 | 0.42 | 0.54 | 0.29 | 0.17 | 0.43 | 0.31 |
| 6 | 107396658 | rs311232 | -0.19 | -0.529 | -0.627 | 0.385 | 0.019 | 0.042 | 0.042 | 0 | 0.74 | 0.85 | 0.84 | 0.89 | 0.81 | 0.79 |
| 6 | 107398769 | rs9386604 | 0.881 | -0.637 | -0.713 | 0.203 | -0.06 | -0.029 | -0.029 | 1.164 | 0.33 | 0.11 | 0.37 | 0.34 | 0.44 | 0.35 |
| 6 | 107399357 | rs9384641 | 0.739 | -0.467 | -0.593 | 0.13 | 0.023 | 0.026 | 0.026 | 0.017 | 0.37 | 0.44 | 0.4 | 0.42 | 0.44 | 0.39 |
| 6 | 107399479 | rs311229 | -0.084 | -0.556 | -0.892 | 0.063 | -0.002 | 0 | 0 | 0 | 0.37 | 0.41 | 0.44 | 0.47 | 0.37 | 0.39 |
| 6 | 107399970 | rs1665912 | -0.067 | -0.558 | -0.893 | 0.062 | -0.002 | 0 | 0 | 0.793 | 0.37 | 0.41 | 0.44 | 0.47 | 0.37 | 0.37 |
| 6 | 107403119 | rs6904836 | 0.794 | -0.416 | -1.204 | 0.04 | 0.019 | 0.036 | 0.036 | 0 | 0.16 | 0.02 | 0.08 | 0.05 | 0.1 | 0.16 |
| 6 | 107408585 | rs311218 | 0.075 | 0.111 | -0.593 | 0.264 | 0.009 | 0.011 | 0.011 | 0 | 0.23 | 0.3 | 0.4 | 0.39 | 0.31 | 0.26 |
| 6 | 107413934 | rs9486523 | -0.539 | 0.052 | -0.763 | 0.136 | -0.026 | -0.018 | -0.018 | 0.28 | 0.15 | 0.18 | 0.04 | 0.08 | 0.06 | 0.14 |
| 6 | 107419041 | rs7762756 | -0.424 | 0.907 | -1.455 | -0.061 | -0.045 | 0.075 | 0.075 | 0 | 0.3 | 0.14 | 0.34 | 0.11 | 0.37 | 0.25 |
| 6 | 107420270 | rs4946810 | 0.537 | 0.7 | -2.22 | 0.24 | -0.007 | 0.004 | 0.004 | 0 | 0.46 | 0.07 | 0.63 | 0.43 | 0.41 | 0.44 |
| 6 | 107420516 | rs4946811 | 0.534 | 0.718 | -2.227 | 0.317 | -0.01 | -0.03 | -0.03 | 0.344 | 0.46 | 0.06 | 0.55 | 0.47 | 0.42 | 0.45 |
| 6 | 107431688 | rs4946812 | -0.986 | 1.794 | -1.108 | 0.4 | 0.016 | -0.004 | -0.004 | 0.379 | 0.15 | 0.16 | 0.24 | 0.15 | 0.3 | 0.17 |
| 6 | 107448680 | rs6918655 | -1.398 | 0.418 | -0.867 | 0.471 | 0.005 | 0.009 | 0.009 | 0 | 0.18 | 0.16 | 0.16 | 0.12 | 0.16 | 0.19 |
| 6 | 107449641 | rs9400094 | -1.512 | 0.107 | -0.077 | 1.215 | -0.055 | 0.091 | 0.091 | 0 | 0.21 | 0.52 | 0.2 | 0.35 | 0.14 | 0.2 |
| 6 | 107460346 | rs4946815 | -1.32 | 0.459 | -0.401 | 0.746 | 0.009 | 0.011 | 0.011 | 0 | 0.14 | 0.09 | 0.11 | 0.09 | 0.14 | 0.18 |
| 6 | 107466630 | rs13202332 | -0.572 | 1.458 | -0.83 | 0.588 | -0.054 | 0.01 | 0.01 | 0 | 0.22 | 0.12 | 0.21 | 0.19 | 0.31 | 0.26 |
| 6 | 107472949 | rs6568468 | 1.283 | 1.084 | 0.153 | 0.435 | 0.02 | -0.017 | -0.017 | 0.882 | 0.6 | 0.55 | 0.61 | 0.69 | 0.56 | 0.57 |
| 6 | 107481461 | rs11759792 | -1.318 | 0.31 | 0.349 | 0.353 | 0.06 | 0.007 | 0.007 | 0 | 0.5 | 0.31 | 0.48 | 0.49 | 0.59 | 0.53 |
| 6 | 107481880 | rs9373923 | -0.491 | 0.125 | 0.141 | 0.173 | 0.026 | -0.049 | -0.049 | 0.282 | 0.19 | 0.07 | 0.18 | 0.26 | 0.14 | 0.25 |
| 6 | 107482956 | rs11964270 | 0.368 | 0.377 | 0.267 | 0.199 | 0.01 | -0.027 | -0.027 | 0 | 0.05 | 0.14 | 0.04 | 0 | 0.08 | 0.02 |
| 6 | 107483340 | rs9373924 | -1.121 | -0.593 | 0.233 | -0.686 | -0.025 | 0.107 | 0.107 | 0.263 | 0.75 | 0.62 | 0.69 | 0.49 | 0.82 | 0.62 |
| 6 | 107484535 | rs6938393 | -0.213 | -0.451 | 0.343 | -0.651 | 0.058 | -0.018 | -0.018 | 0 | 0.29 | 0.42 | 0.28 | 0.23 | 0.38 | 0.26 |
| 6 | 107493418 | rs7774958 | 1.584 | -0.574 | -0.203 | -0.723 | -0.011 | 0.118 | 0.118 | 3.488 | 0.28 | 0.05 | 0.22 | 0.09 | 0.27 | 0.13 |
| 6 | 107501047 | rs6568470 | -1.286 | -0.536 | 0.028 | -0.631 | 0.007 | 0.015 | 0.015 | 1.612 | 0.66 | 0.71 | 0.72 | 0.73 | 0.65 | 0.82 |
| 6 | 107505241 | rs10457161 | 0.115 | -0.459 | -0.259 | -0.414 | 0.017 | -0.018 | -0.018 | 0 | 0.36 | 0.25 | 0.45 | 0.51 | 0.27 | 0.56 |
| 6 | 107508270 | rs12199523 | 0.342 | 1.688 | -0.759 | 0.022 | -0.077 | -0.018 | -0.018 | 0 | 0.14 | 0.01 | 0.14 | 0.13 | 0.23 | 0.09 |
| 6 | 107509052 | rs9386622 | 0.235 | 1.591 | -0.99 | 0.102 | -0.098 | -0.157 | -0.157 | 0.011 | 0.28 | 0.04 | 0.34 | 0.54 | 0.29 | 0.35 |
| 6 | 107517159 | rs11153056 | -0.025 | 1.784 | -0.137 | 0.445 | 0.003 | 0.042 | 0.042 | 0.88 | 0.3 | 0.36 | 0.35 | 0.51 | 0.28 | 0.33 |
| 6 | 107528070 | rs880353 | -0.197 | 2.31 | -0.58 | 0.298 | -0.076 | -0.049 | -0.049 | 0.183 | 0.14 | 0.04 | 0.19 | 0.16 | 0.24 | 0.27 |
| 6 | 107532514 | rs9400105 | 0.108 | 2.153 | 0.136 | 1.121 | 0.002 | 0.005 | 0.005 | 3.653 | 0.27 | 0.28 | 0.17 | 0.37 | 0.26 | 0.26 |
| 6 | 107533160 | rs3734678 | -0.558 | 2.615 | -0.15 | 1.12 | 0.01 | 0.007 | 0.007 | 0 | 0.09 | 0.09 | 0.24 | 0.21 | 0.2 | 0.07 |
| 6 | 107537007 | rs17068076 | 0.425 | 2.499 | -0.166 | 1.116 | -0.035 | -0.052 | -0.052 | 0 | 0.26 | 0.05 | 0.13 | 0.34 | 0.26 | 0.28 |
| 6 | 107538247 | rs10457162 | -0.69 | 2.462 | 0.284 | 1.215 | 0.017 | 0.084 | 0.084 | 0.266 | 0.1 | 0.18 | 0.17 | 0.26 | 0.13 | 0.07 |
| 6 | 107539445 | rs1861419 | -0.752 | 2.426 | 0.341 | 1.197 | 0.018 | 0.092 | 0.092 | 0.61 | 0.11 | 0.2 | 0.17 | 0.26 | 0.13 | 0.07 |
| 6 | 107542087 | rs9386625 | -1.828 | 1.649 | -0.224 | 0.456 | 0.06 | 0.181 | 0.181 | 2.012 | 0.64 | 0.89 | 0.7 | 0.99 | 0.71 | 0.85 |
| 6 | 107549447 | rs9480754 | 0.495 | 1.777 | -0.098 | 0.835 | -0.03 | -0.041 | -0.041 | 2.596 | 0.26 | 0.08 | 0.14 | 0.34 | 0.27 | 0.28 |
| 6 | 107552480 | rs9320208 | -1.052 | 1.952 | 0.864 | 1.164 | 0.09 | 0.156 | 0.156 | 0.412 | 0.15 | 0.39 | 0.34 | 0.38 | 0.22 | 0.28 |
| 6 | 107569957 | rs9486570 | -1.498 | 1.769 | 0.343 | 0.868 | 0.039 | 0.044 | 0.044 | 0.642 | 0.59 | 0.76 | 0.57 | 0.75 | 0.65 | 0.8 |
| 6 | 107573211 | rs2110903 | -2.237 | 1.466 | -0.165 | 0.488 | 0.013 | 0.21 | 0.21 | 1.119 | 0.7 | 0.99 | 0.73 | 1 | 0.73 | 0.86 |
| 6 | 107573570 | rs2110902 | -0.267 | 1.584 | 0.655 | 1.287 | -0.001 | 0.001 | 0.001 | 1.47 | 0.33 | 0.34 | 0.2 | 0.56 | 0.28 | 0.49 |
| 6 | 107592142 | rs10214819 | -0.26 | 1.684 | 0.847 | 1.461 | 0 | 0.002 | 0.002 | 0.635 | 0.17 | 0.22 | 0.12 | 0.21 | 0.22 | 0.22 |
| 6 | 107596774 | rs11967511 | -0.141 | 1.803 | 0.829 | 1.495 | -0.003 | -0.008 | -0.008 | 0.549 | 0.17 | 0.18 | 0.11 | 0.01 | 0.21 | 0.21 |
| 6 | 107611822 | rs9386632 | -0.149 | 1.789 | 0.662 | 1.693 | -0.054 | -0.05 | -0.05 | 0 | 0.08 | 0.01 | 0.26 | 0.23 | 0.18 | 0.09 |
| 6 | 107619620 | rs11751353 | 0.146 | 1.501 | 0.683 | 1.593 | -0.015 | -0.064 | -0.064 | 0 | 0.27 | 0.13 | 0.26 | 0.48 | 0.26 | 0.47 |
| 6 | 107637486 | rs7773295 | 0.584 | 1.453 | 0.601 | 1.612 | -0.036 | -0.09 | -0.09 | 0.019 | 0.23 | 0.02 | 0.1 | 0.32 | 0.22 | 0.29 |
| 6 | 107637582 | rs2500585 | 1.14 | 1.468 | 1.144 | 1.508 | -0.012 | 0.004 | 0.004 | 3.63 | 0.74 | 0.35 | 0.84 | 0.74 | 0.78 | 0.72 |
| 6 | 107639414 | rs2430464 | 0.951 | 1.465 | 1.458 | 1.469 | -0.033 | -0.081 | -0.081 | 2.503 | 0.75 | 0.43 | 0.85 | 0.92 | 0.79 | 0.74 |
| 6 | 107650025 | rs2500578 | 1.761 | 1.577 | 1.116 | 1.491 | -0.012 | -0.01 | -0.01 | 0 | 0.65 | 0.28 | 0.57 | 0.49 | 0.6 | 0.63 |
| 6 | 107652029 | rs9373941 | -0.348 | 1.703 | 1.105 | 1.577 | -0.036 | -0.032 | -0.032 | 0 | 0.09 | 0.04 | 0.26 | 0.24 | 0.18 | 0.09 |
| 6 | 107659857 | rs2500570 | 1.333 | 1.564 | 1.32 | 1.453 | 0.022 | 0.035 | 0.035 | 0.907 | 0.69 | 0.41 | 0.6 | 0.54 | 0.61 | 0.67 |
| 6 | 107671156 | rs6930320 | -1.296 | 1.551 | 1.147 | 1.563 | 0.053 | 0.047 | 0.047 | 0.255 | 0.52 | 0.36 | 0.61 | 0.7 | 0.62 | 0.78 |
| 6 | 107687148 | rs9386636 | -1.68 | 1.813 | 2.075 | 2.556 | 0.035 | 0.081 | 0.081 | 0.027 | 0.11 | 0.22 | 0.21 | 0.29 | 0.15 | 0.16 |
| 6 | 107713321 | rs7745716 | -0.689 | 1.812 | 1.876 | 2.546 | 0 | 0.021 | 0.021 | 0 | 0.07 | 0.09 | 0.19 | 0.23 | 0.14 | 0.12 |
| 6 | 107742215 | rs319090 | 1.272 | 1.553 | 1.843 | 2.524 | 0.005 | -0.009 | -0.009 | 0.019 | 0.69 | 0.66 | 0.69 | 0.5 | 0.78 | 0.39 |
| 6 | 107758124 | rs2353524 | -2.425 | 1.568 | 1.96 | 2.495 | 0.079 | 0.083 | 0.083 | 3.88 | 0.18 | 0.55 | 0.28 | 0.28 | 0.26 | 0.17 |
| 6 | 107763303 | rs319078 | 1.626 | 1.553 | 1.958 | 2.653 | -0.001 | 0.001 | 0.001 | 0 | 0.65 | 0.58 | 0.64 | 0.4 | 0.67 | 0.38 |
| 6 | 107766896 | rs9320215 | -1.327 | 1.582 | 1.322 | 2.798 | -0.012 | 0.006 | 0.006 | 0 | 0.1 | 0.08 | 0.32 | 0.38 | 0.18 | 0.31 |
| 6 | 107767586 | rs319083 | 1.204 | 0.928 | 1.443 | 2.778 | 0.002 | -0.002 | -0.002 | 0 | 0.72 | 0.67 | 0.83 | 0.65 | 0.82 | 0.57 |
| 6 | 107786752 | rs319070 | 2.276 | 0.933 | 1.126 | 2.735 | 0.06 | -0.065 | -0.065 | 1.596 | 0.6 | 0.16 | 0.73 | 0.5 | 0.7 | 0.52 |
| 6 | 107791661 | rs319066 | 0.746 | 0.545 | 1.098 | 2.718 | 0.012 | -0.02 | -0.02 | 0 | 0.88 | 0.9 | 0.94 | 0.69 | 0.91 | 0.78 |
| 6 | 107793039 | rs6911275 | -0.59 | 0.431 | 1.288 | 2.575 | 0.003 | -0.001 | -0.001 | 0 | 0.14 | 0.16 | 0.1 | 0.03 | 0.09 | 0.2 |

| | | | | | | | | | | | | | | | | |
|---|---|---|---|---|---|---|---|---|---|---|---|---|---|---|---|---|
| 6 | 107831841 | rs1321551 | 1.072 | 0.471 | 1.167 | 3.487 | -0.009 | 0.057 | 0.057 | 0 | 0.92 | 0.89 | 0.97 | 0.68 | 0.97 | 0.85 |
| 6 | 107860121 | rs7758767 | -1.139 | -0.045 | 1.439 | 3.539 | -0.008 | 0.108 | 0.108 | 0 | 0.06 | 0.25 | 0.05 | 0.14 | 0.01 | 0.05 |
| 6 | 107865309 | rs13218447 | -1.517 | 0.183 | 0.735 | 3.456 | 0 | 0.041 | 0.041 | 0 | 0.1 | 0.12 | 0.16 | 0.47 | 0.08 | 0.37 |
| 6 | 107868949 | rs9386647 | -1.502 | 0.211 | 0.649 | 3.5 | -0.002 | 0.008 | 0.008 | 0 | 0.1 | 0.08 | 0.16 | 0.56 | 0.09 | 0.35 |
| 6 | 107870546 | rs9373951 | -0.727 | 0.194 | 0.567 | 3.371 | 0.02 | -0.027 | -0.027 | 0 | 0.07 | 0.01 | 0.02 | 0.15 | 0.03 | 0.14 |
| 6 | 107886522 | rs7775895 | -1.069 | -0.233 | 0.94 | 2.522 | -0.019 | -0.038 | -0.038 | 0 | 0.21 | 0.26 | 0.16 | 0.04 | 0.14 | 0.33 |
| 6 | 107899996 | rs12197661 | -0.64 | 0.023 | 0.664 | 2.601 | -0.005 | 0.004 | 0.004 | 0 | 0.09 | 0.04 | 0.05 | 0.05 | 0.11 | 0.05 |
| 6 | 107903654 | rs4425630 | -1.641 | -0.195 | 0.878 | 3.161 | 0.027 | -0.017 | -0.017 | 0.107 | 0.28 | 0.24 | 0.33 | 0.63 | 0.22 | 0.63 |
| 6 | 107904134 | rs12195397 | -0.784 | -0.237 | 1.18 | 2.881 | 0 | -0.008 | -0.008 | 0.056 | 0.1 | 0.14 | 0.06 | 0.05 | 0.11 | 0.05 |
| 6 | 107906652 | rs4440495 | -1.806 | -0.181 | 1.072 | 3.223 | 0.027 | -0.041 | -0.041 | 0 | 0.28 | 0.24 | 0.32 | 0.48 | 0.2 | 0.63 |
| 6 | 107914777 | rs3924727 | -1.413 | 0.034 | 1.675 | 2.369 | -0.044 | -0.057 | -0.057 | 0 | 0.19 | 0.36 | 0.18 | 0.05 | 0.13 | 0.33 |
| 6 | 107918351 | rs9373955 | -1.583 | -0.414 | 2.135 | 3.597 | 0.007 | 0.153 | 0.153 | 0 | 0.13 | 0.23 | 0.1 | 0.43 | 0.13 | 0.3 |
| 6 | 107918382 | rs7748110 | -1.743 | -0.662 | 2.645 | 3.761 | -0.05 | 0.225 | 0.225 | 0.214 | 0.14 | 0.52 | 0.11 | 0.36 | 0.09 | 0.16 |
| 6 | 107921220 | rs2818262 | -2.168 | -0.993 | 2.747 | 3.494 | -0.063 | 0.111 | 0.111 | 0 | 0.24 | 0.51 | 0.27 | 0.57 | 0.17 | 0.56 |
| 6 | 107923284 | rs2818259 | -2.047 | -1.002 | 2.955 | 3.489 | -0.046 | 0.085 | 0.085 | 0 | 0.23 | 0.45 | 0.27 | 0.57 | 0.17 | 0.56 |
| 6 | 107929025 | rs9373956 | -2.465 | -1.087 | 2.939 | 3.195 | -0.023 | -0.014 | -0.014 | 0 | 0.33 | 0.53 | 0.46 | 0.71 | 0.26 | 0.62 |
| 6 | 107933281 | rs9386654 | -1.523 | -1.102 | 2.951 | 3.606 | 0.017 | 0.093 | 0.093 | 0.035 | 0.14 | 0.23 | 0.36 | 0.38 | 0.11 | 0.32 |
| 6 | 107944495 | rs864691 | 2.514 | -1.523 | 2.914 | 3.316 | 0.008 | 0.009 | 0.009 | 0.587 | 0.61 | 0.33 | 0.5 | 0.23 | 0.71 | 0.34 |
| 6 | 107952503 | rs1358414 | -0.496 | -1.487 | 2.982 | 3.49 | -0.003 | 0.002 | 0.002 | 0 | 0.1 | 0.08 | 0.2 | 0.26 | 0.1 | 0.21 |
| 6 | 107957516 | rs699698 | 2.22 | -1.699 | 3.099 | 3.477 | -0.073 | 0.061 | 0.061 | 0.684 | 0.71 | 0.39 | 0.7 | 0.53 | 0.82 | 0.55 |
| 6 | 107960768 | rs846959 | 1.164 | -1.718 | 3.247 | 3.525 | -0.034 | 0.028 | 0.028 | 0.695 | 0.9 | 0.79 | 0.8 | 0.86 | 0.97 | 0.88 |
| 6 | 107961025 | rs3800208 | -1.665 | -1.676 | 3.142 | 3.479 | -0.052 | 0.084 | 0.084 | 0 | 0.21 | 0.38 | 0.12 | 0.34 | 0.15 | 0.36 |
| 6 | 107969113 | rs846964 | -1.705 | -1.659 | 3.127 | 3.435 | -0.07 | 0.12 | 0.12 | 0 | 0.2 | 0.48 | 0.11 | 0.29 | 0.14 | 0.32 |
| 6 | 107975565 | rs3778579 | -0.866 | -1.668 | 3.004 | 3.433 | -0.012 | 0.023 | 0.023 | 0 | 0.16 | 0.25 | 0.09 | 0.25 | 0.14 | 0.3 |
| 6 | 107975732 | rs1728109 | -2.818 | -1.707 | 3.033 | 3.489 | -0.015 | 0.283 | 0.283 | 5 | 0.1 | 0.39 | 0.2 | 0.32 | 0.02 | 0.24 |
| 6 | 107976518 | rs1527874 | 2.235 | -1.737 | 2.704 | 3.267 | -0.009 | -0.002 | -0.002 | 8.027 | 0.7 | 0.3 | 0.7 | 0.43 | 0.83 | 0.45 |
| 6 | 107976756 | rs1527876 | 2.298 | -1.732 | 2.659 | 3.27 | -0.042 | 0.013 | 0.013 | 0.718 | 0.7 | 0.33 | 0.7 | 0.43 | 0.83 | 0.45 |
| 6 | 107985470 | rs761778 | 2.342 | -1.54 | 2.055 | 2.876 | 0.039 | 0.033 | 0.033 | 0.274 | 0.61 | 0.21 | 0.5 | 0.36 | 0.71 | 0.48 |
| 6 | 107989077 | rs846985 | 2.058 | -1.525 | 2.167 | 3.192 | -0.036 | 0.134 | 0.134 | 2.082 | 0.8 | 0.49 | 0.6 | 0.62 | 0.89 | 0.8 |
| 6 | 107989872 | rs9400129 | 1.948 | -1.717 | 2.149 | 3.251 | -0.034 | 0.038 | 0.038 | 2.576 | 0.89 | 0.64 | 0.72 | 0.83 | 0.95 | 0.86 |
| 6 | 107992074 | rs2914 | 2.416 | -1.788 | 1.892 | 3.253 | -0.036 | 0.038 | 0.038 | 0.081 | 0.75 | 0.58 | 0.71 | 0.53 | 0.84 | 0.62 |
| 6 | 107992801 | rs957115 | 1.857 | -1.649 | 1.758 | 2.812 | -0.046 | 0.051 | 0.051 | 0.284 | 0.81 | 0.63 | 0.92 | 0.73 | 0.89 | 0.74 |
| 6 | 107994328 | rs6912306 | -1.256 | -1.48 | 0.918 | 2.577 | 0.009 | -0.029 | -0.029 | 1.679 | 0.22 | 0.22 | 0.08 | 0.26 | 0.19 | 0.24 |
| 6 | 107995121 | rs7754879 | 1.806 | -1.816 | 1.097 | 2.452 | -0.019 | -0.001 | -0.001 | 0.143 | 0.7 | 0.31 | 0.62 | 0.31 | 0.83 | 0.54 |
| 6 | 107998142 | rs846995 | 1.6 | -1.373 | 1.386 | 2.77 | -0.073 | 0.204 | 0.204 | 2.893 | 0.81 | 0.57 | 0.74 | 0.6 | 0.9 | 0.69 |
| 6 | 107998198 | rs846996 | 1.377 | -1.418 | 1.393 | 2.759 | -0.083 | 0.155 | 0.155 | 1.933 | 0.83 | 0.58 | 0.93 | 0.73 | 0.91 | 0.75 |
| 6 | 108000123 | rs9373966 | -1.474 | -1.071 | 0.999 | 2.742 | 0.011 | -0.022 | -0.022 | 0.051 | 0.16 | 0.11 | 0.26 | 0.33 | 0.13 | 0.24 |
| 6 | 108000725 | rs847001 | -1.422 | -0.914 | 1.747 | 2.221 | 0.014 | 0.029 | 0.029 | 0.126 | 0.12 | 0.56 | 0.2 | 0.18 | 0.13 | 0.14 |
| 6 | 108001484 | rs847003 | -2.207 | -1.107 | 2.056 | 2.378 | -0.018 | 0.031 | 0.031 | 0 | 0.35 | 0.52 | 0.47 | 0.41 | 0.3 | 0.4 |
| 6 | 108002078 | rs847005 | -2.186 | -1.126 | 2.34 | 2.346 | 0.006 | -0.004 | -0.004 | 0.162 | 0.37 | 0.7 | 0.5 | 0.55 | 0.31 | 0.4 |
| 6 | 108003003 | rs847007 | -1.516 | -0.836 | 2.754 | 2.563 | -0.01 | 0.058 | 0.058 | 0 | 0.2 | 0.31 | 0.3 | 0.49 | 0.19 | 0.32 |
| 6 | 108006922 | rs1006074 | -1.534 | -0.84 | 2.606 | 2.531 | -0.008 | 0.031 | 0.031 | 0 | 0.2 | 0.29 | 0.3 | 0.36 | 0.17 | 0.32 |
| 6 | 108009760 | rs9400131 | -1.632 | -0.687 | 2.076 | 2.153 | 0.001 | -0.062 | -0.062 | 0 | 0.28 | 0.14 | 0.35 | 0.35 | 0.28 | 0.35 |
| 6 | 108010389 | rs17585366 | -0.593 | -0.734 | 2.237 | 2.204 | -0.002 | -0.004 | -0.004 | 0 | 0.09 | 0.09 | 0.07 | 0.07 | 0.08 | 0.06 |
| 6 | 108010940 | rs7745587 | -1.59 | -0.845 | 2.071 | 2.319 | -0.003 | -0.066 | -0.066 | 0 | 0.27 | 0.13 | 0.35 | 0.35 | 0.28 | 0.35 |
| 6 | 108012831 | rs847015 | -0.761 | -1.238 | 2.308 | 2.453 | 0.01 | 0.247 | 0.247 | 0 | 0.09 | 0.31 | 0.22 | 0.37 | 0.07 | 0.25 |
| 6 | 108013713 | rs699696 | -1.75 | -1.213 | 2.167 | 2.535 | -0.03 | 0.055 | 0.055 | 0 | 0.25 | 0.42 | 0.32 | 0.61 | 0.22 | 0.49 |
| 6 | 108017102 | rs7740153 | -0.588 | -0.995 | 2.014 | 2.558 | 0.001 | 0.099 | 0.099 | 0 | 0.09 | 0.25 | 0.14 | 0.2 | 0.07 | 0.18 |
| 6 | 108019121 | rs846950 | 1.674 | -0.967 | 1.603 | 2.568 | 0.001 | -0.003 | -0.003 | 0 | 0.65 | 0.77 | 0.58 | 0.42 | 0.62 | 0.62 |
| 6 | 108025416 | rs2295279 | -0.831 | -0.992 | 1.2 | 2.746 | -0.026 | -0.075 | -0.075 | 0.314 | 0.22 | 0.04 | 0.15 | 0.21 | 0.24 | 0.23 |
| 6 | 108025645 | rs12216157 | 2.269 | -0.873 | 0.999 | 2.476 | -0.07 | 0.237 | 0.237 | 0.648 | 0.49 | 0.02 | 0.38 | 0.2 | 0.54 | 0.43 |
| 6 | 108026146 | rs1546963 | 0.905 | -0.531 | 1.927 | 2.543 | 0.004 | 0.069 | 0.069 | 0 | 0.91 | 0.83 | 0.92 | 0.73 | 0.91 | 0.94 |
| 6 | 108026317 | rs1546962 | 0.717 | -0.534 | 1.899 | 2.497 | -0.001 | 0.003 | 0.003 | 0 | 0.92 | 0.92 | 0.93 | 0.9 | 0.93 | 0.94 |
| 6 | 108028842 | rs4083443 | 1.102 | -0.543 | 2.068 | 2.509 | 0 | 0.026 | 0.026 | 0.041 | 0.91 | 0.76 | 0.91 | 0.89 | 0.93 | 0.92 |
| 6 | 108029109 | rs3734754 | -1.138 | -0.444 | 1.879 | 2.591 | 0 | -0.017 | -0.017 | 0 | 0.17 | 0.18 | 0.14 | 0.37 | 0.2 | 0.23 |
| 6 | 108029518 | rs1728118 | 1.684 | -0.412 | 2.16 | 2.55 | -0.003 | 0.048 | 0.048 | 0 | 0.74 | 0.63 | 0.79 | 0.51 | 0.72 | 0.68 |
| 6 | 108029900 | rs1619810 | 1.751 | -0.325 | 1.876 | 2.55 | 0 | -0.003 | -0.003 | 0 | 0.73 | 0.69 | 0.79 | 0.51 | 0.71 | 0.68 |
| 6 | 108033181 | rs1728123 | 1.572 | -0.31 | 1.629 | 2.515 | 0.001 | -0.02 | -0.02 | 0 | 0.75 | 0.78 | 0.81 | 0.68 | 0.73 | 0.69 |
| 6 | 108033286 | rs1728125 | 1.572 | -0.31 | 1.629 | 2.515 | 0.001 | -0.02 | -0.02 | 0 | 0.75 | 0.78 | 0.81 | 0.68 | 0.73 | 0.69 |
| 6 | 108034407 | rs1728127 | 1.572 | -0.31 | 1.629 | 2.515 | 0.001 | -0.02 | -0.02 | 0 | 0.75 | 0.78 | 0.81 | 0.68 | 0.73 | 0.69 |
| 6 | 108040154 | rs4083470 | 1.931 | -0.31 | 1.693 | 2.346 | 0.006 | -0.013 | -0.013 | 0.466 | 0.65 | 0.17 | 0.58 | 0.41 | 0.67 | 0.59 |
| 6 | 108040855 | rs12215347 | 2.051 | -0.147 | 1.825 | 2.292 | -0.006 | -0.015 | -0.015 | 0.082 | 0.66 | 0.25 | 0.57 | 0.47 | 0.65 | 0.6 |
| 6 | 108041205 | rs846976 | -1.189 | -0.155 | 2.233 | 2.395 | -0.004 | 0.06 | 0.06 | 0 | 0.19 | 0.35 | 0.14 | 0.29 | 0.19 | 0.23 |
| 6 | 108041677 | rs11751163 | 1.897 | -0.12 | 1.625 | 2.353 | -0.007 | -0.02 | -0.02 | 0.082 | 0.68 | 0.33 | 0.59 | 0.5 | 0.66 | 0.61 |
| 6 | 108041801 | rs11751218 | 1.91 | -0.118 | 1.626 | 2.354 | -0.007 | -0.02 | -0.02 | 0 | 0.68 | 0.32 | 0.59 | 0.51 | 0.66 | 0.61 |
| 6 | 108049042 | rs4946854 | 1.911 | -0.118 | 1.332 | 2.005 | -0.005 | -0.015 | -0.015 | 0.452 | 0.62 | 0.24 | 0.57 | 0.48 | 0.65 | 0.62 |
| 6 | 108050222 | rs4945787 | 0.971 | 0.382 | 1.347 | 1.696 | 0 | 0.017 | 0.017 | 0 | 0.82 | 0.74 | 0.87 | 0.79 | 0.82 | 0.76 |
| 6 | 108053906 | rs1527871 | -1.3 | 0.654 | 1.376 | 1.306 | 0.028 | 0.047 | 0.047 | 0 | 0.13 | 0.26 | 0.22 | 0.21 | 0.15 | 0.14 |
| 6 | 108055170 | rs2049661 | -1.623 | 0.754 | 1.813 | 1.143 | 0.058 | 0.064 | 0.064 | 0 | 0.15 | 0.3 | 0.23 | 0.21 | 0.18 | 0.14 |

| | | | | | | | | | | | | | | | | |
|---|---|---|---|---|---|---|---|---|---|---|---|---|---|---|---|---|
| 6 | 108065135 | rs11759406 | -0.958 | 0.295 | 1.881 | 0.734 | -0.004 | 0.023 | 0.023 | 0 | 0.21 | 0.11 | 0.15 | 0.16 | 0.21 | 0.2 |
| 6 | 108065839 | rs6921876 | -1.477 | 0.287 | 1.917 | 0.803 | -0.009 | -0.007 | -0.007 | 0 | 0.24 | 0.4 | 0.17 | 0.22 | 0.21 | 0.21 |
| 6 | 108068003 | rs6934505 | -0.703 | 0.305 | 1.626 | 0.785 | -0.035 | 0.048 | 0.048 | 0 | 0.19 | 0.02 | 0.14 | 0.1 | 0.19 | 0.17 |
| 6 | 108069833 | rs12664414 | -1.174 | 0.362 | 1.791 | 0.802 | -0.002 | -0.002 | -0.002 | 0 | 0.23 | 0.31 | 0.16 | 0.22 | 0.21 | 0.21 |
| 6 | 108075661 | rs9400138 | -0.837 | 0.336 | 1.554 | 0.701 | 0 | 0.001 | 0.001 | 0 | 0.21 | 0.22 | 0.16 | 0.16 | 0.2 | 0.19 |
| 6 | 108081587 | rs1518524 | 1.118 | 0.296 | 1.433 | 0.35 | -0.026 | -0.093 | -0.093 | 2.947 | 0.75 | 0.42 | 0.8 | 0.9 | 0.8 | 0.81 |
| 6 | 108086370 | rs7772132 | 1.055 | 0.723 | 1.344 | 0.416 | 0.041 | 0.025 | 0.025 | 0.624 | 0.77 | 0.61 | 0.72 | 0.78 | 0.76 | 0.85 |
| 6 | 108088477 | rs4946861 | -1.443 | 0.731 | 2.095 | 0.384 | 0.117 | 0.1 | 0.1 | 0 | 0.14 | 0.48 | 0.28 | 0.19 | 0.17 | 0.14 |
| 6 | 108090454 | rs4391294 | -0.124 | -0.164 | 1.688 | -0.48 | 0.02 | 0.024 | 0.024 | 0.273 | 0.72 | 0.81 | 0.81 | 0.91 | 0.75 | 0.74 |
| 6 | 108093580 | rs6909292 | 1.432 | -0.367 | 1.365 | -0.806 | -0.043 | 0.106 | 0.106 | 0 | 0.47 | 0.1 | 0.46 | 0.73 | 0.53 | 0.58 |
| 6 | 108093657 | rs17528372 | 0.181 | -0.264 | 1.329 | -0.867 | 0.013 | 0.167 | 0.167 | 0.453 | 0.28 | 0.03 | 0.18 | 0.07 | 0.25 | 0.24 |
| 6 | 108094827 | rs2883360 | -0.545 | -0.218 | 1.574 | -0.787 | 0.084 | 0.024 | 0.024 | 0.092 | 0.12 | 0.41 | 0.26 | 0.09 | 0.15 | 0.11 |
| 6 | 108095250 | rs1850654 | -0.423 | -0.213 | 1.562 | -0.784 | 0.081 | 0.025 | 0.025 | 0 | 0.12 | 0.37 | 0.26 | 0.09 | 0.15 | 0.1 |
| 6 | 108102004 | rs938971 | -0.666 | -0.204 | 1.774 | -0.797 | -0.002 | -0.054 | -0.054 | 0 | 0.42 | 0.58 | 0.45 | 0.13 | 0.4 | 0.34 |
| 6 | 108107761 | rs9486692 | -0.731 | -0.153 | 1.859 | -0.636 | -0.007 | 0.03 | 0.03 | 0 | 0.12 | 0.31 | 0.1 | 0.13 | 0.08 | 0.06 |
| 6 | 108111651 | rs11964340 | -0.746 | -0.225 | 1.823 | -0.75 | 0.05 | 0.096 | 0.096 | 0.268 | 0.39 | 0.2 | 0.26 | 0.18 | 0.32 | 0.26 |
| 6 | 108113248 | rs1401171 | 1.542 | -0.261 | 1.738 | -0.789 | -0.063 | 0.093 | 0.093 | 0.084 | 0.45 | 0.07 | 0.45 | 0.71 | 0.52 | 0.6 |
| 6 | 108121953 | rs9320237 | -0.682 | 0.098 | 2.648 | -0.413 | 0.042 | -0.008 | -0.008 | 0 | 0.12 | 0.26 | 0.26 | 0.06 | 0.15 | 0.11 |
| 6 | 108139570 | rs17068819 | -0.652 | -0.075 | 2.42 | -0.458 | 0.005 | 0 | 0 | 0 | 0.11 | 0.11 | 0.08 | 0.12 | 0.07 | 0.03 |
| 6 | 108140137 | rs6568505 | -1.154 | 0.119 | 2.548 | -0.451 | 0.044 | -0.01 | -0.01 | 0 | 0.14 | 0.37 | 0.27 | 0.08 | 0.15 | 0.12 |
| 6 | 108145931 | rs9486730 | -1.472 | 0.087 | 2.13 | -0.466 | 0.022 | -0.014 | -0.014 | 0 | 0.17 | 0.71 | 0.29 | 0.1 | 0.15 | 0.12 |
| 6 | 108147644 | rs4946872 | -1.176 | 0.128 | 1.997 | -0.475 | 0.071 | -0.009 | -0.009 | 0 | 0.15 | 0.58 | 0.29 | 0.08 | 0.15 | 0.12 |
| 6 | 108154872 | rs9400152 | 1.219 | -0.524 | 1.356 | -0.816 | -0.001 | -0.023 | -0.023 | 0.053 | 0.71 | 0.66 | 0.83 | 0.93 | 0.75 | 0.86 |
| 6 | 108161523 | rs10782157 | 1.132 | -0.641 | 2.045 | -0.543 | -0.024 | -0.077 | -0.077 | 0.105 | 0.73 | 0.55 | 0.84 | 0.93 | 0.8 | 0.92 |
| 6 | 108162592 | rs12209269 | 1.393 | -0.691 | 1.107 | -0.986 | -0.026 | 0.069 | 0.069 | 1.181 | 0.44 | 0.1 | 0.64 | 0.7 | 0.55 | 0.7 |
| 6 | 108164951 | rs4945796 | 1.058 | -0.636 | 1.217 | -0.969 | -0.008 | -0.007 | -0.007 | 0 | 0.71 | 0.39 | 0.81 | 0.78 | 0.78 | 0.87 |
| 6 | 108165948 | rs6900051 | 0.373 | -0.559 | 0.843 | -0.773 | -0.009 | 0.11 | 0.11 | 0.218 | 0.24 | 0.01 | 0.15 | 0.07 | 0.23 | 0.18 |
| 6 | 108167542 | rs13192844 | 0.16 | -0.609 | 1.073 | -0.78 | 0.022 | 0.024 | 0.024 | 0 | 0.68 | 0.25 | 0.75 | 0.76 | 0.74 | 0.78 |
| 6 | 108169147 | rs555059 | -1.037 | -0.601 | 0.897 | -0.824 | 0.028 | 0.077 | 0.077 | 0.716 | 0.43 | 0.83 | 0.32 | 0.3 | 0.38 | 0.28 |
| 6 | 108175173 | rs632582 | -1.021 | -0.644 | 0.942 | -0.696 | 0.032 | 0.068 | 0.068 | 1.878 | 0.45 | 0.81 | 0.33 | 0.35 | 0.38 | 0.31 |
| 6 | 108185867 | rs527609 | 0.52 | -0.306 | 1.103 | -0.539 | -0.009 | -0.016 | -0.016 | 0 | 0.86 | 0.92 | 0.91 | 0.71 | 0.86 | 0.9 |
| 6 | 108188033 | rs12528856 | 1.127 | -0.322 | 1.026 | -0.429 | 0.034 | 0.058 | 0.058 | 2.752 | 0.51 | 0.22 | 0.63 | 0.57 | 0.6 | 0.7 |
| 6 | 108189411 | rs6933329 | 0.053 | -0.195 | 0.843 | 0.238 | 0.014 | 0.047 | 0.047 | 0 | 0.08 | 0.16 | 0.06 | 0.13 | 0.1 | 0.29 |
| 6 | 108191007 | rs10872020 | 0.863 | -0.18 | 1.3 | 0.071 | 0.026 | 0.079 | 0.079 | 7.341 | 0.83 | 0.54 | 0.88 | 0.71 | 0.86 | 0.9 |
| 6 | 108213533 | rs663455 | -0.471 | -0.197 | 1.335 | 0.084 | 0.001 | -0.034 | -0.034 | 0 | 0.15 | 0.24 | 0.07 | 0.06 | 0.13 | 0.06 |
| 6 | 108221950 | rs6938487 | 0.269 | -0.188 | 1.321 | 0.094 | -0.011 | -0.017 | -0.017 | 0 | 0.86 | 0.92 | 0.91 | 0.71 | 0.85 | 0.9 |
| 6 | 108233929 | rs675117 | 0.289 | -0.205 | 1.49 | 0.086 | -0.01 | -0.015 | -0.015 | 0 | 0.86 | 0.91 | 0.9 | 0.71 | 0.85 | 0.9 |
| 6 | 108238917 | rs12206846 | 0.976 | -0.275 | 1.45 | 0.026 | 0.029 | 0.047 | 0.047 | 2.385 | 0.52 | 0.29 | 0.65 | 0.57 | 0.59 | 0.7 |
| 6 | 108260459 | rs2064200 | 0.982 | -0.041 | 0.701 | 0.14 | 0.034 | 0.077 | 0.077 | 6.204 | 0.82 | 0.55 | 0.88 | 0.71 | 0.84 | 0.9 |
| 6 | 108261315 | rs2244924 | 1.161 | 0.006 | 0.578 | 0.313 | 0.021 | 0.002 | 0.002 | 0 | 0.77 | 0.68 | 0.86 | 0.78 | 0.75 | 0.65 |
| 6 | 108262363 | rs2143019 | -0.342 | -0.028 | 0.357 | 0.088 | -0.002 | -0.006 | -0.006 | 0 | 0.13 | 0.09 | 0.08 | 0.2 | 0.13 | 0.09 |
| 6 | 108269759 | rs12195281 | -0.169 | 0.001 | 0.234 | -0.112 | 0.003 | 0.101 | 0.101 | 0 | 0.29 | 0.03 | 0.21 | 0.12 | 0.25 | 0.18 |
| 6 | 108281713 | rs12204951 | -0.156 | -0.071 | -0.333 | -0.207 | 0.008 | 0.01 | 0.01 | 0 | 0.31 | 0.28 | 0.24 | 0.13 | 0.26 | 0.19 |
| 6 | 108292031 | rs9384666 | -0.991 | 0.128 | -0.826 | 0.138 | 0 | 0.026 | 0.026 | 0 | 0.2 | 0.12 | 0.1 | 0.13 | 0.15 | 0.17 |
| 6 | 108293757 | rs9486762 | 1.062 | 0.568 | -0.719 | 0.686 | 0.002 | 0.015 | 0.015 | 0 | 0.69 | 0.59 | 0.58 | 0.55 | 0.71 | 0.71 |
| 6 | 108294470 | rs1001145 | -0.603 | 0.52 | -0.923 | 0.471 | -0.047 | 0.08 | 0.08 | 0 | 0.21 | 0.09 | 0.14 | 0.03 | 0.26 | 0.14 |
| 6 | 108300912 | rs11754713 | -0.726 | -0.664 | -0.062 | 1.629 | 0.02 | 0.005 | 0.005 | 0 | 0.22 | 0.13 | 0.11 | 0.19 | 0.14 | 0.14 |
| 6 | 108307246 | rs7748333 | 0.363 | -0.174 | 0.359 | 1.994 | -0.007 | -0.01 | -0.01 | 0 | 0.66 | 0.75 | 0.67 | 0.65 | 0.65 | 0.66 |
| 6 | 108308659 | rs2050042 | 0.948 | -0.406 | 0.743 | 1.964 | 0.013 | 0.006 | 0.006 | 0.093 | 0.51 | 0.32 | 0.48 | 0.58 | 0.54 | 0.56 |
| 6 | 108314167 | rs2355203 | 0.858 | -0.524 | 0.83 | 1.862 | -0.004 | 0.032 | 0.032 | 0.003 | 0.52 | 0.34 | 0.49 | 0.64 | 0.53 | 0.63 |
| 6 | 108317269 | rs7773352 | 0.068 | -0.565 | 0.467 | 1.738 | -0.035 | 0.068 | 0.068 | 0 | 0.65 | 0.79 | 0.63 | 0.85 | 0.58 | 0.72 |
| 6 | 108321035 | rs11153096 | 1.184 | -0.658 | 0.143 | 1.969 | -0.105 | -0.115 | -0.115 | 0 | 0.46 | 0.03 | 0.46 | 0.49 | 0.5 | 0.52 |
| 6 | 108321196 | rs6930921 | -1.334 | -0.372 | 1.177 | 2.21 | 0.051 | 0.193 | 0.193 | 1.739 | 0.06 | 0.43 | 0.04 | 0.1 | 0 | 0.02 |
| 6 | 108323759 | rs17069122 | -0.195 | -0.177 | -0.103 | 2.108 | 0.017 | -0.036 | -0.036 | 0 | 0.06 | 0 | 0.01 | 0.17 | 0.02 | 0.11 |
| 6 | 108325624 | rs17282829 | 0.739 | -0.073 | -0.041 | 1.896 | -0.031 | -0.031 | -0.031 | 0 | 0.56 | 0.17 | 0.51 | 0.5 | 0.58 | 0.56 |
| 6 | 108332351 | rs7770621 | -1.734 | -0.461 | 1.036 | 1.962 | -0.065 | 0.018 | 0.018 | 1.719 | 0.18 | 0.6 | 0.16 | 0.19 | 0.05 | 0.18 |
| 6 | 108335345 | rs7749081 | -1.449 | -0.125 | 0.683 | 2.536 | 0.002 | -0.006 | -0.006 | 3.311 | 0.25 | 0.71 | 0.53 | 0.44 | 0.21 | 0.22 |
| 6 | 108351590 | rs9374002 | -0.365 | -0.226 | 0.634 | 2.673 | -0.047 | -0.045 | -0.045 | 0 | 0.14 | 0.02 | 0.22 | 0.2 | 0.21 | 0.12 |
| 6 | 108352055 | rs9372177 | -1.865 | -0.089 | 1 | 2.899 | -0.001 | 0.009 | 0.009 | 0 | 0.28 | 0.62 | 0.54 | 0.59 | 0.28 | 0.31 |
| 6 | 108354898 | rs9400180 | -1.865 | -0.089 | 1 | 2.899 | -0.001 | 0.009 | 0.009 | 0 | 0.28 | 0.62 | 0.54 | 0.59 | 0.28 | 0.31 |
| 6 | 108357992 | rs2097525 | -0.561 | -0.085 | 1.184 | 2.926 | 0.005 | 0.004 | 0.004 | 0 | 0.15 | 0.12 | 0.24 | 0.25 | 0.24 | 0.12 |
| 6 | 108358328 | rs9374007 | -1.422 | -0.085 | 1.078 | 2.926 | -0.038 | 0.279 | 0.279 | 0 | 0.13 | 0.44 | 0.3 | 0.33 | 0.05 | 0.19 |
| 6 | 108361990 | rs17069204 | -1.877 | -0.087 | 1.055 | 2.921 | -0.001 | 0.009 | 0.009 | 0 | 0.28 | 0.62 | 0.54 | 0.59 | 0.28 | 0.32 |
| 6 | 108364582 | rs12567 | -0.584 | -0.057 | 1.091 | 2.928 | 0.03 | 0.028 | 0.028 | 0 | 0.16 | 0.22 | 0.26 | 0.26 | 0.24 | 0.12 |
| 6 | 108364951 | rs1064346 | -0.584 | -0.087 | 1.088 | 2.928 | 0.029 | 0.029 | 0.029 | 0 | 0.16 | 0.23 | 0.26 | 0.26 | 0.24 | 0.12 |
| 6 | 108365658 | rs9320250 | -1.902 | -0.094 | 0.805 | 2.912 | -0.001 | 0.008 | 0.008 | 0 | 0.28 | 0.62 | 0.54 | 0.59 | 0.28 | 0.32 |
| 6 | 108381347 | rs9374013 | -0.523 | 0.064 | 0.395 | 2.788 | 0.012 | 0.011 | 0.011 | 0 | 0.15 | 0.18 | 0.26 | 0.25 | 0.24 | 0.12 |
| 6 | 108387232 | rs2027876 | -1.139 | 0.073 | 0.365 | 2.698 | 0.004 | 0.011 | 0.011 | 0 | 0.2 | 0.2 | 0.51 | 0.35 | 0.26 | 0.21 |
| 6 | 108393223 | rs6921166 | 0.519 | -0.341 | -0.306 | 2.059 | -0.004 | -0.047 | -0.047 | 0 | 0.85 | 0.99 | 0.96 | 0.82 | 0.88 | 0.87 |

| | | | | | | | | | | | | | | | | |
|---|---|---|---|---|---|---|---|---|---|---|---|---|---|---|---|---|
| 6 | 108407662 | rs2001144 | -1.402 | -0.165 | 0.379 | 2.266 | -0.029 | 0.15 | 0.15 | 0 | 0.14 | 0.62 | 0.33 | 0.38 | 0.07 | 0.2 |
| 6 | 108412714 | rs7769514 | -1.366 | -0.035 | 0.428 | 1.509 | -0.004 | -0.022 | -0.022 | 0 | 0.12 | 0.26 | 0.09 | 0.07 | 0.12 | 0.17 |
| 6 | 108412760 | rs7769535 | -0.754 | -0.781 | -0.66 | 1.453 | 0.054 | -0.005 | -0.005 | 0 | 0.3 | 0.13 | 0.22 | 0.29 | 0.19 | 0.29 |
| 6 | 108415392 | rs4946900 | 1.129 | -0.414 | -0.405 | 1.35 | 0.007 | 0.005 | 0.005 | 0 | 0.81 | 0.84 | 0.9 | 0.86 | 0.84 | 0.86 |
| 6 | 108427424 | rs9374021 | 1.343 | 0.517 | -0.51 | 1.658 | -0.035 | -0.02 | -0.02 | 0 | 0.71 | 0.87 | 0.78 | 0.71 | 0.66 | 0.68 |
| 6 | 108442222 | rs6919324 | -0.955 | 0.385 | 0.514 | 1.686 | 0.017 | 0.017 | 0.017 | 0 | 0.21 | 0.56 | 0.17 | 0.23 | 0.22 | 0.14 |
| 6 | 108446023 | rs17069274 | -0.299 | 0.26 | 0.147 | 1.692 | 0.002 | 0 | 0 | 0 | 0.07 | 0.07 | 0.07 | 0.06 | 0.07 | 0.05 |
| 6 | 108448046 | rs9486815 | -2.059 | -0.627 | 0.104 | 1.92 | 0.003 | -0.004 | -0.004 | 0 | 0.39 | 0.68 | 0.49 | 0.48 | 0.33 | 0.36 |
| 6 | 108449212 | rs4245535 | -0.206 | -0.623 | -0.132 | 1.737 | -0.002 | -0.009 | -0.009 | 0 | 0.07 | 0.02 | 0.07 | 0.06 | 0.07 | 0.05 |
| 6 | 108451893 | rs218294 | -1.931 | -0.656 | 0.084 | 1.816 | 0.02 | -0.014 | -0.014 | 0 | 0.38 | 0.72 | 0.27 | 0.39 | 0.32 | 0.32 |
| 6 | 108454437 | rs218289 | -1.108 | 0.167 | 0.052 | 1.422 | 0.021 | 0.007 | 0.007 | 0 | 0.22 | 0.37 | 0.19 | 0.23 | 0.28 | 0.16 |
| 6 | 108467024 | rs218291 | -0.054 | 0.235 | -1.535 | 0.182 | -0.008 | 0.096 | 0.096 | 0.111 | 0.4 | 0.79 | 0.47 | 0.16 | 0.38 | 0.41 |
| 6 | 108469920 | rs170125 | -0.526 | 0.274 | -2.012 | 0.325 | -0.015 | -0.021 | -0.021 | 1.041 | 0.55 | 0.95 | 0.53 | 0.39 | 0.41 | 0.58 |
| 6 | 108470257 | rs12173906 | -0.465 | 0.262 | -1.086 | 0.681 | 0.001 | 0 | 0 | 11.259 | 0.1 | 0.1 | 0.05 | 0.23 | 0.05 | 0.15 |
| 6 | 108489462 | rs383319 | -0.22 | 0.946 | -1.47 | 0.084 | -0.064 | -0.046 | -0.046 | 1.086 | 0.74 | 0.91 | 0.65 | 0.37 | 0.65 | 0.67 |
| 6 | 108499945 | rs217528 | 0.495 | -0.181 | -0.926 | -0.951 | 0.001 | 0.002 | 0.002 | 0 | 0.46 | 0.46 | 0.38 | 0.25 | 0.5 | 0.3 |
| 6 | 108511004 | rs217534 | -0.743 | -0.317 | -1.291 | -1.126 | -0.183 | -0.098 | -0.098 | 0.208 | 0.71 | 0.98 | 0.61 | 0.39 | 0.59 | 0.72 |
| 6 | 108529085 | rs177717 | 1.151 | -1.443 | -1.328 | -1.794 | -0.045 | 0.097 | 0.097 | 0 | 0.35 | 0.18 | 0.31 | 0.2 | 0.44 | 0.25 |
| 6 | 108535016 | rs579257 | -0.18 | -1.621 | -0.864 | -1.831 | -0.069 | -0.034 | -0.034 | 0 | 0.27 | 0.51 | 0.25 | 0.18 | 0.12 | 0.45 |
| 6 | 108545932 | rs847610 | -0.522 | -1.916 | -0.834 | -1.975 | -0.086 | -0.054 | -0.054 | 0.359 | 0.24 | 0.56 | 0.25 | 0.12 | 0.09 | 0.29 |
| 6 | 108552247 | rs217135 | -1.423 | -1.788 | -0.085 | -1.939 | -0.018 | 0.045 | 0.045 | 0 | 0.64 | 0.74 | 0.69 | 0.8 | 0.55 | 0.75 |
| 6 | 108558280 | rs588409 | -1.568 | -1.965 | -0.389 | -2.462 | -0.008 | -0.016 | -0.016 | 0 | 0.79 | 0.82 | 0.77 | 0.99 | 0.67 | 0.86 |
| 6 | 108561974 | rs657669 | -0.283 | -2.162 | 0.368 | -2.302 | 0.014 | 0.008 | 0.008 | 0.012 | 0.23 | 0.19 | 0.25 | 0.16 | 0.13 | 0.38 |
| 6 | 108571664 | rs635051 | -1.264 | -2.007 | 0.562 | -2.318 | -0.029 | -0.022 | -0.022 | 0.714 | 0.34 | 0.56 | 0.29 | 0.17 | 0.15 | 0.47 |
| 6 | 108602186 | rs488865 | -0.59 | -1.923 | 0.465 | -2.307 | -0.024 | -0.016 | -0.016 | 0 | 0.24 | 0.31 | 0.26 | 0.15 | 0.13 | 0.31 |
| 6 | 108610641 | rs539630 | -1.329 | -2.236 | 0.461 | -2.455 | 0.038 | 0.037 | 0.037 | 0.08 | 0.36 | 0.7 | 0.3 | 0.17 | 0.14 | 0.41 |
| 6 | 108623855 | rs671144 | 0.17 | -2.238 | 0.402 | -2.463 | 0.022 | 0.006 | 0.006 | 0 | 0.16 | 0.08 | 0.21 | 0.15 | 0.11 | 0.29 |
| 6 | 108658974 | rs12528572 | 0.961 | -2.193 | 0.372 | -2.472 | 0.012 | 0.003 | 0.003 | 0 | 0.11 | 0.01 | 0.05 | 0.05 | 0.05 | 0.17 |
| 6 | 108664930 | rs7742646 | -0.915 | -2.168 | 0.45 | -2.457 | -0.006 | 0.001 | 0.001 | 0 | 0.92 | 0.93 | 0.98 | 0.98 | 0.99 | 0.91 |
| 6 | 108687824 | rs10457177 | 1.401 | -1.982 | 0.424 | -2.71 | -0.01 | -0.014 | -0.014 | 0 | 0.21 | 0.18 | 0.23 | 0.01 | 0.34 | 0.13 |
| 6 | 108690156 | rs1537862 | 1.594 | -1.923 | 0.298 | -2.617 | -0.192 | -0.049 | -0.049 | 0 | 0.29 | 0.02 | 0.42 | 0.64 | 0.42 | 0.29 |
| 6 | 108694389 | rs12208637 | 0.807 | -1.895 | 0.508 | -2.639 | -0.016 | 0.06 | 0.06 | 0 | 0.36 | 0.23 | 0.32 | 0.2 | 0.44 | 0.3 |
| 6 | 108713146 | rs11963798 | 0.849 | -2.038 | 0.921 | -2.671 | -0.052 | -0.042 | -0.042 | 0.018 | 0.09 | 0.2 | 0.03 | 0.02 | 0.01 | 0.09 |
| 6 | 108739757 | rs9486880 | 0.792 | -1.952 | 0.823 | -2.624 | -0.021 | 0.064 | 0.064 | 0 | 0.36 | 0.22 | 0.31 | 0.2 | 0.45 | 0.31 |
| 6 | 108767328 | rs11153115 | -1.895 | -1.92 | 0.659 | -2.648 | -0.134 | -0.045 | -0.045 | 0.434 | 0.72 | 0.96 | 0.57 | 0.37 | 0.61 | 0.72 |
| 6 | 108789853 | rs9486893 | 0.288 | -1.646 | 0.566 | -2.725 | 0.019 | -0.023 | -0.023 | 0 | 0.3 | 0.56 | 0.37 | 0.31 | 0.33 | 0.43 |
| 6 | 108791307 | rs12202169 | 0.836 | -1.711 | 0.57 | -2.652 | -0.032 | 0.057 | 0.057 | 0 | 0.36 | 0.24 | 0.32 | 0.2 | 0.48 | 0.33 |
| 6 | 108798498 | rs1332027 | 0.281 | -1.262 | 0.203 | -2.431 | 0.018 | 0.004 | 0.004 | 0 | 0.15 | 0.07 | 0.16 | 0.14 | 0.08 | 0.26 |
| 6 | 108799750 | rs17069590 | 0.872 | -1.198 | 0.197 | -2.423 | -0.015 | -0.025 | -0.025 | 0 | 0.08 | 0.12 | 0.05 | 0 | 0.03 | 0.03 |
| 6 | 108815553 | rs12526579 | -1.506 | -1.213 | 0.111 | -2.468 | -0.055 | 0.041 | 0.041 | 0.911 | 0.69 | 0.85 | 0.56 | 0.3 | 0.58 | 0.67 |
| 6 | 108840870 | rs2274776 | -0.248 | -0.796 | -0.161 | -2.304 | 0 | 0.174 | 0.174 | 0.06 | 0.5 | 0.78 | 0.38 | 0.26 | 0.49 | 0.43 |
| 6 | 108848121 | rs6903501 | -0.798 | -0.629 | -0.141 | -2.022 | -0.012 | 0.017 | 0.017 | 0.43 | 0.62 | 0.68 | 0.52 | 0.38 | 0.56 | 0.67 |
| 6 | 108853493 | rs17041 | -0.297 | -0.348 | 0.01 | -2.174 | -0.002 | 0.037 | 0.037 | 0.007 | 0.48 | 0.44 | 0.46 | 0.25 | 0.49 | 0.45 |
| 6 | 108853606 | rs4946925 | 0.149 | -0.046 | -0.424 | -2.275 | -0.005 | 0.007 | 0.007 | 0 | 0.21 | 0.2 | 0.34 | 0.19 | 0.28 | 0.31 |
| 6 | 108856378 | rs6568547 | -0.033 | -0.682 | -0.034 | -2.302 | -0.03 | -0.115 | -0.115 | 0 | 0.28 | 0.41 | 0.17 | 0.09 | 0.23 | 0.16 |
| 6 | 108858460 | rs9480861 | -0.432 | -0.477 | -0.612 | -2.568 | 0.01 | 0.149 | 0.149 | 0 | 0.55 | 0.77 | 0.29 | 0.15 | 0.43 | 0.43 |
| 6 | 108859593 | rs9374035 | 0.592 | -0.429 | 0.313 | -2.095 | 0.007 | -0.017 | -0.017 | 7.906 | 0.7 | 0.59 | 0.79 | 0.68 | 0.67 | 0.58 |
| 6 | 108865663 | rs6927268 | -0.736 | -1.315 | 0.413 | -2.352 | 0.017 | 0.02 | 0.02 | 0 | 0.29 | 0.23 | 0.15 | 0.27 | 0.24 | 0.37 |
| 6 | 108866570 | rs17069632 | 0.691 | -1.3 | 0.344 | -2.349 | 0.021 | 0.01 | 0.01 | 2.962 | 0.15 | 0.11 | 0.06 | 0 | 0.05 | 0.04 |
| 6 | 108867031 | rs6911407 | -1.139 | -1.558 | 0.664 | -2.524 | 0.052 | 0.105 | 0.105 | 0 | 0.51 | 0.69 | 0.36 | 0.35 | 0.43 | 0.56 |
| 6 | 108875645 | rs9372187 | 0.335 | -1.527 | 0.677 | -2.657 | 0.016 | 0.015 | 0.015 | 0.46 | 0.19 | 0.17 | 0.08 | 0.07 | 0.11 | 0.29 |
| 6 | 108878052 | rs9486902 | 0.814 | -1.621 | 0.943 | -2.726 | -0.003 | -0.052 | -0.052 | 0 | 0.19 | 0.28 | 0.21 | 0.05 | 0.18 | 0.12 |
| 6 | 108885623 | rs1536057 | -0.267 | -1.618 | 0.923 | -2.724 | -0.009 | -0.062 | -0.062 | 0 | 0.29 | 0.42 | 0.27 | 0.21 | 0.32 | 0.2 |
| 6 | 108889822 | rs9372190 | 0.257 | -1.765 | 0.837 | -2.599 | 0.018 | 0.017 | 0.017 | 0.35 | 0.19 | 0.18 | 0.09 | 0.07 | 0.11 | 0.29 |
| 6 | 108913201 | rs13220810 | 0.285 | -0.566 | 1.243 | -1.646 | -0.088 | -0.018 | -0.018 | 0 | 0.14 | 0.01 | 0.19 | 0.1 | 0.21 | 0.16 |
| 6 | 108941393 | rs17598747 | -0.263 | -2.067 | 1.925 | -1.653 | -0.003 | 0.001 | 0.001 | 0 | 0.1 | 0.08 | 0.05 | 0.15 | 0.13 | 0.07 |
| 6 | 108944165 | rs3813498 | 0.652 | -2.14 | 2.173 | -1.744 | -0.034 | -0.073 | -0.073 | 0 | 0.73 | 0.45 | 0.73 | 0.85 | 0.78 | 0.57 |
| 6 | 108951655 | rs9285397 | -0.653 | -2.152 | 2.092 | -1.785 | -0.028 | -0.092 | -0.092 | 0 | 0.25 | 0.42 | 0.23 | 0.07 | 0.19 | 0.18 |
| 6 | 108975562 | rs12203834 | -1.066 | -2.076 | 1.139 | -1.712 | -0.019 | -0.012 | -0.012 | 0 | 0.14 | 0.23 | 0.06 | 0.15 | 0.14 | 0.08 |
| 6 | 108981117 | rs4946933 | 0.579 | -1.706 | 1.184 | -1.51 | -0.012 | 0.002 | 0.002 | 0.119 | 0.94 | 0.88 | 0.96 | 0.93 | 0.96 | 0.75 |
| 6 | 108984067 | rs12196996 | -0.061 | -1.69 | 0.979 | -1.504 | 0.002 | 0.007 | 0.007 | 0 | 0.07 | 0.07 | 0.05 | 0.15 | 0.11 | 0.06 |
| 6 | 108988184 | rs2153960 | 1.451 | -1.749 | 0.786 | -1.57 | 0.044 | 0.119 | 0.119 | 0.686 | 0.59 | 0.07 | 0.66 | 0.7 | 0.64 | 0.49 |
| 6 | 108994826 | rs9398172 | 1.479 | -1.627 | 0.869 | -1.613 | 0.034 | 0.127 | 0.127 | 2.78 | 0.62 | 0.09 | 0.66 | 0.71 | 0.65 | 0.51 |
| 6 | 108996553 | rs3778588 | 0.524 | -1.647 | 1.103 | -1.628 | -0.021 | -0.081 | -0.081 | 0 | 0.75 | 0.38 | 0.76 | 0.93 | 0.81 | 0.82 |
| 6 | 108996963 | rs3800229 | 1.494 | -1.601 | 0.906 | -1.604 | 0.029 | 0.125 | 0.125 | 2.812 | 0.62 | 0.08 | 0.66 | 0.71 | 0.65 | 0.51 |
| 6 | 108998128 | rs3800230 | -0.816 | -1.581 | 0.967 | -1.584 | -0.005 | 0.007 | 0.007 | 0.022 | 0.12 | 0.17 | 0.09 | 0.22 | 0.16 | 0.31 |
| 6 | 108998953 | rs3800232 | -0.816 | -1.581 | 0.967 | -1.584 | -0.005 | 0.007 | 0.007 | 0.022 | 0.12 | 0.17 | 0.09 | 0.22 | 0.16 | 0.3 |
| 6 | 108999287 | rs1935949 | 1.361 | -1.549 | 0.42 | -1.569 | 0.029 | 0.125 | 0.125 | 2.812 | 0.62 | 0.08 | 0.66 | 0.71 | 0.65 | 0.51 |
| 6 | 109007522 | rs1268170 | -1.599 | -1.494 | 0.492 | -1.59 | 0.047 | 0.168 | 0.168 | 0.013 | 0.43 | 0.85 | 0.36 | 0.31 | 0.41 | 0.5 |

| | | | | | | | | | | | | | | | | |
|---|---|---|---|---|---|---|---|---|---|---|---|---|---|---|---|---|
| 6 | 109011327 | rs761779 | -0.112 | -1.522 | 1.105 | -1.622 | -0.011 | -0.121 | -0.121 | 0.247 | 0.2 | 0.46 | 0.21 | 0.05 | 0.18 | 0.11 |
| 6 | 109013930 | rs9486916 | -1.211 | -1.557 | 0.725 | -1.632 | 0.045 | 0.114 | 0.114 | 4.691 | 0.3 | 0.88 | 0.28 | 0.15 | 0.23 | 0.39 |
| 6 | 109017322 | rs1268177 | -1.517 | -1.521 | 0.657 | -1.618 | 0.03 | 0.175 | 0.175 | 0.761 | 0.42 | 0.9 | 0.35 | 0.28 | 0.4 | 0.44 |
| 6 | 109020032 | rs479744 | -1.128 | -1.532 | 0.97 | -1.622 | -0.011 | -0.041 | -0.041 | 0.348 | 0.31 | 0.65 | 0.27 | 0.14 | 0.24 | 0.46 |
| 6 | 109020634 | rs4946938 | -1.506 | -1.754 | 1.61 | -1.131 | -0.061 | -0.061 | -0.061 | 0 | 0.19 | 0.55 | 0.21 | 0.14 | 0.11 | 0.42 |
| 6 | 109024689 | rs517924 | -1.285 | -1.215 | 1.761 | -0.751 | -0.031 | -0.032 | -0.032 | 0 | 0.21 | 0.39 | 0.13 | 0.24 | 0.2 | 0.33 |
| 6 | 109030387 | rs9320268 | -1.466 | -1.273 | 1.659 | -0.577 | -0.034 | -0.017 | -0.017 | 0 | 0.24 | 0.56 | 0.14 | 0.25 | 0.22 | 0.35 |
| 6 | 109031166 | rs556862 | -1.814 | -1.492 | 1.619 | -0.639 | -0.076 | -0.029 | -0.029 | 2.399 | 0.17 | 0.42 | 0.07 | 0.13 | 0.07 | 0.36 |
| 6 | 109032000 | rs564376 | -2.045 | -1.445 | 1.347 | -0.633 | -0.007 | -0.002 | -0.002 | 0 | 0.33 | 0.58 | 0.15 | 0.29 | 0.24 | 0.44 |
| 6 | 109032557 | rs9320269 | -1.659 | -1.069 | 0.971 | -0.482 | -0.055 | -0.051 | -0.051 | 0 | 0.25 | 0.39 | 0.11 | 0.2 | 0.18 | 0.17 |
| 6 | 109032772 | rs10457181 | -0.755 | -0.394 | 0.678 | -0.191 | -0.018 | -0.018 | -0.018 | 0.545 | 0.16 | 0.26 | 0.09 | 0.16 | 0.16 | 0.07 |
| 6 | 109037698 | rs6913023 | -0.225 | 0.311 | 0.053 | -0.169 | -0.001 | -0.008 | -0.008 | 0 | 0.53 | 0.49 | 0.51 | 0.39 | 0.55 | 0.6 |
| 6 | 109038738 | rs9386747 | -1.538 | 0.478 | 0.97 | 0.148 | 0.056 | 0.046 | 0.046 | 0.064 | 0.12 | 0.48 | 0.17 | 0.11 | 0.11 | 0.14 |
| 6 | 109039275 | rs9486931 | -1.441 | 0.554 | 1.217 | 0.068 | 0.142 | 0.017 | 0.017 | 0.109 | 0.07 | 0.45 | 0.07 | 0.01 | 0.09 | 0.05 |
| 6 | 109042459 | rs484754 | -0.516 | 0.698 | -0.231 | 0.379 | -0.104 | -0.099 | -0.099 | 0 | 0.73 | 0.97 | 0.77 | 0.76 | 0.71 | 0.86 |
| 6 | 109051334 | rs6928805 | -0.275 | 0.208 | 0.13 | 0.32 | 0.031 | -0.011 | -0.011 | 0 | 0.47 | 0.82 | 0.61 | 0.55 | 0.41 | 0.54 |
| 6 | 109053056 | rs9384686 | -0.315 | 0.169 | 0.342 | 0.296 | 0.028 | 0.054 | 0.054 | 0.382 | 0.47 | 0.79 | 0.6 | 0.7 | 0.41 | 0.58 |
| 6 | 109053712 | rs2096243 | 0.192 | 0.025 | 0.748 | 0.154 | 0 | 0 | 0 | 0.02 | 0.37 | 0.41 | 0.45 | 0.65 | 0.3 | 0.47 |
| 6 | 109058427 | rs560539 | -0.662 | -0.042 | 0.638 | 0.099 | 0.039 | 0.062 | 0.062 | 1.726 | 0.5 | 0.91 | 0.62 | 0.68 | 0.42 | 0.58 |
| 6 | 109060923 | rs9400242 | -0.064 | -0.057 | 1.203 | 0.083 | -0.008 | -0.005 | -0.005 | 0 | 0.41 | 0.57 | 0.48 | 0.67 | 0.36 | 0.55 |
| 6 | 109061368 | rs578889 | -1.442 | 0.056 | 1.494 | 0.095 | 0.029 | -0.033 | -0.033 | 0.012 | 0.08 | 0.33 | 0.14 | 0.01 | 0.06 | 0.04 |
| 6 | 109061451 | rs579702 | -1.68 | 0.049 | 1.255 | 0.099 | 0.007 | -0.071 | -0.071 | 0 | 0.13 | 0.51 | 0.16 | 0.05 | 0.12 | 0.12 |
| 6 | 109065741 | rs9374042 | -0.342 | -0.102 | 1.244 | 0.211 | 0.001 | 0.002 | 0.002 | 0.158 | 0.44 | 0.44 | 0.59 | 0.64 | 0.41 | 0.59 |
| 6 | 109067998 | rs541040 | -0.762 | -0.028 | 1.074 | 0.279 | 0.004 | 0.021 | 0.021 | 1.582 | 0.54 | 0.74 | 0.67 | 0.64 | 0.45 | 0.66 |
| 6 | 109069491 | rs520324 | -0.605 | 0.05 | 0.955 | 0.297 | 0.004 | 0.01 | 0.01 | 0.65 | 0.52 | 0.58 | 0.55 | 0.64 | 0.44 | 0.66 |
| 6 | 109075953 | rs1268160 | -0.342 | -0.182 | 1.084 | 0.576 | -0.003 | -0.001 | -0.001 | 0 | 0.36 | 0.55 | 0.39 | 0.37 | 0.28 | 0.46 |
| 6 | 109077940 | rs9480877 | -0.282 | -0.221 | 0.725 | 0.673 | 0 | -0.001 | -0.001 | 0.068 | 0.33 | 0.33 | 0.27 | 0.36 | 0.26 | 0.47 |
| 6 | 109080801 | rs571520 | -0.959 | -0.555 | 0.455 | 1.141 | 0.006 | -0.01 | -0.01 | 0 | 0.31 | 0.72 | 0.19 | 0.4 | 0.25 | 0.37 |
| 6 | 109081291 | rs9486948 | -1.472 | -0.55 | 0.981 | 1.109 | 0.08 | 0.07 | 0.07 | 0.775 | 0.06 | 0.5 | 0.04 | 0.02 | 0.03 | 0.06 |
| 6 | 109085636 | rs497961 | -0.523 | -0.582 | 0.499 | 1.269 | 0 | -0.064 | -0.064 | 0 | 0.23 | 0.13 | 0.13 | 0.39 | 0.21 | 0.38 |
| 6 | 109087960 | rs544782 | -1.135 | -0.482 | -0.127 | 1.03 | -0.009 | 0.03 | 0.03 | 0 | 0.65 | 0.91 | 0.59 | 0.73 | 0.67 | 0.87 |
| 6 | 109099027 | rs1146229 | -1.993 | -0.9 | 0.651 | 1.261 | -0.033 | 0.137 | 0.137 | 4.306 | 0.15 | 0.6 | 0.12 | 0.34 | 0.09 | 0.29 |
| 6 | 109107768 | rs560188 | -0.622 | -0.903 | 0.678 | 1.31 | -0.025 | 0.091 | 0.091 | 0.034 | 0.07 | 0.14 | 0.07 | 0.34 | 0.04 | 0.22 |
| 6 | 109115347 | rs472656 | -1.545 | -0.928 | 0.853 | 1.127 | -0.002 | 0.002 | 0.002 | 0.041 | 0.3 | 0.67 | 0.21 | 0.42 | 0.24 | 0.44 |
| 6 | 109116491 | rs1268128 | -1.332 | -0.827 | 0.344 | 0.971 | -0.032 | 0.013 | 0.013 | 0 | 0.64 | 0.91 | 0.6 | 0.7 | 0.65 | 0.85 |
| 6 | 109118557 | rs11961452 | -1.305 | -0.676 | 0.867 | 0.881 | 0.135 | 0.053 | 0.053 | 0.319 | 0.06 | 0.3 | 0.05 | 0.02 | 0.05 | 0.07 |
| 6 | 109123244 | rs953053 | 0.257 | -0.684 | 0.08 | 0.897 | -0.035 | 0.052 | 0.052 | 0 | 0.15 | 0 | 0.07 | 0.06 | 0.14 | 0.16 |
| 6 | 109125822 | rs576546 | -1.212 | -0.71 | 0.143 | 1.028 | -0.01 | 0.042 | 0.042 | 0.05 | 0.64 | 0.88 | 0.59 | 0.74 | 0.65 | 0.85 |
| 6 | 109126624 | rs9374045 | 0.333 | -0.93 | 0.888 | 1.112 | 0.006 | 0.005 | 0.005 | 0 | 0.37 | 0.36 | 0.43 | 0.42 | 0.46 | 0.45 |
| 6 | 109126925 | rs517697 | -1.971 | -1.103 | 1.004 | 1.23 | -0.046 | 0.211 | 0.211 | 2.855 | 0.16 | 0.52 | 0.12 | 0.38 | 0.08 | 0.28 |
| 6 | 109127580 | rs524693 | -1.738 | -1.028 | 1.065 | 1.411 | 0.031 | 0.125 | 0.125 | 6.428 | 0.1 | 0.5 | 0.08 | 0.12 | 0.04 | 0.1 |
| 6 | 109127884 | rs527507 | -1.659 | -1.122 | 0.964 | 1.311 | -0.013 | 0.043 | 0.043 | 6.736 | 0.14 | 0.53 | 0.09 | 0.12 | 0.06 | 0.12 |
| 6 | 109129684 | rs8180632 | 0.762 | -1.063 | 0.426 | 1.256 | -0.054 | -0.063 | -0.063 | 0.902 | 0.31 | 0.15 | 0.39 | 0.33 | 0.36 | 0.3 |
| 6 | 109129939 | rs9372195 | -0.567 | -1.175 | 0.629 | 1.125 | 0.013 | 0.006 | 0.006 | 0 | 0.52 | 0.28 | 0.48 | 0.43 | 0.56 | 0.62 |
| 6 | 109132221 | rs9384688 | 0.395 | -1.007 | 0.807 | 1.115 | -0.015 | -0.035 | -0.035 | 0.54 | 0.36 | 0.27 | 0.41 | 0.41 | 0.42 | 0.34 |
| 6 | 109138491 | rs471422 | -0.946 | -1.089 | 1.118 | 1.14 | 0.003 | 0.285 | 0.285 | 0.417 | 0.12 | 0.45 | 0.11 | 0.35 | 0.07 | 0.22 |
| 6 | 109139055 | rs11153132 | 0.766 | -1.075 | 0.684 | 1.216 | -0.039 | -0.042 | -0.042 | 0 | 0.31 | 0.17 | 0.38 | 0.32 | 0.38 | 0.32 |
| 6 | 109142443 | rs1268125 | -1.727 | -1.15 | 0.518 | 0.595 | -0.041 | 0.102 | 0.102 | 0.01 | 0.67 | 0.93 | 0.62 | 0.83 | 0.68 | 0.86 |
| 6 | 109145305 | rs12209602 | 0.354 | -0.853 | 0.617 | 0.545 | -0.023 | 0.061 | 0.061 | 0 | 0.15 | 0.02 | 0.07 | 0.06 | 0.14 | 0.2 |
| 6 | 109148253 | rs6920522 | 0.092 | -0.894 | 0.664 | 0.651 | -0.058 | -0.065 | -0.065 | 0.063 | 0.44 | 0.14 | 0.45 | 0.44 | 0.53 | 0.48 |
| 6 | 109148861 | rs9486961 | -1.264 | -0.638 | 1.53 | 0.633 | 0.096 | 0.013 | 0.013 | 0 | 0.06 | 0.29 | 0.06 | 0.01 | 0.06 | 0.1 |
| 6 | 109169164 | rs948521 | 0.382 | -1.117 | 1.23 | 0.596 | -0.002 | -0.016 | -0.016 | 0 | 0.2 | 0.24 | 0.11 | 0.17 | 0.19 | 0.24 |
| 6 | 109182944 | rs1268055 | 0.177 | -1.274 | 1.233 | 0.387 | 0 | -0.002 | -0.002 | 0.6 | 0.53 | 0.49 | 0.5 | 0.27 | 0.51 | 0.38 |
| 6 | 109209701 | rs2806369 | -0.483 | -1.006 | 1.272 | 0.812 | 0.105 | 0.054 | 0.054 | 0.273 | 0.87 | 0.67 | 0.86 | 0.79 | 0.79 | 0.83 |
| 6 | 109221846 | rs2806371 | 0.964 | -0.95 | 0.975 | 0.826 | -0.032 | -0.055 | -0.055 | 0.464 | 0.55 | 0.2 | 0.66 | 0.42 | 0.55 | 0.39 |
| 6 | 109237426 | rs2806377 | 0.281 | -0.889 | 0.828 | 0.396 | 0.059 | 0.174 | 0.174 | 0.395 | 0.82 | 0.6 | 0.8 | 0.6 | 0.77 | 0.69 |
| 6 | 109240562 | rs13219800 | 1.748 | -1.127 | 0.621 | 0.171 | -0.111 | 0.072 | 0.072 | 0 | 0.25 | 0.05 | 0.36 | 0.09 | 0.31 | 0.11 |
| 6 | 109243086 | rs1040849 | 0.272 | -1.005 | 0.758 | 0.3 | 0.059 | 0.168 | 0.168 | 0.395 | 0.82 | 0.6 | 0.8 | 0.6 | 0.77 | 0.69 |
| 6 | 109244232 | rs1475055 | 0.998 | -0.885 | 0.533 | 0.077 | -0.004 | -0.133 | -0.133 | 3.588 | 0.27 | 0.01 | 0.28 | 0.33 | 0.24 | 0.27 |
| 6 | 109249965 | rs2798642 | -0.267 | -0.402 | 1.064 | 0.048 | 0.04 | 0.066 | 0.066 | 0.056 | 0.17 | 0.31 | 0.19 | 0.31 | 0.22 | 0.24 |
| 6 | 109259465 | rs6930984 | 0.338 | -0.431 | 0.731 | -0.89 | 0.001 | 0.003 | 0.003 | 0 | 0.38 | 0.46 | 0.47 | 0.58 | 0.38 | 0.4 |
| 6 | 109262787 | rs911478 | 0.048 | -0.489 | -0.511 | -1 | 0.005 | -0.06 | -0.06 | 0 | 0.61 | 0.44 | 0.73 | 0.86 | 0.57 | 0.56 |
| 6 | 109263617 | rs9480897 | -0.631 | -1.171 | 0.314 | -0.208 | 0.05 | 0.069 | 0.069 | 1.852 | 0.05 | 0.33 | 0.03 | 0.07 | 0.03 | 0.05 |
| 6 | 109268050 | rs2798641 | 0.308 | -1.126 | -1.061 | -0.172 | 0.008 | -0.077 | -0.077 | 1.069 | 0.23 | 0.04 | 0.27 | 0.28 | 0.18 | 0.16 |
| 6 | 109270551 | rs2768552 | -0.149 | -0.699 | -1.009 | 0.006 | 0.009 | 0 | 0 | 0 | 0.36 | 0.67 | 0.34 | 0.36 | 0.26 | 0.28 |
| 6 | 109276321 | rs2768548 | 0.451 | -0.428 | -0.776 | -0.538 | 0.003 | 0 | 0 | 0 | 0.67 | 0.69 | 0.85 | 0.82 | 0.73 | 0.64 |
| 6 | 109277165 | rs2806358 | -0.69 | -0.573 | -0.75 | -0.458 | -0.012 | -0.03 | -0.03 | 0 | 0.08 | 0.12 | 0.02 | 0 | 0.03 | 0.02 |
| 6 | 109279105 | rs12195848 | -0.675 | -0.823 | -0.826 | -0.918 | 0.008 | 0.007 | 0.007 | 0 | 0.19 | 0.14 | 0.08 | 0.04 | 0.14 | 0.21 |
| 6 | 109285189 | rs2273669 | -1.149 | -1.065 | -0.196 | -1.088 | -0.028 | -0.075 | -0.075 | 2.43 | 0.22 | 0.35 | 0.09 | 0.08 | 0.16 | 0.25 |

| | | | | | | | | | | | | | | | | |
|---|---|---|---|---|---|---|---|---|---|---|---|---|---|---|---|---|
| 6 | 109292736 | rs1887414 | -1.108 | -1.075 | -0.131 | -1.093 | -0.024 | -0.061 | -0.061 | 0 | 0.22 | 0.31 | 0.1 | 0.08 | 0.16 | 0.26 |
| 6 | 109294752 | rs3734649 | -0.196 | -1.096 | -0.061 | -1.103 | 0.012 | 0.037 | 0.037 | 0 | 0.15 | 0.05 | 0.07 | 0.04 | 0.14 | 0.2 |
| 6 | 109295040 | rs3734651 | -1.188 | -1.095 | -0.06 | -1.103 | -0.024 | -0.061 | -0.061 | 0 | 0.22 | 0.31 | 0.1 | 0.08 | 0.16 | 0.24 |
| 6 | 109302258 | rs12196024 | -0.257 | -1.083 | -0.095 | -1.096 | 0.011 | 0.027 | 0.027 | 0 | 0.16 | 0.09 | 0.08 | 0.04 | 0.14 | 0.2 |
| 6 | 109322554 | rs12214121 | -0.163 | -1.08 | -0.1 | -1.094 | 0.012 | 0.039 | 0.039 | 0 | 0.15 | 0.04 | 0.07 | 0.04 | 0.14 | 0.2 |
| 6 | 109323519 | rs2273668 | -0.322 | -1.079 | -0.137 | -1.094 | -0.009 | 0.056 | 0.056 | 0 | 0.15 | 0.01 | 0.07 | 0.04 | 0.14 | 0.2 |
| 6 | 109337723 | rs12525751 | 0.256 | -0.665 | -0.803 | -1.568 | -0.039 | 0.041 | 0.041 | 0 | 0.18 | 0.07 | 0.18 | 0.07 | 0.23 | 0.19 |
| 6 | 109340558 | rs7757845 | -0.822 | -0.608 | -0.84 | -1.468 | -0.004 | -0.015 | -0.015 | 0 | 0.07 | 0.24 | 0.02 | 0.03 | 0.02 | 0.04 |
| 6 | 109363734 | rs2883970 | 0.011 | -0.628 | -0.987 | -0.79 | 0 | 0.012 | 0.012 | 0 | 0.94 | 0.95 | 0.86 | 0.85 | 0.93 | 0.82 |
| 6 | 109383659 | rs2050310 | -0.532 | -0.655 | -1.382 | -0.263 | -0.072 | -0.009 | -0.009 | 0 | 0.25 | 0.09 | 0.24 | 0.23 | 0.34 | 0.25 |
| 6 | 109410569 | rs6929104 | 0.787 | -0.294 | -1.909 | -0.321 | -0.086 | -0.039 | -0.039 | 0 | 0.28 | 0.01 | 0.2 | 0.24 | 0.33 | 0.16 |
| 6 | 109416586 | rs1106853 | -0.239 | -0.338 | -1.614 | -0.39 | -0.021 | -0.059 | -0.059 | 0 | 0.2 | 0.37 | 0.1 | 0.07 | 0.14 | 0.23 |
| 6 | 109459287 | rs11966104 | 0.251 | -0.341 | -1.615 | -0.414 | -0.013 | -0.046 | -0.046 | 0 | 0.16 | 0.24 | 0.09 | 0.04 | 0.13 | 0.19 |
| 6 | 109461506 | rs1547325 | 0.301 | -0.295 | -1.675 | -0.41 | -0.009 | -0.037 | -0.037 | 0.11 | 0.16 | 0.22 | 0.09 | 0.04 | 0.13 | 0.19 |
| 6 | 109475416 | rs351734 | 0.847 | -0.302 | -1.78 | -0.435 | -0.078 | -0.034 | -0.034 | 0 | 0.28 | 0.01 | 0.2 | 0.24 | 0.33 | 0.13 |
| 6 | 109483826 | rs1998172 | -0.779 | -0.089 | -0.887 | 0.095 | -0.023 | -0.065 | -0.065 | 0.828 | 0.2 | 0.38 | 0.1 | 0.07 | 0.14 | 0.23 |
| 6 | 109497085 | rs3813499 | -0.257 | 0.14 | -1.362 | 0.245 | 0 | -0.008 | -0.008 | 0 | 0.16 | 0.15 | 0.08 | 0.04 | 0.13 | 0.19 |
| 6 | 109499953 | rs9791265 | -0.196 | 0.132 | -1.384 | 0.366 | -0.01 | -0.05 | -0.05 | 0 | 0.31 | 0.21 | 0.4 | 0.54 | 0.31 | 0.43 |
| 6 | 109503976 | rs351739 | 0.077 | -0.008 | -1.015 | 0.414 | 0 | 0 | 0 | 0.012 | 0.64 | 0.37 | 0.78 | 0.7 | 0.67 | 0.75 |
| 6 | 109504086 | rs352849 | -0.573 | 0.339 | -1.443 | 0.624 | -0.017 | -0.015 | -0.015 | 0 | 0.69 | 0.77 | 0.53 | 0.59 | 0.63 | 0.56 |
| 6 | 109510972 | rs7757193 | 0.525 | 0.252 | -1.667 | 0.753 | -0.096 | -0.081 | -0.081 | 0 | 0.39 | 0.03 | 0.56 | 0.41 | 0.38 | 0.49 |
| 6 | 109517706 | rs1761608 | -0.167 | 0.282 | -1.625 | 0.709 | 0.023 | -0.032 | -0.032 | 0 | 0.44 | 0.81 | 0.29 | 0.45 | 0.44 | 0.32 |
| 6 | 109523821 | rs352834 | 1.003 | 0.237 | -1.657 | 0.677 | -0.055 | 0.007 | 0.007 | 0.656 | 0.15 | 0.01 | 0.13 | 0.13 | 0.18 | 0.17 |
| 6 | 109524939 | rs427960 | 0.083 | -0.051 | -0.507 | 0.68 | -0.007 | -0.014 | -0.014 | 1.364 | 0.39 | 0.54 | 0.25 | 0.31 | 0.41 | 0.25 |
| 6 | 109524979 | rs399561 | 0.083 | -0.051 | -0.507 | 0.68 | -0.007 | -0.015 | -0.015 | 0 | 0.39 | 0.53 | 0.25 | 0.31 | 0.41 | 0.25 |
| 6 | 109525045 | rs449850 | 0.083 | -0.051 | -0.507 | 0.701 | -0.007 | -0.016 | -0.016 | 0 | 0.39 | 0.53 | 0.25 | 0.31 | 0.41 | 0.25 |
| 6 | 109527347 | rs191611 | 0.083 | -0.068 | -0.452 | 0.704 | 0.031 | 0.168 | 0.168 | 0 | 0.94 | 0.73 | 0.97 | 0.85 | 0.97 | 0.93 |
| 6 | 109531188 | rs9487008 | 0.235 | -0.579 | -0.67 | 0.315 | -0.056 | -0.056 | -0.056 | 0 | 0.19 | 0.37 | 0.09 | 0.06 | 0.11 | 0.11 |
| 6 | 109548713 | rs59673 | 0.737 | -0.42 | -0.609 | 0.307 | -0.033 | -0.025 | -0.025 | 0 | 0.19 | 0.09 | 0.16 | 0.25 | 0.29 | 0.14 |
| 6 | 109557542 | rs9384701 | 0.983 | -0.377 | -0.687 | 0.291 | -0.084 | -0.073 | -0.073 | 0.047 | 0.18 | 0.01 | 0.15 | 0.24 | 0.27 | 0.13 |
| 6 | 109558471 | rs6919423 | -0.477 | -0.735 | -0.348 | 0.157 | -0.003 | 0.011 | 0.011 | 0 | 0.35 | 0.67 | 0.25 | 0.26 | 0.4 | 0.21 |
| 6 | 109574095 | rs884366 | 1.698 | -0.515 | -0.594 | 0.282 | -0.053 | -0.034 | -0.034 | 0 | 0.36 | 0.07 | 0.41 | 0.39 | 0.32 | 0.51 |
| 6 | 109579723 | rs1113666 | 0.775 | -0.574 | -0.278 | 0.348 | -0.026 | 0.022 | 0.022 | 1.253 | 0.62 | 0.5 | 0.62 | 0.53 | 0.65 | 0.71 |
| 6 | 109583186 | rs1040285 | -0.099 | -0.569 | -0.241 | 0.378 | -0.039 | 0.113 | 0.113 | 1.206 | 0.75 | 0.54 | 0.71 | 0.53 | 0.78 | 0.74 |
| 6 | 109586334 | rs9374069 | -0.11 | -0.487 | -0.157 | 0.4 | 0.005 | 0.057 | 0.057 | 0 | 0.3 | 0.53 | 0.32 | 0.46 | 0.36 | 0.25 |
| 6 | 109586678 | rs932222 | 0.575 | -0.541 | -0.178 | 0.336 | -0.001 | 0.03 | 0.03 | 0 | 0.11 | 0.04 | 0.09 | 0.01 | 0.13 | 0.04 |
| 6 | 109586944 | rs9487019 | -0.003 | -0.557 | -0.076 | 0.344 | 0.003 | -0.048 | -0.048 | 0 | 0.09 | 0.32 | 0.03 | 0 | 0.04 | 0.05 |
| 6 | 109598964 | rs13210693 | -1.444 | -0.546 | -0.24 | 0.362 | -0.031 | -0.014 | -0.014 | 0.162 | 0.5 | 0.89 | 0.46 | 0.48 | 0.53 | 0.34 |
| 6 | 109599263 | rs6910119 | -1.444 | -0.546 | -0.24 | 0.362 | -0.031 | -0.014 | -0.014 | 0 | 0.5 | 0.89 | 0.46 | 0.48 | 0.53 | 0.34 |
| 6 | 109608497 | rs9386791 | -0.863 | -0.53 | -0.227 | 0.142 | 0 | 0.001 | 0.001 | 0 | 0.41 | 0.57 | 0.41 | 0.4 | 0.49 | 0.29 |
| 6 | 109618704 | rs9386796 | -0.112 | -0.554 | -0.179 | 0.144 | 0.002 | -0.009 | -0.009 | 0 | 0.32 | 0.33 | 0.27 | 0.13 | 0.43 | 0.19 |
| 6 | 109619697 | rs9487053 | -1.561 | -0.566 | -0.21 | 0.152 | -0.059 | -0.048 | -0.048 | 0 | 0.51 | 0.92 | 0.45 | 0.4 | 0.53 | 0.34 |
| 6 | 109622494 | rs11153166 | 0.797 | -0.538 | -0.206 | 0.14 | 0.029 | -0.014 | -0.014 | 0 | 0.21 | 0.31 | 0.18 | 0.12 | 0.3 | 0.15 |
| 6 | 109624937 | rs9784763 | -0.698 | -0.564 | -0.739 | 0.143 | 0 | 0.004 | 0.004 | 0 | 0.37 | 0.6 | 0.34 | 0.39 | 0.4 | 0.3 |
| 6 | 109632520 | rs9386797 | -1.419 | -0.502 | -1.373 | 0.192 | -0.055 | -0.041 | -0.041 | 1.185 | 0.51 | 0.91 | 0.45 | 0.4 | 0.53 | 0.33 |
| 6 | 109634828 | rs11966072 | 0.871 | -0.374 | -1.311 | 0.165 | -0.043 | 0.008 | 0.008 | 0 | 0.19 | 0.1 | 0.18 | 0.12 | 0.33 | 0.13 |
| 6 | 109637154 | rs7755798 | -1.404 | -0.288 | -1.568 | 0.193 | -0.038 | -0.012 | -0.012 | 0 | 0.87 | 0.99 | 0.88 | 0.86 | 0.85 | 0.84 |
| 6 | 109640488 | rs11759269 | 1.109 | -0.239 | -1.535 | 0.072 | -0.038 | -0.049 | -0.049 | 0 | 0.43 | 0.11 | 0.43 | 0.46 | 0.35 | 0.55 |
| 6 | 109641411 | rs4946957 | 1.454 | -0.139 | -1.469 | 0.187 | -0.059 | -0.103 | -0.103 | 0 | 0.36 | 0.04 | 0.41 | 0.46 | 0.32 | 0.51 |
| 6 | 109642916 | rs2884036 | 0.032 | 0.248 | -0.388 | 0.06 | 0.045 | -0.056 | -0.056 | 0 | 0.23 | 0.53 | 0.21 | 0.08 | 0.32 | 0.12 |
| 6 | 109647807 | rs1832778 | -0.77 | -0.161 | -0.317 | 0.474 | -0.014 | -0.008 | -0.008 | 0 | 0.16 | 0.23 | 0.11 | 0.14 | 0.12 | 0.11 |
| 6 | 109654132 | rs12529733 | -0.338 | -0.068 | -1.14 | 0.58 | 0.044 | -0.013 | -0.013 | 0 | 0.52 | 0.8 | 0.54 | 0.51 | 0.44 | 0.5 |
| 6 | 109654221 | rs12524502 | 0.219 | -0.46 | -0.278 | 0.521 | -0.009 | -0.03 | -0.03 | 0 | 0.34 | 0.34 | 0.21 | 0.15 | 0.32 | 0.23 |
| 6 | 109668288 | rs2275650 | -0.044 | -0.661 | -0.334 | 0.623 | 0.032 | -0.006 | -0.006 | 0 | 0.37 | 0.17 | 0.31 | 0.35 | 0.35 | 0.39 |
| 6 | 109689431 | rs974034 | -0.986 | -0.699 | 0.206 | 0.598 | 0.045 | -0.001 | -0.001 | 0 | 0.52 | 0.76 | 0.53 | 0.51 | 0.43 | 0.55 |
| 6 | 109689907 | rs11251 | -0.493 | -0.704 | 0.214 | 0.598 | 0.024 | -0.017 | -0.017 | 1.082 | 0.44 | 0.22 | 0.44 | 0.45 | 0.42 | 0.52 |
| 6 | 109692188 | rs9372213 | -1.173 | -0.856 | -0.024 | 1.169 | -0.023 | 0.085 | 0.085 | 0 | 0.57 | 0.87 | 0.56 | 0.68 | 0.47 | 0.57 |
| 6 | 109726739 | rs7754650 | -0.377 | -0.832 | 0.093 | 1.116 | 0.022 | -0.012 | -0.012 | 0.243 | 0.44 | 0.24 | 0.44 | 0.45 | 0.42 | 0.52 |
| 6 | 109736253 | rs13195517 | 0.681 | -0.956 | 0.163 | 1.112 | 0.019 | 0.07 | 0.07 | 0.297 | 0.32 | 0.12 | 0.19 | 0.16 | 0.31 | 0.23 |
| 6 | 109740101 | rs17534632 | 1.404 | -1.284 | -0.219 | 0.74 | -0.02 | 0.126 | 0.126 | 0.145 | 0.2 | 0.01 | 0.14 | 0 | 0.21 | 0.06 |
| 6 | 109742015 | rs9487094 | -0.596 | -1.051 | 0.18 | 1.023 | 0.067 | 0.002 | 0.002 | 0.04 | 0.46 | 0.72 | 0.41 | 0.41 | 0.37 | 0.42 |
| 6 | 109751601 | rs9480957 | -0.036 | -1.043 | 0.461 | 0.989 | 0.017 | 0 | 0 | 0 | 0.38 | 0.25 | 0.32 | 0.35 | 0.35 | 0.39 |
| 6 | 109762236 | rs1048203 | 0.013 | -1.061 | 0.466 | 0.98 | 0.028 | -0.003 | -0.003 | 0 | 0.38 | 0.19 | 0.32 | 0.35 | 0.35 | 0.39 |
| 6 | 109762332 | rs1048197 | 0.753 | -1.037 | 0.377 | 1.028 | -0.003 | 0.038 | 0.038 | 0 | 0.06 | 0.06 | 0.12 | 0.19 | 0.04 | 0.17 |
| 6 | 109762600 | rs749621 | 0.122 | -0.945 | 0.329 | 1.001 | -0.004 | -0.003 | -0.003 | 0 | 0.07 | 0.07 | 0.13 | 0.1 | 0.07 | 0.14 |
| 6 | 109764535 | rs1476387 | -0.985 | -0.968 | 0.353 | 0.976 | 0.048 | -0.003 | -0.003 | 0 | 0.52 | 0.75 | 0.49 | 0.45 | 0.43 | 0.55 |
| 6 | 109768295 | rs910730 | -0.977 | -0.947 | 0.352 | 0.996 | 0.055 | 0 | 0 | 0 | 0.52 | 0.76 | 0.49 | 0.45 | 0.43 | 0.55 |
| 6 | 109771955 | rs7769789 | -1.52 | 0.207 | -0.143 | 0.387 | -0.049 | 0.046 | 0.046 | 0 | 0.92 | 0.99 | 0.87 | 1 | 0.75 | 0.97 |
| 6 | 109775366 | rs4946977 | 0.08 | -0.182 | 0.036 | 0.477 | 0.001 | 0.001 | 0.001 | 0 | 0.08 | 0.12 | 0.13 | 0.1 | 0.07 | 0.14 |

| | | | | | | | | | | | | | | | | |
|---|---|---|---|---|---|---|---|---|---|---|---|---|---|---|---|---|
| 6 | 109775436 | rs2277113 | -0.432 | -0.199 | -0.029 | 0.444 | 0.049 | -0.008 | -0.008 | 0 | 0.52 | 0.76 | 0.49 | 0.46 | 0.43 | 0.55 |
| 6 | 109786980 | rs3734652 | -0.479 | -0.175 | -0.11 | 0.463 | 0.044 | -0.004 | -0.004 | 0.121 | 0.52 | 0.75 | 0.49 | 0.5 | 0.43 | 0.55 |
| 6 | 109797304 | rs2236084 | -0.471 | -0.169 | -0.107 | 0.467 | 0.045 | -0.003 | -0.003 | 0 | 0.52 | 0.75 | 0.49 | 0.5 | 0.43 | 0.55 |
| 6 | 109814411 | rs1054607 | 0.855 | -0.37 | -0.324 | 0.246 | 0.01 | -0.032 | -0.032 | 0 | 0.32 | 0.22 | 0.35 | 0.49 | 0.26 | 0.34 |
| 6 | 109815351 | rs7755941 | 0.542 | -0.389 | -0.049 | 0.219 | 0.012 | 0.001 | 0.001 | 0 | 0.42 | 0.28 | 0.41 | 0.38 | 0.41 | 0.47 |
| 6 | 109817276 | rs11153183 | 0.799 | -0.414 | -0.058 | 0.206 | 0.012 | -0.009 | -0.009 | 0 | 0.4 | 0.17 | 0.41 | 0.38 | 0.41 | 0.47 |
| 6 | 109828474 | rs10457192 | 0.799 | -0.414 | -0.058 | 0.206 | 0.012 | -0.009 | -0.009 | 0 | 0.4 | 0.17 | 0.4 | 0.38 | 0.41 | 0.47 |
| 6 | 109832034 | rs9766863 | 0.802 | -0.553 | 0.016 | 0.318 | 0.022 | -0.031 | -0.031 | 0 | 0.32 | 0.21 | 0.33 | 0.47 | 0.23 | 0.38 |
| 6 | 109849103 | rs6568580 | 0.264 | -0.381 | 0.01 | 0.248 | 0.012 | -0.014 | -0.014 | 0 | 0.72 | 0.32 | 0.73 | 0.85 | 0.65 | 0.85 |
| 6 | 109854243 | rs10782163 | 0.883 | -0.555 | -0.249 | 0.267 | 0.012 | -0.009 | -0.009 | 0 | 0.4 | 0.17 | 0.41 | 0.38 | 0.41 | 0.46 |
| 6 | 109870377 | rs9374097 | 0.231 | -0.565 | -0.239 | 0.281 | 0.013 | -0.014 | -0.014 | 0.056 | 0.72 | 0.32 | 0.73 | 0.85 | 0.64 | 0.85 |
| 6 | 109875361 | rs10434873 | 1.148 | -0.573 | -0.162 | 0.264 | -0.001 | 0.081 | 0.081 | 0 | 0.05 | 0.16 | 0.07 | 0.14 | 0.04 | 0.02 |
| 6 | 109876128 | rs10872041 | 0.235 | -0.565 | -0.316 | 0.286 | 0.013 | -0.014 | -0.014 | 0 | 0.72 | 0.32 | 0.73 | 0.85 | 0.64 | 0.85 |
| 6 | 109885475 | rs10499052 | 1.733 | -0.716 | -0.405 | 0.239 | -0.002 | 0.075 | 0.075 | 0.447 | 0.29 | 0.03 | 0.18 | 0.16 | 0.3 | 0.18 |
| 6 | 109891830 | rs9487156 | 0.144 | -0.703 | -0.385 | 0.245 | 0.013 | -0.014 | -0.014 | 0.036 | 0.72 | 0.32 | 0.73 | 0.85 | 0.64 | 0.85 |
| 6 | 109892465 | rs9374105 | 1.232 | -0.711 | -0.402 | 0.257 | 0.008 | -0.027 | -0.027 | 0.946 | 0.28 | 0.11 | 0.28 | 0.34 | 0.22 | 0.32 |
| 6 | 109892868 | rs9320301 | 1.409 | -0.726 | -0.41 | 0.25 | -0.032 | -0.074 | -0.074 | 0.079 | 0.27 | 0.03 | 0.27 | 0.34 | 0.22 | 0.32 |
| 6 | 109894354 | rs13214717 | 1.49 | -0.717 | -0.375 | 0.249 | 0.019 | 0.077 | 0.077 | 0.171 | 0.31 | 0.09 | 0.2 | 0.16 | 0.31 | 0.21 |
| 6 | 109896308 | rs11153194 | 0.39 | -0.697 | -0.373 | 0.277 | -0.005 | 0.002 | 0.002 | 0.18 | 0.69 | 0.28 | 0.69 | 0.72 | 0.63 | 0.78 |
| 6 | 109918151 | rs12205941 | 1.399 | -0.692 | -0.394 | 0.343 | 0.009 | -0.014 | -0.014 | 0 | 0.06 | 0 | 0.11 | 0.2 | 0.04 | 0.16 |
| 6 | 109946561 | rs12529505 | 2.089 | -0.783 | -0.458 | 0.191 | -0.003 | 0.113 | 0.113 | 1.139 | 0.26 | 0.03 | 0.18 | 0.11 | 0.28 | 0.13 |
| 6 | 109951180 | rs9487176 | 0.238 | -1.079 | -0.272 | 0.303 | 0.013 | -0.014 | -0.014 | 0 | 0.72 | 0.32 | 0.73 | 0.85 | 0.64 | 0.85 |
| 6 | 109953266 | rs12215867 | 0.238 | -1.079 | -0.272 | 0.303 | 0.013 | -0.014 | -0.014 | 0.115 | 0.72 | 0.32 | 0.73 | 0.85 | 0.64 | 0.85 |
| 6 | 109954252 | rs1406957 | 0.238 | -1.079 | -0.272 | 0.303 | 0.013 | -0.014 | -0.014 | 4.126 | 0.72 | 0.32 | 0.73 | 0.85 | 0.64 | 0.85 |
| 6 | 109954518 | rs9374111 | 1.107 | -1.085 | -0.277 | 0.31 | 0.008 | -0.027 | -0.027 | 0 | 0.28 | 0.11 | 0.28 | 0.34 | 0.22 | 0.32 |
| 6 | 109965474 | rs9400310 | 0.275 | -0.938 | -0.142 | 0.31 | 0.013 | -0.014 | -0.014 | 0 | 0.72 | 0.32 | 0.73 | 0.85 | 0.64 | 0.85 |
| 6 | 109987242 | rs9372223 | 1.011 | -1.076 | -0.341 | 0.388 | 0.012 | -0.021 | -0.021 | 0 | 0.28 | 0.11 | 0.28 | 0.34 | 0.22 | 0.32 |
| 6 | 109997529 | rs7767805 | 0.98 | -1.088 | -0.199 | 0.36 | -0.003 | 0.075 | 0.075 | 0 | 0.06 | 0.16 | 0.07 | 0.14 | 0.04 | 0.02 |
| 6 | 110011458 | rs12192268 | 0.777 | -0.943 | -0.306 | 0.468 | 0.011 | -0.002 | -0.002 | 3.198 | 0.4 | 0.16 | 0.4 | 0.34 | 0.41 | 0.46 |
| 6 | 110016656 | rs1096136 | 0.712 | -1.151 | -0.264 | 0.424 | -0.081 | 0.075 | 0.075 | 0.004 | 0.18 | 0.07 | 0.17 | 0.01 | 0.32 | 0.13 |
| 6 | 110017595 | rs1103996 | -1.075 | -1.179 | -0.214 | 0.554 | 0.027 | -0.05 | -0.05 | 0 | 0.68 | 0.83 | 0.67 | 0.53 | 0.77 | 0.61 |
| 6 | 110028962 | rs9386835 | 0.785 | -0.994 | -0.035 | 0.548 | 0.009 | 0.133 | 0.133 | 0 | 0.06 | 0.22 | 0.07 | 0.18 | 0.04 | 0.02 |
| 6 | 110050042 | rs6933962 | -1.514 | -1.093 | -0.4 | 0.609 | 0.002 | -0.017 | -0.017 | 0 | 0.73 | 0.9 | 0.73 | 0.67 | 0.78 | 0.7 |
| 6 | 110051141 | rs2224550 | -1.138 | -1.116 | -0.335 | 0.606 | -0.019 | -0.005 | -0.005 | 0 | 0.41 | 0.76 | 0.52 | 0.53 | 0.47 | 0.47 |
| 6 | 110067773 | rs12206574 | 0.466 | -1.043 | -0.348 | 0.699 | -0.013 | -0.026 | -0.026 | 0.22 | 0.09 | 0.03 | 0.2 | 0.21 | 0.1 | 0.25 |
| 6 | 110068614 | rs12528392 | 0.718 | -1.116 | -0.408 | 0.611 | 0.035 | -0.012 | -0.012 | 0.109 | 0.4 | 0.15 | 0.4 | 0.36 | 0.37 | 0.48 |
| 6 | 110081618 | rs9481003 | -0.375 | -1.015 | -0.019 | 0.364 | -0.018 | 0.249 | 0.249 | 0 | 0.11 | 0.45 | 0.12 | 0.3 | 0.05 | 0.08 |
| 6 | 110092054 | rs9374121 | -0.39 | -0.768 | -0.492 | 0.175 | 0.068 | -0.024 | -0.024 | 0.384 | 0.28 | 0.56 | 0.29 | 0.21 | 0.39 | 0.15 |
| 6 | 110096896 | rs11968978 | 0.893 | -0.917 | -1.082 | 0.242 | 0.047 | 0.09 | 0.09 | 0.119 | 0.31 | 0.08 | 0.2 | 0.14 | 0.28 | 0.23 |
| 6 | 110106234 | rs10499054 | 0.425 | -1.249 | -1.309 | 0.55 | 0.053 | 0.029 | 0.029 | 0 | 0.42 | 0.12 | 0.49 | 0.32 | 0.65 | 0.32 |
| 6 | 110107517 | rs9885672 | -0.607 | -1.919 | -1.511 | 0.833 | 0.066 | -0.027 | -0.027 | 1.751 | 0.35 | 0.77 | 0.26 | 0.41 | 0.16 | 0.48 |
| 6 | 110108855 | rs2492979 | -1.424 | -1.738 | -1.449 | 0.883 | 0.017 | 0.024 | 0.024 | 1.081 | 0.78 | 0.94 | 0.76 | 0.83 | 0.82 | 0.8 |
| 6 | 110117170 | rs4947021 | 0.909 | -1.766 | -1.121 | 1.353 | -0.045 | -0.037 | -0.037 | 0.016 | 0.28 | 0.05 | 0.29 | 0.31 | 0.31 | 0.23 |
| 6 | 110133864 | rs2492977 | -0.948 | -1.522 | -0.615 | 0.89 | -0.052 | 0.086 | 0.086 | 0 | 0.64 | 0.84 | 0.56 | 0.82 | 0.46 | 0.72 |
| 6 | 110137312 | rs4546525 | -0.424 | -1.69 | -0.468 | 1.164 | -0.016 | 0.032 | 0.032 | 0 | 0.58 | 0.72 | 0.51 | 0.69 | 0.45 | 0.61 |
| 6 | 110137431 | rs12527421 | -0.103 | -1.521 | -0.984 | 0.668 | 0.003 | 0.119 | 0.119 | 0 | 0.51 | 0.08 | 0.49 | 0.69 | 0.44 | 0.68 |
| 6 | 110146588 | rs1127775 | -0.026 | -1.513 | -0.98 | 0.673 | 0.002 | 0.118 | 0.118 | 0 | 0.51 | 0.08 | 0.49 | 0.69 | 0.44 | 0.68 |
| 6 | 110156179 | rs7771000 | -0.285 | -1.405 | -0.795 | 0.646 | 0.006 | 0.036 | 0.036 | 0 | 0.53 | 0.35 | 0.5 | 0.7 | 0.44 | 0.68 |
| 6 | 110173831 | rs9487237 | -0.612 | -1.873 | -1.339 | 0.802 | 0.009 | -0.002 | -0.002 | 0 | 0.15 | 0.08 | 0.24 | 0.47 | 0.08 | 0.36 |
| 6 | 110173860 | rs4428539 | 1.382 | -2.225 | -1.789 | 0.481 | 0.004 | 0.1 | 0.1 | 0.073 | 0.23 | 0.02 | 0.19 | 0.06 | 0.2 | 0.05 |
| 6 | 110174645 | rs4601174 | -0.463 | -1.578 | -0.931 | 0.849 | 0 | 0.002 | 0.002 | 0 | 0.6 | 0.59 | 0.53 | 0.73 | 0.41 | 0.69 |
| 6 | 110176543 | rs13209604 | -1.165 | -0.74 | -0.827 | 1.653 | 0.013 | -0.009 | -0.009 | 0.02 | 0.15 | 0.07 | 0.24 | 0.47 | 0.08 | 0.36 |
| 6 | 110189125 | rs9487240 | 0.569 | -0.297 | -1.264 | 0.639 | 0.025 | 0.125 | 0.125 | 2.257 | 0.46 | 0.14 | 0.32 | 0.29 | 0.4 | 0.3 |
| 6 | 110213306 | rs4333457 | -0.577 | -0.559 | -0.111 | 0.504 | 0.014 | 0.026 | 0.026 | 0.88 | 0.5 | 0.54 | 0.64 | 0.71 | 0.6 | 0.69 |
| 6 | 110222925 | rs6903208 | -0.287 | -0.598 | 0.132 | 0.886 | -0.005 | -0.006 | -0.006 | 0 | 0.46 | 0.45 | 0.53 | 0.6 | 0.57 | 0.63 |
| 6 | 110228667 | rs9398226 | 0.315 | -1.06 | -0.297 | 0.568 | 0.023 | 0.005 | 0.005 | 0.531 | 0.16 | 0.07 | 0.05 | 0.16 | 0.1 | 0.24 |
| 6 | 110229633 | rs7740408 | 0.22 | -1.242 | 0.446 | 0.942 | 0.003 | -0.003 | -0.003 | 0 | 0.33 | 0.33 | 0.39 | 0.19 | 0.48 | 0.32 |
| 6 | 110232051 | rs4557564 | -0.229 | -1.234 | 0.261 | 1.043 | -0.025 | 0.094 | 0.094 | 0 | 0.14 | 0.25 | 0.15 | 0.41 | 0.09 | 0.31 |
| 6 | 110239670 | rs6900525 | -0.395 | -1.03 | 0.375 | 0.933 | -0.012 | 0.1 | 0.1 | 0 | 0.14 | 0.21 | 0.23 | 0.51 | 0.1 | 0.3 |
| 6 | 110239704 | rs6900672 | -0.491 | -0.1 | 0.606 | 0.895 | -0.003 | 0.004 | 0.004 | 1.145 | 0.4 | 0.39 | 0.46 | 0.65 | 0.47 | 0.55 |
| 6 | 110241059 | rs9400344 | 0.607 | -0.07 | 0.538 | 0.894 | -0.033 | 0.025 | 0.025 | 7.963 | 0.26 | 0.14 | 0.23 | 0.15 | 0.37 | 0.24 |
| 6 | 110242224 | rs9400345 | -0.293 | -0.019 | 0.453 | 0.451 | -0.007 | 0.012 | 0.012 | 1.115 | 0.4 | 0.33 | 0.45 | 0.65 | 0.47 | 0.55 |
| 6 | 110244389 | rs9400346 | -1.6 | 0.228 | -0.159 | 0.541 | -0.013 | 0.07 | 0.07 | 2.942 | 0.68 | 0.93 | 0.73 | 0.84 | 0.69 | 0.93 |
| 6 | 110245755 | rs4947035 | -0.868 | 0.534 | 0.504 | 0.861 | 0.119 | 0.058 | 0.058 | 0.024 | 0.07 | 0.29 | 0.19 | 0.07 | 0.09 | 0.11 |
| 6 | 110251805 | rs4945848 | -0.681 | 0.172 | -0.144 | 0.612 | 0.004 | -0.004 | -0.004 | 0 | 0.38 | 0.61 | 0.55 | 0.67 | 0.44 | 0.61 |
| 6 | 110255274 | rs9374141 | -0.66 | 0.157 | -0.549 | 0.647 | 0.012 | -0.004 | -0.004 | 0 | 0.39 | 0.49 | 0.45 | 0.61 | 0.43 | 0.61 |
| 6 | 110259941 | rs9372241 | -0.232 | 0.666 | -1.093 | 0.902 | -0.098 | -0.026 | -0.026 | 0 | 0.22 | 0.06 | 0.27 | 0.2 | 0.33 | 0.32 |
| 6 | 110269388 | rs4947041 | -0.685 | -0.445 | -0.632 | 1.389 | 0.012 | 0.011 | 0.011 | 0 | 0.23 | 0.07 | 0.23 | 0.2 | 0.15 | 0.44 |
| 6 | 110273533 | rs4617029 | -1.306 | -0.035 | 0.572 | 1.632 | -0.019 | 0.234 | 0.234 | 7.417 | 0.13 | 0.61 | 0.19 | 0.34 | 0.08 | 0.17 |

| | | | | | | | | | | | | | | | | |
|---|---|---|---|---|---|---|---|---|---|---|---|---|---|---|---|---|
| 6 | 110284860 | rs9386867 | 0.877 | -0.075 | -0.35 | 0.386 | 0.015 | 0.051 | 0.051 | 0.556 | 0.57 | 0.12 | 0.62 | 0.65 | 0.68 | 0.46 |
| 6 | 110285415 | rs2503772 | 0.791 | -0.111 | 0.941 | 0.342 | -0.001 | -0.016 | -0.016 | 0 | 0.63 | 0.49 | 0.65 | 0.65 | 0.68 | 0.5 |
| 6 | 110288776 | rs6909151 | 0.331 | -0.078 | 1.16 | 0.225 | 0.005 | 0.004 | 0.004 | 0 | 0.87 | 0.83 | 0.88 | 0.77 | 0.84 | 0.8 |
| 6 | 110290678 | rs11751776 | -0.93 | -0.133 | 1.335 | 0.167 | -0.034 | -0.066 | -0.066 | 0.01 | 0.23 | 0.38 | 0.24 | 0.13 | 0.18 | 0.33 |
| 6 | 110290740 | rs9372243 | 0.964 | 0.017 | 1.553 | 0.247 | 0.08 | 0.073 | 0.073 | 0 | 0.83 | 0.52 | 0.85 | 0.77 | 0.83 | 0.79 |
| 6 | 110293474 | rs4463297 | 0.495 | 0.209 | 1.436 | 0.398 | 0.094 | 0.104 | 0.104 | 0 | 0.87 | 0.65 | 0.87 | 0.76 | 0.84 | 0.8 |
| 6 | 110296455 | rs9320321 | 1.452 | 0.035 | 0.994 | 0.371 | -0.004 | 0.002 | 0.002 | 1.593 | 0.62 | 0.27 | 0.62 | 0.6 | 0.65 | 0.47 |
| 6 | 110298307 | rs6568622 | 1.097 | 0.323 | 0.936 | 0.544 | 0.021 | 0.018 | 0.018 | 1.363 | 0.75 | 0.39 | 0.82 | 0.71 | 0.75 | 0.69 |
| 6 | 110301661 | rs4354185 | 1.235 | 0.331 | 0.653 | 0.379 | 0.002 | -0.001 | -0.001 | 0 | 0.65 | 0.63 | 0.65 | 0.62 | 0.64 | 0.51 |
| 6 | 110305904 | rs2505039 | -0.547 | -0.011 | -0.785 | -0.929 | -0.071 | 0.128 | 0.128 | 0.34 | 0.62 | 0.95 | 0.49 | 0.84 | 0.44 | 0.56 |
| 6 | 110309036 | rs12526978 | 0.41 | -1.315 | -0.958 | -2.086 | 0 | 0.056 | 0.056 | 0.823 | 0.31 | 0.19 | 0.2 | 0 | 0.35 | 0.11 |
| 6 | 110314783 | rs2501172 | -1.556 | -0.879 | -1.084 | -2.14 | -0.135 | 0.206 | 0.206 | 0.044 | 0.67 | 0.96 | 0.59 | 0.98 | 0.54 | 0.72 |
| 6 | 110320664 | rs12524163 | -0.214 | -1.179 | -0.665 | -2.328 | 0.003 | -0.035 | -0.035 | 0.42 | 0.12 | 0.15 | 0.07 | 0 | 0.16 | 0.06 |
| 6 | 110326686 | rs17542003 | -0.124 | -1.512 | -0.867 | -2.397 | -0.002 | 0.055 | 0.055 | 3.32 | 0.1 | 0.01 | 0.05 | 0.01 | 0.06 | 0.09 |
| 6 | 110334236 | rs9487281 | -1.554 | -1.641 | 0.168 | -2.49 | -0.006 | -0.09 | -0.09 | 0 | 0.27 | 0.61 | 0.23 | 0.02 | 0.22 | 0.16 |
| 6 | 110336743 | rs11757982 | -0.122 | -1.521 | 0.134 | -2.342 | -0.013 | 0.055 | 0.055 | 3.482 | 0.1 | 0 | 0.05 | 0.01 | 0.06 | 0.09 |
| 6 | 110337209 | rs1564078 | -0.842 | -1.675 | 0.611 | -2.482 | -0.006 | -0.108 | -0.108 | 0 | 0.43 | 0.52 | 0.33 | 0.02 | 0.39 | 0.22 |
| 6 | 110337293 | rs1564076 | -0.195 | -1.475 | 0.733 | -2.374 | 0.004 | -0.026 | -0.026 | 0 | 0.32 | 0.31 | 0.27 | 0.01 | 0.34 | 0.13 |
| 6 | 110342651 | rs9487284 | -0.215 | -1.701 | 0.632 | -2.522 | 0.008 | -0.022 | -0.022 | 0 | 0.35 | 0.31 | 0.28 | 0.01 | 0.35 | 0.14 |
| 6 | 110355236 | rs9487293 | -0.014 | -1.432 | 0.716 | -2.382 | 0.002 | 0.054 | 0.054 | 0.164 | 0.3 | 0.15 | 0.18 | 0 | 0.33 | 0.11 |
| 6 | 110356837 | rs6937788 | -0.026 | -1.396 | 0.66 | -2.363 | -0.002 | -0.077 | -0.077 | 0 | 0.31 | 0.33 | 0.19 | 0 | 0.33 | 0.11 |
| 6 | 110357835 | rs9374147 | -0.341 | -1.396 | 0.402 | -1.981 | -0.001 | -0.012 | -0.012 | 0 | 0.39 | 0.38 | 0.22 | 0.15 | 0.34 | 0.16 |
| 6 | 110359250 | rs879134 | -0.21 | -0.987 | 0.747 | -1.249 | 0 | -0.014 | -0.014 | 1.375 | 0.07 | 0.08 | 0.05 | 0 | 0.05 | 0.05 |
| 6 | 110378664 | rs11759874 | -0.777 | -0.926 | 1.287 | -1.23 | 0.058 | -0.021 | -0.021 | 1.06 | 0.08 | 0.29 | 0.07 | 0 | 0.05 | 0.04 |
| 6 | 110385528 | rs9400358 | -1.267 | -1.273 | 1.259 | -1.208 | -0.011 | -0.004 | -0.004 | 0 | 0.16 | 0.19 | 0.08 | 0.15 | 0.08 | 0.12 |
| 6 | 110386342 | rs6912291 | -1.185 | -1.203 | 1.65 | -0.94 | -0.043 | 0.084 | 0.084 | 0 | 0.13 | 0.23 | 0.11 | 0.24 | 0.05 | 0.18 |
| 6 | 110389926 | rs9372248 | -2.615 | -1.323 | 1.91 | -0.976 | -0.012 | 0.147 | 0.147 | 0.473 | 0.13 | 0.37 | 0.12 | 0.25 | 0.08 | 0.2 |
| 6 | 110391085 | rs7747842 | -2.615 | -1.323 | 2.052 | -0.982 | -0.011 | 0.147 | 0.147 | 0.473 | 0.13 | 0.41 | 0.13 | 0.25 | 0.08 | 0.21 |
| 6 | 110391596 | rs7770634 | -0.933 | -1.308 | 1.97 | -0.974 | 0.001 | 0.1 | 0.1 | 0.357 | 0.06 | 0.18 | 0.03 | 0.14 | 0.02 | 0.07 |
| 6 | 110396540 | rs1381688 | -0.933 | -1.308 | 1.97 | -0.974 | 0.001 | 0.1 | 0.1 | 0.357 | 0.06 | 0.18 | 0.03 | 0.14 | 0.02 | 0.07 |
| 6 | 110404180 | rs9386879 | -0.933 | -1.306 | 1.981 | -0.941 | 0.001 | 0.108 | 0.108 | 0.357 | 0.06 | 0.18 | 0.03 | 0.15 | 0.02 | 0.07 |
| 6 | 110408904 | rs12665650 | -0.849 | -1.313 | 2.032 | -0.949 | -0.01 | 0.069 | 0.069 | 0.036 | 0.06 | 0.16 | 0.03 | 0.13 | 0.02 | 0.07 |
| 6 | 110414749 | rs6925392 | -2.084 | -1.689 | 2.044 | -1.133 | -0.031 | 0.034 | 0.034 | 0.785 | 0.19 | 0.37 | 0.17 | 0.23 | 0.1 | 0.23 |
| 6 | 110421361 | rs6568632 | 0.289 | -1.888 | 2.113 | -0.709 | -0.03 | 0.091 | 0.091 | 0 | 0.91 | 0.68 | 0.9 | 0.81 | 0.98 | 0.91 |
| 6 | 110424341 | rs3828927 | 1.886 | -2.036 | 1.514 | -1.156 | 0.079 | -0.042 | -0.042 | 0.03 | 0.55 | 0.05 | 0.6 | 0.58 | 0.68 | 0.62 |
| 6 | 110426046 | rs6940347 | -0.58 | -1.658 | 2.177 | -0.37 | -0.009 | 0.071 | 0.071 | 0.029 | 0.06 | 0.16 | 0.03 | 0.13 | 0.02 | 0.07 |
| 6 | 110447643 | rs4945850 | -2.648 | -1.674 | 2.178 | -0.381 | -0.02 | 0.001 | 0.001 | 2.047 | 0.22 | 0.6 | 0.2 | 0.23 | 0.14 | 0.25 |
| 6 | 110453666 | rs2361500 | -0.303 | -1.675 | 2.188 | -0.381 | -0.007 | 0.014 | 0.014 | 0 | 0.05 | 0.07 | 0.02 | 0.13 | 0.02 | 0.07 |
| 6 | 110468046 | rs6927697 | -2.608 | -1.677 | 2.176 | -0.409 | -0.02 | 0 | 0 | 0.625 | 0.22 | 0.6 | 0.2 | 0.22 | 0.14 | 0.25 |
| 6 | 110474491 | rs12111403 | -0.499 | -1.672 | 2.176 | -0.41 | -0.007 | -0.042 | -0.042 | 0.015 | 0.1 | 0.19 | 0.07 | 0 | 0.06 | 0.04 |
| 6 | 110486412 | rs12191724 | 0.871 | -1.491 | 1.86 | -0.774 | 0.008 | 0.057 | 0.057 | 0 | 0.14 | 0.06 | 0.06 | 0 | 0.11 | 0.05 |
| 6 | 110490570 | rs7750037 | -0.639 | -1.023 | 2.714 | 0.17 | -0.007 | -0.049 | -0.049 | 0 | 0.1 | 0.19 | 0.07 | 0 | 0.06 | 0.04 |
| 6 | 110504680 | rs11757055 | -2.228 | -1.021 | 2.545 | 0.178 | -0.039 | 0.029 | 0.029 | 2.491 | 0.2 | 0.52 | 0.18 | 0.22 | 0.1 | 0.23 |
| 6 | 110516180 | rs11153226 | -2.667 | -1.021 | 2.547 | 0.149 | -0.02 | -0.001 | -0.001 | 0 | 0.22 | 0.6 | 0.2 | 0.22 | 0.14 | 0.25 |
| 6 | 110523342 | rs7775143 | 0.293 | -1.013 | 2.52 | 0.149 | -0.027 | 0.096 | 0.096 | 0 | 0.91 | 0.68 | 0.9 | 0.81 | 0.98 | 0.95 |
| 6 | 110532805 | rs6918148 | -2.26 | -0.84 | 2.541 | 0.005 | -0.039 | 0.029 | 0.029 | 2.491 | 0.2 | 0.52 | 0.18 | 0.22 | 0.1 | 0.23 |
| 6 | 110548240 | rs9487318 | -2.341 | -0.709 | 2.205 | -0.049 | -0.008 | 0.003 | 0.003 | 0.446 | 0.23 | 0.39 | 0.18 | 0.21 | 0.18 | 0.31 |
| 6 | 110555401 | rs9487321 | -2.547 | -0.699 | 2.238 | -0.044 | -0.015 | -0.006 | -0.006 | 0 | 0.24 | 0.54 | 0.19 | 0.21 | 0.18 | 0.31 |
| 6 | 110566053 | rs17071335 | -0.336 | -0.747 | 2.05 | 0.038 | -0.003 | -0.05 | -0.05 | 0 | 0.09 | 0.17 | 0.04 | 0 | 0.05 | 0.05 |
| 6 | 110566328 | rs2334319 | -2.339 | -0.893 | 1.957 | 0.087 | 0.071 | -0.064 | -0.064 | 0.987 | 0.36 | 0.89 | 0.23 | 0.4 | 0.21 | 0.37 |
| 6 | 110570460 | rs9487325 | -2.276 | -1.351 | 2.326 | 0.372 | -0.004 | -0.039 | -0.039 | 0.393 | 0.18 | 0.48 | 0.17 | 0.09 | 0.13 | 0.21 |
| 6 | 110577257 | rs2334323 | -1.786 | -1.424 | 1.833 | 0.244 | 0.093 | -0.01 | -0.01 | 0 | 0.46 | 0.88 | 0.3 | 0.41 | 0.3 | 0.39 |
| 6 | 110577767 | rs17071351 | -2.145 | -0.228 | 2.943 | 0.728 | 0.009 | -0.014 | -0.014 | 0 | 0.14 | 0.24 | 0.15 | 0.09 | 0.13 | 0.21 |
| 6 | 110600193 | rs9487336 | -1.872 | -0.122 | 2.542 | 0.797 | 0.005 | 0.016 | 0.016 | 0 | 0.05 | 0.08 | 0.1 | 0.09 | 0.07 | 0.17 |
| 6 | 110607412 | rs9487342 | -2.03 | -0.535 | 2.733 | 0.732 | -0.023 | -0.087 | -0.087 | 0.004 | 0.13 | 0.57 | 0.08 | 0 | 0.06 | 0.05 |
| 6 | 110620021 | rs2878176 | -1.625 | -0.427 | 2.677 | 0.913 | -0.014 | 0.026 | 0.026 | 0.085 | 0.11 | 0.22 | 0.07 | 0.13 | 0.06 | 0.08 |
| 6 | 110622462 | rs9487358 | -1.969 | -0.464 | 2.821 | 0.996 | -0.02 | -0.007 | -0.007 | 0 | 0.18 | 0.63 | 0.17 | 0.13 | 0.11 | 0.22 |
| 6 | 110628455 | rs12664630 | -2.223 | -0.454 | 2.508 | 1.004 | -0.016 | 0.016 | 0.016 | 2.638 | 0.2 | 0.7 | 0.2 | 0.26 | 0.12 | 0.26 |
| 6 | 110629139 | rs17071424 | -2.419 | -0.464 | 2.421 | 0.999 | 0.01 | -0.009 | -0.009 | 0.558 | 0.22 | 0.81 | 0.21 | 0.26 | 0.12 | 0.26 |
| 6 | 110629590 | rs9487368 | -1.61 | -0.257 | 2.162 | 1.044 | 0.003 | 0.027 | 0.027 | 0.556 | 0.15 | 0.45 | 0.16 | 0.13 | 0.11 | 0.22 |
| 6 | 110641256 | rs9481056 | -1.259 | -0.084 | 1.136 | 1.136 | -0.004 | 0.012 | 0.012 | 0 | 0.14 | 0.33 | 0.15 | 0.14 | 0.1 | 0.26 |
| 6 | 110645562 | rs4947063 | -1.202 | 0.116 | -0.061 | 1.125 | -0.015 | 0.036 | 0.036 | 0.491 | 0.21 | 0.66 | 0.35 | 0.31 | 0.15 | 0.31 |
| 6 | 110655384 | rs9374163 | -0.762 | 0.502 | -0.77 | 1.125 | 0.002 | -0.011 | -0.011 | 0 | 0.29 | 0.19 | 0.27 | 0.34 | 0.28 | 0.25 |
| 6 | 110655782 | rs9386888 | -0.646 | 1.148 | -1.038 | 1.547 | -0.039 | -0.052 | -0.052 | 1.558 | 0.08 | 0.01 | 0.21 | 0.19 | 0.13 | 0.11 |
| 6 | 110655911 | rs9400374 | 1.29 | 1.739 | -0.408 | 1.582 | 0.015 | 0.026 | 0.026 | 0.265 | 0.74 | 0.48 | 0.62 | 0.65 | 0.73 | 0.72 |
| 6 | 110656665 | rs997865 | 0.259 | 1.719 | -1.178 | 1.28 | -0.009 | -0.011 | -0.011 | 0.123 | 0.25 | 0.18 | 0.25 | 0.35 | 0.31 | 0.33 |
| 6 | 110657142 | rs9320326 | 0.82 | 1.983 | -0.707 | 1.359 | -0.017 | -0.034 | -0.034 | 0 | 0.8 | 0.88 | 0.69 | 0.65 | 0.77 | 0.76 |
| 6 | 110659185 | rs976471 | 1.106 | 1.797 | -0.201 | 0.901 | 0.017 | 0.016 | 0.016 | 0 | 0.68 | 0.6 | 0.61 | 0.59 | 0.63 | 0.69 |
| 6 | 110661454 | rs9400375 | -0.429 | 1.733 | -0.089 | 1.121 | 0.008 | 0.005 | 0.005 | 0.924 | 0.39 | 0.34 | 0.35 | 0.36 | 0.34 | 0.42 |

| | | | | | | | | | | | | | | | | |
|---|---|---|---|---|---|---|---|---|---|---|---|---|---|---|---|---|
| 6 | 110662570 | rs2057154 | -0.001 | 1.379 | -0.314 | 0.728 | 0.012 | 0.015 | 0.015 | 0 | 0.35 | 0.28 | 0.3 | 0.28 | 0.3 | 0.34 |
| 6 | 110662915 | rs12524594 | -0.052 | 1.397 | -0.105 | 0.676 | 0.021 | -0.005 | -0.005 | 0 | 0.05 | 0.08 | 0.05 | 0.04 | 0.1 | 0.03 |
| 6 | 110662937 | rs2223972 | 0.391 | 1.47 | -0.392 | 0.659 | -0.077 | -0.084 | -0.084 | 0 | 0.67 | 0.84 | 0.54 | 0.5 | 0.58 | 0.56 |
| 6 | 110664007 | rs7738520 | 0.51 | 1.454 | -0.413 | 0.698 | 0.003 | 0.003 | 0.003 | 0 | 0.31 | 0.28 | 0.28 | 0.28 | 0.27 | 0.33 |
| 6 | 110665556 | rs11963809 | 0.129 | 1.631 | -0.455 | 0.879 | -0.003 | 0 | 0 | 0 | 0.57 | 0.64 | 0.46 | 0.41 | 0.53 | 0.5 |
| 6 | 110669277 | rs9384738 | -0.017 | 1.037 | -0.671 | 0.905 | -0.006 | -0.009 | -0.009 | 2.8 | 0.9 | 0.94 | 0.82 | 0.74 | 0.89 | 0.86 |
| 6 | 110669620 | rs7770648 | 1.379 | 1.014 | -0.748 | 0.204 | -0.016 | 0.029 | 0.029 | 0.389 | 0.27 | 0.04 | 0.18 | 0.17 | 0.21 | 0.19 |
| 6 | 110675112 | rs756487 | 1.079 | 1.167 | -0.279 | 0.248 | -0.007 | -0.016 | -0.016 | 0 | 0.3 | 0.37 | 0.2 | 0.17 | 0.22 | 0.19 |
| 6 | 110680647 | rs7772479 | 0.106 | 0.903 | -0.646 | 0.399 | 0.017 | -0.002 | -0.002 | 0.008 | 0.59 | 0.73 | 0.54 | 0.38 | 0.63 | 0.54 |
| 6 | 110681879 | rs12664330 | 0.197 | 0.94 | -0.682 | 0.413 | 0.004 | 0 | 0 | 0 | 0.58 | 0.63 | 0.53 | 0.38 | 0.63 | 0.54 |
| 6 | 110685093 | rs7757134 | 0.469 | 0.733 | -0.677 | 0.664 | -0.009 | 0.019 | 0.019 | 0 | 0.73 | 0.73 | 0.65 | 0.44 | 0.81 | 0.76 |
| 6 | 110686091 | rs17569742 | -0.344 | 1.749 | -1.119 | 0.281 | -0.075 | -0.023 | -0.023 | 0.256 | 0.17 | 0.02 | 0.24 | 0.14 | 0.26 | 0.3 |
| 6 | 110686893 | rs9487388 | -0.91 | 1.571 | -0.73 | 0.081 | -0.015 | 0.014 | 0.014 | 2.066 | 0.34 | 0.26 | 0.37 | 0.19 | 0.47 | 0.45 |
| 6 | 110690440 | rs9374165 | -0.352 | 1.138 | -0.57 | 0.279 | 0.006 | -0.012 | -0.012 | 0 | 0.42 | 0.5 | 0.41 | 0.34 | 0.52 | 0.46 |
| 6 | 110696370 | rs9374167 | -0.945 | 0.65 | -1.183 | 0.328 | -0.007 | 0.067 | 0.067 | 0.002 | 0.62 | 0.88 | 0.71 | 0.79 | 0.66 | 0.75 |
| 6 | 110706475 | rs9372252 | 0.822 | 1.221 | -1.335 | 0.931 | -0.052 | -0.052 | -0.052 | 0 | 0.09 | 0.01 | 0.09 | 0.17 | 0.16 | 0.07 |
| 6 | 110707820 | rs6568644 | -0.564 | 1.121 | -1.051 | 0.938 | 0.005 | 0.013 | 0.013 | 0.133 | 0.47 | 0.71 | 0.47 | 0.46 | 0.46 | 0.4 |
| 6 | 110710018 | rs964080 | 1.13 | 0.892 | -1.184 | 0.639 | 0.002 | 0.072 | 0.072 | 1.123 | 0.15 | 0.04 | 0.08 | 0 | 0.14 | 0.08 |
| 6 | 110711885 | rs2207357 | -0.116 | 0.947 | -0.464 | 0.743 | -0.001 | 0.001 | 0.001 | 0 | 0.27 | 0.28 | 0.35 | 0.42 | 0.3 | 0.31 |
| 6 | 110716207 | rs9384740 | -0.584 | 1.088 | 0.35 | 0.835 | -0.001 | -0.002 | -0.002 | 0 | 0.46 | 0.54 | 0.46 | 0.43 | 0.46 | 0.42 |
| 6 | 110717493 | rs2057149 | -1.294 | 1.15 | -0.02 | 0.671 | -0.053 | 0.07 | 0.07 | 0.166 | 0.62 | 0.92 | 0.71 | 0.79 | 0.65 | 0.76 |
| 6 | 110724106 | rs9320328 | -1.098 | 1.102 | 0.254 | 0.64 | 0 | -0.027 | -0.027 | 0 | 0.61 | 0.62 | 0.71 | 0.85 | 0.64 | 0.78 |
| 6 | 110724518 | rs1980531 | -0.987 | 1.106 | 0.32 | 0.616 | 0.001 | -0.049 | -0.049 | 0 | 0.58 | 0.52 | 0.69 | 0.84 | 0.63 | 0.76 |
| 6 | 110726236 | rs17492667 | 1.834 | 1.282 | -0.498 | 0.223 | -0.024 | 0.044 | 0.044 | 0.845 | 0.25 | 0.02 | 0.22 | 0.1 | 0.2 | 0.14 |
| 6 | 110727262 | rs11153234 | 0.084 | 1.262 | 0.244 | 0.375 | 0.046 | -0.044 | -0.044 | 0 | 0.87 | 0.75 | 0.95 | 0.95 | 0.86 | 0.93 |
| 6 | 110729270 | rs2073211 | 0.394 | 1.313 | 0.254 | 0.813 | 0.001 | 0.032 | 0.032 | 0.377 | 0.13 | 0.23 | 0.22 | 0.28 | 0.16 | 0.36 |
| 6 | 110729973 | rs2073210 | 0.036 | 1.257 | 0.278 | 0.761 | 0.001 | 0.025 | 0.025 | 0.207 | 0.16 | 0.28 | 0.24 | 0.29 | 0.18 | 0.35 |
| 6 | 110732116 | rs9398235 | 0.492 | 1.285 | 0.271 | 0.831 | 0 | 0.017 | 0.017 | 0.329 | 0.13 | 0.19 | 0.22 | 0.28 | 0.16 | 0.33 |
| 6 | 110732209 | rs9487396 | 0.079 | 1.139 | 0.334 | 0.802 | -0.015 | 0.052 | 0.052 | 0 | 0.17 | 0.25 | 0.1 | 0.35 | 0.14 | 0.09 |
| 6 | 110732314 | rs9481063 | 0.064 | 1.432 | 0.234 | 0.418 | 0.038 | -0.054 | -0.054 | 2.629 | 0.88 | 0.8 | 0.95 | 0.96 | 0.86 | 0.93 |
| 6 | 110732736 | rs9384741 | 0.158 | 1.226 | 0.11 | 0.298 | 0.002 | -0.047 | -0.047 | 0.251 | 0.73 | 0.75 | 0.86 | 0.95 | 0.71 | 0.85 |
| 6 | 110736384 | rs6933664 | -1.086 | 1.087 | 0.311 | 0.991 | -0.001 | 0.039 | 0.039 | 0.393 | 0.32 | 0.57 | 0.3 | 0.47 | 0.32 | 0.54 |
| 6 | 110737115 | rs4945862 | -1.514 | 1.117 | 0.741 | 1.081 | -0.001 | -0.012 | -0.012 | 0 | 0.39 | 0.73 | 0.34 | 0.6 | 0.37 | 0.57 |
| 6 | 110741611 | rs2207356 | -0.372 | 1.092 | 0.818 | 1.166 | -0.002 | 0.014 | 0.014 | 0 | 0.15 | 0.08 | 0.06 | 0.08 | 0.12 | 0.11 |
| 6 | 110741922 | rs12055551 | 0.122 | 1.085 | 0.966 | 1.277 | -0.001 | 0.107 | 0.107 | 0.949 | 0.16 | 0.29 | 0.22 | 0.38 | 0.19 | 0.43 |
| 6 | 110745606 | rs6938431 | -1.067 | 1.084 | 0.541 | 0.899 | 0 | -0.026 | -0.026 | 0 | 0.19 | 0.21 | 0.09 | 0.07 | 0.16 | 0.12 |
| 6 | 110746351 | rs17493602 | 0.903 | 1.035 | 0.127 | 0.755 | 0.004 | 0.017 | 0.017 | 0.961 | 0.12 | 0.05 | 0.06 | 0.05 | 0.09 | 0.06 |
| 6 | 110746526 | rs7756222 | -1.826 | 1.006 | 0.027 | 0.718 | -0.084 | 0.22 | 0.22 | 0.142 | 0.53 | 0.92 | 0.43 | 0.83 | 0.51 | 0.63 |
| 6 | 110746545 | rs9487402 | 0.042 | 0.786 | 0.484 | 0.958 | 0.003 | 0.171 | 0.171 | 0.865 | 0.2 | 0.46 | 0.25 | 0.5 | 0.21 | 0.38 |
| 6 | 110747749 | rs9386898 | 0.779 | 0.77 | 0.627 | 1.184 | 0.018 | 0.073 | 0.073 | 0 | 0.16 | 0.33 | 0.21 | 0.39 | 0.21 | 0.36 |
| 6 | 110748641 | rs3818100 | -0.226 | 0.746 | 0.53 | 1.13 | -0.001 | 0.03 | 0.03 | 0 | 0.27 | 0.41 | 0.27 | 0.45 | 0.29 | 0.42 |
| 6 | 110750148 | rs17578180 | -0.103 | 0.84 | -0.034 | 0.384 | 0.004 | 0.008 | 0.008 | 0 | 0.1 | 0.06 | 0.03 | 0.08 | 0.06 | 0.08 |
| 6 | 110750552 | rs9481067 | -1.307 | 0.906 | -0.046 | 0.29 | 0.019 | 0.243 | 0.243 | 0 | 0.49 | 0.88 | 0.4 | 0.84 | 0.43 | 0.62 |
| 6 | 110757416 | rs221716 | -1.276 | 0.435 | -0.165 | -0.427 | -0.003 | 0.052 | 0.052 | 0.275 | 0.28 | 0.59 | 0.28 | 0.41 | 0.28 | 0.27 |
| 6 | 110758205 | rs221715 | -1.225 | 0.443 | -0.156 | -0.423 | -0.002 | 0.059 | 0.059 | 0.233 | 0.28 | 0.58 | 0.28 | 0.41 | 0.28 | 0.27 |
| 6 | 110761189 | rs13219760 | 1.018 | 0.296 | -0.637 | -0.923 | 0 | 0.005 | 0.005 | 0 | 0.09 | 0.05 | 0.11 | 0.08 | 0.09 | 0.07 |
| 6 | 110763227 | rs221711 | -1.356 | 0.245 | -0.511 | -0.945 | 0.004 | -0.03 | -0.03 | 0 | 0.29 | 0.72 | 0.29 | 0.41 | 0.28 | 0.27 |
| 6 | 110763875 | rs723685 | 0.867 | 0.313 | -0.362 | -0.896 | -0.002 | -0.006 | -0.006 | 0 | 0.09 | 0.1 | 0.11 | 0.08 | 0.09 | 0.06 |
| 6 | 110766308 | rs7768417 | -0.585 | 0.392 | -0.372 | -0.953 | 0.003 | 0.106 | 0.106 | 0 | 0.24 | 0.39 | 0.25 | 0.41 | 0.25 | 0.26 |
| 6 | 110776730 | rs12195708 | -0.701 | 0.033 | -0.532 | -1.062 | -0.012 | -0.019 | -0.019 | 0 | 0.53 | 0.71 | 0.43 | 0.41 | 0.51 | 0.37 |
| 6 | 110778128 | rs714368 | -0.813 | -0.65 | -0.468 | -0.395 | -0.007 | 0.111 | 0.111 | 0 | 0.24 | 0.39 | 0.24 | 0.41 | 0.22 | 0.26 |
| 6 | 110787371 | rs2428192 | 0.741 | -0.878 | -0.768 | -0.539 | 0 | 0.134 | 0.134 | 5.483 | 0.24 | 0.01 | 0.29 | 0.04 | 0.16 | 0.15 |
| 6 | 110790908 | rs7740547 | -1.072 | -0.447 | 0.346 | 0.347 | -0.027 | 0.099 | 0.099 | 0 | 0.25 | 0.39 | 0.23 | 0.41 | 0.19 | 0.24 |
| 6 | 110791310 | rs10214672 | -0.959 | -0.331 | 0.576 | 0.492 | -0.049 | 0.056 | 0.056 | 0.457 | 0.27 | 0.54 | 0.17 | 0.33 | 0.16 | 0.18 |
| 6 | 110792301 | rs2428190 | 1.536 | 0.257 | 1.152 | 0.694 | -0.01 | 0.139 | 0.139 | 0.983 | 0.83 | 0.53 | 0.64 | 0.69 | 0.84 | 0.81 |
| 6 | 110793578 | rs916722 | -1.137 | 0.142 | 1.5 | 0.704 | -0.002 | 0.215 | 0.215 | 0.462 | 0.12 | 0.34 | 0.1 | 0.27 | 0.1 | 0.13 |
| 6 | 110793650 | rs916721 | -1.137 | 0.142 | 1.5 | 0.704 | -0.002 | 0.215 | 0.215 | 0.462 | 0.12 | 0.34 | 0.1 | 0.27 | 0.1 | 0.13 |
| 6 | 110793699 | rs1858928 | -1.137 | 0.142 | 1.5 | 0.704 | -0.002 | 0.215 | 0.215 | 0.462 | 0.12 | 0.34 | 0.1 | 0.27 | 0.1 | 0.13 |
| 6 | 110795117 | rs4543405 | -0.933 | 0.142 | 1.844 | 0.704 | 0.019 | 0.247 | 0.247 | 0 | 0.07 | 0.2 | 0.08 | 0.27 | 0.08 | 0.11 |
| 6 | 110795549 | rs916719 | -1.856 | 0.035 | 2.3 | 0.623 | -0.014 | 0.182 | 0.182 | 0.083 | 0.13 | 0.55 | 0.12 | 0.27 | 0.11 | 0.13 |
| 6 | 110796405 | rs7768422 | -1.644 | 0.069 | 2.334 | 0.629 | 0.049 | 0.35 | 0.35 | 0 | 0.09 | 0.49 | 0.11 | 0.27 | 0.08 | 0.11 |
| 6 | 110799201 | rs6928183 | -0.576 | -0.11 | 1.479 | -0.324 | 0.011 | 0.001 | 0.001 | 0 | 0.71 | 0.79 | 0.79 | 0.97 | 0.76 | 0.84 |
| 6 | 110799548 | rs3806970 | -0.4 | 0.036 | 1.35 | -0.247 | 0.004 | 0.103 | 0.103 | 0 | 0.07 | 0.11 | 0.09 | 0.27 | 0.08 | 0.12 |
| 6 | 110804014 | rs9400393 | 0.071 | 0.753 | 0.52 | -0.32 | -0.06 | -0.065 | -0.065 | 0 | 0.21 | 0.15 | 0.49 | 0.52 | 0.42 | 0.46 |
| 6 | 110807782 | rs2428175 | 1.1 | 0.537 | 0.517 | -0.419 | 0.029 | 0.089 | 0.089 | 4.091 | 0.35 | 0.12 | 0.16 | 0.18 | 0.23 | 0.25 |
| 6 | 110812262 | rs2428169 | -0.035 | 1.215 | 0.504 | -0.515 | -0.006 | 0.005 | 0.005 | 0 | 0.92 | 0.96 | 0.91 | 0.98 | 0.92 | 0.88 |
| 6 | 110813613 | rs2078890 | 0.134 | 1.238 | 0.491 | -0.408 | -0.013 | -0.01 | -0.01 | 0.313 | 0.18 | 0.17 | 0.46 | 0.52 | 0.37 | 0.41 |
| 6 | 110813928 | rs2428168 | 0.478 | 0.555 | 0.665 | -0.256 | 0.012 | -0.011 | -0.011 | 0.055 | 0.42 | 0.35 | 0.26 | 0.45 | 0.31 | 0.36 |
| 6 | 110815878 | rs7759856 | -0.06 | 0.489 | 0.97 | 0.128 | -0.015 | -0.031 | -0.031 | 0 | 0.06 | 0.16 | 0.03 | 0 | 0.03 | 0.02 |

| | | | | | | | | | | | | | | | | |
|---|---|---|---|---|---|---|---|---|---|---|---|---|---|---|---|---|
| 6 | 110819059 | rs6922009 | -1.098 | 0.528 | 1.07 | 0.341 | 0.03 | 0.175 | 0.175 | 0.109 | 0.18 | 0.48 | 0.2 | 0.32 | 0.16 | 0.23 |
| 6 | 110828403 | rs12215879 | 0.949 | 0.42 | 0.413 | -0.235 | 0.027 | 0.022 | 0.022 | 1.485 | 0.29 | 0.19 | 0.16 | 0 | 0.21 | 0.09 |
| 6 | 110837288 | rs9400396 | 0.84 | 0.466 | 0.19 | -0.111 | 0.05 | 0.154 | 0.154 | 0.838 | 0.43 | 0.19 | 0.23 | 0.14 | 0.35 | 0.3 |
| 6 | 110841905 | rs6933588 | 0.073 | 0.568 | 0.606 | -0.145 | 0.01 | -0.029 | -0.029 | 0 | 0.07 | 0.24 | 0.04 | 0 | 0.05 | 0.03 |
| 6 | 110843638 | rs2520573 | 0.443 | 0.974 | 0.034 | 0.316 | -0.03 | -0.028 | -0.028 | 0.015 | 0.31 | 0.18 | 0.46 | 0.43 | 0.37 | 0.43 |
| 6 | 110846248 | rs1989634 | -0.082 | 0.93 | -0.477 | 0.437 | -0.082 | -0.004 | -0.004 | 0 | 0.63 | 0.81 | 0.44 | 0.3 | 0.5 | 0.47 |
| 6 | 110850124 | rs2817781 | 0.357 | 1.047 | -0.506 | 0.499 | -0.055 | -0.098 | -0.098 | 0.011 | 0.31 | 0.11 | 0.49 | 0.54 | 0.39 | 0.44 |
| 6 | 110850515 | rs9386913 | 0.476 | 0.979 | -0.407 | 0.556 | 0.019 | 0.066 | 0.066 | 0.942 | 0.45 | 0.21 | 0.28 | 0.28 | 0.38 | 0.34 |
| 6 | 110851763 | rs2817782 | 0.796 | 0.689 | 0.461 | 1.217 | 0.006 | 0.118 | 0.118 | 0.099 | 0.74 | 0.49 | 0.75 | 0.56 | 0.78 | 0.61 |
| 6 | 110851976 | rs1080261 | 0.049 | 0.521 | -0.111 | 1.424 | -0.001 | -0.013 | -0.013 | 0.23 | 0.16 | 0.13 | 0.16 | 0.31 | 0.21 | 0.13 |
| 6 | 110853510 | rs2023705 | 0.152 | 0.518 | 0.64 | 1.282 | -0.001 | 0.005 | 0.005 | 0 | 0.12 | 0.21 | 0.09 | 0.12 | 0.07 | 0.14 |
| 6 | 110857994 | rs2520571 | 1.216 | 0.38 | 1.209 | 1.631 | 0.002 | 0.045 | 0.045 | 0 | 0.73 | 0.4 | 0.8 | 0.65 | 0.78 | 0.63 |
| 6 | 110865183 | rs739516 | -0.142 | 0.414 | 1.063 | 1.272 | -0.005 | 0.006 | 0.006 | 0 | 0.07 | 0.04 | 0.04 | 0.03 | 0.06 | 0.14 |
| 6 | 110875435 | rs853547 | -1.296 | 0.301 | 0.982 | 1.271 | -0.002 | -0.044 | -0.044 | 0.823 | 0.23 | 0.5 | 0.15 | 0.15 | 0.14 | 0.31 |
| 6 | 110876961 | rs17231299 | -0.739 | 0.389 | 0.237 | 1.282 | 0.001 | -0.022 | -0.022 | 0 | 0.1 | 0.11 | 0.05 | 0.03 | 0.07 | 0.17 |
| 6 | 110883633 | rs2817792 | 0.012 | 0.216 | -0.258 | 1.666 | 0.019 | -0.044 | -0.044 | 0 | 0.78 | 0.88 | 0.84 | 0.62 | 0.81 | 0.81 |
| 6 | 110887169 | rs4464837 | -0.816 | 0.827 | -0.12 | 1.478 | -0.002 | 0 | 0 | 0.519 | 0.42 | 0.43 | 0.59 | 0.56 | 0.58 | 0.62 |
| 6 | 110887577 | rs6932192 | -0.15 | 1.506 | -0.212 | 1.611 | 0 | -0.001 | -0.001 | 2.437 | 0.27 | 0.24 | 0.21 | 0.15 | 0.25 | 0.2 |
| 6 | 110889435 | rs9487444 | -0.159 | 1.519 | -0.227 | 1.618 | 0 | -0.004 | -0.004 | 0 | 0.27 | 0.24 | 0.21 | 0.11 | 0.25 | 0.19 |
| 6 | 110889538 | rs9384748 | -0.366 | 1.79 | -0.319 | 1.822 | -0.091 | -0.102 | -0.102 | 0 | 0.16 | 0.02 | 0.35 | 0.43 | 0.31 | 0.42 |
| 6 | 110907523 | rs6941942 | -1.112 | 0.852 | -0.285 | 1.517 | 0.014 | 0.01 | 0.01 | 2.182 | 0.49 | 0.38 | 0.58 | 0.56 | 0.57 | 0.61 |
| 6 | 110911299 | rs13197089 | -0.994 | 0.854 | 0.388 | 1.688 | -0.001 | -0.001 | -0.001 | 0 | 0.45 | 0.5 | 0.56 | 0.54 | 0.53 | 0.59 |
| 6 | 110912424 | rs9398240 | -0.98 | 0.829 | 0.625 | 1.594 | 0 | -0.018 | -0.018 | 0 | 0.44 | 0.43 | 0.58 | 0.86 | 0.55 | 0.65 |
| 6 | 110924988 | rs2691187 | -0.887 | 0.374 | 0.703 | 1.31 | -0.018 | -0.004 | -0.004 | 1.793 | 0.42 | 0.75 | 0.54 | 0.54 | 0.44 | 0.61 |
| 6 | 110927044 | rs6935101 | -0.382 | 0.835 | 0.812 | 1.72 | 0.003 | -0.003 | -0.003 | 1.374 | 0.29 | 0.39 | 0.45 | 0.22 | 0.32 | 0.25 |
| 6 | 110930564 | rs9386924 | -0.067 | 0.572 | 0.837 | 1.886 | 0.001 | 0.02 | 0.02 | 0 | 0.11 | 0.12 | 0.06 | 0.33 | 0.12 | 0.36 |
| 6 | 110936144 | rs6905041 | -0.144 | 0.258 | 0.91 | 1.376 | -0.017 | -0.012 | -0.012 | 0 | 0.25 | 0.32 | 0.18 | 0.16 | 0.19 | 0.23 |
| 6 | 110950048 | rs13214056 | 0.038 | 0.242 | 0.872 | 1.35 | 0.029 | 0.019 | 0.019 | 0 | 0.23 | 0.13 | 0.15 | 0.16 | 0.18 | 0.22 |
| 6 | 110967879 | rs2691173 | -0.897 | -0.174 | 1.04 | 1.134 | 0.026 | -0.007 | -0.007 | 0 | 0.4 | 0.67 | 0.26 | 0.53 | 0.3 | 0.38 |
| 6 | 110969521 | rs2691172 | -0.024 | -0.126 | 1.33 | 1.198 | -0.04 | 0.07 | 0.07 | 0 | 0.26 | 0.39 | 0.19 | 0.52 | 0.21 | 0.28 |
| 6 | 110972494 | rs2817763 | 0.071 | -0.121 | 1.227 | 1.14 | -0.012 | -0.005 | -0.005 | 0 | 0.25 | 0.31 | 0.17 | 0.19 | 0.19 | 0.21 |
| 6 | 110976628 | rs12201131 | 1.372 | 0.049 | 1.003 | 1.137 | -0.002 | -0.014 | -0.014 | 0 | 0.08 | 0.08 | 0.03 | 0 | 0.06 | 0.01 |
| 6 | 110989662 | rs9386934 | 0.07 | 0.451 | 1.043 | 1.338 | 0.002 | 0.04 | 0.04 | 0 | 0.11 | 0.14 | 0.1 | 0.33 | 0.16 | 0.35 |
| 6 | 111061814 | rs11755241 | 0.143 | 0.166 | 1.406 | 1.221 | 0.014 | 0.006 | 0.006 | 0 | 0.25 | 0.18 | 0.15 | 0.19 | 0.18 | 0.23 |
| 6 | 111081133 | rs9374202 | -0.863 | 0.099 | 1.179 | 1.258 | -0.005 | 0.006 | 0.006 | 0.858 | 0.24 | 0.29 | 0.15 | 0.34 | 0.24 | 0.45 |
| 6 | 111088192 | rs2341860 | -0.78 | -0.015 | 1.139 | 1.04 | 0.026 | -0.026 | -0.026 | 0.452 | 0.79 | 0.97 | 0.87 | 0.81 | 0.84 | 0.79 |
| 6 | 111100687 | rs802658 | -0.955 | 0.147 | 1.273 | 1.128 | -0.013 | 0.02 | 0.02 | 0.785 | 0.25 | 0.35 | 0.18 | 0.68 | 0.26 | 0.52 |
| 6 | 111131329 | rs7771618 | 1.889 | -0.461 | 0.612 | 0.169 | -0.067 | 0.106 | 0.106 | 0.002 | 0.3 | 0.11 | 0.49 | 0.01 | 0.46 | 0.12 |
| 6 | 111135497 | rs3749866 | 0.64 | 0.401 | 0.739 | 0.425 | -0.02 | 0.007 | 0.007 | 0 | 0.06 | 0.02 | 0.29 | 0.01 | 0.16 | 0.03 |
| 6 | 111155400 | rs1279585 | -1.432 | -0.427 | 0.968 | 0.607 | -0.038 | -0.024 | -0.024 | 0 | 0.33 | 0.5 | 0.23 | 0.76 | 0.28 | 0.53 |
| 6 | 111157593 | rs9487538 | 0.893 | -0.268 | 0.859 | 0.509 | 0.005 | -0.027 | -0.027 | 0 | 0.07 | 0.27 | 0.04 | 0.01 | 0.03 | 0.09 |
| 6 | 111161102 | rs802655 | 0.264 | -0.184 | 0.639 | 0.66 | 0 | 0.038 | 0.038 | 0 | 0.11 | 0.14 | 0.1 | 0.33 | 0.14 | 0.34 |
| 6 | 111199032 | rs1008155 | 0.919 | -0.061 | 0.685 | 0.578 | -0.004 | -0.017 | -0.017 | 0.12 | 0.07 | 0.12 | 0.03 | 0.01 | 0.03 | 0.11 |
| 6 | 111200755 | rs1007274 | 0.423 | -0.126 | 0.786 | 0.516 | -0.009 | -0.006 | -0.006 | 0 | 0.25 | 0.26 | 0.16 | 0.21 | 0.2 | 0.26 |
| 6 | 111204046 | rs867374 | 1.16 | -0.075 | 0.618 | 0.685 | -0.014 | 0.04 | 0.04 | 0.229 | 0.75 | 0.6 | 0.83 | 0.35 | 0.75 | 0.53 |
| 6 | 111210420 | rs1027914 | 0.091 | -0.008 | 0.482 | 0.76 | 0 | 0.006 | 0.006 | 0 | 0.83 | 0.84 | 0.86 | 0.37 | 0.78 | 0.64 |
| 6 | 111212283 | rs7768897 | -0.232 | -0.006 | 0.433 | 0.782 | 0.003 | 0.078 | 0.078 | 0 | 0.92 | 0.87 | 0.92 | 0.67 | 0.88 | 0.71 |
| 6 | 111222552 | rs11153258 | -0.376 | 0.049 | 0.437 | 0.806 | 0 | 0.002 | 0.002 | 0 | 0.18 | 0.18 | 0.14 | 0.63 | 0.22 | 0.36 |
| 6 | 111244816 | rs706956 | 0.469 | 0.124 | 0.249 | 0.802 | 0.009 | 0.121 | 0.121 | 0 | 0.08 | 0.19 | 0.09 | 0.33 | 0.13 | 0.29 |
| 6 | 111245606 | rs781494 | -1.472 | -0.063 | 0.314 | 0.702 | -0.031 | 0.04 | 0.04 | 0.372 | 0.28 | 0.53 | 0.19 | 0.65 | 0.25 | 0.49 |
| 6 | 111246020 | rs1279481 | -0.61 | -0.057 | 0.28 | 0.639 | -0.001 | 0.004 | 0.004 | 1.302 | 0.21 | 0.25 | 0.13 | 0.34 | 0.22 | 0.4 |
| 6 | 111248987 | rs1704721 | -0.456 | -0.074 | 0.28 | 0.661 | 0.001 | 0.096 | 0.096 | 0.011 | 0.15 | 0.28 | 0.12 | 0.34 | 0.16 | 0.4 |
| 6 | 111252916 | rs9487557 | 0.598 | -0.116 | 0.217 | 0.224 | 0.003 | -0.003 | -0.003 | 0.734 | 0.11 | 0.12 | 0.04 | 0.01 | 0.06 | 0.03 |
| 6 | 111260923 | rs781499 | -1.239 | 0.184 | 0.168 | 0.12 | -0.009 | 0.194 | 0.194 | 0 | 0.55 | 0.86 | 0.38 | 0.87 | 0.48 | 0.76 |
| 6 | 111277745 | rs12207535 | -0.193 | -0.028 | 0.193 | 0.569 | 0.005 | -0.01 | -0.01 | 0 | 0.17 | 0.18 | 0.07 | 0.02 | 0.09 | 0.15 |
| 6 | 111283592 | rs9374227 | 0.3 | 0.116 | 0.202 | 0.591 | 0.025 | 0.12 | 0.12 | 0 | 0.17 | 0.34 | 0.15 | 0.63 | 0.2 | 0.37 |
| 6 | 111292632 | rs9400437 | 0.3 | 0.116 | 0.202 | 0.591 | 0.025 | 0.12 | 0.12 | 0 | 0.17 | 0.34 | 0.15 | 0.63 | 0.2 | 0.37 |
| 6 | 111299060 | rs9374234 | -1.131 | -0.03 | 0.127 | 0.469 | -0.023 | 0.007 | 0.007 | 0 | 0.34 | 0.53 | 0.22 | 0.66 | 0.29 | 0.52 |
| 6 | 111309591 | rs4947092 | -2.098 | 0.349 | -0.779 | -0.587 | -0.06 | 0.135 | 0.135 | 3.127 | 0.8 | 0.99 | 0.85 | 1 | 0.78 | 0.93 |
| 6 | 111316885 | rs3904925 | -0.044 | 0 | 0.147 | 0.239 | 0.01 | -0.001 | -0.001 | 0 | 0.14 | 0.13 | 0.06 | 0.01 | 0.08 | 0.12 |
| 6 | 111320794 | rs3763195 | 0.312 | 0.171 | 0.086 | 0.454 | 0.005 | 0.021 | 0.021 | 0 | 0.15 | 0.25 | 0.12 | 0.26 | 0.17 | 0.32 |
| 6 | 111324495 | rs6910046 | -0.195 | 0.097 | -0.101 | 0.603 | 0.011 | -0.031 | -0.031 | 0 | 0.16 | 0.1 | 0.1 | 0.47 | 0.13 | 0.15 |
| 6 | 111335687 | rs9372292 | 0.381 | 0.192 | -0.014 | 0.543 | 0.003 | 0.011 | 0.011 | 0 | 0.15 | 0.2 | 0.12 | 0.26 | 0.17 | 0.36 |
| 6 | 111336435 | rs9320354 | -0.166 | 0.096 | 0.232 | 0.401 | -0.007 | 0.061 | 0.061 | 0.536 | 0.16 | 0.22 | 0.11 | 0.49 | 0.13 | 0.14 |
| 6 | 111343394 | rs11751660 | -0.189 | 0.523 | 0.381 | -0.098 | -0.007 | -0.019 | -0.019 | 0.222 | 0.25 | 0.4 | 0.18 | 0.19 | 0.23 | 0.29 |
| 6 | 111352699 | rs4082794 | 0.287 | 0.518 | 0.465 | -0.033 | 0.004 | 0.019 | 0.019 | 0 | 0.16 | 0.27 | 0.12 | 0.26 | 0.16 | 0.37 |
| 6 | 111355890 | rs7748974 | -0.745 | 0.46 | 0.242 | -0.042 | 0.002 | -0.001 | -0.001 | 0 | 0.41 | 0.68 | 0.29 | 0.45 | 0.39 | 0.66 |
| 6 | 111361944 | rs9400461 | -0.371 | 0.3 | 0.003 | -0.311 | -0.01 | -0.009 | -0.009 | 0.172 | 0.3 | 0.39 | 0.29 | 0.79 | 0.25 | 0.55 |
| 6 | 111366465 | rs13202203 | 0.087 | 0.571 | -0.331 | -0.098 | -0.001 | -0.001 | -0.001 | 0 | 0.19 | 0.2 | 0.15 | 0.18 | 0.21 | 0.25 |

| | | | | | | | | | | | | | | | | |
|---|---|---|---|---|---|---|---|---|---|---|---|---|---|---|---|---|
| 6 | 111369781 | rs6920544 | -0.239 | 0.887 | -0.512 | 0.222 | -0.021 | -0.057 | -0.057 | 2.28 | 0.27 | 0.1 | 0.51 | 0.63 | 0.26 | 0.29 |
| 6 | 111374253 | rs3851222 | -0.526 | 0.766 | -0.022 | -0.227 | 0.001 | -0.036 | -0.036 | 0.026 | 0.44 | 0.49 | 0.61 | 0.81 | 0.43 | 0.68 |
| 6 | 111378729 | rs4947099 | -0.532 | 0.789 | -0.215 | -0.216 | 0 | -0.019 | -0.019 | 0.544 | 0.44 | 0.59 | 0.6 | 0.82 | 0.43 | 0.68 |
| 6 | 111381468 | rs6904275 | -0.179 | 0.847 | -0.507 | -0.16 | -0.021 | -0.035 | -0.035 | 0.02 | 0.26 | 0.17 | 0.42 | 0.32 | 0.33 | 0.49 |
| 6 | 111387327 | rs7754456 | 0.359 | 0.458 | -0.748 | -0.705 | 0.006 | 0.057 | 0.057 | 0.226 | 0.31 | 0.23 | 0.22 | 0 | 0.3 | 0.07 |
| 6 | 111390344 | rs6935864 | -0.323 | 0.547 | -0.319 | 0.154 | -0.002 | 0.006 | 0.006 | 0 | 0.27 | 0.28 | 0.41 | 0.36 | 0.25 | 0.48 |
| 6 | 111393061 | rs3904923 | -0.266 | 0.602 | 0.024 | -0.613 | -0.001 | -0.075 | -0.075 | 0 | 0.28 | 0.34 | 0.21 | 0 | 0.29 | 0.07 |
| 6 | 111394421 | rs9320356 | -1.423 | 0.336 | 0.934 | 0.292 | -0.019 | 0.04 | 0.04 | 0 | 0.14 | 0.37 | 0.27 | 0.18 | 0.09 | 0.18 |
| 6 | 111415640 | rs4945876 | -1.653 | 0.917 | 1.68 | 0.19 | 0.076 | 0.027 | 0.027 | 0 | 0.13 | 0.59 | 0.22 | 0.09 | 0.16 | 0.13 |
| 6 | 111424015 | rs9400467 | 0.715 | 1.159 | 1.452 | 0.101 | 0.002 | -0.005 | -0.005 | 0 | 0.73 | 0.71 | 0.66 | 0.75 | 0.63 | 0.76 |
| 6 | 111427286 | rs1022092 | 1.716 | 0.961 | 1.724 | -0.079 | -0.045 | 0.043 | 0.043 | 2.003 | 0.6 | 0.12 | 0.5 | 0.68 | 0.5 | 0.51 |
| 6 | 111458270 | rs4639374 | 0.261 | 0.858 | 2.137 | 0.239 | -0.005 | -0.009 | -0.009 | 0 | 0.1 | 0.07 | 0.1 | 0 | 0.19 | 0.04 |
| 6 | 111497379 | rs395564 | 1.024 | 0.82 | 2.359 | 0.233 | 0 | -0.001 | -0.001 | 0 | 0.81 | 0.8 | 0.76 | 0.74 | 0.82 | 0.66 |
| 6 | 111504655 | rs354529 | 1.961 | 0.848 | 2.57 | 0.212 | 0.018 | -0.027 | -0.027 | 0 | 0.66 | 0.45 | 0.59 | 0.74 | 0.53 | 0.7 |
| 6 | 111510722 | rs1150082 | 1.198 | 0.846 | 2.572 | 0.214 | -0.02 | 0.012 | 0.012 | 0 | 0.8 | 0.58 | 0.74 | 0.75 | 0.82 | 0.78 |
| 6 | 111527696 | rs402014 | 1.813 | 0.721 | 2.397 | 0.278 | 0.033 | -0.041 | -0.041 | 0.044 | 0.66 | 0.48 | 0.59 | 0.74 | 0.53 | 0.7 |
| 6 | 111531412 | rs354545 | 1.884 | 0.753 | 1.915 | 0.272 | -0.014 | 0.018 | 0.018 | 0.347 | 0.62 | 0.28 | 0.59 | 0.69 | 0.52 | 0.65 |
| 6 | 111548813 | rs354532 | 1.736 | 0.496 | 1.598 | 0.485 | -0.018 | 0.008 | 0.008 | 0.04 | 0.58 | 0.27 | 0.49 | 0.57 | 0.5 | 0.63 |
| 6 | 111549162 | rs2297939 | 2.168 | 1.015 | 1.092 | 0.232 | -0.038 | -0.083 | -0.083 | 7.575 | 0.49 | 0.05 | 0.38 | 0.42 | 0.38 | 0.43 |
| 6 | 111558989 | rs7743593 | -1.419 | 1.167 | 1.667 | 0.218 | 0.151 | 0.077 | 0.077 | 0 | 0.09 | 0.39 | 0.22 | 0.08 | 0.15 | 0.15 |
| 6 | 111560946 | rs9487602 | -1.777 | 1.222 | 1.645 | 0.41 | 0.007 | 0.004 | 0.004 | 0.054 | 0.36 | 0.62 | 0.47 | 0.43 | 0.46 | 0.33 |
| 6 | 111565847 | rs411604 | 1.373 | 1.236 | 1.656 | 0.269 | 0.067 | -0.063 | -0.063 | 0 | 0.75 | 0.46 | 0.65 | 0.89 | 0.64 | 0.69 |
| 6 | 111579998 | rs241001 | -0.053 | 0.904 | 1.63 | 0.375 | 0.001 | 0.003 | 0.003 | 0.002 | 0.86 | 0.92 | 0.89 | 0.85 | 0.86 | 0.79 |
| 6 | 111580561 | rs11153277 | 1.986 | 0.831 | 1.495 | 0.62 | 0.036 | -0.109 | -0.109 | 0 | 0.66 | 0.14 | 0.56 | 0.46 | 0.69 | 0.61 |
| 6 | 111589755 | rs240987 | -0.761 | 0.886 | 1.517 | 0.617 | -0.002 | 0.016 | 0.016 | 0 | 0.15 | 0.2 | 0.14 | 0.33 | 0.16 | 0.13 |
| 6 | 111590094 | rs240988 | 0.187 | 0.885 | 1.629 | 0.582 | -0.003 | -0.012 | -0.012 | 0 | 0.85 | 0.73 | 0.88 | 0.85 | 0.86 | 0.79 |
| 6 | 111598246 | rs190246 | -0.657 | 0.916 | 1.388 | 0.653 | -0.002 | 0.013 | 0.013 | 0 | 0.15 | 0.21 | 0.14 | 0.32 | 0.16 | 0.13 |
| 6 | 111620758 | rs465646 | 0.223 | 0.91 | 1.357 | 0.666 | -0.006 | 0.005 | 0.005 | 0 | 0.84 | 0.73 | 0.86 | 0.81 | 0.85 | 0.77 |
| 6 | 111635808 | rs240969 | -1.05 | 0.712 | 1.313 | 0.691 | -0.02 | 0.06 | 0.06 | 0 | 0.18 | 0.37 | 0.2 | 0.36 | 0.16 | 0.19 |
| 6 | 111644524 | rs240965 | 1.31 | 0.692 | 1.344 | 0.562 | -0.001 | -0.006 | -0.006 | 0.472 | 0.67 | 0.42 | 0.64 | 0.77 | 0.66 | 0.69 |
| 6 | 111655530 | rs465969 | -0.605 | 0.592 | 1.222 | 0.475 | -0.001 | -0.003 | -0.003 | 0 | 0.1 | 0.09 | 0.17 | 0.03 | 0.08 | 0.07 |
| 6 | 111691488 | rs459147 | 0.887 | 1.366 | 1.422 | 0.464 | -0.012 | 0.091 | 0.091 | 1.645 | 0.78 | 0.37 | 0.74 | 0.45 | 0.79 | 0.7 |
| 6 | 111695268 | rs455732 | 2.765 | 1.341 | 1.402 | 0.426 | -0.045 | -0.072 | -0.072 | 0 | 0.48 | 0.05 | 0.43 | 0.42 | 0.5 | 0.5 |
| 6 | 111695887 | rs462779 | 0.87 | 1.349 | 1.408 | 0.43 | -0.012 | 0.091 | 0.091 | 1.645 | 0.78 | 0.37 | 0.74 | 0.45 | 0.79 | 0.7 |
| 6 | 111696091 | rs458017 | 0.132 | 1.345 | 1.401 | 0.432 | 0.008 | 0.004 | 0.004 | 0 | 0.08 | 0.04 | 0.07 | 0 | 0.06 | 0.04 |
| 6 | 111696257 | rs461646 | 0.869 | 1.345 | 1.401 | 0.432 | -0.012 | 0.091 | 0.091 | 1.645 | 0.78 | 0.37 | 0.74 | 0.45 | 0.79 | 0.7 |
| 6 | 111699368 | rs455726 | 2.597 | 1.346 | 1.369 | 0.432 | -0.036 | -0.063 | -0.063 | 0 | 0.49 | 0.05 | 0.44 | 0.42 | 0.5 | 0.5 |
| 6 | 111736642 | rs6937734 | 0.784 | 1.279 | 1.715 | 0.496 | -0.01 | 0.162 | 0.162 | 0.854 | 0.79 | 0.44 | 0.75 | 0.45 | 0.79 | 0.7 |
| 6 | 111751853 | rs17090837 | 0.087 | 1.342 | 1.445 | 0.484 | 0.008 | 0.004 | 0.004 | 0 | 0.08 | 0.04 | 0.07 | 0 | 0.06 | 0.04 |
| 6 | 111754314 | rs9487643 | -0.861 | 1.393 | 1.481 | 0.572 | 0.001 | -0.005 | -0.005 | 0 | 0.92 | 0.93 | 0.94 | 0.92 | 0.9 | 0.92 |
| 6 | 111811922 | rs9320364 | 0.772 | 1.362 | 1.188 | 0.211 | -0.002 | -0.055 | -0.055 | 0 | 0.76 | 0.79 | 0.83 | 1 | 0.77 | 0.92 |
| 6 | 111823782 | rs11755082 | 1.474 | 0.959 | 0.736 | 0.29 | -0.025 | -0.023 | -0.023 | 0.144 | 0.08 | 0 | 0.16 | 0.1 | 0.06 | 0.1 |
| 6 | 111843548 | rs9398272 | -0.99 | 0.884 | 1.198 | 0.187 | -0.011 | 0.182 | 0.182 | 1.206 | 0.21 | 0.5 | 0.25 | 0.55 | 0.2 | 0.28 |
| 6 | 111845111 | rs4521634 | 0.963 | 0.633 | 0.922 | -0.014 | -0.011 | -0.096 | -0.096 | 0 | 0.71 | 0.68 | 0.81 | 1 | 0.74 | 0.82 |
| 6 | 111850742 | rs9400481 | -0.876 | 0.754 | 0.866 | 0.253 | -0.008 | 0.131 | 0.131 | 0.355 | 0.2 | 0.42 | 0.24 | 0.55 | 0.2 | 0.28 |
| 6 | 111876985 | rs10484479 | 0.946 | 0.663 | 0.439 | 0.094 | -0.017 | 0.028 | 0.028 | 0 | 0.09 | 0.01 | 0.05 | 0.06 | 0.1 | 0.08 |
| 6 | 111879579 | rs11153297 | 2.284 | 0.658 | 0.211 | -0.076 | -0.075 | -0.095 | -0.095 | 0.53 | 0.46 | 0.06 | 0.47 | 0.42 | 0.51 | 0.49 |
| 6 | 111881268 | rs3777909 | -0.837 | 0.505 | 0.651 | 0.036 | -0.041 | 0.203 | 0.203 | 2.292 | 0.21 | 0.53 | 0.28 | 0.54 | 0.16 | 0.3 |
| 6 | 111897578 | rs2235175 | 1.41 | 1.034 | 0.518 | -0.227 | -0.002 | 0.029 | 0.029 | 0 | 0.52 | 0.29 | 0.58 | 0.56 | 0.5 | 0.63 |
| 6 | 111897852 | rs9320367 | -1.725 | 1.293 | 0.406 | -0.371 | 0.02 | -0.025 | -0.025 | 0 | 0.39 | 0.8 | 0.35 | 0.47 | 0.34 | 0.43 |
| 6 | 111904447 | rs174372 | -0.136 | 1.11 | 0.77 | -0.245 | 0 | 0.109 | 0.109 | 0.642 | 0.8 | 0.57 | 0.82 | 0.55 | 0.81 | 0.73 |
| 6 | 111905066 | rs7766190 | 0.707 | 1.125 | 0.753 | -0.229 | -0.002 | 0.021 | 0.021 | 0 | 0.64 | 0.48 | 0.72 | 0.53 | 0.64 | 0.69 |
| 6 | 111913070 | rs13190932 | 0.017 | 1.136 | 0.541 | -0.293 | 0 | 0.01 | 0.01 | 0 | 0.05 | 0.03 | 0.06 | 0 | 0.07 | 0.03 |
| 6 | 111914248 | rs174382 | 1.628 | 1.119 | 0.657 | -0.369 | 0.007 | -0.018 | -0.018 | 0 | 0.62 | 0.24 | 0.67 | 0.55 | 0.65 | 0.57 |
| 6 | 111914700 | rs174383 | -0.628 | 0.997 | 0.815 | -0.54 | 0.003 | -0.078 | -0.078 | 1.425 | 0.23 | 0.23 | 0.13 | 0 | 0.24 | 0.14 |
| 6 | 111918794 | rs174392 | 0.801 | 1.174 | 0.996 | -0.282 | 0.009 | 0.068 | 0.068 | 4.462 | 0.73 | 0.47 | 0.73 | 0.52 | 0.74 | 0.59 |
| 6 | 111919108 | rs6941014 | 1.668 | 1.153 | 0.938 | -0.357 | -0.006 | 0.007 | 0.007 | 1.144 | 0.53 | 0.2 | 0.56 | 0.52 | 0.56 | 0.53 |
| 6 | 111919334 | rs174396 | 0.357 | 1.137 | 0.943 | -0.363 | -0.009 | 0.048 | 0.048 | 0.708 | 0.15 | 0.04 | 0.09 | 0 | 0.17 | 0.04 |
| 6 | 111919462 | rs174398 | -0.167 | 1.164 | 0.963 | -0.347 | 0.002 | -0.026 | -0.026 | 1.161 | 0.19 | 0.16 | 0.16 | 0.01 | 0.18 | 0.07 |
| 6 | 111922720 | rs13210247 | 0.377 | 0.692 | 0.323 | -0.488 | -0.004 | -0.006 | -0.006 | 0 | 0.12 | 0.18 | 0.17 | 0.06 | 0.08 | 0.07 |
| 6 | 111924410 | rs3777921 | 1.963 | 0.755 | 0.034 | -0.442 | -0.051 | -0.06 | -0.06 | 0 | 0.39 | 0.11 | 0.47 | 0.5 | 0.46 | 0.47 |
| 6 | 111924913 | rs174402 | 0.012 | 0.765 | 0.569 | 0.005 | 0.002 | 0.054 | 0.054 | 1.02 | 0.81 | 0.68 | 0.85 | 0.57 | 0.82 | 0.73 |
| 6 | 111935682 | rs6926543 | 0.926 | 1.27 | 0.05 | -0.268 | 0.006 | 0.017 | 0.017 | 0.438 | 0.62 | 0.44 | 0.61 | 0.46 | 0.64 | 0.61 |
| 6 | 111936578 | rs9487685 | -0.871 | 1.147 | -0.215 | -0.306 | -0.041 | 0.04 | 0.04 | 0.124 | 0.87 | 0.99 | 0.93 | 0.95 | 0.86 | 0.89 |
| 6 | 111940481 | rs4947132 | -1.333 | 1.05 | 0.194 | -0.294 | 0.011 | -0.022 | -0.022 | 0 | 0.1 | 0.26 | 0.13 | 0.02 | 0.09 | 0.12 |
| 6 | 111944006 | rs9387017 | 0.791 | 0.807 | -0.717 | -0.333 | -0.058 | -0.04 | -0.04 | 0 | 0.39 | 0.17 | 0.55 | 0.44 | 0.5 | 0.56 |
| 6 | 111944979 | rs1476043 | 0.818 | 0.819 | -0.725 | -0.323 | -0.062 | -0.046 | -0.046 | 0 | 0.39 | 0.16 | 0.55 | 0.44 | 0.5 | 0.56 |
| 6 | 111949274 | rs9387018 | 0.45 | 0.703 | -0.52 | -0.558 | 0.001 | 0.035 | 0.035 | 5.897 | 0.7 | 0.38 | 0.79 | 0.5 | 0.74 | 0.61 |
| 6 | 111949779 | rs9400484 | 0.774 | 0.388 | -0.612 | -0.444 | -0.02 | -0.044 | -0.044 | 0 | 0.22 | 0.26 | 0.19 | 0.05 | 0.11 | 0.04 |

| | | | | | | | | | | | | | | | | |
|---|---|---|---|---|---|---|---|---|---|---|---|---|---|---|---|---|
| 6 | 111949988 | rs1476044 | 1.039 | 0.716 | -1.251 | -0.271 | -0.076 | -0.091 | -0.091 | 0.166 | 0.33 | 0.08 | 0.45 | 0.39 | 0.42 | 0.44 |
| 6 | 111950649 | rs11153306 | 0.499 | 0.804 | -1.069 | -0.222 | -0.031 | -0.046 | -0.046 | 0 | 0.4 | 0.19 | 0.55 | 0.44 | 0.48 | 0.51 |
| 6 | 111955435 | rs10457242 | 0.313 | 1.006 | -0.499 | 0.149 | 0.02 | 0.047 | 0.047 | 1.391 | 0.26 | 0.45 | 0.22 | 0.38 | 0.28 | 0.15 |
| 6 | 111964747 | rs9481173 | -0.147 | 0.888 | -0.936 | 0.149 | 0.018 | 0.009 | 0.009 | 0 | 0.3 | 0.59 | 0.26 | 0.38 | 0.35 | 0.21 |
| 6 | 111965272 | rs9320373 | -0.087 | 0.936 | -0.914 | 0.163 | 0.021 | 0.012 | 0.012 | 0.054 | 0.29 | 0.59 | 0.25 | 0.38 | 0.34 | 0.2 |
| 6 | 111966134 | rs17072801 | -1.207 | 0.529 | -0.438 | 0.062 | -0.037 | -0.01 | -0.01 | 0 | 0.16 | 0.31 | 0.14 | 0.11 | 0.08 | 0.23 |
| 6 | 111968788 | rs1794818 | 0.122 | 0.967 | -1.249 | -0.026 | -0.047 | 0.137 | 0.137 | 0.17 | 0.59 | 0.88 | 0.74 | 0.8 | 0.59 | 0.45 |
| 6 | 111969489 | rs1794821 | 0.124 | 1.04 | -1.263 | -0.096 | -0.007 | 0.177 | 0.177 | 0.003 | 0.61 | 0.87 | 0.77 | 0.89 | 0.64 | 0.55 |
| 6 | 111973750 | rs1630896 | 0.196 | 1.121 | -1.264 | -0.069 | -0.026 | 0.132 | 0.132 | 0.062 | 0.59 | 0.86 | 0.73 | 0.8 | 0.6 | 0.45 |
| 6 | 111977860 | rs1726131 | -0.059 | 1.079 | -1.311 | -0.121 | -0.037 | 0.181 | 0.181 | 0.023 | 0.64 | 0.93 | 0.77 | 0.89 | 0.65 | 0.56 |
| 6 | 111978744 | rs4947139 | -0.356 | 1.714 | -0.234 | 0.471 | 0.174 | 0.254 | 0.254 | 0.529 | 0.07 | 0.38 | 0.12 | 0.27 | 0.13 | 0.05 |
| 6 | 111982257 | rs6916861 | -0.356 | 1.714 | -0.234 | 0.471 | 0.174 | 0.254 | 0.254 | 0.529 | 0.07 | 0.39 | 0.12 | 0.28 | 0.13 | 0.05 |
| 6 | 111984400 | rs9387025 | 0.658 | 1.466 | -0.199 | 0.592 | -0.043 | 0.038 | 0.038 | 2.556 | 0.49 | 0.13 | 0.54 | 0.55 | 0.49 | 0.38 |
| 6 | 111991321 | rs6919400 | 0.265 | 1.479 | -0.017 | 0.608 | -0.024 | 0.062 | 0.062 | 0.486 | 0.54 | 0.29 | 0.63 | 0.61 | 0.52 | 0.45 |
| 6 | 111996291 | rs13192832 | 1.028 | 1.463 | -0.113 | 0.585 | -0.049 | -0.1 | -0.1 | 0 | 0.46 | 0.04 | 0.5 | 0.46 | 0.44 | 0.34 |
| 6 | 111996419 | rs9374274 | 0.582 | 1.567 | -0.064 | 0.584 | 0.011 | 0.033 | 0.033 | 1.236 | 0.51 | 0.21 | 0.59 | 0.52 | 0.47 | 0.41 |
| 6 | 112003748 | rs706862 | -0.468 | 1.544 | -0.279 | 0.635 | -0.017 | 0.114 | 0.114 | 0.183 | 0.65 | 0.96 | 0.77 | 0.84 | 0.67 | 0.57 |
| 6 | 112005126 | rs2237257 | -0.323 | 1.682 | 0.069 | 0.692 | 0.141 | 0.256 | 0.256 | 0.553 | 0.07 | 0.4 | 0.1 | 0.28 | 0.12 | 0.05 |
| 6 | 112006549 | rs2237259 | 0.865 | 1.181 | -0.167 | 0.687 | -0.011 | -0.027 | -0.027 | 0.172 | 0.47 | 0.16 | 0.51 | 0.46 | 0.45 | 0.35 |
| 6 | 112007280 | rs811470 | 0.159 | 1.196 | -0.1 | 0.469 | -0.003 | 0.026 | 0.026 | 0.316 | 0.55 | 0.39 | 0.65 | 0.61 | 0.53 | 0.47 |
| 6 | 112010019 | rs804192 | 0.353 | 0.836 | -0.242 | 0.433 | -0.003 | -0.008 | -0.008 | 0 | 0.27 | 0.36 | 0.25 | 0.21 | 0.27 | 0.28 |
| 6 | 112015373 | rs2071926 | 0.299 | -0.232 | -0.575 | 0.011 | 0.012 | -0.049 | -0.049 | 0 | 0.52 | 0.08 | 0.59 | 0.47 | 0.59 | 0.43 |
| 6 | 112016261 | rs2071927 | -0.42 | -0.403 | 0.085 | 0.149 | 0.051 | 0.39 | 0.39 | 0.281 | 0.07 | 0.36 | 0.09 | 0.26 | 0.04 | 0.06 |
| 6 | 112020414 | rs809193 | -0.492 | -0.559 | -0.218 | -0.082 | 0.113 | -0.084 | -0.084 | 1.431 | 0.39 | 0.88 | 0.3 | 0.45 | 0.2 | 0.49 |
| 6 | 112024262 | rs3752545 | -0.259 | -0.541 | 0.055 | -0.06 | 0.058 | 0.4 | 0.4 | 0.231 | 0.06 | 0.35 | 0.09 | 0.25 | 0.04 | 0.06 |
| 6 | 112029331 | rs1022646 | 1.009 | -0.647 | 0.011 | -0.195 | 0.07 | 0.084 | 0.084 | 0 | 0.24 | 0.08 | 0.15 | 0.07 | 0.11 | 0.35 |
| 6 | 112029402 | rs1022647 | 0.876 | -0.659 | 0.107 | -0.2 | 0.05 | 0.054 | 0.054 | 0 | 0.25 | 0.13 | 0.15 | 0.07 | 0.12 | 0.35 |
| 6 | 112033370 | rs706882 | -0.985 | -0.659 | 0.137 | -0.195 | 0.028 | 0.061 | 0.061 | 2.174 | 0.09 | 0.38 | 0.06 | 0.13 | 0.04 | 0.08 |
| 6 | 112035002 | rs809192 | -1.075 | -0.585 | 0.068 | -0.219 | -0.047 | 0.006 | 0.006 | 0 | 0.91 | 0.98 | 0.89 | 0.93 | 0.79 | 0.92 |
| 6 | 112037613 | rs706889 | -1.061 | -1.155 | 0.122 | -0.181 | 0.025 | 0.062 | 0.062 | 2.183 | 0.09 | 0.38 | 0.06 | 0.13 | 0.04 | 0.08 |
| 6 | 112041359 | rs706895 | -0.68 | -1.06 | -0.266 | -0.202 | 0.074 | -0.079 | -0.079 | 0 | 0.4 | 0.89 | 0.31 | 0.45 | 0.23 | 0.5 |
| 6 | 112053105 | rs697642 | -2.337 | -1.119 | 0.76 | 0.583 | -0.014 | 0.085 | 0.085 | 7.084 | 0.15 | 0.74 | 0.15 | 0.38 | 0.08 | 0.14 |
| 6 | 112063482 | rs6930230 | -0.639 | -1.263 | 1.427 | 0.45 | 0 | -0.005 | -0.005 | 0 | 0.45 | 0.49 | 0.56 | 0.61 | 0.46 | 0.55 |
| 6 | 112072605 | rs12524424 | -0.366 | -1.266 | 1.34 | 0.446 | 0.016 | 0.028 | 0.028 | 0.253 | 0.05 | 0.24 | 0.04 | 0.01 | 0.01 | 0.04 |
| 6 | 112072692 | rs17072872 | 0.326 | -1.276 | 1.16 | 0.45 | -0.061 | -0.117 | -0.117 | 0.924 | 0.33 | 0.09 | 0.49 | 0.47 | 0.4 | 0.43 |
| 6 | 112075928 | rs6910116 | -0.958 | -1.219 | 1.174 | -0.526 | -0.007 | 0.233 | 0.233 | 0 | 0.49 | 0.8 | 0.64 | 0.86 | 0.5 | 0.61 |
| 6 | 112081938 | rs11964650 | -0.989 | -0.95 | 1.228 | -0.282 | 0.04 | 0.089 | 0.089 | 1.074 | 0.09 | 0.28 | 0.06 | 0.14 | 0.05 | 0.08 |
| 6 | 112089668 | rs9372311 | -0.548 | -1.044 | 0.877 | -0.484 | -0.012 | 0.078 | 0.078 | 0 | 0.44 | 0.67 | 0.61 | 0.86 | 0.49 | 0.53 |
| 6 | 112096345 | rs11963612 | -0.348 | -1.259 | 1.289 | -0.188 | 0.007 | 0.035 | 0.035 | 0 | 0.44 | 0.36 | 0.58 | 0.7 | 0.47 | 0.52 |
| 6 | 112101725 | rs9384805 | -1.082 | -1.286 | 0.867 | -0.328 | -0.026 | 0.129 | 0.129 | 0 | 0.56 | 0.78 | 0.66 | 0.84 | 0.53 | 0.65 |
| 6 | 112121228 | rs2148709 | -2.062 | -1.505 | 1.688 | 0.406 | 0.043 | 0.001 | 0.001 | 2.683 | 0.27 | 0.8 | 0.2 | 0.27 | 0.15 | 0.21 |
| 6 | 112126303 | rs7761230 | -0.585 | -1.556 | 2.323 | 0.514 | -0.045 | 0.017 | 0.017 | 0.085 | 0.15 | 0.33 | 0.12 | 0.2 | 0.1 | 0.07 |
| 6 | 112126339 | rs17072912 | -2.128 | -1.537 | 2.344 | 0.345 | -0.003 | 0.042 | 0.042 | 0.277 | 0.12 | 0.44 | 0.08 | 0.14 | 0.05 | 0.15 |
| 6 | 112126755 | rs1022650 | -1.618 | -1.511 | 2.792 | 0.363 | 0.035 | 0.072 | 0.072 | 0 | 0.08 | 0.26 | 0.05 | 0.12 | 0.04 | 0.1 |
| 6 | 112138882 | rs2182644 | -1.6 | -1.663 | 2.338 | 0.291 | -0.071 | -0.022 | -0.022 | 0.171 | 0.22 | 0.65 | 0.16 | 0.14 | 0.11 | 0.15 |
| 6 | 112152560 | rs1327204 | -1.373 | -1.639 | 2.415 | 0.232 | -0.069 | -0.049 | -0.049 | 0 | 0.2 | 0.43 | 0.11 | 0.12 | 0.11 | 0.17 |
| 6 | 112154200 | rs7761236 | -1.735 | -1.67 | 2.589 | 0.195 | -0.082 | -0.068 | -0.068 | 0 | 0.22 | 0.55 | 0.12 | 0.12 | 0.11 | 0.17 |
| 6 | 112167609 | rs6934113 | -1.677 | -1.693 | 2.742 | 0.299 | 0.017 | 0.048 | 0.048 | 0.91 | 0.08 | 0.28 | 0.07 | 0.12 | 0.05 | 0.09 |
| 6 | 112170222 | rs6568708 | -2.97 | -1.607 | 2.617 | 0.228 | 0.052 | 0.012 | 0.012 | 2.745 | 0.31 | 0.8 | 0.25 | 0.3 | 0.19 | 0.27 |
| 6 | 112175517 | rs9487737 | -0.372 | -1.41 | 2.947 | 0.253 | -0.051 | -0.074 | -0.074 | 0.783 | 0.13 | 0.32 | 0.08 | 0.01 | 0.07 | 0.05 |
| 6 | 112175735 | rs910682 | -2.045 | -1.41 | 2.844 | 0.32 | -0.029 | -0.035 | -0.035 | 0 | 0.24 | 0.64 | 0.18 | 0.14 | 0.16 | 0.16 |
| 6 | 112184621 | rs9320379 | -3.751 | -1.388 | 2.579 | 0.153 | 0.018 | -0.006 | -0.006 | 0 | 0.26 | 0.79 | 0.25 | 0.28 | 0.18 | 0.32 |
| 6 | 112184705 | rs9487738 | -0.857 | -1.431 | 2.762 | 0.038 | 0.007 | -0.011 | -0.011 | 0 | 0.05 | 0.24 | 0.04 | 0.01 | 0.03 | 0.05 |
| 6 | 112195611 | rs1409837 | -0.772 | -1.472 | 2.23 | 0.014 | 0.018 | 0.002 | 0.002 | 0 | 0.16 | 0.1 | 0.07 | 0 | 0.09 | 0.15 |
| 6 | 112203035 | rs9372316 | -2.597 | -1.46 | 2.004 | 0.168 | 0.057 | 0.011 | 0.011 | 0.745 | 0.35 | 0.8 | 0.3 | 0.35 | 0.25 | 0.34 |
| 6 | 112231090 | rs9487747 | -0.113 | -1.209 | 2.176 | 0.289 | -0.011 | -0.027 | -0.027 | 0 | 0.11 | 0.15 | 0.06 | 0 | 0.07 | 0.02 |
| 6 | 112232137 | rs9320383 | -2.492 | -1.293 | 2.517 | 0.321 | -0.005 | -0.001 | -0.001 | 0 | 0.28 | 0.71 | 0.24 | 0.26 | 0.19 | 0.25 |
| 6 | 112232986 | rs9487749 | -2.879 | -1.278 | 2.43 | 0.281 | 0.038 | 0.004 | 0.004 | 1.174 | 0.34 | 0.78 | 0.29 | 0.36 | 0.24 | 0.32 |
| 6 | 112240332 | rs9374302 | -3.649 | -0.504 | 2.888 | 0.823 | 0.003 | -0.005 | -0.005 | 1.575 | 0.24 | 0.72 | 0.24 | 0.36 | 0.17 | 0.3 |
| 6 | 112242154 | rs943436 | -2.139 | -0.476 | 3.248 | 0.92 | 0.032 | 0.223 | 0.223 | 0.307 | 0.1 | 0.3 | 0.15 | 0.36 | 0.09 | 0.13 |
| 6 | 112257478 | rs580900 | -1.853 | -0.493 | 3.195 | 0.917 | 0.016 | 0.212 | 0.212 | 0 | 0.08 | 0.32 | 0.12 | 0.24 | 0.06 | 0.14 |
| 6 | 112259476 | rs17073074 | -1.746 | -0.529 | 3.269 | 0.851 | 0.012 | 0.008 | 0.008 | 0.154 | 0.12 | 0.34 | 0.08 | 0.12 | 0.08 | 0.14 |
| 6 | 112261385 | rs671271 | -3.114 | -0.551 | 2.752 | 0.844 | 0.02 | -0.014 | -0.014 | 0.712 | 0.24 | 0.79 | 0.21 | 0.36 | 0.15 | 0.29 |
| 6 | 112263731 | rs17754910 | 0.325 | 0.002 | 2.28 | 0.544 | -0.047 | 0.052 | 0.052 | 2.771 | 0.11 | 0.03 | 0.1 | 0 | 0.18 | 0.04 |
| 6 | 112281277 | rs9487771 | -2.891 | -0.918 | 2.844 | 1.54 | -0.045 | 0.117 | 0.117 | 1.021 | 0.22 | 0.62 | 0.24 | 0.52 | 0.13 | 0.29 |
| 6 | 112286687 | rs689328 | -2.335 | -0.773 | 2.524 | 1.606 | 0.012 | 0.264 | 0.264 | 0 | 0.09 | 0.45 | 0.13 | 0.29 | 0.06 | 0.18 |
| 6 | 112289582 | rs546913 | -1.964 | -0.677 | 2.322 | 1.663 | 0.025 | 0.268 | 0.268 | 0 | 0.08 | 0.39 | 0.12 | 0.29 | 0.06 | 0.11 |
| 6 | 112294010 | rs9387058 | -1.977 | -0.429 | 1.42 | 1.083 | -0.013 | 0.035 | 0.035 | 0.533 | 0.14 | 0.29 | 0.15 | 0.25 | 0.09 | 0.15 |
| 6 | 112321879 | rs9487783 | -1.791 | -0.294 | 1.485 | 0.607 | -0.01 | 0.038 | 0.038 | 0 | 0.09 | 0.25 | 0.05 | 0.18 | 0.06 | 0.08 |

| | | | | | | | | | | | | | | | | |
|---|---|---|---|---|---|---|---|---|---|---|---|---|---|---|---|---|
| 6 | 112332393 | rs2815787 | -2.216 | -0.301 | 1.489 | 0.446 | -0.013 | -0.004 | -0.004 | 0 | 0.25 | 0.58 | 0.24 | 0.28 | 0.18 | 0.27 |
| 6 | 112338958 | rs1515333 | -2.044 | 0.094 | 0.647 | 0.072 | 0.018 | -0.031 | -0.031 | 0.266 | 0.15 | 0.33 | 0.11 | 0.09 | 0.17 | 0.18 |
| 6 | 112340993 | rs9481218 | -1.398 | -0.305 | 0.263 | 0.022 | -0.006 | -0.042 | -0.042 | 0 | 0.16 | 0.26 | 0.09 | 0.03 | 0.11 | 0.14 |
| 6 | 112342709 | rs1028122 | -1.127 | -0.295 | -0.031 | 0.549 | -0.043 | 0.016 | 0.016 | 0.036 | 0.17 | 0.61 | 0.09 | 0.24 | 0.08 | 0.21 |
| 6 | 112346096 | rs11153327 | -0.643 | 0.31 | -1.452 | 0.26 | -0.059 | 0.026 | 0.026 | 0 | 0.3 | 0.03 | 0.28 | 0.17 | 0.31 | 0.18 |
| 6 | 112350329 | rs925755 | 0.336 | 0.436 | -2.048 | -0.744 | -0.008 | 0.134 | 0.134 | 2.777 | 0.34 | 0.11 | 0.29 | 0.06 | 0.32 | 0.16 |
| 6 | 112351538 | rs6904241 | -0.1 | 0.576 | -1.234 | 0.168 | -0.004 | -0.003 | -0.003 | 0 | 0.62 | 0.62 | 0.53 | 0.45 | 0.52 | 0.49 |
| 6 | 112357513 | rs17073224 | -0.025 | 0.805 | -1.233 | 0.691 | -0.006 | -0.009 | -0.009 | 0 | 0.08 | 0.09 | 0.11 | 0.13 | 0.09 | 0.07 |
| 6 | 112360196 | rs2673307 | 0.441 | 0.304 | -1.312 | 0.836 | 0.052 | 0.045 | 0.045 | 0.967 | 0.5 | 0.3 | 0.44 | 0.44 | 0.42 | 0.4 |
| 6 | 112362670 | rs2815803 | 0.201 | 0.642 | -0.836 | 1.1 | 0.041 | 0.001 | 0.001 | 0 | 0.71 | 0.52 | 0.66 | 0.77 | 0.64 | 0.53 |
| 6 | 112363842 | rs9398296 | 0.326 | -0.14 | -1.356 | 0.787 | 0.04 | -0.026 | -0.026 | 0 | 0.2 | 0.06 | 0.19 | 0.27 | 0.15 | 0.25 |
| 6 | 112367137 | rs4947161 | 0.075 | -0.172 | -1.138 | 0.807 | 0.003 | -0.008 | -0.008 | 0 | 0.68 | 0.35 | 0.63 | 0.79 | 0.63 | 0.59 |
| 6 | 112369882 | rs6900135 | 1.749 | -0.24 | -1.276 | 0.842 | -0.065 | 0.096 | 0.096 | 1.866 | 0.26 | 0.02 | 0.23 | 0.07 | 0.25 | 0.11 |
| 6 | 112370218 | rs4947162 | 1.57 | -0.239 | -1.254 | 0.875 | -0.014 | 0.071 | 0.071 | 1.537 | 0.27 | 0.12 | 0.23 | 0.13 | 0.25 | 0.12 |
| 6 | 112374944 | rs1022313 | 0.969 | -0.183 | -1.159 | 0.995 | -0.006 | -0.021 | -0.021 | 0.666 | 0.2 | 0.13 | 0.22 | 0.34 | 0.23 | 0.13 |
| 6 | 112378555 | rs1230348 | -0.359 | -0.391 | -0.668 | 0.796 | -0.004 | 0.002 | 0.002 | 0 | 0.76 | 0.82 | 0.69 | 0.86 | 0.65 | 0.69 |
| 6 | 112383780 | rs17073268 | 1.039 | -0.496 | -0.21 | 0.844 | -0.001 | 0 | 0 | 0 | 0.21 | 0.18 | 0.22 | 0.35 | 0.24 | 0.13 |
| 6 | 112399750 | rs10456877 | 0.959 | -0.645 | -0.466 | 0.54 | 0.001 | 0.027 | 0.027 | 0.122 | 0.23 | 0.31 | 0.23 | 0.35 | 0.25 | 0.13 |
| 6 | 112402709 | rs9400519 | 0.736 | -0.744 | -0.565 | 0.576 | 0.042 | -0.071 | -0.071 | 0.029 | 0.2 | 0.04 | 0.18 | 0.32 | 0.15 | 0.35 |
| 6 | 112413178 | rs11153336 | 1.307 | -0.63 | -0.432 | 0.439 | 0.002 | 0.012 | 0.012 | 0 | 0.21 | 0.24 | 0.22 | 0.35 | 0.24 | 0.13 |
| 6 | 112421805 | rs916429 | 0.964 | -0.265 | -0.585 | 0.286 | -0.014 | -0.011 | -0.011 | 2.014 | 0.11 | 0 | 0.07 | 0.08 | 0.11 | 0.21 |
| 6 | 112422154 | rs17073353 | 1.301 | -0.279 | -0.287 | 0.424 | 0.002 | 0.013 | 0.013 | 0 | 0.21 | 0.24 | 0.22 | 0.35 | 0.24 | 0.14 |
| 6 | 112422664 | rs2010314 | 0.76 | -0.277 | -0.331 | 0.425 | 0 | 0.042 | 0.042 | 0.238 | 0.24 | 0.42 | 0.23 | 0.35 | 0.24 | 0.14 |
| 6 | 112423810 | rs2213842 | 0.317 | -0.237 | -0.313 | 0.393 | -0.001 | 0.001 | 0.001 | 0 | 0.52 | 0.62 | 0.48 | 0.47 | 0.5 | 0.26 |
| 6 | 112430027 | rs6568718 | 0.785 | -0.23 | -0.332 | 0.387 | 0 | 0.043 | 0.043 | 0.238 | 0.24 | 0.42 | 0.23 | 0.35 | 0.24 | 0.14 |
| 6 | 112431283 | rs12204892 | 1.359 | -0.22 | -0.376 | 0.11 | -0.018 | 0.07 | 0.07 | 1.484 | 0.27 | 0.12 | 0.23 | 0.13 | 0.26 | 0.12 |
| 6 | 112434159 | rs11961355 | 1.254 | -0.166 | -0.25 | 0.245 | 0.003 | 0.018 | 0.018 | 0 | 0.21 | 0.25 | 0.22 | 0.35 | 0.24 | 0.14 |
| 6 | 112435682 | rs6916947 | 0.689 | 0.014 | -0.686 | 0.416 | -0.001 | 0.07 | 0.07 | 0.769 | 0.26 | 0.62 | 0.25 | 0.46 | 0.25 | 0.16 |
| 6 | 112436198 | rs4947169 | -0.418 | -0.097 | -0.736 | 0.373 | -0.002 | -0.003 | -0.003 | 0 | 0.44 | 0.37 | 0.5 | 0.46 | 0.47 | 0.71 |
| 6 | 112438446 | rs1160798 | -1.24 | -0.003 | -0.76 | 0.276 | -0.02 | 0.083 | 0.083 | 1.359 | 0.71 | 0.88 | 0.83 | 0.87 | 0.71 | 0.87 |
| 6 | 112446876 | rs969139 | -0.37 | -0.044 | -0.715 | 0.317 | 0 | -0.001 | -0.001 | 0 | 0.45 | 0.39 | 0.59 | 0.54 | 0.47 | 0.72 |
| 6 | 112456782 | rs12214148 | 1.002 | -0.056 | -0.878 | 0.282 | 0 | 0.023 | 0.023 | 0.055 | 0.24 | 0.29 | 0.23 | 0.36 | 0.24 | 0.16 |
| 6 | 112457390 | rs2032567 | -1.137 | 0.028 | -1.07 | 0.279 | -0.025 | 0.081 | 0.081 | 1.41 | 0.71 | 0.88 | 0.83 | 0.87 | 0.71 | 0.89 |
| 6 | 112460648 | rs2072020 | -0.113 | -0.639 | -0.834 | 0.515 | 0.003 | -0.004 | -0.004 | 0.172 | 0.23 | 0.22 | 0.3 | 0.4 | 0.18 | 0.45 |
| 6 | 112463076 | rs2072026 | -0.477 | 0.444 | -0.569 | 0.887 | 0.016 | -0.007 | -0.007 | 2.478 | 0.59 | 0.65 | 0.6 | 0.54 | 0.64 | 0.45 |
| 6 | 112477628 | rs3798357 | -0.041 | 1.028 | -0.302 | 1.431 | 0.005 | 0.064 | 0.064 | 0.324 | 0.26 | 0.42 | 0.24 | 0.42 | 0.26 | 0.17 |
| 6 | 112480041 | rs3752577 | -0.877 | 1.027 | -0.542 | 1.369 | -0.007 | -0.014 | -0.014 | 0.258 | 0.06 | 0.03 | 0.12 | 0.13 | 0.1 | 0.1 |
| 6 | 112483235 | rs6913656 | 0.893 | 1.42 | -0.132 | 1.736 | 0.028 | 0.064 | 0.064 | 0.686 | 0.75 | 0.67 | 0.62 | 0.57 | 0.65 | 0.54 |
| 6 | 112485798 | rs7766236 | 0.781 | 1.372 | -0.206 | 1.95 | 0.015 | 0.051 | 0.051 | 0 | 0.77 | 0.72 | 0.54 | 0.64 | 0.68 | 0.65 |
| 6 | 112490531 | rs2051649 | 0.564 | 1.165 | 0.643 | 1.985 | 0.03 | 0.077 | 0.077 | 0 | 0.83 | 0.65 | 0.79 | 0.81 | 0.78 | 0.85 |
| 6 | 112493256 | rs9387060 | 1.119 | 0.697 | 0.177 | 1.82 | -0.003 | -0.038 | -0.038 | 0 | 0.6 | 0.23 | 0.58 | 0.55 | 0.59 | 0.57 |
| 6 | 112493872 | rs1050348 | -0.039 | 0.542 | 0.132 | 1.205 | -0.01 | 0.007 | 0.007 | 0 | 0.69 | 0.78 | 0.72 | 0.81 | 0.65 | 0.81 |
| 6 | 112496511 | rs6917763 | 0.652 | 0.411 | 0.789 | 1.127 | -0.033 | -0.045 | -0.045 | 0 | 0.08 | 0.28 | 0.03 | 0 | 0.01 | 0.01 |
| 6 | 112500015 | rs2213839 | -0.069 | 0.427 | 1.109 | 1.136 | 0.001 | -0.002 | -0.002 | 1.925 | 0.69 | 0.71 | 0.72 | 0.81 | 0.65 | 0.81 |
| 6 | 112500650 | rs1158747 | 1.356 | 0.401 | 1.085 | 1.156 | -0.013 | -0.016 | -0.016 | 0 | 0.51 | 0.22 | 0.54 | 0.55 | 0.51 | 0.55 |
| 6 | 112501537 | rs4947172 | -0.388 | 0.399 | 1.183 | 1.156 | 0.016 | 0.069 | 0.069 | 0.168 | 0.09 | 0.2 | 0.14 | 0.26 | 0.12 | 0.26 |
| 6 | 112502989 | rs6907109 | 1.225 | 0.4 | 1.137 | 1.151 | -0.006 | -0.006 | -0.006 | 0 | 0.53 | 0.33 | 0.55 | 0.55 | 0.52 | 0.55 |
| 6 | 112506375 | rs9374309 | 1.403 | 0.347 | 0.947 | 1.334 | -0.033 | -0.013 | -0.013 | 0.178 | 0.51 | 0.12 | 0.53 | 0.51 | 0.51 | 0.54 |
| 6 | 112510818 | rs2072029 | -1.275 | 0.285 | 1.063 | 1.279 | -0.024 | 0.047 | 0.047 | 1.352 | 0.18 | 0.61 | 0.18 | 0.26 | 0.13 | 0.27 |
| 6 | 112511852 | rs2269646 | -1.549 | 0.304 | 1.028 | 1.289 | -0.028 | -0.035 | -0.035 | 0 | 0.49 | 0.89 | 0.47 | 0.45 | 0.47 | 0.46 |
| 6 | 112512905 | rs2072021 | -1.293 | 0.285 | 1.079 | 1.287 | -0.02 | 0.048 | 0.048 | 1.325 | 0.18 | 0.61 | 0.18 | 0.26 | 0.14 | 0.26 |
| 6 | 112514094 | rs2237237 | -0.793 | 0.338 | 1.232 | 1.34 | 0.061 | 0.183 | 0.183 | 1.756 | 0.11 | 0.46 | 0.16 | 0.26 | 0.13 | 0.27 |
| 6 | 112523395 | rs4947177 | -0.029 | -0.142 | 0.432 | 0.809 | 0.009 | -0.013 | -0.013 | 0 | 0.31 | 0.39 | 0.29 | 0.19 | 0.34 | 0.19 |
| 6 | 112524266 | rs9374311 | -0.895 | 0.301 | 0.299 | 1.345 | 0.018 | 0.119 | 0.119 | 0.207 | 0.19 | 0.37 | 0.32 | 0.39 | 0.2 | 0.36 |
| 6 | 112524506 | rs2237244 | -1.233 | -0.001 | -0.625 | 1.227 | -0.021 | 0.066 | 0.066 | 0.475 | 0.56 | 0.94 | 0.75 | 0.62 | 0.61 | 0.67 |
| 6 | 112526101 | rs3777925 | 0.295 | 0.007 | -0.144 | 1.793 | -0.028 | -0.029 | -0.029 | 0.709 | 0.07 | 0.02 | 0.25 | 0.2 | 0.11 | 0.22 |
| 6 | 112526404 | rs7754329 | -0.324 | -0.339 | 0.246 | 1.876 | 0.001 | 0.036 | 0.036 | 0 | 0.92 | 0.91 | 0.86 | 0.86 | 0.91 | 0.95 |
| 6 | 112529268 | rs9487847 | -0.903 | -0.964 | 0.073 | 1.139 | -0.036 | 0.05 | 0.05 | 0.258 | 0.3 | 0.59 | 0.28 | 0.59 | 0.2 | 0.37 |
| 6 | 112529727 | rs2237246 | -0.79 | -0.922 | -0.424 | 1.161 | 0.005 | -0.008 | -0.008 | 0 | 0.31 | 0.72 | 0.29 | 0.58 | 0.2 | 0.36 |
| 6 | 112531660 | rs2282854 | 0.54 | -0.469 | -0.771 | 0.836 | 0.035 | -0.023 | -0.023 | 0.001 | 0.13 | 0.04 | 0.08 | 0.19 | 0.07 | 0.12 |
| 6 | 112535653 | rs9374312 | 0.242 | -0.193 | -1.222 | 0.538 | -0.088 | -0.001 | -0.001 | 0 | 0.34 | 0.12 | 0.42 | 0.28 | 0.46 | 0.39 |
| 6 | 112536990 | rs2213840 | 0.662 | -0.476 | -0.768 | -0.196 | -0.005 | -0.044 | -0.044 | 0 | 0.27 | 0.29 | 0.16 | 0.06 | 0.2 | 0.15 |
| 6 | 112549164 | rs4947179 | 0.19 | -0.414 | -0.967 | 0.143 | 0.018 | -0.012 | -0.012 | 0.055 | 0.44 | 0.17 | 0.67 | 0.42 | 0.59 | 0.59 |
| 6 | 112550015 | rs9487857 | 0.306 | -0.377 | -0.473 | -0.13 | 0 | -0.021 | -0.021 | 0.265 | 0.46 | 0.53 | 0.27 | 0.16 | 0.4 | 0.4 |
| 6 | 112558763 | rs2072024 | 0.251 | 0.199 | -0.521 | 0.508 | -0.036 | -0.056 | -0.056 | 0.409 | 0.19 | 0.08 | 0.5 | 0.41 | 0.28 | 0.37 |
| 6 | 112564517 | rs6926573 | -0.825 | -0.136 | -0.346 | 0.511 | -0.012 | 0.05 | 0.05 | 0.431 | 0.81 | 0.72 | 0.89 | 0.71 | 0.84 | 0.87 |
| 6 | 112578384 | rs11153350 | 1.003 | 0.08 | -1.133 | 0.139 | -0.055 | 0.107 | 0.107 | 0 | 0.24 | 0.09 | 0.17 | 0.01 | 0.32 | 0.21 |
| 6 | 112581415 | rs9481245 | 1.24 | -0.136 | -0.897 | 0.703 | 0.027 | -0.035 | -0.035 | 0 | 0.08 | 0.03 | 0.04 | 0.17 | 0.05 | 0.12 |
| 6 | 112589314 | rs883893 | -0.283 | 0.356 | -0.728 | 0.971 | -0.007 | 0.035 | 0.035 | 0 | 0.91 | 0.93 | 0.64 | 0.59 | 0.86 | 0.78 |

| | | | | | | | | | | | | | | | | |
|---|---|---|---|---|---|---|---|---|---|---|---|---|---|---|---|---|
| 6 | 112590682 | rs17680579 | 1.196 | -0.009 | -0.915 | -0.05 | 0.019 | 0.037 | 0.037 | 1.507 | 0.16 | 0.04 | 0.06 | 0.08 | 0.12 | 0.1 |
| 6 | 112591906 | rs1891472 | 0.429 | 0.081 | -0.588 | -0.328 | -0.01 | 0.007 | 0.007 | 0 | 0.36 | 0.33 | 0.31 | 0.07 | 0.43 | 0.41 |
| 6 | 112596456 | rs1891470 | 0.534 | 0.112 | -0.503 | -0.009 | 0.008 | 0.024 | 0.024 | 0.003 | 0.35 | 0.32 | 0.21 | 0.2 | 0.28 | 0.25 |
| 6 | 112601921 | rs17073644 | 0.01 | -0.021 | -0.548 | 0.852 | -0.024 | -0.036 | -0.036 | 0 | 0.11 | 0.05 | 0.37 | 0.49 | 0.16 | 0.26 |
| 6 | 112603809 | rs6931341 | -0.802 | -0.061 | -0.275 | -0.051 | 0 | -0.021 | -0.021 | 0 | 0.33 | 0.61 | 0.49 | 0.73 | 0.34 | 0.42 |
| 6 | 112605814 | rs12195540 | 0.803 | -0.173 | -0.368 | -0.312 | 0.008 | 0.031 | 0.031 | 2.473 | 0.16 | 0.09 | 0.19 | 0.06 | 0.14 | 0.1 |
| 6 | 112606123 | rs6901099 | 0.046 | -0.249 | 0.069 | -0.245 | 0.016 | -0.006 | -0.006 | 0 | 0.11 | 0.24 | 0.1 | 0.06 | 0.12 | 0.1 |
| 6 | 112606215 | rs6923632 | 0.322 | -0.189 | -0.52 | -0.302 | -0.033 | 0.064 | 0.064 | 0 | 0.34 | 0.13 | 0.48 | 0.23 | 0.42 | 0.34 |
| 6 | 112608146 | rs6935727 | 0.316 | -0.575 | -0.216 | -0.302 | -0.002 | 0.003 | 0.003 | 0 | 0.27 | 0.27 | 0.23 | 0.17 | 0.32 | 0.31 |
| 6 | 112610724 | rs6568729 | 0.082 | -0.76 | -0.191 | -0.546 | -0.003 | 0.007 | 0.007 | 0 | 0.41 | 0.36 | 0.52 | 0.61 | 0.44 | 0.62 |
| 6 | 112612294 | rs9487881 | -0.294 | -0.839 | -0.069 | -0.584 | -0.001 | 0.124 | 0.124 | 0 | 0.47 | 0.71 | 0.54 | 0.77 | 0.47 | 0.69 |
| 6 | 112613737 | rs9487882 | -1.173 | -0.956 | -0.199 | -0.861 | -0.023 | 0.04 | 0.04 | 0 | 0.72 | 0.78 | 0.67 | 0.93 | 0.66 | 0.83 |
| 6 | 112616614 | rs11962065 | 0.088 | -0.707 | -0.512 | -0.648 | 0.002 | 0.002 | 0.002 | 0 | 0.25 | 0.27 | 0.15 | 0.16 | 0.18 | 0.13 |
| 6 | 112621590 | rs3920874 | 0.323 | -0.876 | -0.909 | -0.528 | 0.009 | -0.036 | -0.036 | 0.304 | 0.23 | 0.13 | 0.19 | 0.56 | 0.2 | 0.43 |
| 6 | 112621834 | rs1342593 | -1.359 | -0.997 | -1.337 | -0.953 | -0.129 | 0.171 | 0.171 | 0 | 0.71 | 0.97 | 0.69 | 0.92 | 0.65 | 0.8 |
| 6 | 112622977 | rs12207729 | 0.568 | -0.99 | -1.299 | -0.906 | 0.009 | -0.039 | -0.039 | 0 | 0.23 | 0.12 | 0.19 | 0.57 | 0.2 | 0.43 |
| 6 | 112624037 | rs12209711 | 0.555 | -0.99 | -1.3 | -0.905 | 0.009 | -0.034 | -0.034 | 0 | 0.23 | 0.13 | 0.19 | 0.57 | 0.21 | 0.43 |
| 6 | 112626176 | rs947717 | -1.318 | -0.873 | -1.491 | -1.113 | -0.129 | 0.17 | 0.17 | 0 | 0.71 | 0.97 | 0.69 | 0.92 | 0.65 | 0.8 |
| 6 | 112633545 | rs9320406 | -0.486 | -0.953 | -0.851 | -1.152 | -0.018 | -0.031 | -0.031 | 0 | 0.37 | 0.52 | 0.24 | 0.82 | 0.28 | 0.52 |
| 6 | 112643310 | rs9487887 | 0.802 | -0.803 | -1.473 | -1.004 | 0.005 | -0.055 | -0.055 | 0 | 0.2 | 0.08 | 0.18 | 0.64 | 0.21 | 0.39 |
| 6 | 112644225 | rs17073700 | 0.438 | -0.603 | -1.369 | -0.937 | -0.038 | 0.029 | 0.029 | 0 | 0.12 | 0.21 | 0.04 | 0.17 | 0.06 | 0.14 |
| 6 | 112651206 | rs1935681 | 0.744 | -0.363 | -1.467 | -1.13 | -0.024 | 0.063 | 0.063 | 0 | 0.16 | 0.08 | 0.11 | 0.01 | 0.21 | 0.14 |
| 6 | 112651321 | rs9374318 | 0.954 | -0.79 | -1.009 | -0.491 | -0.042 | 0.003 | 0.003 | 2.608 | 0.16 | 0.05 | 0.32 | 0.15 | 0.17 | 0.11 |
| 6 | 112656121 | rs6927457 | -0.441 | -1.318 | -0.279 | 0.002 | -0.032 | 0.014 | 0.014 | 0 | 0.15 | 0.21 | 0.06 | 0.19 | 0.08 | 0.15 |
| 6 | 112656911 | rs6933749 | -0.533 | -0.853 | -0.783 | -0.5 | 0.049 | 0.068 | 0.068 | 0 | 0.45 | 0.63 | 0.4 | 0.87 | 0.31 | 0.6 |
| 6 | 112661761 | rs9400534 | -0.652 | -0.976 | -0.426 | 0.41 | -0.021 | 0.007 | 0.007 | 0.346 | 0.28 | 0.31 | 0.14 | 0.37 | 0.12 | 0.16 |
| 6 | 112661896 | rs1539403 | 0.785 | -0.892 | -0.397 | -0.019 | 0.071 | 0.006 | 0.006 | 0 | 0.52 | 0.23 | 0.52 | 0.52 | 0.69 | 0.62 |
| 6 | 112668193 | rs9384825 | -0.086 | -1.141 | -0.199 | 0.11 | -0.025 | 0.008 | 0.008 | 0 | 0.19 | 0.27 | 0.14 | 0.3 | 0.08 | 0.17 |
| 6 | 112668922 | rs6931103 | 1.067 | -0.523 | -0.9 | -0.461 | 0.007 | 0.011 | 0.011 | 0.041 | 0.27 | 0.18 | 0.13 | 0.04 | 0.23 | 0.17 |
| 6 | 112670796 | rs12192606 | 1.061 | -0.519 | -0.827 | -0.46 | 0.009 | 0.019 | 0.019 | 0.041 | 0.27 | 0.18 | 0.13 | 0.04 | 0.23 | 0.18 |
| 6 | 112671611 | rs11153361 | 1.05 | -0.524 | -0.765 | -0.462 | 0.003 | -0.016 | -0.016 | 0.512 | 0.27 | 0.21 | 0.13 | 0.04 | 0.23 | 0.18 |
| 6 | 112672155 | rs11153362 | 1.087 | -0.531 | -0.869 | -0.482 | 0.008 | 0.016 | 0.016 | 0 | 0.26 | 0.18 | 0.13 | 0.04 | 0.23 | 0.18 |
| 6 | 112679127 | rs4947185 | -0.159 | -0.328 | -0.54 | -0.119 | -0.031 | -0.03 | -0.03 | 2.336 | 0.68 | 0.77 | 0.72 | 0.56 | 0.52 | 0.57 |
| 6 | 112679304 | rs9400537 | 0.447 | -0.782 | -0.066 | -0.257 | -0.023 | 0.01 | 0.01 | 0.002 | 0.2 | 0.3 | 0.25 | 0.34 | 0.08 | 0.2 |
| 6 | 112680148 | rs10872080 | -0.231 | -0.755 | 0.379 | -0.245 | 0.002 | 0.002 | 0.002 | 0.513 | 0.67 | 0.65 | 0.7 | 0.56 | 0.52 | 0.57 |
| 6 | 112687478 | rs1935679 | -0.327 | -0.747 | 0.278 | -0.244 | -0.031 | -0.031 | -0.031 | 0.054 | 0.67 | 0.76 | 0.72 | 0.56 | 0.52 | 0.56 |
| 6 | 112691284 | rs12527210 | 0.435 | -0.747 | 0.282 | -0.242 | 0.007 | 0.007 | 0.007 | 0.313 | 0.19 | 0.2 | 0.32 | 0.21 | 0.22 | 0.19 |
| 6 | 112693959 | rs12717171 | 0.205 | -0.573 | 0.205 | -0.35 | -0.125 | -0.108 | -0.108 | 0.768 | 0.32 | 0.08 | 0.26 | 0.42 | 0.48 | 0.44 |
| 6 | 112695882 | rs1935676 | -0.347 | -0.705 | 0.413 | -0.203 | 0.037 | -0.023 | -0.023 | 0 | 0.4 | 0.73 | 0.6 | 0.54 | 0.29 | 0.39 |
| 6 | 112703873 | rs1342596 | -0.318 | -0.736 | 0.424 | -0.176 | 0.033 | -0.018 | -0.018 | 0 | 0.39 | 0.73 | 0.6 | 0.55 | 0.29 | 0.39 |
| 6 | 112706267 | rs949578 | 0.214 | -0.923 | 0.502 | -0.119 | -0.006 | 0.037 | 0.037 | 0 | 0.27 | 0.38 | 0.39 | 0.31 | 0.24 | 0.22 |
| 6 | 112710524 | rs9387071 | 0.955 | -0.456 | 0.189 | 0.26 | -0.019 | -0.07 | -0.07 | 2.875 | 0.1 | 0.01 | 0.08 | 0.27 | 0.16 | 0.18 |
| 6 | 112711218 | rs9372328 | 1.382 | -0.434 | 0.186 | -0.122 | 0.008 | 0.128 | 0.128 | 3.253 | 0.3 | 0.09 | 0.14 | 0.07 | 0.25 | 0.19 |
| 6 | 112714501 | rs1935705 | 0.436 | -0.45 | 0.354 | -0.1 | -0.005 | 0.07 | 0.07 | 0 | 0.24 | 0.39 | 0.39 | 0.31 | 0.22 | 0.23 |
| 6 | 112715227 | rs9400542 | -0.206 | -0.482 | 0.374 | -0.117 | -0.095 | -0.055 | -0.055 | 0 | 0.67 | 0.85 | 0.74 | 0.62 | 0.52 | 0.58 |
| 6 | 112717989 | rs9387074 | -0.373 | -0.481 | 0.411 | -0.073 | 0.035 | -0.029 | -0.029 | 1.292 | 0.37 | 0.75 | 0.6 | 0.55 | 0.27 | 0.39 |
| 6 | 112718049 | rs9387075 | 0.645 | -0.48 | 0.377 | -0.088 | -0.041 | 0.05 | 0.05 | 0.112 | 0.13 | 0.35 | 0.21 | 0.25 | 0.05 | 0.16 |
| 6 | 112722413 | rs1108967 | 0.397 | -0.284 | 0.369 | -0.111 | -0.005 | 0.045 | 0.045 | 0 | 0.23 | 0.3 | 0.47 | 0.31 | 0.22 | 0.24 |
| 6 | 112730624 | rs2153882 | -0.246 | 0.692 | 0.22 | -0.114 | -0.105 | -0.096 | -0.096 | 0 | 0.28 | 0.1 | 0.26 | 0.43 | 0.43 | 0.35 |
| 6 | 112755156 | rs7756462 | -0.051 | 1.329 | 1.139 | -0.757 | 0.001 | -0.057 | -0.057 | 0 | 0.59 | 0.6 | 0.68 | 0.92 | 0.51 | 0.73 |
| 6 | 112757968 | rs9487914 | -0.038 | 1.304 | 0.885 | -0.77 | -0.08 | 0.193 | 0.193 | 0.351 | 0.39 | 0.13 | 0.29 | 0.08 | 0.49 | 0.27 |
| 6 | 112760200 | rs1276639 | -0.048 | 1.316 | 0.926 | -0.763 | -0.056 | 0.147 | 0.147 | 0.143 | 0.61 | 0.81 | 0.71 | 0.92 | 0.52 | 0.73 |
| 6 | 112773200 | rs1276628 | -0.044 | 1.319 | 0.979 | -0.762 | -0.039 | 0.1 | 0.1 | 0.119 | 0.61 | 0.77 | 0.71 | 0.92 | 0.53 | 0.73 |
| 6 | 112775377 | rs6934797 | -0.002 | 1.303 | 1.206 | -0.77 | 0.009 | -0.097 | -0.097 | 0 | 0.59 | 0.48 | 0.67 | 0.92 | 0.52 | 0.73 |
| 6 | 112781952 | rs4256466 | 0.08 | 1.32 | 0.879 | -0.773 | 0.005 | -0.088 | -0.088 | 0.696 | 0.58 | 0.53 | 0.61 | 0.91 | 0.51 | 0.71 |
| 6 | 112795161 | rs1150092 | -0.205 | 1.502 | 0.952 | -0.507 | -0.036 | 0.056 | 0.056 | 0.837 | 0.65 | 0.75 | 0.62 | 0.84 | 0.56 | 0.74 |
| 6 | 112799342 | rs9384832 | -0.015 | 1.542 | 0.8 | -0.487 | 0.026 | -0.078 | -0.078 | 0.269 | 0.63 | 0.5 | 0.59 | 0.84 | 0.56 | 0.72 |
| 6 | 112810278 | rs1276584 | 0.206 | 1.656 | 0.719 | -0.341 | 0.008 | -0.049 | -0.049 | 0 | 0.71 | 0.46 | 0.8 | 0.77 | 0.73 | 0.76 |
| 6 | 112818452 | rs1606213 | 0.487 | 2.085 | 1.073 | -0.531 | 0.035 | -0.041 | -0.041 | 0 | 0.69 | 0.54 | 0.54 | 0.79 | 0.58 | 0.69 |
| 6 | 112820230 | rs1276557 | 0.406 | 2.067 | 1.044 | -0.529 | -0.098 | 0.113 | 0.113 | 0.016 | 0.71 | 0.88 | 0.57 | 0.95 | 0.58 | 0.71 |
| 6 | 112847794 | rs1276536 | 0.493 | 2.451 | 0.7 | -0.226 | -0.085 | 0.02 | 0.02 | 0.02 | 0.71 | 0.87 | 0.56 | 0.75 | 0.57 | 0.71 |
| 6 | 112852469 | rs1767965 | 0.442 | 2.451 | 0.718 | -0.226 | -0.017 | 0.007 | 0.007 | 0.005 | 0.7 | 0.76 | 0.54 | 0.75 | 0.57 | 0.7 |
| 6 | 112861788 | rs6926282 | -0.273 | 2.27 | 0.312 | -0.573 | 0 | -0.026 | -0.026 | 0 | 0.1 | 0.13 | 0.05 | 0 | 0.09 | 0.05 |
| 6 | 112876745 | rs1276594 | 0.317 | 2.405 | 0.097 | -0.379 | -0.046 | 0.014 | 0.014 | 0.01 | 0.71 | 0.83 | 0.56 | 0.75 | 0.59 | 0.7 |
| 6 | 112882942 | rs1276535 | -0.217 | 2.351 | -0.376 | -0.529 | -0.102 | 0.085 | 0.085 | 0 | 0.77 | 0.97 | 0.75 | 0.93 | 0.7 | 0.75 |
| 6 | 112891101 | rs11753013 | -0.061 | 2.115 | -0.177 | -0.399 | -0.025 | 0.019 | 0.019 | 0.224 | 0.09 | 0.02 | 0.13 | 0.03 | 0.11 | 0.2 |
| 6 | 112896133 | rs1511667 | 0.071 | 1.658 | -0.085 | -0.148 | 0.001 | -0.003 | -0.003 | 0.014 | 0.68 | 0.66 | 0.7 | 0.54 | 0.72 | 0.76 |
| 6 | 112898350 | rs7738819 | 0.071 | 1.481 | -0.061 | -0.122 | 0.002 | -0.005 | -0.005 | 0.022 | 0.68 | 0.66 | 0.7 | 0.54 | 0.72 | 0.76 |
| 6 | 112909707 | rs2251690 | -0.886 | 1.366 | -0.357 | 0.094 | 0.016 | 0.015 | 0.015 | 0.527 | 0.85 | 0.89 | 0.95 | 0.91 | 0.93 | 0.9 |

| | | | | | | | | | | | | | | | | |
|---|---|---|---|---|---|---|---|---|---|---|---|---|---|---|---|---|
| 6 | 112915138 | rs1511666 | 0.7 | 1.257 | -0.012 | 0.018 | 0.007 | -0.013 | -0.013 | 0 | 0.66 | 0.3 | 0.66 | 0.54 | 0.72 | 0.76 |
| 6 | 112920882 | rs12526710 | 0.913 | 1.236 | -0.551 | -0.26 | -0.028 | 0.035 | 0.035 | 0.737 | 0.08 | 0.01 | 0.19 | 0.01 | 0.1 | 0.05 |
| 6 | 112937551 | rs9398318 | 0.689 | 0.903 | -0.584 | -0.129 | 0.014 | -0.002 | -0.002 | 0.072 | 0.14 | 0 | 0.04 | 0.12 | 0.07 | 0.1 |
| 6 | 112940651 | rs10499064 | 0.144 | 1.374 | -0.755 | -0.621 | -0.017 | 0.021 | 0.021 | 0.215 | 0.14 | 0.07 | 0.17 | 0.01 | 0.17 | 0.24 |
| 6 | 112941943 | rs9372333 | -0.174 | 1.332 | -0.328 | -0.419 | -0.005 | 0.007 | 0.007 | 0 | 0.35 | 0.38 | 0.48 | 0.52 | 0.3 | 0.26 |
| 6 | 112948108 | rs9320411 | -0.706 | 1.817 | -1.072 | -0.551 | -0.083 | -0.017 | -0.017 | 0 | 0.84 | 0.97 | 0.83 | 0.83 | 0.68 | 0.87 |
| 6 | 112955668 | rs12530109 | 0.614 | 1.639 | -0.766 | -0.386 | 0.018 | 0.191 | 0.191 | 0.113 | 0.19 | 0.5 | 0.44 | 0.41 | 0.2 | 0.12 |
| 6 | 112958214 | rs9374351 | -0.779 | 1.76 | -1.304 | -0.455 | -0.083 | -0.017 | -0.017 | 0 | 0.84 | 0.97 | 0.83 | 0.83 | 0.68 | 0.87 |
| 6 | 112961006 | rs9374352 | 0.601 | 1.565 | -1.409 | -0.608 | 0.024 | 0.125 | 0.125 | 0 | 0.54 | 0.81 | 0.61 | 0.74 | 0.45 | 0.52 |
| 6 | 112961019 | rs11759652 | 0.61 | 1.08 | -1.357 | -0.567 | -0.003 | 0.012 | 0.012 | 0.08 | 0.13 | 0.07 | 0.16 | 0.01 | 0.13 | 0.22 |
| 6 | 112961926 | rs1538729 | 1.461 | 0.768 | -1.346 | -0.495 | 0.055 | 0.001 | 0.001 | 0 | 0.33 | 0.13 | 0.15 | 0.31 | 0.25 | 0.4 |
| 6 | 112962393 | rs9372336 | 1.05 | 0.963 | -1.226 | -0.404 | -0.021 | 0.022 | 0.022 | 0.046 | 0.16 | 0.01 | 0.05 | 0.09 | 0.1 | 0.13 |
| 6 | 112963274 | rs9384838 | 1.045 | 0.722 | -1.303 | -0.434 | 0.011 | 0.054 | 0.054 | 0.062 | 0.16 | 0.02 | 0.05 | 0.08 | 0.08 | 0.14 |
| 6 | 112963741 | rs9487935 | 1.051 | 0.215 | -1.468 | -0.419 | 0.021 | -0.006 | -0.006 | 0.049 | 0.28 | 0.2 | 0.2 | 0.33 | 0.19 | 0.39 |
| 6 | 112964352 | rs9387092 | -0.206 | 0.348 | -1.467 | -0.459 | -0.006 | -0.007 | -0.007 | 0 | 0.6 | 0.36 | 0.49 | 0.5 | 0.5 | 0.72 |
| 6 | 112965827 | rs9487939 | 0.422 | 0.093 | -0.938 | 0.02 | 0.016 | -0.004 | -0.004 | 0.073 | 0.27 | 0.2 | 0.2 | 0.3 | 0.19 | 0.35 |
| 6 | 112966440 | rs4947191 | 0.036 | -0.022 | -0.985 | 0.197 | 0.004 | 0 | 0 | 0 | 0.19 | 0.15 | 0.15 | 0.21 | 0.15 | 0.31 |
| 6 | 112967841 | rs4947192 | -0.67 | 0.647 | -0.269 | 0.5 | 0.017 | 0.07 | 0.07 | 0 | 0.27 | 0.42 | 0.29 | 0.43 | 0.31 | 0.46 |
| 6 | 112968423 | rs9487940 | 0.185 | 0.794 | -0.111 | 0.52 | 0.027 | -0.012 | -0.012 | 0 | 0.09 | 0.23 | 0.1 | 0.07 | 0.12 | 0.09 |
| 6 | 112972697 | rs13200368 | 1.185 | 0.69 | -0.581 | 0.294 | 0.008 | 0.042 | 0.042 | 0.173 | 0.21 | 0.05 | 0.1 | 0.11 | 0.15 | 0.09 |
| 6 | 112973229 | rs4947193 | 1.464 | 0.631 | -0.593 | 0.142 | 0.034 | 0.056 | 0.056 | 0.056 | 0.3 | 0.11 | 0.34 | 0.18 | 0.25 | 0.17 |
| 6 | 112975224 | rs6908654 | -0.02 | 0.8 | -0.227 | 0.165 | 0.019 | 0.027 | 0.027 | 3.794 | 0.68 | 0.54 | 0.63 | 0.62 | 0.63 | 0.42 |
| 6 | 112978194 | rs6903991 | -0.006 | 0.898 | -0.165 | 0.165 | 0.007 | 0.009 | 0.009 | 0 | 0.37 | 0.44 | 0.28 | 0.44 | 0.37 | 0.25 |
| 6 | 112990902 | rs1455902 | -0.271 | 0.568 | 0.203 | 0.179 | 0.025 | -0.018 | -0.018 | 0 | 0.91 | 0.8 | 0.9 | 0.94 | 0.88 | 0.91 |
| 6 | 112999076 | rs4945909 | 0.12 | 0.451 | 0.275 | 0.075 | -0.005 | 0.01 | 0.01 | 0 | 0.56 | 0.35 | 0.6 | 0.61 | 0.56 | 0.61 |
| 6 | 113001768 | rs6909472 | 0.639 | 0.513 | 0.062 | 0.058 | -0.005 | -0.033 | -0.033 | 1.807 | 0.17 | 0.1 | 0.18 | 0.34 | 0.2 | 0.37 |
| 6 | 113013481 | rs4947199 | 0.746 | 1.25 | 0.792 | 0.425 | -0.021 | 0.015 | 0.015 | 1.339 | 0.14 | 0.08 | 0.17 | 0.02 | 0.21 | 0.25 |
| 6 | 113017260 | rs6568736 | -0.351 | 1.262 | 0.956 | 0.419 | 0.029 | -0.043 | -0.043 | 0 | 0.79 | 0.73 | 0.73 | 0.92 | 0.68 | 0.66 |
| 6 | 113023402 | rs1378699 | 0.889 | 1.221 | 0.981 | 0.424 | 0.03 | 0.028 | 0.028 | 0 | 0.48 | 0.25 | 0.51 | 0.54 | 0.44 | 0.53 |
| 6 | 113034176 | rs6915011 | 1.987 | 0.934 | 0.697 | 0.325 | -0.013 | -0.022 | -0.022 | 0 | 0.17 | 0.07 | 0.29 | 0.2 | 0.19 | 0.11 |
| 6 | 113047040 | rs11153378 | 0.392 | 0.837 | 0.8 | 0.175 | 0.004 | -0.006 | -0.006 | 0.233 | 0.29 | 0.23 | 0.18 | 0.38 | 0.24 | 0.12 |
| 6 | 113087898 | rs12204764 | 0.503 | 1.122 | 0.943 | 0.406 | 0.015 | 0.067 | 0.067 | 0 | 0.11 | 0.23 | 0.1 | 0.25 | 0.12 | 0.07 |
| 6 | 113092128 | rs4947202 | 0.861 | 1.221 | 0.931 | 0.428 | -0.053 | 0.003 | 0.003 | 0 | 0.65 | 0.21 | 0.6 | 0.62 | 0.55 | 0.59 |
| 6 | 113096276 | rs9374363 | 0.704 | 1.175 | 1.019 | 0.368 | 0.021 | -0.062 | -0.062 | 0 | 0.3 | 0.08 | 0.23 | 0.35 | 0.24 | 0.43 |
| 6 | 113111808 | rs9487982 | -0.447 | 1.687 | 1.712 | 0.242 | 0.052 | 0.008 | 0.008 | 0.009 | 0.09 | 0.34 | 0.12 | 0.09 | 0.11 | 0.09 |
| 6 | 113140667 | rs4593399 | 1.061 | 1.725 | 1.43 | 0.244 | 0.075 | -0.066 | -0.066 | 0 | 0.76 | 0.45 | 0.71 | 0.86 | 0.67 | 0.66 |
| 6 | 113140908 | rs7739145 | 0.571 | 1.258 | 1.261 | 0.666 | 0.088 | 0.031 | 0.031 | 0 | 0.91 | 0.62 | 0.88 | 0.91 | 0.89 | 0.91 |
| 6 | 113149092 | rs9320418 | 0.134 | 1.347 | 1.606 | 0.483 | 0.043 | -0.034 | -0.034 | 0.448 | 0.85 | 0.72 | 0.87 | 0.92 | 0.8 | 0.73 |
| 6 | 113151042 | rs4945917 | 1.245 | 1.705 | 1.69 | 0.346 | -0.04 | 0.016 | 0.016 | 1.568 | 0.73 | 0.19 | 0.75 | 0.77 | 0.65 | 0.62 |
| 6 | 113155367 | rs12212255 | 1.06 | 1.446 | 1.286 | 0.116 | -0.009 | 0.051 | 0.051 | 0 | 0.09 | 0 | 0.07 | 0 | 0.07 | 0.02 |
| 6 | 113162846 | rs9320419 | -0.162 | 1.652 | 1.675 | 0.374 | 0.076 | -0.022 | -0.022 | 0 | 0.08 | 0.25 | 0.1 | 0.03 | 0.13 | 0.1 |
| 6 | 113164380 | rs12524345 | 0.495 | 1.309 | 1.009 | 0.59 | 0.02 | -0.024 | -0.024 | 0 | 0.25 | 0.14 | 0.26 | 0.25 | 0.2 | 0.37 |
| 6 | 113169984 | rs2347389 | -0.115 | 1.334 | 1.157 | 0.548 | -0.003 | -0.005 | -0.005 | 1.932 | 0.32 | 0.25 | 0.34 | 0.28 | 0.33 | 0.46 |
| 6 | 113176057 | rs7766257 | 0.51 | 1.398 | 0.99 | 0.554 | 0.002 | 0.001 | 0.001 | 0 | 0.06 | 0.07 | 0.08 | 0.03 | 0.13 | 0.1 |
| 6 | 113178873 | rs9481295 | 0.313 | 0.962 | 1.276 | 0.594 | -0.011 | 0.008 | 0.008 | 0 | 0.26 | 0.29 | 0.28 | 0.26 | 0.2 | 0.35 |
| 6 | 113179034 | rs9488015 | -0.039 | 0.927 | 1.49 | 0.648 | -0.024 | 0.031 | 0.031 | 0 | 0.3 | 0.55 | 0.3 | 0.38 | 0.21 | 0.36 |
| 6 | 113184692 | rs9320420 | 0.338 | 0.792 | 1.382 | 0.589 | -0.015 | 0.018 | 0.018 | 0 | 0.25 | 0.34 | 0.28 | 0.26 | 0.2 | 0.35 |
| 6 | 113202257 | rs7744587 | -0.022 | 0.919 | 1.225 | 0.642 | -0.038 | 0.064 | 0.064 | 0 | 0.29 | 0.39 | 0.29 | 0.38 | 0.21 | 0.36 |
| 6 | 113205566 | rs9387108 | 1.004 | 0.888 | 0.328 | 0.447 | 0.011 | 0.016 | 0.016 | 0 | 0.32 | 0.14 | 0.2 | 0.31 | 0.29 | 0.14 |
| 6 | 113209728 | rs1542581 | 1.092 | 1.729 | 0.956 | 1.049 | 0.027 | 0.057 | 0.057 | 0.349 | 0.51 | 0.25 | 0.5 | 0.47 | 0.46 | 0.27 |
| 6 | 113213997 | rs17074567 | -0.147 | 1.327 | 0.924 | 1.114 | 0.008 | -0.01 | -0.01 | 3.23 | 0.25 | 0.2 | 0.26 | 0.26 | 0.2 | 0.35 |
| 6 | 113217970 | rs9400576 | 0.146 | 1.41 | 1.192 | 1.116 | -0.036 | 0.078 | 0.078 | 0 | 0.73 | 0.52 | 0.71 | 0.62 | 0.8 | 0.64 |
| 6 | 113219354 | rs9398324 | 0.863 | 2.035 | 1.185 | 1.058 | -0.002 | 0.001 | 0.001 | 0.02 | 0.58 | 0.37 | 0.49 | 0.46 | 0.49 | 0.31 |
| 6 | 113231819 | rs9384850 | 0.198 | 1.105 | 1.218 | 1.114 | -0.035 | 0.056 | 0.056 | 0 | 0.73 | 0.48 | 0.7 | 0.62 | 0.8 | 0.65 |
| 6 | 113234943 | rs11752346 | 0.099 | 1.442 | 1.13 | 1.09 | -0.024 | 0.014 | 0.014 | 0.344 | 0.08 | 0.03 | 0.12 | 0.05 | 0.16 | 0.22 |
| 6 | 113265584 | rs1351469 | -0.3 | 0.777 | 1.163 | 1.276 | -0.012 | 0.017 | 0.017 | 0 | 0.26 | 0.34 | 0.28 | 0.38 | 0.19 | 0.35 |
| 6 | 113270924 | rs1288164 | 0.143 | 0.788 | 1.136 | 1.282 | -0.024 | 0.039 | 0.039 | 0 | 0.74 | 0.62 | 0.72 | 0.62 | 0.81 | 0.65 |
| 6 | 113286947 | rs1288183 | 0.261 | 0.797 | 1.212 | 1.33 | -0.014 | 0.05 | 0.05 | 0 | 0.75 | 0.6 | 0.65 | 0.47 | 0.76 | 0.58 |
| 6 | 113304903 | rs2127965 | 0.538 | 0.223 | 0.924 | 0.555 | 0.05 | 0.111 | 0.111 | 1.372 | 0.49 | 0.23 | 0.5 | 0.37 | 0.62 | 0.47 |
| 6 | 113312142 | rs12205736 | 0.408 | 0.216 | 1.252 | 0.533 | -0.006 | -0.018 | -0.018 | 0 | 0.24 | 0.26 | 0.14 | 0.09 | 0.14 | 0.1 |
| 6 | 113317262 | rs7757130 | 0.291 | 0.702 | 1.013 | 0.78 | -0.003 | 0.001 | 0.001 | 0.021 | 0.07 | 0.04 | 0.04 | 0.06 | 0.08 | 0.15 |
| 6 | 113346306 | rs1481323 | 0.294 | 0.443 | 1.198 | 1.136 | 0.003 | -0.079 | -0.079 | 0 | 0.21 | 0.09 | 0.31 | 0.42 | 0.21 | 0.29 |
| 6 | 113366486 | rs1680101 | -0.258 | 0.424 | 1.184 | 0.951 | 0.001 | -0.004 | -0.004 | 0 | 0.77 | 0.76 | 0.67 | 0.52 | 0.79 | 0.7 |
| 6 | 113367688 | rs1680096 | -0.243 | 0.424 | 1.186 | 0.951 | -0.004 | 0.025 | 0.025 | 0 | 0.77 | 0.72 | 0.67 | 0.52 | 0.79 | 0.7 |
| 6 | 113382012 | rs9387116 | -0.313 | 0.603 | 1.056 | 1.114 | 0 | -0.011 | -0.011 | 0.221 | 0.78 | 0.8 | 0.68 | 0.54 | 0.78 | 0.69 |
| 6 | 113387556 | rs1317552 | 0.361 | 0.888 | 0.816 | 0.466 | -0.004 | 0.026 | 0.026 | 0.27 | 0.33 | 0.25 | 0.41 | 0.27 | 0.36 | 0.4 |
| 6 | 113389114 | rs1112586 | 0.52 | 0.85 | 0.977 | 0.459 | -0.008 | 0.023 | 0.023 | 0.082 | 0.31 | 0.22 | 0.4 | 0.27 | 0.36 | 0.4 |
| 6 | 113393530 | rs6923836 | 0.051 | 0.617 | 1.246 | 0.649 | -0.001 | 0.006 | 0.006 | 0.968 | 0.52 | 0.44 | 0.29 | 0.32 | 0.46 | 0.41 |
| 6 | 113395664 | rs6919954 | 0.461 | 0.726 | 1.246 | 0.457 | -0.002 | -0.003 | -0.003 | 0.702 | 0.21 | 0.18 | 0.13 | 0.25 | 0.23 | 0.3 |

| | | | | | | | | | | | | | | | | |
|---|---|---|---|---|---|---|---|---|---|---|---|---|---|---|---|---|
| 6 | 113395946 | rs6940873 | 0.45 | 0.727 | 1.246 | 0.457 | -0.002 | -0.003 | -0.003 | 0 | 0.21 | 0.18 | 0.13 | 0.25 | 0.23 | 0.3 |
| 6 | 113398162 | rs10457255 | 0.629 | 0.722 | 1.371 | 0.469 | 0 | -0.001 | -0.001 | 1.29 | 0.34 | 0.35 | 0.43 | 0.27 | 0.36 | 0.4 |
| 6 | 113411883 | rs6910441 | 0.377 | 0.738 | 1.399 | 0.5 | -0.002 | -0.005 | -0.005 | 0 | 0.22 | 0.15 | 0.13 | 0.25 | 0.23 | 0.3 |
| 6 | 113414336 | rs1539276 | -0.843 | 0.505 | 1.694 | 0.751 | 0.007 | 0.075 | 0.075 | 0 | 0.85 | 0.79 | 0.72 | 0.59 | 0.82 | 0.8 |
| 6 | 113425130 | rs7770043 | -0.508 | 0.504 | 1.659 | 0.751 | 0.023 | 0.177 | 0.177 | 0 | 0.84 | 0.64 | 0.7 | 0.59 | 0.82 | 0.8 |
| 6 | 113425559 | rs7770743 | 0.559 | 0.468 | 1.47 | 0.691 | 0.004 | 0.002 | 0.002 | 4.361 | 0.3 | 0.26 | 0.16 | 0.07 | 0.22 | 0.11 |
| 6 | 113449502 | rs2502404 | -0.312 | 0.482 | 1.718 | 0.758 | 0.003 | -0.002 | -0.002 | 0 | 0.46 | 0.56 | 0.46 | 0.48 | 0.4 | 0.31 |
| 6 | 113452857 | rs2473979 | 0.029 | 0.431 | 1.675 | 0.793 | 0.002 | 0.032 | 0.032 | 0 | 0.92 | 0.78 | 0.96 | 0.88 | 0.93 | 0.82 |
| 6 | 113463233 | rs12530265 | 0.813 | 0.438 | 1.701 | 0.801 | -0.009 | -0.013 | -0.013 | 0 | 0.14 | 0.1 | 0.27 | 0.4 | 0.18 | 0.19 |
| 6 | 113463252 | rs13218679 | 0.616 | 0.436 | 1.788 | 0.747 | 0.008 | 0.012 | 0.012 | 3.245 | 0.31 | 0.25 | 0.17 | 0.07 | 0.22 | 0.13 |
| 6 | 113464341 | rs6927432 | 0.607 | 0.439 | 1.755 | 0.749 | 0.001 | -0.011 | -0.011 | 0 | 0.31 | 0.28 | 0.17 | 0.07 | 0.22 | 0.13 |
| 6 | 113478468 | rs1321028 | -0.28 | 0.448 | 1.775 | 0.918 | -0.003 | 0.004 | 0.004 | 0.411 | 0.45 | 0.49 | 0.45 | 0.46 | 0.4 | 0.31 |
| 6 | 113479335 | rs2473967 | -0.085 | 0.412 | 1.773 | 0.927 | 0.009 | 0.059 | 0.059 | 0 | 0.92 | 0.77 | 0.96 | 0.86 | 0.93 | 0.82 |
| 6 | 113493636 | rs9400595 | 0.314 | 0.499 | 1.962 | 0.954 | 0.02 | 0.136 | 0.136 | 0 | 0.77 | 0.47 | 0.67 | 0.49 | 0.75 | 0.65 |
| 6 | 113496218 | rs7773871 | 0.861 | 0.471 | 1.588 | 0.859 | -0.066 | 0.013 | 0.013 | 0.106 | 0.45 | 0.14 | 0.49 | 0.43 | 0.52 | 0.51 |
| 6 | 113497970 | rs1321025 | 0.753 | 0.412 | 1.651 | 0.944 | -0.009 | -0.018 | -0.018 | 1.814 | 0.14 | 0.09 | 0.26 | 0.37 | 0.19 | 0.19 |
| 6 | 113502071 | rs1883973 | 0.355 | 0.417 | 1.681 | 0.928 | 0.033 | 0.097 | 0.097 | 0.083 | 0.9 | 0.5 | 0.92 | 0.86 | 0.93 | 0.83 |
| 6 | 113507826 | rs7753145 | -0.371 | 0.367 | 1.297 | 0.817 | 0 | -0.001 | -0.001 | 0.311 | 0.45 | 0.45 | 0.45 | 0.43 | 0.4 | 0.31 |
| 6 | 113509151 | rs12154111 | 0.424 | 0.367 | 1.21 | 0.813 | 0.014 | 0.037 | 0.037 | 4.353 | 0.3 | 0.22 | 0.18 | 0.07 | 0.21 | 0.12 |
| 6 | 113510245 | rs9488095 | 0.468 | 0.421 | 1.071 | 0.853 | 0.009 | 0.069 | 0.069 | 0 | 0.15 | 0.22 | 0.26 | 0.36 | 0.19 | 0.18 |
| 6 | 113512086 | rs9488098 | 0.379 | 0.424 | 0.633 | 0.855 | -0.004 | -0.005 | -0.005 | 0 | 0.15 | 0.12 | 0.26 | 0.37 | 0.19 | 0.19 |
| 6 | 113513574 | rs9488099 | 0.561 | 0.477 | 0.635 | 0.907 | 0.004 | 0.041 | 0.041 | 0 | 0.14 | 0.17 | 0.26 | 0.36 | 0.19 | 0.18 |
| 6 | 113515814 | rs774568 | 0.101 | 0.422 | 0.24 | 0.174 | 0.016 | 0.054 | 0.054 | 5.375 | 0.3 | 0.17 | 0.17 | 0.07 | 0.21 | 0.12 |
| 6 | 113526977 | rs9488105 | 0.344 | 0.276 | 0.141 | 0.805 | 0.012 | -0.006 | -0.006 | 0.211 | 0.57 | 0.39 | 0.66 | 0.54 | 0.64 | 0.58 |
| 6 | 113532598 | rs6918334 | 0.545 | 0.285 | -0.147 | 0.556 | 0.042 | -0.012 | -0.012 | 0 | 0.54 | 0.22 | 0.53 | 0.53 | 0.63 | 0.55 |
| 6 | 113533530 | rs774551 | 0.445 | 0.159 | 0.529 | 0.708 | 0.02 | 0.093 | 0.093 | 0 | 0.93 | 0.75 | 0.94 | 0.9 | 0.96 | 0.88 |
| 6 | 113537529 | rs9374377 | 0.764 | 0.133 | 0.347 | 0.692 | 0.043 | -0.021 | -0.021 | 0 | 0.53 | 0.18 | 0.53 | 0.53 | 0.63 | 0.55 |
| 6 | 113541196 | rs9400601 | 0.13 | -0.214 | 0.31 | 0.44 | 0.019 | 0.018 | 0.018 | 3.493 | 0.71 | 0.8 | 0.87 | 0.88 | 0.85 | 0.77 |
| 6 | 113542698 | rs706922 | 0.359 | -0.203 | -0.136 | 0.397 | 0.054 | 0.081 | 0.081 | 7.007 | 0.23 | 0.07 | 0.08 | 0.02 | 0.11 | 0.1 |
| 6 | 113544965 | rs796495 | 0.646 | -0.225 | 0.042 | 0.459 | 0.017 | 0.019 | 0.019 | 0.229 | 0.6 | 0.33 | 0.56 | 0.63 | 0.68 | 0.69 |
| 6 | 113547561 | rs12191383 | 1.578 | -0.142 | -0.89 | 0.361 | -0.147 | 0.1 | 0.1 | 0 | 0.37 | 0.04 | 0.35 | 0.23 | 0.44 | 0.32 |
| 6 | 113554511 | rs9481326 | -0.333 | -0.182 | 0.211 | 0.448 | 0.016 | 0.044 | 0.044 | 1.977 | 0.15 | 0.28 | 0.32 | 0.34 | 0.2 | 0.21 |
| 6 | 113555945 | rs9320434 | 0.493 | -0.014 | -0.403 | 0.636 | 0.037 | -0.018 | -0.018 | 0 | 0.54 | 0.18 | 0.52 | 0.53 | 0.63 | 0.56 |
| 6 | 113560960 | rs17074931 | -0.522 | 0.674 | -0.188 | 0.882 | 0.002 | 0.004 | 0.004 | 0 | 0.17 | 0.15 | 0.2 | 0.21 | 0.16 | 0.24 |
| 6 | 113562264 | rs774407 | 0.666 | 1.109 | -0.511 | 1.047 | -0.026 | -0.015 | -0.015 | 0.718 | 0.59 | 0.69 | 0.53 | 0.45 | 0.52 | 0.52 |
| 6 | 113562376 | rs774408 | 1.439 | 1.018 | -0.765 | 0.772 | 0.011 | 0.127 | 0.127 | 2.203 | 0.47 | 0.12 | 0.43 | 0.27 | 0.41 | 0.34 |
| 6 | 113564702 | rs13195167 | 0.298 | 1.074 | -0.527 | 0.817 | -0.028 | -0.02 | -0.02 | 0 | 0.06 | 0.01 | 0.05 | 0.09 | 0.09 | 0.18 |
| 6 | 113566803 | rs1491071 | -1.256 | 1.066 | -0.477 | 0.812 | 0 | -0.002 | -0.002 | 0 | 0.26 | 0.72 | 0.25 | 0.4 | 0.23 | 0.36 |
| 6 | 113567100 | rs774410 | 0.172 | 1.06 | 0.234 | 0.934 | 0.013 | 0.01 | 0.01 | 0.96 | 0.71 | 0.6 | 0.71 | 0.71 | 0.64 | 0.8 |
| 6 | 113567992 | rs13208050 | -0.181 | 0.74 | 0.423 | 0.685 | 0 | 0 | 0 | 0 | 0.08 | 0.06 | 0.05 | 0.12 | 0.1 | 0.2 |
| 6 | 113579358 | rs728990 | 0.903 | 1.382 | 0.014 | 0.392 | 0.035 | 0.127 | 0.127 | 3.243 | 0.49 | 0.15 | 0.5 | 0.28 | 0.42 | 0.39 |
| 6 | 113579582 | rs9488119 | -0.491 | 1.306 | 0.896 | 0.449 | 0.015 | 0.015 | 0.015 | 0.644 | 0.12 | 0.33 | 0.08 | 0.12 | 0.11 | 0.2 |
| 6 | 113582427 | rs9488123 | -0.526 | 1.308 | 0.926 | 0.45 | 0.024 | 0.023 | 0.023 | 0.559 | 0.12 | 0.36 | 0.08 | 0.12 | 0.11 | 0.2 |
| 6 | 113589691 | rs774414 | 0.313 | 1.128 | 0.662 | 0.502 | -0.004 | 0.012 | 0.012 | 0 | 0.05 | 0.09 | 0.06 | 0.07 | 0.06 | 0.07 |
| 6 | 113594950 | rs1167969 | -0.894 | 1.381 | 0.502 | 0.582 | -0.027 | -0.038 | -0.038 | 0 | 0.77 | 0.83 | 0.77 | 0.68 | 0.69 | 0.76 |
| 6 | 113595686 | rs12193394 | 1.404 | 1.445 | 0.403 | 0.503 | -0.036 | 0.034 | 0.034 | 0 | 0.06 | 0 | 0.1 | 0 | 0.1 | 0.01 |
| 6 | 113605832 | rs9488132 | -0.725 | 1.27 | 0.827 | 0.789 | 0.013 | 0.022 | 0.022 | 0.429 | 0.27 | 0.49 | 0.25 | 0.37 | 0.28 | 0.38 |
| 6 | 113620484 | rs7768849 | 0.725 | 1.305 | 0.402 | 0.808 | -0.013 | -0.016 | -0.016 | 0 | 0.16 | 0.12 | 0.17 | 0.25 | 0.17 | 0.18 |
| 6 | 113622358 | rs1352751 | 0.464 | 1.194 | 0.366 | 0.759 | -0.039 | -0.055 | -0.055 | 0 | 0.23 | 0.13 | 0.23 | 0.32 | 0.31 | 0.24 |
| 6 | 113636195 | rs11153413 | 1.447 | 1.093 | 0.182 | 0.671 | -0.026 | 0.115 | 0.115 | 3.745 | 0.41 | 0.02 | 0.44 | 0.21 | 0.35 | 0.32 |
| 6 | 113638355 | rs11753543 | 0.465 | 1.337 | 0.355 | 0.822 | -0.053 | -0.071 | -0.071 | 0 | 0.23 | 0.09 | 0.22 | 0.32 | 0.31 | 0.24 |
| 6 | 113641221 | rs9942436 | -0.202 | 1.319 | 0.733 | 0.834 | 0.008 | 0.013 | 0.013 | 0.303 | 0.27 | 0.4 | 0.24 | 0.37 | 0.27 | 0.38 |
| 6 | 113654133 | rs2637534 | -0.58 | 0.813 | 0.695 | 0.7 | 0.016 | 0.026 | 0.026 | 2.356 | 0.5 | 0.6 | 0.48 | 0.69 | 0.58 | 0.61 |
| 6 | 113665288 | rs7765153 | 0.352 | 0.773 | 0.701 | 0.787 | 0.004 | 0.05 | 0.05 | 0 | 0.95 | 0.89 | 0.93 | 0.88 | 0.94 | 0.94 |
| 6 | 113665327 | rs7765175 | 0.989 | 0.781 | 0.596 | 0.653 | 0.008 | 0.134 | 0.134 | 2.188 | 0.42 | 0.1 | 0.45 | 0.21 | 0.38 | 0.36 |
| 6 | 113666000 | rs3851197 | -0.312 | 0.694 | 1.217 | 0.563 | 0.012 | 0.042 | 0.042 | 0.292 | 0.47 | 0.38 | 0.44 | 0.67 | 0.54 | 0.58 |
| 6 | 113667707 | rs1074617 | -0.331 | 0.658 | 1.336 | 0.606 | 0.017 | 0.063 | 0.063 | 0.229 | 0.47 | 0.25 | 0.43 | 0.66 | 0.55 | 0.58 |
| 6 | 113673643 | rs2842826 | 0.217 | 0.67 | 1.332 | 0.614 | 0.004 | 0.06 | 0.06 | 0 | 0.95 | 0.89 | 0.93 | 0.88 | 0.94 | 0.94 |
| 6 | 113680017 | rs9387133 | 0.576 | 1.045 | 1.277 | 0.788 | -0.016 | -0.05 | -0.05 | 0 | 0.2 | 0.1 | 0.17 | 0.31 | 0.24 | 0.19 |
| 6 | 113680516 | rs9400614 | -0.593 | 0.972 | 1.23 | 0.729 | 0.017 | 0.042 | 0.042 | 0.505 | 0.48 | 0.37 | 0.45 | 0.64 | 0.55 | 0.58 |
| 6 | 113682732 | rs7766390 | -0.666 | 0.943 | 0.896 | 0.743 | -0.002 | -0.004 | -0.004 | 0 | 0.48 | 0.54 | 0.46 | 0.65 | 0.56 | 0.58 |
| 6 | 113683756 | rs6937211 | -0.927 | 0.951 | 0.22 | 0.716 | 0.024 | 0.036 | 0.036 | 3.198 | 0.54 | 0.72 | 0.5 | 0.66 | 0.59 | 0.6 |
| 6 | 113683910 | rs1491074 | 0.053 | 1.163 | -0.174 | 0.924 | -0.046 | -0.021 | -0.021 | 0 | 0.26 | 0.13 | 0.26 | 0.33 | 0.32 | 0.34 |
| 6 | 113686661 | rs9320442 | 0.277 | 0.042 | -0.225 | 0.839 | 0.008 | 0.005 | 0.005 | 0 | 0.21 | 0.19 | 0.29 | 0.2 | 0.15 | 0.25 |
| 6 | 113691510 | rs9320443 | 0.868 | 0.198 | -0.91 | 0.683 | -0.145 | 0.033 | 0.033 | 0.394 | 0.41 | 0.07 | 0.36 | 0.33 | 0.49 | 0.41 |
| 6 | 113691659 | rs6910133 | 0.524 | 0.481 | -0.131 | 1.005 | -0.005 | -0.04 | -0.04 | 0 | 0.62 | 0.27 | 0.69 | 0.51 | 0.64 | 0.66 |
| 6 | 113696524 | rs9488159 | 0.512 | 0.487 | -0.127 | 1.013 | -0.007 | -0.041 | -0.041 | 0 | 0.63 | 0.27 | 0.7 | 0.53 | 0.64 | 0.66 |
| 6 | 113700723 | rs509687 | -0.743 | 0.805 | 0.813 | 1.418 | 0.089 | 0.312 | 0.312 | 0.633 | 0.06 | 0.4 | 0.05 | 0.18 | 0.03 | 0.05 |
| 6 | 113703164 | rs515568 | 0.125 | 1.062 | 0.372 | 1.091 | 0.016 | 0.039 | 0.039 | 0 | 0.77 | 0.69 | 0.69 | 0.7 | 0.75 | 0.83 |

| | | | | | | | | | | | | | | | | |
|---|---|---|---|---|---|---|---|---|---|---|---|---|---|---|---|---|
| 6 | 113708819 | rs620611 | -0.584 | 1.014 | 0.907 | 0.837 | 0.037 | 0.077 | 0.077 | 0.162 | 0.15 | 0.39 | 0.09 | 0.15 | 0.12 | 0.1 |
| 6 | 113712036 | rs4945943 | 1.124 | 1.007 | 0.94 | 0.423 | 0.007 | -0.008 | -0.008 | 0 | 0.74 | 0.56 | 0.84 | 0.82 | 0.78 | 0.79 |
| 6 | 113712795 | rs7766374 | 1.305 | 1.207 | 0.331 | -0.012 | 0.029 | 0.006 | 0.006 | 0 | 0.51 | 0.25 | 0.41 | 0.51 | 0.49 | 0.46 |
| 6 | 113716570 | rs6568783 | 0.164 | 0.919 | 0.45 | -0.246 | 0 | -0.004 | -0.004 | 0 | 0.88 | 0.88 | 0.93 | 0.97 | 0.89 | 0.89 |
| 6 | 113727509 | rs9400624 | 1.026 | 0.957 | 0.978 | -0.369 | 0.001 | -0.089 | -0.089 | 0.557 | 0.71 | 0.4 | 0.84 | 0.94 | 0.76 | 0.72 |
| 6 | 113730180 | rs9918379 | 0.391 | 1.037 | 1.12 | -0.323 | 0.021 | -0.029 | -0.029 | 0 | 0.1 | 0.21 | 0.04 | 0.03 | 0.09 | 0.1 |
| 6 | 113732711 | rs9320447 | 0.337 | 0.984 | 1.014 | -0.305 | -0.008 | 0.037 | 0.037 | 0 | 0.86 | 0.92 | 0.92 | 0.96 | 0.86 | 0.83 |
| 6 | 113752301 | rs2049924 | 0.084 | 1.141 | 1.01 | -0.154 | -0.001 | 0.003 | 0.003 | 1.275 | 0.23 | 0.21 | 0.47 | 0.5 | 0.32 | 0.41 |
| 6 | 113759724 | rs2348297 | 0.176 | 1.331 | 0.859 | -0.268 | -0.031 | 0.056 | 0.056 | 0 | 0.85 | 0.95 | 0.91 | 0.96 | 0.84 | 0.81 |
| 6 | 113760063 | rs1033391 | 1.342 | 1.082 | 1.316 | -0.318 | 0.032 | 0.035 | 0.035 | 0.117 | 0.51 | 0.26 | 0.39 | 0.44 | 0.45 | 0.31 |
| 6 | 113760526 | rs1033390 | 1.372 | 1.074 | 1.339 | -0.322 | 0.033 | 0.038 | 0.038 | 0.23 | 0.51 | 0.21 | 0.39 | 0.44 | 0.45 | 0.31 |
| 6 | 113765527 | rs6928402 | -0.016 | 1.068 | 1.594 | -0.287 | 0.031 | 0.108 | 0.108 | 1.507 | 0.25 | 0.47 | 0.49 | 0.49 | 0.32 | 0.42 |
| 6 | 113774396 | rs766330 | 0.61 | 1.082 | 1.525 | -0.293 | 0.002 | 0.001 | 0.001 | 0.733 | 0.63 | 0.63 | 0.44 | 0.47 | 0.54 | 0.41 |
| 6 | 113777858 | rs2142641 | 1.33 | 0.945 | 1.122 | -0.292 | 0.04 | 0.042 | 0.042 | 0.279 | 0.5 | 0.22 | 0.38 | 0.44 | 0.44 | 0.31 |
| 6 | 113780108 | rs4945951 | 0.009 | 1.134 | 1.446 | 0.003 | -0.015 | -0.034 | -0.034 | 0 | 0.23 | 0.16 | 0.48 | 0.49 | 0.33 | 0.42 |
| 6 | 113783774 | rs2179114 | -0.501 | 0.867 | 1.609 | -0.141 | 0.006 | 0.042 | 0.042 | 0.621 | 0.3 | 0.57 | 0.53 | 0.51 | 0.34 | 0.43 |
| 6 | 113786636 | rs10484457 | 1.018 | 0.159 | 0.551 | -0.932 | 0.006 | -0.023 | -0.023 | 0.045 | 0.35 | 0.2 | 0.29 | 0.43 | 0.34 | 0.19 |
| 6 | 113788356 | rs2049923 | -0.437 | 0.37 | 0.276 | -0.831 | -0.031 | 0.155 | 0.155 | 0 | 0.66 | 0.91 | 0.83 | 0.94 | 0.68 | 0.62 |
| 6 | 113792814 | rs12526316 | -0.066 | 0.583 | 0.798 | -0.061 | -0.006 | -0.008 | -0.008 | 0 | 0.24 | 0.2 | 0.49 | 0.5 | 0.33 | 0.42 |
| 6 | 113813639 | rs7769338 | 0.616 | 0.096 | -0.069 | -0.091 | 0.008 | -0.024 | -0.024 | 0 | 0.7 | 0.89 | 0.79 | 0.64 | 0.74 | 0.68 |
| 6 | 113838129 | rs11757557 | 0.042 | 0.374 | 0.273 | -0.27 | -0.041 | 0.029 | 0.029 | 0 | 0.11 | 0.01 | 0.07 | 0.05 | 0.15 | 0.21 |
| 6 | 113869058 | rs2348305 | 0.334 | -0.183 | 0.939 | -0.366 | 0.021 | -0.008 | -0.008 | 0 | 0.13 | 0.02 | 0.05 | 0.17 | 0.08 | 0.11 |
| 6 | 113876688 | rs2127985 | 1.207 | -0.13 | 1.213 | -0.221 | -0.015 | -0.023 | -0.023 | 0 | 0.71 | 0.33 | 0.82 | 0.69 | 0.75 | 0.68 |
| 6 | 113887203 | rs9320463 | -0.178 | -0.004 | 0.929 | -0.308 | -0.046 | 0.019 | 0.019 | 0 | 0.13 | 0.01 | 0.08 | 0.07 | 0.16 | 0.22 |
| 6 | 113893183 | rs4945955 | 0.854 | 0.251 | 1.372 | 0.055 | 0.093 | 0.141 | 0.141 | 0.001 | 0.93 | 0.55 | 0.82 | 0.91 | 0.9 | 0.89 |
| 6 | 113901903 | rs9372366 | -0.992 | 0.358 | 1.215 | -0.19 | -0.001 | -0.013 | -0.013 | 0 | 0.44 | 0.46 | 0.5 | 0.74 | 0.44 | 0.57 |
| 6 | 113910504 | rs4555958 | -0.877 | 0.336 | 1.235 | -0.201 | -0.001 | 0.006 | 0.006 | 0.011 | 0.43 | 0.36 | 0.5 | 0.75 | 0.44 | 0.57 |
| 6 | 113910697 | rs4945957 | 1.244 | 0.338 | 1.058 | -0.218 | -0.007 | 0.154 | 0.154 | 0 | 0.5 | 0.27 | 0.32 | 0.16 | 0.48 | 0.32 |
| 6 | 113920304 | rs1339042 | -0.646 | 0.459 | 0.49 | 0.083 | -0.048 | -0.061 | -0.061 | 0 | 0.29 | 0.09 | 0.32 | 0.34 | 0.32 | 0.42 |
| 6 | 113923883 | rs9400643 | -0.345 | 0.446 | 0.79 | -0.073 | 0.014 | 0.036 | 0.036 | 0 | 0.2 | 0.2 | 0.46 | 0.55 | 0.28 | 0.36 |
| 6 | 113926484 | rs10872099 | -1.16 | 0.434 | 0.653 | -0.213 | -0.022 | 0.086 | 0.086 | 0.234 | 0.44 | 0.26 | 0.63 | 0.82 | 0.51 | 0.67 |
| 6 | 113936188 | rs12190861 | -0.046 | 0.421 | 0.433 | -0.165 | -0.036 | -0.004 | -0.004 | 0.019 | 0.25 | 0.03 | 0.17 | 0.24 | 0.24 | 0.32 |
| 6 | 113940473 | rs9398356 | 0.261 | 0.483 | 0.633 | -0.012 | 0.009 | 0.071 | 0.071 | 0 | 0.14 | 0.12 | 0.3 | 0.46 | 0.19 | 0.24 |
| 6 | 113944431 | rs9384876 | 0.834 | 0.335 | 0.989 | -0.267 | 0 | -0.024 | -0.024 | 0 | 0.54 | 0.48 | 0.34 | 0.17 | 0.48 | 0.33 |
| 6 | 113953169 | rs986543 | 0.662 | 0.147 | 0.532 | -0.509 | -0.001 | -0.014 | -0.014 | 1.294 | 0.42 | 0.46 | 0.31 | 0.24 | 0.44 | 0.31 |
| 6 | 113957581 | rs1416453 | -0.57 | 0.157 | 0.326 | -0.452 | 0 | 0.019 | 0.019 | 0.13 | 0.57 | 0.37 | 0.67 | 0.76 | 0.55 | 0.61 |
| 6 | 113964094 | rs7773293 | 0.513 | 0.749 | 0.642 | 0.038 | 0.01 | 0.124 | 0.124 | 2.715 | 0.21 | 0.31 | 0.34 | 0.45 | 0.19 | 0.25 |
| 6 | 113967632 | rs4945963 | 0.828 | 0.53 | 0.369 | 0.097 | -0.015 | 0.088 | 0.088 | 3.434 | 0.2 | 0.25 | 0.32 | 0.4 | 0.16 | 0.21 |
| 6 | 113971591 | rs6931833 | 0.71 | 0.6 | -0.175 | -0.59 | 0.005 | 0.021 | 0.021 | 0 | 0.29 | 0.26 | 0.15 | 0.2 | 0.24 | 0.25 |
| 6 | 113971691 | rs6932020 | -1.174 | 0.511 | -0.15 | -0.733 | -0.017 | 0.009 | 0.009 | 0.009 | 0.86 | 0.67 | 0.92 | 0.83 | 0.9 | 0.86 |
| 6 | 113972290 | rs9387161 | 0.825 | 0.741 | -0.333 | -0.889 | 0.003 | 0.016 | 0.016 | 2.521 | 0.35 | 0.31 | 0.49 | 0.4 | 0.38 | 0.33 |
| 6 | 113972650 | rs9374411 | 1.011 | 0.853 | -0.759 | -0.639 | -0.048 | -0.02 | -0.02 | 0 | 0.16 | 0.04 | 0.23 | 0.14 | 0.19 | 0.12 |
| 6 | 113977475 | rs13218163 | 0.444 | 0.776 | -0.107 | -0.716 | 0.003 | -0.02 | -0.02 | 0 | 0.65 | 0.43 | 0.72 | 0.74 | 0.61 | 0.57 |
| 6 | 113979801 | rs3862822 | -0.743 | 0.603 | -0.022 | -0.463 | -0.013 | 0.013 | 0.013 | 0 | 0.87 | 0.7 | 0.92 | 0.8 | 0.91 | 0.85 |
| 6 | 113983034 | rs11760159 | -0.316 | 0.865 | -0.246 | -0.583 | 0 | 0 | 0 | 0.477 | 0.58 | 0.49 | 0.43 | 0.49 | 0.55 | 0.63 |
| 6 | 113987376 | rs9374417 | 0.83 | 0.894 | -0.66 | -0.55 | 0 | -0.015 | -0.015 | 0 | 0.27 | 0.19 | 0.41 | 0.48 | 0.26 | 0.24 |
| 6 | 114006061 | rs12213076 | 0.815 | 0.904 | -0.928 | -0.636 | -0.009 | -0.034 | -0.034 | 0 | 0.13 | 0.07 | 0.08 | 0.16 | 0.12 | 0.15 |
| 6 | 114006611 | rs9384880 | 1.007 | 1.093 | -0.446 | -0.526 | 0.009 | -0.066 | -0.066 | 0 | 0.14 | 0.16 | 0.14 | 0 | 0.18 | 0.1 |
| 6 | 114006780 | rs4945968 | 0.874 | 1.118 | -0.481 | -0.51 | -0.015 | -0.084 | -0.084 | 0.24 | 0.26 | 0.1 | 0.39 | 0.47 | 0.25 | 0.23 |
| 6 | 114007818 | rs7761143 | 0.644 | 1.052 | 0.016 | -0.488 | 0.009 | -0.066 | -0.066 | 0.487 | 0.14 | 0.16 | 0.14 | 0 | 0.17 | 0.1 |
| 6 | 114016683 | rs11153434 | 0.019 | 1.28 | -0.29 | -0.214 | -0.005 | 0.004 | 0.004 | 0 | 0.22 | 0.16 | 0.21 | 0.08 | 0.31 | 0.21 |
| 6 | 114020783 | rs9372372 | 1.117 | 1.116 | -0.603 | 0.456 | -0.007 | -0.068 | -0.068 | 0 | 0.19 | 0.06 | 0.27 | 0.38 | 0.17 | 0.25 |
| 6 | 114020940 | rs773682 | -0.235 | 0.964 | -0.981 | -0.085 | 0.005 | 0.024 | 0.024 | 0 | 0.78 | 0.86 | 0.88 | 0.8 | 0.83 | 0.81 |
| 6 | 114021040 | rs773683 | 0.396 | 0.9 | -0.223 | -0.335 | 0.013 | -0.019 | -0.019 | 0 | 0.23 | 0.31 | 0.21 | 0.1 | 0.31 | 0.22 |
| 6 | 114022121 | rs9320467 | 0.472 | 0.809 | -0.496 | -0.245 | -0.001 | 0.001 | 0.001 | 0 | 0.1 | 0.1 | 0.19 | 0.11 | 0.08 | 0.08 |
| 6 | 114027059 | rs9488241 | 0.091 | 0.342 | 0.044 | -0.452 | -0.003 | -0.02 | -0.02 | 0 | 0.28 | 0.31 | 0.23 | 0.1 | 0.31 | 0.24 |
| 6 | 114028139 | rs7774400 | 0.725 | 0.402 | 0.144 | -0.462 | 0 | -0.001 | -0.001 | 2.038 | 0.27 | 0.27 | 0.35 | 0.28 | 0.29 | 0.32 |
| 6 | 114029121 | rs9320468 | 1.025 | 0.475 | -0.165 | -0.382 | -0.071 | -0.049 | -0.049 | 0 | 0.18 | 0.04 | 0.25 | 0.25 | 0.21 | 0.18 |
| 6 | 114030316 | rs9488242 | 1.231 | 0.503 | 0.009 | -0.518 | -0.004 | -0.108 | -0.108 | 0.109 | 0.16 | 0.33 | 0.16 | 0 | 0.19 | 0.12 |
| 6 | 114031275 | rs930563 | -1.086 | 0.441 | -0.218 | -0.518 | -0.017 | -0.092 | -0.092 | 0.262 | 0.7 | 0.9 | 0.8 | 0.53 | 0.72 | 0.64 |
| 6 | 114040930 | rs9400661 | -0.303 | 0.474 | -0.082 | -0.509 | 0.003 | 0.001 | 0.001 | 0.148 | 0.53 | 0.65 | 0.65 | 0.53 | 0.53 | 0.53 |
| 6 | 114040947 | rs9374424 | -0.04 | 0.542 | -0.223 | -0.56 | -0.005 | 0.021 | 0.021 | 0.002 | 0.49 | 0.41 | 0.61 | 0.46 | 0.46 | 0.47 |
| 6 | 114049192 | rs773676 | -0.231 | 0.424 | -0.454 | -0.652 | 0.028 | 0.113 | 0.113 | 0 | 0.47 | 0.76 | 0.51 | 0.23 | 0.44 | 0.41 |
| 6 | 114050439 | rs6936969 | 0.792 | 0.556 | -0.95 | -0.396 | -0.006 | 0.004 | 0.004 | 0 | 0.23 | 0.16 | 0.31 | 0.22 | 0.22 | 0.26 |
| 6 | 114060419 | rs2463314 | -0.206 | 0.531 | -0.81 | -0.496 | 0 | 0.185 | 0.185 | 0 | 0.49 | 0.81 | 0.55 | 0.27 | 0.52 | 0.46 |
| 6 | 114064232 | rs2463310 | 1.502 | 0.941 | -0.672 | -0.018 | 0.007 | -0.067 | -0.067 | 0.204 | 0.18 | 0.07 | 0.12 | 0.36 | 0.15 | 0.2 |
| 6 | 114067127 | rs2030926 | -0.204 | 1.066 | -0.332 | -0.258 | 0.009 | 0.013 | 0.013 | 4.428 | 0.5 | 0.5 | 0.36 | 0.49 | 0.47 | 0.47 |
| 6 | 114077091 | rs9372377 | 0.75 | 0.986 | -0.348 | -0.187 | -0.005 | -0.005 | -0.005 | 0.918 | 0.23 | 0.29 | 0.31 | 0.38 | 0.28 | 0.32 |
| 6 | 114077332 | rs9384882 | 0.742 | 0.912 | -0.185 | -0.308 | -0.011 | 0.007 | 0.007 | 0.799 | 0.33 | 0.33 | 0.26 | 0.35 | 0.3 | 0.25 |

| | | | | | | | | | | | | | | | | |
|---|---|---|---|---|---|---|---|---|---|---|---|---|---|---|---|---|
| 6 | 114080512 | rs12110829 | 1.075 | 0.884 | -0.178 | -0.408 | -0.002 | -0.104 | -0.104 | 0 | 0.18 | 0.26 | 0.15 | 0 | 0.19 | 0.12 |
| 6 | 114082470 | rs9481395 | 1.369 | 1.022 | -0.277 | -0.445 | 0.002 | 0.077 | 0.077 | 0.048 | 0.24 | 0.06 | 0.29 | 0.11 | 0.2 | 0.26 |
| 6 | 114084492 | rs6928789 | -0.003 | 1.473 | 0.095 | 0.083 | 0.018 | 0.007 | 0.007 | 0 | 0.7 | 0.53 | 0.66 | 0.62 | 0.69 | 0.64 |
| 6 | 114087320 | rs10872106 | 0.473 | 1.445 | 0.118 | 0.103 | 0.015 | 0.014 | 0.014 | 0.352 | 0.24 | 0.37 | 0.31 | 0.4 | 0.28 | 0.33 |
| 6 | 114092964 | rs6907551 | 0.113 | 1.483 | 0.105 | 0.125 | -0.003 | 0.007 | 0.007 | 0 | 0.56 | 0.41 | 0.55 | 0.21 | 0.56 | 0.5 |
| 6 | 114099625 | rs7764655 | 0.495 | 1.563 | -0.047 | 0.22 | -0.001 | -0.001 | -0.001 | 0 | 0.23 | 0.29 | 0.3 | 0.36 | 0.27 | 0.32 |
| 6 | 114105634 | rs9372379 | -0.188 | 1.517 | -0.047 | 0.223 | -0.006 | -0.002 | -0.002 | 0.764 | 0.29 | 0.33 | 0.33 | 0.4 | 0.3 | 0.36 |
| 6 | 114111659 | rs1462382 | -0.62 | 1.505 | -0.065 | 0.202 | -0.007 | -0.006 | -0.006 | 0 | 0.39 | 0.43 | 0.4 | 0.5 | 0.44 | 0.46 |
| 6 | 114114057 | rs2615519 | -0.611 | 1.505 | -0.065 | 0.202 | -0.007 | -0.006 | -0.006 | 0.127 | 0.39 | 0.42 | 0.4 | 0.5 | 0.44 | 0.46 |
| 6 | 114126613 | rs17789266 | 0.918 | 1.541 | -0.176 | 0.269 | -0.04 | -0.051 | -0.051 | 0.486 | 0.09 | 0.01 | 0.07 | 0.1 | 0.13 | 0.1 |
| 6 | 114129785 | rs2810179 | -1.09 | 1.411 | -0.132 | 0.261 | 0.024 | -0.051 | -0.051 | 0 | 0.78 | 0.88 | 0.87 | 0.62 | 0.86 | 0.83 |
| 6 | 114130248 | rs963884 | -0.965 | 1.402 | -0.026 | 0.256 | -0.011 | 0.02 | 0.02 | 3.614 | 0.78 | 0.76 | 0.86 | 0.62 | 0.86 | 0.83 |
| 6 | 114141119 | rs9387179 | -1.006 | 1.404 | -0.036 | 0.257 | -0.002 | 0.004 | 0.004 | 3.741 | 0.78 | 0.78 | 0.86 | 0.62 | 0.86 | 0.83 |
| 6 | 114145603 | rs11153450 | 0.54 | 1.544 | -0.043 | 0.224 | 0.003 | 0.005 | 0.005 | 0.168 | 0.23 | 0.3 | 0.29 | 0.36 | 0.24 | 0.29 |
| 6 | 114150024 | rs1031800 | -0.49 | 1.074 | -0.249 | 0.292 | -0.004 | 0.021 | 0.021 | 2.128 | 0.74 | 0.7 | 0.81 | 0.57 | 0.77 | 0.76 |
| 6 | 114150067 | rs2126462 | -0.991 | 0.91 | -0.529 | 0.347 | 0.022 | -0.044 | -0.044 | 0 | 0.78 | 0.87 | 0.87 | 0.62 | 0.86 | 0.83 |
| 6 | 114165133 | rs6918413 | 0.215 | 1.266 | -0.343 | 0.404 | -0.036 | -0.014 | -0.014 | 0.057 | 0.25 | 0.19 | 0.18 | 0.34 | 0.33 | 0.33 |
| 6 | 114168573 | rs7776080 | 0.907 | 1.134 | -0.281 | 0.383 | -0.002 | -0.003 | -0.003 | 0 | 0.55 | 0.36 | 0.68 | 0.48 | 0.49 | 0.48 |
| 6 | 114170427 | rs594930 | -0.353 | 0.684 | -0.308 | 0.274 | -0.017 | -0.038 | -0.038 | 1.607 | 0.09 | 0.28 | 0.04 | 0.04 | 0.03 | 0.04 |
| 6 | 114173871 | rs352081 | -0.148 | 1.151 | -0.916 | 0.542 | -0.025 | -0.047 | -0.047 | 0 | 0.9 | 0.99 | 0.93 | 0.91 | 0.89 | 0.9 |
| 6 | 114185556 | rs9400678 | 0.487 | 1.277 | -1.097 | 0.716 | -0.127 | -0.08 | -0.08 | 0.808 | 0.2 | 0.03 | 0.16 | 0.28 | 0.32 | 0.29 |
| 6 | 114192777 | rs10499076 | -0.384 | 0.934 | -1.059 | 0.479 | -0.07 | -0.001 | -0.001 | 0 | 0.36 | 0.19 | 0.28 | 0.39 | 0.48 | 0.4 |
| 6 | 114194523 | rs11153451 | -0.536 | 0.461 | -1.485 | 0.613 | 0.004 | -0.049 | -0.049 | 0 | 0.59 | 0.23 | 0.41 | 0.55 | 0.6 | 0.64 |
| 6 | 114196489 | rs693834 | 0.537 | 0.497 | -1.128 | 0.599 | -0.004 | 0.009 | 0.009 | 0 | 0.12 | 0.19 | 0.11 | 0.09 | 0.16 | 0.12 |
| 6 | 114202123 | rs515926 | -1.4 | 0.472 | -0.954 | 0.598 | 0.003 | 0.038 | 0.038 | 0 | 0.73 | 0.58 | 0.53 | 0.63 | 0.77 | 0.75 |
| 6 | 114203546 | rs12213550 | 1.016 | 0.553 | -1.228 | 0.515 | -0.108 | -0.081 | -0.081 | 0 | 0.2 | 0.03 | 0.15 | 0.28 | 0.29 | 0.3 |
| 6 | 114206311 | rs503593 | -1.276 | -0.157 | -0.541 | 0.526 | -0.001 | 0.007 | 0.007 | 0 | 0.74 | 0.7 | 0.54 | 0.71 | 0.77 | 0.82 |
| 6 | 114214825 | rs1033804 | 0.236 | -0.409 | -0.463 | 0.383 | 0.053 | 0.014 | 0.014 | 0 | 0.23 | 0.04 | 0.14 | 0.15 | 0.12 | 0.23 |
| 6 | 114215262 | rs352075 | -0.37 | -0.145 | -0.313 | 0.512 | -0.003 | 0.009 | 0.009 | 0 | 0.12 | 0.19 | 0.11 | 0.09 | 0.16 | 0.12 |
| 6 | 114216946 | rs352074 | 0.302 | -0.104 | -0.21 | 0.323 | -0.049 | -0.042 | -0.042 | 0 | 0.7 | 0.53 | 0.8 | 0.84 | 0.81 | 0.74 |
| 6 | 114221955 | rs2758455 | 0.296 | 0.674 | -0.507 | 0.681 | -0.004 | -0.01 | -0.01 | 0 | 0.41 | 0.39 | 0.36 | 0.46 | 0.44 | 0.49 |
| 6 | 114222710 | rs2810160 | -0.153 | 0.699 | -0.688 | 0.768 | -0.015 | 0.134 | 0.134 | 0 | 0.53 | 0.82 | 0.43 | 0.74 | 0.52 | 0.67 |
| 6 | 114225811 | rs352099 | -0.002 | 1.061 | -1.103 | 0.339 | 0 | 0.015 | 0.015 | 0 | 0.81 | 0.87 | 0.79 | 0.93 | 0.78 | 0.85 |
| 6 | 114226057 | rs17075678 | -0.041 | 1.018 | -0.741 | 0.457 | 0 | 0.001 | 0.001 | 0 | 0.11 | 0.09 | 0.05 | 0.28 | 0.1 | 0.16 |
| 6 | 114226540 | rs17075682 | -0.041 | 1.019 | -0.744 | 0.457 | -0.001 | -0.011 | -0.011 | 0 | 0.11 | 0.08 | 0.05 | 0.28 | 0.1 | 0.16 |
| 6 | 114228368 | rs4945507 | 0.946 | 1.111 | -0.308 | 0.395 | -0.002 | 0.002 | 0.002 | 0 | 0.4 | 0.61 | 0.34 | 0.45 | 0.36 | 0.31 |
| 6 | 114228811 | rs517399 | -0.57 | 1.481 | -0.422 | 0.354 | 0.006 | 0.027 | 0.027 | 0 | 0.16 | 0.17 | 0.13 | 0.38 | 0.19 | 0.36 |
| 6 | 114229923 | rs486167 | 0.272 | 1.31 | -0.46 | 0.427 | 0.002 | 0.079 | 0.079 | 0 | 0.55 | 0.75 | 0.46 | 0.76 | 0.54 | 0.67 |
| 6 | 114233582 | rs11758632 | 0.553 | 1.083 | -0.687 | 0.337 | 0.004 | 0.026 | 0.026 | 0 | 0.09 | 0.01 | 0.11 | 0.02 | 0.07 | 0.03 |
| 6 | 114240685 | rs352092 | 0.21 | 1.306 | -0.735 | 0.242 | -0.013 | 0.019 | 0.019 | 0 | 0.91 | 0.94 | 0.75 | 0.92 | 0.84 | 0.85 |
| 6 | 114242226 | rs352091 | 0.456 | 1.32 | -0.19 | 0.1 | 0.045 | 0.021 | 0.021 | 0 | 0.09 | 0.4 | 0.1 | 0.03 | 0.09 | 0.03 |
| 6 | 114242419 | rs17791808 | 0.464 | 1.656 | -0.457 | 0.174 | -0.039 | -0.017 | -0.017 | 0 | 0.08 | 0.01 | 0.1 | 0.08 | 0.12 | 0.11 |
| 6 | 114242899 | rs13212298 | 1.76 | 1.238 | -0.543 | 0.293 | -0.019 | -0.015 | -0.015 | 0.008 | 0.27 | 0.11 | 0.17 | 0.29 | 0.26 | 0.28 |
| 6 | 114255647 | rs17075722 | 0.825 | 1.149 | -0.534 | 0.375 | 0 | 0.023 | 0.023 | 0 | 0.09 | 0.08 | 0.11 | 0.3 | 0.08 | 0.1 |
| 6 | 114260720 | rs3757016 | 0.027 | 1.108 | -0.422 | 0.322 | 0.003 | 0.005 | 0.005 | 0 | 0.55 | 0.4 | 0.52 | 0.7 | 0.57 | 0.63 |
| 6 | 114261852 | rs352068 | 0.522 | 1.08 | -0.797 | 0.419 | -0.036 | 0.017 | 0.017 | 0 | 0.09 | 0.01 | 0.25 | 0.08 | 0.16 | 0.14 |
| 6 | 114265287 | rs9481408 | -0.74 | 1.196 | -0.612 | 0.524 | -0.001 | 0 | 0 | 0.356 | 0.27 | 0.24 | 0.34 | 0.4 | 0.31 | 0.35 |
| 6 | 114273646 | rs10499080 | 0.506 | 0.993 | -0.768 | 0.724 | 0 | 0.028 | 0.028 | 0 | 0.09 | 0.08 | 0.11 | 0.3 | 0.08 | 0.11 |
| 6 | 114285253 | rs2499618 | 0.033 | 1.068 | -0.822 | 0.511 | 0.024 | 0.052 | 0.052 | 0 | 0.36 | 0.19 | 0.38 | 0.28 | 0.34 | 0.35 |
| 6 | 114308159 | rs6920181 | -0.052 | 1.441 | -0.342 | 1 | 0.002 | 0.042 | 0.042 | 0 | 0.1 | 0.14 | 0.12 | 0.28 | 0.09 | 0.11 |
| 6 | 114309018 | rs2281675 | -0.342 | 1.596 | -0.567 | 0.833 | -0.028 | -0.013 | -0.013 | 0 | 0.12 | 0.06 | 0.28 | 0.19 | 0.18 | 0.17 |
| 6 | 114310813 | rs2475628 | -0.452 | 1.273 | -0.374 | 0.579 | 0 | 0.001 | 0.001 | 0 | 0.89 | 0.95 | 0.87 | 0.98 | 0.87 | 0.9 |
| 6 | 114316305 | rs9372385 | 1.022 | 1.446 | -0.417 | 0.535 | 0.014 | -0.013 | -0.013 | 1.384 | 0.43 | 0.25 | 0.29 | 0.44 | 0.36 | 0.53 |
| 6 | 114316404 | rs9374432 | -0.814 | 1.394 | -0.307 | 0.543 | -0.002 | -0.011 | -0.011 | 0.652 | 0.88 | 0.92 | 0.86 | 0.97 | 0.87 | 0.9 |
| 6 | 114319622 | rs2499615 | -0.044 | 1.543 | 0.344 | 0.56 | 0.001 | 0 | 0 | 0 | 0.28 | 0.32 | 0.25 | 0.3 | 0.31 | 0.22 |
| 6 | 114331570 | rs4320400 | 0.4 | 1.332 | 0.341 | 0.511 | -0.005 | 0.013 | 0.013 | 0 | 0.57 | 0.63 | 0.59 | 0.67 | 0.52 | 0.67 |
| 6 | 114346852 | rs9400690 | 1.904 | 1.359 | -0.403 | 0.364 | -0.057 | -0.047 | -0.047 | 0.114 | 0.27 | 0.03 | 0.15 | 0.24 | 0.24 | 0.31 |
| 6 | 114347634 | rs13201675 | -0.11 | 1.39 | -0.044 | 0.582 | 0.001 | -0.023 | -0.023 | 0 | 0.3 | 0.37 | 0.26 | 0.18 | 0.31 | 0.3 |
| 6 | 114348520 | rs11752866 | 0.145 | 1.388 | -0.04 | 0.581 | 0.003 | -0.016 | -0.016 | 0 | 0.28 | 0.32 | 0.25 | 0.18 | 0.31 | 0.21 |
| 6 | 114351627 | rs915203 | -0.468 | 1.422 | 0.227 | 0.743 | 0.003 | -0.01 | -0.01 | 1.927 | 0.4 | 0.52 | 0.52 | 0.43 | 0.44 | 0.45 |
| 6 | 114352617 | rs952111 | 0.591 | 1.439 | 0.033 | 0.762 | -0.009 | 0.006 | 0.006 | 0 | 0.57 | 0.22 | 0.44 | 0.55 | 0.54 | 0.54 |
| 6 | 114352723 | rs12526401 | 0.546 | 1.318 | -0.01 | 0.655 | -0.005 | -0.015 | -0.015 | 0 | 0.09 | 0.06 | 0.26 | 0.25 | 0.13 | 0.15 |
| 6 | 114359294 | rs12198688 | 0.54 | 1.644 | 0.076 | 0.366 | 0.002 | -0.073 | -0.073 | 0.102 | 0.2 | 0.3 | 0.17 | 0.01 | 0.23 | 0.21 |
| 6 | 114361180 | rs11961953 | 0.53 | 1.644 | 0.086 | 0.366 | 0.001 | -0.063 | -0.063 | 0 | 0.2 | 0.28 | 0.17 | 0.01 | 0.23 | 0.2 |
| 6 | 114361189 | rs11961922 | 0.734 | 1.634 | 0.088 | 0.361 | 0.002 | -0.051 | -0.051 | 0.014 | 0.18 | 0.24 | 0.16 | 0 | 0.23 | 0.11 |
| 6 | 114364702 | rs2050389 | -0.41 | 1.607 | 0.278 | 0.439 | 0.007 | -0.032 | -0.032 | 0.21 | 0.31 | 0.49 | 0.44 | 0.26 | 0.36 | 0.37 |
| 6 | 114378146 | rs17075821 | 0.279 | 1.703 | 0.318 | 0.399 | 0.036 | 0.032 | 0.032 | 0 | 0.13 | 0.31 | 0.15 | 0.17 | 0.14 | 0.09 |
| 6 | 114378970 | rs2282237 | -0.17 | 1.78 | -0.172 | 0.388 | -0.006 | -0.019 | -0.019 | 0 | 0.27 | 0.16 | 0.43 | 0.35 | 0.3 | 0.35 |
| 6 | 114383606 | rs875776 | -0.786 | 1.759 | -0.075 | 0.082 | 0.009 | 0.126 | 0.126 | 0 | 0.5 | 0.78 | 0.49 | 0.28 | 0.54 | 0.41 |

| | | | | | | | | | | | | | | | | |
|---|---|---|---|---|---|---|---|---|---|---|---|---|---|---|---|---|
| 6 | 114384171 | rs17189832 | -0.247 | 1.765 | -0.052 | 0.086 | 0.005 | -0.024 | -0.024 | 0 | 0.32 | 0.63 | 0.32 | 0.17 | 0.37 | 0.21 |
| 6 | 114388111 | rs7742134 | -0.276 | 1.685 | -0.575 | -0.066 | -0.002 | 0.01 | 0.01 | 0 | 0.37 | 0.4 | 0.33 | 0.11 | 0.41 | 0.32 |
| 6 | 114388426 | rs1157989 | 0.405 | 1.296 | -1.046 | 0.379 | 0.004 | 0.02 | 0.02 | 0 | 0.18 | 0.12 | 0.17 | 0.11 | 0.18 | 0.21 |
| 6 | 114390653 | rs9488318 | 1.207 | 1.123 | -1.007 | 0.607 | -0.029 | -0.037 | -0.037 | 0 | 0.27 | 0.04 | 0.12 | 0.24 | 0.23 | 0.31 |
| 6 | 114398669 | rs7772117 | -0.118 | 1.301 | -0.098 | -0.057 | -0.007 | 0.011 | 0.011 | 0 | 0.31 | 0.29 | 0.3 | 0.18 | 0.36 | 0.23 |
| 6 | 114407866 | rs9320478 | -0.405 | 1.388 | 0.463 | 0.556 | -0.009 | 0.066 | 0.066 | 0 | 0.86 | 0.78 | 0.87 | 0.77 | 0.9 | 0.87 |
| 6 | 114408925 | rs9488322 | 0.274 | 1.683 | 0.404 | 0.513 | 0.017 | 0.043 | 0.043 | 0 | 0.74 | 0.64 | 0.75 | 0.65 | 0.72 | 0.73 |
| 6 | 114409122 | rs6924016 | 0.408 | 2.002 | 0.215 | 0.268 | 0.038 | -0.008 | -0.008 | 0.025 | 0.16 | 0.3 | 0.2 | 0.18 | 0.22 | 0.17 |
| 6 | 114412037 | rs12211928 | 1.026 | 1.839 | -0.163 | 0.229 | -0.005 | 0.023 | 0.023 | 0 | 0.17 | 0.08 | 0.16 | 0.11 | 0.17 | 0.12 |
| 6 | 114417219 | rs1011910 | 0.846 | 1.901 | -0.191 | 0.276 | -0.015 | -0.028 | -0.028 | 0 | 0.57 | 0.25 | 0.55 | 0.47 | 0.5 | 0.56 |
| 6 | 114422641 | rs12661354 | 0.497 | 1.892 | -0.186 | 0.282 | -0.04 | 0.001 | 0.001 | 0 | 0.11 | 0.03 | 0.11 | 0.13 | 0.18 | 0.14 |
| 6 | 114427189 | rs299416 | -0.823 | 1.649 | -0.009 | 0.402 | -0.006 | 0.034 | 0.034 | 0 | 0.86 | 0.82 | 0.88 | 0.78 | 0.9 | 0.89 |
| 6 | 114435904 | rs1401773 | -0.223 | 1.709 | -0.152 | 0.367 | 0.008 | 0.072 | 0.072 | 0 | 0.8 | 0.63 | 0.8 | 0.66 | 0.79 | 0.78 |
| 6 | 114436933 | rs17794523 | 1.423 | 1.631 | -0.83 | 0.365 | -0.025 | -0.021 | -0.021 | 1.495 | 0.05 | 0.01 | 0.06 | 0.12 | 0.11 | 0.11 |
| 6 | 114437276 | rs2271895 | 1.279 | 1.215 | -0.781 | 0.254 | 0.035 | -0.017 | -0.017 | 0 | 0.38 | 0.11 | 0.38 | 0.36 | 0.33 | 0.36 |
| 6 | 114447603 | rs9488329 | 0.266 | 1.144 | -0.453 | 0.259 | 0.005 | 0.02 | 0.02 | 0 | 0.19 | 0.14 | 0.16 | 0.11 | 0.17 | 0.2 |
| 6 | 114456335 | rs299412 | -0.169 | 1.338 | -0.25 | 0.643 | 0.002 | 0.031 | 0.031 | 0 | 0.86 | 0.82 | 0.87 | 0.78 | 0.88 | 0.88 |
| 6 | 114462618 | rs1022387 | 0.868 | 1.203 | -0.833 | 0.5 | 0.019 | -0.036 | -0.036 | 0 | 0.4 | 0.16 | 0.47 | 0.61 | 0.36 | 0.38 |
| 6 | 114465074 | rs9488335 | 0.615 | 1.6 | -0.415 | 0.329 | 0.013 | 0.012 | 0.012 | 0 | 0.2 | 0.3 | 0.17 | 0.19 | 0.23 | 0.07 |
| 6 | 114467289 | rs872441 | 0.004 | 1.652 | -0.326 | 0.128 | -0.008 | 0.035 | 0.035 | 0.279 | 0.3 | 0.18 | 0.29 | 0.18 | 0.34 | 0.49 |
| 6 | 114475167 | rs884818 | -0.158 | 1.702 | -0.335 | 0.13 | -0.017 | 0.036 | 0.036 | 0.403 | 0.3 | 0.14 | 0.29 | 0.18 | 0.34 | 0.49 |
| 6 | 114484757 | rs7761798 | 0.623 | 1.588 | -0.021 | 0.076 | 0.012 | -0.022 | -0.022 | 0 | 0.07 | 0.12 | 0.05 | 0.01 | 0.07 | 0.03 |
| 6 | 114489321 | rs6908717 | -0.273 | 1.585 | 0.013 | 0.114 | 0.03 | -0.011 | -0.011 | 0.125 | 0.28 | 0.48 | 0.23 | 0.21 | 0.31 | 0.12 |
| 6 | 114492156 | rs4945513 | 0.667 | 1.696 | -0.31 | -0.02 | -0.017 | 0.035 | 0.035 | 0.206 | 0.14 | 0.01 | 0.14 | 0.06 | 0.14 | 0.24 |
| 6 | 114492599 | rs9320482 | -0.566 | 1.214 | -0.417 | 0.02 | 0.002 | 0.009 | 0.009 | 2.755 | 0.55 | 0.63 | 0.43 | 0.45 | 0.52 | 0.29 |
| 6 | 114501492 | rs908936 | 0.439 | 1.244 | -0.768 | 0.071 | -0.014 | 0.097 | 0.097 | 2.902 | 0.44 | 0.22 | 0.56 | 0.67 | 0.48 | 0.7 |
| 6 | 114535977 | rs9320485 | 0.752 | 1.28 | -1.013 | 0.295 | 0.011 | 0.024 | 0.024 | 0 | 0.27 | 0.19 | 0.2 | 0.24 | 0.22 | 0.17 |
| 6 | 114538983 | rs9320486 | 0.064 | 0.986 | -0.999 | 0.331 | -0.001 | 0 | 0 | 0 | 0.45 | 0.46 | 0.36 | 0.45 | 0.44 | 0.23 |
| 6 | 114549540 | rs17193496 | 1.072 | 0.731 | -1.113 | 0.487 | -0.03 | -0.027 | -0.027 | 1.468 | 0.11 | 0 | 0.12 | 0.09 | 0.12 | 0.12 |
| 6 | 114553296 | rs1474171 | 0.81 | 0.438 | -0.819 | 0.293 | 0.023 | 0.013 | 0.013 | 2.215 | 0.23 | 0.15 | 0.2 | 0.25 | 0.13 | 0.14 |
| 6 | 114553493 | rs2492891 | -0.126 | 1.34 | -0.577 | 0.405 | 0.016 | 0.003 | 0.003 | 0 | 0.77 | 0.72 | 0.81 | 0.79 | 0.7 | 0.91 |
| 6 | 114560194 | rs9320488 | 1.165 | 0.629 | -0.662 | 0.123 | 0.002 | 0.001 | 0.001 | 0.106 | 0.06 | 0.03 | 0.05 | 0.01 | 0.05 | 0.01 |
| 6 | 114561617 | rs9320489 | -0.621 | 0.305 | -1.037 | 0.005 | -0.011 | -0.025 | -0.025 | 0.014 | 0.57 | 0.77 | 0.45 | 0.51 | 0.47 | 0.48 |
| 6 | 114563699 | rs6568833 | -0.847 | 0.289 | -1.21 | -0.082 | 0.001 | 0.005 | 0.005 | 0.009 | 0.89 | 0.94 | 0.91 | 0.94 | 0.92 | 0.74 |
| 6 | 114568687 | rs1570437 | -0.11 | -0.418 | -0.582 | 0.104 | -0.014 | 0.016 | 0.016 | 0 | 0.35 | 0.49 | 0.33 | 0.42 | 0.31 | 0.27 |
| 6 | 114572279 | rs1040258 | 0.359 | -0.506 | -0.8 | 0.183 | 0 | 0.028 | 0.028 | 0 | 0.16 | 0.19 | 0.18 | 0.31 | 0.16 | 0.11 |
| 6 | 114579080 | rs1010578 | 1.441 | -0.43 | -1.051 | 0.296 | -0.035 | -0.025 | -0.025 | 0 | 0.08 | 0 | 0.05 | 0.09 | 0.1 | 0.04 |
| 6 | 114579296 | rs9488355 | 0.294 | -0.5 | -0.918 | -0.022 | -0.006 | -0.032 | -0.032 | 0.31 | 0.12 | 0.16 | 0.08 | 0.01 | 0.08 | 0.07 |
| 6 | 114588463 | rs13204144 | -0.291 | -0.532 | -0.837 | 0.03 | 0.007 | -0.016 | -0.016 | 0 | 0.34 | 0.28 | 0.32 | 0.44 | 0.32 | 0.27 |
| 6 | 114588786 | rs9374450 | 1.788 | -0.226 | -1.169 | 0.122 | -0.039 | -0.047 | -0.047 | 0 | 0.09 | 0.01 | 0.27 | 0.32 | 0.13 | 0.11 |
| 6 | 114589360 | rs1535569 | 0.6 | -0.765 | -0.617 | 0.027 | 0.006 | 0.021 | 0.021 | 0 | 0.18 | 0.08 | 0.14 | 0.12 | 0.16 | 0.15 |
| 6 | 114597419 | rs2050613 | -0.337 | -0.265 | -0.128 | 0.029 | 0.009 | 0.003 | 0.003 | 0 | 0.46 | 0.63 | 0.41 | 0.44 | 0.39 | 0.38 |
| 6 | 114598114 | rs17076012 | 1.16 | 0.023 | -0.29 | 0 | -0.004 | -0.035 | -0.035 | 0 | 0.06 | 0.14 | 0.06 | 0.01 | 0.06 | 0.05 |
| 6 | 114600726 | rs2349214 | 0.285 | 0.056 | -0.529 | -0.013 | -0.019 | -0.01 | -0.01 | 0 | 0.1 | 0.02 | 0.04 | 0.08 | 0.09 | 0.1 |
| 6 | 114600745 | rs9400709 | 0.637 | 0.303 | -0.089 | 0.016 | 0.011 | -0.03 | -0.03 | 0 | 0.07 | 0.23 | 0.07 | 0.01 | 0.06 | 0.05 |
| 6 | 114602302 | rs7757785 | -0.179 | 0.551 | -0.715 | 0.275 | 0.003 | 0.001 | 0.001 | 0 | 0.54 | 0.6 | 0.4 | 0.54 | 0.44 | 0.46 |
| 6 | 114612895 | rs785129 | -0.071 | 0.674 | -0.902 | 0.037 | -0.018 | -0.02 | -0.02 | 0 | 0.7 | 0.76 | 0.57 | 0.4 | 0.66 | 0.61 |
| 6 | 114619795 | rs1745580 | -0.218 | 0.645 | -0.172 | 0.022 | 0.003 | 0.003 | 0.003 | 0.188 | 0.7 | 0.68 | 0.57 | 0.4 | 0.66 | 0.61 |
| 6 | 114622214 | rs7770227 | 0.305 | 0.265 | -0.723 | -0.879 | -0.047 | 0.149 | 0.149 | 0 | 0.39 | 0.16 | 0.25 | 0.19 | 0.43 | 0.33 |
| 6 | 114628139 | rs785144 | -0.808 | 0.247 | -0.614 | -0.868 | -0.007 | -0.008 | -0.008 | 0 | 0.7 | 0.71 | 0.57 | 0.4 | 0.66 | 0.61 |
| 6 | 114631425 | rs6927808 | 0.272 | 0.256 | -0.667 | -0.856 | -0.054 | 0.154 | 0.154 | 0 | 0.39 | 0.15 | 0.26 | 0.19 | 0.43 | 0.33 |
| 6 | 114634471 | rs785132 | 0.318 | 0.157 | 0.195 | -0.907 | -0.002 | 0.003 | 0.003 | 0 | 0.4 | 0.39 | 0.27 | 0.19 | 0.43 | 0.42 |
| 6 | 114645237 | rs785143 | -0.977 | 0.074 | 0.09 | -0.944 | -0.024 | -0.019 | -0.019 | 0 | 0.7 | 0.78 | 0.57 | 0.4 | 0.64 | 0.6 |
| 6 | 114653522 | rs1323056 | 0.308 | 0.048 | -0.028 | -1.157 | -0.05 | 0.155 | 0.155 | 0.04 | 0.39 | 0.14 | 0.26 | 0.19 | 0.43 | 0.31 |
| 6 | 114677688 | rs1323046 | -0.274 | 0.249 | 0.252 | -1.331 | -0.004 | -0.049 | -0.049 | 0.231 | 0.4 | 0.5 | 0.39 | 0.17 | 0.37 | 0.33 |
| 6 | 114691511 | rs1415428 | -0.306 | 0.246 | 0.188 | -1.333 | -0.001 | -0.011 | -0.011 | 0 | 0.4 | 0.53 | 0.39 | 0.17 | 0.37 | 0.35 |
| 6 | 114711584 | rs10457277 | 0.834 | 0.054 | -0.062 | -1.486 | 0.009 | -0.089 | -0.089 | 0.056 | 0.17 | 0.27 | 0.13 | 0.01 | 0.18 | 0.04 |
| 6 | 114713323 | rs1033394 | -0.73 | -0.063 | -0.564 | -1.463 | 0 | 0.072 | 0.072 | 0 | 0.71 | 0.86 | 0.82 | 0.81 | 0.73 | 0.74 |
| 6 | 114713988 | rs1033393 | -1.196 | -0.21 | -0.836 | -1.747 | -0.034 | 0.054 | 0.054 | 0.005 | 0.84 | 0.99 | 0.89 | 0.95 | 0.84 | 0.86 |
| 6 | 114714143 | rs1033392 | 0.187 | -0.09 | -0.682 | -1.54 | -0.014 | 0.118 | 0.118 | 0.09 | 0.49 | 0.84 | 0.57 | 0.65 | 0.55 | 0.53 |
| 6 | 114723724 | rs9488390 | 0.191 | -0.105 | 0.004 | -1.481 | -0.02 | -0.046 | -0.046 | 0 | 0.24 | 0.44 | 0.27 | 0.16 | 0.18 | 0.31 |
| 6 | 114733016 | rs10155761 | 0.253 | -0.136 | 0.156 | -1.503 | -0.017 | 0 | 0 | 0.063 | 0.16 | 0.27 | 0.24 | 0.16 | 0.12 | 0.3 |
| 6 | 114740063 | rs9488397 | -0.734 | -0.115 | 0.074 | -1.735 | 0.004 | 0.008 | 0.008 | 0 | 0.33 | 0.62 | 0.37 | 0.17 | 0.31 | 0.34 |
| 6 | 114742697 | rs9320493 | -0.4 | 0.032 | 0.113 | -1.717 | -0.007 | -0.003 | -0.003 | 0 | 0.84 | 0.78 | 0.83 | 0.85 | 0.86 | 0.69 |
| 6 | 114751611 | rs9374461 | -0.331 | 0.074 | 0.123 | -1.66 | -0.006 | -0.002 | -0.002 | 0 | 0.84 | 0.79 | 0.83 | 0.84 | 0.85 | 0.64 |
| 6 | 114762507 | rs12199067 | 1.149 | 0.076 | 0.139 | -1.701 | -0.002 | 0.032 | 0.032 | 0 | 0.15 | 0.06 | 0.1 | 0.01 | 0.18 | 0.04 |
| 6 | 114773600 | rs2142642 | 1.202 | 0.111 | -0.049 | -1.445 | -0.02 | 0 | 0 | 0 | 0.36 | 0.25 | 0.49 | 0.63 | 0.38 | 0.35 |
| 6 | 114791568 | rs4945516 | 0.533 | 0.168 | 0.116 | -1.533 | -0.002 | -0.012 | -0.012 | 0.281 | 0.66 | 0.62 | 0.73 | 0.81 | 0.63 | 0.52 |
| 6 | 114800340 | rs6568841 | 0.62 | 0.151 | 0.274 | -1.541 | -0.003 | -0.014 | -0.014 | 0 | 0.65 | 0.46 | 0.71 | 0.81 | 0.63 | 0.52 |

| | | | | | | | | | | | | | | | | |
|---|---|---|---|---|---|---|---|---|---|---|---|---|---|---|---|---|
| 6 | 114800377 | rs6568842 | -0.295 | 0.147 | 0.048 | -1.558 | -0.006 | 0.002 | 0.002 | 0 | 0.86 | 0.8 | 0.87 | 0.84 | 0.85 | 0.65 |
| 6 | 114811361 | rs12207886 | 1.064 | 0.056 | -0.018 | -1.25 | -0.002 | -0.002 | -0.002 | 0 | 0.15 | 0.1 | 0.1 | 0.01 | 0.17 | 0.07 |
| 6 | 114837030 | rs4946007 | 0.199 | 0.217 | -0.17 | -0.702 | 0.006 | 0.088 | 0.088 | 0 | 0.15 | 0.23 | 0.41 | 0.56 | 0.14 | 0.18 |
| 6 | 114846123 | rs706924 | -0.619 | 0.001 | -0.417 | -0.958 | -0.013 | 0.044 | 0.044 | 0 | 0.62 | 0.77 | 0.74 | 0.71 | 0.57 | 0.48 |
| 6 | 114860722 | rs750377 | -0.439 | -0.438 | 0.227 | -0.655 | -0.043 | -0.069 | -0.069 | 0 | 0.25 | 0.45 | 0.19 | 0.14 | 0.2 | 0.16 |
| 6 | 114865780 | rs17076203 | 0.968 | -0.379 | 0.266 | -0.647 | -0.002 | -0.008 | -0.008 | 0 | 0.1 | 0.11 | 0.1 | 0.1 | 0.12 | 0.1 |
| 6 | 114886995 | rs1041212 | -0.439 | -0.598 | 0.408 | -0.406 | -0.024 | -0.041 | -0.041 | 1.044 | 0.24 | 0.34 | 0.19 | 0.14 | 0.2 | 0.16 |
| 6 | 114889753 | rs1748163 | 0.146 | -0.814 | 0.272 | -0.064 | 0.003 | -0.005 | -0.005 | 0 | 0.36 | 0.31 | 0.33 | 0.4 | 0.3 | 0.42 |
| 6 | 114890253 | rs11968524 | -0.124 | -0.823 | 0.307 | -0.082 | -0.019 | -0.033 | -0.033 | 0.015 | 0.24 | 0.3 | 0.18 | 0.14 | 0.2 | 0.16 |
| 6 | 114904936 | rs7748616 | -0.172 | -0.829 | 0.254 | -0.085 | -0.01 | -0.019 | -0.019 | 0 | 0.24 | 0.28 | 0.18 | 0.14 | 0.2 | 0.16 |
| 6 | 114937265 | rs2222391 | 0.983 | -0.762 | -0.278 | -0.04 | 0.002 | 0.001 | 0.001 | 0.512 | 0.13 | 0.14 | 0.08 | 0.04 | 0.07 | 0.06 |
| 6 | 114943072 | rs9374471 | 0.4 | -0.661 | -0.29 | 0.028 | 0.018 | 0.012 | 0.012 | 0 | 0.5 | 0.37 | 0.58 | 0.56 | 0.62 | 0.52 |
| 6 | 114969280 | rs12190018 | -0.182 | -0.664 | -0.497 | -0.043 | 0.043 | 0.047 | 0.047 | 0 | 0.29 | 0.19 | 0.15 | 0.14 | 0.16 | 0.24 |
| 6 | 114970753 | rs4945521 | 1.076 | -0.656 | -0.51 | -0.034 | -0.049 | -0.057 | -0.057 | 0 | 0.39 | 0.19 | 0.46 | 0.46 | 0.47 | 0.41 |
| 6 | 114973810 | rs3003879 | 0.864 | -0.543 | -0.546 | 0.075 | 0.032 | 0.04 | 0.04 | 0 | 0.2 | 0.29 | 0.26 | 0.3 | 0.23 | 0.25 |
| 6 | 114973962 | rs2947948 | -0.761 | -0.524 | -0.716 | 0.088 | 0.004 | 0.004 | 0.004 | 0 | 0.49 | 0.54 | 0.42 | 0.45 | 0.4 | 0.49 |
| 6 | 114982656 | rs4463291 | -0.688 | -1.23 | -0.922 | -0.263 | 0.071 | 0.103 | 0.103 | 0.241 | 0.3 | 0.12 | 0.15 | 0.12 | 0.17 | 0.21 |
| 6 | 114986804 | rs9372400 | 0.947 | -0.213 | -0.307 | 0.723 | -0.003 | 0.001 | 0.001 | 0 | 0.15 | 0.1 | 0.23 | 0.27 | 0.23 | 0.12 |
| 6 | 114996281 | rs939746 | 0.444 | -0.585 | -0.259 | -0.253 | -0.014 | -0.026 | -0.026 | 0 | 0.16 | 0.22 | 0.13 | 0.1 | 0.15 | 0.1 |
| 6 | 114998125 | rs1521890 | -0.387 | -0.632 | -0.269 | -0.277 | 0.001 | 0.001 | 0.001 | 0 | 0.1 | 0.08 | 0.05 | 0.11 | 0.05 | 0.14 |
| 6 | 114998346 | rs17076291 | 0.566 | -0.629 | -0.307 | -0.276 | 0.006 | 0.006 | 0.006 | 0.067 | 0.16 | 0.13 | 0.13 | 0.1 | 0.15 | 0.1 |
| 6 | 114998487 | rs12201578 | -0.705 | -0.493 | -0.393 | -0.205 | -0.018 | -0.017 | -0.017 | 0 | 0.27 | 0.31 | 0.18 | 0.2 | 0.2 | 0.23 |
| 6 | 115004784 | rs2947945 | 0.267 | -0.146 | -0.746 | -0.447 | 0.012 | 0.029 | 0.029 | 0 | 0.72 | 0.79 | 0.83 | 0.94 | 0.75 | 0.77 |
| 6 | 115005346 | rs11153496 | -0.03 | -0.225 | -0.771 | -0.525 | 0.002 | 0 | 0 | 0 | 0.11 | 0.1 | 0.06 | 0.02 | 0.1 | 0.13 |
| 6 | 115006113 | rs4945522 | 0.267 | -0.171 | -0.863 | -0.569 | 0.004 | -0.002 | -0.002 | 0.133 | 0.29 | 0.54 | 0.32 | 0.28 | 0.26 | 0.35 |
| 6 | 115023523 | rs9400741 | -0.053 | -0.304 | -0.945 | -0.66 | -0.007 | -0.01 | -0.01 | 0 | 0.31 | 0.53 | 0.32 | 0.28 | 0.26 | 0.36 |
| 6 | 115026310 | rs7745314 | 1.488 | -0.289 | -1.286 | -0.482 | -0.065 | -0.022 | -0.022 | 0.017 | 0.37 | 0.16 | 0.49 | 0.59 | 0.47 | 0.41 |
| 6 | 115032573 | rs12204665 | 0.245 | -0.292 | -1.069 | -0.501 | 0.004 | 0.014 | 0.014 | 0 | 0.11 | 0.05 | 0.06 | 0.09 | 0.1 | 0.13 |
| 6 | 115035324 | rs9320509 | -0.63 | -0.312 | -1.07 | -0.517 | 0.012 | 0.032 | 0.032 | 0 | 0.82 | 0.91 | 0.89 | 0.96 | 0.84 | 0.9 |
| 6 | 115036813 | rs9400742 | -0.603 | -0.309 | -1.07 | -0.516 | 0.011 | 0.027 | 0.027 | 0 | 0.81 | 0.89 | 0.89 | 0.96 | 0.84 | 0.9 |
| 6 | 115041835 | rs9374478 | 0.728 | -0.311 | -1.072 | -0.517 | 0.022 | 0.063 | 0.063 | 0 | 0.7 | 0.84 | 0.83 | 0.87 | 0.74 | 0.76 |
| 6 | 115048197 | rs4333451 | -0.604 | -0.311 | -1.076 | -0.517 | 0.011 | 0.027 | 0.027 | 0 | 0.81 | 0.88 | 0.89 | 0.96 | 0.85 | 0.9 |
| 6 | 115049875 | rs7752198 | 0.61 | -0.29 | -1.085 | -0.506 | 0.016 | 0.036 | 0.036 | 0 | 0.7 | 0.8 | 0.82 | 0.87 | 0.74 | 0.76 |
| 6 | 115053726 | rs9387236 | 0.621 | -0.294 | -1.086 | -0.508 | 0.016 | 0.036 | 0.036 | 0 | 0.7 | 0.8 | 0.83 | 0.87 | 0.74 | 0.76 |
| 6 | 115054421 | rs9387237 | 0.611 | -0.288 | -1.084 | -0.505 | 0.016 | 0.036 | 0.036 | 0 | 0.7 | 0.8 | 0.82 | 0.87 | 0.74 | 0.76 |
| 6 | 115058847 | rs4946020 | 0.073 | -0.078 | -0.829 | -0.231 | 0.004 | 0.001 | 0.001 | 0 | 0.81 | 0.85 | 0.89 | 0.96 | 0.84 | 0.9 |
| 6 | 115063621 | rs9387243 | -0.595 | 0.208 | -0.633 | 0.036 | 0.003 | -0.007 | -0.007 | 1.044 | 0.68 | 0.79 | 0.73 | 0.64 | 0.7 | 0.8 |
| 6 | 115068368 | rs9481474 | -0.778 | 0.825 | 0.139 | 0.255 | 0.033 | 0.044 | 0.044 | 0.167 | 0.14 | 0.28 | 0.11 | 0.21 | 0.18 | 0.3 |
| 6 | 115080182 | rs9387247 | -0.187 | 0.581 | 0.253 | 0.266 | -0.001 | 0.087 | 0.087 | 0 | 0.06 | 0.19 | 0.04 | 0.18 | 0.04 | 0.1 |
| 6 | 115087005 | rs9400750 | -0.35 | -0.293 | -0.013 | -0.324 | 0.025 | 0.026 | 0.026 | 0 | 0.85 | 0.94 | 0.94 | 0.91 | 0.93 | 0.82 |
| 6 | 115091389 | rs9488470 | -1.031 | -0.542 | 0.445 | -0.427 | -0.018 | 0.086 | 0.086 | 1.167 | 0.13 | 0.38 | 0.18 | 0.22 | 0.1 | 0.16 |
| 6 | 115093947 | rs11153510 | 1.371 | -0.752 | 0.692 | -0.951 | 0.05 | 0.052 | 0.052 | 0.17 | 0.55 | 0.26 | 0.63 | 0.69 | 0.71 | 0.6 |
| 6 | 115096019 | rs4425620 | 1.358 | -0.751 | 0.707 | -0.953 | 0.052 | 0.053 | 0.053 | 0.143 | 0.55 | 0.26 | 0.63 | 0.69 | 0.71 | 0.6 |
| 6 | 115098954 | rs8180626 | 1.367 | -0.752 | 0.689 | -0.953 | 0.055 | 0.056 | 0.056 | 0.217 | 0.55 | 0.26 | 0.63 | 0.69 | 0.71 | 0.6 |
| 6 | 115106846 | rs9400752 | 1.263 | -0.766 | 0.759 | -0.982 | 0.038 | 0.041 | 0.041 | 0.054 | 0.57 | 0.32 | 0.64 | 0.69 | 0.71 | 0.6 |
| 6 | 115112949 | rs7759732 | 0.782 | -0.311 | 0.17 | -1.186 | 0.002 | -0.002 | -0.002 | 0 | 0.29 | 0.3 | 0.36 | 0.31 | 0.36 | 0.43 |
| 6 | 115137430 | rs9488481 | 0.521 | -0.57 | 0.315 | -0.992 | 0.007 | -0.055 | -0.055 | 2.049 | 0.15 | 0.23 | 0.12 | 0.03 | 0.13 | 0.05 |
| 6 | 115141695 | rs4946038 | 0.455 | -0.682 | 0.352 | -0.995 | -0.049 | -0.007 | -0.007 | 0.012 | 0.72 | 0.57 | 0.76 | 0.72 | 0.84 | 0.66 |
| 6 | 115155076 | rs9400762 | 1.616 | -0.495 | 0.12 | -0.965 | -0.044 | -0.108 | -0.108 | 0 | 0.27 | 0.07 | 0.26 | 0.38 | 0.33 | 0.17 |
| 6 | 115155798 | rs9400763 | 1.251 | -0.977 | 0.783 | -0.578 | -0.013 | 0.011 | 0.011 | 0.161 | 0.72 | 0.47 | 0.7 | 0.72 | 0.78 | 0.79 |
| 6 | 115157476 | rs12173275 | 1.251 | -0.977 | 0.783 | -0.58 | -0.013 | 0.009 | 0.009 | 0.161 | 0.72 | 0.47 | 0.7 | 0.72 | 0.78 | 0.79 |
| 6 | 115168141 | rs9400765 | -0.196 | -1.05 | 0.479 | -0.608 | -0.004 | 0.024 | 0.024 | 0.119 | 0.87 | 0.93 | 0.89 | 0.97 | 0.88 | 0.95 |
| 6 | 115170045 | rs4379324 | 0.066 | -0.837 | 0.602 | -0.497 | 0.034 | 0.106 | 0.106 | 1.474 | 0.72 | 0.89 | 0.84 | 0.91 | 0.81 | 0.77 |
| 6 | 115173667 | rs9320520 | 0.418 | -0.606 | 0.753 | -0.223 | 0.003 | 0.103 | 0.103 | 0 | 0.05 | 0.15 | 0.04 | 0.19 | 0.04 | 0.1 |
| 6 | 115180226 | rs9488492 | -0.817 | -0.729 | 0.651 | -0.495 | -0.016 | -0.029 | -0.029 | 0.005 | 0.38 | 0.45 | 0.31 | 0.16 | 0.24 | 0.3 |
| 6 | 115184297 | rs7743827 | 0.595 | -0.502 | 0.896 | -0.316 | -0.035 | -0.043 | -0.043 | 0 | 0.66 | 0.55 | 0.7 | 0.83 | 0.78 | 0.75 |
| 6 | 115185422 | rs11153521 | -0.166 | 0.916 | 1.464 | -0.011 | 0.017 | 0.138 | 0.138 | 0 | 0.06 | 0.2 | 0.05 | 0.2 | 0.05 | 0.11 |
| 6 | 115191061 | rs7752978 | 0.343 | 1.342 | 1.171 | -0.356 | 0.001 | 0 | 0 | 0 | 0.69 | 0.34 | 0.54 | 0.58 | 0.61 | 0.57 |
| 6 | 115194099 | rs11153523 | -0.023 | 1.329 | 1.418 | -0.312 | 0.014 | 0.145 | 0.145 | 0 | 0.07 | 0.22 | 0.06 | 0.23 | 0.05 | 0.17 |
| 6 | 115212142 | rs9372417 | 1.652 | 1.224 | 1.134 | -0.517 | 0.004 | -0.007 | -0.007 | 0.403 | 0.46 | 0.11 | 0.38 | 0.52 | 0.38 | 0.21 |
| 6 | 115216373 | rs12201099 | 0.408 | 1.247 | 0.827 | -0.512 | -0.019 | 0.015 | 0.015 | 0 | 0.22 | 0.17 | 0.35 | 0.19 | 0.34 | 0.25 |
| 6 | 115230047 | rs4400248 | 0.387 | 1.214 | 0.909 | -0.493 | 0.037 | 0.207 | 0.207 | 0 | 0.92 | 0.66 | 0.89 | 0.77 | 0.95 | 0.83 |
| 6 | 115231965 | rs11153525 | -0.101 | 1.237 | -0.092 | -0.476 | -0.04 | 0.028 | 0.028 | 0 | 0.26 | 0.17 | 0.39 | 0.22 | 0.4 | 0.33 |
| 6 | 115235605 | rs9372420 | 0.68 | 1.214 | -0.088 | -0.527 | -0.004 | -0.002 | -0.002 | 0 | 0.73 | 0.39 | 0.78 | 0.74 | 0.78 | 0.55 |
| 6 | 115265795 | rs17805392 | 0.104 | 1.223 | -0.057 | -0.538 | -0.008 | 0.006 | 0.006 | 0 | 0.23 | 0.24 | 0.34 | 0.19 | 0.34 | 0.26 |
| 6 | 115311251 | rs4434499 | -0.276 | 0.902 | -0.223 | -0.529 | 0.002 | -0.001 | -0.001 | 0.327 | 0.24 | 0.66 | 0.25 | 0.27 | 0.25 | 0.28 |
| 6 | 115313523 | rs4946053 | 0.16 | 1.303 | -0.65 | -0.328 | -0.034 | 0.023 | 0.023 | 0 | 0.22 | 0.15 | 0.35 | 0.19 | 0.38 | 0.26 |
| 6 | 115318098 | rs2350115 | -0.423 | 0.891 | -0.238 | -0.107 | -0.003 | -0.01 | -0.01 | 0 | 0.3 | 0.52 | 0.21 | 0.29 | 0.27 | 0.51 |
| 6 | 115325793 | rs869073 | -0.611 | 1.043 | -1.283 | -0.022 | 0.009 | 0.026 | 0.026 | 3.248 | 0.34 | 0.8 | 0.27 | 0.29 | 0.28 | 0.51 |

| | | | | | | | | | | | | | | | | |
|---|---|---|---|---|---|---|---|---|---|---|---|---|---|---|---|---|
| 6 | 115372264 | rs238562 | -0.436 | 0.883 | -1.626 | 0.148 | 0.058 | -0.033 | -0.033 | 0 | 0.56 | 0.91 | 0.63 | 0.51 | 0.67 | 0.77 |
| 6 | 115377513 | rs17835143 | -0.324 | 1.904 | -1.608 | -0.037 | -0.03 | -0.003 | -0.003 | 0 | 0.19 | 0.13 | 0.36 | 0.26 | 0.33 | 0.19 |
| 6 | 115388972 | rs1431807 | -0.192 | 2.025 | -1.217 | -0.345 | 0.007 | 0.018 | 0.018 | 1.01 | 0.38 | 0.64 | 0.23 | 0.25 | 0.32 | 0.56 |
| 6 | 115514027 | rs35261461 | -0.14 | 1.858 | -0.166 | -0.474 | -0.002 | -0.021 | -0.021 | 0 | 0.36 | 0.45 | 0.28 | 0.25 | 0.35 | 0.53 |
| 6 | 115590427 | rs4946069 | 0.061 | 1.984 | 0.079 | -0.214 | -0.027 | -0.01 | -0.01 | 0.016 | 0.19 | 0.13 | 0.35 | 0.28 | 0.31 | 0.19 |
| 6 | 115595273 | rs1412038 | -0.058 | 1.953 | 0.132 | -0.228 | -0.001 | -0.02 | -0.02 | 0 | 0.65 | 0.55 | 0.72 | 0.75 | 0.65 | 0.51 |
| 6 | 115600669 | rs10456898 | -0.028 | 2.067 | 0.353 | -0.447 | -0.018 | 0.001 | 0.001 | 0.814 | 0.23 | 0.35 | 0.19 | 0.22 | 0.19 | 0.4 |
| 6 | 115605767 | rs1412034 | 1.769 | 2.119 | 0.353 | -0.486 | 0.039 | -0.052 | -0.052 | 2.731 | 0.39 | 0.15 | 0.32 | 0.47 | 0.31 | 0.25 |
| 6 | 115608299 | rs9400815 | 1.563 | 2.11 | 0.365 | -0.493 | 0.036 | -0.046 | -0.046 | 3.448 | 0.43 | 0.17 | 0.36 | 0.47 | 0.34 | 0.26 |
| 6 | 115608842 | rs1548305 | 0.904 | 2.083 | 0.61 | -0.496 | -0.002 | -0.018 | -0.018 | 0 | 0.6 | 0.51 | 0.68 | 0.76 | 0.62 | 0.43 |
| 6 | 115626947 | rs11760145 | -0.679 | 2.199 | 0.55 | -0.506 | -0.017 | -0.014 | -0.014 | 0 | 0.29 | 0.45 | 0.24 | 0.24 | 0.25 | 0.51 |
| 6 | 115650230 | rs13437134 | -0.666 | 2.195 | 0.674 | -0.508 | 0 | 0 | 0 | 0 | 0.3 | 0.67 | 0.26 | 0.24 | 0.25 | 0.52 |
| 6 | 115660938 | rs9374532 | 1.108 | 2.037 | 0.6 | -0.496 | 0.016 | 0.004 | 0.004 | 1.802 | 0.51 | 0.27 | 0.4 | 0.48 | 0.44 | 0.3 |
| 6 | 115673943 | rs9374534 | 1.835 | 1.996 | 0.398 | -0.518 | 0.007 | -0.124 | -0.124 | 0.481 | 0.39 | 0.05 | 0.31 | 0.47 | 0.31 | 0.25 |
| 6 | 115682175 | rs11153556 | -1.363 | 2 | 0.408 | -0.521 | 0.011 | -0.035 | -0.035 | 0.326 | 0.52 | 0.89 | 0.65 | 0.52 | 0.59 | 0.65 |
| 6 | 115685164 | rs4484550 | 0.143 | 2.039 | 0.471 | -0.481 | -0.007 | 0.013 | 0.013 | 0 | 0.91 | 0.95 | 0.96 | 0.99 | 0.9 | 0.89 |
| 6 | 115691951 | rs13211979 | -0.076 | 2.894 | 0.666 | -0.536 | -0.008 | -0.004 | -0.004 | 0 | 0.21 | 0.2 | 0.36 | 0.3 | 0.29 | 0.22 |
| 6 | 115717342 | rs9320533 | -0.225 | 2.56 | 0.555 | -0.488 | -0.009 | 0.009 | 0.009 | 0 | 0.92 | 0.96 | 0.96 | 0.99 | 0.9 | 0.95 |
| 6 | 115756918 | rs4333447 | -0.099 | 2.542 | 0.609 | -0.489 | 0.01 | -0.043 | -0.043 | 0 | 0.92 | 0.89 | 0.96 | 1 | 0.9 | 0.95 |
| 6 | 115779659 | rs7759765 | 1.532 | 2.433 | 0.531 | -0.537 | 0.035 | -0.022 | -0.022 | 1.561 | 0.43 | 0.26 | 0.36 | 0.46 | 0.34 | 0.26 |
| 6 | 115782373 | rs9481563 | -1.032 | 2.314 | 0.589 | -0.497 | -0.003 | -0.027 | -0.027 | 0 | 0.05 | 0.13 | 0.05 | 0 | 0.03 | 0.07 |
| 6 | 115787696 | rs12207938 | -0.427 | 2.355 | 0.704 | -0.585 | -0.011 | -0.003 | -0.003 | 0.033 | 0.24 | 0.39 | 0.19 | 0.23 | 0.2 | 0.39 |
| 6 | 115794381 | rs1158717 | -0.399 | 2.345 | 0.828 | -0.54 | -0.013 | 0.002 | 0.002 | 0.062 | 0.24 | 0.37 | 0.19 | 0.23 | 0.2 | 0.4 |
| 6 | 115812129 | rs10484296 | -1.089 | 2.325 | 0.948 | -0.563 | -0.018 | -0.016 | -0.016 | 0 | 0.3 | 0.56 | 0.24 | 0.23 | 0.22 | 0.47 |
| 6 | 115817157 | rs6912194 | -0.419 | 2.317 | 1.016 | -0.559 | -0.016 | 0.003 | 0.003 | 0.059 | 0.24 | 0.39 | 0.19 | 0.24 | 0.19 | 0.4 |
| 6 | 115817714 | rs9488675 | -0.539 | 2.321 | 0.955 | -0.557 | -0.024 | -0.003 | -0.003 | 0.154 | 0.26 | 0.55 | 0.2 | 0.23 | 0.19 | 0.4 |
| 6 | 115821030 | rs4946087 | -1.653 | 2.444 | 0.846 | -0.6 | 0.03 | -0.056 | -0.056 | 0 | 0.56 | 0.83 | 0.63 | 0.54 | 0.66 | 0.68 |
| 6 | 115821763 | rs9488679 | -0.331 | 2.566 | 0.897 | -0.588 | -0.031 | -0.001 | -0.001 | 0 | 0.24 | 0.17 | 0.38 | 0.3 | 0.37 | 0.23 |
| 6 | 115821926 | rs17076827 | 0.556 | 2.571 | 0.931 | -0.602 | 0.034 | -0.033 | -0.033 | 0 | 0.07 | 0.12 | 0.05 | 0 | 0.1 | 0.05 |
| 6 | 115823786 | rs4946088 | -0.384 | 2.56 | 0.943 | -0.596 | 0.008 | -0.002 | -0.002 | 0.024 | 0.24 | 0.24 | 0.39 | 0.3 | 0.37 | 0.23 |
| 6 | 115834213 | rs9481574 | -0.536 | 2.548 | 0.911 | -0.561 | -0.008 | 0.002 | 0.002 | 0 | 0.26 | 0.22 | 0.39 | 0.3 | 0.37 | 0.29 |
| 6 | 115836618 | rs1467926 | -0.37 | 2.541 | 0.987 | -0.562 | -0.017 | 0.001 | 0.001 | 0.087 | 0.24 | 0.39 | 0.19 | 0.23 | 0.19 | 0.4 |
| 6 | 115844598 | rs9488698 | -0.747 | 2.617 | 0.784 | -0.531 | -0.008 | 0.003 | 0.003 | 0.004 | 0.3 | 0.25 | 0.43 | 0.31 | 0.46 | 0.28 |
| 6 | 115849514 | rs1034460 | -0.324 | 2.571 | 0.744 | -0.511 | -0.051 | -0.009 | -0.009 | 0 | 0.24 | 0.14 | 0.38 | 0.31 | 0.37 | 0.23 |
| 6 | 115875526 | rs985553 | -1.208 | 2.074 | 0.507 | -0.706 | 0.006 | 0.003 | 0.003 | 0.466 | 0.5 | 0.59 | 0.58 | 0.53 | 0.56 | 0.63 |
| 6 | 115886188 | rs12207371 | -2.224 | 2.018 | 0.368 | -0.369 | -0.031 | -0.139 | -0.139 | 3.254 | 0.64 | 0.92 | 0.68 | 0.61 | 0.7 | 0.75 |
| 6 | 115889020 | rs6568908 | -2.224 | 2.018 | 0.368 | -0.369 | -0.031 | -0.139 | -0.139 | 0.286 | 0.64 | 0.92 | 0.68 | 0.61 | 0.7 | 0.75 |
| 6 | 115897699 | rs10223402 | -0.466 | 2.251 | 0.308 | -0.457 | 0.024 | -0.02 | -0.02 | 0.553 | 0.12 | 0.2 | 0.1 | 0.07 | 0.15 | 0.08 |
| 6 | 115905297 | rs9374544 | 1.044 | 2.131 | 0.316 | -0.439 | 0.008 | 0.004 | 0.004 | 0.855 | 0.49 | 0.36 | 0.42 | 0.47 | 0.43 | 0.37 |
| 6 | 115907982 | rs6921093 | 1.022 | 2.207 | 0.326 | -0.42 | 0.009 | 0.003 | 0.003 | 1.82 | 0.51 | 0.36 | 0.41 | 0.47 | 0.43 | 0.37 |
| 6 | 115910044 | rs4128270 | 1.04 | 2.248 | 0.331 | -0.433 | 0.016 | 0.002 | 0.002 | 2.309 | 0.5 | 0.27 | 0.41 | 0.47 | 0.43 | 0.37 |
| 6 | 115916087 | rs2817318 | 0.165 | 2.294 | 0.38 | -0.526 | -0.012 | -0.002 | -0.002 | 0 | 0.93 | 0.94 | 0.89 | 0.79 | 0.88 | 0.76 |
| 6 | 115919941 | rs7755699 | 1.318 | 2.264 | 0.349 | -0.537 | 0.007 | 0.003 | 0.003 | 2.497 | 0.49 | 0.34 | 0.41 | 0.47 | 0.43 | 0.32 |
| 6 | 115922400 | rs9481581 | -0.304 | 2.147 | 0.34 | -0.532 | 0.015 | -0.018 | -0.018 | 0 | 0.11 | 0.19 | 0.09 | 0.07 | 0.13 | 0.07 |
| 6 | 115935290 | rs4320401 | 0.058 | 2.229 | 0.206 | -0.495 | -0.038 | -0.034 | -0.034 | 0 | 0.07 | 0.01 | 0.1 | 0.21 | 0.13 | 0.24 |
| 6 | 115937666 | rs9374553 | 1.451 | 2.139 | 0.241 | -0.618 | 0.006 | 0.029 | 0.029 | 2.904 | 0.62 | 0.19 | 0.7 | 0.7 | 0.67 | 0.53 |
| 6 | 115937730 | rs9400861 | 0.903 | 2.013 | 0.29 | -0.638 | 0.009 | 0.002 | 0.002 | 1.78 | 0.69 | 0.28 | 0.81 | 0.91 | 0.79 | 0.77 |
| 6 | 115944185 | rs9400862 | 1.911 | 2.316 | 0.194 | -0.726 | 0.028 | -0.033 | -0.033 | 0.574 | 0.42 | 0.24 | 0.34 | 0.42 | 0.34 | 0.27 |
| 6 | 115944735 | rs12195710 | 0.38 | 2.22 | 0.161 | -0.762 | -0.043 | -0.021 | -0.021 | 0.357 | 0.19 | 0.11 | 0.35 | 0.29 | 0.3 | 0.19 |
| 6 | 115955544 | rs4946104 | -0.759 | 2.193 | 0.358 | -0.749 | -0.003 | -0.021 | -0.021 | 0.34 | 0.38 | 0.47 | 0.44 | 0.3 | 0.38 | 0.4 |
| 6 | 115967305 | rs7746151 | 0.197 | 2.303 | 0.119 | -0.761 | 0.004 | -0.024 | -0.024 | 0 | 0.8 | 0.77 | 0.84 | 0.93 | 0.77 | 0.88 |
| 6 | 115975295 | rs4266518 | -1.107 | 2.14 | 0.15 | -0.659 | -0.009 | -0.019 | -0.019 | 0 | 0.3 | 0.55 | 0.26 | 0.25 | 0.28 | 0.47 |
| 6 | 115986454 | rs9400867 | -0.74 | 1.647 | -0.714 | -1.192 | -0.024 | 0.035 | 0.035 | 0 | 0.78 | 0.98 | 0.9 | 0.84 | 0.83 | 0.86 |
| 6 | 115995659 | rs1156783 | -0.883 | 2.039 | -0.748 | -1.344 | 0.015 | -0.026 | -0.026 | 0 | 0.31 | 0.39 | 0.44 | 0.23 | 0.38 | 0.33 |
| 6 | 115996511 | rs11153577 | -1.375 | 2.151 | -0.79 | -1.186 | 0.001 | 0 | 0 | 1.385 | 0.38 | 0.41 | 0.57 | 0.45 | 0.56 | 0.57 |
| 6 | 115999839 | rs9320545 | 2.471 | 2.017 | -0.767 | -1.256 | 0.066 | -0.076 | -0.076 | 0 | 0.34 | 0.16 | 0.29 | 0.4 | 0.25 | 0.25 |
| 6 | 116003773 | rs12662480 | -1.137 | 1.688 | -0.711 | -1.289 | 0.021 | -0.05 | -0.05 | 0 | 0.32 | 0.51 | 0.42 | 0.24 | 0.36 | 0.33 |
| 6 | 116004415 | rs17077063 | 0.389 | 1.669 | -0.809 | -1.196 | -0.041 | -0.033 | -0.033 | 0 | 0.07 | 0.01 | 0.14 | 0.21 | 0.17 | 0.24 |
| 6 | 116004497 | rs9374562 | 2.159 | 1.419 | -0.761 | -1.225 | 0.043 | -0.056 | -0.056 | 0 | 0.37 | 0.22 | 0.34 | 0.41 | 0.29 | 0.26 |
| 6 | 116005535 | rs1015157 | -0.615 | 1.606 | -0.559 | -1.304 | 0.01 | -0.021 | -0.021 | 0 | 0.23 | 0.33 | 0.35 | 0.21 | 0.27 | 0.23 |
| 6 | 116006694 | rs6912763 | -0.459 | 1.416 | -0.713 | -0.597 | -0.001 | 0.003 | 0.003 | 0 | 0.39 | 0.46 | 0.35 | 0.43 | 0.38 | 0.41 |
| 6 | 116011927 | rs9372446 | 1.859 | 1.482 | -0.817 | -0.528 | -0.024 | -0.067 | -0.067 | 0.105 | 0.4 | 0.1 | 0.3 | 0.34 | 0.36 | 0.37 |
| 6 | 116011942 | rs10484728 | 0.451 | 1.46 | -0.842 | -0.539 | 0 | 0.021 | 0.021 | 0 | 0.28 | 0.05 | 0.17 | 0.23 | 0.2 | 0.17 |
| 6 | 116014608 | rs9374564 | 2.477 | 1.404 | -0.846 | -0.557 | 0.049 | -0.082 | -0.082 | 0 | 0.34 | 0.06 | 0.24 | 0.32 | 0.25 | 0.26 |
| 6 | 116023522 | rs9387356 | 0.102 | 1.45 | -0.777 | -0.544 | 0.033 | -0.01 | -0.01 | 1.078 | 0.79 | 0.58 | 0.65 | 0.77 | 0.75 | 0.78 |
| 6 | 116030515 | rs9320546 | 2.715 | 1.008 | -1.058 | -0.565 | 0.053 | -0.097 | -0.097 | 0 | 0.29 | 0.06 | 0.21 | 0.31 | 0.21 | 0.18 |
| 6 | 116032842 | rs9387358 | 2.025 | 1.049 | -0.967 | -0.593 | 0.012 | -0.043 | -0.043 | 1.63 | 0.36 | 0.22 | 0.28 | 0.36 | 0.32 | 0.29 |
| 6 | 116040072 | rs12525228 | 0.465 | 0.78 | -0.413 | -0.197 | 0.001 | 0.051 | 0.051 | 0 | 0.28 | 0.05 | 0.16 | 0.2 | 0.21 | 0.17 |
| 6 | 116049370 | rs4946114 | 1.255 | 0.78 | -0.208 | -0.262 | 0.017 | 0.036 | 0.036 | 0 | 0.5 | 0.33 | 0.59 | 0.55 | 0.54 | 0.42 |

| | | | | | | | | | | | | | | | | |
|---|---|---|---|---|---|---|---|---|---|---|---|---|---|---|---|---|
| 6 | 116051927 | rs9488761 | -0.245 | 0.745 | -0.063 | -0.273 | -0.02 | -0.065 | -0.065 | 0 | 0.11 | 0.47 | 0.07 | 0.01 | 0.05 | 0.11 |
| 6 | 116056824 | rs7774310 | 1.813 | 0.899 | -0.1 | -0.195 | -0.027 | -0.05 | -0.05 | 0 | 0.43 | 0.12 | 0.53 | 0.53 | 0.44 | 0.31 |
| 6 | 116063382 | rs7740854 | -0.703 | 0.728 | -0.196 | -0.236 | 0.056 | 0.086 | 0.086 | 0 | 0.46 | 0.65 | 0.25 | 0.22 | 0.3 | 0.37 |
| 6 | 116077361 | rs1935414 | 0.454 | 0.785 | -0.347 | -0.109 | -0.037 | -0.037 | -0.037 | 0 | 0.06 | 0.01 | 0.14 | 0.2 | 0.15 | 0.18 |
| 6 | 116080427 | rs1556614 | 1.804 | 0.776 | -0.36 | -0.112 | -0.065 | -0.052 | -0.052 | 0 | 0.44 | 0.09 | 0.54 | 0.56 | 0.45 | 0.34 |
| 6 | 116085882 | rs6934225 | -1.582 | 0.919 | -0.214 | -0.015 | 0.05 | 0.032 | 0.032 | 0 | 0.12 | 0.21 | 0.2 | 0.22 | 0.25 | 0.28 |
| 6 | 116090265 | rs9968822 | 0.076 | 0.682 | -0.166 | -0.133 | 0 | -0.037 | -0.037 | 0 | 0.08 | 0.13 | 0.06 | 0 | 0.07 | 0.08 |
| 6 | 116090609 | rs3927605 | 1.404 | 0.744 | -0.29 | 0.007 | -0.07 | -0.02 | -0.02 | 0 | 0.45 | 0.08 | 0.58 | 0.58 | 0.54 | 0.4 |
| 6 | 116097510 | rs4946118 | 0.409 | 0.626 | 0.107 | 0.14 | 0.009 | 0.007 | 0.007 | 0 | 0.57 | 0.47 | 0.64 | 0.6 | 0.64 | 0.57 |
| 6 | 116102720 | rs9488775 | -0.494 | 0.624 | -0.14 | 0.127 | -0.005 | -0.004 | -0.004 | 0 | 0.43 | 0.4 | 0.35 | 0.4 | 0.37 | 0.42 |
| 6 | 116105988 | rs2351532 | 0.256 | 1.113 | -0.691 | 0.792 | 0.04 | 0.141 | 0.141 | 4.27 | 0.81 | 0.39 | 0.85 | 0.65 | 0.75 | 0.69 |
| 6 | 116107951 | rs13216090 | 0.571 | 1.047 | -1.421 | 0.667 | 0.008 | 0.043 | 0.043 | 0 | 0.3 | 0.11 | 0.18 | 0.19 | 0.25 | 0.2 |
| 6 | 116115351 | rs1105192 | 0.828 | 1.045 | -1.582 | 0.676 | -0.025 | 0.014 | 0.014 | 0 | 0.28 | 0.05 | 0.15 | 0.19 | 0.23 | 0.21 |
| 6 | 116128667 | rs9320549 | 1.071 | 1.066 | -1.421 | 0.557 | -0.036 | 0.187 | 0.187 | 0 | 0.47 | 0.1 | 0.54 | 0.24 | 0.42 | 0.32 |
| 6 | 116130086 | rs17077265 | -0.207 | 0.492 | -0.929 | 0.559 | 0.006 | -0.007 | -0.007 | 0 | 0.36 | 0.2 | 0.3 | 0.4 | 0.34 | 0.46 |
| 6 | 116131731 | rs6916017 | -0.963 | 0.308 | -0.301 | 0.762 | -0.024 | 0.038 | 0.038 | 0 | 0.27 | 0.5 | 0.4 | 0.36 | 0.23 | 0.43 |
| 6 | 116143788 | rs4946123 | 0.353 | 0.818 | -0.521 | 0.907 | 0.011 | 0.009 | 0.009 | 0.044 | 0.75 | 0.67 | 0.77 | 0.71 | 0.63 | 0.71 |
| 6 | 116147191 | rs10485113 | 0.611 | 0.594 | -0.649 | -0.19 | -0.038 | 0.11 | 0.11 | 0 | 0.22 | 0.04 | 0.18 | 0.02 | 0.22 | 0.07 |
| 6 | 116156793 | rs567564 | 1.318 | 0.35 | -0.453 | -0.157 | -0.01 | -0.02 | -0.02 | 0 | 0.4 | 0.47 | 0.26 | 0.37 | 0.3 | 0.43 |
| 6 | 116169275 | rs9374579 | -0.027 | 0.363 | -0.587 | 0.64 | -0.016 | -0.046 | -0.046 | 0 | 0.28 | 0.13 | 0.44 | 0.28 | 0.3 | 0.28 |
| 6 | 116169307 | rs195482 | -1.707 | 0.369 | -0.616 | 0.295 | 0.01 | 0.025 | 0.025 | 0.591 | 0.62 | 0.74 | 0.76 | 0.64 | 0.7 | 0.59 |
| 6 | 116172203 | rs515413 | -0.106 | 0.643 | -0.621 | -0.076 | 0.048 | -0.031 | -0.031 | 0 | 0.26 | 0.5 | 0.24 | 0.18 | 0.37 | 0.26 |
| 6 | 116172252 | rs7740894 | 0.967 | 0.559 | -1.105 | -0.015 | -0.05 | 0.088 | 0.088 | 0 | 0.18 | 0.01 | 0.15 | 0.02 | 0.2 | 0.03 |
| 6 | 116183352 | rs7751601 | 0.604 | 0.604 | -1.17 | -0.089 | -0.071 | 0.111 | 0.111 | 0 | 0.22 | 0.01 | 0.2 | 0.02 | 0.27 | 0.03 |
| 6 | 116186895 | rs6938448 | 0.051 | 0.41 | -1.226 | 0.115 | -0.027 | 0.025 | 0.025 | 0 | 0.27 | 0.04 | 0.23 | 0.14 | 0.26 | 0.05 |
| 6 | 116192364 | rs195504 | 0.73 | 0.604 | -0.766 | -0.065 | 0.03 | -0.031 | -0.031 | 0 | 0.07 | 0.19 | 0.04 | 0 | 0.06 | 0.04 |
| 6 | 116197988 | rs1416188 | 0.086 | 0.559 | -0.791 | -0.17 | -0.006 | 0.038 | 0.038 | 0 | 0.27 | 0.11 | 0.23 | 0.15 | 0.28 | 0.1 |
| 6 | 116202783 | rs195513 | -0.838 | -0.061 | -0.373 | -0.499 | -0.011 | -0.042 | -0.042 | 0 | 0.41 | 0.5 | 0.29 | 0.16 | 0.36 | 0.27 |
| 6 | 116206371 | rs12203852 | 0.225 | -0.013 | -0.645 | -0.421 | -0.005 | 0.039 | 0.039 | 0 | 0.27 | 0.11 | 0.23 | 0.15 | 0.28 | 0.1 |
| 6 | 116212551 | rs17077343 | 1.383 | -0.046 | -0.7 | -0.35 | 0.025 | -0.023 | -0.023 | 2.898 | 0.06 | 0.02 | 0.03 | 0.13 | 0.01 | 0.02 |
| 6 | 116214243 | rs195468 | -1.427 | 0.261 | -1.053 | -0.281 | -0.032 | -0.087 | -0.087 | 0 | 0.62 | 0.94 | 0.52 | 0.37 | 0.68 | 0.49 |
| 6 | 116234968 | rs195484 | 0.925 | 0.295 | -0.53 | -0.379 | 0.001 | -0.05 | -0.05 | 0 | 0.21 | 0.21 | 0.21 | 0.01 | 0.24 | 0.03 |
| 6 | 116242267 | rs195519 | 1.066 | 0.283 | -0.478 | -0.173 | -0.012 | 0.061 | 0.061 | 0 | 0.06 | 0.14 | 0.2 | 0.14 | 0.04 | 0.03 |
| 6 | 116251893 | rs195529 | -0.583 | 0.217 | -0.512 | -0.529 | -0.002 | -0.02 | -0.02 | 0.707 | 0.4 | 0.4 | 0.43 | 0.16 | 0.37 | 0.22 |
| 6 | 116254141 | rs13216580 | 1.445 | 0.266 | -0.567 | -0.566 | -0.005 | 0 | 0 | 0 | 0.06 | 0.02 | 0.02 | 0 | 0.06 | 0.05 |
| 6 | 116262976 | rs495565 | 0.402 | 0.264 | -0.503 | -0.554 | -0.007 | -0.043 | -0.043 | 0 | 0.59 | 0.52 | 0.56 | 0.84 | 0.63 | 0.78 |
| 6 | 116263181 | rs580396 | 0.402 | 0.264 | -0.503 | -0.554 | -0.007 | -0.043 | -0.043 | 0 | 0.59 | 0.51 | 0.56 | 0.84 | 0.63 | 0.78 |
| 6 | 116263431 | rs1338507 | 1.505 | 0.29 | -0.54 | -0.44 | 0.009 | -0.031 | -0.031 | 0.409 | 0.06 | 0.01 | 0.18 | 0.14 | 0.04 | 0.03 |
| 6 | 116277528 | rs4946130 | 1.505 | 0.29 | -0.54 | -0.44 | 0.009 | -0.031 | -0.031 | 0 | 0.06 | 0.01 | 0.18 | 0.14 | 0.04 | 0.03 |
| 6 | 116277792 | rs17077423 | 1.426 | 0.34 | -0.544 | -0.444 | 0.005 | -0.033 | -0.033 | 1.507 | 0.06 | 0.01 | 0.18 | 0.14 | 0.04 | 0.03 |
| 6 | 116278725 | rs562819 | -0.53 | 0.377 | -0.277 | -0.465 | 0.001 | 0.008 | 0.008 | 0.093 | 0.7 | 0.73 | 0.59 | 0.86 | 0.71 | 0.92 |
| 6 | 116289044 | rs554139 | 0.043 | 0.488 | -0.468 | -0.443 | 0.065 | 0.009 | 0.009 | 0 | 0.23 | 0.46 | 0.23 | 0.2 | 0.35 | 0.28 |
| 6 | 116292489 | rs11752170 | 0.27 | 0.424 | -0.911 | -0.416 | -0.002 | -0.036 | -0.036 | 0.11 | 0.1 | 0.14 | 0.03 | 0.01 | 0.08 | 0.15 |
| 6 | 116292883 | rs3798241 | 1.124 | 0.439 | -0.913 | -0.244 | 0.001 | -0.095 | -0.095 | 0 | 0.36 | 0.06 | 0.33 | 0.64 | 0.28 | 0.5 |
| 6 | 116304866 | rs3777424 | 1.792 | 0.342 | -1.106 | -0.02 | -0.018 | -0.091 | -0.091 | 0 | 0.3 | 0.01 | 0.21 | 0.4 | 0.22 | 0.26 |
| 6 | 116305064 | rs3822858 | 0.198 | 0.537 | -0.772 | -0.163 | -0.001 | 0.018 | 0.018 | 0 | 0.38 | 0.32 | 0.42 | 0.15 | 0.37 | 0.19 |
| 6 | 116306113 | rs9400883 | 0.035 | 0.646 | -0.874 | 0.17 | -0.037 | -0.023 | -0.023 | 0.863 | 0.08 | 0 | 0.14 | 0.31 | 0.1 | 0.25 |
| 6 | 116310336 | rs9320552 | 0.703 | 0.73 | -0.684 | -0.537 | 0 | 0.014 | 0.014 | 0.463 | 0.39 | 0.2 | 0.36 | 0.69 | 0.33 | 0.5 |
| 6 | 116314634 | rs12202641 | -0.031 | 0.214 | -0.185 | -1.353 | 0.007 | 0.009 | 0.009 | 1.373 | 0.44 | 0.39 | 0.43 | 0.05 | 0.41 | 0.26 |
| 6 | 116325142 | rs3756772 | 0.793 | 0.452 | -0.257 | -1.37 | -0.036 | 0.119 | 0.119 | 0.081 | 0.42 | 0.18 | 0.44 | 0.79 | 0.42 | 0.55 |
| 6 | 116355188 | rs12154016 | 0.174 | 0.436 | -0.023 | -1.392 | 0.008 | 0.038 | 0.038 | 0.917 | 0.43 | 0.35 | 0.42 | 0.04 | 0.41 | 0.26 |
| 6 | 116376918 | rs17077492 | 0.581 | -0.086 | -0.08 | -1.558 | -0.01 | -0.049 | -0.049 | 0 | 0.12 | 0.19 | 0.04 | 0.01 | 0.08 | 0.09 |
| 6 | 116386193 | rs4946134 | 0.106 | -0.011 | -0.072 | -1.526 | 0.045 | 0.053 | 0.053 | 0 | 0.86 | 0.54 | 0.86 | 0.82 | 0.83 | 0.81 |
| 6 | 116394809 | rs1338668 | 0.812 | -0.097 | -0.16 | -1.475 | -0.035 | 0.118 | 0.118 | 0.052 | 0.42 | 0.18 | 0.44 | 0.78 | 0.42 | 0.55 |
| 6 | 116406419 | rs17077529 | 1.007 | -0.29 | -0.133 | -1.656 | -0.013 | -0.007 | -0.007 | 0 | 0.08 | 0.09 | 0.17 | 0.03 | 0.04 | 0.07 |
| 6 | 116411005 | rs9688645 | 0.093 | -0.243 | -0.06 | -1.667 | -0.03 | -0.059 | -0.059 | 0 | 0.21 | 0.31 | 0.22 | 0.04 | 0.12 | 0.17 |
| 6 | 116418158 | rs1547030 | -0.559 | -0.2 | -0.131 | -1.618 | 0.021 | -0.045 | -0.045 | 0.829 | 0.64 | 0.53 | 0.66 | 0.81 | 0.55 | 0.76 |
| 6 | 116421127 | rs9400887 | 1.105 | -0.221 | -0.473 | -1.624 | -0.137 | 0.078 | 0.078 | 0.348 | 0.41 | 0.03 | 0.42 | 0.77 | 0.42 | 0.55 |
| 6 | 116421554 | rs7762994 | 0.064 | -0.26 | -0.149 | -1.618 | -0.024 | -0.049 | -0.049 | 0 | 0.21 | 0.29 | 0.22 | 0.04 | 0.13 | 0.21 |
| 6 | 116423299 | rs509002 | -0.771 | 0.217 | 0.487 | -1.405 | 0.039 | -0.135 | -0.135 | 0.002 | 0.72 | 0.6 | 0.73 | 1 | 0.6 | 0.86 |
| 6 | 116430533 | rs482012 | -0.615 | 0.207 | 0.726 | -1.238 | 0.029 | -0.021 | -0.021 | 0 | 0.71 | 0.49 | 0.69 | 0.76 | 0.6 | 0.76 |
| 6 | 116435795 | rs9488836 | 0.472 | 0.228 | 0.769 | -1.215 | 0.039 | -0.033 | -0.033 | 0 | 0.29 | 0.44 | 0.3 | 0.24 | 0.36 | 0.24 |
| 6 | 116440007 | rs1931897 | 1.126 | 0.321 | 0.848 | -1.239 | 0.028 | -0.126 | -0.126 | 0 | 0.24 | 0.35 | 0.21 | 0.01 | 0.27 | 0.13 |
| 6 | 116446576 | rs1064583 | 0.101 | 0.256 | 1.013 | -1.333 | -0.026 | 0.039 | 0.039 | 0.012 | 0.34 | 0.74 | 0.33 | 0.26 | 0.4 | 0.38 |
| 6 | 116451442 | rs2145357 | 1.162 | 0.347 | 1.035 | -1.438 | 0.028 | -0.126 | -0.126 | 0 | 0.23 | 0.35 | 0.21 | 0.01 | 0.27 | 0.13 |
| 6 | 116452203 | rs11965969 | 0.156 | -0.015 | 0.744 | -1.51 | -0.007 | 0.029 | 0.029 | 0 | 0.42 | 0.64 | 0.5 | 0.31 | 0.46 | 0.42 |
| 6 | 116454588 | rs549332 | -1.077 | -0.158 | 0.155 | -1.533 | -0.139 | -0.047 | -0.047 | 0 | 0.61 | 0.98 | 0.58 | 0.35 | 0.57 | 0.5 |
| 6 | 116470561 | rs958738 | 0.211 | -0.042 | 0.615 | -1.425 | -0.005 | 0.056 | 0.056 | 0 | 0.43 | 0.71 | 0.5 | 0.3 | 0.45 | 0.39 |
| 6 | 116486792 | rs11153598 | 0.862 | 0.005 | 0.475 | -1.438 | 0.007 | -0.012 | -0.012 | 0.022 | 0.16 | 0.12 | 0.26 | 0.29 | 0.15 | 0.19 |

| | | | | | | | | | | | | | | | | |
|---|---|---|---|---|---|---|---|---|---|---|---|---|---|---|---|---|
| 6 | 116497317 | rs7755367 | 1.104 | -0.001 | 0.539 | -1.452 | 0.002 | -0.002 | -0.002 | 0 | 0.13 | 0.11 | 0.23 | 0.29 | 0.09 | 0.11 |
| 6 | 116501503 | rs1546943 | 0.207 | 0.008 | 0.492 | -1.471 | -0.004 | 0.05 | 0.05 | 0 | 0.43 | 0.7 | 0.5 | 0.3 | 0.45 | 0.33 |
| 6 | 116508783 | rs11754669 | 1.156 | -0.256 | 0.244 | -1.114 | 0.004 | 0.015 | 0.015 | 1.426 | 0.09 | 0.03 | 0.02 | 0.03 | 0.05 | 0.04 |
| 6 | 116509636 | rs12526817 | 0.116 | -0.143 | 0.438 | -0.976 | -0.007 | 0.012 | 0.012 | 0.022 | 0.18 | 0.25 | 0.29 | 0.3 | 0.18 | 0.26 |
| 6 | 116514046 | rs1204835 | -0.444 | 0.102 | 0.538 | -1.031 | 0.028 | -0.033 | -0.033 | 2.803 | 0.07 | 0.2 | 0.05 | 0.01 | 0.09 | 0.1 |
| 6 | 116514253 | rs1204836 | -1.637 | 0.028 | -0.369 | -0.912 | -0.054 | -0.045 | -0.045 | 0.482 | 0.75 | 0.99 | 0.78 | 0.75 | 0.76 | 0.78 |
| 6 | 116532859 | rs7769748 | 0.006 | 0.001 | 0.229 | -0.997 | -0.001 | -0.026 | -0.026 | 1.517 | 0.07 | 0.26 | 0.05 | 0.01 | 0.05 | 0.06 |
| 6 | 116546924 | rs1204804 | -0.105 | 0.064 | -0.096 | -1.093 | -0.098 | 0.074 | 0.074 | 0 | 0.53 | 0.9 | 0.59 | 0.37 | 0.54 | 0.43 |
| 6 | 116560766 | rs1204817 | 0.048 | -0.407 | 0.002 | -1.135 | -0.022 | 0.012 | 0.012 | 1.757 | 0.25 | 0.59 | 0.35 | 0.36 | 0.22 | 0.32 |
| 6 | 116564388 | rs1204826 | -2.141 | -0.246 | 0.129 | -1.271 | -0.063 | 0.084 | 0.084 | 0 | 0.76 | 0.9 | 0.77 | 1 | 0.64 | 0.9 |
| 6 | 116565725 | rs15679 | 0.884 | -0.395 | 0.472 | -1.239 | -0.024 | 0.039 | 0.039 | 0 | 0.2 | 0.32 | 0.31 | 0.34 | 0.18 | 0.2 |
| 6 | 116570488 | rs1204830 | 0.752 | -0.498 | 0.489 | -1.24 | -0.027 | 0.038 | 0.038 | 0.047 | 0.21 | 0.33 | 0.32 | 0.35 | 0.18 | 0.25 |
| 6 | 116570597 | rs1055814 | 0.167 | -0.511 | 0.419 | -1.292 | -0.018 | 0.078 | 0.078 | 0 | 0.46 | 0.14 | 0.39 | 0.63 | 0.37 | 0.57 |
| 6 | 116571255 | rs6755 | 0.167 | -0.511 | 0.419 | -1.292 | -0.018 | 0.078 | 0.078 | 0 | 0.46 | 0.14 | 0.39 | 0.63 | 0.37 | 0.57 |
| 6 | 116571695 | rs3749893 | 0.327 | -0.517 | 0.41 | -1.296 | -0.021 | 0.076 | 0.076 | 0 | 0.46 | 0.13 | 0.39 | 0.63 | 0.37 | 0.57 |
| 6 | 116572117 | rs11789 | 0.863 | -0.544 | 0.483 | -1.287 | -0.024 | 0.032 | 0.032 | 0.233 | 0.2 | 0.31 | 0.31 | 0.35 | 0.17 | 0.25 |
| 6 | 116573667 | rs9387389 | 0.327 | -0.465 | 0.222 | -1.211 | -0.056 | 0.058 | 0.058 | 0 | 0.45 | 0.1 | 0.38 | 0.63 | 0.38 | 0.57 |
| 6 | 116574455 | rs2232472 | 0.251 | -0.486 | 0.202 | -1.232 | -0.055 | 0.054 | 0.054 | 0 | 0.45 | 0.09 | 0.38 | 0.64 | 0.36 | 0.56 |
| 6 | 116575083 | rs2232470 | -2.411 | -0.433 | 0.235 | -1.228 | -0.116 | 0.136 | 0.136 | 0.023 | 0.78 | 0.96 | 0.79 | 1 | 0.66 | 0.95 |
| 6 | 116587407 | rs1204859 | 1.279 | -0.368 | 0.618 | -1.23 | -0.005 | 0.021 | 0.021 | 0 | 0.18 | 0.22 | 0.31 | 0.31 | 0.22 | 0.14 |
| 6 | 116588069 | rs9400895 | 0.381 | -0.54 | 0.512 | -1.322 | 0.015 | 0.008 | 0.008 | 0 | 0.39 | 0.26 | 0.38 | 0.64 | 0.29 | 0.6 |
| 6 | 116589624 | rs9481617 | -0.034 | -0.529 | 0.474 | -1.299 | -0.028 | -0.081 | -0.081 | 2.997 | 0.19 | 0.39 | 0.09 | 0.04 | 0.13 | 0.16 |
| 6 | 116596247 | rs6568924 | 0.889 | -0.529 | 0.285 | -1.295 | -0.029 | -0.054 | -0.054 | 0 | 0.36 | 0.09 | 0.35 | 0.63 | 0.26 | 0.54 |
| 6 | 116598475 | rs1045182 | 0.699 | -0.511 | 0.311 | -1.285 | 0.006 | -0.014 | -0.014 | 0.06 | 0.36 | 0.16 | 0.36 | 0.63 | 0.26 | 0.54 |
| 6 | 116599665 | rs9400897 | 0.662 | -0.489 | 0.146 | -1.088 | -0.047 | -0.086 | -0.086 | 0 | 0.12 | 0.01 | 0.16 | 0.37 | 0.13 | 0.33 |
| 6 | 116600453 | rs3749894 | 0.164 | -0.591 | 0.271 | -1.136 | -0.016 | -0.046 | -0.046 | 0 | 0.18 | 0.25 | 0.08 | 0.04 | 0.13 | 0.16 |
| 6 | 116601532 | rs6909133 | -0.627 | -0.565 | 0.054 | -1.124 | -0.007 | -0.002 | -0.002 | 0 | 0.37 | 0.53 | 0.39 | 0.35 | 0.35 | 0.3 |
| 6 | 116630334 | rs9387395 | 0.748 | -0.622 | -0.479 | -0.898 | -0.063 | -0.085 | -0.085 | 0 | 0.35 | 0.05 | 0.34 | 0.63 | 0.26 | 0.53 |
| 6 | 116631933 | rs6926908 | 0.101 | -0.665 | -0.008 | -0.927 | -0.037 | -0.091 | -0.091 | 0 | 0.19 | 0.43 | 0.1 | 0.04 | 0.13 | 0.16 |
| 6 | 116634325 | rs11970221 | 0.993 | -0.594 | -0.398 | -0.807 | -0.047 | 0.042 | 0.042 | 0 | 0.4 | 0.17 | 0.52 | 0.31 | 0.57 | 0.19 |
| 6 | 116649483 | rs10782184 | 1.477 | -0.372 | -0.289 | -0.844 | 0.014 | -0.066 | -0.066 | 0 | 0.23 | 0.01 | 0.18 | 0.26 | 0.12 | 0.21 |
| 6 | 116690849 | rs9488914 | 0.124 | 0.679 | -0.1 | -1.326 | -0.006 | -0.063 | -0.063 | 0.28 | 0.55 | 0.48 | 0.57 | 0.94 | 0.42 | 0.72 |
| 6 | 116698931 | rs2213563 | -1.221 | 1.18 | -0.723 | -1.023 | -0.121 | 0.143 | 0.143 | 0 | 0.77 | 0.97 | 0.78 | 1 | 0.65 | 0.95 |
| 6 | 116700809 | rs574871 | -0.934 | 1.135 | -0.589 | -0.992 | -0.022 | -0.001 | -0.001 | 0 | 0.74 | 0.79 | 0.76 | 0.98 | 0.63 | 0.89 |
| 6 | 116703926 | rs3778002 | -0.341 | 0.511 | -0.687 | 0.119 | 0.003 | -0.036 | -0.036 | 0.337 | 0.07 | 0 | 0.05 | 0.18 | 0.05 | 0.18 |
| 6 | 116704697 | rs2858853 | 0.162 | -0.308 | -0.534 | -0.874 | -0.024 | -0.067 | -0.067 | 0 | 0.18 | 0.25 | 0.1 | 0 | 0.1 | 0.1 |
| 6 | 116707299 | rs13203813 | 1.195 | 0.291 | -0.928 | -1.463 | -0.044 | 0.08 | 0.08 | 0 | 0.24 | 0.07 | 0.2 | 0.02 | 0.26 | 0.13 |
| 6 | 116707335 | rs6900245 | 0.784 | 0.021 | -1.09 | -1.061 | 0.008 | -0.03 | -0.03 | 0 | 0.23 | 0.17 | 0.23 | 0.36 | 0.22 | 0.27 |
| 6 | 116708822 | rs519604 | 0.342 | 0.261 | -0.921 | -1.261 | -0.017 | 0.066 | 0.066 | 0.082 | 0.42 | 0.68 | 0.57 | 0.07 | 0.49 | 0.29 |
| 6 | 116717171 | rs473484 | 0.939 | 0.256 | -0.221 | -1.223 | 0.024 | -0.064 | -0.064 | 0 | 0.12 | 0.3 | 0.16 | 0.01 | 0.14 | 0.09 |
| 6 | 116720487 | rs10485183 | 1.629 | 0.107 | -0.374 | -1.201 | -0.037 | 0.07 | 0.07 | 0 | 0.25 | 0.09 | 0.2 | 0.02 | 0.27 | 0.13 |
| 6 | 116726588 | rs3798413 | 0.313 | -0.033 | -0.365 | -1.102 | -0.001 | 0.002 | 0.002 | 0 | 0.34 | 0.38 | 0.39 | 0.38 | 0.35 | 0.31 |
| 6 | 116731075 | rs9488923 | 0.901 | 0.038 | -0.188 | -1.069 | 0.013 | -0.064 | -0.064 | 0 | 0.3 | 0.32 | 0.4 | 0.06 | 0.33 | 0.2 |
| 6 | 116740003 | rs3777995 | -0.018 | -0.068 | -0.45 | -0.92 | 0.008 | -0.029 | -0.029 | 0 | 0.67 | 0.74 | 0.79 | 0.44 | 0.69 | 0.51 |
| 6 | 116740906 | rs9387402 | 2.09 | -0.162 | -0.87 | -0.861 | -0.005 | -0.072 | -0.072 | 0.411 | 0.11 | 0.01 | 0.18 | 0.27 | 0.12 | 0.08 |
| 6 | 116747241 | rs508521 | 0.112 | -0.272 | -0.613 | -0.898 | 0.019 | -0.05 | -0.05 | 0.204 | 0.64 | 0.8 | 0.74 | 0.45 | 0.68 | 0.47 |
| 6 | 116753939 | rs9398433 | 1.577 | -0.279 | -0.913 | -1.059 | -0.095 | -0.06 | -0.06 | 0 | 0.34 | 0.04 | 0.38 | 0.28 | 0.38 | 0.14 |
| 6 | 116753995 | rs549262 | 1.262 | -0.271 | -0.571 | -1.06 | -0.019 | 0.003 | 0.003 | 0 | 0.37 | 0.23 | 0.4 | 0.3 | 0.39 | 0.21 |
| 6 | 116758932 | rs1048386 | 0.009 | -0.315 | -0.275 | -1.109 | 0.002 | -0.016 | -0.016 | 1.711 | 0.28 | 0.36 | 0.33 | 0.15 | 0.3 | 0.26 |
| 6 | 116760990 | rs9488931 | 0.19 | -0.317 | -0.278 | -1.1 | -0.001 | -0.032 | -0.032 | 0 | 0.26 | 0.35 | 0.17 | 0.11 | 0.25 | 0.19 |
| 6 | 116763077 | rs483981 | -0.492 | -0.789 | -0.859 | -1.232 | -0.07 | -0.14 | -0.14 | 0 | 0.82 | 0.98 | 0.85 | 0.47 | 0.75 | 0.64 |
| 6 | 116764563 | rs11153611 | 0.225 | -0.648 | -0.653 | -1.167 | -0.005 | 0.008 | 0.008 | 0 | 0.64 | 0.69 | 0.73 | 0.45 | 0.68 | 0.5 |
| 6 | 116768917 | rs2858829 | 0.677 | 0.184 | -1.15 | -1.199 | 0.057 | -0.007 | -0.007 | 0 | 0.49 | 0.09 | 0.39 | 0.49 | 0.41 | 0.22 |
| 6 | 116769598 | rs17077968 | 1.745 | 0.326 | -1.219 | -1.017 | -0.016 | -0.079 | -0.079 | 1.493 | 0.11 | 0.01 | 0.19 | 0.41 | 0.13 | 0.09 |
| 6 | 116772664 | rs9384978 | 0.197 | 0.701 | -0.827 | -0.98 | -0.054 | -0.062 | -0.062 | 0 | 0.14 | 0.05 | 0.14 | 0.37 | 0.22 | 0.31 |
| 6 | 116774107 | rs9374603 | 0.197 | 0.713 | -0.827 | -0.983 | -0.052 | -0.061 | -0.061 | 0.032 | 0.14 | 0.05 | 0.14 | 0.35 | 0.22 | 0.31 |
| 6 | 116775633 | rs2640882 | 0.169 | 0.576 | -0.494 | -1.207 | 0.003 | -0.028 | -0.028 | 0 | 0.67 | 0.34 | 0.6 | 0.87 | 0.64 | 0.64 |
| 6 | 116787664 | rs2637679 | 0.938 | 0.634 | -0.428 | -1.166 | -0.014 | -0.038 | -0.038 | 0 | 0.17 | 0.2 | 0.04 | 0 | 0.06 | 0.07 |
| 6 | 116788453 | rs2858835 | -0.102 | 1.05 | -0.291 | -1.071 | 0.045 | -0.094 | -0.094 | 0 | 0.77 | 0.49 | 0.78 | 0.89 | 0.73 | 0.73 |
| 6 | 116801573 | rs2637687 | 0.2 | 1.125 | -0.863 | -1.066 | -0.003 | 0.004 | 0.004 | 0 | 0.35 | 0.09 | 0.21 | 0.36 | 0.31 | 0.47 |
| 6 | 116803328 | rs9384981 | 0.816 | 1.029 | -0.842 | -0.7 | -0.061 | -0.145 | -0.145 | 0 | 0.36 | 0.02 | 0.35 | 0.5 | 0.36 | 0.14 |
| 6 | 116821146 | rs7775839 | 0.533 | 0.647 | -0.523 | -0.48 | -0.02 | -0.017 | -0.017 | 0.575 | 0.39 | 0.18 | 0.57 | 0.55 | 0.44 | 0.34 |
| 6 | 116834007 | rs2250263 | -0.307 | 0.605 | -0.098 | -0.888 | 0.03 | -0.06 | -0.06 | 0 | 0.73 | 0.6 | 0.77 | 0.9 | 0.65 | 0.67 |
| 6 | 116837310 | rs3734309 | 0.656 | 0.786 | -0.837 | -0.59 | -0.033 | -0.032 | -0.032 | 0 | 0.39 | 0.12 | 0.56 | 0.55 | 0.44 | 0.34 |
| 6 | 116849485 | rs1741819 | -0.741 | 0.961 | -0.814 | -0.658 | -0.031 | 0.031 | 0.031 | 0 | 0.81 | 0.85 | 0.84 | 0.93 | 0.72 | 0.77 |
| 6 | 116849539 | rs2145335 | 0.363 | 0.888 | -0.863 | -0.265 | -0.039 | -0.04 | -0.04 | 2.516 | 0.39 | 0.11 | 0.51 | 0.55 | 0.44 | 0.33 |
| 6 | 116864570 | rs4142087 | 0.611 | 0.876 | -0.926 | -0.203 | -0.093 | -0.155 | -0.155 | 1.895 | 0.37 | 0.03 | 0.36 | 0.51 | 0.39 | 0.24 |
| 6 | 116866949 | rs17078051 | 0.399 | 0.506 | -0.727 | -0.646 | 0.005 | -0.009 | -0.009 | 0 | 0.21 | 0.15 | 0.15 | 0.35 | 0.17 | 0.3 |
| 6 | 116867514 | rs9374611 | -0.009 | 0.528 | -0.53 | -0.651 | -0.01 | 0.037 | 0.037 | 0 | 0.61 | 0.27 | 0.71 | 0.9 | 0.61 | 0.65 |

| | | | | | | | | | | | | | | | | |
|---|---|---|---|---|---|---|---|---|---|---|---|---|---|---|---|---|
| 6 | 116875427 | rs7762318 | -0.103 | 0.504 | -0.259 | -0.687 | 0.01 | -0.091 | -0.091 | 0 | 0.62 | 0.42 | 0.73 | 0.9 | 0.61 | 0.65 |
| 6 | 116882385 | rs1033722 | 0.752 | 0.533 | -0.475 | -0.647 | -0.039 | -0.04 | -0.04 | 1.154 | 0.39 | 0.11 | 0.51 | 0.55 | 0.44 | 0.33 |
| 6 | 116885557 | rs17078062 | 1.632 | 0.554 | -0.821 | -0.659 | -0.048 | -0.127 | -0.127 | 0.018 | 0.16 | 0.01 | 0.21 | 0.42 | 0.18 | 0.2 |
| 6 | 116887861 | rs1014324 | 0.644 | 0.502 | -0.537 | -0.674 | 0.01 | -0.019 | -0.019 | 0.715 | 0.21 | 0.12 | 0.15 | 0.34 | 0.17 | 0.24 |
| 6 | 116891193 | rs11153615 | 0.414 | 0.292 | -0.066 | -0.812 | 0.001 | 0 | 0 | 0.858 | 0.42 | 0.38 | 0.59 | 0.56 | 0.44 | 0.41 |
| 6 | 116905487 | rs1983721 | 0.519 | 0.16 | 0.282 | -0.879 | 0.001 | 0.003 | 0.003 | 0 | 0.42 | 0.35 | 0.39 | 0.57 | 0.44 | 0.36 |
| 6 | 116912837 | rs10457308 | 0.607 | 0.163 | 0.191 | -0.877 | -0.035 | -0.033 | -0.033 | 0 | 0.4 | 0.1 | 0.37 | 0.55 | 0.43 | 0.34 |
| 6 | 116913417 | rs1998166 | 0.435 | 0.352 | 0.194 | -0.972 | 0.037 | -0.019 | -0.019 | 0 | 0.05 | 0.16 | 0.2 | 0.01 | 0.08 | 0.08 |
| 6 | 116913543 | rs17201530 | 1.976 | 0.295 | 0.06 | -1.023 | -0.037 | 0.039 | 0.039 | 0.184 | 0.19 | 0.02 | 0.14 | 0.09 | 0.19 | 0.04 |
| 6 | 116928857 | rs9400936 | -0.653 | 0.808 | 0.713 | -0.659 | 0.001 | -0.001 | -0.001 | 0.591 | 0.29 | 0.5 | 0.43 | 0.3 | 0.29 | 0.38 |
| 6 | 116937394 | rs11153618 | 0.277 | 0.78 | 0.729 | -0.6 | -0.001 | -0.013 | -0.013 | 0.21 | 0.22 | 0.11 | 0.21 | 0.28 | 0.21 | 0.24 |
| 6 | 116939487 | rs6568953 | -0.887 | 0.581 | 0.634 | -0.762 | -0.006 | -0.008 | -0.008 | 0 | 0.38 | 0.5 | 0.49 | 0.3 | 0.28 | 0.41 |
| 6 | 116950734 | rs6927567 | 0.379 | 0.653 | 0.641 | -0.756 | -0.002 | -0.013 | -0.013 | 0 | 0.22 | 0.11 | 0.21 | 0.29 | 0.22 | 0.24 |
| 6 | 116954069 | rs6925922 | 0.397 | 0.652 | 0.648 | -0.754 | -0.002 | -0.013 | -0.013 | 0 | 0.21 | 0.11 | 0.21 | 0.29 | 0.21 | 0.24 |
| 6 | 116954186 | rs784136 | 0.869 | 0.7 | 0.635 | -0.748 | -0.042 | 0.036 | 0.036 | 0 | 0.2 | 0.12 | 0.12 | 0.12 | 0.27 | 0.24 |
| 6 | 116956489 | rs2295692 | -1 | 0.499 | 0.762 | -0.718 | -0.004 | -0.006 | -0.006 | 0 | 0.39 | 0.5 | 0.49 | 0.31 | 0.29 | 0.42 |
| 6 | 116958653 | rs9384983 | 0.226 | 0.49 | 0.78 | -0.727 | -0.002 | -0.013 | -0.013 | 0 | 0.23 | 0.11 | 0.21 | 0.28 | 0.22 | 0.25 |
| 6 | 116972117 | rs7743637 | -0.975 | 0.511 | 0.76 | -0.71 | -0.004 | -0.006 | -0.006 | 0.857 | 0.39 | 0.5 | 0.49 | 0.31 | 0.29 | 0.43 |
| 6 | 116973182 | rs4946188 | -0.975 | 0.511 | 0.76 | -0.71 | -0.004 | -0.006 | -0.006 | 0 | 0.39 | 0.51 | 0.49 | 0.31 | 0.29 | 0.42 |
| 6 | 116990510 | rs6568960 | -0.997 | 0.475 | 0.872 | -0.688 | 0.016 | 0.011 | 0.011 | 0 | 0.41 | 0.59 | 0.51 | 0.35 | 0.33 | 0.51 |
| 6 | 116992467 | rs9489002 | 1.322 | 0.521 | 0.784 | -0.658 | -0.008 | -0.025 | -0.025 | 0.003 | 0.11 | 0.09 | 0.03 | 0 | 0.03 | 0.02 |
| 6 | 117011184 | rs9481657 | -0.445 | 1.106 | 1.101 | -0.196 | -0.017 | 0.017 | 0.017 | 0 | 0.26 | 0.45 | 0.29 | 0.3 | 0.22 | 0.36 |
| 6 | 117051934 | rs9400957 | 0.104 | 0.871 | 0.692 | -0.197 | 0 | -0.013 | -0.013 | 0.002 | 0.21 | 0.11 | 0.2 | 0.29 | 0.2 | 0.23 |
| 6 | 117053071 | rs598405 | -1.215 | 0.897 | 0.27 | -0.177 | -0.006 | -0.035 | -0.035 | 0 | 0.33 | 0.83 | 0.5 | 0.38 | 0.3 | 0.51 |
| 6 | 117055764 | rs13590 | -0.409 | 0.797 | 0.333 | -0.146 | -0.016 | 0.03 | 0.03 | 0 | 0.24 | 0.45 | 0.29 | 0.3 | 0.2 | 0.34 |
| 6 | 117065528 | rs9398444 | 0.216 | 0.757 | -0.192 | -0.261 | -0.01 | -0.012 | -0.012 | 0 | 0.15 | 0.1 | 0.2 | 0.36 | 0.21 | 0.28 |
| 6 | 117071569 | rs11153626 | 0.539 | 0.692 | -0.542 | -0.241 | -0.002 | 0 | 0 | 0 | 0.2 | 0.21 | 0.12 | 0.15 | 0.27 | 0.31 |
| 6 | 117072651 | rs1336974 | -0.497 | 0.668 | -0.482 | -0.273 | 0 | -0.066 | -0.066 | 0 | 0.32 | 0.48 | 0.21 | 0.15 | 0.31 | 0.37 |
| 6 | 117074628 | rs11153627 | 0.324 | 1.007 | -1.334 | 0.021 | -0.004 | 0.002 | 0.002 | 0 | 0.21 | 0.22 | 0.13 | 0.15 | 0.27 | 0.31 |
| 6 | 117079000 | rs648210 | 0.36 | 0.453 | -1.651 | 0.238 | -0.044 | -0.117 | -0.117 | 0 | 0.46 | 0.03 | 0.38 | 0.46 | 0.43 | 0.26 |
| 6 | 117080851 | rs4946191 | -0.296 | 0.41 | -0.793 | 0.326 | 0.01 | 0 | 0 | 0 | 0.53 | 0.54 | 0.57 | 0.46 | 0.58 | 0.79 |
| 6 | 117081057 | rs648248 | 0.247 | 0.398 | -0.841 | 0.125 | 0.004 | 0.001 | 0.001 | 0 | 0.53 | 0.38 | 0.4 | 0.45 | 0.38 | 0.33 |
| 6 | 117083720 | rs548101 | 0.201 | 0.231 | -0.576 | -0.109 | -0.001 | 0.001 | 0.001 | 0 | 0.56 | 0.55 | 0.58 | 0.51 | 0.43 | 0.33 |
| 6 | 117086378 | rs654128 | 1.741 | -0.56 | -1.198 | -0.644 | 0.013 | 0.079 | 0.079 | 0 | 0.21 | 0.08 | 0.12 | 0.07 | 0.15 | 0.03 |
| 6 | 117091683 | rs686708 | -0.497 | -0.989 | -1.235 | -0.956 | -0.034 | -0.048 | -0.048 | 0 | 0.67 | 0.76 | 0.67 | 0.51 | 0.48 | 0.44 |
| 6 | 117092558 | rs571296 | 0.856 | -1.088 | -1.129 | -1.109 | 0.04 | -0.052 | -0.052 | 2.963 | 0.47 | 0.16 | 0.38 | 0.49 | 0.35 | 0.26 |
| 6 | 117092892 | rs1567700 | 2.282 | -1.136 | -1.178 | -1.09 | -0.029 | -0.097 | -0.097 | 0.018 | 0.13 | 0.01 | 0.21 | 0.39 | 0.13 | 0.1 |
| 6 | 117093645 | rs540136 | -1.17 | 0.065 | -1.268 | -1.182 | -0.004 | -0.001 | -0.001 | 0.021 | 0.8 | 0.78 | 0.81 | 0.54 | 0.74 | 0.65 |
| 6 | 117105519 | rs587771 | -1.08 | 0.216 | -0.826 | -0.876 | 0.035 | 0.075 | 0.075 | 0 | 0.89 | 0.8 | 0.84 | 0.73 | 0.8 | 0.9 |
| 6 | 117113653 | rs615199 | 1.491 | -0.214 | -0.973 | -1.324 | 0.037 | -0.07 | -0.07 | 0 | 0.41 | 0.15 | 0.37 | 0.52 | 0.31 | 0.26 |
| 6 | 117114025 | rs6901250 | 1.453 | -0.224 | -0.655 | -1.329 | 0.042 | -0.039 | -0.039 | 0 | 0.42 | 0.2 | 0.37 | 0.52 | 0.31 | 0.26 |
| 6 | 117114403 | rs6901971 | -1.096 | -0.282 | -0.478 | -1.308 | -0.002 | 0.001 | 0.001 | 0 | 0.79 | 0.77 | 0.78 | 0.55 | 0.73 | 0.61 |
| 6 | 117120786 | rs7768739 | -0.104 | -0.393 | -0.209 | -1.389 | -0.01 | -0.049 | -0.049 | 0.02 | 0.11 | 0.4 | 0.17 | 0.01 | 0.08 | 0.07 |
| 6 | 117123961 | rs17078383 | 0.793 | -0.387 | -0.05 | -1.383 | 0.031 | 0.003 | 0.003 | 0.095 | 0.07 | 0.31 | 0.16 | 0.01 | 0.07 | 0.06 |
| 6 | 117130704 | rs2274911 | -1.143 | -0.356 | -0.332 | -1.369 | -0.002 | 0.001 | 0.001 | 0 | 0.79 | 0.77 | 0.77 | 0.55 | 0.73 | 0.61 |
| 6 | 117147946 | rs6938235 | 1.525 | -0.354 | -0.412 | -1.331 | 0.025 | -0.064 | -0.064 | 0.204 | 0.43 | 0.15 | 0.38 | 0.52 | 0.34 | 0.26 |
| 6 | 117157774 | rs636252 | -0.833 | -0.157 | -0.208 | -1.233 | 0.009 | -0.001 | -0.001 | 0 | 0.46 | 0.4 | 0.45 | 0.47 | 0.58 | 0.68 |
| 6 | 117192899 | rs1996051 | 1.279 | -0.133 | -0.155 | -1.475 | -0.009 | -0.002 | -0.002 | 0 | 0.22 | 0.16 | 0.16 | 0.02 | 0.3 | 0.25 |
| 6 | 117196211 | rs17078438 | 0.483 | -0.096 | 0.328 | -0.739 | 0.02 | -0.004 | -0.004 | 0.351 | 0.06 | 0.23 | 0.15 | 0.01 | 0.07 | 0.07 |
| 6 | 117208926 | rs339329 | -2.083 | -0.264 | -0.258 | -0.923 | 0.013 | 0.072 | 0.072 | 0.109 | 0.81 | 0.95 | 0.88 | 0.93 | 0.85 | 0.98 |
| 6 | 117210052 | rs339331 | 0.458 | 0.368 | 0.082 | -0.515 | 0.001 | 0.001 | 0.001 | 0.007 | 0.21 | 0.24 | 0.26 | 0.4 | 0.31 | 0.41 |
| 6 | 117218693 | rs339346 | -1.575 | -0.042 | -0.016 | -0.759 | 0.058 | 0.108 | 0.108 | 2.59 | 0.69 | 0.84 | 0.84 | 0.89 | 0.8 | 0.87 |
| 6 | 117220071 | rs170951 | -1.575 | -0.042 | 0.06 | -0.759 | 0.043 | 0.069 | 0.069 | 2.24 | 0.69 | 0.79 | 0.83 | 0.89 | 0.8 | 0.87 |
| 6 | 117238480 | rs1321368 | -1.245 | 0.048 | -0.056 | -0.694 | -0.009 | -0.042 | -0.042 | 0 | 0.65 | 0.66 | 0.7 | 0.9 | 0.74 | 0.83 |
| 6 | 117240593 | rs1321366 | -1.173 | 0.528 | 0.417 | -0.081 | 0.046 | 0.052 | 0.052 | 0.28 | 0.27 | 0.49 | 0.35 | 0.42 | 0.32 | 0.47 |
| 6 | 117244374 | rs7770158 | -0.618 | 0.596 | 0.241 | 0.039 | -0.004 | 0.036 | 0.036 | 0.371 | 0.22 | 0.29 | 0.29 | 0.42 | 0.23 | 0.47 |
| 6 | 117246719 | rs4946206 | -0.618 | 0.596 | 0.241 | 0.074 | -0.004 | 0.036 | 0.036 | 0.371 | 0.22 | 0.3 | 0.29 | 0.42 | 0.23 | 0.47 |
| 6 | 117252759 | rs630695 | -0.618 | 0.596 | 0.235 | 0.066 | -0.003 | 0.033 | 0.033 | 0.371 | 0.22 | 0.29 | 0.29 | 0.42 | 0.23 | 0.47 |
| 6 | 117253609 | rs2180621 | 0.516 | 0.851 | 0.051 | -0.071 | -0.026 | 0.026 | 0.026 | 0 | 0.06 | 0.03 | 0.05 | 0 | 0.1 | 0.02 |
| 6 | 117260435 | rs9320588 | 0.939 | 0.61 | -0.099 | -0.698 | -0.01 | -0.043 | -0.043 | 0 | 0.32 | 0.2 | 0.33 | 0.47 | 0.36 | 0.34 |
| 6 | 117267364 | rs1741692 | 1.754 | 0.4 | 0.068 | -0.81 | 0.01 | -0.002 | -0.002 | 0 | 0.34 | 0.3 | 0.2 | 0.1 | 0.25 | 0.1 |
| 6 | 117278289 | rs1406986 | 1.85 | 0.378 | 0.041 | -0.821 | 0.031 | 0.072 | 0.072 | 4.685 | 0.33 | 0.21 | 0.19 | 0.1 | 0.24 | 0.12 |
| 6 | 117279985 | rs1761834 | 0.226 | 0.392 | 0.283 | -0.637 | -0.003 | 0.107 | 0.107 | 0.381 | 0.78 | 0.62 | 0.72 | 0.57 | 0.77 | 0.53 |
| 6 | 117280229 | rs1761833 | 0.44 | 0.413 | 0.379 | -0.968 | 0 | -0.061 | -0.061 | 0 | 0.46 | 0.48 | 0.37 | 0.08 | 0.4 | 0.19 |
| 6 | 117281179 | rs945134 | 0.08 | 0.605 | 0.272 | -0.341 | 0.121 | 0.214 | 0.214 | 0 | 0.89 | 0.63 | 0.81 | 0.73 | 0.8 | 0.86 |
| 6 | 117288499 | rs783642 | 1.23 | 0.52 | 0.052 | -0.58 | 0.041 | 0.053 | 0.053 | 1.943 | 0.4 | 0.29 | 0.21 | 0.13 | 0.26 | 0.27 |
| 6 | 117303710 | rs1591312 | 1.259 | 0.543 | -0.133 | -0.489 | 0.052 | 0.091 | 0.091 | 3.44 | 0.38 | 0.22 | 0.2 | 0.07 | 0.26 | 0.17 |
| 6 | 117320197 | rs955275 | -0.458 | 0.571 | -0.212 | -0.406 | -0.004 | -0.014 | -0.014 | 0.342 | 0.08 | 0.09 | 0.19 | 0.01 | 0.07 | 0.07 |
| 6 | 117327580 | rs1889501 | 0.346 | 0.83 | -0.279 | -0.735 | 0.007 | -0.049 | -0.049 | 0 | 0.66 | 0.64 | 0.54 | 0.13 | 0.52 | 0.45 |

| | | | | | | | | | | | | | | | | |
|---|---|---|---|---|---|---|---|---|---|---|---|---|---|---|---|---|
| 6 | 117344164 | rs13196452 | 0.708 | 1.113 | -0.549 | -0.362 | -0.03 | 0.013 | 0.013 | 0 | 0.22 | 0.1 | 0.12 | 0.18 | 0.21 | 0.34 |
| 6 | 117349993 | rs573837 | 0.372 | 1.617 | -0.151 | 0.034 | -0.092 | -0.158 | -0.158 | 0 | 0.21 | 0.03 | 0.28 | 0.47 | 0.29 | 0.2 |
| 6 | 117350903 | rs9320589 | 0.244 | 0.912 | -0.062 | -0.423 | 0.042 | -0.012 | -0.012 | 1.059 | 0.71 | 0.63 | 0.55 | 0.31 | 0.54 | 0.72 |
| 6 | 117390783 | rs1537612 | 0.039 | 0.851 | -0.774 | 0.011 | 0.025 | 0.075 | 0.075 | 0 | 0.36 | 0.15 | 0.37 | 0.22 | 0.28 | 0.44 |
| 6 | 117394882 | rs666591 | 0.057 | 0.863 | -0.742 | 0.011 | 0.026 | 0.07 | 0.07 | 0 | 0.36 | 0.17 | 0.37 | 0.22 | 0.28 | 0.44 |
| 6 | 117405247 | rs884855 | -0.352 | 0.972 | -0.609 | 0.203 | 0.01 | 0.003 | 0.003 | 0.115 | 0.41 | 0.34 | 0.53 | 0.38 | 0.38 | 0.47 |
| 6 | 117409095 | rs2498930 | -0.337 | 0.972 | -0.624 | 0.203 | 0 | -0.017 | -0.017 | 0 | 0.39 | 0.19 | 0.52 | 0.38 | 0.38 | 0.47 |
| 6 | 117415499 | rs1567252 | -0.315 | 1.048 | -0.577 | 0.366 | -0.017 | -0.016 | -0.016 | 0 | 0.06 | 0.04 | 0.23 | 0.18 | 0.11 | 0.05 |
| 6 | 117423253 | rs2498924 | 0.278 | 1.147 | -0.505 | 0.341 | 0.13 | 0.158 | 0.158 | 0.013 | 0.8 | 0.57 | 0.75 | 0.58 | 0.66 | 0.78 |
| 6 | 117439930 | rs783373 | -0.392 | 0.651 | -0.609 | 0.35 | -0.012 | 0.006 | 0.006 | 0.313 | 0.42 | 0.34 | 0.56 | 0.38 | 0.45 | 0.51 |
| 6 | 117446167 | rs12201798 | -0.303 | 0.678 | -0.75 | 0.385 | -0.019 | -0.021 | -0.021 | 0.259 | 0.06 | 0.04 | 0.16 | 0.16 | 0.12 | 0.06 |
| 6 | 117448820 | rs783371 | -0.508 | 0.585 | -0.696 | 0.353 | -0.011 | 0.06 | 0.06 | 0 | 0.67 | 0.81 | 0.68 | 0.8 | 0.69 | 0.58 |
| 6 | 117459254 | rs9398450 | 0.409 | 0.688 | -0.581 | 0.511 | -0.072 | -0.086 | -0.086 | 0.298 | 0.13 | 0.02 | 0.14 | 0.41 | 0.22 | 0.16 |
| 6 | 117463853 | rs3851205 | -0.032 | 0.494 | -0.567 | 0.389 | 0.02 | 0.014 | 0.014 | 0 | 0.36 | 0.53 | 0.37 | 0.5 | 0.47 | 0.27 |
| 6 | 117467643 | rs12196324 | 0.951 | 0.508 | -0.574 | 0.487 | -0.064 | -0.011 | -0.011 | 0.31 | 0.11 | 0.01 | 0.09 | 0.07 | 0.19 | 0.1 |
| 6 | 117469900 | rs9374634 | 0.135 | 0.299 | -0.457 | 0.593 | 0.02 | 0.015 | 0.015 | 0 | 0.36 | 0.53 | 0.37 | 0.5 | 0.47 | 0.27 |
| 6 | 117481404 | rs560955 | -0.588 | 0.282 | -0.489 | 0.519 | 0.025 | 0.057 | 0.057 | 0.053 | 0.47 | 0.77 | 0.62 | 0.67 | 0.61 | 0.35 |
| 6 | 117484720 | rs783362 | -0.58 | 0.453 | -0.377 | 0.614 | 0.024 | 0.057 | 0.057 | 0.052 | 0.47 | 0.77 | 0.62 | 0.67 | 0.6 | 0.35 |
| 6 | 117487485 | rs7749803 | -0.588 | 0.453 | -0.376 | 0.614 | 0.025 | 0.06 | 0.06 | 0.053 | 0.47 | 0.78 | 0.62 | 0.67 | 0.61 | 0.35 |
| 6 | 117493311 | rs9387452 | 0.149 | 0.465 | -0.339 | 0.64 | 0.018 | 0.011 | 0.011 | 0 | 0.37 | 0.53 | 0.37 | 0.49 | 0.47 | 0.27 |
| 6 | 117495112 | rs9400983 | -0.283 | 0.472 | -0.217 | 0.646 | 0.085 | 0.411 | 0.411 | 0.469 | 0.06 | 0.31 | 0.12 | 0.36 | 0.08 | 0.09 |
| 6 | 117496146 | rs12199802 | -0.591 | 0.397 | -0.918 | 0.67 | 0.01 | 0.038 | 0.038 | 0.073 | 0.07 | 0.11 | 0.16 | 0.16 | 0.12 | 0.06 |
| 6 | 117502777 | rs4946230 | -0.47 | 0.4 | -1.003 | 0.67 | 0.025 | 0.06 | 0.06 | 0.053 | 0.47 | 0.76 | 0.62 | 0.67 | 0.61 | 0.35 |
| 6 | 117510203 | rs6901720 | 0.275 | 0.417 | -0.997 | 0.679 | 0.018 | 0.011 | 0.011 | 0 | 0.37 | 0.53 | 0.37 | 0.49 | 0.47 | 0.27 |
| 6 | 117519509 | rs1534217 | -0.097 | 0.511 | -1.186 | 0.557 | -0.013 | -0.005 | -0.005 | 0 | 0.05 | 0.03 | 0.15 | 0.16 | 0.12 | 0.06 |
| 6 | 117522819 | rs6915129 | -0.397 | 0.488 | -1.194 | 0.532 | 0.023 | 0.058 | 0.058 | 0.004 | 0.46 | 0.77 | 0.62 | 0.67 | 0.6 | 0.33 |
| 6 | 117525530 | rs12191105 | -0.428 | 0.523 | -1.135 | 0.711 | -0.002 | 0.012 | 0.012 | 0 | 0.06 | 0.05 | 0.15 | 0.16 | 0.12 | 0.06 |
| 6 | 117528651 | rs9374640 | 0.078 | 0.448 | -1.222 | 0.241 | 0 | 0 | 0 | 0 | 0.52 | 0.52 | 0.45 | 0.48 | 0.57 | 0.35 |
| 6 | 117532372 | rs1305430 | 0.142 | 2.024 | -1.253 | 0.071 | 0.023 | -0.039 | -0.039 | 0 | 0.1 | 0.13 | 0.1 | 0 | 0.13 | 0.09 |
| 6 | 117536229 | rs1118092 | 0.437 | 1.154 | -1.731 | 0.456 | 0.053 | 0.108 | 0.108 | 0.216 | 0.44 | 0.09 | 0.48 | 0.69 | 0.37 | 0.4 |
| 6 | 117539456 | rs2016523 | 0.467 | 1.184 | -1.704 | 0.471 | 0.056 | 0.107 | 0.107 | 0.336 | 0.44 | 0.11 | 0.49 | 0.69 | 0.37 | 0.39 |
| 6 | 117540701 | rs9372476 | 0.513 | 1.18 | -1.705 | 0.469 | 0.036 | 0.095 | 0.095 | 0.479 | 0.43 | 0.06 | 0.48 | 0.69 | 0.37 | 0.39 |
| 6 | 117542521 | rs9400993 | 0.815 | 1.185 | -1.707 | 0.607 | 0.081 | -0.124 | -0.124 | 2.481 | 0.4 | 0.05 | 0.32 | 0.51 | 0.28 | 0.35 |
| 6 | 117544308 | rs1880781 | 0.754 | 1.187 | -1.706 | 0.608 | 0.093 | -0.089 | -0.089 | 0.058 | 0.41 | 0.1 | 0.33 | 0.51 | 0.28 | 0.35 |
| 6 | 117556500 | rs694859 | 0.039 | 1.207 | -1.668 | -0.44 | -0.064 | -0.006 | -0.006 | 0 | 0.11 | 0.01 | 0.09 | 0.05 | 0.19 | 0.15 |
| 6 | 117560865 | rs620590 | -0.257 | 1.191 | -1.542 | -0.391 | -0.028 | 0.102 | 0.102 | 0 | 0.59 | 0.24 | 0.56 | 0.74 | 0.58 | 0.55 |
| 6 | 117564654 | rs669995 | -0.024 | 1.124 | -1.606 | -0.426 | -0.004 | 0.002 | 0.002 | 0 | 0.16 | 0.19 | 0.1 | 0.06 | 0.2 | 0.16 |
| 6 | 117565453 | rs9400994 | 0.033 | -0.015 | -1.624 | 0.219 | 0.06 | 0.067 | 0.067 | 0.431 | 0.43 | 0.05 | 0.36 | 0.67 | 0.37 | 0.38 |
| 6 | 117573335 | rs398414 | -0.76 | 0.189 | 0.088 | 0.624 | 0.032 | 0.122 | 0.122 | 0 | 0.08 | 0.38 | 0.07 | 0.17 | 0.07 | 0.27 |
| 6 | 117579324 | rs803595 | -0.092 | 0.365 | -0.39 | 0.194 | -0.001 | 0.087 | 0.087 | 0.387 | 0.51 | 0.73 | 0.41 | 0.29 | 0.43 | 0.38 |
| 6 | 117584031 | rs451411 | -0.296 | 0.204 | 0.162 | -0.39 | 0.011 | -0.031 | -0.031 | 0 | 0.29 | 0.51 | 0.28 | 0.19 | 0.26 | 0.24 |
| 6 | 117585553 | rs803596 | 0.154 | 0.397 | 0.718 | -0.505 | 0 | -0.006 | -0.006 | 2.572 | 0.17 | 0.17 | 0.2 | 0.06 | 0.16 | 0.15 |
| 6 | 117587568 | rs706950 | -0.855 | 0.327 | 1.15 | -0.544 | 0.005 | -0.025 | -0.025 | 0 | 0.3 | 0.58 | 0.29 | 0.19 | 0.26 | 0.24 |
| 6 | 117591755 | rs13194610 | -0.382 | 0.189 | 1.096 | -0.426 | 0.027 | 0.03 | 0.03 | 0 | 0.12 | 0.38 | 0.1 | 0.13 | 0.11 | 0.09 |
| 6 | 117594645 | rs3822947 | 0.013 | 0.171 | 0.599 | -0.433 | -0.036 | -0.039 | -0.039 | 0 | 0.09 | 0.01 | 0.06 | 0.13 | 0.1 | 0.1 |
| 6 | 117596993 | rs117083 | -0.583 | 0.309 | 0.659 | -0.418 | -0.002 | -0.004 | -0.004 | 0 | 0.43 | 0.54 | 0.39 | 0.39 | 0.4 | 0.36 |
| 6 | 117599632 | rs441184 | -0.603 | 0.692 | 0.63 | -0.054 | 0 | 0 | 0 | 0 | 0.43 | 0.54 | 0.39 | 0.46 | 0.44 | 0.42 |
| 6 | 117600609 | rs491902 | 0.651 | 0.863 | 0.416 | -0.241 | -0.003 | 0.011 | 0.011 | 0 | 0.52 | 0.32 | 0.57 | 0.54 | 0.51 | 0.52 |
| 6 | 117605749 | rs929122 | 0.055 | 1.033 | 0.264 | -0.247 | 0.032 | -0.084 | -0.084 | 0 | 0.23 | 0.5 | 0.19 | 0.06 | 0.26 | 0.14 |
| 6 | 117605991 | rs13201929 | 0.415 | 1.028 | -0.223 | 0.044 | -0.046 | -0.062 | -0.062 | 0 | 0.11 | 0.01 | 0.08 | 0.17 | 0.12 | 0.1 |
| 6 | 117608318 | rs544047 | -0.278 | 0.695 | -0.634 | -0.033 | -0.001 | 0.004 | 0.004 | 0 | 0.48 | 0.25 | 0.54 | 0.52 | 0.56 | 0.4 |
| 6 | 117609387 | rs929121 | -0.295 | 0.297 | -0.739 | 0.013 | 0.011 | -0.009 | -0.009 | 0 | 0.65 | 0.69 | 0.65 | 0.43 | 0.72 | 0.47 |
| 6 | 117609529 | rs2778 | 0.236 | 0.514 | -1.19 | 0.423 | -0.029 | -0.035 | -0.035 | 0 | 0.36 | 0.1 | 0.42 | 0.39 | 0.38 | 0.23 |
| 6 | 117610618 | rs9489115 | 1.22 | 0.364 | -1.258 | 0.079 | 0.029 | 0.035 | 0.035 | 1.208 | 0.16 | 0.04 | 0.18 | 0.05 | 0.09 | 0.02 |
| 6 | 117619775 | rs3798368 | 0.696 | 0.834 | -1.015 | 0.688 | -0.008 | -0.036 | -0.036 | 0 | 0.32 | 0.13 | 0.4 | 0.41 | 0.34 | 0.29 |
| 6 | 117620139 | rs485768 | -0.067 | 1.052 | -1.063 | 0.783 | -0.056 | -0.038 | -0.038 | 0.04 | 0.15 | 0.03 | 0.16 | 0.16 | 0.22 | 0.15 |
| 6 | 117622188 | rs529156 | -0.067 | 1.052 | -1.046 | 0.783 | -0.045 | -0.029 | -0.029 | 0 | 0.15 | 0.04 | 0.17 | 0.16 | 0.22 | 0.15 |
| 6 | 117622233 | rs529038 | -0.067 | 1.052 | -1.097 | 0.783 | -0.056 | -0.038 | -0.038 | 0.089 | 0.15 | 0.03 | 0.16 | 0.16 | 0.22 | 0.15 |
| 6 | 117624768 | rs172409 | 0.015 | 1.087 | -0.874 | 0.836 | -0.028 | -0.02 | -0.02 | 0 | 0.6 | 0.76 | 0.6 | 0.58 | 0.55 | 0.68 |
| 6 | 117628364 | rs3777955 | 0.128 | 0.878 | -1.044 | 0.248 | -0.013 | -0.034 | -0.034 | 0 | 0.24 | 0.27 | 0.14 | 0.1 | 0.19 | 0.28 |
| 6 | 117628851 | rs507280 | -0.049 | 1.297 | -1.354 | 0.773 | -0.05 | -0.024 | -0.024 | 0 | 0.11 | 0.01 | 0.08 | 0.1 | 0.15 | 0.09 |
| 6 | 117630696 | rs12664183 | 0.335 | 0.878 | -1.356 | 0.848 | 0.001 | -0.005 | -0.005 | 0 | 0.3 | 0.1 | 0.37 | 0.35 | 0.3 | 0.3 |
| 6 | 117631431 | rs552183 | -0.336 | 0.604 | -0.924 | 0.801 | 0 | 0.001 | 0.001 | 0 | 0.15 | 0.13 | 0.11 | 0.12 | 0.15 | 0.08 |
| 6 | 117633847 | rs3777958 | 0.137 | 0.726 | -0.693 | 0.508 | 0.003 | -0.03 | -0.03 | 0 | 0.21 | 0.33 | 0.18 | 0.09 | 0.22 | 0.26 |
| 6 | 117638124 | rs530066 | 0.182 | 0.484 | -0.554 | 0.564 | -0.002 | 0.005 | 0.005 | 0 | 0.16 | 0.11 | 0.12 | 0.15 | 0.16 | 0.18 |
| 6 | 117639104 | rs499992 | -0.404 | 0.577 | -0.291 | 0.597 | 0.002 | -0.005 | -0.005 | 0 | 0.91 | 0.92 | 0.93 | 0.94 | 0.9 | 0.93 |
| 6 | 117639419 | rs497186 | 0.558 | 0.585 | -0.39 | 0.569 | -0.036 | -0.006 | -0.006 | 0.334 | 0.13 | 0.01 | 0.1 | 0.1 | 0.14 | 0.11 |
| 6 | 117640002 | rs9400997 | 1.714 | 0.456 | -0.163 | 0.727 | -0.013 | 0.026 | 0.026 | 0.239 | 0.36 | 0.29 | 0.39 | 0.24 | 0.42 | 0.35 |
| 6 | 117646772 | rs508278 | 0.227 | 0.585 | -0.699 | 1.055 | -0.001 | 0 | 0 | 0 | 0.17 | 0.19 | 0.17 | 0.2 | 0.16 | 0.21 |

| | | | | | | | | | | | | | | | | |
|---|---|---|---|---|---|---|---|---|---|---|---|---|---|---|---|---|
| 6 | 117659005 | rs1000153 | 0.551 | 0.543 | -0.876 | 1.066 | 0 | -0.001 | -0.001 | 0 | 0.19 | 0.16 | 0.15 | 0.1 | 0.2 | 0.26 |
| 6 | 117663047 | rs17079069 | 0.656 | 0.425 | -1.714 | 1.444 | -0.004 | -0.103 | -0.103 | 0 | 0.15 | 0.02 | 0.2 | 0.32 | 0.11 | 0.11 |
| 6 | 117663364 | rs3756891 | 0.429 | 0.79 | -1.424 | 1.243 | -0.01 | 0.011 | 0.011 | 0 | 0.13 | 0.05 | 0.11 | 0.1 | 0.15 | 0.11 |
| 6 | 117666413 | rs3777969 | 0.328 | 0.591 | -1.194 | 1.562 | 0.005 | -0.025 | -0.025 | 0.284 | 0.14 | 0.01 | 0.11 | 0.12 | 0.11 | 0.16 |
| 6 | 117668989 | rs2157535 | 0.272 | 0.981 | -0.563 | 1.772 | 0.001 | -0.019 | -0.019 | 0.767 | 0.78 | 0.85 | 0.73 | 0.57 | 0.79 | 0.84 |
| 6 | 117671962 | rs9398454 | -0.752 | 1.166 | -0.312 | 1.658 | 0.02 | 0.058 | 0.058 | 0 | 0.07 | 0.09 | 0.1 | 0.16 | 0.09 | 0.17 |
| 6 | 117679916 | rs9489138 | 0.569 | 1.154 | -0.746 | 1.314 | -0.03 | -0.017 | -0.017 | 0 | 0.91 | 1 | 0.93 | 0.93 | 0.91 | 0.93 |
| 6 | 117684882 | rs3777971 | -0.094 | 0.877 | -1.043 | 1.297 | 0.002 | -0.001 | -0.001 | 0 | 0.19 | 0.13 | 0.09 | 0.22 | 0.16 | 0.31 |
| 6 | 117684992 | rs1321807 | -0.965 | 0.75 | -0.953 | 1.328 | -0.003 | 0.032 | 0.032 | 0.153 | 0.51 | 0.7 | 0.6 | 0.59 | 0.5 | 0.61 |
| 6 | 117699805 | rs9688879 | 0.595 | 0.763 | -1.114 | 1.335 | -0.025 | 0.001 | 0.001 | 0 | 0.13 | 0.01 | 0.19 | 0.07 | 0.12 | 0.06 |
| 6 | 117708971 | rs9489143 | 1.07 | -0.213 | -1.416 | 0.548 | 0.06 | 0.1 | 0.1 | 12.185 | 0.21 | 0.03 | 0.08 | 0.03 | 0.09 | 0.11 |
| 6 | 117716199 | rs3798384 | -0.035 | -0.276 | -1.416 | 0.634 | 0.007 | -0.011 | -0.011 | 0 | 0.38 | 0.67 | 0.49 | 0.52 | 0.32 | 0.45 |
| 6 | 117718303 | rs9401003 | -0.317 | -0.27 | -0.919 | 0.553 | -0.007 | -0.008 | -0.008 | 0 | 0.37 | 0.58 | 0.3 | 0.27 | 0.25 | 0.27 |
| 6 | 117718813 | rs3798387 | 0.858 | 0.297 | -0.698 | 0.964 | -0.02 | -0.01 | -0.01 | 0 | 0.09 | 0.03 | 0.09 | 0.07 | 0.09 | 0.05 |
| 6 | 117722069 | rs2354342 | 0.365 | 0.232 | -0.497 | 0.842 | 0 | -0.003 | -0.003 | 0.356 | 0.25 | 0.23 | 0.14 | 0.21 | 0.27 | 0.44 |
| 6 | 117724379 | rs2243380 | 0.883 | 0.438 | -0.888 | 0.924 | -0.026 | 0.005 | 0.005 | 0 | 0.08 | 0 | 0.09 | 0.04 | 0.09 | 0.05 |
| 6 | 117724462 | rs2243379 | -0.747 | 0.147 | -1.253 | 0.869 | -0.113 | -0.102 | -0.102 | 0 | 0.56 | 0.93 | 0.52 | 0.56 | 0.49 | 0.72 |
| 6 | 117725448 | rs1998206 | -0.079 | 0.136 | -0.958 | 0.896 | 0.039 | 0.075 | 0.075 | 0 | 0.06 | 0.16 | 0.14 | 0.18 | 0.15 | 0.2 |
| 6 | 117728118 | rs765516 | -0.903 | 0.134 | -0.827 | 0.895 | -0.05 | -0.056 | -0.056 | 0.647 | 0.55 | 0.77 | 0.42 | 0.48 | 0.5 | 0.7 |
| 6 | 117730819 | rs2273601 | 0.141 | 0.079 | -0.662 | 0.82 | -0.007 | -0.008 | -0.008 | 0.022 | 0.39 | 0.51 | 0.19 | 0.27 | 0.26 | 0.48 |
| 6 | 117733717 | rs1407184 | -0.376 | 0.059 | -0.681 | 0.828 | 0.015 | -0.007 | -0.007 | 0 | 0.45 | 0.7 | 0.33 | 0.44 | 0.41 | 0.65 |
| 6 | 117734746 | rs9489152 | -0.659 | 0.1 | -0.638 | 0.916 | 0.002 | 0.045 | 0.045 | 0.011 | 0.52 | 0.59 | 0.3 | 0.31 | 0.45 | 0.6 |
| 6 | 117741446 | rs12332810 | -0.275 | 0.151 | -0.71 | 0.785 | -0.005 | 0.006 | 0.006 | 0 | 0.15 | 0.16 | 0.12 | 0.06 | 0.19 | 0.18 |
| 6 | 117742298 | rs9489160 | 0.324 | 0.177 | -0.59 | 0.901 | 0.004 | -0.009 | -0.009 | 0 | 0.05 | 0.06 | 0.05 | 0 | 0.11 | 0.12 |
| 6 | 117754038 | rs17079166 | -0.119 | 0.173 | -0.564 | 0.899 | 0.03 | -0.028 | -0.028 | 0 | 0.06 | 0.17 | 0.06 | 0.01 | 0.11 | 0.12 |
| 6 | 117759128 | rs9489181 | -0.119 | 0.173 | -0.565 | 0.899 | 0.033 | -0.028 | -0.028 | 0 | 0.06 | 0.17 | 0.06 | 0.01 | 0.11 | 0.12 |
| 6 | 117760579 | rs9372480 | -0.662 | 0.131 | -0.754 | 1.004 | -0.123 | -0.097 | -0.097 | 0 | 0.56 | 0.9 | 0.44 | 0.53 | 0.5 | 0.74 |
| 6 | 117772143 | rs6932793 | -0.535 | 0.203 | -0.595 | 0.812 | -0.007 | 0.006 | 0.006 | 0 | 0.08 | 0.06 | 0.07 | 0.02 | 0.14 | 0.12 |
| 6 | 117785696 | rs6942067 | -0.355 | -0.124 | -0.513 | 0.993 | 0.01 | 0.011 | 0.011 | 0.582 | 0.48 | 0.51 | 0.27 | 0.34 | 0.35 | 0.52 |
| 6 | 117787897 | rs7749229 | -0.153 | -0.129 | -0.502 | 0.991 | 0.002 | 0.002 | 0.002 | 0.367 | 0.45 | 0.5 | 0.25 | 0.32 | 0.32 | 0.51 |
| 6 | 117793798 | rs2104064 | 0.377 | -0.325 | 0.144 | 0.954 | -0.003 | 0.001 | 0.001 | 0 | 0.3 | 0.33 | 0.17 | 0.27 | 0.26 | 0.39 |
| 6 | 117799572 | rs7746536 | 0.164 | -0.416 | 0.283 | 0.898 | 0 | -0.001 | -0.001 | 0 | 0.32 | 0.3 | 0.19 | 0.3 | 0.28 | 0.41 |
| 6 | 117809031 | rs6911915 | -0.554 | -0.312 | 0.287 | 0.837 | -0.01 | -0.021 | -0.021 | 0 | 0.42 | 0.75 | 0.36 | 0.44 | 0.43 | 0.54 |
| 6 | 117814437 | rs13195478 | 0.506 | -0.284 | 0.225 | 0.896 | 0 | -0.001 | -0.001 | 0 | 0.3 | 0.27 | 0.17 | 0.28 | 0.25 | 0.41 |
| 6 | 117816093 | rs4946259 | -1.238 | -0.155 | -0.123 | 0.783 | -0.119 | -0.067 | -0.067 | 0 | 0.59 | 0.92 | 0.55 | 0.61 | 0.54 | 0.74 |
| 6 | 117817555 | rs17574500 | 1.01 | 0.47 | -0.429 | 0.568 | -0.044 | -0.013 | -0.013 | 0 | 0.1 | 0.02 | 0.17 | 0.08 | 0.14 | 0.06 |
| 6 | 117817775 | rs9320605 | 0.962 | 0.45 | -0.226 | 0.395 | 0.025 | 0.043 | 0.043 | 2.269 | 0.15 | 0.04 | 0.07 | 0.04 | 0.08 | 0.09 |
| 6 | 117819718 | rs17574630 | 1.346 | 0.031 | -0.608 | 0.383 | -0.074 | -0.009 | -0.009 | 0.475 | 0.16 | 0 | 0.11 | 0.09 | 0.21 | 0.2 |
| 6 | 117822251 | rs11153664 | -0.351 | 0.163 | -0.318 | 0.473 | -0.011 | -0.048 | -0.048 | 0 | 0.24 | 0.52 | 0.23 | 0.18 | 0.23 | 0.22 |
| 6 | 117822993 | rs4946260 | -1.088 | -0.06 | -0.437 | 0.54 | -0.106 | -0.097 | -0.097 | 0.098 | 0.58 | 0.89 | 0.44 | 0.47 | 0.52 | 0.64 |
| 6 | 117826504 | rs9766914 | 0.185 | -0.116 | -0.349 | 0.118 | 0.017 | -0.057 | -0.057 | 0 | 0.37 | 0.43 | 0.27 | 0.11 | 0.44 | 0.37 |
| 6 | 117831565 | rs9688605 | 0.696 | -0.216 | -0.475 | -0.134 | -0.037 | 0.085 | 0.085 | 0.003 | 0.16 | 0.03 | 0.08 | 0.03 | 0.16 | 0.08 |
| 6 | 117833868 | rs9767548 | -0.379 | -0.286 | -0.259 | 0.158 | 0.053 | 0.156 | 0.156 | 0.852 | 0.05 | 0.35 | 0.06 | 0.09 | 0.01 | 0.04 |
| 6 | 117834444 | rs210610 | -0.559 | -0.256 | -0.297 | 0.229 | -0.013 | -0.019 | -0.019 | 0 | 0.58 | 0.46 | 0.42 | 0.28 | 0.64 | 0.65 |
| 6 | 117844072 | rs210648 | -1.36 | -0.231 | -0.643 | 0.353 | -0.004 | -0.103 | -0.103 | 0 | 0.66 | 0.9 | 0.51 | 0.41 | 0.69 | 0.72 |
| 6 | 117853061 | rs11153668 | -0.784 | -0.249 | -0.355 | 0.292 | -0.011 | -0.01 | -0.01 | 0 | 0.29 | 0.54 | 0.24 | 0.3 | 0.24 | 0.34 |
| 6 | 117854605 | rs210651 | 0.209 | -0.039 | -0.259 | 0.361 | 0.01 | -0.037 | -0.037 | 0 | 0.37 | 0.37 | 0.28 | 0.11 | 0.45 | 0.37 |
| 6 | 117856759 | rs4946263 | -1.148 | -0.049 | 0.232 | 0.609 | 0 | 0.002 | 0.002 | 0.288 | 0.27 | 0.62 | 0.25 | 0.31 | 0.24 | 0.34 |
| 6 | 117857488 | rs3756894 | -0.536 | -0.353 | 0.217 | 1.037 | -0.001 | 0.162 | 0.162 | 0.205 | 0.07 | 0.25 | 0.12 | 0.24 | 0.04 | 0.26 |
| 6 | 117858304 | rs17079343 | -0.062 | -0.627 | 0.067 | 0.895 | -0.052 | 0.044 | 0.044 | 0 | 0.16 | 0.25 | 0.14 | 0.28 | 0.04 | 0.31 |
| 6 | 117859198 | rs9374668 | -0.088 | -0.4 | 0.173 | 1.022 | -0.007 | 0.087 | 0.087 | 0 | 0.08 | 0.18 | 0.26 | 0.24 | 0.09 | 0.12 |
| 6 | 117863175 | rs4946265 | 0.108 | -0.317 | 0.113 | 0.426 | -0.014 | 0.058 | 0.058 | 0.116 | 0.45 | 0.31 | 0.39 | 0.18 | 0.52 | 0.63 |
| 6 | 117868051 | rs389663 | -1.292 | -0.503 | -0.061 | 0.425 | -0.016 | -0.066 | -0.066 | 0 | 0.72 | 0.87 | 0.74 | 0.54 | 0.72 | 0.83 |
| 6 | 117869424 | rs929059 | 0.695 | -0.621 | 0.021 | 0.32 | 0.002 | 0.028 | 0.028 | 0 | 0.16 | 0.09 | 0.08 | 0.03 | 0.13 | 0.07 |
| 6 | 117869850 | rs9489217 | -0.338 | -0.675 | 0.58 | 0.334 | -0.001 | -0.001 | -0.001 | 0 | 0.55 | 0.61 | 0.66 | 0.51 | 0.59 | 0.76 |
| 6 | 117870892 | rs8088 | -0.354 | -0.65 | 0.578 | 0.332 | -0.001 | 0 | 0 | 0 | 0.55 | 0.61 | 0.66 | 0.51 | 0.6 | 0.76 |
| 6 | 117871548 | rs10457315 | 0.697 | -0.672 | 0.372 | 0.3 | -0.02 | 0.09 | 0.09 | 0.101 | 0.44 | 0.26 | 0.39 | 0.18 | 0.52 | 0.64 |
| 6 | 117873991 | rs13202536 | 1.295 | -0.616 | 0.287 | 0.494 | -0.004 | -0.021 | -0.021 | 0 | 0.06 | 0.01 | 0.15 | 0.07 | 0.06 | 0.03 |
| 6 | 117880774 | rs10782187 | -0.903 | -0.494 | 0.537 | 0.489 | -0.007 | 0.027 | 0.027 | 0 | 0.3 | 0.43 | 0.36 | 0.4 | 0.25 | 0.23 |
| 6 | 117882209 | rs1045855 | 0.798 | -0.302 | 0.19 | -0.019 | -0.001 | -0.019 | -0.019 | 0 | 0.16 | 0.16 | 0.12 | 0.01 | 0.2 | 0.16 |
| 6 | 117900944 | rs10484299 | -0.07 | -0.278 | -0.068 | 0.06 | -0.011 | 0.016 | 0.016 | 0 | 0.25 | 0.33 | 0.34 | 0.36 | 0.2 | 0.19 |
| 6 | 117901048 | rs10484298 | 1.652 | -0.328 | -0.232 | -0.01 | -0.08 | 0.045 | 0.045 | 0 | 0.26 | 0.05 | 0.23 | 0.15 | 0.3 | 0.43 |
| 6 | 117901283 | rs210643 | -2.152 | -0.288 | -0.13 | 0.121 | -0.056 | -0.156 | -0.156 | 0 | 0.77 | 0.98 | 0.78 | 0.58 | 0.77 | 0.88 |
| 6 | 117916942 | rs9374670 | -0.728 | -0.238 | 0.031 | 0.175 | -0.011 | 0.025 | 0.025 | 0 | 0.3 | 0.43 | 0.36 | 0.4 | 0.25 | 0.24 |
| 6 | 117923679 | rs3752847 | 0.377 | -0.233 | -0.019 | 0.329 | -0.017 | 0.074 | 0.074 | 0 | 0.1 | 0.24 | 0.24 | 0.21 | 0.07 | 0.13 |
| 6 | 117924602 | rs12200953 | 1.05 | -0.199 | -0.102 | 0.273 | 0.002 | 0.012 | 0.012 | 0 | 0.15 | 0.1 | 0.08 | 0.04 | 0.13 | 0.07 |
| 6 | 117953103 | rs12174680 | -0.346 | -0.186 | -0.017 | 0.106 | -0.003 | 0.082 | 0.082 | 0 | 0.14 | 0.25 | 0.28 | 0.37 | 0.11 | 0.16 |
| 6 | 117955898 | rs4946273 | -0.434 | -0.239 | 0.136 | 0.093 | -0.001 | -0.008 | -0.008 | 0 | 0.54 | 0.56 | 0.59 | 0.81 | 0.47 | 0.36 |
| 6 | 117965710 | rs1967194 | -0.499 | -0.241 | -0.058 | 0.092 | -0.012 | 0.058 | 0.058 | 0.068 | 0.54 | 0.64 | 0.6 | 0.8 | 0.47 | 0.36 |

| | | | | | | | | | | | | | | | | |
|---|---|---|---|---|---|---|---|---|---|---|---|---|---|---|---|---|
| 6 | 117976376 | rs4443534 | 1.797 | -0.187 | -0.898 | 0.107 | -0.009 | -0.076 | -0.076 | 0 | 0.24 | 0.09 | 0.23 | 0.39 | 0.23 | 0.12 |
| 6 | 117988747 | rs11967019 | 0.555 | -0.035 | -0.79 | 0.471 | -0.014 | 0.002 | 0.002 | 0 | 0.08 | 0.18 | 0.16 | 0.07 | 0.05 | 0.06 |
| 6 | 117990153 | rs11153692 | -0.539 | 0.103 | -0.858 | 0.434 | -0.003 | 0.009 | 0.009 | 0 | 0.29 | 0.36 | 0.36 | 0.41 | 0.24 | 0.23 |
| 6 | 117994158 | rs13211542 | 0.705 | 0.356 | -1.386 | 0.197 | -0.074 | 0.036 | 0.036 | 0 | 0.26 | 0.05 | 0.23 | 0.16 | 0.3 | 0.42 |
| 6 | 117998601 | rs4368845 | 0.745 | -0.037 | -1.378 | 0.404 | -0.065 | 0.04 | 0.04 | 0.68 | 0.4 | 0.08 | 0.43 | 0.26 | 0.53 | 0.21 |
| 6 | 117999813 | rs11968450 | 0.144 | -0.257 | -1.124 | 0.608 | 0 | -0.004 | -0.004 | 0.024 | 0.07 | 0.09 | 0.04 | 0 | 0.06 | 0.07 |
| 6 | 118001311 | rs9767278 | 0.144 | -0.257 | -1.124 | 0.608 | 0 | -0.004 | -0.004 | 0 | 0.07 | 0.09 | 0.04 | 0 | 0.06 | 0.07 |
| 6 | 118005198 | rs11751090 | 0.343 | -0.562 | -1.38 | 0.366 | 0.002 | 0.033 | 0.033 | 0 | 0.11 | 0.07 | 0.07 | 0.01 | 0.1 | 0.09 |
| 6 | 118023316 | rs1541317 | 1.159 | -0.543 | -1.449 | 0.405 | -0.116 | -0.001 | -0.001 | 1.318 | 0.39 | 0.03 | 0.43 | 0.26 | 0.54 | 0.22 |
| 6 | 118026126 | rs2498586 | 0.68 | -0.8 | -0.769 | 0.501 | -0.041 | 0.059 | 0.059 | 0 | 0.28 | 0.42 | 0.36 | 0.62 | 0.22 | 0.49 |
| 6 | 118039508 | rs473494 | -1.355 | -0.694 | -1.606 | 0.297 | -0.058 | 0.044 | 0.044 | 0.756 | 0.61 | 0.9 | 0.57 | 0.74 | 0.46 | 0.78 |
| 6 | 118049147 | rs11755703 | 0.27 | -0.619 | -1.579 | 0.482 | 0.006 | 0.001 | 0.001 | 0.495 | 0.12 | 0.09 | 0.05 | 0.03 | 0.06 | 0.06 |
| 6 | 118053656 | rs2501624 | 0.552 | -0.297 | -1.491 | 0.766 | -0.003 | 0.004 | 0.004 | 0 | 0.25 | 0.24 | 0.33 | 0.39 | 0.21 | 0.42 |
| 6 | 118063402 | rs1850624 | 0.409 | 0.423 | -1.562 | 1.036 | 0.02 | 0.001 | 0.001 | 0 | 0.39 | 0.24 | 0.41 | 0.42 | 0.39 | 0.58 |
| 6 | 118076122 | rs636205 | -0.61 | 0.644 | -1.811 | 0.55 | -0.115 | 0.049 | 0.049 | 0 | 0.58 | 0.91 | 0.6 | 0.73 | 0.57 | 0.76 |
| 6 | 118080510 | rs1607369 | 0.058 | 0.535 | -1.503 | 0.461 | -0.001 | -0.008 | -0.008 | 0 | 0.11 | 0.11 | 0.06 | 0.03 | 0.09 | 0.06 |
| 6 | 118091629 | rs2516104 | 0.035 | 0.481 | -1.746 | 0.88 | 0.021 | -0.033 | -0.033 | 0 | 0.4 | 0.24 | 0.42 | 0.53 | 0.4 | 0.64 |
| 6 | 118098501 | rs509931 | -0.174 | 0.88 | -2.046 | 1.078 | -0.068 | -0.152 | -0.152 | 0 | 0.71 | 0.95 | 0.63 | 0.61 | 0.71 | 0.86 |
| 6 | 118103840 | rs622195 | -0.062 | 0.684 | -1.658 | 0.992 | -0.003 | -0.016 | -0.016 | 0 | 0.6 | 0.74 | 0.64 | 0.57 | 0.57 | 0.79 |
| 6 | 118107037 | rs667994 | 0.225 | 0.441 | -1.037 | 0.659 | 0 | -0.055 | -0.055 | 0.67 | 0.11 | 0.42 | 0.06 | 0.02 | 0.05 | 0.06 |
| 6 | 118108940 | rs7742769 | 0.232 | 0.712 | -1.717 | 1.168 | -0.099 | 0.114 | 0.114 | 0.174 | 0.42 | 0.1 | 0.36 | 0.26 | 0.46 | 0.62 |
| 6 | 118113186 | rs7754471 | 0.338 | 0.8 | -1.79 | 0.544 | -0.054 | 0.074 | 0.074 | 0.119 | 0.38 | 0.09 | 0.32 | 0.25 | 0.41 | 0.6 |
| 6 | 118116482 | rs1518358 | -0.017 | 1.035 | -1.264 | 0.902 | -0.002 | -0.01 | -0.01 | 1.949 | 0.41 | 0.32 | 0.51 | 0.43 | 0.39 | 0.4 |
| 6 | 118116823 | rs601136 | 0.061 | 0.641 | -1.19 | 0.684 | -0.016 | -0.022 | -0.022 | 0.39 | 0.57 | 0.74 | 0.43 | 0.44 | 0.56 | 0.71 |
| 6 | 118117141 | rs1518357 | 0.457 | 0.666 | -1.157 | 0.786 | -0.001 | -0.003 | -0.003 | 0 | 0.13 | 0.1 | 0.22 | 0.16 | 0.2 | 0.32 |
| 6 | 118117246 | rs11755280 | 1.019 | 0.667 | -1.403 | 0.579 | -0.054 | -0.038 | -0.038 | 0 | 0.28 | 0.08 | 0.16 | 0.3 | 0.3 | 0.36 |
| 6 | 118117328 | rs1878728 | -0.46 | 0.349 | -0.596 | 0.644 | -0.001 | -0.001 | -0.001 | 0 | 0.41 | 0.36 | 0.51 | 0.28 | 0.34 | 0.29 |
| 6 | 118117602 | rs6903494 | -0.499 | 0.196 | -0.499 | 0.732 | -0.002 | -0.004 | -0.004 | 0.38 | 0.39 | 0.36 | 0.46 | 0.27 | 0.29 | 0.24 |
| 6 | 118136554 | rs9481736 | 0.489 | 0.073 | -0.173 | 0.703 | 0.061 | 0.112 | 0.112 | 0.646 | 0.95 | 0.65 | 0.96 | 0.88 | 0.97 | 0.98 |
| 6 | 118143197 | rs11153705 | 1.015 | 0.073 | -0.543 | 0.701 | -0.156 | -0.02 | -0.02 | 0 | 0.42 | 0.03 | 0.35 | 0.28 | 0.53 | 0.69 |
| 6 | 118148148 | rs2678772 | -0.128 | 0.04 | -0.321 | 0.763 | -0.004 | -0.003 | -0.003 | 0.004 | 0.33 | 0.32 | 0.41 | 0.22 | 0.22 | 0.2 |
| 6 | 118152123 | rs2789007 | 1.208 | -0.053 | -0.401 | 0.667 | 0.001 | 0.047 | 0.047 | 0 | 0.14 | 0.03 | 0.06 | 0 | 0.11 | 0.08 |
| 6 | 118161079 | rs9489262 | 1.254 | -0.042 | -0.336 | 0.67 | -0.051 | -0.027 | -0.027 | 0.544 | 0.38 | 0.12 | 0.35 | 0.34 | 0.48 | 0.66 |
| 6 | 118197118 | rs549115 | -1.425 | -0.026 | -0.26 | 0.326 | -0.005 | -0.012 | -0.012 | 6.174 | 0.82 | 0.88 | 0.86 | 0.7 | 0.79 | 0.93 |
| 6 | 118200950 | rs1885836 | 0.946 | -0.012 | -0.27 | 0.445 | 0.003 | 0.035 | 0.035 | 0 | 0.08 | 0.11 | 0.22 | 0.17 | 0.08 | 0.08 |
| 6 | 118204431 | rs9481742 | 1.192 | 0.051 | -0.451 | 0.465 | -0.051 | -0.08 | -0.08 | 0.738 | 0.38 | 0.13 | 0.45 | 0.43 | 0.48 | 0.66 |
| 6 | 118214732 | rs1923852 | -0.594 | 0.069 | -0.33 | -0.002 | 0.003 | 0.03 | 0.03 | 0 | 0.76 | 0.47 | 0.81 | 0.6 | 0.76 | 0.91 |
| 6 | 118227589 | rs2788997 | -0.549 | 0.087 | -0.168 | 0.022 | 0.064 | 0.049 | 0.049 | 1.831 | 0.06 | 0.41 | 0.05 | 0.07 | 0.03 | 0.02 |
| 6 | 118228527 | rs2788996 | 0.578 | 0.134 | -0.461 | 0.036 | 0.002 | 0.001 | 0.001 | 0 | 0.44 | 0.54 | 0.5 | 0.49 | 0.51 | 0.68 |
| 6 | 118240685 | rs982647 | -0.291 | 0.086 | 0.004 | 0.733 | 0.004 | 0.016 | 0.016 | 0 | 0.75 | 0.47 | 0.81 | 0.61 | 0.75 | 0.91 |
| 6 | 118248247 | rs4946303 | -0.643 | -0.01 | -0.068 | 0.669 | -0.015 | -0.011 | -0.011 | 1.953 | 0.25 | 0.35 | 0.29 | 0.17 | 0.15 | 0.14 |
| 6 | 118249332 | rs1016084 | -0.643 | -0.01 | -0.068 | 0.669 | -0.015 | -0.011 | -0.011 | 0 | 0.25 | 0.35 | 0.29 | 0.17 | 0.15 | 0.14 |
| 6 | 118249495 | rs1016083 | 0.873 | 0.07 | -1.049 | 0.591 | -0.011 | 0.037 | 0.037 | 0 | 0.08 | 0 | 0.03 | 0.02 | 0.04 | 0.06 |
| 6 | 118250697 | rs2355790 | 0.822 | 0.2 | -0.885 | 0.763 | -0.029 | -0.064 | -0.064 | 0 | 0.51 | 0.12 | 0.52 | 0.44 | 0.59 | 0.76 |
| 6 | 118262263 | rs1416421 | -0.454 | 0.315 | -0.002 | 0.554 | 0.004 | 0.016 | 0.016 | 0 | 0.8 | 0.72 | 0.83 | 0.67 | 0.78 | 0.92 |
| 6 | 118263296 | rs7738597 | 0.702 | -0.041 | 0.237 | 0.178 | 0.025 | 0.063 | 0.063 | 0 | 0.59 | 0.21 | 0.72 | 0.59 | 0.67 | 0.84 |
| 6 | 118271405 | rs9489288 | -0.013 | -0.068 | 0.613 | 0.123 | -0.031 | -0.108 | -0.108 | 0.076 | 0.18 | 0.4 | 0.09 | 0.02 | 0.07 | 0.07 |
| 6 | 118281878 | rs5002415 | -0.304 | 0.139 | 0.662 | 0.327 | -0.002 | -0.019 | -0.019 | 0 | 0.28 | 0.38 | 0.37 | 0.17 | 0.29 | 0.38 |
| 6 | 118291417 | rs9387528 | 1.039 | -0.023 | 0.272 | 0.263 | -0.022 | -0.006 | -0.006 | 0 | 0.48 | 0.23 | 0.44 | 0.44 | 0.46 | 0.53 |
| 6 | 118291994 | rs9387530 | 1.368 | 0.014 | 0.144 | 0.316 | -0.029 | -0.016 | -0.016 | 0 | 0.06 | 0.01 | 0.04 | 0.05 | 0.08 | 0.07 |
| 6 | 118300769 | rs1361285 | 0.939 | -0.197 | 0.37 | 0.446 | -0.014 | 0.004 | 0.004 | 0.011 | 0.49 | 0.23 | 0.44 | 0.44 | 0.46 | 0.53 |
| 6 | 118302272 | rs9320622 | 0.724 | -0.191 | 0.39 | 0.449 | 0.001 | 0.013 | 0.013 | 0.047 | 0.52 | 0.27 | 0.45 | 0.47 | 0.48 | 0.54 |
| 6 | 118302310 | rs10499084 | 0.809 | -0.201 | 0.492 | 0.542 | -0.001 | -0.005 | -0.005 | 0 | 0.19 | 0.18 | 0.16 | 0.35 | 0.22 | 0.08 |
| 6 | 118310478 | rs4946312 | 0.476 | -0.145 | 0.572 | 0.491 | -0.005 | -0.03 | -0.03 | 0 | 0.67 | 0.43 | 0.61 | 0.78 | 0.69 | 0.63 |
| 6 | 118320108 | rs9387532 | 0.335 | -0.129 | 0.792 | 0.437 | 0 | -0.022 | -0.022 | 2.085 | 0.7 | 0.64 | 0.62 | 0.82 | 0.69 | 0.64 |
| 6 | 118332156 | rs9385028 | 0.08 | -0.068 | 0.62 | 0.329 | -0.001 | -0.001 | -0.001 | 0 | 0.24 | 0.24 | 0.18 | 0.33 | 0.24 | 0.12 |
| 6 | 118341151 | rs4373391 | 0.29 | -0.246 | 0.457 | 0.465 | -0.001 | 0.008 | 0.008 | 0 | 0.75 | 0.33 | 0.81 | 0.55 | 0.73 | 0.84 |
| 6 | 118342669 | rs4132664 | -0.174 | -0.212 | 0.548 | 0.292 | 0.006 | 0.031 | 0.031 | 0 | 0.22 | 0.62 | 0.18 | 0.36 | 0.26 | 0.15 |
| 6 | 118343461 | rs9489297 | 0.325 | -0.186 | -0.242 | 0.006 | -0.021 | 0.117 | 0.117 | 0 | 0.22 | 0.07 | 0.14 | 0.01 | 0.19 | 0.13 |
| 6 | 118350310 | rs9398478 | 1.108 | -0.19 | -0.178 | 0.228 | -0.016 | -0.029 | -0.029 | 0 | 0.41 | 0.21 | 0.38 | 0.5 | 0.39 | 0.49 |
| 6 | 118352676 | rs9374702 | 1.045 | -0.159 | 0.007 | 0.273 | -0.002 | -0.014 | -0.014 | 0.62 | 0.42 | 0.29 | 0.38 | 0.47 | 0.39 | 0.48 |
| 6 | 118361568 | rs9372495 | 0.83 | -0.344 | 0.642 | 0.928 | 0.002 | -0.021 | -0.021 | 0 | 0.63 | 0.3 | 0.53 | 0.48 | 0.58 | 0.61 |
| 6 | 118365898 | rs9481761 | 0.36 | -0.199 | -0.123 | 1.028 | 0.002 | -0.001 | -0.001 | 0 | 0.14 | 0.13 | 0.3 | 0.14 | 0.15 | 0.24 |
| 6 | 118368339 | rs9320634 | 0.059 | -0.088 | -0.147 | 1.063 | -0.026 | -0.046 | -0.046 | 0.652 | 0.18 | 0.06 | 0.11 | 0.33 | 0.23 | 0.09 |
| 6 | 118372057 | rs11153718 | -0.125 | -0.103 | -0.014 | 0.991 | -0.009 | -0.018 | -0.018 | 1.193 | 0.2 | 0.2 | 0.15 | 0.39 | 0.27 | 0.15 |
| 6 | 118379187 | rs9374704 | -0.252 | -0.09 | -0.091 | 0.998 | 0.002 | 0.008 | 0.008 | 0 | 0.21 | 0.27 | 0.15 | 0.39 | 0.27 | 0.15 |
| 6 | 118396674 | rs7757090 | 0.148 | 0.011 | -0.265 | 0.967 | 0.004 | -0.003 | -0.003 | 0.345 | 0.14 | 0.12 | 0.3 | 0.14 | 0.15 | 0.23 |
| 6 | 118401745 | rs12208433 | -0.123 | 0.195 | 0.149 | 1.474 | 0.018 | 0.032 | 0.032 | 0 | 0.17 | 0.25 | 0.12 | 0.35 | 0.24 | 0.11 |
| 6 | 118411416 | rs6900050 | -0.327 | -0.088 | -0.251 | 0.948 | -0.005 | -0.009 | -0.009 | 0.107 | 0.39 | 0.32 | 0.24 | 0.49 | 0.43 | 0.44 |

| | | | | | | | | | | | | | | | | |
|---|---|---|---|---|---|---|---|---|---|---|---|---|---|---|---|---|
| 6 | 118415013 | rs9374715 | -0.786 | -0.382 | -0.159 | 0.601 | -0.001 | -0.006 | -0.006 | 0.175 | 0.54 | 0.42 | 0.7 | 0.86 | 0.58 | 0.68 |
| 6 | 118424094 | rs4636058 | 0.578 | -0.301 | -0.181 | 0.953 | -0.007 | 0.029 | 0.029 | 0 | 0.15 | 0.18 | 0.41 | 0.36 | 0.15 | 0.24 |
| 6 | 118428025 | rs3923322 | 0.84 | -0.493 | -0.774 | 0.464 | -0.125 | 0.184 | 0.184 | 0.274 | 0.4 | 0.07 | 0.24 | 0.11 | 0.43 | 0.25 |
| 6 | 118428261 | rs4360162 | 0.314 | -0.748 | -0.699 | 0.436 | 0.043 | -0.019 | -0.019 | 0 | 0.59 | 0.24 | 0.39 | 0.46 | 0.68 | 0.35 |
| 6 | 118436112 | rs17079707 | 0.38 | -0.797 | -0.162 | 0.31 | 0.007 | -0.002 | -0.002 | 0 | 0.22 | 0.19 | 0.16 | 0.18 | 0.15 | 0.33 |
| 6 | 118437378 | rs11153722 | 1.118 | -0.739 | -0.364 | 0.267 | -0.089 | 0.199 | 0.199 | 0.204 | 0.4 | 0.14 | 0.25 | 0.07 | 0.43 | 0.25 |
| 6 | 118446510 | rs1334839 | 1.134 | -0.755 | -0.678 | 0.263 | -0.125 | 0.239 | 0.239 | 0.294 | 0.4 | 0.06 | 0.24 | 0.07 | 0.43 | 0.25 |
| 6 | 118457579 | rs10484291 | 0.445 | -0.767 | -0.678 | 0.288 | 0.007 | -0.006 | -0.006 | 0.683 | 0.21 | 0.14 | 0.15 | 0.18 | 0.15 | 0.33 |
| 6 | 118461619 | rs2152076 | 0.418 | -0.731 | -0.634 | 0.403 | 0.026 | 0.037 | 0.037 | 1.919 | 0.61 | 0.27 | 0.4 | 0.25 | 0.58 | 0.58 |
| 6 | 118475798 | rs205966 | -0.306 | -0.074 | -0.754 | 0.833 | -0.001 | 0.004 | 0.004 | 4.172 | 0.81 | 0.85 | 0.81 | 0.48 | 0.76 | 0.91 |
| 6 | 118480795 | rs205959 | -0.464 | -0.179 | -0.844 | 0.237 | -0.004 | -0.02 | -0.02 | 0 | 0.75 | 0.86 | 0.78 | 0.66 | 0.69 | 0.82 |
| 6 | 118482339 | rs1894613 | 0.19 | 0.282 | -0.638 | -0.301 | -0.088 | 0.173 | 0.173 | 0.156 | 0.34 | 0.07 | 0.22 | 0.08 | 0.36 | 0.21 |
| 6 | 118490768 | rs7761235 | -0.28 | 0.188 | -0.14 | 0.407 | -0.069 | -0.047 | -0.047 | 0 | 0.3 | 0.03 | 0.32 | 0.26 | 0.29 | 0.43 |
| 6 | 118496796 | rs205944 | -1.55 | 0.088 | 1.568 | 0.371 | -0.006 | -0.009 | -0.009 | 0 | 0.07 | 0.62 | 0.06 | 0.01 | 0.02 | 0.06 |
| 6 | 118518754 | rs205926 | -1.065 | 0.486 | 0.375 | 0.587 | 0.008 | 0.01 | 0.01 | 0 | 0.08 | 0.26 | 0.07 | 0.05 | 0.06 | 0.14 |
| 6 | 118525699 | rs205924 | 0.495 | 0.828 | -0.461 | 1.018 | -0.015 | -0.055 | -0.055 | 0 | 0.8 | 0.9 | 0.58 | 0.45 | 0.7 | 0.76 |
| 6 | 118527888 | rs1413846 | 0.591 | 0.558 | -0.554 | 0.259 | 0.012 | -0.022 | -0.022 | 0 | 0.68 | 0.79 | 0.5 | 0.35 | 0.64 | 0.58 |
| 6 | 118528018 | rs1413845 | 0.654 | 0.527 | -0.568 | 0.295 | 0.014 | -0.011 | -0.011 | 0 | 0.67 | 0.78 | 0.49 | 0.32 | 0.63 | 0.58 |
| 6 | 118532883 | rs6936121 | 0.836 | 0.204 | 0.551 | -0.052 | -0.011 | 0 | 0 | 0.379 | 0.62 | 0.49 | 0.44 | 0.34 | 0.61 | 0.56 |
| 6 | 118534293 | rs458588 | -0.16 | 0.168 | 0.377 | -0.049 | -0.002 | -0.012 | -0.012 | 0 | 0.06 | 0.1 | 0.04 | 0.02 | 0.06 | 0.14 |
| 6 | 118536165 | rs456384 | -0.672 | -0.101 | 0.136 | -0.306 | 0.003 | 0 | 0 | 2.809 | 0.1 | 0.1 | 0.06 | 0.02 | 0.09 | 0.16 |
| 6 | 118538069 | rs12202829 | -0.083 | -0.04 | 0.182 | -0.227 | 0.007 | -0.067 | -0.067 | 3.507 | 0.26 | 0.11 | 0.46 | 0.63 | 0.3 | 0.28 |
| 6 | 118550873 | rs1034247 | 0.215 | -0.042 | 0.177 | -0.343 | -0.001 | -0.011 | -0.011 | 0 | 0.94 | 0.9 | 0.96 | 0.98 | 0.94 | 0.86 |
| 6 | 118557958 | rs11153725 | 0.489 | 0.336 | 1.136 | -0.494 | -0.005 | -0.006 | -0.006 | 0 | 0.65 | 0.48 | 0.47 | 0.27 | 0.61 | 0.52 |
| 6 | 118559487 | rs2883849 | -0.242 | 0.365 | 1.156 | -0.456 | 0.004 | -0.01 | -0.01 | 0 | 0.29 | 0.25 | 0.48 | 0.69 | 0.33 | 0.37 |
| 6 | 118570514 | rs3901856 | -0.208 | -0.055 | 1.049 | -1.028 | 0.004 | 0.007 | 0.007 | 0.019 | 0.18 | 0.16 | 0.14 | 0.03 | 0.14 | 0.16 |
| 6 | 118576987 | rs169465 | -0.015 | -0.115 | 1.405 | -1.084 | 0.036 | -0.017 | -0.017 | 0.157 | 0.06 | 0.18 | 0.06 | 0.02 | 0.08 | 0.14 |
| 6 | 118588490 | rs731695 | 1.158 | -0.253 | 1.503 | -1.19 | -0.013 | 0.057 | 0.057 | 0.436 | 0.44 | 0.25 | 0.28 | 0.22 | 0.48 | 0.4 |
| 6 | 118591401 | rs6924560 | 0.218 | -0.131 | 1.466 | -1.06 | 0.016 | 0.025 | 0.025 | 0.016 | 0.14 | 0.1 | 0.11 | 0.03 | 0.09 | 0.08 |
| 6 | 118608535 | rs17337518 | 0.751 | 0.355 | 0.809 | -1.634 | -0.013 | 0.013 | 0.013 | 0 | 0.19 | 0.07 | 0.13 | 0.14 | 0.18 | 0.2 |
| 6 | 118610131 | rs444666 | 0.202 | 0.315 | 1.016 | -1.394 | 0.025 | -0.019 | -0.019 | 0 | 0.06 | 0.16 | 0.06 | 0.03 | 0.07 | 0.11 |
| 6 | 118612400 | rs1923282 | -0.686 | 0.174 | 0.961 | -1.491 | 0.003 | -0.013 | -0.013 | 0 | 0.71 | 0.74 | 0.51 | 0.3 | 0.67 | 0.63 |
| 6 | 118612879 | rs430538 | -0.183 | 0.125 | 1.002 | -1.52 | 0.027 | -0.026 | -0.026 | 0 | 0.07 | 0.24 | 0.06 | 0.03 | 0.08 | 0.12 |
| 6 | 118612943 | rs1630266 | 0.603 | 0.076 | 0.793 | -1.524 | 0.015 | 0.034 | 0.034 | 0 | 0.13 | 0.05 | 0.11 | 0.03 | 0.09 | 0.12 |
| 6 | 118613435 | rs399844 | 0.003 | 0.108 | 0.868 | -1.527 | 0.011 | 0.018 | 0.018 | 0.23 | 0.58 | 0.62 | 0.4 | 0.27 | 0.58 | 0.55 |
| 6 | 118615390 | rs1743257 | 0.234 | 0.023 | 0.66 | -1.558 | 0.009 | 0.067 | 0.067 | 0 | 0.19 | 0.05 | 0.15 | 0.03 | 0.14 | 0.14 |
| 6 | 118627503 | rs283046 | 0.162 | 0.044 | 0.619 | -1.663 | 0.037 | 0.056 | 0.056 | 0.29 | 0.55 | 0.71 | 0.38 | 0.27 | 0.59 | 0.54 |
| 6 | 118634133 | rs190756 | -0.944 | 0.066 | 0.622 | -1.637 | 0.005 | -0.012 | -0.012 | 0 | 0.74 | 0.76 | 0.53 | 0.3 | 0.73 | 0.69 |
| 6 | 118634698 | rs11968293 | 0.66 | 0.135 | 0.441 | -1.589 | 0.002 | 0.053 | 0.053 | 0 | 0.46 | 0.3 | 0.3 | 0.24 | 0.5 | 0.42 |
| 6 | 118637042 | rs3734345 | 0.137 | 0.026 | 0.291 | -1.604 | -0.002 | 0.071 | 0.071 | 0 | 0.19 | 0.05 | 0.15 | 0.03 | 0.15 | 0.15 |
| 6 | 118642751 | rs1006932 | -0.485 | 0.537 | 0.506 | -1.502 | 0.003 | 0.004 | 0.004 | 0 | 0.67 | 0.58 | 0.47 | 0.3 | 0.61 | 0.55 |
| 6 | 118653621 | rs6911339 | 0.714 | 0.502 | 0.451 | -1.597 | -0.006 | 0.06 | 0.06 | 0 | 0.46 | 0.27 | 0.28 | 0.24 | 0.44 | 0.39 |
| 6 | 118654836 | rs1743248 | -0.302 | 0.226 | 0.386 | -1.4 | -0.007 | -0.043 | -0.043 | 0 | 0.22 | 0.26 | 0.19 | 0.04 | 0.17 | 0.16 |
| 6 | 118670051 | rs937276 | -0.098 | 0.25 | 0.142 | -1.361 | 0.006 | 0.031 | 0.031 | 0 | 0.2 | 0.14 | 0.18 | 0.07 | 0.16 | 0.16 |
| 6 | 118673075 | rs937277 | -0.453 | 0.336 | 0.292 | -1.381 | -0.004 | 0.005 | 0.005 | 0 | 0.68 | 0.56 | 0.48 | 0.29 | 0.66 | 0.55 |
| 6 | 118680374 | rs11970286 | 0.634 | 0.145 | 0.097 | -1.517 | -0.046 | 0.054 | 0.054 | 0 | 0.41 | 0.2 | 0.22 | 0.22 | 0.46 | 0.36 |
| 6 | 118685043 | rs12665409 | 0.074 | -0.088 | 0.241 | -1.631 | 0.025 | 0.062 | 0.062 | 0 | 0.49 | 0.34 | 0.29 | 0.25 | 0.54 | 0.46 |
| 6 | 118690822 | rs9481784 | -0.237 | -0.29 | 0.882 | -1.757 | 0.058 | 0.028 | 0.028 | 0 | 0.05 | 0.35 | 0.06 | 0.01 | 0.03 | 0.02 |
| 6 | 118692498 | rs7764272 | -0.176 | -0.235 | 0.489 | -1.707 | 0.019 | 0.047 | 0.047 | 0 | 0.55 | 0.69 | 0.35 | 0.26 | 0.56 | 0.48 |
| 6 | 118695049 | rs6903177 | -0.345 | -0.517 | -0.061 | -2.061 | 0.024 | 0.015 | 0.015 | 0.014 | 0.62 | 0.77 | 0.43 | 0.29 | 0.63 | 0.57 |
| 6 | 118700137 | rs17080082 | 0.842 | -0.325 | -0.113 | -1.808 | 0.006 | 0.019 | 0.019 | 0 | 0.12 | 0.04 | 0.1 | 0.03 | 0.09 | 0.1 |
| 6 | 118719413 | rs7758680 | 0.559 | -0.012 | -0.485 | -2.043 | -0.045 | -0.055 | -0.055 | 0 | 0.13 | 0.02 | 0.08 | 0.18 | 0.14 | 0.1 |
| 6 | 118786486 | rs3798420 | 0.07 | -0.342 | 0.311 | -2.317 | 0.023 | 0.053 | 0.053 | 0 | 0.5 | 0.38 | 0.3 | 0.27 | 0.53 | 0.46 |
| 6 | 118794147 | rs7755056 | 1.297 | -0.436 | 0.258 | -2.377 | 0 | 0.003 | 0.003 | 0 | 0.05 | 0.04 | 0.05 | 0.03 | 0.05 | 0.02 |
| 6 | 118803913 | rs9401073 | 1.275 | -0.561 | 0.144 | -2.637 | 0.012 | -0.075 | -0.075 | 0.026 | 0.22 | 0.06 | 0.44 | 0.65 | 0.21 | 0.34 |
| 6 | 118807847 | rs25422 | 0.683 | -0.527 | 0.284 | -2.627 | 0 | -0.002 | -0.002 | 0 | 0.18 | 0.17 | 0.18 | 0.05 | 0.18 | 0.12 |
| 6 | 118819212 | rs12194555 | 0.382 | -0.512 | 0.405 | -2.623 | 0.01 | 0.025 | 0.025 | 0 | 0.51 | 0.56 | 0.31 | 0.27 | 0.53 | 0.46 |
| 6 | 118827836 | rs10484289 | 1.128 | -0.494 | 0.412 | -2.602 | -0.004 | 0.028 | 0.028 | 0.304 | 0.16 | 0.06 | 0.16 | 0.04 | 0.18 | 0.12 |
| 6 | 118840280 | rs12204456 | -0.7 | -0.537 | 0.454 | -2.622 | 0.008 | -0.01 | -0.01 | 0.035 | 0.72 | 0.76 | 0.5 | 0.32 | 0.73 | 0.64 |
| 6 | 118886615 | rs763254 | 0.801 | -0.708 | 0.809 | -2.796 | -0.055 | 0.045 | 0.045 | 0 | 0.4 | 0.21 | 0.23 | 0.25 | 0.46 | 0.34 |
| 6 | 118891582 | rs2181513 | -0.594 | -0.755 | 0.862 | -2.767 | 0.003 | -0.003 | -0.003 | 0 | 0.69 | 0.7 | 0.48 | 0.32 | 0.72 | 0.58 |
| 6 | 118905151 | rs422312 | 1.225 | -0.778 | 0.839 | -2.774 | 0.006 | -0.012 | -0.012 | 0 | 0.23 | 0.2 | 0.45 | 0.65 | 0.21 | 0.34 |
| 6 | 118916627 | rs13220480 | 0.316 | -0.677 | 0.952 | -2.763 | 0.025 | 0.053 | 0.053 | 0 | 0.5 | 0.38 | 0.3 | 0.27 | 0.54 | 0.47 |
| 6 | 118919480 | rs9489456 | -0.712 | -0.68 | 0.822 | -2.767 | 0.003 | -0.003 | -0.003 | 0.156 | 0.69 | 0.7 | 0.48 | 0.32 | 0.72 | 0.58 |
| 6 | 118920285 | rs9489457 | 0.817 | -0.538 | 0.765 | -2.646 | 0.003 | 0.015 | 0.015 | 0.052 | 0.18 | 0.13 | 0.17 | 0.05 | 0.18 | 0.12 |
| 6 | 118961415 | rs12200232 | 0.705 | -0.492 | 0.772 | -2.592 | -0.003 | -0.042 | -0.042 | 0.01 | 0.1 | 0.17 | 0.08 | 0.02 | 0.08 | 0.13 |
| 6 | 118974798 | rs9481842 | -0.364 | -0.604 | 0.906 | -2.693 | -0.02 | -0.119 | -0.119 | 0 | 0.29 | 0.46 | 0.26 | 0.06 | 0.26 | 0.25 |
| 6 | 118976282 | rs17348534 | 1.933 | -0.047 | -0.175 | -2.942 | -0.086 | 0.042 | 0.042 | 0 | 0.19 | 0.01 | 0.09 | 0.05 | 0.22 | 0.1 |
| 6 | 118983924 | rs12210025 | 0.958 | -0.314 | 0.082 | -2.778 | -0.007 | -0.039 | -0.039 | 0.016 | 0.1 | 0.16 | 0.07 | 0.02 | 0.07 | 0.12 |

| | | | | | | | | | | | | | | | | |
|---|---|---|---|---|---|---|---|---|---|---|---|---|---|---|---|---|
| 6 | 118988362 | rs10872167 | 0.399 | -0.34 | -0.11 | -2.708 | -0.048 | 0.04 | 0.04 | 0 | 0.4 | 0.22 | 0.23 | 0.26 | 0.46 | 0.33 |
| 6 | 118998756 | rs2638551 | 0.15 | -0.016 | -0.098 | -2.769 | -0.005 | 0.01 | 0.01 | 0.335 | 0.42 | 0.6 | 0.26 | 0.27 | 0.42 | 0.47 |
| 6 | 119004731 | rs1891715 | 0.517 | -0.149 | -0.38 | -2.963 | -0.078 | 0.138 | 0.138 | 2.414 | 0.27 | 0.03 | 0.23 | 0.03 | 0.32 | 0.13 |
| 6 | 119005785 | rs11754023 | 0.913 | -0.133 | -0.048 | -2.972 | -0.009 | 0.01 | 0.01 | 0.005 | 0.37 | 0.34 | 0.23 | 0.25 | 0.38 | 0.33 |
| 6 | 119009913 | rs6911599 | -0.091 | -0.267 | -0.316 | -3.06 | -0.026 | 0.064 | 0.064 | 0.943 | 0.5 | 0.09 | 0.67 | 0.68 | 0.52 | 0.5 |
| 6 | 119016202 | rs9387596 | 1.446 | 0.035 | -0.092 | -2.857 | 0.032 | -0.073 | -0.073 | 3.721 | 0.23 | 0.06 | 0.43 | 0.66 | 0.21 | 0.38 |
| 6 | 119018502 | rs1771756 | 0.275 | 0.173 | 0.088 | -2.721 | 0.001 | -0.01 | -0.01 | 0 | 0.46 | 0.54 | 0.25 | 0.29 | 0.42 | 0.48 |
| 6 | 119020091 | rs9489504 | -0.037 | -0.077 | 0.182 | -2.216 | 0 | 0 | 0 | 0 | 0.59 | 0.41 | 0.46 | 0.29 | 0.67 | 0.48 |
| 6 | 119031596 | rs794855 | 0.119 | 0.25 | 0.207 | -1.499 | 0.001 | 0.091 | 0.091 | 0 | 0.94 | 0.85 | 0.97 | 0.78 | 0.94 | 0.89 |
| 6 | 119032179 | rs11153771 | 0.464 | 0.519 | 0.064 | -1.611 | 0 | -0.009 | -0.009 | 0 | 0.28 | 0.28 | 0.17 | 0.08 | 0.32 | 0.16 |
| 6 | 119035141 | rs1688609 | 1.081 | 0.327 | 0.042 | -1.849 | 0.006 | 0.045 | 0.045 | 4.492 | 0.31 | 0.21 | 0.25 | 0.08 | 0.31 | 0.24 |
| 6 | 119039118 | rs4142510 | -0.814 | 0.478 | -0.079 | -1.888 | -0.021 | -0.026 | -0.026 | 0 | 0.63 | 0.8 | 0.49 | 0.43 | 0.63 | 0.48 |
| 6 | 119039866 | rs9481852 | 0.979 | 0.454 | 0.678 | -1.479 | 0.004 | -0.013 | -0.013 | 0 | 0.07 | 0.16 | 0.07 | 0.04 | 0.05 | 0.01 |
| 6 | 119040726 | rs1619049 | -1.486 | 0.525 | 0.398 | -1.445 | -0.006 | -0.004 | -0.004 | 0.335 | 0.87 | 0.89 | 0.94 | 1 | 0.88 | 0.9 |
| 6 | 119043898 | rs9489510 | -0.902 | 0.393 | 0.842 | -0.928 | 0.003 | 0.042 | 0.042 | 0.842 | 0.28 | 0.54 | 0.23 | 0.37 | 0.31 | 0.24 |
| 6 | 119044453 | rs10499087 | 0.094 | 0.598 | 0.189 | -0.742 | -0.002 | -0.002 | -0.002 | 0 | 0.21 | 0.2 | 0.12 | 0.32 | 0.26 | 0.17 |
| 6 | 119046017 | rs11153774 | -0.343 | 0.633 | -0.015 | -0.726 | 0.006 | 0.063 | 0.063 | 2.433 | 0.24 | 0.39 | 0.15 | 0.32 | 0.27 | 0.19 |
| 6 | 119048269 | rs9401082 | -0.187 | 0.671 | -0.443 | -0.995 | -0.007 | 0.008 | 0.008 | 0.023 | 0.1 | 0.05 | 0.06 | 0.1 | 0.09 | 0.14 |
| 6 | 119049344 | rs1688615 | 0.956 | 0.64 | -0.423 | -1.668 | -0.005 | 0.056 | 0.056 | 0 | 0.64 | 0.32 | 0.73 | 0.86 | 0.65 | 0.73 |
| 6 | 119051695 | rs6932749 | -0.723 | 0.648 | 0.214 | -1.667 | 0.001 | -0.072 | -0.072 | 1.193 | 0.27 | 0.33 | 0.15 | 0.11 | 0.24 | 0.23 |
| 6 | 119054622 | rs2185559 | -0.278 | 0.644 | 0.281 | -1.355 | 0.035 | -0.024 | -0.024 | 0 | 0.92 | 0.73 | 0.89 | 0.97 | 0.89 | 0.96 |
| 6 | 119055873 | rs794880 | 0.534 | 0.534 | -0.813 | -1.704 | 0.006 | 0.134 | 0.134 | 0 | 0.39 | 0.17 | 0.49 | 0.83 | 0.35 | 0.55 |
| 6 | 119056733 | rs9401084 | -0.086 | 0.685 | -0.613 | -1.475 | 0.002 | -0.004 | -0.004 | 0.248 | 0.11 | 0.14 | 0.07 | 0.11 | 0.12 | 0.13 |
| 6 | 119065289 | rs1417613 | 0.302 | 0.298 | -0.882 | -1.032 | 0.008 | 0.004 | 0.004 | 0 | 0.37 | 0.27 | 0.5 | 0.64 | 0.35 | 0.45 |
| 6 | 119071713 | rs11968468 | 0.282 | 0.41 | -0.984 | -1.333 | 0.003 | 0.1 | 0.1 | 0 | 0.33 | 0.16 | 0.49 | 0.81 | 0.32 | 0.53 |
| 6 | 119076378 | rs6929592 | -0.203 | 0.759 | -0.236 | -1.466 | 0 | -0.027 | -0.027 | 0 | 0.51 | 0.46 | 0.35 | 0.13 | 0.46 | 0.39 |
| 6 | 119079281 | rs7742814 | 0.064 | 0.91 | -0.363 | -1.197 | 0.006 | 0.011 | 0.011 | 0 | 0.4 | 0.35 | 0.29 | 0.25 | 0.34 | 0.32 |
| 6 | 119080379 | rs9489520 | 1.05 | 0.989 | -0.414 | -1.148 | 0.013 | -0.028 | -0.028 | 0.575 | 0.33 | 0.17 | 0.45 | 0.63 | 0.31 | 0.43 |
| 6 | 119091596 | rs17695732 | 0.152 | 1.066 | -0.312 | -0.954 | -0.003 | 0.001 | 0.001 | 0 | 0.08 | 0.09 | 0.1 | 0.09 | 0.11 | 0.14 |
| 6 | 119092253 | rs9489522 | -0.053 | 0.986 | -0.279 | -0.978 | -0.001 | 0.008 | 0.008 | 0 | 0.09 | 0.08 | 0.08 | 0.22 | 0.07 | 0.13 |
| 6 | 119102271 | rs7743045 | -0.891 | 0.895 | 0.233 | -0.927 | 0.002 | 0.005 | 0.005 | 0 | 0.62 | 0.65 | 0.51 | 0.37 | 0.63 | 0.57 |
| 6 | 119110325 | rs2275614 | 0.333 | 0.991 | 0.17 | -0.619 | -0.024 | -0.002 | -0.002 | 0 | 0.07 | 0.03 | 0.08 | 0.09 | 0.09 | 0.15 |
| 6 | 119111339 | rs17080626 | 0.188 | 0.585 | 0.374 | -0.709 | -0.004 | 0.004 | 0.004 | 0.678 | 0.15 | 0.14 | 0.09 | 0.22 | 0.07 | 0.11 |
| 6 | 119113317 | rs9401090 | -0.586 | 0.744 | 0.582 | -0.758 | 0 | 0.001 | 0.001 | 0 | 0.25 | 0.25 | 0.2 | 0.28 | 0.24 | 0.19 |
| 6 | 119130805 | rs445573 | -0.423 | 0.947 | 0.437 | -0.785 | -0.038 | 0.034 | 0.034 | 0 | 0.13 | 0.06 | 0.13 | 0.06 | 0.16 | 0.18 |
| 6 | 119135410 | rs284914 | 0.978 | 0.77 | 0.447 | -0.75 | 0.003 | -0.004 | -0.004 | 0 | 0.12 | 0.1 | 0.05 | 0.2 | 0.04 | 0.09 |
| 6 | 119136759 | rs65685 | 0.366 | 0.832 | 0.435 | -0.757 | -0.031 | 0.004 | 0.004 | 0 | 0.08 | 0.03 | 0.1 | 0.06 | 0.13 | 0.15 |
| 6 | 119218470 | rs7772912 | 0.113 | 0.733 | 0.954 | -0.82 | 0.055 | -0.012 | -0.012 | 0 | 0.11 | 0.3 | 0.15 | 0.07 | 0.16 | 0.07 |
| 6 | 119219245 | rs9489536 | 0.96 | 0.665 | 0.977 | -0.817 | -0.011 | 0.011 | 0.011 | 0.912 | 0.12 | 0.12 | 0.05 | 0.2 | 0.04 | 0.1 |
| 6 | 119220748 | rs9489537 | 0.464 | 0.555 | 1.021 | -0.895 | -0.036 | 0.025 | 0.025 | 0 | 0.15 | 0.21 | 0.06 | 0.21 | 0.05 | 0.13 |
| 6 | 119224275 | rs768995 | 1.085 | 0.583 | 0.992 | -0.872 | 0.003 | -0.003 | -0.003 | 0 | 0.12 | 0.1 | 0.05 | 0.2 | 0.04 | 0.09 |
| 6 | 119224416 | rs11153778 | 0.624 | 0.594 | 1.052 | -0.877 | -0.002 | 0.001 | 0.001 | 0.572 | 0.1 | 0.09 | 0.11 | 0.03 | 0.16 | 0.07 |
| 6 | 119231968 | rs2295709 | -0.397 | 0.59 | 1.019 | -0.929 | -0.01 | 0.034 | 0.034 | 0 | 0.24 | 0.45 | 0.19 | 0.24 | 0.2 | 0.16 |
| 6 | 119240226 | rs988098 | 1.078 | 0.529 | 0.925 | -0.963 | 0 | 0.002 | 0.002 | 0 | 0.59 | 0.4 | 0.67 | 0.69 | 0.64 | 0.63 |
| 6 | 119240928 | rs17506095 | 1.211 | 0.514 | 0.46 | -1.238 | -0.073 | 0.121 | 0.121 | 0 | 0.19 | 0.01 | 0.17 | 0.01 | 0.24 | 0.16 |
| 6 | 119244433 | rs9401098 | 0.898 | 0.731 | 0.585 | -0.348 | -0.05 | -0.06 | -0.06 | 0 | 0.09 | 0 | 0.04 | 0.26 | 0.13 | 0.1 |
| 6 | 119245024 | rs12197931 | 0.946 | 0.378 | 1.166 | -0.931 | 0.003 | -0.003 | -0.003 | 0.38 | 0.12 | 0.1 | 0.05 | 0.2 | 0.04 | 0.09 |
| 6 | 119253682 | rs9320671 | 1.117 | 0.413 | 1.17 | -0.977 | -0.001 | -0.004 | -0.004 | 0.381 | 0.6 | 0.44 | 0.68 | 0.69 | 0.65 | 0.64 |
| 6 | 119254156 | rs6910782 | 1.182 | 0.407 | 1.177 | -0.981 | 0.033 | -0.085 | -0.085 | 0 | 0.72 | 0.53 | 0.73 | 0.89 | 0.68 | 0.73 |
| 6 | 119254373 | rs12191819 | 0.833 | 0.409 | 1.19 | -0.978 | -0.033 | 0.044 | 0.044 | 0.629 | 0.12 | 0.19 | 0.06 | 0.2 | 0.04 | 0.1 |
| 6 | 119256079 | rs1885125 | 0.964 | 0.498 | 0.793 | -0.939 | -0.001 | -0.004 | -0.004 | 0.183 | 0.6 | 0.43 | 0.68 | 0.69 | 0.65 | 0.64 |
| 6 | 119276427 | rs7743694 | 0.636 | 0.639 | 0.243 | -0.823 | -0.018 | 0.044 | 0.044 | 0.071 | 0.83 | 0.91 | 0.86 | 0.93 | 0.85 | 0.82 |
| 6 | 119306412 | rs4946385 | 0.994 | 0.575 | 0.285 | -0.962 | -0.02 | 0.02 | 0.02 | 0 | 0.41 | 0.22 | 0.52 | 0.68 | 0.43 | 0.47 |
| 6 | 119313633 | rs11967889 | 0.074 | 0.48 | 0.853 | -1.046 | -0.001 | -0.004 | -0.004 | 0 | 0.55 | 0.57 | 0.65 | 0.75 | 0.54 | 0.64 |
| 6 | 119322884 | rs9489576 | 0.697 | 0.565 | 0.606 | -1.281 | -0.013 | 0.097 | 0.097 | 0 | 0.22 | 0.06 | 0.17 | 0.01 | 0.24 | 0.19 |
| 6 | 119334129 | rs12213597 | 0.724 | 0.538 | 0.789 | -1.256 | 0.005 | -0.019 | -0.019 | 0.69 | 0.08 | 0.08 | 0.09 | 0 | 0.16 | 0.07 |
| 6 | 119335865 | rs1889561 | -0.491 | 0.493 | 1.247 | -1.319 | 0.011 | -0.077 | -0.077 | 0 | 0.42 | 0.48 | 0.39 | 0.1 | 0.51 | 0.39 |
| 6 | 119340881 | rs3756939 | -0.981 | 0.687 | 1.283 | -1.382 | 0.012 | 0.012 | 0.012 | 0 | 0.49 | 0.56 | 0.47 | 0.1 | 0.58 | 0.42 |
| 6 | 119346010 | rs1775613 | -0.554 | 0.701 | 1.299 | -1.381 | 0.001 | -0.06 | -0.06 | 0.827 | 0.43 | 0.49 | 0.42 | 0.11 | 0.54 | 0.4 |
| 6 | 119346912 | rs6929403 | 1.451 | 0.13 | 1.325 | -1.244 | 0.005 | -0.007 | -0.007 | 0.101 | 0.37 | 0.47 | 0.52 | 0.68 | 0.41 | 0.44 |
| 6 | 119356201 | rs9387629 | 1.401 | -0.581 | 1.48 | -1.179 | -0.054 | 0.069 | 0.069 | 0 | 0.42 | 0.19 | 0.57 | 0.75 | 0.49 | 0.58 |
| 6 | 119359254 | rs12204084 | 0.365 | -0.558 | 1.618 | -1.164 | 0.031 | -0.024 | -0.024 | 0 | 0.09 | 0.15 | 0.11 | 0.03 | 0.16 | 0.07 |
| 6 | 119362155 | rs1342793 | -1.093 | -0.617 | 1.238 | -1.178 | 0.031 | -0.036 | -0.036 | 0.064 | 0.89 | 0.97 | 0.95 | 0.79 | 0.96 | 0.89 |
| 6 | 119369961 | rs4946390 | 0.673 | -0.59 | 1.254 | -1.181 | -0.006 | -0.001 | -0.001 | 0 | 0.72 | 0.39 | 0.79 | 0.76 | 0.76 | 0.79 |
| 6 | 119373912 | rs12526935 | 0.95 | -0.484 | 1.067 | -1.104 | 0.01 | 0.084 | 0.084 | 0 | 0.47 | 0.26 | 0.6 | 0.75 | 0.53 | 0.6 |
| 6 | 119378582 | rs6940495 | 0.331 | -0.556 | 0.953 | -0.994 | -0.003 | -0.027 | -0.027 | 0 | 0.06 | 0.1 | 0.03 | 0 | 0.04 | 0.03 |
| 6 | 119380885 | rs17080799 | -0.444 | -0.5 | 0.948 | -0.982 | -0.001 | -0.017 | -0.017 | 0 | 0.09 | 0.1 | 0.04 | 0 | 0.07 | 0.06 |
| 6 | 119391206 | rs10080387 | -0.095 | -0.584 | 1.237 | -1.018 | -0.001 | -0.053 | -0.053 | 0.058 | 0.26 | 0.28 | 0.2 | 0 | 0.26 | 0.21 |
| 6 | 119397860 | rs1935817 | 1.558 | -0.764 | 1.059 | -0.726 | -0.047 | 0.006 | 0.006 | 0 | 0.42 | 0.16 | 0.32 | 0.68 | 0.5 | 0.53 |

| | | | | | | | | | | | | | | | |
|---|---|---|---|---|---|---|---|---|---|---|---|---|---|---|---|
| 6 | 119400716 | rs1334926 | 0.485 | -0.814 | 1.082 | -0.681 | -0.002 | 0.013 | 0.013 | 0 | 0.22 | 0.15 | 0.18 | 0 | 0.25 | 0.18 |
| 6 | 119407563 | rs12195334 | -0.223 | -0.938 | 1.441 | -0.59 | -0.054 | 0.013 | 0.013 | 0 | 0.19 | 0.26 | 0.07 | 0.21 | 0.07 | 0.19 |
| 6 | 119409529 | rs7749394 | -1.166 | -0.895 | 1.079 | -0.651 | 0.063 | 0.058 | 0.058 | 0 | 0.49 | 0.72 | 0.58 | 0.29 | 0.37 | 0.4 |
| 6 | 119412952 | rs2357161 | -0.821 | -0.932 | 0.996 | -0.696 | -0.09 | 0.022 | 0.022 | 0 | 0.23 | 0.46 | 0.15 | 0.23 | 0.11 | 0.2 |
| 6 | 119425752 | rs17513439 | 0.109 | -0.812 | 0.461 | -0.655 | -0.006 | -0.021 | -0.021 | 0 | 0.05 | 0.08 | 0.02 | 0 | 0.03 | 0.06 |
| 6 | 119425969 | rs4946395 | 0.21 | -0.674 | 0.35 | -0.643 | -0.001 | -0.053 | -0.053 | 0.617 | 0.66 | 0.64 | 0.72 | 0.97 | 0.6 | 0.73 |
| 6 | 119430311 | rs4945630 | 0.668 | -0.408 | 0.143 | -0.401 | 0.002 | 0.003 | 0.003 | 0 | 0.5 | 0.47 | 0.63 | 0.74 | 0.55 | 0.62 |
| 6 | 119433241 | rs6929791 | 0.183 | -0.41 | -0.42 | 0.092 | 0.027 | -0.054 | -0.054 | 0 | 0.13 | 0.04 | 0.07 | 0.27 | 0.07 | 0.11 |
| 6 | 119434014 | rs1360782 | 0.634 | -0.677 | -0.427 | -0.278 | 0.018 | 0.055 | 0.055 | 0 | 0.63 | 0.22 | 0.49 | 0.86 | 0.64 | 0.74 |
| 6 | 119441614 | rs11753851 | 0.323 | -0.522 | -0.591 | -0.338 | -0.035 | 0.069 | 0.069 | 0 | 0.17 | 0.05 | 0.13 | 0.01 | 0.23 | 0.14 |
| 6 | 119443290 | rs676491 | 0.414 | -0.685 | -0.397 | -0.101 | -0.002 | 0.002 | 0.002 | 0 | 0.92 | 0.95 | 0.92 | 1 | 0.89 | 0.94 |
| 6 | 119447352 | rs644640 | 0.896 | -0.754 | 0.153 | 0.032 | -0.013 | 0.029 | 0.029 | 0 | 0.85 | 0.71 | 0.65 | 0.86 | 0.84 | 0.9 |
| 6 | 119449385 | rs1889562 | -0.473 | -0.798 | 0.158 | 0.285 | 0.006 | 0.007 | 0.007 | 0 | 0.41 | 0.37 | 0.28 | 0.28 | 0.36 | 0.35 |
| 6 | 119451257 | rs9320680 | -0.293 | -0.638 | -0.712 | 0.457 | -0.057 | 0.035 | 0.035 | 0 | 0.42 | 0.11 | 0.33 | 0.34 | 0.49 | 0.36 |
| 6 | 119457914 | rs606955 | -0.475 | -0.261 | -0.787 | 0.589 | -0.006 | -0.007 | -0.007 | 0 | 0.9 | 0.92 | 0.69 | 0.9 | 0.87 | 0.92 |
| 6 | 119460716 | rs808031 | -0.341 | -0.511 | 0.32 | 0.758 | 0.006 | 0.072 | 0.072 | 0 | 0.1 | 0.32 | 0.07 | 0.09 | 0.07 | 0.06 |
| 6 | 119460747 | rs9385067 | 0.935 | -0.671 | 0.263 | 0.667 | -0.014 | -0.046 | -0.046 | 0 | 0.34 | 0.22 | 0.28 | 0.55 | 0.32 | 0.5 |
| 6 | 119463018 | rs706983 | 0.773 | -0.678 | 0.519 | 0.686 | 0 | 0.002 | 0.002 | 0 | 0.36 | 0.36 | 0.29 | 0.55 | 0.32 | 0.5 |
| 6 | 119466590 | rs7775389 | 0.686 | -0.611 | 0.428 | 0.548 | -0.001 | 0.01 | 0.01 | 0 | 0.36 | 0.41 | 0.32 | 0.55 | 0.31 | 0.56 |
| 6 | 119467743 | rs808034 | -0.206 | -0.69 | 0.037 | 0.486 | 0.029 | 0.035 | 0.035 | 0.473 | 0.51 | 0.76 | 0.4 | 0.6 | 0.41 | 0.6 |
| 6 | 119469434 | rs1891539 | 0.416 | -0.334 | -0.225 | 0.289 | 0.002 | 0.034 | 0.034 | 0 | 0.35 | 0.22 | 0.43 | 0.2 | 0.35 | 0.19 |
| 6 | 119473094 | rs624829 | 0.997 | -0.239 | -0.243 | 0.649 | -0.005 | -0.018 | -0.018 | 0 | 0.34 | 0.23 | 0.29 | 0.47 | 0.32 | 0.51 |
| 6 | 119494199 | rs4140634 | 0.195 | -0.519 | -0.624 | 0.438 | -0.001 | -0.064 | -0.064 | 1.903 | 0.23 | 0.05 | 0.15 | 0.27 | 0.17 | 0.21 |
| 6 | 119497523 | rs6935104 | -0.369 | -0.545 | -0.263 | 0.271 | 0.002 | -0.034 | -0.034 | 3.707 | 0.68 | 0.44 | 0.55 | 0.77 | 0.66 | 0.79 |
| 6 | 119497565 | rs6916978 | 1.312 | -0.544 | -0.317 | 0.223 | -0.003 | -0.002 | -0.002 | 2.988 | 0.4 | 0.38 | 0.35 | 0.42 | 0.43 | 0.57 |
| 6 | 119497884 | rs6569053 | -0.657 | -0.818 | -0.05 | 0.291 | -0.017 | 0.076 | 0.076 | 0 | 0.23 | 0.53 | 0.2 | 0.3 | 0.2 | 0.2 |
| 6 | 119500239 | rs3798602 | -0.08 | -0.345 | -0.31 | 0.412 | 0 | -0.004 | -0.004 | 1.089 | 0.66 | 0.38 | 0.52 | 0.72 | 0.64 | 0.78 |
| 6 | 119500674 | rs2558 | 1.044 | -0.392 | -0.352 | 0.426 | -0.01 | -0.004 | -0.004 | 0.255 | 0.4 | 0.34 | 0.35 | 0.43 | 0.44 | 0.58 |
| 6 | 119500916 | rs3756943 | 0.322 | -0.592 | -0.739 | 0.666 | 0.015 | -0.051 | -0.051 | 0 | 0.22 | 0.03 | 0.13 | 0.22 | 0.16 | 0.19 |
| 6 | 119501290 | rs3734383 | -0.211 | -0.583 | -0.742 | 0.666 | 0.009 | -0.093 | -0.093 | 0 | 0.26 | 0.04 | 0.17 | 0.28 | 0.2 | 0.2 |
| 6 | 119506043 | rs760784 | 0.24 | -0.627 | -0.477 | 0.542 | 0.024 | -0.009 | -0.009 | 0 | 0.22 | 0.11 | 0.13 | 0.22 | 0.15 | 0.18 |
| 6 | 119508054 | rs3819728 | 0.008 | -0.503 | -0.389 | -0.124 | -0.004 | -0.009 | -0.009 | 0 | 0.69 | 0.65 | 0.56 | 0.78 | 0.72 | 0.77 |
| 6 | 119508207 | rs1012509 | 0.517 | -0.479 | -0.01 | 0.009 | -0.005 | 0.004 | 0.004 | 0 | 0.36 | 0.41 | 0.3 | 0.55 | 0.3 | 0.53 |
| 6 | 119508989 | rs11153794 | 1.012 | -0.777 | 0.429 | -0.04 | -0.031 | -0.026 | -0.026 | 2.328 | 0.73 | 0.37 | 0.61 | 0.83 | 0.83 | 0.83 |
| 6 | 119509003 | rs11153795 | 1.001 | -0.77 | 0.433 | -0.037 | -0.031 | -0.026 | -0.026 | 0 | 0.73 | 0.37 | 0.61 | 0.83 | 0.83 | 0.83 |
| 6 | 119517239 | rs3798603 | -0.186 | -0.81 | 0.685 | -0.058 | -0.002 | 0.001 | 0.001 | 0.184 | 0.87 | 0.85 | 0.7 | 0.84 | 0.9 | 0.89 |
| 6 | 119519092 | rs6917694 | 1.254 | -0.765 | 0.466 | -0.046 | -0.017 | -0.008 | -0.008 | 0 | 0.69 | 0.37 | 0.57 | 0.76 | 0.78 | 0.81 |
| 6 | 119522751 | rs6918295 | -0.018 | -0.628 | 0.475 | 0.09 | -0.02 | 0.013 | 0.013 | 0.216 | 0.87 | 0.75 | 0.7 | 0.84 | 0.92 | 0.9 |
| 6 | 119528316 | rs909715 | -0.287 | -0.661 | 0.124 | -0.068 | -0.021 | -0.052 | -0.052 | 0.007 | 0.15 | 0.19 | 0.07 | 0 | 0.08 | 0.08 |
| 6 | 119528356 | rs2267670 | 0.278 | -0.169 | -0.067 | 0.953 | 0.003 | 0.076 | 0.076 | 0 | 0.1 | 0.12 | 0.14 | 0.36 | 0.16 | 0.08 |
| 6 | 119531053 | rs2267674 | 0.869 | -0.599 | -0.171 | 0.2 | -0.002 | -0.019 | -0.019 | 0 | 0.35 | 0.29 | 0.31 | 0.55 | 0.3 | 0.47 |
| 6 | 119531284 | rs1028326 | -1.366 | -0.843 | -0.237 | 0.028 | 0.01 | -0.014 | -0.014 | 0 | 0.9 | 0.95 | 0.74 | 0.82 | 0.94 | 0.91 |
| 6 | 119531328 | rs1028327 | 0.278 | -0.83 | 0.019 | -0.449 | 0.004 | -0.064 | -0.064 | 0.382 | 0.33 | 0.4 | 0.27 | 0.04 | 0.42 | 0.26 |
| 6 | 119575860 | rs2357163 | 0.365 | -1.215 | -0.025 | -0.327 | -0.039 | 0.001 | 0.001 | 0.048 | 0.69 | 0.62 | 0.64 | 0.7 | 0.81 | 0.78 |
| 6 | 119578310 | rs17080912 | 0.371 | -1.095 | 0.116 | -0.22 | -0.021 | 0.001 | 0.001 | 0 | 0.12 | 0.22 | 0.06 | 0.08 | 0.08 | 0.09 |
| 6 | 119584688 | rs9489638 | 1.297 | -0.911 | -0.107 | -0.22 | -0.014 | -0.045 | -0.045 | 1.932 | 0.34 | 0.17 | 0.29 | 0.55 | 0.3 | 0.44 |
| 6 | 119591307 | rs11962211 | 0.579 | -0.823 | -0.019 | -0.214 | -0.012 | 0.019 | 0.019 | 0.22 | 0.45 | 0.32 | 0.4 | 0.64 | 0.44 | 0.59 |
| 6 | 119596914 | rs6932066 | -0.232 | -1.076 | -0.135 | -0.253 | -0.005 | 0.001 | 0.001 | 0.246 | 0.72 | 0.73 | 0.68 | 0.76 | 0.86 | 0.79 |
| 6 | 119616269 | rs2057554 | 1.165 | -0.778 | -0.333 | -0.097 | -0.006 | -0.039 | -0.039 | 0 | 0.35 | 0.22 | 0.31 | 0.52 | 0.31 | 0.48 |
| 6 | 119633534 | rs3778107 | 0.759 | -0.629 | -0.427 | 0.054 | -0.037 | -0.034 | -0.034 | 0 | 0.09 | 0.01 | 0.09 | 0.11 | 0.13 | 0.11 |
| 6 | 119647135 | rs7741827 | 0.766 | -1.041 | -0.424 | -0.159 | 0.031 | 0.027 | 0.027 | 1.335 | 0.13 | 0.07 | 0.05 | 0 | 0.06 | 0.07 |
| 6 | 119650416 | rs195063 | -0.028 | -1.31 | -0.035 | -0.125 | -0.024 | 0.006 | 0.006 | 0.636 | 0.25 | 0.27 | 0.32 | 0.31 | 0.14 | 0.17 |
| 6 | 119652404 | rs195071 | -0.615 | -1.204 | 0.177 | -0.416 | 0.003 | 0.016 | 0.016 | 0.375 | 0.5 | 0.57 | 0.57 | 0.35 | 0.52 | 0.38 |
| 6 | 119653287 | rs195075 | -0.597 | -1.223 | 0.124 | -0.241 | -0.007 | 0 | 0 | 0 | 0.35 | 0.33 | 0.41 | 0.35 | 0.26 | 0.28 |
| 6 | 119655145 | rs195077 | 0.921 | -0.988 | 0.166 | -0.073 | -0.02 | 0.035 | 0.035 | 0 | 0.13 | 0.17 | 0.27 | 0.23 | 0.07 | 0.1 |
| 6 | 119675453 | rs1935477 | -0.037 | -1.005 | 0.462 | -0.348 | 0.007 | -0.016 | -0.016 | 0 | 0.34 | 0.37 | 0.33 | 0.17 | 0.51 | 0.31 |
| 6 | 119676177 | rs1295392 | -0.676 | -0.641 | 0.26 | -0.745 | 0.017 | -0.002 | -0.002 | 0.325 | 0.52 | 0.73 | 0.6 | 0.47 | 0.56 | 0.4 |
| 6 | 119678981 | rs12206472 | 0.613 | -1.16 | 0.454 | -0.554 | 0.03 | -0.084 | -0.084 | 0 | 0.26 | 0.37 | 0.25 | 0.04 | 0.38 | 0.2 |
| 6 | 119699881 | rs1342316 | -0.541 | -1.432 | 0.081 | -0.496 | -0.014 | 0.046 | 0.046 | 0 | 0.53 | 0.7 | 0.6 | 0.35 | 0.54 | 0.38 |
| 6 | 119704829 | rs9372528 | 0.904 | -1.605 | 0.351 | -0.554 | -0.028 | 0.036 | 0.036 | 0 | 0.14 | 0.24 | 0.27 | 0.22 | 0.07 | 0.1 |
| 6 | 119710923 | rs1342311 | -1.059 | -2.02 | -0.048 | -0.04 | -0.062 | -0.031 | -0.031 | 0 | 0.68 | 0.9 | 0.76 | 0.61 | 0.63 | 0.45 |
| 6 | 119713746 | rs742962 | 1.392 | -2.019 | -0.006 | 0.031 | -0.001 | 0.002 | 0.002 | 0 | 0.16 | 0.2 | 0.16 | 0.25 | 0.09 | 0.07 |
| 6 | 119714337 | rs1342313 | 1.412 | -1.869 | 0.027 | 0.086 | -0.003 | 0.005 | 0.005 | 0 | 0.12 | 0.14 | 0.26 | 0.22 | 0.07 | 0.1 |
| 6 | 119728438 | rs9398511 | 1.412 | -1.869 | 0.045 | 0.086 | -0.01 | 0.015 | 0.015 | 0 | 0.12 | 0.16 | 0.27 | 0.22 | 0.07 | 0.1 |
| 6 | 119729710 | rs9372532 | 1.152 | -1.882 | 0.03 | 0.053 | -0.016 | 0.059 | 0.059 | 0 | 0.15 | 0.24 | 0.3 | 0.3 | 0.11 | 0.12 |
| 6 | 119750565 | rs1342315 | 0.736 | -1.953 | -0.143 | 0.059 | -0.049 | -0.022 | -0.022 | 1.157 | 0.32 | 0.12 | 0.24 | 0.39 | 0.37 | 0.55 |
| 6 | 119756937 | rs12197534 | 0.699 | -1.962 | -0.132 | 0.056 | -0.049 | -0.019 | -0.019 | 1.595 | 0.32 | 0.12 | 0.24 | 0.39 | 0.37 | 0.55 |
| 6 | 119764348 | rs9374796 | -0.577 | -2.029 | 0.117 | -0.034 | 0.003 | -0.028 | -0.028 | 0 | 0.61 | 0.49 | 0.67 | 0.87 | 0.53 | 0.72 |
| 6 | 119767459 | rs9398513 | 0.162 | -2.093 | 0.048 | 0.015 | -0.026 | 0.003 | 0.003 | 0.231 | 0.46 | 0.24 | 0.52 | 0.62 | 0.46 | 0.65 |

| | | | | | | | | | | | | | | | | |
|---|---|---|---|---|---|---|---|---|---|---|---|---|---|---|---|---|
| 6 | 119768643 | rs4945638 | 1.225 | -1.982 | 0.092 | -0.048 | 0.011 | -0.017 | -0.017 | 0 | 0.14 | 0.09 | 0.28 | 0.23 | 0.09 | 0.1 |
| 6 | 119770906 | rs9320690 | -0.498 | -1.953 | 0.165 | -0.131 | 0.002 | -0.025 | -0.025 | 0 | 0.61 | 0.48 | 0.67 | 0.87 | 0.53 | 0.72 |
| 6 | 119771886 | rs2649545 | 0.013 | -1.972 | 0.312 | -0.115 | -0.002 | 0.037 | 0.037 | 2.235 | 0.49 | 0.39 | 0.35 | 0.22 | 0.54 | 0.35 |
| 6 | 119774463 | rs1995224 | 1.037 | -1.731 | 0.266 | 0.09 | -0.016 | 0.028 | 0.028 | 0 | 0.16 | 0.23 | 0.29 | 0.29 | 0.1 | 0.11 |
| 6 | 119780812 | rs9374797 | 0.782 | -1.772 | 0.303 | 0.067 | -0.026 | 0.035 | 0.035 | 0 | 0.17 | 0.29 | 0.3 | 0.29 | 0.1 | 0.13 |
| 6 | 119798030 | rs924397 | -0.155 | -1.818 | 0.141 | -0.132 | -0.018 | 0.004 | 0.004 | 0 | 0.4 | 0.47 | 0.46 | 0.43 | 0.3 | 0.28 |
| 6 | 119798652 | rs10485005 | 0.994 | -1.749 | -0.236 | -0.359 | -0.099 | 0.119 | 0.119 | 0.297 | 0.24 | 0.08 | 0.26 | 0.01 | 0.4 | 0.18 |
| 6 | 119819043 | rs10872180 | -0.102 | -1.917 | -0.069 | -0.187 | -0.031 | 0.012 | 0.012 | 0 | 0.62 | 0.52 | 0.68 | 0.58 | 0.73 | 0.74 |
| 6 | 119820198 | rs11754922 | 1.676 | -1.761 | -0.247 | -0.202 | -0.068 | -0.015 | -0.015 | 0.238 | 0.21 | 0.06 | 0.12 | 0.25 | 0.23 | 0.46 |
| 6 | 119821096 | rs7740164 | 0.73 | -1.808 | -0.111 | -0.195 | -0.013 | -0.034 | -0.034 | 0 | 0.23 | 0.22 | 0.18 | 0.1 | 0.2 | 0.19 |
| 6 | 119826028 | rs17081294 | 0.61 | -1.84 | -0.129 | -0.169 | -0.013 | -0.014 | -0.014 | 0 | 0.24 | 0.22 | 0.18 | 0.15 | 0.2 | 0.21 |
| 6 | 119840553 | rs13210710 | 1.663 | -1.642 | -0.314 | -0.122 | -0.066 | -0.015 | -0.015 | 0.043 | 0.21 | 0.06 | 0.12 | 0.25 | 0.23 | 0.46 |
| 6 | 119841737 | rs12195597 | -0.216 | -1.812 | -0.158 | -0.116 | 0.004 | -0.002 | -0.002 | 0 | 0.6 | 0.55 | 0.58 | 0.68 | 0.53 | 0.74 |
| 6 | 119852918 | rs9401149 | 1.018 | -1.78 | -0.784 | -0.019 | -0.017 | 0.036 | 0.036 | 0 | 0.17 | 0.23 | 0.29 | 0.33 | 0.1 | 0.09 |
| 6 | 119857792 | rs4551205 | 0.882 | -1.302 | -1.085 | -0.132 | 0.004 | -0.003 | -0.003 | 0 | 0.14 | 0.15 | 0.13 | 0.26 | 0.07 | 0.06 |
| 6 | 119864184 | rs9489733 | 0.882 | -1.302 | -1.083 | -0.132 | 0.003 | 0 | 0 | 0.011 | 0.14 | 0.15 | 0.13 | 0.26 | 0.07 | 0.06 |
| 6 | 119878915 | rs6931728 | 0.15 | -0.277 | -0.768 | -0.326 | -0.001 | -0.003 | -0.003 | 1.74 | 0.1 | 0.13 | 0.12 | 0.1 | 0.12 | 0.11 |
| 6 | 119881344 | rs12195793 | 0.281 | 0.073 | -1.047 | -0.311 | -0.119 | 0.132 | 0.132 | 0.137 | 0.22 | 0.05 | 0.25 | 0.01 | 0.39 | 0.17 |
| 6 | 119885481 | rs6905827 | 0.361 | -0.901 | -0.485 | 0.678 | 0.005 | -0.006 | -0.006 | 0.281 | 0.66 | 0.38 | 0.65 | 0.46 | 0.73 | 0.7 |
| 6 | 119889615 | rs6922474 | 1.046 | -0.631 | -1.062 | 0.377 | -0.017 | 0.036 | 0.036 | 0 | 0.23 | 0.11 | 0.25 | 0.19 | 0.24 | 0.44 |
| 6 | 119894743 | rs2016684 | 0.201 | -0.897 | -1.083 | 0.382 | 0.058 | 0.016 | 0.016 | 0 | 0.6 | 0.26 | 0.64 | 0.38 | 0.72 | 0.68 |
| 6 | 119896674 | rs9374805 | 0.017 | -1.09 | -0.252 | 0.251 | -0.001 | 0.152 | 0.152 | 0 | 0.14 | 0.27 | 0.23 | 0.55 | 0.13 | 0.21 |
| 6 | 119902447 | rs6933087 | 1.158 | -0.407 | -0.55 | 0.248 | -0.03 | 0.026 | 0.026 | 0 | 0.28 | 0.15 | 0.33 | 0.18 | 0.35 | 0.17 |
| 6 | 119903523 | rs1603965 | 0.575 | -0.585 | -0.393 | 0.327 | 0.007 | 0.021 | 0.021 | 0 | 0.3 | 0.23 | 0.27 | 0.26 | 0.26 | 0.46 |
| 6 | 119906882 | rs6910329 | 0.609 | -0.455 | -0.42 | 0.366 | -0.046 | 0.058 | 0.058 | 0 | 0.37 | 0.18 | 0.4 | 0.2 | 0.48 | 0.24 |
| 6 | 119915150 | rs12206925 | -0.141 | -0.387 | -0.185 | 0.386 | -0.036 | 0.024 | 0.024 | 0 | 0.35 | 0.47 | 0.33 | 0.55 | 0.26 | 0.3 |
| 6 | 119915332 | rs11962344 | 0.948 | 0.003 | -0.726 | 0.295 | -0.022 | 0.04 | 0.04 | 0 | 0.23 | 0.08 | 0.25 | 0.18 | 0.25 | 0.43 |
| 6 | 119916191 | rs9401158 | 0.823 | 0.036 | -0.682 | 0.839 | -0.017 | -0.041 | -0.041 | 0 | 0.07 | 0 | 0.09 | 0.2 | 0.08 | 0.08 |
| 6 | 119926751 | rs807302 | -1.147 | 0.091 | -0.434 | 0.405 | -0.004 | 0.009 | 0.009 | 0 | 0.79 | 0.81 | 0.75 | 0.87 | 0.75 | 0.56 |
| 6 | 119930874 | rs6905904 | 1.003 | -0.067 | -0.599 | 0.299 | -0.048 | 0.034 | 0.034 | 0.047 | 0.11 | 0.01 | 0.07 | 0.02 | 0.14 | 0.09 |
| 6 | 119935496 | rs9374809 | -0.792 | -0.149 | -0.28 | 0.486 | 0.006 | -0.013 | -0.013 | 0 | 0.73 | 0.65 | 0.71 | 0.8 | 0.68 | 0.56 |
| 6 | 119946363 | rs12212125 | 0.389 | -0.303 | -0.811 | 0.972 | -0.09 | -0.04 | -0.04 | 3.51 | 0.4 | 0.13 | 0.44 | 0.38 | 0.44 | 0.25 |
| 6 | 119949791 | rs2024732 | 0.721 | -0.768 | -0.027 | 1.015 | -0.03 | 0.016 | 0.016 | 0.015 | 0.21 | 0.32 | 0.13 | 0.28 | 0.15 | 0.22 |
| 6 | 119962839 | rs2024731 | 0.87 | -0.642 | -0.517 | 0.839 | -0.047 | -0.018 | -0.018 | 0 | 0.26 | 0.11 | 0.31 | 0.27 | 0.29 | 0.14 |
| 6 | 119965344 | rs17081487 | 0.566 | -0.845 | -0.434 | 1.079 | 0.031 | -0.026 | -0.026 | 0.295 | 0.27 | 0.14 | 0.24 | 0.33 | 0.22 | 0.21 |
| 6 | 119965385 | rs9374813 | 0.928 | -0.726 | -0.601 | 1.166 | -0.047 | -0.018 | -0.018 | 0 | 0.26 | 0.11 | 0.31 | 0.27 | 0.29 | 0.14 |
| 6 | 119968050 | rs9372545 | -0.164 | -0.374 | 0.314 | 0.619 | -0.007 | -0.002 | -0.002 | 0 | 0.58 | 0.64 | 0.62 | 0.6 | 0.53 | 0.36 |
| 6 | 119985036 | rs7757918 | -0.31 | -0.32 | 0.265 | 0.648 | -0.009 | -0.004 | -0.004 | 0.018 | 0.62 | 0.65 | 0.62 | 0.61 | 0.53 | 0.36 |
| 6 | 119990080 | rs1417842 | 0.084 | -0.25 | 0.129 | 0.5 | 0.043 | 0.005 | 0.005 | 0 | 0.5 | 0.59 | 0.53 | 0.57 | 0.35 | 0.31 |
| 6 | 119991684 | rs12198857 | 0.783 | -0.24 | 0.098 | 0.504 | -0.042 | -0.01 | -0.01 | 1.491 | 0.17 | 0.07 | 0.11 | 0.13 | 0.2 | 0.07 |
| 6 | 119993132 | rs9374816 | -0.822 | -0.234 | 0.108 | 0.461 | 0.003 | -0.001 | -0.001 | 1.128 | 0.67 | 0.65 | 0.64 | 0.7 | 0.55 | 0.38 |
| 6 | 119994786 | rs13208384 | 0.45 | 0.041 | 0.271 | 0.758 | -0.042 | 0.036 | 0.036 | 0 | 0.17 | 0.06 | 0.1 | 0.06 | 0.2 | 0.07 |
| 6 | 120006311 | rs2077847 | 0.128 | 0.06 | 0.659 | 0.628 | 0.036 | -0.002 | -0.002 | 0 | 0.18 | 0.36 | 0.14 | 0.12 | 0.21 | 0.07 |
| 6 | 120009445 | rs9481956 | 0.61 | 0.125 | 0.65 | 0.638 | 0.058 | 0.005 | 0.005 | 0 | 0.13 | 0.3 | 0.12 | 0.1 | 0.19 | 0.05 |
| 6 | 120022988 | rs12211253 | 0.139 | 0.053 | 0.622 | 0.542 | 0.022 | -0.04 | -0.04 | 0 | 0.18 | 0.3 | 0.13 | 0.04 | 0.2 | 0.07 |
| 6 | 120025209 | rs721169 | 0.124 | 0.044 | 0.618 | 0.538 | 0.021 | -0.039 | -0.039 | 0 | 0.18 | 0.3 | 0.13 | 0.04 | 0.2 | 0.07 |
| 6 | 120031554 | rs6924981 | 0.662 | -0.396 | 0.47 | 0.363 | 0.059 | -0.032 | -0.032 | 0.028 | 0.41 | 0.18 | 0.45 | 0.57 | 0.32 | 0.35 |
| 6 | 120037070 | rs13193709 | 1.016 | -0.349 | 0.637 | 0.291 | 0.03 | -0.035 | -0.035 | 0.035 | 0.13 | 0.24 | 0.11 | 0.02 | 0.19 | 0.05 |
| 6 | 120038177 | rs706858 | -1.164 | -0.554 | 0.691 | 0.238 | -0.078 | 0.068 | 0.068 | 0.448 | 0.7 | 0.85 | 0.66 | 0.84 | 0.55 | 0.58 |
| 6 | 120041948 | rs4946435 | 0.733 | -0.607 | 0.856 | 0.25 | 0.052 | 0.015 | 0.015 | 0.007 | 0.42 | 0.21 | 0.46 | 0.64 | 0.33 | 0.35 |
| 6 | 120063634 | rs7776137 | -0.253 | -0.558 | 1.003 | 0.286 | 0.013 | -0.018 | -0.018 | 0.248 | 0.6 | 0.51 | 0.6 | 0.68 | 0.53 | 0.42 |
| 6 | 120064600 | rs12196915 | 1.426 | -0.429 | 0.959 | 0.229 | -0.001 | 0 | 0 | 0 | 0.12 | 0.11 | 0.1 | 0.02 | 0.21 | 0.05 |
| 6 | 120064702 | rs9374820 | -0.548 | -0.456 | 1.096 | 0.237 | -0.008 | 0.003 | 0.003 | 0.229 | 0.65 | 0.66 | 0.63 | 0.68 | 0.56 | 0.46 |
| 6 | 120085784 | rs13198646 | 0.977 | -0.437 | 0.896 | 0.234 | -0.069 | 0.056 | 0.056 | 0.446 | 0.3 | 0.15 | 0.34 | 0.18 | 0.42 | 0.41 |
| 6 | 120087630 | rs10872186 | 0.62 | -0.432 | 0.744 | 0.308 | -0.024 | 0.018 | 0.018 | 0 | 0.18 | 0.11 | 0.14 | 0.09 | 0.23 | 0.08 |
| 6 | 120088951 | rs10456925 | 0.124 | -0.471 | 0.736 | 0.302 | -0.026 | 0.027 | 0.027 | 0 | 0.23 | 0.12 | 0.15 | 0.12 | 0.25 | 0.1 |
| 6 | 120093445 | rs13199160 | 0.619 | -0.431 | 1.046 | 0.237 | 0.008 | -0.001 | -0.001 | 0 | 0.33 | 0.38 | 0.38 | 0.32 | 0.42 | 0.54 |
| 6 | 120094482 | rs7739078 | 0.163 | -0.508 | 0.935 | 0.267 | 0.025 | 0.017 | 0.017 | 0 | 0.49 | 0.4 | 0.5 | 0.64 | 0.36 | 0.37 |
| 6 | 120095538 | rs9374827 | 0.672 | -0.482 | 0.581 | 0.263 | 0.061 | -0.04 | -0.04 | 0 | 0.42 | 0.18 | 0.45 | 0.55 | 0.34 | 0.36 |
| 6 | 120101452 | rs17239297 | 1.376 | -0.013 | 0.252 | 0.021 | -0.084 | -0.015 | -0.015 | 0.005 | 0.19 | 0.02 | 0.25 | 0.15 | 0.28 | 0.33 |
| 6 | 120112137 | rs9320707 | 0.271 | -0.656 | 0.217 | -0.425 | -0.011 | 0.064 | 0.064 | 0 | 0.36 | 0.25 | 0.23 | 0.08 | 0.37 | 0.26 |
| 6 | 120112277 | rs9372556 | 0.864 | -0.816 | 0.398 | 0.164 | -0.01 | 0.149 | 0.149 | 0 | 0.09 | 0.11 | 0.25 | 0.41 | 0.08 | 0.09 |
| 6 | 120117845 | rs9489809 | 0.303 | -0.981 | 0.472 | -0.027 | 0 | 0.063 | 0.063 | 0 | 0.55 | 0.34 | 0.48 | 0.21 | 0.6 | 0.63 |
| 6 | 120118220 | rs9489810 | 1.36 | -1.006 | 0.434 | -0.084 | 0.016 | 0.008 | 0.008 | 0 | 0.2 | 0.13 | 0.11 | 0 | 0.16 | 0.18 |
| 6 | 120118349 | rs4946442 | -0.352 | -0.877 | 0.57 | 0.048 | -0.002 | 0.027 | 0.027 | 0 | 0.43 | 0.6 | 0.52 | 0.74 | 0.4 | 0.36 |
| 6 | 120121530 | rs9320713 | 0.389 | -0.771 | 0.608 | 0.321 | -0.007 | 0.249 | 0.249 | 0 | 0.1 | 0.26 | 0.28 | 0.41 | 0.08 | 0.09 |
| 6 | 120122942 | rs1873553 | -0.036 | -0.989 | -0.129 | -0.306 | 0.018 | 0.029 | 0.029 | 0 | 0.5 | 0.75 | 0.4 | 0.44 | 0.44 | 0.65 |
| 6 | 120124035 | rs10457366 | 0.497 | -1.277 | 0.298 | -0.006 | 0.009 | 0.076 | 0.076 | 0 | 0.07 | 0.21 | 0.06 | 0.1 | 0.04 | 0.09 |
| 6 | 120127707 | rs6903627 | 0.088 | -1.465 | 0.118 | -0.111 | -0.053 | -0.01 | -0.01 | 0 | 0.28 | 0.54 | 0.2 | 0.25 | 0.15 | 0.26 |

| | | | | | | | | | | | | | | | | |
|---|---|---|---|---|---|---|---|---|---|---|---|---|---|---|---|---|
| 6 | 120127912 | rs17240135 | 0.956 | -1.427 | 0.034 | -0.149 | -0.047 | 0.023 | 0.023 | 0.7 | 0.31 | 0.2 | 0.34 | 0.24 | 0.42 | 0.44 |
| 6 | 120129674 | rs9372561 | 1.002 | -1.429 | 0.045 | -0.069 | -0.008 | 0.125 | 0.125 | 0 | 0.08 | 0.11 | 0.26 | 0.44 | 0.08 | 0.09 |
| 6 | 120133826 | rs9489815 | 0.257 | -1.438 | 0.064 | -0.097 | -0.019 | 0.037 | 0.037 | 0 | 0.1 | 0.27 | 0.07 | 0.11 | 0.04 | 0.12 |
| 6 | 120134714 | rs7761829 | -0.899 | -1.354 | -0.333 | -0.124 | -0.026 | 0.091 | 0.091 | 0.011 | 0.67 | 0.85 | 0.8 | 0.92 | 0.65 | 0.79 |
| 6 | 120135888 | rs9374834 | 0.993 | -1.198 | -0.32 | 0.002 | -0.005 | 0.1 | 0.1 | 0 | 0.08 | 0.08 | 0.25 | 0.44 | 0.08 | 0.09 |
| 6 | 120141646 | rs2357683 | 0.847 | -1.106 | -0.461 | -0.276 | -0.055 | 0.021 | 0.021 | 0.587 | 0.3 | 0.18 | 0.34 | 0.24 | 0.42 | 0.43 |
| 6 | 120142181 | rs7751642 | -0.841 | -0.947 | -0.423 | -0.505 | -0.025 | 0.056 | 0.056 | 0 | 0.71 | 0.84 | 0.8 | 0.98 | 0.67 | 0.81 |
| 6 | 120143752 | rs7761886 | -0.935 | -0.897 | -0.693 | -0.681 | -0.03 | 0.166 | 0.166 | 0.313 | 0.64 | 0.86 | 0.8 | 0.97 | 0.65 | 0.77 |
| 6 | 120148190 | rs2357684 | -0.467 | -1.024 | -0.658 | -0.658 | -0.013 | 0.143 | 0.143 | 0 | 0.56 | 0.79 | 0.7 | 0.83 | 0.58 | 0.73 |
| 6 | 120148300 | rs4945649 | 0.381 | -1.024 | -0.658 | -0.658 | -0.013 | 0.143 | 0.143 | 0 | 0.44 | 0.21 | 0.3 | 0.17 | 0.42 | 0.27 |
| 6 | 120154449 | rs1330115 | 0.01 | -1.087 | -0.416 | -0.671 | 0.009 | 0.035 | 0.035 | 0 | 0.25 | 0.35 | 0.41 | 0.62 | 0.29 | 0.31 |
| 6 | 120155531 | rs9401181 | -0.669 | -1.142 | -0.332 | -0.727 | 0.001 | 0.011 | 0.011 | 0 | 0.37 | 0.67 | 0.54 | 0.76 | 0.37 | 0.41 |
| 6 | 120160754 | rs9489821 | 0.386 | -1.265 | -0.323 | -0.656 | -0.038 | 0.006 | 0.006 | 0 | 0.16 | 0.3 | 0.09 | 0.14 | 0.08 | 0.12 |
| 6 | 120161947 | rs6928312 | -0.398 | -1.242 | -0.575 | -0.632 | 0.007 | 0.042 | 0.042 | 0 | 0.27 | 0.55 | 0.43 | 0.62 | 0.29 | 0.31 |
| 6 | 120161995 | rs6928479 | -0.836 | -1.151 | -0.611 | -0.632 | -0.001 | 0 | 0 | 1.025 | 0.34 | 0.66 | 0.45 | 0.64 | 0.32 | 0.37 |
| 6 | 120163401 | rs9401182 | 0.767 | -1.235 | -0.743 | -0.631 | -0.028 | -0.045 | -0.045 | 0.004 | 0.14 | 0.06 | 0.33 | 0.49 | 0.23 | 0.2 |
| 6 | 120165025 | rs1330104 | -0.669 | -1.14 | -0.992 | -1.121 | -0.013 | 0.143 | 0.143 | 0 | 0.56 | 0.79 | 0.7 | 0.83 | 0.58 | 0.73 |
| 6 | 120165475 | rs1330106 | 1.035 | -0.773 | -0.965 | -0.73 | -0.01 | 0.015 | 0.015 | 0 | 0.22 | 0.11 | 0.24 | 0.16 | 0.27 | 0.35 |
| 6 | 120177179 | rs11967078 | 1.264 | -0.728 | -1.015 | -0.663 | 0.004 | -0.025 | -0.025 | 0 | 0.09 | 0.04 | 0.1 | 0.14 | 0.07 | 0.05 |
| 6 | 120177484 | rs9489822 | -0.527 | -0.957 | -1.197 | -0.982 | -0.003 | 0.059 | 0.059 | 0 | 0.56 | 0.67 | 0.49 | 0.81 | 0.5 | 0.44 |
| 6 | 120179913 | rs9320717 | 0.117 | -1.002 | -1.198 | -0.981 | 0.006 | 0.006 | 0.006 | 0.108 | 0.42 | 0.63 | 0.38 | 0.61 | 0.43 | 0.4 |
| 6 | 120182610 | rs1329530 | -1.011 | -0.969 | -1.104 | -0.936 | -0.001 | -0.003 | -0.003 | 0 | 0.76 | 0.78 | 0.8 | 0.86 | 0.73 | 0.6 |
| 6 | 120196708 | rs12194033 | 0.071 | -0.961 | -1.072 | -0.899 | -0.006 | -0.004 | -0.004 | 0 | 0.4 | 0.48 | 0.36 | 0.62 | 0.41 | 0.4 |
| 6 | 120200777 | rs4307207 | 1.764 | -0.977 | -1.064 | -0.902 | -0.025 | -0.017 | -0.017 | 0 | 0.07 | 0.14 | 0.03 | 0.02 | 0.02 | 0.01 |
| 6 | 120202190 | rs17636630 | 1.178 | -0.587 | -1.544 | -0.802 | -0.02 | 0.036 | 0.036 | 0 | 0.21 | 0.07 | 0.18 | 0.13 | 0.26 | 0.34 |
| 6 | 120206996 | rs9401188 | 1.14 | -0.46 | -1.592 | -0.06 | -0.045 | -0.061 | -0.061 | 0.461 | 0.1 | 0.01 | 0.06 | 0.19 | 0.13 | 0.13 |
| 6 | 120207151 | rs17636773 | 1.338 | -0.352 | -1.3 | -0.496 | -0.033 | 0.049 | 0.049 | 0 | 0.21 | 0.11 | 0.31 | 0.06 | 0.23 | 0.15 |
| 6 | 120209557 | rs7772450 | 0.089 | -0.402 | -1.272 | -0.507 | 0.019 | 0.028 | 0.028 | 0 | 0.47 | 0.63 | 0.39 | 0.64 | 0.44 | 0.43 |
| 6 | 120213880 | rs2357760 | -0.421 | -0.349 | -1.348 | -0.447 | -0.001 | 0.011 | 0.011 | 0 | 0.68 | 0.75 | 0.71 | 0.72 | 0.67 | 0.58 |
| 6 | 120218447 | rs1873554 | -0.225 | -0.537 | -1.106 | -0.143 | 0.029 | 0.044 | 0.044 | 0 | 0.51 | 0.66 | 0.42 | 0.63 | 0.47 | 0.45 |
| 6 | 120222329 | rs10457371 | 0.012 | -0.743 | -1.023 | -0.381 | 0.01 | 0.011 | 0.011 | 0 | 0.3 | 0.23 | 0.21 | 0.19 | 0.25 | 0.3 |
| 6 | 120229469 | rs1028961 | 1.507 | -0.426 | -1.137 | -0.605 | -0.026 | 0.028 | 0.028 | 0 | 0.2 | 0.12 | 0.35 | 0.07 | 0.22 | 0.08 |
| 6 | 120233977 | rs7773183 | -1.144 | -0.205 | -0.904 | -0.586 | -0.002 | 0.001 | 0.001 | 0 | 0.81 | 0.84 | 0.82 | 0.84 | 0.73 | 0.73 |
| 6 | 120244785 | rs10484965 | 1.202 | -0.589 | -1.05 | -0.857 | -0.02 | -0.069 | -0.069 | 0 | 0.18 | 0.2 | 0.11 | 0 | 0.16 | 0.16 |
| 6 | 120248560 | rs6940289 | -1.588 | -0.33 | -1.177 | -0.804 | -0.073 | -0.017 | -0.017 | 0.337 | 0.83 | 0.99 | 0.85 | 0.84 | 0.78 | 0.75 |
| 6 | 120250121 | rs1023106 | -0.276 | -0.844 | -1.144 | -0.871 | 0 | -0.012 | -0.012 | 0 | 0.66 | 0.71 | 0.73 | 0.85 | 0.6 | 0.56 |
| 6 | 120252180 | rs7775643 | 1.005 | -0.896 | -1.066 | -0.802 | -0.044 | -0.078 | -0.078 | 0.173 | 0.17 | 0.28 | 0.1 | 0.02 | 0.13 | 0.21 |
| 6 | 120252285 | rs9372571 | -0.211 | -0.957 | -1.143 | -0.69 | -0.005 | -0.013 | -0.013 | 2.658 | 0.38 | 0.44 | 0.35 | 0.74 | 0.35 | 0.47 |
| 6 | 120272026 | rs9320725 | 0.551 | -0.996 | -1.158 | -0.612 | -0.005 | 0.063 | 0.063 | 0.081 | 0.26 | 0.45 | 0.24 | 0.56 | 0.26 | 0.4 |
| 6 | 120274398 | rs11750976 | 1.088 | -0.522 | -1.367 | -0.346 | -0.008 | -0.037 | -0.037 | 0.111 | 0.12 | 0.01 | 0.11 | 0.18 | 0.1 | 0.07 |
| 6 | 120277828 | rs7753146 | -0.454 | -0.506 | -1.399 | -0.449 | -0.051 | 0.025 | 0.025 | 0 | 0.7 | 0.89 | 0.71 | 0.69 | 0.63 | 0.71 |
| 6 | 120282927 | rs9401192 | -0.5 | -0.507 | -1.45 | -0.45 | -0.119 | -0.008 | -0.008 | 0 | 0.7 | 0.97 | 0.72 | 0.69 | 0.63 | 0.71 |
| 6 | 120284480 | rs726178 | 1.804 | -0.565 | -1.455 | -0.506 | -0.021 | 0.015 | 0.015 | 0 | 0.22 | 0.17 | 0.36 | 0.11 | 0.22 | 0.09 |
| 6 | 120284989 | rs9401193 | 1.471 | -0.542 | -1.399 | 0.023 | -0.005 | -0.011 | -0.011 | 0.178 | 0.08 | 0.07 | 0.05 | 0.14 | 0.07 | 0.07 |
| 6 | 120301440 | rs7756430 | -0.871 | -0.377 | -1.339 | -0.257 | -0.052 | -0.011 | -0.011 | 0 | 0.78 | 0.9 | 0.76 | 0.69 | 0.67 | 0.73 |
| 6 | 120313023 | rs4946453 | 0.591 | -0.493 | -1.197 | 0.068 | -0.004 | -0.033 | -0.033 | 0 | 0.12 | 0.01 | 0.11 | 0.18 | 0.1 | 0.07 |
| 6 | 120314863 | rs1474780 | 0.752 | -0.147 | -0.687 | 0.064 | -0.008 | 0.007 | 0.007 | 0 | 0.23 | 0.2 | 0.36 | 0.12 | 0.22 | 0.18 |
| 6 | 120342199 | rs6933285 | -0.454 | -0.668 | -0.585 | 0.098 | -0.002 | 0.009 | 0.009 | 0 | 0.57 | 0.59 | 0.54 | 0.76 | 0.59 | 0.68 |
| 6 | 120344175 | rs9285436 | -0.191 | -0.555 | -0.358 | 0.656 | -0.001 | 0.001 | 0.001 | 0 | 0.21 | 0.22 | 0.14 | 0.28 | 0.2 | 0.26 |
| 6 | 120350095 | rs6927020 | -0.77 | -0.747 | -0.04 | 0.352 | -0.025 | -0.028 | -0.028 | 0 | 0.31 | 0.43 | 0.2 | 0.24 | 0.24 | 0.33 |
| 6 | 120352075 | rs9489862 | -0.761 | -0.786 | 0.037 | 0.551 | -0.019 | -0.006 | -0.006 | 0 | 0.31 | 0.4 | 0.2 | 0.32 | 0.24 | 0.33 |
| 6 | 120352426 | rs9489863 | 0.003 | -0.836 | 0.25 | 0.501 | -0.001 | 0 | 0 | 0 | 0.19 | 0.17 | 0.15 | 0.24 | 0.18 | 0.24 |
| 6 | 120359702 | rs6569099 | -0.907 | -1.3 | -0.247 | 0.274 | -0.02 | 0.07 | 0.07 | 0 | 0.6 | 0.77 | 0.59 | 0.83 | 0.58 | 0.67 |
| 6 | 120360717 | rs10456928 | 0.821 | -1.276 | -0.396 | 0.296 | 0.003 | 0.009 | 0.009 | 0 | 0.18 | 0.21 | 0.28 | 0.33 | 0.27 | 0.3 |
| 6 | 120362271 | rs9385111 | -0.032 | -1.245 | -0.481 | 0.05 | -0.018 | -0.012 | -0.012 | 0.492 | 0.27 | 0.37 | 0.19 | 0.24 | 0.22 | 0.3 |
| 6 | 120362309 | rs7761202 | -1.716 | -0.897 | -0.921 | -0.079 | -0.086 | 0.039 | 0.039 | 0 | 0.7 | 0.93 | 0.62 | 0.83 | 0.64 | 0.75 |
| 6 | 120370149 | rs12201870 | 1.815 | -0.939 | -0.912 | -0.072 | -0.082 | 0.012 | 0.012 | 0 | 0.26 | 0.07 | 0.35 | 0.17 | 0.33 | 0.23 |
| 6 | 120375860 | rs6924531 | 1.02 | -1.008 | -0.758 | -0.345 | 0.012 | 0.026 | 0.026 | 0.266 | 0.15 | 0.1 | 0.09 | 0.02 | 0.09 | 0.11 |
| 6 | 120411998 | rs2208884 | -1.986 | -0.75 | -1.149 | 0.055 | -0.075 | -0.001 | -0.001 | 0.001 | 0.73 | 0.91 | 0.66 | 0.75 | 0.68 | 0.77 |
| 6 | 120413143 | rs1040954 | -0.407 | -1.434 | -1.286 | -0.26 | -0.041 | -0.01 | -0.01 | 0 | 0.41 | 0.17 | 0.47 | 0.6 | 0.51 | 0.56 |
| 6 | 120414134 | rs1581748 | 1.217 | -1.557 | -0.926 | -0.339 | -0.012 | -0.04 | -0.04 | 0 | 0.11 | 0.14 | 0.04 | 0 | 0.06 | 0.07 |
| 6 | 120424885 | rs9374863 | 0.055 | -1.643 | -1.209 | -0.119 | 0.005 | -0.03 | -0.03 | 0 | 0.24 | 0.08 | 0.18 | 0.29 | 0.21 | 0.27 |
| 6 | 120428445 | rs2031113 | 0.742 | -1.677 | -0.809 | -0.195 | -0.058 | -0.043 | -0.043 | 0 | 0.19 | 0.49 | 0.09 | 0.08 | 0.09 | 0.11 |
| 6 | 120436965 | rs9385116 | 1.164 | -1.66 | -1.237 | -0.113 | -0.024 | -0.026 | -0.026 | 0 | 0.09 | 0 | 0.07 | 0.1 | 0.11 | 0.04 |
| 6 | 120438850 | rs1475164 | -0.459 | -1.685 | -1.203 | -0.222 | -0.06 | -0.029 | -0.029 | 0.228 | 0.38 | 0.14 | 0.48 | 0.59 | 0.51 | 0.56 |
| 6 | 120442625 | rs11153864 | 1.016 | -1.345 | -1.283 | -0.314 | -0.05 | -0.066 | -0.066 | 0 | 0.17 | 0.06 | 0.3 | 0.31 | 0.3 | 0.29 |
| 6 | 120449389 | rs9387710 | -2.028 | -1.438 | -1.206 | -0.757 | -0.076 | -0.001 | -0.001 | 0 | 0.73 | 0.91 | 0.66 | 0.76 | 0.68 | 0.77 |
| 6 | 120452629 | rs12661222 | 0.561 | -1.461 | -1.064 | -0.686 | -0.008 | -0.013 | -0.013 | 0 | 0.15 | 0.18 | 0.09 | 0.08 | 0.09 | 0.1 |
| 6 | 120460863 | rs262464 | -0.529 | -1.232 | -1.054 | -0.472 | 0 | -0.006 | -0.006 | 0 | 0.47 | 0.53 | 0.53 | 0.67 | 0.54 | 0.6 |

| | | | | | | | | | | | | | | | | |
|---|---|---|---|---|---|---|---|---|---|---|---|---|---|---|---|---|
| 6 | 120468039 | rs12527241 | 0.861 | -0.641 | -1.656 | -0.23 | -0.118 | 0.038 | 0.038 | 0.855 | 0.35 | 0.02 | 0.4 | 0.25 | 0.38 | 0.41 |
| 6 | 120480152 | rs9401215 | 1.235 | -0.567 | -1.175 | -0.091 | -0.056 | 0.021 | 0.021 | 0.186 | 0.29 | 0.13 | 0.35 | 0.24 | 0.34 | 0.25 |
| 6 | 120484764 | rs10484970 | 0.957 | -0.941 | -0.398 | 0.256 | -0.013 | 0.199 | 0.199 | 0 | 0.11 | 0.35 | 0.07 | 0.29 | 0.06 | 0.14 |
| 6 | 120485010 | rs187798 | -0.536 | -1.426 | -0.177 | 0.185 | 0.009 | 0.006 | 0.006 | 0 | 0.88 | 0.89 | 0.94 | 0.94 | 0.94 | 0.93 |
| 6 | 120485517 | rs4272235 | 0.899 | -1.413 | -0.29 | 0.174 | 0.004 | 0.006 | 0.006 | 0 | 0.39 | 0.48 | 0.42 | 0.47 | 0.39 | 0.39 |
| 6 | 120489270 | rs7763073 | 1.622 | -1.328 | -0.42 | 0.196 | -0.056 | 0.068 | 0.068 | 0.188 | 0.29 | 0.13 | 0.34 | 0.18 | 0.34 | 0.25 |
| 6 | 120491715 | rs155156 | 0.295 | -1.1 | -0.398 | 0.271 | -0.032 | -0.014 | -0.014 | 0 | 0.61 | 0.8 | 0.53 | 0.59 | 0.49 | 0.53 |
| 6 | 120503014 | rs9489904 | -0.177 | -1.374 | -0.625 | -0.342 | -0.012 | 0.081 | 0.081 | 0 | 0.26 | 0.08 | 0.2 | 0.08 | 0.24 | 0.2 |
| 6 | 120507545 | rs260320 | -0.41 | -1.305 | -0.119 | 0.084 | -0.035 | 0.002 | 0.002 | 0 | 0.14 | 0.26 | 0.07 | 0.1 | 0.06 | 0.07 |
| 6 | 120508092 | rs260321 | 0.188 | -0.865 | -0.223 | 0.116 | -0.005 | 0.012 | 0.012 | 0 | 0.6 | 0.64 | 0.53 | 0.33 | 0.51 | 0.48 |
| 6 | 120508526 | rs1407801 | 0.367 | -0.844 | 0.049 | 0.255 | -0.009 | 0 | 0 | 0 | 0.09 | 0.12 | 0.05 | 0.06 | 0.04 | 0.07 |
| 6 | 120509191 | rs155162 | 0.177 | -0.972 | 0.15 | 0.465 | 0.011 | -0.01 | -0.01 | 0.907 | 0.55 | 0.55 | 0.49 | 0.52 | 0.52 | 0.53 |
| 6 | 120509979 | rs155164 | -0.245 | -0.817 | 0.412 | 0.707 | -0.021 | 0.099 | 0.099 | 0 | 0.16 | 0.35 | 0.07 | 0.23 | 0.08 | 0.17 |
| 6 | 120516030 | rs261765 | -0.149 | -0.647 | 0.59 | 0.73 | 0.063 | 0.249 | 0.249 | 0 | 0.07 | 0.28 | 0.05 | 0.19 | 0.06 | 0.14 |
| 6 | 120523112 | rs9320737 | 1.16 | -0.575 | 0.409 | 0.566 | -0.074 | 0.022 | 0.022 | 0.141 | 0.3 | 0.14 | 0.37 | 0.29 | 0.38 | 0.28 |
| 6 | 120533642 | rs1968835 | 0.382 | -0.414 | 0.646 | 0.613 | -0.008 | -0.008 | -0.008 | 0 | 0.38 | 0.33 | 0.38 | 0.54 | 0.39 | 0.41 |
| 6 | 120559264 | rs6925376 | 0.418 | -0.036 | 0.537 | 0.935 | 0.009 | 0.095 | 0.095 | 0 | 0.06 | 0.11 | 0.03 | 0.19 | 0.06 | 0.14 |
| 6 | 120559644 | rs6907748 | 1.324 | -0.028 | 0.348 | 0.799 | -0.038 | -0.015 | -0.015 | 0.189 | 0.25 | 0.13 | 0.33 | 0.33 | 0.3 | 0.23 |
| 6 | 120581725 | rs9372587 | 1.329 | 0.085 | -0.299 | 0.829 | -0.035 | -0.011 | -0.011 | 0 | 0.25 | 0.13 | 0.33 | 0.33 | 0.3 | 0.24 |
| 6 | 120590561 | rs9320745 | 0.773 | 0.069 | -0.234 | 0.786 | -0.002 | -0.002 | -0.002 | 0 | 0.32 | 0.29 | 0.37 | 0.52 | 0.36 | 0.38 |
| 6 | 120599100 | rs12216323 | -0.145 | 0.323 | -0.388 | 0.797 | -0.012 | 0.026 | 0.026 | 0 | 0.16 | 0.11 | 0.09 | 0.04 | 0.16 | 0.11 |
| 6 | 120599909 | rs1382521 | -0.191 | 0.012 | -0.32 | 1.396 | -0.003 | -0.015 | -0.015 | 0 | 0.12 | 0.09 | 0.25 | 0.29 | 0.17 | 0.28 |
| 6 | 120606439 | rs9489929 | -0.504 | -0.047 | -0.2 | 0.944 | -0.011 | 0.053 | 0.053 | 0 | 0.1 | 0.14 | 0.07 | 0.19 | 0.08 | 0.15 |
| 6 | 120630456 | rs12211114 | 0.126 | 0.072 | -0.447 | 0.11 | -0.006 | 0.007 | 0.007 | 0.635 | 0.16 | 0.09 | 0.15 | 0 | 0.22 | 0.06 |
| 6 | 120635035 | rs12193636 | 0.126 | 0.082 | -0.447 | 0.11 | -0.005 | 0.007 | 0.007 | 0.635 | 0.16 | 0.09 | 0.15 | 0 | 0.21 | 0.06 |
| 6 | 120636227 | rs1478814 | -1.257 | -0.313 | -0.444 | 0.068 | -0.011 | 0.004 | 0.004 | 0 | 0.64 | 0.72 | 0.62 | 0.54 | 0.66 | 0.61 |
| 6 | 120639044 | rs3013678 | 0.111 | -0.609 | 0.007 | 0.034 | 0.006 | -0.017 | -0.017 | 0 | 0.28 | 0.18 | 0.34 | 0.44 | 0.27 | 0.44 |
| 6 | 120639122 | rs17082113 | 0.49 | -0.636 | 0.126 | -0.016 | -0.006 | 0.02 | 0.02 | 0 | 0.16 | 0.12 | 0.1 | 0.06 | 0.16 | 0.09 |
| 6 | 120641888 | rs1585725 | 0.835 | -0.853 | -0.072 | 0.255 | -0.012 | 0 | 0 | 0 | 0.13 | 0.11 | 0.05 | 0.11 | 0.05 | 0.08 |
| 6 | 120649636 | rs2931096 | -0.755 | -0.813 | 0.003 | 0.199 | 0.007 | 0.005 | 0.005 | 0 | 0.45 | 0.31 | 0.49 | 0.48 | 0.48 | 0.5 |
| 6 | 120653461 | rs1478810 | -1.017 | -0.85 | 0.122 | 0.153 | 0.026 | 0.125 | 0.125 | 0 | 0.5 | 0.7 | 0.34 | 0.2 | 0.45 | 0.25 |
| 6 | 120662422 | rs2927908 | -0.155 | -0.851 | 0.117 | 0.165 | 0.003 | 0.004 | 0.004 | 0.481 | 0.3 | 0.24 | 0.21 | 0.14 | 0.27 | 0.14 |
| 6 | 120675142 | rs2931091 | -1.757 | -0.815 | 0.008 | 0.213 | -0.02 | 0.001 | 0.001 | 0 | 0.65 | 0.75 | 0.62 | 0.54 | 0.66 | 0.61 |
| 6 | 120676158 | rs1486212 | 0.538 | -0.766 | -0.004 | 0.237 | -0.039 | -0.087 | -0.087 | 0 | 0.14 | 0.03 | 0.28 | 0.34 | 0.21 | 0.36 |
| 6 | 120683728 | rs1843599 | 0.5 | -0.753 | 0.072 | 0.244 | -0.005 | -0.005 | -0.005 | 0.3 | 0.14 | 0.08 | 0.29 | 0.34 | 0.21 | 0.36 |
| 6 | 120704438 | rs13194471 | 0.949 | -0.81 | 0.046 | 0.162 | -0.016 | 0.052 | 0.052 | 0.834 | 0.16 | 0.06 | 0.15 | 0 | 0.21 | 0.06 |
| 6 | 120705746 | rs2357906 | -0.881 | -0.562 | 0.545 | 0.383 | 0.007 | 0.02 | 0.02 | 0 | 0.36 | 0.52 | 0.42 | 0.43 | 0.39 | 0.47 |
| 6 | 120706009 | rs9489944 | 0.021 | -0.517 | 0.804 | 0.405 | 0.011 | -0.041 | -0.041 | 0.12 | 0.19 | 0.31 | 0.12 | 0.06 | 0.18 | 0.11 |
| 6 | 120710372 | rs923189 | -0.986 | -0.546 | 0.742 | 0.412 | 0.01 | 0.024 | 0.024 | 0.009 | 0.35 | 0.52 | 0.41 | 0.43 | 0.39 | 0.48 |
| 6 | 120717175 | rs1486216 | -0.803 | -0.526 | 0.715 | 0.414 | 0.019 | 0.042 | 0.042 | 0 | 0.33 | 0.45 | 0.4 | 0.4 | 0.39 | 0.47 |
| 6 | 120732398 | rs17082203 | 0.283 | -0.474 | 0.804 | 0.346 | 0.009 | -0.027 | -0.027 | 0 | 0.16 | 0.27 | 0.11 | 0.06 | 0.16 | 0.09 |
| 6 | 120736857 | rs1352103 | -1.044 | -0.496 | 0.827 | 0.403 | 0.013 | 0.028 | 0.028 | 0 | 0.34 | 0.5 | 0.41 | 0.44 | 0.39 | 0.48 |
| 6 | 120741459 | rs6940560 | 0.601 | -1.067 | 0.54 | 0.279 | 0.005 | 0.006 | 0.006 | 2.553 | 0.25 | 0.29 | 0.21 | 0.12 | 0.22 | 0.15 |
| 6 | 120743193 | rs17082229 | 0.361 | -1.056 | 0.556 | 0.278 | 0.013 | -0.038 | -0.038 | 0 | 0.17 | 0.31 | 0.12 | 0.06 | 0.16 | 0.09 |
| 6 | 120746439 | rs9374908 | 1.995 | -1.116 | 0.44 | 0.285 | -0.116 | -0.007 | -0.007 | 1.217 | 0.27 | 0.04 | 0.32 | 0.33 | 0.34 | 0.3 |
| 6 | 120748778 | rs17082237 | 0.484 | -1.104 | 0.462 | 0.302 | 0.008 | 0.054 | 0.054 | 0 | 0.17 | 0.19 | 0.29 | 0.35 | 0.23 | 0.39 |
| 6 | 120752983 | rs6937194 | -1.198 | -1.071 | 0.267 | 0.293 | -0.009 | 0.016 | 0.016 | 0 | 0.59 | 0.79 | 0.62 | 0.56 | 0.61 | 0.62 |
| 6 | 120761388 | rs9387770 | 1.896 | -0.998 | 0.141 | 0.183 | -0.12 | -0.01 | -0.01 | 0.009 | 0.27 | 0.03 | 0.32 | 0.33 | 0.34 | 0.3 |
| 6 | 120768017 | rs1385961 | -0.489 | -1.023 | 0.381 | 0.124 | 0.01 | -0.026 | -0.026 | 0 | 0.84 | 0.74 | 0.89 | 0.94 | 0.84 | 0.91 |
| 6 | 120778243 | rs9320751 | 0.929 | -1.037 | 0.331 | 0.179 | 0.012 | 0.028 | 0.028 | 1.286 | 0.45 | 0.3 | 0.62 | 0.7 | 0.56 | 0.69 |
| 6 | 120781346 | rs9489956 | 0.403 | -0.951 | -0.039 | 0.143 | 0.008 | 0.013 | 0.013 | 2.539 | 0.24 | 0.26 | 0.21 | 0.12 | 0.22 | 0.14 |
| 6 | 120793375 | rs6923491 | 0.803 | -1.061 | -0.161 | 0.198 | 0.036 | 0.003 | 0.003 | 0 | 0.13 | 0.03 | 0.05 | 0.09 | 0.05 | 0.08 |
| 6 | 120800518 | rs6903715 | 1.593 | -0.881 | -0.227 | 0.137 | -0.069 | 0.019 | 0.019 | 0.062 | 0.28 | 0.12 | 0.32 | 0.33 | 0.34 | 0.3 |
| 6 | 120805969 | rs4534020 | 0.077 | -1.313 | 0.238 | 0.082 | -0.03 | -0.036 | -0.036 | 3.056 | 0.27 | 0.42 | 0.23 | 0.15 | 0.22 | 0.13 |
| 6 | 120807166 | rs9489971 | -0.022 | -1.283 | 0.064 | -0.397 | 0.011 | 0.019 | 0.019 | 2.537 | 0.27 | 0.25 | 0.18 | 0.05 | 0.24 | 0.18 |
| 6 | 120808032 | rs1385966 | -0.277 | -1.407 | 0.137 | -0.436 | -0.05 | -0.046 | -0.046 | 4.226 | 0.74 | 0.55 | 0.86 | 0.86 | 0.83 | 0.84 |
| 6 | 120809500 | rs6569128 | 0.347 | -1.233 | 0.053 | -0.381 | -0.008 | -0.017 | -0.017 | 0.253 | 0.62 | 0.48 | 0.6 | 0.37 | 0.68 | 0.51 |
| 6 | 120811714 | rs6900484 | 0.166 | -0.906 | 0.123 | -0.339 | 0.017 | 0.069 | 0.069 | 1.557 | 0.27 | 0.19 | 0.19 | 0.05 | 0.26 | 0.18 |
| 6 | 120825811 | rs4449663 | 0.558 | -1.042 | 0.408 | -0.335 | -0.018 | -0.061 | -0.061 | 0 | 0.61 | 0.41 | 0.73 | 0.89 | 0.65 | 0.78 |
| 6 | 120834919 | rs4945660 | 0.647 | -1.048 | 0.447 | -0.188 | -0.009 | -0.047 | -0.047 | 0 | 0.14 | 0.07 | 0.26 | 0.47 | 0.15 | 0.29 |
| 6 | 120838127 | rs10155797 | 0.657 | -1.127 | 0.656 | -0.347 | -0.002 | -0.002 | -0.002 | 0 | 0.12 | 0.15 | 0.06 | 0.05 | 0.09 | 0.06 |
| 6 | 120839911 | rs9489974 | 0.304 | -1.072 | 0.57 | -0.344 | 0.013 | 0.056 | 0.056 | 2.137 | 0.25 | 0.19 | 0.19 | 0.05 | 0.26 | 0.17 |
| 6 | 120843771 | rs9489977 | 0.689 | -1.11 | 0.528 | -0.377 | 0.03 | 0.066 | 0.066 | 4.532 | 0.22 | 0.14 | 0.15 | 0.05 | 0.19 | 0.12 |
| 6 | 120850792 | rs7758863 | 0.42 | -1.132 | 0.57 | -0.273 | 0.012 | 0.052 | 0.052 | 2.123 | 0.25 | 0.19 | 0.19 | 0.06 | 0.26 | 0.17 |
| 6 | 120857815 | rs4535586 | 1.167 | -0.943 | 0.558 | -0.198 | -0.012 | 0.058 | 0.058 | 0 | 0.45 | 0.2 | 0.45 | 0.42 | 0.47 | 0.49 |
| 6 | 120859841 | rs4543410 | 0.56 | -0.977 | 0.684 | -0.201 | -0.001 | -0.007 | -0.007 | 0.005 | 0.55 | 0.45 | 0.5 | 0.43 | 0.54 | 0.5 |
| 6 | 120862765 | rs7756874 | -0.115 | -0.937 | 0.882 | -0.302 | -0.014 | -0.015 | -0.015 | 0 | 0.16 | 0.32 | 0.09 | 0.09 | 0.12 | 0.1 |
| 6 | 120863569 | rs1884351 | -0.997 | -0.958 | 0.831 | -0.364 | 0.032 | 0.111 | 0.111 | 1.611 | 0.44 | 0.76 | 0.31 | 0.15 | 0.4 | 0.27 |
| 6 | 120863967 | rs12055391 | -0.169 | -0.949 | 0.852 | -0.361 | -0.011 | -0.103 | -0.103 | 0 | 0.31 | 0.48 | 0.24 | 0.1 | 0.31 | 0.22 |

| | | | | | | | | | | | | | | | | |
|---|---|---|---|---|---|---|---|---|---|---|---|---|---|---|---|---|
| 6 | 120867625 | rs2144462 | 0.107 | -0.938 | 0.456 | -0.34 | -0.026 | -0.026 | -0.026 | 0 | 0.14 | 0.51 | 0.09 | 0.05 | 0.09 | 0.06 |
| 6 | 120868055 | rs2207757 | -0.982 | -0.932 | 0.26 | -0.34 | 0.035 | 0.174 | 0.174 | 1.567 | 0.44 | 0.84 | 0.31 | 0.15 | 0.4 | 0.27 |
| 6 | 120871846 | rs9374946 | 0.033 | -0.75 | 0.19 | -0.26 | 0 | -0.008 | -0.008 | 0 | 0.27 | 0.3 | 0.21 | 0.1 | 0.28 | 0.17 |
| 6 | 120879066 | rs9401296 | -0.823 | -0.501 | 0.045 | 0.198 | 0.008 | -0.019 | -0.019 | 0 | 0.31 | 0.72 | 0.34 | 0.55 | 0.26 | 0.37 |
| 6 | 120889262 | rs12207787 | 0.694 | -0.742 | -0.196 | -0.084 | -0.017 | 0.049 | 0.049 | 0.835 | 0.17 | 0.07 | 0.16 | 0 | 0.26 | 0.1 |
| 6 | 120892140 | rs17082388 | 0.957 | -0.796 | -0.138 | 0.483 | -0.016 | -0.046 | -0.046 | 0 | 0.12 | 0.04 | 0.22 | 0.38 | 0.13 | 0.26 |
| 6 | 120893610 | rs12660166 | -0.732 | -0.845 | 0.429 | 0.424 | -0.002 | 0.011 | 0.011 | 0 | 0.29 | 0.33 | 0.31 | 0.5 | 0.25 | 0.38 |
| 6 | 120894829 | rs6932708 | -0.894 | -0.845 | 0.656 | 0.424 | -0.009 | 0.055 | 0.055 | 0 | 0.3 | 0.46 | 0.32 | 0.5 | 0.25 | 0.38 |
| 6 | 120898758 | rs1203050 | 0.41 | -1.015 | 0.341 | 0.202 | -0.005 | -0.002 | -0.002 | 0.1 | 0.59 | 0.56 | 0.6 | 0.41 | 0.67 | 0.51 |
| 6 | 120909925 | rs12204216 | 1.711 | -0.458 | -0.009 | -0.109 | -0.013 | 0.054 | 0.054 | 0.049 | 0.1 | 0 | 0.05 | 0.03 | 0.07 | 0.02 |
| 6 | 120915342 | rs4945661 | -1.047 | -0.732 | 0.431 | 0.231 | -0.001 | 0 | 0 | 0 | 0.45 | 0.42 | 0.47 | 0.49 | 0.51 | 0.48 |
| 6 | 120920938 | rs1933834 | -0.674 | -0.749 | 0.418 | 0.082 | 0 | -0.007 | -0.007 | 0 | 0.33 | 0.34 | 0.25 | 0.13 | 0.38 | 0.22 |
| 6 | 120929891 | rs1203087 | -0.433 | -0.734 | 0.669 | 0.281 | -0.013 | 0.013 | 0.013 | 0 | 0.18 | 0.53 | 0.1 | 0.12 | 0.12 | 0.11 |
| 6 | 120937827 | rs17082485 | 0.542 | -0.698 | 0.326 | 0.297 | -0.003 | -0.005 | -0.005 | 0 | 0.12 | 0.11 | 0.06 | 0.05 | 0.09 | 0.1 |
| 6 | 120940335 | rs1569882 | -0.155 | -0.698 | 0.276 | 0.297 | -0.008 | 0.008 | 0.008 | 0 | 0.17 | 0.24 | 0.09 | 0.12 | 0.12 | 0.11 |
| 6 | 120943058 | rs1203044 | -0.935 | -0.754 | -0.079 | 0.229 | -0.005 | 0.022 | 0.022 | 0 | 0.85 | 0.99 | 0.92 | 0.93 | 0.89 | 0.92 |
| 6 | 120944006 | rs1203042 | 0.04 | -0.86 | 0.466 | 0.25 | -0.019 | -0.034 | -0.034 | 0 | 0.72 | 0.68 | 0.84 | 0.88 | 0.8 | 0.82 |
| 6 | 120945759 | rs1933846 | 0.727 | -0.853 | 0.178 | 0.269 | 0.055 | 0.005 | 0.005 | 0 | 0.63 | 0.18 | 0.78 | 0.72 | 0.77 | 0.76 |
| 6 | 120951918 | rs12189884 | 0.151 | -0.721 | -0.079 | 0.332 | -0.038 | -0.066 | -0.066 | 0 | 0.28 | 0.09 | 0.36 | 0.37 | 0.4 | 0.37 |
| 6 | 120968061 | rs1320552 | 0.655 | -0.775 | 0.291 | 0.335 | 0.016 | 0.134 | 0.134 | 1.611 | 0.9 | 0.44 | 0.92 | 0.85 | 0.97 | 0.94 |
| 6 | 120978489 | rs1933865 | 0.655 | -0.775 | 0.291 | 0.335 | 0.016 | 0.134 | 0.134 | 1.611 | 0.9 | 0.44 | 0.92 | 0.85 | 0.97 | 0.94 |
| 6 | 120978943 | rs1933862 | 0.655 | -0.775 | 0.291 | 0.335 | 0.016 | 0.134 | 0.134 | 1.611 | 0.9 | 0.44 | 0.92 | 0.85 | 0.97 | 0.94 |
| 6 | 120985483 | rs7740070 | -0.204 | -0.754 | 0.3 | 0.313 | 0.013 | 0.077 | 0.077 | 0.698 | 0.09 | 0.42 | 0.07 | 0.11 | 0.03 | 0.06 |
| 6 | 120987090 | rs12216476 | -1.076 | -0.787 | -0.147 | 0.284 | 0 | -0.001 | -0.001 | 0 | 0.55 | 0.4 | 0.52 | 0.49 | 0.59 | 0.55 |
| 6 | 120989799 | rs1933873 | -1.076 | -0.787 | -0.147 | 0.284 | 0 | -0.001 | -0.001 | 0.231 | 0.55 | 0.4 | 0.52 | 0.49 | 0.59 | 0.55 |
| 6 | 120991959 | rs1933875 | -0.06 | -0.806 | -0.149 | 0.247 | -0.017 | -0.027 | -0.027 | 0 | 0.27 | 0.32 | 0.16 | 0.13 | 0.19 | 0.18 |
| 6 | 120992247 | rs1933877 | -0.06 | -0.774 | -0.149 | 0.263 | -0.016 | -0.026 | -0.026 | 0 | 0.27 | 0.32 | 0.16 | 0.13 | 0.19 | 0.18 |
| 6 | 120992849 | rs1933809 | 0.423 | -0.812 | -0.131 | 0.427 | 0 | 0.02 | 0.02 | 0 | 0.12 | 0.31 | 0.08 | 0.09 | 0.08 | 0.1 |
| 6 | 120996973 | rs12154210 | 0.931 | -0.794 | -0.321 | 0.318 | -0.018 | 0.045 | 0.045 | 1.069 | 0.16 | 0.04 | 0.14 | 0 | 0.26 | 0.1 |
| 6 | 120997216 | rs9387879 | 1.202 | -0.847 | -0.298 | 0.331 | -0.067 | 0.008 | 0.008 | 0 | 0.47 | 0.09 | 0.46 | 0.44 | 0.47 | 0.5 |
| 6 | 120997943 | rs2358048 | -0.096 | -0.713 | -0.224 | 0.451 | -0.005 | 0.185 | 0.185 | 0 | 0.81 | 0.53 | 0.71 | 0.56 | 0.85 | 0.69 |
| 6 | 121006296 | rs1933823 | 0.128 | -0.733 | -0.166 | 0.363 | -0.013 | -0.047 | -0.047 | 0 | 0.76 | 0.54 | 0.79 | 0.86 | 0.73 | 0.85 |
| 6 | 121006692 | rs6907723 | 0.369 | -0.676 | -0.229 | 0.338 | -0.006 | 0.008 | 0.008 | 0 | 0.13 | 0.3 | 0.08 | 0.09 | 0.08 | 0.11 |
| 6 | 121007744 | rs12199690 | 0.6 | -0.49 | -0.086 | 0.508 | -0.018 | 0.045 | 0.045 | 1.069 | 0.16 | 0.04 | 0.14 | 0 | 0.26 | 0.1 |
| 6 | 121009097 | rs13217858 | 0.6 | -0.49 | -0.086 | 0.508 | -0.018 | 0.045 | 0.045 | 1.069 | 0.16 | 0.04 | 0.14 | 0 | 0.26 | 0.1 |
| 6 | 121021878 | rs11963462 | 1.092 | -0.494 | -0.126 | 0.66 | -0.016 | -0.027 | -0.027 | 0.316 | 0.11 | 0.05 | 0.22 | 0.34 | 0.14 | 0.27 |
| 6 | 121023256 | rs1933840 | 0.91 | -0.612 | -0.018 | 0.603 | -0.001 | -0.001 | -0.001 | 0 | 0.63 | 0.37 | 0.56 | 0.52 | 0.57 | 0.58 |
| 6 | 121028021 | rs2358010 | 1.049 | -0.634 | -0.442 | 0.689 | -0.019 | -0.038 | -0.038 | 0.341 | 0.11 | 0.04 | 0.22 | 0.34 | 0.14 | 0.26 |
| 6 | 121058929 | rs4946513 | 0.479 | -0.906 | 0.206 | 0.289 | 0.002 | 0.103 | 0.103 | 0.384 | 0.92 | 0.56 | 0.93 | 0.89 | 0.99 | 0.95 |
| 6 | 121061144 | rs17629983 | 0.423 | -0.637 | -0.795 | 0.118 | -0.01 | 0.012 | 0.012 | 0 | 0.15 | 0.01 | 0.08 | 0.08 | 0.11 | 0.08 |
| 6 | 121079609 | rs13211110 | -0.079 | -0.273 | -0.385 | 0.134 | -0.014 | 0.05 | 0.05 | 0.904 | 0.17 | 0.04 | 0.12 | 0 | 0.25 | 0.09 |
| 6 | 121084577 | rs12194792 | 0.74 | -0.739 | -0.265 | 0.587 | -0.002 | -0.005 | -0.005 | 0.043 | 0.43 | 0.25 | 0.63 | 0.37 | 0.57 | 0.41 |
| 6 | 121093409 | rs9401331 | 0.673 | -1.177 | 0.357 | 0.678 | -0.016 | 0.047 | 0.047 | 0 | 0.24 | 0.36 | 0.18 | 0.39 | 0.15 | 0.31 |
| 6 | 121098457 | rs2817937 | -0.532 | -1.299 | 0.293 | 0.563 | -0.006 | 0.03 | 0.03 | 0 | 0.9 | 0.96 | 0.96 | 0.98 | 0.89 | 0.92 |
| 6 | 121100377 | rs17082790 | 0.359 | -1.317 | 0.176 | 0.828 | -0.002 | -0.032 | -0.032 | 0 | 0.07 | 0.01 | 0.13 | 0.13 | 0.05 | 0.06 |
| 6 | 121102063 | rs9372627 | 1.056 | -1.33 | 0.345 | 0.692 | 0.024 | -0.084 | -0.084 | 0 | 0.22 | 0.07 | 0.17 | 0.39 | 0.14 | 0.3 |
| 6 | 121102335 | rs9320772 | -0.074 | -1.442 | 0.479 | 0.642 | -0.006 | 0 | 0 | 0 | 0.39 | 0.41 | 0.26 | 0.44 | 0.27 | 0.43 |
| 6 | 121106523 | rs2817936 | -0.75 | -1.323 | 0.739 | 0.757 | 0.051 | 0.158 | 0.158 | 0 | 0.06 | 0.24 | 0.14 | 0.14 | 0.03 | 0.04 |
| 6 | 121114974 | rs17082805 | 1.023 | -1.433 | 0.413 | 0.731 | 0.013 | -0.023 | -0.023 | 0.851 | 0.23 | 0.16 | 0.18 | 0.39 | 0.14 | 0.31 |
| 6 | 121117917 | rs10499099 | 0.862 | -1.441 | 0.172 | 0.455 | -0.003 | 0.075 | 0.075 | 0 | 0.15 | 0.01 | 0.07 | 0.03 | 0.12 | 0.13 |
| 6 | 121118144 | rs2817930 | -0.464 | -1.34 | 0.272 | 0.395 | -0.001 | -0.05 | -0.05 | 0 | 0.3 | 0.31 | 0.17 | 0.06 | 0.29 | 0.21 |
| 6 | 121126482 | rs2789074 | -1.729 | -1.171 | 0.046 | 0.633 | -0.073 | 0.038 | 0.038 | 0 | 0.64 | 0.88 | 0.72 | 0.75 | 0.47 | 0.65 |
| 6 | 121147804 | rs2050736 | -0.447 | -1.4 | 0.532 | 0.917 | 0.024 | 0.005 | 0.005 | 0 | 0.46 | 0.69 | 0.53 | 0.6 | 0.31 | 0.51 |
| 6 | 121151328 | rs2789019 | -0.826 | -1.382 | 0.424 | 0.243 | 0.1 | 0.091 | 0.091 | 0 | 0.5 | 0.87 | 0.65 | 0.69 | 0.32 | 0.55 |
| 6 | 121153942 | rs9387896 | 0.863 | -1.472 | 0.419 | 0.456 | 0.032 | -0.05 | -0.05 | 0.058 | 0.23 | 0.13 | 0.36 | 0.43 | 0.12 | 0.31 |
| 6 | 121156480 | rs2817920 | -0.477 | -1.366 | 1.183 | 0.358 | 0.057 | 0 | 0 | 0 | 0.07 | 0.35 | 0.07 | 0 | 0.06 | 0.02 |
| 6 | 121157775 | rs1339013 | -0.734 | -1.514 | 0.982 | 0.254 | 0.041 | 0.012 | 0.012 | 0.532 | 0.49 | 0.6 | 0.62 | 0.6 | 0.32 | 0.52 |
| 6 | 121172873 | rs2789034 | -0.636 | -1.371 | 1.088 | 0.257 | 0.013 | 0.002 | 0.002 | 0 | 0.09 | 0.22 | 0.17 | 0.06 | 0.07 | 0.06 |
| 6 | 121190596 | rs2817949 | -1.948 | -1.211 | 0.581 | -0.161 | -0.098 | -0.023 | -0.023 | 0 | 0.66 | 0.92 | 0.72 | 0.71 | 0.46 | 0.65 |
| 6 | 121196068 | rs2789052 | -1.12 | -1.801 | 0.945 | 0.123 | -0.006 | 0.018 | 0.018 | 0 | 0.12 | 0.4 | 0.2 | 0.11 | 0.08 | 0.1 |
| 6 | 121198620 | rs2789053 | -0.862 | -1.648 | 0.503 | -0.096 | 0.062 | 0.049 | 0.049 | 0 | 0.53 | 0.77 | 0.66 | 0.68 | 0.35 | 0.57 |
| 6 | 121220971 | rs10499101 | -0.501 | -1.231 | 0.64 | -0.247 | -0.006 | 0 | 0 | 0.073 | 0.21 | 0.3 | 0.21 | 0.2 | 0.15 | 0.24 |
| 6 | 121221289 | rs17082894 | 0.301 | -1.216 | 0.693 | -0.294 | -0.02 | -0.018 | -0.018 | 0 | 0.14 | 0.28 | 0.08 | 0.07 | 0.09 | 0.16 |
| 6 | 121225325 | rs4535587 | -1.301 | -1.337 | 1.177 | -0.542 | 0.005 | -0.031 | -0.031 | 0 | 0.15 | 0.48 | 0.11 | 0.06 | 0.14 | 0.1 |
| 6 | 121226570 | rs9490068 | -1.788 | -1.611 | 1.431 | -0.518 | 0.019 | 0.012 | 0.012 | 0.434 | 0.35 | 0.73 | 0.31 | 0.29 | 0.26 | 0.38 |
| 6 | 121236230 | rs9490073 | -0.311 | -1.663 | 1.256 | -0.398 | -0.017 | -0.001 | -0.001 | 0 | 0.2 | 0.25 | 0.2 | 0.23 | 0.12 | 0.28 |
| 6 | 121238247 | rs7748279 | -0.663 | -1.595 | 1.479 | -0.371 | -0.006 | -0.008 | -0.008 | 0.452 | 0.11 | 0.29 | 0.08 | 0.06 | 0.1 | 0.08 |
| 6 | 121252327 | rs4946526 | -1.582 | -1.398 | 1.221 | -0.579 | -0.017 | -0.034 | -0.034 | 0 | 0.34 | 0.58 | 0.51 | 0.26 | 0.27 | 0.38 |
| 6 | 121253636 | rs9320776 | 0.101 | -1.592 | 1.21 | -0.89 | -0.027 | -0.035 | -0.035 | 5.883 | 0.17 | 0.26 | 0.09 | 0.11 | 0.1 | 0.22 |

| | | | | | | | | | | | | | | | | |
|---|---|---|---|---|---|---|---|---|---|---|---|---|---|---|---|---|
| 6 | 121258838 | rs9387908 | 1.029 | -1.634 | 1.293 | -0.865 | 0.021 | -0.005 | -0.005 | 0.091 | 0.85 | 0.65 | 0.58 | 0.86 | 0.85 | 0.86 |
| 6 | 121261915 | rs9320780 | -0.005 | -2.335 | 0.77 | -1.357 | 0.002 | 0.002 | 0.002 | 0 | 0.38 | 0.32 | 0.46 | 0.7 | 0.26 | 0.4 |
| 6 | 121264458 | rs9490091 | 0.513 | -2.403 | 0.726 | -1.404 | -0.041 | -0.038 | -0.038 | 0 | 0.16 | 0.28 | 0.08 | 0.1 | 0.08 | 0.21 |
| 6 | 121264902 | rs7759675 | 0.289 | -2.328 | 0.749 | -1.378 | 0.032 | -0.017 | -0.017 | 0 | 0.93 | 0.68 | 0.94 | 0.99 | 0.94 | 0.94 |
| 6 | 121269693 | rs2358058 | -0.814 | -2.385 | 0.705 | -1.409 | 0.025 | 0.039 | 0.039 | 0 | 0.55 | 0.57 | 0.54 | 0.8 | 0.34 | 0.61 |
| 6 | 121281977 | rs9320782 | 1.003 | -2.335 | 0.891 | -1.367 | 0.014 | 0.065 | 0.065 | 0 | 0.9 | 0.61 | 0.83 | 0.87 | 0.92 | 0.88 |
| 6 | 121284378 | rs1983321 | 0.395 | -2.334 | 0.938 | -1.373 | 0.048 | -0.006 | -0.006 | 1.042 | 0.93 | 0.63 | 0.94 | 0.99 | 0.94 | 0.94 |
| 6 | 121286403 | rs9374988 | 0.085 | -2.442 | 0.918 | -1.427 | 0.027 | 0.023 | 0.023 | 0 | 0.38 | 0.22 | 0.45 | 0.7 | 0.26 | 0.4 |
| 6 | 121288302 | rs12202993 | 1.49 | -2.458 | 0.871 | -1.512 | -0.052 | 0.175 | 0.175 | 0 | 0.35 | 0.12 | 0.3 | 0.07 | 0.58 | 0.28 |
| 6 | 121296521 | rs9490099 | 0.569 | -2.47 | 0.932 | -1.512 | 0.044 | 0.079 | 0.079 | 0 | 0.52 | 0.39 | 0.38 | 0.17 | 0.67 | 0.48 |
| 6 | 121313346 | rs908461 | 0.12 | -2.326 | 1.103 | -1.394 | -0.064 | 0.002 | 0.002 | 0 | 0.8 | 0.57 | 0.79 | 0.76 | 0.91 | 0.74 |
| 6 | 121320685 | rs9320787 | 0.312 | -2.423 | 1.02 | -1.46 | 0.012 | 0.006 | 0.006 | 0.022 | 0.34 | 0.24 | 0.22 | 0.66 | 0.2 | 0.36 |
| 6 | 121328450 | rs10457410 | 1.356 | -2.369 | 0.974 | -1.51 | -0.067 | 0.115 | 0.115 | 0 | 0.37 | 0.09 | 0.4 | 0.19 | 0.59 | 0.3 |
| 6 | 121330661 | rs1878161 | 0.291 | -2.409 | 1.114 | -1.529 | -0.009 | -0.002 | -0.002 | 0.021 | 0.34 | 0.28 | 0.22 | 0.66 | 0.2 | 0.37 |
| 6 | 121334537 | rs9401363 | 0.899 | -2.373 | 1.1 | -1.514 | -0.014 | -0.065 | -0.065 | 0 | 0.71 | 0.37 | 0.62 | 0.85 | 0.79 | 0.67 |
| 6 | 121345011 | rs218839 | -1.59 | -2.369 | 0.994 | -1.519 | -0.075 | 0.201 | 0.201 | 0 | 0.64 | 0.92 | 0.71 | 0.93 | 0.41 | 0.69 |
| 6 | 121345270 | rs6569173 | -0.121 | -2.335 | 1.126 | -1.484 | -0.007 | -0.004 | -0.004 | 0 | 0.21 | 0.26 | 0.42 | 0.25 | 0.14 | 0.27 |
| 6 | 121349899 | rs11968593 | -0.638 | -2.293 | 1.133 | -1.455 | 0.018 | -0.027 | -0.027 | 0 | 0.08 | 0.29 | 0.05 | 0.02 | 0.07 | 0.08 |
| 6 | 121356936 | rs9387915 | 0.348 | -2.591 | 0.992 | -1.461 | -0.024 | 0.004 | 0.004 | 0.072 | 0.32 | 0.35 | 0.23 | 0.61 | 0.22 | 0.37 |
| 6 | 121357796 | rs1935945 | -0.141 | -2.554 | 0.911 | -1.496 | -0.005 | -0.02 | -0.02 | 0 | 0.22 | 0.26 | 0.32 | 0.14 | 0.15 | 0.28 |
| 6 | 121359221 | rs9482115 | -1.43 | -2.436 | 0.74 | -1.458 | -0.026 | 0.135 | 0.135 | 0 | 0.63 | 0.81 | 0.7 | 0.93 | 0.41 | 0.69 |
| 6 | 121369168 | rs6914477 | -0.82 | -2.411 | 0.82 | -1.456 | 0.031 | 0.126 | 0.126 | 1.084 | 0.55 | 0.72 | 0.67 | 0.87 | 0.38 | 0.68 |
| 6 | 121378456 | rs218856 | -0.75 | -2.051 | 0.873 | -1.554 | 0.01 | -0.012 | -0.012 | 0.459 | 0.52 | 0.58 | 0.43 | 0.91 | 0.33 | 0.6 |
| 6 | 121378987 | rs9387916 | 0.551 | -2.132 | 0.728 | -1.472 | 0.046 | -0.061 | -0.061 | 0.17 | 0.31 | 0.04 | 0.2 | 0.63 | 0.21 | 0.32 |
| 6 | 121426900 | rs7751291 | -1.501 | -2.155 | 0.979 | -1.517 | 0.004 | -0.028 | -0.028 | 0.004 | 0.17 | 0.63 | 0.34 | 0.09 | 0.13 | 0.1 |
| 6 | 121441202 | rs905023 | -0.383 | -1.284 | 1.178 | -1.605 | 0.061 | 0.02 | 0.02 | 0 | 0.11 | 0.35 | 0.3 | 0.07 | 0.1 | 0.05 |
| 6 | 121451095 | rs1273728 | -1.635 | -1.299 | 1.054 | -1.579 | 0.003 | 0.016 | 0.016 | 0 | 0.18 | 0.64 | 0.46 | 0.22 | 0.14 | 0.12 |
| 6 | 121456893 | rs9375002 | 0.585 | -1.396 | 0.808 | -1.204 | 0.053 | -0.139 | -0.139 | 1.622 | 0.31 | 0.05 | 0.2 | 0.53 | 0.21 | 0.32 |
| 6 | 121480228 | rs2202065 | 0.153 | -1.307 | 0.859 | -1.177 | 0.086 | -0.048 | -0.048 | 0 | 0.41 | 0.09 | 0.24 | 0.58 | 0.28 | 0.47 |
| 6 | 121491472 | rs1273749 | 0.795 | -1.175 | 1.227 | -1.15 | 0.03 | 0.08 | 0.08 | 0.004 | 0.94 | 0.66 | 0.95 | 0.92 | 0.98 | 0.92 |
| 6 | 121492582 | rs9385187 | 1.378 | -1.172 | 1.195 | -1.172 | -0.006 | -0.005 | -0.005 | 0 | 0.83 | 0.3 | 0.65 | 0.84 | 0.88 | 0.86 |
| 6 | 121493215 | rs9375006 | -0.115 | -1.54 | 0.832 | -1.427 | 0.091 | 0.079 | 0.079 | 0 | 0.45 | 0.16 | 0.38 | 0.7 | 0.32 | 0.51 |
| 6 | 121496479 | rs785400 | 1.063 | -1.259 | 1.222 | -1.374 | -0.023 | 0.085 | 0.085 | 0.595 | 0.88 | 0.44 | 0.91 | 0.86 | 0.95 | 0.9 |
| 6 | 121499838 | rs7766150 | 1.296 | -1.218 | 1.179 | -1.372 | 0.011 | -0.006 | -0.006 | 0 | 0.88 | 0.42 | 0.68 | 0.89 | 0.91 | 0.88 |
| 6 | 121519579 | rs813068 | 0.225 | -1.192 | 1.18 | -1.358 | 0.087 | 0.002 | 0.002 | 0 | 0.06 | 0.23 | 0.27 | 0.02 | 0.07 | 0.04 |
| 6 | 121525517 | rs9320794 | 1.662 | -1.252 | 1.115 | -1.384 | 0.006 | 0.003 | 0.003 | 2.86 | 0.81 | 0.18 | 0.63 | 0.84 | 0.87 | 0.86 |
| 6 | 121526780 | rs17644324 | 1.369 | -1.327 | 0.887 | -1.472 | -0.129 | 0.162 | 0.162 | 0 | 0.37 | 0.03 | 0.28 | 0.11 | 0.57 | 0.35 |
| 6 | 121526898 | rs7749984 | 0.405 | -0.815 | 1.354 | -1.029 | -0.009 | 0.014 | 0.014 | 0.573 | 0.06 | 0.2 | 0.04 | 0.05 | 0.03 | 0.01 |
| 6 | 121542358 | rs4458731 | 0.298 | -0.781 | 1.193 | -0.975 | 0.073 | 0.078 | 0.078 | 0 | 0.93 | 0.79 | 0.63 | 0.85 | 0.92 | 0.95 |
| 6 | 121552020 | rs2389414 | -0.435 | -1.094 | 0.969 | -1.14 | -0.001 | 0.011 | 0.011 | 0.547 | 0.74 | 0.77 | 0.57 | 0.46 | 0.74 | 0.68 |
| 6 | 121577370 | rs7745023 | 1.241 | -0.847 | 1.031 | -1.131 | -0.047 | 0.242 | 0.242 | 1.129 | 0.43 | 0.04 | 0.3 | 0.11 | 0.59 | 0.35 |
| 6 | 121601627 | rs4245508 | -0.285 | -0.806 | 1.182 | -1.086 | 0.078 | 0.079 | 0.079 | 0 | 0.07 | 0.24 | 0.38 | 0.15 | 0.08 | 0.05 |
| 6 | 121629167 | rs9398629 | 0.094 | -0.859 | 0.771 | -1.088 | 0.06 | -0.045 | -0.045 | 0 | 0.35 | 0.09 | 0.21 | 0.59 | 0.26 | 0.47 |
| 6 | 121642851 | rs7767455 | -1.036 | -0.751 | 1.129 | -1.001 | -0.025 | 0.072 | 0.072 | 2.078 | 0.12 | 0.47 | 0.09 | 0.14 | 0.06 | 0.1 |
| 6 | 121644362 | rs9398631 | 1.353 | -0.762 | 0.725 | -0.99 | 0.006 | 0.004 | 0.004 | 2.322 | 0.8 | 0.16 | 0.63 | 0.82 | 0.86 | 0.83 |
| 6 | 121653046 | rs6941089 | -0.183 | -1.04 | 0.92 | -0.669 | -0.012 | 0.037 | 0.037 | 1.935 | 0.07 | 0.31 | 0.06 | 0.05 | 0.04 | 0.01 |
| 6 | 121661170 | rs9398632 | 0.017 | -1.146 | 0.778 | -0.709 | -0.024 | 0.011 | 0.011 | 0.672 | 0.29 | 0.33 | 0.2 | 0.62 | 0.2 | 0.39 |
| 6 | 121669951 | rs9401395 | 0.368 | -0.648 | 1.003 | -0.774 | 0.052 | -0.019 | -0.019 | 0 | 0.35 | 0.13 | 0.22 | 0.59 | 0.26 | 0.46 |
| 6 | 121670666 | rs6919916 | -0.382 | -0.197 | 1.18 | -0.524 | -0.019 | 0.023 | 0.023 | 1.559 | 0.07 | 0.26 | 0.05 | 0.05 | 0.03 | 0.01 |
| 6 | 121671944 | rs9490191 | -0.565 | 0.423 | 0.475 | -0.567 | 0.069 | -0.011 | -0.011 | 0 | 0.11 | 0.25 | 0.12 | 0.09 | 0.18 | 0.08 |
| 6 | 121674586 | rs9320812 | -0.108 | -0.749 | 0.253 | -0.753 | -0.005 | -0.02 | -0.02 | 0 | 0.46 | 0.46 | 0.37 | 0.85 | 0.26 | 0.57 |
| 6 | 121677165 | rs9490199 | -1.019 | -0.252 | 0.359 | -0.377 | -0.026 | 0.073 | 0.073 | 0.724 | 0.16 | 0.63 | 0.23 | 0.29 | 0.12 | 0.14 |
| 6 | 121681930 | rs9372655 | -0.328 | 0.351 | -0.273 | -1.031 | -0.16 | 0.235 | 0.235 | 2.184 | 0.65 | 0.97 | 0.79 | 0.95 | 0.6 | 0.65 |
| 6 | 121683480 | rs9320813 | -1.258 | 0.275 | 1.077 | 0.258 | -0.016 | 0.019 | 0.019 | 0.596 | 0.14 | 0.53 | 0.12 | 0.11 | 0.11 | 0.12 |
| 6 | 121696060 | rs7741391 | -1.233 | 0.305 | 0.967 | 0.284 | -0.014 | 0.025 | 0.025 | 0.373 | 0.14 | 0.54 | 0.12 | 0.11 | 0.11 | 0.12 |
| 6 | 121706199 | rs9482157 | -1.774 | 0.617 | 1.274 | 0.971 | 0.018 | 0.134 | 0.134 | 0.135 | 0.15 | 0.42 | 0.22 | 0.34 | 0.18 | 0.15 |
| 6 | 121707449 | rs9490228 | -1.224 | 0.602 | 1.39 | 1.013 | 0.05 | 0.062 | 0.062 | 0 | 0.09 | 0.3 | 0.08 | 0.13 | 0.13 | 0.13 |
| 6 | 121710583 | rs1555772 | -2.451 | 1.452 | 1.332 | 0.871 | 0.075 | 0.009 | 0.009 | 0 | 0.18 | 0.46 | 0.37 | 0.16 | 0.23 | 0.18 |
| 6 | 121714848 | rs7738656 | -1.838 | 1.469 | 1.292 | 0.936 | 0.049 | 0.144 | 0.144 | 0 | 0.13 | 0.3 | 0.42 | 0.38 | 0.17 | 0.14 |
| 6 | 121727236 | rs6908897 | -2.39 | 1.67 | 1.338 | 0.825 | 0.078 | 0.113 | 0.113 | 0 | 0.18 | 0.37 | 0.47 | 0.38 | 0.25 | 0.16 |
| 6 | 121733669 | rs17083553 | -1.887 | 1.609 | 1.333 | 0.793 | 0.027 | 0.011 | 0.011 | 0 | 0.14 | 0.22 | 0.33 | 0.18 | 0.17 | 0.13 |
| 6 | 121733886 | rs12204978 | 0.538 | 1.716 | 0.147 | -0.314 | -0.085 | 0.191 | 0.191 | 5.019 | 0.3 | 0.03 | 0.15 | 0.04 | 0.3 | 0.26 |
| 6 | 121748542 | rs11154022 | -1.539 | 1.022 | -0.237 | -0.493 | 0.027 | -0.114 | -0.114 | 0.693 | 0.65 | 0.9 | 0.73 | 0.51 | 0.66 | 0.6 |
| 6 | 121750153 | rs1886249 | -0.03 | 0.651 | 0.493 | -0.437 | -0.045 | -0.061 | -0.061 | 0 | 0.17 | 0.39 | 0.1 | 0.06 | 0.11 | 0.16 |
| 6 | 121753943 | rs7740467 | -1.217 | 0.681 | 0.507 | -0.447 | 0.004 | -0.007 | -0.007 | 0 | 0.29 | 0.54 | 0.5 | 0.25 | 0.26 | 0.37 |
| 6 | 121758907 | rs3805787 | -0.199 | 0.281 | -0.696 | -0.039 | -0.039 | -0.088 | -0.088 | 0 | 0.22 | 0.05 | 0.44 | 0.3 | 0.18 | 0.35 |
| 6 | 121759439 | rs11759917 | -0.039 | 0.606 | -0.513 | -0.527 | 0 | 0.033 | 0.033 | 0.071 | 0.27 | 0.2 | 0.13 | 0.04 | 0.26 | 0.25 |
| 6 | 121772983 | rs12204909 | -0.164 | 0.536 | -0.606 | -0.285 | -0.003 | 0.011 | 0.011 | 0 | 0.3 | 0.19 | 0.2 | 0.27 | 0.32 | 0.27 |
| 6 | 121773999 | rs2389541 | -0.755 | 0.285 | -0.396 | -0.413 | 0.007 | 0.044 | 0.044 | 0.023 | 0.35 | 0.79 | 0.24 | 0.27 | 0.33 | 0.27 |

| | | | | | | | | | | | | | | | | |
|---|---|---|---|---|---|---|---|---|---|---|---|---|---|---|---|---|
| 6 | 121774243 | rs12207042 | -0.046 | 0.556 | -0.566 | -0.591 | -0.002 | 0.039 | 0.039 | 0.052 | 0.27 | 0.15 | 0.16 | 0.03 | 0.28 | 0.23 |
| 6 | 121779646 | rs4945688 | -0.699 | 0.667 | -0.638 | 0.055 | -0.003 | -0.039 | -0.039 | 0 | 0.28 | 0.2 | 0.46 | 0.48 | 0.24 | 0.34 |
| 6 | 121790990 | rs6932373 | 0.402 | 0.074 | -0.325 | -0.155 | 0.003 | 0.006 | 0.006 | 0 | 0.59 | 0.37 | 0.74 | 0.62 | 0.61 | 0.71 |
| 6 | 121792093 | rs9490285 | -0.386 | -0.013 | -0.403 | -0.133 | 0.003 | 0.008 | 0.008 | 0 | 0.18 | 0.18 | 0.12 | 0.11 | 0.14 | 0.27 |
| 6 | 121798968 | rs17653941 | -0.06 | 0.056 | -0.845 | -0.222 | -0.074 | -0.089 | -0.089 | 1.747 | 0.14 | 0.01 | 0.12 | 0.29 | 0.2 | 0.11 |
| 6 | 121799206 | rs4946570 | -0.075 | -0.332 | 0.039 | -0.21 | -0.018 | -0.053 | -0.053 | 0 | 0.77 | 0.68 | 0.88 | 0.92 | 0.82 | 0.85 |
| 6 | 121809450 | rs4946572 | 0.795 | -0.334 | 0.126 | -0.177 | 0.006 | 0.009 | 0.009 | 0 | 0.59 | 0.37 | 0.76 | 0.79 | 0.68 | 0.57 |
| 6 | 121811989 | rs1408277 | -0.139 | -0.395 | 0.262 | -0.123 | 0.019 | 0.029 | 0.029 | 0 | 0.8 | 0.86 | 0.91 | 0.92 | 0.86 | 0.85 |
| 6 | 121816879 | rs9490306 | -0.254 | -0.352 | 0.378 | -0.078 | 0.001 | 0.004 | 0.004 | 0.547 | 0.17 | 0.17 | 0.12 | 0.12 | 0.14 | 0.26 |
| 6 | 121817075 | rs6901013 | 0.315 | -0.702 | 0.187 | -0.383 | -0.054 | -0.102 | -0.102 | 0 | 0.68 | 0.51 | 0.85 | 0.91 | 0.82 | 0.79 |
| 6 | 121817108 | rs12201990 | 0.453 | -0.264 | -0.008 | -0.035 | -0.045 | -0.052 | -0.052 | 0 | 0.09 | 0 | 0.07 | 0.28 | 0.14 | 0.06 |
| 6 | 121821077 | rs6925183 | 1.472 | -0.349 | -0.019 | -0.053 | -0.001 | -0.001 | -0.001 | 0 | 0.41 | 0.38 | 0.62 | 0.51 | 0.48 | 0.51 |
| 6 | 121821309 | rs9490308 | 0.074 | -0.459 | -0.21 | -0.09 | 0.002 | 0.012 | 0.012 | 0.012 | 0.16 | 0.09 | 0.11 | 0.11 | 0.13 | 0.19 |
| 6 | 121825381 | rs11154034 | 1.573 | -0.404 | -0.168 | -0.101 | -0.039 | -0.035 | -0.035 | 0 | 0.41 | 0.22 | 0.62 | 0.51 | 0.5 | 0.51 |
| 6 | 121827964 | rs9490310 | 0 | -0.41 | -0.228 | -0.119 | 0.003 | 0.005 | 0.005 | 0 | 0.17 | 0.14 | 0.12 | 0.13 | 0.13 | 0.19 |
| 6 | 121828351 | rs6913516 | 0.797 | -0.408 | -0.145 | -0.121 | 0.049 | 0.075 | 0.075 | 0 | 0.52 | 0.32 | 0.71 | 0.78 | 0.69 | 0.59 |
| 6 | 121832541 | rs9320820 | 0.449 | -0.282 | -0.746 | -0.077 | 0.054 | 0.106 | 0.106 | 0 | 0.54 | 0.69 | 0.72 | 0.78 | 0.67 | 0.6 |
| 6 | 121841447 | rs17083709 | 0.375 | -0.173 | -1.054 | 0.064 | -0.056 | -0.071 | -0.071 | 0 | 0.11 | 0.01 | 0.09 | 0.29 | 0.19 | 0.08 |
| 6 | 121841478 | rs9490315 | 0.168 | -0.237 | -1.04 | -0.226 | 0 | 0.006 | 0.006 | 0 | 0.16 | 0.09 | 0.11 | 0.13 | 0.13 | 0.19 |
| 6 | 121844351 | rs4143301 | -0.106 | 0.056 | -0.74 | 0.021 | 0.039 | 0.076 | 0.076 | 0 | 0.7 | 0.83 | 0.84 | 0.91 | 0.8 | 0.79 |
| 6 | 121850798 | rs12189576 | -1.026 | 0.647 | -0.637 | 0.889 | 0.002 | 0.023 | 0.023 | 0.017 | 0.19 | 0.25 | 0.23 | 0.48 | 0.28 | 0.16 |
| 6 | 121854778 | rs942380 | 0.728 | 0.796 | -0.767 | 1.097 | 0.011 | 0.037 | 0.037 | 0.386 | 0.53 | 0.39 | 0.66 | 0.69 | 0.62 | 0.57 |
| 6 | 121860207 | rs9490318 | -0.333 | 0.722 | -0.749 | 0.862 | -0.001 | -0.005 | -0.005 | 1.446 | 0.17 | 0.22 | 0.15 | 0.16 | 0.14 | 0.17 |
| 6 | 121860377 | rs1324095 | 0.609 | 1.179 | -1.324 | 0.913 | 0.001 | 0.036 | 0.036 | 0 | 0.45 | 0.26 | 0.56 | 0.37 | 0.47 | 0.53 |
| 6 | 121866543 | rs6929781 | 0.838 | 1.021 | -1.039 | 1.287 | -0.007 | 0.01 | 0.01 | 0.566 | 0.41 | 0.27 | 0.59 | 0.38 | 0.46 | 0.55 |
| 6 | 121875648 | rs10457421 | 0.219 | 1.211 | -1.274 | 1.548 | -0.041 | -0.043 | -0.043 | 0 | 0.07 | 0 | 0.1 | 0.22 | 0.14 | 0.04 |
| 6 | 121895061 | rs4946578 | 0.49 | 0.366 | -0.97 | 0.938 | 0.011 | 0.014 | 0.014 | 1.959 | 0.49 | 0.38 | 0.69 | 0.6 | 0.6 | 0.57 |
| 6 | 121900015 | rs9482178 | -0.63 | 0.471 | -1.276 | 1.028 | 0.015 | -0.014 | -0.014 | 0 | 0.34 | 0.21 | 0.2 | 0.38 | 0.26 | 0.27 |
| 6 | 121906297 | rs6906280 | 0.245 | 0.726 | -1.641 | 0.925 | -0.029 | -0.009 | -0.009 | 0.008 | 0.63 | 0.82 | 0.69 | 0.39 | 0.6 | 0.81 |
| 6 | 121907968 | rs9401418 | -1.015 | 0.473 | -1.637 | 1.85 | 0.003 | 0.004 | 0.004 | 0 | 0.1 | 0.1 | 0.08 | 0.38 | 0.07 | 0.08 |
| 6 | 121925056 | rs4523118 | 0.437 | 0.245 | -1.11 | 1.414 | 0 | -0.006 | -0.006 | 0.269 | 0.53 | 0.44 | 0.62 | 0.36 | 0.53 | 0.69 |
| 6 | 121931830 | rs4568483 | 0.379 | 0.204 | -0.923 | 1.392 | -0.001 | 0 | 0 | 0 | 0.54 | 0.5 | 0.62 | 0.36 | 0.53 | 0.69 |
| 6 | 121932660 | rs11759649 | 0.365 | 0.127 | -0.84 | 1.619 | -0.051 | 0.057 | 0.057 | 0.442 | 0.32 | 0.14 | 0.26 | 0.16 | 0.36 | 0.22 |
| 6 | 121933718 | rs12196004 | -0.96 | 0.129 | -0.719 | 1.628 | 0.002 | 0.01 | 0.01 | 0 | 0.13 | 0.13 | 0.09 | 0.48 | 0.1 | 0.08 |
| 6 | 121949958 | rs9401435 | -0.228 | 0.141 | -0.729 | 1.55 | 0 | -0.006 | -0.006 | 0 | 0.65 | 0.63 | 0.71 | 0.84 | 0.64 | 0.78 |
| 6 | 121968356 | rs7772792 | 0.553 | 0.06 | -0.823 | 1.397 | -0.001 | 0.006 | 0.006 | 0.251 | 0.52 | 0.5 | 0.61 | 0.36 | 0.49 | 0.68 |
| 6 | 121986316 | rs10872226 | 1.416 | -0.28 | -0.529 | 0.622 | 0.004 | 0.011 | 0.011 | 0.567 | 0.26 | 0.25 | 0.44 | 0.37 | 0.26 | 0.57 |
| 6 | 121990721 | rs9490336 | 0.522 | -0.459 | -0.717 | 0.411 | 0.003 | 0.005 | 0.005 | 0 | 0.39 | 0.26 | 0.5 | 0.38 | 0.34 | 0.58 |
| 6 | 121995889 | rs7754730 | 0.522 | -0.47 | -0.73 | 0.426 | 0.007 | -0.009 | -0.009 | 0 | 0.38 | 0.42 | 0.55 | 0.38 | 0.41 | 0.66 |
| 6 | 122008684 | rs11757506 | 0.268 | -0.389 | -0.695 | 0.573 | -0.011 | -0.019 | -0.019 | 0 | 0.45 | 0.34 | 0.35 | 0.47 | 0.48 | 0.31 |
| 6 | 122034905 | rs9385212 | -0.144 | -0.413 | -0.759 | 0.57 | 0.004 | -0.005 | -0.005 | 0 | 0.11 | 0.07 | 0.08 | 0.31 | 0.11 | 0.08 |
| 6 | 122043606 | rs6569227 | 0.355 | -0.488 | -0.668 | 0.357 | 0.008 | -0.025 | -0.025 | 0 | 0.09 | 0.09 | 0.09 | 0.01 | 0.15 | 0.1 |
| 6 | 122046232 | rs12055526 | 0.378 | -0.518 | -0.712 | 0.422 | -0.019 | -0.027 | -0.027 | 0 | 0.44 | 0.27 | 0.34 | 0.47 | 0.48 | 0.31 |
| 6 | 122063279 | rs6938429 | 1.511 | -0.463 | -0.541 | 0.209 | -0.007 | -0.034 | -0.034 | 0 | 0.24 | 0.08 | 0.42 | 0.37 | 0.21 | 0.56 |
| 6 | 122064957 | rs4584022 | -0.175 | -0.243 | -0.658 | 0.248 | 0 | 0 | 0 | 0 | 0.5 | 0.46 | 0.31 | 0.38 | 0.46 | 0.67 |
| 6 | 122073750 | rs9385214 | 0.47 | -0.283 | -0.565 | 0.421 | -0.025 | -0.042 | -0.042 | 0 | 0.42 | 0.23 | 0.31 | 0.46 | 0.43 | 0.29 |
| 6 | 122075329 | rs4245517 | -0.302 | -0.297 | -0.476 | 0.406 | 0.001 | -0.006 | -0.006 | 0 | 0.53 | 0.45 | 0.65 | 0.38 | 0.52 | 0.71 |
| 6 | 122120089 | rs7767179 | -0.476 | -0.079 | -0.847 | 1.053 | -0.023 | 0.202 | 0.202 | 2.43 | 0.15 | 0.6 | 0.1 | 0.43 | 0.11 | 0.09 |
| 6 | 122125094 | rs2040424 | -1.186 | -0.251 | -1.14 | 0.908 | -0.065 | -0.11 | -0.11 | 0.626 | 0.74 | 0.99 | 0.54 | 0.65 | 0.75 | 0.4 |
| 6 | 122125427 | rs12208772 | -0.793 | -0.194 | -0.602 | 1.051 | -0.021 | 0.217 | 0.217 | 2.057 | 0.15 | 0.58 | 0.1 | 0.45 | 0.11 | 0.09 |
| 6 | 122188393 | rs12197250 | -0.66 | 0.219 | -1.106 | 0.144 | -0.01 | 0.045 | 0.045 | 0.013 | 0.16 | 0.05 | 0.1 | 0.01 | 0.19 | 0.09 |
| 6 | 122198028 | rs9490354 | -0.446 | 0.218 | -0.977 | 0.54 | -0.025 | -0.009 | -0.009 | 0 | 0.69 | 0.87 | 0.48 | 0.67 | 0.62 | 0.46 |
| 6 | 122198649 | rs9375072 | -0.589 | -0.27 | -0.28 | 1.024 | -0.009 | 0.245 | 0.245 | 3.006 | 0.13 | 0.49 | 0.1 | 0.38 | 0.09 | 0.15 |
| 6 | 122203693 | rs1034443 | -0.393 | -0.225 | 0.252 | 1.056 | 0.003 | 0.251 | 0.251 | 2.153 | 0.12 | 0.37 | 0.09 | 0.42 | 0.08 | 0.14 |
| 6 | 122211520 | rs763075 | -0.42 | -0.413 | -0.164 | 0.744 | -0.037 | 0.035 | 0.035 | 0.843 | 0.6 | 0.86 | 0.42 | 0.71 | 0.54 | 0.44 |
| 6 | 122212926 | rs10872231 | -0.209 | -0.384 | -0.059 | 0.816 | 0.006 | 0.284 | 0.284 | 2.05 | 0.12 | 0.38 | 0.09 | 0.47 | 0.08 | 0.14 |
| 6 | 122213903 | rs11154055 | 0.052 | -0.395 | 0.129 | 0.751 | -0.002 | 0.048 | 0.048 | 1.381 | 0.1 | 0.15 | 0.07 | 0.42 | 0.08 | 0.14 |
| 6 | 122229536 | rs9482193 | -0.118 | -0.616 | -0.344 | 0.383 | -0.005 | 0.066 | 0.066 | 0.398 | 0.49 | 0.86 | 0.39 | 0.65 | 0.49 | 0.42 |
| 6 | 122233342 | rs9398655 | -1.475 | -0.796 | -0.031 | 0.343 | -0.035 | 0.128 | 0.128 | 4.443 | 0.18 | 0.61 | 0.12 | 0.49 | 0.12 | 0.21 |
| 6 | 122246376 | rs4401702 | -1.484 | -0.813 | 0.014 | 0.316 | -0.033 | 0.041 | 0.041 | 3.875 | 0.28 | 0.59 | 0.14 | 0.51 | 0.16 | 0.23 |
| 6 | 122246782 | rs1915559 | -1.282 | -0.845 | 0.019 | 0.303 | -0.041 | 0.045 | 0.045 | 4.834 | 0.26 | 0.59 | 0.13 | 0.44 | 0.13 | 0.22 |
| 6 | 122253894 | rs6420722 | -0.507 | -0.04 | -0.17 | -0.123 | 0 | -0.023 | -0.023 | 0.921 | 0.89 | 0.86 | 0.9 | 1 | 0.85 | 0.94 |
| 6 | 122259154 | rs7747586 | 1.682 | -0.162 | -0.392 | -0.23 | -0.008 | 0.03 | 0.03 | 0 | 0.21 | 0.11 | 0.15 | 0.09 | 0.22 | 0.08 |
| 6 | 122260215 | rs9401471 | 0.177 | -0.27 | 0.077 | 0.291 | -0.002 | 0.174 | 0.174 | 0.933 | 0.1 | 0.2 | 0.11 | 0.46 | 0.09 | 0.23 |
| 6 | 122260679 | rs9401473 | 1.725 | -0.389 | -0.376 | -0.007 | -0.127 | -0.099 | -0.099 | 4.693 | 0.31 | 0.02 | 0.49 | 0.34 | 0.36 | 0.5 |
| 6 | 122265120 | rs1554438 | -1.148 | -0.577 | 0.57 | 0.514 | -0.051 | 0.148 | 0.148 | 0.191 | 0.21 | 0.39 | 0.16 | 0.55 | 0.13 | 0.24 |
| 6 | 122279121 | rs9482203 | 0.666 | -0.625 | 0.428 | 0.685 | -0.006 | -0.017 | -0.017 | 0 | 0.07 | 0.02 | 0.05 | 0.12 | 0.07 | 0.17 |
| 6 | 122287873 | rs6903218 | -1.52 | -0.759 | 0.485 | 0.378 | -0.062 | 0.131 | 0.131 | 0.242 | 0.23 | 0.39 | 0.16 | 0.55 | 0.12 | 0.24 |
| 6 | 122290881 | rs2259338 | -0.183 | -0.552 | 0.767 | 0.331 | 0.027 | -0.059 | -0.059 | 0.132 | 0.87 | 0.76 | 0.89 | 0.99 | 0.82 | 0.88 |

| | | | | | | | | | | | | | | | | |
|---|---|---|---|---|---|---|---|---|---|---|---|---|---|---|---|---|
| 6 | 122293478 | rs1521219 | 1.355 | -0.476 | 0.809 | 0.448 | 0.003 | -0.007 | -0.007 | 0.702 | 0.66 | 0.37 | 0.73 | 0.44 | 0.7 | 0.64 |
| 6 | 122294314 | rs6940106 | -1.167 | -0.531 | 0.837 | 0.559 | -0.018 | 0.039 | 0.039 | 2.471 | 0.29 | 0.42 | 0.22 | 0.44 | 0.23 | 0.26 |
| 6 | 122294945 | rs1521224 | 1.714 | -0.392 | 1.08 | 0.444 | 0.061 | 0.074 | 0.074 | 3.643 | 0.92 | 0.58 | 0.94 | 0.89 | 0.95 | 0.93 |
| 6 | 122296892 | rs2260076 | 0.308 | -0.393 | 1.148 | 0.54 | 0.024 | 0.137 | 0.137 | 0.047 | 0.78 | 0.57 | 0.78 | 0.5 | 0.75 | 0.75 |
| 6 | 122296944 | rs2260077 | 0.509 | -0.404 | 1.266 | 0.537 | 0.009 | 0.098 | 0.098 | 0.779 | 0.78 | 0.38 | 0.78 | 0.57 | 0.78 | 0.75 |
| 6 | 122305408 | rs2679684 | 1.239 | -0.607 | 1.229 | 0.462 | -0.013 | 0.028 | 0.028 | 2.728 | 0.68 | 0.42 | 0.77 | 0.52 | 0.74 | 0.66 |
| 6 | 122313558 | rs2679672 | 1.476 | -0.597 | 1.08 | 0.468 | 0.029 | -0.053 | -0.053 | 0.706 | 0.66 | 0.19 | 0.75 | 0.52 | 0.74 | 0.66 |
| 6 | 122319769 | rs9372673 | 0.056 | -0.47 | 1.152 | 0.473 | -0.017 | 0.08 | 0.08 | 0.661 | 0.19 | 0.28 | 0.17 | 0.43 | 0.15 | 0.24 |
| 6 | 122323908 | rs1521242 | 1.204 | -0.51 | 0.601 | 0.432 | 0.028 | -0.048 | -0.048 | 0 | 0.64 | 0.18 | 0.73 | 0.45 | 0.72 | 0.66 |
| 6 | 122330339 | rs7751455 | 0.428 | -0.432 | 0.638 | 0.556 | 0.015 | 0.103 | 0.103 | 1.172 | 0.76 | 0.47 | 0.81 | 0.57 | 0.77 | 0.67 |
| 6 | 122331465 | rs12526782 | -0.511 | -0.528 | 0.758 | 0.337 | -0.036 | -0.05 | -0.05 | 0 | 0.15 | 0.46 | 0.11 | 0.08 | 0.08 | 0.08 |
| 6 | 122332598 | rs7766484 | -0.07 | -0.001 | 0.469 | 0.445 | 0.054 | 0.159 | 0.159 | 0.419 | 0.77 | 0.5 | 0.79 | 0.51 | 0.72 | 0.74 |
| 6 | 122333898 | rs7772387 | 1.485 | 0.248 | -0.703 | 0.18 | -0.042 | 0.028 | 0.028 | 0 | 0.19 | 0.01 | 0.13 | 0.08 | 0.2 | 0.07 |
| 6 | 122342787 | rs12214218 | 1.225 | 0.357 | -0.659 | 0.304 | -0.041 | -0.028 | -0.028 | 0 | 0.3 | 0.07 | 0.47 | 0.32 | 0.33 | 0.5 |
| 6 | 122346667 | rs13210237 | 0.25 | 0.173 | -0.357 | 0.046 | -0.003 | -0.021 | -0.021 | 0 | 0.34 | 0.35 | 0.23 | 0.1 | 0.31 | 0.15 |
| 6 | 122357985 | rs9375094 | 0.085 | 0.263 | -0.494 | 0.892 | 0.009 | 0.006 | 0.006 | 0 | 0.05 | 0.01 | 0.03 | 0.22 | 0.02 | 0.03 |
| 6 | 122363499 | rs1474226 | 0.968 | -0.044 | -1.128 | 0.786 | 0.018 | 0.016 | 0.016 | 0.037 | 0.53 | 0.12 | 0.6 | 0.35 | 0.63 | 0.56 |
| 6 | 122363734 | rs1357056 | -0.576 | -0.107 | -0.477 | 0.212 | -0.016 | -0.073 | -0.073 | 0 | 0.12 | 0.55 | 0.07 | 0 | 0.07 | 0.04 |
| 6 | 122367334 | rs2679699 | 0.049 | 0.23 | -0.378 | 0.706 | -0.001 | 0 | 0 | 0 | 0.71 | 0.7 | 0.74 | 0.63 | 0.76 | 0.64 |
| 6 | 122372524 | rs9385230 | -1.034 | 0.572 | -0.175 | 0.724 | 0.001 | 0.035 | 0.035 | 0 | 0.08 | 0.06 | 0.07 | 0.27 | 0.08 | 0.04 |
| 6 | 122374344 | rs7742105 | -1.279 | 0.485 | 0.058 | 0.238 | -0.002 | -0.005 | -0.005 | 1.498 | 0.27 | 0.38 | 0.26 | 0.27 | 0.25 | 0.13 |
| 6 | 122374469 | rs2684257 | -1.15 | 0.201 | 0.128 | 0.32 | -0.016 | 0.012 | 0.012 | 0 | 0.24 | 0.37 | 0.23 | 0.27 | 0.21 | 0.12 |
| 6 | 122377265 | rs9320853 | -1.095 | 0.215 | 0.551 | 0.403 | 0.056 | 0.046 | 0.046 | 0 | 0.08 | 0.41 | 0.07 | 0.07 | 0.06 | 0.04 |
| 6 | 122377707 | rs2684256 | -1.641 | -0.332 | 0.454 | 0.123 | -0.031 | 0.021 | 0.021 | 0 | 0.23 | 0.59 | 0.16 | 0.28 | 0.15 | 0.13 |
| 6 | 122379099 | rs9388041 | -0.496 | 0.157 | 0.315 | 0.018 | 0.002 | -0.006 | -0.006 | 0 | 0.16 | 0.12 | 0.11 | 0.21 | 0.16 | 0.09 |
| 6 | 122386599 | rs2816109 | 0.205 | 0.283 | -0.167 | -0.134 | 0 | -0.006 | -0.006 | 0 | 0.55 | 0.56 | 0.43 | 0.63 | 0.51 | 0.55 |
| 6 | 122388268 | rs2816106 | -0.19 | 0.54 | -0.107 | -0.444 | -0.013 | -0.004 | -0.004 | 0 | 0.29 | 0.3 | 0.44 | 0.33 | 0.26 | 0.37 |
| 6 | 122388851 | rs1402538 | 1.299 | 0.477 | -0.133 | -0.253 | -0.009 | 0.002 | 0.002 | 0 | 0.43 | 0.55 | 0.34 | 0.39 | 0.35 | 0.47 |
| 6 | 122396987 | rs2816095 | -0.384 | 1.218 | 0.072 | -0.138 | 0.098 | -0.038 | -0.038 | 1.225 | 0.84 | 0.54 | 0.84 | 0.89 | 0.79 | 0.91 |
| 6 | 122410187 | rs9401492 | 0.3 | 1.368 | -0.148 | -0.052 | -0.058 | -0.051 | -0.051 | 0 | 0.13 | 0.02 | 0.1 | 0.21 | 0.18 | 0.08 |
| 6 | 122414296 | rs2606604 | -0.635 | 1.293 | 0.062 | -0.119 | 0.083 | -0.006 | -0.006 | 0.437 | 0.87 | 0.72 | 0.87 | 0.89 | 0.8 | 0.91 |
| 6 | 122428444 | rs720204 | -0.299 | 0.589 | -0.269 | -0.479 | 0.003 | 0.003 | 0.003 | 0.114 | 0.36 | 0.3 | 0.47 | 0.28 | 0.3 | 0.4 |
| 6 | 122428475 | rs720205 | -0.63 | 0.918 | -0.035 | -0.349 | 0.071 | 0.005 | 0.005 | 0.107 | 0.87 | 0.71 | 0.87 | 0.86 | 0.8 | 0.9 |
| 6 | 122433229 | rs11969110 | 1.086 | 0.756 | -0.277 | -0.507 | -0.036 | 0.046 | 0.046 | 0.025 | 0.08 | 0 | 0.1 | 0.01 | 0.12 | 0.09 |
| 6 | 122433255 | rs2606645 | 0.473 | 0.809 | -0.085 | -0.288 | 0.056 | 0.015 | 0.015 | 0 | 0.71 | 0.39 | 0.73 | 0.64 | 0.62 | 0.82 |
| 6 | 122454702 | rs9401494 | 0.404 | 0.929 | 0.125 | -0.273 | -0.017 | -0.014 | -0.014 | 0.219 | 0.14 | 0.09 | 0.12 | 0.22 | 0.19 | 0.08 |
| 6 | 122455798 | rs1456705 | -0.134 | 0.122 | -0.281 | -0.662 | 0.017 | 0.018 | 0.018 | 0.324 | 0.43 | 0.32 | 0.5 | 0.29 | 0.33 | 0.51 |
| 6 | 122482716 | rs2078231 | 1.477 | 0.236 | -0.394 | -0.632 | 0 | 0 | 0 | 0 | 0.29 | 0.27 | 0.23 | 0.38 | 0.27 | 0.32 |
| 6 | 122493544 | rs9401495 | 0.994 | 0.369 | -0.512 | -0.633 | -0.053 | -0.064 | -0.064 | 0 | 0.12 | 0.01 | 0.11 | 0.22 | 0.17 | 0.07 |
| 6 | 122505593 | rs17084326 | 1.8 | 0.238 | -0.465 | -0.598 | 0 | -0.082 | -0.082 | 2.965 | 0.27 | 0.05 | 0.21 | 0.35 | 0.27 | 0.31 |
| 6 | 122532273 | rs12662844 | 1.798 | 0.241 | -0.416 | -0.453 | 0.003 | -0.061 | -0.061 | 0.024 | 0.27 | 0.08 | 0.21 | 0.33 | 0.27 | 0.31 |
| 6 | 122537218 | rs12201864 | 0.738 | 0.281 | -0.421 | -0.468 | -0.028 | -0.023 | -0.023 | 0 | 0.14 | 0.07 | 0.13 | 0.22 | 0.19 | 0.08 |
| 6 | 122537787 | rs6905707 | 1.711 | 0.247 | -0.352 | -0.486 | 0.001 | -0.002 | -0.002 | 0 | 0.27 | 0.21 | 0.22 | 0.33 | 0.27 | 0.31 |
| 6 | 122545849 | rs9385236 | 0.656 | 0.35 | -0.337 | -0.421 | -0.013 | -0.01 | -0.01 | 0 | 0.14 | 0.1 | 0.14 | 0.22 | 0.19 | 0.08 |
| 6 | 122570412 | rs17084371 | 0.714 | -0.121 | -0.125 | -0.44 | 0.004 | -0.004 | -0.004 | 0 | 0.17 | 0.15 | 0.31 | 0.25 | 0.1 | 0.34 |
| 6 | 122572618 | rs9632500 | 1.087 | 0.214 | -0.059 | -0.307 | -0.008 | 0.002 | 0.002 | 0.012 | 0.13 | 0.11 | 0.12 | 0.13 | 0.2 | 0.1 |
| 6 | 122581103 | rs7765736 | 0.799 | 0.178 | 0.157 | -0.286 | 0.026 | 0.03 | 0.03 | 0 | 0.15 | 0.24 | 0.14 | 0.22 | 0.18 | 0.07 |
| 6 | 122581156 | rs11756232 | 0.396 | -0.15 | 0.137 | -0.476 | 0.002 | 0.001 | 0.001 | 0 | 0.25 | 0.25 | 0.38 | 0.25 | 0.19 | 0.31 |
| 6 | 122585720 | rs1927869 | 1.271 | -0.032 | -0.001 | -0.364 | -0.012 | -0.06 | -0.06 | 0 | 0.1 | 0.01 | 0.29 | 0.25 | 0.09 | 0.26 |
| 6 | 122586107 | rs1379087 | -0.155 | -0.008 | 0.103 | -0.374 | 0.002 | 0.016 | 0.016 | 0 | 0.26 | 0.27 | 0.43 | 0.47 | 0.27 | 0.33 |
| 6 | 122588540 | rs699402 | 0.604 | 0.006 | -0.046 | -0.429 | 0 | 0.023 | 0.023 | 0.171 | 0.17 | 0.1 | 0.12 | 0.03 | 0.15 | 0.24 |
| 6 | 122605510 | rs699413 | 0.006 | 0.417 | 0.248 | -0.1 | 0.034 | 0.009 | 0.009 | 0 | 0.72 | 0.66 | 0.74 | 0.65 | 0.62 | 0.84 |
| 6 | 122629902 | rs9375112 | -0.3 | 0.365 | 0.365 | 0.195 | 0.053 | 0.038 | 0.038 | 0 | 0.3 | 0.54 | 0.28 | 0.42 | 0.39 | 0.18 |
| 6 | 122650980 | rs17084411 | 0.738 | 0.214 | 0.365 | 0.229 | -0.004 | 0.01 | 0.01 | 0 | 0.12 | 0.15 | 0.3 | 0.25 | 0.09 | 0.26 |
| 6 | 122661084 | rs1028901 | 0.295 | 0.85 | 0.502 | 0.511 | 0.018 | 0.021 | 0.021 | 0 | 0.15 | 0.2 | 0.13 | 0.23 | 0.18 | 0.07 |
| 6 | 122661338 | rs7751410 | 0.642 | 0.678 | 0.468 | 0.329 | -0.011 | -0.013 | -0.013 | 0 | 0.41 | 0.32 | 0.39 | 0.45 | 0.47 | 0.35 |
| 6 | 122663662 | rs9372690 | -0.564 | 0.773 | 0.509 | 0.311 | 0.069 | 0.054 | 0.054 | 0 | 0.29 | 0.46 | 0.27 | 0.43 | 0.38 | 0.18 |
| 6 | 122663943 | rs12212486 | 0.427 | 0.772 | 0.528 | 0.308 | 0.036 | 0.042 | 0.042 | 0 | 0.16 | 0.29 | 0.14 | 0.23 | 0.18 | 0.07 |
| 6 | 122666394 | rs1339532 | 0.824 | 0.798 | 0.298 | 0.329 | 0.004 | 0.001 | 0.001 | 0.009 | 0.13 | 0.16 | 0.13 | 0.2 | 0.2 | 0.11 |
| 6 | 122675030 | rs9401513 | 0.178 | 0.709 | 0.403 | 0.285 | 0.028 | 0.031 | 0.031 | 0 | 0.17 | 0.33 | 0.14 | 0.23 | 0.18 | 0.07 |
| 6 | 122677364 | rs1610023 | 1.232 | 0.651 | 0.242 | 0.395 | 0.006 | -0.01 | -0.01 | 0 | 0.28 | 0.19 | 0.22 | 0.31 | 0.26 | 0.27 |
| 6 | 122683423 | rs2316669 | -0.355 | 0.581 | 0.203 | 0.395 | 0.014 | 0.012 | 0.012 | 0.111 | 0.58 | 0.65 | 0.5 | 0.67 | 0.65 | 0.44 |
| 6 | 122687584 | rs2094574 | -0.365 | 0.619 | 0.175 | 0.465 | 0.024 | 0.025 | 0.025 | 0.085 | 0.58 | 0.67 | 0.49 | 0.68 | 0.64 | 0.4 |
| 6 | 122691294 | rs9320866 | 0.212 | 0.763 | 0.304 | 0.646 | 0.029 | 0.028 | 0.028 | 0 | 0.17 | 0.33 | 0.14 | 0.21 | 0.18 | 0.07 |
| 6 | 122693552 | rs7744565 | -0.311 | 0.696 | 0.151 | 0.567 | 0.024 | 0.025 | 0.025 | 0.085 | 0.58 | 0.67 | 0.49 | 0.68 | 0.64 | 0.4 |
| 6 | 122708203 | rs11154078 | 0.378 | 0.885 | 0.021 | 0.718 | 0.042 | 0.02 | 0.02 | 0.105 | 0.14 | 0.26 | 0.14 | 0.21 | 0.2 | 0.07 |
| 6 | 122709054 | rs1416733 | -0.19 | 0.763 | -0.028 | 0.598 | 0.007 | 0.017 | 0.017 | 0.285 | 0.55 | 0.61 | 0.47 | 0.69 | 0.62 | 0.4 |
| 6 | 122709984 | rs530928 | 0.228 | 0.738 | -0.114 | 0.549 | 0.014 | 0.017 | 0.017 | 0.629 | 0.43 | 0.37 | 0.47 | 0.32 | 0.31 | 0.58 |
| 6 | 122712377 | rs9490444 | 1.149 | 0.719 | -0.628 | 0.672 | -0.051 | -0.043 | -0.043 | 0 | 0.28 | 0.06 | 0.26 | 0.27 | 0.34 | 0.28 |

| | | | | | | | | | | | | | | | | |
|---|---|---|---|---|---|---|---|---|---|---|---|---|---|---|---|---|
| 6 | 122721510 | rs1741820 | 0.68 | 0.772 | -0.578 | 0.996 | 0.019 | 0.023 | 0.023 | 0 | 0.63 | 0.53 | 0.61 | 0.45 | 0.56 | 0.72 |
| 6 | 122745009 | rs556439 | 0.012 | 0.78 | -0.726 | 1.079 | 0.044 | 0.035 | 0.035 | 0 | 0.78 | 0.65 | 0.74 | 0.7 | 0.71 | 0.8 |
| 6 | 122752447 | rs12192360 | 0.512 | 0.761 | -0.693 | 1.076 | 0.003 | -0.032 | -0.032 | 0 | 0.15 | 0.1 | 0.12 | 0.25 | 0.15 | 0.08 |
| 6 | 122752471 | rs12205759 | 0.371 | 0.769 | -0.691 | 1.076 | 0 | -0.03 | -0.03 | 0 | 0.15 | 0.08 | 0.13 | 0.2 | 0.15 | 0.07 |
| 6 | 122759415 | rs577838 | -0.886 | 0.8 | -0.659 | 1.048 | 0.091 | 0.031 | 0.031 | 0.682 | 0.87 | 0.69 | 0.87 | 0.84 | 0.8 | 0.95 |
| 6 | 122763602 | rs9388074 | 0.374 | 0.736 | -0.74 | 1.052 | 0 | -0.036 | -0.036 | 6.648 | 0.14 | 0.08 | 0.12 | 0.25 | 0.15 | 0.08 |
| 6 | 122765026 | rs13371 | -0.874 | 0.806 | -0.654 | 1.019 | 0.091 | 0.031 | 0.031 | 0.682 | 0.87 | 0.69 | 0.87 | 0.84 | 0.8 | 0.95 |
| 6 | 122765777 | rs11154080 | 1.627 | 0.795 | -0.896 | 0.774 | -0.018 | 0.028 | 0.028 | 0.178 | 0.06 | 0.01 | 0.03 | 0 | 0.07 | 0.05 |
| 6 | 122765905 | rs487098 | 0.731 | 0.775 | -0.792 | 1.019 | 0.013 | 0.017 | 0.017 | 0 | 0.64 | 0.55 | 0.61 | 0.45 | 0.56 | 0.72 |
| 6 | 122765948 | rs12524 | 0.775 | 0.768 | -0.778 | 1.004 | 0.019 | 0.025 | 0.025 | 0 | 0.63 | 0.49 | 0.61 | 0.45 | 0.56 | 0.72 |
| 6 | 122786706 | rs9385246 | 0.404 | 0.698 | -0.833 | 1.003 | 0 | -0.036 | -0.036 | 0.06 | 0.14 | 0.08 | 0.12 | 0.25 | 0.15 | 0.08 |
| 6 | 122787063 | rs13437168 | 0.784 | 0.706 | -0.812 | 0.996 | 0.019 | 0.025 | 0.025 | 0 | 0.63 | 0.49 | 0.61 | 0.45 | 0.56 | 0.72 |
| 6 | 122792090 | rs17201852 | 1.444 | 0.291 | -1.263 | 0.815 | -0.04 | 0.002 | 0.002 | 0.45 | 0.25 | 0.03 | 0.15 | 0.15 | 0.26 | 0.24 |
| 6 | 122798376 | rs9490459 | 0.299 | 0.479 | -1.027 | 0.949 | 0.003 | -0.03 | -0.03 | 0 | 0.15 | 0.1 | 0.12 | 0.25 | 0.15 | 0.08 |
| 6 | 122801699 | rs10872238 | 0.342 | 0.477 | -1.041 | 0.949 | 0 | -0.038 | -0.038 | 0 | 0.15 | 0.08 | 0.13 | 0.25 | 0.15 | 0.08 |
| 6 | 122807318 | rs2039784 | 0.477 | 0.455 | -1.058 | 0.956 | 0.002 | -0.04 | -0.04 | 0 | 0.15 | 0.08 | 0.13 | 0.25 | 0.15 | 0.08 |
| 6 | 122809599 | rs7767114 | 0.795 | 0.401 | -0.973 | 1.064 | 0.019 | 0.026 | 0.026 | 0 | 0.64 | 0.5 | 0.61 | 0.45 | 0.57 | 0.72 |
| 6 | 122829471 | rs155464 | 0.182 | 0.407 | -1.087 | 1.074 | 0.04 | 0.052 | 0.052 | 0 | 0.73 | 0.53 | 0.75 | 0.59 | 0.66 | 0.87 |
| 6 | 122855421 | rs1367490 | 0.272 | 0.424 | -1.213 | 1.059 | -0.008 | -0.029 | -0.029 | 0.13 | 0.09 | 0.01 | 0.13 | 0.14 | 0.1 | 0.15 |
| 6 | 122869514 | rs9320878 | 0.301 | 0.502 | -1.116 | 0.905 | 0.002 | -0.022 | -0.022 | 0 | 0.14 | 0.13 | 0.13 | 0.26 | 0.15 | 0.07 |
| 6 | 122870132 | rs197680 | -0.765 | 0.706 | -0.985 | 0.768 | -0.007 | 0 | 0 | 0 | 0.88 | 0.91 | 0.89 | 0.98 | 0.81 | 0.96 |
| 6 | 122874842 | rs197684 | -0.703 | 0.725 | -0.924 | 0.843 | 0.07 | 0.021 | 0.021 | 1.065 | 0.88 | 0.76 | 0.88 | 0.84 | 0.82 | 0.95 |
| 6 | 122910337 | rs6930161 | 0.87 | 0.878 | -1.399 | 0.905 | -0.081 | -0.087 | -0.087 | 0 | 0.3 | 0.09 | 0.28 | 0.33 | 0.39 | 0.19 |
| 6 | 122913123 | rs1430816 | -1.18 | 1.101 | -1.154 | 0.874 | 0.115 | 0.018 | 0.018 | 0 | 0.9 | 0.67 | 0.88 | 0.9 | 0.84 | 0.97 |
| 6 | 122914320 | rs2566837 | 0.159 | 0.835 | -1.378 | 0.834 | -0.001 | -0.002 | -0.002 | 0 | 0.49 | 0.46 | 0.5 | 0.31 | 0.37 | 0.59 |
| 6 | 122918372 | rs4554346 | 0.151 | 0.823 | -2.087 | 0.689 | 0.007 | -0.045 | -0.045 | 0.676 | 0.1 | 0.02 | 0.1 | 0.21 | 0.08 | 0.19 |
| 6 | 122927060 | rs13191198 | 0.238 | 0.853 | -2.318 | 0.562 | -0.023 | -0.017 | -0.017 | 12.96 | 0.44 | 0.1 | 0.34 | 0.38 | 0.43 | 0.4 |
| 6 | 122928126 | rs9482260 | 0.998 | 1.691 | -1.527 | 0.67 | 0.028 | -0.04 | -0.04 | 0 | 0.09 | 0.14 | 0.11 | 0 | 0.15 | 0.02 |
| 6 | 122935161 | rs9388103 | 0.729 | 1.384 | -1.674 | 1.18 | -0.031 | -0.045 | -0.045 | 3.036 | 0.08 | 0.02 | 0.09 | 0.18 | 0.12 | 0.06 |
| 6 | 122937369 | rs17675201 | 1.05 | 1.278 | -1.617 | 0.767 | -0.057 | -0.021 | -0.021 | 0 | 0.21 | 0.06 | 0.18 | 0.15 | 0.28 | 0.11 |
| 6 | 122941354 | rs9385262 | 0.238 | 0.945 | -1.15 | 1.025 | 0.002 | 0.011 | 0.011 | 0.374 | 0.09 | 0.09 | 0.09 | 0.13 | 0.11 | 0.09 |
| 6 | 122941551 | rs9398679 | 0.099 | 1 | -1.461 | 1.033 | -0.013 | -0.064 | -0.064 | 0.226 | 0.22 | 0.09 | 0.21 | 0.37 | 0.29 | 0.18 |
| 6 | 122945140 | rs9388105 | -0.358 | 0.538 | -1.57 | 0.121 | 0 | 0 | 0 | 0 | 0.63 | 0.39 | 0.5 | 0.4 | 0.71 | 0.31 |
| 6 | 122949909 | rs6914480 | 0.658 | 0.514 | -1.584 | 0.038 | -0.018 | 0.075 | 0.075 | 0.024 | 0.4 | 0.27 | 0.29 | 0.03 | 0.42 | 0.13 |
| 6 | 122950770 | rs2815584 | 0.126 | 0.465 | -1.222 | 0.326 | -0.008 | 0.016 | 0.016 | 1.843 | 0.37 | 0.55 | 0.5 | 0.57 | 0.29 | 0.68 |
| 6 | 122965282 | rs2566830 | 0.112 | 0.353 | -1.268 | 0.27 | -0.006 | 0.024 | 0.024 | 1.227 | 0.35 | 0.55 | 0.5 | 0.57 | 0.29 | 0.68 |
| 6 | 122972905 | rs2815600 | 0.155 | 0.403 | -0.944 | 0.483 | -0.016 | 0.024 | 0.024 | 2.008 | 0.36 | 0.46 | 0.49 | 0.48 | 0.29 | 0.66 |
| 6 | 122978422 | rs2316962 | 0.974 | 0.515 | -1.334 | -0.044 | -0.016 | 0 | 0 | 0.02 | 0.31 | 0.26 | 0.24 | 0 | 0.35 | 0.06 |
| 6 | 122980559 | rs17052989 | 0.598 | 0.391 | -0.974 | 0.147 | -0.006 | 0.101 | 0.101 | 0.952 | 0.12 | 0.21 | 0.29 | 0.28 | 0.08 | 0.39 |
| 6 | 122990817 | rs9320886 | -0.002 | 0.606 | -1.772 | -0.135 | 0.029 | 0.004 | 0.004 | 1.404 | 0.53 | 0.35 | 0.45 | 0.39 | 0.65 | 0.25 |
| 6 | 122992957 | rs17676223 | 1.457 | 0.643 | -1.915 | -0.423 | -0.023 | -0.073 | -0.073 | 0 | 0.14 | 0.01 | 0.13 | 0.24 | 0.19 | 0.09 |
| 6 | 122998592 | rs7769704 | 0.665 | 0.773 | -1.441 | -0.719 | -0.028 | 0.029 | 0.029 | 0 | 0.29 | 0.17 | 0.23 | 0 | 0.35 | 0.06 |
| 6 | 123006095 | rs7741881 | -0.153 | 0.801 | -0.961 | 0.302 | -0.015 | -0.005 | -0.005 | 0.376 | 0.34 | 0.33 | 0.31 | 0.32 | 0.38 | 0.19 |
| 6 | 123014708 | rs7771724 | -0.309 | 0.755 | -1.341 | 0.293 | 0.009 | 0.016 | 0.016 | 0.221 | 0.46 | 0.34 | 0.41 | 0.54 | 0.53 | 0.3 |
| 6 | 123018092 | rs9490525 | 1.002 | 0.928 | -1.394 | 0.054 | -0.012 | -0.016 | -0.016 | 0 | 0.2 | 0.24 | 0.17 | 0 | 0.26 | 0.05 |
| 6 | 123020680 | rs9401572 | -0.953 | 0.353 | -1.373 | -0.398 | -0.022 | 0.02 | 0.02 | 0.445 | 0.69 | 0.45 | 0.57 | 0.56 | 0.77 | 0.4 |
| 6 | 123022496 | rs4382328 | 0.462 | 0.338 | -1.415 | -0.424 | -0.013 | 0.081 | 0.081 | 0.294 | 0.44 | 0.33 | 0.33 | 0.03 | 0.49 | 0.17 |
| 6 | 123027077 | rs9401575 | 0.624 | 0.252 | -1.424 | -0.198 | -0.008 | -0.083 | -0.083 | 3.208 | 0.25 | 0.1 | 0.24 | 0.44 | 0.27 | 0.22 |
| 6 | 123029669 | rs13218313 | 1.002 | -0.355 | -1.034 | 0.039 | 0.007 | 0.064 | 0.064 | 0 | 0.23 | 0.08 | 0.15 | 0.01 | 0.24 | 0.11 |
| 6 | 123031749 | rs9490531 | 0.753 | -0.765 | -0.735 | 0.264 | -0.023 | 0.027 | 0.027 | 2.579 | 0.21 | 0.43 | 0.33 | 0.25 | 0.17 | 0.45 |
| 6 | 123032317 | rs4406270 | 0.162 | -1.087 | -1.113 | 0.115 | 0.044 | 0.063 | 0.063 | 0.061 | 0.35 | 0.09 | 0.26 | 0.27 | 0.29 | 0.3 |
| 6 | 123038540 | rs3813492 | -0.53 | -0.847 | -0.741 | -0.039 | 0.011 | 0.009 | 0.009 | 2.273 | 0.41 | 0.59 | 0.56 | 0.65 | 0.32 | 0.69 |
| 6 | 123039932 | rs12197460 | 0.885 | -0.815 | -0.386 | -0.001 | 0.021 | 0.014 | 0.014 | 1.418 | 0.22 | 0.36 | 0.36 | 0.26 | 0.25 | 0.45 |
| 6 | 123042792 | rs12209418 | 0.553 | -0.308 | -0.641 | -0.512 | 0.006 | 0.077 | 0.077 | 0 | 0.26 | 0.09 | 0.17 | 0.1 | 0.27 | 0.15 |
| 6 | 123046832 | rs1132630 | 0.109 | -0.389 | -0.62 | -0.503 | 0.01 | 0.009 | 0.009 | 1.64 | 0.41 | 0.58 | 0.56 | 0.65 | 0.32 | 0.69 |
| 6 | 123047016 | rs1132643 | 0.459 | 0.289 | 0.131 | 0.356 | -0.125 | -0.012 | -0.012 | 0 | 0.29 | 0.1 | 0.24 | 0.23 | 0.39 | 0.14 |
| 6 | 123049011 | rs6569275 | 0.188 | 0.319 | 0.215 | 0.354 | -0.03 | 0.014 | 0.014 | 3.063 | 0.33 | 0.26 | 0.27 | 0.25 | 0.41 | 0.16 |
| 6 | 123052266 | rs7761403 | -0.194 | 0.266 | 0.186 | 0.539 | -0.021 | 0.005 | 0.005 | 2.379 | 0.37 | 0.31 | 0.3 | 0.35 | 0.46 | 0.19 |
| 6 | 123058132 | rs12204690 | 0.245 | 0.269 | 0.24 | 0.619 | -0.016 | -0.014 | -0.014 | 0 | 0.57 | 0.61 | 0.62 | 0.53 | 0.52 | 0.75 |
| 6 | 123059586 | rs9388125 | 0.128 | 0.284 | -0.203 | 0.648 | -0.103 | -0.072 | -0.072 | 0 | 0.35 | 0.12 | 0.31 | 0.37 | 0.42 | 0.2 |
| 6 | 123071112 | rs6913003 | 0.563 | 0.412 | -0.056 | 0.711 | -0.006 | 0.015 | 0.015 | 0 | 0.94 | 0.85 | 0.95 | 0.9 | 0.95 | 0.94 |
| 6 | 123072973 | rs11963361 | 0.227 | 0.455 | 0.116 | 0.708 | -0.001 | -0.001 | -0.001 | 0.871 | 0.57 | 0.46 | 0.61 | 0.54 | 0.51 | 0.75 |
| 6 | 123074621 | rs9401589 | -0.021 | 0.782 | -0.318 | 0.894 | -0.007 | -0.003 | -0.003 | 0 | 0.34 | 0.28 | 0.31 | 0.35 | 0.42 | 0.18 |
| 6 | 123075986 | rs9388133 | 1.044 | 0.622 | 0.157 | 1.065 | -0.012 | 0.035 | 0.035 | 0.294 | 0.83 | 0.58 | 0.82 | 0.8 | 0.85 | 0.88 |
| 6 | 123077625 | rs9490545 | 0.061 | 0.618 | -0.372 | 0.757 | 0.013 | 0.039 | 0.039 | 0.22 | 0.41 | 0.23 | 0.41 | 0.32 | 0.62 | 0.37 |
| 6 | 123078637 | rs9490546 | -0.093 | 1.307 | -0.5 | 0.644 | -0.022 | 0.004 | 0.004 | 0 | 0.17 | 0.11 | 0.2 | 0.19 | 0.24 | 0.17 |
| 6 | 123087429 | rs936806 | -0.011 | 0.476 | -0.781 | 0.096 | -0.013 | 0.195 | 0.195 | 0 | 0.47 | 0.23 | 0.33 | 0.21 | 0.46 | 0.31 |
| 6 | 123089365 | rs1485649 | -0.339 | 0.323 | -0.647 | 0.429 | -0.041 | -0.021 | -0.021 | 0 | 0.8 | 0.43 | 0.85 | 0.85 | 0.9 | 0.83 |
| 6 | 123091132 | rs7742826 | 1.222 | 0.357 | -0.693 | 0.487 | -0.051 | 0.04 | 0.04 | 0 | 0.25 | 0.1 | 0.21 | 0.12 | 0.38 | 0.19 |

| | | | | | | | | | | | | | | | | |
|---|---|---|---|---|---|---|---|---|---|---|---|---|---|---|---|---|
| 6 | 123092085 | rs9385270 | 0.643 | -0.073 | -0.577 | 0.7 | -0.001 | 0.054 | 0.054 | 0 | 0.1 | 0.09 | 0.31 | 0.52 | 0.08 | 0.34 |
| 6 | 123095188 | rs7764703 | -0.22 | -0.078 | -0.506 | 0.035 | -0.051 | -0.019 | -0.019 | 0 | 0.8 | 0.42 | 0.85 | 0.85 | 0.91 | 0.84 |
| 6 | 123095881 | rs13214367 | 0.234 | 0.516 | -0.879 | 0.301 | 0.008 | 0.131 | 0.131 | 0 | 0.29 | 0.06 | 0.13 | 0.05 | 0.23 | 0.14 |
| 6 | 123101647 | rs2243372 | 0.041 | 0.718 | -0.423 | 0.696 | 0.002 | 0.01 | 0.01 | 0 | 0.54 | 0.55 | 0.66 | 0.7 | 0.57 | 0.7 |
| 6 | 123104307 | rs9490550 | 0.334 | 0.894 | -0.222 | 1.131 | 0.005 | -0.005 | -0.005 | 0.474 | 0.36 | 0.35 | 0.32 | 0.28 | 0.43 | 0.26 |
| 6 | 123104703 | rs9401596 | 0.893 | 0.851 | -0.169 | 1.327 | -0.002 | 0.005 | 0.005 | 0 | 0.82 | 0.73 | 0.82 | 0.75 | 0.83 | 0.83 |
| 6 | 123107697 | rs9482286 | -0.976 | 1.051 | -0.24 | 0.894 | 0 | -0.009 | -0.009 | 0 | 0.22 | 0.33 | 0.34 | 0.24 | 0.21 | 0.22 |
| 6 | 123109596 | rs6904500 | 0.3 | 1.441 | -0.781 | 1.164 | -0.017 | -0.04 | -0.04 | 1.482 | 0.75 | 0.84 | 0.68 | 0.53 | 0.7 | 0.64 |
| 6 | 123113273 | rs9490551 | -0.617 | 1.133 | -0.06 | 0.704 | 0.022 | -0.036 | -0.036 | 1.137 | 0.08 | 0.25 | 0.06 | 0 | 0.06 | 0.02 |
| 6 | 123118500 | rs9375185 | -0.331 | 1.35 | -0.831 | 0.787 | -0.016 | -0.054 | -0.054 | 0 | 0.14 | 0.05 | 0.21 | 0.33 | 0.18 | 0.04 |
| 6 | 123122385 | rs9372718 | -0.267 | 1.363 | -0.923 | 0.855 | -0.046 | -0.094 | -0.094 | 0.008 | 0.13 | 0.01 | 0.21 | 0.34 | 0.18 | 0.05 |
| 6 | 123124386 | rs2046446 | 0.866 | 1.985 | 0.202 | 0.09 | 0.187 | 0.021 | 0.021 | 0 | 0.93 | 0.68 | 0.92 | 1 | 0.87 | 0.96 |
| 6 | 123127597 | rs13192569 | 0.302 | 0.969 | -0.956 | -0.205 | 0.022 | 0.007 | 0.007 | 0 | 0.21 | 0.17 | 0.08 | 0.17 | 0.14 | 0.23 |
| 6 | 123129159 | rs11751996 | 0.143 | 1.115 | -1.551 | -0.922 | -0.027 | 0.088 | 0.088 | 0 | 0.32 | 0.13 | 0.26 | 0.13 | 0.32 | 0.25 |
| 6 | 123131515 | rs9375186 | 0.181 | 0.837 | -1.562 | -1.209 | -0.003 | 0.169 | 0.169 | 0 | 0.56 | 0.84 | 0.61 | 0.87 | 0.46 | 0.72 |
| 6 | 123139281 | rs7738249 | -0.14 | 0.884 | -1.782 | -1.122 | -0.078 | 0.155 | 0.155 | 0 | 0.39 | 0.12 | 0.34 | 0.14 | 0.45 | 0.26 |
| 6 | 123148664 | rs11154116 | 0.108 | 1.034 | -1.939 | -1.141 | -0.025 | 0.104 | 0.104 | 0 | 0.32 | 0.11 | 0.25 | 0.15 | 0.31 | 0.25 |
| 6 | 123154371 | rs7451567 | 0 | 1.362 | -1.482 | -0.753 | -0.019 | 0.018 | 0.018 | 0 | 0.95 | 0.99 | 0.95 | 1 | 0.88 | 0.98 |
| 6 | 123156684 | rs6920848 | 0.078 | 0.831 | -1.28 | -0.384 | -0.001 | -0.01 | -0.01 | 4.721 | 0.23 | 0.19 | 0.18 | 0.25 | 0.2 | 0.19 |
| 6 | 123167378 | rs12332942 | -0.154 | 0.393 | -0.513 | -1.033 | -0.025 | -0.119 | -0.119 | 2.911 | 0.18 | 0.4 | 0.12 | 0.04 | 0.14 | 0.12 |
| 6 | 123172683 | rs4962262 | -0.3 | 0.551 | -0.749 | -0.55 | 0.007 | 0.037 | 0.037 | 0 | 0.17 | 0.28 | 0.39 | 0.65 | 0.17 | 0.34 |
| 6 | 123176725 | rs6935233 | -0.217 | 0.4 | -1.221 | -1.033 | 0 | -0.013 | -0.013 | 0 | 0.63 | 0.39 | 0.76 | 0.85 | 0.66 | 0.67 |
| 6 | 123182193 | rs7759311 | 0.192 | 0.269 | -1.391 | -0.726 | 0.006 | -0.008 | -0.008 | 0 | 0.79 | 0.81 | 0.63 | 0.34 | 0.84 | 0.62 |
| 6 | 123183020 | rs6569294 | 0.469 | 0.593 | -1.627 | -0.947 | -0.057 | 0.105 | 0.105 | 0 | 0.33 | 0.13 | 0.32 | 0.13 | 0.44 | 0.2 |
| 6 | 123189265 | rs9490566 | 0.014 | 0.53 | -1.727 | -0.97 | -0.037 | 0.155 | 0.155 | 0.215 | 0.45 | 0.18 | 0.36 | 0.21 | 0.5 | 0.34 |
| 6 | 123189470 | rs9490568 | 0.293 | 0.516 | -1.814 | -0.993 | -0.045 | 0.178 | 0.178 | 1.047 | 0.41 | 0.12 | 0.32 | 0.15 | 0.45 | 0.26 |
| 6 | 123190840 | rs7740530 | -0.019 | 0.967 | -1.8 | -0.982 | -0.062 | 0.144 | 0.144 | 0.714 | 0.43 | 0.16 | 0.36 | 0.21 | 0.53 | 0.32 |
| 6 | 123196340 | rs9401605 | -0.139 | 0.864 | -1.441 | 0.012 | -0.011 | -0.021 | -0.021 | 0.41 | 0.35 | 0.32 | 0.39 | 0.46 | 0.4 | 0.25 |
| 6 | 123200118 | rs9401609 | 0.221 | 1.369 | -1.393 | -0.029 | 0.007 | 0.029 | 0.029 | 0.432 | 0.23 | 0.31 | 0.34 | 0.41 | 0.32 | 0.14 |
| 6 | 123206331 | rs7758880 | -0.583 | 0.693 | -1.116 | -0.434 | 0.003 | -0.005 | -0.005 | 0.002 | 0.7 | 0.68 | 0.79 | 0.77 | 0.69 | 0.79 |
| 6 | 123209819 | rs7382626 | 0.572 | 0.955 | -0.881 | -0.032 | -0.002 | 0.001 | 0.001 | 0.326 | 0.22 | 0.24 | 0.33 | 0.41 | 0.32 | 0.14 |
| 6 | 123218858 | rs7449946 | 0.353 | 1.254 | -0.476 | -0.59 | 0.004 | -0.038 | -0.038 | 0 | 0.61 | 0.59 | 0.79 | 0.88 | 0.63 | 0.56 |
| 6 | 123222617 | rs9490584 | 0.187 | 1.404 | -0.354 | -0.672 | 0.038 | -0.021 | -0.021 | 0.578 | 0.84 | 0.77 | 0.87 | 0.92 | 0.75 | 0.78 |
| 6 | 123223696 | rs9482300 | 0.808 | 1.425 | -0.41 | -0.377 | -0.006 | -0.005 | -0.005 | 0.598 | 0.3 | 0.35 | 0.4 | 0.33 | 0.3 | 0.42 |
| 6 | 123224427 | rs9388153 | 0.093 | 1.975 | 0.21 | 0.318 | -0.002 | 0.007 | 0.007 | 0.174 | 0.35 | 0.42 | 0.44 | 0.55 | 0.37 | 0.18 |
| 6 | 123225410 | rs9482302 | 0.163 | 2.16 | 0.168 | 0.261 | 0.005 | -0.004 | -0.004 | 0 | 0.61 | 0.57 | 0.53 | 0.39 | 0.51 | 0.67 |
| 6 | 123226577 | rs9398707 | -0.014 | 1.567 | 0.244 | 0.339 | 0.031 | -0.01 | -0.01 | 0.049 | 0.84 | 0.71 | 0.9 | 0.91 | 0.83 | 0.77 |
| 6 | 123233105 | rs9398709 | 0.705 | 1.56 | 0.312 | 0.498 | 0.001 | -0.009 | -0.009 | 0 | 0.56 | 0.52 | 0.44 | 0.71 | 0.52 | 0.63 |
| 6 | 123236315 | rs4962257 | 0.737 | 1.839 | 0.316 | 0.492 | -0.006 | 0.05 | 0.05 | 0 | 0.22 | 0.07 | 0.18 | 0.08 | 0.21 | 0.07 |
| 6 | 123236650 | rs1601450 | 0.292 | 1.838 | 0.362 | 0.492 | -0.001 | -0.01 | -0.01 | 0.056 | 0.26 | 0.24 | 0.22 | 0.09 | 0.27 | 0.1 |
| 6 | 123238525 | rs9482308 | 0.292 | 1.838 | 0.362 | 0.51 | -0.001 | -0.009 | -0.009 | 0 | 0.26 | 0.24 | 0.23 | 0.09 | 0.27 | 0.1 |
| 6 | 123238676 | rs7753475 | -0.306 | 1.853 | 0.487 | 0.648 | -0.018 | -0.015 | -0.015 | 0.029 | 0.84 | 0.9 | 0.68 | 0.79 | 0.78 | 0.73 |
| 6 | 123241452 | rs1905855 | -0.456 | 1.282 | 0.443 | 0.777 | -0.019 | 0.213 | 0.213 | 0.594 | 0.17 | 0.43 | 0.22 | 0.47 | 0.14 | 0.15 |
| 6 | 123244145 | rs724469 | 0.021 | 1.294 | 0.389 | 0.814 | 0.007 | 0.357 | 0.357 | 2.407 | 0.11 | 0.42 | 0.21 | 0.48 | 0.09 | 0.1 |
| 6 | 123245633 | rs7452973 | 1.606 | 1.358 | -0.442 | 0.623 | 0.028 | 0.105 | 0.105 | 0 | 0.4 | 0.21 | 0.23 | 0.23 | 0.37 | 0.5 |
| 6 | 123245782 | rs7453004 | -0.82 | 1.467 | -0.423 | 0.621 | 0.013 | 0.002 | 0.002 | 0.012 | 0.43 | 0.71 | 0.46 | 0.57 | 0.42 | 0.25 |
| 6 | 123251303 | rs13200297 | 0.636 | 1.437 | -0.514 | 0.576 | -0.026 | 0.053 | 0.053 | 0 | 0.23 | 0.08 | 0.2 | 0.08 | 0.26 | 0.09 |
| 6 | 123253629 | rs12205678 | 1.646 | 1.467 | -0.496 | 0.583 | 0.022 | 0.106 | 0.106 | 0 | 0.41 | 0.21 | 0.24 | 0.21 | 0.38 | 0.5 |
| 6 | 123254662 | rs6938524 | 1.199 | 1.399 | -0.121 | 1.013 | -0.003 | 0.018 | 0.018 | 0.48 | 0.63 | 0.3 | 0.44 | 0.3 | 0.65 | 0.59 |
| 6 | 123268345 | rs6569313 | 1.19 | 1.379 | -0.174 | 1 | -0.006 | 0.022 | 0.022 | 0.459 | 0.64 | 0.29 | 0.44 | 0.28 | 0.64 | 0.59 |
| 6 | 123286751 | rs4896501 | 0.07 | 1.712 | -0.484 | 1.098 | -0.03 | -0.028 | -0.028 | 0 | 0.16 | 0.09 | 0.31 | 0.24 | 0.2 | 0.24 |
| 6 | 123296698 | rs7754344 | 0.81 | 1.002 | 0.058 | -0.111 | -0.006 | -0.046 | -0.046 | 0 | 0.43 | 0.46 | 0.26 | 0.17 | 0.39 | 0.54 |
| 6 | 123296853 | rs12662473 | 0.873 | 1.06 | -0.057 | 0.456 | 0.01 | -0.031 | -0.031 | 0 | 0.11 | 0.05 | 0.26 | 0.37 | 0.06 | 0.16 |
| 6 | 123305899 | rs9385280 | -0.475 | 1.266 | 0.254 | 0.305 | 0.029 | 0.12 | 0.12 | 0.005 | 0.19 | 0.34 | 0.22 | 0.37 | 0.22 | 0.21 |
| 6 | 123305965 | rs9372726 | 0.861 | 0.441 | -0.045 | 0.444 | 0.01 | -0.031 | -0.031 | 0.838 | 0.11 | 0.05 | 0.26 | 0.37 | 0.06 | 0.16 |
| 6 | 123319382 | rs11154131 | 0.246 | 0.606 | -0.001 | -0.837 | -0.035 | 0.035 | 0.035 | 0 | 0.22 | 0.15 | 0.23 | 0.09 | 0.31 | 0.08 |
| 6 | 123323407 | rs7762002 | -0.173 | 0.632 | 0.074 | -0.801 | -0.031 | 0.046 | 0.046 | 0 | 0.74 | 0.83 | 0.74 | 0.91 | 0.68 | 0.9 |
| 6 | 123344094 | rs7741544 | 1.326 | 0.557 | 0.099 | -0.989 | -0.003 | -0.047 | -0.047 | 0 | 0.41 | 0.45 | 0.24 | 0.2 | 0.38 | 0.49 |
| 6 | 123346362 | rs12660210 | 0.16 | 0.572 | -0.14 | -0.95 | -0.003 | -0.001 | -0.001 | 0.064 | 0.25 | 0.25 | 0.46 | 0.7 | 0.22 | 0.36 |
| 6 | 123352977 | rs9401632 | -0.217 | 0.804 | -0.777 | -0.787 | -0.036 | 0.054 | 0.054 | 0 | 0.74 | 0.85 | 0.75 | 0.92 | 0.68 | 0.91 |
| 6 | 123364258 | rs7774522 | -0.694 | 0.263 | -0.187 | -0.185 | -0.02 | -0.029 | -0.029 | 0.807 | 0.32 | 0.44 | 0.49 | 0.7 | 0.27 | 0.39 |
| 6 | 123378134 | rs9388176 | 1.026 | 0.504 | -1.176 | -0.344 | -0.01 | 0.038 | 0.038 | 0 | 0.73 | 0.81 | 0.77 | 0.88 | 0.69 | 0.74 |
| 6 | 123385245 | rs9482329 | -0.501 | 0.442 | -1.15 | -0.392 | -0.009 | -0.028 | -0.028 | 4.246 | 0.12 | 0.21 | 0.06 | 0.02 | 0.1 | 0.18 |
| 6 | 123386063 | rs7768952 | 1.559 | 0.166 | -1.345 | -0.503 | -0.08 | 0.212 | 0.212 | 0.075 | 0.45 | 0.09 | 0.26 | 0.19 | 0.48 | 0.5 |
| 6 | 123390699 | rs7757286 | 0.099 | 0.298 | -1.003 | -0.296 | -0.031 | 0.005 | 0.005 | 0 | 0.85 | 0.95 | 0.83 | 0.92 | 0.8 | 0.92 |
| 6 | 123391750 | rs4618543 | -0.817 | -0.052 | -0.688 | -0.361 | -0.014 | -0.045 | -0.045 | 0 | 0.13 | 0.24 | 0.07 | 0.04 | 0.1 | 0.18 |
| 6 | 123395743 | rs12198275 | 1.112 | 0.186 | -1.402 | -0.471 | -0.1 | 0.196 | 0.196 | 0.023 | 0.41 | 0.09 | 0.29 | 0.16 | 0.49 | 0.31 |
| 6 | 123400798 | rs6939697 | -0.609 | -0.284 | -1.254 | -0.254 | 0.062 | 0.06 | 0.06 | 0.191 | 0.38 | 0.79 | 0.36 | 0.75 | 0.25 | 0.48 |
| 6 | 123404591 | rs9401635 | -0.627 | -0.274 | -1.259 | -0.228 | 0.056 | 0.071 | 0.071 | 0.295 | 0.39 | 0.79 | 0.36 | 0.8 | 0.25 | 0.48 |

| | | | | | | | | | | | | | | | | |
|---|---|---|---|---|---|---|---|---|---|---|---|---|---|---|---|---|
| 6 | 123414078 | rs9375219 | -0.46 | -0.178 | -1.193 | -0.174 | 0.058 | 0.028 | 0.028 | 2.637 | 0.39 | 0.77 | 0.33 | 0.72 | 0.24 | 0.41 |
| 6 | 123417732 | rs11962240 | 0.027 | 0.022 | -1.223 | -0.075 | 0.03 | -0.021 | -0.021 | 0.049 | 0.3 | 0.17 | 0.29 | 0.68 | 0.22 | 0.39 |
| 6 | 123426679 | rs12200173 | 0.319 | 0.355 | -1.177 | -0.109 | -0.018 | 0.013 | 0.013 | 0 | 0.81 | 0.89 | 0.83 | 0.91 | 0.77 | 0.72 |
| 6 | 123434838 | rs4624888 | 0.273 | 0.346 | -1.113 | -0.118 | -0.001 | 0 | 0 | 0 | 0.8 | 0.81 | 0.8 | 0.86 | 0.75 | 0.7 |
| 6 | 123440754 | rs9401637 | -0.155 | 0.325 | -0.86 | 0.229 | 0.023 | -0.043 | -0.043 | 0 | 0.27 | 0.12 | 0.26 | 0.67 | 0.2 | 0.37 |
| 6 | 123470555 | rs9375224 | -1.217 | 1.347 | -0.118 | 0.312 | 0.002 | -0.003 | -0.003 | 0 | 0.19 | 0.19 | 0.12 | 0.1 | 0.23 | 0.27 |
| 6 | 123479459 | rs9375226 | -1.042 | 1.623 | -0.332 | 0.192 | -0.044 | 0.034 | 0.034 | 0.271 | 0.18 | 0.15 | 0.15 | 0.1 | 0.25 | 0.2 |
| 6 | 123479765 | rs9375227 | -1.198 | 1.699 | -0.476 | -0.014 | -0.056 | 0.063 | 0.063 | 0.076 | 0.22 | 0.13 | 0.17 | 0.08 | 0.29 | 0.22 |
| 6 | 123480590 | rs9482343 | -1.507 | 1.539 | 0.49 | 0.156 | 0.018 | -0.031 | -0.031 | 0.035 | 0.32 | 0.61 | 0.3 | 0.17 | 0.37 | 0.31 |
| 6 | 123481911 | rs6916837 | -0.205 | 1.207 | -0.049 | 0.808 | -0.042 | -0.141 | -0.141 | 8.771 | 0.82 | 0.98 | 0.86 | 0.52 | 0.85 | 0.69 |
| 6 | 123483042 | rs4896709 | 1.376 | 1.304 | 0.247 | 0.672 | -0.027 | 0.042 | 0.042 | 0.758 | 0.54 | 0.22 | 0.58 | 0.36 | 0.55 | 0.39 |
| 6 | 123491487 | rs13201756 | -0.708 | 1.463 | 0.728 | 0.451 | -0.031 | 0.042 | 0.042 | 0.089 | 0.22 | 0.12 | 0.19 | 0.03 | 0.28 | 0.22 |
| 6 | 123511016 | rs7741907 | -1.064 | 1.445 | 1.367 | 0.51 | 0.025 | -0.025 | -0.025 | 0 | 0.24 | 0.32 | 0.24 | 0.13 | 0.29 | 0.24 |
| 6 | 123529456 | rs13192453 | -0.896 | 1.199 | 0.902 | 0.112 | -0.035 | 0.062 | 0.062 | 0 | 0.25 | 0.12 | 0.21 | 0.04 | 0.29 | 0.22 |
| 6 | 123537837 | rs7743126 | 0.937 | 0.858 | 1.188 | 0.577 | 0.016 | 0.008 | 0.008 | 1.14 | 0.52 | 0.65 | 0.59 | 0.37 | 0.44 | 0.34 |
| 6 | 123539684 | rs361508 | -0.508 | 1.122 | 0.928 | 0.724 | -0.033 | -0.141 | -0.141 | 0 | 0.78 | 0.98 | 0.83 | 0.52 | 0.8 | 0.63 |
| 6 | 123549399 | rs6569329 | -0.516 | 1.073 | 1.021 | 0.714 | -0.03 | -0.141 | -0.141 | 0 | 0.78 | 0.98 | 0.83 | 0.52 | 0.8 | 0.64 |
| 6 | 123549610 | rs4591876 | 1.091 | 1.05 | 1.267 | 0.755 | 0.022 | -0.004 | -0.004 | 2.594 | 0.53 | 0.78 | 0.6 | 0.41 | 0.44 | 0.26 |
| 6 | 123549991 | rs7741771 | 1.345 | 1.096 | 0.912 | 0.711 | 0.027 | 0.033 | 0.033 | 1.578 | 0.49 | 0.35 | 0.57 | 0.33 | 0.44 | 0.25 |
| 6 | 123552306 | rs10457444 | 1.47 | 1.097 | 0.527 | 0.711 | 0.025 | 0.033 | 0.033 | 1.799 | 0.48 | 0.23 | 0.57 | 0.33 | 0.44 | 0.25 |
| 6 | 123556891 | rs9388193 | -0.784 | 1.546 | 0.773 | 0.25 | 0.007 | -0.016 | -0.016 | 1.767 | 0.25 | 0.27 | 0.23 | 0.06 | 0.36 | 0.28 |
| 6 | 123571228 | rs4345414 | 1.12 | 1.412 | 0.247 | 0.503 | -0.016 | 0.006 | 0.006 | 0.218 | 0.7 | 0.33 | 0.78 | 0.76 | 0.7 | 0.61 |
| 6 | 123590126 | rs2195729 | 0.293 | 1.784 | 0.285 | 0.338 | 0.019 | -0.001 | -0.001 | 0 | 0.88 | 0.86 | 0.9 | 0.88 | 0.82 | 0.84 |
| 6 | 123591424 | rs11757138 | -0.219 | 1.773 | 0.414 | 0.303 | 0.008 | -0.023 | -0.023 | 4.573 | 0.14 | 0.23 | 0.1 | 0.08 | 0.13 | 0.22 |
| 6 | 123596860 | rs12206156 | 0.857 | 1.191 | -0.535 | 0.638 | 0.012 | 0.069 | 0.069 | 0.084 | 0.5 | 0.32 | 0.58 | 0.29 | 0.52 | 0.26 |
| 6 | 123600707 | rs1346920 | 0.461 | 1.085 | -0.596 | 0.608 | 0.018 | -0.111 | -0.111 | 4.65 | 0.23 | 0.09 | 0.21 | 0.51 | 0.19 | 0.37 |
| 6 | 123620716 | rs7745242 | 0.845 | 1.119 | -0.621 | 0.436 | 0.02 | 0.093 | 0.093 | 0.078 | 0.5 | 0.32 | 0.58 | 0.25 | 0.51 | 0.25 |
| 6 | 123627146 | rs720203 | 0.597 | 0.541 | -1.31 | 0.716 | -0.005 | -0.003 | -0.003 | 0 | 0.11 | 0.02 | 0.27 | 0.09 | 0.13 | 0.13 |
| 6 | 123628287 | rs9320926 | -0.05 | 0.605 | -0.973 | 1.042 | 0.019 | -0.098 | -0.098 | 0 | 0.24 | 0.08 | 0.21 | 0.64 | 0.19 | 0.37 |
| 6 | 123639051 | rs9375243 | 0.063 | 0.658 | -0.998 | 0.727 | 0.013 | -0.087 | -0.087 | 0 | 0.24 | 0.07 | 0.25 | 0.63 | 0.18 | 0.43 |
| 6 | 123642389 | rs7746138 | 0.085 | 1.045 | -0.62 | 0.115 | 0.007 | -0.027 | -0.027 | 2.033 | 0.16 | 0.2 | 0.1 | 0.08 | 0.15 | 0.24 |
| 6 | 123646414 | rs17679624 | 1.093 | 0.787 | -1.28 | -0.331 | -0.008 | 0.013 | 0.013 | 0.739 | 0.07 | 0 | 0.05 | 0.03 | 0.06 | 0.02 |
| 6 | 123649770 | rs1019862 | -0.982 | 1.417 | -0.515 | -0.655 | 0.009 | -0.023 | -0.023 | 0 | 0.3 | 0.58 | 0.23 | 0.16 | 0.31 | 0.35 |
| 6 | 123658680 | rs12216080 | 1.178 | 1.51 | -0.444 | -0.938 | 0.011 | 0.038 | 0.038 | 0.364 | 0.41 | 0.35 | 0.49 | 0.14 | 0.43 | 0.16 |
| 6 | 123664207 | rs7769239 | 0.589 | 1.46 | -0.462 | -0.951 | 0.006 | 0.082 | 0.082 | 0.117 | 0.51 | 0.35 | 0.59 | 0.21 | 0.6 | 0.2 |
| 6 | 123668347 | rs9375249 | 0.391 | 1.462 | -0.337 | -1.372 | 0.015 | -0.032 | -0.032 | 0 | 0.75 | 0.43 | 0.83 | 0.83 | 0.77 | 0.62 |
| 6 | 123669127 | rs1933899 | 0.924 | 1.333 | -0.314 | -1.337 | 0.007 | -0.009 | -0.009 | 0.195 | 0.65 | 0.42 | 0.72 | 0.75 | 0.61 | 0.58 |
| 6 | 123672656 | rs3850235 | -0.637 | 1.301 | -0.335 | -1.426 | -0.007 | 0.005 | 0.005 | 1.588 | 0.94 | 0.97 | 0.95 | 0.99 | 0.92 | 0.94 |
| 6 | 123683028 | rs9320930 | -0.547 | 1.11 | -0.066 | -1.533 | 0.013 | 0.125 | 0.125 | 2.209 | 0.5 | 0.28 | 0.45 | 0.78 | 0.49 | 0.74 |
| 6 | 123688832 | rs12202709 | 1.163 | 0.815 | -0.169 | -1.682 | 0.001 | 0.086 | 0.086 | 0.228 | 0.37 | 0.24 | 0.46 | 0.11 | 0.42 | 0.16 |
| 6 | 123690604 | rs9320931 | -0.197 | 0.888 | -0.132 | -1.582 | 0.01 | 0.014 | 0.014 | 0.993 | 0.39 | 0.32 | 0.36 | 0.7 | 0.3 | 0.5 |
| 6 | 123696766 | rs17737379 | -0.369 | 0.658 | -0.174 | -1.798 | -0.018 | -0.058 | -0.058 | 0 | 0.18 | 0.31 | 0.12 | 0.07 | 0.13 | 0.05 |
| 6 | 123697583 | rs2163447 | -1.17 | 0.66 | -0.493 | -2.065 | -0.032 | 0.056 | 0.056 | 0.956 | 0.9 | 0.99 | 0.91 | 0.99 | 0.86 | 0.93 |
| 6 | 123699042 | rs6901953 | 1.27 | 0.708 | -0.422 | -2.186 | 0 | -0.002 | -0.002 | 0.045 | 0.34 | 0.34 | 0.43 | 0.1 | 0.37 | 0.15 |
| 6 | 123700559 | rs6569341 | 1.27 | 0.708 | -0.418 | -2.186 | 0.001 | 0.004 | 0.004 | 0 | 0.34 | 0.33 | 0.43 | 0.1 | 0.37 | 0.15 |
| 6 | 123702642 | rs9482382 | 0.132 | 0.701 | -0.351 | -2.185 | -0.005 | -0.02 | -0.02 | 0 | 0.17 | 0.21 | 0.12 | 0.07 | 0.13 | 0.05 |
| 6 | 123703983 | rs2043148 | 0.419 | 0.705 | -0.228 | -2.19 | -0.009 | 0.033 | 0.033 | 0.201 | 0.37 | 0.26 | 0.35 | 0.82 | 0.36 | 0.7 |
| 6 | 123705062 | rs7752950 | -1.012 | 0.702 | -0.196 | -2.17 | -0.008 | 0.014 | 0.014 | 1.124 | 0.89 | 0.95 | 0.91 | 0.91 | 0.86 | 0.89 |
| 6 | 123705434 | rs7773653 | -0.017 | 0.667 | -0.169 | -2.139 | 0.002 | 0.039 | 0.039 | 0.011 | 0.53 | 0.66 | 0.56 | 0.17 | 0.5 | 0.19 |
| 6 | 123707732 | rs1431284 | 1.192 | 0.665 | -0.175 | -2.142 | -0.003 | -0.06 | -0.06 | 0 | 0.35 | 0.44 | 0.43 | 0.1 | 0.37 | 0.15 |
| 6 | 123711840 | rs17737857 | -0.032 | 0.79 | -0.234 | -2.14 | 0.009 | 0.085 | 0.085 | 0.047 | 0.53 | 0.72 | 0.56 | 0.25 | 0.48 | 0.23 |
| 6 | 123724482 | rs873460 | 0.899 | 0.723 | -0.478 | -2.18 | 0.002 | 0.045 | 0.045 | 0.33 | 0.37 | 0.3 | 0.44 | 0.1 | 0.4 | 0.16 |
| 6 | 123728539 | rs1560465 | 0.572 | 0.702 | -0.459 | -2.179 | -0.007 | -0.023 | -0.023 | 0 | 0.13 | 0.2 | 0.1 | 0.07 | 0.11 | 0.04 |
| 6 | 123731285 | rs9388238 | 1.23 | 0.718 | -0.567 | -2.172 | 0.004 | -0.069 | -0.069 | 0 | 0.25 | 0.07 | 0.25 | 0.62 | 0.19 | 0.46 |
| 6 | 123735555 | rs1933896 | 0.621 | 0.887 | -0.422 | -2.158 | 0.003 | 0.001 | 0.001 | 0.015 | 0.39 | 0.38 | 0.45 | 0.1 | 0.4 | 0.16 |
| 6 | 123739731 | rs12665088 | 0.5 | 1.04 | -0.671 | -2.321 | 0.031 | 0.06 | 0.06 | 2.913 | 0.44 | 0.3 | 0.38 | 0.69 | 0.36 | 0.5 |
| 6 | 123742101 | rs11154157 | 0.937 | 1.076 | -0.662 | -2.282 | 0 | 0.064 | 0.064 | 0.515 | 0.31 | 0.22 | 0.4 | 0.1 | 0.33 | 0.16 |
| 6 | 123744048 | rs4320388 | 1.128 | 1.137 | -0.595 | -2.27 | 0.073 | 0.013 | 0.013 | 0 | 0.11 | 0.21 | 0.1 | 0.12 | 0.17 | 0.23 |
| 6 | 123749752 | rs1343962 | 0.373 | 1.088 | -0.677 | -2.274 | -0.016 | 0.026 | 0.026 | 0 | 0.41 | 0.64 | 0.5 | 0.32 | 0.47 | 0.41 |
| 6 | 123750519 | rs12208822 | 0.854 | 1.121 | -0.726 | -2.456 | 0.002 | 0.026 | 0.026 | 0.423 | 0.32 | 0.28 | 0.4 | 0.1 | 0.33 | 0.16 |
| 6 | 123754142 | rs9401671 | -0.539 | 1.265 | -0.667 | -2.442 | 0 | -0.036 | -0.036 | 0.015 | 0.56 | 0.6 | 0.49 | 0.87 | 0.5 | 0.71 |
| 6 | 123754480 | rs12203141 | 0.233 | 1.212 | -0.567 | -2.274 | 0.004 | -0.067 | -0.067 | 0 | 0.13 | 0.33 | 0.1 | 0.02 | 0.1 | 0.06 |
| 6 | 123760725 | rs13196705 | -0.495 | 0.973 | -0.714 | -2.319 | 0.002 | -0.034 | -0.034 | 0 | 0.3 | 0.39 | 0.18 | 0.14 | 0.27 | 0.4 |
| 6 | 123769149 | rs9490756 | 0.318 | 0.871 | -0.692 | -2.16 | 0.006 | -0.065 | -0.065 | 1.046 | 0.13 | 0.31 | 0.1 | 0.02 | 0.1 | 0.06 |
| 6 | 123776501 | rs9482391 | 0.437 | 1.068 | -1.023 | -2.072 | -0.01 | 0.009 | 0.009 | 0 | 0.18 | 0.14 | 0.09 | 0.13 | 0.19 | 0.3 |
| 6 | 123779012 | rs9401672 | 1.057 | 0.983 | -1.427 | -2.251 | -0.029 | -0.09 | -0.09 | 0 | 0.23 | 0.07 | 0.26 | 0.62 | 0.22 | 0.3 |
| 6 | 123783377 | rs9490772 | 0.539 | 0.871 | -1.215 | -2.008 | -0.01 | 0.008 | 0.008 | 0 | 0.18 | 0.14 | 0.1 | 0.14 | 0.19 | 0.3 |
| 6 | 123788232 | rs4571590 | -0.774 | 1.013 | -1.121 | -2.006 | 0.003 | 0.048 | 0.048 | 2.057 | 0.86 | 0.97 | 0.91 | 1 | 0.86 | 0.94 |
| 6 | 123810308 | rs2153134 | 1.076 | 1.02 | -0.942 | -2.726 | -0.003 | 0.001 | 0.001 | 0.142 | 0.14 | 0.13 | 0.32 | 0.13 | 0.22 | 0.07 |

| | | | | | | | | | | | | | | | | |
|---|---|---|---|---|---|---|---|---|---|---|---|---|---|---|---|---|
| 6 | 123814633 | rs17739564 | 0.946 | 0.873 | -0.778 | -2.561 | -0.06 | 0.041 | 0.041 | 0 | 0.1 | 0 | 0.06 | 0.03 | 0.15 | 0.24 |
| 6 | 123815838 | rs4897255 | 0.191 | 0.787 | -0.516 | -2.728 | 0.001 | -0.024 | -0.024 | 1.316 | 0.45 | 0.46 | 0.61 | 0.24 | 0.51 | 0.25 |
| 6 | 123818871 | rs9320939 | 0.38 | 0.581 | -1.009 | -2.61 | -0.02 | 0.061 | 0.061 | 0.5 | 0.42 | 0.29 | 0.57 | 0.27 | 0.5 | 0.44 |
| 6 | 123819847 | rs9320940 | -0.348 | 0.434 | -0.919 | -2.562 | -0.004 | -0.011 | -0.011 | 0 | 0.81 | 0.73 | 0.86 | 0.88 | 0.86 | 0.84 |
| 6 | 123829676 | rs9398732 | 0.933 | 0.468 | -0.904 | -2.54 | -0.037 | 0.044 | 0.044 | 0.274 | 0.26 | 0.41 | 0.21 | 0.6 | 0.21 | 0.35 |
| 6 | 123835147 | rs4612191 | -0.548 | 0.562 | -1.061 | -2.526 | -0.001 | -0.006 | -0.006 | 0 | 0.83 | 0.79 | 0.87 | 0.88 | 0.86 | 0.84 |
| 6 | 123838281 | rs17689345 | 0.548 | 1.054 | -1.379 | -1.987 | -0.01 | 0.039 | 0.039 | 0.916 | 0.13 | 0.03 | 0.09 | 0 | 0.15 | 0.06 |
| 6 | 123846449 | rs4897271 | 0.435 | 0.725 | -1.257 | -1.849 | -0.004 | 0.016 | 0.016 | 0.701 | 0.31 | 0.26 | 0.5 | 0.24 | 0.36 | 0.19 |
| 6 | 123850194 | rs9490795 | 0.052 | -0.022 | -0.699 | -1.223 | 0.011 | 0.01 | 0.01 | 6.538 | 0.57 | 0.39 | 0.66 | 0.37 | 0.66 | 0.51 |
| 6 | 123858425 | rs7771989 | -0.312 | 0.243 | -0.444 | -1.261 | 0 | 0.013 | 0.013 | 0 | 0.06 | 0.15 | 0.04 | 0.05 | 0.03 | 0.02 |
| 6 | 123861461 | rs13212480 | -0.109 | 0.125 | -0.648 | -1.957 | 0.006 | -0.079 | -0.079 | 0 | 0.44 | 0.51 | 0.32 | 0.08 | 0.53 | 0.42 |
| 6 | 123865515 | rs7349870 | -0.651 | 0.101 | -0.949 | -1.96 | -0.016 | 0.231 | 0.231 | 0 | 0.53 | 0.85 | 0.4 | 0.14 | 0.58 | 0.45 |
| 6 | 123868992 | rs6905496 | -0.209 | 0.254 | -0.892 | -1.914 | -0.004 | 0.057 | 0.057 | 0 | 0.46 | 0.6 | 0.35 | 0.1 | 0.56 | 0.43 |
| 6 | 123885639 | rs6569352 | 0.592 | 0.574 | -1.422 | -1.482 | -0.017 | 0.006 | 0.006 | 0 | 0.92 | 0.99 | 0.96 | 0.95 | 0.91 | 0.94 |
| 6 | 123886231 | rs4897302 | 0.506 | 0.448 | -1.572 | -1.66 | -0.021 | 0.112 | 0.112 | 0 | 0.37 | 0.17 | 0.58 | 0.84 | 0.36 | 0.48 |
| 6 | 123888182 | rs1998262 | 0.702 | 0.437 | -1.177 | -1.324 | 0.004 | 0.017 | 0.017 | 0 | 0.75 | 0.85 | 0.88 | 0.9 | 0.76 | 0.72 |
| 6 | 123892001 | rs2145735 | 0.155 | 0.606 | -1.292 | -1.323 | -0.006 | 0.119 | 0.119 | 0 | 0.43 | 0.26 | 0.59 | 0.83 | 0.36 | 0.54 |
| 6 | 123892432 | rs1407220 | 0.553 | 1.626 | -0.444 | -1.786 | 0.046 | -0.079 | -0.079 | 0 | 0.81 | 0.64 | 0.84 | 0.94 | 0.79 | 0.86 |
| 6 | 123893417 | rs11757798 | 0.246 | 1.506 | -0.555 | -1.375 | -0.006 | 0.025 | 0.025 | 0 | 0.37 | 0.53 | 0.34 | 0.48 | 0.33 | 0.38 |
| 6 | 123893646 | rs9385304 | -0.08 | 1.534 | -0.748 | -1.627 | -0.001 | -0.002 | -0.002 | 0 | 0.53 | 0.47 | 0.58 | 0.53 | 0.56 | 0.51 |
| 6 | 123897071 | rs9388260 | 0.169 | 1.934 | -0.187 | -1.546 | 0.084 | -0.006 | -0.006 | 0 | 0.06 | 0.15 | 0.05 | 0.03 | 0.12 | 0.07 |
| 6 | 123897553 | rs4897305 | -0.318 | 1.664 | 0.057 | -1.794 | 0.001 | -0.026 | -0.026 | 0.232 | 0.84 | 0.88 | 0.9 | 1 | 0.87 | 0.89 |
| 6 | 123897712 | rs1407221 | 0.174 | 1.33 | -0.018 | -2.102 | 0.001 | 0.022 | 0.022 | 0 | 0.51 | 0.67 | 0.48 | 0.44 | 0.5 | 0.54 |
| 6 | 123898505 | rs9375268 | 0.06 | 1.295 | -0.001 | -1.999 | 0.152 | 0.025 | 0.025 | 0 | 0.08 | 0.27 | 0.07 | 0.05 | 0.13 | 0.1 |
| 6 | 123900022 | rs6922576 | 0.846 | 1.199 | -0.496 | -1.922 | 0.013 | -0.045 | -0.045 | 0.015 | 0.28 | 0.16 | 0.38 | 0.52 | 0.26 | 0.28 |
| 6 | 123902541 | rs1321084 | 0.071 | 1.078 | -0.741 | -1.817 | -0.005 | 0.025 | 0.025 | 0 | 0.47 | 0.59 | 0.52 | 0.59 | 0.47 | 0.51 |
| 6 | 123909246 | rs6903080 | 0.162 | 1.415 | -0.82 | -1.451 | -0.014 | 0.001 | 0.001 | 0.003 | 0.39 | 0.36 | 0.41 | 0.38 | 0.45 | 0.38 |
| 6 | 123912806 | rs13191900 | 0.119 | 1.529 | -1.194 | -1.002 | -0.07 | 0.02 | 0.02 | 0.317 | 0.14 | 0.02 | 0.14 | 0.07 | 0.25 | 0.1 |
| 6 | 123912911 | rs9398738 | 0.165 | 0.857 | -1.175 | -0.396 | -0.031 | -0.105 | -0.105 | 1.512 | 0.25 | 0.06 | 0.39 | 0.57 | 0.27 | 0.27 |
| 6 | 123913242 | rs9490824 | -0.584 | 0.804 | 0.412 | 0.658 | 0.016 | 0.051 | 0.051 | 0 | 0.29 | 0.39 | 0.35 | 0.42 | 0.33 | 0.36 |
| 6 | 123917565 | rs1590246 | 1.361 | 0.026 | 1.588 | 1.217 | -0.003 | 0.013 | 0.013 | 0.178 | 0.9 | 0.69 | 0.94 | 0.92 | 0.9 | 0.89 |
| 6 | 123924927 | rs7760343 | 1.07 | -0.238 | 1.504 | 1.31 | 0.007 | 0.026 | 0.026 | 0 | 0.57 | 0.4 | 0.67 | 0.26 | 0.56 | 0.48 |
| 6 | 123925257 | rs1535353 | 1.732 | -0.243 | 2.081 | 1.645 | -0.018 | 0.046 | 0.046 | 0 | 0.81 | 0.63 | 0.88 | 0.74 | 0.81 | 0.82 |
| 6 | 123939553 | rs9388272 | -0.353 | -0.361 | 2.207 | 1.748 | -0.001 | 0.016 | 0.016 | 0 | 0.24 | 0.22 | 0.2 | 0.44 | 0.24 | 0.37 |
| 6 | 123941672 | rs1570186 | -1.505 | -0.413 | 2.281 | 1.714 | 0.04 | -0.03 | -0.03 | 0.358 | 0.38 | 0.86 | 0.29 | 0.62 | 0.31 | 0.43 |
| 6 | 123945300 | rs1321874 | -1.317 | -0.4 | 2.334 | 1.718 | 0.034 | -0.028 | -0.028 | 1.661 | 0.36 | 0.84 | 0.29 | 0.62 | 0.31 | 0.43 |
| 6 | 123950488 | rs9375280 | -0.299 | -0.324 | 2.076 | 1.775 | 0.002 | 0.041 | 0.041 | 0 | 0.25 | 0.29 | 0.22 | 0.44 | 0.26 | 0.37 |
| 6 | 123955723 | rs7774820 | -1.783 | -0.425 | 2.136 | 1.457 | -0.059 | 0.065 | 0.065 | 3.605 | 0.13 | 0.62 | 0.07 | 0.18 | 0.05 | 0.06 |
| 6 | 123955967 | rs11757453 | -1.082 | -0.388 | 2.099 | 1.193 | -0.028 | -0.045 | -0.045 | 0.032 | 0.09 | 0.24 | 0.04 | 0.01 | 0.05 | 0.03 |
| 6 | 123958493 | rs3813325 | -1.626 | 0.068 | 2.166 | 1.463 | -0.007 | 0.025 | 0.025 | 0.424 | 0.07 | 0.16 | 0.06 | 0.13 | 0.06 | 0.03 |
| 6 | 123965497 | rs1853157 | -0.348 | 0.13 | 2.18 | 1.466 | 0.009 | 0.064 | 0.064 | 0 | 0.24 | 0.3 | 0.43 | 0.55 | 0.27 | 0.38 |
| 6 | 123970838 | rs1853155 | -0.43 | 0.096 | 1.885 | 1.296 | 0.01 | 0.099 | 0.099 | 0 | 0.26 | 0.39 | 0.46 | 0.56 | 0.28 | 0.39 |
| 6 | 123975313 | rs2169092 | 1.383 | 0.049 | 1.04 | 0.917 | 0 | 0.055 | 0.055 | 1.201 | 0.53 | 0.06 | 0.42 | 0.31 | 0.6 | 0.43 |
| 6 | 123983066 | rs979537 | -1.035 | 0.466 | 1.565 | 1.34 | 0.001 | 0.003 | 0.003 | 0.832 | 0.36 | 0.65 | 0.54 | 0.7 | 0.35 | 0.5 |
| 6 | 124003483 | rs7774699 | -2.197 | 0.552 | 2.549 | 1.555 | 0.078 | 0.176 | 0.176 | 1.431 | 0.06 | 0.42 | 0.04 | 0.07 | 0.02 | 0.02 |
| 6 | 124008919 | rs9482426 | -2.073 | 0.534 | 2.937 | 1.508 | 0.095 | 0.066 | 0.066 | 2.658 | 0.05 | 0.46 | 0.05 | 0 | 0.02 | 0.01 |
| 6 | 124012601 | rs2611428 | 2.68 | 0.442 | 2.864 | 1.406 | -0.025 | -0.017 | -0.017 | 0.061 | 0.81 | 0.28 | 0.86 | 0.9 | 0.88 | 0.87 |
| 6 | 124023039 | rs2611432 | 2.506 | 0.486 | 2.651 | 1.43 | -0.041 | -0.001 | -0.001 | 0 | 0.86 | 0.31 | 0.91 | 0.9 | 0.92 | 0.88 |
| 6 | 124032062 | rs4897436 | -0.787 | 0.179 | 2.412 | 2.046 | -0.021 | -0.011 | -0.011 | 0.012 | 0.07 | 0.01 | 0.15 | 0.64 | 0.07 | 0.29 |
| 6 | 124039569 | rs11963671 | -0.418 | 0.098 | 2.66 | 0.924 | -0.016 | 0.027 | 0.027 | 0.062 | 0.06 | 0.02 | 0.04 | 0 | 0.08 | 0.02 |
| 6 | 124059254 | rs9482443 | -2.761 | -0.157 | 2.457 | 1.205 | -0.026 | -0.006 | -0.006 | 0 | 0.14 | 0.69 | 0.11 | 0.08 | 0.1 | 0.05 |
| 6 | 124081147 | rs9490917 | -1.663 | -0.135 | 2.6 | 1.202 | 0.019 | -0.029 | -0.029 | 0 | 0.06 | 0.22 | 0.07 | 0 | 0.06 | 0.01 |
| 6 | 124083410 | rs9490919 | -1.884 | -0.133 | 2.601 | 1.261 | 0.006 | 0.01 | 0.01 | 0.452 | 0.07 | 0.24 | 0.07 | 0.05 | 0.06 | 0.03 |
| 6 | 124089172 | rs7751806 | -2.213 | -0.119 | 1.885 | 1.245 | -0.019 | 0.031 | 0.031 | 7.624 | 0.13 | 0.63 | 0.11 | 0.11 | 0.1 | 0.05 |
| 6 | 124095959 | rs3861460 | -2.105 | -0.063 | 1.267 | 1.273 | -0.018 | 0.011 | 0.011 | 9.999 | 0.13 | 0.76 | 0.11 | 0.11 | 0.1 | 0.04 |
| 6 | 124133445 | rs9490938 | -1.73 | -0.774 | 0.673 | 1.377 | 0.002 | -0.008 | -0.008 | 1.791 | 0.27 | 0.77 | 0.29 | 0.43 | 0.11 | 0.46 |
| 6 | 124137674 | rs7769832 | -1.845 | -0.638 | 1.52 | 0.854 | -0.02 | -0.048 | -0.048 | 0.816 | 0.1 | 0.59 | 0.07 | 0.01 | 0.05 | 0.03 |
| 6 | 124141597 | rs12207431 | -1.17 | -0.575 | 1.568 | 0.896 | 0.007 | -0.012 | -0.012 | 0 | 0.05 | 0.2 | 0.04 | 0.01 | 0.05 | 0.02 |
| 6 | 124142979 | rs928529 | -1.674 | -0.867 | 1.248 | 1.337 | -0.053 | 0.072 | 0.072 | 1.831 | 0.23 | 0.61 | 0.28 | 0.56 | 0.13 | 0.4 |
| 6 | 124144456 | rs9490950 | 1.284 | -0.654 | 1.187 | 1.074 | 0.009 | -0.048 | -0.048 | 1.773 | 0.92 | 0.79 | 0.92 | 1 | 0.91 | 0.92 |
| 6 | 124144516 | rs9490951 | -1.645 | -0.659 | 1.149 | 1.279 | -0.052 | 0.067 | 0.067 | 1.932 | 0.23 | 0.61 | 0.28 | 0.59 | 0.13 | 0.4 |
| 6 | 124149869 | rs9388284 | -0.008 | -0.234 | -0.631 | 0.397 | -0.041 | -0.009 | -0.009 | 0 | 0.46 | 0.11 | 0.43 | 0.6 | 0.48 | 0.53 |
| 6 | 124150058 | rs9401703 | 0.295 | 0.122 | -1.705 | -0.455 | -0.076 | -0.054 | -0.054 | 0 | 0.46 | 0.06 | 0.46 | 0.55 | 0.49 | 0.48 |
| 6 | 124150232 | rs9372759 | 0.339 | 0.374 | -1.711 | -0.64 | -0.042 | 0.003 | 0.003 | 5.41 | 0.48 | 0.15 | 0.45 | 0.55 | 0.51 | 0.47 |
| 6 | 124156176 | rs12207255 | -0.17 | 0.442 | -0.548 | -0.903 | -0.001 | -0.007 | -0.007 | 0 | 0.22 | 0.2 | 0.13 | 0.06 | 0.2 | 0.21 |
| 6 | 124156824 | rs9375285 | 0.85 | 0.219 | -1.11 | -0.652 | -0.053 | -0.035 | -0.035 | 0.148 | 0.45 | 0.13 | 0.43 | 0.53 | 0.51 | 0.49 |
| 6 | 124159438 | rs1490352 | -0.169 | 0.265 | -0.86 | -0.57 | 0.001 | 0.015 | 0.015 | 0 | 0.74 | 0.7 | 0.64 | 0.63 | 0.78 | 0.77 |
| 6 | 124160157 | rs6569365 | -0.53 | 0.312 | -1.443 | -0.438 | -0.03 | 0.082 | 0.082 | 0.254 | 0.49 | 0.84 | 0.59 | 0.35 | 0.47 | 0.47 |
| 6 | 124163015 | rs6936906 | 0.154 | 0.411 | -0.775 | 0.116 | -0.003 | 0.007 | 0.007 | 0 | 0.56 | 0.56 | 0.52 | 0.71 | 0.55 | 0.6 |

| | | | | | | | | | | | | | | | | |
|---|---|---|---|---|---|---|---|---|---|---|---|---|---|---|---|---|
| 6 | 124166282 | rs7759596 | 0.173 | 0.234 | -0.456 | 0.106 | -0.002 | 0.012 | 0.012 | 0 | 0.55 | 0.39 | 0.48 | 0.73 | 0.55 | 0.61 |
| 6 | 124174336 | rs17749667 | 0.084 | 0.346 | -0.876 | 0.517 | -0.016 | -0.004 | -0.004 | 0 | 0.28 | 0.09 | 0.28 | 0.28 | 0.32 | 0.26 |
| 6 | 124178280 | rs6569368 | -0.226 | 0.407 | -0.119 | 0.355 | 0.005 | -0.015 | -0.015 | 0 | 0.35 | 0.55 | 0.35 | 0.29 | 0.38 | 0.22 |
| 6 | 124180018 | rs6932634 | -0.929 | 0.131 | -0.173 | 0.331 | -0.003 | 0.007 | 0.007 | 0 | 0.64 | 0.68 | 0.72 | 0.76 | 0.64 | 0.59 |
| 6 | 124181267 | rs6939381 | -0.955 | 0.106 | -0.237 | 0.306 | -0.037 | 0.029 | 0.029 | 0 | 0.65 | 0.85 | 0.72 | 0.76 | 0.63 | 0.59 |
| 6 | 124181361 | rs6939886 | 0.413 | -0.241 | -0.25 | 0.205 | 0.003 | 0.012 | 0.012 | 0.377 | 0.61 | 0.29 | 0.62 | 0.68 | 0.61 | 0.75 |
| 6 | 124182990 | rs6907969 | 0.075 | -0.42 | 0.003 | 0.23 | -0.016 | 0.031 | 0.031 | 0 | 0.34 | 0.41 | 0.35 | 0.64 | 0.26 | 0.46 |
| 6 | 124183703 | rs17693403 | 0.104 | -0.346 | -0.523 | -0.008 | -0.034 | 0.066 | 0.066 | 0 | 0.27 | 0.08 | 0.27 | 0.1 | 0.31 | 0.14 |
| 6 | 124187922 | rs4512241 | -1.193 | -0.352 | -0.469 | -0.143 | -0.039 | -0.012 | -0.012 | 0 | 0.64 | 0.88 | 0.74 | 0.73 | 0.63 | 0.61 |
| 6 | 124189423 | rs2318064 | 0.473 | -0.311 | -0.534 | -0.027 | -0.021 | -0.082 | -0.082 | 0 | 0.25 | 0.07 | 0.36 | 0.49 | 0.23 | 0.36 |
| 6 | 124190888 | rs6924068 | 0.146 | -0.145 | -0.477 | -0.131 | -0.038 | 0.065 | 0.065 | 0 | 0.26 | 0.1 | 0.3 | 0.09 | 0.35 | 0.18 |
| 6 | 124195220 | rs9388286 | 0.157 | -0.231 | -0.098 | 0.093 | -0.005 | -0.058 | -0.058 | 0 | 0.27 | 0.11 | 0.36 | 0.5 | 0.22 | 0.37 |
| 6 | 124200765 | rs9388287 | -1.06 | -0.157 | 0.345 | -0.078 | -0.002 | -0.005 | -0.005 | 3.434 | 0.52 | 0.51 | 0.68 | 0.7 | 0.56 | 0.56 |
| 6 | 124201150 | rs2130589 | 0.019 | -0.154 | 0.459 | -0.103 | -0.014 | 0.018 | 0.018 | 0 | 0.21 | 0.14 | 0.27 | 0.08 | 0.31 | 0.19 |
| 6 | 124202712 | rs1352678 | -0.829 | -0.5 | 0.528 | 0.113 | 0.001 | 0.012 | 0.012 | 0 | 0.53 | 0.59 | 0.68 | 0.69 | 0.55 | 0.56 |
| 6 | 124206586 | rs9388288 | -0.467 | -0.521 | -0.214 | 0.134 | -0.006 | -0.004 | -0.004 | 0 | 0.48 | 0.25 | 0.63 | 0.58 | 0.53 | 0.56 |
| 6 | 124207776 | rs2873615 | -0.279 | -0.445 | -0.915 | 0.078 | -0.006 | -0.066 | -0.066 | 0 | 0.39 | 0.15 | 0.46 | 0.62 | 0.39 | 0.44 |
| 6 | 124217985 | rs4897581 | -0.011 | -0.505 | -1.111 | -0.005 | -0.047 | -0.056 | -0.056 | 0.043 | 0.34 | 0.09 | 0.44 | 0.51 | 0.37 | 0.43 |
| 6 | 124221118 | rs9482487 | -1.202 | -0.198 | -0.453 | 0.158 | 0 | -0.001 | -0.001 | 0 | 0.2 | 0.71 | 0.25 | 0.19 | 0.17 | 0.15 |
| 6 | 124222153 | rs1008742 | -1.203 | -0.221 | -0.463 | 0.145 | 0 | -0.001 | -0.001 | 0 | 0.2 | 0.7 | 0.25 | 0.19 | 0.17 | 0.15 |
| 6 | 124235689 | rs11755174 | -0.477 | -0.531 | -0.876 | 0.021 | -0.025 | -0.039 | -0.039 | 0 | 0.34 | 0.21 | 0.46 | 0.67 | 0.38 | 0.46 |
| 6 | 124236221 | rs1490359 | -0.569 | -0.734 | -0.638 | -0.07 | 0.005 | -0.005 | -0.005 | 0.149 | 0.4 | 0.33 | 0.46 | 0.65 | 0.38 | 0.46 |
| 6 | 124236284 | rs12530266 | -0.43 | -0.756 | -0.637 | -0.082 | 0.007 | -0.003 | -0.003 | 0 | 0.4 | 0.36 | 0.46 | 0.65 | 0.38 | 0.46 |
| 6 | 124238179 | rs12192551 | 0.278 | -0.929 | -0.822 | 0.023 | -0.114 | -0.142 | -0.142 | 0 | 0.28 | 0.02 | 0.42 | 0.53 | 0.35 | 0.42 |
| 6 | 124238200 | rs584050 | 1.473 | -0.919 | -0.866 | -0.225 | -0.015 | 0.022 | 0.022 | 0 | 0.4 | 0.11 | 0.24 | 0.26 | 0.37 | 0.39 |
| 6 | 124241003 | rs235665 | 1.27 | -0.846 | -0.695 | -0.174 | 0.001 | 0.003 | 0.003 | 0 | 0.43 | 0.52 | 0.28 | 0.26 | 0.38 | 0.41 |
| 6 | 124241640 | rs9401711 | -0.389 | -0.781 | -0.902 | -0.061 | -0.015 | 0.04 | 0.04 | 0.244 | 0.05 | 0.14 | 0.02 | 0.12 | 0.01 | 0.03 |
| 6 | 124243131 | rs11755255 | 0.128 | -0.738 | -0.906 | -0.061 | -0.059 | -0.085 | -0.085 | 0 | 0.29 | 0.08 | 0.42 | 0.53 | 0.35 | 0.42 |
| 6 | 124244564 | rs13192934 | 0.196 | -0.699 | -0.886 | -0.04 | -0.061 | -0.086 | -0.086 | 0 | 0.29 | 0.08 | 0.42 | 0.53 | 0.35 | 0.42 |
| 6 | 124250159 | rs9401713 | -0.166 | -0.352 | -1.056 | -0.167 | -0.049 | -0.07 | -0.07 | 0 | 0.35 | 0.15 | 0.44 | 0.65 | 0.36 | 0.44 |
| 6 | 124253062 | rs235698 | 0.42 | -0.172 | -0.891 | -0.188 | -0.006 | 0.008 | 0.008 | 0 | 0.8 | 0.84 | 0.73 | 0.92 | 0.74 | 0.87 |
| 6 | 124253962 | rs173105 | 0.19 | -0.122 | -0.844 | -0.124 | 0.004 | -0.003 | -0.003 | 1.287 | 0.85 | 0.88 | 0.75 | 0.92 | 0.76 | 0.88 |
| 6 | 124254836 | rs235694 | 0.941 | -0.3 | -0.811 | -0.085 | 0.022 | 0.036 | 0.036 | 0 | 0.49 | 0.59 | 0.29 | 0.28 | 0.39 | 0.46 |
| 6 | 124255182 | rs235692 | 1.041 | -0.278 | -0.864 | -0.06 | 0.022 | 0.022 | 0.022 | 0 | 0.46 | 0.58 | 0.28 | 0.28 | 0.34 | 0.41 |
| 6 | 124270952 | rs9388291 | 0.085 | -0.266 | -1.127 | 0.051 | -0.109 | -0.162 | -0.162 | 0 | 0.29 | 0.02 | 0.43 | 0.63 | 0.35 | 0.39 |
| 6 | 124271440 | rs235667 | -0.041 | -0.085 | -1.17 | -0.137 | -0.098 | 0.025 | 0.025 | 0 | 0.82 | 0.99 | 0.73 | 0.92 | 0.69 | 0.82 |
| 6 | 124278812 | rs4895951 | 1.041 | -0.577 | -1.091 | -0.016 | 0.001 | -0.024 | -0.024 | 1.998 | 0.37 | 0.11 | 0.34 | 0.28 | 0.28 | 0.39 |
| 6 | 124284365 | rs1267993 | 0.65 | -0.6 | -0.634 | -0.03 | 0.002 | 0.002 | 0.002 | 0 | 0.46 | 0.47 | 0.39 | 0.28 | 0.3 | 0.4 |
| 6 | 124286626 | rs1267992 | 0.929 | -0.319 | -0.963 | 0.141 | 0.029 | 0.017 | 0.017 | 0 | 0.4 | 0.3 | 0.35 | 0.28 | 0.29 | 0.4 |
| 6 | 124289841 | rs7753102 | -0.786 | -0.031 | -0.843 | 0.841 | -0.019 | 0.035 | 0.035 | 0 | 0.06 | 0.12 | 0.03 | 0.17 | 0.02 | 0.09 |
| 6 | 124291308 | rs9401715 | -0.134 | 0.516 | -1.13 | 0.714 | -0.026 | -0.064 | -0.064 | 0 | 0.27 | 0.15 | 0.22 | 0.46 | 0.29 | 0.3 |
| 6 | 124298718 | rs9320967 | -0.569 | 0.501 | -0.674 | 0.414 | -0.004 | 0.025 | 0.025 | 2.2 | 0.31 | 0.78 | 0.23 | 0.18 | 0.33 | 0.17 |
| 6 | 124304371 | rs13217719 | 0.544 | 0.531 | -0.471 | 0.694 | 0.009 | 0.017 | 0.017 | 0 | 0.44 | 0.31 | 0.54 | 0.33 | 0.38 | 0.48 |
| 6 | 124306595 | rs9375303 | 0.177 | 0.708 | -1.219 | 0.963 | -0.082 | -0.114 | -0.114 | 0 | 0.28 | 0.02 | 0.15 | 0.37 | 0.28 | 0.32 |
| 6 | 124310248 | rs1397199 | -0.276 | 0.855 | -0.667 | 0.859 | -0.024 | 0.014 | 0.014 | 0 | 0.78 | 0.86 | 0.79 | 0.84 | 0.65 | 0.82 |
| 6 | 124310564 | rs6901490 | 0.428 | 0.736 | -0.345 | 0.618 | 0.029 | -0.005 | -0.005 | 1.105 | 0.45 | 0.66 | 0.6 | 0.46 | 0.34 | 0.48 |
| 6 | 124313842 | rs1355283 | -0.69 | 0.571 | -0.209 | 0.235 | 0.01 | 0.04 | 0.04 | 0 | 0.46 | 0.79 | 0.61 | 0.63 | 0.4 | 0.49 |
| 6 | 124315768 | rs7742884 | -0.216 | 0.4 | 0.159 | 0.098 | -0.003 | -0.002 | -0.002 | 0 | 0.34 | 0.65 | 0.43 | 0.62 | 0.37 | 0.36 |
| 6 | 124317132 | rs7748718 | -0.855 | 0.531 | 0.061 | 0.023 | -0.01 | -0.03 | -0.03 | 0 | 0.12 | 0.12 | 0.19 | 0.01 | 0.05 | 0.14 |
| 6 | 124318522 | rs7754719 | 0.095 | 0.67 | 0.518 | 0.474 | 0.022 | 0.035 | 0.035 | 0 | 0.28 | 0.42 | 0.38 | 0.42 | 0.31 | 0.28 |
| 6 | 124328531 | rs3861386 | 0.34 | 0.67 | 0.51 | 0.179 | -0.016 | -0.031 | -0.031 | 0 | 0.25 | 0.2 | 0.34 | 0.4 | 0.27 | 0.27 |
| 6 | 124329017 | rs1104940 | 0.404 | 0.525 | 0.283 | -0.214 | -0.001 | -0.003 | -0.003 | 0 | 0.57 | 0.5 | 0.4 | 0.59 | 0.6 | 0.58 |
| 6 | 124329313 | rs1510295 | 0.173 | 0.518 | 0.17 | -0.169 | 0.011 | 0.021 | 0.021 | 0 | 0.83 | 0.92 | 0.77 | 0.99 | 0.87 | 0.84 |
| 6 | 124332293 | rs4363051 | -0.337 | 0.843 | 0.404 | 0.302 | 0.001 | 0.008 | 0.008 | 0 | 0.36 | 0.52 | 0.42 | 0.6 | 0.34 | 0.35 |
| 6 | 124333162 | rs9320971 | -0.49 | 0.824 | 0.932 | 0.231 | 0.027 | 0.211 | 0.211 | 0 | 0.05 | 0.27 | 0.04 | 0.2 | 0.04 | 0.07 |
| 6 | 124337383 | rs17627947 | -1.079 | 1.228 | 0.448 | -1.078 | 0 | -0.047 | -0.047 | 0 | 0.23 | 0.21 | 0.27 | 0.01 | 0.18 | 0.15 |
| 6 | 124342195 | rs6935830 | -1.228 | 0.886 | 0.83 | -0.956 | -0.023 | -0.161 | -0.161 | 0 | 0.25 | 0.43 | 0.3 | 0.01 | 0.19 | 0.2 |
| 6 | 124346434 | rs6905913 | -0.955 | 0.314 | 0.83 | -0.906 | -0.034 | -0.113 | -0.113 | 0 | 0.23 | 0.26 | 0.26 | 0.01 | 0.16 | 0.2 |
| 6 | 124355206 | rs234481 | -0.395 | 0.539 | 0.418 | -0.003 | -0.003 | -0.01 | -0.01 | 0.271 | 0.24 | 0.22 | 0.2 | 0.35 | 0.32 | 0.25 |
| 6 | 124357337 | rs9491054 | -1.284 | 0.348 | 0.851 | -0.218 | 0.019 | 0.004 | 0.004 | 3.264 | 0.07 | 0.47 | 0.05 | 0 | 0.01 | 0.02 |
| 6 | 124361146 | rs234478 | 0.344 | 0.786 | -0.312 | -0.449 | -0.011 | 0.105 | 0.105 | 0.054 | 0.54 | 0.32 | 0.54 | 0.79 | 0.5 | 0.56 |
| 6 | 124371914 | rs1510292 | -0.839 | 0.875 | 0.467 | -0.39 | 0.005 | -0.05 | -0.05 | 0.578 | 0.1 | 0.22 | 0.07 | 0.02 | 0.08 | 0.13 |
| 6 | 124376164 | rs2064687 | -1.772 | 1.244 | 1.324 | 0.397 | 0.028 | 0.058 | 0.058 | 0 | 0.17 | 0.41 | 0.13 | 0.27 | 0.18 | 0.26 |
| 6 | 124376272 | rs1510293 | 1.193 | 1.294 | 0.899 | 0.325 | -0.011 | 0.052 | 0.052 | 0 | 0.39 | 0.18 | 0.24 | 0.28 | 0.43 | 0.4 |
| 6 | 124379742 | rs9320974 | 1.154 | 1.309 | 0.894 | 0.422 | -0.028 | 0.046 | 0.046 | 0 | 0.39 | 0.18 | 0.26 | 0.31 | 0.47 | 0.42 |
| 6 | 124380321 | rs7765184 | -1.312 | 1.088 | 1.017 | 0.315 | -0.021 | 0.026 | 0.026 | 0.026 | 0.21 | 0.41 | 0.11 | 0.25 | 0.13 | 0.25 |
| 6 | 124388163 | rs6902462 | -0.482 | 1.188 | 0.943 | 0.295 | 0.014 | 0.058 | 0.058 | 0 | 0.14 | 0.32 | 0.1 | 0.27 | 0.14 | 0.25 |
| 6 | 124392114 | rs1858234 | -0.215 | 1.138 | 1.051 | 0.251 | -0.012 | -0.042 | -0.042 | 0.404 | 0.62 | 0.78 | 0.39 | 0.54 | 0.6 | 0.67 |
| 6 | 124396364 | rs6932669 | -1.416 | 1.339 | 1.192 | 0.233 | -0.009 | 0.018 | 0.018 | 0 | 0.23 | 0.57 | 0.13 | 0.26 | 0.17 | 0.25 |

| | | | | | | | | | | | | | | | | |
|---|---|---|---|---|---|---|---|---|---|---|---|---|---|---|---|---|
| 6 | 124398002 | rs17086577 | -0.969 | 1.458 | 0.81 | 0.671 | 0.013 | 0.109 | 0.109 | 0 | 0.05 | 0.13 | 0.03 | 0.23 | 0.05 | 0.12 |
| 6 | 124401214 | rs2048338 | 0.947 | 1.325 | 0.536 | 0.371 | -0.006 | 0.061 | 0.061 | 0 | 0.39 | 0.19 | 0.25 | 0.24 | 0.42 | 0.41 |
| 6 | 124404539 | rs9320979 | -1.51 | 1.489 | 0.8 | 0.869 | 0.034 | 0.058 | 0.058 | 0 | 0.06 | 0.12 | 0.05 | 0.2 | 0.1 | 0.14 |
| 6 | 124405409 | rs9375306 | -1.682 | 1.233 | 1.038 | 0.705 | 0.049 | 0.063 | 0.063 | 0.046 | 0.07 | 0.29 | 0.05 | 0.12 | 0.06 | 0.12 |
| 6 | 124416847 | rs9388300 | -0.521 | 1.1 | 0.684 | 0.905 | -0.002 | 0.054 | 0.054 | 0 | 0.19 | 0.25 | 0.4 | 0.47 | 0.17 | 0.2 |
| 6 | 124421114 | rs173067 | 0.939 | 1.508 | 0.32 | 0.231 | -0.022 | 0.041 | 0.041 | 0 | 0.66 | 0.82 | 0.76 | 0.77 | 0.67 | 0.67 |
| 6 | 124434003 | rs163283 | 1.699 | 1.295 | 0.569 | 0.059 | 0.011 | 0.048 | 0.048 | 0 | 0.46 | 0.27 | 0.32 | 0.31 | 0.48 | 0.48 |
| 6 | 124441233 | rs12204100 | -0.456 | 1.417 | 0.531 | -0.105 | -0.055 | 0.147 | 0.147 | 0 | 0.24 | 0.03 | 0.15 | 0.03 | 0.22 | 0.22 |
| 6 | 124442286 | rs594664 | 2.365 | 1.538 | 0.546 | -0.09 | 0.006 | 0.027 | 0.027 | 0 | 0.41 | 0.27 | 0.26 | 0.29 | 0.43 | 0.44 |
| 6 | 124443660 | rs645673 | 1.374 | 1.003 | 0.539 | -0.376 | -0.023 | 0.05 | 0.05 | 0 | 0.65 | 0.78 | 0.73 | 0.78 | 0.65 | 0.7 |
| 6 | 124454316 | rs11758638 | 0.664 | 1.062 | 0.417 | -0.448 | -0.027 | 0.06 | 0.06 | 0 | 0.1 | 0 | 0.06 | 0 | 0.09 | 0.03 |
| 6 | 124461446 | rs163286 | 0.43 | 0.988 | 0.515 | -0.33 | 0.001 | -0.002 | -0.002 | 1.189 | 0.87 | 0.87 | 0.91 | 0.87 | 0.84 | 0.88 |
| 6 | 124468892 | rs9375309 | 1.18 | 0.98 | 0.48 | -0.361 | -0.01 | 0.031 | 0.031 | 0 | 0.63 | 0.65 | 0.7 | 0.77 | 0.64 | 0.65 |
| 6 | 124474595 | rs802252 | 1.562 | 0.364 | 1.2 | -0.115 | -0.011 | -0.015 | -0.015 | 0 | 0.83 | 0.54 | 0.84 | 0.86 | 0.89 | 0.91 |
| 6 | 124476771 | rs802262 | 0.629 | 0.377 | 1.228 | -0.111 | 0.003 | -0.028 | -0.028 | 0.478 | 0.89 | 0.73 | 0.92 | 0.94 | 0.91 | 0.95 |
| 6 | 124481519 | rs802272 | 1.267 | 0.374 | 1.11 | -0.104 | -0.003 | -0.003 | -0.003 | 0 | 0.86 | 0.8 | 0.87 | 0.86 | 0.89 | 0.9 |
| 6 | 124481616 | rs802273 | 1.873 | 0.344 | 1.29 | -0.116 | -0.015 | -0.012 | -0.012 | 0 | 0.83 | 0.45 | 0.83 | 0.84 | 0.89 | 0.86 |
| 6 | 124483459 | rs802274 | 2.03 | 0.327 | 1.369 | -0.129 | -0.021 | -0.022 | -0.022 | 0 | 0.82 | 0.38 | 0.83 | 0.85 | 0.89 | 0.87 |
| 6 | 124496256 | rs2758826 | 0.378 | 0.198 | 1.095 | -0.049 | -0.002 | 0.008 | 0.008 | 0 | 0.91 | 0.93 | 0.93 | 0.93 | 0.91 | 0.95 |
| 6 | 124500739 | rs9398750 | -0.273 | -0.08 | 1.004 | 0.422 | -0.006 | 0.003 | 0.003 | 0.134 | 0.22 | 0.25 | 0.22 | 0.3 | 0.15 | 0.18 |
| 6 | 124501118 | rs802295 | 1.686 | -0.081 | 0.454 | 0.404 | 0.05 | -0.018 | -0.018 | 0 | 0.6 | 0.25 | 0.61 | 0.54 | 0.74 | 0.68 |
| 6 | 124511758 | rs6908839 | -1.291 | 0.306 | 0.974 | 0.899 | 0.039 | 0.097 | 0.097 | 0 | 0.07 | 0.39 | 0.09 | 0.08 | 0.02 | 0.05 |
| 6 | 124521622 | rs802276 | 1.122 | 0.603 | 0.741 | 0.887 | -0.004 | 0.008 | 0.008 | 0 | 0.7 | 0.61 | 0.69 | 0.65 | 0.74 | 0.75 |
| 6 | 124543314 | rs9482518 | 0.201 | 0.622 | 0.737 | 0.977 | 0.002 | 0.021 | 0.021 | 0.01 | 0.81 | 0.73 | 0.78 | 0.73 | 0.83 | 0.85 |
| 6 | 124571367 | rs2689887 | -1.583 | 0.803 | -0.977 | 0.917 | -0.071 | -0.1 | -0.1 | 3.031 | 0.55 | 0.96 | 0.52 | 0.52 | 0.49 | 0.45 |
| 6 | 124579645 | rs2689886 | -1.625 | 0.42 | 0.187 | 1.34 | 0.009 | -0.002 | -0.002 | 0.126 | 0.38 | 0.78 | 0.39 | 0.4 | 0.31 | 0.26 |
| 6 | 124580035 | rs1523977 | -1.049 | 0.439 | 0.204 | 1.332 | -0.002 | 0.002 | 0.002 | 0.09 | 0.28 | 0.69 | 0.33 | 0.34 | 0.24 | 0.21 |
| 6 | 124582245 | rs2753138 | -0.988 | 0.331 | 0.204 | 1.33 | -0.003 | 0.006 | 0.006 | 0.207 | 0.28 | 0.66 | 0.32 | 0.34 | 0.24 | 0.18 |
| 6 | 124584708 | rs11758916 | -0.505 | 0.337 | 0.157 | 1.332 | 0.003 | 0.006 | 0.006 | 0 | 0.09 | 0.07 | 0.06 | 0.06 | 0.07 | 0.05 |
| 6 | 124584845 | rs2689898 | -0.871 | 0.356 | 0.11 | 1.321 | -0.001 | 0.021 | 0.021 | 0.262 | 0.27 | 0.56 | 0.31 | 0.34 | 0.24 | 0.18 |
| 6 | 124588039 | rs1973961 | -1.017 | 0.53 | -0.678 | 1.212 | 0.01 | -0.009 | -0.009 | 0 | 0.38 | 0.78 | 0.39 | 0.42 | 0.31 | 0.26 |
| 6 | 124603390 | rs802497 | -1.455 | 0.269 | -0.663 | 1.357 | 0.021 | -0.028 | -0.028 | 0.389 | 0.38 | 0.88 | 0.25 | 0.51 | 0.31 | 0.33 |
| 6 | 124603670 | rs713215 | -1.064 | 0.309 | 0.384 | 1.431 | 0.004 | 0.041 | 0.041 | 0.082 | 0.18 | 0.26 | 0.13 | 0.27 | 0.18 | 0.13 |
| 6 | 124606520 | rs7771967 | -0.591 | 0.433 | 0.523 | 1.55 | 0.004 | 0.046 | 0.046 | 0.312 | 0.11 | 0.17 | 0.07 | 0.27 | 0.12 | 0.11 |
| 6 | 124608621 | rs2753147 | 0.394 | 0.324 | 1.052 | 1.099 | -0.002 | -0.062 | -0.062 | 0.106 | 0.87 | 0.7 | 0.9 | 0.97 | 0.86 | 0.91 |
| 6 | 124610320 | rs802500 | -2.381 | 0.516 | 1.129 | 1.869 | 0.029 | -0.045 | -0.045 | 0 | 0.41 | 0.94 | 0.36 | 0.56 | 0.31 | 0.35 |
| 6 | 124612578 | rs2689869 | -2.074 | 0.234 | 1.923 | 2.164 | -0.029 | 0.009 | 0.009 | 1.188 | 0.23 | 0.52 | 0.15 | 0.22 | 0.17 | 0.18 |
| 6 | 124613921 | rs2626115 | -2.299 | 0.233 | 2.092 | 2.175 | -0.014 | 0.02 | 0.02 | 1.85 | 0.24 | 0.66 | 0.17 | 0.31 | 0.18 | 0.18 |
| 6 | 124616517 | rs11753035 | -0.956 | 0.228 | 2.088 | 2.177 | 0.007 | 0.078 | 0.078 | 0.079 | 0.11 | 0.2 | 0.07 | 0.27 | 0.12 | 0.1 |
| 6 | 124617031 | rs2753159 | -2.017 | 0.145 | 2.17 | 2.124 | -0.043 | -0.077 | -0.077 | 1.593 | 0.15 | 0.66 | 0.11 | 0.03 | 0.07 | 0.08 |
| 6 | 124617746 | rs4513814 | -0.797 | 0.148 | 2.223 | 2.146 | -0.001 | 0.012 | 0.012 | 0.019 | 0.11 | 0.11 | 0.06 | 0.19 | 0.1 | 0.1 |
| 6 | 124619270 | rs2753161 | -1.808 | 0.072 | 2.267 | 1.991 | -0.03 | -0.079 | -0.079 | 0 | 0.14 | 0.54 | 0.1 | 0.03 | 0.06 | 0.08 |
| 6 | 124619294 | rs9491108 | -0.948 | 0.089 | 2.069 | 2.045 | 0 | 0.066 | 0.066 | 0.079 | 0.11 | 0.2 | 0.07 | 0.25 | 0.11 | 0.1 |
| 6 | 124619846 | rs2753162 | -2.436 | 0.1 | 1.578 | 2.053 | 0.028 | -0.015 | -0.015 | 0 | 0.26 | 0.84 | 0.18 | 0.28 | 0.18 | 0.18 |
| 6 | 124627115 | rs1851780 | -2.597 | 0.088 | 1.423 | 2.025 | 0.077 | -0.091 | -0.091 | 0 | 0.35 | 0.91 | 0.3 | 0.53 | 0.2 | 0.31 |
| 6 | 124632478 | rs6921885 | -2.144 | 0.283 | 2.009 | 2.219 | -0.052 | 0.177 | 0.177 | 1.41 | 0.14 | 0.47 | 0.16 | 0.29 | 0.05 | 0.21 |
| 6 | 124632800 | rs802504 | -1.013 | 0.343 | 1.945 | 2.065 | 0.014 | -0.02 | -0.02 | 0.084 | 0.05 | 0.15 | 0.03 | 0 | 0.04 | 0.02 |
| 6 | 124633499 | rs11751387 | -0.417 | 0.35 | 1.804 | 2.368 | -0.031 | -0.053 | -0.053 | 0 | 0.08 | 0 | 0.07 | 0.18 | 0.11 | 0.11 |
| 6 | 124635249 | rs2208345 | -2.209 | 0.332 | 2.062 | 2.389 | -0.022 | 0.061 | 0.061 | 2.25 | 0.24 | 0.57 | 0.23 | 0.46 | 0.15 | 0.3 |
| 6 | 124635641 | rs9482533 | -1.533 | 0.228 | 1.554 | 2.188 | -0.024 | 0.13 | 0.13 | 0 | 0.1 | 0.25 | 0.13 | 0.23 | 0.03 | 0.15 |
| 6 | 124637664 | rs9491113 | -1.885 | 0.087 | 1.186 | 2.242 | -0.022 | 0.032 | 0.032 | 0 | 0.15 | 0.22 | 0.15 | 0.23 | 0.08 | 0.15 |
| 6 | 124641194 | rs9375319 | -0.9 | 0.025 | 1.045 | 2.277 | 0.009 | -0.003 | -0.003 | 0 | 0.08 | 0.04 | 0.12 | 0.23 | 0.03 | 0.15 |
| 6 | 124643000 | rs2753172 | -2.099 | 0.348 | 0.294 | 2.189 | -0.006 | -0.104 | -0.104 | 0 | 0.33 | 0.92 | 0.29 | 0.54 | 0.25 | 0.31 |
| 6 | 124646119 | rs6899511 | -1.268 | 0.288 | 0.555 | 1.978 | -0.031 | 0.04 | 0.04 | 0 | 0.15 | 0.22 | 0.13 | 0.25 | 0.07 | 0.14 |
| 6 | 124650048 | rs9401729 | -1.31 | 0.624 | 0.418 | 1.749 | -0.002 | 0.024 | 0.024 | 1.474 | 0.3 | 0.38 | 0.26 | 0.58 | 0.27 | 0.33 |
| 6 | 124650870 | rs2626128 | -0.461 | 0.332 | 0.153 | 1.392 | -0.003 | 0.006 | 0.006 | 0.017 | 0.42 | 0.34 | 0.42 | 0.38 | 0.48 | 0.37 |
| 6 | 124651358 | rs2626130 | -1.294 | -0.104 | 0.485 | 1.398 | -0.002 | 0.001 | 0.001 | 0.895 | 0.39 | 0.69 | 0.38 | 0.45 | 0.4 | 0.37 |
| 6 | 124651861 | rs11154212 | 0.017 | 0.322 | 0.152 | 0.948 | -0.052 | 0.101 | 0.101 | 0 | 0.19 | 0.02 | 0.23 | 0.03 | 0.19 | 0.18 |
| 6 | 124652232 | rs2626131 | 0.837 | 0.237 | 0.716 | 1.552 | -0.002 | 0 | 0 | 0.035 | 0.64 | 0.71 | 0.6 | 0.69 | 0.6 | 0.66 |
| 6 | 124656470 | rs4516960 | -0.433 | 0.072 | 1.486 | 1.473 | 0 | 0.017 | 0.017 | 0.76 | 0.1 | 0.31 | 0.16 | 0.09 | 0.09 | 0.07 |
| 6 | 124671329 | rs12195743 | 0.154 | 0.039 | 1.484 | 1.515 | -0.007 | 0.006 | 0.006 | 0 | 0.2 | 0.12 | 0.29 | 0.15 | 0.18 | 0.17 |
| 6 | 124673700 | rs12201181 | -0.011 | 0.032 | 1.523 | 1.493 | 0 | 0.011 | 0.011 | 0 | 0.22 | 0.15 | 0.24 | 0.12 | 0.18 | 0.17 |
| 6 | 124677633 | rs4478425 | 1.07 | -0.045 | 1.851 | 1.563 | -0.003 | 0.009 | 0.009 | 0 | 0.77 | 0.63 | 0.75 | 0.75 | 0.73 | 0.77 |
| 6 | 124681503 | rs12204812 | -1.085 | 0.018 | 1.724 | 1.415 | -0.003 | -0.026 | -0.026 | 0 | 0.34 | 0.49 | 0.42 | 0.21 | 0.32 | 0.27 |
| 6 | 124685182 | rs1407529 | 1.604 | -0.109 | 1.78 | 1.455 | 0.001 | 0.004 | 0.004 | 0 | 0.49 | 0.45 | 0.43 | 0.5 | 0.45 | 0.52 |
| 6 | 124687473 | rs9401741 | 2.273 | -0.103 | 1.766 | 1.457 | -0.033 | -0.022 | -0.022 | 0 | 0.4 | 0.12 | 0.33 | 0.43 | 0.38 | 0.45 |
| 6 | 124688755 | rs911959 | -0.285 | -0.068 | 1.938 | 1.47 | -0.002 | -0.007 | -0.007 | 0.373 | 0.25 | 0.29 | 0.32 | 0.15 | 0.22 | 0.2 |
| 6 | 124688857 | rs911960 | 0.041 | -0.098 | 1.919 | 1.458 | -0.014 | -0.029 | -0.029 | 0 | 0.87 | 0.96 | 0.85 | 0.81 | 0.81 | 0.84 |
| 6 | 124690371 | rs980658 | -0.288 | -0.072 | 2.408 | 1.358 | -0.001 | -0.004 | -0.004 | 0 | 0.25 | 0.25 | 0.31 | 0.16 | 0.22 | 0.2 |

| | | | | | | | | | | | | | | | | |
|---|---|---|---|---|---|---|---|---|---|---|---|---|---|---|---|---|
| 6 | 124691352 | rs9491141 | -0.708 | -0.07 | 2.401 | 1.359 | 0.013 | 0.069 | 0.069 | 0.453 | 0.11 | 0.4 | 0.12 | 0.16 | 0.1 | 0.07 |
| 6 | 124691424 | rs6938403 | 0.256 | -0.059 | 2.361 | 1.35 | -0.004 | -0.007 | -0.007 | 4.745 | 0.08 | 0.12 | 0.09 | 0.06 | 0.08 | 0.07 |
| 6 | 124691506 | rs2146288 | -0.26 | -0.05 | 2.28 | 1.341 | -0.005 | 0.016 | 0.016 | 0 | 0.12 | 0.19 | 0.1 | 0.16 | 0.11 | 0.12 |
| 6 | 124693042 | rs11755888 | -0.165 | -0.018 | 2.015 | 1.316 | -0.001 | 0.001 | 0.001 | 0 | 0.24 | 0.2 | 0.31 | 0.07 | 0.22 | 0.2 |
| 6 | 124695544 | rs4510696 | -1.547 | -0.013 | 1.997 | 1.383 | -0.008 | 0.038 | 0.038 | 0.56 | 0.22 | 0.56 | 0.21 | 0.32 | 0.21 | 0.2 |
| 6 | 124698568 | rs10457447 | -0.067 | 0.085 | 1.776 | 1.249 | -0.014 | 0.059 | 0.059 | 0 | 0.22 | 0.08 | 0.3 | 0.06 | 0.22 | 0.2 |
| 6 | 124699827 | rs918913 | 0.946 | 0.014 | 2.028 | 1.458 | 0 | 0.007 | 0.007 | 0 | 0.76 | 0.72 | 0.75 | 0.63 | 0.69 | 0.72 |
| 6 | 124700720 | rs9491143 | 1.054 | -0.111 | 2.28 | 1.526 | -0.003 | 0.006 | 0.006 | 0.101 | 0.53 | 0.59 | 0.44 | 0.47 | 0.45 | 0.51 |
| 6 | 124701564 | rs7751845 | 0.497 | -0.111 | 2.134 | 1.489 | -0.045 | -0.038 | -0.038 | 0.061 | 0.59 | 0.8 | 0.48 | 0.57 | 0.5 | 0.59 |
| 6 | 124703501 | rs990802 | -1.818 | 0.083 | 2.427 | 1.49 | 0.011 | 0.056 | 0.056 | 0.068 | 0.08 | 0.27 | 0.04 | 0.1 | 0.07 | 0.05 |
| 6 | 124704731 | rs9320988 | -1.974 | 0.073 | 2.61 | 1.737 | -0.008 | 0.062 | 0.062 | 0 | 0.16 | 0.45 | 0.15 | 0.16 | 0.14 | 0.17 |
| 6 | 124705214 | rs6937991 | -2.272 | 0.054 | 2.505 | 1.727 | -0.022 | 0.045 | 0.045 | 0.053 | 0.17 | 0.53 | 0.15 | 0.16 | 0.14 | 0.17 |
| 6 | 124706331 | rs17051777 | -0.696 | 0.162 | 1.985 | 1.614 | 0.007 | -0.004 | -0.004 | 0.846 | 0.1 | 0.19 | 0.11 | 0.07 | 0.11 | 0.13 |
| 6 | 124708702 | rs12055404 | 1.412 | -0.161 | 1.432 | 1.735 | -0.031 | 0.062 | 0.062 | 0.872 | 0.45 | 0.17 | 0.36 | 0.36 | 0.42 | 0.52 |
| 6 | 124708814 | rs9491150 | -1.225 | -0.091 | 1.874 | 1.848 | 0 | 0.028 | 0.028 | 0.068 | 0.07 | 0.17 | 0.09 | 0.11 | 0.07 | 0.08 |
| 6 | 124713527 | rs332611 | 0.135 | -0.142 | 1.555 | 2.079 | -0.054 | 0.009 | 0.009 | 0 | 0.71 | 0.91 | 0.65 | 0.78 | 0.65 | 0.74 |
| 6 | 124715205 | rs9401743 | 1.033 | -0.106 | 1.869 | 1.932 | -0.002 | 0.006 | 0.006 | 0 | 0.52 | 0.68 | 0.42 | 0.45 | 0.45 | 0.56 |
| 6 | 124715576 | rs9385331 | 1.824 | 0.014 | 1.119 | 1.76 | -0.033 | 0.046 | 0.046 | 0.444 | 0.42 | 0.11 | 0.35 | 0.35 | 0.38 | 0.51 |
| 6 | 124734727 | rs558725 | -0.783 | -0.093 | 1.384 | 1.569 | 0 | 0.002 | 0.002 | 0 | 0.19 | 0.19 | 0.19 | 0.25 | 0.21 | 0.13 |
| 6 | 124738540 | rs491104 | -0.591 | -0.087 | 1.353 | 1.565 | -0.022 | -0.031 | -0.031 | 3 | 0.17 | 0.06 | 0.18 | 0.25 | 0.2 | 0.13 |
| 6 | 124739792 | rs523240 | -1.724 | -0.15 | 2.136 | 1.8 | 0.013 | 0.077 | 0.077 | 0 | 0.25 | 0.45 | 0.24 | 0.33 | 0.26 | 0.19 |
| 6 | 124743507 | rs531930 | -0.21 | -0.278 | 1.847 | 1.79 | 0.001 | 0.03 | 0.03 | 0.666 | 0.74 | 0.87 | 0.75 | 0.85 | 0.75 | 0.81 |
| 6 | 124745863 | rs2325944 | 1.959 | -0.417 | 1.556 | 1.646 | -0.069 | -0.04 | -0.04 | 0.009 | 0.45 | 0.13 | 0.49 | 0.52 | 0.48 | 0.59 |
| 6 | 124761137 | rs592322 | -1.252 | -0.528 | 2.008 | 1.723 | -0.009 | 0.004 | 0.004 | 0 | 0.22 | 0.3 | 0.2 | 0.25 | 0.19 | 0.14 |
| 6 | 124763669 | rs9766943 | 1.493 | -0.622 | 1.748 | 1.777 | -0.02 | 0.026 | 0.026 | 0.05 | 0.75 | 0.39 | 0.8 | 0.68 | 0.81 | 0.8 |
| 6 | 124764631 | rs9401746 | 0.252 | -0.654 | 1.467 | 1.744 | -0.013 | -0.072 | -0.072 | 0.091 | 0.86 | 0.98 | 0.86 | 0.74 | 0.88 | 0.91 |
| 6 | 124768132 | rs11154239 | 1.368 | -0.634 | 1.634 | 1.809 | -0.005 | 0.007 | 0.007 | 0.695 | 0.75 | 0.32 | 0.79 | 0.65 | 0.82 | 0.83 |
| 6 | 124785449 | rs579554 | -1.529 | -0.557 | 1.479 | 1.69 | -0.006 | -0.006 | -0.006 | 0 | 0.26 | 0.67 | 0.25 | 0.26 | 0.24 | 0.14 |
| 6 | 124794887 | rs688197 | 0.221 | -0.703 | 1.116 | 1.43 | 0.01 | 0.016 | 0.016 | 0.01 | 0.72 | 0.82 | 0.74 | 0.86 | 0.75 | 0.77 |
| 6 | 124805874 | rs7767478 | -0.654 | -0.746 | 1.247 | 1.574 | -0.029 | -0.001 | -0.001 | 0 | 0.31 | 0.45 | 0.28 | 0.22 | 0.26 | 0.25 |
| 6 | 124807814 | rs587706 | -1.117 | -1.586 | 0.294 | 1.162 | 0.018 | 0.007 | 0.007 | 0 | 0.41 | 0.34 | 0.47 | 0.64 | 0.35 | 0.4 |
| 6 | 124808189 | rs1549355 | -0.717 | -1.621 | -0.104 | 1.568 | 0.014 | -0.06 | -0.06 | 0 | 0.23 | 0.16 | 0.26 | 0.43 | 0.19 | 0.25 |
| 6 | 124809084 | rs1572471 | -0.69 | -1.46 | -0.251 | 1.529 | -0.025 | -0.053 | -0.053 | 0.57 | 0.35 | 0.23 | 0.41 | 0.44 | 0.47 | 0.38 |
| 6 | 124811874 | rs519862 | 0.053 | -1.639 | -0.891 | 1.208 | 0.033 | -0.074 | -0.074 | 0 | 0.25 | 0.08 | 0.22 | 0.35 | 0.16 | 0.23 |
| 6 | 124817317 | rs9388339 | 0.157 | -1.377 | -0.069 | 1.174 | -0.006 | 0.002 | 0.002 | 0.527 | 0.8 | 0.79 | 0.78 | 0.72 | 0.86 | 0.88 |
| 6 | 124827707 | rs582940 | 0.859 | -1.154 | 0.563 | 0.769 | -0.001 | 0.037 | 0.037 | 0 | 0.66 | 0.73 | 0.76 | 0.83 | 0.61 | 0.76 |
| 6 | 124829945 | rs539252 | 1.62 | -1.049 | 0.738 | 0.83 | 0.007 | -0.011 | -0.011 | 0.689 | 0.53 | 0.34 | 0.51 | 0.54 | 0.51 | 0.55 |
| 6 | 124837391 | rs7753567 | -0.428 | -0.995 | 0.431 | 0.844 | -0.068 | 0.05 | 0.05 | 0 | 0.29 | 0.07 | 0.21 | 0.16 | 0.39 | 0.23 |
| 6 | 124838090 | rs1842129 | 1.586 | -1.056 | 0.587 | 0.755 | -0.013 | -0.053 | -0.053 | 0 | 0.53 | 0.15 | 0.5 | 0.55 | 0.51 | 0.57 |
| 6 | 124841336 | rs643002 | 1.643 | -1.188 | 1.114 | 0.294 | -0.044 | -0.062 | -0.062 | 0 | 0.9 | 0.47 | 0.94 | 0.99 | 0.99 | 0.94 |
| 6 | 124857257 | rs589665 | 1.236 | -1.009 | 0.32 | 0.349 | 0.037 | 0.03 | 0.03 | 0.353 | 0.56 | 0.17 | 0.57 | 0.67 | 0.6 | 0.64 |
| 6 | 124859468 | rs2054466 | 1.402 | -1.029 | 0.853 | 0.068 | -0.007 | -0.071 | -0.071 | 0.173 | 0.89 | 0.72 | 0.94 | 1 | 0.93 | 0.98 |
| 6 | 124863962 | rs535492 | -1.18 | -0.749 | 0.172 | -0.006 | -0.001 | -0.007 | -0.007 | 0.01 | 0.44 | 0.52 | 0.41 | 0.3 | 0.44 | 0.24 |
| 6 | 124865155 | rs670842 | 0.069 | -0.49 | -0.311 | 0.081 | -0.08 | -0.065 | -0.065 | 0 | 0.14 | 0.01 | 0.2 | 0.25 | 0.24 | 0.15 |
| 6 | 124865450 | rs7770974 | -0.903 | -1.21 | 0.159 | -0.355 | -0.013 | -0.054 | -0.054 | 0.462 | 0.16 | 0.22 | 0.1 | 0 | 0.11 | 0.1 |
| 6 | 124867530 | rs575595 | 1.947 | -1.627 | 0.626 | -0.57 | -0.001 | -0.052 | -0.052 | 4.773 | 0.71 | 0.31 | 0.82 | 1 | 0.82 | 0.87 |
| 6 | 124867958 | rs991884 | 2.116 | -1.629 | 0.211 | -0.688 | -0.037 | 0.131 | 0.131 | 0.265 | 0.45 | 0.17 | 0.57 | 0.75 | 0.54 | 0.66 |
| 6 | 124869624 | rs7768294 | 0.545 | -1.631 | 0.427 | -0.683 | -0.011 | 0.033 | 0.033 | 0 | 0.65 | 0.77 | 0.69 | 0.74 | 0.62 | 0.78 |
| 6 | 124873435 | rs7747665 | -0.664 | -1.597 | 0.246 | -0.708 | -0.012 | -0.049 | -0.049 | 0.81 | 0.15 | 0.21 | 0.08 | 0 | 0.09 | 0.09 |
| 6 | 124876943 | rs971927 | 1.62 | -1.514 | -0.035 | -0.643 | -0.021 | 0.159 | 0.159 | 0.206 | 0.47 | 0.19 | 0.58 | 0.76 | 0.55 | 0.67 |
| 6 | 124879373 | rs530822 | -0.328 | -1.121 | 0.38 | -0.331 | -0.01 | 0.009 | 0.009 | 0 | 0.27 | 0.18 | 0.38 | 0.27 | 0.33 | 0.21 |
| 6 | 124881787 | rs802691 | 1.069 | -1.332 | 0.961 | -0.157 | -0.022 | -0.081 | -0.081 | 0 | 0.88 | 0.67 | 0.91 | 1 | 0.91 | 0.95 |
| 6 | 124882382 | rs527307 | 1.011 | -1.4 | 0.701 | 0.073 | -0.011 | -0.012 | -0.012 | 0 | 0.6 | 0.51 | 0.54 | 0.68 | 0.63 | 0.65 |
| 6 | 124887359 | rs17052010 | -1.556 | -1.388 | 1.446 | 0.39 | -0.004 | -0.072 | -0.072 | 0 | 0.12 | 0.4 | 0.11 | 0 | 0.09 | 0.06 |
| 6 | 124889636 | rs612146 | 2.311 | -1.886 | 1.139 | 0.777 | 0.093 | 0.001 | 0.001 | 0.837 | 0.61 | 0.18 | 0.62 | 0.56 | 0.71 | 0.57 |
| 6 | 124890305 | rs6905287 | -1.213 | -1.683 | 1.192 | 0.703 | 0.022 | -0.008 | -0.008 | 0 | 0.14 | 0.06 | 0.18 | 0.2 | 0.11 | 0.22 |
| 6 | 124890704 | rs9491192 | -0.944 | -1.448 | 1.553 | 0.668 | 0.029 | -0.026 | -0.026 | 0 | 0.07 | 0.24 | 0.08 | 0 | 0.07 | 0.05 |
| 6 | 124894966 | rs9491194 | -1.463 | -1.498 | 1.456 | 0.632 | 0.034 | -0.06 | -0.06 | 0.026 | 0.11 | 0.49 | 0.12 | 0 | 0.1 | 0.06 |
| 6 | 124896353 | rs611601 | 0.804 | -1.69 | 1.077 | 0.705 | -0.017 | 0.002 | 0.002 | 0 | 0.88 | 0.85 | 0.93 | 0.83 | 0.93 | 0.81 |
| 6 | 124896441 | rs575793 | 0.365 | -1.626 | 0.725 | 1.087 | -0.024 | -0.031 | -0.031 | 0.326 | 0.19 | 0.07 | 0.24 | 0.2 | 0.2 | 0.08 |
| 6 | 124896527 | rs576646 | 1.976 | -1.768 | 0.623 | 1.298 | 0.015 | 0.004 | 0.004 | 0.116 | 0.48 | 0.08 | 0.51 | 0.34 | 0.58 | 0.54 |
| 6 | 124897878 | rs520478 | 1.038 | -1.773 | 0.797 | 1.378 | 0.033 | -0.026 | -0.026 | 0 | 0.6 | 0.72 | 0.66 | 0.52 | 0.68 | 0.61 |
| 6 | 124900701 | rs2325854 | -1.019 | -1.669 | 1.207 | 1.378 | -0.025 | -0.013 | -0.013 | 0 | 0.16 | 0.25 | 0.12 | 0.16 | 0.13 | 0.23 |
| 6 | 124907009 | rs649757 | 0.738 | -1.578 | 1.567 | 1.48 | 0.067 | 0.192 | 0.192 | 0.048 | 0.95 | 0.71 | 0.93 | 0.81 | 0.96 | 0.97 |
| 6 | 124910388 | rs637547 | 1.804 | -1.668 | 1.671 | 1.6 | -0.011 | 0.103 | 0.103 | 0.631 | 0.86 | 0.59 | 0.88 | 0.72 | 0.9 | 0.88 |
| 6 | 124910739 | rs635837 | 0.268 | -1.735 | 1.522 | 1.564 | -0.008 | -0.002 | -0.002 | 0.833 | 0.2 | 0.11 | 0.22 | 0.18 | 0.2 | 0.07 |
| 6 | 124911534 | rs1871329 | -0.711 | -1.754 | 1.652 | 1.63 | 0.004 | -0.004 | -0.004 | 0.171 | 0.3 | 0.27 | 0.26 | 0.3 | 0.25 | 0.17 |
| 6 | 124911565 | rs641498 | 1.868 | -1.812 | 1.665 | 1.618 | 0.039 | 0.024 | 0.024 | 0 | 0.51 | 0.32 | 0.57 | 0.36 | 0.59 | 0.58 |
| 6 | 124912456 | rs1382648 | -0.936 | -1.558 | 1.395 | 1.728 | -0.017 | -0.005 | -0.005 | 0.302 | 0.15 | 0.22 | 0.2 | 0.16 | 0.12 | 0.24 |

| | | | | | | | | | | | | | | | | |
|---|---|---|---|---|---|---|---|---|---|---|---|---|---|---|---|---|
| 6 | 124917584 | rs10499128 | -0.348 | -1.446 | 1.008 | 1.793 | -0.002 | 0.001 | 0.001 | 0 | 0.06 | 0.06 | 0.02 | 0.09 | 0.04 | 0.1 |
| 6 | 124919697 | rs630683 | 1.204 | -1.45 | 1.028 | 1.627 | -0.014 | 0.062 | 0.062 | 0 | 0.9 | 0.82 | 0.93 | 0.72 | 0.93 | 0.87 |
| 6 | 124923560 | rs1382647 | 1.652 | -1.143 | 0.266 | 1.115 | -0.038 | -0.01 | -0.01 | 0 | 0.4 | 0.23 | 0.49 | 0.36 | 0.47 | 0.45 |
| 6 | 124923968 | rs9491199 | -1.023 | -1.289 | 0.046 | 1.305 | 0.023 | -0.01 | -0.01 | 0 | 0.15 | 0.09 | 0.18 | 0.19 | 0.08 | 0.26 |
| 6 | 124934207 | rs10499129 | -1.216 | -1.205 | 0.072 | 1.245 | -0.003 | -0.001 | -0.001 | 0 | 0.16 | 0.2 | 0.16 | 0.17 | 0.09 | 0.25 |
| 6 | 124935890 | rs17708334 | -0.745 | -0.913 | -0.267 | 1.393 | 0.027 | 0.002 | 0.002 | 0 | 0.14 | 0.09 | 0.09 | 0.16 | 0.08 | 0.22 |
| 6 | 124937930 | rs9375345 | -0.959 | -0.375 | 0.038 | 2.026 | -0.01 | 0.105 | 0.105 | 1.013 | 0.08 | 0.1 | 0.06 | 0.41 | 0.05 | 0.17 |
| 6 | 124937974 | rs9401758 | -0.563 | 0.138 | -0.123 | 0.796 | -0.001 | -0.013 | -0.013 | 0.153 | 0.19 | 0.23 | 0.23 | 0.14 | 0.18 | 0.08 |
| 6 | 124941785 | rs9491204 | -0.376 | 0.204 | -0.608 | 0.821 | -0.001 | 0.008 | 0.008 | 0.062 | 0.18 | 0.15 | 0.13 | 0.13 | 0.18 | 0.07 |
| 6 | 124945120 | rs497817 | -0.533 | 0.073 | -0.689 | 0.814 | -0.002 | 0.006 | 0.006 | 0.09 | 0.19 | 0.15 | 0.23 | 0.15 | 0.2 | 0.08 |
| 6 | 124950825 | rs505959 | 0.8 | -0.168 | -0.891 | 1.051 | 0.008 | -0.025 | -0.025 | 0 | 0.8 | 0.89 | 0.71 | 0.71 | 0.82 | 0.75 |
| 6 | 124951063 | rs504008 | 1.181 | -1.267 | -0.423 | 0.753 | 0.011 | 0.001 | 0.001 | 1.389 | 0.61 | 0.35 | 0.62 | 0.63 | 0.75 | 0.63 |
| 6 | 124965290 | rs6932501 | -1.126 | -1.343 | -0.159 | 0.707 | -0.014 | 0.001 | 0.001 | 0.38 | 0.29 | 0.35 | 0.32 | 0.32 | 0.23 | 0.29 |
| 6 | 124975861 | rs2619076 | -0.03 | -0.849 | -1.34 | 0.211 | 0.017 | -0.004 | -0.004 | 0 | 0.58 | 0.19 | 0.58 | 0.41 | 0.6 | 0.5 |
| 6 | 124978387 | rs781486 | -0.045 | -0.821 | -1.046 | 0.286 | -0.011 | -0.01 | -0.01 | 0.498 | 0.81 | 0.74 | 0.91 | 0.86 | 0.88 | 0.82 |
| 6 | 124978932 | rs10872289 | 0.616 | -0.736 | -1.318 | 0.157 | -0.005 | -0.001 | -0.001 | 0 | 0.36 | 0.48 | 0.39 | 0.35 | 0.36 | 0.3 |
| 6 | 124979689 | rs1479039 | -0.641 | -0.751 | -1.304 | 0.174 | -0.005 | -0.004 | -0.004 | 1.513 | 0.81 | 0.78 | 0.91 | 0.85 | 0.88 | 0.82 |
| 6 | 124980834 | rs1677933 | 0.269 | -0.846 | -1.438 | 0.181 | -0.041 | -0.002 | -0.002 | 0.021 | 0.44 | 0.17 | 0.51 | 0.43 | 0.48 | 0.47 |
| 6 | 124982187 | rs6928014 | -0.066 | -0.39 | -1.687 | 0.077 | -0.001 | 0.005 | 0.005 | 0 | 0.26 | 0.27 | 0.29 | 0.39 | 0.25 | 0.36 |
| 6 | 124984982 | rs2325913 | -0.426 | -0.669 | -1.649 | 0.061 | -0.045 | 0.052 | 0.052 | 0.277 | 0.61 | 0.77 | 0.65 | 0.74 | 0.58 | 0.65 |
| 6 | 124990342 | rs9491210 | 1.009 | -0.672 | -1.572 | 0.077 | 0.004 | 0.011 | 0.011 | 0 | 0.33 | 0.48 | 0.37 | 0.34 | 0.36 | 0.26 |
| 6 | 124990380 | rs781474 | -0.48 | -0.505 | -1.615 | 0.147 | 0.012 | -0.004 | -0.004 | 0 | 0.49 | 0.27 | 0.54 | 0.51 | 0.52 | 0.52 |
| 6 | 125001170 | rs1479047 | -0.028 | -0.412 | -1.177 | 0.519 | -0.006 | 0.028 | 0.028 | 0 | 0.21 | 0.26 | 0.29 | 0.39 | 0.22 | 0.37 |
| 6 | 125011951 | rs7774956 | -0.858 | -0.212 | -1.12 | 0.495 | -0.022 | 0.03 | 0.03 | 0.06 | 0.63 | 0.73 | 0.65 | 0.74 | 0.6 | 0.65 |
| 6 | 125018981 | rs6908208 | -0.627 | -0.395 | -0.952 | 1.035 | -0.015 | 0.051 | 0.051 | 0 | 0.25 | 0.31 | 0.29 | 0.5 | 0.21 | 0.36 |
| 6 | 125023343 | rs4540279 | -0.371 | -0.415 | -1.018 | 1.014 | -0.012 | 0.039 | 0.039 | 0.3 | 0.22 | 0.3 | 0.28 | 0.39 | 0.19 | 0.35 |
| 6 | 125027223 | rs2325911 | 0.141 | -0.404 | -1.271 | 1.076 | 0.003 | -0.007 | -0.007 | 1.232 | 0.17 | 0.15 | 0.25 | 0.35 | 0.15 | 0.25 |
| 6 | 125035966 | rs715944 | -0.031 | -0.417 | -1.292 | 0.948 | -0.001 | -0.003 | -0.003 | 0.851 | 0.43 | 0.57 | 0.41 | 0.33 | 0.44 | 0.38 |
| 6 | 125037288 | rs17767102 | -0.432 | -0.578 | -1.326 | 0.818 | 0.01 | -0.016 | -0.016 | 0.036 | 0.07 | 0.01 | 0.04 | 0.09 | 0.06 | 0.09 |
| 6 | 125038952 | rs6569393 | -0.154 | -0.326 | -1.382 | 0.868 | 0 | -0.001 | -0.001 | 0 | 0.46 | 0.57 | 0.55 | 0.44 | 0.48 | 0.41 |
| 6 | 125053277 | rs11154260 | -0.727 | -0.465 | -1.504 | 1.067 | 0.024 | -0.059 | -0.059 | 2.052 | 0.2 | 0.09 | 0.25 | 0.43 | 0.17 | 0.26 |
| 6 | 125054180 | rs10782227 | 0.119 | -0.244 | -1.484 | 1.022 | -0.001 | 0 | 0 | 0.447 | 0.44 | 0.55 | 0.54 | 0.44 | 0.44 | 0.41 |
| 6 | 125055529 | rs870583 | 0.098 | -0.25 | -1.48 | 1.029 | 0.013 | -0.041 | -0.041 | 0 | 0.13 | 0.06 | 0.21 | 0.33 | 0.11 | 0.17 |
| 6 | 125060716 | rs1415766 | -0.558 | -0.035 | -1.567 | 0.931 | -0.005 | 0.008 | 0.008 | 0 | 0.57 | 0.61 | 0.69 | 0.72 | 0.56 | 0.59 |
| 6 | 125063042 | rs17711154 | 0.547 | -0.025 | -1.697 | 0.926 | -0.015 | 0.029 | 0.029 | 0 | 0.41 | 0.26 | 0.32 | 0.29 | 0.47 | 0.43 |
| 6 | 125066144 | rs756281 | 0.087 | 0.073 | -2.095 | 0.698 | -0.093 | -0.078 | -0.078 | 0.21 | 0.36 | 0.02 | 0.46 | 0.35 | 0.4 | 0.34 |
| 6 | 125078183 | rs1337871 | 0.285 | -0.175 | -0.929 | 0.91 | 0.016 | -0.01 | -0.01 | 0 | 0.62 | 0.34 | 0.59 | 0.48 | 0.71 | 0.55 |
| 6 | 125080405 | rs13213832 | 0.215 | 0.563 | -1.309 | 0.731 | -0.051 | -0.029 | -0.029 | 0.286 | 0.18 | 0.03 | 0.16 | 0.18 | 0.2 | 0.09 |
| 6 | 125082498 | rs9372792 | -0.538 | 0.146 | -1.027 | 1.185 | 0 | 0.025 | 0.025 | 0 | 0.14 | 0.14 | 0.16 | 0.39 | 0.12 | 0.18 |
| 6 | 125085648 | rs7754584 | -0.354 | 0.013 | -0.734 | 0.404 | 0.002 | 0.004 | 0.004 | 0 | 0.26 | 0.2 | 0.32 | 0.22 | 0.24 | 0.29 |
| 6 | 125087972 | rs9385345 | -0.466 | -0.56 | -0.345 | -0.102 | -0.031 | -0.046 | -0.046 | 0 | 0.9 | 0.79 | 0.94 | 1 | 0.97 | 0.97 |
| 6 | 125096524 | rs11154263 | -0.481 | -0.769 | -0.657 | -0.28 | -0.022 | -0.032 | -0.032 | 0 | 0.29 | 0.19 | 0.33 | 0.56 | 0.31 | 0.43 |
| 6 | 125097062 | rs9482574 | 0.961 | -1.527 | -0.679 | -0.86 | -0.052 | -0.103 | -0.103 | 0 | 0.19 | 0.35 | 0.16 | 0.02 | 0.11 | 0.13 |
| 6 | 125097427 | rs1415769 | 1.105 | -1.547 | -0.792 | -0.863 | -0.062 | -0.02 | -0.02 | 0.676 | 0.41 | 0.12 | 0.45 | 0.42 | 0.54 | 0.41 |
| 6 | 125098980 | rs2065973 | 1.266 | -1.413 | -0.738 | -0.83 | -0.041 | -0.012 | -0.012 | 0.15 | 0.11 | 0.02 | 0.21 | 0.08 | 0.14 | 0.22 |
| 6 | 125107056 | rs2786910 | 0.765 | -1.277 | -0.512 | -0.639 | -0.043 | -0.007 | -0.007 | 0 | 0.4 | 0.19 | 0.45 | 0.41 | 0.53 | 0.4 |
| 6 | 125108969 | rs9321006 | 0.401 | -1.17 | -0.426 | -0.637 | 0 | 0.019 | 0.019 | 0.468 | 0.18 | 0.18 | 0.11 | 0.51 | 0.18 | 0.22 |
| 6 | 125110808 | rs1998413 | 0.764 | -0.911 | -0.519 | -0.471 | -0.089 | -0.053 | -0.053 | 0 | 0.33 | 0.11 | 0.4 | 0.4 | 0.47 | 0.35 |
| 6 | 125111409 | rs9401773 | -0.066 | -0.898 | -0.277 | -0.29 | -0.001 | -0.001 | -0.001 | 0 | 0.51 | 0.53 | 0.5 | 0.59 | 0.45 | 0.61 |
| 6 | 125116245 | rs1337868 | -0.087 | -0.883 | 0.231 | -0.031 | -0.028 | -0.052 | -0.052 | 2.867 | 0.9 | 0.63 | 0.93 | 1 | 0.97 | 0.97 |
| 6 | 125122731 | rs9375372 | -0.76 | -0.277 | -0.499 | 0.503 | -0.005 | -0.032 | -0.032 | 0.694 | 0.22 | 0.15 | 0.26 | 0.58 | 0.22 | 0.39 |
| 6 | 125134393 | rs9385352 | -0.573 | -0.276 | -0.37 | 0.72 | -0.006 | 0.045 | 0.045 | 0 | 0.08 | 0.06 | 0.06 | 0.52 | 0.08 | 0.15 |
| 6 | 125135997 | rs9401776 | 1.276 | -0.683 | -0.317 | 0.019 | 0.001 | -0.014 | -0.014 | 0 | 0.65 | 0.72 | 0.7 | 0.45 | 0.66 | 0.51 |
| 6 | 125143279 | rs6569399 | -0.64 | -1.129 | -0.17 | -0.396 | -0.012 | -0.059 | -0.059 | 0 | 0.15 | 0.22 | 0.07 | 0 | 0.1 | 0.05 |
| 6 | 125144352 | rs2025251 | 1.827 | -2.22 | -0.895 | -0.962 | -0.083 | -0.136 | -0.136 | 0 | 0.39 | 0.04 | 0.48 | 0.44 | 0.57 | 0.46 |
| 6 | 125146971 | rs9388381 | -0.471 | -2.014 | 0.062 | -0.776 | 0.045 | 0.408 | 0.408 | 0.028 | 0.12 | 0.43 | 0.09 | 0.52 | 0.09 | 0.17 |
| 6 | 125147538 | rs1923728 | 0.861 | -1.942 | -0.134 | -0.986 | 0 | -0.037 | -0.037 | 0 | 0.64 | 0.65 | 0.72 | 0.94 | 0.71 | 0.64 |
| 6 | 125156449 | rs11154271 | -0.153 | -1.582 | -0.532 | -0.764 | 0.008 | 0.047 | 0.047 | 0.815 | 0.78 | 0.92 | 0.77 | 0.94 | 0.79 | 0.72 |
| 6 | 125168198 | rs1591705 | 1.441 | -1.05 | -0.288 | -0.56 | -0.015 | -0.007 | -0.007 | 0 | 0.43 | 0.29 | 0.46 | 0.39 | 0.54 | 0.48 |
| 6 | 125172488 | rs1409294 | 0.618 | -0.247 | -0.203 | -0.645 | -0.01 | -0.008 | -0.008 | 0 | 0.58 | 0.67 | 0.45 | 0.44 | 0.49 | 0.52 |
| 6 | 125175526 | rs881112 | -0.776 | -0.24 | -0.331 | -0.736 | -0.027 | 0.052 | 0.052 | 0 | 0.84 | 0.97 | 0.78 | 0.94 | 0.81 | 0.76 |
| 6 | 125178368 | rs1885693 | -0.252 | -0.357 | 0.119 | -0.688 | 0.015 | -0.047 | -0.047 | 0.116 | 0.09 | 0.3 | 0.06 | 0 | 0.06 | 0.02 |
| 6 | 125181691 | rs1326156 | 1.446 | -0.142 | -0.083 | -0.896 | -0.01 | -0.067 | -0.067 | 0 | 0.4 | 0.16 | 0.33 | 0.44 | 0.38 | 0.47 |
| 6 | 125201001 | rs781736 | 0.208 | -0.007 | -0.256 | -0.825 | -0.07 | -0.098 | -0.098 | 0 | 0.33 | 0.1 | 0.5 | 0.56 | 0.45 | 0.46 |
| 6 | 125202367 | rs12211560 | -0.685 | -0.113 | 0.683 | -1.122 | 0.005 | -0.048 | -0.048 | 0 | 0.09 | 0.22 | 0.05 | 0 | 0.05 | 0.02 |
| 6 | 125203438 | rs781738 | 0.785 | 0.061 | 0.307 | -0.726 | -0.011 | -0.102 | -0.102 | 0.748 | 0.84 | 0.68 | 0.9 | 1 | 0.89 | 0.95 |
| 6 | 125220250 | rs781720 | 0.457 | 0.254 | 0.121 | -0.375 | -0.003 | -0.007 | -0.007 | 0 | 0.67 | 0.36 | 0.68 | 0.83 | 0.71 | 0.74 |
| 6 | 125227328 | rs781728 | -0.148 | -0.174 | 0.626 | -0.556 | -0.012 | -0.141 | -0.141 | 0 | 0.77 | 0.43 | 0.81 | 1 | 0.78 | 0.91 |
| 6 | 125229093 | rs781732 | 1.047 | -0.319 | 0.46 | -0.542 | 0.059 | 0.051 | 0.051 | 2.862 | 0.66 | 0.12 | 0.69 | 0.82 | 0.75 | 0.78 |

| | | | | | | | | | | | | | | | | |
|---|---|---|---|---|---|---|---|---|---|---|---|---|---|---|---|---|
| 6 | 125232333 | rs781739 | 0.428 | 0.437 | -0.169 | -0.152 | -0.098 | -0.138 | -0.138 | 0 | 0.19 | 0.01 | 0.28 | 0.41 | 0.29 | 0.26 |
| 6 | 125244265 | rs781762 | 1.605 | 0.458 | -0.157 | -0.28 | -0.009 | 0.029 | 0.029 | 0 | 0.06 | 0 | 0.04 | 0 | 0.07 | 0.01 |
| 6 | 125250416 | rs6910745 | 0.425 | 0.373 | 0.125 | -0.208 | 0.001 | 0.001 | 0.001 | 0.352 | 0.68 | 0.32 | 0.71 | 0.82 | 0.75 | 0.83 |
| 6 | 125251374 | rs6916777 | 0.411 | 0.659 | 0.44 | -0.16 | 0.006 | 0.005 | 0.005 | 0 | 0.61 | 0.48 | 0.54 | 0.61 | 0.52 | 0.67 |
| 6 | 125252521 | rs4896595 | -0.389 | -0.016 | 0.719 | -0.215 | -0.007 | -0.008 | -0.008 | 0.174 | 0.73 | 0.7 | 0.75 | 0.84 | 0.76 | 0.83 |
| 6 | 125253532 | rs1889436 | 0.309 | -0.34 | 0.045 | -0.662 | 0.002 | 0 | 0 | 0 | 0.16 | 0.17 | 0.06 | 0.1 | 0.11 | 0.08 |
| 6 | 125255208 | rs962641 | 0.095 | -0.131 | 0.117 | -0.526 | 0 | -0.002 | -0.002 | 0 | 0.54 | 0.56 | 0.47 | 0.44 | 0.5 | 0.59 |
| 6 | 125276801 | rs12663635 | 1.343 | 0.08 | -0.693 | -0.436 | -0.135 | -0.046 | -0.046 | 0 | 0.34 | 0.02 | 0.37 | 0.34 | 0.36 | 0.52 |
| 6 | 125281126 | rs4896619 | -0.009 | -0.091 | -0.272 | -0.5 | -0.004 | -0.018 | -0.018 | 0 | 0.8 | 0.84 | 0.9 | 0.89 | 0.84 | 0.88 |
| 6 | 125298036 | rs7761012 | 0.296 | -0.163 | -0.159 | -0.511 | -0.031 | -0.06 | -0.06 | 0 | 0.78 | 0.68 | 0.89 | 0.88 | 0.84 | 0.88 |
| 6 | 125303483 | rs1413594 | -0.017 | -0.104 | -0.533 | -0.484 | -0.003 | -0.013 | -0.013 | 0 | 0.8 | 0.86 | 0.91 | 0.89 | 0.84 | 0.87 |
| 6 | 125311779 | rs2325762 | 0.356 | 0.04 | -0.568 | -0.097 | -0.025 | -0.066 | -0.066 | 0 | 0.2 | 0.1 | 0.28 | 0.4 | 0.27 | 0.25 |
| 6 | 125319983 | rs9491274 | 0.312 | -0.146 | -0.468 | -0.308 | -0.003 | -0.033 | -0.033 | 0 | 0.27 | 0.37 | 0.26 | 0.13 | 0.25 | 0.19 |
| 6 | 125324404 | rs9388397 | 0.195 | 0.461 | -1.35 | 0.176 | -0.036 | -0.078 | -0.078 | 0 | 0.23 | 0.08 | 0.31 | 0.43 | 0.3 | 0.28 |
| 6 | 125329613 | rs1931657 | -0.073 | 0.354 | -1.379 | 0.122 | -0.011 | -0.001 | -0.001 | 0 | 0.93 | 0.91 | 0.96 | 0.88 | 0.95 | 0.95 |
| 6 | 125332811 | rs9388398 | -0.189 | 0.465 | -1.702 | 0.149 | 0.057 | 0.143 | 0.143 | 0 | 0.52 | 0.13 | 0.64 | 0.75 | 0.61 | 0.71 |
| 6 | 125333797 | rs9388399 | -0.621 | 0.648 | -0.875 | 0.69 | -0.037 | -0.086 | -0.086 | 0 | 0.23 | 0.08 | 0.3 | 0.47 | 0.3 | 0.28 |
| 6 | 125344138 | rs4895621 | 1.12 | -0.182 | -0.048 | 0.076 | -0.04 | 0.056 | 0.056 | 0 | 0.89 | 0.63 | 0.93 | 0.86 | 0.97 | 0.89 |
| 6 | 125350432 | rs2759263 | 0.497 | -0.14 | -0.708 | -0.625 | 0.005 | 0.004 | 0.004 | 0 | 0.63 | 0.41 | 0.76 | 0.87 | 0.7 | 0.85 |
| 6 | 125354154 | rs9375383 | 0.64 | 0.321 | -1.36 | -0.339 | -0.017 | -0.012 | -0.012 | 1.198 | 0.46 | 0.26 | 0.61 | 0.52 | 0.45 | 0.57 |
| 6 | 125359388 | rs6911261 | -0.031 | 0.651 | -1.532 | -0.249 | -0.045 | 0.088 | 0.088 | 0 | 0.86 | 0.99 | 0.89 | 1 | 0.84 | 0.97 |
| 6 | 125365295 | rs482014 | 0.438 | 0.757 | -1.163 | 0.051 | -0.004 | -0.006 | -0.006 | 0 | 0.42 | 0.31 | 0.57 | 0.52 | 0.4 | 0.56 |
| 6 | 125368709 | rs1830651 | -0.469 | 0.626 | -1.412 | 0.597 | -0.006 | -0.065 | -0.065 | 0 | 0.32 | 0.12 | 0.26 | 0.37 | 0.37 | 0.3 |
| 6 | 125369674 | rs12664303 | -0.478 | 0.633 | -1.487 | 0.578 | -0.014 | -0.095 | -0.095 | 0 | 0.32 | 0.08 | 0.26 | 0.4 | 0.38 | 0.31 |
| 6 | 125370626 | rs576419 | -0.43 | 0.766 | -0.716 | 0.118 | -0.005 | 0.012 | 0.012 | 0 | 0.44 | 0.35 | 0.41 | 0.34 | 0.5 | 0.42 |
| 6 | 125370852 | rs11154288 | -1.277 | 0.553 | 0.254 | -0.057 | 0.045 | 0.001 | 0.001 | 0.234 | 0.12 | 0.46 | 0.08 | 0.06 | 0.09 | 0.18 |
| 6 | 125374466 | rs567448 | -0.028 | 0.18 | 0.217 | -0.812 | -0.006 | 0.065 | 0.065 | 0.777 | 0.61 | 0.76 | 0.39 | 0.28 | 0.61 | 0.5 |
| 6 | 125376102 | rs497277 | 0.269 | -0.198 | 0.518 | -1.392 | 0.025 | -0.05 | -0.05 | 0 | 0.35 | 0.46 | 0.27 | 0.21 | 0.43 | 0.26 |
| 6 | 125383146 | rs11154290 | 0.364 | -0.515 | 0.354 | -1.703 | -0.078 | 0.092 | 0.092 | 0.383 | 0.33 | 0.11 | 0.24 | 0.18 | 0.44 | 0.29 |
| 6 | 125385470 | rs9385362 | -0.137 | -0.787 | 1.407 | -1.789 | -0.007 | 0.081 | 0.081 | 0 | 0.44 | 0.26 | 0.29 | 0.27 | 0.51 | 0.42 |
| 6 | 125387475 | rs554999 | -1.234 | -0.817 | 1.278 | -1.808 | -0.045 | 0.028 | 0.028 | 1.239 | 0.68 | 0.94 | 0.43 | 0.3 | 0.66 | 0.58 |
| 6 | 125396993 | rs10485341 | -0.223 | -0.907 | 1.516 | -1.955 | 0 | -0.011 | -0.011 | 0 | 0.52 | 0.5 | 0.33 | 0.25 | 0.52 | 0.45 |
| 6 | 125398702 | rs9491304 | 0.485 | -0.894 | 1.462 | -1.954 | 0.003 | 0 | 0 | 0.009 | 0.14 | 0.12 | 0.06 | 0.02 | 0.11 | 0.1 |
| 6 | 125402711 | rs492800 | -0.49 | -0.876 | 1.194 | -1.983 | 0.007 | 0.08 | 0.08 | 0 | 0.55 | 0.7 | 0.38 | 0.27 | 0.58 | 0.45 |
| 6 | 125403279 | rs1322633 | 0.756 | -0.98 | 0.754 | -2.048 | -0.17 | 0.114 | 0.114 | 0.682 | 0.35 | 0.03 | 0.26 | 0.17 | 0.46 | 0.24 |
| 6 | 125403394 | rs1322634 | -0.232 | -1.068 | 1.031 | -1.973 | 0.018 | 0.091 | 0.091 | 0 | 0.52 | 0.34 | 0.34 | 0.27 | 0.58 | 0.46 |
| 6 | 125416400 | rs539794 | -1.436 | -1.164 | 1.287 | -1.458 | 0.053 | 0.094 | 0.094 | 3.634 | 0.35 | 0.8 | 0.18 | 0.12 | 0.22 | 0.32 |
| 6 | 125418472 | rs10046333 | -1.363 | -0.711 | 1.096 | -0.837 | -0.03 | -0.054 | -0.054 | 0 | 0.19 | 0.34 | 0.09 | 0.1 | 0.1 | 0.21 |
| 6 | 125427388 | rs6569415 | 0.124 | -0.044 | 0.232 | 0.048 | -0.032 | -0.034 | -0.034 | 0 | 0.64 | 0.8 | 0.56 | 0.55 | 0.58 | 0.6 |
| 6 | 125433716 | rs7451644 | 0.701 | -0.596 | 0.741 | -0.083 | -0.013 | -0.012 | -0.012 | 0.221 | 0.43 | 0.35 | 0.47 | 0.49 | 0.48 | 0.47 |
| 6 | 125434358 | rs12200663 | 1.069 | -0.58 | 0.404 | -0.167 | -0.061 | -0.078 | -0.078 | 0 | 0.36 | 0.17 | 0.42 | 0.52 | 0.45 | 0.37 |
| 6 | 125437307 | rs579493 | -1.306 | -0.59 | 1.443 | -0.129 | 0.017 | -0.04 | -0.04 | 0.531 | 0.09 | 0.38 | 0.08 | 0 | 0.05 | 0.12 |
| 6 | 125461810 | rs11154296 | 0.406 | -0.392 | 0.343 | 0.483 | -0.013 | 0.03 | 0.03 | 0.252 | 0.57 | 0.28 | 0.47 | 0.66 | 0.54 | 0.51 |
| 6 | 125469657 | rs9482603 | 0.03 | -0.471 | 1.13 | 0.493 | 0.005 | -0.023 | -0.023 | 0 | 0.06 | 0.15 | 0.04 | 0 | 0.05 | 0.02 |
| 6 | 125470012 | rs9491317 | -1.224 | -0.003 | 1.346 | 0.636 | 0.09 | 0.042 | 0.042 | 0.034 | 0.1 | 0.36 | 0.1 | 0.12 | 0.16 | 0.11 |
| 6 | 125470996 | rs13192443 | -1.206 | -0.074 | 1.278 | 0.638 | 0.065 | 0.047 | 0.047 | 0 | 0.07 | 0.21 | 0.06 | 0.12 | 0.12 | 0.09 |
| 6 | 125471950 | rs7749760 | -1.27 | -0.115 | 1.29 | 1.539 | 0.077 | 0.267 | 0.267 | 0 | 0.07 | 0.23 | 0.08 | 0.36 | 0.13 | 0.09 |
| 6 | 125475645 | rs557339 | -1.028 | -1.423 | 0.815 | 1.128 | 0.01 | 0.023 | 0.023 | 0.239 | 0.57 | 0.79 | 0.61 | 0.64 | 0.43 | 0.56 |
| 6 | 125476616 | rs476436 | -1.028 | -1.423 | 0.815 | 1.128 | 0.01 | 0.023 | 0.023 | 0.239 | 0.57 | 0.79 | 0.61 | 0.63 | 0.43 | 0.56 |
| 6 | 125476813 | rs478265 | -0.928 | -1.529 | 0.991 | 1.043 | 0.008 | 0.012 | 0.012 | 0.477 | 0.54 | 0.66 | 0.57 | 0.3 | 0.4 | 0.49 |
| 6 | 125479803 | rs9321023 | 0.357 | -1.512 | 0.753 | 1.219 | -0.006 | -0.023 | -0.023 | 0.146 | 0.05 | 0.1 | 0.04 | 0 | 0.04 | 0.02 |
| 6 | 125481306 | rs514751 | -0.506 | -1.879 | 0.809 | 0.932 | 0.024 | 0.019 | 0.019 | 0 | 0.4 | 0.66 | 0.49 | 0.27 | 0.3 | 0.39 |
| 6 | 125500112 | rs1741748 | 0.053 | -1.986 | 1.032 | 0.84 | -0.009 | -0.001 | -0.001 | 0 | 0.31 | 0.34 | 0.46 | 0.26 | 0.26 | 0.35 |
| 6 | 125500202 | rs17052811 | -1.251 | -2.202 | 1.052 | 0.917 | -0.017 | 0.017 | 0.017 | 0 | 0.23 | 0.26 | 0.15 | 0.38 | 0.16 | 0.17 |
| 6 | 125500534 | rs564373 | -1.989 | -2.186 | 0.792 | 0.908 | -0.053 | -0.019 | -0.019 | 0 | 0.62 | 0.95 | 0.66 | 0.63 | 0.46 | 0.55 |
| 6 | 125501314 | rs482263 | -0.696 | -2.214 | 0.895 | 0.887 | 0.047 | 0.039 | 0.039 | 0 | 0.42 | 0.73 | 0.51 | 0.26 | 0.3 | 0.39 |
| 6 | 125502382 | rs17052816 | -1.007 | -1.996 | 0.965 | 1.031 | 0 | 0 | 0 | 0 | 0.2 | 0.22 | 0.15 | 0.38 | 0.16 | 0.16 |
| 6 | 125508443 | rs506824 | -0.825 | -1.832 | 0.826 | 0.709 | 0.025 | 0.03 | 0.03 | 0.1 | 0.46 | 0.36 | 0.56 | 0.29 | 0.39 | 0.44 |
| 6 | 125509459 | rs1555373 | -0.891 | -1.654 | 0.455 | 0.63 | -0.012 | -0.007 | -0.007 | 0 | 0.22 | 0.28 | 0.19 | 0.14 | 0.18 | 0.27 |
| 6 | 125510922 | rs481706 | -1.676 | -1.967 | -0.186 | 1.012 | -0.078 | -0.025 | -0.025 | 0.776 | 0.63 | 0.92 | 0.66 | 0.61 | 0.48 | 0.59 |
| 6 | 125521338 | rs554004 | -0.86 | -2.203 | 0.511 | 1.167 | -0.02 | 0.034 | 0.034 | 0 | 0.19 | 0.33 | 0.11 | 0.28 | 0.13 | 0.14 |
| 6 | 125522271 | rs6917198 | 0.017 | -2.499 | 0.755 | 1.137 | 0.003 | -0.005 | -0.005 | 0 | 0.3 | 0.29 | 0.42 | 0.29 | 0.27 | 0.39 |
| 6 | 125524718 | rs474688 | -2.236 | -2.585 | 0.49 | 1.053 | -0.013 | 0.012 | 0.012 | 0.141 | 0.62 | 0.66 | 0.64 | 0.66 | 0.49 | 0.61 |
| 6 | 125528577 | rs13192014 | 0.073 | -2.639 | 0.357 | 1.069 | -0.035 | -0.085 | -0.085 | 0 | 0.28 | 0.04 | 0.41 | 0.29 | 0.27 | 0.4 |
| 6 | 125528974 | rs495191 | -1.556 | -2.473 | 0.988 | 1.06 | -0.035 | 0.02 | 0.02 | 0.003 | 0.28 | 0.41 | 0.19 | 0.38 | 0.18 | 0.22 |
| 6 | 125531454 | rs12110572 | -0.621 | -2.293 | 0.916 | 1.062 | 0 | -0.022 | -0.022 | 0.228 | 0.05 | 0.14 | 0.03 | 0 | 0.03 | 0 |
| 6 | 125541862 | rs3823222 | -0.717 | -2.231 | 0.903 | 1.127 | -0.011 | 0.018 | 0.018 | 0 | 0.18 | 0.26 | 0.09 | 0.28 | 0.13 | 0.14 |
| 6 | 125551895 | rs6916559 | -0.54 | -1.954 | 0.853 | 1.182 | -0.002 | 0.026 | 0.026 | 0 | 0.16 | 0.25 | 0.11 | 0.3 | 0.13 | 0.14 |
| 6 | 125553176 | rs6923152 | -1.171 | -1.884 | 0.858 | 1.028 | -0.007 | -0.04 | -0.04 | 0 | 0.07 | 0.17 | 0.06 | 0 | 0.05 | 0.01 |

| | | | | | | | | | | | | | | | | |
|---|---|---|---|---|---|---|---|---|---|---|---|---|---|---|---|---|
| 6 | 125567330 | rs9321026 | -1.992 | -2.187 | 0.412 | 1 | 0.082 | -0.024 | -0.024 | 0 | 0.39 | 0.89 | 0.29 | 0.38 | 0.25 | 0.23 |
| 6 | 125569047 | rs2273560 | -0.4 | -2.194 | 0.573 | 1.009 | -0.001 | 0.024 | 0.024 | 0 | 0.16 | 0.26 | 0.11 | 0.29 | 0.13 | 0.15 |
| 6 | 125569863 | rs3823219 | -2.579 | -2.428 | 0.2 | 0.859 | -0.132 | -0.005 | -0.005 | 0 | 0.64 | 0.94 | 0.7 | 0.67 | 0.5 | 0.65 |
| 6 | 125572026 | rs3806997 | -0.838 | -2.382 | 0.452 | 0.895 | 0 | 0.052 | 0.052 | 0 | 0.2 | 0.39 | 0.14 | 0.31 | 0.16 | 0.16 |
| 6 | 125574509 | rs3799732 | -1.218 | -2.407 | 0.784 | 1.043 | 0 | 0.001 | 0.001 | 0.051 | 0.39 | 0.39 | 0.52 | 0.6 | 0.38 | 0.56 |
| 6 | 125574696 | rs2275623 | -2.218 | -2.456 | 0.499 | 1.054 | -0.082 | 0.023 | 0.023 | 0.875 | 0.59 | 0.94 | 0.68 | 0.67 | 0.47 | 0.62 |
| 6 | 125583672 | rs2024971 | -1.252 | -2.228 | 1.06 | 1.116 | -0.007 | -0.004 | -0.004 | 0 | 0.4 | 0.54 | 0.53 | 0.61 | 0.35 | 0.57 |
| 6 | 125585103 | rs1160495 | -0.773 | -2.217 | 1.149 | 1.092 | -0.009 | 0.005 | 0.005 | 1.225 | 0.29 | 0.32 | 0.44 | 0.3 | 0.25 | 0.4 |
| 6 | 125585976 | rs9321030 | 0.131 | -2.178 | 0.771 | 1.095 | -0.023 | -0.1 | -0.1 | 0 | 0.2 | 0.02 | 0.37 | 0.3 | 0.19 | 0.39 |
| 6 | 125587961 | rs9482620 | -1.277 | -2.173 | 1.21 | 0.955 | -0.006 | -0.054 | -0.054 | 0 | 0.1 | 0.25 | 0.08 | 0 | 0.06 | 0.01 |
| 6 | 125596957 | rs1040803 | -2.245 | -2.375 | 0.749 | 1.064 | -0.103 | 0.064 | 0.064 | 0.995 | 0.59 | 0.96 | 0.69 | 0.7 | 0.47 | 0.63 |
| 6 | 125609217 | rs3778451 | -1.384 | -2.08 | 1.372 | 1.16 | 0.016 | 0.16 | 0.16 | 0.089 | 0.13 | 0.42 | 0.32 | 0.29 | 0.12 | 0.16 |
| 6 | 125618969 | rs17053023 | -0.64 | -2.049 | 1.002 | 1.217 | 0.005 | 0.048 | 0.048 | 0 | 0.09 | 0.15 | 0.28 | 0.26 | 0.11 | 0.16 |
| 6 | 125623810 | rs12523848 | -2.043 | -2.03 | 0.765 | 1.095 | 0.031 | -0.064 | -0.064 | 0 | 0.38 | 0.93 | 0.52 | 0.39 | 0.28 | 0.24 |
| 6 | 125630380 | rs9491381 | -1.162 | -1.859 | 1.305 | 0.689 | 0.012 | 0.047 | 0.047 | 5.269 | 0.07 | 0.39 | 0.06 | 0.05 | 0.01 | 0.01 |
| 6 | 125635050 | rs17053047 | -0.324 | -1.875 | 0.775 | 0.605 | -0.018 | -0.048 | -0.048 | 0.324 | 0.11 | 0.19 | 0.05 | 0 | 0.06 | 0.04 |
| 6 | 125635692 | rs9491388 | -1.557 | -1.963 | 0.554 | 0.655 | 0.002 | 0.001 | 0.001 | 5.942 | 0.35 | 0.72 | 0.26 | 0.16 | 0.26 | 0.15 |
| 6 | 125636350 | rs6935961 | -1.583 | -1.826 | 0.188 | 0.919 | 0.03 | 0.031 | 0.031 | 1.462 | 0.51 | 0.81 | 0.44 | 0.38 | 0.45 | 0.48 |
| 6 | 125645991 | rs9482644 | -1.317 | -1.996 | 0.125 | 1.056 | 0.022 | 0.033 | 0.033 | 0.135 | 0.43 | 0.74 | 0.63 | 0.64 | 0.4 | 0.51 |
| 6 | 125647124 | rs1341200 | -1.073 | -2.145 | 0.5 | 1.087 | -0.02 | 0.223 | 0.223 | 1.451 | 0.18 | 0.48 | 0.35 | 0.4 | 0.14 | 0.12 |
| 6 | 125648345 | rs9388416 | -0.495 | -1.733 | 0.103 | 1.01 | 0.001 | 0.001 | 0.001 | 0 | 0.26 | 0.28 | 0.28 | 0.24 | 0.26 | 0.39 |
| 6 | 125658409 | rs9388421 | -1.005 | -1.478 | -0.429 | 0.275 | -0.01 | 0.007 | 0.007 | 0 | 0.61 | 0.71 | 0.64 | 0.62 | 0.46 | 0.57 |
| 6 | 125672037 | rs9482645 | -0.999 | -1.119 | -0.463 | 0.438 | 0.006 | 0.013 | 0.013 | 0 | 0.56 | 0.71 | 0.62 | 0.59 | 0.48 | 0.58 |
| 6 | 125673963 | rs9482647 | -0.276 | -0.933 | -0.968 | -0.177 | -0.006 | -0.003 | -0.003 | 0 | 0.21 | 0.27 | 0.2 | 0.14 | 0.2 | 0.14 |
| 6 | 125677053 | rs13208476 | -0.617 | -0.731 | -1.519 | -0.811 | 0.007 | 0.031 | 0.031 | 0.04 | 0.24 | 0.01 | 0.12 | 0.21 | 0.19 | 0.35 |
| 6 | 125678114 | rs1547786 | 0.217 | -0.227 | -0.996 | -1.197 | 0 | 0.001 | 0.001 | 0 | 0.51 | 0.53 | 0.62 | 0.62 | 0.54 | 0.54 |
| 6 | 125682719 | rs7756600 | -1.491 | 0.243 | -0.956 | -2.29 | -0.043 | 0.214 | 0.214 | 0.073 | 0.48 | 0.88 | 0.31 | 0.15 | 0.39 | 0.36 |
| 6 | 125683552 | rs17797323 | -0.616 | 0.313 | -1.037 | -2.374 | -0.059 | 0.071 | 0.071 | 0.191 | 0.2 | 0.01 | 0.12 | 0.03 | 0.18 | 0.21 |
| 6 | 125690300 | rs6931700 | -0.811 | -0.422 | 0.209 | -2.221 | -0.022 | -0.043 | -0.043 | 0.289 | 0.27 | 0.31 | 0.11 | 0.04 | 0.13 | 0.22 |
| 6 | 125692100 | rs17053086 | -0.696 | -0.325 | 0.558 | -2.227 | 0.04 | 0.032 | 0.032 | 0.006 | 0.1 | 0.36 | 0.08 | 0.05 | 0.08 | 0.02 |
| 6 | 125693818 | rs10485334 | -1.558 | -0.282 | 0.573 | -2.274 | 0.063 | -0.013 | -0.013 | 0.003 | 0.16 | 0.44 | 0.14 | 0.1 | 0.18 | 0.12 |
| 6 | 125694182 | rs11969350 | -0.668 | -0.266 | 0.542 | -2.272 | 0.048 | 0.016 | 0.016 | 0.032 | 0.1 | 0.37 | 0.08 | 0.04 | 0.08 | 0.02 |
| 6 | 125694227 | rs9491396 | -1.642 | -0.184 | -0.379 | -2.166 | 0.03 | 0.177 | 0.177 | 0.115 | 0.45 | 0.87 | 0.25 | 0.13 | 0.31 | 0.33 |
| 6 | 125697350 | rs1260742 | 1.493 | -0.186 | -0.38 | -2.166 | 0.031 | 0.179 | 0.179 | 0.269 | 0.55 | 0.12 | 0.75 | 0.87 | 0.69 | 0.67 |
| 6 | 125698805 | rs10456957 | -1.001 | 0.349 | 0.395 | -1.86 | 0.011 | -0.011 | -0.011 | 0 | 0.07 | 0.11 | 0.06 | 0.05 | 0.1 | 0.11 |
| 6 | 125724359 | rs1353202 | 1.066 | -0.609 | 0.787 | -2.164 | -0.045 | -0.084 | -0.084 | 0.075 | 0.79 | 0.61 | 0.85 | 0.92 | 0.88 | 0.8 |
| 6 | 125731930 | rs9321038 | 1.103 | -0.625 | 1.001 | -2.172 | -0.055 | -0.103 | -0.103 | 0 | 0.79 | 0.51 | 0.85 | 0.92 | 0.88 | 0.8 |
| 6 | 125735639 | rs6933106 | 1.106 | -0.632 | 1.016 | -2.175 | -0.055 | -0.102 | -0.102 | 0 | 0.79 | 0.51 | 0.85 | 0.92 | 0.88 | 0.8 |
| 6 | 125748384 | rs984977 | 1.235 | -0.677 | 0.298 | -2.98 | 0.042 | 0.201 | 0.201 | 1.692 | 0.53 | 0.2 | 0.68 | 0.92 | 0.61 | 0.74 |
| 6 | 125761549 | rs17741928 | -0.034 | -0.752 | -0.003 | -2.957 | -0.056 | 0.139 | 0.139 | 1.68 | 0.22 | 0.01 | 0.14 | 0 | 0.26 | 0.07 |
| 6 | 125763141 | rs6904555 | -0.793 | -0.96 | 0.722 | -2.929 | -0.02 | -0.044 | -0.044 | 0 | 0.11 | 0.26 | 0.05 | 0.03 | 0.06 | 0.07 |
| 6 | 125763226 | rs1494137 | 1.091 | -0.978 | 1.077 | -2.941 | -0.023 | -0.058 | -0.058 | 0.472 | 0.87 | 0.63 | 0.94 | 0.97 | 0.94 | 0.92 |
| 6 | 125763730 | rs10085328 | -0.531 | -0.625 | 0.953 | -2.756 | 0.007 | 0.028 | 0.028 | 4.87 | 0.11 | 0.03 | 0.14 | 0.05 | 0.09 | 0.13 |
| 6 | 125776333 | rs1353197 | 0.809 | -0.627 | 0.821 | -2.82 | 0.026 | 0.059 | 0.059 | 1.043 | 0.6 | 0.26 | 0.69 | 0.94 | 0.66 | 0.8 |
| 6 | 125782201 | rs9385374 | 0.951 | -0.655 | 0.958 | -2.83 | 0.012 | -0.018 | -0.018 | 0 | 0.92 | 0.4 | 0.96 | 1 | 0.97 | 0.99 |
| 6 | 125784867 | rs6930486 | -0.715 | -0.263 | -0.943 | -2.579 | 0.004 | 0.129 | 0.129 | 0.159 | 0.32 | 0.12 | 0.26 | 0.06 | 0.31 | 0.19 |
| 6 | 125791136 | rs9385377 | 0.506 | -0.107 | -1.426 | -2.887 | -0.052 | 0.287 | 0.287 | 0.254 | 0.53 | 0.12 | 0.66 | 0.9 | 0.55 | 0.67 |
| 6 | 125797437 | rs4895744 | 0.325 | -0.447 | -0.613 | -3.301 | -0.013 | -0.058 | -0.058 | 0 | 0.64 | 0.44 | 0.8 | 0.92 | 0.76 | 0.71 |
| 6 | 125797983 | rs4895745 | 1.108 | -0.246 | -0.645 | -3.334 | -0.034 | 0.318 | 0.318 | 0.386 | 0.51 | 0.12 | 0.66 | 0.92 | 0.55 | 0.64 |
| 6 | 125798475 | rs931546 | -0.002 | -1.068 | -0.146 | -3.269 | 0.005 | -0.145 | -0.145 | 0 | 0.8 | 0.57 | 0.87 | 0.99 | 0.84 | 0.83 |
| 6 | 125806422 | rs2326186 | 1.03 | -1.204 | -0.168 | -3.322 | -0.023 | -0.064 | -0.064 | 0.014 | 0.67 | 0.42 | 0.84 | 0.92 | 0.83 | 0.74 |
| 6 | 125808193 | rs17053272 | -0.573 | -1.596 | 0.062 | -3.275 | -0.005 | -0.002 | -0.002 | 0 | 0.22 | 0.21 | 0.1 | 0.07 | 0.09 | 0.16 |
| 6 | 125815808 | rs1353200 | 0.416 | -0.732 | -0.649 | -3.469 | -0.052 | 0.177 | 0.177 | 0 | 0.41 | 0.1 | 0.63 | 0.84 | 0.55 | 0.53 |
| 6 | 125822889 | rs1494145 | 0.407 | -0.706 | -0.839 | -3.463 | -0.048 | 0.188 | 0.188 | 0 | 0.41 | 0.09 | 0.66 | 0.84 | 0.56 | 0.54 |
| 6 | 125847454 | rs9375400 | -0.239 | -1.055 | 0.393 | -3.884 | -0.004 | -0.12 | -0.12 | 0 | 0.67 | 0.52 | 0.73 | 0.94 | 0.72 | 0.78 |
| 6 | 125856917 | rs1353201 | 0.912 | -0.291 | -0.452 | -4.369 | -0.083 | 0.072 | 0.072 | 0 | 0.32 | 0.05 | 0.5 | 0.84 | 0.4 | 0.48 |
| 6 | 125861535 | rs2326146 | 0.387 | -1.123 | -0.166 | -4.438 | 0.038 | 0.217 | 0.217 | 0 | 0.45 | 0.15 | 0.63 | 0.9 | 0.64 | 0.58 |
| 6 | 125861936 | rs17053352 | 0.552 | -1.065 | 0.514 | -4.461 | 0.001 | -0.103 | -0.103 | 4.002 | 0.29 | 0.43 | 0.24 | 0.04 | 0.22 | 0.17 |
| 6 | 125866064 | rs1353203 | 0.523 | -1.172 | -0.137 | -4.558 | 0.004 | 0.258 | 0.258 | 0.739 | 0.45 | 0.12 | 0.62 | 0.9 | 0.63 | 0.59 |
| 6 | 125866455 | rs9321042 | 0.35 | -1.672 | 0.039 | -4.55 | -0.002 | -0.11 | -0.11 | 0.004 | 0.34 | 0.51 | 0.32 | 0.04 | 0.26 | 0.2 |
| 6 | 125867237 | rs9321045 | 0.624 | -1.982 | -0.61 | -4.503 | -0.038 | 0.152 | 0.152 | 3.389 | 0.39 | 0.06 | 0.63 | 0.85 | 0.63 | 0.54 |
| 6 | 125867395 | rs9321046 | 0.241 | -2.084 | -0.551 | -4.54 | 0.068 | 0.21 | 0.21 | 1.485 | 0.46 | 0.15 | 0.66 | 0.86 | 0.69 | 0.56 |
| 6 | 125895248 | rs10457464 | -0.625 | -2.019 | 0.256 | -4.525 | -0.007 | -0.076 | -0.076 | 0 | 0.68 | 0.62 | 0.75 | 0.96 | 0.77 | 0.79 |
| 6 | 125914993 | rs11964411 | -0.29 | -1.875 | 0.014 | -4.409 | -0.028 | -0.053 | -0.053 | 0 | 0.09 | 0.2 | 0.03 | 0.02 | 0.05 | 0.05 |
| 6 | 125921642 | rs4262206 | 0.405 | -1.901 | -0.486 | -4.464 | 0.094 | 0.272 | 0.272 | 3.405 | 0.54 | 0.09 | 0.78 | 0.87 | 0.74 | 0.73 |
| 6 | 125940188 | rs6925158 | 0.856 | -1.843 | -0.8 | -4.444 | -0.1 | 0.05 | 0.05 | 1.534 | 0.33 | 0.06 | 0.58 | 0.79 | 0.59 | 0.47 |
| 6 | 125944755 | rs2326188 | 0.137 | -1.917 | -0.357 | -4.432 | 0.043 | 0.075 | 0.075 | 1.565 | 0.56 | 0.3 | 0.79 | 0.87 | 0.74 | 0.73 |
| 6 | 125985540 | rs4897150 | -0.207 | -0.949 | -0.162 | -3.446 | 0.02 | 0.039 | 0.039 | 0 | 0.48 | 0.43 | 0.6 | 0.9 | 0.58 | 0.63 |
| 6 | 125988575 | rs11964098 | 0.43 | -0.847 | -0.155 | -3.538 | 0.009 | -0.008 | -0.008 | 0.001 | 0.23 | 0.16 | 0.1 | 0 | 0.15 | 0.06 |

| | | | | | | | | | | | | | | | | |
|---|---|---|---|---|---|---|---|---|---|---|---|---|---|---|---|---|
| 6 | 125997436 | rs1832979 | 0.41 | -0.414 | -0.32 | -3.162 | -0.028 | 0.107 | 0.107 | 0.154 | 0.39 | 0.22 | 0.54 | 0.83 | 0.55 | 0.58 |
| 6 | 126007620 | rs2747714 | -0.484 | -0.411 | -0.253 | -3.187 | -0.006 | 0.038 | 0.038 | 0.221 | 0.6 | 0.63 | 0.45 | 0.13 | 0.44 | 0.42 |
| 6 | 126022602 | rs1777220 | 0.144 | -0.67 | 0.03 | -3.21 | -0.002 | -0.032 | -0.032 | 0 | 0.52 | 0.42 | 0.39 | 0.13 | 0.41 | 0.37 |
| 6 | 126029043 | rs1268092 | -0.508 | -0.618 | 0.02 | -3.08 | -0.028 | 0.15 | 0.15 | 1.071 | 0.59 | 0.88 | 0.41 | 0.17 | 0.41 | 0.42 |
| 6 | 126036184 | rs1343121 | -0.203 | -0.582 | 0.154 | -3.081 | 0 | -0.015 | -0.015 | 0.1 | 0.55 | 0.5 | 0.37 | 0.14 | 0.4 | 0.41 |
| 6 | 126052729 | rs4895799 | 0.267 | -0.077 | -0.17 | -2.957 | -0.007 | 0.051 | 0.051 | 0 | 0.46 | 0.42 | 0.6 | 0.78 | 0.57 | 0.43 |
| 6 | 126057748 | rs9398785 | -0.453 | -0.115 | -0.193 | -3.006 | 0.005 | 0.011 | 0.011 | 0 | 0.57 | 0.47 | 0.67 | 0.78 | 0.67 | 0.49 |
| 6 | 126068001 | rs980014 | 0.709 | -0.226 | -0.131 | -2.632 | -0.094 | 0.065 | 0.065 | 0 | 0.39 | 0.11 | 0.54 | 0.73 | 0.51 | 0.41 |
| 6 | 126069702 | rs1343116 | -0.364 | -0.314 | -0.062 | -2.646 | 0.015 | 0.043 | 0.043 | 0 | 0.53 | 0.44 | 0.61 | 0.77 | 0.6 | 0.48 |
| 6 | 126079968 | rs9321054 | 0.75 | -0.296 | -0.141 | -2.647 | -0.084 | 0.062 | 0.062 | 0 | 0.38 | 0.12 | 0.45 | 0.73 | 0.5 | 0.39 |
| 6 | 126081319 | rs3734637 | -0.345 | -0.343 | 0.04 | -2.656 | 0.017 | 0.047 | 0.047 | 0 | 0.52 | 0.43 | 0.53 | 0.76 | 0.6 | 0.46 |
| 6 | 126081612 | rs6940469 | -0.326 | -0.315 | 0.072 | -2.624 | 0 | -0.014 | -0.014 | 0 | 0.12 | 0.18 | 0.06 | 0.04 | 0.09 | 0.07 |
| 6 | 126090277 | rs1159974 | 0.711 | -0.302 | 0.05 | -2.625 | -0.06 | 0.058 | 0.058 | 0 | 0.38 | 0.17 | 0.45 | 0.73 | 0.5 | 0.39 |
| 6 | 126099246 | rs1739384 | -0.483 | -0.408 | 0.853 | -2.71 | 0.008 | 0.023 | 0.023 | 0 | 0.54 | 0.59 | 0.54 | 0.77 | 0.59 | 0.46 |
| 6 | 126106016 | rs1268071 | -0.796 | -0.242 | 1.04 | -2.262 | 0.007 | -0.04 | -0.04 | 0 | 0.14 | 0.26 | 0.08 | 0.04 | 0.1 | 0.07 |
| 6 | 126113822 | rs1268087 | -0.997 | -0.237 | 0.906 | -2.237 | 0.005 | -0.055 | -0.055 | 0.042 | 0.15 | 0.42 | 0.09 | 0.04 | 0.1 | 0.07 |
| 6 | 126122564 | rs11967627 | 0.428 | -0.379 | 0.589 | -2.39 | 0.006 | -0.012 | -0.012 | 0 | 0.16 | 0.11 | 0.06 | 0 | 0.11 | 0.04 |
| 6 | 126133389 | rs600709 | 0.93 | -0.268 | 0.744 | -2.161 | -0.002 | -0.113 | -0.113 | 0.031 | 0.67 | 0.52 | 0.81 | 0.92 | 0.71 | 0.84 |
| 6 | 126134936 | rs549438 | 1.352 | -0.524 | 0.314 | -2.248 | 0.014 | 0.184 | 0.184 | 3.645 | 0.61 | 0.11 | 0.78 | 0.88 | 0.7 | 0.83 |
| 6 | 126142576 | rs644096 | 0.751 | -0.409 | 0.733 | -2.231 | 0.001 | -0.084 | -0.084 | 1.81 | 0.68 | 0.57 | 0.81 | 0.92 | 0.71 | 0.84 |
| 6 | 126146711 | rs622936 | 0.819 | -0.349 | -0.16 | -2.436 | 0.004 | 0.055 | 0.055 | 0.116 | 0.29 | 0.12 | 0.47 | 0.2 | 0.3 | 0.45 |
| 6 | 126156943 | rs9321057 | 1.298 | -0.463 | -0.402 | -1.642 | -0.138 | -0.162 | -0.162 | 0 | 0.31 | 0.03 | 0.32 | 0.51 | 0.39 | 0.38 |
| 6 | 126163537 | rs674859 | 0.773 | -0.598 | 0.4 | -1.696 | 0.015 | 0.032 | 0.032 | 0 | 0.32 | 0.24 | 0.47 | 0.23 | 0.3 | 0.46 |
| 6 | 126163599 | rs17708107 | 0.33 | -0.571 | 0.207 | -1.797 | -0.039 | 0.044 | 0.044 | 0 | 0.09 | 0.01 | 0.05 | 0 | 0.1 | 0.06 |
| 6 | 126165003 | rs584032 | -0.231 | -0.583 | 0.378 | -1.885 | -0.001 | -0.014 | -0.014 | 0 | 0.45 | 0.53 | 0.55 | 0.25 | 0.42 | 0.52 |
| 6 | 126165465 | rs481233 | -0.191 | -0.77 | 0.563 | -1.82 | 0.022 | 0.179 | 0.179 | 0.584 | 0.92 | 0.69 | 0.93 | 0.76 | 0.91 | 0.93 |
| 6 | 126174747 | rs6569439 | -1.035 | -0.573 | 0.078 | -1.763 | 0.002 | 0.033 | 0.033 | 0.056 | 0.71 | 0.84 | 0.53 | 0.76 | 0.68 | 0.54 |
| 6 | 126181956 | rs9401851 | 0.622 | -0.465 | -0.101 | -1.703 | -0.085 | -0.08 | -0.08 | 0 | 0.4 | 0.08 | 0.37 | 0.51 | 0.49 | 0.43 |
| 6 | 126190894 | rs7754833 | 1.163 | -0.489 | -0.142 | -1.651 | -0.156 | -0.165 | -0.165 | 0 | 0.32 | 0.03 | 0.41 | 0.53 | 0.4 | 0.37 |
| 6 | 126195500 | rs2456110 | -1.132 | -0.577 | -0.055 | -1.751 | 0.015 | 0.024 | 0.024 | 3.401 | 0.39 | 0.89 | 0.22 | 0.31 | 0.31 | 0.16 |
| 6 | 126197049 | rs10080352 | -0.404 | -0.608 | 0.221 | -1.785 | 0.011 | -0.043 | -0.043 | 0.222 | 0.11 | 0.19 | 0.06 | 0 | 0.1 | 0.04 |
| 6 | 126199862 | rs1570383 | 0.01 | -0.644 | 0.587 | -1.791 | -0.026 | 0.005 | 0.005 | 0.388 | 0.24 | 0.44 | 0.13 | 0.25 | 0.19 | 0.11 |
| 6 | 126209245 | rs6934080 | -0.895 | -0.634 | 0.478 | -1.79 | -0.002 | -0.007 | -0.007 | 0 | 0.35 | 0.68 | 0.2 | 0.25 | 0.29 | 0.15 |
| 6 | 126210395 | rs6919947 | 0.161 | -0.741 | 0.382 | -1.698 | 0.002 | 0.003 | 0.003 | 0 | 0.45 | 0.4 | 0.49 | 0.59 | 0.52 | 0.42 |
| 6 | 126218600 | rs6941158 | -0.959 | -0.474 | -0.231 | -1.53 | -0.003 | 0.062 | 0.062 | 2 | 0.72 | 0.97 | 0.79 | 0.83 | 0.74 | 0.55 |
| 6 | 126223944 | rs4897160 | 0.682 | -0.546 | -0.055 | -1.614 | -0.025 | -0.012 | -0.012 | 0.016 | 0.45 | 0.22 | 0.61 | 0.52 | 0.48 | 0.41 |
| 6 | 126230445 | rs6911452 | -0.978 | -0.641 | -0.191 | -1.61 | 0.015 | 0.064 | 0.064 | 1.508 | 0.71 | 0.92 | 0.76 | 0.83 | 0.7 | 0.54 |
| 6 | 126232063 | rs9401855 | -0.272 | -0.592 | 0.16 | -1.641 | 0.004 | 0.02 | 0.02 | 0.351 | 0.62 | 0.75 | 0.7 | 0.77 | 0.61 | 0.49 |
| 6 | 126238310 | rs715880 | 0.114 | -0.586 | 0.527 | -1.627 | 0.008 | -0.001 | -0.001 | 0 | 0.56 | 0.45 | 0.66 | 0.53 | 0.57 | 0.46 |
| 6 | 126244682 | rs1322755 | -0.656 | -0.36 | 0.616 | -1.53 | 0.048 | 0.063 | 0.063 | 0 | 0.54 | 0.75 | 0.71 | 0.77 | 0.66 | 0.48 |
| 6 | 126245644 | rs9482716 | 0.06 | 0 | 0.583 | -1.307 | 0.003 | 0.071 | 0.071 | 0 | 0.12 | 0.27 | 0.08 | 0.24 | 0.12 | 0.07 |
| 6 | 126249914 | rs1567 | 0.043 | 0.007 | 0.543 | -1.304 | 0.001 | 0.062 | 0.062 | 0 | 0.12 | 0.26 | 0.08 | 0.24 | 0.12 | 0.07 |
| 6 | 126255034 | rs10872303 | -0.137 | 0.3 | 0.353 | -1.632 | 0 | -0.004 | -0.004 | 0 | 0.4 | 0.45 | 0.63 | 0.61 | 0.54 | 0.4 |
| 6 | 126256410 | rs11751595 | 0.401 | 0.359 | 0.048 | -1.364 | 0 | -0.001 | -0.001 | 0 | 0.31 | 0.3 | 0.58 | 0.61 | 0.45 | 0.37 |
| 6 | 126271503 | rs9398792 | 0.461 | 0.008 | -0.297 | -2.388 | -0.033 | 0.128 | 0.128 | 0 | 0.42 | 0.2 | 0.65 | 0.95 | 0.5 | 0.63 |
| 6 | 126288023 | rs3757212 | -0.198 | -0.286 | -0.18 | -2.505 | -0.006 | 0.087 | 0.087 | 0 | 0.66 | 0.78 | 0.79 | 0.96 | 0.65 | 0.73 |
| 6 | 126300445 | rs6909567 | -0.59 | -0.541 | 0.676 | -2.565 | 0.061 | 0.057 | 0.057 | 5.414 | 0.05 | 0.4 | 0.05 | 0 | 0 | 0 |
| 6 | 126307962 | rs6569448 | -0.274 | -0.49 | 0.485 | -2.608 | -0.008 | 0.087 | 0.087 | 0 | 0.66 | 0.78 | 0.78 | 0.96 | 0.64 | 0.73 |
| 6 | 126331545 | rs10485179 | 0.08 | -0.509 | 0.553 | -2.619 | 0.005 | -0.011 | -0.011 | 0 | 0.6 | 0.31 | 0.73 | 0.96 | 0.64 | 0.73 |
| 6 | 126352051 | rs10872306 | 0.48 | -0.37 | 0.661 | -2.28 | -0.038 | 0.142 | 0.142 | 0 | 0.42 | 0.19 | 0.64 | 0.93 | 0.47 | 0.58 |
| 6 | 126356389 | rs9375421 | -0.117 | -0.413 | 0.526 | -2.307 | -0.015 | 0.229 | 0.229 | 0 | 0.5 | 0.75 | 0.69 | 0.95 | 0.48 | 0.59 |
| 6 | 126373765 | rs1465104 | 0.551 | -0.369 | 0.469 | -2.33 | -0.063 | 0.188 | 0.188 | 0 | 0.42 | 0.14 | 0.63 | 0.93 | 0.47 | 0.58 |
| 6 | 126392441 | rs17053767 | -1.914 | -0.365 | 1.569 | -1.519 | -0.024 | -0.036 | -0.036 | 0 | 0.08 | 0.63 | 0.07 | 0 | 0.01 | 0 |
| 6 | 126402418 | rs2186061 | -1.094 | -0.373 | 2.014 | -1.303 | 0.026 | 0.097 | 0.097 | 0.235 | 0.05 | 0.23 | 0.06 | 0.11 | 0.04 | 0.03 |
| 6 | 126404590 | rs17053778 | -1.956 | -0.376 | 1.425 | -1.293 | -0.029 | 0.03 | 0.03 | 6.479 | 0.1 | 0.61 | 0.09 | 0.11 | 0.04 | 0.03 |
| 6 | 126413814 | rs6939492 | -2.077 | -0.409 | 1.255 | -1.324 | -0.037 | -0.002 | -0.002 | 0.872 | 0.11 | 0.69 | 0.1 | 0.11 | 0.04 | 0.03 |
| 6 | 126437248 | rs10499133 | -1.728 | -0.238 | 1.036 | -1.384 | -0.019 | -0.009 | -0.009 | 0.579 | 0.13 | 0.64 | 0.1 | 0.11 | 0.07 | 0.05 |
| 6 | 126444684 | rs6569454 | -0.948 | 0.103 | 1.055 | -1.889 | 0.026 | -0.089 | -0.089 | 0 | 0.17 | 0.36 | 0.15 | 0.02 | 0.18 | 0.24 |
| 6 | 126445961 | rs6569455 | -0.963 | 0.235 | 1.051 | -1.894 | 0.033 | -0.089 | -0.089 | 0 | 0.17 | 0.36 | 0.15 | 0.03 | 0.19 | 0.24 |
| 6 | 126449652 | rs9321060 | -0.121 | -0.288 | 0.299 | -2.625 | 0.015 | 0.077 | 0.077 | 0 | 0.33 | 0.18 | 0.2 | 0.08 | 0.31 | 0.15 |
| 6 | 126471447 | rs9388472 | -0.235 | -0.513 | 0.397 | -2.624 | 0.011 | 0.019 | 0.019 | 0 | 0.28 | 0.22 | 0.13 | 0.09 | 0.23 | 0.1 |
| 6 | 126472811 | rs12198445 | -0.891 | -0.536 | 0.451 | -2.791 | 0.003 | -0.012 | -0.012 | 0.365 | 0.47 | 0.46 | 0.29 | 0.11 | 0.46 | 0.35 |
| 6 | 126481408 | rs1330674 | -0.546 | -0.243 | 0.707 | -2.515 | -0.008 | -0.104 | -0.104 | 0 | 0.27 | 0.6 | 0.19 | 0.09 | 0.27 | 0.08 |
| 6 | 126491310 | rs6935937 | -0.835 | 0.027 | 0.817 | -2.273 | 0.012 | -0.138 | -0.138 | 0.022 | 0.26 | 0.45 | 0.2 | 0.02 | 0.28 | 0.27 |
| 6 | 126522821 | rs9491587 | -1.73 | -0.217 | 1.432 | -1.775 | 0.03 | 0.027 | 0.027 | 5.801 | 0.05 | 0.4 | 0.03 | 0 | 0 | 0 |
| 6 | 126524030 | rs9491589 | -1.8 | -0.224 | 1.512 | -1.778 | 0.032 | 0.029 | 0.029 | 6.383 | 0.06 | 0.42 | 0.03 | 0 | 0 | 0 |
| 6 | 126573579 | rs6569457 | -0.187 | 0.216 | 0.73 | -0.838 | -0.007 | -0.021 | -0.021 | 0 | 0.16 | 0.12 | 0.32 | 0.54 | 0.17 | 0.31 |
| 6 | 126594267 | rs643394 | -0.42 | 0.193 | 1.307 | -0.787 | -0.009 | 0.237 | 0.237 | 0.956 | 0.16 | 0.41 | 0.32 | 0.53 | 0.13 | 0.3 |
| 6 | 126599758 | rs1455736 | -0.112 | 0.208 | 1.107 | -0.769 | 0.002 | 0.004 | 0.004 | 1.882 | 0.14 | 0.12 | 0.3 | 0.52 | 0.12 | 0.29 |

| | | | | | | | | | | | | | | | | |
|---|---|---|---|---|---|---|---|---|---|---|---|---|---|---|---|---|
| 6 | 126601960 | rs9388477 | -0.354 | 0.221 | 1.102 | -0.785 | -0.002 | 0.012 | 0.012 | 1.321 | 0.16 | 0.18 | 0.32 | 0.53 | 0.17 | 0.3 |
| 6 | 126609124 | rs607779 | 0.049 | 0.106 | 0.475 | -1.99 | 0.008 | -0.05 | -0.05 | 0 | 0.47 | 0.41 | 0.62 | 0.97 | 0.38 | 0.57 |
| 6 | 126638124 | rs9372837 | 0.184 | 0.407 | 0.21 | -2.476 | 0.005 | -0.11 | -0.11 | 0 | 0.6 | 0.43 | 0.68 | 0.98 | 0.49 | 0.65 |
| 6 | 126642219 | rs647463 | -0.992 | 1.03 | -0.277 | -2.806 | -0.054 | 0.086 | 0.086 | 0 | 0.84 | 0.98 | 0.91 | 1 | 0.79 | 0.98 |
| 6 | 126655243 | rs1378579 | 0.9 | 0.889 | -0.195 | -2.823 | -0.055 | 0.226 | 0.226 | 2.857 | 0.57 | 0.12 | 0.66 | 0.98 | 0.49 | 0.65 |
| 6 | 126659043 | rs6919397 | 0.722 | 0.729 | 0.061 | -2.911 | 0 | -0.069 | -0.069 | 0 | 0.58 | 0.4 | 0.69 | 0.97 | 0.45 | 0.65 |
| 6 | 126661154 | rs9388486 | 0.858 | 0.729 | 0.036 | -2.911 | 0.004 | 0.111 | 0.111 | 0.969 | 0.57 | 0.21 | 0.67 | 0.97 | 0.45 | 0.65 |
| 6 | 126661858 | rs9375435 | 0.822 | 0.718 | 0.011 | -2.94 | -0.002 | 0.109 | 0.109 | 0.956 | 0.58 | 0.21 | 0.68 | 0.97 | 0.45 | 0.64 |
| 6 | 126664333 | rs17754780 | 0.83 | 0.722 | 0.027 | -2.898 | 0 | 0.116 | 0.116 | 1.048 | 0.58 | 0.2 | 0.68 | 0.97 | 0.45 | 0.65 |
| 6 | 126674354 | rs2045258 | 0.878 | 0.701 | -0.001 | -2.918 | -0.003 | 0.168 | 0.168 | 1.309 | 0.57 | 0.17 | 0.67 | 0.97 | 0.45 | 0.66 |
| 6 | 126686407 | rs12202204 | 0.114 | 0.76 | 0.484 | -3.031 | -0.038 | 0.038 | 0.038 | 0 | 0.17 | 0.06 | 0.17 | 0.02 | 0.29 | 0.31 |
| 6 | 126698719 | rs9388489 | 1.074 | 0.813 | 0.418 | -3.002 | -0.004 | 0.166 | 0.166 | 1.353 | 0.57 | 0.17 | 0.67 | 0.97 | 0.45 | 0.65 |
| 6 | 126708470 | rs7755075 | -0.927 | 0.793 | 1.023 | -2.993 | 0.012 | -0.084 | -0.084 | 0 | 0.82 | 0.71 | 0.9 | 1 | 0.79 | 0.98 |
| 6 | 126744087 | rs9321065 | 0.962 | 0.765 | 0.842 | -2.978 | -0.003 | 0.035 | 0.035 | 0.424 | 0.58 | 0.26 | 0.68 | 0.97 | 0.45 | 0.65 |
| 6 | 126802598 | rs7753563 | -0.199 | 0.691 | 1.089 | -3.025 | 0.025 | -0.135 | -0.135 | 0 | 0.77 | 0.57 | 0.86 | 1 | 0.74 | 0.96 |
| 6 | 126822635 | rs9372840 | 0.949 | 0.563 | 0.706 | -3.023 | -0.001 | 0.035 | 0.035 | 0.198 | 0.57 | 0.29 | 0.68 | 0.97 | 0.45 | 0.65 |
| 6 | 126845380 | rs1337736 | 1.12 | 0.318 | 0.428 | -2.179 | -0.001 | 0.019 | 0.019 | 0 | 0.2 | 0.21 | 0.26 | 0.52 | 0.2 | 0.33 |
| 6 | 126877523 | rs9401888 | 0.755 | 0.034 | -0.037 | -2.447 | -0.01 | 0.223 | 0.223 | 2.373 | 0.54 | 0.16 | 0.67 | 0.97 | 0.48 | 0.69 |
| 6 | 126897969 | rs3861455 | 0.755 | 0.034 | -0.037 | -2.447 | -0.01 | 0.223 | 0.223 | 2.373 | 0.54 | 0.16 | 0.67 | 0.97 | 0.48 | 0.69 |
| 6 | 126903011 | rs9388501 | 0.756 | 0.008 | 0.239 | -2.46 | 0 | -0.043 | -0.043 | 0 | 0.55 | 0.4 | 0.69 | 0.97 | 0.48 | 0.69 |
| 6 | 126936235 | rs7738836 | 0.65 | 0.026 | 0.26 | -2.442 | -0.001 | 0.033 | 0.033 | 1.504 | 0.55 | 0.32 | 0.69 | 0.97 | 0.48 | 0.67 |
| 6 | 126964510 | rs4273712 | 1.136 | 0.12 | 0.217 | -2.233 | 0.037 | -0.118 | -0.118 | 0 | 0.36 | 0.05 | 0.43 | 0.44 | 0.28 | 0.35 |
| 6 | 127011064 | rs2746187 | 0.502 | 0.396 | 0.317 | -2.376 | -0.002 | -0.06 | -0.06 | 0 | 0.24 | 0.29 | 0.11 | 0 | 0.23 | 0.03 |
| 6 | 127046153 | rs853983 | -1.15 | 0.433 | -0.314 | -2.401 | -0.026 | 0.233 | 0.233 | 4.728 | 0.46 | 0.84 | 0.32 | 0.03 | 0.51 | 0.31 |
| 6 | 127079341 | rs1262554 | -0.016 | 0.253 | 0.064 | -1.952 | -0.008 | -0.143 | -0.143 | 0 | 0.31 | 0.44 | 0.16 | 0 | 0.28 | 0.04 |
| 6 | 127094587 | rs4895816 | 1.014 | 0.497 | -0.175 | -1.152 | -0.001 | 0.071 | 0.071 | 0 | 0.21 | 0.32 | 0.27 | 0.54 | 0.22 | 0.35 |
| 6 | 127105529 | rs9398816 | 0.614 | 0.421 | -0.584 | -0.932 | -0.045 | -0.082 | -0.082 | 0.537 | 0.39 | 0.11 | 0.36 | 0.49 | 0.45 | 0.42 |
| 6 | 127147704 | rs9375459 | 0.762 | 0.475 | -0.41 | -0.937 | -0.021 | -0.049 | -0.049 | 0.121 | 0.4 | 0.22 | 0.37 | 0.49 | 0.45 | 0.43 |
| 6 | 127164572 | rs17563605 | -0.301 | 1.068 | -0.491 | -1.718 | -0.048 | 0.047 | 0.047 | 0 | 0.13 | 0.03 | 0.18 | 0.02 | 0.25 | 0.25 |
| 6 | 127208090 | rs2225642 | 0.83 | 0.903 | -0.408 | -1.501 | -0.038 | -0.001 | -0.001 | 0.341 | 0.42 | 0.27 | 0.43 | 0.55 | 0.49 | 0.43 |
| 6 | 127208390 | rs4897193 | 0.695 | 0.938 | -0.063 | -1.444 | -0.004 | 0.034 | 0.034 | 0 | 0.45 | 0.35 | 0.45 | 0.66 | 0.52 | 0.39 |
| 6 | 127210093 | rs2151522 | 1.367 | 0.92 | -0.149 | -1.362 | -0.012 | 0.004 | 0.004 | 0.051 | 0.34 | 0.2 | 0.36 | 0.64 | 0.41 | 0.42 |
| 6 | 127212478 | rs17054192 | 0.254 | -0.369 | 0.449 | -1.496 | 0.008 | -0.083 | -0.083 | 0 | 0.13 | 0.24 | 0.06 | 0 | 0.1 | 0.07 |
| 6 | 127221252 | rs2027033 | 1.333 | -0.351 | 0.378 | -1.281 | -0.017 | 0.015 | 0.015 | 0 | 0.38 | 0.34 | 0.41 | 0.67 | 0.48 | 0.43 |
| 6 | 127230163 | rs9388539 | 0.243 | -0.701 | 0.626 | -1.124 | -0.016 | 0.008 | 0.008 | 0.009 | 0.2 | 0.25 | 0.28 | 0.27 | 0.12 | 0.24 |
| 6 | 127262500 | rs9398824 | 1.013 | -0.016 | 0.447 | -0.747 | -0.052 | 0.02 | 0.02 | 0 | 0.35 | 0.19 | 0.39 | 0.69 | 0.45 | 0.43 |
| 6 | 127263275 | rs1343658 | 0.566 | -0.047 | 0.451 | -0.766 | 0.009 | -0.014 | -0.014 | 0 | 0.39 | 0.49 | 0.47 | 0.69 | 0.46 | 0.45 |
| 6 | 127276881 | rs10872310 | 0.623 | 0.024 | 0.367 | -0.738 | 0.014 | -0.017 | -0.017 | 0.324 | 0.39 | 0.47 | 0.42 | 0.69 | 0.45 | 0.44 |
| 6 | 127284568 | rs1930958 | 0.613 | -0.009 | 0.361 | -0.9 | 0.015 | -0.032 | -0.032 | 0 | 0.39 | 0.51 | 0.42 | 0.7 | 0.46 | 0.42 |
| 6 | 127284605 | rs1930959 | 0.989 | 0.023 | 0.387 | -0.886 | -0.018 | 0.018 | 0.018 | 0 | 0.36 | 0.31 | 0.4 | 0.7 | 0.45 | 0.4 |
| 6 | 127298394 | rs9401928 | 0.649 | -0.032 | 0.1 | -0.923 | -0.018 | 0.035 | 0.035 | 0 | 0.39 | 0.31 | 0.42 | 0.71 | 0.47 | 0.41 |
| 6 | 127311647 | rs1014054 | 0.689 | 0.217 | 0.099 | -0.777 | 0.003 | -0.006 | -0.006 | 0 | 0.38 | 0.41 | 0.41 | 0.7 | 0.45 | 0.42 |
| 6 | 127337699 | rs1034207 | 0.079 | 0.68 | -0.169 | -0.315 | 0 | 0.016 | 0.016 | 1.179 | 0.92 | 0.96 | 0.96 | 0.95 | 0.94 | 0.92 |
| 6 | 127380482 | rs6902885 | 1.258 | 0.468 | -0.077 | -0.31 | 0.008 | 0.006 | 0.006 | 0 | 0.45 | 0.37 | 0.38 | 0.59 | 0.42 | 0.46 |
| 6 | 127381956 | rs2745359 | -0.255 | 0.511 | -0.248 | -0.333 | 0.014 | 0.017 | 0.017 | 0.657 | 0.12 | 0.06 | 0.06 | 0.04 | 0.06 | 0.02 |
| 6 | 127392977 | rs9321069 | 1.305 | 0.476 | -0.135 | -0.288 | 0.005 | 0.003 | 0.003 | 0 | 0.44 | 0.37 | 0.38 | 0.59 | 0.42 | 0.45 |
| 6 | 127400325 | rs2800718 | -1.093 | 0.26 | -0.008 | -0.292 | 0.003 | -0.029 | -0.029 | 0.042 | 0.25 | 0.24 | 0.2 | 0.12 | 0.25 | 0.18 |
| 6 | 127409882 | rs4580892 | -0.47 | 0.463 | -0.056 | -0.075 | -0.004 | 0.024 | 0.024 | 0 | 0.29 | 0.37 | 0.4 | 0.43 | 0.28 | 0.34 |
| 6 | 127412728 | rs12192275 | -0.505 | 0.91 | -0.041 | 0.052 | 0.027 | -0.013 | -0.013 | 0.039 | 0.09 | 0.1 | 0.11 | 0.05 | 0.2 | 0.13 |
| 6 | 127421733 | rs1936790 | -1.052 | 0.45 | -0.049 | 0.091 | -0.019 | 0.004 | 0.004 | 0 | 0.43 | 0.49 | 0.44 | 0.5 | 0.29 | 0.35 |
| 6 | 127423055 | rs1936792 | -0.154 | 0.394 | 0.094 | 0.042 | -0.001 | 0.007 | 0.007 | 0.827 | 0.25 | 0.27 | 0.38 | 0.43 | 0.24 | 0.29 |
| 6 | 127432657 | rs1936797 | -0.923 | 0.355 | -0.093 | -0.015 | -0.01 | 0.016 | 0.016 | 0.583 | 0.34 | 0.41 | 0.4 | 0.44 | 0.29 | 0.36 |
| 6 | 127438642 | rs10872311 | -0.782 | 0.283 | -0.154 | -0.054 | -0.007 | 0.018 | 0.018 | 0.531 | 0.32 | 0.38 | 0.38 | 0.45 | 0.25 | 0.32 |
| 6 | 127455138 | rs7766106 | -0.652 | 0.186 | -0.157 | -0.111 | -0.011 | -0.011 | -0.011 | 0 | 0.4 | 0.33 | 0.48 | 0.5 | 0.46 | 0.46 |
| 6 | 127456122 | rs9491697 | -0.15 | 0.234 | -0.121 | -0.058 | -0.002 | -0.002 | -0.002 | 1.361 | 0.32 | 0.29 | 0.46 | 0.51 | 0.42 | 0.4 |
| 6 | 127460591 | rs1892170 | 0.115 | 0.086 | -0.091 | 0.002 | 0.004 | 0.003 | 0.003 | 1.551 | 0.09 | 0.05 | 0.09 | 0.08 | 0.01 | 0.01 |
| 6 | 127461682 | rs1936802 | -1.519 | 0.052 | 0.071 | -0.103 | 0 | 0.003 | 0.003 | 0 | 0.54 | 0.58 | 0.6 | 0.58 | 0.51 | 0.53 |
| 6 | 127476516 | rs1892172 | -0.932 | 0.276 | -0.088 | -0.074 | -0.001 | -0.008 | -0.008 | 0 | 0.46 | 0.6 | 0.52 | 0.49 | 0.5 | 0.52 |
| 6 | 127481154 | rs4644087 | -0.952 | 0.313 | -0.449 | -0.056 | 0 | -0.003 | -0.003 | 0 | 0.46 | 0.56 | 0.52 | 0.49 | 0.5 | 0.52 |
| 6 | 127483308 | rs13202608 | 0.203 | 0.628 | -0.71 | -0.019 | -0.028 | 0.026 | 0.026 | 0.075 | 0.06 | 0.01 | 0.06 | 0 | 0.1 | 0.03 |
| 6 | 127507403 | rs11759578 | 0.126 | -0.01 | -0.56 | 0.163 | 0.002 | 0.03 | 0.03 | 0 | 0.07 | 0.02 | 0.02 | 0 | 0.05 | 0.06 |
| 6 | 127509157 | rs7745274 | -0.997 | 0.199 | -0.461 | 0.291 | 0.002 | 0.004 | 0.004 | 0 | 0.54 | 0.59 | 0.6 | 0.57 | 0.51 | 0.53 |
| 6 | 127519234 | rs10457487 | -0.614 | 1.334 | -0.812 | 0.854 | -0.017 | -0.017 | -0.017 | 1.125 | 0.34 | 0.28 | 0.47 | 0.48 | 0.45 | 0.38 |
| 6 | 127553539 | rs6569476 | 0.249 | 1.061 | -1.002 | 0.475 | -0.011 | -0.01 | -0.01 | 0 | 0.77 | 0.84 | 0.79 | 0.79 | 0.74 | 0.58 |
| 6 | 127574044 | rs9388553 | 0.544 | 0.777 | -0.282 | 0.552 | 0.001 | 0.008 | 0.008 | 0 | 0.69 | 0.66 | 0.75 | 0.7 | 0.7 | 0.55 |
| 6 | 127587086 | rs877102 | 0.864 | -0.058 | -0.431 | 0.223 | -0.008 | 0 | 0 | 0 | 0.62 | 0.58 | 0.73 | 0.69 | 0.7 | 0.5 |
| 6 | 127596985 | rs12210968 | 0.922 | 0.165 | 0.318 | 0.397 | 0.01 | 0.062 | 0.062 | 0 | 0.57 | 0.32 | 0.44 | 0.1 | 0.66 | 0.45 |
| 6 | 127600630 | rs2180341 | -0.245 | 0.133 | 0.409 | 0.439 | 0.005 | -0.009 | -0.009 | 0.364 | 0.73 | 0.7 | 0.77 | 0.8 | 0.72 | 0.59 |
| 6 | 127605898 | rs4897207 | -0.245 | 0.133 | 0.409 | 0.439 | 0.005 | -0.009 | -0.009 | 0 | 0.73 | 0.7 | 0.77 | 0.8 | 0.72 | 0.59 |

| | | | | | | | | | | | | | | | | |
|---|---|---|---|---|---|---|---|---|---|---|---|---|---|---|---|---|
| 6 | 127641960 | rs3823042 | 0.415 | 0.917 | 0.023 | -0.251 | -0.056 | 0.009 | 0.009 | 0 | 0.09 | 0.01 | 0.15 | 0.04 | 0.19 | 0.13 |
| 6 | 127649319 | rs3823043 | 0.754 | 0.429 | 0.035 | -0.64 | -0.002 | -0.005 | -0.005 | 1.886 | 0.07 | 0.09 | 0.05 | 0.05 | 0.07 | 0.1 |
| 6 | 127649350 | rs11961994 | 0.446 | 0.433 | 0.007 | -0.602 | 0.01 | 0.062 | 0.062 | 0 | 0.57 | 0.32 | 0.44 | 0.1 | 0.66 | 0.46 |
| 6 | 127657099 | rs13194280 | 1.018 | -0.333 | -0.547 | -1.305 | 0.035 | 0.063 | 0.063 | 4.181 | 0.13 | 0.03 | 0.09 | 0 | 0.11 | 0.08 |
| 6 | 127663116 | rs3798859 | 0.247 | 0.08 | -0.223 | -0.964 | 0.01 | 0.061 | 0.061 | 0 | 0.57 | 0.32 | 0.44 | 0.1 | 0.66 | 0.46 |
| 6 | 127669744 | rs9398840 | -1.044 | 0.142 | -0.056 | -0.904 | 0.001 | -0.003 | -0.003 | 1.583 | 0.73 | 0.72 | 0.77 | 0.8 | 0.72 | 0.59 |
| 6 | 127685893 | rs6900074 | -0.724 | 0.113 | -0.079 | -0.919 | -0.004 | -0.017 | -0.017 | 0.033 | 0.69 | 0.62 | 0.76 | 0.8 | 0.72 | 0.59 |
| 6 | 127690694 | rs9388568 | -0.734 | 0.116 | -0.078 | -0.917 | -0.004 | -0.017 | -0.017 | 0 | 0.7 | 0.62 | 0.76 | 0.8 | 0.72 | 0.59 |
| 6 | 127719062 | rs9401955 | -1.581 | 0.218 | 0.015 | -0.818 | -0.024 | 0.106 | 0.106 | 0.561 | 0.18 | 0.39 | 0.35 | 0.69 | 0.1 | 0.13 |
| 6 | 127723584 | rs9401957 | -1.275 | 0.181 | 0.059 | -0.704 | -0.03 | 0.129 | 0.129 | 0.195 | 0.16 | 0.37 | 0.33 | 0.69 | 0.06 | 0.13 |
| 6 | 127726196 | rs6569487 | 0.075 | 0.245 | -0.113 | -0.846 | -0.004 | -0.002 | -0.002 | 0 | 0.56 | 0.56 | 0.69 | 0.64 | 0.63 | 0.4 |
| 6 | 127740939 | rs4523101 | -0.006 | 0.538 | -0.251 | -1.259 | 0.002 | -0.063 | -0.063 | 0.291 | 0.57 | 0.6 | 0.41 | 0.07 | 0.56 | 0.4 |
| 6 | 127764305 | rs17205908 | 1.149 | 0.19 | -0.837 | -1.635 | -0.061 | 0.215 | 0.215 | 0.185 | 0.3 | 0.01 | 0.16 | 0 | 0.3 | 0.15 |
| 6 | 127768472 | rs2236026 | -0.102 | 0.258 | -0.61 | -1.162 | -0.023 | -0.01 | -0.01 | 0 | 0.08 | 0.02 | 0.3 | 0.68 | 0.09 | 0.08 |
| 6 | 127768762 | rs6902288 | 0.067 | 0.193 | -0.385 | -1.257 | 0.004 | -0.082 | -0.082 | 0.254 | 0.57 | 0.58 | 0.42 | 0.07 | 0.56 | 0.4 |
| 6 | 127771452 | rs3734447 | -0.105 | 0.234 | -0.347 | -1.236 | 0.01 | -0.086 | -0.086 | 0 | 0.59 | 0.58 | 0.42 | 0.07 | 0.56 | 0.42 |
| 6 | 127775166 | rs9375513 | -0.198 | 0.288 | -0.458 | -1.153 | -0.023 | -0.015 | -0.015 | 0.132 | 0.09 | 0.03 | 0.3 | 0.69 | 0.09 | 0.08 |
| 6 | 127777516 | rs7738385 | -0.695 | 0.165 | -0.312 | -1.251 | 0.004 | -0.005 | -0.005 | 0 | 0.68 | 0.66 | 0.72 | 0.76 | 0.64 | 0.5 |
| 6 | 127787878 | rs6910292 | 0.815 | 0.394 | -0.397 | -1.385 | 0.003 | 0.054 | 0.054 | 0 | 0.36 | 0.21 | 0.21 | 0 | 0.35 | 0.17 |
| 6 | 127797179 | rs3734448 | -0.916 | 0.596 | -0.245 | -0.895 | -0.003 | -0.003 | -0.003 | 0.16 | 0.23 | 0.2 | 0.45 | 0.74 | 0.24 | 0.32 |
| 6 | 127797374 | rs3734450 | -0.715 | 0.629 | -0.215 | -0.867 | -0.012 | -0.013 | -0.013 | 0 | 0.2 | 0.13 | 0.44 | 0.74 | 0.24 | 0.29 |
| 6 | 127801055 | rs12211079 | 0.718 | 0.783 | -0.271 | -0.791 | -0.041 | -0.021 | -0.021 | 0.018 | 0.07 | 0.01 | 0.2 | 0.07 | 0.14 | 0.16 |
| 6 | 127805089 | rs1512449 | -0.505 | 0.287 | 0.257 | -1.04 | -0.001 | 0.008 | 0.008 | 0 | 0.63 | 0.7 | 0.7 | 0.76 | 0.59 | 0.49 |
| 6 | 127808959 | rs13220247 | -1.149 | 0.126 | 0.482 | -0.41 | -0.016 | 0.129 | 0.129 | 0.166 | 0.17 | 0.38 | 0.28 | 0.67 | 0.1 | 0.14 |
| 6 | 127827468 | rs13206423 | -0.475 | -0.094 | 0.017 | -0.343 | 0.006 | -0.002 | -0.002 | 0 | 0.1 | 0.11 | 0.03 | 0.1 | 0.05 | 0.05 |
| 6 | 127829926 | rs553228 | -0.088 | -0.138 | 0.442 | -0.313 | -0.007 | 0.007 | 0.007 | 0 | 0.77 | 0.64 | 0.81 | 0.75 | 0.81 | 0.54 |
| 6 | 127830319 | rs577906 | 1.063 | -0.409 | 0.383 | -0.662 | 0.044 | 0.02 | 0.02 | 0.063 | 0.54 | 0.31 | 0.71 | 0.62 | 0.67 | 0.44 |
| 6 | 127834542 | rs6903353 | -0.097 | -0.654 | 0.366 | -0.66 | 0.006 | -0.002 | -0.002 | 0.069 | 0.1 | 0.11 | 0.03 | 0.1 | 0.05 | 0.05 |
| 6 | 127839809 | rs1858424 | -0.313 | -0.546 | 0.293 | -1.33 | -0.017 | 0.064 | 0.064 | 0 | 0.38 | 0.21 | 0.63 | 0.9 | 0.42 | 0.57 |
| 6 | 127842630 | rs17054787 | 0.628 | -0.508 | 0.049 | -1.131 | 0.011 | -0.027 | -0.027 | 0 | 0.14 | 0.06 | 0.11 | 0.22 | 0.12 | 0.32 |
| 6 | 127843776 | rs9482815 | -0.856 | -1.014 | 0.353 | -1.655 | -0.013 | -0.014 | -0.014 | 0 | 0.22 | 0.28 | 0.1 | 0.09 | 0.12 | 0.22 |
| 6 | 127850966 | rs1512454 | -1.119 | -0.669 | 0.31 | -1.655 | -0.013 | -0.005 | -0.005 | 0.096 | 0.58 | 0.66 | 0.73 | 0.99 | 0.51 | 0.77 |
| 6 | 127857752 | rs4897215 | -1.938 | -0.697 | -0.072 | -1.705 | -0.156 | 0.246 | 0.246 | 0 | 0.67 | 0.98 | 0.79 | 1 | 0.56 | 0.82 |
| 6 | 127866950 | rs6931244 | -0.566 | -0.653 | 0.316 | -1.527 | -0.022 | -0.023 | -0.023 | 0 | 0.06 | 0.2 | 0.03 | 0.01 | 0.02 | 0.02 |
| 6 | 127868931 | rs7739336 | -0.608 | -0.655 | 0.315 | -1.528 | -0.023 | -0.024 | -0.024 | 0 | 0.06 | 0.2 | 0.03 | 0.01 | 0.02 | 0.02 |
| 6 | 127871194 | rs1606331 | -1.744 | -0.559 | -0.112 | -1.622 | -0.103 | 0.2 | 0.2 | 0 | 0.67 | 0.93 | 0.79 | 1 | 0.56 | 0.82 |
| 6 | 127887455 | rs3891518 | 0.227 | -0.933 | -0.187 | -1.125 | -0.011 | -0.001 | -0.001 | 0 | 0.15 | 0.11 | 0.1 | 0.2 | 0.06 | 0.25 |
| 6 | 127888292 | rs7746884 | -1.906 | -0.737 | -0.417 | -1.713 | -0.128 | 0.168 | 0.168 | 0.015 | 0.69 | 0.93 | 0.78 | 1 | 0.52 | 0.81 |
| 6 | 127895753 | rs1374983 | -1.793 | -0.786 | -0.306 | -1.729 | -0.087 | 0.113 | 0.113 | 0 | 0.68 | 0.87 | 0.78 | 1 | 0.52 | 0.81 |
| 6 | 127898305 | rs9388590 | -0.331 | -0.922 | -0.395 | -1.81 | 0.03 | 0.1 | 0.1 | 0 | 0.44 | 0.24 | 0.61 | 0.93 | 0.37 | 0.55 |
| 6 | 127903054 | rs1080436 | -1.712 | -0.455 | -0.585 | -1.576 | -0.123 | 0.155 | 0.155 | 0.208 | 0.68 | 0.93 | 0.78 | 1 | 0.51 | 0.8 |
| 6 | 127909720 | rs6569504 | -1.723 | -0.178 | -0.862 | -1.652 | -0.077 | 0.174 | 0.174 | 0.735 | 0.7 | 0.95 | 0.81 | 1 | 0.63 | 0.8 |
| 6 | 127909977 | rs2121881 | -0.49 | -0.389 | -0.037 | -1.09 | 0.003 | 0 | 0 | 0 | 0.44 | 0.5 | 0.54 | 0.56 | 0.53 | 0.43 |
| 6 | 127910550 | rs721508 | 0.645 | 0.006 | -0.22 | -1.121 | -0.022 | -0.029 | -0.029 | 0 | 0.31 | 0.17 | 0.45 | 0.62 | 0.4 | 0.26 |
| 6 | 127913413 | rs2166835 | -1.766 | -0.894 | -0.277 | -1.367 | -0.082 | 0.105 | 0.105 | 0.662 | 0.7 | 0.91 | 0.78 | 1 | 0.54 | 0.81 |
| 6 | 127920925 | rs4897222 | -0.391 | -1.328 | 0.22 | -1.322 | -0.007 | -0.009 | -0.009 | 0 | 0.44 | 0.48 | 0.49 | 0.68 | 0.39 | 0.38 |
| 6 | 127923526 | rs960104 | 0.753 | -1.372 | 0.263 | -1.37 | 0.002 | -0.017 | -0.017 | 0 | 0.3 | 0.19 | 0.42 | 0.62 | 0.3 | 0.24 |
| 6 | 127929801 | rs1448063 | -0.397 | -1.32 | 0.109 | -1.367 | -0.001 | -0.001 | -0.001 | 0 | 0.44 | 0.44 | 0.48 | 0.68 | 0.39 | 0.38 |
| 6 | 127935511 | rs7744028 | -1.861 | -1.275 | -0.304 | -1.364 | -0.097 | 0.131 | 0.131 | 1.466 | 0.67 | 0.9 | 0.75 | 1 | 0.5 | 0.76 |
| 6 | 127936007 | rs11154421 | -0.394 | -1.033 | -0.148 | -1.079 | -0.006 | -0.009 | -0.009 | 0.36 | 0.44 | 0.53 | 0.49 | 0.68 | 0.39 | 0.37 |
| 6 | 127953106 | rs10080285 | 0.075 | -0.938 | -0.117 | -0.64 | -0.028 | -0.037 | -0.037 | 0 | 0.08 | 0.16 | 0.02 | 0 | 0.01 | 0.07 |
| 6 | 127953990 | rs2922248 | 0.399 | -0.826 | -0.059 | -0.437 | -0.022 | 0.015 | 0.015 | 0 | 0.75 | 0.68 | 0.74 | 0.68 | 0.86 | 0.58 |
| 6 | 127954614 | rs2922249 | 0.54 | -0.056 | -0.218 | -0.624 | 0.003 | -0.04 | -0.04 | 0 | 0.38 | 0.34 | 0.3 | 0.13 | 0.53 | 0.36 |
| 6 | 127965850 | rs7758223 | 0.566 | 0.112 | -0.528 | 0.01 | 0.011 | 0.016 | 0.016 | 0 | 0.05 | 0.01 | 0.14 | 0.31 | 0.03 | 0.27 |
| 6 | 127977160 | rs579177 | 0.528 | 0.404 | -0.398 | -1.292 | -0.024 | -0.001 | -0.001 | 0.763 | 0.36 | 0.25 | 0.24 | 0 | 0.52 | 0.18 |
| 6 | 128013896 | rs674751 | 0.985 | 0.598 | -0.285 | -1.362 | -0.094 | 0.123 | 0.123 | 0.853 | 0.3 | 0.11 | 0.2 | 0 | 0.44 | 0.16 |
| 6 | 128023698 | rs498692 | 0.077 | 0.661 | -0.31 | -1.268 | -0.004 | 0.014 | 0.014 | 0 | 0.4 | 0.33 | 0.28 | 0.09 | 0.57 | 0.2 |
| 6 | 128025366 | rs12201026 | 1.085 | 0.676 | -0.43 | -1.259 | -0.114 | 0.147 | 0.147 | 0.941 | 0.3 | 0.1 | 0.2 | 0 | 0.44 | 0.16 |
| 6 | 128035339 | rs525267 | 0.714 | 0.668 | -0.42 | -1.256 | -0.083 | 0.125 | 0.125 | 0.565 | 0.33 | 0.13 | 0.24 | 0.09 | 0.49 | 0.18 |
| 6 | 128040676 | rs11758347 | -0.341 | 0.673 | -0.455 | -1.234 | 0.006 | 0.032 | 0.032 | 0 | 0.06 | 0.01 | 0.03 | 0 | 0.07 | 0.02 |
| 6 | 128042115 | rs527206 | 0.26 | 0.643 | -0.406 | -1.256 | -0.005 | 0.031 | 0.031 | 0 | 0.4 | 0.32 | 0.29 | 0.1 | 0.57 | 0.2 |
| 6 | 128047943 | rs486235 | 1.073 | 0.752 | -0.713 | -1.313 | -0.125 | 0.161 | 0.161 | 1.286 | 0.29 | 0.06 | 0.19 | 0 | 0.44 | 0.16 |
| 6 | 128051957 | rs7776202 | -0.224 | 0.418 | -0.034 | -1.45 | 0.001 | -0.044 | -0.044 | 0 | 0.54 | 0.57 | 0.69 | 0.94 | 0.42 | 0.8 |
| 6 | 128056719 | rs6940280 | -1.238 | 0.62 | 0.215 | -1.365 | 0.019 | -0.031 | -0.031 | 0 | 0.13 | 0.2 | 0.09 | 0.06 | 0.15 | 0.05 |
| 6 | 128062827 | rs9491858 | 0.071 | 0.721 | 0.181 | -1.628 | 0 | -0.085 | -0.085 | 0.424 | 0.43 | 0.48 | 0.3 | 0.06 | 0.53 | 0.19 |
| 6 | 128064363 | rs9491859 | -0.132 | 0.628 | 0.386 | -1.704 | 0.032 | 0.026 | 0.026 | 0 | 0.48 | 0.6 | 0.37 | 0.06 | 0.63 | 0.24 |
| 6 | 128121143 | rs12153861 | -1.315 | 0.616 | 0.118 | -1.272 | -0.008 | -0.033 | -0.033 | 0.224 | 0.09 | 0.13 | 0.03 | 0.01 | 0.05 | 0.04 |
| 6 | 128129380 | rs920992 | -1.178 | 0.627 | 0.135 | -1.267 | -0.003 | -0.02 | -0.02 | 0 | 0.08 | 0.11 | 0.03 | 0.01 | 0.05 | 0.04 |
| 6 | 128135898 | rs12214570 | -0.608 | 0.814 | -0.031 | -1.358 | 0 | -0.009 | -0.009 | 0.452 | 0.07 | 0.04 | 0.03 | 0.01 | 0.05 | 0.04 |

| | | | | | | | | | | | | | | | | |
|---|---|---|---|---|---|---|---|---|---|---|---|---|---|---|---|---|
| 6 | 128139197 | rs12663768 | -0.079 | 0.944 | 0.132 | -1.309 | 0.045 | 0.003 | 0.003 | 0 | 0.05 | 0.14 | 0.05 | 0.03 | 0.09 | 0.02 |
| 6 | 128147046 | rs12525945 | 0.139 | 0.208 | -0.005 | -0.851 | 0.007 | -0.005 | -0.005 | 0 | 0.15 | 0.13 | 0.2 | 0.37 | 0.08 | 0.41 |
| 6 | 128148815 | rs6901443 | -0.19 | 0.032 | 0.006 | -1.302 | 0.007 | 0.028 | 0.028 | 0 | 0.48 | 0.64 | 0.58 | 0.62 | 0.49 | 0.43 |
| 6 | 128149241 | rs6922159 | -1.009 | 0.057 | -0.033 | -1.308 | -0.036 | 0.055 | 0.055 | 0 | 0.63 | 0.76 | 0.78 | 0.99 | 0.56 | 0.84 |
| 6 | 128153284 | rs270002 | -2.144 | 0.005 | -0.17 | -1.349 | -0.13 | 0.161 | 0.161 | 0.942 | 0.74 | 0.97 | 0.82 | 1 | 0.61 | 0.87 |
| 6 | 128155948 | rs7767438 | 0.367 | 0.014 | 0.033 | -1.196 | -0.008 | -0.002 | -0.002 | 0 | 0.43 | 0.51 | 0.52 | 0.59 | 0.4 | 0.41 |
| 6 | 128162928 | rs4243492 | -1.404 | 0.81 | -0.291 | -0.977 | -0.014 | 0.089 | 0.089 | 0 | 0.69 | 0.88 | 0.82 | 0.99 | 0.68 | 0.88 |
| 6 | 128173126 | rs270028 | -0.446 | 0.971 | -0.347 | -0.979 | 0.016 | 0.062 | 0.062 | 0 | 0.57 | 0.31 | 0.67 | 0.83 | 0.62 | 0.66 |
| 6 | 128175722 | rs10484716 | 0.219 | 1.054 | -0.443 | -0.856 | -0.006 | -0.029 | -0.029 | 0.288 | 0.18 | 0.14 | 0.27 | 0.39 | 0.19 | 0.47 |
| 6 | 128184738 | rs270041 | 0.609 | 1.101 | -0.451 | -0.885 | -0.013 | -0.013 | -0.013 | 0.396 | 0.39 | 0.2 | 0.4 | 0.45 | 0.43 | 0.2 |
| 6 | 128187461 | rs11752561 | 1.163 | 1.078 | -0.817 | -1.144 | -0.042 | 0.044 | 0.044 | 0 | 0.08 | 0.02 | 0.04 | 0 | 0.11 | 0.14 |
| 6 | 128189717 | rs270050 | -1.68 | 0.327 | -1.273 | -1.556 | -0.038 | 0.067 | 0.067 | 0 | 0.8 | 0.97 | 0.83 | 0.95 | 0.77 | 0.88 |
| 6 | 128194473 | rs12194062 | 2.029 | 0.058 | -1.285 | -1.664 | -0.056 | 0.147 | 0.147 | 1.187 | 0.24 | 0.03 | 0.14 | 0 | 0.29 | 0.11 |
| 6 | 128207306 | rs270017 | 0.128 | -0.175 | -0.723 | -1.534 | 0.011 | 0.1 | 0.1 | 0 | 0.53 | 0.65 | 0.62 | 0.79 | 0.43 | 0.57 |
| 6 | 128211078 | rs156061 | 1.267 | -0.03 | -0.856 | -1.654 | -0.023 | 0.1 | 0.1 | 0 | 0.19 | 0.03 | 0.17 | 0.01 | 0.21 | 0.12 |
| 6 | 128214882 | rs156060 | -0.153 | -0.232 | -0.554 | -1.38 | 0.013 | 0.113 | 0.113 | 0 | 0.47 | 0.35 | 0.38 | 0.21 | 0.56 | 0.43 |
| 6 | 128222626 | rs156065 | -0.039 | -0.244 | -0.492 | -1.387 | -0.004 | 0.156 | 0.156 | 0 | 0.46 | 0.22 | 0.37 | 0.21 | 0.56 | 0.41 |
| 6 | 128223935 | rs12525721 | -0.023 | -0.273 | -0.14 | -1.005 | -0.026 | 0.13 | 0.13 | 0 | 0.13 | 0.23 | 0.23 | 0.37 | 0.09 | 0.36 |
| 6 | 128246717 | rs802713 | -0.796 | -0.434 | -0.427 | -1.122 | 0.003 | 0 | 0 | 0 | 0.6 | 0.46 | 0.6 | 0.59 | 0.67 | 0.77 |
| 6 | 128255965 | rs9402006 | 0.221 | -0.292 | -0.668 | -0.99 | -0.002 | 0.026 | 0.026 | 0 | 0.12 | 0.13 | 0.21 | 0.37 | 0.07 | 0.35 |
| 6 | 128262515 | rs802753 | -0.959 | -0.426 | -0.491 | -1.074 | -0.006 | -0.015 | -0.015 | 0 | 0.65 | 0.81 | 0.65 | 0.59 | 0.67 | 0.77 |
| 6 | 128265919 | rs802750 | -0.049 | -0.103 | -0.681 | -1.727 | -0.044 | 0.145 | 0.145 | 0.075 | 0.4 | 0.2 | 0.29 | 0.15 | 0.43 | 0.37 |
| 6 | 128266148 | rs802749 | -0.808 | -0.159 | -0.615 | -1.641 | -0.009 | 0.127 | 0.127 | 0 | 0.93 | 0.89 | 0.81 | 0.59 | 0.95 | 0.71 |
| 6 | 128268374 | rs11751484 | 1.477 | -0.242 | -0.71 | -1.685 | -0.033 | -0.069 | -0.069 | 0.017 | 0.27 | 0.04 | 0.18 | 0.33 | 0.26 | 0.17 |
| 6 | 128270067 | rs9491889 | 0.91 | -0.371 | -0.363 | -1.174 | -0.036 | 0.056 | 0.056 | 0.849 | 0.12 | 0.07 | 0.06 | 0 | 0.14 | 0.13 |
| 6 | 128272324 | rs802744 | -0.492 | -0.34 | -0.26 | -1.147 | -0.025 | 0.118 | 0.118 | 0.189 | 0.4 | 0.26 | 0.31 | 0.15 | 0.43 | 0.37 |
| 6 | 128275171 | rs802741 | -0.401 | -0.276 | -0.243 | -1.06 | 0.001 | 0.072 | 0.072 | 0 | 0.92 | 0.92 | 0.79 | 0.59 | 0.92 | 0.71 |
| 6 | 128278798 | rs802734 | -0.097 | -0.203 | -0.596 | -1.192 | -0.009 | 0.057 | 0.057 | 0 | 0.28 | 0.11 | 0.24 | 0.15 | 0.29 | 0.24 |
| 6 | 128281861 | rs802726 | -0.632 | -0.062 | -0.524 | -1.13 | -0.032 | 0.105 | 0.105 | 0.594 | 0.39 | 0.26 | 0.31 | 0.15 | 0.42 | 0.34 |
| 6 | 128284771 | rs802721 | -0.712 | -0.067 | -0.411 | -1.122 | -0.001 | -0.055 | -0.055 | 0 | 0.41 | 0.52 | 0.31 | 0.16 | 0.4 | 0.34 |
| 6 | 128287848 | rs761332 | 0.575 | -0.081 | -1.466 | -1.069 | -0.004 | -0.012 | -0.012 | 0 | 0.13 | 0.16 | 0.08 | 0 | 0.14 | 0.13 |
| 6 | 128289511 | rs802718 | 0.129 | -0.239 | -1.453 | -0.296 | 0.01 | -0.007 | -0.007 | 0 | 0.88 | 0.89 | 0.79 | 0.45 | 0.9 | 0.64 |
| 6 | 128291199 | rs3190930 | 0.033 | -0.1 | -1.389 | -0.652 | -0.007 | 0.046 | 0.046 | 0 | 0.2 | 0.1 | 0.2 | 0.02 | 0.23 | 0.13 |
| 6 | 128299370 | rs3901172 | 0.932 | -0.399 | -1.128 | 0.287 | 0.01 | -0.033 | -0.033 | 0 | 0.85 | 0.88 | 0.8 | 0.37 | 0.87 | 0.58 |
| 6 | 128300949 | rs9491896 | 0.244 | -0.184 | -1.173 | 0.203 | -0.005 | -0.011 | -0.011 | 0 | 0.1 | 0.05 | 0.04 | 0.3 | 0.09 | 0.11 |
| 6 | 128305888 | rs9402011 | 0.316 | -0.152 | -1.235 | -0.077 | -0.004 | -0.004 | -0.004 | 4.187 | 0.22 | 0.23 | 0.38 | 0.45 | 0.29 | 0.27 |
| 6 | 128306769 | rs6919966 | -0.845 | -0.3 | -0.975 | -0.109 | 0.004 | 0.013 | 0.013 | 0 | 0.4 | 0.47 | 0.46 | 0.51 | 0.41 | 0.4 |
| 6 | 128307725 | rs6905941 | 0.545 | -0.218 | -1.41 | -0.087 | -0.105 | -0.005 | -0.005 | 0 | 0.2 | 0.01 | 0.39 | 0.14 | 0.32 | 0.22 |
| 6 | 128311140 | rs9375543 | -0.545 | -0.047 | -1.28 | -0.35 | 0 | 0 | 0 | 0.017 | 0.35 | 0.34 | 0.46 | 0.45 | 0.41 | 0.33 |
| 6 | 128319352 | rs2024557 | 0.38 | 0.045 | -1.259 | -0.294 | -0.002 | -0.003 | -0.003 | 0 | 0.62 | 0.63 | 0.54 | 0.54 | 0.59 | 0.67 |
| 6 | 128320912 | rs9398863 | -0.244 | 0.384 | -1.189 | -0.138 | -0.002 | -0.033 | -0.033 | 0 | 0.86 | 0.9 | 0.95 | 1 | 0.9 | 0.94 |
| 6 | 128324241 | rs717851 | 0.255 | 0.333 | -1.193 | -0.155 | 0 | 0.008 | 0.008 | 0 | 0.58 | 0.53 | 0.54 | 0.48 | 0.59 | 0.59 |
| 6 | 128325577 | rs6569519 | 0.356 | 0.896 | -1.538 | -0.5 | -0.005 | -0.051 | -0.051 | 2.873 | 0.86 | 0.84 | 0.95 | 1 | 0.9 | 0.94 |
| 6 | 128334236 | rs2326671 | -0.465 | 1.378 | -1.716 | 0.214 | -0.116 | 0.157 | 0.157 | 0 | 0.3 | 0.03 | 0.25 | 0.05 | 0.34 | 0.17 |
| 6 | 128343763 | rs7776358 | -1.348 | 1.624 | 0.099 | 0.346 | 0.08 | -0.128 | -0.128 | 0.043 | 0.19 | 0.5 | 0.16 | 0.01 | 0.22 | 0.09 |
| 6 | 128343842 | rs10456973 | 0.244 | 1.097 | -0.146 | 0.008 | 0.017 | 0.034 | 0.034 | 0.134 | 0.6 | 0.31 | 0.45 | 0.19 | 0.66 | 0.54 |
| 6 | 128351143 | rs9482858 | -1.181 | 1.413 | 1.078 | 0.192 | 0.053 | -0.059 | -0.059 | 0.221 | 0.1 | 0.19 | 0.1 | 0 | 0.14 | 0.01 |
| 6 | 128367589 | rs9482868 | 1.985 | 1.318 | 1.443 | 0.384 | 0.051 | -0.048 | -0.048 | 0 | 0.84 | 0.54 | 0.88 | 0.95 | 0.83 | 0.94 |
| 6 | 128391766 | rs7757761 | 1.207 | 1.411 | 1.361 | 0.356 | 0.046 | -0.011 | -0.011 | 0.849 | 0.9 | 0.78 | 0.89 | 0.95 | 0.85 | 0.98 |
| 6 | 128394889 | rs4544930 | 1.115 | 1.649 | 1.139 | 0.23 | 0.021 | -0.045 | -0.045 | 0.035 | 0.87 | 0.82 | 0.89 | 0.99 | 0.83 | 0.98 |
| 6 | 128397007 | rs6912749 | 1.587 | 1.53 | 1.125 | 0.207 | 0.055 | -0.06 | -0.06 | 1.694 | 0.82 | 0.44 | 0.85 | 0.95 | 0.8 | 0.97 |
| 6 | 128403745 | rs17828130 | 0.666 | 1.057 | -0.035 | 0.184 | 0 | 0.032 | 0.032 | 0 | 0.55 | 0.12 | 0.65 | 0.59 | 0.64 | 0.57 |
| 6 | 128442857 | rs4072328 | -0.707 | 0.944 | 0.71 | 0.398 | -0.008 | -0.015 | -0.015 | 0.337 | 0.08 | 0.24 | 0.04 | 0.04 | 0.05 | 0.04 |
| 6 | 128450019 | rs4498385 | -1.081 | 1.318 | 0.627 | 0.316 | 0.013 | -0.023 | -0.023 | 0.21 | 0.11 | 0.24 | 0.07 | 0.04 | 0.12 | 0.07 |
| 6 | 128461788 | rs9491913 | -1.513 | 0.904 | 1.365 | 0.783 | -0.016 | 0.237 | 0.237 | 2.286 | 0.13 | 0.56 | 0.13 | 0.38 | 0.06 | 0.06 |
| 6 | 128530307 | rs1012049 | -0.953 | 1.004 | 0.573 | -0.004 | 0.027 | -0.011 | -0.011 | 0.028 | 0.36 | 0.89 | 0.28 | 0.36 | 0.28 | 0.35 |
| 6 | 128547177 | rs12528715 | -0.503 | 0.902 | 1.286 | 0.254 | -0.002 | -0.017 | -0.017 | 0 | 0.09 | 0.09 | 0.03 | 0.03 | 0.04 | 0.15 |
| 6 | 128566321 | rs9784880 | -1.661 | 0.999 | 1.674 | 0.382 | 0.008 | 0.33 | 0.33 | 0.365 | 0.06 | 0.33 | 0.09 | 0.33 | 0.02 | 0.03 |
| 6 | 128593134 | rs6923988 | -0.525 | 1.365 | 0.861 | -0.595 | -0.019 | 0.033 | 0.033 | 0 | 0.13 | 0.06 | 0.11 | 0 | 0.19 | 0.04 |
| 6 | 128609148 | rs11966128 | -1.783 | 1.114 | 1.441 | -0.41 | -0.001 | -0.059 | -0.059 | 0 | 0.11 | 0.32 | 0.06 | 0.01 | 0.07 | 0.05 |
| 6 | 128611211 | rs17055564 | -1.468 | 1.106 | 1.538 | -0.414 | -0.001 | -0.02 | -0.02 | 0.081 | 0.09 | 0.11 | 0.04 | 0.01 | 0.07 | 0.05 |
| 6 | 128615092 | rs17352289 | -0.794 | 0.786 | 1.659 | -0.691 | 0.011 | 0.013 | 0.013 | 0 | 0.15 | 0.09 | 0.04 | 0.04 | 0.07 | 0.13 |
| 6 | 128626631 | rs17456385 | -1.259 | 0.838 | 1.652 | -0.557 | 0 | -0.003 | -0.003 | 0 | 0.08 | 0.06 | 0.03 | 0.01 | 0.06 | 0.05 |
| 6 | 128632340 | rs17055612 | -1.566 | 0.857 | 1.946 | -0.545 | -0.002 | -0.038 | -0.038 | 0 | 0.09 | 0.14 | 0.05 | 0.01 | 0.07 | 0.05 |
| 6 | 128636055 | rs17055628 | -1.242 | 0.857 | 1.936 | -0.545 | -0.001 | -0.026 | -0.026 | 0 | 0.08 | 0.1 | 0.03 | 0.01 | 0.06 | 0.05 |
| 6 | 128641017 | rs7773248 | -1.556 | 0.854 | 2.046 | -0.546 | 0.007 | -0.05 | -0.05 | 1.232 | 0.1 | 0.28 | 0.06 | 0.01 | 0.06 | 0.05 |
| 6 | 128675821 | rs9491939 | 2.463 | 0.484 | 1.881 | 0.005 | 0.008 | -0.011 | -0.011 | 0 | 0.81 | 0.2 | 0.85 | 0.64 | 0.91 | 0.88 |
| 6 | 128691353 | rs7748155 | 2.266 | 0.943 | 2.233 | -0.19 | 0.025 | -0.12 | -0.12 | 0.207 | 0.83 | 0.5 | 0.87 | 0.99 | 0.84 | 0.87 |
| 6 | 128709493 | rs953549 | 2.335 | 0.581 | 2.227 | 0.111 | -0.005 | 0.006 | 0.006 | 12.388 | 0.84 | 0.21 | 0.85 | 0.65 | 0.92 | 0.88 |

| | | | | | | | | | | | | | | | | |
|---|---|---|---|---|---|---|---|---|---|---|---|---|---|---|---|---|
| 6 | 128740387 | rs3822945 | -2.266 | 0.879 | 1.335 | -0.134 | 0.032 | -0.017 | -0.017 | 0 | 0.36 | 0.9 | 0.22 | 0.4 | 0.24 | 0.31 |
| 6 | 128741476 | rs9398869 | -0.638 | 0.964 | 1.41 | -0.751 | -0.003 | 0.046 | 0.046 | 0 | 0.15 | 0.02 | 0.05 | 0.06 | 0.09 | 0.14 |
| 6 | 128745590 | rs17055854 | -1.63 | 1.155 | 1.919 | -0.419 | 0.027 | -0.054 | -0.054 | 0.006 | 0.1 | 0.45 | 0.07 | 0.01 | 0.06 | 0.05 |
| 6 | 128760481 | rs7741248 | -2.019 | 1.051 | 1.453 | -0.225 | 0.013 | -0.063 | -0.063 | 0.092 | 0.11 | 0.43 | 0.06 | 0.01 | 0.05 | 0.14 |
| 6 | 128769272 | rs1935129 | -1.963 | 1.253 | 1.781 | -0.2 | 0.043 | -0.025 | -0.025 | 0.44 | 0.07 | 0.25 | 0.03 | 0.01 | 0.05 | 0.06 |
| 6 | 128782919 | rs6914064 | -1.504 | 1.373 | 0.857 | 0.024 | -0.059 | 0.018 | 0.018 | 0.249 | 0.45 | 0.9 | 0.43 | 0.69 | 0.39 | 0.41 |
| 6 | 128790787 | rs9491960 | -1.848 | 1.522 | 1.759 | -0.102 | 0.039 | -0.115 | -0.115 | 0.002 | 0.19 | 0.35 | 0.13 | 0.01 | 0.23 | 0.18 |
| 6 | 128795887 | rs17053280 | -1.933 | 1.487 | 2.023 | -0.12 | 0.047 | -0.13 | -0.13 | 0 | 0.19 | 0.39 | 0.13 | 0.01 | 0.22 | 0.18 |
| 6 | 128813085 | rs17460940 | -0.777 | 1.643 | 1.816 | -0.041 | -0.037 | 0.042 | 0.042 | 0.064 | 0.1 | 0.02 | 0.09 | 0 | 0.17 | 0.1 |
| 6 | 128814111 | rs9482888 | -1.053 | 1.511 | 1.826 | -0.144 | 0.004 | -0.039 | -0.039 | 0.973 | 0.08 | 0.17 | 0.03 | 0 | 0.04 | 0.03 |
| 6 | 128815679 | rs13196432 | -2.346 | 1.506 | 2.173 | -0.002 | 0.103 | -0.001 | -0.001 | 0 | 0.08 | 0.34 | 0.04 | 0.02 | 0.06 | 0.09 |
| 6 | 128829399 | rs6938035 | -1.586 | 1.156 | 1.736 | 0.495 | 0 | 0.004 | 0.004 | 0.062 | 0.37 | 0.73 | 0.49 | 0.7 | 0.32 | 0.41 |
| 6 | 128832425 | rs17364090 | -2.043 | 1.916 | 2.077 | 0.654 | 0.014 | -0.026 | -0.026 | 0 | 0.09 | 0.14 | 0.08 | 0 | 0.16 | 0.09 |
| 6 | 128833290 | rs17364118 | -2.06 | 1.912 | 2.228 | 0.652 | 0.034 | -0.041 | -0.041 | 1.732 | 0.09 | 0.17 | 0.08 | 0 | 0.16 | 0.09 |
| 6 | 128839470 | rs6923822 | -2.896 | 1.931 | 2.613 | 0.718 | 0.135 | -0.052 | -0.052 | 0.018 | 0.11 | 0.33 | 0.08 | 0.02 | 0.15 | 0.12 |
| 6 | 128850216 | rs10085247 | -2.365 | 1.907 | 2.462 | 0.753 | 0.068 | -0.038 | -0.038 | 0 | 0.08 | 0.18 | 0.07 | 0 | 0.14 | 0.09 |
| 6 | 128879393 | rs9492001 | -2.861 | 1.9 | 3.269 | 0.729 | 0.185 | -0.029 | -0.029 | 0.042 | 0.1 | 0.48 | 0.08 | 0.03 | 0.13 | 0.13 |
| 6 | 128900663 | rs1341585 | -2.054 | 1.935 | 3.139 | 0.744 | 0.138 | 0.012 | 0.012 | 0.892 | 0.06 | 0.3 | 0.06 | 0 | 0.09 | 0.05 |
| 6 | 128904306 | rs2326702 | -2.329 | 1.999 | 3.115 | 0.747 | 0.261 | 0.02 | 0.02 | 0.632 | 0.07 | 0.39 | 0.08 | 0 | 0.13 | 0.06 |
| 6 | 128929094 | rs9482905 | -1.989 | 2.15 | 2.682 | 1.543 | 0.033 | 0.219 | 0.219 | 0 | 0.06 | 0.24 | 0.21 | 0.31 | 0.06 | 0.06 |
| 6 | 128938905 | rs12208509 | -0.794 | 2.409 | 2.257 | 1.21 | -0.005 | 0.009 | 0.009 | 0 | 0.07 | 0.06 | 0.05 | 0 | 0.09 | 0.05 |
| 6 | 128941251 | rs6932252 | -2.358 | 2.302 | 2.535 | 1.324 | 0.167 | 0.212 | 0.212 | 0.005 | 0.11 | 0.43 | 0.27 | 0.31 | 0.17 | 0.11 |
| 6 | 128951005 | rs17463514 | -0.91 | 1.733 | 1.491 | 0.838 | -0.033 | 0.038 | 0.038 | 1.924 | 0.12 | 0.04 | 0.11 | 0.03 | 0.17 | 0.26 |
| 6 | 128955282 | rs9372897 | -1.864 | 1.72 | 2.124 | 1.213 | 0.031 | 0.024 | 0.024 | 2.913 | 0.19 | 0.35 | 0.25 | 0.26 | 0.21 | 0.36 |
| 6 | 128963710 | rs12201837 | -1.999 | 1.603 | 2.022 | 0.651 | 0.018 | 0.002 | 0.002 | 1.349 | 0.23 | 0.38 | 0.27 | 0.27 | 0.24 | 0.35 |
| 6 | 128965521 | rs17056165 | -0.94 | 1.29 | 2.111 | 0.362 | -0.013 | -0.03 | -0.03 | 1.364 | 0.06 | 0.11 | 0.05 | 0 | 0.03 | 0.05 |
| 6 | 128973435 | rs6569539 | -2.236 | 0.496 | 2.249 | -0.241 | -0.004 | -0.032 | -0.032 | 0 | 0.35 | 0.53 | 0.32 | 0.27 | 0.3 | 0.41 |
| 6 | 128980678 | rs542925 | -1.534 | 0.681 | 2.26 | -0.057 | 0.015 | 0.04 | 0.04 | 0.515 | 0.22 | 0.31 | 0.37 | 0.59 | 0.25 | 0.36 |
| 6 | 128981829 | rs471145 | -2.75 | 0.625 | 2.265 | -0.093 | 0.009 | -0.003 | -0.003 | 0 | 0.41 | 0.8 | 0.49 | 0.59 | 0.34 | 0.46 |
| 6 | 128989302 | rs6933943 | 2.783 | 0.484 | 1.444 | -0.641 | -0.104 | 0.041 | 0.041 | 0 | 0.43 | 0.09 | 0.35 | 0.28 | 0.5 | 0.36 |
| 6 | 128994200 | rs6901011 | 0.826 | 0.768 | 1.231 | -1.113 | -0.026 | 0.035 | 0.035 | 0 | 0.09 | 0.04 | 0.1 | 0 | 0.11 | 0.03 |
| 6 | 128994522 | rs6921491 | 0.319 | 0.66 | 1.712 | -0.816 | -0.001 | -0.022 | -0.022 | 0 | 0.11 | 0.1 | 0.04 | 0 | 0.05 | 0.05 |
| 6 | 128994545 | rs7758040 | -0.873 | 0.652 | 1.76 | -0.822 | -0.008 | -0.045 | -0.045 | 0 | 0.17 | 0.19 | 0.1 | 0 | 0.09 | 0.1 |
| 6 | 128998864 | rs7766952 | 0.316 | 0.664 | 1.689 | -0.81 | -0.001 | -0.022 | -0.022 | 0 | 0.11 | 0.1 | 0.04 | 0 | 0.05 | 0.05 |
| 6 | 129003449 | rs1396297 | -1.355 | 0.726 | 1.796 | -0.764 | 0.029 | 0.088 | 0.088 | 0.43 | 0.22 | 0.4 | 0.38 | 0.57 | 0.25 | 0.36 |
| 6 | 129009593 | rs9402063 | -1.514 | 0.719 | 1.802 | -0.768 | 0.018 | 0.06 | 0.06 | 0.017 | 0.23 | 0.39 | 0.39 | 0.57 | 0.26 | 0.37 |
| 6 | 129022203 | rs506953 | -2.382 | 0.665 | 1.301 | -0.856 | -0.009 | -0.069 | -0.069 | 0 | 0.42 | 0.91 | 0.5 | 0.56 | 0.35 | 0.47 |
| 6 | 129039345 | rs7754832 | -2.052 | 0.014 | 1.986 | -0.649 | -0.024 | 0.067 | 0.067 | 0 | 0.26 | 0.62 | 0.37 | 0.52 | 0.18 | 0.21 |
| 6 | 129048640 | rs9492055 | -1.228 | -0.047 | 1.807 | -0.857 | -0.044 | -0.127 | -0.127 | 0 | 0.2 | 0.48 | 0.13 | 0.01 | 0.11 | 0.13 |
| 6 | 129051683 | rs2326743 | -1.392 | -0.02 | 1.607 | -0.661 | 0.028 | 0.386 | 0.386 | 0.005 | 0.09 | 0.36 | 0.26 | 0.52 | 0.08 | 0.08 |
| 6 | 129059014 | rs9482919 | -0.79 | -0.056 | 0.985 | -0.841 | 0.017 | 0.021 | 0.021 | 0 | 0.17 | 0.1 | 0.09 | 0.01 | 0.1 | 0.13 |
| 6 | 129059113 | rs774284 | -2.372 | -0.032 | 0.681 | -0.769 | 0.051 | -0.111 | -0.111 | 0 | 0.3 | 0.88 | 0.39 | 0.52 | 0.18 | 0.21 |
| 6 | 129062588 | rs774280 | -0.044 | -0.481 | 0.548 | -2.532 | -0.034 | 0.071 | 0.071 | 0 | 0.71 | 0.89 | 0.71 | 0.85 | 0.66 | 0.7 |
| 6 | 129072536 | rs9482920 | 1.263 | -0.75 | 1.261 | -2.223 | -0.057 | -0.133 | -0.133 | 1.72 | 0.8 | 0.5 | 0.88 | 0.99 | 0.9 | 0.87 |
| 6 | 129093272 | rs9372901 | 1.306 | -0.679 | 0.622 | -2.325 | 0.026 | 0.174 | 0.174 | 0 | 0.46 | 0.14 | 0.57 | 0.8 | 0.6 | 0.59 |
| 6 | 129094378 | rs9402072 | 1.584 | -0.647 | 0.607 | -2.334 | -0.051 | 0.128 | 0.128 | 0 | 0.41 | 0.11 | 0.47 | 0.79 | 0.53 | 0.52 |
| 6 | 129114003 | rs2661908 | -1.699 | -0.943 | 1.221 | -2.355 | -0.031 | -0.089 | -0.089 | 0 | 0.21 | 0.63 | 0.11 | 0.01 | 0.11 | 0.11 |
| 6 | 129125184 | rs884557 | -0.456 | -0.864 | 1.682 | -2.289 | 0.026 | -0.037 | -0.037 | 0 | 0.11 | 0.46 | 0.06 | 0.03 | 0.07 | 0.05 |
| 6 | 129138471 | rs9482936 | -1.725 | -0.957 | 1.992 | -2.355 | -0.052 | -0.087 | -0.087 | 0 | 0.13 | 0.39 | 0.07 | 0 | 0.04 | 0.06 |
| 6 | 129139283 | rs2661897 | 0.999 | -1.096 | 1.865 | -2.518 | 0.016 | 0.046 | 0.046 | 0 | 0.52 | 0.39 | 0.52 | 0.77 | 0.59 | 0.58 |
| 6 | 129141297 | rs6569558 | -0.005 | -1.048 | 1.412 | -2.802 | 0.006 | 0.066 | 0.066 | 0 | 0.68 | 0.83 | 0.71 | 0.84 | 0.69 | 0.71 |
| 6 | 129141326 | rs6569559 | 1.192 | -1.043 | 1.195 | -2.808 | 0.005 | -0.002 | -0.002 | 0 | 0.51 | 0.51 | 0.62 | 0.83 | 0.58 | 0.56 |
| 6 | 129149507 | rs6902060 | 1.464 | -0.949 | 0.957 | -2.736 | 0.029 | 0.216 | 0.216 | 0.505 | 0.47 | 0.16 | 0.53 | 0.83 | 0.58 | 0.58 |
| 6 | 129156907 | rs6917682 | -1.271 | -0.775 | 1.188 | -2.12 | -0.027 | -0.05 | -0.05 | 0 | 0.09 | 0.18 | 0.05 | 0 | 0.04 | 0.1 |
| 6 | 129167449 | rs9492129 | -0.966 | -0.588 | 1.019 | -1.971 | -0.002 | -0.047 | -0.047 | 0.865 | 0.1 | 0.3 | 0.04 | 0.01 | 0.06 | 0.06 |
| 6 | 129169909 | rs4487603 | -0.89 | -0.274 | 0.624 | -2.103 | 0.007 | 0.181 | 0.181 | 2.365 | 0.46 | 0.84 | 0.39 | 0.17 | 0.44 | 0.38 |
| 6 | 129173136 | rs6569567 | -1.372 | -0.258 | 0.525 | -2.095 | -0.042 | 0.235 | 0.235 | 1.546 | 0.52 | 0.88 | 0.42 | 0.17 | 0.5 | 0.44 |
| 6 | 129174561 | rs4532461 | -1.104 | -0.085 | 0.639 | -1.905 | 0.007 | 0.096 | 0.096 | 1.3 | 0.45 | 0.7 | 0.37 | 0.13 | 0.44 | 0.38 |
| 6 | 129175472 | rs4074861 | -0.622 | 0.556 | 0.141 | -1.534 | -0.001 | -0.017 | -0.017 | 0 | 0.34 | 0.34 | 0.31 | 0.13 | 0.42 | 0.35 |
| 6 | 129185534 | rs17056587 | -0.662 | 0.246 | 0.49 | -0.993 | -0.014 | -0.029 | -0.029 | 1.054 | 0.06 | 0.12 | 0.05 | 0 | 0.02 | 0.01 |
| 6 | 129188482 | rs4512245 | -1.576 | 0.082 | 0.588 | -1.799 | 0.011 | 0.027 | 0.027 | 0.02 | 0.37 | 0.67 | 0.24 | 0.1 | 0.33 | 0.28 |
| 6 | 129190200 | rs10457510 | -0.487 | -0.054 | 0.681 | -1.693 | -0.001 | 0.003 | 0.003 | 0 | 0.11 | 0.08 | 0.09 | 0.11 | 0.13 | 0.06 |
| 6 | 129201417 | rs13202274 | -0.291 | -0.098 | 0.88 | -1.786 | 0.002 | -0.006 | -0.006 | 0 | 0.1 | 0.1 | 0.05 | 0 | 0.11 | 0.11 |
| 6 | 129201696 | rs9372909 | -0.48 | 0.006 | 0.819 | -1.723 | -0.003 | 0.004 | 0.004 | 0 | 0.11 | 0.08 | 0.1 | 0.1 | 0.13 | 0.06 |
| 6 | 129219050 | rs4285344 | -0.6 | -0.167 | 1.005 | -1.834 | -0.003 | -0.035 | -0.035 | 0.009 | 0.12 | 0.13 | 0.05 | 0 | 0.11 | 0.1 |
| 6 | 129219490 | rs12208997 | -1.202 | -0.302 | 1.57 | -1.732 | 0.001 | -0.011 | -0.011 | 0 | 0.16 | 0.3 | 0.12 | 0.14 | 0.17 | 0.13 |
| 6 | 129222295 | rs4551190 | -2.6 | -0.238 | 1.903 | -1.79 | 0.033 | 0.089 | 0.089 | 0 | 0.37 | 0.8 | 0.24 | 0.16 | 0.3 | 0.28 |
| 6 | 129235633 | rs9482954 | -1.873 | -0.474 | 2.081 | -1.726 | -0.041 | -0.158 | -0.158 | 0 | 0.21 | 0.5 | 0.12 | 0 | 0.14 | 0.15 |
| 6 | 129236954 | rs9482955 | -1.012 | -0.451 | 2.172 | -1.678 | -0.006 | -0.038 | -0.038 | 0 | 0.08 | 0.24 | 0.06 | 0 | 0.03 | 0.05 |

| | | | | | | | | | | | | | | | | |
|---|---|---|---|---|---|---|---|---|---|---|---|---|---|---|---|---|
| 6 | 129266168 | rs4897284 | 0.334 | -0.239 | 1.729 | -1.774 | -0.001 | 0.004 | 0.004 | 0 | 0.86 | 0.83 | 0.89 | 0.83 | 0.85 | 0.93 |
| 6 | 129280105 | rs4897285 | 1.674 | -0.196 | 1.221 | -1.755 | 0.032 | 0.082 | 0.082 | 0 | 0.62 | 0.21 | 0.76 | 0.84 | 0.7 | 0.73 |
| 6 | 129302280 | rs11154452 | -1.047 | -0.512 | 2.053 | -1.857 | -0.032 | -0.111 | -0.111 | 0.058 | 0.14 | 0.45 | 0.08 | 0 | 0.07 | 0.16 |
| 6 | 129312758 | rs4594973 | 1.403 | -0.47 | 1.213 | -2.504 | 0.01 | 0.236 | 0.236 | 0 | 0.57 | 0.15 | 0.72 | 0.96 | 0.67 | 0.67 |
| 6 | 129314221 | rs9285464 | -1.175 | -0.546 | 1.913 | -2.593 | 0.003 | -0.105 | -0.105 | 0.441 | 0.32 | 0.41 | 0.21 | 0.01 | 0.29 | 0.22 |
| 6 | 129329546 | rs6569573 | -1.402 | -0.553 | 2.229 | -2.601 | -0.001 | -0.104 | -0.104 | 0.093 | 0.34 | 0.63 | 0.24 | 0.01 | 0.28 | 0.22 |
| 6 | 129330021 | rs10499146 | -0.427 | -0.498 | 2.368 | -2.644 | 0.001 | -0.059 | -0.059 | 0 | 0.16 | 0.21 | 0.07 | 0.01 | 0.15 | 0.18 |
| 6 | 129344023 | rs4895840 | 0.978 | -0.523 | 2.233 | -2.616 | -0.028 | -0.011 | -0.011 | 0 | 0.9 | 0.8 | 0.97 | 0.97 | 0.97 | 0.89 |
| 6 | 129344194 | rs4895841 | 1.808 | -0.521 | 2.192 | -2.66 | 0.02 | 0.079 | 0.079 | 0 | 0.57 | 0.34 | 0.75 | 0.96 | 0.68 | 0.66 |
| 6 | 129346877 | rs9375609 | 1.876 | -0.476 | 1.801 | -2.637 | 0.003 | 0.285 | 0.285 | 0 | 0.56 | 0.12 | 0.72 | 0.96 | 0.68 | 0.67 |
| 6 | 129349030 | rs4576277 | 1.868 | -0.476 | 1.806 | -2.637 | 0.014 | 0.262 | 0.262 | 0 | 0.56 | 0.16 | 0.72 | 0.96 | 0.68 | 0.67 |
| 6 | 129351484 | rs4076090 | 1.577 | -0.329 | 2.006 | -2.562 | 0 | 0.048 | 0.048 | 0 | 0.63 | 0.26 | 0.76 | 0.96 | 0.69 | 0.67 |
| 6 | 129364595 | rs10457513 | -0.487 | -0.266 | 2.033 | -2.548 | 0 | -0.032 | -0.032 | 0 | 0.16 | 0.18 | 0.07 | 0.01 | 0.15 | 0.18 |
| 6 | 129367953 | rs9402095 | 1.8 | -0.238 | 1.686 | -2.662 | -0.026 | 0.223 | 0.223 | 0.065 | 0.51 | 0.15 | 0.67 | 0.83 | 0.56 | 0.63 |
| 6 | 129368803 | rs11154461 | -1.058 | -0.221 | 1.81 | -2.572 | -0.005 | -0.05 | -0.05 | 0.213 | 0.29 | 0.34 | 0.17 | 0.13 | 0.28 | 0.23 |
| 6 | 129371533 | rs9402098 | 0.136 | -0.175 | 1.749 | -2.502 | -0.007 | 0.002 | 0.002 | 0 | 0.95 | 0.89 | 0.98 | 0.97 | 0.97 | 0.89 |
| 6 | 129383680 | rs12190465 | -1.044 | -0.13 | 1.842 | -2.553 | -0.004 | -0.035 | -0.035 | 3.262 | 0.29 | 0.32 | 0.17 | 0.14 | 0.28 | 0.23 |
| 6 | 129383986 | rs10456976 | -0.902 | -0.155 | 1.977 | -2.561 | -0.009 | -0.067 | -0.067 | 0 | 0.3 | 0.4 | 0.17 | 0.14 | 0.28 | 0.23 |
| 6 | 129390242 | rs9482978 | -0.495 | -0.057 | 1.839 | -2.342 | 0.036 | -0.116 | -0.116 | 0.007 | 0.16 | 0.45 | 0.16 | 0 | 0.13 | 0.04 |
| 6 | 129391246 | rs17782500 | -0.06 | -0.025 | 1.2 | -2.321 | -0.008 | 0.002 | 0.002 | 0 | 0.13 | 0.1 | 0.13 | 0 | 0.13 | 0.04 |
| 6 | 129394663 | rs13216682 | 1.132 | 0.069 | 0.591 | -2.638 | -0.036 | 0.083 | 0.083 | 0.361 | 0.15 | 0.04 | 0.12 | 0 | 0.2 | 0.04 |
| 6 | 129408902 | rs4569989 | -0.675 | -0.289 | 1.516 | -1.933 | -0.01 | -0.027 | -0.027 | 0 | 0.19 | 0.45 | 0.13 | 0.13 | 0.15 | 0.06 |
| 6 | 129421876 | rs7771065 | -1.535 | -0.564 | 1.771 | -1.73 | -0.022 | -0.048 | -0.048 | 0 | 0.27 | 0.43 | 0.16 | 0.17 | 0.2 | 0.22 |
| 6 | 129422405 | rs7751685 | -1.158 | -0.48 | 2.126 | -1.676 | 0.039 | 0.081 | 0.081 | 0 | 0.05 | 0.21 | 0.05 | 0.13 | 0.06 | 0.1 |
| 6 | 129425564 | rs9388686 | 1.709 | -0.483 | 2.247 | -1.816 | -0.008 | -0.023 | -0.023 | 0.079 | 0.75 | 0.35 | 0.85 | 0.84 | 0.82 | 0.74 |
| 6 | 129427422 | rs9492234 | -1.553 | -0.539 | 2.31 | -1.243 | 0.027 | 0.034 | 0.034 | 0 | 0.22 | 0.4 | 0.14 | 0.26 | 0.21 | 0.25 |
| 6 | 129435888 | rs899350 | 1.869 | -0.575 | 2.26 | -1.345 | -0.002 | -0.02 | -0.02 | 0 | 0.73 | 0.36 | 0.82 | 0.83 | 0.78 | 0.75 |
| 6 | 129442344 | rs2448012 | 1.747 | -0.532 | 2.277 | -1.322 | 0.002 | -0.021 | -0.021 | 0 | 0.75 | 0.35 | 0.84 | 0.84 | 0.79 | 0.75 |
| 6 | 129466298 | rs3778141 | -1.557 | -0.544 | 2.339 | -1.323 | 0.028 | -0.056 | -0.056 | 0 | 0.22 | 0.41 | 0.14 | 0.14 | 0.21 | 0.25 |
| 6 | 129473791 | rs17056866 | -1.015 | -0.656 | 1.713 | 0.134 | 0.003 | -0.067 | -0.067 | 0 | 0.11 | 0.01 | 0.18 | 0.51 | 0.06 | 0.15 |
| 6 | 129483375 | rs17056876 | -1.217 | -0.641 | 2.513 | -0.082 | -0.013 | -0.011 | -0.011 | 0.089 | 0.07 | 0.17 | 0.03 | 0.04 | 0.02 | 0.08 |
| 6 | 129505485 | rs1478808 | -1.915 | -0.78 | 2.476 | -0.248 | 0.004 | -0.12 | -0.12 | 0 | 0.24 | 0.53 | 0.14 | 0.04 | 0.2 | 0.15 |
| 6 | 129505832 | rs9492262 | -2.628 | -1.016 | 1.986 | -0.52 | 0.021 | 0.104 | 0.104 | 0 | 0.38 | 0.79 | 0.22 | 0.17 | 0.28 | 0.32 |
| 6 | 129509207 | rs2876021 | -2.15 | -0.978 | 2.027 | -0.474 | 0.003 | 0.013 | 0.013 | 0.05 | 0.28 | 0.71 | 0.17 | 0.17 | 0.22 | 0.25 |
| 6 | 129519697 | rs12215704 | -1.942 | -0.958 | 2.195 | -0.504 | -0.016 | -0.131 | -0.131 | 0 | 0.31 | 0.45 | 0.17 | 0.03 | 0.22 | 0.2 |
| 6 | 129519893 | rs12195187 | -0.006 | -0.895 | 1.949 | -0.466 | 0 | -0.012 | -0.012 | 0 | 0.08 | 0.06 | 0.04 | 0 | 0.06 | 0.07 |
| 6 | 129525473 | rs7766548 | -2.438 | -0.953 | 2.034 | -0.393 | -0.004 | -0.014 | -0.014 | 0 | 0.35 | 0.61 | 0.2 | 0.17 | 0.24 | 0.31 |
| 6 | 129533619 | rs1478806 | -0.619 | -0.968 | 2.032 | 0.959 | -0.009 | -0.038 | -0.038 | 0.088 | 0.15 | 0.05 | 0.11 | 0.32 | 0.13 | 0.11 |
| 6 | 129537876 | rs265360 | 2.109 | -0.968 | 2.061 | 1.032 | 0.001 | -0.002 | -0.002 | 0 | 0.66 | 0.34 | 0.76 | 0.53 | 0.77 | 0.73 |
| 6 | 129540114 | rs265354 | 1.694 | -0.894 | 1.842 | 1.02 | 0.045 | 0.194 | 0.194 | 2.361 | 0.95 | 0.71 | 0.96 | 0.89 | 0.99 | 0.93 |
| 6 | 129551233 | rs265332 | 2.048 | -0.918 | 1.16 | 1.03 | 0.02 | -0.084 | -0.084 | 0 | 0.65 | 0.15 | 0.7 | 0.53 | 0.75 | 0.72 |
| 6 | 129554948 | rs10447440 | 0.234 | -0.37 | 0.878 | 1.344 | -0.056 | -0.066 | -0.066 | 0.001 | 0.16 | 0.01 | 0.21 | 0.22 | 0.16 | 0.2 |
| 6 | 129561490 | rs265379 | 1.572 | -0.625 | 2.39 | 1.742 | -0.015 | 0.068 | 0.068 | 0.003 | 0.76 | 0.48 | 0.76 | 0.37 | 0.83 | 0.75 |
| 6 | 129564944 | rs265387 | 1.933 | -0.59 | 1.778 | 1.517 | -0.003 | 0.009 | 0.009 | 0 | 0.7 | 0.4 | 0.71 | 0.36 | 0.76 | 0.71 |
| 6 | 129570213 | rs7751571 | -1.283 | -0.226 | 1.815 | 1.413 | 0.003 | -0.026 | -0.026 | 6.779 | 0.17 | 0.23 | 0.11 | 0.08 | 0.15 | 0.24 |
| 6 | 129571330 | rs3816665 | -1.82 | -0.293 | 2.413 | 1.378 | 0.001 | -0.069 | -0.069 | 0 | 0.19 | 0.6 | 0.13 | 0.08 | 0.16 | 0.24 |
| 6 | 129574015 | rs7754620 | -1.797 | -0.268 | 2.423 | 1.396 | 0.003 | -0.065 | -0.065 | 1.327 | 0.19 | 0.59 | 0.13 | 0.11 | 0.16 | 0.25 |
| 6 | 129580545 | rs265401 | -2.508 | -0.512 | 1.99 | 1.751 | 0.023 | 0.035 | 0.035 | 0 | 0.35 | 0.79 | 0.38 | 0.76 | 0.28 | 0.39 |
| 6 | 129583473 | rs3823006 | -1.37 | -0.274 | 1.664 | 1.657 | 0.003 | -0.032 | -0.032 | 0.56 | 0.18 | 0.26 | 0.11 | 0.1 | 0.16 | 0.24 |
| 6 | 129590111 | rs3778119 | -1.269 | -0.275 | 1.367 | 1.641 | -0.004 | 0.007 | 0.007 | 0.012 | 0.18 | 0.2 | 0.12 | 0.11 | 0.17 | 0.26 |
| 6 | 129591515 | rs187851 | -0.42 | -0.318 | 1.005 | 2.22 | 0.012 | -0.037 | -0.037 | 0 | 0.1 | 0.01 | 0.1 | 0.44 | 0.09 | 0.05 |
| 6 | 129592284 | rs2448011 | -1.015 | -0.317 | 1.078 | 1.654 | 0.014 | -0.028 | -0.028 | 0 | 0.14 | 0.07 | 0.23 | 0.64 | 0.11 | 0.14 |
| 6 | 129594759 | rs265395 | 1.474 | -0.106 | 1.464 | 1.101 | 0.015 | -0.079 | -0.079 | 0.011 | 0.8 | 0.58 | 0.87 | 0.87 | 0.83 | 0.73 |
| 6 | 129604372 | rs2326794 | 1.58 | -0.458 | 1.187 | 1.632 | -0.001 | 0 | 0 | 0 | 0.69 | 0.34 | 0.76 | 0.55 | 0.76 | 0.74 |
| 6 | 129605056 | rs902370 | 1.998 | -0.436 | 1.127 | 1.75 | 0.039 | -0.105 | -0.105 | 0 | 0.65 | 0.16 | 0.62 | 0.49 | 0.74 | 0.68 |
| 6 | 129612273 | rs1994620 | 1.392 | -0.471 | 1.547 | 1.672 | -0.032 | 0.082 | 0.082 | 0 | 0.73 | 0.5 | 0.76 | 0.53 | 0.77 | 0.71 |
| 6 | 129612563 | rs1015845 | 2.007 | -0.479 | 1.41 | 1.506 | 0.011 | -0.015 | -0.015 | 0 | 0.66 | 0.32 | 0.62 | 0.46 | 0.74 | 0.65 |
| 6 | 129616516 | rs2101575 | 1.324 | -0.366 | 1.486 | 1.202 | 0.003 | -0.052 | -0.052 | 0.587 | 0.82 | 0.62 | 0.88 | 0.88 | 0.83 | 0.77 |
| 6 | 129619410 | rs12206487 | -0.488 | -0.465 | 1.062 | 1.599 | 0.015 | -0.019 | -0.019 | 0.79 | 0.11 | 0.1 | 0.24 | 0.42 | 0.08 | 0.1 |
| 6 | 129620506 | rs12208401 | 1.192 | -0.077 | 0.401 | 0.879 | -0.056 | 0.104 | 0.104 | 0 | 0.56 | 0.11 | 0.56 | 0.71 | 0.54 | 0.51 |
| 6 | 129620833 | rs6926104 | -1.215 | -0.154 | 1.107 | 0.798 | 0.012 | -0.038 | -0.038 | 0.224 | 0.34 | 0.52 | 0.38 | 0.16 | 0.42 | 0.37 |
| 6 | 129621144 | rs9385485 | -1.326 | -0.142 | 1.11 | 0.846 | -0.028 | 0.063 | 0.063 | 0.807 | 0.2 | 0.38 | 0.28 | 0.54 | 0.12 | 0.23 |
| 6 | 129630354 | rs9321159 | 2.162 | -0.032 | 1.068 | 0.828 | -0.046 | 0.069 | 0.069 | 0 | 0.46 | 0.09 | 0.34 | 0.31 | 0.46 | 0.41 |
| 6 | 129635800 | rs2306942 | -0.823 | -0.158 | 1.101 | 0.789 | -0.015 | 0.02 | 0.02 | 0.223 | 0.08 | 0.22 | 0.06 | 0.11 | 0.04 | 0.12 |
| 6 | 129637915 | rs1490392 | 1.399 | -0.11 | 1.134 | 0.794 | -0.007 | 0.002 | 0.002 | 0 | 0.55 | 0.43 | 0.4 | 0.43 | 0.5 | 0.53 |
| 6 | 129640389 | rs2451686 | 0.468 | 0.173 | 1.23 | 0.393 | 0.003 | -0.009 | -0.009 | 0.779 | 0.93 | 0.93 | 0.94 | 0.96 | 0.89 | 0.92 |
| 6 | 129642470 | rs4642505 | -0.762 | 0.134 | 1.584 | 0.376 | -0.004 | 0.001 | 0.001 | 0 | 0.26 | 0.24 | 0.27 | 0.1 | 0.31 | 0.29 |
| 6 | 129644548 | rs17057149 | -0.439 | 0.225 | 1.827 | 0.283 | -0.002 | -0.024 | -0.024 | 0.116 | 0.12 | 0.15 | 0.18 | 0.05 | 0.09 | 0.12 |
| 6 | 129648704 | rs1994619 | 1.188 | 0.229 | 1.818 | 0.284 | 0.003 | -0.038 | -0.038 | 0 | 0.82 | 0.77 | 0.76 | 0.92 | 0.8 | 0.8 |

| | | | | | | | | | | | | | | | | |
|---|---|---|---|---|---|---|---|---|---|---|---|---|---|---|---|---|
| 6 | 129653602 | rs1472057 | 1.49 | 0.182 | 1.802 | 0.411 | -0.022 | 0.047 | 0.047 | 0 | 0.74 | 0.43 | 0.63 | 0.39 | 0.77 | 0.69 |
| 6 | 129659104 | rs9372924 | 2.477 | 0.172 | 1.728 | 0.362 | -0.045 | 0.138 | 0.138 | 0 | 0.46 | 0.1 | 0.34 | 0.27 | 0.46 | 0.4 |
| 6 | 129660222 | rs2451679 | 0.119 | -0.166 | 1.766 | 0.373 | -0.036 | 0.037 | 0.037 | 0.023 | 0.15 | 0.05 | 0.11 | 0.05 | 0.22 | 0.17 |
| 6 | 129661369 | rs11154471 | -1.181 | -0.379 | 1.857 | 0.354 | -0.002 | -0.056 | -0.056 | 0 | 0.27 | 0.31 | 0.27 | 0.09 | 0.25 | 0.28 |
| 6 | 129662357 | rs1873329 | -1.122 | -0.379 | 1.828 | 0.354 | 0 | 0.036 | 0.036 | 0 | 0.27 | 0.2 | 0.25 | 0.09 | 0.25 | 0.28 |
| 6 | 129663271 | rs1387918 | 2.418 | -0.384 | 1.77 | 0.371 | -0.015 | 0.11 | 0.11 | 0 | 0.47 | 0.11 | 0.36 | 0.3 | 0.52 | 0.41 |
| 6 | 129666012 | rs11154473 | -0.935 | -0.282 | 1.745 | 0.476 | -0.008 | 0.033 | 0.033 | 0 | 0.25 | 0.18 | 0.24 | 0.15 | 0.26 | 0.28 |
| 6 | 129666380 | rs7766742 | 2.465 | -0.287 | 1.545 | 0.464 | -0.09 | 0.183 | 0.183 | 0 | 0.47 | 0.05 | 0.35 | 0.22 | 0.51 | 0.41 |
| 6 | 129670548 | rs17057158 | -0.232 | -0.244 | 1.535 | 0.314 | 0.003 | 0.003 | 0.003 | 0 | 0.12 | 0.11 | 0.15 | 0.05 | 0.07 | 0.12 |
| 6 | 129671311 | rs9398901 | 1.466 | -0.249 | 1.597 | 0.312 | -0.006 | 0.012 | 0.012 | 0 | 0.67 | 0.74 | 0.67 | 0.88 | 0.63 | 0.63 |
| 6 | 129673136 | rs17799083 | 0.182 | -0.244 | 1.579 | 0.317 | -0.03 | 0.049 | 0.049 | 0 | 0.14 | 0.04 | 0.1 | 0.03 | 0.19 | 0.16 |
| 6 | 129674757 | rs3813367 | -0.55 | -0.35 | 1.612 | 0.383 | -0.01 | 0.017 | 0.017 | 0.006 | 0.07 | 0.16 | 0.05 | 0.11 | 0.04 | 0.12 |
| 6 | 129678863 | rs7453910 | 1.161 | -0.347 | 1.595 | 0.37 | 0.001 | -0.035 | -0.035 | 0 | 0.81 | 0.78 | 0.77 | 0.91 | 0.82 | 0.79 |
| 6 | 129686150 | rs9388696 | -1.557 | -0.373 | 1.625 | 0.375 | -0.007 | -0.002 | -0.002 | 0 | 0.33 | 0.62 | 0.41 | 0.63 | 0.31 | 0.39 |
| 6 | 129689145 | rs4296900 | 1.912 | -0.664 | 1.6 | 0.741 | -0.008 | -0.031 | -0.031 | 0 | 0.74 | 0.52 | 0.73 | 0.79 | 0.78 | 0.68 |
| 6 | 129698081 | rs9402119 | 1.588 | -0.559 | 1.386 | 0.81 | 0.012 | 0.025 | 0.025 | 0 | 0.86 | 0.72 | 0.88 | 0.83 | 0.85 | 0.79 |
| 6 | 129699599 | rs7356971 | -0.104 | -0.702 | 0.793 | 0.753 | -0.001 | 0.057 | 0.057 | 0 | 0.11 | 0.14 | 0.23 | 0.46 | 0.08 | 0.1 |
| 6 | 129700264 | rs6904763 | -1.035 | -0.335 | 0.445 | -0.035 | -0.005 | -0.026 | -0.026 | 0.326 | 0.15 | 0.17 | 0.19 | 0.05 | 0.1 | 0.12 |
| 6 | 129700998 | rs17057200 | -0.679 | -0.219 | 0.029 | 0.069 | -0.006 | 0.016 | 0.016 | 0 | 0.06 | 0.1 | 0.04 | 0.11 | 0.03 | 0.12 |
| 6 | 129702793 | rs6899448 | 1.187 | -0.202 | -0.006 | 0.081 | -0.001 | 0 | 0 | 0 | 0.86 | 0.79 | 0.88 | 0.85 | 0.85 | 0.79 |
| 6 | 129707118 | rs7770270 | -1.061 | -0.121 | -0.327 | 0.026 | 0.01 | 0.023 | 0.023 | 0 | 0.15 | 0.13 | 0.17 | 0.05 | 0.1 | 0.12 |
| 6 | 129715289 | rs2437095 | 0.551 | 0.218 | -0.287 | 0.079 | -0.003 | 0.01 | 0.01 | 0 | 0.83 | 0.84 | 0.78 | 0.92 | 0.8 | 0.74 |
| 6 | 129715459 | rs17057233 | -0.243 | 0.099 | -0.281 | 0.029 | 0 | 0.008 | 0.008 | 0.028 | 0.1 | 0.06 | 0.06 | 0.08 | 0.08 | 0.13 |
| 6 | 129715775 | rs2437093 | 0.673 | -0.265 | 0.268 | 0.629 | -0.018 | 0.032 | 0.032 | 0 | 0.71 | 0.57 | 0.55 | 0.46 | 0.75 | 0.64 |
| 6 | 129715822 | rs6914003 | 0.161 | -0.706 | -0.895 | -0.192 | 0.028 | 0.059 | 0.059 | 0 | 0.33 | 0.21 | 0.12 | 0.08 | 0.21 | 0.24 |
| 6 | 129716209 | rs2437092 | 0.163 | -0.556 | -0.517 | 0.028 | -0.019 | 0.043 | 0.043 | 0 | 0.73 | 0.45 | 0.53 | 0.46 | 0.75 | 0.64 |
| 6 | 129717246 | rs2437090 | 0.5 | 0.156 | -1.045 | 0.472 | -0.107 | -0.033 | -0.033 | 2.079 | 0.34 | 0.1 | 0.35 | 0.36 | 0.49 | 0.34 |
| 6 | 129720863 | rs2437086 | -0.232 | 0.141 | -0.688 | 0.873 | 0.006 | 0.013 | 0.013 | 0.329 | 0.55 | 0.71 | 0.54 | 0.64 | 0.59 | 0.6 |
| 6 | 129722389 | rs3749877 | 0.008 | 0.386 | -0.176 | 1.26 | -0.007 | -0.002 | -0.002 | 0 | 0.56 | 0.66 | 0.47 | 0.55 | 0.46 | 0.52 |
| 6 | 129722425 | rs3749878 | 0.016 | 0.384 | -0.176 | 1.248 | -0.007 | -0.003 | -0.003 | 0.344 | 0.56 | 0.66 | 0.47 | 0.55 | 0.46 | 0.52 |
| 6 | 129724894 | rs2326817 | 0.009 | 0.373 | -0.175 | 1.259 | -0.005 | -0.004 | -0.004 | 0 | 0.56 | 0.66 | 0.47 | 0.54 | 0.46 | 0.52 |
| 6 | 129734028 | rs4143750 | 0.049 | 0.393 | -0.02 | 1.203 | -0.001 | -0.008 | -0.008 | 0.432 | 0.54 | 0.51 | 0.41 | 0.36 | 0.45 | 0.46 |
| 6 | 129739884 | rs2326805 | 0.109 | 0.378 | -0.035 | 1.192 | 0.002 | 0.01 | 0.01 | 0 | 0.53 | 0.43 | 0.4 | 0.36 | 0.45 | 0.46 |
| 6 | 129762112 | rs2297738 | -0.641 | 0.43 | -0.243 | 1.279 | -0.001 | 0.002 | 0.002 | 0 | 0.15 | 0.15 | 0.23 | 0.22 | 0.16 | 0.17 |
| 6 | 129768893 | rs1414738 | -0.717 | 0.441 | -0.138 | 1.283 | 0.001 | 0.019 | 0.019 | 0 | 0.13 | 0.17 | 0.22 | 0.22 | 0.13 | 0.16 |
| 6 | 129783642 | rs7755729 | 0.676 | -0.086 | -0.275 | 1.057 | 0 | 0 | 0 | 0.014 | 0.46 | 0.42 | 0.39 | 0.38 | 0.45 | 0.44 |
| 6 | 129785391 | rs1414736 | -0.348 | -0.154 | -0.351 | 1.194 | -0.059 | -0.069 | -0.069 | 1.581 | 0.62 | 0.86 | 0.63 | 0.55 | 0.58 | 0.61 |
| 6 | 129788658 | rs6917532 | 0.659 | -0.149 | 0.021 | 1.105 | 0 | 0 | 0 | 0.175 | 0.46 | 0.49 | 0.39 | 0.33 | 0.46 | 0.44 |
| 6 | 129793971 | rs6905196 | 0.18 | -0.056 | -0.217 | 1.167 | -0.019 | 0.047 | 0.047 | 0 | 0.56 | 0.79 | 0.42 | 0.31 | 0.42 | 0.44 |
| 6 | 129798084 | rs2571577 | -1.025 | -0.234 | -0.008 | 0.911 | 0.001 | -0.02 | -0.02 | 0 | 0.22 | 0.2 | 0.29 | 0.3 | 0.2 | 0.37 |
| 6 | 129799682 | rs2784895 | 1.527 | -0.134 | 0.597 | 0.791 | 0 | 0.016 | 0.016 | 0 | 0.71 | 0.53 | 0.66 | 0.69 | 0.72 | 0.6 |
| 6 | 129808776 | rs7740552 | 1.545 | -0.158 | 0.586 | 0.952 | -0.004 | 0.019 | 0.019 | 0 | 0.69 | 0.41 | 0.62 | 0.62 | 0.72 | 0.6 |
| 6 | 129808901 | rs7740749 | 1.534 | -0.153 | 0.586 | 0.883 | -0.005 | 0.004 | 0.004 | 0 | 0.69 | 0.41 | 0.65 | 0.7 | 0.72 | 0.6 |
| 6 | 129813053 | rs2244008 | -1.217 | 0.014 | 0.351 | 0.819 | -0.002 | 0.012 | 0.012 | 0 | 0.06 | 0.12 | 0.07 | 0.13 | 0.07 | 0.15 |
| 6 | 129814998 | rs10499160 | -0.54 | -0.3 | -0.029 | 0.518 | -0.021 | 0.075 | 0.075 | 0.007 | 0.2 | 0.06 | 0.22 | 0.05 | 0.18 | 0.16 |
| 6 | 129822837 | rs9321170 | 1.148 | -0.298 | 0.195 | 0.623 | -0.006 | 0.006 | 0.006 | 0 | 0.67 | 0.42 | 0.63 | 0.64 | 0.72 | 0.66 |
| 6 | 129827022 | rs11154483 | 1.125 | -0.332 | 0.174 | 0.431 | -0.003 | -0.001 | -0.001 | 0 | 0.64 | 0.41 | 0.63 | 0.69 | 0.69 | 0.65 |
| 6 | 129833023 | rs2571574 | 0.733 | -0.663 | -0.118 | 0.399 | 0.015 | 0.001 | 0.001 | 0 | 0.7 | 0.76 | 0.71 | 0.75 | 0.77 | 0.68 |
| 6 | 129834205 | rs2571575 | 0.661 | -0.781 | 0.084 | 0.366 | -0.018 | -0.003 | -0.003 | 2.249 | 0.69 | 0.59 | 0.77 | 0.77 | 0.79 | 0.67 |
| 6 | 129834224 | rs2571576 | 1.085 | -0.827 | -0.242 | 0.031 | -0.006 | -0.007 | -0.007 | 0 | 0.57 | 0.5 | 0.63 | 0.74 | 0.66 | 0.62 |
| 6 | 129834629 | rs12205363 | 0.107 | -0.518 | -0.718 | 0.111 | -0.019 | 0.035 | 0.035 | 0 | 0.07 | 0.01 | 0.04 | 0 | 0.08 | 0.02 |
| 6 | 129835814 | rs2297740 | 0.077 | -1.389 | -0.8 | -0.017 | -0.061 | 0.017 | 0.017 | 0 | 0.72 | 0.54 | 0.81 | 0.73 | 0.85 | 0.68 |
| 6 | 129836336 | rs13208666 | 1.334 | -1.187 | -1.278 | 0.013 | -0.063 | -0.063 | -0.063 | 0 | 0.31 | 0.18 | 0.44 | 0.58 | 0.45 | 0.43 |
| 6 | 129838002 | rs6939307 | 0.245 | -1.124 | -1.19 | -0.432 | 0.001 | 0.002 | 0.002 | 0 | 0.3 | 0.32 | 0.2 | 0.03 | 0.29 | 0.17 |
| 6 | 129844019 | rs17809707 | -0.514 | -1.15 | -1.191 | -0.38 | 0.004 | 0.039 | 0.039 | 0 | 0.43 | 0.43 | 0.4 | 0.15 | 0.43 | 0.27 |
| 6 | 129844740 | rs12214392 | 0.187 | -0.996 | -1.238 | -0.327 | -0.012 | 0.095 | 0.095 | 0 | 0.29 | 0.2 | 0.21 | 0.04 | 0.31 | 0.18 |
| 6 | 129849509 | rs875128 | 0.715 | -0.244 | -0.825 | 0.466 | -0.04 | -0.027 | -0.027 | 0.641 | 0.29 | 0.2 | 0.33 | 0.66 | 0.45 | 0.41 |
| 6 | 129850455 | rs2571595 | 0.764 | -0.014 | -0.326 | 1.337 | 0.011 | 0.136 | 0.136 | 0 | 0.82 | 0.65 | 0.81 | 0.68 | 0.82 | 0.61 |
| 6 | 129854355 | rs977608 | -1.132 | 0.029 | -1.034 | 0.824 | 0.027 | 0.036 | 0.036 | 0.087 | 0.19 | 0.1 | 0.15 | 0.09 | 0.11 | 0.16 |
| 6 | 129856375 | rs9385497 | -0.916 | 0.056 | -0.94 | 0.904 | 0.018 | 0.001 | 0.001 | 0.038 | 0.18 | 0.1 | 0.15 | 0.13 | 0.11 | 0.16 |
| 6 | 129861392 | rs2275215 | -1.151 | 0.526 | -0.307 | 1.498 | 0.001 | 0.064 | 0.064 | 2.455 | 0.25 | 0.55 | 0.24 | 0.39 | 0.26 | 0.51 |
| 6 | 129862792 | rs2571591 | 0.843 | 0.616 | 0.415 | 2.458 | 0.023 | 0.117 | 0.117 | 1.025 | 0.83 | 0.69 | 0.77 | 0.5 | 0.75 | 0.62 |
| 6 | 129865823 | rs2571590 | 0.859 | 0.615 | 0.414 | 2.557 | 0.022 | 0.132 | 0.132 | 1.077 | 0.83 | 0.69 | 0.78 | 0.55 | 0.75 | 0.61 |
| 6 | 129866241 | rs2784890 | 0.472 | 0.639 | -0.618 | 1.693 | -0.053 | 0.096 | 0.096 | 0 | 0.41 | 0.21 | 0.49 | 0.3 | 0.43 | 0.43 |
| 6 | 129869267 | rs7760943 | -0.602 | 0.655 | -0.084 | 2.456 | 0.02 | 0.122 | 0.122 | 0 | 0.15 | 0.24 | 0.29 | 0.47 | 0.21 | 0.23 |
| 6 | 129872966 | rs10457525 | -0.673 | 0.809 | 0.206 | 2.579 | 0.017 | 0.046 | 0.046 | 0 | 0.15 | 0.34 | 0.26 | 0.2 | 0.17 | 0.19 |
| 6 | 129876535 | rs9402140 | 0.401 | 0.784 | -0.077 | 2.009 | -0.016 | -0.031 | -0.031 | 0 | 0.33 | 0.27 | 0.3 | 0.5 | 0.36 | 0.45 |
| 6 | 129896501 | rs10457526 | -0.541 | 0.489 | 0.249 | 2.184 | 0 | 0.011 | 0.011 | 1.107 | 0.16 | 0.17 | 0.37 | 0.24 | 0.2 | 0.33 |
| 6 | 129907587 | rs6935162 | 0.68 | 0.627 | 0.002 | 2.019 | -0.016 | 0.015 | 0.015 | 0 | 0.28 | 0.21 | 0.24 | 0.29 | 0.31 | 0.43 |

| | | | | | | | | | | | | | | | | |
|---|---|---|---|---|---|---|---|---|---|---|---|---|---|---|---|---|
| 6 | 129913087 | rs4504494 | -1.205 | 0.077 | 0.52 | 2.139 | -0.062 | 0.194 | 0.194 | 1.543 | 0.16 | 0.53 | 0.32 | 0.3 | 0.05 | 0.16 |
| 6 | 129918239 | rs7758025 | 0.694 | -0.37 | -0.001 | 1.359 | -0.013 | -0.016 | -0.016 | 0 | 0.22 | 0.27 | 0.14 | 0.07 | 0.15 | 0.06 |
| 6 | 129918743 | rs9385502 | 0.381 | -0.056 | -0.286 | 1.438 | 0.004 | 0.017 | 0.017 | 0 | 0.08 | 0.09 | 0.11 | 0.26 | 0.16 | 0.06 |
| 6 | 129920784 | rs9402145 | 0.54 | -0.029 | -0.378 | 0.918 | -0.01 | -0.007 | -0.007 | 0 | 0.07 | 0.07 | 0.15 | 0.13 | 0.16 | 0.06 |
| 6 | 129921216 | rs9321174 | -0.179 | -0.072 | -0.325 | 0.688 | -0.005 | 0.013 | 0.013 | 0 | 0.29 | 0.22 | 0.33 | 0.13 | 0.36 | 0.19 |
| 6 | 129925003 | rs12530181 | 0.658 | -0.075 | -0.501 | 0.629 | 0.008 | 0.057 | 0.057 | 0 | 0.22 | 0.07 | 0.17 | 0 | 0.21 | 0.13 |
| 6 | 129927014 | rs3813366 | -1.013 | -0.16 | -0.452 | 0.857 | 0.014 | -0.02 | -0.02 | 0 | 0.44 | 0.71 | 0.5 | 0.45 | 0.41 | 0.37 |
| 6 | 129935610 | rs13205216 | 0.77 | -0.336 | -0.426 | 0.358 | 0.006 | 0.009 | 0.009 | 0 | 0.19 | 0.13 | 0.12 | 0.05 | 0.15 | 0.04 |
| 6 | 129942817 | rs9375638 | 0.003 | -0.463 | -1.009 | 0.012 | -0.051 | 0.054 | 0.054 | 0 | 0.26 | 0.11 | 0.2 | 0.17 | 0.3 | 0.2 |
| 6 | 129952001 | rs9375644 | -0.799 | -0.599 | -0.895 | 0.135 | -0.012 | 0.021 | 0.021 | 0 | 0.55 | 0.74 | 0.43 | 0.36 | 0.48 | 0.37 |
| 6 | 129955487 | rs4509146 | 0.087 | -0.616 | -1.02 | -0.631 | -0.044 | 0.075 | 0.075 | 0.029 | 0.31 | 0.13 | 0.27 | 0.18 | 0.34 | 0.23 |
| 6 | 129959097 | rs9402155 | 0.411 | -0.63 | -1.077 | -0.658 | -0.055 | 0.005 | 0.005 | 0 | 0.18 | 0.08 | 0.19 | 0.17 | 0.25 | 0.19 |
| 6 | 129961249 | rs9483050 | -0.255 | -0.696 | -0.623 | -0.713 | 0.002 | 0.003 | 0.003 | 0 | 0.27 | 0.26 | 0.25 | 0.11 | 0.22 | 0.16 |
| 6 | 129962131 | rs4473880 | -0.11 | -0.324 | -0.563 | -0.491 | -0.001 | -0.006 | -0.006 | 0 | 0.06 | 0.07 | 0.12 | 0 | 0.08 | 0.09 |
| 6 | 129962633 | rs10499163 | -0.502 | -0.471 | -0.377 | -0.464 | -0.008 | 0 | 0 | 0 | 0.28 | 0.26 | 0.22 | 0.27 | 0.33 | 0.26 |
| 6 | 129966405 | rs12216321 | 0.16 | -0.603 | -0.18 | -0.279 | -0.015 | 0.015 | 0.015 | 0 | 0.1 | 0.16 | 0.03 | 0.1 | 0.07 | 0.08 |
| 6 | 129970962 | rs9398913 | -0.786 | -0.818 | -0.13 | -0.66 | 0 | 0.007 | 0.007 | 1.259 | 0.47 | 0.66 | 0.41 | 0.41 | 0.43 | 0.37 |
| 6 | 129971544 | rs4479964 | -0.261 | -0.846 | 0.035 | -0.672 | 0 | 0 | 0 | 1.471 | 0.41 | 0.36 | 0.28 | 0.37 | 0.37 | 0.31 |
| 6 | 129971929 | rs9388721 | 0.995 | -0.93 | -0.111 | -0.738 | -0.046 | 0.005 | 0.005 | 0 | 0.22 | 0.07 | 0.2 | 0.2 | 0.24 | 0.21 |
| 6 | 129973247 | rs4487604 | 0.282 | -0.93 | -0.047 | -0.743 | -0.003 | 0.003 | 0.003 | 0 | 0.28 | 0.32 | 0.33 | 0.24 | 0.3 | 0.27 |
| 6 | 129974695 | rs9402157 | 0.653 | -0.93 | -0.228 | -0.74 | -0.019 | 0.019 | 0.019 | 0 | 0.24 | 0.18 | 0.22 | 0.21 | 0.24 | 0.2 |
| 6 | 129974846 | rs12192896 | 0.436 | -1.025 | -0.129 | -0.701 | -0.013 | 0.02 | 0.02 | 0 | 0.17 | 0.22 | 0.07 | 0.17 | 0.13 | 0.1 |
| 6 | 129978268 | rs4337949 | -1.348 | -1.044 | -0.125 | -0.755 | 0.007 | 0.011 | 0.011 | 1.08 | 0.49 | 0.68 | 0.42 | 0.41 | 0.43 | 0.37 |
| 6 | 129978396 | rs4499954 | -0.064 | -1.112 | -0.154 | -0.623 | -0.01 | 0.024 | 0.024 | 0 | 0.3 | 0.29 | 0.33 | 0.25 | 0.3 | 0.27 |
| 6 | 129982285 | rs9388722 | -1.325 | -1.595 | -0.335 | -0.927 | -0.012 | -0.006 | -0.006 | 0 | 0.6 | 0.72 | 0.46 | 0.42 | 0.47 | 0.39 |
| 6 | 129982914 | rs13209334 | 0.348 | -1.534 | -0.609 | -0.932 | -0.04 | 0.13 | 0.13 | 0 | 0.38 | 0.07 | 0.29 | 0.2 | 0.35 | 0.25 |
| 6 | 129987492 | rs13204501 | 1.03 | -1.418 | -0.59 | -0.814 | -0.063 | -0.012 | -0.012 | 0 | 0.23 | 0.07 | 0.21 | 0.23 | 0.27 | 0.22 |
| 6 | 129989626 | rs4379317 | -1.235 | -1.782 | -0.485 | -0.997 | 0.021 | 0.009 | 0.009 | 0 | 0.62 | 0.49 | 0.46 | 0.4 | 0.49 | 0.46 |
| 6 | 129993700 | rs12195164 | 0.884 | -1.909 | -0.409 | -1.005 | -0.033 | 0.007 | 0.007 | 0 | 0.18 | 0.23 | 0.06 | 0.17 | 0.07 | 0.15 |
| 6 | 129996025 | rs17057571 | -0.533 | -1.817 | -0.636 | -0.964 | 0.019 | 0.017 | 0.017 | 0.47 | 0.42 | 0.63 | 0.26 | 0.21 | 0.22 | 0.25 |
| 6 | 129997515 | rs9388723 | 0.112 | -0.878 | -0.959 | -0.768 | -0.026 | 0.125 | 0.125 | 0 | 0.42 | 0.18 | 0.39 | 0.23 | 0.41 | 0.31 |
| 6 | 130005667 | rs6920979 | 0.008 | -0.962 | -0.465 | -0.57 | -0.005 | -0.004 | -0.004 | 0.011 | 0.46 | 0.46 | 0.32 | 0.25 | 0.31 | 0.49 |
| 6 | 130008445 | rs12660691 | 0.191 | -1.561 | -0.279 | -0.913 | -0.063 | -0.004 | -0.004 | 0 | 0.24 | 0.35 | 0.26 | 0.18 | 0.11 | 0.2 |
| 6 | 130008932 | rs2051632 | 0.021 | -0.543 | -0.699 | -0.305 | 0 | 0 | 0 | 0 | 0.5 | 0.49 | 0.56 | 0.46 | 0.48 | 0.46 |
| 6 | 130010372 | rs9375656 | -0.898 | -0.454 | -0.636 | -0.26 | 0.024 | 0.1 | 0.1 | 0.089 | 0.77 | 0.6 | 0.7 | 0.52 | 0.72 | 0.79 |
| 6 | 130012373 | rs11759328 | 1.342 | -0.475 | -0.819 | -0.338 | -0.001 | 0.056 | 0.056 | 0 | 0.21 | 0.09 | 0.13 | 0.05 | 0.19 | 0.11 |
| 6 | 130015331 | rs1894642 | -0.083 | -0.496 | -0.532 | 0.197 | -0.001 | -0.005 | -0.005 | 0 | 0.48 | 0.59 | 0.56 | 0.65 | 0.56 | 0.55 |
| 6 | 130018030 | rs17467757 | 1.462 | -0.078 | -0.889 | 0.227 | -0.079 | -0.012 | -0.012 | 0 | 0.17 | 0.02 | 0.17 | 0.17 | 0.26 | 0.2 |
| 6 | 130020347 | rs763132 | 0.653 | -0.121 | -0.816 | 0.273 | -0.057 | -0.048 | -0.048 | 0 | 0.22 | 0.14 | 0.27 | 0.37 | 0.38 | 0.33 |
| 6 | 130020811 | rs2187896 | 0.266 | -0.311 | -0.562 | 0.173 | -0.045 | 0.014 | 0.014 | 0 | 0.28 | 0.47 | 0.3 | 0.27 | 0.18 | 0.22 |
| 6 | 130021128 | rs2157460 | 1.074 | -0.368 | -0.538 | 0.183 | -0.047 | 0.011 | 0.011 | 0 | 0.19 | 0.26 | 0.16 | 0.18 | 0.08 | 0.16 |
| 6 | 130022390 | rs2097506 | -1.243 | -0.307 | -0.718 | 0.011 | 0.001 | 0.011 | 0.011 | 0 | 0.78 | 0.76 | 0.81 | 0.7 | 0.82 | 0.88 |
| 6 | 130023213 | rs12529067 | 1.086 | -0.294 | -0.737 | 0.005 | -0.002 | -0.002 | -0.002 | 0 | 0.08 | 0.09 | 0.13 | 0.09 | 0.1 | 0.06 |
| 6 | 130024200 | rs1023480 | 0.498 | -0.352 | -0.805 | -0.036 | -0.024 | 0.137 | 0.137 | 0 | 0.45 | 0.17 | 0.41 | 0.24 | 0.53 | 0.53 |
| 6 | 130024629 | rs6917887 | -0.13 | -0.458 | -0.511 | 0.112 | -0.024 | 0.041 | 0.041 | 0 | 0.33 | 0.58 | 0.4 | 0.46 | 0.29 | 0.34 |
| 6 | 130028542 | rs11154494 | 0.682 | -0.445 | -0.486 | 0.049 | -0.004 | -0.034 | -0.034 | 0 | 0.23 | 0.29 | 0.16 | 0.07 | 0.21 | 0.24 |
| 6 | 130029839 | rs2068685 | 0.542 | -0.496 | -0.538 | 0.378 | -0.022 | 0.002 | 0.002 | 0 | 0.2 | 0.21 | 0.16 | 0.19 | 0.08 | 0.17 |
| 6 | 130031215 | rs3752536 | -0.556 | 0.235 | -0.527 | 0.559 | 0 | 0.003 | 0.003 | 0 | 0.81 | 0.83 | 0.82 | 0.71 | 0.84 | 0.87 |
| 6 | 130033866 | rs9385512 | -0.716 | 0.021 | -0.42 | -0.211 | 0.008 | -0.07 | -0.07 | 0.05 | 0.78 | 0.68 | 0.84 | 0.94 | 0.79 | 0.75 |
| 6 | 130035812 | rs9388726 | 0.545 | 0.055 | -0.62 | 0.291 | 0 | 0.028 | 0.028 | 0 | 0.06 | 0.08 | 0.02 | 0.26 | 0.04 | 0.04 |
| 6 | 130035924 | rs11154496 | 0.111 | 0 | -0.971 | -0.112 | -0.018 | 0.043 | 0.043 | 0 | 0.18 | 0.08 | 0.18 | 0.06 | 0.23 | 0.09 |
| 6 | 130036632 | rs17468526 | 1.069 | -0.087 | -0.881 | -0.248 | 0.003 | -0.02 | -0.02 | 0 | 0.19 | 0.22 | 0.14 | 0.07 | 0.21 | 0.26 |
| 6 | 130036813 | rs12203891 | 0.137 | 0.006 | -0.845 | -0.145 | -0.014 | 0.035 | 0.035 | 0 | 0.18 | 0.09 | 0.19 | 0.06 | 0.23 | 0.09 |
| 6 | 130037874 | rs7768656 | -0.411 | -0.018 | -0.737 | 0.129 | 0.005 | 0.077 | 0.077 | 0 | 0.9 | 0.76 | 0.9 | 0.77 | 0.92 | 0.88 |
| 6 | 130037920 | rs12205840 | 1.298 | -0.225 | -1.968 | 0.006 | -0.084 | -0.101 | -0.101 | 0 | 0.25 | 0.06 | 0.3 | 0.35 | 0.33 | 0.29 |
| 6 | 130038005 | rs1984408 | 0.045 | -0.123 | -1.589 | -0.048 | 0.007 | 0.029 | 0.029 | 0 | 0.33 | 0.28 | 0.31 | 0.1 | 0.3 | 0.39 |
| 6 | 130045677 | rs9398918 | 0.449 | 0.133 | -1.804 | 0 | 0.001 | -0.003 | -0.003 | 0 | 0.21 | 0.16 | 0.14 | 0.26 | 0.23 | 0.12 |
| 6 | 130048450 | rs17384497 | 0.261 | 0.126 | -1.811 | -0.224 | 0.025 | 0.025 | 0.025 | 0 | 0.15 | 0.04 | 0.16 | 0.04 | 0.12 | 0.16 |
| 6 | 130049171 | rs17384593 | 0.126 | 0.253 | -1.695 | -0.245 | -0.004 | 0.004 | 0.004 | 0 | 0.2 | 0.18 | 0.14 | 0.06 | 0.2 | 0.25 |
| 6 | 130050610 | rs7753327 | -0.138 | 0.347 | -1.352 | 0.178 | -0.001 | 0.042 | 0.042 | 0 | 0.43 | 0.26 | 0.43 | 0.32 | 0.41 | 0.53 |
| 6 | 130055734 | rs6914728 | 0.267 | -0.212 | -1.222 | -0.049 | 0.019 | 0.021 | 0.021 | 1.983 | 0.71 | 0.81 | 0.83 | 0.84 | 0.82 | 0.83 |
| 6 | 130056581 | rs17469847 | 0.056 | 0.425 | -1.479 | -0.624 | -0.014 | 0.027 | 0.027 | 0.01 | 0.2 | 0.15 | 0.16 | 0.06 | 0.23 | 0.16 |
| 6 | 130059070 | rs376632 | -0.211 | 0.06 | -1.7 | 0.302 | 0.006 | 0.006 | 0.006 | 1.47 | 0.76 | 0.82 | 0.87 | 0.84 | 0.87 | 0.84 |
| 6 | 130062717 | rs208869 | 0.122 | 0.235 | -1.481 | 0.01 | 0 | 0.005 | 0.005 | 0 | 0.43 | 0.51 | 0.53 | 0.51 | 0.48 | 0.53 |
| 6 | 130062855 | rs9492364 | -0.46 | 0.17 | -1.134 | -0.581 | -0.006 | -0.021 | -0.021 | 0 | 0.3 | 0.38 | 0.22 | 0.17 | 0.25 | 0.2 |
| 6 | 130063709 | rs10484284 | 0.131 | 0.094 | -1.076 | -0.566 | 0 | 0.001 | 0.001 | 0 | 0.21 | 0.24 | 0.17 | 0.16 | 0.2 | 0.14 |
| 6 | 130064670 | rs10484283 | 0.711 | 0.141 | -0.953 | -0.549 | -0.012 | -0.025 | -0.025 | 0.048 | 0.1 | 0.2 | 0.07 | 0.04 | 0.07 | 0.05 |
| 6 | 130064727 | rs9492365 | 0.711 | 0.141 | -0.965 | -0.549 | -0.012 | -0.023 | -0.023 | 0 | 0.1 | 0.19 | 0.07 | 0.04 | 0.08 | 0.06 |
| 6 | 130066078 | rs10484282 | 0.47 | 0.428 | -1.463 | -0.901 | -0.054 | 0.05 | 0.05 | 0 | 0.25 | 0.07 | 0.19 | 0.15 | 0.31 | 0.15 |

| | | | | | | | | | | | | | | | | |
|---|---|---|---|---|---|---|---|---|---|---|---|---|---|---|---|---|
| 6 | 130066736 | rs9402168 | -0.226 | 0.088 | -1.125 | -0.94 | -0.001 | 0.005 | 0.005 | 0 | 0.41 | 0.58 | 0.39 | 0.5 | 0.35 | 0.44 |
| 6 | 130068065 | rs6933604 | 0.751 | 0.761 | -0.716 | -0.17 | 0.006 | 0.106 | 0.106 | 0 | 0.11 | 0.2 | 0.2 | 0.39 | 0.09 | 0.2 |
| 6 | 130071629 | rs172318 | 1.203 | 0.823 | -0.767 | -0.309 | 0.006 | 0.018 | 0.018 | 0 | 0.2 | 0.22 | 0.36 | 0.35 | 0.28 | 0.37 |
| 6 | 130071823 | rs9388729 | -0.028 | 0.804 | -0.721 | -0.288 | -0.019 | -0.04 | -0.04 | 0.047 | 0.15 | 0.06 | 0.18 | 0.27 | 0.14 | 0.17 |
| 6 | 130071841 | rs9375662 | 1.037 | 0.763 | -0.78 | -0.206 | -0.042 | -0.051 | -0.051 | 0 | 0.09 | 0 | 0.13 | 0.27 | 0.09 | 0.12 |
| 6 | 130072556 | rs208865 | -1.189 | 0.635 | -0.731 | -0.356 | -0.01 | -0.038 | -0.038 | 0 | 0.85 | 0.87 | 0.95 | 1 | 0.94 | 0.97 |
| 6 | 130072861 | rs208867 | 0.646 | 0.836 | -0.899 | -0.238 | 0.01 | 0.015 | 0.015 | 0 | 0.26 | 0.29 | 0.41 | 0.35 | 0.33 | 0.42 |
| 6 | 130073291 | rs182320 | 0.183 | 0.678 | -0.673 | -0.641 | -0.002 | 0 | 0 | 0 | 0.34 | 0.29 | 0.54 | 0.65 | 0.41 | 0.53 |
| 6 | 130073727 | rs208868 | -0.581 | 0.537 | -0.475 | -0.473 | -0.06 | 0.046 | 0.046 | 0.714 | 0.83 | 0.64 | 0.87 | 0.77 | 0.94 | 0.9 |
| 6 | 130075541 | rs433351 | 0.362 | 1.648 | -0.948 | -0.978 | -0.054 | 0.205 | 0.205 | 0 | 0.39 | 0.12 | 0.27 | 0.04 | 0.5 | 0.44 |
| 6 | 130078071 | rs6569626 | 0.068 | 1.872 | -0.501 | -0.321 | 0.015 | 0.105 | 0.105 | 0 | 0.11 | 0.2 | 0.21 | 0.32 | 0.15 | 0.11 |
| 6 | 130081673 | rs17057830 | 0.648 | 1.793 | -0.48 | -0.492 | -0.005 | -0.022 | -0.022 | 0.015 | 0.16 | 0.22 | 0.13 | 0.07 | 0.12 | 0.16 |
| 6 | 130082061 | rs4897336 | 0.695 | 1.837 | -0.667 | -0.207 | -0.001 | 0.036 | 0.036 | 0 | 0.13 | 0.17 | 0.26 | 0.54 | 0.09 | 0.18 |
| 6 | 130082258 | rs12195685 | 0.575 | 2.323 | -0.888 | -1.498 | -0.037 | 0.02 | 0.02 | 0 | 0.09 | 0 | 0.06 | 0.06 | 0.08 | 0.09 |
| 6 | 130084368 | rs208872 | -0.656 | 2.655 | -0.276 | -0.981 | 0.086 | 0.068 | 0.068 | 0 | 0.87 | 0.71 | 0.74 | 0.78 | 0.76 | 0.77 |
| 6 | 130086447 | rs17057836 | 0.428 | 0.908 | -1.199 | -2.122 | 0.002 | 0.002 | 0.002 | 0 | 0.18 | 0.12 | 0.11 | 0 | 0.15 | 0.1 |
| 6 | 130086475 | rs899860 | -0.336 | 1.147 | -1.233 | -1.469 | -0.005 | 0.085 | 0.085 | 0 | 0.81 | 0.96 | 0.87 | 0.99 | 0.76 | 0.69 |
| 6 | 130088262 | rs6905254 | -0.593 | 1.469 | -0.561 | -0.911 | 0.006 | -0.041 | -0.041 | 1.879 | 0.39 | 0.38 | 0.32 | 0.13 | 0.43 | 0.36 |
| 6 | 130088924 | rs11754070 | -0.534 | 1.772 | -0.609 | -0.461 | -0.011 | 0.008 | 0.008 | 1.194 | 0.12 | 0.1 | 0.1 | 0.03 | 0.21 | 0.11 |
| 6 | 130090404 | rs17057865 | -0.838 | 1.114 | 0.147 | -0.51 | 0.024 | -0.003 | -0.003 | 0.015 | 0.07 | 0.27 | 0.1 | 0.02 | 0.05 | 0.1 |
| 6 | 130091555 | rs7759860 | -0.74 | 0.979 | -0.158 | 0.098 | -0.028 | -0.045 | -0.045 | 0 | 0.21 | 0.09 | 0.29 | 0.31 | 0.24 | 0.2 |
| 6 | 130091761 | rs9385515 | -0.982 | 0.365 | -0.662 | -0.371 | -0.017 | -0.018 | -0.018 | 0 | 0.18 | 0.27 | 0.24 | 0.14 | 0.14 | 0.23 |
| 6 | 130092334 | rs4897339 | -1.428 | -0.016 | -1.112 | -1.342 | -0.046 | 0.085 | 0.085 | 0 | 0.6 | 0.8 | 0.57 | 0.29 | 0.5 | 0.48 |
| 6 | 130092485 | rs4895856 | -0.161 | -0.177 | -0.915 | -1.374 | -0.008 | 0 | 0 | 0 | 0.16 | 0.16 | 0.24 | 0.17 | 0.13 | 0.22 |
| 6 | 130098771 | rs6931600 | -1.478 | -0.154 | -0.754 | -1.41 | -0.037 | 0.075 | 0.075 | 0 | 0.6 | 0.78 | 0.58 | 0.28 | 0.5 | 0.48 |
| 6 | 130101130 | rs12202100 | 1.007 | -0.522 | -0.075 | -1.692 | -0.023 | -0.048 | -0.048 | 0.629 | 0.11 | 0.18 | 0.07 | 0 | 0.04 | 0.05 |
| 6 | 130101822 | rs6927242 | -2.036 | -0.742 | -0.729 | -1.761 | -0.046 | 0.067 | 0.067 | 0 | 0.68 | 0.85 | 0.77 | 0.86 | 0.57 | 0.79 |
| 6 | 130101879 | rs10484280 | 0.777 | -0.688 | -0.798 | -1.823 | -0.062 | 0.101 | 0.101 | 0 | 0.17 | 0.01 | 0.1 | 0.03 | 0.19 | 0.09 |
| 6 | 130102959 | rs9321180 | -0.028 | -0.594 | -0.554 | -1.74 | 0.006 | -0.066 | -0.066 | 0 | 0.31 | 0.42 | 0.26 | 0.11 | 0.32 | 0.2 |
| 6 | 130113882 | rs9321183 | -0.903 | -0.544 | -0.878 | -1.553 | -0.023 | 0.06 | 0.06 | 0 | 0.59 | 0.75 | 0.57 | 0.27 | 0.49 | 0.45 |
| 6 | 130116227 | rs11968591 | 1.491 | -0.388 | -1.044 | -1.572 | -0.043 | 0.051 | 0.051 | 0 | 0.28 | 0.15 | 0.22 | 0.14 | 0.39 | 0.2 |
| 6 | 130117403 | rs4895858 | 0.375 | -0.927 | -0.932 | -1.054 | 0.012 | -0.077 | -0.077 | 0 | 0.79 | 0.93 | 0.73 | 0.43 | 0.8 | 0.59 |
| 6 | 130121834 | rs1480519 | 0.057 | -0.938 | -0.819 | -1.239 | 0.008 | -0.021 | -0.021 | 1.806 | 0.85 | 0.89 | 0.7 | 0.29 | 0.84 | 0.78 |
| 6 | 130122482 | rs11154504 | -0.017 | -0.88 | -0.652 | -1.397 | 0.007 | -0.001 | -0.001 | 4.588 | 0.09 | 0.1 | 0.03 | 0 | 0.05 | 0.21 |
| 6 | 130135714 | rs11967794 | -0.151 | -1.032 | -0.439 | -1.215 | 0.01 | 0 | 0 | 0 | 0.11 | 0.14 | 0.04 | 0 | 0.04 | 0.23 |
| 6 | 130140120 | rs17473162 | 0.805 | -0.993 | -0.455 | -1.222 | -0.015 | 0.013 | 0.013 | 0 | 0.05 | 0.03 | 0.06 | 0 | 0.11 | 0.03 |
| 6 | 130142606 | rs17057971 | -0.359 | -1.09 | -0.387 | -1.229 | 0.005 | -0.01 | -0.01 | 0 | 0.12 | 0.17 | 0.05 | 0 | 0.06 | 0.25 |
| 6 | 130147261 | rs9402178 | 0.254 | -1.247 | -0.738 | -1.467 | 0.019 | -0.004 | -0.004 | 0 | 0.21 | 0.08 | 0.11 | 0.22 | 0.14 | 0.23 |
| 6 | 130150861 | rs12206334 | 1.501 | -1.742 | -0.643 | -1.842 | -0.015 | 0.047 | 0.047 | 0 | 0.34 | 0.22 | 0.31 | 0.14 | 0.57 | 0.24 |
| 6 | 130151533 | rs4624895 | -0.215 | -2.006 | -0.035 | -1.961 | -0.016 | 0.056 | 0.056 | 0 | 0.79 | 0.66 | 0.51 | 0.36 | 0.77 | 0.72 |
| 6 | 130152440 | rs9372948 | 0.721 | -2.049 | -0.094 | -1.997 | 0.005 | -0.012 | -0.012 | 0 | 0.21 | 0.06 | 0.12 | 0.22 | 0.16 | 0.24 |
| 6 | 130152452 | rs9375666 | 0.721 | -2.049 | -0.094 | -1.997 | 0.005 | -0.012 | -0.012 | 0.109 | 0.21 | 0.06 | 0.12 | 0.22 | 0.16 | 0.24 |
| 6 | 130152479 | rs7758540 | 0.433 | -2.065 | -0.025 | -2.014 | 0.001 | -0.008 | -0.008 | 0.806 | 0.58 | 0.61 | 0.39 | 0.14 | 0.62 | 0.47 |
| 6 | 130152520 | rs7776426 | 2.233 | -2.011 | -0.055 | -2.029 | -0.042 | 0.033 | 0.033 | 0.044 | 0.3 | 0.18 | 0.25 | 0.14 | 0.46 | 0.21 |
| 6 | 130154667 | rs9492393 | -0.972 | -2.013 | -0.033 | -2.025 | 0.025 | 0.037 | 0.037 | 0 | 0.5 | 0.49 | 0.27 | 0.23 | 0.32 | 0.5 |
| 6 | 130154686 | rs4629709 | -0.203 | -1.939 | 0.021 | -1.884 | -0.016 | 0.061 | 0.061 | 0 | 0.79 | 0.67 | 0.51 | 0.36 | 0.77 | 0.71 |
| 6 | 130163525 | rs17396328 | 0.51 | -1.928 | -0.053 | -1.873 | 0.011 | -0.008 | -0.008 | 1.339 | 0.21 | 0.07 | 0.12 | 0.22 | 0.16 | 0.24 |
| 6 | 130165115 | rs9492397 | 1.268 | -1.907 | 0.046 | -1.901 | -0.039 | -0.071 | -0.071 | 0 | 0.13 | 0.22 | 0.05 | 0 | 0.04 | 0.05 |
| 6 | 130168701 | rs9483068 | 0.541 | -1.827 | -0.028 | -1.859 | 0.022 | 0.05 | 0.05 | 0 | 0.57 | 0.65 | 0.41 | 0.14 | 0.62 | 0.47 |
| 6 | 130170928 | rs10872348 | -0.117 | -1.631 | -0.028 | -1.732 | -0.008 | 0.022 | 0.022 | 0 | 0.79 | 0.73 | 0.51 | 0.34 | 0.77 | 0.71 |
| 6 | 130171667 | rs9402186 | 0.004 | -1.569 | -0.171 | -1.747 | -0.005 | 0.015 | 0.015 | 0 | 0.24 | 0.2 | 0.18 | 0.23 | 0.23 | 0.23 |
| 6 | 130174927 | rs9398926 | 0.344 | -1.552 | -0.071 | -1.684 | 0.016 | -0.003 | -0.003 | 0.137 | 0.2 | 0.15 | 0.12 | 0.23 | 0.14 | 0.22 |
| 6 | 130175138 | rs9321194 | -0.41 | -1.662 | 0.064 | -1.774 | -0.022 | -0.046 | -0.046 | 0 | 0.38 | 0.49 | 0.24 | 0.23 | 0.27 | 0.27 |
| 6 | 130175635 | rs1383318 | 0.035 | -1.463 | -0.324 | -1.443 | 0.017 | 0.011 | 0.011 | 0 | 0.11 | 0.1 | 0.03 | 0 | 0.05 | 0.23 |
| 6 | 130179253 | rs2876063 | 0.581 | -1.725 | -0.204 | -1.46 | 0.03 | 0.035 | 0.035 | 0.004 | 0.55 | 0.42 | 0.39 | 0.37 | 0.61 | 0.63 |
| 6 | 130179710 | rs11154513 | 0.842 | -1.767 | -0.301 | -1.556 | 0.033 | 0.057 | 0.057 | 0 | 0.52 | 0.4 | 0.4 | 0.33 | 0.63 | 0.36 |
| 6 | 130187499 | rs9492408 | 0.765 | -2.092 | 0.086 | -1.769 | -0.029 | -0.057 | -0.057 | 2.539 | 0.15 | 0.21 | 0.06 | 0 | 0.04 | 0.06 |
| 6 | 130188359 | rs17397741 | 0.344 | -2.022 | -0.308 | -1.702 | -0.001 | -0.008 | -0.008 | 0.737 | 0.16 | 0.18 | 0.23 | 0.07 | 0.14 | 0.18 |
| 6 | 130190531 | rs17755837 | 0.746 | -2.069 | -0.289 | -1.988 | -0.008 | 0.005 | 0.005 | 0 | 0.12 | 0.08 | 0.14 | 0.03 | 0.18 | 0.11 |
| 6 | 130190985 | rs2127706 | 0.33 | -1.992 | -0.324 | -1.82 | -0.001 | -0.018 | -0.018 | 0 | 0.16 | 0.18 | 0.23 | 0.06 | 0.14 | 0.18 |
| 6 | 130194204 | rs6939565 | 0.966 | -2.085 | -0.198 | -1.834 | -0.007 | 0.026 | 0.026 | 0 | 0.42 | 0.37 | 0.46 | 0.72 | 0.54 | 0.39 |
| 6 | 130194422 | rs11752628 | 0.56 | -1.652 | -0.461 | -1.276 | -0.002 | 0.074 | 0.074 | 0 | 0.1 | 0.11 | 0.18 | 0.48 | 0.08 | 0.19 |
| 6 | 130194876 | rs10499165 | 1.674 | -1.636 | -0.182 | -1.439 | -0.006 | 0.002 | 0.002 | 0 | 0.31 | 0.31 | 0.28 | 0.25 | 0.46 | 0.2 |
| 6 | 130195659 | rs10499166 | 1.795 | -1.622 | -0.406 | -1.431 | -0.039 | 0.005 | 0.005 | 0 | 0.28 | 0.21 | 0.27 | 0.25 | 0.46 | 0.24 |
| 6 | 130205142 | rs17058133 | 0.062 | -1.695 | -0.466 | -1.429 | 0.014 | 0.003 | 0.003 | 0 | 0.15 | 0.12 | 0.07 | 0 | 0.09 | 0.22 |
| 6 | 130214112 | rs1572572 | -0.647 | -1.273 | -0.621 | -1.244 | -0.011 | -0.002 | -0.002 | 1.585 | 0.86 | 0.9 | 0.79 | 0.86 | 0.76 | 0.85 |
| 6 | 130217473 | rs9375675 | -0.199 | -1.564 | -0.487 | -0.962 | -0.003 | 0.074 | 0.074 | 0 | 0.13 | 0.18 | 0.32 | 0.53 | 0.1 | 0.27 |
| 6 | 130222234 | rs11752675 | 0.731 | -0.766 | -0.96 | -1.122 | -0.03 | -0.035 | -0.035 | 0.144 | 0.23 | 0.19 | 0.31 | 0.53 | 0.32 | 0.29 |
| 6 | 130231608 | rs12660061 | 0.726 | -0.755 | -0.985 | -1.115 | -0.02 | -0.024 | -0.024 | 0.255 | 0.23 | 0.21 | 0.31 | 0.53 | 0.32 | 0.29 |

| | | | | | | | | | | | | | | | | |
|---|---|---|---|---|---|---|---|---|---|---|---|---|---|---|---|---|
| 6 | 130232043 | rs9375677 | 0.256 | -0.805 | -1.056 | -1.194 | -0.025 | -0.02 | -0.02 | 0 | 0.28 | 0.25 | 0.45 | 0.59 | 0.35 | 0.38 |
| 6 | 130236932 | rs7759830 | -0.223 | -1.045 | -1.055 | -1.336 | 0.002 | 0.051 | 0.051 | 0.106 | 0.6 | 0.67 | 0.37 | 0.18 | 0.44 | 0.46 |
| 6 | 130238237 | rs9492417 | 0.617 | -1.328 | -0.971 | -1.585 | -0.011 | -0.065 | -0.065 | 0 | 0.36 | 0.32 | 0.21 | 0.04 | 0.21 | 0.34 |
| 6 | 130239391 | rs9398929 | -0.013 | -1.321 | -0.971 | -1.58 | -0.001 | -0.004 | -0.004 | 0 | 0.48 | 0.58 | 0.61 | 0.73 | 0.57 | 0.49 |
| 6 | 130242858 | rs7753901 | -1.661 | -1.26 | -0.948 | -1.443 | -0.012 | -0.016 | -0.016 | 0 | 0.84 | 0.9 | 0.81 | 0.77 | 0.79 | 0.83 |
| 6 | 130251793 | rs2039683 | 1.73 | -1.086 | -0.858 | -1.347 | 0.002 | -0.035 | -0.035 | 0 | 0.14 | 0.15 | 0.11 | 0 | 0.22 | 0.09 |
| 6 | 130253378 | rs9402199 | -0.428 | -1.762 | -0.717 | -1.195 | -0.012 | -0.02 | -0.02 | 0 | 0.56 | 0.59 | 0.67 | 0.73 | 0.65 | 0.55 |
| 6 | 130254385 | rs6909737 | 0.479 | -1.284 | -1.369 | -0.98 | -0.05 | -0.037 | -0.037 | 0.151 | 0.26 | 0.21 | 0.46 | 0.59 | 0.39 | 0.37 |
| 6 | 130255964 | rs12524191 | 1.031 | -1.06 | -1.694 | -0.707 | -0.007 | -0.013 | -0.013 | 1.813 | 0.09 | 0.03 | 0.17 | 0.38 | 0.08 | 0.18 |
| 6 | 130256258 | rs9388752 | -0.818 | -1.44 | -1.675 | -1.528 | 0.004 | 0.012 | 0.012 | 0 | 0.69 | 0.95 | 0.77 | 0.76 | 0.74 | 0.78 |
| 6 | 130258638 | rs12197285 | 0.918 | -0.849 | -1.049 | -0.975 | -0.003 | -0.003 | -0.003 | 0 | 0.21 | 0.15 | 0.15 | 0 | 0.24 | 0.13 |
| 6 | 130262763 | rs1890143 | -0.274 | 0.387 | -0.616 | 0.105 | 0.045 | 0.039 | 0.039 | 0 | 0.75 | 0.71 | 0.66 | 0.46 | 0.62 | 0.72 |
| 6 | 130263591 | rs9375683 | -0.892 | 0.339 | -0.892 | 0.263 | 0.002 | -0.007 | -0.007 | 0 | 0.21 | 0.17 | 0.32 | 0.27 | 0.22 | 0.26 |
| 6 | 130267753 | rs1538185 | -0.175 | 0.701 | -0.804 | 0.572 | -0.003 | -0.012 | -0.012 | 0.093 | 0.15 | 0.11 | 0.18 | 0.24 | 0.19 | 0.16 |
| 6 | 130274198 | rs9385528 | -0.101 | 0.663 | -0.878 | 0.292 | -0.044 | -0.009 | -0.009 | 0 | 0.64 | 0.81 | 0.63 | 0.61 | 0.54 | 0.58 |
| 6 | 130275975 | rs11757973 | 0.239 | -0.09 | -0.683 | -0.565 | -0.004 | -0.032 | -0.032 | 0 | 0.22 | 0.23 | 0.14 | 0.02 | 0.15 | 0.17 |
| 6 | 130285951 | rs6931642 | 0.245 | -0.296 | -0.215 | -0.69 | 0.015 | -0.048 | -0.048 | 0 | 0.12 | 0.31 | 0.1 | 0.02 | 0.1 | 0.12 |
| 6 | 130287785 | rs957185 | -0.034 | -0.642 | -0.269 | -1.214 | 0 | -0.046 | -0.046 | 0 | 0.42 | 0.6 | 0.39 | 0.04 | 0.42 | 0.39 |
| 6 | 130290443 | rs1890142 | 0.918 | -0.705 | -0.073 | -1.318 | -0.007 | 0.021 | 0.021 | 0 | 0.26 | 0.16 | 0.16 | 0.01 | 0.28 | 0.15 |
| 6 | 130292225 | rs9402203 | 0.623 | -0.792 | -0.102 | -1.07 | -0.002 | 0.128 | 0.128 | 0 | 0.08 | 0.18 | 0.19 | 0.56 | 0.08 | 0.13 |
| 6 | 130295827 | rs1577910 | -1.192 | -0.85 | -0.19 | -1.15 | -0.034 | -0.018 | -0.018 | 0 | 0.63 | 0.81 | 0.63 | 0.58 | 0.54 | 0.61 |
| 6 | 130302794 | rs10499171 | 0.672 | -0.93 | 0.052 | -1.12 | 0.024 | -0.039 | -0.039 | 0 | 0.11 | 0.34 | 0.11 | 0.03 | 0.1 | 0.12 |
| 6 | 130304725 | rs9385529 | 0.041 | -0.883 | -0.309 | -0.909 | 0.014 | -0.021 | -0.021 | 0.076 | 0.22 | 0.24 | 0.24 | 0.53 | 0.13 | 0.23 |
| 6 | 130309435 | rs9321202 | -0.701 | -0.723 | -0.344 | -0.883 | 0.019 | 0.009 | 0.009 | 0 | 0.51 | 0.42 | 0.52 | 0.54 | 0.44 | 0.5 |
| 6 | 130310518 | rs9321203 | -0.68 | -0.623 | -0.012 | -0.308 | -0.013 | 0.022 | 0.022 | 0 | 0.25 | 0.35 | 0.37 | 0.53 | 0.16 | 0.35 |
| 6 | 130311090 | rs4897358 | -0.744 | -0.581 | -0.13 | -0.325 | -0.002 | 0.004 | 0.004 | 0 | 0.26 | 0.31 | 0.39 | 0.54 | 0.19 | 0.36 |
| 6 | 130311831 | rs9321204 | 0.26 | -0.153 | -0.163 | -0.988 | -0.003 | 0.013 | 0.013 | 0 | 0.58 | 0.65 | 0.61 | 0.78 | 0.6 | 0.47 |
| 6 | 130312067 | rs7747276 | 0.738 | 0.362 | 0.425 | -0.969 | 0.007 | 0.016 | 0.016 | 0 | 0.49 | 0.49 | 0.47 | 0.64 | 0.44 | 0.37 |
| 6 | 130320223 | rs1544144 | 0.56 | 0.698 | 0.567 | -0.984 | -0.028 | 0.039 | 0.039 | 0.19 | 0.62 | 0.7 | 0.69 | 0.81 | 0.55 | 0.58 |
| 6 | 130321244 | rs945889 | -0.297 | 0.739 | 1.502 | -0.352 | -0.023 | 0.123 | 0.123 | 0.275 | 0.13 | 0.47 | 0.15 | 0.22 | 0.08 | 0.11 |
| 6 | 130322357 | rs6569644 | -0.152 | 1.722 | 1.057 | -0.21 | -0.026 | 0.012 | 0.012 | 0 | 0.83 | 0.9 | 0.81 | 0.89 | 0.73 | 0.68 |
| 6 | 130328399 | rs13203469 | 1.463 | 1.701 | 0.999 | -0.275 | -0.074 | 0.032 | 0.032 | 0.213 | 0.62 | 0.13 | 0.54 | 0.73 | 0.6 | 0.48 |
| 6 | 130328639 | rs6904762 | 0.569 | 1.744 | 1.37 | -0.299 | -0.002 | -0.004 | -0.004 | 0 | 0.74 | 0.75 | 0.67 | 0.88 | 0.68 | 0.53 |
| 6 | 130331093 | rs7754627 | -0.096 | 1.802 | 1.457 | -0.228 | 0.015 | 0.007 | 0.007 | 0 | 0.07 | 0.14 | 0.06 | 0.08 | 0.09 | 0.05 |
| 6 | 130334365 | rs1337775 | 1.351 | 1.792 | 1.386 | -0.257 | -0.022 | 0.001 | 0.001 | 0 | 0.7 | 0.27 | 0.56 | 0.73 | 0.62 | 0.51 |
| 6 | 130335109 | rs1112847 | 1.702 | 1.771 | 1.058 | -0.209 | -0.054 | -0.023 | -0.023 | 0 | 0.54 | 0.14 | 0.45 | 0.57 | 0.5 | 0.37 |
| 6 | 130337266 | rs6569647 | -1.475 | 1.65 | 1.24 | -0.064 | 0.005 | -0.008 | -0.008 | 0 | 0.27 | 0.74 | 0.24 | 0.33 | 0.21 | 0.19 |
| 6 | 130345835 | rs1415701 | -1.012 | 2.058 | 1.415 | 0.174 | 0.012 | 0.052 | 0.052 | 0 | 0.26 | 0.41 | 0.32 | 0.38 | 0.28 | 0.26 |
| 6 | 130349119 | rs6569648 | -0.15 | 2.307 | 0.907 | 0.065 | -0.073 | 0.024 | 0.024 | 0 | 0.88 | 0.99 | 0.88 | 0.95 | 0.78 | 0.88 |
| 6 | 130349475 | rs12530176 | 1.801 | 2.399 | 0.836 | -0.023 | -0.028 | -0.042 | -0.042 | 0 | 0.53 | 0.1 | 0.39 | 0.55 | 0.43 | 0.38 |
| 6 | 130354855 | rs9388766 | 1.299 | 2.628 | 1.876 | 0.044 | 0.052 | -0.05 | -0.05 | 0.767 | 0.78 | 0.45 | 0.7 | 0.85 | 0.7 | 0.59 |
| 6 | 130368508 | rs6928313 | 1.884 | 2.603 | 1.23 | 0.04 | -0.125 | 0.038 | 0.038 | 0.188 | 0.63 | 0.11 | 0.55 | 0.76 | 0.55 | 0.53 |
| 6 | 130374102 | rs9388768 | 1.713 | 2.411 | 1.282 | 0.059 | -0.046 | 0.024 | 0.024 | 0.18 | 0.65 | 0.11 | 0.61 | 0.76 | 0.65 | 0.54 |
| 6 | 130379954 | rs9388769 | 1.862 | 2.433 | 1.277 | -0.02 | -0.022 | 0.031 | 0.031 | 0.051 | 0.66 | 0.17 | 0.62 | 0.76 | 0.65 | 0.55 |
| 6 | 130395240 | rs12528442 | -0.405 | 2.399 | 1.717 | 0.068 | 0.004 | 0.023 | 0.023 | 0 | 0.1 | 0.2 | 0.18 | 0.11 | 0.08 | 0.1 |
| 6 | 130397238 | rs9321207 | 2.326 | 2.58 | 1.618 | 0.061 | -0.079 | -0.025 | -0.025 | 0.007 | 0.64 | 0.14 | 0.57 | 0.65 | 0.58 | 0.5 |
| 6 | 130397515 | rs4548027 | -1.43 | 2.535 | 2.254 | 0 | 0.027 | -0.113 | -0.113 | 0 | 0.19 | 0.41 | 0.17 | 0.03 | 0.24 | 0.14 |
| 6 | 130410069 | rs6569654 | -2.213 | 2.396 | 2.05 | -0.393 | -0.018 | 0.035 | 0.035 | 0 | 0.32 | 0.7 | 0.4 | 0.13 | 0.4 | 0.24 |
| 6 | 130416949 | rs9483083 | -1.035 | 2.336 | 1.914 | -0.379 | -0.021 | 0.064 | 0.064 | 0 | 0.17 | 0.06 | 0.12 | 0.03 | 0.23 | 0.14 |
| 6 | 130419555 | rs7755589 | -1.516 | 2.33 | 2.119 | -0.382 | 0.002 | -0.003 | -0.003 | 0 | 0.2 | 0.22 | 0.29 | 0.13 | 0.28 | 0.21 |
| 6 | 130429016 | rs9388772 | 2.361 | 2.913 | 1.902 | -0.306 | -0.115 | 0.044 | 0.044 | 0.075 | 0.63 | 0.14 | 0.54 | 0.76 | 0.51 | 0.52 |
| 6 | 130432420 | rs9402216 | 2.366 | 3.021 | 1.951 | -0.391 | -0.119 | 0.053 | 0.053 | 0.172 | 0.64 | 0.13 | 0.54 | 0.76 | 0.52 | 0.5 |
| 6 | 130433290 | rs9321208 | -1.11 | 3.288 | 1.788 | -0.351 | -0.069 | 0.07 | 0.07 | 0 | 0.16 | 0.04 | 0.17 | 0.03 | 0.32 | 0.14 |
| 6 | 130441464 | rs11154522 | -1.724 | 3.21 | 1.571 | -0.281 | 0.029 | -0.016 | -0.016 | 0.836 | 0.21 | 0.27 | 0.34 | 0.13 | 0.36 | 0.22 |
| 6 | 130449280 | rs12194709 | -1.152 | 3.265 | 1.417 | -0.277 | 0.039 | -0.002 | -0.002 | 0 | 0.12 | 0.22 | 0.23 | 0.1 | 0.2 | 0.15 |
| 6 | 130453842 | rs3777428 | -0.429 | 2.792 | 0.932 | -0.216 | -0.023 | -0.004 | -0.004 | 0 | 0.13 | 0.33 | 0.17 | 0.1 | 0.06 | 0.12 |
| 6 | 130454535 | rs13207727 | -0.569 | 2.799 | 0.94 | -0.207 | 0.034 | -0.013 | -0.013 | 0 | 0.14 | 0.33 | 0.23 | 0.1 | 0.16 | 0.12 |
| 6 | 130456938 | rs6933511 | 1.219 | 2.821 | -0.229 | 0.066 | -0.006 | 0.003 | 0.003 | 0 | 0.81 | 0.82 | 0.81 | 0.86 | 0.67 | 0.62 |
| 6 | 130465401 | rs11154524 | -0.466 | 2.531 | -0.331 | 0.015 | 0.001 | 0.03 | 0.03 | 0 | 0.06 | 0 | 0.03 | 0 | 0.06 | 0.01 |
| 6 | 130466676 | rs4897370 | -0.527 | 2.87 | -0.606 | 0.305 | -0.016 | -0.058 | -0.058 | 0 | 0.27 | 0.13 | 0.29 | 0.51 | 0.28 | 0.26 |
| 6 | 130471063 | rs2095452 | -0.945 | 2.647 | -0.394 | 0 | -0.003 | -0.011 | -0.011 | 0 | 0.37 | 0.31 | 0.49 | 0.54 | 0.38 | 0.42 |
| 6 | 130472920 | rs870816 | -0.337 | 2.589 | -0.458 | -0.06 | -0.039 | -0.062 | -0.062 | 0 | 0.28 | 0.12 | 0.32 | 0.46 | 0.33 | 0.26 |
| 6 | 130474597 | rs1932106 | 0.056 | 2.098 | -0.516 | -0.738 | 0 | -0.006 | -0.006 | 0.28 | 0.67 | 0.64 | 0.56 | 0.82 | 0.59 | 0.67 |
| 6 | 130479049 | rs1889625 | 1.386 | 1.506 | -0.047 | -0.684 | 0.029 | -0.032 | -0.032 | 0 | 0.85 | 0.73 | 0.71 | 0.94 | 0.77 | 0.77 |
| 6 | 130483222 | rs9321214 | 1.054 | 1.49 | -0.028 | -0.665 | 0.008 | -0.023 | -0.023 | 0 | 0.87 | 0.76 | 0.75 | 0.94 | 0.83 | 0.78 |
| 6 | 130487689 | rs6939344 | 0.648 | 1.448 | 0.129 | -0.678 | -0.008 | -0.043 | -0.043 | 0.907 | 0.91 | 0.79 | 0.79 | 0.98 | 0.92 | 0.85 |
| 6 | 130488626 | rs1547334 | 1.121 | 1.462 | 0.164 | -0.701 | 0.018 | -0.054 | -0.054 | 0 | 0.88 | 0.75 | 0.75 | 0.98 | 0.85 | 0.85 |
| 6 | 130490603 | rs9492471 | -2.427 | 1.449 | 0.83 | -0.751 | 0.153 | 0.004 | 0.004 | 0.454 | 0.11 | 0.47 | 0.12 | 0.04 | 0.19 | 0.08 |
| 6 | 130490969 | rs9492472 | -1.863 | 1.432 | 0.845 | -0.753 | 0.1 | 0.035 | 0.035 | 0 | 0.09 | 0.43 | 0.08 | 0.04 | 0.12 | 0.09 |

| | | | | | | | | | | | | | | | | |
|---|---|---|---|---|---|---|---|---|---|---|---|---|---|---|---|---|
| 6 | 130499476 | rs1034263 | -1.871 | 1.364 | 0.81 | -0.823 | 0.077 | -0.011 | -0.011 | 0 | 0.05 | 0.28 | 0.06 | 0 | 0.07 | 0.01 |
| 6 | 130499861 | rs1034262 | 1.121 | 1.248 | 0.244 | -0.717 | 0.043 | -0.08 | -0.08 | 0 | 0.85 | 0.59 | 0.75 | 0.98 | 0.8 | 0.81 |
| 6 | 130503306 | rs9402219 | 0.394 | 0.951 | -0.67 | -0.655 | -0.008 | -0.012 | -0.012 | 1.021 | 0.56 | 0.68 | 0.47 | 0.49 | 0.55 | 0.56 |
| 6 | 130506382 | rs7749850 | -0.017 | 0.339 | -1.22 | -1.248 | -0.004 | 0.056 | 0.056 | 0 | 0.48 | 0.41 | 0.36 | 0.09 | 0.51 | 0.41 |
| 6 | 130507274 | rs13216555 | 0.74 | 0.465 | -1.02 | -0.875 | -0.001 | -0.005 | -0.005 | 0 | 0.47 | 0.41 | 0.42 | 0.4 | 0.5 | 0.54 |
| 6 | 130511560 | rs6900507 | -0.238 | 0.209 | -1.296 | -1.088 | 0.005 | 0.008 | 0.008 | 5.142 | 0.21 | 0.21 | 0.12 | 0.1 | 0.16 | 0.16 |
| 6 | 130512136 | rs12197003 | 0.372 | 0.321 | -1.461 | -0.978 | -0.027 | -0.048 | -0.048 | 0 | 0.23 | 0.19 | 0.35 | 0.47 | 0.28 | 0.21 |
| 6 | 130520672 | rs1932115 | 1.101 | -0.143 | -1.81 | -1.468 | 0.002 | -0.126 | -0.126 | 0 | 0.33 | 0.5 | 0.25 | 0 | 0.38 | 0.19 |
| 6 | 130521794 | rs7768884 | -1.033 | 0.765 | -1.499 | -0.76 | -0.006 | -0.087 | -0.087 | 2.628 | 0.79 | 0.92 | 0.72 | 0.59 | 0.81 | 0.59 |
| 6 | 130530815 | rs1932100 | -0.945 | 0.831 | -1.464 | -0.759 | 0 | -0.06 | -0.06 | 0 | 0.78 | 0.87 | 0.71 | 0.59 | 0.81 | 0.6 |
| 6 | 130535420 | rs4897377 | -0.475 | 0.617 | -1.449 | -0.622 | -0.004 | -0.012 | -0.012 | 0 | 0.73 | 0.67 | 0.66 | 0.59 | 0.73 | 0.58 |
| 6 | 130537005 | rs2326951 | -0.369 | 0.73 | -0.981 | 0.142 | -0.047 | -0.082 | -0.082 | 0 | 0.22 | 0.07 | 0.34 | 0.47 | 0.26 | 0.21 |
| 6 | 130544761 | rs4897379 | -1.5 | 0.321 | -0.177 | -0.386 | -0.014 | -0.052 | -0.052 | 0 | 0.26 | 0.57 | 0.21 | 0.13 | 0.19 | 0.22 |
| 6 | 130549070 | rs6569667 | -1.295 | 0.425 | 0.149 | -0.473 | -0.022 | -0.076 | -0.076 | 0 | 0.17 | 0.74 | 0.11 | 0 | 0.09 | 0.06 |
| 6 | 130549114 | rs6569668 | -0.321 | 0.382 | 0.515 | -0.465 | -0.008 | -0.059 | -0.059 | 0 | 0.11 | 0.45 | 0.05 | 0 | 0.03 | 0.04 |
| 6 | 130549801 | rs10447418 | -1.604 | 0.367 | 0.088 | -0.243 | -0.003 | -0.008 | -0.008 | 0.028 | 0.24 | 0.2 | 0.34 | 0.48 | 0.28 | 0.39 |
| 6 | 130550421 | rs10485399 | -0.256 | 0.269 | 0.104 | -0.203 | 0.01 | -0.014 | -0.014 | 0 | 0.09 | 0.05 | 0.17 | 0.28 | 0.07 | 0.13 |
| 6 | 130550964 | rs6927351 | -1.021 | 0.274 | 0.12 | -0.226 | -0.003 | 0.029 | 0.029 | 0 | 0.14 | 0.16 | 0.21 | 0.35 | 0.13 | 0.21 |
| 6 | 130554677 | rs6908798 | -1.075 | 0.277 | 0.022 | -0.225 | 0.003 | -0.02 | -0.02 | 0 | 0.14 | 0.1 | 0.2 | 0.35 | 0.13 | 0.21 |
| 6 | 130567718 | rs7759336 | -1.491 | 0.181 | 0.071 | -0.435 | 0.019 | 0.073 | 0.073 | 1.232 | 0.25 | 0.86 | 0.23 | 0.14 | 0.19 | 0.21 |
| 6 | 130570115 | rs2024974 | -2.194 | 0.283 | 0.163 | -0.391 | -0.027 | -0.037 | -0.037 | 0 | 0.41 | 0.94 | 0.45 | 0.49 | 0.37 | 0.45 |
| 6 | 130576615 | rs2326946 | -2.218 | 0.25 | 0.27 | -0.373 | 0.005 | 0 | 0 | 0 | 0.39 | 0.65 | 0.42 | 0.49 | 0.37 | 0.48 |
| 6 | 130577677 | rs962011 | -2.105 | 0.256 | 0.233 | -0.37 | 0.01 | 0 | 0 | 0 | 0.4 | 0.78 | 0.44 | 0.49 | 0.37 | 0.48 |
| 6 | 130601380 | rs12212688 | -2.105 | 0.176 | 0.354 | -0.3 | -0.001 | -0.003 | -0.003 | 0.582 | 0.29 | 0.31 | 0.39 | 0.49 | 0.33 | 0.44 |
| 6 | 130607703 | rs12333039 | -2.093 | 0.201 | 0.393 | -0.347 | 0.003 | 0.009 | 0.009 | 0 | 0.29 | 0.33 | 0.38 | 0.47 | 0.33 | 0.39 |
| 6 | 130611364 | rs10457548 | -0.435 | 0.21 | 0.407 | -0.375 | -0.003 | 0.004 | 0.004 | 0 | 0.12 | 0.09 | 0.14 | 0.12 | 0.14 | 0.11 |
| 6 | 130617329 | rs12528700 | -1.106 | 0.195 | 0.459 | -0.294 | -0.008 | -0.038 | -0.038 | 0 | 0.15 | 0.08 | 0.24 | 0.35 | 0.18 | 0.27 |
| 6 | 130627814 | rs6569673 | -2.212 | 0.095 | 0.443 | -0.362 | 0.016 | -0.009 | -0.009 | 0 | 0.28 | 0.77 | 0.31 | 0.36 | 0.21 | 0.32 |
| 6 | 130633537 | rs7760869 | -1.068 | 0.132 | 0.542 | -0.341 | -0.006 | 0.037 | 0.037 | 0 | 0.15 | 0.19 | 0.21 | 0.36 | 0.14 | 0.29 |
| 6 | 130638922 | rs10872361 | 0.005 | 0.184 | 0.502 | -0.394 | -0.007 | 0.008 | 0.008 | 0 | 0.1 | 0.06 | 0.13 | 0.12 | 0.12 | 0.11 |
| 6 | 130639471 | rs6928618 | -1.575 | 0.146 | 0.715 | -0.424 | -0.04 | 0.056 | 0.056 | 0.103 | 0.23 | 0.45 | 0.24 | 0.35 | 0.17 | 0.31 |
| 6 | 130653601 | rs4895881 | -2.312 | 0.135 | 0.624 | -0.438 | 0.006 | -0.002 | -0.002 | 0.32 | 0.37 | 0.79 | 0.44 | 0.47 | 0.34 | 0.44 |
| 6 | 130657718 | rs4897413 | -2.35 | 0.133 | 0.643 | -0.439 | 0.002 | -0.01 | -0.01 | 0.635 | 0.38 | 0.84 | 0.44 | 0.47 | 0.34 | 0.44 |
| 6 | 130665389 | rs6916087 | -2.151 | 0.137 | 0.64 | -0.437 | 0.001 | -0.002 | -0.002 | 0.048 | 0.36 | 0.65 | 0.42 | 0.47 | 0.35 | 0.42 |
| 6 | 130673672 | rs6914528 | -1.107 | 0.133 | 0.638 | -0.439 | 0.014 | -0.095 | -0.095 | 0.669 | 0.21 | 0.51 | 0.21 | 0.12 | 0.2 | 0.14 |
| 6 | 130675996 | rs6917875 | -2.58 | 0.12 | 0.635 | -0.443 | -0.002 | -0.009 | -0.009 | 0.042 | 0.39 | 0.87 | 0.44 | 0.47 | 0.35 | 0.44 |
| 6 | 130681201 | rs9483116 | -1.112 | 0.135 | 0.652 | -0.437 | 0.014 | -0.095 | -0.095 | 0 | 0.21 | 0.51 | 0.21 | 0.12 | 0.2 | 0.14 |
| 6 | 130702701 | rs3850236 | -1.092 | 0.068 | 0.867 | -0.472 | 0.012 | -0.116 | -0.116 | 0 | 0.21 | 0.49 | 0.22 | 0.07 | 0.2 | 0.14 |
| 6 | 130704268 | rs11154574 | -1.211 | 0.063 | 0.83 | -0.464 | 0.002 | 0.001 | 0.001 | 0 | 0.15 | 0.17 | 0.18 | 0.34 | 0.13 | 0.3 |
| 6 | 130723467 | rs1398817 | -1.604 | 0.025 | 0.835 | -0.525 | -0.002 | -0.037 | -0.037 | 0 | 0.25 | 0.7 | 0.25 | 0.08 | 0.2 | 0.15 |
| 6 | 130737708 | rs6919251 | -1.218 | 0.01 | 0.798 | -0.483 | 0.006 | -0.016 | -0.016 | 0 | 0.15 | 0.12 | 0.18 | 0.34 | 0.14 | 0.29 |
| 6 | 130739501 | rs12190331 | -1.178 | 0.027 | 0.817 | -0.48 | 0 | 0.022 | 0.022 | 0 | 0.15 | 0.16 | 0.2 | 0.34 | 0.17 | 0.28 |
| 6 | 130747461 | rs9492587 | -1.953 | 0.002 | 0.796 | -0.53 | 0.003 | 0.036 | 0.036 | 0.576 | 0.26 | 0.79 | 0.27 | 0.14 | 0.21 | 0.15 |
| 6 | 130752428 | rs10155668 | -1.248 | -0.044 | 0.847 | -0.545 | -0.006 | -0.097 | -0.097 | 0 | 0.22 | 0.54 | 0.22 | 0.13 | 0.17 | 0.14 |
| 6 | 130754217 | rs7738424 | -1.393 | 0.017 | 0.848 | -0.545 | -0.009 | -0.082 | -0.082 | 0 | 0.23 | 0.56 | 0.23 | 0.13 | 0.18 | 0.15 |
| 6 | 130756091 | rs10485239 | 0.075 | -0.014 | 0.804 | -0.484 | 0.011 | -0.051 | -0.051 | 0.861 | 0.08 | 0.01 | 0.13 | 0.27 | 0.06 | 0.17 |
| 6 | 130757127 | rs17059002 | -1.351 | -0.002 | 0.821 | -0.487 | -0.006 | 0.02 | 0.02 | 0 | 0.15 | 0.17 | 0.18 | 0.34 | 0.14 | 0.29 |
| 6 | 130761957 | rs3813359 | -1.955 | -0.008 | 0.823 | -0.491 | -0.033 | 0.074 | 0.074 | 0 | 0.19 | 0.37 | 0.2 | 0.34 | 0.14 | 0.29 |
| 6 | 130763281 | rs9492594 | -0.304 | -0.021 | 0.88 | -0.506 | -0.006 | -0.032 | -0.032 | 0 | 0.09 | 0.41 | 0.04 | 0 | 0.02 | 0.03 |
| 6 | 130767104 | rs4555944 | -2.344 | 0.094 | 0.634 | -0.489 | 0.011 | 0 | 0 | 0.96 | 0.37 | 0.71 | 0.4 | 0.47 | 0.32 | 0.44 |
| 6 | 130769858 | rs17059066 | -1.475 | 0.162 | 0.577 | -0.525 | -0.002 | -0.005 | -0.005 | 0.212 | 0.07 | 0.13 | 0.04 | 0.07 | 0.07 | 0.12 |
| 6 | 130771125 | rs2209151 | -2.087 | 0.156 | 0.454 | -0.536 | 0.013 | 0.068 | 0.068 | 1.148 | 0.31 | 0.79 | 0.28 | 0.2 | 0.25 | 0.28 |
| 6 | 130772152 | rs6901603 | -1.592 | 0.185 | 0.387 | -0.355 | 0.022 | -0.002 | -0.002 | 0 | 0.36 | 0.84 | 0.39 | 0.41 | 0.31 | 0.42 |
| 6 | 130773815 | rs13212605 | 0.616 | 0.596 | 0.178 | -1.241 | -0.006 | 0.025 | 0.025 | 0 | 0.3 | 0.21 | 0.24 | 0 | 0.34 | 0.19 |
| 6 | 130778955 | rs11154583 | -1.362 | 0.655 | 0.419 | -0.789 | 0.002 | 0.001 | 0.001 | 0 | 0.05 | 0.07 | 0.06 | 0.07 | 0.08 | 0.12 |
| 6 | 130781452 | rs9285466 | -1.432 | 0.544 | 0.594 | -0.605 | 0.013 | 0.05 | 0.05 | 0.019 | 0.06 | 0.19 | 0.05 | 0.1 | 0.06 | 0.14 |
| 6 | 130783882 | rs12208365 | -0.995 | 0.739 | 0.467 | -0.689 | 0.005 | 0.008 | 0.008 | 0 | 0.06 | 0.11 | 0.04 | 0.07 | 0.09 | 0.14 |
| 6 | 130784063 | rs13218932 | -0.145 | 0.519 | 0.747 | -1.291 | 0.006 | 0.07 | 0.07 | 2.677 | 0.48 | 0.31 | 0.41 | 0.13 | 0.44 | 0.34 |
| 6 | 130784336 | rs13203169 | 0.654 | 0.401 | 0.777 | -1.376 | -0.003 | -0.025 | -0.025 | 0 | 0.33 | 0.29 | 0.25 | 0.06 | 0.36 | 0.23 |
| 6 | 130787077 | rs6927777 | -0.138 | 0.401 | 0.796 | -1.365 | -0.001 | 0.031 | 0.031 | 0.081 | 0.46 | 0.31 | 0.41 | 0.07 | 0.46 | 0.33 |
| 6 | 130787438 | rs11154584 | -0.535 | 0.348 | 0.689 | -1.304 | 0.01 | 0.083 | 0.083 | 0 | 0.52 | 0.33 | 0.48 | 0.14 | 0.57 | 0.48 |
| 6 | 130787776 | rs11751990 | -0.18 | 0.442 | 0.102 | -0.917 | 0.005 | 0.038 | 0.038 | 3.099 | 0.16 | 0.05 | 0.18 | 0.06 | 0.12 | 0.11 |
| 6 | 130789541 | rs9492602 | -0.317 | 0.173 | 0.347 | -0.948 | 0.004 | -0.066 | -0.066 | 0.522 | 0.43 | 0.5 | 0.38 | 0.15 | 0.47 | 0.41 |
| 6 | 130789807 | rs9483132 | -2.485 | -0.339 | 1.04 | 0.548 | 0.001 | -0.019 | -0.019 | 0.353 | 0.32 | 0.72 | 0.47 | 0.49 | 0.28 | 0.51 |
| 6 | 130791847 | rs9375715 | -1.181 | -0.384 | 0.736 | 0.529 | -0.008 | -0.003 | -0.003 | 0.558 | 0.08 | 0.05 | 0.08 | 0.24 | 0.13 | 0.15 |
| 6 | 130792769 | rs17059127 | -0.428 | -0.453 | 0.759 | 0.495 | 0.032 | 0.042 | 0.042 | 10.319 | 0.15 | 0.06 | 0.16 | 0.06 | 0.06 | 0.15 |
| 6 | 130796396 | rs9492605 | -2.259 | -0.454 | 0.96 | 0.526 | 0.047 | 0.135 | 0.135 | 0.33 | 0.13 | 0.45 | 0.18 | 0.29 | 0.14 | 0.19 |
| 6 | 130805694 | rs12197132 | -1.186 | -0.629 | 0.875 | 0.543 | -0.005 | -0.002 | -0.002 | 0 | 0.08 | 0.06 | 0.08 | 0.24 | 0.13 | 0.19 |
| 6 | 130810954 | rs4897427 | -2.299 | -0.624 | 0.82 | 0.562 | -0.034 | 0.008 | 0.008 | 2.302 | 0.3 | 0.52 | 0.35 | 0.37 | 0.21 | 0.35 |

| | | | | | | | | | | | | | | | | |
|---|---|---|---|---|---|---|---|---|---|---|---|---|---|---|---|---|
| 6 | 130818014 | rs2876086 | -1.847 | -0.598 | 0.394 | 0.552 | -0.019 | 0.002 | 0.002 | 2.436 | 0.26 | 0.39 | 0.26 | 0.31 | 0.19 | 0.34 |
| 6 | 130824446 | rs11962865 | -1.049 | -0.657 | -0.33 | 0.324 | 0.017 | 0.017 | 0.017 | 6.904 | 0.2 | 0.15 | 0.18 | 0.06 | 0.07 | 0.16 |
| 6 | 130827482 | rs9402234 | -1.525 | -0.133 | -0.005 | 1.157 | 0.01 | 0.072 | 0.072 | 0 | 0.11 | 0.22 | 0.16 | 0.34 | 0.11 | 0.15 |
| 6 | 130829447 | rs1203356 | -0.621 | -1.143 | -0.701 | 1.554 | 0.016 | -0.122 | -0.122 | 0.009 | 0.2 | 0.02 | 0.32 | 0.46 | 0.18 | 0.17 |
| 6 | 130832922 | rs17059166 | -0.819 | -1.726 | 0.337 | 0.861 | -0.028 | 0.037 | 0.037 | 0 | 0.26 | 0.53 | 0.22 | 0.34 | 0.19 | 0.49 |
| 6 | 130839788 | rs873146 | -0.661 | -1.827 | 1.062 | 0.702 | -0.006 | -0.035 | -0.035 | 0.373 | 0.15 | 0.33 | 0.13 | 0.09 | 0.14 | 0.18 |
| 6 | 130841206 | rs7771642 | -0.661 | -1.78 | 1.061 | 0.702 | -0.002 | -0.035 | -0.035 | 0 | 0.15 | 0.33 | 0.13 | 0.11 | 0.14 | 0.18 |
| 6 | 130842360 | rs6899931 | -1.573 | -1.731 | 0.988 | 0.637 | -0.023 | -0.091 | -0.091 | 0.423 | 0.25 | 0.5 | 0.18 | 0.11 | 0.2 | 0.24 |
| 6 | 130844791 | rs9492632 | -1.615 | -1.721 | 1.102 | 0.674 | -0.044 | -0.091 | -0.091 | 0 | 0.22 | 0.48 | 0.15 | 0.11 | 0.13 | 0.23 |
| 6 | 130845421 | rs9375723 | -0.839 | -2.579 | 0.856 | 0.747 | -0.007 | -0.003 | -0.003 | 0 | 0.39 | 0.45 | 0.33 | 0.67 | 0.31 | 0.42 |
| 6 | 130847255 | rs9492633 | -0.62 | -2.59 | 0.857 | 0.774 | 0.003 | 0 | 0 | 0.194 | 0.37 | 0.38 | 0.32 | 0.66 | 0.3 | 0.42 |
| 6 | 130852623 | rs7768204 | -0.686 | -2.628 | 1.147 | 0.904 | -0.036 | -0.035 | -0.035 | 0 | 0.14 | 0.31 | 0.1 | 0.09 | 0.07 | 0.17 |
| 6 | 130854536 | rs9492635 | 0.548 | -2.508 | 1.148 | 0.96 | 0 | -0.009 | -0.009 | 0 | 0.07 | 0.12 | 0.03 | 0 | 0.05 | 0.05 |
| 6 | 130859449 | rs9483138 | -0.88 | -2.523 | 1.215 | 0.967 | -0.006 | -0.005 | -0.005 | 0 | 0.41 | 0.51 | 0.38 | 0.74 | 0.32 | 0.54 |
| 6 | 130868480 | rs1933770 | -1.463 | -2.512 | 1.115 | 0.972 | 0.057 | 0.082 | 0.082 | 0.026 | 0.51 | 0.71 | 0.43 | 0.76 | 0.38 | 0.59 |
| 6 | 130868894 | rs17059212 | 0.205 | -2.479 | 1.107 | 1.056 | -0.014 | -0.135 | -0.135 | 0.059 | 0.26 | 0.02 | 0.27 | 0.4 | 0.25 | 0.2 |
| 6 | 130871599 | rs1203345 | -1.69 | -2.823 | 0.745 | 0.621 | -0.01 | 0.017 | 0.017 | 0 | 0.64 | 0.75 | 0.46 | 0.76 | 0.45 | 0.7 |
| 6 | 130876090 | rs6914156 | 0.214 | -2.492 | 0.217 | -0.032 | -0.012 | -0.028 | -0.028 | 0 | 0.75 | 0.74 | 0.88 | 0.91 | 0.85 | 0.73 |
| 6 | 130886683 | rs9492647 | 0.37 | -2.121 | -0.142 | 0.062 | -0.004 | -0.029 | -0.029 | 0 | 0.08 | 0.15 | 0.04 | 0 | 0.05 | 0.05 |
| 6 | 130887916 | rs6932172 | 1.248 | -2.122 | -0.376 | 0.249 | -0.013 | -0.023 | -0.023 | 0.036 | 0.37 | 0.29 | 0.53 | 0.49 | 0.54 | 0.48 |
| 6 | 130892429 | rs7451336 | 0.127 | -2.008 | -0.802 | 0.215 | 0.001 | -0.094 | -0.094 | 0.595 | 0.27 | 0.03 | 0.27 | 0.36 | 0.24 | 0.2 |
| 6 | 130894602 | rs1316537 | 0.317 | -1.286 | -0.115 | 0.763 | 0 | 0 | 0 | 0.266 | 0.45 | 0.54 | 0.6 | 0.61 | 0.57 | 0.59 |
| 6 | 130910342 | rs9321242 | -1.015 | -1.287 | -0.084 | 1.241 | -0.004 | 0.012 | 0.012 | 0 | 0.08 | 0.09 | 0.1 | 0.19 | 0.05 | 0.12 |
| 6 | 130913854 | rs7745661 | 0.321 | -1.633 | -0.652 | 0.962 | 0.017 | 0.018 | 0.018 | 0 | 0.64 | 0.84 | 0.65 | 0.7 | 0.66 | 0.72 |
| 6 | 130916665 | rs11154595 | -0.281 | -1.589 | -0.397 | 1.951 | 0.006 | -0.017 | -0.017 | 0.124 | 0.09 | 0.04 | 0.16 | 0.19 | 0.07 | 0.11 |
| 6 | 130918607 | rs9375749 | -0.984 | -1.786 | -0.029 | 1.79 | -0.007 | 0.005 | 0.005 | 0 | 0.16 | 0.2 | 0.2 | 0.19 | 0.1 | 0.14 |
| 6 | 130921300 | rs4144213 | -1.695 | -2.153 | 0.306 | 1.95 | -0.023 | 0.023 | 0.023 | 0 | 0.28 | 0.64 | 0.36 | 0.53 | 0.17 | 0.51 |
| 6 | 130922149 | rs9483149 | 1.059 | -1.76 | 0.907 | 2.332 | -0.02 | 0.064 | 0.064 | 0 | 0.86 | 0.73 | 0.83 | 0.73 | 0.89 | 0.62 |
| 6 | 130923327 | rs9388813 | -1.456 | -1.55 | 0.728 | 2.644 | -0.021 | 0.138 | 0.138 | 0 | 0.11 | 0.23 | 0.23 | 0.36 | 0.07 | 0.19 |
| 6 | 130926413 | rs4897445 | -1.323 | -1.408 | 0.583 | 1.91 | -0.026 | 0.119 | 0.119 | 0 | 0.15 | 0.29 | 0.22 | 0.46 | 0.09 | 0.47 |
| 6 | 130936249 | rs4897446 | -0.931 | -1.404 | 0.398 | 1.412 | -0.028 | 0.014 | 0.014 | 1.081 | 0.36 | 0.5 | 0.41 | 0.62 | 0.29 | 0.55 |
| 6 | 130936406 | rs6903457 | 0.114 | -1.53 | 0.479 | 1.095 | 0.012 | 0.018 | 0.018 | 0.598 | 0.15 | 0.07 | 0.09 | 0 | 0.14 | 0.08 |
| 6 | 130936495 | rs4144214 | -0.633 | -1.4 | 0.254 | 0.767 | -0.01 | -0.038 | -0.038 | 0 | 0.35 | 0.53 | 0.31 | 0.2 | 0.33 | 0.15 |
| 6 | 130941233 | rs4895890 | -0.432 | -1.554 | 0.597 | 1.277 | -0.017 | 0.091 | 0.091 | 0 | 0.08 | 0.19 | 0.1 | 0.22 | 0.03 | 0.12 |
| 6 | 130943247 | rs6924980 | 0.453 | -1.524 | 0.907 | 0.974 | -0.029 | -0.01 | -0.01 | 0.523 | 0.16 | 0.3 | 0.15 | 0.15 | 0.12 | 0.09 |
| 6 | 130953757 | rs9321245 | -0.176 | -1.49 | 0.503 | 1.202 | -0.004 | -0.039 | -0.039 | 0.009 | 0.09 | 0.03 | 0.13 | 0.26 | 0.08 | 0.3 |
| 6 | 130966801 | rs2136055 | -1.46 | -1.68 | 0.495 | 1.205 | 0.036 | 0.031 | 0.031 | 0 | 0.41 | 0.7 | 0.45 | 0.73 | 0.3 | 0.62 |
| 6 | 130967588 | rs9492683 | -1.062 | -2.048 | 0.569 | 1.04 | -0.011 | 0.042 | 0.042 | 2.592 | 0.22 | 0.29 | 0.21 | 0.47 | 0.15 | 0.25 |
| 6 | 130968384 | rs4897447 | -0.457 | -1.944 | 0.641 | 0.962 | -0.014 | 0.075 | 0.075 | 0 | 0.16 | 0.29 | 0.16 | 0.32 | 0.1 | 0.23 |
| 6 | 130969493 | rs13218604 | 0.357 | -1.699 | 0.253 | 0.743 | -0.009 | 0.02 | 0.02 | 0 | 0.18 | 0.12 | 0.16 | 0.05 | 0.2 | 0.06 |
| 6 | 130970281 | rs2326972 | 0.481 | -1.479 | 0.29 | 1.067 | -0.023 | -0.074 | -0.074 | 0.089 | 0.14 | 0.24 | 0.1 | 0 | 0.11 | 0.08 |
| 6 | 130977969 | rs9375754 | -0.532 | -1.071 | -0.301 | 1.507 | -0.002 | -0.018 | -0.018 | 0 | 0.07 | 0.03 | 0.09 | 0.26 | 0.06 | 0.29 |
| 6 | 130984945 | rs2020202 | -0.901 | -1.648 | -0.837 | 1.14 | 0.019 | -0.125 | -0.125 | 0 | 0.28 | 0.06 | 0.2 | 0.57 | 0.2 | 0.43 |
| 6 | 130992155 | rs9388825 | -1.448 | -1.604 | -0.877 | 0.537 | -0.01 | 0.09 | 0.09 | 4.888 | 0.56 | 0.79 | 0.53 | 0.79 | 0.48 | 0.65 |
| 6 | 130995644 | rs11754539 | 0.21 | -1.434 | 0.175 | 0.392 | -0.007 | -0.076 | -0.076 | 0 | 0.24 | 0.27 | 0.18 | 0 | 0.25 | 0.14 |
| 6 | 131003355 | rs7752280 | -0.543 | -1.361 | 0.725 | 0.551 | 0.02 | 0.165 | 0.165 | 0 | 0.05 | 0.24 | 0.12 | 0.21 | 0.03 | 0.08 |
| 6 | 131004998 | rs9388831 | -1.399 | -1.402 | 0.643 | 0.419 | 0.009 | 0.089 | 0.089 | 4.797 | 0.55 | 0.77 | 0.53 | 0.79 | 0.48 | 0.66 |
| 6 | 131018295 | rs9402269 | 0.287 | -0.438 | 0.968 | 0.564 | 0.033 | -0.014 | -0.014 | 0 | 0.13 | 0.21 | 0.08 | 0.09 | 0.18 | 0.19 |
| 6 | 131021812 | rs994211 | -0.218 | -0.745 | 0.914 | 1.149 | 0.005 | -0.019 | -0.019 | 0 | 0.17 | 0.15 | 0.18 | 0.28 | 0.17 | 0.18 |
| 6 | 131030583 | rs9483167 | -1.142 | -0.452 | 1.14 | 0.957 | 0.045 | 0.021 | 0.021 | 0 | 0.27 | 0.47 | 0.36 | 0.32 | 0.3 | 0.33 |
| 6 | 131032549 | rs9388833 | -0.655 | -0.49 | 1.142 | 1.035 | 0.007 | 0.042 | 0.042 | 0 | 0.14 | 0.15 | 0.25 | 0.36 | 0.14 | 0.23 |
| 6 | 131035958 | rs9483170 | -1.336 | -0.699 | 1.042 | 0.809 | 0.001 | 0.004 | 0.004 | 2.962 | 0.5 | 0.56 | 0.39 | 0.34 | 0.51 | 0.49 |
| 6 | 131040736 | rs12191580 | 0.736 | -0.584 | 0.928 | 0.822 | 0.016 | -0.007 | -0.007 | 0.504 | 0.13 | 0.15 | 0.08 | 0.08 | 0.18 | 0.19 |
| 6 | 131044450 | rs1396250 | -1.253 | -0.69 | 1.024 | 0.823 | -0.004 | 0.006 | 0.006 | 4.276 | 0.46 | 0.57 | 0.47 | 0.44 | 0.49 | 0.55 |
| 6 | 131044773 | rs1848410 | -0.886 | -0.765 | 1.112 | 0.844 | 0.007 | 0.008 | 0.008 | 0 | 0.31 | 0.37 | 0.26 | 0.34 | 0.35 | 0.37 |
| 6 | 131044799 | rs2221651 | 0.669 | -0.795 | 1.113 | 0.868 | -0.011 | -0.003 | -0.003 | 0.14 | 0.2 | 0.25 | 0.18 | 0.19 | 0.17 | 0.14 |
| 6 | 131046822 | rs4897451 | -1.742 | -0.853 | 1.111 | 0.708 | 0 | -0.022 | -0.022 | 0.255 | 0.45 | 0.42 | 0.56 | 0.88 | 0.5 | 0.62 |
| 6 | 131046888 | rs4897452 | -0.022 | -0.842 | 1.061 | 0.727 | -0.037 | 0.03 | 0.03 | 0 | 0.1 | 0.02 | 0.2 | 0.05 | 0.11 | 0.13 |
| 6 | 131047443 | rs1497883 | -2.573 | -0.781 | 1.061 | 0.779 | -0.004 | 0.039 | 0.039 | 5.011 | 0.64 | 0.71 | 0.69 | 0.88 | 0.66 | 0.75 |
| 6 | 131054125 | rs10457556 | -1.417 | -0.729 | 1.005 | 0.817 | -0.007 | -0.002 | -0.002 | 0 | 0.41 | 0.36 | 0.51 | 0.58 | 0.47 | 0.51 |
| 6 | 131063056 | rs9402276 | 0.786 | -0.708 | 1.037 | 0.724 | 0.039 | -0.001 | -0.001 | 0 | 0.12 | 0.21 | 0.07 | 0.09 | 0.18 | 0.2 |
| 6 | 131063500 | rs4549635 | 1.341 | -1.039 | 1.127 | 0.699 | 0.004 | -0.016 | -0.016 | 0 | 0.16 | 0.17 | 0.12 | 0 | 0.16 | 0.13 |
| 6 | 131063724 | rs4897458 | -0.533 | -1.072 | 1.04 | 0.804 | 0.007 | -0.004 | -0.004 | 0 | 0.19 | 0.15 | 0.18 | 0.22 | 0.17 | 0.16 |
| 6 | 131074746 | rs1497890 | -0.275 | -1.08 | 0.895 | 0.799 | 0.016 | -0.021 | -0.021 | 0 | 0.18 | 0.09 | 0.18 | 0.21 | 0.17 | 0.16 |
| 6 | 131077027 | rs1876545 | -0.987 | -1.13 | 0.913 | 0.914 | 0.001 | -0.004 | -0.004 | 0 | 0.44 | 0.54 | 0.52 | 0.72 | 0.49 | 0.59 |
| 6 | 131080130 | rs6933145 | -2.536 | -1.105 | 0.817 | 0.761 | -0.011 | 0.084 | 0.084 | 3.303 | 0.64 | 0.77 | 0.7 | 0.93 | 0.66 | 0.75 |
| 6 | 131084057 | rs4448122 | -1.178 | -1.109 | 0.832 | 0.771 | -0.014 | 0.053 | 0.053 | 1.338 | 0.47 | 0.71 | 0.54 | 0.72 | 0.49 | 0.6 |
| 6 | 131089467 | rs4897466 | -0.421 | -1.082 | 0.514 | 0.727 | -0.012 | 0.022 | 0.022 | 0.395 | 0.37 | 0.33 | 0.39 | 0.14 | 0.45 | 0.46 |
| 6 | 131092501 | rs761836 | -2.41 | -0.887 | -0.083 | 0.796 | -0.006 | 0.044 | 0.044 | 3.8 | 0.65 | 0.74 | 0.71 | 0.93 | 0.66 | 0.75 |

| | | | | | | | | | | | | | | | | |
|---|---|---|---|---|---|---|---|---|---|---|---|---|---|---|---|---|
| 6 | 131095294 | rs11154605 | -0.065 | -0.697 | -0.568 | 1.271 | -0.02 | -0.07 | -0.07 | 0 | 0.15 | 0.01 | 0.15 | 0.21 | 0.16 | 0.16 |
| 6 | 131097518 | rs9375768 | -0.835 | -0.697 | -0.504 | 0.867 | 0.01 | -0.02 | -0.02 | 0 | 0.2 | 0.07 | 0.26 | 0.79 | 0.2 | 0.3 |
| 6 | 131100306 | rs2326985 | -0.961 | -0.691 | -0.431 | 0.831 | 0.026 | 0.032 | 0.032 | 0.239 | 0.38 | 0.25 | 0.38 | 0.8 | 0.36 | 0.43 |
| 6 | 131103572 | rs11154606 | -0.708 | -0.282 | -0.597 | 1.291 | -0.011 | -0.124 | -0.124 | 0 | 0.17 | 0.03 | 0.22 | 0.48 | 0.17 | 0.21 |
| 6 | 131107930 | rs7748328 | -0.694 | 0.057 | -0.148 | 0.583 | 0.002 | -0.007 | -0.007 | 0.07 | 0.23 | 0.2 | 0.29 | 0.1 | 0.28 | 0.32 |
| 6 | 131110370 | rs7741731 | -1.409 | -0.758 | 0.227 | 0.886 | -0.03 | 0.024 | 0.024 | 0 | 0.23 | 0.41 | 0.2 | 0.3 | 0.18 | 0.26 |
| 6 | 131112116 | rs12527906 | 1.173 | -0.402 | -0.418 | 0.126 | -0.037 | 0.075 | 0.075 | 2.386 | 0.36 | 0.17 | 0.36 | 0.18 | 0.41 | 0.38 |
| 6 | 131112865 | rs17059616 | 0.09 | -0.579 | -0.063 | 0.125 | -0.029 | -0.03 | -0.03 | 0 | 0.06 | 0.15 | 0.02 | 0 | 0 | 0.03 |
| 6 | 131114386 | rs761841 | 0.069 | -0.413 | 0.181 | 0.13 | 0.006 | 0.01 | 0.01 | 1.212 | 0.53 | 0.44 | 0.59 | 0.25 | 0.59 | 0.55 |
| 6 | 131121992 | rs12215315 | -2.133 | -0.503 | 0.551 | 0.525 | -0.036 | 0.063 | 0.063 | 1.041 | 0.22 | 0.62 | 0.25 | 0.37 | 0.14 | 0.26 |
| 6 | 131128370 | rs7761842 | -2.304 | -0.476 | 0.412 | 0.383 | 0.009 | -0.031 | -0.031 | 0 | 0.36 | 0.77 | 0.44 | 0.43 | 0.3 | 0.4 |
| 6 | 131131650 | rs11758228 | -0.195 | -0.222 | 0.665 | 0.364 | 0 | 0 | 0 | 0.057 | 0.06 | 0.04 | 0.1 | 0.09 | 0.08 | 0.16 |
| 6 | 131131934 | rs7744295 | -0.882 | -0.253 | 0.812 | 0.273 | 0.004 | -0.012 | -0.012 | 0.822 | 0.15 | 0.18 | 0.2 | 0.08 | 0.16 | 0.14 |
| 6 | 131135714 | rs9385562 | -0.444 | -0.22 | 0.893 | 0.694 | -0.016 | 0.191 | 0.191 | 0 | 0.06 | 0.2 | 0.15 | 0.41 | 0.04 | 0.08 |
| 6 | 131136313 | rs7758259 | 1.688 | 0.031 | 0.518 | 0.292 | -0.031 | 0.049 | 0.049 | 0.924 | 0.33 | 0.16 | 0.28 | 0.18 | 0.4 | 0.38 |
| 6 | 131145052 | rs1114644 | -0.22 | 0.172 | 0.515 | -0.246 | 0.015 | -0.007 | -0.007 | 0 | 0.59 | 0.72 | 0.69 | 0.99 | 0.69 | 0.82 |
| 6 | 131148426 | rs6940163 | -0.137 | 0.117 | 0.503 | -0.196 | -0.015 | -0.011 | -0.011 | 0.085 | 0.06 | 0.08 | 0.04 | 0.01 | 0.02 | 0.05 |
| 6 | 131149795 | rs6908154 | -0.48 | 0.053 | 0.985 | -0.051 | -0.006 | -0.006 | -0.006 | 0 | 0.62 | 0.59 | 0.67 | 0.74 | 0.71 | 0.84 |
| 6 | 131149902 | rs6908659 | -0.313 | 0.05 | 1.066 | -0.045 | -0.012 | -0.012 | -0.012 | 0 | 0.62 | 0.54 | 0.66 | 0.75 | 0.71 | 0.84 |
| 6 | 131150127 | rs9388858 | -0.489 | 0.054 | 1.011 | -0.042 | 0.011 | 0.011 | 0.011 | 0 | 0.62 | 0.63 | 0.67 | 0.75 | 0.71 | 0.84 |
| 6 | 131152851 | rs2281490 | -1.622 | 0.284 | 0.884 | -0.065 | 0.013 | 0.024 | 0.024 | 0.528 | 0.26 | 0.37 | 0.4 | 0.68 | 0.31 | 0.51 |
| 6 | 131159590 | rs9385564 | 2.021 | 0.175 | 0.712 | -0.017 | -0.028 | -0.065 | -0.065 | 1.469 | 0.32 | 0.09 | 0.34 | 0.46 | 0.39 | 0.36 |
| 6 | 131160876 | rs6536 | -0.878 | 0.155 | 0.858 | -0.097 | 0.01 | 0.108 | 0.108 | 0.178 | 0.66 | 0.88 | 0.74 | 1 | 0.71 | 0.87 |
| 6 | 131161504 | rs1043596 | -2.047 | 0.147 | 0.977 | -0.05 | 0.007 | -0.006 | -0.006 | 0 | 0.31 | 0.57 | 0.33 | 0.28 | 0.3 | 0.48 |
| 6 | 131163067 | rs17059683 | -0.703 | 0.285 | 1.016 | -0.073 | 0.048 | 0.056 | 0.056 | 0 | 0.05 | 0.17 | 0.12 | 0.13 | 0.06 | 0.12 |
| 6 | 131163098 | rs6935267 | -1.547 | 0.024 | 0.689 | -0.382 | 0.016 | 0.022 | 0.022 | 0 | 0.77 | 0.89 | 0.81 | 1 | 0.82 | 0.89 |
| 6 | 131163660 | rs12194612 | -2.203 | 0.319 | 1.003 | -0.049 | 0.012 | -0.005 | -0.005 | 0 | 0.3 | 0.47 | 0.32 | 0.28 | 0.3 | 0.48 |
| 6 | 131170273 | rs9402292 | 1.289 | 0.311 | 0.847 | -0.09 | 0 | -0.046 | -0.046 | 1.445 | 0.37 | 0.17 | 0.35 | 0.48 | 0.39 | 0.4 |
| 6 | 131175997 | rs17831009 | -1.055 | 0.397 | 0.825 | -0.374 | 0.008 | 0.003 | 0.003 | 0 | 0.13 | 0.17 | 0.14 | 0.19 | 0.17 | 0.33 |
| 6 | 131177645 | rs2184408 | -0.891 | 0.342 | 0.716 | -0.448 | 0.009 | 0.113 | 0.113 | 0.178 | 0.66 | 0.88 | 0.74 | 1 | 0.71 | 0.87 |
| 6 | 131183678 | rs933064 | -1.882 | 0.245 | 1.16 | -0.103 | 0.026 | 0.364 | 0.364 | 0.031 | 0.09 | 0.45 | 0.12 | 0.36 | 0.05 | 0.14 |
| 6 | 131184770 | rs915172 | -2.426 | 0.248 | 1.189 | -0.257 | 0.009 | -0.001 | -0.001 | 0 | 0.3 | 0.59 | 0.37 | 0.29 | 0.3 | 0.48 |
| 6 | 131184914 | rs915171 | -1.371 | 0.297 | 1.147 | -0.284 | 0.028 | -0.016 | -0.016 | 0.609 | 0.21 | 0.39 | 0.27 | 0.15 | 0.27 | 0.32 |
| 6 | 131189874 | rs10457562 | 0.194 | 0.329 | 1.004 | -0.274 | -0.043 | -0.008 | -0.008 | 0 | 0.07 | 0 | 0.1 | 0.06 | 0.14 | 0.21 |
| 6 | 131190838 | rs17059736 | -0.465 | 0.218 | 1.115 | -0.467 | 0.006 | -0.009 | -0.009 | 0.244 | 0.13 | 0.25 | 0.21 | 0.09 | 0.13 | 0.11 |
| 6 | 131209404 | rs6938586 | -1.755 | 0.149 | 1.087 | -0.455 | -0.016 | -0.001 | -0.001 | 0 | 0.25 | 0.6 | 0.37 | 0.23 | 0.19 | 0.33 |
| 6 | 131215271 | rs4897473 | 2.022 | 0.214 | 0.805 | -0.656 | -0.074 | 0.141 | 0.141 | 1.686 | 0.29 | 0.05 | 0.27 | 0.05 | 0.36 | 0.29 |
| 6 | 131221902 | rs2068147 | 1.187 | -0.084 | 0.962 | -0.555 | -0.048 | -0.052 | -0.052 | 0 | 0.4 | 0.18 | 0.4 | 0.52 | 0.51 | 0.51 |
| 6 | 131234008 | rs1334689 | -0.909 | -0.258 | 1.048 | -0.588 | 0.03 | 0.109 | 0.109 | 0.581 | 0.67 | 0.89 | 0.79 | 1 | 0.73 | 0.88 |
| 6 | 131241971 | rs4357159 | -1.672 | -0.246 | 1.448 | -0.516 | -0.006 | 0.077 | 0.077 | 0.854 | 0.24 | 0.66 | 0.37 | 0.47 | 0.21 | 0.32 |
| 6 | 131250223 | rs3822863 | 1.313 | -0.236 | 1.261 | -0.532 | -0.089 | -0.11 | -0.11 | 0 | 0.39 | 0.08 | 0.4 | 0.52 | 0.53 | 0.52 |
| 6 | 131253973 | rs7766004 | -0.78 | -0.187 | 1.34 | -0.515 | 0.017 | -0.007 | -0.007 | 0 | 0.67 | 0.79 | 0.78 | 1 | 0.73 | 0.88 |
| 6 | 131266757 | rs3777464 | -0.717 | -0.036 | 1.321 | -0.116 | 0.008 | 0.116 | 0.116 | 0.009 | 0.12 | 0.23 | 0.21 | 0.34 | 0.11 | 0.12 |
| 6 | 131278669 | rs12202345 | -0.271 | 0.003 | 1.501 | -0.195 | 0.018 | -0.019 | -0.019 | 0 | 0.07 | 0.33 | 0.08 | 0 | 0.06 | 0.08 |
| 6 | 131282210 | rs6941712 | -1.157 | 0.079 | 1.391 | -0.042 | 0.027 | 0.203 | 0.203 | 0.01 | 0.14 | 0.29 | 0.29 | 0.47 | 0.14 | 0.23 |
| 6 | 131294299 | rs12529894 | -0.05 | 0.166 | 1.144 | -0.242 | 0.009 | 0.004 | 0.004 | 0 | 0.08 | 0.12 | 0.2 | 0.09 | 0.09 | 0.09 |
| 6 | 131304233 | rs9321265 | -1.12 | 0.106 | 1.162 | -0.299 | 0.023 | 0.043 | 0.043 | 0.101 | 0.16 | 0.31 | 0.31 | 0.22 | 0.16 | 0.2 |
| 6 | 131306828 | rs1977578 | -1.123 | -0.142 | 0.626 | -0.721 | 0.017 | 0.106 | 0.106 | 0.164 | 0.73 | 0.94 | 0.82 | 1 | 0.77 | 0.9 |
| 6 | 131308551 | rs2327006 | -0.666 | -0.001 | 0.857 | -0.439 | 0.006 | -0.007 | -0.007 | 0 | 0.12 | 0.21 | 0.23 | 0.08 | 0.13 | 0.09 |
| 6 | 131313416 | rs9375791 | -1.903 | 0.144 | 0.964 | -0.562 | 0.009 | -0.044 | -0.044 | 3.909 | 0.33 | 0.48 | 0.42 | 0.8 | 0.36 | 0.47 |
| 6 | 131320232 | rs6905151 | -0.884 | 0.125 | 0.943 | -0.338 | -0.002 | -0.011 | -0.011 | 0 | 0.2 | 0.17 | 0.2 | 0.45 | 0.22 | 0.36 |
| 6 | 131327956 | rs6921207 | -1.989 | 0.107 | 0.675 | -0.402 | -0.001 | -0.008 | -0.008 | 0.181 | 0.36 | 0.65 | 0.45 | 0.54 | 0.37 | 0.44 |
| 6 | 131332532 | rs9375795 | -0.236 | 0.125 | 0.565 | -0.314 | -0.049 | -0.08 | -0.08 | 0 | 0.12 | 0.01 | 0.15 | 0.47 | 0.16 | 0.25 |
| 6 | 131335324 | rs1888959 | -0.48 | -0.131 | 0.452 | -0.634 | 0.006 | 0 | 0 | 0 | 0.63 | 0.65 | 0.7 | 0.61 | 0.75 | 0.75 |
| 6 | 131339257 | rs11969461 | 0.055 | -0.08 | 0.439 | -0.759 | 0.002 | -0.001 | -0.001 | 0 | 0.1 | 0.09 | 0.21 | 0.09 | 0.15 | 0.08 |
| 6 | 131345133 | rs6906478 | 1.51 | -0.11 | 0.384 | -0.862 | -0.011 | 0.031 | 0.031 | 0.836 | 0.34 | 0.3 | 0.29 | 0.06 | 0.38 | 0.31 |
| 6 | 131345397 | rs9388876 | -0.083 | -0.118 | 0.304 | -0.812 | -0.049 | -0.079 | -0.079 | 1.33 | 0.12 | 0.01 | 0.15 | 0.45 | 0.16 | 0.25 |
| 6 | 131345664 | rs2007339 | -1.155 | -0.162 | 0.372 | -0.811 | 0.032 | 0.087 | 0.087 | 0.243 | 0.68 | 0.89 | 0.74 | 0.87 | 0.76 | 0.78 |
| 6 | 131348212 | rs4895903 | 2.012 | -0.165 | 0.374 | -0.803 | -0.088 | 0.136 | 0.136 | 2.16 | 0.29 | 0.07 | 0.26 | 0.05 | 0.38 | 0.31 |
| 6 | 131349617 | rs12203760 | -0.824 | -0.165 | 0.582 | -0.749 | 0.005 | 0.088 | 0.088 | 0 | 0.21 | 0.34 | 0.21 | 0.45 | 0.22 | 0.36 |
| 6 | 131371748 | rs7765757 | -1.087 | -0.124 | 0.346 | -0.825 | 0.032 | 0.107 | 0.107 | 0.123 | 0.68 | 0.92 | 0.75 | 0.87 | 0.76 | 0.78 |
| 6 | 131372135 | rs13437326 | 1.479 | 0.087 | 0.442 | -0.756 | -0.004 | -0.003 | -0.003 | 0 | 0.34 | 0.33 | 0.3 | 0.06 | 0.38 | 0.31 |
| 6 | 131394554 | rs7752642 | -1.357 | 0.4 | 0.21 | -0.883 | 0.009 | -0.022 | -0.022 | 0.733 | 0.37 | 0.55 | 0.48 | 0.81 | 0.44 | 0.49 |
| 6 | 131402416 | rs9492803 | 1.225 | 0.729 | 0.213 | -0.805 | -0.004 | -0.005 | -0.005 | 0 | 0.34 | 0.45 | 0.31 | 0.29 | 0.36 | 0.4 |
| 6 | 131402814 | rs13208443 | 0.731 | 1.391 | 0.093 | -0.346 | -0.02 | 0.035 | 0.035 | 0 | 0.1 | 0.03 | 0.08 | 0 | 0.12 | 0.04 |
| 6 | 131404668 | rs9398976 | -0.094 | 1.089 | 0.15 | -0.175 | -0.045 | -0.03 | -0.03 | 0 | 0.09 | 0.02 | 0.14 | 0.47 | 0.15 | 0.23 |
| 6 | 131405423 | rs11154614 | -0.165 | 0.883 | 0.331 | -0.368 | 0.008 | -0.014 | -0.014 | 0 | 0.19 | 0.23 | 0.27 | 0.1 | 0.24 | 0.13 |
| 6 | 131422545 | rs6915839 | 0.714 | 0.765 | 0.175 | -0.284 | -0.004 | -0.002 | -0.002 | 11.518 | 0.46 | 0.52 | 0.43 | 0.42 | 0.42 | 0.53 |
| 6 | 131422563 | rs6915845 | -0.107 | 0.43 | -0.363 | -0.159 | -0.007 | -0.047 | -0.047 | 1.362 | 0.67 | 0.86 | 0.7 | 0.52 | 0.65 | 0.65 |

| | | | | | | | | | | | | | | | | |
|---|---|---|---|---|---|---|---|---|---|---|---|---|---|---|---|---|
| 6 | 131427176 | rs1919453 | -0.068 | 0.491 | -0.458 | -0.126 | -0.009 | -0.052 | -0.052 | 0 | 0.67 | 0.85 | 0.71 | 0.53 | 0.65 | 0.65 |
| 6 | 131433715 | rs509716 | -0.887 | 0.495 | -0.681 | -0.397 | 0.032 | 0.117 | 0.117 | 0 | 0.8 | 0.97 | 0.92 | 0.99 | 0.89 | 0.87 |
| 6 | 131434031 | rs12212943 | 0.057 | 1.055 | -0.549 | -0.185 | -0.039 | 0.029 | 0.029 | 0 | 0.07 | 0.01 | 0.11 | 0.01 | 0.17 | 0.19 |
| 6 | 131436888 | rs7776083 | 0.262 | 1.373 | -0.383 | -0.385 | 0.002 | -0.019 | -0.019 | 0 | 0.19 | 0.18 | 0.26 | 0.05 | 0.23 | 0.12 |
| 6 | 131438063 | rs643459 | -0.828 | 1.234 | -0.07 | -0.041 | 0.027 | 0.022 | 0.022 | 0 | 0.31 | 0.35 | 0.48 | 0.47 | 0.47 | 0.38 |
| 6 | 131443843 | rs516473 | -0.641 | 1.357 | -0.32 | -0.802 | 0.028 | -0.008 | -0.008 | 0 | 0.32 | 0.44 | 0.46 | 0.29 | 0.44 | 0.44 |
| 6 | 131451026 | rs600306 | -0.165 | 1.024 | -0.371 | -1.255 | -0.018 | -0.071 | -0.071 | 0.231 | 0.73 | 0.66 | 0.83 | 0.98 | 0.86 | 0.86 |
| 6 | 131451112 | rs543363 | -0.648 | 1.993 | -0.613 | -0.378 | -0.017 | 0.002 | 0.002 | 0 | 0.06 | 0.01 | 0.15 | 0.08 | 0.18 | 0.2 |
| 6 | 131459907 | rs516545 | -0.378 | 1.247 | -0.437 | -0.892 | -0.003 | -0.067 | -0.067 | 4.14 | 0.26 | 0.31 | 0.26 | 0.04 | 0.28 | 0.19 |
| 6 | 131465988 | rs12206629 | 0.01 | 1.765 | -1.312 | -1.3 | -0.063 | 0.018 | 0.018 | 3.856 | 0.19 | 0.06 | 0.11 | 0.11 | 0.27 | 0.23 |
| 6 | 131469461 | rs497082 | -0.784 | 1.077 | -1.686 | -1.586 | -0.011 | 0.003 | 0.003 | 0.062 | 0.64 | 0.85 | 0.48 | 0.29 | 0.63 | 0.4 |
| 6 | 131476528 | rs11965549 | 0.88 | 0.562 | -1.078 | -2.105 | 0.005 | -0.019 | -0.019 | 0 | 0.19 | 0.18 | 0.1 | 0 | 0.1 | 0.1 |
| 6 | 131481670 | rs1569196 | 0.856 | 0.79 | -1.461 | -2.225 | 0.006 | -0.002 | -0.002 | 0 | 0.24 | 0.15 | 0.32 | 0.75 | 0.24 | 0.48 |
| 6 | 131482127 | rs7772245 | 0.005 | 0.967 | -1.12 | -2.32 | -0.007 | 0.041 | 0.041 | 0.829 | 0.6 | 0.32 | 0.51 | 0.77 | 0.56 | 0.77 |
| 6 | 131489100 | rs1886073 | -0.241 | 0.859 | -0.624 | -2.439 | 0.008 | -0.031 | -0.031 | 0 | 0.65 | 0.47 | 0.55 | 0.79 | 0.59 | 0.8 |
| 6 | 131489690 | rs9385571 | 0.814 | 0.899 | -0.726 | -2.591 | 0 | 0.026 | 0.026 | 0.142 | 0.35 | 0.21 | 0.4 | 0.76 | 0.36 | 0.49 |
| 6 | 131490472 | rs1408027 | 0.202 | 0.952 | -0.742 | -2.591 | 0 | 0.003 | 0.003 | 0.024 | 0.53 | 0.45 | 0.5 | 0.76 | 0.45 | 0.55 |
| 6 | 131498603 | rs1534625 | 0.08 | 0.89 | -0.341 | -2.267 | 0 | -0.018 | -0.018 | 0 | 0.48 | 0.44 | 0.52 | 0.94 | 0.5 | 0.74 |
| 6 | 131502771 | rs1746469 | -0.651 | 0.818 | -0.144 | -1.568 | -0.002 | 0.029 | 0.029 | 0 | 0.28 | 0.16 | 0.3 | 0.21 | 0.32 | 0.21 |
| 6 | 131504851 | rs1746471 | -0.637 | 0.744 | -0.033 | -1.706 | 0 | 0.038 | 0.038 | 0 | 0.31 | 0.21 | 0.24 | 0.05 | 0.35 | 0.23 |
| 6 | 131509096 | rs1209159 | -0.637 | 0.744 | -0.037 | -1.706 | 0 | 0.045 | 0.045 | 0 | 0.31 | 0.2 | 0.23 | 0.05 | 0.35 | 0.23 |
| 6 | 131512245 | rs1190799 | -0.565 | 0.757 | -0.056 | -1.699 | -0.005 | 0.077 | 0.077 | 0 | 0.3 | 0.15 | 0.23 | 0.05 | 0.35 | 0.23 |
| 6 | 131517901 | rs1190792 | 0.163 | 1.332 | -0.084 | -1.554 | -0.015 | -0.013 | -0.013 | 0.187 | 0.94 | 0.95 | 0.79 | 0.91 | 0.82 | 0.9 |
| 6 | 131520655 | rs1190788 | -0.663 | 0.7 | -0.046 | -1.612 | -0.008 | 0.086 | 0.086 | 0 | 0.3 | 0.14 | 0.23 | 0.05 | 0.35 | 0.23 |
| 6 | 131520994 | rs3903759 | -0.045 | 0.736 | -0.147 | -1.686 | -0.003 | 0.011 | 0.011 | 0.176 | 0.09 | 0.06 | 0.07 | 0.05 | 0.14 | 0.14 |
| 6 | 131521704 | rs1190813 | 0.119 | 0.638 | -0.161 | -1.667 | -0.04 | 0.128 | 0.128 | 0.082 | 0.21 | 0.02 | 0.14 | 0 | 0.21 | 0.09 |
| 6 | 131547341 | rs11154621 | -0.923 | 0.193 | 0.999 | -1.873 | -0.006 | -0.083 | -0.083 | 1.96 | 0.37 | 0.42 | 0.25 | 0.06 | 0.35 | 0.25 |
| 6 | 131549060 | rs10484763 | -0.825 | 0.178 | 0.838 | -1.834 | 0.013 | 0.063 | 0.063 | 0 | 0.34 | 0.2 | 0.33 | 0.21 | 0.34 | 0.25 |
| 6 | 131554291 | rs1885640 | -1.308 | 0.095 | 0.931 | -1.854 | 0.001 | 0.008 | 0.008 | 0 | 0.46 | 0.58 | 0.57 | 0.31 | 0.49 | 0.36 |
| 6 | 131563577 | rs4629710 | 0.046 | 0.116 | 0.489 | -1.858 | -0.01 | 0.155 | 0.155 | 0 | 0.24 | 0.02 | 0.15 | 0 | 0.2 | 0.1 |
| 6 | 131573433 | rs3777483 | -1.425 | -0.162 | 0.855 | -1.9 | 0.018 | 0.106 | 0.106 | 0 | 0.52 | 0.76 | 0.59 | 0.31 | 0.5 | 0.37 |
| 6 | 131589506 | rs6924196 | -0.666 | 0.13 | 0.643 | -1.716 | -0.008 | -0.013 | -0.013 | 0 | 0.17 | 0.14 | 0.38 | 0.3 | 0.28 | 0.24 |
| 6 | 131600188 | rs10484765 | -0.825 | -0.029 | 0.797 | -1.738 | -0.001 | -0.025 | -0.025 | 0 | 0.44 | 0.52 | 0.57 | 0.26 | 0.5 | 0.35 |
| 6 | 131613602 | rs6569737 | -1.14 | 0.139 | 0.121 | -1.634 | -0.006 | 0.12 | 0.12 | 0 | 0.54 | 0.71 | 0.5 | 0.11 | 0.52 | 0.39 |
| 6 | 131622339 | rs2876142 | -0.865 | -0.039 | 0.552 | -1.714 | -0.021 | -0.045 | -0.045 | 0.292 | 0.21 | 0.35 | 0.25 | 0.09 | 0.17 | 0.18 |
| 6 | 131622480 | rs2104209 | -1.264 | -0.04 | -0.263 | -1.653 | -0.081 | 0.109 | 0.109 | 0 | 0.57 | 0.88 | 0.58 | 0.24 | 0.53 | 0.41 |
| 6 | 131637690 | rs9483246 | -1.389 | -0.257 | 0.572 | -1.711 | -0.008 | -0.021 | -0.021 | 0 | 0.3 | 0.44 | 0.41 | 0.23 | 0.3 | 0.29 |
| 6 | 131638681 | rs9492908 | -0.702 | -0.159 | 0.525 | -1.676 | 0 | 0 | 0 | 0 | 0.19 | 0.18 | 0.25 | 0.08 | 0.18 | 0.18 |
| 6 | 131671671 | rs2146829 | -1.563 | -0.345 | -0.035 | -2.146 | -0.107 | 0.127 | 0.127 | 0.702 | 0.56 | 0.94 | 0.61 | 0.21 | 0.52 | 0.43 |
| 6 | 131671687 | rs9483252 | -1.01 | -0.178 | 0.955 | -1.667 | -0.015 | -0.044 | -0.044 | 0 | 0.22 | 0.34 | 0.27 | 0.07 | 0.18 | 0.18 |
| 6 | 131671763 | rs9483253 | -1.023 | -0.182 | 0.975 | -1.668 | -0.016 | -0.047 | -0.047 | 0.202 | 0.22 | 0.34 | 0.27 | 0.06 | 0.18 | 0.18 |
| 6 | 131673070 | rs4897509 | -1.037 | -0.182 | 0.986 | -1.668 | -0.017 | -0.045 | -0.045 | 0.232 | 0.22 | 0.36 | 0.27 | 0.07 | 0.18 | 0.18 |
| 6 | 131675201 | rs9483254 | -0.804 | -0.166 | 1.023 | -1.675 | -0.002 | -0.009 | -0.009 | 0 | 0.21 | 0.23 | 0.27 | 0.07 | 0.19 | 0.18 |
| 6 | 131684994 | rs9483259 | -0.806 | -0.515 | 1.108 | -1.639 | -0.049 | -0.094 | -0.094 | 0.059 | 0.14 | 0.36 | 0.05 | 0 | 0.04 | 0.03 |
| 6 | 131691964 | rs6939316 | -1.219 | 0.1 | 1.071 | -1.305 | 0.1 | 0.156 | 0.156 | 0 | 0.06 | 0.24 | 0.1 | 0.16 | 0.11 | 0.05 |
| 6 | 131698944 | rs11965227 | -0.404 | -0.055 | 0.034 | -1.916 | -0.072 | 0.116 | 0.116 | 1.873 | 0.29 | 0.06 | 0.39 | 0.09 | 0.37 | 0.32 |
| 6 | 131703293 | rs942150 | -1.616 | -0.731 | -0.381 | -1.949 | -0.138 | 0.03 | 0.03 | 0.662 | 0.56 | 0.96 | 0.61 | 0.3 | 0.53 | 0.41 |
| 6 | 131707088 | rs6916044 | -1.135 | -0.702 | 0.565 | -1.801 | -0.001 | -0.004 | -0.004 | 0.043 | 0.46 | 0.56 | 0.52 | 0.27 | 0.49 | 0.37 |
| 6 | 131709068 | rs12210559 | 0.095 | -0.647 | 0.508 | -1.838 | -0.066 | 0.092 | 0.092 | 1.72 | 0.26 | 0.06 | 0.35 | 0.09 | 0.36 | 0.28 |
| 6 | 131709941 | rs2031205 | -0.782 | -0.656 | 0.586 | -1.812 | -0.023 | 0.013 | 0.013 | 0 | 0.17 | 0.51 | 0.12 | 0.17 | 0.11 | 0.06 |
| 6 | 131712051 | rs2143774 | -0.319 | -0.591 | 0.121 | -1.831 | -0.049 | 0.08 | 0.08 | 0.587 | 0.75 | 0.93 | 0.68 | 0.91 | 0.67 | 0.73 |
| 6 | 131719021 | rs6933890 | -0.743 | -0.612 | 0.499 | -1.558 | -0.024 | 0.006 | 0.006 | 0 | 0.15 | 0.3 | 0.09 | 0.17 | 0.09 | 0.05 |
| 6 | 131721876 | rs4897517 | 1.188 | -0.484 | -1.141 | -1.798 | -0.105 | -0.108 | -0.108 | 0.002 | 0.31 | 0.03 | 0.41 | 0.62 | 0.38 | 0.53 |
| 6 | 131723249 | rs9492955 | -0.621 | -0.483 | -0.22 | -1.749 | 0.015 | 0.064 | 0.064 | 0 | 0.1 | 0.27 | 0.1 | 0.17 | 0.11 | 0.04 |
| 6 | 131728258 | rs4487609 | 0.458 | -0.75 | -0.273 | -2.094 | 0.007 | -0.012 | -0.012 | 0 | 0.62 | 0.49 | 0.54 | 0.67 | 0.53 | 0.65 |
| 6 | 131731609 | rs9483277 | -0.477 | -0.687 | -0.067 | -1.896 | 0.046 | 0.142 | 0.142 | 0 | 0.07 | 0.23 | 0.1 | 0.17 | 0.08 | 0.02 |
| 6 | 131737302 | rs1033540 | -0.194 | -0.788 | -0.961 | -2.183 | -0.028 | 0.038 | 0.038 | 1.166 | 0.27 | 0.12 | 0.33 | 0.16 | 0.33 | 0.35 |
| 6 | 131737654 | rs722027 | -0.173 | -0.991 | -0.889 | -2.229 | -0.011 | -0.079 | -0.079 | 0 | 0.84 | 0.66 | 0.89 | 0.99 | 0.88 | 0.95 |
| 6 | 131738514 | rs11154637 | 0.868 | -0.997 | -1.48 | -1.83 | -0.04 | 0.019 | 0.019 | 2.663 | 0.48 | 0.11 | 0.44 | 0.64 | 0.46 | 0.59 |
| 6 | 131746742 | rs12215468 | 1.187 | -0.764 | -1.37 | -1.595 | -0.055 | -0.011 | -0.011 | 0 | 0.24 | 0.11 | 0.39 | 0.74 | 0.38 | 0.51 |
| 6 | 131754930 | rs2179771 | -0.539 | -0.404 | -0.661 | -1.263 | 0.03 | 0.057 | 0.057 | 0 | 0.06 | 0.12 | 0.08 | 0.15 | 0.1 | 0.04 |
| 6 | 131761677 | rs2078265 | -1.073 | -1.273 | -1.526 | -1.94 | 0.035 | 0.122 | 0.122 | 1.155 | 0.39 | 0.91 | 0.32 | 0.16 | 0.3 | 0.13 |
| 6 | 131763378 | rs2608903 | -2.031 | -1.392 | -1.863 | -2.09 | -0.13 | -0.074 | -0.074 | 0 | 0.7 | 0.98 | 0.64 | 0.29 | 0.63 | 0.44 |
| 6 | 131770803 | rs2608905 | 0.482 | -1.232 | -1.276 | -1.495 | -0.076 | -0.045 | -0.045 | 0.19 | 0.16 | 0.01 | 0.28 | 0.19 | 0.25 | 0.27 |
| 6 | 131780576 | rs17060292 | 0.576 | -1.961 | -0.072 | -1.427 | -0.009 | -0.047 | -0.047 | 0.148 | 0.1 | 0.16 | 0.07 | 0 | 0.07 | 0.03 |
| 6 | 131783628 | rs12717182 | 0.589 | -2.108 | -0.25 | -1.505 | -0.006 | 0.057 | 0.057 | 0 | 0.12 | 0.01 | 0.09 | 0.01 | 0.11 | 0.06 |
| 6 | 131789736 | rs2608935 | 0.926 | -1.853 | 0.629 | -1.184 | 0.014 | 0.009 | 0.009 | 0 | 0.82 | 0.51 | 0.82 | 0.84 | 0.83 | 0.93 |
| 6 | 131802325 | rs2807276 | -1.06 | -2.069 | -0.634 | -1.164 | 0.073 | 0.058 | 0.058 | 0 | 0.47 | 0.94 | 0.5 | 0.32 | 0.36 | 0.34 |
| 6 | 131807510 | rs2249821 | -1.065 | -2.067 | -0.634 | -1.163 | 0.074 | 0.059 | 0.059 | 0.019 | 0.47 | 0.94 | 0.5 | 0.32 | 0.36 | 0.34 |

| | | | | | | | | | | | | | | | | |
|---|---|---|---|---|---|---|---|---|---|---|---|---|---|---|---|---|
| 6 | 131808591 | rs9493000 | -0.848 | -1.99 | -0.082 | -1.129 | 0.012 | 0.094 | 0.094 | 0 | 0.07 | 0.2 | 0.09 | 0.2 | 0.06 | 0.04 |
| 6 | 131809436 | rs3861462 | -1.092 | -2.302 | 0.355 | -1.378 | -0.038 | 0.008 | 0.008 | 0.004 | 0.22 | 0.48 | 0.16 | 0.2 | 0.13 | 0.08 |
| 6 | 131809920 | rs2807278 | -0.417 | -2.437 | 0.281 | -1.484 | -0.01 | -0.034 | -0.034 | 0 | 0.35 | 0.4 | 0.38 | 0.12 | 0.3 | 0.29 |
| 6 | 131809958 | rs2608968 | -1.608 | -2.136 | -0.175 | -1.313 | -0.045 | 0.007 | 0.007 | 0.161 | 0.57 | 0.96 | 0.56 | 0.32 | 0.42 | 0.37 |
| 6 | 131814278 | rs2246281 | -0.494 | -2.14 | 0.436 | -1.304 | 0.01 | 0.036 | 0.036 | 1.065 | 0.19 | 0.34 | 0.35 | 0.3 | 0.23 | 0.26 |
| 6 | 131814574 | rs2807292 | -1.158 | -2.344 | 0.111 | -1.409 | 0.02 | 0.001 | 0.001 | 1.23 | 0.31 | 0.76 | 0.41 | 0.3 | 0.25 | 0.28 |
| 6 | 131843741 | rs9493012 | -0.745 | -2.483 | 0.262 | -1.614 | -0.054 | -0.146 | -0.146 | 0 | 0.32 | 0.49 | 0.18 | 0.01 | 0.19 | 0.11 |
| 6 | 131845152 | rs2781641 | -0.078 | -1.989 | 0.257 | -1.061 | -0.011 | 0.019 | 0.019 | 1.104 | 0.14 | 0.08 | 0.27 | 0.11 | 0.18 | 0.21 |
| 6 | 131857907 | rs9493022 | -0.252 | -2.025 | 0.27 | -1.086 | -0.016 | -0.066 | -0.066 | 0.739 | 0.16 | 0.3 | 0.11 | 0.01 | 0.13 | 0.09 |
| 6 | 131858779 | rs2749928 | -1.783 | -2.004 | -0.372 | -1.013 | 0.064 | 0.069 | 0.069 | 0 | 0.47 | 0.96 | 0.51 | 0.33 | 0.35 | 0.35 |
| 6 | 131876943 | rs2781651 | -1.478 | -1.762 | 0.338 | -0.663 | 0.035 | 0.011 | 0.011 | 0.553 | 0.31 | 0.76 | 0.4 | 0.31 | 0.23 | 0.28 |
| 6 | 131879281 | rs2608975 | -0.938 | -1.66 | 0.77 | -0.61 | -0.007 | 0.016 | 0.016 | 0.227 | 0.22 | 0.32 | 0.34 | 0.31 | 0.22 | 0.26 |
| 6 | 131879805 | rs3895535 | -0.074 | -1.327 | 0.187 | -0.484 | -0.042 | -0.007 | -0.007 | 0 | 0.13 | 0.01 | 0.25 | 0.13 | 0.17 | 0.21 |
| 6 | 131883790 | rs17060390 | -0.593 | -1.43 | 0.926 | -0.389 | 0.012 | 0.142 | 0.142 | 1.321 | 0.09 | 0.33 | 0.1 | 0.17 | 0.05 | 0.02 |
| 6 | 131888147 | rs2608937 | -0.224 | -1.34 | 0.542 | -0.349 | -0.019 | -0.002 | -0.002 | 0 | 0.13 | 0.04 | 0.25 | 0.12 | 0.17 | 0.22 |
| 6 | 131893463 | rs2608897 | -1.352 | -1.746 | 0.315 | -0.761 | 0.043 | 0.039 | 0.039 | 0 | 0.44 | 0.71 | 0.44 | 0.32 | 0.32 | 0.31 |
| 6 | 131895144 | rs2781667 | -1.413 | -1.763 | 0.269 | -0.779 | 0.072 | 0.067 | 0.067 | 0 | 0.44 | 0.77 | 0.48 | 0.32 | 0.32 | 0.32 |
| 6 | 131897278 | rs2781668 | -0.051 | -1.37 | 0.41 | -0.266 | 0 | 0.004 | 0.004 | 0.315 | 0.12 | 0.11 | 0.25 | 0.25 | 0.17 | 0.21 |
| 6 | 131898208 | rs2246012 | -0.062 | -1.369 | 0.478 | 0.246 | 0.004 | 0.043 | 0.043 | 0.314 | 0.12 | 0.12 | 0.25 | 0.34 | 0.17 | 0.21 |
| 6 | 131904719 | rs17599586 | -0.366 | -1.685 | 0.367 | -0.207 | 0.022 | 0.029 | 0.029 | 0 | 0.19 | 0.12 | 0.09 | 0.01 | 0.13 | 0.08 |
| 6 | 131905717 | rs17184300 | -0.425 | -1.71 | 0.368 | -0.219 | 0.023 | 0.028 | 0.028 | 0 | 0.2 | 0.11 | 0.09 | 0.01 | 0.13 | 0.09 |
| 6 | 131950233 | rs3756784 | -1.144 | -2.3 | -0.565 | -0.349 | 0.052 | -0.002 | -0.002 | 0.016 | 0.34 | 0.76 | 0.34 | 0.32 | 0.2 | 0.3 |
| 6 | 131954879 | rs1206684 | 1.663 | -0.613 | -1.551 | -0.485 | -0.1 | -0.177 | -0.177 | 0 | 0.32 | 0.02 | 0.34 | 0.51 | 0.37 | 0.48 |
| 6 | 131958485 | rs9375819 | -0.162 | 0.173 | -0.836 | -0.264 | -0.099 | 0.105 | 0.105 | 1.905 | 0.83 | 0.99 | 0.84 | 1 | 0.74 | 0.88 |
| 6 | 131960898 | rs12663047 | -0.913 | -0.149 | 0.247 | 0.442 | 0.006 | 0.069 | 0.069 | 1.206 | 0.15 | 0.23 | 0.27 | 0.31 | 0.16 | 0.26 |
| 6 | 131986057 | rs12207523 | -1.315 | -0.185 | -0.028 | 0.719 | -0.006 | 0.049 | 0.049 | 0 | 0.06 | 0.05 | 0.15 | 0.32 | 0.06 | 0.18 |
| 6 | 131992549 | rs10080319 | -1.246 | -0.528 | 0.892 | 0.492 | 0.008 | -0.004 | -0.004 | 1.03 | 0.06 | 0.28 | 0.04 | 0 | 0.02 | 0.03 |
| 6 | 131992714 | rs2066889 | -0.102 | -1.589 | 1.217 | -0.001 | -0.053 | -0.067 | -0.067 | 0 | 0.15 | 0.21 | 0.04 | 0 | 0.02 | 0.02 |
| 6 | 131995313 | rs9493048 | -1.812 | -1.587 | 1.086 | 0.03 | -0.063 | 0.032 | 0.032 | 0 | 0.27 | 0.54 | 0.23 | 0.33 | 0.1 | 0.24 |
| 6 | 131999582 | rs3883882 | -0.377 | -1.296 | 0.133 | -0.777 | -0.14 | 0.176 | 0.176 | 1.222 | 0.72 | 0.99 | 0.78 | 0.97 | 0.61 | 0.8 |
| 6 | 132005325 | rs9493050 | -1.331 | -1.646 | 0.575 | -0.581 | -0.061 | 0.038 | 0.038 | 0 | 0.26 | 0.52 | 0.23 | 0.32 | 0.1 | 0.23 |
| 6 | 132005531 | rs12210900 | -0.692 | -1.566 | 0.251 | -0.539 | -0.003 | 0.055 | 0.055 | 0 | 0.06 | 0.05 | 0.15 | 0.32 | 0.06 | 0.18 |
| 6 | 132012886 | rs6901900 | -1.609 | -1.833 | 0.33 | -0.679 | 0.105 | -0.028 | -0.028 | 0 | 0.33 | 0.83 | 0.28 | 0.33 | 0.11 | 0.26 |
| 6 | 132038303 | rs12206403 | -0.748 | -1.589 | 0.377 | -0.542 | -0.003 | 0.01 | 0.01 | 0.014 | 0.06 | 0.02 | 0.21 | 0.32 | 0.05 | 0.2 |
| 6 | 132043056 | rs2270089 | -1.393 | -1.706 | 0.891 | -0.609 | -0.019 | 0.002 | 0.002 | 0 | 0.28 | 0.68 | 0.3 | 0.32 | 0.08 | 0.21 |
| 6 | 132049450 | rs3800198 | -1.151 | -1.646 | 1.122 | -0.587 | 0.01 | 0.214 | 0.214 | 0.221 | 0.08 | 0.22 | 0.23 | 0.32 | 0.05 | 0.2 |
| 6 | 132051395 | rs9483334 | -1.1 | -1.571 | 0.499 | -0.555 | -0.018 | 0.002 | 0.002 | 0 | 0.28 | 0.68 | 0.3 | 0.32 | 0.09 | 0.21 |
| 6 | 132055621 | rs365475 | 1.098 | -0.495 | -0.331 | -0.321 | -0.12 | -0.025 | -0.025 | 1.304 | 0.4 | 0.09 | 0.45 | 0.64 | 0.5 | 0.54 |
| 6 | 132061420 | rs17601580 | -0.344 | -0.361 | 0.398 | -0.198 | -0.017 | 0.104 | 0.104 | 0.7 | 0.17 | 0.01 | 0.09 | 0.01 | 0.2 | 0.16 |
| 6 | 132066501 | rs7759504 | -1.458 | -0.446 | 0.762 | 0.712 | -0.007 | 0.191 | 0.191 | 0 | 0.07 | 0.19 | 0.24 | 0.45 | 0.04 | 0.16 |
| 6 | 132068237 | rs11754555 | -1.397 | -0.503 | 1.408 | 0.374 | -0.013 | -0.142 | -0.142 | 0.2 | 0.31 | 0.36 | 0.18 | 0.01 | 0.29 | 0.24 |
| 6 | 132071193 | rs4383836 | -1.107 | -0.491 | 1.447 | 0.375 | -0.01 | -0.127 | -0.127 | 0 | 0.24 | 0.33 | 0.16 | 0.01 | 0.25 | 0.23 |
| 6 | 132076328 | rs9493088 | -2.272 | -0.574 | 1.757 | 0.483 | -0.016 | 0.12 | 0.12 | 0 | 0.22 | 0.44 | 0.33 | 0.47 | 0.13 | 0.23 |
| 6 | 132086493 | rs17060705 | -0.524 | -0.506 | 1.53 | 0.721 | -0.004 | -0.011 | -0.011 | 0 | 0.08 | 0.05 | 0.14 | 0.2 | 0.07 | 0.18 |
| 6 | 132089007 | rs6911803 | -0.663 | -0.465 | 1.682 | 0.656 | -0.005 | -0.05 | -0.05 | 0 | 0.25 | 0.26 | 0.15 | 0.05 | 0.26 | 0.4 |
| 6 | 132093249 | rs12524612 | -0.545 | -0.399 | 0.929 | 0.709 | -0.004 | -0.013 | -0.013 | 0 | 0.08 | 0.06 | 0.14 | 0.2 | 0.07 | 0.18 |
| 6 | 132095002 | rs3844006 | -0.799 | -0.408 | 1.078 | 0.659 | 0.005 | 0.012 | 0.012 | 0.032 | 0.27 | 0.21 | 0.15 | 0.09 | 0.26 | 0.4 |
| 6 | 132099761 | rs7773292 | -1.84 | -0.26 | 0.665 | 0.83 | 0.03 | 0.035 | 0.035 | 2.784 | 0.48 | 0.79 | 0.51 | 0.53 | 0.41 | 0.65 |
| 6 | 132100610 | rs7758525 | -1.251 | -0.247 | 0.783 | 0.711 | -0.011 | -0.02 | -0.02 | 0.026 | 0.38 | 0.55 | 0.31 | 0.28 | 0.33 | 0.58 |
| 6 | 132112533 | rs3850253 | -1.44 | -0.392 | -0.022 | 0.553 | 0.025 | 0.004 | 0.004 | 2.62 | 0.43 | 0.7 | 0.5 | 0.53 | 0.37 | 0.61 |
| 6 | 132113403 | rs9493099 | -0.665 | -0.394 | -0.498 | 0.549 | 0.006 | -0.078 | -0.078 | 0.179 | 0.31 | 0.1 | 0.41 | 0.5 | 0.31 | 0.58 |
| 6 | 132119366 | rs6925433 | -1.089 | -0.137 | -0.456 | 0.721 | 0.05 | 0.005 | 0.005 | 2.524 | 0.46 | 0.8 | 0.51 | 0.52 | 0.38 | 0.62 |
| 6 | 132124599 | rs9385584 | 0.772 | 0.246 | -0.28 | 0.118 | 0.009 | 0.071 | 0.071 | 0 | 0.5 | 0.33 | 0.43 | 0.75 | 0.5 | 0.41 |
| 6 | 132126320 | rs6935458 | -1.049 | -0.261 | 0.346 | 0.479 | -0.029 | 0.123 | 0.123 | 0.656 | 0.2 | 0.47 | 0.21 | 0.59 | 0.11 | 0.12 |
| 6 | 132127293 | rs11154644 | 0.678 | 0.119 | 0.097 | 0.013 | -0.039 | 0.045 | 0.045 | 5.42 | 0.67 | 0.77 | 0.54 | 0.8 | 0.56 | 0.45 |
| 6 | 132128564 | rs1800949 | -0.11 | 0.148 | 0.011 | 0.132 | -0.082 | -0.044 | -0.044 | 0 | 0.13 | 0.01 | 0.22 | 0.18 | 0.22 | 0.13 |
| 6 | 132131601 | rs766592 | 0.854 | -0.05 | 0.029 | -0.027 | -0.059 | 0.051 | 0.051 | 3.966 | 0.71 | 0.88 | 0.65 | 0.82 | 0.61 | 0.74 |
| 6 | 132133116 | rs6569759 | 0.894 | -0.344 | 0.216 | -0.333 | -0.031 | 0.067 | 0.067 | 5.193 | 0.57 | 0.75 | 0.52 | 0.73 | 0.49 | 0.49 |
| 6 | 132138338 | rs9493105 | 0.955 | -0.338 | 0.14 | -0.315 | 0.015 | 0.006 | 0.006 | 0 | 0.13 | 0.03 | 0.1 | 0.09 | 0.11 | 0.25 |
| 6 | 132140203 | rs9398995 | 1.825 | -0.378 | 0.418 | 0.163 | -0.008 | -0.022 | -0.022 | 0 | 0.39 | 0.19 | 0.37 | 0.48 | 0.39 | 0.36 |
| 6 | 132146520 | rs9375830 | -0.696 | -0.626 | 0.707 | 0.347 | -0.027 | 0.036 | 0.036 | 0.344 | 0.3 | 0.54 | 0.27 | 0.35 | 0.22 | 0.34 |
| 6 | 132150439 | rs2021966 | 1.701 | -0.446 | 0.549 | -0.756 | -0.013 | 0.047 | 0.047 | 1.357 | 0.42 | 0.28 | 0.5 | 0.7 | 0.45 | 0.47 |
| 6 | 132151105 | rs7771841 | -0.934 | -1.36 | 1.43 | -0.16 | 0.009 | 0.053 | 0.053 | 0.054 | 0.09 | 0.23 | 0.12 | 0.19 | 0.09 | 0.14 |
| 6 | 132153152 | rs9372999 | -0.141 | -1.32 | 1.484 | -0.405 | -0.02 | -0.024 | -0.024 | 0 | 0.17 | 0.26 | 0.11 | 0.09 | 0.14 | 0.26 |
| 6 | 132153207 | rs858338 | -0.878 | -1.338 | 1.2 | -0.5 | 0.007 | 0.006 | 0.006 | 0 | 0.23 | 0.13 | 0.15 | 0.01 | 0.21 | 0.15 |
| 6 | 132157149 | rs7775386 | -1.526 | -1.408 | 1.541 | -0.016 | 0.052 | 0.06 | 0.06 | 0 | 0.13 | 0.53 | 0.16 | 0.17 | 0.13 | 0.17 |
| 6 | 132160455 | rs858341 | -1.518 | -1.575 | 1.433 | -0.124 | 0.015 | 0.035 | 0.035 | 0 | 0.48 | 0.64 | 0.38 | 0.29 | 0.41 | 0.56 |
| 6 | 132160643 | rs858342 | -0.679 | -1.573 | 1.263 | -0.146 | 0.041 | 0.142 | 0.142 | 0 | 0.34 | 0.11 | 0.22 | 0.12 | 0.28 | 0.39 |
| 6 | 132163617 | rs858345 | -1.425 | -1.575 | 1.355 | 0.022 | 0.026 | 0.052 | 0.052 | 0 | 0.49 | 0.68 | 0.45 | 0.37 | 0.42 | 0.57 |

| | | | | | | | | | | | | | | | | |
|---|---|---|---|---|---|---|---|---|---|---|---|---|---|---|---|---|
| 6 | 132167290 | rs6926970 | -1.914 | -1.607 | 0.906 | -0.512 | 0.037 | 0.045 | 0.045 | 1.291 | 0.25 | 0.79 | 0.18 | 0.1 | 0.13 | 0.18 |
| 6 | 132167676 | rs9402348 | 0.011 | -1.646 | 0.728 | -0.523 | -0.064 | 0.048 | 0.048 | 0.599 | 0.28 | 0.01 | 0.27 | 0.18 | 0.29 | 0.37 |
| 6 | 132167977 | rs997509 | -0.795 | -1.632 | 2.056 | 0.013 | -0.001 | -0.017 | -0.017 | 0 | 0.1 | 0.11 | 0.02 | 0 | 0.04 | 0.03 |
| 6 | 132168997 | rs7773477 | -1.313 | -1.631 | 2.17 | 0.016 | -0.027 | -0.067 | -0.067 | 0 | 0.13 | 0.25 | 0.05 | 0.01 | 0.06 | 0.09 |
| 6 | 132175059 | rs7749493 | -2.277 | -1.549 | 1.972 | 0.067 | 0.022 | 0.025 | 0.025 | 0.862 | 0.25 | 0.78 | 0.18 | 0.1 | 0.13 | 0.14 |
| 6 | 132193395 | rs7769712 | -1.263 | -0.83 | 1.57 | 0.735 | 0.031 | 0.086 | 0.086 | 0.221 | 0.07 | 0.3 | 0.1 | 0.1 | 0.06 | 0.09 |
| 6 | 132203432 | rs7768480 | -1.649 | -1.021 | 1.102 | 0.774 | -0.045 | 0.034 | 0.034 | 0 | 0.16 | 0.54 | 0.16 | 0.14 | 0.08 | 0.12 |
| 6 | 132211534 | rs1804025 | -1.219 | -1.129 | 1.313 | 0.426 | 0.005 | -0.023 | -0.023 | 0 | 0.06 | 0.24 | 0.03 | 0 | 0.03 | 0.03 |
| 6 | 132211763 | rs1044548 | -1.739 | -1.123 | 1.256 | 0.431 | -0.006 | -0.06 | -0.06 | 0 | 0.1 | 0.34 | 0.06 | 0 | 0.05 | 0.03 |
| 6 | 132211814 | rs1044558 | -1.739 | -1.123 | 1.256 | 0.431 | -0.006 | -0.06 | -0.06 | 0 | 0.1 | 0.34 | 0.06 | 0 | 0.05 | 0.03 |
| 6 | 132212694 | rs7754561 | -1.006 | -2.606 | 0.369 | 0.591 | 0.216 | 0.046 | 0.046 | 0.586 | 0.5 | 0.87 | 0.5 | 0.55 | 0.27 | 0.46 |
| 6 | 132212752 | rs11154647 | -1.493 | -1.741 | 0.776 | 0.49 | -0.006 | -0.061 | -0.061 | 0.277 | 0.1 | 0.35 | 0.06 | 0 | 0.05 | 0.03 |
| 6 | 132217687 | rs7753048 | -1.414 | -1.708 | 1.132 | 0.64 | -0.019 | 0.025 | 0.025 | 0 | 0.14 | 0.33 | 0.13 | 0.12 | 0.12 | 0.17 |
| 6 | 132228577 | rs6569765 | -1.195 | -2.107 | 1.012 | 1.037 | 0.074 | 0.037 | 0.037 | 0 | 0.5 | 0.73 | 0.47 | 0.54 | 0.37 | 0.57 |
| 6 | 132228769 | rs6907728 | -0.394 | -2.466 | 1.078 | 1.365 | -0.004 | 0.081 | 0.081 | 0.157 | 0.13 | 0.33 | 0.14 | 0.24 | 0.1 | 0.08 |
| 6 | 132234136 | rs4897553 | -0.763 | -2.516 | 0.755 | 1.389 | 0.007 | 0.002 | 0.002 | 0 | 0.36 | 0.31 | 0.32 | 0.29 | 0.27 | 0.48 |
| 6 | 132234716 | rs9493133 | -1.087 | -2.072 | 0.873 | 1.249 | -0.023 | 0.085 | 0.085 | 1.519 | 0.17 | 0.54 | 0.18 | 0.24 | 0.12 | 0.11 |
| 6 | 132238279 | rs9373002 | -0.277 | -2.102 | -0.279 | 1.33 | 0.009 | -0.019 | -0.019 | 3.188 | 0.27 | 0.11 | 0.31 | 0.29 | 0.26 | 0.51 |
| 6 | 132243683 | rs9375851 | -1.016 | -2.02 | 0.217 | 1.176 | -0.011 | 0.001 | 0.001 | 0 | 0.43 | 0.54 | 0.51 | 0.52 | 0.38 | 0.62 |
| 6 | 132247361 | rs2151533 | -0.581 | -1.895 | -0.417 | 1.261 | 0.018 | -0.061 | -0.061 | 1.467 | 0.36 | 0.2 | 0.48 | 0.52 | 0.36 | 0.57 |
| 6 | 132250141 | rs1029122 | -0.919 | -1.942 | -1.278 | 0.964 | -0.079 | -0.087 | -0.087 | 0 | 0.6 | 0.96 | 0.49 | 0.57 | 0.43 | 0.63 |
| 6 | 132254064 | rs6910279 | -0.481 | -0.21 | -1.045 | 0.755 | 0.033 | 0.073 | 0.073 | 0.89 | 0.86 | 0.8 | 0.7 | 0.57 | 0.8 | 0.76 |
| 6 | 132254785 | rs12200211 | -0.161 | -0.6 | -1.32 | -0.071 | 0.009 | -0.042 | -0.042 | 0 | 0.12 | 0.21 | 0.14 | 0 | 0.15 | 0.03 |
| 6 | 132257444 | rs6940184 | 0.014 | -1.035 | -1.934 | -0.05 | 0.092 | -0.011 | -0.011 | 0.098 | 0.37 | 0.2 | 0.17 | 0.33 | 0.22 | 0.55 |
| 6 | 132262991 | rs4897554 | -0.799 | 0.168 | -1.96 | -0.725 | -0.098 | 0.147 | 0.147 | 0 | 0.7 | 0.9 | 0.76 | 1 | 0.59 | 0.81 |
| 6 | 132268964 | rs9399005 | -0.181 | 0.122 | -1.881 | 0.342 | -0.045 | -0.105 | -0.105 | 0 | 0.24 | 0.06 | 0.29 | 0.48 | 0.3 | 0.33 |
| 6 | 132274730 | rs2151532 | 0.126 | 0.236 | -1.351 | 0.511 | 0.001 | -0.013 | -0.013 | 0 | 0.88 | 0.89 | 0.79 | 0.77 | 0.92 | 0.88 |
| 6 | 132275437 | rs928501 | -0.691 | 0.147 | -0.654 | 0.188 | -0.011 | -0.043 | -0.043 | 0 | 0.34 | 0.43 | 0.22 | 0.11 | 0.27 | 0.21 |
| 6 | 132277333 | rs7768619 | 0.219 | 0.464 | -0.563 | 0.57 | -0.01 | -0.004 | -0.004 | 0 | 0.13 | 0.06 | 0.26 | 0.45 | 0.22 | 0.28 |
| 6 | 132277462 | rs9388949 | -1.351 | 0.141 | -0.183 | 0.302 | 0.001 | 0.001 | 0.001 | 1.703 | 0.48 | 0.55 | 0.49 | 0.56 | 0.49 | 0.51 |
| 6 | 132287349 | rs12663962 | 0.043 | 0.358 | -0.451 | 0.465 | -0.062 | -0.098 | -0.098 | 0.653 | 0.15 | 0.02 | 0.23 | 0.45 | 0.25 | 0.24 |
| 6 | 132289157 | rs9493157 | -0.621 | 0.348 | 0.246 | 0.468 | -0.027 | 0.004 | 0.004 | 0 | 0.32 | 0.51 | 0.33 | 0.31 | 0.23 | 0.26 |
| 6 | 132298141 | rs928505 | -1.179 | 0.409 | 0.057 | 0.321 | -0.027 | 0.081 | 0.081 | 0 | 0.43 | 0.21 | 0.55 | 0.76 | 0.49 | 0.51 |
| 6 | 132303676 | rs11154656 | -0.035 | 0.726 | -0.041 | 0.132 | -0.034 | -0.063 | -0.063 | 0 | 0.14 | 0.04 | 0.23 | 0.43 | 0.24 | 0.25 |
| 6 | 132305683 | rs233908 | -0.933 | 0.563 | 0.444 | -0.002 | 0.003 | 0.028 | 0.028 | 0 | 0.42 | 0.59 | 0.28 | 0.19 | 0.35 | 0.37 |
| 6 | 132309046 | rs233912 | -1.909 | 0.552 | 0.157 | -0.007 | -0.006 | -0.005 | -0.005 | 1.645 | 0.58 | 0.85 | 0.55 | 0.62 | 0.59 | 0.62 |
| 6 | 132314595 | rs9402381 | 0.223 | 0.663 | 0.303 | 0.208 | -0.035 | -0.06 | -0.06 | 0 | 0.13 | 0.04 | 0.22 | 0.41 | 0.23 | 0.26 |
| 6 | 132316561 | rs170881 | -1.715 | 0.571 | 0.121 | 0.001 | -0.043 | -0.025 | -0.025 | 0.445 | 0.4 | 0.81 | 0.43 | 0.59 | 0.42 | 0.41 |
| 6 | 132319670 | rs233940 | -0.765 | 0.569 | 0.83 | -0.052 | 0.011 | -0.013 | -0.013 | 0 | 0.23 | 0.38 | 0.16 | 0.18 | 0.19 | 0.14 |
| 6 | 132319751 | rs2027035 | -0.406 | 0.521 | 0.905 | -0.093 | -0.011 | 0.027 | 0.027 | 0 | 0.14 | 0.24 | 0.09 | 0.18 | 0.1 | 0.1 |
| 6 | 132329574 | rs511587 | 0.704 | 0.712 | 0.616 | 0.022 | -0.005 | -0.007 | -0.007 | 0.881 | 0.09 | 0.08 | 0.06 | 0 | 0.08 | 0.04 |
| 6 | 132339176 | rs9483372 | -0.225 | 0.585 | 0.714 | 0.068 | 0.008 | 0.048 | 0.048 | 0.019 | 0.17 | 0.3 | 0.22 | 0.41 | 0.26 | 0.29 |
| 6 | 132342745 | rs233928 | -1.023 | 0.567 | 0.996 | 0.034 | 0.024 | 0.038 | 0.038 | 0 | 0.27 | 0.36 | 0.29 | 0.41 | 0.34 | 0.33 |
| 6 | 132344140 | rs233930 | -0.816 | 0.595 | 0.76 | 0.05 | -0.019 | -0.027 | -0.027 | 0 | 0.24 | 0.17 | 0.28 | 0.42 | 0.34 | 0.32 |
| 6 | 132363507 | rs975515 | -2.437 | 0.506 | 0.432 | -0.058 | -0.046 | -0.037 | -0.037 | 2.761 | 0.61 | 0.91 | 0.54 | 0.65 | 0.62 | 0.63 |
| 6 | 132364521 | rs1206737 | -0.714 | 0.553 | 0.926 | -0.079 | -0.005 | -0.007 | -0.007 | 0.018 | 0.21 | 0.36 | 0.15 | 0.17 | 0.16 | 0.14 |
| 6 | 132368241 | rs1206738 | 0.058 | 0.718 | 0.664 | 0.065 | -0.011 | -0.025 | -0.025 | 0 | 0.17 | 0.1 | 0.21 | 0.43 | 0.27 | 0.28 |
| 6 | 132373986 | rs2327194 | -1.455 | 0.652 | 0.791 | -0.057 | 0.003 | 0.026 | 0.026 | 0 | 0.39 | 0.71 | 0.27 | 0.22 | 0.3 | 0.29 |
| 6 | 132375483 | rs4895927 | -2.731 | 0.708 | 0.757 | -0.065 | -0.065 | -0.053 | -0.053 | 0 | 0.61 | 0.95 | 0.54 | 0.65 | 0.62 | 0.62 |
| 6 | 132376200 | rs9493174 | -1.348 | 0.704 | 0.803 | -0.071 | 0.005 | 0.027 | 0.027 | 0 | 0.39 | 0.72 | 0.27 | 0.22 | 0.3 | 0.3 |
| 6 | 132380046 | rs9321321 | -1.575 | 0.618 | 1.041 | -0.091 | -0.029 | -0.019 | -0.019 | 0 | 0.42 | 0.77 | 0.43 | 0.5 | 0.5 | 0.49 |
| 6 | 132380559 | rs11154658 | -0.088 | 0.614 | 1.02 | -0.085 | -0.003 | -0.001 | -0.001 | 0 | 0.19 | 0.18 | 0.12 | 0.15 | 0.12 | 0.13 |
| 6 | 132380668 | rs9483377 | -0.589 | 0.634 | 1.033 | -0.088 | 0.015 | 0.049 | 0.049 | 0.015 | 0.22 | 0.26 | 0.27 | 0.43 | 0.32 | 0.33 |
| 6 | 132381115 | rs9385594 | -1.402 | 0.625 | 1.04 | -0.102 | -0.005 | -0.004 | -0.004 | 0 | 0.39 | 0.36 | 0.39 | 0.5 | 0.5 | 0.49 |
| 6 | 132381855 | rs7754913 | -0.125 | 0.611 | 1.035 | -0.102 | -0.013 | -0.003 | -0.003 | 0 | 0.2 | 0.35 | 0.12 | 0.16 | 0.12 | 0.13 |
| 6 | 132383261 | rs11961001 | 0.173 | 0.611 | 0.482 | -0.079 | -0.041 | 0.052 | 0.052 | 0 | 0.17 | 0.04 | 0.12 | 0.06 | 0.18 | 0.17 |
| 6 | 132383953 | rs9373010 | 0.197 | 0.613 | 0.514 | -0.064 | -0.014 | -0.03 | -0.03 | 0 | 0.16 | 0.08 | 0.21 | 0.43 | 0.27 | 0.28 |
| 6 | 132397902 | rs11962845 | 0.111 | 0.659 | 0.14 | -0.457 | -0.041 | 0.052 | 0.052 | 0 | 0.17 | 0.04 | 0.12 | 0.06 | 0.19 | 0.17 |
| 6 | 132404255 | rs7770134 | -0.476 | 0.797 | 0.108 | -0.426 | 0.001 | -0.014 | -0.014 | 0.076 | 0.23 | 0.22 | 0.19 | 0.07 | 0.25 | 0.22 |
| 6 | 132408304 | rs969522 | -0.478 | 0.802 | 0.169 | -0.426 | 0.006 | -0.032 | -0.032 | 0 | 0.23 | 0.27 | 0.2 | 0.07 | 0.25 | 0.22 |
| 6 | 132424554 | rs11154661 | -0.053 | 0.582 | 0.042 | -0.373 | 0.002 | -0.001 | -0.001 | 0 | 0.18 | 0.13 | 0.1 | 0.16 | 0.11 | 0.13 |
| 6 | 132425242 | rs6935514 | -1.889 | 0.51 | -0.265 | -0.033 | -0.055 | -0.125 | -0.125 | 0 | 0.62 | 0.95 | 0.55 | 0.53 | 0.63 | 0.62 |
| 6 | 132429885 | rs731033 | -0.64 | 0.622 | 0.532 | -0.254 | -0.005 | -0.04 | -0.04 | 0 | 0.17 | 0.2 | 0.12 | 0.05 | 0.14 | 0.19 |
| 6 | 132434060 | rs997607 | -0.53 | 0.457 | 0.404 | -0.213 | -0.007 | -0.024 | -0.024 | 0 | 0.16 | 0.17 | 0.1 | 0.05 | 0.12 | 0.19 |
| 6 | 132444234 | rs2064086 | -1.325 | 0.351 | 0.016 | -0.19 | 0.003 | 0.048 | 0.048 | 0 | 0.44 | 0.79 | 0.33 | 0.31 | 0.35 | 0.34 |
| 6 | 132450342 | rs9483384 | 0.021 | 0.305 | 0.123 | -0.2 | -0.01 | -0.008 | -0.008 | 0 | 0.13 | 0.15 | 0.05 | 0.07 | 0.08 | 0.15 |
| 6 | 132451644 | rs1358771 | -0.448 | 0.268 | 0.013 | -0.232 | 0.002 | -0.001 | -0.001 | 1.68 | 0.27 | 0.27 | 0.24 | 0.23 | 0.26 | 0.18 |
| 6 | 132459667 | rs6909982 | -1.08 | 0.387 | -0.132 | -0.149 | 0.004 | 0.028 | 0.028 | 0 | 0.43 | 0.66 | 0.32 | 0.31 | 0.35 | 0.34 |
| 6 | 132460724 | rs9483387 | -1.274 | 0.407 | -0.607 | -0.136 | 0 | 0.055 | 0.055 | 0 | 0.45 | 0.83 | 0.33 | 0.31 | 0.35 | 0.34 |

| | | | | | | | | | | | | | | | | |
|---|---|---|---|---|---|---|---|---|---|---|---|---|---|---|---|---|
| 6 | 132466452 | rs1321267 | -1.399 | 0.358 | -0.801 | -0.098 | -0.028 | 0.021 | 0.021 | 0 | 0.54 | 0.82 | 0.45 | 0.33 | 0.51 | 0.42 |
| 6 | 132476741 | rs9493202 | -0.799 | -0.595 | -0.429 | -0.016 | 0.044 | 0.029 | 0.029 | 3.704 | 0.06 | 0.55 | 0.05 | 0 | 0.01 | 0 |
| 6 | 132482287 | rs728371 | 0.041 | -0.252 | -1.193 | -0.429 | -0.074 | 0.094 | 0.094 | 0.003 | 0.21 | 0.08 | 0.21 | 0.01 | 0.33 | 0.21 |
| 6 | 132483263 | rs12661294 | -0.217 | -0.695 | -1.096 | 0.017 | 0.039 | 0.012 | 0.012 | 0 | 0.15 | 0.01 | 0.02 | 0.08 | 0.07 | 0.17 |
| 6 | 132500493 | rs12529280 | 0.611 | -0.192 | -0.797 | 0.542 | -0.032 | -0.023 | -0.023 | 0 | 0.07 | 0.01 | 0.1 | 0.24 | 0.08 | 0.05 |
| 6 | 132507974 | rs7761015 | -1.159 | -0.364 | -0.526 | 0.011 | -0.027 | -0.039 | -0.039 | 0 | 0.08 | 0.66 | 0.08 | 0 | 0.01 | 0 |
| 6 | 132511836 | rs9493231 | -0.896 | -0.326 | -0.388 | 0.041 | 0.012 | 0.003 | 0.003 | 5.959 | 0.07 | 0.57 | 0.07 | 0 | 0.01 | 0 |
| 6 | 132515568 | rs10456994 | 0.517 | -0.502 | -0.79 | 0.112 | -0.036 | 0.135 | 0.135 | 0.005 | 0.53 | 0.16 | 0.53 | 0.76 | 0.49 | 0.51 |
| 6 | 132516995 | rs9375863 | 0.082 | -0.096 | -1.161 | 0.35 | -0.014 | 0.023 | 0.023 | 0.033 | 0.6 | 0.67 | 0.59 | 0.76 | 0.49 | 0.52 |
| 6 | 132530470 | rs12211717 | 0.868 | 0.246 | -1.707 | 0.77 | -0.098 | -0.102 | -0.102 | 0.126 | 0.31 | 0.04 | 0.41 | 0.4 | 0.33 | 0.33 |
| 6 | 132533363 | rs3927419 | -1.038 | -0.467 | -1.458 | 0.951 | 0.019 | 0.081 | 0.081 | 0.233 | 0.47 | 0.78 | 0.49 | 0.67 | 0.38 | 0.66 |
| 6 | 132534652 | rs11154667 | -0.615 | -0.826 | -0.321 | 1.32 | -0.007 | 0.193 | 0.193 | 0 | 0.15 | 0.27 | 0.16 | 0.39 | 0.12 | 0.19 |
| 6 | 132537406 | rs7756650 | -0.163 | -1.01 | 0.044 | 0.973 | 0 | 0 | 0 | 0 | 0.15 | 0.12 | 0.12 | 0.18 | 0.19 | 0.28 |
| 6 | 132537834 | rs7760956 | -1.714 | -1.097 | 0.347 | 0.929 | -0.001 | -0.011 | -0.011 | 0.884 | 0.35 | 0.67 | 0.32 | 0.61 | 0.32 | 0.47 |
| 6 | 132539383 | rs17061077 | -0.674 | -1.057 | 0.384 | 0.92 | -0.01 | -0.007 | -0.007 | 0.648 | 0.12 | 0.14 | 0.06 | 0.05 | 0.06 | 0.19 |
| 6 | 132541408 | rs7739499 | -2.094 | -1.159 | 0.562 | 0.873 | 0.023 | 0.059 | 0.059 | 0.286 | 0.47 | 0.69 | 0.37 | 0.66 | 0.38 | 0.66 |
| 6 | 132557000 | rs1998073 | -1.106 | -1.297 | 1.452 | 1.101 | 0.005 | -0.018 | -0.018 | 0 | 0.25 | 0.29 | 0.15 | 0.23 | 0.25 | 0.4 |
| 6 | 132557900 | rs11154669 | -1.103 | -1.322 | 1.37 | 1.099 | -0.019 | -0.136 | -0.136 | 0.104 | 0.16 | 0.04 | 0.16 | 0.43 | 0.13 | 0.25 |
| 6 | 132560968 | rs1998075 | -0.256 | -1.52 | 1.483 | 1.092 | 0.004 | -0.003 | -0.003 | 0 | 0.17 | 0.17 | 0.14 | 0.18 | 0.2 | 0.27 |
| 6 | 132561805 | rs7738716 | -1.39 | -1.461 | 1.464 | 1.122 | -0.003 | -0.039 | -0.039 | 0.03 | 0.28 | 0.46 | 0.19 | 0.23 | 0.25 | 0.4 |
| 6 | 132567078 | rs9321334 | -1.593 | -1.463 | 1.146 | 1.355 | 0 | 0 | 0 | 0 | 0.31 | 0.62 | 0.21 | 0.27 | 0.25 | 0.4 |
| 6 | 132571889 | rs12528986 | 0.593 | -1.456 | 0.773 | 1.433 | -0.017 | -0.003 | -0.003 | 0 | 0.07 | 0.02 | 0.12 | 0.34 | 0.06 | 0.07 |
| 6 | 132593091 | rs7762331 | -2.532 | -1.636 | 0.341 | 1.144 | -0.178 | 0.017 | 0.017 | 0.617 | 0.62 | 0.98 | 0.58 | 0.79 | 0.52 | 0.71 |
| 6 | 132598642 | rs7739007 | -0.703 | -1.411 | 1.284 | 1.349 | -0.003 | -0.001 | -0.001 | 0.991 | 0.08 | 0.07 | 0.02 | 0.08 | 0.05 | 0.13 |
| 6 | 132603042 | rs1983568 | -1.139 | -1.412 | 1.427 | 1.582 | -0.001 | 0.045 | 0.045 | 0 | 0.13 | 0.1 | 0.2 | 0.38 | 0.1 | 0.23 |
| 6 | 132611762 | rs9388971 | -0.655 | -1.506 | 1.483 | 1.305 | 0.014 | -0.008 | -0.008 | 0 | 0.17 | 0.23 | 0.14 | 0.21 | 0.22 | 0.28 |
| 6 | 132617072 | rs6902875 | -2.185 | -1.604 | 1.152 | 1.28 | -0.03 | -0.01 | -0.01 | 0 | 0.33 | 0.9 | 0.24 | 0.3 | 0.28 | 0.4 |
| 6 | 132617469 | rs6903616 | -1.47 | -1.695 | 1.65 | 1.344 | -0.036 | 0.042 | 0.042 | 0.198 | 0.11 | 0.33 | 0.04 | 0.09 | 0.06 | 0.12 |
| 6 | 132623184 | rs4897574 | -1.958 | -1.642 | 1.367 | 1.329 | 0.012 | 0.028 | 0.028 | 1.89 | 0.41 | 0.73 | 0.38 | 0.68 | 0.34 | 0.61 |
| 6 | 132635480 | rs6917708 | -0.725 | -1.596 | 0.958 | 1.596 | -0.012 | 0.005 | 0.005 | 0 | 0.05 | 0.08 | 0.01 | 0.05 | 0.03 | 0.06 |
| 6 | 132642770 | rs2235434 | -1.147 | -1.535 | 0.069 | 1.361 | 0.025 | -0.073 | -0.073 | 0.017 | 0.16 | 0.02 | 0.11 | 0.29 | 0.13 | 0.42 |
| 6 | 132643117 | rs589756 | -0.41 | -1.231 | 0.269 | 0.948 | 0.018 | -0.037 | -0.037 | 0 | 0.14 | 0.17 | 0.14 | 0 | 0.19 | 0.11 |
| 6 | 132644035 | rs604196 | -1.72 | -1.252 | -0.233 | 1.213 | -0.015 | 0.036 | 0.036 | 1.929 | 0.41 | 0.85 | 0.45 | 0.66 | 0.35 | 0.6 |
| 6 | 132647962 | rs10484483 | -1.073 | -1.878 | 0.504 | 1.681 | -0.009 | 0.022 | 0.022 | 0 | 0.1 | 0.09 | 0.08 | 0.2 | 0.09 | 0.27 |
| 6 | 132655551 | rs6909136 | -2.032 | -2.104 | -0.06 | 1.153 | 0.081 | 0.016 | 0.016 | 0 | 0.33 | 0.83 | 0.23 | 0.26 | 0.21 | 0.39 |
| 6 | 132681877 | rs13196312 | -1.283 | -2.164 | 0.392 | 1.229 | -0.013 | -0.002 | -0.002 | 0 | 0.23 | 0.25 | 0.16 | 0.22 | 0.17 | 0.32 |
| 6 | 132683762 | rs17061202 | -1.469 | -2.158 | 0.457 | 1.243 | -0.04 | -0.006 | -0.006 | 0 | 0.25 | 0.36 | 0.17 | 0.23 | 0.17 | 0.32 |
| 6 | 132690239 | rs9375877 | -1.424 | -2.154 | 0.404 | 1.029 | -0.051 | 0.017 | 0.017 | 0 | 0.26 | 0.4 | 0.27 | 0.67 | 0.17 | 0.34 |
| 6 | 132694751 | rs6569792 | -1.738 | -2.013 | -0.66 | 0.82 | 0.102 | 0.07 | 0.07 | 0.179 | 0.39 | 0.9 | 0.38 | 0.73 | 0.27 | 0.46 |
| 6 | 132696016 | rs3734744 | -1.34 | -2.127 | -0.176 | 1.225 | -0.017 | 0.004 | 0.004 | 0.009 | 0.21 | 0.69 | 0.18 | 0.26 | 0.15 | 0.29 |
| 6 | 132711200 | rs2275394 | -1.498 | -2.501 | -0.814 | 0.563 | 0.125 | -0.003 | -0.003 | 0 | 0.35 | 0.89 | 0.33 | 0.66 | 0.2 | 0.36 |
| 6 | 132716562 | rs1577995 | -0.546 | -1.529 | -0.847 | -0.049 | -0.029 | 0.072 | 0.072 | 0.163 | 0.21 | 0.07 | 0.11 | 0 | 0.21 | 0.17 |
| 6 | 132718231 | rs7745230 | -1.504 | -1.962 | -1.369 | -0.557 | -0.066 | 0.14 | 0.14 | 0.523 | 0.6 | 0.91 | 0.51 | 0.87 | 0.46 | 0.61 |
| 6 | 132724116 | rs6900501 | -0.806 | -2.255 | -0.726 | -0.026 | -0.018 | 0.02 | 0.02 | 0 | 0.26 | 0.71 | 0.24 | 0.44 | 0.15 | 0.16 |
| 6 | 132724298 | rs6901304 | -1.285 | -2.327 | -0.769 | -0.159 | 0.095 | 0.105 | 0.105 | 0 | 0.37 | 0.81 | 0.38 | 0.87 | 0.24 | 0.38 |
| 6 | 132730947 | rs1416100 | -0.45 | -1.993 | -0.195 | -0.094 | -0.001 | -0.007 | -0.007 | 1.321 | 0.26 | 0.26 | 0.16 | 0 | 0.24 | 0.24 |
| 6 | 132733418 | rs6923916 | -0.635 | -1.966 | 0.135 | 0.019 | -0.068 | 0.193 | 0.193 | 0 | 0.2 | 0.43 | 0.2 | 0.42 | 0.12 | 0.07 |
| 6 | 132749484 | rs7750935 | -1.122 | -1.902 | -0.004 | 0.106 | 0.033 | -0.003 | -0.003 | 0 | 0.45 | 0.7 | 0.35 | 0.42 | 0.35 | 0.3 |
| 6 | 132749585 | rs7754992 | -1.235 | -1.894 | 0.012 | 0.092 | 0.028 | -0.002 | -0.002 | 4.488 | 0.45 | 0.68 | 0.34 | 0.42 | 0.35 | 0.3 |
| 6 | 132749870 | rs9483446 | -0.246 | -1.843 | 0.878 | 0.634 | -0.029 | -0.039 | -0.039 | 0 | 0.08 | 0.16 | 0.04 | 0 | 0.04 | 0.03 |
| 6 | 132757467 | rs9388996 | -1.011 | -1.849 | 0.617 | 0.608 | -0.059 | -0.049 | -0.049 | 0 | 0.21 | 0.39 | 0.26 | 0.12 | 0.13 | 0.18 |
| 6 | 132762095 | rs9385608 | -1.147 | -1.835 | -0.134 | 0.615 | 0.017 | -0.012 | -0.012 | 1.639 | 0.31 | 0.72 | 0.4 | 0.54 | 0.17 | 0.2 |
| 6 | 132765669 | rs17794361 | -1.205 | -1.185 | 0.111 | 0.475 | -0.013 | -0.009 | -0.009 | 0.098 | 0.21 | 0.33 | 0.15 | 0.22 | 0.17 | 0.24 |
| 6 | 132766737 | rs10499186 | -0.103 | -1.13 | -0.605 | -0.157 | 0.015 | 0.046 | 0.046 | 2.989 | 0.2 | 0.13 | 0.13 | 0.08 | 0.15 | 0.12 |
| 6 | 132780167 | rs1044083 | -1.072 | 0.068 | 0.142 | 0.481 | 0.007 | 0.005 | 0.005 | 0.277 | 0.3 | 0.3 | 0.21 | 0.16 | 0.21 | 0.25 |
| 6 | 132781339 | rs3183732 | 0.608 | -0.163 | -0.185 | 0.009 | -0.006 | 0.007 | 0.007 | 0 | 0.31 | 0.22 | 0.42 | 0.69 | 0.31 | 0.47 |
| 6 | 132783865 | rs9483454 | 0.644 | -0.311 | 0.091 | -0.128 | 0.002 | -0.005 | -0.005 | 0 | 0.27 | 0.21 | 0.25 | 0.55 | 0.18 | 0.4 |
| 6 | 132788323 | rs6935785 | -1.586 | -0.104 | 0.516 | -0.574 | -0.024 | 0.046 | 0.046 | 0 | 0.69 | 0.8 | 0.67 | 0.85 | 0.56 | 0.79 |
| 6 | 132797275 | rs1416391 | 0.38 | -0.088 | 1.271 | -0.465 | -0.024 | -0.071 | -0.071 | 0 | 0.27 | 0.47 | 0.19 | 0.16 | 0.2 | 0.25 |
| 6 | 132797923 | rs17061324 | 1.148 | -0.05 | 1.112 | -0.437 | 0.011 | 0.047 | 0.047 | 2.485 | 0.2 | 0.12 | 0.12 | 0.1 | 0.15 | 0.12 |
| 6 | 132799544 | rs9493329 | 0.718 | -0.081 | 1.131 | -0.393 | 0.012 | -0.02 | -0.02 | 0.353 | 0.28 | 0.2 | 0.28 | 0.55 | 0.21 | 0.4 |
| 6 | 132799913 | rs13205427 | 0.465 | -0.045 | 1.144 | -0.667 | -0.004 | 0.007 | 0.007 | 0 | 0.31 | 0.25 | 0.43 | 0.71 | 0.31 | 0.47 |
| 6 | 132801875 | rs1002799 | 0.562 | -0.209 | 1.116 | -0.708 | -0.013 | 0.001 | 0.001 | 0 | 0.29 | 0.3 | 0.27 | 0.65 | 0.21 | 0.39 |
| 6 | 132805152 | rs9483463 | 0.485 | -0.173 | 1.248 | -0.681 | -0.03 | 0.004 | 0.004 | 0 | 0.3 | 0.38 | 0.27 | 0.65 | 0.21 | 0.38 |
| 6 | 132814037 | rs2788942 | -0.207 | 0.494 | 1.012 | -0.429 | -0.014 | 0.01 | 0.01 | 0 | 0.69 | 0.73 | 0.59 | 0.66 | 0.62 | 0.61 |
| 6 | 132818354 | rs2788940 | 0.723 | 0.576 | 1.069 | -0.188 | 0.046 | 0.023 | 0.023 | 0 | 0.87 | 0.79 | 0.73 | 0.8 | 0.81 | 0.73 |
| 6 | 132818540 | rs2842889 | 0.723 | 0.603 | 1.044 | -0.185 | 0.095 | 0.001 | 0.001 | 0 | 0.91 | 0.76 | 0.8 | 0.93 | 0.86 | 0.85 |
| 6 | 132821407 | rs9493336 | 0.392 | -0.085 | 0.444 | -0.071 | 0.013 | -0.033 | -0.033 | 0 | 0.27 | 0.14 | 0.22 | 0.54 | 0.21 | 0.37 |
| 6 | 132825982 | rs7758170 | 0.73 | 0.168 | 0.671 | -0.758 | 0.022 | 0.093 | 0.093 | 0 | 0.05 | 0.18 | 0.06 | 0.15 | 0.05 | 0.12 |

| | | | | | | | | | | | | | | | | |
|---|---|---|---|---|---|---|---|---|---|---|---|---|---|---|---|---|
| 6 | 132830944 | rs9493341 | 0.135 | -0.175 | -0.116 | -1.079 | 0.039 | 0.033 | 0.033 | 1.967 | 0.46 | 0.24 | 0.41 | 0.61 | 0.36 | 0.5 |
| 6 | 132836473 | rs7745902 | -0.299 | -0.091 | 0.198 | -1.54 | -0.022 | -0.035 | -0.035 | 0 | 0.31 | 0.43 | 0.31 | 0.79 | 0.25 | 0.51 |
| 6 | 132842396 | rs1591811 | -0.52 | 0.35 | 0.281 | -1.749 | 0.005 | -0.022 | -0.022 | 0.118 | 0.67 | 0.66 | 0.64 | 0.87 | 0.65 | 0.77 |
| 6 | 132845366 | rs2153177 | 0.629 | 0.137 | 0.26 | -1.789 | -0.026 | 0.029 | 0.029 | 0.084 | 0.34 | 0.22 | 0.44 | 0.23 | 0.43 | 0.29 |
| 6 | 132846134 | rs6936245 | 0.222 | 0.013 | 0.507 | -1.674 | -0.008 | -0.018 | -0.018 | 0 | 0.05 | 0.25 | 0.02 | 0 | 0.01 | 0.03 |
| 6 | 132854027 | rs2066076 | 0.184 | 0.151 | 0.295 | -1.608 | 0.003 | -0.004 | -0.004 | 0 | 0.28 | 0.24 | 0.17 | 0.64 | 0.2 | 0.42 |
| 6 | 132854616 | rs9493366 | 0.056 | 0.196 | 0.256 | -1.595 | -0.039 | 0.024 | 0.024 | 0 | 0.3 | 0.34 | 0.18 | 0.64 | 0.21 | 0.42 |
| 6 | 132857687 | rs17061369 | 0.285 | 0.597 | 0.08 | -1.536 | 0.009 | -0.019 | -0.019 | 0.666 | 0.08 | 0.1 | 0.06 | 0.02 | 0.09 | 0.07 |
| 6 | 132859437 | rs9402420 | 0.187 | 0.65 | 0.077 | -1.539 | 0.01 | -0.02 | -0.02 | 0 | 0.08 | 0.1 | 0.06 | 0.02 | 0.1 | 0.09 |
| 6 | 132860260 | rs9389004 | 0.239 | 0.251 | 0.099 | -1.511 | -0.005 | 0.012 | 0.012 | 0 | 0.06 | 0.01 | 0.03 | 0.02 | 0.05 | 0.04 |
| 6 | 132860575 | rs6930707 | 0.278 | 0.291 | 0.122 | -1.489 | 0.016 | -0.014 | -0.014 | 0 | 0.06 | 0.1 | 0.05 | 0.02 | 0.07 | 0.06 |
| 6 | 132862349 | rs2153174 | 0.428 | 0.172 | 0.168 | -1.823 | 0.004 | -0.013 | -0.013 | 0 | 0.39 | 0.46 | 0.49 | 0.23 | 0.47 | 0.32 |
| 6 | 132864548 | rs9321348 | -1.244 | 0.06 | 0.14 | -1.878 | -0.012 | 0.053 | 0.053 | 0 | 0.74 | 0.85 | 0.7 | 0.88 | 0.74 | 0.81 |
| 6 | 132867959 | rs9389005 | -0.248 | 0.249 | 0.481 | -1.899 | 0.049 | -0.008 | -0.008 | 0 | 0.08 | 0.23 | 0.06 | 0.02 | 0.07 | 0.06 |
| 6 | 132871653 | rs1933988 | 0.028 | 0.329 | 0.402 | -2.1 | 0.063 | -0.035 | -0.035 | 0 | 0.26 | 0.44 | 0.43 | 0.15 | 0.41 | 0.28 |
| 6 | 132871758 | rs2840835 | -0.659 | 0.266 | 0.351 | -2.135 | 0.002 | 0.128 | 0.128 | 0.267 | 0.47 | 0.75 | 0.55 | 0.24 | 0.54 | 0.38 |
| 6 | 132874814 | rs8192627 | 0.747 | 0.317 | 0.181 | -2.044 | -0.003 | 0.002 | 0.002 | 0 | 0.06 | 0.03 | 0.04 | 0.02 | 0.07 | 0.06 |
| 6 | 132878322 | rs11963639 | 1.2 | 0.29 | 0.114 | -1.986 | 0.003 | 0.003 | 0.003 | 2.887 | 0.19 | 0.16 | 0.12 | 0.09 | 0.14 | 0.12 |
| 6 | 132879258 | rs2327212 | 1.389 | 0.297 | 0.048 | -1.979 | 0.022 | 0.04 | 0.04 | 3.032 | 0.18 | 0.07 | 0.1 | 0.09 | 0.12 | 0.1 |
| 6 | 132880608 | rs17061429 | 1.167 | 0.294 | 0.107 | -1.98 | -0.018 | -0.046 | -0.046 | 0 | 0.2 | 0.3 | 0.14 | 0.09 | 0.14 | 0.12 |
| 6 | 132880648 | rs2255071 | 0.504 | 0.615 | -0.769 | -1.823 | 0.026 | -0.011 | -0.011 | 0 | 0.2 | 0.3 | 0.37 | 0.13 | 0.34 | 0.22 |
| 6 | 132883583 | rs2840837 | 0.949 | 0.661 | -1.094 | -2.149 | -0.003 | -0.002 | -0.002 | 0 | 0.17 | 0.17 | 0.36 | 0.04 | 0.33 | 0.19 |
| 6 | 132884701 | rs1361281 | -1.043 | -0.148 | -1.1 | -1.615 | 0 | -0.041 | -0.041 | 0.19 | 0.55 | 0.44 | 0.34 | 0.86 | 0.44 | 0.57 |
| 6 | 132885210 | rs11968522 | -0.168 | -0.292 | -1.324 | -1.718 | -0.023 | 0.113 | 0.113 | 2.497 | 0.29 | 0.06 | 0.16 | 0.1 | 0.24 | 0.15 |
| 6 | 132885345 | rs9321350 | -0.077 | -0.231 | -0.291 | -0.902 | 0.014 | -0.066 | -0.066 | 0 | 0.21 | 0.04 | 0.14 | 0.76 | 0.17 | 0.4 |
| 6 | 132888831 | rs9493381 | -0.798 | 0.017 | 0.03 | -0.795 | -0.009 | -0.044 | -0.044 | 0 | 0.37 | 0.46 | 0.3 | 0.79 | 0.31 | 0.48 |
| 6 | 132890341 | rs9389007 | -0.751 | 0.324 | -1.276 | -0.641 | -0.004 | 0.058 | 0.058 | 0 | 0.34 | 0.16 | 0.27 | 0.79 | 0.31 | 0.48 |
| 6 | 132892253 | rs8192624 | -0.927 | 0.383 | -0.857 | -0.354 | -0.005 | 0.019 | 0.019 | 0 | 0.1 | 0.05 | 0.06 | 0.02 | 0.09 | 0.08 |
| 6 | 132893712 | rs4305745 | -0.193 | 0.063 | -0.688 | -0.357 | 0 | -0.001 | -0.001 | 0 | 0.58 | 0.61 | 0.47 | 0.41 | 0.54 | 0.64 |
| 6 | 132897029 | rs6924201 | -0.256 | -0.029 | -0.909 | -0.31 | 0.013 | -0.016 | -0.016 | 1.949 | 0.18 | 0.35 | 0.27 | 0.18 | 0.18 | 0.12 |
| 6 | 132897200 | rs6904438 | -0.194 | -0.93 | -0.86 | -0.959 | 0.024 | 0.077 | 0.077 | 0 | 0.52 | 0.65 | 0.6 | 0.3 | 0.57 | 0.39 |
| 6 | 132897858 | rs7765655 | -1.454 | -0.9 | -0.659 | -0.925 | -0.018 | -0.07 | -0.07 | 0.017 | 0.75 | 0.86 | 0.73 | 0.59 | 0.73 | 0.77 |
| 6 | 132899017 | rs17194904 | 0.66 | -0.98 | -0.093 | -0.982 | -0.001 | 0 | 0 | 0.362 | 0.12 | 0.1 | 0.22 | 0.16 | 0.16 | 0.09 |
| 6 | 132899531 | rs6937423 | -0.45 | -0.996 | 0.346 | -1.008 | 0.059 | 0.022 | 0.022 | 0 | 0.05 | 0.24 | 0.04 | 0.01 | 0.03 | 0.03 |
| 6 | 132901735 | rs12526035 | 0.542 | -0.964 | 0.218 | -0.949 | 0 | -0.001 | -0.001 | 0.632 | 0.12 | 0.12 | 0.22 | 0.16 | 0.16 | 0.09 |
| 6 | 132903972 | rs12528813 | 0.836 | -0.984 | 0.155 | -0.945 | -0.04 | 0.007 | 0.007 | 0 | 0.11 | 0.01 | 0.2 | 0.06 | 0.16 | 0.08 |
| 6 | 132905187 | rs9321354 | -0.691 | -1.667 | 0.239 | -1.242 | -0.037 | -0.004 | -0.004 | 0 | 0.27 | 0.39 | 0.25 | 0.32 | 0.14 | 0.43 |
| 6 | 132907913 | rs4897591 | -1.082 | -1.715 | 0.239 | -1.249 | -0.01 | -0.002 | -0.002 | 0.368 | 0.43 | 0.48 | 0.48 | 0.49 | 0.3 | 0.53 |
| 6 | 132908879 | rs4458717 | 0.157 | -1.77 | 0.258 | -1.281 | 0.001 | 0.002 | 0.002 | 0 | 0.2 | 0.21 | 0.29 | 0.17 | 0.2 | 0.13 |
| 6 | 132910612 | rs3813355 | -1.484 | -1.708 | 0.382 | -1.264 | 0 | 0.002 | 0.002 | 0.114 | 0.69 | 0.73 | 0.68 | 0.56 | 0.67 | 0.72 |
| 6 | 132910634 | rs3813354 | -0.535 | -1.445 | 0.78 | -1.027 | -0.003 | -0.002 | -0.002 | 0 | 0.21 | 0.19 | 0.19 | 0.32 | 0.1 | 0.39 |
| 6 | 132912612 | rs4605911 | -0.797 | -1.491 | 0.682 | -1.068 | -0.06 | 0.011 | 0.011 | 0.047 | 0.21 | 0.34 | 0.21 | 0.31 | 0.08 | 0.38 |
| 6 | 132913582 | rs17061481 | -0.465 | -1.511 | 0.621 | -1.116 | -0.007 | 0.015 | 0.015 | 0 | 0.13 | 0.23 | 0.11 | 0.26 | 0.08 | 0.29 |
| 6 | 132915931 | rs9389009 | -0.682 | -1.508 | 0.637 | -1.196 | 0 | -0.002 | -0.002 | 0 | 0.43 | 0.6 | 0.32 | 0.31 | 0.46 | 0.4 |
| 6 | 132917612 | rs4305747 | -1.273 | -1.493 | 0.665 | -1.188 | -0.045 | 0.056 | 0.056 | 0.112 | 0.55 | 0.85 | 0.46 | 0.39 | 0.49 | 0.61 |
| 6 | 132919550 | rs12197997 | 0.952 | -0.909 | 0.759 | -0.902 | -0.042 | 0.043 | 0.043 | 1.112 | 0.21 | 0.1 | 0.15 | 0.05 | 0.32 | 0.16 |
| 6 | 132920150 | rs17215186 | 0.478 | -0.892 | 0.538 | -0.835 | 0.016 | -0.043 | -0.043 | 0 | 0.26 | 0.23 | 0.18 | 0.05 | 0.36 | 0.18 |
| 6 | 132921520 | rs17061497 | -0.415 | -0.898 | 0.552 | -0.821 | -0.013 | 0.045 | 0.045 | 0.161 | 0.13 | 0.32 | 0.11 | 0.26 | 0.08 | 0.21 |
| 6 | 132921834 | rs12203326 | 0.241 | -0.279 | 0.412 | -0.884 | 0.02 | -0.046 | -0.046 | 0 | 0.25 | 0.23 | 0.17 | 0.05 | 0.35 | 0.17 |
| 6 | 132925613 | rs6917098 | -0.405 | -0.095 | 0.001 | -0.746 | 0.017 | -0.105 | -0.105 | 1.359 | 0.35 | 0.43 | 0.27 | 0.08 | 0.39 | 0.32 |
| 6 | 132929016 | rs12200377 | -1.299 | -0.363 | 0.148 | 0.041 | -0.023 | 0.032 | 0.032 | 0 | 0.15 | 0.6 | 0.15 | 0.14 | 0.09 | 0.16 |
| 6 | 132929084 | rs12200430 | -0.857 | -0.127 | -0.828 | -0.321 | -0.009 | -0.008 | -0.008 | 0.052 | 0.45 | 0.76 | 0.52 | 0.53 | 0.48 | 0.58 |
| 6 | 132929147 | rs9493399 | -0.788 | -0.25 | -0.667 | -0.476 | -0.015 | 0 | 0 | 0 | 0.14 | 0.23 | 0.15 | 0.2 | 0.11 | 0.19 |
| 6 | 132929276 | rs9399036 | -0.826 | -0.383 | -1.658 | 0.099 | -0.035 | 0.062 | 0.062 | 0.078 | 0.59 | 0.84 | 0.52 | 0.66 | 0.55 | 0.57 |
| 6 | 132930198 | rs17061508 | -0.063 | -0.613 | -1.298 | 0.302 | 0.001 | -0.001 | -0.001 | 0.502 | 0.06 | 0.06 | 0.07 | 0.12 | 0.03 | 0.09 |
| 6 | 132930324 | rs7759004 | -0.321 | -0.895 | -0.405 | 0.309 | -0.028 | 0.064 | 0.064 | 1.64 | 0.1 | 0.56 | 0.1 | 0.09 | 0.03 | 0.09 |
| 6 | 132931359 | rs1081076 | 0.553 | -0.929 | -0.348 | 0.684 | 0.004 | 0.052 | 0.052 | 0 | 0.06 | 0.19 | 0.08 | 0.08 | 0.03 | 0.04 |
| 6 | 132933476 | rs9483484 | -0.795 | -0.78 | -0.161 | 0.58 | -0.007 | 0.023 | 0.023 | 1.149 | 0.13 | 0.61 | 0.17 | 0.1 | 0.07 | 0.11 |
| 6 | 132934131 | rs13220366 | 1.1 | -0.766 | -0.723 | 0.35 | -0.054 | 0.085 | 0.085 | 0 | 0.22 | 0.02 | 0.12 | 0.05 | 0.22 | 0.15 |
| 6 | 132934579 | rs9389011 | 0.602 | -0.699 | -0.94 | 1.164 | -0.063 | -0.086 | -0.086 | 0 | 0.15 | 0.02 | 0.18 | 0.3 | 0.24 | 0.2 |
| 6 | 132940293 | rs4451148 | -0.321 | -0.89 | -0.833 | 1.414 | -0.016 | -0.112 | -0.112 | 0.587 | 0.23 | 0.09 | 0.26 | 0.6 | 0.28 | 0.35 |
| 6 | 132941616 | rs9389012 | -1.008 | -0.966 | -1.189 | 1.351 | 0.006 | 0.059 | 0.059 | 0.035 | 0.51 | 0.26 | 0.43 | 0.73 | 0.52 | 0.56 |
| 6 | 132942718 | rs11968252 | 1.287 | -0.886 | -1.339 | 1.121 | -0.056 | 0.082 | 0.082 | 0.015 | 0.21 | 0.02 | 0.12 | 0.05 | 0.22 | 0.11 |
| 6 | 132942953 | rs9321355 | -0.571 | -0.951 | -1.12 | 1.264 | -0.035 | 0.023 | 0.023 | 2.158 | 0.13 | 0.71 | 0.18 | 0.16 | 0.04 | 0.12 |
| 6 | 132943233 | rs9321356 | -0.814 | -0.917 | -1.103 | 1.222 | -0.034 | 0.048 | 0.048 | 0.872 | 0.16 | 0.7 | 0.18 | 0.35 | 0.06 | 0.2 |
| 6 | 132943436 | rs9389015 | -1.063 | -0.945 | -1.364 | 0.947 | -0.067 | 0.149 | 0.149 | 0.407 | 0.57 | 0.86 | 0.54 | 0.85 | 0.53 | 0.63 |
| 6 | 132944523 | rs9385619 | -0.926 | -1.35 | -1.446 | 0.097 | -0.051 | -0.054 | -0.054 | 0.065 | 0.34 | 0.19 | 0.36 | 0.49 | 0.44 | 0.46 |
| 6 | 132944997 | rs4129256 | -0.302 | -1.237 | -1.488 | 0.183 | -0.022 | 0.016 | 0.016 | 0 | 0.2 | 0.07 | 0.18 | 0.17 | 0.2 | 0.21 |
| 6 | 132951025 | rs7756164 | 0.47 | -1.345 | -1.265 | 0.247 | 0.021 | 0.281 | 0.281 | 0.471 | 0.08 | 0.53 | 0.11 | 0.26 | 0.03 | 0.1 |

| | | | | | | | | | | | | | | | | |
|---|---|---|---|---|---|---|---|---|---|---|---|---|---|---|---|---|
| 6 | 132951889 | rs10782248 | -1.014 | -1.673 | -1.535 | -0.013 | 0.048 | 0.207 | 0.207 | 0 | 0.5 | 0.8 | 0.48 | 0.86 | 0.43 | 0.53 |
| 6 | 132951990 | rs4549637 | 0.28 | -1.648 | -1.58 | 0.015 | -0.059 | -0.093 | -0.093 | 0.062 | 0.31 | 0.08 | 0.24 | 0.34 | 0.32 | 0.28 |
| 6 | 132956202 | rs4345419 | 1.741 | -1.6 | -1.742 | -0.123 | 0.046 | 0.099 | 0.099 | 0 | 0.17 | 0.06 | 0.08 | 0.02 | 0.08 | 0.04 |
| 6 | 132963041 | rs6569819 | -1.767 | -1.34 | -1.63 | -0.007 | -0.072 | 0.142 | 0.142 | 0 | 0.65 | 0.92 | 0.6 | 0.89 | 0.58 | 0.66 |
| 6 | 132963053 | rs9389020 | -0.588 | -1.011 | -2.041 | 0.347 | -0.009 | -0.012 | -0.012 | 0 | 0.2 | 0.16 | 0.25 | 0.46 | 0.29 | 0.33 |
| 6 | 132966279 | rs8192620 | 0.303 | -1.014 | -2.043 | 0.365 | -0.011 | -0.012 | -0.012 | 0 | 0.14 | 0.1 | 0.17 | 0.32 | 0.24 | 0.24 |
| 6 | 132966348 | rs8192619 | -0.114 | -0.946 | -1.994 | 0.178 | -0.001 | 0.003 | 0.003 | 0 | 0.05 | 0.05 | 0.07 | 0.14 | 0.05 | 0.09 |
| 6 | 132982421 | rs7762377 | 0.479 | -0.787 | -2.062 | 0.247 | -0.066 | -0.084 | -0.084 | 0 | 0.13 | 0.02 | 0.15 | 0.34 | 0.21 | 0.23 |
| 6 | 132983625 | rs17061564 | -0.049 | -1.081 | -1.863 | 0.16 | 0.003 | -0.002 | -0.002 | 0 | 0.4 | 0.37 | 0.38 | 0.59 | 0.39 | 0.4 |
| 6 | 132992748 | rs1016503 | -0.584 | -0.896 | -1.769 | 0.011 | -0.049 | 0.045 | 0.045 | 0 | 0.85 | 0.99 | 0.89 | 0.95 | 0.84 | 0.87 |
| 6 | 132996866 | rs2840808 | 0.412 | -0.772 | -2.223 | 0.076 | -0.006 | 0.021 | 0.021 | 0 | 0.44 | 0.33 | 0.54 | 0.35 | 0.46 | 0.48 |
| 6 | 132999572 | rs4897599 | -0.277 | -0.495 | -2.035 | 0.37 | -0.005 | 0.108 | 0.108 | 0 | 0.09 | 0.21 | 0.13 | 0.23 | 0.07 | 0.13 |
| 6 | 133002269 | rs2064312 | 0.265 | -0.511 | -2.732 | 0.097 | -0.025 | -0.045 | -0.045 | 0.348 | 0.35 | 0.09 | 0.25 | 0.35 | 0.35 | 0.26 |
| 6 | 133002505 | rs2745425 | 1.419 | 0.063 | -2.844 | -0.002 | -0.038 | 0.037 | 0.037 | 0 | 0.07 | 0 | 0.05 | 0 | 0.13 | 0.02 |
| 6 | 133003344 | rs2745426 | 0.769 | -0.264 | -2.684 | 0.088 | 0.019 | 0.039 | 0.039 | 0.382 | 0.25 | 0.61 | 0.39 | 0.35 | 0.29 | 0.25 |
| 6 | 133012322 | rs3798800 | 0.821 | -0.164 | -2.859 | -0.04 | -0.088 | -0.09 | -0.09 | 0 | 0.13 | 0.02 | 0.16 | 0.34 | 0.28 | 0.19 |
| 6 | 133014623 | rs742516 | 0.323 | -0.361 | -2.922 | -0.121 | 0 | 0.005 | 0.005 | 0 | 0.34 | 0.66 | 0.57 | 0.58 | 0.36 | 0.45 |
| 6 | 133014962 | rs12665089 | 1.514 | -0.344 | -3.218 | -0.052 | 0.026 | -0.031 | -0.031 | 0.402 | 0.06 | 0.01 | 0.16 | 0.1 | 0.03 | 0.15 |
| 6 | 133014991 | rs909975 | 0.77 | -0.309 | -3.172 | -0.045 | -0.007 | -0.019 | -0.019 | 0 | 0.28 | 0.75 | 0.42 | 0.48 | 0.34 | 0.3 |
| 6 | 133015606 | rs6930032 | 0.382 | 0.34 | -2.842 | -0.125 | -0.043 | -0.046 | -0.046 | 0.517 | 0.16 | 0.08 | 0.19 | 0.47 | 0.32 | 0.25 |
| 6 | 133018701 | rs17604042 | 1.033 | -0.887 | -2.534 | -0.009 | 0.056 | 0.099 | 0.099 | 0 | 0.21 | 0.04 | 0.09 | 0.01 | 0.13 | 0.07 |
| 6 | 133019203 | rs2223613 | 0.02 | -0.655 | -1.514 | 0.6 | -0.001 | 0.012 | 0.012 | 0.13 | 0.31 | 0.65 | 0.55 | 0.46 | 0.32 | 0.4 |
| 6 | 133024183 | rs6936833 | -0.068 | -0.573 | -2.04 | 0.629 | 0.004 | -0.037 | -0.037 | 0.106 | 0.32 | 0.78 | 0.56 | 0.45 | 0.32 | 0.4 |
| 6 | 133030492 | rs2300077 | -0.116 | -0.173 | -2.327 | 0.812 | 0.03 | -0.03 | -0.03 | 0 | 0.39 | 0.86 | 0.57 | 0.39 | 0.34 | 0.52 |
| 6 | 133032668 | rs3798793 | -0.705 | -0.189 | -2.419 | 0.768 | 0.043 | -0.051 | -0.051 | 0 | 0.49 | 0.92 | 0.63 | 0.52 | 0.41 | 0.6 |
| 6 | 133032759 | rs3798792 | -0.413 | 0.137 | -1.872 | 0.718 | 0.028 | -0.055 | -0.055 | 0 | 0.38 | 0.83 | 0.57 | 0.47 | 0.34 | 0.52 |
| 6 | 133035098 | rs2294757 | -0.456 | 0.11 | -2.204 | 0.358 | -0.005 | -0.034 | -0.034 | 0 | 0.39 | 0.9 | 0.57 | 0.38 | 0.37 | 0.54 |
| 6 | 133037861 | rs6569827 | -0.113 | -0.046 | -1.754 | 0.176 | -0.047 | -0.034 | -0.034 | 0 | 0.64 | 0.84 | 0.69 | 0.41 | 0.53 | 0.53 |
| 6 | 133039321 | rs6569828 | -0.109 | 0.127 | -1.657 | 0.243 | -0.022 | 0.045 | 0.045 | 0.101 | 0.64 | 0.84 | 0.67 | 0.28 | 0.49 | 0.51 |
| 6 | 133045040 | rs9483495 | -0.954 | -0.093 | -0.039 | 0.133 | 0.111 | 0.232 | 0.232 | 1.616 | 0.06 | 0.4 | 0.07 | 0.13 | 0.05 | 0.02 |
| 6 | 133045812 | rs2294758 | 0.526 | -1.149 | 0.008 | -0.672 | 0.024 | 0.04 | 0.04 | 0 | 0.36 | 0.24 | 0.43 | 0.23 | 0.34 | 0.35 |
| 6 | 133052616 | rs4895943 | -0.721 | -1.46 | -0.175 | -0.862 | -0.089 | -0.03 | -0.03 | 0 | 0.61 | 0.92 | 0.7 | 0.39 | 0.53 | 0.54 |
| 6 | 133055170 | rs6942306 | -0.669 | -1.471 | -0.146 | -0.902 | -0.02 | 0.107 | 0.107 | 0.122 | 0.6 | 0.85 | 0.66 | 0.26 | 0.48 | 0.52 |
| 6 | 133061091 | rs6569843 | -0.137 | -1.625 | 0.06 | -0.742 | -0.004 | -0.011 | -0.011 | 0.923 | 0.47 | 0.48 | 0.45 | 0.23 | 0.39 | 0.46 |
| 6 | 133065049 | rs12211125 | 0.843 | -1.711 | -0.697 | -0.8 | -0.017 | 0.031 | 0.031 | 0 | 0.06 | 0 | 0.12 | 0 | 0.09 | 0.01 |
| 6 | 133067782 | rs2267952 | -0.414 | -1.605 | -0.693 | -0.649 | 0.054 | 0.058 | 0.058 | 0.127 | 0.52 | 0.82 | 0.65 | 0.37 | 0.45 | 0.5 |
| 6 | 133076896 | rs9483499 | -0.273 | -1.679 | -0.684 | -0.722 | -0.046 | -0.017 | -0.017 | 3.642 | 0.2 | 0.57 | 0.23 | 0.13 | 0.14 | 0.17 |
| 6 | 133077063 | rs1883613 | 0.111 | -1.149 | -0.553 | -0.659 | 0.015 | 0.027 | 0.027 | 0 | 0.35 | 0.26 | 0.44 | 0.24 | 0.34 | 0.36 |
| 6 | 133085116 | rs7738071 | -1.06 | -1.249 | 0.289 | -0.707 | -0.05 | -0.108 | -0.108 | 0 | 0.19 | 0.38 | 0.09 | 0.08 | 0.13 | 0.13 |
| 6 | 133091293 | rs998630 | -1.071 | -1.41 | 1.367 | -0.772 | 0.004 | -0.01 | -0.01 | 0.458 | 0.07 | 0.28 | 0.03 | 0 | 0.03 | 0.05 |
| 6 | 133100128 | rs7773338 | -0.786 | -0.878 | 0.209 | -1.361 | -0.172 | 0.051 | 0.051 | 1.095 | 0.76 | 0.98 | 0.84 | 0.92 | 0.6 | 0.83 |
| 6 | 133105120 | rs6926101 | -1.402 | -0.999 | 1.047 | -1.364 | -0.016 | -0.052 | -0.052 | 0 | 0.1 | 0.53 | 0.06 | 0.03 | 0.06 | 0.11 |
| 6 | 133107989 | rs761125 | 0.685 | -1.787 | 0.809 | -1.05 | 0.035 | 0.016 | 0.016 | 0.028 | 0.36 | 0.25 | 0.53 | 0.22 | 0.3 | 0.34 |
| 6 | 133116664 | rs6934190 | -0.512 | -1.739 | 1.522 | -1.008 | -0.003 | -0.033 | -0.033 | 0 | 0.08 | 0.22 | 0.04 | 0.03 | 0.06 | 0.11 |
| 6 | 133122831 | rs12204027 | -1.003 | -1.793 | 1.511 | -1.01 | -0.055 | -0.113 | -0.113 | 0 | 0.2 | 0.41 | 0.1 | 0.08 | 0.13 | 0.18 |
| 6 | 133126220 | rs5006218 | -1.132 | -1.794 | 1.421 | -1.01 | -0.031 | -0.065 | -0.065 | 0.022 | 0.21 | 0.55 | 0.11 | 0.08 | 0.13 | 0.18 |
| 6 | 133132500 | rs12211425 | 0.319 | -1.756 | 1.318 | -0.987 | -0.029 | -0.053 | -0.053 | 0 | 0.12 | 0.2 | 0.06 | 0.06 | 0.08 | 0.07 |
| 6 | 133135807 | rs9493450 | -0.557 | -1.636 | 1.051 | -0.901 | 0.007 | 0.028 | 0.028 | 0.201 | 0.57 | 0.78 | 0.63 | 0.31 | 0.43 | 0.54 |
| 6 | 133138010 | rs9389034 | -0.111 | -1.641 | 1.004 | -0.894 | 0.027 | 0.027 | 0.027 | 0 | 0.49 | 0.57 | 0.6 | 0.28 | 0.38 | 0.42 |
| 6 | 133150887 | rs755895 | -0.513 | -1.702 | 0.939 | -0.911 | 0.013 | 0.027 | 0.027 | 0.09 | 0.57 | 0.67 | 0.6 | 0.3 | 0.42 | 0.55 |
| 6 | 133155584 | rs9493458 | 0.124 | -1.782 | 0.857 | -0.947 | 0.068 | 0.084 | 0.084 | 0 | 0.46 | 0.22 | 0.55 | 0.27 | 0.36 | 0.45 |
| 6 | 133164442 | rs9402461 | 0.217 | -1.92 | 1.096 | -1.041 | 0.01 | 0.017 | 0.017 | 0.182 | 0.42 | 0.42 | 0.55 | 0.24 | 0.35 | 0.45 |
| 6 | 133168582 | rs4897617 | -0.58 | -1.999 | 1.504 | -1.047 | -0.002 | -0.004 | -0.004 | 0 | 0.57 | 0.52 | 0.6 | 0.3 | 0.43 | 0.55 |
| 6 | 133171832 | rs9389037 | 0.326 | -1.992 | 1.477 | -1.028 | 0.023 | 0.045 | 0.045 | 0.088 | 0.4 | 0.26 | 0.52 | 0.22 | 0.35 | 0.43 |
| 6 | 133178712 | rs12209781 | -0.143 | -1.979 | 1.547 | -1.019 | -0.017 | -0.028 | -0.028 | 0 | 0.15 | 0.14 | 0.06 | 0.05 | 0.08 | 0.1 |
| 6 | 133186168 | rs12193474 | -1.023 | -1.981 | 1.492 | -1.026 | -0.066 | -0.115 | -0.115 | 0.217 | 0.2 | 0.34 | 0.09 | 0.06 | 0.12 | 0.14 |
| 6 | 133188860 | rs1408064 | -0.755 | -1.981 | 1.488 | -1.022 | 0.001 | 0.03 | 0.03 | 0 | 0.59 | 0.77 | 0.62 | 0.3 | 0.44 | 0.55 |
| 6 | 133195740 | rs9402467 | 0.664 | -1.838 | 1.176 | -0.977 | 0.023 | -0.006 | -0.006 | 0 | 0.3 | 0.26 | 0.47 | 0.24 | 0.26 | 0.36 |
| 6 | 133207105 | rs9375928 | 0.604 | -1.831 | 1.159 | -0.976 | 0.021 | -0.007 | -0.007 | 0 | 0.29 | 0.24 | 0.45 | 0.21 | 0.26 | 0.34 |
| 6 | 133219461 | rs525153 | 0.491 | -2.034 | 1.02 | -0.918 | 0.026 | -0.023 | -0.023 | 0.109 | 0.3 | 0.22 | 0.45 | 0.25 | 0.26 | 0.3 |
| 6 | 133221231 | rs411639 | -0.14 | -2.04 | 1.017 | -0.938 | 0.001 | 0.001 | 0.001 | 1.795 | 0.39 | 0.58 | 0.49 | 0.25 | 0.31 | 0.37 |
| 6 | 133228839 | rs421500 | 0.505 | -2.034 | 0.831 | -0.937 | 0.105 | 0.064 | 0.064 | 0.42 | 0.5 | 0.26 | 0.43 | 0.66 | 0.61 | 0.52 |
| 6 | 133236406 | rs271118 | 0.219 | -1.651 | 0.82 | -0.716 | -0.003 | 0.091 | 0.091 | 0.491 | 0.46 | 0.2 | 0.26 | 0.27 | 0.54 | 0.41 |
| 6 | 133237051 | rs9375930 | 0.517 | -1.535 | 0.928 | -0.095 | -0.001 | 0.028 | 0.028 | 0 | 0.14 | 0.18 | 0.19 | 0.44 | 0.12 | 0.13 |
| 6 | 133237557 | rs271119 | 1.11 | -1.534 | 0.687 | -0.146 | -0.042 | 0.055 | 0.055 | 0.568 | 0.33 | 0.16 | 0.18 | 0.14 | 0.41 | 0.28 |
| 6 | 133242963 | rs4897620 | 0.131 | -1.05 | 1.463 | 0.504 | 0 | 0.082 | 0.082 | 0.01 | 0.14 | 0.24 | 0.19 | 0.45 | 0.12 | 0.13 |
| 6 | 133243774 | rs11154703 | -0.575 | -1.109 | 1.491 | 0.491 | -0.005 | 0.178 | 0.178 | 0.848 | 0.19 | 0.41 | 0.23 | 0.59 | 0.17 | 0.24 |
| 6 | 133256495 | rs6912746 | -0.588 | -1.125 | 1.342 | 0.475 | -0.019 | 0.132 | 0.132 | 0.444 | 0.2 | 0.41 | 0.22 | 0.45 | 0.17 | 0.19 |
| 6 | 133258002 | rs17062020 | -1.033 | -1.103 | 1.342 | 0.485 | -0.007 | 0.043 | 0.043 | 0.104 | 0.07 | 0.12 | 0.03 | 0.14 | 0.07 | 0.13 |

| | | | | | | | | | | | | | | | | |
|---|---|---|---|---|---|---|---|---|---|---|---|---|---|---|---|---|
| 6 | 133260475 | rs9375933 | 0.12 | -1.007 | 1.141 | 0.335 | -0.006 | 0.129 | 0.129 | 0.132 | 0.15 | 0.3 | 0.2 | 0.45 | 0.12 | 0.13 |
| 6 | 133267403 | rs6904471 | -0.948 | -0.886 | 0.659 | -0.592 | 0.006 | 0.108 | 0.108 | 9.754 | 0.54 | 0.74 | 0.69 | 0.86 | 0.48 | 0.67 |
| 6 | 133271320 | rs271151 | -0.321 | -0.59 | 0.734 | -0.276 | -0.002 | -0.018 | -0.018 | 0 | 0.45 | 0.51 | 0.64 | 0.71 | 0.48 | 0.57 |
| 6 | 133275728 | rs271156 | -0.239 | -0.6 | 0.949 | -0.151 | 0 | 0 | 0 | 4.105 | 0.41 | 0.45 | 0.62 | 0.71 | 0.43 | 0.5 |
| 6 | 133279564 | rs271137 | -0.858 | -0.503 | 0.59 | -0.326 | 0.035 | 0.165 | 0.165 | 7.579 | 0.5 | 0.75 | 0.7 | 0.84 | 0.46 | 0.62 |
| 6 | 133292440 | rs378273 | 0.058 | -0.715 | -0.06 | -0.038 | -0.054 | -0.11 | -0.11 | 0 | 0.31 | 0.04 | 0.47 | 0.59 | 0.32 | 0.34 |
| 6 | 133294795 | rs885488 | 0.094 | -0.334 | 0.069 | -0.291 | -0.024 | 0.009 | 0.009 | 0 | 0.1 | 0.01 | 0.06 | 0.03 | 0.12 | 0.16 |
| 6 | 133323872 | rs271167 | -0.691 | -0.578 | 0.878 | -0.285 | 0.006 | 0.039 | 0.039 | 3.088 | 0.52 | 0.43 | 0.6 | 0.7 | 0.5 | 0.59 |
| 6 | 133340139 | rs2926349 | -1.199 | 1.01 | 0.95 | -0.434 | 0.015 | 0.158 | 0.158 | 3.342 | 0.63 | 0.91 | 0.79 | 0.87 | 0.66 | 0.79 |
| 6 | 133355708 | rs1459239 | -0.611 | 0.807 | 0.968 | -0.366 | 0 | -0.003 | -0.003 | 0 | 0.52 | 0.53 | 0.62 | 0.7 | 0.5 | 0.59 |
| 6 | 133366486 | rs4571580 | 0.32 | 0.75 | 0.594 | -0.347 | 0.021 | 0.114 | 0.114 | 1.996 | 0.51 | 0.78 | 0.45 | 0.3 | 0.51 | 0.41 |
| 6 | 133396241 | rs7766945 | -1.769 | 0.92 | 1.647 | 0.195 | 0.006 | -0.001 | -0.001 | 0 | 0.18 | 0.76 | 0.19 | 0.22 | 0.18 | 0.22 |
| 6 | 133396480 | rs4487594 | -0.749 | 1.226 | 1.333 | 0.254 | -0.032 | 0.028 | 0.028 | 0 | 0.94 | 0.99 | 0.94 | 1 | 0.89 | 0.99 |
| 6 | 133416156 | rs9493515 | -1.343 | 0.993 | 1.199 | 0.456 | 0.018 | 0.005 | 0.005 | 4.194 | 0.14 | 0.77 | 0.18 | 0.21 | 0.17 | 0.12 |
| 6 | 133421492 | rs12192605 | 0.362 | 1.285 | 0.808 | 0.592 | 0.014 | -0.004 | -0.004 | 0 | 0.06 | 0.05 | 0.1 | 0.16 | 0.16 | 0.08 |
| 6 | 133438126 | rs7764524 | -1.028 | 0.575 | 1.332 | 1.187 | -0.022 | -0.006 | -0.006 | 0.002 | 0.35 | 0.78 | 0.48 | 0.64 | 0.33 | 0.39 |
| 6 | 133447476 | rs9399051 | -1.15 | 0.441 | 1.479 | 1.084 | -0.001 | -0.001 | -0.001 | 0.029 | 0.36 | 0.65 | 0.47 | 0.64 | 0.33 | 0.39 |
| 6 | 133447768 | rs9389049 | 0.048 | 0.444 | 1.049 | 1.464 | 0 | -0.063 | -0.063 | 0 | 0.19 | 0.09 | 0.33 | 0.34 | 0.16 | 0.22 |
| 6 | 133450298 | rs6569867 | -1.408 | 0.479 | 0.889 | 1.365 | -0.028 | -0.006 | -0.006 | 0.143 | 0.36 | 0.79 | 0.48 | 0.64 | 0.36 | 0.39 |
| 6 | 133454738 | rs6913444 | -1.334 | 0.623 | 1.2 | 1.405 | -0.002 | -0.001 | -0.001 | 0 | 0.18 | 0.79 | 0.17 | 0.26 | 0.18 | 0.17 |
| 6 | 133459633 | rs6932057 | -1.198 | 0.631 | 1.261 | 1.432 | 0.015 | 0.029 | 0.029 | 1.564 | 0.17 | 0.74 | 0.17 | 0.26 | 0.18 | 0.17 |
| 6 | 133459694 | rs6931767 | -0.775 | 0.848 | 0.713 | 1.454 | -0.052 | -0.038 | -0.038 | 0.036 | 0.89 | 0.99 | 0.8 | 0.82 | 0.83 | 0.88 |
| 6 | 133474673 | rs1832392 | 0.309 | 0.675 | 1.143 | 1.272 | 0.04 | 0.015 | 0.015 | 0 | 0.09 | 0.12 | 0.11 | 0.16 | 0.17 | 0.1 |
| 6 | 133476778 | rs6927764 | -1.502 | 0.628 | 0.781 | 1.264 | -0.003 | -0.003 | -0.003 | 0 | 0.18 | 0.8 | 0.17 | 0.26 | 0.18 | 0.17 |
| 6 | 133479661 | rs12196256 | 0.53 | 0.765 | 0.573 | 1.253 | -0.052 | -0.044 | -0.044 | 0.892 | 0.08 | 0.01 | 0.09 | 0.17 | 0.17 | 0.09 |
| 6 | 133481191 | rs531156 | -0.246 | 0.238 | -0.198 | 0.676 | -0.13 | -0.139 | -0.139 | 0.442 | 0.42 | 0.07 | 0.64 | 0.55 | 0.52 | 0.55 |
| 6 | 133490022 | rs7767378 | -0.816 | -0.024 | 0.461 | 0.287 | 0.045 | -0.016 | -0.016 | 0 | 0.2 | 0.51 | 0.17 | 0.18 | 0.2 | 0.2 |
| 6 | 133495524 | rs546329 | -1.104 | -0.303 | -0.737 | 0.011 | -0.009 | 0.097 | 0.097 | 1.44 | 0.61 | 0.89 | 0.79 | 0.81 | 0.62 | 0.68 |
| 6 | 133500908 | rs968274 | -0.076 | -0.214 | -0.529 | -0.005 | -0.037 | -0.054 | -0.054 | 0.097 | 0.1 | 0.23 | 0.05 | 0.01 | 0.02 | 0.05 |
| 6 | 133502198 | rs9402484 | -0.638 | 0.008 | -0.602 | 0.163 | 0.005 | -0.001 | -0.001 | 0 | 0.19 | 0.24 | 0.16 | 0.18 | 0.2 | 0.14 |
| 6 | 133517005 | rs509685 | -1.542 | -0.061 | -1.047 | 0.159 | -0.038 | 0.08 | 0.08 | 5.525 | 0.7 | 0.94 | 0.84 | 0.86 | 0.72 | 0.83 |
| 6 | 133523516 | rs12215534 | 0.883 | 0.521 | -0.782 | 0.62 | -0.048 | -0.046 | -0.046 | 0 | 0.08 | 0.01 | 0.1 | 0.18 | 0.17 | 0.09 |
| 6 | 133523789 | rs477094 | 0.905 | 0.418 | -1.021 | 0.774 | -0.043 | -0.04 | -0.04 | 0.628 | 0.11 | 0.02 | 0.21 | 0.17 | 0.17 | 0.15 |
| 6 | 133542649 | rs492771 | -1.632 | 0.502 | -1.029 | 0.613 | -0.054 | -0.06 | -0.06 | 0 | 0.89 | 1 | 0.91 | 0.82 | 0.85 | 0.96 |
| 6 | 133546476 | rs2145349 | 0.086 | 0.01 | -0.503 | 0.811 | 0.012 | -0.06 | -0.06 | 0 | 0.25 | 0.18 | 0.4 | 0.44 | 0.25 | 0.37 |
| 6 | 133546689 | rs2145351 | -0.273 | -0.059 | -0.235 | 0.776 | 0.041 | -0.061 | -0.061 | 0.78 | 0.31 | 0.15 | 0.39 | 0.44 | 0.26 | 0.42 |
| 6 | 133547481 | rs1361147 | -1.128 | -0.046 | -0.207 | 0.736 | -0.005 | 0.01 | 0.01 | 0.056 | 0.44 | 0.37 | 0.54 | 0.61 | 0.45 | 0.51 |
| 6 | 133550176 | rs1338012 | -1.135 | -0.045 | -0.15 | 0.718 | -0.005 | 0.01 | 0.01 | 0.047 | 0.44 | 0.39 | 0.54 | 0.61 | 0.45 | 0.51 |
| 6 | 133550279 | rs1338013 | -0.239 | -0.022 | -0.235 | 0.722 | 0.039 | -0.082 | -0.082 | 0 | 0.31 | 0.13 | 0.39 | 0.44 | 0.26 | 0.42 |
| 6 | 133552737 | rs10485235 | 0.1 | -0.008 | -0.335 | 0.75 | 0.042 | 0.018 | 0.018 | 0 | 0.12 | 0.21 | 0.15 | 0.17 | 0.19 | 0.09 |
| 6 | 133556249 | rs9389058 | 0.471 | 0.027 | -0.556 | 0.751 | 0 | -0.004 | -0.004 | 0 | 0.1 | 0.11 | 0.13 | 0.17 | 0.19 | 0.09 |
| 6 | 133561646 | rs3813346 | -0.97 | -0.001 | -0.173 | 0.692 | -0.001 | -0.002 | -0.002 | 0 | 0.45 | 0.55 | 0.55 | 0.61 | 0.45 | 0.51 |
| 6 | 133573447 | rs722488 | 0.33 | 0.196 | -0.612 | 1.161 | -0.031 | -0.124 | -0.124 | 0.012 | 0.17 | 0.01 | 0.33 | 0.41 | 0.15 | 0.29 |
| 6 | 133576444 | rs561355 | 0.524 | 0.185 | -0.568 | 1.033 | -0.01 | -0.017 | -0.017 | 0 | 0.13 | 0.09 | 0.21 | 0.2 | 0.17 | 0.14 |
| 6 | 133578733 | rs973714 | -0.868 | -0.089 | 0.029 | 0.725 | -0.013 | 0.011 | 0.011 | 1.304 | 0.4 | 0.47 | 0.47 | 0.48 | 0.35 | 0.56 |
| 6 | 133586425 | rs7745129 | -1.54 | 0.019 | -0.271 | 0.6 | -0.006 | 0.082 | 0.082 | 1.306 | 0.5 | 0.8 | 0.61 | 0.65 | 0.5 | 0.63 |
| 6 | 133589583 | rs3777799 | -0.482 | 0.058 | 0.447 | 0.674 | 0.058 | -0.01 | -0.01 | 0.702 | 0.19 | 0.39 | 0.16 | 0.16 | 0.21 | 0.15 |
| 6 | 133595684 | rs3734277 | 0.091 | -0.221 | 0.025 | 0.731 | 0.002 | -0.052 | -0.052 | 0 | 0.2 | 0.15 | 0.36 | 0.46 | 0.19 | 0.33 |
| 6 | 133606617 | rs9385643 | -1.501 | 0.403 | 0.071 | 0.788 | -0.02 | 0.088 | 0.088 | 3.084 | 0.51 | 0.84 | 0.6 | 0.66 | 0.47 | 0.65 |
| 6 | 133607305 | rs17062270 | 0.099 | 0.518 | 0.337 | 1.231 | 0 | -0.021 | -0.021 | 0 | 0.18 | 0.14 | 0.34 | 0.43 | 0.15 | 0.28 |
| 6 | 133610598 | rs211432 | -0.707 | 0.312 | 1.069 | 0.868 | -0.001 | -0.034 | -0.034 | 0 | 0.3 | 0.44 | 0.29 | 0.18 | 0.33 | 0.29 |
| 6 | 133611341 | rs2224382 | -0.953 | 0.808 | 1.305 | 1.328 | 0.029 | 0.038 | 0.038 | 0.817 | 0.33 | 0.45 | 0.43 | 0.45 | 0.36 | 0.45 |
| 6 | 133612992 | rs211436 | -0.624 | 0.366 | 1.239 | 0.994 | 0.01 | 0.022 | 0.022 | 0 | 0.25 | 0.48 | 0.33 | 0.25 | 0.25 | 0.26 |
| 6 | 133614576 | rs433294 | -1.129 | 0.409 | 0.655 | 0.943 | 0 | 0.075 | 0.075 | 0 | 0.45 | 0.76 | 0.64 | 0.69 | 0.4 | 0.54 |
| 6 | 133621996 | rs9373046 | 0.531 | 0.489 | 0.85 | 1.146 | 0.007 | 0.001 | 0.001 | 0 | 0.09 | 0.13 | 0.13 | 0.15 | 0.16 | 0.09 |
| 6 | 133624199 | rs9373048 | -0.368 | 0.444 | 1.053 | 1.112 | 0.033 | -0.002 | -0.002 | 0 | 0.15 | 0.29 | 0.16 | 0.16 | 0.19 | 0.15 |
| 6 | 133625980 | rs11154722 | 0.251 | 0.27 | 0.873 | 1.249 | 0.005 | -0.03 | -0.03 | 0 | 0.16 | 0.11 | 0.27 | 0.44 | 0.14 | 0.26 |
| 6 | 133631190 | rs950374 | -0.391 | 0.242 | 1.14 | 1.253 | -0.034 | 0.158 | 0.158 | 0.081 | 0.21 | 0.39 | 0.31 | 0.5 | 0.14 | 0.35 |
| 6 | 133632333 | rs6941879 | -0.583 | 0.396 | 0.756 | 1.057 | -0.022 | -0.015 | -0.015 | 0.056 | 0.85 | 0.91 | 0.72 | 0.81 | 0.79 | 0.87 |
| 6 | 133633350 | rs1343867 | -1.38 | 0.337 | 0.765 | 0.918 | 0.003 | 0.032 | 0.032 | 0.672 | 0.36 | 0.7 | 0.25 | 0.19 | 0.31 | 0.38 |
| 6 | 133635272 | rs1336527 | -0.467 | 0.47 | 0.411 | 0.738 | -0.017 | -0.007 | -0.007 | 0 | 0.84 | 0.88 | 0.72 | 0.81 | 0.79 | 0.87 |
| 6 | 133639892 | rs17062339 | 1.721 | 0.448 | 0.343 | 0.666 | 0.013 | 0.074 | 0.074 | 0.243 | 0.42 | 0.27 | 0.23 | 0.15 | 0.39 | 0.3 |
| 6 | 133640448 | rs6937974 | 0.441 | 0.464 | 0.426 | 0.694 | -0.002 | -0.022 | -0.022 | 0 | 0.65 | 0.5 | 0.42 | 0.3 | 0.66 | 0.53 |
| 6 | 133642013 | rs12523816 | 2.403 | 0.423 | 0.416 | 0.657 | -0.05 | 0.12 | 0.12 | 1.373 | 0.35 | 0.06 | 0.2 | 0.15 | 0.35 | 0.23 |
| 6 | 133642918 | rs12524250 | 1.262 | 0.419 | 0.58 | 0.818 | 0.006 | 0.021 | 0.021 | 0.034 | 0.52 | 0.4 | 0.36 | 0.3 | 0.55 | 0.39 |
| 6 | 133648156 | rs17062356 | -0.125 | 0.126 | 0.967 | 1.16 | 0.005 | -0.005 | -0.005 | 0 | 0.12 | 0.1 | 0.06 | 0 | 0.11 | 0.14 |
| 6 | 133654340 | rs6924629 | -0.639 | 0.094 | 0.914 | 1.372 | -0.047 | 0.155 | 0.155 | 0.468 | 0.22 | 0.51 | 0.31 | 0.51 | 0.14 | 0.34 |
| 6 | 133658299 | rs745091 | -0.792 | 0.487 | 0.734 | 0.954 | 0.007 | -0.012 | -0.012 | 0 | 0.21 | 0.21 | 0.18 | 0.15 | 0.26 | 0.23 |
| 6 | 133675770 | rs2184783 | -0.751 | 0.146 | 1.221 | 1.679 | -0.036 | 0.006 | 0.006 | 0.476 | 0.31 | 0.48 | 0.36 | 0.28 | 0.25 | 0.37 |

| | | | | | | | | | | | | | | | | |
|---|---|---|---|---|---|---|---|---|---|---|---|---|---|---|---|---|
| 6 | 133678108 | rs9373051 | -0.924 | 0.238 | 1.042 | 1.985 | 0.002 | 0.039 | 0.039 | 0 | 0.14 | 0.2 | 0.12 | 0.35 | 0.15 | 0.17 |
| 6 | 133683963 | rs212831 | 0.999 | 0.191 | 0.697 | 1.959 | 0.016 | -0.06 | -0.06 | 0 | 0.6 | 0.15 | 0.52 | 0.56 | 0.63 | 0.54 |
| 6 | 133685363 | rs7766140 | 0.365 | 0.233 | 0.656 | 2.004 | 0.006 | 0 | 0 | 0 | 0.06 | 0.04 | 0.01 | 0.19 | 0.03 | 0.08 |
| 6 | 133687403 | rs211629 | 0.553 | 0.18 | 1.025 | 2.004 | -0.031 | 0.013 | 0.013 | 0.487 | 0.71 | 0.52 | 0.63 | 0.72 | 0.74 | 0.65 |
| 6 | 133692366 | rs10484669 | 0.931 | 0.321 | 0.75 | 2.063 | -0.033 | -0.032 | -0.032 | 0 | 0.05 | 0 | 0.08 | 0.16 | 0.11 | 0.08 |
| 6 | 133694338 | rs211609 | 1.325 | 0.282 | 0.876 | 1.967 | 0.028 | 0.036 | 0.036 | 2.147 | 0.55 | 0.29 | 0.52 | 0.37 | 0.6 | 0.46 |
| 6 | 133698217 | rs211605 | 1.49 | 0.178 | 0.846 | 2.11 | -0.015 | 0.046 | 0.046 | 0 | 0.76 | 0.39 | 0.73 | 0.6 | 0.81 | 0.76 |
| 6 | 133699845 | rs211604 | 0.425 | 0.091 | 0.498 | 1.677 | -0.011 | 0.031 | 0.031 | 0 | 0.92 | 0.76 | 0.85 | 0.95 | 0.95 | 0.92 |
| 6 | 133701158 | rs9493613 | -0.084 | 0.276 | -0.363 | 1.643 | -0.011 | -0.042 | -0.042 | 0 | 0.21 | 0.06 | 0.21 | 0.23 | 0.21 | 0.3 |
| 6 | 133702425 | rs159407 | 1.343 | 1.144 | -0.179 | 1.937 | -0.092 | -0.028 | -0.028 | 0.42 | 0.58 | 0.12 | 0.4 | 0.39 | 0.59 | 0.53 |
| 6 | 133703325 | rs2235485 | -0.445 | 1.55 | -0.317 | 2.228 | -0.068 | -0.083 | -0.083 | 0 | 0.13 | 0.01 | 0.21 | 0.28 | 0.15 | 0.22 |
| 6 | 133705590 | rs159410 | -1.684 | 1.518 | 0.657 | 2.325 | 0.033 | 0.033 | 0.033 | 0 | 0.31 | 0.41 | 0.44 | 0.52 | 0.38 | 0.33 |
| 6 | 133710214 | rs159415 | 1.633 | 0.802 | 0.829 | 2.647 | -0.015 | -0.057 | -0.057 | 1.202 | 0.72 | 0.2 | 0.56 | 0.62 | 0.72 | 0.76 |
| 6 | 133711721 | rs7760570 | -0.822 | 0.949 | 1.014 | 2.649 | 0.049 | 0.202 | 0.202 | 0.206 | 0.1 | 0.33 | 0.2 | 0.31 | 0.11 | 0.17 |
| 6 | 133714231 | rs159417 | 2.455 | 0.544 | 0.883 | 2.624 | 0.013 | -0.042 | -0.042 | 0.048 | 0.65 | 0.22 | 0.51 | 0.5 | 0.73 | 0.61 |
| 6 | 133715210 | rs3777811 | -0.286 | 0.754 | 0.841 | 2.641 | -0.029 | -0.057 | -0.057 | 0 | 0.09 | 0.01 | 0.16 | 0.22 | 0.09 | 0.18 |
| 6 | 133716549 | rs455704 | 0.373 | 0.732 | 0.94 | 2.573 | -0.026 | 0.031 | 0.031 | 0 | 0.08 | 0.03 | 0.06 | 0 | 0.11 | 0.01 |
| 6 | 133725974 | rs296424 | 1.484 | 0.638 | 1.238 | 2.688 | -0.01 | 0.084 | 0.084 | 0 | 0.78 | 0.44 | 0.64 | 0.61 | 0.82 | 0.82 |
| 6 | 133726502 | rs11968392 | -0.106 | 0.603 | 1.074 | 2.719 | -0.013 | 0.186 | 0.186 | 0 | 0.06 | 0.11 | 0.13 | 0.31 | 0.02 | 0.11 |
| 6 | 133727421 | rs296429 | 1.646 | 0.694 | 1.087 | 2.292 | -0.002 | 0.02 | 0.02 | 0 | 0.69 | 0.38 | 0.5 | 0.46 | 0.73 | 0.67 |
| 6 | 133728758 | rs211621 | 0.778 | 0.482 | 1.429 | 1.881 | 0.038 | 0.113 | 0.113 | 0 | 0.93 | 0.67 | 0.85 | 0.92 | 0.94 | 0.94 |
| 6 | 133732908 | rs211598 | 1.259 | 0.147 | 2.126 | 1.973 | -0.005 | 0.033 | 0.033 | 0.195 | 0.9 | 0.68 | 0.83 | 0.91 | 0.91 | 0.92 |
| 6 | 133733988 | rs9483581 | -0.647 | 0.174 | 1.685 | 2.162 | -0.019 | -0.032 | -0.032 | 3.988 | 0.12 | 0.01 | 0.21 | 0.12 | 0.1 | 0.18 |
| 6 | 133743221 | rs212785 | 1.564 | 0.014 | 1.925 | 2.683 | -0.027 | 0.113 | 0.113 | 0 | 0.86 | 0.67 | 0.73 | 0.68 | 0.9 | 0.87 |
| 6 | 133748381 | rs9385647 | -1.764 | -0.109 | 1.056 | 2.128 | 0.013 | 0.06 | 0.06 | 0 | 0.44 | 0.67 | 0.58 | 0.78 | 0.36 | 0.5 |
| 6 | 133755207 | rs9375963 | -0.054 | -0.122 | 1.243 | 2.163 | 0.002 | 0.011 | 0.011 | 0 | 0.18 | 0.15 | 0.2 | 0.35 | 0.21 | 0.23 |
| 6 | 133756500 | rs3777860 | -0.736 | -0.117 | 1.433 | 2.178 | -0.027 | 0.163 | 0.163 | 0 | 0.13 | 0.35 | 0.14 | 0.31 | 0.06 | 0.09 |
| 6 | 133763905 | rs6907872 | -1.278 | 0.137 | 0.82 | 2.065 | -0.001 | 0 | 0 | 0 | 0.3 | 0.28 | 0.44 | 0.46 | 0.31 | 0.4 |
| 6 | 133766431 | rs9389078 | 0.044 | 0.042 | 0.648 | 2.059 | -0.036 | -0.069 | -0.069 | 2.16 | 0.16 | 0.03 | 0.18 | 0.34 | 0.2 | 0.25 |
| 6 | 133768525 | rs17062561 | -0.058 | 0.024 | 0.61 | 2.081 | -0.004 | -0.058 | -0.058 | 0 | 0.09 | 0 | 0.12 | 0.24 | 0.05 | 0.05 |
| 6 | 133776112 | rs10872406 | -1.479 | 0.028 | 0.782 | 2.097 | -0.002 | 0.023 | 0.023 | 0 | 0.29 | 0.32 | 0.33 | 0.58 | 0.26 | 0.29 |
| 6 | 133776122 | rs11154727 | -0.208 | 0.154 | 0.442 | 1.918 | -0.006 | -0.01 | -0.01 | 0 | 0.16 | 0.09 | 0.2 | 0.34 | 0.21 | 0.24 |
| 6 | 133789728 | rs9493627 | -0.186 | -0.883 | 0.178 | 0.999 | -0.011 | 0.002 | 0.002 | 0.51 | 0.4 | 0.57 | 0.34 | 0.34 | 0.31 | 0.41 |
| 6 | 133790251 | rs212765 | 0.49 | -0.817 | 0.497 | 1.023 | 0 | -0.003 | -0.003 | 0 | 0.42 | 0.54 | 0.46 | 0.45 | 0.44 | 0.27 |
| 6 | 133790391 | rs1022514 | 0.144 | -0.68 | 0.161 | 1.006 | -0.009 | 0.002 | 0.002 | 0.498 | 0.4 | 0.57 | 0.34 | 0.35 | 0.31 | 0.41 |
| 6 | 133790493 | rs212764 | -0.62 | -0.224 | 0.166 | 1.233 | -0.071 | -0.041 | -0.041 | 0 | 0.79 | 0.97 | 0.79 | 0.73 | 0.75 | 0.66 |
| 6 | 133817456 | rs568911 | 0.425 | -0.253 | 0.125 | 0.954 | -0.001 | -0.002 | -0.002 | 0 | 0.29 | 0.3 | 0.37 | 0.38 | 0.28 | 0.22 |
| 6 | 133821417 | rs3777878 | -0.533 | -0.34 | 0.65 | 1.127 | 0.029 | 0.394 | 0.394 | 0 | 0.07 | 0.37 | 0.16 | 0.28 | 0.02 | 0.08 |
| 6 | 133827354 | rs2277083 | -0.414 | -0.203 | 0.009 | 0.68 | -0.024 | 0.039 | 0.039 | 0 | 0.68 | 0.71 | 0.69 | 0.82 | 0.57 | 0.81 |
| 6 | 133836188 | rs11154730 | 0.36 | -0.653 | -0.174 | 0.522 | -0.017 | 0.008 | 0.008 | 0 | 0.36 | 0.55 | 0.3 | 0.35 | 0.24 | 0.38 |
| 6 | 133847876 | rs2327351 | 0.234 | -0.912 | -0.704 | 0.192 | -0.027 | -0.066 | -0.066 | 0.102 | 0.17 | 0.1 | 0.35 | 0.38 | 0.28 | 0.27 |
| 6 | 133849966 | rs3734279 | 0.438 | -0.801 | -0.562 | -0.376 | -0.013 | 0.052 | 0.052 | 0 | 0.41 | 0.2 | 0.53 | 0.69 | 0.4 | 0.64 |
| 6 | 133851377 | rs9483586 | -1.316 | -1.316 | -0.595 | -0.238 | -0.09 | 0.02 | 0.02 | 0 | 0.76 | 0.96 | 0.79 | 0.82 | 0.73 | 0.82 |
| 6 | 133852051 | rs17053541 | -0.348 | -2.107 | 0.451 | -0.077 | -0.038 | -0.036 | -0.036 | 0 | 0.24 | 0.52 | 0.14 | 0.13 | 0.14 | 0.15 |
| 6 | 133857604 | rs6934616 | -0.226 | -2.061 | 0.235 | -0.052 | -0.05 | -0.048 | -0.048 | 0 | 0.24 | 0.38 | 0.13 | 0.13 | 0.14 | 0.15 |
| 6 | 133863172 | rs9493636 | -0.657 | -1.619 | -0.063 | -0.271 | -0.032 | 0.013 | 0.013 | 0.009 | 0.34 | 0.52 | 0.51 | 0.52 | 0.32 | 0.38 |
| 6 | 133864250 | rs4895970 | -0.784 | -1.533 | -0.571 | -0.472 | 0.037 | 0.027 | 0.027 | 0.793 | 0.49 | 0.35 | 0.53 | 0.58 | 0.43 | 0.48 |
| 6 | 133865464 | rs9389085 | 0.45 | -1.666 | -1.196 | -0.391 | 0.019 | -0.102 | -0.102 | 0 | 0.17 | 0.05 | 0.34 | 0.53 | 0.13 | 0.32 |
| 6 | 133866326 | rs17062664 | -0.148 | -1.183 | -0.57 | -0.516 | -0.008 | -0.047 | -0.047 | 0 | 0.2 | 0.2 | 0.13 | 0.05 | 0.15 | 0.13 |
| 6 | 133869809 | rs4437491 | -0.365 | -1.166 | -0.937 | -0.478 | 0.042 | 0.108 | 0.108 | 3.048 | 0.52 | 0.2 | 0.59 | 0.64 | 0.5 | 0.51 |
| 6 | 133871529 | rs4424102 | -1.171 | -1.767 | -0.409 | -0.951 | 0.013 | -0.009 | -0.009 | 1.227 | 0.25 | 0.8 | 0.2 | 0.03 | 0.15 | 0.11 |
| 6 | 133872201 | rs7759024 | -0.756 | -1.774 | 0.596 | -0.933 | 0.016 | -0.001 | -0.001 | 0 | 0.06 | 0.3 | 0.06 | 0 | 0.02 | 0.01 |
| 6 | 133873486 | rs9285474 | -0.7 | -1.805 | 0.542 | -0.978 | -0.029 | -0.128 | -0.128 | 0 | 0.19 | 0.45 | 0.15 | 0.03 | 0.15 | 0.12 |
| 6 | 133885723 | rs6569884 | -1.553 | -1.28 | -0.105 | -0.911 | -0.01 | 0.054 | 0.054 | 2.209 | 0.46 | 0.92 | 0.38 | 0.35 | 0.36 | 0.45 |
| 6 | 133886140 | rs9483590 | -0.745 | -1.244 | 0.527 | -0.965 | -0.043 | -0.095 | -0.095 | 0 | 0.21 | 0.47 | 0.14 | 0.08 | 0.14 | 0.13 |
| 6 | 133890195 | rs10046269 | -1.21 | -0.98 | 0.757 | -0.74 | 0.05 | 0.105 | 0.105 | 5.567 | 0.06 | 0.53 | 0.05 | 0.05 | 0 | 0 |
| 6 | 133892991 | rs9321406 | -1.007 | -1.109 | 0.11 | -0.868 | -0.001 | -0.015 | -0.015 | 0 | 0.25 | 0.72 | 0.19 | 0.07 | 0.13 | 0.11 |
| 6 | 133897107 | rs1319087 | -1.191 | -0.764 | 0.049 | -0.613 | 0.005 | 0.01 | 0.01 | 0.852 | 0.39 | 0.66 | 0.39 | 0.4 | 0.31 | 0.36 |
| 6 | 133901157 | rs12664733 | -0.11 | -0.769 | -0.205 | -0.557 | 0.014 | -0.016 | -0.016 | 6.183 | 0.22 | 0.1 | 0.09 | 0.25 | 0.12 | 0.18 |
| 6 | 133903856 | rs6902621 | -0.847 | -0.714 | 0.251 | -0.713 | 0.007 | -0.009 | -0.009 | 0 | 0.2 | 0.67 | 0.3 | 0.18 | 0.18 | 0.29 |
| 6 | 133907283 | rs1878551 | -0.761 | -0.787 | 0.516 | -0.751 | 0.002 | 0.001 | 0.001 | 3.038 | 0.28 | 0.68 | 0.13 | 0.25 | 0.13 | 0.18 |
| 6 | 133909759 | rs9373062 | 0.04 | -0.652 | 0.316 | -0.682 | -0.07 | -0.046 | -0.046 | 0 | 0.13 | 0.01 | 0.25 | 0.18 | 0.18 | 0.27 |
| 6 | 133919776 | rs2636601 | 0.341 | -0.617 | 1.02 | -0.74 | 0.04 | 0.001 | 0.001 | 0 | 0.84 | 0.7 | 0.72 | 0.82 | 0.82 | 0.72 |
| 6 | 133920563 | rs2677818 | 0.185 | -0.596 | 0.921 | -0.729 | 0.034 | 0.005 | 0.005 | 0 | 0.86 | 0.78 | 0.73 | 0.82 | 0.82 | 0.72 |
| 6 | 133923265 | rs2256702 | 1.176 | -0.728 | 0.434 | -0.825 | 0.004 | 0.024 | 0.024 | 3.408 | 0.55 | 0.22 | 0.53 | 0.57 | 0.63 | 0.52 |
| 6 | 133937427 | rs2636591 | 0.953 | -0.83 | 0.583 | -0.885 | 0 | 0.011 | 0.011 | 2.69 | 0.58 | 0.33 | 0.61 | 0.6 | 0.68 | 0.53 |
| 6 | 133946416 | rs2636623 | 1.342 | -0.852 | 0.497 | -0.878 | 0.002 | 0.039 | 0.039 | 2.607 | 0.53 | 0.19 | 0.53 | 0.59 | 0.64 | 0.49 |
| 6 | 133955119 | rs2636627 | 0.713 | -0.911 | 1.179 | -0.885 | 0.005 | 0.006 | 0.006 | 0.187 | 0.79 | 0.55 | 0.68 | 0.78 | 0.82 | 0.66 |
| 6 | 133956851 | rs4410743 | 1.259 | -0.312 | -0.294 | -1.806 | -0.098 | -0.133 | -0.133 | 0 | 0.3 | 0.02 | 0.38 | 0.58 | 0.37 | 0.32 |

| | | | | | | | | | | | | | | | | |
|---|---|---|---|---|---|---|---|---|---|---|---|---|---|---|---|---|
| 6 | 133958612 | rs12207040 | 0.235 | -0.377 | 0.134 | -1.809 | 0.002 | 0.007 | 0.007 | 1.192 | 0.25 | 0.22 | 0.16 | 0.17 | 0.18 | 0.17 |
| 6 | 133962467 | rs4895979 | 0.966 | -0.299 | -0.093 | -1.717 | -0.011 | 0.058 | 0.058 | 2.946 | 0.56 | 0.19 | 0.53 | 0.64 | 0.64 | 0.5 |
| 6 | 133968194 | rs7772627 | 0.442 | -0.846 | 0.335 | -1.738 | -0.026 | -0.015 | -0.015 | 0 | 0.09 | 0.3 | 0.04 | 0.04 | 0.02 | 0.04 |
| 6 | 133968745 | rs6569890 | -1.094 | -0.898 | 0.045 | -1.858 | 0.027 | 0.031 | 0.031 | 1.017 | 0.35 | 0.7 | 0.25 | 0.28 | 0.21 | 0.23 |
| 6 | 133980726 | rs6941397 | 0.91 | -0.858 | -0.522 | -1.89 | -0.004 | 0.01 | 0.01 | 1.377 | 0.19 | 0.14 | 0.09 | 0.16 | 0.14 | 0.15 |
| 6 | 133988652 | rs6914525 | -0.247 | -0.644 | -0.756 | -1.809 | 0.013 | 0.02 | 0.02 | 0.71 | 0.31 | 0.67 | 0.16 | 0.2 | 0.16 | 0.19 |
| 6 | 133995787 | rs9375980 | 0.919 | -0.06 | -1.431 | -1.597 | -0.124 | -0.124 | -0.124 | 0 | 0.28 | 0.03 | 0.43 | 0.61 | 0.41 | 0.33 |
| 6 | 133998430 | rs9321411 | -0.888 | -0.128 | -0.802 | -1.428 | 0.046 | 0.125 | 0.125 | 1.9 | 0.38 | 0.87 | 0.27 | 0.21 | 0.22 | 0.2 |
| 6 | 134013272 | rs9493698 | -1.098 | -0.065 | -0.874 | -1.423 | 0.029 | 0.117 | 0.117 | 2.891 | 0.4 | 0.92 | 0.29 | 0.23 | 0.23 | 0.21 |
| 6 | 134021034 | rs2017324 | 0.431 | 0.001 | -0.726 | -1.409 | -0.019 | 0.029 | 0.029 | 0.217 | 0.26 | 0.09 | 0.1 | 0.21 | 0.18 | 0.16 |
| 6 | 134038999 | rs10457611 | 0.473 | 0.256 | -0.983 | -1.45 | -0.05 | 0.03 | 0.03 | 0 | 0.14 | 0.01 | 0.19 | 0.06 | 0.15 | 0.29 |
| 6 | 134048274 | rs6921832 | -0.45 | 0.173 | -0.982 | -1.498 | 0.024 | 0.08 | 0.08 | 2.515 | 0.36 | 0.79 | 0.27 | 0.19 | 0.22 | 0.2 |
| 6 | 134069785 | rs12196724 | 0.469 | 0.333 | -1.233 | -1.353 | -0.05 | 0.033 | 0.033 | 0 | 0.14 | 0.01 | 0.19 | 0.06 | 0.15 | 0.26 |
| 6 | 134089019 | rs1999756 | -0.597 | 0.03 | -1.312 | -1.41 | 0.05 | 0.113 | 0.113 | 2.522 | 0.36 | 0.89 | 0.27 | 0.21 | 0.2 | 0.19 |
| 6 | 134090307 | rs12201533 | 0.297 | 0.327 | -0.535 | -1.129 | 0.005 | -0.003 | -0.003 | 0 | 0.18 | 0.23 | 0.23 | 0.15 | 0.18 | 0.26 |
| 6 | 134090759 | rs4895986 | -0.679 | 0.323 | -0.746 | -1.24 | 0.006 | 0.086 | 0.086 | 0 | 0.48 | 0.72 | 0.5 | 0.33 | 0.44 | 0.42 |
| 6 | 134091859 | rs10485156 | 0.375 | -0.32 | -0.921 | -1.078 | -0.04 | -0.011 | -0.011 | 0 | 0.2 | 0.02 | 0.08 | 0.19 | 0.14 | 0.16 |
| 6 | 134091966 | rs1208228 | -1.006 | -0.285 | -1.085 | -1.248 | -0.03 | 0.096 | 0.096 | 0.529 | 0.83 | 0.98 | 0.89 | 1 | 0.79 | 0.93 |
| 6 | 134094349 | rs9389095 | 0.421 | -0.322 | -0.524 | -1.183 | -0.002 | -0.023 | -0.023 | 0.33 | 0.55 | 0.57 | 0.69 | 0.78 | 0.61 | 0.74 |
| 6 | 134100937 | rs1208242 | -0.607 | -0.395 | -0.562 | -0.903 | 0 | -0.032 | -0.032 | 0.573 | 0.82 | 0.85 | 0.87 | 1 | 0.77 | 0.93 |
| 6 | 134105003 | rs12196907 | 0.871 | -0.378 | -0.682 | -0.401 | 0 | 0.001 | 0.001 | 0 | 0.26 | 0.26 | 0.46 | 0.36 | 0.29 | 0.37 |
| 6 | 134108701 | rs1208260 | -0.76 | -0.505 | -0.504 | -0.195 | -0.005 | -0.004 | -0.004 | 0 | 0.84 | 0.89 | 0.88 | 1 | 0.77 | 0.93 |
| 6 | 134123832 | rs11154745 | 0.745 | -0.825 | -0.494 | -0.177 | -0.008 | -0.008 | -0.008 | 0 | 0.32 | 0.26 | 0.53 | 0.36 | 0.33 | 0.44 |
| 6 | 134129501 | rs1208248 | 0.395 | -1.183 | -0.57 | -0.261 | 0.023 | 0.032 | 0.032 | 0.229 | 0.55 | 0.35 | 0.67 | 0.72 | 0.64 | 0.72 |
| 6 | 134131421 | rs1208252 | -1.035 | -1.178 | -0.728 | -0.338 | -0.008 | 0.041 | 0.041 | 0 | 0.84 | 0.91 | 0.88 | 1 | 0.79 | 0.93 |
| 6 | 134132238 | rs2209044 | 0.562 | -1.356 | -0.756 | -0.374 | 0.042 | 0.022 | 0.022 | 0.296 | 0.22 | 0.04 | 0.09 | 0.22 | 0.14 | 0.2 |
| 6 | 134132516 | rs1208254 | -0.035 | -0.816 | -0.341 | -0.355 | -0.017 | -0.084 | -0.084 | 0 | 0.78 | 0.37 | 0.76 | 0.94 | 0.78 | 0.92 |
| 6 | 134141309 | rs1041714 | 0.672 | -0.406 | -1.664 | -0.164 | -0.064 | -0.084 | -0.084 | 0.181 | 0.22 | 0.06 | 0.14 | 0.33 | 0.31 | 0.22 |
| 6 | 134142660 | rs942132 | -0.439 | -0.559 | -1.227 | -0.747 | -0.009 | 0.046 | 0.046 | 0 | 0.84 | 0.91 | 0.88 | 1 | 0.79 | 0.94 |
| 6 | 134151297 | rs12193973 | 0.566 | -0.336 | -1.378 | -0.26 | -0.068 | -0.054 | -0.054 | 0 | 0.28 | 0.06 | 0.44 | 0.44 | 0.32 | 0.44 |
| 6 | 134152181 | rs1746015 | -0.774 | -0.763 | -1.926 | -0.657 | -0.013 | 0.154 | 0.154 | 4.433 | 0.77 | 0.98 | 0.87 | 1 | 0.75 | 0.92 |
| 6 | 134158800 | rs1208285 | 0.623 | -1.162 | -2.091 | -0.656 | -0.107 | -0.139 | -0.139 | 2.089 | 0.24 | 0.03 | 0.22 | 0.48 | 0.35 | 0.29 |
| 6 | 134170048 | rs1208276 | 0.128 | -1.384 | -1.452 | -0.829 | 0.006 | 0.042 | 0.042 | 0 | 0.13 | 0.04 | 0.06 | 0.02 | 0.11 | 0.11 |
| 6 | 134171132 | rs1535616 | 0.875 | -1.263 | -0.614 | -0.793 | 0.005 | 0.011 | 0.011 | 0 | 0.32 | 0.37 | 0.45 | 0.5 | 0.29 | 0.44 |
| 6 | 134172950 | rs1126392 | 0.808 | -1.335 | -0.287 | -0.833 | -0.022 | -0.041 | -0.041 | 0 | 0.25 | 0.17 | 0.33 | 0.42 | 0.35 | 0.28 |
| 6 | 134178405 | rs9375987 | 0.807 | -1.342 | -0.231 | -0.836 | -0.07 | -0.122 | -0.122 | 0 | 0.25 | 0.08 | 0.31 | 0.42 | 0.35 | 0.28 |
| 6 | 134196674 | rs12524865 | 0.883 | -1.29 | -0.123 | -0.858 | 0.014 | 0.035 | 0.035 | 0 | 0.32 | 0.46 | 0.46 | 0.5 | 0.29 | 0.43 |
| 6 | 134205092 | rs969282 | -0.131 | -1.136 | -0.225 | -0.788 | -0.059 | -0.071 | -0.071 | 2.755 | 0.64 | 0.83 | 0.62 | 0.55 | 0.55 | 0.63 |
| 6 | 134209837 | rs2327429 | 1.255 | -1.252 | -0.221 | -0.776 | -0.065 | -0.117 | -0.117 | 0 | 0.27 | 0.06 | 0.44 | 0.53 | 0.3 | 0.45 |
| 6 | 134210037 | rs2327430 | 0.597 | -0.878 | 0.365 | -0.833 | 0.019 | 0.013 | 0.013 | 0.167 | 0.57 | 0.39 | 0.76 | 0.94 | 0.65 | 0.73 |
| 6 | 134211922 | rs2327431 | 0.154 | -0.445 | -0.443 | -0.645 | 0.002 | 0.002 | 0.002 | 0 | 0.88 | 0.75 | 0.93 | 0.85 | 0.91 | 0.92 |
| 6 | 134216475 | rs9389114 | -0.137 | -0.388 | -1.149 | -0.616 | -0.001 | -0.001 | -0.001 | 0 | 0.89 | 0.87 | 0.96 | 0.85 | 0.91 | 0.92 |
| 6 | 134216994 | rs17481786 | 0.546 | -0.453 | -1.3 | -0.725 | 0.025 | 0.148 | 0.148 | 8.35 | 0.22 | 0.02 | 0.1 | 0 | 0.18 | 0.06 |
| 6 | 134217699 | rs7774514 | 0.023 | -0.256 | -1.122 | -0.541 | -0.002 | -0.002 | -0.002 | 0 | 0.89 | 0.85 | 0.95 | 0.85 | 0.91 | 0.92 |
| 6 | 134229319 | rs229919 | -0.089 | -0.198 | -0.841 | -0.607 | 0.004 | 0.169 | 0.169 | 5.55 | 0.72 | 0.98 | 0.86 | 0.97 | 0.75 | 0.88 |
| 6 | 134230714 | rs151106 | 0.249 | -0.2 | -0.745 | -0.605 | 0.008 | -0.013 | -0.013 | 5.157 | 0.68 | 0.7 | 0.83 | 0.96 | 0.75 | 0.86 |
| 6 | 134232194 | rs9399080 | 0.453 | 0.165 | -1.598 | -0.355 | -0.101 | -0.121 | -0.121 | 0 | 0.18 | 0.02 | 0.27 | 0.28 | 0.29 | 0.18 |
| 6 | 134234787 | rs229925 | 0.436 | -0.838 | -0.829 | -1.119 | 0.033 | 0.145 | 0.145 | 5.294 | 0.72 | 0.94 | 0.86 | 0.97 | 0.75 | 0.88 |
| 6 | 134259753 | rs9483632 | -1.316 | -0.484 | 0.378 | -0.375 | -0.035 | 0.036 | 0.036 | 0 | 0.13 | 0.39 | 0.2 | 0.15 | 0.09 | 0.09 |
| 6 | 134266699 | rs162192 | 0.132 | -0.031 | -0.359 | -0.844 | -0.127 | 0.123 | 0.123 | 3.402 | 0.35 | 0.07 | 0.32 | 0.16 | 0.5 | 0.23 |
| 6 | 134268822 | rs228867 | -0.559 | -0.08 | 0.297 | -0.893 | 0.008 | 0.041 | 0.041 | 0.235 | 0.47 | 0.46 | 0.44 | 0.17 | 0.6 | 0.29 |
| 6 | 134287736 | rs6902440 | -0.811 | -0.098 | 0.408 | -0.885 | -0.038 | -0.067 | -0.067 | 0 | 0.13 | 0.3 | 0.13 | 0.04 | 0.09 | 0.07 |
| 6 | 134288098 | rs9373073 | 0.198 | -0.101 | -0.392 | -0.903 | -0.061 | 0.102 | 0.102 | 2.276 | 0.36 | 0.16 | 0.33 | 0.16 | 0.52 | 0.23 |
| 6 | 134292257 | rs9399083 | -0.817 | -0.136 | -0.659 | -0.903 | 0.022 | 0.072 | 0.072 | 0 | 0.52 | 0.75 | 0.54 | 0.3 | 0.6 | 0.31 |
| 6 | 134304023 | rs2076583 | 0.228 | 0.024 | -0.807 | -0.872 | -0.123 | 0.123 | 0.123 | 3.154 | 0.35 | 0.07 | 0.32 | 0.15 | 0.5 | 0.22 |
| 6 | 134310874 | rs1385066 | 0.188 | -0.268 | -0.801 | -0.894 | -0.093 | 0.139 | 0.139 | 2.876 | 0.36 | 0.07 | 0.34 | 0.15 | 0.54 | 0.23 |
| 6 | 134314963 | rs9493790 | -1.385 | -0.773 | 0.32 | -0.424 | -0.064 | 0.009 | 0.009 | 0.71 | 0.16 | 0.67 | 0.21 | 0.15 | 0.08 | 0.09 |
| 6 | 134317730 | rs229908 | 0.496 | -0.374 | 0.154 | -1.241 | -0.008 | -0.122 | -0.122 | 0.47 | 0.68 | 0.56 | 0.77 | 0.89 | 0.71 | 0.56 |
| 6 | 134318961 | rs229906 | 0.616 | -0.797 | 0.09 | -1.785 | -0.002 | -0.044 | -0.044 | 0.953 | 0.58 | 0.58 | 0.76 | 0.88 | 0.66 | 0.54 |
| 6 | 134319474 | rs229904 | 0.217 | -0.595 | -0.291 | -1.599 | -0.011 | 0.069 | 0.069 | 0.23 | 0.76 | 0.88 | 0.83 | 0.91 | 0.75 | 0.6 |
| 6 | 134325029 | rs9376000 | -0.138 | 0.094 | -0.196 | -1.115 | 0.005 | 0.001 | 0.001 | 0 | 0.12 | 0.11 | 0.22 | 0.18 | 0.16 | 0.11 |
| 6 | 134325409 | rs127171 | -0.119 | -0.228 | -0.388 | -1.546 | -0.032 | 0.096 | 0.096 | 0 | 0.23 | 0.08 | 0.17 | 0.05 | 0.23 | 0.33 |
| 6 | 134326106 | rs47137 | -1.154 | -0.854 | 0.215 | -1.355 | 0.002 | -0.005 | -0.005 | 0 | 0.43 | 0.43 | 0.44 | 0.36 | 0.41 | 0.46 |
| 6 | 134327863 | rs9376001 | 0.528 | -0.942 | 0.561 | -0.834 | -0.024 | 0.055 | 0.055 | 9.944 | 0.09 | 0.19 | 0.03 | 0.14 | 0.02 | 0.02 |
| 6 | 134329997 | rs2811675 | -1.645 | -1.485 | 0.691 | -0.154 | -0.014 | 0.012 | 0.012 | 1.954 | 0.54 | 0.72 | 0.47 | 0.55 | 0.43 | 0.51 |
| 6 | 134330160 | rs12176172 | 0.082 | -1.081 | 0.089 | -1.311 | 0.016 | 0.006 | 0.006 | 0 | 0.12 | 0.15 | 0.22 | 0.19 | 0.16 | 0.12 |
| 6 | 134330274 | rs2627239 | -0.377 | -1.385 | -0.217 | -1.315 | -0.001 | 0.001 | 0.001 | 1.02 | 0.27 | 0.25 | 0.2 | 0.17 | 0.22 | 0.33 |
| 6 | 134333273 | rs9373076 | -0.47 | -1.084 | -0.082 | -1.47 | 0 | 0.014 | 0.014 | 2.688 | 0.19 | 0.19 | 0.2 | 0.32 | 0.16 | 0.12 |
| 6 | 134340501 | rs9389128 | -1.126 | -1.263 | -0.373 | -1.548 | -0.031 | 0.035 | 0.035 | 3.32 | 0.29 | 0.43 | 0.26 | 0.41 | 0.19 | 0.12 |

| | | | | | | | | | | | | | | | | |
|---|---|---|---|---|---|---|---|---|---|---|---|---|---|---|---|---|
| 6 | 134343797 | rs11154757 | -0.428 | -1.238 | -0.417 | -1.533 | -0.001 | 0.005 | 0.005 | 2.688 | 0.19 | 0.17 | 0.21 | 0.32 | 0.16 | 0.11 |
| 6 | 134344468 | rs2627232 | -1.065 | -1.037 | -0.732 | -1.569 | 0.002 | 0.006 | 0.006 | 2.027 | 0.48 | 0.46 | 0.42 | 0.42 | 0.4 | 0.43 |
| 6 | 134345361 | rs1564774 | -0.155 | -0.636 | -1.346 | -1.24 | -0.007 | 0.027 | 0.027 | 0 | 0.22 | 0.27 | 0.24 | 0.42 | 0.17 | 0.11 |
| 6 | 134347008 | rs2492303 | -0.218 | -0.41 | -1.049 | -0.827 | -0.011 | -0.004 | -0.004 | 0 | 0.39 | 0.24 | 0.4 | 0.43 | 0.4 | 0.44 |
| 6 | 134362607 | rs2811679 | -0.143 | 0.105 | -1.622 | -0.858 | -0.041 | 0.006 | 0.006 | 0.294 | 0.62 | 0.79 | 0.53 | 0.36 | 0.51 | 0.55 |
| 6 | 134365043 | rs2254235 | -0.52 | 0.549 | -1.459 | -1.371 | -0.046 | 0.064 | 0.064 | 0.033 | 0.63 | 0.82 | 0.52 | 0.28 | 0.52 | 0.67 |
| 6 | 134368953 | rs965652 | 0.05 | -0.037 | -1.356 | -1.719 | -0.019 | 0.076 | 0.076 | 0.51 | 0.44 | 0.31 | 0.39 | 0.26 | 0.46 | 0.59 |
| 6 | 134369424 | rs2811686 | 0.322 | 0.86 | -1.72 | -1.353 | -0.025 | 0.049 | 0.049 | 0 | 0.22 | 0.11 | 0.17 | 0.11 | 0.25 | 0.42 |
| 6 | 134374673 | rs2627231 | 0.004 | 1.111 | -1.278 | -0.444 | -0.031 | 0.056 | 0.056 | 0.165 | 0.66 | 0.85 | 0.56 | 0.28 | 0.57 | 0.67 |
| 6 | 134391373 | rs7738638 | -0.982 | 1.047 | -0.068 | -0.309 | 0.028 | -0.061 | -0.061 | 0.688 | 0.15 | 0.46 | 0.33 | 0.1 | 0.15 | 0.21 |
| 6 | 134392815 | rs9389139 | -1.101 | 1.055 | 0.165 | -0.333 | 0.038 | -0.048 | -0.048 | 0 | 0.14 | 0.44 | 0.32 | 0.1 | 0.15 | 0.22 |
| 6 | 134413537 | rs13215360 | -1.388 | 1.148 | 0.262 | -0.615 | 0.001 | -0.041 | -0.041 | 0 | 0.17 | 0.27 | 0.3 | 0.1 | 0.17 | 0.22 |
| 6 | 134416205 | rs9483651 | -0.099 | 1.635 | -0.285 | -1.118 | -0.011 | 0.024 | 0.024 | 0 | 0.27 | 0.2 | 0.24 | 0.14 | 0.33 | 0.16 |
| 6 | 134417322 | rs6908581 | 0.537 | 1.994 | -0.837 | -0.482 | -0.009 | -0.006 | -0.006 | 0.012 | 0.73 | 0.73 | 0.44 | 0.24 | 0.68 | 0.51 |
| 6 | 134420012 | rs6569926 | 1.083 | 1.845 | -0.744 | -0.506 | -0.008 | -0.021 | -0.021 | 0 | 0.45 | 0.48 | 0.2 | 0.1 | 0.35 | 0.35 |
| 6 | 134426215 | rs7766159 | 0.674 | 2.022 | -0.958 | 0.568 | -0.005 | -0.007 | -0.007 | 0 | 0.72 | 0.66 | 0.61 | 0.47 | 0.66 | 0.55 |
| 6 | 134427657 | rs6921560 | 0.009 | 1.674 | -1.073 | 0.688 | -0.031 | -0.041 | -0.041 | 0 | 0.26 | 0.09 | 0.19 | 0.22 | 0.29 | 0.15 |
| 6 | 134430223 | rs7749507 | -0.959 | 0.684 | -0.923 | 0.445 | -0.001 | 0.013 | 0.013 | 0.102 | 0.29 | 0.26 | 0.41 | 0.28 | 0.26 | 0.25 |
| 6 | 134431870 | rs7741722 | -1.433 | 0.282 | -1.697 | 0.324 | -0.154 | 0.015 | 0.015 | 0.967 | 0.59 | 0.98 | 0.71 | 0.75 | 0.56 | 0.55 |
| 6 | 134434375 | rs17063374 | -0.726 | 0.113 | -1.072 | 0.534 | -0.019 | -0.006 | -0.006 | 0 | 0.16 | 0.28 | 0.11 | 0.14 | 0.1 | 0.19 |
| 6 | 134445276 | rs4243446 | -1.161 | -0.007 | -0.703 | 0.319 | 0.014 | 0.073 | 0.073 | 2.118 | 0.48 | 0.8 | 0.61 | 0.71 | 0.48 | 0.44 |
| 6 | 134445763 | rs17063391 | -0.596 | 0.042 | -0.054 | 0.497 | 0.004 | 0.005 | 0.005 | 0 | 0.23 | 0.26 | 0.39 | 0.31 | 0.28 | 0.22 |
| 6 | 134445774 | rs11963187 | -0.632 | -0.122 | -0.033 | 0.574 | -0.014 | -0.05 | -0.05 | 0 | 0.15 | 0.27 | 0.1 | 0.04 | 0.1 | 0.12 |
| 6 | 134445802 | rs11968128 | -0.632 | -0.122 | -0.033 | 0.574 | -0.014 | -0.05 | -0.05 | 0 | 0.15 | 0.26 | 0.1 | 0.04 | 0.1 | 0.12 |
| 6 | 134447959 | rs12662210 | -0.015 | 0.057 | -0.559 | 0.816 | 0.001 | -0.047 | -0.047 | 1.021 | 0.17 | 0.09 | 0.17 | 0.4 | 0.2 | 0.2 |
| 6 | 134465900 | rs9321428 | -0.365 | -0.126 | -0.569 | 0.256 | -0.003 | 0.011 | 0.011 | 0 | 0.21 | 0.31 | 0.3 | 0.3 | 0.19 | 0.17 |
| 6 | 134466473 | rs4645436 | -0.142 | 0.044 | -1.042 | 0.281 | -0.047 | -0.125 | -0.125 | 0 | 0.22 | 0.02 | 0.15 | 0.35 | 0.29 | 0.25 |
| 6 | 134466561 | rs4318899 | -0.885 | 0.002 | -1.018 | 0.143 | -0.014 | -0.042 | -0.042 | 0 | 0.3 | 0.2 | 0.27 | 0.35 | 0.41 | 0.28 |
| 6 | 134466694 | rs17063429 | 0.223 | 0.064 | -1.23 | 0.144 | -0.003 | -0.003 | -0.003 | 0 | 0.08 | 0.09 | 0.09 | 0 | 0.13 | 0.03 |
| 6 | 134467724 | rs9493844 | 0 | 0.057 | -1.228 | 0.186 | 0.015 | 0.079 | 0.079 | 0 | 0.06 | 0.13 | 0.14 | 0.21 | 0.07 | 0.13 |
| 6 | 134468423 | rs2811690 | -1.153 | -0.174 | -2.251 | 0.166 | 0.007 | -0.015 | -0.015 | 0 | 0.36 | 0.22 | 0.28 | 0.35 | 0.44 | 0.35 |
| 6 | 134469171 | rs2811692 | 1.452 | -0.218 | -2.765 | 0.02 | -0.098 | -0.127 | -0.127 | 1.037 | 0.39 | 0.06 | 0.38 | 0.41 | 0.36 | 0.44 |
| 6 | 134469853 | rs17234055 | -0.948 | -0.286 | -1.484 | 0.387 | 0.003 | 0.035 | 0.035 | 0.672 | 0.13 | 0.08 | 0.1 | 0 | 0.15 | 0.11 |
| 6 | 134474280 | rs1626029 | -0.682 | 0.191 | -1.187 | 0.718 | -0.009 | 0.024 | 0.024 | 0 | 0.13 | 0.13 | 0.13 | 0 | 0.19 | 0.05 |
| 6 | 134481965 | rs2758145 | 0.695 | 0.152 | -0.92 | 1.636 | 0.002 | -0.069 | -0.069 | 0 | 0.82 | 0.93 | 0.86 | 0.65 | 0.78 | 0.78 |
| 6 | 134488960 | rs9402571 | -1.082 | 0.217 | -0.314 | 1.484 | -0.004 | 0.052 | 0.052 | 0 | 0.2 | 0.28 | 0.15 | 0.38 | 0.2 | 0.24 |
| 6 | 134493397 | rs1057293 | -0.947 | 0.456 | -0.319 | 1.162 | 0 | 0.011 | 0.011 | 0 | 0.07 | 0.07 | 0.13 | 0.22 | 0.08 | 0.08 |
| 6 | 134493947 | rs1743966 | -1.667 | 0.148 | 0.288 | 0.685 | -0.003 | 0.01 | 0.01 | 0 | 0.19 | 0.33 | 0.24 | 0.26 | 0.2 | 0.1 |
| 6 | 134499854 | rs1763527 | 0.549 | -0.3 | -0.138 | -0.287 | -0.006 | 0.045 | 0.045 | 0 | 0.41 | 0.31 | 0.39 | 0.16 | 0.5 | 0.35 |
| 6 | 134502227 | rs1763528 | -0.598 | -0.699 | -0.72 | -0.294 | 0.035 | -0.03 | -0.03 | 0.479 | 0.61 | 0.94 | 0.66 | 0.46 | 0.68 | 0.45 |
| 6 | 134504204 | rs1075427 | 0.735 | -0.838 | -0.656 | -0.476 | -0.07 | 0.167 | 0.167 | 0 | 0.37 | 0.13 | 0.37 | 0.16 | 0.51 | 0.35 |
| 6 | 134509254 | rs1114707 | -0.27 | -0.687 | -0.103 | -0.316 | 0.042 | 0.041 | 0.041 | 0 | 0.55 | 0.71 | 0.6 | 0.45 | 0.64 | 0.43 |
| 6 | 134516019 | rs9373085 | 0.152 | -0.593 | -0.716 | -0.269 | -0.016 | -0.039 | -0.039 | 0.29 | 0.37 | 0.07 | 0.39 | 0.57 | 0.34 | 0.52 |
| 6 | 134526205 | rs6569934 | 0.147 | -0.547 | 0.071 | -0.587 | 0.011 | 0.019 | 0.019 | 0.27 | 0.55 | 0.61 | 0.45 | 0.18 | 0.61 | 0.43 |
| 6 | 134530697 | rs9493857 | 2.256 | 0.242 | 0.153 | -0.153 | 0.023 | -0.017 | -0.017 | 0 | 0.78 | 0.1 | 0.58 | 0.73 | 0.84 | 0.92 |
| 6 | 134535159 | rs9493858 | -1.122 | 0.707 | 0.646 | -0.374 | -0.005 | -0.075 | -0.075 | 0.203 | 0.16 | 0.35 | 0.13 | 0.03 | 0.14 | 0.07 |
| 6 | 134539179 | rs4896028 | 2.033 | 0.685 | 0.479 | -0.379 | 0.005 | 0.004 | 0.004 | 0.006 | 0.71 | 0.27 | 0.56 | 0.72 | 0.74 | 0.88 |
| 6 | 134544733 | rs1763500 | 0.787 | 0.75 | 0.118 | -0.296 | -0.003 | -0.04 | -0.04 | 0 | 0.93 | 0.76 | 0.92 | 1 | 0.95 | 1 |
| 6 | 134557207 | rs1763510 | 1.723 | 1.772 | -0.67 | -0.369 | -0.087 | 0.007 | 0.007 | 1.04 | 0.63 | 0.06 | 0.43 | 0.69 | 0.63 | 0.81 |
| 6 | 134559089 | rs9376020 | 0.288 | 2.158 | -1.163 | -0.076 | -0.077 | -0.098 | -0.098 | 0.396 | 0.13 | 0.01 | 0.14 | 0.46 | 0.24 | 0.18 |
| 6 | 134560021 | rs1743940 | 1.56 | 2.177 | -0.048 | -0.009 | 0.065 | 0.066 | 0.066 | 0 | 0.86 | 0.63 | 0.66 | 0.74 | 0.82 | 0.94 |
| 6 | 134560518 | rs1743939 | 0.774 | 1.986 | -0.072 | -0.158 | 0.056 | -0.062 | -0.062 | 0 | 0.9 | 0.66 | 0.89 | 1 | 0.88 | 0.98 |
| 6 | 134564866 | rs9493871 | -0.829 | 1.615 | -0.599 | -0.01 | 0.032 | -0.021 | -0.021 | 2.221 | 0.06 | 0.21 | 0.13 | 0.02 | 0.07 | 0.07 |
| 6 | 134568921 | rs734738 | 1.174 | 2.048 | -1.521 | 0.058 | -0.138 | 0.001 | 0.001 | 0.548 | 0.62 | 0.06 | 0.39 | 0.68 | 0.57 | 0.79 |
| 6 | 134570482 | rs1763523 | 1.027 | 1.711 | -1.793 | 0.684 | -0.114 | -0.001 | -0.001 | 1.188 | 0.44 | 0.07 | 0.42 | 0.43 | 0.51 | 0.55 |
| 6 | 134571178 | rs9373086 | -0.578 | 1.421 | -1.386 | 1.489 | -0.001 | 0.009 | 0.009 | 0 | 0.09 | 0.03 | 0.15 | 0.37 | 0.09 | 0.13 |
| 6 | 134574934 | rs4896032 | 1.219 | 1.614 | -0.314 | 1.448 | 0 | 0 | 0 | 0 | 0.57 | 0.41 | 0.51 | 0.54 | 0.63 | 0.69 |
| 6 | 134578667 | rs1743948 | 1.091 | 1.498 | -0.436 | 1.323 | -0.015 | 0.023 | 0.023 | 0 | 0.67 | 0.45 | 0.68 | 0.63 | 0.73 | 0.82 |
| 6 | 134578920 | rs17063576 | -0.347 | 1.49 | -0.489 | 1.303 | -0.017 | -0.011 | -0.011 | 0 | 0.11 | 0.07 | 0.19 | 0.08 | 0.09 | 0.13 |
| 6 | 134581016 | rs1981093 | -0.965 | 1.41 | 0.333 | 1.443 | 0 | 0.077 | 0.077 | 0 | 0.1 | 0.25 | 0.19 | 0.16 | 0.09 | 0.11 |
| 6 | 134581118 | rs1763505 | 1.208 | 1.429 | 0.261 | 1.45 | -0.004 | 0.012 | 0.012 | 0.089 | 0.64 | 0.59 | 0.59 | 0.55 | 0.71 | 0.71 |
| 6 | 134582426 | rs4896036 | -2.125 | 1.507 | 0.15 | 1.17 | 0.039 | 0.114 | 0.114 | 0 | 0.49 | 0.74 | 0.62 | 0.6 | 0.4 | 0.4 |
| 6 | 134599215 | rs9493883 | -1.786 | 1.548 | 0.441 | 1.608 | -0.016 | 0.008 | 0.008 | 0.155 | 0.28 | 0.33 | 0.45 | 0.63 | 0.22 | 0.29 |
| 6 | 134602454 | rs9376026 | -1.934 | 2.382 | 0.423 | 1.418 | 0.01 | -0.015 | -0.015 | 1.056 | 0.31 | 0.38 | 0.52 | 0.8 | 0.38 | 0.36 |
| 6 | 134614761 | rs9389154 | -0.341 | 2.592 | 0.685 | 1.668 | 0.004 | -0.025 | -0.025 | 0.54 | 0.12 | 0.05 | 0.15 | 0.45 | 0.09 | 0.12 |
| 6 | 134615030 | rs12202855 | -1.709 | 2.518 | 0.751 | 1.126 | 0.044 | -0.065 | -0.065 | 0 | 0.12 | 0.28 | 0.14 | 0.01 | 0.17 | 0.08 |
| 6 | 134620673 | rs6569937 | -2.405 | 2.556 | 1.044 | 1.14 | 0.019 | -0.078 | -0.078 | 0 | 0.37 | 0.46 | 0.66 | 0.9 | 0.49 | 0.37 |
| 6 | 134623840 | rs9493891 | -1.101 | 2.177 | 0.959 | 1.2 | 0.007 | -0.036 | -0.036 | 0.114 | 0.08 | 0.21 | 0.09 | 0.01 | 0.1 | 0.06 |
| 6 | 134625104 | rs6909876 | -2.4 | 2.257 | 1.202 | 1.189 | 0.024 | -0.096 | -0.096 | 0.014 | 0.36 | 0.46 | 0.66 | 0.89 | 0.46 | 0.37 |

| | | | | | | | | | | | | | | | | |
|---|---|---|---|---|---|---|---|---|---|---|---|---|---|---|---|---|
| 6 | 134627600 | rs13216187 | -0.423 | 2.093 | 1.205 | 1.61 | -0.041 | -0.075 | -0.075 | 0 | 0.13 | 0.03 | 0.27 | 0.51 | 0.19 | 0.14 |
| 6 | 134630783 | rs9493897 | -0.765 | 1.998 | 1.257 | 1.372 | -0.004 | 0.071 | 0.071 | 0 | 0.1 | 0.16 | 0.24 | 0.3 | 0.09 | 0.08 |
| 6 | 134632345 | rs9399091 | -1.54 | 2.031 | 1.218 | 1.871 | -0.075 | -0.127 | -0.127 | 0 | 0.19 | 0.05 | 0.34 | 0.5 | 0.28 | 0.22 |
| 6 | 134633308 | rs7762476 | -2.077 | 1.221 | 1.865 | 1.181 | 0.01 | -0.008 | -0.008 | 0 | 0.08 | 0.27 | 0.11 | 0.06 | 0.06 | 0.09 |
| 6 | 134666311 | rs7749856 | -2.168 | 1.227 | 2.161 | 1.345 | 0.018 | 0.237 | 0.237 | 0 | 0.11 | 0.48 | 0.29 | 0.31 | 0.09 | 0.21 |
| 6 | 134669083 | rs2269566 | -1.683 | 1.199 | 2.264 | 1.114 | 0.106 | 0.077 | 0.077 | 0 | 0.05 | 0.34 | 0.05 | 0.01 | 0.04 | 0.06 |
| 6 | 134674130 | rs727981 | -2.426 | 1.323 | 1.913 | 1.16 | -0.001 | 0.012 | 0.012 | 0.934 | 0.17 | 0.69 | 0.34 | 0.28 | 0.15 | 0.27 |
| 6 | 134684613 | rs6908972 | -1.168 | 1.407 | 1.235 | 0.684 | 0.013 | -0.025 | -0.025 | 0.41 | 0.05 | 0.12 | 0.04 | 0 | 0.07 | 0.01 |
| 6 | 134685508 | rs9493917 | -2.077 | 1.992 | 1.279 | 0.02 | 0.035 | -0.118 | -0.118 | 0.193 | 0.3 | 0.52 | 0.3 | 0.05 | 0.38 | 0.26 |
| 6 | 134686860 | rs10485350 | -1.232 | 1.925 | 0.79 | 0.028 | -0.002 | -0.013 | -0.013 | 0 | 0.22 | 0.18 | 0.24 | 0.05 | 0.31 | 0.25 |
| 6 | 134690745 | rs12199570 | -1.21 | 1.903 | 0.691 | 0.035 | -0.011 | 0.013 | 0.013 | 0 | 0.22 | 0.15 | 0.24 | 0.05 | 0.3 | 0.24 |
| 6 | 134692578 | rs6569941 | -1.978 | 1.649 | 0.042 | 0.41 | -0.141 | -0.042 | -0.042 | 0 | 0.41 | 0.9 | 0.47 | 0.43 | 0.48 | 0.39 |
| 6 | 134693503 | rs12210842 | -0.599 | 1.754 | 0.18 | -0.448 | -0.009 | 0.021 | 0.021 | 0 | 0.1 | 0.05 | 0.1 | 0.05 | 0.12 | 0.12 |
| 6 | 134703671 | rs11969046 | -1.746 | 1.317 | 0.213 | -0.495 | -0.044 | 0.174 | 0.174 | 3.124 | 0.38 | 0.87 | 0.38 | 0.08 | 0.43 | 0.29 |
| 6 | 134704195 | rs11154765 | -1.658 | 1.15 | 0.087 | -0.643 | -0.041 | 0.21 | 0.21 | 1.36 | 0.4 | 0.87 | 0.37 | 0.07 | 0.43 | 0.27 |
| 6 | 134706333 | rs6911285 | -1.006 | 0.936 | 0.922 | -0.531 | 0.015 | 0.006 | 0.006 | 4.621 | 0.06 | 0.32 | 0.03 | 0 | 0 | 0 |
| 6 | 134709696 | rs6934484 | -1.187 | 0.921 | 0.532 | -0.539 | 0.011 | 0.001 | 0.001 | 7.272 | 0.07 | 0.51 | 0.05 | 0 | 0 | 0 |
| 6 | 134715654 | rs6918011 | -1.267 | 1.299 | -0.763 | -0.658 | -0.033 | 0.167 | 0.167 | 1.79 | 0.39 | 0.84 | 0.36 | 0.04 | 0.44 | 0.27 |
| 6 | 134720067 | rs9402592 | 0.694 | 0.485 | -1.472 | -0.11 | -0.076 | -0.023 | -0.023 | 0 | 0.42 | 0.08 | 0.6 | 0.55 | 0.5 | 0.58 |
| 6 | 134721958 | rs6932621 | 0.526 | -0.575 | -0.49 | -0.043 | -0.04 | -0.03 | -0.03 | 0 | 0.67 | 0.58 | 0.65 | 0.85 | 0.79 | 0.8 |
| 6 | 134724334 | rs9493929 | -1.379 | -0.717 | 0.169 | -0.067 | -0.044 | -0.06 | -0.06 | 0 | 0.35 | 0.52 | 0.2 | 0.16 | 0.22 | 0.27 |
| 6 | 134727620 | rs736476 | -1.514 | 0.004 | -0.991 | 0.34 | -0.094 | 0.029 | 0.029 | 0.605 | 0.47 | 0.94 | 0.64 | 0.57 | 0.45 | 0.32 |
| 6 | 134730255 | rs6569943 | -0.748 | -0.984 | -0.015 | -0.26 | -0.066 | -0.092 | -0.092 | 1.053 | 0.14 | 0.37 | 0.08 | 0 | 0.04 | 0.09 |
| 6 | 134732534 | rs9493942 | -1.33 | -0.982 | -0.691 | -0.21 | 0.12 | 0.113 | 0.113 | 1.843 | 0.35 | 0.89 | 0.22 | 0.16 | 0.17 | 0.18 |
| 6 | 134733728 | rs2327484 | -1.462 | -0.937 | -0.564 | -0.186 | 0.094 | 0.089 | 0.089 | 0.885 | 0.34 | 0.82 | 0.22 | 0.16 | 0.17 | 0.18 |
| 6 | 134736804 | rs7773254 | -0.865 | -0.383 | 0.095 | -0.01 | -0.032 | -0.023 | -0.023 | 0 | 0.23 | 0.6 | 0.16 | 0.16 | 0.14 | 0.09 |
| 6 | 134737409 | rs7754232 | 0.452 | 0.217 | -0.192 | 0.424 | -0.008 | -0.007 | -0.007 | 0 | 0.79 | 0.76 | 0.51 | 0.48 | 0.68 | 0.79 |
| 6 | 134737675 | rs4896043 | -1.129 | -0.244 | 0.136 | 0.394 | -0.001 | -0.007 | -0.007 | 0 | 0.29 | 0.28 | 0.43 | 0.45 | 0.34 | 0.26 |
| 6 | 134738088 | rs4896044 | -1.147 | -0.906 | 0.037 | -0.269 | 0.034 | 0.084 | 0.084 | 0.359 | 0.51 | 0.74 | 0.65 | 0.69 | 0.42 | 0.48 |
| 6 | 134740464 | rs9385688 | -1.141 | -1.034 | 0.533 | -0.652 | -0.023 | 0.039 | 0.039 | 1.163 | 0.05 | 0.23 | 0.04 | 0.07 | 0 | 0.06 |
| 6 | 134740564 | rs9373094 | -1.315 | -1.392 | 0.776 | -0.868 | -0.01 | -0.002 | -0.002 | 0 | 0.27 | 0.72 | 0.18 | 0.22 | 0.13 | 0.21 |
| 6 | 134740669 | rs7774624 | 0.104 | -1.47 | 1.126 | -0.929 | -0.013 | -0.083 | -0.083 | 1.752 | 0.13 | 0.25 | 0.08 | 0 | 0.09 | 0.07 |
| 6 | 134740760 | rs6940101 | -1.277 | -1.392 | 0.99 | -0.814 | -0.042 | 0.091 | 0.091 | 1.478 | 0.13 | 0.39 | 0.08 | 0.22 | 0.05 | 0.14 |
| 6 | 134744184 | rs9493950 | -0.33 | -1.185 | 0.55 | -0.745 | -0.016 | -0.055 | -0.055 | 0 | 0.75 | 0.89 | 0.51 | 0.45 | 0.72 | 0.78 |
| 6 | 134744212 | rs9389172 | -0.769 | -1.16 | 0.587 | -1.203 | -0.01 | 0.008 | 0.008 | 0 | 0.17 | 0.28 | 0.1 | 0.22 | 0.12 | 0.14 |
| 6 | 134746878 | rs6902048 | 0.913 | -1.127 | -0.017 | -1.84 | -0.011 | 0.008 | 0.008 | 0 | 0.39 | 0.35 | 0.31 | 0.13 | 0.47 | 0.37 |
| 6 | 134752626 | rs7772299 | -0.422 | -0.437 | -0.037 | -1.374 | -0.025 | -0.02 | -0.02 | 0 | 0.33 | 0.2 | 0.56 | 0.58 | 0.36 | 0.29 |
| 6 | 134752930 | rs2024692 | -0.074 | -0.526 | -0.176 | -1.563 | -0.006 | -0.005 | -0.005 | 0.259 | 0.83 | 0.84 | 0.92 | 0.8 | 0.85 | 0.86 |
| 6 | 134755476 | rs12663404 | 0.361 | -0.318 | -0.294 | -1.412 | -0.056 | -0.109 | -0.109 | 0 | 0.23 | 0.05 | 0.49 | 0.61 | 0.23 | 0.22 |
| 6 | 134757127 | rs4896046 | 0.193 | -0.358 | -0.259 | -1.393 | -0.039 | -0.095 | -0.095 | 0 | 0.25 | 0.06 | 0.5 | 0.6 | 0.23 | 0.22 |
| 6 | 134766051 | rs6910410 | -0.455 | -0.317 | 0.152 | -1.339 | -0.007 | 0.005 | 0.005 | 2.597 | 0.28 | 0.42 | 0.46 | 0.67 | 0.24 | 0.34 |
| 6 | 134774577 | rs2327478 | 0.48 | -0.306 | 0.187 | -1.293 | -0.002 | -0.003 | -0.003 | 0 | 0.21 | 0.16 | 0.41 | 0.57 | 0.23 | 0.26 |
| 6 | 134775505 | rs2295199 | 1.467 | -0.403 | 0.114 | -1.536 | -0.033 | 0.053 | 0.053 | 1.073 | 0.42 | 0.19 | 0.35 | 0.32 | 0.46 | 0.46 |
| 6 | 134778896 | rs6936551 | -0.594 | -0.297 | 0.628 | -0.846 | -0.006 | 0.011 | 0.011 | 2.569 | 0.27 | 0.4 | 0.45 | 0.68 | 0.23 | 0.34 |
| 6 | 134780275 | rs7738980 | -0.356 | -0.215 | 0.433 | -0.852 | 0 | 0.004 | 0.004 | 1.258 | 0.25 | 0.28 | 0.43 | 0.71 | 0.23 | 0.34 |
| 6 | 134786163 | rs9402608 | -1.127 | -0.258 | 0.17 | -0.783 | -0.012 | 0.111 | 0.111 | 0.477 | 0.09 | 0.39 | 0.06 | 0.16 | 0.03 | 0.13 |
| 6 | 134787054 | rs2876252 | 1.044 | -0.203 | -0.11 | -1.007 | 0.011 | 0.104 | 0.104 | 3.365 | 0.53 | 0.17 | 0.43 | 0.32 | 0.6 | 0.57 |
| 6 | 134789041 | rs17063789 | -0.647 | 0.073 | -0.204 | -1.194 | -0.009 | 0.112 | 0.112 | 4.408 | 0.2 | 0.45 | 0.21 | 0.41 | 0.15 | 0.23 |
| 6 | 134790138 | rs11154772 | 0.351 | 0.622 | -0.318 | -0.925 | 0.074 | 0.145 | 0.145 | 0.48 | 0.2 | 0.41 | 0.46 | 0.55 | 0.26 | 0.21 |
| 6 | 134790368 | rs4896055 | 0.21 | 0.583 | -0.358 | -0.92 | 0.02 | 0.137 | 0.137 | 0.734 | 0.22 | 0.45 | 0.44 | 0.55 | 0.21 | 0.2 |
| 6 | 134791801 | rs2024690 | 0.221 | 0.197 | -0.356 | -0.98 | 0.061 | 0.141 | 0.141 | 0.756 | 0.22 | 0.45 | 0.46 | 0.52 | 0.26 | 0.21 |
| 6 | 134792784 | rs2327477 | -0.736 | 0.442 | -1.243 | -1.16 | -0.031 | 0.096 | 0.096 | 0 | 0.85 | 0.99 | 0.92 | 1 | 0.84 | 0.91 |
| 6 | 134796043 | rs4896060 | -0.905 | 0.722 | -0.399 | -0.512 | -0.002 | -0.004 | -0.004 | 0 | 0.34 | 0.37 | 0.59 | 0.84 | 0.34 | 0.41 |
| 6 | 134796909 | rs6908662 | -0.796 | 0.843 | -0.771 | -0.654 | -0.012 | 0.031 | 0.031 | 0.023 | 0.33 | 0.27 | 0.56 | 0.88 | 0.35 | 0.42 |
| 6 | 134804441 | rs9376046 | -0.305 | 0.292 | -0.464 | -0.028 | -0.006 | 0.038 | 0.038 | 4.521 | 0.12 | 0.15 | 0.14 | 0.29 | 0.07 | 0.17 |
| 6 | 134809080 | rs4895410 | -0.501 | 0.433 | -0.626 | -1.681 | 0.022 | 0.012 | 0.012 | 0.004 | 0.5 | 0.45 | 0.73 | 0.95 | 0.59 | 0.61 |
| 6 | 134817135 | rs2144219 | -0.351 | 0.677 | -0.76 | -1.83 | 0.003 | 0.003 | 0.003 | 1.553 | 0.23 | 0.21 | 0.13 | 0.08 | 0.2 | 0.28 |
| 6 | 134818314 | rs6929591 | -0.066 | 0.846 | -0.937 | -2.191 | -0.072 | 0.063 | 0.063 | 0.479 | 0.25 | 0.15 | 0.57 | 0.86 | 0.39 | 0.33 |
| 6 | 134821469 | rs4895411 | -0.098 | 1.182 | -0.943 | -2.294 | -0.033 | 0.036 | 0.036 | 0.698 | 0.22 | 0.2 | 0.56 | 0.84 | 0.31 | 0.29 |
| 6 | 134822574 | rs6935757 | -0.073 | -0.68 | -1.197 | -3.097 | 0.032 | 0.066 | 0.066 | 4.092 | 0.43 | 0.65 | 0.22 | 0.1 | 0.32 | 0.41 |
| 6 | 134831205 | rs9321460 | 0.42 | -1.422 | -0.933 | -3.247 | 0.013 | 0.008 | 0.008 | 1.043 | 0.31 | 0.26 | 0.15 | 0.07 | 0.23 | 0.23 |
| 6 | 134831486 | rs4896067 | 0.871 | -1.731 | -1.46 | -3.469 | -0.139 | -0.019 | -0.019 | 0 | 0.22 | 0.04 | 0.58 | 0.75 | 0.4 | 0.38 |
| 6 | 134835139 | rs6934912 | 0.266 | -2.181 | -1.19 | -3.577 | 0.008 | -0.01 | -0.01 | 0.138 | 0.36 | 0.34 | 0.17 | 0.07 | 0.27 | 0.34 |
| 6 | 134840948 | rs6938621 | -0.231 | -2.413 | -1.012 | -3.737 | -0.014 | -0.099 | -0.099 | 0.363 | 0.72 | 0.42 | 0.84 | 0.94 | 0.79 | 0.85 |
| 6 | 134845350 | rs6920474 | 0.843 | -2.151 | -1.211 | -3.509 | 0.02 | 0.035 | 0.035 | 0.824 | 0.31 | 0.22 | 0.14 | 0.13 | 0.23 | 0.26 |
| 6 | 134855869 | rs6901993 | -0.893 | -1.777 | -1.06 | -3.383 | -0.017 | 0.024 | 0.024 | 0 | 0.85 | 0.93 | 0.89 | 1 | 0.83 | 0.94 |
| 6 | 134856353 | rs9389191 | -0.896 | -1.511 | -0.563 | -2.991 | 0.034 | 0.134 | 0.134 | 0 | 0.56 | 0.75 | 0.78 | 0.93 | 0.63 | 0.69 |
| 6 | 134858347 | rs7752190 | -0.305 | -1.242 | -0.177 | -2.651 | 0.003 | -0.016 | -0.016 | 0.481 | 0.38 | 0.39 | 0.7 | 0.76 | 0.54 | 0.53 |
| 6 | 134873987 | rs13199020 | -0.246 | -0.643 | -0.677 | -2.477 | -0.037 | 0.047 | 0.047 | 0 | 0.33 | 0.25 | 0.65 | 0.85 | 0.47 | 0.45 |

| | | | | | | | | | | | | | | | | |
|---|---|---|---|---|---|---|---|---|---|---|---|---|---|---|---|---|
| 6 | 134877551 | rs7739403 | 0.215 | -1.256 | -0.003 | -1.84 | -0.011 | 0.041 | 0.041 | 0 | 0.06 | 0.19 | 0.03 | 0.11 | 0.03 | 0.03 |
| 6 | 134884553 | rs6569954 | -0.416 | -1.212 | -0.191 | -2.198 | 0.014 | 0.005 | 0.005 | 0 | 0.56 | 0.45 | 0.73 | 0.92 | 0.63 | 0.67 |
| 6 | 134887047 | rs228420 | -1.547 | -1.202 | 0.597 | -1.206 | -0.063 | -0.109 | -0.109 | 0 | 0.23 | 0.51 | 0.1 | 0.07 | 0.12 | 0.24 |
| 6 | 134888788 | rs228423 | -1.054 | -0.67 | 0.627 | -0.638 | -0.023 | -0.018 | -0.018 | 0 | 0.09 | 0.29 | 0.05 | 0.05 | 0.04 | 0.06 |
| 6 | 134904563 | rs4895415 | -0.633 | -1.364 | 0.433 | -0.395 | -0.029 | 0.005 | 0.005 | 0 | 0.25 | 0.29 | 0.12 | 0.21 | 0.14 | 0.18 |
| 6 | 134919354 | rs4896079 | 0.015 | -1.459 | 0.177 | -0.372 | -0.001 | -0.008 | -0.008 | 0.001 | 0.17 | 0.02 | 0.07 | 0.1 | 0.12 | 0.11 |
| 6 | 134922729 | rs2327457 | -0.138 | -1.556 | 0.133 | -0.421 | 0.004 | -0.003 | -0.003 | 0 | 0.18 | 0.02 | 0.07 | 0.1 | 0.12 | 0.11 |
| 6 | 134938220 | rs7754115 | -0.822 | -2.02 | -1.018 | -1.355 | -0.079 | -0.039 | -0.039 | 0 | 0.62 | 0.96 | 0.33 | 0.25 | 0.38 | 0.4 |
| 6 | 134943460 | rs7743660 | -0.016 | -2.285 | -0.136 | -1.523 | -0.02 | -0.03 | -0.03 | 0 | 0.38 | 0.5 | 0.18 | 0.15 | 0.26 | 0.28 |
| 6 | 134948543 | rs4438975 | -1.084 | -2.02 | 0.091 | -1.213 | -0.016 | 0.135 | 0.135 | 0.588 | 0.08 | 0.27 | 0.04 | 0.14 | 0.02 | 0.05 |
| 6 | 134969201 | rs9494013 | -0.78 | -1.917 | -0.633 | -1.499 | -0.021 | 0.066 | 0.066 | 0 | 0.61 | 0.84 | 0.34 | 0.22 | 0.37 | 0.49 |
| 6 | 134973208 | rs4896086 | 0.466 | -1.996 | -0.786 | -1.537 | 0.017 | 0.052 | 0.052 | 0.196 | 0.19 | 0.02 | 0.09 | 0.07 | 0.1 | 0.12 |
| 6 | 134979007 | rs11964024 | 0.905 | -2.254 | -0.52 | -1.742 | -0.02 | 0.117 | 0.117 | 0.053 | 0.19 | 0.02 | 0.09 | 0 | 0.17 | 0.18 |
| 6 | 134982427 | rs9494019 | -0.651 | -2.222 | -0.083 | -1.683 | -0.023 | -0.002 | -0.002 | 0 | 0.15 | 0.26 | 0.08 | 0.12 | 0.1 | 0.18 |
| 6 | 134988449 | rs4571581 | -0.339 | -1.388 | 0.345 | -1.634 | -0.003 | -0.022 | -0.022 | 0 | 0.31 | 0.65 | 0.3 | 0.16 | 0.32 | 0.35 |
| 6 | 134994331 | rs9494022 | -1.09 | -1.33 | 0.673 | -1.57 | 0.044 | 0.059 | 0.059 | 0.071 | 0.07 | 0.29 | 0.13 | 0.12 | 0.11 | 0.17 |
| 6 | 134998188 | rs7747509 | -0.851 | -1.389 | 0.232 | -1.776 | 0.016 | 0.069 | 0.069 | 0 | 0.53 | 0.7 | 0.39 | 0.22 | 0.4 | 0.47 |
| 6 | 135005295 | rs7765392 | 0.644 | -1.238 | 0.74 | -1.459 | 0.007 | 0.012 | 0.012 | 0 | 0.19 | 0.12 | 0.11 | 0 | 0.17 | 0.16 |
| 6 | 135005715 | rs4617006 | -0.174 | -1.245 | 0.906 | -1.431 | -0.039 | -0.071 | -0.071 | 0.675 | 0.27 | 0.54 | 0.17 | 0.11 | 0.21 | 0.26 |
| 6 | 135011576 | rs6569962 | 0.047 | -1.389 | 0.694 | -1.753 | -0.02 | -0.104 | -0.104 | 0 | 0.3 | 0.39 | 0.16 | 0 | 0.21 | 0.16 |
| 6 | 135013954 | rs9483738 | -0.606 | -0.882 | -0.055 | -0.915 | -0.006 | -0.031 | -0.031 | 1.754 | 0.54 | 0.86 | 0.45 | 0.44 | 0.47 | 0.42 |
| 6 | 135014966 | rs6914876 | 0.427 | -0.741 | 0.059 | -0.973 | -0.036 | -0.146 | -0.146 | 0 | 0.3 | 0.53 | 0.17 | 0 | 0.19 | 0.16 |
| 6 | 135022253 | rs9765870 | 0.36 | 0.376 | -1.183 | 0.075 | -0.045 | -0.097 | -0.097 | 0 | 0.73 | 0.86 | 0.5 | 0.48 | 0.63 | 0.68 |
| 6 | 135022619 | rs12205010 | -0.26 | 0.383 | -0.851 | 0.168 | -0.011 | -0.022 | -0.022 | 1.647 | 0.05 | 0.01 | 0.04 | 0.16 | 0.08 | 0.06 |
| 6 | 135025753 | rs9389206 | 0.221 | 0.334 | -0.733 | -0.079 | 0.046 | 0.002 | 0.002 | 2.6 | 0.46 | 0.3 | 0.35 | 0.44 | 0.36 | 0.45 |
| 6 | 135027967 | rs1321613 | 0.866 | 0.413 | -0.997 | -0.118 | -0.007 | 0.021 | 0.021 | 0 | 0.07 | 0.03 | 0.06 | 0.05 | 0.07 | 0.08 |
| 6 | 135030045 | rs7757871 | -0.117 | 0.496 | -0.574 | -0.161 | 0.005 | -0.004 | -0.004 | 0 | 0.55 | 0.39 | 0.41 | 0.5 | 0.47 | 0.53 |
| 6 | 135033315 | rs7754149 | -0.376 | 1.351 | -0.293 | -0.126 | 0.06 | -0.023 | -0.023 | 0 | 0.84 | 0.71 | 0.71 | 0.92 | 0.74 | 0.85 |
| 6 | 135043731 | rs9376055 | 0.792 | 1.105 | -0.688 | -0.2 | -0.04 | -0.084 | -0.084 | 2.426 | 0.14 | 0.01 | 0.19 | 0.28 | 0.17 | 0.21 |
| 6 | 135044376 | rs9376056 | 0.354 | 0.99 | -0.444 | -0.302 | -0.026 | -0.045 | -0.045 | 0 | 0.37 | 0.23 | 0.36 | 0.5 | 0.38 | 0.4 |
| 6 | 135045825 | rs1884952 | -0.189 | 1.389 | 0.202 | -0.172 | 0.04 | -0.017 | -0.017 | 0.628 | 0.85 | 0.75 | 0.72 | 0.91 | 0.74 | 0.85 |
| 6 | 135052032 | rs9376058 | -0.893 | 1.012 | -0.224 | -0.118 | -0.01 | -0.003 | -0.003 | 0 | 0.41 | 0.68 | 0.49 | 0.41 | 0.46 | 0.43 |
| 6 | 135052046 | rs9376059 | -0.381 | 1.024 | 0.044 | -0.085 | -0.007 | -0.003 | -0.003 | 0 | 0.3 | 0.43 | 0.31 | 0.33 | 0.29 | 0.33 |
| 6 | 135056757 | rs1407107 | 0.836 | 1.246 | -0.232 | 0.07 | 0.019 | 0.008 | 0.008 | 0 | 0.52 | 0.32 | 0.53 | 0.6 | 0.45 | 0.45 |
| 6 | 135059694 | rs7775195 | 0.459 | 1.122 | -0.349 | 0.145 | 0.004 | -0.008 | -0.008 | 0 | 0.16 | 0.15 | 0.2 | 0.28 | 0.17 | 0.21 |
| 6 | 135063873 | rs6922476 | 1.168 | 1.124 | -0.154 | 0.09 | -0.024 | -0.022 | -0.022 | 0.272 | 0.4 | 0.48 | 0.38 | 0.32 | 0.28 | 0.25 |
| 6 | 135071826 | rs2327534 | -0.579 | 1.915 | -0.061 | 0.508 | -0.01 | -0.015 | -0.015 | 0 | 0.84 | 0.86 | 0.81 | 0.78 | 0.79 | 0.73 |
| 6 | 135081569 | rs9385703 | 0.317 | 1.837 | 0.034 | 0.445 | 0.034 | 0.024 | 0.024 | 0 | 0.67 | 0.51 | 0.67 | 0.67 | 0.61 | 0.59 |
| 6 | 135089554 | rs958747 | 0.157 | 2.066 | 0.028 | 0.91 | 0.006 | 0 | 0 | 0 | 0.83 | 0.84 | 0.77 | 0.91 | 0.66 | 0.8 |
| 6 | 135104930 | rs9402651 | 0.206 | 2.118 | -0.493 | 1.31 | 0.015 | -0.051 | -0.051 | 0 | 0.1 | 0.03 | 0.13 | 0.28 | 0.07 | 0.16 |
| 6 | 135117042 | rs9373112 | 0.712 | 2.479 | -0.363 | 0.809 | -0.04 | -0.044 | -0.044 | 0 | 0.1 | 0.03 | 0.28 | 0.26 | 0.18 | 0.08 |
| 6 | 135121662 | rs4895424 | 0.623 | 1.826 | 0.143 | 0.002 | 0.093 | -0.061 | -0.061 | 0 | 0.78 | 0.53 | 0.72 | 0.91 | 0.67 | 0.8 |
| 6 | 135128858 | rs9483751 | -0.399 | 1.105 | -0.328 | 0.561 | -0.002 | 0.003 | 0.003 | 0 | 0.37 | 0.51 | 0.56 | 0.62 | 0.38 | 0.38 |
| 6 | 135134270 | rs12202002 | 0.533 | 0.352 | -0.86 | 0.181 | 0.057 | 0.053 | 0.053 | 2.576 | 0.33 | 0.12 | 0.1 | 0.2 | 0.19 | 0.31 |
| 6 | 135135717 | rs9376067 | 0.37 | 0.939 | -0.211 | 0.572 | -0.001 | 0 | 0 | 3.478 | 0.57 | 0.44 | 0.48 | 0.47 | 0.51 | 0.65 |
| 6 | 135138360 | rs9402653 | 0.576 | 0.784 | 0.529 | 0.464 | 0 | -0.001 | -0.001 | 0 | 0.56 | 0.39 | 0.46 | 0.47 | 0.53 | 0.58 |
| 6 | 135138990 | rs2049953 | 0.074 | 1.109 | 0.362 | 0.612 | -0.003 | 0.011 | 0.011 | 0 | 0.75 | 0.84 | 0.81 | 0.82 | 0.72 | 0.72 |
| 6 | 135142218 | rs2179170 | 0.01 | 1.389 | 0.754 | 0.873 | 0.025 | 0.093 | 0.093 | 0 | 0.16 | 0.3 | 0.33 | 0.35 | 0.19 | 0.13 |
| 6 | 135142809 | rs11753042 | 0.128 | 1.313 | 0.346 | 0.395 | -0.006 | 0.012 | 0.012 | 0 | 0.07 | 0.02 | 0.05 | 0.06 | 0.09 | 0.07 |
| 6 | 135143748 | rs7744034 | 0.994 | 0.969 | 0.749 | 0.085 | -0.002 | -0.018 | -0.018 | 0 | 0.63 | 0.41 | 0.71 | 0.76 | 0.66 | 0.64 |
| 6 | 135144622 | rs11754258 | 0.193 | 0.908 | 0.65 | 0.078 | 0.001 | 0.012 | 0.012 | 0 | 0.08 | 0.03 | 0.05 | 0.06 | 0.09 | 0.11 |
| 6 | 135150133 | rs2876265 | 1.763 | 1.078 | 0.532 | 0.39 | -0.024 | -0.017 | -0.017 | 8.749 | 0.46 | 0.09 | 0.44 | 0.44 | 0.41 | 0.5 |
| 6 | 135150488 | rs2876266 | -0.163 | 1.012 | 0.708 | 0.396 | -0.022 | 0.049 | 0.049 | 0 | 0.78 | 0.9 | 0.82 | 0.86 | 0.75 | 0.77 |
| 6 | 135150550 | rs9321476 | -1.377 | 0.998 | 0.908 | 0.584 | 0.003 | -0.032 | -0.032 | 0 | 0.31 | 0.8 | 0.39 | 0.42 | 0.33 | 0.27 |
| 6 | 135160309 | rs11752655 | -0.952 | 0.981 | 1.052 | 0.572 | -0.011 | 0.003 | 0.003 | 1.657 | 0.13 | 0.46 | 0.11 | 0.09 | 0.09 | 0.13 |
| 6 | 135168474 | rs9385707 | 0.07 | 1.125 | 1.011 | 0.685 | -0.006 | -0.009 | -0.009 | 0.005 | 0.16 | 0.16 | 0.25 | 0.33 | 0.24 | 0.14 |
| 6 | 135172345 | rs1883225 | 0.289 | 1.145 | 1.314 | 0.445 | 0.005 | -0.04 | -0.04 | 0 | 0.77 | 0.65 | 0.81 | 0.85 | 0.77 | 0.7 |
| 6 | 135172880 | rs9494090 | -0.246 | 0.854 | 2.075 | 1.056 | 0.006 | 0.042 | 0.042 | 0.827 | 0.16 | 0.21 | 0.25 | 0.34 | 0.24 | 0.09 |
| 6 | 135176176 | rs1040806 | -0.894 | 0.772 | 2.171 | 1.057 | 0.019 | 0.158 | 0.158 | 1.487 | 0.19 | 0.49 | 0.29 | 0.35 | 0.24 | 0.09 |
| 6 | 135178852 | rs9373114 | -1.645 | 0.91 | 1.866 | 0.981 | 0 | 0.024 | 0.024 | 2.095 | 0.28 | 0.36 | 0.32 | 0.41 | 0.31 | 0.34 |
| 6 | 135182281 | rs909627 | -2.089 | 0.88 | 1.91 | 0.918 | 0.01 | -0.04 | -0.04 | 3.979 | 0.41 | 0.79 | 0.53 | 0.54 | 0.41 | 0.47 |
| 6 | 135183233 | rs742103 | -1.296 | 0.801 | 1.709 | 0.938 | -0.01 | 0.008 | 0.008 | 0 | 0.12 | 0.32 | 0.11 | 0.11 | 0.08 | 0.25 |
| 6 | 135187044 | rs17776494 | -0.888 | 0.836 | 1.586 | 0.956 | -0.007 | 0.006 | 0.006 | 0 | 0.11 | 0.2 | 0.09 | 0.11 | 0.07 | 0.25 |
| 6 | 135192075 | rs9385708 | -0.195 | 0.945 | 1.232 | 1.03 | -0.021 | -0.039 | -0.039 | 0 | 0.15 | 0.07 | 0.22 | 0.28 | 0.24 | 0.08 |
| 6 | 135198116 | rs9402661 | -1.863 | 0.891 | 1.274 | 0.912 | 0.012 | -0.06 | -0.06 | 0 | 0.41 | 0.86 | 0.53 | 0.53 | 0.41 | 0.48 |
| 6 | 135199287 | rs9483768 | -1.16 | 0.847 | 1.842 | 0.889 | 0.014 | 0.022 | 0.022 | 0.006 | 0.08 | 0.27 | 0.1 | 0.08 | 0.07 | 0.12 |
| 6 | 135199596 | rs9494101 | -1.16 | 0.847 | 1.842 | 0.889 | 0.014 | 0.022 | 0.022 | 0.006 | 0.08 | 0.27 | 0.1 | 0.08 | 0.07 | 0.12 |
| 6 | 135209883 | rs7759527 | -1.454 | 0.862 | 1.974 | 0.893 | 0.052 | 0.043 | 0.043 | 0.466 | 0.09 | 0.43 | 0.11 | 0.08 | 0.08 | 0.15 |
| 6 | 135210182 | rs962189 | -1.999 | 0.83 | 1.419 | 0.863 | 0.004 | 0.003 | 0.003 | 2.275 | 0.21 | 0.79 | 0.32 | 0.21 | 0.17 | 0.27 |

| | | | | | | | | | | | | | | | | |
|---|---|---|---|---|---|---|---|---|---|---|---|---|---|---|---|---|
| 6 | 135231513 | rs2281038 | 1.554 | 1.188 | 1.321 | 1.085 | -0.01 | -0.016 | -0.016 | 0.095 | 0.61 | 0.28 | 0.46 | 0.45 | 0.55 | 0.54 |
| 6 | 135233909 | rs9321479 | 1.136 | 1.189 | 1.804 | 1.093 | 0.013 | 0.042 | 0.042 | 0.344 | 0.71 | 0.42 | 0.61 | 0.59 | 0.62 | 0.74 |
| 6 | 135235534 | rs9494115 | -1.445 | 1.018 | 2.299 | 0.991 | -0.02 | -0.006 | -0.006 | 1.87 | 0.14 | 0.51 | 0.2 | 0.13 | 0.07 | 0.19 |
| 6 | 135237158 | rs6569979 | -1.588 | 0.998 | 2.453 | 0.98 | -0.029 | -0.016 | -0.016 | 0 | 0.14 | 0.51 | 0.2 | 0.13 | 0.07 | 0.19 |
| 6 | 135237686 | rs4646880 | 1.573 | 1.075 | 2.277 | 0.978 | -0.01 | -0.025 | -0.025 | 0 | 0.6 | 0.2 | 0.45 | 0.47 | 0.55 | 0.54 |
| 6 | 135246294 | rs728030 | 0.839 | 0.874 | 2.118 | 1.247 | 0.013 | 0.01 | 0.01 | 0.352 | 0.75 | 0.71 | 0.72 | 0.76 | 0.65 | 0.71 |
| 6 | 135250530 | rs2064101 | 0.857 | 0.858 | 2.249 | 1.307 | -0.024 | -0.017 | -0.017 | 0 | 0.76 | 0.83 | 0.74 | 0.76 | 0.65 | 0.72 |
| 6 | 135252195 | rs2294319 | -0.905 | 0.775 | 2.599 | 1.496 | -0.022 | 0.11 | 0.11 | 0 | 0.11 | 0.31 | 0.22 | 0.29 | 0.06 | 0.19 |
| 6 | 135253451 | rs2072825 | 0.801 | 0.782 | 2.514 | 1.516 | -0.004 | -0.003 | -0.003 | 0 | 0.75 | 0.75 | 0.73 | 0.76 | 0.64 | 0.72 |
| 6 | 135253603 | rs2072827 | 2.258 | 0.602 | 1.89 | 1.023 | -0.117 | -0.068 | -0.068 | 0 | 0.49 | 0.03 | 0.41 | 0.47 | 0.47 | 0.46 |
| 6 | 135258010 | rs2235263 | -1.152 | 0.501 | 2.39 | 1.159 | -0.033 | 0.123 | 0.123 | 0 | 0.13 | 0.37 | 0.24 | 0.29 | 0.06 | 0.2 |
| 6 | 135258942 | rs2142725 | 0.8 | 0.653 | 2.355 | 1.092 | 0.02 | 0.018 | 0.018 | 0.026 | 0.75 | 0.62 | 0.72 | 0.76 | 0.65 | 0.71 |
| 6 | 135259366 | rs1022491 | 1.128 | 0.651 | 2.399 | 1.075 | 0.022 | 0.013 | 0.013 | 0.147 | 0.73 | 0.47 | 0.71 | 0.76 | 0.65 | 0.71 |
| 6 | 135263557 | rs3813342 | -0.193 | 0.507 | 2.207 | 1.108 | 0.002 | -0.05 | -0.05 | 0.361 | 0.08 | 0 | 0.2 | 0.29 | 0.06 | 0.2 |
| 6 | 135264294 | rs6919309 | -0.929 | 0.481 | 2.57 | 1.099 | -0.025 | 0.086 | 0.086 | 0 | 0.11 | 0.28 | 0.22 | 0.28 | 0.06 | 0.2 |
| 6 | 135265617 | rs4646871 | -1.825 | 0.552 | 2.642 | 1.083 | -0.001 | 0.009 | 0.009 | 0.685 | 0.37 | 0.62 | 0.5 | 0.53 | 0.41 | 0.49 |
| 6 | 135271886 | rs4646869 | -0.547 | 0.446 | 2.468 | 1.017 | 0.008 | -0.008 | -0.008 | 2.156 | 0.1 | 0.09 | 0.2 | 0.28 | 0.07 | 0.21 |
| 6 | 135271946 | rs4646868 | -1.73 | 0.434 | 2.578 | 1.011 | 0.002 | 0.014 | 0.014 | 0.355 | 0.36 | 0.43 | 0.48 | 0.53 | 0.42 | 0.5 |
| 6 | 135272477 | rs4267967 | -2.038 | 0.405 | 2.595 | 0.996 | -0.002 | 0.014 | 0.014 | 0.534 | 0.38 | 0.51 | 0.49 | 0.53 | 0.42 | 0.51 |
| 6 | 135275485 | rs4289677 | -0.713 | 0.432 | 2.633 | 0.906 | -0.022 | -0.015 | -0.015 | 0 | 0.23 | 0.18 | 0.24 | 0.24 | 0.35 | 0.29 |
| 6 | 135276065 | rs4896118 | -0.806 | 0.438 | 2.567 | 0.909 | 0.011 | 0.006 | 0.006 | 0.198 | 0.24 | 0.27 | 0.26 | 0.24 | 0.35 | 0.29 |
| 6 | 135282225 | rs7739722 | -1.057 | 0.465 | 2.538 | 0.91 | 0.014 | 0.008 | 0.008 | 0.022 | 0.26 | 0.33 | 0.27 | 0.24 | 0.35 | 0.29 |
| 6 | 135282656 | rs9389248 | -1.822 | 0.417 | 2.572 | 1.013 | 0.001 | 0.01 | 0.01 | 0.983 | 0.37 | 0.42 | 0.47 | 0.54 | 0.44 | 0.51 |
| 6 | 135283348 | rs4475342 | -2.139 | 0.928 | 2.534 | 0.881 | -0.011 | -0.008 | -0.008 | 0 | 0.43 | 0.73 | 0.55 | 0.54 | 0.49 | 0.53 |
| 6 | 135285148 | rs9399129 | -1.807 | 0.925 | 2.443 | 0.879 | 0.006 | 0.005 | 0.005 | 1.813 | 0.4 | 0.42 | 0.51 | 0.54 | 0.49 | 0.53 |
| 6 | 135305304 | rs12525425 | -0.715 | 0.921 | 2.314 | 0.891 | -0.001 | -0.004 | -0.004 | 0.017 | 0.14 | 0.12 | 0.26 | 0.29 | 0.13 | 0.25 |
| 6 | 135315398 | rs9376080 | -0.941 | 0.848 | 2.272 | 0.853 | 0 | -0.001 | -0.001 | 0.322 | 0.15 | 0.15 | 0.26 | 0.29 | 0.13 | 0.25 |
| 6 | 135315570 | rs6909975 | -2.174 | 0.858 | 2.217 | 0.863 | -0.019 | -0.013 | -0.013 | 1.155 | 0.43 | 0.79 | 0.55 | 0.54 | 0.49 | 0.53 |
| 6 | 135334600 | rs1014021 | -2.001 | 0.884 | 1.949 | 0.877 | -0.003 | -0.002 | -0.002 | 0 | 0.42 | 0.6 | 0.52 | 0.53 | 0.49 | 0.53 |
| 6 | 135349756 | rs4896128 | -0.64 | 0.931 | 1.207 | 0.867 | -0.007 | -0.005 | -0.005 | 1.534 | 0.23 | 0.19 | 0.23 | 0.24 | 0.36 | 0.28 |
| 6 | 135368331 | rs9389260 | -0.74 | 0.961 | 0.632 | 0.9 | -0.01 | -0.037 | -0.037 | 0 | 0.13 | 0.06 | 0.25 | 0.29 | 0.13 | 0.25 |
| 6 | 135383995 | rs2327582 | 1.981 | 1.116 | -0.021 | 0.645 | -0.035 | -0.123 | -0.123 | 0 | 0.42 | 0.03 | 0.37 | 0.46 | 0.36 | 0.34 |
| 6 | 135390836 | rs1547247 | -0.302 | 1.195 | 0.679 | 0.96 | -0.064 | -0.018 | -0.018 | 0 | 0.15 | 0.03 | 0.16 | 0.11 | 0.29 | 0.09 |
| 6 | 135395548 | rs13220662 | -1.852 | 1.097 | 1.193 | 1.211 | 0.003 | 0.003 | 0.003 | 0.42 | 0.27 | 0.67 | 0.26 | 0.26 | 0.32 | 0.27 |
| 6 | 135411228 | rs9376090 | -0.375 | 1.268 | 0.681 | 1.347 | -0.071 | -0.084 | -0.084 | 0 | 0.13 | 0.02 | 0.14 | 0.25 | 0.26 | 0.11 |
| 6 | 135414793 | rs9494139 | -0.982 | 1.252 | 0.496 | 1.394 | -0.014 | -0.028 | -0.028 | 0 | 0.15 | 0.08 | 0.24 | 0.29 | 0.17 | 0.28 |
| 6 | 135419018 | rs9399137 | -0.406 | 1.432 | 0.669 | 1.308 | -0.049 | -0.058 | -0.058 | 0 | 0.13 | 0.04 | 0.15 | 0.24 | 0.26 | 0.11 |
| 6 | 135423412 | rs1074849 | -1.049 | 1.222 | 0.829 | 1.201 | -0.02 | -0.036 | -0.036 | 1.582 | 0.19 | 0.12 | 0.27 | 0.33 | 0.24 | 0.38 |
| 6 | 135424203 | rs6930223 | -2.225 | 1.217 | 1.453 | 1.2 | -0.035 | -0.006 | -0.006 | 0.303 | 0.41 | 0.78 | 0.54 | 0.59 | 0.53 | 0.49 |
| 6 | 135426573 | rs4895441 | -0.453 | 1.187 | 1.597 | 1.199 | -0.03 | -0.034 | -0.034 | 0 | 0.14 | 0.08 | 0.18 | 0.26 | 0.27 | 0.1 |
| 6 | 135427144 | rs9376092 | -0.584 | 1.181 | 1.895 | 1.196 | 0.021 | 0.03 | 0.03 | 0.058 | 0.15 | 0.22 | 0.2 | 0.26 | 0.28 | 0.11 |
| 6 | 135427159 | rs9389269 | -0.431 | 1.183 | 1.839 | 1.197 | -0.041 | -0.046 | -0.046 | 0.493 | 0.14 | 0.05 | 0.18 | 0.26 | 0.27 | 0.1 |
| 6 | 135431318 | rs6920211 | -1.452 | 0.83 | 2.256 | 1.195 | 0.012 | 0.062 | 0.062 | 0 | 0.19 | 0.43 | 0.24 | 0.26 | 0.25 | 0.11 |
| 6 | 135432552 | rs9494145 | -0.4 | 0.825 | 1.961 | 0.864 | -0.02 | -0.04 | -0.04 | 0.042 | 0.13 | 0.06 | 0.16 | 0.24 | 0.22 | 0.09 |
| 6 | 135435171 | rs2223385 | -1.339 | 0.927 | 2.143 | 0.793 | 0.005 | 0.018 | 0.018 | 1.108 | 0.22 | 0.32 | 0.29 | 0.34 | 0.25 | 0.39 |
| 6 | 135439263 | rs2026937 | -2.255 | 0.415 | 1.566 | 0.488 | 0.017 | 0.042 | 0.042 | 0.53 | 0.49 | 0.78 | 0.53 | 0.61 | 0.5 | 0.54 |
| 6 | 135447773 | rs9321485 | -1.566 | 0.509 | 2.155 | 0.075 | 0.008 | 0.035 | 0.035 | 1.083 | 0.23 | 0.39 | 0.29 | 0.37 | 0.25 | 0.4 |
| 6 | 135451580 | rs1569534 | -0.941 | 0.441 | 1.411 | 0.055 | -0.009 | -0.022 | -0.022 | 0 | 0.22 | 0.13 | 0.27 | 0.36 | 0.24 | 0.4 |
| 6 | 135463947 | rs2050019 | -1.211 | 0.435 | 1.489 | -0.028 | 0 | 0.009 | 0.009 | 0 | 0.25 | 0.36 | 0.31 | 0.36 | 0.24 | 0.4 |
| 6 | 135465474 | rs17064262 | -0.77 | 0.326 | 1.099 | -0.17 | -0.011 | -0.016 | -0.016 | 0.053 | 0.15 | 0.21 | 0.17 | 0.1 | 0.16 | 0.15 |
| 6 | 135468891 | rs9494155 | -1.787 | 0.402 | 0.561 | -0.168 | 0.036 | 0.014 | 0.014 | 0.52 | 0.44 | 0.84 | 0.51 | 0.46 | 0.41 | 0.59 |
| 6 | 135482566 | rs6569999 | -1.726 | 0.567 | 0.696 | -0.025 | 0.017 | -0.013 | -0.013 | 0 | 0.4 | 0.77 | 0.48 | 0.5 | 0.39 | 0.56 |
| 6 | 135487614 | rs7757054 | -0.76 | 0.85 | 0.776 | 0.112 | 0.005 | 0.004 | 0.004 | 0 | 0.11 | 0.07 | 0.13 | 0.09 | 0.14 | 0.16 |
| 6 | 135490898 | rs9483798 | -1.146 | 0.808 | 1.2 | 0.034 | 0.004 | 0.06 | 0.06 | 0.394 | 0.24 | 0.61 | 0.34 | 0.41 | 0.22 | 0.35 |
| 6 | 135498201 | rs1320960 | -1.747 | 0.831 | 1.176 | 0.014 | 0.011 | -0.029 | -0.029 | 0 | 0.36 | 0.76 | 0.48 | 0.49 | 0.37 | 0.55 |
| 6 | 135515448 | rs11154794 | -0.989 | 0.613 | 1.315 | 0.064 | -0.01 | -0.009 | -0.009 | 0.899 | 0.11 | 0.23 | 0.13 | 0.08 | 0.11 | 0.15 |
| 6 | 135522939 | rs743589 | -0.954 | 1.078 | 0.652 | -0.256 | 0.004 | 0.001 | 0.001 | 0 | 0.37 | 0.61 | 0.45 | 0.43 | 0.41 | 0.42 |
| 6 | 135525396 | rs3819409 | -1.029 | 1.091 | 0.557 | -0.248 | -0.002 | -0.001 | -0.001 | 0 | 0.38 | 0.62 | 0.47 | 0.43 | 0.42 | 0.43 |
| 6 | 135535240 | rs210943 | -0.535 | 1.119 | 0.612 | -0.213 | 0.009 | 0.008 | 0.008 | 0 | 0.3 | 0.36 | 0.42 | 0.43 | 0.38 | 0.35 |
| 6 | 135535447 | rs210942 | -0.519 | 1.139 | 0.621 | -0.203 | 0.009 | 0.008 | 0.008 | 0 | 0.3 | 0.36 | 0.42 | 0.43 | 0.38 | 0.35 |
| 6 | 135536496 | rs210941 | -1.254 | 1.186 | -0.08 | -0.189 | -0.023 | -0.014 | -0.014 | 0 | 0.45 | 0.8 | 0.56 | 0.51 | 0.5 | 0.54 |
| 6 | 135540380 | rs210937 | -1.387 | 1.706 | -0.249 | -0.119 | -0.079 | -0.064 | -0.064 | 1.389 | 0.48 | 0.95 | 0.4 | 0.51 | 0.5 | 0.52 |
| 6 | 135541194 | rs6923895 | 0.521 | 1.845 | 0.092 | -0.166 | -0.002 | -0.016 | -0.016 | 0 | 0.15 | 0.15 | 0.15 | 0.06 | 0.18 | 0.09 |
| 6 | 135541395 | rs9399143 | -0.762 | 2.102 | 0.099 | -0.172 | -0.03 | -0.07 | -0.07 | 0.263 | 0.22 | 0.11 | 0.1 | 0.45 | 0.22 | 0.39 |
| 6 | 135545460 | rs9321496 | -1.634 | 1.87 | -0.044 | -0.198 | -0.081 | -0.059 | -0.059 | 0.277 | 0.5 | 0.95 | 0.4 | 0.51 | 0.5 | 0.52 |
| 6 | 135550343 | rs12196516 | -1.102 | 1.911 | 1.148 | 0.416 | 0.001 | 0.004 | 0.004 | 0 | 0.22 | 0.18 | 0.08 | 0.16 | 0.18 | 0.37 |
| 6 | 135555913 | rs4895445 | -1.603 | 2.026 | 1.727 | 0.38 | 0.107 | 0.111 | 0.111 | 0 | 0.12 | 0.36 | 0.2 | 0.24 | 0.19 | 0.08 |
| 6 | 135559005 | rs9483803 | -1.747 | 2.04 | 1.806 | 0.395 | 0.142 | 0.151 | 0.151 | 0 | 0.13 | 0.46 | 0.21 | 0.24 | 0.19 | 0.08 |
| 6 | 135566943 | rs4896134 | -1.754 | 1.35 | 1.093 | 0.383 | -0.043 | -0.003 | -0.003 | 0 | 0.36 | 0.82 | 0.32 | 0.35 | 0.39 | 0.41 |

| | | | | | | | | | | | | | | | | |
|---|---|---|---|---|---|---|---|---|---|---|---|---|---|---|---|---|
| 6 | 135606565 | rs1052502 | -1.95 | 0.407 | 1.739 | 0.622 | 0.012 | 0.003 | 0.003 | 7.702 | 0.13 | 0.84 | 0.15 | 0.13 | 0.06 | 0.14 |
| 6 | 135617601 | rs7766656 | -1.829 | 0.411 | 1.936 | 0.628 | -0.007 | -0.005 | -0.005 | 0 | 0.11 | 0.78 | 0.15 | 0.07 | 0.06 | 0.13 |
| 6 | 135619996 | rs9285480 | -1.746 | 0.391 | 2.175 | 0.631 | -0.024 | -0.041 | -0.041 | 0 | 0.1 | 0.69 | 0.11 | 0.01 | 0.03 | 0.11 |
| 6 | 135622819 | rs4896142 | -1.786 | 0.494 | 0.803 | 0.66 | -0.054 | -0.092 | -0.092 | 0 | 0.43 | 0.95 | 0.46 | 0.42 | 0.43 | 0.46 |
| 6 | 135623833 | rs2207000 | -1.793 | 0.643 | 1.511 | 0.689 | -0.022 | -0.053 | -0.053 | 0 | 0.25 | 0.52 | 0.19 | 0.11 | 0.21 | 0.13 |
| 6 | 135627369 | rs6935146 | 1.253 | 0.659 | 1.304 | 1.037 | -0.032 | -0.026 | -0.026 | 0 | 0.67 | 0.78 | 0.61 | 0.62 | 0.58 | 0.54 |
| 6 | 135634006 | rs9402702 | 2.094 | 0.54 | 1.032 | 0.949 | -0.12 | -0.099 | -0.099 | 0 | 0.54 | 0.04 | 0.5 | 0.56 | 0.51 | 0.49 |
| 6 | 135639644 | rs6914831 | 1.559 | 0.559 | 2.008 | 0.992 | 0.006 | 0.004 | 0.004 | 0.042 | 0.64 | 0.62 | 0.61 | 0.63 | 0.57 | 0.54 |
| 6 | 135642756 | rs2064430 | 1.774 | 0.58 | 2.14 | 0.998 | 0.008 | 0.005 | 0.005 | 0 | 0.62 | 0.45 | 0.58 | 0.62 | 0.56 | 0.53 |
| 6 | 135696597 | rs2746432 | 1.639 | 0.678 | 2.09 | 1.049 | -0.012 | -0.009 | -0.009 | 0 | 0.66 | 0.63 | 0.58 | 0.63 | 0.58 | 0.55 |
| 6 | 135721285 | rs13208164 | -0.756 | 0.989 | 1.688 | 1.156 | -0.084 | -0.043 | -0.043 | 0 | 0.17 | 0.01 | 0.14 | 0.15 | 0.27 | 0.19 |
| 6 | 135732194 | rs17778438 | -0.043 | 0.63 | 1.414 | 1.602 | -0.019 | -0.054 | -0.054 | 0.364 | 0.1 | 0.01 | 0.06 | 0.25 | 0.11 | 0.07 |
| 6 | 135739062 | rs2614290 | 2.008 | 0.716 | 2.174 | 1.624 | 0.112 | 0.087 | 0.087 | 3.573 | 0.93 | 0.5 | 0.89 | 0.97 | 0.96 | 0.94 |
| 6 | 135739355 | rs11154801 | -1.357 | 0.712 | 1.981 | 1.572 | -0.084 | -0.061 | -0.061 | 0.099 | 0.32 | 0.08 | 0.33 | 0.32 | 0.37 | 0.41 |
| 6 | 135749202 | rs1547079 | 1.867 | 0.663 | 2.332 | 1.501 | 0.078 | 0.049 | 0.049 | 4.806 | 0.92 | 0.5 | 0.89 | 0.97 | 0.94 | 0.94 |
| 6 | 135757022 | rs1535435 | 2.743 | 0.606 | 2.392 | 1.516 | -0.019 | -0.008 | -0.008 | 0 | 0.88 | 0.3 | 0.87 | 0.9 | 0.93 | 0.9 |
| 6 | 135765129 | rs717120 | -0.355 | 0.654 | 1.834 | 1.558 | -0.061 | -0.036 | -0.036 | 0 | 0.17 | 0.01 | 0.11 | 0.14 | 0.22 | 0.2 |
| 6 | 135793706 | rs6908428 | -1.49 | 0.573 | 2.146 | 1.567 | -0.018 | -0.007 | -0.007 | 0 | 0.33 | 0.2 | 0.35 | 0.31 | 0.37 | 0.41 |
| 6 | 135829796 | rs4896151 | 1.698 | 0.526 | 2.283 | 1.519 | 0.044 | 0.026 | 0.026 | 2.71 | 0.92 | 0.63 | 0.9 | 0.96 | 0.95 | 0.94 |
| 6 | 135848223 | rs9389295 | 2.17 | 0.506 | 1.997 | 1.545 | -0.101 | -0.072 | -0.072 | 0.073 | 0.54 | 0.05 | 0.52 | 0.59 | 0.57 | 0.5 |
| 6 | 135851573 | rs9494266 | -2.533 | 0.5 | 2.042 | 1.538 | -0.011 | -0.007 | -0.007 | 0 | 0.13 | 0.8 | 0.15 | 0.1 | 0.07 | 0.09 |
| 6 | 135870778 | rs9402717 | 2.31 | 0.52 | 1.803 | 1.527 | -0.106 | -0.062 | -0.062 | 0.093 | 0.53 | 0.04 | 0.5 | 0.59 | 0.53 | 0.49 |
| 6 | 135880480 | rs9494274 | -2.144 | 0.487 | 1.962 | 1.651 | 0.003 | -0.011 | -0.011 | 0 | 0.1 | 0.7 | 0.13 | 0.04 | 0.05 | 0.06 |
| 6 | 135893723 | rs13208505 | -1.471 | 0.487 | 1.907 | 1.665 | -0.035 | -0.012 | -0.012 | 0 | 0.33 | 0.14 | 0.34 | 0.32 | 0.36 | 0.41 |
| 6 | 135897935 | rs10484771 | 0.929 | 0.257 | 1.877 | 1.556 | -0.015 | -0.031 | -0.031 | 0 | 0.06 | 0 | 0.04 | 0.24 | 0.07 | 0.05 |
| 6 | 135899344 | rs2092556 | 2.027 | 0.209 | 2.313 | 1.089 | 0.046 | 0.025 | 0.025 | 6.051 | 0.9 | 0.46 | 0.88 | 0.96 | 0.95 | 0.94 |
| 6 | 135921139 | rs9385729 | -2.287 | -0.092 | 2.234 | 1.209 | -0.001 | 0.002 | 0.002 | 0.82 | 0.39 | 0.7 | 0.35 | 0.29 | 0.37 | 0.44 |
| 6 | 135926863 | rs4896156 | -2.407 | -0.114 | 2.213 | 1.194 | -0.003 | 0.034 | 0.034 | 0.924 | 0.43 | 0.85 | 0.4 | 0.29 | 0.39 | 0.46 |
| 6 | 135927526 | rs6941396 | -1.42 | -0.082 | 2.028 | 1.192 | -0.015 | 0.004 | 0.004 | 0.004 | 0.29 | 0.13 | 0.23 | 0.24 | 0.31 | 0.36 |
| 6 | 135933005 | rs2143681 | -2.247 | -0.064 | 1.901 | 1.31 | -0.003 | 0.002 | 0.002 | 0.839 | 0.4 | 0.76 | 0.36 | 0.31 | 0.37 | 0.44 |
| 6 | 135940481 | rs1883818 | -0.512 | -0.078 | 1.958 | 1.347 | -0.002 | 0.027 | 0.027 | 0 | 0.05 | 0.18 | 0.05 | 0.06 | 0.04 | 0.05 |
| 6 | 135952009 | rs9402729 | -1.9 | 0.228 | 1.689 | 1.247 | -0.012 | -0.006 | -0.006 | 0.194 | 0.37 | 0.39 | 0.32 | 0.35 | 0.4 | 0.48 |
| 6 | 135957359 | rs13201869 | -0.559 | 0.297 | 1.344 | 1.452 | -0.058 | -0.061 | -0.061 | 0 | 0.14 | 0.01 | 0.08 | 0.18 | 0.17 | 0.17 |
| 6 | 135957792 | rs6902485 | -2.263 | -0.031 | 1.816 | 1.124 | -0.025 | 0.02 | 0.02 | 6.967 | 0.12 | 0.61 | 0.15 | 0.05 | 0.06 | 0.07 |
| 6 | 135967964 | rs9494312 | -1.321 | -0.026 | 1.584 | 1.113 | -0.005 | -0.011 | -0.011 | 0 | 0.37 | 0.53 | 0.31 | 0.24 | 0.29 | 0.44 |
| 6 | 135983115 | rs12210389 | -1.259 | -0.018 | 1.719 | 1.85 | -0.003 | 0.005 | 0.005 | 0 | 0.36 | 0.54 | 0.31 | 0.38 | 0.31 | 0.41 |
| 6 | 136001034 | rs9402743 | -1.883 | -0.124 | 1.857 | 1.676 | 0.006 | -0.027 | -0.027 | 0.448 | 0.4 | 0.81 | 0.49 | 0.45 | 0.34 | 0.47 |
| 6 | 136008608 | rs9494332 | -1.014 | -0.149 | 2.272 | 1.712 | 0.002 | 0.012 | 0.012 | 0 | 0.32 | 0.28 | 0.4 | 0.39 | 0.31 | 0.41 |
| 6 | 136015672 | rs1569719 | 1.41 | -0.395 | 1.249 | 1.073 | -0.051 | 0.139 | 0.139 | 2.104 | 0.28 | 0.03 | 0.28 | 0.05 | 0.29 | 0.19 |
| 6 | 136018274 | rs4129778 | -2.914 | -0.636 | 2.972 | 1.986 | -0.005 | 0.079 | 0.079 | 8.163 | 0.1 | 0.53 | 0.1 | 0.05 | 0.04 | 0.07 |
| 6 | 136019147 | rs12202212 | -1.217 | -0.637 | 2.637 | 2.03 | -0.034 | -0.077 | -0.077 | 1.159 | 0.3 | 0.09 | 0.35 | 0.39 | 0.29 | 0.41 |
| 6 | 136053695 | rs11154819 | -0.859 | -0.828 | 2.507 | 2.111 | -0.032 | -0.038 | -0.038 | 0 | 0.07 | 0.01 | 0.11 | 0.17 | 0.08 | 0.11 |
| 6 | 136056234 | rs1475069 | -0.951 | -0.854 | 2.574 | 1.717 | -0.071 | -0.129 | -0.129 | 0.098 | 0.28 | 0.06 | 0.37 | 0.45 | 0.27 | 0.42 |
| 6 | 136059177 | rs2038551 | -1.019 | -0.858 | 2.632 | 1.715 | -0.043 | -0.092 | -0.092 | 0.02 | 0.28 | 0.09 | 0.38 | 0.45 | 0.27 | 0.42 |
| 6 | 136059423 | rs911507 | -2.37 | -0.911 | 2.686 | 1.684 | 0.027 | -0.054 | -0.054 | 0.092 | 0.44 | 0.9 | 0.5 | 0.45 | 0.3 | 0.5 |
| 6 | 136064441 | rs12201044 | -1.083 | -0.883 | 2.871 | 1.727 | 0 | -0.017 | -0.017 | 0 | 0.3 | 0.29 | 0.38 | 0.4 | 0.25 | 0.37 |
| 6 | 136065411 | rs7739804 | -2.021 | -0.886 | 2.838 | 1.722 | 0.001 | 0.054 | 0.054 | 0 | 0.07 | 0.23 | 0.08 | 0.06 | 0.05 | 0.12 |
| 6 | 136073927 | rs12216021 | -0.954 | -0.859 | 2.478 | 1.724 | -0.025 | -0.086 | -0.086 | 0 | 0.28 | 0.11 | 0.35 | 0.4 | 0.25 | 0.37 |
| 6 | 136080464 | rs9399161 | -1.907 | -0.868 | 2.755 | 1.712 | 0.007 | -0.005 | -0.005 | 0.278 | 0.4 | 0.66 | 0.47 | 0.45 | 0.3 | 0.45 |
| 6 | 136107630 | rs981580 | -1.028 | -0.909 | 2.708 | 1.714 | -0.032 | -0.104 | -0.104 | 0 | 0.28 | 0.13 | 0.35 | 0.4 | 0.25 | 0.36 |
| 6 | 136116013 | rs9494390 | -2.091 | -0.812 | 2.736 | 1.716 | 0.039 | 0.001 | 0.001 | 0.402 | 0.45 | 0.79 | 0.49 | 0.46 | 0.32 | 0.49 |
| 6 | 136119672 | rs1342240 | -1.693 | -1.004 | 2.841 | 1.594 | 0.086 | 0.155 | 0.155 | 0 | 0.05 | 0.26 | 0.09 | 0.05 | 0.04 | 0.07 |
| 6 | 136121559 | rs7767870 | -3.221 | -1.061 | 2.776 | 1.644 | -0.024 | -0.001 | -0.001 | 0 | 0.2 | 0.74 | 0.27 | 0.13 | 0.1 | 0.18 |
| 6 | 136127782 | rs7776097 | -1.577 | -1.046 | 2.592 | 1.615 | -0.017 | 0.003 | 0.003 | 0 | 0.07 | 0.2 | 0.13 | 0.06 | 0.02 | 0.04 |
| 6 | 136131756 | rs927330 | -1.79 | -1.166 | 2.788 | 1.667 | -0.006 | 0.003 | 0.003 | 0 | 0.4 | 0.48 | 0.43 | 0.41 | 0.26 | 0.41 |
| 6 | 136131888 | rs2027516 | -1.499 | -1.181 | 2.833 | 1.719 | -0.011 | 0.016 | 0.016 | 0 | 0.37 | 0.62 | 0.44 | 0.41 | 0.26 | 0.38 |
| 6 | 136132370 | rs1885271 | -1.811 | -1.228 | 2.834 | 1.554 | -0.018 | 0.021 | 0.021 | 0 | 0.1 | 0.24 | 0.08 | 0.05 | 0.05 | 0.13 |
| 6 | 136133245 | rs9399168 | -1.16 | -1.216 | 2.786 | 1.56 | 0.017 | 0.055 | 0.055 | 0 | 0.06 | 0.18 | 0.07 | 0.05 | 0.04 | 0.1 |
| 6 | 136133324 | rs7765591 | -1.767 | -1.212 | 2.766 | 1.46 | -0.003 | 0.079 | 0.079 | 0.146 | 0.08 | 0.42 | 0.09 | 0.07 | 0.02 | 0.04 |
| 6 | 136133659 | rs947583 | -1.101 | -1.19 | 2.172 | 1.438 | -0.001 | 0.002 | 0.002 | 0 | 0.3 | 0.28 | 0.35 | 0.34 | 0.23 | 0.36 |
| 6 | 136136872 | rs6570045 | -1.84 | -1.031 | 1.326 | 1.543 | 0.061 | -0.005 | -0.005 | 2.169 | 0.47 | 0.86 | 0.52 | 0.49 | 0.3 | 0.52 |
| 6 | 136139013 | rs6913708 | -1.393 | -0.846 | 1.466 | 1.384 | -0.018 | 0.05 | 0.05 | 0 | 0.05 | 0.2 | 0.06 | 0.11 | 0.02 | 0.02 |
| 6 | 136145479 | rs11154825 | -0.6 | -0.895 | 0.939 | 1.347 | -0.012 | -0.067 | -0.067 | 1.959 | 0.23 | 0.09 | 0.3 | 0.33 | 0.2 | 0.34 |
| 6 | 136146508 | rs9376154 | -1.397 | -0.907 | 1.765 | 1.456 | -0.037 | 0.053 | 0.053 | 0 | 0.14 | 0.37 | 0.14 | 0.17 | 0.08 | 0.18 |
| 6 | 136150544 | rs1342237 | -1.561 | -0.921 | 1.468 | 1.449 | -0.069 | 0.044 | 0.044 | 0 | 0.18 | 0.55 | 0.15 | 0.17 | 0.08 | 0.18 |
| 6 | 136163201 | rs2009129 | -1.457 | -0.853 | 0.762 | 1.275 | 0.052 | -0.024 | -0.024 | 0.163 | 0.39 | 0.79 | 0.42 | 0.5 | 0.23 | 0.45 |
| 6 | 136185184 | rs9402765 | -1.143 | -0.785 | 1.232 | 1.183 | -0.037 | 0.023 | 0.023 | 0 | 0.13 | 0.33 | 0.14 | 0.12 | 0.06 | 0.1 |
| 6 | 136198594 | rs6570048 | 0.704 | -0.524 | -0.016 | 1.084 | -0.045 | 0.02 | 0.02 | 0 | 0.44 | 0.11 | 0.45 | 0.44 | 0.51 | 0.45 |
| 6 | 136209082 | rs7767574 | -1.318 | -0.689 | 0.953 | 0.906 | 0.006 | 0.003 | 0.003 | 3.347 | 0.08 | 0.39 | 0.05 | 0 | 0.04 | 0.05 |

| | | | | | | | | | | | | | | | | |
|---|---|---|---|---|---|---|---|---|---|---|---|---|---|---|---|---|
| 6 | 136213506 | rs1557142 | -0.669 | -0.876 | 0.313 | 1.047 | -0.058 | 0.068 | 0.068 | 0 | 0.71 | 0.89 | 0.78 | 0.86 | 0.65 | 0.67 |
| 6 | 136217299 | rs6570053 | -0.521 | -0.746 | 0.757 | 1.42 | -0.041 | 0.048 | 0.048 | 0 | 0.67 | 0.86 | 0.71 | 0.8 | 0.61 | 0.61 |
| 6 | 136217496 | rs7744517 | -0.691 | -0.612 | 0.741 | 1.505 | -0.003 | 0.008 | 0.008 | 0 | 0.09 | 0.11 | 0.09 | 0.1 | 0.08 | 0.07 |
| 6 | 136218110 | rs9402774 | 0.742 | -0.395 | 0.569 | 1.352 | 0 | -0.007 | -0.007 | 0 | 0.44 | 0.34 | 0.51 | 0.59 | 0.43 | 0.48 |
| 6 | 136220416 | rs4382284 | 0.134 | -0.171 | 0.488 | 1.23 | 0.003 | 0.004 | 0.004 | 0 | 0.55 | 0.64 | 0.55 | 0.66 | 0.4 | 0.5 |
| 6 | 136222577 | rs9494412 | -0.647 | 0.643 | -0.057 | 1.519 | 0.022 | 0.022 | 0.022 | 0 | 0.08 | 0.13 | 0.11 | 0.1 | 0.13 | 0.09 |
| 6 | 136222807 | rs10872448 | 0.349 | 0.271 | -0.545 | 1.367 | -0.046 | -0.016 | -0.016 | 2.306 | 0.41 | 0.12 | 0.44 | 0.46 | 0.41 | 0.43 |
| 6 | 136224650 | rs9389342 | -1.29 | -0.122 | 0.391 | 0.651 | -0.024 | -0.046 | -0.046 | 0 | 0.12 | 0.24 | 0.06 | 0.02 | 0.06 | 0.12 |
| 6 | 136226163 | rs9483889 | -0.895 | -0.825 | 0.794 | -0.098 | -0.035 | -0.129 | -0.129 | 0 | 0.18 | 0.41 | 0.12 | 0 | 0.12 | 0.15 |
| 6 | 136227558 | rs7752169 | -0.797 | -0.499 | -0.111 | 0.079 | -0.001 | -0.001 | -0.001 | 0.179 | 0.58 | 0.57 | 0.5 | 0.47 | 0.54 | 0.56 |
| 6 | 136228219 | rs2327672 | 0.288 | -0.049 | -0.204 | 0.128 | 0.012 | 0.066 | 0.066 | 0 | 0.12 | 0.15 | 0.22 | 0.34 | 0.15 | 0.27 |
| 6 | 136230385 | rs6570057 | -0.267 | -0.434 | -0.409 | 0.048 | -0.059 | -0.077 | -0.077 | 0 | 0.23 | 0.44 | 0.15 | 0.08 | 0.14 | 0.2 |
| 6 | 136232680 | rs6926933 | -0.96 | -0.317 | -0.6 | -0.176 | -0.034 | -0.104 | -0.104 | 0 | 0.3 | 0.52 | 0.2 | 0.07 | 0.2 | 0.3 |
| 6 | 136234351 | rs7751423 | -0.882 | -0.442 | -0.552 | -0.168 | -0.053 | -0.11 | -0.11 | 0 | 0.29 | 0.49 | 0.19 | 0.07 | 0.19 | 0.28 |
| 6 | 136235568 | rs12208105 | 0.415 | -0.474 | -0.665 | -0.026 | -0.031 | -0.08 | -0.08 | 0 | 0.36 | 0.1 | 0.38 | 0.47 | 0.41 | 0.38 |
| 6 | 136236826 | rs6899330 | -1.461 | -0.365 | -0.727 | -0.081 | -0.043 | 0.072 | 0.072 | 0.051 | 0.7 | 0.83 | 0.71 | 0.85 | 0.65 | 0.84 |
| 6 | 136239179 | rs1891362 | 0.759 | -0.366 | -0.625 | 0.194 | -0.03 | -0.054 | -0.054 | 0 | 0.32 | 0.11 | 0.35 | 0.38 | 0.38 | 0.3 |
| 6 | 136239856 | rs4499949 | -1.222 | -0.547 | -0.358 | 0.404 | 0.014 | -0.008 | -0.008 | 0 | 0.38 | 0.64 | 0.36 | 0.47 | 0.28 | 0.54 |
| 6 | 136244220 | rs6919578 | -0.313 | -0.359 | -0.46 | 0.538 | -0.001 | 0.018 | 0.018 | 0 | 0.14 | 0.14 | 0.2 | 0.4 | 0.13 | 0.28 |
| 6 | 136246471 | rs9494421 | -0.871 | -0.482 | -0.427 | 0.281 | -0.023 | -0.069 | -0.069 | 0 | 0.33 | 0.48 | 0.25 | 0.21 | 0.27 | 0.49 |
| 6 | 136247928 | rs11154829 | 0.721 | -0.643 | -1.225 | 0.789 | -0.063 | -0.117 | -0.117 | 0 | 0.23 | 0.01 | 0.3 | 0.31 | 0.28 | 0.28 |
| 6 | 136260580 | rs17248872 | -0.231 | 0.291 | -1.147 | 0.374 | -0.049 | 0.001 | 0.001 | 0 | 0.17 | 0.05 | 0.11 | 0.2 | 0.24 | 0.29 |
| 6 | 136261407 | rs6916391 | -1.142 | -0.425 | -0.931 | -0.058 | 0.004 | -0.004 | -0.004 | 0 | 0.78 | 0.72 | 0.74 | 0.86 | 0.71 | 0.81 |
| 6 | 136263815 | rs2876317 | 0.042 | -0.616 | -0.607 | -0.095 | 0 | 0.004 | 0.004 | 0 | 0.6 | 0.58 | 0.63 | 0.65 | 0.49 | 0.52 |
| 6 | 136270587 | rs1342242 | 0.413 | -1.275 | -0.234 | -0.053 | -0.003 | 0.001 | 0.001 | 0.481 | 0.39 | 0.43 | 0.52 | 0.47 | 0.38 | 0.35 |
| 6 | 136271963 | rs6907337 | 0.376 | -1.186 | -0.397 | -0.083 | -0.069 | -0.028 | -0.028 | 0.937 | 0.17 | 0.05 | 0.17 | 0.21 | 0.24 | 0.28 |
| 6 | 136276087 | rs10872455 | 0.639 | -1.248 | -0.139 | -0.098 | 0.018 | -0.023 | -0.023 | 0 | 0.38 | 0.3 | 0.51 | 0.48 | 0.38 | 0.35 |
| 6 | 136295854 | rs7761961 | 0.568 | -1.357 | -0.037 | -0.02 | -0.007 | 0.007 | 0.007 | 0.141 | 0.41 | 0.53 | 0.55 | 0.47 | 0.41 | 0.36 |
| 6 | 136297906 | rs4133053 | 1.569 | -1.45 | -0.289 | -0.12 | -0.097 | 0.06 | 0.06 | 0 | 0.2 | 0.01 | 0.16 | 0.09 | 0.25 | 0.15 |
| 6 | 136303107 | rs9399176 | 0.59 | -1.259 | 0.009 | 0.043 | 0.005 | 0.028 | 0.028 | 0 | 0.4 | 0.27 | 0.27 | 0.23 | 0.35 | 0.31 |
| 6 | 136306350 | rs4075416 | 0.188 | -1.415 | 0.064 | 0.162 | 0.009 | 0.007 | 0.007 | 0.176 | 0.43 | 0.65 | 0.56 | 0.55 | 0.41 | 0.41 |
| 6 | 136311863 | rs4254999 | 0.808 | -1.512 | -0.601 | 0.24 | -0.068 | -0.183 | -0.183 | 0 | 0.34 | 0.03 | 0.39 | 0.45 | 0.37 | 0.35 |
| 6 | 136333229 | rs9402782 | 0.818 | -1.501 | -0.592 | 0.21 | -0.064 | -0.16 | -0.16 | 0 | 0.34 | 0.03 | 0.39 | 0.39 | 0.37 | 0.34 |
| 6 | 136340592 | rs4333432 | -0.334 | -1.046 | -0.283 | 0.431 | -0.042 | -0.032 | -0.032 | 0 | 0.85 | 0.95 | 0.86 | 0.79 | 0.83 | 0.79 |
| 6 | 136344062 | rs9389361 | 0.555 | -1.089 | 0.201 | 0.461 | -0.048 | -0.15 | -0.15 | 0 | 0.3 | 0.05 | 0.4 | 0.4 | 0.37 | 0.34 |
| 6 | 136349307 | rs6916194 | -0.247 | -1.252 | 0.354 | 0.51 | -0.008 | 0.012 | 0.012 | 0 | 0.37 | 0.56 | 0.56 | 0.51 | 0.4 | 0.4 |
| 6 | 136351726 | rs9389363 | -0.179 | -1.078 | -0.426 | 0.378 | 0.012 | -0.004 | -0.004 | 0 | 0.37 | 0.3 | 0.44 | 0.57 | 0.38 | 0.39 |
| 6 | 136359446 | rs9376173 | 0.207 | -0.394 | -0.696 | 0.495 | 0.01 | -0.046 | -0.046 | 0.161 | 0.31 | 0.21 | 0.42 | 0.52 | 0.33 | 0.16 |
| 6 | 136365636 | rs3813341 | 0.553 | 0.197 | 0.124 | -0.214 | 0.003 | -0.023 | -0.023 | 0.867 | 0.32 | 0.35 | 0.21 | 0.16 | 0.34 | 0.21 |
| 6 | 136366397 | rs13212385 | -1.248 | 0.165 | 0.66 | 0.535 | -0.003 | -0.075 | -0.075 | 3.84 | 0.14 | 0.25 | 0.12 | 0 | 0.13 | 0.09 |
| 6 | 136368978 | rs7747433 | -1.28 | -0.059 | 0.056 | 1.296 | -0.019 | 0.031 | 0.031 | 0.167 | 0.21 | 0.6 | 0.34 | 0.25 | 0.13 | 0.28 |
| 6 | 136370658 | rs6921041 | -0.147 | 0.312 | -0.314 | 0.988 | -0.019 | 0.004 | 0.004 | 0 | 0.39 | 0.66 | 0.51 | 0.42 | 0.44 | 0.53 |
| 6 | 136380147 | rs1383393 | -0.421 | -0.084 | -0.979 | 1.006 | -0.007 | 0.053 | 0.053 | 0 | 0.61 | 0.78 | 0.67 | 0.78 | 0.62 | 0.67 |
| 6 | 136392020 | rs899894 | 0.14 | -0.102 | -0.687 | 1.071 | -0.008 | 0.007 | 0.007 | 0 | 0.38 | 0.73 | 0.43 | 0.37 | 0.39 | 0.52 |
| 6 | 136394895 | rs1013020 | 0.289 | -0.171 | 0.012 | 1.035 | 0.001 | -0.001 | -0.001 | 0 | 0.37 | 0.59 | 0.4 | 0.36 | 0.39 | 0.51 |
| 6 | 136410563 | rs7770239 | -0.645 | -0.488 | 0.936 | 0.733 | -0.001 | -0.019 | -0.019 | 0.067 | 0.07 | 0.32 | 0.03 | 0 | 0.02 | 0.06 |
| 6 | 136437533 | rs1480645 | -1.161 | -0.436 | 0.079 | 0.694 | -0.1 | 0.058 | 0.058 | 0 | 0.74 | 0.99 | 0.85 | 0.91 | 0.7 | 0.76 |
| 6 | 136450147 | rs3799386 | 0.694 | 0.002 | 0.292 | 1.085 | -0.11 | -0.104 | -0.104 | 0 | 0.27 | 0.02 | 0.37 | 0.33 | 0.34 | 0.44 |
| 6 | 136464760 | rs7770469 | -0.112 | 0.072 | 0.863 | 1.102 | 0.002 | -0.002 | -0.002 | 0.016 | 0.37 | 0.64 | 0.45 | 0.42 | 0.36 | 0.5 |
| 6 | 136471707 | rs1471545 | -0.058 | -0.1 | 0.8 | 1.271 | -0.003 | -0.02 | -0.02 | 0 | 0.38 | 0.25 | 0.53 | 0.55 | 0.4 | 0.47 |
| 6 | 136471800 | rs1471546 | -0.24 | -0.042 | 0.563 | 1.306 | 0.004 | -0.004 | -0.004 | 0.043 | 0.42 | 0.63 | 0.56 | 0.56 | 0.4 | 0.47 |
| 6 | 136475578 | rs1871693 | -0.051 | 0.214 | 0.372 | 1.191 | -0.028 | -0.08 | -0.08 | 0.19 | 0.26 | 0.09 | 0.29 | 0.38 | 0.32 | 0.21 |
| 6 | 136501828 | rs487278 | 0.657 | 0.14 | 1.274 | 0.975 | -0.03 | -0.003 | -0.003 | 0.191 | 0.66 | 0.12 | 0.5 | 0.59 | 0.63 | 0.68 |
| 6 | 136506199 | rs7746642 | -1.589 | -0.398 | 2.166 | 0.62 | 0.032 | 0.028 | 0.028 | 5.008 | 0.06 | 0.49 | 0.05 | 0.02 | 0 | 0 |
| 6 | 136516257 | rs7753890 | -2.567 | -0.631 | 2.3 | 0.741 | -0.001 | 0 | 0 | 0 | 0.12 | 0.88 | 0.12 | 0.15 | 0.02 | 0.12 |
| 6 | 136525585 | rs11965553 | -2.215 | -0.547 | 2.56 | 0.392 | -0.027 | -0.03 | -0.03 | 0 | 0.08 | 0.66 | 0.08 | 0.01 | 0.01 | 0.05 |
| 6 | 136538031 | rs9494473 | -1.206 | -0.607 | 2.591 | 0.398 | -0.036 | -0.015 | -0.015 | 0.538 | 0.19 | 0.4 | 0.13 | 0.2 | 0.12 | 0.22 |
| 6 | 136548432 | rs9494484 | -2.395 | -0.426 | 2.898 | 0.353 | -0.036 | -0.039 | -0.039 | 0.709 | 0.08 | 0.68 | 0.09 | 0 | 0 | 0.01 |
| 6 | 136552086 | rs6570075 | -2.377 | -0.426 | 2.914 | 0.353 | -0.033 | -0.036 | -0.036 | 0.346 | 0.08 | 0.67 | 0.09 | 0 | 0 | 0.01 |
| 6 | 136559847 | rs9494492 | -1.07 | -0.005 | 2.578 | 0.088 | 0.038 | -0.062 | -0.062 | 0 | 0.14 | 0.32 | 0.13 | 0.05 | 0.18 | 0.11 |
| 6 | 136564395 | rs6908463 | -0.137 | -0.732 | 1.875 | 0.439 | 0.007 | 0.047 | 0.047 | 9.608 | 0.07 | 0.43 | 0.07 | 0.07 | 0.01 | 0.04 |
| 6 | 136571465 | rs797553 | -0.552 | -0.834 | 1.479 | 0.28 | -0.03 | -0.033 | -0.033 | 0 | 0.16 | 0.28 | 0.12 | 0.13 | 0.14 | 0.12 |
| 6 | 136572311 | rs797554 | -1.539 | -1.166 | 1.952 | 0.411 | -0.062 | -0.031 | -0.031 | 0 | 0.23 | 0.53 | 0.15 | 0.21 | 0.15 | 0.17 |
| 6 | 136595752 | rs811753 | -0.759 | -1.35 | 0.248 | -0.236 | -0.013 | 0.043 | 0.043 | 1.611 | 0.76 | 0.97 | 0.84 | 0.84 | 0.77 | 0.87 |
| 6 | 136645817 | rs6934308 | -0.55 | -1.49 | -0.123 | 0.854 | -0.022 | -0.128 | -0.128 | 0.653 | 0.7 | 0.96 | 0.68 | 0.6 | 0.71 | 0.82 |
| 6 | 136648627 | rs6931974 | -0.01 | -1.783 | 0.199 | 1.449 | -0.016 | 0.07 | 0.07 | 0 | 0.17 | 0.24 | 0.27 | 0.45 | 0.11 | 0.16 |
| 6 | 136650494 | rs2142921 | -0.105 | -1.652 | 0.745 | 1.154 | -0.001 | -0.001 | -0.001 | 0 | 0.86 | 0.83 | 0.89 | 0.85 | 0.85 | 0.87 |
| 6 | 136655206 | rs4896207 | -1.581 | -1.954 | 1.377 | 1.321 | -0.058 | 0.049 | 0.049 | 14.64 | 0.11 | 0.61 | 0.22 | 0.2 | 0.01 | 0.05 |
| 6 | 136663008 | rs2038082 | -0.13 | -1.874 | 0.93 | 1.186 | -0.003 | -0.003 | -0.003 | 0 | 0.86 | 0.83 | 0.9 | 0.85 | 0.85 | 0.87 |

| | | | | | | | | | | | | | | | | |
|---|---|---|---|---|---|---|---|---|---|---|---|---|---|---|---|---|
| 6 | 136689360 | rs2038077 | 0.721 | -2.067 | 1.104 | 1.414 | -0.037 | 0.054 | 0.054 | 0.102 | 0.78 | 0.61 | 0.76 | 0.6 | 0.84 | 0.84 |
| 6 | 136695212 | rs3778297 | -0.373 | -2.115 | 1.109 | 1.393 | -0.038 | 0.104 | 0.104 | 4.445 | 0.09 | 0.25 | 0.18 | 0.28 | 0.01 | 0.04 |
| 6 | 136743135 | rs2076193 | 0.499 | -2.606 | 0.534 | 0.859 | -0.014 | -0.001 | -0.001 | 1.156 | 0.62 | 0.56 | 0.65 | 0.57 | 0.71 | 0.8 |
| 6 | 136748317 | rs3823167 | 0.629 | -1.381 | 0.972 | 1.325 | -0.009 | -0.007 | -0.007 | 0 | 0.81 | 0.74 | 0.84 | 0.82 | 0.85 | 0.86 |
| 6 | 136751118 | rs997139 | 0.927 | -1.384 | 0.977 | 1.328 | -0.045 | 0.068 | 0.068 | 0 | 0.77 | 0.57 | 0.72 | 0.57 | 0.84 | 0.83 |
| 6 | 136753218 | rs3799419 | 0.603 | -0.868 | 0.893 | 1.839 | -0.191 | -0.067 | -0.067 | 0.266 | 0.28 | 0.03 | 0.35 | 0.26 | 0.41 | 0.25 |
| 6 | 136765459 | rs3778307 | 1.25 | -0.933 | 1.721 | 2.291 | -0.04 | 0.134 | 0.134 | 0 | 0.81 | 0.54 | 0.71 | 0.56 | 0.85 | 0.84 |
| 6 | 136774165 | rs4896210 | 0.186 | -0.876 | 1.417 | 2.197 | -0.001 | -0.001 | -0.001 | 0.26 | 0.87 | 0.84 | 0.9 | 0.85 | 0.85 | 0.88 |
| 6 | 136781280 | rs926370 | 0.228 | -0.86 | 1.445 | 2.318 | -0.006 | 0.005 | 0.005 | 0 | 0.87 | 0.81 | 0.89 | 0.81 | 0.85 | 0.88 |
| 6 | 136812315 | rs2181096 | 1.495 | -0.569 | 1.92 | 2.421 | -0.03 | 0.063 | 0.063 | 0.845 | 0.77 | 0.43 | 0.68 | 0.59 | 0.81 | 0.85 |
| 6 | 136817488 | rs9399183 | -1.929 | -0.883 | 2.008 | 2.468 | 0.024 | -0.02 | -0.02 | 0 | 0.24 | 0.82 | 0.33 | 0.41 | 0.16 | 0.15 |
| 6 | 136839188 | rs3799453 | -0.04 | -0.656 | 1.657 | 2.674 | 0.004 | 0.007 | 0.007 | 0 | 0.12 | 0.06 | 0.09 | 0.14 | 0.15 | 0.11 |
| 6 | 136839738 | rs3799458 | -2.126 | -0.56 | 1.827 | 2.723 | 0.035 | 0.105 | 0.105 | 0.633 | 0.07 | 0.37 | 0.12 | 0.12 | 0.01 | 0.02 |
| 6 | 136860731 | rs13191834 | 0.025 | -0.432 | 0.82 | 2.678 | -0.035 | -0.047 | -0.047 | 0 | 0.11 | 0.01 | 0.12 | 0.15 | 0.12 | 0.34 |
| 6 | 136877044 | rs28715589 | -1.256 | -0.393 | 1.485 | 2.816 | -0.018 | 0.064 | 0.064 | 0 | 0.17 | 0.31 | 0.26 | 0.42 | 0.14 | 0.13 |
| 6 | 136893813 | rs9389406 | 0.065 | -0.338 | 1.434 | 2.816 | -0.016 | -0.013 | -0.013 | 0 | 0.1 | 0.04 | 0.08 | 0.16 | 0.12 | 0.1 |
| 6 | 136898341 | rs9389407 | -0.546 | -0.352 | 1.494 | 2.86 | -0.011 | 0.007 | 0.007 | 0 | 0.13 | 0.21 | 0.11 | 0.18 | 0.12 | 0.1 |
| 6 | 136904235 | rs6570084 | -1.179 | -0.335 | 1.477 | 2.885 | -0.017 | 0.089 | 0.089 | 0 | 0.16 | 0.3 | 0.26 | 0.43 | 0.13 | 0.13 |
| 6 | 136907588 | rs9389409 | 0.044 | -0.342 | 1.461 | 2.894 | -0.002 | -0.031 | -0.031 | 0 | 0.1 | 0.05 | 0.16 | 0.33 | 0.13 | 0.11 |
| 6 | 136915573 | rs2282857 | -2.081 | -0.319 | 1.568 | 2.765 | 0.003 | 0.233 | 0.233 | 0 | 0.06 | 0.29 | 0.18 | 0.25 | 0.01 | 0.03 |
| 6 | 136918854 | rs17796516 | -1.783 | -0.29 | 1.513 | 2.945 | -0.003 | 0.274 | 0.274 | 0 | 0.06 | 0.22 | 0.19 | 0.3 | 0.01 | 0.03 |
| 6 | 136929935 | rs12164028 | -1.857 | -0.357 | 1.656 | 3.037 | 0 | 0.322 | 0.322 | 0 | 0.07 | 0.29 | 0.2 | 0.31 | 0.01 | 0.04 |
| 6 | 136934748 | rs9389413 | -1.284 | -0.369 | 1.628 | 2.949 | -0.009 | 0.059 | 0.059 | 0 | 0.06 | 0.17 | 0.09 | 0.09 | 0.01 | 0.02 |
| 6 | 136939818 | rs9321564 | -1.257 | -0.337 | 1.595 | 2.807 | -0.027 | 0.134 | 0.134 | 0 | 0.22 | 0.59 | 0.31 | 0.49 | 0.17 | 0.17 |
| 6 | 136952930 | rs2237268 | -0.521 | 0.552 | 0.45 | 1.955 | -0.069 | 0.04 | 0.04 | 0 | 0.68 | 0.87 | 0.71 | 0.76 | 0.61 | 0.78 |
| 6 | 136960278 | rs6917580 | -0.156 | 0.651 | 0.537 | 1.775 | 0.009 | 0.006 | 0.006 | 0 | 0.62 | 0.49 | 0.56 | 0.6 | 0.61 | 0.74 |
| 6 | 136963493 | rs9494547 | -1.477 | 0.133 | 0.985 | 1.995 | -0.011 | 0.162 | 0.162 | 0.762 | 0.12 | 0.57 | 0.14 | 0.24 | 0.04 | 0.08 |
| 6 | 136963890 | rs9376219 | 0.568 | 0.216 | 0.011 | 1.579 | 0.02 | 0.097 | 0.097 | 0 | 0.36 | 0.17 | 0.22 | 0.15 | 0.32 | 0.2 |
| 6 | 136966741 | rs2237270 | -1.041 | 0.566 | 0.697 | 2.308 | 0.01 | 0.128 | 0.128 | 0 | 0.17 | 0.33 | 0.25 | 0.39 | 0.17 | 0.26 |
| 6 | 136968541 | rs2237271 | -1.299 | 0.544 | 0.735 | 2.382 | 0.003 | 0.164 | 0.164 | 0 | 0.18 | 0.36 | 0.25 | 0.45 | 0.17 | 0.26 |
| 6 | 136973316 | rs9402838 | -0.944 | 0.673 | 0.316 | 2.106 | -0.153 | -0.005 | -0.005 | 0.149 | 0.66 | 0.97 | 0.73 | 0.73 | 0.59 | 0.83 |
| 6 | 136974283 | rs6904753 | -1.718 | 0.623 | 1.382 | 2.628 | -0.001 | 0.149 | 0.149 | 0.019 | 0.19 | 0.4 | 0.27 | 0.55 | 0.17 | 0.3 |
| 6 | 136974623 | rs17796708 | -0.013 | 0.823 | 0.701 | 2.359 | -0.021 | -0.034 | -0.034 | 0.093 | 0.08 | 0.01 | 0.12 | 0.21 | 0.13 | 0.19 |
| 6 | 136980798 | rs6914406 | -0.859 | 0.704 | 0.822 | 2.35 | -0.002 | 0.253 | 0.253 | 0 | 0.07 | 0.27 | 0.14 | 0.29 | 0.02 | 0.08 |
| 6 | 136985219 | rs7773600 | -0.752 | 0.944 | 0.1 | 1.696 | 0.043 | 0.259 | 0.259 | 0 | 0.17 | 0.46 | 0.29 | 0.49 | 0.18 | 0.31 |
| 6 | 136989035 | rs13195420 | 0.056 | 0.642 | -0.943 | 0.627 | 0.018 | 0.126 | 0.126 | 0.002 | 0.42 | 0.17 | 0.38 | 0.17 | 0.41 | 0.21 |
| 6 | 136990283 | rs3765259 | -0.443 | 0.471 | -1.001 | 0.678 | -0.016 | -0.009 | -0.009 | 0 | 0.45 | 0.28 | 0.58 | 0.46 | 0.49 | 0.54 |
| 6 | 136992379 | rs9376221 | -0.477 | 0.313 | 0.261 | 0.723 | -0.021 | -0.007 | -0.007 | 1.06 | 0.16 | 0.31 | 0.14 | 0.14 | 0.1 | 0.06 |
| 6 | 136995210 | rs9402839 | -1.282 | -0.066 | 0.736 | 0.597 | -0.051 | -0.033 | -0.033 | 0.486 | 0.23 | 0.47 | 0.19 | 0.17 | 0.14 | 0.09 |
| 6 | 136997619 | rs4363056 | -1.27 | -0.017 | -0.254 | 0.568 | -0.109 | -0.048 | -0.048 | 0.638 | 0.65 | 0.96 | 0.68 | 0.68 | 0.62 | 0.72 |
| 6 | 137001106 | rs9494555 | -0.717 | 0.088 | 0.644 | 0.588 | -0.04 | -0.02 | -0.02 | 0.426 | 0.18 | 0.41 | 0.15 | 0.14 | 0.1 | 0.06 |
| 6 | 137017749 | rs1009709 | -0.416 | 0.852 | 1.166 | 0.809 | -0.003 | 0.007 | 0.007 | 0 | 0.53 | 0.58 | 0.65 | 0.58 | 0.49 | 0.67 |
| 6 | 137021524 | rs11154880 | -0.235 | 0.807 | 1.35 | 0.872 | 0 | 0.001 | 0.001 | 0 | 0.49 | 0.43 | 0.63 | 0.54 | 0.49 | 0.66 |
| 6 | 137022722 | rs4243449 | -0.112 | 0.569 | 1.49 | 0.856 | 0.006 | -0.001 | -0.001 | 0 | 0.48 | 0.3 | 0.6 | 0.53 | 0.47 | 0.65 |
| 6 | 137026266 | rs9321567 | -0.547 | 0.658 | 1.354 | 0.809 | -0.017 | 0.029 | 0.029 | 0 | 0.54 | 0.8 | 0.65 | 0.58 | 0.47 | 0.67 |
| 6 | 137034464 | rs911182 | 0.438 | 0.674 | 1.364 | 0.797 | -0.015 | 0.031 | 0.031 | 0 | 0.46 | 0.2 | 0.35 | 0.42 | 0.53 | 0.34 |
| 6 | 137042403 | rs9389429 | 0.455 | 0.427 | 1.318 | 0.83 | -0.036 | 0.014 | 0.014 | 0 | 0.45 | 0.16 | 0.34 | 0.43 | 0.49 | 0.39 |
| 6 | 137049452 | rs7740264 | -1.507 | 0.534 | 1.433 | 0.754 | 0.052 | 0.048 | 0.048 | 2.34 | 0.05 | 0.54 | 0.05 | 0 | 0 | 0 |
| 6 | 137056229 | rs6911217 | 0.086 | 0.517 | 1.38 | 0.764 | -0.001 | 0.006 | 0.006 | 0 | 0.51 | 0.62 | 0.38 | 0.46 | 0.49 | 0.41 |
| 6 | 137071655 | rs6570098 | 0.001 | 0.657 | 0.512 | 0.793 | -0.004 | 0.018 | 0.018 | 0 | 0.52 | 0.65 | 0.37 | 0.43 | 0.49 | 0.41 |
| 6 | 137079542 | rs4896219 | 0.232 | 0.744 | 0.29 | 0.677 | -0.02 | 0 | 0 | 0 | 0.49 | 0.13 | 0.36 | 0.43 | 0.52 | 0.43 |
| 6 | 137083138 | rs932589 | 0.111 | 0.635 | 1.255 | 0.707 | 0.032 | 0.014 | 0.014 | 0 | 0.51 | 0.61 | 0.35 | 0.43 | 0.43 | 0.44 |
| 6 | 137101726 | rs1570054 | -0.526 | 0.694 | 1.868 | 0.785 | 0.028 | 0.02 | 0.02 | 0 | 0.53 | 0.65 | 0.68 | 0.6 | 0.6 | 0.61 |
| 6 | 137108757 | rs6912539 | -0.66 | 0.684 | 1.821 | 0.794 | 0.049 | -0.004 | -0.004 | 0 | 0.55 | 0.78 | 0.69 | 0.57 | 0.6 | 0.55 |
| 6 | 137135004 | rs9494573 | -0.19 | 0.718 | 2.319 | 0.954 | 0.01 | 0.003 | 0.003 | 0 | 0.47 | 0.37 | 0.63 | 0.55 | 0.56 | 0.6 |
| 6 | 137140454 | rs6919857 | -0.556 | 0.734 | 2.21 | 0.984 | 0.063 | -0.029 | -0.029 | 0.013 | 0.52 | 0.78 | 0.69 | 0.52 | 0.6 | 0.55 |
| 6 | 137142946 | rs3757169 | -0.179 | 0.737 | 2.554 | 0.96 | 0.018 | -0.008 | -0.008 | 0 | 0.47 | 0.38 | 0.64 | 0.52 | 0.56 | 0.55 |
| 6 | 137150543 | rs7764028 | -0.524 | 0.732 | 2.347 | 0.951 | 0.043 | 0.013 | 0.013 | 0 | 0.52 | 0.7 | 0.68 | 0.55 | 0.6 | 0.61 |
| 6 | 137161671 | rs7740484 | -0.644 | 0.738 | 2.153 | 0.868 | 0.046 | 0.006 | 0.006 | 0 | 0.54 | 0.7 | 0.64 | 0.56 | 0.61 | 0.62 |
| 6 | 137179387 | rs2012700 | -0.639 | 0.73 | 2.15 | 0.874 | 0.046 | 0.004 | 0.004 | 0 | 0.54 | 0.71 | 0.64 | 0.55 | 0.61 | 0.62 |
| 6 | 137205972 | rs1342643 | 0.01 | 1.24 | 1.915 | 1.021 | 0.001 | 0 | 0 | 0 | 0.05 | 0.05 | 0.15 | 0.05 | 0.1 | 0.05 |
| 6 | 137207304 | rs1342645 | -0.015 | 1.229 | 2.383 | 1.021 | -0.033 | -0.001 | -0.001 | 0.523 | 0.43 | 0.23 | 0.43 | 0.47 | 0.47 | 0.52 |
| 6 | 137209591 | rs2327766 | 0.636 | 1.326 | 2.051 | 1.152 | -0.019 | -0.042 | -0.042 | 0.018 | 0.13 | 0.01 | 0.14 | 0.22 | 0.14 | 0.24 |
| 6 | 137222671 | rs7757102 | -0.583 | 1.248 | 2.055 | 0.978 | 0.021 | 0.001 | 0.001 | 0 | 0.51 | 0.59 | 0.52 | 0.51 | 0.58 | 0.6 |
| 6 | 137231671 | rs9389449 | -0.562 | 1.165 | 1.863 | 1.018 | 0.005 | -0.001 | -0.001 | 0 | 0.5 | 0.47 | 0.49 | 0.52 | 0.55 | 0.59 |
| 6 | 137234970 | rs1050803 | -0.511 | 1.157 | 1.859 | 1.014 | 0.004 | -0.001 | -0.001 | 0 | 0.49 | 0.47 | 0.49 | 0.52 | 0.55 | 0.59 |
| 6 | 137241328 | rs9494600 | -0.501 | 1.149 | 1.497 | 1.047 | 0.05 | 0.016 | 0.016 | 0 | 0.51 | 0.3 | 0.61 | 0.55 | 0.61 | 0.59 |
| 6 | 137253162 | rs6570115 | -0.64 | 0.711 | 1.069 | 0.906 | 0.008 | 0.013 | 0.013 | 0 | 0.53 | 0.64 | 0.51 | 0.61 | 0.57 | 0.59 |
| 6 | 137256419 | rs6570116 | -0.719 | 0.567 | 0.441 | 0.804 | -0.012 | -0.026 | -0.026 | 0 | 0.61 | 0.87 | 0.6 | 0.61 | 0.61 | 0.7 |

| | | | | | | | | | | | | | | | | |
|---|---|---|---|---|---|---|---|---|---|---|---|---|---|---|---|---|
| 6 | 137259725 | rs12191772 | 0.388 | 0.463 | 0.258 | 0.504 | -0.033 | -0.056 | -0.056 | 0 | 0.14 | 0.01 | 0.12 | 0.25 | 0.19 | 0.2 |
| 6 | 137263940 | rs6570118 | -0.916 | 0.374 | 0.856 | 0.509 | -0.025 | -0.059 | -0.059 | 0 | 0.64 | 0.83 | 0.58 | 0.53 | 0.58 | 0.71 |
| 6 | 137264165 | rs7750197 | 0.216 | 0.096 | 1.027 | 0.378 | -0.008 | 0.003 | 0.003 | 0 | 0.42 | 0.39 | 0.42 | 0.38 | 0.5 | 0.37 |
| 6 | 137266341 | rs1891486 | -1.152 | 0.166 | 0.227 | 0.296 | -0.053 | -0.097 | -0.097 | 0.248 | 0.65 | 0.9 | 0.58 | 0.53 | 0.58 | 0.71 |
| 6 | 137268106 | rs7743308 | 0.093 | 0.221 | 0.254 | 0.42 | -0.011 | -0.001 | -0.001 | 0.093 | 0.42 | 0.45 | 0.43 | 0.44 | 0.5 | 0.38 |
| 6 | 137271663 | rs6928748 | -0.833 | -0.086 | 0.172 | 0.169 | -0.042 | -0.044 | -0.044 | 0 | 0.19 | 0.24 | 0.14 | 0.09 | 0.07 | 0.29 |
| 6 | 137282117 | rs4895471 | 0.434 | 0.505 | -0.701 | 0.087 | 0.014 | 0.053 | 0.053 | 0.535 | 0.57 | 0.34 | 0.69 | 0.65 | 0.61 | 0.53 |
| 6 | 137288483 | rs1935743 | 0.179 | 0.472 | -0.472 | -0.123 | -0.009 | -0.027 | -0.027 | 0 | 0.69 | 0.65 | 0.79 | 0.81 | 0.73 | 0.61 |
| 6 | 137290613 | rs6936698 | 0.726 | 0.671 | -1.489 | 0.199 | -0.011 | -0.011 | -0.011 | 1.301 | 0.26 | 0.06 | 0.29 | 0.18 | 0.19 | 0.18 |
| 6 | 137294656 | rs10872465 | 0.683 | 0.996 | -1 | 0.192 | -0.012 | -0.002 | -0.002 | 1.296 | 0.53 | 0.25 | 0.49 | 0.43 | 0.49 | 0.46 |
| 6 | 137295402 | rs2153893 | -0.561 | 1.233 | -0.212 | 0.716 | 0.027 | 0.111 | 0.111 | 0.38 | 0.18 | 0.3 | 0.3 | 0.39 | 0.25 | 0.29 |
| 6 | 137306563 | rs6935119 | 0.622 | 1.876 | -1.062 | 0.235 | -0.052 | 0.008 | 0.008 | 0 | 0.78 | 0.86 | 0.81 | 0.85 | 0.68 | 0.82 |
| 6 | 137307421 | rs9385780 | 0.473 | 1.141 | -2.122 | -0.378 | -0.026 | 0.123 | 0.123 | 2.734 | 0.45 | 0.08 | 0.5 | 0.8 | 0.46 | 0.52 |
| 6 | 137316830 | rs10484795 | 0.388 | 0.916 | -2.697 | -0.585 | -0.073 | 0.074 | 0.074 | 0.309 | 0.2 | 0.02 | 0.18 | 0.12 | 0.25 | 0.18 |
| 6 | 137323213 | rs1342642 | 0.414 | 0.893 | -1.566 | -1.41 | 0.006 | -0.022 | -0.022 | 0 | 0.25 | 0.24 | 0.18 | 0.07 | 0.26 | 0.23 |
| 6 | 137324655 | rs1744061 | 1.225 | 0.56 | -1.883 | -1.592 | -0.074 | 0.008 | 0.008 | 0 | 0.21 | 0.01 | 0.23 | 0.1 | 0.27 | 0.12 |
| 6 | 137326103 | rs1626680 | -0.357 | 0.473 | -1.263 | -1.617 | -0.003 | 0.036 | 0.036 | 1.865 | 0.69 | 0.28 | 0.77 | 0.93 | 0.73 | 0.77 |
| 6 | 137333718 | rs2010614 | -0.27 | 0.592 | -0.83 | -1.461 | 0.022 | 0.008 | 0.008 | 2.55 | 0.93 | 0.45 | 0.94 | 1 | 0.99 | 1 |
| 6 | 137333851 | rs9402865 | 0.647 | 0.576 | -1.64 | -1.498 | -0.046 | 0.092 | 0.092 | 0.306 | 0.42 | 0.08 | 0.5 | 0.8 | 0.44 | 0.49 |
| 6 | 137341012 | rs11755272 | 0.803 | 0.877 | -1.641 | -1.481 | -0.114 | 0.075 | 0.075 | 0.847 | 0.23 | 0.02 | 0.22 | 0.14 | 0.32 | 0.18 |
| 6 | 137343720 | rs1167846 | 0.031 | -0.224 | -1.331 | -1.519 | 0.027 | 0.099 | 0.099 | 2.344 | 0.67 | 0.16 | 0.76 | 0.93 | 0.76 | 0.78 |
| 6 | 137349025 | rs1167849 | 0.519 | -0.12 | -1.586 | -1.422 | 0.025 | 0.185 | 0.185 | 1.648 | 0.61 | 0.08 | 0.73 | 0.9 | 0.73 | 0.56 |
| 6 | 137352519 | rs276504 | -0.635 | -0.601 | -1.737 | -0.65 | 0.048 | -0.037 | -0.037 | 0 | 0.53 | 0.84 | 0.42 | 0.45 | 0.35 | 0.63 |
| 6 | 137361292 | rs9376243 | 1.151 | -0.596 | -1.776 | -0.566 | 0.037 | -0.1 | -0.1 | 3.578 | 0.13 | 0 | 0.13 | 0.34 | 0.06 | 0.18 |
| 6 | 137361670 | rs276511 | 0.348 | -0.867 | -1.39 | -0.929 | -0.001 | -0.007 | -0.007 | 0.105 | 0.27 | 0.3 | 0.17 | 0.08 | 0.25 | 0.37 |
| 6 | 137363835 | rs3799487 | 0.965 | -0.286 | -1.887 | -0.693 | -0.138 | -0.12 | -0.12 | 4.453 | 0.34 | 0.03 | 0.42 | 0.41 | 0.45 | 0.24 |
| 6 | 137378863 | rs12214209 | 0.863 | -0.314 | -1.532 | -0.738 | 0.015 | -0.018 | -0.018 | 0.302 | 0.14 | 0.11 | 0.1 | 0.23 | 0.06 | 0.14 |
| 6 | 137379548 | rs276583 | -0.498 | 0.117 | -1.456 | -0.699 | -0.044 | 0.022 | 0.022 | 0.114 | 0.59 | 0.82 | 0.44 | 0.33 | 0.48 | 0.68 |
| 6 | 137401777 | rs276550 | -0.533 | -0.161 | -1.339 | -0.641 | -0.049 | 0.018 | 0.018 | 0.482 | 0.6 | 0.84 | 0.44 | 0.33 | 0.49 | 0.68 |
| 6 | 137408632 | rs9376250 | -0.006 | -0.646 | -0.5 | -0.245 | -0.021 | 0.016 | 0.016 | 0 | 0.17 | 0.21 | 0.12 | 0.21 | 0.09 | 0.36 |
| 6 | 137415146 | rs276555 | 0.799 | -0.705 | -1.285 | -0.656 | -0.078 | -0.097 | -0.097 | 0.306 | 0.37 | 0.12 | 0.36 | 0.52 | 0.55 | 0.35 |
| 6 | 137422058 | rs276567 | -0.149 | -0.994 | -0.559 | -0.682 | -0.026 | 0.007 | 0.007 | 0 | 0.79 | 0.72 | 0.83 | 0.81 | 0.87 | 0.79 |
| 6 | 137422342 | rs9402870 | 0.131 | -1.732 | -0.975 | -1.258 | 0.06 | -0.001 | -0.001 | 1.711 | 0.38 | 0.12 | 0.33 | 0.33 | 0.31 | 0.35 |
| 6 | 137422752 | rs276568 | 0.958 | -1.744 | -0.54 | -1.259 | 0.079 | 0.059 | 0.059 | 0 | 0.51 | 0.37 | 0.59 | 0.67 | 0.65 | 0.62 |
| 6 | 137423779 | rs17066063 | 0.703 | -1.582 | -0.32 | -1.177 | -0.005 | -0.012 | -0.012 | 0.022 | 0.26 | 0.27 | 0.23 | 0.14 | 0.23 | 0.16 |
| 6 | 137430021 | rs4896232 | -0.733 | -1.562 | -0.358 | -1.171 | 0.043 | 0.029 | 0.029 | 0 | 0.46 | 0.57 | 0.39 | 0.34 | 0.34 | 0.37 |
| 6 | 137433986 | rs276573 | 1.364 | -1.547 | -0.485 | -1.264 | -0.04 | -0.042 | -0.042 | 0 | 0.34 | 0.26 | 0.38 | 0.54 | 0.53 | 0.51 |
| 6 | 137439066 | rs9376258 | 1.332 | -1.547 | -0.717 | -1.575 | -0.003 | -0.079 | -0.079 | 0 | 0.13 | 0 | 0.1 | 0.24 | 0.08 | 0.16 |
| 6 | 137443682 | rs10484797 | 1.369 | -1.346 | -0.817 | -1.702 | -0.004 | 0.025 | 0.025 | 0 | 0.18 | 0.07 | 0.22 | 0.12 | 0.12 | 0.12 |
| 6 | 137444054 | rs4895477 | 0.615 | -1.028 | -0.362 | -1.556 | -0.034 | 0.036 | 0.036 | 0 | 0.2 | 0.31 | 0.15 | 0.24 | 0.11 | 0.18 |
| 6 | 137452537 | rs6912319 | -1.21 | -0.821 | -0.463 | -1.436 | 0.009 | 0.01 | 0.01 | 0 | 0.65 | 0.57 | 0.6 | 0.5 | 0.46 | 0.65 |
| 6 | 137454459 | rs72761 | -0.733 | -0.081 | -0.501 | -1.567 | 0 | -0.004 | -0.004 | 0 | 0.58 | 0.62 | 0.59 | 0.34 | 0.53 | 0.6 |
| 6 | 137464218 | rs276467 | -0.097 | 0.549 | -0.76 | -1.23 | -0.027 | 0.038 | 0.038 | 0 | 0.23 | 0.1 | 0.11 | 0.18 | 0.23 | 0.26 |
| 6 | 137464916 | rs202565 | -0.44 | 0.271 | -0.216 | -0.707 | -0.034 | 0.074 | 0.074 | 0 | 0.88 | 0.98 | 0.8 | 1 | 0.83 | 0.93 |
| 6 | 137465074 | rs202566 | -0.44 | 0.271 | -0.216 | -0.696 | -0.034 | 0.068 | 0.068 | 0 | 0.88 | 0.98 | 0.8 | 1 | 0.83 | 0.93 |
| 6 | 137466087 | rs28366 | -0.323 | 0.319 | -0.664 | -0.713 | -0.034 | 0.079 | 0.079 | 0 | 0.87 | 0.98 | 0.79 | 1 | 0.82 | 0.92 |
| 6 | 137470756 | rs10484798 | -0.082 | 0.753 | 0.082 | 0.202 | 0.011 | 0.09 | 0.09 | 0 | 0.17 | 0.27 | 0.27 | 0.4 | 0.21 | 0.12 |
| 6 | 137475858 | rs7774349 | -1.434 | 0.618 | -0.167 | -0.13 | -0.059 | 0.105 | 0.105 | 1.177 | 0.58 | 0.91 | 0.49 | 0.73 | 0.48 | 0.66 |
| 6 | 137478424 | rs9494674 | -0.869 | 0.815 | 0.407 | 0.219 | 0.022 | -0.028 | -0.028 | 0 | 0.07 | 0.27 | 0.09 | 0 | 0.07 | 0.04 |
| 6 | 137482982 | rs11154915 | 0.207 | 0.718 | 0.324 | 0.389 | 0.022 | 0.036 | 0.036 | 0 | 0.19 | 0.07 | 0.09 | 0.15 | 0.17 | 0.12 |
| 6 | 137483258 | rs1040622 | 0.177 | 0.714 | 0.332 | 0.387 | 0.023 | 0.035 | 0.035 | 0 | 0.2 | 0.07 | 0.09 | 0.15 | 0.17 | 0.12 |
| 6 | 137495707 | rs742931 | -1.156 | 0.848 | 0.503 | 0.556 | 0.028 | 0.035 | 0.035 | 1.515 | 0.45 | 0.76 | 0.48 | 0.61 | 0.41 | 0.56 |
| 6 | 137500786 | rs7749054 | 0.445 | 0.926 | -0.294 | 0.589 | 0.016 | 0.13 | 0.13 | 0 | 0.15 | 0.27 | 0.26 | 0.4 | 0.17 | 0.11 |
| 6 | 137501535 | rs1891596 | -0.786 | 1.044 | -1.415 | 0.521 | 0.035 | 0.051 | 0.051 | 1.259 | 0.46 | 0.84 | 0.48 | 0.61 | 0.41 | 0.56 |
| 6 | 137515233 | rs12215787 | -0.257 | 0.886 | -0.921 | 0.425 | -0.009 | -0.02 | -0.02 | 0 | 0.24 | 0.32 | 0.17 | 0.19 | 0.21 | 0.36 |
| 6 | 137519780 | rs3799488 | 0.278 | 0.822 | -1.247 | 0.676 | -0.046 | -0.058 | -0.058 | 0 | 0.09 | 0.01 | 0.14 | 0.26 | 0.13 | 0.11 |
| 6 | 137531031 | rs9376267 | -0.257 | 0.695 | -1.266 | 0.367 | -0.026 | -0.068 | -0.068 | 0.734 | 0.25 | 0.14 | 0.24 | 0.46 | 0.26 | 0.56 |
| 6 | 137532751 | rs9376268 | 0.278 | 0.726 | -1.412 | 0.379 | -0.059 | -0.094 | -0.094 | 0 | 0.21 | 0.07 | 0.21 | 0.4 | 0.25 | 0.25 |
| 6 | 137539810 | rs10457655 | 0.13 | 0.704 | -1.112 | 0.279 | -0.027 | -0.04 | -0.04 | 0 | 0.23 | 0.29 | 0.14 | 0.11 | 0.15 | 0.08 |
| 6 | 137540370 | rs7749390 | -0.423 | 1.005 | -0.971 | 0.444 | 0.003 | 0.001 | 0.001 | 0 | 0.5 | 0.5 | 0.36 | 0.59 | 0.39 | 0.63 |
| 6 | 137541075 | rs1327474 | -0.51 | 0.821 | -2.198 | -0.412 | -0.097 | 0.197 | 0.197 | 0 | 0.65 | 0.96 | 0.67 | 0.92 | 0.54 | 0.79 |
| 6 | 137546106 | rs12529779 | -0.478 | 0.496 | -1.056 | 0.81 | -0.001 | -0.004 | -0.004 | 0 | 0.2 | 0.05 | 0.18 | 0.18 | 0.2 | 0.1 |
| 6 | 137547038 | rs608914 | 0.432 | 0.796 | -0.05 | 1.264 | 0.056 | 0.07 | 0.07 | 1.419 | 0.77 | 0.58 | 0.7 | 0.35 | 0.71 | 0.6 |
| 6 | 137551037 | rs641029 | 0.248 | -0.161 | 0.08 | 1.374 | -0.003 | -0.021 | -0.021 | 0.417 | 0.88 | 0.95 | 0.87 | 0.76 | 0.84 | 0.84 |
| 6 | 137578168 | rs2807899 | 0.632 | -1.218 | 0.114 | 1.453 | 0.037 | -0.008 | -0.008 | 0.078 | 0.58 | 0.68 | 0.62 | 0.53 | 0.72 | 0.51 |
| 6 | 137580222 | rs1539495 | 0.173 | -1.268 | 0.082 | 1.451 | 0.035 | 0 | 0 | 0 | 0.26 | 0.19 | 0.24 | 0.21 | 0.11 | 0.15 |
| 6 | 137585425 | rs2807893 | 1.081 | -1.434 | 0.312 | 1.527 | 0.022 | 0.039 | 0.039 | 0 | 0.48 | 0.58 | 0.48 | 0.22 | 0.61 | 0.52 |
| 6 | 137586045 | rs12176587 | -0.734 | -1.432 | 0.312 | 1.519 | -0.001 | -0.002 | -0.002 | 0 | 0.39 | 0.4 | 0.39 | 0.54 | 0.23 | 0.33 |
| 6 | 137588858 | rs2797679 | 1.056 | -1.319 | 0.378 | 1.296 | 0.006 | 0.006 | 0.006 | 0 | 0.47 | 0.54 | 0.48 | 0.2 | 0.6 | 0.33 |

| | | | | | | | | | | | | | | | | |
|---|---|---|---|---|---|---|---|---|---|---|---|---|---|---|---|---|
| 6 | 137593756 | rs6906644 | -1 | -1.346 | 0.63 | 1.223 | -0.005 | 0.047 | 0.047 | 0.322 | 0.2 | 0.42 | 0.25 | 0.35 | 0.18 | 0.2 |
| 6 | 137603442 | rs4896251 | -0.292 | -1.227 | 0.237 | 1.213 | -0.008 | 0 | 0 | 0 | 0.26 | 0.26 | 0.23 | 0.23 | 0.11 | 0.16 |
| 6 | 137605627 | rs9321590 | -0.635 | -0.918 | 0.242 | 0.866 | -0.008 | 0 | 0 | 0.847 | 0.43 | 0.53 | 0.48 | 0.58 | 0.27 | 0.35 |
| 6 | 137609488 | rs4896253 | -1.064 | -0.515 | 0.077 | 0.574 | -0.01 | 0.015 | 0.015 | 0 | 0.09 | 0 | 0.09 | 0.05 | 0.07 | 0.19 |
| 6 | 137610097 | rs9321591 | -1.436 | -0.578 | 0.618 | 0.805 | -0.018 | 0.061 | 0.061 | 0 | 0.14 | 0.28 | 0.18 | 0.28 | 0.09 | 0.21 |
| 6 | 137617136 | rs1760084 | 0.622 | -0.36 | 0.715 | 0.226 | -0.006 | -0.016 | -0.016 | 0 | 0.84 | 0.6 | 0.83 | 0.9 | 0.84 | 0.89 |
| 6 | 137617284 | rs17066290 | -0.767 | -0.382 | 0.873 | -0.285 | -0.002 | -0.045 | -0.045 | 0 | 0.11 | 0.17 | 0.15 | 0.03 | 0.11 | 0.1 |
| 6 | 137617340 | rs9402884 | 1.466 | -0.37 | 0.55 | -0.693 | 0.005 | 0.015 | 0.015 | 0 | 0.59 | 0.31 | 0.61 | 0.72 | 0.61 | 0.67 |
| 6 | 137619278 | rs963386 | -1.366 | -0.502 | 0.29 | -1.01 | -0.006 | -0.094 | -0.094 | 0 | 0.29 | 0.48 | 0.25 | 0.05 | 0.3 | 0.13 |
| 6 | 137625920 | rs9484011 | 0.27 | -0.469 | 0.256 | -0.922 | -0.001 | 0 | 0 | 2.826 | 0.11 | 0.1 | 0.12 | 0.05 | 0.13 | 0.05 |
| 6 | 137631222 | rs7747814 | 0.107 | -0.924 | -0.793 | -2.056 | -0.036 | 0.2 | 0.2 | 2.524 | 0.53 | 0.83 | 0.62 | 0.87 | 0.52 | 0.74 |
| 6 | 137635497 | rs3850205 | -0.447 | -0.933 | -0.844 | -2.057 | -0.067 | 0.17 | 0.17 | 8.101 | 0.64 | 0.93 | 0.74 | 0.92 | 0.64 | 0.79 |
| 6 | 137640173 | rs6909860 | 0.343 | -0.956 | -0.83 | -2.106 | -0.042 | 0.16 | 0.16 | 5.601 | 0.34 | 0.1 | 0.26 | 0.02 | 0.33 | 0.18 |
| 6 | 137641316 | rs6570146 | 0.03 | -1.08 | -0.38 | -2.083 | 0 | -0.006 | -0.006 | 0 | 0.57 | 0.6 | 0.61 | 0.89 | 0.6 | 0.74 |
| 6 | 137650094 | rs1145965 | 1.462 | -0.593 | -1.089 | -1.835 | -0.1 | -0.131 | -0.131 | 0 | 0.19 | 0.01 | 0.25 | 0.52 | 0.28 | 0.32 |
| 6 | 137656235 | rs4896258 | -0.757 | -0.713 | -0.413 | -1.696 | -0.026 | -0.01 | -0.01 | 0 | 0.88 | 0.97 | 0.88 | 0.91 | 0.85 | 0.92 |
| 6 | 137664402 | rs3914898 | 0.264 | -0.729 | -0.325 | -1.706 | -0.008 | -0.013 | -0.013 | 0 | 0.81 | 0.7 | 0.76 | 0.86 | 0.81 | 0.87 |
| 6 | 137664631 | rs6570147 | 0.415 | -0.595 | -0.015 | -1.713 | -0.001 | 0 | 0 | 0 | 0.38 | 0.4 | 0.45 | 0.69 | 0.42 | 0.64 |
| 6 | 137666088 | rs4895484 | 0.401 | -0.44 | -0.809 | -1.575 | -0.008 | -0.003 | -0.003 | 0.058 | 0.05 | 0.15 | 0.12 | 0.05 | 0.05 | 0.05 |
| 6 | 137670808 | rs9321593 | 0.43 | -0.308 | -1.438 | -1.549 | -0.006 | 0.115 | 0.115 | 3.863 | 0.4 | 0.19 | 0.29 | 0.02 | 0.37 | 0.21 |
| 6 | 137674040 | rs1022400 | -0.177 | 0.534 | -1.385 | -0.682 | -0.081 | 0.039 | 0.039 | 5.718 | 0.35 | 0.14 | 0.4 | 0.7 | 0.42 | 0.61 |
| 6 | 137681617 | rs13208135 | -0.926 | 0.824 | -0.85 | -0.713 | -0.001 | -0.023 | -0.023 | 0 | 0.14 | 0.12 | 0.09 | 0.02 | 0.15 | 0.24 |
| 6 | 137682677 | rs958791 | -0.21 | 0.402 | -1.375 | -0.623 | 0.01 | -0.009 | -0.009 | 0 | 0.29 | 0.22 | 0.44 | 0.67 | 0.26 | 0.26 |
| 6 | 137692864 | rs1591208 | -0.08 | 0.476 | -1.289 | 1.005 | 0.003 | -0.016 | -0.016 | 0 | 0.26 | 0.2 | 0.27 | 0.41 | 0.25 | 0.26 |
| 6 | 137693167 | rs9376273 | -0.574 | 0.633 | -1.515 | 0.529 | -0.036 | -0.062 | -0.062 | 1.123 | 0.29 | 0.16 | 0.37 | 0.41 | 0.36 | 0.29 |
| 6 | 137695321 | rs12662095 | 0.43 | 0.722 | -1.43 | 0.617 | 0.004 | -0.016 | -0.016 | 0 | 0.26 | 0.2 | 0.27 | 0.41 | 0.25 | 0.25 |
| 6 | 137698614 | rs4896263 | -0.477 | 0.603 | -0.843 | 0.74 | -0.001 | 0.002 | 0.002 | 0 | 0.25 | 0.25 | 0.33 | 0.45 | 0.34 | 0.28 |
| 6 | 137710766 | rs12526230 | 0.047 | 0.813 | -0.79 | 0.68 | 0.013 | -0.048 | -0.048 | 0 | 0.14 | 0.19 | 0.16 | 0.02 | 0.16 | 0.22 |
| 6 | 137712235 | rs9389496 | 0.419 | 0.794 | -0.796 | 0.974 | -0.002 | 0.047 | 0.047 | 0 | 0.11 | 0.14 | 0.18 | 0.45 | 0.18 | 0.05 |
| 6 | 137712485 | rs9321598 | -1.164 | 0.692 | -0.545 | 0.511 | 0.002 | -0.003 | -0.003 | 0 | 0.51 | 0.53 | 0.7 | 0.83 | 0.62 | 0.68 |
| 6 | 137724238 | rs9389498 | 0.661 | 0.805 | -0.525 | 0.07 | 0.006 | 0.007 | 0.007 | 0 | 0.12 | 0.2 | 0.1 | 0.18 | 0.18 | 0.04 |
| 6 | 137731441 | rs9321604 | -1.3 | 0.655 | 0.182 | 0.002 | 0.009 | -0.005 | -0.005 | 0 | 0.56 | 0.67 | 0.66 | 0.54 | 0.64 | 0.72 |
| 6 | 137735758 | rs737467 | 0.758 | 0.75 | -0.109 | 0.028 | -0.007 | -0.01 | -0.01 | 0.189 | 0.2 | 0.09 | 0.24 | 0.3 | 0.22 | 0.23 |
| 6 | 137739024 | rs13204631 | 0.457 | 0.708 | 0.014 | 0.044 | 0.004 | 0.002 | 0.002 | 0 | 0.13 | 0.2 | 0.11 | 0.17 | 0.2 | 0.06 |
| 6 | 137742167 | rs7756465 | -0.707 | 0.313 | 0.052 | -0.264 | -0.005 | -0.055 | -0.055 | 0 | 0.27 | 0.57 | 0.34 | 0.07 | 0.22 | 0.44 |
| 6 | 137743424 | rs1022501 | -0.424 | 0.268 | 0.407 | -0.305 | 0.069 | -0.084 | -0.084 | 0 | 0.16 | 0.38 | 0.3 | 0.03 | 0.2 | 0.29 |
| 6 | 137745877 | rs12212451 | -0.318 | 0.09 | 0.097 | -0.341 | 0.009 | 0.008 | 0.008 | 0.045 | 0.25 | 0.21 | 0.33 | 0.25 | 0.19 | 0.32 |
| 6 | 137746446 | rs11756814 | 0.256 | 0.14 | -0.083 | -0.389 | 0.01 | 0.001 | 0.001 | 0 | 0.21 | 0.26 | 0.18 | 0.23 | 0.31 | 0.12 |
| 6 | 137746585 | rs6570156 | -1.049 | 0.097 | -0.072 | -0.414 | 0.014 | 0.035 | 0.035 | 1.602 | 0.39 | 0.68 | 0.53 | 0.32 | 0.34 | 0.59 |
| 6 | 137747699 | rs1333623 | -1.097 | 0.114 | -0.138 | -0.504 | 0.036 | 0.157 | 0.157 | 0.411 | 0.47 | 0.74 | 0.55 | 0.2 | 0.42 | 0.54 |
| 6 | 137747828 | rs9402895 | 0.077 | -0.476 | -0.713 | -1.137 | -0.004 | 0.003 | 0.003 | 0 | 0.3 | 0.23 | 0.21 | 0.28 | 0.31 | 0.26 |
| 6 | 137748849 | rs6929588 | -0.897 | -0.396 | -0.572 | -1.043 | 0.044 | 0.202 | 0.202 | 1.176 | 0.49 | 0.83 | 0.55 | 0.25 | 0.43 | 0.62 |
| 6 | 137749301 | rs12527578 | 1.048 | -0.392 | -0.608 | -1.045 | 0.004 | -0.006 | -0.006 | 0.449 | 0.09 | 0.11 | 0.14 | 0.02 | 0.14 | 0.18 |
| 6 | 137753946 | rs9484036 | -0.127 | -0.525 | -0.317 | -1.05 | -0.046 | -0.084 | -0.084 | 0 | 0.28 | 0.43 | 0.33 | 0.17 | 0.19 | 0.32 |
| 6 | 137759063 | rs9385795 | -0.597 | -0.475 | -0.347 | -1.036 | 0.026 | -0.005 | -0.005 | 0 | 0.33 | 0.56 | 0.35 | 0.32 | 0.46 | 0.39 |
| 6 | 137768325 | rs7773301 | -0.373 | -0.471 | -0.319 | -1.034 | 0.052 | -0.019 | -0.019 | 0 | 0.31 | 0.48 | 0.33 | 0.27 | 0.46 | 0.31 |
| 6 | 137774574 | rs9484045 | -0.088 | -0.535 | -0.399 | -1.047 | -0.03 | -0.058 | -0.058 | 0.184 | 0.27 | 0.38 | 0.32 | 0.17 | 0.19 | 0.32 |
| 6 | 137786289 | rs7775153 | -0.299 | -0.428 | -0.324 | -0.952 | -0.037 | -0.062 | -0.062 | 0 | 0.38 | 0.55 | 0.39 | 0.22 | 0.28 | 0.44 |
| 6 | 137786979 | rs13205934 | 1.042 | -0.219 | -0.651 | -0.887 | -0.02 | 0.017 | 0.017 | 0 | 0.09 | 0.04 | 0.14 | 0.02 | 0.14 | 0.18 |
| 6 | 137792213 | rs7769121 | -2.029 | -0.138 | -0.878 | -0.678 | -0.066 | -0.15 | -0.15 | 2.25 | 0.79 | 0.98 | 0.81 | 0.63 | 0.78 | 0.84 |
| 6 | 137798044 | rs6907990 | -1.353 | 0.147 | -0.341 | -0.669 | -0.013 | -0.035 | -0.035 | 0 | 0.61 | 0.89 | 0.66 | 0.5 | 0.65 | 0.71 |
| 6 | 137799725 | rs7756356 | -0.18 | -0.44 | 0.305 | -0.104 | -0.065 | -0.074 | -0.074 | 0 | 0.13 | 0.4 | 0.08 | 0 | 0.02 | 0.07 |
| 6 | 137801164 | rs1360606 | -0.307 | -0.388 | -0.323 | -0.151 | 0.009 | 0.012 | 0.012 | 0 | 0.2 | 0.18 | 0.27 | 0.18 | 0.17 | 0.26 |
| 6 | 137801589 | rs12198924 | -0.673 | -0.445 | -0.203 | -0.103 | -0.015 | -0.02 | -0.02 | 2.778 | 0.3 | 0.21 | 0.37 | 0.44 | 0.37 | 0.42 |
| 6 | 137801687 | rs12191965 | -0.673 | -0.445 | -0.203 | -0.103 | -0.015 | -0.02 | -0.02 | 0 | 0.3 | 0.21 | 0.37 | 0.44 | 0.37 | 0.42 |
| 6 | 137804081 | rs10457664 | -0.544 | -0.417 | -0.319 | -0.089 | -0.004 | -0.005 | -0.005 | 0.083 | 0.3 | 0.3 | 0.37 | 0.44 | 0.37 | 0.42 |
| 6 | 137806717 | rs1000041 | -0.201 | -0.746 | -0.231 | 0.016 | -0.034 | -0.05 | -0.05 | 0.26 | 0.1 | 0.18 | 0.04 | 0 | 0.02 | 0.07 |
| 6 | 137812161 | rs6918078 | -0.549 | -0.241 | -0.533 | 0.403 | -0.027 | 0.003 | 0.003 | 0.075 | 0.09 | 0.17 | 0.11 | 0.16 | 0.03 | 0.13 |
| 6 | 137815948 | rs6909958 | 0.011 | 0.218 | -1.409 | 0.705 | -0.054 | -0.06 | -0.06 | 0.109 | 0.63 | 0.9 | 0.63 | 0.57 | 0.5 | 0.56 |
| 6 | 137822718 | rs7767115 | -0.106 | -0.112 | -0.795 | 1.087 | 0.007 | 0.064 | 0.064 | 0.282 | 0.25 | 0.52 | 0.39 | 0.59 | 0.29 | 0.14 |
| 6 | 137823134 | rs2179298 | 0.617 | -0.83 | -1.016 | 0.692 | -0.003 | -0.002 | -0.002 | 0 | 0.11 | 0.13 | 0.11 | 0.08 | 0.09 | 0.15 |
| 6 | 137825631 | rs13200244 | 0.741 | -0.865 | -0.821 | 1.121 | -0.02 | -0.061 | -0.061 | 1.241 | 0.18 | 0.06 | 0.24 | 0.27 | 0.24 | 0.08 |
| 6 | 137826868 | rs9494794 | 0.831 | -1.296 | -1.051 | 0.519 | -0.008 | -0.061 | -0.061 | 0 | 0.31 | 0.33 | 0.2 | 0 | 0.15 | 0.26 |
| 6 | 137828235 | rs1000738 | -1.335 | -1.186 | -0.379 | 0.731 | -0.173 | -0.004 | -0.004 | 0.009 | 0.69 | 0.97 | 0.7 | 0.77 | 0.54 | 0.61 |
| 6 | 137841810 | rs4896271 | 1.18 | -2.027 | -0.848 | 0.11 | 0.05 | 0.141 | 0.141 | 1.349 | 0.21 | 0.01 | 0.15 | 0 | 0.11 | 0.2 |
| 6 | 137842399 | rs7775535 | -1.418 | -1.588 | -0.849 | 0.268 | -0.13 | -0.044 | -0.044 | 0 | 0.69 | 0.97 | 0.69 | 0.67 | 0.48 | 0.6 |
| 6 | 137843002 | rs4896272 | -0.142 | -1.696 | -0.075 | 0.106 | 0.058 | 0.065 | 0.065 | 0.222 | 0.43 | 0.34 | 0.3 | 0.1 | 0.26 | 0.42 |
| 6 | 137846731 | rs6570174 | 0.476 | -1.787 | 0.024 | 0.594 | 0.028 | -0.037 | -0.037 | 0.818 | 0.22 | 0.11 | 0.26 | 0.27 | 0.16 | 0.11 |
| 6 | 137848642 | rs9402902 | 0.153 | -2.07 | 0.351 | 0.666 | -0.054 | 0.056 | 0.056 | 0 | 0.2 | 0.36 | 0.18 | 0.31 | 0.15 | 0.22 |

| | | | | | | | | | | | | | | | | |
|---|---|---|---|---|---|---|---|---|---|---|---|---|---|---|---|---|
| 6 | 137850196 | rs423680 | 0.541 | -0.639 | 0.13 | 0.264 | -0.006 | -0.012 | -0.012 | 0 | 0.64 | 0.31 | 0.59 | 0.41 | 0.59 | 0.59 |
| 6 | 137853460 | rs996243 | -0.195 | -0.555 | 0.198 | 0.296 | 0.089 | 0.006 | 0.006 | 0.501 | 0.85 | 0.67 | 0.77 | 0.81 | 0.74 | 0.86 |
| 6 | 137855509 | rs391233 | -0.591 | -0.39 | 0.198 | 0.249 | 0.107 | -0.044 | -0.044 | 0 | 0.88 | 0.69 | 0.86 | 0.96 | 0.77 | 0.93 |
| 6 | 137856957 | rs4896278 | 0.443 | -0.505 | 0.178 | 0.25 | -0.001 | -0.007 | -0.007 | 0.943 | 0.67 | 0.32 | 0.68 | 0.56 | 0.62 | 0.65 |
| 6 | 137857927 | rs12110715 | 0.635 | -0.624 | 0.126 | 0.171 | 0.012 | -0.076 | -0.076 | 0 | 0.24 | 0.1 | 0.32 | 0.41 | 0.19 | 0.16 |
| 6 | 137861948 | rs203698 | 0.971 | -0.674 | 0.074 | 0.177 | -0.027 | 0.134 | 0.134 | 0.873 | 0.43 | 0.22 | 0.35 | 0.15 | 0.44 | 0.49 |
| 6 | 137867448 | rs203691 | 0.032 | -0.548 | -0.179 | 0.297 | 0.007 | 0.011 | 0.011 | 0 | 0.64 | 0.59 | 0.54 | 0.51 | 0.59 | 0.71 |
| 6 | 137870673 | rs6910797 | 0.092 | -0.556 | -0.362 | 0.625 | 0.022 | -0.031 | -0.031 | 0 | 0.19 | 0.15 | 0.15 | 0.36 | 0.15 | 0.22 |
| 6 | 137873084 | rs9689119 | -0.264 | 0.043 | -0.062 | 0.444 | 0.032 | 0.002 | 0.002 | 0.082 | 0.15 | 0.22 | 0.2 | 0.22 | 0.24 | 0.19 |
| 6 | 137873221 | rs12209996 | 0.364 | -0.302 | -0.358 | 0.664 | 0.008 | -0.01 | -0.01 | 0 | 0.16 | 0.15 | 0.16 | 0.38 | 0.17 | 0.3 |
| 6 | 137873321 | rs1412925 | -0.373 | -0.45 | -0.505 | 0.121 | 0.02 | 0.002 | 0.002 | 0.34 | 0.6 | 0.55 | 0.51 | 0.44 | 0.56 | 0.72 |
| 6 | 137874408 | rs9321619 | 0.09 | -0.051 | -1.293 | -1.04 | -0.01 | 0.051 | 0.051 | 2.391 | 0.42 | 0.22 | 0.47 | 0.63 | 0.49 | 0.57 |
| 6 | 137889729 | rs650434 | -0.85 | -0.142 | -1.382 | -1.491 | 0.001 | 0.093 | 0.093 | 0 | 0.7 | 0.87 | 0.8 | 0.83 | 0.78 | 0.75 |
| 6 | 137889913 | rs662100 | 0.139 | -0.285 | -1.36 | -1.798 | -0.001 | 0.034 | 0.034 | 0 | 0.43 | 0.34 | 0.48 | 0.09 | 0.47 | 0.39 |
| 6 | 137891497 | rs526676 | 0.684 | -0.442 | -1.393 | -1.92 | -0.012 | 0.038 | 0.038 | 0.211 | 0.34 | 0.24 | 0.36 | 0.04 | 0.4 | 0.27 |
| 6 | 137893893 | rs603183 | -0.298 | -0.567 | -1.299 | -1.7 | 0.004 | 0.006 | 0.006 | 0 | 0.59 | 0.4 | 0.68 | 0.67 | 0.75 | 0.63 |
| 6 | 137899792 | rs558015 | -0.687 | -0.506 | -0.923 | -2.333 | -0.001 | -0.01 | -0.01 | 0.421 | 0.42 | 0.54 | 0.49 | 0.29 | 0.43 | 0.34 |
| 6 | 137901443 | rs6570177 | 0.758 | -0.525 | -0.977 | -2.291 | -0.011 | 0.028 | 0.028 | 0 | 0.09 | 0.21 | 0.09 | 0.16 | 0.03 | 0.12 |
| 6 | 137902440 | rs491335 | 0.143 | -0.451 | -1.701 | -2.325 | -0.004 | 0.006 | 0.006 | 0 | 0.28 | 0.25 | 0.35 | 0.09 | 0.39 | 0.14 |
| 6 | 137903756 | rs11970302 | 1.087 | -0.396 | -1.963 | -2.286 | 0.013 | 0.03 | 0.03 | 0.106 | 0.11 | 0.05 | 0.11 | 0.01 | 0.13 | 0.07 |
| 6 | 137906476 | rs2175890 | 1.57 | -0.232 | -2.216 | -2.336 | 0.003 | 0.073 | 0.073 | 0 | 0.16 | 0.03 | 0.17 | 0 | 0.14 | 0.24 |
| 6 | 137911899 | rs659512 | -0.542 | 0.302 | -1.862 | -1.883 | -0.079 | 0.061 | 0.061 | 0 | 0.61 | 0.87 | 0.69 | 0.3 | 0.57 | 0.58 |
| 6 | 137920217 | rs526077 | -1.234 | 0.414 | -1.912 | -1.628 | -0.052 | 0.027 | 0.027 | 0 | 0.78 | 0.97 | 0.89 | 0.83 | 0.81 | 0.8 |
| 6 | 137922827 | rs646807 | 0.607 | 0.682 | -1.195 | -1.076 | 0.002 | -0.001 | -0.001 | 0 | 0.15 | 0.17 | 0.19 | 0.09 | 0.22 | 0.07 |
| 6 | 137923390 | rs9376287 | 0.857 | 0.521 | -1.244 | -0.734 | -0.035 | -0.075 | -0.075 | 0 | 0.16 | 0.06 | 0.26 | 0.51 | 0.22 | 0.22 |
| 6 | 137924218 | rs10080841 | 0.174 | 0.292 | -0.948 | -1.329 | -0.003 | -0.035 | -0.035 | 0 | 0.09 | 0.12 | 0.12 | 0 | 0.14 | 0.06 |
| 6 | 137924662 | rs695151 | -0.545 | -0.126 | -0.86 | -2.103 | 0.001 | -0.016 | -0.016 | 0 | 0.56 | 0.59 | 0.69 | 0.89 | 0.64 | 0.61 |
| 6 | 137926346 | rs596014 | 1.181 | 0.045 | -1.093 | -1.986 | -0.074 | -0.036 | -0.036 | 0 | 0.15 | 0.02 | 0.17 | 0.17 | 0.21 | 0.08 |
| 6 | 137929810 | rs569420 | 0.451 | -0.483 | -0.858 | -2.312 | -0.055 | 0.123 | 0.123 | 0 | 0.39 | 0.16 | 0.35 | 0.17 | 0.43 | 0.27 |
| 6 | 137930302 | rs6917072 | 0.314 | -0.501 | -0.546 | -1.819 | -0.021 | 0.029 | 0.029 | 0 | 0.11 | 0.27 | 0.1 | 0.18 | 0.05 | 0.16 |
| 6 | 137933374 | rs608242 | -1.189 | -0.207 | -1.12 | -1.871 | 0.01 | 0.036 | 0.036 | 0 | 0.78 | 0.87 | 0.9 | 0.92 | 0.86 | 0.8 |
| 6 | 137933924 | rs610803 | -1.271 | 0.18 | -1.038 | -1.628 | -0.022 | -0.044 | -0.044 | 0 | 0.85 | 0.91 | 0.72 | 0.48 | 0.76 | 0.78 |
| 6 | 137934634 | rs17066624 | 0.432 | -0.033 | -0.837 | -2.112 | -0.012 | -0.09 | -0.09 | 0.093 | 0.21 | 0.26 | 0.16 | 0 | 0.18 | 0.14 |
| 6 | 137936396 | rs556742 | -0.378 | 0.326 | -0.791 | -1.815 | -0.012 | 0.045 | 0.045 | 0.101 | 0.55 | 0.33 | 0.47 | 0.32 | 0.51 | 0.45 |
| 6 | 137940936 | rs495610 | -0.32 | 0.28 | -0.831 | -2.242 | 0 | -0.019 | -0.019 | 0.294 | 0.47 | 0.51 | 0.52 | 0.21 | 0.53 | 0.27 |
| 6 | 137941096 | rs13209759 | 0.94 | 0.508 | -0.902 | -2.284 | -0.029 | 0.028 | 0.028 | 0 | 0.06 | 0 | 0.09 | 0 | 0.13 | 0.06 |
| 6 | 137945924 | rs6570180 | 0.033 | 0.481 | -0.869 | -2.316 | 0.013 | -0.079 | -0.079 | 0 | 0.2 | 0.24 | 0.24 | 0.01 | 0.28 | 0.16 |
| 6 | 137947988 | rs487438 | -1.374 | 0.438 | -0.934 | -2.34 | 0.006 | -0.003 | -0.003 | 0.081 | 0.68 | 0.61 | 0.62 | 0.29 | 0.66 | 0.56 |
| 6 | 137948434 | rs636107 | 0.932 | -0.211 | -1.25 | -2.168 | -0.027 | 0.02 | 0.02 | 1.663 | 0.25 | 0.09 | 0.22 | 0.17 | 0.25 | 0.1 |
| 6 | 137951047 | rs473892 | -0.915 | -0.185 | -1.2 | -2.133 | 0.02 | -0.023 | -0.023 | 0 | 0.62 | 0.51 | 0.48 | 0.25 | 0.52 | 0.51 |
| 6 | 137951850 | rs556008 | -0.405 | 0.361 | -1.165 | -1.832 | 0 | 0 | 0 | 0 | 0.4 | 0.39 | 0.37 | 0.17 | 0.39 | 0.2 |
| 6 | 137955032 | rs6941060 | -0.148 | 0.414 | -0.63 | -1.382 | -0.002 | -0.079 | -0.079 | 0.126 | 0.14 | 0.23 | 0.14 | 0 | 0.14 | 0.1 |
| 6 | 137956333 | rs9688558 | 0.223 | 0.422 | -1.235 | -0.987 | 0.004 | 0.025 | 0.025 | 0 | 0.16 | 0.07 | 0.09 | 0.08 | 0.13 | 0.23 |
| 6 | 137959235 | rs6933404 | -0.278 | 0.724 | -0.972 | -0.923 | -0.005 | 0.004 | 0.004 | 0 | 0.13 | 0.09 | 0.1 | 0 | 0.17 | 0.11 |
| 6 | 137959455 | rs631204 | 0.125 | 0.744 | -0.932 | -1.215 | 0.005 | 0.016 | 0.016 | 0 | 0.5 | 0.59 | 0.48 | 0.33 | 0.42 | 0.4 |
| 6 | 137960364 | rs566097 | 0.938 | 0.628 | -0.936 | -1.588 | 0.002 | 0.006 | 0.006 | 0.55 | 0.18 | 0.37 | 0.15 | 0.23 | 0.13 | 0.11 |
| 6 | 137965418 | rs13207033 | 0.48 | 0.571 | -1.101 | -1.594 | -0.005 | 0.108 | 0.108 | 0 | 0.3 | 0.14 | 0.28 | 0.09 | 0.28 | 0.28 |
| 6 | 137967214 | rs13192841 | 0.514 | 0.567 | -1.103 | -1.603 | -0.011 | 0.111 | 0.111 | 0.014 | 0.29 | 0.13 | 0.28 | 0.09 | 0.28 | 0.28 |
| 6 | 137967252 | rs12527282 | 0.471 | 0.572 | -1.102 | -1.6 | -0.005 | 0.108 | 0.108 | 0 | 0.3 | 0.14 | 0.28 | 0.09 | 0.28 | 0.28 |
| 6 | 137973068 | rs2327832 | 0.45 | 0.642 | -0.991 | -1.497 | -0.004 | 0 | 0 | 0 | 0.13 | 0.09 | 0.1 | 0 | 0.17 | 0.11 |
| 6 | 137974450 | rs583816 | -0.455 | 0.617 | -1.163 | -1.107 | 0.005 | 0.005 | 0.005 | 0 | 0.6 | 0.59 | 0.54 | 0.49 | 0.52 | 0.52 |
| 6 | 137984935 | rs498549 | 0.126 | 0.544 | -1.087 | -1.293 | 0 | -0.003 | -0.003 | 0 | 0.49 | 0.54 | 0.44 | 0.32 | 0.43 | 0.4 |
| 6 | 137987387 | rs10447442 | 0.376 | 0.596 | -1.083 | -1.273 | -0.001 | -0.019 | -0.019 | 0.031 | 0.11 | 0.08 | 0.06 | 0.15 | 0.09 | 0.12 |
| 6 | 137993049 | rs536331 | 0.131 | 0.574 | -1.077 | -1.291 | 0 | -0.002 | -0.002 | 0 | 0.49 | 0.55 | 0.44 | 0.32 | 0.42 | 0.37 |
| 6 | 138001115 | rs6921233 | 0.107 | 0.585 | -0.94 | -1.29 | 0.002 | 0.002 | 0.002 | 0 | 0.4 | 0.53 | 0.47 | 0.52 | 0.47 | 0.43 |
| 6 | 138006504 | rs6920220 | 0.836 | 0.649 | -0.961 | -1.332 | -0.004 | -0.006 | -0.006 | 0.455 | 0.12 | 0.1 | 0.1 | 0 | 0.17 | 0.1 |
| 6 | 138007530 | rs636393 | -0.249 | 0.57 | -1.051 | -1.433 | 0.01 | 0.141 | 0.141 | 0.28 | 0.57 | 0.8 | 0.64 | 0.82 | 0.6 | 0.51 |
| 6 | 138008124 | rs1518048 | 0.429 | 0.621 | -0.949 | -1.337 | 0.012 | 0.013 | 0.013 | 0 | 0.37 | 0.46 | 0.45 | 0.52 | 0.46 | 0.4 |
| 6 | 138018439 | rs6917441 | 0.958 | 0.514 | -0.694 | -1.045 | 0.015 | 0.088 | 0.088 | 0 | 0.25 | 0.35 | 0.35 | 0.52 | 0.29 | 0.3 |
| 6 | 138018675 | rs10499195 | 0.714 | 0.44 | -1.008 | -1.169 | -0.036 | 0.2 | 0.2 | 0.117 | 0.41 | 0.18 | 0.38 | 0.03 | 0.45 | 0.38 |
| 6 | 138020871 | rs640353 | 1.266 | 0.427 | -0.744 | -1.079 | -0.001 | 0.011 | 0.011 | 0.023 | 0.16 | 0.18 | 0.13 | 0.29 | 0.13 | 0.1 |
| 6 | 138021613 | rs4896297 | 0.14 | 0.407 | -0.834 | -1.296 | 0 | -0.024 | -0.024 | 0.042 | 0.12 | 0.07 | 0.07 | 0.16 | 0.1 | 0.12 |
| 6 | 138022540 | rs647108 | -0.003 | 0.544 | -0.667 | -1.12 | -0.005 | 0.088 | 0.088 | 0.449 | 0.44 | 0.66 | 0.49 | 0.81 | 0.43 | 0.41 |
| 6 | 138023549 | rs597846 | 0.111 | 0.328 | -0.208 | -0.604 | -0.002 | -0.013 | -0.013 | 0.22 | 0.19 | 0.24 | 0.16 | 0.09 | 0.18 | 0.09 |
| 6 | 138025552 | rs585985 | -0.113 | 0.23 | -0.586 | -0.945 | -0.007 | 0.022 | 0.022 | 0 | 0.9 | 0.94 | 0.9 | 0.98 | 0.91 | 0.84 |
| 6 | 138026272 | rs8180657 | 1.026 | 0.342 | -0.586 | -1.119 | -0.001 | 0.023 | 0.023 | 0.001 | 0.34 | 0.37 | 0.29 | 0.5 | 0.34 | 0.34 |
| 6 | 138027115 | rs4432993 | 1.107 | 0.32 | -0.49 | -1.13 | 0 | 0.011 | 0.011 | 0 | 0.33 | 0.32 | 0.28 | 0.5 | 0.34 | 0.34 |
| 6 | 138027662 | rs6901613 | 0.638 | 0.253 | -0.593 | -1.103 | 0.001 | -0.002 | -0.002 | 0.08 | 0.63 | 0.66 | 0.72 | 0.55 | 0.69 | 0.66 |
| 6 | 138028152 | rs633010 | 0.199 | 0.157 | -0.624 | -0.874 | 0.009 | -0.03 | -0.03 | 0 | 0.24 | 0.15 | 0.16 | 0.43 | 0.2 | 0.17 |

| | | | | | | | | | | | | | | | | |
|---|---|---|---|---|---|---|---|---|---|---|---|---|---|---|---|---|
| 6 | 138029909 | rs604927 | -0.168 | 0.306 | -0.745 | -1.209 | -0.03 | -0.023 | -0.023 | 2.188 | 0.88 | 0.99 | 0.89 | 0.91 | 0.88 | 0.88 |
| 6 | 138033233 | rs651645 | 0.239 | 0.162 | -0.46 | -0.752 | -0.001 | 0.042 | 0.042 | 0.43 | 0.56 | 0.66 | 0.46 | 0.29 | 0.51 | 0.56 |
| 6 | 138037366 | rs1518047 | 0.586 | 0.264 | -0.159 | -1.201 | -0.001 | -0.098 | -0.098 | 3.597 | 0.18 | 0.33 | 0.14 | 0.01 | 0.13 | 0.1 |
| 6 | 138046665 | rs9494850 | 0.046 | 0.323 | -0.591 | -1.16 | -0.01 | 0.063 | 0.063 | 2.149 | 0.34 | 0.27 | 0.42 | 0.07 | 0.42 | 0.31 |
| 6 | 138047994 | rs6570184 | -0.331 | 0.31 | -0.525 | -1.12 | 0.009 | 0.168 | 0.168 | 0.895 | 0.49 | 0.27 | 0.49 | 0.07 | 0.54 | 0.34 |
| 6 | 138052680 | rs643746 | 0.576 | -0.015 | 0.169 | -1.17 | 0.003 | 0.039 | 0.039 | 0.288 | 0.43 | 0.38 | 0.36 | 0.84 | 0.42 | 0.53 |
| 6 | 138066687 | rs6924473 | -0.133 | 0.029 | -0.112 | -1.19 | -0.023 | -0.032 | -0.032 | 0.197 | 0.85 | 0.97 | 0.87 | 0.86 | 0.85 | 0.83 |
| 6 | 138070058 | rs1890370 | -0.08 | -0.088 | -0.216 | -1.461 | -0.001 | 0.051 | 0.051 | 0.105 | 0.72 | 0.84 | 0.82 | 0.88 | 0.75 | 0.84 |
| 6 | 138071287 | rs999638 | 0.597 | -0.01 | -0.084 | -1.47 | 0 | -0.014 | -0.014 | 0.322 | 0.44 | 0.45 | 0.43 | 0.84 | 0.52 | 0.34 |
| 6 | 138075443 | rs11961697 | 0.847 | 0.074 | -0.344 | -1.557 | 0 | 0.006 | 0.006 | 0.458 | 0.39 | 0.36 | 0.35 | 0.82 | 0.43 | 0.32 |
| 6 | 138076153 | rs6927210 | 0.418 | 0.051 | -0.459 | -1.567 | -0.008 | 0.099 | 0.099 | 0.25 | 0.47 | 0.68 | 0.51 | 0.84 | 0.53 | 0.41 |
| 6 | 138082762 | rs946227 | 1.667 | -0.513 | -1.094 | -1.386 | -0.122 | -0.056 | -0.056 | 0 | 0.24 | 0.03 | 0.2 | 0.24 | 0.38 | 0.17 |
| 6 | 138085196 | rs7753364 | 0.592 | 0.007 | -0.647 | -1.756 | 0.002 | -0.024 | -0.024 | 0.02 | 0.45 | 0.5 | 0.45 | 0.82 | 0.53 | 0.41 |
| 6 | 138085248 | rs7753394 | 0.551 | 0.013 | -0.779 | -1.753 | -0.003 | 0.009 | 0.009 | 0.164 | 0.45 | 0.42 | 0.44 | 0.82 | 0.53 | 0.36 |
| 6 | 138099740 | rs3866716 | 1.442 | -0.25 | -0.966 | -1.861 | -0.057 | 0.076 | 0.076 | 4.125 | 0.13 | 0.01 | 0.25 | 0.01 | 0.17 | 0.16 |
| 6 | 138100695 | rs2683065 | 0.209 | -0.642 | -0.866 | -1.909 | -0.013 | -0.056 | -0.056 | 0 | 0.59 | 0.5 | 0.73 | 0.86 | 0.7 | 0.56 |
| 6 | 138102476 | rs10782267 | 0.757 | -0.078 | -0.942 | -1.756 | -0.082 | 0.131 | 0.131 | 3.251 | 0.21 | 0.01 | 0.29 | 0.01 | 0.26 | 0.39 |
| 6 | 138104861 | rs1416172 | -0.179 | -0.019 | -0.344 | -1.363 | 0.003 | 0.027 | 0.027 | 0 | 0.86 | 0.71 | 0.81 | 0.84 | 0.88 | 0.84 |
| 6 | 138106955 | rs6570189 | -0.57 | 0.073 | -0.832 | -1.322 | -0.032 | -0.039 | -0.039 | 1.649 | 0.87 | 0.99 | 0.89 | 0.88 | 0.88 | 0.89 |
| 6 | 138115266 | rs654039 | 0.155 | 0.113 | -0.689 | -1.065 | 0.001 | 0.018 | 0.018 | 0.176 | 0.44 | 0.4 | 0.42 | 0.83 | 0.48 | 0.38 |
| 6 | 138125829 | rs9385808 | 1.059 | 0.122 | -0.997 | -0.023 | -0.038 | -0.081 | -0.081 | 0 | 0.12 | 0.01 | 0.16 | 0.3 | 0.16 | 0.07 |
| 6 | 138127204 | rs675640 | 0.06 | -0.001 | -0.338 | -1.027 | 0.001 | -0.053 | -0.053 | 0 | 0.91 | 0.86 | 0.92 | 0.98 | 0.91 | 0.89 |
| 6 | 138128750 | rs17066886 | -0.166 | -0.407 | -0.123 | -1.209 | -0.029 | -0.073 | -0.073 | 0 | 0.12 | 0.37 | 0.09 | 0.01 | 0.05 | 0.08 |
| 6 | 138130479 | rs619796 | -0.461 | -0.154 | -1.335 | -1.134 | -0.029 | -0.028 | -0.028 | 0.464 | 0.88 | 0.98 | 0.89 | 0.92 | 0.86 | 0.92 |
| 6 | 138132445 | rs9494868 | -0.404 | -0.26 | -1.337 | -1.308 | 0.012 | -0.001 | -0.001 | 0.69 | 0.3 | 0.63 | 0.23 | 0.03 | 0.22 | 0.21 |
| 6 | 138137551 | rs679670 | 0.625 | -0.116 | -1.417 | -1.165 | 0.036 | 0.147 | 0.147 | 0.051 | 0.58 | 0.28 | 0.68 | 0.89 | 0.64 | 0.71 |
| 6 | 138142754 | rs694764 | 0.229 | -0.527 | -1.039 | -0.932 | 0.004 | 0.004 | 0.004 | 2.428 | 0.76 | 0.28 | 0.8 | 0.89 | 0.86 | 0.89 |
| 6 | 138144173 | rs6923167 | 1.322 | -0.613 | -0.35 | -0.986 | -0.014 | -0.048 | -0.048 | 1.873 | 0.08 | 0.22 | 0.04 | 0 | 0.03 | 0 |
| 6 | 138146505 | rs657597 | 0.896 | -0.389 | -1.226 | -1.431 | 0.012 | 0.11 | 0.11 | 4.514 | 0.6 | 0.22 | 0.62 | 0.86 | 0.7 | 0.72 |
| 6 | 138147048 | rs670369 | 0.225 | -0.411 | -1.163 | -1.406 | 0.008 | 0.01 | 0.01 | 1.99 | 0.73 | 0.29 | 0.83 | 0.9 | 0.84 | 0.85 |
| 6 | 138150434 | rs2788287 | 1.174 | -0.475 | -1.852 | -0.739 | 0.001 | -0.126 | -0.126 | 0 | 0.19 | 0.03 | 0.2 | 0.48 | 0.18 | 0.25 |
| 6 | 138156425 | rs17779870 | 1.296 | -0.183 | -1.617 | -0.857 | -0.05 | 0.044 | 0.044 | 7.456 | 0.13 | 0.01 | 0.16 | 0.03 | 0.13 | 0.1 |
| 6 | 138157037 | rs9389536 | 1.105 | -0.091 | -1.818 | -0.583 | -0.016 | -0.019 | -0.019 | 0.02 | 0.08 | 0.01 | 0.24 | 0.46 | 0.09 | 0.24 |
| 6 | 138161013 | rs6918329 | -0.432 | -0.679 | -1.859 | -1.956 | 0.037 | 0.221 | 0.221 | 3.234 | 0.45 | 0.83 | 0.4 | 0.1 | 0.31 | 0.26 |
| 6 | 138172383 | rs600144 | 1.336 | 0.099 | -1.069 | -1.097 | -0.043 | -0.03 | -0.03 | 0 | 0.17 | 0.02 | 0.16 | 0.17 | 0.23 | 0.06 |
| 6 | 138179136 | rs11970361 | 0.855 | -0.19 | -0.17 | -1.663 | -0.005 | -0.035 | -0.035 | 0.237 | 0.1 | 0.12 | 0.07 | 0.01 | 0.06 | 0.04 |
| 6 | 138192062 | rs5029933 | 0.964 | -0.194 | -0.182 | -1.654 | -0.004 | -0.037 | -0.037 | 0 | 0.09 | 0.09 | 0.03 | 0 | 0.06 | 0.04 |
| 6 | 138192761 | rs719150 | -0.086 | -0.282 | 0.462 | -1.682 | -0.015 | -0.032 | -0.032 | 0 | 0.14 | 0.52 | 0.15 | 0.06 | 0.09 | 0.1 |
| 6 | 138195693 | rs643177 | -0.192 | -0.376 | 0.347 | -1.848 | 0.001 | -0.021 | -0.021 | 0 | 0.71 | 0.64 | 0.72 | 0.91 | 0.73 | 0.75 |
| 6 | 138199417 | rs610604 | -0.012 | -0.497 | -0.631 | -1.516 | 0.001 | 0.039 | 0.039 | 4.128 | 0.59 | 0.32 | 0.61 | 0.9 | 0.66 | 0.67 |
| 6 | 138222510 | rs9321634 | -0.374 | -0.389 | -1.762 | -1.409 | -0.02 | 0.196 | 0.196 | 6.034 | 0.45 | 0.9 | 0.43 | 0.15 | 0.36 | 0.37 |
| 6 | 138248841 | rs7755566 | 0.006 | -0.098 | -0.774 | -0.935 | -0.001 | -0.011 | -0.011 | 1.125 | 0.39 | 0.41 | 0.39 | 0.21 | 0.58 | 0.24 |
| 6 | 138253479 | rs9389541 | 0.829 | -0.56 | -1.315 | -1.179 | 0.014 | 0.046 | 0.046 | 0 | 0.17 | 0.05 | 0.17 | 0.08 | 0.15 | 0.13 |
| 6 | 138260037 | rs9373201 | -0.191 | -0.828 | -1.097 | -1.407 | 0.051 | 0.085 | 0.085 | 1.347 | 0.52 | 0.72 | 0.44 | 0.27 | 0.61 | 0.41 |
| 6 | 138265348 | rs6941603 | -0.354 | -0.792 | -1.2 | -1.513 | 0.044 | 0.099 | 0.099 | 1.982 | 0.53 | 0.86 | 0.46 | 0.3 | 0.61 | 0.47 |
| 6 | 138266248 | rs10872479 | 0.68 | -0.472 | -1.269 | -1.712 | -0.098 | 0.044 | 0.044 | 1.215 | 0.31 | 0.12 | 0.3 | 0.18 | 0.47 | 0.17 |
| 6 | 138266431 | rs10499198 | 1.253 | 0.025 | -1.343 | -1.726 | -0.078 | -0.04 | -0.04 | 0 | 0.18 | 0.03 | 0.2 | 0.18 | 0.31 | 0.08 |
| 6 | 138266685 | rs9321637 | 0.161 | -0.241 | -0.77 | -1.6 | 0.058 | -0.018 | -0.018 | 0 | 0.11 | 0.37 | 0.1 | 0.03 | 0.11 | 0.08 |
| 6 | 138275769 | rs6899975 | 0.635 | -0.115 | -1.008 | -1.264 | 0.008 | 0.001 | 0.001 | 0 | 0.21 | 0.29 | 0.31 | 0.23 | 0.37 | 0.14 |
| 6 | 138282375 | rs4388301 | -0.722 | -0.385 | -1.48 | -1.406 | 0.013 | 0.009 | 0.009 | 2.893 | 0.58 | 0.76 | 0.58 | 0.38 | 0.63 | 0.56 |
| 6 | 138283265 | rs9376307 | -0.218 | -0.569 | -2.301 | -1.232 | 0 | -0.007 | -0.007 | 0 | 0.34 | 0.32 | 0.24 | 0.15 | 0.26 | 0.42 |
| 6 | 138285462 | rs9373202 | 0.769 | -0.05 | -2.189 | -0.901 | -0.015 | 0.04 | 0.04 | 1.656 | 0.2 | 0.04 | 0.18 | 0.1 | 0.19 | 0.17 |
| 6 | 138287560 | rs4895501 | 0.046 | 0.286 | -2.017 | -0.552 | -0.003 | -0.004 | -0.004 | 0 | 0.56 | 0.47 | 0.52 | 0.62 | 0.41 | 0.63 |
| 6 | 138288536 | rs11154948 | 0.753 | 0.992 | -2.946 | -0.279 | -0.097 | -0.037 | -0.037 | 0.971 | 0.17 | 0.03 | 0.22 | 0.18 | 0.34 | 0.08 |
| 6 | 138291360 | rs4454154 | -0.618 | 0.411 | -2.278 | 0.166 | 0.045 | 0.033 | 0.033 | 1.925 | 0.48 | 0.25 | 0.48 | 0.56 | 0.38 | 0.43 |
| 6 | 138299093 | rs9399229 | -0.673 | 1.012 | -1.904 | -0.308 | -0.027 | 0.022 | 0.022 | 1.552 | 0.22 | 0.09 | 0.25 | 0.14 | 0.26 | 0.21 |
| 6 | 138304169 | rs7768419 | -0.593 | 0.655 | -1.618 | -0.08 | -0.006 | 0.002 | 0.002 | 0 | 0.08 | 0.04 | 0.03 | 0.08 | 0.07 | 0.09 |
| 6 | 138304615 | rs11154954 | -0.197 | 0.149 | -2.33 | -0.213 | 0.067 | -0.058 | -0.058 | 0.002 | 0.58 | 0.88 | 0.74 | 0.45 | 0.71 | 0.63 |
| 6 | 138315199 | rs6925237 | -0.444 | 0.444 | -1.815 | -0.616 | -0.009 | 0.002 | 0.002 | 0 | 0.08 | 0.05 | 0.04 | 0.08 | 0.08 | 0.05 |
| 6 | 138325071 | rs4341022 | -0.134 | 0.083 | -2.334 | -1.585 | 0.04 | 0.15 | 0.15 | 0 | 0.39 | 0.18 | 0.24 | 0.17 | 0.29 | 0.35 |
| 6 | 138326557 | rs4479955 | 0.343 | -0.237 | -2.574 | -1.872 | 0.028 | 0.042 | 0.042 | 0.743 | 0.22 | 0.15 | 0.08 | 0.16 | 0.14 | 0.26 |
| 6 | 138326693 | rs9402935 | -0.387 | -0.355 | -1.977 | -2.587 | -0.01 | -0.111 | -0.111 | 0 | 0.8 | 0.66 | 0.85 | 1 | 0.83 | 0.93 |
| 6 | 138328450 | rs9321643 | 0.109 | -0.536 | -1.851 | -2.731 | 0.024 | 0.083 | 0.083 | 6.161 | 0.29 | 0.19 | 0.15 | 0.15 | 0.23 | 0.29 |
| 6 | 138329630 | rs6908176 | -0.833 | -0.402 | -1.711 | -2.66 | -0.013 | -0.064 | -0.064 | 0 | 0.89 | 0.79 | 0.93 | 0.98 | 0.92 | 0.96 |
| 6 | 138330626 | rs9389547 | 0.151 | -0.525 | -1.793 | -2.725 | 0.02 | 0.022 | 0.022 | 0.476 | 0.57 | 0.62 | 0.78 | 0.83 | 0.69 | 0.67 |
| 6 | 138335037 | rs9376314 | 0.861 | -0.075 | -1.516 | -2.208 | 0.027 | -0.038 | -0.038 | 0.248 | 0.32 | 0.42 | 0.5 | 0.69 | 0.46 | 0.52 |
| 6 | 138335260 | rs12207754 | 0.134 | -0.269 | -1.626 | -1.949 | 0.036 | 0.195 | 0.195 | 0 | 0.39 | 0.11 | 0.23 | 0.15 | 0.28 | 0.34 |
| 6 | 138335296 | rs9373206 | 0.124 | 0.375 | -1.144 | -1.283 | 0.005 | -0.024 | -0.024 | 0.144 | 0.41 | 0.55 | 0.59 | 0.69 | 0.54 | 0.55 |
| 6 | 138337903 | rs12203234 | -0.813 | 0.686 | -0.836 | -0.802 | 0.001 | -0.01 | -0.01 | 0.23 | 0.19 | 0.23 | 0.28 | 0.12 | 0.19 | 0.25 |

| | | | | | | | | | | | | | | | | |
|---|---|---|---|---|---|---|---|---|---|---|---|---|---|---|---|---|
| 6 | 138338207 | rs6570205 | -0.394 | 0.443 | -0.659 | -0.235 | 0.016 | -0.001 | -0.001 | 0.005 | 0.48 | 0.58 | 0.53 | 0.51 | 0.6 | 0.28 |
| 6 | 138338340 | rs6905679 | -0.369 | 0.859 | -0.101 | -0.059 | 0.002 | -0.004 | -0.004 | 0 | 0.26 | 0.46 | 0.22 | 0.15 | 0.19 | 0.14 |
| 6 | 138340676 | rs4464817 | -0.258 | 0.547 | -0.946 | -0.419 | -0.008 | 0.106 | 0.106 | 0 | 0.74 | 0.93 | 0.79 | 0.87 | 0.74 | 0.8 |
| 6 | 138341562 | rs4624886 | 0.212 | 0.895 | -0.353 | 0.053 | 0 | 0.047 | 0.047 | 0.08 | 0.23 | 0.28 | 0.29 | 0.36 | 0.21 | 0.29 |
| 6 | 138343019 | rs9389549 | 0.531 | 0.8 | -0.126 | -0.161 | 0.04 | -0.04 | -0.04 | 0 | 0.77 | 0.52 | 0.68 | 0.86 | 0.72 | 0.75 |
| 6 | 138345526 | rs9321645 | -0.168 | 0.652 | 0.05 | -0.004 | 0.046 | 0.07 | 0.07 | 0.472 | 0.09 | 0.33 | 0.18 | 0.12 | 0.1 | 0.13 |
| 6 | 138348968 | rs9373209 | 0.715 | 0.444 | -0.24 | 0.118 | -0.006 | 0.001 | 0.001 | 0 | 0.06 | 0.06 | 0.02 | 0.11 | 0.05 | 0.04 |
| 6 | 138351878 | rs2473092 | 0.332 | 0.648 | -0.209 | 0.147 | -0.02 | 0.094 | 0.094 | 0.844 | 0.2 | 0.29 | 0.24 | 0.35 | 0.14 | 0.29 |
| 6 | 138354180 | rs4895508 | 0.136 | 0.607 | -0.269 | 0.147 | 0.007 | 0.009 | 0.009 | 0 | 0.64 | 0.65 | 0.59 | 0.64 | 0.62 | 0.59 |
| 6 | 138361317 | rs13214334 | 0.995 | 0.544 | -1.175 | -0.002 | 0.041 | 0.025 | 0.025 | 0.227 | 0.13 | 0.01 | 0.09 | 0.1 | 0.08 | 0.08 |
| 6 | 138363064 | rs7762658 | -0.325 | 0.739 | -0.774 | 0.21 | 0.049 | 0.085 | 0.085 | 0.188 | 0.09 | 0.24 | 0.27 | 0.24 | 0.18 | 0.05 |
| 6 | 138365571 | rs12212617 | -0.106 | 0.732 | -0.896 | 0.394 | -0.002 | 0.045 | 0.045 | 0.122 | 0.11 | 0.09 | 0.22 | 0.44 | 0.11 | 0.28 |
| 6 | 138366274 | rs13213035 | -0.73 | 1.061 | -1.183 | -0.333 | -0.032 | 0.026 | 0.026 | 0 | 0.88 | 0.94 | 0.85 | 0.99 | 0.75 | 0.87 |
| 6 | 138366807 | rs17613612 | 1.814 | 0.158 | -1.784 | -0.865 | -0.072 | 0.113 | 0.113 | 0 | 0.28 | 0.03 | 0.14 | 0.06 | 0.25 | 0.18 |
| 6 | 138367590 | rs9376319 | 1.337 | 0.038 | -1.745 | -0.792 | 0.002 | -0.02 | -0.02 | 0 | 0.07 | 0 | 0.01 | 0.11 | 0.05 | 0.04 |
| 6 | 138374999 | rs6420764 | 0.032 | 0.002 | -1.529 | -0.888 | 0 | 0 | 0 | 0 | 0.29 | 0.34 | 0.5 | 0.74 | 0.32 | 0.39 |
| 6 | 138385481 | rs9402941 | 1.082 | -0.109 | -1.628 | -0.961 | -0.028 | 0.122 | 0.122 | 0 | 0.36 | 0.09 | 0.16 | 0.18 | 0.3 | 0.23 |
| 6 | 138391562 | rs9321647 | 0.421 | -0.281 | -1.129 | -1.113 | 0.001 | 0 | 0 | 0 | 0.52 | 0.43 | 0.29 | 0.22 | 0.43 | 0.32 |
| 6 | 138393028 | rs10440882 | 0.68 | -0.347 | -0.919 | -0.881 | -0.014 | -0.063 | -0.063 | 0 | 0.16 | 0.33 | 0.13 | 0.04 | 0.13 | 0.09 |
| 6 | 138393314 | rs643571 | 1.925 | -0.417 | -1.149 | -0.917 | 0.044 | 0.085 | 0.085 | 0.02 | 0.15 | 0.01 | 0.1 | 0.05 | 0.09 | 0.11 |
| 6 | 138404048 | rs9484111 | -0.25 | -0.318 | -0.969 | -0.857 | -0.03 | -0.049 | -0.049 | 0 | 0.23 | 0.37 | 0.15 | 0.17 | 0.18 | 0.13 |
| 6 | 138406100 | rs7749355 | 1.146 | -0.042 | -1.234 | -0.31 | -0.005 | 0.223 | 0.223 | 0 | 0.05 | 0.09 | 0.17 | 0.43 | 0.04 | 0.26 |
| 6 | 138406328 | rs480631 | -0.35 | 0.092 | -1.194 | -0.478 | 0.023 | 0.023 | 0.023 | 0.023 | 0.47 | 0.65 | 0.52 | 0.45 | 0.44 | 0.29 |
| 6 | 138409379 | rs681144 | 0.375 | 0.08 | -1.224 | -0.075 | 0.021 | -0.004 | -0.004 | 0 | 0.46 | 0.23 | 0.29 | 0.52 | 0.37 | 0.53 |
| 6 | 138411514 | rs514552 | -0.348 | 0.99 | -0.718 | -0.527 | 0.004 | 0.02 | 0.02 | 0 | 0.49 | 0.46 | 0.68 | 0.87 | 0.53 | 0.6 |
| 6 | 138413269 | rs648396 | -0.282 | 1.645 | -0.237 | -0.236 | 0.017 | 0.001 | 0.001 | 0 | 0.66 | 0.55 | 0.53 | 0.72 | 0.58 | 0.77 |
| 6 | 138414945 | rs2473097 | 0.355 | 1.314 | -0.108 | -1.474 | 0.022 | -0.044 | -0.044 | 0 | 0.14 | 0.18 | 0.12 | 0.02 | 0.21 | 0.18 |
| 6 | 138415087 | rs666769 | -0.693 | 1.351 | -0.199 | -1.517 | -0.005 | -0.127 | -0.127 | 0 | 0.67 | 0.49 | 0.73 | 0.94 | 0.66 | 0.81 |
| 6 | 138415246 | rs2473098 | 0.284 | 1.349 | -0.236 | -1.364 | 0.033 | -0.066 | -0.066 | 0 | 0.15 | 0.27 | 0.13 | 0.02 | 0.21 | 0.18 |
| 6 | 138416343 | rs808443 | -0.53 | 1.296 | -0.33 | -1.097 | 0.002 | -0.083 | -0.083 | 0.072 | 0.68 | 0.59 | 0.74 | 0.94 | 0.67 | 0.8 |
| 6 | 138417118 | rs2484066 | 0.139 | 1.28 | -0.102 | -1.158 | 0.001 | -0.045 | -0.045 | 0 | 0.43 | 0.54 | 0.35 | 0.08 | 0.52 | 0.29 |
| 6 | 138417301 | rs4896313 | -0.176 | 1.108 | -0.546 | -0.492 | -0.034 | -0.076 | -0.076 | 1.844 | 0.14 | 0.03 | 0.33 | 0.7 | 0.15 | 0.31 |
| 6 | 138417415 | rs2484067 | 0.507 | 1.17 | 0.064 | -0.899 | -0.008 | 0.077 | 0.077 | 0.74 | 0.44 | 0.32 | 0.32 | 0.06 | 0.48 | 0.25 |
| 6 | 138418015 | rs9494937 | -1.241 | 0.694 | 0.806 | -0.078 | -0.001 | -0.003 | -0.003 | 0 | 0.42 | 0.54 | 0.46 | 0.7 | 0.38 | 0.59 |
| 6 | 138419207 | rs2327871 | -1.336 | 0.649 | 0.906 | -0.165 | 0.014 | 0.044 | 0.044 | 1.553 | 0.25 | 0.43 | 0.35 | 0.7 | 0.27 | 0.43 |
| 6 | 138425860 | rs9494940 | -1.234 | 0.724 | 0.983 | -0.019 | 0.071 | 0.322 | 0.322 | 1.071 | 0.07 | 0.26 | 0.26 | 0.59 | 0.13 | 0.11 |
| 6 | 138426378 | rs952802 | -1.601 | 0.723 | 0.912 | -0.079 | -0.002 | -0.007 | -0.007 | 0 | 0.43 | 0.53 | 0.46 | 0.71 | 0.38 | 0.6 |
| 6 | 138431353 | rs9494941 | -1.556 | 0.814 | 0.907 | -0.047 | -0.003 | -0.011 | -0.011 | 0 | 0.42 | 0.52 | 0.42 | 0.72 | 0.38 | 0.57 |
| 6 | 138438723 | rs1537621 | -0.912 | 0.789 | 0.776 | -0.066 | 0.023 | 0.086 | 0.086 | 0 | 0.35 | 0.2 | 0.16 | 0.19 | 0.26 | 0.46 |
| 6 | 138440838 | rs1931867 | -1.548 | 0.799 | 0.855 | -0.04 | -0.002 | -0.016 | -0.016 | 0 | 0.42 | 0.5 | 0.42 | 0.79 | 0.39 | 0.57 |
| 6 | 138444360 | rs10457667 | 0.643 | 0.926 | 0.494 | -0.205 | -0.057 | 0.048 | 0.048 | 0.06 | 0.09 | 0.01 | 0.1 | 0.01 | 0.18 | 0.06 |
| 6 | 138444930 | rs6922466 | -1.306 | -0.172 | 1.028 | 0.352 | 0.013 | 0.107 | 0.107 | 0.024 | 0.34 | 0.14 | 0.15 | 0.18 | 0.25 | 0.46 |
| 6 | 138448062 | rs12660547 | -1.336 | -0.19 | 1.075 | 0.391 | 0.02 | 0.089 | 0.089 | 0.004 | 0.36 | 0.21 | 0.18 | 0.19 | 0.28 | 0.44 |
| 6 | 138449555 | rs7773257 | -0.77 | -0.114 | 0.439 | 0.427 | 0.007 | 0.015 | 0.015 | 0.244 | 0.19 | 0.11 | 0.1 | 0.17 | 0.14 | 0.29 |
| 6 | 138450069 | rs6920846 | -1.416 | -0.079 | 0.446 | 0.399 | 0.006 | 0.032 | 0.032 | 0.002 | 0.38 | 0.22 | 0.38 | 0.74 | 0.33 | 0.48 |
| 6 | 138451274 | rs4896318 | -1.441 | -0.078 | 0.471 | 0.399 | 0.007 | 0.022 | 0.022 | 0 | 0.39 | 0.27 | 0.38 | 0.74 | 0.33 | 0.48 |
| 6 | 138459129 | rs9389562 | -1.393 | 0.012 | 0.28 | 0.1 | 0.003 | 0.06 | 0.06 | 0.021 | 0.39 | 0.2 | 0.38 | 0.78 | 0.34 | 0.49 |
| 6 | 138461875 | rs9402947 | -1.294 | 0.041 | 0.195 | -0.15 | -0.001 | 0.089 | 0.089 | 0 | 0.37 | 0.18 | 0.41 | 0.82 | 0.34 | 0.48 |
| 6 | 138465225 | rs9389565 | 0.296 | -0.049 | -0.03 | -0.391 | -0.027 | 0.092 | 0.092 | 0 | 0.16 | 0 | 0.07 | 0.02 | 0.14 | 0.14 |
| 6 | 138465341 | rs11154962 | -0.071 | -0.077 | -0.106 | -0.313 | -0.017 | -0.026 | -0.026 | 0.234 | 0.14 | 0 | 0.06 | 0.18 | 0.11 | 0.25 |
| 6 | 138465435 | rs17067151 | -0.111 | 0.076 | 0.332 | -0.257 | 0.003 | -0.004 | -0.004 | 0.094 | 0.18 | 0.13 | 0.09 | 0.02 | 0.14 | 0.15 |
| 6 | 138467413 | rs9402950 | -0.768 | 0.033 | 0.062 | -0.132 | 0.002 | 0.06 | 0.06 | 0.006 | 0.31 | 0.08 | 0.13 | 0.22 | 0.24 | 0.27 |
| 6 | 138475527 | rs714935 | 0.247 | 0.145 | 0.351 | 0.133 | 0.009 | -0.02 | -0.02 | 0 | 0.14 | 0.04 | 0.06 | 0.19 | 0.1 | 0.2 |
| 6 | 138477031 | rs1883468 | -0.994 | 0.296 | 0.967 | 0.252 | 0.008 | 0.083 | 0.083 | 0 | 0.46 | 0.63 | 0.49 | 0.87 | 0.43 | 0.44 |
| 6 | 138479242 | rs9385824 | -0.527 | 0.287 | 0.96 | 0.737 | -0.005 | 0.012 | 0.012 | 0 | 0.06 | 0.02 | 0.03 | 0.02 | 0.06 | 0.05 |
| 6 | 138484929 | rs4896325 | -2.188 | -0.629 | 1.814 | 1.408 | -0.02 | 0.2 | 0.2 | 0.542 | 0.19 | 0.54 | 0.29 | 0.45 | 0.16 | 0.12 |
| 6 | 138490354 | rs4896326 | -1.358 | -0.573 | 1.916 | 1.569 | 0.003 | 0.253 | 0.253 | 0 | 0.06 | 0.16 | 0.17 | 0.37 | 0.07 | 0.08 |
| 6 | 138493350 | rs6925058 | -1.717 | -0.665 | 2.703 | 1.517 | 0.008 | -0.027 | -0.027 | 0 | 0.09 | 0.34 | 0.07 | 0.02 | 0.07 | 0.04 |
| 6 | 138496694 | rs1013054 | -2.612 | -0.672 | 2.635 | 1.544 | -0.015 | 0.274 | 0.274 | 0.269 | 0.16 | 0.51 | 0.24 | 0.38 | 0.13 | 0.12 |
| 6 | 138509838 | rs1008206 | -2.115 | -0.694 | 2.534 | 1.564 | -0.014 | 0.143 | 0.143 | 0 | 0.12 | 0.2 | 0.21 | 0.39 | 0.1 | 0.12 |
| 6 | 138514589 | rs4896329 | -2.815 | -0.732 | 2.813 | 1.553 | -0.026 | 0.185 | 0.185 | 0.753 | 0.17 | 0.58 | 0.25 | 0.4 | 0.13 | 0.12 |
| 6 | 138519929 | rs1509683 | -2.754 | -0.726 | 2.75 | 1.487 | -0.023 | 0.221 | 0.221 | 0.462 | 0.16 | 0.51 | 0.23 | 0.38 | 0.13 | 0.12 |
| 6 | 138525847 | rs6903101 | -2.067 | -0.733 | 2.684 | 1.259 | -0.006 | -0.044 | -0.044 | 0 | 0.1 | 0.49 | 0.08 | 0 | 0.05 | 0.04 |
| 6 | 138531936 | rs9402956 | -1.456 | -0.626 | 2.6 | 1.637 | 0.027 | 0.336 | 0.336 | 0 | 0.06 | 0.23 | 0.16 | 0.36 | 0.06 | 0.05 |
| 6 | 138535935 | rs9494967 | -1.474 | -0.691 | 2.932 | 1.58 | 0.029 | 0.016 | 0.016 | 0 | 0.05 | 0.27 | 0.04 | 0.01 | 0.03 | 0.06 |
| 6 | 138537432 | rs9402958 | -1.167 | -0.691 | 2.927 | 1.587 | 0.015 | 0.281 | 0.281 | 0 | 0.05 | 0.15 | 0.15 | 0.36 | 0.06 | 0.05 |
| 6 | 138537859 | rs3813338 | -1.167 | -0.692 | 2.95 | 1.587 | 0.015 | 0.281 | 0.281 | 0 | 0.05 | 0.15 | 0.15 | 0.36 | 0.06 | 0.05 |
| 6 | 138539556 | rs11758517 | -1.167 | -0.689 | 2.882 | 1.588 | 0.015 | 0.281 | 0.281 | 0 | 0.05 | 0.15 | 0.16 | 0.36 | 0.06 | 0.05 |
| 6 | 138539960 | rs7768738 | -1.854 | -0.681 | 2.392 | 1.532 | 0.026 | 0.4 | 0.4 | 0 | 0.1 | 0.41 | 0.19 | 0.38 | 0.09 | 0.11 |

| | | | | | | | | | | | | | | | | |
|---|---|---|---|---|---|---|---|---|---|---|---|---|---|---|---|---|
| 6 | 138540924 | rs760972 | -1.334 | -1.496 | 1.293 | 1.073 | 0.034 | 0.005 | 0.005 | 0 | 0.38 | 0.69 | 0.33 | 0.61 | 0.23 | 0.45 |
| 6 | 138566893 | rs9402966 | -1.948 | -0.79 | 1.685 | 1.5 | -0.01 | 0.256 | 0.256 | 1.045 | 0.12 | 0.59 | 0.22 | 0.37 | 0.09 | 0.11 |
| 6 | 138570001 | rs9389574 | -1.743 | -0.728 | 1.541 | 1.513 | 0.008 | 0.345 | 0.345 | 0.486 | 0.11 | 0.52 | 0.23 | 0.38 | 0.09 | 0.08 |
| 6 | 138591166 | rs203146 | -0.357 | -0.622 | 0.029 | 0.203 | 0.006 | 0.006 | 0.006 | 0.89 | 0.4 | 0.38 | 0.42 | 0.42 | 0.3 | 0.54 |
| 6 | 138599285 | rs203138 | -0.543 | -0.721 | 0.033 | 0.16 | 0.012 | 0.017 | 0.017 | 0.488 | 0.43 | 0.4 | 0.42 | 0.39 | 0.3 | 0.55 |
| 6 | 138606252 | rs203136 | -0.751 | -0.81 | 0.261 | 0.023 | 0.041 | 0.074 | 0.074 | 3.569 | 0.47 | 0.86 | 0.5 | 0.43 | 0.35 | 0.62 |
| 6 | 138611312 | rs9376339 | -0.508 | -1.268 | 0.227 | 0.505 | -0.012 | 0.071 | 0.071 | 0 | 0.22 | 0.32 | 0.22 | 0.5 | 0.18 | 0.23 |
| 6 | 138612441 | rs17067346 | 0.05 | -1.702 | -0.229 | 0.344 | 0.036 | -0.013 | -0.013 | 0 | 0.14 | 0.06 | 0.22 | 0.2 | 0.05 | 0.24 |
| 6 | 138615148 | rs203145 | 0.34 | -2.006 | -0.462 | 0.03 | 0.006 | 0.012 | 0.012 | 0 | 0.16 | 0.07 | 0.14 | 0.16 | 0.14 | 0.16 |
| 6 | 138616344 | rs17067360 | -0.031 | -2.046 | -0.022 | 0.375 | 0.046 | -0.015 | -0.015 | 0.544 | 0.14 | 0.05 | 0.22 | 0.19 | 0.05 | 0.27 |
| 6 | 138617294 | rs4896339 | -1.122 | -2.086 | 0.021 | 0.372 | 0.065 | -0.035 | -0.035 | 0 | 0.32 | 0.1 | 0.39 | 0.64 | 0.2 | 0.43 |
| 6 | 138619511 | rs2076265 | -0.911 | -1.967 | 0.419 | 0.404 | -0.028 | 0.098 | 0.098 | 0 | 0.24 | 0.35 | 0.2 | 0.46 | 0.16 | 0.19 |
| 6 | 138628723 | rs2272892 | -0.677 | -1.832 | 0.131 | 0.478 | -0.013 | 0.066 | 0.066 | 0 | 0.21 | 0.28 | 0.18 | 0.46 | 0.16 | 0.16 |
| 6 | 138628807 | rs2272891 | -0.677 | -1.832 | 0.131 | 0.478 | -0.013 | 0.066 | 0.066 | 0 | 0.21 | 0.28 | 0.18 | 0.46 | 0.16 | 0.16 |
| 6 | 138630513 | rs9376340 | -0.101 | -1.794 | -0.226 | 0.499 | -0.012 | -0.131 | -0.131 | 0 | 0.17 | 0.02 | 0.15 | 0.46 | 0.15 | 0.16 |
| 6 | 138634579 | rs3817832 | -0.893 | -1.903 | 0.307 | -0.118 | -0.031 | -0.004 | -0.004 | 0.742 | 0.17 | 0.26 | 0.32 | 0.2 | 0.08 | 0.27 |
| 6 | 138642177 | rs3736707 | -0.819 | -1.715 | 0.122 | -0.118 | -0.01 | -0.002 | -0.002 | 0 | 0.18 | 0.22 | 0.27 | 0.2 | 0.11 | 0.33 |
| 6 | 138655028 | rs4895520 | -1.275 | -1.573 | 0.64 | -0.249 | -0.039 | -0.018 | -0.018 | 0 | 0.15 | 0.62 | 0.15 | 0.12 | 0.06 | 0.05 |
| 6 | 138656512 | rs203142 | 0.384 | -1.966 | -0.162 | -0.791 | -0.024 | 0.063 | 0.063 | 0.017 | 0.18 | 0.02 | 0.16 | 0.04 | 0.15 | 0.15 |
| 6 | 138657683 | rs6932923 | -1.101 | -1.921 | 0.032 | -0.728 | -0.039 | -0.018 | -0.018 | 0 | 0.16 | 0.66 | 0.15 | 0.12 | 0.06 | 0.05 |
| 6 | 138662396 | rs11964805 | -0.015 | -1.93 | 0.139 | -0.984 | 0.001 | -0.001 | -0.001 | 0 | 0.08 | 0.08 | 0.06 | 0.09 | 0.03 | 0.01 |
| 6 | 138663385 | rs7740144 | -1.45 | -2.001 | 1.169 | -1.026 | -0.034 | -0.02 | -0.02 | 1.338 | 0.15 | 0.44 | 0.13 | 0.09 | 0.06 | 0.05 |
| 6 | 138674754 | rs203133 | -1.317 | -2.04 | 1.158 | -1.159 | 0.037 | 0.095 | 0.095 | 0.048 | 0.41 | 0.76 | 0.38 | 0.15 | 0.29 | 0.22 |
| 6 | 138683525 | rs17067480 | -0.734 | -1.951 | 0.974 | -1.104 | -0.036 | 0.005 | 0.005 | 1.115 | 0.11 | 0.36 | 0.09 | 0.09 | 0.03 | 0.01 |
| 6 | 138696398 | rs203139 | -1.679 | -2.401 | -0.264 | -0.994 | -0.103 | -0.122 | -0.122 | 0 | 0.64 | 0.94 | 0.65 | 0.61 | 0.44 | 0.53 |
| 6 | 138715607 | rs388676 | 0.446 | -1.945 | 0.52 | -1.639 | -0.024 | 0.031 | 0.031 | 0.127 | 0.19 | 0.02 | 0.16 | 0.08 | 0.16 | 0.17 |
| 6 | 138723962 | rs4896353 | -1.903 | -1.569 | 1.128 | -1.289 | -0.07 | -0.048 | -0.048 | 0 | 0.17 | 0.49 | 0.14 | 0.09 | 0.05 | 0.07 |
| 6 | 138734029 | rs2076279 | -0.336 | -1.238 | 0.566 | -1.059 | 0.007 | -0.002 | -0.002 | 0 | 0.08 | 0.05 | 0.06 | 0.1 | 0.03 | 0.01 |
| 6 | 138735409 | rs9321658 | -1.11 | -1.021 | 0.85 | -1.072 | 0.021 | -0.046 | -0.046 | 0.014 | 0.08 | 0.28 | 0.11 | 0 | 0.08 | 0.05 |
| 6 | 138740439 | rs6570235 | -1.036 | -1.021 | 0.651 | -1.244 | -0.013 | -0.035 | -0.035 | 0.668 | 0.23 | 0.35 | 0.14 | 0.1 | 0.14 | 0.06 |
| 6 | 138742172 | rs12208658 | 1.344 | -0.436 | 0.07 | -1.979 | -0.051 | 0.041 | 0.041 | 0 | 0.21 | 0.07 | 0.17 | 0.07 | 0.32 | 0.22 |
| 6 | 138742249 | rs7776384 | 1.091 | -0.814 | 0.695 | -1.693 | 0.021 | 0.043 | 0.043 | 0 | 0.48 | 0.34 | 0.64 | 0.9 | 0.67 | 0.73 |
| 6 | 138747618 | rs4895523 | -1.17 | -0.492 | 0.912 | -1.318 | 0.01 | -0.071 | -0.071 | 0 | 0.12 | 0.44 | 0.1 | 0.02 | 0.06 | 0.03 |
| 6 | 138753337 | rs2039121 | 0.644 | -0.464 | 0.717 | -1.366 | -0.006 | -0.045 | -0.045 | 0 | 0.63 | 0.57 | 0.72 | 0.9 | 0.74 | 0.8 |
| 6 | 138754323 | rs2327891 | -0.124 | -0.272 | -0.15 | -1.185 | -0.004 | 0.103 | 0.103 | 0 | 0.74 | 0.95 | 0.82 | 0.93 | 0.79 | 0.83 |
| 6 | 138764089 | rs9968931 | -1.147 | 0.389 | 1.015 | -0.524 | -0.038 | -0.133 | -0.133 | 0 | 0.23 | 0.62 | 0.18 | 0.02 | 0.13 | 0.04 |
| 6 | 138771952 | rs10499200 | -1.032 | 1.28 | 0.626 | 0.125 | -0.001 | -0.021 | -0.021 | 0 | 0.12 | 0.13 | 0.12 | 0 | 0.17 | 0.09 |
| 6 | 138775667 | rs17726669 | -1.466 | 1.843 | 0.509 | 0.056 | -0.064 | 0.056 | 0.056 | 0 | 0.15 | 0.1 | 0.17 | 0 | 0.29 | 0.1 |
| 6 | 138791986 | rs12529521 | -1.158 | 0.959 | 0.838 | 1.127 | 0.007 | 0.021 | 0.021 | 0 | 0.16 | 0.22 | 0.35 | 0.29 | 0.2 | 0.16 |
| 6 | 138794628 | rs7765573 | -1.193 | 1.003 | 0.881 | 1.062 | 0.018 | 0.05 | 0.05 | 0 | 0.16 | 0.24 | 0.4 | 0.32 | 0.22 | 0.17 |
| 6 | 138807577 | rs6937832 | -1.185 | 0.449 | 0.981 | 0.67 | -0.007 | -0.024 | -0.024 | 0.325 | 0.1 | 0.29 | 0.08 | 0.04 | 0.07 | 0.01 |
| 6 | 138818837 | rs4896370 | 0.169 | 0.643 | 0.148 | 1.016 | -0.003 | -0.005 | -0.005 | 0.343 | 0.61 | 0.63 | 0.53 | 0.52 | 0.62 | 0.47 |
| 6 | 138819510 | rs7746234 | 0.522 | 0.292 | 0.317 | 0.892 | 0 | 0 | 0 | 0.692 | 0.53 | 0.45 | 0.48 | 0.5 | 0.59 | 0.46 |
| 6 | 138820324 | rs10457670 | -0.035 | 0.243 | -0.445 | 0.982 | -0.059 | -0.044 | -0.044 | 0.388 | 0.29 | 0.09 | 0.38 | 0.38 | 0.33 | 0.44 |
| 6 | 138821708 | rs12527125 | 0.62 | -0.366 | -0.349 | 0.989 | 0.001 | 0.041 | 0.041 | 1.385 | 0.5 | 0.19 | 0.45 | 0.43 | 0.58 | 0.44 |
| 6 | 138822629 | rs9389585 | -0.955 | -0.971 | -0.611 | 1.825 | 0.013 | -0.043 | -0.043 | 3.101 | 0.13 | 0.07 | 0.09 | 0.34 | 0.12 | 0.19 |
| 6 | 138823548 | rs9389586 | -0.284 | -1.718 | -1.477 | 1.634 | 0.085 | -0.045 | -0.045 | 0.791 | 0.39 | 0.08 | 0.27 | 0.41 | 0.29 | 0.32 |
| 6 | 138823991 | rs12193890 | -0.078 | -1.859 | -1.528 | 1.629 | 0.068 | -0.074 | -0.074 | 0.117 | 0.35 | 0.07 | 0.26 | 0.42 | 0.28 | 0.28 |
| 6 | 138824122 | rs9495093 | -0.771 | -2.206 | -0.167 | 2.045 | -0.034 | 0.36 | 0.36 | 0.686 | 0.11 | 0.5 | 0.28 | 0.37 | 0.04 | 0.19 |
| 6 | 138825687 | rs7754416 | -1.359 | -1.751 | 0.549 | 1.703 | 0.048 | 0.077 | 0.077 | 0.059 | 0.12 | 0.39 | 0.17 | 0.16 | 0.14 | 0.12 |
| 6 | 138830566 | rs6922771 | -0.961 | -2.329 | 0.221 | 0.584 | -0.021 | 0.023 | 0.023 | 1.439 | 0.69 | 0.73 | 0.7 | 0.94 | 0.51 | 0.64 |
| 6 | 138830872 | rs9402978 | 0.07 | -2.408 | 0.624 | 0.614 | 0.036 | 0.057 | 0.057 | 1.842 | 0.57 | 0.33 | 0.51 | 0.85 | 0.37 | 0.57 |
| 6 | 138833114 | rs9402979 | 0.704 | -2.288 | 0.542 | 0.998 | 0.057 | -0.131 | -0.131 | 0 | 0.39 | 0.05 | 0.33 | 0.47 | 0.27 | 0.27 |
| 6 | 138834164 | rs6570241 | -1.159 | -1.825 | 0.905 | 1.164 | 0.064 | 0.004 | 0.004 | 0.001 | 0.1 | 0.43 | 0.11 | 0.06 | 0.08 | 0.07 |
| 6 | 138836972 | rs4074123 | -2.372 | -1.862 | 0.978 | 1.179 | -0.011 | 0.03 | 0.03 | 0.547 | 0.15 | 0.58 | 0.2 | 0.17 | 0.12 | 0.18 |
| 6 | 138841157 | rs6570242 | 0.703 | -1.942 | 0.671 | 1.03 | 0.079 | -0.043 | -0.043 | 0 | 0.46 | 0.11 | 0.37 | 0.46 | 0.34 | 0.27 |
| 6 | 138846307 | rs9484171 | -0.682 | -1.775 | 0.994 | 0.635 | 0.079 | 0.052 | 0.052 | 0 | 0.07 | 0.36 | 0.07 | 0.06 | 0.05 | 0.03 |
| 6 | 138857296 | rs3923724 | -0.531 | -1.931 | 0.717 | 0.671 | -0.029 | 0.018 | 0.018 | 1.553 | 0.63 | 0.78 | 0.62 | 0.65 | 0.46 | 0.43 |
| 6 | 138864637 | rs9399243 | -0.296 | -1.887 | 1.178 | 0.452 | 0 | 0 | 0 | 0 | 0.58 | 0.58 | 0.55 | 0.61 | 0.44 | 0.46 |
| 6 | 138865324 | rs4895532 | -0.046 | -2.052 | 1.1 | 0.465 | 0.029 | 0.027 | 0.027 | 1.419 | 0.54 | 0.32 | 0.48 | 0.59 | 0.41 | 0.4 |
| 6 | 138872074 | rs4131177 | 0.225 | -2.1 | 1.035 | 0.608 | 0.04 | 0.004 | 0.004 | 1.574 | 0.5 | 0.52 | 0.5 | 0.49 | 0.32 | 0.27 |
| 6 | 138872645 | rs4481452 | 0.507 | -2.042 | 0.496 | 0.639 | 0.129 | -0.025 | -0.025 | 0.054 | 0.47 | 0.16 | 0.47 | 0.49 | 0.32 | 0.27 |
| 6 | 138874805 | rs6914360 | -1.295 | -1.471 | 0.999 | 0.671 | -0.001 | 0.003 | 0.003 | 0 | 0.09 | 0.1 | 0.08 | 0.11 | 0.09 | 0.14 |
| 6 | 138875426 | rs4498379 | -0.064 | -1.725 | 1.402 | 0.432 | -0.016 | -0.025 | -0.025 | 0 | 0.12 | 0.43 | 0.07 | 0.03 | 0.06 | 0.02 |
| 6 | 138876411 | rs9495126 | -1.93 | -1.823 | 1.314 | 0.383 | 0.034 | 0.021 | 0.021 | 1.379 | 0.24 | 0.79 | 0.17 | 0.14 | 0.15 | 0.17 |
| 6 | 138877865 | rs6901012 | -1.03 | -1.683 | 1.263 | 0.443 | -0.001 | 0.003 | 0.003 | 0 | 0.09 | 0.09 | 0.07 | 0.11 | 0.09 | 0.14 |
| 6 | 138893460 | rs7774918 | -0.533 | -1.761 | 1.903 | 0.289 | -0.026 | -0.077 | -0.077 | 0 | 0.14 | 0.5 | 0.08 | 0.02 | 0.07 | 0.03 |
| 6 | 138898188 | rs9495139 | -0.606 | -1.653 | 1.96 | 0.39 | -0.009 | -0.037 | -0.037 | 0.174 | 0.15 | 0.38 | 0.12 | 0.07 | 0.1 | 0.09 |
| 6 | 138899640 | rs7754723 | -1.625 | -1.792 | 1.779 | 0.478 | -0.005 | -0.002 | -0.002 | 0 | 0.24 | 0.68 | 0.15 | 0.19 | 0.15 | 0.15 |

| | | | | | | | | | | | | | | | | |
|---|---|---|---|---|---|---|---|---|---|---|---|---|---|---|---|---|
| 6 | 138900935 | rs9495144 | -0.396 | -1.799 | 1.799 | 0.393 | -0.027 | -0.061 | -0.061 | 0 | 0.14 | 0.38 | 0.07 | 0.04 | 0.07 | 0.03 |
| 6 | 138902869 | rs7751302 | -1.928 | -1.799 | 1.4 | 0.431 | 0.022 | 0.011 | 0.011 | 1.64 | 0.27 | 0.75 | 0.22 | 0.2 | 0.2 | 0.25 |
| 6 | 138905857 | rs4576272 | -1.064 | -1.692 | 1.668 | 0.502 | 0.025 | -0.062 | -0.062 | 0 | 0.11 | 0.43 | 0.09 | 0.01 | 0.08 | 0.02 |
| 6 | 138909661 | rs6923972 | -0.478 | -2.536 | 0.779 | 0.301 | 0.121 | 0.056 | 0.056 | 0.695 | 0.46 | 0.79 | 0.55 | 0.58 | 0.34 | 0.48 |
| 6 | 138909908 | rs4629702 | 1.031 | -2.397 | 0.59 | 0.325 | 0.048 | -0.142 | -0.142 | 0 | 0.28 | 0.03 | 0.34 | 0.39 | 0.21 | 0.28 |
| 6 | 138914474 | rs4445072 | 0.042 | -2.36 | 0.804 | 0.282 | -0.031 | 0.008 | 0.008 | 0.873 | 0.39 | 0.45 | 0.42 | 0.4 | 0.28 | 0.31 |
| 6 | 138929076 | rs4896377 | 0.026 | -1.971 | 0.046 | -0.324 | 0.071 | 0.107 | 0.107 | 0.677 | 0.54 | 0.23 | 0.51 | 0.78 | 0.37 | 0.6 |
| 6 | 138929434 | rs9484186 | 0.092 | -0.93 | 0.092 | -0.171 | -0.001 | 0.001 | 0.001 | 0 | 0.24 | 0.27 | 0.2 | 0.23 | 0.23 | 0.33 |
| 6 | 138933559 | rs4896378 | -0.813 | -0.936 | 0.07 | -0.276 | 0.003 | 0.012 | 0.012 | 0.024 | 0.36 | 0.78 | 0.34 | 0.32 | 0.32 | 0.43 |
| 6 | 138934878 | rs9399248 | 0.899 | -1.264 | 0.035 | 0.023 | 0 | -0.058 | -0.058 | 1.838 | 0.1 | 0.01 | 0.07 | 0.27 | 0.07 | 0.17 |
| 6 | 138939675 | rs9495159 | -0.578 | -1.191 | 0.58 | -0.042 | 0.038 | 0.023 | 0.023 | 0 | 0.1 | 0.36 | 0.11 | 0.08 | 0.1 | 0.1 |
| 6 | 138943139 | rs9389597 | -1.723 | -0.656 | 0.057 | -0.213 | -0.033 | 0.031 | 0.031 | 1.128 | 0.78 | 0.85 | 0.82 | 0.97 | 0.68 | 0.91 |
| 6 | 138943931 | rs10085225 | -0.492 | -0.905 | 0.937 | 0.443 | -0.016 | 0.015 | 0.015 | 0 | 0.17 | 0.28 | 0.12 | 0.28 | 0.11 | 0.18 |
| 6 | 138944142 | rs6928475 | -1.625 | -0.857 | 0.543 | 0.395 | 0.017 | -0.002 | -0.002 | 0 | 0.5 | 0.75 | 0.49 | 0.6 | 0.46 | 0.62 |
| 6 | 138945826 | rs12190796 | -0.368 | -0.19 | 0.597 | 0.161 | -0.019 | 0.027 | 0.027 | 0 | 0.2 | 0.11 | 0.17 | 0.16 | 0.22 | 0.26 |
| 6 | 138966214 | rs6570248 | -1.159 | -0.226 | 0.723 | 0.227 | 0.006 | 0.013 | 0.013 | 0 | 0.27 | 0.5 | 0.29 | 0.31 | 0.27 | 0.41 |
| 6 | 138967387 | rs12197358 | -0.386 | -0.23 | 0.731 | 0.224 | -0.017 | 0.009 | 0.009 | 0 | 0.21 | 0.11 | 0.18 | 0.18 | 0.22 | 0.26 |
| 6 | 138970367 | rs6570251 | -1.166 | -0.235 | 0.705 | 0.203 | 0.006 | 0.013 | 0.013 | 0 | 0.27 | 0.49 | 0.29 | 0.31 | 0.27 | 0.41 |
| 6 | 138984235 | rs7769368 | -1.314 | -0.146 | 0.888 | 0.511 | 0.027 | 0.067 | 0.067 | 0.151 | 0.11 | 0.46 | 0.15 | 0.14 | 0.08 | 0.16 |
| 6 | 138994934 | rs12196991 | -0.53 | -0.208 | -0.772 | 0.089 | -0.015 | 0.062 | 0.062 | 0.928 | 0.22 | 0.04 | 0.16 | 0.16 | 0.21 | 0.34 |
| 6 | 138996154 | rs4896381 | -1.156 | -0.384 | -1.416 | -0.355 | -0.101 | 0.164 | 0.164 | 4.284 | 0.63 | 0.93 | 0.69 | 0.92 | 0.51 | 0.84 |
| 6 | 138998697 | rs6570261 | -1.147 | -0.349 | -1.051 | -0.372 | -0.056 | 0.121 | 0.121 | 4.33 | 0.62 | 0.84 | 0.68 | 0.92 | 0.52 | 0.85 |
| 6 | 139006406 | rs9376361 | -0.652 | -0.64 | -0.648 | 0.371 | 0.007 | -0.004 | -0.004 | 0 | 0.42 | 0.67 | 0.5 | 0.54 | 0.39 | 0.68 |
| 6 | 139013015 | rs9321669 | -0.02 | -0.839 | 0.35 | 0.34 | -0.057 | 0.089 | 0.089 | 0 | 0.2 | 0.47 | 0.18 | 0.38 | 0.14 | 0.12 |
| 6 | 139017780 | rs4348337 | -0.345 | -0.962 | 0.44 | 0.722 | -0.001 | -0.004 | -0.004 | 0 | 0.15 | 0.17 | 0.27 | 0.22 | 0.15 | 0.14 |
| 6 | 139018132 | rs1129180 | -0.211 | -0.807 | 0.421 | 0.677 | -0.02 | 0.012 | 0.012 | 0.503 | 0.25 | 0.48 | 0.23 | 0.33 | 0.22 | 0.52 |
| 6 | 139019029 | rs6901919 | -0.082 | -0.926 | -0.094 | 0.693 | -0.006 | 0.003 | 0.003 | 0.82 | 0.24 | 0.22 | 0.21 | 0.32 | 0.21 | 0.52 |
| 6 | 139023312 | rs4598095 | -0.816 | -1.069 | -0.287 | 0.681 | 0.01 | -0.009 | -0.009 | 0.462 | 0.28 | 0.69 | 0.23 | 0.44 | 0.2 | 0.25 |
| 6 | 139025468 | rs4896387 | -1.633 | -0.648 | -0.944 | 0.346 | -0.137 | 0.155 | 0.155 | 2.57 | 0.63 | 0.94 | 0.68 | 0.92 | 0.53 | 0.84 |
| 6 | 139026203 | rs6912737 | -1.487 | -1.208 | -0.87 | 0.357 | -0.052 | 0.167 | 0.167 | 0.775 | 0.61 | 0.84 | 0.65 | 0.92 | 0.47 | 0.83 |
| 6 | 139034727 | rs4895538 | -0.866 | -0.934 | -0.724 | 0.63 | -0.001 | -0.001 | -0.001 | 1.495 | 0.46 | 0.42 | 0.37 | 0.63 | 0.38 | 0.61 |
| 6 | 139035790 | rs4288220 | 0.185 | -0.919 | -1.478 | 0.65 | 0.024 | 0.041 | 0.041 | 0 | 0.09 | 0.02 | 0.05 | 0 | 0.05 | 0.02 |
| 6 | 139042338 | rs4896391 | -0.341 | -0.903 | -1.503 | 1.029 | 0.03 | -0.098 | -0.098 | 0 | 0.18 | 0.03 | 0.13 | 0.41 | 0.13 | 0.13 |
| 6 | 139044135 | rs4896392 | -0.801 | -0.936 | -1.372 | 0.335 | 0.09 | 0.064 | 0.064 | 0 | 0.44 | 0.78 | 0.47 | 0.72 | 0.32 | 0.53 |
| 6 | 139050948 | rs10457675 | 0.584 | 0.02 | -2.049 | -0.237 | -0.072 | 0.058 | 0.058 | 4.521 | 0.33 | 0.04 | 0.22 | 0.11 | 0.32 | 0.1 |
| 6 | 139051062 | rs4434486 | -0.113 | 0.381 | -2.277 | -0.702 | -0.012 | 0.07 | 0.07 | 0 | 0.83 | 0.91 | 0.89 | 1 | 0.86 | 0.92 |
| 6 | 139056007 | rs4895539 | 0.217 | 0.165 | -2.409 | -0.489 | -0.001 | -0.039 | -0.039 | 0.003 | 0.27 | 0.09 | 0.47 | 0.68 | 0.29 | 0.58 |
| 6 | 139062907 | rs7749041 | 0.367 | -0.442 | -1.592 | -1.121 | 0.017 | -0.126 | -0.126 | 0.531 | 0.68 | 0.55 | 0.73 | 0.9 | 0.69 | 0.73 |
| 6 | 139075341 | rs4538729 | 0.107 | -0.296 | -1.601 | -1.401 | -0.008 | 0.178 | 0.178 | 0.114 | 0.48 | 0.28 | 0.6 | 0.8 | 0.5 | 0.74 |
| 6 | 139083742 | rs6911335 | -0.36 | -0.273 | -1.491 | -1.607 | 0.001 | 0.155 | 0.155 | 0.98 | 0.58 | 0.79 | 0.73 | 0.89 | 0.57 | 0.78 |
| 6 | 139094355 | rs2273509 | 1.317 | -0.347 | -1.123 | -0.638 | -0.026 | -0.086 | -0.086 | 0 | 0.18 | 0.08 | 0.22 | 0.41 | 0.18 | 0.33 |
| 6 | 139095721 | rs9376369 | 0.307 | -0.342 | -1.018 | -0.662 | -0.007 | -0.062 | -0.062 | 0 | 0.27 | 0.16 | 0.42 | 0.6 | 0.28 | 0.53 |
| 6 | 139096620 | rs9403000 | 0.405 | -0.298 | -1.03 | -0.628 | -0.03 | -0.083 | -0.083 | 0 | 0.26 | 0.1 | 0.42 | 0.62 | 0.28 | 0.61 |
| 6 | 139102366 | rs6927344 | -0.156 | -0.185 | 0.301 | -0.379 | 0.031 | 0.041 | 0.041 | 0 | 0.05 | 0.14 | 0.03 | 0.12 | 0.04 | 0.11 |
| 6 | 139110345 | rs11155003 | -0.04 | -0.274 | 0.102 | -0.454 | -0.005 | 0.004 | 0.004 | 0.1 | 0.31 | 0.52 | 0.23 | 0.29 | 0.23 | 0.22 |
| 6 | 139114163 | rs6756 | -0.177 | -0.309 | -0.066 | -0.622 | -0.006 | -0.013 | -0.013 | 0 | 0.34 | 0.47 | 0.26 | 0.27 | 0.28 | 0.12 |
| 6 | 139115518 | rs1474962 | -0.087 | -0.198 | -0.495 | -0.447 | 0 | -0.001 | -0.001 | 0 | 0.3 | 0.65 | 0.25 | 0.27 | 0.23 | 0.14 |
| 6 | 139122240 | rs10457678 | 0.735 | -0.194 | -0.968 | -0.485 | -0.054 | -0.046 | -0.046 | 0.396 | 0.24 | 0.04 | 0.25 | 0.17 | 0.25 | 0.09 |
| 6 | 139122341 | rs9321674 | 0.725 | -0.194 | -0.968 | -0.485 | -0.054 | -0.046 | -0.046 | 0 | 0.24 | 0.04 | 0.25 | 0.17 | 0.25 | 0.09 |
| 6 | 139125140 | rs722653 | -0.143 | -0.182 | -0.864 | -0.968 | -0.012 | 0.067 | 0.067 | 0.294 | 0.55 | 0.74 | 0.59 | 0.73 | 0.5 | 0.72 |
| 6 | 139130213 | rs911081 | -0.773 | -0.193 | -0.8 | -0.94 | -0.008 | 0.007 | 0.007 | 0 | 0.79 | 0.86 | 0.84 | 0.89 | 0.75 | 0.78 |
| 6 | 139131062 | rs9373223 | 1.093 | -0.199 | -0.786 | -0.696 | 0.001 | -0.032 | -0.032 | 0 | 0.33 | 0.17 | 0.31 | 0.42 | 0.3 | 0.34 |
| 6 | 139131118 | rs9376370 | -0.569 | -0.126 | -0.349 | -0.335 | 0.007 | 0.111 | 0.111 | 2.301 | 0.18 | 0.5 | 0.28 | 0.31 | 0.21 | 0.38 |
| 6 | 139132840 | rs12664876 | 0.283 | -0.217 | -1.092 | -0.194 | -0.001 | -0.089 | -0.089 | 0 | 0.3 | 0.13 | 0.45 | 0.4 | 0.32 | 0.28 |
| 6 | 139134112 | rs746864 | 0.038 | -0.334 | -1.025 | -0.097 | -0.003 | -0.005 | -0.005 | 0 | 0.24 | 0.3 | 0.25 | 0.29 | 0.27 | 0.19 |
| 6 | 139134754 | rs4895548 | 0.382 | -0.403 | -0.67 | -0.109 | -0.002 | -0.002 | -0.002 | 0 | 0.44 | 0.48 | 0.48 | 0.61 | 0.38 | 0.65 |
| 6 | 139135077 | rs9389619 | 0.303 | -0.377 | -1.119 | -0.122 | -0.005 | 0.012 | 0.012 | 0 | 0.35 | 0.27 | 0.32 | 0.29 | 0.36 | 0.21 |
| 6 | 139135418 | rs9389620 | 1.326 | -0.306 | -1.45 | -0.197 | -0.016 | -0.061 | -0.061 | 3.819 | 0.26 | 0.05 | 0.14 | 0.35 | 0.21 | 0.44 |
| 6 | 139139274 | rs7751893 | -0.545 | 0.069 | -0.756 | -0.163 | -0.021 | 0.036 | 0.036 | 0 | 0.78 | 0.85 | 0.79 | 0.9 | 0.72 | 0.84 |
| 6 | 139141137 | rs9321677 | 0.515 | 0.159 | -0.295 | 0.186 | 0.005 | 0.02 | 0.02 | 0 | 0.7 | 0.62 | 0.54 | 0.63 | 0.66 | 0.64 |
| 6 | 139142909 | rs12193993 | 0.876 | 0.529 | -1.015 | 0.358 | -0.06 | 0.066 | 0.066 | 0 | 0.24 | 0.05 | 0.27 | 0.09 | 0.28 | 0.11 |
| 6 | 139142941 | rs998035 | -0.256 | 0.493 | -0.667 | 0.395 | -0.001 | -0.001 | -0.001 | 0.648 | 0.34 | 0.33 | 0.5 | 0.35 | 0.38 | 0.31 |
| 6 | 139143007 | rs6938729 | 0.374 | 0.402 | -0.957 | 0.429 | 0.011 | -0.015 | -0.015 | 0 | 0.08 | 0.02 | 0.19 | 0.26 | 0.08 | 0.2 |
| 6 | 139143088 | rs7773177 | 0.787 | 0.403 | -0.924 | 0.412 | -0.02 | 0.055 | 0.055 | 0 | 0.25 | 0.13 | 0.28 | 0.09 | 0.28 | 0.11 |
| 6 | 139148695 | rs17067940 | 0.332 | 0.474 | -1.159 | 0.468 | 0.004 | -0.016 | -0.016 | 0 | 0.08 | 0.02 | 0.2 | 0.26 | 0.09 | 0.2 |
| 6 | 139151784 | rs11155012 | 0.685 | 0.686 | -1.08 | -0.068 | -0.042 | 0.068 | 0.068 | 0 | 0.24 | 0.09 | 0.27 | 0.09 | 0.28 | 0.11 |
| 6 | 139158192 | rs4896409 | -0.096 | 0.304 | -0.832 | 0.043 | -0.001 | 0.006 | 0.006 | 0 | 0.59 | 0.58 | 0.5 | 0.8 | 0.46 | 0.75 |
| 6 | 139161261 | rs9376375 | -0.075 | 0.087 | -0.966 | 0.206 | 0.006 | 0.009 | 0.009 | 0 | 0.42 | 0.6 | 0.27 | 0.39 | 0.35 | 0.56 |
| 6 | 139167579 | rs9399258 | 0.612 | 0.667 | -1.192 | 0.399 | -0.031 | -0.062 | -0.062 | 0 | 0.62 | 0.22 | 0.44 | 0.54 | 0.6 | 0.65 |

| | | | | | | | | | | | | | | | | |
|---|---|---|---|---|---|---|---|---|---|---|---|---|---|---|---|---|
| 6 | 139186074 | rs1730387 | -0.388 | 0.756 | -0.981 | -0.119 | 0.021 | 0.065 | 0.065 | 0 | 0.47 | 0.63 | 0.7 | 0.22 | 0.59 | 0.36 |
| 6 | 139187231 | rs12213988 | 0.484 | 1.118 | -1.116 | -0.181 | -0.023 | 0.048 | 0.048 | 0 | 0.27 | 0.16 | 0.4 | 0.12 | 0.35 | 0.2 |
| 6 | 139192374 | rs1188852 | -0.903 | 0.891 | -1.276 | 0.031 | 0.025 | 0.008 | 0.008 | 2.111 | 0.55 | 0.87 | 0.73 | 0.38 | 0.63 | 0.46 |
| 6 | 139193603 | rs10872499 | -0.077 | 1.052 | -0.817 | -0.216 | -0.005 | 0.013 | 0.013 | 0 | 0.29 | 0.22 | 0.42 | 0.12 | 0.35 | 0.2 |
| 6 | 139196861 | rs6937315 | 0.328 | 0.443 | -0.708 | -0.016 | 0.021 | 0.018 | 0.018 | 0 | 0.45 | 0.6 | 0.34 | 0.36 | 0.35 | 0.54 |
| 6 | 139199890 | rs6933108 | 0.373 | -0.019 | -0.687 | -0.279 | -0.011 | -0.011 | -0.011 | 0 | 0.39 | 0.5 | 0.24 | 0.32 | 0.32 | 0.5 |
| 6 | 139200330 | rs1188857 | 0.801 | 0.918 | -1.202 | -0.77 | -0.088 | 0.149 | 0.149 | 0 | 0.22 | 0.02 | 0.18 | 0 | 0.27 | 0.1 |
| 6 | 139201909 | rs6902027 | -0.6 | 0.181 | -0.282 | -0.451 | 0.001 | -0.046 | -0.046 | 0 | 0.16 | 0.36 | 0.21 | 0.1 | 0.15 | 0.1 |
| 6 | 139205386 | rs9376382 | 0.293 | -0.417 | -0.362 | -0.126 | 0.017 | 0.009 | 0.009 | 0 | 0.5 | 0.45 | 0.28 | 0.66 | 0.29 | 0.61 |
| 6 | 139213267 | rs1188838 | 0.723 | 0.102 | -0.391 | -0.047 | -0.114 | 0.094 | 0.094 | 0 | 0.3 | 0.06 | 0.54 | 0.11 | 0.44 | 0.26 |
| 6 | 139213276 | rs11962409 | -0.105 | 0.055 | -0.146 | -0.016 | -0.014 | 0.222 | 0.222 | 0 | 0.07 | 0.3 | 0.04 | 0.22 | 0.02 | 0.05 |
| 6 | 139221850 | rs1188873 | -0.735 | -0.254 | -0.128 | -0.051 | 0.009 | -0.04 | -0.04 | 0 | 0.75 | 0.87 | 0.83 | 0.64 | 0.77 | 0.74 |
| 6 | 139225925 | rs12619 | 0.298 | 0.024 | -0.343 | 0.058 | -0.001 | -0.001 | -0.001 | 0 | 0.37 | 0.39 | 0.58 | 0.33 | 0.46 | 0.31 |
| 6 | 139229872 | rs1044418 | 1.063 | -0.056 | -0.319 | 0.05 | 0.027 | -0.033 | -0.033 | 0 | 0.08 | 0.17 | 0.09 | 0.03 | 0.14 | 0.05 |
| 6 | 139241332 | rs9495285 | 1.063 | -0.051 | -0.337 | 0.052 | 0.027 | -0.033 | -0.033 | 0 | 0.08 | 0.17 | 0.09 | 0.03 | 0.14 | 0.06 |
| 6 | 139247076 | rs3763188 | 1.576 | -0.281 | -0.548 | -0.001 | 0.026 | -0.022 | -0.022 | 0 | 0.25 | 0.07 | 0.13 | 0.28 | 0.17 | 0.37 |
| 6 | 139248639 | rs1188880 | 0.499 | 0.307 | -0.395 | -0.117 | -0.113 | 0.091 | 0.091 | 0 | 0.3 | 0.09 | 0.55 | 0.11 | 0.45 | 0.26 |
| 6 | 139277883 | rs953229 | -0.035 | 0.266 | -0.268 | -0.016 | -0.024 | 0.155 | 0.155 | 0 | 0.06 | 0.21 | 0.03 | 0.22 | 0.02 | 0.05 |
| 6 | 139285630 | rs9376387 | 0.398 | 0.307 | -0.527 | -0.07 | -0.002 | -0.002 | -0.002 | 1.176 | 0.55 | 0.45 | 0.32 | 0.64 | 0.39 | 0.63 |
| 6 | 139294201 | rs6937040 | -1.156 | 0.361 | -0.626 | -0.266 | -0.125 | 0.167 | 0.167 | 0 | 0.73 | 0.96 | 0.76 | 1 | 0.63 | 0.85 |
| 6 | 139294798 | rs11962973 | -0.626 | -0.856 | -0.035 | 0.397 | 0.017 | 0.129 | 0.129 | 0.443 | 0.15 | 0.47 | 0.13 | 0.25 | 0.16 | 0.11 |
| 6 | 139319717 | rs9403023 | 0.193 | -1.155 | -1.135 | -0.164 | 0.114 | -0.027 | -0.027 | 0 | 0.28 | 0.05 | 0.09 | 0.27 | 0.11 | 0.38 |
| 6 | 139319971 | rs7761041 | -1.131 | -0.187 | -1.111 | -0.3 | -0.125 | 0.056 | 0.056 | 0.103 | 0.77 | 0.94 | 0.43 | 0.88 | 0.57 | 0.78 |
| 6 | 139325711 | rs9376393 | -0.432 | -0.682 | -0.736 | -0.285 | 0.001 | 0.014 | 0.014 | 0.051 | 0.65 | 0.69 | 0.35 | 0.83 | 0.4 | 0.71 |
| 6 | 139328660 | rs2498635 | 0.487 | -0.937 | -0.15 | 0.06 | -0.024 | 0.019 | 0.019 | 0.189 | 0.34 | 0.42 | 0.21 | 0.57 | 0.28 | 0.31 |
| 6 | 139328765 | rs12664486 | -0.188 | -0.673 | -0.646 | 0.13 | 0.029 | 0.022 | 0.022 | 0 | 0.46 | 0.68 | 0.32 | 0.61 | 0.43 | 0.39 |
| 6 | 139338617 | rs9484223 | 0.314 | -0.664 | -0.13 | 0.635 | -0.028 | 0.022 | 0.022 | 0 | 0.14 | 0.15 | 0.08 | 0.26 | 0.05 | 0.33 |
| 6 | 139339423 | rs9399268 | 0.316 | -0.705 | 0.065 | 0.626 | -0.05 | 0.052 | 0.052 | 0 | 0.16 | 0.25 | 0.1 | 0.27 | 0.06 | 0.35 |
| 6 | 139348484 | rs4391279 | -0.454 | -0.398 | -0.048 | 0.364 | 0.003 | 0.011 | 0.011 | 0 | 0.49 | 0.55 | 0.51 | 0.51 | 0.54 | 0.61 |
| 6 | 139355158 | rs3813331 | 0.308 | -1.697 | 0.025 | -0.1 | -0.001 | -0.001 | -0.001 | 0 | 0.07 | 0.07 | 0.08 | 0.06 | 0.1 | 0.06 |
| 6 | 139357615 | rs6918066 | -0.026 | -2.087 | 0.217 | -0.616 | -0.014 | 0.017 | 0.017 | 0.211 | 0.35 | 0.69 | 0.44 | 0.12 | 0.51 | 0.22 |
| 6 | 139358723 | rs7760083 | 0.534 | -2.426 | -0.05 | -0.449 | 0.003 | -0.004 | -0.004 | 0 | 0.14 | 0.11 | 0.13 | 0.19 | 0.05 | 0.28 |
| 6 | 139361937 | rs2504377 | 0.129 | -2.247 | 0.065 | -0.621 | 0.003 | 0.085 | 0.085 | 0.648 | 0.61 | 0.26 | 0.53 | 0.91 | 0.43 | 0.76 |
| 6 | 139363102 | rs6906235 | 0.721 | -1.844 | 0.51 | -0.494 | 0.071 | -0.08 | -0.08 | 0 | 0.25 | 0.35 | 0.32 | 0.05 | 0.39 | 0.15 |
| 6 | 139365230 | rs9495324 | 0.832 | -1.724 | 0.553 | -0.535 | 0.098 | -0.095 | -0.095 | 0.018 | 0.24 | 0.42 | 0.29 | 0.05 | 0.36 | 0.14 |
| 6 | 139369543 | rs7769010 | 0.862 | -2.043 | -0.098 | -0.139 | 0.063 | 0.029 | 0.029 | 0 | 0.42 | 0.12 | 0.41 | 0.72 | 0.37 | 0.5 |
| 6 | 139379566 | rs2498646 | 1.677 | -2.958 | 1.063 | 0.51 | 0.033 | 0.028 | 0.028 | 0.137 | 0.62 | 0.15 | 0.66 | 0.75 | 0.72 | 0.64 |
| 6 | 139391491 | rs2498680 | 1.016 | -2.785 | 1.105 | 0.568 | -0.022 | -0.041 | -0.041 | 0 | 0.71 | 0.73 | 0.76 | 0.91 | 0.8 | 0.66 |
| 6 | 139401474 | rs10872502 | 1.84 | -2.356 | 0.798 | 0.774 | 0.019 | -0.01 | -0.01 | 0.006 | 0.76 | 0.21 | 0.72 | 0.75 | 0.85 | 0.76 |
| 6 | 139403783 | rs2473500 | 0.571 | -2.172 | 0.896 | 0.694 | -0.017 | -0.038 | -0.038 | 0 | 0.87 | 0.8 | 0.86 | 0.94 | 0.92 | 0.87 |
| 6 | 139411399 | rs4896431 | 2.403 | -2.32 | 0.432 | 1.066 | -0.001 | -0.018 | -0.018 | 0 | 0.6 | 0.32 | 0.64 | 0.6 | 0.59 | 0.56 |
| 6 | 139412610 | rs9385849 | -0.443 | -2.28 | 0.417 | 1.1 | -0.011 | -0.015 | -0.015 | 0 | 0.06 | 0.04 | 0.04 | 0.17 | 0.07 | 0.1 |
| 6 | 139416226 | rs17068331 | -0.947 | -2.286 | 0.513 | 1.123 | 0.01 | 0.007 | 0.007 | 0.892 | 0.2 | 0.32 | 0.16 | 0.18 | 0.24 | 0.24 |
| 6 | 139418007 | rs9403037 | 1.958 | -2.241 | 0.265 | 1.141 | -0.001 | -0.008 | -0.008 | 0 | 0.61 | 0.39 | 0.65 | 0.6 | 0.59 | 0.56 |
| 6 | 139418576 | rs9495342 | -0.127 | -1.322 | -0.419 | 1.391 | -0.02 | -0.019 | -0.019 | 0 | 0.12 | 0.05 | 0.1 | 0.12 | 0.17 | 0.15 |
| 6 | 139420730 | rs2498652 | -1.537 | -1.208 | -0.006 | 1.246 | 0.045 | 0.083 | 0.083 | 0 | 0.09 | 0.2 | 0.07 | 0.18 | 0.14 | 0.12 |
| 6 | 139430596 | rs2498667 | -1.426 | -1.316 | -1.121 | 0.675 | 0.036 | 0.066 | 0.066 | 0.106 | 0.13 | 0.25 | 0.13 | 0.26 | 0.26 | 0.29 |
| 6 | 139431557 | rs10485229 | 0.698 | -1.399 | -1.307 | 0.727 | -0.019 | -0.077 | -0.077 | 0 | 0.11 | 0.01 | 0.14 | 0.41 | 0.09 | 0.14 |
| 6 | 139435577 | rs2473528 | -1.974 | -0.868 | -0.973 | 0.003 | 0.019 | 0.066 | 0.066 | 0.196 | 0.17 | 0.29 | 0.34 | 0.35 | 0.27 | 0.41 |
| 6 | 139437827 | rs17068343 | -1.362 | -1.062 | -0.843 | -0.983 | -0.008 | 0.021 | 0.021 | 0 | 0.08 | 0.11 | 0.23 | 0.16 | 0.05 | 0.16 |
| 6 | 139470762 | rs3777677 | -0.121 | -1.284 | -1.035 | -1.205 | 0.02 | 0.141 | 0.141 | 5.166 | 0.57 | 0.8 | 0.69 | 0.22 | 0.66 | 0.58 |
| 6 | 139497680 | rs3777671 | -0.416 | -1.242 | -0.958 | -1.165 | 0 | 0.025 | 0.025 | 0 | 0.06 | 0.13 | 0.04 | 0.06 | 0.04 | 0.03 |
| 6 | 139500135 | rs7745259 | 0.512 | -1.215 | -1.097 | -1.184 | 0.016 | 0.04 | 0.04 | 3.916 | 0.48 | 0.58 | 0.46 | 0.06 | 0.6 | 0.34 |
| 6 | 139500742 | rs3777667 | -1.476 | -0.914 | 0.164 | 0.408 | -0.009 | 0.094 | 0.094 | 0 | 0.08 | 0.21 | 0.23 | 0.16 | 0.04 | 0.16 |
| 6 | 139513381 | rs6932100 | -1.684 | -1.022 | 0.158 | 0.55 | -0.021 | 0.057 | 0.057 | 0 | 0.09 | 0.22 | 0.23 | 0.15 | 0.05 | 0.21 |
| 6 | 139518546 | rs7765885 | -1.442 | -0.962 | 0.08 | 0.353 | -0.015 | 0.035 | 0.035 | 0 | 0.09 | 0.22 | 0.18 | 0.12 | 0.04 | 0.23 |
| 6 | 139540103 | rs4596498 | 0.409 | -1.075 | -0.681 | -0.101 | 0.037 | 0.045 | 0.045 | 1.505 | 0.7 | 0.87 | 0.81 | 0.85 | 0.78 | 0.8 |
| 6 | 139549046 | rs7770356 | 0.493 | -1.144 | -0.386 | -0.188 | -0.007 | -0.012 | -0.012 | 0 | 0.56 | 0.45 | 0.72 | 0.81 | 0.68 | 0.7 |
| 6 | 139559885 | rs4896446 | 0.384 | -1.264 | -0.51 | 0.065 | 0.037 | 0.022 | 0.022 | 0.118 | 0.58 | 0.38 | 0.64 | 0.7 | 0.67 | 0.7 |
| 6 | 139563622 | rs9321711 | -1.006 | -0.977 | -0.551 | -0.49 | 0.001 | -0.002 | -0.002 | 0 | 0.13 | 0.09 | 0.05 | 0 | 0.1 | 0.03 |
| 6 | 139575852 | rs4534002 | -1.246 | -0.569 | -1.414 | -0.832 | -0.042 | 0.123 | 0.123 | 0.233 | 0.42 | 0.9 | 0.32 | 0.17 | 0.36 | 0.36 |
| 6 | 139576736 | rs17068451 | -1.115 | -0.591 | -0.083 | -0.951 | -0.017 | -0.088 | -0.088 | 0.339 | 0.21 | 0.55 | 0.13 | 0.03 | 0.15 | 0.06 |
| 6 | 139579009 | rs943651 | -0.522 | -0.655 | -0.037 | -0.95 | 0.017 | -0.022 | -0.022 | 0 | 0.2 | 0.33 | 0.19 | 0.11 | 0.21 | 0.29 |
| 6 | 139580792 | rs6903798 | -1.451 | -0.977 | -0.921 | -0.851 | -0.102 | 0.124 | 0.124 | 1.513 | 0.58 | 0.96 | 0.61 | 0.76 | 0.55 | 0.69 |
| 6 | 139590045 | rs726794 | -1.478 | -0.667 | -0.525 | -0.767 | -0.001 | 0.046 | 0.046 | 0.935 | 0.32 | 0.88 | 0.27 | 0.17 | 0.26 | 0.31 |
| 6 | 139593772 | rs6930395 | -0.599 | -0.556 | 0.395 | -0.53 | 0.013 | -0.003 | -0.003 | 0.888 | 0.15 | 0.27 | 0.15 | 0.12 | 0.17 | 0.26 |
| 6 | 139599209 | rs11965233 | -0.527 | -0.713 | 0.393 | -0.61 | 0.009 | -0.008 | -0.008 | 0 | 0.16 | 0.29 | 0.15 | 0.12 | 0.16 | 0.26 |
| 6 | 139600127 | rs9495409 | -0.829 | -0.743 | 0.892 | -0.592 | 0.036 | 0.012 | 0.012 | 0 | 0.14 | 0.37 | 0.15 | 0.11 | 0.14 | 0.25 |
| 6 | 139602317 | rs9495410 | -1.072 | -0.847 | 0.663 | -0.578 | 0 | -0.019 | -0.019 | 0 | 0.28 | 0.42 | 0.27 | 0.23 | 0.26 | 0.36 |

| | | | | | | | | | | | | | | | | |
|---|---|---|---|---|---|---|---|---|---|---|---|---|---|---|---|---|
| 6 | 139613017 | rs7746727 | -1.376 | -0.866 | -0.22 | -0.545 | 0.022 | 0.015 | 0.015 | 0.18 | 0.34 | 0.83 | 0.32 | 0.29 | 0.26 | 0.34 |
| 6 | 139617753 | rs9376406 | 0.056 | -0.623 | -0.496 | -0.267 | 0.033 | 0.013 | 0.013 | 0.076 | 0.45 | 0.27 | 0.57 | 0.56 | 0.63 | 0.5 |
| 6 | 139620056 | rs7756020 | -0.152 | -0.703 | 0.283 | -0.262 | 0.007 | 0.004 | 0.004 | 0 | 0.48 | 0.54 | 0.61 | 0.56 | 0.63 | 0.5 |
| 6 | 139622545 | rs3861395 | -0.484 | -0.895 | -0.178 | 0.266 | 0.063 | 0.212 | 0.212 | 0.657 | 0.52 | 0.79 | 0.78 | 0.81 | 0.65 | 0.58 |
| 6 | 139625074 | rs9385856 | -0.513 | -0.575 | -0.193 | 0.557 | -0.036 | -0.058 | -0.058 | 0.575 | 0.6 | 0.84 | 0.5 | 0.53 | 0.43 | 0.52 |
| 6 | 139625649 | rs9376408 | 1.331 | -0.96 | -0.519 | 0.271 | 0.008 | 0.021 | 0.021 | 7.317 | 0.3 | 0.01 | 0.12 | 0.16 | 0.2 | 0.17 |
| 6 | 139627411 | rs9495423 | 0.375 | -0.397 | 0.358 | 0.631 | 0.02 | 0.038 | 0.038 | 1.232 | 0.49 | 0.46 | 0.22 | 0.26 | 0.33 | 0.42 |
| 6 | 139630138 | rs6925036 | -1.555 | -0.094 | 0.455 | 1.139 | -0.021 | 0.129 | 0.129 | 1.869 | 0.12 | 0.57 | 0.29 | 0.28 | 0.1 | 0.1 |
| 6 | 139637172 | rs7770507 | 0.21 | 0.736 | -0.445 | 0.936 | -0.008 | -0.01 | -0.01 | 0.076 | 0.64 | 0.65 | 0.48 | 0.47 | 0.46 | 0.66 |
| 6 | 139637874 | rs7775616 | 0.727 | 1.152 | -0.717 | 1.197 | -0.034 | -0.049 | -0.049 | 0 | 0.92 | 1 | 0.87 | 0.69 | 0.87 | 0.88 |
| 6 | 139645295 | rs9484261 | -0.867 | 1.656 | -0.223 | -0.222 | 0.063 | -0.127 | -0.127 | 0 | 0.23 | 0.57 | 0.27 | 0.01 | 0.32 | 0.3 |
| 6 | 139645671 | rs13199562 | -0.346 | 2.03 | -0.051 | -0.172 | 0.031 | -0.047 | -0.047 | 0.307 | 0.11 | 0.2 | 0.11 | 0 | 0.18 | 0.13 |
| 6 | 139647006 | rs7757846 | -0.235 | 1.358 | -1.028 | -0.239 | 0.007 | 0.02 | 0.02 | 0 | 0.6 | 0.76 | 0.55 | 0.29 | 0.57 | 0.53 |
| 6 | 139651545 | rs11155050 | -0.272 | 0.827 | -1.387 | -1.263 | 0.03 | -0.009 | -0.009 | 0.475 | 0.66 | 0.86 | 0.68 | 0.68 | 0.71 | 0.7 |
| 6 | 139652888 | rs11754546 | -0.477 | 0.269 | -0.631 | -2.673 | 0.002 | -0.049 | -0.049 | 0.493 | 0.45 | 0.57 | 0.34 | 0.01 | 0.39 | 0.27 |
| 6 | 139660012 | rs11155053 | -0.406 | 0.363 | -0.322 | -1.986 | 0.025 | -0.045 | -0.045 | 0.56 | 0.09 | 0.14 | 0.08 | 0 | 0.11 | 0.04 |
| 6 | 139660091 | rs6925308 | -0.433 | -0.341 | -0.65 | -2.181 | 0.017 | 0.004 | 0.004 | 2.613 | 0.26 | 0.22 | 0.19 | 0 | 0.17 | 0.13 |
| 6 | 139663885 | rs7744152 | 0.271 | 0.09 | -0.507 | -1.677 | -0.025 | -0.008 | -0.008 | 0.742 | 0.35 | 0.26 | 0.42 | 0.69 | 0.35 | 0.53 |
| 6 | 139670131 | rs4243461 | 0.085 | 0.082 | 0.43 | -1.484 | 0.008 | 0.01 | 0.01 | 1.013 | 0.74 | 0.7 | 0.64 | 0.71 | 0.65 | 0.64 |
| 6 | 139675479 | rs9495450 | 1.034 | -0.147 | 0.203 | -1.67 | -0.025 | 0.055 | 0.055 | 0 | 0.47 | 0.22 | 0.55 | 0.68 | 0.45 | 0.56 |
| 6 | 139683605 | rs10872514 | -1.127 | -0.739 | 0.541 | -2.522 | 0.047 | 0.231 | 0.231 | 0 | 0.47 | 0.81 | 0.36 | 0.03 | 0.42 | 0.24 |
| 6 | 139686322 | rs6901049 | -0.226 | -0.7 | 0.84 | -2.526 | 0.004 | -0.066 | -0.066 | 0 | 0.21 | 0.31 | 0.21 | 0.03 | 0.23 | 0.16 |
| 6 | 139688287 | rs6913402 | -1.017 | -0.599 | 0.143 | -2.478 | 0.04 | 0.194 | 0.194 | 0 | 0.46 | 0.78 | 0.36 | 0.03 | 0.42 | 0.24 |
| 6 | 139691315 | rs4428519 | 0.19 | -0.456 | -0.225 | -2.366 | -0.045 | 0.045 | 0.045 | 0.641 | 0.16 | 0.02 | 0.15 | 0.03 | 0.2 | 0.15 |
| 6 | 139693834 | rs1131431 | -0.491 | -0.505 | -0.071 | -2.361 | -0.02 | -0.098 | -0.098 | 0 | 0.29 | 0.57 | 0.16 | 0 | 0.2 | 0.1 |
| 6 | 139695061 | rs1131400 | -0.367 | 0.132 | -0.287 | -1.339 | -0.009 | 0.017 | 0.017 | 1.096 | 0.16 | 0.11 | 0.16 | 0.03 | 0.19 | 0.15 |
| 6 | 139706222 | rs6927712 | -0.712 | -0.451 | -0.675 | -1.123 | 0.008 | -0.025 | -0.025 | 0 | 0.23 | 0.17 | 0.08 | 0 | 0.1 | 0.08 |
| 6 | 139716715 | rs10457687 | -0.797 | 0.711 | -1.244 | -1.584 | -0.054 | 0.102 | 0.102 | 0 | 0.23 | 0.06 | 0.1 | 0.01 | 0.24 | 0.11 |
| 6 | 139716941 | rs9373237 | -0.193 | 0.733 | -1.821 | -0.917 | -0.004 | -0.045 | -0.045 | 0 | 0.25 | 0.16 | 0.29 | 0.4 | 0.24 | 0.29 |
| 6 | 139721842 | rs11155061 | -0.844 | 0.546 | -0.573 | -0.805 | 0 | 0 | 0 | 0.009 | 0.49 | 0.52 | 0.41 | 0.4 | 0.44 | 0.38 |
| 6 | 139731038 | rs11155065 | -0.747 | 0.735 | -0.564 | -1.047 | -0.004 | 0 | 0 | 0.163 | 0.52 | 0.53 | 0.56 | 0.43 | 0.54 | 0.48 |
| 6 | 139742496 | rs9321717 | -0.15 | 0.108 | -0.464 | -0.85 | -0.008 | -0.006 | -0.006 | 0 | 0.58 | 0.55 | 0.57 | 0.61 | 0.65 | 0.52 |
| 6 | 139742788 | rs887780 | -0.298 | -0.471 | -0.245 | -0.998 | -0.002 | 0 | 0 | 0.016 | 0.72 | 0.74 | 0.67 | 0.66 | 0.72 | 0.57 |
| 6 | 139744364 | rs2041620 | 0.293 | 0.113 | -0.654 | -1.786 | 0.006 | 0.002 | 0.002 | 0 | 0.31 | 0.62 | 0.44 | 0.64 | 0.35 | 0.4 |
| 6 | 139745739 | rs1019237 | -0.795 | -0.005 | -0.673 | -1.725 | 0.002 | 0 | 0 | 1.967 | 0.72 | 0.78 | 0.68 | 0.7 | 0.72 | 0.59 |
| 6 | 139750875 | rs9495470 | -0.219 | -0.58 | -0.189 | -1.917 | -0.053 | -0.039 | -0.039 | 0.025 | 0.15 | 0.28 | 0.09 | 0.06 | 0.07 | 0.06 |
| 6 | 139757582 | rs10085329 | -0.749 | -0.486 | -0.431 | -0.505 | -0.03 | 0.042 | 0.042 | 2.325 | 0.23 | 0.66 | 0.29 | 0.43 | 0.14 | 0.25 |
| 6 | 139758486 | rs12529425 | 0.762 | -0.024 | -0.825 | -0.507 | -0.096 | -0.106 | -0.106 | 0.016 | 0.22 | 0.03 | 0.21 | 0.31 | 0.29 | 0.16 |
| 6 | 139759144 | rs9403072 | 0.766 | -0.027 | -0.827 | -0.508 | -0.096 | -0.106 | -0.106 | 0 | 0.22 | 0.03 | 0.21 | 0.31 | 0.29 | 0.16 |
| 6 | 139760195 | rs3010311 | -0.17 | -0.25 | -0.408 | -0.731 | 0.005 | 0.007 | 0.007 | 0 | 0.57 | 0.48 | 0.56 | 0.69 | 0.63 | 0.5 |
| 6 | 139760858 | rs9385862 | 0.39 | -0.121 | -0.138 | -0.516 | 0 | 0.182 | 0.182 | 0 | 0.05 | 0.11 | 0.18 | 0.36 | 0.06 | 0.19 |
| 6 | 139762442 | rs3010304 | -0.919 | -0.468 | 0.521 | -0.794 | -0.002 | 0.008 | 0.008 | 0 | 0.72 | 0.77 | 0.64 | 0.72 | 0.69 | 0.56 |
| 6 | 139763531 | rs2908498 | -0.073 | -0.323 | 0.273 | -0.886 | 0.008 | 0.038 | 0.038 | 0 | 0.54 | 0.3 | 0.37 | 0.33 | 0.53 | 0.28 |
| 6 | 139767988 | rs9389685 | 1.408 | -0.192 | 0.183 | -0.459 | -0.047 | -0.054 | -0.054 | 0 | 0.1 | 0.04 | 0.14 | 0.27 | 0.16 | 0.05 |
| 6 | 139771895 | rs9495484 | 0.64 | -0.352 | 0.179 | -0.583 | -0.024 | -0.027 | -0.027 | 0 | 0.16 | 0.17 | 0.09 | 0.06 | 0.09 | 0.08 |
| 6 | 139774005 | rs4896456 | 0.454 | -0.319 | -0.068 | -0.461 | 0.005 | 0.08 | 0.08 | 0 | 0.15 | 0.24 | 0.23 | 0.32 | 0.16 | 0.27 |
| 6 | 139774853 | rs3010301 | 1.179 | -0.091 | -0.619 | -0.563 | -0.04 | 0.129 | 0.129 | 0.212 | 0.28 | 0.08 | 0.15 | 0.07 | 0.27 | 0.15 |
| 6 | 139783488 | rs2908501 | -1.205 | -0.268 | -0.797 | -0.355 | -0.007 | 0.007 | 0.007 | 0 | 0.72 | 0.82 | 0.65 | 0.71 | 0.7 | 0.56 |
| 6 | 139787721 | rs2908504 | 0.62 | -0.1 | -1.046 | -0.534 | -0.082 | -0.001 | -0.001 | 0 | 0.4 | 0.14 | 0.31 | 0.35 | 0.46 | 0.21 |
| 6 | 139788803 | rs4410751 | -0.28 | -0.165 | -0.976 | -0.867 | -0.023 | 0.021 | 0.021 | 0 | 0.44 | 0.2 | 0.4 | 0.38 | 0.46 | 0.3 |
| 6 | 139791745 | rs2110183 | -0.21 | -0.066 | -1.137 | -1.007 | -0.006 | -0.004 | -0.004 | 1.193 | 0.47 | 0.47 | 0.58 | 0.35 | 0.57 | 0.4 |
| 6 | 139794302 | rs3010297 | 0.289 | 0.305 | -0.93 | -1.104 | 0.037 | -0.081 | -0.081 | 0.316 | 0.24 | 0.37 | 0.37 | 0.05 | 0.32 | 0.3 |
| 6 | 139801295 | rs9376419 | 1.248 | -0.036 | -1.405 | -0.701 | -0.02 | -0.086 | -0.086 | 1.763 | 0.17 | 0.02 | 0.16 | 0.35 | 0.18 | 0.27 |
| 6 | 139802205 | rs2080112 | -0.125 | -0.22 | -0.574 | -1.196 | 0.001 | -0.116 | -0.116 | 0 | 0.29 | 0.53 | 0.33 | 0.05 | 0.33 | 0.25 |
| 6 | 139803189 | rs3010291 | -0.903 | -0.413 | -0.847 | -0.382 | 0.021 | -0.006 | -0.006 | 0.037 | 0.81 | 0.87 | 0.85 | 0.74 | 0.86 | 0.75 |
| 6 | 139803301 | rs3010290 | -0.09 | 0.001 | -0.743 | -0.462 | -0.003 | -0.008 | -0.008 | 1.165 | 0.63 | 0.38 | 0.72 | 0.7 | 0.68 | 0.63 |
| 6 | 139806747 | rs7766444 | 0.585 | 0.384 | -1.233 | -0.006 | -0.066 | -0.072 | -0.072 | 0 | 0.23 | 0.06 | 0.33 | 0.31 | 0.3 | 0.18 |
| 6 | 139807600 | rs6916887 | -0.826 | -0.136 | -1.581 | -0.457 | -0.026 | 0.13 | 0.13 | 0 | 0.59 | 0.86 | 0.57 | 0.78 | 0.59 | 0.46 |
| 6 | 139808000 | rs4895561 | 0.636 | -0.366 | -1.328 | -0.532 | -0.012 | -0.043 | -0.043 | 0.195 | 0.19 | 0.13 | 0.14 | 0.43 | 0.23 | 0.22 |
| 6 | 139809258 | rs11970753 | 1.327 | -0.558 | -0.937 | -0.603 | -0.025 | -0.037 | -0.037 | 0 | 0.07 | 0.09 | 0.02 | 0 | 0.03 | 0.01 |
| 6 | 139811689 | rs9495497 | -0.292 | -0.582 | -0.752 | -0.609 | -0.019 | 0.066 | 0.066 | 0 | 0.32 | 0.43 | 0.21 | 0.46 | 0.3 | 0.29 |
| 6 | 139817115 | rs9376422 | -0.532 | -0.457 | -1.005 | -0.492 | 0.009 | 0.089 | 0.089 | 0 | 0.46 | 0.82 | 0.52 | 0.7 | 0.47 | 0.35 |
| 6 | 139820898 | rs3010286 | -1.47 | -0.389 | -1.236 | -0.742 | 0.011 | 0.037 | 0.037 | 0 | 0.73 | 0.85 | 0.77 | 0.8 | 0.78 | 0.65 |
| 6 | 139820958 | rs2908511 | 0.392 | -0.419 | -0.543 | -0.449 | 0.005 | -0.083 | -0.083 | 0.184 | 0.19 | 0.4 | 0.24 | 0.02 | 0.19 | 0.2 |
| 6 | 139821622 | rs2908513 | -1.266 | -0.37 | -0.612 | -0.423 | -0.003 | 0.058 | 0.058 | 0 | 0.6 | 0.83 | 0.72 | 0.73 | 0.66 | 0.56 |
| 6 | 139821672 | rs2908514 | 0.364 | -0.445 | -0.061 | -0.37 | 0.001 | -0.036 | -0.036 | 0.021 | 0.17 | 0.25 | 0.22 | 0.02 | 0.19 | 0.2 |
| 6 | 139822327 | rs4895563 | -0.692 | -0.362 | -0.143 | -0.02 | -0.041 | 0.167 | 0.167 | 0.095 | 0.19 | 0.36 | 0.16 | 0.4 | 0.17 | 0.19 |
| 6 | 139827777 | rs633417 | -0.647 | -0.297 | -0.606 | -1.045 | -0.003 | -0.042 | -0.042 | 0.631 | 0.29 | 0.34 | 0.28 | 0.06 | 0.27 | 0.27 |
| 6 | 139829917 | rs17513809 | 0.923 | -0.289 | -1.327 | -0.88 | -0.077 | -0.058 | -0.058 | 0 | 0.26 | 0.01 | 0.13 | 0.25 | 0.2 | 0.37 |

| | | | | | | | | | | | | | | | | |
|---|---|---|---|---|---|---|---|---|---|---|---|---|---|---|---|---|
| 6 | 139834012 | rs632057 | -0.348 | -0.446 | -1.112 | -1.276 | 0.007 | -0.012 | -0.012 | 0 | 0.66 | 0.43 | 0.6 | 0.32 | 0.61 | 0.75 |
| 6 | 139835689 | rs668459 | -0.248 | -0.296 | -1.241 | -1.283 | -0.004 | 0.005 | 0.005 | 0.053 | 0.63 | 0.3 | 0.56 | 0.32 | 0.57 | 0.74 |
| 6 | 139837171 | rs12205386 | 0.401 | -0.257 | -1.314 | -1.304 | 0.005 | -0.011 | -0.011 | 0.448 | 0.32 | 0.23 | 0.4 | 0.04 | 0.33 | 0.32 |
| 6 | 139839423 | rs643381 | 0.764 | 0.023 | -1.171 | -0.349 | -0.045 | 0.112 | 0.112 | 0 | 0.52 | 0.09 | 0.49 | 0.29 | 0.51 | 0.68 |
| 6 | 139843787 | rs4896469 | -0.511 | 1.134 | -0.339 | 0.687 | 0.045 | 0.042 | 0.042 | 0 | 0.22 | 0.32 | 0.28 | 0.64 | 0.34 | 0.23 |
| 6 | 139845923 | rs9321725 | -0.746 | 0.713 | -0.968 | 0.754 | -0.019 | -0.035 | -0.035 | 0 | 0.35 | 0.77 | 0.45 | 0.62 | 0.4 | 0.39 |
| 6 | 139846381 | rs11155074 | -0.191 | 0.947 | -0.917 | 1.434 | -0.021 | -0.026 | -0.026 | 0.841 | 0.08 | 0.02 | 0.08 | 0.27 | 0.13 | 0.12 |
| 6 | 139846416 | rs9484282 | -0.649 | 0.391 | -0.282 | 0.933 | -0.007 | 0.005 | 0.005 | 0 | 0.36 | 0.45 | 0.5 | 0.44 | 0.33 | 0.3 |
| 6 | 139847721 | rs7759066 | 0.301 | 0.187 | -0.517 | 0.493 | -0.001 | -0.002 | -0.002 | 0 | 0.22 | 0.16 | 0.16 | 0.11 | 0.13 | 0.11 |
| 6 | 139851661 | rs6903764 | -0.003 | 0.369 | -0.261 | 0.446 | -0.022 | 0.04 | 0.04 | 0 | 0.29 | 0.48 | 0.45 | 0.44 | 0.26 | 0.26 |
| 6 | 139853301 | rs6911238 | -0.735 | 0.603 | -1.04 | 0.397 | 0.023 | -0.029 | -0.029 | 0 | 0.53 | 0.82 | 0.66 | 0.54 | 0.39 | 0.37 |
| 6 | 139858293 | rs10484820 | 0.137 | 0.823 | -1.114 | 0.519 | -0.048 | -0.055 | -0.055 | 0 | 0.11 | 0.02 | 0.11 | 0.28 | 0.2 | 0.13 |
| 6 | 139862246 | rs6913354 | -1.282 | 0.568 | -0.951 | 0.212 | -0.031 | 0.103 | 0.103 | 0.679 | 0.65 | 0.84 | 0.75 | 0.83 | 0.59 | 0.5 |
| 6 | 139864959 | rs13209699 | -0.653 | 0.484 | -0.844 | 0.527 | -0.005 | -0.032 | -0.032 | 0 | 0.41 | 0.24 | 0.3 | 0.4 | 0.36 | 0.26 |
| 6 | 139867807 | rs9389694 | 0.022 | -0.083 | -1.041 | 0.472 | 0.011 | 0.044 | 0.044 | 4.349 | 0.2 | 0.03 | 0.16 | 0.07 | 0.12 | 0.08 |
| 6 | 139871202 | rs4896470 | -0.29 | -0.186 | -0.646 | 0.606 | -0.001 | 0.023 | 0.023 | 0 | 0.18 | 0.2 | 0.35 | 0.43 | 0.18 | 0.23 |
| 6 | 139871653 | rs9403090 | -1.152 | 0.173 | -0.771 | 0.068 | -0.018 | 0.071 | 0.071 | 2.21 | 0.63 | 0.72 | 0.71 | 0.83 | 0.53 | 0.54 |
| 6 | 139878255 | rs4895566 | 0.113 | -0.476 | -0.711 | 0.142 | 0.015 | -0.022 | -0.022 | 0 | 0.85 | 0.91 | 0.91 | 0.7 | 0.89 | 0.91 |
| 6 | 139892016 | rs7750201 | -0.626 | -0.314 | -0.613 | 0.068 | 0.001 | 0.008 | 0.008 | 0.64 | 0.48 | 0.63 | 0.61 | 0.54 | 0.41 | 0.4 |
| 6 | 139896146 | rs9399283 | 0.74 | -0.742 | -0.424 | 0.156 | -0.028 | -0.027 | -0.027 | 1.17 | 0.59 | 0.51 | 0.52 | 0.25 | 0.71 | 0.75 |
| 6 | 139898067 | rs2057034 | -0.176 | -0.857 | -0.638 | 0.308 | 0.036 | -0.034 | -0.034 | 0 | 0.81 | 0.94 | 0.9 | 0.74 | 0.88 | 0.9 |
| 6 | 139898472 | rs910069 | 0.916 | -0.958 | -0.497 | 0.36 | 0.024 | 0.003 | 0.003 | 5.197 | 0.65 | 0.71 | 0.65 | 0.65 | 0.8 | 0.84 |
| 6 | 139898858 | rs9495528 | -0.384 | -0.617 | -0.184 | 0.49 | -0.031 | -0.023 | -0.023 | 0.809 | 0.16 | 0.22 | 0.26 | 0.09 | 0.08 | 0.06 |
| 6 | 139899272 | rs1033503 | -0.953 | -0.622 | -0.546 | 0.37 | -0.007 | 0.015 | 0.015 | 2.587 | 0.31 | 0.34 | 0.4 | 0.49 | 0.22 | 0.26 |
| 6 | 139902326 | rs7765883 | -0.765 | 0.056 | -0.413 | 0.242 | 0.006 | 0.001 | 0.001 | 0 | 0.12 | 0.15 | 0.12 | 0.15 | 0.17 | 0.11 |
| 6 | 139902376 | rs11751645 | -0.356 | -0.314 | -0.089 | 0.219 | -0.01 | 0.034 | 0.034 | 0 | 0.15 | 0.24 | 0.08 | 0.18 | 0.1 | 0.35 |
| 6 | 139902770 | rs9321730 | -0.342 | -0.771 | -0.867 | -0.594 | 0.055 | -0.008 | -0.008 | 1.071 | 0.61 | 0.9 | 0.56 | 0.34 | 0.74 | 0.64 |
| 6 | 139902907 | rs9321731 | 1.393 | -0.805 | -0.659 | -0.709 | 0.014 | -0.099 | -0.099 | 0.283 | 0.31 | 0.38 | 0.34 | 0 | 0.46 | 0.17 |
| 6 | 139904892 | rs12664553 | 0.095 | -0.899 | -0.785 | -0.273 | 0.005 | -0.012 | -0.012 | 0 | 0.15 | 0.09 | 0.07 | 0.18 | 0.1 | 0.35 |
| 6 | 139911322 | rs727098 | -0.142 | -1.083 | -0.968 | -0.223 | 0.059 | -0.001 | -0.001 | 0 | 0.61 | 0.9 | 0.57 | 0.34 | 0.74 | 0.64 |
| 6 | 139916724 | rs9495535 | 0.546 | -1.295 | -0.565 | -0.505 | 0.036 | 0.047 | 0.047 | 0 | 0.51 | 0.45 | 0.46 | 0.16 | 0.67 | 0.3 |
| 6 | 139919937 | rs9495538 | 1.169 | -1.225 | -0.614 | -0.779 | -0.028 | 0.056 | 0.056 | 0 | 0.37 | 0.28 | 0.37 | 0.08 | 0.54 | 0.21 |
| 6 | 139927204 | rs9484295 | 0.503 | -1.33 | -0.531 | -0.678 | 0.066 | 0.088 | 0.088 | 0 | 0.51 | 0.39 | 0.67 | 0.16 | 0.71 | 0.32 |
| 6 | 139928089 | rs12525587 | 0.294 | -1.013 | -0.231 | -0.299 | 0.022 | -0.035 | -0.035 | 0 | 0.09 | 0.02 | 0.04 | 0.17 | 0.06 | 0.32 |
| 6 | 139958370 | rs4895574 | 0.254 | -1.263 | -0.266 | 0.133 | -0.008 | 0.003 | 0.003 | 0 | 0.59 | 0.53 | 0.61 | 0.5 | 0.68 | 0.51 |
| 6 | 139962127 | rs11751597 | 0.023 | -1.296 | -0.562 | 0.241 | 0.002 | -0.164 | -0.164 | 0.067 | 0.26 | 0.03 | 0.31 | 0.51 | 0.21 | 0.43 |
| 6 | 139962517 | rs7765897 | -1.452 | -1.338 | -0.507 | 0.045 | 0.006 | 0.006 | 0.006 | 0 | 0.44 | 0.61 | 0.5 | 0.6 | 0.42 | 0.54 |
| 6 | 139963500 | rs9484299 | -1.128 | -1.728 | -0.254 | -0.158 | 0.014 | -0.002 | -0.002 | 0 | 0.43 | 0.56 | 0.46 | 0.57 | 0.38 | 0.53 |
| 6 | 139963957 | rs9321743 | -0.823 | -1.432 | -0.297 | 0.027 | -0.004 | 0.016 | 0.016 | 0 | 0.37 | 0.4 | 0.43 | 0.57 | 0.35 | 0.52 |
| 6 | 139966136 | rs9403098 | 0.031 | -1.286 | 0.112 | 0.051 | 0.022 | -0.016 | -0.016 | 0 | 0.11 | 0.18 | 0.11 | 0.07 | 0.15 | 0.09 |
| 6 | 139966673 | rs9321744 | -0.795 | -1.198 | -0.024 | 0.031 | 0.003 | 0.008 | 0.008 | 0 | 0.37 | 0.37 | 0.42 | 0.57 | 0.37 | 0.52 |
| 6 | 139967247 | rs17069144 | -0.308 | -1.358 | -0.329 | -0.124 | -0.008 | -0.023 | -0.023 | 0 | 0.07 | 0.12 | 0.05 | 0.02 | 0.06 | 0.02 |
| 6 | 139971166 | rs17069153 | 0.12 | -1.374 | -0.691 | 0.126 | -0.013 | -0.15 | -0.15 | 0.348 | 0.26 | 0.02 | 0.31 | 0.41 | 0.21 | 0.45 |
| 6 | 139972909 | rs11757727 | 0.12 | -1.374 | -0.691 | 0.126 | -0.013 | -0.15 | -0.15 | 3.705 | 0.26 | 0.02 | 0.32 | 0.41 | 0.21 | 0.44 |
| 6 | 139972987 | rs9495557 | -1.377 | -1.306 | -0.507 | 0.064 | 0.02 | -0.01 | -0.01 | 1.283 | 0.47 | 0.73 | 0.51 | 0.52 | 0.41 | 0.55 |
| 6 | 139976184 | rs9376443 | -0.391 | -1.062 | -0.314 | 0.036 | 0.006 | -0.007 | -0.007 | 0 | 0.12 | 0.16 | 0.1 | 0.08 | 0.14 | 0.09 |
| 6 | 139977660 | rs2327977 | -1.344 | -2.227 | -0.935 | -0.161 | 0.087 | 0.013 | 0.013 | 0 | 0.37 | 0.82 | 0.32 | 0.67 | 0.23 | 0.5 |
| 6 | 139981124 | rs6937998 | -1.329 | -1.903 | -0.51 | -0.346 | 0.017 | 0.001 | 0.001 | 0.078 | 0.24 | 0.74 | 0.19 | 0.27 | 0.16 | 0.36 |
| 6 | 139981210 | rs9399289 | 0.327 | -1.909 | -0.49 | -0.297 | 0.039 | -0.046 | -0.046 | 0.241 | 0.13 | 0.03 | 0.06 | 0.26 | 0.09 | 0.35 |
| 6 | 139984667 | rs9321747 | -0.476 | -1.96 | 0.138 | -0.363 | -0.072 | 0.039 | 0.039 | 0 | 0.18 | 0.35 | 0.11 | 0.24 | 0.12 | 0.34 |
| 6 | 139989016 | rs9403101 | -0.415 | -2.012 | 0.128 | -0.386 | -0.069 | 0.062 | 0.062 | 0 | 0.17 | 0.36 | 0.11 | 0.24 | 0.12 | 0.34 |
| 6 | 139990860 | rs9373243 | 0.562 | -2.011 | -0.195 | -0.385 | 0.029 | -0.064 | -0.064 | 0 | 0.11 | 0.02 | 0.06 | 0.24 | 0.09 | 0.34 |
| 6 | 139995603 | rs7768976 | -1.72 | -2.047 | -0.15 | -0.439 | 0.024 | 0.007 | 0.007 | 0.153 | 0.26 | 0.75 | 0.2 | 0.25 | 0.16 | 0.35 |
| 6 | 139998244 | rs6904272 | -0.402 | -1.418 | -0.007 | -0.498 | -0.005 | -0.036 | -0.036 | 0 | 0.07 | 0.19 | 0.06 | 0 | 0.04 | 0.01 |
| 6 | 140003853 | rs9403105 | -0.711 | -1.221 | -0.082 | -1.057 | 0.002 | -0.04 | -0.04 | 6.148 | 0.52 | 0.56 | 0.58 | 0.91 | 0.45 | 0.7 |
| 6 | 140016409 | rs9376452 | 0.146 | -1.343 | -0.351 | -1.187 | -0.02 | -0.018 | -0.018 | 3.119 | 0.31 | 0.34 | 0.21 | 0.27 | 0.27 | 0.44 |
| 6 | 140017022 | rs6912263 | -2.053 | -1.353 | -0.631 | -1.358 | -0.032 | 0.041 | 0.041 | 0 | 0.73 | 0.8 | 0.72 | 0.91 | 0.65 | 0.82 |
| 6 | 140017757 | rs13216266 | 0.233 | -1.401 | -0.657 | -1.362 | 0.013 | 0.024 | 0.024 | 0 | 0.37 | 0.22 | 0.47 | 0.64 | 0.34 | 0.38 |
| 6 | 140027475 | rs10499212 | 1.461 | -1.142 | -0.86 | -1.058 | 0.04 | -0.036 | -0.036 | 0 | 0.12 | 0.01 | 0.03 | 0.18 | 0.03 | 0.26 |
| 6 | 140033899 | rs1936212 | -2.032 | -0.948 | -0.843 | -1.167 | -0.144 | 0.125 | 0.125 | 0 | 0.76 | 0.98 | 0.72 | 0.93 | 0.64 | 0.82 |
| 6 | 140068985 | rs9385876 | 1.117 | -1.101 | -0.691 | -0.936 | 0.018 | -0.048 | -0.048 | 0 | 0.16 | 0.06 | 0.1 | 0.31 | 0.11 | 0.32 |
| 6 | 140081223 | rs9403115 | -0.148 | -1.072 | -0.483 | -1.147 | -0.009 | -0.001 | -0.001 | 0.777 | 0.3 | 0.59 | 0.22 | 0.31 | 0.25 | 0.38 |
| 6 | 140084137 | rs11969282 | 0.233 | -1.054 | -0.898 | -0.584 | 0.008 | 0.01 | 0.01 | 0 | 0.32 | 0.22 | 0.35 | 0.3 | 0.29 | 0.28 |
| 6 | 140102938 | rs6911905 | -0.265 | -0.964 | -0.68 | -0.011 | -0.006 | -0.004 | -0.004 | 0.027 | 0.32 | 0.34 | 0.18 | 0.3 | 0.26 | 0.39 |
| 6 | 140118907 | rs9403122 | 0.257 | -0.511 | -0.952 | 0.395 | -0.008 | -0.061 | -0.061 | 0.227 | 0.1 | 0 | 0.22 | 0.32 | 0.1 | 0.17 |
| 6 | 140124470 | rs3015438 | -0.119 | -0.399 | -0.592 | 0.067 | 0.001 | -0.024 | -0.024 | 0.167 | 0.94 | 0.87 | 0.97 | 1 | 0.94 | 0.98 |
| 6 | 140135009 | rs9399294 | 0.427 | -0.981 | -0.717 | 0.013 | 0.016 | -0.012 | -0.012 | 0 | 0.22 | 0.15 | 0.12 | 0.31 | 0.14 | 0.29 |
| 6 | 140136192 | rs3001014 | -0.209 | -1.522 | -0.465 | -0.203 | -0.016 | 0.006 | 0.006 | 1.269 | 0.78 | 0.65 | 0.84 | 0.75 | 0.8 | 0.79 |
| 6 | 140136531 | rs7776108 | 0.211 | -1.488 | -0.21 | -0.35 | -0.011 | 0.002 | 0.002 | 0.002 | 0.27 | 0.27 | 0.16 | 0.31 | 0.2 | 0.37 |

| | | | | | | | | | | | | | | | | |
|---|---|---|---|---|---|---|---|---|---|---|---|---|---|---|---|---|
| 6 | 140140431 | rs17754188 | 0.777 | -1.423 | -0.252 | -0.337 | 0.018 | -0.026 | -0.026 | 1.867 | 0.05 | 0.12 | 0.04 | 0 | 0.06 | 0.08 |
| 6 | 140144453 | rs3001017 | 0.998 | -1.396 | -0.595 | -0.311 | 0.04 | 0.003 | 0.003 | 0.056 | 0.51 | 0.36 | 0.68 | 0.44 | 0.6 | 0.43 |
| 6 | 140144919 | rs17069360 | 0.706 | -1.404 | -1.254 | 0.027 | 0.006 | -0.034 | -0.034 | 0 | 0.11 | 0.04 | 0.21 | 0.32 | 0.1 | 0.17 |
| 6 | 140150368 | rs17069373 | 0.605 | -1.217 | -1.119 | 0.178 | 0.022 | -0.019 | -0.019 | 0 | 0.21 | 0.12 | 0.12 | 0.31 | 0.14 | 0.29 |
| 6 | 140152588 | rs12525319 | 0.441 | -0.569 | -1.146 | -0.05 | -0.042 | -0.018 | -0.018 | 0.025 | 0.09 | 0 | 0.12 | 0.07 | 0.12 | 0.07 |
| 6 | 140154211 | rs9389724 | -0.258 | -1.078 | -0.907 | -0.059 | 0.033 | -0.022 | -0.022 | 0 | 0.37 | 0.28 | 0.23 | 0.56 | 0.28 | 0.46 |
| 6 | 140155090 | rs4349829 | -0.83 | -0.616 | -1.18 | -0.378 | -0.002 | 0.03 | 0.03 | 0 | 0.47 | 0.66 | 0.63 | 0.63 | 0.5 | 0.51 |
| 6 | 140156942 | rs6570376 | -0.009 | 0.139 | -1.195 | -0.459 | 0.021 | -0.042 | -0.042 | 0 | 0.61 | 0.79 | 0.76 | 0.44 | 0.66 | 0.52 |
| 6 | 140158828 | rs9389726 | -0.028 | 0.194 | -1.255 | -0.806 | 0.007 | -0.047 | -0.047 | 0 | 0.3 | 0.2 | 0.4 | 0.41 | 0.34 | 0.3 |
| 6 | 140159178 | rs4143943 | -0.051 | 0.399 | -1.313 | -0.688 | 0.007 | -0.016 | -0.016 | 0.297 | 0.58 | 0.7 | 0.63 | 0.43 | 0.63 | 0.47 |
| 6 | 140164286 | rs4401680 | -1.039 | 0.463 | -1.528 | -0.641 | 0.018 | -0.01 | -0.01 | 0 | 0.72 | 0.89 | 0.86 | 0.68 | 0.77 | 0.68 |
| 6 | 140167314 | rs12203040 | 1.608 | 0.65 | -1.537 | -0.681 | -0.119 | 0.166 | 0.166 | 0 | 0.24 | 0.02 | 0.17 | 0.02 | 0.28 | 0.16 |
| 6 | 140171922 | rs9399295 | 0.217 | 0.593 | -0.667 | -0.036 | 0 | -0.003 | -0.003 | 0 | 0.59 | 0.65 | 0.61 | 0.43 | 0.61 | 0.46 |
| 6 | 140172490 | rs7750105 | -0.309 | 0.617 | -0.748 | -0.113 | -0.007 | 0 | 0 | 5.737 | 0.9 | 0.92 | 0.85 | 1 | 0.91 | 0.93 |
| 6 | 140173634 | rs3946063 | -0.264 | 0.617 | -0.748 | -0.113 | -0.007 | 0.001 | 0.001 | 5.093 | 0.9 | 0.92 | 0.85 | 1 | 0.91 | 0.93 |
| 6 | 140173790 | rs9385883 | 0.952 | 0.524 | -0.755 | -0.032 | 0.018 | -0.021 | -0.021 | 0 | 0.22 | 0.12 | 0.13 | 0.32 | 0.17 | 0.34 |
| 6 | 140181109 | rs9403127 | 0.043 | 0.546 | -0.731 | -0.078 | 0.005 | -0.014 | -0.014 | 0 | 0.32 | 0.22 | 0.24 | 0.58 | 0.29 | 0.49 |
| 6 | 140181699 | rs6570383 | -1.289 | 0.98 | -0.404 | -0.169 | -0.014 | 0.035 | 0.035 | 17.068 | 0.93 | 0.99 | 0.98 | 1 | 0.94 | 0.98 |
| 6 | 140182546 | rs9376471 | 0.816 | 1.001 | -0.21 | -0.166 | 0.024 | 0.015 | 0.015 | 0.77 | 0.8 | 0.75 | 0.74 | 0.75 | 0.79 | 0.79 |
| 6 | 140183615 | rs9373252 | 0.379 | 0.963 | -0.601 | -0.069 | 0.001 | 0.002 | 0.002 | 2.118 | 0.83 | 0.87 | 0.87 | 0.75 | 0.82 | 0.84 |
| 6 | 140183693 | rs12111348 | -0.063 | 0.809 | -0.467 | 0.072 | 0.011 | 0.049 | 0.049 | 0 | 0.1 | 0.12 | 0.1 | 0.25 | 0.12 | 0.14 |
| 6 | 140190858 | rs9484321 | 0.145 | 0.828 | -0.695 | -0.218 | -0.001 | 0 | 0 | 0 | 0.53 | 0.63 | 0.57 | 0.42 | 0.55 | 0.43 |
| 6 | 140194149 | rs12524738 | 0.248 | 0.366 | -1.16 | -0.256 | 0.019 | 0.011 | 0.011 | 0.045 | 0.14 | 0.05 | 0.09 | 0.1 | 0.11 | 0.08 |
| 6 | 140194296 | rs6938814 | 0.99 | 0.262 | -1.179 | -0.296 | 0.012 | -0.084 | -0.084 | 0 | 0.23 | 0.11 | 0.13 | 0.51 | 0.19 | 0.37 |
| 6 | 140194763 | rs9373254 | 1.184 | 0.574 | -1.019 | -0.616 | -0.016 | 0.092 | 0.092 | 0 | 0.26 | 0.13 | 0.15 | 0.05 | 0.25 | 0.18 |
| 6 | 140197079 | rs4896495 | -0.015 | 0.403 | -0.488 | -0.4 | 0.007 | -0.008 | -0.008 | 0 | 0.42 | 0.55 | 0.44 | 0.53 | 0.36 | 0.56 |
| 6 | 140204777 | rs9376478 | 1.102 | 0.348 | -0.449 | -0.454 | 0.015 | -0.065 | -0.065 | 0 | 0.27 | 0.12 | 0.18 | 0.47 | 0.22 | 0.39 |
| 6 | 140205618 | rs9376479 | 0.939 | 0.448 | -0.553 | -0.644 | 0.004 | 0.074 | 0.074 | 0 | 0.33 | 0.22 | 0.18 | 0.06 | 0.29 | 0.2 |
| 6 | 140206760 | rs4895576 | -0.093 | 0.578 | -0.658 | -0.548 | 0.054 | 0.024 | 0.024 | 0.008 | 0.06 | 0.25 | 0.21 | 0.06 | 0.07 | 0.11 |
| 6 | 140214597 | rs642162 | -0.289 | 0.617 | -0.908 | -0.517 | -0.03 | -0.054 | -0.054 | 2.275 | 0.76 | 0.83 | 0.63 | 0.59 | 0.65 | 0.76 |
| 6 | 140219414 | rs12215417 | 1.636 | 0.533 | -0.991 | -0.645 | -0.091 | 0.133 | 0.133 | 0.085 | 0.25 | 0.02 | 0.14 | 0.06 | 0.27 | 0.19 |
| 6 | 140223597 | rs12197457 | 0.683 | 0.786 | -0.711 | -0.465 | -0.001 | -0.028 | -0.028 | 0 | 0.37 | 0.37 | 0.38 | 0.13 | 0.36 | 0.3 |
| 6 | 140227929 | rs591269 | 0.099 | 0.662 | -0.747 | -0.419 | -0.054 | -0.078 | -0.078 | 0 | 0.71 | 0.82 | 0.46 | 0.52 | 0.59 | 0.65 |
| 6 | 140241272 | rs627705 | 0.248 | 0.624 | -0.723 | -0.105 | -0.025 | 0.021 | 0.021 | 0.221 | 0.35 | 0.46 | 0.25 | 0.46 | 0.27 | 0.42 |
| 6 | 140251292 | rs586246 | -0.056 | 1.081 | -1.403 | 0.155 | -0.001 | -0.049 | -0.049 | 0 | 0.77 | 0.86 | 0.7 | 0.64 | 0.73 | 0.81 |
| 6 | 140261867 | rs609488 | 0.118 | 1.159 | -1.395 | 0.143 | -0.013 | -0.051 | -0.051 | 0 | 0.75 | 0.83 | 0.53 | 0.6 | 0.7 | 0.76 |
| 6 | 140263141 | rs1028471 | 0.449 | 1.332 | -1.406 | 0.514 | -0.038 | -0.021 | -0.021 | 2.412 | 0.08 | 0.01 | 0.09 | 0.11 | 0.12 | 0.09 |
| 6 | 140280398 | rs646695 | 0.295 | 0.81 | -1.193 | 0.322 | 0.009 | -0.046 | -0.046 | 0 | 0.28 | 0.13 | 0.2 | 0.41 | 0.24 | 0.41 |
| 6 | 140291158 | rs1933743 | 0.099 | 0.934 | -1.321 | 0.147 | -0.01 | -0.04 | -0.04 | 0.116 | 0.62 | 0.82 | 0.61 | 0.52 | 0.66 | 0.51 |
| 6 | 140303401 | rs4895579 | -0.577 | 0.92 | -1.379 | 0.135 | 0.013 | -0.045 | -0.045 | 0 | 0.72 | 0.86 | 0.8 | 0.58 | 0.76 | 0.58 |
| 6 | 140313718 | rs1416288 | -0.523 | 0.707 | -1.465 | 0.048 | 0.028 | -0.035 | -0.035 | 4.204 | 0.69 | 0.87 | 0.8 | 0.61 | 0.76 | 0.59 |
| 6 | 140325871 | rs1933741 | 0.56 | 0.749 | -0.473 | 0.013 | 0.001 | -0.011 | -0.011 | 0 | 0.1 | 0.11 | 0.2 | 0.04 | 0.1 | 0.07 |
| 6 | 140337034 | rs17812384 | 0.577 | 0.75 | -0.44 | 0.014 | -0.012 | 0.019 | 0.019 | 4.891 | 0.1 | 0.05 | 0.19 | 0.04 | 0.1 | 0.07 |
| 6 | 140337443 | rs6921044 | -0.754 | 0.717 | -1.048 | -0.535 | -0.041 | 0.004 | 0.004 | 0.508 | 0.41 | 0.83 | 0.66 | 0.56 | 0.47 | 0.4 |
| 6 | 140353664 | rs4243464 | -0.89 | 0.63 | -1.43 | -0.521 | -0.044 | 0.012 | 0.012 | 0.747 | 0.71 | 0.99 | 0.81 | 0.78 | 0.73 | 0.65 |
| 6 | 140362365 | rs11755397 | 1.01 | 0.77 | -1.181 | -0.404 | -0.012 | 0.018 | 0.018 | 4.891 | 0.1 | 0.05 | 0.19 | 0.04 | 0.1 | 0.07 |
| 6 | 140387945 | rs1408295 | -0.912 | 0.796 | -1.21 | -0.422 | -0.011 | 0.053 | 0.053 | 5.498 | 0.91 | 0.98 | 0.83 | 0.99 | 0.92 | 0.93 |
| 6 | 140391106 | rs1324130 | 0.475 | 0.789 | -1.405 | -0.239 | -0.015 | -0.062 | -0.062 | 0 | 0.14 | 0.07 | 0.28 | 0.41 | 0.21 | 0.16 |
| 6 | 140406272 | rs477737 | -0.976 | 0.734 | -0.587 | -0.308 | -0.006 | 0.012 | 0.012 | 5.48 | 0.89 | 0.92 | 0.81 | 0.96 | 0.9 | 0.91 |
| 6 | 140444057 | rs505897 | 0.949 | 0.7 | -0.643 | -0.374 | -0.044 | 0.009 | 0.009 | 1.027 | 0.59 | 0.16 | 0.4 | 0.44 | 0.53 | 0.6 |
| 6 | 140463013 | rs7761894 | 0.349 | 0.689 | -0.092 | -0.312 | -0.002 | 0.01 | 0.01 | 0 | 0.15 | 0.14 | 0.23 | 0.39 | 0.21 | 0.16 |
| 6 | 140475571 | rs2328026 | 0.793 | 0.676 | 0.518 | -0.379 | 0.007 | -0.003 | -0.003 | 0 | 0.12 | 0.18 | 0.12 | 0.13 | 0.16 | 0.11 |
| 6 | 140476088 | rs201285 | -2.194 | 0.451 | 0.114 | -0.611 | -0.081 | 0.146 | 0.146 | 0 | 0.75 | 0.98 | 0.87 | 0.94 | 0.74 | 0.83 |
| 6 | 140565473 | rs6570390 | -0.718 | 0.032 | 0.304 | -0.711 | 0.045 | -0.01 | -0.01 | 0 | 0.63 | 0.86 | 0.71 | 0.58 | 0.72 | 0.52 |
| 6 | 140614699 | rs9389785 | 0.457 | 0.09 | 0.263 | -0.789 | -0.002 | 0.002 | 0.002 | 0 | 0.15 | 0.12 | 0.24 | 0.39 | 0.21 | 0.21 |
| 6 | 140636631 | rs1157002 | -1.062 | -0.224 | 0.309 | -0.97 | 0.005 | 0.015 | 0.015 | 5.896 | 0.9 | 0.94 | 0.92 | 0.96 | 0.93 | 0.95 |
| 6 | 140676761 | rs9376517 | 0.653 | -0.283 | 0.279 | -0.924 | 0.047 | -0.031 | -0.031 | 0.076 | 0.34 | 0.16 | 0.27 | 0.42 | 0.25 | 0.44 |
| 6 | 140681277 | rs11965946 | 0.898 | -0.153 | 0.379 | -0.948 | 0.01 | -0.004 | -0.004 | 0 | 0.12 | 0.17 | 0.12 | 0.13 | 0.15 | 0.12 |
| 6 | 140717053 | rs17069951 | 0.874 | -0.324 | 0.394 | -0.923 | 0.029 | -0.007 | -0.007 | 0 | 0.06 | 0.27 | 0.07 | 0.01 | 0.03 | 0.07 |
| 6 | 140730784 | rs1856756 | 0.714 | -0.322 | -0.371 | -0.875 | 0.023 | -0.049 | -0.049 | 0 | 0.34 | 0.1 | 0.27 | 0.42 | 0.26 | 0.43 |
| 6 | 140732105 | rs13213875 | 0.295 | -0.064 | -0.465 | -0.779 | -0.015 | -0.013 | -0.013 | 0.347 | 0.15 | 0.1 | 0.34 | 0.39 | 0.24 | 0.21 |
| 6 | 140734969 | rs12205549 | 2.011 | 0.238 | -0.807 | -0.964 | -0.076 | 0.147 | 0.147 | 0 | 0.22 | 0.02 | 0.11 | 0.02 | 0.24 | 0.13 |
| 6 | 140779173 | rs13210978 | 0.464 | 0.255 | -0.821 | -0.507 | -0.058 | -0.081 | -0.081 | 0.007 | 0.13 | 0.03 | 0.26 | 0.37 | 0.23 | 0.17 |
| 6 | 140780824 | rs3850223 | 1.02 | 0.203 | -1.095 | -0.672 | -0.029 | -0.124 | -0.124 | 0 | 0.32 | 0.04 | 0.36 | 0.45 | 0.27 | 0.41 |
| 6 | 140785977 | rs17069970 | 0.916 | 0.166 | -0.862 | -0.779 | 0.009 | -0.003 | -0.003 | 0 | 0.11 | 0.16 | 0.12 | 0.12 | 0.15 | 0.11 |
| 6 | 140794431 | rs17069974 | 0.755 | 0.048 | -0.713 | -0.808 | -0.002 | -0.031 | -0.031 | 0.305 | 0.06 | 0.17 | 0.06 | 0 | 0.03 | 0.08 |
| 6 | 140801497 | rs9403183 | 1.089 | 0.099 | -1.339 | -0.59 | -0.047 | -0.017 | -0.017 | 0 | 0.29 | 0.02 | 0.3 | 0.23 | 0.26 | 0.34 |
| 6 | 140814202 | rs11754117 | 1.33 | 0.158 | -1.195 | -0.562 | 0 | 0.045 | 0.045 | 4.069 | 0.1 | 0.03 | 0.04 | 0.04 | 0.08 | 0.08 |
| 6 | 140849702 | rs9376523 | 0.861 | 0.047 | -1.247 | -0.371 | -0.046 | -0.043 | -0.043 | 0.243 | 0.55 | 0.12 | 0.49 | 0.52 | 0.52 | 0.54 |

| | | | | | | | | | | | | | | | | |
|---|---|---|---|---|---|---|---|---|---|---|---|---|---|---|---|---|
| 6 | 140863146 | rs9321772 | 0.983 | 0.049 | -1.481 | -0.389 | -0.082 | -0.054 | -0.054 | 0 | 0.53 | 0.06 | 0.47 | 0.46 | 0.51 | 0.53 |
| 6 | 140864558 | rs1414831 | -0.55 | 0.173 | -1.241 | -0.263 | -0.001 | 0.007 | 0.007 | 0.425 | 0.8 | 0.81 | 0.85 | 0.82 | 0.77 | 0.81 |
| 6 | 140878731 | rs9389808 | 0.897 | -0.282 | -1.1 | -0.314 | -0.06 | -0.042 | -0.042 | 0 | 0.28 | 0.01 | 0.3 | 0.24 | 0.26 | 0.34 |
| 6 | 140882872 | rs9389809 | 0.674 | -0.304 | -1.058 | -0.472 | -0.036 | -0.115 | -0.115 | 0 | 0.32 | 0.05 | 0.37 | 0.41 | 0.28 | 0.41 |
| 6 | 140902819 | rs1414840 | -0.173 | -0.32 | -0.018 | 0.009 | 0.007 | 0 | 0 | 0 | 0.64 | 0.56 | 0.6 | 0.65 | 0.58 | 0.65 |
| 6 | 140911389 | rs13212274 | 1.724 | -0.354 | -0.332 | -0.132 | -0.071 | 0.134 | 0.134 | 0 | 0.22 | 0.02 | 0.11 | 0.02 | 0.25 | 0.12 |
| 6 | 140931910 | rs6570402 | -0.21 | -0.326 | -0.081 | 0.055 | -0.057 | -0.028 | -0.028 | 0 | 0.66 | 0.85 | 0.62 | 0.64 | 0.57 | 0.68 |
| 6 | 140933310 | rs1953635 | 1.11 | -0.221 | -0.086 | -0.794 | -0.014 | -0.012 | -0.012 | 0 | 0.45 | 0.12 | 0.39 | 0.43 | 0.43 | 0.47 |
| 6 | 140948427 | rs1577654 | 0.085 | 0.071 | -0.141 | -0.61 | 0.002 | 0.004 | 0.004 | 0 | 0.63 | 0.57 | 0.57 | 0.56 | 0.57 | 0.68 |
| 6 | 140958047 | rs1336370 | -0.624 | 0.205 | -0.349 | -0.576 | -0.038 | -0.083 | -0.083 | 0 | 0.75 | 0.9 | 0.64 | 0.63 | 0.67 | 0.76 |
| 6 | 140988341 | rs4394230 | -0.007 | 0.234 | -0.191 | -0.572 | -0.017 | -0.029 | -0.029 | 0.148 | 0.64 | 0.72 | 0.57 | 0.56 | 0.57 | 0.67 |
| 6 | 140995321 | rs6570404 | 0.206 | 0.215 | -0.076 | -0.581 | 0.001 | 0.003 | 0.003 | 0 | 0.62 | 0.56 | 0.57 | 0.56 | 0.57 | 0.67 |
| 6 | 141001420 | rs6901977 | -1.083 | 0.095 | -0.264 | -0.608 | -0.003 | -0.005 | -0.005 | 0 | 0.78 | 0.82 | 0.64 | 0.63 | 0.7 | 0.76 |
| 6 | 141015758 | rs4565324 | 0.094 | 0.092 | -0.25 | -0.571 | -0.003 | -0.01 | -0.01 | 0.283 | 0.63 | 0.65 | 0.57 | 0.56 | 0.57 | 0.67 |
| 6 | 141024733 | rs12194318 | 1.756 | 0.257 | -0.806 | -0.65 | -0.056 | 0.096 | 0.096 | 0 | 0.2 | 0.01 | 0.13 | 0.05 | 0.23 | 0.1 |
| 6 | 141036372 | rs2460952 | -0.118 | 0.457 | -0.045 | -0.252 | 0.005 | 0.012 | 0.012 | 0 | 0.67 | 0.67 | 0.58 | 0.57 | 0.59 | 0.67 |
| 6 | 141045961 | rs1521148 | -1.109 | 0.546 | -0.237 | -0.192 | -0.034 | -0.07 | -0.07 | 0 | 0.8 | 0.95 | 0.66 | 0.65 | 0.7 | 0.76 |
| 6 | 141061418 | rs6901618 | -0.687 | 0.257 | -0.014 | -0.746 | -0.015 | 0.022 | 0.022 | 1.612 | 0.56 | 0.7 | 0.43 | 0.41 | 0.51 | 0.46 |
| 6 | 141087211 | rs988434 | -0.21 | 0.063 | -0.092 | -0.783 | 0.033 | 0.049 | 0.049 | 0.003 | 0.44 | 0.74 | 0.37 | 0.34 | 0.36 | 0.37 |
| 6 | 141100043 | rs985072 | -0.091 | 0.069 | -0.075 | -0.779 | 0.026 | 0.041 | 0.041 | 0 | 0.43 | 0.72 | 0.37 | 0.34 | 0.35 | 0.37 |
| 6 | 141114515 | rs594433 | 0.193 | 0.205 | -0.034 | -0.703 | 0.044 | 0.076 | 0.076 | 0 | 0.53 | 0.25 | 0.61 | 0.66 | 0.6 | 0.61 |
| 6 | 141133003 | rs582203 | 0.266 | 0.048 | 0.126 | -0.628 | 0.057 | 0.083 | 0.083 | 0 | 0.53 | 0.25 | 0.6 | 0.66 | 0.61 | 0.64 |
| 6 | 141151470 | rs671124 | 0.788 | 0.13 | -0.148 | -0.665 | -0.069 | 0.023 | 0.023 | 4.095 | 0.43 | 0.11 | 0.56 | 0.59 | 0.5 | 0.55 |
| 6 | 141153358 | rs672301 | -0.14 | 0.082 | 0.319 | -0.748 | 0.012 | 0.011 | 0.011 | 0 | 0.58 | 0.41 | 0.65 | 0.67 | 0.66 | 0.72 |
| 6 | 141171423 | rs685606 | -0.437 | 0.067 | 0.579 | -0.865 | 0.019 | 0.045 | 0.045 | 0 | 0.63 | 0.7 | 0.76 | 0.78 | 0.66 | 0.8 |
| 6 | 141171561 | rs17070386 | -0.244 | 0.226 | -0.012 | -0.882 | 0.016 | -0.035 | -0.035 | 0.569 | 0.15 | 0.24 | 0.09 | 0.07 | 0.15 | 0.17 |
| 6 | 141174222 | rs661704 | -1.5 | 0.115 | -0.43 | -0.927 | -0.035 | 0.117 | 0.117 | 0 | 0.77 | 0.96 | 0.86 | 0.95 | 0.74 | 0.9 |
| 6 | 141177470 | rs11758974 | 0.699 | 0.263 | -0.286 | -0.884 | -0.034 | 0.01 | 0.01 | 3.595 | 0.09 | 0.01 | 0.04 | 0.07 | 0.11 | 0.08 |
| 6 | 141182922 | rs651226 | 0.814 | 0.245 | -0.187 | -0.785 | -0.048 | 0.048 | 0.048 | 3.088 | 0.43 | 0.16 | 0.59 | 0.64 | 0.51 | 0.55 |
| 6 | 141185228 | rs849294 | 0.163 | 0.073 | 0.247 | -0.475 | 0.007 | 0.011 | 0.011 | 0 | 0.59 | 0.44 | 0.68 | 0.71 | 0.64 | 0.71 |
| 6 | 141214296 | rs1856384 | -0.714 | 0.094 | 0.312 | -0.249 | -0.011 | 0.003 | 0.003 | 0 | 0.12 | 0.21 | 0.11 | 0.18 | 0.1 | 0.1 |
| 6 | 141222516 | rs1970349 | -0.275 | 0.334 | 0.109 | -0.097 | 0 | -0.007 | -0.007 | 0 | 0.09 | 0.13 | 0.08 | 0.19 | 0.07 | 0.08 |
| 6 | 141233620 | rs6914898 | 0.499 | 0.32 | 0.08 | -0.564 | 0.002 | -0.044 | -0.044 | 0 | 0.3 | 0.24 | 0.18 | 0 | 0.3 | 0.19 |
| 6 | 141244330 | rs9389832 | 0.977 | 0.27 | -0.422 | -0.414 | -0.09 | -0.053 | -0.053 | 0 | 0.27 | 0.03 | 0.32 | 0.29 | 0.31 | 0.27 |
| 6 | 141244368 | rs6924310 | 0.169 | 0.167 | -0.033 | -0.547 | -0.001 | 0.028 | 0.028 | 0 | 0.38 | 0.26 | 0.37 | 0.29 | 0.35 | 0.39 |
| 6 | 141259938 | rs10499222 | 0.573 | 0.203 | -0.223 | -0.51 | 0.002 | -0.01 | -0.01 | 0.706 | 0.09 | 0.1 | 0.03 | 0 | 0.04 | 0.12 |
| 6 | 141267021 | rs11155146 | 0.759 | 0.258 | -0.259 | -0.318 | 0 | -0.051 | -0.051 | 0.238 | 0.3 | 0.24 | 0.18 | 0 | 0.29 | 0.19 |
| 6 | 141270660 | rs9389835 | 0.365 | 0.437 | -0.283 | 0.056 | -0.005 | -0.039 | -0.039 | 0.128 | 0.39 | 0.45 | 0.26 | 0.2 | 0.37 | 0.27 |
| 6 | 141272231 | rs9389836 | 0.009 | 0.432 | -0.594 | 0.617 | -0.045 | -0.021 | -0.021 | 0 | 0.28 | 0.07 | 0.32 | 0.29 | 0.31 | 0.27 |
| 6 | 141288687 | rs9376553 | 0.282 | -0.249 | -0.395 | 0.596 | 0.033 | 0.006 | 0.006 | 0 | 0.63 | 0.81 | 0.62 | 0.39 | 0.7 | 0.47 |
| 6 | 141301526 | rs1832449 | -0.468 | -0.576 | -0.593 | 1.106 | -0.036 | -0.1 | -0.1 | 0 | 0.79 | 0.94 | 0.68 | 0.64 | 0.77 | 0.74 |
| 6 | 141319040 | rs12175481 | -0.191 | -0.684 | -0.027 | 0.924 | 0.007 | 0.021 | 0.021 | 0 | 0.09 | 0.21 | 0.09 | 0.15 | 0.06 | 0.08 |
| 6 | 141325406 | rs9495873 | -0.275 | -0.711 | -0.049 | 0.947 | 0.026 | -0.025 | -0.025 | 1.283 | 0.16 | 0.09 | 0.06 | 0.25 | 0.07 | 0.28 |
| 6 | 141332731 | rs6906941 | 0.489 | -0.647 | -0.027 | 0.849 | 0.031 | 0.004 | 0.004 | 0 | 0.64 | 0.8 | 0.62 | 0.39 | 0.7 | 0.47 |
| 6 | 141357672 | rs9321797 | 0.681 | -0.753 | 0.837 | 0.775 | -0.014 | -0.007 | -0.007 | 0 | 0.62 | 0.56 | 0.54 | 0.39 | 0.7 | 0.47 |
| 6 | 141364472 | rs7751709 | 0.852 | -0.74 | 0.765 | 0.754 | 0.014 | 0.043 | 0.043 | 0 | 0.57 | 0.67 | 0.49 | 0.24 | 0.65 | 0.4 |
| 6 | 141371851 | rs1832446 | 1.276 | -0.426 | -0.243 | -0.028 | -0.092 | 0.014 | 0.014 | 0 | 0.33 | 0.02 | 0.25 | 0.24 | 0.36 | 0.28 |
| 6 | 141399812 | rs1819632 | 0.407 | -0.635 | 0.612 | 0.211 | 0 | 0 | 0 | 0 | 0.62 | 0.62 | 0.54 | 0.39 | 0.7 | 0.46 |
| 6 | 141409979 | rs9495898 | 0.11 | -0.311 | 0.255 | 0.452 | 0.003 | -0.009 | -0.009 | 0.221 | 0.06 | 0.03 | 0.04 | 0.08 | 0.05 | 0.1 |
| 6 | 141428255 | rs9389870 | -0.474 | -0.312 | 0.284 | 0.419 | 0.001 | 0.018 | 0.018 | 0 | 0.12 | 0.16 | 0.12 | 0.24 | 0.11 | 0.18 |
| 6 | 141442825 | rs4385332 | -0.424 | -0.342 | 0.069 | 0.57 | -0.005 | -0.01 | -0.01 | 1.701 | 0.8 | 0.85 | 0.59 | 0.62 | 0.77 | 0.72 |
| 6 | 141453145 | rs7738388 | 0.914 | -0.393 | 0.043 | 0.792 | -0.012 | -0.01 | -0.01 | 0.276 | 0.31 | 0.24 | 0.28 | 0.35 | 0.34 | 0.26 |
| 6 | 141453781 | rs9376578 | -0.914 | -1.574 | 0.749 | 0.55 | -0.144 | -0.023 | -0.023 | 0.554 | 0.2 | 0.52 | 0.09 | 0.11 | 0.04 | 0.23 |
| 6 | 141457722 | rs1037441 | 0.058 | -1.234 | 1.082 | 0.981 | 0 | 0.109 | 0.109 | 0 | 0.24 | 0.41 | 0.35 | 0.43 | 0.24 | 0.28 |
| 6 | 141461775 | rs9399348 | 0.056 | -1.346 | 1.225 | 0.851 | -0.067 | 0.024 | 0.024 | 0 | 0.12 | 0.29 | 0.04 | 0.1 | 0.03 | 0.21 |
| 6 | 141462594 | rs7763002 | -0.087 | -1.349 | 1.285 | 0.845 | -0.071 | 0.058 | 0.058 | 0.001 | 0.12 | 0.4 | 0.05 | 0.1 | 0.03 | 0.23 |
| 6 | 141473363 | rs1452991 | 1.174 | -1.117 | 0.509 | 0.666 | -0.035 | 0.034 | 0.034 | 0 | 0.37 | 0.24 | 0.3 | 0.28 | 0.39 | 0.26 |
| 6 | 141488481 | rs1452992 | -0.624 | -1.173 | 0.734 | 0.819 | 0.001 | -0.001 | -0.001 | 0 | 0.37 | 0.56 | 0.44 | 0.6 | 0.28 | 0.52 |
| 6 | 141519035 | rs2328195 | -0.048 | -0.943 | 0.844 | 0.94 | 0.001 | 0.103 | 0.103 | 0 | 0.25 | 0.37 | 0.39 | 0.45 | 0.24 | 0.3 |
| 6 | 141529979 | rs9495923 | 0.109 | -0.956 | 0.912 | 0.933 | -0.003 | -0.042 | -0.042 | 0.016 | 0.24 | 0.14 | 0.36 | 0.45 | 0.24 | 0.3 |
| 6 | 141530140 | rs9403259 | -0.835 | -1.314 | 1.141 | 0.852 | 0.035 | 0.002 | 0.002 | 0 | 0.38 | 0.65 | 0.44 | 0.62 | 0.28 | 0.52 |
| 6 | 141562759 | rs1452989 | 0.306 | -1.293 | 1.071 | 0.641 | -0.002 | -0.004 | -0.004 | 0.144 | 0.62 | 0.63 | 0.69 | 0.71 | 0.63 | 0.54 |
| 6 | 141567407 | rs2045017 | 0.345 | -1.247 | 1.041 | 0.663 | -0.007 | -0.066 | -0.066 | 3.53 | 0.24 | 0.11 | 0.35 | 0.45 | 0.23 | 0.3 |
| 6 | 141582182 | rs9495937 | 0.468 | -1.263 | 1.364 | 0.607 | 0.002 | 0.003 | 0.003 | 0 | 0.5 | 0.56 | 0.36 | 0.42 | 0.44 | 0.48 |
| 6 | 141587036 | rs9376596 | -0.111 | -1.179 | 1.468 | 0.603 | -0.058 | 0.02 | 0.02 | 0 | 0.13 | 0.29 | 0.06 | 0.17 | 0.04 | 0.23 |
| 6 | 141596752 | rs9385947 | 0.083 | -1.185 | 1.417 | 0.607 | -0.04 | 0.009 | 0.009 | 0 | 0.12 | 0.18 | 0.05 | 0.17 | 0.04 | 0.23 |
| 6 | 141602679 | rs9321807 | 0.083 | -1.185 | 1.417 | 0.628 | -0.04 | 0.011 | 0.011 | 0 | 0.12 | 0.18 | 0.05 | 0.17 | 0.04 | 0.23 |
| 6 | 141624854 | rs2328202 | 0.55 | -1.215 | 1.48 | 0.614 | 0.004 | 0.007 | 0.007 | 1.983 | 0.6 | 0.38 | 0.66 | 0.72 | 0.63 | 0.54 |
| 6 | 141634266 | rs9376605 | -0.545 | -1.15 | 1.449 | 0.586 | -0.002 | -0.004 | -0.004 | 0 | 0.74 | 0.78 | 0.72 | 0.89 | 0.66 | 0.77 |

| | | | | | | | | | | | | | | | | |
|---|---|---|---|---|---|---|---|---|---|---|---|---|---|---|---|---|
| 6 | 141656138 | rs1335301 | 0.943 | -1.204 | 0.933 | 1.025 | -0.054 | -0.083 | -0.083 | 0 | 0.16 | 0.04 | 0.3 | 0.28 | 0.22 | 0.26 |
| 6 | 141667778 | rs9403287 | -0.023 | -1.272 | 1.172 | 0.761 | -0.046 | 0.01 | 0.01 | 0 | 0.13 | 0.2 | 0.05 | 0.17 | 0.04 | 0.22 |
| 6 | 141671732 | rs9376609 | 1.277 | -1.088 | 1.07 | 0.581 | -0.008 | 0.01 | 0.01 | 0 | 0.36 | 0.35 | 0.33 | 0.26 | 0.39 | 0.24 |
| 6 | 141672901 | rs9403288 | 1.405 | -1.088 | 0.484 | 0.581 | -0.103 | 0.002 | 0.002 | 0.715 | 0.34 | 0.1 | 0.3 | 0.26 | 0.38 | 0.24 |
| 6 | 141683547 | rs9373305 | 1.437 | -1.199 | 0.515 | 0.58 | -0.071 | 0.027 | 0.027 | 0.183 | 0.35 | 0.16 | 0.32 | 0.26 | 0.41 | 0.25 |
| 6 | 141718812 | rs6932353 | 0.812 | -1.555 | 0.554 | 0.345 | 0.042 | 0.031 | 0.031 | 0 | 0.59 | 0.31 | 0.69 | 0.31 | 0.76 | 0.51 |
| 6 | 141720314 | rs9403302 | 0.177 | -1.648 | 0.905 | 0.467 | -0.071 | 0.015 | 0.015 | 0 | 0.86 | 0.65 | 0.92 | 0.86 | 0.96 | 0.8 |
| 6 | 141728211 | rs2184529 | 0.36 | -1.648 | 0.703 | 0.476 | -0.001 | -0.079 | -0.079 | 0.168 | 0.21 | 0.09 | 0.21 | 0.55 | 0.18 | 0.29 |
| 6 | 141740918 | rs1414123 | 0.552 | -1.477 | 0.367 | 0.267 | -0.021 | 0.046 | 0.046 | 0 | 0.25 | 0.18 | 0.36 | 0.12 | 0.35 | 0.21 |
| 6 | 141747538 | rs1335295 | 0.765 | -1.569 | 0.402 | 0.293 | -0.093 | -0.025 | -0.025 | 0 | 0.81 | 0.4 | 0.89 | 0.87 | 0.95 | 0.81 |
| 6 | 141748163 | rs1029180 | 0.291 | -1.466 | 0.43 | 0.34 | -0.083 | 0.013 | 0.013 | 0.007 | 0.85 | 0.53 | 0.92 | 0.87 | 0.96 | 0.81 |
| 6 | 141753243 | rs11962370 | 1.46 | -1.123 | -0.159 | 0.255 | -0.035 | 0.04 | 0.04 | 0 | 0.34 | 0.21 | 0.33 | 0.18 | 0.42 | 0.24 |
| 6 | 141764919 | rs12212395 | -0.094 | -0.981 | -0.295 | 0.272 | -0.024 | 0.078 | 0.078 | 0.149 | 0.28 | 0.17 | 0.36 | 0.12 | 0.36 | 0.24 |
| 6 | 141787794 | rs4386830 | 0.624 | -0.89 | -0.186 | 0.366 | 0.018 | 0.008 | 0.008 | 0 | 0.43 | 0.3 | 0.29 | 0.36 | 0.32 | 0.42 |
| 6 | 141793547 | rs4473876 | 0.529 | -1.181 | -0.211 | 0.466 | -0.012 | 0.003 | 0.003 | 0 | 0.09 | 0.12 | 0.04 | 0.17 | 0.02 | 0.14 |
| 6 | 141799534 | rs13201721 | 0.732 | -1.013 | -0.574 | 0.806 | -0.057 | -0.102 | -0.102 | 0 | 0.19 | 0.06 | 0.21 | 0.37 | 0.27 | 0.24 |
| 6 | 141804428 | rs6905051 | 1.55 | -0.93 | -0.455 | 0.661 | 0.008 | 0.021 | 0.021 | 0 | 0.33 | 0.19 | 0.26 | 0.18 | 0.3 | 0.28 |
| 6 | 141809473 | rs9376624 | 0.174 | -0.753 | 0.087 | 0.47 | 0.018 | -0.03 | -0.03 | 1.948 | 0.63 | 0.49 | 0.5 | 0.72 | 0.58 | 0.66 |
| 6 | 141827882 | rs6570464 | -0.584 | -0.58 | 0.318 | 0.189 | 0.013 | -0.072 | -0.072 | 0 | 0.78 | 0.69 | 0.67 | 0.94 | 0.76 | 0.82 |
| 6 | 141848595 | rs12204830 | -0.112 | -0.686 | 0.458 | 0.284 | 0.002 | 0.003 | 0.003 | 0 | 0.15 | 0.17 | 0.14 | 0.24 | 0.17 | 0.13 |
| 6 | 141857115 | rs6904106 | 0.608 | -0.979 | 0.257 | 0.439 | -0.014 | -0.049 | -0.049 | 0.013 | 0.23 | 0.15 | 0.23 | 0.37 | 0.28 | 0.27 |
| 6 | 141860331 | rs9376628 | 1.7 | -1.207 | 0.146 | 0.312 | 0.014 | -0.002 | -0.002 | 0 | 0.36 | 0.17 | 0.25 | 0.34 | 0.31 | 0.43 |
| 6 | 141869622 | rs12204767 | 0.366 | -1.301 | -0.272 | 0.208 | -0.011 | -0.018 | -0.018 | 0 | 0.16 | 0.1 | 0.13 | 0.24 | 0.17 | 0.13 |
| 6 | 141869843 | rs2225906 | -1.033 | -1.029 | -0.157 | 0.19 | -0.02 | 0.053 | 0.053 | 0 | 0.8 | 0.92 | 0.69 | 0.94 | 0.76 | 0.82 |
| 6 | 141882519 | rs13208315 | 0.655 | -1.138 | -0.146 | 0.622 | -0.049 | -0.062 | -0.062 | 0 | 0.22 | 0.07 | 0.23 | 0.31 | 0.28 | 0.26 |
| 6 | 141894535 | rs1999876 | 1.271 | -1.443 | -0.195 | 0.108 | -0.008 | -0.005 | -0.005 | 0 | 0.5 | 0.14 | 0.42 | 0.52 | 0.53 | 0.48 |
| 6 | 141905766 | rs1933283 | -0.7 | -1.46 | 0.135 | 0.101 | 0.009 | -0.027 | -0.027 | 0.41 | 0.76 | 0.74 | 0.66 | 0.89 | 0.71 | 0.79 |
| 6 | 141919054 | rs9496093 | -0.76 | -1.447 | 0.233 | 0.174 | -0.02 | 0.096 | 0.096 | 0.015 | 0.21 | 0.53 | 0.23 | 0.36 | 0.17 | 0.27 |
| 6 | 141919745 | rs17178941 | 0.582 | -1.487 | -0.006 | 0.234 | -0.056 | -0.075 | -0.075 | 0 | 0.26 | 0.06 | 0.22 | 0.34 | 0.3 | 0.24 |
| 6 | 141937741 | rs9496101 | -0.792 | -1.191 | 0.584 | 0.551 | -0.019 | 0.077 | 0.077 | 0 | 0.21 | 0.42 | 0.18 | 0.35 | 0.17 | 0.27 |
| 6 | 141951995 | rs9496108 | 1.448 | -1.114 | 0.455 | 0.641 | -0.013 | 0.016 | 0.016 | 0 | 0.06 | 0.08 | 0.04 | 0.16 | 0.02 | 0.18 |
| 6 | 141964290 | rs1415909 | -0.92 | -1.139 | 0.55 | 0.626 | -0.014 | 0.041 | 0.041 | 0 | 0.21 | 0.33 | 0.16 | 0.35 | 0.18 | 0.31 |
| 6 | 141966721 | rs9389940 | 0.287 | -1.119 | 0.349 | 0.496 | -0.01 | -0.011 | -0.011 | 0.713 | 0.34 | 0.11 | 0.29 | 0.35 | 0.35 | 0.29 |
| 6 | 141988466 | rs1338103 | -0.71 | -0.464 | 0.688 | 0.77 | -0.005 | 0.062 | 0.062 | 0 | 0.18 | 0.29 | 0.15 | 0.35 | 0.17 | 0.27 |
| 6 | 141992359 | rs4513816 | -0.832 | -0.178 | 0.367 | 0.653 | -0.006 | -0.004 | -0.004 | 0.072 | 0.78 | 0.83 | 0.66 | 0.74 | 0.75 | 0.66 |
| 6 | 141994355 | rs1577948 | -0.626 | -0.103 | 0.516 | 0.676 | 0.032 | 0.07 | 0.07 | 0 | 0.13 | 0.29 | 0.11 | 0.19 | 0.15 | 0.09 |
| 6 | 142021799 | rs9321832 | -0.066 | -0.141 | 0.033 | 0.638 | -0.074 | -0.067 | -0.067 | 0 | 0.36 | 0.13 | 0.4 | 0.55 | 0.48 | 0.36 |
| 6 | 142021883 | rs9376653 | -0.412 | -0.237 | 0.115 | 0.583 | -0.039 | -0.03 | -0.03 | 0 | 0.4 | 0.22 | 0.42 | 0.56 | 0.49 | 0.38 |
| 6 | 142024307 | rs6936180 | -0.333 | -0.268 | 0.13 | 0.56 | -0.016 | -0.026 | -0.026 | 0 | 0.09 | 0.12 | 0.03 | 0 | 0.03 | 0.03 |
| 6 | 142024463 | rs9321833 | 0.083 | -0.554 | 0.034 | 0.141 | 0.02 | 0.011 | 0.011 | 1.614 | 0.07 | 0.38 | 0.04 | 0 | 0.01 | 0.02 |
| 6 | 142027493 | rs973118 | -0.348 | 0.055 | -0.174 | 0.517 | -0.02 | -0.013 | -0.013 | 0 | 0.41 | 0.3 | 0.43 | 0.55 | 0.5 | 0.36 |
| 6 | 142028607 | rs975858 | -1.345 | 0.101 | -0.51 | 0.533 | -0.029 | -0.052 | -0.052 | 0 | 0.64 | 0.56 | 0.76 | 0.83 | 0.75 | 0.71 |
| 6 | 142032704 | rs4896551 | -0.025 | 0.417 | -1.02 | 0.868 | -0.021 | -0.021 | -0.021 | 0 | 0.22 | 0.1 | 0.33 | 0.28 | 0.25 | 0.34 |
| 6 | 142037511 | rs9373323 | -0.575 | 0.079 | -0.73 | 0.808 | -0.001 | 0 | 0 | 0.067 | 0.29 | 0.29 | 0.25 | 0.3 | 0.27 | 0.24 |
| 6 | 142038365 | rs6913816 | -1.183 | -0.583 | -1.824 | 0.068 | -0.07 | 0.08 | 0.08 | 0 | 0.66 | 0.99 | 0.76 | 0.84 | 0.7 | 0.65 |
| 6 | 142038634 | rs9484549 | 0.023 | -0.394 | -1.485 | -0.054 | -0.001 | 0 | 0 | 0.03 | 0.1 | 0.1 | 0.09 | 0.05 | 0.18 | 0.07 |
| 6 | 142039815 | rs12209540 | -0.89 | -0.714 | -1.023 | 0.198 | -0.003 | -0.002 | -0.002 | 0 | 0.4 | 0.63 | 0.44 | 0.33 | 0.34 | 0.39 |
| 6 | 142041859 | rs2024658 | 0.441 | -0.825 | -0.906 | 0.145 | 0.045 | 0.038 | 0.038 | 0 | 0.21 | 0.13 | 0.27 | 0.13 | 0.11 | 0.26 |
| 6 | 142047297 | rs6913823 | 0.373 | -0.868 | -0.27 | 0.127 | -0.019 | -0.015 | -0.015 | 2.909 | 0.21 | 0.25 | 0.27 | 0.13 | 0.11 | 0.26 |
| 6 | 142061431 | rs6903549 | -0.078 | -0.508 | 0.403 | 0.27 | 0.009 | -0.006 | -0.006 | 0 | 0.11 | 0.13 | 0.09 | 0.06 | 0.18 | 0.07 |
| 6 | 142076325 | rs12191465 | 0.162 | -0.491 | 0.501 | 0.247 | 0.001 | 0.034 | 0.034 | 0 | 0.06 | 0.11 | 0.06 | 0.16 | 0.05 | 0.07 |
| 6 | 142093922 | rs9496139 | -1.422 | -0.778 | 0.427 | 0.057 | -0.008 | -0.021 | -0.021 | 0.07 | 0.34 | 0.64 | 0.31 | 0.2 | 0.28 | 0.24 |
| 6 | 142112455 | rs9484553 | -0.517 | -0.679 | 0.149 | -0.191 | -0.001 | 0.001 | 0.001 | 0 | 0.19 | 0.16 | 0.14 | 0.05 | 0.22 | 0.09 |
| 6 | 142130009 | rs9496146 | -0.519 | -0.721 | -0.034 | -0.214 | -0.003 | 0.04 | 0.04 | 0 | 0.19 | 0.13 | 0.13 | 0.05 | 0.22 | 0.09 |
| 6 | 142164431 | rs9496163 | -0.711 | -0.76 | 0.201 | -0.25 | 0.001 | -0.084 | -0.084 | 0.343 | 0.22 | 0.42 | 0.15 | 0.05 | 0.23 | 0.09 |
| 6 | 142169481 | rs9389952 | 0.224 | -0.763 | -0.389 | -0.169 | -0.087 | -0.108 | -0.108 | 0 | 0.18 | 0.01 | 0.13 | 0.32 | 0.23 | 0.22 |
| 6 | 142169787 | rs17071172 | 0.638 | -0.995 | 0.022 | -0.946 | -0.021 | -0.055 | -0.055 | 10.442 | 0.12 | 0.18 | 0.12 | 0 | 0.05 | 0.08 |
| 6 | 142174962 | rs910325 | -0.781 | -1.107 | -0.127 | -0.86 | 0 | -0.003 | -0.003 | 0 | 0.44 | 0.53 | 0.48 | 0.69 | 0.42 | 0.46 |
| 6 | 142175291 | rs1884055 | -1.941 | -1.036 | -0.182 | -0.852 | -0.031 | 0.035 | 0.035 | 0 | 0.65 | 0.9 | 0.62 | 0.73 | 0.64 | 0.56 |
| 6 | 142177674 | rs7753254 | -1.02 | -1.05 | -0.051 | -0.677 | 0.004 | 0.024 | 0.024 | 0.12 | 0.47 | 0.62 | 0.5 | 0.69 | 0.42 | 0.48 |
| 6 | 142188406 | rs11155211 | -0.259 | -1.557 | 0.09 | -1.064 | -0.059 | -0.103 | -0.103 | 0.177 | 0.19 | 0.3 | 0.13 | 0 | 0.05 | 0.09 |
| 6 | 142188711 | rs9484567 | -0.126 | -1.374 | -0.352 | -1.217 | -0.023 | 0.052 | 0.052 | 0 | 0.18 | 0.07 | 0.12 | 0.01 | 0.22 | 0.12 |
| 6 | 142191527 | rs12202271 | -0.194 | -1.445 | -0.066 | -1.017 | -0.034 | -0.07 | -0.07 | 1.511 | 0.18 | 0.25 | 0.11 | 0 | 0.05 | 0.09 |
| 6 | 142200163 | rs10457728 | -0.028 | -0.856 | 0.122 | -0.789 | -0.018 | 0.078 | 0.078 | 0.009 | 0.24 | 0.1 | 0.2 | 0.01 | 0.25 | 0.22 |
| 6 | 142219217 | rs7758295 | -1.048 | -1.087 | 0.233 | -0.438 | 0.01 | 0.013 | 0.013 | 0 | 0.48 | 0.57 | 0.36 | 0.29 | 0.37 | 0.4 |
| 6 | 142232392 | rs17071231 | 0.624 | -1.053 | 0.235 | -0.312 | 0.035 | -0.054 | -0.054 | 2.376 | 0.16 | 0.09 | 0.11 | 0.28 | 0.1 | 0.14 |
| 6 | 142236407 | rs9389954 | 0.026 | -0.789 | 0.191 | -0.098 | 0.019 | 0.083 | 0.083 | 0 | 0.14 | 0.17 | 0.2 | 0.43 | 0.17 | 0.23 |
| 6 | 142245981 | rs9376660 | -0.444 | -0.772 | 0.165 | -0.233 | 0.001 | 0.005 | 0.005 | 0.031 | 0.31 | 0.26 | 0.31 | 0.75 | 0.27 | 0.38 |
| 6 | 142253976 | rs4072064 | -1.411 | -0.658 | 0.17 | -0.246 | -0.002 | 0.005 | 0.005 | 0 | 0.59 | 0.64 | 0.55 | 0.73 | 0.54 | 0.58 |

| | | | | | | | | | | | | | | | | |
|---|---|---|---|---|---|---|---|---|---|---|---|---|---|---|---|---|
| 6 | 142259172 | rs12197645 | 0.518 | -0.24 | 0.054 | -0.019 | -0.038 | 0.085 | 0.085 | 0 | 0.21 | 0.06 | 0.19 | 0.01 | 0.24 | 0.2 |
| 6 | 142267503 | rs2784940 | -0.567 | -0.257 | 0.055 | 0.037 | -0.018 | -0.018 | -0.018 | 0.237 | 0.34 | 0.25 | 0.41 | 0.45 | 0.41 | 0.41 |
| 6 | 142276494 | rs1577619 | 0.398 | -0.168 | -0.77 | -0.004 | -0.015 | 0.029 | 0.029 | 0.01 | 0.21 | 0.12 | 0.2 | 0.01 | 0.24 | 0.19 |
| 6 | 142277769 | rs1414743 | 0.512 | -0.221 | -1.102 | 0.042 | -0.029 | 0.016 | 0.016 | 1.203 | 0.21 | 0.23 | 0.13 | 0.28 | 0.11 | 0.2 |
| 6 | 142278682 | rs1006257 | 0.281 | 0.227 | -2.086 | -0.557 | -0.047 | 0.109 | 0.109 | 0 | 0.21 | 0.04 | 0.2 | 0.01 | 0.25 | 0.19 |
| 6 | 142283198 | rs2784945 | -0.237 | 0.699 | -1.402 | 0.246 | -0.025 | 0.072 | 0.072 | 0 | 0.59 | 0.76 | 0.55 | 0.73 | 0.53 | 0.61 |
| 6 | 142287111 | rs7772396 | -0.779 | 1.206 | -1.47 | 0.735 | -0.062 | -0.121 | -0.121 | 0 | 0.25 | 0.12 | 0.35 | 0.48 | 0.36 | 0.31 |
| 6 | 142306066 | rs17054023 | -0.835 | 0.664 | 0.376 | 0.438 | -0.026 | 0.016 | 0.016 | 2.373 | 0.06 | 0.18 | 0.03 | 0.05 | 0 | 0.28 |
| 6 | 142322456 | rs9385984 | -0.93 | 1.323 | 0.091 | 0.312 | -0.032 | -0.041 | -0.041 | 1.125 | 0.17 | 0.09 | 0.29 | 0.68 | 0.24 | 0.16 |
| 6 | 142328285 | rs4596499 | -0.594 | 0.702 | -0.091 | -0.201 | 0.022 | 0.016 | 0.016 | 0 | 0.9 | 0.95 | 0.97 | 1 | 0.96 | 0.93 |
| 6 | 142332081 | rs9389968 | -1.135 | 1.185 | 0.564 | 0.059 | 0.006 | 0.01 | 0.01 | 0.255 | 0.18 | 0.21 | 0.3 | 0.68 | 0.24 | 0.16 |
| 6 | 142343409 | rs7773151 | -1.062 | 1.213 | 0.457 | -0.108 | -0.016 | -0.015 | -0.015 | 0 | 0.19 | 0.14 | 0.3 | 0.71 | 0.25 | 0.18 |
| 6 | 142346745 | rs7764878 | -0.279 | 1.408 | 0.889 | -0.769 | -0.001 | -0.038 | -0.038 | 0.512 | 0.22 | 0.24 | 0.17 | 0.06 | 0.22 | 0.38 |
| 6 | 142349998 | rs9496231 | -0.597 | 1.11 | 1.087 | -0.826 | -0.009 | -0.09 | -0.09 | 0 | 0.25 | 0.45 | 0.2 | 0.06 | 0.22 | 0.38 |
| 6 | 142359772 | rs12212141 | -0.355 | 1.001 | 0.973 | -0.877 | 0 | -0.008 | -0.008 | 0 | 0.24 | 0.22 | 0.17 | 0.06 | 0.23 | 0.33 |
| 6 | 142377095 | rs9484591 | -1.394 | 1.339 | 1.456 | -0.652 | 0.015 | -0.026 | -0.026 | 0 | 0.4 | 0.49 | 0.53 | 0.74 | 0.47 | 0.44 |
| 6 | 142389382 | rs9376669 | -1.209 | 1.343 | 1.523 | -0.652 | 0.001 | -0.004 | -0.004 | 0 | 0.4 | 0.62 | 0.55 | 0.74 | 0.47 | 0.44 |
| 6 | 142398726 | rs6902780 | -0.856 | 1.4 | 1.466 | -0.563 | 0.024 | 0.035 | 0.035 | 0 | 0.2 | 0.27 | 0.36 | 0.7 | 0.24 | 0.2 |
| 6 | 142455731 | rs9376674 | 0.984 | 1.43 | 0.292 | -0.763 | -0.064 | -0.039 | -0.039 | 0 | 0.16 | 0.01 | 0.09 | 0.16 | 0.2 | 0.21 |
| 6 | 142461644 | rs225616 | 0.934 | 1.532 | 0.873 | -0.674 | 0.017 | -0.011 | -0.011 | 0.052 | 0.63 | 0.56 | 0.48 | 0.28 | 0.51 | 0.55 |
| 6 | 142523865 | rs1989181 | -0.477 | 1.69 | 0.903 | -0.448 | 0.005 | 0.03 | 0.03 | 0.147 | 0.13 | 0.15 | 0.32 | 0.67 | 0.19 | 0.16 |
| 6 | 142531037 | rs225699 | 1.01 | 1.678 | 0.744 | -0.535 | 0.009 | -0.005 | -0.005 | 0.28 | 0.65 | 0.58 | 0.5 | 0.29 | 0.57 | 0.55 |
| 6 | 142545280 | rs225712 | -0.014 | 1.238 | -0.163 | -0.551 | -0.061 | 0.016 | 0.016 | 0 | 0.8 | 0.97 | 0.76 | 0.92 | 0.76 | 0.71 |
| 6 | 142548099 | rs225717 | -0.206 | 1.24 | -0.085 | -0.55 | -0.024 | 0.03 | 0.03 | 0 | 0.8 | 0.91 | 0.75 | 0.92 | 0.76 | 0.73 |
| 6 | 142557418 | rs225735 | -0.112 | 1.218 | -0.064 | -0.565 | -0.004 | 0.009 | 0.009 | 0 | 0.79 | 0.82 | 0.73 | 0.92 | 0.76 | 0.73 |
| 6 | 142564217 | rs9399396 | -0.792 | 1.325 | -0.254 | -0.195 | -0.008 | -0.003 | -0.003 | 0 | 0.13 | 0.12 | 0.32 | 0.64 | 0.22 | 0.16 |
| 6 | 142568305 | rs2782527 | 0.562 | 1.407 | -0.325 | 0.486 | 0.006 | 0.213 | 0.213 | 0 | 0.95 | 0.89 | 0.83 | 0.46 | 0.93 | 0.93 |
| 6 | 142568452 | rs1931986 | -0.716 | 0.91 | 0.061 | 0.205 | 0.009 | 0.03 | 0.03 | 0 | 0.16 | 0.26 | 0.29 | 0.18 | 0.17 | 0.13 |
| 6 | 142569314 | rs1572265 | 0.214 | 1.189 | -0.749 | -0.052 | -0.019 | -0.038 | -0.038 | 0.406 | 0.74 | 0.81 | 0.6 | 0.39 | 0.68 | 0.54 |
| 6 | 142580952 | rs7745188 | -0.551 | 0.803 | -0.402 | 0.063 | 0.018 | -0.009 | -0.009 | 0 | 0.33 | 0.42 | 0.44 | 0.24 | 0.38 | 0.42 |
| 6 | 142583426 | rs1953319 | -0.841 | 0.464 | -1.697 | -0.392 | 0.123 | -0.048 | -0.048 | 0 | 0.55 | 0.84 | 0.7 | 0.46 | 0.74 | 0.74 |
| 6 | 142586578 | rs1931989 | 0.349 | 0.767 | -1.432 | 0.107 | 0.03 | 0.002 | 0.002 | 0 | 0.71 | 0.61 | 0.59 | 0.24 | 0.57 | 0.61 |
| 6 | 142587324 | rs9321861 | 0.524 | 0.913 | -2.618 | -0.311 | -0.052 | -0.007 | -0.007 | 0 | 0.18 | 0.07 | 0.13 | 0.21 | 0.31 | 0.23 |
| 6 | 142600563 | rs1319584 | -0.573 | 0.299 | -2.399 | -0.301 | -0.066 | 0.093 | 0.093 | 1.857 | 0.38 | 0.16 | 0.33 | 0.18 | 0.48 | 0.46 |
| 6 | 142607176 | rs9484615 | -0.77 | 0.57 | -2.802 | -0.067 | -0.059 | 0.148 | 0.148 | 2.301 | 0.48 | 0.91 | 0.49 | 0.25 | 0.5 | 0.48 |
| 6 | 142621947 | rs1342059 | 0.232 | 0.463 | -1.936 | 0.17 | -0.01 | 0.029 | 0.029 | 0 | 0.06 | 0 | 0.03 | 0 | 0.06 | 0.02 |
| 6 | 142624280 | rs17790314 | -0.429 | 0.619 | -1.792 | 0.422 | -0.008 | 0.002 | 0.002 | 0 | 0.08 | 0.07 | 0.09 | 0.1 | 0.12 | 0.08 |
| 6 | 142626699 | rs9376683 | -0.736 | 0.055 | -1.796 | 0.998 | 0.015 | -0.011 | -0.011 | 0 | 0.12 | 0.1 | 0.22 | 0.65 | 0.06 | 0.13 |
| 6 | 142629135 | rs9376684 | -0.736 | 0.024 | -1.878 | 0.979 | 0.025 | -0.028 | -0.028 | 0.891 | 0.12 | 0.05 | 0.22 | 0.65 | 0.06 | 0.13 |
| 6 | 142633744 | rs9389983 | -0.474 | 0.446 | -2.554 | 0.211 | -0.16 | 0.064 | 0.064 | 0 | 0.59 | 0.98 | 0.67 | 0.8 | 0.54 | 0.65 |
| 6 | 142641556 | rs9385993 | -0.6 | -0.142 | -2.164 | 0.912 | 0.018 | -0.023 | -0.023 | 0 | 0.13 | 0.07 | 0.12 | 0.27 | 0.08 | 0.07 |
| 6 | 142655490 | rs2039987 | -1.481 | -0.118 | -1.686 | 0.514 | 0 | 0 | 0 | 0.219 | 0.31 | 0.3 | 0.32 | 0.35 | 0.31 | 0.3 |
| 6 | 142658162 | rs2050157 | -1.853 | -0.717 | -0.689 | 0.437 | 0.019 | 0.008 | 0.008 | 0.182 | 0.39 | 0.72 | 0.37 | 0.4 | 0.32 | 0.31 |
| 6 | 142679572 | rs6570507 | -1.73 | -0.807 | -0.825 | 0.414 | 0.027 | -0.032 | -0.032 | 4.051 | 0.4 | 0.96 | 0.41 | 0.4 | 0.32 | 0.31 |
| 6 | 142691549 | rs11155242 | -0.441 | -0.824 | -0.767 | 0.41 | 0.01 | -0.027 | -0.027 | 0 | 0.18 | 0.23 | 0.2 | 0.09 | 0.23 | 0.23 |
| 6 | 142703877 | rs4896582 | -1.953 | -0.998 | -0.775 | 0.97 | 0.039 | 0.043 | 0.043 | 0.36 | 0.43 | 0.96 | 0.52 | 0.75 | 0.33 | 0.34 |
| 6 | 142709601 | rs9403384 | -0.115 | -0.775 | -0.601 | 0.72 | 0.022 | -0.027 | -0.027 | 0 | 0.13 | 0.07 | 0.12 | 0.26 | 0.08 | 0.07 |
| 6 | 142750516 | rs3817928 | -0.514 | -0.625 | -0.407 | 0.667 | 0.003 | -0.013 | -0.013 | 1.491 | 0.18 | 0.21 | 0.2 | 0.09 | 0.23 | 0.23 |
| 6 | 142755024 | rs6570510 | -1.579 | -0.926 | -0.24 | 0.593 | 0.019 | 0.01 | 0.01 | 0.015 | 0.42 | 0.67 | 0.37 | 0.39 | 0.32 | 0.31 |
| 6 | 142767633 | rs3748069 | -1.579 | -0.925 | 0.051 | 0.59 | 0.018 | 0.01 | 0.01 | 0.015 | 0.42 | 0.67 | 0.38 | 0.39 | 0.32 | 0.31 |
| 6 | 142773504 | rs12529186 | 0.552 | -0.138 | -0.557 | 0.137 | -0.082 | -0.056 | -0.056 | 0 | 0.13 | 0.01 | 0.15 | 0.15 | 0.25 | 0.16 |
| 6 | 142777064 | rs7776375 | -1.494 | -1.289 | -0.069 | -0.166 | 0.019 | 0.01 | 0.01 | 0.015 | 0.42 | 0.67 | 0.38 | 0.39 | 0.32 | 0.31 |
| 6 | 142811590 | rs10214911 | -0.462 | -1.333 | 0.008 | -0.212 | 0.006 | -0.004 | -0.004 | 0 | 0.18 | 0.17 | 0.14 | 0.26 | 0.09 | 0.07 |
| 6 | 142818428 | rs17071819 | -0.278 | -1.291 | -0.23 | -0.212 | 0.026 | -0.011 | -0.011 | 0 | 0.17 | 0.04 | 0.14 | 0.22 | 0.1 | 0.07 |
| 6 | 142839498 | rs262121 | -1.584 | -1.323 | 0.065 | -0.273 | 0.069 | 0.049 | 0.049 | 0 | 0.42 | 0.85 | 0.4 | 0.36 | 0.32 | 0.34 |
| 6 | 142846565 | rs155259 | -1.513 | -1.308 | 0.169 | -0.254 | 0.016 | 0.012 | 0.012 | 0 | 0.42 | 0.65 | 0.37 | 0.36 | 0.32 | 0.34 |
| 6 | 142846707 | rs2328339 | -0.616 | -1.352 | 0.365 | -0.272 | -0.062 | 0.015 | 0.015 | 1.738 | 0.21 | 0.43 | 0.18 | 0.26 | 0.1 | 0.09 |
| 6 | 142853486 | rs262130 | -0.781 | -0.736 | -0.061 | -0.299 | 0 | -0.003 | -0.003 | 0.577 | 0.2 | 0.21 | 0.19 | 0.1 | 0.22 | 0.25 |
| 6 | 142879589 | rs605790 | 0.832 | -1.063 | -0.803 | -0.57 | 0.011 | 0.182 | 0.182 | 0 | 0.5 | 0.15 | 0.62 | 0.7 | 0.55 | 0.56 |
| 6 | 142879693 | rs702293 | 0.099 | -1.585 | -0.254 | -0.966 | -0.001 | 0.001 | 0.001 | 0 | 0.77 | 0.8 | 0.8 | 0.78 | 0.71 | 0.85 |
| 6 | 142882955 | rs263114 | 1.126 | -1.748 | 0.866 | -1.047 | -0.001 | -0.024 | -0.024 | 0 | 0.57 | 0.49 | 0.7 | 0.78 | 0.58 | 0.59 |
| 6 | 142884774 | rs865141 | 0.235 | -1.84 | 1.073 | -1.097 | -0.003 | -0.03 | -0.03 | 0 | 0.07 | 0.09 | 0.04 | 0 | 0.09 | 0.01 |
| 6 | 142888515 | rs146250 | 0.053 | -1.836 | 1.203 | -1.101 | 0.003 | -0.045 | -0.045 | 1.398 | 0.08 | 0.17 | 0.05 | 0 | 0.09 | 0.01 |
| 6 | 142896655 | rs263164 | -0.982 | -1.711 | 1.127 | -1.103 | -0.021 | -0.098 | -0.098 | 0 | 0.22 | 0.3 | 0.11 | 0 | 0.19 | 0.27 |
| 6 | 142907515 | rs263156 | 2.217 | -2.367 | 0.632 | -1.173 | -0.075 | 0.046 | 0.046 | 0 | 0.45 | 0.04 | 0.62 | 0.66 | 0.55 | 0.56 |
| 6 | 142911971 | rs12201257 | 0.414 | -2.325 | 1.182 | -1.113 | -0.012 | 0.028 | 0.028 | 0 | 0.06 | 0.2 | 0.05 | 0.08 | 0.03 | 0.02 |
| 6 | 142921498 | rs13217677 | 0.139 | -2.332 | 1.094 | -1.132 | 0 | -0.001 | -0.001 | 0 | 0.21 | 0.19 | 0.18 | 0.21 | 0.23 | 0.14 |
| 6 | 142922061 | rs2077836 | -2.412 | -2.738 | 0.65 | -1.04 | -0.14 | -0.083 | -0.083 | 0 | 0.61 | 0.96 | 0.42 | 0.39 | 0.48 | 0.43 |
| 6 | 142936816 | rs17071903 | 0.41 | -2.43 | 1.187 | -0.798 | -0.014 | 0.007 | 0.007 | 0 | 0.07 | 0.07 | 0.06 | 0.05 | 0.02 | 0.03 |

| | | | | | | | | | | | | | | | | |
|---|---|---|---|---|---|---|---|---|---|---|---|---|---|---|---|---|
| 6 | 142946384 | rs12191946 | 0.161 | -2.469 | 1.319 | -0.822 | -0.014 | 0.001 | 0.001 | 0 | 0.08 | 0.12 | 0.04 | 0.08 | 0.03 | 0.02 |
| 6 | 142949260 | rs9373355 | -1.705 | -2.681 | 1.342 | -0.882 | -0.006 | 0.004 | 0.004 | 0.056 | 0.36 | 0.61 | 0.3 | 0.39 | 0.29 | 0.21 |
| 6 | 142959329 | rs9403402 | -1.344 | -2.595 | 1.502 | -0.973 | -0.041 | -0.012 | -0.012 | 0.557 | 0.19 | 0.36 | 0.11 | 0.1 | 0.12 | 0.06 |
| 6 | 142964625 | rs6570521 | -1.682 | -2.706 | 1.686 | -0.937 | 0.001 | -0.001 | -0.001 | 0.084 | 0.32 | 0.65 | 0.3 | 0.27 | 0.35 | 0.18 |
| 6 | 142965039 | rs6928084 | -2.156 | -2.745 | 1.7 | -0.982 | 0.005 | 0.008 | 0.008 | 0 | 0.4 | 0.72 | 0.34 | 0.35 | 0.38 | 0.2 |
| 6 | 142969334 | rs7757435 | -1.527 | -2.984 | 1.623 | -1.213 | -0.075 | -0.043 | -0.043 | 0 | 0.24 | 0.51 | 0.14 | 0.1 | 0.14 | 0.1 |
| 6 | 142969570 | rs9390016 | -2.074 | -2.923 | 1.389 | -1.238 | 0.01 | 0.022 | 0.022 | 0 | 0.44 | 0.71 | 0.36 | 0.34 | 0.41 | 0.24 |
| 6 | 142973369 | rs9403404 | -0.489 | -2.918 | 1.597 | -1.224 | -0.022 | -0.025 | -0.025 | 0 | 0.15 | 0.24 | 0.07 | 0.05 | 0.11 | 0.06 |
| 6 | 142975097 | rs6936988 | -1.464 | -2.915 | 1.152 | -1.204 | -0.057 | -0.033 | -0.033 | 0 | 0.24 | 0.59 | 0.16 | 0.1 | 0.14 | 0.1 |
| 6 | 142976691 | rs7771119 | 0.162 | -2.762 | 0.979 | -1.242 | -0.028 | -0.009 | -0.009 | 0.534 | 0.17 | 0.12 | 0.19 | 0.24 | 0.27 | 0.14 |
| 6 | 142990718 | rs1396898 | -1.754 | -3.002 | 0.95 | -1.441 | 0.01 | 0.121 | 0.121 | 0 | 0.51 | 0.72 | 0.37 | 0.24 | 0.48 | 0.42 |
| 6 | 142991072 | rs13206398 | 0.253 | -2.895 | 0.577 | -1.384 | -0.012 | -0.046 | -0.046 | 0 | 0.17 | 0.19 | 0.06 | 0 | 0.09 | 0.22 |
| 6 | 142993151 | rs7753052 | -1.91 | -2.586 | -0.1 | -1.121 | -0.072 | 0.035 | 0.035 | 0 | 0.6 | 0.86 | 0.41 | 0.3 | 0.5 | 0.47 |
| 6 | 142998843 | rs11155260 | 0.049 | -2.054 | -0.362 | -0.622 | -0.028 | -0.029 | -0.029 | 0 | 0.16 | 0.09 | 0.18 | 0.27 | 0.23 | 0.09 |
| 6 | 143007165 | rs11155262 | -0.294 | -2.031 | -0.459 | -0.791 | -0.02 | 0.005 | 0.005 | 1.495 | 0.26 | 0.12 | 0.21 | 0.26 | 0.26 | 0.11 |
| 6 | 143008964 | rs7775764 | -1.65 | -2.092 | -0.423 | -0.862 | -0.031 | 0.017 | 0.017 | 0 | 0.6 | 0.74 | 0.42 | 0.33 | 0.49 | 0.52 |
| 6 | 143012781 | rs597531 | 1.559 | -2.268 | -0.183 | -0.466 | 0.058 | 0.051 | 0.051 | 0.346 | 0.5 | 0.32 | 0.57 | 0.61 | 0.61 | 0.46 |
| 6 | 143014234 | rs614081 | 1.523 | -2.262 | -0.587 | -0.462 | 0.074 | 0.064 | 0.064 | 0.629 | 0.49 | 0.22 | 0.57 | 0.6 | 0.61 | 0.46 |
| 6 | 143015405 | rs3861401 | -0.438 | -2.326 | -0.798 | -0.115 | 0.032 | 0.046 | 0.046 | 0 | 0.15 | 0.07 | 0.05 | 0.02 | 0.09 | 0.25 |
| 6 | 143020207 | rs6570524 | 0.523 | -1.745 | -0.321 | 0.251 | -0.034 | 0.024 | 0.024 | 0 | 0.89 | 0.74 | 0.93 | 0.88 | 0.96 | 0.91 |
| 6 | 143020752 | rs12200517 | -0.566 | -1.093 | -1.082 | 0.419 | -0.003 | -0.026 | -0.026 | 0 | 0.13 | 0.06 | 0.21 | 0.27 | 0.14 | 0.13 |
| 6 | 143031228 | rs478884 | 0.836 | -1.409 | -1.376 | 1.095 | -0.014 | -0.094 | -0.094 | 0 | 0.85 | 1 | 0.96 | 0.73 | 0.9 | 0.68 |
| 6 | 143031271 | rs6902684 | -0.835 | -2.867 | -1.362 | 0.757 | 0.024 | 0.002 | 0.002 | 3.263 | 0.28 | 0.71 | 0.26 | 0.3 | 0.11 | 0.28 |
| 6 | 143031519 | rs502984 | 0.739 | -1.524 | -2.277 | 0.203 | -0.094 | 0.043 | 0.043 | 0.254 | 0.35 | 0.07 | 0.56 | 0.24 | 0.45 | 0.32 |
| 6 | 143032521 | rs533183 | 0.959 | -1.313 | -2.01 | 0.289 | -0.101 | 0.01 | 0.01 | 0.165 | 0.3 | 0.08 | 0.44 | 0.24 | 0.43 | 0.3 |
| 6 | 143033454 | rs600030 | 0.446 | -1.272 | -1.026 | 0.475 | 0.025 | 0.02 | 0.02 | 0 | 0.93 | 0.83 | 0.94 | 0.91 | 0.91 | 0.91 |
| 6 | 143039141 | rs534230 | 0.236 | -1.874 | -0.661 | 0.115 | -0.003 | -0.007 | -0.007 | 0 | 0.6 | 0.47 | 0.68 | 0.8 | 0.69 | 0.78 |
| 6 | 143044071 | rs574975 | -0.195 | -1.468 | -0.844 | 0.147 | 0.007 | 0.019 | 0.019 | 1.018 | 0.25 | 0.33 | 0.26 | 0.57 | 0.29 | 0.51 |
| 6 | 143051130 | rs653458 | 0.419 | -1.974 | -0.099 | -0.175 | -0.019 | -0.01 | -0.01 | 0 | 0.8 | 0.7 | 0.84 | 0.9 | 0.92 | 0.92 |
| 6 | 143051810 | rs639954 | 0.304 | -1.866 | -0.068 | 0.218 | 0.052 | 0.089 | 0.089 | 0.713 | 0.17 | 0.21 | 0.2 | 0.44 | 0.26 | 0.4 |
| 6 | 143056556 | rs9390022 | 1.724 | -2.007 | -0.182 | 0.12 | -0.048 | 0.035 | 0.035 | 0 | 0.3 | 0.11 | 0.4 | 0.22 | 0.4 | 0.25 |
| 6 | 143057067 | rs4895598 | 0.866 | -2.105 | -0.091 | 0.109 | 0.005 | -0.009 | -0.009 | 0.9 | 0.42 | 0.43 | 0.5 | 0.36 | 0.5 | 0.34 |
| 6 | 143057547 | rs12526857 | 0.314 | -2.134 | -0.254 | 0.089 | 0.005 | 0.001 | 0.001 | 0 | 0.1 | 0.1 | 0.08 | 0.09 | 0.1 | 0.08 |
| 6 | 143058408 | rs538289 | -0.098 | -2.187 | -0.256 | 0.067 | 0.008 | 0.008 | 0.008 | 2.432 | 0.6 | 0.61 | 0.68 | 0.8 | 0.76 | 0.74 |
| 6 | 143060440 | rs17071978 | -0.019 | -1.714 | -0.5 | 0.147 | -0.004 | 0.001 | 0.001 | 0 | 0.11 | 0.16 | 0.08 | 0.13 | 0.1 | 0.08 |
| 6 | 143062803 | rs197490 | 0.018 | -1.775 | -0.763 | 0.023 | 0.003 | 0.002 | 0.002 | 0 | 0.38 | 0.53 | 0.27 | 0.44 | 0.38 | 0.48 |
| 6 | 143069315 | rs197482 | -1.658 | -1.351 | -0.757 | -0.762 | -0.07 | 0.061 | 0.061 | 0.042 | 0.72 | 0.91 | 0.71 | 0.83 | 0.59 | 0.75 |
| 6 | 143069864 | rs578343 | 0.034 | -1.518 | -0.816 | -0.791 | 0.015 | 0.011 | 0.011 | 0.644 | 0.14 | 0.07 | 0.14 | 0.06 | 0.1 | 0.08 |
| 6 | 143070763 | rs6930478 | 1.026 | -1.645 | -0.629 | -0.943 | -0.04 | -0.119 | -0.119 | 0 | 0.25 | 0.34 | 0.1 | 0.01 | 0.14 | 0.09 |
| 6 | 143072350 | rs576420 | -0.016 | -1.226 | -1.063 | -0.691 | 0.008 | 0.011 | 0.011 | 0.716 | 0.14 | 0.08 | 0.16 | 0.06 | 0.11 | 0.08 |
| 6 | 143073041 | rs3383 | -1.335 | -0.795 | -1.048 | -0.444 | -0.046 | 0.055 | 0.055 | 0 | 0.72 | 0.87 | 0.72 | 0.83 | 0.6 | 0.75 |
| 6 | 143074700 | rs169098 | -1.335 | -0.772 | -1.047 | -0.443 | -0.045 | 0.051 | 0.051 | 0.02 | 0.72 | 0.87 | 0.72 | 0.83 | 0.61 | 0.78 |
| 6 | 143076543 | rs12524093 | 0.23 | -0.506 | -1.283 | -0.242 | -0.03 | -0.023 | -0.023 | 0 | 0.05 | 0.01 | 0.07 | 0.09 | 0.12 | 0.13 |
| 6 | 143077232 | rs197470 | 0.069 | -0.226 | -1.165 | -0.205 | 0.006 | 0.007 | 0.007 | 0 | 0.52 | 0.74 | 0.53 | 0.55 | 0.45 | 0.38 |
| 6 | 143077289 | rs197469 | -0.671 | -0.008 | -2.023 | -0.088 | -0.11 | 0.004 | 0.004 | 0.072 | 0.66 | 0.93 | 0.64 | 0.76 | 0.58 | 0.72 |
| 6 | 143080039 | rs197465 | -0.5 | -0.177 | -1.507 | 0.177 | -0.009 | -0.056 | -0.056 | 0.006 | 0.24 | 0.16 | 0.35 | 0.49 | 0.26 | 0.25 |
| 6 | 143083063 | rs638880 | -0.227 | -0.517 | -1.112 | -0.423 | 0.041 | 0.068 | 0.068 | 0.908 | 0.42 | 0.34 | 0.29 | 0.09 | 0.31 | 0.16 |
| 6 | 143086596 | rs197458 | 0.224 | -0.069 | -1.147 | 0.154 | 0.02 | 0.009 | 0.009 | 0.46 | 0.59 | 0.65 | 0.51 | 0.54 | 0.66 | 0.58 |
| 6 | 143087003 | rs197457 | 0.098 | -0.153 | -1.382 | 0.112 | -0.014 | -0.054 | -0.054 | 0 | 0.89 | 0.98 | 0.69 | 0.63 | 0.89 | 0.84 |
| 6 | 143091263 | rs109836 | -0.319 | -0.21 | -1.387 | 0.051 | 0.009 | 0.035 | 0.035 | 0 | 0.93 | 0.99 | 0.97 | 1 | 0.97 | 0.95 |
| 6 | 143097493 | rs197508 | 0.797 | -0.324 | -0.432 | -0.239 | -0.003 | -0.003 | -0.003 | 0 | 0.56 | 0.57 | 0.48 | 0.53 | 0.6 | 0.56 |
| 6 | 143101562 | rs84578 | 0.824 | -0.355 | -0.424 | -0.135 | 0.005 | -0.032 | -0.032 | 0 | 0.78 | 0.61 | 0.59 | 0.77 | 0.8 | 0.79 |
| 6 | 143101663 | rs126343 | 0.627 | -0.341 | -0.797 | -0.224 | -0.025 | -0.07 | -0.07 | 0 | 0.81 | 0.96 | 0.62 | 0.64 | 0.8 | 0.79 |
| 6 | 143115636 | rs491414 | 0.705 | -0.562 | -0.565 | -0.247 | 0.009 | 0.001 | 0.001 | 0 | 0.62 | 0.35 | 0.49 | 0.58 | 0.68 | 0.66 |
| 6 | 143118715 | rs12525269 | 0.152 | -0.56 | -0.732 | -0.259 | -0.011 | 0.021 | 0.021 | 0 | 0.06 | 0 | 0.03 | 0.01 | 0.07 | 0.04 |
| 6 | 143130226 | rs512808 | 0.769 | -0.639 | -0.431 | -0.267 | 0.025 | 0.002 | 0.002 | 0 | 0.61 | 0.25 | 0.47 | 0.58 | 0.68 | 0.66 |
| 6 | 143137188 | rs12190656 | 0.346 | -0.653 | -0.551 | -0.332 | 0.002 | 0.078 | 0.078 | 0 | 0.18 | 0.06 | 0.1 | 0.04 | 0.18 | 0.16 |
| 6 | 143142313 | rs9791312 | 1.088 | 0.312 | -1.221 | -0.093 | -0.096 | -0.088 | -0.088 | 0 | 0.19 | 0.05 | 0.25 | 0.28 | 0.33 | 0.39 |
| 6 | 143144885 | rs926372 | -0.347 | -0.104 | -1.006 | 0.055 | 0.003 | -0.017 | -0.017 | 1.135 | 0.81 | 0.83 | 0.61 | 0.61 | 0.86 | 0.83 |
| 6 | 143149300 | rs493985 | -0.299 | -0.417 | -0.651 | -0.112 | -0.005 | 0.005 | 0.005 | 1.41 | 0.77 | 0.77 | 0.63 | 0.6 | 0.87 | 0.84 |
| 6 | 143150043 | rs9373363 | 0.831 | -0.661 | -0.765 | -0.477 | 0.008 | 0.014 | 0.014 | 0.082 | 0.21 | 0.29 | 0.18 | 0.28 | 0.25 | 0.2 |
| 6 | 143152086 | rs198663 | -0.334 | -0.056 | -1.44 | -0.498 | 0.005 | 0.05 | 0.05 | 0.09 | 0.52 | 0.7 | 0.38 | 0.33 | 0.54 | 0.45 |
| 6 | 143165203 | rs12203103 | 1.093 | 0.804 | -1.935 | -0.703 | -0.074 | 0.101 | 0.101 | 0 | 0.17 | 0.01 | 0.13 | 0.02 | 0.23 | 0.16 |
| 6 | 143166851 | rs198652 | 0.282 | 0.527 | -1.372 | -0.578 | 0.169 | 0.087 | 0.087 | 2.796 | 0.06 | 0.48 | 0.07 | 0.04 | 0.07 | 0.02 |
| 6 | 143172013 | rs728589 | 0.829 | 0.341 | -1.769 | -0.565 | 0.013 | -0.001 | -0.001 | 0.255 | 0.21 | 0.15 | 0.16 | 0.2 | 0.2 | 0.2 |
| 6 | 143175960 | rs12190352 | 0.77 | 0.614 | -2.085 | -0.658 | -0.088 | 0.157 | 0.157 | 0 | 0.25 | 0.02 | 0.15 | 0.03 | 0.28 | 0.24 |
| 6 | 143185683 | rs765875 | 0.161 | 0.489 | -1.896 | -0.201 | -0.001 | 0.004 | 0.004 | 1.314 | 0.46 | 0.24 | 0.39 | 0.53 | 0.5 | 0.57 |
| 6 | 143193344 | rs9390025 | 0.493 | 0.587 | -1.62 | 0.395 | -0.032 | -0.066 | -0.066 | 0 | 0.24 | 0.08 | 0.24 | 0.35 | 0.3 | 0.38 |
| 6 | 143200547 | rs198683 | 0.258 | 0.388 | -1.806 | 0.144 | 0.001 | 0.065 | 0.065 | 0 | 0.55 | 0.63 | 0.35 | 0.04 | 0.56 | 0.4 |

| | | | | | | | | | | | | | | | | |
|---|---|---|---|---|---|---|---|---|---|---|---|---|---|---|---|---|
| 6 | 143205435 | rs11155266 | -0.316 | 0.337 | -1.597 | 0.693 | 0.004 | -0.037 | -0.037 | 0.386 | 0.19 | 0.13 | 0.36 | 0.42 | 0.15 | 0.15 |
| 6 | 143213127 | rs12201410 | -0.233 | 0.496 | -2.013 | 0.176 | 0 | 0.108 | 0.108 | 0 | 0.48 | 0.18 | 0.45 | 0.31 | 0.42 | 0.33 |
| 6 | 143214829 | rs12205042 | -0.049 | 0.21 | -2.395 | -0.288 | -0.016 | 0.159 | 0.159 | 0 | 0.29 | 0.09 | 0.18 | 0.07 | 0.24 | 0.23 |
| 6 | 143220180 | rs9496475 | -1.247 | 0.129 | -1.829 | -0.107 | 0.045 | 0.069 | 0.069 | 0 | 0.69 | 0.6 | 0.56 | 0.51 | 0.58 | 0.51 |
| 6 | 143220643 | rs7749585 | 0.68 | -0.079 | -1.632 | 0.104 | -0.014 | 0.003 | 0.003 | 3.293 | 0.07 | 0.34 | 0.04 | 0.03 | 0.01 | 0.03 |
| 6 | 143224510 | rs9390030 | 1.276 | 0.351 | -2.218 | 0.179 | -0.019 | -0.042 | -0.042 | 0 | 0.11 | 0.01 | 0.07 | 0.17 | 0.09 | 0.09 |
| 6 | 143231149 | rs9403421 | 1.276 | 0.351 | -2.218 | 0.179 | -0.019 | -0.042 | -0.042 | 0.277 | 0.11 | 0.02 | 0.07 | 0.17 | 0.09 | 0.09 |
| 6 | 143235252 | rs6923298 | -0.151 | 0.502 | -1.907 | 0.526 | 0.001 | 0.017 | 0.017 | 0.408 | 0.36 | 0.17 | 0.39 | 0.32 | 0.33 | 0.28 |
| 6 | 143256102 | rs9496482 | -0.338 | 0.888 | -1.296 | -0.083 | 0.004 | 0.009 | 0.009 | 0 | 0.28 | 0.29 | 0.17 | 0.09 | 0.22 | 0.19 |
| 6 | 143260600 | rs6940880 | -0.329 | 0.861 | -1.289 | -0.153 | 0.005 | 0.008 | 0.008 | 0 | 0.28 | 0.27 | 0.17 | 0.08 | 0.22 | 0.19 |
| 6 | 143270216 | rs2095369 | 0.266 | 1.065 | -0.5 | 0.749 | 0.01 | 0.039 | 0.039 | 0.022 | 0.2 | 0.25 | 0.39 | 0.46 | 0.28 | 0.37 |
| 6 | 143270247 | rs1413627 | 0.442 | 1.024 | -0.082 | 0.943 | 0.057 | 0.106 | 0.106 | 0 | 0.93 | 0.76 | 0.92 | 0.85 | 0.93 | 0.85 |
| 6 | 143272689 | rs9386015 | 0.508 | 1.011 | 0.015 | 0.936 | 0.079 | 0.139 | 0.139 | 0 | 0.93 | 0.71 | 0.91 | 0.85 | 0.93 | 0.84 |
| 6 | 143274232 | rs6925124 | 0.551 | 0.681 | -0.386 | 0.683 | 0.001 | -0.009 | -0.009 | 5.497 | 0.58 | 0.25 | 0.53 | 0.39 | 0.6 | 0.53 |
| 6 | 143275398 | rs9496486 | 0.606 | 0.812 | -0.399 | 0.736 | -0.002 | -0.025 | -0.025 | 0 | 0.6 | 0.35 | 0.55 | 0.46 | 0.6 | 0.58 |
| 6 | 143282725 | rs6923804 | -0.029 | 0.937 | -0.792 | 0.494 | -0.004 | -0.008 | -0.008 | 0 | 0.31 | 0.27 | 0.4 | 0.45 | 0.36 | 0.43 |
| 6 | 143288285 | rs1041656 | -0.305 | 0.755 | -0.583 | -0.269 | 0.007 | -0.011 | -0.011 | 0.281 | 0.64 | 0.55 | 0.49 | 0.23 | 0.58 | 0.45 |
| 6 | 143289361 | rs1041654 | -0.311 | 0.316 | -0.66 | -0.803 | -0.001 | -0.052 | -0.052 | 0 | 0.44 | 0.41 | 0.35 | 0.04 | 0.41 | 0.33 |
| 6 | 143297660 | rs6920570 | -0.399 | 0.65 | -0.333 | -0.27 | 0 | 0.047 | 0.047 | 0 | 0.15 | 0.27 | 0.23 | 0.24 | 0.16 | 0.24 |
| 6 | 143304760 | rs1413628 | 0.097 | 0.363 | -0.662 | -1.001 | 0.004 | -0.097 | -0.097 | 0.035 | 0.4 | 0.44 | 0.27 | 0.08 | 0.4 | 0.17 |
| 6 | 143307904 | rs7764982 | 0.283 | 0.263 | -0.244 | -1.076 | -0.001 | -0.012 | -0.012 | 0.014 | 0.26 | 0.27 | 0.19 | 0.09 | 0.23 | 0.09 |
| 6 | 143308879 | rs9321880 | -0.968 | 0.033 | 0.255 | -0.428 | -0.004 | -0.015 | -0.015 | 0 | 0.54 | 0.75 | 0.61 | 0.42 | 0.57 | 0.38 |
| 6 | 143310827 | rs1360738 | -0.483 | -0.011 | 0.704 | -0.457 | -0.007 | 0.041 | 0.041 | 0 | 0.32 | 0.52 | 0.43 | 0.42 | 0.35 | 0.31 |
| 6 | 143311522 | rs2184360 | -0.372 | -0.03 | 0.66 | -0.438 | 0.003 | 0.049 | 0.049 | 0.044 | 0.25 | 0.39 | 0.37 | 0.32 | 0.29 | 0.24 |
| 6 | 143311994 | rs6570541 | -0.837 | -0.04 | 0.685 | -0.483 | -0.02 | -0.004 | -0.004 | 1.906 | 0.47 | 0.72 | 0.57 | 0.42 | 0.51 | 0.33 |
| 6 | 143312232 | rs9321883 | -0.475 | 0.091 | 0.423 | -0.092 | 0.002 | 0.036 | 0.036 | 0 | 0.24 | 0.3 | 0.38 | 0.35 | 0.29 | 0.23 |
| 6 | 143320871 | rs4896611 | -0.674 | 0.094 | 0.381 | -0.1 | -0.002 | 0.108 | 0.108 | 0.018 | 0.26 | 0.45 | 0.4 | 0.42 | 0.3 | 0.27 |
| 6 | 143327325 | rs199255 | 0.093 | 0.217 | 0.151 | -0.116 | 0.001 | 0.01 | 0.01 | 0.01 | 0.78 | 0.76 | 0.65 | 0.76 | 0.71 | 0.81 |
| 6 | 143327839 | rs9496500 | 0.509 | 0.411 | -0.069 | -0.248 | 0.002 | 0.007 | 0.007 | 0 | 0.6 | 0.57 | 0.51 | 0.54 | 0.54 | 0.53 |
| 6 | 143329123 | rs199256 | -0.049 | 0.246 | 0.004 | -0.073 | -0.001 | -0.001 | -0.001 | 0 | 0.76 | 0.77 | 0.64 | 0.65 | 0.73 | 0.64 |
| 6 | 143332403 | rs9496502 | 0.193 | 0.286 | 0.434 | -0.006 | 0.044 | -0.021 | -0.021 | 0.328 | 0.05 | 0.16 | 0.06 | 0 | 0.06 | 0.02 |
| 6 | 143332882 | rs199259 | 0.078 | 0.251 | 0.022 | -0.13 | 0 | -0.001 | -0.001 | 0.202 | 0.26 | 0.18 | 0.23 | 0.2 | 0.32 | 0.17 |
| 6 | 143333813 | rs2050003 | 0.828 | -0.099 | -0.505 | 0.09 | 0.048 | -0.095 | -0.095 | 0 | 0.2 | 0 | 0.07 | 0.29 | 0.08 | 0.14 |
| 6 | 143334993 | rs199248 | 0.512 | 0.931 | 0.475 | 0.188 | -0.01 | 0.003 | 0.003 | 0 | 0.08 | 0.04 | 0.09 | 0 | 0.16 | 0.04 |
| 6 | 143337546 | rs199245 | -1.342 | 0.909 | 0.517 | 0.221 | -0.002 | -0.001 | -0.001 | 0 | 0.2 | 0.72 | 0.23 | 0.22 | 0.25 | 0.13 |
| 6 | 143340590 | rs9321886 | 0.454 | 0.642 | 0.45 | 0.284 | -0.005 | -0.018 | -0.018 | 0 | 0.19 | 0.21 | 0.15 | 0.13 | 0.17 | 0.1 |
| 6 | 143340668 | rs199244 | -0.676 | 0.167 | 0.341 | 0.556 | -0.051 | 0.061 | 0.061 | 0.251 | 0.4 | 0.72 | 0.35 | 0.27 | 0.47 | 0.3 |
| 6 | 143340858 | rs199243 | -0.132 | 0.255 | 0.066 | 0.835 | 0.02 | -0.019 | -0.019 | 0 | 0.31 | 0.34 | 0.26 | 0.22 | 0.4 | 0.18 |
| 6 | 143346704 | rs9321887 | -1.323 | -0.5 | 0.017 | 1.675 | -0.014 | 0.051 | 0.051 | 0.503 | 0.14 | 0.2 | 0.23 | 0.49 | 0.08 | 0.37 |
| 6 | 143359013 | rs12664681 | -0.797 | -0.491 | -0.029 | 1.799 | 0.029 | -0.031 | -0.031 | 0.329 | 0.11 | 0.06 | 0.03 | 0.41 | 0.07 | 0.28 |
| 6 | 143382110 | rs7774011 | -1.627 | -0.682 | 0.167 | 1.662 | -0.028 | 0.041 | 0.041 | 2.659 | 0.21 | 0.3 | 0.26 | 0.5 | 0.11 | 0.37 |
| 6 | 143384258 | rs3804527 | -1.706 | -0.679 | 0.039 | 1.638 | -0.047 | 0.065 | 0.065 | 5.041 | 0.22 | 0.35 | 0.27 | 0.5 | 0.11 | 0.37 |
| 6 | 143400092 | rs3804528 | -0.752 | -0.626 | -0.271 | 1.835 | 0.024 | -0.019 | -0.019 | 0.007 | 0.11 | 0.08 | 0.03 | 0.42 | 0.04 | 0.27 |
| 6 | 143428150 | rs6919707 | -1.413 | -1.137 | 1.01 | 1.64 | 0.054 | 0.099 | 0.099 | 3.577 | 0.06 | 0.46 | 0.05 | 0.05 | 0.01 | 0.05 |
| 6 | 143437284 | rs7738387 | -2.409 | -1.395 | 1.136 | 1.563 | -0.074 | 0.048 | 0.048 | 9.5 | 0.23 | 0.65 | 0.1 | 0.47 | 0.06 | 0.34 |
| 6 | 143437481 | rs6900049 | -1.773 | -1.371 | 1.107 | 1.575 | -0.033 | 0.029 | 0.029 | 2.24 | 0.18 | 0.27 | 0.06 | 0.47 | 0.05 | 0.34 |
| 6 | 143438246 | rs17590859 | -0.11 | -1.292 | 0.927 | 1.754 | 0.04 | -0.035 | -0.035 | 0 | 0.08 | 0 | 0.02 | 0.37 | 0.03 | 0.21 |
| 6 | 143480314 | rs6570550 | 0.059 | -0.968 | 0.142 | 0.67 | -0.015 | 0.044 | 0.044 | 0.01 | 0.46 | 0.73 | 0.68 | 0.61 | 0.47 | 0.42 |
| 6 | 143480743 | rs6908441 | 0.417 | -0.92 | 0.159 | 0.82 | -0.001 | -0.007 | -0.007 | 0 | 0.32 | 0.36 | 0.38 | 0.15 | 0.31 | 0.29 |
| 6 | 143481592 | rs9403452 | 0.078 | -1.029 | -0.053 | 0.928 | -0.051 | -0.066 | -0.066 | 0.662 | 0.37 | 0.17 | 0.49 | 0.44 | 0.47 | 0.39 |
| 6 | 143481743 | rs4895601 | -0.52 | -1.031 | -0.04 | 0.887 | -0.018 | 0.017 | 0.017 | 0.041 | 0.46 | 0.76 | 0.71 | 0.58 | 0.49 | 0.46 |
| 6 | 143488566 | rs1573992 | 0.702 | -0.828 | -0.012 | 1.107 | -0.017 | 0.049 | 0.049 | 0.799 | 0.27 | 0.17 | 0.2 | 0.07 | 0.29 | 0.26 |
| 6 | 143502969 | rs17629067 | -1.086 | -0.501 | -0.05 | 1.49 | 0.006 | -0.011 | -0.011 | 0.057 | 0.17 | 0.14 | 0.05 | 0.4 | 0.06 | 0.31 |
| 6 | 143519095 | rs7772833 | 0.003 | 0.593 | -0.012 | 1.083 | -0.001 | -0.01 | -0.01 | 0.045 | 0.13 | 0.15 | 0.09 | 0.01 | 0.17 | 0.13 |
| 6 | 143521442 | rs7745568 | -1.055 | 0.444 | 0.475 | 1.583 | -0.003 | 0.064 | 0.064 | 0 | 0.21 | 0.31 | 0.35 | 0.48 | 0.22 | 0.25 |
| 6 | 143539299 | rs11758699 | -1.287 | 1.018 | 0.149 | 1.705 | 0.007 | 0.023 | 0.023 | 0 | 0.19 | 0.23 | 0.39 | 0.31 | 0.24 | 0.15 |
| 6 | 143543360 | rs9403466 | -0.783 | 1.584 | 0.02 | 1.716 | -0.017 | -0.017 | -0.017 | 0 | 0.07 | 0.06 | 0.06 | 0.19 | 0.12 | 0.18 |
| 6 | 143544740 | rs7740762 | -0.879 | 1.374 | 0.671 | 1.309 | 0.032 | -0.052 | -0.052 | 0.811 | 0.1 | 0.3 | 0.09 | 0 | 0.09 | 0.1 |
| 6 | 143548316 | rs9399430 | -1.604 | 1.314 | 0.559 | 1.76 | 0.051 | 0.119 | 0.119 | 0 | 0.11 | 0.29 | 0.34 | 0.4 | 0.14 | 0.25 |
| 6 | 143555358 | rs12191886 | -1.49 | 0.553 | 0.326 | 1.392 | 0.001 | 0.001 | 0.001 | 0 | 0.28 | 0.28 | 0.42 | 0.71 | 0.24 | 0.45 |
| 6 | 143556934 | rs6910775 | -1.184 | 0.53 | 0.615 | 1.836 | 0 | 0.128 | 0.128 | 0 | 0.21 | 0.37 | 0.42 | 0.39 | 0.21 | 0.19 |
| 6 | 143562004 | rs4342440 | -1.957 | 0.158 | 0.101 | 0.863 | 0.025 | 0.169 | 0.169 | 1.484 | 0.49 | 0.69 | 0.57 | 0.86 | 0.43 | 0.7 |
| 6 | 143565218 | rs986589 | -0.826 | 0.07 | -0.192 | 1.083 | -0.068 | -0.135 | -0.135 | 0.622 | 0.28 | 0.08 | 0.3 | 0.44 | 0.35 | 0.29 |
| 6 | 143567039 | rs12207891 | 0.091 | -0.037 | -0.11 | 1.137 | 0.012 | -0.068 | -0.068 | 0.131 | 0.11 | 0 | 0.03 | 0.33 | 0.05 | 0.33 |
| 6 | 143585300 | rs17530287 | -0.059 | 0.14 | 0.055 | 0.581 | -0.044 | 0.031 | 0.031 | 0.707 | 0.08 | 0.01 | 0.06 | 0.03 | 0.11 | 0.11 |
| 6 | 143586007 | rs892713 | -0.945 | 0.118 | -0.084 | 0.576 | -0.024 | -0.023 | -0.023 | 1.528 | 0.27 | 0.03 | 0.25 | 0.71 | 0.26 | 0.45 |
| 6 | 143588399 | rs11757344 | -0.586 | 0.112 | -0.138 | 0.573 | -0.065 | -0.063 | -0.063 | 0 | 0.22 | 0.02 | 0.24 | 0.71 | 0.26 | 0.45 |
| 6 | 143588803 | rs17072414 | -0.256 | 0.194 | 0.149 | 0.626 | 0.008 | -0.019 | -0.019 | 0 | 0.09 | 0.14 | 0.07 | 0.03 | 0.11 | 0.11 |
| 6 | 143589210 | rs9390073 | -0.204 | 0.168 | -0.021 | 0.621 | -0.021 | 0.016 | 0.016 | 0.566 | 0.12 | 0.04 | 0.07 | 0.08 | 0.14 | 0.18 |

| | | | | | | | | | | | | | | | | |
|---|---|---|---|---|---|---|---|---|---|---|---|---|---|---|---|---|
| 6 | 143592386 | rs1570350 | -2.305 | -0.093 | -0.198 | 0.418 | -0.124 | 0.228 | 0.228 | 0.503 | 0.55 | 0.93 | 0.44 | 0.88 | 0.46 | 0.77 |
| 6 | 143594364 | rs9285499 | -1.417 | -0.092 | 0.486 | 0.544 | -0.01 | -0.009 | -0.009 | 0 | 0.31 | 0.55 | 0.27 | 0.76 | 0.22 | 0.46 |
| 6 | 143596413 | rs9373384 | -2.289 | -0.072 | 0.659 | 0.479 | 0.004 | -0.04 | -0.04 | 0 | 0.24 | 0.5 | 0.53 | 0.19 | 0.24 | 0.29 |
| 6 | 143598107 | rs9390075 | 0.541 | -0.136 | 0.577 | 0.882 | -0.053 | -0.088 | -0.088 | 0 | 0.12 | 0.01 | 0.09 | 0.35 | 0.16 | 0.15 |
| 6 | 143601132 | rs11155287 | -1.522 | -0.185 | 0.787 | 0.849 | 0.009 | 0.03 | 0.03 | 0 | 0.31 | 0.19 | 0.18 | 0.75 | 0.26 | 0.49 |
| 6 | 143602840 | rs13205655 | -0.863 | -0.162 | 0.936 | 0.831 | 0.006 | -0.008 | -0.008 | 0 | 0.14 | 0.2 | 0.37 | 0.11 | 0.18 | 0.2 |
| 6 | 143604514 | rs9321898 | -2.084 | -0.191 | 0.935 | 0.82 | -0.001 | -0.033 | -0.033 | 0 | 0.42 | 0.56 | 0.61 | 0.85 | 0.39 | 0.65 |
| 6 | 143604645 | rs11155288 | -1.825 | -0.111 | 0.919 | 0.807 | -0.01 | -0.02 | -0.02 | 0 | 0.32 | 0.33 | 0.51 | 0.48 | 0.38 | 0.41 |
| 6 | 143609638 | rs9484719 | 0.043 | -0.084 | 0.918 | 0.919 | 0.011 | -0.006 | -0.006 | 0 | 0.12 | 0.08 | 0.03 | 0.38 | 0.05 | 0.28 |
| 6 | 143610748 | rs9403481 | 0.838 | 0.004 | 0.906 | 0.711 | -0.027 | 0.007 | 0.007 | 0.544 | 0.06 | 0 | 0.03 | 0.04 | 0.08 | 0.11 |
| 6 | 143614813 | rs11755477 | -1.183 | -0.218 | 0.99 | 0.825 | -0.018 | 0.07 | 0.07 | 0.205 | 0.22 | 0.54 | 0.25 | 0.37 | 0.17 | 0.2 |
| 6 | 143616718 | rs9496565 | -1.725 | -0.221 | 0.791 | 0.824 | -0.004 | 0.007 | 0.007 | 0.986 | 0.28 | 0.66 | 0.3 | 0.41 | 0.22 | 0.26 |
| 6 | 143617983 | rs9376740 | -1.004 | 0.197 | 0.646 | 0.946 | 0.009 | -0.001 | -0.001 | 0 | 0.21 | 0.26 | 0.13 | 0.18 | 0.25 | 0.29 |
| 6 | 143621631 | rs11754065 | 0.079 | 0.121 | 0.346 | 1.102 | -0.02 | -0.058 | -0.058 | 0 | 0.14 | 0.05 | 0.1 | 0.32 | 0.16 | 0.16 |
| 6 | 143635012 | rs10457047 | 0.287 | 0.069 | 0.365 | 0.834 | 0.019 | -0.034 | -0.034 | 0 | 0.11 | 0.03 | 0.03 | 0.31 | 0.05 | 0.26 |
| 6 | 143647931 | rs9376744 | -0.923 | 0.046 | 0.31 | 0.445 | -0.055 | -0.061 | -0.061 | 0 | 0.28 | 0.12 | 0.25 | 0.58 | 0.32 | 0.36 |
| 6 | 143651969 | rs9403484 | -0.552 | 0.123 | 0.307 | 0.498 | -0.045 | -0.035 | -0.035 | 0 | 0.15 | 0.06 | 0.33 | 0.22 | 0.2 | 0.22 |
| 6 | 143654625 | rs2303386 | -1.864 | 0.117 | 0.68 | 0.358 | -0.008 | -0.012 | -0.012 | 0 | 0.35 | 0.4 | 0.56 | 0.55 | 0.37 | 0.61 |
| 6 | 143657292 | rs9376745 | -2.707 | 0.025 | 0.754 | 0.188 | 0.002 | 0.012 | 0.012 | 0.003 | 0.61 | 0.74 | 0.71 | 1 | 0.56 | 0.89 |
| 6 | 143658576 | rs1885650 | -0.695 | 0.029 | 0.444 | 0.317 | -0.008 | -0.026 | -0.026 | 0 | 0.19 | 0.11 | 0.12 | 0.29 | 0.17 | 0.4 |
| 6 | 143659578 | rs17072445 | -1.151 | -0.032 | 0.773 | 0.315 | -0.066 | -0.023 | -0.023 | 0 | 0.12 | 0.57 | 0.09 | 0.07 | 0.02 | 0.09 |
| 6 | 143660432 | rs12527382 | 0.181 | 0.018 | 0.685 | 0.315 | -0.002 | -0.001 | -0.001 | 0.06 | 0.08 | 0.06 | 0.05 | 0.03 | 0.08 | 0.13 |
| 6 | 143661166 | rs9908 | -1.243 | -0.009 | 0.691 | 0.407 | 0 | 0.001 | 0.001 | 2.351 | 0.27 | 0.7 | 0.18 | 0.46 | 0.19 | 0.27 |
| 6 | 143669993 | rs9399441 | -0.892 | 0.092 | 0.434 | 0.41 | -0.001 | 0 | 0 | 0.378 | 0.2 | 0.19 | 0.36 | 0.23 | 0.21 | 0.24 |
| 6 | 143686864 | rs1345990 | -1.209 | 0.08 | 0.428 | 0.245 | -0.009 | 0.002 | 0.002 | 0.196 | 0.36 | 0.29 | 0.46 | 0.63 | 0.37 | 0.43 |
| 6 | 143691393 | rs4896651 | -1.99 | 0.016 | 0.416 | 0.156 | 0.007 | 0.115 | 0.115 | 1.359 | 0.46 | 0.84 | 0.62 | 0.73 | 0.4 | 0.52 |
| 6 | 143703478 | rs9496607 | -0.063 | -0.104 | 0.651 | 0.202 | -0.028 | 0.019 | 0.019 | 0.349 | 0.07 | 0.32 | 0.05 | 0.06 | 0.02 | 0.08 |
| 6 | 143703613 | rs223597 | 0.396 | -0.054 | 0.136 | 0.169 | -0.015 | -0.024 | -0.024 | 0.072 | 0.83 | 0.93 | 0.88 | 0.75 | 0.83 | 0.63 |
| 6 | 143706420 | rs223607 | 1.952 | 0.367 | 0.131 | 0.103 | -0.035 | -0.053 | -0.053 | 0 | 0.52 | 0.12 | 0.34 | 0.43 | 0.58 | 0.3 |
| 6 | 143707820 | rs6942294 | -0.46 | 0.436 | 0.57 | 0.185 | -0.026 | 0.038 | 0.038 | 1.506 | 0.09 | 0.52 | 0.16 | 0.1 | 0.02 | 0.09 |
| 6 | 143711698 | rs17072527 | -0.081 | 1.363 | -0.877 | 0.598 | -0.055 | -0.015 | -0.015 | 0 | 0.13 | 0.02 | 0.08 | 0.12 | 0.18 | 0.16 |
| 6 | 143713705 | rs4896653 | -0.644 | 1.084 | -0.8 | 0.436 | -0.013 | -0.04 | -0.04 | 0 | 0.22 | 0.16 | 0.14 | 0.48 | 0.24 | 0.23 |
| 6 | 143714240 | rs223612 | 0.674 | 1.217 | -0.823 | 0.776 | -0.021 | -0.031 | -0.031 | 0 | 0.72 | 0.83 | 0.56 | 0.58 | 0.69 | 0.54 |
| 6 | 143720467 | rs7757142 | -0.862 | 0.982 | -0.878 | 1.167 | -0.017 | -0.066 | -0.066 | 0 | 0.19 | 0.08 | 0.13 | 0.5 | 0.21 | 0.25 |
| 6 | 143721486 | rs2473452 | 0.522 | 0.983 | -0.874 | 0.122 | 0.002 | 0.067 | 0.067 | 1.685 | 0.53 | 0.39 | 0.28 | 0.24 | 0.42 | 0.34 |
| 6 | 143721737 | rs223587 | -0.414 | 1.105 | -0.314 | 0.623 | 0.049 | -0.033 | -0.033 | 0.236 | 0.23 | 0.32 | 0.24 | 0.13 | 0.37 | 0.21 |
| 6 | 143722512 | rs223588 | -0.715 | 0.656 | -1.083 | 0.445 | -0.007 | 0.024 | 0.024 | 0 | 0.42 | 0.59 | 0.31 | 0.32 | 0.48 | 0.41 |
| 6 | 143722679 | rs9390088 | -0.604 | 0.297 | -1.281 | 1.441 | -0.002 | -0.061 | -0.061 | 9.551 | 0.1 | 0.01 | 0.23 | 0.24 | 0.05 | 0.22 |
| 6 | 143725481 | rs223592 | -0.461 | 0.266 | -1.327 | 1.276 | 0.016 | -0.051 | -0.051 | 0 | 0.16 | 0.03 | 0.05 | 0.26 | 0.09 | 0.21 |
| 6 | 143725635 | rs742380 | -1.151 | 0.788 | -0.693 | 1.782 | 0.009 | 0.081 | 0.081 | 0 | 0.2 | 0.26 | 0.3 | 0.57 | 0.27 | 0.29 |
| 6 | 143725670 | rs760954 | -1.17 | -0.147 | -0.233 | 0.907 | -0.009 | -0.015 | -0.015 | 2.089 | 0.36 | 0.49 | 0.3 | 0.8 | 0.26 | 0.41 |
| 6 | 143725775 | rs9484743 | -1.155 | -0.1 | 0.491 | 1.33 | -0.039 | 0.096 | 0.096 | 0 | 0.15 | 0.41 | 0.26 | 0.24 | 0.05 | 0.19 |
| 6 | 143726697 | rs9376748 | 0.037 | 0.493 | -0.447 | 1.068 | -0.037 | -0.02 | -0.02 | 0 | 0.16 | 0.04 | 0.12 | 0.15 | 0.19 | 0.18 |
| 6 | 143727457 | rs9376749 | -0.99 | 0.344 | -0.574 | 1.127 | -0.026 | -0.074 | -0.074 | 0.087 | 0.28 | 0.05 | 0.35 | 0.32 | 0.25 | 0.36 |
| 6 | 143727723 | rs926451 | -0.398 | 0.506 | 0.367 | 1.619 | 0.075 | 0.391 | 0.391 | 0 | 0.07 | 0.29 | 0.14 | 0.39 | 0.09 | 0.11 |
| 6 | 143735762 | rs2496590 | 0.598 | 0.459 | -0.009 | 0.206 | -0.012 | 0.028 | 0.028 | 2.892 | 0.58 | 0.66 | 0.6 | 0.18 | 0.56 | 0.36 |
| 6 | 143747270 | rs1061220 | 0.61 | 0.418 | -0.028 | 0.185 | -0.012 | 0.03 | 0.03 | 3.463 | 0.58 | 0.67 | 0.59 | 0.18 | 0.54 | 0.35 |
| 6 | 143748722 | rs3088203 | 0.636 | 0.419 | -0.028 | 0.185 | -0.012 | 0.032 | 0.032 | 3.34 | 0.58 | 0.66 | 0.59 | 0.18 | 0.54 | 0.35 |
| 6 | 143749295 | rs1065355 | 0.26 | 0.149 | 0.266 | -0.142 | -0.003 | -0.005 | -0.005 | 0 | 0.2 | 0.27 | 0.16 | 0.16 | 0.21 | 0.17 |
| 6 | 143754112 | rs4896656 | 0.606 | 0.118 | 0.51 | -0.132 | 0 | -0.014 | -0.014 | 0.165 | 0.57 | 0.57 | 0.58 | 0.19 | 0.54 | 0.36 |
| 6 | 143765778 | rs6929191 | 0.485 | 0.132 | 0.423 | -0.124 | -0.012 | 0.032 | 0.032 | 3.432 | 0.58 | 0.66 | 0.59 | 0.19 | 0.54 | 0.36 |
| 6 | 143772311 | rs2272924 | -0.183 | 0.028 | 0.463 | -0.09 | -0.014 | -0.013 | -0.013 | 0 | 0.22 | 0.39 | 0.28 | 0.18 | 0.23 | 0.23 |
| 6 | 143778954 | rs3804538 | 0.015 | 0.012 | 0.389 | -0.123 | 0.004 | 0.007 | 0.007 | 0 | 0.22 | 0.2 | 0.15 | 0.16 | 0.23 | 0.19 |
| 6 | 143785485 | rs161070 | 0.338 | 0.008 | 0.071 | 0.144 | -0.007 | -0.037 | -0.037 | 0.077 | 0.13 | 0.01 | 0.12 | 0.18 | 0.11 | 0.15 |
| 6 | 143796333 | rs161064 | 1.337 | 0.258 | 0.672 | 0.399 | -0.015 | 0.02 | 0.02 | 0.994 | 0.4 | 0.25 | 0.43 | 0.32 | 0.42 | 0.24 |
| 6 | 143802490 | rs161061 | -0.178 | 0.279 | 0.691 | 0.5 | -0.013 | -0.037 | -0.037 | 0 | 0.66 | 0.8 | 0.75 | 0.52 | 0.67 | 0.49 |
| 6 | 143806551 | rs161057 | 0.33 | 0.507 | 0.821 | 0.776 | 0.04 | 0.185 | 0.185 | 0 | 0.06 | 0.16 | 0.15 | 0.31 | 0.11 | 0.12 |
| 6 | 143810909 | rs10809 | 0.524 | 0.452 | 0.351 | 0.679 | -0.038 | -0.035 | -0.035 | 0.631 | 0.17 | 0.01 | 0.13 | 0.16 | 0.21 | 0.16 |
| 6 | 143816859 | rs8161 | -1.147 | 0.373 | 0.519 | 0.644 | 0.002 | 0.103 | 0.103 | 0 | 0.29 | 0.48 | 0.44 | 0.48 | 0.34 | 0.36 |
| 6 | 143823112 | rs3762001 | -0.03 | 0.297 | 0.254 | 0.441 | 0.005 | 0.006 | 0.006 | 0 | 0.2 | 0.12 | 0.15 | 0.17 | 0.22 | 0.19 |
| 6 | 143825104 | rs11155297 | 1.715 | -0.009 | -0.989 | -0.042 | -0.096 | 0.181 | 0.181 | 3.731 | 0.27 | 0.01 | 0.13 | 0 | 0.27 | 0.06 |
| 6 | 143827102 | rs1038390 | -0.063 | -0.489 | -0.107 | 1.023 | -0.018 | -0.03 | -0.03 | 0 | 0.19 | 0.01 | 0.12 | 0.18 | 0.21 | 0.12 |
| 6 | 143833126 | rs4896663 | -1.343 | -0.23 | -0.896 | 1.262 | 0.032 | -0.038 | -0.038 | 0 | 0.39 | 0.78 | 0.6 | 0.52 | 0.38 | 0.35 |
| 6 | 143834669 | rs573119 | -0.407 | 0.685 | -0.581 | 1.713 | 0.001 | 0.034 | 0.034 | 0 | 0.08 | 0.08 | 0.15 | 0.29 | 0.14 | 0.17 |
| 6 | 143838433 | rs9390098 | 0.893 | 0.613 | -0.716 | 1.563 | -0.015 | -0.029 | -0.029 | 0 | 0.86 | 0.88 | 0.6 | 0.66 | 0.82 | 0.78 |
| 6 | 143842099 | rs9496652 | 0.532 | 0.276 | -0.752 | 1.613 | -0.007 | -0.026 | -0.026 | 0 | 0.87 | 0.89 | 0.61 | 0.67 | 0.82 | 0.78 |
| 6 | 143842182 | rs9496654 | 1.156 | 0.026 | -0.262 | 1.72 | 0.003 | 0.053 | 0.053 | 0.107 | 0.71 | 0.64 | 0.51 | 0.49 | 0.69 | 0.48 |
| 6 | 143844171 | rs6913701 | -0.276 | -0.598 | -1.231 | 0.842 | 0.049 | -0.035 | -0.035 | 0 | 0.42 | 0.81 | 0.32 | 0.48 | 0.31 | 0.37 |
| 6 | 143846791 | rs2128977 | -0.958 | -0.488 | -1.19 | 0.884 | -0.048 | 0.045 | 0.045 | 0.372 | 0.57 | 0.85 | 0.51 | 0.65 | 0.44 | 0.71 |

| | | | | | | | | | | | | | | | | |
|---|---|---|---|---|---|---|---|---|---|---|---|---|---|---|---|---|
| 6 | 143848143 | rs17072752 | -0.036 | -0.895 | -1.125 | 1.278 | 0.024 | -0.066 | -0.066 | 1.976 | 0.12 | 0.01 | 0.1 | 0.26 | 0.08 | 0.19 |
| 6 | 143851294 | rs9403498 | -0.746 | -1.109 | -0.64 | 1.083 | -0.001 | 0.017 | 0.017 | 0.06 | 0.28 | 0.27 | 0.23 | 0.46 | 0.27 | 0.26 |
| 6 | 143853438 | rs9403499 | -0.441 | -1.119 | -0.605 | 1.089 | 0.001 | -0.003 | -0.003 | 0 | 0.26 | 0.21 | 0.21 | 0.46 | 0.23 | 0.25 |
| 6 | 143860599 | rs1466533 | -0.227 | -1.539 | -0.134 | 1.06 | -0.047 | 0 | 0 | 0 | 0.27 | 0.38 | 0.26 | 0.24 | 0.14 | 0.42 |
| 6 | 143861900 | rs1970352 | -0.607 | -1.474 | 0.279 | 1.188 | -0.01 | -0.001 | -0.001 | 0 | 0.41 | 0.59 | 0.38 | 0.37 | 0.22 | 0.54 |
| 6 | 143863044 | rs4896665 | -0.266 | -0.601 | 0.668 | 1.221 | 0.016 | 0.035 | 0.035 | 0 | 0.18 | 0.29 | 0.13 | 0.3 | 0.19 | 0.17 |
| 6 | 143863992 | rs9390103 | -0.238 | -0.594 | 0.64 | 1.225 | 0.013 | 0.029 | 0.029 | 0 | 0.18 | 0.26 | 0.13 | 0.3 | 0.19 | 0.17 |
| 6 | 143867336 | rs9484764 | 0.637 | -0.793 | 1.048 | 1.026 | -0.016 | 0.001 | 0.001 | 0 | 0.61 | 0.53 | 0.42 | 0.39 | 0.67 | 0.51 |
| 6 | 143868242 | rs9376755 | 0.119 | -0.75 | 0.949 | 0.854 | -0.017 | -0.009 | -0.009 | 0 | 0.26 | 0.15 | 0.15 | 0.28 | 0.28 | 0.37 |
| 6 | 143872563 | rs12530338 | 0.548 | -0.784 | 0.862 | 0.85 | -0.006 | 0 | 0 | 0 | 0.61 | 0.44 | 0.4 | 0.39 | 0.67 | 0.52 |
| 6 | 143875758 | rs12529563 | 0.411 | -0.758 | 0.69 | 0.669 | 0.002 | 0.016 | 0.016 | 0 | 0.22 | 0.12 | 0.12 | 0.18 | 0.21 | 0.3 |
| 6 | 143890039 | rs10979 | 0.506 | -0.944 | 0.945 | 0.734 | -0.001 | 0.004 | 0.004 | 0 | 0.65 | 0.61 | 0.47 | 0.39 | 0.64 | 0.56 |
| 6 | 143892771 | rs9403511 | 0.01 | -0.976 | 1.015 | 0.679 | 0.014 | -0.011 | -0.011 | 0.215 | 0.32 | 0.25 | 0.23 | 0.38 | 0.25 | 0.43 |
| 6 | 143896343 | rs9496677 | 1.556 | -0.698 | 1.032 | 0.569 | -0.003 | -0.049 | -0.049 | 0 | 0.33 | 0.3 | 0.24 | 0.01 | 0.38 | 0.13 |
| 6 | 143903156 | rs11155304 | -0.362 | -0.656 | 0.878 | 0.78 | 0.005 | 0.002 | 0.002 | 0 | 0.05 | 0.04 | 0.06 | 0.07 | 0.09 | 0.11 |
| 6 | 143904692 | rs4142501 | -0.153 | -0.892 | 1.167 | 0.709 | -0.004 | 0.001 | 0.001 | 0.414 | 0.34 | 0.38 | 0.25 | 0.4 | 0.26 | 0.45 |
| 6 | 143906340 | rs10782290 | -0.24 | -0.818 | 1.153 | 0.653 | -0.002 | 0.014 | 0.014 | 0.139 | 0.73 | 0.69 | 0.53 | 0.45 | 0.73 | 0.68 |
| 6 | 143906672 | rs13194572 | 0.296 | -0.888 | 1.179 | 0.628 | 0.026 | 0.048 | 0.048 | 0.028 | 0.28 | 0.14 | 0.18 | 0.15 | 0.22 | 0.35 |
| 6 | 143908013 | rs9403515 | -0.285 | -0.842 | 1.204 | 0.821 | -0.001 | 0.01 | 0.01 | 0.139 | 0.73 | 0.7 | 0.55 | 0.55 | 0.73 | 0.7 |
| 6 | 143914596 | rs10484802 | 1.188 | -0.846 | 1.179 | 0.81 | 0.009 | -0.002 | -0.002 | 0.013 | 0.39 | 0.47 | 0.31 | 0.35 | 0.42 | 0.24 |
| 6 | 143915677 | rs10484803 | 0.147 | -0.877 | 0.96 | 0.606 | 0.029 | 0.055 | 0.055 | 0.129 | 0.29 | 0.14 | 0.2 | 0.16 | 0.24 | 0.36 |
| 6 | 143918718 | rs12527367 | 0.804 | -0.718 | 1.191 | 0.584 | 0 | -0.012 | -0.012 | 0.055 | 0.62 | 0.4 | 0.44 | 0.26 | 0.62 | 0.49 |
| 6 | 143920811 | rs9496698 | 1.161 | -0.59 | 1.161 | 0.54 | -0.008 | 0.006 | 0.006 | 0 | 0.38 | 0.39 | 0.33 | 0.14 | 0.48 | 0.24 |
| 6 | 143925374 | rs12216096 | 1.65 | -0.71 | 0.926 | 0.604 | -0.037 | 0.076 | 0.076 | 0 | 0.33 | 0.17 | 0.24 | 0.11 | 0.37 | 0.14 |
| 6 | 143928437 | rs9376769 | 1.152 | -0.728 | 1.194 | 0.807 | -0.032 | 0.041 | 0.041 | 0 | 0.39 | 0.22 | 0.28 | 0.23 | 0.44 | 0.23 |
| 6 | 143929874 | rs1338535 | 0.422 | -0.934 | 1.429 | 0.642 | -0.007 | -0.075 | -0.075 | 0 | 0.25 | 0.4 | 0.17 | 0.09 | 0.25 | 0.3 |
| 6 | 143930768 | rs10484804 | -0.219 | -0.906 | 1.161 | 0.837 | 0.001 | -0.074 | -0.074 | 0 | 0.74 | 0.84 | 0.6 | 0.53 | 0.76 | 0.67 |
| 6 | 143934587 | rs17368239 | 1.937 | -0.859 | 1.17 | 0.366 | -0.079 | 0.173 | 0.173 | 0.318 | 0.33 | 0.06 | 0.21 | 0.01 | 0.37 | 0.12 |
| 6 | 143934705 | rs9376770 | -0.4 | -0.984 | 1.011 | 0.555 | -0.004 | -0.003 | -0.003 | 0 | 0.86 | 0.85 | 0.63 | 0.52 | 0.83 | 0.69 |
| 6 | 143937943 | rs12193494 | -0.024 | -1.074 | 1.274 | 0.322 | 0.011 | 0.041 | 0.041 | 0 | 0.34 | 0.28 | 0.18 | 0.09 | 0.31 | 0.32 |
| 6 | 143940112 | rs9373395 | -0.45 | -0.952 | 1.314 | 0.525 | 0.015 | 0.181 | 0.181 | 0.035 | 0.08 | 0.22 | 0.08 | 0.29 | 0.07 | 0.1 |
| 6 | 143940767 | rs17072830 | 0.19 | -1.006 | 1.267 | 0.441 | -0.014 | 0.003 | 0.003 | 0 | 0.11 | 0.16 | 0.13 | 0.11 | 0.07 | 0.14 |
| 6 | 143943314 | rs9390123 | -0.068 | -1.072 | 1.012 | 0.358 | -0.004 | -0.009 | -0.009 | 0 | 0.64 | 0.67 | 0.48 | 0.42 | 0.58 | 0.41 |
| 6 | 143945634 | rs9496711 | 0.785 | -1.044 | 0.373 | 0.376 | 0.015 | 0.019 | 0.019 | 0.44 | 0.13 | 0.09 | 0.04 | 0 | 0.07 | 0.02 |
| 6 | 143948472 | rs9399452 | -0.19 | -0.793 | 0.435 | 0.628 | 0.009 | 0.063 | 0.063 | 0 | 0.07 | 0.15 | 0.06 | 0.2 | 0.06 | 0.09 |
| 6 | 143952725 | rs10484805 | 0.324 | -0.814 | 0.16 | 0.541 | 0.016 | -0.003 | -0.003 | 0 | 0.11 | 0.02 | 0.12 | 0.12 | 0.07 | 0.13 |
| 6 | 143955378 | rs12215937 | 0.97 | -0.825 | 0.198 | 0.538 | -0.044 | 0.123 | 0.123 | 0.012 | 0.44 | 0.14 | 0.35 | 0.23 | 0.46 | 0.28 |
| 6 | 143959214 | rs1890403 | -0.289 | -0.545 | 0.308 | 1.083 | -0.032 | -0.032 | -0.032 | 0 | 0.9 | 0.98 | 0.78 | 0.55 | 0.84 | 0.81 |
| 6 | 143961610 | rs6931801 | -0.703 | -1.237 | 0.896 | 1.045 | -0.014 | 0.044 | 0.044 | 0 | 0.18 | 0.31 | 0.21 | 0.31 | 0.14 | 0.29 |
| 6 | 143967030 | rs7766091 | -1.23 | -1.26 | 0.809 | 0.786 | 0 | 0 | 0 | 0 | 0.53 | 0.45 | 0.39 | 0.46 | 0.48 | 0.57 |
| 6 | 143968766 | rs11155314 | -0.213 | -1.262 | 0.469 | 0.625 | 0.007 | 0.04 | 0.04 | 0 | 0.35 | 0.21 | 0.18 | 0.18 | 0.34 | 0.29 |
| 6 | 143982187 | rs7741998 | -1.101 | -1.182 | 0.644 | 0.724 | 0.001 | 0.033 | 0.033 | 0 | 0.52 | 0.36 | 0.36 | 0.32 | 0.48 | 0.6 |
| 6 | 143994199 | rs9403521 | -0.225 | -1.192 | 0.401 | 0.669 | 0.002 | -0.031 | -0.031 | 0 | 0.16 | 0.11 | 0.17 | 0.36 | 0.15 | 0.31 |
| 6 | 144000856 | rs7760178 | -0.898 | -1.272 | 0.344 | 0.563 | 0.016 | -0.023 | -0.023 | 0 | 0.46 | 0.25 | 0.34 | 0.48 | 0.44 | 0.56 |
| 6 | 144004981 | rs9403523 | -0.628 | -1.329 | 0.672 | 0.659 | -0.011 | 0.076 | 0.076 | 0 | 0.19 | 0.33 | 0.2 | 0.38 | 0.15 | 0.32 |
| 6 | 144005795 | rs9496729 | -1.454 | -1.356 | 1.084 | 0.626 | -0.001 | -0.002 | -0.002 | 0.08 | 0.53 | 0.56 | 0.39 | 0.49 | 0.48 | 0.61 |
| 6 | 144005949 | rs9484792 | -0.382 | -1.338 | 0.987 | 0.598 | 0.004 | 0.035 | 0.035 | 0 | 0.35 | 0.24 | 0.19 | 0.1 | 0.33 | 0.29 |
| 6 | 144011546 | rs9496733 | -1.446 | -1.346 | 1.07 | 0.661 | -0.017 | -0.022 | -0.022 | 0.45 | 0.54 | 0.72 | 0.4 | 0.49 | 0.48 | 0.61 |
| 6 | 144015633 | rs6907742 | -0.667 | -1.386 | 1.094 | 0.542 | -0.017 | -0.071 | -0.071 | 0 | 0.17 | 0.37 | 0.07 | 0.04 | 0.11 | 0.13 |
| 6 | 144017250 | rs910419 | -1.255 | -1.334 | 0.383 | 0.615 | -0.015 | -0.013 | -0.013 | 0.051 | 0.55 | 0.7 | 0.4 | 0.49 | 0.49 | 0.61 |
| 6 | 144017778 | rs1884172 | 0.698 | -1.274 | -0.051 | 0.595 | -0.041 | 0.047 | 0.047 | 0 | 0.2 | 0.04 | 0.22 | 0.09 | 0.24 | 0.25 |
| 6 | 144020900 | rs9496739 | -1.021 | -1.398 | 0.308 | 0.729 | 0.048 | -0.012 | -0.012 | 0 | 0.36 | 0.77 | 0.32 | 0.41 | 0.23 | 0.37 |
| 6 | 144025295 | rs7739094 | -1.085 | -0.876 | 0.405 | 0.599 | 0.006 | 0.019 | 0.019 | 0 | 0.62 | 0.55 | 0.45 | 0.46 | 0.62 | 0.64 |
| 6 | 144025782 | rs9321935 | -0.732 | -0.972 | 0.043 | 0.598 | -0.002 | -0.003 | -0.003 | 0 | 0.54 | 0.35 | 0.38 | 0.45 | 0.49 | 0.58 |
| 6 | 144028153 | rs4896668 | -0.398 | -1.333 | -0.384 | 1.031 | 0.024 | -0.15 | -0.15 | 0.191 | 0.26 | 0.06 | 0.2 | 0.52 | 0.19 | 0.33 |
| 6 | 144030493 | rs2488099 | -0.172 | -1.107 | 0.25 | 0.962 | 0.002 | -0.064 | -0.064 | 0 | 0.36 | 0.52 | 0.35 | 0.16 | 0.42 | 0.36 |
| 6 | 144031193 | rs2488100 | -0.388 | -1.598 | -0.507 | 0.824 | 0.006 | 0.113 | 0.113 | 0 | 0.51 | 0.16 | 0.37 | 0.33 | 0.56 | 0.44 |
| 6 | 144031408 | rs10457752 | 0.37 | -1.472 | -0.774 | 1.3 | 0.007 | -0.01 | -0.01 | 0 | 0.16 | 0.11 | 0.15 | 0.36 | 0.09 | 0.25 |
| 6 | 144032317 | rs2488101 | -1.275 | -1.281 | -0.683 | 1.083 | -0.047 | 0.014 | 0.014 | 0.361 | 0.64 | 0.83 | 0.59 | 0.71 | 0.63 | 0.69 |
| 6 | 144033992 | rs9496751 | -0.569 | -1.517 | -0.551 | 1.211 | -0.015 | 0.019 | 0.019 | 0 | 0.28 | 0.34 | 0.37 | 0.47 | 0.2 | 0.38 |
| 6 | 144034848 | rs1015339 | -1.494 | -1.194 | -0.965 | 1.165 | -0.119 | 0.018 | 0.018 | 0.858 | 0.68 | 0.91 | 0.71 | 0.8 | 0.61 | 0.8 |
| 6 | 144035357 | rs1015340 | -0.965 | -0.345 | -0.558 | 0.972 | -0.001 | 0.009 | 0.009 | 0 | 0.48 | 0.43 | 0.4 | 0.46 | 0.52 | 0.59 |
| 6 | 144035464 | rs1015341 | -0.883 | -0.322 | -0.999 | 0.973 | -0.004 | 0.011 | 0.011 | 0 | 0.47 | 0.38 | 0.41 | 0.46 | 0.53 | 0.59 |
| 6 | 144036214 | rs12662850 | 0.043 | -0.343 | -1.229 | 0.883 | -0.01 | -0.007 | -0.007 | 1.169 | 0.12 | 0.05 | 0.07 | 0.13 | 0.12 | 0.18 |
| 6 | 144036993 | rs733399 | 0.427 | -0.357 | -1.384 | 0.775 | -0.052 | 0.043 | 0.043 | 0 | 0.22 | 0.03 | 0.25 | 0.13 | 0.26 | 0.24 |
| 6 | 144038185 | rs7769992 | -0.609 | -0.395 | -1.02 | 1.041 | -0.007 | 0.115 | 0.115 | 4.556 | 0.07 | 0.64 | 0.05 | 0.12 | 0.01 | 0.05 |
| 6 | 144041537 | rs9403525 | -1.137 | -0.104 | -1.323 | 0.912 | -0.123 | -0.119 | -0.119 | 0 | 0.54 | 0.94 | 0.46 | 0.54 | 0.54 | 0.62 |
| 6 | 144041704 | rs6915752 | -1.104 | -0.066 | -1.285 | 0.931 | -0.073 | -0.071 | -0.071 | 0.677 | 0.54 | 0.89 | 0.46 | 0.54 | 0.54 | 0.62 |
| 6 | 144042059 | rs6915880 | -0.552 | 0.556 | -0.888 | 0.395 | -0.005 | 0.015 | 0.015 | 0 | 0.36 | 0.28 | 0.34 | 0.3 | 0.42 | 0.41 |

| | | | | | | | | | | | | | | | | |
|---|---|---|---|---|---|---|---|---|---|---|---|---|---|---|---|---|
| 6 | 144047833 | rs9403527 | -0.684 | 0.482 | -1.268 | 0.171 | 0.016 | 0.044 | 0.044 | 0 | 0.35 | 0.18 | 0.28 | 0.24 | 0.33 | 0.35 |
| 6 | 144048080 | rs9376784 | -0.717 | 1.037 | -1.351 | -0.018 | -0.054 | 0.064 | 0.064 | 0 | 0.37 | 0.14 | 0.32 | 0.24 | 0.42 | 0.33 |
| 6 | 144048476 | rs12209142 | -0.63 | 1.065 | -0.567 | -0.423 | 0.012 | -0.069 | -0.069 | 0 | 0.11 | 0.23 | 0.06 | 0 | 0.07 | 0.17 |
| 6 | 144053496 | rs9386046 | -0.895 | 0.855 | -0.032 | -0.614 | 0.01 | 0.066 | 0.066 | 0 | 0.44 | 0.7 | 0.38 | 0.24 | 0.39 | 0.41 |
| 6 | 144056863 | rs7738300 | -0.635 | 1.047 | -0.062 | -0.009 | 0.004 | 0.023 | 0.023 | 0 | 0.22 | 0.3 | 0.29 | 0.36 | 0.25 | 0.31 |
| 6 | 144056981 | rs9390131 | -0.158 | 1.157 | -0.658 | -0.26 | -0.082 | 0.031 | 0.031 | 0.484 | 0.16 | 0.02 | 0.11 | 0.1 | 0.22 | 0.29 |
| 6 | 144064167 | rs12208670 | -0.28 | 0.793 | -0.488 | -0.031 | -0.072 | -0.023 | -0.023 | 0 | 0.27 | 0.09 | 0.32 | 0.3 | 0.38 | 0.35 |
| 6 | 144065262 | rs9403528 | -0.835 | 0.746 | -0.289 | -0.057 | -0.045 | -0.013 | -0.013 | 0 | 0.33 | 0.24 | 0.38 | 0.4 | 0.45 | 0.49 |
| 6 | 144068641 | rs11962678 | -0.697 | 0.582 | 0.279 | -0.06 | 0.02 | 0.003 | 0.003 | 0 | 0.33 | 0.41 | 0.36 | 0.37 | 0.38 | 0.42 |
| 6 | 144072118 | rs12207527 | -0.277 | 0.643 | 0.072 | -0.023 | -0.069 | -0.039 | -0.039 | 0.177 | 0.28 | 0.09 | 0.32 | 0.34 | 0.38 | 0.36 |
| 6 | 144079788 | rs6931825 | -0.571 | 0.778 | 0.215 | -0.06 | 0.003 | -0.015 | -0.015 | 0 | 0.11 | 0.18 | 0.08 | 0.07 | 0.1 | 0.16 |
| 6 | 144079854 | rs6931865 | -1.746 | 0.638 | -0.105 | -0.107 | 0.004 | -0.032 | -0.032 | 0 | 0.63 | 0.73 | 0.66 | 0.6 | 0.66 | 0.72 |
| 6 | 144081609 | rs2073214 | 0.14 | 0.558 | -0.056 | -0.044 | -0.017 | -0.023 | -0.023 | 0 | 0.21 | 0.29 | 0.24 | 0.14 | 0.17 | 0.13 |
| 6 | 144086394 | rs9496767 | -0.577 | 0.625 | 0.057 | 0.192 | 0.003 | 0.008 | 0.008 | 0.34 | 0.49 | 0.39 | 0.55 | 0.46 | 0.55 | 0.53 |
| 6 | 144094948 | rs2295202 | 0.78 | 0.537 | -0.295 | 0.54 | -0.009 | 0.11 | 0.11 | 4.093 | 0.15 | 0.22 | 0.2 | 0.41 | 0.1 | 0.18 |
| 6 | 144100865 | rs6570581 | -0.574 | 0.844 | -0.638 | 0.466 | 0.045 | 0.003 | 0.003 | 1.617 | 0.53 | 0.28 | 0.54 | 0.62 | 0.66 | 0.65 |
| 6 | 144103420 | rs4624889 | -0.257 | 1.457 | -1.106 | 1.059 | -0.01 | -0.06 | -0.06 | 0 | 0.14 | 0.05 | 0.06 | 0.27 | 0.13 | 0.21 |
| 6 | 144105403 | rs7742964 | -0.373 | 1.715 | -0.965 | 1.196 | -0.03 | -0.028 | -0.028 | 0 | 0.19 | 0.11 | 0.3 | 0.31 | 0.29 | 0.28 |
| 6 | 144111346 | rs9496771 | -1.516 | 1.413 | -0.248 | 0.972 | 0.016 | 0.018 | 0.018 | 2.748 | 0.52 | 0.54 | 0.61 | 0.67 | 0.63 | 0.63 |
| 6 | 144116036 | rs2328516 | -0.737 | 1.857 | 0.158 | 1.32 | 0.047 | 0.026 | 0.026 | 0 | 0.19 | 0.3 | 0.39 | 0.27 | 0.29 | 0.27 |
| 6 | 144116086 | rs2876572 | -1.965 | 1.628 | 0.255 | 1.343 | -0.003 | -0.004 | -0.004 | 0.31 | 0.49 | 0.54 | 0.67 | 0.69 | 0.63 | 0.66 |
| 6 | 144125035 | rs9386048 | -0.955 | 1.63 | 0.166 | 1.425 | -0.04 | -0.069 | -0.069 | 0.067 | 0.28 | 0.1 | 0.17 | 0.38 | 0.33 | 0.39 |
| 6 | 144125687 | rs761626 | 0.015 | 1.644 | 0.018 | 1.488 | -0.024 | -0.09 | -0.09 | 0 | 0.15 | 0.01 | 0.07 | 0.31 | 0.14 | 0.26 |
| 6 | 144128686 | rs9376791 | -1.893 | 1.851 | 0.137 | 1.488 | 0.001 | 0.024 | 0.024 | 4.452 | 0.35 | 0.46 | 0.31 | 0.46 | 0.36 | 0.48 |
| 6 | 144129632 | rs7752960 | 0.225 | 1.884 | 0 | 1.435 | -0.057 | -0.037 | -0.037 | 0 | 0.15 | 0.06 | 0.37 | 0.22 | 0.26 | 0.18 |
| 6 | 144130089 | rs9386049 | -0.275 | 1.621 | 0.305 | 1.36 | 0.003 | -0.009 | -0.009 | 0 | 0.14 | 0.13 | 0.12 | 0.06 | 0.18 | 0.13 |
| 6 | 144133948 | rs6920228 | -1.581 | 1.581 | 0.457 | 1.385 | -0.001 | 0.017 | 0.017 | 4.053 | 0.36 | 0.54 | 0.23 | 0.45 | 0.36 | 0.48 |
| 6 | 144136099 | rs9376793 | -1.067 | 1.479 | -0.083 | 1.357 | 0 | -0.026 | -0.026 | 0 | 0.27 | 0.16 | 0.18 | 0.37 | 0.29 | 0.37 |
| 6 | 144139385 | rs9390143 | -1.631 | 1.598 | 0.31 | 1.112 | 0.018 | -0.001 | -0.001 | 2.37 | 0.34 | 0.54 | 0.6 | 0.34 | 0.38 | 0.34 |
| 6 | 144139564 | rs926794 | -0.7 | 1.477 | 0.378 | 1.173 | 0.017 | 0.028 | 0.028 | 0 | 0.18 | 0.3 | 0.49 | 0.26 | 0.23 | 0.18 |
| 6 | 144139851 | rs13204629 | -1.577 | 0.515 | 0.314 | 1.456 | 0.017 | -0.023 | -0.023 | 0.291 | 0.24 | 0.19 | 0.12 | 0.44 | 0.14 | 0.31 |
| 6 | 144141545 | rs9496773 | -1.481 | 0.585 | 0.504 | 0.997 | -0.021 | 0.036 | 0.036 | 0.141 | 0.31 | 0.45 | 0.35 | 0.44 | 0.24 | 0.39 |
| 6 | 144145797 | rs1082 | -1.472 | 0.635 | 0.505 | 0.998 | -0.02 | 0.036 | 0.036 | 0.13 | 0.31 | 0.44 | 0.35 | 0.44 | 0.24 | 0.39 |
| 6 | 144150609 | rs1072385 | -0.9 | 0.652 | 0.522 | 1.028 | 0.004 | -0.007 | -0.007 | 0.142 | 0.22 | 0.23 | 0.12 | 0.35 | 0.18 | 0.2 |
| 6 | 144153002 | rs12197107 | -0.862 | 0.657 | 0.475 | 1.027 | 0.011 | -0.043 | -0.043 | 0 | 0.22 | 0.13 | 0.11 | 0.35 | 0.18 | 0.21 |
| 6 | 144154634 | rs9496777 | -1.55 | 0.584 | 0.513 | 0.948 | -0.028 | 0.047 | 0.047 | 0.028 | 0.3 | 0.48 | 0.35 | 0.43 | 0.23 | 0.38 |
| 6 | 144156309 | rs10440888 | -1.29 | 0.613 | 0.526 | 0.963 | -0.017 | 0.081 | 0.081 | 0 | 0.26 | 0.48 | 0.35 | 0.43 | 0.23 | 0.37 |
| 6 | 144159200 | rs17614442 | -0.509 | 0.749 | 0.144 | 0.651 | 0.002 | 0 | 0 | 0 | 0.05 | 0.03 | 0.03 | 0.05 | 0.04 | 0.15 |
| 6 | 144160280 | rs9386051 | 1.725 | 0.96 | 0.078 | 0.259 | -0.009 | 0.034 | 0.034 | 0.343 | 0.35 | 0.19 | 0.22 | 0.2 | 0.38 | 0.3 |
| 6 | 144167477 | rs2277078 | -1.25 | 0.91 | -0.131 | 0.426 | 0.001 | -0.006 | -0.006 | 0.106 | 0.26 | 0.29 | 0.2 | 0.49 | 0.22 | 0.39 |
| 6 | 144172082 | rs4896678 | -1.54 | 0.908 | -0.028 | 0.424 | -0.006 | 0.029 | 0.029 | 1.546 | 0.28 | 0.42 | 0.31 | 0.51 | 0.24 | 0.43 |
| 6 | 144181619 | rs7454959 | -0.503 | 0.913 | -0.062 | 0.365 | -0.001 | -0.001 | -0.001 | 0 | 0.06 | 0.07 | 0.02 | 0 | 0.05 | 0.09 |
| 6 | 144185414 | rs6918444 | -0.386 | 0.978 | -0.089 | 0.222 | -0.029 | 0.014 | 0.014 | 0 | 0.7 | 0.88 | 0.56 | 0.7 | 0.66 | 0.72 |
| 6 | 144187628 | rs11155329 | -0.829 | 0.465 | 0.124 | 0.532 | 0.001 | -0.04 | -0.04 | 0 | 0.2 | 0.18 | 0.16 | 0.41 | 0.19 | 0.29 |
| 6 | 144188805 | rs2328508 | -1.665 | 0.488 | 0.106 | 0.46 | 0 | 0 | 0 | 0 | 0.29 | 0.35 | 0.3 | 0.51 | 0.25 | 0.43 |
| 6 | 144204610 | rs4896684 | -1.304 | 0.646 | -0.064 | 0.283 | 0 | -0.004 | -0.004 | 0.268 | 0.07 | 0.16 | 0.12 | 0.04 | 0.07 | 0.09 |
| 6 | 144216973 | rs761541 | -1.409 | 0.537 | 0.342 | 0.187 | 0.032 | 0.002 | 0.002 | 2.386 | 0.1 | 0.49 | 0.17 | 0.04 | 0.06 | 0.06 |
| 6 | 144220404 | rs7764938 | 0.147 | 0.56 | 0.05 | 0.003 | 0.009 | 0.03 | 0.03 | 0 | 0.23 | 0.13 | 0.33 | 0.17 | 0.2 | 0.25 |
| 6 | 144222930 | rs9496805 | -0.953 | 0.549 | -0.135 | 0.162 | 0.02 | -0.037 | -0.037 | 0 | 0.08 | 0.34 | 0.06 | 0 | 0.04 | 0.03 |
| 6 | 144257444 | rs12528289 | -0.321 | 0.806 | -0.569 | -0.016 | 0.011 | -0.039 | -0.039 | 0 | 0.35 | 0.42 | 0.43 | 0.25 | 0.32 | 0.51 |
| 6 | 144258244 | rs6570592 | -1.037 | 0.69 | -0.924 | -0.294 | -0.004 | 0.098 | 0.098 | 0.404 | 0.52 | 0.68 | 0.52 | 0.26 | 0.45 | 0.58 |
| 6 | 144258619 | rs1884087 | -1.972 | 0.65 | -1.159 | -0.074 | -0.011 | 0.015 | 0.015 | 0.257 | 0.67 | 0.79 | 0.65 | 0.76 | 0.57 | 0.81 |
| 6 | 144259554 | rs3736746 | -0.911 | 0.661 | -1.123 | -0.353 | -0.001 | 0.11 | 0.11 | 0.297 | 0.51 | 0.68 | 0.52 | 0.26 | 0.45 | 0.56 |
| 6 | 144261480 | rs1049846 | -0.012 | 0.785 | -1.095 | -0.216 | 0.011 | -0.027 | -0.027 | 0.046 | 0.33 | 0.41 | 0.42 | 0.25 | 0.32 | 0.48 |
| 6 | 144264366 | rs9403539 | 0.641 | 0.422 | -1.065 | -0.005 | -0.008 | -0.06 | -0.06 | 2.67 | 0.87 | 0.75 | 0.92 | 1 | 0.89 | 0.94 |
| 6 | 144268094 | rs6930726 | -0.449 | 0.687 | -1.534 | 0.173 | -0.001 | -0.003 | -0.003 | 0 | 0.48 | 0.47 | 0.55 | 0.69 | 0.46 | 0.6 |
| 6 | 144268652 | rs2076682 | -0.036 | 0.726 | -2.595 | 0.555 | -0.002 | -0.126 | -0.126 | 0 | 0.16 | 0 | 0.11 | 0.44 | 0.13 | 0.23 |
| 6 | 144268954 | rs2076683 | 0.463 | 1.081 | -2.043 | 0.239 | -0.043 | 0.035 | 0.035 | 0 | 0.37 | 0.14 | 0.36 | 0.26 | 0.43 | 0.32 |
| 6 | 144272474 | rs7770529 | 0.224 | 1.283 | -2.32 | 0.057 | -0.159 | 0.028 | 0.028 | 0 | 0.71 | 0.97 | 0.78 | 0.78 | 0.65 | 0.61 |
| 6 | 144274210 | rs12528876 | -0.009 | -0.03 | -1.896 | -0.083 | -0.018 | -0.012 | -0.012 | 0 | 0.24 | 0.28 | 0.36 | 0.17 | 0.18 | 0.2 |
| 6 | 144277568 | rs9321951 | 0.222 | -0.158 | -2.187 | 0.36 | 0.009 | -0.001 | -0.001 | 0 | 0.61 | 0.69 | 0.61 | 0.62 | 0.68 | 0.54 |
| 6 | 144278095 | rs2268447 | 0.262 | -0.391 | -1.432 | 0.241 | -0.02 | -0.003 | -0.003 | 3.597 | 0.58 | 0.47 | 0.59 | 0.62 | 0.68 | 0.54 |
| 6 | 144280799 | rs3819812 | 0.203 | -0.393 | -1.198 | 0.301 | -0.024 | -0.006 | -0.006 | 3.57 | 0.59 | 0.49 | 0.59 | 0.65 | 0.67 | 0.58 |
| 6 | 144281385 | rs7770000 | 1.22 | -0.207 | -1.714 | 0.076 | -0.04 | 0.066 | 0.066 | 0 | 0.15 | 0.02 | 0.12 | 0.03 | 0.21 | 0.11 |
| 6 | 144285159 | rs2268450 | 0.696 | -0.151 | -1.818 | 1.015 | -0.059 | -0.047 | -0.047 | 0.002 | 0.17 | 0.04 | 0.21 | 0.23 | 0.26 | 0.2 |
| 6 | 144287218 | rs9373409 | 0.985 | -0.294 | -1.114 | 0.882 | -0.02 | 0.018 | 0.018 | 0 | 0.39 | 0.27 | 0.31 | 0.3 | 0.39 | 0.38 |
| 6 | 144288550 | rs17073273 | -0.021 | -0.603 | -1.44 | 0.848 | -0.077 | -0.16 | -0.16 | 0 | 0.25 | 0.05 | 0.27 | 0.52 | 0.33 | 0.39 |
| 6 | 144289461 | rs2273323 | 0.965 | -0.541 | -1.399 | 0.716 | -0.008 | -0.052 | -0.052 | 0 | 0.09 | 0.01 | 0.07 | 0.17 | 0.08 | 0.2 |
| 6 | 144291150 | rs13218225 | 1.624 | 0.333 | -1.774 | -0.008 | -0.05 | 0.056 | 0.056 | 2.679 | 0.1 | 0.01 | 0.11 | 0 | 0.13 | 0.02 |

| | | | | | | | | | | | | | | | | |
|---|---|---|---|---|---|---|---|---|---|---|---|---|---|---|---|---|
| 6 | 144291182 | rs9496829 | -0.617 | -0.548 | -1.177 | 0.223 | -0.044 | -0.071 | -0.071 | 0 | 0.68 | 0.53 | 0.63 | 0.86 | 0.75 | 0.79 |
| 6 | 144292587 | rs9403540 | 1.453 | -0.694 | -1.297 | 0.186 | -0.177 | -0.036 | -0.036 | 0 | 0.36 | 0.03 | 0.29 | 0.35 | 0.41 | 0.39 |
| 6 | 144304727 | rs9321955 | -0.516 | -0.663 | -0.959 | 0.064 | -0.053 | -0.066 | -0.066 | 1.601 | 0.71 | 0.61 | 0.65 | 0.87 | 0.8 | 0.8 |
| 6 | 144305568 | rs9403542 | 0.23 | -0.624 | -1.027 | 0.096 | 0.045 | 0.05 | 0.05 | 0 | 0.56 | 0.28 | 0.51 | 0.72 | 0.6 | 0.7 |
| 6 | 144318433 | rs7752440 | -0.594 | -0.802 | -0.988 | 0 | -0.02 | -0.027 | -0.027 | 0 | 0.76 | 0.72 | 0.66 | 0.87 | 0.8 | 0.82 |
| 6 | 144318998 | rs4552753 | -0.174 | -0.793 | -0.773 | -0.318 | -0.012 | -0.056 | -0.056 | 0 | 0.11 | 0.21 | 0.04 | 0 | 0.07 | 0.02 |
| 6 | 144319407 | rs6570599 | -0.122 | -0.684 | -1.115 | -0.051 | 0.012 | -0.031 | -0.031 | 5.052 | 0.67 | 0.77 | 0.49 | 0.49 | 0.66 | 0.52 |
| 6 | 144323572 | rs2328537 | 0.892 | -0.307 | -1.167 | -0.235 | -0.009 | 0.014 | 0.014 | 0 | 0.39 | 0.35 | 0.33 | 0.33 | 0.4 | 0.44 |
| 6 | 144331491 | rs9321957 | 1.231 | -0.198 | -0.883 | -0.124 | -0.002 | 0.011 | 0.011 | 0 | 0.36 | 0.26 | 0.26 | 0.34 | 0.33 | 0.4 |
| 6 | 144334195 | rs6937531 | 0.261 | -0.157 | -0.779 | -0.105 | -0.006 | 0.008 | 0.008 | 0 | 0.23 | 0.17 | 0.33 | 0.17 | 0.25 | 0.13 |
| 6 | 144337886 | rs2207851 | 0.385 | -0.427 | -0.717 | -0.04 | 0.001 | -0.006 | -0.006 | 0 | 0.59 | 0.43 | 0.6 | 0.5 | 0.58 | 0.53 |
| 6 | 144344172 | rs9484838 | -0.08 | -1.193 | -0.133 | 0.554 | -0.017 | 0.024 | 0.024 | 0 | 0.81 | 0.81 | 0.79 | 0.63 | 0.85 | 0.76 |
| 6 | 144358117 | rs9321958 | 1.407 | -0.907 | -0.701 | 0.22 | -0.158 | -0.025 | -0.025 | 0.407 | 0.39 | 0.03 | 0.27 | 0.34 | 0.44 | 0.48 |
| 6 | 144381928 | rs10457757 | -0.694 | -0.96 | 0.03 | 0.27 | 0.017 | 0.063 | 0.063 | 0 | 0.38 | 0.28 | 0.43 | 0.17 | 0.33 | 0.28 |
| 6 | 144382294 | rs2328542 | -1.858 | -1.265 | 0.086 | 0.206 | -0.026 | 0.008 | 0.008 | 1.057 | 0.64 | 0.76 | 0.68 | 0.65 | 0.49 | 0.55 |
| 6 | 144382983 | rs9484840 | -0.08 | -1.312 | 0.26 | 0.451 | -0.061 | 0.095 | 0.095 | 0 | 0.26 | 0.49 | 0.26 | 0.49 | 0.15 | 0.26 |
| 6 | 144395930 | rs12204640 | 0.508 | -1.024 | -0.019 | 0.137 | 0 | -0.008 | -0.008 | 0 | 0.07 | 0.08 | 0.05 | 0 | 0.08 | 0.02 |
| 6 | 144402945 | rs6929066 | -0.176 | -1.342 | -0.04 | -0.001 | 0.026 | 0.045 | 0.045 | 0 | 0.29 | 0.15 | 0.3 | 0.15 | 0.23 | 0.26 |
| 6 | 144411489 | rs9496856 | -0.967 | -1.556 | 0.473 | -0.161 | -0.019 | -0.065 | -0.065 | 0 | 0.28 | 0.52 | 0.17 | 0.13 | 0.24 | 0.2 |
| 6 | 144412067 | rs9496857 | -1.741 | -1.336 | 0.512 | 0.184 | 0.015 | 0.04 | 0.04 | 0 | 0.45 | 0.8 | 0.47 | 0.29 | 0.41 | 0.29 |
| 6 | 144412107 | rs11155342 | 0.173 | -1.257 | 0.203 | 0.21 | 0.003 | -0.066 | -0.066 | 0 | 0.11 | 0.3 | 0.09 | 0 | 0.09 | 0.03 |
| 6 | 144413470 | rs7752460 | 0.23 | -0.974 | -0.509 | 0.434 | 0.005 | 0.004 | 0.004 | 0 | 0.17 | 0.12 | 0.07 | 0.13 | 0.15 | 0.19 |
| 6 | 144417730 | rs12199296 | 0.029 | -1.144 | -0.528 | 0.495 | -0.041 | -0.007 | -0.007 | 0 | 0.31 | 0.41 | 0.28 | 0.33 | 0.23 | 0.28 |
| 6 | 144419525 | rs9321961 | -1.893 | -0.628 | -0.544 | 0.514 | -0.171 | -0.097 | -0.097 | 0 | 0.7 | 0.97 | 0.72 | 0.61 | 0.58 | 0.63 |
| 6 | 144419647 | rs12203051 | -0.597 | -0.86 | -0.331 | 0.498 | 0.011 | 0.003 | 0.003 | 0.003 | 0.48 | 0.53 | 0.35 | 0.45 | 0.37 | 0.47 |
| 6 | 144425331 | rs2328543 | -1.634 | -0.548 | -1.129 | 0.595 | -0.143 | -0.082 | -0.082 | 1.081 | 0.68 | 0.95 | 0.7 | 0.58 | 0.56 | 0.56 |
| 6 | 144426021 | rs4896697 | -1.077 | -1.237 | -1.144 | 0.631 | -0.019 | -0.015 | -0.015 | 1.421 | 0.54 | 0.9 | 0.47 | 0.56 | 0.45 | 0.45 |
| 6 | 144428387 | rs4895631 | 0.105 | -1.32 | -0.929 | 0.909 | -0.006 | -0.014 | -0.014 | 0.057 | 0.26 | 0.15 | 0.22 | 0.3 | 0.25 | 0.33 |
| 6 | 144433502 | rs7747271 | -2.198 | -1.101 | -1.15 | 1.169 | -0.131 | 0.026 | 0.026 | 1.34 | 0.66 | 0.94 | 0.73 | 0.7 | 0.6 | 0.56 |
| 6 | 144434024 | rs6935203 | 0.211 | -1.331 | -0.506 | 1.381 | -0.006 | 0.015 | 0.015 | 0 | 0.11 | 0.21 | 0.13 | 0.14 | 0.12 | 0.09 |
| 6 | 144434871 | rs9403548 | 0.129 | -1.404 | -0.207 | 1.348 | -0.003 | -0.004 | -0.004 | 0 | 0.22 | 0.22 | 0.31 | 0.13 | 0.21 | 0.11 |
| 6 | 144444514 | rs6570606 | -0.077 | -1.112 | -0.215 | 1.079 | 0.022 | -0.018 | -0.018 | 0.689 | 0.25 | 0.15 | 0.18 | 0.31 | 0.17 | 0.34 |
| 6 | 144446518 | rs7760281 | -0.692 | -0.671 | 0.461 | 1.046 | -0.001 | 0.003 | 0.003 | 0 | 0.36 | 0.31 | 0.46 | 0.28 | 0.43 | 0.23 |
| 6 | 144454797 | rs6570607 | -1.349 | -1.42 | 1.359 | 1.449 | -0.002 | 0.011 | 0.011 | 0 | 0.26 | 0.57 | 0.2 | 0.32 | 0.25 | 0.1 |
| 6 | 144456712 | rs10457052 | 0.161 | -1.445 | 0.721 | 1.481 | -0.048 | -0.022 | -0.022 | 0 | 0.23 | 0.01 | 0.23 | 0.16 | 0.21 | 0.29 |
| 6 | 144457899 | rs12208207 | -0.455 | -1.283 | 1.723 | 1.706 | 0.016 | -0.009 | -0.009 | 0 | 0.17 | 0.03 | 0.08 | 0.16 | 0.14 | 0.08 |
| 6 | 144459961 | rs6910108 | -0.35 | -1.337 | 1.897 | 1.752 | -0.04 | 0.002 | 0.002 | 0 | 0.08 | 0.18 | 0.03 | 0.1 | 0.03 | 0.08 |
| 6 | 144460762 | rs6916177 | -0.939 | -1.198 | 1.633 | 1.78 | 0.004 | -0.014 | -0.014 | 0 | 0.23 | 0.28 | 0.16 | 0.16 | 0.25 | 0.09 |
| 6 | 144465219 | rs9496878 | -1.05 | -1.363 | 1.843 | 2.058 | -0.064 | 0.126 | 0.126 | 0 | 0.12 | 0.37 | 0.07 | 0.26 | 0.05 | 0.09 |
| 6 | 144491331 | rs2328565 | -1.543 | -0.958 | 1.45 | 2.871 | -0.017 | 0.296 | 0.296 | 0 | 0.07 | 0.22 | 0.05 | 0.45 | 0.06 | 0.17 |
| 6 | 144495433 | rs4896702 | -1.486 | -0.958 | 1.432 | 2.873 | -0.015 | 0.298 | 0.298 | 0 | 0.07 | 0.22 | 0.05 | 0.44 | 0.06 | 0.17 |
| 6 | 144497577 | rs4896703 | -0.911 | -1.189 | 0.344 | 2.23 | -0.013 | 0.085 | 0.085 | 0 | 0.56 | 0.95 | 0.45 | 0.7 | 0.42 | 0.44 |
| 6 | 144499144 | rs4895634 | -0.299 | -0.916 | 0.682 | 1.8 | 0.022 | 0.079 | 0.079 | 0.509 | 0.43 | 0.29 | 0.35 | 0.14 | 0.35 | 0.25 |
| 6 | 144508698 | rs3734228 | -1.028 | -0.407 | 0.796 | 1.98 | 0.012 | 0.219 | 0.219 | 0 | 0.06 | 0.28 | 0.05 | 0.23 | 0.06 | 0.11 |
| 6 | 144510926 | rs3734227 | -0.349 | -0.312 | -0.142 | 1.427 | -0.002 | -0.01 | -0.01 | 0 | 0.35 | 0.17 | 0.23 | 0.36 | 0.3 | 0.3 |
| 6 | 144519150 | rs4896706 | -0.36 | -0.535 | -0.38 | 0.797 | 0.01 | 0.032 | 0.032 | 0.476 | 0.41 | 0.19 | 0.44 | 0.7 | 0.39 | 0.45 |
| 6 | 144522587 | rs3822904 | -0.478 | -0.556 | -0.438 | 0.786 | -0.033 | -0.003 | -0.003 | 0.769 | 0.4 | 0.07 | 0.44 | 0.7 | 0.39 | 0.45 |
| 6 | 144530533 | rs4286803 | -0.983 | -1.243 | 0.29 | 0.5 | 0.079 | 0.144 | 0.144 | 0.2 | 0.49 | 0.73 | 0.5 | 0.82 | 0.38 | 0.46 |
| 6 | 144537937 | rs9376812 | -0.862 | -1.319 | 0.549 | 0.509 | 0.043 | 0.113 | 0.113 | 0 | 0.56 | 0.77 | 0.52 | 0.8 | 0.39 | 0.57 |
| 6 | 144544807 | rs9403554 | -0.267 | -1.304 | 0.546 | 0.567 | 0.053 | 0.033 | 0.033 | 3.026 | 0.46 | 0.12 | 0.44 | 0.68 | 0.39 | 0.54 |
| 6 | 144550345 | rs1547622 | -0.03 | -1.45 | 0.393 | 0.443 | 0.009 | -0.011 | -0.011 | 0 | 0.41 | 0.08 | 0.42 | 0.68 | 0.37 | 0.55 |
| 6 | 144554543 | rs1322450 | -0.182 | -1.466 | 0.746 | 0.436 | 0.017 | 0.023 | 0.023 | 3.784 | 0.43 | 0.35 | 0.44 | 0.72 | 0.37 | 0.56 |
| 6 | 144556657 | rs7767952 | -0.942 | -1.286 | 0.04 | 0.286 | 0.018 | 0.14 | 0.14 | 0.87 | 0.57 | 0.9 | 0.54 | 0.83 | 0.41 | 0.58 |
| 6 | 144585022 | rs7748053 | -1.541 | -1.457 | 0.856 | 0.511 | -0.064 | 0.019 | 0.019 | 1.33 | 0.13 | 0.46 | 0.07 | 0.1 | 0.04 | 0.01 |
| 6 | 144586746 | rs7767688 | -0.348 | -1.365 | 0.392 | 0.456 | 0.035 | 0.03 | 0.03 | 0 | 0.45 | 0.35 | 0.4 | 0.71 | 0.36 | 0.59 |
| 6 | 144588926 | rs2502627 | -0.536 | -1.283 | 0.402 | 0.358 | 0.04 | 0.048 | 0.048 | 0 | 0.48 | 0.41 | 0.42 | 0.77 | 0.36 | 0.6 |
| 6 | 144597047 | rs658562 | -0.791 | -0.483 | 0.213 | 1.005 | 0.012 | 0.024 | 0.024 | 0 | 0.52 | 0.69 | 0.5 | 0.4 | 0.39 | 0.56 |
| 6 | 144600884 | rs583788 | -0.764 | -0.492 | 0.524 | 1.06 | 0.01 | 0.012 | 0.012 | 0 | 0.52 | 0.64 | 0.49 | 0.41 | 0.38 | 0.56 |
| 6 | 144607646 | rs588629 | -0.179 | -0.822 | 0.056 | 0.853 | -0.023 | -0.028 | -0.028 | 0 | 0.27 | 0.08 | 0.3 | 0.26 | 0.22 | 0.43 |
| 6 | 144608800 | rs664541 | -1.52 | -1.01 | 0.145 | 1.296 | 0.013 | -0.026 | -0.026 | 0.434 | 0.53 | 0.8 | 0.52 | 0.5 | 0.38 | 0.58 |
| 6 | 144610302 | rs637232 | -0.858 | -1.158 | 1.019 | 1.227 | -0.009 | -0.006 | -0.006 | 0.138 | 0.41 | 0.49 | 0.41 | 0.37 | 0.3 | 0.49 |
| 6 | 144610379 | rs615755 | -1.727 | -1.138 | 0.859 | 1.204 | 0.025 | -0.025 | -0.025 | 0 | 0.53 | 0.81 | 0.51 | 0.5 | 0.37 | 0.58 |
| 6 | 144610433 | rs636408 | -0.038 | -1.145 | 0.86 | 1.201 | -0.018 | -0.023 | -0.023 | 0 | 0.27 | 0.08 | 0.31 | 0.26 | 0.22 | 0.43 |
| 6 | 144613822 | rs672468 | -1.722 | -1.13 | 0.886 | 1.18 | 0.014 | -0.023 | -0.023 | 0.49 | 0.53 | 0.81 | 0.52 | 0.49 | 0.38 | 0.58 |
| 6 | 144613989 | rs589892 | -1.722 | -1.13 | 0.886 | 1.18 | 0.014 | -0.023 | -0.023 | 0 | 0.53 | 0.81 | 0.52 | 0.49 | 0.38 | 0.58 |
| 6 | 144616939 | rs2473133 | -1.693 | -1.201 | 0.975 | 1.134 | 0.038 | 0.006 | 0.006 | 0.759 | 0.27 | 0.85 | 0.22 | 0.24 | 0.16 | 0.15 |
| 6 | 144618167 | rs1322447 | -1.431 | -1.201 | 1.249 | 1.134 | 0.005 | 0 | 0 | 0.033 | 0.26 | 0.67 | 0.2 | 0.24 | 0.15 | 0.16 |
| 6 | 144619178 | rs2473137 | -0.252 | -1.213 | 1.429 | 1.138 | -0.004 | -0.049 | -0.049 | 0.944 | 0.13 | 0.24 | 0.09 | 0.06 | 0.08 | 0.06 |
| 6 | 144619627 | rs1474630 | -0.882 | -1.21 | 1.385 | 1.179 | -0.005 | 0.126 | 0.126 | 0.062 | 0.12 | 0.43 | 0.12 | 0.19 | 0.08 | 0.1 |

| | | | | | | | | | | | | | | | | |
|---|---|---|---|---|---|---|---|---|---|---|---|---|---|---|---|---|
| 6 | 144620145 | rs2473138 | -0.655 | -1.22 | 1.762 | 1.182 | -0.01 | 0.084 | 0.084 | 0 | 0.15 | 0.31 | 0.12 | 0.24 | 0.07 | 0.08 |
| 6 | 144622964 | rs2473147 | -1.656 | -1.24 | 1.448 | 1.045 | 0.021 | 0.032 | 0.032 | 0.917 | 0.27 | 0.77 | 0.2 | 0.06 | 0.15 | 0.13 |
| 6 | 144624144 | rs675261 | -0.843 | -1.181 | 1.448 | 1.172 | 0.012 | 0.044 | 0.044 | 0 | 0.39 | 0.21 | 0.37 | 0.28 | 0.29 | 0.47 |
| 6 | 144632199 | rs926397 | -0.918 | -1.218 | 1.894 | 1.327 | -0.008 | 0.005 | 0.005 | 0 | 0.41 | 0.51 | 0.42 | 0.46 | 0.29 | 0.49 |
| 6 | 144633601 | rs2473153 | -1.718 | -1.183 | 1.801 | 1.327 | -0.002 | -0.062 | -0.062 | 1.062 | 0.53 | 0.92 | 0.52 | 0.46 | 0.37 | 0.56 |
| 6 | 144637332 | rs2502634 | 0.149 | -1.116 | 2.087 | 1.439 | -0.001 | -0.019 | -0.019 | 0 | 0.1 | 0.12 | 0.06 | 0.03 | 0.07 | 0.05 |
| 6 | 144638157 | rs6903169 | -1.725 | -1.123 | 2.381 | 1.489 | 0.079 | 0.389 | 0.389 | 3.805 | 0.05 | 0.51 | 0.08 | 0.19 | 0.01 | 0.02 |
| 6 | 144641358 | rs2982239 | 0.071 | -0.957 | 2.3 | 1.303 | 0.002 | -0.019 | -0.019 | 0 | 0.1 | 0.13 | 0.08 | 0.02 | 0.09 | 0.05 |
| 6 | 144656653 | rs760855 | -0.788 | -1.339 | 1.762 | 1.264 | -0.036 | -0.024 | -0.024 | 0 | 0.3 | 0.07 | 0.35 | 0.26 | 0.27 | 0.39 |
| 6 | 144665357 | rs6917261 | -1.865 | -1.343 | 2.11 | 1.421 | 0.011 | 0.204 | 0.204 | 5.568 | 0.07 | 0.6 | 0.09 | 0.16 | 0.01 | 0.02 |
| 6 | 144671771 | rs9321970 | 1.59 | -1.414 | 1.806 | 1.336 | 0.041 | -0.017 | -0.017 | 0 | 0.53 | 0.09 | 0.49 | 0.58 | 0.65 | 0.49 |
| 6 | 144675091 | rs6922608 | -1.785 | -1.421 | 1.975 | 1.456 | 0.013 | 0.201 | 0.201 | 5.758 | 0.06 | 0.6 | 0.09 | 0.16 | 0.01 | 0.02 |
| 6 | 144686774 | rs7765923 | 0.375 | -1.324 | 1.441 | 1.798 | -0.015 | -0.045 | -0.045 | 2.319 | 0.69 | 0.92 | 0.65 | 0.68 | 0.75 | 0.62 |
| 6 | 144699731 | rs6904422 | 0.453 | -1.294 | 1.398 | 1.742 | -0.02 | -0.034 | -0.034 | 0 | 0.68 | 0.92 | 0.64 | 0.68 | 0.73 | 0.62 |
| 6 | 144702116 | rs9496948 | -1.972 | -1.092 | 1.915 | 1.633 | 0.063 | 0.152 | 0.152 | 2.502 | 0.07 | 0.49 | 0.12 | 0.13 | 0.06 | 0.03 |
| 6 | 144709028 | rs7745285 | -1.986 | -1.092 | 1.716 | 1.511 | 0.036 | 0.098 | 0.098 | 0.102 | 0.1 | 0.49 | 0.27 | 0.13 | 0.09 | 0.05 |
| 6 | 144714689 | rs11965388 | -0.799 | -1.126 | 1.753 | 1.386 | -0.006 | -0.06 | -0.06 | 0.016 | 0.11 | 0.21 | 0.21 | 0.01 | 0.11 | 0.04 |
| 6 | 144737670 | rs4896716 | 0.161 | -1.043 | 0.012 | 1.634 | -0.013 | -0.03 | -0.03 | 0.097 | 0.12 | 0.09 | 0.2 | 0.21 | 0.1 | 0.08 |
| 6 | 144740319 | rs9321974 | 1.37 | -1.166 | -0.68 | 1.222 | -0.078 | -0.067 | -0.067 | 1.703 | 0.41 | 0.06 | 0.3 | 0.46 | 0.53 | 0.43 |
| 6 | 144747242 | rs7356801 | -0.307 | -0.755 | 0.411 | 1.334 | 0.027 | 0.046 | 0.046 | 0.076 | 0.13 | 0.24 | 0.23 | 0.19 | 0.12 | 0.08 |
| 6 | 144749359 | rs6570628 | 0.626 | -0.994 | 0.072 | 1.218 | -0.011 | 0.005 | 0.005 | 0 | 0.92 | 0.86 | 0.95 | 0.92 | 0.94 | 0.85 |
| 6 | 144761383 | rs4282425 | -1.099 | -0.892 | 0.353 | 0.918 | -0.019 | -0.003 | -0.003 | 0 | 0.32 | 0.59 | 0.48 | 0.29 | 0.22 | 0.21 |
| 6 | 144771845 | rs4895643 | -0.577 | -0.367 | 0.06 | 0.767 | -0.011 | 0.026 | 0.026 | 0 | 0.37 | 0.49 | 0.51 | 0.53 | 0.31 | 0.34 |
| 6 | 144790022 | rs6904280 | -1.442 | -0.791 | 0.739 | 0.329 | -0.022 | -0.029 | -0.029 | 0.229 | 0.15 | 0.63 | 0.13 | 0.12 | 0.07 | 0.03 |
| 6 | 144806691 | rs9496982 | -1 | -1.384 | 0.169 | -0.156 | 0.039 | 0.057 | 0.057 | 0.713 | 0.37 | 0.73 | 0.22 | 0.14 | 0.23 | 0.34 |
| 6 | 144814016 | rs6912724 | -0.837 | -1.368 | 0.861 | -0.283 | 0.007 | 0.004 | 0.004 | 0 | 0.34 | 0.34 | 0.2 | 0.08 | 0.25 | 0.33 |
| 6 | 144832834 | rs7739143 | -1.06 | -1.47 | 0.99 | -0.212 | 0.038 | 0.064 | 0.064 | 0.402 | 0.49 | 0.62 | 0.4 | 0.26 | 0.36 | 0.55 |
| 6 | 144843858 | rs3811099 | -0.093 | -1.515 | 0.669 | -0.223 | -0.018 | -0.022 | -0.022 | 0 | 0.2 | 0.03 | 0.27 | 0.2 | 0.18 | 0.24 |
| 6 | 144852201 | rs12204734 | 0.369 | -1.548 | 0.649 | -0.25 | -0.019 | 0.077 | 0.077 | 0 | 0.25 | 0.01 | 0.1 | 0.06 | 0.2 | 0.31 |
| 6 | 144854707 | rs12661566 | -0.68 | -1.633 | 0.694 | 0.179 | -0.004 | 0.189 | 0.189 | 0.122 | 0.07 | 0.11 | 0.1 | 0.53 | 0.05 | 0.13 |
| 6 | 144857413 | rs12190718 | 0.464 | -1.647 | 0.793 | 0.123 | 0.017 | 0.061 | 0.061 | 0 | 0.25 | 0.09 | 0.11 | 0.06 | 0.2 | 0.31 |
| 6 | 144861946 | rs10499231 | -1.334 | -1.673 | 0.816 | 0.094 | 0.002 | 0.183 | 0.183 | 2.119 | 0.55 | 0.8 | 0.52 | 0.85 | 0.41 | 0.67 |
| 6 | 144866705 | rs7770106 | -1.509 | -1.643 | 0.868 | 0.103 | 0.01 | 0.173 | 0.173 | 1.526 | 0.55 | 0.77 | 0.52 | 0.85 | 0.41 | 0.67 |
| 6 | 144869785 | rs1534443 | -0.267 | -1.648 | 1.638 | 0.139 | -0.012 | 0.011 | 0.011 | 0 | 0.32 | 0.33 | 0.21 | 0.6 | 0.25 | 0.44 |
| 6 | 144872051 | rs1005974 | -0.31 | -1.583 | 1.758 | 0.222 | -0.018 | -0.019 | -0.019 | 0 | 0.22 | 0.36 | 0.29 | 0.2 | 0.16 | 0.24 |
| 6 | 144878337 | rs2293536 | -1.004 | -1.443 | 1.79 | 0.332 | 0.012 | 0.351 | 0.351 | 0.137 | 0.07 | 0.21 | 0.11 | 0.53 | 0.05 | 0.13 |
| 6 | 144891032 | rs9376824 | -1.608 | -1.448 | 1.909 | 0.323 | 0.022 | 0.371 | 0.371 | 1.39 | 0.1 | 0.32 | 0.13 | 0.53 | 0.07 | 0.14 |
| 6 | 144907979 | rs6915747 | -1.421 | -1.545 | 1.405 | 0.125 | 0.008 | 0.177 | 0.177 | 1.532 | 0.55 | 0.81 | 0.53 | 0.84 | 0.41 | 0.67 |
| 6 | 144909029 | rs1918751 | -0.803 | -1.585 | 1.373 | 0.225 | -0.004 | 0.157 | 0.157 | 0.128 | 0.07 | 0.09 | 0.1 | 0.53 | 0.05 | 0.13 |
| 6 | 144910079 | rs9390173 | -0.988 | -1.607 | 1.498 | 0.147 | 0.055 | 0.131 | 0.131 | 0.627 | 0.5 | 0.25 | 0.47 | 0.78 | 0.39 | 0.66 |
| 6 | 144926230 | rs1554816 | -0.666 | -1.669 | 1.383 | -0.002 | -0.03 | 0 | 0 | 0.003 | 0.25 | 0.47 | 0.31 | 0.26 | 0.17 | 0.32 |
| 6 | 144937933 | rs12200263 | 0.537 | -1.617 | 0.45 | -0.095 | -0.02 | 0.073 | 0.073 | 0 | 0.24 | 0.02 | 0.1 | 0.06 | 0.2 | 0.31 |
| 6 | 144943157 | rs9390180 | 0.064 | -1.533 | 0.595 | 0.151 | -0.008 | -0.035 | -0.035 | 0 | 0.19 | 0.03 | 0.29 | 0.22 | 0.15 | 0.29 |
| 6 | 144952491 | rs11155364 | -1.25 | -1.443 | 0.529 | 0.146 | -0.014 | -0.038 | -0.038 | 0 | 0.11 | 0.03 | 0.11 | 0.52 | 0.13 | 0.2 |
| 6 | 144958656 | rs10434890 | -0.515 | -1.718 | 1.068 | -0.156 | 0.018 | 0.016 | 0.016 | 0.021 | 0.44 | 0.32 | 0.41 | 0.34 | 0.35 | 0.45 |
| 6 | 144965205 | rs6929795 | -1.53 | -1.722 | 1.353 | -0.187 | -0.02 | 0.106 | 0.106 | 1.097 | 0.59 | 0.73 | 0.56 | 0.86 | 0.48 | 0.64 |
| 6 | 144967564 | rs9390184 | -0.934 | -1.616 | 1.553 | -0.052 | 0.016 | 0.231 | 0.231 | 0.034 | 0.09 | 0.22 | 0.12 | 0.51 | 0.12 | 0.17 |
| 6 | 144968195 | rs7753246 | -0.278 | -1.648 | 1.398 | -0.124 | -0.001 | -0.001 | -0.001 | 0 | 0.23 | 0.19 | 0.31 | 0.24 | 0.17 | 0.21 |
| 6 | 144980551 | rs9390185 | 0.493 | -1.666 | 0.954 | -0.08 | -0.004 | -0.055 | -0.055 | 0 | 0.16 | 0.02 | 0.23 | 0.24 | 0.1 | 0.16 |
| 6 | 144982326 | rs4896735 | -1.562 | -1.713 | 1.067 | -0.143 | -0.038 | 0.163 | 0.163 | 1.45 | 0.59 | 0.81 | 0.57 | 0.87 | 0.48 | 0.64 |
| 6 | 144988922 | rs12663692 | -0.484 | -1.908 | 1.278 | 0.43 | -0.005 | -0.001 | -0.001 | 0.886 | 0.21 | 0.21 | 0.3 | 0.24 | 0.12 | 0.18 |
| 6 | 144998088 | rs9386076 | -1.305 | -1.953 | 1.02 | 0.472 | -0.002 | 0.064 | 0.064 | 0.037 | 0.12 | 0.15 | 0.13 | 0.56 | 0.13 | 0.2 |
| 6 | 144999763 | rs9376837 | -1.269 | -1.951 | 0.993 | 0.439 | -0.001 | 0.072 | 0.072 | 0.011 | 0.12 | 0.13 | 0.13 | 0.5 | 0.14 | 0.2 |
| 6 | 145001253 | rs7742705 | -1.785 | -1.981 | 0.972 | 0.416 | -0.027 | -0.042 | -0.042 | 0 | 0.34 | 0.52 | 0.44 | 0.8 | 0.25 | 0.38 |
| 6 | 145001887 | rs11155370 | -1.449 | -1.535 | 0.196 | 0.974 | 0 | 0.019 | 0.019 | 0.005 | 0.11 | 0.07 | 0.11 | 0.5 | 0.13 | 0.2 |
| 6 | 145012869 | rs9376841 | -0.153 | -1.885 | -0.204 | 0.818 | 0.004 | -0.117 | -0.117 | 0.196 | 0.16 | 0.02 | 0.1 | 0.48 | 0.11 | 0.25 |
| 6 | 145025611 | rs9376844 | -0.158 | -1.892 | -0.207 | 0.82 | 0.004 | -0.118 | -0.118 | 0 | 0.16 | 0.02 | 0.1 | 0.48 | 0.11 | 0.25 |
| 6 | 145028992 | rs9390200 | -1.526 | -2.627 | -0.424 | 0.491 | 0.141 | 0.157 | 0.157 | 0.193 | 0.5 | 0.85 | 0.64 | 0.78 | 0.29 | 0.53 |
| 6 | 145030284 | rs9321987 | -1.279 | -2.7 | -0.453 | 0.542 | 0.08 | 0.031 | 0.031 | 0.867 | 0.34 | 0.84 | 0.54 | 0.32 | 0.2 | 0.29 |
| 6 | 145031860 | rs9373424 | -0.094 | -2.23 | -0.211 | 0.819 | 0.016 | -0.062 | -0.062 | 0.116 | 0.17 | 0.07 | 0.11 | 0.46 | 0.1 | 0.24 |
| 6 | 145033615 | rs13204138 | -0.672 | -2.3 | 0.226 | 0.606 | 0.003 | -0.004 | -0.004 | 0.17 | 0.05 | 0.07 | 0.05 | 0.05 | 0.06 | 0.05 |
| 6 | 145043655 | rs13208243 | 0.103 | -2.501 | 0.629 | 1.133 | 0.028 | -0.058 | -0.058 | 0 | 0.16 | 0.07 | 0.11 | 0.46 | 0.08 | 0.27 |
| 6 | 145054953 | rs12525007 | -0.14 | -2.617 | 0.52 | 1.031 | 0.031 | -0.042 | -0.042 | 0 | 0.14 | 0.04 | 0.38 | 0.25 | 0.1 | 0.12 |
| 6 | 145062080 | rs4419685 | -0.916 | -2.716 | 1.285 | 1 | -0.043 | 0.034 | 0.034 | 0 | 0.19 | 0.33 | 0.44 | 0.28 | 0.11 | 0.17 |
| 6 | 145073793 | rs4131501 | -0.269 | -2.745 | 1.691 | 0.973 | -0.023 | 0.167 | 0.167 | 0 | 0.21 | 0.45 | 0.17 | 0.51 | 0.14 | 0.31 |
| 6 | 145078694 | rs6927442 | -1.193 | -2.792 | 1.606 | 0.87 | 0.025 | 0.012 | 0.012 | 0.564 | 0.35 | 0.3 | 0.53 | 0.74 | 0.2 | 0.43 |
| 6 | 145096340 | rs4243474 | -1.65 | -2.982 | 1.735 | 0.826 | 0.097 | 0.107 | 0.107 | 0.384 | 0.42 | 0.78 | 0.59 | 0.82 | 0.21 | 0.44 |
| 6 | 145098212 | rs4272231 | -1.65 | -2.982 | 1.735 | 0.841 | 0.097 | 0.107 | 0.107 | 0.384 | 0.42 | 0.78 | 0.59 | 0.82 | 0.2 | 0.44 |
| 6 | 145100672 | rs6901062 | -1.063 | -2.916 | 1.765 | 0.913 | 0.041 | -0.001 | -0.001 | 0.28 | 0.32 | 0.24 | 0.51 | 0.71 | 0.18 | 0.38 |

| | | | | | | | | | | | | | | | | |
|---|---|---|---|---|---|---|---|---|---|---|---|---|---|---|---|---|
| 6 | 145105354 | rs6926377 | -1.288 | -3.324 | 1.025 | 0.537 | 0.069 | -0.032 | -0.032 | 0 | 0.41 | 0.77 | 0.56 | 0.46 | 0.31 | 0.38 |
| 6 | 145114935 | rs4243475 | -1.757 | -3.335 | 1.02 | 0.578 | 0.019 | 0.137 | 0.137 | 0 | 0.58 | 0.79 | 0.66 | 0.89 | 0.41 | 0.64 |
| 6 | 145118264 | rs4266507 | 0.172 | -3.34 | 1.215 | 0.659 | -0.031 | 0.017 | 0.017 | 0 | 0.16 | 0.21 | 0.42 | 0.25 | 0.09 | 0.12 |
| 6 | 145120256 | rs7748226 | 0.29 | -3.314 | 0.803 | 0.672 | 0.006 | -0.019 | -0.019 | 0 | 0.19 | 0.06 | 0.1 | 0.19 | 0.2 | 0.26 |
| 6 | 145122139 | rs17074210 | 0.267 | -3.336 | 0.84 | 0.675 | 0.036 | -0.064 | -0.064 | 1.182 | 0.14 | 0.02 | 0.38 | 0.25 | 0.09 | 0.12 |
| 6 | 145130578 | rs7759754 | 0.779 | -3.305 | 0.878 | 0.552 | 0.01 | 0.027 | 0.027 | 2.718 | 0.16 | 0.04 | 0.07 | 0.06 | 0.15 | 0.17 |
| 6 | 145131707 | rs9321993 | 0.042 | -3.191 | 0.918 | 0.552 | -0.025 | 0.079 | 0.079 | 0 | 0.21 | 0.31 | 0.13 | 0.45 | 0.11 | 0.26 |
| 6 | 145135148 | rs4895655 | -0.845 | -3.013 | 0.434 | 0.704 | -0.006 | 0.002 | 0.002 | 0 | 0.34 | 0.37 | 0.49 | 0.44 | 0.24 | 0.34 |
| 6 | 145148111 | rs2297846 | -0.122 | -3.279 | 0.455 | 1.295 | 0.018 | -0.052 | -0.052 | 0 | 0.19 | 0.1 | 0.12 | 0.44 | 0.1 | 0.26 |
| 6 | 145159917 | rs7754727 | -1.948 | -2.778 | 1.038 | 1.408 | 0.091 | 0.153 | 0.153 | 0 | 0.09 | 0.38 | 0.1 | 0.17 | 0.12 | 0.14 |
| 6 | 145162001 | rs4895657 | -2.238 | -3.262 | 0.346 | 1.069 | -0.02 | 0.344 | 0.344 | 0 | 0.54 | 0.98 | 0.68 | 0.95 | 0.37 | 0.62 |
| 6 | 145162225 | rs1111808 | -1.009 | -3.297 | 0.482 | 1.151 | -0.007 | 0.042 | 0.042 | 0 | 0.11 | 0.21 | 0.08 | 0.19 | 0.11 | 0.16 |
| 6 | 145162404 | rs4895658 | -0.685 | -3.366 | 0.499 | 1.307 | 0.001 | -0.003 | -0.003 | 0 | 0.18 | 0.18 | 0.43 | 0.3 | 0.14 | 0.19 |
| 6 | 145165362 | rs10499238 | -0.081 | -3.131 | 0.591 | 0.917 | 0.005 | 0 | 0 | 0 | 0.08 | 0.05 | 0.03 | 0.05 | 0.06 | 0.09 |
| 6 | 145168455 | rs9390235 | -1.038 | -3.133 | 0.629 | 0.94 | -0.007 | 0.04 | 0.04 | 0 | 0.11 | 0.2 | 0.08 | 0.19 | 0.11 | 0.16 |
| 6 | 145169607 | rs6903007 | -0.822 | -3.134 | 0.626 | 0.94 | 0.004 | -0.006 | -0.006 | 0 | 0.1 | 0.05 | 0.06 | 0.1 | 0.11 | 0.16 |
| 6 | 145170020 | rs9376865 | -2.229 | -3.193 | 0.881 | 0.887 | 0.047 | 0.224 | 0.224 | 0 | 0.54 | 0.82 | 0.67 | 0.95 | 0.37 | 0.62 |
| 6 | 145174098 | rs10192 | -0.954 | -3.21 | 0.9 | 0.917 | -0.058 | -0.007 | -0.007 | 0.23 | 0.37 | 0.45 | 0.53 | 0.72 | 0.19 | 0.39 |
| 6 | 145174832 | rs9390238 | -1.922 | -3.202 | 0.886 | 0.874 | 0.082 | 0.111 | 0.111 | 0 | 0.49 | 0.68 | 0.62 | 0.91 | 0.3 | 0.56 |
| 6 | 145184504 | rs4321838 | -1.071 | -3.144 | 0.893 | 0.824 | -0.005 | 0.001 | 0.001 | 0 | 0.12 | 0.16 | 0.08 | 0.1 | 0.12 | 0.16 |
| 6 | 145185269 | rs1933752 | -0.956 | -3.16 | 0.903 | 0.812 | -0.027 | -0.021 | -0.021 | 0 | 0.4 | 0.46 | 0.56 | 0.76 | 0.25 | 0.45 |
| 6 | 145185780 | rs6936275 | -0.395 | -3.082 | 0.713 | 0.35 | -0.009 | -0.001 | -0.001 | 0 | 0.09 | 0.16 | 0.05 | 0.06 | 0.06 | 0.09 |
| 6 | 145187154 | rs2153112 | -1.804 | -2.881 | 0.32 | 0.31 | 0.046 | 0.225 | 0.225 | 0 | 0.54 | 0.82 | 0.67 | 0.94 | 0.37 | 0.62 |
| 6 | 145201896 | rs11968590 | -0.924 | -3.024 | 1.791 | 2.082 | -0.018 | 0.023 | 0.023 | 0 | 0.27 | 0.3 | 0.18 | 0.46 | 0.14 | 0.34 |
| 6 | 145204339 | rs17805722 | -0.273 | -3.163 | 1.712 | 1.935 | 0.004 | 0.008 | 0.008 | 0 | 0.08 | 0.04 | 0.21 | 0 | 0.09 | 0.03 |
| 6 | 145206430 | rs9399515 | -0.691 | -3.2 | 1.866 | 2.143 | 0.025 | -0.026 | -0.026 | 0 | 0.26 | 0.2 | 0.17 | 0.37 | 0.14 | 0.35 |
| 6 | 145207006 | rs9403622 | -1.7 | -3.244 | 1.803 | 2.066 | -0.041 | 0.021 | 0.021 | 0 | 0.38 | 0.46 | 0.4 | 0.46 | 0.24 | 0.39 |
| 6 | 145208507 | rs17074404 | -2.17 | -3.234 | 1.841 | 2.441 | -0.041 | 0.106 | 0.106 | 0.003 | 0.17 | 0.51 | 0.3 | 0.25 | 0.11 | 0.17 |
| 6 | 145209041 | rs12175421 | -1.874 | -3.21 | 1.846 | 2.345 | -0.018 | 0.005 | 0.005 | 0 | 0.4 | 0.55 | 0.41 | 0.5 | 0.25 | 0.44 |
| 6 | 145241719 | rs7769065 | -0.652 | -3.202 | 1.812 | 2.441 | 0.052 | -0.048 | -0.048 | 0.461 | 0.26 | 0.09 | 0.15 | 0.35 | 0.14 | 0.34 |
| 6 | 145243339 | rs2153110 | -2.026 | -3.241 | 1.819 | 2.39 | 0.03 | -0.006 | -0.006 | 1.135 | 0.42 | 0.66 | 0.42 | 0.52 | 0.25 | 0.45 |
| 6 | 145253946 | rs2050719 | -0.599 | -3.274 | 1.865 | 2.435 | 0.03 | -0.025 | -0.025 | 0 | 0.25 | 0.16 | 0.14 | 0.36 | 0.09 | 0.34 |
| 6 | 145257393 | rs13216871 | -1.377 | -3.291 | 1.879 | 2.416 | -0.047 | 0.023 | 0.023 | 0 | 0.32 | 0.58 | 0.18 | 0.54 | 0.12 | 0.41 |
| 6 | 145258506 | rs6570663 | -1.201 | -3.222 | 1.879 | 2.389 | -0.075 | 0.057 | 0.057 | 0 | 0.29 | 0.53 | 0.18 | 0.52 | 0.11 | 0.34 |
| 6 | 145266524 | rs7757828 | -1.924 | -3.206 | 1.909 | 2.364 | 0.039 | 0 | 0 | 0 | 0.41 | 0.67 | 0.43 | 0.56 | 0.22 | 0.46 |
| 6 | 145267460 | rs6904538 | -1.346 | -3.2 | 1.927 | 2.368 | -0.024 | 0.009 | 0.009 | 0 | 0.32 | 0.62 | 0.21 | 0.56 | 0.13 | 0.41 |
| 6 | 145276644 | rs9373437 | -2.154 | -3.423 | 1.441 | 2.356 | 0.156 | 0.185 | 0.185 | 0 | 0.45 | 0.86 | 0.59 | 0.83 | 0.24 | 0.49 |
| 6 | 145282984 | rs17819123 | -0.446 | -3.397 | 1.531 | 1.988 | 0.008 | 0.01 | 0.01 | 0 | 0.1 | 0.05 | 0.12 | 0.01 | 0.07 | 0.04 |
| 6 | 145283881 | rs9373438 | -0.579 | -3.393 | 1.479 | 2.076 | 0.059 | -0.063 | -0.063 | 0.277 | 0.24 | 0.07 | 0.14 | 0.36 | 0.09 | 0.34 |
| 6 | 145286368 | rs1856416 | -2.166 | -3.471 | 1.297 | 1.891 | 0.159 | -0.054 | -0.054 | 0.372 | 0.45 | 0.9 | 0.36 | 0.54 | 0.21 | 0.46 |
| 6 | 145288265 | rs7760387 | -1.016 | -3.202 | 1.127 | 1.993 | -0.054 | 0.041 | 0.041 | 0 | 0.3 | 0.57 | 0.2 | 0.53 | 0.12 | 0.41 |
| 6 | 145307190 | rs4896766 | -1.959 | -3.142 | -0.655 | 1.97 | 0.149 | -0.064 | -0.064 | 0 | 0.44 | 0.89 | 0.36 | 0.53 | 0.21 | 0.47 |
| 6 | 145313726 | rs9399525 | -0.332 | -2.55 | -0.425 | 2.068 | 0.031 | -0.075 | -0.075 | 0 | 0.18 | 0.08 | 0.1 | 0.42 | 0.1 | 0.26 |
| 6 | 145317932 | rs949772 | -0.815 | -2.417 | -0.418 | 1.945 | 0.002 | -0.003 | -0.003 | 0.013 | 0.11 | 0.08 | 0.13 | 0.15 | 0.08 | 0.07 |
| 6 | 145319640 | rs9399527 | -0.536 | -2.203 | -0.596 | 1.854 | 0.008 | -0.06 | -0.06 | 0.031 | 0.19 | 0.02 | 0.13 | 0.26 | 0.12 | 0.22 |
| 6 | 145329620 | rs6913592 | -1.111 | -2.062 | -0.721 | 1.568 | 0.057 | 0.033 | 0.033 | 0.276 | 0.47 | 0.88 | 0.51 | 0.66 | 0.37 | 0.52 |
| 6 | 145331760 | rs9376889 | -0.072 | -2.328 | -0.132 | 1.889 | 0.002 | -0.006 | -0.006 | 0.362 | 0.22 | 0.15 | 0.27 | 0.44 | 0.18 | 0.36 |
| 6 | 145333573 | rs1890472 | -1.394 | -2.386 | 0.459 | 1.836 | 0.033 | 0.018 | 0.018 | 0.166 | 0.43 | 0.68 | 0.49 | 0.66 | 0.36 | 0.49 |
| 6 | 145334972 | rs9390273 | -1.288 | -2.702 | 0.631 | 1.517 | 0.023 | 0.036 | 0.036 | 0.199 | 0.43 | 0.3 | 0.44 | 0.72 | 0.35 | 0.54 |
| 6 | 145335490 | rs12528747 | -0.667 | -2.715 | 0.658 | 1.511 | 0 | 0.01 | 0.01 | 0 | 0.13 | 0.15 | 0.14 | 0.09 | 0.15 | 0.06 |
| 6 | 145336871 | rs9376890 | -2.044 | -2.716 | 0.659 | 1.504 | 0.041 | 0.228 | 0.228 | 0.049 | 0.53 | 0.87 | 0.53 | 0.85 | 0.39 | 0.65 |
| 6 | 145342352 | rs1933927 | -1.95 | -2.726 | 1.024 | 1.499 | 0.057 | 0.187 | 0.187 | 0 | 0.52 | 0.78 | 0.52 | 0.85 | 0.39 | 0.65 |
| 6 | 145342763 | rs2069128 | -1.107 | -2.736 | 1.053 | 1.532 | -0.005 | -0.002 | -0.002 | 0.156 | 0.17 | 0.29 | 0.16 | 0.18 | 0.16 | 0.08 |
| 6 | 145347127 | rs9497171 | -2.024 | -2.727 | 1.088 | 1.524 | 0.052 | 0.2 | 0.2 | 0.081 | 0.53 | 0.84 | 0.53 | 0.85 | 0.39 | 0.65 |
| 6 | 145348881 | rs2328629 | -0.605 | -2.653 | 1.36 | 1.658 | -0.014 | 0.006 | 0.006 | 0 | 0.3 | 0.29 | 0.3 | 0.62 | 0.2 | 0.49 |
| 6 | 145360102 | rs17074746 | -0.843 | -2.476 | 1.305 | 1.628 | -0.03 | 0.026 | 0.026 | 0 | 0.1 | 0.23 | 0.05 | 0.14 | 0.06 | 0.01 |
| 6 | 145361484 | rs946316 | -0.545 | -2.502 | 0.862 | 1.639 | 0.011 | -0.077 | -0.077 | 0.41 | 0.32 | 0.15 | 0.46 | 0.45 | 0.29 | 0.39 |
| 6 | 145364350 | rs7753282 | -0.219 | -2.507 | 0.887 | 1.724 | -0.047 | 0.124 | 0.124 | 0.027 | 0.1 | 0.19 | 0.05 | 0.3 | 0.03 | 0.17 |
| 6 | 145365169 | rs7740371 | -1.62 | -2.574 | 0.818 | 1.637 | 0.111 | 0.2 | 0.2 | 0.167 | 0.51 | 0.86 | 0.6 | 0.79 | 0.34 | 0.65 |
| 6 | 145370722 | rs2153143 | 0.192 | -2.303 | 0.625 | 1.811 | 0 | -0.067 | -0.067 | 0.04 | 0.27 | 0.1 | 0.43 | 0.35 | 0.24 | 0.41 |
| 6 | 145374694 | rs4896774 | 0.157 | -2.219 | 0.635 | 1.84 | -0.004 | -0.056 | -0.056 | 0.394 | 0.27 | 0.11 | 0.43 | 0.35 | 0.25 | 0.41 |
| 6 | 145377731 | rs521141 | -0.329 | -2.224 | 0.632 | 1.762 | -0.004 | -0.099 | -0.099 | 0 | 0.73 | 0.89 | 0.57 | 0.54 | 0.74 | 0.59 |
| 6 | 145378208 | rs13199258 | -0.615 | -2.015 | 0.904 | 1.794 | -0.052 | 0.096 | 0.096 | 0.183 | 0.11 | 0.19 | 0.05 | 0.23 | 0.03 | 0.2 |
| 6 | 145389949 | rs6913807 | -0.693 | -1.705 | 0.53 | 1.308 | -0.013 | 0.024 | 0.024 | 0 | 0.11 | 0.07 | 0.14 | 0.06 | 0.14 | 0.07 |
| 6 | 145399182 | rs635627 | 1.619 | -1.9 | 0.852 | 1.291 | -0.005 | 0.036 | 0.036 | 0 | 0.42 | 0.35 | 0.47 | 0.22 | 0.53 | 0.26 |
| 6 | 145405137 | rs6920213 | -0.324 | -1.86 | 0.605 | 0.648 | 0.003 | 0.019 | 0.019 | 0 | 0.09 | 0.07 | 0.06 | 0.06 | 0.08 | 0.05 |
| 6 | 145421935 | rs556577 | 1.231 | -1.98 | 0.871 | 0.765 | 0.003 | 0.007 | 0.007 | 0 | 0.48 | 0.57 | 0.51 | 0.21 | 0.56 | 0.25 |
| 6 | 145422462 | rs4141953 | -0.131 | -2.025 | 0.91 | 0.925 | -0.016 | 0.028 | 0.028 | 0.086 | 0.1 | 0.11 | 0.05 | 0.24 | 0.06 | 0.19 |
| 6 | 145433457 | rs649933 | 1.672 | -2.025 | 1.03 | 0.906 | -0.002 | -0.005 | -0.005 | 1.322 | 0.43 | 0.4 | 0.48 | 0.09 | 0.53 | 0.24 |

| | | | | | | | | | | | | | | | | |
|---|---|---|---|---|---|---|---|---|---|---|---|---|---|---|---|---|
| 6 | 145433861 | rs9403663 | -1.183 | -2.018 | 0.944 | 0.925 | -0.026 | 0.02 | 0.02 | 0.375 | 0.2 | 0.29 | 0.12 | 0.29 | 0.13 | 0.24 |
| 6 | 145444461 | rs577911 | -1.278 | -2.009 | 0.995 | 1.082 | 0.027 | 0.089 | 0.089 | 0.352 | 0.52 | 0.71 | 0.5 | 0.81 | 0.42 | 0.67 |
| 6 | 145457784 | rs7738467 | 0.239 | -1.557 | -0.351 | 0.917 | 0.017 | -0.001 | -0.001 | 0 | 0.06 | 0.12 | 0.12 | 0.04 | 0.08 | 0.12 |
| 6 | 145467487 | rs9497193 | -0.914 | -1.585 | -0.311 | 0.916 | -0.015 | -0.014 | -0.014 | 0.063 | 0.12 | 0.25 | 0.14 | 0.07 | 0.1 | 0.11 |
| 6 | 145472281 | rs644583 | -0.526 | -1.145 | -0.804 | 1.136 | -0.004 | -0.028 | -0.028 | 0 | 0.34 | 0.24 | 0.4 | 0.5 | 0.34 | 0.53 |
| 6 | 145477959 | rs9376900 | 0.463 | -1.093 | -1.33 | 1.175 | 0.008 | -0.036 | -0.036 | 0 | 0.07 | 0 | 0.03 | 0.1 | 0.05 | 0.09 |
| 6 | 145482735 | rs510667 | 0.066 | -0.988 | -1.27 | 1.203 | -0.069 | -0.187 | -0.187 | 1.948 | 0.27 | 0.01 | 0.28 | 0.47 | 0.27 | 0.42 |
| 6 | 145488159 | rs7757238 | -0.143 | -0.817 | -1.013 | 1.196 | 0.024 | -0.004 | -0.004 | 0 | 0.15 | 0.13 | 0.08 | 0.15 | 0.1 | 0.13 |
| 6 | 145497217 | rs9497196 | -0.433 | -0.039 | -0.732 | 1.366 | -0.001 | -0.005 | -0.005 | 0 | 0.35 | 0.3 | 0.4 | 0.5 | 0.34 | 0.52 |
| 6 | 145518231 | rs7768714 | -0.111 | 1.229 | -1.48 | 1.247 | -0.102 | -0.068 | -0.068 | 0.117 | 0.22 | 0.05 | 0.17 | 0.26 | 0.33 | 0.29 |
| 6 | 145518501 | rs544254 | -0.143 | 0.627 | -1.665 | 1.07 | 0.001 | -0.019 | -0.019 | 2.059 | 0.68 | 0.73 | 0.53 | 0.55 | 0.67 | 0.62 |
| 6 | 145521889 | rs2179285 | 0.527 | -0.459 | -2.032 | 0.185 | 0.046 | -0.028 | -0.028 | 0 | 0.29 | 0.01 | 0.25 | 0.24 | 0.19 | 0.22 |
| 6 | 145524534 | rs12190534 | 0.503 | 0.07 | -2.117 | -0.013 | 0.053 | 0.027 | 0.027 | 0.061 | 0.31 | 0.05 | 0.28 | 0.23 | 0.23 | 0.22 |
| 6 | 145528057 | rs10484702 | -0.223 | 0.357 | -2.098 | 0.38 | -0.021 | -0.002 | -0.002 | 0 | 0.62 | 0.29 | 0.69 | 0.64 | 0.5 | 0.61 |
| 6 | 145534679 | rs1884955 | 0.535 | 0.252 | -1.898 | 0.465 | 0.015 | -0.03 | -0.03 | 0 | 0.29 | 0.2 | 0.51 | 0.44 | 0.23 | 0.35 |
| 6 | 145534850 | rs491131 | 0.083 | 0.553 | -1.399 | 0.368 | -0.001 | -0.008 | -0.008 | 0.283 | 0.35 | 0.29 | 0.57 | 0.44 | 0.35 | 0.46 |
| 6 | 145539000 | rs12198531 | 0.617 | 0.544 | -1.442 | 0.365 | 0.02 | -0.038 | -0.038 | 0 | 0.29 | 0.18 | 0.4 | 0.41 | 0.23 | 0.35 |
| 6 | 145539467 | rs12206002 | 0.8 | 0.536 | -1.484 | 0.364 | 0.036 | 0.032 | 0.032 | 0 | 0.33 | 0.05 | 0.29 | 0.23 | 0.26 | 0.24 |
| 6 | 145539545 | rs475017 | 0.154 | 0.606 | -1.397 | 0.378 | 0.006 | -0.015 | -0.015 | 0.85 | 0.06 | 0.06 | 0.06 | 0 | 0.12 | 0.11 |
| 6 | 145546952 | rs1033425 | -0.07 | 0.545 | -0.969 | 0.387 | 0.014 | -0.004 | -0.004 | 0 | 0.38 | 0.7 | 0.5 | 0.41 | 0.34 | 0.36 |
| 6 | 145549551 | rs547175 | -1.528 | 0.534 | -0.995 | 0.441 | -0.06 | -0.05 | -0.05 | 0 | 0.79 | 0.94 | 0.85 | 0.71 | 0.71 | 0.72 |
| 6 | 145549781 | rs4896785 | 0.076 | 0.489 | -0.372 | 0.432 | -0.005 | 0.005 | 0.005 | 0 | 0.37 | 0.55 | 0.59 | 0.44 | 0.34 | 0.36 |
| 6 | 145554027 | rs7744166 | -0.167 | 0.372 | -0.424 | 0.454 | -0.004 | -0.004 | -0.004 | 0.123 | 0.39 | 0.64 | 0.61 | 0.42 | 0.41 | 0.36 |
| 6 | 145559759 | rs7747556 | 0.333 | 0.322 | -0.346 | 0.42 | 0.078 | -0.028 | -0.028 | 0 | 0.08 | 0.35 | 0.08 | 0 | 0.11 | 0.01 |
| 6 | 145563233 | rs2753088 | -1.519 | 0.405 | -0.742 | 0.472 | -0.094 | -0.081 | -0.081 | 0.292 | 0.79 | 0.96 | 0.84 | 0.69 | 0.71 | 0.71 |
| 6 | 145564733 | rs17074886 | 1.132 | 0.018 | -1.06 | 0.448 | 0.012 | -0.09 | -0.09 | 0 | 0.15 | 0.02 | 0.28 | 0.31 | 0.12 | 0.11 |
| 6 | 145568161 | rs11966141 | 0.049 | 0.081 | -0.98 | 0.418 | -0.002 | -0.039 | -0.039 | 0 | 0.3 | 0.19 | 0.51 | 0.43 | 0.3 | 0.35 |
| 6 | 145569264 | rs12212061 | 0.016 | 0.08 | -0.963 | 0.418 | 0 | -0.029 | -0.029 | 0.012 | 0.31 | 0.21 | 0.51 | 0.43 | 0.3 | 0.35 |
| 6 | 145572515 | rs9399540 | 0.703 | 0.066 | -0.905 | 0.402 | 0.006 | -0.001 | -0.001 | 0 | 0.07 | 0.09 | 0.05 | 0.06 | 0.1 | 0.01 |
| 6 | 145575145 | rs4896789 | -0.541 | 0.06 | -0.872 | 0.395 | -0.002 | -0.005 | -0.005 | 0 | 0.38 | 0.36 | 0.57 | 0.49 | 0.4 | 0.36 |
| 6 | 145582842 | rs579734 | 1.194 | -0.153 | -1.109 | 0.343 | 0.028 | 0.04 | 0.04 | 0 | 0.32 | 0.06 | 0.18 | 0.21 | 0.26 | 0.22 |
| 6 | 145584096 | rs502530 | -0.048 | -0.193 | -1.028 | 0.303 | -0.013 | -0.033 | -0.033 | 0.184 | 0.07 | 0.12 | 0.05 | 0 | 0.05 | 0.12 |
| 6 | 145585481 | rs2038473 | 1.232 | -0.197 | -1.045 | 0.304 | 0.028 | 0.039 | 0.039 | 0 | 0.32 | 0.06 | 0.19 | 0.21 | 0.26 | 0.22 |
| 6 | 145586348 | rs17074939 | 0.492 | -0.241 | -1.039 | 0.36 | 0.024 | -0.039 | -0.039 | 0.604 | 0.25 | 0.15 | 0.44 | 0.37 | 0.2 | 0.29 |
| 6 | 145586542 | rs566661 | 0.241 | -0.245 | -0.996 | 0.332 | 0.037 | 0.068 | 0.068 | 0 | 0.45 | 0.28 | 0.28 | 0.27 | 0.41 | 0.36 |
| 6 | 145587688 | rs543831 | -0.763 | -0.21 | -0.853 | 0.332 | -0.003 | -0.03 | -0.03 | 0 | 0.14 | 0.24 | 0.1 | 0.06 | 0.15 | 0.14 |
| 6 | 145587882 | rs11970013 | 0.904 | -0.416 | -1.009 | 0.517 | 0.014 | -0.015 | -0.015 | 0 | 0.19 | 0.14 | 0.32 | 0.28 | 0.15 | 0.11 |
| 6 | 145588073 | rs1415808 | -0.399 | -0.781 | -1.06 | 0.177 | 0.005 | -0.04 | -0.04 | 0 | 0.14 | 0.31 | 0.1 | 0.06 | 0.16 | 0.03 |
| 6 | 145590770 | rs511290 | 0.548 | -0.885 | -1.172 | 0.163 | 0.018 | 0.007 | 0.007 | 0 | 0.35 | 0.31 | 0.21 | 0.27 | 0.23 | 0.25 |
| 6 | 145591148 | rs17373628 | 0.912 | -0.768 | -1.13 | 0.231 | -0.012 | -0.019 | -0.019 | 0 | 0.05 | 0.01 | 0.12 | 0.06 | 0.04 | 0.14 |
| 6 | 145592220 | rs545242 | -0.046 | -0.768 | -1.124 | 0.204 | 0.038 | 0.042 | 0.042 | 0 | 0.45 | 0.33 | 0.28 | 0.28 | 0.39 | 0.27 |
| 6 | 145593806 | rs491794 | -0.269 | -0.605 | -1.135 | 0.305 | -0.005 | -0.03 | -0.03 | 0 | 0.7 | 0.79 | 0.47 | 0.54 | 0.68 | 0.66 |
| 6 | 145601560 | rs1321529 | 1.15 | -0.243 | -0.88 | 0.431 | 0.07 | 0.022 | 0.022 | 0 | 0.25 | 0.41 | 0.2 | 0.28 | 0.29 | 0.39 |
| 6 | 145602159 | rs501447 | 0.276 | -0.319 | -0.901 | 0.367 | -0.039 | 0.002 | 0.002 | 0.262 | 0.35 | 0.71 | 0.29 | 0.33 | 0.43 | 0.43 |
| 6 | 145603116 | rs494020 | 0.935 | -0.333 | -0.952 | 0.382 | -0.014 | 0.013 | 0.013 | 0.252 | 0.06 | 0.06 | 0.07 | 0 | 0.13 | 0.01 |
| 6 | 145604408 | rs9403680 | 0.717 | -0.313 | -1.003 | 0.326 | 0.027 | 0.009 | 0.009 | 0 | 0.3 | 0.51 | 0.24 | 0.33 | 0.34 | 0.44 |
| 6 | 145605868 | rs9376913 | 0.48 | -0.281 | -1.081 | 0.378 | 0.042 | 0.008 | 0.008 | 0 | 0.33 | 0.42 | 0.36 | 0.33 | 0.37 | 0.57 |
| 6 | 145609950 | rs581088 | 0.107 | -0.856 | -1.469 | 0.235 | 0.03 | -0.035 | -0.035 | 0 | 0.61 | 0.86 | 0.48 | 0.56 | 0.66 | 0.64 |
| 6 | 145617145 | rs13191080 | 0.875 | -0.778 | -1.467 | 0.173 | -0.073 | 0.024 | 0.024 | 0 | 0.44 | 0.08 | 0.3 | 0.32 | 0.48 | 0.55 |
| 6 | 145623921 | rs552965 | 0.914 | -0.343 | -0.909 | 0.194 | 0.052 | -0.008 | -0.008 | 0 | 0.95 | 0.87 | 0.93 | 0.97 | 0.89 | 0.93 |
| 6 | 145626208 | rs9386110 | -1.351 | -0.376 | -0.765 | 0.217 | 0.09 | 0.049 | 0.049 | 0.074 | 0.07 | 0.29 | 0.07 | 0.08 | 0.12 | 0.07 |
| 6 | 145632202 | rs17074985 | -0.21 | -0.56 | -1.038 | 0.319 | -0.015 | -0.015 | -0.015 | 0 | 0.08 | 0.03 | 0.15 | 0.06 | 0.08 | 0.18 |
| 6 | 145648326 | rs581129 | 0.255 | -1.017 | -0.4 | 0.432 | 0.01 | -0.111 | -0.111 | 0 | 0.28 | 0.07 | 0.37 | 0.6 | 0.25 | 0.25 |
| 6 | 145652638 | rs534037 | 0.195 | -1.166 | -0.225 | 0.234 | 0.042 | 0.006 | 0.006 | 0.146 | 0.64 | 0.22 | 0.56 | 0.33 | 0.71 | 0.75 |
| 6 | 145667005 | rs9386112 | 0.284 | -1.107 | -0.225 | -0.114 | -0.004 | -0.041 | -0.041 | 0 | 0.39 | 0.14 | 0.45 | 0.63 | 0.38 | 0.32 |
| 6 | 145667105 | rs4895672 | 0.206 | -0.745 | 0.717 | -0.044 | 0.023 | -0.02 | -0.02 | 0 | 0.93 | 0.71 | 0.96 | 1 | 0.93 | 1 |
| 6 | 145670357 | rs4538734 | 0.695 | -0.563 | 0.754 | -0.104 | 0.055 | -0.04 | -0.04 | 0 | 0.91 | 0.68 | 0.92 | 1 | 0.87 | 0.99 |
| 6 | 145671077 | rs1856275 | 0.889 | 0.025 | -0.159 | -0.59 | -0.047 | 0.12 | 0.12 | 5.326 | 0.5 | 0.03 | 0.52 | 0.7 | 0.45 | 0.39 |
| 6 | 145677803 | rs9390307 | 0.618 | -0.002 | 0.588 | -0.621 | 0.008 | 0.053 | 0.053 | 3.366 | 0.54 | 0.36 | 0.55 | 0.72 | 0.45 | 0.46 |
| 6 | 145689259 | rs6902581 | 0.532 | -0.01 | 1.117 | -0.688 | -0.01 | -0.064 | -0.064 | 2.236 | 0.75 | 0.66 | 0.79 | 0.94 | 0.79 | 0.82 |
| 6 | 145691464 | rs1832378 | 0.938 | -0.1 | 1.1 | -0.801 | 0.005 | 0.06 | 0.06 | 1.468 | 0.53 | 0.34 | 0.57 | 0.71 | 0.47 | 0.44 |
| 6 | 145703134 | rs1970317 | 0.424 | -0.139 | 1.018 | -0.854 | 0.002 | 0.001 | 0.001 | 0 | 0.16 | 0.13 | 0.15 | 0.06 | 0.09 | 0.17 |
| 6 | 145704346 | rs6908878 | -0.098 | -0.067 | 1.142 | -0.836 | 0.031 | -0.041 | -0.041 | 0 | 0.92 | 0.79 | 0.94 | 1 | 0.88 | 0.99 |
| 6 | 145709371 | rs12664698 | 1.061 | -0.506 | 1.21 | -0.78 | -0.017 | 0.006 | 0.006 | 1.496 | 0.19 | 0.13 | 0.2 | 0.17 | 0.3 | 0.31 |
| 6 | 145714664 | rs12208709 | 0.511 | -0.52 | 1.206 | -0.789 | 0.001 | 0 | 0 | 0.068 | 0.16 | 0.12 | 0.15 | 0.06 | 0.09 | 0.17 |
| 6 | 145715513 | rs9376924 | 1.084 | -0.523 | 1.251 | -0.79 | -0.04 | 0.012 | 0.012 | 1.259 | 0.19 | 0.1 | 0.2 | 0.17 | 0.31 | 0.31 |
| 6 | 145720470 | rs12215687 | 0.499 | -0.511 | 1.036 | -0.753 | 0.001 | 0 | 0 | 0.068 | 0.16 | 0.12 | 0.14 | 0.06 | 0.09 | 0.17 |
| 6 | 145729196 | rs9322025 | -0.379 | -0.441 | 0.937 | -0.751 | -0.016 | 0.027 | 0.027 | 0.085 | 0.37 | 0.24 | 0.36 | 0.26 | 0.41 | 0.56 |
| 6 | 145732167 | rs7773601 | -0.873 | -0.692 | 1.077 | -0.804 | 0.012 | -0.023 | -0.023 | 0 | 0.94 | 0.84 | 0.97 | 1 | 0.93 | 1 |

| | | | | | | | | | | | | | | | | |
|---|---|---|---|---|---|---|---|---|---|---|---|---|---|---|---|---|
| 6 | 145734928 | rs11155418 | 0.106 | -0.707 | 1.003 | -0.812 | 0.012 | 0.016 | 0.016 | 0 | 0.18 | 0.1 | 0.16 | 0.08 | 0.11 | 0.25 |
| 6 | 145759193 | rs9403701 | 0.406 | -0.626 | 1.012 | -0.791 | -0.01 | -0.032 | -0.032 | 0 | 0.76 | 0.75 | 0.81 | 0.92 | 0.81 | 0.73 |
| 6 | 145765450 | rs1118772 | -0.89 | -0.559 | 0.959 | -0.738 | 0.013 | -0.022 | -0.022 | 0 | 0.95 | 0.85 | 0.98 | 1 | 0.94 | 1 |
| 6 | 145769507 | rs1509205 | 0.366 | -0.656 | 1.181 | -0.788 | -0.023 | -0.056 | -0.056 | 2.909 | 0.75 | 0.68 | 0.8 | 0.92 | 0.81 | 0.73 |
| 6 | 145773164 | rs9399547 | 1.226 | -0.603 | 1.315 | -0.827 | -0.001 | -0.003 | -0.003 | 0.297 | 0.68 | 0.37 | 0.78 | 0.84 | 0.75 | 0.69 |
| 6 | 145774684 | rs2136319 | 1.226 | -0.588 | 1.217 | -0.827 | -0.001 | -0.008 | -0.008 | 2.233 | 0.68 | 0.37 | 0.78 | 0.88 | 0.75 | 0.69 |
| 6 | 145784805 | rs867056 | -0.524 | -0.45 | 1.008 | -0.726 | -0.018 | 0.017 | 0.017 | 0.154 | 0.77 | 0.83 | 0.83 | 0.86 | 0.68 | 0.59 |
| 6 | 145800915 | rs6910793 | -0.516 | -0.54 | 1.176 | -0.778 | 0.042 | 0.08 | 0.08 | 0.196 | 0.06 | 0.36 | 0.06 | 0.07 | 0.03 | 0.02 |
| 6 | 145801020 | rs398408 | -0.516 | -0.617 | 1.238 | -0.838 | -0.012 | 0.033 | 0.033 | 0 | 0.38 | 0.25 | 0.23 | 0.17 | 0.4 | 0.54 |
| 6 | 145802833 | rs365515 | -1.811 | -0.794 | 1.128 | -0.119 | -0.006 | -0.009 | -0.009 | 0 | 0.5 | 0.75 | 0.45 | 0.51 | 0.48 | 0.62 |
| 6 | 145804384 | rs12197582 | 0.744 | -0.768 | 0.899 | -0.111 | 0.021 | 0.036 | 0.036 | 0 | 0.14 | 0.04 | 0.06 | 0.03 | 0.08 | 0.13 |
| 6 | 145807214 | rs9403708 | -0.195 | -0.714 | 0.908 | -0.076 | -0.018 | 0.134 | 0.134 | 0 | 0.06 | 0.14 | 0.15 | 0.28 | 0.05 | 0.06 |
| 6 | 145809983 | rs406868 | 0.469 | -0.696 | 0.885 | 0.024 | -0.017 | 0.005 | 0.005 | 0.22 | 0.23 | 0.17 | 0.17 | 0.21 | 0.32 | 0.41 |
| 6 | 145810849 | rs399516 | -0.911 | -0.642 | 1.021 | 0.151 | 0.001 | -0.004 | -0.004 | 0 | 0.43 | 0.41 | 0.35 | 0.43 | 0.44 | 0.59 |
| 6 | 145834745 | rs378937 | -0.402 | -0.626 | 0.986 | 0.088 | -0.013 | 0.028 | 0.028 | 0 | 0.39 | 0.25 | 0.26 | 0.26 | 0.42 | 0.57 |
| 6 | 145841024 | rs371744 | -1.289 | -0.607 | 0.929 | 0.085 | 0 | -0.002 | -0.002 | 0 | 0.48 | 0.46 | 0.4 | 0.45 | 0.48 | 0.63 |
| 6 | 145841515 | rs398060 | -1.515 | -0.602 | 0.803 | 0.106 | -0.005 | -0.005 | -0.005 | 1.397 | 0.5 | 0.75 | 0.44 | 0.51 | 0.48 | 0.63 |
| 6 | 145872147 | rs12526956 | -0.224 | -0.523 | 0.782 | -0.008 | -0.011 | 0.036 | 0.036 | 0 | 0.08 | 0.13 | 0.14 | 0.19 | 0.06 | 0.06 |
| 6 | 145907007 | rs9403729 | -0.239 | -0.556 | 0.55 | -0.107 | -0.02 | 0.041 | 0.041 | 0 | 0.37 | 0.23 | 0.23 | 0.23 | 0.4 | 0.55 |
| 6 | 145932576 | rs951143 | -1.525 | -0.49 | 0.312 | -0.041 | -0.013 | 0.006 | 0.006 | 1.136 | 0.5 | 0.83 | 0.45 | 0.49 | 0.48 | 0.63 |
| 6 | 145945778 | rs9399559 | -0.737 | -0.371 | 0.546 | -0.195 | 0 | -0.004 | -0.004 | 0 | 0.41 | 0.62 | 0.3 | 0.29 | 0.42 | 0.57 |
| 6 | 145955305 | rs9497380 | -0.754 | -0.442 | 0.833 | -0.23 | 0 | -0.006 | -0.006 | 0 | 0.41 | 0.6 | 0.29 | 0.29 | 0.42 | 0.57 |
| 6 | 146004677 | rs702316 | -0.92 | -0.386 | 0.781 | -0.437 | -0.028 | -0.006 | -0.006 | 0 | 0.82 | 0.91 | 0.84 | 0.79 | 0.71 | 0.57 |
| 6 | 146014168 | rs4896837 | 0.486 | -0.397 | 0.816 | -0.393 | -0.006 | 0.016 | 0.016 | 0 | 0.61 | 0.67 | 0.76 | 0.77 | 0.59 | 0.45 |
| 6 | 146025662 | rs6930154 | 0.756 | -0.388 | 0.758 | -0.389 | 0.003 | -0.024 | -0.024 | 0 | 0.59 | 0.44 | 0.71 | 0.76 | 0.58 | 0.44 |
| 6 | 146056848 | rs2235481 | -0.371 | -0.269 | 0.588 | -0.326 | -0.012 | 0.004 | 0.004 | 0 | 0.77 | 0.82 | 0.83 | 0.79 | 0.67 | 0.56 |
| 6 | 146064889 | rs1415745 | 1.122 | -0.383 | -0.101 | -0.887 | -0.055 | 0.1 | 0.1 | 1.058 | 0.17 | 0.01 | 0.12 | 0 | 0.2 | 0.08 |
| 6 | 146118225 | rs12199362 | -0.209 | -1.207 | 0.483 | -0.691 | -0.032 | 0.049 | 0.049 | 0 | 0.36 | 0.2 | 0.23 | 0.23 | 0.41 | 0.51 |
| 6 | 146122137 | rs12200607 | 1.213 | -1.247 | 0.422 | -0.729 | 0.012 | 0.047 | 0.047 | 0 | 0.12 | 0.01 | 0.06 | 0.03 | 0.08 | 0.11 |
| 6 | 146123536 | rs7739735 | -0.804 | -1.189 | 0.559 | -0.675 | 0 | -0.003 | -0.003 | 0 | 0.41 | 0.61 | 0.3 | 0.29 | 0.43 | 0.53 |
| 6 | 146126419 | rs9373475 | -0.265 | -1.187 | 0.535 | -0.677 | -0.019 | 0.036 | 0.036 | 0 | 0.37 | 0.25 | 0.23 | 0.23 | 0.41 | 0.51 |
| 6 | 146145678 | rs9390358 | -0.436 | -1.153 | 0.729 | -0.654 | -0.004 | 0.005 | 0.005 | 0 | 0.38 | 0.35 | 0.25 | 0.29 | 0.41 | 0.52 |
| 6 | 146207465 | rs1062068 | -0.354 | -1.205 | 0.684 | -0.735 | -0.012 | 0.044 | 0.044 | 0 | 0.4 | 0.27 | 0.24 | 0.26 | 0.41 | 0.52 |
| 6 | 146207563 | rs1062067 | -0.354 | -1.205 | 0.684 | -0.734 | -0.012 | 0.028 | 0.028 | 0 | 0.4 | 0.27 | 0.24 | 0.31 | 0.41 | 0.52 |
| 6 | 146207657 | rs2256998 | -0.354 | -1.205 | 0.684 | -0.734 | -0.012 | 0.028 | 0.028 | 0 | 0.4 | 0.27 | 0.24 | 0.31 | 0.41 | 0.52 |
| 6 | 146225271 | rs9497424 | 0.544 | -1.211 | 0.726 | -0.737 | 0 | -0.005 | -0.005 | 0.076 | 0.57 | 0.44 | 0.72 | 0.69 | 0.59 | 0.48 |
| 6 | 146234644 | rs9497426 | -1.643 | -1.1 | 0.729 | -0.807 | -0.048 | -0.028 | -0.028 | 0 | 0.87 | 0.96 | 0.86 | 0.81 | 0.75 | 0.76 |
| 6 | 146285204 | rs10223507 | 0.552 | -1.417 | 0.592 | -0.724 | -0.001 | -0.002 | -0.002 | 0.024 | 0.57 | 0.42 | 0.72 | 0.74 | 0.59 | 0.48 |
| 6 | 146304603 | rs2777475 | -1.564 | -1.435 | 0.732 | -0.468 | -0.045 | -0.04 | -0.04 | 0 | 0.62 | 0.83 | 0.46 | 0.5 | 0.51 | 0.6 |
| 6 | 146368041 | rs9403765 | -0.712 | -1.453 | 0.621 | -0.456 | 0.003 | 0.012 | 0.012 | 0.365 | 0.5 | 0.5 | 0.29 | 0.31 | 0.43 | 0.52 |
| 6 | 146371074 | rs960385 | 0.362 | -1.519 | 0.573 | -0.514 | -0.019 | 0.029 | 0.029 | 0 | 0.31 | 0.17 | 0.18 | 0.21 | 0.34 | 0.42 |
| 6 | 146394655 | rs969694 | -0.801 | -1.363 | 0.761 | -0.33 | 0 | 0 | 0 | 0.758 | 0.52 | 0.5 | 0.29 | 0.44 | 0.43 | 0.53 |
| 6 | 146396699 | rs10872583 | -0.594 | -1.373 | 0.625 | -0.329 | 0.015 | 0.053 | 0.053 | 0.369 | 0.5 | 0.28 | 0.27 | 0.39 | 0.43 | 0.52 |
| 6 | 146406294 | rs1474871 | -0.878 | -1.284 | 0.606 | -0.403 | 0 | 0.006 | 0.006 | 0.702 | 0.54 | 0.57 | 0.3 | 0.32 | 0.44 | 0.53 |
| 6 | 146413758 | rs6913797 | -0.698 | -1.261 | 0.459 | -0.423 | 0.006 | 0.045 | 0.045 | 0.229 | 0.52 | 0.35 | 0.28 | 0.31 | 0.44 | 0.53 |
| 6 | 146447172 | rs4896857 | -0.767 | -0.764 | 0.613 | -0.202 | 0.051 | 0.069 | 0.069 | 0.026 | 0.08 | 0.33 | 0.18 | 0.07 | 0.07 | 0.06 |
| 6 | 146475319 | rs4629704 | -1.594 | -0.935 | 0.754 | -0.243 | -0.011 | 0.009 | 0.009 | 1.159 | 0.14 | 0.7 | 0.22 | 0.14 | 0.11 | 0.09 |
| 6 | 146488167 | rs4551188 | -1.041 | -0.927 | 0.88 | -0.285 | 0.042 | -0.018 | -0.018 | 0.082 | 0.12 | 0.42 | 0.1 | 0.07 | 0.1 | 0.08 |
| 6 | 146503040 | rs13218303 | 0.977 | -1.185 | 0.248 | -0.458 | 0.032 | 0.07 | 0.07 | 0.017 | 0.18 | 0.05 | 0.07 | 0.06 | 0.1 | 0.14 |
| 6 | 146504240 | rs9497474 | -1.33 | -1.012 | 0.411 | -0.502 | -0.006 | 0.013 | 0.013 | 0.769 | 0.13 | 0.67 | 0.21 | 0.13 | 0.11 | 0.09 |
| 6 | 146510447 | rs2206517 | -1.33 | -1.012 | 0.411 | -0.502 | -0.006 | 0.013 | 0.013 | 0.769 | 0.13 | 0.67 | 0.21 | 0.13 | 0.11 | 0.09 |
| 6 | 146516537 | rs9386147 | 1.093 | -0.795 | -0.311 | -0.399 | -0.074 | -0.075 | -0.075 | 1.243 | 0.16 | 0.02 | 0.17 | 0.3 | 0.27 | 0.28 |
| 6 | 146535519 | rs9322052 | 0.175 | -0.737 | 0.062 | -0.505 | 0.045 | 0.052 | 0.052 | 0 | 0.06 | 0.33 | 0.05 | 0.06 | 0.04 | 0.03 |
| 6 | 146543192 | rs2056994 | 0.175 | -0.737 | 0.062 | -0.505 | 0.045 | 0.052 | 0.052 | 0 | 0.06 | 0.33 | 0.05 | 0.05 | 0.04 | 0.03 |
| 6 | 146551363 | rs9485058 | -1.017 | -0.76 | 0.111 | -0.501 | -0.016 | 0.056 | 0.056 | 1.574 | 0.1 | 0.67 | 0.2 | 0.13 | 0.06 | 0.06 |
| 6 | 146554978 | rs12210031 | 0.464 | -0.946 | -0.035 | -0.67 | 0.029 | 0.099 | 0.099 | 0 | 0.23 | 0.09 | 0.09 | 0 | 0.15 | 0.22 |
| 6 | 146559200 | rs2223587 | -0.97 | -0.884 | -0.017 | -0.615 | -0.018 | 0.049 | 0.049 | 1.574 | 0.1 | 0.68 | 0.2 | 0.13 | 0.06 | 0.06 |
| 6 | 146563922 | rs4272232 | -2.065 | -1.117 | -0.676 | -0.516 | -0.16 | -0.175 | -0.175 | 1.38 | 0.69 | 0.99 | 0.53 | 0.54 | 0.57 | 0.64 |
| 6 | 146571933 | rs6900247 | 0.09 | -0.554 | 0.065 | -0.127 | 0.05 | 0.058 | 0.058 | 0 | 0.06 | 0.33 | 0.06 | 0.05 | 0.04 | 0.03 |
| 6 | 146578751 | rs11155459 | 0.472 | -0.602 | -1.093 | -0.143 | -0.079 | -0.095 | -0.095 | 0 | 0.18 | 0.02 | 0.19 | 0.33 | 0.29 | 0.23 |
| 6 | 146615383 | rs12200797 | 0.114 | -0.738 | -1.363 | -0.388 | 0.02 | 0.103 | 0.103 | 0.336 | 0.22 | 0.07 | 0.08 | 0 | 0.15 | 0.22 |
| 6 | 146620665 | rs1555084 | -0.933 | -0.679 | -1.343 | -0.347 | 0.052 | 0.076 | 0.076 | 2.674 | 0.33 | 0.85 | 0.3 | 0.13 | 0.21 | 0.28 |
| 6 | 146633818 | rs362962 | -0.916 | -0.662 | -1.335 | -0.338 | 0.051 | 0.075 | 0.075 | 2.548 | 0.33 | 0.85 | 0.3 | 0.13 | 0.21 | 0.28 |
| 6 | 146652354 | rs362949 | -1.116 | 0.04 | -0.494 | 0.633 | -0.021 | 0.033 | 0.033 | 1.562 | 0.1 | 0.72 | 0.21 | 0.13 | 0.06 | 0.06 |
| 6 | 146665424 | rs10872587 | -0.367 | 0.466 | -1.405 | 0.631 | -0.082 | -0.093 | -0.093 | 1.571 | 0.25 | 0.04 | 0.22 | 0.38 | 0.35 | 0.2 |
| 6 | 146665616 | rs1983634 | -0.576 | -0.32 | -1.103 | -0.443 | 0.004 | -0.011 | -0.011 | 0 | 0.39 | 0.43 | 0.21 | 0.11 | 0.26 | 0.3 |
| 6 | 146665672 | rs1983635 | -0.254 | 0.152 | -1.774 | -0.481 | -0.044 | 0.146 | 0.146 | 0 | 0.32 | 0.03 | 0.22 | 0.09 | 0.28 | 0.31 |
| 6 | 146669045 | rs56566 | -0.342 | -0.085 | -0.822 | -0.094 | -0.019 | -0.054 | -0.054 | 0 | 0.34 | 0.42 | 0.23 | 0.17 | 0.24 | 0.31 |
| 6 | 146672103 | rs362871 | -1.324 | 0.16 | -1.031 | 0.498 | -0.002 | -0.001 | -0.001 | 0.01 | 0.66 | 0.66 | 0.46 | 0.66 | 0.54 | 0.67 |

| | | | | | | | | | | | | | | | | |
|---|---|---|---|---|---|---|---|---|---|---|---|---|---|---|---|---|
| 6 | 146672674 | rs2268661 | -0.52 | -0.035 | -0.829 | 1.186 | -0.012 | 0.01 | 0.01 | 0 | 0.24 | 0.07 | 0.17 | 0.23 | 0.24 | 0.29 |
| 6 | 146677281 | rs3819832 | -0.52 | 0.02 | -0.781 | 1.067 | -0.013 | 0.009 | 0.009 | 0 | 0.24 | 0.07 | 0.17 | 0.23 | 0.24 | 0.29 |
| 6 | 146679735 | rs362839 | -0.713 | -0.186 | 0.135 | 1.47 | -0.016 | 0.055 | 0.055 | 0 | 0.25 | 0.41 | 0.27 | 0.34 | 0.22 | 0.24 |
| 6 | 146679840 | rs362840 | -0.159 | -0.26 | 0.463 | 1.212 | -0.006 | -0.024 | -0.024 | 0.269 | 0.19 | 0.24 | 0.13 | 0.1 | 0.17 | 0.13 |
| 6 | 146680449 | rs362842 | -1.218 | -0.026 | 0.88 | 1.523 | 0.008 | 0.109 | 0.109 | 3.322 | 0.11 | 0.49 | 0.09 | 0.16 | 0.08 | 0.11 |
| 6 | 146680907 | rs362845 | -1.344 | -0.103 | 0.768 | 1.653 | -0.019 | 0.092 | 0.092 | 0.829 | 0.15 | 0.4 | 0.11 | 0.29 | 0.12 | 0.17 |
| 6 | 146690260 | rs9485078 | -0.948 | -0.665 | 0.041 | 1.312 | -0.001 | 0 | 0 | 0 | 0.55 | 0.54 | 0.33 | 0.56 | 0.41 | 0.41 |
| 6 | 146698225 | rs6930314 | -0.894 | -0.465 | 0.074 | 0.975 | -0.012 | 0.067 | 0.067 | 0.065 | 0.14 | 0.28 | 0.1 | 0.29 | 0.12 | 0.17 |
| 6 | 146700087 | rs12195183 | 0.186 | -0.443 | -0.119 | 0.973 | 0.021 | 0.125 | 0.125 | 1.139 | 0.35 | 0.09 | 0.21 | 0.11 | 0.29 | 0.18 |
| 6 | 146712761 | rs362852 | -0.465 | -0.187 | -0.008 | 1.141 | -0.001 | -0.001 | -0.001 | 0 | 0.5 | 0.49 | 0.32 | 0.4 | 0.41 | 0.36 |
| 6 | 146719883 | rs362856 | -1.287 | -0.71 | 0.886 | 1.404 | -0.028 | 0.235 | 0.235 | 1.693 | 0.12 | 0.46 | 0.07 | 0.27 | 0.05 | 0.08 |
| 6 | 146720966 | rs2235875 | -1.797 | -0.894 | 1.133 | 1.313 | -0.053 | 0.073 | 0.073 | 4.656 | 0.15 | 0.65 | 0.09 | 0.27 | 0.05 | 0.08 |
| 6 | 146721766 | rs9399577 | -0.454 | -0.871 | 0.944 | 1.271 | -0.003 | 0.022 | 0.022 | 0.47 | 0.08 | 0.05 | 0.04 | 0.2 | 0.04 | 0.07 |
| 6 | 146726569 | rs7755078 | 0.134 | -0.775 | 0.39 | 0.904 | -0.067 | 0.182 | 0.182 | 0.36 | 0.38 | 0.05 | 0.28 | 0.12 | 0.38 | 0.23 |
| 6 | 146729216 | rs1033534 | -1.884 | -0.822 | 0.117 | 1.099 | -0.169 | -0.012 | -0.012 | 0 | 0.72 | 0.98 | 0.51 | 0.81 | 0.57 | 0.72 |
| 6 | 146733790 | rs3804300 | -1.584 | -0.607 | 0.257 | 0.543 | -0.105 | -0.093 | -0.093 | 0 | 0.67 | 0.95 | 0.49 | 0.6 | 0.55 | 0.66 |
| 6 | 146739590 | rs12190176 | 0.296 | -0.61 | 0.226 | 0.412 | -0.073 | 0.181 | 0.181 | 0.33 | 0.37 | 0.05 | 0.28 | 0.1 | 0.38 | 0.23 |
| 6 | 146743903 | rs2024589 | -0.662 | -0.392 | 0.883 | 0.95 | -0.002 | 0.005 | 0.005 | 0 | 0.2 | 0.24 | 0.13 | 0.42 | 0.12 | 0.4 |
| 6 | 146748607 | rs9386151 | -0.9 | -0.039 | 1.005 | 0.814 | -0.001 | 0 | 0 | 0 | 0.54 | 0.49 | 0.46 | 0.55 | 0.54 | 0.63 |
| 6 | 146752460 | rs726714 | -1.516 | -0.082 | 1.037 | 0.813 | -0.062 | -0.072 | -0.072 | 0 | 0.63 | 0.94 | 0.51 | 0.61 | 0.58 | 0.66 |
| 6 | 146754127 | rs362817 | -0.86 | -0.068 | 1.297 | 0.818 | -0.002 | -0.012 | -0.012 | 0 | 0.56 | 0.72 | 0.47 | 0.55 | 0.55 | 0.63 |
| 6 | 146755140 | rs2942 | -0.882 | -0.098 | 1.266 | 0.808 | -0.001 | 0 | 0 | 0 | 0.54 | 0.49 | 0.46 | 0.55 | 0.55 | 0.63 |
| 6 | 146757211 | rs3804293 | -1.267 | -0.084 | 1.177 | 0.811 | -0.01 | -0.015 | -0.015 | 0 | 0.61 | 0.8 | 0.5 | 0.61 | 0.58 | 0.67 |
| 6 | 146758542 | rs362822 | -0.958 | -0.065 | 0.844 | 0.785 | -0.001 | 0.003 | 0.003 | 0 | 0.55 | 0.51 | 0.46 | 0.55 | 0.55 | 0.64 |
| 6 | 146759364 | rs9376993 | -0.926 | -0.062 | 0.839 | 0.786 | -0.001 | 0 | 0 | 0 | 0.55 | 0.49 | 0.46 | 0.55 | 0.55 | 0.64 |
| 6 | 146766138 | rs13191599 | -0.986 | -0.527 | 0.798 | 1.007 | -0.007 | 0.005 | 0.005 | 0 | 0.43 | 0.47 | 0.35 | 0.54 | 0.35 | 0.59 |
| 6 | 146785984 | rs9403788 | -0.429 | -0.655 | 0.671 | 0.954 | -0.021 | -0.122 | -0.122 | 0 | 0.31 | 0.07 | 0.25 | 0.52 | 0.27 | 0.45 |
| 6 | 146794697 | rs846410 | -1.171 | -0.639 | 0.65 | 0.953 | -0.007 | 0.006 | 0.006 | 0 | 0.42 | 0.45 | 0.35 | 0.54 | 0.34 | 0.6 |
| 6 | 146797757 | rs17076083 | -0.077 | -0.468 | -0.049 | 0.972 | -0.01 | -0.041 | -0.041 | 0.105 | 0.11 | 0.23 | 0.08 | 0.04 | 0.07 | 0.15 |
| 6 | 146809655 | rs2025157 | -1.299 | -0.357 | -0.792 | 1.056 | 0.026 | -0.038 | -0.038 | 0 | 0.5 | 0.94 | 0.39 | 0.58 | 0.36 | 0.63 |
| 6 | 146816019 | rs846411 | -0.085 | -0.164 | -0.688 | 1.057 | -0.062 | -0.014 | -0.014 | 0 | 0.19 | 0.01 | 0.17 | 0.15 | 0.18 | 0.2 |
| 6 | 146823438 | rs4896869 | -0.4 | -0.383 | -0.49 | 1.187 | -0.03 | -0.021 | -0.021 | 0.54 | 0.11 | 0.02 | 0.13 | 0.19 | 0.13 | 0.19 |
| 6 | 146824515 | rs9322064 | 1.121 | -0.658 | -0.365 | 1.47 | -0.005 | -0.068 | -0.068 | 0 | 0.76 | 0.83 | 0.78 | 0.42 | 0.79 | 0.54 |
| 6 | 146826657 | rs1570282 | 2.072 | -1.046 | -0.045 | 1.445 | 0.055 | 0.068 | 0.068 | 0.012 | 0.53 | 0.19 | 0.54 | 0.26 | 0.69 | 0.34 |
| 6 | 146827281 | rs4896870 | -0.151 | -0.931 | 0.364 | 1.355 | -0.013 | -0.071 | -0.071 | 0.395 | 0.15 | 0.26 | 0.19 | 0 | 0.09 | 0.14 |
| 6 | 146828922 | rs1358902 | 1.126 | -0.757 | 0.2 | 1.639 | -0.001 | 0.012 | 0.012 | 0 | 0.74 | 0.67 | 0.77 | 0.38 | 0.79 | 0.52 |
| 6 | 146849283 | rs9497566 | -0.084 | -0.609 | -0.426 | 1.752 | -0.093 | -0.089 | -0.089 | 0.183 | 0.82 | 0.99 | 0.81 | 0.65 | 0.79 | 0.78 |
| 6 | 146849677 | rs719787 | 0.442 | -0.587 | 0.084 | 1.443 | -0.001 | -0.046 | -0.046 | 0.099 | 0.65 | 0.74 | 0.73 | 0.42 | 0.72 | 0.59 |
| 6 | 146855097 | rs11155474 | 1.312 | -0.77 | 0.075 | 1.351 | 0.042 | 0.047 | 0.047 | 1.67 | 0.52 | 0.05 | 0.52 | 0.28 | 0.68 | 0.43 |
| 6 | 146868048 | rs3804288 | 0.193 | -0.755 | 0.018 | 1.064 | 0 | -0.019 | -0.019 | 0.25 | 0.65 | 0.75 | 0.73 | 0.36 | 0.72 | 0.6 |
| 6 | 146872685 | rs11155475 | 0.948 | -0.29 | -0.323 | 0.585 | 0.017 | 0.121 | 0.121 | 1.463 | 0.52 | 0.04 | 0.52 | 0.22 | 0.68 | 0.43 |
| 6 | 146873614 | rs9377004 | 0.805 | -0.22 | -0.02 | 0.621 | 0.018 | 0.034 | 0.034 | 0.25 | 0.53 | 0.38 | 0.54 | 0.22 | 0.68 | 0.43 |
| 6 | 146900032 | rs9386159 | 0.989 | 0.268 | -1.103 | 0.287 | -0.085 | 0.25 | 0.25 | 0.157 | 0.47 | 0.05 | 0.53 | 0.08 | 0.53 | 0.32 |
| 6 | 146900616 | rs7775840 | 0.431 | 0.316 | -0.597 | 0.672 | 0.015 | 0.04 | 0.04 | 0 | 0.93 | 0.86 | 0.94 | 0.87 | 0.93 | 0.89 |
| 6 | 146906650 | rs9485099 | 0.098 | -0.007 | 0.171 | 0.989 | 0.001 | -0.023 | -0.023 | 0 | 0.31 | 0.36 | 0.25 | 0.29 | 0.3 | 0.31 |
| 6 | 146910111 | rs12662795 | -0.631 | -0.086 | 0.447 | 1.288 | 0.013 | -0.01 | -0.01 | 0.019 | 0.13 | 0.15 | 0.1 | 0.5 | 0.09 | 0.26 |
| 6 | 146911430 | rs9377007 | 0.682 | 0.051 | 0.475 | 1.234 | 0.026 | 0.034 | 0.034 | 0.033 | 0.92 | 0.85 | 0.93 | 0.87 | 0.92 | 0.88 |
| 6 | 146918950 | rs2275606 | -0.202 | 0.006 | 0.331 | 1.93 | 0.004 | 0.009 | 0.009 | 0 | 0.08 | 0.1 | 0.06 | 0.27 | 0.07 | 0.16 |
| 6 | 146919485 | rs9377009 | 1.802 | -0.087 | 0.562 | 1.712 | -0.026 | 0.042 | 0.042 | 0.093 | 0.77 | 0.45 | 0.77 | 0.37 | 0.82 | 0.63 |
| 6 | 146929483 | rs9497578 | -0.509 | 0.042 | 0.73 | 1.794 | -0.004 | 0.057 | 0.057 | 0 | 0.1 | 0.2 | 0.08 | 0.24 | 0.08 | 0.15 |
| 6 | 146929596 | rs1575376 | 1.504 | 0.134 | 0.228 | 1.358 | -0.06 | 0.193 | 0.193 | 2.4 | 0.45 | 0.07 | 0.51 | 0.1 | 0.46 | 0.31 |
| 6 | 146934383 | rs9390399 | 0.215 | 0.158 | 0.556 | 1.608 | 0.005 | 0.01 | 0.01 | 0 | 0.59 | 0.51 | 0.64 | 0.39 | 0.54 | 0.49 |
| 6 | 146940316 | rs9386161 | 0.163 | -0.584 | -0.055 | 0.956 | 0 | -0.001 | -0.001 | 0.237 | 0.51 | 0.48 | 0.58 | 0.4 | 0.57 | 0.48 |
| 6 | 146944056 | rs9497581 | 0.528 | -0.375 | -0.129 | 0.682 | -0.009 | -0.015 | -0.015 | 0 | 0.17 | 0.13 | 0.15 | 0.25 | 0.22 | 0.17 |
| 6 | 146944800 | rs7746671 | -0.046 | -0.377 | -0.191 | 0.682 | 0.003 | 0.017 | 0.017 | 3.727 | 0.22 | 0.42 | 0.19 | 0.25 | 0.22 | 0.18 |
| 6 | 146953108 | rs9390403 | 0.264 | -0.377 | -0.238 | 0.734 | 0.012 | 0.04 | 0.04 | 0.103 | 0.47 | 0.2 | 0.53 | 0.33 | 0.56 | 0.35 |
| 6 | 146953914 | rs17826169 | 0.126 | -0.308 | -0.578 | 0.742 | 0 | 0.019 | 0.019 | 0 | 0.49 | 0.37 | 0.48 | 0.28 | 0.46 | 0.44 |
| 6 | 146957941 | rs957547 | 0.654 | -0.286 | -1.335 | 0.22 | -0.039 | 0.159 | 0.159 | 0 | 0.39 | 0.1 | 0.39 | 0.09 | 0.37 | 0.31 |
| 6 | 146968702 | rs4895700 | 1.028 | 0.052 | -1.239 | 0.403 | -0.07 | 0.091 | 0.091 | 0 | 0.31 | 0.05 | 0.38 | 0.09 | 0.33 | 0.27 |
| 6 | 146971848 | rs7768539 | -0.421 | 0.331 | -0.515 | 0.473 | -0.012 | 0.02 | 0.02 | 0 | 0.75 | 0.85 | 0.69 | 0.73 | 0.74 | 0.72 |
| 6 | 146972068 | rs912000 | -0.141 | 0.186 | -0.373 | 0.568 | -0.002 | -0.02 | -0.02 | 0 | 0.7 | 0.75 | 0.66 | 0.53 | 0.68 | 0.62 |
| 6 | 146972237 | rs9390409 | 0.071 | 0.261 | -0.393 | 0.669 | -0.001 | 0 | 0 | 0.154 | 0.62 | 0.59 | 0.59 | 0.45 | 0.6 | 0.53 |
| 6 | 146973073 | rs9390411 | 0.075 | 0.072 | -0.763 | 0.459 | 0.003 | -0.009 | -0.009 | 0 | 0.75 | 0.74 | 0.66 | 0.71 | 0.73 | 0.82 |
| 6 | 146973453 | rs9322069 | 0 | 0.053 | -0.998 | 0.393 | 0.017 | -0.075 | -0.075 | 0 | 0.73 | 0.92 | 0.84 | 0.69 | 0.71 | 0.83 |
| 6 | 146981247 | rs973572 | 0.124 | 0.431 | -1.326 | -0.003 | -0.074 | -0.119 | -0.119 | 0.01 | 0.33 | 0.11 | 0.36 | 0.52 | 0.38 | 0.47 |
| 6 | 146986428 | rs9390412 | -0.016 | 0.297 | -0.871 | -0.072 | -0.008 | -0.022 | -0.022 | 0 | 0.36 | 0.35 | 0.39 | 0.54 | 0.38 | 0.48 |
| 6 | 146991045 | rs2226067 | -0.363 | 0.243 | -1.108 | -0.036 | 0.018 | -0.041 | -0.041 | 0 | 0.73 | 0.88 | 0.84 | 0.72 | 0.71 | 0.86 |
| 6 | 146991066 | rs9386162 | 0.524 | 0.641 | -0.949 | 0.43 | -0.021 | -0.104 | -0.104 | 0.683 | 0.15 | 0.01 | 0.12 | 0.38 | 0.15 | 0.22 |
| 6 | 146995026 | rs4895701 | 0.691 | 0.695 | -0.54 | -0.444 | -0.004 | -0.081 | -0.081 | 0 | 0.27 | 0.51 | 0.34 | 0.13 | 0.21 | 0.28 |

| | | | | | | | | | | | | | | | | |
|---|---|---|---|---|---|---|---|---|---|---|---|---|---|---|---|---|
| 6 | 146998463 | rs4576267 | 0.683 | 0.841 | -0.77 | -0.564 | -0.04 | 0.041 | 0.041 | 0.015 | 0.14 | 0.05 | 0.28 | 0.08 | 0.2 | 0.23 |
| 6 | 146998804 | rs6907896 | 0.475 | 0.367 | -0.27 | -0.222 | -0.001 | -0.11 | -0.11 | 0.526 | 0.27 | 0.5 | 0.35 | 0.09 | 0.21 | 0.26 |
| 6 | 147001409 | rs1417588 | 0.64 | 0.349 | -0.369 | -0.231 | 0.003 | -0.08 | -0.08 | 3.367 | 0.26 | 0.35 | 0.33 | 0.09 | 0.2 | 0.26 |
| 6 | 147002401 | rs9377017 | -0.314 | 0.406 | -0.831 | -0.105 | -0.031 | -0.039 | -0.039 | 0 | 0.65 | 0.97 | 0.83 | 0.75 | 0.64 | 0.82 |
| 6 | 147006845 | rs12195808 | 0.163 | 0.592 | -0.576 | 0.026 | -0.001 | -0.012 | -0.012 | 0.268 | 0.29 | 0.65 | 0.36 | 0.13 | 0.21 | 0.29 |
| 6 | 147013320 | rs9497607 | 0.39 | 0.78 | -0.894 | 0.105 | 0.007 | -0.075 | -0.075 | 0.14 | 0.27 | 0.43 | 0.34 | 0.12 | 0.23 | 0.28 |
| 6 | 147015780 | rs4129313 | 0.166 | 1.133 | -1.5 | 0.824 | -0.037 | -0.027 | -0.027 | 0 | 0.61 | 0.81 | 0.56 | 0.38 | 0.54 | 0.45 |
| 6 | 147019755 | rs6905055 | -0.609 | 1.283 | -1.233 | 1.193 | -0.011 | -0.03 | -0.03 | 0 | 0.3 | 0.15 | 0.34 | 0.46 | 0.38 | 0.32 |
| 6 | 147034592 | rs2225136 | -0.656 | 1.496 | -1.726 | 1.288 | -0.094 | -0.05 | -0.05 | 0.009 | 0.21 | 0.01 | 0.16 | 0.2 | 0.26 | 0.21 |
| 6 | 147035035 | rs6901516 | -0.376 | 1.596 | -1.134 | 1.818 | 0.033 | 0.103 | 0.103 | 0.318 | 0.13 | 0.32 | 0.17 | 0.32 | 0.19 | 0.17 |
| 6 | 147039724 | rs9497613 | -1.345 | 0.845 | -0.455 | 1.297 | -0.02 | -0.01 | -0.01 | 0.951 | 0.44 | 0.71 | 0.37 | 0.58 | 0.49 | 0.46 |
| 6 | 147039821 | rs9497614 | -0.895 | 0.749 | 0.006 | 1.339 | -0.006 | -0.008 | -0.008 | 0.716 | 0.35 | 0.32 | 0.32 | 0.51 | 0.45 | 0.38 |
| 6 | 147041059 | rs913308 | -0.831 | 0.209 | -0.232 | 0.902 | -0.012 | -0.007 | -0.007 | 0 | 0.43 | 0.35 | 0.36 | 0.47 | 0.49 | 0.42 |
| 6 | 147041608 | rs4896885 | -0.943 | 0.133 | -0.83 | 0.162 | -0.064 | 0.014 | 0.014 | 0 | 0.82 | 0.95 | 0.8 | 0.87 | 0.73 | 0.88 |
| 6 | 147047341 | rs3747754 | -0.057 | -0.203 | -0.578 | 0.347 | 0.003 | 0.004 | 0.004 | 0 | 0.28 | 0.08 | 0.16 | 0.2 | 0.26 | 0.21 |
| 6 | 147048743 | rs1327670 | -0.9 | -0.166 | -0.076 | 0.558 | 0.031 | 0.039 | 0.039 | 0.985 | 0.46 | 0.72 | 0.45 | 0.63 | 0.43 | 0.44 |
| 6 | 147052588 | rs11155481 | -1.293 | -0.285 | -0.176 | 0.345 | -0.048 | 0.033 | 0.033 | 0.653 | 0.56 | 0.86 | 0.5 | 0.64 | 0.48 | 0.5 |
| 6 | 147053632 | rs4896886 | 0.308 | -0.139 | 0.263 | 0.459 | 0.012 | 0.113 | 0.113 | 0 | 0.13 | 0.19 | 0.25 | 0.43 | 0.16 | 0.23 |
| 6 | 147062220 | rs1327666 | -0.802 | -0.148 | 0.07 | 0.234 | 0.024 | 0.018 | 0.018 | 1.571 | 0.44 | 0.69 | 0.45 | 0.61 | 0.43 | 0.44 |
| 6 | 147066010 | rs9322073 | -0.72 | -0.153 | -0.159 | 0.224 | 0.031 | 0.028 | 0.028 | 1.77 | 0.45 | 0.76 | 0.34 | 0.61 | 0.43 | 0.44 |
| 6 | 147091074 | rs17535206 | 0.171 | -0.093 | -0.277 | 0.219 | -0.002 | 0.027 | 0.027 | 0.949 | 0.1 | 0.07 | 0.03 | 0 | 0.05 | 0.06 |
| 6 | 147094975 | rs259417 | 1.194 | -0.115 | -0.011 | 0.233 | -0.005 | 0.033 | 0.033 | 0.138 | 0.46 | 0.23 | 0.63 | 0.38 | 0.53 | 0.5 |
| 6 | 147098991 | rs4896888 | 1.179 | 0.085 | 0.094 | 0.099 | -0.003 | 0.003 | 0.003 | 0 | 0.44 | 0.42 | 0.63 | 0.4 | 0.46 | 0.48 |
| 6 | 147103416 | rs259405 | 0.087 | 0.197 | -0.331 | 0.219 | -0.013 | -0.042 | -0.042 | 0 | 0.8 | 0.85 | 0.85 | 0.61 | 0.79 | 0.78 |
| 6 | 147104187 | rs259404 | 0.051 | 0.286 | -0.205 | -0.369 | 0.025 | -0.033 | -0.033 | 0.038 | 0.06 | 0.14 | 0.04 | 0.01 | 0.09 | 0.09 |
| 6 | 147106841 | rs259391 | 0.476 | 0.222 | -0.967 | -0.282 | 0.011 | 0.013 | 0.013 | 0 | 0.52 | 0.55 | 0.69 | 0.41 | 0.57 | 0.56 |
| 6 | 147109823 | rs259395 | -0.566 | 1.01 | -0.91 | -0.105 | 0.012 | 0.036 | 0.036 | 0.454 | 0.29 | 0.42 | 0.26 | 0.6 | 0.36 | 0.33 |
| 6 | 147115291 | rs259401 | 0.322 | 0.779 | -1.95 | -0.311 | -0.029 | -0.047 | -0.047 | 0.61 | 0.67 | 0.83 | 0.73 | 0.45 | 0.62 | 0.55 |
| 6 | 147121763 | rs638144 | -0.199 | 0.691 | -1.974 | -0.517 | -0.113 | -0.008 | -0.008 | 0.084 | 0.68 | 0.97 | 0.69 | 0.71 | 0.68 | 0.64 |
| 6 | 147126145 | rs259387 | -0.073 | 0.531 | -0.575 | -0.943 | 0.007 | -0.062 | -0.062 | 0 | 0.21 | 0.31 | 0.14 | 0.04 | 0.24 | 0.16 |
| 6 | 147126874 | rs9390422 | 0.256 | 0.493 | -1.134 | -1.106 | -0.037 | -0.032 | -0.032 | 0 | 0.24 | 0.18 | 0.47 | 0.69 | 0.35 | 0.41 |
| 6 | 147134202 | rs259371 | -0.982 | -0.533 | -1.646 | -1.622 | -0.136 | 0.084 | 0.084 | 1.363 | 0.66 | 0.98 | 0.68 | 0.82 | 0.65 | 0.66 |
| 6 | 147144142 | rs9377031 | 0.403 | -0.44 | -0.713 | -1.595 | 0.013 | 0.006 | 0.006 | 0.969 | 0.25 | 0.33 | 0.45 | 0.69 | 0.36 | 0.42 |
| 6 | 147144192 | rs259368 | 0.906 | -0.663 | -0.571 | -1.715 | -0.002 | 0.054 | 0.054 | 0 | 0.18 | 0.06 | 0.09 | 0.04 | 0.19 | 0.11 |
| 6 | 147158227 | rs1924695 | -1.095 | -0.694 | 0.155 | -1.74 | -0.035 | 0.064 | 0.064 | 1.946 | 0.65 | 0.85 | 0.64 | 0.82 | 0.64 | 0.65 |
| 6 | 147162249 | rs1324681 | 0.47 | -0.456 | 0.262 | -1.521 | -0.048 | -0.035 | -0.035 | 0 | 0.24 | 0.17 | 0.45 | 0.68 | 0.36 | 0.43 |
| 6 | 147178335 | rs9497651 | 0.254 | -0.696 | 0.897 | -1.619 | -0.028 | -0.111 | -0.111 | 0 | 0.22 | 0.54 | 0.13 | 0.04 | 0.19 | 0.11 |
| 6 | 147179022 | rs9399586 | 0.587 | -0.571 | 0.383 | -1.623 | -0.087 | -0.065 | -0.065 | 0 | 0.22 | 0.06 | 0.44 | 0.69 | 0.36 | 0.46 |
| 6 | 147182339 | rs9322079 | -0.32 | -0.741 | 0.645 | -1.645 | 0.001 | -0.015 | -0.015 | 0 | 0.46 | 0.48 | 0.56 | 0.74 | 0.56 | 0.57 |
| 6 | 147185851 | rs2274170 | -0.286 | -0.651 | 0.327 | -1.691 | 0.015 | -0.021 | -0.021 | 0 | 0.4 | 0.51 | 0.54 | 0.71 | 0.51 | 0.5 |
| 6 | 147187237 | rs6937015 | -1.251 | -0.865 | -0.365 | -1.915 | -0.12 | 0.081 | 0.081 | 0.912 | 0.65 | 0.98 | 0.67 | 0.85 | 0.65 | 0.67 |
| 6 | 147188206 | rs1924694 | -0.438 | -0.949 | -0.257 | -2 | 0.07 | 0.109 | 0.109 | 0.919 | 0.43 | 0.74 | 0.22 | 0.11 | 0.3 | 0.21 |
| 6 | 147189967 | rs9390434 | -0.193 | -0.935 | 0.096 | -2.053 | -0.003 | 0.005 | 0.005 | 0.468 | 0.39 | 0.41 | 0.53 | 0.75 | 0.45 | 0.46 |
| 6 | 147195422 | rs12215283 | 1.152 | -0.926 | 0.118 | -2.047 | -0.002 | -0.017 | -0.017 | 0 | 0.17 | 0.16 | 0.1 | 0.03 | 0.18 | 0.11 |
| 6 | 147200786 | rs1324684 | -0.431 | -0.967 | 0.124 | -2.054 | 0.002 | -0.025 | -0.025 | 0 | 0.43 | 0.58 | 0.54 | 0.8 | 0.47 | 0.5 |
| 6 | 147202796 | rs1122570 | -0.641 | -0.946 | 0.086 | -2.043 | 0.005 | 0.058 | 0.058 | 0 | 0.56 | 0.66 | 0.63 | 0.78 | 0.63 | 0.57 |
| 6 | 147203404 | rs878617 | 1.052 | -0.937 | 0.015 | -2.035 | -0.002 | -0.011 | -0.011 | 0 | 0.18 | 0.18 | 0.1 | 0.09 | 0.19 | 0.11 |
| 6 | 147210677 | rs1022877 | -0.435 | -0.497 | 0.248 | -1.725 | 0.005 | -0.017 | -0.017 | 0 | 0.39 | 0.46 | 0.54 | 0.79 | 0.46 | 0.47 |
| 6 | 147216278 | rs927672 | 0.223 | 0.086 | -0.457 | -1.519 | -0.001 | -0.002 | -0.002 | 0 | 0.7 | 0.71 | 0.66 | 0.83 | 0.66 | 0.6 |
| 6 | 147229457 | rs17076418 | -1.255 | 0.223 | -0.434 | -0.902 | -0.03 | -0.042 | -0.042 | 0.031 | 0.08 | 0.27 | 0.03 | 0.01 | 0.03 | 0.1 |
| 6 | 147238475 | rs1998641 | -0.295 | 0.545 | -0.867 | -0.625 | -0.024 | -0.02 | -0.02 | 0.352 | 0.25 | 0.17 | 0.46 | 0.67 | 0.39 | 0.49 |
| 6 | 147243237 | rs9390441 | 0.257 | 0.547 | -1.051 | -1.074 | -0.009 | 0.04 | 0.04 | 0 | 0.69 | 0.8 | 0.78 | 0.9 | 0.68 | 0.64 |
| 6 | 147243419 | rs4896896 | 0.26 | 0.355 | -2.142 | -0.064 | -0.013 | -0.01 | -0.01 | 0.029 | 0.53 | 0.12 | 0.45 | 0.45 | 0.57 | 0.51 |
| 6 | 147250951 | rs9322081 | 0.048 | -0.315 | -0.82 | 0.27 | 0 | -0.002 | -0.002 | 0.044 | 0.46 | 0.45 | 0.54 | 0.6 | 0.51 | 0.49 |
| 6 | 147259868 | rs11155494 | -0.458 | -0.393 | -0.604 | 0.613 | 0.007 | 0.01 | 0.01 | 0 | 0.5 | 0.57 | 0.56 | 0.48 | 0.55 | 0.45 |
| 6 | 147268277 | rs7742400 | 0.323 | -0.439 | -0.291 | 0.562 | -0.005 | 0 | 0 | 0 | 0.42 | 0.38 | 0.53 | 0.4 | 0.51 | 0.39 |
| 6 | 147271295 | rs9403837 | 0.186 | -0.542 | -0.384 | 0.528 | -0.062 | -0.036 | -0.036 | 0 | 0.43 | 0.15 | 0.5 | 0.48 | 0.53 | 0.42 |
| 6 | 147276458 | rs4452660 | -0.5 | -0.531 | -0.34 | 0.612 | 0.004 | 0.015 | 0.015 | 0 | 0.5 | 0.53 | 0.57 | 0.68 | 0.55 | 0.54 |
| 6 | 147288120 | rs4143743 | -0.567 | -0.46 | -0.385 | 0.667 | 0.013 | -0.031 | -0.031 | 0.004 | 0.08 | 0.03 | 0.04 | 0.28 | 0.06 | 0.15 |
| 6 | 147289349 | rs1529013 | -0.774 | -0.545 | -0.371 | 0.567 | 0.002 | 0.026 | 0.026 | 0 | 0.56 | 0.64 | 0.6 | 0.8 | 0.59 | 0.65 |
| 6 | 147296394 | rs1112833 | -0.717 | -0.721 | -0.003 | 0.309 | -0.02 | 0.006 | 0.006 | 0.009 | 0.53 | 0.78 | 0.45 | 0.61 | 0.54 | 0.53 |
| 6 | 147301862 | rs9485136 | 0.626 | -0.796 | 0.041 | 0.278 | -0.059 | -0.054 | -0.054 | 0.146 | 0.37 | 0.12 | 0.34 | 0.45 | 0.43 | 0.39 |
| 6 | 147302374 | rs7775644 | -0.135 | -0.758 | 0.011 | 0.31 | -0.001 | -0.004 | -0.004 | 0 | 0.46 | 0.64 | 0.4 | 0.53 | 0.48 | 0.43 |
| 6 | 147306600 | rs17076524 | -1.164 | -0.746 | 0.017 | 0.301 | 0.001 | -0.01 | -0.01 | 0 | 0.09 | 0.08 | 0.06 | 0.14 | 0.1 | 0.12 |
| 6 | 147311453 | rs10457807 | 0.155 | -0.739 | 0.073 | 0.317 | -0.001 | -0.001 | -0.001 | 0 | 0.42 | 0.57 | 0.38 | 0.45 | 0.44 | 0.39 |
| 6 | 147326634 | rs1406537 | -0.266 | -0.779 | -0.051 | 0.223 | 0.002 | -0.012 | -0.012 | 0 | 0.47 | 0.72 | 0.42 | 0.53 | 0.48 | 0.47 |
| 6 | 147332567 | rs9399593 | -0.53 | -0.769 | -0.141 | 0.218 | -0.008 | 0.011 | 0.011 | 0.006 | 0.51 | 0.77 | 0.44 | 0.61 | 0.52 | 0.52 |
| 6 | 147351302 | rs1529017 | 0.406 | -0.71 | 0.559 | 0.393 | -0.057 | -0.063 | -0.063 | 0 | 0.31 | 0.14 | 0.32 | 0.45 | 0.4 | 0.42 |
| 6 | 147354068 | rs6570785 | -0.961 | -0.807 | 0.631 | 0.263 | -0.004 | -0.004 | -0.004 | 5.28 | 0.49 | 0.76 | 0.43 | 0.54 | 0.48 | 0.52 |

| | | | | | | | | | | | | | | | | |
|---|---|---|---|---|---|---|---|---|---|---|---|---|---|---|---|---|
| 6 | 147354092 | rs6570786 | 0.113 | -0.813 | 0.694 | 0.26 | 0.014 | 0.022 | 0.022 | 0 | 0.36 | 0.5 | 0.36 | 0.45 | 0.4 | 0.43 |
| 6 | 147357238 | rs6570787 | -1.103 | -0.824 | 0.512 | 0.208 | -0.011 | 0.014 | 0.014 | 0 | 0.52 | 0.78 | 0.45 | 0.62 | 0.52 | 0.56 |
| 6 | 147357913 | rs17076578 | -1.253 | -0.646 | 0.628 | 0.49 | -0.003 | -0.003 | -0.003 | 0 | 0.12 | 0.14 | 0.06 | 0.08 | 0.08 | 0.08 |
| 6 | 147363661 | rs4143744 | 0.042 | -0.758 | 0.618 | 0.477 | -0.031 | -0.054 | -0.054 | 0.039 | 0.38 | 0.19 | 0.39 | 0.55 | 0.49 | 0.52 |
| 6 | 147366538 | rs11969474 | -1.176 | -0.537 | 1.16 | 0.871 | 0.004 | 0.072 | 0.072 | 0.538 | 0.06 | 0.22 | 0.05 | 0.14 | 0.06 | 0.09 |
| 6 | 147368934 | rs6907758 | -1.353 | -0.559 | 1.178 | 0.852 | -0.022 | 0.02 | 0.02 | 0.159 | 0.11 | 0.3 | 0.07 | 0.14 | 0.07 | 0.1 |
| 6 | 147373917 | rs9403843 | 0.305 | -0.711 | 0.865 | 0.783 | -0.042 | -0.045 | -0.045 | 1.886 | 0.43 | 0.17 | 0.31 | 0.51 | 0.49 | 0.51 |
| 6 | 147394522 | rs6910250 | -0.283 | -0.558 | 1.386 | 0.887 | -0.001 | 0 | 0 | 0 | 0.55 | 0.49 | 0.38 | 0.65 | 0.57 | 0.62 |
| 6 | 147406342 | rs9386178 | -0.564 | -0.577 | 1.338 | 0.706 | -0.028 | 0.032 | 0.032 | 0 | 0.59 | 0.74 | 0.42 | 0.71 | 0.56 | 0.64 |
| 6 | 147408639 | rs9322090 | 0.187 | -0.641 | 1.154 | 0.726 | -0.039 | 0.002 | 0.002 | 0 | 0.48 | 0.17 | 0.33 | 0.58 | 0.51 | 0.56 |
| 6 | 147409180 | rs12664682 | -0.98 | -0.526 | 1.597 | 0.772 | -0.016 | 0.044 | 0.044 | 0.045 | 0.07 | 0.2 | 0.05 | 0.13 | 0.05 | 0.09 |
| 6 | 147409235 | rs12661753 | -1.452 | -0.527 | 1.6 | 0.771 | -0.012 | 0.068 | 0.068 | 0.369 | 0.08 | 0.29 | 0.05 | 0.13 | 0.05 | 0.09 |
| 6 | 147414032 | rs7771711 | -0.771 | -0.51 | 0.908 | 0.78 | -0.015 | 0.026 | 0.026 | 0 | 0.06 | 0.13 | 0.04 | 0.13 | 0.05 | 0.09 |
| 6 | 147435953 | rs9485144 | -1.576 | -0.522 | 0.484 | 0.767 | -0.046 | 0.018 | 0.018 | 5.534 | 0.12 | 0.67 | 0.09 | 0.14 | 0.05 | 0.09 |
| 6 | 147463439 | rs9485147 | -1.128 | -0.374 | 0.445 | 0.722 | -0.018 | 0.049 | 0.049 | 2.338 | 0.1 | 0.51 | 0.08 | 0.12 | 0.06 | 0.09 |
| 6 | 147471730 | rs2256216 | -0.386 | 0.038 | -0.488 | 0.615 | -0.03 | 0.02 | 0.02 | 0 | 0.59 | 0.79 | 0.4 | 0.7 | 0.56 | 0.66 |
| 6 | 147486533 | rs649544 | -0.682 | 0.365 | 0.118 | 0.737 | 0.006 | 0.07 | 0.07 | 0.717 | 0.53 | 0.72 | 0.54 | 0.76 | 0.58 | 0.55 |
| 6 | 147490398 | rs1660690 | -0.162 | 0.431 | 0.345 | 1.438 | -0.03 | -0.008 | -0.008 | 0.458 | 0.39 | 0.25 | 0.37 | 0.44 | 0.5 | 0.46 |
| 6 | 147492742 | rs693573 | -0.191 | 0.404 | 0.341 | 1.467 | -0.041 | -0.014 | -0.014 | 0 | 0.39 | 0.22 | 0.36 | 0.44 | 0.49 | 0.45 |
| 6 | 147500484 | rs694625 | -0.319 | 0.388 | 0.644 | 1.472 | -0.014 | -0.003 | -0.003 | 0 | 0.43 | 0.31 | 0.4 | 0.47 | 0.56 | 0.51 |
| 6 | 147501129 | rs2818868 | -0.423 | 0.28 | 0.976 | 1.444 | -0.014 | 0.052 | 0.052 | 0.027 | 0.44 | 0.32 | 0.47 | 0.77 | 0.53 | 0.48 |
| 6 | 147501449 | rs1112062 | 0.525 | 0.651 | 0.472 | 1.899 | -0.103 | -0.143 | -0.143 | 0 | 0.2 | 0.02 | 0.25 | 0.38 | 0.31 | 0.24 |
| 6 | 147503799 | rs598549 | -1.091 | 0.327 | 1.161 | 1.543 | 0 | 0 | 0 | 0.86 | 0.5 | 0.56 | 0.52 | 0.72 | 0.6 | 0.49 |
| 6 | 147511721 | rs485434 | -0.3 | 0.207 | 1.121 | 1.638 | -0.05 | 0.011 | 0.011 | 0.054 | 0.4 | 0.19 | 0.38 | 0.43 | 0.54 | 0.42 |
| 6 | 147535305 | rs1221638 | -0.908 | 0.167 | 1.736 | 1.514 | -0.001 | 0.021 | 0.021 | 0 | 0.47 | 0.64 | 0.44 | 0.7 | 0.49 | 0.49 |
| 6 | 147539165 | rs1765028 | -0.53 | 0.081 | 1.568 | 1.553 | -0.028 | -0.042 | -0.042 | 0 | 0.42 | 0.22 | 0.38 | 0.54 | 0.47 | 0.46 |
| 6 | 147549602 | rs657155 | -0.668 | -0.114 | 1.714 | 1.502 | -0.001 | -0.001 | -0.001 | 0 | 0.45 | 0.55 | 0.43 | 0.43 | 0.48 | 0.43 |
| 6 | 147566285 | rs2119490 | -0.217 | -0.085 | 1.713 | 1.514 | -0.067 | -0.027 | -0.027 | 0.183 | 0.38 | 0.06 | 0.33 | 0.42 | 0.43 | 0.4 |
| 6 | 147570723 | rs608147 | -0.373 | -0.057 | 1.706 | 1.516 | -0.017 | 0.008 | 0.008 | 0 | 0.4 | 0.19 | 0.34 | 0.43 | 0.43 | 0.4 |
| 6 | 147620401 | rs2850177 | -0.643 | 0.107 | 1.668 | 1.207 | 0 | 0.006 | 0.006 | 0 | 0.43 | 0.43 | 0.4 | 0.7 | 0.44 | 0.49 |
| 6 | 147635547 | rs1039085 | 0.453 | 0.015 | 1.662 | 1.207 | 0 | 0.007 | 0.007 | 0 | 0.57 | 0.57 | 0.6 | 0.31 | 0.57 | 0.57 |
| 6 | 147639534 | rs9373522 | 1.096 | 0.014 | 1.182 | 1.056 | -0.001 | 0.089 | 0.089 | 2.414 | 0.48 | 0.22 | 0.49 | 0.29 | 0.49 | 0.45 |
| 6 | 147670086 | rs12191720 | 1.089 | 0.047 | 0.608 | 1.118 | -0.009 | 0.14 | 0.14 | 2.422 | 0.5 | 0.11 | 0.48 | 0.28 | 0.5 | 0.52 |
| 6 | 147680359 | rs9390459 | 0.619 | 0.455 | 0.457 | 1.426 | -0.001 | 0.001 | 0.001 | 0.005 | 0.58 | 0.57 | 0.56 | 0.29 | 0.54 | 0.56 |
| 6 | 147685480 | rs6924763 | -0.464 | 0.543 | 0.331 | 1.49 | -0.015 | -0.004 | -0.004 | 0 | 0.85 | 0.98 | 0.89 | 0.82 | 0.82 | 0.87 |
| 6 | 147699123 | rs2328813 | 0.18 | 0.363 | 0.663 | 1.434 | -0.002 | 0.031 | 0.031 | 0.031 | 0.58 | 0.66 | 0.6 | 0.29 | 0.56 | 0.53 |
| 6 | 147701133 | rs9373523 | -0.116 | 0.382 | 0.575 | 1.41 | -0.016 | 0.005 | 0.005 | 0.026 | 0.62 | 0.76 | 0.66 | 0.58 | 0.57 | 0.61 |
| 6 | 147709180 | rs7739314 | 0.262 | 0.513 | 0.26 | 0.988 | -0.001 | -0.014 | -0.014 | 0 | 0.56 | 0.51 | 0.55 | 0.29 | 0.52 | 0.55 |
| 6 | 147723577 | rs582437 | -0.255 | -0.312 | -0.511 | 1.155 | 0.05 | 0.041 | 0.041 | 0 | 0.54 | 0.88 | 0.51 | 0.56 | 0.4 | 0.47 |
| 6 | 147730324 | rs593289 | 0.142 | -0.681 | 0.376 | 0.859 | 0.002 | 0.002 | 0.002 | 1.76 | 0.44 | 0.41 | 0.41 | 0.27 | 0.31 | 0.42 |
| 6 | 147738146 | rs642527 | 0.079 | -0.613 | 0.48 | 0.874 | -0.013 | -0.013 | -0.013 | 0 | 0.47 | 0.49 | 0.43 | 0.27 | 0.33 | 0.43 |
| 6 | 147740456 | rs547161 | 1.515 | -0.283 | 1.26 | 1.571 | 0.022 | 0.292 | 0.292 | 1.82 | 0.92 | 0.61 | 0.86 | 0.75 | 0.94 | 0.87 |
| 6 | 147748240 | rs478495 | 1.276 | -0.778 | 1.153 | 1.253 | -0.047 | 0.121 | 0.121 | 2.312 | 0.83 | 0.52 | 0.81 | 0.75 | 0.89 | 0.84 |
| 6 | 147759111 | rs377609 | 1.566 | -0.7 | 1.185 | 1.313 | 0.025 | 0.318 | 0.318 | 5.146 | 0.91 | 0.55 | 0.85 | 0.76 | 0.94 | 0.87 |
| 6 | 147764267 | rs325105 | 0.791 | -0.509 | -0.273 | 1.121 | -0.171 | -0.061 | -0.061 | 0.158 | 0.28 | 0.02 | 0.29 | 0.32 | 0.42 | 0.28 |
| 6 | 147767235 | rs485486 | -0.532 | -0.819 | -0.086 | 1.221 | 0 | -0.029 | -0.029 | 0 | 0.78 | 0.88 | 0.72 | 0.74 | 0.79 | 0.71 |
| 6 | 147780366 | rs10447451 | 0.436 | -1.054 | -0.385 | 0.912 | -0.014 | 0.016 | 0.016 | 0 | 0.22 | 0.05 | 0.2 | 0.18 | 0.23 | 0.2 |
| 6 | 147790322 | rs635513 | 0.15 | -1.161 | 0.241 | 1.106 | 0.006 | 0 | 0 | 0 | 0.09 | 0.06 | 0.05 | 0 | 0.06 | 0.04 |
| 6 | 147799122 | rs587008 | 0.917 | -1.303 | 0.797 | 1.249 | 0.039 | -0.021 | -0.021 | 0.271 | 0.6 | 0.24 | 0.61 | 0.52 | 0.73 | 0.62 |
| 6 | 147813016 | rs389674 | 0.539 | -1.307 | 1.503 | 1.314 | -0.029 | -0.033 | -0.033 | 0.545 | 0.64 | 0.47 | 0.67 | 0.77 | 0.73 | 0.64 |
| 6 | 147814584 | rs453032 | 0.897 | -1.317 | 1.305 | 1.346 | 0.048 | -0.04 | -0.04 | 1.22 | 0.61 | 0.21 | 0.6 | 0.57 | 0.73 | 0.61 |
| 6 | 147816995 | rs452577 | 0.184 | -1.049 | 1.586 | 1.082 | -0.028 | -0.023 | -0.023 | 0 | 0.73 | 0.51 | 0.75 | 0.81 | 0.8 | 0.71 |
| 6 | 147819598 | rs325118 | -0.054 | -1.041 | 1.578 | 1.077 | -0.019 | -0.016 | -0.016 | 0.071 | 0.75 | 0.56 | 0.75 | 0.81 | 0.8 | 0.7 |
| 6 | 147822074 | rs954818 | -0.433 | -1.044 | 1.561 | 0.993 | -0.009 | -0.006 | -0.006 | 0 | 0.26 | 0.26 | 0.28 | 0.26 | 0.2 | 0.3 |
| 6 | 147822152 | rs586354 | -1.212 | -1.211 | 1.753 | 0.884 | 0.014 | 0.023 | 0.023 | 0.986 | 0.39 | 0.63 | 0.39 | 0.27 | 0.3 | 0.37 |
| 6 | 147824569 | rs473720 | 0.031 | -0.763 | 1.617 | 1.01 | 0.001 | -0.002 | -0.002 | 0 | 0.23 | 0.25 | 0.22 | 0.23 | 0.2 | 0.3 |
| 6 | 147832444 | rs1333712 | 0.447 | -0.667 | 1.005 | 1.21 | -0.022 | -0.05 | -0.05 | 0.067 | 0.19 | 0.03 | 0.2 | 0.22 | 0.2 | 0.17 |
| 6 | 147834827 | rs494395 | -1.266 | -0.287 | 2.042 | 1.277 | 0 | -0.002 | -0.002 | 0 | 0.34 | 0.63 | 0.33 | 0.24 | 0.27 | 0.37 |
| 6 | 147836218 | rs472577 | -1.35 | -0.258 | 1.623 | 1.336 | 0.006 | -0.052 | -0.052 | 0.052 | 0.36 | 0.82 | 0.34 | 0.39 | 0.27 | 0.41 |
| 6 | 147839691 | rs536996 | -1.607 | -0.311 | 0.98 | 1.327 | 0.006 | -0.074 | -0.074 | 0 | 0.39 | 0.83 | 0.44 | 0.45 | 0.32 | 0.43 |
| 6 | 147840595 | rs669830 | -0.911 | -0.013 | 0.654 | 0.953 | 0.005 | -0.036 | -0.036 | 0.097 | 0.3 | 0.41 | 0.32 | 0.21 | 0.26 | 0.35 |
| 6 | 147847523 | rs480456 | -1.395 | 0.429 | 0.21 | 1.335 | -0.001 | 0.009 | 0.009 | 0 | 0.3 | 0.66 | 0.4 | 0.41 | 0.28 | 0.38 |
| 6 | 147847718 | rs765153 | -0.777 | 0.833 | -0.148 | 0.88 | 0.002 | 0 | 0 | 0.379 | 0.16 | 0.14 | 0.3 | 0.22 | 0.22 | 0.3 |
| 6 | 147851291 | rs1333715 | 0.042 | 0.371 | -0.347 | -0.02 | 0.009 | 0.015 | 0.015 | 0 | 0.58 | 0.59 | 0.77 | 0.89 | 0.67 | 0.69 |
| 6 | 147851974 | rs12203200 | -0.606 | 0.851 | -0.287 | 0.99 | -0.001 | 0.009 | 0.009 | 0 | 0.13 | 0.13 | 0.15 | 0.29 | 0.15 | 0.3 |
| 6 | 147852697 | rs9377056 | -0.887 | 0.315 | -0.015 | 0.508 | -0.024 | 0.01 | 0.01 | 0 | 0.34 | 0.42 | 0.26 | 0.39 | 0.27 | 0.38 |
| 6 | 147859406 | rs9403860 | -0.255 | 0.152 | 0.241 | 0.035 | -0.001 | 0.012 | 0.012 | 0 | 0.54 | 0.57 | 0.59 | 0.7 | 0.51 | 0.51 |
| 6 | 147864300 | rs6900377 | -0.27 | -0.121 | 0.408 | 0.096 | 0.023 | 0.086 | 0.086 | 0 | 0.56 | 0.33 | 0.74 | 0.93 | 0.63 | 0.65 |
| 6 | 147866616 | rs1473691 | -0.918 | -0.244 | 0.393 | -0.12 | -0.001 | 0.156 | 0.156 | 0 | 0.65 | 0.88 | 0.81 | 1 | 0.68 | 0.77 |

| | | | | | | | | | | | | | | | | |
|---|---|---|---|---|---|---|---|---|---|---|---|---|---|---|---|---|
| 6 | 147872216 | rs702328 | -0.47 | -0.643 | 0.395 | 0.506 | -0.007 | 0.008 | 0.008 | 0 | 0.35 | 0.37 | 0.45 | 0.53 | 0.3 | 0.43 |
| 6 | 147876038 | rs702335 | -0.304 | -0.9 | 0.454 | 0.828 | -0.013 | 0.007 | 0.007 | 0 | 0.32 | 0.37 | 0.41 | 0.59 | 0.24 | 0.39 |
| 6 | 147879709 | rs4896924 | 0.577 | -0.261 | 0.307 | 0.637 | -0.046 | -0.019 | -0.019 | 0 | 0.25 | 0.1 | 0.32 | 0.31 | 0.37 | 0.28 |
| 6 | 147881001 | rs702336 | -0.507 | -0.839 | 0.728 | 0.594 | 0 | 0.023 | 0.023 | 0 | 0.5 | 0.61 | 0.5 | 0.63 | 0.54 | 0.45 |
| 6 | 147881171 | rs702337 | 0 | -1.002 | 0.646 | 0.547 | -0.005 | 0.004 | 0.004 | 0 | 0.43 | 0.41 | 0.44 | 0.63 | 0.52 | 0.39 |
| 6 | 147885803 | rs702341 | -0.735 | -1.007 | 0.777 | 0.692 | -0.011 | 0.176 | 0.176 | 0.464 | 0.13 | 0.32 | 0.28 | 0.31 | 0.1 | 0.24 |
| 6 | 147886592 | rs702343 | -0.706 | -1.07 | 0.709 | 0.486 | -0.016 | -0.026 | -0.026 | 1.197 | 0.07 | 0.23 | 0.04 | 0.01 | 0.03 | 0.02 |
| 6 | 147887218 | rs702346 | -1.468 | -1.084 | 0.164 | 0.164 | -0.128 | 0.226 | 0.226 | 0 | 0.64 | 0.97 | 0.79 | 0.97 | 0.62 | 0.76 |
| 6 | 147888022 | rs702349 | -1.469 | -1.09 | 0.16 | 0.161 | -0.128 | 0.226 | 0.226 | 0 | 0.64 | 0.96 | 0.78 | 0.97 | 0.62 | 0.76 |
| 6 | 147888368 | rs9399608 | 0.311 | -0.95 | 0.522 | 0.724 | 0.007 | -0.015 | -0.015 | 0 | 0.08 | 0.04 | 0.25 | 0.34 | 0.07 | 0.28 |
| 6 | 147888494 | rs702350 | -1.045 | -0.897 | 0.8 | 0.593 | -0.011 | -0.036 | -0.036 | 1.65 | 0.08 | 0.27 | 0.05 | 0 | 0.03 | 0.01 |
| 6 | 147888927 | rs702351 | -0.921 | -0.836 | 0.746 | 0.48 | -0.001 | 0.009 | 0.009 | 0 | 0.33 | 0.57 | 0.39 | 0.38 | 0.37 | 0.27 |
| 6 | 147891505 | rs844589 | -0.978 | -0.915 | 0.386 | 0.442 | -0.003 | 0.008 | 0.008 | 0 | 0.34 | 0.58 | 0.39 | 0.39 | 0.37 | 0.28 |
| 6 | 147894916 | rs970285 | -0.35 | -1.081 | 0.265 | 0.638 | 0.004 | 0.006 | 0.006 | 0 | 0.26 | 0.31 | 0.36 | 0.38 | 0.35 | 0.25 |
| 6 | 147898558 | rs844626 | -0.381 | -1.295 | 0.316 | 0.222 | -0.009 | -0.028 | -0.028 | 0 | 0.07 | 0.13 | 0.03 | 0 | 0.03 | 0.02 |
| 6 | 147901135 | rs10484687 | 0.518 | -1.042 | -0.266 | -0.15 | -0.051 | 0.01 | 0.01 | 0 | 0.13 | 0.02 | 0.07 | 0.04 | 0.14 | 0.16 |
| 6 | 147902467 | rs12191000 | -1.803 | -1.27 | -0.153 | -0.32 | -0.002 | -0.025 | -0.025 | 0 | 0.63 | 0.58 | 0.74 | 0.76 | 0.63 | 0.77 |
| 6 | 147908995 | rs844562 | -1.801 | -1.407 | 0.002 | -0.234 | -0.001 | -0.001 | -0.001 | 0 | 0.62 | 0.65 | 0.6 | 0.46 | 0.6 | 0.53 |
| 6 | 147910426 | rs844565 | -1.316 | -1.473 | 0.022 | -0.307 | -0.008 | 0.007 | 0.007 | 0 | 0.37 | 0.52 | 0.41 | 0.4 | 0.31 | 0.51 |
| 6 | 147912444 | rs844569 | -2.106 | -1.62 | -0.087 | -0.25 | -0.001 | -0.002 | -0.002 | 0 | 0.64 | 0.61 | 0.76 | 0.71 | 0.65 | 0.76 |
| 6 | 147912801 | rs11970117 | 0.943 | -1.416 | -0.106 | -0.147 | -0.012 | -0.012 | -0.012 | 0 | 0.13 | 0.23 | 0.25 | 0.13 | 0.09 | 0.11 |
| 6 | 147913541 | rs13218203 | 1.202 | -1.248 | 0.058 | -0.095 | -0.013 | 0.089 | 0.089 | 0 | 0.08 | 0.12 | 0.19 | 0.3 | 0.07 | 0.2 |
| 6 | 147915466 | rs1944645 | -2.298 | -1.208 | -0.227 | -0.127 | -0.025 | 0.021 | 0.021 | 0.016 | 0.69 | 0.85 | 0.79 | 0.77 | 0.67 | 0.77 |
| 6 | 147916111 | rs11963682 | 0.873 | -1.11 | -0.234 | -0.109 | -0.037 | 0.02 | 0.02 | 0 | 0.13 | 0.04 | 0.07 | 0.04 | 0.14 | 0.15 |
| 6 | 147916971 | rs6911479 | 0.391 | -1.146 | -0.184 | -0.124 | 0.004 | -0.028 | -0.028 | 0 | 0.16 | 0.24 | 0.09 | 0.04 | 0.15 | 0.16 |
| 6 | 147917452 | rs7750010 | 2.775 | -1.112 | -0.816 | -0.103 | -0.022 | 0.004 | 0.004 | 0 | 0.06 | 0 | 0.09 | 0.05 | 0.06 | 0.06 |
| 6 | 147918240 | rs17387507 | 0.929 | -1.105 | -0.81 | -0.101 | -0.046 | 0.009 | 0.009 | 0 | 0.13 | 0.02 | 0.07 | 0.04 | 0.14 | 0.15 |
| 6 | 147919087 | rs2204299 | 0.779 | -1.078 | -0.74 | -0.057 | -0.079 | -0.09 | -0.09 | 0 | 0.29 | 0.05 | 0.35 | 0.37 | 0.35 | 0.34 |
| 6 | 147920283 | rs12204941 | 0.81 | -1.067 | -0.735 | -0.052 | -0.08 | -0.09 | -0.09 | 0.394 | 0.28 | 0.05 | 0.36 | 0.37 | 0.35 | 0.33 |
| 6 | 147920983 | rs844607 | -2.291 | -0.986 | -0.712 | -0.015 | -0.032 | 0.019 | 0.019 | 0.047 | 0.7 | 0.88 | 0.79 | 0.77 | 0.66 | 0.78 |
| 6 | 147922185 | rs4896928 | 0.111 | -0.815 | -0.41 | 0.221 | -0.001 | -0.005 | -0.005 | 0.071 | 0.33 | 0.29 | 0.38 | 0.38 | 0.35 | 0.33 |
| 6 | 147927031 | rs9377061 | -0.312 | -0.817 | -0.409 | 0.024 | -0.005 | 0.007 | 0.007 | 0 | 0.39 | 0.32 | 0.47 | 0.4 | 0.41 | 0.39 |
| 6 | 147928723 | rs7766811 | 0.027 | -0.786 | -0.306 | 0.076 | -0.002 | 0.001 | 0.001 | 0 | 0.33 | 0.29 | 0.38 | 0.35 | 0.36 | 0.34 |
| 6 | 147933349 | rs12662247 | -0.38 | -0.79 | 0.093 | 0.121 | -0.044 | 0.181 | 0.181 | 0 | 0.16 | 0.45 | 0.25 | 0.32 | 0.09 | 0.23 |
| 6 | 147934662 | rs9497819 | -0.403 | -0.727 | -0.238 | -0.406 | -0.02 | -0.054 | -0.054 | 0.651 | 0.2 | 0.63 | 0.12 | 0.04 | 0.15 | 0.15 |
| 6 | 147934947 | rs9377063 | -0.557 | -0.18 | -0.441 | -0.501 | 0.05 | -0.088 | -0.088 | 4.703 | 0.3 | 0.48 | 0.27 | 0.1 | 0.4 | 0.22 |
| 6 | 147934995 | rs9377064 | -0.279 | -0.439 | -0.281 | -0.352 | 0.011 | -0.075 | -0.075 | 4.046 | 0.24 | 0.46 | 0.21 | 0.1 | 0.23 | 0.2 |
| 6 | 147935016 | rs9403864 | -0.837 | -0.991 | -0.44 | -0.019 | -0.001 | -0.003 | -0.003 | 0.008 | 0.54 | 0.5 | 0.57 | 0.43 | 0.55 | 0.52 |
| 6 | 147939736 | rs9497822 | -0.291 | -0.49 | -0.774 | 0.028 | -0.016 | 0.004 | 0.004 | 0.1 | 0.44 | 0.28 | 0.55 | 0.61 | 0.46 | 0.57 |
| 6 | 147941453 | rs9497824 | -0.269 | -0.455 | -0.823 | 0.046 | -0.021 | 0.001 | 0.001 | 0.069 | 0.44 | 0.26 | 0.55 | 0.61 | 0.46 | 0.57 |
| 6 | 147945346 | rs11155511 | 0.118 | -0.693 | -0.649 | 0.058 | 0.004 | 0.073 | 0.073 | 0 | 0.48 | 0.7 | 0.45 | 0.33 | 0.5 | 0.44 |
| 6 | 147948040 | rs9497829 | -0.093 | -0.726 | -0.734 | 0.401 | 0.004 | -0.005 | -0.005 | 0.013 | 0.11 | 0.15 | 0.22 | 0.28 | 0.09 | 0.3 |
| 6 | 147949628 | rs719454 | 0.15 | -0.935 | -0.407 | 0.152 | 0.001 | 0.037 | 0.037 | 0 | 0.53 | 0.37 | 0.44 | 0.32 | 0.53 | 0.49 |
| 6 | 147950909 | rs6935781 | 0.172 | -0.94 | -0.405 | 0.149 | 0.002 | 0.037 | 0.037 | 0 | 0.53 | 0.37 | 0.44 | 0.32 | 0.53 | 0.49 |
| 6 | 147955887 | rs6933618 | -0.178 | -1.018 | -0.567 | 0.15 | 0 | -0.032 | -0.032 | 0.001 | 0.59 | 0.25 | 0.63 | 0.57 | 0.61 | 0.74 |
| 6 | 147956348 | rs4896932 | 0.224 | -1.224 | -0.797 | 0.027 | 0.012 | -0.02 | -0.02 | 0.416 | 0.53 | 0.19 | 0.6 | 0.57 | 0.58 | 0.68 |
| 6 | 147956771 | rs17388532 | 0.381 | -0.903 | -0.912 | -0.143 | 0.009 | -0.036 | -0.036 | 0 | 0.13 | 0.16 | 0.09 | 0.01 | 0.16 | 0.13 |
| 6 | 147958767 | rs17388649 | 1.701 | -0.911 | -1.587 | -0.105 | -0.095 | -0.041 | -0.041 | 0.022 | 0.29 | 0.03 | 0.3 | 0.3 | 0.32 | 0.29 |
| 6 | 147968492 | rs844595 | -1.077 | -1.133 | -1.455 | 0.028 | -0.005 | -0.002 | -0.002 | 0 | 0.71 | 0.7 | 0.76 | 0.75 | 0.67 | 0.83 |
| 6 | 147976024 | rs12525574 | 0.569 | -0.877 | -2.346 | 0.215 | 0.007 | -0.055 | -0.055 | 0 | 0.09 | 0.01 | 0.2 | 0.26 | 0.09 | 0.25 |
| 6 | 147976121 | rs1944802 | 0.843 | -0.774 | -2.319 | 0.004 | -0.049 | 0.062 | 0.062 | 0.019 | 0.46 | 0.1 | 0.47 | 0.38 | 0.52 | 0.48 |
| 6 | 147977738 | rs10457815 | 1.51 | -0.572 | -2.497 | 0.156 | -0.091 | -0.021 | -0.021 | 1.027 | 0.32 | 0.03 | 0.32 | 0.3 | 0.35 | 0.3 |
| 6 | 147985603 | rs9390488 | -0.526 | -0.543 | -2.013 | 0.203 | 0.007 | -0.008 | -0.008 | 0 | 0.44 | 0.28 | 0.45 | 0.51 | 0.44 | 0.48 |
| 6 | 147986024 | rs9399613 | 0.296 | 0.17 | -2.129 | 0.452 | -0.059 | -0.052 | -0.052 | 0 | 0.25 | 0.07 | 0.2 | 0.27 | 0.31 | 0.17 |
| 6 | 147987530 | rs1523965 | 0.835 | 0.004 | -1.956 | 0.124 | 0.007 | -0.022 | -0.022 | 0 | 0.6 | 0.11 | 0.6 | 0.59 | 0.65 | 0.56 |
| 6 | 147987946 | rs9377073 | -0.975 | -0.262 | -1.503 | 0.407 | 0.001 | -0.019 | -0.019 | 0.601 | 0.14 | 0.1 | 0.25 | 0.29 | 0.15 | 0.26 |
| 6 | 147988029 | rs9322103 | 1.148 | -0.437 | -0.876 | -0.032 | -0.012 | -0.051 | -0.051 | 0 | 0.84 | 0.32 | 0.87 | 0.95 | 0.89 | 0.89 |
| 6 | 147988841 | rs9373532 | 0.656 | -0.645 | -1.229 | -0.027 | -0.008 | 0.001 | 0.001 | 0 | 0.54 | 0.3 | 0.6 | 0.56 | 0.53 | 0.51 |
| 6 | 147992200 | rs7745338 | -0.318 | -1.026 | -1.462 | -0.23 | -0.037 | -0.038 | -0.038 | 0.702 | 0.13 | 0.24 | 0.05 | 0.05 | 0.04 | 0.11 |
| 6 | 147993054 | rs9390491 | 1.258 | -0.757 | -1.865 | -0.103 | -0.168 | -0.164 | -0.164 | 0.262 | 0.41 | 0.03 | 0.55 | 0.48 | 0.49 | 0.41 |
| 6 | 147994707 | rs4896938 | -0.401 | -0.123 | -1.01 | 0.229 | 0.01 | -0.021 | -0.021 | 0 | 0.1 | 0.05 | 0.12 | 0.2 | 0.08 | 0.22 |
| 6 | 148002142 | rs1608530 | -0.703 | 0.238 | -0.425 | -0.287 | -0.002 | -0.055 | -0.055 | 0.729 | 0.14 | 0.16 | 0.07 | 0 | 0.12 | 0.08 |
| 6 | 148007144 | rs9497839 | 0.462 | 0.628 | -0.148 | 0.203 | 0.029 | 0.02 | 0.02 | 0 | 0.79 | 0.64 | 0.8 | 0.75 | 0.74 | 0.85 |
| 6 | 148021701 | rs9485215 | -0.943 | -0.44 | -0.647 | -0.697 | 0.039 | 0.052 | 0.052 | 1.885 | 0.33 | 0.75 | 0.17 | 0.04 | 0.17 | 0.2 |
| 6 | 148021909 | rs17077261 | -0.373 | -0.553 | -0.225 | -0.747 | -0.004 | -0.039 | -0.039 | 0 | 0.15 | 0.16 | 0.05 | 0 | 0.08 | 0.05 |
| 6 | 148024669 | rs6914269 | -0.185 | -0.033 | -0.863 | -0.281 | -0.06 | -0.107 | -0.107 | 0 | 0.78 | 0.93 | 0.73 | 0.45 | 0.72 | 0.69 |
| 6 | 148024843 | rs9322106 | -0.087 | -0.031 | -0.634 | -0.392 | 0.003 | -0.016 | -0.016 | 0 | 0.14 | 0.12 | 0.04 | 0 | 0.08 | 0.05 |
| 6 | 148034286 | rs9485217 | -0.59 | -0.124 | 0.445 | -0.225 | 0.01 | 0 | 0 | 0.087 | 0.1 | 0.39 | 0.06 | 0.04 | 0.04 | 0.06 |
| 6 | 148034869 | rs3908266 | 0.477 | -0.265 | 0.47 | -0.294 | 0.002 | 0.009 | 0.009 | 0 | 0.59 | 0.67 | 0.65 | 0.41 | 0.64 | 0.57 |

| | | | | | | | | | | | | | | | | |
|---|---|---|---|---|---|---|---|---|---|---|---|---|---|---|---|---|
| 6 | 148037791 | rs7764867 | 1.432 | -0.144 | 0.216 | -0.478 | -0.043 | 0.022 | 0.022 | 0.78 | 0.39 | 0.18 | 0.5 | 0.35 | 0.49 | 0.46 |
| 6 | 148044515 | rs3908265 | -0.907 | -0.324 | 0.864 | -0.494 | -0.012 | -0.124 | -0.124 | 1.762 | 0.16 | 0.49 | 0.11 | 0 | 0.1 | 0.1 |
| 6 | 148045181 | rs9322108 | 0.961 | 0.163 | 1.046 | -0.477 | 0.001 | 0.011 | 0.011 | 1.462 | 0.52 | 0.43 | 0.59 | 0.37 | 0.56 | 0.52 |
| 6 | 148048684 | rs9485219 | 0.342 | -0.001 | 0.922 | -0.535 | 0 | -0.008 | -0.008 | 0 | 0.05 | 0.07 | 0.01 | 0 | 0.03 | 0.05 |
| 6 | 148049016 | rs9497852 | -0.909 | 0.037 | 1.328 | -0.716 | -0.018 | -0.137 | -0.137 | 0.109 | 0.19 | 0.48 | 0.08 | 0.02 | 0.13 | 0.1 |
| 6 | 148049229 | rs4122006 | -0.133 | 0.451 | 1.099 | -0.183 | -0.041 | -0.086 | -0.086 | 0.012 | 0.77 | 0.88 | 0.68 | 0.51 | 0.71 | 0.7 |
| 6 | 148053041 | rs9377077 | -0.016 | 0.346 | 1.156 | -0.175 | -0.004 | -0.008 | -0.008 | 0 | 0.77 | 0.77 | 0.68 | 0.51 | 0.73 | 0.7 |
| 6 | 148053476 | rs12210064 | 0.599 | 0.425 | 0.998 | -0.256 | -0.003 | 0.008 | 0.008 | 3.06 | 0.11 | 0.06 | 0.03 | 0 | 0.09 | 0.05 |
| 6 | 148056480 | rs9399619 | 1.653 | 0.309 | 0.585 | -0.168 | -0.077 | 0.029 | 0.029 | 4.793 | 0.47 | 0.09 | 0.52 | 0.42 | 0.52 | 0.46 |
| 6 | 148062400 | rs2282605 | 1.127 | 0.27 | 0.691 | -0.204 | 0.003 | 0.015 | 0.015 | 1.398 | 0.55 | 0.37 | 0.59 | 0.4 | 0.59 | 0.5 |
| 6 | 148066394 | rs12214565 | 0.627 | 0.43 | -0.142 | -0.343 | -0.008 | 0.008 | 0.008 | 2.706 | 0.11 | 0.06 | 0.04 | 0 | 0.11 | 0.06 |
| 6 | 148074155 | rs17077306 | 0.101 | 0.406 | -0.27 | -0.199 | -0.008 | -0.024 | -0.024 | 0 | 0.06 | 0.12 | 0.01 | 0 | 0.03 | 0.05 |
| 6 | 148074336 | rs11760126 | 0.578 | 0.524 | -1.033 | -0.527 | -0.019 | 0.018 | 0.018 | 0.092 | 0.16 | 0.01 | 0.16 | 0.14 | 0.16 | 0.2 |
| 6 | 148086791 | rs9377079 | 1.153 | 0.484 | -0.464 | 0.126 | -0.074 | -0.032 | -0.032 | 0.092 | 0.47 | 0.08 | 0.53 | 0.5 | 0.52 | 0.47 |
| 6 | 148103855 | rs17077323 | -0.102 | 0.553 | 0.181 | 0.658 | -0.004 | 0.095 | 0.095 | 0 | 0.09 | 0.14 | 0.18 | 0.34 | 0.09 | 0.17 |
| 6 | 148105936 | rs4121999 | -1.189 | 0.382 | 0.552 | 0.27 | -0.013 | 0.003 | 0.003 | 0 | 0.37 | 0.52 | 0.31 | 0.34 | 0.29 | 0.32 |
| 6 | 148108890 | rs12662237 | -1.152 | 0.397 | 0.479 | 0.221 | -0.008 | 0.005 | 0.005 | 0.127 | 0.37 | 0.53 | 0.32 | 0.35 | 0.32 | 0.3 |
| 6 | 148126416 | rs949073 | -1.343 | 0.449 | 0.596 | 0.178 | 0.013 | -0.004 | -0.004 | 1.187 | 0.41 | 0.71 | 0.36 | 0.39 | 0.34 | 0.35 |
| 6 | 148127362 | rs6936905 | -0.418 | 0.432 | 0.687 | -0.138 | -0.007 | -0.043 | -0.043 | 0 | 0.08 | 0.25 | 0.06 | 0.01 | 0.05 | 0.02 |
| 6 | 148128141 | rs12195214 | 0.062 | 0.465 | 0.58 | -0.13 | 0 | -0.032 | -0.032 | 0 | 0.19 | 0.17 | 0.08 | 0 | 0.15 | 0.09 |
| 6 | 148138449 | rs9386196 | -0.628 | 0.788 | 0.019 | -0.179 | -0.077 | -0.024 | -0.024 | 0 | 0.85 | 0.99 | 0.72 | 0.85 | 0.75 | 0.88 |
| 6 | 148141501 | rs17077431 | -1.57 | 0.147 | 0.999 | 0.083 | -0.016 | -0.025 | -0.025 | 0 | 0.14 | 0.29 | 0.1 | 0.08 | 0.11 | 0.14 |
| 6 | 148145648 | rs12195694 | -0.905 | 0.143 | 1.043 | 0.026 | 0.006 | 0.004 | 0.004 | 0 | 0.25 | 0.2 | 0.11 | 0.03 | 0.2 | 0.16 |
| 6 | 148152077 | rs10484683 | -0.566 | 0.231 | 0.892 | 0.16 | 0.007 | 0.051 | 0.051 | 0.476 | 0.26 | 0.15 | 0.13 | 0.03 | 0.21 | 0.16 |
| 6 | 148159028 | rs1528868 | -1.133 | 0.299 | 0.512 | 0.508 | -0.006 | 0.095 | 0.095 | 0 | 0.14 | 0.25 | 0.22 | 0.4 | 0.13 | 0.27 |
| 6 | 148160266 | rs9485225 | -1.047 | 0.434 | 0.191 | 0.583 | 0.018 | -0.006 | -0.006 | 0 | 0.41 | 0.75 | 0.36 | 0.39 | 0.34 | 0.39 |
| 6 | 148162328 | rs2294805 | -0.732 | 0.659 | 0.307 | 1.507 | 0.001 | 0.11 | 0.11 | 0 | 0.09 | 0.15 | 0.19 | 0.34 | 0.09 | 0.17 |
| 6 | 148180666 | rs4084217 | 0.23 | 0.791 | -0.567 | 0.281 | -0.008 | 0.037 | 0.037 | 0.314 | 0.64 | 0.75 | 0.64 | 0.76 | 0.56 | 0.74 |
| 6 | 148182196 | rs3861425 | 0.284 | 0.771 | -1.119 | 0.664 | -0.042 | -0.024 | -0.024 | 0.611 | 0.37 | 0.1 | 0.4 | 0.37 | 0.41 | 0.31 |
| 6 | 148183190 | rs2294806 | -0.094 | 0.488 | -1.66 | 0.58 | 0.032 | -0.045 | -0.045 | 1.366 | 0.59 | 0.19 | 0.6 | 0.47 | 0.65 | 0.45 |
| 6 | 148183894 | rs3887007 | 0.973 | 0.524 | -1.662 | 0.612 | -0.043 | -0.014 | -0.014 | 0 | 0.37 | 0.12 | 0.4 | 0.35 | 0.41 | 0.31 |
| 6 | 148184099 | rs3861426 | -0.273 | 0.262 | -0.904 | 0.249 | -0.001 | -0.006 | -0.006 | 0 | 0.28 | 0.23 | 0.19 | 0.07 | 0.26 | 0.13 |
| 6 | 148185587 | rs500110 | -0.513 | -0.088 | -0.738 | 0.664 | -0.008 | -0.02 | -0.02 | 0 | 0.82 | 0.9 | 0.84 | 0.72 | 0.8 | 0.64 |
| 6 | 148186670 | rs531173 | 0.132 | -0.17 | -0.003 | 0.621 | -0.001 | 0 | 0 | 0.345 | 0.74 | 0.72 | 0.78 | 0.69 | 0.75 | 0.57 |
| 6 | 148188229 | rs510748 | -0.757 | -0.263 | 0.829 | 0.703 | -0.024 | 0.035 | 0.035 | 0.26 | 0.17 | 0.49 | 0.23 | 0.18 | 0.13 | 0.16 |
| 6 | 148192919 | rs678691 | 0.161 | -0.21 | 0.855 | 0.747 | 0.006 | 0.074 | 0.074 | 4.638 | 0.09 | 0.38 | 0.14 | 0.13 | 0.08 | 0.11 |
| 6 | 148193929 | rs13198106 | 1.587 | -0.052 | 0.254 | 0.759 | -0.043 | -0.059 | -0.059 | 0 | 0.36 | 0.11 | 0.4 | 0.44 | 0.39 | 0.3 |
| 6 | 148202273 | rs614002 | -1.641 | -0.141 | 0.289 | 1.065 | -0.048 | 0.028 | 0.028 | 1.101 | 0.64 | 0.86 | 0.69 | 0.67 | 0.59 | 0.7 |
| 6 | 148205415 | rs9377083 | 0.759 | -0.202 | 0.197 | 1.135 | 0.012 | -0.041 | -0.041 | 0 | 0.15 | 0.06 | 0.14 | 0.32 | 0.12 | 0.33 |
| 6 | 148205420 | rs508902 | 0.007 | -0.257 | 0.396 | 1.079 | 0.002 | -0.012 | -0.012 | 0 | 0.16 | 0.15 | 0.3 | 0.24 | 0.16 | 0.16 |
| 6 | 148205955 | rs671555 | -1.516 | -0.221 | 0.403 | 1.058 | -0.026 | 0.036 | 0.036 | 0.686 | 0.6 | 0.78 | 0.66 | 0.68 | 0.54 | 0.68 |
| 6 | 148207211 | rs9322110 | -1.136 | -0.244 | 0.184 | 0.913 | -0.009 | 0.009 | 0.009 | 0 | 0.52 | 0.33 | 0.51 | 0.44 | 0.49 | 0.63 |
| 6 | 148207621 | rs10457061 | 0.009 | -0.183 | 0.128 | 0.742 | -0.007 | 0.041 | 0.041 | 0 | 0.27 | 0.2 | 0.21 | 0.11 | 0.27 | 0.19 |
| 6 | 148213642 | rs3895168 | 0.285 | -0.258 | 0.266 | 1.336 | -0.001 | 0.007 | 0.007 | 0 | 0.16 | 0.18 | 0.15 | 0.33 | 0.13 | 0.33 |
| 6 | 148215688 | rs4385334 | 0.418 | -0.135 | 0.002 | 1.169 | -0.064 | 0.027 | 0.027 | 0 | 0.18 | 0.02 | 0.16 | 0.08 | 0.21 | 0.18 |
| 6 | 148216298 | rs2876669 | 0.232 | -0.217 | 0.55 | 1.117 | 0.001 | -0.009 | -0.009 | 0 | 0.2 | 0.21 | 0.18 | 0.08 | 0.21 | 0.18 |
| 6 | 148219940 | rs474377 | -0.742 | -0.372 | 0.71 | 1.174 | -0.033 | 0.034 | 0.034 | 5.512 | 0.25 | 0.58 | 0.36 | 0.3 | 0.22 | 0.19 |
| 6 | 148221015 | rs600222 | -1.314 | -0.387 | 0.669 | 1.146 | 0.018 | 0.01 | 0.01 | 6.463 | 0.45 | 0.81 | 0.53 | 0.38 | 0.43 | 0.38 |
| 6 | 148221417 | rs17077550 | 0.376 | -0.35 | 0.655 | 1.053 | 0.002 | -0.014 | -0.014 | 0 | 0.2 | 0.22 | 0.18 | 0.09 | 0.21 | 0.18 |
| 6 | 148222844 | rs624015 | -1.695 | -0.281 | 0.35 | 1.085 | -0.062 | 0.082 | 0.082 | 5.333 | 0.58 | 0.88 | 0.68 | 0.72 | 0.56 | 0.7 |
| 6 | 148229629 | rs618688 | -0.986 | -0.329 | 0.71 | 1.036 | 0.009 | 0.01 | 0.01 | 7.64 | 0.43 | 0.67 | 0.52 | 0.37 | 0.4 | 0.38 |
| 6 | 148235498 | rs12110586 | -0.206 | -0.187 | 0.812 | 0.895 | 0.077 | -0.039 | -0.039 | 0 | 0.07 | 0.22 | 0.06 | 0 | 0.1 | 0.04 |
| 6 | 148236385 | rs526847 | -1.449 | -0.428 | 0.335 | 1.12 | -0.005 | -0.039 | -0.039 | 0.25 | 0.41 | 0.81 | 0.48 | 0.52 | 0.38 | 0.58 |
| 6 | 148242514 | rs10080530 | 0.318 | -0.391 | 0.317 | 1.051 | 0.032 | -0.029 | -0.029 | 0 | 0.06 | 0.12 | 0.05 | 0 | 0.1 | 0.04 |
| 6 | 148247765 | rs589143 | -1.094 | -0.435 | 0.366 | 1.087 | 0 | -0.001 | -0.001 | 0 | 0.28 | 0.65 | 0.39 | 0.28 | 0.28 | 0.24 |
| 6 | 148251243 | rs3915135 | -0.254 | -0.319 | -0.043 | 1.153 | 0.022 | -0.036 | -0.036 | 0 | 0.28 | 0.15 | 0.21 | 0.36 | 0.23 | 0.48 |
| 6 | 148253856 | rs17647552 | 0.622 | -0.088 | -0.072 | 1.142 | -0.024 | 0.057 | 0.057 | 1.751 | 0.1 | 0.01 | 0.16 | 0 | 0.1 | 0.1 |
| 6 | 148255082 | rs17647618 | 0.607 | -0.005 | 0.003 | 1.178 | -0.002 | 0.001 | 0.001 | 0 | 0.05 | 0.04 | 0.05 | 0 | 0.1 | 0.04 |
| 6 | 148258033 | rs646839 | -0.451 | -0.231 | 0.145 | 1.192 | -0.029 | 0.027 | 0.027 | 2.653 | 0.21 | 0.37 | 0.32 | 0.27 | 0.18 | 0.21 |
| 6 | 148259210 | rs9390510 | 0.596 | -0.248 | 0.111 | 1.232 | -0.006 | -0.038 | -0.038 | 0 | 0.12 | 0.04 | 0.08 | 0.25 | 0.1 | 0.34 |
| 6 | 148262501 | rs533034 | -0.872 | -0.17 | 0.127 | 0.888 | -0.013 | 0.024 | 0.024 | 0.31 | 0.33 | 0.4 | 0.4 | 0.53 | 0.28 | 0.55 |
| 6 | 148266755 | rs13217257 | 0.475 | 0.259 | -0.148 | 0.585 | -0.007 | 0.004 | 0.004 | 0 | 0.05 | 0.03 | 0.04 | 0 | 0.09 | 0.03 |
| 6 | 148270210 | rs12524732 | 0.015 | 0.122 | 0.188 | 0.458 | -0.008 | -0.055 | -0.055 | 0 | 0.14 | 0.19 | 0.07 | 0 | 0.11 | 0.15 |
| 6 | 148270789 | rs12525177 | 0.048 | 0.125 | -0.098 | 0.46 | 0.006 | 0.045 | 0.045 | 1.354 | 0.13 | 0.06 | 0.07 | 0 | 0.11 | 0.14 |
| 6 | 148270822 | rs17582242 | 0.447 | 0.302 | -0.049 | 0.569 | -0.005 | 0.004 | 0.004 | 0 | 0.05 | 0.04 | 0.05 | 0 | 0.1 | 0.04 |
| 6 | 148274611 | rs6904457 | 0.182 | -0.136 | 0.198 | 0.775 | -0.017 | 0.002 | 0.002 | 6.359 | 0.8 | 0.72 | 0.75 | 0.79 | 0.82 | 0.82 |
| 6 | 148274761 | rs3861444 | 0.378 | -0.164 | 0.234 | 0.77 | -0.015 | -0.009 | -0.009 | 0.058 | 0.14 | 0.24 | 0.13 | 0.1 | 0.12 | 0.13 |
| 6 | 148276339 | rs11155530 | 2.085 | 0.305 | -0.105 | 0.426 | -0.023 | -0.011 | -0.011 | 0 | 0.36 | 0.13 | 0.3 | 0.38 | 0.36 | 0.28 |
| 6 | 148283379 | rs6899879 | -0.172 | 0.164 | 0.19 | 0.221 | 0.033 | -0.146 | -0.146 | 0 | 0.22 | 0.45 | 0.13 | 0 | 0.21 | 0.17 |

| | | | | | | | | | | | | | | | | |
|---|---|---|---|---|---|---|---|---|---|---|---|---|---|---|---|---|
| 6 | 148284197 | rs9322113 | 1.208 | 0.108 | 0.154 | 0.2 | 0.021 | 0.114 | 0.114 | 4.254 | 0.54 | 0.2 | 0.55 | 0.78 | 0.58 | 0.63 |
| 6 | 148284529 | rs3903597 | -0.181 | 0.14 | 0.165 | 0.204 | 0.032 | -0.148 | -0.148 | 0 | 0.22 | 0.47 | 0.14 | 0 | 0.21 | 0.17 |
| 6 | 148286285 | rs11155534 | 0.415 | 0.15 | 0.09 | 0.322 | 0.002 | -0.002 | -0.002 | 0.052 | 0.65 | 0.65 | 0.5 | 0.4 | 0.71 | 0.48 |
| 6 | 148289312 | rs12660201 | 0.672 | 0.231 | -0.041 | 0.398 | 0.002 | -0.052 | -0.052 | 0 | 0.14 | 0.04 | 0.2 | 0.38 | 0.11 | 0.34 |
| 6 | 148291002 | rs3903594 | -0.748 | 0.22 | 0.125 | 0.277 | 0.008 | 0.012 | 0.012 | 1.189 | 0.36 | 0.51 | 0.34 | 0.37 | 0.32 | 0.5 |
| 6 | 148292041 | rs9322114 | 0.341 | 0.393 | 0.023 | 0.319 | -0.025 | -0.079 | -0.079 | 0.205 | 0.17 | 0.04 | 0.24 | 0.38 | 0.21 | 0.34 |
| 6 | 148292162 | rs10484678 | 2.356 | 0.299 | 0.023 | 0.312 | -0.015 | 0.001 | 0.001 | 1.469 | 0.37 | 0.18 | 0.31 | 0.38 | 0.37 | 0.29 |
| 6 | 148295038 | rs9791215 | 0.975 | 0.194 | 0.314 | 0.232 | -0.004 | -0.016 | -0.016 | 0 | 0.68 | 0.32 | 0.61 | 0.76 | 0.69 | 0.78 |
| 6 | 148297083 | rs17649567 | 0.739 | 0.14 | 0.218 | 0.251 | -0.001 | -0.02 | -0.02 | 0.004 | 0.14 | 0.12 | 0.06 | 0 | 0.12 | 0.14 |
| 6 | 148297485 | rs11969459 | -0.794 | 0.107 | 0.193 | 0.268 | 0.013 | -0.026 | -0.026 | 0 | 0.94 | 0.98 | 0.83 | 0.88 | 0.94 | 0.95 |
| 6 | 148297601 | rs2328836 | 0.07 | 0.118 | 0.275 | 0.26 | -0.023 | -0.005 | -0.005 | 0.286 | 0.76 | 0.68 | 0.68 | 0.76 | 0.79 | 0.81 |
| 6 | 148297991 | rs17077821 | 0.68 | 0.275 | -0.091 | 0.331 | 0.001 | -0.055 | -0.055 | 0 | 0.14 | 0.04 | 0.2 | 0.37 | 0.11 | 0.34 |
| 6 | 148298888 | rs6901528 | 0.528 | 0.276 | 0.017 | 0.04 | -0.002 | -0.024 | -0.024 | 0 | 0.14 | 0.13 | 0.06 | 0 | 0.11 | 0.14 |
| 6 | 148300251 | rs3861449 | -0.79 | 0.31 | 0.055 | 0.048 | -0.001 | -0.051 | -0.051 | 0.267 | 0.26 | 0.67 | 0.22 | 0.12 | 0.25 | 0.17 |
| 6 | 148302445 | rs3903592 | 0.12 | 0.245 | -1.033 | -0.164 | -0.005 | 0.059 | 0.059 | 0 | 0.51 | 0.19 | 0.48 | 0.62 | 0.52 | 0.61 |
| 6 | 148303941 | rs11756401 | 0.623 | 0.594 | -1.181 | 0.126 | -0.065 | -0.086 | -0.086 | 0 | 0.28 | 0.04 | 0.3 | 0.32 | 0.28 | 0.44 |
| 6 | 148305952 | rs4122017 | -0.012 | 0.545 | -1.13 | 0.059 | -0.027 | 0.055 | 0.055 | 0 | 0.51 | 0.18 | 0.47 | 0.61 | 0.51 | 0.61 |
| 6 | 148306502 | rs12193181 | -0.244 | 0.625 | -1.172 | 0.04 | 0.008 | 0.044 | 0.044 | 0 | 0.54 | 0.18 | 0.51 | 0.62 | 0.6 | 0.61 |
| 6 | 148307343 | rs7755750 | -0.53 | 0.657 | -0.923 | -0.026 | 0.023 | 0.004 | 0.004 | 0 | 0.62 | 0.3 | 0.56 | 0.61 | 0.69 | 0.64 |
| 6 | 148307918 | rs6570808 | -1.938 | 0.199 | -0.918 | -0.096 | 0.01 | -0.001 | -0.001 | 0 | 0.9 | 0.94 | 0.83 | 0.89 | 0.93 | 0.92 |
| 6 | 148308874 | rs17077838 | 0.394 | 0.457 | -0.419 | -0.163 | 0.029 | -0.01 | -0.01 | 0.072 | 0.29 | 0.43 | 0.29 | 0.28 | 0.32 | 0.28 |
| 6 | 148310261 | rs9377090 | 1.074 | 0.235 | -0.629 | 0.033 | 0.005 | -0.013 | -0.013 | 0 | 0.24 | 0.15 | 0.18 | 0.29 | 0.22 | 0.16 |
| 6 | 148312897 | rs12662092 | 1.39 | 0.224 | -0.376 | -0.131 | -0.003 | -0.007 | -0.007 | 0 | 0.18 | 0.12 | 0.2 | 0.28 | 0.18 | 0.24 |
| 6 | 148315090 | rs9403906 | -0.803 | 0.165 | -0.644 | -0.73 | -0.001 | -0.003 | -0.003 | 0.766 | 0.81 | 0.84 | 0.64 | 0.85 | 0.81 | 0.74 |
| 6 | 148315259 | rs9377092 | 0.732 | 0.101 | -1.069 | -1.2 | -0.019 | 0.168 | 0.168 | 0 | 0.45 | 0.18 | 0.42 | 0.84 | 0.44 | 0.59 |
| 6 | 148319277 | rs9390519 | 0.625 | -0.29 | -0.423 | -0.964 | -0.001 | 0.002 | 0.002 | 0 | 0.47 | 0.46 | 0.29 | 0.25 | 0.46 | 0.22 |
| 6 | 148321738 | rs6933891 | -0.181 | -0.157 | -0.357 | -1.082 | 0.003 | -0.044 | -0.044 | 0 | 0.72 | 0.62 | 0.57 | 0.84 | 0.69 | 0.73 |
| 6 | 148326813 | rs7767522 | 0.344 | -0.13 | -0.396 | -1.009 | 0.019 | 0.053 | 0.053 | 0 | 0.5 | 0.43 | 0.36 | 0.23 | 0.57 | 0.24 |
| 6 | 148328367 | rs1404484 | 0.321 | -0.587 | -0.386 | -0.942 | -0.013 | 0.019 | 0.019 | 0 | 0.3 | 0.4 | 0.27 | 0.62 | 0.24 | 0.49 |
| 6 | 148333774 | rs12524192 | 1.225 | -0.466 | -0.567 | -1.184 | -0.04 | 0.019 | 0.019 | 0 | 0.29 | 0.19 | 0.28 | 0.22 | 0.39 | 0.22 |
| 6 | 148338152 | rs4335008 | 0.223 | -0.689 | -0.481 | -1.256 | 0.007 | 0.01 | 0.01 | 0 | 0.51 | 0.47 | 0.4 | 0.23 | 0.63 | 0.26 |
| 6 | 148338672 | rs2272450 | 1.043 | -0.624 | -0.488 | -1.233 | -0.046 | 0.024 | 0.024 | 0 | 0.33 | 0.21 | 0.32 | 0.23 | 0.49 | 0.23 |
| 6 | 148340422 | rs4896955 | 0.232 | -0.781 | -0.039 | -0.868 | -0.011 | -0.028 | -0.028 | 0 | 0.63 | 0.62 | 0.59 | 0.85 | 0.74 | 0.72 |
| 6 | 148341758 | rs1881826 | -0.852 | -0.771 | 0.067 | -0.843 | -0.004 | -0.003 | -0.003 | 0 | 0.82 | 0.82 | 0.67 | 0.85 | 0.88 | 0.76 |
| 6 | 148348007 | rs4626446 | 0.829 | -0.756 | 0.059 | -0.858 | 0.003 | -0.025 | -0.025 | 4.385 | 0.18 | 0.19 | 0.08 | 0 | 0.14 | 0.03 |
| 6 | 148349012 | rs4895729 | -0.154 | -0.924 | 0.174 | -0.646 | -0.026 | 0.164 | 0.164 | 0 | 0.18 | 0.33 | 0.18 | 0.53 | 0.13 | 0.41 |
| 6 | 148350275 | rs9322119 | 0.705 | -0.853 | 0.113 | -0.633 | 0.003 | 0 | 0 | 0 | 0.63 | 0.4 | 0.47 | 0.32 | 0.74 | 0.43 |
| 6 | 148350620 | rs12174997 | 0.38 | -0.655 | 0.025 | -0.492 | -0.001 | 0.107 | 0.107 | 0 | 0.08 | 0.1 | 0.12 | 0.53 | 0.08 | 0.3 |
| 6 | 148354753 | rs9390528 | -0.962 | -0.648 | 0.071 | -0.52 | 0.007 | -0.013 | -0.013 | 0 | 0.4 | 0.66 | 0.31 | 0.51 | 0.37 | 0.44 |
| 6 | 148357629 | rs9399632 | 0.377 | -0.292 | 0.014 | -0.327 | -0.006 | 0.002 | 0.002 | 0 | 0.08 | 0.01 | 0.11 | 0.53 | 0.08 | 0.3 |
| 6 | 148359153 | rs9390531 | 0.183 | -0.287 | 0.088 | -0.325 | -0.003 | 0.138 | 0.138 | 0 | 0.09 | 0.16 | 0.12 | 0.53 | 0.08 | 0.3 |
| 6 | 148373852 | rs12665328 | -0.333 | -0.35 | 0.097 | -0.131 | -0.012 | 0.089 | 0.089 | 0 | 0.14 | 0.23 | 0.13 | 0.44 | 0.1 | 0.24 |
| 6 | 148374346 | rs9403922 | 0.158 | -0.442 | 0.107 | -0.886 | -0.001 | -0.014 | -0.014 | 0 | 0.59 | 0.58 | 0.53 | 0.84 | 0.62 | 0.7 |
| 6 | 148375031 | rs7759893 | 0.624 | -0.321 | 0.063 | -0.806 | 0.005 | -0.001 | -0.001 | 0 | 0.52 | 0.49 | 0.49 | 0.84 | 0.58 | 0.62 |
| 6 | 148377085 | rs12661581 | 0.555 | -0.442 | 0.107 | -0.686 | -0.002 | 0.102 | 0.102 | 0 | 0.09 | 0.13 | 0.12 | 0.53 | 0.08 | 0.29 |
| 6 | 148386997 | rs9497926 | 0.753 | -0.157 | 0.061 | -0.896 | 0.024 | 0.019 | 0.019 | 0 | 0.46 | 0.39 | 0.42 | 0.32 | 0.6 | 0.33 |
| 6 | 148398001 | rs1934797 | 1.073 | 0.349 | 0.194 | -0.818 | 0 | -0.023 | -0.023 | 0 | 0.12 | 0.11 | 0.04 | 0 | 0.12 | 0.02 |
| 6 | 148398847 | rs12665723 | -0.229 | -0.112 | 0.235 | -0.495 | 0.012 | -0.031 | -0.031 | 0 | 0.28 | 0.19 | 0.31 | 0.55 | 0.22 | 0.49 |
| 6 | 148403405 | rs1007475 | 0.959 | 0.522 | 0.311 | -0.436 | -0.007 | 0.06 | 0.06 | 0 | 0.33 | 0.19 | 0.21 | 0.11 | 0.29 | 0.27 |
| 6 | 148407655 | rs9497930 | 0.41 | 0.6 | 0.832 | -0.537 | 0 | -0.001 | -0.001 | 0 | 0.68 | 0.69 | 0.48 | 0.34 | 0.68 | 0.51 |
| 6 | 148409331 | rs7764264 | 0.059 | 0.738 | 0.899 | -0.53 | 0.007 | -0.022 | -0.022 | 0 | 0.9 | 0.89 | 0.93 | 0.99 | 0.85 | 0.9 |
| 6 | 148410391 | rs2328849 | -0.461 | 0.724 | 0.902 | -0.523 | -0.007 | -0.004 | -0.004 | 0 | 0.93 | 0.89 | 0.81 | 0.94 | 0.95 | 0.9 |
| 6 | 148426922 | rs9377106 | 0.504 | 0.774 | 0.388 | -0.47 | 0.004 | -0.118 | -0.118 | 0.69 | 0.21 | 0.05 | 0.27 | 0.59 | 0.16 | 0.4 |
| 6 | 148435295 | rs6936689 | 1.077 | 0.816 | -0.584 | -0.53 | -0.011 | 0.049 | 0.049 | 0 | 0.31 | 0.19 | 0.2 | 0.06 | 0.29 | 0.26 |
| 6 | 148439462 | rs9373551 | 0.691 | 1.255 | 0.028 | 0.185 | -0.007 | 0.032 | 0.032 | 0 | 0.52 | 0.32 | 0.48 | 0.65 | 0.45 | 0.64 |
| 6 | 148439839 | rs4896965 | 0.188 | 1.37 | -0.703 | 0.493 | -0.009 | 0.009 | 0.009 | 0 | 0.68 | 0.81 | 0.74 | 0.72 | 0.62 | 0.76 |
| 6 | 148447884 | rs7765680 | 0.494 | 1.843 | -0.196 | 0.506 | 0.05 | -0.023 | -0.023 | 0 | 0.73 | 0.48 | 0.73 | 0.8 | 0.62 | 0.76 |
| 6 | 148462046 | rs13216119 | 0.903 | 1.876 | -0.915 | 0.565 | 0.043 | -0.023 | -0.023 | 0.3 | 0.39 | 0.18 | 0.47 | 0.52 | 0.28 | 0.58 |
| 6 | 148462967 | rs11155553 | -0.024 | 1.777 | -0.439 | 0.416 | 0 | 0.001 | 0.001 | 0 | 0.35 | 0.35 | 0.26 | 0.27 | 0.34 | 0.18 |
| 6 | 148465078 | rs9377111 | 0.173 | 1.691 | -0.72 | 0.474 | -0.032 | 0.004 | 0.004 | 0 | 0.21 | 0.1 | 0.19 | 0.17 | 0.29 | 0.22 |
| 6 | 148467702 | rs9377112 | 0.224 | 1.412 | -1.043 | 0.584 | 0.008 | -0.033 | -0.033 | 0 | 0.22 | 0.11 | 0.16 | 0.34 | 0.2 | 0.15 |
| 6 | 148468876 | rs9485250 | 0.933 | 1.328 | -1.101 | 0.569 | 0.037 | -0.088 | -0.088 | 0 | 0.38 | 0.11 | 0.47 | 0.49 | 0.28 | 0.59 |
| 6 | 148469586 | rs12524329 | 0.612 | 1.529 | -0.751 | 0.598 | -0.006 | 0.007 | 0.007 | 0 | 0.08 | 0.06 | 0.06 | 0 | 0.1 | 0.01 |
| 6 | 148469844 | rs9322127 | 0.785 | 1.767 | -0.935 | 0.188 | 0.006 | 0.034 | 0.034 | 0 | 0.51 | 0.23 | 0.49 | 0.55 | 0.48 | 0.66 |
| 6 | 148470301 | rs12173665 | 0.052 | 1.643 | -0.884 | 0.451 | 0.003 | -0.021 | -0.021 | 0.595 | 0.2 | 0.13 | 0.27 | 0.33 | 0.22 | 0.18 |
| 6 | 148470431 | rs2153543 | 0.375 | 2.024 | -0.647 | 0.348 | 0.007 | -0.014 | -0.014 | 0 | 0.7 | 0.35 | 0.76 | 0.84 | 0.67 | 0.79 |
| 6 | 148470854 | rs6570818 | 0.073 | 1.981 | -0.689 | 0.321 | 0.006 | -0.021 | -0.021 | 0 | 0.2 | 0.12 | 0.26 | 0.33 | 0.21 | 0.18 |
| 6 | 148471747 | rs1934809 | 0.533 | 1.623 | -0.692 | 0.374 | 0.018 | 0.023 | 0.023 | 0.394 | 0.49 | 0.31 | 0.35 | 0.4 | 0.44 | 0.52 |
| 6 | 148471975 | rs12209762 | 0.503 | 1.639 | -0.463 | 0.382 | 0.008 | 0.01 | 0.01 | 0.368 | 0.5 | 0.39 | 0.36 | 0.4 | 0.44 | 0.53 |

| | | | | | | | | | | | | | | | | |
|---|---|---|---|---|---|---|---|---|---|---|---|---|---|---|---|---|
| 6 | 148475673 | rs9322128 | -0.616 | 1.251 | -0.669 | 0.427 | 0.007 | 0.008 | 0.008 | 1.122 | 0.48 | 0.66 | 0.61 | 0.57 | 0.48 | 0.45 |
| 6 | 148486349 | rs9403930 | -0.198 | 0.64 | -1.001 | 0.486 | 0.01 | -0.049 | -0.049 | 0 | 0.35 | 0.14 | 0.28 | 0.51 | 0.31 | 0.31 |
| 6 | 148486753 | rs11155555 | -0.062 | 0.613 | -0.638 | 0.413 | 0.013 | -0.001 | -0.001 | 0 | 0.43 | 0.32 | 0.42 | 0.48 | 0.37 | 0.32 |
| 6 | 148491574 | rs6570819 | -0.501 | 0.493 | -0.69 | 0.539 | -0.018 | -0.02 | -0.02 | 0 | 0.63 | 0.76 | 0.58 | 0.6 | 0.58 | 0.45 |
| 6 | 148495280 | rs9497952 | -0.057 | 0.366 | -0.601 | 0.638 | 0 | -0.001 | -0.001 | 0.439 | 0.05 | 0.05 | 0.03 | 0.06 | 0.01 | 0.03 |
| 6 | 148508148 | rs6920454 | -0.313 | 0.178 | -0.181 | 0.458 | 0.002 | -0.001 | -0.001 | 0 | 0.49 | 0.54 | 0.52 | 0.53 | 0.47 | 0.44 |
| 6 | 148511237 | rs1934804 | -0.467 | 0.025 | -0.018 | 0.42 | 0.018 | 0.01 | 0.01 | 0 | 0.5 | 0.66 | 0.6 | 0.55 | 0.42 | 0.38 |
| 6 | 148513761 | rs9497963 | -0.961 | 0.137 | 0.901 | 0.323 | 0.041 | -0.05 | -0.05 | 0 | 0.17 | 0.5 | 0.17 | 0.07 | 0.14 | 0.12 |
| 6 | 148517345 | rs6570821 | -1.171 | 0.029 | 0.382 | 0.28 | -0.04 | -0.061 | -0.061 | 0 | 0.64 | 0.81 | 0.53 | 0.54 | 0.58 | 0.47 |
| 6 | 148519573 | rs2328856 | 0.306 | 0.093 | 0.429 | 0.241 | 0.002 | -0.025 | -0.025 | 0 | 0.06 | 0.08 | 0.06 | 0 | 0.07 | 0.06 |
| 6 | 148530140 | rs885344 | -0.74 | -0.01 | 0.314 | 0.152 | -0.002 | -0.018 | -0.018 | 0.567 | 0.31 | 0.65 | 0.25 | 0.13 | 0.28 | 0.17 |
| 6 | 148533303 | rs9377117 | 0.343 | 0.12 | 0.128 | 0.393 | 0.004 | -0.033 | -0.033 | 1.386 | 0.3 | 0.14 | 0.28 | 0.4 | 0.28 | 0.3 |
| 6 | 148537736 | rs7756362 | 0.702 | 0.111 | 0.202 | 0.567 | 0.001 | -0.004 | -0.004 | 0.002 | 0.25 | 0.16 | 0.24 | 0.33 | 0.25 | 0.28 |
| 6 | 148540356 | rs17078169 | 0.41 | 0.231 | -0.161 | 0.774 | -0.032 | -0.029 | -0.029 | 0 | 0.09 | 0.02 | 0.08 | 0.14 | 0.1 | 0.08 |
| 6 | 148542357 | rs9377118 | 0.788 | 0.189 | 0.015 | 0.743 | 0.001 | -0.028 | -0.028 | 0.099 | 0.24 | 0.1 | 0.23 | 0.33 | 0.25 | 0.28 |
| 6 | 148542743 | rs9322131 | 0.404 | -0.038 | 0.067 | 0.402 | -0.017 | -0.033 | -0.033 | 0 | 0.19 | 0.26 | 0.11 | 0.07 | 0.13 | 0.06 |
| 6 | 148542967 | rs9322133 | -0.263 | -0.032 | -0.028 | 0.344 | 0.003 | -0.01 | -0.01 | 0 | 0.25 | 0.3 | 0.17 | 0.07 | 0.2 | 0.11 |
| 6 | 148543861 | rs6570822 | -0.739 | 0.007 | -0.216 | 0.832 | 0 | 0.096 | 0.096 | 0 | 0.51 | 0.66 | 0.6 | 0.74 | 0.45 | 0.53 |
| 6 | 148545817 | rs6570824 | -1.08 | 0.129 | -0.629 | 1.016 | -0.047 | 0.071 | 0.071 | 0 | 0.65 | 0.8 | 0.69 | 0.83 | 0.59 | 0.58 |
| 6 | 148547065 | rs12195413 | 1.161 | 0.194 | -1.063 | 1.677 | -0.029 | -0.055 | -0.055 | 0 | 0.12 | 0.01 | 0.08 | 0.17 | 0.11 | 0.15 |
| 6 | 148549835 | rs1538893 | -0.112 | 0.308 | -0.807 | 1.176 | 0.012 | -0.035 | -0.035 | 0 | 0.33 | 0.11 | 0.31 | 0.39 | 0.32 | 0.35 |
| 6 | 148553550 | rs7753842 | 0.673 | 0.59 | -0.909 | 0.599 | -0.001 | -0.003 | -0.003 | 0 | 0.23 | 0.22 | 0.11 | 0.14 | 0.21 | 0.23 |
| 6 | 148553992 | rs6899613 | -0.093 | 1.043 | -0.687 | 1.226 | 0.016 | 0.007 | 0.007 | 0 | 0.83 | 0.81 | 0.77 | 0.86 | 0.77 | 0.66 |
| 6 | 148556084 | rs7758765 | -0.02 | 0.762 | -1.376 | 1.002 | 0.014 | -0.029 | -0.029 | 0.893 | 0.27 | 0.1 | 0.24 | 0.36 | 0.24 | 0.28 |
| 6 | 148560857 | rs7767314 | 0.155 | 1.403 | -1.051 | 0.667 | -0.035 | 0.025 | 0.025 | 0 | 0.86 | 0.96 | 0.79 | 0.95 | 0.81 | 0.78 |
| 6 | 148562917 | rs9497980 | 0.124 | 0.518 | -0.304 | 1.575 | -0.036 | 0.091 | 0.091 | 0 | 0.15 | 0.26 | 0.08 | 0.36 | 0.07 | 0.19 |
| 6 | 148565442 | rs7769910 | 0.238 | 1.087 | -0.093 | 1.464 | -0.007 | 0.033 | 0.033 | 0 | 0.16 | 0.22 | 0.12 | 0.38 | 0.12 | 0.18 |
| 6 | 148567159 | rs6914370 | -0.345 | 1.065 | -0.165 | 1.575 | -0.008 | 0.036 | 0.036 | 0 | 0.21 | 0.29 | 0.27 | 0.42 | 0.16 | 0.26 |
| 6 | 148567462 | rs6914932 | -0.146 | 1.077 | -0.198 | 1.567 | -0.003 | 0.013 | 0.013 | 0.058 | 0.19 | 0.22 | 0.16 | 0.37 | 0.16 | 0.22 |
| 6 | 148569583 | rs10499249 | 0.708 | 1.451 | -0.541 | 2.006 | -0.026 | -0.028 | -0.028 | 0.638 | 0.06 | 0 | 0.05 | 0.18 | 0.06 | 0.06 |
| 6 | 148575687 | rs4896978 | 0.094 | 1.442 | -0.613 | 1.688 | -0.004 | -0.005 | -0.005 | 1.665 | 0.86 | 0.88 | 0.69 | 0.76 | 0.8 | 0.8 |
| 6 | 148582765 | rs9390549 | -0.2 | 0.936 | -0.071 | 1.114 | -0.01 | 0.012 | 0.012 | 0 | 0.3 | 0.35 | 0.23 | 0.42 | 0.28 | 0.29 |
| 6 | 148584115 | rs17078231 | 1.271 | 1.015 | -0.212 | 1.073 | -0.026 | -0.023 | -0.023 | 0.797 | 0.06 | 0 | 0.05 | 0.06 | 0.05 | 0.05 |
| 6 | 148585295 | rs9390550 | 0.87 | 0.947 | -0.231 | 1.017 | -0.012 | 0.001 | 0.001 | 0 | 0.08 | 0.01 | 0.04 | 0.06 | 0.1 | 0.05 |
| 6 | 148587096 | rs10499250 | 1.663 | 0.761 | -0.442 | 0.2 | -0.016 | 0.06 | 0.06 | 0 | 0.26 | 0.13 | 0.17 | 0.09 | 0.26 | 0.3 |
| 6 | 148588469 | rs9497985 | -0.041 | 0.733 | -0.278 | 0.245 | -0.009 | 0.024 | 0.024 | 0 | 0.61 | 0.69 | 0.5 | 0.25 | 0.57 | 0.55 |
| 6 | 148591116 | rs9373554 | 0.261 | 0.711 | -0.23 | 0.065 | -0.004 | 0.052 | 0.052 | 0 | 0.52 | 0.38 | 0.43 | 0.22 | 0.48 | 0.54 |
| 6 | 148591999 | rs7752482 | 1.912 | 0.604 | -0.103 | 0.482 | -0.007 | 0.059 | 0.059 | 0 | 0.16 | 0.26 | 0.15 | 0.36 | 0.14 | 0.2 |
| 6 | 148593153 | rs6901639 | 0.337 | 0.565 | -0.152 | 0.472 | -0.002 | -0.001 | -0.001 | 0 | 0.12 | 0.14 | 0.19 | 0.15 | 0.14 | 0.11 |
| 6 | 148606036 | rs9399642 | 0.005 | 0.633 | -0.42 | 0.282 | -0.009 | 0.006 | 0.006 | 0 | 0.58 | 0.69 | 0.52 | 0.35 | 0.54 | 0.55 |
| 6 | 148608485 | rs9322139 | 0.751 | 0.744 | -0.688 | 0.291 | -0.016 | 0.019 | 0.019 | 0 | 0.37 | 0.53 | 0.3 | 0.46 | 0.36 | 0.3 |
| 6 | 148609400 | rs6925130 | -0.05 | 0.961 | -0.9 | 0.095 | -0.025 | -0.02 | -0.02 | 0 | 0.19 | 0.09 | 0.31 | 0.24 | 0.27 | 0.22 |
| 6 | 148610348 | rs2493923 | -0.153 | 0.734 | -0.7 | 0.093 | 0.014 | 0.012 | 0.012 | 0 | 0.49 | 0.64 | 0.45 | 0.58 | 0.43 | 0.42 |
| 6 | 148613771 | rs4434491 | 0.069 | 0.776 | -0.79 | -0.064 | 0.009 | 0.017 | 0.017 | 0 | 0.46 | 0.68 | 0.44 | 0.63 | 0.43 | 0.46 |
| 6 | 148614267 | rs2493926 | 0.417 | 0.819 | -0.607 | 0.514 | 0.006 | 0.019 | 0.019 | 0 | 0.14 | 0.19 | 0.17 | 0.27 | 0.12 | 0.12 |
| 6 | 148617199 | rs4896987 | 0.316 | 0.484 | -0.655 | 0.28 | 0.001 | 0 | 0 | 0 | 0.43 | 0.58 | 0.33 | 0.39 | 0.4 | 0.39 |
| 6 | 148621018 | rs6570836 | -0.486 | 0.766 | -0.734 | 0.264 | -0.003 | 0.001 | 0.001 | 0 | 0.58 | 0.64 | 0.66 | 0.64 | 0.62 | 0.6 |
| 6 | 148621930 | rs7763618 | 0.07 | 0.71 | -0.723 | 0.625 | -0.002 | -0.002 | -0.002 | 0.23 | 0.77 | 0.78 | 0.64 | 0.69 | 0.71 | 0.74 |
| 6 | 148628009 | rs9390554 | 0.555 | 0.061 | -0.791 | 0.248 | -0.026 | 0.008 | 0.008 | 0 | 0.28 | 0.51 | 0.2 | 0.31 | 0.27 | 0.24 |
| 6 | 148630465 | rs6902928 | 0.299 | 0.32 | -0.883 | -0.282 | 0 | -0.008 | -0.008 | 0 | 0.31 | 0.29 | 0.24 | 0.09 | 0.25 | 0.37 |
| 6 | 148633283 | rs6920755 | -0.108 | 0.51 | -0.869 | -0.167 | 0.01 | 0.008 | 0.008 | 1.223 | 0.41 | 0.36 | 0.42 | 0.34 | 0.37 | 0.45 |
| 6 | 148634783 | rs13195974 | -0.205 | 0.548 | -1.054 | -0.151 | 0.023 | 0 | 0 | 0 | 0.4 | 0.19 | 0.43 | 0.42 | 0.39 | 0.46 |
| 6 | 148638111 | rs6918668 | 0.935 | 0.502 | -0.6 | -0.54 | -0.011 | -0.04 | -0.04 | 0 | 0.32 | 0.39 | 0.21 | 0.19 | 0.26 | 0.28 |
| 6 | 148638158 | rs4445079 | 0.049 | 1.126 | -0.379 | -0.225 | 0.026 | -0.033 | -0.033 | 0 | 0.06 | 0.13 | 0.07 | 0 | 0.09 | 0.01 |
| 6 | 148643960 | rs9390556 | 0.436 | 0.82 | -0.318 | -0.175 | 0.004 | -0.05 | -0.05 | 0.405 | 0.81 | 0.64 | 0.77 | 0.9 | 0.82 | 0.71 |
| 6 | 148663110 | rs9377127 | -0.317 | 0.805 | -0.784 | 0.091 | -0.002 | -0.06 | -0.06 | 0.64 | 0.83 | 0.96 | 0.77 | 0.73 | 0.82 | 0.84 |
| 6 | 148667310 | rs2493913 | 0.583 | 1.081 | -0.478 | 0.235 | 0.019 | 0.07 | 0.07 | 0 | 0.78 | 0.67 | 0.67 | 0.59 | 0.73 | 0.71 |
| 6 | 148670925 | rs17078259 | 1.228 | 1.167 | -0.769 | 0.339 | -0.03 | 0.035 | 0.035 | 0 | 0.05 | 0.01 | 0.13 | 0.31 | 0.08 | 0.06 |
| 6 | 148671794 | rs9403946 | 0.914 | 0.981 | -1.009 | -0.257 | -0.041 | 0.048 | 0.048 | 2.192 | 0.21 | 0.09 | 0.15 | 0.12 | 0.28 | 0.12 |
| 6 | 148673778 | rs2495954 | -0.262 | 0.955 | -0.665 | -0.098 | 0.015 | -0.001 | -0.001 | 0 | 0.92 | 0.88 | 0.94 | 0.91 | 0.89 | 0.88 |
| 6 | 148678866 | rs17078272 | 1.329 | 0.416 | -0.921 | -0.56 | -0.019 | -0.012 | -0.012 | 0 | 0.26 | 0.31 | 0.22 | 0.19 | 0.17 | 0.36 |
| 6 | 148679552 | rs4407729 | 1.172 | 0.719 | -0.705 | -0.201 | -0.001 | -0.005 | -0.005 | 0 | 0.46 | 0.49 | 0.36 | 0.3 | 0.44 | 0.48 |
| 6 | 148679737 | rs2495956 | 0.354 | 0.745 | -0.726 | -0.101 | 0.045 | 0.133 | 0.133 | 0 | 0.77 | 0.58 | 0.66 | 0.59 | 0.73 | 0.71 |
| 6 | 148679852 | rs6570845 | 0.206 | 0.186 | -1.106 | -0.19 | 0 | -0.002 | -0.002 | 0 | 0.69 | 0.91 | 0.71 | 0.71 | 0.71 | 0.76 |
| 6 | 148682423 | rs6570846 | -0.609 | 0.266 | -0.508 | -0.792 | -0.001 | -0.057 | -0.057 | 0.139 | 0.31 | 0.3 | 0.27 | 0.04 | 0.23 | 0.28 |
| 6 | 148690764 | rs4537167 | -0.51 | 0.507 | -0.301 | 0.02 | 0.03 | 0.076 | 0.076 | 0.124 | 0.78 | 0.62 | 0.83 | 0.59 | 0.71 | 0.83 |
| 6 | 148695175 | rs17078277 | 0.015 | -0.377 | -0.463 | 0.04 | -0.002 | -0.001 | -0.001 | 0 | 0.17 | 0.08 | 0.19 | 0.2 | 0.07 | 0.23 |
| 6 | 148698507 | rs9485288 | 1.244 | 0.069 | -0.747 | -0.037 | 0 | 0.002 | 0.002 | 0 | 0.33 | 0.34 | 0.42 | 0.26 | 0.33 | 0.4 |
| 6 | 148702262 | rs4897002 | -0.808 | 0.394 | -0.796 | -0.058 | 0.06 | 0.15 | 0.15 | 0.066 | 0.77 | 0.58 | 0.82 | 0.55 | 0.69 | 0.82 |

| | | | | | | | | | | | | | | | | |
|---|---|---|---|---|---|---|---|---|---|---|---|---|---|---|---|---|
| 6 | 148704612 | rs4895752 | 0.237 | 0.353 | -0.939 | -0.111 | 0.025 | 0.03 | 0.03 | 2.414 | 0.53 | 0.37 | 0.62 | 0.42 | 0.43 | 0.64 |
| 6 | 148707493 | rs10457832 | 0.994 | 0.68 | -1.368 | -0.771 | 0.001 | 0.069 | 0.069 | 0 | 0.36 | 0.24 | 0.42 | 0.21 | 0.36 | 0.39 |
| 6 | 148710800 | rs9498028 | -0.649 | 0.932 | -0.931 | -0.126 | -0.005 | 0.006 | 0.006 | 2.366 | 0.25 | 0.21 | 0.21 | 0.13 | 0.27 | 0.2 |
| 6 | 148710896 | rs9498029 | -0.714 | 0.986 | -0.996 | -0.147 | -0.006 | 0.007 | 0.007 | 2.274 | 0.25 | 0.21 | 0.21 | 0.13 | 0.26 | 0.19 |
| 6 | 148712650 | rs12196375 | -0.443 | 0.966 | -0.651 | -0.095 | -0.001 | -0.034 | -0.034 | 0 | 0.16 | 0.18 | 0.18 | 0.02 | 0.15 | 0.29 |
| 6 | 148721578 | rs12527170 | -0.173 | 1.239 | -0.931 | 0.357 | -0.041 | -0.024 | -0.024 | 0 | 0.11 | 0.01 | 0.23 | 0.14 | 0.12 | 0.07 |
| 6 | 148726746 | rs9373560 | -0.871 | 1.578 | -0.242 | 0.387 | 0.025 | 0.004 | 0.004 | 1.598 | 0.3 | 0.39 | 0.38 | 0.33 | 0.34 | 0.2 |
| 6 | 148731834 | rs17078341 | 0.453 | 1.555 | -0.794 | 0.641 | 0.004 | -0.021 | -0.021 | 0 | 0.21 | 0.15 | 0.16 | 0.29 | 0.2 | 0.09 |
| 6 | 148734174 | rs11155566 | -0.304 | 1.426 | -0.196 | 0.513 | -0.003 | -0.001 | -0.001 | 0 | 0.27 | 0.17 | 0.34 | 0.23 | 0.28 | 0.18 |
| 6 | 148736448 | rs9390567 | 0.117 | 1.39 | 0.377 | 0.509 | 0.026 | 0.013 | 0.013 | 0 | 0.13 | 0.27 | 0.26 | 0.13 | 0.13 | 0.07 |
| 6 | 148737230 | rs6910915 | 0.582 | 1.422 | 0.233 | 0.513 | 0.002 | 0 | 0 | 0 | 0.11 | 0.08 | 0.24 | 0.12 | 0.13 | 0.07 |
| 6 | 148744408 | rs11754081 | 0.652 | 1.15 | 0.236 | -0.138 | 0.002 | -0.013 | -0.013 | 0 | 0.17 | 0.12 | 0.11 | 0.01 | 0.15 | 0.33 |
| 6 | 148749393 | rs1768326 | 0.353 | 1.015 | 0.381 | -0.154 | -0.009 | -0.01 | -0.01 | 1.314 | 0.18 | 0.34 | 0.12 | 0.1 | 0.14 | 0.1 |
| 6 | 148752678 | rs909980 | 1.378 | 1.133 | 0.221 | -0.031 | 0 | -0.002 | -0.002 | 1.957 | 0.28 | 0.33 | 0.33 | 0.37 | 0.26 | 0.36 |
| 6 | 148759707 | rs4895754 | -0.931 | 1.265 | 0.374 | -0.065 | 0.006 | 0.005 | 0.005 | 0 | 0.88 | 0.84 | 0.75 | 0.85 | 0.87 | 0.93 |
| 6 | 148760518 | rs9390569 | 0.392 | 1.219 | 0.044 | -0.157 | 0 | -0.031 | -0.031 | 0 | 0.19 | 0.15 | 0.11 | 0.01 | 0.15 | 0.35 |
| 6 | 148761603 | rs9498038 | 0.694 | 1.245 | 0.082 | 0.178 | 0.004 | 0.002 | 0.002 | 0 | 0.46 | 0.41 | 0.43 | 0.4 | 0.41 | 0.71 |
| 6 | 148767836 | rs9386227 | -0.26 | 1.621 | -0.304 | 0.526 | -0.052 | -0.048 | -0.048 | 0 | 0.87 | 0.99 | 0.74 | 0.76 | 0.81 | 0.9 |
| 6 | 148770108 | rs7763869 | 0.725 | 1.572 | -0.194 | 0.419 | 0 | 0.002 | 0.002 | 3.426 | 0.17 | 0.16 | 0.16 | 0.27 | 0.22 | 0.1 |
| 6 | 148770505 | rs7746800 | -0.082 | 1.466 | -0.052 | 0.359 | 0.004 | 0.005 | 0.005 | 2.68 | 0.25 | 0.33 | 0.22 | 0.36 | 0.32 | 0.15 |
| 6 | 148771965 | rs1022544 | -0.942 | 1.33 | -0.133 | 0.4 | -0.025 | -0.035 | -0.035 | 0.381 | 0.89 | 0.98 | 0.77 | 0.83 | 0.87 | 0.93 |
| 6 | 148772470 | rs6941174 | -0.396 | 1.166 | -0.218 | 0.15 | -0.011 | -0.002 | -0.002 | 0 | 0.4 | 0.36 | 0.37 | 0.42 | 0.51 | 0.26 |
| 6 | 148772589 | rs9377135 | 0.568 | 0.799 | 0.138 | -0.137 | -0.008 | 0.178 | 0.178 | 0.096 | 0.16 | 0.38 | 0.16 | 0.34 | 0.11 | 0.14 |
| 6 | 148772998 | rs9377136 | -0.468 | 0.704 | 0.05 | -0.004 | 0.013 | 0.021 | 0.021 | 0 | 0.52 | 0.46 | 0.5 | 0.7 | 0.61 | 0.39 |
| 6 | 148773318 | rs9399648 | -0.45 | 0.784 | -0.093 | 0.207 | -0.009 | -0.019 | -0.019 | 0 | 0.35 | 0.25 | 0.31 | 0.48 | 0.38 | 0.43 |
| 6 | 148773594 | rs7746540 | -0.28 | 0.867 | -0.181 | 0.181 | 0 | 0.003 | 0.003 | 0 | 0.61 | 0.55 | 0.52 | 0.42 | 0.58 | 0.53 |
| 6 | 148776295 | rs2328901 | 0.633 | 1.206 | -0.704 | -0.753 | -0.001 | 0.048 | 0.048 | 0 | 0.36 | 0.25 | 0.29 | 0.08 | 0.32 | 0.3 |
| 6 | 148776487 | rs17641840 | 1.561 | 1.406 | -1.149 | -0.69 | -0.032 | 0.035 | 0.035 | 2.522 | 0.09 | 0.01 | 0.07 | 0.05 | 0.12 | 0.03 |
| 6 | 148778680 | rs9322150 | 0.935 | 1.432 | -1.349 | -0.205 | -0.03 | -0.092 | -0.092 | 0.171 | 0.14 | 0.03 | 0.2 | 0.51 | 0.17 | 0.13 |
| 6 | 148780913 | rs7756365 | -1.085 | 1.376 | -1.202 | -0.399 | 0 | -0.001 | -0.001 | 0 | 0.87 | 0.9 | 0.86 | 0.89 | 0.86 | 0.93 |
| 6 | 148781825 | rs9399650 | -0.106 | 1.419 | -1.19 | -0.384 | -0.001 | 0.011 | 0.011 | 0 | 0.28 | 0.18 | 0.31 | 0.68 | 0.3 | 0.32 |
| 6 | 148785054 | rs9377141 | -0.581 | 1.464 | -0.793 | -0.482 | 0 | -0.013 | -0.013 | 0 | 0.4 | 0.42 | 0.4 | 0.74 | 0.44 | 0.38 |
| 6 | 148785423 | rs12664363 | 0.488 | 1.398 | -1.21 | 0.018 | -0.009 | -0.046 | -0.046 | 0 | 0.13 | 0.08 | 0.1 | 0.18 | 0.14 | 0.18 |
| 6 | 148787137 | rs12210132 | 0.543 | 1.668 | -0.901 | -0.125 | -0.031 | 0.025 | 0.025 | 0 | 0.1 | 0.03 | 0.08 | 0.07 | 0.14 | 0.16 |
| 6 | 148787159 | rs12527089 | -0.016 | 1.06 | -0.401 | 0.367 | -0.009 | 0.023 | 0.023 | 0.124 | 0.05 | 0.01 | 0.12 | 0.02 | 0.04 | 0.16 |
| 6 | 148787467 | rs7743102 | -0.881 | 1.086 | -0.114 | 0.299 | 0.012 | -0.12 | -0.12 | 0 | 0.22 | 0.43 | 0.24 | 0.09 | 0.21 | 0.24 |
| 6 | 148788006 | rs6930337 | -0.584 | 0.745 | -0.346 | 0.497 | 0.003 | -0.06 | -0.06 | 0.281 | 0.11 | 0.31 | 0.15 | 0.02 | 0.06 | 0.2 |
| 6 | 148788604 | rs1474706 | -0.155 | 0.982 | -1.195 | 1.064 | -0.006 | -0.008 | -0.008 | 0 | 0.88 | 0.96 | 0.86 | 0.88 | 0.87 | 0.93 |
| 6 | 148789171 | rs4897015 | 0.314 | 0.974 | -1.129 | 1.225 | -0.001 | 0.014 | 0.014 | 0 | 0.85 | 0.83 | 0.82 | 0.73 | 0.84 | 0.85 |
| 6 | 148789370 | rs17642099 | 0.619 | 1.052 | -1.23 | 0.807 | -0.031 | 0.026 | 0.026 | 4.282 | 0.08 | 0.01 | 0.07 | 0.07 | 0.12 | 0.02 |
| 6 | 148790330 | rs6570854 | 0.326 | 0.721 | -1.211 | 1.32 | -0.005 | -0.09 | -0.09 | 0 | 0.82 | 0.97 | 0.72 | 0.38 | 0.82 | 0.82 |
| 6 | 148792617 | rs1883625 | -0.744 | 0.641 | -0.224 | 0.88 | -0.012 | -0.028 | -0.028 | 0 | 0.17 | 0.4 | 0.1 | 0.14 | 0.14 | 0.21 |
| 6 | 148792852 | rs2294776 | -1.268 | 0.405 | -0.123 | 2.001 | -0.022 | 0.133 | 0.133 | 0.191 | 0.23 | 0.43 | 0.17 | 0.56 | 0.2 | 0.39 |
| 6 | 148794249 | rs7760319 | -0.478 | 0.825 | -0.333 | 1.034 | 0.003 | -0.01 | -0.01 | 0 | 0.7 | 0.7 | 0.53 | 0.84 | 0.63 | 0.7 |
| 6 | 148795472 | rs2294777 | 1.338 | 0.767 | -0.79 | 0.718 | -0.017 | 0.028 | 0.028 | 1.693 | 0.09 | 0.01 | 0.06 | 0.03 | 0.1 | 0.04 |
| 6 | 148797824 | rs9485292 | -0.646 | 0.825 | -0.587 | 0.78 | -0.02 | -0.002 | -0.002 | 0 | 0.73 | 0.82 | 0.65 | 0.75 | 0.66 | 0.73 |
| 6 | 148803550 | rs9322151 | -0.712 | 0.775 | -0.697 | 0.73 | -0.07 | -0.02 | -0.02 | 0 | 0.74 | 0.94 | 0.65 | 0.75 | 0.66 | 0.72 |
| 6 | 148804893 | rs6928016 | 0.077 | 0.892 | -0.277 | 0.42 | 0.019 | -0.09 | -0.09 | 2.428 | 0.16 | 0.37 | 0.23 | 0.03 | 0.16 | 0.11 |
| 6 | 148807213 | rs11961740 | 0 | 0.854 | -0.112 | 0.541 | -0.009 | -0.026 | -0.026 | 0.519 | 0.17 | 0.32 | 0.09 | 0.15 | 0.14 | 0.22 |
| 6 | 148815558 | rs1624157 | 0.611 | 0.831 | -0.044 | 0.479 | 0.001 | -0.019 | -0.019 | 0 | 0.06 | 0.03 | 0.03 | 0 | 0.07 | 0.04 |
| 6 | 148820512 | rs9390576 | -0.426 | 0.68 | 0.138 | 1.269 | -0.003 | 0.132 | 0.132 | 0 | 0.08 | 0.16 | 0.24 | 0.45 | 0.08 | 0.17 |
| 6 | 148821379 | rs12214798 | 1.474 | 0.605 | -0.151 | 1.106 | -0.026 | 0.049 | 0.049 | 0.307 | 0.34 | 0.05 | 0.19 | 0.14 | 0.3 | 0.26 |
| 6 | 148823562 | rs6931504 | 0.226 | 0.637 | 0.084 | 1.063 | -0.002 | 0.019 | 0.019 | 0 | 0.54 | 0.35 | 0.51 | 0.62 | 0.48 | 0.5 |
| 6 | 148823914 | rs6936843 | -0.022 | 0.968 | 0.027 | 1.066 | 0 | 0 | 0 | 0 | 0.63 | 0.63 | 0.41 | 0.32 | 0.61 | 0.52 |
| 6 | 148824840 | rs7455026 | 1.404 | 1.286 | -0.483 | 1.05 | -0.026 | 0.023 | 0.023 | 1.872 | 0.09 | 0.01 | 0.06 | 0.03 | 0.11 | 0.04 |
| 6 | 148825030 | rs1618036 | -0.353 | 1.285 | -0.149 | 1.272 | 0.017 | -0.021 | -0.021 | 0 | 0.17 | 0.32 | 0.1 | 0.15 | 0.17 | 0.22 |
| 6 | 148827807 | rs1147865 | -0.143 | 1.191 | -0.627 | 1.475 | -0.009 | 0.02 | 0.02 | 0 | 0.72 | 0.86 | 0.56 | 0.77 | 0.68 | 0.73 |
| 6 | 148829305 | rs6570859 | 0.366 | 1.252 | -0.289 | 1.152 | -0.001 | -0.014 | -0.014 | 0 | 0.62 | 0.57 | 0.66 | 0.7 | 0.64 | 0.57 |
| 6 | 148829441 | rs6935952 | 0.472 | 1.216 | -0.431 | 0.958 | 0.001 | 0 | 0 | 0 | 0.58 | 0.59 | 0.72 | 0.61 | 0.59 | 0.5 |
| 6 | 148829474 | rs1147864 | 0.392 | 1.38 | -0.703 | 0.549 | -0.001 | 0.022 | 0.022 | 0.215 | 0.66 | 0.69 | 0.68 | 0.86 | 0.67 | 0.78 |
| 6 | 148829910 | rs9377143 | -0.79 | 1.852 | -0.678 | 1.642 | 0.012 | 0.07 | 0.07 | 0 | 0.1 | 0.15 | 0.11 | 0.45 | 0.18 | 0.28 |
| 6 | 148831153 | rs11751252 | 0.705 | 1.599 | -1.212 | 0.967 | -0.024 | 0.032 | 0.032 | 0.714 | 0.11 | 0.01 | 0.08 | 0.05 | 0.12 | 0.08 |
| 6 | 148831252 | rs1147862 | 1.071 | 1.426 | -1.693 | 0.426 | 0.006 | 0.045 | 0.045 | 0.892 | 0.3 | 0.09 | 0.2 | 0.13 | 0.29 | 0.22 |
| 6 | 148835416 | rs1057793 | 0.499 | 1.45 | -1.62 | 0.779 | -0.009 | -0.021 | -0.021 | 0 | 0.17 | 0.1 | 0.22 | 0.32 | 0.22 | 0.12 |
| 6 | 148835939 | rs1147861 | -0.577 | 1.13 | -1.006 | 0.431 | 0.002 | 0.004 | 0.004 | 0 | 0.73 | 0.76 | 0.67 | 0.83 | 0.81 | 0.66 |
| 6 | 148838112 | rs1147858 | -0.722 | 1.047 | -1.399 | 0.489 | 0.01 | 0 | 0 | 0 | 0.76 | 0.89 | 0.69 | 0.81 | 0.83 | 0.72 |
| 6 | 148841752 | rs1147857 | -0.024 | 1.174 | -1.523 | 0.472 | -0.016 | 0.103 | 0.103 | 0 | 0.52 | 0.24 | 0.46 | 0.69 | 0.54 | 0.47 |
| 6 | 148844451 | rs1147855 | 0.609 | 0.611 | -1.576 | 0.588 | 0.033 | 0.031 | 0.031 | 1.751 | 0.28 | 0.06 | 0.14 | 0.13 | 0.22 | 0.2 |
| 6 | 148848919 | rs2110198 | 0.865 | 0.774 | -0.861 | 1.177 | -0.004 | 0.034 | 0.034 | 0 | 0.88 | 0.77 | 0.83 | 0.78 | 0.89 | 0.84 |

| | | | | | | | | | | | | | | | | |
|---|---|---|---|---|---|---|---|---|---|---|---|---|---|---|---|---|
| 6 | 148849562 | rs2328897 | 0.435 | 0.515 | -1.307 | 0.966 | 0.002 | -0.039 | -0.039 | 0 | 0.71 | 0.84 | 0.75 | 0.62 | 0.73 | 0.72 |
| 6 | 148849648 | rs1147854 | -0.357 | 0.7 | -1.014 | 0.632 | 0.007 | 0.011 | 0.011 | 0.082 | 0.51 | 0.57 | 0.53 | 0.66 | 0.61 | 0.46 |
| 6 | 148852325 | rs2294783 | 1.417 | 1.188 | -1.432 | 0.505 | -0.033 | 0.024 | 0.024 | 0 | 0.1 | 0.01 | 0.07 | 0.05 | 0.11 | 0.05 |
| 6 | 148852391 | rs2294784 | -0.017 | 1.165 | -1.23 | 0.601 | 0.005 | 0.012 | 0.012 | 0 | 0.15 | 0.22 | 0.13 | 0.15 | 0.17 | 0.15 |
| 6 | 148853867 | rs1465307 | 0.149 | 0.994 | -1.563 | 0.571 | 0.003 | 0.002 | 0.002 | 0 | 0.47 | 0.41 | 0.52 | 0.45 | 0.46 | 0.59 |
| 6 | 148854984 | rs2294787 | 1.192 | 1.08 | -1.738 | 0.591 | -0.025 | 0.024 | 0.024 | 0 | 0.1 | 0.01 | 0.07 | 0.05 | 0.1 | 0.05 |
| 6 | 148855462 | rs759106 | 0.323 | 1.008 | -1.557 | 1.066 | -0.011 | -0.042 | -0.042 | 0.276 | 0.93 | 1 | 0.89 | 0.77 | 0.93 | 0.87 |
| 6 | 148859288 | rs9498061 | -0.408 | 1.483 | -1.504 | 0.591 | -0.004 | 0.003 | 0.003 | 0 | 0.17 | 0.17 | 0.13 | 0.11 | 0.2 | 0.16 |
| 6 | 148859387 | rs1569655 | 0.595 | 1.31 | -1.998 | 0.407 | 0.047 | -0.02 | -0.02 | 0.496 | 0.43 | 0.17 | 0.52 | 0.42 | 0.38 | 0.36 |
| 6 | 148860133 | rs9689438 | 0.557 | 1.71 | -1.951 | 0.07 | -0.019 | 0.016 | 0.016 | 1.818 | 0.27 | 0.2 | 0.32 | 0.17 | 0.31 | 0.33 |
| 6 | 148861716 | rs6909677 | 0.04 | 1.712 | -1.788 | 0.192 | -0.001 | 0.001 | 0.001 | 0 | 0.12 | 0.1 | 0.12 | 0.1 | 0.15 | 0.11 |
| 6 | 148864499 | rs208697 | 0.72 | 1.275 | -2.154 | -0.165 | -0.011 | 0.108 | 0.108 | 2.76 | 0.25 | 0 | 0.11 | 0.05 | 0.2 | 0.13 |
| 6 | 148865257 | rs208696 | 0.644 | 1.889 | -1.676 | 0.318 | -0.014 | 0.004 | 0.004 | 0 | 0.65 | 0.7 | 0.62 | 0.36 | 0.57 | 0.64 |
| 6 | 148869370 | rs17643290 | 0.862 | 2.124 | -1.663 | 0.229 | -0.058 | 0.04 | 0.04 | 0.012 | 0.1 | 0.01 | 0.1 | 0.04 | 0.18 | 0.06 |
| 6 | 148869910 | rs8641 | -0.104 | 1.941 | -1.712 | 0.557 | 0.002 | -0.003 | -0.003 | 0 | 0.77 | 0.79 | 0.74 | 0.69 | 0.77 | 0.79 |
| 6 | 148871389 | rs9485296 | 0.807 | 1.959 | -1.673 | 0.561 | 0.001 | 0.026 | 0.026 | 0 | 0.17 | 0.08 | 0.15 | 0.11 | 0.18 | 0.15 |
| 6 | 148874974 | rs208739 | 0.899 | 1.89 | -1.481 | 0.646 | -0.003 | -0.023 | -0.023 | 0 | 0.21 | 0.11 | 0.16 | 0.41 | 0.19 | 0.21 |
| 6 | 148882824 | rs208729 | 0.77 | 2.104 | -1.086 | 0.301 | -0.007 | 0.013 | 0.013 | 0 | 0.58 | 0.63 | 0.58 | 0.7 | 0.5 | 0.63 |
| 6 | 148883305 | rs83099 | 0.279 | 2.09 | -1.052 | 0.355 | -0.003 | -0.004 | -0.004 | 0.168 | 0.69 | 0.67 | 0.68 | 0.75 | 0.69 | 0.69 |
| 6 | 148891724 | rs208707 | 1.156 | 2.168 | -0.256 | 1.136 | 0.037 | 0.038 | 0.038 | 0.068 | 0.81 | 0.68 | 0.76 | 0.73 | 0.73 | 0.86 |
| 6 | 148895566 | rs6928856 | -1.149 | 1.703 | -0.529 | 1.059 | 0.002 | 0.037 | 0.037 | 0.043 | 0.07 | 0.16 | 0.12 | 0.12 | 0.07 | 0.11 |
| 6 | 148895973 | rs10484601 | -0.646 | 1.884 | -0.602 | 1.051 | 0.003 | 0.01 | 0.01 | 1.106 | 0.15 | 0.21 | 0.19 | 0.24 | 0.18 | 0.12 |
| 6 | 148897510 | rs208704 | -0.834 | 2.097 | -1.164 | 1.16 | -0.038 | -0.052 | -0.052 | 0 | 0.23 | 0.1 | 0.26 | 0.37 | 0.34 | 0.15 |
| 6 | 148897782 | rs208703 | -0.826 | 1.528 | -1.331 | 0.611 | 0.042 | 0.139 | 0.139 | 0 | 0.55 | 0.8 | 0.65 | 0.83 | 0.66 | 0.62 |
| 6 | 148898485 | rs208702 | -0.939 | 1.541 | -0.927 | 0.789 | 0.048 | 0.041 | 0.041 | 0 | 0.5 | 0.66 | 0.63 | 0.65 | 0.65 | 0.42 |
| 6 | 148901519 | rs12209297 | 0.241 | 1.651 | -1.206 | 0.987 | -0.05 | 0.001 | 0.001 | 0.156 | 0.11 | 0.01 | 0.08 | 0.09 | 0.17 | 0.05 |
| 6 | 148902151 | rs208699 | -0.675 | 1.414 | -0.766 | 0.868 | -0.003 | -0.037 | -0.037 | 0 | 0.64 | 0.61 | 0.65 | 0.83 | 0.69 | 0.73 |
| 6 | 148914294 | rs9498066 | -0.737 | 2.285 | -0.797 | 1.582 | -0.013 | 0.015 | 0.015 | 0.782 | 0.19 | 0.17 | 0.28 | 0.07 | 0.27 | 0.17 |
| 6 | 148916365 | rs1535509 | -1.075 | 1.43 | -0.566 | 2.34 | 0.003 | 0.05 | 0.05 | 0 | 0.07 | 0.05 | 0.08 | 0.31 | 0.11 | 0.04 |
| 6 | 148919901 | rs9377145 | -1.102 | 1.244 | -0.244 | 1.427 | 0.034 | 0.108 | 0.108 | 3.143 | 0.22 | 0.49 | 0.18 | 0.38 | 0.24 | 0.22 |
| 6 | 148920164 | rs6909404 | -0.033 | 1.085 | -1.109 | -0.201 | 0.003 | 0.017 | 0.017 | 0 | 0.58 | 0.68 | 0.52 | 0.99 | 0.59 | 0.75 |
| 6 | 148920352 | rs9390586 | -0.894 | 0.895 | -1.779 | 1.015 | -0.025 | -0.092 | -0.092 | 1.472 | 0.16 | 0.01 | 0.14 | 0.39 | 0.18 | 0.11 |
| 6 | 148921825 | rs9377146 | 0.063 | 1.084 | -2.261 | 1.043 | -0.141 | -0.177 | -0.177 | 0.021 | 0.31 | 0.02 | 0.29 | 0.38 | 0.34 | 0.18 |
| 6 | 148924798 | rs1570363 | -0.597 | 0.384 | -1.728 | 1.048 | 0.013 | -0.016 | -0.016 | 0 | 0.29 | 0.28 | 0.4 | 0.45 | 0.26 | 0.44 |
| 6 | 148928031 | rs9373570 | -0.892 | 0.921 | -0.985 | 0.831 | 0 | 0.019 | 0.019 | 0.592 | 0.4 | 0.52 | 0.54 | 0.56 | 0.45 | 0.53 |
| 6 | 148929075 | rs1475144 | -1.1 | 0.927 | -0.775 | 0.834 | 0 | 0.016 | 0.016 | 0.572 | 0.4 | 0.54 | 0.55 | 0.56 | 0.45 | 0.53 |
| 6 | 148929867 | rs9390588 | -0.501 | 0.817 | -0.517 | 0.796 | 0.097 | 0.012 | 0.012 | 0 | 0.11 | 0.31 | 0.15 | 0.12 | 0.19 | 0.09 |
| 6 | 148934993 | rs911979 | -0.279 | 0.616 | -0.72 | 1.036 | 0.011 | -0.007 | -0.007 | 0 | 0.23 | 0.18 | 0.38 | 0.3 | 0.2 | 0.35 |
| 6 | 148935711 | rs9498070 | 0.199 | 0.579 | -0.758 | 1.028 | -0.009 | 0.028 | 0.028 | 0 | 0.05 | 0.09 | 0.03 | 0.15 | 0.04 | 0.08 |
| 6 | 148937464 | rs874841 | -1.519 | 0.377 | -0.927 | 0.684 | 0.006 | -0.045 | -0.045 | 1.29 | 0.66 | 0.89 | 0.7 | 0.66 | 0.71 | 0.81 |
| 6 | 148940600 | rs6928140 | 0.589 | 0.503 | -0.572 | 0.444 | 0.024 | -0.057 | -0.057 | 0 | 0.24 | 0.34 | 0.15 | 0.1 | 0.27 | 0.25 |
| 6 | 148940808 | rs13193931 | 0.582 | 0.661 | -1.169 | 0.465 | -0.042 | 0.016 | 0.016 | 0 | 0.07 | 0.01 | 0.11 | 0.05 | 0.16 | 0.05 |
| 6 | 148943174 | rs9403969 | 0.999 | 0.398 | -0.943 | 0.461 | -0.058 | 0.059 | 0.059 | 0 | 0.22 | 0.08 | 0.13 | 0.09 | 0.27 | 0.25 |
| 6 | 148946393 | rs2065147 | 0.256 | 0.317 | -0.984 | 0.781 | 0.006 | -0.052 | -0.052 | 0 | 0.2 | 0.11 | 0.35 | 0.4 | 0.21 | 0.34 |
| 6 | 148952353 | rs4897023 | -0.677 | 0.793 | -0.779 | 0.43 | -0.063 | 0.066 | 0.066 | 0 | 0.76 | 0.94 | 0.86 | 0.86 | 0.74 | 0.73 |
| 6 | 148971363 | rs6929864 | -0.254 | 0.945 | -0.498 | 0.616 | -0.005 | -0.005 | -0.005 | 0 | 0.18 | 0.13 | 0.21 | 0.41 | 0.22 | 0.3 |
| 6 | 148979117 | rs7449871 | -0.917 | 0.768 | -0.365 | 0.446 | -0.01 | 0.019 | 0.019 | 0 | 0.74 | 0.82 | 0.84 | 0.91 | 0.69 | 0.77 |
| 6 | 148981394 | rs7451498 | -0.458 | 0.884 | -0.413 | 0.404 | -0.035 | 0.038 | 0.038 | 0 | 0.7 | 0.81 | 0.75 | 0.89 | 0.55 | 0.74 |
| 6 | 148996235 | rs11754224 | 0.329 | 1.254 | -0.415 | 0.654 | -0.043 | -0.081 | -0.081 | 0 | 0.12 | 0.01 | 0.08 | 0.36 | 0.14 | 0.22 |
| 6 | 148997061 | rs9498107 | 1.247 | 0.838 | -0.229 | 0.479 | 0.052 | -0.009 | -0.009 | 0.163 | 0.39 | 0.13 | 0.48 | 0.42 | 0.29 | 0.28 |
| 6 | 149013861 | rs11155587 | 1.35 | 1.026 | -0.273 | 0.421 | 0.062 | -0.014 | -0.014 | 0 | 0.41 | 0.14 | 0.48 | 0.44 | 0.29 | 0.29 |
| 6 | 149015365 | rs9377154 | -0.234 | 1.173 | -0.18 | 0.503 | -0.003 | 0.009 | 0.009 | 0.007 | 0.17 | 0.15 | 0.21 | 0.41 | 0.23 | 0.3 |
| 6 | 149016528 | rs9498131 | -0.511 | 1.134 | 0.301 | 0.468 | -0.005 | -0.064 | -0.064 | 0.015 | 0.67 | 0.49 | 0.8 | 0.87 | 0.68 | 0.68 |
| 6 | 149016692 | rs7453519 | 0.674 | 0.919 | 0.175 | 0.476 | 0.018 | 0.037 | 0.037 | 0.073 | 0.5 | 0.35 | 0.6 | 0.46 | 0.45 | 0.38 |
| 6 | 149027366 | rs11155589 | 0.775 | 1.119 | 0.135 | 0.475 | 0.028 | 0.03 | 0.03 | 0.391 | 0.52 | 0.37 | 0.53 | 0.43 | 0.34 | 0.4 |
| 6 | 149027559 | rs6918447 | -0.864 | 1.376 | -0.095 | 0.537 | 0.018 | 0.04 | 0.04 | 0 | 0.2 | 0.25 | 0.3 | 0.45 | 0.35 | 0.33 |
| 6 | 149038080 | rs7356870 | 0.361 | 1.219 | 0.235 | -0.306 | -0.021 | 0.033 | 0.033 | 0 | 0.65 | 0.71 | 0.73 | 0.86 | 0.53 | 0.62 |
| 6 | 149042800 | rs10457834 | 0.42 | 0.682 | 0.404 | -0.422 | 0 | 0.009 | 0.009 | 0 | 0.65 | 0.36 | 0.81 | 0.88 | 0.67 | 0.64 |
| 6 | 149044264 | rs4897043 | -0.41 | 0.787 | 0.174 | -0.419 | -0.037 | 0.061 | 0.061 | 0 | 0.74 | 0.91 | 0.85 | 0.88 | 0.69 | 0.72 |
| 6 | 149048092 | rs4130936 | -1.298 | 0.751 | 0.901 | -0.038 | 0.047 | 0 | 0 | 0 | 0.08 | 0.41 | 0.03 | 0 | 0.03 | 0.09 |
| 6 | 149053030 | rs9377155 | -1.235 | 1.117 | 0.705 | 0.145 | 0.048 | 0.043 | 0.043 | 0.325 | 0.12 | 0.38 | 0.3 | 0.2 | 0.12 | 0.33 |
| 6 | 149058808 | rs7764228 | 0.011 | 0.954 | -0.707 | -0.181 | -0.078 | 0.027 | 0.027 | 0 | 0.53 | 0.85 | 0.44 | 0.37 | 0.54 | 0.54 |
| 6 | 149066468 | rs6920109 | 0.034 | 0.727 | -0.307 | -0.262 | 0.002 | -0.005 | -0.005 | 0 | 0.7 | 0.71 | 0.6 | 0.44 | 0.74 | 0.7 |
| 6 | 149074229 | rs9498147 | -0.554 | 1.708 | -0.101 | -0.347 | 0.012 | 0.001 | 0.001 | 0 | 0.38 | 0.5 | 0.42 | 0.42 | 0.44 | 0.47 |
| 6 | 149080301 | rs11751562 | -0.331 | 1.718 | 0.194 | -0.375 | 0.008 | -0.001 | -0.001 | 0 | 0.36 | 0.32 | 0.38 | 0.36 | 0.41 | 0.45 |
| 6 | 149082853 | rs12189648 | 1.21 | 1.239 | -0.285 | -0.843 | -0.007 | 0.1 | 0.1 | 0.008 | 0.31 | 0.15 | 0.2 | 0.06 | 0.36 | 0.24 |
| 6 | 149083115 | rs7760963 | -0.491 | 1.837 | 0.432 | -0.207 | 0.048 | 0.013 | 0.013 | 0 | 0.27 | 0.36 | 0.27 | 0.28 | 0.35 | 0.36 |
| 6 | 149087254 | rs7754504 | -1.076 | 1.856 | 0.455 | -0.269 | -0.041 | 0.023 | 0.023 | 0.042 | 0.37 | 0.67 | 0.37 | 0.28 | 0.46 | 0.41 |
| 6 | 149087712 | rs17645399 | 0.401 | 2.053 | 0.226 | -0.282 | -0.01 | 0.037 | 0.037 | 0 | 0.16 | 0.1 | 0.12 | 0.03 | 0.17 | 0.18 |

| | | | | | | | | | | | | | | | | |
|---|---|---|---|---|---|---|---|---|---|---|---|---|---|---|---|---|
| 6 | 149091694 | rs17645564 | -0.071 | 2.074 | 0.312 | -0.268 | 0.04 | -0.039 | -0.039 | 0.128 | 0.07 | 0.16 | 0.09 | 0 | 0.1 | 0.05 |
| 6 | 149092757 | rs6570896 | -0.816 | 2.11 | 0.118 | -0.17 | 0.049 | 0.01 | 0.01 | 0 | 0.28 | 0.44 | 0.25 | 0.25 | 0.32 | 0.33 |
| 6 | 149093372 | rs6933887 | 0.052 | 1.775 | -0.142 | 0.029 | 0.107 | 0.161 | 0.161 | 0 | 0.11 | 0.31 | 0.13 | 0.21 | 0.15 | 0.16 |
| 6 | 149099378 | rs6570897 | -0.296 | 0.929 | -0.732 | 0.187 | 0.02 | -0.01 | -0.01 | 0.759 | 0.34 | 0.36 | 0.32 | 0.24 | 0.42 | 0.39 |
| 6 | 149105738 | rs9386234 | -0.27 | 0.538 | -1.041 | 1.237 | 0.016 | -0.128 | -0.128 | 0 | 0.16 | 0.01 | 0.19 | 0.43 | 0.09 | 0.12 |
| 6 | 149112756 | rs7753810 | -0.849 | 0.842 | -0.049 | 0.038 | 0.001 | -0.069 | -0.069 | 0 | 0.27 | 0.48 | 0.37 | 0.12 | 0.31 | 0.24 |
| 6 | 149112958 | rs9390608 | -0.976 | 0.694 | -0.57 | 1.275 | -0.018 | -0.048 | -0.048 | 2.412 | 0.1 | 0.04 | 0.05 | 0.5 | 0.1 | 0.08 |
| 6 | 149113186 | rs9390610 | -0.898 | 0.563 | -0.416 | 1.819 | 0 | 0.016 | 0.016 | 0 | 0.09 | 0.05 | 0.05 | 0.54 | 0.1 | 0.06 |
| 6 | 149113918 | rs17085135 | -0.969 | -0.013 | 0.186 | 1.956 | -0.018 | 0.013 | 0.013 | 0 | 0.27 | 0.35 | 0.19 | 0.31 | 0.18 | 0.3 |
| 6 | 149114811 | rs9390611 | -1.009 | 0.223 | 0.107 | 1.745 | -0.022 | -0.022 | -0.022 | 0 | 0.11 | 0.06 | 0.14 | 0.58 | 0.26 | 0.11 |
| 6 | 149114996 | rs11753679 | 0.326 | -0.126 | -0.261 | 0.911 | 0.006 | 0.009 | 0.009 | 0 | 0.14 | 0.1 | 0.24 | 0.18 | 0.11 | 0.31 |
| 6 | 149117485 | rs4897050 | -1.302 | 0.063 | -0.495 | -0.452 | -0.001 | -0.043 | -0.043 | 0.034 | 0.79 | 0.84 | 0.89 | 1 | 0.81 | 0.87 |
| 6 | 149118099 | rs9485331 | -0.348 | 0.024 | -0.687 | 0.037 | -0.017 | -0.04 | -0.04 | 0.042 | 0.24 | 0.15 | 0.37 | 0.62 | 0.36 | 0.41 |
| 6 | 149118926 | rs4897051 | 0.062 | -0.53 | -0.38 | 0.019 | -0.02 | -0.013 | -0.013 | 0.168 | 0.14 | 0.18 | 0.12 | 0.08 | 0.07 | 0.2 |
| 6 | 149120342 | rs9390612 | -0.155 | 0.057 | -0.469 | 0.373 | -0.001 | 0.009 | 0.009 | 0.001 | 0.1 | 0.06 | 0.06 | 0.25 | 0.11 | 0.09 |
| 6 | 149123419 | rs9498164 | -0.046 | -0.054 | -0.863 | -0.069 | -0.063 | -0.064 | -0.064 | 0 | 0.35 | 0.17 | 0.46 | 0.45 | 0.47 | 0.36 |
| 6 | 149124034 | rs6570904 | -0.468 | -0.115 | -1.157 | -0.87 | -0.058 | 0.064 | 0.064 | 0.054 | 0.81 | 0.93 | 0.85 | 1 | 0.74 | 0.91 |
| 6 | 149125116 | rs9390613 | 0.457 | -0.584 | -0.792 | -0.96 | -0.044 | 0.053 | 0.053 | 0 | 0.26 | 0.5 | 0.28 | 0.41 | 0.19 | 0.29 |
| 6 | 149126723 | rs7751854 | 0.063 | -0.595 | -0.809 | -1.184 | 0 | -0.062 | -0.062 | 0 | 0.61 | 0.45 | 0.6 | 0.9 | 0.61 | 0.66 |
| 6 | 149131352 | rs900475 | -0.917 | -0.254 | -1.087 | -1.101 | -0.035 | 0.022 | 0.022 | 0.138 | 0.81 | 0.87 | 0.82 | 0.98 | 0.67 | 0.87 |
| 6 | 149135814 | rs11155599 | 0.322 | -0.309 | -0.876 | -1.122 | -0.007 | -0.079 | -0.079 | 0.26 | 0.25 | 0.33 | 0.3 | 0.1 | 0.24 | 0.26 |
| 6 | 149141278 | rs7761991 | -0.932 | -0.005 | -1.31 | -0.959 | -0.065 | 0.065 | 0.065 | 0.209 | 0.82 | 0.94 | 0.82 | 0.98 | 0.67 | 0.87 |
| 6 | 149156611 | rs17663626 | 0.608 | 0.927 | -0.801 | -0.171 | 0.007 | -0.018 | -0.018 | 0 | 0.06 | 0.09 | 0.06 | 0 | 0.1 | 0.01 |
| 6 | 149160809 | rs2128332 | -1.128 | 0.169 | -0.814 | 0.087 | -0.001 | -0.033 | -0.033 | 0 | 0.54 | 0.4 | 0.67 | 0.94 | 0.52 | 0.63 |
| 6 | 149161504 | rs2128333 | 0.322 | 0.419 | -1.263 | 0.61 | -0.099 | -0.144 | -0.144 | 0 | 0.25 | 0.03 | 0.39 | 0.52 | 0.3 | 0.34 |
| 6 | 149163503 | rs12529264 | 0.028 | 0.381 | -1.212 | 0.402 | -0.021 | -0.045 | -0.045 | 0 | 0.19 | 0.21 | 0.23 | 0.1 | 0.14 | 0.25 |
| 6 | 149166697 | rs9377166 | -0.843 | 1.084 | -1.394 | 1.359 | -0.088 | 0.126 | 0.126 | 0 | 0.32 | 0.09 | 0.42 | 0.84 | 0.35 | 0.39 |
| 6 | 149168205 | rs9403990 | -0.4 | 1.15 | -1.469 | 1.503 | -0.074 | -0.041 | -0.041 | 0 | 0.26 | 0.06 | 0.39 | 0.66 | 0.3 | 0.34 |
| 6 | 149175496 | rs9377167 | -0.991 | 0.636 | -0.895 | 1.763 | 0.001 | -0.029 | -0.029 | 0.008 | 0.08 | 0.02 | 0.04 | 0.33 | 0.07 | 0.06 |
| 6 | 149186342 | rs9283119 | -0.918 | 0.919 | -0.69 | 1.226 | -0.005 | -0.026 | -0.026 | 0.405 | 0.24 | 0.15 | 0.35 | 0.56 | 0.22 | 0.31 |
| 6 | 149195423 | rs11965507 | -1.069 | 1.009 | 0.19 | 0.586 | 0.112 | -0.01 | -0.01 | 0 | 0.06 | 0.24 | 0.13 | 0.01 | 0.11 | 0.05 |
| 6 | 149200719 | rs1384670 | -0.368 | 0.295 | 0.055 | 0.373 | 0.015 | -0.007 | -0.007 | 0 | 0.31 | 0.24 | 0.25 | 0.41 | 0.21 | 0.45 |
| 6 | 149202876 | rs9485337 | -1.144 | 0.432 | 0.323 | 0.582 | 0.076 | -0.004 | -0.004 | 0.955 | 0.09 | 0.3 | 0.18 | 0.07 | 0.14 | 0.08 |
| 6 | 149205617 | rs9377172 | -0.533 | 0.05 | -0.05 | 0.717 | -0.011 | 0.054 | 0.054 | 0 | 0.11 | 0.15 | 0.2 | 0.37 | 0.08 | 0.14 |
| 6 | 149210734 | rs9498175 | 0.647 | 0.296 | -0.715 | 0.29 | -0.035 | -0.088 | -0.088 | 0 | 0.31 | 0.14 | 0.4 | 0.48 | 0.39 | 0.37 |
| 6 | 149214810 | rs1351698 | -0.381 | -0.314 | 0.063 | 0.199 | -0.036 | -0.067 | -0.067 | 0 | 0.28 | 0.36 | 0.12 | 0.12 | 0.18 | 0.35 |
| 6 | 149216304 | rs4897066 | -0.888 | 0.067 | -0.323 | 0.37 | 0.021 | 0.022 | 0.022 | 0.18 | 0.46 | 0.74 | 0.46 | 0.47 | 0.39 | 0.54 |
| 6 | 149216375 | rs10080691 | -0.932 | 0.502 | 0.335 | 1.012 | 0.002 | 0.046 | 0.046 | 0 | 0.24 | 0.42 | 0.32 | 0.37 | 0.23 | 0.32 |
| 6 | 149221294 | rs1871921 | 0.291 | 0.225 | -0.548 | 0.705 | -0.007 | 0.177 | 0.177 | 0.738 | 0.36 | 0.09 | 0.25 | 0.1 | 0.34 | 0.3 |
| 6 | 149222646 | rs7757423 | 0.042 | 0.507 | -0.314 | 0.642 | 0 | -0.004 | -0.004 | 0.327 | 0.67 | 0.57 | 0.69 | 0.54 | 0.68 | 0.63 |
| 6 | 149224628 | rs12192351 | 0.166 | 0.867 | 0.241 | 0.548 | -0.004 | 0 | 0 | 0 | 0.32 | 0.26 | 0.4 | 0.1 | 0.43 | 0.26 |
| 6 | 149228093 | rs4897070 | -0.235 | 0.862 | -0.154 | 1.334 | -0.006 | -0.01 | -0.01 | 0 | 0.82 | 0.76 | 0.84 | 0.72 | 0.8 | 0.83 |
| 6 | 149228992 | rs6570908 | 0.122 | 0.858 | -0.229 | 0.994 | 0 | 0.005 | 0.005 | 0.151 | 0.78 | 0.71 | 0.77 | 0.36 | 0.79 | 0.68 |
| 6 | 149230849 | rs13215793 | 0.237 | 1.079 | -0.658 | 0.542 | -0.014 | 0.024 | 0.024 | 0 | 0.15 | 0.11 | 0.12 | 0.05 | 0.16 | 0.1 |
| 6 | 149233373 | rs6908004 | 0.772 | 0.928 | -0.632 | 0.234 | 0.007 | 0.108 | 0.108 | 1.078 | 0.36 | 0.16 | 0.26 | 0.11 | 0.34 | 0.27 |
| 6 | 149233995 | rs9403998 | 0.562 | 0.95 | -0.43 | 0.666 | 0 | 0.057 | 0.057 | 0 | 0.94 | 0.89 | 0.97 | 0.81 | 0.96 | 0.91 |
| 6 | 149234031 | rs9404000 | 0.266 | 1.097 | -0.529 | 1.049 | -0.006 | -0.043 | -0.043 | 1.258 | 0.11 | 0.03 | 0.18 | 0.43 | 0.11 | 0.28 |
| 6 | 149234163 | rs17665513 | 1.09 | 1.109 | -0.522 | 0.991 | 0.011 | 0.107 | 0.107 | 0.347 | 0.35 | 0.16 | 0.25 | 0.11 | 0.33 | 0.26 |
| 6 | 149234271 | rs9399670 | 0.389 | 1.062 | -0.507 | 1.004 | 0.007 | -0.004 | -0.004 | 0.101 | 0.47 | 0.2 | 0.44 | 0.53 | 0.45 | 0.55 |
| 6 | 149234718 | rs17721845 | 1.158 | 0.949 | -0.402 | 0.939 | 0.021 | 0.061 | 0.061 | 0.357 | 0.35 | 0.22 | 0.25 | 0.11 | 0.28 | 0.26 |
| 6 | 149237105 | rs1471677 | 0.419 | 1.007 | -0.166 | 0.828 | 0.002 | 0.065 | 0.065 | 0 | 0.95 | 0.89 | 0.97 | 0.81 | 0.95 | 0.93 |
| 6 | 149237490 | rs9390633 | 0.574 | 0.972 | -0.354 | 0.685 | 0.034 | 0.002 | 0.002 | 0.549 | 0.47 | 0.25 | 0.43 | 0.53 | 0.39 | 0.54 |
| 6 | 149238522 | rs9498176 | -0.191 | 1.103 | -0.614 | 0.183 | 0 | -0.015 | -0.015 | 0 | 0.06 | 0.07 | 0.05 | 0 | 0.06 | 0.02 |
| 6 | 149239806 | rs9498177 | -0.158 | 1.007 | -0.983 | -0.113 | 0.001 | 0.09 | 0.09 | 0 | 0.25 | 0.08 | 0.17 | 0.06 | 0.21 | 0.13 |
| 6 | 149240393 | rs12201109 | -0.118 | 1.433 | 0.002 | 0.432 | 0.025 | -0.002 | -0.002 | 0 | 0.21 | 0.28 | 0.28 | 0.13 | 0.32 | 0.21 |
| 6 | 149246402 | rs11155607 | -0.707 | 1.068 | 0.362 | 0.526 | 0.004 | 0.058 | 0.058 | 0 | 0.2 | 0.5 | 0.25 | 0.2 | 0.19 | 0.18 |
| 6 | 149246525 | rs1026900 | 0.395 | 0.687 | -0.49 | 0.312 | 0.018 | 0.002 | 0.002 | 0 | 0.4 | 0.3 | 0.33 | 0.41 | 0.35 | 0.44 |
| 6 | 149252323 | rs9404007 | 0.21 | 0.633 | -0.492 | 0.285 | 0.015 | 0.001 | 0.001 | 0 | 0.4 | 0.31 | 0.33 | 0.41 | 0.35 | 0.43 |
| 6 | 149252715 | rs10499254 | 0.456 | 0.593 | -0.896 | 0.363 | 0 | -0.001 | -0.001 | 0 | 0.07 | 0.07 | 0.13 | 0.1 | 0.06 | 0.13 |
| 6 | 149255811 | rs17080152 | -0.057 | 0.408 | -0.956 | 0.212 | -0.009 | 0.006 | 0.006 | 1.01 | 0.11 | 0.09 | 0.09 | 0 | 0.14 | 0.02 |
| 6 | 149258535 | rs17080218 | 1.325 | 0.014 | -1.181 | 0.351 | 0 | -0.033 | -0.033 | 0.114 | 0.18 | 0.04 | 0.11 | 0.16 | 0.15 | 0.18 |
| 6 | 149259321 | rs979429 | -0.547 | 0.388 | -0.351 | 0.489 | -0.002 | 0.004 | 0.004 | 0 | 0.54 | 0.48 | 0.6 | 0.54 | 0.5 | 0.63 |
| 6 | 149264090 | rs2449392 | 0.891 | 0.67 | -0.512 | 0.498 | -0.028 | -0.027 | -0.027 | 0 | 0.16 | 0.11 | 0.26 | 0.31 | 0.25 | 0.2 |
| 6 | 149265407 | rs9373582 | -0.137 | 0.646 | -0.652 | 0.574 | 0.009 | -0.001 | -0.001 | 0 | 0.13 | 0.25 | 0.1 | 0.15 | 0.14 | 0.17 |
| 6 | 149266798 | rs6570913 | 0.722 | 0.642 | -0.662 | 0.566 | 0.032 | -0.044 | -0.044 | 0 | 0.41 | 0.23 | 0.44 | 0.39 | 0.35 | 0.45 |
| 6 | 149267234 | rs9390643 | 0.213 | 0.691 | -0.628 | 0.547 | 0.019 | 0.021 | 0.021 | 1.858 | 0.09 | 0.2 | 0.07 | 0.16 | 0.11 | 0.16 |
| 6 | 149269890 | rs17728703 | -0.055 | 0.874 | -0.824 | 0.184 | -0.003 | -0.009 | -0.009 | 0 | 0.11 | 0.09 | 0.09 | 0 | 0.14 | 0.02 |
| 6 | 149270829 | rs13219957 | 1.241 | 0.899 | -0.968 | 0.261 | -0.009 | -0.021 | -0.021 | 0.055 | 0.11 | 0.04 | 0.18 | 0.11 | 0.13 | 0.12 |
| 6 | 149271125 | rs1156561 | 0.882 | 1.206 | -0.645 | -0.041 | 0.026 | 0.008 | 0.008 | 0 | 0.43 | 0.34 | 0.45 | 0.54 | 0.36 | 0.61 |

| | | | | | | | | | | | | | | | | |
|---|---|---|---|---|---|---|---|---|---|---|---|---|---|---|---|---|
| 6 | 149271254 | rs2500511 | -0.785 | 1.651 | -1.307 | -0.03 | -0.063 | 0.097 | 0.097 | 0 | 0.75 | 0.99 | 0.84 | 0.92 | 0.75 | 0.86 |
| 6 | 149273772 | rs2170534 | -0.26 | 0.854 | -0.696 | 0.315 | -0.012 | 0.019 | 0.019 | 0 | 0.57 | 0.68 | 0.56 | 0.62 | 0.5 | 0.66 |
| 6 | 149274164 | rs17728886 | -0.53 | 1.14 | -0.401 | 0.545 | 0.002 | -0.013 | -0.013 | 0 | 0.07 | 0.1 | 0.07 | 0 | 0.11 | 0.02 |
| 6 | 149279188 | rs9386240 | -0.699 | 0.62 | -0.02 | 1.041 | -0.011 | 0.06 | 0.06 | 0 | 0.21 | 0.35 | 0.33 | 0.29 | 0.2 | 0.23 |
| 6 | 149283811 | rs17729143 | 0.344 | 0.475 | -1.057 | 0.87 | -0.031 | -0.025 | -0.025 | 0.85 | 0.18 | 0.01 | 0.11 | 0.11 | 0.16 | 0.1 |
| 6 | 149285138 | rs6933251 | 0.158 | 0.364 | -0.915 | 0.815 | 0.014 | 0.016 | 0.016 | 0 | 0.41 | 0.34 | 0.26 | 0.33 | 0.36 | 0.34 |
| 6 | 149285719 | rs3734377 | 0.13 | 0.322 | -0.935 | 0.816 | 0.015 | 0.016 | 0.016 | 0 | 0.42 | 0.34 | 0.27 | 0.34 | 0.36 | 0.34 |
| 6 | 149287703 | rs2500513 | -0.645 | 0.535 | -1.259 | 0.551 | -0.012 | -0.041 | -0.041 | 0.029 | 0.75 | 0.9 | 0.75 | 0.67 | 0.75 | 0.71 |
| 6 | 149295702 | rs9404010 | 0.946 | 0.586 | -1.291 | 0.415 | -0.016 | 0.006 | 0.006 | 0 | 0.36 | 0.16 | 0.25 | 0.32 | 0.37 | 0.33 |
| 6 | 149297245 | rs2500515 | 0.072 | 0.429 | -0.105 | 0.335 | -0.007 | -0.002 | -0.002 | 0 | 0.33 | 0.26 | 0.43 | 0.35 | 0.36 | 0.37 |
| 6 | 149300485 | rs12204964 | 0.866 | 0.418 | 0.047 | 0.306 | -0.008 | 0.07 | 0.07 | 0 | 0.39 | 0.18 | 0.26 | 0.25 | 0.39 | 0.34 |
| 6 | 149300868 | rs2449381 | -0.495 | 0.039 | -0.071 | 0.674 | -0.016 | -0.029 | -0.029 | 0 | 0.77 | 0.91 | 0.79 | 0.73 | 0.75 | 0.75 |
| 6 | 149301135 | rs922832 | 0.117 | 0.108 | 0.209 | 0.664 | -0.001 | -0.005 | -0.005 | 0 | 0.34 | 0.3 | 0.48 | 0.48 | 0.34 | 0.42 |
| 6 | 149304906 | rs17729786 | 1.058 | 0.162 | -0.041 | 0.592 | -0.003 | -0.004 | -0.004 | 0 | 0.09 | 0 | 0.04 | 0.08 | 0.07 | 0.06 |
| 6 | 149305513 | rs2486405 | -0.272 | 0.088 | 0.161 | 0.475 | 0.028 | -0.023 | -0.023 | 0 | 0.3 | 0.38 | 0.34 | 0.26 | 0.4 | 0.32 |
| 6 | 149307684 | rs7741054 | -0.489 | 0.079 | 0.24 | 0.632 | 0.034 | -0.015 | -0.015 | 0.034 | 0.2 | 0.5 | 0.23 | 0.19 | 0.24 | 0.18 |
| 6 | 149308950 | rs1564601 | 0.963 | -0.129 | -0.168 | 0.57 | -0.008 | 0.019 | 0.019 | 0 | 0.4 | 0.21 | 0.27 | 0.33 | 0.41 | 0.39 |
| 6 | 149310677 | rs2500522 | 0.191 | -0.116 | 0.079 | 0.868 | 0.001 | -0.013 | -0.013 | 0 | 0.34 | 0.26 | 0.52 | 0.49 | 0.33 | 0.35 |
| 6 | 149312417 | rs2449385 | 0.133 | -0.221 | 0.326 | 1.087 | -0.014 | -0.021 | -0.021 | 0 | 0.34 | 0.42 | 0.32 | 0.25 | 0.29 | 0.33 |
| 6 | 149314780 | rs9390648 | 1.035 | -0.149 | 0.294 | 0.956 | 0.003 | 0.051 | 0.051 | 0 | 0.41 | 0.25 | 0.22 | 0.27 | 0.41 | 0.41 |
| 6 | 149315885 | rs2500526 | 0.273 | -0.576 | 0.342 | 0.86 | -0.062 | 0.095 | 0.095 | 0 | 0.76 | 0.57 | 0.57 | 0.59 | 0.85 | 0.74 |
| 6 | 149322058 | rs9390650 | 0.516 | -0.217 | 0.611 | 0.772 | 0.002 | 0.055 | 0.055 | 0 | 0.43 | 0.32 | 0.24 | 0.27 | 0.44 | 0.41 |
| 6 | 149327223 | rs6911677 | -0.058 | -0.21 | 0.601 | 0.888 | 0.011 | 0.001 | 0.001 | 0 | 0.18 | 0.19 | 0.2 | 0.19 | 0.26 | 0.2 |
| 6 | 149328196 | rs12214448 | 0.838 | -0.194 | 0.521 | 0.829 | -0.007 | 0.045 | 0.045 | 0 | 0.4 | 0.25 | 0.23 | 0.27 | 0.42 | 0.4 |
| 6 | 149329220 | rs2486420 | 0.906 | -0.224 | 0.594 | 0.859 | -0.01 | -0.003 | -0.003 | 0 | 0.32 | 0.36 | 0.3 | 0.29 | 0.26 | 0.33 |
| 6 | 149329489 | rs2500536 | 0.982 | 0.032 | 0.467 | 0.978 | 0.007 | 0.024 | 0.024 | 0 | 0.72 | 0.67 | 0.54 | 0.56 | 0.68 | 0.73 |
| 6 | 149336354 | rs2500548 | 0.391 | 0.086 | 0.34 | 1.085 | 0.003 | 0.004 | 0.004 | 0 | 0.91 | 0.91 | 0.76 | 0.75 | 0.94 | 0.93 |
| 6 | 149337165 | rs12201653 | -0.383 | -0.037 | 0.484 | 0.962 | 0.018 | 0.01 | 0.01 | 0 | 0.19 | 0.22 | 0.28 | 0.22 | 0.27 | 0.17 |
| 6 | 149337489 | rs9390652 | 0.695 | -0.326 | 0.124 | 0.637 | -0.022 | 0.096 | 0.096 | 0 | 0.37 | 0.16 | 0.22 | 0.17 | 0.4 | 0.32 |
| 6 | 149339960 | rs3798592 | 1.412 | -0.335 | -0.069 | 0.518 | -0.002 | 0.003 | 0.003 | 0 | 0.15 | 0.1 | 0.09 | 0.15 | 0.15 | 0.19 |
| 6 | 149344832 | rs2500549 | 0.846 | -0.585 | 0.484 | 0.757 | 0.024 | 0.017 | 0.017 | 0 | 0.32 | 0.14 | 0.22 | 0.27 | 0.25 | 0.39 |
| 6 | 149345952 | rs10499252 | 0.6 | -0.173 | 0.265 | 0.527 | -0.037 | -0.023 | -0.023 | 0 | 0.06 | 0 | 0.13 | 0.08 | 0.13 | 0.09 |
| 6 | 149348504 | rs2340802 | -0.237 | -0.614 | 0.543 | 0.41 | 0.045 | -0.019 | -0.019 | 0 | 0.59 | 0.79 | 0.54 | 0.39 | 0.68 | 0.47 |
| 6 | 149351381 | rs9377188 | -0.588 | -0.654 | 0.424 | 0.401 | 0.039 | -0.075 | -0.075 | 0.536 | 0.64 | 0.87 | 0.56 | 0.47 | 0.71 | 0.55 |
| 6 | 149354307 | rs9399678 | -0.635 | -0.599 | 0.438 | 0.436 | 0.034 | -0.079 | -0.079 | 0.156 | 0.65 | 0.89 | 0.56 | 0.47 | 0.71 | 0.56 |
| 6 | 149357384 | rs6570921 | 0.811 | -0.324 | 0.717 | 0.27 | -0.009 | 0.038 | 0.038 | 0 | 0.38 | 0.26 | 0.23 | 0.18 | 0.41 | 0.32 |
| 6 | 149361111 | rs6936176 | -0.26 | -0.273 | 1.129 | 0.023 | 0.045 | 0.031 | 0.031 | 0 | 0.55 | 0.68 | 0.52 | 0.28 | 0.65 | 0.47 |
| 6 | 149366064 | rs2153252 | 0.887 | -0.361 | 1.323 | -0.046 | -0.001 | -0.001 | -0.001 | 0.239 | 0.35 | 0.29 | 0.21 | 0.07 | 0.35 | 0.31 |
| 6 | 149370158 | rs6915298 | -0.539 | -0.364 | 1.386 | -0.054 | 0.029 | 0.039 | 0.039 | 0 | 0.57 | 0.87 | 0.52 | 0.28 | 0.63 | 0.47 |
| 6 | 149371138 | rs12202158 | 0.827 | -0.352 | 1.424 | -0.053 | -0.003 | 0.041 | 0.041 | 0 | 0.35 | 0.21 | 0.2 | 0.07 | 0.35 | 0.32 |
| 6 | 149374484 | rs9322164 | -0.195 | -0.293 | 1.119 | 0.016 | -0.012 | -0.009 | -0.009 | 0 | 0.2 | 0.19 | 0.29 | 0.28 | 0.27 | 0.24 |
| 6 | 149388387 | rs2340798 | -0.534 | -0.188 | 0.878 | 0.079 | 0.033 | 0.008 | 0.008 | 0 | 0.58 | 0.82 | 0.5 | 0.36 | 0.63 | 0.54 |
| 6 | 149391605 | rs9485353 | 0.846 | -0.156 | 0.947 | 0.016 | -0.001 | -0.06 | -0.06 | 0 | 0.36 | 0.38 | 0.21 | 0.07 | 0.36 | 0.32 |
| 6 | 149398249 | rs6939103 | 0.091 | -0.009 | 0.685 | 0.148 | 0.05 | 0.023 | 0.023 | 0 | 0.18 | 0.3 | 0.29 | 0.2 | 0.26 | 0.15 |
| 6 | 149401466 | rs2486409 | 0.954 | -0.156 | 0.474 | 0.327 | -0.023 | -0.056 | -0.056 | 0 | 0.33 | 0.03 | 0.24 | 0.36 | 0.31 | 0.4 |
| 6 | 149405218 | rs2153251 | 0.819 | 0.095 | 0.668 | 0.186 | 0 | -0.032 | -0.032 | 0 | 0.37 | 0.36 | 0.24 | 0.07 | 0.37 | 0.32 |
| 6 | 149411165 | rs11755147 | 0.087 | 0.086 | 0.518 | 0.261 | 0.051 | 0.033 | 0.033 | 0 | 0.18 | 0.34 | 0.27 | 0.2 | 0.23 | 0.14 |
| 6 | 149411689 | rs7761119 | -0.475 | -0.046 | 0.04 | 0.3 | 0.003 | 0.04 | 0.04 | 0 | 0.58 | 0.92 | 0.53 | 0.27 | 0.63 | 0.46 |
| 6 | 149416054 | rs1934225 | 0.288 | 0.188 | 0.283 | 0.43 | 0.037 | 0.027 | 0.027 | 0 | 0.17 | 0.31 | 0.26 | 0.19 | 0.22 | 0.14 |
| 6 | 149417022 | rs1934226 | -0.117 | 0.123 | -0.167 | 0.536 | -0.014 | -0.045 | -0.045 | 0.085 | 0.92 | 0.99 | 0.77 | 0.73 | 0.93 | 0.94 |
| 6 | 149418310 | rs9498214 | 0.481 | 0.069 | 0.172 | -0.601 | 0 | 0.005 | 0.005 | 0 | 0.39 | 0.33 | 0.25 | 0.08 | 0.4 | 0.33 |
| 6 | 149423228 | rs4897080 | 0.298 | 0.352 | -0.612 | -0.445 | 0.026 | 0.016 | 0.016 | 0 | 0.17 | 0.26 | 0.26 | 0.2 | 0.23 | 0.14 |
| 6 | 149429546 | rs6902452 | -0.352 | 0.088 | -1.034 | -0.2 | -0.028 | -0.042 | -0.042 | 0 | 0.61 | 0.96 | 0.54 | 0.38 | 0.64 | 0.54 |
| 6 | 149435003 | rs3935945 | -0.08 | 0.443 | -0.114 | -0.045 | 0.041 | 0.022 | 0.022 | 0 | 0.22 | 0.41 | 0.3 | 0.22 | 0.25 | 0.16 |
| 6 | 149435111 | rs7772697 | 0.047 | 0.123 | -0.448 | -0.577 | -0.002 | -0.023 | -0.023 | 0 | 0.42 | 0.54 | 0.28 | 0.18 | 0.4 | 0.45 |
| 6 | 149436538 | rs4897081 | -0.667 | 0.201 | 0.356 | -0.456 | 0.01 | -0.007 | -0.007 | 0 | 0.34 | 0.56 | 0.39 | 0.28 | 0.4 | 0.39 |
| 6 | 149437584 | rs6900527 | 0.513 | 0.214 | 0.507 | -0.478 | 0 | -0.005 | -0.005 | 0.118 | 0.16 | 0.18 | 0.12 | 0.04 | 0.19 | 0.17 |
| 6 | 149442245 | rs7761698 | -0.011 | 0.202 | 0.646 | -0.478 | -0.006 | -0.023 | -0.023 | 0 | 0.2 | 0.3 | 0.13 | 0.1 | 0.2 | 0.26 |
| 6 | 149454963 | rs12191264 | -0.646 | 0.241 | 0.634 | -0.481 | 0.004 | -0.005 | -0.005 | 0 | 0.34 | 0.39 | 0.38 | 0.21 | 0.39 | 0.31 |
| 6 | 149460966 | rs4357160 | 0.618 | 0.295 | 0.56 | -0.638 | 0.008 | -0.003 | -0.003 | 0 | 0.16 | 0.21 | 0.2 | 0.15 | 0.19 | 0.14 |
| 6 | 149469655 | rs6570947 | -1.088 | -0.185 | 0.162 | -1.014 | -0.007 | 0.027 | 0.027 | 0 | 0.61 | 0.84 | 0.5 | 0.32 | 0.59 | 0.61 |
| 6 | 149471139 | rs6919716 | 0.859 | -0.191 | 0.159 | -1.065 | 0.001 | 0.02 | 0.02 | 0.574 | 0.24 | 0.15 | 0.15 | 0.07 | 0.19 | 0.22 |
| 6 | 149472075 | rs2069314 | -0.492 | -0.058 | 0.298 | -1.005 | 0.009 | -0.038 | -0.038 | 0 | 0.36 | 0.54 | 0.34 | 0.18 | 0.39 | 0.32 |
| 6 | 149486333 | rs6926365 | -0.577 | -0.384 | 0.363 | -0.976 | 0.002 | -0.002 | -0.002 | 0 | 0.45 | 0.56 | 0.29 | 0.12 | 0.4 | 0.43 |
| 6 | 149488513 | rs13214977 | 0.217 | -0.424 | -0.214 | -0.793 | 0 | -0.018 | -0.018 | 0.052 | 0.26 | 0.24 | 0.16 | 0.07 | 0.2 | 0.23 |
| 6 | 149493632 | rs7746208 | -0.366 | -0.479 | -0.641 | -0.536 | 0.063 | 0.13 | 0.13 | 0 | 0.48 | 0.77 | 0.26 | 0.25 | 0.33 | 0.48 |
| 6 | 149495480 | rs12200379 | 0.224 | -0.705 | -0.88 | -0.662 | 0.004 | 0.032 | 0.032 | 0 | 0.15 | 0.05 | 0.08 | 0.03 | 0.11 | 0.21 |
| 6 | 149498383 | rs7751665 | -0.178 | -0.787 | 0.075 | -0.689 | -0.022 | -0.029 | -0.029 | 0 | 0.18 | 0.33 | 0.1 | 0.11 | 0.12 | 0.26 |
| 6 | 149499799 | rs6925848 | 0.234 | -0.721 | 0.293 | -0.617 | -0.026 | -0.056 | -0.056 | 0 | 0.3 | 0.47 | 0.17 | 0.22 | 0.22 | 0.27 |

| | | | | | | | | | | | | | | | | |
|---|---|---|---|---|---|---|---|---|---|---|---|---|---|---|---|---|
| 6 | 149504792 | rs7451236 | -0.877 | -0.081 | -0.8 | 0.205 | -0.093 | -0.1 | -0.1 | 0.196 | 0.73 | 0.98 | 0.74 | 0.65 | 0.66 | 0.69 |
| 6 | 149513519 | rs7766009 | -0.315 | -0.292 | -0.417 | -0.076 | -0.011 | -0.011 | -0.011 | 0 | 0.7 | 0.78 | 0.55 | 0.39 | 0.58 | 0.69 |
| 6 | 149517720 | rs11155632 | 0.932 | -0.463 | -0.967 | -0.215 | -0.006 | -0.025 | -0.025 | 0 | 0.1 | 0 | 0.21 | 0.1 | 0.1 | 0.13 |
| 6 | 149518083 | rs6570952 | -0.446 | -0.757 | -0.511 | -0.51 | 0.016 | 0.02 | 0.02 | 0 | 0.45 | 0.6 | 0.41 | 0.27 | 0.34 | 0.52 |
| 6 | 149523760 | rs565357 | -0.874 | -0.635 | -0.791 | -0.412 | -0.075 | -0.074 | -0.074 | 0 | 0.72 | 0.89 | 0.56 | 0.39 | 0.57 | 0.69 |
| 6 | 149526345 | rs558902 | -0.061 | -0.583 | -0.482 | -0.473 | -0.005 | 0.042 | 0.042 | 0 | 0.62 | 0.76 | 0.59 | 0.77 | 0.59 | 0.67 |
| 6 | 149527462 | rs9377195 | 0.106 | -0.655 | -0.249 | -0.447 | -0.003 | 0.006 | 0.006 | 0 | 0.35 | 0.52 | 0.44 | 0.65 | 0.35 | 0.5 |
| 6 | 149533147 | rs659119 | 1.134 | -0.287 | -0.56 | -0.489 | -0.013 | 0.058 | 0.058 | 0 | 0.25 | 0.09 | 0.14 | 0.12 | 0.23 | 0.15 |
| 6 | 149536923 | rs17725371 | 1.108 | -0.325 | -0.38 | 0.129 | -0.03 | -0.039 | -0.039 | 0 | 0.21 | 0.08 | 0.39 | 0.33 | 0.3 | 0.24 |
| 6 | 149539773 | rs2272901 | -0.213 | -0.715 | -0.134 | 0.253 | -0.005 | -0.068 | -0.068 | 0 | 0.29 | 0.11 | 0.17 | 0.41 | 0.26 | 0.27 |
| 6 | 149546620 | rs12201953 | -0.123 | -0.515 | 0.404 | -0.056 | -0.025 | -0.03 | -0.03 | 0 | 0.07 | 0.14 | 0.01 | 0.02 | 0.02 | 0.16 |
| 6 | 149552517 | rs6910470 | -0.15 | -0.553 | 0.591 | -0.021 | -0.024 | -0.019 | -0.019 | 0 | 0.1 | 0.24 | 0.04 | 0.05 | 0.04 | 0.05 |
| 6 | 149552587 | rs926408 | -0.007 | -0.72 | 0.028 | -0.19 | 0.013 | 0.043 | 0.043 | 2.021 | 0.47 | 0.29 | 0.49 | 0.36 | 0.45 | 0.39 |
| 6 | 149557858 | rs7750814 | 0.828 | -0.303 | 0.339 | -0.024 | 0.018 | 0.045 | 0.045 | 0 | 0.17 | 0.26 | 0.28 | 0.3 | 0.21 | 0.24 |
| 6 | 149560966 | rs7748981 | 1.077 | -0.525 | 0.27 | -0.14 | -0.002 | 0 | 0 | 0.189 | 0.27 | 0.26 | 0.43 | 0.28 | 0.34 | 0.25 |
| 6 | 149562839 | rs6570956 | -0.42 | -0.69 | 0.13 | -0.368 | 0.015 | 0.062 | 0.062 | 0 | 0.72 | 0.84 | 0.86 | 0.96 | 0.74 | 0.7 |
| 6 | 149565631 | rs9399682 | 1.004 | 0.038 | 0.399 | 0.225 | -0.012 | -0.018 | -0.018 | 2.48 | 0.15 | 0.12 | 0.26 | 0.3 | 0.2 | 0.22 |
| 6 | 149566825 | rs6935566 | 0.276 | -0.024 | 0.387 | 0.209 | 0.005 | 0.005 | 0.005 | 0 | 0.42 | 0.49 | 0.7 | 0.58 | 0.55 | 0.48 |
| 6 | 149573828 | rs4897097 | 0.977 | 0.11 | -0.296 | 0.21 | -0.073 | -0.023 | -0.023 | 0 | 0.24 | 0.04 | 0.41 | 0.28 | 0.34 | 0.25 |
| 6 | 149577079 | rs7758080 | -0.357 | -0.452 | 0.662 | 0.411 | -0.015 | 0.006 | 0.006 | 0 | 0.38 | 0.43 | 0.26 | 0.4 | 0.25 | 0.31 |
| 6 | 149580983 | rs11968300 | 0.082 | -0.631 | 0.901 | 0.369 | 0.008 | -0.02 | -0.02 | 0.469 | 0.29 | 0.17 | 0.21 | 0.37 | 0.21 | 0.23 |
| 6 | 149581692 | rs4897101 | 0.862 | -0.729 | 1.251 | 0.327 | 0.002 | 0.007 | 0.007 | 0 | 0.14 | 0.16 | 0.24 | 0.24 | 0.18 | 0.23 |
| 6 | 149582909 | rs6937786 | -0.058 | -0.694 | 1.05 | 0.335 | -0.005 | 0.019 | 0.019 | 0 | 0.23 | 0.47 | 0.29 | 0.27 | 0.22 | 0.28 |
| 6 | 149585510 | rs6570957 | -0.728 | -0.611 | 0.668 | 0.176 | 0.016 | 0.051 | 0.051 | 0.132 | 0.51 | 0.63 | 0.49 | 0.7 | 0.4 | 0.49 |
| 6 | 149587364 | rs6908247 | -0.728 | -0.611 | 0.66 | 0.176 | 0.015 | 0.047 | 0.047 | 0.132 | 0.51 | 0.63 | 0.5 | 0.7 | 0.4 | 0.5 |
| 6 | 149594286 | rs9285518 | -0.508 | -0.656 | 0.424 | 0.165 | 0.027 | 0.052 | 0.052 | 1.413 | 0.46 | 0.76 | 0.49 | 0.66 | 0.36 | 0.44 |
| 6 | 149596746 | rs4897104 | -0.742 | -0.667 | 0.422 | 0.142 | 0.02 | 0.083 | 0.083 | 0.046 | 0.51 | 0.72 | 0.5 | 0.7 | 0.4 | 0.5 |
| 6 | 149597217 | rs9404025 | -0.435 | -0.589 | 0.317 | 0.185 | 0.006 | 0.013 | 0.013 | 0.63 | 0.44 | 0.62 | 0.47 | 0.66 | 0.36 | 0.44 |
| 6 | 149600310 | rs6904464 | -0.908 | -0.628 | -0.384 | 0.364 | 0.012 | 0.08 | 0.08 | 1.728 | 0.55 | 0.87 | 0.5 | 0.7 | 0.38 | 0.5 |
| 6 | 149605182 | rs9498283 | -0.487 | -0.498 | 0.064 | 0.907 | 0.002 | -0.003 | -0.003 | 0.395 | 0.28 | 0.2 | 0.19 | 0.42 | 0.16 | 0.22 |
| 6 | 149614718 | rs610707 | -0.963 | -0.191 | 0.093 | 1.003 | 0.025 | -0.011 | -0.011 | 0.209 | 0.39 | 0.71 | 0.29 | 0.44 | 0.29 | 0.34 |
| 6 | 149622136 | rs613276 | -0.8 | -0.322 | 0.855 | 0.935 | -0.015 | 0.018 | 0.018 | 0 | 0.34 | 0.41 | 0.26 | 0.44 | 0.29 | 0.34 |
| 6 | 149625169 | rs563373 | -1.043 | -0.35 | 0.653 | 0.921 | 0.041 | -0.025 | -0.025 | 0 | 0.38 | 0.73 | 0.28 | 0.44 | 0.29 | 0.34 |
| 6 | 149625612 | rs597144 | -0.714 | -0.328 | 0.582 | 0.932 | -0.013 | 0.012 | 0.012 | 0.027 | 0.35 | 0.55 | 0.27 | 0.44 | 0.29 | 0.34 |
| 6 | 149628277 | rs661838 | -1.054 | 0.076 | -0.356 | 1.025 | 0.019 | -0.017 | -0.017 | 0 | 0.46 | 0.89 | 0.38 | 0.57 | 0.36 | 0.5 |
| 6 | 149658978 | rs504985 | -1.576 | 0.955 | 0.776 | 1.11 | 0.005 | -0.011 | -0.011 | 0 | 0.36 | 0.32 | 0.44 | 0.64 | 0.35 | 0.39 |
| 6 | 149685617 | rs506268 | -1.955 | 0.824 | 1.814 | 0.95 | -0.006 | 0.001 | 0.001 | 0.422 | 0.4 | 0.57 | 0.47 | 0.64 | 0.37 | 0.49 |
| 6 | 149688090 | rs6911411 | -0.536 | 0.714 | 1.701 | 0.894 | 0 | -0.024 | -0.024 | 0.318 | 0.2 | 0.22 | 0.17 | 0.02 | 0.21 | 0.11 |
| 6 | 149691355 | rs6570963 | 2.08 | 0.703 | 2.028 | 0.886 | -0.014 | 0.043 | 0.043 | 0.111 | 0.56 | 0.27 | 0.4 | 0.26 | 0.52 | 0.41 |
| 6 | 149692935 | rs9285519 | -1.777 | 0.693 | 2.12 | 0.89 | -0.004 | 0.004 | 0.004 | 0.164 | 0.36 | 0.44 | 0.46 | 0.64 | 0.35 | 0.39 |
| 6 | 149705635 | rs11155647 | -0.933 | 0.629 | 1.952 | 1.044 | 0 | 0.006 | 0.006 | 0.076 | 0.25 | 0.24 | 0.19 | 0.09 | 0.25 | 0.19 |
| 6 | 149718650 | rs237028 | -2.274 | 0.591 | 1.827 | 1.028 | -0.021 | 0.072 | 0.072 | 0.468 | 0.45 | 0.77 | 0.6 | 0.74 | 0.48 | 0.6 |
| 6 | 149730846 | rs652921 | -0.168 | 0.657 | 1.756 | 1.107 | 0.004 | -0.015 | -0.015 | 1.378 | 0.1 | 0.06 | 0.25 | 0.55 | 0.09 | 0.2 |
| 6 | 149731904 | rs2744434 | -0.237 | 0.657 | 1.756 | 1.107 | 0.004 | -0.018 | -0.018 | 0 | 0.1 | 0.06 | 0.25 | 0.55 | 0.09 | 0.2 |
| 6 | 149743871 | rs2744432 | -1.373 | 0.631 | 1.986 | 1.122 | 0 | 0.034 | 0.034 | 0.506 | 0.29 | 0.36 | 0.42 | 0.56 | 0.29 | 0.28 |
| 6 | 149743930 | rs237018 | -2.328 | 0.489 | 1.868 | 0.989 | -0.024 | 0.097 | 0.097 | 0.558 | 0.46 | 0.81 | 0.61 | 0.74 | 0.49 | 0.61 |
| 6 | 149748411 | rs674919 | -0.236 | 0.581 | 1.827 | 1.103 | -0.005 | 0.086 | 0.086 | 0 | 0.1 | 0.17 | 0.27 | 0.42 | 0.1 | 0.2 |
| 6 | 149750334 | rs237004 | -1.704 | 0.581 | 1.876 | 0.355 | 0.032 | -0.05 | -0.05 | 0.153 | 0.3 | 0.51 | 0.3 | 0.11 | 0.34 | 0.33 |
| 6 | 149756355 | rs9498349 | 0.208 | 0.378 | 1.636 | 0.349 | -0.019 | -0.045 | -0.045 | 0 | 0.13 | 0.25 | 0.09 | 0.01 | 0.08 | 0.08 |
| 6 | 149762537 | rs2179367 | -1.883 | 0.213 | 1.64 | 0.475 | 0.027 | 0.054 | 0.054 | 0.882 | 0.42 | 0.73 | 0.6 | 0.78 | 0.37 | 0.51 |
| 6 | 149762851 | rs9498354 | -0.29 | 0.611 | 1.63 | 0.524 | 0.009 | -0.079 | -0.079 | 2.465 | 0.21 | 0.34 | 0.18 | 0.02 | 0.21 | 0.11 |
| 6 | 149767105 | rs12192159 | -0.222 | 0.666 | 1.251 | 0.56 | 0.001 | -0.026 | -0.026 | 0 | 0.19 | 0.23 | 0.17 | 0.02 | 0.21 | 0.11 |
| 6 | 149782908 | rs376757 | -0.604 | 0.915 | 0.988 | 0.622 | 0.077 | -0.033 | -0.033 | 0 | 0.26 | 0.49 | 0.25 | 0.11 | 0.35 | 0.21 |
| 6 | 149795154 | rs376269 | -0.668 | 0.791 | 0.786 | 0.533 | 0.059 | -0.035 | -0.035 | 0.001 | 0.28 | 0.54 | 0.27 | 0.12 | 0.38 | 0.23 |
| 6 | 149798835 | rs656461 | -0.965 | 0.6 | 0.186 | 0.962 | 0.011 | -0.044 | -0.044 | 0 | 0.16 | 0.07 | 0.38 | 0.76 | 0.14 | 0.29 |
| 6 | 149802179 | rs2789503 | -0.845 | 0.467 | 0.094 | 0.554 | 0 | 0 | 0 | 0 | 0.54 | 0.5 | 0.46 | 0.53 | 0.49 | 0.57 |
| 6 | 149814762 | rs2341551 | -1.17 | -0.61 | 0.365 | -0.065 | -0.031 | -0.017 | -0.017 | 0.25 | 0.31 | 0.64 | 0.49 | 0.77 | 0.18 | 0.4 |
| 6 | 149825264 | rs4895792 | -0.797 | -0.405 | -0.312 | -0.557 | -0.053 | 0.026 | 0.026 | 0 | 0.86 | 1 | 0.91 | 0.94 | 0.87 | 0.8 |
| 6 | 149845117 | rs17087579 | 0.225 | -0.971 | 0.676 | -0.508 | -0.047 | -0.062 | -0.062 | 0 | 0.11 | 0.25 | 0.02 | 0 | 0.02 | 0.1 |
| 6 | 149845650 | rs10485097 | -0.202 | -1.058 | 0.757 | -0.545 | 0.011 | -0.021 | -0.021 | 0 | 0.18 | 0.09 | 0.44 | 0.77 | 0.16 | 0.3 |
| 6 | 149855742 | rs761617 | -1.671 | -1.07 | 0.951 | -0.561 | -0.044 | -0.025 | -0.025 | 0 | 0.3 | 0.44 | 0.46 | 0.78 | 0.18 | 0.4 |
| 6 | 149861981 | rs11961781 | 0.163 | -0.957 | 1.016 | -0.422 | -0.047 | -0.062 | -0.062 | 0 | 0.11 | 0.25 | 0.02 | 0 | 0.02 | 0.1 |
| 6 | 149896318 | rs4870139 | -0.12 | -0.52 | 0.66 | -0.45 | -0.012 | 0.028 | 0.028 | 0.092 | 0.31 | 0.14 | 0.6 | 0.82 | 0.35 | 0.39 |
| 6 | 149906712 | rs9285521 | -0.461 | -0.598 | 1.051 | -0.746 | -0.001 | -0.005 | -0.005 | 0.163 | 0.34 | 0.33 | 0.62 | 0.84 | 0.36 | 0.38 |
| 6 | 149913867 | rs11155662 | -0.218 | -0.557 | 0.997 | -0.726 | -0.011 | 0.043 | 0.043 | 0 | 0.3 | 0.14 | 0.6 | 0.82 | 0.34 | 0.38 |
| 6 | 149915053 | rs3734295 | -0.403 | -0.568 | 1.333 | -0.733 | -0.007 | -0.039 | -0.039 | 0 | 0.33 | 0.45 | 0.62 | 0.82 | 0.34 | 0.38 |
| 6 | 149943594 | rs10782310 | -0.424 | -0.616 | 1.45 | -0.697 | -0.003 | -0.02 | -0.02 | 0 | 0.36 | 0.59 | 0.64 | 0.82 | 0.36 | 0.39 |
| 6 | 149972087 | rs12529698 | -0.376 | -0.622 | 1.496 | -0.7 | -0.002 | -0.035 | -0.035 | 0 | 0.34 | 0.42 | 0.62 | 0.82 | 0.36 | 0.38 |
| 6 | 149979416 | rs1125 | -0.031 | -0.671 | 1.426 | 0.097 | -0.021 | -0.071 | -0.071 | 0 | 0.29 | 0.13 | 0.49 | 0.52 | 0.35 | 0.27 |

| | | | | | | | | | | | | | | | | |
|---|---|---|---|---|---|---|---|---|---|---|---|---|---|---|---|---|
| 6 | 149982417 | rs9800736 | -0.407 | -0.674 | 1.438 | -0.09 | -0.002 | -0.039 | -0.039 | 0 | 0.34 | 0.42 | 0.62 | 0.82 | 0.36 | 0.38 |
| 6 | 149983216 | rs3924871 | -0.443 | -0.683 | 1.471 | -0.095 | -0.005 | -0.031 | -0.031 | 0 | 0.36 | 0.57 | 0.63 | 0.82 | 0.36 | 0.38 |
| 6 | 150004779 | rs3798761 | -0.435 | -0.655 | 1.503 | -0.091 | -0.006 | -0.03 | -0.03 | 0 | 0.36 | 0.58 | 0.64 | 0.82 | 0.36 | 0.38 |
| 6 | 150032247 | rs11155674 | -0.409 | -0.68 | 1.51 | -0.094 | -0.004 | -0.03 | -0.03 | 0.812 | 0.36 | 0.57 | 0.63 | 0.82 | 0.36 | 0.38 |
| 6 | 150039148 | rs12174349 | -0.422 | -0.673 | 1.521 | -0.09 | -0.005 | -0.033 | -0.033 | 0 | 0.36 | 0.57 | 0.63 | 0.82 | 0.36 | 0.38 |
| 6 | 150046219 | rs9479094 | -0.422 | -0.67 | 1.485 | -0.088 | -0.005 | -0.033 | -0.033 | 0 | 0.36 | 0.57 | 0.63 | 0.82 | 0.36 | 0.38 |
| 6 | 150049261 | rs17088269 | 0.595 | -0.89 | 1.652 | -0.198 | -0.039 | -0.037 | -0.037 | 0 | 0.09 | 0.21 | 0.02 | 0.01 | 0.01 | 0.11 |
| 6 | 150050676 | rs12526675 | -0.098 | -0.866 | 1.79 | -0.194 | -0.003 | 0.022 | 0.022 | 0 | 0.32 | 0.24 | 0.61 | 0.82 | 0.36 | 0.38 |
| 6 | 150061959 | rs12212415 | 2.253 | -0.676 | 1.143 | -0.549 | -0.096 | 0.113 | 0.113 | 0 | 0.28 | 0.01 | 0.18 | 0.05 | 0.34 | 0.14 |
| 6 | 150067675 | rs10872646 | -0.648 | -1.279 | 1.913 | 0.164 | -0.003 | -0.03 | -0.03 | 0 | 0.35 | 0.54 | 0.64 | 0.82 | 0.36 | 0.38 |
| 6 | 150072028 | rs12528279 | -0.648 | -1.279 | 1.913 | 0.164 | -0.003 | -0.03 | -0.03 | 3.366 | 0.35 | 0.54 | 0.64 | 0.82 | 0.36 | 0.38 |
| 6 | 150073891 | rs10782316 | -0.648 | -1.279 | 1.913 | 0.164 | -0.003 | -0.03 | -0.03 | 0 | 0.35 | 0.54 | 0.64 | 0.82 | 0.36 | 0.38 |
| 6 | 150145001 | rs9371201 | -0.163 | -1.218 | 1.176 | 0.375 | -0.019 | -0.077 | -0.077 | 0.433 | 0.29 | 0.13 | 0.49 | 0.51 | 0.35 | 0.28 |
| 6 | 150146216 | rs9478848 | -0.855 | -1.168 | 1.096 | -0.012 | 0.045 | 0.118 | 0.118 | 0.061 | 0.45 | 0.76 | 0.65 | 0.83 | 0.37 | 0.48 |
| 6 | 150166569 | rs1108889 | -0.058 | -0.982 | 0.838 | 0.063 | -0.034 | 0.056 | 0.056 | 0 | 0.31 | 0.1 | 0.49 | 0.84 | 0.35 | 0.39 |
| 6 | 150169932 | rs7769115 | -0.322 | -0.991 | 1.222 | 0.079 | 0.001 | 0.028 | 0.028 | 0 | 0.33 | 0.26 | 0.51 | 0.84 | 0.35 | 0.39 |
| 6 | 150174507 | rs6899595 | -0.306 | -1.094 | 1.738 | 0.019 | -0.047 | -0.04 | -0.04 | 0.37 | 0.11 | 0.42 | 0.03 | 0.01 | 0.01 | 0.14 |
| 6 | 150177132 | rs4869750 | -0.933 | -1.056 | 1.416 | 0.085 | 0.038 | 0.089 | 0.089 | 0.008 | 0.44 | 0.73 | 0.55 | 0.85 | 0.36 | 0.52 |
| 6 | 150182629 | rs1537590 | -0.622 | -0.936 | 1.392 | 0.612 | 0.002 | 0.177 | 0.177 | 0.021 | 0.1 | 0.22 | 0.08 | 0.47 | 0.1 | 0.18 |
| 6 | 150187873 | rs1999632 | 0.535 | -1.015 | 1.158 | 0.2 | -0.037 | -0.09 | -0.09 | 0 | 0.23 | 0.04 | 0.36 | 0.35 | 0.25 | 0.23 |
| 6 | 150195384 | rs6557133 | -0.731 | -1.172 | 1.507 | 0.055 | 0 | -0.032 | -0.032 | 0 | 0.39 | 0.41 | 0.53 | 0.89 | 0.42 | 0.47 |
| 6 | 150198491 | rs9322227 | -0.606 | -1.149 | 1.341 | 0.079 | -0.003 | 0.003 | 0.003 | 0.259 | 0.13 | 0.11 | 0.12 | 0.55 | 0.16 | 0.25 |
| 6 | 150201720 | rs10457852 | 0.646 | -1.129 | 1.127 | 0.061 | -0.05 | -0.088 | -0.088 | 0 | 0.23 | 0.04 | 0.37 | 0.34 | 0.26 | 0.23 |
| 6 | 150205095 | rs7742993 | -0.688 | -1.151 | 1.54 | -0.026 | -0.001 | 0 | 0 | 0.505 | 0.39 | 0.38 | 0.5 | 0.89 | 0.42 | 0.48 |
| 6 | 150210681 | rs9371533 | -1.242 | -1.3 | 1.768 | -0.139 | 0.033 | 0.244 | 0.244 | 1.212 | 0.51 | 0.84 | 0.54 | 0.9 | 0.43 | 0.62 |
| 6 | 150210723 | rs6925151 | 0.101 | -0.55 | 2.205 | 0.459 | 0 | 0.024 | 0.024 | 0 | 0.26 | 0.39 | 0.39 | 0.33 | 0.26 | 0.22 |
| 6 | 150210802 | rs6933882 | -0.965 | -0.546 | 2.164 | 0.46 | 0.001 | -0.054 | -0.054 | 1.835 | 0.39 | 0.46 | 0.51 | 0.89 | 0.42 | 0.48 |
| 6 | 150211100 | rs3798763 | 0.425 | -0.54 | 1.89 | 0.465 | -0.05 | -0.083 | -0.083 | 0 | 0.23 | 0.04 | 0.37 | 0.33 | 0.26 | 0.22 |
| 6 | 150212003 | rs9383583 | -1.015 | -0.519 | 1.878 | 0.488 | 0.001 | 0.097 | 0.097 | 0.581 | 0.15 | 0.22 | 0.12 | 0.55 | 0.16 | 0.26 |
| 6 | 150217360 | rs9371544 | -0.844 | -0.527 | 1.89 | 0.351 | -0.003 | 0.036 | 0.036 | 0.584 | 0.39 | 0.33 | 0.49 | 0.87 | 0.42 | 0.48 |
| 6 | 150226059 | rs9383940 | -0.896 | -0.461 | 1.763 | 0.469 | 0.004 | 0.089 | 0.089 | 1.016 | 0.42 | 0.74 | 0.53 | 0.85 | 0.41 | 0.47 |
| 6 | 150232879 | rs4458704 | 0.71 | -0.455 | 1.72 | 0.447 | 0.004 | 0.073 | 0.073 | 0.941 | 0.59 | 0.3 | 0.47 | 0.15 | 0.59 | 0.53 |
| 6 | 150242165 | rs6927913 | -1.045 | -0.417 | 1.555 | 0.617 | 0.019 | 0.315 | 0.315 | 0.528 | 0.16 | 0.43 | 0.13 | 0.54 | 0.15 | 0.24 |
| 6 | 150245110 | rs4869767 | -0.3 | -0.609 | 0.823 | 0.608 | -0.005 | -0.013 | -0.013 | 0.012 | 0.76 | 0.82 | 0.63 | 0.68 | 0.74 | 0.78 |
| 6 | 150245203 | rs4869768 | -0.291 | -0.606 | 0.825 | 0.61 | -0.005 | -0.013 | -0.013 | 0.876 | 0.76 | 0.82 | 0.63 | 0.68 | 0.74 | 0.78 |
| 6 | 150246386 | rs17079048 | 0.355 | -1.273 | 1.001 | 0.155 | -0.007 | -0.044 | -0.044 | 3.535 | 0.46 | 0.45 | 0.22 | 0.1 | 0.35 | 0.39 |
| 6 | 150247274 | rs7762285 | 0.489 | -0.698 | 0.791 | 0.475 | -0.01 | -0.032 | -0.032 | 1.248 | 0.81 | 0.75 | 0.92 | 0.89 | 0.85 | 0.76 |
| 6 | 150251382 | rs4283898 | 0.086 | -0.598 | 0.985 | 1.225 | -0.001 | 0.004 | 0.004 | 0 | 0.58 | 0.51 | 0.29 | 0.59 | 0.48 | 0.49 |
| 6 | 150256609 | rs4504486 | -0.876 | -0.912 | 1.343 | 1.819 | 0.005 | 0.047 | 0.047 | 0 | 0.07 | 0.04 | 0.11 | 0.33 | 0.03 | 0.16 |
| 6 | 150261361 | rs9479184 | -0.624 | -1.076 | 1.518 | 1.856 | -0.021 | 0.049 | 0.049 | 0.099 | 0.11 | 0.15 | 0.05 | 0.38 | 0.05 | 0.06 |
| 6 | 150271944 | rs12524576 | -1.555 | -0.834 | 1.193 | 1.549 | -0.001 | 0.038 | 0.038 | 0 | 0.2 | 0.23 | 0.38 | 0.36 | 0.17 | 0.28 |
| 6 | 150272532 | rs6557200 | -1.482 | -0.751 | 0.222 | 1.535 | 0.004 | -0.009 | -0.009 | 0 | 0.25 | 0.75 | 0.42 | 0.37 | 0.18 | 0.29 |
| 6 | 150284435 | rs4425606 | -1.378 | -0.625 | 0.042 | 1.504 | 0.009 | -0.024 | -0.024 | 0 | 0.23 | 0.79 | 0.24 | 0.36 | 0.15 | 0.24 |
| 6 | 150298842 | rs1853665 | -1.024 | 0.038 | 0.132 | 0.983 | -0.025 | 0.029 | 0.029 | 0.185 | 0.25 | 0.5 | 0.32 | 0.36 | 0.18 | 0.18 |
| 6 | 150303710 | rs7773464 | 0.226 | 0.519 | -0.293 | 0.655 | 0.001 | -0.003 | -0.003 | 0.077 | 0.58 | 0.55 | 0.54 | 0.59 | 0.51 | 0.65 |
| 6 | 150318351 | rs9383633 | -0.149 | -0.073 | -1.413 | 0.173 | -0.006 | 0.062 | 0.062 | 0 | 0.43 | 0.13 | 0.28 | 0.26 | 0.42 | 0.38 |
| 6 | 150318754 | rs6922844 | -0.849 | -0.363 | -0.766 | 1.114 | -0.014 | 0.125 | 0.125 | 0 | 0.08 | 0.18 | 0.09 | 0.22 | 0.04 | 0.09 |
| 6 | 150330061 | rs2343266 | 0.816 | 0.08 | -1.183 | 1 | -0.01 | -0.033 | -0.033 | 1.5 | 0.83 | 0.85 | 0.72 | 0.47 | 0.79 | 0.71 |
| 6 | 150338065 | rs9479405 | 0.739 | -0.482 | -0.682 | 0.274 | -0.002 | 0.075 | 0.075 | 0 | 0.48 | 0.38 | 0.3 | 0.24 | 0.44 | 0.37 |
| 6 | 150340035 | rs6913561 | -0.352 | -0.473 | -0.101 | 0.245 | 0.007 | 0.015 | 0.015 | 0 | 0.44 | 0.36 | 0.47 | 0.23 | 0.41 | 0.4 |
| 6 | 150344374 | rs11757186 | 1.077 | -0.593 | 0.112 | 0.111 | -0.036 | 0.114 | 0.114 | 0 | 0.42 | 0.24 | 0.29 | 0.24 | 0.44 | 0.37 |
| 6 | 150349132 | rs12209388 | 0.499 | -0.889 | -0.336 | -0.071 | -0.059 | -0.014 | -0.014 | 0 | 0.12 | 0.01 | 0.18 | 0.06 | 0.19 | 0.13 |
| 6 | 150350099 | rs13209192 | 1.251 | -0.996 | -0.007 | -0.085 | 0.001 | -0.03 | -0.03 | 0 | 0.45 | 0.54 | 0.4 | 0.24 | 0.45 | 0.4 |
| 6 | 150355441 | rs1413901 | 0.131 | -1.013 | 0.034 | -0.02 | 0.012 | 0.115 | 0.115 | 0 | 0.9 | 0.85 | 0.87 | 0.48 | 0.86 | 0.79 |
| 6 | 150358012 | rs9479482 | 1.606 | -1.098 | 0.17 | -0.14 | -0.061 | 0.14 | 0.14 | 0 | 0.41 | 0.19 | 0.29 | 0.24 | 0.44 | 0.37 |
| 6 | 150359199 | rs562425 | 1.4 | -1.437 | -0.008 | -0.865 | -0.093 | 0.085 | 0.085 | 0 | 0.33 | 0.17 | 0.28 | 0.21 | 0.41 | 0.25 |
| 6 | 150364009 | rs644866 | 0.97 | -1.503 | 0.765 | 0.378 | -0.02 | 0.086 | 0.086 | 0 | 0.75 | 0.48 | 0.67 | 0.5 | 0.8 | 0.69 |
| 6 | 150366143 | rs6557262 | 0.1 | -1.478 | 0.586 | 0.179 | -0.008 | -0.002 | -0.002 | 0 | 0.18 | 0.16 | 0.3 | 0.16 | 0.25 | 0.28 |
| 6 | 150368264 | rs11155700 | -0.227 | -0.315 | 0.907 | 0.167 | 0.036 | 0.006 | 0.006 | 0 | 0.2 | 0.45 | 0.32 | 0.16 | 0.25 | 0.28 |
| 6 | 150369345 | rs4870236 | -0.524 | -0.663 | 0.804 | -0.06 | -0.026 | -0.027 | -0.027 | 0 | 0.31 | 0.57 | 0.29 | 0.09 | 0.17 | 0.2 |
| 6 | 150370552 | rs17054300 | -0.411 | -0.11 | 0.749 | -0.014 | 0.003 | 0.004 | 0.004 | 0 | 0.28 | 0.26 | 0.21 | 0.09 | 0.23 | 0.17 |
| 6 | 150376318 | rs11155703 | 0.502 | -0.118 | 0.378 | 0.13 | -0.033 | -0.018 | -0.018 | 0.165 | 0.18 | 0.26 | 0.19 | 0.12 | 0.12 | 0.13 |
| 6 | 150382573 | rs4869814 | -0.121 | 0.21 | -0.437 | 1.096 | 0 | 0.019 | 0.019 | 0 | 0.91 | 0.9 | 0.82 | 0.71 | 0.88 | 0.88 |
| 6 | 150385730 | rs912558 | 1.778 | 0.203 | -0.694 | 0.893 | 0.008 | 0.02 | 0.02 | 0 | 0.43 | 0.29 | 0.37 | 0.41 | 0.36 | 0.43 |
| 6 | 150387746 | rs12202737 | 0.026 | 0.607 | -1.07 | 1.003 | -0.048 | -0.044 | -0.044 | 0 | 0.18 | 0.05 | 0.23 | 0.21 | 0.29 | 0.28 |
| 6 | 150404286 | rs12197847 | 0.112 | 0.742 | -1.088 | 0.817 | -0.053 | -0.028 | -0.028 | 0 | 0.17 | 0.05 | 0.23 | 0.17 | 0.27 | 0.25 |
| 6 | 150407607 | rs17510968 | 1.318 | 0.183 | -1.37 | 0.587 | 0.02 | 0.072 | 0.072 | 0 | 0.16 | 0.01 | 0.07 | 0.02 | 0.1 | 0.1 |
| 6 | 150408096 | rs789830 | -0.523 | 0.507 | -1.168 | 1.065 | -0.023 | -0.012 | -0.012 | 0 | 0.92 | 0.99 | 0.83 | 0.6 | 0.88 | 0.87 |
| 6 | 150409072 | rs789824 | 1.661 | 0.313 | -0.664 | 0.833 | -0.001 | -0.006 | -0.006 | 0 | 0.41 | 0.61 | 0.27 | 0.22 | 0.34 | 0.33 |

| | | | | | | | | | | | | | | | | |
|---|---|---|---|---|---|---|---|---|---|---|---|---|---|---|---|---|
| 6 | 150410910 | rs1953320 | 0.379 | 0.414 | -0.342 | 1.603 | 0.002 | 0.052 | 0.052 | 0.03 | 0.87 | 0.88 | 0.79 | 0.46 | 0.84 | 0.78 |
| 6 | 150412077 | rs17079296 | 0.161 | 0.241 | -0.337 | 1.608 | 0.004 | -0.031 | -0.031 | 1.651 | 0.14 | 0.06 | 0.15 | 0.57 | 0.13 | 0.17 |
| 6 | 150412758 | rs6933642 | -0.945 | 0.245 | 0.324 | 0.873 | -0.017 | 0.085 | 0.085 | 0.003 | 0.25 | 0.49 | 0.29 | 0.62 | 0.23 | 0.32 |
| 6 | 150414026 | rs9479642 | -1.114 | 0.326 | 0.557 | 0.513 | 0.003 | 0.138 | 0.138 | 2.099 | 0.17 | 0.64 | 0.24 | 0.48 | 0.17 | 0.19 |
| 6 | 150416202 | rs9371711 | -1.042 | 0.453 | 0.162 | -0.375 | 0.002 | 0.009 | 0.009 | 0.143 | 0.35 | 0.65 | 0.34 | 0.82 | 0.32 | 0.43 |
| 6 | 150416731 | rs9479658 | 0.37 | 0.587 | 0.148 | -0.616 | 0.002 | -0.033 | -0.033 | 0 | 0.11 | 0.12 | 0.07 | 0 | 0.12 | 0.05 |
| 6 | 150421248 | rs7758305 | -0.993 | 0.739 | -0.258 | -0.961 | -0.002 | 0.036 | 0.036 | 0.737 | 0.77 | 0.85 | 0.69 | 0.89 | 0.76 | 0.74 |
| 6 | 150422657 | rs9397629 | 0.334 | 0.974 | 0.016 | -0.634 | -0.022 | 0.019 | 0.019 | 0 | 0.1 | 0.05 | 0.07 | 0.05 | 0.14 | 0.14 |
| 6 | 150426272 | rs13205318 | 1.46 | 0.886 | -0.01 | -0.74 | -0.009 | 0.107 | 0.107 | 0 | 0.27 | 0.08 | 0.18 | 0.02 | 0.25 | 0.2 |
| 6 | 150429697 | rs13218442 | 1.432 | 0.907 | 0.04 | -0.741 | -0.002 | 0.081 | 0.081 | 0 | 0.27 | 0.12 | 0.18 | 0.02 | 0.24 | 0.22 |
| 6 | 150434824 | rs9478465 | -0.941 | 0.825 | 0.435 | -0.622 | 0.018 | 0.008 | 0.008 | 0 | 0.22 | 0.41 | 0.18 | 0.25 | 0.24 | 0.25 |
| 6 | 150437749 | rs520028 | 0.092 | 0.918 | 0.275 | -0.475 | -0.001 | 0.001 | 0.001 | 0 | 0.47 | 0.52 | 0.36 | 0.27 | 0.47 | 0.46 |
| 6 | 150444811 | rs9479734 | -1.057 | 0.847 | 0.354 | -0.1 | 0.024 | 0.118 | 0.118 | 0 | 0.19 | 0.36 | 0.2 | 0.62 | 0.22 | 0.28 |
| 6 | 150447142 | rs7743003 | 0.269 | 0.82 | 0.075 | -0.308 | 0 | 0.004 | 0.004 | 0 | 0.45 | 0.55 | 0.33 | 0.26 | 0.42 | 0.43 |
| 6 | 150448358 | rs6916591 | -0.288 | 0.7 | -0.587 | -0.569 | -0.04 | 0.179 | 0.179 | 0 | 0.54 | 0.82 | 0.46 | 0.77 | 0.49 | 0.55 |
| 6 | 150449865 | rs4870302 | 0.099 | 0.694 | -0.371 | -0.238 | 0.003 | 0.006 | 0.006 | 0 | 0.1 | 0.05 | 0.14 | 0.61 | 0.1 | 0.23 |
| 6 | 150449951 | rs4870303 | -0.595 | 0.652 | -0.836 | -0.648 | -0.063 | 0.109 | 0.109 | 0 | 0.67 | 0.87 | 0.54 | 0.87 | 0.59 | 0.62 |
| 6 | 150450034 | rs6922684 | -1.173 | 0.621 | 0.776 | 0.041 | 0.01 | 0.154 | 0.154 | 0.019 | 0.19 | 0.48 | 0.21 | 0.61 | 0.18 | 0.25 |
| 6 | 150451024 | rs7769185 | -0.729 | 0.674 | 0.456 | -0.354 | 0.008 | 0.021 | 0.021 | 0 | 0.4 | 0.7 | 0.31 | 0.72 | 0.35 | 0.35 |
| 6 | 150451873 | rs9479738 | 0.591 | 0.783 | 0.837 | -0.343 | 0.047 | -0.023 | -0.023 | 1.323 | 0.06 | 0.22 | 0.04 | 0 | 0.08 | 0.04 |
| 6 | 150452268 | rs1107836 | -0.723 | 0.626 | 1.064 | -0.554 | 0.007 | 0.033 | 0.033 | 0 | 0.38 | 0.3 | 0.36 | 0.86 | 0.31 | 0.37 |
| 6 | 150452759 | rs2050388 | -0.594 | 0.605 | 1.252 | -0.479 | -0.014 | -0.006 | -0.006 | 0 | 0.35 | 0.5 | 0.28 | 0.72 | 0.27 | 0.29 |
| 6 | 150454520 | rs2065715 | 0.505 | 0.899 | 0.865 | -0.778 | 0.002 | 0.057 | 0.057 | 0 | 0.52 | 0.67 | 0.44 | 0.25 | 0.47 | 0.4 |
| 6 | 150463883 | rs35455053 | -0.077 | 0.775 | 0.633 | -0.419 | 0 | 0.002 | 0.002 | 0 | 0.18 | 0.16 | 0.19 | 0.6 | 0.17 | 0.23 |
| 6 | 150465300 | rs1889551 | -0.178 | 0.795 | 0.45 | -0.478 | 0.017 | 0.034 | 0.034 | 0 | 0.1 | 0.24 | 0.14 | 0.14 | 0.12 | 0.1 |
| 6 | 150467100 | rs9371775 | -1.009 | 0.762 | 0.113 | -0.797 | 0.011 | 0.1 | 0.1 | 0 | 0.47 | 0.68 | 0.42 | 0.86 | 0.39 | 0.44 |
| 6 | 150467903 | rs9479762 | 0.987 | 0.855 | -0.184 | -0.491 | 0.009 | 0.038 | 0.038 | 0 | 0.16 | 0.04 | 0.07 | 0.12 | 0.1 | 0.1 |
| 6 | 150471263 | rs1408505 | 0.562 | 1.098 | 0.078 | -0.572 | 0.008 | -0.026 | -0.026 | 0 | 0.16 | 0.19 | 0.31 | 0.03 | 0.23 | 0.33 |
| 6 | 150471594 | rs1408504 | 0.301 | 1.097 | 0.374 | 0.103 | 0.006 | -0.016 | -0.016 | 0 | 0.64 | 0.53 | 0.52 | 0.67 | 0.61 | 0.53 |
| 6 | 150472250 | rs9885654 | 0.974 | 0.841 | -0.172 | -0.772 | -0.024 | 0.091 | 0.091 | 0 | 0.34 | 0.18 | 0.24 | 0.02 | 0.39 | 0.23 |
| 6 | 150473849 | rs1889552 | -1.252 | 0.642 | -0.496 | -0.451 | -0.018 | 0.087 | 0.087 | 0 | 0.78 | 0.98 | 0.89 | 0.88 | 0.82 | 0.91 |
| 6 | 150477700 | rs6557340 | 0.084 | 0.672 | -0.205 | -0.011 | -0.013 | 0.013 | 0.013 | 0 | 0.23 | 0.17 | 0.34 | 0.13 | 0.28 | 0.4 |
| 6 | 150477907 | rs13199127 | 0.073 | 0.646 | -0.159 | 0.079 | -0.004 | 0.069 | 0.069 | 0 | 0.13 | 0.14 | 0.15 | 0.47 | 0.11 | 0.17 |
| 6 | 150483531 | rs2343536 | -0.719 | 0.763 | -0.407 | -0.499 | 0 | -0.014 | -0.014 | 0.041 | 0.71 | 0.68 | 0.79 | 0.87 | 0.72 | 0.86 |
| 6 | 150483910 | rs9371787 | 0.199 | 1.047 | -0.585 | -0.22 | 0.003 | 0.026 | 0.026 | 0 | 0.34 | 0.24 | 0.29 | 0.19 | 0.33 | 0.25 |
| 6 | 150490917 | rs9397708 | -0.723 | 1.115 | -0.31 | 0.91 | 0 | -0.005 | -0.005 | 0.084 | 0.05 | 0.03 | 0.03 | 0.17 | 0.05 | 0.06 |
| 6 | 150494069 | rs1886359 | -0.438 | 1.428 | -0.128 | 0.442 | 0.001 | -0.001 | -0.001 | 0 | 0.16 | 0.2 | 0.37 | 0.15 | 0.25 | 0.37 |
| 6 | 150494873 | rs2057557 | -0.421 | 1.098 | -0.24 | 0.495 | -0.006 | -0.001 | -0.001 | 0 | 0.14 | 0.14 | 0.4 | 0.15 | 0.17 | 0.35 |
| 6 | 150496965 | rs6918437 | -0.926 | 0.724 | 0.041 | 0.205 | 0.003 | -0.044 | -0.044 | 0.676 | 0.36 | 0.49 | 0.54 | 0.79 | 0.35 | 0.62 |
| 6 | 150498828 | rs9371803 | -0.784 | 0.693 | 0.226 | 0.818 | 0.002 | 0.03 | 0.03 | 0 | 0.24 | 0.4 | 0.39 | 0.34 | 0.23 | 0.44 |
| 6 | 150501505 | rs11155710 | -0.143 | 0.519 | 1.083 | 0.94 | 0.01 | 0.032 | 0.032 | 0 | 0.09 | 0.2 | 0.06 | 0.11 | 0.08 | 0.08 |
| 6 | 150502047 | rs9397722 | -0.293 | 0.48 | 0.101 | 0.186 | -0.027 | 0.017 | 0.017 | 0 | 0.56 | 0.86 | 0.65 | 0.42 | 0.54 | 0.7 |
| 6 | 150506330 | rs6905409 | 0.337 | 0.419 | 0.129 | 0.136 | 0 | -0.024 | -0.024 | 0 | 0.44 | 0.54 | 0.58 | 0.27 | 0.46 | 0.6 |
| 6 | 150506561 | rs11155711 | 0.268 | 0.488 | -0.068 | 0.278 | -0.01 | -0.04 | -0.04 | 0.336 | 0.13 | 0.05 | 0.16 | 0.43 | 0.13 | 0.18 |
| 6 | 150511399 | rs9397194 | -0.355 | 0.145 | 0.171 | -0.413 | -0.027 | 0.081 | 0.081 | 0 | 0.16 | 0.44 | 0.14 | 0.2 | 0.09 | 0.16 |
| 6 | 150512533 | rs12215988 | 0.156 | 0.388 | -0.266 | -0.222 | -0.007 | 0.063 | 0.063 | 0 | 0.52 | 0.68 | 0.56 | 0.31 | 0.51 | 0.63 |
| 6 | 150521594 | rs910534 | -0.32 | 0.769 | -0.47 | -0.528 | 0.001 | -0.011 | -0.011 | 0 | 0.41 | 0.44 | 0.49 | 0.29 | 0.44 | 0.46 |
| 6 | 150521638 | rs975211 | 0.203 | 0.671 | -1.154 | -0.722 | -0.038 | 0.106 | 0.106 | 0 | 0.25 | 0.06 | 0.17 | 0.01 | 0.28 | 0.06 |
| 6 | 150525198 | rs7773289 | 0.505 | 0.257 | -0.878 | -0.25 | -0.004 | 0.017 | 0.017 | 0 | 0.85 | 0.91 | 0.92 | 0.87 | 0.86 | 0.86 |
| 6 | 150525260 | rs2281438 | 0.754 | -0.209 | -0.829 | -0.041 | 0.023 | 0.015 | 0.015 | 0.028 | 0.48 | 0.36 | 0.62 | 0.46 | 0.58 | 0.61 |
| 6 | 150525413 | rs2281437 | -1.074 | -0.838 | -0.286 | -0.015 | -0.03 | 0.054 | 0.054 | 0.24 | 0.18 | 0.61 | 0.16 | 0.35 | 0.08 | 0.14 |
| 6 | 150525540 | rs2281436 | 0.933 | -0.492 | -0.045 | -0.327 | -0.006 | -0.017 | -0.017 | 0 | 0.67 | 0.64 | 0.77 | 0.77 | 0.72 | 0.77 |
| 6 | 150525791 | rs11759558 | -0.72 | -0.244 | -0.045 | -0.359 | 0.001 | -0.052 | -0.052 | 0.437 | 0.19 | 0.21 | 0.14 | 0.01 | 0.2 | 0.12 |
| 6 | 150532152 | rs877906 | -0.178 | -1.188 | -0.066 | -0.3 | 0.021 | 0.019 | 0.019 | 0 | 0.47 | 0.64 | 0.27 | 0.3 | 0.29 | 0.26 |
| 6 | 150532221 | rs877905 | -0.504 | -1.236 | 0.106 | -0.398 | 0.009 | 0.017 | 0.017 | 0 | 0.57 | 0.67 | 0.33 | 0.3 | 0.37 | 0.3 |
| 6 | 150532488 | rs877907 | -0.575 | -1.242 | -0.031 | -0.39 | 0.018 | 0.032 | 0.032 | 0.022 | 0.58 | 0.76 | 0.34 | 0.3 | 0.37 | 0.3 |
| 6 | 150535437 | rs9478585 | -0.814 | -1.177 | 0.346 | -0.11 | -0.027 | 0.029 | 0.029 | 0 | 0.1 | 0.15 | 0.16 | 0.19 | 0.07 | 0.1 |
| 6 | 150535702 | rs926930 | -0.702 | -1.167 | 0.205 | -0.104 | -0.005 | 0.001 | 0.001 | 0 | 0.09 | 0.07 | 0.15 | 0.19 | 0.07 | 0.09 |
| 6 | 150538654 | rs9885695 | -0.765 | -1.527 | 0.843 | -0.491 | -0.023 | -0.137 | -0.137 | 0.328 | 0.18 | 0.47 | 0.11 | 0.01 | 0.14 | 0.1 |
| 6 | 150538665 | rs9885696 | -1.72 | -1.622 | 0.788 | -0.538 | -0.047 | -0.059 | -0.059 | 0 | 0.28 | 0.48 | 0.28 | 0.19 | 0.21 | 0.19 |
| 6 | 150547144 | rs7741317 | -1.061 | -1.564 | 0.729 | -0.443 | -0.03 | 0.014 | 0.014 | 0 | 0.12 | 0.23 | 0.18 | 0.19 | 0.08 | 0.1 |
| 6 | 150557220 | rs6936990 | -1.61 | -1.252 | 1.009 | -0.557 | -0.033 | -0.01 | -0.01 | 0 | 0.17 | 0.35 | 0.12 | 0.18 | 0.12 | 0.13 |
| 6 | 150557525 | rs9384264 | -0.633 | -1.227 | 0.923 | -0.534 | 0.001 | -0.002 | -0.002 | 0.102 | 0.1 | 0.1 | 0.06 | 0.17 | 0.08 | 0.08 |
| 6 | 150557735 | rs12526741 | 0.279 | -1.162 | 1.375 | 0.019 | 0.003 | 0.021 | 0.021 | 0 | 0.11 | 0.19 | 0.17 | 0.16 | 0.11 | 0.12 |
| 6 | 150558985 | rs4870386 | -1.289 | -1.707 | 0.731 | -0.093 | -0.038 | -0.07 | -0.07 | 0 | 0.62 | 0.86 | 0.47 | 0.51 | 0.43 | 0.48 |
| 6 | 150561826 | rs4870389 | -1.12 | -1.64 | 0.784 | -0.059 | -0.02 | -0.047 | -0.047 | 0.35 | 0.6 | 0.81 | 0.46 | 0.53 | 0.44 | 0.48 |
| 6 | 150564916 | rs1086206 | -0.535 | -1.698 | 0.893 | -0.078 | 0.041 | 0.041 | 0.041 | 0.009 | 0.49 | 0.7 | 0.39 | 0.41 | 0.35 | 0.4 |
| 6 | 150570867 | rs3734729 | -0.331 | -1.708 | 0.892 | -0.091 | 0 | -0.001 | -0.001 | 0 | 0.08 | 0.06 | 0.05 | 0.1 | 0.05 | 0.07 |
| 6 | 150571265 | rs1151 | -1.385 | -1.776 | 1.069 | -0.096 | -0.025 | 0.017 | 0.017 | 0 | 0.25 | 0.51 | 0.27 | 0.27 | 0.22 | 0.24 |

| | | | | | | | | | | | | | | | | |
|---|---|---|---|---|---|---|---|---|---|---|---|---|---|---|---|---|
| 6 | 150572208 | rs1211947 | -1.215 | -1.639 | 0.615 | -0.066 | -0.044 | -0.076 | -0.076 | 0 | 0.61 | 0.88 | 0.48 | 0.53 | 0.44 | 0.48 |
| 6 | 150575200 | rs11155716 | -0.923 | -1.34 | 0.913 | -0.021 | -0.025 | 0.002 | 0.002 | 0 | 0.11 | 0.18 | 0.08 | 0.11 | 0.08 | 0.07 |
| 6 | 150579911 | rs789498 | -1.446 | -1.719 | 0.298 | -0.263 | 0.06 | 0.062 | 0.062 | 0 | 0.44 | 0.82 | 0.42 | 0.33 | 0.33 | 0.34 |
| 6 | 150582038 | rs9480096 | -0.924 | -1.788 | 0.611 | -0.323 | -0.019 | -0.02 | -0.02 | 0.036 | 0.28 | 0.32 | 0.19 | 0.16 | 0.17 | 0.17 |
| 6 | 150584385 | rs789501 | -0.2 | -1.733 | 0.334 | -0.562 | 0 | -0.024 | -0.024 | 0.003 | 0.25 | 0.21 | 0.13 | 0.05 | 0.22 | 0.15 |
| 6 | 150584654 | rs789503 | -1.089 | -1.325 | 0.023 | -0.1 | -0.017 | 0.016 | 0.016 | 0.239 | 0.55 | 0.83 | 0.43 | 0.4 | 0.48 | 0.51 |
| 6 | 150589471 | rs12212378 | -0.793 | -1.507 | 0.204 | -0.055 | 0.003 | 0.022 | 0.022 | 0.591 | 0.52 | 0.52 | 0.28 | 0.27 | 0.46 | 0.46 |
| 6 | 150590784 | rs7741814 | 0.1 | -1.616 | -0.487 | -0.268 | 0.015 | 0.083 | 0.083 | 0 | 0.36 | 0.13 | 0.17 | 0.22 | 0.32 | 0.37 |
| 6 | 150591429 | rs11760085 | -1.298 | -1.511 | -0.382 | -0.091 | 0.001 | -0.001 | -0.001 | 0 | 0.24 | 0.75 | 0.27 | 0.2 | 0.18 | 0.17 |
| 6 | 150594363 | rs789481 | -1.169 | -1.551 | -0.387 | -0.189 | -0.005 | 0.007 | 0.007 | 1.958 | 0.54 | 0.84 | 0.41 | 0.42 | 0.45 | 0.5 |
| 6 | 150596143 | rs789483 | 0.033 | -0.911 | -0.968 | 0.117 | -0.014 | 0.03 | 0.03 | 0 | 0.56 | 0.32 | 0.44 | 0.31 | 0.52 | 0.6 |
| 6 | 150599561 | rs789490 | -0.779 | -0.592 | -1.001 | 0.084 | -0.024 | -0.028 | -0.028 | 0 | 0.67 | 0.82 | 0.63 | 0.5 | 0.59 | 0.7 |
| 6 | 150601041 | rs9384331 | 0.517 | -0.702 | -0.966 | 0.662 | -0.021 | -0.046 | -0.046 | 0.373 | 0.18 | 0.05 | 0.25 | 0.21 | 0.15 | 0.28 |
| 6 | 150605469 | rs12215647 | -0.323 | -0.596 | -0.463 | 0.212 | -0.004 | 0.008 | 0.008 | 0.023 | 0.35 | 0.37 | 0.36 | 0.48 | 0.28 | 0.37 |
| 6 | 150606341 | rs6928831 | -0.466 | -0.686 | 0.029 | 0.164 | -0.011 | 0.011 | 0.011 | 0 | 0.37 | 0.52 | 0.47 | 0.47 | 0.3 | 0.49 |
| 6 | 150608138 | rs9397821 | -0.093 | -0.725 | -0.245 | -0.216 | 0.001 | 0.041 | 0.041 | 0 | 0.49 | 0.37 | 0.44 | 0.69 | 0.5 | 0.49 |
| 6 | 150610615 | rs9480154 | 0.633 | -0.644 | -0.672 | -0.1 | 0.014 | -0.016 | -0.016 | 0.347 | 0.15 | 0.1 | 0.06 | 0.27 | 0.07 | 0.16 |
| 6 | 150611385 | rs4391275 | 0.812 | -0.374 | -0.61 | -0.095 | -0.076 | -0.058 | -0.058 | 0 | 0.29 | 0.12 | 0.34 | 0.4 | 0.4 | 0.28 |
| 6 | 150611573 | rs4407726 | -0.451 | 0.117 | -0.158 | -0.706 | -0.001 | 0.068 | 0.068 | 0.375 | 0.72 | 0.81 | 0.85 | 0.92 | 0.73 | 0.82 |
| 6 | 150615876 | rs1556879 | -0.631 | 0.367 | -0.031 | -0.782 | 0.01 | -0.144 | -0.144 | 0 | 0.68 | 0.55 | 0.8 | 0.97 | 0.69 | 0.78 |
| 6 | 150617997 | rs7356994 | 0.46 | -0.767 | -0.535 | -0.589 | 0.001 | -0.001 | -0.001 | 1.716 | 0.28 | 0.25 | 0.33 | 0.73 | 0.29 | 0.35 |
| 6 | 150618750 | rs11751092 | 1.031 | -0.712 | -1.227 | -0.581 | 0.028 | 0.002 | 0.002 | 0 | 0.14 | 0.01 | 0.17 | 0.11 | 0.07 | 0.11 |
| 6 | 150625469 | rs9371388 | 0.166 | -0.517 | -0.747 | -0.275 | -0.012 | 0.01 | 0.01 | 0.015 | 0.28 | 0.49 | 0.33 | 0.72 | 0.28 | 0.35 |
| 6 | 150625572 | rs6904151 | -0.147 | -0.153 | -0.97 | -0.103 | 0.003 | 0.036 | 0.036 | 0 | 0.21 | 0.16 | 0.13 | 0.04 | 0.21 | 0.21 |
| 6 | 150634383 | rs9397264 | -0.386 | -0.213 | -0.8 | 0.771 | -0.01 | -0.006 | -0.006 | 0 | 0.44 | 0.42 | 0.57 | 0.71 | 0.36 | 0.56 |
| 6 | 150637477 | rs11155719 | 0.162 | 0.372 | -0.529 | 1.364 | 0.001 | 0 | 0 | 0 | 0.6 | 0.59 | 0.52 | 0.44 | 0.53 | 0.61 |
| 6 | 150639079 | rs4870466 | -1.219 | -0.372 | -0.512 | 2.017 | 0.009 | -0.053 | -0.053 | 0 | 0.2 | 0.11 | 0.29 | 0.55 | 0.19 | 0.36 |
| 6 | 150644030 | rs12527319 | -0.723 | -0.562 | 0.21 | 2.596 | -0.007 | 0.02 | 0.02 | 0 | 0.06 | 0.08 | 0.14 | 0.16 | 0.03 | 0.04 |
| 6 | 150649726 | rs9397277 | -1.263 | -1.561 | 0.113 | 1.715 | 0.079 | -0.016 | -0.016 | 0 | 0.46 | 0.74 | 0.54 | 0.59 | 0.37 | 0.56 |
| 6 | 150657006 | rs9397886 | -1.771 | -1.288 | 0.471 | 1.959 | 0.024 | -0.034 | -0.034 | 0.119 | 0.34 | 0.74 | 0.42 | 0.54 | 0.27 | 0.51 |
| 6 | 150658401 | rs884678 | -1.046 | -1.349 | 0.67 | 1.929 | -0.009 | 0.018 | 0.018 | 0 | 0.17 | 0.23 | 0.19 | 0.32 | 0.16 | 0.3 |
| 6 | 150659958 | rs9371955 | -1.681 | -1.402 | 0.815 | 1.903 | -0.013 | 0.012 | 0.012 | 0.232 | 0.22 | 0.29 | 0.22 | 0.34 | 0.18 | 0.39 |
| 6 | 150660611 | rs4869889 | -1.407 | -1.343 | 0.775 | 1.887 | 0.017 | -0.017 | -0.017 | 0 | 0.1 | 0.08 | 0.16 | 0.19 | 0.06 | 0.16 |
| 6 | 150664867 | rs4869892 | -0.166 | -1.312 | 0.858 | 1.902 | 0.008 | -0.01 | -0.01 | 0 | 0.05 | 0.03 | 0.13 | 0.17 | 0.03 | 0.03 |
| 6 | 150665278 | rs7770283 | -1.358 | -1.347 | 0.937 | 1.918 | -0.007 | -0.002 | -0.002 | 0 | 0.08 | 0.16 | 0.09 | 0.06 | 0.04 | 0.13 |
| 6 | 150665752 | rs9397900 | -1.078 | -1.39 | 1.187 | 1.93 | -0.008 | 0.022 | 0.022 | 0 | 0.16 | 0.21 | 0.16 | 0.27 | 0.16 | 0.28 |
| 6 | 150667739 | rs1000674 | -1.512 | -1.325 | 1.029 | 1.912 | -0.01 | 0.015 | 0.015 | 0 | 0.1 | 0.19 | 0.19 | 0.21 | 0.05 | 0.1 |
| 6 | 150670899 | rs654694 | 1.966 | -1.296 | 1.079 | 1.919 | -0.03 | 0.048 | 0.048 | 0.017 | 0.71 | 0.54 | 0.62 | 0.49 | 0.77 | 0.57 |
| 6 | 150673177 | rs623882 | 1.446 | -1.242 | 1.057 | 1.825 | -0.025 | 0.046 | 0.046 | 0 | 0.79 | 0.59 | 0.77 | 0.67 | 0.84 | 0.66 |
| 6 | 150674258 | rs643603 | 1.661 | -1.06 | 0.895 | 1.93 | 0.012 | 0.007 | 0.007 | 0.091 | 0.63 | 0.3 | 0.58 | 0.33 | 0.69 | 0.44 |
| 6 | 150675830 | rs688015 | 1.393 | -0.978 | 1.186 | 2.032 | -0.001 | 0.168 | 0.168 | 0 | 0.89 | 0.65 | 0.83 | 0.67 | 0.9 | 0.76 |
| 6 | 150676502 | rs623742 | 1.557 | -1.061 | 0.929 | 1.938 | 0.025 | 0.012 | 0.012 | 0 | 0.6 | 0.21 | 0.55 | 0.37 | 0.66 | 0.41 |
| 6 | 150676716 | rs622842 | 1.672 | -0.801 | 1.554 | 1.951 | 0.006 | 0.222 | 0.222 | 0 | 0.91 | 0.59 | 0.85 | 0.68 | 0.94 | 0.83 |
| 6 | 150680274 | rs683620 | 1.6 | 0.355 | 0.982 | 1.216 | 0.018 | -0.02 | -0.02 | 0.514 | 0.9 | 0.84 | 0.89 | 0.98 | 0.86 | 0.87 |
| 6 | 150688664 | rs678182 | 1.471 | 0.398 | 1.142 | 1.238 | 0.045 | -0.024 | -0.024 | 0 | 0.92 | 0.85 | 0.9 | 0.98 | 0.86 | 0.87 |
| 6 | 150692480 | rs620102 | 1.401 | 0.322 | 1.105 | 1.2 | 0 | 0.002 | 0.002 | 0 | 0.92 | 0.92 | 0.9 | 0.98 | 0.86 | 0.87 |
| 6 | 150698674 | rs588036 | 1.216 | -0.496 | 0.333 | 0.599 | -0.053 | 0.161 | 0.161 | 0 | 0.45 | 0.19 | 0.37 | 0.22 | 0.52 | 0.27 |
| 6 | 150698839 | rs12211918 | -0.522 | -0.428 | 0.321 | 0.583 | 0.02 | -0.004 | -0.004 | 0 | 0.06 | 0.08 | 0.08 | 0.02 | 0.13 | 0.13 |
| 6 | 150698920 | rs12211979 | -0.322 | -0.426 | 0.291 | 0.585 | -0.03 | 0.008 | 0.008 | 0 | 0.05 | 0 | 0.08 | 0.02 | 0.13 | 0.13 |
| 6 | 150701927 | rs674026 | 0.784 | -0.813 | 0.954 | 1.186 | -0.007 | -0.009 | -0.009 | 0 | 0.56 | 0.67 | 0.53 | 0.54 | 0.55 | 0.4 |
| 6 | 150704465 | rs608175 | 0.819 | -0.846 | 0.977 | 1.176 | -0.005 | -0.006 | -0.006 | 0.021 | 0.56 | 0.66 | 0.52 | 0.53 | 0.55 | 0.4 |
| 6 | 150713782 | rs2076292 | -0.519 | -0.808 | 1.35 | 1.326 | 0.008 | 0.246 | 0.246 | 0 | 0.09 | 0.41 | 0.15 | 0.31 | 0.05 | 0.14 |
| 6 | 150715666 | rs2076289 | -0.67 | -0.81 | 1.326 | 1.325 | 0.001 | 0.238 | 0.238 | 0 | 0.1 | 0.47 | 0.15 | 0.31 | 0.05 | 0.14 |
| 6 | 150719404 | rs2076285 | -0.688 | -0.825 | 1.337 | 1.337 | -0.002 | 0.238 | 0.238 | 0 | 0.1 | 0.45 | 0.16 | 0.32 | 0.05 | 0.15 |
| 6 | 150719613 | rs2076284 | -0.688 | -0.825 | 1.337 | 1.337 | -0.002 | 0.238 | 0.238 | 0 | 0.1 | 0.45 | 0.15 | 0.32 | 0.05 | 0.15 |
| 6 | 150719710 | rs675804 | 1.509 | -1.134 | 1.075 | 1.378 | 0.026 | 0.118 | 0.118 | 0 | 0.48 | 0.21 | 0.39 | 0.25 | 0.58 | 0.29 |
| 6 | 150720439 | rs672605 | 0.477 | -1.23 | 0.984 | 1.81 | -0.015 | -0.05 | -0.05 | 5.556 | 0.92 | 1 | 0.85 | 0.79 | 0.93 | 0.77 |
| 6 | 150720571 | rs601223 | 0.997 | -1.251 | 1.693 | 1.799 | 0.006 | 0.085 | 0.085 | 0 | 0.9 | 0.72 | 0.83 | 0.79 | 0.93 | 0.77 |
| 6 | 150720908 | rs660153 | 1.801 | -1.286 | 1.673 | 1.797 | -0.04 | 0.194 | 0.194 | 0 | 0.81 | 0.55 | 0.7 | 0.47 | 0.88 | 0.62 |
| 6 | 150721229 | rs587713 | 2.076 | -2.314 | 1.102 | 1.218 | -0.014 | 0.006 | 0.006 | 0.726 | 0.65 | 0.43 | 0.64 | 0.43 | 0.8 | 0.6 |
| 6 | 150721401 | rs657972 | 2.137 | -1.72 | 1.408 | 1.473 | -0.006 | 0.056 | 0.056 | 0 | 0.71 | 0.48 | 0.59 | 0.42 | 0.75 | 0.58 |
| 6 | 150721430 | rs1416481 | -0.126 | -1.751 | 1.845 | 1.508 | -0.002 | 0.204 | 0.204 | 0 | 0.09 | 0.26 | 0.14 | 0.33 | 0.06 | 0.15 |
| 6 | 150721445 | rs6911086 | -1.275 | -1.752 | 1.563 | 1.509 | -0.027 | 0.103 | 0.103 | 0.031 | 0.18 | 0.51 | 0.21 | 0.37 | 0.12 | 0.16 |
| 6 | 150721882 | rs12208588 | -1.231 | -1.742 | 1.612 | 1.542 | -0.025 | 0.165 | 0.165 | 0.011 | 0.18 | 0.44 | 0.29 | 0.39 | 0.12 | 0.17 |
| 6 | 150722115 | rs6933986 | 0.87 | -1.726 | 1.573 | 1.512 | -0.001 | -0.001 | -0.001 | 0.217 | 0.07 | 0.05 | 0.04 | 0.04 | 0.07 | 0.02 |
| 6 | 150722210 | rs11758050 | -0.164 | -1.704 | 1.407 | 1.499 | 0.005 | 0.001 | 0.001 | 0 | 0.23 | 0.16 | 0.14 | 0.18 | 0.21 | 0.25 |
| 6 | 150722772 | rs2211106 | -0.186 | -1.769 | 1.388 | 1.5 | 0.006 | -0.004 | -0.004 | 0 | 0.23 | 0.17 | 0.14 | 0.22 | 0.21 | 0.25 |
| 6 | 150723616 | rs7763368 | -1.23 | -1.616 | 1.481 | 1.577 | -0.022 | 0.164 | 0.164 | 0.007 | 0.18 | 0.46 | 0.29 | 0.38 | 0.12 | 0.16 |
| 6 | 150723812 | rs7768510 | -1.23 | -1.616 | 1.481 | 1.577 | -0.022 | 0.164 | 0.164 | 0.007 | 0.18 | 0.46 | 0.29 | 0.38 | 0.12 | 0.16 |

| | | | | | | | | | | | | | | | | |
|---|---|---|---|---|---|---|---|---|---|---|---|---|---|---|---|---|
| 6 | 150724581 | rs9372012 | 0.077 | -1.581 | 1.095 | 1.643 | -0.01 | 0.058 | 0.058 | 0 | 0.08 | 0.08 | 0.21 | 0.34 | 0.05 | 0.14 |
| 6 | 150724853 | rs3734737 | 0.015 | -1.686 | 0.784 | 1.407 | 0 | -0.007 | -0.007 | 0 | 0.2 | 0.12 | 0.13 | 0.19 | 0.21 | 0.16 |
| 6 | 150725278 | rs956440 | -0.386 | -1.661 | 0.869 | 1.388 | 0.002 | -0.001 | -0.001 | 0 | 0.25 | 0.21 | 0.16 | 0.22 | 0.23 | 0.26 |
| 6 | 150726042 | rs805996 | 1.85 | -1.626 | 0.509 | 0.439 | -0.027 | 0.174 | 0.174 | 0 | 0.48 | 0.16 | 0.36 | 0.2 | 0.53 | 0.23 |
| 6 | 150726097 | rs13197942 | -0.268 | -1.592 | 0.663 | 0.509 | 0.004 | -0.004 | -0.004 | 0 | 0.07 | 0.08 | 0.08 | 0.02 | 0.1 | 0.11 |
| 6 | 150726746 | rs2008273 | -0.178 | -1.757 | 0.655 | 0.608 | 0.022 | -0.058 | -0.058 | 1.099 | 0.24 | 0.16 | 0.31 | 0.57 | 0.2 | 0.36 |
| 6 | 150727036 | rs1883395 | -1.044 | -1.774 | 0.876 | 0.579 | 0 | -0.002 | -0.002 | 0 | 0.34 | 0.35 | 0.42 | 0.6 | 0.34 | 0.52 |
| 6 | 150727505 | rs649744 | 0.748 | -1.696 | 1.169 | 0.584 | -0.008 | 0.06 | 0.06 | 0 | 0.89 | 0.71 | 0.83 | 0.82 | 0.91 | 0.76 |
| 6 | 150727790 | rs648460 | 0.734 | -1.691 | 1.171 | 0.587 | -0.008 | 0.06 | 0.06 | 0 | 0.89 | 0.71 | 0.83 | 0.82 | 0.91 | 0.76 |
| 6 | 150729878 | rs9480384 | -0.134 | -1.625 | 0.731 | 0.305 | -0.013 | -0.015 | -0.015 | 0 | 0.13 | 0.22 | 0.11 | 0.04 | 0.1 | 0.02 |
| 6 | 150731232 | rs638990 | 1.742 | -1.638 | 0.706 | 0.381 | 0.004 | -0.02 | -0.02 | 0 | 0.68 | 0.52 | 0.75 | 0.81 | 0.66 | 0.59 |
| 6 | 150731655 | rs590179 | 1.795 | -1.638 | 0.133 | 0.143 | -0.089 | 0.202 | 0.202 | 0 | 0.43 | 0.13 | 0.31 | 0.16 | 0.46 | 0.22 |
| 6 | 150732960 | rs9480387 | -0.427 | -1.35 | 0.961 | 0.581 | -0.006 | -0.012 | -0.012 | 0.06 | 0.12 | 0.32 | 0.1 | 0.04 | 0.1 | 0.05 |
| 6 | 150734256 | rs665670 | 1.345 | -0.929 | 0.662 | 0.71 | 0.011 | 0.055 | 0.055 | 0.119 | 0.89 | 0.64 | 0.92 | 0.88 | 0.91 | 0.79 |
| 6 | 150740285 | rs13208297 | -0.332 | -0.48 | -0.618 | 0.518 | 0.025 | -0.02 | -0.02 | 3.081 | 0.35 | 0.17 | 0.38 | 0.4 | 0.29 | 0.33 |
| 6 | 150741436 | rs652800 | 0.264 | 0.651 | -1.408 | -0.372 | -0.07 | 0.103 | 0.103 | 0.022 | 0.27 | 0.09 | 0.19 | 0.08 | 0.36 | 0.15 |
| 6 | 150744204 | rs609421 | 1.085 | 0.976 | -1.56 | -0.47 | -0.022 | 0.005 | 0.005 | 0 | 0.06 | 0 | 0.02 | 0.03 | 0.07 | 0.02 |
| 6 | 150744837 | rs4869908 | 1.248 | 0.654 | -1.155 | -0.652 | 0.005 | 0.024 | 0.024 | 0 | 0.25 | 0.34 | 0.35 | 0.5 | 0.28 | 0.41 |
| 6 | 150744999 | rs9384509 | 0.639 | 0.978 | -1.492 | -0.764 | -0.041 | 0.086 | 0.086 | 0.582 | 0.2 | 0.05 | 0.13 | 0.04 | 0.22 | 0.12 |
| 6 | 150747497 | rs810958 | -0.58 | 0.891 | -1.091 | -0.746 | 0.041 | 0.04 | 0.04 | 0 | 0.57 | 0.82 | 0.56 | 0.6 | 0.64 | 0.6 |
| 6 | 150747577 | rs909785 | -1.375 | 0.238 | -1.366 | -0.862 | 0.029 | 0.081 | 0.081 | 0.078 | 0.78 | 0.98 | 0.92 | 0.88 | 0.86 | 0.82 |
| 6 | 150748147 | rs594747 | 0.091 | 0.867 | -0.298 | -0.363 | 0.025 | -0.017 | -0.017 | 0 | 0.11 | 0.28 | 0.07 | 0.07 | 0.14 | 0.06 |
| 6 | 150751874 | rs7769234 | 0.757 | 0.744 | -0.276 | -0.099 | 0.021 | 0.088 | 0.088 | 0 | 0.27 | 0.48 | 0.35 | 0.51 | 0.29 | 0.41 |
| 6 | 150754157 | rs661005 | -0.116 | 0.212 | -0.728 | 0.056 | 0.065 | 0.054 | 0.054 | 0 | 0.57 | 0.77 | 0.66 | 0.65 | 0.7 | 0.61 |
| 6 | 150756506 | rs7753635 | 0.644 | 1.115 | 0.52 | 1.598 | 0.051 | 0.15 | 0.15 | 0 | 0.85 | 0.6 | 0.65 | 0.65 | 0.78 | 0.7 |
| 6 | 150761290 | rs12528277 | 0.213 | 0.906 | 0.129 | 0.878 | -0.018 | 0.035 | 0.035 | 0 | 0.43 | 0.34 | 0.38 | 0.25 | 0.44 | 0.39 |
| 6 | 150761928 | rs7751747 | -0.014 | 0.835 | 0.148 | 0.73 | -0.013 | 0.017 | 0.017 | 0 | 0.86 | 0.89 | 0.88 | 0.96 | 0.76 | 0.9 |
| 6 | 150764310 | rs2050047 | -0.228 | 0.783 | 0.625 | 0.798 | 0.029 | 0.088 | 0.088 | 0 | 0.5 | 0.69 | 0.65 | 0.7 | 0.51 | 0.64 |
| 6 | 150765473 | rs4870521 | -0.238 | 0.777 | 0.512 | 0.758 | 0.027 | 0.128 | 0.128 | 0 | 0.51 | 0.8 | 0.66 | 0.72 | 0.51 | 0.66 |
| 6 | 150765510 | rs17800315 | 1.076 | 0.744 | 0.286 | 0.753 | 0.056 | 0.056 | 0.056 | 0 | 0.35 | 0.06 | 0.22 | 0.24 | 0.25 | 0.24 |
| 6 | 150766205 | rs720920 | 0.251 | 0.861 | 0.327 | 0.778 | -0.018 | 0.036 | 0.036 | 0 | 0.43 | 0.34 | 0.38 | 0.25 | 0.44 | 0.39 |
| 6 | 150770756 | rs1003240 | 0.316 | 1.083 | 0.135 | 0.639 | -0.002 | 0 | 0 | 0 | 0.26 | 0.25 | 0.28 | 0.22 | 0.29 | 0.29 |
| 6 | 150771106 | rs1013663 | 0.108 | 1.107 | 0.051 | 0.641 | 0.001 | -0.001 | -0.001 | 0 | 0.27 | 0.31 | 0.29 | 0.22 | 0.3 | 0.29 |
| 6 | 150772453 | rs742315 | 0.849 | 1.047 | 0.12 | 0.231 | 0.01 | 0.015 | 0.015 | 0 | 0.43 | 0.36 | 0.27 | 0.28 | 0.35 | 0.3 |
| 6 | 150774227 | rs17676258 | 0.873 | 1.313 | -0.045 | 0.271 | -0.026 | 0.027 | 0.027 | 0 | 0.07 | 0.02 | 0.13 | 0.01 | 0.15 | 0.04 |
| 6 | 150775678 | rs9384533 | 1.199 | 1.433 | 0.171 | 0.147 | -0.001 | -0.011 | -0.011 | 0.289 | 0.15 | 0.14 | 0.09 | 0.03 | 0.14 | 0.1 |
| 6 | 150776864 | rs3860828 | 1.077 | 1.042 | 0.159 | 0.089 | 0.039 | 0.043 | 0.043 | 0 | 0.38 | 0.22 | 0.22 | 0.26 | 0.26 | 0.24 |
| 6 | 150776874 | rs4134337 | 1.864 | 0.797 | -0.125 | -0.108 | 0.04 | 0.022 | 0.022 | 0.001 | 0.27 | 0.14 | 0.17 | 0.22 | 0.15 | 0.17 |
| 6 | 150778742 | rs11754699 | 0.963 | 0.84 | -0.025 | -0.108 | 0.043 | 0.052 | 0.052 | 0 | 0.38 | 0.19 | 0.21 | 0.26 | 0.26 | 0.24 |
| 6 | 150779197 | rs4870524 | 0.6 | 0.993 | 0.086 | 0.036 | 0.004 | 0.052 | 0.052 | 0 | 0.53 | 0.33 | 0.3 | 0.28 | 0.4 | 0.34 |
| 6 | 150786457 | rs9372048 | -0.474 | 1.101 | 0.332 | 0.369 | -0.001 | -0.035 | -0.035 | 0 | 0.72 | 0.82 | 0.74 | 0.66 | 0.71 | 0.73 |
| 6 | 150786882 | rs13210742 | 1.307 | 1.003 | 0.326 | 0.356 | 0.02 | 0.061 | 0.061 | 0 | 0.43 | 0.34 | 0.26 | 0.22 | 0.3 | 0.27 |
| 6 | 150790291 | rs9372051 | -0.01 | 1.068 | 0.19 | 0.469 | -0.004 | -0.008 | -0.008 | 0 | 0.22 | 0.22 | 0.35 | 0.43 | 0.27 | 0.41 |
| 6 | 150790679 | rs4074888 | -0.615 | 1.102 | 0.076 | 0.362 | -0.002 | -0.023 | -0.023 | 0 | 0.77 | 0.83 | 0.76 | 0.66 | 0.75 | 0.72 |
| 6 | 150792757 | rs12209935 | 1.376 | 1.23 | 0.071 | 0.437 | -0.01 | 0.003 | 0.003 | 0 | 0.1 | 0.04 | 0.05 | 0.08 | 0.11 | 0.1 |
| 6 | 150799494 | rs12213006 | 1.376 | 1.23 | 0.071 | 0.416 | -0.01 | 0.006 | 0.006 | 0 | 0.1 | 0.04 | 0.05 | 0.07 | 0.11 | 0.1 |
| 6 | 150799903 | rs9480461 | -0.306 | 1.2 | 0.342 | 0.491 | 0.013 | 0.069 | 0.069 | 0 | 0.16 | 0.25 | 0.3 | 0.36 | 0.18 | 0.32 |
| 6 | 150801064 | rs6911714 | -0.949 | 1.32 | 0.042 | 0.667 | -0.01 | -0.013 | -0.013 | 0 | 0.92 | 0.97 | 0.93 | 0.93 | 0.9 | 0.95 |
| 6 | 150802064 | rs9398008 | 0.208 | 1.356 | 0.24 | 0.586 | 0.016 | 0.069 | 0.069 | 0.318 | 0.11 | 0.2 | 0.3 | 0.36 | 0.16 | 0.33 |
| 6 | 150807451 | rs4367393 | -0.842 | 1.062 | -0.066 | 0.252 | 0.041 | 0.034 | 0.034 | 0.098 | 0.51 | 0.68 | 0.59 | 0.45 | 0.61 | 0.56 |
| 6 | 150811384 | rs9384545 | 1.389 | 1.301 | 0.081 | 0.05 | 0.002 | -0.028 | -0.028 | 0 | 0.16 | 0.18 | 0.1 | 0.02 | 0.16 | 0.09 |
| 6 | 150812668 | rs6908890 | -0.868 | 1.161 | 0.263 | 0.144 | 0.036 | 0.034 | 0.034 | 0.144 | 0.51 | 0.66 | 0.61 | 0.42 | 0.62 | 0.54 |
| 6 | 150816326 | rs17079796 | 1.437 | 0.994 | -0.003 | 0.358 | 0.032 | 0.013 | 0.013 | 0.471 | 0.34 | 0.19 | 0.25 | 0.29 | 0.23 | 0.2 |
| 6 | 150819398 | rs9480467 | -0.501 | 1.097 | 0.264 | -0.068 | -0.015 | -0.001 | -0.001 | 0 | 0.35 | 0.25 | 0.5 | 0.37 | 0.45 | 0.52 |
| 6 | 150820346 | rs12205882 | 0.514 | 1.06 | 0.565 | -0.093 | 0 | -0.032 | -0.032 | 0 | 0.27 | 0.28 | 0.15 | 0.05 | 0.27 | 0.18 |
| 6 | 150823321 | rs6938615 | -1.492 | 1.021 | 0.596 | -0.061 | -0.007 | 0.003 | 0.003 | 0.012 | 0.63 | 0.56 | 0.63 | 0.43 | 0.71 | 0.7 |
| 6 | 150829932 | rs4131417 | -0.497 | 0.877 | 0.452 | -0.01 | 0.002 | -0.007 | -0.007 | 0 | 0.29 | 0.23 | 0.35 | 0.37 | 0.3 | 0.49 |
| 6 | 150835885 | rs7764084 | -1.176 | 0.919 | 0.333 | 0.117 | -0.014 | 0.018 | 0.018 | 0 | 0.58 | 0.74 | 0.52 | 0.38 | 0.59 | 0.67 |
| 6 | 150843677 | rs9478758 | -0.546 | 0.941 | 0.412 | 0.189 | 0 | -0.004 | -0.004 | 0 | 0.29 | 0.23 | 0.36 | 0.36 | 0.31 | 0.49 |
| 6 | 150851876 | rs9688466 | 1.361 | 0.909 | 0.452 | 0.029 | 0.008 | -0.042 | -0.042 | 0 | 0.16 | 0.22 | 0.1 | 0.02 | 0.18 | 0.08 |
| 6 | 150855864 | rs12195563 | 0.105 | 0.635 | 0.341 | 0.025 | -0.003 | -0.094 | -0.094 | 0 | 0.34 | 0.4 | 0.16 | 0.06 | 0.31 | 0.18 |
| 6 | 150860123 | rs7341280 | -0.097 | 0.759 | -0.372 | 0.183 | -0.007 | -0.034 | -0.034 | 0 | 0.23 | 0.12 | 0.34 | 0.36 | 0.28 | 0.49 |
| 6 | 150868443 | rs1492574 | -0.929 | 0.754 | -0.268 | 0.216 | 0.009 | 0.021 | 0.021 | 0.017 | 0.54 | 0.66 | 0.52 | 0.42 | 0.59 | 0.64 |
| 6 | 150869099 | rs9371462 | -0.429 | 0.601 | -0.487 | 0.139 | -0.014 | -0.023 | -0.023 | 0.061 | 0.79 | 0.81 | 0.62 | 0.69 | 0.72 | 0.81 |
| 6 | 150873111 | rs7747720 | -0.517 | 0.241 | -0.502 | 0.234 | 0.006 | 0.035 | 0.035 | 0.849 | 0.52 | 0.58 | 0.49 | 0.35 | 0.56 | 0.63 |
| 6 | 150873233 | rs7758730 | 0.05 | 0.263 | -0.454 | 0.13 | -0.004 | -0.015 | -0.015 | 0 | 0.23 | 0.14 | 0.34 | 0.3 | 0.27 | 0.48 |
| 6 | 150873623 | rs9383831 | 1.727 | 0.324 | -0.312 | 0.277 | 0.039 | -0.008 | -0.008 | 0.741 | 0.26 | 0.16 | 0.12 | 0.34 | 0.14 | 0.17 |
| 6 | 150873766 | rs7740233 | -0.641 | 0.634 | -0.108 | 0.606 | 0.007 | -0.004 | -0.004 | 0.513 | 0.51 | 0.58 | 0.49 | 0.47 | 0.56 | 0.65 |
| 6 | 150880499 | rs9384561 | -0.884 | 0.89 | -0.325 | 0.998 | -0.003 | -0.008 | -0.008 | 0 | 0.18 | 0.14 | 0.31 | 0.3 | 0.24 | 0.45 |

| | | | | | | | | | | | | | | | | |
|---|---|---|---|---|---|---|---|---|---|---|---|---|---|---|---|---|
| 6 | 150881887 | rs9478760 | -0.209 | 1.106 | -0.681 | 1.242 | -0.037 | -0.054 | -0.054 | 0.059 | 0.1 | 0.02 | 0.28 | 0.29 | 0.16 | 0.3 |
| 6 | 150890623 | rs1122537 | 0.077 | 0.793 | -0.897 | 0.605 | -0.015 | 0.063 | 0.063 | 0 | 0.17 | 0.04 | 0.11 | 0.02 | 0.17 | 0.1 |
| 6 | 150891162 | rs9397336 | -0.436 | 0.81 | 0.067 | 0.966 | 0.048 | 0.13 | 0.13 | 0 | 0.13 | 0.32 | 0.28 | 0.28 | 0.2 | 0.32 |
| 6 | 150892649 | rs9372073 | 0.183 | 0.919 | 0.257 | 1.077 | 0.026 | 0.034 | 0.034 | 0 | 0.12 | 0.26 | 0.24 | 0.18 | 0.16 | 0.09 |
| 6 | 150894274 | rs9480482 | 0.497 | 0.815 | -0.103 | 0.664 | -0.002 | -0.002 | -0.002 | 0.273 | 0.13 | 0.11 | 0.14 | 0.02 | 0.17 | 0.15 |
| 6 | 150894638 | rs9322266 | -1.597 | 0.714 | -0.462 | 1.056 | 0.014 | 0.071 | 0.071 | 0 | 0.53 | 0.78 | 0.69 | 0.62 | 0.64 | 0.63 |
| 6 | 150894902 | rs9372074 | 0.302 | 0.76 | -0.213 | 1.133 | 0.004 | 0.097 | 0.097 | 0 | 0.09 | 0.17 | 0.17 | 0.27 | 0.13 | 0.21 |
| 6 | 150897232 | rs9322267 | -1.377 | 0.787 | -0.806 | 0.927 | -0.007 | 0.088 | 0.088 | 0 | 0.54 | 0.87 | 0.7 | 0.64 | 0.63 | 0.62 |
| 6 | 150900125 | rs7742124 | 0.278 | 0.926 | -0.277 | 1.015 | -0.001 | -0.009 | -0.009 | 0 | 0.16 | 0.2 | 0.12 | 0.1 | 0.17 | 0.09 |
| 6 | 150908613 | rs1492576 | -0.683 | 0.727 | -0.689 | 1.336 | -0.001 | -0.011 | -0.011 | 0 | 0.72 | 0.8 | 0.72 | 0.67 | 0.7 | 0.73 |
| 6 | 150911634 | rs9322269 | 0.652 | 0.969 | -1.245 | 1.347 | -0.001 | 0.022 | 0.022 | 0 | 0.31 | 0.14 | 0.19 | 0.22 | 0.25 | 0.27 |
| 6 | 150913528 | rs9397338 | 0.517 | 0.974 | -1.209 | 1.143 | -0.002 | -0.058 | -0.058 | 0 | 0.32 | 0.17 | 0.38 | 0.42 | 0.27 | 0.26 |
| 6 | 150915881 | rs7750629 | -0.26 | 0.792 | -0.775 | 0.653 | -0.013 | 0.034 | 0.034 | 0.029 | 0.37 | 0.25 | 0.31 | 0.27 | 0.43 | 0.43 |
| 6 | 150923115 | rs3050 | -0.204 | 0.443 | -0.352 | 0.805 | 0.013 | 0.017 | 0.017 | 0 | 0.73 | 0.78 | 0.83 | 0.82 | 0.82 | 0.8 |
| 6 | 150924617 | rs6904488 | 0.055 | 0.342 | 0.281 | 1.055 | -0.033 | -0.01 | -0.01 | 0 | 0.71 | 0.59 | 0.68 | 0.69 | 0.81 | 0.78 |
| 6 | 150925777 | rs10484776 | 0.286 | 0.593 | 0.325 | 1.305 | -0.001 | 0.001 | 0.001 | 0 | 0.07 | 0.05 | 0.07 | 0.16 | 0.13 | 0.19 |
| 6 | 150926499 | rs2047001 | -0.06 | 0.406 | 0.267 | 1.121 | -0.033 | -0.01 | -0.01 | 0.152 | 0.71 | 0.59 | 0.68 | 0.69 | 0.81 | 0.78 |
| 6 | 150931131 | rs1492577 | 1.27 | 0.562 | -0.351 | 0.735 | 0.004 | 0.079 | 0.079 | 0.108 | 0.31 | 0.1 | 0.17 | 0.13 | 0.26 | 0.16 |
| 6 | 150935556 | rs6557547 | 0.165 | 0.785 | 0.42 | 1.426 | 0 | -0.001 | -0.001 | 0 | 0.52 | 0.57 | 0.62 | 0.48 | 0.48 | 0.37 |
| 6 | 150935797 | rs11754524 | -0.311 | 1.112 | 0.336 | 1.628 | 0.022 | 0.031 | 0.031 | 0 | 0.21 | 0.24 | 0.24 | 0.35 | 0.31 | 0.37 |
| 6 | 150936716 | rs4870536 | 0.164 | 0.765 | -0.108 | 1.601 | 0.001 | -0.002 | -0.002 | 0.063 | 0.14 | 0.18 | 0.21 | 0.2 | 0.13 | 0.09 |
| 6 | 150947524 | rs12527099 | 0.424 | 0.899 | -0.553 | 1.436 | 0 | -0.001 | -0.001 | 0 | 0.12 | 0.08 | 0.21 | 0.21 | 0.13 | 0.09 |
| 6 | 150947644 | rs17422247 | 0.417 | 0.844 | -0.706 | 1.178 | -0.016 | -0.028 | -0.028 | 0.642 | 0.07 | 0 | 0.02 | 0.07 | 0.05 | 0.07 |
| 6 | 150948810 | rs11155745 | 0.757 | 0.699 | -0.654 | 0.998 | -0.046 | 0.044 | 0.044 | 0 | 0.32 | 0.08 | 0.27 | 0.17 | 0.33 | 0.23 |
| 6 | 150952507 | rs11758926 | -0.022 | 0.464 | 0.223 | 1.08 | 0.001 | -0.006 | -0.006 | 0.023 | 0.1 | 0.1 | 0.09 | 0.12 | 0.14 | 0.24 |
| 6 | 150952769 | rs6557548 | -0.402 | 0.125 | 0.329 | 1.299 | 0.003 | -0.009 | -0.009 | 0 | 0.89 | 0.92 | 0.93 | 0.85 | 0.92 | 0.9 |
| 6 | 150953184 | rs4323318 | -1.655 | 0.116 | 1.079 | 1.143 | 0.036 | 0.115 | 0.115 | 0.135 | 0.11 | 0.47 | 0.25 | 0.16 | 0.08 | 0.13 |
| 6 | 150959792 | rs9384570 | -0.888 | 0.223 | 0.301 | 1.14 | -0.007 | 0.005 | 0.005 | 1.347 | 0.21 | 0.17 | 0.18 | 0.17 | 0.23 | 0.28 |
| 6 | 150967130 | rs6557549 | -1.344 | -0.215 | 0.708 | 1.151 | -0.028 | 0.012 | 0.012 | 0.158 | 0.23 | 0.42 | 0.22 | 0.26 | 0.16 | 0.28 |
| 6 | 150967919 | rs17080052 | -1.219 | -0.051 | 0.554 | 1.288 | -0.015 | 0.088 | 0.088 | 0.043 | 0.06 | 0.2 | 0.14 | 0.13 | 0 | 0.06 |
| 6 | 150983318 | rs4577811 | -0.899 | -0.234 | 0.194 | 1.386 | 0 | 0 | 0 | 0.14 | 0.18 | 0.18 | 0.07 | 0.31 | 0.15 | 0.24 |
| 6 | 150987367 | rs12525903 | -0.838 | -0.273 | 0.634 | 1.437 | -0.003 | 0.002 | 0.002 | 0 | 0.17 | 0.22 | 0.07 | 0.25 | 0.13 | 0.14 |
| 6 | 150987778 | rs9480512 | -0.906 | 0.065 | 0.408 | 1.501 | 0.007 | 0.008 | 0.008 | 0.001 | 0.06 | 0.07 | 0.12 | 0.08 | 0.1 | 0.2 |
| 6 | 150988903 | rs7349946 | -1.517 | -0.179 | -0.446 | 1.257 | -0.004 | 0.019 | 0.019 | 0.188 | 0.7 | 0.9 | 0.73 | 0.74 | 0.7 | 0.8 |
| 6 | 150997440 | rs17080093 | -0.09 | -1.287 | 0.232 | 1.044 | -0.03 | -0.014 | -0.014 | 0.014 | 0.14 | 0.22 | 0.2 | 0.12 | 0.08 | 0.17 |
| 6 | 151000532 | rs9480526 | -0.275 | -1.208 | 0.256 | 1.082 | -0.002 | -0.003 | -0.003 | 0 | 0.16 | 0.19 | 0.31 | 0.14 | 0.16 | 0.27 |
| 6 | 151004289 | rs6900518 | -0.473 | -1.368 | 0.858 | 1.12 | -0.027 | 0.071 | 0.071 | 0.655 | 0.18 | 0.48 | 0.08 | 0.28 | 0.13 | 0.14 |
| 6 | 151006924 | rs7738394 | -1.333 | -1.363 | 0.962 | 1.127 | -0.01 | 0.008 | 0.008 | 1.594 | 0.33 | 0.57 | 0.38 | 0.43 | 0.29 | 0.41 |
| 6 | 151010614 | rs9478783 | -1.173 | -1.468 | 1.165 | 1.2 | -0.029 | 0.03 | 0.03 | 0.699 | 0.22 | 0.57 | 0.33 | 0.32 | 0.17 | 0.37 |
| 6 | 151015996 | rs275351 | -0.835 | -1.29 | 0.675 | 1.075 | -0.004 | -0.007 | -0.007 | 0.042 | 0.55 | 0.38 | 0.51 | 0.5 | 0.48 | 0.55 |
| 6 | 151018420 | rs6921304 | 0.346 | -1.339 | 0.586 | 1.19 | 0.001 | 0 | 0 | 0 | 0.83 | 0.82 | 0.71 | 0.86 | 0.87 | 0.62 |
| 6 | 151024280 | rs1494309 | -0.114 | -1.356 | 0.531 | 1.17 | -0.028 | -0.014 | -0.014 | 0 | 0.42 | 0.2 | 0.32 | 0.41 | 0.43 | 0.38 |
| 6 | 151025628 | rs9480551 | -0.892 | -1.207 | 1.323 | 1.099 | 0.078 | 0.181 | 0.181 | 1.727 | 0.07 | 0.33 | 0.11 | 0.19 | 0.07 | 0.14 |
| 6 | 151026401 | rs6940976 | 0.685 | -1.279 | 1.486 | 1.048 | -0.028 | 0.034 | 0.034 | 0 | 0.82 | 0.53 | 0.65 | 0.8 | 0.86 | 0.69 |
| 6 | 151026886 | rs9478787 | -0.567 | -1.19 | 0.852 | 0.898 | 0.048 | 0.067 | 0.067 | 0 | 0.06 | 0.15 | 0.16 | 0.1 | 0.08 | 0.19 |
| 6 | 151027008 | rs7776166 | 0.528 | -1.078 | 0.86 | 1.081 | -0.06 | 0.037 | 0.037 | 0 | 0.83 | 0.5 | 0.78 | 0.76 | 0.92 | 0.78 |
| 6 | 151031607 | rs6905228 | 0.526 | -1.079 | 0.894 | 0.987 | -0.059 | 0.006 | 0.006 | 0 | 0.83 | 0.5 | 0.78 | 0.8 | 0.92 | 0.79 |
| 6 | 151032785 | rs7769973 | 0.934 | -0.948 | 0.477 | 0.892 | -0.01 | 0 | 0 | 0 | 0.44 | 0.3 | 0.55 | 0.52 | 0.5 | 0.46 |
| 6 | 151035800 | rs4272224 | -0.162 | -0.788 | 0.468 | 0.994 | 0.02 | 0.009 | 0.009 | 0.085 | 0.24 | 0.34 | 0.3 | 0.26 | 0.33 | 0.44 |
| 6 | 151036450 | rs4449641 | 0.991 | -0.781 | 0.182 | 0.786 | 0.003 | 0.008 | 0.008 | 0.467 | 0.51 | 0.38 | 0.47 | 0.44 | 0.51 | 0.36 |
| 6 | 151048139 | rs6919370 | 1.416 | -1.701 | -0.019 | 0.827 | 0.048 | -0.035 | -0.035 | 0.259 | 0.62 | 0.27 | 0.57 | 0.49 | 0.77 | 0.55 |
| 6 | 151048986 | rs6901930 | -0.84 | -1.503 | 0.236 | 0.927 | 0.017 | 0.171 | 0.171 | 0.049 | 0.12 | 0.38 | 0.07 | 0.28 | 0.09 | 0.12 |
| 6 | 151058043 | rs9478132 | 0.335 | -1.713 | -0.356 | 0.628 | 0.021 | 0.017 | 0.017 | 0.08 | 0.83 | 0.91 | 0.81 | 0.91 | 0.94 | 0.84 |
| 6 | 151067274 | rs13198996 | -0.89 | -0.779 | 0.038 | 0.998 | -0.002 | -0.003 | -0.003 | 0.073 | 0.15 | 0.15 | 0.25 | 0.23 | 0.16 | 0.2 |
| 6 | 151072347 | rs9478809 | -0.976 | -0.915 | 0.186 | 0.839 | -0.055 | -0.055 | -0.055 | 3.305 | 0.21 | 0.58 | 0.12 | 0.12 | 0.1 | 0.09 |
| 6 | 151075128 | rs4869945 | -1.636 | -0.922 | 0.224 | 0.937 | 0.014 | -0.003 | -0.003 | 0 | 0.27 | 0.73 | 0.3 | 0.28 | 0.18 | 0.24 |
| 6 | 151076883 | rs9918380 | -0.662 | -0.761 | 0.114 | 0.907 | 0.018 | 0.067 | 0.067 | 0 | 0.06 | 0.11 | 0.17 | 0.15 | 0.07 | 0.15 |
| 6 | 151083111 | rs13211253 | -1.74 | -1.373 | 0.155 | 1.004 | 0.011 | -0.012 | -0.012 | 0 | 0.33 | 0.67 | 0.3 | 0.52 | 0.18 | 0.29 |
| 6 | 151084927 | rs7772605 | -1.432 | -1.168 | -0.032 | 0.516 | -0.066 | 0.005 | 0.005 | 3.004 | 0.29 | 0.53 | 0.27 | 0.37 | 0.17 | 0.17 |
| 6 | 151096419 | rs742636 | -0.476 | -0.374 | -0.71 | 0.071 | 0.013 | 0.014 | 0.014 | 0 | 0.14 | 0.09 | 0.08 | 0.05 | 0.09 | 0.08 |
| 6 | 151110620 | rs9371472 | -0.004 | 0.425 | -1.144 | -0.153 | -0.04 | 0.063 | 0.063 | 0.002 | 0.71 | 0.85 | 0.63 | 0.9 | 0.65 | 0.66 |
| 6 | 151112333 | rs9371474 | 0.861 | 0.066 | -1.72 | 0.154 | -0.035 | -0.065 | -0.065 | 0.066 | 0.32 | 0.1 | 0.29 | 0.38 | 0.3 | 0.33 |
| 6 | 151118271 | rs2006958 | 0.304 | -0.191 | -1.111 | 0.307 | -0.058 | -0.034 | -0.034 | 0 | 0.43 | 0.15 | 0.38 | 0.44 | 0.49 | 0.48 |
| 6 | 151119123 | rs6557097 | -0.798 | -0.299 | -1.334 | 0.173 | -0.089 | 0.149 | 0.149 | 1.176 | 0.68 | 0.93 | 0.62 | 0.92 | 0.62 | 0.68 |
| 6 | 151120686 | rs9383848 | -0.243 | -0.114 | -0.949 | 0.233 | -0.011 | 0.206 | 0.206 | 0 | 0.49 | 0.8 | 0.52 | 0.8 | 0.48 | 0.56 |
| 6 | 151121196 | rs4869947 | 0.934 | -0.049 | -0.231 | 0.3 | -0.007 | 0.032 | 0.032 | 0.272 | 0.29 | 0.34 | 0.27 | 0.39 | 0.24 | 0.27 |
| 6 | 151121915 | rs2073061 | -0.466 | -0.04 | -0.285 | 0.281 | -0.011 | 0.207 | 0.207 | 0 | 0.49 | 0.8 | 0.52 | 0.8 | 0.48 | 0.56 |
| 6 | 151122094 | rs2073060 | 0.724 | 0.091 | -0.477 | 0.354 | -0.042 | -0.021 | -0.021 | 0.395 | 0.08 | 0.01 | 0.07 | 0.08 | 0.16 | 0.16 |
| 6 | 151123155 | rs9371198 | 1.084 | 0.126 | -0.09 | 0.464 | 0 | -0.024 | -0.024 | 0 | 0.26 | 0.12 | 0.22 | 0.36 | 0.24 | 0.21 |

| | | | | | | | | | | | | | | | | |
|---|---|---|---|---|---|---|---|---|---|---|---|---|---|---|---|---|
| 6 | 151123175 | rs9397024 | 0.117 | 0.155 | -0.021 | 0.468 | -0.005 | -0.005 | -0.005 | 0 | 0.37 | 0.59 | 0.39 | 0.7 | 0.3 | 0.34 |
| 6 | 151124914 | rs2167 | 0.117 | 0.155 | -0.021 | 0.468 | -0.005 | -0.005 | -0.005 | 2.09 | 0.37 | 0.6 | 0.39 | 0.7 | 0.3 | 0.34 |
| 6 | 151125251 | rs10046456 | 0.282 | 0.185 | 0.044 | 0.663 | -0.001 | 0.085 | 0.085 | 0 | 0.08 | 0.2 | 0.04 | 0.2 | 0.07 | 0.09 |
| 6 | 151125896 | rs712218 | -0.401 | 0.214 | -0.334 | 0.398 | 0.004 | 0 | 0 | 0 | 0.58 | 0.46 | 0.46 | 0.58 | 0.53 | 0.55 |
| 6 | 151126585 | rs7754056 | 0.328 | 0.282 | -0.499 | 0.755 | 0.005 | -0.022 | -0.022 | 0 | 0.25 | 0.14 | 0.22 | 0.36 | 0.22 | 0.2 |
| 6 | 151132928 | rs712213 | -0.382 | 0.296 | -1.308 | 0.036 | 0.009 | 0.054 | 0.054 | 0.074 | 0.35 | 0.2 | 0.2 | 0.18 | 0.32 | 0.14 |
| 6 | 151135576 | rs17080350 | -1.068 | 0.39 | -1.202 | 0.352 | 0 | 0.009 | 0.009 | 0.007 | 0.11 | 0.15 | 0.06 | 0.17 | 0.11 | 0.07 |
| 6 | 151144242 | rs1178409 | 0.038 | 0.662 | -1.8 | -0.446 | -0.036 | 0.132 | 0.132 | 0.278 | 0.25 | 0.04 | 0.13 | 0.01 | 0.25 | 0.1 |
| 6 | 151145278 | rs761264 | -0.497 | 0.246 | -1.736 | -0.866 | -0.033 | -0.032 | -0.032 | 0 | 0.72 | 0.66 | 0.81 | 0.82 | 0.81 | 0.85 |
| 6 | 151149077 | rs803413 | 0.754 | 0.264 | -1.703 | -0.911 | -0.016 | 0.118 | 0.118 | 0.046 | 0.26 | 0.11 | 0.15 | 0.01 | 0.25 | 0.1 |
| 6 | 151149833 | rs17426727 | 0.236 | 1.1 | -1.901 | -0.748 | -0.022 | -0.013 | -0.013 | 0 | 0.07 | 0.03 | 0.21 | 0.1 | 0.14 | 0.17 |
| 6 | 151152059 | rs2073065 | -0.309 | 0.93 | -1.92 | -0.734 | -0.013 | -0.009 | -0.009 | 0 | 0.09 | 0.07 | 0.2 | 0.13 | 0.13 | 0.2 |
| 6 | 151152257 | rs926624 | 1.252 | 0.794 | -2.026 | -0.433 | -0.033 | -0.056 | -0.056 | 0 | 0.21 | 0.12 | 0.33 | 0.38 | 0.31 | 0.27 |
| 6 | 151152656 | rs803411 | -0.296 | 0.638 | -1.929 | -0.538 | -0.013 | -0.001 | -0.001 | 0.155 | 0.61 | 0.61 | 0.72 | 0.54 | 0.71 | 0.64 |
| 6 | 151155341 | rs6917461 | 1.279 | 0.638 | -2.042 | -0.527 | -0.033 | -0.055 | -0.055 | 0 | 0.21 | 0.12 | 0.33 | 0.38 | 0.31 | 0.27 |
| 6 | 151156327 | rs803407 | -0.326 | 0.534 | -2.326 | -0.653 | 0.036 | -0.002 | -0.002 | 0.12 | 0.71 | 0.96 | 0.84 | 0.72 | 0.8 | 0.74 |
| 6 | 151158446 | rs803403 | 1.121 | 0.943 | -1.981 | -0.306 | -0.015 | 0.12 | 0.12 | 0.017 | 0.24 | 0.09 | 0.14 | 0.01 | 0.22 | 0.09 |
| 6 | 151159630 | rs803401 | 0.145 | 0.625 | -1.387 | 0.429 | -0.009 | 0.002 | 0.002 | 0 | 0.62 | 0.6 | 0.72 | 0.52 | 0.69 | 0.64 |
| 6 | 151163115 | rs17427389 | -1.369 | 1.52 | -1.35 | 1.164 | -0.018 | -0.011 | -0.011 | 0 | 0.09 | 0.04 | 0.23 | 0.13 | 0.15 | 0.22 |
| 6 | 151164764 | rs9397027 | -0.691 | 1.625 | -1.289 | 1.306 | -0.012 | -0.05 | -0.05 | 0.002 | 0.1 | 0.02 | 0.09 | 0.28 | 0.1 | 0.21 |
| 6 | 151168602 | rs10484779 | -0.849 | 1.741 | -0.837 | 0.868 | -0.015 | -0.013 | -0.013 | 0.738 | 0.06 | 0.03 | 0.22 | 0.11 | 0.12 | 0.15 |
| 6 | 151188317 | rs9322288 | -0.712 | 1.42 | -0.962 | 0.698 | -0.021 | -0.011 | -0.011 | 0.211 | 0.45 | 0.78 | 0.53 | 0.44 | 0.45 | 0.42 |
| 6 | 151193451 | rs11757561 | -0.89 | 1.767 | -0.15 | 0.538 | 0.038 | 0.201 | 0.201 | 0.248 | 0.17 | 0.4 | 0.33 | 0.45 | 0.2 | 0.25 |
| 6 | 151196754 | rs13194204 | -0.123 | 1.993 | -0.225 | 0.83 | -0.025 | 0.033 | 0.033 | 0 | 0.06 | 0.01 | 0.09 | 0 | 0.09 | 0.08 |
| 6 | 151197731 | rs2073066 | -0.233 | 1.922 | -0.133 | 0.694 | 0.006 | 0.007 | 0.007 | 0.022 | 0.23 | 0.1 | 0.12 | 0.23 | 0.2 | 0.26 |
| 6 | 151198895 | rs2073064 | -0.656 | 1.745 | 0.274 | 0.691 | 0.034 | 0.135 | 0.135 | 0.103 | 0.11 | 0.34 | 0.27 | 0.21 | 0.11 | 0.07 |
| 6 | 151199071 | rs2073063 | -1.114 | 1.651 | -0.111 | 0.699 | -0.003 | 0 | 0 | 0 | 0.45 | 0.59 | 0.53 | 0.45 | 0.47 | 0.43 |
| 6 | 151209997 | rs4869953 | -1.089 | 1.584 | -0.188 | 0.757 | 0.002 | -0.004 | -0.004 | 0 | 0.47 | 0.63 | 0.59 | 0.49 | 0.53 | 0.45 |
| 6 | 151213211 | rs17349743 | -1.141 | 1.835 | -0.417 | 0.626 | -0.005 | -0.003 | -0.003 | 0 | 0.2 | 0.21 | 0.46 | 0.22 | 0.3 | 0.2 |
| 6 | 151215594 | rs803422 | -0.236 | 1.742 | -1.624 | -0.757 | -0.056 | 0.163 | 0.163 | 0.526 | 0.72 | 0.95 | 0.85 | 1 | 0.7 | 0.89 |
| 6 | 151217919 | rs17080461 | 0.168 | 1.52 | -1.8 | 0.035 | -0.003 | -0.013 | -0.013 | 0.71 | 0.09 | 0.07 | 0.21 | 0.27 | 0.15 | 0.25 |
| 6 | 151219414 | rs803420 | -0.345 | 1.413 | -2.371 | -0.616 | -0.1 | 0.145 | 0.145 | 1.094 | 0.59 | 0.92 | 0.77 | 0.84 | 0.58 | 0.77 |
| 6 | 151223680 | rs997429 | 0.484 | 1.148 | -1.917 | -0.65 | -0.004 | 0.006 | 0.006 | 0.928 | 0.19 | 0.25 | 0.13 | 0.18 | 0.17 | 0.23 |
| 6 | 151227086 | rs711879 | -0.052 | 1.091 | -2.655 | -0.615 | -0.044 | -0.1 | -0.1 | 0 | 0.23 | 0.1 | 0.54 | 0.62 | 0.34 | 0.46 |
| 6 | 151228287 | rs712208 | 0.346 | 1.063 | -2.673 | -0.636 | 0.005 | -0.086 | -0.086 | 0 | 0.19 | 0.09 | 0.31 | 0.62 | 0.22 | 0.44 |
| 6 | 151229012 | rs12195069 | 1.084 | 0.838 | -2.583 | -0.907 | 0 | -0.012 | -0.012 | 0 | 0.07 | 0.07 | 0.05 | 0 | 0.07 | 0.02 |
| 6 | 151230837 | rs1771845 | -0.11 | 1.142 | -2.655 | -0.662 | -0.062 | -0.101 | -0.101 | 0.253 | 0.25 | 0.1 | 0.58 | 0.62 | 0.39 | 0.47 |
| 6 | 151236287 | rs175860 | -0.405 | 0.587 | -2.522 | -0.731 | -0.026 | -0.023 | -0.023 | 0 | 0.53 | 0.82 | 0.53 | 0.43 | 0.47 | 0.53 |
| 6 | 151243234 | rs2295084 | 0.132 | -0.466 | -1.911 | -1.411 | 0 | 0 | 0 | 0 | 0.23 | 0.2 | 0.22 | 0.24 | 0.14 | 0.22 |
| 6 | 151243700 | rs9478858 | 0.325 | -0.29 | -1.404 | -0.868 | -0.031 | -0.04 | -0.04 | 0 | 0.09 | 0.19 | 0.06 | 0.03 | 0.05 | 0.11 |
| 6 | 151244271 | rs6557101 | 0.273 | -0.289 | -1.414 | -0.868 | -0.032 | -0.041 | -0.041 | 0.635 | 0.09 | 0.19 | 0.06 | 0.03 | 0.05 | 0.11 |
| 6 | 151245314 | rs175853 | -0.958 | -0.074 | -1.841 | -0.822 | -0.015 | 0.096 | 0.096 | 0.733 | 0.61 | 0.9 | 0.71 | 0.79 | 0.66 | 0.79 |
| 6 | 151245988 | rs803456 | -0.427 | 0.787 | -1.3 | -0.131 | -0.068 | 0.013 | 0.013 | 1.72 | 0.33 | 0.21 | 0.49 | 0.72 | 0.5 | 0.56 |
| 6 | 151246984 | rs12199747 | -0.528 | 0.125 | -0.561 | -0.105 | -0.032 | -0.04 | -0.04 | 0.061 | 0.09 | 0.18 | 0.05 | 0.03 | 0.05 | 0.11 |
| 6 | 151247439 | rs803455 | 0.528 | 0.307 | -0.929 | 0.215 | -0.011 | -0.029 | -0.029 | 0.049 | 0.91 | 0.98 | 0.86 | 0.88 | 0.93 | 0.96 |
| 6 | 151248028 | rs4869700 | -1.087 | 0.504 | -0.748 | -0.067 | -0.001 | 0.005 | 0.005 | 0 | 0.19 | 0.17 | 0.33 | 0.16 | 0.18 | 0.32 |
| 6 | 151250567 | rs4869954 | -1.244 | 0.795 | -0.495 | -0.812 | -0.002 | 0.042 | 0.042 | 0.968 | 0.33 | 0.72 | 0.37 | 0.07 | 0.34 | 0.41 |
| 6 | 151252917 | rs6902664 | -0.008 | 0.739 | 0.348 | -0.87 | 0.02 | -0.064 | -0.064 | 0 | 0.16 | 0.25 | 0.12 | 0.02 | 0.2 | 0.22 |
| 6 | 151252962 | rs6922248 | -1.513 | 0.71 | 0.791 | -0.914 | -0.002 | -0.07 | -0.07 | 0 | 0.3 | 0.61 | 0.32 | 0.03 | 0.27 | 0.29 |
| 6 | 151253382 | rs1474787 | -1.329 | 0.708 | 0.791 | -0.915 | -0.002 | -0.069 | -0.069 | 0 | 0.3 | 0.62 | 0.32 | 0.03 | 0.27 | 0.29 |
| 6 | 151254080 | rs11155758 | -1.092 | 0.686 | 0.948 | -0.894 | 0.002 | -0.091 | -0.091 | 0 | 0.24 | 0.59 | 0.3 | 0.04 | 0.23 | 0.27 |
| 6 | 151255129 | rs6919680 | -0.168 | 0.337 | 1.045 | -0.109 | -0.012 | 0.248 | 0.248 | 0 | 0.11 | 0.27 | 0.13 | 0.51 | 0.07 | 0.27 |
| 6 | 151255434 | rs2096066 | -1.786 | 0.337 | 1.041 | -0.126 | 0.016 | -0.03 | -0.03 | 0 | 0.4 | 0.74 | 0.45 | 0.55 | 0.33 | 0.56 |
| 6 | 151255774 | rs4869955 | -0.384 | 0.332 | 1.135 | -0.179 | -0.018 | -0.045 | -0.045 | 1.267 | 0.11 | 0.18 | 0.17 | 0.01 | 0.05 | 0.06 |
| 6 | 151256680 | rs6557103 | 0.285 | 0.339 | 1.138 | -0.193 | 0.023 | -0.062 | -0.062 | 0 | 0.16 | 0.25 | 0.13 | 0.02 | 0.2 | 0.22 |
| 6 | 151257871 | rs742832 | -0.663 | 0.278 | 1.087 | -0.169 | -0.026 | -0.062 | -0.062 | 0 | 0.12 | 0.22 | 0.18 | 0.01 | 0.05 | 0.06 |
| 6 | 151258770 | rs742829 | -1.823 | 0.271 | 1.13 | -0.082 | -0.059 | 0.14 | 0.14 | 0 | 0.24 | 0.53 | 0.32 | 0.53 | 0.14 | 0.34 |
| 6 | 151260473 | rs11155761 | -0.967 | 0.462 | 0.926 | -0.257 | 0 | -0.042 | -0.042 | 0 | 0.31 | 0.65 | 0.34 | 0.03 | 0.28 | 0.29 |
| 6 | 151261539 | rs4869959 | -0.671 | 0.566 | 0.649 | -0.189 | 0.009 | -0.124 | -0.124 | 0 | 0.26 | 0.38 | 0.31 | 0.02 | 0.27 | 0.28 |
| 6 | 151264778 | rs803447 | -0.712 | 0.409 | -0.577 | -0.515 | -0.053 | -0.061 | -0.061 | 0.008 | 0.63 | 0.83 | 0.61 | 0.57 | 0.56 | 0.74 |
| 6 | 151265214 | rs803446 | 0.686 | -0.048 | 0.107 | -1.268 | -0.018 | -0.09 | -0.09 | 0.99 | 0.22 | 0.26 | 0.14 | 0.01 | 0.18 | 0.07 |
| 6 | 151266717 | rs803444 | 0.033 | 0.662 | 0.016 | -1.048 | 0.003 | 0.019 | 0.019 | 1.141 | 0.3 | 0.32 | 0.3 | 0.5 | 0.29 | 0.51 |
| 6 | 151270495 | rs803470 | 0.489 | 0.817 | 0.218 | -1.209 | 0.031 | -0.049 | -0.049 | 0 | 0.87 | 0.63 | 0.8 | 0.96 | 0.87 | 0.82 |
| 6 | 151271356 | rs1555179 | 0.956 | 0.743 | -0.215 | -1.463 | -0.036 | -0.081 | -0.081 | 0 | 0.31 | 0.18 | 0.35 | 0.44 | 0.38 | 0.22 |
| 6 | 151278172 | rs1738586 | 0.059 | 0.733 | 0.159 | -1.311 | -0.003 | -0.012 | -0.012 | 0 | 0.29 | 0.22 | 0.26 | 0.51 | 0.28 | 0.5 |
| 6 | 151279026 | rs1738583 | 0.865 | 0.479 | 0.636 | -1.184 | 0.02 | -0.058 | -0.058 | 0 | 0.86 | 0.65 | 0.8 | 0.97 | 0.87 | 0.8 |
| 6 | 151281567 | rs1771798 | 0.874 | 0.568 | 0.913 | -0.949 | 0.054 | 0.063 | 0.063 | 0.047 | 0.91 | 0.69 | 0.93 | 0.88 | 0.9 | 0.81 |
| 6 | 151281668 | rs7739248 | 0.071 | 0.407 | 0.539 | -1.616 | -0.014 | -0.071 | -0.071 | 0 | 0.23 | 0.3 | 0.14 | 0.01 | 0.2 | 0.08 |
| 6 | 151282731 | rs1738579 | 0.234 | 0.135 | 0.108 | -1.641 | -0.006 | -0.006 | -0.006 | 0 | 0.56 | 0.5 | 0.52 | 0.37 | 0.57 | 0.62 |

| | | | | | | | | | | | | | | | | |
|---|---|---|---|---|---|---|---|---|---|---|---|---|---|---|---|---|
| 6 | 151286224 | rs1627462 | 0.169 | -0.115 | 0.067 | -1.727 | 0 | 0 | 0 | 0 | 0.61 | 0.34 | 0.43 | 0.36 | 0.67 | 0.66 |
| 6 | 151288991 | rs563440 | -0.545 | -0.151 | 0.092 | -1.682 | -0.022 | 0.11 | 0.11 | 0.368 | 0.2 | 0.36 | 0.32 | 0.53 | 0.16 | 0.16 |
| 6 | 151289442 | rs558936 | -1.078 | -0.129 | -0.346 | -1.726 | 0.017 | 0.068 | 0.068 | 0 | 0.76 | 0.87 | 0.85 | 0.9 | 0.75 | 0.79 |
| 6 | 151294238 | rs561882 | 0.802 | -0.241 | -0.448 | -1.632 | -0.007 | 0.059 | 0.059 | 0 | 0.1 | 0 | 0.06 | 0 | 0.11 | 0.08 |
| 6 | 151300056 | rs569110 | -0.538 | -0.276 | -0.333 | -1.663 | 0.044 | 0.031 | 0.031 | 0.801 | 0.46 | 0.71 | 0.62 | 0.63 | 0.4 | 0.37 |
| 6 | 151301256 | rs9371487 | 0.048 | -0.28 | -0.257 | -1.641 | -0.026 | 0.002 | 0.002 | 0 | 0.34 | 0.59 | 0.4 | 0.63 | 0.27 | 0.28 |
| 6 | 151301951 | rs7752450 | 0.749 | -0.268 | -0.482 | -1.565 | 0.001 | 0.074 | 0.074 | 0 | 0.07 | 0.05 | 0.25 | 0.52 | 0.05 | 0.08 |
| 6 | 151303497 | rs4242276 | 0.262 | -0.374 | -0.432 | -1.621 | -0.038 | -0.063 | -0.063 | 0 | 0.32 | 0.58 | 0.31 | 0.1 | 0.23 | 0.21 |
| 6 | 151308395 | rs579283 | -0.655 | -0.105 | -0.542 | -1.52 | 0.054 | 0.046 | 0.046 | 1.248 | 0.5 | 0.73 | 0.63 | 0.64 | 0.41 | 0.4 |
| 6 | 151310274 | rs519861 | -1.169 | 0.207 | -0.792 | -1.796 | 0 | -0.039 | -0.039 | 0.503 | 0.82 | 0.85 | 0.88 | 0.99 | 0.83 | 0.79 |
| 6 | 151313367 | rs9968875 | 0.649 | 0.637 | -0.768 | -1.365 | 0.003 | 0 | 0 | 0 | 0.08 | 0.04 | 0.07 | 0.03 | 0.12 | 0.05 |
| 6 | 151315241 | rs803503 | -0.953 | -0.172 | -0.593 | -1.206 | -0.028 | 0.037 | 0.037 | 0.348 | 0.29 | 0.61 | 0.5 | 0.54 | 0.2 | 0.2 |
| 6 | 151317192 | rs538017 | -0.767 | -0.126 | -0.631 | -1.236 | -0.013 | 0.041 | 0.041 | 0.832 | 0.27 | 0.63 | 0.51 | 0.54 | 0.23 | 0.21 |
| 6 | 151318039 | rs9478162 | 0.895 | -0.504 | -0.741 | -1.628 | 0.03 | 0.056 | 0.056 | 0 | 0.22 | 0.14 | 0.13 | 0.09 | 0.21 | 0.2 |
| 6 | 151318079 | rs10457867 | 1.308 | -0.003 | -0.756 | -1.447 | -0.074 | -0.057 | -0.057 | 0 | 0.17 | 0.03 | 0.14 | 0.22 | 0.24 | 0.3 |
| 6 | 151319827 | rs561060 | -0.704 | -0.968 | 0.189 | -1.568 | 0.032 | 0.026 | 0.026 | 0.617 | 0.45 | 0.69 | 0.39 | 0.1 | 0.36 | 0.32 |
| 6 | 151322448 | rs9478908 | -0.449 | -0.46 | -0.176 | -0.832 | 0.047 | 0.072 | 0.072 | 0 | 0.3 | 0.22 | 0.15 | 0.13 | 0.22 | 0.23 |
| 6 | 151323664 | rs9478911 | -0.298 | -0.434 | -0.162 | -0.816 | -0.008 | -0.02 | -0.02 | 1.093 | 0.3 | 0.35 | 0.16 | 0.13 | 0.22 | 0.23 |
| 6 | 151327033 | rs7770982 | -0.912 | 0.095 | 0.291 | -0.333 | 0.004 | -0.055 | -0.055 | 2.822 | 0.13 | 0.24 | 0.06 | 0.03 | 0.1 | 0.06 |
| 6 | 151327345 | rs530971 | -0.672 | -0.386 | 0.595 | -0.55 | -0.017 | -0.034 | -0.034 | 0.387 | 0.07 | 0.14 | 0.04 | 0.01 | 0.06 | 0.06 |
| 6 | 151327735 | rs484994 | -1.423 | -0.454 | 0.725 | -0.585 | -0.027 | -0.055 | -0.055 | 1.265 | 0.1 | 0.33 | 0.04 | 0.01 | 0.06 | 0.06 |
| 6 | 151327848 | rs505358 | -1.762 | -0.556 | 0.027 | -0.019 | 0.023 | -0.062 | -0.062 | 0 | 0.4 | 0.92 | 0.45 | 0.58 | 0.32 | 0.22 |
| 6 | 151327909 | rs505206 | -1.088 | -0.45 | 0.34 | -0.005 | -0.026 | -0.054 | -0.054 | 0.013 | 0.1 | 0.33 | 0.05 | 0.01 | 0.06 | 0.07 |
| 6 | 151327929 | rs505148 | -1.088 | -0.45 | 0.34 | -0.005 | -0.026 | -0.054 | -0.054 | 0 | 0.1 | 0.33 | 0.04 | 0.01 | 0.05 | 0.07 |
| 6 | 151327946 | rs17080689 | -0.16 | -0.346 | 0.615 | 0.029 | 0.001 | -0.025 | -0.025 | 0.426 | 0.11 | 0.18 | 0.07 | 0.04 | 0.09 | 0.08 |
| 6 | 151328122 | rs503348 | -0.442 | -0.386 | 0.43 | 0.18 | -0.017 | -0.034 | -0.034 | 0 | 0.08 | 0.13 | 0.03 | 0.01 | 0.05 | 0.07 |
| 6 | 151330686 | rs526586 | -0.922 | -0.613 | -0.376 | 0.37 | 0.055 | 0.064 | 0.064 | 0 | 0.5 | 0.84 | 0.47 | 0.6 | 0.41 | 0.24 |
| 6 | 151332186 | rs7760043 | -0.033 | -0.795 | 0.782 | -0.166 | -0.007 | -0.144 | -0.144 | 0 | 0.27 | 0.45 | 0.2 | 0.01 | 0.27 | 0.08 |
| 6 | 151338793 | rs9397373 | -0.329 | -0.873 | 1.016 | 0.21 | -0.005 | 0.02 | 0.02 | 0.635 | 0.32 | 0.49 | 0.32 | 0.6 | 0.33 | 0.15 |
| 6 | 151340454 | rs17080705 | 0.305 | -0.843 | 0.505 | 0.057 | -0.001 | -0.012 | -0.012 | 0.095 | 0.25 | 0.2 | 0.19 | 0.01 | 0.27 | 0.08 |
| 6 | 151341298 | rs472703 | -0.926 | -0.967 | 0.693 | 0.044 | -0.042 | 0.024 | 0.024 | 0.394 | 0.25 | 0.55 | 0.27 | 0.68 | 0.14 | 0.14 |
| 6 | 151343977 | rs1079660 | 0.256 | -0.616 | 0.215 | -0.069 | -0.001 | -0.016 | -0.016 | 0 | 0.25 | 0.22 | 0.19 | 0.01 | 0.27 | 0.08 |
| 6 | 151358299 | rs9478918 | 0.176 | -0.712 | -0.346 | 0.15 | 0.034 | 0.066 | 0.066 | 0 | 0.79 | 0.96 | 0.82 | 0.94 | 0.87 | 0.88 |
| 6 | 151362659 | rs9322299 | 0.529 | -0.835 | -0.226 | -0.08 | -0.002 | -0.001 | -0.001 | 0 | 0.65 | 0.66 | 0.68 | 0.34 | 0.76 | 0.76 |
| 6 | 151370204 | rs11155770 | -0.041 | -0.301 | -0.917 | 0.141 | -0.034 | 0.004 | 0.004 | 0.073 | 0.51 | 0.87 | 0.45 | 0.43 | 0.47 | 0.68 |
| 6 | 151371751 | rs6928026 | -0.391 | -0.842 | -0.863 | -0.035 | 0.012 | 0.018 | 0.018 | 0 | 0.51 | 0.7 | 0.49 | 0.56 | 0.48 | 0.43 |
| 6 | 151372383 | rs6914261 | -0.449 | -0.666 | -0.876 | -0.117 | -0.006 | 0.039 | 0.039 | 0 | 0.52 | 0.7 | 0.6 | 0.63 | 0.52 | 0.45 |
| 6 | 151379664 | rs9478934 | 0.15 | -0.641 | -0.325 | -0.101 | -0.023 | -0.052 | -0.052 | 0 | 0.11 | 0.24 | 0.06 | 0 | 0.06 | 0.06 |
| 6 | 151387215 | rs1570190 | -0.126 | -0.378 | -0.456 | -0.059 | 0 | 0.001 | 0.001 | 0 | 0.12 | 0.18 | 0.17 | 0.06 | 0.11 | 0.22 |
| 6 | 151388060 | rs17354394 | 0.059 | -0.418 | -0.451 | -0.053 | -0.001 | -0.001 | -0.001 | 0 | 0.11 | 0.18 | 0.07 | 0.06 | 0.07 | 0.21 |
| 6 | 151388867 | rs4869713 | -0.276 | -0.48 | -0.968 | -0.312 | 0.002 | 0.092 | 0.092 | 0 | 0.52 | 0.21 | 0.6 | 0.7 | 0.58 | 0.45 |
| 6 | 151389129 | rs12211869 | 0.679 | -0.437 | -1.053 | -0.425 | -0.051 | -0.005 | -0.005 | 0 | 0.29 | 0.1 | 0.24 | 0.24 | 0.34 | 0.53 |
| 6 | 151391321 | rs9322301 | 0.448 | -0.365 | -1.173 | -0.301 | -0.106 | -0.004 | -0.004 | 0 | 0.42 | 0.06 | 0.45 | 0.71 | 0.47 | 0.36 |
| 6 | 151392209 | rs6923486 | -0.828 | -0.467 | -0.428 | -0.343 | -0.034 | -0.067 | -0.067 | 0.84 | 0.75 | 0.36 | 0.8 | 0.93 | 0.85 | 0.9 |
| 6 | 151392273 | rs6923669 | 1.209 | -0.273 | -0.763 | -0.277 | -0.037 | -0.035 | -0.035 | 0.106 | 0.1 | 0.02 | 0.26 | 0.54 | 0.13 | 0.14 |
| 6 | 151396691 | rs9397032 | -0.007 | -0.299 | -0.551 | -0.156 | -0.094 | 0.073 | 0.073 | 0.216 | 0.48 | 0.1 | 0.5 | 0.72 | 0.54 | 0.41 |
| 6 | 151399891 | rs730489 | -0.942 | -0.294 | -0.299 | -0.202 | -0.024 | -0.078 | -0.078 | 0 | 0.74 | 0.39 | 0.71 | 0.94 | 0.81 | 0.89 |
| 6 | 151403148 | rs9371494 | 0.096 | -0.322 | -0.434 | -0.276 | -0.001 | -0.008 | -0.008 | 0.642 | 0.62 | 0.35 | 0.63 | 0.79 | 0.68 | 0.7 |
| 6 | 151404303 | rs11155773 | 1.118 | -0.211 | -0.26 | -0.16 | 0.001 | -0.001 | -0.001 | 0 | 0.26 | 0.21 | 0.22 | 0.19 | 0.28 | 0.34 |
| 6 | 151405391 | rs7742684 | 0.051 | -0.243 | 0.129 | -0.179 | 0.004 | -0.006 | -0.006 | 0 | 0.15 | 0.22 | 0.19 | 0.14 | 0.18 | 0.2 |
| 6 | 151407272 | rs3849793 | 0.847 | -0.068 | -0.126 | 0.034 | -0.057 | -0.096 | -0.096 | 0.452 | 0.35 | 0.1 | 0.4 | 0.57 | 0.4 | 0.22 |
| 6 | 151407472 | rs17080757 | -0.058 | 0.003 | 0.224 | 0.155 | 0.001 | 0.002 | 0.002 | 0 | 0.34 | 0.26 | 0.26 | 0.4 | 0.34 | 0.62 |
| 6 | 151409027 | rs12216229 | 0.779 | 0.108 | 0.227 | -0.032 | 0 | 0 | 0 | 0 | 0.25 | 0.19 | 0.22 | 0.17 | 0.28 | 0.34 |
| 6 | 151416939 | rs3849795 | -0.772 | -0.307 | 0.553 | -0.041 | -0.01 | 0.08 | 0.08 | 0.206 | 0.13 | 0.43 | 0.05 | 0.18 | 0.08 | 0.29 |
| 6 | 151418320 | rs11753794 | 0.021 | -0.239 | 0.046 | 0.145 | 0.023 | 0.031 | 0.031 | 0.891 | 0.42 | 0.25 | 0.32 | 0.34 | 0.39 | 0.35 |
| 6 | 151419806 | rs13199827 | 0.69 | -0.239 | -0.091 | 0.139 | 0.015 | 0.047 | 0.047 | 0.295 | 0.32 | 0.16 | 0.28 | 0.2 | 0.32 | 0.16 |
| 6 | 151422414 | rs13212150 | 0.85 | -0.236 | -0.089 | -0.134 | 0.013 | 0.068 | 0.068 | 0.336 | 0.32 | 0.16 | 0.28 | 0.15 | 0.32 | 0.16 |
| 6 | 151422750 | rs7646 | 1.338 | -0.034 | -0.241 | -0.273 | -0.015 | 0.018 | 0.018 | 0 | 0.21 | 0.12 | 0.23 | 0.09 | 0.28 | 0.23 |
| 6 | 151422955 | rs1047665 | 0.719 | -0.128 | -0.799 | -0.11 | -0.014 | -0.039 | -0.039 | 0 | 0.08 | 0 | 0.03 | 0.14 | 0.07 | 0.18 |
| 6 | 151423606 | rs6912269 | -0.2 | -0.165 | -0.679 | 0.036 | 0.031 | 0.026 | 0.026 | 0.153 | 0.48 | 0.35 | 0.33 | 0.35 | 0.39 | 0.49 |
| 6 | 151427385 | rs6934983 | 0.727 | -0.231 | -0.311 | 0.44 | -0.047 | 0.009 | 0.009 | 0.466 | 0.08 | 0.19 | 0.02 | 0.06 | 0 | 0.14 |
| 6 | 151427734 | rs6557111 | -0.694 | 0.453 | -1.018 | 0.594 | -0.014 | -0.046 | -0.046 | 0.155 | 0.73 | 0.83 | 0.68 | 0.44 | 0.72 | 0.74 |
| 6 | 151430051 | rs12661373 | -0.149 | 0.129 | -0.963 | 0.026 | 0.012 | 0.038 | 0.038 | 1.257 | 0.26 | 0.15 | 0.25 | 0.14 | 0.26 | 0.14 |
| 6 | 151432485 | rs4869988 | -0.139 | 0.585 | -0.355 | 0.224 | -0.001 | -0.003 | -0.003 | 0 | 0.48 | 0.46 | 0.56 | 0.34 | 0.6 | 0.45 |
| 6 | 151433994 | rs7772926 | 0.29 | 0.917 | -0.468 | -0.124 | 0.017 | -0.086 | -0.086 | 0 | 0.36 | 0.41 | 0.43 | 0.13 | 0.44 | 0.29 |
| 6 | 151435955 | rs6923696 | -0.515 | 0.535 | -0.611 | 0.337 | 0.008 | -0.02 | -0.02 | 1.396 | 0.57 | 0.86 | 0.63 | 0.48 | 0.65 | 0.47 |
| 6 | 151439698 | rs9478170 | -0.364 | 0.264 | 0.009 | 0.297 | -0.023 | 0.007 | 0.007 | 1.286 | 0.62 | 0.5 | 0.64 | 0.46 | 0.73 | 0.49 |
| 6 | 151441417 | rs4869989 | -0.386 | 0.332 | 0.063 | 0.318 | -0.017 | 0.018 | 0.018 | 1.163 | 0.63 | 0.51 | 0.64 | 0.5 | 0.73 | 0.49 |
| 6 | 151443655 | rs11758761 | 0.577 | 0.368 | -0.152 | 0.152 | -0.036 | 0.013 | 0.013 | 0.221 | 0.2 | 0.12 | 0.15 | 0.18 | 0.24 | 0.15 |

| | | | | | | | | | | | | | | | | |
|---|---|---|---|---|---|---|---|---|---|---|---|---|---|---|---|---|
| 6 | 151446475 | rs11753902 | 0.597 | 0.383 | -0.24 | 0.159 | -0.062 | -0.004 | -0.004 | 0.001 | 0.19 | 0.04 | 0.14 | 0.18 | 0.24 | 0.15 |
| 6 | 151450188 | rs7760165 | -1.035 | 0.132 | 0.664 | 0.61 | 0.084 | 0.044 | 0.044 | 0.724 | 0.06 | 0.46 | 0.05 | 0 | 0.06 | 0.01 |
| 6 | 151452358 | rs7771304 | -1.142 | 0.156 | 0.648 | 0.625 | 0.017 | 0.037 | 0.037 | 0.422 | 0.12 | 0.51 | 0.21 | 0.12 | 0.1 | 0.06 |
| 6 | 151455122 | rs6911282 | -0.906 | 0.05 | -0.352 | 0.869 | -0.023 | -0.079 | -0.079 | 0.45 | 0.84 | 0.99 | 0.89 | 0.79 | 0.87 | 0.71 |
| 6 | 151459830 | rs6557113 | 0.234 | 0.048 | -0.257 | 0.769 | 0.018 | 0.011 | 0.011 | 1.571 | 0.62 | 0.82 | 0.63 | 0.38 | 0.72 | 0.44 |
| 6 | 151461931 | rs12660161 | 0.549 | -0.11 | -0.334 | 0.902 | -0.017 | -0.107 | -0.107 | 3.353 | 0.17 | 0.02 | 0.21 | 0.31 | 0.12 | 0.13 |
| 6 | 151461965 | rs12660147 | 0.621 | -0.077 | -0.355 | 0.88 | -0.027 | -0.118 | -0.118 | 0 | 0.17 | 0.01 | 0.2 | 0.32 | 0.12 | 0.13 |
| 6 | 151462646 | rs9383557 | 0.285 | 0.228 | 0.302 | 0.629 | -0.004 | -0.006 | -0.006 | 0 | 0.71 | 0.37 | 0.7 | 0.75 | 0.73 | 0.65 |
| 6 | 151467182 | rs6924562 | 0.657 | 0.494 | -0.054 | 0.067 | 0 | -0.013 | -0.013 | 0.01 | 0.48 | 0.5 | 0.32 | 0.26 | 0.51 | 0.36 |
| 6 | 151467445 | rs4869991 | -0.108 | 0.907 | -0.487 | 0.577 | -0.023 | -0.083 | -0.083 | 1.265 | 0.78 | 0.94 | 0.76 | 0.62 | 0.78 | 0.65 |
| 6 | 151469255 | rs6909007 | -0.924 | 0.861 | 0.424 | -0.075 | 0.044 | -0.039 | -0.039 | 0.072 | 0.1 | 0.4 | 0.19 | 0 | 0.08 | 0.05 |
| 6 | 151472251 | rs942382 | 0.111 | 1.158 | -0.575 | -0.381 | -0.034 | 0.058 | 0.058 | 0.019 | 0.66 | 0.77 | 0.75 | 0.79 | 0.58 | 0.62 |
| 6 | 151473330 | rs4869993 | -0.676 | 1.034 | -0.285 | 0.05 | 0.013 | 0.029 | 0.029 | 0.26 | 0.28 | 0.35 | 0.31 | 0.4 | 0.33 | 0.31 |
| 6 | 151476147 | rs10085219 | -0.301 | 0.979 | -0.321 | -0.311 | -0.001 | 0.004 | 0.004 | 0 | 0.07 | 0.1 | 0.16 | 0 | 0.07 | 0.07 |
| 6 | 151481007 | rs6934196 | -0.47 | 0.182 | -0.919 | -1.286 | 0.023 | 0.058 | 0.058 | 0.335 | 0.71 | 0.88 | 0.9 | 0.92 | 0.8 | 0.77 |
| 6 | 151481364 | rs752104 | 0.256 | -0.157 | -0.36 | -1.452 | -0.017 | -0.076 | -0.076 | 0 | 0.59 | 0.51 | 0.71 | 0.9 | 0.73 | 0.7 |
| 6 | 151485501 | rs2346892 | -0.645 | -0.136 | -0.87 | -1.415 | 0.028 | 0.068 | 0.068 | 0.623 | 0.7 | 0.87 | 0.9 | 0.92 | 0.8 | 0.76 |
| 6 | 151488316 | rs9478967 | 0.056 | -0.184 | -0.811 | -1.448 | 0.032 | 0.083 | 0.083 | 0 | 0.63 | 0.8 | 0.74 | 0.92 | 0.73 | 0.7 |
| 6 | 151488718 | rs9478180 | -0.179 | -0.287 | -0.375 | -1.019 | -0.002 | -0.004 | -0.004 | 0 | 0.25 | 0.65 | 0.31 | 0.31 | 0.21 | 0.23 |
| 6 | 151495387 | rs9322304 | 0.215 | -0.244 | -0.728 | -1.065 | -0.007 | 0.028 | 0.028 | 0 | 0.08 | 0.05 | 0.17 | 0 | 0.08 | 0.07 |
| 6 | 151496340 | rs1535831 | -0.593 | -0.672 | -0.76 | -1.01 | 0.025 | 0.025 | 0.025 | 0 | 0.42 | 0.71 | 0.51 | 0.35 | 0.34 | 0.37 |
| 6 | 151497673 | rs9478183 | 0.538 | -0.588 | -1.15 | -0.958 | 0.023 | 0.088 | 0.088 | 0 | 0.17 | 0.06 | 0.2 | 0 | 0.14 | 0.1 |
| 6 | 151501035 | rs752543 | -0.615 | -0.331 | -1.436 | -1.014 | -0.139 | -0.076 | -0.076 | 0 | 0.61 | 0.94 | 0.59 | 0.42 | 0.48 | 0.46 |
| 6 | 151503261 | rs6918015 | -0.092 | -0.126 | -0.105 | -0.417 | 0.008 | 0.044 | 0.044 | 0 | 0.23 | 0.48 | 0.3 | 0.34 | 0.2 | 0.26 |
| 6 | 151513858 | rs9478984 | -0.57 | 0.219 | -0.316 | 0.609 | -0.007 | 0.026 | 0.026 | 0 | 0.45 | 0.44 | 0.52 | 0.72 | 0.46 | 0.67 |
| 6 | 151527056 | rs9478986 | 0.175 | 0.252 | -0.493 | 0.92 | 0.002 | -0.016 | -0.016 | 0 | 0.3 | 0.31 | 0.31 | 0.44 | 0.27 | 0.31 |
| 6 | 151527565 | rs1890136 | 0.083 | 0.295 | -0.483 | 0.923 | -0.002 | 0.004 | 0.004 | 0 | 0.32 | 0.43 | 0.32 | 0.43 | 0.27 | 0.35 |
| 6 | 151527772 | rs4869999 | -0.563 | 0.567 | -0.212 | 1.133 | 0.006 | 0.038 | 0.038 | 0 | 0.15 | 0.21 | 0.19 | 0.28 | 0.17 | 0.34 |
| 6 | 151542187 | rs9478186 | -0.077 | 1.096 | -1.144 | 1.348 | -0.067 | -0.004 | -0.004 | 0 | 0.29 | 0.11 | 0.41 | 0.26 | 0.37 | 0.3 |
| 6 | 151546061 | rs9322306 | -0.504 | -0.08 | -0.771 | 1.373 | -0.016 | 0.005 | 0.005 | 1.082 | 0.65 | 0.64 | 0.67 | 0.56 | 0.71 | 0.66 |
| 6 | 151563575 | rs6557123 | 0.292 | 0.483 | -1.311 | 1.335 | -0.07 | 0.016 | 0.016 | 0 | 0.32 | 0.12 | 0.39 | 0.19 | 0.37 | 0.33 |
| 6 | 151569243 | rs4870002 | -0.399 | 0.079 | -0.889 | 0.977 | 0.009 | 0.016 | 0.016 | 0 | 0.5 | 0.59 | 0.73 | 0.64 | 0.57 | 0.59 |
| 6 | 151572302 | rs10499265 | 0.759 | -0.293 | -1.322 | 0.844 | 0.03 | 0.077 | 0.077 | 2.193 | 0.14 | 0.04 | 0.05 | 0.01 | 0.09 | 0.02 |
| 6 | 151572494 | rs9285530 | 0.465 | -0.079 | -1.459 | 1.12 | -0.144 | -0.014 | -0.014 | 0 | 0.36 | 0.07 | 0.39 | 0.25 | 0.4 | 0.39 |
| 6 | 151575250 | rs7764018 | -0.345 | -0.345 | -0.672 | 0.471 | -0.004 | -0.039 | -0.039 | 0 | 0.63 | 0.61 | 0.73 | 0.87 | 0.62 | 0.79 |
| 6 | 151575848 | rs17080949 | 0.31 | -0.229 | -1.228 | 0.583 | 0 | -0.035 | -0.035 | 0.06 | 0.12 | 0.08 | 0.28 | 0.25 | 0.08 | 0.21 |
| 6 | 151575870 | rs12202922 | -0.197 | -0.681 | -1.327 | -0.054 | 0.085 | 0.221 | 0.221 | 8.743 | 0.49 | 0.23 | 0.72 | 0.97 | 0.65 | 0.83 |
| 6 | 151575944 | rs17080959 | 0.614 | -0.671 | -0.996 | 0.301 | 0.001 | 0 | 0 | 0 | 0.18 | 0.19 | 0.13 | 0.15 | 0.16 | 0.06 |
| 6 | 151577270 | rs7742224 | -0.211 | -0.196 | -1.129 | 0.349 | -0.107 | -0.079 | -0.079 | 0 | 0.6 | 0.86 | 0.49 | 0.55 | 0.52 | 0.39 |
| 6 | 151579432 | rs4869723 | 0.077 | -0.354 | -0.704 | 0.165 | 0.005 | 0.002 | 0.002 | 6.16 | 0.53 | 0.6 | 0.43 | 0.55 | 0.48 | 0.37 |
| 6 | 151582095 | rs9478188 | 0.143 | 0.007 | -0.743 | 0.295 | -0.004 | -0.022 | -0.022 | 0 | 0.06 | 0.08 | 0.03 | 0.01 | 0.03 | 0.04 |
| 6 | 151582632 | rs9383562 | -0.626 | -0.091 | -1.119 | 0.207 | -0.127 | 0.039 | 0.039 | 0.206 | 0.7 | 0.93 | 0.76 | 0.82 | 0.59 | 0.61 |
| 6 | 151583977 | rs7382659 | 0.55 | -0.58 | -1.291 | -0.142 | 0.039 | -0.049 | -0.049 | 0.457 | 0.26 | 0.07 | 0.3 | 0.25 | 0.17 | 0.22 |
| 6 | 151583994 | rs11961573 | -0.922 | -0.168 | -0.737 | 0.185 | -0.002 | 0.016 | 0.016 | 0 | 0.11 | 0.21 | 0.07 | 0.17 | 0.09 | 0.19 |
| 6 | 151587799 | rs11967865 | 0.019 | -0.213 | -0.83 | 0.152 | -0.003 | -0.025 | -0.025 | 0.357 | 0.06 | 0.08 | 0.03 | 0.01 | 0.04 | 0.02 |
| 6 | 151592001 | rs9383875 | -0.825 | -0.008 | -0.871 | 0.46 | 0.007 | 0.024 | 0.024 | 0 | 0.1 | 0.13 | 0.12 | 0.2 | 0.16 | 0.06 |
| 6 | 151592056 | rs926557 | -0.363 | -0.297 | -0.742 | 0.096 | 0.045 | -0.013 | -0.013 | 0.083 | 0.45 | 0.65 | 0.51 | 0.49 | 0.35 | 0.34 |
| 6 | 151594564 | rs2257102 | -0.779 | -0.059 | -0.807 | 0.049 | -0.046 | -0.021 | -0.021 | 3.101 | 0.68 | 0.83 | 0.64 | 0.64 | 0.58 | 0.56 |
| 6 | 151596949 | rs6938445 | 0.215 | -0.215 | -0.697 | -0.02 | 0.013 | -0.015 | -0.015 | 1.051 | 0.49 | 0.21 | 0.49 | 0.47 | 0.46 | 0.35 |
| 6 | 151597937 | rs11966225 | 1.139 | -0.521 | -0.799 | -0.418 | -0.006 | 0.044 | 0.044 | 0.03 | 0.21 | 0.08 | 0.1 | 0.02 | 0.21 | 0.07 |
| 6 | 151598107 | rs6557129 | 0.805 | -0.871 | -0.325 | -0.44 | -0.007 | -0.029 | -0.029 | 0 | 0.08 | 0.13 | 0.06 | 0 | 0.06 | 0.05 |
| 6 | 151599211 | rs17081014 | 1.248 | -0.851 | -0.463 | -0.244 | 0.052 | -0.05 | -0.05 | 0 | 0.23 | 0.06 | 0.3 | 0.26 | 0.13 | 0.24 |
| 6 | 151601387 | rs4431446 | 0.585 | -0.755 | -0.362 | -0.285 | -0.009 | 0.012 | 0.012 | 0 | 0.26 | 0.17 | 0.21 | 0.21 | 0.31 | 0.11 |
| 6 | 151606123 | rs9383877 | -0.098 | -0.747 | -0.449 | -0.186 | -0.007 | 0.093 | 0.093 | 0.832 | 0.54 | 0.3 | 0.61 | 0.79 | 0.5 | 0.57 |
| 6 | 151607502 | rs7760914 | 0.87 | -0.76 | -0.446 | -0.081 | 0.036 | -0.084 | -0.084 | 0 | 0.27 | 0.1 | 0.4 | 0.57 | 0.19 | 0.46 |
| 6 | 151608694 | rs12174672 | 0.155 | -0.742 | -0.312 | -0.06 | -0.007 | -0.009 | -0.009 | 0 | 0.33 | 0.27 | 0.32 | 0.63 | 0.39 | 0.44 |
| 6 | 151612694 | rs12195473 | 1.187 | -0.543 | -0.689 | -0.115 | -0.141 | 0.042 | 0.042 | 0.609 | 0.25 | 0.02 | 0.22 | 0.1 | 0.37 | 0.22 |
| 6 | 151616291 | rs9383878 | 0.754 | -0.749 | -0.352 | -0.024 | -0.03 | -0.02 | -0.02 | 0.892 | 0.31 | 0.21 | 0.35 | 0.41 | 0.44 | 0.43 |
| 6 | 151617320 | rs9322313 | 0.016 | -0.723 | -0.4 | 0.026 | 0.011 | 0.021 | 0.021 | 0.477 | 0.56 | 0.29 | 0.52 | 0.38 | 0.41 | 0.52 |
| 6 | 151618235 | rs9479007 | 0.224 | -0.725 | -0.391 | -0.336 | 0.003 | 0.172 | 0.172 | 0.117 | 0.52 | 0.21 | 0.62 | 0.89 | 0.49 | 0.69 |
| 6 | 151618879 | rs1474719 | -0.171 | -0.733 | -0.178 | -0.315 | 0.027 | -0.025 | -0.025 | 0.269 | 0.93 | 0.73 | 0.94 | 1 | 0.95 | 0.98 |
| 6 | 151619282 | rs10155709 | 0.411 | -0.733 | -0.358 | -0.296 | -0.011 | -0.046 | -0.046 | 0.276 | 0.1 | 0.18 | 0.06 | 0 | 0.05 | 0.06 |
| 6 | 151619432 | rs3818832 | 0.782 | -0.714 | -0.202 | -0.031 | -0.004 | -0.03 | -0.03 | 0 | 0.25 | 0.16 | 0.21 | 0.45 | 0.24 | 0.33 |
| 6 | 151620494 | rs9479008 | 0.566 | -0.555 | -0.503 | 0.05 | -0.003 | 0.025 | 0.025 | 0 | 0.28 | 0.35 | 0.22 | 0.44 | 0.25 | 0.33 |
| 6 | 151621492 | rs926559 | 0.277 | 0.024 | -0.463 | -0.598 | 0.018 | 0.027 | 0.027 | 0.021 | 0.53 | 0.39 | 0.58 | 0.69 | 0.4 | 0.65 |
| 6 | 151621611 | rs2251681 | 1.09 | 0.325 | -0.644 | 0.174 | 0.028 | 0.097 | 0.097 | 0 | 0.9 | 0.75 | 0.89 | 0.77 | 0.86 | 0.95 |
| 6 | 151626671 | rs2473609 | -0.259 | -0.149 | -1.588 | -0.621 | 0.054 | 0.081 | 0.081 | 0.355 | 0.77 | 0.93 | 0.77 | 1 | 0.85 | 0.83 |
| 6 | 151627562 | rs2294794 | -0.227 | 1.36 | -1.808 | -0.443 | -0.12 | 0.077 | 0.077 | 0.538 | 0.24 | 0.09 | 0.35 | 0.06 | 0.44 | 0.24 |
| 6 | 151629865 | rs9397389 | -0.052 | 1.055 | -1.059 | 0.103 | 0.016 | 0.013 | 0.013 | 0.145 | 0.27 | 0.43 | 0.37 | 0.28 | 0.26 | 0.1 |

| | | | | | | | | | | | | | | | | |
|---|---|---|---|---|---|---|---|---|---|---|---|---|---|---|---|---|
| 6 | 151632427 | rs4870010 | -0.266 | 1.81 | -1.598 | -0.014 | -0.055 | 0.047 | 0.047 | 1.083 | 0.16 | 0.05 | 0.22 | 0.05 | 0.28 | 0.16 |
| 6 | 151632899 | rs926561 | 0.083 | 1.128 | -1.944 | 0.599 | 0.006 | -0.102 | -0.102 | 0 | 0.25 | 0.1 | 0.28 | 0.52 | 0.21 | 0.34 |
| 6 | 151632968 | rs7740243 | -0.241 | 1.374 | -1.635 | 0.111 | -0.02 | 0.062 | 0.062 | 0.343 | 0.25 | 0.1 | 0.25 | 0.06 | 0.27 | 0.12 |
| 6 | 151634078 | rs6908030 | -1.027 | 0.758 | -0.48 | 0.563 | 0.004 | -0.043 | -0.043 | 1.434 | 0.31 | 0.44 | 0.36 | 0.86 | 0.33 | 0.5 |
| 6 | 151635626 | rs9383882 | -1.176 | 0.643 | -0.738 | 2.182 | 0.003 | 0.044 | 0.044 | 0.096 | 0.06 | 0.06 | 0.15 | 0.39 | 0.05 | 0.09 |
| 6 | 151639599 | rs12210237 | -0.763 | 0.573 | -0.585 | 1.797 | -0.012 | -0.028 | -0.028 | 1.457 | 0.15 | 0.09 | 0.11 | 0.36 | 0.23 | 0.25 |
| 6 | 151641001 | rs12205375 | -1.286 | 0.196 | -0.211 | 1.103 | 0.007 | -0.027 | -0.027 | 0.462 | 0.07 | 0.15 | 0.06 | 0.01 | 0.08 | 0.14 |
| 6 | 151642006 | rs2347436 | -1.267 | -0.375 | -0.128 | 1.532 | -0.002 | 0.001 | 0.001 | 0 | 0.28 | 0.31 | 0.33 | 0.35 | 0.23 | 0.31 |
| 6 | 151643910 | rs2485542 | -1.652 | -0.612 | 0.64 | 1.849 | 0.003 | 0.183 | 0.183 | 0.38 | 0.13 | 0.38 | 0.22 | 0.34 | 0.11 | 0.23 |
| 6 | 151645569 | rs2248861 | -1.083 | -1.585 | -0.793 | 1.11 | -0.032 | 0.056 | 0.056 | 3.803 | 0.57 | 0.95 | 0.49 | 0.76 | 0.38 | 0.66 |
| 6 | 151654269 | rs2626356 | -0.269 | -2.266 | -1.111 | 0.736 | 0.041 | -0.016 | -0.016 | 0.151 | 0.16 | 0.02 | 0.14 | 0.14 | 0.06 | 0.15 |
| 6 | 151655001 | rs7749357 | -0.142 | -3.029 | -0.377 | 0.29 | -0.079 | -0.145 | -0.145 | 0 | 0.26 | 0.45 | 0.11 | 0.01 | 0.1 | 0.16 |
| 6 | 151658189 | rs11155784 | 0.469 | -3.161 | -0.518 | 0.225 | 0.05 | 0.095 | 0.095 | 0 | 0.2 | 0.04 | 0.07 | 0.01 | 0.09 | 0.15 |
| 6 | 151659478 | rs222575 | 0.469 | -3.278 | -0.246 | 0.672 | -0.002 | -0.008 | -0.008 | 0 | 0.23 | 0.24 | 0.24 | 0.42 | 0.22 | 0.36 |
| 6 | 151660477 | rs222577 | -1.855 | -3.288 | -0.297 | 0.663 | -0.075 | -0.113 | -0.113 | 0.354 | 0.63 | 0.94 | 0.47 | 0.58 | 0.34 | 0.62 |
| 6 | 151663289 | rs756009 | -0.322 | -3.359 | 0.117 | 0.624 | -0.064 | -0.107 | -0.107 | 0 | 0.26 | 0.56 | 0.11 | 0.01 | 0.08 | 0.13 |
| 6 | 151663560 | rs900653 | 0.533 | -3.16 | 0.226 | 0.756 | 0.004 | -0.004 | -0.004 | 0 | 0.63 | 0.63 | 0.64 | 0.43 | 0.74 | 0.52 |
| 6 | 151670172 | rs3734799 | 0.664 | -3.148 | 0.493 | 0.885 | 0.004 | -0.005 | -0.005 | 0 | 0.63 | 0.36 | 0.63 | 0.46 | 0.76 | 0.59 |
| 6 | 151670897 | rs900654 | -0.518 | -3.12 | 0.214 | 0.84 | -0.001 | -0.006 | -0.006 | 0 | 0.78 | 0.77 | 0.78 | 0.59 | 0.78 | 0.64 |
| 6 | 151672812 | rs3734797 | 1.09 | -3.123 | 0.155 | 0.802 | 0.029 | -0.009 | -0.009 | 0 | 0.1 | 0.02 | 0.11 | 0.13 | 0.02 | 0.03 |
| 6 | 151673589 | rs12201388 | 0.222 | -3.321 | 0.13 | 0.661 | 0.03 | 0.021 | 0.021 | 1.207 | 0.23 | 0.15 | 0.08 | 0.01 | 0.09 | 0.17 |
| 6 | 151674326 | rs3823310 | 1.457 | -3.305 | 0.437 | 0.674 | 0.023 | -0.012 | -0.012 | 0.269 | 0.41 | 0.22 | 0.55 | 0.44 | 0.67 | 0.43 |
| 6 | 151674593 | rs3734795 | 0.257 | -3.275 | 0.082 | 0.537 | -0.064 | 0.023 | 0.023 | 3.275 | 0.13 | 0.35 | 0.14 | 0.14 | 0.02 | 0.03 |
| 6 | 151677359 | rs1042167 | 0.06 | -2.816 | 0.003 | 0.419 | 0.077 | 0.114 | 0.114 | 1.282 | 0.23 | 0.06 | 0.08 | 0.01 | 0.09 | 0.18 |
| 6 | 151686905 | rs11155787 | 0.837 | -1.994 | 0.361 | 0.95 | 0.001 | 0.004 | 0.004 | 3.903 | 0.39 | 0.35 | 0.54 | 0.28 | 0.64 | 0.37 |
| 6 | 151696304 | rs6557131 | -1.033 | -1.965 | 0.961 | 1.434 | 0.009 | 0.119 | 0.119 | 0.101 | 0.06 | 0.25 | 0.03 | 0.16 | 0.04 | 0.05 |
| 6 | 151705334 | rs7775640 | -1.603 | -2.388 | 0.711 | 1.237 | -0.012 | 0.003 | 0.003 | 0.008 | 0.33 | 0.54 | 0.36 | 0.7 | 0.23 | 0.38 |
| 6 | 151706663 | rs3849797 | -0.477 | -2.509 | 0.604 | 1.491 | -0.004 | 0.002 | 0.002 | 0 | 0.15 | 0.02 | 0.17 | 0.13 | 0.12 | 0.08 |
| 6 | 151715105 | rs10002 | -1.852 | -2.455 | 0.71 | 1.469 | 0.001 | -0.001 | -0.001 | 0 | 0.33 | 0.63 | 0.37 | 0.71 | 0.23 | 0.37 |
| 6 | 151721749 | rs7771156 | -2.209 | -3.122 | 0.341 | 1.02 | 0.099 | 0.108 | 0.108 | 2.221 | 0.49 | 0.75 | 0.45 | 0.72 | 0.34 | 0.53 |
| 6 | 151726829 | rs3757315 | -1.696 | -2.904 | 0.621 | 1.16 | 0.002 | -0.001 | -0.001 | 0.073 | 0.33 | 0.63 | 0.37 | 0.7 | 0.23 | 0.37 |
| 6 | 151730784 | rs9478202 | -1.116 | -2.884 | 1.088 | 1.409 | -0.032 | 0.194 | 0.194 | 1.037 | 0.17 | 0.5 | 0.16 | 0.4 | 0.07 | 0.26 |
| 6 | 151730802 | rs9383889 | -1.668 | -3.104 | 1.344 | 1.439 | -0.041 | 0.046 | 0.046 | 4.065 | 0.31 | 0.52 | 0.22 | 0.54 | 0.19 | 0.42 |
| 6 | 151743948 | rs9478203 | -0.589 | -2.982 | 1.605 | 1.175 | 0.045 | 0.031 | 0.031 | 1.424 | 0.05 | 0.37 | 0.04 | 0 | 0.01 | 0.02 |
| 6 | 151754273 | rs6904364 | -1.799 | -2.911 | 1.192 | 0.965 | 0.003 | 0.003 | 0.003 | 1.486 | 0.47 | 0.46 | 0.43 | 0.74 | 0.41 | 0.49 |
| 6 | 151759192 | rs3800274 | -1.517 | -3.059 | 1.205 | 0.913 | -0.016 | 0.013 | 0.013 | 0.159 | 0.32 | 0.48 | 0.35 | 0.73 | 0.28 | 0.38 |
| 6 | 151759630 | rs9322321 | -1.43 | -3.099 | 1.246 | 1.196 | 0.006 | -0.001 | -0.001 | 0 | 0.35 | 0.36 | 0.3 | 0.35 | 0.28 | 0.28 |
| 6 | 151765296 | rs3823311 | -0.896 | -3.067 | 1.271 | 1.231 | -0.035 | -0.021 | -0.021 | 5.235 | 0.22 | 0.39 | 0.13 | 0.17 | 0.17 | 0.2 |
| 6 | 151768495 | rs9383896 | -0.407 | -3.044 | 1.166 | 1.279 | 0.004 | 0.05 | 0.05 | 0 | 0.12 | 0.16 | 0.13 | 0.39 | 0.11 | 0.22 |
| 6 | 151773504 | rs13201167 | -2.491 | -3.124 | 1.05 | 1.255 | 0.033 | 0.11 | 0.11 | 5.284 | 0.53 | 0.88 | 0.47 | 0.77 | 0.41 | 0.5 |
| 6 | 151774339 | rs3757316 | -2.053 | -3.138 | 1.133 | 1.265 | 0.05 | 0.009 | 0.009 | 1.323 | 0.47 | 0.69 | 0.42 | 0.61 | 0.36 | 0.42 |
| 6 | 151779064 | rs6916340 | -2.044 | -3.137 | 1.129 | 1.265 | 0.051 | 0.009 | 0.009 | 1.259 | 0.47 | 0.7 | 0.42 | 0.61 | 0.36 | 0.42 |
| 6 | 151779157 | rs6911936 | 0.147 | -3.118 | 0.893 | 1.19 | 0.017 | 0.034 | 0.034 | 0 | 0.15 | 0.09 | 0.08 | 0.02 | 0.12 | 0.12 |
| 6 | 151786177 | rs6931104 | -2.335 | -3.023 | 0.848 | 1.248 | 0.033 | 0.11 | 0.11 | 5.284 | 0.53 | 0.88 | 0.47 | 0.77 | 0.41 | 0.5 |
| 6 | 151791737 | rs13202913 | -0.562 | -2.744 | 1.232 | 1.535 | -0.004 | 0.002 | 0.002 | 0 | 0.12 | 0.11 | 0.14 | 0.53 | 0.12 | 0.23 |
| 6 | 151793095 | rs9478207 | -0.906 | -2.847 | 1.615 | 1.262 | -0.046 | -0.144 | -0.144 | 0 | 0.21 | 0.41 | 0.12 | 0.01 | 0.14 | 0.15 |
| 6 | 151799924 | rs13198669 | -0.756 | -2.924 | 1.214 | 1.348 | -0.005 | -0.038 | -0.038 | 0 | 0.15 | 0.09 | 0.16 | 0.43 | 0.12 | 0.22 |
| 6 | 151806338 | rs9479050 | -0.722 | -2.89 | 1.486 | 1.057 | -0.041 | -0.101 | -0.101 | 0 | 0.19 | 0.34 | 0.1 | 0.01 | 0.12 | 0.13 |
| 6 | 151816094 | rs1971255 | -0.756 | -2.89 | 1.465 | 1.057 | -0.045 | -0.108 | -0.108 | 0.004 | 0.19 | 0.35 | 0.1 | 0.01 | 0.12 | 0.13 |
| 6 | 151817992 | rs17081325 | -0.675 | -2.881 | 1.619 | 1.097 | -0.007 | -0.034 | -0.034 | 0.137 | 0.19 | 0.21 | 0.09 | 0.01 | 0.12 | 0.13 |
| 6 | 151824449 | rs12173791 | -0.843 | -3.03 | 1.361 | 1.227 | -0.028 | 0.116 | 0.116 | 0 | 0.13 | 0.23 | 0.17 | 0.36 | 0.08 | 0.19 |
| 6 | 151830769 | rs4870024 | -1.62 | -3.01 | 0.782 | 1.227 | -0.015 | 0.003 | 0.003 | 0.095 | 0.33 | 0.63 | 0.28 | 0.38 | 0.2 | 0.32 |
| 6 | 151832668 | rs9371528 | -0.986 | -2.808 | 0.709 | 1.469 | -0.002 | 0.004 | 0.004 | 0 | 0.21 | 0.23 | 0.16 | 0.41 | 0.14 | 0.22 |
| 6 | 151837570 | rs17081341 | 0.052 | -2.504 | 0.36 | 1.204 | 0.008 | 0 | 0 | 0.018 | 0.08 | 0.07 | 0.14 | 0.2 | 0.04 | 0.12 |
| 6 | 151847967 | rs9479055 | 1.669 | -1.964 | 0.026 | 0.508 | 0.02 | 0.134 | 0.134 | 0.81 | 0.47 | 0.14 | 0.48 | 0.2 | 0.6 | 0.44 |
| 6 | 151849040 | rs17688135 | 0.072 | -1.915 | 0.183 | 0.506 | 0.001 | -0.004 | -0.004 | 0.429 | 0.05 | 0.08 | 0.1 | 0.07 | 0.08 | 0.08 |
| 6 | 151851124 | rs6900089 | 0.034 | -1.917 | 0.229 | 0.505 | 0.002 | -0.006 | -0.006 | 0 | 0.06 | 0.09 | 0.1 | 0.07 | 0.08 | 0.08 |
| 6 | 151852258 | rs6933202 | -0.144 | -2.093 | 0.344 | 0.37 | -0.002 | 0.022 | 0.022 | 0 | 0.88 | 0.96 | 0.94 | 1 | 0.86 | 0.89 |
| 6 | 151853291 | rs6912538 | -1.292 | -2.222 | 0.6 | 0.399 | 0.01 | 0.005 | 0.005 | 3.062 | 0.35 | 0.67 | 0.35 | 0.72 | 0.18 | 0.36 |
| 6 | 151853949 | rs9479060 | -0.924 | -1.894 | 1.385 | 0.78 | -0.051 | 0.052 | 0.052 | 3.719 | 0.14 | 0.28 | 0.05 | 0.21 | 0.04 | 0.11 |
| 6 | 151855488 | rs6557144 | 0.826 | -1.85 | 1.146 | 0.778 | 0.017 | -0.011 | -0.011 | 0 | 0.78 | 0.92 | 0.91 | 0.79 | 0.82 | 0.8 |
| 6 | 151857370 | rs1532401 | 2.043 | -1.89 | 1.33 | 0.675 | 0.028 | 0.035 | 0.035 | 0.288 | 0.63 | 0.22 | 0.75 | 0.71 | 0.72 | 0.7 |
| 6 | 151857594 | rs6931664 | 2.466 | -2.239 | 1.08 | 0.538 | -0.003 | 0.117 | 0.117 | 3.435 | 0.46 | 0.13 | 0.47 | 0.21 | 0.59 | 0.43 |
| 6 | 151859314 | rs4870034 | 1.878 | -2.263 | 1.11 | 0.511 | 0.025 | 0.032 | 0.032 | 0.243 | 0.63 | 0.28 | 0.75 | 0.71 | 0.72 | 0.68 |
| 6 | 151860110 | rs1856859 | 0.107 | -2.253 | 1.1 | 0.516 | 0.012 | -0.07 | -0.07 | 0 | 0.17 | 0.09 | 0.29 | 0.5 | 0.13 | 0.27 |
| 6 | 151862142 | rs7769700 | 0.128 | -2.234 | 1.149 | 0.473 | -0.006 | 0.007 | 0.007 | 0 | 0.06 | 0.03 | 0.09 | 0.07 | 0.08 | 0.07 |
| 6 | 151862228 | rs4463269 | 1.312 | -2.246 | 1.285 | 0.464 | -0.004 | 0.005 | 0.005 | 0 | 0.71 | 0.87 | 0.81 | 0.72 | 0.73 | 0.7 |
| 6 | 151865531 | rs3757317 | -0.668 | -2.259 | 1.257 | 0.467 | -0.022 | -0.056 | -0.056 | 4.326 | 0.16 | 0.03 | 0.12 | 0.27 | 0.14 | 0.18 |
| 6 | 151867730 | rs9479066 | -0.56 | -2.252 | 1.693 | 0.573 | 0.046 | 0.114 | 0.114 | 1.253 | 0.06 | 0.4 | 0.04 | 0.04 | 0 | 0.02 |

| | | | | | | | | | | | | | | | | |
|---|---|---|---|---|---|---|---|---|---|---|---|---|---|---|---|---|
| 6 | 151868129 | rs1023939 | 1.922 | -2.251 | 1.462 | 0.57 | 0.025 | 0.014 | 0.014 | 0.225 | 0.63 | 0.29 | 0.75 | 0.68 | 0.72 | 0.68 |
| 6 | 151868736 | rs9371534 | 1.922 | -2.251 | 1.443 | 0.57 | 0.026 | 0.014 | 0.014 | 0.225 | 0.63 | 0.28 | 0.75 | 0.68 | 0.72 | 0.68 |
| 6 | 151868835 | rs9371536 | 1.922 | -2.251 | 1.443 | 0.57 | 0.026 | 0.014 | 0.014 | 0.225 | 0.63 | 0.28 | 0.75 | 0.68 | 0.72 | 0.68 |
| 6 | 151869219 | rs9383923 | 0.815 | -2.255 | 1.536 | 0.56 | -0.033 | -0.101 | -0.101 | 0.196 | 0.8 | 0.41 | 0.88 | 0.99 | 0.87 | 0.87 |
| 6 | 151869645 | rs3734803 | -0.729 | -2.133 | 1.144 | 0.602 | -0.006 | -0.024 | -0.024 | 0 | 0.16 | 0.06 | 0.12 | 0.27 | 0.14 | 0.18 |
| 6 | 151873833 | rs7767221 | -0.109 | -2.055 | 1.069 | 0.661 | 0 | 0.029 | 0.029 | 0 | 0.87 | 0.94 | 0.93 | 1 | 0.87 | 0.89 |
| 6 | 151873896 | rs13213582 | -0.14 | -2.031 | 1.111 | 0.796 | 0.014 | -0.046 | -0.046 | 3.601 | 0.18 | 0.15 | 0.29 | 0.5 | 0.13 | 0.26 |
| 6 | 151877049 | rs7754143 | -0.303 | -1.901 | 1.114 | 0.755 | 0.013 | 0.07 | 0.07 | 0.41 | 0.05 | 0.27 | 0.03 | 0.04 | 0 | 0.02 |
| 6 | 151887150 | rs958283 | -0.081 | -1.887 | 0.663 | 0.79 | 0.007 | -0.017 | -0.017 | 0.38 | 0.17 | 0.15 | 0.29 | 0.47 | 0.13 | 0.26 |
| 6 | 151891022 | rs12205762 | -0.296 | -1.905 | 0.48 | 0.369 | 0.004 | -0.015 | -0.015 | 0 | 0.07 | 0.09 | 0.09 | 0.07 | 0.08 | 0.07 |
| 6 | 151892135 | rs4869737 | 1.434 | -2.002 | 0.295 | 0.299 | -0.018 | -0.026 | -0.026 | 1.218 | 0.64 | 0.54 | 0.76 | 0.71 | 0.72 | 0.68 |
| 6 | 151894505 | rs953767 | -0.035 | -1.654 | -0.241 | 0.336 | -0.022 | 0.067 | 0.067 | 0 | 0.88 | 0.99 | 0.94 | 1 | 0.88 | 0.89 |
| 6 | 151898506 | rs1871859 | -0.426 | -1.591 | 0.208 | 0.863 | 0.013 | -0.034 | -0.034 | 0 | 0.22 | 0.21 | 0.3 | 0.52 | 0.16 | 0.26 |
| 6 | 151900238 | rs9478223 | -1.136 | -1.481 | 0.245 | 0.475 | 0.019 | -0.027 | -0.027 | 0.022 | 0.1 | 0.3 | 0.11 | 0.07 | 0.08 | 0.07 |
| 6 | 151900811 | rs6913493 | -0.902 | -1.388 | 0.291 | 0.52 | 0.033 | -0.015 | -0.015 | 0.384 | 0.09 | 0.3 | 0.11 | 0.08 | 0.08 | 0.1 |
| 6 | 151901409 | rs4870044 | -1.001 | -1.328 | -0.815 | 0.537 | 0.048 | 0.086 | 0.086 | 2.311 | 0.42 | 0.78 | 0.46 | 0.81 | 0.29 | 0.47 |
| 6 | 151906293 | rs4869741 | -1.033 | -1.315 | -0.947 | 0.544 | 0.054 | 0.099 | 0.099 | 2.773 | 0.43 | 0.8 | 0.47 | 0.81 | 0.29 | 0.48 |
| 6 | 151914113 | rs3757318 | -0.548 | -1.009 | -0.824 | 1.092 | -0.006 | -0.014 | -0.014 | 0 | 0.07 | 0.02 | 0.05 | 0.26 | 0.08 | 0.04 |
| 6 | 151914143 | rs17081463 | -1.087 | -1.18 | -0.559 | 0.899 | -0.006 | -0.007 | -0.007 | 0 | 0.08 | 0.12 | 0.04 | 0.08 | 0.05 | 0.11 |
| 6 | 151918856 | rs12662670 | -0.723 | -1.272 | -0.765 | 1.045 | 0.003 | -0.029 | -0.029 | 0.034 | 0.09 | 0.04 | 0.06 | 0.31 | 0.08 | 0.05 |
| 6 | 151919936 | rs9397066 | -1.49 | -1.975 | -1.341 | 0.642 | 0.094 | 0.174 | 0.174 | 0.707 | 0.43 | 0.87 | 0.49 | 0.82 | 0.28 | 0.35 |
| 6 | 151920772 | rs1340874 | -1.127 | -1.967 | -1.265 | 0.695 | 0.095 | 0.068 | 0.068 | 1.111 | 0.37 | 0.84 | 0.47 | 0.74 | 0.23 | 0.24 |
| 6 | 151920913 | rs9383587 | -1.127 | -1.967 | -1.265 | 0.695 | 0.095 | 0.068 | 0.068 | 1.111 | 0.37 | 0.84 | 0.47 | 0.74 | 0.23 | 0.24 |
| 6 | 151928342 | rs7746854 | 1.479 | -1.52 | -1.661 | 0.301 | 0.015 | 0.23 | 0.23 | 1.604 | 0.47 | 0.12 | 0.42 | 0.17 | 0.59 | 0.49 |
| 6 | 151929942 | rs6557158 | 0.471 | -1.502 | -0.634 | 0.43 | -0.022 | 0.028 | 0.028 | 1.344 | 0.63 | 0.55 | 0.56 | 0.54 | 0.7 | 0.64 |
| 6 | 151930249 | rs12663827 | -0.54 | -0.962 | -0.825 | 0.704 | 0.004 | -0.025 | -0.025 | 0 | 0.09 | 0.04 | 0.06 | 0.3 | 0.08 | 0.05 |
| 6 | 151932778 | rs1023940 | -0.735 | -0.646 | -1.403 | 0.23 | -0.067 | -0.08 | -0.08 | 0 | 0.64 | 0.86 | 0.42 | 0.48 | 0.48 | 0.5 |
| 6 | 151933175 | rs1038304 | -0.721 | -0.607 | -1.362 | 0.221 | -0.063 | -0.073 | -0.073 | 0 | 0.64 | 0.85 | 0.42 | 0.48 | 0.49 | 0.5 |
| 6 | 151936677 | rs6929137 | -0.376 | -1.495 | -0.706 | -0.421 | -0.03 | -0.014 | -0.014 | 0 | 0.41 | 0.5 | 0.25 | 0.33 | 0.3 | 0.3 |
| 6 | 151939181 | rs3734804 | -1.282 | -1.383 | -0.461 | -0.545 | -0.037 | -0.012 | -0.012 | 0 | 0.63 | 0.76 | 0.38 | 0.43 | 0.5 | 0.47 |
| 6 | 151939350 | rs3734805 | 0.295 | -1.667 | -0.571 | -0.772 | 0.02 | -0.049 | -0.049 | 1.725 | 0.11 | 0.03 | 0.06 | 0.32 | 0.08 | 0.05 |
| 6 | 151939560 | rs6932260 | -1.544 | -1.668 | -0.571 | -0.774 | -0.037 | -0.013 | -0.013 | 0 | 0.63 | 0.76 | 0.38 | 0.43 | 0.5 | 0.47 |
| 6 | 151939627 | rs6904261 | 0.31 | -1.804 | 0.032 | -0.85 | 0.016 | 0.031 | 0.031 | 0 | 0.26 | 0.17 | 0.15 | 0.02 | 0.21 | 0.25 |
| 6 | 151939770 | rs6932603 | -1.602 | -1.786 | 0.014 | -0.837 | -0.036 | -0.012 | -0.012 | 0 | 0.63 | 0.76 | 0.38 | 0.43 | 0.51 | 0.47 |
| 6 | 151939848 | rs9383935 | 0.316 | -1.776 | 0.01 | -0.819 | 0.02 | -0.049 | -0.049 | 0 | 0.11 | 0.03 | 0.06 | 0.31 | 0.08 | 0.05 |
| 6 | 151940260 | rs9383589 | 0.316 | -1.776 | -0.011 | -0.819 | 0.019 | -0.062 | -0.062 | 0 | 0.11 | 0.02 | 0.06 | 0.31 | 0.08 | 0.05 |
| 6 | 151941461 | rs3734806 | -0.59 | -1.85 | 0.056 | -0.869 | -0.03 | -0.014 | -0.014 | 0 | 0.4 | 0.5 | 0.25 | 0.34 | 0.3 | 0.3 |
| 6 | 151942194 | rs3757322 | -0.807 | -1.864 | 0.049 | -0.876 | -0.03 | -0.014 | -0.014 | 0 | 0.4 | 0.5 | 0.25 | 0.34 | 0.3 | 0.3 |
| 6 | 151943014 | rs9479087 | 0.232 | -1.811 | 0.065 | -0.876 | -0.003 | -0.04 | -0.04 | 0.052 | 0.23 | 0.28 | 0.13 | 0.09 | 0.21 | 0.17 |
| 6 | 151946658 | rs7751941 | 0.62 | -1.977 | -0.068 | -0.987 | 0.006 | 0.018 | 0.018 | 0 | 0.21 | 0.16 | 0.14 | 0.02 | 0.19 | 0.23 |
| 6 | 151947068 | rs7752591 | -1.575 | -1.933 | -0.112 | -0.899 | -0.032 | -0.012 | -0.012 | 0 | 0.62 | 0.74 | 0.37 | 0.43 | 0.51 | 0.48 |
| 6 | 151949806 | rs6913578 | -0.925 | -1.984 | 0.094 | -0.772 | -0.029 | -0.01 | -0.01 | 0 | 0.39 | 0.5 | 0.23 | 0.34 | 0.29 | 0.31 |
| 6 | 151950235 | rs6557161 | -0.866 | -1.917 | -0.099 | -0.762 | 0.025 | 0.008 | 0.008 | 0 | 0.4 | 0.66 | 0.26 | 0.36 | 0.31 | 0.32 |
| 6 | 151952890 | rs6908732 | -0.623 | -1.73 | -0.403 | -0.948 | -0.044 | 0.129 | 0.129 | 0 | 0.74 | 0.96 | 0.83 | 0.91 | 0.73 | 0.75 |
| 6 | 151953859 | rs9397068 | 0.458 | -1.778 | -0.404 | -0.944 | 0.027 | -0.058 | -0.058 | 2.736 | 0.11 | 0.02 | 0.05 | 0.31 | 0.07 | 0.06 |
| 6 | 151954127 | rs6900157 | -0.797 | -1.653 | -0.331 | -0.89 | 0.043 | 0.015 | 0.015 | 0 | 0.41 | 0.7 | 0.26 | 0.36 | 0.31 | 0.32 |
| 6 | 151962202 | rs852003 | -1.516 | -1.394 | -0.088 | -0.629 | -0.031 | -0.019 | -0.019 | 0 | 0.67 | 0.74 | 0.43 | 0.43 | 0.57 | 0.57 |
| 6 | 151974422 | rs10484919 | -0.334 | -1.644 | 0.45 | -0.404 | -0.046 | 0.113 | 0.113 | 0 | 0.15 | 0.37 | 0.08 | 0.3 | 0.08 | 0.09 |
| 6 | 151977473 | rs851971 | 0.056 | -1.885 | -0.047 | -0.401 | -0.04 | -0.143 | -0.143 | 0 | 0.33 | 0.04 | 0.34 | 0.49 | 0.32 | 0.39 |
| 6 | 151978439 | rs712219 | -2.023 | -1.801 | 0.034 | -0.68 | -0.1 | 0.086 | 0.086 | 0 | 0.7 | 0.9 | 0.63 | 0.86 | 0.59 | 0.69 |
| 6 | 151985550 | rs851997 | -0.805 | -1.839 | 0.156 | -0.639 | 0.017 | 0.035 | 0.035 | 0 | 0.47 | 0.41 | 0.43 | 0.79 | 0.4 | 0.48 |
| 6 | 151986003 | rs9397442 | -0.843 | -1.858 | 0.255 | -0.486 | -0.051 | 0.028 | 0.028 | 0 | 0.19 | 0.37 | 0.27 | 0.72 | 0.12 | 0.22 |
| 6 | 151987806 | rs7745737 | -0.874 | -1.798 | 0.64 | -0.178 | -0.047 | 0.154 | 0.154 | 0 | 0.2 | 0.44 | 0.12 | 0.37 | 0.14 | 0.14 |
| 6 | 151998105 | rs1293942 | 0.16 | -1.691 | 0.272 | -0.243 | -0.02 | -0.072 | -0.072 | 0 | 0.32 | 0.12 | 0.34 | 0.44 | 0.32 | 0.4 |
| 6 | 151998263 | rs980281 | -0.816 | -1.853 | 0.739 | -0.414 | -0.002 | 0 | 0 | 0.874 | 0.54 | 0.57 | 0.57 | 0.51 | 0.46 | 0.61 |
| 6 | 151999091 | rs1293940 | -1.064 | -1.827 | 0.599 | -0.39 | -0.015 | -0.016 | -0.016 | 0 | 0.59 | 0.69 | 0.59 | 0.52 | 0.51 | 0.66 |
| 6 | 152002932 | rs6557164 | 0.288 | -1.573 | 0.991 | -0.39 | -0.017 | -0.072 | -0.072 | 0 | 0.16 | 0.38 | 0.14 | 0.03 | 0.14 | 0.15 |
| 6 | 152005534 | rs851995 | -0.795 | -1.288 | 0.54 | -0.386 | -0.007 | 0 | 0 | 0 | 0.59 | 0.62 | 0.4 | 0.15 | 0.54 | 0.52 |
| 6 | 152006011 | rs851993 | -1.749 | -1.17 | -0.05 | -0.412 | -0.098 | 0.093 | 0.093 | 2.164 | 0.71 | 0.92 | 0.65 | 0.86 | 0.61 | 0.72 |
| 6 | 152006178 | rs9383939 | 0.391 | -0.921 | -0.02 | -0.019 | -0.012 | 0.162 | 0.162 | 0 | 0.08 | 0.14 | 0.05 | 0.37 | 0.06 | 0.06 |
| 6 | 152006581 | rs851991 | -0.311 | -0.878 | 0.034 | -0.592 | 0.011 | 0.115 | 0.115 | 0 | 0.5 | 0.35 | 0.35 | 0.11 | 0.49 | 0.43 |
| 6 | 152007889 | rs851987 | 0.345 | -0.415 | 0.3 | -0.045 | -0.001 | 0.045 | 0.045 | 0 | 0.71 | 0.55 | 0.54 | 0.61 | 0.71 | 0.65 |
| 6 | 152008780 | rs3020331 | 0.114 | 0.909 | 0.393 | -0.427 | -0.003 | -0.003 | -0.003 | 1.754 | 0.28 | 0.28 | 0.37 | 0.1 | 0.45 | 0.21 |
| 6 | 152009638 | rs2941740 | 0.096 | 0.967 | 0.616 | -0.425 | 0.027 | -0.032 | -0.032 | 0 | 0.28 | 0.35 | 0.37 | 0.1 | 0.44 | 0.21 |
| 6 | 152010254 | rs3020333 | -0.06 | 0.944 | 1.045 | -0.327 | 0.084 | -0.061 | -0.061 | 0 | 0.29 | 0.53 | 0.43 | 0.15 | 0.47 | 0.22 |
| 6 | 152012956 | rs3020334 | -0.025 | 0.92 | 1.114 | -0.366 | 0.053 | -0.059 | -0.059 | 0 | 0.29 | 0.42 | 0.38 | 0.1 | 0.44 | 0.21 |
| 6 | 152017691 | rs1293936 | -0.051 | 1.331 | 1.211 | -0.334 | 0.087 | -0.062 | -0.062 | 0 | 0.28 | 0.56 | 0.41 | 0.12 | 0.46 | 0.22 |
| 6 | 152024415 | rs851983 | 0.224 | 1.3 | 0.782 | -0.326 | -0.014 | 0.01 | 0.01 | 0.01 | 0.26 | 0.25 | 0.36 | 0.1 | 0.42 | 0.21 |
| 6 | 152024985 | rs851982 | 0.206 | 1.319 | 0.784 | -0.327 | -0.047 | 0.037 | 0.037 | 0 | 0.25 | 0.18 | 0.34 | 0.1 | 0.42 | 0.21 |

| | | | | | | | | | | | | | | | | |
|---|---|---|---|---|---|---|---|---|---|---|---|---|---|---|---|---|
| 6 | 152040291 | rs12525163 | -0.483 | 1.841 | 0.921 | -0.161 | 0.041 | 0.004 | 0.004 | 0.196 | 0.12 | 0.22 | 0.26 | 0.12 | 0.27 | 0.16 |
| 6 | 152042260 | rs10484921 | -0.248 | 1.913 | 0.892 | -0.181 | 0.021 | -0.003 | -0.003 | 0 | 0.11 | 0.17 | 0.23 | 0.1 | 0.24 | 0.15 |
| 6 | 152048022 | rs1159327 | -0.85 | 1.841 | 1.117 | -0.067 | 0.105 | 0.061 | 0.061 | 1.457 | 0.14 | 0.44 | 0.31 | 0.16 | 0.26 | 0.15 |
| 6 | 152054363 | rs3020343 | -0.501 | 1.284 | -0.085 | -0.424 | 0.023 | 0.08 | 0.08 | 0.077 | 0.44 | 0.25 | 0.53 | 0.2 | 0.63 | 0.37 |
| 6 | 152054374 | rs12195741 | 0.546 | 1.388 | -0.216 | -0.484 | -0.12 | 0.155 | 0.155 | 0.118 | 0.31 | 0.02 | 0.24 | 0.03 | 0.36 | 0.22 |
| 6 | 152057914 | rs3020348 | -0.285 | 0.528 | 0.184 | -0.22 | 0.021 | 0.07 | 0.07 | 0 | 0.45 | 0.29 | 0.54 | 0.2 | 0.63 | 0.39 |
| 6 | 152059563 | rs2982552 | -0.079 | 0.475 | 0.155 | -0.245 | -0.007 | 0.141 | 0.141 | 0.002 | 0.43 | 0.18 | 0.49 | 0.14 | 0.6 | 0.38 |
| 6 | 152061210 | rs2982551 | -0.408 | 0.288 | 0.922 | -0.332 | 0.019 | 0.09 | 0.09 | 0.289 | 0.47 | 0.67 | 0.54 | 0.17 | 0.6 | 0.38 |
| 6 | 152065886 | rs3020306 | 0.052 | 0.165 | 0.565 | -0.098 | 0.01 | 0.1 | 0.1 | 0 | 0.46 | 0.18 | 0.44 | 0.21 | 0.6 | 0.36 |
| 6 | 152068364 | rs1999805 | -0.288 | 0.137 | 0.594 | -0.084 | 0.007 | 0.011 | 0.011 | 0 | 0.48 | 0.56 | 0.49 | 0.24 | 0.63 | 0.37 |
| 6 | 152082369 | rs1336981 | -0.951 | -0.403 | -0.075 | 0.272 | 0.015 | -0.049 | -0.049 | 0 | 0.64 | 0.86 | 0.74 | 0.52 | 0.72 | 0.7 |
| 6 | 152085275 | rs1890010 | -0.335 | -0.915 | -0.355 | -0.336 | 0.009 | -0.001 | -0.001 | 1.229 | 0.61 | 0.66 | 0.73 | 0.95 | 0.72 | 0.64 |
| 6 | 152089768 | rs2485209 | 0.009 | -1.132 | -0.443 | -0.336 | 0.031 | 0.029 | 0.029 | 0.972 | 0.51 | 0.71 | 0.52 | 0.52 | 0.61 | 0.64 |
| 6 | 152090045 | rs6939257 | -0.352 | -1.34 | -0.058 | 0.053 | -0.04 | 0.054 | 0.054 | 0 | 0.24 | 0.55 | 0.34 | 0.74 | 0.17 | 0.25 |
| 6 | 152090324 | rs6939683 | 0.914 | -1.311 | -0.35 | 0.058 | -0.004 | 0.02 | 0.02 | 0 | 0.17 | 0.19 | 0.26 | 0.44 | 0.13 | 0.11 |
| 6 | 152090707 | rs2504063 | 0.506 | -1.109 | -1 | -0.087 | 0.017 | 0.054 | 0.054 | 0.068 | 0.46 | 0.33 | 0.46 | 0.18 | 0.63 | 0.4 |
| 6 | 152092749 | rs4870053 | -0.56 | -1.078 | -0.985 | 0.074 | 0.005 | 0.001 | 0.001 | 0 | 0.33 | 0.7 | 0.39 | 0.78 | 0.22 | 0.4 |
| 6 | 152095167 | rs2504065 | 0.442 | -0.796 | -0.979 | -0.096 | -0.002 | -0.008 | -0.008 | 0 | 0.44 | 0.37 | 0.42 | 0.43 | 0.57 | 0.46 |
| 6 | 152095332 | rs2248586 | -0.532 | -0.443 | -0.97 | -0.12 | 0.007 | 0.06 | 0.06 | 0.27 | 0.69 | 0.85 | 0.77 | 0.87 | 0.71 | 0.67 |
| 6 | 152103792 | rs1285057 | -0.524 | 0.174 | -0.508 | 0.685 | 0.035 | 0.07 | 0.07 | 0.28 | 0.57 | 0.75 | 0.58 | 0.72 | 0.68 | 0.58 |
| 6 | 152110943 | rs543650 | 0.09 | 0.747 | 0.131 | 1.774 | -0.012 | -0.009 | -0.009 | 0 | 0.65 | 0.8 | 0.64 | 0.59 | 0.58 | 0.68 |
| 6 | 152116655 | rs17081685 | -0.263 | 0.692 | 0.585 | 2.091 | 0.013 | 0.147 | 0.147 | 0.013 | 0.09 | 0.24 | 0.29 | 0.33 | 0.08 | 0.12 |
| 6 | 152125444 | rs488133 | -0.601 | 0.87 | 0.444 | 1.738 | -0.105 | 0.111 | 0.111 | 0 | 0.74 | 0.98 | 0.79 | 0.95 | 0.67 | 0.79 |
| 6 | 152129077 | rs2077647 | -0.2 | 0.297 | 1.533 | 2.494 | -0.001 | -0.001 | -0.001 | 0 | 0.5 | 0.52 | 0.47 | 0.38 | 0.44 | 0.51 |
| 6 | 152130918 | rs532010 | -0.136 | 0.258 | 1.556 | 2.458 | 0.007 | 0.004 | 0.004 | 0.001 | 0.48 | 0.44 | 0.43 | 0.38 | 0.34 | 0.48 |
| 6 | 152148870 | rs7759411 | -0.357 | 0.213 | 1.563 | 1.655 | -0.031 | 0.025 | 0.025 | 0 | 0.07 | 0.17 | 0.03 | 0.12 | 0.02 | 0.17 |
| 6 | 152149435 | rs11155813 | -1.23 | 0.256 | 1.66 | 1.728 | 0.017 | 0.193 | 0.193 | 0 | 0.12 | 0.42 | 0.26 | 0.3 | 0.12 | 0.24 |
| 6 | 152156197 | rs827423 | 0.193 | 0.401 | 0.922 | 1.468 | 0.008 | 0.017 | 0.017 | 0 | 0.49 | 0.34 | 0.68 | 0.63 | 0.55 | 0.5 |
| 6 | 152157881 | rs6902771 | -0.343 | 0.549 | 0.608 | 1.383 | 0.005 | 0.005 | 0.005 | 0 | 0.49 | 0.5 | 0.34 | 0.4 | 0.43 | 0.39 |
| 6 | 152177663 | rs827419 | 0.324 | -0.345 | -0.413 | 1.008 | -0.04 | 0.209 | 0.209 | 0.034 | 0.36 | 0.07 | 0.29 | 0.02 | 0.36 | 0.19 |
| 6 | 152182315 | rs1514348 | -0.115 | -0.157 | -0.222 | 1.388 | 0.018 | -0.008 | -0.008 | 0.01 | 0.46 | 0.68 | 0.38 | 0.48 | 0.39 | 0.42 |
| 6 | 152183551 | rs1643821 | -0.107 | -0.154 | -0.326 | 1.372 | 0.013 | -0.007 | -0.007 | 0 | 0.46 | 0.66 | 0.38 | 0.48 | 0.39 | 0.42 |
| 6 | 152199359 | rs11155819 | 0.291 | 0.23 | -0.842 | 0.908 | -0.048 | 0.078 | 0.078 | 1.402 | 0.31 | 0.1 | 0.25 | 0.18 | 0.33 | 0.21 |
| 6 | 152200129 | rs9322335 | 0.701 | 0.243 | -0.117 | 1.539 | 0.004 | 0.073 | 0.073 | 0 | 0.77 | 0.69 | 0.67 | 0.52 | 0.75 | 0.61 |
| 6 | 152200430 | rs9322336 | 0.632 | 0.254 | -0.765 | 1.775 | -0.06 | -0.134 | -0.134 | 1.382 | 0.81 | 0.99 | 0.71 | 0.63 | 0.78 | 0.61 |
| 6 | 152203104 | rs6557170 | 0.65 | 0.301 | -0.695 | 1.589 | -0.007 | -0.109 | -0.109 | 0 | 0.79 | 0.94 | 0.72 | 0.52 | 0.77 | 0.61 |
| 6 | 152204210 | rs11155820 | 0.866 | 0.038 | -1.372 | 0.819 | -0.095 | 0.215 | 0.215 | 1.399 | 0.33 | 0.02 | 0.23 | 0.02 | 0.32 | 0.27 |
| 6 | 152212508 | rs7761846 | -0.89 | -0.347 | 0.322 | 1.398 | 0.054 | 0.049 | 0.049 | 0 | 0.1 | 0.38 | 0.24 | 0.12 | 0.07 | 0.11 |
| 6 | 152234593 | rs6557171 | 1.618 | -0.389 | 0.616 | 1.431 | -0.012 | 0.023 | 0.023 | 0 | 0.67 | 0.49 | 0.49 | 0.4 | 0.7 | 0.45 |
| 6 | 152241150 | rs988328 | -0.669 | -0.626 | 0.601 | 1.641 | 0.013 | -0.087 | -0.087 | 0.166 | 0.18 | 0.06 | 0.22 | 0.45 | 0.16 | 0.28 |
| 6 | 152260206 | rs6912184 | 0.962 | -0.688 | 0.624 | 1.51 | 0.035 | -0.081 | -0.081 | 0.713 | 0.74 | 0.88 | 0.73 | 0.5 | 0.79 | 0.57 |
| 6 | 152260668 | rs4583998 | 1.671 | -0.709 | 0.775 | 1.453 | 0.008 | -0.005 | -0.005 | 0 | 0.63 | 0.39 | 0.48 | 0.39 | 0.71 | 0.45 |
| 6 | 152266377 | rs3020410 | 2.029 | -0.83 | 0.772 | 1.543 | 0 | 0 | 0 | 0.029 | 0.87 | 0.88 | 0.9 | 0.88 | 0.89 | 0.73 |
| 6 | 152266468 | rs3003917 | 1.108 | -0.98 | 1.107 | 1.713 | -0.036 | 0.011 | 0.011 | 0 | 0.75 | 0.63 | 0.7 | 0.31 | 0.83 | 0.55 |
| 6 | 152268483 | rs3020424 | 1.35 | -0.844 | 0.814 | 1.571 | 0.002 | -0.004 | -0.004 | 3.768 | 0.67 | 0.68 | 0.5 | 0.39 | 0.73 | 0.48 |
| 6 | 152279514 | rs3003921 | 0.817 | -0.789 | 0.622 | 1.445 | -0.02 | 0.028 | 0.028 | 0 | 0.75 | 0.71 | 0.75 | 0.58 | 0.83 | 0.57 |
| 6 | 152283044 | rs3020401 | 1.689 | -0.774 | 0.506 | 1.31 | 0.037 | 0.001 | 0.001 | 0 | 0.61 | 0.3 | 0.51 | 0.39 | 0.71 | 0.42 |
| 6 | 152283458 | rs985191 | -0.846 | -0.563 | 0.843 | 1.287 | 0.015 | -0.012 | -0.012 | 1.592 | 0.12 | 0.26 | 0.24 | 0.11 | 0.12 | 0.14 |
| 6 | 152284458 | rs3003925 | 0.927 | -0.721 | 0.723 | 1.481 | 0.008 | -0.023 | -0.023 | 0 | 0.8 | 0.85 | 0.77 | 0.66 | 0.83 | 0.62 |
| 6 | 152285122 | rs2982688 | 1.806 | -0.641 | 1.168 | 1.593 | -0.008 | 0.003 | 0.003 | 0.71 | 0.62 | 0.47 | 0.7 | 0.38 | 0.69 | 0.45 |
| 6 | 152285152 | rs2982689 | 0.932 | -0.679 | 1.284 | 1.614 | 0 | 0.015 | 0.015 | 0.648 | 0.82 | 0.84 | 0.83 | 0.55 | 0.84 | 0.59 |
| 6 | 152286705 | rs985695 | -1.164 | -0.696 | 1.425 | 1.572 | -0.008 | -0.042 | -0.042 | 0 | 0.2 | 0.45 | 0.15 | 0.17 | 0.15 | 0.14 |
| 6 | 152287295 | rs2347869 | 1.963 | -0.771 | 1.303 | 1.579 | 0.042 | 0.045 | 0.045 | 0.15 | 0.58 | 0.15 | 0.64 | 0.3 | 0.68 | 0.41 |
| 6 | 152292660 | rs2347871 | 1.791 | -0.78 | 1.36 | 1.581 | 0.03 | -0.003 | -0.003 | 0.319 | 0.6 | 0.34 | 0.65 | 0.39 | 0.69 | 0.44 |
| 6 | 152292672 | rs2347872 | 0.422 | -0.792 | 1.366 | 1.631 | 0.017 | -0.049 | -0.049 | 0 | 0.85 | 0.93 | 0.85 | 0.66 | 0.87 | 0.66 |
| 6 | 152293905 | rs3020325 | 1.495 | -0.702 | 1.383 | 1.325 | 0.027 | -0.002 | -0.002 | 0 | 0.6 | 0.36 | 0.65 | 0.39 | 0.69 | 0.44 |
| 6 | 152298435 | rs2982683 | 0.813 | -0.366 | 0.914 | 1.017 | -0.098 | 0.033 | 0.033 | 0.364 | 0.29 | 0.02 | 0.35 | 0.16 | 0.31 | 0.28 |
| 6 | 152303437 | rs728524 | -1.362 | -0.211 | 2.12 | 1.792 | 0.034 | 0.197 | 0.197 | 0 | 0.06 | 0.27 | 0.05 | 0.16 | 0.03 | 0.12 |
| 6 | 152304328 | rs9397463 | -1.431 | -0.302 | 2.282 | 1.751 | 0.003 | -0.016 | -0.016 | 0 | 0.13 | 0.25 | 0.1 | 0.11 | 0.11 | 0.14 |
| 6 | 152305047 | rs926777 | 2.075 | -0.424 | 2.087 | 1.802 | -0.025 | 0.049 | 0.049 | 1.077 | 0.73 | 0.41 | 0.76 | 0.49 | 0.8 | 0.61 |
| 6 | 152306204 | rs2982684 | 0.827 | -0.412 | 2.189 | 1.849 | -0.032 | 0.113 | 0.113 | 0.769 | 0.87 | 0.8 | 0.89 | 0.6 | 0.92 | 0.76 |
| 6 | 152306346 | rs9371236 | -1.374 | -0.254 | 1.767 | 1.778 | 0.037 | 0.209 | 0.209 | 0 | 0.06 | 0.27 | 0.05 | 0.16 | 0.03 | 0.12 |
| 6 | 152307261 | rs3020407 | 2.032 | -0.155 | 1.082 | 1.51 | 0.052 | -0.018 | -0.018 | 0.305 | 0.63 | 0.13 | 0.66 | 0.37 | 0.73 | 0.44 |
| 6 | 152307706 | rs2144025 | 1.415 | -0.147 | 1.981 | 1.717 | -0.028 | 0.235 | 0.235 | 0 | 0.83 | 0.51 | 0.81 | 0.46 | 0.86 | 0.72 |
| 6 | 152309132 | rs7743290 | -2.154 | -0.053 | 2.201 | 1.511 | -0.036 | 0.036 | 0.036 | 0 | 0.27 | 0.41 | 0.15 | 0.41 | 0.21 | 0.31 |
| 6 | 152313146 | rs9340941 | -1.388 | 0.021 | 2.102 | 1.528 | 0 | 0.004 | 0.004 | 0 | 0.07 | 0.05 | 0.04 | 0.12 | 0.05 | 0.17 |
| 6 | 152313718 | rs9340944 | -1.97 | -0.029 | 2.12 | 1.494 | 0.002 | 0.007 | 0.007 | 0 | 0.17 | 0.11 | 0.07 | 0.07 | 0.14 | 0.19 |
| 6 | 152322885 | rs722208 | -2.761 | -0.063 | 2.333 | 1.512 | 0.008 | -0.009 | -0.009 | 0 | 0.37 | 0.66 | 0.21 | 0.54 | 0.27 | 0.52 |
| 6 | 152328484 | rs13216134 | -0.544 | -0.172 | 2.399 | 1.526 | -0.012 | 0.044 | 0.044 | 0 | 0.12 | 0.13 | 0.07 | 0.34 | 0.1 | 0.19 |

| | | | | | | | | | | | | | | | | |
|---|---|---|---|---|---|---|---|---|---|---|---|---|---|---|---|---|
| 6 | 152328616 | rs1569788 | -2.735 | -0.159 | 2.309 | 1.53 | 0.014 | -0.014 | -0.014 | 0 | 0.37 | 0.67 | 0.21 | 0.54 | 0.27 | 0.51 |
| 6 | 152330673 | rs9340958 | -0.701 | -0.176 | 1.964 | 1.471 | 0.001 | 0.003 | 0.003 | 0 | 0.11 | 0.07 | 0.04 | 0.06 | 0.08 | 0.08 |
| 6 | 152333104 | rs13203975 | -0.965 | -0.15 | 1.83 | 1.474 | -0.029 | 0.099 | 0.099 | 0 | 0.14 | 0.29 | 0.1 | 0.34 | 0.11 | 0.21 |
| 6 | 152333945 | rs9340978 | -0.118 | 0.009 | 1.054 | 1.239 | -0.013 | 0.037 | 0.037 | 0 | 0.08 | 0.04 | 0.03 | 0 | 0.07 | 0.01 |
| 6 | 152345162 | rs3020418 | -2.327 | 0.314 | 1.199 | 1.353 | 0.003 | -0.006 | -0.006 | 0 | 0.34 | 0.68 | 0.19 | 0.45 | 0.28 | 0.42 |
| 6 | 152348901 | rs9478265 | -0.342 | 0.39 | 1.101 | 1.383 | -0.008 | -0.028 | -0.028 | 0 | 0.07 | 0.02 | 0.05 | 0.13 | 0.06 | 0.14 |
| 6 | 152356270 | rs6941835 | -0.017 | 0.359 | 1.366 | 1.404 | -0.013 | 0.087 | 0.087 | 0 | 0.09 | 0.15 | 0.07 | 0.33 | 0.08 | 0.18 |
| 6 | 152358940 | rs3020434 | -1.531 | 0.343 | 1.51 | 1.375 | -0.007 | 0.006 | 0.006 | 0 | 0.18 | 0.12 | 0.09 | 0.1 | 0.18 | 0.19 |
| 6 | 152367993 | rs3020365 | -2.542 | 0.324 | 1.52 | 1.39 | 0 | -0.024 | -0.024 | 0 | 0.41 | 0.74 | 0.25 | 0.6 | 0.33 | 0.55 |
| 6 | 152368758 | rs3020366 | -2.699 | 0.327 | 1.597 | 1.408 | 0.001 | -0.012 | -0.012 | 0 | 0.4 | 0.6 | 0.24 | 0.59 | 0.34 | 0.56 |
| 6 | 152371190 | rs3020368 | -0.74 | 0.372 | 1.465 | 1.204 | -0.001 | -0.007 | -0.007 | 0.244 | 0.12 | 0.13 | 0.06 | 0.09 | 0.09 | 0.09 |
| 6 | 152376475 | rs6932864 | -1.145 | 0.284 | 1.494 | 1.24 | -0.053 | 0.162 | 0.162 | 0 | 0.16 | 0.46 | 0.11 | 0.37 | 0.1 | 0.23 |
| 6 | 152382311 | rs2273206 | -1.152 | 0.286 | 1.494 | 1.24 | -0.054 | 0.16 | 0.16 | 0 | 0.16 | 0.46 | 0.11 | 0.37 | 0.1 | 0.24 |
| 6 | 152382325 | rs2273207 | -0.042 | 0.306 | 1.401 | 1.253 | -0.012 | 0.07 | 0.07 | 0 | 0.09 | 0.14 | 0.07 | 0.33 | 0.08 | 0.18 |
| 6 | 152382382 | rs2207396 | -1.999 | 0.278 | 1.124 | 1.216 | 0.003 | -0.005 | -0.005 | 0 | 0.25 | 0.27 | 0.13 | 0.2 | 0.24 | 0.35 |
| 6 | 152387664 | rs3778082 | -0.905 | 0.312 | 0.648 | 1.241 | -0.046 | 0.153 | 0.153 | 0 | 0.16 | 0.47 | 0.11 | 0.36 | 0.11 | 0.21 |
| 6 | 152389968 | rs3020375 | -2.137 | 0.44 | -0.222 | 1.193 | 0.006 | -0.04 | -0.04 | 0 | 0.42 | 0.77 | 0.25 | 0.59 | 0.36 | 0.59 |
| 6 | 152392561 | rs12199102 | 0.516 | 1.216 | -0.647 | 0.905 | -0.02 | 0.032 | 0.032 | 0 | 0.07 | 0 | 0.04 | 0 | 0.08 | 0.02 |
| 6 | 152412137 | rs3020382 | -1.847 | 1.168 | -0.289 | 1.221 | -0.016 | 0.02 | 0.02 | 0 | 0.17 | 0.11 | 0.09 | 0.11 | 0.17 | 0.2 |
| 6 | 152414992 | rs2982900 | -0.579 | 1.192 | -0.614 | 1.356 | -0.001 | 0.004 | 0.004 | 0 | 0.09 | 0.02 | 0.03 | 0.1 | 0.06 | 0.07 |
| 6 | 152418575 | rs3778099 | -0.908 | 0.544 | 0.047 | 1.998 | -0.013 | 0.096 | 0.096 | 0 | 0.11 | 0.22 | 0.09 | 0.36 | 0.1 | 0.19 |
| 6 | 152420095 | rs2228480 | -0.931 | 0.957 | -0.15 | 1.84 | 0.001 | 0.005 | 0.005 | 0 | 0.11 | 0.15 | 0.24 | 0.2 | 0.18 | 0.21 |
| 6 | 152421854 | rs3798758 | -1.393 | 0.638 | 0.162 | 2.842 | -0.014 | 0.106 | 0.106 | 0 | 0.05 | 0.14 | 0.12 | 0.26 | 0.03 | 0.12 |
| 6 | 152425582 | rs2813544 | 0.03 | 1.313 | 0.617 | 2.518 | 0.041 | -0.022 | -0.022 | 0.141 | 0.14 | 0.22 | 0.2 | 0.13 | 0.23 | 0.1 |
| 6 | 152432902 | rs910416 | -1.153 | 1.302 | 0.579 | 2.379 | 0.004 | 0 | 0 | 0 | 0.42 | 0.52 | 0.5 | 0.57 | 0.55 | 0.49 |
| 6 | 152436158 | rs9383963 | -1.21 | 1.27 | 0.629 | 2.417 | -0.012 | -0.006 | -0.006 | 0 | 0.14 | 0.11 | 0.2 | 0.36 | 0.23 | 0.29 |
| 6 | 152437016 | rs2747652 | -1.518 | 1.175 | 0.862 | 2.352 | 0.003 | -0.001 | -0.001 | 0 | 0.43 | 0.51 | 0.53 | 0.58 | 0.55 | 0.52 |
| 6 | 152438103 | rs7450824 | -1.099 | 1.19 | 0.79 | 2.356 | -0.034 | -0.041 | -0.041 | 1.069 | 0.13 | 0.08 | 0.2 | 0.36 | 0.23 | 0.27 |
| 6 | 152441587 | rs2813550 | -1.582 | 0.586 | -0.331 | 1.833 | -0.069 | 0.032 | 0.032 | 0 | 0.76 | 0.98 | 0.87 | 0.88 | 0.78 | 0.86 |
| 6 | 152442821 | rs2813558 | 1.179 | 0.529 | -0.18 | 1.841 | 0.035 | -0.033 | -0.033 | 0 | 0.17 | 0.29 | 0.2 | 0.13 | 0.22 | 0.11 |
| 6 | 152443389 | rs12681 | 1.165 | 0.534 | -0.179 | 1.842 | 0.034 | -0.034 | -0.034 | 0 | 0.17 | 0.29 | 0.2 | 0.13 | 0.22 | 0.11 |
| 6 | 152443468 | rs2250122 | 0.944 | 0.212 | 0.034 | 1.683 | 0.034 | -0.006 | -0.006 | 0.118 | 0.29 | 0.1 | 0.3 | 0.31 | 0.22 | 0.36 |
| 6 | 152443761 | rs2295192 | 0.615 | 0.108 | -0.188 | 1.691 | 0.009 | -0.003 | -0.003 | 0.225 | 0.08 | 0.06 | 0.06 | 0.07 | 0.04 | 0.08 |
| 6 | 152449729 | rs2673781 | 0.253 | 0.184 | 0.007 | 1.724 | -0.035 | -0.005 | -0.005 | 0 | 0.38 | 0.5 | 0.41 | 0.38 | 0.27 | 0.4 |
| 6 | 152450763 | rs2459110 | -0.127 | 0.242 | -0.037 | 1.719 | -0.002 | -0.001 | -0.001 | 0 | 0.44 | 0.47 | 0.46 | 0.44 | 0.42 | 0.52 |
| 6 | 152453094 | rs2295193 | -0.957 | 0.254 | -0.173 | 1.568 | -0.021 | 0.032 | 0.032 | 0 | 0.6 | 0.72 | 0.66 | 0.74 | 0.55 | 0.76 |
| 6 | 152456475 | rs3818109 | -0.205 | 0.082 | -0.077 | 1.398 | 0.029 | 0.032 | 0.032 | 0 | 0.47 | 0.7 | 0.42 | 0.38 | 0.39 | 0.53 |
| 6 | 152457028 | rs6934016 | -0.035 | 0.194 | -0.191 | 1.37 | 0.017 | -0.003 | -0.003 | 0 | 0.08 | 0.11 | 0.06 | 0.06 | 0.15 | 0.1 |
| 6 | 152459143 | rs9322361 | 1.393 | 0.254 | -0.417 | 1.329 | -0.058 | 0.027 | 0.027 | 0.032 | 0.14 | 0.01 | 0.09 | 0.04 | 0.18 | 0.04 |
| 6 | 152463488 | rs10484863 | -0.143 | -0.08 | 0.043 | 1.915 | -0.016 | -0.038 | -0.038 | 1.641 | 0.08 | 0.01 | 0.14 | 0.3 | 0.08 | 0.16 |
| 6 | 152464367 | rs2747660 | 0.29 | -0.201 | 0.248 | 1.511 | -0.027 | -0.016 | -0.016 | 1.926 | 0.33 | 0.55 | 0.46 | 0.29 | 0.26 | 0.4 |
| 6 | 152464839 | rs2256135 | -1.307 | -0.094 | 0.033 | 1.531 | -0.035 | 0.035 | 0.035 | 0 | 0.6 | 0.78 | 0.71 | 0.69 | 0.54 | 0.7 |
| 6 | 152466582 | rs2813487 | -0.196 | -0.311 | -0.39 | 1.413 | -0.014 | -0.034 | -0.034 | 0 | 0.41 | 0.47 | 0.51 | 0.21 | 0.33 | 0.4 |
| 6 | 152466674 | rs2747662 | -0.126 | -0.225 | -0.304 | 1.236 | 0.026 | 0.025 | 0.025 | 0 | 0.41 | 0.34 | 0.51 | 0.3 | 0.33 | 0.41 |
| 6 | 152469188 | rs2252755 | -0.299 | -0.211 | -0.324 | 1.243 | -0.009 | -0.008 | -0.008 | 0 | 0.42 | 0.46 | 0.51 | 0.3 | 0.33 | 0.41 |
| 6 | 152469331 | rs2252748 | 0.095 | -0.215 | -0.354 | 1.239 | 0.025 | 0.02 | 0.02 | 0 | 0.37 | 0.31 | 0.46 | 0.3 | 0.28 | 0.4 |
| 6 | 152470752 | rs17082236 | -0.71 | -0.145 | -0.198 | 1.506 | -0.045 | -0.074 | -0.074 | 0 | 0.12 | 0.33 | 0.09 | 0 | 0.06 | 0.02 |
| 6 | 152473181 | rs910415 | 0.205 | 0.449 | -0.981 | 1.863 | -0.011 | -0.024 | -0.024 | 0.681 | 0.06 | 0.06 | 0.05 | 0 | 0.05 | 0.02 |
| 6 | 152474790 | rs9397084 | -1.215 | 0.929 | -1.121 | 2.346 | -0.016 | -0.044 | -0.044 | 0 | 0.15 | 0.07 | 0.22 | 0.43 | 0.2 | 0.28 |
| 6 | 152476648 | rs2635475 | -0.297 | 0.856 | -1.474 | 1.594 | -0.078 | -0.074 | -0.074 | 0.272 | 0.41 | 0.08 | 0.47 | 0.49 | 0.45 | 0.54 |
| 6 | 152478820 | rs9397085 | -1.087 | 0.41 | -0.233 | 1.554 | -0.019 | 0.026 | 0.026 | 0 | 0.26 | 0.6 | 0.15 | 0.33 | 0.22 | 0.36 |
| 6 | 152479587 | rs17082253 | -0.585 | 0.727 | -0.003 | 0.792 | -0.005 | -0.072 | -0.072 | 0 | 0.2 | 0.39 | 0.11 | 0.08 | 0.17 | 0.1 |
| 6 | 152484288 | rs2635474 | 0.399 | 0.938 | -0.449 | 1.06 | 0.001 | 0.002 | 0.002 | 0 | 0.66 | 0.7 | 0.77 | 0.7 | 0.69 | 0.8 |
| 6 | 152484763 | rs4870079 | -0.179 | 0.946 | -0.306 | 1.129 | 0.005 | -0.003 | -0.003 | 0.119 | 0.31 | 0.54 | 0.26 | 0.29 | 0.32 | 0.55 |
| 6 | 152486279 | rs6901280 | -1.274 | 0.892 | -0.385 | 2.034 | -0.001 | 0.04 | 0.04 | 0 | 0.11 | 0.14 | 0.23 | 0.45 | 0.15 | 0.23 |
| 6 | 152486372 | rs12213919 | -0.815 | 1.089 | -0.125 | 1.302 | 0.009 | -0.016 | -0.016 | 0 | 0.1 | 0.19 | 0.21 | 0.08 | 0.17 | 0.11 |
| 6 | 152486648 | rs2635473 | 0.784 | 0.707 | 0.115 | 1.052 | -0.003 | -0.004 | -0.004 | 0 | 0.67 | 0.43 | 0.57 | 0.33 | 0.7 | 0.44 |
| 6 | 152487003 | rs2635472 | -0.02 | 0.375 | -0.607 | 0.174 | 0.005 | 0.098 | 0.098 | 2.193 | 0.34 | 0.19 | 0.22 | 0.12 | 0.33 | 0.16 |
| 6 | 152487184 | rs2635471 | -0.271 | 0.434 | 0.115 | 1.131 | 0.001 | 0.158 | 0.158 | 0 | 0.88 | 0.8 | 0.85 | 0.6 | 0.88 | 0.67 |
| 6 | 152487830 | rs2813497 | -0.121 | 0.355 | 0.367 | 0.37 | -0.003 | -0.042 | -0.042 | 0 | 0.55 | 0.52 | 0.7 | 0.88 | 0.58 | 0.8 |
| 6 | 152487926 | rs17082273 | 0.297 | 0.426 | -0.054 | 0.4 | -0.039 | -0.025 | -0.025 | 0 | 0.07 | 0 | 0.18 | 0.11 | 0.1 | 0.16 |
| 6 | 152488577 | rs17082275 | 0.106 | 0.383 | 0.01 | 0.393 | -0.011 | -0.003 | -0.003 | 0 | 0.08 | 0.03 | 0.18 | 0.11 | 0.09 | 0.16 |
| 6 | 152489037 | rs997389 | 1.575 | 0.174 | -0.09 | 0.391 | -0.1 | 0.072 | 0.072 | 0 | 0.3 | 0.04 | 0.35 | 0.14 | 0.34 | 0.21 |
| 6 | 152489038 | rs997769 | -0.372 | 0.166 | 0.115 | 0.335 | 0.017 | 0.056 | 0.056 | 0.552 | 0.56 | 0.64 | 0.72 | 0.88 | 0.58 | 0.8 |
| 6 | 152489913 | rs2635447 | 0.068 | 0.089 | 0.155 | 0.615 | 0 | -0.016 | -0.016 | 0 | 0.46 | 0.5 | 0.53 | 0.76 | 0.48 | 0.62 |
| 6 | 152490251 | rs9479232 | 1.032 | 0.441 | -0.818 | 0.646 | -0.1 | 0.044 | 0.044 | 0 | 0.3 | 0.04 | 0.35 | 0.16 | 0.34 | 0.21 |
| 6 | 152491710 | rs2813498 | -0.69 | -0.104 | 0.159 | 1.265 | -0.027 | 0.113 | 0.113 | 0.119 | 0.22 | 0.42 | 0.18 | 0.55 | 0.15 | 0.44 |
| 6 | 152492517 | rs2763031 | -0.527 | -0.299 | -0.123 | 1.22 | 0.001 | 0.001 | 0.001 | 0 | 0.45 | 0.51 | 0.38 | 0.59 | 0.31 | 0.61 |
| 6 | 152493465 | rs2635469 | -0.634 | -0.386 | 0.195 | 1.496 | 0.049 | 0.02 | 0.02 | 0 | 0.46 | 0.71 | 0.42 | 0.64 | 0.34 | 0.64 |

| | | | | | | | | | | | | | | | | |
|---|---|---|---|---|---|---|---|---|---|---|---|---|---|---|---|---|
| 6 | 152493719 | rs2635468 | -0.754 | -0.467 | 0.568 | 1.739 | -0.008 | 0.098 | 0.098 | 0 | 0.09 | 0.23 | 0.17 | 0.28 | 0.05 | 0.23 |
| 6 | 152495811 | rs2813503 | -1.05 | -0.479 | 0.667 | 1.711 | 0.006 | 0.122 | 0.122 | 0.041 | 0.1 | 0.3 | 0.18 | 0.28 | 0.07 | 0.23 |
| 6 | 152495843 | rs2813504 | -0.846 | -0.537 | 0.67 | 1.704 | 0.069 | 0.005 | 0.005 | 0 | 0.47 | 0.77 | 0.43 | 0.58 | 0.34 | 0.62 |
| 6 | 152498606 | rs1000864 | -0.026 | -0.509 | 0.599 | 1.677 | -0.02 | 0.004 | 0.004 | 0 | 0.27 | 0.36 | 0.18 | 0.28 | 0.2 | 0.38 |
| 6 | 152503969 | rs2813510 | 1.036 | 0.196 | -0.116 | 1.266 | -0.078 | -0.102 | -0.102 | 0 | 0.17 | 0.01 | 0.3 | 0.21 | 0.19 | 0.16 |
| 6 | 152511420 | rs2813526 | 0.098 | 0.569 | 0.171 | 1.128 | -0.008 | -0.003 | -0.003 | 0 | 0.35 | 0.48 | 0.27 | 0.34 | 0.34 | 0.44 |
| 6 | 152515258 | rs7772542 | -0.002 | 0.824 | 0.003 | 1.097 | 0.002 | 0.01 | 0.01 | 0 | 0.06 | 0.08 | 0.18 | 0.18 | 0.09 | 0.19 |
| 6 | 152515301 | rs4870081 | -0.556 | 0.715 | 0.043 | 0.852 | 0 | 0.001 | 0.001 | 0 | 0.66 | 0.74 | 0.76 | 0.67 | 0.66 | 0.62 |
| 6 | 152516441 | rs2813532 | -0.97 | 0.899 | 0.03 | 0.71 | 0.011 | -0.016 | -0.016 | 0 | 0.12 | 0.29 | 0.11 | 0.11 | 0.14 | 0.12 |
| 6 | 152517503 | rs9479258 | 0.055 | 0.947 | -0.146 | 0.666 | 0.008 | -0.034 | -0.034 | 0.233 | 0.06 | 0.15 | 0.04 | 0 | 0.08 | 0.01 |
| 6 | 152520162 | rs2813535 | -0.012 | 0.876 | -0.448 | 0.522 | 0.002 | 0.016 | 0.016 | 0 | 0.28 | 0.21 | 0.19 | 0.14 | 0.28 | 0.14 |
| 6 | 152521955 | rs6900213 | 0.872 | 0.993 | -1.011 | 0.868 | -0.024 | -0.071 | -0.071 | 0 | 0.17 | 0.03 | 0.26 | 0.22 | 0.13 | 0.16 |
| 6 | 152522222 | rs2635442 | 0.777 | 0.992 | -0.953 | 0.753 | -0.008 | 0.033 | 0.033 | 0 | 0.22 | 0.06 | 0.15 | 0.14 | 0.21 | 0.14 |
| 6 | 152522812 | rs2673776 | -0.785 | 0.839 | -0.288 | 0.931 | 0.001 | -0.004 | -0.004 | 0.082 | 0.63 | 0.61 | 0.72 | 0.63 | 0.59 | 0.62 |
| 6 | 152523550 | rs2253407 | -0.471 | 1.062 | -0.622 | 1.304 | -0.021 | 0.032 | 0.032 | 1.474 | 0.48 | 0.2 | 0.6 | 0.57 | 0.46 | 0.49 |
| 6 | 152526387 | rs17082323 | -0.313 | 0.755 | -0.562 | 1.386 | 0.001 | 0.008 | 0.008 | 0 | 0.07 | 0.08 | 0.18 | 0.13 | 0.08 | 0.21 |
| 6 | 152528757 | rs2763026 | -0.405 | 0.59 | -0.858 | 1.033 | 0.01 | 0.025 | 0.025 | 0 | 0.79 | 0.9 | 0.86 | 0.87 | 0.83 | 0.87 |
| 6 | 152529260 | rs3798756 | 0.327 | 0.456 | -0.942 | 1.054 | 0.017 | -0.022 | -0.022 | 0.366 | 0.2 | 0.11 | 0.24 | 0.21 | 0.13 | 0.13 |
| 6 | 152532222 | rs2763028 | 0.258 | 0.958 | -0.268 | 0.978 | -0.001 | 0.004 | 0.004 | 0 | 0.76 | 0.74 | 0.8 | 0.72 | 0.78 | 0.84 |
| 6 | 152542548 | rs9383976 | -0.712 | 1.41 | -0.145 | 1.451 | 0.01 | 0.055 | 0.055 | 0.377 | 0.16 | 0.3 | 0.25 | 0.54 | 0.21 | 0.38 |
| 6 | 152542868 | rs9397493 | -1.185 | 1.103 | -0.168 | 0.574 | 0 | -0.027 | -0.027 | 0 | 0.42 | 0.5 | 0.65 | 0.75 | 0.42 | 0.61 |
| 6 | 152542962 | rs9383977 | -1.085 | 1.053 | -0.444 | 0.478 | 0.002 | -0.026 | -0.026 | 0 | 0.33 | 0.22 | 0.41 | 0.55 | 0.32 | 0.51 |
| 6 | 152544371 | rs7382254 | -0.031 | 1.041 | -0.509 | 0.553 | -0.036 | -0.043 | -0.043 | 0 | 0.07 | 0.02 | 0.22 | 0.2 | 0.11 | 0.1 |
| 6 | 152547151 | rs1359323 | -1.407 | 0.879 | 0.32 | 0.324 | -0.003 | 0.011 | 0.011 | 0 | 0.41 | 0.4 | 0.64 | 0.74 | 0.43 | 0.6 |
| 6 | 152548027 | rs17803505 | 0.277 | 0.767 | 0.127 | 0.22 | -0.017 | 0.07 | 0.07 | 0.049 | 0.12 | 0.01 | 0.17 | 0.01 | 0.1 | 0.11 |
| 6 | 152550483 | rs4142055 | -1.932 | 1.088 | 0.219 | -0.474 | 0.004 | 0.003 | 0.003 | 0 | 0.54 | 0.58 | 0.75 | 0.78 | 0.61 | 0.66 |
| 6 | 152554396 | rs12110479 | 0.308 | 1.108 | 0.034 | -0.519 | 0.04 | -0.049 | -0.049 | 1.027 | 0.14 | 0.24 | 0.11 | 0.05 | 0.19 | 0.06 |
| 6 | 152555112 | rs2296254 | -1.053 | 0.716 | -0.504 | -0.189 | -0.027 | -0.015 | -0.015 | 0 | 0.25 | 0.05 | 0.4 | 0.24 | 0.22 | 0.24 |
| 6 | 152561737 | rs9322364 | -1.08 | 0.15 | -0.761 | -0.835 | 0.036 | 0.18 | 0.18 | 0 | 0.53 | 0.76 | 0.65 | 0.88 | 0.58 | 0.65 |
| 6 | 152565897 | rs3888239 | -0.748 | 0.011 | -0.462 | -0.773 | 0.004 | -0.022 | -0.022 | 0 | 0.4 | 0.57 | 0.58 | 0.75 | 0.53 | 0.55 |
| 6 | 152567047 | rs6901631 | -0.252 | -0.335 | -0.646 | -1.052 | -0.019 | -0.021 | -0.021 | 0.846 | 0.26 | 0.3 | 0.23 | 0.14 | 0.14 | 0.18 |
| 6 | 152567445 | rs6940845 | -1.889 | -0.307 | -0.679 | -1.014 | 0.004 | 0.101 | 0.101 | 0.15 | 0.66 | 0.85 | 0.81 | 0.89 | 0.67 | 0.73 |
| 6 | 152570415 | rs6557210 | -1.25 | 0.021 | -0.617 | -0.76 | 0.018 | 0.048 | 0.048 | 0.075 | 0.53 | 0.65 | 0.63 | 0.88 | 0.59 | 0.64 |
| 6 | 152570971 | rs7757670 | 0.532 | 0.02 | -0.629 | -0.664 | 0.017 | 0.002 | 0.002 | 0 | 0.12 | 0.07 | 0.05 | 0.13 | 0.05 | 0.09 |
| 6 | 152577752 | rs17082389 | 0.568 | 0.021 | -0.65 | -0.664 | 0.024 | 0.001 | 0.001 | 0 | 0.12 | 0.06 | 0.05 | 0.13 | 0.05 | 0.09 |
| 6 | 152578782 | rs9371247 | 0.19 | 0.263 | -0.727 | 0.417 | -0.044 | -0.031 | -0.031 | 0.174 | 0.08 | 0.02 | 0.25 | 0.46 | 0.15 | 0.2 |
| 6 | 152585454 | rs4870090 | -1.161 | 0.124 | -0.662 | 0.269 | 0.02 | 0.051 | 0.051 | 0.068 | 0.52 | 0.64 | 0.63 | 0.83 | 0.59 | 0.64 |
| 6 | 152591432 | rs9397496 | 0.364 | 0.45 | -1.08 | 0.586 | -0.032 | 0.046 | 0.046 | 0 | 0.05 | 0.02 | 0.24 | 0.46 | 0.13 | 0.15 |
| 6 | 152592751 | rs633891 | -0.319 | 0.414 | -1.209 | 0.131 | -0.056 | -0.044 | -0.044 | 0 | 0.3 | 0.17 | 0.31 | 0.33 | 0.4 | 0.51 |
| 6 | 152593067 | rs6904737 | -0.067 | 0.524 | -0.948 | 0.113 | -0.002 | -0.013 | -0.013 | 0 | 0.13 | 0.08 | 0.13 | 0.2 | 0.13 | 0.13 |
| 6 | 152611876 | rs2788381 | -0.046 | 0.254 | -1.168 | -0.486 | -0.03 | -0.111 | -0.111 | 0.003 | 0.7 | 0.5 | 0.81 | 0.92 | 0.76 | 0.82 |
| 6 | 152618420 | rs1408458 | -0.063 | 0.518 | -1.547 | -0.03 | -0.036 | -0.026 | -0.026 | 0 | 0.31 | 0.46 | 0.25 | 0.24 | 0.24 | 0.39 |
| 6 | 152623532 | rs1408459 | -0.668 | 0.879 | -2.229 | 0.561 | -0.024 | -0.035 | -0.035 | 0.002 | 0.81 | 0.84 | 0.64 | 0.49 | 0.7 | 0.78 |
| 6 | 152625963 | rs11752464 | 0.228 | 0.428 | -2.579 | -0.527 | 0.048 | 0.154 | 0.154 | 0.39 | 0.32 | 0.01 | 0.21 | 0.1 | 0.2 | 0.28 |
| 6 | 152629586 | rs9371581 | 0.124 | 1.35 | -1.505 | 0.235 | -0.025 | -0.006 | -0.006 | 0 | 0.36 | 0.32 | 0.4 | 0.65 | 0.55 | 0.39 |
| 6 | 152629617 | rs9383985 | 0.535 | 1.381 | -2.127 | 0.366 | -0.059 | -0.14 | -0.14 | 1.446 | 0.27 | 0.05 | 0.26 | 0.6 | 0.33 | 0.35 |
| 6 | 152629815 | rs4112833 | -0.042 | 0.535 | -1.672 | -0.309 | -0.015 | 0.082 | 0.082 | 0 | 0.78 | 0.94 | 0.89 | 0.98 | 0.81 | 0.78 |
| 6 | 152632750 | rs9397501 | 0.243 | 0.308 | -1.562 | -0.287 | 0.025 | 0.079 | 0.079 | 0 | 0.72 | 0.88 | 0.89 | 0.93 | 0.82 | 0.71 |
| 6 | 152632823 | rs9371582 | -0.249 | 0.059 | -1.29 | 0.164 | 0.003 | 0.001 | 0.001 | 0 | 0.22 | 0.2 | 0.15 | 0.18 | 0.17 | 0.2 |
| 6 | 152635806 | rs9371583 | -0.965 | -0.238 | -1.233 | 1.036 | 0.011 | -0.024 | -0.024 | 1.917 | 0.15 | 0.19 | 0.19 | 0.66 | 0.14 | 0.23 |
| 6 | 152635846 | rs9383986 | -0.308 | -0.229 | -1.26 | 1.318 | 0.01 | -0.033 | -0.033 | 0 | 0.1 | 0.05 | 0.15 | 0.53 | 0.11 | 0.2 |
| 6 | 152638209 | rs1890100 | -0.362 | -0.129 | -1.784 | 0.386 | -0.063 | 0.126 | 0.126 | 0 | 0.73 | 0.93 | 0.85 | 0.96 | 0.69 | 0.84 |
| 6 | 152640248 | rs2306915 | -0.49 | -0.128 | -1.613 | 0.478 | 0.002 | -0.005 | -0.005 | 0.047 | 0.13 | 0.1 | 0.17 | 0.55 | 0.14 | 0.25 |
| 6 | 152640525 | rs9397095 | 0.625 | -0.329 | -0.799 | 0.35 | -0.004 | -0.003 | -0.003 | 0 | 0.58 | 0.65 | 0.66 | 0.41 | 0.55 | 0.58 |
| 6 | 152641656 | rs7741183 | -0.943 | -0.483 | -0.773 | 0.742 | -0.009 | 0.069 | 0.069 | 0 | 0.06 | 0.17 | 0.06 | 0.09 | 0.02 | 0.06 |
| 6 | 152643365 | rs9322367 | -0.137 | -0.342 | -0.852 | 0.633 | -0.031 | 0.045 | 0.045 | 0 | 0.69 | 0.78 | 0.81 | 0.82 | 0.67 | 0.8 |
| 6 | 152644111 | rs7738189 | 0.708 | -0.466 | -0.797 | 0.591 | -0.02 | 0.068 | 0.068 | 0 | 0.56 | 0.67 | 0.64 | 0.26 | 0.53 | 0.55 |
| 6 | 152648927 | rs6557213 | 0.808 | -0.328 | -0.441 | 0.722 | -0.001 | -0.017 | -0.017 | 0.364 | 0.45 | 0.54 | 0.63 | 0.13 | 0.44 | 0.35 |
| 6 | 152654160 | rs959024 | -0.519 | -0.29 | -0.587 | 1.309 | -0.003 | 0.06 | 0.06 | 0 | 0.06 | 0.04 | 0.07 | 0.54 | 0.07 | 0.16 |
| 6 | 152656215 | rs1999829 | -0.303 | -0.476 | -0.689 | 0.842 | 0.038 | 0.122 | 0.122 | 0 | 0.66 | 0.91 | 0.71 | 0.86 | 0.74 | 0.61 |
| 6 | 152658142 | rs9479297 | -0.37 | -0.486 | -0.704 | 0.837 | 0.017 | 0.128 | 0.128 | 0 | 0.66 | 0.96 | 0.71 | 0.86 | 0.74 | 0.61 |
| 6 | 152658216 | rs9479299 | 1.592 | -0.695 | -0.581 | 0.571 | -0.048 | 0.075 | 0.075 | 0.345 | 0.34 | 0.1 | 0.47 | 0.13 | 0.39 | 0.22 |
| 6 | 152658974 | rs9478316 | 0.671 | -0.902 | -0.12 | 0.406 | 0.003 | 0.003 | 0.003 | 0 | 0.49 | 0.51 | 0.6 | 0.6 | 0.63 | 0.39 |
| 6 | 152660837 | rs9371587 | -1.088 | -0.893 | 0.06 | 0.445 | -0.008 | 0.023 | 0.023 | 0 | 0.22 | 0.48 | 0.16 | 0.27 | 0.21 | 0.19 |
| 6 | 152662127 | rs9371589 | 1.033 | -0.67 | 0.081 | 0.553 | -0.003 | -0.001 | -0.001 | 0.017 | 0.41 | 0.37 | 0.53 | 0.57 | 0.48 | 0.35 |
| 6 | 152663276 | rs7759171 | 1.027 | -0.738 | 0.116 | 0.588 | 0 | -0.001 | -0.001 | 0 | 0.41 | 0.39 | 0.53 | 0.56 | 0.48 | 0.38 |
| 6 | 152665261 | rs4645434 | 0.488 | -1.232 | 0.657 | 0.619 | 0.003 | 0.004 | 0.004 | 0 | 0.52 | 0.41 | 0.62 | 0.65 | 0.6 | 0.51 |
| 6 | 152671475 | rs9478320 | 0.404 | -1.1 | 0.621 | 0.926 | 0.018 | 0.016 | 0.016 | 0 | 0.5 | 0.58 | 0.63 | 0.62 | 0.6 | 0.52 |
| 6 | 152672540 | rs4632900 | -0.19 | -1.085 | 0.639 | 0.763 | 0.02 | -0.068 | -0.068 | 0 | 0.13 | 0.21 | 0.08 | 0 | 0.16 | 0.09 |

| | | | | | | | | | | | | | | | | |
|---|---|---|---|---|---|---|---|---|---|---|---|---|---|---|---|---|
| 6 | 152673147 | rs6908392 | 0.362 | -1.001 | 0.649 | 0.829 | 0.002 | 0.002 | 0.002 | 0 | 0.5 | 0.51 | 0.62 | 0.62 | 0.6 | 0.52 |
| 6 | 152679087 | rs9478325 | -0.168 | -0.978 | 0.652 | 0.694 | 0.04 | -0.074 | -0.074 | 0 | 0.12 | 0.28 | 0.08 | 0 | 0.16 | 0.09 |
| 6 | 152683413 | rs4407724 | -0.262 | -1.079 | 0.67 | 1.052 | -0.001 | -0.003 | -0.003 | 0 | 0.57 | 0.58 | 0.69 | 0.79 | 0.68 | 0.59 |
| 6 | 152689195 | rs9371592 | 0.28 | -0.336 | 0.978 | 1.216 | -0.019 | -0.02 | -0.02 | 0 | 0.4 | 0.23 | 0.51 | 0.46 | 0.53 | 0.4 |
| 6 | 152691944 | rs9371595 | -0.241 | -0.344 | 0.893 | 1.264 | 0.017 | 0.063 | 0.063 | 0 | 0.07 | 0.19 | 0.09 | 0.17 | 0.1 | 0.07 |
| 6 | 152692103 | rs9383620 | -0.028 | -0.3 | 0.595 | 1.286 | -0.004 | -0.002 | -0.002 | 0.01 | 0.07 | 0.07 | 0.07 | 0.17 | 0.1 | 0.07 |
| 6 | 152694184 | rs6913579 | 0.117 | -0.082 | 0.826 | 1 | 0.002 | 0.002 | 0.002 | 0.015 | 0.41 | 0.44 | 0.54 | 0.46 | 0.53 | 0.41 |
| 6 | 152697706 | rs214955 | -0.136 | 0.12 | -0.126 | 1.068 | 0 | 0.001 | 0.001 | 0 | 0.58 | 0.55 | 0.47 | 0.54 | 0.47 | 0.59 |
| 6 | 152702821 | rs214959 | -0.286 | 0.028 | -0.836 | 1.17 | -0.136 | -0.058 | -0.058 | 0 | 0.66 | 0.95 | 0.5 | 0.64 | 0.54 | 0.65 |
| 6 | 152704907 | rs1933689 | -0.925 | -0.453 | 0.352 | 1.367 | -0.044 | 0.157 | 0.157 | 0.669 | 0.1 | 0.39 | 0.07 | 0.21 | 0.03 | 0.17 |
| 6 | 152708310 | rs214950 | 0.392 | -0.81 | -0.095 | 1.217 | 0.037 | -0.041 | -0.041 | 6.212 | 0.31 | 0.15 | 0.16 | 0.4 | 0.22 | 0.32 |
| 6 | 152711002 | rs214942 | -0.33 | -0.574 | 0.41 | 1.165 | 0.042 | 0.071 | 0.071 | 0.59 | 0.45 | 0.73 | 0.31 | 0.75 | 0.37 | 0.47 |
| 6 | 152711045 | rs9479307 | 0.425 | -0.579 | 0.855 | 1.211 | 0.005 | 0.042 | 0.042 | 0.052 | 0.1 | 0.15 | 0.12 | 0.35 | 0.15 | 0.15 |
| 6 | 152712302 | rs9942542 | 0.326 | -0.345 | 0.535 | 0.833 | -0.002 | 0.005 | 0.005 | 0 | 0.27 | 0.25 | 0.3 | 0.2 | 0.31 | 0.29 |
| 6 | 152714446 | rs20585 | -0.213 | -0.561 | 1.142 | 1.05 | 0.01 | 0.015 | 0.015 | 0.775 | 0.46 | 0.38 | 0.3 | 0.75 | 0.39 | 0.48 |
| 6 | 152718491 | rs550685 | -0.572 | -0.635 | 1.031 | 0.992 | 0.067 | 0.139 | 0.139 | 0.966 | 0.51 | 0.83 | 0.35 | 0.75 | 0.43 | 0.51 |
| 6 | 152738335 | rs7738528 | -0.76 | -0.466 | 1.509 | 1.247 | -0.001 | 0.073 | 0.073 | 0 | 0.17 | 0.4 | 0.16 | 0.36 | 0.2 | 0.21 |
| 6 | 152738345 | rs214989 | 0.8 | -0.616 | 1.334 | 1.27 | 0.038 | 0.016 | 0.016 | 0.096 | 0.3 | 0.15 | 0.14 | 0.23 | 0.19 | 0.29 |
| 6 | 152752324 | rs525210 | -0.637 | -0.597 | 1.365 | 1.199 | 0.06 | 0.114 | 0.114 | 0.067 | 0.51 | 0.81 | 0.34 | 0.75 | 0.42 | 0.53 |
| 6 | 152755628 | rs215006 | 0.454 | -0.556 | 1.693 | 1.299 | -0.007 | 0.001 | 0.001 | 0.047 | 0.34 | 0.37 | 0.17 | 0.38 | 0.21 | 0.33 |
| 6 | 152762098 | rs6915736 | -0.192 | -0.392 | 1.615 | 1.397 | 0.045 | 0.208 | 0.208 | 0 | 0.12 | 0.39 | 0.14 | 0.37 | 0.15 | 0.16 |
| 6 | 152765201 | rs12526147 | -0.105 | -0.42 | 1.523 | 1.401 | 0.018 | 0.142 | 0.142 | 0 | 0.12 | 0.34 | 0.13 | 0.37 | 0.15 | 0.16 |
| 6 | 152766186 | rs214993 | -0.564 | -0.345 | 1.355 | 0.882 | 0.042 | 0.093 | 0.093 | 0.39 | 0.51 | 0.77 | 0.37 | 0.75 | 0.45 | 0.53 |
| 6 | 152766766 | rs214992 | 0.832 | -0.457 | 1.238 | 0.918 | 0.028 | -0.021 | -0.021 | 0.398 | 0.3 | 0.15 | 0.19 | 0.34 | 0.22 | 0.29 |
| 6 | 152771849 | rs17082701 | 0.495 | -0.407 | 1.207 | 0.778 | -0.007 | -0.023 | -0.023 | 0 | 0.06 | 0.07 | 0.04 | 0 | 0.07 | 0.05 |
| 6 | 152772264 | rs214976 | -0.652 | -0.461 | 1.267 | 0.758 | 0.051 | 0.122 | 0.122 | 0.377 | 0.52 | 0.81 | 0.37 | 0.75 | 0.45 | 0.52 |
| 6 | 152772494 | rs9397105 | 0.391 | -0.494 | 1.238 | 0.813 | -0.035 | -0.073 | -0.073 | 0.349 | 0.11 | 0.01 | 0.08 | 0.43 | 0.15 | 0.19 |
| 6 | 152774033 | rs70018 | -0.677 | -0.486 | 1.243 | 0.817 | 0.051 | 0.121 | 0.121 | 0.42 | 0.52 | 0.81 | 0.37 | 0.75 | 0.45 | 0.52 |
| 6 | 152778598 | rs214966 | 0.85 | -0.561 | 1.269 | 0.772 | 0.045 | 0.022 | 0.022 | 0 | 0.29 | 0.13 | 0.17 | 0.22 | 0.19 | 0.26 |
| 6 | 152790533 | rs9371600 | -0.92 | -0.335 | 1.544 | 0.871 | -0.008 | 0.066 | 0.066 | 0 | 0.2 | 0.35 | 0.15 | 0.43 | 0.23 | 0.24 |
| 6 | 152797065 | rs9397510 | -0.334 | -0.36 | 1.482 | 0.901 | 0.009 | 0.103 | 0.103 | 0 | 0.15 | 0.31 | 0.13 | 0.43 | 0.18 | 0.19 |
| 6 | 152806563 | rs9397513 | -0.614 | -0.405 | 1.493 | 0.69 | -0.001 | 0.048 | 0.048 | 0 | 0.2 | 0.31 | 0.17 | 0.45 | 0.25 | 0.24 |
| 6 | 152831067 | rs2623971 | 0.71 | -0.934 | 1.436 | 0.664 | 0.04 | 0.025 | 0.025 | 0.478 | 0.3 | 0.14 | 0.18 | 0.27 | 0.2 | 0.28 |
| 6 | 152831248 | rs17842543 | 0.452 | -0.867 | 1.444 | 0.621 | -0.007 | -0.033 | -0.033 | 0 | 0.07 | 0.13 | 0.05 | 0 | 0.07 | 0.05 |
| 6 | 152836660 | rs9371605 | -0.059 | -0.503 | 1.216 | 0.394 | -0.008 | -0.008 | -0.008 | 0 | 0.23 | 0.22 | 0.2 | 0.45 | 0.38 | 0.22 |
| 6 | 152836721 | rs9384004 | 0.128 | -0.5 | 1.015 | 0.396 | -0.034 | -0.043 | -0.043 | 0 | 0.21 | 0.17 | 0.2 | 0.45 | 0.38 | 0.22 |
| 6 | 152841248 | rs2623958 | -0.572 | -0.526 | 0.74 | 0.438 | -0.017 | 0.039 | 0.039 | 0.062 | 0.61 | 0.79 | 0.46 | 0.74 | 0.65 | 0.56 |
| 6 | 152842018 | rs2623981 | -1.366 | -0.415 | 0.376 | 0.167 | -0.113 | 0.09 | 0.09 | 0.634 | 0.78 | 0.99 | 0.65 | 0.95 | 0.73 | 0.79 |
| 6 | 152853674 | rs2623966 | 0.247 | -0.465 | 1.074 | 0.605 | 0.042 | 0.002 | 0.002 | 0.762 | 0.29 | 0.13 | 0.18 | 0.3 | 0.2 | 0.28 |
| 6 | 152857887 | rs2496124 | 0.198 | -0.467 | 1.055 | 0.605 | -0.007 | -0.001 | -0.001 | 0.19 | 0.3 | 0.22 | 0.19 | 0.29 | 0.2 | 0.28 |
| 6 | 152869726 | rs2695261 | 0.323 | -0.147 | -0.512 | 0.338 | 0.047 | 0.061 | 0.061 | 0.782 | 0.29 | 0.1 | 0.17 | 0.15 | 0.2 | 0.24 |
| 6 | 152872272 | rs9397523 | -0.643 | 0.575 | 0.119 | 0.866 | 0.034 | 0.056 | 0.056 | 0 | 0.31 | 0.53 | 0.25 | 0.47 | 0.4 | 0.24 |
| 6 | 152872911 | rs9322378 | -0.596 | 0.295 | -0.084 | 0.448 | -0.001 | -0.001 | -0.001 | 0.066 | 0.48 | 0.46 | 0.64 | 0.57 | 0.53 | 0.35 |
| 6 | 152873957 | rs9322379 | -0.504 | 0.818 | 0.812 | 0.752 | 0.033 | 0.129 | 0.129 | 0 | 0.2 | 0.46 | 0.16 | 0.59 | 0.19 | 0.21 |
| 6 | 152875862 | rs9383626 | -0.009 | 0.947 | 0.837 | 0.978 | -0.008 | -0.01 | -0.01 | 0 | 0.14 | 0.11 | 0.12 | 0.46 | 0.14 | 0.13 |
| 6 | 152877077 | rs1554783 | 0.735 | 0.795 | 0.349 | -0.018 | -0.002 | -0.002 | -0.002 | 0 | 0.3 | 0.29 | 0.23 | 0.19 | 0.23 | 0.31 |
| 6 | 152879190 | rs12201313 | 0.248 | 0.948 | 0.006 | -0.231 | 0.013 | 0.014 | 0.014 | 0.218 | 0.49 | 0.57 | 0.57 | 0.2 | 0.4 | 0.41 |
| 6 | 152887445 | rs2623964 | 0.015 | 1.257 | -0.001 | -0.256 | -0.011 | 0.031 | 0.031 | 0.405 | 0.42 | 0.28 | 0.34 | 0.79 | 0.44 | 0.54 |
| 6 | 152892738 | rs818442 | 0.021 | 0.689 | 0.474 | -0.314 | 0.015 | -0.008 | -0.008 | 0 | 0.37 | 0.39 | 0.32 | 0.33 | 0.44 | 0.43 |
| 6 | 152893252 | rs818441 | 0.466 | 0.496 | -0.144 | -0.593 | -0.091 | 0.035 | 0.035 | 0.712 | 0.31 | 0.07 | 0.27 | 0.21 | 0.39 | 0.35 |
| 6 | 152896974 | rs12528046 | 0.242 | -0.338 | 0.527 | -0.087 | -0.02 | -0.065 | -0.065 | 0.494 | 0.2 | 0.32 | 0.34 | 0.01 | 0.17 | 0.1 |
| 6 | 152902786 | rs17779152 | 0.418 | -0.247 | -0.277 | -0.043 | 0.008 | -0.002 | -0.002 | 0 | 0.19 | 0.16 | 0.32 | 0.01 | 0.17 | 0.1 |
| 6 | 152915796 | rs818452 | 0 | 0.171 | -0.722 | 0.427 | -0.006 | -0.046 | -0.046 | 0 | 0.07 | 0 | 0.02 | 0.18 | 0.06 | 0.17 |
| 6 | 152923465 | rs860173 | -0.278 | -0.279 | -0.578 | 0.121 | 0.017 | 0 | 0 | 0 | 0.71 | 0.77 | 0.58 | 0.77 | 0.74 | 0.69 |
| 6 | 152924673 | rs818451 | 0.189 | -0.245 | -0.115 | 0.163 | -0.018 | -0.002 | -0.002 | 0 | 0.61 | 0.54 | 0.53 | 0.67 | 0.64 | 0.59 |
| 6 | 152926517 | rs2695253 | 0.395 | -0.279 | -0.606 | -0.535 | -0.034 | -0.015 | -0.015 | 0 | 0.37 | 0.2 | 0.43 | 0.63 | 0.48 | 0.38 |
| 6 | 152927749 | rs6909384 | 0.503 | -0.039 | -0.31 | -0.265 | 0.001 | -0.002 | -0.002 | 0 | 0.1 | 0.2 | 0.06 | 0.1 | 0.1 | 0.1 |
| 6 | 152937865 | rs1028651 | -0.522 | -0.552 | -0.392 | -0.432 | -0.001 | -0.002 | -0.002 | 0 | 0.71 | 0.72 | 0.62 | 0.81 | 0.68 | 0.65 |
| 6 | 152939439 | rs1744387 | -0.522 | -0.552 | -0.392 | -0.432 | -0.001 | -0.002 | -0.002 | 0 | 0.71 | 0.72 | 0.62 | 0.81 | 0.68 | 0.65 |
| 6 | 152939886 | rs1631293 | -0.288 | -0.637 | -0.052 | -0.393 | -0.017 | 0.018 | 0.018 | 0 | 0.67 | 0.56 | 0.53 | 0.67 | 0.66 | 0.59 |
| 6 | 152940466 | rs1744385 | -0.564 | -0.623 | -0.182 | -0.459 | -0.001 | -0.002 | -0.002 | 0 | 0.71 | 0.72 | 0.62 | 0.81 | 0.68 | 0.65 |
| 6 | 152941053 | rs1407490 | -0.375 | -1.227 | -0.645 | -1.019 | 0.017 | 0.052 | 0.052 | 0.019 | 0.54 | 0.64 | 0.61 | 0.81 | 0.55 | 0.55 |
| 6 | 152942015 | rs6940651 | 0.691 | -1.314 | -0.44 | -1.049 | -0.02 | -0.118 | -0.118 | 0 | 0.23 | 0.38 | 0.11 | 0.04 | 0.15 | 0.15 |
| 6 | 152946514 | rs1631457 | -0.676 | -1.219 | -0.184 | -0.712 | -0.008 | 0.008 | 0.008 | 0 | 0.68 | 0.65 | 0.61 | 0.68 | 0.68 | 0.57 |
| 6 | 152963826 | rs1830642 | -0.89 | -1.242 | -0.099 | -0.707 | -0.005 | -0.003 | -0.003 | 1.285 | 0.72 | 0.7 | 0.66 | 0.78 | 0.71 | 0.64 |
| 6 | 152974550 | rs2258974 | -2.103 | -1.349 | -0.282 | -0.714 | 0.018 | 0.007 | 0.007 | 0 | 0.83 | 0.9 | 0.76 | 0.91 | 0.81 | 0.72 |
| 6 | 152979834 | rs9383636 | 0.341 | -0.98 | -0.406 | -0.984 | -0.08 | -0.033 | -0.033 | 0.146 | 0.37 | 0.13 | 0.44 | 0.62 | 0.43 | 0.32 |
| 6 | 152982768 | rs9384022 | 1.22 | -0.119 | -0.749 | -0.262 | -0.05 | -0.097 | -0.097 | 0 | 0.16 | 0.01 | 0.09 | 0.31 | 0.17 | 0.05 |
| 6 | 152985508 | rs2758767 | -0.198 | -0.445 | -0.45 | -0.49 | 0.011 | 0.025 | 0.025 | 0.082 | 0.64 | 0.33 | 0.72 | 0.77 | 0.72 | 0.67 |

| | | | | | | | | | | | | | | | | |
|---|---|---|---|---|---|---|---|---|---|---|---|---|---|---|---|---|
| 6 | 153000482 | rs9285533 | -0.202 | -0.34 | -0.455 | -0.415 | 0.005 | -0.013 | -0.013 | 0 | 0.41 | 0.44 | 0.54 | 0.76 | 0.44 | 0.38 |
| 6 | 153003011 | rs1923545 | 0.259 | -0.3 | -0.653 | -0.403 | -0.114 | -0.064 | -0.064 | 0 | 0.36 | 0.09 | 0.4 | 0.59 | 0.42 | 0.31 |
| 6 | 153006016 | rs2800625 | 1.041 | -0.414 | -0.473 | -0.445 | 0.004 | -0.003 | -0.003 | 0 | 0.33 | 0.3 | 0.39 | 0.33 | 0.32 | 0.43 |
| 6 | 153015245 | rs2800641 | -0.572 | -0.358 | -0.499 | -0.613 | -0.022 | -0.153 | -0.153 | 0 | 0.69 | 0.5 | 0.83 | 0.95 | 0.73 | 0.75 |
| 6 | 153021809 | rs1322524 | -0.109 | -0.313 | -0.565 | -0.538 | 0.011 | -0.023 | -0.023 | 0 | 0.4 | 0.49 | 0.54 | 0.77 | 0.44 | 0.39 |
| 6 | 153025827 | rs2758787 | -0.461 | -0.334 | -0.53 | -0.619 | -0.012 | -0.102 | -0.102 | 0.558 | 0.68 | 0.6 | 0.83 | 0.95 | 0.73 | 0.75 |
| 6 | 153027903 | rs2244217 | -0.518 | 0.007 | -0.48 | -0.601 | -0.021 | -0.151 | -0.151 | 0.367 | 0.69 | 0.52 | 0.84 | 0.95 | 0.73 | 0.75 |
| 6 | 153034155 | rs2800637 | 0.747 | -0.06 | -0.565 | -0.536 | -0.009 | 0.004 | 0.004 | 0 | 0.33 | 0.39 | 0.4 | 0.33 | 0.32 | 0.43 |
| 6 | 153034248 | rs2758789 | 1.587 | -0.04 | -0.812 | -0.56 | 0.01 | 0 | 0 | 0 | 0.28 | 0.1 | 0.3 | 0.19 | 0.29 | 0.36 |
| 6 | 153043035 | rs17710008 | 1.143 | 0.419 | -0.842 | -0.599 | -0.061 | -0.006 | -0.006 | 0.277 | 0.13 | 0.01 | 0.24 | 0.09 | 0.16 | 0.12 |
| 6 | 153045227 | rs6934838 | -0.014 | 0.348 | 0.188 | -0.381 | -0.007 | -0.022 | -0.022 | 0 | 0.64 | 0.6 | 0.8 | 0.8 | 0.7 | 0.62 |
| 6 | 153050768 | rs9397554 | 0.166 | 0.45 | 0.142 | -0.298 | 0.017 | 0.034 | 0.034 | 0.63 | 0.62 | 0.34 | 0.75 | 0.76 | 0.7 | 0.62 |
| 6 | 153055554 | rs9322396 | 0.309 | 0.469 | 0.13 | -0.221 | 0.045 | 0.043 | 0.043 | 0.658 | 0.58 | 0.21 | 0.66 | 0.62 | 0.67 | 0.55 |
| 6 | 153062947 | rs1282448 | 0.016 | 0.423 | 0.625 | -0.547 | -0.035 | -0.009 | -0.009 | 0.564 | 0.18 | 0.39 | 0.17 | 0.18 | 0.13 | 0.22 |
| 6 | 153063431 | rs637572 | 0.001 | 0.423 | 0.422 | -0.547 | -0.031 | -0.011 | -0.011 | 0 | 0.19 | 0.35 | 0.16 | 0.18 | 0.13 | 0.22 |
| 6 | 153066078 | rs680314 | -1.194 | 0.567 | -0.041 | -0.576 | 0.057 | 0.023 | 0.023 | 0 | 0.66 | 0.88 | 0.64 | 0.8 | 0.69 | 0.62 |
| 6 | 153066298 | rs647749 | -0.218 | 0.635 | 0.486 | -0.614 | 0.003 | -0.006 | -0.006 | 0 | 0.46 | 0.38 | 0.28 | 0.54 | 0.42 | 0.41 |
| 6 | 153066369 | rs13211784 | 0.597 | 0.634 | 0.315 | -0.757 | 0 | -0.002 | -0.002 | 0 | 0.18 | 0.19 | 0.11 | 0.12 | 0.14 | 0.19 |
| 6 | 153066802 | rs17710848 | 0.035 | 0.763 | 0.346 | -0.542 | 0.026 | -0.032 | -0.032 | 0 | 0.06 | 0.18 | 0.06 | 0 | 0.08 | 0.02 |
| 6 | 153066908 | rs2756117 | 0.988 | 0.544 | 0.166 | -0.018 | -0.038 | 0.002 | 0.002 | 0 | 0.07 | 0.01 | 0.06 | 0.42 | 0.13 | 0.1 |
| 6 | 153069342 | rs17083011 | 0.025 | 0.372 | 0.472 | -0.372 | -0.024 | -0.092 | -0.092 | 0 | 0.21 | 0.52 | 0.14 | 0.1 | 0.15 | 0.18 |
| 6 | 153072696 | rs3799142 | 0.183 | 0.053 | 0.42 | -1.127 | -0.01 | -0.032 | -0.032 | 0.134 | 0.2 | 0.29 | 0.11 | 0.1 | 0.14 | 0.15 |
| 6 | 153074322 | rs3823082 | -0.251 | 0.201 | -0.122 | -1.034 | -0.031 | -0.09 | -0.09 | 0 | 0.34 | 0.58 | 0.19 | 0.11 | 0.19 | 0.21 |
| 6 | 153076789 | rs12201030 | 0.088 | 0.957 | -0.203 | -0.012 | -0.027 | -0.035 | -0.035 | 0 | 0.09 | 0.01 | 0.1 | 0.11 | 0.1 | 0.07 |
| 6 | 153080061 | rs688136 | 0.133 | 0.882 | 0.094 | -0.444 | 0.001 | 0.001 | 0.001 | 0 | 0.43 | 0.57 | 0.31 | 0.19 | 0.37 | 0.34 |
| 6 | 153082369 | rs627728 | 0.331 | 0.978 | -0.602 | -0.575 | 0.002 | 0.028 | 0.028 | 0 | 0.38 | 0.3 | 0.26 | 0.13 | 0.33 | 0.29 |
| 6 | 153085234 | rs671330 | 0.082 | 1.024 | -0.744 | 0.126 | -0.003 | 0.041 | 0.041 | 0 | 0.7 | 0.81 | 0.6 | 0.83 | 0.66 | 0.64 |
| 6 | 153094557 | rs606445 | 0.912 | 1.127 | 0.232 | 0.67 | 0.043 | 0.065 | 0.065 | 0 | 0.85 | 0.59 | 0.79 | 0.83 | 0.81 | 0.91 |
| 6 | 153096002 | rs4870141 | -0.763 | 1.03 | -0.143 | 0.762 | -0.004 | -0.033 | -0.033 | 0 | 0.11 | 0.03 | 0.15 | 0.17 | 0.09 | 0.11 |
| 6 | 153096503 | rs1410489 | -0.358 | 1.797 | 0.138 | 0.415 | 0.023 | -0.012 | -0.012 | 0 | 0.24 | 0.34 | 0.39 | 0.25 | 0.34 | 0.41 |
| 6 | 153099969 | rs596362 | 0.44 | 1.511 | -0.029 | -0.821 | -0.007 | 0.051 | 0.051 | 0 | 0.53 | 0.4 | 0.54 | 0.11 | 0.51 | 0.55 |
| 6 | 153101244 | rs612400 | 1.189 | 1.302 | -0.351 | -1.409 | -0.005 | -0.003 | -0.003 | 0 | 0.17 | 0.13 | 0.14 | 0 | 0.21 | 0.11 |
| 6 | 153103035 | rs4870142 | 0.19 | 1.5 | 0.257 | -1.254 | 0 | -0.005 | -0.005 | 0 | 0.42 | 0.44 | 0.46 | 0.27 | 0.42 | 0.47 |
| 6 | 153105851 | rs311349 | 1.395 | 1.236 | 0.21 | -0.419 | -0.032 | -0.115 | -0.115 | 0.102 | 0.22 | 0.02 | 0.16 | 0.31 | 0.22 | 0.13 |
| 6 | 153106182 | rs7740619 | 0.829 | 1.244 | 0.238 | -0.445 | 0.003 | -0.026 | -0.026 | 2.262 | 0.26 | 0.33 | 0.15 | 0.16 | 0.25 | 0.1 |
| 6 | 153106506 | rs12194558 | 1.342 | 1.246 | -0.043 | -0.404 | -0.003 | 0.02 | 0.02 | 0 | 0.16 | 0.1 | 0.3 | 0.11 | 0.18 | 0.36 |
| 6 | 153107539 | rs506140 | -0.247 | 1.18 | 0.035 | -0.4 | 0.011 | -0.024 | -0.024 | 0.08 | 0.61 | 0.73 | 0.64 | 0.52 | 0.65 | 0.7 |
| 6 | 153111532 | rs311352 | -0.985 | 1.188 | -0.236 | -0.522 | -0.039 | -0.059 | -0.059 | 0 | 0.9 | 1 | 0.84 | 0.83 | 0.89 | 0.88 |
| 6 | 153119212 | rs7767039 | 0.563 | 1.26 | -0.26 | -0.492 | 0.002 | 0.001 | 0.001 | 0 | 0.07 | 0.09 | 0.07 | 0.06 | 0.15 | 0.03 |
| 6 | 153120813 | rs620598 | 0.893 | 1.093 | -0.08 | -0.515 | 0.011 | 0.035 | 0.035 | 0 | 0.2 | 0.31 | 0.18 | 0.28 | 0.22 | 0.24 |
| 6 | 153120904 | rs311334 | 1.421 | 1.228 | -0.742 | -0.399 | -0.006 | -0.129 | -0.129 | 0 | 0.2 | 0.02 | 0.14 | 0.34 | 0.2 | 0.13 |
| 6 | 153123215 | rs9479396 | 0.997 | 1.558 | -0.149 | -0.367 | 0.006 | -0.002 | -0.002 | 0.154 | 0.22 | 0.19 | 0.13 | 0.26 | 0.13 | 0.2 |
| 6 | 153124301 | rs578512 | 0.599 | 1.642 | -0.566 | -0.041 | 0.001 | 0.001 | 0.001 | 0 | 0.4 | 0.55 | 0.28 | 0.28 | 0.33 | 0.31 |
| 6 | 153137096 | rs311330 | 1.183 | 2.117 | -0.546 | 0.181 | -0.018 | -0.002 | -0.002 | 1.874 | 0.78 | 0.8 | 0.74 | 0.77 | 0.69 | 0.8 |
| 6 | 153137621 | rs311326 | 0.598 | 1.175 | -0.516 | 0.393 | -0.008 | -0.009 | -0.009 | 0.069 | 0.85 | 0.88 | 0.81 | 0.83 | 0.85 | 0.81 |
| 6 | 153143932 | rs670307 | -0.462 | 0.741 | -0.278 | -0.299 | -0.005 | -0.079 | -0.079 | 0 | 0.18 | 0.23 | 0.09 | 0 | 0.17 | 0.1 |
| 6 | 153143973 | rs17083085 | -0.372 | 1.328 | -1.208 | 0.671 | -0.058 | -0.103 | -0.103 | 0.064 | 0.16 | 0.02 | 0.23 | 0.32 | 0.22 | 0.16 |
| 6 | 153146484 | rs484558 | -0.291 | 1.317 | -1.12 | 0.778 | 0 | 0.002 | 0.002 | 0 | 0.79 | 0.77 | 0.76 | 0.61 | 0.78 | 0.85 |
| 6 | 153151179 | rs13205491 | 0.146 | 1.041 | -0.699 | -0.058 | -0.007 | -0.082 | -0.082 | 0.078 | 0.28 | 0.38 | 0.35 | 0.06 | 0.21 | 0.2 |
| 6 | 153171417 | rs192846 | 0.247 | 1.15 | -0.902 | -0.115 | -0.002 | 0.075 | 0.075 | 0.057 | 0.52 | 0.68 | 0.57 | 0.22 | 0.51 | 0.33 |
| 6 | 153173999 | rs9384046 | 0.594 | 1.196 | -0.981 | 0.415 | -0.002 | -0.002 | -0.002 | 0 | 0.12 | 0.09 | 0.13 | 0.24 | 0.17 | 0.12 |
| 6 | 153176559 | rs592896 | 0.172 | 0.714 | -0.838 | -0.149 | 0.001 | -0.007 | -0.007 | 0 | 0.32 | 0.31 | 0.49 | 0.18 | 0.34 | 0.24 |
| 6 | 153180096 | rs1359964 | 1.723 | 0.649 | -0.368 | -0.297 | 0.002 | -0.011 | -0.011 | 0 | 0.2 | 0.17 | 0.06 | 0.04 | 0.16 | 0.09 |
| 6 | 153186742 | rs9397564 | 0.221 | 0.646 | -0.611 | -0.122 | -0.003 | 0.005 | 0.005 | 0 | 0.36 | 0.45 | 0.24 | 0.4 | 0.37 | 0.52 |
| 6 | 153188447 | rs603992 | -1.274 | 0.636 | -0.737 | -0.056 | 0.003 | -0.029 | -0.029 | 0 | 0.68 | 0.76 | 0.73 | 0.55 | 0.71 | 0.76 |
| 6 | 153189269 | rs694975 | 1.428 | 0.507 | -0.678 | -0.517 | -0.045 | 0.008 | 0.008 | 0.046 | 0.16 | 0.01 | 0.19 | 0.08 | 0.2 | 0.1 |
| 6 | 153200563 | rs604150 | -0.944 | 0.164 | -0.885 | -0.248 | -0.064 | -0.118 | -0.118 | 0.538 | 0.75 | 0.94 | 0.71 | 0.54 | 0.64 | 0.79 |
| 6 | 153200869 | rs602823 | -0.418 | 0.095 | -0.702 | -0.179 | 0.008 | 0.035 | 0.035 | 0.334 | 0.51 | 0.67 | 0.62 | 0.36 | 0.44 | 0.39 |
| 6 | 153208857 | rs575592 | -0.747 | 0.093 | -0.704 | -0.184 | -0.009 | -0.004 | -0.004 | 0 | 0.55 | 0.67 | 0.64 | 0.5 | 0.48 | 0.7 |
| 6 | 153209197 | rs9371645 | 1.438 | 0.044 | -0.665 | -0.218 | -0.013 | -0.002 | -0.002 | 0 | 0.18 | 0.24 | 0.12 | 0.22 | 0.09 | 0.15 |
| 6 | 153215035 | rs570343 | 0.614 | 0.098 | -0.685 | -0.326 | -0.003 | 0.004 | 0.004 | 0 | 0.23 | 0.17 | 0.24 | 0.11 | 0.26 | 0.14 |
| 6 | 153218261 | rs656610 | -0.238 | 0.07 | -0.665 | -0.274 | 0.004 | -0.034 | -0.034 | 0 | 0.34 | 0.43 | 0.5 | 0.14 | 0.35 | 0.24 |
| 6 | 153224190 | rs584258 | -0.294 | -0.113 | -0.546 | -0.126 | -0.002 | -0.004 | -0.004 | 0 | 0.33 | 0.43 | 0.48 | 0.28 | 0.34 | 0.53 |
| 6 | 153226106 | rs539107 | -0.448 | 0.026 | -0.632 | -0.084 | 0.02 | 0.002 | 0.002 | 1.694 | 0.51 | 0.67 | 0.6 | 0.5 | 0.42 | 0.68 |
| 6 | 153242271 | rs6912840 | 0.976 | 0.203 | -0.552 | -0.044 | 0.013 | -0.001 | -0.001 | 0 | 0.06 | 0.14 | 0.23 | 0.03 | 0.06 | 0.06 |
| 6 | 153250976 | rs6939590 | -0.791 | 0.391 | -0.973 | -0.06 | -0.126 | -0.175 | -0.175 | 0 | 0.73 | 0.97 | 0.7 | 0.58 | 0.62 | 0.8 |
| 6 | 153253547 | rs6909370 | 1.099 | 0.206 | -0.784 | -0.287 | -0.004 | 0.016 | 0.016 | 0.078 | 0.18 | 0.09 | 0.21 | 0.08 | 0.22 | 0.1 |
| 6 | 153257363 | rs9479442 | 1.118 | 0.271 | -0.262 | -0.143 | -0.004 | -0.075 | -0.075 | 0.006 | 0.2 | 0.35 | 0.09 | 0.07 | 0.17 | 0.12 |
| 6 | 153258023 | rs4869773 | 0.104 | 0.121 | -0.444 | -0.134 | 0.008 | 0.014 | 0.014 | 2.451 | 0.48 | 0.43 | 0.43 | 0.44 | 0.41 | 0.62 |

| | | | | | | | | | | | | | | | |
|---|---|---|---|---|---|---|---|---|---|---|---|---|---|---|---|
| 6 | 153258346 | rs6923712 | 0.527 | 0.025 | -0.558 | -0.284 | 0.003 | 0.01 | 0.01 | 0.054 | 0.26 | 0.22 | 0.12 | 0.18 | 0.21 | 0.43 |
| 6 | 153259384 | rs9479445 | 1.401 | 0.148 | -0.275 | -0.343 | 0.006 | -0.022 | -0.022 | 3.147 | 0.17 | 0.19 | 0.08 | 0.07 | 0.19 | 0.12 |
| 6 | 153262157 | rs11969689 | 0.258 | -0.015 | -0.277 | -0.518 | -0.004 | -0.037 | -0.037 | 0.067 | 0.38 | 0.4 | 0.33 | 0.12 | 0.37 | 0.24 |
| 6 | 153267724 | rs2144358 | -0.819 | 0.248 | 0.116 | 0.113 | -0.104 | -0.16 | -0.16 | 0.02 | 0.74 | 0.95 | 0.72 | 0.57 | 0.66 | 0.8 |
| 6 | 153268295 | rs7454210 | 0.237 | 0.22 | 0.206 | 0.092 | 0.009 | -0.037 | -0.037 | 0.029 | 0.24 | 0.4 | 0.28 | 0.12 | 0.25 | 0.18 |
| 6 | 153271852 | rs10484865 | 1.204 | 0.255 | 0.183 | 0.102 | 0.007 | -0.001 | -0.001 | 1.353 | 0.05 | 0.15 | 0.2 | 0.02 | 0.05 | 0.06 |
| 6 | 153273743 | rs6900526 | 0.14 | 0.212 | -0.17 | 0.13 | 0.027 | -0.008 | -0.008 | 1.227 | 0.53 | 0.79 | 0.61 | 0.51 | 0.43 | 0.59 |
| 6 | 153274321 | rs9285536 | 1.536 | 0.273 | -0.182 | 0.075 | -0.036 | 0.04 | 0.04 | 0 | 0.15 | 0.05 | 0.07 | 0.07 | 0.19 | 0.08 |
| 6 | 153276423 | rs9479453 | 0.482 | -0.1 | -0.013 | 0.133 | 0.01 | 0.013 | 0.013 | 0 | 0.07 | 0.15 | 0.05 | 0.13 | 0.09 | 0.22 |
| 6 | 153280912 | rs4869774 | -0.768 | -0.156 | 0.1 | -0.037 | 0.004 | 0.006 | 0.006 | 0 | 0.14 | 0.32 | 0.25 | 0.15 | 0.14 | 0.29 |
| 6 | 153281015 | rs4869775 | 1.24 | -0.158 | 0.106 | -0.039 | -0.003 | -0.007 | -0.007 | 0 | 0.05 | 0.08 | 0.19 | 0.02 | 0.05 | 0.07 |
| 6 | 153285650 | rs4870155 | -0.772 | -0.153 | 0.102 | -0.011 | 0.004 | 0.01 | 0.01 | 0 | 0.14 | 0.32 | 0.25 | 0.16 | 0.15 | 0.28 |
| 6 | 153290888 | rs874302 | 1.271 | -0.276 | 0.081 | -0.026 | -0.003 | -0.004 | -0.004 | 0.072 | 0.38 | 0.38 | 0.36 | 0.32 | 0.29 | 0.26 |
| 6 | 153291277 | rs9397573 | 1.821 | 0.127 | -0.527 | 0.072 | -0.091 | 0.011 | 0.011 | 0.001 | 0.15 | 0.01 | 0.11 | 0.07 | 0.24 | 0.1 |
| 6 | 153292712 | rs1997960 | -0.636 | 0.069 | -0.513 | 0.123 | -0.086 | -0.166 | -0.166 | 0 | 0.73 | 0.98 | 0.74 | 0.6 | 0.68 | 0.79 |
| 6 | 153296370 | rs7763565 | 0.733 | 0.379 | -0.037 | -0.015 | 0 | -0.008 | -0.008 | 0 | 0.06 | 0.16 | 0.2 | 0.02 | 0.05 | 0.06 |
| 6 | 153301542 | rs9479476 | 0.378 | 0.389 | -0.046 | -0.001 | 0.01 | 0.018 | 0.018 | 0 | 0.07 | 0.15 | 0.05 | 0.13 | 0.09 | 0.22 |
| 6 | 153301614 | rs2207609 | -0.55 | 0.407 | -0.019 | -0.03 | 0.023 | -0.017 | -0.017 | 0 | 0.29 | 0.37 | 0.37 | 0.22 | 0.38 | 0.38 |
| 6 | 153306005 | rs3757066 | -0.641 | 0.395 | -0.213 | 0.027 | 0.002 | 0.002 | 0.002 | 0 | 0.14 | 0.24 | 0.3 | 0.15 | 0.17 | 0.29 |
| 6 | 153307483 | rs9384060 | 1.394 | 0.564 | -0.389 | 0.031 | -0.047 | 0.026 | 0.026 | 0 | 0.16 | 0.06 | 0.11 | 0.07 | 0.23 | 0.1 |
| 6 | 153308312 | rs12189965 | -0.061 | 0.466 | -0.485 | 0.076 | 0.001 | 0.004 | 0.004 | 0 | 0.09 | 0.14 | 0.11 | 0.13 | 0.11 | 0.23 |
| 6 | 153313534 | rs11756873 | 1.198 | 0.591 | -0.68 | 0.046 | -0.09 | 0.005 | 0.005 | 0.279 | 0.15 | 0.01 | 0.1 | 0.07 | 0.24 | 0.09 |
| 6 | 153313832 | rs766127 | 0.028 | -0.244 | -0.33 | 0.07 | 0.004 | -0.01 | -0.01 | 0 | 0.66 | 0.71 | 0.73 | 0.54 | 0.7 | 0.64 |
| 6 | 153315617 | rs3799135 | -0.495 | -0.524 | -0.352 | -0.122 | -0.009 | -0.01 | -0.01 | 0 | 0.18 | 0.27 | 0.3 | 0.15 | 0.16 | 0.32 |
| 6 | 153315921 | rs3799133 | -1.644 | -0.53 | -0.534 | 0.422 | -0.053 | -0.05 | -0.05 | 0.048 | 0.86 | 0.99 | 0.88 | 0.83 | 0.85 | 0.91 |
| 6 | 153333328 | rs650018 | 0.852 | -0.331 | -0.186 | 0.073 | 0.003 | 0.008 | 0.008 | 0 | 0.49 | 0.44 | 0.43 | 0.39 | 0.53 | 0.32 |
| 6 | 153335954 | rs567425 | -0.289 | -0.194 | -0.244 | 0.25 | 0.003 | -0.012 | -0.012 | 0 | 0.68 | 0.71 | 0.73 | 0.56 | 0.7 | 0.75 |
| 6 | 153345185 | rs672170 | -0.228 | 0.209 | -0.277 | 0.429 | 0.017 | -0.039 | -0.039 | 0 | 0.62 | 0.77 | 0.55 | 0.55 | 0.65 | 0.68 |
| 6 | 153348742 | rs598623 | 0.071 | 0.198 | -0.251 | 0.506 | -0.005 | -0.008 | -0.008 | 0.015 | 0.45 | 0.47 | 0.43 | 0.46 | 0.41 | 0.57 |
| 6 | 153349379 | rs595797 | 0.328 | 0.268 | -0.552 | 0.457 | -0.008 | -0.006 | -0.006 | 0 | 0.35 | 0.42 | 0.34 | 0.37 | 0.34 | 0.34 |
| 6 | 153349446 | rs3799125 | 0.651 | 1.002 | -1.204 | 0.515 | -0.042 | -0.027 | -0.027 | 0 | 0.08 | 0.03 | 0.09 | 0.11 | 0.2 | 0.1 |
| 6 | 153349813 | rs583297 | 0.377 | -0.283 | -1.253 | 0.161 | 0.017 | 0.035 | 0.035 | 0 | 0.41 | 0.24 | 0.39 | 0.28 | 0.4 | 0.37 |
| 6 | 153352156 | rs1292829 | -0.162 | -0.544 | -1.345 | 0.015 | 0.029 | -0.019 | -0.019 | 0 | 0.8 | 0.99 | 0.86 | 0.85 | 0.88 | 0.84 |
| 6 | 153352474 | rs7774354 | 1.424 | -1.122 | -1.215 | -0.445 | -0.027 | -0.041 | -0.041 | 0 | 0.19 | 0.3 | 0.07 | 0.05 | 0.11 | 0.07 |
| 6 | 153353126 | rs9479484 | 0.451 | -1.161 | -1.08 | -0.551 | -0.001 | -0.029 | -0.029 | 0 | 0.07 | 0.1 | 0.05 | 0 | 0.08 | 0.02 |
| 6 | 153353459 | rs1292830 | 0.228 | -0.687 | -1.072 | -0.588 | 0.002 | 0.019 | 0.019 | 1.5 | 0.47 | 0.54 | 0.64 | 0.75 | 0.49 | 0.66 |
| 6 | 153354211 | rs2057181 | 0.332 | -0.053 | -0.588 | -0.439 | 0.024 | -0.058 | -0.058 | 0 | 0.33 | 0.44 | 0.2 | 0.1 | 0.39 | 0.18 |
| 6 | 153365384 | rs2295232 | 0.26 | -0.282 | -0.406 | 0.288 | 0.054 | 0.016 | 0.016 | 0.006 | 0.49 | 0.59 | 0.38 | 0.64 | 0.29 | 0.52 |
| 6 | 153367910 | rs4599660 | 0.285 | -0.309 | 0.07 | 0.287 | 0.015 | 0.004 | 0.004 | 0 | 0.49 | 0.56 | 0.37 | 0.63 | 0.29 | 0.52 |
| 6 | 153368195 | rs9478383 | 0.988 | -0.714 | 0.365 | 0.711 | 0.007 | 0.082 | 0.082 | 0 | 0.1 | 0.18 | 0.1 | 0.3 | 0.09 | 0.08 |
| 6 | 153371183 | rs9479488 | 0.012 | 0.238 | 0.322 | 0.81 | -0.002 | 0.001 | 0.001 | 0 | 0.61 | 0.65 | 0.5 | 0.7 | 0.57 | 0.61 |
| 6 | 153377562 | rs872371 | 1.057 | 0.332 | -0.052 | 0.801 | -0.054 | -0.044 | -0.044 | 0.504 | 0.18 | 0.02 | 0.28 | 0.14 | 0.22 | 0.25 |
| 6 | 153383590 | rs9397581 | 1.494 | 0.031 | 0.025 | 0.644 | 0.015 | -0.003 | -0.003 | 0.013 | 0.37 | 0.32 | 0.21 | 0.36 | 0.25 | 0.18 |
| 6 | 153384196 | rs1933262 | 0.701 | 0.341 | 0.331 | 0.501 | 0.119 | -0.037 | -0.037 | 0 | 0.14 | 0.31 | 0.16 | 0.07 | 0.27 | 0.09 |
| 6 | 153394728 | rs4385321 | 0.569 | 0.03 | 0.274 | 0.616 | -0.003 | -0.002 | -0.002 | 0 | 0.53 | 0.51 | 0.4 | 0.67 | 0.35 | 0.59 |
| 6 | 153397729 | rs7743048 | -0.923 | 0.046 | 0.051 | 0.453 | -0.029 | -0.056 | -0.056 | 0 | 0.86 | 0.99 | 0.85 | 0.84 | 0.87 | 0.89 |
| 6 | 153397958 | rs9479495 | 0.879 | 0.286 | 0.155 | 0.438 | 0.012 | -0.011 | -0.011 | 0 | 0.16 | 0.12 | 0.16 | 0.05 | 0.33 | 0.12 |
| 6 | 153398700 | rs17083510 | -0.635 | 0.005 | 0.214 | 0.488 | 0.003 | 0.005 | 0.005 | 0 | 0.46 | 0.45 | 0.57 | 0.3 | 0.59 | 0.38 |
| 6 | 153402841 | rs6920313 | 1.396 | -0.068 | 0.238 | 0.514 | 0.025 | -0.014 | -0.014 | 0 | 0.41 | 0.33 | 0.23 | 0.43 | 0.31 | 0.26 |
| 6 | 153404825 | rs9479504 | 0.285 | -0.061 | 0.293 | 0.512 | -0.025 | 0.022 | 0.022 | 0 | 0.31 | 0.26 | 0.43 | 0.13 | 0.5 | 0.3 |
| 6 | 153405328 | rs10499270 | 1.041 | -0.048 | 0.053 | 0.528 | -0.001 | -0.062 | -0.062 | 0.064 | 0.13 | 0.01 | 0.13 | 0.17 | 0.08 | 0.09 |
| 6 | 153406345 | rs17083512 | -0.233 | -0.025 | 0.222 | 0.603 | -0.013 | 0.093 | 0.093 | 0 | 0.05 | 0.17 | 0.02 | 0.24 | 0.01 | 0.29 |
| 6 | 153406657 | rs687508 | -0.564 | -0.012 | 0.174 | 0.599 | 0.01 | 0.026 | 0.026 | 0 | 0.44 | 0.35 | 0.57 | 0.3 | 0.59 | 0.38 |
| 6 | 153412476 | rs3910736 | 1.161 | 0.258 | 0.353 | 0.151 | -0.006 | -0.002 | -0.002 | 2.72 | 0.45 | 0.44 | 0.25 | 0.65 | 0.31 | 0.54 |
| 6 | 153413309 | rs7740777 | 0.402 | 0.335 | 0.396 | 0.126 | -0.006 | -0.034 | -0.034 | 0.059 | 0.6 | 0.52 | 0.39 | 0.84 | 0.4 | 0.63 |
| 6 | 153423571 | rs1281971 | 0.723 | 0.471 | 0.242 | 0.15 | -0.002 | 0.001 | 0.001 | 0 | 0.08 | 0.06 | 0.17 | 0.03 | 0.1 | 0.07 |
| 6 | 153427265 | rs9479509 | 1.466 | 0.35 | 0.027 | 0.269 | 0.02 | -0.008 | -0.008 | 0 | 0.4 | 0.34 | 0.24 | 0.58 | 0.3 | 0.51 |
| 6 | 153441079 | rs1933488 | 0.549 | 0.439 | 0.095 | -0.232 | -0.006 | -0.04 | -0.04 | 0 | 0.57 | 0.48 | 0.39 | 0.84 | 0.4 | 0.63 |
| 6 | 153443159 | rs6557273 | 1.417 | 0.502 | -0.147 | -0.206 | 0.024 | 0.007 | 0.007 | 0 | 0.41 | 0.33 | 0.25 | 0.66 | 0.31 | 0.51 |
| 6 | 153450489 | rs611499 | -0.08 | 0.593 | -0.215 | -0.186 | -0.035 | 0.035 | 0.035 | 0 | 0.67 | 0.78 | 0.58 | 0.87 | 0.5 | 0.7 |
| 6 | 153453987 | rs12198739 | 0.181 | 0.609 | -0.145 | 0.186 | -0.015 | -0.057 | -0.057 | 0 | 0.14 | 0.01 | 0.14 | 0.18 | 0.09 | 0.12 |
| 6 | 153457680 | rs1281958 | -0.701 | 0.823 | -0.025 | 0.043 | 0.062 | -0.105 | -0.105 | 1.135 | 0.28 | 0.54 | 0.37 | 0.06 | 0.43 | 0.18 |
| 6 | 153463915 | rs683039 | -1.137 | 0.752 | -0.545 | 0.174 | -0.003 | 0.048 | 0.048 | 0 | 0.44 | 0.67 | 0.51 | 0.24 | 0.52 | 0.31 |
| 6 | 153479726 | rs12191018 | 0.05 | 0.808 | -0.787 | 0.644 | -0.012 | -0.082 | -0.082 | 0.193 | 0.14 | 0.01 | 0.13 | 0.23 | 0.09 | 0.11 |
| 6 | 153483603 | rs1124016 | -1.315 | 0.809 | -0.505 | 0.184 | -0.01 | 0.04 | 0.04 | 0.319 | 0.43 | 0.76 | 0.5 | 0.29 | 0.51 | 0.25 |
| 6 | 153488244 | rs1281988 | -1.049 | 1.09 | -0.104 | 0.077 | 0 | -0.007 | -0.007 | 0 | 0.29 | 0.72 | 0.37 | 0.09 | 0.42 | 0.15 |
| 6 | 153496654 | rs4869781 | -1.014 | 0.978 | 0.04 | 0.061 | 0.019 | -0.055 | -0.055 | 0 | 0.3 | 0.61 | 0.35 | 0.09 | 0.37 | 0.14 |
| 6 | 153503386 | rs9478395 | 0.29 | 1.659 | -0.505 | 0.259 | -0.073 | 0.032 | 0.032 | 0.064 | 0.11 | 0.01 | 0.1 | 0.04 | 0.25 | 0.06 |
| 6 | 153504165 | rs553072 | -0.713 | 1.288 | 0.189 | 0.406 | 0.009 | -0.012 | -0.012 | 3.784 | 0.14 | 0.29 | 0.33 | 0.09 | 0.14 | 0.12 |

| | | | | | | | | | | | | | | | | |
|---|---|---|---|---|---|---|---|---|---|---|---|---|---|---|---|---|
| 6 | 153504188 | rs11753918 | -0.739 | 1.527 | 0.239 | 0.601 | 0.004 | -0.006 | -0.006 | 0 | 0.14 | 0.18 | 0.08 | 0.15 | 0.15 | 0.35 |
| 6 | 153504519 | rs12153830 | -0.112 | 1.002 | -0.159 | 0.756 | 0.003 | -0.02 | -0.02 | 0.006 | 0.14 | 0.06 | 0.15 | 0.21 | 0.1 | 0.1 |
| 6 | 153515011 | rs315428 | 1.035 | 0.787 | 0.085 | 1.118 | -0.009 | 0.034 | 0.034 | 0.001 | 0.7 | 0.41 | 0.77 | 0.46 | 0.8 | 0.49 |
| 6 | 153515792 | rs9384084 | 0.43 | 0.723 | -0.108 | 1.433 | 0.006 | -0.043 | -0.043 | 0.012 | 0.07 | 0.01 | 0.24 | 0.17 | 0.04 | 0.06 |
| 6 | 153517909 | rs13191030 | 1.399 | 0.64 | -0.624 | 0.731 | -0.005 | 0.019 | 0.019 | 0 | 0.34 | 0.13 | 0.31 | 0.25 | 0.36 | 0.27 |
| 6 | 153518604 | rs17724757 | 1.262 | 0.621 | -0.584 | 0.722 | 0.004 | 0.023 | 0.023 | 0.105 | 0.35 | 0.17 | 0.32 | 0.24 | 0.36 | 0.27 |
| 6 | 153520113 | rs316299 | -0.446 | 0.808 | -0.106 | 0.994 | -0.007 | 0.102 | 0.102 | 0.019 | 0.19 | 0.49 | 0.1 | 0.34 | 0.13 | 0.39 |
| 6 | 153524411 | rs315414 | -0.776 | 1.008 | -0.136 | 1.024 | 0.002 | -0.01 | -0.01 | 0 | 0.89 | 0.9 | 0.87 | 0.79 | 0.92 | 0.88 |
| 6 | 153526941 | rs521222 | 0.248 | 1.097 | -0.191 | 0.948 | -0.022 | 0.042 | 0.042 | 0 | 0.32 | 0.23 | 0.44 | 0.2 | 0.42 | 0.2 |
| 6 | 153528141 | rs317077 | 0.363 | 1.034 | -0.099 | 0.964 | -0.012 | 0.014 | 0.014 | 0.073 | 0.56 | 0.59 | 0.42 | 0.6 | 0.5 | 0.69 |
| 6 | 153529163 | rs685341 | -0.11 | 1.369 | 0.282 | 1.111 | -0.003 | -0.001 | -0.001 | 0.09 | 0.37 | 0.41 | 0.45 | 0.39 | 0.35 | 0.43 |
| 6 | 153529372 | rs567323 | 0.073 | 1.431 | 0.347 | 1.126 | 0.001 | -0.011 | -0.011 | 0.532 | 0.35 | 0.43 | 0.47 | 0.28 | 0.39 | 0.19 |
| 6 | 153529919 | rs9478397 | 1.19 | 1.417 | 0.283 | 0.863 | 0.03 | 0.008 | 0.008 | 0 | 0.43 | 0.29 | 0.34 | 0.35 | 0.36 | 0.37 |
| 6 | 153532572 | rs315409 | 0.128 | 1.388 | 0.368 | 0.772 | 0.017 | 0.01 | 0.01 | 0.066 | 0.84 | 0.68 | 0.82 | 0.82 | 0.84 | 0.83 |
| 6 | 153539218 | rs1871400 | -0.18 | 1.354 | 0.275 | 0.714 | -0.001 | 0 | 0 | 0 | 0.34 | 0.36 | 0.43 | 0.33 | 0.35 | 0.39 |
| 6 | 153540142 | rs2155 | 1.008 | 1.626 | 0.242 | 0.835 | -0.028 | -0.027 | -0.027 | 0 | 0.06 | 0 | 0.03 | 0.12 | 0.09 | 0.07 |
| 6 | 153541661 | rs7744062 | 0.528 | 1.099 | 0.388 | 0.507 | -0.012 | -0.004 | -0.004 | 0 | 0.16 | 0.04 | 0.07 | 0.16 | 0.19 | 0.28 |
| 6 | 153543995 | rs317105 | 1.377 | 1.002 | -0.424 | 0.516 | -0.027 | -0.021 | -0.021 | 0.194 | 0.47 | 0.18 | 0.37 | 0.46 | 0.48 | 0.46 |
| 6 | 153546315 | rs317104 | 0.543 | 1.046 | 0.224 | 0.598 | 0.003 | 0.009 | 0.009 | 0 | 0.72 | 0.77 | 0.74 | 0.77 | 0.78 | 0.78 |
| 6 | 153549673 | rs315408 | 0.352 | 0.963 | 0.483 | 0.393 | -0.001 | -0.045 | -0.045 | 0 | 0.16 | 0.18 | 0.11 | 0.01 | 0.14 | 0.08 |
| 6 | 153550147 | rs9479561 | -1.096 | 1.058 | 0.833 | 1.176 | 0.096 | 0.124 | 0.124 | 0 | 0.09 | 0.54 | 0.3 | 0.14 | 0.11 | 0.03 |
| 6 | 153551654 | rs1962749 | 0.546 | 0.855 | 0.081 | 1.226 | 0 | 0.012 | 0.012 | 0 | 0.9 | 0.91 | 0.84 | 0.77 | 0.91 | 0.89 |
| 6 | 153559877 | rs6929221 | -0.937 | 0.669 | 0.609 | 1.264 | 0.003 | 0.058 | 0.058 | 0 | 0.18 | 0.41 | 0.19 | 0.29 | 0.16 | 0.27 |
| 6 | 153566855 | rs7755493 | 0.854 | 0.652 | -0.262 | 0.855 | -0.067 | -0.062 | -0.062 | 0 | 0.34 | 0.07 | 0.3 | 0.29 | 0.39 | 0.4 |
| 6 | 153572513 | rs7762618 | 0.247 | 0.473 | -0.215 | 0.565 | -0.001 | -0.001 | -0.001 | 0.002 | 0.55 | 0.4 | 0.63 | 0.37 | 0.59 | 0.54 |
| 6 | 153576121 | rs11155901 | -0.873 | 0.317 | -0.032 | 0.761 | -0.027 | 0.072 | 0.072 | 0 | 0.23 | 0.59 | 0.2 | 0.37 | 0.17 | 0.3 |
| 6 | 153576772 | rs6908893 | -0.485 | 0.351 | 0.242 | 0.743 | -0.01 | 0.059 | 0.059 | 0 | 0.58 | 0.81 | 0.53 | 0.66 | 0.6 | 0.7 |
| 6 | 153577020 | rs9384087 | 0.172 | 0.378 | 0.328 | 0.773 | -0.031 | 0.152 | 0.152 | 0 | 0.11 | 0.36 | 0.1 | 0.28 | 0.06 | 0.22 |
| 6 | 153577503 | rs6905569 | 0.895 | 0.72 | 0.016 | 0.593 | -0.01 | -0.004 | -0.004 | 0 | 0.35 | 0.22 | 0.33 | 0.29 | 0.43 | 0.41 |
| 6 | 153578655 | rs17083816 | -0.606 | 0.769 | -0.093 | 0.463 | 0.003 | -0.001 | -0.001 | 1.61 | 0.08 | 0.15 | 0.07 | 0.09 | 0.07 | 0.06 |
| 6 | 153586473 | rs7771485 | -0.459 | 0.739 | -0.006 | 0.2 | 0.016 | -0.033 | -0.033 | 0 | 0.22 | 0.61 | 0.18 | 0.09 | 0.21 | 0.17 |
| 6 | 153587328 | rs9383655 | 1.029 | 0.531 | -0.673 | 0.178 | -0.006 | -0.011 | -0.011 | 0 | 0.2 | 0.06 | 0.13 | 0.27 | 0.19 | 0.28 |
| 6 | 153588258 | rs1321602 | -1.573 | 0.306 | -1.011 | 0.241 | -0.059 | -0.016 | -0.016 | 0 | 0.78 | 0.98 | 0.9 | 0.77 | 0.79 | 0.85 |
| 6 | 153590578 | rs7768368 | -0.36 | 0.234 | -0.035 | 0.326 | 0.068 | 0.063 | 0.063 | 1.949 | 0.07 | 0.49 | 0.07 | 0.02 | 0.04 | 0.03 |
| 6 | 153592043 | rs7761260 | -1.116 | 0.032 | -0.217 | 0.701 | -0.03 | 0.063 | 0.063 | 0.473 | 0.18 | 0.62 | 0.15 | 0.25 | 0.12 | 0.15 |
| 6 | 153594954 | rs17083833 | 0.646 | 0.132 | -0.537 | 0.816 | 0.01 | 0.005 | 0.005 | 0 | 0.08 | 0.04 | 0.06 | 0.23 | 0.05 | 0.1 |
| 6 | 153600685 | rs9397597 | 0.09 | 0.327 | -0.548 | 0.046 | -0.02 | -0.009 | -0.009 | 0.044 | 0.38 | 0.22 | 0.27 | 0.59 | 0.36 | 0.49 |
| 6 | 153602202 | rs965500 | -0.497 | 0.245 | -0.264 | -0.071 | -0.006 | 0.06 | 0.06 | 0 | 0.5 | 0.78 | 0.37 | 0.59 | 0.45 | 0.54 |
| 6 | 153620144 | rs1159927 | 0.039 | 0.243 | 0.277 | -0.052 | -0.003 | 0.002 | 0.002 | 0 | 0.4 | 0.33 | 0.28 | 0.59 | 0.35 | 0.5 |
| 6 | 153627062 | rs9371687 | 0.031 | 0.493 | 0.318 | -0.18 | -0.001 | 0.003 | 0.003 | 0 | 0.29 | 0.25 | 0.53 | 0.18 | 0.34 | 0.3 |
| 6 | 153632118 | rs1479438 | -1.391 | 0.558 | -0.447 | -0.145 | -0.028 | 0.046 | 0.046 | 0 | 0.54 | 0.85 | 0.72 | 0.29 | 0.6 | 0.48 |
| 6 | 153632737 | rs9397143 | -1.407 | 0.585 | -0.641 | -0.145 | -0.044 | 0.034 | 0.034 | 0 | 0.55 | 0.92 | 0.73 | 0.29 | 0.6 | 0.48 |
| 6 | 153633163 | rs9371689 | -0.328 | 0.88 | -0.862 | -0.26 | -0.027 | -0.012 | -0.012 | 0.019 | 0.25 | 0.11 | 0.28 | 0.21 | 0.26 | 0.23 |
| 6 | 153637046 | rs9397612 | -0.719 | 0.897 | -0.887 | -0.065 | -0.003 | 0.038 | 0.038 | 0 | 0.46 | 0.68 | 0.67 | 0.27 | 0.5 | 0.43 |
| 6 | 153637126 | rs9371690 | -0.865 | 0.745 | -1.071 | 0.028 | 0.013 | 0.09 | 0.09 | 0 | 0.51 | 0.75 | 0.71 | 0.27 | 0.57 | 0.47 |
| 6 | 153640499 | rs12192110 | -0.457 | 0.674 | -1.061 | 0.477 | -0.01 | 0.006 | 0.006 | 0 | 0.24 | 0.12 | 0.32 | 0.19 | 0.33 | 0.27 |
| 6 | 153653153 | rs2503797 | -0.157 | 0.628 | -1.276 | 0.688 | -0.085 | 0.049 | 0.049 | 0.759 | 0.55 | 0.9 | 0.6 | 0.61 | 0.51 | 0.6 |
| 6 | 153664462 | rs4870193 | 0.3 | 0.952 | -1.13 | 0.965 | 0.001 | 0.002 | 0.002 | 0.369 | 0.08 | 0.07 | 0.13 | 0.18 | 0.18 | 0.17 |
| 6 | 153664718 | rs17083928 | -0.364 | 0.736 | -0.252 | 0.262 | 0.013 | -0.017 | -0.017 | 0 | 0.09 | 0.32 | 0.27 | 0.02 | 0.07 | 0.06 |
| 6 | 153666260 | rs2185276 | -0.493 | 0.702 | -0.372 | 0.309 | -0.075 | 0.073 | 0.073 | 0.704 | 0.54 | 0.89 | 0.59 | 0.63 | 0.5 | 0.6 |
| 6 | 153671375 | rs1038555 | 0.365 | 0.631 | 0.309 | 0.314 | 0.002 | 0.011 | 0.011 | 0 | 0.42 | 0.31 | 0.29 | 0.6 | 0.41 | 0.53 |
| 6 | 153672328 | rs11969504 | -0.275 | 0.61 | 0.348 | 0.284 | 0.023 | 0.002 | 0.002 | 0 | 0.05 | 0.28 | 0.03 | 0 | 0.03 | 0.01 |
| 6 | 153678811 | rs12665473 | 0.777 | 1.074 | -0.401 | 0.156 | -0.031 | 0.025 | 0.025 | 0.149 | 0.16 | 0.06 | 0.15 | 0.13 | 0.21 | 0.16 |
| 6 | 153678935 | rs1684802 | -0.099 | 0.687 | -0.301 | 0.104 | 0.004 | 0.013 | 0.013 | 1.307 | 0.6 | 0.69 | 0.49 | 0.73 | 0.64 | 0.7 |
| 6 | 153688995 | rs1221923 | 1.192 | 0.525 | -0.903 | 0.201 | -0.029 | -0.016 | -0.016 | 0 | 0.27 | 0.02 | 0.12 | 0.25 | 0.21 | 0.2 |
| 6 | 153690928 | rs1221930 | 0.195 | 0.538 | 0.016 | -0.031 | 0.007 | 0.04 | 0.04 | 2.307 | 0.63 | 0.75 | 0.51 | 0.82 | 0.67 | 0.71 |
| 6 | 153693962 | rs2129345 | 0.039 | 0.883 | -0.043 | -0.03 | 0.002 | 0.033 | 0.033 | 3.865 | 0.28 | 0.15 | 0.16 | 0.13 | 0.25 | 0.22 |
| 6 | 153694667 | rs4870194 | -0.703 | 0.952 | 0.362 | 0.537 | 0.009 | 0.044 | 0.044 | 0.4 | 0.05 | 0.2 | 0.25 | 0.07 | 0.05 | 0.05 |
| 6 | 153694723 | rs17083960 | -1.237 | 0.976 | 0.374 | 0.447 | 0.018 | 0.03 | 0.03 | 0.558 | 0.07 | 0.22 | 0.27 | 0.06 | 0.08 | 0.06 |
| 6 | 153695725 | rs1386208 | -0.824 | 0.962 | -0.06 | 0.501 | -0.003 | 0.016 | 0.016 | 0 | 0.06 | 0.18 | 0.25 | 0.06 | 0.05 | 0.05 |
| 6 | 153700256 | rs9791224 | -0.745 | 1.231 | -0.455 | 0.169 | -0.005 | 0.018 | 0.018 | 0 | 0.21 | 0.14 | 0.17 | 0.14 | 0.23 | 0.2 |
| 6 | 153701034 | rs901418 | -1.023 | 1.587 | 0.03 | 0.208 | 0.033 | -0.031 | -0.031 | 0 | 0.23 | 0.38 | 0.21 | 0.18 | 0.26 | 0.14 |
| 6 | 153701111 | rs2790090 | -0.28 | 1.551 | -0.41 | 0.55 | -0.032 | -0.01 | -0.01 | 0.487 | 0.33 | 0.29 | 0.39 | 0.41 | 0.51 | 0.31 |
| 6 | 153704179 | rs17083982 | -0.577 | 0.731 | -0.194 | 0.41 | 0 | 0 | 0 | 0 | 0.19 | 0.18 | 0.31 | 0.13 | 0.13 | 0.17 |
| 6 | 153704558 | rs7738326 | -1.367 | 0.801 | 0.058 | 0.669 | -0.003 | 0.004 | 0.004 | 0.035 | 0.36 | 0.45 | 0.45 | 0.45 | 0.32 | 0.4 |
| 6 | 153707246 | rs6922464 | -1.025 | 1.131 | 0.127 | 0.945 | 0.024 | 0.075 | 0.075 | 3.135 | 0.19 | 0.44 | 0.15 | 0.32 | 0.18 | 0.22 |
| 6 | 153714440 | rs6923536 | -0.065 | 1.045 | -0.695 | 0.596 | 0 | 0.009 | 0.009 | 0.021 | 0.52 | 0.68 | 0.38 | 0.45 | 0.52 | 0.41 |
| 6 | 153724545 | rs1221934 | 0.313 | 1.055 | -1.422 | 0.082 | -0.013 | -0.056 | -0.056 | 0 | 0.36 | 0.1 | 0.26 | 0.58 | 0.35 | 0.37 |
| 6 | 153724684 | rs1237546 | -0.575 | 0.878 | -1.634 | -0.542 | 0.043 | 0.072 | 0.072 | 0 | 0.69 | 0.83 | 0.81 | 0.88 | 0.8 | 0.6 |

| | | | | | | | | | | | | | | | | |
|---|---|---|---|---|---|---|---|---|---|---|---|---|---|---|---|---|
| 6 | 153736996 | rs9479596 | 0.177 | 1.451 | -1.354 | -0.473 | 0.015 | -0.018 | -0.018 | 1.091 | 0.06 | 0.09 | 0.08 | 0.01 | 0.14 | 0.04 |
| 6 | 153737895 | rs961846 | 0.097 | 1.445 | -1.323 | -0.476 | 0.019 | -0.022 | -0.022 | 2.074 | 0.06 | 0.1 | 0.08 | 0.01 | 0.14 | 0.04 |
| 6 | 153742076 | rs270289 | 0.825 | 1.423 | -1.481 | -0.254 | -0.031 | -0.073 | -0.073 | 0.398 | 0.35 | 0.12 | 0.27 | 0.56 | 0.36 | 0.4 |
| 6 | 153742913 | rs4869792 | -0.081 | 1.603 | -0.472 | 0.179 | 0.014 | 0.018 | 0.018 | 0 | 0.21 | 0.28 | 0.41 | 0.29 | 0.24 | 0.16 |
| 6 | 153748051 | rs2054263 | -0.068 | 1.624 | -0.472 | 0.172 | 0.013 | 0.018 | 0.018 | 0 | 0.21 | 0.27 | 0.41 | 0.29 | 0.24 | 0.16 |
| 6 | 153753285 | rs270301 | 0.482 | 1.115 | -0.214 | -0.503 | -0.004 | -0.027 | -0.027 | 0 | 0.62 | 0.38 | 0.76 | 0.85 | 0.67 | 0.65 |
| 6 | 153754313 | rs7758257 | -1.293 | 1.327 | 0.143 | -0.486 | -0.005 | -0.036 | -0.036 | 2.977 | 0.3 | 0.55 | 0.26 | 0.2 | 0.28 | 0.24 |
| 6 | 153754860 | rs270304 | 1.382 | 2.01 | 0.303 | -0.572 | 0.011 | -0.03 | -0.03 | 0 | 0.61 | 0.4 | 0.65 | 0.76 | 0.58 | 0.63 |
| 6 | 153757434 | rs12110552 | -0.817 | 1.635 | 0.561 | -0.659 | 0.081 | -0.031 | -0.031 | 0 | 0.09 | 0.45 | 0.07 | 0.01 | 0.1 | 0.02 |
| 6 | 153757736 | rs270309 | 1.004 | 1.481 | 0.246 | -0.597 | 0.036 | 0.019 | 0.019 | 0 | 0.72 | 0.85 | 0.8 | 0.77 | 0.83 | 0.67 |
| 6 | 153769973 | rs11752411 | -1.535 | 2.138 | 1.073 | -0.689 | 0.008 | -0.032 | -0.032 | 0 | 0.23 | 0.32 | 0.18 | 0.15 | 0.24 | 0.22 |
| 6 | 153770054 | rs270311 | 2.462 | 2.565 | 1.4 | -0.434 | 0.014 | -0.011 | -0.011 | 0 | 0.66 | 0.36 | 0.67 | 0.76 | 0.55 | 0.65 |
| 6 | 153771612 | rs7758954 | -1.496 | 2.181 | 1.287 | -0.383 | 0.008 | -0.031 | -0.031 | 0.308 | 0.22 | 0.32 | 0.18 | 0.15 | 0.24 | 0.22 |
| 6 | 153775310 | rs7761788 | -1.522 | 2.182 | 1.293 | -0.383 | 0.008 | -0.033 | -0.033 | 0.01 | 0.23 | 0.33 | 0.18 | 0.15 | 0.24 | 0.22 |
| 6 | 153782560 | rs12526771 | -0.377 | 2.179 | 1.27 | -0.336 | -0.001 | -0.002 | -0.002 | 0.175 | 0.1 | 0.11 | 0.1 | 0.14 | 0.09 | 0.2 |
| 6 | 153798767 | rs270320 | 1.455 | 2.269 | 1.52 | -0.359 | 0.031 | -0.024 | -0.024 | 0 | 0.84 | 0.7 | 0.89 | 0.91 | 0.79 | 0.86 |
| 6 | 153802064 | rs13208348 | -0.515 | 2.263 | 1.676 | -0.369 | 0.012 | -0.035 | -0.035 | 0 | 0.09 | 0.13 | 0.06 | 0.01 | 0.15 | 0.03 |
| 6 | 153803254 | rs270314 | 1.997 | 2.188 | 1.793 | -0.385 | 0.019 | -0.021 | -0.021 | 0.022 | 0.72 | 0.51 | 0.79 | 0.77 | 0.69 | 0.66 |
| 6 | 153822736 | rs270255 | 2.265 | 2.177 | 1.591 | -0.367 | 0.003 | -0.014 | -0.014 | 0.71 | 0.7 | 0.37 | 0.75 | 0.78 | 0.69 | 0.66 |
| 6 | 153827582 | rs541247 | 0.59 | 2.231 | 1.612 | -0.311 | 0 | 0.004 | 0.004 | 0 | 0.9 | 0.88 | 0.89 | 0.85 | 0.91 | 0.8 |
| 6 | 153828648 | rs270276 | -1.721 | 2.695 | 1.876 | -0.337 | 0.164 | 0.031 | 0.031 | 0 | 0.12 | 0.4 | 0.18 | 0.09 | 0.21 | 0.14 |
| 6 | 153830536 | rs270274 | 0.128 | 2.69 | 1.876 | -0.337 | -0.006 | -0.009 | -0.009 | 0 | 0.05 | 0.03 | 0.05 | 0.08 | 0.07 | 0.11 |
| 6 | 153833568 | rs270271 | 2.247 | 2.679 | 1.875 | -0.345 | 0.069 | 0.015 | 0.015 | 0 | 0.78 | 0.46 | 0.72 | 0.76 | 0.7 | 0.66 |
| 6 | 153844303 | rs654894 | 1.996 | 2.808 | 1.773 | 0.343 | 0.119 | 0.125 | 0.125 | 0 | 0.82 | 0.53 | 0.7 | 0.68 | 0.71 | 0.74 |
| 6 | 153847756 | rs17084250 | -1.465 | 2.814 | 1.861 | 0.338 | 0.156 | -0.01 | -0.01 | 0 | 0.07 | 0.28 | 0.12 | 0.01 | 0.15 | 0.02 |
| 6 | 153848527 | rs581094 | 0.531 | 2.956 | 1.291 | 0.434 | -0.008 | -0.012 | -0.012 | 0 | 0.9 | 0.92 | 0.83 | 0.69 | 0.86 | 0.76 |
| 6 | 153851594 | rs11756366 | 1.917 | 2.449 | 1.113 | 0.169 | 0.011 | 0.007 | 0.007 | 0.099 | 0.66 | 0.63 | 0.62 | 0.59 | 0.49 | 0.58 |
| 6 | 153855165 | rs9371705 | 1.527 | 2.129 | 0.735 | 0.225 | -0.005 | -0.012 | -0.012 | 0 | 0.69 | 0.73 | 0.71 | 0.61 | 0.63 | 0.6 |
| 6 | 153857936 | rs559901 | 1.131 | 2.126 | 0.711 | 0.223 | -0.01 | -0.019 | -0.019 | 0 | 0.75 | 0.84 | 0.76 | 0.68 | 0.7 | 0.71 |
| 6 | 153858614 | rs1674726 | 1.195 | 1.989 | 0.703 | 0.209 | -0.006 | -0.013 | -0.013 | 0 | 0.69 | 0.73 | 0.71 | 0.61 | 0.63 | 0.6 |
| 6 | 153863909 | rs598761 | -0.95 | 1.588 | 0.307 | -0.254 | -0.004 | -0.005 | -0.005 | 0 | 0.41 | 0.51 | 0.29 | 0.27 | 0.38 | 0.27 |
| 6 | 153864063 | rs1674729 | 1.413 | 1.565 | 0.267 | -0.201 | 0.007 | 0.007 | 0.007 | 0.168 | 0.5 | 0.46 | 0.61 | 0.59 | 0.53 | 0.53 |
| 6 | 153865421 | rs1590194 | 0.925 | 1.618 | 0.257 | -0.192 | -0.007 | 0.003 | 0.003 | 0 | 0.59 | 0.52 | 0.64 | 0.6 | 0.58 | 0.57 |
| 6 | 153865679 | rs627056 | 0.329 | 1.154 | 0.101 | -0.12 | -0.004 | -0.001 | -0.001 | 0 | 0.88 | 0.91 | 0.88 | 0.87 | 0.88 | 0.76 |
| 6 | 153865890 | rs2219512 | 0.829 | 1.33 | 0.055 | -0.12 | 0 | -0.001 | -0.001 | 0 | 0.57 | 0.56 | 0.57 | 0.58 | 0.45 | 0.55 |
| 6 | 153875899 | rs476003 | 0.775 | 1.66 | -0.242 | 0.093 | 0.015 | 0.014 | 0.014 | 0 | 0.89 | 0.9 | 0.85 | 0.8 | 0.83 | 0.8 |
| 6 | 153880741 | rs1154029 | 1.185 | 1.108 | -0.515 | 0.018 | 0 | 0.001 | 0.001 | 0 | 0.51 | 0.45 | 0.39 | 0.52 | 0.54 | 0.4 |
| 6 | 153881380 | rs576496 | -0.822 | 0.443 | -0.457 | 0.129 | 0.005 | 0.004 | 0.004 | 0 | 0.38 | 0.41 | 0.45 | 0.27 | 0.29 | 0.4 |
| 6 | 153882337 | rs9384111 | -0.797 | 0.819 | -0.888 | 1.013 | -0.059 | -0.093 | -0.093 | 0 | 0.13 | 0.02 | 0.36 | 0.27 | 0.15 | 0.39 |
| 6 | 153902816 | rs1937834 | 0.434 | 0.38 | -0.529 | 0.444 | 0.001 | 0.004 | 0.004 | 0 | 0.84 | 0.89 | 0.81 | 0.77 | 0.83 | 0.91 |
| 6 | 153915549 | rs4489167 | -1.03 | -0.657 | -0.584 | -0.458 | 0.05 | 0.071 | 0.071 | 0.955 | 0.29 | 0.8 | 0.2 | 0.01 | 0.17 | 0.19 |
| 6 | 153917304 | rs4493751 | -0.927 | -0.68 | -0.056 | -0.468 | -0.013 | -0.062 | -0.062 | 0 | 0.27 | 0.68 | 0.19 | 0.01 | 0.17 | 0.13 |
| 6 | 153924485 | rs3935975 | -1.219 | -0.746 | -0.256 | -0.501 | 0.074 | 0.152 | 0.152 | 0.645 | 0.3 | 0.9 | 0.22 | 0.02 | 0.18 | 0.13 |
| 6 | 153924566 | rs3935974 | -1.219 | -0.746 | -0.256 | -0.501 | 0.074 | 0.152 | 0.152 | 0.645 | 0.3 | 0.9 | 0.21 | 0.02 | 0.18 | 0.13 |
| 6 | 153926453 | rs7754521 | -0.792 | -0.708 | 0.106 | -0.482 | -0.044 | -0.131 | -0.131 | 0 | 0.26 | 0.54 | 0.18 | 0.01 | 0.17 | 0.12 |
| 6 | 153928396 | rs9479657 | -0.842 | -0.836 | -0.147 | -0.474 | 0.042 | 0.033 | 0.033 | 0.848 | 0.21 | 0.85 | 0.14 | 0.01 | 0.08 | 0.03 |
| 6 | 153930390 | rs7757781 | -0.776 | -0.654 | -0.186 | 0.357 | 0.005 | -0.002 | -0.002 | 0 | 0.3 | 0.68 | 0.28 | 0.33 | 0.17 | 0.34 |
| 6 | 153932450 | rs11753844 | -0.776 | -0.654 | -0.186 | 0.357 | 0.005 | -0.002 | -0.002 | 0.373 | 0.3 | 0.68 | 0.28 | 0.33 | 0.18 | 0.34 |
| 6 | 153945567 | rs9479664 | 0.054 | -0.608 | -0.277 | 0.255 | -0.003 | 0.007 | 0.007 | 0 | 0.08 | 0.03 | 0.07 | 0.01 | 0.1 | 0.1 |
| 6 | 153955406 | rs4870208 | -0.661 | -0.693 | 0.014 | 0.257 | -0.035 | 0.02 | 0.02 | 0 | 0.28 | 0.55 | 0.27 | 0.33 | 0.18 | 0.34 |
| 6 | 153955693 | rs13218904 | 0.985 | -0.415 | -1.063 | 0.356 | -0.065 | -0.119 | -0.119 | 0 | 0.2 | 0.02 | 0.16 | 0.33 | 0.21 | 0.25 |
| 6 | 153966047 | rs12664772 | -1.466 | -0.602 | -0.576 | 0.33 | 0.046 | -0.012 | -0.012 | 0 | 0.4 | 0.91 | 0.37 | 0.34 | 0.28 | 0.45 |
| 6 | 153977548 | rs9478457 | -1.266 | -0.48 | -0.157 | 0.378 | 0.022 | 0.006 | 0.006 | 0.006 | 0.38 | 0.72 | 0.35 | 0.34 | 0.28 | 0.43 |
| 6 | 153978001 | rs7759898 | -0.752 | -0.396 | -0.006 | 0.643 | -0.04 | 0.026 | 0.026 | 0 | 0.28 | 0.49 | 0.26 | 0.33 | 0.18 | 0.32 |
| 6 | 153985502 | rs4870213 | -0.646 | -0.311 | 0.318 | 0.729 | -0.015 | 0.013 | 0.013 | 0.013 | 0.25 | 0.38 | 0.21 | 0.28 | 0.17 | 0.26 |
| 6 | 153990281 | rs11155929 | -1.02 | -0.369 | 0.46 | 0.694 | -0.039 | 0.017 | 0.017 | 0.01 | 0.28 | 0.54 | 0.23 | 0.28 | 0.17 | 0.27 |
| 6 | 153999020 | rs9371714 | 0.008 | -0.426 | 0.208 | 0.662 | 0 | -0.002 | -0.002 | 0 | 0.13 | 0.13 | 0.14 | 0.23 | 0.16 | 0.13 |
| 6 | 154000377 | rs6906652 | -0.287 | -0.247 | 0.096 | 0.405 | 0.004 | 0.011 | 0.011 | 0 | 0.51 | 0.56 | 0.48 | 0.68 | 0.45 | 0.52 |
| 6 | 154005563 | rs9384125 | 0.391 | -0.198 | 0.05 | 0.471 | 0 | -0.002 | -0.002 | 0.102 | 0.13 | 0.13 | 0.14 | 0.23 | 0.16 | 0.13 |
| 6 | 154005613 | rs2186140 | -0.171 | -0.101 | -0.031 | 0.06 | 0 | 0 | 0 | 0 | 0.54 | 0.56 | 0.68 | 0.59 | 0.47 | 0.59 |
| 6 | 154005736 | rs6922133 | 1.164 | -0.098 | -0.082 | 0.049 | -0.003 | -0.016 | -0.016 | 0 | 0.11 | 0.07 | 0.14 | 0.23 | 0.16 | 0.11 |
| 6 | 154011024 | rs2000371 | -1.077 | -0.073 | -0.107 | 0.009 | -0.069 | -0.022 | -0.022 | 0 | 0.68 | 0.93 | 0.79 | 0.7 | 0.63 | 0.66 |
| 6 | 154011615 | rs9371718 | 0.768 | -0.173 | -0.06 | 0.005 | 0 | 0 | 0 | 0 | 0.13 | 0.14 | 0.14 | 0.23 | 0.16 | 0.13 |
| 6 | 154017197 | rs1937600 | 0.901 | -0.292 | 0.066 | -0.088 | 0.006 | 0.028 | 0.028 | 0.446 | 0.15 | 0.1 | 0.3 | 0.02 | 0.17 | 0.1 |
| 6 | 154022718 | rs9397637 | -0.449 | -0.35 | 0.117 | -0.125 | -0.024 | -0.03 | -0.03 | 0 | 0.31 | 0.5 | 0.44 | 0.26 | 0.33 | 0.26 |
| 6 | 154025836 | rs1937594 | 1.762 | -0.362 | 0.007 | -0.114 | 0.012 | -0.05 | -0.05 | 1.07 | 0.23 | 0.08 | 0.23 | 0.34 | 0.15 | 0.34 |
| 6 | 154033449 | rs1954623 | -1.713 | -0.018 | -0.029 | -0.116 | -0.002 | -0.047 | -0.047 | 7.459 | 0.75 | 0.91 | 0.76 | 0.66 | 0.83 | 0.66 |
| 6 | 154036895 | rs1937590 | 0.5 | -0.286 | 0.21 | 0.117 | -0.005 | 0.002 | 0.002 | 0 | 0.12 | 0.12 | 0.09 | 0.1 | 0.16 | 0.07 |
| 6 | 154042101 | rs17084425 | 0.501 | -0.189 | -0.126 | 0.1 | -0.003 | 0.009 | 0.009 | 0 | 0.14 | 0.23 | 0.14 | 0.23 | 0.16 | 0.15 |

| | | | | | | | | | | | | | | | | |
|---|---|---|---|---|---|---|---|---|---|---|---|---|---|---|---|---|
| 6 | 154046247 | rs6939855 | -1.058 | -0.111 | -0.264 | 0.083 | -0.068 | -0.021 | -0.021 | 0 | 0.68 | 0.93 | 0.79 | 0.7 | 0.63 | 0.65 |
| 6 | 154050494 | rs9397645 | 0.63 | -0.403 | -0.051 | 0.133 | 0.002 | -0.013 | -0.013 | 0 | 0.13 | 0.1 | 0.12 | 0.22 | 0.16 | 0.13 |
| 6 | 154055754 | rs1316368 | 0.201 | -0.407 | 0.022 | 0.348 | 0.008 | 0.034 | 0.034 | 0.01 | 0.18 | 0.35 | 0.18 | 0.27 | 0.2 | 0.2 |
| 6 | 154057114 | rs2096411 | 1.125 | -0.382 | -0.045 | 0.344 | -0.01 | -0.032 | -0.032 | 2.418 | 0.12 | 0.04 | 0.12 | 0.22 | 0.18 | 0.11 |
| 6 | 154058060 | rs10457894 | 1.807 | -0.448 | -0.118 | 0.296 | 0.035 | -0.033 | -0.033 | 0.945 | 0.23 | 0.08 | 0.21 | 0.29 | 0.14 | 0.34 |
| 6 | 154059970 | rs1937588 | -0.402 | -0.245 | -0.039 | 0.132 | -0.006 | -0.03 | -0.03 | 0 | 0.33 | 0.47 | 0.48 | 0.25 | 0.38 | 0.26 |
| 6 | 154076643 | rs1937580 | -0.51 | -0.25 | 0.03 | 0.105 | -0.055 | -0.037 | -0.037 | 0 | 0.61 | 0.82 | 0.76 | 0.56 | 0.54 | 0.67 |
| 6 | 154082228 | rs1937645 | 0.414 | -0.253 | -0.06 | 0.183 | -0.01 | 0.007 | 0.007 | 0.012 | 0.42 | 0.5 | 0.4 | 0.54 | 0.34 | 0.55 |
| 6 | 154093266 | rs6557316 | -0.51 | -0.234 | -0.435 | 0.144 | 0.001 | -0.025 | -0.025 | 0 | 0.33 | 0.71 | 0.32 | 0.49 | 0.38 | 0.33 |
| 6 | 154099619 | rs1892361 | -0.395 | -0.234 | -0.396 | 0.147 | -0.001 | 0.001 | 0.001 | 0.029 | 0.32 | 0.64 | 0.32 | 0.49 | 0.38 | 0.31 |
| 6 | 154100603 | rs9371307 | -0.176 | -0.195 | -0.663 | 0.078 | -0.037 | 0.068 | 0.068 | 0 | 0.57 | 0.79 | 0.53 | 0.76 | 0.52 | 0.67 |
| 6 | 154105011 | rs1937631 | -1.052 | 0.077 | -0.635 | 0.273 | -0.076 | -0.025 | -0.025 | 0.236 | 0.73 | 0.92 | 0.88 | 0.78 | 0.72 | 0.81 |
| 6 | 154111389 | rs9397655 | 0.097 | 0.061 | -0.287 | 0.294 | 0.006 | -0.007 | -0.007 | 0 | 0.53 | 0.36 | 0.67 | 0.53 | 0.49 | 0.59 |
| 6 | 154112042 | rs1892359 | 0.442 | -0.23 | -0.181 | 0.285 | -0.004 | -0.004 | -0.004 | 3.117 | 0.42 | 0.44 | 0.56 | 0.29 | 0.31 | 0.41 |
| 6 | 154112263 | rs1892356 | 0.542 | -0.203 | -0.174 | 0.295 | -0.006 | -0.005 | -0.005 | 0.96 | 0.41 | 0.44 | 0.55 | 0.29 | 0.29 | 0.39 |
| 6 | 154116716 | rs1748293 | 0.348 | -0.16 | -0.259 | 0.107 | 0.004 | 0.024 | 0.024 | 0 | 0.66 | 0.52 | 0.67 | 0.52 | 0.59 | 0.62 |
| 6 | 154121293 | rs1781633 | -0.408 | -0.121 | -0.111 | 0.055 | -0.001 | 0.003 | 0.003 | 0.042 | 0.79 | 0.7 | 0.81 | 0.76 | 0.77 | 0.78 |
| 6 | 154126171 | rs1781624 | 1.237 | -0.08 | -0.305 | 0.134 | -0.117 | -0.102 | -0.102 | 0.334 | 0.23 | 0.03 | 0.12 | 0.27 | 0.29 | 0.22 |
| 6 | 154126730 | rs9397657 | 0.777 | 0.002 | -0.097 | 0.057 | -0.003 | -0.007 | -0.007 | 0 | 0.13 | 0.09 | 0.13 | 0.23 | 0.19 | 0.17 |
| 6 | 154126738 | rs17084501 | 0.417 | -0.093 | -0.049 | -0.161 | 0.009 | -0.065 | -0.065 | 0 | 0.14 | 0.25 | 0.33 | 0.01 | 0.18 | 0.05 |
| 6 | 154135968 | rs1781619 | -0.076 | -0.576 | 0.08 | 0.419 | -0.001 | -0.001 | -0.001 | 0 | 0.79 | 0.73 | 0.82 | 0.78 | 0.77 | 0.79 |
| 6 | 154140199 | rs10485262 | 1.005 | -1.272 | 0.191 | 0.758 | 0.034 | -0.014 | -0.014 | 1.771 | 0.23 | 0.14 | 0.2 | 0.3 | 0.09 | 0.3 |
| 6 | 154155918 | rs2350402 | -0.75 | -1.009 | 0 | 0.808 | 0.018 | 0.028 | 0.028 | 0.236 | 0.57 | 0.83 | 0.49 | 0.71 | 0.42 | 0.72 |
| 6 | 154159710 | rs11752884 | 0.543 | -1.045 | 0.007 | 0.849 | -0.001 | -0.001 | -0.001 | 0.041 | 0.13 | 0.1 | 0.13 | 0.2 | 0.18 | 0.16 |
| 6 | 154165415 | rs7762407 | 0.868 | -1.115 | 0.074 | 0.851 | -0.006 | -0.001 | -0.001 | 0 | 0.28 | 0.26 | 0.28 | 0.29 | 0.16 | 0.34 |
| 6 | 154168881 | rs9371753 | -0.656 | -1.108 | 0.121 | 0.875 | -0.026 | 0.058 | 0.058 | 0.065 | 0.27 | 0.5 | 0.2 | 0.42 | 0.27 | 0.34 |
| 6 | 154171882 | rs9479721 | 0.809 | -1.056 | 0.015 | 0.901 | -0.005 | -0.001 | -0.001 | 0 | 0.29 | 0.26 | 0.28 | 0.3 | 0.16 | 0.34 |
| 6 | 154173845 | rs4869813 | -0.028 | -0.431 | -0.233 | 0.901 | -0.032 | 0.063 | 0.063 | 0 | 0.12 | 0.29 | 0.07 | 0.23 | 0.08 | 0.17 |
| 6 | 154182625 | rs6927084 | -0.182 | -0.034 | -0.976 | 0.516 | 0.017 | 0.03 | 0.03 | 0.341 | 0.57 | 0.81 | 0.48 | 0.72 | 0.42 | 0.72 |
| 6 | 154192175 | rs2065139 | -0.726 | -0.123 | 0.081 | 0.594 | -0.024 | 0.028 | 0.028 | 0.151 | 0.14 | 0.41 | 0.08 | 0.15 | 0.11 | 0.17 |
| 6 | 154198820 | rs689219 | -0.451 | 0.39 | -0.428 | 0.567 | -0.014 | -0.008 | -0.008 | 0.295 | 0.82 | 0.94 | 0.93 | 0.84 | 0.8 | 0.81 |
| 6 | 154199581 | rs7739612 | 0.244 | 0.968 | -1.075 | 0.312 | -0.055 | -0.007 | -0.007 | 0 | 0.14 | 0.03 | 0.1 | 0.08 | 0.16 | 0.08 |
| 6 | 154204182 | rs11155944 | 0.228 | 0.348 | -1.123 | 0.574 | -0.001 | -0.009 | -0.009 | 0.018 | 0.23 | 0.23 | 0.43 | 0.41 | 0.25 | 0.4 |
| 6 | 154213973 | rs9371762 | 0.249 | 0.414 | -1.156 | 0.628 | -0.001 | -0.008 | -0.008 | 0 | 0.23 | 0.23 | 0.43 | 0.42 | 0.25 | 0.39 |
| 6 | 154219177 | rs9384159 | 0.392 | 0.415 | -1.17 | 0.626 | 0.008 | -0.032 | -0.032 | 0 | 0.2 | 0.1 | 0.4 | 0.35 | 0.21 | 0.38 |
| 6 | 154221214 | rs581564 | 0.09 | 0.117 | -0.589 | 0.319 | 0 | -0.004 | -0.004 | 0 | 0.55 | 0.41 | 0.48 | 0.43 | 0.54 | 0.41 |
| 6 | 154237443 | rs12202611 | -0.286 | -0.482 | -0.116 | 0.459 | 0.015 | -0.027 | -0.027 | 0 | 0.12 | 0.35 | 0.24 | 0.06 | 0.09 | 0.04 |
| 6 | 154237937 | rs4870255 | 0.993 | -0.493 | -0.305 | 0.441 | 0.072 | -0.061 | -0.061 | 0.098 | 0.26 | 0.06 | 0.26 | 0.28 | 0.18 | 0.22 |
| 6 | 154263279 | rs282096 | 0.542 | 0.477 | 0.06 | 0.69 | -0.001 | 0.036 | 0.036 | 0.232 | 0.64 | 0.49 | 0.44 | 0.44 | 0.61 | 0.48 |
| 6 | 154266565 | rs7745695 | -0.637 | 0.53 | 0.5 | 0.74 | -0.016 | 0.033 | 0.033 | 0 | 0.34 | 0.44 | 0.52 | 0.49 | 0.35 | 0.51 |
| 6 | 154266709 | rs9397673 | 0.037 | 0.525 | 0.356 | 0.913 | 0.008 | -0.007 | -0.007 | 0.381 | 0.23 | 0.17 | 0.47 | 0.34 | 0.25 | 0.42 |
| 6 | 154266737 | rs7745892 | -1.092 | 0.485 | 0.354 | 0.888 | -0.002 | 0.02 | 0.02 | 0.125 | 0.36 | 0.53 | 0.56 | 0.56 | 0.39 | 0.52 |
| 6 | 154267147 | rs2169411 | 0.771 | 0.424 | 0.247 | 0.836 | -0.004 | -0.021 | -0.021 | 0 | 0.16 | 0.09 | 0.28 | 0.33 | 0.2 | 0.41 |
| 6 | 154269402 | rs654359 | 0.793 | 0.086 | -0.414 | 0.24 | 0.003 | 0.003 | 0.003 | 0 | 0.4 | 0.45 | 0.28 | 0.37 | 0.38 | 0.37 |
| 6 | 154271235 | rs7744543 | 1.063 | 0.026 | -0.392 | 0.217 | -0.024 | -0.04 | -0.04 | 0 | 0.08 | 0.13 | 0.19 | 0 | 0.05 | 0.02 |
| 6 | 154274843 | rs574350 | 0.429 | 0.041 | -0.515 | 0.167 | 0.001 | 0.002 | 0.002 | 0 | 0.45 | 0.56 | 0.38 | 0.43 | 0.49 | 0.49 |
| 6 | 154278203 | rs613515 | -0.712 | 0.168 | -0.949 | -0.03 | 0.003 | 0.023 | 0.023 | 0.014 | 0.68 | 0.9 | 0.86 | 0.77 | 0.74 | 0.82 |
| 6 | 154282201 | rs546536 | -0.733 | 0.167 | -0.95 | -0.031 | 0.001 | 0.021 | 0.021 | 0.02 | 0.69 | 0.9 | 0.86 | 0.77 | 0.74 | 0.82 |
| 6 | 154283838 | rs11759340 | 1.134 | 0.155 | -0.918 | 0.003 | -0.009 | -0.018 | -0.018 | 0 | 0.07 | 0.03 | 0.04 | 0.16 | 0.1 | 0.08 |
| 6 | 154288032 | rs6900805 | -0.778 | 0.189 | -0.925 | -0.045 | 0.004 | 0.021 | 0.021 | 0.011 | 0.69 | 0.9 | 0.86 | 0.77 | 0.74 | 0.83 |
| 6 | 154294803 | rs10457900 | -0.084 | 0.591 | -0.441 | 0.095 | -0.015 | -0.004 | -0.004 | 0.26 | 0.11 | 0.24 | 0.21 | 0.07 | 0.09 | 0.07 |
| 6 | 154294917 | rs10457901 | -0.084 | 0.591 | -0.441 | 0.095 | -0.015 | -0.004 | -0.004 | 0 | 0.11 | 0.24 | 0.21 | 0.07 | 0.09 | 0.07 |
| 6 | 154297720 | rs10457902 | -1.039 | 0.701 | -0.503 | 0.154 | 0.063 | 0.047 | 0.047 | 0 | 0.08 | 0.21 | 0.12 | 0.13 | 0.14 | 0.08 |
| 6 | 154307735 | rs9371769 | -0.303 | 0.577 | -0.708 | 0.582 | -0.013 | -0.034 | -0.034 | 0 | 0.27 | 0.17 | 0.57 | 0.43 | 0.36 | 0.46 |
| 6 | 154309594 | rs1675903 | -0.487 | 0.564 | -0.65 | 0.715 | -0.002 | 0.014 | 0.014 | 0 | 0.89 | 0.9 | 0.94 | 0.71 | 0.87 | 0.84 |
| 6 | 154312116 | rs13220748 | 1.897 | 0.305 | -1.043 | 0.416 | -0.015 | 0.075 | 0.075 | 0 | 0.18 | 0 | 0.07 | 0.05 | 0.12 | 0.06 |
| 6 | 154314916 | rs6914262 | -0.944 | 0.417 | -0.146 | 0.856 | 0.014 | 0.053 | 0.053 | 0.105 | 0.31 | 0.43 | 0.6 | 0.5 | 0.4 | 0.53 |
| 6 | 154315367 | rs1675902 | -0.555 | 0.422 | -0.236 | 0.879 | -0.021 | -0.044 | -0.044 | 0 | 0.92 | 0.99 | 0.96 | 0.78 | 0.9 | 0.91 |
| 6 | 154319434 | rs1116706 | 0.444 | 0.394 | -0.193 | 0.674 | 0.001 | 0.022 | 0.022 | 0 | 0.17 | 0.18 | 0.29 | 0.38 | 0.2 | 0.41 |
| 6 | 154319802 | rs4870261 | -0.066 | 0.388 | -0.219 | 0.516 | -0.007 | 0.018 | 0.018 | 0 | 0.11 | 0.08 | 0.29 | 0.06 | 0.16 | 0.05 |
| 6 | 154323909 | rs9478495 | -0.445 | 0.355 | -0.119 | 0.499 | -0.001 | -0.026 | -0.026 | 0 | 0.13 | 0.23 | 0.3 | 0.06 | 0.16 | 0.05 |
| 6 | 154326053 | rs1304725 | 1.08 | 0.355 | -0.196 | 0.469 | 0.002 | -0.011 | -0.011 | 0 | 0.6 | 0.46 | 0.35 | 0.28 | 0.5 | 0.38 |
| 6 | 154335150 | rs712244 | -0.075 | 0.369 | -0.544 | 0.671 | -0.08 | -0.166 | -0.166 | 0 | 0.22 | 0.02 | 0.31 | 0.58 | 0.3 | 0.5 |
| 6 | 154360797 | rs1799971 | 0.457 | 0.531 | -0.207 | 0.704 | -0.041 | -0.096 | -0.096 | 3.516 | 0.14 | 0.01 | 0.2 | 0.39 | 0.16 | 0.42 |
| 6 | 154368786 | rs509544 | -0.797 | 0.614 | 0.128 | 0.356 | -0.012 | 0.02 | 0.02 | 0 | 0.13 | 0.1 | 0.31 | 0.05 | 0.2 | 0.06 |
| 6 | 154368998 | rs4870266 | 0.511 | 0.295 | -0.305 | 0.048 | 0.011 | -0.058 | -0.058 | 0 | 0.16 | 0.01 | 0.08 | 0.19 | 0.07 | 0.19 |
| 6 | 154372692 | rs499796 | -1.22 | 0.672 | 0.582 | -0.377 | 0.029 | -0.055 | -0.055 | 0 | 0.16 | 0.39 | 0.34 | 0.05 | 0.21 | 0.06 |
| 6 | 154377925 | rs3778146 | -0.82 | 0.737 | 0.467 | -0.339 | 0.016 | 0 | 0 | 0 | 0.06 | 0.11 | 0.13 | 0.05 | 0.15 | 0.03 |
| 6 | 154393138 | rs589046 | -1.721 | 0.701 | 0.635 | -0.341 | 0.021 | -0.005 | -0.005 | 0 | 0.2 | 0.55 | 0.36 | 0.14 | 0.24 | 0.12 |

| | | | | | | | | | | | | | | | | |
|---|---|---|---|---|---|---|---|---|---|---|---|---|---|---|---|---|
| 6 | 154393884 | rs3778153 | -0.645 | 0.945 | 0.163 | -0.237 | 0.026 | 0 | 0 | 0.229 | 0.06 | 0.1 | 0.13 | 0.05 | 0.15 | 0.03 |
| 6 | 154407967 | rs563649 | -0.338 | 0.497 | 0.196 | -0.167 | -0.005 | 0 | 0 | 0 | 0.11 | 0.11 | 0.21 | 0.09 | 0.09 | 0.08 |
| 6 | 154414446 | rs540825 | -1.68 | 0.918 | -0.189 | -0.824 | -0.052 | 0.125 | 0.125 | 0.057 | 0.75 | 0.95 | 0.83 | 0.95 | 0.77 | 0.87 |
| 6 | 154414563 | rs675026 | -1.204 | 0.907 | 0.104 | -0.83 | -0.016 | 0.028 | 0.028 | 0 | 0.71 | 0.73 | 0.73 | 0.9 | 0.68 | 0.85 |
| 6 | 154414573 | rs562859 | -1.199 | 0.858 | 0.377 | -0.855 | 0.012 | -0.047 | -0.047 | 0 | 0.7 | 0.63 | 0.72 | 0.9 | 0.68 | 0.85 |
| 6 | 154415434 | rs660756 | -1.191 | 0.84 | 0.374 | -0.859 | 0.011 | -0.046 | -0.046 | 0 | 0.7 | 0.63 | 0.72 | 0.9 | 0.68 | 0.85 |
| 6 | 154417745 | rs12203621 | 0.222 | 0.701 | 0.136 | -0.891 | -0.002 | 0.001 | 0.001 | 0 | 0.08 | 0.08 | 0.02 | 0.1 | 0.05 | 0.07 |
| 6 | 154422744 | rs647192 | -1.735 | 0.759 | 0.052 | -0.887 | -0.04 | 0.108 | 0.108 | 0 | 0.74 | 0.93 | 0.83 | 0.93 | 0.77 | 0.87 |
| 6 | 154424321 | rs9322447 | 0.143 | 0.665 | 0.304 | -1.057 | -0.004 | 0.054 | 0.054 | 0.087 | 0.47 | 0.35 | 0.45 | 0.8 | 0.54 | 0.67 |
| 6 | 154428537 | rs650825 | 1.244 | 1.148 | 0.491 | -0.845 | -0.004 | 0.022 | 0.022 | 0 | 0.26 | 0.14 | 0.19 | 0.08 | 0.23 | 0.13 |
| 6 | 154441864 | rs558948 | 1.219 | 1.045 | 0.006 | -0.852 | -0.008 | 0.057 | 0.057 | 0.044 | 0.27 | 0.13 | 0.18 | 0.07 | 0.23 | 0.14 |
| 6 | 154445215 | rs10485058 | 0.621 | 1.072 | -0.096 | -0.749 | -0.008 | 0.004 | 0.004 | 0 | 0.07 | 0.04 | 0.08 | 0.04 | 0.16 | 0.05 |
| 6 | 154458207 | rs2010884 | 0.333 | 0.937 | -0.429 | -0.37 | 0 | -0.046 | -0.046 | 0 | 0.23 | 0.11 | 0.16 | 0.37 | 0.23 | 0.18 |
| 6 | 154458856 | rs655059 | 0.746 | 0.589 | -0.403 | -0.393 | 0.012 | 0.042 | 0.042 | 0 | 0.08 | 0.01 | 0.23 | 0 | 0.06 | 0.1 |
| 6 | 154461123 | rs13203628 | 0.913 | 1.081 | -0.108 | -0.178 | 0 | -0.04 | -0.04 | 0.322 | 0.29 | 0.34 | 0.2 | 0.07 | 0.24 | 0.13 |
| 6 | 154461274 | rs12660296 | 0.909 | 1.083 | -0.107 | -0.177 | 0 | -0.038 | -0.038 | 0 | 0.29 | 0.34 | 0.2 | 0.07 | 0.24 | 0.13 |
| 6 | 154462086 | rs9397687 | -0.361 | 0.765 | -0.411 | 0.286 | -0.001 | -0.001 | -0.001 | 0 | 0.39 | 0.38 | 0.44 | 0.4 | 0.44 | 0.25 |
| 6 | 154462378 | rs518596 | -0.305 | 0.769 | -0.633 | 0.083 | 0.005 | 0.027 | 0.027 | 0 | 0.57 | 0.37 | 0.65 | 0.82 | 0.64 | 0.81 |
| 6 | 154463054 | rs6557337 | -0.225 | 0.856 | -0.624 | 0.625 | -0.01 | -0.003 | -0.003 | 0 | 0.43 | 0.51 | 0.38 | 0.38 | 0.37 | 0.54 |
| 6 | 154463968 | rs483481 | 0.194 | 1.194 | -0.901 | 0.537 | 0.002 | 0.053 | 0.053 | 0.583 | 0.46 | 0.32 | 0.5 | 0.27 | 0.44 | 0.3 |
| 6 | 154464330 | rs569284 | -1.282 | 1.014 | -0.159 | 0.676 | -0.005 | 0.014 | 0.014 | 0 | 0.07 | 0.15 | 0.13 | 0.08 | 0.05 | 0.11 |
| 6 | 154465199 | rs12207811 | -0.579 | 1.038 | -0.331 | 1.518 | 0.001 | 0.049 | 0.049 | 0 | 0.08 | 0.09 | 0.06 | 0.42 | 0.08 | 0.27 |
| 6 | 154466818 | rs4869817 | 0.13 | 1.101 | -0.13 | 0.566 | 0 | -0.001 | -0.001 | 1.067 | 0.48 | 0.46 | 0.42 | 0.33 | 0.51 | 0.31 |
| 6 | 154470217 | rs9397178 | -0.017 | 1.179 | -0.051 | 0.717 | 0 | 0.061 | 0.061 | 0 | 0.08 | 0.1 | 0.06 | 0.42 | 0.08 | 0.27 |
| 6 | 154472327 | rs4870268 | 0.189 | 1.527 | 0.411 | 0.405 | -0.001 | 0.002 | 0.002 | 0.398 | 0.44 | 0.43 | 0.32 | 0.3 | 0.46 | 0.19 |
| 6 | 154474871 | rs9383693 | -0.17 | 1.689 | 0.429 | 0.327 | -0.001 | -0.001 | -0.001 | 1.002 | 0.53 | 0.53 | 0.39 | 0.49 | 0.56 | 0.41 |
| 6 | 154476528 | rs9479767 | -0.145 | 1.594 | 0.579 | 0.233 | -0.003 | -0.002 | -0.002 | 0 | 0.53 | 0.51 | 0.39 | 0.49 | 0.56 | 0.41 |
| 6 | 154477913 | rs17277929 | 1.188 | 1.32 | 0.231 | -0.143 | 0.019 | 0.062 | 0.062 | 3.84 | 0.19 | 0.03 | 0.1 | 0.06 | 0.15 | 0.07 |
| 6 | 154478440 | rs2236256 | -0.468 | 1.349 | 0.329 | -0.046 | -0.003 | -0.002 | -0.002 | 0.174 | 0.53 | 0.51 | 0.39 | 0.49 | 0.56 | 0.41 |
| 6 | 154478893 | rs2236259 | -0.532 | 1.359 | 0.216 | -0.04 | 0.002 | -0.012 | -0.012 | 0 | 0.56 | 0.75 | 0.42 | 0.49 | 0.57 | 0.41 |
| 6 | 154483146 | rs9479769 | 0.175 | 1.324 | 0.157 | -0.001 | 0 | 0.009 | 0.009 | 0.447 | 0.44 | 0.41 | 0.31 | 0.3 | 0.44 | 0.2 |
| 6 | 154485640 | rs12199124 | -0.125 | 1.209 | 0.147 | 0.229 | 0.002 | 0.037 | 0.037 | 0 | 0.1 | 0.06 | 0.07 | 0.41 | 0.08 | 0.26 |
| 6 | 154487421 | rs2281617 | -0.765 | 1.418 | 0.586 | 0.07 | 0 | 0.12 | 0.12 | 0 | 0.16 | 0.27 | 0.12 | 0.41 | 0.14 | 0.29 |
| 6 | 154490658 | rs6938037 | -1.235 | 1.518 | 0.4 | -0.164 | -0.017 | 0.154 | 0.154 | 0 | 0.64 | 0.88 | 0.48 | 0.9 | 0.65 | 0.67 |
| 6 | 154491137 | rs12208947 | -0.468 | 1.483 | 0.473 | -0.138 | 0 | 0.051 | 0.051 | 0 | 0.15 | 0.16 | 0.11 | 0.41 | 0.14 | 0.29 |
| 6 | 154492854 | rs7745499 | -0.861 | 1.521 | 0.595 | -0.316 | -0.012 | 0.076 | 0.076 | 0 | 0.6 | 0.8 | 0.43 | 0.71 | 0.59 | 0.47 |
| 6 | 154500139 | rs13437608 | 0.089 | 1.461 | 0.817 | -0.291 | 0.013 | -0.027 | -0.027 | 0 | 0.05 | 0.16 | 0.04 | 0 | 0.06 | 0.02 |
| 6 | 154501125 | rs17278409 | 0.347 | 1.415 | 0.699 | -0.408 | -0.002 | -0.019 | -0.019 | 0 | 0.27 | 0.27 | 0.16 | 0.06 | 0.22 | 0.11 |
| 6 | 154503512 | rs11965988 | -0.098 | 1.486 | 1.464 | 0.033 | 0.057 | -0.008 | -0.008 | 0 | 0.05 | 0.2 | 0.05 | 0 | 0.06 | 0.02 |
| 6 | 154513198 | rs4626436 | -0.431 | 1.471 | 1.448 | 0.018 | 0.022 | -0.039 | -0.039 | 0.256 | 0.08 | 0.25 | 0.06 | 0 | 0.06 | 0.04 |
| 6 | 154517262 | rs11970612 | -0.149 | 1.514 | 1.406 | 0.025 | 0.068 | -0.001 | -0.001 | 0 | 0.05 | 0.23 | 0.05 | 0 | 0.06 | 0.02 |
| 6 | 154522661 | rs9322451 | -1.214 | 1.513 | 1.288 | 0.247 | 0.016 | 0.08 | 0.08 | 0.536 | 0.18 | 0.26 | 0.15 | 0.61 | 0.2 | 0.49 |
| 6 | 154522740 | rs1852629 | -0.209 | 1.363 | 1.152 | -0.081 | 0 | -0.016 | -0.016 | 0.009 | 0.57 | 0.57 | 0.41 | 0.89 | 0.59 | 0.64 |
| 6 | 154530980 | rs6913456 | -0.111 | 1.19 | 0.963 | 0.141 | 0.009 | -0.004 | -0.004 | 0.039 | 0.57 | 0.65 | 0.42 | 0.49 | 0.62 | 0.44 |
| 6 | 154534412 | rs790252 | 0.81 | 0.898 | 0.735 | -0.14 | 0.014 | 0.058 | 0.058 | 0 | 0.25 | 0.13 | 0.45 | 0.1 | 0.22 | 0.26 |
| 6 | 154535578 | rs9478526 | -0.083 | 0.983 | 0.985 | -0.108 | 0.021 | -0.024 | -0.024 | 0.78 | 0.05 | 0.16 | 0.04 | 0 | 0.07 | 0.02 |
| 6 | 154540306 | rs790262 | -1.03 | 0.983 | 1.07 | -0.109 | 0.021 | 0.101 | 0.101 | 0 | 0.5 | 0.71 | 0.63 | 0.32 | 0.46 | 0.52 |
| 6 | 154542491 | rs2272381 | -1.128 | 1.043 | 0.965 | -0.097 | -0.001 | 0 | 0 | 0 | 0.18 | 0.23 | 0.12 | 0.21 | 0.17 | 0.23 |
| 6 | 154548429 | rs2103277 | -0.012 | 0.893 | 0.75 | -0.173 | 0.015 | 0.051 | 0.051 | 0 | 0.35 | 0.26 | 0.51 | 0.1 | 0.33 | 0.28 |
| 6 | 154550853 | rs9397697 | 0.8 | 0.854 | 0.682 | -0.119 | 0.017 | 0.068 | 0.068 | 0 | 0.26 | 0.09 | 0.47 | 0.09 | 0.23 | 0.28 |
| 6 | 154551118 | rs12212773 | 0.258 | 1.072 | 0.641 | 0.188 | -0.008 | -0.002 | -0.002 | 0.298 | 0.09 | 0.03 | 0.05 | 0.44 | 0.07 | 0.28 |
| 6 | 154560337 | rs6935927 | -0.484 | 1.041 | 1.042 | -0.624 | 0.001 | -0.005 | -0.005 | 0 | 0.44 | 0.43 | 0.61 | 0.07 | 0.43 | 0.41 |
| 6 | 154563653 | rs9479796 | 1.275 | 0.999 | 1.157 | -0.533 | -0.001 | -0.02 | -0.02 | 0 | 0.41 | 0.44 | 0.28 | 0.27 | 0.43 | 0.19 |
| 6 | 154565335 | rs4314511 | 0.25 | 1.022 | 0.923 | -0.448 | -0.009 | -0.028 | -0.028 | 0 | 0.86 | 0.91 | 0.9 | 0.34 | 0.86 | 0.6 |
| 6 | 154567384 | rs9383697 | 0.495 | 1.042 | 0.651 | -0.665 | 0.004 | 0.003 | 0.003 | 0 | 0.29 | 0.25 | 0.49 | 0.07 | 0.26 | 0.35 |
| 6 | 154567676 | rs7759388 | -0.336 | 1.109 | 0.373 | -0.618 | -0.016 | 0.034 | 0.034 | 0 | 0.15 | 0.09 | 0.11 | 0 | 0.17 | 0.06 |
| 6 | 154574830 | rs9479800 | 1.291 | 1.064 | 0.667 | -0.502 | -0.005 | 0.031 | 0.031 | 0.439 | 0.4 | 0.27 | 0.27 | 0.27 | 0.43 | 0.19 |
| 6 | 154578345 | rs13202830 | 0.405 | 0.97 | 0.575 | -0.561 | 0.006 | 0.016 | 0.016 | 0 | 0.3 | 0.27 | 0.49 | 0.07 | 0.27 | 0.35 |
| 6 | 154581997 | rs2350877 | -0.954 | 1.146 | 0.765 | -0.439 | -0.003 | 0.007 | 0.007 | 0.239 | 0.57 | 0.6 | 0.7 | 0.73 | 0.53 | 0.79 |
| 6 | 154582669 | rs9322455 | 1.285 | 1.022 | 0.35 | -0.625 | -0.029 | 0.022 | 0.022 | 0.201 | 0.35 | 0.17 | 0.23 | 0.27 | 0.38 | 0.18 |
| 6 | 154583206 | rs11966376 | 0.068 | 1.017 | 0.596 | -0.724 | 0.014 | -0.042 | -0.042 | 0 | 0.08 | 0.16 | 0.06 | 0 | 0.07 | 0.04 |
| 6 | 154583379 | rs7761411 | 0.664 | 1.024 | 0.121 | -0.791 | -0.024 | 0.062 | 0.062 | 1.081 | 0.45 | 0.28 | 0.31 | 0.27 | 0.5 | 0.21 |
| 6 | 154590432 | rs7766597 | -1.504 | 0.802 | -0.035 | -0.394 | 0.002 | 0.001 | 0.001 | 0 | 0.38 | 0.67 | 0.27 | 0.67 | 0.36 | 0.52 |
| 6 | 154597660 | rs17085260 | -0.54 | 0.682 | -0.204 | 0.23 | -0.001 | 0.022 | 0.022 | 0 | 0.18 | 0.17 | 0.11 | 0.58 | 0.17 | 0.39 |
| 6 | 154600367 | rs9479807 | 0.5 | 0.705 | -0.965 | -0.173 | -0.011 | 0.027 | 0.027 | 0 | 0.19 | 0.04 | 0.43 | 0.15 | 0.19 | 0.28 |
| 6 | 154601755 | rs1547588 | -1.286 | 0.879 | -1.216 | -0.204 | -0.088 | 0.045 | 0.045 | 0.499 | 0.58 | 0.92 | 0.72 | 0.73 | 0.54 | 0.79 |
| 6 | 154603498 | rs13204400 | -0.174 | 0.694 | -0.474 | 0.499 | 0.006 | 0.097 | 0.097 | 0 | 0.11 | 0.16 | 0.08 | 0.42 | 0.12 | 0.24 |
| 6 | 154609173 | rs7753809 | -0.53 | 0.761 | -1.06 | -0.409 | -0.007 | 0.073 | 0.073 | 0 | 0.28 | 0.11 | 0.48 | 0.13 | 0.3 | 0.27 |
| 6 | 154614426 | rs4118099 | -0.72 | 0.865 | -1.785 | -0.508 | -0.108 | -0.005 | -0.005 | 0.223 | 0.71 | 0.94 | 0.49 | 0.72 | 0.61 | 0.73 |

| | | | | | | | | | | | | | | | | |
|---|---|---|---|---|---|---|---|---|---|---|---|---|---|---|---|---|
| 6 | 154616419 | rs10872705 | -0.188 | 0.417 | -1.869 | -0.805 | -0.058 | 0.179 | 0.179 | 0.288 | 0.57 | 0.86 | 0.51 | 0.85 | 0.52 | 0.82 |
| 6 | 154618934 | rs13193677 | -0.025 | 0.233 | -1.299 | -0.68 | 0 | -0.022 | -0.022 | 0 | 0.12 | 0.13 | 0.05 | 0 | 0.12 | 0.03 |
| 6 | 154622914 | rs1406056 | -1.526 | 0.253 | -1.662 | -0.693 | -0.114 | 0.027 | 0.027 | 0 | 0.73 | 0.98 | 0.83 | 0.85 | 0.73 | 0.89 |
| 6 | 154625192 | rs1534446 | -0.068 | -0.309 | -1.229 | -0.08 | -0.013 | 0.081 | 0.081 | 2.959 | 0.15 | 0.3 | 0.16 | 0.32 | 0.1 | 0.33 |
| 6 | 154627294 | rs2882586 | -0.972 | -0.699 | -0.951 | -0.327 | 0 | -0.045 | -0.045 | 0.238 | 0.62 | 0.57 | 0.76 | 0.81 | 0.68 | 0.79 |
| 6 | 154629080 | rs7739525 | -0.566 | -0.698 | -1.074 | -0.251 | -0.009 | 0.023 | 0.023 | 2.845 | 0.21 | 0.27 | 0.17 | 0.36 | 0.14 | 0.43 |
| 6 | 154630541 | rs883273 | 0.794 | -0.318 | -1.062 | -0.117 | 0.038 | 0.059 | 0.059 | 0 | 0.27 | 0.46 | 0.37 | 0.37 | 0.31 | 0.31 |
| 6 | 154637393 | rs1293927 | -0.22 | -0.075 | -1.653 | -0.939 | -0.012 | 0.033 | 0.033 | 0.274 | 0.81 | 0.88 | 0.74 | 0.88 | 0.72 | 0.86 |
| 6 | 154640746 | rs1293928 | 0.663 | -0.018 | -1.695 | -0.876 | -0.007 | 0.044 | 0.044 | 0 | 0.36 | 0.27 | 0.22 | 0.21 | 0.32 | 0.21 |
| 6 | 154652025 | rs12665293 | -0.408 | 0.262 | -1.82 | -0.104 | 0.012 | 0 | 0 | 0.276 | 0.13 | 0.07 | 0.21 | 0.16 | 0.08 | 0.36 |
| 6 | 154656520 | rs2092417 | -0.576 | 0.381 | -1.846 | -0.082 | 0.007 | 0.001 | 0.001 | 1.936 | 0.2 | 0.11 | 0.25 | 0.24 | 0.17 | 0.46 |
| 6 | 154658031 | rs9371790 | 0.25 | 0.9 | -1.828 | 0.417 | -0.097 | -0.122 | -0.122 | 0 | 0.22 | 0.01 | 0.38 | 0.65 | 0.29 | 0.33 |
| 6 | 154668463 | rs2499640 | 0.594 | 0.636 | -1.77 | -0.207 | -0.048 | 0.117 | 0.117 | 0.148 | 0.43 | 0.14 | 0.35 | 0.2 | 0.42 | 0.29 |
| 6 | 154669628 | rs2038155 | -0.625 | 0.754 | -1.017 | -0.063 | -0.002 | 0.016 | 0.016 | 0.044 | 0.75 | 0.71 | 0.63 | 0.32 | 0.74 | 0.61 |
| 6 | 154669917 | rs6935066 | 0.121 | 0.704 | -0.792 | -0.198 | 0.012 | -0.06 | -0.06 | 0 | 0.17 | 0.28 | 0.12 | 0.01 | 0.23 | 0.05 |
| 6 | 154675915 | rs2499649 | -0.011 | 0.572 | -1.318 | 0.493 | 0.007 | -0.02 | -0.02 | 0 | 0.38 | 0.2 | 0.44 | 0.38 | 0.28 | 0.32 |
| 6 | 154677723 | rs742517 | -0.145 | 0.66 | -1.243 | 0.513 | -0.009 | -0.004 | -0.004 | 0.497 | 0.4 | 0.22 | 0.45 | 0.35 | 0.34 | 0.32 |
| 6 | 154681326 | rs2473548 | -0.451 | 0.518 | -0.901 | 0.33 | -0.003 | 0.004 | 0.004 | 0 | 0.36 | 0.3 | 0.32 | 0.3 | 0.33 | 0.29 |
| 6 | 154684845 | rs7765522 | -0.056 | -0.42 | -0.334 | 0.62 | -0.001 | -0.001 | -0.001 | 0 | 0.32 | 0.32 | 0.27 | 0.35 | 0.33 | 0.38 |
| 6 | 154685454 | rs761130 | -1.139 | -0.849 | -0.716 | 0.58 | -0.065 | 0.151 | 0.151 | 0 | 0.56 | 0.87 | 0.62 | 0.74 | 0.5 | 0.57 |
| 6 | 154690654 | rs12193925 | -0.813 | -0.769 | -0.066 | 0.726 | -0.024 | 0.091 | 0.091 | 0.004 | 0.24 | 0.39 | 0.34 | 0.4 | 0.18 | 0.19 |
| 6 | 154691448 | rs11968059 | -0.816 | -0.961 | -0.4 | 0.568 | 0.001 | 0.006 | 0.006 | 0 | 0.14 | 0.09 | 0.29 | 0.14 | 0.12 | 0.2 |
| 6 | 154694179 | rs9384206 | -0.114 | -0.751 | -0.003 | 0.9 | -0.073 | 0.267 | 0.267 | 0.031 | 0.16 | 0.5 | 0.09 | 0.38 | 0.08 | 0.11 |
| 6 | 154699068 | rs9479825 | 0.221 | -0.553 | -0.213 | 0.452 | -0.001 | 0.001 | 0.001 | 0 | 0.29 | 0.31 | 0.27 | 0.33 | 0.32 | 0.36 |
| 6 | 154701025 | rs4870281 | -1.035 | -0.572 | -0.821 | 0.33 | -0.058 | 0.209 | 0.209 | 0.337 | 0.57 | 0.9 | 0.63 | 0.86 | 0.51 | 0.7 |
| 6 | 154705702 | rs9397714 | -0.784 | -1.061 | 0.103 | 1.212 | -0.06 | 0.11 | 0.11 | 0.167 | 0.18 | 0.29 | 0.08 | 0.49 | 0.1 | 0.24 |
| 6 | 154707256 | rs9479827 | -0.402 | -0.483 | 0.226 | 0.924 | -0.043 | -0.013 | -0.013 | 2.234 | 0.25 | 0.49 | 0.15 | 0.25 | 0.15 | 0.18 |
| 6 | 154708633 | rs12526737 | 0.082 | 0.1 | -0.013 | 1.366 | -0.001 | -0.004 | -0.004 | 0.915 | 0.2 | 0.23 | 0.18 | 0.38 | 0.17 | 0.23 |
| 6 | 154715911 | rs1933685 | -0.656 | 0.347 | -0.076 | 0.95 | -0.001 | -0.002 | -0.002 | 0.013 | 0.11 | 0.07 | 0.27 | 0.13 | 0.09 | 0.26 |
| 6 | 154716112 | rs1361218 | -0.88 | 0.182 | -0.305 | 0.43 | -0.002 | 0.097 | 0.097 | 0 | 0.54 | 0.71 | 0.67 | 0.88 | 0.44 | 0.64 |
| 6 | 154717442 | rs9383705 | -1.01 | 0.248 | -0.593 | 0.496 | -0.026 | 0.123 | 0.123 | 0 | 0.59 | 0.81 | 0.72 | 0.88 | 0.51 | 0.65 |
| 6 | 154719380 | rs3812243 | -0.714 | 0.212 | -0.203 | 1.596 | -0.008 | 0.044 | 0.044 | 0 | 0.08 | 0.13 | 0.11 | 0.28 | 0.03 | 0.26 |
| 6 | 154727408 | rs9397716 | -0.511 | 0.345 | -0.268 | 1.381 | -0.002 | -0.017 | -0.017 | 0 | 0.2 | 0.13 | 0.22 | 0.28 | 0.22 | 0.13 |
| 6 | 154730651 | rs9384211 | -0.64 | 0.245 | 0.149 | 1.408 | -0.005 | 0.005 | 0.005 | 0 | 0.21 | 0.23 | 0.23 | 0.22 | 0.17 | 0.14 |
| 6 | 154733074 | rs7745077 | 0.144 | 0.244 | -0.257 | 0.525 | -0.004 | 0.034 | 0.034 | 0 | 0.48 | 0.65 | 0.39 | 0.64 | 0.48 | 0.46 |
| 6 | 154733843 | rs11752945 | 0.666 | 0.352 | 0.286 | 0.357 | -0.006 | -0.003 | -0.003 | 0 | 0.35 | 0.27 | 0.31 | 0.59 | 0.34 | 0.43 |
| 6 | 154736749 | rs7752017 | 0.712 | 0.131 | 0.759 | 0.71 | 0.03 | -0.041 | -0.041 | 0 | 0.33 | 0.46 | 0.24 | 0.23 | 0.42 | 0.2 |
| 6 | 154736783 | rs7750888 | -1.392 | -0.068 | 0.566 | 0.652 | -0.003 | 0.02 | 0.02 | 0 | 0.7 | 0.77 | 0.61 | 0.86 | 0.65 | 0.6 |
| 6 | 154739219 | rs9383708 | 0.114 | -0.049 | 0.706 | 0.851 | -0.015 | 0.125 | 0.125 | 0 | 0.09 | 0.14 | 0.13 | 0.4 | 0.03 | 0.26 |
| 6 | 154739357 | rs12528745 | 0.371 | 0.118 | 0.834 | 0.301 | 0.013 | -0.034 | -0.034 | 0 | 0.35 | 0.56 | 0.24 | 0.22 | 0.4 | 0.22 |
| 6 | 154740655 | rs9384215 | -1.295 | 0.179 | 0.904 | 0.82 | 0.015 | 0.038 | 0.038 | 0 | 0.14 | 0.26 | 0.2 | 0.27 | 0.15 | 0.16 |
| 6 | 154740770 | rs9397190 | -0.939 | 0.163 | 0.636 | 0.826 | -0.002 | 0.021 | 0.021 | 0 | 0.13 | 0.16 | 0.18 | 0.25 | 0.11 | 0.15 |
| 6 | 154743525 | rs7771159 | 0.946 | 0.136 | 0.63 | 0.62 | -0.003 | 0.014 | 0.014 | 0 | 0.34 | 0.29 | 0.23 | 0.21 | 0.35 | 0.21 |
| 6 | 154748630 | rs9322464 | 0.493 | 0.127 | 0.65 | 0.558 | -0.016 | -0.002 | -0.002 | 0 | 0.4 | 0.28 | 0.38 | 0.39 | 0.42 | 0.31 |
| 6 | 154749087 | rs9397717 | -0.133 | 0.107 | 0.41 | 0.56 | 0 | 0 | 0 | 1.419 | 0.5 | 0.51 | 0.43 | 0.41 | 0.51 | 0.42 |
| 6 | 154750505 | rs11155973 | -0.124 | -0.094 | 0.399 | 0.621 | -0.055 | 0.131 | 0.131 | 0 | 0.2 | 0.38 | 0.19 | 0.45 | 0.09 | 0.26 |
| 6 | 154758463 | rs4869823 | -1.216 | 0.081 | -0.528 | 0.422 | -0.085 | 0.104 | 0.104 | 0 | 0.71 | 0.96 | 0.63 | 0.88 | 0.59 | 0.67 |
| 6 | 154758890 | rs12332787 | -0.106 | 0.541 | 0.037 | 0.526 | 0.001 | -0.02 | -0.02 | 0.194 | 0.07 | 0.07 | 0.04 | 0 | 0.08 | 0.06 |
| 6 | 154759822 | rs9479855 | -0.575 | 0.487 | 0.031 | 0.616 | 0.001 | 0.009 | 0.009 | 0 | 0.14 | 0.08 | 0.08 | 0.07 | 0.12 | 0.08 |
| 6 | 154760264 | rs9371342 | -1.007 | 0.566 | 0.504 | 0.834 | 0.11 | 0.309 | 0.309 | 0.364 | 0.06 | 0.57 | 0.14 | 0.24 | 0.04 | 0.1 |
| 6 | 154761064 | rs6910929 | 0.489 | 0.577 | -0.389 | 0.096 | -0.022 | 0.098 | 0.098 | 0 | 0.32 | 0.13 | 0.22 | 0.12 | 0.33 | 0.24 |
| 6 | 154762524 | rs6916438 | -1.368 | 0.421 | 0.872 | 0.54 | 0.067 | 0.019 | 0.019 | 0.216 | 0.06 | 0.44 | 0.05 | 0 | 0.03 | 0.04 |
| 6 | 154762980 | rs9479857 | -1.067 | 0.474 | 0.585 | 0.855 | 0.002 | 0.003 | 0.003 | 0 | 0.42 | 0.6 | 0.59 | 0.62 | 0.36 | 0.54 |
| 6 | 154763401 | rs6557351 | -1.024 | 0.464 | 0.344 | 1.296 | -0.004 | 0.003 | 0.003 | 0 | 0.28 | 0.62 | 0.36 | 0.29 | 0.24 | 0.28 |
| 6 | 154768778 | rs9322466 | -1.271 | 0.463 | 0.301 | 1.541 | -0.003 | 0.031 | 0.031 | 0 | 0.29 | 0.64 | 0.4 | 0.47 | 0.25 | 0.4 |
| 6 | 154771277 | rs9322467 | -0.965 | 0.641 | -0.715 | 0.412 | -0.037 | 0.095 | 0.095 | 0 | 0.72 | 0.86 | 0.76 | 0.9 | 0.67 | 0.8 |
| 6 | 154774050 | rs4870289 | 0 | 0.77 | -0.528 | 0.467 | 0 | 0.009 | 0.009 | 0 | 0.34 | 0.26 | 0.22 | 0.3 | 0.31 | 0.39 |
| 6 | 154782097 | rs10782333 | -0.767 | 0.941 | -0.849 | 0.501 | -0.015 | 0.043 | 0.043 | 0 | 0.65 | 0.77 | 0.59 | 0.76 | 0.62 | 0.66 |
| 6 | 154783783 | rs6557357 | 0.057 | 0.634 | -0.01 | 0.877 | 0.012 | 0.047 | 0.047 | 0 | 0.28 | 0.44 | 0.37 | 0.37 | 0.31 | 0.22 |
| 6 | 154784600 | rs7742597 | -0.918 | 0.562 | -0.64 | 0.67 | -0.013 | -0.006 | -0.006 | 0 | 0.47 | 0.85 | 0.45 | 0.52 | 0.43 | 0.43 |
| 6 | 154787639 | rs4870294 | -0.736 | 0.395 | -0.095 | 0.405 | -0.011 | -0.019 | -0.019 | 0.815 | 0.19 | 0.31 | 0.09 | 0.13 | 0.12 | 0.22 |
| 6 | 154797178 | rs4398752 | 0.118 | 0.302 | -1.305 | 0.465 | -0.02 | -0.048 | -0.048 | 0.673 | 0.28 | 0.02 | 0.17 | 0.25 | 0.24 | 0.24 |
| 6 | 154800018 | rs12194277 | -0.405 | 0.468 | -0.87 | 0.307 | -0.004 | 0.01 | 0.01 | 0 | 0.53 | 0.41 | 0.56 | 0.58 | 0.52 | 0.42 |
| 6 | 154801450 | rs11155976 | 0.211 | 0.585 | -1.088 | 0.417 | 0.001 | 0.081 | 0.081 | 0 | 0.33 | 0.1 | 0.19 | 0.19 | 0.31 | 0.24 |
| 6 | 154801853 | rs9479877 | -0.441 | 0.498 | -0.879 | 0.281 | -0.006 | 0.013 | 0.013 | 0 | 0.54 | 0.36 | 0.58 | 0.55 | 0.52 | 0.43 |
| 6 | 154803883 | rs946236 | -0.131 | 0.578 | -0.628 | 0.844 | 0.002 | -0.053 | -0.053 | 0.76 | 0.19 | 0.09 | 0.36 | 0.37 | 0.2 | 0.2 |
| 6 | 154804650 | rs4352688 | -0.665 | 0.773 | -0.117 | 0.503 | -0.002 | 0.02 | 0.02 | 0 | 0.34 | 0.48 | 0.44 | 0.55 | 0.35 | 0.41 |
| 6 | 154806742 | rs7762516 | -0.051 | 1.07 | 0.164 | 0.006 | 0.023 | -0.051 | -0.051 | 0 | 0.26 | 0.5 | 0.19 | 0.12 | 0.27 | 0.21 |
| 6 | 154812444 | rs13217240 | -0.722 | 0.785 | -0.049 | 0.099 | 0.002 | 0.01 | 0.01 | 0 | 0.28 | 0.2 | 0.37 | 0.24 | 0.27 | 0.24 |

| | | | | | | | | | | | | | | | | |
|---|---|---|---|---|---|---|---|---|---|---|---|---|---|---|---|---|
| 6 | 154815891 | rs6911971 | -1.099 | 0.706 | 0.685 | -0.167 | 0.001 | -0.026 | -0.026 | 0 | 0.36 | 0.47 | 0.36 | 0.29 | 0.35 | 0.3 |
| 6 | 154817844 | rs6914034 | 0.283 | 0.691 | 0.434 | -0.098 | -0.003 | 0.007 | 0.007 | 0 | 0.1 | 0.06 | 0.22 | 0.07 | 0.14 | 0.05 |
| 6 | 154818897 | rs6557361 | -0.532 | 0.684 | 1.098 | -0.165 | 0.03 | -0.041 | -0.041 | 0 | 0.13 | 0.39 | 0.08 | 0.05 | 0.1 | 0.09 |
| 6 | 154819484 | rs6942272 | -0.715 | 0.671 | 1.069 | -0.142 | -0.001 | -0.033 | -0.033 | 0 | 0.21 | 0.29 | 0.29 | 0.12 | 0.22 | 0.14 |
| 6 | 154823016 | rs10872708 | 0.013 | 0.653 | 0.594 | -0.091 | -0.039 | -0.04 | -0.04 | 0 | 0.11 | 0 | 0.07 | 0.16 | 0.11 | 0.1 |
| 6 | 154825116 | rs1538340 | 0.904 | 0.405 | 0.792 | -0.307 | -0.005 | 0.039 | 0.039 | 0 | 0.39 | 0.29 | 0.27 | 0.23 | 0.38 | 0.31 |
| 6 | 154826359 | rs11967674 | -1.32 | 0.097 | 2.07 | -0.191 | 0.004 | -0.073 | -0.073 | 0 | 0.16 | 0.35 | 0.11 | 0.04 | 0.14 | 0.08 |
| 6 | 154827861 | rs7768635 | -0.503 | 0.01 | 2.219 | -0.091 | 0.004 | -0.013 | -0.013 | 0.496 | 0.15 | 0.26 | 0.14 | 0.12 | 0.19 | 0.08 |
| 6 | 154837628 | rs871070 | -1.911 | -0.219 | 2.217 | -0.206 | -0.006 | -0.02 | -0.02 | 0.309 | 0.32 | 0.6 | 0.32 | 0.19 | 0.29 | 0.15 |
| 6 | 154844905 | rs9918406 | -0.384 | -0.133 | 2.278 | 0.002 | -0.001 | 0.038 | 0.038 | 0 | 0.14 | 0.34 | 0.13 | 0.18 | 0.15 | 0.08 |
| 6 | 154854851 | rs12524025 | -0.797 | 0.149 | 1.861 | -0.183 | 0.015 | -0.028 | -0.028 | 0.062 | 0.09 | 0.18 | 0.07 | 0.06 | 0.1 | 0.06 |
| 6 | 154876504 | rs17085526 | -1.979 | -0.076 | 1.748 | 0.168 | 0.01 | 0.004 | 0.004 | 0 | 0.37 | 0.65 | 0.34 | 0.38 | 0.32 | 0.18 |
| 6 | 154876716 | rs17085528 | -0.763 | -0.019 | 1.776 | 0.242 | -0.003 | -0.007 | -0.007 | 0.491 | 0.18 | 0.35 | 0.24 | 0.15 | 0.19 | 0.1 |
| 6 | 154879757 | rs11155982 | -0.571 | 0.181 | 1.066 | 0.292 | 0.01 | 0.032 | 0.032 | 0 | 0.08 | 0.15 | 0.06 | 0.18 | 0.11 | 0.06 |
| 6 | 154881835 | rs9784771 | -0.864 | 0.07 | 1.187 | 0.199 | -0.019 | -0.041 | -0.041 | 0 | 0.12 | 0.16 | 0.07 | 0.05 | 0.05 | 0.03 |
| 6 | 154891396 | rs12524781 | -0.728 | 0.353 | 0.933 | 0.431 | 0.011 | 0.029 | 0.029 | 0 | 0.08 | 0.14 | 0.06 | 0.18 | 0.12 | 0.06 |
| 6 | 154899702 | rs1022404 | -0.379 | 0.269 | 0.895 | 0.303 | -0.001 | 0.003 | 0.003 | 0 | 0.16 | 0.18 | 0.2 | 0.27 | 0.19 | 0.13 |
| 6 | 154927883 | rs2351313 | -0.885 | 0.375 | 0.933 | 0.119 | -0.002 | 0.008 | 0.008 | 0 | 0.23 | 0.37 | 0.25 | 0.2 | 0.23 | 0.12 |
| 6 | 154929308 | rs1533746 | 0.176 | 0.402 | 0.628 | 0.138 | -0.022 | -0.063 | -0.063 | 1.737 | 0.15 | 0.02 | 0.17 | 0.19 | 0.17 | 0.11 |
| 6 | 154933171 | rs11155986 | -1.245 | 0.318 | 0.9 | 0.111 | -0.005 | 0.003 | 0.003 | 0 | 0.25 | 0.39 | 0.28 | 0.2 | 0.25 | 0.13 |
| 6 | 154934989 | rs9479902 | -0.976 | 0.323 | 0.91 | 0.116 | -0.001 | 0.007 | 0.007 | 0 | 0.23 | 0.34 | 0.26 | 0.2 | 0.24 | 0.13 |
| 6 | 154942072 | rs11155993 | -0.572 | 0.33 | 0.787 | 0.123 | 0 | -0.002 | -0.002 | 0.338 | 0.2 | 0.2 | 0.23 | 0.19 | 0.22 | 0.11 |
| 6 | 154946930 | rs1996737 | -0.029 | 0.343 | 0.808 | 0.171 | -0.004 | 0.009 | 0.009 | 0 | 0.16 | 0.19 | 0.19 | 0.19 | 0.16 | 0.11 |
| 6 | 154958156 | rs7766057 | -0.152 | 0.541 | 0.46 | 0.412 | -0.006 | -0.025 | -0.025 | 0 | 0.2 | 0.15 | 0.35 | 0.46 | 0.25 | 0.37 |
| 6 | 154961462 | rs7745756 | -0.727 | -0.043 | 0.363 | -0.318 | -0.005 | 0.048 | 0.048 | 0 | 0.17 | 0.04 | 0.11 | 0.14 | 0.14 | 0.1 |
| 6 | 154970556 | rs9479923 | -1.025 | 0.044 | 0.474 | -0.266 | -0.025 | -0.073 | -0.073 | 0.262 | 0.22 | 0.43 | 0.16 | 0.15 | 0.17 | 0.11 |
| 6 | 154975467 | rs3126042 | -0.439 | 0.518 | 0.172 | -0.178 | -0.002 | -0.015 | -0.015 | 0.007 | 0.26 | 0.32 | 0.14 | 0.25 | 0.22 | 0.32 |
| 6 | 154976670 | rs9322473 | -0.467 | 0.124 | 0.218 | 0.094 | -0.036 | 0.076 | 0.076 | 0.007 | 0.23 | 0.36 | 0.39 | 0.53 | 0.18 | 0.38 |
| 6 | 154981639 | rs6910245 | -1.076 | 0.813 | -0.261 | -0.249 | 0.005 | -0.021 | -0.021 | 2.058 | 0.33 | 0.73 | 0.5 | 0.57 | 0.33 | 0.41 |
| 6 | 154986571 | rs9371822 | -0.557 | 0.964 | -0.231 | -0.256 | -0.021 | 0.075 | 0.075 | 0.007 | 0.25 | 0.39 | 0.4 | 0.55 | 0.23 | 0.38 |
| 6 | 154986664 | rs9397738 | -0.566 | 0.998 | -0.572 | -0.464 | -0.014 | -0.004 | -0.004 | 0 | 0.84 | 0.93 | 0.91 | 0.81 | 0.85 | 0.76 |
| 6 | 154991843 | rs4870304 | 0.945 | 0.802 | -0.699 | -0.529 | -0.009 | 0.04 | 0.04 | 0 | 0.55 | 0.36 | 0.68 | 0.76 | 0.57 | 0.56 |
| 6 | 154995031 | rs11756851 | 1.633 | 0.722 | -0.914 | -0.51 | -0.116 | 0.04 | 0.04 | 0 | 0.39 | 0.05 | 0.35 | 0.25 | 0.45 | 0.26 |
| 6 | 154996850 | rs10499306 | 0.052 | 0.395 | -0.465 | -0.15 | -0.034 | 0.141 | 0.141 | 0.073 | 0.17 | 0.31 | 0.33 | 0.52 | 0.14 | 0.32 |
| 6 | 154997553 | rs7738774 | -1.36 | 0.881 | -1.215 | -0.453 | -0.012 | 0.02 | 0.02 | 0 | 0.1 | 0.05 | 0.12 | 0.05 | 0.12 | 0.09 |
| 6 | 155001410 | rs6420139 | 0.892 | 0.794 | -1.172 | -0.427 | -0.001 | 0.062 | 0.062 | 0 | 0.65 | 0.77 | 0.81 | 0.84 | 0.69 | 0.63 |
| 6 | 155011339 | rs9479939 | -0.419 | 0.619 | -0.989 | -0.239 | -0.019 | -0.033 | -0.033 | 0 | 0.1 | 0.15 | 0.06 | 0 | 0.06 | 0.05 |
| 6 | 155012182 | rs9397196 | -0.647 | 0.93 | -1.178 | -0.204 | -0.04 | -0.008 | -0.008 | 0 | 0.18 | 0.1 | 0.17 | 0.22 | 0.2 | 0.29 |
| 6 | 155012313 | rs9397197 | -1.131 | 1.007 | -1.122 | -0.114 | -0.004 | 0.015 | 0.015 | 0 | 0.28 | 0.25 | 0.23 | 0.22 | 0.25 | 0.33 |
| 6 | 155019223 | rs9397199 | -0.711 | 1.254 | -1.285 | 0.651 | -0.015 | -0.048 | -0.048 | 8.191 | 0.2 | 0.14 | 0.34 | 0.51 | 0.24 | 0.31 |
| 6 | 155022713 | rs7754964 | -1.471 | 0.761 | -1.862 | -0.128 | -0.117 | 0.119 | 0.119 | 1.128 | 0.62 | 0.93 | 0.59 | 0.84 | 0.53 | 0.72 |
| 6 | 155072320 | rs4869834 | -0.828 | 0.529 | -1.478 | 0.118 | 0.016 | -0.028 | -0.028 | 0.052 | 0.27 | 0.16 | 0.37 | 0.63 | 0.24 | 0.29 |
| 6 | 155092199 | rs9479954 | -1.406 | 0.229 | 0.2 | -0.255 | -0.001 | -0.032 | -0.032 | 8.838 | 0.38 | 0.39 | 0.48 | 0.9 | 0.35 | 0.56 |
| 6 | 155094764 | rs7744421 | -2.168 | 0.007 | 0.324 | -0.402 | 0 | 0.219 | 0.219 | 0 | 0.52 | 0.82 | 0.57 | 0.91 | 0.45 | 0.66 |
| 6 | 155096462 | rs10872710 | -2.25 | -0.03 | 0.22 | -0.472 | -0.116 | 0.283 | 0.283 | 0.058 | 0.56 | 0.94 | 0.59 | 0.91 | 0.48 | 0.7 |
| 6 | 155111359 | rs9397749 | -2.232 | -0.023 | 0.253 | -0.469 | -0.06 | 0.239 | 0.239 | 0.059 | 0.56 | 0.87 | 0.59 | 0.91 | 0.48 | 0.7 |
| 6 | 155113232 | rs3800003 | -0.503 | -0.063 | 0.367 | -0.507 | 0.01 | -0.016 | -0.016 | 0 | 0.25 | 0.2 | 0.36 | 0.63 | 0.21 | 0.31 |
| 6 | 155138942 | rs2351846 | 0.338 | -0.098 | 0.093 | -0.017 | -0.009 | -0.03 | -0.03 | 1.532 | 0.13 | 0.07 | 0.29 | 0.44 | 0.16 | 0.2 |
| 6 | 155145017 | rs12523716 | 2.093 | -0.023 | 0.047 | -0.119 | -0.012 | 0.245 | 0.245 | 0.047 | 0.47 | 0.15 | 0.43 | 0.09 | 0.55 | 0.3 |
| 6 | 155156221 | rs1980602 | 2.2 | -0.001 | -0.021 | -0.092 | -0.061 | 0.238 | 0.238 | 0.038 | 0.44 | 0.12 | 0.41 | 0.09 | 0.52 | 0.3 |
| 6 | 155160436 | rs17085709 | 0.212 | -0.03 | 0.087 | 0.003 | -0.014 | -0.034 | -0.034 | 0 | 0.1 | 0.03 | 0.1 | 0.26 | 0.11 | 0.25 |
| 6 | 155162121 | rs12199781 | 1.996 | 0.238 | 0.04 | -0.176 | -0.054 | 0.246 | 0.246 | 0.37 | 0.45 | 0.12 | 0.42 | 0.09 | 0.54 | 0.31 |
| 6 | 155165469 | rs2351843 | 0.078 | -0.078 | -0.287 | -0.241 | -0.004 | -0.025 | -0.025 | 0.084 | 0.25 | 0.21 | 0.41 | 0.42 | 0.26 | 0.26 |
| 6 | 155175477 | rs9384247 | -0.023 | -0.106 | -0.281 | -0.25 | -0.006 | -0.041 | -0.041 | 0 | 0.26 | 0.19 | 0.41 | 0.43 | 0.26 | 0.26 |
| 6 | 155182333 | rs764419 | 0.881 | -0.202 | -0.363 | -0.233 | -0.065 | -0.089 | -0.089 | 0 | 0.14 | 0.03 | 0.36 | 0.41 | 0.2 | 0.26 |
| 6 | 155182564 | rs764420 | -0.602 | -0.3 | -0.158 | -0.388 | -0.001 | 0 | 0 | 0 | 0.25 | 0.21 | 0.27 | 0.28 | 0.2 | 0.3 |
| 6 | 155185451 | rs9479968 | 0.633 | -0.285 | -0.22 | -0.398 | -0.009 | -0.042 | -0.042 | 0 | 0.25 | 0.18 | 0.41 | 0.42 | 0.25 | 0.28 |
| 6 | 155194880 | rs17085762 | -1.02 | -0.615 | 0.087 | -0.3 | -0.024 | 0.05 | 0.05 | 0 | 0.28 | 0.35 | 0.3 | 0.49 | 0.21 | 0.34 |
| 6 | 155200674 | rs7763655 | -0.878 | -0.47 | 0.231 | -0.402 | -0.007 | 0.013 | 0.013 | 0 | 0.23 | 0.29 | 0.25 | 0.26 | 0.2 | 0.28 |
| 6 | 155206501 | rs6902018 | 0.138 | -0.196 | 0.023 | -0.111 | 0.001 | 0.006 | 0.006 | 0 | 0.64 | 0.48 | 0.53 | 0.39 | 0.65 | 0.6 |
| 6 | 155216846 | rs6557379 | 0.433 | -0.038 | 0.035 | -0.077 | 0.002 | -0.002 | -0.002 | 0 | 0.6 | 0.51 | 0.49 | 0.34 | 0.57 | 0.55 |
| 6 | 155218117 | rs7760792 | 0.901 | 0.039 | -0.369 | -0.599 | 0.009 | 0.133 | 0.133 | 0 | 0.37 | 0.15 | 0.22 | 0.11 | 0.36 | 0.28 |
| 6 | 155219158 | rs7753068 | 0.901 | 0.039 | -0.369 | -0.599 | 0.009 | 0.133 | 0.133 | 0 | 0.37 | 0.15 | 0.22 | 0.11 | 0.36 | 0.28 |
| 6 | 155221600 | rs6557383 | 0.305 | 0.366 | -0.304 | -0.147 | 0.006 | 0.007 | 0.007 | 0 | 0.48 | 0.52 | 0.29 | 0.3 | 0.4 | 0.39 |
| 6 | 155225638 | rs7747703 | -0.59 | 0.559 | -0.336 | -0.111 | 0.001 | 0.023 | 0.023 | 2.75 | 0.18 | 0.53 | 0.13 | 0.18 | 0.17 | 0.1 |
| 6 | 155225683 | rs911193 | -0.274 | 0.745 | -1.188 | -0.283 | -0.02 | 0.036 | 0.036 | 0 | 0.58 | 0.74 | 0.36 | 0.32 | 0.51 | 0.51 |
| 6 | 155229566 | rs1123928 | 0.628 | 0.845 | -1.374 | -0.217 | -0.057 | 0.082 | 0.082 | 0 | 0.32 | 0.05 | 0.18 | 0.15 | 0.29 | 0.31 |
| 6 | 155232421 | rs1555374 | -1.124 | 1.172 | 0.053 | -0.008 | 0.063 | 0.057 | 0.057 | 0 | 0.12 | 0.61 | 0.12 | 0.13 | 0.15 | 0.24 |
| 6 | 155235078 | rs9479977 | 0.228 | 0.949 | -0.08 | -0.443 | 0 | 0.007 | 0.007 | 0 | 0.54 | 0.42 | 0.29 | 0.26 | 0.43 | 0.38 |

| | | | | | | | | | | | | | | | | |
|---|---|---|---|---|---|---|---|---|---|---|---|---|---|---|---|---|
| 6 | 155235217 | rs6557385 | -0.182 | 0.984 | -0.041 | -0.012 | 0.008 | 0.012 | 0.012 | 0 | 0.36 | 0.6 | 0.38 | 0.49 | 0.44 | 0.42 |
| 6 | 155235639 | rs9371835 | -0.054 | 1.062 | -0.114 | -0.024 | 0.018 | 0.04 | 0.04 | 0 | 0.33 | 0.48 | 0.35 | 0.47 | 0.4 | 0.31 |
| 6 | 155244584 | rs9322484 | -0.626 | 0.871 | -1.273 | -0.358 | -0.069 | -0.028 | -0.028 | 0 | 0.56 | 0.93 | 0.47 | 0.56 | 0.53 | 0.55 |
| 6 | 155247355 | rs7768412 | -0.034 | 0.756 | -0.544 | -0.539 | 0.002 | 0.03 | 0.03 | 0 | 0.55 | 0.66 | 0.33 | 0.3 | 0.42 | 0.49 |
| 6 | 155248159 | rs9478593 | 0.795 | 0.831 | -0.721 | -0.797 | -0.015 | 0.042 | 0.042 | 0 | 0.36 | 0.24 | 0.22 | 0.21 | 0.36 | 0.32 |
| 6 | 155248596 | rs1983785 | -0.146 | 1.795 | -0.494 | -0.24 | -0.035 | 0 | 0 | 0 | 0.92 | 0.97 | 0.87 | 0.56 | 0.84 | 0.8 |
| 6 | 155249485 | rs17085806 | -1.035 | 0.8 | -0.079 | -1.196 | 0.02 | -0.004 | -0.004 | 0 | 0.14 | 0.45 | 0.11 | 0.09 | 0.08 | 0.14 |
| 6 | 155251481 | rs7752042 | 0.151 | 0.985 | -0.66 | -1.259 | 0.015 | 0.126 | 0.126 | 0 | 0.5 | 0.3 | 0.27 | 0.2 | 0.44 | 0.34 |
| 6 | 155253299 | rs6925269 | 0.112 | 1.004 | -0.466 | -0.804 | 0.005 | 0.014 | 0.014 | 0.12 | 0.53 | 0.57 | 0.28 | 0.23 | 0.41 | 0.37 |
| 6 | 155255049 | rs9322486 | 0.348 | 1.606 | -0.244 | -1.133 | 0.017 | -0.062 | -0.062 | 0 | 0.8 | 0.76 | 0.8 | 1 | 0.68 | 0.88 |
| 6 | 155255431 | rs9322487 | -0.206 | 1.433 | -0.397 | -0.735 | 0.002 | -0.012 | -0.012 | 0 | 0.1 | 0.06 | 0.08 | 0.23 | 0.08 | 0.23 |
| 6 | 155255820 | rs9371837 | 0.151 | 1.052 | -0.375 | -1.674 | -0.005 | -0.017 | -0.017 | 1.672 | 0.41 | 0.47 | 0.27 | 0.26 | 0.35 | 0.45 |
| 6 | 155256289 | rs9397210 | -0.132 | 1.429 | 0.118 | -1.095 | -0.011 | -0.001 | -0.001 | 0 | 0.82 | 0.89 | 0.82 | 1 | 0.73 | 0.89 |
| 6 | 155256667 | rs9397754 | -0.413 | 1.277 | 0.354 | -1.001 | -0.009 | 0.052 | 0.052 | 0 | 0.12 | 0.22 | 0.1 | 0.25 | 0.08 | 0.31 |
| 6 | 155270087 | rs9371840 | 0.095 | 1.395 | 0.18 | -1.239 | 0.033 | -0.115 | -0.115 | 1.502 | 0.79 | 0.65 | 0.79 | 0.99 | 0.72 | 0.81 |
| 6 | 155271178 | rs7745637 | 0.544 | 1.424 | -0.031 | -0.886 | -0.004 | 0.002 | 0.002 | 0 | 0.71 | 0.77 | 0.72 | 0.77 | 0.64 | 0.6 |
| 6 | 155271211 | rs9384260 | 0.139 | 0.82 | -0.326 | -0.996 | -0.004 | -0.001 | -0.001 | 0 | 0.26 | 0.21 | 0.49 | 0.7 | 0.27 | 0.33 |
| 6 | 155271952 | rs6557389 | -0.694 | 0.627 | -0.57 | -1.001 | 0.003 | 0.017 | 0.017 | 0 | 0.29 | 0.72 | 0.17 | 0.14 | 0.24 | 0.22 |
| 6 | 155273364 | rs4870322 | -0.061 | 0.759 | -0.639 | -0.326 | -0.024 | 0.029 | 0.029 | 0 | 0.63 | 0.72 | 0.63 | 0.26 | 0.52 | 0.37 |
| 6 | 155274063 | rs6925622 | -0.261 | 0.073 | -0.542 | 0.095 | -0.023 | -0.101 | -0.101 | 0 | 0.17 | 0.52 | 0.1 | 0.05 | 0.17 | 0.17 |
| 6 | 155279785 | rs6934056 | -0.521 | 0.753 | -1.702 | 1 | -0.106 | -0.16 | -0.16 | 0 | 0.76 | 0.99 | 0.83 | 0.56 | 0.73 | 0.59 |
| 6 | 155281493 | rs4870327 | 0.155 | 0.658 | -1.421 | 1.119 | -0.043 | -0.046 | -0.046 | 0 | 0.65 | 0.82 | 0.76 | 0.56 | 0.59 | 0.58 |
| 6 | 155295590 | rs11156007 | -0.413 | 0.33 | -1.023 | 1.353 | -0.018 | 0.085 | 0.085 | 0 | 0.17 | 0.3 | 0.41 | 0.34 | 0.15 | 0.18 |
| 6 | 155301046 | rs7745332 | 0.032 | 0.452 | -1.825 | 1.007 | -0.005 | -0.092 | -0.092 | 0 | 0.72 | 0.87 | 0.84 | 0.51 | 0.77 | 0.63 |
| 6 | 155310510 | rs4484532 | 0.162 | 0.36 | -1.76 | 0.293 | 0.029 | -0.065 | -0.065 | 0.362 | 0.31 | 0.08 | 0.22 | 0.38 | 0.24 | 0.35 |
| 6 | 155310743 | rs2351542 | -0.51 | 1.389 | -1.423 | -0.764 | -0.056 | 0.092 | 0.092 | 0 | 0.33 | 0.2 | 0.48 | 0.1 | 0.44 | 0.16 |
| 6 | 155312633 | rs4869840 | -0.598 | 1.353 | -1.062 | -0.821 | -0.02 | 0.022 | 0.022 | 0 | 0.36 | 0.3 | 0.53 | 0.12 | 0.49 | 0.18 |
| 6 | 155314787 | rs6916651 | 0.398 | 0.753 | -1.091 | -0.094 | -0.006 | -0.028 | -0.028 | 0.4 | 0.82 | 0.89 | 0.67 | 0.63 | 0.79 | 0.61 |
| 6 | 155317769 | rs9479998 | -0.285 | 0.444 | -0.481 | -0.333 | -0.008 | -0.065 | -0.065 | 4.329 | 0.13 | 0.26 | 0.14 | 0.02 | 0.1 | 0.06 |
| 6 | 155319155 | rs9322494 | -0.318 | 0.683 | -0.901 | -0.189 | 0.004 | -0.028 | -0.028 | 0 | 0.21 | 0.11 | 0.34 | 0.37 | 0.23 | 0.37 |
| 6 | 155325444 | rs12216118 | -0.32 | 1.063 | -0.335 | -0.347 | -0.012 | 0.002 | 0.002 | 0 | 0.16 | 0.11 | 0.08 | 0.14 | 0.15 | 0.09 |
| 6 | 155329941 | rs12210329 | -0.28 | 1.084 | -0.359 | -0.348 | -0.019 | -0.002 | -0.002 | 0.001 | 0.15 | 0.1 | 0.09 | 0.16 | 0.17 | 0.09 |
| 6 | 155330107 | rs9480006 | -1.038 | 0.095 | 0.433 | -0.546 | -0.027 | -0.111 | -0.111 | 1.103 | 0.21 | 0.41 | 0.18 | 0.03 | 0.15 | 0.13 |
| 6 | 155337220 | rs2498491 | -1.228 | 0.27 | 0.59 | -0.137 | -0.004 | -0.048 | -0.048 | 2.408 | 0.15 | 0.43 | 0.32 | 0.08 | 0.14 | 0.13 |
| 6 | 155345181 | rs7746011 | 1.157 | 0.066 | 0.185 | -0.002 | 0.004 | 0.001 | 0.001 | 0 | 0.92 | 0.94 | 0.75 | 0.9 | 0.93 | 0.82 |
| 6 | 155354681 | rs7756226 | 0.737 | -0.279 | 0.309 | -1.244 | -0.003 | -0.04 | -0.04 | 0.256 | 0.61 | 0.6 | 0.6 | 0.88 | 0.71 | 0.7 |
| 6 | 155359196 | rs13200965 | -0.264 | -0.032 | 0.374 | -0.586 | 0.007 | 0.052 | 0.052 | 0 | 0.3 | 0.17 | 0.14 | 0.18 | 0.23 | 0.15 |
| 6 | 155359624 | rs4484531 | 1.008 | -0.146 | 0.631 | 0.247 | -0.012 | -0.004 | -0.004 | 2.827 | 0.44 | 0.32 | 0.45 | 0.43 | 0.53 | 0.38 |
| 6 | 155362547 | rs4476844 | 0.244 | -0.205 | 0.769 | 0.391 | 0.003 | 0.01 | 0.01 | 0 | 0.31 | 0.28 | 0.19 | 0.23 | 0.25 | 0.18 |
| 6 | 155365412 | rs9371850 | 1.067 | -0.345 | 0.609 | 0.587 | -0.017 | 0.031 | 0.031 | 0 | 0.42 | 0.26 | 0.4 | 0.36 | 0.53 | 0.35 |
| 6 | 155366098 | rs7762675 | 1.106 | -0.349 | 0.62 | 0.926 | -0.015 | 0.066 | 0.066 | 0 | 0.42 | 0.27 | 0.4 | 0.21 | 0.52 | 0.34 |
| 6 | 155367203 | rs6557391 | -0.026 | -0.667 | 0.031 | 0.873 | 0.03 | 0.048 | 0.048 | 0 | 0.8 | 0.94 | 0.69 | 0.9 | 0.85 | 0.72 |
| 6 | 155368063 | rs7744915 | 0.317 | -0.544 | 0.359 | 1.027 | -0.002 | -0.054 | -0.054 | 0 | 0.16 | 0.06 | 0.12 | 0.32 | 0.16 | 0.26 |
| 6 | 155371513 | rs9322495 | -0.285 | -0.564 | 0.846 | 0.835 | -0.002 | -0.023 | -0.023 | 0.03 | 0.37 | 0.29 | 0.24 | 0.5 | 0.35 | 0.41 |
| 6 | 155372284 | rs7766864 | 1.005 | -0.008 | 0.307 | 0.484 | 0 | 0.04 | 0.04 | 0 | 0.46 | 0.32 | 0.42 | 0.22 | 0.47 | 0.29 |
| 6 | 155372751 | rs9322496 | 0.634 | 0.253 | -0.046 | 0.997 | -0.023 | -0.059 | -0.059 | 0 | 0.87 | 0.95 | 0.74 | 0.77 | 0.85 | 0.72 |
| 6 | 155375896 | rs2062411 | -0.01 | -0.057 | -0.006 | 0.443 | -0.012 | -0.02 | -0.02 | 0.036 | 0.33 | 0.46 | 0.37 | 0.3 | 0.26 | 0.19 |
| 6 | 155375989 | rs7774757 | 0.725 | -0.02 | -0.013 | -0.245 | -0.002 | -0.019 | -0.019 | 0.074 | 0.41 | 0.35 | 0.29 | 0.06 | 0.44 | 0.26 |
| 6 | 155376270 | rs9397770 | 0.658 | -0.346 | -0.099 | -0.132 | 0.031 | 0.058 | 0.058 | 0 | 0.21 | 0.08 | 0.1 | 0.1 | 0.17 | 0.17 |
| 6 | 155379737 | rs9397772 | -0.312 | -0.683 | 0.088 | -0.276 | -0.033 | -0.042 | -0.042 | 1.603 | 0.15 | 0.27 | 0.06 | 0.07 | 0.08 | 0.12 |
| 6 | 155382807 | rs9322497 | 1.22 | -0.138 | 0.104 | -0.644 | -0.062 | 0.057 | 0.057 | 0 | 0.4 | 0.16 | 0.25 | 0.27 | 0.4 | 0.29 |
| 6 | 155383198 | rs11156008 | -0.02 | 0.004 | 0.321 | 0.234 | -0.001 | -0.001 | -0.001 | 0 | 0.64 | 0.74 | 0.52 | 0.62 | 0.55 | 0.58 |
| 6 | 155387526 | rs6900877 | 0.963 | 0.034 | 0.018 | 0.183 | -0.038 | 0.1 | 0.1 | 0 | 0.52 | 0.18 | 0.28 | 0.35 | 0.48 | 0.41 |
| 6 | 155389334 | rs17085911 | 0.858 | -0.055 | 0.083 | 0.538 | -0.001 | -0.001 | -0.001 | 0 | 0.07 | 0.05 | 0.04 | 0.21 | 0.06 | 0.16 |
| 6 | 155392569 | rs6935629 | 1.057 | 0.16 | 0.075 | 0.21 | -0.005 | 0.009 | 0.009 | 0 | 0.48 | 0.41 | 0.45 | 0.54 | 0.46 | 0.41 |
| 6 | 155392692 | rs6931203 | 0.19 | 0.435 | 0.45 | 0.337 | -0.003 | -0.024 | -0.024 | 0 | 0.17 | 0.19 | 0.07 | 0.04 | 0.11 | 0.19 |
| 6 | 155395055 | rs9384280 | 0.315 | 0.721 | 0.258 | 0.604 | -0.004 | 0.002 | 0.002 | 0 | 0.17 | 0.1 | 0.34 | 0.44 | 0.24 | 0.36 |
| 6 | 155404129 | rs6901517 | 0.482 | 0.707 | 0.344 | -0.022 | -0.009 | -0.061 | -0.061 | 0.555 | 0.16 | 0.23 | 0.08 | 0.04 | 0.12 | 0.18 |
| 6 | 155407208 | rs9322498 | 1.133 | 0.84 | -0.212 | 0.108 | -0.048 | 0.081 | 0.081 | 0.379 | 0.4 | 0.11 | 0.24 | 0.28 | 0.39 | 0.23 |
| 6 | 155410710 | rs12207246 | 0.265 | 1.003 | 0.07 | -0.153 | 0.002 | -0.036 | -0.036 | 0 | 0.07 | 0.14 | 0.25 | 0 | 0.08 | 0.09 |
| 6 | 155411270 | rs9384283 | 0.877 | 0.997 | -0.063 | 0.333 | -0.021 | -0.056 | -0.056 | 0 | 0.09 | 0 | 0.05 | 0.31 | 0.07 | 0.26 |
| 6 | 155411606 | rs6915661 | -0.196 | 1.548 | 0.133 | -0.14 | 0.043 | -0.027 | -0.027 | 0 | 0.08 | 0.16 | 0.44 | 0.02 | 0.14 | 0.1 |
| 6 | 155412553 | rs4869845 | 0.412 | 1.379 | 0.109 | -0.159 | 0.01 | 0.088 | 0.088 | 0 | 0.49 | 0.3 | 0.67 | 0.28 | 0.5 | 0.33 |
| 6 | 155417066 | rs13203300 | 1.088 | 1.417 | -0.137 | -0.155 | -0.003 | 0.078 | 0.078 | 0.373 | 0.39 | 0.09 | 0.24 | 0.28 | 0.36 | 0.21 |
| 6 | 155418824 | rs11156010 | 0.124 | 1.041 | 0.182 | -0.019 | -0.01 | 0.114 | 0.114 | 0 | 0.55 | 0.85 | 0.72 | 0.3 | 0.52 | 0.32 |
| 6 | 155420691 | rs10223883 | 0.986 | 1.077 | 0.194 | -0.345 | -0.008 | -0.008 | -0.008 | 0 | 0.34 | 0.3 | 0.57 | 0.04 | 0.37 | 0.25 |
| 6 | 155422393 | rs6557400 | 0.097 | 1.12 | 0.271 | -0.19 | 0.011 | 0.09 | 0.09 | 0 | 0.54 | 0.75 | 0.71 | 0.27 | 0.52 | 0.31 |
| 6 | 155424250 | rs9478613 | 0.301 | 1.168 | 0.131 | -0.181 | -0.001 | 0.017 | 0.017 | 0 | 0.51 | 0.61 | 0.54 | 0.28 | 0.48 | 0.3 |
| 6 | 155427624 | rs6942152 | -0.552 | 1.211 | -0.182 | 0.215 | 0.001 | 0.035 | 0.035 | 0 | 0.25 | 0.36 | 0.17 | 0.35 | 0.24 | 0.46 |

| | | | | | | | | | | | | | | | | |
|---|---|---|---|---|---|---|---|---|---|---|---|---|---|---|---|---|
| 6 | 155428222 | rs7757550 | 1.019 | 0.673 | -0.968 | 0.479 | 0.002 | -0.017 | -0.017 | 0 | 0.15 | 0.11 | 0.1 | 0.24 | 0.12 | 0.06 |
| 6 | 155437978 | rs9383743 | -0.205 | 0.855 | -1.222 | 0.421 | -0.038 | -0.11 | -0.11 | 0.701 | 0.22 | 0.07 | 0.12 | 0.36 | 0.2 | 0.44 |
| 6 | 155439372 | rs4870356 | -0.886 | 0.705 | -1.033 | 0.218 | 0.01 | 0.023 | 0.023 | 0.885 | 0.42 | 0.72 | 0.4 | 0.64 | 0.32 | 0.5 |
| 6 | 155442052 | rs6557403 | -0.844 | 0.67 | -1.048 | 0.23 | 0.013 | 0.017 | 0.017 | 0.921 | 0.42 | 0.72 | 0.25 | 0.63 | 0.32 | 0.5 |
| 6 | 155444724 | rs9322500 | -0.851 | 0.717 | -1.015 | 0.231 | 0.018 | 0.034 | 0.034 | 1.153 | 0.42 | 0.78 | 0.39 | 0.65 | 0.3 | 0.5 |
| 6 | 155449531 | rs9480063 | 1.07 | 1.037 | -1.582 | -0.368 | -0.084 | 0.197 | 0.197 | 0.63 | 0.33 | 0.04 | 0.43 | 0.02 | 0.37 | 0.26 |
| 6 | 155451352 | rs931312 | -0.16 | 1.071 | -1.297 | 0.176 | -0.015 | -0.058 | -0.058 | 0 | 0.21 | 0.07 | 0.25 | 0.3 | 0.18 | 0.44 |
| 6 | 155456335 | rs9478616 | 0.855 | 1.202 | -1.284 | -0.304 | -0.082 | 0.17 | 0.17 | 0.983 | 0.3 | 0.03 | 0.43 | 0.03 | 0.34 | 0.24 |
| 6 | 155456818 | rs9397228 | -0.441 | 1.116 | -0.164 | 0.184 | -0.017 | 0.037 | 0.037 | 0 | 0.7 | 0.51 | 0.78 | 0.57 | 0.66 | 0.73 |
| 6 | 155460725 | rs13219130 | 0.268 | 1.195 | -0.249 | 0.156 | -0.006 | 0.015 | 0.015 | 0 | 0.14 | 0.1 | 0.19 | 0.05 | 0.12 | 0.17 |
| 6 | 155461595 | rs7759406 | 0.208 | 1.204 | -0.222 | 0.227 | -0.002 | 0.001 | 0.001 | 0 | 0.56 | 0.44 | 0.59 | 0.52 | 0.54 | 0.56 |
| 6 | 155472539 | rs2352150 | 0.221 | 1.301 | -0.705 | 0.232 | -0.006 | -0.001 | -0.001 | 2.267 | 0.54 | 0.28 | 0.56 | 0.52 | 0.51 | 0.52 |
| 6 | 155473551 | rs3805832 | 0.641 | 1.189 | -0.613 | 0.691 | -0.001 | 0.005 | 0.005 | 0 | 0.06 | 0.05 | 0.04 | 0.24 | 0.07 | 0.04 |
| 6 | 155485535 | rs1032140 | 0.013 | 1.31 | -0.282 | 0.321 | 0 | 0.009 | 0.009 | 0 | 0.66 | 0.36 | 0.75 | 0.56 | 0.6 | 0.7 |
| 6 | 155485554 | rs1032141 | 0.006 | 1.268 | -0.279 | 0.325 | -0.001 | 0.009 | 0.009 | 0 | 0.66 | 0.36 | 0.75 | 0.56 | 0.61 | 0.7 |
| 6 | 155491391 | rs998318 | 0.319 | 1.564 | -0.648 | -0.088 | -0.08 | 0.16 | 0.16 | 1.589 | 0.28 | 0.02 | 0.4 | 0.03 | 0.33 | 0.24 |
| 6 | 155503116 | rs7770051 | -0.318 | 1.404 | -0.058 | 0.315 | 0.002 | 0.003 | 0.003 | 0 | 0.57 | 0.57 | 0.63 | 0.53 | 0.64 | 0.58 |
| 6 | 155503151 | rs7756965 | -0.561 | 1.24 | -0.945 | 0.358 | 0.02 | 0.002 | 0.002 | 0.006 | 0.62 | 0.89 | 0.71 | 0.58 | 0.7 | 0.66 |
| 6 | 155504960 | rs6905473 | -0.328 | 1.219 | -0.563 | 0.67 | -0.012 | -0.002 | -0.002 | 0 | 0.44 | 0.26 | 0.5 | 0.48 | 0.48 | 0.56 |
| 6 | 155509242 | rs9371870 | -0.238 | 1.339 | -0.889 | 0.581 | -0.024 | -0.041 | -0.041 | 0.136 | 0.34 | 0.17 | 0.45 | 0.41 | 0.37 | 0.46 |
| 6 | 155511901 | rs9480081 | 0.719 | 1.319 | -0.022 | 0.069 | 0.02 | -0.067 | -0.067 | 0 | 0.88 | 0.72 | 0.93 | 1 | 0.87 | 0.96 |
| 6 | 155514059 | rs9478621 | 1.185 | 1.304 | -0.291 | -0.017 | -0.001 | 0.004 | 0.004 | 0.065 | 0.57 | 0.39 | 0.51 | 0.75 | 0.53 | 0.65 |
| 6 | 155515255 | rs11759031 | -0.541 | 1.324 | -0.302 | 0.358 | -0.002 | -0.021 | -0.021 | 0 | 0.17 | 0.06 | 0.08 | 0.24 | 0.2 | 0.23 |
| 6 | 155516499 | rs17086041 | -1.471 | 1.402 | 0.073 | 0.165 | 0.007 | -0.029 | -0.029 | 0 | 0.33 | 0.56 | 0.55 | 0.21 | 0.36 | 0.33 |
| 6 | 155517672 | rs6919431 | -0.691 | 0.832 | -0.47 | 1.032 | 0.01 | -0.096 | -0.096 | 1.568 | 0.23 | 0.11 | 0.18 | 0.49 | 0.25 | 0.26 |
| 6 | 155519888 | rs17086046 | -0.528 | 0.919 | 0.201 | 0.945 | -0.002 | 0.062 | 0.062 | 0 | 0.08 | 0.22 | 0.15 | 0.2 | 0.05 | 0.14 |
| 6 | 155530844 | rs9384296 | -1.09 | 1.606 | -0.166 | 0.976 | -0.001 | 0.007 | 0.007 | 0 | 0.15 | 0.17 | 0.17 | 0.36 | 0.12 | 0.22 |
| 6 | 155532689 | rs1485754 | -0.626 | 1.693 | 0.007 | 0.548 | -0.002 | 0.027 | 0.027 | 0 | 0.29 | 0.6 | 0.56 | 0.53 | 0.29 | 0.27 |
| 6 | 155536225 | rs9397235 | -0.809 | 1.708 | -0.095 | 0.551 | 0 | 0.007 | 0.007 | 0 | 0.15 | 0.16 | 0.08 | 0.35 | 0.13 | 0.23 |
| 6 | 155536606 | rs9480091 | -0.524 | 1.87 | 0.627 | 0.387 | -0.003 | 0.023 | 0.023 | 0 | 0.29 | 0.32 | 0.58 | 0.51 | 0.32 | 0.32 |
| 6 | 155537311 | rs9384297 | -0.93 | 1.794 | 0.368 | 0.612 | -0.005 | -0.005 | -0.005 | 1.111 | 0.15 | 0.13 | 0.08 | 0.35 | 0.14 | 0.26 |
| 6 | 155539010 | rs9383745 | 0.01 | 1.614 | 0.186 | 0.546 | -0.005 | -0.038 | -0.038 | 0 | 0.07 | 0 | 0.04 | 0.2 | 0.05 | 0.16 |
| 6 | 155544003 | rs9397793 | -1.361 | 1.63 | 0.57 | 0.046 | 0.002 | -0.033 | -0.033 | 0 | 0.38 | 0.56 | 0.63 | 0.87 | 0.34 | 0.45 |
| 6 | 155544150 | rs6557409 | -0.154 | 2.061 | 0.102 | -0.174 | -0.023 | 0.022 | 0.022 | 0 | 0.94 | 0.97 | 0.92 | 1 | 0.88 | 0.96 |
| 6 | 155544257 | rs6557410 | 0.653 | 2.206 | 0.538 | 0.067 | -0.015 | -0.017 | -0.017 | 0 | 0.73 | 0.75 | 0.39 | 0.48 | 0.65 | 0.71 |
| 6 | 155544397 | rs4143683 | -0.115 | 1.932 | 0.3 | 0.151 | -0.022 | -0.062 | -0.062 | 0 | 0.2 | 0.13 | 0.52 | 0.52 | 0.23 | 0.25 |
| 6 | 155545107 | rs9384302 | -0.012 | 1.362 | 0.546 | 0.047 | 0.003 | -0.008 | -0.008 | 0 | 0.07 | 0.07 | 0.04 | 0.2 | 0.05 | 0.08 |
| 6 | 155551221 | rs6930662 | 1.32 | 1.421 | 0.598 | -0.162 | -0.002 | -0.002 | -0.002 | 0 | 0.7 | 0.33 | 0.73 | 0.57 | 0.67 | 0.67 |
| 6 | 155556469 | rs2882936 | 0.039 | 1.295 | 0.334 | -0.943 | -0.001 | -0.016 | -0.016 | 0 | 0.25 | 0.17 | 0.5 | 0.62 | 0.24 | 0.34 |
| 6 | 155561796 | rs11751128 | 1.74 | 0.595 | -0.628 | -1.795 | -0.07 | 0.156 | 0.156 | 0 | 0.24 | 0.01 | 0.15 | 0 | 0.28 | 0.07 |
| 6 | 155568881 | rs3011906 | 0.369 | 0.369 | -0.023 | -1.854 | 0.001 | 0.004 | 0.004 | 0 | 0.57 | 0.6 | 0.3 | 0.19 | 0.48 | 0.38 |
| 6 | 155569862 | rs3011903 | -0.691 | 0.453 | 0.027 | -1.853 | -0.002 | 0.002 | 0.002 | 0 | 0.74 | 0.69 | 0.48 | 0.35 | 0.75 | 0.57 |
| 6 | 155571336 | rs3011901 | 1.403 | 0.04 | 0.187 | -2.052 | 0.019 | 0.131 | 0.131 | 0 | 0.36 | 0.12 | 0.19 | 0.01 | 0.32 | 0.25 |
| 6 | 155572854 | rs1108371 | 0.54 | 0.116 | 0.152 | -1.983 | 0.001 | -0.001 | -0.001 | 0 | 0.27 | 0.26 | 0.52 | 0.66 | 0.25 | 0.39 |
| 6 | 155574427 | rs2274825 | 1.138 | 0.079 | 0.261 | -1.986 | 0 | -0.001 | -0.001 | 0 | 0.07 | 0.09 | 0.04 | 0.18 | 0.05 | 0.07 |
| 6 | 155581024 | rs1408756 | -0.043 | 0.292 | 0.317 | -2.021 | -0.007 | 0.007 | 0.007 | 0 | 0.37 | 0.35 | 0.62 | 0.8 | 0.41 | 0.59 |
| 6 | 155585154 | rs11966646 | 1.084 | 0.301 | 0.337 | -2.036 | 0.014 | -0.019 | -0.019 | 3.403 | 0.06 | 0.06 | 0.08 | 0 | 0.11 | 0.04 |
| 6 | 155585278 | rs2985700 | -0.526 | 0.282 | 0.445 | -2.033 | -0.001 | -0.059 | -0.059 | 0 | 0.45 | 0.49 | 0.66 | 0.97 | 0.46 | 0.66 |
| 6 | 155585325 | rs9480098 | 0.481 | 0.273 | 0.419 | -1.995 | -0.001 | 0 | 0 | 0 | 0.28 | 0.3 | 0.53 | 0.64 | 0.26 | 0.45 |
| 6 | 155586131 | rs1041098 | -2.258 | 0.475 | 0.046 | -2.183 | -0.019 | 0.14 | 0.14 | 0 | 0.78 | 0.99 | 0.9 | 1 | 0.81 | 0.94 |
| 6 | 155596700 | rs1014664 | 0.555 | 0.892 | 1.223 | -1.449 | -0.003 | 0.02 | 0.02 | 0 | 0.08 | 0.17 | 0.06 | 0.18 | 0.06 | 0.13 |
| 6 | 155597147 | rs1555774 | 0.367 | 0.907 | 1.146 | -1.457 | 0.001 | -0.001 | -0.001 | 0.569 | 0.27 | 0.26 | 0.52 | 0.65 | 0.25 | 0.39 |
| 6 | 155606325 | rs324356 | -1.14 | 0.939 | 1.191 | -1.478 | -0.015 | 0.118 | 0.118 | 0 | 0.53 | 0.68 | 0.71 | 0.83 | 0.53 | 0.6 |
| 6 | 155615348 | rs12529366 | 0.758 | 0.947 | 1.205 | -1.449 | 0.019 | -0.019 | -0.019 | 0 | 0.06 | 0.06 | 0.08 | 0 | 0.13 | 0.04 |
| 6 | 155615728 | rs162965 | -1.706 | 0.922 | 1.219 | -1.523 | 0.021 | 0.056 | 0.056 | 0 | 0.63 | 0.73 | 0.8 | 0.99 | 0.68 | 0.75 |
| 6 | 155621019 | rs950994 | -0.277 | 0.934 | 1.282 | -1.45 | -0.02 | -0.023 | -0.023 | 0 | 0.39 | 0.46 | 0.56 | 0.66 | 0.33 | 0.43 |
| 6 | 155625104 | rs9371371 | -0.994 | 0.901 | 1.232 | -1.45 | 0.021 | 0.064 | 0.064 | 0 | 0.52 | 0.34 | 0.72 | 0.81 | 0.57 | 0.7 |
| 6 | 155630653 | rs9384310 | 0.758 | 0.687 | 1.183 | -1.387 | -0.059 | -0.032 | -0.032 | 0 | 0.09 | 0.01 | 0.08 | 0.14 | 0.15 | 0.21 |
| 6 | 155632762 | rs56237 | -0.077 | 0.596 | 1.516 | -1.244 | 0.018 | 0.035 | 0.035 | 0 | 0.1 | 0.28 | 0.07 | 0.17 | 0.09 | 0.12 |
| 6 | 155632844 | rs721101 | 0.796 | 0.538 | 1.447 | -1.323 | -0.04 | 0.026 | 0.026 | 0 | 0.18 | 0.05 | 0.13 | 0.14 | 0.24 | 0.26 |
| 6 | 155632938 | rs162984 | -1.289 | 0.015 | 1.601 | -1.151 | 0.008 | 0.01 | 0.01 | 0 | 0.61 | 0.64 | 0.78 | 0.98 | 0.65 | 0.83 |
| 6 | 155635624 | rs1334688 | -0.649 | 0.046 | 1.224 | -1.12 | 0.019 | 0.055 | 0.055 | 0 | 0.52 | 0.36 | 0.72 | 0.81 | 0.59 | 0.7 |
| 6 | 155643247 | rs4585573 | 0.132 | 0.343 | 1.371 | -0.667 | -0.03 | 0.036 | 0.036 | 0 | 0.25 | 0.12 | 0.22 | 0.14 | 0.33 | 0.3 |
| 6 | 155643733 | rs17086159 | -0.691 | -0.25 | 2.475 | 0.527 | -0.001 | 0.005 | 0.005 | 0 | 0.09 | 0.16 | 0.04 | 0.17 | 0.08 | 0.07 |
| 6 | 155647047 | rs9397238 | -0.792 | -0.176 | 1.873 | 0.796 | -0.019 | 0.003 | 0.003 | 0 | 0.22 | 0.28 | 0.48 | 0.69 | 0.18 | 0.39 |
| 6 | 155652382 | rs114266 | -0.949 | 0.286 | 1.96 | 1.308 | -0.004 | 0.083 | 0.083 | 0 | 0.08 | 0.18 | 0.03 | 0.16 | 0.05 | 0.07 |
| 6 | 155661820 | rs335405 | -2.037 | 0.37 | 2.558 | 1.82 | 0.012 | 0.091 | 0.091 | 0 | 0.23 | 0.46 | 0.44 | 0.43 | 0.24 | 0.33 |
| 6 | 155661931 | rs9322513 | -1.794 | 0.377 | 2.823 | 1.847 | 0 | 0.039 | 0.039 | 0 | 0.21 | 0.26 | 0.42 | 0.39 | 0.2 | 0.32 |
| 6 | 155683997 | rs7741784 | -0.629 | 0.304 | 2.894 | 1.701 | -0.008 | 0.033 | 0.033 | 0 | 0.07 | 0.19 | 0.03 | 0.13 | 0.03 | 0.05 |

| | | | | | | | | | | | | | | | | |
|---|---|---|---|---|---|---|---|---|---|---|---|---|---|---|---|---|
| 6 | 155692133 | rs9480123 | -2.149 | 0.303 | 2.972 | 1.746 | 0.006 | 0.118 | 0.118 | 0 | 0.23 | 0.44 | 0.43 | 0.47 | 0.23 | 0.34 |
| 6 | 155697906 | rs9322520 | -1.147 | 0.302 | 2.898 | 1.743 | 0.008 | 0.037 | 0.037 | 0 | 0.15 | 0.25 | 0.39 | 0.26 | 0.17 | 0.28 |
| 6 | 155699732 | rs9383760 | -2.015 | 0.299 | 2.944 | 1.752 | 0.004 | 0.076 | 0.076 | 0 | 0.22 | 0.32 | 0.42 | 0.47 | 0.23 | 0.35 |
| 6 | 155700624 | rs9397818 | -1.324 | 0.295 | 2.782 | 1.696 | -0.001 | 0.013 | 0.013 | 0 | 0.1 | 0.12 | 0.04 | 0.21 | 0.08 | 0.07 |
| 6 | 155704119 | rs17086231 | -0.382 | 0.302 | 2.961 | 1.99 | -0.013 | -0.019 | -0.019 | 0 | 0.08 | 0.03 | 0.44 | 0.24 | 0.12 | 0.16 |
| 6 | 155705018 | rs7759871 | -1.2 | 0.271 | 3.306 | 1.893 | 0.029 | 0.07 | 0.07 | 0 | 0.15 | 0.38 | 0.38 | 0.22 | 0.16 | 0.25 |
| 6 | 155722119 | rs163003 | -2.231 | 0.277 | 3.396 | 2.023 | -0.001 | 0.002 | 0.002 | 0.064 | 0.23 | 0.72 | 0.28 | 0.4 | 0.21 | 0.31 |
| 6 | 155728232 | rs12524299 | -1.043 | 0.281 | 3.473 | 1.933 | 0.002 | 0.005 | 0.005 | 0 | 0.14 | 0.17 | 0.22 | 0.19 | 0.15 | 0.23 |
| 6 | 155733557 | rs17086266 | -1.471 | 0.321 | 3.074 | 1.898 | -0.001 | 0.002 | 0.002 | 0 | 0.17 | 0.28 | 0.23 | 0.2 | 0.17 | 0.23 |
| 6 | 155733636 | rs11759841 | -1.862 | 0.326 | 2.816 | 1.884 | 0.007 | 0.094 | 0.094 | 0 | 0.2 | 0.46 | 0.26 | 0.33 | 0.19 | 0.28 |
| 6 | 155740285 | rs231946 | -0.966 | 0.588 | 0.984 | 0.889 | 0.026 | 0.04 | 0.04 | 0 | 0.09 | 0.33 | 0.03 | 0.09 | 0.07 | 0.05 |
| 6 | 155740528 | rs2294680 | -1.195 | 0.971 | -0.063 | 0.78 | 0.005 | 0.004 | 0.004 | 0 | 0.15 | 0.15 | 0.23 | 0.2 | 0.18 | 0.22 |
| 6 | 155743245 | rs13207865 | -0.41 | 0.877 | -0.457 | -0.048 | -0.001 | -0.019 | -0.019 | 0 | 0.26 | 0.21 | 0.25 | 0.07 | 0.27 | 0.17 |
| 6 | 155748212 | rs2281253 | -0.734 | 0.275 | -0.947 | 0.442 | 0.005 | 0.011 | 0.011 | 0 | 0.36 | 0.23 | 0.35 | 0.74 | 0.34 | 0.34 |
| 6 | 155749404 | rs231951 | -0.327 | 0.832 | -1.447 | -0.551 | -0.036 | 0.044 | 0.044 | 0 | 0.86 | 0.97 | 0.88 | 0.96 | 0.84 | 0.93 |
| 6 | 155750035 | rs231954 | 0.078 | 0.637 | -1.309 | -0.461 | -0.013 | 0.071 | 0.071 | 0 | 0.58 | 0.75 | 0.63 | 0.91 | 0.57 | 0.79 |
| 6 | 155754323 | rs231957 | 0.28 | 0.844 | -1.364 | -0.747 | -0.019 | 0.026 | 0.026 | 0 | 0.17 | 0.1 | 0.13 | 0.06 | 0.24 | 0.14 |
| 6 | 155755404 | rs231958 | -0.681 | 0.434 | -0.532 | -0.17 | 0.001 | 0 | 0 | 0 | 0.36 | 0.55 | 0.29 | 0.58 | 0.39 | 0.38 |
| 6 | 155758610 | rs2235668 | -0.195 | 0.02 | -0.414 | -0.664 | -0.022 | 0.011 | 0.011 | 0 | 0.23 | 0.35 | 0.18 | 0.34 | 0.15 | 0.23 |
| 6 | 155759200 | rs12191893 | 0.368 | 0.454 | -0.719 | -0.896 | -0.037 | 0.06 | 0.06 | 0 | 0.21 | 0.08 | 0.15 | 0.09 | 0.28 | 0.17 |
| 6 | 155759778 | rs9322524 | -0.078 | 0.56 | -0.083 | -0.822 | 0.02 | -0.059 | -0.059 | 0 | 0.13 | 0.2 | 0.12 | 0.02 | 0.14 | 0.21 |
| 6 | 155760195 | rs2235670 | -0.011 | 0.053 | -0.037 | -0.335 | -0.017 | 0.023 | 0.023 | 0.177 | 0.16 | 0.24 | 0.16 | 0.33 | 0.12 | 0.19 |
| 6 | 155761246 | rs3749930 | 0.043 | 0.39 | -0.249 | 0.091 | -0.004 | 0.134 | 0.134 | 0 | 0.06 | 0.07 | 0.19 | 0.48 | 0.05 | 0.23 |
| 6 | 155766362 | rs6919626 | -0.026 | 0.807 | -0.301 | -0.484 | 0.005 | 0.01 | 0.01 | 0 | 0.63 | 0.59 | 0.51 | 0.49 | 0.64 | 0.65 |
| 6 | 155766687 | rs6935359 | 0.522 | 0.391 | -1.032 | -0.566 | 0.036 | 0 | 0 | 0 | 0.35 | 0.22 | 0.29 | 0.41 | 0.27 | 0.32 |
| 6 | 155771008 | rs2235674 | 0.082 | 1.511 | -1.089 | -0.101 | -0.049 | -0.053 | -0.053 | 0 | 0.18 | 0.1 | 0.39 | 0.53 | 0.34 | 0.4 |
| 6 | 155771597 | rs12661812 | -0.342 | 0.806 | -1.012 | -0.412 | 0.002 | 0.004 | 0.004 | 0.07 | 0.09 | 0.09 | 0.06 | 0.02 | 0.08 | 0.09 |
| 6 | 155771929 | rs2235675 | -0.613 | 0.857 | -1.424 | -0.12 | -0.042 | -0.073 | -0.073 | 0 | 0.71 | 0.87 | 0.51 | 0.55 | 0.64 | 0.65 |
| 6 | 155772039 | rs9478652 | -0.877 | 0.419 | -0.347 | -0.358 | 0.031 | -0.05 | -0.05 | 0 | 0.2 | 0.45 | 0.11 | 0.1 | 0.22 | 0.16 |
| 6 | 155775606 | rs13203407 | 1.404 | -0.011 | -0.619 | -0.427 | 0.016 | 0.01 | 0.01 | 0 | 0.15 | 0.11 | 0.23 | 0.12 | 0.1 | 0.12 |
| 6 | 155775980 | rs12195525 | 1.408 | -0.015 | -0.613 | -0.429 | 0.014 | 0.009 | 0.009 | 0 | 0.15 | 0.11 | 0.23 | 0.12 | 0.1 | 0.12 |
| 6 | 155778870 | rs9478654 | -0.115 | -0.237 | -0.749 | -0.748 | -0.019 | 0.032 | 0.032 | 0.867 | 0.22 | 0.14 | 0.12 | 0.06 | 0.26 | 0.3 |
| 6 | 155779534 | rs6935922 | -0.935 | -0.337 | -0.431 | -0.793 | -0.003 | -0.032 | -0.032 | 0.934 | 0.39 | 0.56 | 0.36 | 0.18 | 0.37 | 0.42 |
| 6 | 155791824 | rs9322525 | -0.6 | -0.373 | -0.829 | -0.728 | -0.007 | -0.014 | -0.014 | 0.312 | 0.28 | 0.17 | 0.16 | 0.39 | 0.28 | 0.43 |
| 6 | 155796830 | rs10457095 | -1.486 | -0.35 | -0.618 | -0.752 | 0.034 | 0.033 | 0.033 | 0.173 | 0.5 | 0.72 | 0.44 | 0.51 | 0.43 | 0.57 |
| 6 | 155804649 | rs2181069 | 1.43 | -0.235 | -0.812 | -0.471 | 0.013 | -0.005 | -0.005 | 0 | 0.16 | 0.07 | 0.23 | 0.15 | 0.13 | 0.14 |
| 6 | 155805281 | rs7738860 | -1.632 | -0.256 | -0.869 | -0.197 | -0.005 | -0.041 | -0.041 | 0.889 | 0.55 | 0.85 | 0.46 | 0.59 | 0.46 | 0.65 |
| 6 | 155806127 | rs9384333 | 0.314 | -0.227 | -0.608 | -0.225 | -0.004 | -0.006 | -0.006 | 0 | 0.15 | 0.15 | 0.07 | 0.41 | 0.18 | 0.35 |
| 6 | 155808279 | rs7763526 | -0.817 | -0.396 | -0.347 | -0.611 | 0.023 | 0.03 | 0.03 | 0.914 | 0.4 | 0.68 | 0.38 | 0.18 | 0.28 | 0.29 |
| 6 | 155813894 | rs4869856 | -0.594 | -0.294 | -0.293 | -0.61 | 0.028 | 0.041 | 0.041 | 0.031 | 0.37 | 0.75 | 0.37 | 0.16 | 0.27 | 0.26 |
| 6 | 155828491 | rs2145982 | -0.372 | -0.15 | -0.879 | -0.652 | 0.042 | 0.064 | 0.064 | 0.082 | 0.36 | 0.79 | 0.37 | 0.14 | 0.26 | 0.24 |
| 6 | 155832225 | rs12199201 | 1.289 | -0.276 | -1.027 | -0.736 | 0.023 | -0.005 | -0.005 | 0 | 0.15 | 0.04 | 0.25 | 0.13 | 0.11 | 0.13 |
| 6 | 155836065 | rs4869857 | -1.616 | 0.025 | -1.272 | -0.159 | -0.029 | -0.062 | -0.062 | 0 | 0.55 | 0.89 | 0.47 | 0.57 | 0.47 | 0.64 |
| 6 | 155840770 | rs9371892 | 0.288 | 0.026 | -1.266 | -0.109 | -0.051 | -0.098 | -0.098 | 0 | 0.18 | 0.04 | 0.09 | 0.43 | 0.21 | 0.4 |
| 6 | 155847549 | rs4869859 | -1.616 | 0.017 | -1.248 | -0.118 | -0.027 | -0.049 | -0.049 | 0 | 0.55 | 0.88 | 0.47 | 0.56 | 0.47 | 0.64 |
| 6 | 155863106 | rs10872716 | -0.473 | -0.103 | -1.146 | -0.512 | 0.042 | 0.065 | 0.065 | 0.034 | 0.36 | 0.78 | 0.37 | 0.14 | 0.25 | 0.24 |
| 6 | 155870828 | rs7752075 | -1.286 | -0.005 | -1.167 | -0.309 | -0.04 | -0.002 | -0.002 | 0.01 | 0.56 | 0.84 | 0.61 | 0.49 | 0.5 | 0.71 |
| 6 | 155872642 | rs9480156 | -1.1 | -0.077 | -0.505 | -0.034 | -0.029 | 0.03 | 0.03 | 0.748 | 0.22 | 0.56 | 0.19 | 0.35 | 0.17 | 0.41 |
| 6 | 155872975 | rs12190086 | 1.322 | 0.337 | -0.935 | -0.173 | -0.045 | -0.007 | -0.007 | 0 | 0.16 | 0.02 | 0.37 | 0.13 | 0.19 | 0.13 |
| 6 | 155878027 | rs12195819 | 1.296 | 0.281 | -0.942 | -0.205 | -0.068 | 0.004 | 0.004 | 0 | 0.17 | 0.01 | 0.25 | 0.13 | 0.2 | 0.13 |
| 6 | 155896513 | rs9480160 | 0.348 | 0.215 | -0.964 | -0.038 | -0.02 | -0.064 | -0.064 | 0 | 0.11 | 0.03 | 0.06 | 0.26 | 0.11 | 0.41 |
| 6 | 155896760 | rs9371895 | 0.372 | 0.18 | -0.957 | -0.399 | -0.001 | 0.04 | 0.04 | 0 | 0.13 | 0.1 | 0.31 | 0.55 | 0.16 | 0.16 |
| 6 | 155897444 | rs12663818 | 0.389 | 0.172 | -0.92 | -0.388 | -0.002 | 0.056 | 0.056 | 0 | 0.12 | 0.15 | 0.29 | 0.56 | 0.13 | 0.15 |
| 6 | 155898417 | rs4242289 | 0.014 | -0.031 | -0.599 | -0.55 | 0 | 0 | 0 | 0 | 0.63 | 0.68 | 0.66 | 0.57 | 0.65 | 0.42 |
| 6 | 155899723 | rs11156030 | 1.409 | 0.492 | -1.06 | -0.956 | -0.036 | 0.049 | 0.049 | 0.023 | 0.18 | 0.06 | 0.15 | 0.01 | 0.24 | 0.07 |
| 6 | 155911026 | rs11755068 | -0.281 | -1.144 | -0.299 | -0.009 | 0.003 | -0.007 | -0.007 | 0.058 | 0.13 | 0.1 | 0.14 | 0.05 | 0.17 | 0.08 |
| 6 | 155912676 | rs9371379 | 0.816 | -1.547 | -0.694 | -0.142 | -0.001 | 0.155 | 0.155 | 0 | 0.53 | 0.15 | 0.53 | 0.22 | 0.53 | 0.48 |
| 6 | 155915224 | rs9371900 | 1.055 | -1.91 | 0.7 | -0.299 | -0.001 | -0.014 | -0.014 | 0 | 0.51 | 0.49 | 0.52 | 0.17 | 0.5 | 0.36 |
| 6 | 155920891 | rs6557436 | -0.263 | -1.982 | 0.825 | -0.23 | 0 | 0 | 0 | 0 | 0.73 | 0.76 | 0.7 | 0.23 | 0.71 | 0.54 |
| 6 | 155921475 | rs9480167 | -0.887 | -1.971 | 0.868 | -0.219 | 0 | -0.026 | -0.026 | 0.019 | 0.19 | 0.22 | 0.17 | 0.05 | 0.2 | 0.16 |
| 6 | 155935243 | rs1909301 | -0.784 | -1.978 | 0.715 | -0.022 | 0 | -0.017 | -0.017 | 0 | 0.18 | 0.17 | 0.17 | 0.05 | 0.2 | 0.12 |
| 6 | 155938820 | rs9478667 | -0.899 | -1.954 | 0.705 | -0.02 | 0.001 | -0.028 | -0.028 | 0 | 0.19 | 0.22 | 0.17 | 0.05 | 0.2 | 0.12 |
| 6 | 155949243 | rs9397828 | 0.788 | -2.033 | 0.716 | -0.005 | -0.004 | 0.014 | 0.014 | 1.447 | 0.57 | 0.61 | 0.55 | 0.58 | 0.52 | 0.5 |
| 6 | 155950001 | rs9480176 | -0.886 | -1.971 | 0.755 | -0.002 | 0 | -0.028 | -0.028 | 0.283 | 0.19 | 0.22 | 0.17 | 0.05 | 0.2 | 0.14 |
| 6 | 155959814 | rs1498636 | -0.533 | -1.952 | 0.607 | 0.035 | -0.014 | -0.036 | -0.036 | 0.001 | 0.76 | 0.82 | 0.72 | 0.63 | 0.73 | 0.63 |
| 6 | 155970936 | rs6913088 | 0.919 | -2.018 | 1.25 | 0.036 | -0.001 | 0.014 | 0.014 | 0.235 | 0.54 | 0.35 | 0.53 | 0.58 | 0.51 | 0.5 |
| 6 | 155974396 | rs9371382 | 0.528 | -2.001 | 1.105 | -0.005 | 0 | -0.001 | -0.001 | 0 | 0.59 | 0.37 | 0.56 | 0.61 | 0.55 | 0.57 |
| 6 | 155982996 | rs9371910 | -1.119 | -2.007 | 1.513 | 0.207 | -0.018 | 0.179 | 0.179 | 0 | 0.13 | 0.33 | 0.23 | 0.58 | 0.08 | 0.32 |
| 6 | 155987825 | rs9478671 | -0.812 | -1.918 | 0.924 | 0.081 | -0.005 | 0.01 | 0.01 | 0 | 0.17 | 0.14 | 0.14 | 0.04 | 0.2 | 0.13 |

| | | | | | | | | | | | | | | | | |
|---|---|---|---|---|---|---|---|---|---|---|---|---|---|---|---|---|
| 6 | 155994436 | rs9397831 | 0.915 | -1.858 | 1.068 | 0.31 | 0.017 | -0.007 | -0.007 | 0 | 0.57 | 0.31 | 0.44 | 0.41 | 0.59 | 0.34 |
| 6 | 155998045 | rs12662240 | -0.304 | -1.727 | 0.858 | 0.44 | 0.024 | 0.131 | 0.131 | 0 | 0.09 | 0.34 | 0.22 | 0.15 | 0.08 | 0.26 |
| 6 | 155998741 | rs7739122 | 0.227 | -1.76 | 0.855 | 0.441 | -0.001 | -0.036 | -0.036 | 0 | 0.7 | 0.86 | 0.68 | 0.59 | 0.68 | 0.63 |
| 6 | 155999212 | rs1498623 | -0.327 | -1.766 | 0.85 | 0.441 | -0.012 | -0.045 | -0.045 | 0 | 0.76 | 0.88 | 0.71 | 0.63 | 0.72 | 0.66 |
| 6 | 155999495 | rs17086570 | -0.297 | -1.743 | 0.883 | 0.409 | 0.024 | 0.131 | 0.131 | 0 | 0.09 | 0.34 | 0.22 | 0.15 | 0.08 | 0.26 |
| 6 | 155999789 | rs6908070 | 0.724 | -1.792 | 0.724 | 0.336 | -0.032 | 0.025 | 0.025 | 0 | 0.59 | 0.79 | 0.57 | 0.58 | 0.53 | 0.58 |
| 6 | 156000576 | rs1391656 | 0.673 | -1.593 | 0.856 | 0.481 | -0.038 | 0.004 | 0.004 | 0 | 0.62 | 0.8 | 0.58 | 0.59 | 0.56 | 0.59 |
| 6 | 156002324 | rs9371386 | 1.377 | -1.508 | 0.784 | 0.504 | 0 | 0.009 | 0.009 | 0 | 0.51 | 0.26 | 0.34 | 0.43 | 0.47 | 0.3 |
| 6 | 156005810 | rs17753470 | -0.306 | -1.185 | 0.466 | 0.465 | -0.003 | -0.014 | -0.014 | 0 | 0.13 | 0.16 | 0.12 | 0.01 | 0.16 | 0.1 |
| 6 | 156007516 | rs2133282 | -0.384 | -1.239 | 0.189 | 0.642 | -0.039 | 0.008 | 0.008 | 0 | 0.8 | 0.94 | 0.76 | 0.8 | 0.75 | 0.8 |
| 6 | 156012352 | rs12663232 | -1.048 | -1.227 | 0.645 | 0.933 | 0.004 | -0.013 | -0.013 | 0 | 0.24 | 0.49 | 0.34 | 0.16 | 0.24 | 0.35 |
| 6 | 156018291 | rs4407725 | -0.174 | -1.049 | 0.229 | 1.02 | -0.004 | -0.025 | -0.025 | 0 | 0.77 | 0.82 | 0.68 | 0.39 | 0.72 | 0.73 |
| 6 | 156021432 | rs10457919 | 0.053 | -1.24 | 0.459 | 0.062 | -0.011 | -0.008 | -0.008 | 0 | 0.34 | 0.17 | 0.5 | 0.37 | 0.4 | 0.43 |
| 6 | 156032199 | rs9322533 | 0.07 | -1.146 | 0.519 | 0.109 | -0.001 | 0 | 0 | 0 | 0.36 | 0.3 | 0.53 | 0.37 | 0.41 | 0.43 |
| 6 | 156050029 | rs4256444 | 0.734 | -1.234 | 0.716 | -0.267 | -0.093 | 0.21 | 0.21 | 0.538 | 0.39 | 0.11 | 0.32 | 0.08 | 0.43 | 0.25 |
| 6 | 156056334 | rs9480197 | 0.164 | -1.186 | 1.215 | -0.221 | 0.014 | -0.038 | -0.038 | 0 | 0.07 | 0.13 | 0.03 | 0 | 0.07 | 0.03 |
| 6 | 156063738 | rs2647709 | 0.485 | -1.129 | 1.179 | -0.079 | 0.003 | -0.001 | -0.001 | 0 | 0.79 | 0.85 | 0.88 | 0.48 | 0.88 | 0.71 |
| 6 | 156066601 | rs9371389 | 0.702 | -1.137 | 1.467 | -0.083 | -0.046 | 0.121 | 0.121 | 0 | 0.78 | 0.68 | 0.86 | 0.48 | 0.88 | 0.71 |
| 6 | 156068182 | rs716434 | -0.073 | -1.119 | 1.322 | -0.099 | 0.003 | 0.014 | 0.014 | 0.006 | 0.87 | 0.93 | 0.92 | 0.48 | 0.9 | 0.79 |
| 6 | 156080059 | rs2817430 | -0.905 | -1.094 | 1.257 | -0.119 | -0.062 | 0.146 | 0.146 | 0 | 0.24 | 0.43 | 0.14 | 0.51 | 0.12 | 0.29 |
| 6 | 156081507 | rs11156041 | 0.135 | -0.364 | 1.353 | 0.151 | -0.002 | 0.004 | 0.004 | 0 | 0.36 | 0.44 | 0.53 | 0.41 | 0.44 | 0.46 |
| 6 | 156084508 | rs12207602 | 1.295 | -0.218 | 1.862 | 1.262 | -0.034 | 0.137 | 0.137 | 0.167 | 0.78 | 0.52 | 0.8 | 0.48 | 0.86 | 0.73 |
| 6 | 156087233 | rs11965880 | -2.358 | 0.23 | 2.055 | 0.459 | 0.064 | 0.048 | 0.048 | 0.377 | 0.08 | 0.53 | 0.08 | 0.05 | 0.05 | 0.03 |
| 6 | 156088762 | rs6557446 | -1.677 | 0.232 | 2.09 | 0.435 | 0.098 | 0.132 | 0.132 | 0.359 | 0.05 | 0.45 | 0.06 | 0.04 | 0.02 | 0.02 |
| 6 | 156099690 | rs9480200 | -2.266 | 0.21 | 1.726 | 0.421 | 0.027 | 0.008 | 0.008 | 0.339 | 0.1 | 0.56 | 0.09 | 0.05 | 0.05 | 0.05 |
| 6 | 156101388 | rs11967053 | -1.266 | 0.422 | 0.951 | 0.382 | 0.076 | 0.025 | 0.025 | 0 | 0.05 | 0.25 | 0.04 | 0.01 | 0.04 | 0.03 |
| 6 | 156107830 | rs9480206 | -0.648 | 0.659 | 0.153 | 0.142 | 0 | -0.014 | -0.014 | 0 | 0.32 | 0.42 | 0.29 | 0.18 | 0.3 | 0.26 |
| 6 | 156109579 | rs4329126 | 0.957 | 0.908 | 0.571 | 0.738 | 0.02 | 0.097 | 0.097 | 0 | 0.78 | 0.6 | 0.57 | 0.64 | 0.78 | 0.75 |
| 6 | 156118550 | rs7749754 | 0.966 | 0.647 | 0.6 | 0.812 | 0.002 | -0.003 | -0.003 | 0 | 0.88 | 0.88 | 0.91 | 0.82 | 0.94 | 0.96 |
| 6 | 156121031 | rs7761161 | 1.012 | 0.616 | 0.586 | 0.9 | 0.003 | -0.004 | -0.004 | 0 | 0.88 | 0.88 | 0.91 | 0.8 | 0.94 | 0.96 |
| 6 | 156122600 | rs4424096 | 1.392 | 0.72 | 0.574 | 1.073 | 0.021 | 0.124 | 0.124 | 0 | 0.77 | 0.55 | 0.57 | 0.54 | 0.77 | 0.74 |
| 6 | 156126988 | rs4128674 | -0.196 | 0.411 | -0.136 | 1.156 | 0.009 | -0.062 | -0.062 | 0 | 0.12 | 0.04 | 0.16 | 0.38 | 0.1 | 0.26 |
| 6 | 156131953 | rs9480214 | 0.574 | 0.595 | -0.209 | -0.296 | 0.001 | -0.022 | -0.022 | 1.016 | 0.37 | 0.3 | 0.16 | 0.05 | 0.35 | 0.25 |
| 6 | 156132330 | rs9397843 | -0.637 | 0.869 | 0.083 | 0.397 | 0.04 | 0.159 | 0.159 | 0 | 0.19 | 0.36 | 0.32 | 0.44 | 0.27 | 0.39 |
| 6 | 156133888 | rs10499299 | -0.041 | 0.86 | 0.13 | 0.105 | 0.015 | -0.013 | -0.013 | 0 | 0.09 | 0.16 | 0.14 | 0.06 | 0.13 | 0.1 |
| 6 | 156136688 | rs13213139 | 0.565 | 0.949 | 0.126 | -0.508 | -0.002 | -0.056 | -0.056 | 0.019 | 0.21 | 0.22 | 0.09 | 0.06 | 0.21 | 0.13 |
| 6 | 156139323 | rs4869870 | 0.109 | 0.986 | 0.458 | -0.128 | 0.03 | -0.049 | -0.049 | 0.853 | 0.08 | 0.21 | 0.1 | 0 | 0.11 | 0.03 |
| 6 | 156140443 | rs4599663 | -0.2 | 1.091 | 0.244 | 0.575 | 0.001 | 0.011 | 0.011 | 0 | 0.34 | 0.43 | 0.42 | 0.56 | 0.32 | 0.37 |
| 6 | 156141694 | rs4382275 | -0.186 | 1.078 | 0.153 | 0.579 | -0.001 | -0.011 | -0.011 | 0 | 0.34 | 0.33 | 0.39 | 0.5 | 0.3 | 0.38 |
| 6 | 156142928 | rs9383776 | -0.362 | 1.055 | 0.186 | 0.524 | -0.008 | -0.001 | -0.001 | 0 | 0.42 | 0.39 | 0.51 | 0.57 | 0.43 | 0.43 |
| 6 | 156149244 | rs4455683 | -0.494 | 0.794 | -0.245 | 0.544 | 0.001 | 0.07 | 0.07 | 0 | 0.12 | 0.19 | 0.21 | 0.27 | 0.12 | 0.3 |
| 6 | 156152890 | rs4260759 | -1.125 | 0.896 | -0.232 | 0.764 | 0 | 0.002 | 0.002 | 0 | 0.51 | 0.64 | 0.67 | 0.5 | 0.49 | 0.51 |
| 6 | 156153561 | rs9371928 | -0.945 | 0.8 | -0.09 | 0.801 | 0.008 | 0.023 | 0.023 | 0 | 0.19 | 0.27 | 0.3 | 0.24 | 0.23 | 0.33 |
| 6 | 156162006 | rs9480221 | -0.436 | 0.697 | -0.215 | 0.366 | -0.004 | 0.002 | 0.002 | 0 | 0.28 | 0.3 | 0.42 | 0.67 | 0.3 | 0.4 |
| 6 | 156162032 | rs9371932 | -0.421 | 0.555 | -0.346 | 0.227 | 0.001 | -0.014 | -0.014 | 0.675 | 0.07 | 0.04 | 0.15 | 0.13 | 0.05 | 0.21 |
| 6 | 156168371 | rs7755229 | 0.424 | -0.04 | -0.536 | -0.712 | 0.018 | 0.027 | 0.027 | 0.205 | 0.24 | 0.16 | 0.1 | 0.05 | 0.19 | 0.12 |
| 6 | 156172676 | rs4502946 | 0.254 | 0.294 | -0.456 | -0.615 | -0.002 | 0 | 0 | 0 | 0.27 | 0.3 | 0.42 | 0.67 | 0.31 | 0.4 |
| 6 | 156173871 | rs13191915 | -0.408 | 0.115 | 0.354 | -0.752 | -0.008 | -0.041 | -0.041 | 1.902 | 0.34 | 0.42 | 0.34 | 0.2 | 0.33 | 0.41 |
| 6 | 156175005 | rs6917282 | -1.423 | -0.004 | 0.403 | -0.811 | -0.007 | 0.122 | 0.122 | 0 | 0.63 | 0.85 | 0.76 | 0.84 | 0.63 | 0.8 |
| 6 | 156177108 | rs4449642 | -0.929 | -0.006 | 0.784 | -0.625 | 0.011 | 0.091 | 0.091 | 0 | 0.11 | 0.34 | 0.15 | 0.13 | 0.06 | 0.28 |
| 6 | 156178985 | rs11968187 | -0.791 | -0.143 | 0.252 | -0.873 | 0.026 | 0.03 | 0.03 | 0 | 0.38 | 0.63 | 0.3 | 0.19 | 0.27 | 0.42 |
| 6 | 156191577 | rs4455684 | -0.071 | -0.032 | 0.354 | -0.995 | 0 | -0.043 | -0.043 | 0 | 0.4 | 0.48 | 0.6 | 0.82 | 0.4 | 0.71 |
| 6 | 156198168 | rs6557456 | -0.051 | 0.036 | 0.676 | -0.938 | 0.02 | -0.047 | -0.047 | 0 | 0.37 | 0.55 | 0.54 | 0.71 | 0.44 | 0.48 |
| 6 | 156198945 | rs9480227 | 0.119 | 0.036 | 0.656 | -0.889 | -0.002 | 0.008 | 0.008 | 0 | 0.34 | 0.35 | 0.45 | 0.71 | 0.36 | 0.46 |
| 6 | 156199192 | rs12665588 | 1.012 | 0.377 | 0.603 | 0.227 | 0.003 | 0.088 | 0.088 | 0 | 0.08 | 0.08 | 0.09 | 0.4 | 0.11 | 0.17 |
| 6 | 156200568 | rs9784833 | 0.426 | 0.191 | 0.529 | -0.246 | 0 | 0.003 | 0.003 | 0 | 0.17 | 0.12 | 0.08 | 0.05 | 0.16 | 0.08 |
| 6 | 156201766 | rs6941564 | -0.78 | -0.005 | 0.498 | -0.207 | -0.005 | 0.019 | 0.019 | 0 | 0.58 | 0.57 | 0.53 | 0.72 | 0.55 | 0.51 |
| 6 | 156207989 | rs4495282 | -1.854 | -0.129 | 0.463 | -0.296 | -0.002 | 0.008 | 0.008 | 0 | 0.69 | 0.79 | 0.8 | 0.87 | 0.68 | 0.84 |
| 6 | 156209335 | rs9384389 | -0.665 | -0.114 | 0.497 | -0.264 | -0.002 | 0.084 | 0.084 | 0 | 0.51 | 0.71 | 0.72 | 0.82 | 0.52 | 0.75 |
| 6 | 156214101 | rs12174424 | 0.503 | -0.109 | 0.435 | -0.043 | 0.017 | -0.009 | -0.009 | 0 | 0.08 | 0.06 | 0.14 | 0.12 | 0.04 | 0.24 |
| 6 | 156230765 | rs9384393 | 0.055 | -0.134 | 0.466 | 0.012 | -0.002 | -0.003 | -0.003 | 0.071 | 0.4 | 0.34 | 0.54 | 0.59 | 0.43 | 0.44 |
| 6 | 156233006 | rs6932698 | 0.899 | -0.109 | 0.421 | 0.052 | 0 | 0.022 | 0.022 | 0 | 0.13 | 0.07 | 0.06 | 0.04 | 0.14 | 0.09 |
| 6 | 156234522 | rs9371942 | -0.504 | -0.135 | 0.683 | 0.346 | -0.027 | 0.013 | 0.013 | 0 | 0.13 | 0.23 | 0.09 | 0.19 | 0.04 | 0.25 |
| 6 | 156237804 | rs9397861 | -0.651 | -0.156 | 0.678 | 0.346 | -0.038 | 0.067 | 0.067 | 0 | 0.15 | 0.3 | 0.15 | 0.27 | 0.05 | 0.3 |
| 6 | 156240505 | rs17086839 | -0.058 | -0.1 | 0.556 | 0.361 | -0.006 | 0.006 | 0.006 | 0 | 0.11 | 0.15 | 0.09 | 0.21 | 0.04 | 0.25 |
| 6 | 156241520 | rs9384394 | -0.11 | -0.052 | 0.553 | 0.365 | -0.005 | 0.005 | 0.005 | 0 | 0.12 | 0.15 | 0.09 | 0.22 | 0.04 | 0.29 |
| 6 | 156254710 | rs10457924 | 1.023 | 0.034 | 0.116 | 0.527 | 0 | 0.031 | 0.031 | 0 | 0.09 | 0.08 | 0.07 | 0.32 | 0.1 | 0.15 |
| 6 | 156258961 | rs7771021 | 0.768 | 0.195 | -0.162 | -0.51 | 0.001 | 0.003 | 0.003 | 0 | 0.1 | 0.08 | 0.04 | 0.04 | 0.09 | 0.04 |
| 6 | 156259621 | rs9371407 | -1.068 | 0.134 | -0.063 | -0.526 | -0.049 | 0.059 | 0.059 | 0 | 0.24 | 0.44 | 0.21 | 0.27 | 0.15 | 0.37 |

| | | | | | | | | | | | | | | | | |
|---|---|---|---|---|---|---|---|---|---|---|---|---|---|---|---|---|
| 6 | 156261012 | rs6920311 | -1.5 | 0.021 | -0.593 | -0.52 | -0.094 | 0.197 | 0.197 | 0 | 0.66 | 0.98 | 0.79 | 0.91 | 0.65 | 0.79 |
| 6 | 156264656 | rs11156067 | 0.281 | 0.121 | -0.244 | -0.278 | 0 | 0 | 0 | 0 | 0.4 | 0.41 | 0.54 | 0.59 | 0.43 | 0.42 |
| 6 | 156265573 | rs9384399 | 0.218 | 0.117 | -0.3 | -0.281 | 0.001 | 0.001 | 0.001 | 0 | 0.41 | 0.45 | 0.55 | 0.61 | 0.43 | 0.42 |
| 6 | 156266096 | rs9371408 | -0.102 | 0.14 | -0.597 | -0.248 | 0.03 | -0.04 | -0.04 | 0 | 0.11 | 0.04 | 0.09 | 0.17 | 0.04 | 0.28 |
| 6 | 156277741 | rs9480235 | -0.078 | 0.643 | -0.314 | -0.159 | -0.002 | -0.004 | -0.004 | 0.007 | 0.39 | 0.35 | 0.53 | 0.59 | 0.42 | 0.37 |
| 6 | 156284136 | rs9480236 | -0.47 | 0.742 | -0.473 | -0.027 | 0.005 | -0.007 | -0.007 | 0 | 0.44 | 0.51 | 0.54 | 0.62 | 0.47 | 0.44 |
| 6 | 156285173 | rs9371946 | -0.247 | 0.827 | -1.201 | -0.336 | 0.01 | 0.017 | 0.017 | 0 | 0.46 | 0.42 | 0.43 | 0.34 | 0.47 | 0.45 |
| 6 | 156285880 | rs6906938 | -0.284 | 0.874 | -1.199 | -0.274 | -0.01 | 0.036 | 0.036 | 0 | 0.46 | 0.43 | 0.46 | 0.3 | 0.54 | 0.39 |
| 6 | 156286410 | rs9371947 | -0.293 | 0.855 | -0.902 | -0.483 | -0.023 | 0.098 | 0.098 | 0.048 | 0.58 | 0.73 | 0.73 | 0.87 | 0.54 | 0.74 |
| 6 | 156286540 | rs9397869 | -0.021 | 0.286 | -0.521 | 0.092 | -0.004 | 0.002 | 0.002 | 0.738 | 0.28 | 0.33 | 0.31 | 0.29 | 0.28 | 0.27 |
| 6 | 156290380 | rs9383788 | -0.018 | -0.144 | -0.112 | 0.232 | 0.001 | -0.002 | -0.002 | 0.155 | 0.4 | 0.44 | 0.42 | 0.33 | 0.47 | 0.35 |
| 6 | 156293797 | rs4591861 | 0.426 | -0.756 | -0.406 | 0.156 | 0.029 | 0.005 | 0.005 | 0 | 0.17 | 0.11 | 0.22 | 0.17 | 0.09 | 0.08 |
| 6 | 156299960 | rs4363050 | -0.691 | -0.752 | -0.18 | 0.346 | -0.005 | -0.003 | -0.003 | 0 | 0.7 | 0.74 | 0.74 | 0.64 | 0.68 | 0.53 |
| 6 | 156305117 | rs4496814 | -0.49 | -1.077 | 0.011 | 0.301 | -0.014 | -0.01 | -0.01 | 0.267 | 0.35 | 0.6 | 0.33 | 0.22 | 0.21 | 0.15 |
| 6 | 156314920 | rs9371951 | 0.356 | -0.705 | 0.126 | 0.479 | -0.006 | -0.005 | -0.005 | 0.015 | 0.4 | 0.61 | 0.43 | 0.4 | 0.47 | 0.38 |
| 6 | 156315056 | rs9384411 | -0.222 | -0.821 | 0.056 | 0.425 | -0.004 | 0.008 | 0.008 | 0 | 0.54 | 0.74 | 0.67 | 0.53 | 0.57 | 0.45 |
| 6 | 156319368 | rs4869879 | 0.316 | -0.844 | 0.147 | 0.533 | 0.007 | 0 | 0 | 0 | 0.15 | 0.16 | 0.24 | 0.14 | 0.1 | 0.07 |
| 6 | 156326130 | rs6931064 | -0.141 | -0.829 | 0.081 | 0.563 | -0.001 | 0.003 | 0.003 | 0 | 0.44 | 0.41 | 0.32 | 0.47 | 0.42 | 0.55 |
| 6 | 156335489 | rs1449679 | -0.959 | -1.013 | -0.004 | 0.493 | -0.036 | -0.032 | -0.032 | 0.153 | 0.64 | 0.81 | 0.58 | 0.61 | 0.52 | 0.64 |
| 6 | 156335857 | rs1449678 | -0.223 | -0.916 | -0.24 | 0.382 | 0.005 | 0.014 | 0.014 | 0 | 0.47 | 0.35 | 0.32 | 0.48 | 0.43 | 0.56 |
| 6 | 156337304 | rs17086954 | 0.771 | -0.797 | -0.791 | 0.472 | -0.082 | -0.085 | -0.085 | 0 | 0.23 | 0.04 | 0.23 | 0.39 | 0.3 | 0.46 |
| 6 | 156337865 | rs6928730 | -0.007 | -1.106 | -0.302 | 0.381 | -0.001 | 0.001 | 0.001 | 0.014 | 0.38 | 0.55 | 0.37 | 0.41 | 0.39 | 0.54 |
| 6 | 156339112 | rs9285544 | -0.359 | -1.207 | -0.691 | 0.338 | -0.038 | -0.037 | -0.037 | 0 | 0.34 | 0.44 | 0.15 | 0.19 | 0.2 | 0.12 |
| 6 | 156341409 | rs1449676 | 1.25 | -1.084 | -1.642 | 0.431 | 0.015 | -0.002 | -0.002 | 0 | 0.11 | 0.06 | 0.06 | 0.1 | 0.07 | 0.03 |
| 6 | 156345146 | rs1449672 | -0.75 | -1.079 | -1.683 | 0.5 | -0.019 | -0.025 | -0.025 | 0 | 0.64 | 0.7 | 0.47 | 0.56 | 0.55 | 0.63 |
| 6 | 156345213 | rs13218955 | 0.139 | -0.602 | -2.036 | 0.857 | -0.086 | -0.064 | -0.064 | 0 | 0.3 | 0.06 | 0.31 | 0.37 | 0.35 | 0.5 |
| 6 | 156348083 | rs4870421 | -0.544 | -0.576 | -2.057 | 0.657 | -0.059 | -0.05 | -0.05 | 0 | 0.47 | 0.16 | 0.41 | 0.51 | 0.48 | 0.6 |
| 6 | 156348805 | rs9397891 | -0.063 | 0.119 | -1.688 | 0.847 | -0.099 | -0.093 | -0.093 | 0.105 | 0.31 | 0.06 | 0.48 | 0.64 | 0.37 | 0.55 |
| 6 | 156350450 | rs4870425 | 0.397 | -0.115 | -1.302 | 0.249 | 0.039 | 0.048 | 0.048 | 0 | 0.58 | 0.27 | 0.58 | 0.78 | 0.71 | 0.65 |
| 6 | 156351816 | rs1349239 | 0.049 | 0.287 | -1.337 | -0.428 | -0.008 | 0.035 | 0.035 | 0.192 | 0.71 | 0.81 | 0.81 | 0.89 | 0.77 | 0.91 |
| 6 | 156353867 | rs6909234 | 1.09 | 0.421 | -0.827 | -0.349 | -0.001 | -0.005 | -0.005 | 0.187 | 0.46 | 0.48 | 0.58 | 0.64 | 0.48 | 0.55 |
| 6 | 156355252 | rs13194470 | -0.058 | 0.529 | -1.309 | -0.499 | -0.027 | 0.013 | 0.013 | 0.459 | 0.19 | 0.06 | 0.13 | 0.19 | 0.23 | 0.17 |
| 6 | 156363648 | rs9322557 | 0.097 | 0.617 | -0.881 | -0.355 | -0.006 | 0.043 | 0.043 | 0.185 | 0.71 | 0.82 | 0.81 | 0.91 | 0.78 | 0.91 |
| 6 | 156365973 | rs9322560 | -0.18 | 0.781 | -0.718 | -0.207 | -0.001 | -0.005 | -0.005 | 0 | 0.86 | 0.88 | 0.93 | 0.96 | 0.88 | 0.94 |
| 6 | 156369514 | rs6935860 | -0.074 | 1.065 | -0.668 | -0.152 | -0.001 | -0.003 | -0.003 | 0 | 0.43 | 0.45 | 0.36 | 0.45 | 0.45 | 0.59 |
| 6 | 156377582 | rs1584535 | -0.297 | 1.159 | -0.502 | 0.052 | 0.001 | -0.001 | -0.001 | 0 | 0.56 | 0.56 | 0.63 | 0.49 | 0.55 | 0.42 |
| 6 | 156377806 | rs9397895 | 0.603 | 1.051 | -0.709 | 0.231 | 0.001 | -0.003 | -0.003 | 11.508 | 0.18 | 0.18 | 0.12 | 0.06 | 0.1 | 0.04 |
| 6 | 156386980 | rs6557469 | -0.005 | 1.431 | -0.786 | 0.651 | -0.031 | -0.071 | -0.071 | 0.234 | 0.31 | 0.16 | 0.48 | 0.42 | 0.46 | 0.33 |
| 6 | 156392232 | rs9480253 | -0.278 | 1.404 | 0.099 | 0.542 | 0.068 | 0.025 | 0.025 | 0 | 0.17 | 0.53 | 0.16 | 0.19 | 0.24 | 0.18 |
| 6 | 156393579 | rs17185162 | 0.763 | 0.941 | -0.473 | 0.935 | 0.014 | -0.02 | -0.02 | 3.347 | 0.16 | 0.09 | 0.31 | 0.21 | 0.09 | 0.05 |
| 6 | 156395008 | rs2354391 | 0.791 | 0.936 | -0.52 | 0.943 | 0.023 | -0.033 | -0.033 | 0.053 | 0.16 | 0.07 | 0.31 | 0.2 | 0.09 | 0.05 |
| 6 | 156400939 | rs9397897 | 0.022 | 1.651 | -1.093 | 1.055 | -0.068 | -0.05 | -0.05 | 0 | 0.13 | 0.01 | 0.11 | 0.22 | 0.19 | 0.16 |
| 6 | 156404902 | rs1375714 | -0.201 | 0.455 | -1.077 | 0.392 | 0.029 | 0.011 | 0.011 | 1.023 | 0.5 | 0.63 | 0.68 | 0.49 | 0.44 | 0.43 |
| 6 | 156405387 | rs7751239 | 1.618 | 0.573 | -1.218 | 0.431 | 0.004 | 0.011 | 0.011 | 0.046 | 0.1 | 0.05 | 0.08 | 0 | 0.12 | 0.03 |
| 6 | 156410838 | rs9480255 | 0.086 | 0.717 | -1.416 | 0.193 | -0.039 | -0.041 | -0.041 | 0 | 0.34 | 0.2 | 0.21 | 0.51 | 0.38 | 0.44 |
| 6 | 156418695 | rs13191871 | 0.573 | 0.142 | -0.649 | 0.493 | -0.015 | 0.009 | 0.009 | 0 | 0.2 | 0.19 | 0.22 | 0.27 | 0.12 | 0.33 |
| 6 | 156421907 | rs9371410 | 0.255 | 0.682 | -0.988 | 0.632 | -0.027 | -0.023 | -0.023 | 0 | 0.15 | 0.08 | 0.12 | 0.25 | 0.17 | 0.29 |
| 6 | 156425884 | rs11156079 | 0.03 | 0.647 | -1.077 | 0.506 | -0.035 | -0.011 | -0.011 | 0 | 0.19 | 0.1 | 0.12 | 0.24 | 0.22 | 0.33 |
| 6 | 156427957 | rs9480261 | 1.155 | 0.586 | -0.588 | 0.728 | 0.019 | 0.096 | 0.096 | 0 | 0.2 | 0.37 | 0.37 | 0.36 | 0.2 | 0.15 |
| 6 | 156432407 | rs10499290 | 0.765 | 0.522 | -1.216 | 0.869 | -0.017 | -0.047 | -0.047 | 1.54 | 0.13 | 0.01 | 0.12 | 0.12 | 0.16 | 0.15 |
| 6 | 156434933 | rs6925781 | -1.47 | 0.581 | -1.364 | 0.603 | -0.099 | 0.022 | 0.022 | 0 | 0.77 | 0.99 | 0.87 | 0.88 | 0.73 | 0.8 |
| 6 | 156435767 | rs6905034 | 0.091 | 0.371 | -0.876 | 0.596 | -0.004 | 0.005 | 0.005 | 0 | 0.23 | 0.19 | 0.13 | 0.22 | 0.19 | 0.37 |
| 6 | 156439521 | rs11156080 | 0.179 | 0.402 | -0.293 | 0.827 | 0.004 | 0.039 | 0.039 | 0 | 0.28 | 0.49 | 0.44 | 0.36 | 0.3 | 0.18 |
| 6 | 156446501 | rs17186394 | 0.308 | 0.547 | -0.41 | 0.71 | 0 | -0.001 | -0.001 | 0.018 | 0.27 | 0.26 | 0.43 | 0.2 | 0.32 | 0.1 |
| 6 | 156447429 | rs7746475 | -1.277 | 0.755 | -0.897 | 0.631 | -0.096 | 0.028 | 0.028 | 0 | 0.78 | 0.99 | 0.88 | 0.88 | 0.73 | 0.83 |
| 6 | 156447991 | rs2122304 | 0.948 | 0.725 | -0.086 | 1.308 | 0.01 | 0.02 | 0.02 | 0 | 0.16 | 0.21 | 0.33 | 0.2 | 0.17 | 0.07 |
| 6 | 156449758 | rs930670 | -0.114 | 0.471 | 0.285 | 1.128 | -0.017 | 0.048 | 0.048 | 0 | 0.26 | 0.49 | 0.22 | 0.42 | 0.2 | 0.44 |
| 6 | 156449765 | rs12209400 | 0.421 | 0.763 | 0.366 | 1.293 | 0.008 | -0.026 | -0.026 | 0 | 0.2 | 0.26 | 0.18 | 0.07 | 0.2 | 0.1 |
| 6 | 156455669 | rs2354393 | 0.076 | 0.66 | 0.354 | 1.333 | -0.015 | 0.014 | 0.014 | 0 | 0.31 | 0.5 | 0.23 | 0.44 | 0.24 | 0.47 |
| 6 | 156458742 | rs4394225 | 0.047 | 0.541 | 0.348 | 1.346 | -0.013 | 0.002 | 0.002 | 0 | 0.31 | 0.52 | 0.23 | 0.42 | 0.24 | 0.47 |
| 6 | 156461430 | rs11962960 | 1.3 | 0.581 | 0.165 | 1.096 | -0.006 | -0.006 | -0.006 | 0 | 0.13 | 0.16 | 0.13 | 0.05 | 0.07 | 0.02 |
| 6 | 156463321 | rs6914206 | -0.939 | 0.679 | -0.017 | 1.287 | -0.025 | 0.004 | 0.004 | 0 | 0.48 | 0.82 | 0.46 | 0.6 | 0.44 | 0.54 |
| 6 | 156463939 | rs6914961 | -1.122 | 0.679 | -0.013 | 1.271 | 0.002 | 0.01 | 0.01 | 0 | 0.52 | 0.84 | 0.47 | 0.62 | 0.46 | 0.57 |
| 6 | 156467842 | rs6557473 | -0.217 | 0.343 | 0.182 | 1.202 | -0.003 | -0.008 | -0.008 | 0 | 0.4 | 0.46 | 0.33 | 0.31 | 0.33 | 0.41 |
| 6 | 156469089 | rs1037287 | 0.084 | 0.457 | -0.582 | 1.098 | -0.012 | 0.013 | 0.013 | 0 | 0.28 | 0.18 | 0.2 | 0.26 | 0.26 | 0.4 |
| 6 | 156474355 | rs875642 | -0.138 | 0.504 | -0.501 | 1.029 | 0.002 | 0.066 | 0.066 | 0 | 0.4 | 0.21 | 0.27 | 0.27 | 0.31 | 0.43 |
| 6 | 156475282 | rs12201446 | -0.42 | 0.545 | -0.126 | 0.951 | 0.005 | -0.01 | -0.01 | 0 | 0.53 | 0.69 | 0.38 | 0.57 | 0.38 | 0.58 |
| 6 | 156476113 | rs7767863 | -0.362 | 1.011 | 0.091 | 1.096 | 0 | 0.015 | 0.015 | 0 | 0.47 | 0.74 | 0.37 | 0.57 | 0.43 | 0.61 |
| 6 | 156477121 | rs6557474 | -0.766 | 1.251 | 0.172 | 1.251 | 0 | 0.001 | 0.001 | 0 | 0.69 | 0.83 | 0.54 | 0.71 | 0.7 | 0.67 |

| | | | | | | | | | | | | | | | | |
|---|---|---|---|---|---|---|---|---|---|---|---|---|---|---|---|---|
| 6 | 156485290 | rs6936091 | -0.438 | 1.408 | -0.299 | 1.345 | -0.007 | 0.002 | 0.002 | 0 | 0.65 | 0.64 | 0.5 | 0.69 | 0.57 | 0.65 |
| 6 | 156490528 | rs7749940 | 0.286 | 1.402 | -0.395 | 1.126 | 0.013 | 0.002 | 0.002 | 0.077 | 0.33 | 0.45 | 0.22 | 0.36 | 0.33 | 0.3 |
| 6 | 156492674 | rs9397906 | 0.163 | 1.518 | -0.685 | 1.172 | 0 | 0.067 | 0.067 | 0.157 | 0.13 | 0.24 | 0.1 | 0.31 | 0.13 | 0.23 |
| 6 | 156492897 | rs9371966 | -1.043 | 1.341 | -0.719 | 1.077 | -0.004 | -0.008 | -0.008 | 0 | 0.69 | 0.86 | 0.49 | 0.69 | 0.69 | 0.67 |
| 6 | 156496460 | rs7742923 | 0.869 | 1.358 | -0.715 | 1.064 | -0.034 | 0.074 | 0.074 | 0.648 | 0.27 | 0.06 | 0.14 | 0.11 | 0.21 | 0.2 |
| 6 | 156502670 | rs6557479 | -0.283 | 1.394 | -0.674 | 1.111 | -0.012 | 0.002 | 0.002 | 0 | 0.41 | 0.69 | 0.34 | 0.38 | 0.48 | 0.38 |
| 6 | 156504495 | rs10499286 | 0.382 | 1.128 | -0.569 | 1.142 | -0.028 | -0.124 | -0.124 | 0.077 | 0.27 | 0.44 | 0.19 | 0.06 | 0.24 | 0.13 |
| 6 | 156507643 | rs6557480 | -0.161 | 1.389 | -1.031 | 0.989 | -0.006 | -0.007 | -0.007 | 0 | 0.69 | 0.87 | 0.5 | 0.69 | 0.7 | 0.67 |
| 6 | 156517298 | rs12214175 | -0.859 | 1.815 | -0.369 | 1.169 | 0.024 | -0.002 | -0.002 | 0 | 0.08 | 0.15 | 0.14 | 0.06 | 0.16 | 0.08 |
| 6 | 156517374 | rs4486031 | -0.33 | 2.366 | -0.711 | 1.933 | -0.053 | -0.059 | -0.059 | 0 | 0.09 | 0.01 | 0.09 | 0.27 | 0.16 | 0.22 |
| 6 | 156519131 | rs6557482 | -0.547 | 1.961 | -0.357 | 1.657 | 0 | 0.004 | 0.004 | 0 | 0.39 | 0.57 | 0.35 | 0.44 | 0.39 | 0.4 |
| 6 | 156520101 | rs9480284 | -0.034 | 1.828 | -0.476 | 1.693 | -0.028 | 0.055 | 0.055 | 0 | 0.29 | 0.17 | 0.22 | 0.13 | 0.29 | 0.2 |
| 6 | 156520159 | rs9371972 | 0.427 | 1.913 | -0.496 | 1.755 | -0.031 | -0.047 | -0.047 | 0 | 0.08 | 0.01 | 0.07 | 0.26 | 0.09 | 0.22 |
| 6 | 156520533 | rs4573097 | -0.51 | 1.977 | -0.311 | 1.747 | -0.023 | -0.05 | -0.05 | 0 | 0.35 | 0.49 | 0.26 | 0.18 | 0.24 | 0.24 |
| 6 | 156527067 | rs6934898 | -0.649 | 1.947 | -0.109 | 1.853 | 0.122 | 0.207 | 0.207 | 0.064 | 0.13 | 0.39 | 0.15 | 0.31 | 0.23 | 0.25 |
| 6 | 156529762 | rs4562153 | -0.871 | 1.81 | -0.574 | 1.459 | -0.03 | -0.005 | -0.005 | 0 | 0.43 | 0.73 | 0.55 | 0.55 | 0.53 | 0.61 |
| 6 | 156540417 | rs2820406 | -0.224 | 1.78 | -0.353 | 1.501 | 0.009 | 0.022 | 0.022 | 0 | 0.32 | 0.4 | 0.41 | 0.48 | 0.37 | 0.48 |
| 6 | 156543930 | rs6918330 | -0.982 | 1.752 | -0.453 | 1.428 | -0.013 | 0.182 | 0.182 | 0 | 0.51 | 0.91 | 0.49 | 0.75 | 0.56 | 0.63 |
| 6 | 156546947 | rs12203183 | 1.252 | 1.816 | -0.426 | 1.214 | 0.001 | 0.066 | 0.066 | 0 | 0.21 | 0.05 | 0.09 | 0.07 | 0.12 | 0.12 |
| 6 | 156549168 | rs2818055 | -1.207 | 1.942 | -0.115 | 1.186 | 0.051 | -0.041 | -0.041 | 0 | 0.17 | 0.62 | 0.25 | 0.07 | 0.33 | 0.15 |
| 6 | 156549813 | rs9384441 | -0.202 | 1.902 | 0.17 | 1.314 | -0.009 | 0.018 | 0.018 | 0 | 0.35 | 0.62 | 0.26 | 0.44 | 0.23 | 0.39 |
| 6 | 156550400 | rs7760889 | -0.855 | 1.941 | 0.141 | 1.316 | -0.017 | -0.025 | -0.025 | 0.332 | 0.46 | 0.87 | 0.43 | 0.5 | 0.42 | 0.52 |
| 6 | 156550589 | rs7763778 | -0.737 | 1.979 | 0.264 | 1.398 | 0.036 | 0.305 | 0.305 | 0.354 | 0.14 | 0.5 | 0.15 | 0.37 | 0.1 | 0.29 |
| 6 | 156553950 | rs2820397 | -1.549 | 2 | 0.186 | 1.54 | 0.003 | 0.003 | 0.003 | 0.086 | 0.25 | 0.67 | 0.33 | 0.35 | 0.4 | 0.4 |
| 6 | 156555020 | rs17087225 | -0.526 | 1.999 | 0.166 | 1.507 | 0.035 | -0.019 | -0.019 | 0 | 0.12 | 0.23 | 0.18 | 0.07 | 0.2 | 0.13 |
| 6 | 156560746 | rs9384443 | 0.329 | 1.967 | -0.044 | 1.579 | -0.047 | -0.057 | -0.057 | 0 | 0.09 | 0.01 | 0.09 | 0.24 | 0.14 | 0.23 |
| 6 | 156561221 | rs2818050 | -1.474 | 2.002 | 0.017 | 1.56 | 0.024 | 0.026 | 0.026 | 0.061 | 0.26 | 0.63 | 0.36 | 0.36 | 0.4 | 0.43 |
| 6 | 156566142 | rs2820409 | -1.122 | 1.88 | -0.53 | 1.539 | -0.019 | -0.062 | -0.062 | 0.026 | 0.52 | 0.94 | 0.49 | 0.47 | 0.57 | 0.55 |
| 6 | 156568138 | rs9371978 | 0.006 | 1.96 | -0.351 | 1.585 | -0.011 | -0.011 | -0.011 | 0 | 0.11 | 0.09 | 0.11 | 0.22 | 0.16 | 0.26 |
| 6 | 156572211 | rs12205158 | -0.345 | 1.711 | 0.117 | 1.304 | 0.146 | 0.075 | 0.075 | 0.116 | 0.05 | 0.37 | 0.06 | 0.05 | 0.11 | 0.03 |
| 6 | 156573593 | rs7755697 | 1.091 | 1.595 | -0.473 | 1.279 | -0.01 | 0.029 | 0.029 | 0 | 0.21 | 0.01 | 0.08 | 0.09 | 0.13 | 0.12 |
| 6 | 156574933 | rs9397917 | -0.628 | 1.581 | -0.278 | 1.365 | 0.002 | 0.027 | 0.027 | 0 | 0.42 | 0.29 | 0.53 | 0.71 | 0.42 | 0.67 |
| 6 | 156578396 | rs9371979 | 0.291 | 1.644 | -0.962 | 1.631 | -0.03 | -0.064 | -0.064 | 0 | 0.1 | 0.01 | 0.05 | 0.25 | 0.09 | 0.26 |
| 6 | 156578620 | rs10499281 | 0.68 | 1.737 | -0.942 | 1.616 | -0.002 | -0.087 | -0.087 | 0 | 0.2 | 0.07 | 0.31 | 0.4 | 0.16 | 0.32 |
| 6 | 156580748 | rs17090905 | -0.734 | 1.818 | -0.692 | 1.598 | 0.028 | 0.004 | 0.004 | 0 | 0.05 | 0.15 | 0.16 | 0.04 | 0.12 | 0.07 |
| 6 | 156581815 | rs4382278 | -0.317 | 1.783 | -0.546 | 1.753 | -0.01 | 0.105 | 0.105 | 0.113 | 0.22 | 0.56 | 0.2 | 0.58 | 0.18 | 0.41 |
| 6 | 156582195 | rs12661043 | 0.83 | 1.789 | -0.652 | 1.72 | 0.002 | -0.085 | -0.085 | 0 | 0.2 | 0.05 | 0.31 | 0.4 | 0.16 | 0.32 |
| 6 | 156586155 | rs6917784 | -0.69 | 1.988 | 0.002 | 1.497 | 0.005 | 0.017 | 0.017 | 0.23 | 0.22 | 0.62 | 0.19 | 0.29 | 0.24 | 0.33 |
| 6 | 156586753 | rs9478696 | 1 | 1.703 | -0.377 | 1.355 | -0.037 | -0.036 | -0.036 | 0.216 | 0.5 | 0.08 | 0.56 | 0.51 | 0.47 | 0.46 |
| 6 | 156588729 | rs6925021 | 1.061 | 1.96 | 0.622 | 1.548 | 0.018 | 0.026 | 0.026 | 0.077 | 0.76 | 0.44 | 0.8 | 0.7 | 0.73 | 0.66 |
| 6 | 156590382 | rs9480298 | -1.375 | 1.529 | 0.429 | 1.143 | 0.056 | -0.01 | -0.01 | 2.193 | 0.1 | 0.22 | 0.18 | 0.08 | 0.19 | 0.11 |
| 6 | 156612353 | rs4870441 | -0.379 | 1.046 | -0.531 | 1.114 | -0.054 | 0.113 | 0.113 | 0 | 0.27 | 0.04 | 0.23 | 0.09 | 0.32 | 0.3 |
| 6 | 156614578 | rs6910494 | -0.632 | 0.818 | -0.545 | 0.992 | -0.002 | -0.003 | -0.003 | 0 | 0.52 | 0.65 | 0.7 | 0.48 | 0.56 | 0.45 |
| 6 | 156617792 | rs6918956 | -0.297 | 0.563 | 0.177 | 0.914 | -0.004 | 0.004 | 0.004 | 0 | 0.37 | 0.58 | 0.28 | 0.5 | 0.36 | 0.51 |
| 6 | 156618923 | rs11759982 | 0.695 | 0.666 | -0.607 | 0.184 | -0.027 | 0.058 | 0.058 | 0 | 0.21 | 0.09 | 0.16 | 0.08 | 0.18 | 0.19 |
| 6 | 156620324 | rs9480304 | -0.008 | 0.689 | -0.086 | 0.52 | -0.003 | 0.023 | 0.023 | 0 | 0.37 | 0.48 | 0.28 | 0.5 | 0.38 | 0.52 |
| 6 | 156628874 | rs9397922 | 0.064 | 0.662 | -0.059 | 0.547 | -0.002 | 0.021 | 0.021 | 0 | 0.36 | 0.41 | 0.28 | 0.47 | 0.37 | 0.52 |
| 6 | 156641429 | rs7739135 | -0.344 | 0.834 | -0.238 | -0.275 | 0 | -0.008 | -0.008 | 0 | 0.41 | 0.44 | 0.59 | 0.7 | 0.39 | 0.5 |
| 6 | 156657842 | rs12213468 | 0.164 | 0.474 | -0.596 | -0.623 | -0.021 | 0.065 | 0.065 | 0.2 | 0.27 | 0.07 | 0.2 | 0.09 | 0.32 | 0.25 |
| 6 | 156671119 | rs7762328 | -0.349 | 0.127 | -0.017 | 0.172 | 0.001 | -0.006 | -0.006 | 0 | 0.17 | 0.09 | 0.12 | 0.17 | 0.13 | 0.19 |
| 6 | 156675495 | rs9480313 | 0.249 | 0.147 | -0.102 | 0.158 | 0.007 | -0.022 | -0.022 | 1.116 | 0.1 | 0.02 | 0.08 | 0.1 | 0.07 | 0.07 |
| 6 | 156677813 | rs12204446 | 1.084 | 0.327 | -0.236 | 0.159 | -0.039 | 0.046 | 0.046 | 0 | 0.15 | 0.02 | 0.15 | 0.04 | 0.19 | 0.08 |
| 6 | 156681169 | rs9384456 | -1.022 | 0.289 | -0.338 | 0.139 | -0.01 | 0.081 | 0.081 | 0.307 | 0.87 | 1 | 0.93 | 1 | 0.88 | 0.96 |
| 6 | 156690480 | rs17087341 | -0.703 | 0.329 | 0.489 | 0.828 | 0.002 | 0.001 | 0.001 | 0.73 | 0.21 | 0.19 | 0.13 | 0.18 | 0.14 | 0.19 |
| 6 | 156691015 | rs12199342 | 0.816 | 0.319 | 0.453 | 0.846 | -0.051 | 0.027 | 0.027 | 0 | 0.14 | 0.02 | 0.16 | 0.05 | 0.2 | 0.08 |
| 6 | 156692804 | rs9371995 | -0.168 | 0.13 | 1.056 | 0.78 | 0.011 | 0.072 | 0.072 | 0 | 0.24 | 0.48 | 0.46 | 0.55 | 0.26 | 0.3 |
| 6 | 156695781 | rs9322573 | -1.19 | -0.099 | 1.148 | 0.696 | -0.009 | 0.081 | 0.081 | 0 | 0.45 | 0.73 | 0.62 | 0.73 | 0.43 | 0.49 |
| 6 | 156697455 | rs10457099 | -0.008 | -0.062 | 0.81 | 0.776 | 0.02 | 0.041 | 0.041 | 0 | 0.23 | 0.35 | 0.47 | 0.54 | 0.3 | 0.3 |
| 6 | 156697681 | rs1377973 | -0.911 | -0.024 | 0.175 | 1.047 | 0.002 | -0.004 | -0.004 | 0 | 0.21 | 0.21 | 0.33 | 0.32 | 0.15 | 0.2 |
| 6 | 156700816 | rs2819083 | -0.244 | 0.077 | 0.229 | 0.077 | -0.021 | -0.017 | -0.017 | 1.122 | 0.27 | 0.19 | 0.38 | 0.59 | 0.37 | 0.35 |
| 6 | 156700918 | rs12201190 | -0.639 | -0.341 | 0.242 | -0.177 | 0.013 | 0.015 | 0.015 | 0 | 0.23 | 0.19 | 0.11 | 0.13 | 0.17 | 0.2 |
| 6 | 156705461 | rs11156105 | 0.398 | -0.077 | 0.601 | -0.008 | 0.012 | -0.004 | -0.004 | 1.41 | 0.65 | 0.57 | 0.58 | 0.37 | 0.59 | 0.65 |
| 6 | 156706749 | rs1349890 | 0.181 | -0.043 | 0.312 | 0.017 | -0.036 | -0.002 | -0.002 | 0.683 | 0.67 | 0.8 | 0.59 | 0.37 | 0.59 | 0.65 |
| 6 | 156711543 | rs11156106 | 0.406 | -0.069 | 0.498 | -0.054 | 0.009 | 0 | 0 | 0 | 0.65 | 0.59 | 0.57 | 0.4 | 0.59 | 0.65 |
| 6 | 156712682 | rs13214940 | 0.271 | -0.155 | -0.14 | -0.097 | -0.003 | 0.002 | 0.002 | 0 | 0.65 | 0.66 | 0.57 | 0.37 | 0.58 | 0.65 |
| 6 | 156713154 | rs4870448 | 0.172 | -0.172 | -0.161 | -0.082 | -0.017 | 0.004 | 0.004 | 0 | 0.67 | 0.73 | 0.58 | 0.37 | 0.57 | 0.65 |
| 6 | 156714899 | rs4429960 | 0.799 | -0.328 | 0.182 | 0.038 | 0 | -0.092 | -0.092 | 0 | 0.44 | 0.52 | 0.3 | 0.1 | 0.42 | 0.41 |
| 6 | 156715899 | rs2123987 | -0.112 | -0.232 | -0.6 | 0.146 | -0.039 | 0.073 | 0.073 | 0 | 0.44 | 0.19 | 0.48 | 0.3 | 0.47 | 0.33 |
| 6 | 156717565 | rs1078299 | 0.565 | -0.148 | -0.546 | 0.401 | -0.012 | 0.168 | 0.168 | 0 | 0.88 | 0.74 | 0.78 | 0.45 | 0.89 | 0.72 |

| | | | | | | | | | | | | | | | | |
|---|---|---|---|---|---|---|---|---|---|---|---|---|---|---|---|---|
| 6 | 156719068 | rs13218460 | 0.725 | -0.08 | -1.13 | 0.147 | 0.011 | 0.123 | 0.123 | 0 | 0.47 | 0.68 | 0.32 | 0.13 | 0.42 | 0.46 |
| 6 | 156720114 | rs2603450 | 0.316 | -0.11 | -1.306 | 0.34 | -0.006 | 0.092 | 0.092 | 0.01 | 0.89 | 0.83 | 0.78 | 0.46 | 0.88 | 0.73 |
| 6 | 156721621 | rs2354988 | 0.694 | -0.101 | -1.325 | 0.258 | 0.011 | 0.139 | 0.139 | 0.017 | 0.48 | 0.69 | 0.33 | 0.14 | 0.42 | 0.46 |
| 6 | 156722390 | rs1902064 | -0.073 | 0.081 | -1.641 | 0.371 | -0.077 | 0.015 | 0.015 | 0 | 0.41 | 0.12 | 0.46 | 0.32 | 0.46 | 0.27 |
| 6 | 156730538 | rs11156108 | -0.572 | 0.073 | -0.938 | 0.907 | -0.008 | 0.271 | 0.271 | 0.883 | 0.1 | 0.27 | 0.14 | 0.55 | 0.05 | 0.21 |
| 6 | 156731149 | rs9384464 | -0.572 | 0.073 | -0.938 | 0.907 | -0.008 | 0.271 | 0.271 | 0.883 | 0.1 | 0.27 | 0.14 | 0.55 | 0.05 | 0.21 |
| 6 | 156735886 | rs2819088 | -0.529 | -0.201 | -2.31 | -0.117 | -0.059 | 0.199 | 0.199 | 0 | 0.55 | 0.93 | 0.7 | 0.86 | 0.52 | 0.59 |
| 6 | 156751703 | rs2603430 | 0.567 | 0.485 | -1.617 | 1.089 | -0.003 | -0.031 | -0.031 | 0.338 | 0.84 | 0.91 | 0.83 | 0.52 | 0.83 | 0.68 |
| 6 | 156761338 | rs980814 | 0.218 | 0.287 | -1.973 | 0.295 | -0.044 | 0.146 | 0.146 | 0.022 | 0.47 | 0.13 | 0.52 | 0.22 | 0.47 | 0.43 |
| 6 | 156763643 | rs2444785 | -0.804 | 0.512 | -1.161 | 0.71 | 0.005 | 0.02 | 0.02 | 0.026 | 0.4 | 0.65 | 0.35 | 0.35 | 0.4 | 0.43 |
| 6 | 156764075 | rs2603433 | 0.31 | 0.041 | -1.181 | 0.819 | 0.005 | -0.006 | -0.006 | 0 | 0.89 | 0.84 | 0.9 | 0.89 | 0.86 | 0.92 |
| 6 | 156765127 | rs6557499 | -0.516 | 0.02 | -0.742 | 1.623 | 0.007 | -0.005 | -0.005 | 0 | 0.37 | 0.64 | 0.31 | 0.48 | 0.31 | 0.46 |
| 6 | 156766474 | rs2045138 | -0.749 | -0.074 | -0.867 | 1.947 | 0.054 | -0.046 | -0.046 | 0 | 0.39 | 0.8 | 0.37 | 0.58 | 0.31 | 0.51 |
| 6 | 156769759 | rs9371430 | -0.345 | -0.064 | -0.116 | 1.976 | -0.062 | 0.009 | 0.009 | 0 | 0.31 | 0.46 | 0.23 | 0.38 | 0.21 | 0.43 |
| 6 | 156778660 | rs10782344 | -0.482 | 0.005 | -0.12 | 1.946 | -0.036 | 0.034 | 0.034 | 0 | 0.33 | 0.46 | 0.25 | 0.49 | 0.25 | 0.43 |
| 6 | 156782481 | rs2493698 | -0.021 | 0.131 | -0.218 | 1.905 | -0.015 | 0.012 | 0.012 | 1.467 | 0.92 | 0.99 | 0.93 | 0.97 | 0.91 | 0.93 |
| 6 | 156785323 | rs2444783 | 0.627 | -0.001 | 0.17 | 1.856 | 0.004 | 0.001 | 0.001 | 0 | 0.87 | 0.83 | 0.85 | 0.85 | 0.83 | 0.87 |
| 6 | 156787460 | rs6931416 | -0.866 | -0.173 | 0.277 | 1.899 | -0.028 | 0.004 | 0.004 | 0 | 0.33 | 0.46 | 0.27 | 0.39 | 0.29 | 0.45 |
| 6 | 156788415 | rs6920956 | -0.784 | -0.282 | 0.29 | 1.944 | -0.023 | 0.025 | 0.025 | 0 | 0.31 | 0.52 | 0.2 | 0.45 | 0.26 | 0.43 |
| 6 | 156792873 | rs9372001 | -1.481 | -0.645 | 0.842 | 2.013 | 0.026 | -0.03 | -0.03 | 0 | 0.3 | 0.74 | 0.14 | 0.51 | 0.18 | 0.44 |
| 6 | 156794133 | rs1017643 | 1.797 | -0.74 | 0.625 | 1.957 | 0.034 | -0.108 | -0.108 | 0.081 | 0.57 | 0.05 | 0.62 | 0.37 | 0.68 | 0.45 |
| 6 | 156797257 | rs7739058 | -0.728 | -0.739 | 1.155 | 1.992 | -0.068 | 0.04 | 0.04 | 0 | 0.23 | 0.35 | 0.07 | 0.37 | 0.1 | 0.36 |
| 6 | 156801192 | rs958386 | 1.827 | -0.715 | 0.902 | 1.951 | 0.066 | -0.064 | -0.064 | 0 | 0.57 | 0.08 | 0.63 | 0.36 | 0.68 | 0.44 |
| 6 | 156809036 | rs2819066 | 1.628 | -0.778 | 1.119 | 2.009 | 0.07 | -0.035 | -0.035 | 0 | 0.63 | 0.21 | 0.67 | 0.36 | 0.75 | 0.47 |
| 6 | 156817434 | rs2444787 | 1.28 | -0.475 | 1.072 | 1.389 | -0.007 | 0.008 | 0.008 | 0.081 | 0.84 | 0.75 | 0.79 | 0.79 | 0.85 | 0.84 |
| 6 | 156819939 | rs2250759 | 0.736 | -0.482 | 1.073 | 1.294 | 0.001 | -0.009 | -0.009 | 0.155 | 0.87 | 0.81 | 0.8 | 0.9 | 0.86 | 0.88 |
| 6 | 156826343 | rs2493242 | 1.231 | -0.455 | 1.128 | 1.313 | -0.003 | 0.002 | 0.002 | 0.184 | 0.84 | 0.75 | 0.8 | 0.81 | 0.85 | 0.84 |
| 6 | 156833278 | rs2817431 | 0.622 | -0.402 | 1.03 | 1.091 | 0.002 | -0.004 | -0.004 | 0.213 | 0.87 | 0.85 | 0.81 | 0.9 | 0.85 | 0.89 |
| 6 | 156834043 | rs6557501 | -1.631 | -0.331 | 1.764 | 1.221 | 0.004 | 0.068 | 0.068 | 0 | 0.18 | 0.53 | 0.2 | 0.26 | 0.15 | 0.25 |
| 6 | 156842462 | rs2493701 | 1.928 | -0.561 | 1.094 | 1.23 | 0.007 | -0.099 | -0.099 | 1.951 | 0.58 | 0.05 | 0.67 | 0.51 | 0.69 | 0.47 |
| 6 | 156844950 | rs1327184 | -1.779 | 0.174 | 1.764 | 1.336 | 0.038 | 0.004 | 0.004 | 0 | 0.2 | 0.5 | 0.23 | 0.18 | 0.2 | 0.18 |
| 6 | 156846366 | rs13201016 | -1.2 | -0.258 | 1.455 | 1.461 | -0.002 | 0.039 | 0.039 | 0 | 0.18 | 0.4 | 0.21 | 0.23 | 0.16 | 0.25 |
| 6 | 156849322 | rs9478709 | -0.501 | -0.204 | 0.572 | 1.326 | -0.026 | 0.032 | 0.032 | 0 | 0.07 | 0.11 | 0.01 | 0.18 | 0.02 | 0.18 |
| 6 | 156858484 | rs9478712 | -1.654 | 0.087 | -0.224 | 1.035 | -0.02 | -0.066 | -0.066 | 0 | 0.36 | 0.96 | 0.3 | 0.38 | 0.28 | 0.49 |
| 6 | 156861144 | rs9480344 | -1.168 | 0.027 | 0.816 | 0.989 | -0.02 | -0.04 | -0.04 | 0 | 0.24 | 0.49 | 0.15 | 0.18 | 0.17 | 0.25 |
| 6 | 156864146 | rs11753286 | -1.943 | 0.189 | 0.188 | 0.77 | -0.029 | -0.013 | -0.013 | 0 | 0.35 | 0.95 | 0.33 | 0.33 | 0.29 | 0.48 |
| 6 | 156866619 | rs9480346 | -1.843 | 0.233 | 0.4 | 0.792 | 0.009 | 0.019 | 0.019 | 0 | 0.34 | 0.86 | 0.32 | 0.33 | 0.28 | 0.47 |
| 6 | 156866882 | rs11156113 | -1.899 | 0.29 | -0.128 | 0.48 | -0.027 | 0.044 | 0.044 | 0 | 0.39 | 0.92 | 0.34 | 0.34 | 0.32 | 0.48 |
| 6 | 156867121 | rs7750601 | -0.475 | 1.244 | -0.058 | 0.509 | -0.017 | -0.015 | -0.015 | 0.022 | 0.08 | 0.05 | 0.12 | 0.14 | 0.17 | 0.18 |
| 6 | 156868092 | rs288977 | -1.628 | 1.459 | 0.551 | 0.588 | 0.042 | 0.056 | 0.056 | 0.136 | 0.17 | 0.31 | 0.18 | 0.3 | 0.24 | 0.28 |
| 6 | 156868110 | rs288976 | -1.918 | 1.334 | 0.408 | 0.533 | -0.018 | 0.006 | 0.006 | 0 | 0.4 | 0.74 | 0.36 | 0.41 | 0.41 | 0.55 |
| 6 | 156868655 | rs9480347 | -1.046 | 1.954 | 0.097 | 0.882 | -0.001 | -0.002 | -0.002 | 0 | 0.1 | 0.08 | 0.06 | 0.13 | 0.1 | 0.15 |
| 6 | 156873144 | rs9480349 | -0.613 | 1.698 | 0.121 | 0.423 | -0.012 | 0.007 | 0.007 | 0 | 0.14 | 0.12 | 0.18 | 0.1 | 0.17 | 0.14 |
| 6 | 156874734 | rs288968 | -0.45 | 1.133 | -0.638 | -0.423 | 0.012 | 0.083 | 0.083 | 0.135 | 0.84 | 0.95 | 0.91 | 0.99 | 0.88 | 0.91 |
| 6 | 156878678 | rs288975 | -1.17 | 0.724 | -0.219 | -0.72 | 0.001 | -0.045 | -0.045 | 0 | 0.2 | 0.64 | 0.13 | 0.11 | 0.18 | 0.21 |
| 6 | 156882687 | rs288959 | -0.782 | 0.759 | 0.216 | -0.705 | 0.006 | -0.022 | -0.022 | 0.041 | 0.16 | 0.21 | 0.11 | 0.1 | 0.17 | 0.19 |
| 6 | 156893548 | rs288956 | 0.6 | 1.077 | 0.5 | -0.281 | -0.002 | -0.014 | -0.014 | 0 | 0.57 | 0.48 | 0.42 | 0.69 | 0.58 | 0.66 |
| 6 | 156897183 | rs2767577 | -1.235 | 1.065 | 0.608 | -0.281 | 0.088 | 0.12 | 0.12 | 0.35 | 0.13 | 0.36 | 0.37 | 0.29 | 0.23 | 0.25 |
| 6 | 156904256 | rs1998930 | 1.133 | 1.008 | -0.126 | -0.301 | 0.038 | 0.065 | 0.065 | 0 | 0.48 | 0.13 | 0.3 | 0.6 | 0.39 | 0.52 |
| 6 | 156907168 | rs9480356 | 0.161 | 1.111 | 0.312 | 0.293 | -0.027 | 0.041 | 0.041 | 0 | 0.11 | 0.05 | 0.12 | 0 | 0.13 | 0.03 |
| 6 | 156910285 | rs2819080 | -1.161 | 1.237 | 1.041 | 0.15 | 0.141 | 0.04 | 0.04 | 0 | 0.1 | 0.36 | 0.14 | 0.12 | 0.2 | 0.15 |
| 6 | 156913012 | rs2817461 | -1.205 | 1.269 | 0.297 | 0.168 | -0.045 | 0.011 | 0.011 | 0 | 0.2 | 0.15 | 0.25 | 0.19 | 0.32 | 0.24 |
| 6 | 156916754 | rs2767582 | -0.891 | 1.102 | 0.348 | 0.397 | -0.01 | -0.006 | -0.006 | 0 | 0.09 | 0.07 | 0.11 | 0.16 | 0.21 | 0.16 |
| 6 | 156916766 | rs9384478 | -0.725 | 0.576 | 0.085 | 0.961 | 0.025 | -0.05 | -0.05 | 0 | 0.13 | 0.05 | 0.06 | 0.24 | 0.07 | 0.14 |
| 6 | 156921064 | rs1022984 | -1.203 | 0.659 | 0.273 | 0.91 | -0.021 | -0.021 | -0.021 | 0 | 0.41 | 0.33 | 0.37 | 0.56 | 0.5 | 0.44 |
| 6 | 156922277 | rs913209 | -1.537 | 0.579 | 0.716 | 1.052 | -0.001 | 0.001 | 0.001 | 0 | 0.45 | 0.54 | 0.4 | 0.61 | 0.51 | 0.46 |
| 6 | 156922747 | rs2767584 | -1.254 | 0.664 | 0.687 | 0.971 | -0.013 | -0.014 | -0.014 | 0 | 0.41 | 0.33 | 0.37 | 0.56 | 0.5 | 0.45 |
| 6 | 156942543 | rs9480365 | -0.723 | 0.764 | 0.59 | 0.912 | 0.025 | -0.025 | -0.025 | 2.493 | 0.16 | 0.27 | 0.13 | 0.14 | 0.2 | 0.17 |
| 6 | 156949912 | rs11156119 | 1.372 | 0.574 | 0.709 | 0.636 | 0.005 | 0.002 | 0.002 | 0 | 0.57 | 0.44 | 0.59 | 0.44 | 0.52 | 0.51 |
| 6 | 156954218 | rs1246172 | -0.331 | 0.546 | 0.165 | 0.688 | -0.042 | 0.018 | 0.018 | 0 | 0.81 | 0.92 | 0.8 | 0.84 | 0.69 | 0.85 |
| 6 | 156954566 | rs7740342 | 1.138 | 0.575 | 0.208 | 0.679 | -0.008 | -0.007 | -0.007 | 0 | 0.56 | 0.69 | 0.6 | 0.44 | 0.46 | 0.51 |
| 6 | 156961391 | rs4870475 | -0.335 | 0.511 | -0.439 | 0.638 | -0.017 | 0.021 | 0.021 | 0 | 0.79 | 0.83 | 0.79 | 0.85 | 0.68 | 0.85 |
| 6 | 156964319 | rs17087661 | -0.368 | 0.223 | -0.636 | 0.925 | 0.017 | -0.052 | -0.052 | 0.005 | 0.12 | 0.03 | 0.11 | 0.25 | 0.05 | 0.19 |
| 6 | 156966162 | rs1246176 | 0.855 | 0.14 | -0.279 | 0.739 | -0.001 | -0.01 | -0.01 | 0 | 0.47 | 0.26 | 0.53 | 0.42 | 0.41 | 0.54 |
| 6 | 156971688 | rs9397956 | -0.314 | 0.101 | -0.115 | 0.89 | -0.023 | -0.03 | -0.03 | 1.108 | 0.17 | 0.04 | 0.09 | 0.21 | 0.14 | 0.25 |
| 6 | 156974859 | rs6916489 | -1.092 | -0.3 | 0.742 | 0.953 | -0.054 | 0.082 | 0.082 | 0 | 0.11 | 0.37 | 0.12 | 0.26 | 0.05 | 0.12 |
| 6 | 156976650 | rs6919504 | -0.827 | -0.235 | 0.73 | 0.923 | -0.024 | 0.052 | 0.052 | 0 | 0.09 | 0.33 | 0.08 | 0.16 | 0.05 | 0.09 |
| 6 | 156994250 | rs34352212 | -0.526 | -0.464 | 0.623 | 0.811 | 0.007 | -0.006 | -0.006 | 0 | 0.22 | 0.17 | 0.14 | 0.29 | 0.14 | 0.28 |
| 6 | 157003689 | rs7768451 | -0.517 | -0.438 | 0.453 | 0.794 | -0.021 | 0.006 | 0.006 | 0 | 0.08 | 0.14 | 0.07 | 0.16 | 0.05 | 0.09 |

| | | | | | | | | | | | | | | | | |
|---|---|---|---|---|---|---|---|---|---|---|---|---|---|---|---|---|
| 6 | 157009381 | rs9384488 | -1.619 | -0.413 | -0.097 | 0.711 | 0.048 | -0.068 | -0.068 | 0 | 0.44 | 0.86 | 0.31 | 0.55 | 0.3 | 0.52 |
| 6 | 157020791 | rs9480379 | -1.391 | -0.454 | 0.273 | 0.653 | -0.035 | 0.012 | 0.012 | 0 | 0.33 | 0.47 | 0.22 | 0.45 | 0.2 | 0.43 |
| 6 | 157022885 | rs10499278 | -1.087 | -0.04 | 0.411 | 0.851 | -0.032 | 0.066 | 0.066 | 0 | 0.09 | 0.26 | 0.08 | 0.17 | 0.05 | 0.09 |
| 6 | 157042018 | rs9397313 | -0.158 | 0.488 | -0.61 | 0.468 | -0.004 | 0.05 | 0.05 | 0.025 | 0.47 | 0.71 | 0.35 | 0.38 | 0.43 | 0.46 |
| 6 | 157048637 | rs1537583 | -0.303 | 0.592 | -0.43 | 0.732 | 0.004 | 0 | 0 | 0 | 0.4 | 0.36 | 0.19 | 0.37 | 0.32 | 0.33 |
| 6 | 157059428 | rs6934980 | 0.106 | 0.756 | -0.181 | 0.769 | -0.005 | -0.006 | -0.006 | 0 | 0.58 | 0.46 | 0.77 | 0.67 | 0.64 | 0.65 |
| 6 | 157064953 | rs2151884 | 0.569 | 1.394 | -0.729 | 0.88 | -0.004 | 0.029 | 0.029 | 0 | 0.4 | 0.22 | 0.55 | 0.38 | 0.42 | 0.39 |
| 6 | 157066200 | rs4870480 | -0.402 | 1.463 | -0.407 | 0.979 | 0.006 | 0.006 | 0.006 | 0 | 0.14 | 0.14 | 0.2 | 0.19 | 0.19 | 0.21 |
| 6 | 157072815 | rs9372014 | 0.001 | 0.915 | -0.89 | 0.515 | 0.043 | -0.003 | -0.003 | 0 | 0.36 | 0.12 | 0.23 | 0.34 | 0.28 | 0.37 |
| 6 | 157072952 | rs10457935 | -0.347 | 1.728 | -0.216 | 0.13 | 0.038 | -0.03 | -0.03 | 0 | 0.09 | 0.12 | 0.15 | 0.02 | 0.19 | 0.1 |
| 6 | 157075575 | rs4869890 | -1.086 | 1.151 | 0.451 | -0.03 | 0.002 | 0.024 | 0.024 | 0 | 0.1 | 0.38 | 0.06 | 0.08 | 0.06 | 0.08 |
| 6 | 157089510 | rs4870483 | -1.268 | 1.441 | 0.612 | 0.014 | 0.007 | -0.006 | -0.006 | 0 | 0.17 | 0.18 | 0.2 | 0.09 | 0.23 | 0.14 |
| 6 | 157103832 | rs6934734 | -1.666 | 1.342 | 1.044 | -0.035 | 0.026 | -0.038 | -0.038 | 0 | 0.2 | 0.53 | 0.22 | 0.09 | 0.23 | 0.14 |
| 6 | 157108163 | rs6927409 | -2.081 | 1.059 | 0.386 | -0.151 | -0.01 | 0.007 | 0.007 | 0.655 | 0.37 | 0.77 | 0.37 | 0.38 | 0.35 | 0.47 |
| 6 | 157113617 | rs9397968 | 0.452 | 1.138 | 0.262 | 0.061 | -0.011 | -0.007 | -0.007 | 4.213 | 0.07 | 0.05 | 0.09 | 0.26 | 0.07 | 0.24 |
| 6 | 157127825 | rs4870490 | -0.413 | 0.865 | 0.279 | 0.025 | 0.002 | -0.006 | -0.006 | 0.493 | 0.07 | 0.05 | 0.04 | 0.09 | 0.06 | 0.09 |
| 6 | 157130042 | rs9397970 | 0.305 | 0.833 | 0.326 | 0.003 | 0 | 0.015 | 0.015 | 0 | 0.08 | 0.11 | 0.1 | 0.27 | 0.05 | 0.24 |
| 6 | 157135476 | rs542224 | 1.141 | 0.341 | -0.38 | -1.307 | 0.016 | 0.081 | 0.081 | 0 | 0.23 | 0.08 | 0.1 | 0 | 0.17 | 0.11 |
| 6 | 157136631 | rs1709065 | -1.191 | 0.175 | -0.156 | -1.374 | -0.041 | 0.002 | 0.002 | 0 | 0.61 | 0.79 | 0.36 | 0.41 | 0.46 | 0.58 |
| 6 | 157144362 | rs182308 | 1.341 | 0.111 | 0.058 | -1.488 | 0.007 | 0.069 | 0.069 | 0 | 0.25 | 0.11 | 0.1 | 0 | 0.21 | 0.12 |
| 6 | 157149930 | rs9480393 | 0.819 | 0.298 | 0.051 | -1.39 | -0.001 | 0.004 | 0.004 | 0 | 0.1 | 0.1 | 0.12 | 0.32 | 0.07 | 0.25 |
| 6 | 157150230 | rs12206233 | 1.017 | 0.619 | -0.165 | -1.519 | -0.044 | 0.043 | 0.043 | 0 | 0.08 | 0.01 | 0.15 | 0 | 0.18 | 0.04 |
| 6 | 157161580 | rs287905 | -1.269 | 0.26 | 0.236 | -1.576 | -0.011 | -0.002 | -0.002 | 0.57 | 0.65 | 0.7 | 0.48 | 0.4 | 0.61 | 0.59 |
| 6 | 157177969 | rs287914 | -1.427 | 0.292 | 0.426 | -1.589 | 0 | 0 | 0 | 0 | 0.67 | 0.73 | 0.5 | 0.44 | 0.63 | 0.61 |
| 6 | 157183559 | rs9397976 | -1.516 | 0.301 | 0.389 | -1.583 | -0.016 | -0.02 | -0.02 | 0 | 0.68 | 0.75 | 0.5 | 0.44 | 0.63 | 0.65 |
| 6 | 157188134 | rs9372022 | -1.21 | 0.618 | 0.493 | -1.794 | -0.008 | -0.025 | -0.025 | 0 | 0.29 | 0.44 | 0.21 | 0.32 | 0.2 | 0.39 |
| 6 | 157188892 | rs288550 | -1.316 | 0.441 | 0.047 | -1.923 | -0.022 | 0.012 | 0.012 | 0 | 0.59 | 0.84 | 0.44 | 0.42 | 0.54 | 0.54 |
| 6 | 157193250 | rs9397978 | -0.755 | 0.503 | 0.282 | -1.88 | 0.012 | 0.014 | 0.014 | 0 | 0.26 | 0.31 | 0.32 | 0.41 | 0.32 | 0.39 |
| 6 | 157196760 | rs6905191 | 0.1 | 0.502 | 0.298 | -1.859 | 0.022 | 0.005 | 0.005 | 0 | 0.13 | 0.21 | 0.18 | 0.09 | 0.22 | 0.1 |
| 6 | 157196974 | rs10485205 | 0.452 | 0.5 | 0.278 | -1.852 | -0.005 | -0.011 | -0.011 | 0 | 0.13 | 0.1 | 0.14 | 0.32 | 0.1 | 0.26 |
| 6 | 157197802 | rs913971 | -1.84 | 0.431 | 0.245 | -1.966 | -0.03 | 0.018 | 0.018 | 0.044 | 0.42 | 0.83 | 0.37 | 0.41 | 0.41 | 0.46 |
| 6 | 157199844 | rs17266366 | -0.284 | 0.399 | 0.239 | -1.953 | 0.001 | -0.006 | -0.006 | 1.72 | 0.23 | 0.23 | 0.21 | 0.09 | 0.3 | 0.15 |
| 6 | 157201931 | rs158258 | 1.033 | 0.339 | 0.29 | -2.045 | -0.004 | -0.053 | -0.053 | 0 | 0.24 | 0.29 | 0.11 | 0.01 | 0.17 | 0.12 |
| 6 | 157203063 | rs287873 | -1.688 | 0.427 | 0.138 | -1.84 | -0.024 | -0.014 | -0.014 | 0 | 0.62 | 0.88 | 0.46 | 0.42 | 0.58 | 0.56 |
| 6 | 157216039 | rs184074 | -1.04 | 0.771 | 0.08 | -0.973 | -0.007 | -0.011 | -0.011 | 0 | 0.71 | 0.78 | 0.7 | 0.67 | 0.57 | 0.73 |
| 6 | 157216166 | rs9372026 | -0.237 | 1.22 | 1.119 | -1.026 | 0.05 | 0.042 | 0.042 | 0 | 0.1 | 0.26 | 0.12 | 0.09 | 0.18 | 0.1 |
| 6 | 157216200 | rs9384510 | 0.445 | 1.257 | 0.865 | -0.918 | -0.006 | -0.005 | -0.005 | 0 | 0.11 | 0.1 | 0.13 | 0.34 | 0.11 | 0.27 |
| 6 | 157218902 | rs167007 | -1.455 | 1.054 | 1.268 | -1.564 | 0.004 | 0.034 | 0.034 | 0 | 0.48 | 0.6 | 0.31 | 0.1 | 0.45 | 0.32 |
| 6 | 157219303 | rs9384511 | -0.24 | 1.052 | 0.898 | -1.521 | 0.003 | -0.001 | -0.001 | 0.116 | 0.13 | 0.13 | 0.17 | 0.09 | 0.2 | 0.12 |
| 6 | 157223707 | rs287945 | -0.559 | 0.99 | 0.347 | -1.498 | 0.008 | 0.017 | 0.017 | 0.34 | 0.34 | 0.32 | 0.14 | 0.01 | 0.25 | 0.19 |
| 6 | 157232519 | rs4870497 | -0.899 | 1.027 | 0.211 | -1.168 | 0 | 0.003 | 0.003 | 0.097 | 0.24 | 0.77 | 0.23 | 0.11 | 0.28 | 0.19 |
| 6 | 157242653 | rs284422 | -0.211 | 1.033 | -0.536 | -1.128 | 0.009 | 0.064 | 0.064 | 0 | 0.26 | 0.19 | 0.1 | 0 | 0.21 | 0.12 |
| 6 | 157250189 | rs284430 | -1.501 | 1.216 | -0.017 | -0.593 | -0.002 | 0.013 | 0.013 | 0 | 0.35 | 0.68 | 0.27 | 0.1 | 0.37 | 0.27 |
| 6 | 157251223 | rs284431 | -0.859 | 1.235 | 0.067 | -0.428 | 0.003 | -0.043 | -0.043 | 0 | 0.14 | 0.24 | 0.06 | 0.02 | 0.09 | 0.12 |
| 6 | 157255918 | rs284418 | -1.02 | 0.953 | -0.855 | 0.007 | -0.097 | 0.097 | 0.097 | 0 | 0.54 | 0.91 | 0.51 | 0.68 | 0.52 | 0.58 |
| 6 | 157262028 | rs6906983 | -0.956 | 0.834 | 0.515 | 0.593 | -0.01 | 0.021 | 0.021 | 0 | 0.14 | 0.24 | 0.1 | 0.21 | 0.12 | 0.08 |
| 6 | 157264716 | rs2236421 | -1.574 | 0.746 | 1.145 | 0.715 | -0.031 | 0.148 | 0.148 | 0 | 0.18 | 0.52 | 0.12 | 0.3 | 0.13 | 0.14 |
| 6 | 157265752 | rs10485203 | -1.156 | 0.896 | 0.498 | 0.415 | -0.009 | 0.03 | 0.03 | 0 | 0.14 | 0.25 | 0.1 | 0.24 | 0.13 | 0.09 |
| 6 | 157266505 | rs4870499 | -0.852 | 1.304 | 0.075 | 0.033 | -0.027 | 0.027 | 0.027 | 0 | 0.17 | 0.1 | 0.19 | 0 | 0.26 | 0.09 |
| 6 | 157274799 | rs1466648 | -1.467 | 1.271 | 0.271 | 0.175 | -0.004 | 0.002 | 0.002 | 0.85 | 0.24 | 0.78 | 0.34 | 0.22 | 0.31 | 0.15 |
| 6 | 157287299 | rs12208834 | -0.743 | 1.496 | 0.024 | 0.248 | -0.036 | -0.017 | -0.017 | 0 | 0.18 | 0.11 | 0.28 | 0.23 | 0.3 | 0.15 |
| 6 | 157297182 | rs2356043 | 0.199 | 0.903 | 0.348 | 0.291 | -0.004 | 0.011 | 0.011 | 0 | 0.23 | 0.2 | 0.2 | 0.42 | 0.18 | 0.41 |
| 6 | 157312750 | rs9372033 | 0.363 | 0.822 | 0.766 | 0.215 | -0.034 | 0.089 | 0.089 | 0 | 0.21 | 0.35 | 0.2 | 0.42 | 0.16 | 0.39 |
| 6 | 157318748 | rs6557516 | -1.116 | 0.969 | -0.037 | -0.454 | -0.02 | -0.024 | -0.024 | 0 | 0.66 | 0.98 | 0.64 | 0.75 | 0.75 | 0.66 |
| 6 | 157320902 | rs12214873 | 0.487 | 1.314 | 0.084 | -0.479 | -0.016 | 0.007 | 0.007 | 0 | 0.11 | 0.07 | 0.15 | 0 | 0.22 | 0.06 |
| 6 | 157321061 | rs10485201 | -0.084 | 1.306 | 0.32 | -0.457 | 0.002 | 0 | 0 | 0 | 0.19 | 0.21 | 0.29 | 0.23 | 0.3 | 0.15 |
| 6 | 157334519 | rs9478745 | 0.839 | 1.38 | 0.115 | -0.592 | -0.022 | 0.028 | 0.028 | 0 | 0.19 | 0.13 | 0.12 | 0.11 | 0.22 | 0.13 |
| 6 | 157337988 | rs6909426 | -0.218 | 1.351 | 0.505 | -0.508 | 0.004 | -0.007 | -0.007 | 0 | 0.41 | 0.45 | 0.49 | 0.65 | 0.48 | 0.55 |
| 6 | 157338328 | rs9285548 | -0.095 | 1.335 | 0.399 | -0.504 | -0.014 | 0.018 | 0.018 | 0 | 0.4 | 0.26 | 0.47 | 0.67 | 0.48 | 0.55 |
| 6 | 157352528 | rs11753090 | 1.025 | 1.326 | 0.243 | -0.454 | 0.001 | -0.001 | -0.001 | 0 | 0.22 | 0.17 | 0.2 | 0.41 | 0.18 | 0.4 |
| 6 | 157357091 | rs2207227 | -0.393 | 1.536 | 0.224 | -0.232 | 0.082 | 0.023 | 0.023 | 0.033 | 0.2 | 0.47 | 0.3 | 0.23 | 0.3 | 0.15 |
| 6 | 157368518 | rs9322596 | -0.161 | 1.591 | -0.296 | -0.227 | -0.009 | -0.003 | -0.003 | 0.635 | 0.18 | 0.18 | 0.29 | 0.23 | 0.3 | 0.15 |
| 6 | 157376672 | rs12211179 | 0.935 | 1.421 | -0.705 | -0.338 | -0.036 | -0.065 | -0.065 | 0 | 0.26 | 0.01 | 0.33 | 0.24 | 0.22 | 0.26 |
| 6 | 157388615 | rs9480424 | 0.481 | 1.441 | -0.716 | -0.46 | -0.012 | -0.006 | -0.006 | 0 | 0.07 | 0.09 | 0.05 | 0.03 | 0.03 | 0.06 |
| 6 | 157389895 | rs7770280 | 0.616 | 1.928 | -1.113 | -0.61 | -0.03 | 0.037 | 0.037 | 0 | 0.17 | 0.06 | 0.12 | 0.07 | 0.22 | 0.12 |
| 6 | 157389909 | rs7755878 | -0.307 | 1.929 | -1.214 | -0.322 | -0.027 | 0.029 | 0.029 | 0.01 | 0.4 | 0.76 | 0.44 | 0.35 | 0.48 | 0.35 |
| 6 | 157390781 | rs13216156 | 0.899 | 1.845 | -1.539 | 0.232 | -0.037 | -0.028 | -0.028 | 0 | 0.17 | 0 | 0.06 | 0.14 | 0.14 | 0.13 |
| 6 | 157394217 | rs2207229 | -0.356 | 1.51 | -1.231 | 0.441 | -0.086 | -0.08 | -0.08 | 0 | 0.71 | 0.98 | 0.65 | 0.73 | 0.73 | 0.74 |
| 6 | 157398175 | rs17088067 | -0.297 | 1.604 | 0.197 | 0.602 | 0.052 | -0.057 | -0.057 | 0 | 0.16 | 0.37 | 0.12 | 0.07 | 0.19 | 0.11 |

| | | | | | | | | | | | | | | | | |
|---|---|---|---|---|---|---|---|---|---|---|---|---|---|---|---|---|
| 6 | 157405761 | rs3734440 | 0.763 | 1.613 | -0.105 | 0.863 | -0.015 | -0.007 | -0.007 | 0.453 | 0.58 | 0.27 | 0.63 | 0.59 | 0.52 | 0.61 |
| 6 | 157405930 | rs3734441 | 0.777 | 1.577 | -0.12 | 0.871 | -0.014 | -0.009 | -0.009 | 0 | 0.58 | 0.28 | 0.63 | 0.57 | 0.52 | 0.58 |
| 6 | 157413820 | rs9397995 | 0.937 | 1.773 | -0.56 | 0.194 | -0.006 | -0.004 | -0.004 | 0 | 0.54 | 0.27 | 0.39 | 0.53 | 0.53 | 0.54 |
| 6 | 157416680 | rs767860 | 0.654 | 1.386 | -0.353 | 0.186 | -0.001 | -0.002 | -0.002 | 0 | 0.6 | 0.33 | 0.44 | 0.46 | 0.45 | 0.62 |
| 6 | 157417484 | rs6916789 | 0.613 | 1.089 | -0.367 | 0.129 | 0 | 0.005 | 0.005 | 0 | 0.51 | 0.4 | 0.5 | 0.6 | 0.57 | 0.53 |
| 6 | 158022695 | rs16900291 | -0.277 | 0.726 | -0.31 | 0.184 | -0.006 | -0.019 | -0.019 | 0 | 0.31 | 0.43 | 0.4 | 0.21 | 0.27 | 0.26 |
| 6 | 158022745 | rs4708866 | -0.386 | 0.795 | -1.433 | 0.404 | -0.059 | 0.002 | 0.002 | 0 | 0.65 | 0.9 | 0.64 | 0.76 | 0.55 | 0.64 |
| 6 | 158029872 | rs34162221 | 0.3 | 0.44 | -1.039 | 0.495 | 0.006 | 0.002 | 0.002 | 0.69 | 0.28 | 0.23 | 0.14 | 0.18 | 0.19 | 0.2 |
| 6 | 158030998 | rs34192579 | -0.303 | 0.622 | -1.195 | 0.366 | -0.011 | 0.005 | 0.005 | 0 | 0.72 | 0.84 | 0.65 | 0.76 | 0.66 | 0.7 |
| 6 | 158034980 | rs3933566 | 0.041 | 0.196 | -0.923 | 0.537 | 0.004 | -0.004 | -0.004 | 0.061 | 0.16 | 0.16 | 0.13 | 0.27 | 0.11 | 0.16 |
| 6 | 158069399 | rs602890 | -0.088 | 0.685 | -1.276 | -0.188 | -0.06 | 0.099 | 0.099 | 0 | 0.35 | 0.21 | 0.61 | 0.11 | 0.51 | 0.42 |
| 6 | 158070192 | rs9355850 | 0.222 | -0.072 | -0.54 | 0.146 | -0.02 | 0.028 | 0.028 | 0 | 0.28 | 0.34 | 0.15 | 0.59 | 0.16 | 0.3 |
| 6 | 158070339 | rs587437 | 0.2 | -0.064 | -0.482 | -0.137 | 0.012 | -0.064 | -0.064 | 0.629 | 0.34 | 0.49 | 0.38 | 0.11 | 0.4 | 0.35 |
| 6 | 158070812 | rs586141 | -0.371 | 0.053 | -0.158 | -0.069 | -0.011 | 0.021 | 0.021 | 0 | 0.6 | 0.69 | 0.52 | 0.7 | 0.56 | 0.65 |
| 6 | 158077053 | rs9355853 | 0.342 | -0.1 | 0.503 | 0.134 | -0.008 | 0.01 | 0.01 | 0 | 0.28 | 0.32 | 0.13 | 0.54 | 0.16 | 0.31 |
| 6 | 158077359 | rs634718 | -0.46 | -0.06 | 0.292 | -0.278 | -0.006 | -0.007 | -0.007 | 0 | 0.65 | 0.7 | 0.53 | 0.64 | 0.62 | 0.65 |
| 6 | 158078716 | rs618368 | 0.375 | -0.048 | 0.413 | -0.208 | 0.029 | -0.095 | -0.095 | 1.804 | 0.32 | 0.43 | 0.37 | 0.1 | 0.4 | 0.33 |
| 6 | 158080044 | rs602237 | -0.963 | -0.007 | 0.106 | -0.323 | -0.009 | -0.019 | -0.019 | 0 | 0.66 | 0.84 | 0.76 | 0.65 | 0.68 | 0.74 |
| 6 | 158080876 | rs628389 | 0.659 | 0.01 | 0.026 | -0.315 | -0.047 | 0.074 | 0.074 | 0 | 0.29 | 0.13 | 0.35 | 0.1 | 0.39 | 0.33 |
| 6 | 158085284 | rs9365242 | 0.669 | -0.097 | 0.009 | -0.279 | 0.039 | -0.076 | -0.076 | 0 | 0.25 | 0.14 | 0.14 | 0.54 | 0.18 | 0.29 |
| 6 | 158086912 | rs625880 | -0.732 | -0.077 | 0.253 | -0.294 | -0.002 | -0.003 | -0.003 | 0 | 0.65 | 0.78 | 0.75 | 0.69 | 0.68 | 0.75 |
| 6 | 158088225 | rs12055716 | 0.414 | -0.198 | 0.235 | 0.002 | 0.028 | -0.06 | -0.06 | 0 | 0.27 | 0.17 | 0.15 | 0.56 | 0.2 | 0.31 |
| 6 | 158088737 | rs6927266 | -0.742 | 0.295 | 0.479 | -0.112 | 0.018 | -0.036 | -0.036 | 0 | 0.09 | 0.3 | 0.25 | 0.02 | 0.09 | 0.08 |
| 6 | 158090541 | rs672764 | -0.513 | 0.604 | -0.074 | -0.432 | 0.001 | -0.025 | -0.025 | 1.691 | 0.28 | 0.64 | 0.42 | 0.09 | 0.32 | 0.2 |
| 6 | 158090743 | rs9295140 | 0.631 | 0.267 | -0.633 | -0.192 | 0.022 | -0.061 | -0.061 | 0.023 | 0.28 | 0.19 | 0.17 | 0.53 | 0.22 | 0.29 |
| 6 | 158091108 | rs2038586 | 0.442 | 0.522 | -0.605 | -0.028 | 0 | -0.001 | -0.001 | 0 | 0.14 | 0.1 | 0.17 | 0.12 | 0.15 | 0.23 |
| 6 | 158092638 | rs688181 | -0.887 | 0.616 | -0.626 | 0.056 | -0.031 | 0.002 | 0.002 | 0 | 0.69 | 0.86 | 0.79 | 0.72 | 0.68 | 0.72 |
| 6 | 158092980 | rs9355333 | 0.494 | -0.098 | -0.733 | 0.158 | 0.065 | -0.074 | -0.074 | 0 | 0.35 | 0.18 | 0.21 | 0.51 | 0.25 | 0.31 |
| 6 | 158096328 | rs3749839 | -0.703 | 0.732 | -0.421 | 0.433 | -0.105 | -0.06 | -0.06 | 0 | 0.69 | 0.96 | 0.77 | 0.71 | 0.67 | 0.77 |
| 6 | 158097475 | rs629218 | -0.655 | 0.733 | -0.089 | 0.452 | -0.037 | -0.033 | -0.033 | 0 | 0.69 | 0.86 | 0.76 | 0.66 | 0.67 | 0.76 |
| 6 | 158099736 | rs9347482 | -0.274 | 0.706 | 0.585 | 0.437 | -0.001 | -0.001 | -0.001 | 0 | 0.64 | 0.72 | 0.74 | 0.58 | 0.63 | 0.68 |
| 6 | 158101005 | rs13205348 | 0.325 | 0.687 | 0.544 | 0.272 | -0.002 | 0.012 | 0.012 | 0 | 0.13 | 0.12 | 0.27 | 0.07 | 0.15 | 0.17 |
| 6 | 158101468 | rs941888 | -0.836 | 0.698 | 0.413 | 0.293 | -0.049 | -0.021 | -0.021 | 0 | 0.69 | 0.9 | 0.76 | 0.69 | 0.67 | 0.76 |
| 6 | 158101911 | rs675658 | -0.408 | 0.732 | 0.671 | 0.046 | -0.002 | -0.003 | -0.003 | 0 | 0.65 | 0.71 | 0.74 | 0.58 | 0.63 | 0.67 |
| 6 | 158102714 | rs877676 | 0.242 | 0.741 | 0.246 | 0.005 | 0 | -0.004 | -0.004 | 0 | 0.13 | 0.16 | 0.29 | 0.07 | 0.17 | 0.18 |
| 6 | 158106863 | rs585635 | 0.23 | 0.554 | 0.07 | 0.13 | 0.01 | 0.009 | 0.009 | 0 | 0.5 | 0.36 | 0.44 | 0.51 | 0.48 | 0.51 |
| 6 | 158115457 | rs9355891 | 0.194 | 0.557 | 0.065 | 0.114 | 0.004 | 0.006 | 0.006 | 0 | 0.49 | 0.29 | 0.44 | 0.51 | 0.48 | 0.43 |
| 6 | 158115787 | rs672786 | 0.091 | 0.432 | 0.084 | 0.133 | 0.011 | 0.011 | 0.011 | 0 | 0.5 | 0.37 | 0.46 | 0.5 | 0.49 | 0.43 |
| 6 | 158117108 | rs683735 | -0.469 | 0.622 | -0.367 | 0.085 | -0.064 | -0.064 | -0.064 | 0 | 0.67 | 0.9 | 0.75 | 0.62 | 0.61 | 0.66 |
| 6 | 158117808 | rs6920371 | 0.337 | 0.596 | -0.113 | -0.403 | 0 | -0.019 | -0.019 | 0 | 0.19 | 0.18 | 0.29 | 0.06 | 0.2 | 0.21 |
| 6 | 158118344 | rs590906 | 0.343 | 0.409 | -0.197 | -0.526 | 0.003 | -0.161 | -0.161 | 0.546 | 0.28 | 0.48 | 0.23 | 0 | 0.3 | 0.16 |
| 6 | 158131946 | rs9458285 | 0.509 | 0.396 | -0.794 | 0.187 | 0.037 | -0.08 | -0.08 | 0.536 | 0.21 | 0.09 | 0.16 | 0.49 | 0.13 | 0.31 |
| 6 | 158132992 | rs7766023 | -0.086 | 0.583 | -1.085 | -0.014 | -0.077 | 0.163 | 0.163 | 0 | 0.52 | 0.89 | 0.63 | 0.23 | 0.53 | 0.46 |
| 6 | 158137474 | rs7765247 | -0.267 | 0.376 | -0.536 | 0.017 | 0.025 | 0.002 | 0.002 | 0 | 0.5 | 0.42 | 0.58 | 0.54 | 0.41 | 0.56 |
| 6 | 158138616 | rs9458301 | -0.287 | 0.315 | -0.482 | -0.084 | 0.012 | 0.021 | 0.021 | 0 | 0.5 | 0.47 | 0.6 | 0.71 | 0.44 | 0.62 |
| 6 | 158139618 | rs930125 | 0.14 | 0.446 | -0.798 | -0.308 | -0.002 | -0.047 | -0.047 | 0.002 | 0.28 | 0.33 | 0.41 | 0.06 | 0.29 | 0.22 |
| 6 | 158150982 | rs599898 | -0.747 | 0.781 | -1.408 | -0.089 | -0.125 | -0.06 | -0.06 | 0 | 0.73 | 0.98 | 0.82 | 0.74 | 0.68 | 0.79 |
| 6 | 158161866 | rs659978 | 0.811 | 0.387 | -0.955 | -0.308 | 0.005 | -0.08 | -0.08 | 0 | 0.22 | 0.33 | 0.18 | 0 | 0.23 | 0.2 |
| 6 | 158163207 | rs595436 | -0.088 | 0.409 | -1.481 | -0.261 | 0.013 | 0.001 | 0.001 | 0 | 0.49 | 0.63 | 0.39 | 0.52 | 0.4 | 0.52 |
| 6 | 158171888 | rs4709555 | 0.559 | 0.882 | -2.515 | -0.194 | -0.031 | 0.075 | 0.075 | 0 | 0.23 | 0.11 | 0.39 | 0.07 | 0.32 | 0.23 |
| 6 | 158178348 | rs9347573 | -0.177 | 0.698 | -2.258 | 0.859 | 0.035 | -0.063 | -0.063 | 0 | 0.25 | 0.12 | 0.19 | 0.48 | 0.17 | 0.31 |
| 6 | 158179374 | rs597619 | -0.232 | 0.838 | -2.604 | 0.86 | -0.071 | -0.059 | -0.059 | 0 | 0.76 | 0.95 | 0.83 | 0.74 | 0.73 | 0.82 |
| 6 | 158180026 | rs9347574 | -0.634 | 0.634 | -2.18 | 1.217 | 0.034 | -0.07 | -0.07 | 0.683 | 0.27 | 0.19 | 0.2 | 0.51 | 0.18 | 0.31 |
| 6 | 158184853 | rs1864500 | -0.57 | 1.274 | -2.28 | 1.37 | -0.046 | -0.139 | -0.139 | 0.968 | 0.24 | 0.08 | 0.23 | 0.49 | 0.25 | 0.27 |
| 6 | 158190377 | rs1753196 | -0.493 | 1.251 | -1.818 | 1.577 | -0.065 | -0.106 | -0.106 | 0.538 | 0.21 | 0.05 | 0.14 | 0.35 | 0.25 | 0.23 |
| 6 | 158193244 | rs16900421 | -1.203 | 0.788 | -1.16 | 2.093 | -0.004 | 0.059 | 0.059 | 0.1 | 0.14 | 0.16 | 0.16 | 0.5 | 0.1 | 0.23 |
| 6 | 158193664 | rs352193 | 0.323 | 0.957 | -1.306 | 1.387 | -0.016 | 0.028 | 0.028 | 0 | 0.57 | 0.72 | 0.69 | 0.61 | 0.51 | 0.46 |
| 6 | 158193725 | rs1034058 | 0.215 | 0.681 | -0.826 | 0.244 | -0.014 | 0.045 | 0.045 | 0 | 0.6 | 0.74 | 0.71 | 0.73 | 0.57 | 0.55 |
| 6 | 158194153 | rs168455 | -1.019 | 0.161 | 0.237 | 0.248 | 0.031 | 0.203 | 0.203 | 0.466 | 0.15 | 0.5 | 0.14 | 0.37 | 0.15 | 0.33 |
| 6 | 158194819 | rs2770113 | -0.59 | -0.197 | -0.185 | -0.702 | -0.006 | 0.03 | 0.03 | 1.465 | 0.55 | 0.61 | 0.41 | 0.77 | 0.5 | 0.58 |
| 6 | 158195695 | rs928008 | -0.14 | -0.787 | -0.052 | -0.924 | -0.021 | 0.094 | 0.094 | 0.001 | 0.54 | 0.71 | 0.64 | 0.72 | 0.51 | 0.51 |
| 6 | 158197326 | rs240642 | 0.084 | -0.906 | 0.102 | -0.98 | -0.01 | 0.005 | 0.005 | 0 | 0.52 | 0.71 | 0.61 | 0.49 | 0.49 | 0.66 |
| 6 | 158198680 | rs9365376 | -0.475 | -1.219 | 0.116 | -1.073 | -0.039 | 0.077 | 0.077 | 0 | 0.19 | 0.31 | 0.13 | 0.51 | 0.09 | 0.15 |
| 6 | 158201121 | rs11753617 | 0.716 | -0.955 | -0.135 | -1.433 | 0.009 | -0.025 | -0.025 | 0 | 0.18 | 0.33 | 0.15 | 0.12 | 0.19 | 0.2 |
| 6 | 158201642 | rs13217265 | 0.771 | -0.952 | -0.153 | -1.43 | 0.01 | -0.023 | -0.023 | 0.501 | 0.18 | 0.32 | 0.15 | 0.12 | 0.19 | 0.17 |
| 6 | 158206901 | rs2025994 | -0.482 | -0.687 | -0.878 | -1.115 | -0.003 | -0.002 | -0.002 | 0 | 0.19 | 0.18 | 0.11 | 0.77 | 0.08 | 0.36 |
| 6 | 158207928 | rs2803358 | -0.427 | -0.135 | -2.069 | -1.154 | 0.114 | 0.283 | 0.283 | 0.825 | 0.47 | 0.92 | 0.33 | 0.9 | 0.3 | 0.61 |
| 6 | 158208870 | rs12192757 | 1.482 | -0.062 | -2.094 | -1.136 | -0.077 | 0.06 | 0.06 | 1.956 | 0.23 | 0.02 | 0.39 | 0.1 | 0.27 | 0.2 |
| 6 | 158209347 | rs2770111 | -0.46 | 0.335 | -1.719 | -0.964 | -0.111 | 0.165 | 0.165 | 0 | 0.71 | 0.94 | 0.73 | 1 | 0.59 | 0.8 |

| | | | | | | | | | | | | | | | | |
|---|---|---|---|---|---|---|---|---|---|---|---|---|---|---|---|---|
| 6 | 158209783 | rs2770110 | -0.057 | 0.318 | -0.996 | 0.084 | -0.018 | -0.053 | -0.053 | 0 | 0.37 | 0.08 | 0.48 | 0.59 | 0.34 | 0.3 |
| 6 | 158216126 | rs2803362 | 0.647 | 0.532 | 0.061 | 0.126 | -0.011 | -0.015 | -0.015 | 0.026 | 0.63 | 0.54 | 0.63 | 0.3 | 0.69 | 0.53 |
| 6 | 158216859 | rs2803363 | 0.286 | 0.785 | -0.227 | -0.57 | 0 | 0.005 | 0.005 | 0 | 0.74 | 0.83 | 0.8 | 0.99 | 0.76 | 0.91 |
| 6 | 158222240 | rs9458540 | -0.904 | 1.338 | 0.622 | -0.217 | 0.098 | -0.013 | -0.013 | 0 | 0.06 | 0.23 | 0.06 | 0 | 0.11 | 0.01 |
| 6 | 158223703 | rs9456774 | -1.868 | 1.4 | 1.459 | -0.209 | 0.079 | -0.013 | -0.013 | 0 | 0.13 | 0.53 | 0.16 | 0.13 | 0.18 | 0.13 |
| 6 | 158240876 | rs6934158 | -1.623 | 1.215 | 1.554 | -0.615 | 0.04 | -0.11 | -0.11 | 0 | 0.16 | 0.38 | 0.1 | 0 | 0.19 | 0.13 |
| 6 | 158242573 | rs13202017 | -1.301 | 1.214 | 1.707 | -0.615 | 0.008 | -0.045 | -0.045 | 0 | 0.14 | 0.17 | 0.1 | 0 | 0.19 | 0.13 |
| 6 | 158249111 | rs1359984 | -2.058 | 1.195 | 1.728 | -0.603 | 0.033 | -0.062 | -0.062 | 0.485 | 0.21 | 0.52 | 0.18 | 0.14 | 0.26 | 0.24 |
| 6 | 158262162 | rs9356062 | -0.778 | 1.199 | 1.753 | -0.463 | 0.04 | 0.082 | 0.082 | 0 | 0.05 | 0.23 | 0.07 | 0.14 | 0.06 | 0.11 |
| 6 | 158264260 | rs2770104 | -1.083 | 1.389 | 1.127 | -0.372 | 0.021 | 0.011 | 0.011 | 0.174 | 0.5 | 0.66 | 0.4 | 0.47 | 0.54 | 0.49 |
| 6 | 158274398 | rs2803353 | 0.069 | 1.651 | 0.816 | -0.387 | -0.049 | -0.095 | -0.095 | 2.213 | 0.29 | 0.03 | 0.21 | 0.33 | 0.27 | 0.25 |
| 6 | 158297540 | rs9295212 | 0.034 | 1.694 | 1.122 | -0.466 | 0.025 | -0.021 | -0.021 | 0 | 0.06 | 0.09 | 0.05 | 0 | 0.12 | 0.01 |
| 6 | 158304083 | rs7741141 | -1.553 | 1.942 | 1.137 | -0.489 | 0.051 | -0.044 | -0.044 | 0 | 0.2 | 0.38 | 0.19 | 0.16 | 0.26 | 0.16 |
| 6 | 158304915 | rs2746217 | 0.951 | 1.461 | 0.087 | -0.534 | -0.034 | 0 | 0 | 0 | 0.25 | 0.01 | 0.37 | 0.14 | 0.25 | 0.19 |
| 6 | 158307549 | rs1883823 | -1.472 | 1.468 | 0.804 | -0.288 | 0.049 | -0.06 | -0.06 | 0.203 | 0.2 | 0.48 | 0.17 | 0.14 | 0.26 | 0.24 |
| 6 | 158310790 | rs1033528 | -1.545 | 1.441 | 0.782 | -0.302 | 0.034 | -0.069 | -0.069 | 0 | 0.21 | 0.47 | 0.17 | 0.14 | 0.26 | 0.26 |
| 6 | 158313916 | rs9458784 | -0.783 | 1.485 | 0.559 | -0.271 | 0.024 | -0.076 | -0.076 | 0 | 0.14 | 0.26 | 0.09 | 0 | 0.18 | 0.13 |
| 6 | 158327269 | rs9365567 | -1.37 | 1.469 | 0.555 | -0.263 | 0.014 | -0.035 | -0.035 | 0 | 0.2 | 0.29 | 0.14 | 0.14 | 0.25 | 0.25 |
| 6 | 158330765 | rs3211067 | -1.234 | 1.52 | 0.562 | -0.26 | 0.007 | -0.016 | -0.016 | 0 | 0.19 | 0.22 | 0.14 | 0.14 | 0.25 | 0.25 |
| 6 | 158357201 | rs724897 | -1.356 | 1.418 | 0.586 | -0.198 | 0.015 | -0.036 | -0.036 | 0 | 0.2 | 0.29 | 0.14 | 0.14 | 0.25 | 0.25 |
| 6 | 158362686 | rs9458900 | -0.825 | 1.412 | 0.578 | -0.224 | 0.005 | -0.057 | -0.057 | 0 | 0.16 | 0.2 | 0.09 | 0 | 0.19 | 0.16 |
| 6 | 158364714 | rs1043863 | -0.913 | 1.436 | 0.701 | -0.224 | 0.02 | -0.092 | -0.092 | 0 | 0.16 | 0.29 | 0.1 | 0 | 0.2 | 0.16 |
| 6 | 158365035 | rs577 | -1.282 | 1.419 | 0.621 | -0.15 | 0.03 | -0.045 | -0.045 | 0.044 | 0.2 | 0.36 | 0.15 | 0.15 | 0.26 | 0.25 |
| 6 | 158373208 | rs9356166 | -1.291 | 1.505 | 0.79 | -0.088 | 0.1 | 0.012 | 0.012 | 0 | 0.13 | 0.42 | 0.13 | 0.12 | 0.19 | 0.13 |
| 6 | 158376269 | rs6931002 | -0.454 | 1.334 | -0.132 | -0.365 | -0.035 | 0.015 | 0.015 | 0.078 | 0.68 | 0.9 | 0.54 | 0.74 | 0.62 | 0.65 |
| 6 | 158381509 | rs11755838 | 0.39 | 1.188 | -0.402 | -0.445 | -0.039 | -0.007 | -0.007 | 0.112 | 0.3 | 0.09 | 0.23 | 0.29 | 0.31 | 0.22 |
| 6 | 158382074 | rs1118242 | 0.494 | 0.898 | -0.218 | -0.288 | -0.009 | 0.019 | 0.019 | 0.283 | 0.29 | 0.65 | 0.26 | 0.43 | 0.21 | 0.35 |
| 6 | 158387117 | rs9295253 | -0.822 | 1.134 | -0.568 | -0.229 | -0.012 | -0.003 | -0.003 | 0 | 0.69 | 0.91 | 0.55 | 0.72 | 0.65 | 0.65 |
| 6 | 158387610 | rs9654570 | 0.586 | 1.079 | -0.281 | -0.246 | -0.015 | 0.043 | 0.043 | 0 | 0.28 | 0.4 | 0.25 | 0.42 | 0.21 | 0.37 |
| 6 | 158389741 | rs9458975 | -0.366 | 1.026 | -0.158 | -0.542 | 0.002 | -0.007 | -0.007 | 0.008 | 0.43 | 0.54 | 0.31 | 0.29 | 0.43 | 0.28 |
| 6 | 158397559 | rs9456931 | -0.698 | 0.796 | -0.281 | -0.743 | 0.001 | -0.03 | -0.03 | 0 | 0.25 | 0.34 | 0.19 | 0.17 | 0.24 | 0.27 |
| 6 | 158399002 | rs2025641 | -0.265 | 0.949 | -0.582 | -1.093 | 0 | -0.011 | -0.011 | 0 | 0.42 | 0.41 | 0.3 | 0.16 | 0.4 | 0.27 |
| 6 | 158399449 | rs4263608 | 0.497 | 0.747 | -0.789 | -1.127 | -0.004 | 0.055 | 0.055 | 0 | 0.2 | 0.06 | 0.17 | 0.13 | 0.18 | 0.18 |
| 6 | 158410377 | rs7772395 | 0.389 | 0.768 | -0.482 | -1.166 | 0.007 | -0.072 | -0.072 | 0.026 | 0.21 | 0.33 | 0.14 | 0.02 | 0.22 | 0.08 |
| 6 | 158411364 | rs9459056 | -1.071 | 0.933 | -1.327 | -0.836 | -0.018 | -0.001 | -0.001 | 0 | 0.58 | 0.83 | 0.46 | 0.45 | 0.56 | 0.58 |
| 6 | 158412184 | rs10455934 | -0.866 | 0.756 | -1.047 | -0.686 | 0.017 | 0.059 | 0.059 | 0 | 0.49 | 0.78 | 0.38 | 0.45 | 0.42 | 0.53 |
| 6 | 158412519 | rs10455936 | -0.034 | 0.636 | -0.742 | -0.562 | 0.003 | -0.005 | -0.005 | 0.098 | 0.27 | 0.23 | 0.23 | 0.42 | 0.2 | 0.45 |
| 6 | 158412832 | rs10945973 | -0.385 | 0.668 | -1.513 | -0.495 | -0.028 | 0.011 | 0.011 | 0 | 0.58 | 0.79 | 0.43 | 0.45 | 0.51 | 0.55 |
| 6 | 158420388 | rs12198470 | 0.078 | 0.839 | -1.218 | 0.051 | -0.014 | -0.023 | -0.023 | 0 | 0.05 | 0.01 | 0.06 | 0.14 | 0.06 | 0.25 |
| 6 | 158421274 | rs6904665 | -0.747 | 1.036 | -1.473 | 0.397 | -0.016 | -0.024 | -0.024 | 0.605 | 0.34 | 0.75 | 0.3 | 0.4 | 0.34 | 0.43 |
| 6 | 158422080 | rs12202135 | -0.214 | 1.061 | -1.552 | 0.475 | -0.031 | -0.039 | -0.039 | 0 | 0.12 | 0.02 | 0.11 | 0.18 | 0.13 | 0.3 |
| 6 | 158424693 | rs9459093 | -0.332 | 0.814 | -1.816 | 0.235 | -0.002 | -0.012 | -0.012 | 0 | 0.47 | 0.81 | 0.38 | 0.47 | 0.43 | 0.33 |
| 6 | 158431736 | rs1998791 | -0.69 | 1.037 | -1.845 | 0.235 | -0.046 | -0.074 | -0.074 | 0.217 | 0.76 | 0.89 | 0.54 | 0.62 | 0.7 | 0.69 |
| 6 | 158435572 | rs9365723 | 0.174 | 0.882 | -0.815 | 0.128 | 0.016 | 0.032 | 0.032 | 0.266 | 0.67 | 0.45 | 0.45 | 0.46 | 0.63 | 0.45 |
| 6 | 158435623 | rs9365724 | -0.508 | 0.868 | -0.402 | 0.117 | 0.014 | 0.036 | 0.036 | 0 | 0.73 | 0.61 | 0.51 | 0.6 | 0.7 | 0.7 |
| 6 | 158439029 | rs881751 | 0.324 | 0.217 | -0.293 | -0.226 | 0.008 | 0.008 | 0.008 | 0.917 | 0.5 | 0.52 | 0.34 | 0.26 | 0.4 | 0.29 |
| 6 | 158443106 | rs9459154 | -1.07 | 0.231 | -0.068 | -0.069 | 0.088 | 0.142 | 0.142 | 0 | 0.09 | 0.45 | 0.1 | 0.14 | 0.08 | 0.24 |
| 6 | 158450937 | rs7768038 | 0.313 | -0.238 | -0.071 | -0.048 | 0.032 | 0.043 | 0.043 | 3.236 | 0.48 | 0.41 | 0.31 | 0.27 | 0.36 | 0.29 |
| 6 | 158454874 | rs2295893 | 0.236 | -0.214 | -0.188 | -0.031 | 0.103 | 0.157 | 0.157 | 1.913 | 0.5 | 0.75 | 0.32 | 0.27 | 0.36 | 0.29 |
| 6 | 158455055 | rs2295894 | 0.297 | -0.164 | -0.271 | -0.005 | 0.004 | 0.002 | 0.002 | 1.851 | 0.48 | 0.48 | 0.29 | 0.27 | 0.36 | 0.29 |
| 6 | 158462506 | rs1750043 | 0.072 | -0.066 | -0.663 | 0.221 | -0.001 | -0.001 | -0.001 | 0.891 | 0.56 | 0.62 | 0.4 | 0.51 | 0.45 | 0.64 |
| 6 | 158467599 | rs16900563 | -0.692 | 0.094 | -0.275 | 0.689 | 0.014 | 0.109 | 0.109 | 0.615 | 0.08 | 0.24 | 0.06 | 0.24 | 0.04 | 0.32 |
| 6 | 158467930 | rs750997 | -0.292 | 0.352 | -0.939 | 0.897 | -0.061 | -0.12 | -0.12 | 1.216 | 0.83 | 0.99 | 0.52 | 0.64 | 0.79 | 0.89 |
| 6 | 158467989 | rs2181190 | 0.621 | 0.136 | -0.804 | 0.845 | -0.019 | -0.06 | -0.06 | 2.512 | 0.16 | 0.02 | 0.12 | 0.24 | 0.17 | 0.24 |
| 6 | 158472445 | rs2502620 | -0.582 | 0.434 | 0.14 | 0.973 | 0.002 | 0.002 | 0.002 | 3.768 | 0.41 | 0.56 | 0.37 | 0.48 | 0.49 | 0.63 |
| 6 | 158473387 | rs2502618 | -0.175 | 0.494 | 0.11 | 0.86 | 0.007 | -0.007 | -0.007 | 0 | 0.34 | 0.34 | 0.22 | 0.33 | 0.41 | 0.48 |
| 6 | 158480144 | rs1750034 | 0.279 | 0.615 | -0.164 | 1.2 | 0.012 | 0.095 | 0.095 | 0.155 | 0.93 | 0.8 | 0.86 | 0.85 | 0.92 | 0.86 |
| 6 | 158487143 | rs1009014 | -0.228 | 0.634 | -0.755 | 0.678 | -0.003 | 0.016 | 0.016 | 0 | 0.36 | 0.31 | 0.22 | 0.31 | 0.38 | 0.46 |
| 6 | 158489240 | rs1750047 | -0.369 | 1.133 | -1.314 | 0.695 | -0.023 | 0 | 0 | 0 | 0.31 | 0.27 | 0.49 | 0.3 | 0.41 | 0.29 |
| 6 | 158490779 | rs751873 | -0.472 | 0.512 | -0.761 | 0.468 | 0.009 | -0.006 | -0.006 | 0 | 0.41 | 0.34 | 0.52 | 0.45 | 0.34 | 0.4 |
| 6 | 158493701 | rs4333441 | 0.272 | 0.266 | -0.504 | 0.33 | -0.02 | 0.001 | 0.001 | 0 | 0.22 | 0.28 | 0.12 | 0.25 | 0.15 | 0.27 |
| 6 | 158494775 | rs6455990 | 0.218 | 0.695 | -0.315 | 0.217 | -0.001 | -0.006 | -0.006 | 0 | 0.72 | 0.69 | 0.54 | 0.87 | 0.72 | 0.8 |
| 6 | 158495863 | rs12203105 | -0.319 | 0.708 | -0.448 | 0.224 | -0.008 | 0.019 | 0.019 | 0 | 0.27 | 0.26 | 0.45 | 0.13 | 0.28 | 0.2 |
| 6 | 158496856 | rs1744178 | -0.008 | -0.028 | -1.049 | -0.752 | 0.041 | 0.08 | 0.08 | 0.037 | 0.39 | 0.27 | 0.2 | 0.01 | 0.29 | 0.2 |
| 6 | 158497717 | rs2296508 | 0.064 | 0.38 | -1.53 | -0.426 | -0.015 | 0.017 | 0.017 | 0.66 | 0.52 | 0.76 | 0.35 | 0.59 | 0.51 | 0.55 |
| 6 | 158498519 | rs1750041 | -0.262 | 0.549 | -1.532 | -0.605 | -0.019 | 0.021 | 0.021 | 0.045 | 0.92 | 0.95 | 0.58 | 0.6 | 0.85 | 0.83 |
| 6 | 158498555 | rs1750040 | 1.374 | 0.578 | -1.561 | -0.653 | 0.009 | -0.022 | -0.022 | 0 | 0.06 | 0.11 | 0.05 | 0 | 0.1 | 0.08 |
| 6 | 158499302 | rs2296507 | 0.277 | 0.532 | -1.396 | -0.683 | 0.003 | 0.027 | 0.027 | 0.513 | 0.5 | 0.67 | 0.35 | 0.58 | 0.51 | 0.55 |
| 6 | 158501843 | rs350292 | 1.026 | 0.541 | -1.382 | -0.683 | 0.02 | -0.037 | -0.037 | 0.867 | 0.07 | 0.18 | 0.06 | 0 | 0.1 | 0.08 |

| | | | | | | | | | | | | | | | | |
|---|---|---|---|---|---|---|---|---|---|---|---|---|---|---|---|---|
| 6 | 158501951 | rs1885543 | 0.3 | 0.511 | -1.398 | -0.691 | 0.006 | 0.031 | 0.031 | 0.385 | 0.5 | 0.66 | 0.35 | 0.58 | 0.51 | 0.55 |
| 6 | 158504365 | rs1744175 | -0.225 | 0.485 | -1.49 | -0.672 | 0.007 | -0.045 | -0.045 | 0.1 | 0.57 | 0.87 | 0.42 | 0.58 | 0.61 | 0.62 |
| 6 | 158505088 | rs1744173 | -0.232 | 0.763 | -1.272 | -0.392 | -0.034 | -0.007 | -0.007 | 0 | 0.92 | 0.98 | 0.59 | 0.59 | 0.85 | 0.83 |
| 6 | 158505921 | rs17489860 | 0.66 | 0.61 | -1.137 | -0.593 | 0.034 | -0.045 | -0.045 | 0 | 0.08 | 0.23 | 0.06 | 0 | 0.1 | 0.08 |
| 6 | 158509750 | rs350289 | -0.482 | 0.741 | -1.148 | -0.369 | -0.032 | 0 | 0 | 0 | 0.93 | 0.98 | 0.59 | 0.61 | 0.87 | 0.85 |
| 6 | 158513586 | rs2477809 | -0.034 | 0.554 | -1.046 | -0.695 | 0.002 | 0.023 | 0.023 | 0.765 | 0.49 | 0.62 | 0.36 | 0.57 | 0.51 | 0.56 |
| 6 | 158517308 | rs2502601 | -0.042 | 0.68 | -1.028 | -0.7 | 0.001 | 0.019 | 0.019 | 0.807 | 0.49 | 0.61 | 0.36 | 0.57 | 0.51 | 0.61 |
| 6 | 158517451 | rs2502600 | -0.075 | 0.673 | -1.009 | -0.704 | 0.001 | 0.023 | 0.023 | 0.707 | 0.5 | 0.66 | 0.36 | 0.57 | 0.51 | 0.61 |
| 6 | 158518507 | rs1048394 | -0.04 | 0.674 | -1.01 | -0.704 | 0.001 | 0.019 | 0.019 | 0.807 | 0.49 | 0.61 | 0.36 | 0.57 | 0.51 | 0.61 |
| 6 | 158518639 | rs17490133 | 1.265 | 0.432 | -1.242 | -0.947 | 0 | 0.195 | 0.195 | 1.159 | 0.28 | 0.02 | 0.14 | 0.01 | 0.22 | 0.14 |
| 6 | 158526159 | rs2475559 | 0.085 | 0.429 | -1.185 | -0.916 | -0.001 | 0.025 | 0.025 | 0.744 | 0.5 | 0.68 | 0.36 | 0.57 | 0.51 | 0.61 |
| 6 | 158526581 | rs1358940 | -0.629 | 0.459 | -1.192 | -0.943 | 0 | -0.027 | -0.027 | 0 | 0.86 | 0.86 | 0.93 | 1 | 0.87 | 0.9 |
| 6 | 158530037 | rs9365886 | 0.68 | 0.461 | -1.205 | -0.941 | -0.063 | -0.133 | -0.133 | 0 | 0.36 | 0.04 | 0.56 | 0.43 | 0.36 | 0.29 |
| 6 | 158531168 | rs2502607 | 0.976 | 0.917 | -1.185 | -1.166 | -0.001 | -0.003 | -0.003 | 0 | 0.3 | 0.29 | 0.17 | 0.27 | 0.3 | 0.38 |
| 6 | 158532187 | rs2502606 | 0.586 | 0.933 | -1.191 | -1.245 | 0.038 | 0.011 | 0.011 | 2.24 | 0.24 | 0.31 | 0.23 | 0.3 | 0.35 | 0.27 |
| 6 | 158555641 | rs1951543 | -0.808 | 0.824 | -1.216 | -1.201 | 0 | -0.034 | -0.034 | 0.324 | 0.86 | 0.85 | 0.93 | 1 | 0.87 | 0.9 |
| 6 | 158559176 | rs9459503 | 1.402 | 0.445 | -1.578 | -1.251 | 0.01 | -0.052 | -0.052 | 0 | 0.13 | 0.01 | 0.04 | 0.14 | 0.08 | 0.08 |
| 6 | 158566296 | rs13202019 | 1.735 | 0.665 | -1.584 | -1.28 | 0.013 | 0.196 | 0.196 | 1.123 | 0.28 | 0.02 | 0.14 | 0.01 | 0.22 | 0.14 |
| 6 | 158570802 | rs9348053 | 1.11 | 0.759 | -1.646 | -1.197 | -0.047 | -0.135 | -0.135 | 0 | 0.36 | 0.04 | 0.56 | 0.42 | 0.36 | 0.29 |
| 6 | 158571501 | rs6929274 | -0.588 | 1.244 | -1.049 | -0.869 | 0 | -0.003 | -0.003 | 0 | 0.7 | 0.71 | 0.82 | 0.73 | 0.69 | 0.65 |
| 6 | 158572975 | rs11756587 | 1.702 | 2.009 | -0.844 | -0.48 | -0.029 | -0.002 | -0.002 | 0.414 | 0.06 | 0.01 | 0.08 | 0.3 | 0.13 | 0.24 |
| 6 | 158584691 | rs10946113 | 0.539 | 1.474 | -0.654 | -0.454 | -0.038 | -0.063 | -0.063 | 0 | 0.11 | 0.02 | 0.42 | 0.42 | 0.14 | 0.15 |
| 6 | 158586682 | rs890458 | -0.837 | 1.551 | -0.684 | -0.482 | -0.047 | -0.029 | -0.029 | 0 | 0.92 | 1 | 0.94 | 0.88 | 0.85 | 0.89 |
| 6 | 158588702 | rs9365947 | -0.877 | 0.641 | -0.259 | -0.291 | 0.004 | 0.019 | 0.019 | 0 | 0.23 | 0.35 | 0.51 | 0.42 | 0.26 | 0.25 |
| 6 | 158598474 | rs9348087 | -1.025 | 0.634 | -0.341 | -0.296 | 0.024 | 0.1 | 0.1 | 0 | 0.24 | 0.44 | 0.51 | 0.42 | 0.26 | 0.25 |
| 6 | 158605689 | rs10946136 | 0.29 | 0.64 | -0.475 | -0.293 | -0.037 | -0.072 | -0.072 | 0 | 0.11 | 0.03 | 0.43 | 0.42 | 0.14 | 0.15 |
| 6 | 158606615 | rs9459620 | -0.943 | 0.623 | -0.447 | -0.291 | 0.014 | 0.077 | 0.077 | 0 | 0.25 | 0.6 | 0.52 | 0.42 | 0.26 | 0.25 |
| 6 | 158620608 | rs2363781 | -0.032 | 0.534 | -0.696 | -0.704 | -0.008 | -0.036 | -0.036 | 0 | 0.13 | 0.11 | 0.44 | 0.44 | 0.14 | 0.18 |
| 6 | 158622125 | rs1124275 | 0.198 | 0.548 | -0.247 | -0.85 | 0.009 | -0.053 | -0.053 | 0 | 0.11 | 0.18 | 0.08 | 0 | 0.12 | 0.1 |
| 6 | 158622391 | rs662185 | -0.324 | 0.491 | -0.484 | -1.045 | 0 | -0.001 | -0.001 | 0.463 | 0.55 | 0.59 | 0.7 | 0.71 | 0.58 | 0.6 |
| 6 | 158624814 | rs644637 | 0.154 | 0.356 | 0.078 | -0.717 | -0.005 | -0.135 | -0.135 | 0 | 0.65 | 0.6 | 0.8 | 1 | 0.66 | 0.82 |
| 6 | 158626332 | rs11962291 | -1.07 | 0.597 | 0.729 | 0.533 | 0.036 | -0.088 | -0.088 | 0.01 | 0.13 | 0.33 | 0.1 | 0 | 0.13 | 0.1 |
| 6 | 158628384 | rs911532 | -2.011 | 0.702 | 0.806 | 0.927 | 0.03 | 0.116 | 0.116 | 0 | 0.23 | 0.49 | 0.44 | 0.55 | 0.24 | 0.32 |
| 6 | 158634175 | rs6909925 | -1.302 | 0.669 | 0.742 | 1.137 | 0.039 | 0.343 | 0.343 | 0 | 0.1 | 0.39 | 0.34 | 0.41 | 0.09 | 0.16 |
| 6 | 158634878 | rs9347131 | -0.877 | 0.701 | 0.284 | 1.153 | 0.003 | 0.118 | 0.118 | 0 | 0.08 | 0.15 | 0.33 | 0.39 | 0.09 | 0.16 |
| 6 | 158638232 | rs7769220 | -1.062 | 0.706 | 0.397 | 1.023 | 0.034 | 0.035 | 0.035 | 0 | 0.17 | 0.42 | 0.11 | 0.14 | 0.15 | 0.15 |
| 6 | 158643526 | rs9348153 | -1.262 | 0.996 | 0.499 | 1.223 | 0.013 | 0.263 | 0.263 | 0 | 0.09 | 0.26 | 0.34 | 0.41 | 0.08 | 0.17 |
| 6 | 158644859 | rs7755193 | -1.135 | 0.862 | 0.106 | 0.641 | 0.071 | 0.052 | 0.052 | 0 | 0.14 | 0.43 | 0.11 | 0.13 | 0.15 | 0.15 |
| 6 | 158649353 | rs4710078 | -0.325 | 0.468 | -1.121 | 0.752 | 0.009 | -0.026 | -0.026 | 0.334 | 0.18 | 0.18 | 0.36 | 0.55 | 0.16 | 0.28 |
| 6 | 158651395 | rs6929952 | -0.987 | 0.597 | -1.349 | 0.412 | 0 | 0.004 | 0.004 | 0.329 | 0.4 | 0.34 | 0.51 | 0.57 | 0.44 | 0.38 |
| 6 | 158655732 | rs9356514 | -0.711 | 0.669 | -1.426 | 0.45 | -0.006 | -0.014 | -0.014 | 0 | 0.36 | 0.22 | 0.46 | 0.56 | 0.4 | 0.36 |
| 6 | 158687103 | rs9366083 | -0.624 | 0.602 | -1.417 | 0.594 | -0.047 | -0.091 | -0.091 | 0 | 0.36 | 0.1 | 0.46 | 0.46 | 0.42 | 0.32 |
| 6 | 158689753 | rs9355613 | -0.624 | 0.602 | -1.417 | 0.594 | -0.047 | -0.091 | -0.091 | 0 | 0.36 | 0.1 | 0.46 | 0.46 | 0.42 | 0.32 |
| 6 | 158693267 | rs166391 | 0.107 | 0.458 | -1.537 | 0.504 | 0.001 | -0.002 | -0.002 | 0 | 0.08 | 0.09 | 0.04 | 0.1 | 0.04 | 0.08 |
| 6 | 158698463 | rs4709150 | -0.747 | 0.465 | -1.497 | 0.485 | -0.013 | -0.028 | -0.028 | 0.805 | 0.4 | 0.26 | 0.51 | 0.46 | 0.46 | 0.33 |
| 6 | 158704735 | rs9356571 | -0.736 | 0.466 | -1.497 | 0.485 | -0.014 | -0.029 | -0.029 | 0 | 0.4 | 0.26 | 0.51 | 0.46 | 0.46 | 0.33 |
| 6 | 158708822 | rs9348250 | 0.266 | 0.591 | -1.515 | 0.574 | -0.007 | -0.111 | -0.111 | 3.652 | 0.16 | 0.03 | 0.34 | 0.45 | 0.15 | 0.23 |
| 6 | 158711855 | rs705930 | -0.906 | 0.803 | -0.933 | -0.015 | -0.001 | 0.012 | 0.012 | 0 | 0.33 | 0.33 | 0.21 | 0.11 | 0.35 | 0.18 |
| 6 | 158732602 | rs3812099 | 0.819 | 0.53 | -0.626 | 0.357 | -0.02 | 0.006 | 0.006 | 0 | 0.87 | 0.75 | 0.91 | 0.9 | 0.92 | 0.92 |
| 6 | 158732819 | rs9456285 | -0.112 | 0.814 | -0.916 | -0.106 | -0.035 | 0.063 | 0.063 | 0 | 0.19 | 0.06 | 0.12 | 0.01 | 0.27 | 0.09 |
| 6 | 158733310 | rs7741924 | 0.59 | 0.801 | -0.902 | -0.099 | -0.02 | 0.005 | 0.005 | 0 | 0.87 | 0.74 | 0.91 | 0.9 | 0.92 | 0.92 |
| 6 | 158733887 | rs4478433 | 0.566 | 0.795 | -0.897 | -0.102 | -0.018 | 0.006 | 0.006 | 0 | 0.88 | 0.75 | 0.92 | 0.9 | 0.92 | 0.93 |
| 6 | 158734275 | rs4433000 | 0.938 | 0.679 | -0.949 | -0.144 | -0.022 | -0.001 | -0.001 | 1.602 | 0.86 | 0.74 | 0.91 | 0.9 | 0.92 | 0.93 |
| 6 | 158734876 | rs9347227 | 0.39 | 0.48 | -1.558 | 0.455 | -0.011 | -0.113 | -0.113 | 0.119 | 0.16 | 0.03 | 0.34 | 0.44 | 0.15 | 0.23 |
| 6 | 158740218 | rs10945541 | 0.738 | 0.51 | -1.423 | 0.407 | 0 | -0.01 | -0.01 | 0 | 0.68 | 0.66 | 0.78 | 0.89 | 0.65 | 0.84 |
| 6 | 158750050 | rs3196816 | 0.274 | 0.485 | -1.39 | 0.478 | -0.021 | 0.012 | 0.012 | 0 | 0.87 | 0.72 | 0.9 | 0.9 | 0.92 | 0.93 |
| 6 | 158750330 | rs7767022 | 0.909 | 0.573 | -1.598 | 0.418 | 0.01 | -0.018 | -0.018 | 0 | 0.69 | 0.71 | 0.56 | 0.48 | 0.75 | 0.7 |
| 6 | 158754879 | rs9456291 | 0.238 | 0.544 | -1.528 | 0.211 | -0.023 | 0.009 | 0.009 | 0 | 0.88 | 0.73 | 0.91 | 0.9 | 0.92 | 0.93 |
| 6 | 158756832 | rs6455580 | 0.219 | 0.535 | -1.51 | 0.206 | -0.018 | 0.009 | 0.009 | 0 | 0.88 | 0.74 | 0.91 | 0.9 | 0.92 | 0.93 |
| 6 | 158804647 | rs827977 | 0.541 | 0.761 | -1.705 | 0.497 | 0.022 | -0.004 | -0.004 | 0 | 0.72 | 0.8 | 0.62 | 0.34 | 0.76 | 0.67 |
| 6 | 158806047 | rs705932 | 1.298 | 1.075 | -1.784 | 1.033 | 0.001 | 0.007 | 0.007 | 0 | 0.48 | 0.65 | 0.45 | 0.45 | 0.43 | 0.57 |
| 6 | 158811727 | rs705935 | 0.88 | 1.041 | -1.746 | 1.134 | 0.007 | -0.009 | -0.009 | 0.446 | 0.67 | 0.68 | 0.58 | 0.44 | 0.72 | 0.67 |
| 6 | 158812361 | rs705936 | 0.235 | 0.874 | -1.835 | 1.02 | -0.01 | -0.002 | -0.002 | 0 | 0.88 | 0.85 | 0.96 | 0.93 | 0.93 | 0.93 |
| 6 | 158816875 | rs697505 | 0.807 | 1.343 | -2.543 | 0.634 | -0.076 | -0.038 | -0.038 | 0.178 | 0.11 | 0.01 | 0.26 | 0.14 | 0.23 | 0.3 |
| 6 | 158822682 | rs1329346 | 0.444 | 1.132 | -2.385 | 0.507 | 0.021 | -0.069 | -0.069 | 0 | 0.18 | 0.05 | 0.34 | 0.59 | 0.16 | 0.26 |
| 6 | 158830239 | rs341168 | 0.177 | 1.535 | -2.624 | -0.089 | -0.13 | 0.163 | 0.163 | 0 | 0.25 | 0.04 | 0.18 | 0.01 | 0.37 | 0.11 |
| 6 | 158832854 | rs341160 | 0.874 | 1.314 | -1.926 | 0.019 | 0.002 | -0.002 | -0.002 | 0.324 | 0.44 | 0.59 | 0.42 | 0.45 | 0.38 | 0.56 |
| 6 | 158863830 | rs705960 | 0.111 | 1.06 | -1.465 | -0.236 | -0.015 | -0.003 | -0.003 | 0 | 0.89 | 0.81 | 0.95 | 0.92 | 0.91 | 0.92 |
| 6 | 158870081 | rs705956 | 0.12 | 1.283 | -1.965 | -0.529 | -0.001 | -0.001 | -0.001 | 0.132 | 0.62 | 0.78 | 0.53 | 0.4 | 0.59 | 0.65 |

| | | | | | | | | | | | | | | | | |
|---|---|---|---|---|---|---|---|---|---|---|---|---|---|---|---|---|
| 6 | 158887729 | rs598665 | 0.636 | 1.324 | -1.949 | -0.654 | 0.009 | 0.009 | 0.009 | 0 | 0.4 | 0.66 | 0.4 | 0.36 | 0.31 | 0.54 |
| 6 | 158897452 | rs600752 | -0.096 | 1.142 | -1.607 | -0.357 | -0.003 | -0.005 | -0.005 | 0 | 0.09 | 0.15 | 0.04 | 0.07 | 0.11 | 0.06 |
| 6 | 158909773 | rs6928393 | 0.764 | 1.259 | -1.705 | -0.549 | 0 | 0.01 | 0.01 | 0.005 | 0.49 | 0.53 | 0.48 | 0.36 | 0.45 | 0.59 |
| 6 | 158910491 | rs2274884 | 0.281 | 1.467 | -1.825 | -0.634 | -0.007 | -0.003 | -0.003 | 0.518 | 0.6 | 0.78 | 0.52 | 0.41 | 0.56 | 0.66 |
| 6 | 158914825 | rs17504313 | 0.443 | 0.938 | -1.996 | -0.847 | -0.005 | 0.011 | 0.011 | 0.505 | 0.11 | 0.08 | 0.05 | 0.05 | 0.13 | 0.09 |
| 6 | 158916972 | rs1041566 | 0.443 | 0.912 | -2.051 | -0.844 | -0.052 | -0.142 | -0.142 | 1.662 | 0.29 | 0.04 | 0.42 | 0.53 | 0.31 | 0.27 |
| 6 | 158921724 | rs7753885 | 1.102 | 0.84 | -1.852 | -0.873 | -0.004 | 0.018 | 0.018 | 0.01 | 0.4 | 0.65 | 0.42 | 0.34 | 0.39 | 0.5 |
| 6 | 158924595 | rs3749853 | -0.533 | 1.034 | -1.419 | -0.679 | 0.009 | -0.003 | -0.003 | 0 | 0.15 | 0.17 | 0.1 | 0.14 | 0.19 | 0.11 |
| 6 | 158929489 | rs3749854 | 0.342 | 0.982 | -1.568 | -1.037 | -0.023 | -0.035 | -0.035 | 0 | 0.31 | 0.14 | 0.45 | 0.56 | 0.35 | 0.29 |
| 6 | 158930696 | rs6918518 | 0.535 | 0.957 | -1.553 | -1.186 | -0.002 | 0.044 | 0.044 | 0 | 0.51 | 0.63 | 0.45 | 0.4 | 0.49 | 0.57 |
| 6 | 158930744 | rs9347237 | 0.346 | 0.966 | -1.802 | -1.025 | -0.019 | -0.041 | -0.041 | 0 | 0.31 | 0.16 | 0.45 | 0.54 | 0.35 | 0.3 |
| 6 | 158935739 | rs6916832 | 0.681 | 1.006 | -1.702 | -0.63 | -0.003 | -0.011 | -0.011 | 0 | 0.41 | 0.46 | 0.41 | 0.37 | 0.37 | 0.51 |
| 6 | 158941011 | rs1866896 | -0.137 | 1.282 | -2.551 | -0.277 | -0.116 | -0.118 | -0.118 | 3.784 | 0.63 | 0.97 | 0.51 | 0.46 | 0.61 | 0.65 |
| 6 | 158957104 | rs9456318 | -0.508 | 1.295 | -1.575 | -0.53 | -0.021 | 0.041 | 0.041 | 0 | 0.18 | 0.1 | 0.08 | 0.01 | 0.23 | 0.11 |
| 6 | 158966332 | rs2033970 | 0.397 | 1.072 | -1.382 | -0.278 | 0 | -0.002 | -0.002 | 0 | 0.94 | 0.97 | 0.97 | 1 | 0.91 | 0.97 |
| 6 | 158967384 | rs2083587 | 0.265 | 0.975 | -1.755 | -0.148 | -0.025 | -0.027 | -0.027 | 0 | 0.3 | 0.13 | 0.46 | 0.57 | 0.31 | 0.29 |
| 6 | 158969177 | rs10447366 | 0.173 | 0.919 | -1.62 | 0.087 | 0.001 | -0.037 | -0.037 | 0 | 0.31 | 0.16 | 0.44 | 0.56 | 0.29 | 0.3 |
| 6 | 158973475 | rs2362575 | -0.323 | 0.685 | -1.155 | -0.614 | -0.01 | 0.019 | 0.019 | 0.799 | 0.61 | 0.72 | 0.63 | 0.7 | 0.56 | 0.46 |
| 6 | 158976559 | rs7746250 | 0.073 | 0.455 | -0.341 | -0.831 | -0.001 | 0.004 | 0.004 | 0 | 0.28 | 0.26 | 0.16 | 0.14 | 0.28 | 0.22 |
| 6 | 158980351 | rs9457403 | 0.107 | 0.179 | 0.402 | -1.188 | 0.015 | 0.026 | 0.026 | 0 | 0.44 | 0.64 | 0.52 | 0.72 | 0.33 | 0.4 |
| 6 | 158982201 | rs7754383 | 0.451 | 0.047 | 0.708 | -1.309 | -0.02 | -0.066 | -0.066 | 0 | 0.13 | 0.25 | 0.07 | 0 | 0.07 | 0.1 |
| 6 | 158982921 | rs750618 | -0.577 | 0.162 | 0.328 | -1.569 | -0.041 | 0.042 | 0.042 | 0 | 0.63 | 0.76 | 0.64 | 0.73 | 0.53 | 0.51 |
| 6 | 158998253 | rs7764967 | 0.994 | -0.092 | 0.55 | -1.351 | -0.001 | 0 | 0 | 0 | 0.32 | 0.34 | 0.42 | 0.63 | 0.26 | 0.27 |
| 6 | 158998571 | rs10223384 | -0.643 | 0.316 | 0.351 | -1.723 | 0.006 | -0.076 | -0.076 | 3.747 | 0.4 | 0.45 | 0.32 | 0.1 | 0.44 | 0.28 |
| 6 | 159007004 | rs4709221 | -0.917 | 0.355 | 0.259 | -1.756 | -0.009 | 0.001 | 0.001 | 0.56 | 0.73 | 0.84 | 0.75 | 0.73 | 0.7 | 0.55 |
| 6 | 159010379 | rs1545217 | 0.072 | 0.009 | 0.694 | -1.598 | 0.031 | 0.085 | 0.085 | 0.58 | 0.49 | 0.69 | 0.57 | 0.73 | 0.41 | 0.43 |
| 6 | 159012810 | rs6925536 | -0.589 | 0.162 | 0.598 | -1.597 | -0.044 | 0.051 | 0.051 | 0.002 | 0.63 | 0.82 | 0.67 | 0.73 | 0.58 | 0.53 |
| 6 | 159012832 | rs9457409 | -0.507 | 0.122 | 0.758 | -1.631 | 0.006 | -0.135 | -0.135 | 0 | 0.28 | 0.38 | 0.21 | 0.01 | 0.32 | 0.2 |
| 6 | 159014026 | rs9457410 | 0.57 | 0.078 | 0.804 | -1.582 | 0 | -0.075 | -0.075 | 0.005 | 0.14 | 0.19 | 0.11 | 0 | 0.13 | 0.1 |
| 6 | 159016076 | rs1036741 | 0.758 | 0.232 | 0.724 | -1.528 | -0.022 | -0.016 | -0.016 | 0.019 | 0.35 | 0.48 | 0.47 | 0.72 | 0.25 | 0.33 |
| 6 | 159032670 | rs9457413 | -0.429 | 0.402 | 0.653 | -1.454 | -0.011 | 0.013 | 0.013 | 0 | 0.59 | 0.69 | 0.63 | 0.65 | 0.57 | 0.48 |
| 6 | 159045294 | rs6915032 | 0.131 | 0.459 | 0.5 | -1.248 | 0 | 0.002 | 0.002 | 0 | 0.46 | 0.49 | 0.51 | 0.64 | 0.43 | 0.38 |
| 6 | 159047536 | rs16900886 | 0.95 | 0.469 | 0.244 | -1.274 | 0.006 | -0.007 | -0.007 | 0 | 0.31 | 0.25 | 0.41 | 0.64 | 0.26 | 0.28 |
| 6 | 159049367 | rs4708800 | 0.95 | 0.5 | 0.244 | -1.274 | 0.005 | -0.007 | -0.007 | 0 | 0.31 | 0.25 | 0.41 | 0.64 | 0.26 | 0.29 |
| 6 | 159055282 | rs3199561 | 0.334 | 0.528 | 0.115 | -1.247 | -0.002 | -0.005 | -0.005 | 0.001 | 0.13 | 0.12 | 0.1 | 0.01 | 0.18 | 0.1 |
| 6 | 159068714 | rs13198609 | 0.445 | 0.546 | -0.108 | -1.233 | -0.016 | 0.028 | 0.028 | 0 | 0.13 | 0.06 | 0.09 | 0.01 | 0.2 | 0.1 |
| 6 | 159069879 | rs2165450 | 0.814 | 0.475 | -0.069 | -1.035 | 0.007 | -0.026 | -0.026 | 0 | 0.3 | 0.14 | 0.42 | 0.65 | 0.25 | 0.28 |
| 6 | 159081429 | rs4709242 | 0.836 | 0.479 | -0.04 | -1.079 | 0.012 | -0.014 | -0.014 | 0 | 0.31 | 0.24 | 0.42 | 0.65 | 0.24 | 0.28 |
| 6 | 159085005 | rs317795 | 0.328 | 0.678 | -0.593 | -1.426 | -0.031 | 0.067 | 0.067 | 0.05 | 0.13 | 0.02 | 0.07 | 0 | 0.15 | 0.13 |
| 6 | 159085981 | rs16900918 | 0.11 | 0.725 | -0.029 | -1.322 | 0.006 | -0.024 | -0.024 | 0 | 0.16 | 0.26 | 0.14 | 0.08 | 0.2 | 0.13 |
| 6 | 159087492 | rs317798 | -0.167 | 0.566 | -0.385 | -1.302 | -0.026 | 0.005 | 0.005 | 0 | 0.7 | 0.83 | 0.7 | 0.73 | 0.63 | 0.55 |
| 6 | 159091331 | rs317802 | -0.29 | 0.161 | -0.989 | -1.935 | 0.024 | 0.194 | 0.194 | 0 | 0.53 | 0.8 | 0.66 | 0.88 | 0.38 | 0.65 |
| 6 | 159095699 | rs2583077 | 0.064 | -0.721 | -0.78 | -2.576 | 0.04 | 0.035 | 0.035 | 0.717 | 0.36 | 0.68 | 0.14 | 0.09 | 0.12 | 0.25 |
| 6 | 159101293 | rs317789 | -0.548 | -0.542 | -0.822 | -2.458 | 0.098 | 0.165 | 0.165 | 0 | 0.49 | 0.79 | 0.33 | 0.21 | 0.28 | 0.53 |
| 6 | 159104769 | rs10452575 | 0.814 | -0.952 | -0.123 | -2.408 | -0.047 | -0.102 | -0.102 | 0.314 | 0.24 | 0.33 | 0.08 | 0.01 | 0.08 | 0.11 |
| 6 | 159111946 | rs6913508 | 0.727 | -0.975 | 0.184 | -2.42 | -0.06 | -0.125 | -0.125 | 1.522 | 0.23 | 0.37 | 0.09 | 0.01 | 0.08 | 0.13 |
| 6 | 159117018 | rs6905421 | -0.171 | -0.706 | -0.032 | -2.662 | 0.016 | -0.037 | -0.037 | 1.488 | 0.37 | 0.49 | 0.36 | 0.21 | 0.46 | 0.51 |
| 6 | 159118978 | rs9457437 | -0.206 | -0.726 | -0.323 | -2.683 | -0.008 | 0.02 | 0.02 | 1.662 | 0.4 | 0.67 | 0.37 | 0.22 | 0.47 | 0.53 |
| 6 | 159119811 | rs9456340 | -0.743 | -0.384 | -0.464 | -2.443 | -0.031 | 0.028 | 0.028 | 0 | 0.68 | 0.81 | 0.48 | 0.22 | 0.58 | 0.63 |
| 6 | 159123590 | rs6930297 | 0.114 | -0.346 | -0.157 | -2.355 | -0.002 | -0.004 | -0.004 | 0.768 | 0.3 | 0.28 | 0.2 | 0.11 | 0.15 | 0.34 |
| 6 | 159123826 | rs6919936 | 0.217 | -0.175 | -0.004 | -2.436 | -0.034 | -0.064 | -0.064 | 0.149 | 0.3 | 0.42 | 0.21 | 0.12 | 0.17 | 0.37 |
| 6 | 159128675 | rs13206612 | 0.609 | 0.331 | -0.799 | -2.881 | -0.058 | 0.072 | 0.072 | 0 | 0.14 | 0.04 | 0.14 | 0.01 | 0.26 | 0.08 |
| 6 | 159128772 | rs7758665 | 0.401 | 0.306 | -0.61 | -2.893 | 0.01 | -0.043 | -0.043 | 0.028 | 0.15 | 0.18 | 0.16 | 0.01 | 0.26 | 0.08 |
| 6 | 159129551 | rs9456341 | 0.594 | -0.098 | -1.052 | -3.132 | 0.034 | 0.118 | 0.118 | 0 | 0.27 | 0.09 | 0.18 | 0.07 | 0.17 | 0.38 |
| 6 | 159135863 | rs13214253 | -0.417 | 0.302 | -0.538 | -2.901 | 0.027 | -0.103 | -0.103 | 3.563 | 0.23 | 0.35 | 0.21 | 0.01 | 0.32 | 0.14 |
| 6 | 159145196 | rs7751477 | 0.511 | 0.203 | -1.092 | -2.717 | -0.049 | 0.1 | 0.1 | 0 | 0.2 | 0.04 | 0.16 | 0.01 | 0.27 | 0.11 |
| 6 | 159146871 | rs6455600 | -1.123 | 0.2 | -1.236 | -2.619 | -0.093 | 0.071 | 0.071 | 0 | 0.69 | 0.93 | 0.47 | 0.24 | 0.6 | 0.68 |
| 6 | 159150338 | rs9457456 | -0.504 | 0.021 | -0.875 | -2.653 | -0.02 | 0.044 | 0.044 | 0 | 0.6 | 0.72 | 0.41 | 0.23 | 0.53 | 0.65 |
| 6 | 159155998 | rs2129209 | 0.456 | 0.03 | -0.263 | -2.684 | -0.005 | -0.011 | -0.011 | 0.537 | 0.22 | 0.31 | 0.19 | 0.21 | 0.18 | 0.44 |
| 6 | 159161428 | rs2171210 | 0.474 | -0.129 | -0.604 | -3.171 | -0.02 | 0.098 | 0.098 | 0 | 0.25 | 0.08 | 0.15 | 0.01 | 0.27 | 0.15 |
| 6 | 159166329 | rs6933401 | -0.462 | -0.298 | -0.227 | -3.184 | -0.001 | -0.008 | -0.008 | 0.285 | 0.51 | 0.57 | 0.31 | 0.12 | 0.49 | 0.46 |
| 6 | 159166883 | rs6455601 | 0.203 | -0.384 | -0.276 | -3.191 | -0.005 | -0.078 | -0.078 | 3.203 | 0.27 | 0.45 | 0.16 | 0.11 | 0.25 | 0.31 |
| 6 | 159171342 | rs3123093 | -0.631 | -0.211 | -0.458 | -2.911 | -0.011 | 0.061 | 0.061 | 0 | 0.6 | 0.74 | 0.35 | 0.18 | 0.56 | 0.55 |
| 6 | 159183351 | rs9347251 | 0.065 | 0.108 | -0.338 | -2.994 | 0.004 | -0.004 | -0.004 | 0.268 | 0.28 | 0.3 | 0.27 | 0.2 | 0.47 | 0.43 |
| 6 | 159183562 | rs2171209 | -1.002 | -0.693 | -0.548 | -3.372 | 0.099 | 0.072 | 0.072 | 0 | 0.66 | 0.88 | 0.47 | 0.24 | 0.79 | 0.67 |
| 6 | 159184507 | rs2291386 | 0.137 | -0.811 | -0.072 | -3.434 | -0.026 | -0.033 | -0.033 | 0 | 0.1 | 0.31 | 0.03 | 0.03 | 0.04 | 0.04 |
| 6 | 159188282 | rs9364994 | 1.939 | -0.478 | -0.304 | -3.516 | -0.044 | 0.063 | 0.063 | 0 | 0.16 | 0.02 | 0.1 | 0.07 | 0.19 | 0.29 |
| 6 | 159191533 | rs12201279 | 0.188 | -0.21 | 0.012 | -3.432 | 0.108 | 0.013 | 0.013 | 0 | 0.1 | 0.19 | 0.13 | 0.13 | 0.25 | 0.15 |
| 6 | 159194788 | rs753017 | -0.77 | -0.917 | -0.116 | -3.508 | -0.032 | -0.015 | -0.015 | 7.843 | 0.63 | 0.58 | 0.48 | 0.28 | 0.82 | 0.67 |

| | | | | | | | | | | | | | | | | |
|---|---|---|---|---|---|---|---|---|---|---|---|---|---|---|---|---|
| 6 | 159196603 | rs9456352 | -1.173 | -0.632 | -0.089 | -3.543 | 0.101 | 0.071 | 0.071 | 0 | 0.66 | 0.93 | 0.47 | 0.23 | 0.78 | 0.67 |
| 6 | 159198730 | rs3102971 | -0.113 | -0.573 | -0.049 | -3.663 | 0.003 | 0 | 0 | 0 | 0.4 | 0.38 | 0.35 | 0.21 | 0.6 | 0.51 |
| 6 | 159205668 | rs3103003 | -0.151 | -0.587 | -0.032 | -3.663 | -0.001 | -0.01 | -0.01 | 0.473 | 0.41 | 0.41 | 0.36 | 0.21 | 0.6 | 0.51 |
| 6 | 159208889 | rs729892 | 1.365 | -0.678 | 0.007 | -3.695 | -0.001 | -0.027 | -0.027 | 0.051 | 0.15 | 0.15 | 0.08 | 0.01 | 0.14 | 0.12 |
| 6 | 159210126 | rs3123116 | 0.794 | -0.539 | -0.071 | -3.726 | 0.001 | -0.036 | -0.036 | 0 | 0.13 | 0.11 | 0.09 | 0.01 | 0.13 | 0.07 |
| 6 | 159215408 | rs6455604 | 0.478 | -0.542 | -0.07 | -3.731 | -0.008 | 0.005 | 0.005 | 0.003 | 0.28 | 0.28 | 0.26 | 0.19 | 0.47 | 0.44 |
| 6 | 159217558 | rs7760064 | 0.212 | -0.615 | -0.042 | -3.759 | -0.049 | -0.144 | -0.144 | 0 | 0.25 | 0.46 | 0.11 | 0.03 | 0.18 | 0.16 |
| 6 | 159220659 | rs3102993 | 0.796 | -0.545 | -0.165 | -3.728 | 0.001 | -0.036 | -0.036 | 0 | 0.13 | 0.11 | 0.09 | 0.01 | 0.13 | 0.07 |
| 6 | 159220863 | rs3102992 | -0.136 | -0.548 | -0.166 | -3.731 | -0.001 | -0.01 | -0.01 | 0.018 | 0.4 | 0.39 | 0.35 | 0.21 | 0.6 | 0.51 |
| 6 | 159221547 | rs6455605 | 0.408 | -0.408 | -0.291 | -3.633 | -0.008 | 0.005 | 0.005 | 0.007 | 0.27 | 0.28 | 0.26 | 0.19 | 0.47 | 0.44 |
| 6 | 159247859 | rs1471836 | -1.273 | -0.607 | -0.63 | -3.502 | 0.091 | 0.058 | 0.058 | 0 | 0.67 | 0.93 | 0.47 | 0.23 | 0.79 | 0.67 |
| 6 | 159256346 | rs9347260 | 1.05 | -0.196 | -0.562 | -3.45 | -0.013 | 0.029 | 0.029 | 0 | 0.2 | 0.15 | 0.15 | 0.06 | 0.25 | 0.31 |
| 6 | 159257004 | rs7746304 | 0.828 | -0.641 | -0.263 | -3.339 | -0.008 | -0.059 | -0.059 | 0 | 0.17 | 0.2 | 0.07 | 0 | 0.15 | 0.12 |
| 6 | 159262953 | rs3127158 | 0.962 | -0.431 | -0.325 | -3.275 | -0.001 | -0.042 | -0.042 | 0 | 0.1 | 0.11 | 0.08 | 0.01 | 0.11 | 0.02 |
| 6 | 159264974 | rs6924755 | -0.273 | -0.15 | -0.372 | -3.076 | -0.003 | 0.053 | 0.053 | 0 | 0.37 | 0.26 | 0.29 | 0.22 | 0.55 | 0.55 |
| 6 | 159270650 | rs1994351 | 0.444 | -0.663 | -0.27 | -2.863 | 0.037 | -0.133 | -0.133 | 0.273 | 0.33 | 0.41 | 0.24 | 0.02 | 0.46 | 0.18 |
| 6 | 159270803 | rs2129210 | 0.297 | -0.592 | -0.201 | -2.767 | 0.002 | -0.018 | -0.018 | 0 | 0.25 | 0.36 | 0.22 | 0.19 | 0.27 | 0.43 |
| 6 | 159270945 | rs2129211 | -0.343 | -0.616 | -0.524 | -2.74 | -0.009 | -0.032 | -0.032 | 0 | 0.36 | 0.47 | 0.24 | 0.22 | 0.33 | 0.53 |
| 6 | 159277326 | rs7746144 | -0.021 | -0.702 | -0.931 | -2.786 | -0.008 | -0.059 | -0.059 | 0 | 0.54 | 0.51 | 0.39 | 0.09 | 0.66 | 0.52 |
| 6 | 159279274 | rs4397254 | 0.215 | -0.489 | -1.67 | -2.617 | -0.004 | -0.023 | -0.023 | 0 | 0.13 | 0.14 | 0.05 | 0.03 | 0.09 | 0.14 |
| 6 | 159282859 | rs10945563 | 0.786 | 0.12 | -2.288 | -2.472 | 0.001 | -0.04 | -0.04 | 0 | 0.25 | 0.25 | 0.16 | 0.01 | 0.35 | 0.17 |
| 6 | 159299216 | rs9457490 | -0.407 | 0.391 | -2.709 | -1.248 | -0.014 | -0.061 | -0.061 | 0.161 | 0.76 | 0.96 | 0.48 | 0.32 | 0.77 | 0.73 |
| 6 | 159308110 | rs9347268 | -0.064 | 0.566 | -2.353 | -0.822 | -0.013 | 0.018 | 0.018 | 0 | 0.19 | 0.04 | 0.14 | 0.14 | 0.21 | 0.29 |
| 6 | 159316430 | rs9457507 | 0.095 | 0.185 | -2.268 | -0.643 | 0.028 | -0.006 | -0.006 | 0 | 0.65 | 0.86 | 0.44 | 0.34 | 0.7 | 0.62 |
| 6 | 159317483 | rs9457508 | 0.514 | 0.395 | -2.483 | -0.632 | 0.002 | 0.035 | 0.035 | 0 | 0.6 | 0.89 | 0.44 | 0.31 | 0.64 | 0.51 |
| 6 | 159317512 | rs9457509 | 0.207 | 0.297 | -1.987 | -0.103 | 0.021 | -0.07 | -0.07 | 0 | 0.73 | 0.93 | 0.49 | 0.37 | 0.77 | 0.67 |
| 6 | 159324086 | rs9355684 | -0.587 | 0.647 | -1.532 | -0.164 | 0 | 0.01 | 0.01 | 0 | 0.19 | 0.19 | 0.1 | 0.11 | 0.22 | 0.15 |
| 6 | 159325462 | rs9457511 | 0.663 | 1.234 | -2.281 | -0.573 | 0.032 | 0.154 | 0.154 | 0.01 | 0.52 | 0.82 | 0.35 | 0.31 | 0.39 | 0.55 |
| 6 | 159326830 | rs12200537 | 1.078 | 0.605 | -1.335 | -1.099 | 0.005 | -0.02 | -0.02 | 0.145 | 0.36 | 0.33 | 0.26 | 0.04 | 0.29 | 0.33 |
| 6 | 159328353 | rs9457512 | 0.612 | 0.633 | -0.848 | -1.118 | 0.002 | 0.001 | 0.001 | 0 | 0.46 | 0.56 | 0.31 | 0.12 | 0.42 | 0.41 |
| 6 | 159331667 | rs11965281 | -1.032 | 0.391 | 0.137 | -0.488 | -0.001 | 0.015 | 0.015 | 0 | 0.13 | 0.18 | 0.07 | 0.28 | 0.15 | 0.22 |
| 6 | 159335251 | rs9457516 | -1.063 | 0.321 | 0.592 | -0.49 | 0.037 | 0.19 | 0.19 | 0 | 0.09 | 0.44 | 0.06 | 0.25 | 0.06 | 0.17 |
| 6 | 159336324 | rs9346772 | -1.827 | 0.534 | 0.575 | -0.447 | 0.021 | 0.048 | 0.048 | 0.462 | 0.2 | 0.62 | 0.14 | 0.33 | 0.22 | 0.29 |
| 6 | 159338019 | rs7738389 | -1.537 | 0.072 | 0.721 | -0.72 | 0.005 | 0.009 | 0.009 | 0.836 | 0.39 | 0.64 | 0.22 | 0.31 | 0.33 | 0.34 |
| 6 | 159340025 | rs9365007 | -1.733 | 0.077 | 0.676 | -0.943 | 0.026 | 0.102 | 0.102 | 0.246 | 0.52 | 0.7 | 0.64 | 0.9 | 0.44 | 0.6 |
| 6 | 159340672 | rs9365009 | -1.168 | 0.159 | 0.878 | -0.936 | -0.029 | -0.031 | -0.031 | 0.793 | 0.41 | 0.49 | 0.56 | 0.83 | 0.27 | 0.51 |
| 6 | 159340892 | rs7774277 | -0.425 | 0.308 | 0.897 | -0.905 | 0.023 | -0.003 | -0.003 | 0 | 0.11 | 0.22 | 0.08 | 0.07 | 0.17 | 0.09 |
| 6 | 159343610 | rs3923900 | -1.543 | 0.375 | 0.904 | -0.826 | 0.023 | 0.095 | 0.095 | 0.116 | 0.51 | 0.68 | 0.63 | 0.8 | 0.44 | 0.61 |
| 6 | 159347415 | rs9355688 | -1.162 | 0.629 | 0.747 | -0.238 | -0.008 | 0.038 | 0.038 | 0 | 0.41 | 0.28 | 0.59 | 0.67 | 0.42 | 0.43 |
| 6 | 159369287 | rs9457535 | -1.614 | 0.487 | 1.304 | 0.573 | -0.036 | 0.135 | 0.135 | 0 | 0.2 | 0.41 | 0.4 | 0.39 | 0.12 | 0.24 |
| 6 | 159375207 | rs12529684 | -0.554 | 0.501 | 1.114 | 0.571 | 0.014 | -0.024 | -0.024 | 0 | 0.12 | 0.06 | 0.34 | 0.39 | 0.08 | 0.21 |
| 6 | 159398633 | rs3756988 | -1.065 | 0.501 | 1.117 | 0.57 | 0.005 | -0.044 | -0.044 | 0 | 0.14 | 0.07 | 0.37 | 0.39 | 0.1 | 0.22 |
| 6 | 159398700 | rs3756987 | -1.065 | 0.501 | 1.117 | 0.57 | 0.005 | -0.044 | -0.044 | 0.813 | 0.14 | 0.07 | 0.37 | 0.39 | 0.1 | 0.22 |
| 6 | 159399348 | rs768994 | -0.536 | 0.493 | 1.119 | 0.574 | 0.015 | -0.028 | -0.028 | 0.234 | 0.12 | 0.05 | 0.34 | 0.39 | 0.08 | 0.21 |
| 6 | 159414899 | rs16889320 | -0.556 | 0.494 | 1.119 | 0.574 | 0.014 | -0.024 | -0.024 | 0 | 0.12 | 0.06 | 0.34 | 0.39 | 0.08 | 0.21 |
| 6 | 159416444 | rs2092540 | -0.216 | 0.531 | 1.004 | 0.495 | -0.06 | 0.053 | 0.053 | 0.084 | 0.79 | 0.92 | 0.88 | 0.91 | 0.72 | 0.85 |
| 6 | 159420147 | rs7773373 | -1.858 | 0.426 | 1.358 | 0.518 | -0.048 | 0.135 | 0.135 | 0 | 0.21 | 0.47 | 0.4 | 0.39 | 0.12 | 0.24 |
| 6 | 159422031 | rs12205296 | -0.811 | 0.226 | 1.25 | 0.565 | 0.008 | -0.04 | -0.04 | 0 | 0.13 | 0.06 | 0.36 | 0.39 | 0.09 | 0.22 |
| 6 | 159423454 | rs9365019 | -0.223 | 0.375 | 1.402 | 0.519 | -0.004 | 0.01 | 0.01 | 0.994 | 0.23 | 0.24 | 0.13 | 0.09 | 0.25 | 0.15 |
| 6 | 159425707 | rs2016588 | 1.441 | 0.226 | 1.275 | 0.596 | 0.004 | -0.02 | -0.02 | 0 | 0.58 | 0.3 | 0.51 | 0.54 | 0.62 | 0.67 |
| 6 | 159438349 | rs3117156 | 0.538 | 0.298 | 1.355 | 0.523 | 0.013 | -0.073 | -0.073 | 0 | 0.72 | 0.57 | 0.84 | 0.91 | 0.7 | 0.84 |
| 6 | 159453337 | rs4449655 | -1.329 | 0.074 | 1.059 | 0.777 | -0.01 | 0.06 | 0.06 | 0 | 0.15 | 0.16 | 0.33 | 0.38 | 0.1 | 0.19 |
| 6 | 159456266 | rs4709266 | -1.345 | 0.064 | 1.053 | 0.771 | -0.01 | 0.059 | 0.059 | 0 | 0.15 | 0.16 | 0.33 | 0.38 | 0.1 | 0.19 |
| 6 | 159456375 | rs4709267 | -1.328 | 0.068 | 1.055 | 0.773 | -0.01 | 0.06 | 0.06 | 0 | 0.15 | 0.16 | 0.33 | 0.38 | 0.1 | 0.19 |
| 6 | 159461931 | rs2073138 | -0.985 | 0.04 | 1.055 | 0.777 | -0.023 | 0.09 | 0.09 | 0 | 0.13 | 0.15 | 0.3 | 0.38 | 0.08 | 0.17 |
| 6 | 159463452 | rs926657 | -1.253 | 0.245 | 1.112 | 0.381 | 0.069 | 0.035 | 0.035 | 0 | 0.08 | 0.45 | 0.05 | 0.03 | 0.06 | 0.04 |
| 6 | 159463964 | rs9295089 | -1.385 | 0.412 | 1.034 | 0.294 | 0.082 | -0.032 | -0.032 | 0 | 0.12 | 0.45 | 0.09 | 0.03 | 0.1 | 0.05 |
| 6 | 159465977 | rs1738074 | 1.235 | 0.249 | 0.292 | 0.631 | -0.006 | 0.044 | 0.044 | 0 | 0.53 | 0.3 | 0.45 | 0.45 | 0.57 | 0.62 |
| 6 | 159467753 | rs16889434 | -0.152 | 0.229 | 0.282 | 0.675 | 0.009 | -0.031 | -0.031 | 0 | 0.09 | 0.01 | 0.32 | 0.45 | 0.06 | 0.16 |
| 6 | 159469574 | rs182429 | 1.18 | 0.295 | 0.331 | 0.518 | -0.008 | 0.044 | 0.044 | 0 | 0.53 | 0.3 | 0.45 | 0.45 | 0.56 | 0.63 |
| 6 | 159475468 | rs2073137 | -0.537 | 0.241 | 0.265 | 0.94 | 0.006 | -0.036 | -0.036 | 0 | 0.09 | 0.01 | 0.32 | 0.46 | 0.06 | 0.16 |
| 6 | 159478308 | rs9457545 | -1.061 | 0.138 | 0.595 | 0.943 | -0.011 | 0.11 | 0.11 | 0 | 0.08 | 0.11 | 0.32 | 0.46 | 0.03 | 0.21 |
| 6 | 159484776 | rs169858 | 0.359 | 0.12 | 0.644 | 0.243 | -0.021 | 0.046 | 0.046 | 0.079 | 0.71 | 0.83 | 0.82 | 0.93 | 0.69 | 0.82 |
| 6 | 159490436 | rs212388 | 1.144 | -0.43 | 0.859 | 0.031 | 0.022 | 0.076 | 0.076 | 0 | 0.5 | 0.29 | 0.42 | 0.44 | 0.6 | 0.54 |
| 6 | 159498306 | rs1572527 | -1.356 | -0.35 | 1.339 | 0.359 | -0.004 | 0.323 | 0.323 | 0 | 0.08 | 0.3 | 0.37 | 0.5 | 0.03 | 0.21 |
| 6 | 159501256 | rs3103418 | -0.808 | -0.337 | 1.247 | 0.372 | -0.007 | 0.134 | 0.134 | 0 | 0.06 | 0.09 | 0.36 | 0.49 | 0.03 | 0.22 |
| 6 | 159506600 | rs2451258 | 0.156 | -0.581 | 1.023 | -0.015 | -0.059 | 0.196 | 0.196 | 0.006 | 0.63 | 0.91 | 0.83 | 0.94 | 0.65 | 0.78 |
| 6 | 159507027 | rs3127211 | -0.724 | -0.591 | 1.091 | 0.102 | -0.008 | 0.13 | 0.13 | 0 | 0.07 | 0.09 | 0.37 | 0.5 | 0.03 | 0.22 |
| 6 | 159507348 | rs9355697 | 0.523 | -0.69 | 0.92 | -0.284 | 0.001 | -0.006 | -0.006 | 2.382 | 0.13 | 0.15 | 0.1 | 0.1 | 0.12 | 0.05 |

| | | | | | | | | | | | | | | | | |
|---|---|---|---|---|---|---|---|---|---|---|---|---|---|---|---|---|
| 6 | 159511488 | rs3127214 | -0.736 | -0.683 | 0.648 | -0.222 | -0.015 | 0.173 | 0.173 | 0 | 0.07 | 0.16 | 0.37 | 0.49 | 0.03 | 0.22 |
| 6 | 159515077 | rs2451279 | -0.162 | -0.627 | 0.198 | -0.596 | -0.049 | 0.164 | 0.164 | 0.006 | 0.64 | 0.87 | 0.82 | 0.94 | 0.64 | 0.77 |
| 6 | 159515270 | rs575998 | 1.142 | -0.59 | 0.302 | -0.416 | -0.014 | 0.033 | 0.033 | 0 | 0.38 | 0.27 | 0.31 | 0.35 | 0.48 | 0.49 |
| 6 | 159516770 | rs755617 | 1.183 | -0.425 | 0.195 | -0.692 | -0.014 | 0.052 | 0.052 | 0 | 0.36 | 0.25 | 0.29 | 0.27 | 0.45 | 0.44 |
| 6 | 159517059 | rs12209018 | 0.562 | -0.422 | -0.283 | -0.84 | -0.006 | 0.115 | 0.115 | 0.014 | 0.19 | 0.03 | 0.08 | 0 | 0.15 | 0.1 |
| 6 | 159517672 | rs3103413 | -0.98 | -0.231 | 0.217 | -0.177 | -0.019 | 0.296 | 0.296 | 0 | 0.09 | 0.3 | 0.33 | 0.5 | 0.03 | 0.2 |
| 6 | 159521128 | rs2451241 | 1.28 | -0.146 | -0.65 | -0.66 | -0.006 | 0.026 | 0.026 | 0 | 0.36 | 0.3 | 0.29 | 0.28 | 0.43 | 0.43 |
| 6 | 159526689 | rs2485368 | -0.424 | 0.153 | -0.758 | -0.753 | -0.054 | 0.102 | 0.102 | 0 | 0.81 | 0.98 | 0.9 | 0.98 | 0.79 | 0.84 |
| 6 | 159528254 | rs7772013 | -0.374 | 0.155 | -0.526 | -0.669 | -0.003 | 0.006 | 0.006 | 0 | 0.23 | 0.6 | 0.26 | 0.21 | 0.2 | 0.14 |
| 6 | 159538494 | rs7759630 | 0.206 | 0.153 | -0.09 | -1.234 | -0.004 | -0.15 | -0.15 | 0.012 | 0.2 | 0.46 | 0.11 | 0 | 0.16 | 0.1 |
| 6 | 159539213 | rs644165 | 0.279 | 0.304 | -0.538 | -0.392 | -0.008 | 0.047 | 0.047 | 0 | 0.37 | 0.21 | 0.27 | 0.33 | 0.42 | 0.46 |
| 6 | 159549389 | rs7765276 | -0.619 | 0.733 | -0.179 | -1.058 | -0.006 | 0.004 | 0.004 | 0.062 | 0.26 | 0.24 | 0.21 | 0.11 | 0.32 | 0.27 |
| 6 | 159550853 | rs2451244 | 0.608 | 0.7 | -0.419 | -1.166 | 0.006 | 0.085 | 0.085 | 0 | 0.54 | 0.67 | 0.44 | 0.76 | 0.48 | 0.59 |
| 6 | 159556357 | rs2485328 | -0.504 | 0.655 | -0.189 | -0.72 | 0.004 | 0 | 0 | 0.015 | 0.39 | 0.48 | 0.59 | 0.62 | 0.46 | 0.54 |
| 6 | 159572922 | rs694704 | -0.916 | 0.615 | -0.166 | -0.803 | 0.027 | 0.044 | 0.044 | 0 | 0.59 | 0.81 | 0.73 | 0.77 | 0.65 | 0.66 |
| 6 | 159580324 | rs945773 | 0.002 | 0.281 | -0.096 | -0.801 | -0.061 | -0.005 | -0.005 | 0 | 0.43 | 0.15 | 0.59 | 0.62 | 0.48 | 0.55 |
| 6 | 159585941 | rs295340 | -0.283 | 0.472 | -0.063 | -0.687 | 0 | -0.002 | -0.002 | 0.35 | 0.48 | 0.54 | 0.66 | 0.62 | 0.48 | 0.55 |
| 6 | 159589126 | rs295335 | -0.132 | 0.503 | -0.039 | -0.645 | 0 | -0.002 | -0.002 | 0.01 | 0.46 | 0.5 | 0.64 | 0.62 | 0.46 | 0.55 |
| 6 | 159603958 | rs294879 | 0.363 | 0.61 | -0.414 | -0.746 | -0.001 | 0.072 | 0.072 | 0 | 0.46 | 0.26 | 0.32 | 0.22 | 0.49 | 0.41 |
| 6 | 159604352 | rs294878 | -1.334 | 0.976 | 0.335 | -0.205 | 0.018 | 0.022 | 0.022 | 0 | 0.18 | 0.54 | 0.15 | 0.17 | 0.22 | 0.12 |
| 6 | 159604988 | rs9457562 | -1.228 | 1.529 | 0.237 | -0.589 | 0.02 | -0.043 | -0.043 | 0 | 0.12 | 0.2 | 0.09 | 0.01 | 0.14 | 0.07 |
| 6 | 159606873 | rs294877 | -1.364 | 1.057 | -0.222 | -0.263 | 0.02 | 0.025 | 0.025 | 0 | 0.38 | 0.81 | 0.39 | 0.64 | 0.31 | 0.4 |
| 6 | 159608041 | rs12210751 | -0.563 | 0.875 | 0.526 | -0.334 | -0.004 | -0.041 | -0.041 | 5.133 | 0.15 | 0.12 | 0.08 | 0 | 0.1 | 0.08 |
| 6 | 159611599 | rs1810984 | -1.737 | 0.841 | 0.02 | -0.729 | 0.016 | 0.061 | 0.061 | 0 | 0.46 | 0.81 | 0.43 | 0.63 | 0.38 | 0.43 |
| 6 | 159626072 | rs294918 | -0.782 | 1.032 | 1.054 | -0.549 | 0.022 | 0.072 | 0.072 | 0.077 | 0.18 | 0.37 | 0.28 | 0.63 | 0.23 | 0.3 |
| 6 | 159627030 | rs12203580 | -0.521 | 0.956 | 1.292 | -0.657 | -0.006 | -0.066 | -0.066 | 0.428 | 0.19 | 0.2 | 0.11 | 0 | 0.12 | 0.08 |
| 6 | 159628978 | rs294915 | -1.723 | 0.961 | 1.375 | -0.639 | 0 | 0.002 | 0.002 | 0 | 0.37 | 0.65 | 0.39 | 0.64 | 0.35 | 0.38 |
| 6 | 159630244 | rs446504 | 0.125 | 0.929 | 1.487 | -0.679 | 0.019 | -0.019 | -0.019 | 0 | 0.06 | 0.27 | 0.1 | 0 | 0.05 | 0.06 |
| 6 | 159633600 | rs449602 | -1.661 | 0.932 | 1.497 | 0.175 | -0.012 | -0.013 | -0.013 | 1.641 | 0.29 | 0.58 | 0.25 | 0.3 | 0.22 | 0.19 |
| 6 | 159634608 | rs12526577 | 1.95 | 1.018 | 0.829 | -0.225 | -0.048 | 0.111 | 0.111 | 0 | 0.37 | 0.05 | 0.28 | 0.24 | 0.43 | 0.34 |
| 6 | 159635095 | rs552986 | -0.157 | 1.076 | 1.417 | 0.278 | 0.008 | 0.118 | 0.118 | 0 | 0.08 | 0.13 | 0.15 | 0.34 | 0.13 | 0.19 |
| 6 | 159639482 | rs381568 | -1.583 | 1.086 | 1.475 | 0.292 | -0.003 | -0.004 | -0.004 | 0.925 | 0.3 | 0.6 | 0.25 | 0.32 | 0.26 | 0.2 |
| 6 | 159640565 | rs419626 | -2.026 | 0.989 | 1.5 | 0.141 | 0.001 | 0.015 | 0.015 | 0 | 0.38 | 0.71 | 0.39 | 0.65 | 0.36 | 0.38 |
| 6 | 159642501 | rs399369 | -1.808 | 1.011 | 1.64 | 0.236 | 0 | 0.001 | 0.001 | 0 | 0.36 | 0.69 | 0.38 | 0.64 | 0.35 | 0.39 |
| 6 | 159660105 | rs433256 | 0.657 | 1.146 | 1.387 | 0.692 | -0.001 | 0.127 | 0.127 | 0 | 0.89 | 0.71 | 0.86 | 0.63 | 0.91 | 0.87 |
| 6 | 159662136 | rs537693 | -0.659 | 1.198 | 1.01 | 0.839 | 0.005 | 0.094 | 0.094 | 0 | 0.09 | 0.14 | 0.17 | 0.35 | 0.14 | 0.28 |
| 6 | 159667227 | rs294913 | 0.641 | 1.189 | 1.095 | 0.83 | 0 | 0.112 | 0.112 | 0 | 0.91 | 0.81 | 0.87 | 0.64 | 0.91 | 0.87 |
| 6 | 159667961 | rs294911 | 0.42 | 1.319 | 0.883 | 0.473 | -0.052 | 0.065 | 0.065 | 0 | 0.25 | 0.04 | 0.32 | 0.1 | 0.24 | 0.2 |
| 6 | 159672542 | rs550443 | -0.635 | 0.983 | 1.812 | 1.138 | 0.032 | 0.172 | 0.172 | 0 | 0.09 | 0.19 | 0.18 | 0.34 | 0.14 | 0.25 |
| 6 | 159672937 | rs294898 | 1.674 | 0.872 | 1.947 | 0.968 | 0.006 | 0.037 | 0.037 | 0.27 | 0.75 | 0.38 | 0.67 | 0.66 | 0.75 | 0.62 |
| 6 | 159673608 | rs500550 | -0.558 | 0.864 | 1.885 | 1.039 | 0.033 | 0.175 | 0.175 | 0 | 0.09 | 0.19 | 0.18 | 0.34 | 0.14 | 0.25 |
| 6 | 159692405 | rs3814448 | 0.034 | 0.798 | 1.698 | 0.98 | 0.021 | 0.263 | 0.263 | 0 | 0.05 | 0.2 | 0.04 | 0.36 | 0.04 | 0.07 |
| 6 | 159692596 | rs419009 | 0.436 | 0.809 | 1.676 | 0.902 | 0 | -0.029 | -0.029 | 0.367 | 0.87 | 0.87 | 0.94 | 0.99 | 0.89 | 0.88 |
| 6 | 159703373 | rs625284 | 1.266 | 0.813 | 1.53 | 0.934 | -0.008 | 0.036 | 0.036 | 0.424 | 0.79 | 0.36 | 0.81 | 0.64 | 0.84 | 0.8 |
| 6 | 159714412 | rs2501175 | 1.23 | 0.821 | 1.603 | 0.938 | -0.008 | 0.053 | 0.053 | 0.443 | 0.79 | 0.4 | 0.82 | 0.63 | 0.84 | 0.8 |
| 6 | 159716957 | rs578217 | 0.334 | 0.855 | 1.237 | 0.754 | -0.006 | 0.013 | 0.013 | 0 | 0.87 | 0.91 | 0.88 | 0.99 | 0.88 | 0.87 |
| 6 | 159723661 | rs10455858 | -0.646 | 0.829 | 1.332 | 1.007 | 0.026 | 0.156 | 0.156 | 0 | 0.09 | 0.21 | 0.18 | 0.33 | 0.14 | 0.25 |
| 6 | 159726375 | rs9364510 | 0.03 | 0.751 | 1.609 | 0.981 | 0.017 | 0.263 | 0.263 | 0 | 0.06 | 0.21 | 0.04 | 0.37 | 0.04 | 0.07 |
| 6 | 159727848 | rs12204649 | -0.416 | 0.778 | 1.555 | 1.029 | 0.027 | 0.168 | 0.168 | 0 | 0.09 | 0.21 | 0.18 | 0.34 | 0.14 | 0.26 |
| 6 | 159733694 | rs905684 | 0.455 | 0.881 | 1.73 | 0.781 | 0.008 | -0.053 | -0.053 | 0 | 0.86 | 0.82 | 0.87 | 1 | 0.87 | 0.87 |
| 6 | 159735202 | rs875276 | 1.284 | 0.711 | 1.977 | 0.809 | -0.003 | 0.028 | 0.028 | 0.424 | 0.78 | 0.35 | 0.8 | 0.66 | 0.82 | 0.8 |
| 6 | 159756737 | rs9365046 | 0.041 | 0.775 | 1.445 | 0.857 | 0.039 | 0.327 | 0.327 | 0 | 0.06 | 0.27 | 0.05 | 0.34 | 0.04 | 0.07 |
| 6 | 159771467 | rs4709305 | 1.199 | 0.713 | -0.228 | 0.303 | -0.011 | 0.078 | 0.078 | 0 | 0.36 | 0.08 | 0.25 | 0.2 | 0.33 | 0.26 |
| 6 | 159779084 | rs1339326 | 0.224 | 1.265 | -0.413 | 0.219 | -0.003 | -0.017 | -0.017 | 1.203 | 0.87 | 0.87 | 0.87 | 1 | 0.88 | 0.87 |
| 6 | 159781191 | rs9365049 | 0.591 | 1.284 | -0.763 | 0.247 | -0.093 | 0.099 | 0.099 | 0 | 0.29 | 0.01 | 0.35 | 0.09 | 0.33 | 0.24 |
| 6 | 159781278 | rs10945608 | -0.684 | 1.25 | 0.605 | 1.37 | 0.028 | 0.179 | 0.179 | 0 | 0.08 | 0.21 | 0.18 | 0.38 | 0.14 | 0.26 |
| 6 | 159785533 | rs9365050 | 1.588 | 1.266 | 0.502 | 1.278 | 0.021 | 0.057 | 0.057 | 0 | 0.79 | 0.66 | 0.54 | 0.6 | 0.74 | 0.61 |
| 6 | 159785799 | rs946457 | 0.84 | 0.901 | -0.229 | 0.246 | -0.02 | 0.028 | 0.028 | 0 | 0.87 | 0.95 | 0.88 | 1 | 0.88 | 0.87 |
| 6 | 159795332 | rs6455633 | -1.077 | 0.3 | -0.004 | 0.793 | 0.004 | -0.036 | -0.036 | 0.021 | 0.37 | 0.78 | 0.33 | 0.44 | 0.36 | 0.26 |
| 6 | 159801024 | rs1544167 | -0.932 | -0.226 | 1.132 | 0.917 | -0.004 | 0.011 | 0.011 | 0 | 0.27 | 0.3 | 0.23 | 0.33 | 0.21 | 0.21 |
| 6 | 159803460 | rs4709313 | 0.239 | -0.42 | 0.645 | 1.169 | -0.012 | -0.065 | -0.065 | 0.872 | 0.15 | 0.04 | 0.24 | 0.41 | 0.16 | 0.35 |
| 6 | 159805250 | rs9355721 | -0.592 | -0.426 | 1.214 | 0.899 | -0.014 | 0.213 | 0.213 | 0 | 0.06 | 0.2 | 0.04 | 0.28 | 0.02 | 0.05 |
| 6 | 159807939 | rs9365056 | -1.481 | -0.427 | 1.09 | 0.826 | -0.002 | 0.002 | 0.002 | 0 | 0.26 | 0.75 | 0.24 | 0.31 | 0.16 | 0.18 |
| 6 | 159814581 | rs12214847 | -0.461 | -0.227 | 0.871 | 1.016 | -0.008 | 0.116 | 0.116 | 0 | 0.07 | 0.16 | 0.06 | 0.26 | 0.06 | 0.07 |
| 6 | 159817482 | rs3823037 | -1.699 | -0.652 | 0.53 | 0.947 | 0.016 | -0.003 | -0.003 | 0 | 0.47 | 0.91 | 0.41 | 0.52 | 0.37 | 0.56 |
| 6 | 159817950 | rs11752900 | 0.21 | -0.6 | 0.547 | 0.952 | -0.004 | -0.037 | -0.037 | 0.041 | 0.18 | 0.05 | 0.15 | 0.23 | 0.17 | 0.37 |
| 6 | 159822140 | rs9295098 | -1.603 | -0.631 | 0.797 | 0.93 | 0.023 | 0.011 | 0.011 | 0 | 0.45 | 0.8 | 0.4 | 0.53 | 0.37 | 0.56 |
| 6 | 159833206 | rs9456400 | -0.518 | -0.614 | 0.556 | 0.962 | -0.008 | 0.013 | 0.013 | 0 | 0.15 | 0.09 | 0.16 | 0.08 | 0.14 | 0.13 |
| 6 | 159834759 | rs12524603 | 0.081 | -0.602 | 0.524 | 0.943 | -0.014 | -0.043 | -0.043 | 0 | 0.19 | 0.04 | 0.15 | 0.23 | 0.18 | 0.38 |

| | | | | | | | | | | | | | | | | |
|---|---|---|---|---|---|---|---|---|---|---|---|---|---|---|---|---|
| 6 | 159837795 | rs1361325 | -1.439 | -0.418 | 0.856 | 1.019 | -0.001 | -0.013 | -0.013 | 0 | 0.21 | 0.18 | 0.21 | 0.3 | 0.19 | 0.19 |
| 6 | 159843841 | rs9364516 | -0.478 | -0.143 | 0.861 | 1.048 | -0.003 | 0.047 | 0.047 | 0 | 0.07 | 0.09 | 0.06 | 0.22 | 0.06 | 0.07 |
| 6 | 159852780 | rs9457631 | -0.138 | 0.297 | 0.244 | 0.277 | 0.06 | -0.031 | -0.031 | 0.419 | 0.06 | 0.15 | 0.12 | 0 | 0.1 | 0.05 |
| 6 | 159854623 | rs7758822 | -1.104 | -0.307 | -0.387 | 0.542 | 0.039 | -0.059 | -0.059 | 1.002 | 0.19 | 0.12 | 0.13 | 0.49 | 0.12 | 0.35 |
| 6 | 159856361 | rs4708823 | 0.666 | -0.001 | -0.049 | 0.383 | 0.002 | 0.012 | 0.012 | 0.069 | 0.65 | 0.44 | 0.67 | 0.52 | 0.67 | 0.49 |
| 6 | 159857289 | rs4709321 | -0.274 | 0.301 | -0.685 | -0.133 | 0.004 | 0.015 | 0.015 | 0 | 0.46 | 0.38 | 0.51 | 0.62 | 0.38 | 0.48 |
| 6 | 159857915 | rs4709322 | -0.144 | 0.675 | -1.185 | -0.855 | 0.004 | 0.164 | 0.164 | 0 | 0.49 | 0.8 | 0.55 | 0.73 | 0.56 | 0.64 |
| 6 | 159860514 | rs9457636 | -0.433 | 0.902 | -0.716 | -0.333 | -0.01 | 0.002 | 0.002 | 0 | 0.08 | 0.06 | 0.13 | 0 | 0.12 | 0.04 |
| 6 | 159861286 | rs6455639 | -0.137 | 0.603 | -0.64 | -0.598 | 0.001 | 0.004 | 0.004 | 0 | 0.24 | 0.2 | 0.29 | 0.25 | 0.19 | 0.11 |
| 6 | 159862025 | rs2066105 | -1.059 | 0.299 | 0.051 | -0.807 | -0.002 | -0.078 | -0.078 | 0 | 0.17 | 0.3 | 0.18 | 0 | 0.16 | 0.08 |
| 6 | 159862832 | rs9457637 | -1.429 | 1.122 | 0.377 | -0.675 | 0.065 | -0.01 | -0.01 | 0.115 | 0.14 | 0.28 | 0.33 | 0.15 | 0.22 | 0.16 |
| 6 | 159865757 | rs1890582 | -0.643 | 0.092 | 0.884 | -0.19 | -0.048 | 0.042 | 0.042 | 1.049 | 0.25 | 0.39 | 0.34 | 0.42 | 0.17 | 0.18 |
| 6 | 159866409 | rs4709326 | 0.32 | 0.552 | 0.373 | -0.798 | -0.007 | -0.024 | -0.024 | 0 | 0.62 | 0.53 | 0.71 | 0.71 | 0.67 | 0.74 |
| 6 | 159868031 | rs4521619 | 0.962 | 0.815 | 0.39 | -0.864 | 0.004 | 0.021 | 0.021 | 0 | 0.52 | 0.35 | 0.44 | 0.56 | 0.54 | 0.63 |
| 6 | 159868795 | rs6455640 | 0.486 | 0.716 | 0.325 | -0.813 | -0.005 | -0.005 | -0.005 | 0 | 0.59 | 0.49 | 0.54 | 0.56 | 0.63 | 0.67 |
| 6 | 159874165 | rs1977751 | 0.343 | 0.631 | 0.374 | -0.885 | -0.003 | -0.007 | -0.007 | 0 | 0.61 | 0.43 | 0.7 | 0.71 | 0.67 | 0.74 |
| 6 | 159876028 | rs6926782 | 1.424 | 0.206 | -0.492 | -0.887 | -0.01 | 0.083 | 0.083 | 0 | 0.29 | 0.03 | 0.2 | 0.12 | 0.32 | 0.23 |
| 6 | 159881056 | rs1551214 | 0.477 | 0.432 | 0.111 | -0.516 | 0.008 | 0.007 | 0.007 | 0 | 0.57 | 0.36 | 0.53 | 0.56 | 0.63 | 0.67 |
| 6 | 159886303 | rs2880107 | -0.677 | 0.963 | -0.257 | -0.666 | -0.073 | 0.079 | 0.079 | 0.189 | 0.86 | 0.97 | 0.88 | 0.99 | 0.8 | 0.86 |
| 6 | 159893829 | rs4708831 | -0.679 | 0.355 | 0.342 | -0.724 | -0.029 | -0.086 | -0.086 | 0 | 0.28 | 0.46 | 0.33 | 0.1 | 0.23 | 0.17 |
| 6 | 159895128 | rs13215914 | -0.73 | 0.342 | 0.528 | -0.73 | -0.035 | -0.099 | -0.099 | 0 | 0.28 | 0.5 | 0.33 | 0.1 | 0.23 | 0.17 |
| 6 | 159896460 | rs6919753 | -0.497 | 0.279 | 0.702 | -0.788 | 0.002 | -0.006 | -0.006 | 0 | 0.13 | 0.4 | 0.27 | 0.1 | 0.11 | 0.07 |
| 6 | 159898528 | rs1901548 | 0.561 | 0.272 | 0.7 | -0.802 | -0.001 | -0.014 | -0.014 | 0 | 0.07 | 0.1 | 0.1 | 0 | 0.08 | 0.03 |
| 6 | 159899298 | rs16890402 | 0.315 | -0.079 | 0.985 | -0.291 | -0.025 | 0.148 | 0.148 | 0 | 0.06 | 0.19 | 0.11 | 0.26 | 0 | 0.03 |
| 6 | 159903581 | rs6929061 | 0.684 | -0.017 | 0.768 | -0.345 | 0.015 | 0.03 | 0.03 | 0 | 0.15 | 0.06 | 0.06 | 0 | 0.11 | 0.1 |
| 6 | 159904320 | rs720694 | -1.581 | 0.563 | 0.796 | -0.361 | -0.012 | 0.017 | 0.017 | 0.791 | 0.4 | 0.83 | 0.49 | 0.42 | 0.34 | 0.28 |
| 6 | 159912337 | rs192138 | -0.358 | 0.652 | 1.02 | -0.494 | -0.017 | -0.074 | -0.074 | 0 | 0.83 | 0.7 | 0.93 | 1 | 0.89 | 0.9 |
| 6 | 159915825 | rs240863 | -0.385 | 0.672 | 0.517 | -0.452 | -0.019 | -0.077 | -0.077 | 1.562 | 0.83 | 0.79 | 0.93 | 1 | 0.89 | 0.9 |
| 6 | 159922541 | rs240832 | 0.582 | 0.882 | -0.299 | -0.5 | -0.078 | 0.115 | 0.115 | 1.119 | 0.2 | 0.02 | 0.15 | 0.01 | 0.25 | 0.2 |
| 6 | 159926784 | rs6455651 | -0.798 | 0.918 | 0.529 | 1.118 | 0.085 | 0.275 | 0.275 | 0 | 0.11 | 0.34 | 0.18 | 0.49 | 0.12 | 0.13 |
| 6 | 159926996 | rs7772414 | -1.063 | 0.806 | 0.561 | 1.292 | 0.07 | 0.303 | 0.303 | 0 | 0.13 | 0.37 | 0.19 | 0.53 | 0.12 | 0.13 |
| 6 | 159927175 | rs7771621 | -0.789 | 0.951 | 0.653 | 1.164 | 0.057 | 0.106 | 0.106 | 0 | 0.11 | 0.36 | 0.06 | 0.18 | 0.08 | 0.11 |
| 6 | 159927218 | rs7754221 | -1.005 | 0.907 | 0.517 | 1.115 | 0.078 | 0.169 | 0.169 | 0 | 0.13 | 0.38 | 0.1 | 0.32 | 0.12 | 0.13 |
| 6 | 159927599 | rs6455652 | -0.112 | 1.241 | -0.061 | 0.568 | -0.001 | -0.025 | -0.025 | 0.281 | 0.44 | 0.53 | 0.43 | 0.81 | 0.42 | 0.53 |
| 6 | 159928324 | rs2341786 | 0.702 | 1.416 | -0.621 | 0.288 | -0.001 | -0.021 | -0.021 | 0.283 | 0.09 | 0.1 | 0.06 | 0 | 0.09 | 0.04 |
| 6 | 159931105 | rs9355736 | 0.419 | 1.341 | -1.093 | 0.573 | 0.021 | -0.037 | -0.037 | 0 | 0.37 | 0.23 | 0.52 | 0.53 | 0.37 | 0.5 |
| 6 | 159933238 | rs13206227 | 0.611 | 1.597 | -1.644 | 0.219 | -0.022 | 0.225 | 0.225 | 0 | 0.39 | 0.05 | 0.25 | 0.03 | 0.38 | 0.33 |
| 6 | 159942291 | rs6455655 | -0.319 | 1.613 | -1.283 | 1.073 | -0.008 | -0.116 | -0.116 | 0.243 | 0.67 | 0.93 | 0.66 | 0.56 | 0.67 | 0.53 |
| 6 | 159944996 | rs409359 | -0.338 | 1.705 | -0.841 | 0.451 | 0.013 | 0.038 | 0.038 | 0 | 0.52 | 0.27 | 0.49 | 0.66 | 0.54 | 0.61 |
| 6 | 159946240 | rs10080577 | 0.014 | 1.734 | -0.528 | 0.488 | -0.016 | 0.015 | 0.015 | 0 | 0.1 | 0.05 | 0.06 | 0 | 0.1 | 0.09 |
| 6 | 159946955 | rs9457666 | 0.014 | 1.734 | -0.528 | 0.488 | -0.016 | 0.015 | 0.015 | 0 | 0.1 | 0.05 | 0.06 | 0 | 0.1 | 0.09 |
| 6 | 159947896 | rs9295103 | -0.349 | 1.609 | -0.014 | 0.439 | 0.013 | -0.063 | -0.063 | 0 | 0.12 | 0.26 | 0.09 | 0 | 0.1 | 0.09 |
| 6 | 159949340 | rs378512 | -0.711 | 1.232 | -0.09 | 0.84 | 0.002 | 0 | 0 | 0 | 0.33 | 0.27 | 0.36 | 0.68 | 0.29 | 0.51 |
| 6 | 159951508 | rs686168 | -1.448 | 1.59 | 0.71 | 1.237 | 0.039 | 0.086 | 0.086 | 0.002 | 0.09 | 0.36 | 0.25 | 0.14 | 0.1 | 0.1 |
| 6 | 159952432 | rs10945617 | -1.395 | 2.174 | 0.914 | 1.745 | 0.047 | 0.059 | 0.059 | 0 | 0.15 | 0.29 | 0.16 | 0.21 | 0.2 | 0.19 |
| 6 | 159952609 | rs10945618 | -1.124 | 2.177 | 0.895 | 1.794 | 0.045 | 0.073 | 0.073 | 0 | 0.13 | 0.25 | 0.13 | 0.22 | 0.16 | 0.17 |
| 6 | 159953147 | rs372440 | -1.113 | 1.592 | 0.316 | 1.652 | -0.024 | -0.001 | -0.001 | 0 | 0.36 | 0.23 | 0.46 | 0.55 | 0.4 | 0.42 |
| 6 | 159953434 | rs398132 | -1.241 | 1.595 | 0.506 | 1.231 | 0.009 | 0.03 | 0.03 | 0 | 0.44 | 0.35 | 0.55 | 0.66 | 0.55 | 0.54 |
| 6 | 159953548 | rs413634 | -2.233 | 1.737 | 1.433 | 1.233 | 0.015 | -0.022 | -0.022 | 1.094 | 0.17 | 0.41 | 0.34 | 0.09 | 0.21 | 0.14 |
| 6 | 159953928 | rs391045 | -2.071 | 1.343 | 1.111 | 1.267 | 0.036 | 0.007 | 0.007 | 0.12 | 0.53 | 0.88 | 0.72 | 0.55 | 0.57 | 0.56 |
| 6 | 159954027 | rs380609 | -1.244 | 1.535 | 1.109 | 0.742 | -0.013 | -0.007 | -0.007 | 0 | 0.35 | 0.37 | 0.54 | 0.44 | 0.43 | 0.39 |
| 6 | 159959673 | rs555627 | -0.459 | 1.067 | 0.764 | 0.595 | -0.026 | -0.023 | -0.023 | 0 | 0.09 | 0.01 | 0.17 | 0.07 | 0.08 | 0.06 |
| 6 | 159959831 | rs9347325 | -0.329 | 1.197 | 1.004 | 0.604 | -0.003 | -0.001 | -0.001 | 0 | 0.25 | 0.24 | 0.34 | 0.35 | 0.34 | 0.31 |
| 6 | 159962019 | rs240853 | -1.613 | 1.112 | 1.291 | 0.56 | 0.002 | -0.002 | -0.002 | 0 | 0.42 | 0.61 | 0.56 | 0.42 | 0.44 | 0.38 |
| 6 | 159962962 | rs240852 | -2.044 | 1.246 | 1.359 | 0.512 | 0.023 | 0.039 | 0.039 | 0.036 | 0.49 | 0.72 | 0.63 | 0.42 | 0.51 | 0.5 |
| 6 | 159963386 | rs4543399 | -0.906 | 1.147 | 1.237 | 0.481 | 0.013 | -0.007 | -0.007 | 0 | 0.32 | 0.42 | 0.38 | 0.35 | 0.35 | 0.31 |
| 6 | 159966964 | rs240844 | -1.025 | 1.378 | 1.141 | 0.512 | 0.01 | -0.011 | -0.011 | 0 | 0.12 | 0.21 | 0.22 | 0.07 | 0.12 | 0.12 |
| 6 | 159977925 | rs240895 | -0.521 | 1.173 | 0.926 | 0.616 | -0.002 | 0.004 | 0.004 | 0 | 0.1 | 0.1 | 0.17 | 0.07 | 0.08 | 0.08 |
| 6 | 159978469 | rs9365078 | -0.333 | 1.272 | 0.999 | 0.678 | -0.002 | -0.001 | -0.001 | 0 | 0.24 | 0.2 | 0.34 | 0.34 | 0.34 | 0.26 |
| 6 | 159982326 | rs9364527 | -0.404 | 1.305 | 0.86 | 0.63 | 0.01 | 0.004 | 0.004 | 0 | 0.25 | 0.25 | 0.34 | 0.36 | 0.35 | 0.26 |
| 6 | 159984005 | rs240880 | -1.246 | 1.286 | 0.899 | 0.585 | -0.001 | -0.017 | -0.017 | 1.769 | 0.21 | 0.28 | 0.25 | 0.05 | 0.16 | 0.2 |
| 6 | 159985424 | rs240877 | -1.577 | 1.089 | 0.638 | 0.68 | 0.012 | 0.034 | 0.034 | 0.502 | 0.47 | 0.7 | 0.62 | 0.41 | 0.52 | 0.46 |
| 6 | 159985856 | rs240874 | -1.301 | 0.576 | 0.489 | 0.329 | -0.026 | -0.054 | -0.054 | 0 | 0.18 | 0.33 | 0.22 | 0.05 | 0.1 | 0.08 |
| 6 | 159987048 | rs240873 | -0.987 | 1.209 | 0.186 | 0.847 | 0.024 | -0.005 | -0.005 | 0 | 0.34 | 0.54 | 0.4 | 0.61 | 0.37 | 0.37 |
| 6 | 159988748 | rs9456423 | -0.89 | 0.763 | -0.023 | 0.921 | 0.001 | -0.012 | -0.012 | 0 | 0.43 | 0.62 | 0.26 | 0.47 | 0.35 | 0.42 |
| 6 | 159991061 | rs6941197 | -0.778 | 0.716 | 0.305 | 0.892 | -0.001 | -0.001 | -0.001 | 0 | 0.4 | 0.56 | 0.26 | 0.54 | 0.35 | 0.43 |
| 6 | 159991083 | rs12190682 | -0.272 | 0.749 | 0.919 | 1.001 | 0.005 | 0.028 | 0.028 | 0 | 0.13 | 0.25 | 0.11 | 0.17 | 0.12 | 0.21 |
| 6 | 159993192 | rs6942219 | -1.452 | 0.707 | 1.037 | 0.995 | -0.007 | -0.018 | -0.018 | 0.018 | 0.42 | 0.74 | 0.27 | 0.54 | 0.36 | 0.43 |
| 6 | 159998878 | rs7755526 | -0.187 | 0.586 | 1.134 | 0.838 | 0.003 | 0.008 | 0.008 | 0 | 0.21 | 0.19 | 0.12 | 0.37 | 0.23 | 0.23 |

| | | | | | | | | | | | | | | | | |
|---|---|---|---|---|---|---|---|---|---|---|---|---|---|---|---|---|
| 6 | 159999186 | rs10945621 | -0.295 | 0.61 | 1.123 | 0.898 | 0.004 | 0.03 | 0.03 | 0 | 0.14 | 0.25 | 0.11 | 0.18 | 0.12 | 0.21 |
| 6 | 160005397 | rs9457676 | -0.822 | 0.609 | 1.191 | 0.896 | 0.003 | 0.012 | 0.012 | 0 | 0.27 | 0.26 | 0.15 | 0.37 | 0.24 | 0.23 |
| 6 | 160006485 | rs1452132 | -1.459 | 0.606 | 1.205 | 0.896 | 0.001 | 0.001 | 0.001 | 0 | 0.4 | 0.5 | 0.26 | 0.55 | 0.36 | 0.43 |
| 6 | 160011716 | rs9456431 | -1.433 | 0.632 | 1.219 | 0.849 | 0.007 | 0.011 | 0.011 | 0 | 0.38 | 0.49 | 0.25 | 0.44 | 0.36 | 0.42 |
| 6 | 160013592 | rs1817115 | 0.091 | 0.653 | 1.092 | 0.856 | 0.021 | -0.004 | -0.004 | 0 | 0.11 | 0.23 | 0.1 | 0.07 | 0.12 | 0.19 |
| 6 | 160017361 | rs12205349 | 0.117 | 0.644 | 1.072 | 0.853 | 0.02 | -0.005 | -0.005 | 0 | 0.11 | 0.22 | 0.1 | 0.07 | 0.12 | 0.2 |
| 6 | 160024243 | rs1551220 | -0.284 | 0.618 | 0.994 | 0.933 | 0.013 | 0.044 | 0.044 | 0.003 | 0.17 | 0.23 | 0.11 | 0.37 | 0.2 | 0.23 |
| 6 | 160025456 | rs9456433 | -0.418 | 0.838 | 1.074 | 0.804 | 0.051 | 0.076 | 0.076 | 0 | 0.18 | 0.3 | 0.15 | 0.38 | 0.27 | 0.22 |
| 6 | 160025471 | rs12525670 | -0.843 | 0.69 | 1.173 | 0.606 | -0.023 | -0.02 | -0.02 | 0 | 0.25 | 0.43 | 0.16 | 0.17 | 0.16 | 0.23 |
| 6 | 160030008 | rs9355741 | 1.622 | 0.669 | 0.426 | 0.121 | 0.049 | 0.065 | 0.065 | 0 | 0.4 | 0.19 | 0.41 | 0.32 | 0.3 | 0.32 |
| 6 | 160031215 | rs4382297 | -1.355 | 0.455 | 0.738 | 0.117 | -0.019 | 0.013 | 0.013 | 0 | 0.46 | 0.64 | 0.3 | 0.57 | 0.43 | 0.46 |
| 6 | 160037460 | rs12190302 | 0.222 | 0.543 | 0.892 | 0.151 | 0.018 | -0.038 | -0.038 | 0 | 0.17 | 0.37 | 0.12 | 0.06 | 0.15 | 0.22 |
| 6 | 160038263 | rs4368835 | -1.165 | 0.534 | 0.876 | 0.16 | -0.017 | -0.012 | -0.012 | 0 | 0.42 | 0.64 | 0.3 | 0.46 | 0.43 | 0.44 |
| 6 | 160044559 | rs12193610 | 0.364 | 0.537 | 0.766 | 0.138 | 0.008 | -0.023 | -0.023 | 0 | 0.16 | 0.24 | 0.11 | 0.06 | 0.15 | 0.19 |
| 6 | 160045332 | rs4708838 | -1.238 | 0.441 | 0.363 | -0.022 | -0.037 | 0.01 | 0.01 | 0 | 0.45 | 0.77 | 0.31 | 0.58 | 0.43 | 0.43 |
| 6 | 160050415 | rs4708839 | -0.43 | 0.847 | 0.45 | -0.144 | 0.048 | 0.079 | 0.079 | 0 | 0.29 | 0.44 | 0.23 | 0.48 | 0.36 | 0.33 |
| 6 | 160054518 | rs1570319 | -0.918 | 0.603 | 0.668 | -0.699 | 0.016 | 0.016 | 0.016 | 0 | 0.08 | 0.15 | 0.04 | 0.1 | 0.06 | 0.07 |
| 6 | 160061229 | rs7754295 | -1.018 | 0.601 | 0.665 | -0.7 | 0.02 | 0.021 | 0.021 | 0 | 0.08 | 0.19 | 0.04 | 0.1 | 0.06 | 0.07 |
| 6 | 160063852 | rs2842997 | 0.611 | 0.399 | 0.103 | -1.232 | 0.005 | 0.164 | 0.164 | 0 | 0.46 | 0.23 | 0.56 | 0.14 | 0.46 | 0.51 |
| 6 | 160067287 | rs2758317 | -0.021 | 0.493 | 0.122 | -1.151 | 0 | -0.012 | -0.012 | 0 | 0.54 | 0.52 | 0.61 | 0.23 | 0.52 | 0.58 |
| 6 | 160069037 | rs2842994 | 0.479 | 0.297 | -0.018 | -1.523 | -0.001 | -0.026 | -0.026 | 0 | 0.24 | 0.26 | 0.23 | 0.1 | 0.16 | 0.2 |
| 6 | 160075188 | rs2758322 | -0.166 | 0.747 | 0.58 | -0.686 | 0.057 | 0.243 | 0.243 | 0 | 0.83 | 0.53 | 0.86 | 0.54 | 0.78 | 0.82 |
| 6 | 160075197 | rs2842989 | 0.835 | 0.771 | 0.541 | -0.899 | 0.011 | 0.07 | 0.07 | 0 | 0.46 | 0.35 | 0.57 | 0.12 | 0.45 | 0.48 |
| 6 | 160085434 | rs2842985 | -0.54 | 0.769 | 0.697 | -0.902 | 0.025 | 0.124 | 0.124 | 0 | 0.85 | 0.76 | 0.88 | 0.54 | 0.79 | 0.85 |
| 6 | 160085841 | rs9365085 | 0.607 | 0.635 | 0.506 | -0.977 | -0.006 | -0.047 | -0.047 | 0 | 0.26 | 0.15 | 0.24 | 0.42 | 0.27 | 0.28 |
| 6 | 160091560 | rs732498 | -0.235 | 0.643 | 0.259 | -0.967 | -0.004 | -0.027 | -0.027 | 0 | 0.37 | 0.18 | 0.29 | 0.42 | 0.32 | 0.35 |
| 6 | 160098222 | rs4342445 | 0.604 | 0.644 | 0.259 | -0.989 | -0.006 | -0.047 | -0.047 | 0 | 0.26 | 0.14 | 0.24 | 0.42 | 0.27 | 0.28 |
| 6 | 160099260 | rs6917589 | 0.604 | 0.644 | 0.259 | -0.989 | -0.006 | -0.047 | -0.047 | 0 | 0.26 | 0.15 | 0.24 | 0.42 | 0.27 | 0.28 |
| 6 | 160101320 | rs5746151 | 0.203 | 0.642 | 0.258 | -0.993 | 0.002 | 0.015 | 0.015 | 0 | 0.1 | 0.03 | 0.05 | 0 | 0.06 | 0.07 |
| 6 | 160103084 | rs5746136 | -0.024 | 0.47 | 0.394 | -0.847 | -0.004 | -0.03 | -0.03 | 2.813 | 0.37 | 0.18 | 0.29 | 0.42 | 0.33 | 0.35 |
| 6 | 160105070 | rs2758331 | 0.789 | 0.512 | 0.212 | -0.908 | -0.006 | 0.208 | 0.208 | 0 | 0.45 | 0.19 | 0.56 | 0.12 | 0.45 | 0.47 |
| 6 | 160106125 | rs2855116 | 0.789 | 0.512 | 0.212 | -0.908 | -0.006 | 0.208 | 0.208 | 0 | 0.45 | 0.19 | 0.56 | 0.12 | 0.45 | 0.47 |
| 6 | 160113872 | rs4880 | 0.642 | 0.728 | 0.115 | -0.762 | 0.008 | 0.027 | 0.027 | 0 | 0.48 | 0.42 | 0.58 | 0.13 | 0.47 | 0.51 |
| 6 | 160120799 | rs6912979 | 0.277 | 1.179 | -0.024 | -0.387 | -0.005 | -0.019 | -0.019 | 0 | 0.3 | 0.22 | 0.26 | 0.43 | 0.31 | 0.31 |
| 6 | 160122921 | rs2758352 | 0.062 | 1.233 | 0.095 | 0.383 | 0.03 | 0.131 | 0.131 | 0 | 0.85 | 0.68 | 0.87 | 0.59 | 0.79 | 0.85 |
| 6 | 160133957 | rs752779 | 0.062 | 1.233 | 0.086 | 0.383 | 0.028 | 0.127 | 0.127 | 0 | 0.85 | 0.68 | 0.87 | 0.59 | 0.79 | 0.85 |
| 6 | 160135577 | rs9355747 | 0.277 | 1.189 | -0.002 | -0.383 | -0.004 | -0.014 | -0.014 | 0.401 | 0.3 | 0.24 | 0.26 | 0.43 | 0.3 | 0.31 |
| 6 | 160137687 | rs4516970 | 0.227 | 1.124 | 0.251 | -0.61 | 0.009 | -0.029 | -0.029 | 0 | 0.05 | 0.12 | 0.02 | 0 | 0.03 | 0.07 |
| 6 | 160139961 | rs4709363 | 0.41 | 1.117 | 0.302 | -0.607 | -0.004 | -0.016 | -0.016 | 0 | 0.3 | 0.23 | 0.26 | 0.43 | 0.3 | 0.31 |
| 6 | 160147507 | rs1048021 | 0.387 | 1.083 | 0.294 | -0.624 | -0.006 | -0.022 | -0.022 | 0.09 | 0.31 | 0.22 | 0.26 | 0.43 | 0.3 | 0.31 |
| 6 | 160147630 | rs1048019 | 0.3 | 1.007 | 0.517 | -0.646 | 0.007 | 0.039 | 0.039 | 0 | 0.32 | 0.4 | 0.27 | 0.43 | 0.3 | 0.31 |
| 6 | 160159034 | rs2842974 | -0.47 | 0.858 | 0.759 | -0.402 | 0.025 | 0.118 | 0.118 | 0 | 0.85 | 0.75 | 0.87 | 0.59 | 0.79 | 0.85 |
| 6 | 160159221 | rs6925350 | 0.665 | 0.999 | 0.728 | -0.615 | 0 | -0.013 | -0.013 | 0 | 0.21 | 0.17 | 0.22 | 0.43 | 0.21 | 0.29 |
| 6 | 160161225 | rs4709364 | 0.665 | 0.999 | 0.762 | -0.615 | 0 | 0.005 | 0.005 | 0.061 | 0.21 | 0.22 | 0.22 | 0.43 | 0.21 | 0.29 |
| 6 | 160174303 | rs3822849 | -0.222 | 0.994 | 0.615 | -0.618 | -0.007 | 0.055 | 0.055 | 0.493 | 0.28 | 0.48 | 0.25 | 0.43 | 0.24 | 0.36 |
| 6 | 160174456 | rs1440 | 0.636 | 0.981 | 0.48 | -0.655 | -0.04 | 0.178 | 0.178 | 0.062 | 0.57 | 0.19 | 0.62 | 0.16 | 0.55 | 0.49 |
| 6 | 160182096 | rs927450 | 0.901 | 0.599 | 0.308 | -0.851 | -0.021 | 0.255 | 0.255 | 0 | 0.51 | 0.17 | 0.6 | 0.16 | 0.54 | 0.49 |
| 6 | 160188891 | rs9347340 | 0.542 | 0.587 | 0.48 | -0.816 | 0.001 | 0.082 | 0.082 | 0 | 0.74 | 0.53 | 0.83 | 0.59 | 0.76 | 0.78 |
| 6 | 160196343 | rs25683 | 0.943 | 0.583 | 0.483 | -0.821 | -0.019 | 0.257 | 0.257 | 0 | 0.51 | 0.17 | 0.6 | 0.16 | 0.55 | 0.49 |
| 6 | 160198395 | rs3465 | 1.205 | 0.531 | -0.038 | -1.332 | -0.053 | 0.163 | 0.163 | 0 | 0.34 | 0.14 | 0.35 | 0.1 | 0.38 | 0.28 |
| 6 | 160199687 | rs15982 | 0.495 | 0.814 | 0.327 | -0.788 | 0.001 | 0.082 | 0.082 | 0 | 0.74 | 0.53 | 0.83 | 0.59 | 0.76 | 0.78 |
| 6 | 160200026 | rs4832 | 0.828 | 0.826 | 0.351 | -0.789 | 0 | -0.015 | -0.015 | 0.04 | 0.21 | 0.17 | 0.22 | 0.43 | 0.21 | 0.29 |
| 6 | 160202860 | rs4235925 | -0.547 | 0.816 | 0.436 | -0.8 | 0.018 | 0.099 | 0.099 | 0 | 0.85 | 0.78 | 0.88 | 0.59 | 0.8 | 0.85 |
| 6 | 160204127 | rs4709368 | 0.465 | 0.854 | 0.235 | -0.781 | 0.001 | 0.082 | 0.082 | 0 | 0.74 | 0.53 | 0.83 | 0.59 | 0.76 | 0.78 |
| 6 | 160206631 | rs3818299 | 0.744 | 0.882 | 0.186 | -0.767 | -0.002 | 0.156 | 0.156 | 0 | 0.52 | 0.26 | 0.61 | 0.16 | 0.54 | 0.49 |
| 6 | 160207570 | rs4709369 | -1.024 | 1.014 | 0.147 | -0.771 | -0.046 | -0.077 | -0.077 | 0 | 0.89 | 0.98 | 0.92 | 0.59 | 0.86 | 0.86 |
| 6 | 160211039 | rs6929124 | 0.472 | 1.006 | 0.137 | -0.79 | 0.002 | -0.03 | -0.03 | 0 | 0.24 | 0.18 | 0.23 | 0.42 | 0.24 | 0.3 |
| 6 | 160211728 | rs11757681 | 0.408 | 1.132 | -0.197 | -1.068 | -0.001 | 0.186 | 0.186 | 0 | 0.5 | 0.25 | 0.6 | 0.16 | 0.51 | 0.49 |
| 6 | 160218006 | rs6908294 | 0.098 | 1.123 | -0.224 | -1.065 | -0.002 | -0.053 | -0.053 | 0 | 0.55 | 0.51 | 0.62 | 0.16 | 0.54 | 0.55 |
| 6 | 160219337 | rs4739 | 0.314 | 1.124 | -0.993 | -0.869 | 0 | 0.148 | 0.148 | 0 | 0.5 | 0.3 | 0.6 | 0.16 | 0.51 | 0.49 |
| 6 | 160219640 | rs13202793 | 0.314 | 1.124 | -0.993 | -0.869 | 0 | 0.148 | 0.148 | 0 | 0.5 | 0.3 | 0.6 | 0.16 | 0.51 | 0.49 |
| 6 | 160237489 | rs2342480 | 0.262 | 1.513 | -1.043 | -0.003 | -0.038 | -0.047 | -0.047 | 0.56 | 0.77 | 0.86 | 0.84 | 0.61 | 0.67 | 0.79 |
| 6 | 160238709 | rs2038819 | 0.776 | 1.342 | -1.247 | -0.724 | -0.001 | 0.065 | 0.065 | 0 | 0.3 | 0.18 | 0.28 | 0.11 | 0.28 | 0.28 |
| 6 | 160239268 | rs876380 | 0.009 | 1.095 | -1.138 | -0.772 | -0.018 | -0.028 | -0.028 | 0 | 0.73 | 0.83 | 0.82 | 0.62 | 0.68 | 0.79 |
| 6 | 160242028 | rs4708850 | 0.495 | 1.099 | -1.051 | -0.897 | -0.004 | 0.052 | 0.052 | 0 | 0.28 | 0.15 | 0.27 | 0.12 | 0.28 | 0.33 |
| 6 | 160247249 | rs6902339 | -0.062 | 1.262 | -0.644 | -1.184 | -0.01 | 0.034 | 0.034 | 0.004 | 0.41 | 0.38 | 0.3 | 0.1 | 0.53 | 0.46 |
| 6 | 160248090 | rs4709380 | 0.355 | 0.634 | -0.449 | -0.914 | 0.017 | 0.049 | 0.049 | 0 | 0.38 | 0.22 | 0.3 | 0.18 | 0.3 | 0.36 |
| 6 | 160249368 | rs11965350 | -0.066 | 0.439 | 0.633 | 0.075 | -0.016 | 0.091 | 0.091 | 0.541 | 0.13 | 0.47 | 0.1 | 0.2 | 0.08 | 0.07 |
| 6 | 160251955 | rs9457732 | -1.216 | 0.818 | 0.371 | 0.169 | 0.005 | -0.013 | -0.013 | 0 | 0.29 | 0.53 | 0.39 | 0.7 | 0.32 | 0.37 |

| | | | | | | | | | | | | | | | | |
|---|---|---|---|---|---|---|---|---|---|---|---|---|---|---|---|---|
| 6 | 160284771 | rs9365107 | 1.279 | 0.99 | 0.713 | -0.106 | 0.012 | 0.014 | 0.014 | 0.457 | 0.74 | 0.56 | 0.57 | 0.31 | 0.65 | 0.67 |
| 6 | 160286188 | rs6924573 | 0.485 | 1.007 | 0.738 | -0.145 | 0.005 | 0.03 | 0.03 | 0 | 0.87 | 0.82 | 0.67 | 0.32 | 0.83 | 0.8 |
| 6 | 160295377 | rs7767760 | 1.607 | 1.012 | 0.741 | -0.197 | 0.007 | 0.009 | 0.009 | 0.215 | 0.76 | 0.62 | 0.57 | 0.29 | 0.68 | 0.7 |
| 6 | 160299505 | rs9365111 | 0.558 | 1.032 | 0.836 | -0.128 | 0.009 | 0.053 | 0.053 | 0 | 0.86 | 0.76 | 0.67 | 0.31 | 0.83 | 0.8 |
| 6 | 160301358 | rs6925813 | 0.373 | 1.042 | 0.795 | -0.286 | -0.001 | -0.001 | -0.001 | 0.109 | 0.86 | 0.87 | 0.68 | 0.21 | 0.83 | 0.8 |
| 6 | 160311752 | rs220735 | -1.679 | 0.892 | 0.716 | -0.46 | 0.002 | -0.031 | -0.031 | 0 | 0.27 | 0.44 | 0.44 | 0.81 | 0.33 | 0.28 |
| 6 | 160323406 | rs220729 | -0.981 | 0.654 | 0.38 | -0.487 | 0.005 | -0.001 | -0.001 | 0 | 0.21 | 0.15 | 0.35 | 0.78 | 0.23 | 0.25 |
| 6 | 160326567 | rs220723 | -0.242 | 1.23 | -0.307 | -1.053 | -0.073 | 0.012 | 0.012 | 0 | 0.76 | 0.98 | 0.66 | 0.83 | 0.73 | 0.65 |
| 6 | 160328620 | rs220721 | -0.753 | 1.343 | 0.737 | -0.234 | 0 | 0.025 | 0.025 | 0.051 | 0.19 | 0.29 | 0.34 | 0.67 | 0.24 | 0.17 |
| 6 | 160329423 | rs170219 | -1.638 | 1.349 | 0.893 | -0.397 | -0.001 | 0.001 | 0.001 | 0.613 | 0.28 | 0.6 | 0.43 | 0.7 | 0.34 | 0.28 |
| 6 | 160344699 | rs6455670 | -0.827 | 1.073 | 0.796 | -0.285 | 0.002 | 0.043 | 0.043 | 0.139 | 0.21 | 0.4 | 0.34 | 0.67 | 0.25 | 0.18 |
| 6 | 160345027 | rs9355765 | -0.17 | 1.097 | 0.878 | -0.317 | 0.023 | -0.005 | -0.005 | 0 | 0.06 | 0.16 | 0.07 | 0.03 | 0.09 | 0.09 |
| 6 | 160352497 | rs1983633 | 0.021 | 1.131 | 0.275 | -0.299 | -0.004 | 0.008 | 0.008 | 0 | 0.05 | 0.04 | 0.06 | 0.03 | 0.09 | 0.09 |
| 6 | 160357037 | rs1040574 | -1.709 | 1.021 | 0.408 | -0.287 | -0.009 | -0.003 | -0.003 | 0.946 | 0.3 | 0.49 | 0.42 | 0.71 | 0.34 | 0.28 |
| 6 | 160370811 | rs2013600 | -1.291 | 1.008 | 0.52 | -0.275 | -0.005 | 0.011 | 0.011 | 0.069 | 0.25 | 0.33 | 0.35 | 0.66 | 0.25 | 0.18 |
| 6 | 160384403 | rs6934383 | -1.202 | 1.067 | 0.47 | 0.165 | -0.014 | 0.171 | 0.171 | 0 | 0.15 | 0.37 | 0.28 | 0.46 | 0.17 | 0.16 |
| 6 | 160389147 | rs8191687 | 0.791 | 1.076 | 0.43 | 0.008 | -0.006 | -0.018 | -0.018 | 0 | 0.05 | 0.07 | 0.01 | 0 | 0.01 | 0.01 |
| 6 | 160394247 | rs3798207 | -0.658 | 1.15 | 0.286 | 0.109 | -0.004 | 0.07 | 0.07 | 0 | 0.13 | 0.23 | 0.27 | 0.48 | 0.16 | 0.16 |
| 6 | 160395632 | rs9457795 | -1.595 | 1.116 | 0.179 | -0.111 | 0 | 0 | 0 | 0 | 0.28 | 0.32 | 0.42 | 0.7 | 0.34 | 0.26 |
| 6 | 160397713 | rs645851 | -1.202 | 1.528 | -1.029 | -0.751 | -0.056 | 0.125 | 0.125 | 0 | 0.81 | 0.99 | 0.86 | 1 | 0.81 | 0.87 |
| 6 | 160398330 | rs9456490 | -1.15 | 1.816 | -0.372 | 0.077 | -0.001 | -0.021 | -0.021 | 0 | 0.17 | 0.18 | 0.3 | 0.5 | 0.24 | 0.25 |
| 6 | 160402705 | rs6917747 | -0.41 | 1.88 | -0.433 | -0.059 | -0.027 | 0.034 | 0.034 | 0 | 0.09 | 0.05 | 0.06 | 0.03 | 0.14 | 0.13 |
| 6 | 160403788 | rs3798201 | -0.336 | 1.772 | -0.073 | 0.212 | -0.013 | 0.142 | 0.142 | 0 | 0.12 | 0.22 | 0.24 | 0.46 | 0.13 | 0.13 |
| 6 | 160404832 | rs635551 | 0.457 | 1.967 | -0.445 | -0.421 | -0.039 | 0.026 | 0.026 | 0 | 0.1 | 0.01 | 0.1 | 0.04 | 0.16 | 0.13 |
| 6 | 160415341 | rs6455678 | -0.768 | 1.538 | 0.276 | -0.048 | -0.008 | 0.097 | 0.097 | 0 | 0.15 | 0.26 | 0.26 | 0.58 | 0.16 | 0.25 |
| 6 | 160422785 | rs9295119 | -0.729 | 1.543 | 0.182 | -0.046 | -0.006 | 0.065 | 0.065 | 0 | 0.15 | 0.23 | 0.26 | 0.58 | 0.16 | 0.25 |
| 6 | 160425247 | rs3822843 | -0.682 | 1.529 | 0.19 | -0.047 | -0.011 | 0.11 | 0.11 | 0 | 0.15 | 0.28 | 0.27 | 0.57 | 0.16 | 0.25 |
| 6 | 160428858 | rs6455679 | -0.682 | 1.529 | 0.183 | -0.047 | -0.011 | 0.11 | 0.11 | 0 | 0.15 | 0.28 | 0.27 | 0.57 | 0.16 | 0.25 |
| 6 | 160429883 | rs3734181 | -0.753 | 1.626 | -0.001 | -0.109 | -0.001 | -0.001 | -0.001 | 0 | 0.23 | 0.23 | 0.3 | 0.57 | 0.25 | 0.27 |
| 6 | 160430656 | rs1003737 | -0.947 | 1.512 | -0.19 | -0.5 | 0.012 | -0.034 | -0.034 | 0 | 0.37 | 0.53 | 0.46 | 0.72 | 0.45 | 0.47 |
| 6 | 160435569 | rs3798189 | -0.738 | 1.434 | -0.288 | -1.003 | 0 | -0.051 | -0.051 | 0 | 0.46 | 0.51 | 0.44 | 0.84 | 0.49 | 0.61 |
| 6 | 160435929 | rs3798188 | 0.023 | 1.576 | -0.252 | -0.469 | 0.002 | -0.001 | -0.001 | 0 | 0.17 | 0.13 | 0.28 | 0.53 | 0.18 | 0.24 |
| 6 | 160437347 | rs9347380 | -0.292 | 1.659 | -0.116 | -0.472 | -0.016 | -0.003 | -0.003 | 0 | 0.27 | 0.24 | 0.14 | 0.31 | 0.29 | 0.35 |
| 6 | 160440172 | rs9347381 | 0.317 | 1.571 | -0.236 | -0.43 | -0.001 | -0.005 | -0.005 | 0 | 0.15 | 0.09 | 0.05 | 0.21 | 0.14 | 0.25 |
| 6 | 160443390 | rs9456496 | 0.413 | 1.482 | -0.229 | -0.421 | 0.022 | 0.042 | 0.042 | 0 | 0.12 | 0.03 | 0.1 | 0.02 | 0.1 | 0.09 |
| 6 | 160443861 | rs9457811 | 0.265 | 1.547 | -0.176 | -0.384 | -0.001 | -0.001 | -0.001 | 0 | 0.16 | 0.14 | 0.06 | 0.21 | 0.15 | 0.27 |
| 6 | 160447988 | rs6455680 | -1.396 | 1.153 | -0.438 | -1.245 | -0.006 | -0.034 | -0.034 | 0 | 0.65 | 0.6 | 0.73 | 0.83 | 0.65 | 0.69 |
| 6 | 160448462 | rs2297356 | 0.609 | 1.004 | -0.563 | -0.773 | -0.035 | -0.05 | -0.05 | 0 | 0.14 | 0.02 | 0.05 | 0.22 | 0.15 | 0.2 |
| 6 | 160453561 | rs4709390 | -0.159 | 0.882 | -0.6 | -0.861 | 0.02 | -0.055 | -0.055 | 0 | 0.21 | 0.1 | 0.3 | 0.53 | 0.2 | 0.26 |
| 6 | 160455254 | rs2297357 | -0.107 | 0.88 | -0.607 | -0.862 | 0.019 | -0.057 | -0.057 | 0.208 | 0.21 | 0.09 | 0.29 | 0.53 | 0.19 | 0.26 |
| 6 | 160456163 | rs1867348 | 0.671 | 1.04 | -0.548 | -0.69 | -0.018 | -0.017 | -0.017 | 0.058 | 0.08 | 0.01 | 0.19 | 0.51 | 0.1 | 0.17 |
| 6 | 160461085 | rs3777414 | 0.065 | 1.152 | -0.239 | -0.926 | 0.005 | 0.028 | 0.028 | 0 | 0.36 | 0.25 | 0.35 | 0.75 | 0.35 | 0.47 |
| 6 | 160461334 | rs8191772 | -0.05 | 1.231 | -0.402 | -0.886 | -0.003 | -0.016 | -0.016 | 0 | 0.38 | 0.37 | 0.35 | 0.75 | 0.35 | 0.47 |
| 6 | 160465054 | rs687088 | -0.195 | 1.139 | -0.755 | -0.533 | 0 | -0.002 | -0.002 | 0 | 0.72 | 0.77 | 0.79 | 0.67 | 0.66 | 0.72 |
| 6 | 160465291 | rs688359 | -0.224 | 1.177 | -0.756 | -0.623 | -0.002 | -0.005 | -0.005 | 0 | 0.61 | 0.41 | 0.54 | 0.34 | 0.62 | 0.57 |
| 6 | 160466103 | rs8191794 | 0.233 | 1.286 | -1.007 | -0.562 | -0.014 | 0.037 | 0.037 | 0 | 0.07 | 0 | 0.04 | 0.01 | 0.07 | 0.14 |
| 6 | 160467890 | rs416572 | 0.066 | 2.476 | -1.351 | -0.635 | -0.111 | -0.017 | -0.017 | 0.127 | 0.85 | 0.99 | 0.84 | 0.85 | 0.71 | 0.84 |
| 6 | 160468278 | rs998075 | 0.79 | 1.46 | -0.55 | -0.617 | -0.001 | 0.032 | 0.032 | 0 | 0.55 | 0.64 | 0.49 | 0.69 | 0.48 | 0.53 |
| 6 | 160469149 | rs648253 | -1.123 | 1.622 | 0.104 | 0.171 | 0.003 | 0.027 | 0.027 | 0 | 0.5 | 0.63 | 0.64 | 0.64 | 0.48 | 0.58 |
| 6 | 160472032 | rs4709393 | 1.592 | 1.341 | -0.156 | -0.374 | 0.007 | -0.002 | -0.002 | 0.588 | 0.36 | 0.26 | 0.2 | 0.37 | 0.29 | 0.31 |
| 6 | 160476267 | rs384167 | -0.705 | 1.688 | -0.368 | -0.499 | -0.042 | -0.037 | -0.037 | 0.943 | 0.93 | 0.99 | 0.72 | 0.84 | 0.81 | 0.84 |
| 6 | 160477885 | rs8191824 | 1.055 | 1.249 | 0.108 | -0.641 | -0.012 | -0.043 | -0.043 | 0.725 | 0.17 | 0.16 | 0.08 | 0.01 | 0.08 | 0.18 |
| 6 | 160478789 | rs4709395 | 1.332 | 1.316 | 0.149 | -0.512 | 0.046 | 0.017 | 0.017 | 0.206 | 0.34 | 0.13 | 0.18 | 0.27 | 0.28 | 0.23 |
| 6 | 160479478 | rs8191829 | 0.398 | 1.391 | 0.386 | -0.484 | 0.012 | 0.04 | 0.04 | 0.792 | 0.19 | 0.33 | 0.29 | 0.34 | 0.21 | 0.21 |
| 6 | 160479882 | rs2274849 | 0.443 | 1.438 | 0.385 | -0.485 | 0.011 | 0.042 | 0.042 | 0.821 | 0.18 | 0.33 | 0.29 | 0.34 | 0.2 | 0.21 |
| 6 | 160481537 | rs7746102 | 0.184 | 1.439 | 0.396 | -0.483 | 0.009 | 0.038 | 0.038 | 0.495 | 0.2 | 0.36 | 0.29 | 0.34 | 0.21 | 0.21 |
| 6 | 160490314 | rs920812 | 0.861 | 1.223 | 0.482 | -0.749 | 0.018 | 0.012 | 0.012 | 0 | 0.43 | 0.31 | 0.25 | 0.35 | 0.33 | 0.36 |
| 6 | 160491741 | rs3798180 | 0.282 | 1.21 | 0.573 | -0.741 | 0.023 | 0.04 | 0.04 | 0 | 0.5 | 0.61 | 0.44 | 0.69 | 0.38 | 0.48 |
| 6 | 160494521 | rs2282140 | 1.854 | 1.082 | 0.555 | -0.707 | 0.031 | -0.002 | -0.002 | 0.504 | 0.27 | 0.14 | 0.12 | 0.26 | 0.16 | 0.18 |
| 6 | 160495055 | rs8191890 | 1.408 | 1.197 | 0.756 | -0.689 | 0.001 | 0.013 | 0.013 | 0 | 0.07 | 0.08 | 0.06 | 0.14 | 0.13 | 0.06 |
| 6 | 160496055 | rs3777406 | 1.006 | 1.208 | 0.765 | -0.554 | 0.002 | -0.009 | -0.009 | 3.752 | 0.16 | 0.2 | 0.14 | 0.1 | 0.17 | 0.19 |
| 6 | 160503656 | rs9457827 | -1.139 | 1.005 | 1.092 | -0.269 | 0.024 | 0.399 | 0.399 | 0.153 | 0.07 | 0.31 | 0.2 | 0.43 | 0.05 | 0.19 |
| 6 | 160506755 | rs13220323 | 1.343 | 1.001 | 1.001 | -0.295 | 0.045 | -0.004 | -0.004 | 0 | 0.28 | 0.12 | 0.12 | 0.26 | 0.16 | 0.18 |
| 6 | 160508172 | rs3798176 | -0.1 | 1.029 | 0.998 | -0.32 | 0.02 | 0.036 | 0.036 | 0 | 0.49 | 0.6 | 0.44 | 0.69 | 0.37 | 0.47 |
| 6 | 160511354 | rs1130861 | 1.331 | 1.019 | 0.974 | -0.314 | 0.027 | 0 | 0 | 0 | 0.28 | 0.15 | 0.12 | 0.26 | 0.16 | 0.18 |
| 6 | 160514541 | rs1888727 | -0.058 | 1.13 | 0.976 | -0.276 | 0.025 | 0.044 | 0.044 | 0 | 0.49 | 0.61 | 0.44 | 0.7 | 0.37 | 0.48 |
| 6 | 160517481 | rs1803989 | -1.002 | 1.698 | 1.453 | 0.38 | 0.028 | 0.383 | 0.383 | 0.116 | 0.06 | 0.26 | 0.2 | 0.42 | 0.05 | 0.19 |
| 6 | 160523709 | rs2297367 | 0.756 | 0.983 | 1.183 | -0.226 | -0.012 | -0.01 | -0.01 | 0.763 | 0.06 | 0.1 | 0.07 | 0.04 | 0.05 | 0.09 |
| 6 | 160527180 | rs1050015 | 0.863 | 1.095 | 1.013 | -0.396 | -0.021 | 0.031 | 0.031 | 0.075 | 0.13 | 0.08 | 0.11 | 0 | 0.16 | 0.12 |

| | | | | | | | | | | | | | | | | |
|---|---|---|---|---|---|---|---|---|---|---|---|---|---|---|---|---|
| 6 | 160528057 | rs7753051 | -0.256 | 1.019 | 1.391 | -0.202 | 0.017 | 0.167 | 0.167 | 0.903 | 0.19 | 0.46 | 0.32 | 0.51 | 0.21 | 0.29 |
| 6 | 160530626 | rs1479355 | -0.251 | 1.023 | 1.372 | -0.204 | 0.019 | 0.164 | 0.164 | 0.84 | 0.19 | 0.45 | 0.31 | 0.51 | 0.21 | 0.29 |
| 6 | 160538921 | rs456598 | 1.045 | 0.782 | 1.128 | -0.326 | -0.019 | 0.051 | 0.051 | 0.017 | 0.13 | 0.04 | 0.25 | 0 | 0.14 | 0.12 |
| 6 | 160539046 | rs4709399 | -0.717 | 0.871 | 1.38 | -0.114 | -0.009 | 0.086 | 0.086 | 1.876 | 0.26 | 0.55 | 0.39 | 0.56 | 0.25 | 0.37 |
| 6 | 160543123 | rs1867351 | 0.083 | 0.924 | 1.219 | -0.03 | 0.008 | 0.072 | 0.072 | 0.332 | 0.17 | 0.33 | 0.29 | 0.39 | 0.18 | 0.2 |
| 6 | 160543562 | rs461473 | 1.611 | 0.986 | 0.633 | -0.204 | 0.006 | 0.038 | 0.038 | 0.29 | 0.13 | 0.03 | 0.07 | 0 | 0.09 | 0.01 |
| 6 | 160548433 | rs3822841 | 0.029 | 1.81 | 1.213 | 0.695 | -0.016 | -0.049 | -0.049 | 0 | 0.16 | 0.12 | 0.28 | 0.39 | 0.18 | 0.2 |
| 6 | 160552803 | rs3798174 | 0.339 | 1.882 | 1.25 | 0.633 | 0.001 | -0.016 | -0.016 | 4.663 | 0.07 | 0.09 | 0.09 | 0.04 | 0.08 | 0.09 |
| 6 | 160553007 | rs4646275 | -0.29 | 1.611 | 1.235 | 0.64 | -0.009 | 0.183 | 0.183 | 0 | 0.09 | 0.15 | 0.25 | 0.41 | 0.08 | 0.17 |
| 6 | 160556022 | rs3798173 | 0.048 | 1.767 | 0.757 | 0.726 | -0.018 | -0.048 | -0.048 | 0 | 0.16 | 0.12 | 0.28 | 0.38 | 0.18 | 0.2 |
| 6 | 160560845 | rs628031 | 0.095 | 1.614 | 0.488 | 0.456 | -0.015 | 0.007 | 0.007 | 0.376 | 0.72 | 0.73 | 0.78 | 0.74 | 0.59 | 0.61 |
| 6 | 160560933 | rs9457843 | 0.545 | 1.171 | 0.226 | 0.653 | 0 | -0.015 | -0.015 | 0 | 0.11 | 0.16 | 0.08 | 0 | 0.11 | 0.08 |
| 6 | 160563520 | rs7766568 | 0.138 | 0.957 | 0.34 | 0.25 | -0.012 | -0.018 | -0.018 | 0 | 0.07 | 0.11 | 0.08 | 0.04 | 0.06 | 0.06 |
| 6 | 160567928 | rs3798167 | 1.06 | 0.921 | 0.274 | 0.096 | 0.048 | -0.004 | -0.004 | 0 | 0.35 | 0.14 | 0.15 | 0.31 | 0.2 | 0.21 |
| 6 | 160568896 | rs654993 | -1.001 | 1.651 | 0.974 | 0.23 | -0.003 | -0.003 | -0.003 | 0.002 | 0.55 | 0.66 | 0.71 | 0.52 | 0.55 | 0.63 |
| 6 | 160570172 | rs9295124 | 0.798 | 1.518 | 1.02 | 0.439 | 0 | 0.005 | 0.005 | 0 | 0.56 | 0.59 | 0.29 | 0.34 | 0.4 | 0.36 |
| 6 | 160572866 | rs622342 | 0.51 | 1.93 | 0.99 | 0.542 | -0.008 | 0.005 | 0.005 | 0.592 | 0.81 | 0.82 | 0.6 | 0.85 | 0.62 | 0.75 |
| 6 | 160574535 | rs650284 | 0.48 | 1.973 | 1.014 | 0.555 | -0.004 | 0.003 | 0.003 | 0.485 | 0.81 | 0.82 | 0.6 | 0.85 | 0.62 | 0.75 |
| 6 | 160574802 | rs1443844 | 1.051 | 1.894 | 1.022 | 0.333 | 0.001 | 0.003 | 0.003 | 0 | 0.53 | 0.55 | 0.29 | 0.35 | 0.4 | 0.36 |
| 6 | 160575202 | rs3798164 | 1.055 | 1.942 | 0.936 | 0.097 | 0.002 | 0.005 | 0.005 | 0 | 0.53 | 0.53 | 0.28 | 0.35 | 0.39 | 0.36 |
| 6 | 160575366 | rs11753995 | 0.342 | 2.046 | 0.911 | 0.068 | -0.048 | 0.078 | 0.078 | 0 | 0.13 | 0.01 | 0.26 | 0.01 | 0.16 | 0.12 |
| 6 | 160575985 | rs2297374 | 1.045 | 2.079 | 0.948 | 0.112 | 0.004 | 0.006 | 0.006 | 0 | 0.52 | 0.53 | 0.27 | 0.34 | 0.38 | 0.36 |
| 6 | 160578271 | rs9295125 | 1.025 | 2.308 | 0.139 | 0.375 | -0.003 | -0.006 | -0.006 | 2.087 | 0.53 | 0.51 | 0.28 | 0.35 | 0.39 | 0.36 |
| 6 | 160578860 | rs1564348 | -0.014 | 0.614 | -0.034 | 0.29 | -0.002 | -0.02 | -0.02 | 0 | 0.14 | 0.11 | 0.27 | 0 | 0.16 | 0.12 |
| 6 | 160578914 | rs609468 | -0.805 | 0.289 | 0.217 | 1.094 | -0.019 | 0.035 | 0.035 | 0 | 0.28 | 0.36 | 0.32 | 0.52 | 0.22 | 0.39 |
| 6 | 160579527 | rs622591 | -0.732 | 0.364 | -0.166 | 1.133 | -0.002 | 0.004 | 0.004 | 0 | 0.28 | 0.29 | 0.32 | 0.52 | 0.22 | 0.39 |
| 6 | 160581502 | rs4646283 | -0.807 | 1.325 | -0.459 | 0.71 | -0.011 | 0.01 | 0.01 | 0.18 | 0.09 | 0.06 | 0.1 | 0.04 | 0.14 | 0.15 |
| 6 | 160581911 | rs6455682 | -1.08 | 1.282 | -0.183 | 1.979 | 0.002 | 0.161 | 0.161 | 0.002 | 0.08 | 0.12 | 0.09 | 0.37 | 0.09 | 0.12 |
| 6 | 160582340 | rs9456505 | -0.855 | 0.855 | 0.155 | 1.777 | -0.007 | 0.012 | 0.012 | 0 | 0.1 | 0.13 | 0.27 | 0.22 | 0.08 | 0.25 |
| 6 | 160583048 | rs2083867 | 0.747 | 1.011 | -0.577 | 1.039 | 0.003 | 0.023 | 0.023 | 0 | 0.58 | 0.71 | 0.46 | 0.62 | 0.62 | 0.44 |
| 6 | 160583904 | rs7744238 | 0.744 | 1.064 | -0.776 | 0.819 | 0.004 | 0.028 | 0.028 | 0 | 0.58 | 0.74 | 0.47 | 0.62 | 0.61 | 0.45 |
| 6 | 160584853 | rs3101826 | 1.348 | 1.13 | -0.162 | -0.148 | 0 | -0.064 | -0.064 | 0.598 | 0.35 | 0.31 | 0.21 | 0 | 0.35 | 0.1 |
| 6 | 160587521 | rs1871389 | 0.459 | 0.995 | -0.081 | -0.096 | 0.002 | 0.005 | 0.005 | 0 | 0.56 | 0.4 | 0.44 | 0.62 | 0.61 | 0.45 |
| 6 | 160589491 | rs7773429 | 0.14 | 1.134 | -0.365 | -0.076 | -0.056 | -0.098 | -0.098 | 2.048 | 0.18 | 0.04 | 0.2 | 0.61 | 0.23 | 0.34 |
| 6 | 160598486 | rs791195 | -0.342 | 0.899 | -0.211 | -0.111 | -0.016 | 0.055 | 0.055 | 0 | 0.68 | 0.88 | 0.73 | 0.85 | 0.69 | 0.7 |
| 6 | 160610124 | rs791183 | 0.579 | 0.932 | -0.058 | 0.203 | 0.007 | 0.004 | 0.004 | 0 | 0.55 | 0.38 | 0.43 | 0.52 | 0.61 | 0.45 |
| 6 | 160622637 | rs476235 | 1.421 | 0.838 | -0.193 | -0.372 | 0 | -0.066 | -0.066 | 0 | 0.35 | 0.33 | 0.21 | 0.01 | 0.35 | 0.12 |
| 6 | 160622708 | rs476064 | 0.016 | 0.681 | 0.063 | 0.223 | -0.019 | 0.075 | 0.075 | 0 | 0.72 | 0.9 | 0.75 | 0.89 | 0.72 | 0.7 |
| 6 | 160625978 | rs2619268 | 1.151 | 0.311 | 0.457 | 0.631 | -0.024 | 0.02 | 0.02 | 0 | 0.66 | 0.39 | 0.59 | 0.49 | 0.76 | 0.51 |
| 6 | 160635886 | rs10945656 | -0.896 | 1.332 | 0.676 | 0.185 | 0.023 | -0.012 | -0.012 | 0.544 | 0.07 | 0.19 | 0.07 | 0.04 | 0.12 | 0.09 |
| 6 | 160637844 | rs3127592 | -0.876 | 1.269 | 1.03 | 0.287 | 0.026 | -0.011 | -0.011 | 1.384 | 0.07 | 0.19 | 0.07 | 0.04 | 0.12 | 0.09 |
| 6 | 160637890 | rs694812 | 0.538 | 1.297 | 1.298 | 0.135 | 0.012 | 0.004 | 0.004 | 0 | 0.88 | 0.7 | 0.91 | 0.87 | 0.89 | 0.85 |
| 6 | 160638076 | rs3103353 | -0.775 | 1.301 | 1.278 | 0.15 | 0.026 | -0.008 | -0.008 | 2.692 | 0.07 | 0.19 | 0.07 | 0.05 | 0.12 | 0.09 |
| 6 | 160638725 | rs3103352 | -0.742 | 1.301 | 1.28 | 0.135 | 0.025 | -0.011 | -0.011 | 0.495 | 0.07 | 0.19 | 0.07 | 0.04 | 0.12 | 0.09 |
| 6 | 160638745 | rs2450974 | 0.773 | 1.282 | 1.36 | 0.128 | 0.011 | 0.002 | 0.002 | 0 | 0.87 | 0.67 | 0.91 | 0.87 | 0.89 | 0.85 |
| 6 | 160640494 | rs10755577 | -1.576 | 1.278 | 1.341 | 0.131 | 0.033 | 0.048 | 0.048 | 0 | 0.1 | 0.32 | 0.09 | 0.14 | 0.12 | 0.09 |
| 6 | 160642528 | rs316000 | 1.282 | 1.284 | 1.332 | 0.121 | 0.034 | -0.011 | -0.011 | 0 | 0.82 | 0.58 | 0.86 | 0.83 | 0.77 | 0.77 |
| 6 | 160645832 | rs316003 | 1.523 | 1.438 | 1.348 | 0.131 | 0.031 | 0.009 | 0.009 | 0.279 | 0.8 | 0.42 | 0.84 | 0.8 | 0.77 | 0.77 |
| 6 | 160646173 | rs316004 | 1.386 | 1.436 | 1.404 | 0.13 | 0.044 | 0.019 | 0.019 | 0.064 | 0.81 | 0.48 | 0.84 | 0.8 | 0.77 | 0.77 |
| 6 | 160646280 | rs316005 | 1.519 | 1.458 | 1.392 | 0.13 | 0.028 | 0.009 | 0.009 | 0.279 | 0.8 | 0.43 | 0.84 | 0.8 | 0.77 | 0.77 |
| 6 | 160646612 | rs479078 | 1.492 | 1.419 | 1.399 | 0.117 | 0.013 | -0.003 | -0.003 | 1.929 | 0.79 | 0.36 | 0.84 | 0.8 | 0.77 | 0.78 |
| 6 | 160652858 | rs315978 | -0.028 | 1.444 | 1.436 | 0.114 | 0.014 | -0.037 | -0.037 | 0.675 | 0.9 | 0.77 | 0.92 | 0.96 | 0.88 | 0.85 |
| 6 | 160659723 | rs315984 | 1.561 | 1.43 | 1.453 | 0.135 | 0.023 | 0.006 | 0.006 | 0.228 | 0.8 | 0.4 | 0.83 | 0.8 | 0.77 | 0.77 |
| 6 | 160660694 | rs315988 | 1.561 | 1.43 | 1.453 | 0.135 | 0.023 | 0.006 | 0.006 | 0.228 | 0.8 | 0.4 | 0.83 | 0.8 | 0.77 | 0.77 |
| 6 | 160662143 | rs315990 | 1.395 | 1.43 | 1.449 | 0.135 | 0.018 | 0.003 | 0.003 | 0.422 | 0.8 | 0.37 | 0.83 | 0.8 | 0.77 | 0.77 |
| 6 | 160665461 | rs315995 | 1.561 | 1.441 | 1.33 | 0.141 | 0.023 | 0.006 | 0.006 | 0.228 | 0.8 | 0.4 | 0.83 | 0.8 | 0.77 | 0.77 |
| 6 | 160668041 | rs316021 | 1.532 | 1.326 | 1.186 | 0.478 | 0.039 | 0.054 | 0.054 | 0 | 0.8 | 0.44 | 0.83 | 0.76 | 0.76 | 0.78 |
| 6 | 160668389 | rs2279463 | -1.036 | 1.33 | 1.241 | 0.385 | 0.012 | 0 | 0 | 0 | 0.09 | 0.19 | 0.07 | 0.08 | 0.13 | 0.08 |
| 6 | 160669081 | rs316020 | -0.008 | 1.334 | 1.126 | 0.393 | 0.009 | -0.026 | -0.026 | 0 | 0.91 | 0.82 | 0.91 | 0.96 | 0.89 | 0.87 |
| 6 | 160669718 | rs3912161 | -0.257 | 1.396 | 0.889 | 0.536 | -0.025 | -0.024 | -0.024 | 0.088 | 0.08 | 0 | 0.1 | 0.11 | 0.07 | 0.13 |
| 6 | 160670282 | rs316019 | 0.078 | 1.35 | 1.13 | 0.383 | 0.008 | 0.019 | 0.019 | 0 | 0.91 | 0.81 | 0.91 | 0.86 | 0.89 | 0.87 |
| 6 | 160670494 | rs316018 | 1.58 | 1.375 | 1.078 | 0.393 | 0.025 | 0.026 | 0.026 | 0 | 0.8 | 0.53 | 0.83 | 0.78 | 0.76 | 0.79 |
| 6 | 160671500 | rs316016 | 1.457 | 1.381 | 1.007 | 0.396 | 0.022 | 0.023 | 0.023 | 0.033 | 0.79 | 0.5 | 0.83 | 0.78 | 0.76 | 0.79 |
| 6 | 160672264 | rs316015 | 1.486 | 1.378 | 1.01 | 0.394 | 0.025 | 0.022 | 0.022 | 0.063 | 0.79 | 0.44 | 0.83 | 0.78 | 0.76 | 0.79 |
| 6 | 160672625 | rs316013 | 1.457 | 1.379 | 1.009 | 0.393 | 0.026 | 0.023 | 0.023 | 0.033 | 0.79 | 0.5 | 0.83 | 0.78 | 0.76 | 0.79 |
| 6 | 160673884 | rs316012 | 1.514 | 1.378 | 0.984 | 0.393 | 0.021 | 0.019 | 0.019 | 0.057 | 0.79 | 0.41 | 0.83 | 0.78 | 0.76 | 0.79 |
| 6 | 160675764 | rs316009 | -0.042 | 1.364 | 0.91 | 0.262 | -0.001 | -0.002 | -0.002 | 0 | 0.92 | 0.9 | 0.93 | 0.96 | 0.89 | 0.87 |
| 6 | 160676408 | rs316007 | 1.076 | 1.339 | 0.774 | 0.292 | 0.028 | -0.054 | -0.054 | 0 | 0.79 | 0.48 | 0.83 | 0.87 | 0.76 | 0.8 |
| 6 | 160678493 | rs638360 | 1.085 | 1.315 | 0.768 | 0.292 | 0.024 | -0.055 | -0.055 | 0 | 0.79 | 0.48 | 0.84 | 0.87 | 0.76 | 0.79 |

| | | | | | | | | | | | | | | | | |
|---|---|---|---|---|---|---|---|---|---|---|---|---|---|---|---|---|
| 6 | 160679400 | rs624249 | 1.252 | 1.161 | -0.067 | 0.076 | 0.029 | 0.093 | 0.093 | 0.176 | 0.45 | 0.27 | 0.27 | 0.15 | 0.39 | 0.25 |
| 6 | 160681393 | rs3127573 | -0.169 | 1.391 | 0.372 | 0.373 | 0.018 | 0.013 | 0.013 | 0 | 0.06 | 0.14 | 0.06 | 0.1 | 0.12 | 0.07 |
| 6 | 160682218 | rs3127572 | -0.667 | 1.129 | 0.722 | 0.256 | 0.004 | -0.002 | -0.002 | 0.217 | 0.11 | 0.23 | 0.07 | 0.1 | 0.13 | 0.08 |
| 6 | 160682237 | rs316024 | -0.105 | 1.212 | 0.583 | 0.263 | 0 | -0.024 | -0.024 | 0 | 0.31 | 0.23 | 0.54 | 0.48 | 0.32 | 0.36 |
| 6 | 160682897 | rs7757997 | 0.097 | 1.168 | 0.708 | 0.198 | -0.014 | 0.087 | 0.087 | 0.024 | 0.06 | 0.19 | 0.06 | 0.18 | 0.04 | 0.18 |
| 6 | 160683269 | rs492453 | 1.892 | 1.409 | 0.758 | -0.048 | 0.012 | 0.023 | 0.023 | 0 | 0.35 | 0.24 | 0.21 | 0.07 | 0.29 | 0.13 |
| 6 | 160683381 | rs316025 | 0.411 | 1.61 | 0.737 | -0.052 | -0.01 | 0.008 | 0.008 | 0 | 0.78 | 0.82 | 0.84 | 0.83 | 0.74 | 0.75 |
| 6 | 160684370 | rs316026 | -0.404 | 1.617 | 0.619 | -0.019 | -0.001 | -0.001 | -0.001 | 0 | 0.38 | 0.38 | 0.59 | 0.49 | 0.4 | 0.45 |
| 6 | 160686668 | rs316029 | -0.983 | 1.698 | 0.571 | 0.019 | -0.012 | 0.005 | 0.005 | 0 | 0.89 | 0.92 | 0.89 | 0.92 | 0.86 | 0.83 |
| 6 | 160687866 | rs316030 | -0.044 | 1.708 | 0.609 | -0.012 | -0.016 | -0.034 | -0.034 | 3.259 | 0.35 | 0.23 | 0.55 | 0.49 | 0.38 | 0.45 |
| 6 | 160689558 | rs7757336 | -0.827 | 1.697 | 0.671 | -0.018 | 0.021 | 0.016 | 0.016 | 0.238 | 0.16 | 0.41 | 0.12 | 0.18 | 0.16 | 0.08 |
| 6 | 160689730 | rs316033 | -0.085 | 1.672 | 0.446 | 0.033 | -0.009 | -0.02 | -0.02 | 0 | 0.35 | 0.27 | 0.56 | 0.57 | 0.38 | 0.45 |
| 6 | 160692694 | rs492315 | 1.686 | 1.309 | 0.482 | 0.108 | -0.001 | -0.007 | -0.007 | 2.584 | 0.37 | 0.33 | 0.21 | 0.16 | 0.29 | 0.12 |
| 6 | 160692719 | rs614564 | 1.403 | 1.298 | 0.49 | 0.139 | 0.001 | 0 | 0 | 0 | 0.41 | 0.36 | 0.25 | 0.34 | 0.33 | 0.3 |
| 6 | 160693107 | rs3119311 | 0.315 | 1.328 | 0.54 | 0.111 | 0.008 | 0.023 | 0.023 | 0 | 0.1 | 0.2 | 0.07 | 0.18 | 0.13 | 0.08 |
| 6 | 160694081 | rs598135 | 0.79 | 1.38 | 0.345 | 0.204 | 0.009 | 0.005 | 0.005 | 0 | 0.5 | 0.54 | 0.32 | 0.44 | 0.46 | 0.38 |
| 6 | 160697628 | rs532440 | -0.318 | 1.593 | -0.025 | -0.077 | -0.002 | -0.004 | -0.004 | 0 | 0.3 | 0.27 | 0.44 | 0.67 | 0.3 | 0.55 |
| 6 | 160697762 | rs533452 | 0.196 | 1.848 | -0.315 | -0.215 | -0.017 | 0.028 | 0.028 | 0 | 0.72 | 0.77 | 0.68 | 0.92 | 0.65 | 0.74 |
| 6 | 160699534 | rs2619276 | 0.275 | 1.792 | -0.249 | 0.009 | -0.003 | -0.001 | -0.001 | 0 | 0.2 | 0.18 | 0.24 | 0.19 | 0.25 | 0.28 |
| 6 | 160699698 | rs578560 | 0.133 | 2.116 | -0.436 | 0.111 | -0.068 | -0.03 | -0.03 | 2.284 | 0.1 | 0.01 | 0.16 | 0.1 | 0.17 | 0.13 |
| 6 | 160700301 | rs3119312 | 1.577 | 1.36 | -0.032 | -0.002 | -0.003 | -0.006 | -0.006 | 0 | 0.36 | 0.33 | 0.21 | 0.2 | 0.32 | 0.15 |
| 6 | 160700441 | rs474617 | 0.329 | 1.17 | 0.071 | 0.226 | -0.003 | -0.002 | -0.002 | 0 | 0.61 | 0.56 | 0.5 | 0.33 | 0.66 | 0.45 |
| 6 | 160705154 | rs568567 | -0.874 | 1.069 | 0.096 | 0.261 | 0.007 | -0.009 | -0.009 | 0 | 0.79 | 0.81 | 0.7 | 0.55 | 0.85 | 0.71 |
| 6 | 160706661 | rs1685986 | 0.898 | 1.371 | -0.076 | -0.069 | -0.002 | 0.054 | 0.054 | 0 | 0.4 | 0.27 | 0.25 | 0.15 | 0.41 | 0.17 |
| 6 | 160714425 | rs368239 | 1.732 | 1.328 | -0.32 | -0.239 | 0.029 | 0.129 | 0.129 | 0 | 0.31 | 0.07 | 0.17 | 0.07 | 0.26 | 0.1 |
| 6 | 160716361 | rs415317 | 0.505 | 1.422 | -0.131 | -0.247 | -0.004 | 0.076 | 0.076 | 0 | 0.52 | 0.31 | 0.43 | 0.24 | 0.51 | 0.39 |
| 6 | 160719593 | rs368123 | -0.146 | 0.933 | -0.37 | -0.184 | -0.007 | 0.004 | 0.004 | 0 | 0.61 | 0.55 | 0.51 | 0.4 | 0.66 | 0.46 |
| 6 | 160720997 | rs448033 | 1.504 | 0.85 | -0.507 | -0.26 | 0.038 | 0.113 | 0.113 | 0 | 0.32 | 0.11 | 0.18 | 0.07 | 0.26 | 0.1 |
| 6 | 160741106 | rs446291 | 1.639 | 0.895 | -0.642 | -0.219 | 0.027 | 0.119 | 0.119 | 0 | 0.3 | 0.07 | 0.17 | 0.08 | 0.26 | 0.1 |
| 6 | 160742239 | rs393271 | 0.187 | 0.84 | -0.204 | 0.206 | -0.007 | 0.006 | 0.006 | 0 | 0.61 | 0.55 | 0.51 | 0.43 | 0.66 | 0.45 |
| 6 | 160756037 | rs316170 | 0.064 | 0.64 | -0.313 | 0.135 | 0.015 | -0.009 | -0.009 | 0 | 0.59 | 0.65 | 0.51 | 0.43 | 0.65 | 0.46 |
| 6 | 160756729 | rs2481031 | -0.265 | 0.964 | -0.111 | -0.118 | 0.076 | -0.035 | -0.035 | 0 | 0.25 | 0.39 | 0.3 | 0.15 | 0.35 | 0.15 |
| 6 | 160760078 | rs316174 | 0.215 | 1.414 | -0.038 | -0.23 | -0.002 | 0.021 | 0.021 | 0 | 0.53 | 0.57 | 0.46 | 0.36 | 0.51 | 0.39 |
| 6 | 160765029 | rs572149 | 1.056 | 1.239 | 0.351 | -0.188 | -0.021 | -0.002 | -0.002 | 0 | 0.37 | 0.43 | 0.24 | 0.36 | 0.3 | 0.31 |
| 6 | 160765589 | rs637575 | 1.055 | 1.235 | 0.349 | -0.158 | -0.021 | -0.001 | -0.001 | 0.018 | 0.37 | 0.43 | 0.24 | 0.36 | 0.3 | 0.31 |
| 6 | 160767183 | rs569919 | 1.576 | 1.204 | 0.125 | -0.305 | 0.028 | -0.001 | -0.001 | 0 | 0.31 | 0.12 | 0.19 | 0.28 | 0.27 | 0.3 |
| 6 | 160770312 | rs474513 | 0.29 | 1.08 | 0.268 | -0.561 | -0.001 | 0.034 | 0.034 | 0 | 0.49 | 0.57 | 0.45 | 0.29 | 0.48 | 0.36 |
| 6 | 160770360 | rs539298 | 0.29 | 1.08 | 0.268 | -0.561 | -0.001 | 0.034 | 0.034 | 0 | 0.49 | 0.57 | 0.45 | 0.29 | 0.48 | 0.36 |
| 6 | 160770918 | rs501470 | 0.29 | 1.07 | 0.263 | -0.347 | -0.001 | 0.034 | 0.034 | 0 | 0.49 | 0.57 | 0.45 | 0.29 | 0.48 | 0.36 |
| 6 | 160771618 | rs2484241 | 1.103 | 0.812 | 0.211 | -0.433 | -0.062 | 0.062 | 0.062 | 0 | 0.11 | 0.02 | 0.2 | 0 | 0.21 | 0.08 |
| 6 | 160774486 | rs487152 | 0.26 | 0.78 | 0.248 | -0.386 | -0.001 | 0.035 | 0.035 | 0 | 0.49 | 0.57 | 0.45 | 0.29 | 0.48 | 0.36 |
| 6 | 160774928 | rs512077 | -0.414 | 0.245 | -0.124 | -0.347 | -0.032 | -0.021 | -0.021 | 0.197 | 0.72 | 0.65 | 0.68 | 0.83 | 0.84 | 0.69 |
| 6 | 160775652 | rs519118 | 0.405 | 0.482 | 0.093 | -0.153 | 0 | 0.031 | 0.031 | 0 | 0.49 | 0.56 | 0.45 | 0.29 | 0.48 | 0.36 |
| 6 | 160776325 | rs675162 | 0.328 | 0.476 | 0.314 | -0.102 | 0.002 | 0.035 | 0.035 | 0 | 0.51 | 0.63 | 0.46 | 0.37 | 0.49 | 0.36 |
| 6 | 160778639 | rs394487 | 1.081 | 0.452 | 0.404 | -0.108 | -0.018 | -0.012 | -0.012 | 0 | 0.37 | 0.55 | 0.26 | 0.3 | 0.27 | 0.26 |
| 6 | 160779134 | rs641990 | 0.257 | 0.461 | 0.377 | -0.149 | -0.003 | 0.027 | 0.027 | 0 | 0.51 | 0.62 | 0.47 | 0.38 | 0.5 | 0.35 |
| 6 | 160780912 | rs394468 | 1.789 | 0.411 | 0.426 | -0.175 | 0.012 | 0.004 | 0.004 | 0 | 0.31 | 0.28 | 0.22 | 0.3 | 0.27 | 0.26 |
| 6 | 160781666 | rs2504934 | 0.564 | 0.505 | 0.304 | -0.158 | -0.022 | 0.012 | 0.012 | 0 | 0.14 | 0.1 | 0.21 | 0 | 0.23 | 0.08 |
| 6 | 160781733 | rs377551 | 1.174 | 0.456 | 0.344 | -0.143 | -0.022 | -0.014 | -0.014 | 0 | 0.37 | 0.54 | 0.26 | 0.3 | 0.28 | 0.26 |
| 6 | 160784500 | rs2665354 | -0.446 | 0.851 | 0.137 | -0.139 | -0.024 | 0.018 | 0.018 | 0 | 0.86 | 0.91 | 0.78 | 0.92 | 0.77 | 0.92 |
| 6 | 160784836 | rs2665353 | -0.446 | 0.851 | 0.137 | -0.139 | -0.024 | 0.018 | 0.018 | 0 | 0.86 | 0.91 | 0.78 | 0.92 | 0.77 | 0.92 |
| 6 | 160785976 | rs2457556 | -0.446 | 0.851 | 0.137 | -0.163 | -0.024 | 0.014 | 0.014 | 0 | 0.86 | 0.91 | 0.79 | 1 | 0.77 | 0.92 |
| 6 | 160786342 | rs437865 | -0.446 | 0.851 | 0.137 | -0.139 | -0.024 | 0.018 | 0.018 | 0 | 0.86 | 0.91 | 0.78 | 0.92 | 0.77 | 0.92 |
| 6 | 160789296 | rs402219 | 1.613 | 0.318 | 0.099 | 0.124 | 0.031 | -0.022 | -0.022 | 0 | 0.34 | 0.18 | 0.42 | 0.38 | 0.28 | 0.41 |
| 6 | 160796058 | rs1018234 | 1.25 | 0.194 | 0.722 | 0.058 | -0.025 | 0 | 0 | 0 | 0.38 | 0.45 | 0.26 | 0.36 | 0.28 | 0.41 |
| 6 | 160796565 | rs316244 | 1.706 | 0.194 | 0.72 | 0.058 | 0.029 | -0.016 | -0.016 | 0.171 | 0.34 | 0.17 | 0.22 | 0.36 | 0.27 | 0.41 |
| 6 | 160797625 | rs6905958 | -0.326 | 0.217 | 0.793 | 0.001 | -0.002 | 0.001 | 0.001 | 0 | 0.46 | 0.38 | 0.53 | 0.57 | 0.5 | 0.51 |
| 6 | 160802239 | rs3123634 | -0.29 | 0.196 | 0.783 | 0.025 | -0.003 | 0.001 | 0.001 | 0 | 0.46 | 0.37 | 0.52 | 0.57 | 0.5 | 0.51 |
| 6 | 160805669 | rs1510229 | 1.741 | 0.185 | 0.827 | 0.007 | 0.03 | -0.015 | -0.015 | 0 | 0.34 | 0.18 | 0.22 | 0.36 | 0.27 | 0.41 |
| 6 | 160807161 | rs1317652 | -0.287 | 0.22 | 0.797 | 0.007 | -0.002 | 0.001 | 0.001 | 0 | 0.46 | 0.37 | 0.52 | 0.57 | 0.5 | 0.51 |
| 6 | 160807432 | rs884742 | -0.289 | 0.222 | 0.8 | 0.01 | -0.002 | 0.001 | 0.001 | 0 | 0.46 | 0.37 | 0.52 | 0.57 | 0.5 | 0.51 |
| 6 | 160808395 | rs394352 | 1.777 | 0.242 | 0.793 | 0.013 | 0.027 | -0.016 | -0.016 | 0.166 | 0.34 | 0.17 | 0.22 | 0.36 | 0.27 | 0.41 |
| 6 | 160808447 | rs2221750 | 0.115 | 0.304 | 0.574 | -0.034 | -0.019 | -0.022 | -0.022 | 0 | 0.3 | 0.35 | 0.34 | 0.24 | 0.21 | 0.23 |
| 6 | 160808901 | rs7740824 | 0.774 | 0.311 | 0.568 | -0.034 | -0.1 | -0.109 | -0.109 | 0 | 0.16 | 0.02 | 0.19 | 0.33 | 0.28 | 0.28 |
| 6 | 160810720 | rs3105748 | -0.622 | 0.313 | 0.579 | -0.033 | -0.002 | 0.001 | 0.001 | 0 | 0.46 | 0.37 | 0.52 | 0.57 | 0.5 | 0.51 |
| 6 | 160811154 | rs440962 | 2.095 | 0.312 | 0.579 | -0.034 | 0.028 | -0.016 | -0.016 | 0 | 0.34 | 0.17 | 0.22 | 0.36 | 0.27 | 0.41 |
| 6 | 160812262 | rs1567441 | 2.795 | -0.15 | 0.288 | -0.401 | -0.012 | 0.162 | 0.162 | 0 | 0.24 | 0.01 | 0.17 | 0 | 0.22 | 0.06 |
| 6 | 160813756 | rs446926 | 1.486 | 0.154 | 0.541 | -0.068 | -0.033 | 0 | 0 | 0 | 0.38 | 0.48 | 0.26 | 0.36 | 0.28 | 0.41 |
| 6 | 160815710 | rs10499314 | 0.325 | 0.116 | 1.146 | -0.083 | 0.014 | 0.016 | 0.016 | 0 | 0.29 | 0.16 | 0.33 | 0.24 | 0.22 | 0.23 |

| | | | | | | | | | | | | | | | | |
|---|---|---|---|---|---|---|---|---|---|---|---|---|---|---|---|---|
| 6 | 160818263 | rs1510225 | 2.049 | 0.124 | 0.752 | -0.077 | 0.028 | -0.016 | -0.016 | 0.834 | 0.33 | 0.17 | 0.22 | 0.36 | 0.27 | 0.41 |
| 6 | 160820978 | rs3105749 | -0.63 | 0.139 | 0.706 | -0.07 | -0.002 | 0.001 | 0.001 | 0 | 0.46 | 0.37 | 0.52 | 0.57 | 0.5 | 0.51 |
| 6 | 160826095 | rs1605964 | -0.608 | 0.145 | 0.712 | -0.064 | -0.002 | 0.001 | 0.001 | 0 | 0.46 | 0.37 | 0.52 | 0.57 | 0.5 | 0.51 |
| 6 | 160833468 | rs7745775 | 0.177 | 0.209 | 0.677 | -0.009 | -0.018 | -0.022 | -0.022 | 0.658 | 0.3 | 0.35 | 0.34 | 0.24 | 0.22 | 0.23 |
| 6 | 160833664 | rs9364554 | 0.803 | 0.287 | 0.444 | -0.038 | -0.089 | -0.1 | -0.1 | 0 | 0.16 | 0.02 | 0.19 | 0.33 | 0.28 | 0.29 |
| 6 | 160834515 | rs12194182 | 0.789 | 0.29 | 0.445 | -0.037 | -0.091 | -0.1 | -0.1 | 0 | 0.16 | 0.02 | 0.19 | 0.33 | 0.28 | 0.29 |
| 6 | 160834828 | rs2457571 | 0.34 | 0.26 | 0.431 | -0.057 | -0.152 | -0.082 | -0.082 | 0 | 0.7 | 0.92 | 0.61 | 0.67 | 0.5 | 0.63 |
| 6 | 160838361 | rs3106166 | -0.803 | 0.13 | 0.378 | 0.039 | 0.007 | 0.031 | 0.031 | 0 | 0.49 | 0.75 | 0.56 | 0.59 | 0.46 | 0.48 |
| 6 | 160840252 | rs7758229 | 0.378 | 0.138 | 0.221 | 0.125 | -0.106 | -0.061 | -0.061 | 0 | 0.2 | 0.05 | 0.21 | 0.25 | 0.32 | 0.29 |
| 6 | 160842211 | rs9456537 | 0.25 | 0.058 | 0.617 | 0.164 | -0.015 | -0.004 | -0.004 | 0 | 0.27 | 0.34 | 0.32 | 0.24 | 0.18 | 0.21 |
| 6 | 160842537 | rs3123636 | 0.376 | 0.16 | 0.406 | 0.188 | -0.116 | -0.069 | -0.069 | 0 | 0.2 | 0.03 | 0.21 | 0.25 | 0.32 | 0.29 |
| 6 | 160848167 | rs2665357 | -1.082 | 0.061 | 0.535 | 0.074 | -0.004 | 0.005 | 0.005 | 0 | 0.54 | 0.75 | 0.59 | 0.56 | 0.52 | 0.5 |
| 6 | 160850273 | rs3106164 | 0.452 | 0.24 | 0.478 | 0.156 | -0.125 | -0.109 | -0.109 | 0.016 | 0.2 | 0.01 | 0.22 | 0.33 | 0.34 | 0.28 |
| 6 | 160850911 | rs2661834 | 1.625 | 0.132 | 0.633 | -0.031 | 0.047 | 0.004 | 0.004 | 0 | 0.38 | 0.2 | 0.23 | 0.36 | 0.27 | 0.41 |
| 6 | 160851766 | rs376563 | -0.972 | 0.119 | 0.64 | -0.04 | 0.009 | 0.062 | 0.062 | 0 | 0.51 | 0.76 | 0.59 | 0.64 | 0.54 | 0.5 |
| 6 | 160853286 | rs388170 | 0.497 | 0.153 | 0.537 | -0.011 | -0.125 | -0.14 | -0.14 | 0.346 | 0.2 | 0.01 | 0.22 | 0.42 | 0.34 | 0.29 |
| 6 | 160853393 | rs9456538 | 0.03 | 0.129 | 0.792 | -0.028 | -0.029 | -0.014 | -0.014 | 0 | 0.28 | 0.4 | 0.33 | 0.21 | 0.18 | 0.21 |
| 6 | 160853929 | rs4709432 | 0.336 | 0.135 | 0.786 | -0.025 | -0.021 | -0.005 | -0.005 | 0.017 | 0.25 | 0.37 | 0.32 | 0.22 | 0.18 | 0.21 |
| 6 | 160853995 | rs2504928 | -1.423 | 0.319 | 0.594 | 0.028 | -0.057 | 0.058 | 0.058 | 0 | 0.89 | 0.98 | 0.83 | 1 | 0.81 | 0.92 |
| 6 | 160854138 | rs2457577 | -1.397 | 0.311 | 0.592 | 0.024 | -0.057 | 0.059 | 0.059 | 0 | 0.89 | 0.98 | 0.83 | 1 | 0.81 | 0.92 |
| 6 | 160856843 | rs9365164 | 0.136 | 0.308 | 0.703 | 0.129 | -0.027 | -0.03 | -0.03 | 1.661 | 0.22 | 0.19 | 0.25 | 0.41 | 0.35 | 0.29 |
| 6 | 160858188 | rs2292334 | 0.021 | 0.266 | 0.672 | -0.192 | -0.066 | -0.089 | -0.089 | 0.045 | 0.23 | 0.1 | 0.46 | 0.46 | 0.35 | 0.29 |
| 6 | 160860154 | rs2063345 | 1.892 | 0.087 | 0.688 | -0.3 | 0.049 | -0.027 | -0.027 | 0 | 0.36 | 0.12 | 0.22 | 0.43 | 0.27 | 0.42 |
| 6 | 160860430 | rs2504927 | 0.615 | 0.087 | 0.877 | -0.306 | 0.002 | 0.007 | 0.007 | 0 | 0.6 | 0.49 | 0.31 | 0.53 | 0.43 | 0.62 |
| 6 | 160863108 | rs2174914 | 1.898 | 0.073 | 0.719 | -0.297 | 0.049 | -0.025 | -0.025 | 0 | 0.36 | 0.12 | 0.22 | 0.42 | 0.27 | 0.42 |
| 6 | 160863532 | rs2048327 | 0.473 | 0.125 | 0.607 | -0.289 | -0.123 | -0.149 | -0.149 | 0 | 0.21 | 0.03 | 0.45 | 0.46 | 0.35 | 0.29 |
| 6 | 160868327 | rs2457575 | 1.895 | 0.095 | 0.596 | -0.306 | 0.05 | -0.03 | -0.03 | 0 | 0.36 | 0.11 | 0.23 | 0.43 | 0.27 | 0.42 |
| 6 | 160868668 | rs3918286 | 0.494 | 0.121 | 0.538 | -0.304 | -0.123 | -0.149 | -0.149 | 0 | 0.21 | 0.03 | 0.45 | 0.46 | 0.35 | 0.29 |
| 6 | 160868701 | rs2457574 | 0.694 | 0.102 | 0.643 | -0.321 | 0.001 | 0.003 | 0.003 | 0 | 0.59 | 0.41 | 0.31 | 0.54 | 0.44 | 0.62 |
| 6 | 160869091 | rs2457573 | -1.149 | 0.116 | 0.636 | -0.314 | -0.012 | -0.001 | -0.001 | 0 | 0.88 | 0.89 | 0.82 | 1 | 0.81 | 0.92 |
| 6 | 160870476 | rs2941383 | -1.171 | 0.116 | 0.636 | -0.314 | -0.012 | -0.002 | -0.002 | 0 | 0.88 | 0.89 | 0.82 | 1 | 0.81 | 0.92 |
| 6 | 160872151 | rs1810126 | 0.431 | 0.119 | 0.605 | -0.303 | -0.096 | -0.12 | -0.12 | 0 | 0.21 | 0.07 | 0.45 | 0.46 | 0.35 | 0.29 |
| 6 | 160876290 | rs2504926 | 0.686 | 0.089 | 0.6 | -0.27 | 0 | 0.003 | 0.003 | 0 | 0.59 | 0.41 | 0.31 | 0.54 | 0.44 | 0.62 |
| 6 | 160876356 | rs7744658 | 0.522 | 0.081 | 0.717 | -0.278 | 0 | 0 | 0 | 0 | 0.23 | 0.24 | 0.08 | 0.11 | 0.16 | 0.21 |
| 6 | 160876394 | rs2504925 | -0.627 | 0.656 | 0.637 | 0.103 | -0.003 | -0.02 | -0.02 | 1.005 | 0.87 | 0.86 | 0.81 | 1 | 0.79 | 0.91 |
| 6 | 160877640 | rs2457564 | 0.854 | 0.712 | 0.313 | 0.212 | 0 | 0.003 | 0.003 | 0 | 0.59 | 0.41 | 0.31 | 0.46 | 0.43 | 0.61 |
| 6 | 160882029 | rs2063347 | 0.283 | 0.911 | -0.069 | 0.14 | -0.125 | -0.149 | -0.149 | 0 | 0.21 | 0.03 | 0.45 | 0.46 | 0.36 | 0.28 |
| 6 | 160892241 | rs3127602 | 0.496 | 1.088 | 0.143 | -0.657 | 0.004 | -0.014 | -0.014 | 0 | 0.31 | 0.31 | 0.15 | 0.11 | 0.32 | 0.22 |
| 6 | 160896431 | rs2136675 | 1.813 | 0.97 | 0.143 | -0.629 | 0.028 | -0.072 | -0.072 | 0.007 | 0.3 | 0.12 | 0.45 | 0.54 | 0.27 | 0.42 |
| 6 | 160897872 | rs2504920 | -0.042 | 0.953 | 0.281 | -0.766 | 0.003 | -0.085 | -0.085 | 0.78 | 0.6 | 0.58 | 0.67 | 0.96 | 0.53 | 0.75 |
| 6 | 160898831 | rs6914575 | 0.069 | 1.278 | 0.307 | -0.507 | 0.003 | -0.068 | -0.068 | 3.13 | 0.22 | 0.06 | 0.18 | 0.35 | 0.2 | 0.28 |
| 6 | 160902269 | rs9346817 | 0.1 | 1.344 | 0.339 | -0.625 | 0 | -0.065 | -0.065 | 0.925 | 0.21 | 0.06 | 0.17 | 0.34 | 0.19 | 0.27 |
| 6 | 160903178 | rs9355803 | 0.1 | 1.344 | 0.339 | -0.625 | 0 | -0.065 | -0.065 | 0 | 0.21 | 0.06 | 0.17 | 0.34 | 0.19 | 0.27 |
| 6 | 160903622 | rs9355804 | 0.1 | 1.344 | 0.339 | -0.625 | 0 | -0.065 | -0.065 | 0 | 0.21 | 0.06 | 0.17 | 0.34 | 0.19 | 0.27 |
| 6 | 160906086 | rs3123629 | 0.979 | 1.389 | 0.422 | -0.682 | 0.003 | -0.013 | -0.013 | 0 | 0.31 | 0.31 | 0.15 | 0.11 | 0.32 | 0.22 |
| 6 | 160910899 | rs10945673 | 0.348 | 1.468 | 0.443 | -0.616 | -0.02 | -0.084 | -0.084 | 0 | 0.19 | 0.04 | 0.16 | 0.34 | 0.19 | 0.27 |
| 6 | 160911053 | rs3123630 | -1.151 | 1.508 | 0.367 | -0.639 | -0.066 | 0.062 | 0.062 | 0 | 0.89 | 0.99 | 0.83 | 0.99 | 0.81 | 0.92 |
| 6 | 160914460 | rs9346818 | 0.905 | 1.515 | 0.396 | -0.624 | 0.004 | -0.009 | -0.009 | 0 | 0.61 | 0.43 | 0.6 | 0.66 | 0.59 | 0.64 |
| 6 | 160921566 | rs9457930 | -0.13 | 1.318 | 0.183 | -0.249 | 0.038 | 0.193 | 0.193 | 0 | 0.49 | 0.18 | 0.62 | 0.89 | 0.47 | 0.7 |
| 6 | 160940120 | rs9365168 | -0.615 | 1.511 | -0.117 | -0.028 | -0.016 | 0.012 | 0.012 | 0 | 0.64 | 0.61 | 0.5 | 0.45 | 0.71 | 0.58 |
| 6 | 160946449 | rs3124787 | -1.268 | 1.556 | -0.29 | -0.046 | 0.019 | -0.081 | -0.081 | 0 | 0.68 | 0.89 | 0.55 | 0.45 | 0.7 | 0.54 |
| 6 | 160957114 | rs7449650 | -0.459 | 1.594 | -0.221 | 0.019 | -0.028 | 0.016 | 0.016 | 0 | 0.61 | 0.55 | 0.5 | 0.45 | 0.69 | 0.54 |
| 6 | 160970963 | rs7761293 | -0.105 | 0.793 | -0.654 | -0.269 | -0.008 | 0.162 | 0.162 | 0 | 0.46 | 0.81 | 0.37 | 0.11 | 0.49 | 0.28 |
| 6 | 160981736 | rs9365171 | -1.415 | 1.016 | 0.331 | 0.631 | 0.007 | 0.004 | 0.004 | 0 | 0.34 | 0.41 | 0.26 | 0.35 | 0.37 | 0.34 |
| 6 | 161019866 | rs6926458 | -1.048 | 1.05 | 0.249 | 0.612 | -0.026 | -0.084 | -0.084 | 1.789 | 0.23 | 0.09 | 0.15 | 0.43 | 0.26 | 0.27 |
| 6 | 161029728 | rs13202636 | -1.057 | 1.025 | 0.453 | 0.6 | -0.025 | -0.085 | -0.085 | 0 | 0.23 | 0.09 | 0.15 | 0.43 | 0.26 | 0.27 |
| 6 | 161032481 | rs7761377 | -1.931 | 0.988 | 1.161 | 0.524 | -0.006 | -0.009 | -0.009 | 0 | 0.39 | 0.65 | 0.3 | 0.44 | 0.41 | 0.35 |
| 6 | 161069941 | rs10945682 | -1.717 | 0.883 | 1.218 | 0.544 | 0.01 | 0.015 | 0.015 | 0 | 0.35 | 0.58 | 0.27 | 0.43 | 0.4 | 0.35 |
| 6 | 161081842 | rs1321196 | 1.309 | 0.99 | 0.804 | 0.604 | 0.027 | 0.038 | 0.038 | 0 | 0.68 | 0.54 | 0.74 | 0.57 | 0.6 | 0.65 |
| 6 | 161082695 | rs1367211 | 1.281 | 1.03 | 0.8 | 0.053 | 0.012 | -0.051 | -0.051 | 0.446 | 0.71 | 0.48 | 0.8 | 0.81 | 0.71 | 0.72 |
| 6 | 161090520 | rs1358754 | -0.353 | 1.15 | 0.612 | -0.112 | 0.016 | -0.04 | -0.04 | 2.974 | 0.09 | 0.14 | 0.1 | 0.01 | 0.14 | 0.07 |
| 6 | 161108536 | rs6935921 | 0.708 | 0.993 | 0.681 | 0.084 | -0.007 | 0.03 | 0.03 | 0.314 | 0.7 | 0.56 | 0.5 | 0.36 | 0.73 | 0.51 |
| 6 | 161111113 | rs7767705 | -0.669 | 1.138 | 0.264 | 0.056 | 0.012 | 0.002 | 0.002 | 0 | 0.22 | 0.41 | 0.19 | 0.18 | 0.25 | 0.17 |
| 6 | 161115245 | rs7772437 | 1.312 | 1.074 | 0.163 | 0.001 | -0.001 | -0.006 | -0.006 | 0.792 | 0.53 | 0.43 | 0.34 | 0.18 | 0.49 | 0.41 |
| 6 | 161118577 | rs9458005 | -0.471 | 0.935 | -0.682 | 0.15 | -0.008 | -0.005 | -0.005 | 0 | 0.19 | 0.16 | 0.16 | 0.18 | 0.24 | 0.17 |
| 6 | 161121958 | rs4252050 | 0.38 | 0.714 | -0.681 | 0.506 | -0.003 | -0.07 | -0.07 | 0.234 | 0.77 | 0.9 | 0.59 | 0.36 | 0.81 | 0.58 |
| 6 | 161128264 | rs4252072 | 0.59 | 0.483 | -0.482 | -0.907 | 0.003 | -0.048 | -0.048 | 0 | 0.32 | 0.28 | 0.2 | 0 | 0.3 | 0.08 |
| 6 | 161136294 | rs9295131 | 0.595 | 0.47 | -0.366 | -0.685 | 0 | -0.002 | -0.002 | 0.811 | 0.63 | 0.38 | 0.46 | 0.45 | 0.68 | 0.48 |
| 6 | 161137518 | rs4252107 | 1.804 | 0.455 | -0.326 | -0.74 | -0.018 | 0.094 | 0.094 | 0 | 0.25 | 0.08 | 0.17 | 0 | 0.29 | 0.08 |

| | | | | | | | | | | | | | | | | |
|---|---|---|---|---|---|---|---|---|---|---|---|---|---|---|---|---|
| 6 | 161137663 | rs4252109 | 1.669 | 0.424 | -0.332 | -0.756 | -0.003 | 0.033 | 0.033 | 0 | 0.26 | 0.14 | 0.17 | 0 | 0.29 | 0.08 |
| 6 | 161137779 | rs14224 | -0.419 | 0.112 | 0.578 | -0.888 | -0.002 | -0.001 | -0.001 | 2.352 | 0.48 | 0.55 | 0.33 | 0.46 | 0.4 | 0.44 |
| 6 | 161137990 | rs783147 | 0.736 | 0.431 | 0.768 | -0.956 | -0.011 | -0.004 | -0.004 | 0.034 | 0.44 | 0.18 | 0.57 | 0.52 | 0.45 | 0.4 |
| 6 | 161139480 | rs1130656 | 0.412 | 0.414 | 0.735 | -0.953 | -0.01 | -0.035 | -0.035 | 0 | 0.38 | 0.24 | 0.28 | 0.46 | 0.39 | 0.4 |
| 6 | 161139502 | rs2295368 | 0.46 | 0.415 | 0.736 | -0.952 | -0.012 | -0.036 | -0.036 | 0.029 | 0.37 | 0.24 | 0.28 | 0.46 | 0.39 | 0.4 |
| 6 | 161139857 | rs13231 | 1.896 | 0.454 | 0.779 | -1.154 | -0.003 | 0.031 | 0.031 | 0 | 0.26 | 0.15 | 0.17 | 0 | 0.29 | 0.08 |
| 6 | 161143190 | rs4252114 | 0.864 | 0.438 | 0.856 | -1.171 | -0.019 | -0.146 | -0.146 | 1.997 | 0.37 | 0.49 | 0.23 | 0 | 0.31 | 0.12 |
| 6 | 161143233 | rs4252116 | -0.497 | 0.422 | 0.965 | -1.179 | -0.053 | -0.06 | -0.06 | 0 | 0.1 | 0.3 | 0.05 | 0 | 0.01 | 0.04 |
| 6 | 161143376 | rs4252117 | 1.946 | 0.42 | 0.969 | -1.181 | -0.025 | 0.111 | 0.111 | 0 | 0.26 | 0.07 | 0.17 | 0 | 0.3 | 0.08 |
| 6 | 161143608 | rs4252120 | 1.87 | 0.409 | 1.05 | -1.186 | -0.006 | 0.047 | 0.047 | 0 | 0.26 | 0.15 | 0.18 | 0 | 0.3 | 0.08 |
| 6 | 161143634 | rs4252121 | -0.496 | 0.409 | 1.051 | -1.185 | -0.053 | -0.06 | -0.06 | 0 | 0.1 | 0.3 | 0.05 | 0 | 0.01 | 0.04 |
| 6 | 161144729 | rs1853025 | -0.496 | 0.409 | 1.051 | -1.185 | -0.053 | -0.06 | -0.06 | 0 | 0.1 | 0.3 | 0.05 | 0 | 0.01 | 0.04 |
| 6 | 161146704 | rs1897108 | 0.903 | 0.567 | 1.02 | -1.112 | -0.014 | -0.128 | -0.128 | 0 | 0.36 | 0.45 | 0.23 | 0 | 0.31 | 0.12 |
| 6 | 161152294 | rs4252126 | 1.947 | 0.568 | 1.187 | -1.111 | -0.023 | 0.107 | 0.107 | 0 | 0.26 | 0.08 | 0.17 | 0 | 0.3 | 0.08 |
| 6 | 161152449 | rs783145 | -0.978 | 0.388 | 0.98 | -1.405 | -0.047 | 0.176 | 0.176 | 0 | 0.6 | 0.83 | 0.72 | 0.98 | 0.51 | 0.73 |
| 6 | 161153080 | rs4252130 | 2.139 | 0.448 | 1.011 | -1.412 | -0.023 | 0.107 | 0.107 | 0 | 0.26 | 0.07 | 0.17 | 0 | 0.3 | 0.08 |
| 6 | 161153990 | rs813641 | 0.309 | 0.44 | 1.021 | -1.248 | -0.001 | -0.009 | -0.009 | 0 | 0.18 | 0.11 | 0.4 | 0.49 | 0.15 | 0.32 |
| 6 | 161154232 | rs4252135 | 2.091 | 0.495 | 0.871 | -1.468 | -0.025 | 0.111 | 0.111 | 0 | 0.26 | 0.07 | 0.17 | 0 | 0.3 | 0.08 |
| 6 | 161154955 | rs1317026 | -0.924 | 0.705 | 0.966 | -1.362 | -0.035 | -0.069 | -0.069 | 0 | 0.11 | 0.42 | 0.06 | 0 | 0.02 | 0.04 |
| 6 | 161160505 | rs4252155 | -0.577 | 0.595 | 0.918 | -1.359 | -0.053 | -0.06 | -0.06 | 0.197 | 0.1 | 0.31 | 0.05 | 0 | 0.01 | 0.04 |
| 6 | 161160644 | rs4252159 | 0.976 | 0.696 | 0.566 | -1.185 | -0.023 | -0.02 | -0.02 | 0.549 | 0.1 | 0.01 | 0.07 | 0.09 | 0.09 | 0.06 |
| 6 | 161161559 | rs4252165 | 2.054 | 0.652 | 0.481 | -1.328 | -0.006 | 0.041 | 0.041 | 0 | 0.26 | 0.15 | 0.18 | 0 | 0.3 | 0.08 |
| 6 | 161162406 | rs4252170 | 0.871 | 0.705 | 0.422 | -1.173 | -0.022 | -0.017 | -0.017 | 0 | 0.11 | 0.02 | 0.07 | 0.09 | 0.09 | 0.06 |
| 6 | 161168548 | rs783182 | -0.718 | 0.904 | 0.799 | -1.365 | 0 | -0.085 | -0.085 | 0.779 | 0.55 | 0.45 | 0.68 | 0.98 | 0.5 | 0.71 |
| 6 | 161170835 | rs783176 | 0.041 | 0.47 | 0.546 | -1.549 | -0.003 | -0.07 | -0.07 | 0.063 | 0.23 | 0.05 | 0.19 | 0.43 | 0.17 | 0.33 |
| 6 | 161173946 | rs11060 | -0.84 | 0.438 | 0.606 | -1.604 | 0 | -0.084 | -0.084 | 0.604 | 0.55 | 0.53 | 0.71 | 0.98 | 0.52 | 0.71 |
| 6 | 161177842 | rs783165 | -0.966 | 0.37 | 0.606 | -1.623 | -0.001 | -0.1 | -0.1 | 5.212 | 0.55 | 0.47 | 0.69 | 0.98 | 0.52 | 0.71 |
| 6 | 161186889 | rs7450979 | 0.283 | 0.382 | 0.738 | -1.576 | 0 | 0 | 0 | 0 | 0.31 | 0.3 | 0.54 | 0.58 | 0.35 | 0.38 |
| 6 | 161187044 | rs2465836 | 0.044 | 0.403 | 0.602 | -1.553 | 0.015 | -0.043 | -0.043 | 0 | 0.23 | 0.07 | 0.14 | 0.4 | 0.17 | 0.33 |
| 6 | 161190199 | rs1247558 | -0.969 | 0.388 | 0.674 | -1.608 | 0 | -0.081 | -0.081 | 0 | 0.55 | 0.53 | 0.7 | 0.98 | 0.51 | 0.71 |
| 6 | 161192189 | rs1652500 | 0.083 | 0.473 | 0.529 | -1.521 | -0.008 | 0.006 | 0.006 | 0 | 0.85 | 0.86 | 0.88 | 0.98 | 0.78 | 0.82 |
| 6 | 161196800 | rs1782627 | 0.945 | 0.341 | 0.616 | -1.543 | 0.01 | 0.028 | 0.028 | 0 | 0.48 | 0.39 | 0.31 | 0.4 | 0.43 | 0.4 |
| 6 | 161196966 | rs1621801 | -1.37 | 0.292 | 0.791 | -1.47 | -0.003 | -0.041 | -0.041 | 0 | 0.6 | 0.6 | 0.72 | 0.98 | 0.52 | 0.74 |
| 6 | 161201960 | rs13199812 | -0.281 | 0.34 | 0.821 | -1.432 | -0.001 | -0.008 | -0.008 | 0 | 0.37 | 0.48 | 0.57 | 0.58 | 0.35 | 0.42 |
| 6 | 161203756 | rs1247559 | -0.1 | 0.378 | 0.508 | -1.349 | 0.013 | -0.024 | -0.024 | 0 | 0.23 | 0.13 | 0.15 | 0.4 | 0.17 | 0.32 |
| 6 | 161213047 | rs9458036 | -0.086 | 0.267 | 0.718 | -1.307 | 0.009 | 0.003 | 0.003 | 0 | 0.33 | 0.38 | 0.56 | 0.58 | 0.36 | 0.42 |
| 6 | 161216608 | rs2064712 | 0.605 | 0.753 | 0.272 | -1.628 | -0.03 | 0.005 | 0.005 | 0 | 0.09 | 0.02 | 0.09 | 0.14 | 0.15 | 0.15 |
| 6 | 161231177 | rs783156 | -0.967 | 0.214 | 0.579 | -2.292 | -0.002 | -0.025 | -0.025 | 0 | 0.55 | 0.59 | 0.69 | 0.98 | 0.51 | 0.72 |
| 6 | 161232224 | rs9458040 | 0.195 | 0.246 | 0.449 | -2.198 | 0.002 | -0.001 | -0.001 | 0 | 0.31 | 0.31 | 0.52 | 0.63 | 0.35 | 0.4 |
| 6 | 161234047 | rs813021 | -0.078 | 0.383 | 0.353 | -1.968 | 0.018 | -0.033 | -0.033 | 1.107 | 0.22 | 0.06 | 0.16 | 0.35 | 0.15 | 0.31 |
| 6 | 161246303 | rs1629794 | -1.306 | 0.279 | 0.366 | -2.118 | -0.036 | 0.101 | 0.101 | 0 | 0.63 | 0.79 | 0.75 | 0.98 | 0.54 | 0.73 |
| 6 | 161250259 | rs9355839 | 0.703 | 0.271 | 0.255 | -2.052 | -0.006 | -0.043 | -0.043 | 0 | 0.15 | 0.02 | 0.12 | 0.17 | 0.1 | 0.07 |
| 6 | 161254403 | rs1652471 | -1.249 | 0.311 | 0.385 | -1.991 | -0.015 | 0.016 | 0.016 | 0 | 0.63 | 0.7 | 0.74 | 0.98 | 0.54 | 0.72 |
| 6 | 161257418 | rs9347445 | 0.159 | 0.225 | 0.375 | -1.966 | -0.006 | 0.007 | 0.007 | 0 | 0.36 | 0.26 | 0.55 | 0.62 | 0.38 | 0.45 |
| 6 | 161260530 | rs2489952 | -0.34 | 0.366 | 0.43 | -1.909 | -0.017 | 0.003 | 0.003 | 0 | 0.27 | 0.37 | 0.19 | 0.36 | 0.16 | 0.28 |
| 6 | 161267335 | rs9355841 | -0.864 | 0.644 | -0.19 | -1.962 | -0.005 | -0.028 | -0.028 | 0 | 0.63 | 0.65 | 0.73 | 0.98 | 0.54 | 0.76 |
| 6 | 161280379 | rs1247513 | -0.051 | 0.798 | -0.153 | -1.901 | 0.014 | -0.016 | -0.016 | 0 | 0.1 | 0.15 | 0.09 | 0.02 | 0.2 | 0.16 |
| 6 | 161281175 | rs2465850 | -0.802 | 0.615 | -0.137 | -1.748 | -0.006 | -0.012 | -0.012 | 0 | 0.6 | 0.65 | 0.72 | 0.98 | 0.53 | 0.73 |
| 6 | 161292913 | rs2489944 | -0.134 | 0.253 | -0.522 | -1.21 | 0.01 | 0.011 | 0.011 | 0.036 | 0.21 | 0.09 | 0.15 | 0.16 | 0.15 | 0.17 |
| 6 | 161305715 | rs1247517 | -0.247 | 0.76 | -0.442 | -1.024 | -0.058 | -0.009 | -0.009 | 0 | 0.29 | 0.13 | 0.52 | 0.65 | 0.42 | 0.43 |
| 6 | 161315130 | rs9346841 | -0.401 | -0.258 | 0.25 | -1.116 | -0.092 | -0.045 | -0.045 | 0 | 0.27 | 0.51 | 0.13 | 0.22 | 0.14 | 0.25 |
| 6 | 161318009 | rs1937470 | 1.225 | -0.053 | 0.547 | -1.677 | -0.011 | -0.104 | -0.104 | 0 | 0.3 | 0.32 | 0.17 | 0 | 0.24 | 0.1 |
| 6 | 161319672 | rs11970528 | 0.754 | 0.239 | 0.489 | -1.522 | 0.001 | -0.02 | -0.02 | 0 | 0.41 | 0.41 | 0.31 | 0.17 | 0.41 | 0.26 |
| 6 | 161323288 | rs9365233 | -0.941 | 0.639 | 0.13 | -1.364 | -0.017 | -0.04 | -0.04 | 0 | 0.77 | 0.85 | 0.51 | 0.49 | 0.73 | 0.69 |
| 6 | 161326650 | rs1247330 | -0.848 | 0.77 | 0.712 | -1.589 | 0.033 | -0.031 | -0.031 | 0 | 0.27 | 0.4 | 0.23 | 0.18 | 0.33 | 0.31 |
| 6 | 161333781 | rs1247320 | 0.429 | 0.82 | 0.695 | -1.549 | 0.021 | 0.012 | 0.012 | 0 | 0.13 | 0.21 | 0.16 | 0.16 | 0.18 | 0.16 |
| 6 | 161343052 | rs11963931 | 0.784 | 0.721 | 0.476 | -1.738 | 0.005 | 0.04 | 0.04 | 0 | 0.4 | 0.21 | 0.22 | 0.02 | 0.38 | 0.25 |
| 6 | 161344717 | rs1962358 | 0.279 | 0.688 | 0.599 | -1.697 | -0.01 | 0.014 | 0.014 | 0 | 0.6 | 0.35 | 0.34 | 0.3 | 0.55 | 0.53 |
| 6 | 161348069 | rs935181 | 1.47 | 0.643 | 0.473 | -1.733 | -0.016 | -0.127 | -0.127 | 0 | 0.29 | 0.39 | 0.16 | 0 | 0.23 | 0.11 |
| 6 | 161349326 | rs7756550 | 0.828 | 0.449 | -0.248 | -1.61 | 0.044 | 0.063 | 0.063 | 0 | 0.19 | 0.07 | 0.09 | 0.06 | 0.14 | 0.11 |
| 6 | 161358808 | rs591676 | 0.458 | 1.147 | -0.256 | -1.261 | 0.023 | 0.017 | 0.017 | 0 | 0.12 | 0.21 | 0.13 | 0.16 | 0.15 | 0.16 |
| 6 | 161360378 | rs673319 | -1.413 | 0.646 | -1.034 | -1.596 | -0.005 | 0.061 | 0.061 | 0 | 0.73 | 0.88 | 0.83 | 1 | 0.74 | 0.89 |
| 6 | 161362678 | rs1530395 | -0.569 | 0.46 | -0.567 | -0.32 | 0.038 | 0.017 | 0.017 | 0 | 0.28 | 0.2 | 0.15 | 0.29 | 0.21 | 0.31 |
| 6 | 161363705 | rs11752957 | -0.634 | 1.509 | -0.093 | 0.164 | 0.006 | -0.019 | -0.019 | 0 | 0.06 | 0.07 | 0.04 | 0.01 | 0.1 | 0.11 |
| 6 | 161368268 | rs9458066 | -0.832 | 1.175 | -0.015 | 0.27 | 0 | -0.017 | -0.017 | 0.112 | 0.06 | 0.11 | 0.04 | 0.02 | 0.04 | 0.08 |
| 6 | 161377557 | rs1247336 | -1.892 | 2.261 | 0.791 | 0.655 | 0.076 | 0.029 | 0.029 | 0 | 0.17 | 0.6 | 0.2 | 0.22 | 0.27 | 0.22 |
| 6 | 161378192 | rs590884 | -1.867 | 2.1 | 0.17 | 0.118 | -0.029 | 0.021 | 0.021 | 0 | 0.34 | 0.72 | 0.53 | 0.25 | 0.39 | 0.35 |
| 6 | 161383479 | rs604924 | -1.717 | 1.632 | 0.404 | 0.937 | 0.004 | 0.004 | 0.004 | 0 | 0.4 | 0.6 | 0.27 | 0.68 | 0.33 | 0.41 |
| 6 | 161385757 | rs7760467 | -0.9 | 1.86 | -0.156 | 1.157 | -0.041 | -0.094 | -0.094 | 0.002 | 0.21 | 0.09 | 0.14 | 0.55 | 0.23 | 0.2 |

| | | | | | | | | | | | | | | | | |
|---|---|---|---|---|---|---|---|---|---|---|---|---|---|---|---|---|
| 6 | 161385816 | rs675293 | -1.724 | 1.828 | 0.588 | 1.195 | -0.02 | 0.034 | 0.034 | 1.139 | 0.34 | 0.3 | 0.27 | 0.76 | 0.36 | 0.35 |
| 6 | 161386772 | rs2099938 | -0.744 | 1.736 | 0.576 | 1.214 | 0.03 | -0.078 | -0.078 | 1.388 | 0.2 | 0.1 | 0.1 | 0.55 | 0.15 | 0.2 |
| 6 | 161391546 | rs685388 | -0.44 | 1.731 | 0.645 | 1.183 | 0.008 | 0.017 | 0.017 | 0 | 0.14 | 0.22 | 0.13 | 0.22 | 0.14 | 0.15 |
| 6 | 161393311 | rs657030 | -1.922 | 1.469 | 1.13 | 0.993 | -0.002 | 0.003 | 0.003 | 1.377 | 0.43 | 0.63 | 0.36 | 0.76 | 0.45 | 0.38 |
| 6 | 161395857 | rs9355861 | 0.049 | 1.368 | 0.697 | 1.183 | 0.044 | -0.065 | -0.065 | 0 | 0.12 | 0.05 | 0.05 | 0.4 | 0.04 | 0.12 |
| 6 | 161396138 | rs7766870 | -0.027 | 1.502 | 0.822 | 1.072 | -0.003 | 0.003 | 0.003 | 0 | 0.06 | 0.04 | 0.06 | 0 | 0.1 | 0.04 |
| 6 | 161396158 | rs617514 | -2.078 | 1.474 | 0.385 | 0.506 | -0.002 | 0.155 | 0.155 | 0 | 0.6 | 0.88 | 0.45 | 0.84 | 0.59 | 0.56 |
| 6 | 161406871 | rs9458073 | -0.418 | 1.767 | 0.939 | 0.647 | 0.11 | 0.06 | 0.06 | 0 | 0.09 | 0.21 | 0.11 | 0.13 | 0.19 | 0.06 |
| 6 | 161407819 | rs673468 | -1.409 | 1.621 | 1.038 | 0.804 | 0.001 | -0.001 | -0.001 | 3.041 | 0.4 | 0.62 | 0.28 | 0.42 | 0.3 | 0.3 |
| 6 | 161409387 | rs602837 | -1.538 | 1.711 | 1.089 | 0.802 | 0.009 | -0.006 | -0.006 | 0 | 0.45 | 0.63 | 0.31 | 0.48 | 0.36 | 0.36 |
| 6 | 161411973 | rs678290 | -1.963 | 1.545 | 0.83 | 0.604 | -0.036 | 0.195 | 0.195 | 0 | 0.56 | 0.94 | 0.43 | 0.82 | 0.54 | 0.53 |
| 6 | 161418642 | rs625977 | 0.967 | 1.379 | 1.144 | 1.187 | -0.058 | 0.037 | 0.037 | 0 | 0.68 | 0.49 | 0.86 | 0.56 | 0.84 | 0.7 |
| 6 | 161424682 | rs606817 | 1.001 | 1.293 | 1.11 | 1.163 | -0.058 | 0.038 | 0.038 | 0 | 0.68 | 0.5 | 0.86 | 0.54 | 0.84 | 0.7 |
| 6 | 161430468 | rs644110 | 0.015 | 1.212 | 1.303 | 1.268 | -0.006 | 0.013 | 0.013 | 0.617 | 0.85 | 0.8 | 0.95 | 0.63 | 0.95 | 0.86 |
| 6 | 161433981 | rs4709485 | 1.402 | 1.211 | 1.28 | 1.151 | -0.017 | -0.004 | -0.004 | 0 | 0.61 | 0.43 | 0.78 | 0.61 | 0.78 | 0.71 |
| 6 | 161449576 | rs13200744 | 0.299 | 1.216 | 1.147 | 1.325 | 0.044 | -0.055 | -0.055 | 3.501 | 0.13 | 0.07 | 0.04 | 0.33 | 0.04 | 0.14 |
| 6 | 161454462 | rs9365245 | 1.181 | 1.234 | 1.255 | 1.078 | -0.017 | -0.14 | -0.14 | 0 | 0.72 | 0.45 | 0.77 | 0.97 | 0.76 | 0.84 |
| 6 | 161460937 | rs1562026 | 0.033 | 1.245 | 1.184 | 1.027 | -0.004 | -0.01 | -0.01 | 0 | 0.86 | 0.87 | 0.84 | 0.99 | 0.82 | 0.92 |
| 6 | 161466180 | rs2314379 | 2.474 | 1.146 | 1.123 | 0.977 | -0.021 | 0.173 | 0.173 | 0 | 0.43 | 0.08 | 0.55 | 0.19 | 0.44 | 0.46 |
| 6 | 161468817 | rs1822520 | 1.388 | 1.063 | 1.068 | 0.947 | -0.015 | -0.118 | -0.118 | 0 | 0.7 | 0.41 | 0.76 | 0.97 | 0.75 | 0.84 |
| 6 | 161477664 | rs9365246 | -1.153 | 0.953 | 1.004 | 1.474 | -0.089 | 0.006 | 0.006 | 0.064 | 0.27 | 0.47 | 0.11 | 0.33 | 0.11 | 0.21 |
| 6 | 161478446 | rs1001808 | 1.499 | 0.97 | 1 | 1.516 | -0.019 | -0.015 | -0.015 | 0 | 0.61 | 0.43 | 0.77 | 0.72 | 0.78 | 0.72 |
| 6 | 161482494 | rs12665456 | 0.287 | 0.963 | 0.906 | 1.595 | 0.044 | -0.044 | -0.044 | 0.399 | 0.13 | 0.07 | 0.04 | 0.26 | 0.04 | 0.13 |
| 6 | 161487794 | rs4709487 | -1.6 | 1.062 | 0.935 | 1.573 | -0.048 | 0.02 | 0.02 | 0 | 0.34 | 0.48 | 0.15 | 0.63 | 0.17 | 0.4 |
| 6 | 161502094 | rs9347479 | 0.122 | 2.044 | 0.532 | 0.88 | -0.062 | 0.115 | 0.115 | 0 | 0.8 | 0.96 | 0.78 | 1 | 0.75 | 0.91 |
| 6 | 161521680 | rs11758794 | 0.02 | 1.831 | 1.288 | 1.291 | -0.018 | -0.027 | -0.027 | 0.559 | 0.12 | 0.13 | 0.05 | 0.02 | 0.05 | 0.08 |
| 6 | 161522365 | rs4709491 | -0.376 | 1.855 | 1.336 | 1.274 | 0.008 | -0.042 | -0.042 | 0 | 0.89 | 0.84 | 0.87 | 0.99 | 0.89 | 0.94 |
| 6 | 161522478 | rs9458121 | 0.136 | 1.837 | 1.283 | 1.302 | -0.017 | -0.039 | -0.039 | 0 | 0.11 | 0.18 | 0.04 | 0.02 | 0.06 | 0.08 |
| 6 | 161529678 | rs2272197 | -1.047 | 2.098 | 0.942 | 1.562 | -0.018 | -0.016 | -0.016 | 0 | 0.1 | 0.07 | 0.1 | 0.26 | 0.17 | 0.21 |
| 6 | 161538250 | rs1488 | 1.091 | 2.135 | 0.913 | 1.536 | 0.038 | -0.006 | -0.006 | 0 | 0.69 | 0.51 | 0.48 | 0.75 | 0.65 | 0.82 |
| 6 | 161541909 | rs12527150 | 0.847 | 1.884 | 0.894 | 1.098 | -0.024 | -0.131 | -0.131 | 0 | 0.73 | 0.43 | 0.8 | 0.97 | 0.83 | 0.88 |
| 6 | 161544198 | rs9347480 | -0.771 | 2.154 | 0.488 | 1.685 | -0.055 | -0.052 | -0.052 | 0.198 | 0.08 | 0.01 | 0.1 | 0.26 | 0.16 | 0.21 |
| 6 | 161550999 | rs2293286 | -2.277 | 1.926 | 0.993 | 1.74 | 0.002 | 0.013 | 0.013 | 1.608 | 0.2 | 0.76 | 0.32 | 0.27 | 0.19 | 0.29 |
| 6 | 161551024 | rs2293287 | -1.277 | 1.77 | 1.83 | 1.768 | 0.082 | 0.19 | 0.19 | 0.078 | 0.08 | 0.33 | 0.15 | 0.16 | 0.09 | 0.17 |
| 6 | 161555840 | rs10945713 | 1.645 | 1.468 | 0.711 | 0.834 | -0.007 | 0.115 | 0.115 | 0 | 0.39 | 0.11 | 0.31 | 0.22 | 0.41 | 0.39 |
| 6 | 161556159 | rs3798225 | -1.461 | 1.325 | 1.092 | 0.684 | -0.006 | 0.002 | 0.002 | 0 | 0.32 | 0.68 | 0.55 | 0.23 | 0.41 | 0.22 |
| 6 | 161557662 | rs3734462 | 0.175 | 1.524 | 0.859 | 0.757 | -0.013 | -0.012 | -0.012 | 0 | 0.05 | 0.02 | 0.33 | 0.18 | 0.19 | 0.06 |
| 6 | 161558036 | rs7768457 | -0.924 | 1.411 | 1.236 | 0.656 | 0.111 | 0.114 | 0.114 | 0 | 0.17 | 0.44 | 0.4 | 0.22 | 0.25 | 0.15 |
| 6 | 161559807 | rs2314190 | -1.327 | 0.82 | 1.345 | -0.154 | -0.034 | -0.1 | -0.1 | 0 | 0.23 | 0.58 | 0.18 | 0.04 | 0.15 | 0.15 |
| 6 | 161561256 | rs1807426 | -2.021 | 0.808 | 1.346 | -0.145 | 0.057 | 0.028 | 0.028 | 0 | 0.08 | 0.45 | 0.11 | 0.03 | 0.1 | 0.09 |
| 6 | 161564104 | rs10945718 | -1.435 | 0.775 | 1.328 | -0.204 | -0.037 | -0.062 | -0.062 | 0 | 0.14 | 0.45 | 0.13 | 0.03 | 0.11 | 0.1 |
| 6 | 161565158 | rs12191615 | -0.067 | 0.805 | 1.536 | -0.211 | 0.007 | -0.058 | -0.058 | 0.297 | 0.12 | 0.23 | 0.1 | 0.01 | 0.1 | 0.07 |
| 6 | 161566405 | rs1880367 | -1.798 | 0.79 | 1.522 | -0.223 | -0.026 | -0.121 | -0.121 | 0 | 0.25 | 0.54 | 0.21 | 0.04 | 0.21 | 0.16 |
| 6 | 161566909 | rs6455707 | -1.256 | 0.786 | 1.606 | -0.172 | -0.02 | -0.07 | -0.07 | 0.183 | 0.2 | 0.35 | 0.14 | 0.04 | 0.13 | 0.14 |
| 6 | 161567333 | rs2277091 | 2.64 | 0.758 | 0.707 | -0.228 | -0.031 | 0.161 | 0.161 | 1.218 | 0.41 | 0.05 | 0.29 | 0.21 | 0.42 | 0.4 |
| 6 | 161567968 | rs1001421 | -1.541 | 0.511 | 1.375 | 0.007 | -0.011 | -0.046 | -0.046 | 2.924 | 0.2 | 0.27 | 0.15 | 0.04 | 0.14 | 0.15 |
| 6 | 161571151 | rs11965825 | -0.51 | 0.633 | 1.097 | 0.034 | 0.008 | -0.072 | -0.072 | 0 | 0.13 | 0.34 | 0.11 | 0.01 | 0.11 | 0.07 |
| 6 | 161576790 | rs3798920 | -0.532 | 0.36 | 0.542 | 0.037 | 0.037 | 0.041 | 0.041 | 0 | 0.16 | 0.08 | 0.06 | 0.03 | 0.07 | 0.14 |
| 6 | 161582535 | rs6934457 | -1.982 | 0.162 | 1.128 | -0.138 | -0.012 | -0.055 | -0.055 | 0.729 | 0.26 | 0.64 | 0.22 | 0.04 | 0.21 | 0.18 |
| 6 | 161586840 | rs729986 | -0.572 | -0.497 | 0.651 | 0.039 | -0.013 | 0.006 | 0.006 | 0.564 | 0.21 | 0.21 | 0.29 | 0.27 | 0.1 | 0.26 |
| 6 | 161589477 | rs7754843 | -1.812 | 0.002 | 0.866 | -0.178 | -0.01 | -0.098 | -0.098 | 0 | 0.25 | 0.43 | 0.22 | 0.04 | 0.25 | 0.17 |
| 6 | 161589595 | rs12208953 | 0.642 | 0.114 | 0.185 | -0.016 | -0.041 | 0.043 | 0.043 | 0 | 0.09 | 0.01 | 0.09 | 0.01 | 0.11 | 0.06 |
| 6 | 161592100 | rs1125640 | -0.421 | 0.262 | 0.098 | 0.095 | -0.028 | -0.018 | -0.018 | 0 | 0.94 | 1 | 0.96 | 0.92 | 0.93 | 0.91 |
| 6 | 161601836 | rs3798925 | -0.465 | 0.344 | 0.327 | 0.105 | 0.001 | 0 | 0 | 0 | 0.1 | 0.11 | 0.05 | 0.04 | 0.06 | 0.11 |
| 6 | 161603980 | rs12203149 | 0.477 | 0.618 | 0.284 | 0.178 | -0.04 | 0.045 | 0.045 | 0 | 0.09 | 0.01 | 0.1 | 0.01 | 0.11 | 0.06 |
| 6 | 161609016 | rs2064721 | 2.381 | 0.606 | -0.248 | -0.072 | 0.016 | 0.131 | 0.131 | 0 | 0.37 | 0.09 | 0.26 | 0.2 | 0.36 | 0.37 |
| 6 | 161609789 | rs746731 | 1.021 | 1.102 | 0.442 | 0.098 | 0 | -0.044 | -0.044 | 2.207 | 0.79 | 0.51 | 0.83 | 0.9 | 0.78 | 0.81 |
| 6 | 161614799 | rs11756131 | -1.343 | 1.144 | 0.373 | 0.103 | 0.026 | 0.019 | 0.019 | 0.037 | 0.25 | 0.39 | 0.23 | 0.66 | 0.29 | 0.32 |
| 6 | 161619810 | rs7769321 | -0.055 | 1.208 | 0.51 | 0.088 | -0.029 | -0.022 | -0.022 | 0.17 | 0.89 | 0.98 | 0.9 | 0.9 | 0.87 | 0.82 |
| 6 | 161619821 | rs7770491 | -0.601 | 0.904 | 0.82 | 0.146 | -0.012 | -0.077 | -0.077 | 0 | 0.12 | 0.52 | 0.08 | 0 | 0.09 | 0.02 |
| 6 | 161622023 | rs12204872 | 0.326 | 1.05 | 0.453 | 0.46 | -0.003 | -0.001 | -0.001 | 0.006 | 0.11 | 0.08 | 0.09 | 0.01 | 0.11 | 0.09 |
| 6 | 161622991 | rs3778227 | -0.582 | 0.854 | 0.759 | 1.062 | -0.049 | 0.086 | 0.086 | 0 | 0.16 | 0.34 | 0.3 | 0.31 | 0.05 | 0.23 |
| 6 | 161624149 | rs4709501 | -1.511 | 1.016 | 0.56 | 0.913 | -0.021 | 0.014 | 0.014 | 0 | 0.39 | 0.52 | 0.47 | 0.41 | 0.27 | 0.49 |
| 6 | 161626204 | rs9458150 | -1.929 | 1.347 | 1.546 | 1.207 | 0.07 | 0.094 | 0.094 | 0.083 | 0.23 | 0.45 | 0.23 | 0.4 | 0.33 | 0.14 |
| 6 | 161628111 | rs9456641 | -1.275 | 1.578 | 1.41 | 1.598 | 0.025 | 0.05 | 0.05 | 0 | 0.13 | 0.14 | 0.17 | 0.41 | 0.24 | 0.13 |
| 6 | 161639071 | rs3778228 | -0.955 | 1.583 | 0.818 | 1.711 | -0.01 | 0.009 | 0.009 | 0 | 0.15 | 0.19 | 0.3 | 0.28 | 0.08 | 0.16 |
| 6 | 161641480 | rs9458157 | -0.675 | 2.274 | 0.744 | 1.535 | 0.025 | 0 | 0 | 0 | 0.05 | 0.04 | 0.11 | 0.14 | 0.17 | 0.07 |
| 6 | 161644540 | rs3778229 | -1.154 | 1.402 | 1.015 | 1.388 | -0.014 | -0.006 | -0.006 | 0 | 0.37 | 0.27 | 0.33 | 0.33 | 0.4 | 0.34 |
| 6 | 161647962 | rs3798937 | -1.04 | 1.539 | 1.471 | 1.399 | -0.001 | 0.021 | 0.021 | 0 | 0.22 | 0.34 | 0.16 | 0.26 | 0.18 | 0.15 |

| | | | | | | | | | | | | | | | | |
|---|---|---|---|---|---|---|---|---|---|---|---|---|---|---|---|---|
| 6 | 161651368 | rs7738367 | -1.654 | 1.255 | 1.422 | 0.678 | -0.009 | 0.02 | 0.02 | 0 | 0.42 | 0.64 | 0.33 | 0.23 | 0.4 | 0.33 |
| 6 | 161655357 | rs3798939 | -0.284 | 1.298 | 1.046 | 0.345 | 0.005 | -0.018 | -0.018 | 0 | 0.18 | 0.17 | 0.16 | 0.05 | 0.23 | 0.19 |
| 6 | 161658435 | rs6455713 | -1.119 | 0.937 | 0.945 | 0.611 | -0.008 | -0.049 | -0.049 | 0.16 | 0.22 | 0.38 | 0.11 | 0.06 | 0.14 | 0.12 |
| 6 | 161665523 | rs3757025 | -0.425 | 0.881 | 0.716 | 1.343 | 0.001 | 0.065 | 0.065 | 0 | 0.06 | 0.07 | 0.04 | 0.3 | 0.08 | 0.07 |
| 6 | 161672443 | rs1884480 | -1.324 | 0.736 | 1.279 | 1.355 | 0.016 | 0.265 | 0.265 | 0 | 0.1 | 0.41 | 0.09 | 0.33 | 0.08 | 0.08 |
| 6 | 161675491 | rs3798943 | 1.242 | 0.708 | 0.855 | 1.146 | -0.001 | -0.001 | -0.001 | 0 | 0.55 | 0.44 | 0.66 | 0.43 | 0.54 | 0.6 |
| 6 | 161675950 | rs6911180 | -1.814 | 0.293 | 0.685 | 0.284 | -0.015 | -0.021 | -0.021 | 0.121 | 0.58 | 0.74 | 0.6 | 0.5 | 0.51 | 0.42 |
| 6 | 161677404 | rs3757030 | -1.935 | 0.251 | 0.662 | 0.294 | -0.035 | -0.044 | -0.044 | 0 | 0.59 | 0.82 | 0.61 | 0.51 | 0.51 | 0.43 |
| 6 | 161677634 | rs3757031 | -1.032 | 0.301 | 0.727 | 0.377 | 0.005 | 0.017 | 0.017 | 0 | 0.29 | 0.65 | 0.47 | 0.44 | 0.27 | 0.29 |
| 6 | 161678751 | rs9458173 | -0.031 | 0.409 | 0.409 | 0.607 | 0.043 | 0.246 | 0.246 | 0 | 0.06 | 0.24 | 0.07 | 0.24 | 0.05 | 0.06 |
| 6 | 161679188 | rs12662114 | -0.843 | 0.394 | -0.042 | 0.417 | -0.02 | 0.316 | 0.316 | 0 | 0.12 | 0.48 | 0.32 | 0.39 | 0.06 | 0.12 |
| 6 | 161680844 | rs3823058 | -0.202 | 0.566 | -1.364 | 0.134 | -0.028 | 0.04 | 0.04 | 0 | 0.19 | 0.02 | 0.09 | 0.11 | 0.15 | 0.14 |
| 6 | 161683974 | rs6926382 | -1.278 | 1.045 | -0.951 | 0.543 | -0.046 | -0.057 | -0.057 | 0 | 0.59 | 0.87 | 0.62 | 0.53 | 0.51 | 0.43 |
| 6 | 161690121 | rs3798955 | -0.12 | 0.943 | -0.804 | 0.857 | 0.001 | 0.003 | 0.003 | 0.096 | 0.07 | 0.02 | 0.03 | 0.22 | 0.05 | 0.05 |
| 6 | 161692286 | rs747866 | -0.804 | 1.076 | -1.054 | 0.61 | -0.022 | 0.001 | 0.001 | 0 | 0.51 | 0.8 | 0.57 | 0.52 | 0.41 | 0.39 |
| 6 | 161697400 | rs3757037 | -1.025 | 1.152 | -0.797 | 0.767 | -0.003 | -0.011 | -0.011 | 0 | 0.37 | 0.79 | 0.3 | 0.41 | 0.33 | 0.26 |
| 6 | 161698258 | rs1321852 | -0.708 | 0.733 | -0.477 | 0.795 | 0.015 | 0.037 | 0.037 | 0 | 0.16 | 0.34 | 0.12 | 0.21 | 0.16 | 0.07 |
| 6 | 161702270 | rs374255 | -0.351 | 0.448 | -1.836 | 0.678 | 0.034 | -0.079 | -0.079 | 0.867 | 0.56 | 0.9 | 0.44 | 0.54 | 0.55 | 0.46 |
| 6 | 161707465 | rs1077281 | 0.44 | 0.218 | -1.252 | 0.4 | -0.011 | -0.002 | -0.002 | 0 | 0.74 | 0.34 | 0.64 | 0.72 | 0.79 | 0.82 |
| 6 | 161712035 | rs1080085 | 0.087 | 1.225 | -1.85 | -0.435 | -0.039 | 0.028 | 0.028 | 0 | 0.19 | 0.17 | 0.2 | 0 | 0.31 | 0.15 |
| 6 | 161712880 | rs9347486 | 0.629 | 0.345 | -2.597 | -0.486 | -0.036 | 0.072 | 0.072 | 0 | 0.3 | 0.03 | 0.23 | 0.13 | 0.24 | 0.25 |
| 6 | 161713967 | rs9364590 | 0.115 | 0.272 | -1.78 | 0.33 | 0 | 0.001 | 0.001 | 0 | 0.24 | 0.2 | 0.22 | 0.57 | 0.24 | 0.43 |
| 6 | 161722057 | rs7760006 | 0.536 | 0.478 | -1.131 | 0.741 | -0.022 | 0.03 | 0.03 | 0 | 0.83 | 0.72 | 0.78 | 0.75 | 0.89 | 0.82 |
| 6 | 161723705 | rs11758285 | -0.57 | 1.2 | -2.226 | 0.463 | -0.089 | -0.155 | -0.155 | 2.17 | 0.25 | 0.01 | 0.22 | 0.43 | 0.35 | 0.18 |
| 6 | 161725297 | rs1407211 | 0.372 | 1.122 | -2.1 | -0.441 | -0.028 | 0.027 | 0.027 | 0 | 0.62 | 0.82 | 0.52 | 0.36 | 0.58 | 0.53 |
| 6 | 161725838 | rs6455718 | -0.227 | 1.227 | -2.164 | -0.34 | -0.06 | -0.071 | -0.071 | 0 | 0.77 | 0.95 | 0.6 | 0.46 | 0.69 | 0.81 |
| 6 | 161731010 | rs9347493 | -0.26 | 0.174 | -1.588 | -1.208 | -0.014 | -0.022 | -0.022 | 1.074 | 0.4 | 0.52 | 0.37 | 0.31 | 0.4 | 0.3 |
| 6 | 161746170 | rs1079951 | 0.332 | 0.236 | -1.725 | -1.182 | -0.028 | -0.029 | -0.029 | 0.098 | 0.25 | 0.06 | 0.3 | 0.3 | 0.28 | 0.24 |
| 6 | 161751279 | rs9365281 | 0.078 | 0.1 | -1.618 | -1.639 | 0.017 | 0.032 | 0.032 | 0 | 0.43 | 0.23 | 0.38 | 0.38 | 0.37 | 0.55 |
| 6 | 161757885 | rs9458222 | 0.464 | 0.301 | -0.985 | -2.749 | -0.011 | 0.03 | 0.03 | 0 | 0.13 | 0.04 | 0.1 | 0 | 0.15 | 0.07 |
| 6 | 161766391 | rs9458225 | -1.204 | 0.288 | -0.771 | -2.961 | -0.005 | -0.07 | -0.07 | 3.369 | 0.44 | 0.56 | 0.31 | 0.08 | 0.45 | 0.54 |
| 6 | 161769835 | rs3734464 | 0.49 | 0.258 | -0.864 | -2.964 | 0.014 | 0.02 | 0.02 | 0 | 0.09 | 0.05 | 0.06 | 0.02 | 0.1 | 0.07 |
| 6 | 161772739 | rs9458229 | -0.851 | 0.384 | -0.738 | -2.882 | -0.003 | -0.119 | -0.119 | 0 | 0.36 | 0.52 | 0.25 | 0.07 | 0.35 | 0.47 |
| 6 | 161772949 | rs3798964 | -0.383 | 0.295 | -0.485 | -2.65 | -0.007 | -0.082 | -0.082 | 0 | 0.22 | 0.4 | 0.14 | 0.06 | 0.21 | 0.3 |
| 6 | 161779706 | rs7752498 | -0.802 | 0.482 | -0.65 | -2.711 | -0.002 | -0.067 | -0.067 | 0.031 | 0.44 | 0.57 | 0.31 | 0.07 | 0.46 | 0.55 |
| 6 | 161782938 | rs2145573 | -0.718 | 0.428 | 0.025 | -2.621 | -0.002 | -0.133 | -0.133 | 0.142 | 0.28 | 0.54 | 0.24 | 0.02 | 0.32 | 0.12 |
| 6 | 161793319 | rs7766508 | -1.108 | 0.519 | -0.233 | -2.691 | 0 | -0.034 | -0.034 | 0 | 0.34 | 0.39 | 0.28 | 0.11 | 0.39 | 0.37 |
| 6 | 161802448 | rs12215325 | -1.102 | 0.472 | -0.363 | -2.734 | 0.003 | -0.041 | -0.041 | 0.047 | 0.13 | 0.16 | 0.15 | 0 | 0.2 | 0.1 |
| 6 | 161803952 | rs3890730 | -1.273 | 0.481 | -0.445 | -2.694 | 0.003 | -0.007 | -0.007 | 0 | 0.17 | 0.17 | 0.27 | 0.12 | 0.27 | 0.29 |
| 6 | 161804054 | rs9458240 | 0.753 | 0.435 | -0.543 | -2.799 | -0.007 | 0.02 | 0.02 | 0.104 | 0.12 | 0.07 | 0.09 | 0 | 0.13 | 0.04 |
| 6 | 161806542 | rs9355894 | -0.859 | 0.589 | -0.849 | -2.708 | -0.003 | -0.017 | -0.017 | 1.238 | 0.13 | 0.17 | 0.15 | 0.08 | 0.17 | 0.26 |
| 6 | 161806657 | rs17641094 | -0.755 | 0.593 | -0.928 | -2.694 | -0.011 | 0.012 | 0.012 | 0 | 0.07 | 0.03 | 0.13 | 0.03 | 0.11 | 0.06 |
| 6 | 161806740 | rs17641124 | 0.847 | 1.319 | -1.288 | -2.754 | -0.049 | -0.029 | -0.029 | 0 | 0.09 | 0.02 | 0.24 | 0.15 | 0.18 | 0.1 |
| 6 | 161808668 | rs4315992 | 0.313 | 1.179 | -0.997 | -3.118 | -0.009 | -0.018 | -0.018 | 0 | 0.07 | 0 | 0.07 | 0.09 | 0.09 | 0.22 |
| 6 | 161810282 | rs12199584 | -0.326 | 1.052 | -1.167 | -3.363 | -0.02 | 0.043 | 0.043 | 0 | 0.13 | 0.04 | 0.13 | 0.03 | 0.18 | 0.09 |
| 6 | 161810908 | rs4131770 | -0.961 | 0.577 | -1.259 | -3.531 | -0.019 | 0.088 | 0.088 | 0.014 | 0.73 | 0.87 | 0.8 | 0.94 | 0.76 | 0.76 |
| 6 | 161812626 | rs11964284 | -0.716 | 0.825 | -1.014 | -3.208 | 0.001 | 0.002 | 0.002 | 0 | 0.13 | 0.14 | 0.15 | 0.08 | 0.17 | 0.26 |
| 6 | 161815583 | rs3924680 | 0.266 | 0.878 | -1.06 | -3.153 | -0.001 | -0.017 | -0.017 | 0.25 | 0.05 | 0.06 | 0.09 | 0 | 0.08 | 0.04 |
| 6 | 161817168 | rs9365285 | -0.622 | 0.766 | -1.311 | -3.288 | -0.023 | 0.097 | 0.097 | 0.335 | 0.72 | 0.88 | 0.79 | 0.93 | 0.74 | 0.7 |
| 6 | 161824515 | rs9347505 | -0.704 | 0.431 | -1.047 | -2.645 | -0.001 | -0.005 | -0.005 | 0.024 | 0.17 | 0.17 | 0.24 | 0.08 | 0.21 | 0.25 |
| 6 | 161830541 | rs9355347 | 0.371 | 0.12 | -0.857 | -2.457 | -0.003 | 0.034 | 0.034 | 0 | 0.3 | 0.17 | 0.38 | 0.85 | 0.26 | 0.35 |
| 6 | 161837789 | rs13202365 | -0.616 | -0.135 | -0.329 | -2.083 | 0.015 | -0.021 | -0.021 | 0 | 0.09 | 0.25 | 0.13 | 0.03 | 0.13 | 0.07 |
| 6 | 161839781 | rs4709517 | -0.102 | -0.309 | -0.983 | -2.135 | -0.003 | -0.027 | -0.027 | 0.044 | 0.44 | 0.54 | 0.5 | 0.81 | 0.4 | 0.42 |
| 6 | 161841791 | rs4454114 | -0.101 | 0.364 | -1.411 | -1.819 | -0.018 | 0.179 | 0.179 | 0 | 0.46 | 0.77 | 0.36 | 0.08 | 0.5 | 0.35 |
| 6 | 161847010 | rs9355898 | -0.03 | -0.335 | -1.026 | -1.495 | 0.033 | 0.036 | 0.036 | 0 | 0.48 | 0.61 | 0.43 | 0.82 | 0.3 | 0.55 |
| 6 | 161847281 | rs9458246 | -0.846 | -0.268 | -1.13 | -2.25 | 0.007 | 0 | 0 | 0.2 | 0.15 | 0.08 | 0.11 | 0 | 0.12 | 0.03 |
| 6 | 161855785 | rs9347511 | 0.013 | -0.007 | -1.026 | -2.746 | -0.002 | -0.008 | -0.008 | 0 | 0.55 | 0.47 | 0.65 | 0.84 | 0.55 | 0.66 |
| 6 | 161859628 | rs12526387 | 0.229 | -0.491 | -1.033 | -2.899 | 0.005 | 0.011 | 0.011 | 0 | 0.07 | 0.04 | 0.04 | 0.01 | 0.04 | 0.09 |
| 6 | 161859655 | rs6909832 | -0.715 | -0.741 | -0.92 | -3.149 | -0.004 | -0.045 | -0.045 | 0 | 0.27 | 0.38 | 0.2 | 0.06 | 0.26 | 0.19 |
| 6 | 161860169 | rs9458252 | -0.831 | -0.377 | -0.816 | -3.753 | 0.002 | -0.028 | -0.028 | 1.828 | 0.61 | 0.61 | 0.4 | 0.1 | 0.57 | 0.38 |
| 6 | 161863510 | rs9458254 | 1.027 | -0.349 | -0.95 | -3.653 | -0.004 | -0.008 | -0.008 | 6.122 | 0.16 | 0.18 | 0.12 | 0.06 | 0.13 | 0.08 |
| 6 | 161863770 | rs9458255 | 1.045 | -0.357 | -0.794 | -3.679 | -0.007 | -0.041 | -0.041 | 0 | 0.16 | 0.19 | 0.12 | 0.01 | 0.12 | 0.06 |
| 6 | 161865232 | rs11752647 | -0.356 | -0.099 | -1.184 | -3.856 | -0.009 | 0.048 | 0.048 | 0 | 0.28 | 0.13 | 0.21 | 0.01 | 0.32 | 0.2 |
| 6 | 161866296 | rs6904671 | -0.419 | 0.426 | -1.035 | -3.461 | -0.007 | 0.019 | 0.019 | 0 | 0.3 | 0.23 | 0.26 | 0.13 | 0.36 | 0.19 |
| 6 | 161867419 | rs12154057 | -0.39 | 0.539 | -0.614 | -2.634 | 0.002 | 0.015 | 0.015 | 0 | 0.46 | 0.57 | 0.28 | 0.16 | 0.4 | 0.29 |
| 6 | 161869692 | rs17647823 | -0.821 | 0.805 | -0.552 | -1.144 | -0.005 | 0.014 | 0.014 | 0 | 0.2 | 0.14 | 0.13 | 0.01 | 0.21 | 0.16 |
| 6 | 161871513 | rs9456674 | -1.347 | 0.651 | 0.215 | -0.236 | 0.02 | 0.026 | 0.026 | 0 | 0.1 | 0.21 | 0.08 | 0.12 | 0.12 | 0.07 |
| 6 | 161874448 | rs11968360 | -1.595 | 0.854 | 0.228 | -0.421 | 0.051 | -0.024 | -0.024 | 0 | 0.07 | 0.19 | 0.07 | 0.04 | 0.11 | 0.03 |
| 6 | 161882221 | rs9346856 | -0.84 | 0.139 | -0.32 | -0.336 | 0.003 | 0.004 | 0.004 | 0 | 0.08 | 0.06 | 0.04 | 0.04 | 0.03 | 0.03 |

| | | | | | | | | | | | | | | | | |
|---|---|---|---|---|---|---|---|---|---|---|---|---|---|---|---|---|
| 6 | 161884630 | rs1886237 | -0.365 | 0.169 | -1.739 | -0.342 | -0.122 | 0.009 | 0.009 | 0 | 0.58 | 0.97 | 0.54 | 0.34 | 0.52 | 0.36 |
| 6 | 161885595 | rs2209249 | -0.509 | -0.416 | -0.346 | -0.726 | -0.037 | -0.102 | -0.102 | 0 | 0.16 | 0.35 | 0.11 | 0.02 | 0.09 | 0.07 |
| 6 | 161886037 | rs16892547 | -1.343 | -0.478 | 0.248 | -0.77 | -0.008 | -0.037 | -0.037 | 0 | 0.26 | 0.68 | 0.17 | 0.06 | 0.13 | 0.1 |
| 6 | 161888411 | rs6903203 | -1.182 | -0.495 | 0.592 | -0.75 | -0.056 | -0.11 | -0.11 | 0 | 0.25 | 0.55 | 0.17 | 0.06 | 0.11 | 0.09 |
| 6 | 161891198 | rs9458273 | -1.545 | -0.791 | 0.348 | -0.468 | 0.039 | 0.043 | 0.043 | 0.646 | 0.31 | 0.78 | 0.38 | 0.17 | 0.21 | 0.24 |
| 6 | 161891388 | rs6906519 | -0.016 | 0.28 | -0.078 | 0.14 | -0.059 | -0.066 | -0.066 | 0 | 0.9 | 0.99 | 0.79 | 0.76 | 0.84 | 0.78 |
| 6 | 161891485 | rs9458274 | 0.29 | 0.011 | 0.49 | -0.017 | -0.018 | -0.023 | -0.023 | 0 | 0.06 | 0.12 | 0.05 | 0.02 | 0.01 | 0 |
| 6 | 161894974 | rs16892570 | -0.805 | -0.024 | 1.139 | -0.077 | 0.025 | -0.036 | -0.036 | 0 | 0.09 | 0.34 | 0.05 | 0 | 0.04 | 0.06 |
| 6 | 161898721 | rs9458277 | -0.942 | -0.014 | 1.435 | -0.1 | -0.003 | 0.043 | 0.043 | 0 | 0.09 | 0.25 | 0.15 | 0.14 | 0.08 | 0.19 |
| 6 | 161899244 | rs9456676 | -1.375 | -0.101 | 1.532 | -0.17 | -0.028 | -0.002 | -0.002 | 0 | 0.16 | 0.35 | 0.19 | 0.14 | 0.13 | 0.2 |
| 6 | 161900199 | rs4709531 | 1.304 | 0.633 | 1.322 | -0.046 | 0.043 | 0.024 | 0.024 | 0.003 | 0.78 | 0.4 | 0.67 | 0.7 | 0.73 | 0.72 |
| 6 | 161901679 | rs7774267 | -0.724 | 0.672 | 1.474 | -0.143 | 0.007 | -0.046 | -0.046 | 0 | 0.08 | 0.17 | 0.04 | 0 | 0.06 | 0.09 |
| 6 | 161906107 | rs9364599 | 1.911 | 0.673 | 1.037 | 0.426 | -0.119 | -0.146 | -0.146 | 0 | 0.58 | 0.04 | 0.43 | 0.51 | 0.58 | 0.51 |
| 6 | 161909253 | rs9355349 | 1.855 | 0.678 | 1.345 | 0.435 | -0.035 | -0.057 | -0.057 | 0 | 0.59 | 0.16 | 0.43 | 0.51 | 0.58 | 0.51 |
| 6 | 161909909 | rs766313 | 0.538 | 0.872 | 1.414 | 0.59 | 0.073 | 0.134 | 0.134 | 0 | 0.87 | 0.73 | 0.71 | 0.67 | 0.78 | 0.79 |
| 6 | 161914659 | rs555637 | 0.971 | 0.602 | 0.2 | -0.082 | -0.096 | -0.054 | -0.054 | 0 | 0.34 | 0.02 | 0.32 | 0.64 | 0.35 | 0.47 |
| 6 | 161916480 | rs9458289 | -0.415 | 0.684 | 0.595 | -0.197 | 0.037 | 0.215 | 0.215 | 2.913 | 0.51 | 0.82 | 0.43 | 0.14 | 0.44 | 0.29 |
| 6 | 161921295 | rs11969944 | 0.118 | 0.676 | 0.928 | -0.196 | 0.002 | 0.015 | 0.015 | 0 | 0.07 | 0.12 | 0.11 | 0.11 | 0.08 | 0.16 |
| 6 | 161923155 | rs577876 | 0.301 | 0.542 | 1.491 | 0.282 | 0.021 | 0.028 | 0.028 | 4.52 | 0.5 | 0.28 | 0.32 | 0.53 | 0.43 | 0.53 |
| 6 | 161923538 | rs693282 | 0.86 | 0.68 | 1.856 | 0.235 | 0.055 | 0.079 | 0.079 | 0 | 0.84 | 0.66 | 0.62 | 0.66 | 0.75 | 0.76 |
| 6 | 161929039 | rs7769885 | 0.149 | 0.605 | 1.735 | 0.166 | 0.006 | -0.017 | -0.017 | 0 | 0.05 | 0.17 | 0.07 | 0 | 0.05 | 0.01 |
| 6 | 161933935 | rs16892673 | -0.239 | 0.644 | 1.629 | 0.133 | -0.006 | -0.045 | -0.045 | 0 | 0.11 | 0.17 | 0.04 | 0 | 0.07 | 0.07 |
| 6 | 161935166 | rs517010 | 0.628 | 0.866 | 1.185 | -0.071 | -0.006 | 0.002 | 0.002 | 3.591 | 0.39 | 0.17 | 0.36 | 0.6 | 0.38 | 0.58 |
| 6 | 161935928 | rs545310 | -0.085 | 0.947 | 1.293 | 0.026 | 0.011 | 0.031 | 0.031 | 0 | 0.9 | 0.86 | 0.69 | 0.65 | 0.81 | 0.77 |
| 6 | 161936234 | rs548060 | 0.824 | 0.923 | 1.358 | 0.008 | 0.012 | 0.018 | 0.018 | 0 | 0.79 | 0.67 | 0.66 | 0.65 | 0.74 | 0.72 |
| 6 | 161938219 | rs569463 | 1.167 | 0.949 | 1.109 | -0.039 | 0.009 | 0.004 | 0.004 | 0 | 0.72 | 0.64 | 0.61 | 0.63 | 0.66 | 0.65 |
| 6 | 161939571 | rs492267 | -0.082 | 0.93 | 1.099 | -0.031 | 0.011 | 0.03 | 0.03 | 0 | 0.9 | 0.86 | 0.69 | 0.65 | 0.81 | 0.77 |
| 6 | 161939647 | rs11965303 | 0.015 | 0.881 | 1.072 | -0.062 | -0.012 | -0.047 | -0.047 | 0 | 0.11 | 0.19 | 0.04 | 0 | 0.06 | 0.05 |
| 6 | 161942019 | rs2223768 | -0.562 | 0.892 | 1.068 | -0.065 | -0.002 | -0.03 | -0.03 | 0 | 0.18 | 0.22 | 0.08 | 0.03 | 0.13 | 0.13 |
| 6 | 161942405 | rs510849 | 1.137 | 1.071 | 1.062 | -0.033 | 0.005 | 0.004 | 0.004 | 0 | 0.72 | 0.64 | 0.64 | 0.63 | 0.69 | 0.66 |
| 6 | 161944019 | rs12209107 | 1.542 | 1.134 | 1.263 | -0.127 | 0.006 | -0.001 | -0.001 | 0.135 | 0.26 | 0.15 | 0.16 | 0 | 0.23 | 0.06 |
| 6 | 161949490 | rs496759 | -0.076 | 1.327 | 1.163 | -0.056 | 0.005 | 0.031 | 0.031 | 0 | 0.9 | 0.87 | 0.71 | 0.66 | 0.84 | 0.78 |
| 6 | 161950447 | rs6455733 | 0.585 | 1.215 | 1.371 | -0.095 | -0.001 | -0.021 | -0.021 | 2.043 | 0.05 | 0.13 | 0.06 | 0 | 0.05 | 0.01 |
| 6 | 161952887 | rs480557 | 0.016 | 1.191 | 1.628 | -0.157 | 0.001 | 0.017 | 0.017 | 0.596 | 0.52 | 0.37 | 0.49 | 0.63 | 0.52 | 0.67 |
| 6 | 161953285 | rs505555 | -0.132 | 1.253 | 1.499 | -0.061 | 0.011 | 0.035 | 0.035 | 0 | 0.9 | 0.87 | 0.72 | 0.66 | 0.81 | 0.78 |
| 6 | 161954107 | rs12189751 | 1.569 | 1.084 | 1.311 | -0.131 | 0.008 | 0.065 | 0.065 | 4.075 | 0.25 | 0.11 | 0.16 | 0 | 0.23 | 0.06 |
| 6 | 161955143 | rs16892736 | 0.017 | 1.071 | 1.241 | 0.002 | 0.018 | -0.008 | -0.008 | 2.422 | 0.06 | 0.2 | 0.07 | 0 | 0.05 | 0.01 |
| 6 | 161956675 | rs574165 | -0.393 | 1.165 | 0.827 | 0.095 | 0 | -0.001 | -0.001 | 0 | 0.58 | 0.4 | 0.49 | 0.64 | 0.56 | 0.74 |
| 6 | 161958165 | rs6928997 | 0.925 | 0.959 | 0.069 | 0.354 | -0.029 | -0.141 | -0.141 | 0 | 0.32 | 0.06 | 0.25 | 0.48 | 0.31 | 0.42 |
| 6 | 161958898 | rs475221 | 0.149 | 1.308 | 0.673 | 0.324 | -0.001 | 0.018 | 0.018 | 5.176 | 0.45 | 0.48 | 0.48 | 0.23 | 0.5 | 0.33 |
| 6 | 161960215 | rs530128 | 0.22 | 1.273 | 0.397 | 0.355 | 0.001 | 0.003 | 0.003 | 0 | 0.08 | 0.12 | 0.1 | 0.11 | 0.09 | 0.08 |
| 6 | 161962121 | rs568670 | 0.46 | 1.259 | 0.667 | 0.579 | 0.026 | 0.024 | 0.024 | 0 | 0.78 | 0.6 | 0.61 | 0.69 | 0.74 | 0.75 |
| 6 | 161963415 | rs527960 | 0.583 | 0.835 | 0.264 | 0.303 | 0.003 | 0.036 | 0.036 | 0.628 | 0.67 | 0.46 | 0.49 | 0.56 | 0.67 | 0.51 |
| 6 | 161963892 | rs12198566 | 1.024 | 0.875 | -0.606 | 0.147 | -0.004 | 0.062 | 0.062 | 6.354 | 0.33 | 0.19 | 0.29 | 0.05 | 0.35 | 0.19 |
| 6 | 161967901 | rs12526148 | 0.347 | 0.845 | -0.576 | 0.772 | -0.031 | -0.098 | -0.098 | 0.046 | 0.3 | 0.07 | 0.19 | 0.45 | 0.31 | 0.35 |
| 6 | 161971398 | rs9355906 | 0.289 | 0.278 | -0.697 | 0.686 | 0.023 | -0.116 | -0.116 | 0.014 | 0.28 | 0.05 | 0.14 | 0.48 | 0.21 | 0.33 |
| 6 | 161971598 | rs9355907 | 0.088 | -0.345 | -0.749 | 0.362 | 0.072 | -0.065 | -0.065 | 0.139 | 0.25 | 0.11 | 0.11 | 0.56 | 0.11 | 0.27 |
| 6 | 161972186 | rs6910938 | 0.276 | -0.285 | -0.331 | -0.614 | -0.004 | 0.006 | 0.006 | 0.173 | 0.59 | 0.6 | 0.41 | 0.67 | 0.59 | 0.43 |
| 6 | 161974956 | rs9364602 | 0.896 | -0.324 | -0.115 | -0.825 | 0.022 | 0.042 | 0.042 | 1.074 | 0.47 | 0.38 | 0.3 | 0.68 | 0.44 | 0.36 |
| 6 | 161975378 | rs9346868 | 0.453 | -0.278 | 0.025 | -0.852 | 0.001 | -0.008 | -0.008 | 0 | 0.57 | 0.46 | 0.36 | 0.69 | 0.53 | 0.44 |
| 6 | 161976974 | rs9355908 | 1.029 | -0.337 | -0.282 | -0.66 | -0.013 | -0.117 | -0.117 | 2.554 | 0.22 | 0.05 | 0.12 | 0.59 | 0.21 | 0.3 |
| 6 | 161978943 | rs6921987 | 0.775 | -0.255 | 0.073 | -0.62 | 0.032 | 0.044 | 0.044 | 1.151 | 0.47 | 0.33 | 0.29 | 0.63 | 0.44 | 0.36 |
| 6 | 161979264 | rs6901951 | 0.251 | -0.189 | 0.242 | -0.989 | -0.017 | -0.052 | -0.052 | 0 | 0.18 | 0.23 | 0.11 | 0.04 | 0.13 | 0.1 |
| 6 | 161984377 | rs6900355 | 0.26 | -0.326 | 0.577 | -1.311 | 0.004 | -0.027 | -0.027 | 0.009 | 0.52 | 0.59 | 0.48 | 0.12 | 0.45 | 0.36 |
| 6 | 161987050 | rs9365300 | 0.215 | -0.445 | 0.937 | -1.014 | -0.014 | 0.011 | 0.011 | 3.226 | 0.31 | 0.6 | 0.21 | 0.57 | 0.27 | 0.31 |
| 6 | 161988513 | rs9456684 | 0.533 | -0.622 | 0.557 | -1.263 | -0.001 | -0.001 | -0.001 | 0 | 0.56 | 0.55 | 0.42 | 0.71 | 0.47 | 0.53 |
| 6 | 161988767 | rs9355351 | 0.222 | -0.636 | 0.323 | -1.229 | -0.041 | -0.007 | -0.007 | 0 | 0.31 | 0.51 | 0.26 | 0.71 | 0.25 | 0.5 |
| 6 | 161989903 | rs885782 | 0.569 | -0.438 | -0.273 | -1.046 | 0.003 | 0 | 0 | 1.017 | 0.32 | 0.23 | 0.19 | 0.67 | 0.31 | 0.37 |
| 6 | 161990339 | rs10945756 | 1.379 | -0.575 | -0.852 | -1.211 | 0.023 | 0.039 | 0.039 | 7.972 | 0.27 | 0.17 | 0.17 | 0 | 0.22 | 0.04 |
| 6 | 161996370 | rs7755681 | 0.442 | 0.06 | -0.682 | -0.412 | -0.051 | 0.041 | 0.041 | 0 | 0.65 | 0.78 | 0.48 | 0.72 | 0.6 | 0.57 |
| 6 | 162006836 | rs9364605 | 0.675 | 0.007 | -0.988 | -0.184 | -0.01 | -0.073 | -0.073 | 0.175 | 0.21 | 0.08 | 0.1 | 0.55 | 0.21 | 0.32 |
| 6 | 162010329 | rs6927285 | 0.856 | -0.063 | -0.733 | -0.247 | -0.022 | 0.057 | 0.057 | 0 | 0.58 | 0.22 | 0.43 | 0.64 | 0.59 | 0.51 |
| 6 | 162019444 | rs9458312 | 0.182 | 0.314 | -1.101 | -0.058 | -0.097 | 0.023 | 0.023 | 0.097 | 0.71 | 0.95 | 0.59 | 0.75 | 0.66 | 0.63 |
| 6 | 162038638 | rs7750988 | 0.058 | 0.221 | -0.559 | 0.456 | -0.001 | -0.015 | -0.015 | 0.784 | 0.22 | 0.16 | 0.11 | 0.56 | 0.23 | 0.29 |
| 6 | 162039061 | rs6455737 | -0.022 | 0.156 | -1.424 | -0.071 | -0.111 | 0.006 | 0.006 | 0.708 | 0.68 | 0.95 | 0.46 | 0.73 | 0.62 | 0.54 |
| 6 | 162039711 | rs12210797 | 1.137 | 0.14 | -0.462 | -0.265 | 0.002 | 0.007 | 0.007 | 1.537 | 0.42 | 0.35 | 0.34 | 0.13 | 0.41 | 0.23 |
| 6 | 162042839 | rs9365308 | -0.174 | 0.052 | -0.503 | 0.344 | -0.011 | -0.058 | -0.058 | 0 | 0.22 | 0.08 | 0.13 | 0.6 | 0.25 | 0.35 |
| 6 | 162043005 | rs9365309 | -0.663 | -0.247 | -0.397 | 0.812 | 0.022 | -0.065 | -0.065 | 0 | 0.21 | 0.1 | 0.13 | 0.51 | 0.16 | 0.32 |
| 6 | 162050324 | rs2144208 | -0.712 | -0.058 | 0.755 | 0.657 | 0.002 | -0.001 | -0.001 | 0 | 0.2 | 0.2 | 0.11 | 0.21 | 0.21 | 0.16 |

| | | | | | | | | | | | | | | | | |
|---|---|---|---|---|---|---|---|---|---|---|---|---|---|---|---|---|
| 6 | 162060468 | rs6455744 | -0.339 | 0.173 | 0.898 | 0.07 | -0.039 | -0.023 | -0.023 | 0 | 0.64 | 0.85 | 0.4 | 0.45 | 0.58 | 0.49 |
| 6 | 162061459 | rs6455745 | -1.318 | -0.155 | 0.85 | -0.186 | 0.014 | 0.008 | 0.008 | 0 | 0.33 | 0.75 | 0.28 | 0.33 | 0.3 | 0.31 |
| 6 | 162063500 | rs7761805 | -0.193 | -0.127 | 1.02 | -0.236 | 0 | 0 | 0 | 0.339 | 0.26 | 0.23 | 0.18 | 0.22 | 0.3 | 0.19 |
| 6 | 162065367 | rs10945761 | 1.911 | -0.189 | 0.924 | -0.287 | -0.018 | 0.127 | 0.127 | 0 | 0.31 | 0.11 | 0.17 | 0.12 | 0.29 | 0.19 |
| 6 | 162065816 | rs6914754 | -0.098 | -0.151 | 0.782 | -0.235 | -0.006 | 0.008 | 0.008 | 0 | 0.25 | 0.21 | 0.18 | 0.21 | 0.3 | 0.18 |
| 6 | 162066421 | rs9647610 | 1.801 | -0.148 | 0.386 | -0.217 | -0.015 | 0.078 | 0.078 | 0 | 0.31 | 0.12 | 0.17 | 0.18 | 0.29 | 0.2 |
| 6 | 162068432 | rs10945764 | -0.847 | -0.158 | 0.284 | -0.211 | -0.085 | -0.07 | -0.07 | 0 | 0.7 | 0.97 | 0.46 | 0.39 | 0.67 | 0.5 |
| 6 | 162068565 | rs10945765 | -1.567 | -0.35 | 0.797 | -0.132 | -0.014 | 0.088 | 0.088 | 0 | 0.39 | 0.86 | 0.3 | 0.27 | 0.38 | 0.31 |
| 6 | 162069753 | rs6939092 | -0.978 | -0.398 | 1.284 | -0.166 | -0.005 | 0.023 | 0.023 | 0 | 0.29 | 0.69 | 0.21 | 0.22 | 0.31 | 0.19 |
| 6 | 162074015 | rs16892840 | -0.539 | -0.216 | 1.305 | -0.077 | -0.01 | -0.013 | -0.013 | 0 | 0.09 | 0.17 | 0.08 | 0.05 | 0.07 | 0.11 |
| 6 | 162076539 | rs11756024 | -0.683 | -0.206 | 1.384 | -0.054 | 0.018 | -0.022 | -0.022 | 0 | 0.25 | 0.33 | 0.2 | 0.22 | 0.31 | 0.18 |
| 6 | 162077619 | rs6908057 | -1.543 | -0.358 | 1.131 | -0.07 | 0.012 | 0.023 | 0.023 | 0 | 0.36 | 0.77 | 0.26 | 0.31 | 0.34 | 0.31 |
| 6 | 162079189 | rs7757239 | -0.014 | -0.733 | 1.262 | 0.001 | -0.016 | -0.031 | -0.031 | 0 | 0.68 | 0.63 | 0.55 | 0.8 | 0.8 | 0.69 |
| 6 | 162081478 | rs7749966 | 0.995 | -0.667 | 1.054 | 0.177 | 0.017 | 0.024 | 0.024 | 0 | 0.48 | 0.31 | 0.35 | 0.51 | 0.58 | 0.44 |
| 6 | 162081763 | rs9365311 | 0.985 | -0.679 | 1.052 | 0.193 | 0.016 | 0.019 | 0.019 | 0 | 0.47 | 0.31 | 0.35 | 0.51 | 0.58 | 0.45 |
| 6 | 162082773 | rs12528063 | 0.215 | -0.618 | 0.782 | 0.217 | -0.008 | 0 | 0 | 0 | 0.17 | 0.11 | 0.19 | 0.21 | 0.23 | 0.15 |
| 6 | 162084808 | rs12210817 | 1.505 | -0.62 | 0.573 | 0.249 | -0.08 | -0.079 | -0.079 | 3.98 | 0.39 | 0.16 | 0.29 | 0.44 | 0.5 | 0.38 |
| 6 | 162086509 | rs9458331 | -1.064 | -0.235 | 0.628 | 0.073 | 0.018 | -0.028 | -0.028 | 0 | 0.07 | 0.25 | 0.06 | 0 | 0.06 | 0.02 |
| 6 | 162090188 | rs910417 | 0.072 | -0.153 | -0.064 | 0.194 | -0.003 | 0.001 | 0.001 | 0 | 0.82 | 0.84 | 0.8 | 0.8 | 0.77 | 0.85 |
| 6 | 162090981 | rs16888746 | 0.498 | -0.344 | -0.482 | 0.178 | 0.01 | -0.015 | -0.015 | 0 | 0.23 | 0.02 | 0.37 | 0.2 | 0.12 | 0.25 |
| 6 | 162092964 | rs9347531 | 0.697 | -0.142 | 0.299 | 0.708 | 0.027 | 0.07 | 0.07 | 0 | 0.76 | 0.51 | 0.76 | 0.69 | 0.72 | 0.85 |
| 6 | 162093021 | rs9355357 | 0.499 | 0.176 | 0.191 | -0.642 | 0 | 0.011 | 0.011 | 0 | 0.48 | 0.44 | 0.34 | 0.61 | 0.55 | 0.61 |
| 6 | 162093941 | rs9347532 | 0.972 | -0.278 | -0.107 | -0.726 | -0.024 | 0.007 | 0.007 | 4.097 | 0.42 | 0.24 | 0.3 | 0.57 | 0.49 | 0.51 |
| 6 | 162094867 | rs4708918 | -0.915 | -1.291 | 0.18 | -0.705 | 0.012 | 0.012 | 0.012 | 0 | 0.48 | 0.52 | 0.51 | 0.33 | 0.31 | 0.52 |
| 6 | 162095089 | rs4708919 | 0.044 | -1.014 | 0.218 | -0.579 | 0.022 | 0.033 | 0.033 | 1.185 | 0.32 | 0.12 | 0.42 | 0.26 | 0.21 | 0.33 |
| 6 | 162095455 | rs4708920 | -0.434 | -0.948 | 0.286 | -0.554 | -0.007 | -0.008 | -0.008 | 0.29 | 0.4 | 0.4 | 0.48 | 0.29 | 0.26 | 0.39 |
| 6 | 162096342 | rs16892910 | 0.274 | -0.519 | -0.511 | -0.391 | 0.017 | 0.026 | 0.026 | 0.359 | 0.27 | 0.06 | 0.39 | 0.21 | 0.17 | 0.25 |
| 6 | 162114707 | rs9365319 | 0.282 | -0.323 | 0.145 | -0.134 | 0 | 0.001 | 0.001 | 0 | 0.88 | 0.82 | 0.88 | 0.84 | 0.84 | 0.86 |
| 6 | 162114809 | rs9364608 | 0.532 | -0.245 | 0.135 | -0.243 | 0.007 | 0 | 0 | 0 | 0.4 | 0.29 | 0.51 | 0.54 | 0.57 | 0.35 |
| 6 | 162119957 | rs9456701 | 0.034 | -0.528 | 0.011 | -0.149 | 0.04 | 0.015 | 0.015 | 0 | 0.54 | 0.35 | 0.57 | 0.58 | 0.7 | 0.55 |
| 6 | 162128543 | rs6921226 | 0.034 | 0.414 | 0.346 | -0.412 | 0.004 | -0.016 | -0.016 | 0 | 0.57 | 0.46 | 0.66 | 0.3 | 0.42 | 0.52 |
| 6 | 162133154 | rs6455762 | -0.21 | 0.535 | 0.505 | -0.473 | -0.015 | 0.016 | 0.016 | 0.092 | 0.6 | 0.65 | 0.7 | 0.29 | 0.44 | 0.53 |
| 6 | 162136058 | rs11962671 | -1.422 | 0.522 | 1.225 | -0.258 | -0.008 | -0.044 | -0.044 | 0 | 0.14 | 0.65 | 0.11 | 0.04 | 0.12 | 0.12 |
| 6 | 162136159 | rs11962721 | -1.01 | 0.466 | 1.394 | -0.259 | -0.001 | -0.041 | -0.041 | 0 | 0.12 | 0.53 | 0.08 | 0.03 | 0.09 | 0.06 |
| 6 | 162141896 | rs1018462 | -1.271 | 0.41 | 0.261 | -0.377 | 0 | 0.016 | 0.016 | 0.007 | 0.35 | 0.75 | 0.28 | 0.22 | 0.34 | 0.31 |
| 6 | 162144016 | rs6455765 | -0.46 | 0.817 | 0.397 | -0.058 | -0.008 | -0.005 | -0.005 | 3.798 | 0.19 | 0.13 | 0.17 | 0.18 | 0.22 | 0.21 |
| 6 | 162157656 | rs4708926 | -0.07 | 0.9 | -0.245 | -0.394 | 0.009 | 0.091 | 0.091 | 0 | 0.52 | 0.68 | 0.5 | 0.78 | 0.5 | 0.59 |
| 6 | 162157860 | rs4708928 | 0.663 | 0.816 | 0.01 | 0.171 | -0.002 | -0.001 | -0.001 | 0 | 0.31 | 0.27 | 0.32 | 0.27 | 0.25 | 0.28 |
| 6 | 162158268 | rs2982903 | 0.815 | 1.075 | -0.629 | -0.248 | -0.043 | 0.069 | 0.069 | 0.036 | 0.27 | 0.02 | 0.18 | 0.12 | 0.22 | 0.25 |
| 6 | 162158580 | rs2180025 | -0.031 | 1.079 | -0.304 | -0.183 | 0.017 | 0.05 | 0.05 | 0 | 0.48 | 0.58 | 0.47 | 0.79 | 0.45 | 0.5 |
| 6 | 162159011 | rs964863 | 0.106 | 1.211 | -0.419 | 0.19 | 0.002 | 0.032 | 0.032 | 0 | 0.16 | 0.17 | 0.14 | 0.52 | 0.2 | 0.23 |
| 6 | 162161666 | rs1884158 | 0.154 | 1.265 | -0.64 | -0.159 | -0.006 | 0.01 | 0.01 | 0 | 0.23 | 0.22 | 0.33 | 0.08 | 0.29 | 0.16 |
| 6 | 162164735 | rs3019450 | -0.239 | 1.308 | -0.572 | -0.138 | 0.014 | -0.044 | -0.044 | 0.021 | 0.76 | 0.62 | 0.65 | 0.92 | 0.69 | 0.84 |
| 6 | 162165362 | rs16892988 | -0.237 | 1.282 | -0.915 | 0.606 | 0.004 | 0.093 | 0.093 | 0 | 0.13 | 0.18 | 0.11 | 0.57 | 0.16 | 0.28 |
| 6 | 162168836 | rs3019444 | -0.154 | 1.093 | -1.427 | -0.147 | -0.044 | 0.081 | 0.081 | 0.271 | 0.28 | 0.02 | 0.17 | 0.12 | 0.21 | 0.24 |
| 6 | 162169823 | rs16893004 | -0.43 | 1.244 | -0.676 | 0.133 | 0.023 | -0.019 | -0.019 | 0 | 0.06 | 0.15 | 0.06 | 0.02 | 0.08 | 0.08 |
| 6 | 162170039 | rs1569836 | -0.375 | 0.314 | -1.098 | 0.191 | 0.007 | 0.003 | 0.003 | 0 | 0.35 | 0.35 | 0.43 | 0.76 | 0.22 | 0.45 |
| 6 | 162170141 | rs9355359 | 0.065 | 0.571 | -1.704 | 0.517 | -0.101 | -0.038 | -0.038 | 0 | 0.59 | 0.92 | 0.43 | 0.46 | 0.49 | 0.44 |
| 6 | 162172718 | rs7760443 | -0.913 | 0.222 | -0.384 | 0.883 | 0.001 | 0.07 | 0.07 | 0.295 | 0.19 | 0.36 | 0.16 | 0.4 | 0.18 | 0.31 |
| 6 | 162182126 | rs9355931 | 0.641 | -0.001 | -0.078 | 0.276 | 0.001 | -0.007 | -0.007 | 1.325 | 0.56 | 0.51 | 0.48 | 0.58 | 0.56 | 0.47 |
| 6 | 162183907 | rs6455771 | 0.394 | 0.105 | 0.204 | 0.068 | 0.014 | -0.047 | -0.047 | 0 | 0.89 | 0.77 | 0.73 | 0.97 | 0.9 | 0.93 |
| 6 | 162192856 | rs3016563 | 1.094 | -0.464 | -0.2 | 0.205 | -0.002 | 0.037 | 0.037 | 0.295 | 0.51 | 0.08 | 0.46 | 0.56 | 0.59 | 0.37 |
| 6 | 162193485 | rs9458364 | -0.601 | -0.567 | 0.413 | 0.219 | -0.007 | -0.012 | -0.012 | 0 | 0.4 | 0.45 | 0.28 | 0.38 | 0.37 | 0.55 |
| 6 | 162199157 | rs12208431 | -0.641 | -0.568 | 0.561 | 0.243 | -0.005 | -0.008 | -0.008 | 0.119 | 0.4 | 0.52 | 0.26 | 0.4 | 0.37 | 0.55 |
| 6 | 162202943 | rs11968578 | -0.651 | -0.46 | 0.597 | -0.367 | -0.001 | -0.025 | -0.025 | 0 | 0.37 | 0.62 | 0.27 | 0.22 | 0.38 | 0.55 |
| 6 | 162204946 | rs3016558 | 1.398 | -0.663 | 1.176 | -0.169 | -0.032 | -0.055 | -0.055 | 0 | 0.85 | 0.66 | 0.93 | 0.94 | 0.93 | 0.91 |
| 6 | 162205731 | rs3019436 | 1.531 | -0.636 | 1.174 | -0.156 | -0.038 | -0.067 | -0.067 | 0 | 0.84 | 0.49 | 0.92 | 0.94 | 0.92 | 0.91 |
| 6 | 162209619 | rs3019435 | 1.636 | -0.622 | 0.911 | -0.149 | -0.041 | -0.069 | -0.069 | 2.26 | 0.83 | 0.49 | 0.91 | 0.94 | 0.92 | 0.87 |
| 6 | 162225428 | rs3019424 | 1.279 | -0.436 | 0.649 | 0.003 | -0.012 | -0.031 | -0.031 | 0.984 | 0.89 | 0.59 | 0.95 | 0.97 | 0.97 | 0.89 |
| 6 | 162234333 | rs7739802 | 0.174 | -0.254 | -0.118 | -0.114 | 0.011 | 0.027 | 0.027 | 0 | 0.49 | 0.58 | 0.44 | 0.36 | 0.53 | 0.46 |
| 6 | 162236075 | rs3016539 | -0.125 | -0.483 | -0.601 | -0.096 | 0.036 | 0.039 | 0.039 | 0 | 0.82 | 0.92 | 0.93 | 0.91 | 0.88 | 0.76 |
| 6 | 162236682 | rs3016538 | 0.458 | 0.061 | -0.637 | -0.392 | -0.014 | 0.014 | 0.014 | 0 | 0.2 | 0.14 | 0.22 | 0.15 | 0.26 | 0.23 |
| 6 | 162236866 | rs1001092 | -0.136 | -0.094 | -0.244 | -0.471 | 0.001 | 0.001 | 0.001 | 0 | 0.51 | 0.47 | 0.69 | 0.71 | 0.59 | 0.59 |
| 6 | 162237644 | rs3016537 | -1.201 | -0.033 | -0.143 | -0.472 | 0.003 | -0.036 | -0.036 | 1.26 | 0.7 | 0.55 | 0.78 | 0.79 | 0.71 | 0.77 |
| 6 | 162237949 | rs3016536 | -0.709 | -0.131 | -0.309 | -0.012 | 0.022 | 0.058 | 0.058 | 0 | 0.49 | 0.35 | 0.56 | 0.69 | 0.45 | 0.57 |
| 6 | 162238518 | rs3019438 | -1.054 | -0.092 | -0.333 | 0.655 | 0.043 | 0.145 | 0.145 | 0 | 0.07 | 0.24 | 0.09 | 0.2 | 0.07 | 0.15 |
| 6 | 162242404 | rs1016085 | 0.934 | -0.138 | -0.523 | 0.555 | -0.015 | -0.111 | -0.111 | 0 | 0.22 | 0.04 | 0.39 | 0.4 | 0.24 | 0.24 |
| 6 | 162244325 | rs9458392 | 0.489 | -0.098 | -0.439 | 0.496 | 0.017 | 0.071 | 0.071 | 0 | 0.51 | 0.65 | 0.43 | 0.31 | 0.52 | 0.42 |
| 6 | 162245686 | rs2022993 | 0.049 | 0.344 | 0.214 | 0.974 | -0.003 | 0.049 | 0.049 | 0 | 0.68 | 0.51 | 0.54 | 0.53 | 0.66 | 0.7 |

| | | | | | | | | | | | | | | | | |
|---|---|---|---|---|---|---|---|---|---|---|---|---|---|---|---|---|
| 6 | 162248169 | rs2022991 | -0.066 | 0.539 | -0.298 | 1.09 | -0.001 | 0.015 | 0.015 | 0 | 0.69 | 0.65 | 0.55 | 0.54 | 0.66 | 0.7 |
| 6 | 162250522 | rs4280941 | 1.483 | 0.647 | 0.004 | 1.237 | 0.005 | 0.095 | 0.095 | 0 | 0.91 | 0.73 | 0.91 | 0.78 | 0.93 | 0.72 |
| 6 | 162252602 | rs9347543 | -0.1 | 0.494 | -0.652 | 1.177 | 0.01 | -0.045 | -0.045 | 0 | 0.29 | 0.15 | 0.22 | 0.42 | 0.24 | 0.21 |
| 6 | 162254146 | rs10155686 | -0.681 | 1.395 | -0.022 | 0.627 | 0.019 | -0.006 | -0.006 | 0 | 0.26 | 0.28 | 0.23 | 0.27 | 0.36 | 0.4 |
| 6 | 162254546 | rs7738279 | -1.191 | 1.528 | 0.121 | 0.335 | 0.025 | -0.048 | -0.048 | 3.329 | 0.1 | 0.15 | 0.1 | 0.01 | 0.14 | 0.12 |
| 6 | 162254823 | rs9347544 | 1.521 | 1.025 | 0.577 | 0.351 | 0.021 | -0.027 | -0.027 | 0 | 0.89 | 0.7 | 0.94 | 0.96 | 0.89 | 0.86 |
| 6 | 162255535 | rs9458393 | -0.23 | 0.427 | 0.056 | 0.637 | 0.026 | 0.037 | 0.037 | 0 | 0.48 | 0.71 | 0.38 | 0.39 | 0.39 | 0.49 |
| 6 | 162256288 | rs9458397 | -0.131 | 0.431 | -0.004 | 0.715 | 0.01 | 0.005 | 0.005 | 1.034 | 0.48 | 0.58 | 0.35 | 0.45 | 0.4 | 0.55 |
| 6 | 162264031 | rs6922518 | 0.103 | 0.028 | 0.017 | 0.606 | 0.01 | 0.024 | 0.024 | 0 | 0.47 | 0.33 | 0.38 | 0.64 | 0.44 | 0.56 |
| 6 | 162266376 | rs12111115 | 0.749 | -0.102 | -0.645 | 0.693 | -0.034 | 0.024 | 0.024 | 0.871 | 0.27 | 0.11 | 0.25 | 0.24 | 0.34 | 0.36 |
| 6 | 162267179 | rs7755755 | -0.485 | 0.077 | -0.249 | 0.465 | 0.005 | -0.025 | -0.025 | 0 | 0.56 | 0.56 | 0.46 | 0.73 | 0.54 | 0.61 |
| 6 | 162270070 | rs10945782 | 0.08 | 0.35 | -0.394 | 0.476 | 0 | 0.005 | 0.005 | 0.402 | 0.46 | 0.56 | 0.45 | 0.29 | 0.44 | 0.32 |
| 6 | 162270741 | rs3798218 | -0.077 | 0.059 | -0.14 | 0.705 | -0.001 | -0.01 | -0.01 | 0 | 0.55 | 0.53 | 0.43 | 0.66 | 0.51 | 0.63 |
| 6 | 162270981 | rs6926642 | -0.221 | 0.104 | -0.106 | 0.76 | 0.001 | -0.011 | -0.011 | 0 | 0.55 | 0.59 | 0.46 | 0.74 | 0.51 | 0.61 |
| 6 | 162274042 | rs10945783 | 0.482 | 0.143 | -0.678 | 0.941 | 0.008 | -0.011 | -0.011 | 0 | 0.17 | 0.1 | 0.18 | 0.29 | 0.09 | 0.16 |
| 6 | 162275397 | rs12665316 | 0.919 | 0.211 | -0.667 | 0.598 | -0.021 | 0.057 | 0.057 | 0.582 | 0.32 | 0.12 | 0.23 | 0.22 | 0.37 | 0.41 |
| 6 | 162279763 | rs13205685 | 0.992 | 0.057 | -0.649 | 0.627 | -0.004 | 0.048 | 0.048 | 0.452 | 0.31 | 0.15 | 0.22 | 0.22 | 0.35 | 0.39 |
| 6 | 162281168 | rs12524695 | 0.42 | 0.494 | -0.232 | 0.306 | 0.034 | 0.059 | 0.059 | 0 | 0.49 | 0.33 | 0.4 | 0.63 | 0.44 | 0.49 |
| 6 | 162283824 | rs9689943 | -0.529 | 0.56 | -0.269 | 0.105 | -0.005 | 0.013 | 0.013 | 4.248 | 0.45 | 0.56 | 0.47 | 0.24 | 0.48 | 0.39 |
| 6 | 162291767 | rs9295174 | -0.624 | 0.229 | -0.059 | 0.663 | 0.002 | 0.036 | 0.036 | 0 | 0.29 | 0.42 | 0.39 | 0.42 | 0.27 | 0.31 |
| 6 | 162292450 | rs9365333 | -0.738 | 0.284 | -0.217 | 0.672 | 0 | 0.015 | 0.015 | 0 | 0.3 | 0.35 | 0.4 | 0.43 | 0.27 | 0.31 |
| 6 | 162296941 | rs9347551 | -0.21 | 0.45 | -0.319 | 0.808 | -0.035 | -0.083 | -0.083 | 0 | 0.2 | 0.05 | 0.28 | 0.41 | 0.22 | 0.28 |
| 6 | 162298351 | rs6909000 | 0.316 | 0.245 | -0.305 | 0.136 | -0.001 | -0.01 | -0.01 | 0 | 0.67 | 0.63 | 0.53 | 0.22 | 0.71 | 0.51 |
| 6 | 162300908 | rs9456721 | 0.513 | 0.184 | -0.391 | 0.024 | 0.012 | 0.101 | 0.101 | 0 | 0.46 | 0.25 | 0.42 | 0.23 | 0.55 | 0.38 |
| 6 | 162302567 | rs768150 | -0.976 | 0.856 | 0.459 | 0.945 | 0.034 | 0.045 | 0.045 | 0.043 | 0.28 | 0.49 | 0.41 | 0.46 | 0.33 | 0.4 |
| 6 | 162305402 | rs9347553 | -1.134 | 0.723 | 0.603 | 1.168 | -0.02 | 0.001 | 0.001 | 0 | 0.4 | 0.7 | 0.47 | 0.6 | 0.43 | 0.54 |
| 6 | 162307273 | rs9364621 | -0.826 | 0.841 | 0.892 | 1.264 | 0.056 | 0.061 | 0.061 | 0 | 0.31 | 0.52 | 0.44 | 0.51 | 0.39 | 0.4 |
| 6 | 162308434 | rs9355947 | -1.425 | 0.529 | 0.243 | 1.093 | -0.001 | 0.064 | 0.064 | 0 | 0.47 | 0.76 | 0.53 | 0.68 | 0.48 | 0.66 |
| 6 | 162310496 | rs9355950 | -0.429 | 0.745 | -0.101 | 1.382 | -0.024 | -0.029 | -0.029 | 0.044 | 0.22 | 0.2 | 0.39 | 0.47 | 0.32 | 0.29 |
| 6 | 162314554 | rs9347555 | -1.168 | 0.695 | -0.452 | 1.192 | -0.043 | 0.006 | 0.006 | 0.011 | 0.4 | 0.83 | 0.47 | 0.63 | 0.37 | 0.54 |
| 6 | 162317097 | rs9355954 | -0.306 | 1.133 | -0.469 | 1.176 | -0.059 | -0.013 | -0.013 | 0 | 0.18 | 0.07 | 0.35 | 0.26 | 0.28 | 0.27 |
| 6 | 162317269 | rs9458413 | -0.902 | 0.096 | 0.251 | 1.036 | -0.068 | 0.01 | 0.01 | 0 | 0.21 | 0.43 | 0.12 | 0.18 | 0.12 | 0.29 |
| 6 | 162319145 | rs9365344 | 0.085 | 0.4 | -0.47 | 0.755 | 0.012 | 0.013 | 0.013 | 0 | 0.47 | 0.32 | 0.47 | 0.58 | 0.41 | 0.36 |
| 6 | 162321377 | rs9355371 | -0.164 | 0.284 | -0.69 | 0.492 | -0.004 | 0.007 | 0.007 | 0.014 | 0.53 | 0.62 | 0.37 | 0.58 | 0.49 | 0.42 |
| 6 | 162322332 | rs9355958 | 0.642 | -0.091 | -1.099 | 0.13 | -0.03 | 0.064 | 0.064 | 0 | 0.25 | 0.1 | 0.18 | 0.11 | 0.31 | 0.14 |
| 6 | 162324657 | rs12214649 | 1.155 | 0.313 | -1.559 | 0.137 | -0.06 | -0.019 | -0.019 | 0 | 0.18 | 0.01 | 0.15 | 0.14 | 0.21 | 0.19 |
| 6 | 162329287 | rs6899760 | 0.532 | -0.106 | -1.075 | -0.215 | -0.003 | -0.07 | -0.07 | 1.171 | 0.4 | 0.48 | 0.28 | 0.11 | 0.42 | 0.16 |
| 6 | 162335467 | rs9347564 | 1.577 | -0.225 | -0.963 | 0.07 | -0.008 | 0 | 0 | 0 | 0.1 | 0.11 | 0.08 | 0.1 | 0.04 | 0.16 |
| 6 | 162336354 | rs9355374 | 0.635 | -0.202 | -0.764 | 0.037 | -0.003 | -0.057 | -0.057 | 0.008 | 0.4 | 0.44 | 0.27 | 0.11 | 0.42 | 0.16 |
| 6 | 162339060 | rs12198926 | 1.719 | 0.052 | -1.12 | -0.143 | -0.04 | 0.056 | 0.056 | 0 | 0.11 | 0 | 0.21 | 0.01 | 0.13 | 0.11 |
| 6 | 162339286 | rs9364628 | -0.069 | 0.016 | -0.797 | 0.002 | 0.012 | 0.094 | 0.094 | 0.676 | 0.52 | 0.66 | 0.36 | 0.21 | 0.46 | 0.32 |
| 6 | 162340950 | rs9365352 | 0.409 | 0.049 | -0.589 | -0.07 | -0.001 | -0.065 | -0.065 | 0.165 | 0.4 | 0.43 | 0.27 | 0.11 | 0.42 | 0.16 |
| 6 | 162344712 | rs9458419 | -0.03 | 0.449 | -0.279 | 0.257 | -0.012 | -0.007 | -0.007 | 0 | 0.57 | 0.71 | 0.47 | 0.56 | 0.55 | 0.48 |
| 6 | 162349894 | rs13194166 | 0.972 | 0.706 | -0.998 | 0.42 | -0.07 | -0.028 | -0.028 | 0 | 0.18 | 0.01 | 0.16 | 0.15 | 0.21 | 0.19 |
| 6 | 162365901 | rs1790022 | -0.542 | 0.057 | -0.077 | 0.238 | 0.001 | 0 | 0 | 0 | 0.59 | 0.62 | 0.45 | 0.55 | 0.54 | 0.52 |
| 6 | 162368885 | rs9458424 | -0.065 | 0.367 | -0.22 | 0.004 | 0.003 | -0.042 | -0.042 | 1.528 | 0.08 | 0.16 | 0.05 | 0 | 0.09 | 0.01 |
| 6 | 162371705 | rs6902304 | 0.311 | 0.396 | -0.355 | 0.019 | 0.001 | -0.025 | -0.025 | 0.02 | 0.06 | 0.09 | 0.04 | 0 | 0.09 | 0.01 |
| 6 | 162375909 | rs1784597 | -0.642 | 0.546 | -0.676 | 0.043 | -0.012 | -0.007 | -0.007 | 0 | 0.29 | 0.18 | 0.25 | 0.34 | 0.41 | 0.29 |
| 6 | 162376946 | rs9458425 | 0.682 | 0.289 | -0.918 | -0.179 | 0.011 | 0.015 | 0.015 | 0 | 0.22 | 0.09 | 0.13 | 0.15 | 0.14 | 0.25 |
| 6 | 162376982 | rs1624390 | 0.197 | 0.518 | -0.805 | -0.1 | -0.018 | -0.029 | -0.029 | 0 | 0.23 | 0.11 | 0.21 | 0.34 | 0.32 | 0.29 |
| 6 | 162382734 | rs1784594 | -0.59 | 0.373 | -0.319 | -0.328 | -0.001 | -0.008 | -0.008 | 0 | 0.41 | 0.33 | 0.32 | 0.45 | 0.4 | 0.53 |
| 6 | 162392902 | rs11964796 | -0.411 | 0.338 | -0.268 | -0.446 | -0.01 | -0.032 | -0.032 | 0 | 0.29 | 0.4 | 0.19 | 0.16 | 0.24 | 0.27 |
| 6 | 162396518 | rs1789995 | -0.674 | 0.349 | -0.314 | -0.428 | -0.006 | -0.026 | -0.026 | 0 | 0.39 | 0.25 | 0.3 | 0.44 | 0.4 | 0.54 |
| 6 | 162396659 | rs1789994 | 0.05 | 0.36 | -0.323 | -0.405 | -0.007 | -0.014 | -0.014 | 0 | 0.19 | 0.14 | 0.19 | 0.29 | 0.26 | 0.28 |
| 6 | 162398976 | rs1789993 | -1.277 | 0.243 | -0.136 | -0.432 | -0.001 | -0.001 | -0.001 | 0.153 | 0.46 | 0.49 | 0.36 | 0.44 | 0.5 | 0.56 |
| 6 | 162401412 | rs1784608 | 0.326 | 0.285 | -0.186 | -0.396 | -0.043 | -0.064 | -0.064 | 0.945 | 0.17 | 0.03 | 0.17 | 0.29 | 0.26 | 0.28 |
| 6 | 162405853 | rs1784607 | -0.719 | 0.251 | -0.074 | -0.412 | 0.02 | 0 | 0 | 0 | 0.26 | 0.38 | 0.25 | 0.28 | 0.36 | 0.29 |
| 6 | 162406957 | rs1789990 | -1.201 | 0.283 | -0.016 | -0.451 | -0.001 | 0.006 | 0.006 | 0 | 0.47 | 0.58 | 0.38 | 0.43 | 0.51 | 0.53 |
| 6 | 162426517 | rs9458430 | 0.399 | 0.45 | -0.022 | -0.545 | 0.001 | 0.001 | 0.001 | 0 | 0.59 | 0.55 | 0.66 | 0.58 | 0.56 | 0.48 |
| 6 | 162428289 | rs9347577 | -0.121 | 0.561 | -0.475 | -0.491 | -0.068 | -0.075 | -0.075 | 0 | 0.82 | 0.97 | 0.82 | 0.73 | 0.73 | 0.75 |
| 6 | 162436969 | rs13201582 | 1.437 | 0.455 | -0.763 | -1.025 | 0 | 0.08 | 0.08 | 0 | 0.29 | 0.13 | 0.18 | 0.04 | 0.29 | 0.22 |
| 6 | 162437387 | rs2156745 | 1.437 | 0.589 | -0.763 | -1.025 | -0.01 | 0.08 | 0.08 | 0 | 0.29 | 0.13 | 0.19 | 0.04 | 0.3 | 0.22 |
| 6 | 162437697 | rs1006650 | 0.274 | 0.793 | -0.484 | -0.791 | -0.003 | 0 | 0 | 0 | 0.77 | 0.78 | 0.73 | 0.75 | 0.65 | 0.76 |
| 6 | 162440687 | rs1893895 | 0.495 | 0.601 | -0.606 | -0.549 | -0.002 | 0.084 | 0.084 | 0 | 0.24 | 0.39 | 0.4 | 0.52 | 0.17 | 0.25 |
| 6 | 162440758 | rs9347579 | 0.013 | 0.788 | -0.935 | -0.273 | -0.047 | -0.046 | -0.046 | 0.11 | 0.09 | 0.01 | 0.16 | 0.18 | 0.18 | 0.15 |
| 6 | 162445941 | rs2849570 | 0.383 | 0.622 | -0.952 | -0.434 | -0.034 | -0.049 | -0.049 | 0 | 0.83 | 0.98 | 0.78 | 0.79 | 0.75 | 0.76 |
| 6 | 162447490 | rs1954917 | 0.558 | 0.514 | -0.702 | -0.632 | 0.004 | 0.015 | 0.015 | 0 | 0.59 | 0.7 | 0.61 | 0.65 | 0.56 | 0.47 |
| 6 | 162447612 | rs1954916 | 0.44 | 0.098 | -0.697 | -1.042 | 0 | 0.025 | 0.025 | 0 | 0.21 | 0.12 | 0.15 | 0.15 | 0.17 | 0.31 |
| 6 | 162450471 | rs2849567 | -1.052 | 0.094 | -0.776 | -1.109 | -0.005 | -0.047 | -0.047 | 0 | 0.91 | 1 | 0.86 | 0.86 | 0.92 | 0.78 |

| | | | | | | | | | | | | | | | | |
|---|---|---|---|---|---|---|---|---|---|---|---|---|---|---|---|---|
| 6 | 162452035 | rs884193 | 0.074 | 0.154 | 0.139 | -1.079 | -0.01 | 0.021 | 0.021 | 0 | 0.08 | 0.26 | 0.06 | 0.09 | 0.07 | 0.06 |
| 6 | 162459664 | rs2849564 | 0.677 | 0.05 | -0.222 | -1.682 | -0.002 | 0.028 | 0.028 | 0 | 0.42 | 0.18 | 0.39 | 0.67 | 0.37 | 0.46 |
| 6 | 162460484 | rs9458437 | 1.215 | 0.077 | -0.565 | -1.861 | -0.089 | 0.048 | 0.048 | 0 | 0.17 | 0.02 | 0.29 | 0.07 | 0.31 | 0.16 |
| 6 | 162461329 | rs6911067 | -0.513 | 0.014 | 0.564 | -1.5 | -0.018 | -0.095 | -0.095 | 0.441 | 0.14 | 0.36 | 0.1 | 0 | 0.12 | 0.04 |
| 6 | 162465203 | rs9458439 | 0.706 | 0.159 | 0.077 | -1.418 | -0.025 | 0.04 | 0.04 | 0 | 0.41 | 0.25 | 0.49 | 0.26 | 0.49 | 0.47 |
| 6 | 162470896 | rs2023048 | -1.551 | 0.179 | 0.146 | -1.33 | 0.011 | 0.018 | 0.018 | 0 | 0.78 | 0.86 | 0.93 | 0.88 | 0.86 | 0.81 |
| 6 | 162472025 | rs2187198 | 0.355 | 0.219 | 0.122 | -1.335 | 0 | 0.045 | 0.045 | 0 | 0.48 | 0.35 | 0.54 | 0.26 | 0.57 | 0.52 |
| 6 | 162474893 | rs9456734 | 0.638 | 0.386 | 0.177 | -1.023 | -0.005 | 0.001 | 0.001 | 0 | 0.17 | 0.15 | 0.3 | 0.54 | 0.17 | 0.24 |
| 6 | 162475434 | rs9365364 | -0.701 | 0.207 | 0.629 | -1.033 | -0.02 | -0.043 | -0.043 | 0 | 0.72 | 0.52 | 0.88 | 0.81 | 0.78 | 0.75 |
| 6 | 162489533 | rs4709566 | -1.259 | 0.396 | 0.146 | -1.144 | 0 | 0.02 | 0.02 | 0 | 0.77 | 0.85 | 0.92 | 0.88 | 0.8 | 0.78 |
| 6 | 162491042 | rs7767694 | -0.124 | 0.446 | 0.273 | -1.254 | 0.003 | 0.054 | 0.054 | 0 | 0.27 | 0.44 | 0.36 | 0.61 | 0.24 | 0.24 |
| 6 | 162492316 | rs6900778 | -0.051 | 0.463 | 0.308 | -1.251 | 0.004 | 0.042 | 0.042 | 0 | 0.27 | 0.4 | 0.35 | 0.61 | 0.23 | 0.24 |
| 6 | 162493682 | rs9458448 | 0.408 | 0.53 | -0.243 | -1.062 | -0.02 | 0.044 | 0.044 | 0 | 0.4 | 0.21 | 0.5 | 0.27 | 0.47 | 0.46 |
| 6 | 162507690 | rs1893551 | -1.207 | 0.674 | -0.456 | -1.104 | -0.05 | 0.11 | 0.11 | 0.384 | 0.68 | 0.91 | 0.74 | 0.86 | 0.72 | 0.73 |
| 6 | 162510974 | rs6908394 | 0.472 | 0.512 | -0.059 | -1.179 | 0.003 | 0.026 | 0.026 | 0.873 | 0.35 | 0.26 | 0.46 | 0.23 | 0.4 | 0.42 |
| 6 | 162511254 | rs9347586 | -0.549 | 0.45 | 0.583 | -1.218 | -0.021 | 0.028 | 0.028 | 0 | 0.29 | 0.43 | 0.25 | 0.63 | 0.27 | 0.27 |
| 6 | 162513967 | rs9346902 | -0.342 | 0.183 | 0.816 | -1.166 | -0.003 | 0.019 | 0.019 | 0.56 | 0.58 | 0.59 | 0.65 | 0.79 | 0.6 | 0.68 |
| 6 | 162514407 | rs7759298 | 0.091 | 0.146 | 0.806 | -1.042 | -0.009 | 0.001 | 0.001 | 0 | 0.07 | 0.17 | 0.07 | 0.09 | 0.08 | 0.01 |
| 6 | 162519165 | rs2187204 | -0.639 | -0.046 | 0.606 | -1.231 | 0.003 | 0.065 | 0.065 | 0 | 0.61 | 0.68 | 0.68 | 0.83 | 0.69 | 0.66 |
| 6 | 162527631 | rs1954925 | -0.29 | -0.034 | 0.758 | -1.134 | -0.003 | -0.014 | -0.014 | 0 | 0.54 | 0.42 | 0.62 | 0.75 | 0.64 | 0.65 |
| 6 | 162530070 | rs9458459 | 0.372 | -0.075 | 0.941 | -1.131 | -0.024 | -0.017 | -0.017 | 0 | 0.08 | 0.17 | 0.06 | 0.04 | 0.04 | 0.07 |
| 6 | 162530432 | rs7761505 | -0.406 | 0.094 | 0.774 | -1.052 | -0.001 | -0.006 | -0.006 | 0 | 0.55 | 0.48 | 0.62 | 0.72 | 0.63 | 0.65 |
| 6 | 162534642 | rs6939051 | 0.27 | 0.171 | 0.094 | -1.101 | 0.003 | 0.013 | 0.013 | 0.007 | 0.45 | 0.56 | 0.38 | 0.25 | 0.36 | 0.31 |
| 6 | 162538351 | rs7756400 | 1.1 | 0.176 | -0.106 | -1.123 | 0.001 | 0.031 | 0.031 | 0.229 | 0.31 | 0.3 | 0.26 | 0.18 | 0.24 | 0.23 |
| 6 | 162539650 | rs9456745 | 1.248 | 0.328 | -0.583 | -1.204 | -0.107 | 0.075 | 0.075 | 0 | 0.21 | 0.02 | 0.31 | 0.05 | 0.33 | 0.18 |
| 6 | 162541706 | rs876145 | 0.148 | 0.229 | -0.219 | -0.962 | 0.001 | 0 | 0 | 0 | 0.49 | 0.58 | 0.4 | 0.24 | 0.4 | 0.35 |
| 6 | 162546042 | rs9355982 | 0.389 | 0.182 | -0.433 | -0.983 | 0.005 | 0.015 | 0.015 | 0 | 0.46 | 0.49 | 0.39 | 0.21 | 0.37 | 0.31 |
| 6 | 162551099 | rs2023051 | -0.121 | -0.246 | -0.609 | -0.238 | -0.008 | 0.167 | 0.167 | 0 | 0.87 | 0.83 | 0.89 | 0.49 | 0.91 | 0.79 |
| 6 | 162559158 | rs12661355 | 0.571 | -0.188 | -0.656 | -0.269 | -0.022 | -0.06 | -0.06 | 0.608 | 0.11 | 0.01 | 0.11 | 0.2 | 0.12 | 0.23 |
| 6 | 162559180 | rs7773019 | -0.565 | -0.11 | -0.052 | -0.55 | -0.015 | -0.042 | -0.042 | 0 | 0.14 | 0.26 | 0.14 | 0.04 | 0.12 | 0.08 |
| 6 | 162565718 | rs4605857 | -0.101 | 0.217 | -0.323 | -0.355 | 0.002 | 0.064 | 0.064 | 0 | 0.51 | 0.71 | 0.45 | 0.26 | 0.43 | 0.37 |
| 6 | 162568613 | rs12202992 | -0.087 | -0.366 | -0.411 | 0.026 | 0.04 | 0.128 | 0.128 | 0.049 | 0.43 | 0.23 | 0.28 | 0.23 | 0.36 | 0.37 |
| 6 | 162571263 | rs713053 | 0.815 | -0.445 | -0.654 | 0.821 | -0.118 | 0.034 | 0.034 | 1.834 | 0.41 | 0.06 | 0.53 | 0.26 | 0.53 | 0.38 |
| 6 | 162577258 | rs2849609 | 0.774 | -0.614 | 0.744 | 0.995 | -0.007 | 0.069 | 0.069 | 0.58 | 0.88 | 0.42 | 0.89 | 0.86 | 0.92 | 0.86 |
| 6 | 162587892 | rs9347594 | 0.681 | -0.666 | -0.124 | 0.774 | -0.11 | 0.035 | 0.035 | 1.01 | 0.44 | 0.03 | 0.54 | 0.26 | 0.56 | 0.41 |
| 6 | 162592807 | rs10945799 | 0.241 | -0.508 | 0.124 | 0.739 | 0.025 | 0.151 | 0.151 | 0.157 | 0.41 | 0.14 | 0.25 | 0.19 | 0.35 | 0.36 |
| 6 | 162597463 | rs12202396 | 0.525 | -0.431 | 0.594 | 1.03 | 0.014 | 0.045 | 0.045 | 2.604 | 0.44 | 0.2 | 0.57 | 0.3 | 0.58 | 0.39 |
| 6 | 162601116 | rs6935949 | 0.46 | -0.433 | 0.702 | 1.138 | 0.019 | 0.007 | 0.007 | 1.185 | 0.46 | 0.33 | 0.59 | 0.4 | 0.59 | 0.44 |
| 6 | 162601721 | rs1954948 | -0.218 | -0.421 | 0.303 | 0.793 | 0.05 | 0.092 | 0.092 | 0.808 | 0.48 | 0.37 | 0.3 | 0.23 | 0.38 | 0.49 |
| 6 | 162605266 | rs9355987 | 0.379 | -0.504 | -0.207 | 0.951 | 0.062 | 0.071 | 0.071 | 2.743 | 0.47 | 0.17 | 0.6 | 0.32 | 0.64 | 0.48 |
| 6 | 162605330 | rs7765100 | -0.738 | -0.435 | 0.478 | 0.971 | -0.017 | 0.035 | 0.035 | 0 | 0.26 | 0.61 | 0.28 | 0.33 | 0.2 | 0.12 |
| 6 | 162606080 | rs9458476 | -1.303 | -0.131 | 0.467 | 0.485 | 0.006 | -0.05 | -0.05 | 1.217 | 0.09 | 0.22 | 0.04 | 0 | 0.05 | 0.02 |
| 6 | 162612003 | rs9355988 | 0.173 | 0.292 | -0.722 | 1.075 | -0.084 | 0.105 | 0.105 | 0 | 0.62 | 0.91 | 0.69 | 0.78 | 0.52 | 0.7 |
| 6 | 162614973 | rs2023075 | 1.108 | 0.389 | -0.943 | 0.834 | -0.008 | 0.036 | 0.036 | 0 | 0.87 | 0.95 | 0.93 | 0.99 | 0.85 | 0.91 |
| 6 | 162615606 | rs12529283 | -0.262 | 0.106 | -1.254 | 0.559 | -0.021 | 0.085 | 0.085 | 0.019 | 0.45 | 0.8 | 0.67 | 0.69 | 0.46 | 0.44 |
| 6 | 162617643 | rs9456751 | 0.857 | 0.103 | -1.431 | 0.567 | -0.006 | 0.101 | 0.101 | 0.163 | 0.44 | 0.12 | 0.26 | 0.29 | 0.4 | 0.49 |
| 6 | 162619277 | rs10945802 | 0.701 | 0.151 | -1.455 | 0.624 | -0.052 | 0.046 | 0.046 | 0 | 0.28 | 0.08 | 0.26 | 0.21 | 0.36 | 0.19 |
| 6 | 162621262 | rs9355387 | 0.303 | 0.162 | -1.369 | 0.717 | -0.015 | -0.041 | -0.041 | 0.708 | 0.09 | 0.02 | 0.26 | 0.12 | 0.07 | 0.14 |
| 6 | 162622828 | rs10945803 | 0.489 | 0.353 | -1.788 | 0.48 | -0.079 | 0.075 | 0.075 | 0 | 0.34 | 0.09 | 0.29 | 0.23 | 0.46 | 0.2 |
| 6 | 162624646 | rs9458481 | 0.733 | 0.321 | -1.739 | 0.367 | -0.001 | 0.057 | 0.057 | 0 | 0.39 | 0.12 | 0.23 | 0.29 | 0.32 | 0.48 |
| 6 | 162627290 | rs9458486 | 0.74 | 0.557 | -2.098 | -0.364 | -0.064 | 0.077 | 0.077 | 0 | 0.23 | 0.01 | 0.11 | 0.1 | 0.21 | 0.18 |
| 6 | 162632171 | rs2023073 | -0.639 | 0.685 | -0.893 | 0.018 | 0.015 | 0.005 | 0.005 | 0 | 0.16 | 0.32 | 0.32 | 0.12 | 0.16 | 0.15 |
| 6 | 162633118 | rs2023071 | 0.161 | 0.83 | -1.803 | -0.104 | -0.113 | 0.064 | 0.064 | 0 | 0.33 | 0.07 | 0.29 | 0.23 | 0.46 | 0.2 |
| 6 | 162633618 | rs4546464 | -0.088 | 0.55 | -1.632 | -0.06 | -0.09 | 0.1 | 0.1 | 0 | 0.38 | 0.14 | 0.56 | 0.21 | 0.52 | 0.31 |
| 6 | 162634337 | rs2187213 | -0.074 | -0.118 | -0.978 | 0.292 | 0.017 | 0.016 | 0.016 | 0 | 0.45 | 0.6 | 0.43 | 0.68 | 0.35 | 0.57 |
| 6 | 162642724 | rs9346906 | -0.897 | -0.688 | -1.012 | 0.948 | 0.034 | -0.054 | -0.054 | 0 | 0.24 | 0.17 | 0.23 | 0.58 | 0.12 | 0.35 |
| 6 | 162646769 | rs12664568 | -0.406 | -0.825 | -0.528 | 0.673 | -0.004 | 0.008 | 0.008 | 0 | 0.13 | 0.19 | 0.13 | 0.18 | 0.11 | 0.3 |
| 6 | 162646892 | rs9365388 | -1.502 | -0.735 | 0.396 | 0.537 | 0.01 | 0.019 | 0.019 | 0 | 0.41 | 0.62 | 0.4 | 0.79 | 0.3 | 0.68 |
| 6 | 162653609 | rs10945805 | -0.28 | -0.687 | 0.398 | 0.567 | -0.003 | 0.004 | 0.004 | 0 | 0.13 | 0.17 | 0.13 | 0.18 | 0.11 | 0.3 |
| 6 | 162660066 | rs12203066 | -1.086 | -0.821 | 0.507 | 1.11 | -0.008 | 0.041 | 0.041 | 0 | 0.18 | 0.31 | 0.15 | 0.3 | 0.14 | 0.41 |
| 6 | 162660989 | rs2096982 | -1.749 | -1.037 | -0.476 | 0.195 | -0.188 | 0.171 | 0.171 | 0 | 0.66 | 0.98 | 0.68 | 0.93 | 0.51 | 0.82 |
| 6 | 162661990 | rs951196 | -1.41 | -1.153 | 0.704 | 0.507 | -0.012 | -0.013 | -0.013 | 0.366 | 0.4 | 0.58 | 0.37 | 0.78 | 0.25 | 0.68 |
| 6 | 162664364 | rs2023068 | -0.806 | -1.237 | 0.846 | 0.496 | -0.003 | 0 | 0 | 0 | 0.33 | 0.34 | 0.34 | 0.66 | 0.21 | 0.56 |
| 6 | 162666206 | rs9346908 | -0.426 | -1.199 | 0.685 | 0.453 | -0.008 | 0.021 | 0.021 | 0 | 0.06 | 0.12 | 0.03 | 0.1 | 0.04 | 0.1 |
| 6 | 162667011 | rs1954939 | -1.607 | -1.214 | 0.8 | 0.453 | 0.047 | 0.17 | 0.17 | 0 | 0.45 | 0.81 | 0.43 | 0.86 | 0.34 | 0.69 |
| 6 | 162668889 | rs6922813 | -0.748 | -1.075 | 0.868 | 0.485 | -0.011 | 0.028 | 0.028 | 0 | 0.07 | 0.15 | 0.03 | 0.12 | 0.04 | 0.12 |
| 6 | 162671390 | rs7770788 | -0.817 | -1.281 | 0.751 | 0.469 | -0.038 | -0.065 | -0.065 | 0 | 0.25 | 0.42 | 0.17 | 0.13 | 0.16 | 0.39 |
| 6 | 162674093 | rs6921358 | -0.868 | -1.277 | 0.762 | 0.471 | -0.041 | -0.068 | -0.068 | 0.646 | 0.25 | 0.44 | 0.17 | 0.13 | 0.16 | 0.42 |
| 6 | 162683876 | rs7744798 | -1.525 | -1.217 | 0.402 | 0.606 | 0.021 | -0.037 | -0.037 | 0 | 0.36 | 0.82 | 0.27 | 0.38 | 0.28 | 0.53 |

| | | | | | | | | | | | | | | | | |
|---|---|---|---|---|---|---|---|---|---|---|---|---|---|---|---|---|
| 6 | 162687814 | rs6930862 | 0.38 | -1.209 | 0.578 | 0.513 | 0 | 0 | 0 | 0.885 | 0.11 | 0.14 | 0.09 | 0.13 | 0.11 | 0.29 |
| 6 | 162690022 | rs6942198 | -1.353 | -1.137 | 0.735 | 0.61 | -0.005 | 0 | 0 | 0.013 | 0.25 | 0.69 | 0.15 | 0.2 | 0.18 | 0.24 |
| 6 | 162690634 | rs1954954 | -0.945 | -1.08 | 0.566 | 0.626 | -0.025 | -0.023 | -0.023 | 2.718 | 0.21 | 0.46 | 0.12 | 0.11 | 0.14 | 0.2 |
| 6 | 162697914 | rs9347608 | 0.687 | -0.977 | -0.147 | 0.998 | 0.012 | -0.042 | -0.042 | 0 | 0.1 | 0.01 | 0.16 | 0.49 | 0.05 | 0.17 |
| 6 | 162709948 | rs9364646 | -0.731 | -1.328 | 0.26 | 0.688 | -0.031 | 0.041 | 0.041 | 0.773 | 0.26 | 0.36 | 0.25 | 0.52 | 0.1 | 0.28 |
| 6 | 162713427 | rs9458499 | 0.223 | -1.474 | -0.24 | -0.034 | -0.023 | -0.011 | -0.011 | 0 | 0.35 | 0.53 | 0.37 | 0.28 | 0.26 | 0.48 |
| 6 | 162716541 | rs952388 | -0.947 | -0.73 | -0.763 | -0.126 | -0.059 | 0.156 | 0.156 | 0 | 0.65 | 0.9 | 0.63 | 0.92 | 0.42 | 0.81 |
| 6 | 162724247 | rs4392700 | -0.834 | -0.631 | -0.447 | 0.561 | 0.001 | -0.001 | -0.001 | 0.035 | 0.25 | 0.67 | 0.28 | 0.55 | 0.17 | 0.28 |
| 6 | 162729381 | rs4709595 | 0.557 | 0.001 | -1.271 | -0.23 | -0.064 | 0.149 | 0.149 | 0 | 0.53 | 0.11 | 0.45 | 0.24 | 0.5 | 0.48 |
| 6 | 162733752 | rs7752706 | 0.118 | -0.457 | -1.075 | -0.426 | -0.018 | 0.083 | 0.083 | 0 | 0.52 | 0.74 | 0.5 | 0.29 | 0.51 | 0.51 |
| 6 | 162736336 | rs6455807 | -1.291 | -0.486 | -1.232 | -0.148 | -0.06 | 0.003 | 0.003 | 0.242 | 0.8 | 0.9 | 0.74 | 0.8 | 0.67 | 0.8 |
| 6 | 162736476 | rs7760647 | -0.323 | -0.72 | -0.329 | -0.148 | 0.017 | -0.004 | -0.004 | 0 | 0.7 | 0.57 | 0.69 | 0.73 | 0.62 | 0.71 |
| 6 | 162737217 | rs9458512 | -0.085 | -0.786 | -0.292 | -0.322 | -0.034 | 0.041 | 0.041 | 0.485 | 0.33 | 0.46 | 0.26 | 0.53 | 0.17 | 0.3 |
| 6 | 162738705 | rs9458515 | 1.106 | -0.756 | -0.382 | -0.317 | -0.075 | 0.126 | 0.126 | 0 | 0.37 | 0.11 | 0.43 | 0.18 | 0.46 | 0.41 |
| 6 | 162738804 | rs7756134 | 0.17 | -0.655 | -0.114 | -0.146 | 0.002 | 0.037 | 0.037 | 0 | 0.07 | 0.15 | 0.03 | 0.06 | 0.04 | 0.06 |
| 6 | 162741184 | rs9365401 | -0.635 | -0.69 | -0.07 | -0.164 | 0.012 | 0.004 | 0.004 | 0 | 0.68 | 0.62 | 0.55 | 0.32 | 0.6 | 0.61 |
| 6 | 162746359 | rs6930880 | -0.499 | -0.688 | -0.073 | -0.121 | 0.009 | 0.01 | 0.01 | 0 | 0.67 | 0.62 | 0.55 | 0.38 | 0.6 | 0.6 |
| 6 | 162748555 | rs9458524 | 0.384 | -0.738 | -0.296 | -0.424 | -0.006 | 0.029 | 0.029 | 0 | 0.54 | 0.37 | 0.47 | 0.25 | 0.5 | 0.52 |
| 6 | 162750426 | rs9347621 | 1.117 | 0.094 | -1.306 | -0.574 | 0.011 | 0.045 | 0.045 | 0 | 0.3 | 0.12 | 0.21 | 0.2 | 0.33 | 0.35 |
| 6 | 162751907 | rs1954950 | -1.433 | 0.449 | -0.878 | -0.435 | -0.062 | 0.047 | 0.047 | 0.165 | 0.8 | 0.89 | 0.73 | 0.89 | 0.67 | 0.8 |
| 6 | 162752050 | rs2000753 | -1.419 | 0.449 | -0.878 | -0.435 | -0.062 | 0.047 | 0.047 | 0.202 | 0.8 | 0.89 | 0.73 | 0.89 | 0.67 | 0.8 |
| 6 | 162767682 | rs6935149 | 0.208 | 0.208 | -0.792 | -0.421 | -0.043 | 0.054 | 0.054 | 0 | 0.39 | 0.3 | 0.38 | 0.24 | 0.45 | 0.3 |
| 6 | 162771688 | rs9456769 | 0.293 | 0.396 | 0.04 | -0.721 | -0.001 | -0.107 | -0.107 | 0.012 | 0.29 | 0.41 | 0.18 | 0.04 | 0.25 | 0.12 |
| 6 | 162779151 | rs10945816 | 0.46 | 0.211 | 0.036 | -0.455 | 0.002 | 0.011 | 0.011 | 0 | 0.27 | 0.32 | 0.4 | 0.64 | 0.32 | 0.36 |
| 6 | 162789197 | rs7768039 | 0.799 | 0.199 | -0.004 | -0.452 | 0.005 | 0.007 | 0.007 | 0 | 0.22 | 0.3 | 0.25 | 0.3 | 0.26 | 0.19 |
| 6 | 162789300 | rs7769367 | -0.31 | 0.134 | 0.316 | -0.525 | -0.001 | -0.002 | -0.002 | 0 | 0.49 | 0.52 | 0.42 | 0.33 | 0.51 | 0.3 |
| 6 | 162790478 | rs6455813 | -1.659 | 0.102 | 0.011 | -0.567 | -0.057 | 0.097 | 0.097 | 0 | 0.64 | 0.93 | 0.61 | 0.82 | 0.58 | 0.56 |
| 6 | 162792115 | rs10806757 | 0.397 | 0.204 | 0.134 | -0.542 | 0.009 | 0.016 | 0.016 | 0 | 0.28 | 0.42 | 0.41 | 0.68 | 0.32 | 0.36 |
| 6 | 162792379 | rs9365409 | 1.917 | 0.073 | -0.732 | -0.933 | -0.029 | 0.039 | 0.039 | 0.001 | 0.17 | 0.01 | 0.1 | 0.08 | 0.17 | 0.28 |
| 6 | 162794024 | rs7752854 | 1.349 | 0.246 | -0.23 | -0.494 | 0.013 | -0.012 | -0.012 | 0 | 0.18 | 0.32 | 0.24 | 0.19 | 0.25 | 0.15 |
| 6 | 162795058 | rs9365410 | 0.539 | -0.132 | -0.073 | -0.289 | -0.026 | 0.267 | 0.267 | 1.936 | 0.15 | 0.37 | 0.19 | 0.51 | 0.07 | 0.27 |
| 6 | 162799117 | rs1008676 | 0.662 | 0.145 | -0.479 | -0.264 | -0.005 | 0.039 | 0.039 | 0.011 | 0.27 | 0.17 | 0.16 | 0.04 | 0.24 | 0.11 |
| 6 | 162800335 | rs4709602 | 1.242 | 0.317 | -0.14 | 0.646 | -0.009 | -0.031 | -0.031 | 0 | 0.09 | 0.01 | 0.16 | 0.43 | 0.07 | 0.19 |
| 6 | 162800665 | rs11759064 | 0.832 | 0.341 | 0.058 | 0.091 | 0.004 | -0.002 | -0.002 | 0 | 0.15 | 0.3 | 0.21 | 0.2 | 0.18 | 0.14 |
| 6 | 162803457 | rs2056909 | 1.335 | -0.091 | -0.339 | 0.867 | 0.015 | -0.02 | -0.02 | 0 | 0.08 | 0.02 | 0.16 | 0.37 | 0.05 | 0.18 |
| 6 | 162803645 | rs2142527 | -0.67 | -0.09 | -0.023 | 0.245 | 0.003 | 0.021 | 0.021 | 0 | 0.47 | 0.54 | 0.53 | 0.72 | 0.46 | 0.51 |
| 6 | 162804504 | rs9456772 | -0.664 | -0.31 | 0.064 | -0.006 | 0.015 | 0.031 | 0.031 | 0.003 | 0.42 | 0.7 | 0.46 | 0.68 | 0.36 | 0.47 |
| 6 | 162809009 | rs10945825 | 0.932 | -0.227 | 0.51 | -0.016 | 0.003 | -0.012 | -0.012 | 0.228 | 0.2 | 0.26 | 0.23 | 0.19 | 0.24 | 0.15 |
| 6 | 162811902 | rs10945826 | 1.736 | -0.251 | 0.35 | -0.029 | -0.038 | -0.005 | -0.005 | 0.708 | 0.16 | 0.07 | 0.21 | 0.19 | 0.24 | 0.15 |
| 6 | 162812916 | rs12665151 | 1.672 | -0.577 | 0.375 | 0.12 | 0.015 | 0.007 | 0.007 | 0 | 0.08 | 0.04 | 0.15 | 0.38 | 0.05 | 0.18 |
| 6 | 162820114 | rs9356012 | 0.462 | -0.488 | 0.559 | -0.025 | 0.004 | 0.004 | 0.004 | 0 | 0.08 | 0.07 | 0.05 | 0.1 | 0.06 | 0.13 |
| 6 | 162828829 | rs9364654 | 0.053 | -0.491 | 0.67 | -0.026 | 0.017 | 0.028 | 0.028 | 0 | 0.09 | 0.23 | 0.07 | 0.12 | 0.06 | 0.14 |
| 6 | 162831336 | rs6455824 | -1.464 | -0.711 | -0.154 | -0.269 | -0.032 | 0.038 | 0.038 | 0 | 0.7 | 0.99 | 0.84 | 0.83 | 0.76 | 0.68 |
| 6 | 162835388 | rs6937352 | -1.092 | -0.599 | 0.68 | -0.069 | -0.007 | 0.077 | 0.077 | 0 | 0.52 | 0.81 | 0.56 | 0.68 | 0.47 | 0.52 |
| 6 | 162836296 | rs13191310 | -1.034 | -0.538 | 0.687 | -0.038 | 0.007 | 0.078 | 0.078 | 0 | 0.51 | 0.77 | 0.56 | 0.68 | 0.47 | 0.51 |
| 6 | 162838024 | rs6911768 | -1.106 | -0.534 | 0.683 | -0.037 | -0.001 | 0.086 | 0.086 | 0 | 0.52 | 0.79 | 0.56 | 0.69 | 0.47 | 0.56 |
| 6 | 162841417 | rs9365417 | -1.042 | -0.53 | 0.769 | -0.033 | 0.01 | 0.045 | 0.045 | 0 | 0.5 | 0.61 | 0.54 | 0.67 | 0.47 | 0.52 |
| 6 | 162841979 | rs10806758 | -0.499 | -0.545 | 0.894 | -0.03 | 0.002 | -0.003 | -0.003 | 0 | 0.4 | 0.33 | 0.47 | 0.57 | 0.4 | 0.38 |
| 6 | 162847889 | rs9347636 | 0.077 | -0.527 | 0.88 | -0.02 | 0.022 | 0.023 | 0.023 | 0 | 0.1 | 0.28 | 0.07 | 0.1 | 0.07 | 0.14 |
| 6 | 162850909 | rs10945830 | -0.433 | -0.545 | 0.874 | -0.028 | -0.006 | 0.002 | 0.002 | 0 | 0.41 | 0.47 | 0.49 | 0.57 | 0.4 | 0.38 |
| 6 | 162851519 | rs7773233 | -1.033 | -0.589 | 0.879 | -0.003 | -0.029 | 0.054 | 0.054 | 0 | 0.31 | 0.53 | 0.33 | 0.52 | 0.23 | 0.43 |
| 6 | 162851541 | rs9458548 | 0.024 | -0.408 | 0.968 | 0.039 | 0.023 | 0.02 | 0.02 | 0 | 0.1 | 0.27 | 0.06 | 0.1 | 0.07 | 0.14 |
| 6 | 162855426 | rs13215957 | -0.509 | -0.403 | 0.643 | 0.032 | -0.001 | 0.004 | 0.004 | 0 | 0.33 | 0.68 | 0.31 | 0.28 | 0.31 | 0.34 |
| 6 | 162864839 | rs10455909 | 0.716 | -0.237 | 0.557 | 0.148 | 0.006 | -0.024 | -0.024 | 2.072 | 0.21 | 0.27 | 0.24 | 0.16 | 0.24 | 0.17 |
| 6 | 162866295 | rs10945834 | -0.285 | -0.279 | -0.158 | 0.235 | -0.001 | 0.019 | 0.019 | 0 | 0.41 | 0.63 | 0.4 | 0.28 | 0.41 | 0.39 |
| 6 | 162868012 | rs6933733 | -0.54 | -0.079 | -0.653 | 0.056 | 0.018 | 0.088 | 0.088 | 0 | 0.5 | 0.76 | 0.56 | 0.68 | 0.46 | 0.57 |
| 6 | 162871034 | rs2142524 | 0.439 | 0.032 | -0.687 | -0.299 | -0.014 | 0.031 | 0.031 | 0 | 0.33 | 0.23 | 0.31 | 0.19 | 0.4 | 0.2 |
| 6 | 162871297 | rs6914080 | -0.568 | -0.521 | -0.005 | 0.163 | 0.005 | 0.024 | 0.024 | 0.066 | 0.41 | 0.68 | 0.49 | 0.72 | 0.36 | 0.53 |
| 6 | 162872974 | rs9346914 | 0.207 | -0.446 | -0.531 | 0.14 | -0.004 | 0.03 | 0.03 | 0 | 0.2 | 0.1 | 0.19 | 0.1 | 0.19 | 0.23 |
| 6 | 162874140 | rs9458560 | -0.339 | -0.637 | -0.278 | 0.536 | 0.02 | -0.026 | -0.026 | 1.361 | 0.32 | 0.73 | 0.39 | 0.57 | 0.28 | 0.32 |
| 6 | 162875706 | rs4709612 | -0.372 | -0.455 | -0.117 | 0.446 | -0.009 | 0.02 | 0.02 | 0 | 0.33 | 0.4 | 0.39 | 0.57 | 0.34 | 0.34 |
| 6 | 162878407 | rs12661450 | -0.624 | -0.532 | -0.015 | 0.836 | -0.004 | 0.279 | 0.279 | 0.613 | 0.14 | 0.43 | 0.18 | 0.38 | 0.11 | 0.19 |
| 6 | 162878694 | rs9347645 | -0.636 | -0.37 | -1.02 | 0.709 | 0.003 | -0.016 | -0.016 | 0 | 0.15 | 0.08 | 0.19 | 0.36 | 0.17 | 0.28 |
| 6 | 162880494 | rs2056907 | -0.755 | -0.274 | -1.56 | 0.097 | -0.179 | -0.036 | -0.036 | 0 | 0.64 | 0.97 | 0.63 | 0.65 | 0.55 | 0.64 |
| 6 | 162881462 | rs7451668 | -0.187 | 0.058 | -0.828 | 0.628 | -0.004 | 0.049 | 0.049 | 0 | 0.14 | 0.17 | 0.18 | 0.3 | 0.1 | 0.2 |
| 6 | 162883484 | rs12174214 | 0.268 | -0.169 | -0.763 | 0.597 | -0.036 | -0.012 | -0.012 | 0.001 | 0.21 | 0.45 | 0.18 | 0.15 | 0.16 | 0.2 |
| 6 | 162884946 | rs1893116 | -0.033 | 0.479 | -1.207 | 0.872 | -0.02 | 0.043 | 0.043 | 0 | 0.62 | 0.8 | 0.65 | 0.69 | 0.62 | 0.66 |
| 6 | 162886686 | rs2186815 | -0.185 | 0.486 | -0.783 | 1.378 | -0.015 | 0.122 | 0.122 | 0.442 | 0.08 | 0.13 | 0.15 | 0.3 | 0.05 | 0.17 |
| 6 | 162888522 | rs4709619 | 0.101 | 1.056 | -1.727 | 0.04 | -0.15 | 0.134 | 0.134 | 0 | 0.7 | 0.98 | 0.7 | 0.92 | 0.62 | 0.82 |

| | | | | | | | | | | | | | | | | |
|---|---|---|---|---|---|---|---|---|---|---|---|---|---|---|---|---|
| 6 | 162889975 | rs7454474 | -0.601 | 0.773 | -1.251 | 0.125 | 0.01 | -0.007 | -0.007 | 0.405 | 0.54 | 0.69 | 0.48 | 0.46 | 0.47 | 0.46 |
| 6 | 162904165 | rs4493732 | -0.662 | 0.993 | -1.222 | 0.092 | -0.008 | -0.007 | -0.007 | 0 | 0.55 | 0.71 | 0.5 | 0.52 | 0.53 | 0.48 |
| 6 | 162906595 | rs1893114 | -0.677 | 0.986 | -1.225 | 0.113 | -0.008 | -0.009 | -0.009 | 0 | 0.55 | 0.71 | 0.5 | 0.52 | 0.53 | 0.48 |
| 6 | 162912514 | rs7740031 | 0.36 | 0.623 | -0.589 | -0.319 | 0.026 | -0.089 | -0.089 | 0.813 | 0.17 | 0.42 | 0.18 | 0.03 | 0.2 | 0.1 |
| 6 | 162914986 | rs9346917 | -0.327 | 0.863 | -0.93 | -0.243 | 0.018 | -0.036 | -0.036 | 0 | 0.37 | 0.24 | 0.32 | 0.48 | 0.35 | 0.36 |
| 6 | 162917942 | rs1893107 | -0.73 | 0.76 | -0.461 | -0.111 | 0 | 0 | 0 | 0 | 0.53 | 0.55 | 0.5 | 0.51 | 0.54 | 0.46 |
| 6 | 162925492 | rs12524235 | -0.022 | 0.735 | -1.233 | -0.395 | -0.026 | 0.091 | 0.091 | 0 | 0.28 | 0.16 | 0.26 | 0.01 | 0.38 | 0.16 |
| 6 | 162927803 | rs2023001 | -0.136 | 0.62 | -1.093 | -0.404 | -0.005 | 0.024 | 0.024 | 0 | 0.3 | 0.27 | 0.26 | 0.03 | 0.38 | 0.17 |
| 6 | 162928898 | rs6930136 | -0.531 | 0.347 | -0.802 | 0.555 | 0.013 | -0.052 | -0.052 | 0.055 | 0.22 | 0.16 | 0.21 | 0.45 | 0.16 | 0.28 |
| 6 | 162933253 | rs2022999 | -0.134 | 0.386 | -0.878 | 0.304 | 0.003 | -0.03 | -0.03 | 0 | 0.23 | 0.16 | 0.23 | 0.5 | 0.21 | 0.25 |
| 6 | 162934144 | rs7764218 | -0.522 | 0.446 | -0.859 | 0.124 | 0 | -0.007 | -0.007 | 0 | 0.08 | 0.11 | 0.09 | 0.05 | 0.06 | 0.19 |
| 6 | 162935431 | rs9458572 | 0.209 | 0.037 | -0.678 | -0.177 | -0.01 | 0.001 | 0.001 | 1.046 | 0.08 | 0.14 | 0.06 | 0.07 | 0.05 | 0.03 |
| 6 | 162936958 | rs6925670 | -0.731 | -0.127 | -1.399 | -0.895 | 0.018 | 0.021 | 0.021 | 0 | 0.47 | 0.27 | 0.41 | 0.54 | 0.41 | 0.54 |
| 6 | 162943061 | rs6908330 | 0.11 | -0.116 | -0.722 | -0.877 | 0.011 | 0.217 | 0.217 | 0.972 | 0.85 | 0.55 | 0.85 | 0.57 | 0.89 | 0.93 |
| 6 | 162945539 | rs6904956 | -0.241 | -0.145 | -0.64 | -0.921 | 0 | -0.024 | -0.024 | 0 | 0.1 | 0.18 | 0.1 | 0.03 | 0.08 | 0.2 |
| 6 | 162946697 | rs2846508 | 0.81 | -0.144 | -0.603 | -0.908 | -0.01 | 0.074 | 0.074 | 0.478 | 0.75 | 0.46 | 0.75 | 0.53 | 0.82 | 0.73 |
| 6 | 162957095 | rs13202905 | -0.142 | 0.109 | -0.93 | -1.31 | -0.006 | 0.059 | 0.059 | 0.004 | 0.23 | 0.1 | 0.2 | 0.03 | 0.23 | 0.31 |
| 6 | 162957575 | rs2846470 | 1.963 | 0.084 | -1.103 | -1.48 | -0.054 | 0.149 | 0.149 | 0.367 | 0.27 | 0.08 | 0.24 | 0.01 | 0.36 | 0.15 |
| 6 | 162962262 | rs2846468 | 0.974 | 0.078 | -1.083 | -1.491 | -0.033 | 0.245 | 0.245 | 1.813 | 0.43 | 0.13 | 0.33 | 0.02 | 0.49 | 0.25 |
| 6 | 162962828 | rs2803073 | -1.151 | 0.002 | -0.971 | -1.294 | -0.01 | -0.004 | -0.004 | 0 | 0.75 | 0.86 | 0.69 | 0.25 | 0.74 | 0.72 |
| 6 | 162966852 | rs13193473 | 0.845 | -0.069 | -1.008 | -1.213 | 0.009 | 0.057 | 0.057 | 0 | 0.16 | 0.04 | 0.09 | 0 | 0.13 | 0.09 |
| 6 | 162967096 | rs6930532 | -0.568 | -0.005 | -0.704 | -1.049 | -0.023 | 0.005 | 0.005 | 0.12 | 0.22 | 0.49 | 0.23 | 0.21 | 0.15 | 0.23 |
| 6 | 162969924 | rs957906 | 1.159 | 0.079 | -0.956 | -0.961 | -0.022 | -0.107 | -0.107 | 0.037 | 0.2 | 0.03 | 0.21 | 0.51 | 0.2 | 0.26 |
| 6 | 162970471 | rs7740421 | -0.035 | 0.09 | -0.915 | -1.008 | -0.001 | -0.019 | -0.019 | 0 | 0.1 | 0.14 | 0.1 | 0.03 | 0.08 | 0.2 |
| 6 | 162972240 | rs9458583 | -1.429 | 0.172 | -0.968 | -1.155 | -0.007 | 0.013 | 0.013 | 0 | 0.67 | 0.66 | 0.65 | 0.74 | 0.59 | 0.82 |
| 6 | 162976142 | rs9365460 | 0.313 | 0.032 | -1.106 | -1.331 | 0.03 | -0.049 | -0.049 | 0 | 0.36 | 0.15 | 0.29 | 0.5 | 0.29 | 0.35 |
| 6 | 162979056 | rs13207671 | 0.62 | -0.01 | -1.069 | -1.271 | 0.02 | 0.078 | 0.078 | 0 | 0.19 | 0.04 | 0.1 | 0 | 0.15 | 0.1 |
| 6 | 162980498 | rs9347666 | 0.092 | 0.009 | -1.066 | -1.273 | 0.026 | -0.042 | -0.042 | 0 | 0.4 | 0.16 | 0.32 | 0.5 | 0.35 | 0.37 |
| 6 | 162992704 | rs4709641 | 1.415 | 0.005 | -1.069 | -1.269 | -0.039 | -0.139 | -0.139 | 0.106 | 0.2 | 0.04 | 0.21 | 0.49 | 0.2 | 0.26 |
| 6 | 162993672 | rs6906849 | 0.109 | 0.138 | -0.867 | -1.274 | 0.003 | 0.034 | 0.034 | 0 | 0.18 | 0.39 | 0.22 | 0.22 | 0.14 | 0.24 |
| 6 | 162994422 | rs2803062 | 1.941 | 0.288 | -0.871 | -1.286 | -0.033 | 0.13 | 0.13 | 0.482 | 0.27 | 0.1 | 0.24 | 0.01 | 0.35 | 0.15 |
| 6 | 162995680 | rs9456798 | -0.943 | 0.136 | -0.833 | -1.163 | 0 | -0.003 | -0.003 | 2.336 | 0.58 | 0.55 | 0.54 | 0.71 | 0.49 | 0.61 |
| 6 | 162996263 | rs9458611 | -0.649 | 0.113 | -0.824 | -1.244 | -0.018 | -0.04 | -0.04 | 0.447 | 0.37 | 0.43 | 0.32 | 0.21 | 0.28 | 0.33 |
| 6 | 163007394 | rs9347669 | -1.305 | 0.131 | -0.94 | -1.335 | -0.011 | 0.021 | 0.021 | 0.41 | 0.63 | 0.69 | 0.64 | 0.95 | 0.54 | 0.63 |
| 6 | 163023054 | rs9364664 | 1.014 | 0.227 | -0.833 | -1.157 | 0 | -0.066 | -0.066 | 0.091 | 0.22 | 0.12 | 0.22 | 0.5 | 0.2 | 0.27 |
| 6 | 163029097 | rs2803054 | 0.469 | 0.189 | -0.918 | -1.322 | -0.001 | 0.096 | 0.096 | 0.199 | 0.54 | 0.33 | 0.37 | 0.04 | 0.61 | 0.47 |
| 6 | 163036511 | rs1893119 | 0.131 | 0.247 | -1.141 | -1.124 | -0.008 | 0.018 | 0.018 | 0 | 0.54 | 0.78 | 0.66 | 0.47 | 0.55 | 0.37 |
| 6 | 163039045 | rs2846530 | 0.44 | 0.258 | -0.982 | -1.115 | 0.006 | 0.093 | 0.093 | 0 | 0.5 | 0.67 | 0.57 | 0.18 | 0.5 | 0.34 |
| 6 | 163046601 | rs9356053 | 1.263 | 0.184 | -1.112 | -1.136 | -0.043 | -0.146 | -0.146 | 0 | 0.21 | 0.02 | 0.22 | 0.48 | 0.22 | 0.27 |
| 6 | 163050848 | rs6911406 | -0.053 | 0.041 | -1.051 | -1.303 | -0.018 | -0.007 | -0.007 | 0.17 | 0.22 | 0.4 | 0.31 | 0.17 | 0.15 | 0.19 |
| 6 | 163059427 | rs2846568 | -0.718 | 0.084 | -1.096 | -1.321 | -0.001 | -0.001 | -0.001 | 0.128 | 0.92 | 0.93 | 0.98 | 0.95 | 0.89 | 0.72 |
| 6 | 163059714 | rs2846566 | -0.118 | 0.053 | -1.115 | -1.357 | -0.008 | 0.06 | 0.06 | 0 | 0.77 | 0.89 | 0.89 | 0.95 | 0.77 | 0.65 |
| 6 | 163068845 | rs2803103 | 0.168 | 0.113 | -1.114 | -1.359 | 0.002 | 0.017 | 0.017 | 0 | 0.54 | 0.78 | 0.67 | 0.46 | 0.57 | 0.37 |
| 6 | 163081812 | rs13202339 | 0.847 | -0.031 | -1.113 | -1.318 | -0.001 | 0.081 | 0.081 | 0 | 0.16 | 0.04 | 0.09 | 0 | 0.12 | 0.08 |
| 6 | 163088783 | rs2156258 | 0.13 | -0.036 | -1.113 | -1.335 | -0.006 | 0.003 | 0.003 | 0 | 0.55 | 0.78 | 0.72 | 0.48 | 0.55 | 0.36 |
| 6 | 163100595 | rs2803086 | 1.631 | -0.079 | -1.131 | -1.372 | -0.048 | 0.029 | 0.029 | 0 | 0.3 | 0.15 | 0.33 | 0.29 | 0.4 | 0.16 |
| 6 | 163122220 | rs1893537 | 1.516 | -0.076 | -1.147 | -1.325 | -0.043 | -0.144 | -0.144 | 0 | 0.2 | 0.02 | 0.17 | 0.5 | 0.21 | 0.27 |
| 6 | 163126601 | rs10945858 | -0.031 | -0.092 | -1.087 | -1.342 | -0.016 | 0.007 | 0.007 | 0 | 0.22 | 0.4 | 0.36 | 0.19 | 0.16 | 0.2 |
| 6 | 163132529 | rs10806763 | 1.509 | -0.075 | -1.119 | -1.326 | -0.043 | -0.143 | -0.143 | 0 | 0.2 | 0.02 | 0.17 | 0.5 | 0.21 | 0.27 |
| 6 | 163132822 | rs13192618 | 0.734 | -0.15 | -1.13 | -1.37 | -0.002 | 0.085 | 0.085 | 0 | 0.16 | 0.04 | 0.14 | 0 | 0.13 | 0.08 |
| 6 | 163134248 | rs2849518 | -0.013 | -0.138 | -1.102 | -1.369 | 0.001 | -0.001 | -0.001 | 0 | 0.75 | 0.8 | 0.89 | 0.97 | 0.77 | 0.63 |
| 6 | 163144683 | rs9347682 | -0.207 | -0.142 | -1.122 | -1.363 | -0.006 | -0.028 | -0.028 | 0 | 0.42 | 0.47 | 0.54 | 0.68 | 0.37 | 0.46 |
| 6 | 163149497 | rs2276201 | -0.863 | -0.078 | -1.173 | -1.334 | 0.046 | 0.08 | 0.08 | 0 | 0.41 | 0.73 | 0.48 | 0.2 | 0.28 | 0.29 |
| 6 | 163151399 | rs9356058 | -1.812 | -0.073 | -1.191 | -1.322 | -0.052 | 0.032 | 0.032 | 0.003 | 0.7 | 0.86 | 0.67 | 0.71 | 0.6 | 0.84 |
| 6 | 163151824 | rs9347684 | 1.328 | -0.045 | -1.207 | -1.334 | -0.032 | -0.137 | -0.137 | 1.487 | 0.21 | 0.05 | 0.16 | 0.49 | 0.2 | 0.25 |
| 6 | 163153238 | rs6455848 | -0.134 | -0.148 | -0.915 | -1.315 | -0.024 | 0.004 | 0.004 | 0 | 0.23 | 0.42 | 0.36 | 0.19 | 0.16 | 0.2 |
| 6 | 163159187 | rs13195186 | -1.385 | -0.144 | -0.933 | -1.303 | 0.023 | 0.155 | 0.155 | 0 | 0.49 | 0.81 | 0.5 | 0.24 | 0.39 | 0.57 |
| 6 | 163162350 | rs7764309 | -0.401 | -0.079 | -0.868 | -1.269 | -0.002 | -0.139 | -0.139 | 3.963 | 0.27 | 0.38 | 0.15 | 0.03 | 0.25 | 0.37 |
| 6 | 163171970 | rs9347686 | -0.413 | -0.001 | -1.029 | -1.218 | -0.034 | -0.017 | -0.017 | 0 | 0.45 | 0.16 | 0.29 | 0.52 | 0.45 | 0.63 |
| 6 | 163176151 | rs7744306 | 0.34 | 0.015 | -1.07 | -1.212 | -0.042 | -0.13 | -0.13 | 0.201 | 0.36 | 0.06 | 0.26 | 0.5 | 0.32 | 0.35 |
| 6 | 163190044 | rs10806765 | 1.365 | -0.025 | -1.078 | -1.216 | -0.037 | -0.139 | -0.139 | 0.175 | 0.21 | 0.02 | 0.17 | 0.5 | 0.2 | 0.27 |
| 6 | 163192038 | rs11961035 | -0.155 | -0.121 | -0.87 | -1.301 | -0.028 | -0.001 | -0.001 | 1.441 | 0.23 | 0.36 | 0.35 | 0.19 | 0.14 | 0.19 |
| 6 | 163201379 | rs7759501 | 0.644 | -0.208 | -0.64 | -1.35 | -0.007 | 0.073 | 0.073 | 0.071 | 0.11 | 0.28 | 0.12 | 0.15 | 0.06 | 0.05 |
| 6 | 163203092 | rs13212746 | -0.021 | -0.101 | -0.84 | -1.34 | -0.015 | 0.085 | 0.085 | 0 | 0.24 | 0.12 | 0.12 | 0.02 | 0.25 | 0.37 |
| 6 | 163203212 | rs11966414 | -0.319 | -0.042 | -0.714 | -1.192 | -0.036 | -0.003 | -0.003 | 0.39 | 0.23 | 0.41 | 0.35 | 0.18 | 0.14 | 0.19 |
| 6 | 163212299 | rs10806767 | -1.422 | 0.177 | -1.021 | -1.074 | -0.039 | 0.029 | 0.029 | 0.004 | 0.69 | 0.8 | 0.66 | 0.69 | 0.59 | 0.84 |
| 6 | 163213454 | rs1333955 | -0.634 | 0.079 | -0.981 | -1.043 | -0.032 | 0.054 | 0.054 | 0.222 | 0.6 | 0.69 | 0.63 | 0.68 | 0.46 | 0.55 |
| 6 | 163219891 | rs1041632 | -1.428 | -0.245 | -0.966 | -0.655 | -0.007 | -0.019 | -0.019 | 0 | 0.55 | 0.75 | 0.6 | 0.52 | 0.45 | 0.57 |
| 6 | 163220087 | rs1041634 | -1.442 | -0.25 | -0.955 | -0.657 | -0.013 | -0.035 | -0.035 | 0 | 0.55 | 0.78 | 0.59 | 0.52 | 0.45 | 0.58 |

| | | | | | | | | | | | | | | | | |
|---|---|---|---|---|---|---|---|---|---|---|---|---|---|---|---|---|
| 6 | 163221792 | rs4458655 | -0.392 | -0.503 | -0.689 | -0.719 | -0.038 | -0.033 | -0.033 | 0.113 | 0.22 | 0.4 | 0.38 | 0.14 | 0.13 | 0.17 |
| 6 | 163231084 | rs9364672 | -1.531 | -0.637 | -0.766 | -0.769 | 0.014 | -0.004 | -0.004 | 7.042 | 0.39 | 0.69 | 0.49 | 0.44 | 0.31 | 0.51 |
| 6 | 163232332 | rs1514338 | -1.072 | -0.605 | -0.773 | -0.773 | -0.001 | -0.005 | -0.005 | 0 | 0.32 | 0.61 | 0.42 | 0.17 | 0.26 | 0.48 |
| 6 | 163239119 | rs9356065 | 0.201 | -0.547 | -0.807 | -0.695 | -0.006 | -0.011 | -0.011 | 2.133 | 0.06 | 0.06 | 0.07 | 0.27 | 0.05 | 0.02 |
| 6 | 163248589 | rs6917499 | -0.707 | -0.599 | -0.777 | -0.762 | -0.043 | -0.021 | -0.021 | 0.752 | 0.25 | 0.47 | 0.35 | 0.21 | 0.14 | 0.17 |
| 6 | 163249395 | rs11963648 | -0.726 | -0.511 | -0.822 | -0.752 | -0.024 | -0.02 | -0.02 | 0 | 0.26 | 0.49 | 0.35 | 0.22 | 0.16 | 0.17 |
| 6 | 163250778 | rs949902 | 0.129 | -0.549 | -1.004 | -0.433 | -0.015 | 0.027 | 0.027 | 0 | 0.12 | 0.02 | 0.21 | 0.08 | 0.11 | 0.14 |
| 6 | 163259260 | rs868708 | -1.441 | -0.51 | -1.051 | -0.397 | 0.05 | 0.004 | 0.004 | 0.311 | 0.49 | 0.69 | 0.56 | 0.56 | 0.34 | 0.34 |
| 6 | 163270143 | rs13202482 | 0.797 | -0.519 | -1.177 | -0.641 | -0.018 | 0.022 | 0.022 | 0 | 0.07 | 0.02 | 0.02 | 0.02 | 0.1 | 0.28 |
| 6 | 163273193 | rs9456814 | -1.432 | -0.518 | -1.107 | -0.573 | 0.022 | 0.089 | 0.089 | 0.023 | 0.52 | 0.78 | 0.49 | 0.33 | 0.4 | 0.6 |
| 6 | 163273326 | rs9456815 | -1.114 | -0.187 | -0.677 | -0.142 | -0.027 | -0.011 | -0.011 | 0 | 0.25 | 0.43 | 0.34 | 0.23 | 0.16 | 0.17 |
| 6 | 163274306 | rs12665063 | 0.044 | 0.206 | -1.345 | 0.273 | -0.014 | -0.041 | -0.041 | 0 | 0.07 | 0.01 | 0.11 | 0.15 | 0.06 | 0.03 |
| 6 | 163279979 | rs9346937 | -0.211 | 0.364 | -2.181 | 0.032 | -0.1 | -0.111 | -0.111 | 0.554 | 0.63 | 0.96 | 0.67 | 0.47 | 0.62 | 0.58 |
| 6 | 163280204 | rs4709655 | 0.118 | 0.016 | -2.279 | -0.336 | 0.011 | 0.018 | 0.018 | 0 | 0.16 | 0.13 | 0.12 | 0.06 | 0.13 | 0.1 |
| 6 | 163284038 | rs1399221 | -0.259 | 0.575 | -2.705 | -0.492 | -0.134 | -0.08 | -0.08 | 1.42 | 0.6 | 0.96 | 0.68 | 0.49 | 0.61 | 0.52 |
| 6 | 163286352 | rs9346938 | -0.706 | 0.155 | -2.046 | -0.79 | 0.013 | 0.001 | 0.001 | 0 | 0.37 | 0.35 | 0.37 | 0.4 | 0.29 | 0.41 |
| 6 | 163289036 | rs13196605 | -0.378 | 0.375 | -1.222 | -0.966 | 0.047 | -0.032 | -0.032 | 0 | 0.08 | 0.24 | 0.04 | 0.02 | 0.1 | 0.05 |
| 6 | 163289067 | rs7760425 | -1.365 | 0.489 | -1.262 | -0.98 | -0.107 | -0.097 | -0.097 | 1.146 | 0.67 | 0.95 | 0.69 | 0.48 | 0.63 | 0.56 |
| 6 | 163291300 | rs6936963 | 0.071 | 0.634 | -0.924 | -0.871 | 0.052 | -0.022 | -0.022 | 0 | 0.07 | 0.25 | 0.04 | 0 | 0.08 | 0.03 |
| 6 | 163291953 | rs11968371 | 1.65 | 0.447 | -0.572 | -0.941 | -0.004 | -0.003 | -0.003 | 0.459 | 0.21 | 0.16 | 0.25 | 0.07 | 0.26 | 0.1 |
| 6 | 163296879 | rs6940541 | -0.179 | 0.263 | -0.508 | -0.867 | -0.013 | 0.058 | 0.058 | 0 | 0.13 | 0.2 | 0.21 | 0.34 | 0.09 | 0.24 |
| 6 | 163301044 | rs9458662 | -0.265 | 0.546 | -0.884 | -0.539 | -0.006 | 0.017 | 0.017 | 0.065 | 0.6 | 0.35 | 0.6 | 0.76 | 0.54 | 0.77 |
| 6 | 163304290 | rs6455854 | 0.481 | 0.368 | -1.301 | -0.854 | -0.01 | 0.21 | 0.21 | 0 | 0.43 | 0.14 | 0.4 | 0.13 | 0.46 | 0.27 |
| 6 | 163305426 | rs9295203 | 0.353 | 0.254 | -1.204 | -0.825 | 0.022 | 0.057 | 0.057 | 0.005 | 0.24 | 0.12 | 0.16 | 0.06 | 0.21 | 0.18 |
| 6 | 163307736 | rs13202023 | -0.247 | 0.149 | -0.505 | -0.495 | -0.027 | 0.195 | 0.195 | 0.016 | 0.15 | 0.47 | 0.24 | 0.36 | 0.1 | 0.24 |
| 6 | 163309796 | rs2874453 | -0.167 | 0.16 | -0.389 | -0.615 | -0.008 | -0.05 | -0.05 | 0.089 | 0.31 | 0.35 | 0.2 | 0.07 | 0.27 | 0.21 |
| 6 | 163312690 | rs9347700 | 0.081 | 0.213 | -0.226 | -0.426 | -0.016 | 0.124 | 0.124 | 0 | 0.13 | 0.28 | 0.22 | 0.36 | 0.09 | 0.24 |
| 6 | 163312845 | rs9365505 | -0.284 | 0.427 | -0.323 | -0.424 | -0.003 | -0.001 | -0.001 | 0 | 0.58 | 0.59 | 0.64 | 0.5 | 0.56 | 0.52 |
| 6 | 163316412 | rs9364679 | 1.537 | 0.416 | -0.707 | -0.345 | -0.11 | -0.108 | -0.108 | 0.158 | 0.27 | 0.03 | 0.33 | 0.35 | 0.31 | 0.4 |
| 6 | 163317274 | rs9356075 | -0.825 | 0.463 | -0.656 | -0.32 | -0.03 | -0.06 | -0.06 | 0 | 0.9 | 0.99 | 0.88 | 0.85 | 0.88 | 0.91 |
| 6 | 163318480 | rs1877771 | -1.083 | 0.445 | -0.201 | -0.629 | -0.047 | -0.025 | -0.025 | 0 | 0.64 | 0.82 | 0.55 | 0.48 | 0.57 | 0.5 |
| 6 | 163319108 | rs908018 | -0.93 | 0.436 | -0.15 | -0.634 | -0.033 | -0.017 | -0.017 | 0.044 | 0.63 | 0.78 | 0.54 | 0.5 | 0.57 | 0.5 |
| 6 | 163320074 | rs7738903 | -0.005 | 0.278 | 0.393 | -0.319 | 0.012 | 0.012 | 0.012 | 0 | 0.24 | 0.18 | 0.16 | 0.07 | 0.2 | 0.17 |
| 6 | 163320125 | rs7738930 | -0.273 | 0.204 | 0.594 | -0.212 | -0.036 | 0.168 | 0.168 | 0.197 | 0.1 | 0.27 | 0.09 | 0.35 | 0.02 | 0.18 |
| 6 | 163321232 | rs13211224 | -0.283 | 0.485 | 0.34 | -0.303 | 0.014 | -0.032 | -0.032 | 0 | 0.07 | 0.12 | 0.04 | 0 | 0.08 | 0.04 |
| 6 | 163324842 | rs915013 | 0.709 | 0.681 | 0.12 | -0.296 | -0.003 | 0.001 | 0.001 | 0 | 0.62 | 0.59 | 0.69 | 0.55 | 0.67 | 0.53 |
| 6 | 163326692 | rs9295206 | -1.52 | 0.386 | -0.066 | -0.25 | 0.044 | -0.062 | -0.062 | 0 | 0.46 | 0.92 | 0.41 | 0.52 | 0.36 | 0.5 |
| 6 | 163328595 | rs1399222 | -0.615 | 0.506 | 0.433 | -0.212 | 0 | -0.041 | -0.041 | 0 | 0.08 | 0.14 | 0.04 | 0 | 0.07 | 0.04 |
| 6 | 163332533 | rs2048697 | -0.904 | 0.532 | 0.582 | 0.052 | 0.069 | 0.088 | 0.088 | 0.899 | 0.06 | 0.35 | 0.11 | 0.09 | 0.04 | 0.07 |
| 6 | 163333585 | rs1828819 | -0.676 | 0.378 | 0.2 | -0.007 | -0.002 | 0 | 0 | 0.119 | 0.32 | 0.32 | 0.26 | 0.44 | 0.26 | 0.39 |
| 6 | 163339332 | rs9365506 | 1.238 | 0.962 | 0.017 | -0.008 | -0.025 | -0.008 | -0.008 | 0 | 0.54 | 0.27 | 0.52 | 0.5 | 0.53 | 0.46 |
| 6 | 163340367 | rs2151652 | 0.213 | 1.121 | 0.363 | 0.048 | -0.007 | 0.014 | 0.014 | 0 | 0.85 | 0.88 | 0.84 | 0.96 | 0.82 | 0.8 |
| 6 | 163354979 | rs1588787 | 1.193 | 1.058 | 0.807 | 0.14 | 0.015 | 0.018 | 0.018 | 0 | 0.7 | 0.39 | 0.49 | 0.52 | 0.59 | 0.63 |
| 6 | 163361196 | rs9347707 | 0.449 | 1.114 | 1.202 | 0.299 | -0.011 | -0.002 | -0.002 | 0 | 0.78 | 0.8 | 0.63 | 0.78 | 0.65 | 0.74 |
| 6 | 163371058 | rs9347712 | 1.279 | 1.066 | 1.747 | 0.346 | 0.08 | 0.267 | 0.267 | 0.365 | 0.95 | 0.74 | 0.88 | 0.74 | 0.94 | 0.9 |
| 6 | 163376777 | rs4593343 | 1.11 | 1.057 | 1.68 | 0.273 | 0.047 | 0.053 | 0.053 | 0 | 0.72 | 0.57 | 0.51 | 0.52 | 0.6 | 0.64 |
| 6 | 163378724 | rs7769707 | 1.344 | 1.047 | 1.269 | 0.31 | -0.011 | -0.013 | -0.013 | 5.355 | 0.69 | 0.31 | 0.49 | 0.51 | 0.59 | 0.64 |
| 6 | 163387377 | rs9458689 | -1.711 | 0.722 | 1.422 | 0.539 | 0.077 | 0.26 | 0.26 | 0.269 | 0.07 | 0.4 | 0.12 | 0.26 | 0.06 | 0.11 |
| 6 | 163395063 | rs4329088 | 0.674 | 0.405 | 0.577 | 0.129 | -0.019 | 0.029 | 0.029 | 0 | 0.55 | 0.28 | 0.42 | 0.57 | 0.52 | 0.6 |
| 6 | 163401767 | rs6909464 | 0.837 | -0.027 | 0.515 | 0.231 | 0.038 | -0.015 | -0.015 | 1.741 | 0.44 | 0.2 | 0.34 | 0.48 | 0.37 | 0.45 |
| 6 | 163420626 | rs4709670 | 0.003 | -0.14 | 0.568 | -0.169 | 0.046 | 0.138 | 0.138 | 0.74 | 0.54 | 0.83 | 0.47 | 0.7 | 0.39 | 0.55 |
| 6 | 163429010 | rs1324965 | 0.808 | -0.129 | 0.537 | -0.101 | 0.039 | 0.075 | 0.075 | 0.036 | 0.45 | 0.14 | 0.41 | 0.63 | 0.37 | 0.5 |
| 6 | 163430436 | rs2025546 | 0.095 | 0.18 | 1.067 | -0.214 | -0.002 | -0.018 | -0.018 | 0 | 0.56 | 0.39 | 0.55 | 0.92 | 0.51 | 0.59 |
| 6 | 163435051 | rs6924858 | -0.912 | 0.357 | 1.658 | 0.23 | 0.022 | 0.079 | 0.079 | 0 | 0.13 | 0.33 | 0.14 | 0.29 | 0.15 | 0.09 |
| 6 | 163448750 | rs1408710 | -0.466 | 0.349 | 1.581 | 0.172 | 0.016 | 0.063 | 0.063 | 0 | 0.12 | 0.25 | 0.14 | 0.3 | 0.15 | 0.13 |
| 6 | 163450269 | rs9347717 | 0.186 | 0.027 | 1.302 | -0.286 | -0.013 | -0.016 | -0.016 | 0 | 0.4 | 0.52 | 0.37 | 0.66 | 0.29 | 0.48 |
| 6 | 163455782 | rs3939989 | -0.608 | 0.07 | 0.806 | -0.292 | 0.015 | 0.256 | 0.256 | 0 | 0.52 | 0.83 | 0.51 | 0.96 | 0.43 | 0.62 |
| 6 | 163459795 | rs1536034 | 0.241 | -0.002 | 0.765 | -0.185 | -0.005 | -0.006 | -0.006 | 0 | 0.4 | 0.58 | 0.36 | 0.66 | 0.28 | 0.48 |
| 6 | 163475289 | rs575885 | -0.63 | 0.468 | -0.006 | -0.411 | -0.088 | 0.27 | 0.27 | 0 | 0.56 | 0.89 | 0.52 | 0.94 | 0.48 | 0.67 |
| 6 | 163476794 | rs547694 | -0.608 | 0.41 | 0.064 | -0.43 | -0.066 | 0.262 | 0.262 | 0 | 0.55 | 0.87 | 0.52 | 0.96 | 0.47 | 0.67 |
| 6 | 163477104 | rs571644 | -0.684 | 0.378 | 0.51 | -0.44 | -0.011 | 0.065 | 0.065 | 0 | 0.54 | 0.69 | 0.5 | 0.94 | 0.46 | 0.67 |
| 6 | 163486937 | rs561440 | 0.131 | 0.417 | 0.301 | 0.251 | 0.007 | 0.005 | 0.005 | 0 | 0.37 | 0.18 | 0.35 | 0.61 | 0.3 | 0.49 |
| 6 | 163489668 | rs2763983 | 0.31 | 0.255 | 0.195 | 0.395 | 0.002 | -0.033 | -0.033 | 0.039 | 0.23 | 0.13 | 0.27 | 0.62 | 0.19 | 0.45 |
| 6 | 163496289 | rs1575472 | 0.393 | 0.207 | 0.063 | 1.106 | -0.038 | -0.025 | -0.025 | 0 | 0.09 | 0.02 | 0.16 | 0.35 | 0.13 | 0.24 |
| 6 | 163507051 | rs573070 | -0.481 | 0.078 | 0.154 | 1.293 | 0.014 | -0.049 | -0.049 | 0 | 0.28 | 0.18 | 0.34 | 0.46 | 0.22 | 0.43 |
| 6 | 163507487 | rs520741 | -0.511 | 0.042 | 0.406 | 1.328 | -0.02 | 0.051 | 0.051 | 0 | 0.3 | 0.41 | 0.38 | 0.54 | 0.22 | 0.43 |
| 6 | 163521368 | rs548705 | 0.012 | 0.223 | -0.295 | 1.297 | -0.032 | -0.047 | -0.047 | 0.599 | 0.24 | 0.1 | 0.14 | 0.28 | 0.25 | 0.38 |
| 6 | 163527859 | rs11964879 | 0.919 | 0.254 | -0.276 | 0.786 | 0.002 | 0.021 | 0.021 | 0.106 | 0.12 | 0.06 | 0.06 | 0 | 0.1 | 0.04 |
| 6 | 163528806 | rs577398 | -0.985 | -0.018 | 0.086 | 0.802 | 0.02 | -0.016 | -0.016 | 0 | 0.46 | 0.69 | 0.48 | 0.54 | 0.38 | 0.47 |

| | | | | | | | | | | | | | | | | |
|---|---|---|---|---|---|---|---|---|---|---|---|---|---|---|---|---|
| 6 | 163529076 | rs535479 | -0.972 | -0.027 | 0.32 | 0.818 | 0.007 | -0.005 | -0.005 | 0 | 0.44 | 0.56 | 0.34 | 0.47 | 0.37 | 0.47 |
| 6 | 163530425 | rs568619 | -0.107 | 0.188 | 0.635 | 1.111 | -0.023 | -0.099 | -0.099 | 0.536 | 0.27 | 0.11 | 0.25 | 0.47 | 0.27 | 0.42 |
| 6 | 163530507 | rs524763 | -0.861 | 0.214 | 0.522 | 1.126 | 0.006 | -0.02 | -0.02 | 0 | 0.34 | 0.63 | 0.42 | 0.54 | 0.28 | 0.44 |
| 6 | 163536607 | rs520273 | -0.504 | 0.658 | -0.428 | 0.884 | -0.054 | -0.039 | -0.039 | 0.192 | 0.75 | 0.91 | 0.72 | 0.77 | 0.65 | 0.67 |
| 6 | 163538017 | rs9295211 | -1.153 | 0.384 | 0.373 | 0.818 | 0.086 | 0.14 | 0.14 | 2.548 | 0.05 | 0.41 | 0.16 | 0.07 | 0.01 | 0.01 |
| 6 | 163543699 | rs536980 | -0.446 | 0.779 | -1.085 | 1.071 | -0.062 | -0.083 | -0.083 | 0 | 0.24 | 0.04 | 0.28 | 0.27 | 0.24 | 0.36 |
| 6 | 163544705 | rs6908157 | 0.291 | 0.433 | -0.902 | 1.017 | 0.006 | -0.009 | -0.009 | 0 | 0.13 | 0.08 | 0.18 | 0.23 | 0.09 | 0.07 |
| 6 | 163546499 | rs2476467 | -0.706 | -0.155 | -0.37 | 0.967 | -0.013 | -0.001 | -0.001 | 0 | 0.32 | 0.56 | 0.28 | 0.25 | 0.18 | 0.37 |
| 6 | 163546636 | rs6455864 | -0.899 | -0.096 | -0.684 | 0.951 | 0.006 | -0.007 | -0.007 | 0 | 0.29 | 0.66 | 0.26 | 0.35 | 0.24 | 0.23 |
| 6 | 163547053 | rs2476468 | 0.342 | 0.556 | -0.9 | 0.333 | -0.012 | -0.007 | -0.007 | 1.199 | 0.66 | 0.34 | 0.64 | 0.37 | 0.61 | 0.62 |
| 6 | 163547203 | rs2874514 | 1.532 | 0.835 | -1.269 | 0.087 | -0.063 | 0.243 | 0.243 | 0 | 0.32 | 0.04 | 0.24 | 0 | 0.35 | 0.22 |
| 6 | 163549871 | rs554427 | -0.585 | 1.19 | -0.961 | 0.414 | 0.009 | 0.023 | 0.023 | 0 | 0.2 | 0.26 | 0.32 | 0.47 | 0.28 | 0.23 |
| 6 | 163550269 | rs558017 | -0.469 | 1.189 | -0.955 | 0.413 | 0.019 | 0.042 | 0.042 | 0 | 0.2 | 0.3 | 0.32 | 0.47 | 0.28 | 0.23 |
| 6 | 163551376 | rs504844 | -0.771 | 1.211 | -1.168 | 0.49 | -0.017 | -0.027 | -0.027 | 0 | 0.83 | 0.89 | 0.82 | 0.72 | 0.8 | 0.81 |
| 6 | 163551992 | rs498657 | 0.659 | 1.156 | -1.167 | 0.351 | -0.017 | -0.07 | -0.07 | 0 | 0.18 | 0.06 | 0.18 | 0.25 | 0.16 | 0.33 |
| 6 | 163553313 | rs520929 | -0.775 | 1.178 | -1.209 | 0.317 | -0.014 | -0.022 | -0.022 | 0 | 0.86 | 0.89 | 0.82 | 0.7 | 0.81 | 0.82 |
| 6 | 163553748 | rs16894545 | 0.458 | 1.047 | -1.097 | 0.111 | 0.001 | 0 | 0 | 0 | 0.13 | 0.15 | 0.07 | 0.11 | 0.15 | 0.11 |
| 6 | 163554041 | rs566126 | -0.475 | 0.939 | -1.177 | -0.167 | -0.017 | 0.098 | 0.098 | 0 | 0.4 | 0.79 | 0.35 | 0.21 | 0.39 | 0.25 |
| 6 | 163555157 | rs483222 | 0.818 | 0.914 | -1.133 | 0.069 | 0.012 | -0.04 | -0.04 | 0.252 | 0.18 | 0.07 | 0.19 | 0.26 | 0.12 | 0.33 |
| 6 | 163555732 | rs13209731 | 0.329 | 1.065 | -0.92 | -0.105 | 0.005 | 0 | 0 | 0 | 0.13 | 0.15 | 0.07 | 0.11 | 0.18 | 0.11 |
| 6 | 163559907 | rs1033579 | 0.703 | 0.894 | -0.153 | -0.219 | -0.041 | -0.153 | -0.153 | 0 | 0.22 | 0.44 | 0.14 | 0 | 0.15 | 0.04 |
| 6 | 163563674 | rs547434 | -0.091 | 1.04 | -0.412 | -0.112 | 0 | 0.006 | 0.006 | 0 | 0.34 | 0.41 | 0.35 | 0.57 | 0.31 | 0.53 |
| 6 | 163566467 | rs1205662 | -1.592 | 1.046 | -0.709 | -0.13 | -0.059 | 0.021 | 0.021 | 0 | 0.71 | 0.98 | 0.71 | 0.78 | 0.68 | 0.74 |
| 6 | 163568075 | rs1205665 | -0.181 | 1.076 | -0.4 | -0.017 | -0.002 | -0.002 | -0.002 | 0 | 0.34 | 0.33 | 0.35 | 0.57 | 0.33 | 0.55 |
| 6 | 163569272 | rs10455920 | 1.074 | 0.904 | -0.616 | -0.031 | 0.012 | 0.058 | 0.058 | 0 | 0.18 | 0.08 | 0.12 | 0 | 0.15 | 0.05 |
| 6 | 163573298 | rs1408705 | -0.462 | 0.987 | -0.325 | 0.272 | 0.029 | 0.014 | 0.014 | 0.026 | 0.05 | 0.15 | 0.15 | 0.11 | 0.07 | 0.07 |
| 6 | 163574674 | rs13215511 | 0.709 | 1.03 | -0.551 | 0.585 | 0.003 | 0.032 | 0.032 | 0 | 0.11 | 0.12 | 0.15 | 0.31 | 0.14 | 0.15 |
| 6 | 163575022 | rs9346951 | 0.879 | 0.663 | -0.799 | 0.5 | 0.024 | -0.041 | -0.041 | 0.035 | 0.16 | 0.06 | 0.17 | 0.25 | 0.09 | 0.31 |
| 6 | 163576022 | rs10945874 | 1.052 | 0.714 | -0.602 | 0.522 | 0.019 | -0.042 | -0.042 | 0 | 0.15 | 0.06 | 0.16 | 0.25 | 0.09 | 0.31 |
| 6 | 163576293 | rs2747688 | -1.397 | 0.874 | -0.871 | -0.045 | -0.059 | 0.017 | 0.017 | 0 | 0.7 | 0.98 | 0.71 | 0.77 | 0.69 | 0.74 |
| 6 | 163577178 | rs7742985 | -0.284 | 0.798 | 0.27 | -0.259 | 0.036 | -0.017 | -0.017 | 0 | 0.18 | 0.39 | 0.11 | 0.12 | 0.23 | 0.16 |
| 6 | 163577921 | rs12197662 | -0.359 | 0.688 | 0.621 | -0.439 | -0.002 | -0.032 | -0.032 | 0.208 | 0.36 | 0.39 | 0.22 | 0.12 | 0.36 | 0.21 |
| 6 | 163578666 | rs10945875 | 0.002 | 0.387 | 0.773 | -0.32 | -0.046 | -0.101 | -0.101 | 0 | 0.24 | 0.46 | 0.26 | 0.09 | 0.17 | 0.13 |
| 6 | 163579586 | rs6932546 | 0.59 | 0.545 | 0.024 | -0.239 | 0.004 | 0.004 | 0.004 | 0 | 0.2 | 0.16 | 0.24 | 0.09 | 0.16 | 0.11 |
| 6 | 163583121 | rs12197200 | 1.095 | 0.658 | -0.339 | -0.292 | 0.005 | 0.057 | 0.057 | 0 | 0.16 | 0.05 | 0.12 | 0 | 0.15 | 0.04 |
| 6 | 163583286 | rs2747690 | 0.55 | 0.729 | -0.258 | 0.357 | 0.008 | -0.029 | -0.029 | 2.431 | 0.17 | 0.04 | 0.18 | 0.23 | 0.12 | 0.22 |
| 6 | 163584973 | rs2747691 | 0.025 | 0.839 | -0.566 | 0.086 | -0.023 | 0 | 0 | 0.329 | 0.23 | 0.05 | 0.16 | 0.22 | 0.24 | 0.32 |
| 6 | 163586569 | rs4709679 | 0.092 | 1.22 | 0.323 | 0.962 | 0.019 | 0.082 | 0.082 | 0 | 0.16 | 0.24 | 0.24 | 0.43 | 0.22 | 0.24 |
| 6 | 163590256 | rs2235993 | -0.833 | 1.187 | 0.47 | 0.77 | 0.002 | 0.001 | 0.001 | 0 | 0.34 | 0.41 | 0.35 | 0.57 | 0.44 | 0.44 |
| 6 | 163592235 | rs2763986 | 1.242 | 0.899 | -0.204 | 0.46 | -0.02 | 0.107 | 0.107 | 0 | 0.39 | 0.09 | 0.36 | 0.23 | 0.39 | 0.29 |
| 6 | 163597103 | rs2747695 | 0.534 | 0.865 | 0.169 | 0.506 | 0.005 | 0.039 | 0.039 | 0 | 0.53 | 0.32 | 0.46 | 0.35 | 0.56 | 0.41 |
| 6 | 163597536 | rs761621 | -0.825 | 0.737 | -0.288 | 0.467 | 0.012 | 0.003 | 0.003 | 0 | 0.81 | 0.95 | 0.85 | 0.86 | 0.88 | 0.8 |
| 6 | 163600767 | rs2985668 | 1.504 | 0.762 | -0.289 | 0.487 | -0.026 | 0.118 | 0.118 | 0 | 0.41 | 0.13 | 0.38 | 0.23 | 0.44 | 0.29 |
| 6 | 163600801 | rs4709680 | 0.29 | 1.099 | -0.027 | 0.653 | 0.004 | 0.047 | 0.047 | 0 | 0.21 | 0.28 | 0.36 | 0.47 | 0.26 | 0.31 |
| 6 | 163603470 | rs2763994 | 0.094 | 1.083 | -0.255 | 0.633 | -0.027 | -0.001 | -0.001 | 0 | 0.7 | 0.83 | 0.77 | 0.75 | 0.71 | 0.66 |
| 6 | 163608555 | rs13198883 | 0.384 | 1.167 | -0.044 | 0.542 | -0.027 | -0.017 | -0.017 | 0 | 0.1 | 0.05 | 0.06 | 0.11 | 0.14 | 0.12 |
| 6 | 163609693 | rs13217038 | 0.327 | 1.155 | -0.023 | 0.557 | -0.016 | -0.01 | -0.01 | 0.722 | 0.1 | 0.06 | 0.06 | 0.11 | 0.14 | 0.13 |
| 6 | 163610096 | rs1008295 | 0.232 | 1.198 | -0.005 | 0.884 | 0.023 | 0.073 | 0.073 | 0 | 0.22 | 0.37 | 0.37 | 0.38 | 0.28 | 0.3 |
| 6 | 163610526 | rs7759538 | -0.87 | 0.943 | 0.042 | 1.007 | 0.036 | 0.015 | 0.015 | 0 | 0.14 | 0.33 | 0.08 | 0.11 | 0.15 | 0.13 |
| 6 | 163611793 | rs6907521 | 0.226 | 0.666 | -0.112 | 1.059 | -0.022 | 0.004 | 0.004 | 0 | 0.76 | 0.63 | 0.81 | 0.76 | 0.85 | 0.81 |
| 6 | 163615962 | rs9456839 | -0.485 | 1.044 | -0.503 | 1.225 | -0.024 | -0.034 | -0.034 | 0 | 0.1 | 0.02 | 0.07 | 0.11 | 0.12 | 0.13 |
| 6 | 163620616 | rs9456841 | -1.277 | 1.541 | 0.166 | 1.77 | 0.071 | 0.142 | 0.142 | 0 | 0.24 | 0.49 | 0.42 | 0.49 | 0.3 | 0.3 |
| 6 | 163622117 | rs9458747 | -0.034 | 1.987 | -0.128 | 1.855 | -0.021 | -0.022 | -0.022 | 0 | 0.06 | 0.03 | 0.06 | 0.12 | 0.09 | 0.08 |
| 6 | 163622658 | rs1022611 | -1.411 | 1.468 | 0.751 | 1.56 | 0.058 | 0.127 | 0.127 | 0.013 | 0.26 | 0.52 | 0.41 | 0.5 | 0.3 | 0.36 |
| 6 | 163623619 | rs9458749 | -1.465 | 1.332 | 0.582 | 1.523 | -0.022 | 0.018 | 0.018 | 0 | 0.05 | 0.11 | 0.01 | 0.08 | 0.01 | 0.07 |
| 6 | 163623703 | rs9458750 | -1.66 | 1.346 | 0.484 | 2.038 | 0.028 | 0.059 | 0.059 | 0.198 | 0.27 | 0.62 | 0.36 | 0.5 | 0.32 | 0.32 |
| 6 | 163623977 | rs1122682 | -0.508 | 1.429 | 0.289 | 1.93 | -0.012 | -0.009 | -0.009 | 2.109 | 0.16 | 0.17 | 0.32 | 0.42 | 0.3 | 0.22 |
| 6 | 163625641 | rs763598 | -0.852 | 0.547 | -0.777 | 1.164 | -0.066 | 0.017 | 0.017 | 0.008 | 0.67 | 0.97 | 0.71 | 0.76 | 0.65 | 0.69 |
| 6 | 163629787 | rs9458753 | -1.554 | 1.173 | 0.364 | 1.439 | 0.034 | 0.035 | 0.035 | 0.165 | 0.25 | 0.67 | 0.36 | 0.43 | 0.34 | 0.31 |
| 6 | 163630086 | rs13206290 | -0.542 | 1.672 | -0.402 | 1.697 | -0.004 | -0.004 | -0.004 | 0 | 0.16 | 0.17 | 0.32 | 0.42 | 0.34 | 0.23 |
| 6 | 163631801 | rs10945881 | -0.468 | 0.839 | -0.917 | 1.299 | -0.077 | -0.1 | -0.1 | 0.252 | 0.29 | 0.07 | 0.41 | 0.6 | 0.29 | 0.36 |
| 6 | 163631995 | rs2143071 | -0.846 | 0.219 | -0.463 | 0.505 | -0.019 | -0.031 | -0.031 | 0 | 0.64 | 0.52 | 0.72 | 0.77 | 0.63 | 0.79 |
| 6 | 163635880 | rs6906182 | -1.29 | 0.203 | -0.786 | 0.558 | -0.039 | 0.007 | 0.007 | 0.005 | 0.73 | 0.96 | 0.77 | 0.81 | 0.65 | 0.81 |
| 6 | 163638267 | rs16888819 | 0.143 | 0.091 | -0.33 | 0.477 | 0.012 | 0.048 | 0.048 | 0 | 0.14 | 0.31 | 0.18 | 0.2 | 0.15 | 0.22 |
| 6 | 163640489 | rs12213157 | 0.57 | -0.15 | 0.053 | 0.28 | -0.009 | -0.092 | -0.092 | 0 | 0.23 | 0.39 | 0.16 | 0.04 | 0.2 | 0.2 |
| 6 | 163642205 | rs4709683 | -0.002 | -0.084 | -0.408 | 0.385 | -0.021 | -0.048 | -0.048 | 0 | 0.35 | 0.21 | 0.42 | 0.57 | 0.3 | 0.38 |
| 6 | 163647830 | rs12210929 | -0.36 | 0.078 | -0.397 | 0.696 | -0.002 | -0.002 | -0.002 | 0 | 0.57 | 0.57 | 0.57 | 0.6 | 0.5 | 0.58 |
| 6 | 163650431 | rs6916948 | -0.501 | 0.223 | -1.121 | 0.753 | -0.008 | -0.001 | -0.001 | 0 | 0.6 | 0.73 | 0.59 | 0.62 | 0.5 | 0.65 |
| 6 | 163651283 | rs6922754 | -0.393 | 0.219 | -1.115 | 0.456 | -0.009 | -0.005 | -0.005 | 0 | 0.6 | 0.73 | 0.41 | 0.4 | 0.5 | 0.64 |

| | | | | | | | | | | | | | | | | |
|---|---|---|---|---|---|---|---|---|---|---|---|---|---|---|---|---|
| 6 | 163661513 | rs6908925 | -0.321 | -0.005 | -1.215 | 0.233 | 0.013 | 0.018 | 0.018 | 0 | 0.25 | 0.13 | 0.15 | 0.23 | 0.17 | 0.3 |
| 6 | 163662002 | rs10945884 | -1.059 | 0.016 | -1.697 | 0.202 | -0.119 | -0.104 | -0.104 | 0 | 0.63 | 0.91 | 0.5 | 0.49 | 0.48 | 0.7 |
| 6 | 163663237 | rs11964522 | -0.047 | -0.315 | -1.069 | 0.392 | 0.002 | 0.005 | 0.005 | 0.401 | 0.37 | 0.68 | 0.34 | 0.25 | 0.32 | 0.4 |
| 6 | 163665303 | rs16894692 | -0.926 | -0.441 | -0.769 | 1.401 | -0.011 | 0.168 | 0.168 | 0 | 0.05 | 0.2 | 0.11 | 0.23 | 0.01 | 0.12 |
| 6 | 163666336 | rs2294458 | -0.432 | -0.177 | -1.363 | 0.704 | 0.035 | -0.014 | -0.014 | 0.012 | 0.17 | 0.06 | 0.11 | 0.21 | 0.09 | 0.22 |
| 6 | 163673166 | rs9295214 | -0.038 | 0.289 | -1.704 | 0.616 | 0.027 | 0.017 | 0.017 | 0 | 0.46 | 0.81 | 0.46 | 0.4 | 0.42 | 0.45 |
| 6 | 163677027 | rs1555064 | -0.376 | 0.457 | -1.593 | 0.789 | 0 | -0.017 | -0.017 | 0 | 0.69 | 0.78 | 0.6 | 0.63 | 0.72 | 0.67 |
| 6 | 163683522 | rs7760002 | -0.116 | 0.331 | -0.905 | 0.454 | -0.006 | 0.007 | 0.007 | 0.463 | 0.34 | 0.32 | 0.24 | 0.24 | 0.39 | 0.36 |
| 6 | 163690429 | rs4470828 | -0.186 | 0.123 | -0.374 | 0.527 | 0.007 | -0.005 | -0.005 | 0 | 0.35 | 0.6 | 0.24 | 0.34 | 0.39 | 0.43 |
| 6 | 163697439 | rs7750165 | -0.545 | -0.085 | -0.245 | 0.391 | -0.011 | 0.024 | 0.024 | 0 | 0.42 | 0.69 | 0.28 | 0.37 | 0.44 | 0.47 |
| 6 | 163703866 | rs6922089 | -1.266 | -0.053 | -0.611 | 0.508 | -0.033 | -0.081 | -0.081 | 0.221 | 0.73 | 0.93 | 0.66 | 0.66 | 0.71 | 0.67 |
| 6 | 163704008 | rs233 | 0.924 | -0.183 | -0.564 | 0.382 | -0.01 | -0.023 | -0.023 | 0 | 0.27 | 0.11 | 0.37 | 0.29 | 0.26 | 0.21 |
| 6 | 163705708 | rs9458774 | -0.564 | -0.174 | -0.544 | 0.633 | 0.001 | -0.066 | -0.066 | 0 | 0.66 | 0.9 | 0.62 | 0.62 | 0.66 | 0.63 |
| 6 | 163706213 | rs13200025 | -0.105 | 0.008 | -0.244 | 0.388 | 0 | 0 | 0 | 0.414 | 0.54 | 0.51 | 0.53 | 0.49 | 0.5 | 0.46 |
| 6 | 163709810 | rs10945886 | 0.713 | 0.287 | -0.356 | 0.363 | -0.037 | 0.033 | 0.033 | 0.384 | 0.13 | 0.05 | 0.1 | 0.06 | 0.2 | 0.15 |
| 6 | 163710165 | rs6916146 | 0.905 | 0.177 | -0.29 | 0.422 | -0.003 | 0.006 | 0.006 | 0 | 0.19 | 0.29 | 0.11 | 0.2 | 0.19 | 0.18 |
| 6 | 163710391 | rs13202719 | 0.914 | 0.104 | -0.943 | 0.462 | -0.017 | -0.027 | -0.027 | 0.664 | 0.28 | 0.1 | 0.38 | 0.28 | 0.27 | 0.21 |
| 6 | 163711677 | rs10455921 | 0.933 | 0.132 | -0.756 | 0.581 | -0.034 | -0.081 | -0.081 | 0 | 0.26 | 0.07 | 0.34 | 0.35 | 0.3 | 0.34 |
| 6 | 163713283 | rs7768540 | -0.655 | -0.051 | -0.352 | 0.544 | -0.058 | -0.011 | -0.011 | 0 | 0.14 | 0.35 | 0.05 | 0.12 | 0.05 | 0.08 |
| 6 | 163719115 | rs6927758 | 0.199 | 0.551 | -1.149 | 0.753 | 0.002 | -0.002 | -0.002 | 0.193 | 0.46 | 0.57 | 0.56 | 0.48 | 0.44 | 0.41 |
| 6 | 163719619 | rs742333 | 0.067 | 0.866 | -1.333 | 0.851 | 0.01 | 0.016 | 0.016 | 0.105 | 0.47 | 0.36 | 0.43 | 0.38 | 0.45 | 0.42 |
| 6 | 163719877 | rs6455889 | -0.536 | 0.999 | -1.844 | 0.804 | -0.036 | 0.019 | 0.019 | 0.374 | 0.34 | 0.85 | 0.29 | 0.29 | 0.35 | 0.27 |
| 6 | 163721073 | rs12526892 | -0.947 | 1.161 | -1.738 | 1.37 | -0.062 | -0.127 | -0.127 | 0 | 0.18 | 0.03 | 0.36 | 0.39 | 0.3 | 0.24 |
| 6 | 163721184 | rs9458781 | 0.924 | 1.3 | -1.113 | 0.893 | 0.002 | 0.004 | 0.004 | 0.174 | 0.55 | 0.48 | 0.39 | 0.5 | 0.46 | 0.46 |
| 6 | 163722268 | rs4709696 | -1.307 | 1.139 | 0.285 | 0.678 | 0.065 | 0.169 | 0.169 | 0.785 | 0.12 | 0.45 | 0.26 | 0.25 | 0.15 | 0.19 |
| 6 | 163722764 | rs4709698 | -2.202 | 1.011 | 0.174 | 0.271 | -0.018 | 0.002 | 0.002 | 1.654 | 0.4 | 0.78 | 0.52 | 0.44 | 0.45 | 0.52 |
| 6 | 163723545 | rs10455922 | -1.155 | 0.981 | 0.289 | 0.203 | -0.001 | -0.082 | -0.082 | 0 | 0.2 | 0.24 | 0.19 | 0.02 | 0.18 | 0.14 |
| 6 | 163723916 | rs4708987 | -2.261 | 1.008 | 0.191 | 0.227 | -0.01 | 0.03 | 0.03 | 0.876 | 0.33 | 0.76 | 0.45 | 0.28 | 0.33 | 0.35 |
| 6 | 163725498 | rs4708990 | -2.396 | 1.013 | 0.174 | 0.371 | -0.031 | -0.046 | -0.046 | 0 | 0.4 | 0.82 | 0.52 | 0.48 | 0.45 | 0.52 |
| 6 | 163726229 | rs11759691 | -1.019 | 0.752 | 0.471 | 0.235 | 0.071 | 0.23 | 0.23 | 1.686 | 0.1 | 0.42 | 0.25 | 0.24 | 0.12 | 0.13 |
| 6 | 163726426 | rs3799339 | -2.121 | 0.775 | 0.222 | 0.207 | 0.004 | 0.001 | 0.001 | 1.271 | 0.25 | 0.71 | 0.39 | 0.27 | 0.24 | 0.29 |
| 6 | 163730006 | rs10945890 | -1.318 | 0.922 | 0.347 | 0.465 | 0.029 | 0.128 | 0.128 | 0.184 | 0.15 | 0.36 | 0.31 | 0.34 | 0.23 | 0.23 |
| 6 | 163730334 | rs11757855 | -0.245 | 0.711 | -0.17 | -0.289 | -0.008 | -0.015 | -0.015 | 0.957 | 0.07 | 0.05 | 0.06 | 0.09 | 0.11 | 0.11 |
| 6 | 163730672 | rs11753570 | 0.006 | 0.735 | -0.21 | -0.275 | -0.019 | -0.025 | -0.025 | 0 | 0.07 | 0.01 | 0.06 | 0.09 | 0.11 | 0.13 |
| 6 | 163731502 | rs12197625 | -1.075 | 0.517 | 0.005 | -0.391 | -0.027 | -0.065 | -0.065 | 0.457 | 0.14 | 0.26 | 0.14 | 0.02 | 0.09 | 0.1 |
| 6 | 163731915 | rs11759099 | -0.441 | 1.082 | -0.358 | 0.07 | -0.039 | -0.038 | -0.038 | 0 | 0.08 | 0.01 | 0.09 | 0.15 | 0.13 | 0.11 |
| 6 | 163732939 | rs12529659 | -0.617 | 0.236 | 0.093 | -0.684 | -0.028 | -0.099 | -0.099 | 0 | 0.2 | 0.35 | 0.09 | 0.02 | 0.13 | 0.13 |
| 6 | 163733168 | rs12211969 | -0.891 | 0.273 | -0.334 | -0.821 | 0.007 | 0.013 | 0.013 | 0 | 0.18 | 0.14 | 0.19 | 0.06 | 0.18 | 0.11 |
| 6 | 163733438 | rs9365548 | 0.424 | 0.201 | -0.385 | -0.559 | 0.046 | 0.027 | 0.027 | 0 | 0.44 | 0.28 | 0.31 | 0.67 | 0.32 | 0.51 |
| 6 | 163733462 | rs7743292 | -1.316 | 0.986 | 0.29 | -0.436 | 0.015 | -0.09 | -0.09 | 0 | 0.15 | 0.58 | 0.07 | 0.04 | 0.11 | 0.08 |
| 6 | 163734204 | rs9346956 | 0.447 | 1.389 | -0.356 | 0.21 | 0.005 | 0.001 | 0.001 | 0 | 0.65 | 0.39 | 0.48 | 0.6 | 0.57 | 0.66 |
| 6 | 163736204 | rs9306 | -0.155 | 1.188 | -0.747 | 0.026 | -0.009 | 0.027 | 0.027 | 0.399 | 0.42 | 0.69 | 0.48 | 0.67 | 0.42 | 0.57 |
| 6 | 163736502 | rs6903758 | -0.101 | 1.274 | -0.603 | -0.128 | 0.01 | 0.02 | 0.02 | 0.779 | 0.39 | 0.24 | 0.42 | 0.67 | 0.35 | 0.51 |
| 6 | 163739362 | rs4709701 | -0.385 | 1.506 | -1.099 | -0.156 | -0.043 | -0.043 | -0.043 | 0.985 | 0.84 | 0.97 | 0.83 | 0.83 | 0.82 | 0.9 |
| 6 | 163747722 | rs2321514 | 0.292 | 1.194 | -0.636 | 0.328 | 0 | 0.003 | 0.003 | 0 | 0.66 | 0.67 | 0.71 | 0.45 | 0.66 | 0.68 |
| 6 | 163748325 | rs9458797 | 0.247 | 0.776 | -0.816 | 0.189 | 0.006 | -0.021 | -0.021 | 0 | 0.23 | 0.18 | 0.33 | 0.28 | 0.21 | 0.3 |
| 6 | 163748659 | rs6925566 | 0.508 | 0.804 | -0.871 | -0.303 | -0.011 | 0.023 | 0.023 | 0 | 0.18 | 0.09 | 0.25 | 0.13 | 0.22 | 0.09 |
| 6 | 163749552 | rs13211088 | 0.508 | 0.603 | -0.68 | -0.387 | 0 | 0.001 | 0.001 | 0 | 0.17 | 0.16 | 0.23 | 0.13 | 0.19 | 0.07 |
| 6 | 163758941 | rs4709704 | 0.233 | 0.713 | -0.467 | 0.07 | 0.001 | 0.006 | 0.006 | 0 | 0.18 | 0.16 | 0.14 | 0.36 | 0.2 | 0.24 |
| 6 | 163761520 | rs4709708 | 0.243 | 0.633 | -0.565 | 0.14 | -0.006 | -0.008 | -0.008 | 0 | 0.18 | 0.15 | 0.14 | 0.28 | 0.19 | 0.22 |
| 6 | 163764137 | rs11752187 | 0.257 | 0.23 | -1.205 | -0.622 | -0.035 | 0.092 | 0.092 | 0 | 0.38 | 0.25 | 0.44 | 0.13 | 0.47 | 0.29 |
| 6 | 163764804 | rs10945893 | -0.619 | -0.252 | -1.162 | -0.684 | -0.014 | -0.057 | -0.057 | 0.455 | 0.66 | 0.83 | 0.58 | 0.54 | 0.66 | 0.66 |
| 6 | 163767824 | rs4370339 | 0.527 | 0.075 | -0.788 | -1.182 | -0.002 | -0.076 | -0.076 | 0 | 0.35 | 0.42 | 0.4 | 0.14 | 0.39 | 0.24 |
| 6 | 163768368 | rs13202030 | 0.483 | -0.056 | -1.469 | -1.024 | -0.07 | -0.005 | -0.005 | 0.861 | 0.3 | 0.09 | 0.33 | 0.2 | 0.32 | 0.21 |
| 6 | 163769090 | rs9347738 | 0.618 | -0.584 | -1.056 | -0.314 | -0.012 | -0.086 | -0.086 | 0.532 | 0.22 | 0.11 | 0.33 | 0.38 | 0.21 | 0.23 |
| 6 | 163772096 | rs13203110 | 0.56 | -0.601 | -0.802 | -0.528 | -0.001 | -0.021 | -0.021 | 0 | 0.36 | 0.58 | 0.4 | 0.14 | 0.34 | 0.21 |
| 6 | 163772350 | rs16894832 | 0.995 | -0.595 | -1.001 | -0.668 | -0.026 | -0.071 | -0.071 | 0 | 0.27 | 0.1 | 0.16 | 0.41 | 0.33 | 0.2 |
| 6 | 163773393 | rs9347742 | -0.403 | -0.58 | -0.911 | -0.713 | -0.008 | 0.007 | 0.007 | 0 | 0.91 | 0.92 | 0.91 | 0.97 | 0.88 | 0.86 |
| 6 | 163773687 | rs10945895 | 0.491 | -0.706 | -0.537 | -0.741 | -0.003 | 0.004 | 0.004 | 0 | 0.6 | 0.47 | 0.51 | 0.51 | 0.66 | 0.42 |
| 6 | 163773871 | rs9364686 | 0.006 | -0.324 | -0.654 | -0.519 | -0.018 | 0.103 | 0.103 | 0 | 0.54 | 0.3 | 0.53 | 0.87 | 0.54 | 0.65 |
| 6 | 163774147 | rs4140499 | 0.634 | -0.515 | -0.657 | -0.769 | 0.003 | -0.022 | -0.022 | 0 | 0.29 | 0.2 | 0.15 | 0.44 | 0.3 | 0.42 |
| 6 | 163776851 | rs735751 | 0.138 | 0.06 | -1.142 | -0.697 | 0.002 | -0.005 | -0.005 | 0 | 0.9 | 0.89 | 0.92 | 0.97 | 0.89 | 0.86 |
| 6 | 163777215 | rs735752 | 0.353 | -0.044 | -1.133 | -0.693 | -0.019 | 0.001 | 0.001 | 0 | 0.36 | 0.45 | 0.19 | 0.41 | 0.32 | 0.43 |
| 6 | 163779672 | rs9355427 | 0.012 | -0.135 | -1.057 | -0.238 | -0.003 | 0 | 0 | 0 | 0.94 | 0.94 | 0.93 | 0.98 | 0.91 | 0.9 |
| 6 | 163782827 | rs13220178 | 0.695 | -0.296 | -1.196 | -0.278 | -0.016 | -0.079 | -0.079 | 0 | 0.7 | 0.87 | 0.56 | 0.51 | 0.68 | 0.67 |
| 6 | 163783291 | rs1764001 | -0.464 | -0.35 | -1.235 | 0.002 | -0.017 | -0.134 | -0.134 | 0 | 0.22 | 0.06 | 0.36 | 0.47 | 0.22 | 0.22 |
| 6 | 163788191 | rs1028355 | -0.579 | -0.375 | -1.256 | -0.045 | -0.011 | -0.138 | -0.138 | 0 | 0.23 | 0.07 | 0.36 | 0.49 | 0.22 | 0.37 |
| 6 | 163789838 | rs1737317 | 0.138 | -0.457 | -0.899 | -1.544 | 0.006 | 0.068 | 0.068 | 0 | 0.66 | 0.78 | 0.8 | 0.96 | 0.72 | 0.76 |
| 6 | 163792420 | rs1629251 | -0.435 | -0.581 | -0.431 | -1.615 | 0.001 | -0.003 | -0.003 | 0.009 | 0.74 | 0.83 | 0.82 | 1 | 0.77 | 0.86 |

| | | | | | | | | | | | | | | | | |
|---|---|---|---|---|---|---|---|---|---|---|---|---|---|---|---|---|
| 6 | 163794506 | rs2744422 | -0.548 | -1.085 | 0.121 | -1.825 | -0.038 | -0.042 | -0.042 | 0.016 | 0.3 | 0.49 | 0.39 | 0.2 | 0.2 | 0.27 |
| 6 | 163794927 | rs2064225 | -0.296 | -1.107 | 0.331 | -2.132 | 0 | -0.028 | -0.028 | 0 | 0.63 | 0.67 | 0.76 | 0.94 | 0.7 | 0.8 |
| 6 | 163796510 | rs1737319 | 0.343 | -1.151 | 0.454 | -2.196 | 0.002 | -0.003 | -0.003 | 0 | 0.19 | 0.21 | 0.39 | 0.26 | 0.15 | 0.17 |
| 6 | 163797775 | rs1737323 | 0.253 | -1.202 | 0.636 | -2.262 | -0.002 | -0.094 | -0.094 | 0 | 0.53 | 0.53 | 0.74 | 0.94 | 0.64 | 0.7 |
| 6 | 163801031 | rs4709714 | 0.138 | -1.087 | 0.68 | -2.179 | 0.006 | 0.013 | 0.013 | 0 | 0.51 | 0.52 | 0.41 | 0.73 | 0.59 | 0.67 |
| 6 | 163805742 | rs16894934 | 0.231 | -0.985 | 0.699 | -2.131 | -0.017 | 0.008 | 0.008 | 0 | 0.2 | 0.28 | 0.39 | 0.25 | 0.15 | 0.18 |
| 6 | 163817911 | rs9364687 | 1.197 | -0.931 | 0.085 | -2.217 | -0.162 | -0.03 | -0.03 | 0 | 0.3 | 0.02 | 0.31 | 0.73 | 0.46 | 0.52 |
| 6 | 163844392 | rs2759386 | -0.118 | -1.056 | 0.746 | -1.834 | 0.003 | 0.004 | 0.004 | 0 | 0.91 | 0.92 | 0.97 | 0.96 | 0.92 | 0.91 |
| 6 | 163863551 | rs9458828 | 0.023 | -1.087 | 1.198 | -1.895 | 0.002 | -0.031 | -0.031 | 0.216 | 0.09 | 0.12 | 0.06 | 0 | 0.07 | 0.05 |
| 6 | 163873745 | rs813640 | -0.157 | -1.09 | 1.54 | -2.108 | -0.003 | -0.115 | -0.115 | 1.306 | 0.36 | 0.51 | 0.23 | 0 | 0.3 | 0.2 |
| 6 | 163887376 | rs1744925 | -0.865 | -1.082 | 1.152 | -2.101 | 0.001 | -0.023 | -0.023 | 0 | 0.74 | 0.78 | 0.84 | 1 | 0.78 | 0.86 |
| 6 | 163891894 | rs1744926 | -0.315 | -1.124 | 1.325 | -2.141 | -0.008 | -0.115 | -0.115 | 0 | 0.69 | 0.45 | 0.8 | 1 | 0.76 | 0.82 |
| 6 | 163892317 | rs1737604 | -0.966 | -1.113 | 1.23 | -2.135 | -0.008 | 0.07 | 0.07 | 0 | 0.76 | 0.89 | 0.85 | 1 | 0.78 | 0.86 |
| 6 | 163895307 | rs1230321 | 0.069 | -1.108 | 0.95 | -2.141 | -0.001 | -0.084 | -0.084 | 0.021 | 0.65 | 0.64 | 0.78 | 1 | 0.71 | 0.8 |
| 6 | 163907885 | rs1016502 | -0.958 | -1.088 | 0.933 | -2.128 | -0.058 | 0.166 | 0.166 | 0 | 0.76 | 0.98 | 0.85 | 1 | 0.78 | 0.86 |
| 6 | 163910325 | rs6931903 | -0.356 | -1.19 | 0.929 | -2.152 | 0.003 | -0.005 | -0.005 | 0.113 | 0.17 | 0.16 | 0.2 | 0.23 | 0.14 | 0.16 |
| 6 | 163920573 | rs2038148 | -1.018 | -1.179 | 0.754 | -2.106 | 0.001 | -0.015 | -0.015 | 0 | 0.74 | 0.78 | 0.84 | 0.99 | 0.78 | 0.86 |
| 6 | 163929340 | rs9364689 | 0.818 | -0.855 | 0.284 | -1.893 | -0.169 | 0.041 | 0.041 | 0 | 0.39 | 0.03 | 0.55 | 0.73 | 0.54 | 0.56 |
| 6 | 163963084 | rs2294703 | -0.011 | -0.636 | 0.069 | -1.578 | 0.006 | -0.008 | -0.008 | 1.149 | 0.12 | 0.15 | 0.17 | 0.23 | 0.08 | 0.16 |
| 6 | 163975823 | rs11964059 | 0.265 | -0.557 | 0.07 | -1.578 | 0.021 | -0.031 | -0.031 | 0.027 | 0.06 | 0.17 | 0.04 | 0 | 0.05 | 0.02 |
| 6 | 163981387 | rs2143358 | 0.067 | -0.52 | -0.968 | -1.267 | 0.009 | -0.013 | -0.013 | 0 | 0.12 | 0.1 | 0.17 | 0.23 | 0.08 | 0.16 |
| 6 | 163995737 | rs2294698 | 0.108 | 0.019 | -1.034 | -0.935 | 0.022 | 0.104 | 0.104 | 0 | 0.47 | 0.23 | 0.6 | 0.73 | 0.59 | 0.57 |
| 6 | 164007423 | rs4709724 | -0.071 | 0.423 | -1.352 | -0.985 | -0.022 | 0.161 | 0.161 | 0 | 0.54 | 0.81 | 0.66 | 0.87 | 0.55 | 0.75 |
| 6 | 164008408 | rs4709725 | 0.486 | 0.466 | -1.887 | 0.125 | 0.009 | -0.102 | -0.102 | 0.372 | 0.3 | 0.08 | 0.43 | 0.43 | 0.28 | 0.34 |
| 6 | 164010321 | rs4709726 | 0.245 | 0.863 | -1.27 | -0.605 | 0.026 | 0.024 | 0.024 | 1.251 | 0.63 | 0.54 | 0.67 | 0.51 | 0.53 | 0.57 |
| 6 | 164012052 | rs7761240 | 0.264 | 1.043 | -0.305 | -1.014 | 0.004 | 0.011 | 0.011 | 0 | 0.5 | 0.54 | 0.59 | 0.76 | 0.48 | 0.44 |
| 6 | 164015506 | rs12202352 | 0.371 | 1.507 | 0.088 | -1.207 | 0.015 | 0.022 | 0.022 | 0.02 | 0.12 | 0.12 | 0.15 | 0.19 | 0.18 | 0.09 |
| 6 | 164017390 | rs9295225 | 0.236 | 1.418 | 0.739 | -1.388 | -0.002 | 0.044 | 0.044 | 0 | 0.35 | 0.22 | 0.26 | 0.06 | 0.31 | 0.26 |
| 6 | 164021962 | rs4529277 | -0.126 | 1.343 | 0.73 | -1.282 | 0.02 | 0.026 | 0.026 | 0.021 | 0.14 | 0.21 | 0.16 | 0.19 | 0.19 | 0.09 |
| 6 | 164023708 | rs11756061 | -0.654 | 0.59 | 0.39 | -1.659 | 0.018 | 0.058 | 0.058 | 0 | 0.26 | 0.43 | 0.27 | 0.4 | 0.29 | 0.29 |
| 6 | 164027544 | rs10945909 | -0.934 | 0.692 | 0.455 | -1.755 | -0.003 | -0.007 | -0.007 | 0 | 0.6 | 0.7 | 0.51 | 0.46 | 0.54 | 0.52 |
| 6 | 164027663 | rs17592412 | 0.777 | 0.08 | 0.426 | -1.651 | -0.023 | 0.009 | 0.009 | 0.632 | 0.1 | 0.15 | 0.03 | 0.18 | 0.03 | 0.17 |
| 6 | 164027758 | rs9365584 | 0.303 | 0.377 | 0.198 | -1.781 | 0.006 | 0.004 | 0.004 | 0.02 | 0.34 | 0.29 | 0.23 | 0.06 | 0.25 | 0.24 |
| 6 | 164030045 | rs6935776 | 0.202 | 0.35 | 0.012 | -1.825 | 0.006 | 0.001 | 0.001 | 0.258 | 0.35 | 0.3 | 0.21 | 0.05 | 0.26 | 0.27 |
| 6 | 164031621 | rs7756185 | 0.322 | 0.516 | 0.214 | -1.783 | 0.01 | 0.01 | 0.01 | 0 | 0.49 | 0.56 | 0.51 | 0.7 | 0.37 | 0.52 |
| 6 | 164033649 | rs9355432 | 0.276 | 0.713 | -0.106 | -1.648 | -0.003 | -0.003 | -0.003 | 0 | 0.08 | 0.13 | 0.03 | 0.09 | 0.05 | 0.13 |
| 6 | 164037047 | rs4708999 | 0.217 | 0.955 | -0.011 | -1.549 | 0.012 | 0.001 | 0.001 | 0.404 | 0.8 | 0.8 | 0.71 | 0.75 | 0.62 | 0.74 |
| 6 | 164043825 | rs6933936 | -0.539 | 0.21 | 0.285 | -2.405 | 0.019 | 0.029 | 0.029 | 0.206 | 0.43 | 0.59 | 0.24 | 0.11 | 0.26 | 0.3 |
| 6 | 164050359 | rs1983789 | -0.705 | 0.338 | 0.396 | -2.345 | 0.033 | 0.066 | 0.066 | 0.435 | 0.45 | 0.63 | 0.24 | 0.12 | 0.27 | 0.32 |
| 6 | 164054933 | rs6455910 | 0.194 | 0.626 | 0.248 | -2.295 | -0.002 | 0.014 | 0.014 | 0 | 0.27 | 0.16 | 0.19 | 0 | 0.22 | 0.11 |
| 6 | 164057051 | rs7754222 | 0.287 | 0.766 | 0.065 | -2.148 | -0.008 | 0.002 | 0.002 | 0 | 0.91 | 0.91 | 0.87 | 1 | 0.8 | 0.88 |
| 6 | 164059096 | rs7744302 | 0.143 | 0.799 | 0.013 | -2.132 | -0.034 | 0.032 | 0.032 | 0 | 0.92 | 0.97 | 0.88 | 1 | 0.8 | 0.88 |
| 6 | 164059136 | rs7744325 | -0.011 | 0.678 | -0.183 | -2.457 | 0.018 | 0.099 | 0.099 | 0.231 | 0.32 | 0.15 | 0.18 | 0.01 | 0.23 | 0.15 |
| 6 | 164059156 | rs7744332 | 1.175 | -0.072 | 0.221 | -2.231 | 0.008 | -0.025 | -0.025 | 0 | 0.8 | 0.33 | 0.81 | 0.91 | 0.75 | 0.74 |
| 6 | 164060523 | rs7750406 | -0.838 | -0.757 | 0.692 | -2.033 | -0.012 | 0.021 | 0.021 | 0.319 | 0.09 | 0.24 | 0.05 | 0.11 | 0.04 | 0.14 |
| 6 | 164061846 | rs13207959 | -0.257 | -0.544 | 0.123 | -2.378 | 0.031 | 0.096 | 0.096 | 0.322 | 0.3 | 0.15 | 0.12 | 0.01 | 0.2 | 0.13 |
| 6 | 164063263 | rs13191187 | -0.374 | -0.607 | 0.257 | -2.359 | 0.011 | 0.001 | 0.001 | 0 | 0.32 | 0.28 | 0.15 | 0.01 | 0.23 | 0.16 |
| 6 | 164065525 | rs9364696 | 1.353 | -0.521 | -0.168 | -2.28 | -0.102 | 0.145 | 0.145 | 0.195 | 0.43 | 0.11 | 0.46 | 0.77 | 0.51 | 0.48 |
| 6 | 164066386 | rs7755199 | -0.316 | -0.604 | 0.418 | -2.277 | -0.012 | -0.023 | -0.023 | 3.325 | 0.35 | 0.44 | 0.21 | 0.23 | 0.23 | 0.36 |
| 6 | 164067316 | rs9458875 | 0.043 | -0.596 | 0.338 | -2.311 | 0.018 | 0.027 | 0.027 | 0 | 0.3 | 0.27 | 0.17 | 0.09 | 0.19 | 0.22 |
| 6 | 164071406 | rs12191684 | -0.028 | -0.592 | 0.302 | -2.315 | 0.034 | 0.097 | 0.097 | 0.081 | 0.29 | 0.15 | 0.16 | 0.08 | 0.19 | 0.22 |
| 6 | 164071823 | rs1322888 | -0.904 | -0.534 | 0.718 | -2.368 | -0.001 | -0.048 | -0.048 | 0 | 0.45 | 0.49 | 0.47 | 0.09 | 0.43 | 0.37 |
| 6 | 164073368 | rs9458878 | 1.105 | -0.569 | 0.618 | -2.403 | 0.008 | 0.13 | 0.13 | 0.802 | 0.48 | 0.28 | 0.49 | 0.91 | 0.54 | 0.6 |
| 6 | 164074842 | rs1407658 | 0.991 | -0.772 | 0.975 | -2.069 | -0.018 | -0.069 | -0.069 | 2.751 | 0.89 | 0.58 | 0.95 | 1 | 0.96 | 0.97 |
| 6 | 164079396 | rs9347774 | 1.517 | -0.769 | -0.109 | -1.995 | -0.035 | 0.137 | 0.137 | 0.434 | 0.46 | 0.12 | 0.47 | 0.71 | 0.54 | 0.47 |
| 6 | 164082190 | rs7773362 | 0.783 | -0.123 | 0.422 | -2.141 | 0.005 | -0.037 | -0.037 | 0.067 | 0.91 | 0.52 | 0.95 | 1 | 0.97 | 0.97 |
| 6 | 164084671 | rs4709737 | 0.946 | -0.049 | -0.209 | -2.139 | 0.027 | -0.116 | -0.116 | 0.069 | 0.8 | 0.47 | 0.66 | 1 | 0.73 | 0.81 |
| 6 | 164087931 | rs11755805 | -0.01 | -0.152 | -0.881 | -2.183 | 0.013 | 0.04 | 0.04 | 0.224 | 0.25 | 0.16 | 0.17 | 0.16 | 0.18 | 0.22 |
| 6 | 164088863 | rs4709738 | 0.885 | 0.189 | -1.392 | -1.957 | -0.006 | 0.195 | 0.195 | 0.323 | 0.54 | 0.18 | 0.48 | 0.83 | 0.55 | 0.59 |
| 6 | 164089110 | rs12191767 | -0.915 | 0.941 | -0.819 | -0.993 | -0.001 | -0.017 | -0.017 | 0 | 0.12 | 0.12 | 0.29 | 0 | 0.24 | 0.16 |
| 6 | 164091657 | rs1407662 | 0.528 | -0.292 | -1.151 | -1.141 | 0.044 | 0.062 | 0.062 | 0 | 0.74 | 0.88 | 0.68 | 0.93 | 0.84 | 0.83 |
| 6 | 164092291 | rs4709741 | -0.84 | 0.062 | -0.899 | -0.7 | -0.001 | -0.001 | -0.001 | 0 | 0.1 | 0.11 | 0.24 | 0.09 | 0.09 | 0.14 |
| 6 | 164095511 | rs4709742 | -0.771 | 0.322 | -1.593 | -0.077 | -0.012 | -0.065 | -0.065 | 1.512 | 0.3 | 0.19 | 0.39 | 0.52 | 0.28 | 0.53 |
| 6 | 164097101 | rs1475161 | 1.029 | 0.476 | -1.15 | -0.799 | -0.006 | -0.078 | -0.078 | 0 | 0.4 | 0.46 | 0.26 | 0.07 | 0.37 | 0.24 |
| 6 | 164104828 | rs9458886 | 1.194 | 0.106 | -0.796 | -1.33 | -0.01 | 0.019 | 0.019 | 0 | 0.29 | 0.22 | 0.22 | 0.01 | 0.33 | 0.14 |
| 6 | 164105984 | rs4709745 | -0.269 | -0.064 | -0.417 | -1.131 | 0.007 | -0.017 | -0.017 | 0.983 | 0.38 | 0.29 | 0.42 | 0.53 | 0.3 | 0.57 |
| 6 | 164106739 | rs9365597 | -0.458 | -0.048 | -0.135 | -0.926 | 0 | 0.004 | 0.004 | 0 | 0.71 | 0.73 | 0.66 | 0.6 | 0.66 | 0.75 |
| 6 | 164109051 | rs9346970 | 0.964 | 0.077 | -0.307 | -0.843 | -0.018 | -0.083 | -0.083 | 0 | 0.19 | 0.07 | 0.12 | 0.44 | 0.15 | 0.39 |
| 6 | 164109438 | rs9356135 | 1.002 | 0.059 | -0.312 | -0.853 | -0.023 | -0.09 | -0.09 | 0.714 | 0.18 | 0.05 | 0.12 | 0.44 | 0.15 | 0.39 |

| | | | | | | | | | | | | | | | | |
|---|---|---|---|---|---|---|---|---|---|---|---|---|---|---|---|---|
| 6 | 164111266 | rs750810 | -0.17 | -0.011 | 0.532 | -0.898 | -0.028 | -0.019 | -0.019 | 1.014 | 0.12 | 0.31 | 0.05 | 0.06 | 0.05 | 0.11 |
| 6 | 164111361 | rs750811 | -0.507 | -0.162 | 0.723 | -0.988 | -0.014 | 0.02 | 0.02 | 0.082 | 0.34 | 0.55 | 0.2 | 0.5 | 0.22 | 0.49 |
| 6 | 164133150 | rs9295227 | -0.431 | 0.035 | 0.605 | -0.909 | -0.005 | -0.016 | -0.016 | 1.213 | 0.72 | 0.8 | 0.68 | 0.6 | 0.66 | 0.74 |
| 6 | 164142124 | rs6931774 | -0.641 | -0.556 | 0.133 | -1.464 | 0 | 0.027 | 0.027 | 0 | 0.54 | 0.61 | 0.32 | 0.79 | 0.42 | 0.55 |
| 6 | 164143918 | rs7766259 | -0.317 | -0.505 | 0.201 | -1.437 | -0.001 | -0.005 | -0.005 | 0.518 | 0.93 | 0.93 | 0.82 | 0.95 | 0.89 | 0.84 |
| 6 | 164149859 | rs6904842 | 1.028 | -0.544 | 0.317 | -1.516 | -0.002 | -0.003 | -0.003 | 0.167 | 0.37 | 0.34 | 0.49 | 0.16 | 0.48 | 0.31 |
| 6 | 164153758 | rs10046169 | 0.689 | -0.63 | 0.51 | -1.439 | 0.007 | 0.025 | 0.025 | 0 | 0.19 | 0.21 | 0.15 | 0.34 | 0.23 | 0.1 |
| 6 | 164155654 | rs9347781 | 0.482 | -0.577 | 0.403 | -1.405 | 0.055 | -0.103 | -0.103 | 0 | 0.31 | 0.09 | 0.16 | 0.5 | 0.18 | 0.44 |
| 6 | 164158276 | rs7775785 | -0.464 | -0.333 | 0.801 | -1.719 | -0.007 | 0.021 | 0.021 | 0 | 0.56 | 0.58 | 0.33 | 0.86 | 0.46 | 0.62 |
| 6 | 164163461 | rs4709750 | 0.701 | -0.176 | 0.591 | -1.69 | 0.005 | -0.026 | -0.026 | 0 | 0.85 | 0.49 | 0.78 | 0.89 | 0.85 | 0.77 |
| 6 | 164164233 | rs9458896 | 1.087 | -0.131 | 0.559 | -1.686 | -0.043 | 0.097 | 0.097 | 0.99 | 0.34 | 0.19 | 0.47 | 0.1 | 0.43 | 0.23 |
| 6 | 164164404 | rs7741441 | 0.081 | -0.121 | 0.651 | -1.627 | 0.005 | -0.039 | -0.039 | 0.322 | 0.91 | 0.77 | 0.81 | 0.96 | 0.89 | 0.85 |
| 6 | 164166908 | rs9365603 | -0.004 | -0.117 | 0.348 | -1.625 | -0.002 | -0.032 | -0.032 | 0 | 0.91 | 0.84 | 0.81 | 0.96 | 0.89 | 0.85 |
| 6 | 164169311 | rs6455917 | 0.293 | -0.161 | 0.257 | -1.628 | -0.044 | -0.075 | -0.075 | 0 | 0.78 | 0.55 | 0.77 | 0.88 | 0.84 | 0.73 |
| 6 | 164169511 | rs6918609 | 0.07 | -0.135 | -0.025 | -1.292 | -0.003 | 0.033 | 0.033 | 0 | 0.44 | 0.37 | 0.24 | 0.73 | 0.4 | 0.49 |
| 6 | 164173328 | rs4709755 | 0.954 | 0.08 | -0.298 | -1.285 | 0.014 | 0.084 | 0.084 | 0.356 | 0.34 | 0.13 | 0.41 | 0.15 | 0.34 | 0.24 |
| 6 | 164173448 | rs6926801 | 0.213 | 0.021 | -0.363 | -1.391 | -0.002 | 0.004 | 0.004 | 0 | 0.4 | 0.39 | 0.32 | 0.73 | 0.47 | 0.47 |
| 6 | 164174617 | rs6932044 | 0.796 | -0.023 | 0.476 | -1.163 | -0.043 | -0.089 | -0.089 | 0.249 | 0.74 | 0.54 | 0.73 | 0.88 | 0.81 | 0.71 |
| 6 | 164186315 | rs12199352 | -0.763 | 0.177 | -0.036 | -0.408 | 0.001 | -0.002 | -0.002 | 0 | 0.3 | 0.32 | 0.23 | 0.3 | 0.23 | 0.18 |
| 6 | 164186677 | rs10945919 | -1.255 | -0.026 | 0.313 | -0.641 | -0.009 | 0.001 | 0.001 | 0 | 0.42 | 0.45 | 0.28 | 0.37 | 0.3 | 0.28 |
| 6 | 164187884 | rs1998491 | -0.192 | 0.556 | 0.757 | -0.752 | 0.02 | -0.035 | -0.035 | 0 | 0.94 | 0.89 | 0.94 | 0.98 | 0.89 | 0.89 |
| 6 | 164190919 | rs6455920 | -0.736 | 0.292 | 0.661 | -0.801 | 0 | -0.046 | -0.046 | 0 | 0.23 | 0.28 | 0.14 | 0.06 | 0.19 | 0.13 |
| 6 | 164193413 | rs9356147 | 1.083 | -0.039 | 0.903 | -1.053 | 0.001 | -0.003 | -0.003 | 0.537 | 0.64 | 0.5 | 0.68 | 0.69 | 0.63 | 0.72 |
| 6 | 164193518 | rs9356148 | 1.307 | -0.078 | 0.751 | -1.046 | 0 | 0.001 | 0.001 | 0.357 | 0.57 | 0.35 | 0.65 | 0.64 | 0.59 | 0.63 |
| 6 | 164194292 | rs7742306 | 0.052 | 0.26 | 0.415 | -0.937 | -0.007 | -0.032 | -0.032 | 0.169 | 0.23 | 0.15 | 0.17 | 0.29 | 0.25 | 0.15 |
| 6 | 164205230 | rs6938579 | 0.028 | 0.261 | 0.403 | -0.963 | -0.017 | -0.038 | -0.038 | 0 | 0.23 | 0.15 | 0.17 | 0.31 | 0.27 | 0.15 |
| 6 | 164211973 | rs1111705 | 0.644 | 0.215 | 0.783 | -1.125 | -0.005 | -0.005 | -0.005 | 1.636 | 0.65 | 0.7 | 0.76 | 0.63 | 0.6 | 0.66 |
| 6 | 164214737 | rs9365610 | -0.154 | 0.244 | 0.763 | -1.109 | 0.004 | -0.017 | -0.017 | 0 | 0.88 | 0.88 | 0.93 | 0.92 | 0.86 | 0.81 |
| 6 | 164236262 | rs9347792 | 1.735 | -0.077 | 0.671 | -1.343 | -0.053 | -0.035 | -0.035 | 0 | 0.43 | 0.14 | 0.62 | 0.63 | 0.49 | 0.54 |
| 6 | 164237237 | rs10945924 | -0.366 | 0.006 | 0.877 | -1.29 | -0.004 | -0.006 | -0.006 | 1.119 | 0.77 | 0.8 | 0.82 | 0.71 | 0.73 | 0.85 |
| 6 | 164239275 | rs9355439 | 0.785 | -0.03 | 1.191 | -1.33 | 0.015 | 0.01 | 0.01 | 0 | 0.62 | 0.52 | 0.72 | 0.63 | 0.56 | 0.65 |
| 6 | 164241690 | rs1040641 | 0.835 | -0.036 | 1.539 | -1.335 | -0.001 | -0.001 | -0.001 | 0 | 0.61 | 0.41 | 0.72 | 0.62 | 0.56 | 0.65 |
| 6 | 164254001 | rs2321977 | -0.103 | -0.046 | 1.146 | -1.317 | 0.003 | 0.009 | 0.009 | 0 | 0.73 | 0.68 | 0.79 | 0.71 | 0.69 | 0.85 |
| 6 | 164259420 | rs9355442 | 1.589 | -0.111 | 0.877 | -1.332 | -0.008 | 0.004 | 0.004 | 1.185 | 0.43 | 0.26 | 0.6 | 0.63 | 0.45 | 0.53 |
| 6 | 164260935 | rs7752540 | 0.109 | -0.05 | 0.804 | -1.282 | -0.013 | -0.038 | -0.038 | 0 | 0.07 | 0.22 | 0.05 | 0 | 0.03 | 0.05 |
| 6 | 164264584 | rs17441835 | 0.016 | 0.012 | 0.503 | -1.239 | -0.004 | -0.033 | -0.033 | 0 | 0.12 | 0.12 | 0.08 | 0 | 0.08 | 0.07 |
| 6 | 164280686 | rs9347799 | 1.403 | 0.017 | 0.151 | -1.169 | -0.017 | -0.037 | -0.037 | 0 | 0.39 | 0.21 | 0.55 | 0.56 | 0.45 | 0.48 |
| 6 | 164284054 | rs6938128 | -1.42 | -0.049 | 0.526 | -1.099 | -0.003 | -0.024 | -0.024 | 0 | 0.34 | 0.61 | 0.25 | 0.14 | 0.24 | 0.34 |
| 6 | 164289040 | rs6903346 | -0.57 | -0.156 | 0.865 | -1.055 | -0.035 | -0.088 | -0.088 | 0.018 | 0.14 | 0.31 | 0.07 | 0 | 0.05 | 0.06 |
| 6 | 164289220 | rs6925698 | -1.311 | -0.124 | 0.999 | -1.21 | -0.019 | -0.089 | -0.089 | 0 | 0.3 | 0.43 | 0.22 | 0.13 | 0.21 | 0.33 |
| 6 | 164289249 | rs2073219 | 1.502 | 0.071 | 0.65 | -1.36 | -0.001 | -0.037 | -0.037 | 3.377 | 0.4 | 0.15 | 0.56 | 0.58 | 0.42 | 0.44 |
| 6 | 164291168 | rs9356161 | 0.623 | 0.017 | 0.82 | -1.407 | 0.001 | -0.009 | -0.009 | 0.005 | 0.54 | 0.47 | 0.67 | 0.69 | 0.5 | 0.63 |
| 6 | 164292513 | rs9355443 | 0.092 | 0.069 | 0.901 | -1.367 | 0 | 0 | 0 | 0.021 | 0.66 | 0.63 | 0.75 | 0.69 | 0.55 | 0.69 |
| 6 | 164297041 | rs9295235 | -0.621 | 0.176 | 0.867 | -1.409 | -0.001 | -0.048 | -0.048 | 0 | 0.19 | 0.39 | 0.14 | 0.12 | 0.12 | 0.23 |
| 6 | 164298459 | rs4709786 | 0.015 | 0.236 | 0.707 | -1.428 | -0.015 | -0.009 | -0.009 | 0 | 0.63 | 0.72 | 0.69 | 0.6 | 0.54 | 0.64 |
| 6 | 164299483 | rs9295239 | -0.903 | 0.246 | 0.733 | -1.399 | 0.006 | -0.077 | -0.077 | 0.434 | 0.24 | 0.41 | 0.19 | 0.14 | 0.22 | 0.37 |
| 6 | 164299928 | rs9456887 | 0.101 | 0.243 | 0.755 | -1.403 | 0.007 | -0.056 | -0.056 | 0.415 | 0.1 | 0.27 | 0.06 | 0.03 | 0.1 | 0.16 |
| 6 | 164302394 | rs9458917 | -1.322 | 0.198 | 0.481 | -1.447 | 0.002 | -0.006 | -0.006 | 0.32 | 0.37 | 0.59 | 0.26 | 0.14 | 0.28 | 0.44 |
| 6 | 164307139 | rs9295241 | 0.955 | 0.202 | 0.427 | -1.433 | 0.04 | 0.014 | 0.014 | 0 | 0.48 | 0.33 | 0.61 | 0.58 | 0.42 | 0.43 |
| 6 | 164311616 | rs6914455 | -1.057 | 0.271 | 0.155 | -1.354 | 0.005 | 0.011 | 0.011 | 0.688 | 0.37 | 0.63 | 0.27 | 0.14 | 0.28 | 0.44 |
| 6 | 164314825 | rs9346980 | 1.5 | 0.311 | -0.452 | -1.225 | 0.021 | -0.053 | -0.053 | 0.058 | 0.36 | 0.17 | 0.53 | 0.57 | 0.37 | 0.37 |
| 6 | 164314976 | rs6933409 | -0.409 | 0.6 | -0.31 | -0.847 | 0.01 | -0.061 | -0.061 | 0 | 0.11 | 0.34 | 0.06 | 0.03 | 0.1 | 0.16 |
| 6 | 164316375 | rs4709792 | -0.126 | 0.705 | -1.312 | -0.627 | -0.136 | -0.114 | -0.114 | 0 | 0.75 | 0.98 | 0.81 | 0.72 | 0.65 | 0.81 |
| 6 | 164319963 | rs16895510 | -0.294 | 0.373 | -0.699 | -1.254 | -0.031 | -0.138 | -0.138 | 1.438 | 0.3 | 0.58 | 0.15 | 0.01 | 0.19 | 0.19 |
| 6 | 164322346 | rs9456899 | -0.458 | 0.669 | -1.59 | -0.91 | -0.018 | 0.035 | 0.035 | 0 | 0.59 | 0.72 | 0.34 | 0.29 | 0.46 | 0.55 |
| 6 | 164322735 | rs13215377 | -0.804 | 0.708 | -1.052 | -0.64 | 0.02 | 0.025 | 0.025 | 0.029 | 0.14 | 0.28 | 0.1 | 0.18 | 0.16 | 0.18 |
| 6 | 164322839 | rs12199159 | -0.277 | 0.602 | -1.647 | -0.951 | 0.021 | 0.036 | 0.036 | 0 | 0.43 | 0.68 | 0.23 | 0.05 | 0.33 | 0.34 |
| 6 | 164324188 | rs2038299 | 0.443 | 0.971 | -2.253 | 0.187 | -0.062 | -0.084 | -0.084 | 1.013 | 0.55 | 0.07 | 0.38 | 0.52 | 0.48 | 0.41 |
| 6 | 164324219 | rs9458932 | -0.574 | 1.526 | -1.323 | 0.514 | 0.019 | 0.02 | 0.02 | 0 | 0.11 | 0.13 | 0.25 | 0.16 | 0.21 | 0.18 |
| 6 | 164325395 | rs9355445 | -0.079 | 0.595 | -1.512 | 0.201 | 0.02 | -0.009 | -0.009 | 0 | 0.25 | 0.1 | 0.11 | 0.25 | 0.2 | 0.29 |
| 6 | 164325492 | rs11755754 | 0.218 | 0.911 | -1.806 | -0.143 | -0.049 | 0.053 | 0.053 | 0 | 0.31 | 0.11 | 0.22 | 0.15 | 0.4 | 0.41 |
| 6 | 164330598 | rs16895531 | -0.816 | 0.771 | -1 | 0.327 | -0.006 | 0.036 | 0.036 | 0 | 0.06 | 0.17 | 0.06 | 0.08 | 0.04 | 0.13 |
| 6 | 164340663 | rs9355447 | 0.076 | 0.862 | -0.557 | 0.12 | -0.012 | 0.001 | 0.001 | 0.019 | 0.61 | 0.62 | 0.32 | 0.34 | 0.54 | 0.6 |
| 6 | 164341220 | rs6921880 | -0.311 | 0.786 | -0.717 | 0.058 | -0.026 | -0.033 | -0.033 | 0 | 0.65 | 0.69 | 0.37 | 0.38 | 0.58 | 0.73 |
| 6 | 164343775 | rs9356171 | 0.297 | 0.777 | -0.44 | -0.394 | -0.014 | 0.013 | 0.013 | 0 | 0.32 | 0.25 | 0.23 | 0.16 | 0.44 | 0.46 |
| 6 | 164347626 | rs1040644 | -0.209 | 0.558 | -0.501 | -0.447 | -0.013 | 0.003 | 0.003 | 0 | 0.56 | 0.47 | 0.37 | 0.44 | 0.64 | 0.63 |
| 6 | 164350087 | rs2322177 | 0.883 | 1.069 | -0.215 | -0.585 | 0.001 | -0.049 | -0.049 | 0 | 0.29 | 0.33 | 0.16 | 0.05 | 0.28 | 0.27 |
| 6 | 164351521 | rs6928944 | 0.603 | 1.151 | -0.701 | -0.425 | -0.034 | -0.104 | -0.104 | 5.266 | 0.21 | 0.02 | 0.12 | 0.29 | 0.19 | 0.16 |
| 6 | 164353061 | rs12205431 | -0.347 | 1.158 | -0.047 | 0.076 | 0.053 | 0.021 | 0.021 | 0.166 | 0.05 | 0.08 | 0.09 | 0.11 | 0.17 | 0.22 |

| | | | | | | | | | | | | | | | | |
|---|---|---|---|---|---|---|---|---|---|---|---|---|---|---|---|---|
| 6 | 164354268 | rs6936893 | 0.562 | 1.056 | -0.176 | 0.037 | -0.005 | 0.051 | 0.051 | 0 | 0.49 | 0.27 | 0.27 | 0.33 | 0.47 | 0.43 |
| 6 | 164355784 | rs9647611 | 0.096 | 0.928 | -0.98 | -0.01 | -0.014 | -0.032 | -0.032 | 0 | 0.13 | 0.13 | 0.02 | 0 | 0.04 | 0.09 |
| 6 | 164359009 | rs9295251 | 0.133 | 1.515 | -1.053 | 0.306 | -0.003 | 0.007 | 0.007 | 0 | 0.55 | 0.57 | 0.3 | 0.36 | 0.51 | 0.45 |
| 6 | 164359880 | rs875772 | -1.268 | 1.491 | -1.09 | 0.166 | 0.015 | -0.011 | -0.011 | 1.823 | 0.1 | 0.11 | 0.11 | 0.06 | 0.15 | 0.19 |
| 6 | 164360657 | rs6907523 | -0.315 | 0.633 | -2.187 | 0.198 | -0.157 | -0.031 | -0.031 | 0 | 0.61 | 0.97 | 0.57 | 0.41 | 0.5 | 0.56 |
| 6 | 164361544 | rs7775629 | -0.392 | 0.091 | -1.659 | 0.163 | 0.003 | 0.027 | 0.027 | 0 | 0.44 | 0.7 | 0.27 | 0.41 | 0.39 | 0.42 |
| 6 | 164362709 | rs4709025 | -0.736 | 0.4 | -0.474 | -0.079 | 0.105 | -0.024 | -0.024 | 0 | 0.1 | 0.31 | 0.12 | 0.04 | 0.14 | 0.08 |
| 6 | 164362833 | rs4709026 | -1.064 | -0.172 | -0.483 | 0.05 | -0.039 | -0.038 | -0.038 | 0.907 | 0.27 | 0.45 | 0.17 | 0.19 | 0.17 | 0.29 |
| 6 | 164363807 | rs2322178 | -0.064 | -0.569 | -1.079 | 0.028 | -0.002 | -0.013 | -0.013 | 0 | 0.12 | 0.01 | 0.06 | 0.1 | 0.1 | 0.13 |
| 6 | 164366131 | rs7748452 | -0.332 | -0.673 | -1.085 | 0.187 | 0.003 | -0.053 | -0.053 | 0 | 0.12 | 0.04 | 0.06 | 0.24 | 0.1 | 0.2 |
| 6 | 164368087 | rs9458956 | 0.42 | -0.645 | -0.708 | 0.152 | 0.005 | 0.005 | 0.005 | 0 | 0.08 | 0.12 | 0.05 | 0.16 | 0.1 | 0.09 |
| 6 | 164369451 | rs6910079 | -1.682 | -0.862 | -0.986 | -0.008 | -0.04 | -0.021 | -0.021 | 6.075 | 0.55 | 0.8 | 0.45 | 0.59 | 0.51 | 0.59 |
| 6 | 164372151 | rs7743762 | -0.319 | -0.486 | -0.329 | 0.288 | -0.002 | -0.002 | -0.002 | 0 | 0.16 | 0.15 | 0.22 | 0.16 | 0.13 | 0.18 |
| 6 | 164372312 | rs9355450 | -0.339 | -0.52 | -0.343 | 0.265 | -0.03 | -0.046 | -0.046 | 0.619 | 0.15 | 0.07 | 0.11 | 0.29 | 0.19 | 0.24 |
| 6 | 164373180 | rs725759 | -1.3 | -0.568 | 0.063 | 0.27 | -0.004 | -0.008 | -0.008 | 0 | 0.28 | 0.23 | 0.21 | 0.43 | 0.31 | 0.37 |
| 6 | 164373654 | rs2078446 | -0.897 | -0.127 | -0.684 | 0.314 | -0.005 | -0.013 | -0.013 | 0 | 0.36 | 0.29 | 0.39 | 0.47 | 0.36 | 0.45 |
| 6 | 164377679 | rs9458957 | -1.251 | 0.011 | -1.612 | 0.299 | -0.008 | -0.012 | -0.012 | 0 | 0.55 | 0.71 | 0.32 | 0.57 | 0.47 | 0.54 |
| 6 | 164378685 | rs4379268 | -0.759 | 0.422 | -1.527 | 0.569 | -0.041 | -0.045 | -0.045 | 0.095 | 0.19 | 0.03 | 0.09 | 0.3 | 0.19 | 0.2 |
| 6 | 164382899 | rs9356174 | -0.922 | 0.313 | -1.452 | 0.437 | -0.009 | -0.011 | -0.011 | 0 | 0.32 | 0.11 | 0.26 | 0.35 | 0.3 | 0.36 |
| 6 | 164383521 | rs12660673 | -0.817 | 0.473 | -0.849 | 0.03 | 0.004 | -0.016 | -0.016 | 0 | 0.26 | 0.32 | 0.24 | 0.2 | 0.29 | 0.21 |
| 6 | 164384219 | rs9456909 | -0.702 | -0.135 | -0.326 | 0.215 | -0.031 | -0.024 | -0.024 | 0 | 0.3 | 0.36 | 0.21 | 0.24 | 0.2 | 0.28 |
| 6 | 164388079 | rs1865916 | -0.68 | 0.396 | -0.167 | 0.275 | 0.005 | 0.001 | 0.001 | 0 | 0.31 | 0.35 | 0.35 | 0.34 | 0.31 | 0.28 |
| 6 | 164391762 | rs7754718 | -0.682 | 0.248 | 0.171 | 0.134 | 0.02 | 0.005 | 0.005 | 0 | 0.23 | 0.36 | 0.33 | 0.24 | 0.24 | 0.18 |
| 6 | 164402861 | rs4709802 | -0.627 | 0.111 | 0.935 | 0.016 | 0.048 | -0.003 | -0.003 | 0.433 | 0.23 | 0.44 | 0.31 | 0.23 | 0.23 | 0.17 |
| 6 | 164407062 | rs1756 | -1.319 | -0.2 | 0.945 | -0.118 | -0.002 | 0.004 | 0.004 | 0 | 0.3 | 0.31 | 0.33 | 0.25 | 0.27 | 0.21 |
| 6 | 164407297 | rs12191111 | -1.127 | -0.234 | 1.039 | -0.135 | -0.01 | 0.011 | 0.011 | 0 | 0.28 | 0.22 | 0.33 | 0.25 | 0.27 | 0.2 |
| 6 | 164412742 | rs617616 | -1.602 | -0.38 | 1.653 | -0.171 | 0.008 | 0.034 | 0.034 | 0.02 | 0.34 | 0.77 | 0.37 | 0.25 | 0.28 | 0.22 |
| 6 | 164414940 | rs587114 | -0.88 | -0.093 | 2.007 | -0.095 | 0.04 | -0.004 | -0.004 | 1.218 | 0.23 | 0.49 | 0.32 | 0.24 | 0.24 | 0.2 |
| 6 | 164416580 | rs493605 | -0.853 | -0.105 | 1.834 | -0.093 | 0.025 | -0.005 | -0.005 | 0.362 | 0.23 | 0.6 | 0.33 | 0.24 | 0.24 | 0.19 |
| 6 | 164421002 | rs205982 | -1.175 | -0.346 | 1.048 | -0.239 | -0.024 | -0.019 | -0.019 | 0.094 | 0.18 | 0.68 | 0.13 | 0.09 | 0.1 | 0.15 |
| 6 | 164423425 | rs1005541 | 0.385 | 0.315 | 0.853 | 0.025 | -0.01 | 0.006 | 0.006 | 0 | 0.8 | 0.82 | 0.85 | 0.81 | 0.77 | 0.82 |
| 6 | 164426165 | rs206001 | 0.282 | 0.323 | 0.857 | 0.029 | -0.011 | 0.008 | 0.008 | 0 | 0.8 | 0.82 | 0.85 | 0.81 | 0.77 | 0.81 |
| 6 | 164426765 | rs206003 | 0.22 | 0.346 | 0.794 | 0.041 | -0.026 | 0.014 | 0.014 | 0 | 0.8 | 0.88 | 0.85 | 0.82 | 0.77 | 0.81 |
| 6 | 164434628 | rs388305 | 0.27 | 0.38 | 0.858 | 0.06 | 0.02 | -0.012 | -0.012 | 0.696 | 0.79 | 0.7 | 0.84 | 0.81 | 0.77 | 0.83 |
| 6 | 164440149 | rs4709810 | 0.803 | 0.257 | 0.85 | -0.269 | 0.017 | -0.055 | -0.055 | 0 | 0.67 | 0.53 | 0.72 | 0.84 | 0.64 | 0.75 |
| 6 | 164441635 | rs4709816 | 1.133 | 0.286 | -0.104 | 0.104 | -0.018 | -0.099 | -0.099 | 0 | 0.28 | 0.15 | 0.42 | 0.49 | 0.3 | 0.4 |
| 6 | 164442269 | rs9458967 | 0.284 | 0.178 | 0.169 | 0.181 | 0.012 | 0.01 | 0.01 | 0 | 0.38 | 0.27 | 0.32 | 0.32 | 0.35 | 0.37 |
| 6 | 164445349 | rs6455932 | 1.104 | 0.344 | 0.239 | 0.2 | -0.008 | -0.069 | -0.069 | 0 | 0.28 | 0.16 | 0.41 | 0.49 | 0.3 | 0.39 |
| 6 | 164445664 | rs85750 | -0.02 | 0.362 | 0.572 | -0.078 | 0 | -0.004 | -0.004 | 0 | 0.76 | 0.78 | 0.79 | 0.84 | 0.75 | 0.81 |
| 6 | 164447167 | rs205989 | -0.184 | 0.398 | 0.189 | -0.141 | -0.043 | 0.066 | 0.066 | 0 | 0.77 | 0.92 | 0.8 | 0.88 | 0.75 | 0.81 |
| 6 | 164451546 | rs9365646 | 0.36 | 0.367 | 0.585 | 0.504 | -0.001 | -0.001 | -0.001 | 4.87 | 0.37 | 0.34 | 0.29 | 0.37 | 0.33 | 0.47 |
| 6 | 164452203 | rs9355453 | 0.317 | 0.337 | 0.54 | 0.499 | -0.002 | 0 | 0 | 0.231 | 0.37 | 0.36 | 0.29 | 0.38 | 0.33 | 0.47 |
| 6 | 164456716 | rs2064793 | -0.415 | 0.435 | 0.623 | 0.716 | 0.001 | -0.002 | -0.002 | 0 | 0.53 | 0.6 | 0.6 | 0.5 | 0.53 | 0.45 |
| 6 | 164459012 | rs9458972 | 0.513 | 0.389 | 0.657 | 0.435 | 0.008 | -0.007 | -0.007 | 0.597 | 0.36 | 0.26 | 0.51 | 0.37 | 0.37 | 0.32 |
| 6 | 164461976 | rs12208802 | 1.233 | 0.443 | 0.55 | 0.496 | 0.01 | -0.042 | -0.042 | 1.269 | 0.28 | 0.12 | 0.25 | 0.34 | 0.22 | 0.37 |
| 6 | 164463355 | rs4709819 | 0.041 | 0.493 | 0.666 | 0.435 | 0.01 | -0.003 | -0.003 | 0 | 0.43 | 0.37 | 0.53 | 0.42 | 0.43 | 0.38 |
| 6 | 164466497 | rs2064792 | -0.097 | 0.473 | 0.685 | 0.445 | 0.003 | 0 | 0 | 0 | 0.45 | 0.41 | 0.55 | 0.42 | 0.48 | 0.39 |
| 6 | 164485356 | rs392649 | 0.681 | 0.47 | 0.449 | 0.47 | 0.015 | -0.016 | -0.016 | 0 | 0.33 | 0.18 | 0.28 | 0.35 | 0.28 | 0.44 |
| 6 | 164492126 | rs1436857 | 0.937 | 0.522 | 0.295 | 0.464 | -0.005 | -0.042 | -0.042 | 0 | 0.33 | 0.1 | 0.28 | 0.35 | 0.29 | 0.44 |
| 6 | 164502228 | rs183275 | 0.012 | 0.249 | -0.043 | 0.135 | 0.018 | -0.017 | -0.017 | 0 | 0.33 | 0.74 | 0.5 | 0.36 | 0.3 | 0.25 |
| 6 | 164502635 | rs206730 | -0.99 | 0.372 | -0.216 | 0.08 | -0.083 | -0.021 | -0.021 | 0 | 0.81 | 0.96 | 0.86 | 0.77 | 0.76 | 0.83 |
| 6 | 164502943 | rs3888819 | 0.603 | 0.218 | -0.233 | -0.033 | 0.013 | 0 | 0 | 0 | 0.44 | 0.24 | 0.34 | 0.42 | 0.4 | 0.52 |
| 6 | 164503129 | rs10945951 | 1.637 | -0.1 | -0.663 | -0.264 | -0.058 | -0.057 | -0.057 | 1.155 | 0.3 | 0.05 | 0.29 | 0.32 | 0.32 | 0.42 |
| 6 | 164503657 | rs206731 | 0.754 | -0.298 | 0.166 | -0.292 | -0.063 | 0.08 | 0.08 | 0 | 0.21 | 0.35 | 0.32 | 0.36 | 0.11 | 0.18 |
| 6 | 164504114 | rs206735 | -0.338 | -0.348 | 0.045 | -0.314 | -0.005 | -0.001 | -0.001 | 0 | 0.33 | 0.63 | 0.39 | 0.35 | 0.21 | 0.26 |
| 6 | 164507352 | rs10455932 | 0.657 | 0.056 | 0.009 | -0.465 | 0.006 | 0.005 | 0.005 | 0 | 0.45 | 0.38 | 0.41 | 0.42 | 0.45 | 0.52 |
| 6 | 164508276 | rs206738 | 0.757 | -0.304 | -0.298 | -0.049 | 0.015 | 0.016 | 0.016 | 0 | 0.48 | 0.6 | 0.47 | 0.42 | 0.42 | 0.55 |
| 6 | 164509315 | rs2033705 | 2.539 | -0.388 | -0.911 | -0.125 | -0.01 | -0.085 | -0.085 | 0 | 0.24 | 0.04 | 0.3 | 0.36 | 0.19 | 0.36 |
| 6 | 164509450 | rs12333017 | 0.301 | -0.133 | -0.284 | -0.274 | 0 | -0.012 | -0.012 | 0 | 0.2 | 0.21 | 0.11 | 0.05 | 0.24 | 0.16 |
| 6 | 164512178 | rs172272 | 0.775 | -0.231 | -0.23 | -0.221 | 0.005 | 0.016 | 0.016 | 0 | 0.09 | 0.05 | 0.03 | 0.01 | 0.07 | 0.1 |
| 6 | 164518318 | rs10945955 | -0.575 | -0.188 | 0.022 | -0.189 | 0.005 | -0.053 | -0.053 | 0.051 | 0.22 | 0.36 | 0.22 | 0.08 | 0.28 | 0.17 |
| 6 | 164520503 | rs933251 | 0.932 | -0.145 | -0.112 | 0.077 | 0.01 | 0.008 | 0.008 | 0 | 0.51 | 0.54 | 0.48 | 0.61 | 0.39 | 0.58 |
| 6 | 164523846 | rs9456927 | -0.492 | -0.15 | -0.007 | 0.077 | 0.001 | -0.043 | -0.043 | 0 | 0.22 | 0.33 | 0.12 | 0.07 | 0.26 | 0.17 |
| 6 | 164523901 | rs2322278 | 1.983 | -0.379 | 0.023 | 0.061 | 0.016 | -0.047 | -0.047 | 0 | 0.32 | 0.16 | 0.48 | 0.47 | 0.29 | 0.4 |
| 6 | 164525967 | rs13205991 | -0.6 | 0.09 | 0.332 | 0.039 | 0.003 | -0.044 | -0.044 | 0 | 0.21 | 0.32 | 0.12 | 0.07 | 0.26 | 0.17 |
| 6 | 164528095 | rs206701 | -1.183 | 0.117 | 0.42 | 0.016 | -0.003 | -0.083 | -0.083 | 0 | 0.28 | 0.54 | 0.15 | 0.08 | 0.28 | 0.24 |
| 6 | 164528407 | rs9456928 | 0.813 | 0.107 | 0.769 | 0.074 | 0.02 | -0.022 | -0.022 | 0.024 | 0.11 | 0.23 | 0.08 | 0.05 | 0.18 | 0.1 |
| 6 | 164528619 | rs206704 | 0.061 | 0.203 | 0.49 | 0.187 | -0.062 | -0.043 | -0.043 | 0 | 0.66 | 0.88 | 0.56 | 0.6 | 0.62 | 0.66 |
| 6 | 164529381 | rs3752534 | 1.158 | 0.159 | -0.313 | 0.28 | 0.027 | 0.013 | 0.013 | 0 | 0.5 | 0.35 | 0.46 | 0.55 | 0.44 | 0.56 |

| | | | | | | | | | | | | | | | | |
|---|---|---|---|---|---|---|---|---|---|---|---|---|---|---|---|---|
| 6 | 164533751 | rs9458989 | 0.77 | 0.276 | -0.392 | 0.265 | 0 | -0.007 | -0.007 | 0 | 0.11 | 0.15 | 0.06 | 0.05 | 0.12 | 0.1 |
| 6 | 164534582 | rs4709823 | 0.555 | 0.678 | -0.625 | 0.744 | -0.002 | -0.023 | -0.023 | 0 | 0.14 | 0.07 | 0.28 | 0.27 | 0.13 | 0.14 |
| 6 | 164538607 | rs206686 | -1.07 | 0.863 | -0.518 | 0.612 | -0.004 | -0.002 | -0.002 | 0 | 0.39 | 0.68 | 0.38 | 0.42 | 0.47 | 0.26 |
| 6 | 164539435 | rs9458994 | 0.622 | 1.24 | -0.224 | 0.57 | 0.028 | -0.024 | -0.024 | 0 | 0.07 | 0.11 | 0.16 | 0 | 0.16 | 0.04 |
| 6 | 164539696 | rs206687 | -0.961 | 1.001 | -0.507 | 0.424 | -0.005 | -0.014 | -0.014 | 0.314 | 0.56 | 0.76 | 0.49 | 0.55 | 0.61 | 0.45 |
| 6 | 164540137 | rs9458995 | 0.081 | 1.105 | -1.039 | 0.309 | 0.001 | 0.019 | 0.019 | 0 | 0.29 | 0.21 | 0.31 | 0.13 | 0.34 | 0.28 |
| 6 | 164541324 | rs9458996 | -0.6 | 1.213 | -0.264 | 0.875 | -0.014 | -0.015 | -0.015 | 0 | 0.23 | 0.18 | 0.16 | 0.33 | 0.28 | 0.15 |
| 6 | 164549712 | rs9347819 | -0.235 | 0.652 | 0.131 | 0.795 | -0.011 | 0.034 | 0.034 | 0 | 0.07 | 0.12 | 0.1 | 0.15 | 0.05 | 0.14 |
| 6 | 164551891 | rs206716 | -0.413 | 0.729 | -0.6 | 0.67 | -0.111 | -0.159 | -0.159 | 0 | 0.64 | 0.94 | 0.57 | 0.51 | 0.61 | 0.8 |
| 6 | 164557489 | rs7746403 | 0.478 | 0.986 | -0.18 | 0.382 | -0.005 | -0.006 | -0.006 | 0.793 | 0.11 | 0.08 | 0.08 | 0 | 0.18 | 0.05 |
| 6 | 164561709 | rs9356186 | -0.69 | 1.021 | 0.091 | 0.496 | 0.019 | -0.01 | -0.01 | 0.905 | 0.65 | 0.78 | 0.6 | 0.59 | 0.72 | 0.81 |
| 6 | 164567872 | rs17550239 | 0.598 | 1.137 | -0.155 | 0.508 | -0.019 | 0.018 | 0.018 | 0 | 0.11 | 0.05 | 0.1 | 0 | 0.2 | 0.06 |
| 6 | 164568038 | rs807926 | -0.087 | 0.991 | 0.02 | 0.478 | -0.002 | 0.022 | 0.022 | 0 | 0.47 | 0.64 | 0.44 | 0.42 | 0.54 | 0.54 |
| 6 | 164569414 | rs1359484 | 1.575 | 0.49 | -0.135 | 0.566 | -0.018 | 0.033 | 0.033 | 0.227 | 0.23 | 0.13 | 0.19 | 0.16 | 0.25 | 0.33 |
| 6 | 164571038 | rs1325410 | 0.778 | -0.806 | -0.438 | 0.334 | 0.004 | -0.004 | -0.004 | 0 | 0.16 | 0.18 | 0.15 | 0.18 | 0.12 | 0.24 |
| 6 | 164576988 | rs807921 | -1.141 | -0.487 | -0.512 | 0.148 | -0.03 | -0.093 | -0.093 | 1.505 | 0.76 | 0.98 | 0.68 | 0.65 | 0.83 | 0.83 |
| 6 | 164577878 | rs9346995 | 0.827 | 0.242 | 0.307 | 0.684 | -0.037 | -0.005 | -0.005 | 0 | 0.24 | 0.16 | 0.22 | 0.25 | 0.34 | 0.34 |
| 6 | 164579132 | rs7748431 | -0.75 | -0.287 | 1.327 | 0.803 | 0.014 | -0.062 | -0.062 | 0 | 0.12 | 0.21 | 0.08 | 0 | 0.15 | 0.05 |
| 6 | 164584877 | rs807914 | -1.456 | -0.713 | 0.885 | 1.388 | -0.008 | 0.001 | 0.001 | 0.457 | 0.28 | 0.61 | 0.28 | 0.33 | 0.2 | 0.39 |
| 6 | 164588002 | rs807903 | -1.465 | -0.701 | 0.142 | 1.395 | -0.007 | 0.001 | 0.001 | 0.471 | 0.28 | 0.61 | 0.28 | 0.33 | 0.2 | 0.39 |
| 6 | 164589045 | rs929099 | -0.935 | -0.125 | -0.767 | 1.495 | 0.014 | -0.024 | -0.024 | 0.35 | 0.2 | 0.14 | 0.23 | 0.37 | 0.15 | 0.35 |
| 6 | 164596778 | rs9459017 | -0.798 | 0.041 | -0.535 | 1.154 | -0.006 | 0.005 | 0.005 | 0 | 0.13 | 0.16 | 0.11 | 0.17 | 0.12 | 0.09 |
| 6 | 164597168 | rs11970080 | -0.951 | -0.006 | -0.104 | 2.225 | -0.002 | 0.153 | 0.153 | 0 | 0.09 | 0.15 | 0.18 | 0.36 | 0.08 | 0.12 |
| 6 | 164598220 | rs11757902 | 0.472 | 0.267 | -1.064 | 0.756 | -0.053 | 0.016 | 0.016 | 0.408 | 0.38 | 0.12 | 0.41 | 0.39 | 0.43 | 0.27 |
| 6 | 164599359 | rs4709827 | 0.553 | 0.605 | -1.261 | 0.529 | -0.046 | -0.023 | -0.023 | 1.539 | 0.33 | 0.09 | 0.38 | 0.39 | 0.34 | 0.25 |
| 6 | 164602642 | rs973215 | 0.408 | 1.855 | -0.788 | 0.155 | -0.023 | 0.024 | 0.024 | 0 | 0.93 | 0.96 | 0.9 | 1 | 0.81 | 0.95 |
| 6 | 164604869 | rs807898 | 0.756 | 1.137 | -1.234 | -0.363 | 0.006 | 0.082 | 0.082 | 1.869 | 0.52 | 0.11 | 0.42 | 0.41 | 0.4 | 0.42 |
| 6 | 164610887 | rs807894 | 0.357 | 1.011 | -1.175 | -0.282 | -0.001 | -0.005 | -0.005 | 0 | 0.5 | 0.21 | 0.54 | 0.58 | 0.57 | 0.4 |
| 6 | 164611239 | rs733336 | 0.646 | 0.917 | -0.264 | -0.164 | 0.013 | -0.019 | -0.019 | 0 | 0.14 | 0.27 | 0.14 | 0.09 | 0.13 | 0.25 |
| 6 | 164612480 | rs807891 | 1.615 | 1.19 | -1.023 | -0.6 | -0.037 | -0.086 | -0.086 | 0 | 0.16 | 0.01 | 0.12 | 0.28 | 0.17 | 0.15 |
| 6 | 164612793 | rs12528404 | 1.44 | 1.095 | -0.788 | -0.535 | 0.053 | 0.078 | 0.078 | 3.029 | 0.19 | 0.06 | 0.04 | 0.02 | 0.07 | 0.13 |
| 6 | 164612818 | rs807888 | -0.127 | 1.302 | -0.728 | -0.057 | -0.031 | -0.002 | -0.002 | 0 | 0.92 | 0.97 | 0.84 | 0.94 | 0.85 | 0.93 |
| 6 | 164613647 | rs807887 | -0.17 | 1.259 | -0.43 | -0.062 | 0.054 | -0.025 | -0.025 | 0 | 0.78 | 0.61 | 0.69 | 0.85 | 0.73 | 0.69 |
| 6 | 164616757 | rs807882 | 0.478 | 0.938 | -0.405 | -0.032 | 0.059 | 0.033 | 0.033 | 1.976 | 0.48 | 0.21 | 0.4 | 0.6 | 0.35 | 0.39 |
| 6 | 164621949 | rs2764650 | 0.059 | 0.966 | -0.265 | -0.098 | 0.01 | 0.011 | 0.011 | 0.451 | 0.68 | 0.42 | 0.51 | 0.4 | 0.58 | 0.69 |
| 6 | 164628975 | rs10484505 | 1.159 | 0.994 | -0.339 | -0.187 | -0.067 | -0.08 | -0.08 | 0.02 | 0.26 | 0.02 | 0.29 | 0.34 | 0.28 | 0.2 |
| 6 | 164629089 | rs10484504 | 0.616 | 0.991 | -0.329 | -0.239 | 0.006 | -0.039 | -0.039 | 0.663 | 0.23 | 0.27 | 0.17 | 0.04 | 0.24 | 0.36 |
| 6 | 164631462 | rs807911 | 0.231 | 1.59 | -0.641 | 0.18 | 0.022 | 0.008 | 0.008 | 0.095 | 0.71 | 0.59 | 0.58 | 0.67 | 0.59 | 0.75 |
| 6 | 164632576 | rs807907 | 0.627 | 1.168 | -0.864 | 0.228 | -0.044 | -0.054 | -0.054 | 0 | 0.26 | 0.03 | 0.29 | 0.32 | 0.27 | 0.2 |
| 6 | 164634589 | rs6917331 | 0.85 | 1.686 | 0.004 | 0.171 | 0.027 | -0.002 | -0.002 | 0 | 0.71 | 0.61 | 0.57 | 0.69 | 0.58 | 0.74 |
| 6 | 164634781 | rs6941584 | 0.818 | 1.937 | -0.261 | 0.087 | 0.003 | 0 | 0 | 0 | 0.82 | 0.84 | 0.68 | 0.84 | 0.68 | 0.84 |
| 6 | 164636164 | rs10447371 | -0.834 | 1.69 | -0.252 | -0.744 | 0.013 | -0.111 | -0.111 | 0 | 0.34 | 0.51 | 0.33 | 0.06 | 0.41 | 0.21 |
| 6 | 164642517 | rs11755623 | -0.204 | 1.236 | 0.29 | 0.298 | 0.002 | 0.145 | 0.145 | 0 | 0.09 | 0.21 | 0.04 | 0.28 | 0.06 | 0.12 |
| 6 | 164642757 | rs10945966 | -0.255 | 1.226 | 0.353 | 0.293 | 0.004 | 0.16 | 0.16 | 0 | 0.09 | 0.24 | 0.05 | 0.28 | 0.06 | 0.12 |
| 6 | 164647755 | rs9459032 | -0.144 | 0.871 | 1.003 | 0.444 | -0.013 | 0.198 | 0.198 | 0.004 | 0.17 | 0.46 | 0.21 | 0.43 | 0.12 | 0.21 |
| 6 | 164655660 | rs942734 | 1.555 | 0.604 | -0.746 | -0.336 | 0.054 | -0.049 | -0.049 | 0 | 0.3 | 0.07 | 0.16 | 0.4 | 0.19 | 0.32 |
| 6 | 164657235 | rs4709832 | -1.062 | 1.18 | -0.264 | -0.789 | 0.021 | 0.062 | 0.062 | 1.026 | 0.37 | 0.84 | 0.44 | 0.22 | 0.33 | 0.26 |
| 6 | 164657797 | rs6930968 | -0.277 | 1.631 | 0.389 | -0.713 | 0.005 | 0.008 | 0.008 | 0 | 0.11 | 0.24 | 0.07 | 0.08 | 0.09 | 0.11 |
| 6 | 164659258 | rs13203846 | 0.215 | 1.59 | 0.604 | -0.694 | 0.017 | 0.005 | 0.005 | 0.35 | 0.13 | 0.25 | 0.16 | 0.09 | 0.19 | 0.12 |
| 6 | 164663686 | rs6915442 | -1.096 | 1.763 | 0.199 | -0.987 | 0.013 | -0.05 | -0.05 | 0 | 0.74 | 0.86 | 0.77 | 0.35 | 0.75 | 0.72 |
| 6 | 164665485 | rs942731 | -1.843 | 2.104 | 0.255 | -1.171 | 0.009 | -0.001 | -0.001 | 0 | 0.39 | 0.6 | 0.57 | 0.34 | 0.45 | 0.41 |
| 6 | 164666479 | rs9456942 | -1.263 | 2.12 | 0.3 | -1.175 | 0.031 | 0.028 | 0.028 | 0.705 | 0.28 | 0.4 | 0.43 | 0.33 | 0.3 | 0.34 |
| 6 | 164667024 | rs9346998 | 0.682 | 2.149 | 0.12 | -1.004 | 0 | -0.005 | -0.005 | 0 | 0.11 | 0.08 | 0.27 | 0.25 | 0.08 | 0.13 |
| 6 | 164670331 | rs1408915 | -0.692 | 2.277 | 0.071 | -0.975 | 0.086 | 0.025 | 0.025 | 0 | 0.08 | 0.17 | 0.1 | 0.08 | 0.15 | 0.14 |
| 6 | 164672324 | rs11961044 | 0.528 | 2.318 | -0.232 | -0.961 | -0.001 | 0.017 | 0.017 | 0 | 0.12 | 0.15 | 0.28 | 0.25 | 0.07 | 0.13 |
| 6 | 164673314 | rs10214423 | 0.373 | 2.183 | -0.424 | -0.565 | 0.036 | -0.012 | -0.012 | 0 | 0.05 | 0.06 | 0.07 | 0.01 | 0.11 | 0.1 |
| 6 | 164675307 | rs1359486 | -0.495 | 1.929 | -0.526 | -0.553 | 0.003 | -0.008 | -0.008 | 0 | 0.72 | 0.84 | 0.74 | 0.28 | 0.71 | 0.66 |
| 6 | 164683715 | rs794108 | 0.797 | 2.259 | 0.065 | 0.374 | 0.021 | 0 | 0 | 0 | 0.62 | 0.49 | 0.44 | 0.65 | 0.56 | 0.58 |
| 6 | 164688107 | rs9459051 | -1.311 | 2.691 | 0.06 | -0.045 | 0.068 | -0.021 | -0.021 | 0.119 | 0.06 | 0.12 | 0.08 | 0.01 | 0.12 | 0.11 |
| 6 | 164691459 | rs942727 | -1.541 | 2.007 | 1.358 | 0.377 | 0.152 | 0.275 | 0.275 | 0.015 | 0.05 | 0.39 | 0.05 | 0.13 | 0.05 | 0.09 |
| 6 | 164693724 | rs2791524 | -0.816 | 2.394 | 0.843 | 0.043 | 0.012 | -0.013 | -0.013 | 0 | 0.36 | 0.42 | 0.39 | 0.71 | 0.43 | 0.53 |
| 6 | 164693961 | rs2791526 | -0.264 | 2.179 | 0.489 | -0.287 | 0.011 | 0.068 | 0.068 | 0 | 0.17 | 0.27 | 0.32 | 0.36 | 0.17 | 0.21 |
| 6 | 164698486 | rs1340083 | 0.909 | 2.287 | 0.403 | -0.518 | 0.085 | 0.087 | 0.087 | 0 | 0.91 | 0.7 | 0.92 | 0.86 | 0.87 | 0.87 |
| 6 | 164703988 | rs794132 | -0.464 | 2.287 | 0.401 | -0.52 | 0.004 | -0.011 | -0.011 | 0 | 0.95 | 0.92 | 0.96 | 0.99 | 0.92 | 0.98 |
| 6 | 164704293 | rs1361429 | 1.12 | 2.58 | 0.368 | -0.732 | 0.019 | 0.02 | 0.02 | 0 | 0.64 | 0.51 | 0.39 | 0.55 | 0.5 | 0.57 |
| 6 | 164711772 | rs12525909 | 1.349 | 2.681 | 0.313 | -0.768 | 0.001 | 0.003 | 0.003 | 0 | 0.61 | 0.57 | 0.39 | 0.55 | 0.49 | 0.57 |
| 6 | 164713980 | rs705779 | -0.481 | 2.677 | 0.206 | -0.752 | 0.009 | -0.018 | -0.018 | 0 | 0.95 | 0.9 | 0.96 | 0.99 | 0.92 | 0.98 |
| 6 | 164715165 | rs942723 | 0.573 | 2.674 | -0.002 | -0.797 | -0.008 | 0.076 | 0.076 | 0 | 0.17 | 0.24 | 0.12 | 0.51 | 0.13 | 0.28 |
| 6 | 164716390 | rs4709836 | -0.087 | 3.051 | -0.412 | -0.782 | -0.018 | -0.03 | -0.03 | 0 | 0.19 | 0.11 | 0.45 | 0.3 | 0.2 | 0.28 |

| | | | | | | | | | | | | | | | | |
|---|---|---|---|---|---|---|---|---|---|---|---|---|---|---|---|---|
| 6 | 164717284 | rs9365676 | -0.148 | 2.156 | -0.269 | -0.624 | -0.069 | -0.016 | -0.016 | 0 | 0.26 | 0.12 | 0.53 | 0.3 | 0.36 | 0.29 |
| 6 | 164721435 | rs9295263 | -0.178 | 2.058 | -0.319 | -0.657 | 0.055 | 0.07 | 0.07 | 0 | 0.26 | 0.42 | 0.54 | 0.44 | 0.37 | 0.39 |
| 6 | 164721947 | rs7759775 | 0.794 | 2.042 | -0.339 | -0.678 | 0.001 | 0.019 | 0.019 | 0 | 0.16 | 0.16 | 0.11 | 0.51 | 0.13 | 0.29 |
| 6 | 164722031 | rs1886675 | 1.245 | 1.676 | -1.134 | -0.932 | 0.05 | 0.178 | 0.178 | 0 | 0.39 | 0.14 | 0.22 | 0.04 | 0.34 | 0.29 |
| 6 | 164724047 | rs794114 | -0.92 | 2.082 | -1.106 | -0.138 | 0.057 | 0.062 | 0.062 | 0.009 | 0.29 | 0.46 | 0.54 | 0.43 | 0.39 | 0.39 |
| 6 | 164734449 | rs7750881 | 1.433 | 1.78 | -1.514 | -0.114 | 0.001 | 0 | 0 | 0.042 | 0.49 | 0.5 | 0.35 | 0.56 | 0.4 | 0.58 |
| 6 | 164752315 | rs1340076 | -0.194 | 1.914 | -1.718 | 0.289 | -0.047 | -0.038 | -0.038 | 0.12 | 0.22 | 0.12 | 0.45 | 0.29 | 0.31 | 0.23 |
| 6 | 164752476 | rs9459066 | -0.662 | 0.967 | -1.235 | 0.429 | 0.029 | 0.147 | 0.147 | 0 | 0.08 | 0.3 | 0.05 | 0.15 | 0.06 | 0.11 |
| 6 | 164753144 | rs1934602 | 0.717 | 1.211 | -2.332 | -0.004 | -0.062 | 0.09 | 0.09 | 0 | 0.85 | 0.97 | 0.89 | 0.99 | 0.81 | 0.94 |
| 6 | 164756364 | rs1340079 | -0.384 | 0.66 | -1.873 | 0.845 | 0.025 | 0.004 | 0.004 | 7.07 | 0.47 | 0.87 | 0.55 | 0.52 | 0.35 | 0.42 |
| 6 | 164766124 | rs9347838 | -1.024 | 0.788 | -1.299 | 1.355 | -0.003 | -0.016 | -0.016 | 0 | 0.16 | 0.09 | 0.28 | 0.27 | 0.13 | 0.25 |
| 6 | 164766576 | rs793979 | -1.053 | 0.484 | -0.933 | 1.878 | -0.004 | 0.008 | 0.008 | 4.811 | 0.19 | 0.71 | 0.18 | 0.22 | 0.12 | 0.28 |
| 6 | 164766717 | rs6930276 | -1.808 | 0.395 | -0.031 | 3.074 | -0.001 | 0.241 | 0.241 | 0 | 0.08 | 0.2 | 0.11 | 0.39 | 0.06 | 0.15 |
| 6 | 164773524 | rs9654571 | -0.96 | 0.311 | 0.003 | 2.478 | 0 | -0.01 | -0.01 | 0 | 0.05 | 0.11 | 0.04 | 0.03 | 0.05 | 0.03 |
| 6 | 164783891 | rs1496118 | -1.466 | 1.121 | 0.673 | 2.032 | 0.012 | -0.051 | -0.051 | 0.553 | 0.26 | 0.57 | 0.21 | 0.15 | 0.27 | 0.25 |
| 6 | 164790822 | rs2101878 | -0.878 | 0.735 | -0.464 | 1.808 | -0.022 | -0.043 | -0.043 | 0 | 0.61 | 0.85 | 0.57 | 0.51 | 0.55 | 0.64 |
| 6 | 164798906 | rs793981 | 0.932 | 0.638 | 0.028 | 1.974 | 0.011 | 0.085 | 0.085 | 0 | 0.16 | 0.28 | 0.29 | 0.28 | 0.2 | 0.15 |
| 6 | 164810702 | rs12215802 | 0.308 | 0.183 | -0.454 | 1.218 | -0.064 | -0.067 | -0.067 | 0 | 0.26 | 0.12 | 0.38 | 0.41 | 0.33 | 0.46 |
| 6 | 164816435 | rs12333136 | 0.233 | 0.345 | -0.445 | 1.191 | -0.067 | -0.06 | -0.06 | 0 | 0.27 | 0.13 | 0.39 | 0.42 | 0.35 | 0.46 |
| 6 | 164818834 | rs9356206 | 0.437 | 0.317 | -0.212 | 1.149 | 0.002 | 0.008 | 0.008 | 0 | 0.06 | 0.15 | 0.04 | 0.1 | 0.06 | 0.08 |
| 6 | 164823252 | rs1832651 | -1.046 | 0.033 | -0.472 | 0.815 | -0.01 | 0.035 | 0.035 | 0 | 0.68 | 0.64 | 0.6 | 0.53 | 0.66 | 0.74 |
| 6 | 164825347 | rs9356209 | -0.71 | -0.198 | -0.229 | 0.996 | 0 | 0 | 0 | 0.331 | 0.42 | 0.49 | 0.48 | 0.52 | 0.41 | 0.54 |
| 6 | 164827880 | rs9356210 | -0.429 | -0.16 | -0.403 | 1.016 | -0.031 | -0.028 | -0.028 | 0 | 0.37 | 0.29 | 0.45 | 0.52 | 0.41 | 0.54 |
| 6 | 164831395 | rs9365699 | -0.398 | -0.137 | -0.671 | 1.028 | -0.047 | -0.043 | -0.043 | 0 | 0.36 | 0.24 | 0.45 | 0.52 | 0.41 | 0.54 |
| 6 | 164834193 | rs1340080 | 0.9 | 0.089 | -0.636 | 1.14 | -0.005 | -0.038 | -0.038 | 2.134 | 0.15 | 0.06 | 0.26 | 0.37 | 0.12 | 0.29 |
| 6 | 164838366 | rs2132434 | 0.248 | 0.265 | -0.649 | 0.718 | -0.042 | -0.051 | -0.051 | 0 | 0.24 | 0.14 | 0.37 | 0.42 | 0.3 | 0.45 |
| 6 | 164851966 | rs1832649 | 0.02 | 0.569 | -0.63 | 0.666 | -0.033 | -0.037 | -0.037 | 0 | 0.27 | 0.16 | 0.39 | 0.43 | 0.33 | 0.45 |
| 6 | 164852232 | rs1417056 | -0.001 | 0.571 | -0.618 | 0.668 | -0.03 | -0.033 | -0.033 | 4.438 | 0.27 | 0.18 | 0.39 | 0.42 | 0.33 | 0.45 |
| 6 | 164871044 | rs1496117 | 1.036 | 0.56 | -0.435 | 0.872 | -0.005 | -0.02 | -0.02 | 0.969 | 0.15 | 0.09 | 0.28 | 0.38 | 0.14 | 0.28 |
| 6 | 164884849 | rs9459108 | 0.888 | 0.558 | -0.4 | 0.889 | 0.002 | 0.027 | 0.027 | 0.137 | 0.16 | 0.16 | 0.28 | 0.37 | 0.14 | 0.3 |
| 6 | 164888366 | rs2029870 | 0.3 | 0.542 | -0.409 | 0.871 | 0.001 | 0.029 | 0.029 | 0.281 | 0.2 | 0.29 | 0.32 | 0.4 | 0.19 | 0.33 |
| 6 | 164914089 | rs10455943 | -0.056 | 0.481 | -0.864 | 0.982 | -0.004 | -0.066 | -0.066 | 0 | 0.28 | 0.13 | 0.3 | 0.47 | 0.26 | 0.35 |
| 6 | 164914812 | rs9459122 | 0.148 | 0.923 | -0.93 | 0.766 | -0.014 | 0.014 | 0.014 | 0 | 0.17 | 0.14 | 0.15 | 0.06 | 0.24 | 0.18 |
| 6 | 164920393 | rs9456968 | 0.967 | 0.706 | -0.909 | 0.816 | -0.026 | -0.044 | -0.044 | 0 | 0.15 | 0.04 | 0.23 | 0.29 | 0.15 | 0.23 |
| 6 | 164925854 | rs11752324 | 0.684 | 0.404 | -0.475 | 0.322 | -0.017 | -0.045 | -0.045 | 0 | 0.11 | 0.17 | 0.05 | 0 | 0.09 | 0.08 |
| 6 | 164926983 | rs6903174 | 0.294 | 0.954 | -0.485 | 0.819 | -0.02 | 0 | 0 | 0 | 0.55 | 0.72 | 0.54 | 0.43 | 0.53 | 0.47 |
| 6 | 164928141 | rs9347865 | 0.059 | 1.588 | -0.334 | 1.054 | 0.014 | 0.004 | 0.004 | 0 | 0.14 | 0.14 | 0.14 | 0.16 | 0.24 | 0.18 |
| 6 | 164930105 | rs11963489 | 0.71 | 0.929 | -0.208 | 0.949 | 0 | 0 | 0 | 0.148 | 0.66 | 0.31 | 0.57 | 0.3 | 0.75 | 0.59 |
| 6 | 164931002 | rs9356221 | 0.735 | 0.809 | -0.224 | 1.378 | -0.01 | -0.034 | -0.034 | 0.963 | 0.14 | 0.1 | 0.22 | 0.32 | 0.14 | 0.24 |
| 6 | 164938541 | rs9459128 | 0.062 | 0.553 | -0.177 | 1.275 | 0.006 | -0.022 | -0.022 | 0 | 0.05 | 0.03 | 0.07 | 0.14 | 0.03 | 0.11 |
| 6 | 164945340 | rs1909056 | -0.384 | 0.566 | 0.198 | 1.344 | 0.011 | 0.03 | 0.03 | 0 | 0.82 | 0.76 | 0.71 | 0.38 | 0.79 | 0.74 |
| 6 | 164946941 | rs10945998 | 1.648 | 0.517 | -0.121 | 0.312 | 0.009 | 0.202 | 0.202 | 2.098 | 0.39 | 0.08 | 0.34 | 0.04 | 0.37 | 0.23 |
| 6 | 164949755 | rs6907115 | 0.266 | 0.492 | 0.862 | 0.801 | 0.019 | -0.057 | -0.057 | 0.116 | 0.11 | 0.25 | 0.11 | 0.01 | 0.16 | 0.17 |
| 6 | 164953220 | rs10946001 | 0.906 | 0.671 | 0.42 | 1.057 | 0.003 | 0.055 | 0.055 | 0 | 0.7 | 0.55 | 0.62 | 0.48 | 0.75 | 0.64 |
| 6 | 164968016 | rs1874710 | 0.783 | 0.132 | 0.052 | -0.156 | -0.041 | -0.142 | -0.142 | 0 | 0.77 | 0.53 | 0.75 | 0.98 | 0.82 | 0.77 |
| 6 | 164988133 | rs7760143 | -0.055 | 0.61 | -0.768 | 0.452 | -0.025 | -0.032 | -0.032 | 0 | 0.3 | 0.2 | 0.59 | 0.39 | 0.41 | 0.47 |
| 6 | 164988737 | rs2132411 | 0.037 | 0.176 | -0.738 | -0.559 | 0.007 | -0.016 | -0.016 | 0.264 | 0.3 | 0.31 | 0.18 | 0.61 | 0.27 | 0.4 |
| 6 | 164989332 | rs2172966 | 0.241 | -0.238 | -0.085 | -0.576 | 0.031 | 0.022 | 0.022 | 1.533 | 0.94 | 0.66 | 0.96 | 1 | 0.99 | 1 |
| 6 | 164990438 | rs9365729 | 0.02 | -0.171 | -0.247 | -0.62 | 0 | 0.001 | 0.001 | 0 | 0.62 | 0.44 | 0.37 | 0.61 | 0.56 | 0.52 |
| 6 | 164994771 | rs9347874 | 0.676 | -0.258 | -0.745 | -0.166 | -0.032 | -0.001 | -0.001 | 0 | 0.8 | 0.41 | 0.87 | 0.82 | 0.87 | 0.88 |
| 6 | 164998556 | rs9347876 | 0.51 | -0.509 | -1.027 | -0.335 | -0.017 | 0.028 | 0.028 | 0 | 0.47 | 0.27 | 0.27 | 0.44 | 0.45 | 0.37 |
| 6 | 165005155 | rs6455953 | 0.608 | -0.558 | -1.028 | -0.038 | -0.042 | -0.014 | -0.014 | 0 | 0.33 | 0.14 | 0.56 | 0.31 | 0.43 | 0.51 |
| 6 | 165015261 | rs9347879 | 0.527 | -0.539 | -1.031 | -0.044 | -0.027 | 0.029 | 0.029 | 0 | 0.47 | 0.25 | 0.28 | 0.44 | 0.46 | 0.37 |
| 6 | 165015966 | rs9459153 | 0.567 | -0.526 | -1.222 | -0.057 | -0.085 | 0.014 | 0.014 | 0.27 | 0.27 | 0.06 | 0.31 | 0.23 | 0.37 | 0.28 |
| 6 | 165016007 | rs2055261 | 0.568 | -0.607 | -1.392 | -0.202 | 0.007 | 0.047 | 0.047 | 0 | 0.5 | 0.4 | 0.56 | 0.64 | 0.5 | 0.58 |
| 6 | 165016833 | rs2047282 | 0.419 | -0.248 | -1.594 | -0.126 | -0.085 | 0.079 | 0.079 | 0.305 | 0.27 | 0.05 | 0.27 | 0.17 | 0.37 | 0.28 |
| 6 | 165034661 | rs9347882 | 0.316 | -0.332 | -1.802 | -0.458 | 0.024 | 0.112 | 0.112 | 0 | 0.37 | 0.2 | 0.44 | 0.17 | 0.34 | 0.28 |
| 6 | 165038961 | rs7776144 | 0.859 | 0.187 | -1.753 | -0.151 | -0.009 | -0.01 | -0.01 | 0.186 | 0.15 | 0.12 | 0.13 | 0.37 | 0.19 | 0.11 |
| 6 | 165040373 | rs11968290 | -0.214 | -0.581 | -1.129 | -0.77 | -0.019 | 0.043 | 0.043 | 0 | 0.27 | 0.46 | 0.37 | 0.36 | 0.22 | 0.26 |
| 6 | 165045334 | rs727811 | 0.11 | -0.177 | -1.382 | -0.327 | 0.026 | 0.044 | 0.044 | 0.109 | 0.46 | 0.2 | 0.46 | 0.64 | 0.58 | 0.64 |
| 6 | 165059966 | rs6919146 | 0.176 | -0.371 | -0.861 | -0.379 | 0 | -0.005 | -0.005 | 0 | 0.59 | 0.6 | 0.65 | 0.81 | 0.64 | 0.62 |
| 6 | 165060975 | rs9456987 | 0.085 | 0.117 | -1.006 | 0.498 | 0.007 | -0.016 | -0.016 | 0 | 0.13 | 0.04 | 0.15 | 0.14 | 0.11 | 0.3 |
| 6 | 165062425 | rs9356236 | 0.653 | -0.041 | -0.587 | 0.46 | -0.013 | -0.017 | -0.017 | 0.02 | 0.78 | 0.4 | 0.78 | 0.88 | 0.78 | 0.9 |
| 6 | 165063209 | rs8180617 | 0.453 | 0.903 | -1.01 | 1.234 | -0.052 | -0.066 | -0.066 | 0 | 0.16 | 0.05 | 0.12 | 0.35 | 0.25 | 0.19 |
| 6 | 165066871 | rs4709871 | 0.725 | 0.602 | -0.303 | 0.924 | 0.001 | -0.002 | -0.002 | 0 | 0.56 | 0.49 | 0.6 | 0.37 | 0.49 | 0.44 |
| 6 | 165072478 | rs9364750 | 0.769 | 0.58 | -0.266 | 1.01 | -0.006 | 0.024 | 0.024 | 0 | 0.56 | 0.4 | 0.6 | 0.29 | 0.49 | 0.44 |
| 6 | 165072761 | rs9347892 | -0.518 | 0.474 | -0.278 | 1.075 | -0.006 | 0.041 | 0.041 | 7.381 | 0.06 | 0.08 | 0.31 | 0.2 | 0.03 | 0.06 |
| 6 | 165093175 | rs1496007 | 0.327 | 0.712 | -0.153 | 1.139 | 0.005 | 0.025 | 0.025 | 0 | 0.18 | 0.2 | 0.14 | 0.57 | 0.25 | 0.33 |
| 6 | 165095352 | rs9347899 | 0.282 | 0.355 | -0.23 | 0.895 | -0.02 | -0.039 | -0.039 | 0 | 0.67 | 0.46 | 0.7 | 0.84 | 0.71 | 0.63 |

| | | | | | | | | | | | | | | | | |
|---|---|---|---|---|---|---|---|---|---|---|---|---|---|---|---|---|
| 6 | 165115007 | rs903791 | -0.065 | 0.272 | -0.134 | 0.902 | -0.06 | -0.039 | -0.039 | 0 | 0.75 | 0.51 | 0.86 | 0.83 | 0.86 | 0.69 |
| 6 | 165164603 | rs1511088 | 1.196 | 0.529 | -0.223 | 0.943 | 0.012 | 0.077 | 0.077 | 0 | 0.48 | 0.31 | 0.62 | 0.27 | 0.42 | 0.29 |
| 6 | 165170398 | rs1012175 | 1.744 | 0.545 | -0.499 | 0.789 | -0.009 | 0.169 | 0.169 | 0.541 | 0.42 | 0.19 | 0.31 | 0.07 | 0.38 | 0.22 |
| 6 | 165173588 | rs9356265 | 0.503 | 1.213 | -0.198 | 1.419 | -0.079 | -0.106 | -0.106 | 0 | 0.16 | 0.03 | 0.14 | 0.46 | 0.27 | 0.23 |
| 6 | 165177313 | rs7747058 | 0.104 | 1.045 | -0.11 | 1.275 | -0.027 | -0.028 | -0.028 | 0 | 0.75 | 0.79 | 0.76 | 0.43 | 0.57 | 0.62 |
| 6 | 165182669 | rs9456996 | -0.632 | 0.423 | 0.643 | 1.176 | -0.003 | 0.005 | 0.005 | 0 | 0.27 | 0.3 | 0.4 | 0.32 | 0.21 | 0.44 |
| 6 | 165188223 | rs6905430 | 0.333 | 0.493 | 0.843 | 1.133 | 0.022 | -0.029 | -0.029 | 0 | 0.83 | 0.5 | 0.8 | 0.89 | 0.79 | 0.91 |
| 6 | 165189841 | rs11752397 | 1.201 | 0.625 | 0.69 | 1.204 | 0.002 | -0.007 | -0.007 | 0 | 0.57 | 0.21 | 0.4 | 0.57 | 0.59 | 0.48 |
| 6 | 165193140 | rs10946027 | 1.212 | 0.627 | 0.671 | 1.008 | 0.003 | -0.031 | -0.031 | 0 | 0.57 | 0.2 | 0.4 | 0.49 | 0.58 | 0.47 |
| 6 | 165208598 | rs907223 | -0.403 | 0.577 | 0.843 | 0.976 | 0.007 | -0.01 | -0.01 | 0.018 | 0.85 | 0.77 | 0.83 | 0.89 | 0.79 | 0.92 |
| 6 | 165209942 | rs9356274 | 0.185 | 0.578 | 0.826 | 0.977 | 0.009 | -0.03 | -0.03 | 0 | 0.82 | 0.59 | 0.81 | 0.89 | 0.79 | 0.92 |
| 6 | 165222329 | rs10806806 | 1.198 | 0.675 | 0.236 | 1.032 | 0.003 | -0.031 | -0.031 | 0 | 0.57 | 0.2 | 0.4 | 0.49 | 0.58 | 0.47 |
| 6 | 165230761 | rs1511087 | -0.918 | 0.523 | 0.335 | 1.201 | -0.015 | 0.039 | 0.039 | 0 | 0.28 | 0.58 | 0.43 | 0.39 | 0.21 | 0.44 |
| 6 | 165237939 | rs9356281 | -0.903 | 0.542 | 0.314 | 1.211 | -0.015 | 0.042 | 0.042 | 0 | 0.28 | 0.58 | 0.43 | 0.4 | 0.21 | 0.44 |
| 6 | 165262895 | rs9365785 | -0.89 | 1.246 | 0.079 | 1.449 | 0.032 | 0.011 | 0.011 | 0.103 | 0.31 | 0.5 | 0.25 | 0.62 | 0.44 | 0.39 |
| 6 | 165273510 | rs7356835 | 0.177 | 0.537 | 0.114 | 1.022 | -0.01 | 0.015 | 0.015 | 0 | 0.7 | 0.69 | 0.49 | 0.58 | 0.7 | 0.61 |
| 6 | 165274092 | rs7356875 | 0.257 | 0.549 | 0.15 | 1.019 | -0.006 | 0.021 | 0.021 | 0 | 0.7 | 0.67 | 0.47 | 0.42 | 0.68 | 0.55 |
| 6 | 165276034 | rs1912672 | 0.283 | -0.051 | -0.292 | 0.452 | -0.006 | -0.001 | -0.001 | 0 | 0.61 | 0.6 | 0.63 | 0.59 | 0.59 | 0.55 |
| 6 | 165277087 | rs6911539 | 2.003 | -0.24 | -0.363 | 0.224 | 0.036 | 0.042 | 0.042 | 0 | 0.31 | 0.13 | 0.24 | 0.06 | 0.16 | 0.17 |
| 6 | 165278401 | rs9459222 | 1.043 | -0.049 | -0.449 | 0.276 | 0 | 0.015 | 0.015 | 0.351 | 0.16 | 0.14 | 0.07 | 0.03 | 0.19 | 0.1 |
| 6 | 165279451 | rs1912675 | 0.753 | 0.75 | -0.412 | 0.305 | 0 | 0.004 | 0.004 | 0.077 | 0.61 | 0.39 | 0.44 | 0.5 | 0.62 | 0.52 |
| 6 | 165284996 | rs7759442 | -0.37 | 0.752 | -0.408 | 0.34 | -0.039 | 0.046 | 0.046 | 0.263 | 0.29 | 0.23 | 0.15 | 0.21 | 0.34 | 0.21 |
| 6 | 165287626 | rs6936556 | -0.485 | 0.731 | -0.38 | 0.363 | -0.017 | 0.02 | 0.02 | 0 | 0.76 | 0.73 | 0.53 | 0.68 | 0.77 | 0.63 |
| 6 | 165295240 | rs9347937 | 1.399 | 0.764 | -0.356 | 0.292 | 0.003 | 0.015 | 0.015 | 0 | 0.44 | 0.33 | 0.35 | 0.3 | 0.4 | 0.37 |
| 6 | 165296363 | rs4380732 | -0.455 | 0.759 | -0.359 | 0.369 | -0.019 | 0.015 | 0.015 | 0 | 0.76 | 0.71 | 0.57 | 0.69 | 0.77 | 0.63 |
| 6 | 165298267 | rs7767429 | 0.281 | 0.768 | -0.369 | 0.369 | 0.005 | -0.004 | -0.004 | 0.6 | 0.14 | 0.21 | 0.08 | 0.1 | 0.15 | 0.11 |
| 6 | 165298415 | rs7742715 | -0.408 | 0.594 | -0.382 | 0.394 | -0.01 | -0.008 | -0.008 | 0 | 0.32 | 0.39 | 0.22 | 0.39 | 0.37 | 0.26 |
| 6 | 165298534 | rs4394186 | 0.962 | 0.705 | -0.726 | 0.515 | 0.003 | 0.015 | 0.015 | 0 | 0.44 | 0.33 | 0.35 | 0.3 | 0.4 | 0.37 |
| 6 | 165309797 | rs7764525 | 0.769 | 0.838 | -0.892 | 0.267 | 0.005 | 0.007 | 0.007 | 0.127 | 0.48 | 0.41 | 0.44 | 0.54 | 0.52 | 0.49 |
| 6 | 165312013 | rs12530062 | 0.803 | 0.649 | -1.383 | 0.193 | 0.023 | 0.055 | 0.055 | 2.205 | 0.16 | 0.1 | 0.07 | 0.03 | 0.14 | 0.08 |
| 6 | 165314143 | rs9365808 | 0.792 | 0.635 | -1.402 | 0.507 | 0.015 | -0.041 | -0.041 | 1.763 | 0.09 | 0.02 | 0.3 | 0.34 | 0.07 | 0.35 |
| 6 | 165314534 | rs911725 | 0.965 | 0.559 | -1.598 | 0.17 | 0.099 | -0.009 | -0.009 | 0.533 | 0.42 | 0.07 | 0.59 | 0.6 | 0.26 | 0.54 |
| 6 | 165316048 | rs6455970 | -0.675 | 1.263 | -0.826 | -0.177 | 0.113 | -0.074 | -0.074 | 0 | 0.19 | 0.48 | 0.13 | 0.1 | 0.22 | 0.14 |
| 6 | 165316941 | rs10455948 | -0.171 | 0.876 | -0.808 | -0.042 | 0.061 | -0.053 | -0.053 | 0 | 0.15 | 0.45 | 0.09 | 0.07 | 0.12 | 0.04 |
| 6 | 165317841 | rs1227989 | -0.36 | 0.568 | -0.729 | 0.414 | 0 | -0.004 | -0.004 | 0 | 0.37 | 0.42 | 0.48 | 0.66 | 0.44 | 0.62 |
| 6 | 165321314 | rs1227991 | 0.452 | 0.427 | -0.935 | 0.284 | -0.002 | 0 | 0 | 0 | 0.49 | 0.3 | 0.61 | 0.44 | 0.47 | 0.45 |
| 6 | 165323041 | rs1116435 | -0.138 | 0.325 | -0.524 | 0.259 | 0.005 | -0.02 | -0.02 | 0 | 0.74 | 0.67 | 0.8 | 0.86 | 0.69 | 0.75 |
| 6 | 165324358 | rs996301 | -0.026 | 0.101 | -0.263 | 0.504 | -0.019 | 0.025 | 0.025 | 0 | 0.19 | 0.26 | 0.15 | 0.3 | 0.17 | 0.13 |
| 6 | 165326296 | rs752398 | -0.399 | 0.12 | -0.235 | 0.532 | -0.027 | 0.044 | 0.044 | 0 | 0.22 | 0.33 | 0.17 | 0.42 | 0.19 | 0.28 |
| 6 | 165327219 | rs6910737 | 1.416 | -0.15 | -0.827 | -0.032 | -0.048 | 0.148 | 0.148 | 0 | 0.25 | 0.03 | 0.27 | 0.02 | 0.27 | 0.17 |
| 6 | 165327502 | rs6934720 | 0.408 | -0.054 | -0.675 | 0.092 | 0.001 | 0.044 | 0.044 | 0 | 0.18 | 0.31 | 0.14 | 0.3 | 0.18 | 0.14 |
| 6 | 165329095 | rs16897235 | 0.408 | -0.044 | -0.657 | 0.015 | 0.001 | 0.091 | 0.091 | 0 | 0.18 | 0.31 | 0.14 | 0.37 | 0.18 | 0.14 |
| 6 | 165329230 | rs6924121 | -0.133 | -0.142 | -0.708 | -0.182 | 0.004 | -0.006 | -0.006 | 0.019 | 0.62 | 0.63 | 0.58 | 0.57 | 0.68 | 0.52 |
| 6 | 165329794 | rs6929521 | 0.88 | -0.106 | -0.51 | -0.043 | 0.007 | -0.019 | -0.019 | 0 | 0.13 | 0.11 | 0.14 | 0.05 | 0.19 | 0.07 |
| 6 | 165332233 | rs6921473 | -0.556 | -0.231 | -0.485 | 0.266 | 0 | 0.001 | 0.001 | 0 | 0.73 | 0.71 | 0.64 | 0.6 | 0.8 | 0.71 |
| 6 | 165332257 | rs12189587 | 0.69 | -0.418 | -0.448 | 0.335 | -0.001 | 0 | 0 | 0.003 | 0.22 | 0.25 | 0.33 | 0.35 | 0.1 | 0.25 |
| 6 | 165332880 | rs7772470 | 0.092 | -0.1 | -0.304 | 0.554 | -0.021 | 0.206 | 0.206 | 1.504 | 0.09 | 0.26 | 0.08 | 0.29 | 0.03 | 0.07 |
| 6 | 165343557 | rs2093442 | 0.928 | -0.214 | -0.97 | 0.012 | -0.02 | 0.109 | 0.109 | 0 | 0.3 | 0.08 | 0.27 | 0.13 | 0.31 | 0.32 |
| 6 | 165346371 | rs6920410 | 0.013 | -0.651 | -0.766 | 0.011 | -0.042 | -0.006 | -0.006 | 0 | 0.31 | 0.61 | 0.43 | 0.65 | 0.13 | 0.32 |
| 6 | 165355350 | rs9347950 | 0.217 | -0.728 | -0.537 | 0.237 | -0.042 | 0.218 | 0.218 | 0.385 | 0.1 | 0.3 | 0.09 | 0.29 | 0.02 | 0.04 |
| 6 | 165357645 | rs13215619 | 0.728 | -0.652 | -0.713 | 0.017 | -0.005 | 0.041 | 0.041 | 0 | 0.12 | 0.07 | 0.06 | 0.03 | 0.12 | 0.18 |
| 6 | 165358162 | rs6917857 | -0.102 | -0.678 | -0.564 | 0.227 | -0.042 | -0.006 | -0.006 | 0 | 0.31 | 0.62 | 0.43 | 0.65 | 0.13 | 0.32 |
| 6 | 165368163 | rs9355519 | 0.384 | -0.605 | -0.508 | 0.441 | -0.035 | 0.154 | 0.154 | 2.111 | 0.09 | 0.29 | 0.05 | 0.2 | 0.02 | 0.06 |
| 6 | 165369127 | rs9365832 | 0.944 | -0.614 | -0.809 | 0.295 | 0.015 | -0.056 | -0.056 | 0.338 | 0.2 | 0.1 | 0.32 | 0.35 | 0.1 | 0.26 |
| 6 | 165377627 | rs1570277 | -0.001 | -0.216 | -0.928 | -0.022 | -0.001 | -0.002 | -0.002 | 0 | 0.41 | 0.64 | 0.49 | 0.73 | 0.28 | 0.39 |
| 6 | 165377997 | rs6455978 | 0.389 | 0.284 | -0.618 | 0.366 | 0.002 | -0.009 | -0.009 | 0 | 0.72 | 0.74 | 0.77 | 0.87 | 0.6 | 0.72 |
| 6 | 165381837 | rs7749037 | -0.116 | 0.169 | -0.699 | 0.956 | -0.002 | -0.003 | -0.003 | 0.301 | 0.95 | 0.96 | 0.97 | 0.95 | 0.93 | 0.96 |
| 6 | 165394580 | rs10946054 | -0.428 | -0.205 | -0.743 | 1.087 | 0.008 | -0.012 | -0.012 | 0.001 | 0.85 | 0.89 | 0.7 | 0.66 | 0.93 | 0.77 |
| 6 | 165395250 | rs7745328 | -0.891 | -0.879 | -0.265 | 1.038 | -0.051 | 0.148 | 0.148 | 1.083 | 0.19 | 0.48 | 0.15 | 0.36 | 0.13 | 0.22 |
| 6 | 165396461 | rs11754642 | 1.12 | -0.603 | -0.623 | 0.771 | -0.024 | 0.002 | 0.002 | 1.477 | 0.09 | 0.03 | 0.06 | 0.09 | 0.15 | 0.07 |
| 6 | 165396894 | rs7772966 | -0.269 | -0.671 | -0.292 | 0.87 | -0.001 | 0.021 | 0.021 | 0 | 0.21 | 0.26 | 0.17 | 0.33 | 0.25 | 0.15 |
| 6 | 165397843 | rs9347055 | -0.269 | -0.671 | -0.292 | 0.87 | -0.001 | 0.021 | 0.021 | 0 | 0.21 | 0.26 | 0.17 | 0.33 | 0.25 | 0.15 |
| 6 | 165398137 | rs9355525 | -0.269 | -0.671 | -0.292 | 0.87 | -0.001 | 0.021 | 0.021 | 0 | 0.21 | 0.26 | 0.17 | 0.33 | 0.25 | 0.15 |
| 6 | 165401154 | rs9459275 | 1.379 | -0.427 | -0.823 | 0.29 | -0.089 | 0.292 | 0.292 | 0 | 0.43 | 0.07 | 0.34 | 0.06 | 0.45 | 0.39 |
| 6 | 165405678 | rs911727 | 1.012 | -0.46 | -0.638 | 0.53 | 0.01 | -0.015 | -0.015 | 0 | 0.15 | 0.12 | 0.3 | 0.34 | 0.08 | 0.23 |
| 6 | 165407250 | rs13209326 | 0.73 | -0.381 | -0.718 | 0.149 | -0.001 | -0.001 | -0.001 | 0 | 0.07 | 0.09 | 0.04 | 0.04 | 0.09 | 0.18 |
| 6 | 165410311 | rs9459277 | 1.866 | -0.482 | -0.985 | -0.161 | -0.09 | 0.232 | 0.232 | 0 | 0.33 | 0.03 | 0.29 | 0.02 | 0.33 | 0.21 |
| 6 | 165410382 | rs9457028 | 0.083 | -0.365 | -0.526 | 0.231 | -0.002 | -0.012 | -0.012 | 0 | 0.08 | 0.12 | 0.05 | 0.04 | 0.09 | 0.18 |
| 6 | 165412500 | rs9365855 | 0.308 | -0.376 | -0.523 | 0.491 | 0.003 | -0.04 | -0.04 | 0.431 | 0.1 | 0.04 | 0.05 | 0.16 | 0.1 | 0.07 |

| | | | | | | | | | | | | | | | | |
|---|---|---|---|---|---|---|---|---|---|---|---|---|---|---|---|---|
| 6 | 165412717 | rs16897299 | 1.171 | -0.374 | -0.356 | 0.378 | 0.025 | -0.022 | -0.022 | 0 | 0.11 | 0.11 | 0.13 | 0.07 | 0.19 | 0.06 |
| 6 | 165413895 | rs9356334 | 0.791 | -0.803 | -0.564 | 0.606 | 0.022 | -0.02 | -0.02 | 0.014 | 0.06 | 0.03 | 0.03 | 0.1 | 0.02 | 0.04 |
| 6 | 165425189 | rs1322201 | 0.295 | -0.733 | -0.346 | 0.568 | 0.001 | 0.083 | 0.083 | 0 | 0.06 | 0.25 | 0.03 | 0.13 | 0.04 | 0.14 |
| 6 | 165427920 | rs6937822 | 0.081 | -0.649 | -0.901 | 0.603 | 0.006 | -0.011 | -0.011 | 3.167 | 0.88 | 0.92 | 0.89 | 0.75 | 0.9 | 0.92 |
| 6 | 165434315 | rs395044 | -0.078 | -0.648 | -0.957 | 0.615 | 0.003 | -0.001 | -0.001 | 0.061 | 0.11 | 0.09 | 0.11 | 0.24 | 0.09 | 0.08 |
| 6 | 165439363 | rs227484 | -1.379 | -0.75 | -1.264 | 0.212 | -0.04 | 0.213 | 0.213 | 0 | 0.63 | 0.89 | 0.69 | 0.99 | 0.65 | 0.79 |
| 6 | 165441064 | rs1322197 | 0.712 | -0.736 | -1.159 | 0.927 | 0.011 | -0.014 | -0.014 | 0 | 0.16 | 0.16 | 0.29 | 0.28 | 0.08 | 0.23 |
| 6 | 165442362 | rs9355531 | -0.266 | -0.62 | -1.374 | 0.634 | 0.011 | -0.018 | -0.018 | 0 | 0.13 | 0.06 | 0.12 | 0.28 | 0.1 | 0.08 |
| 6 | 165443820 | rs979515 | -0.913 | -0.406 | -1.16 | 0.244 | -0.002 | 0.102 | 0.102 | 0 | 0.59 | 0.74 | 0.65 | 0.94 | 0.61 | 0.65 |
| 6 | 165445066 | rs2066897 | -0.721 | -0.388 | -1.104 | 1.151 | 0.013 | -0.019 | -0.019 | 0 | 0.15 | 0.08 | 0.12 | 0.33 | 0.1 | 0.15 |
| 6 | 165447942 | rs6913534 | 0.203 | 0.352 | -1.635 | 0.095 | -0.066 | 0.159 | 0.159 | 0 | 0.42 | 0.2 | 0.25 | 0.12 | 0.47 | 0.26 |
| 6 | 165451901 | rs760812 | 0.395 | 0.357 | -2.408 | 0.323 | 0.015 | -0.021 | -0.021 | 0 | 0.56 | 0.11 | 0.61 | 0.41 | 0.65 | 0.56 |
| 6 | 165452463 | rs227465 | 0.428 | 0.169 | -1.311 | 0.159 | -0.029 | 0.134 | 0.134 | 0.564 | 0.21 | 0.38 | 0.18 | 0.48 | 0.15 | 0.31 |
| 6 | 165453907 | rs1923382 | -0.616 | 0.117 | -0.693 | -0.04 | 0.02 | -0.03 | -0.03 | 0 | 0.88 | 0.73 | 0.85 | 0.96 | 0.88 | 0.94 |
| 6 | 165454688 | rs13219159 | 1.972 | 0.55 | -0.848 | -0.468 | -0.044 | 0.102 | 0.102 | 0 | 0.28 | 0.13 | 0.17 | 0 | 0.33 | 0.2 |
| 6 | 165455420 | rs10455803 | -0.111 | 0.407 | -0.862 | -0.072 | 0.003 | 0.004 | 0.004 | 0 | 0.62 | 0.69 | 0.44 | 0.68 | 0.62 | 0.57 |
| 6 | 165458279 | rs9364788 | 0.832 | 0.45 | -1.137 | -0.026 | -0.04 | -0.078 | -0.078 | 0 | 0.27 | 0.03 | 0.41 | 0.28 | 0.26 | 0.37 |
| 6 | 165466890 | rs9347983 | 0.387 | 1.098 | -0.69 | -0.096 | 0.03 | 0.095 | 0.095 | 0 | 0.16 | 0.23 | 0.21 | 0.52 | 0.24 | 0.33 |
| 6 | 165467714 | rs9295300 | 1.834 | 1.078 | -0.875 | -0.25 | -0.023 | 0.107 | 0.107 | 0 | 0.29 | 0.12 | 0.16 | 0 | 0.31 | 0.18 |
| 6 | 165469641 | rs227457 | -0.651 | 0.811 | -0.605 | 0.623 | 0.006 | 0.015 | 0.015 | 0 | 0.44 | 0.56 | 0.57 | 0.77 | 0.4 | 0.56 |
| 6 | 165472153 | rs1883340 | -0.007 | 0.91 | -0.182 | 0.809 | 0.009 | -0.026 | -0.026 | 0 | 0.88 | 0.79 | 0.86 | 0.97 | 0.89 | 0.94 |
| 6 | 165478051 | rs227455 | -1.113 | 0.806 | -0.038 | 1.037 | 0.013 | 0.148 | 0.148 | 2.253 | 0.15 | 0.25 | 0.13 | 0.47 | 0.14 | 0.21 |
| 6 | 165484164 | rs10452574 | 0.34 | 0.596 | -0.262 | 0.277 | -0.001 | -0.002 | -0.002 | 0.058 | 0.16 | 0.17 | 0.23 | 0.11 | 0.08 | 0.06 |
| 6 | 165487131 | rs1322193 | 1.305 | 0.798 | -0.634 | -0.106 | -0.032 | 0.058 | 0.058 | 0 | 0.39 | 0.13 | 0.27 | 0.23 | 0.47 | 0.25 |
| 6 | 165487496 | rs2223376 | 0.708 | 0.498 | -0.671 | -0.081 | -0.001 | -0.001 | -0.001 | 0 | 0.16 | 0.17 | 0.23 | 0.11 | 0.08 | 0.06 |
| 6 | 165489704 | rs17490175 | 1.092 | 0.882 | -0.459 | -0.104 | -0.028 | 0.052 | 0.052 | 0 | 0.42 | 0.21 | 0.3 | 0.24 | 0.53 | 0.28 |
| 6 | 165491551 | rs1923380 | 1.01 | 0.375 | -0.419 | -0.167 | -0.012 | -0.009 | -0.009 | 0 | 0.65 | 0.57 | 0.66 | 0.69 | 0.76 | 0.8 |
| 6 | 165493183 | rs2743961 | -1.104 | 0.503 | -0.884 | -0.507 | -0.002 | -0.063 | -0.063 | 0.441 | 0.36 | 0.91 | 0.4 | 0.41 | 0.28 | 0.29 |
| 6 | 165494322 | rs9356344 | 0.093 | 0.454 | -0.697 | -0.175 | -0.077 | -0.153 | -0.153 | 0 | 0.23 | 0.02 | 0.37 | 0.5 | 0.3 | 0.47 |
| 6 | 165496918 | rs526725 | -0.701 | 0.208 | 0.236 | -0.274 | -0.004 | -0.005 | -0.005 | 0 | 0.42 | 0.41 | 0.51 | 0.83 | 0.52 | 0.69 |
| 6 | 165501693 | rs544603 | -1.175 | 0.226 | -0.289 | -0.278 | 0.01 | 0.133 | 0.133 | 0 | 0.49 | 0.71 | 0.55 | 0.83 | 0.55 | 0.69 |
| 6 | 165501975 | rs568444 | -0.332 | 0.818 | -0.818 | 0.095 | -0.071 | -0.035 | -0.035 | 0.751 | 0.32 | 0.13 | 0.45 | 0.6 | 0.45 | 0.58 |
| 6 | 165504192 | rs2675741 | 0.236 | 0.873 | -0.752 | 0.118 | -0.05 | -0.019 | -0.019 | 0 | 0.67 | 0.83 | 0.55 | 0.39 | 0.55 | 0.42 |
| 6 | 165508286 | rs2758937 | -1.182 | 0.418 | 0.454 | 0.574 | 0.005 | 0.109 | 0.109 | 0.216 | 0.17 | 0.55 | 0.11 | 0.34 | 0.17 | 0.27 |
| 6 | 165510101 | rs1923607 | -0.938 | 0.257 | -0.645 | -0.721 | -0.048 | 0.234 | 0.234 | 0 | 0.68 | 0.98 | 0.81 | 0.98 | 0.72 | 0.83 |
| 6 | 165511920 | rs9347988 | -1.259 | 0.048 | 0.898 | 0.491 | -0.003 | -0.004 | -0.004 | 0 | 0.22 | 0.31 | 0.35 | 0.25 | 0.2 | 0.19 |
| 6 | 165520199 | rs9457038 | -0.023 | 0.02 | 0.998 | 0.221 | -0.001 | -0.016 | -0.016 | 0 | 0.09 | 0.07 | 0.06 | 0 | 0.09 | 0.03 |
| 6 | 165520298 | rs2675733 | -0.871 | -0.013 | 1.241 | 0.452 | 0.002 | 0.044 | 0.044 | 0 | 0.09 | 0.17 | 0.27 | 0.27 | 0.1 | 0.1 |
| 6 | 165522144 | rs2758932 | -2.282 | -0.024 | 1 | 0.781 | -0.028 | -0.024 | -0.024 | 0 | 0.49 | 0.88 | 0.52 | 0.57 | 0.46 | 0.43 |
| 6 | 165523008 | rs7747916 | -0.761 | -0.007 | 1.647 | 0.901 | -0.019 | 0.165 | 0.165 | 0.015 | 0.1 | 0.36 | 0.06 | 0.25 | 0.05 | 0.09 |
| 6 | 165528860 | rs2758933 | -1.807 | 0.088 | 1.435 | 0.56 | 0.017 | -0.003 | -0.003 | 0 | 0.29 | 0.42 | 0.4 | 0.32 | 0.32 | 0.31 |
| 6 | 165530602 | rs7762850 | -1.627 | -0.205 | 1.079 | 0.833 | -0.042 | 0.039 | 0.039 | 0 | 0.18 | 0.43 | 0.26 | 0.26 | 0.1 | 0.14 |
| 6 | 165533455 | rs13215765 | -1.406 | -0.11 | 0.9 | 0.859 | -0.014 | 0.023 | 0.023 | 0 | 0.14 | 0.08 | 0.09 | 0.05 | 0.17 | 0.15 |
| 6 | 165535286 | rs7759656 | -0.154 | -0.097 | 0.664 | 0.958 | -0.007 | 0.019 | 0.019 | 0.01 | 0.06 | 0.13 | 0.16 | 0.14 | 0.02 | 0.05 |
| 6 | 165543352 | rs9364791 | -0.179 | -0.096 | 0.608 | 0.906 | -0.007 | -0.023 | -0.023 | 0 | 0.06 | 0.01 | 0.07 | 0.13 | 0.06 | 0.09 |
| 6 | 165557550 | rs894240 | -2.365 | -0.102 | 0.59 | 0.92 | -0.006 | -0.063 | -0.063 | 0.299 | 0.44 | 0.86 | 0.45 | 0.54 | 0.38 | 0.37 |
| 6 | 165558993 | rs1441899 | -2.385 | -0.114 | 0.316 | 0.926 | -0.02 | -0.07 | -0.07 | 0 | 0.45 | 0.92 | 0.45 | 0.57 | 0.38 | 0.37 |
| 6 | 165560336 | rs6926303 | -2.059 | 0.416 | 0.83 | 1.715 | -0.043 | 0.178 | 0.178 | 0.008 | 0.22 | 0.48 | 0.29 | 0.52 | 0.14 | 0.18 |
| 6 | 165564573 | rs957967 | -2.024 | 0.455 | 0.755 | 1.659 | 0.002 | 0.054 | 0.054 | 0 | 0.31 | 0.46 | 0.37 | 0.55 | 0.3 | 0.35 |
| 6 | 165566852 | rs2758944 | -2.127 | 0.478 | 0.518 | 1.634 | -0.002 | 0.052 | 0.052 | 0 | 0.32 | 0.46 | 0.37 | 0.56 | 0.29 | 0.35 |
| 6 | 165569308 | rs2675723 | -1.902 | 0.826 | 0.31 | 1.825 | 0.012 | 0.085 | 0.085 | 0 | 0.27 | 0.41 | 0.36 | 0.54 | 0.27 | 0.34 |
| 6 | 165579317 | rs7769494 | -0.92 | 0.614 | 0.365 | 0.886 | -0.002 | -0.012 | -0.012 | 0 | 0.16 | 0.15 | 0.26 | 0.31 | 0.16 | 0.28 |
| 6 | 165581006 | rs506879 | -1.121 | 0.676 | 0.079 | 0.95 | 0.004 | 0.012 | 0.012 | 0 | 0.25 | 0.4 | 0.32 | 0.31 | 0.24 | 0.3 |
| 6 | 165588018 | rs1764047 | 0.037 | 0.661 | -0.682 | 0.956 | 0.005 | -0.015 | -0.015 | 0.122 | 0.59 | 0.73 | 0.74 | 0.48 | 0.62 | 0.72 |
| 6 | 165588626 | rs12660977 | 0.22 | 0.772 | -1.352 | 1.518 | -0.01 | -0.091 | -0.091 | 0 | 0.18 | 0.04 | 0.15 | 0.49 | 0.16 | 0.29 |
| 6 | 165588629 | rs6913287 | -0.4 | 0.792 | -1.766 | 1.122 | -0.024 | 0.009 | 0.009 | 0 | 0.09 | 0.01 | 0.2 | 0.08 | 0.09 | 0.08 |
| 6 | 165590516 | rs1744509 | -0.244 | 1.381 | -2.435 | 0.735 | -0.052 | 0.019 | 0.019 | 0.013 | 0.59 | 0.82 | 0.62 | 0.39 | 0.53 | 0.33 |
| 6 | 165591182 | rs1899358 | 0.377 | 1.561 | -3.037 | 0.013 | -0.067 | 0.075 | 0.075 | 0 | 0.12 | 0.01 | 0.12 | 0 | 0.24 | 0.1 |
| 6 | 165591630 | rs2583822 | 0.56 | 0.686 | -2.115 | -0.38 | -0.031 | -0.015 | -0.015 | 0 | 0.11 | 0.28 | 0.06 | 0.08 | 0.04 | 0.06 |
| 6 | 165594793 | rs6923393 | -0.379 | 1.272 | -2.285 | -0.773 | -0.096 | 0.11 | 0.11 | 0 | 0.24 | 0.04 | 0.35 | 0.08 | 0.36 | 0.2 |
| 6 | 165597348 | rs2583824 | -0.118 | 1.129 | -1.927 | -0.796 | 0.032 | 0.01 | 0.01 | 2.278 | 0.35 | 0.7 | 0.27 | 0.31 | 0.17 | 0.14 |
| 6 | 165597904 | rs1613866 | -0.116 | 1.117 | -1.932 | -0.802 | 0.032 | 0.009 | 0.009 | 2.249 | 0.35 | 0.7 | 0.27 | 0.31 | 0.17 | 0.14 |
| 6 | 165598461 | rs1764038 | -1.965 | 0.966 | -2.211 | -0.961 | -0.017 | -0.116 | -0.116 | 0.014 | 0.73 | 0.95 | 0.79 | 0.56 | 0.76 | 0.68 |
| 6 | 165599178 | rs12525918 | 0.823 | 0.991 | -2.184 | -0.969 | -0.006 | -0.005 | -0.005 | 0.226 | 0.14 | 0.15 | 0.17 | 0.17 | 0.23 | 0.35 |
| 6 | 165600056 | rs1441894 | -1.054 | 0.916 | -2.181 | -0.94 | -0.047 | -0.113 | -0.113 | 0.339 | 0.62 | 0.94 | 0.57 | 0.48 | 0.64 | 0.58 |
| 6 | 165601421 | rs1441896 | 0.898 | 0.696 | -1.917 | -0.961 | -0.047 | 0.007 | 0.007 | 2.996 | 0.24 | 0.47 | 0.22 | 0.23 | 0.13 | 0.08 |
| 6 | 165606180 | rs2839869 | 0.408 | 0.918 | -2.032 | -0.874 | -0.029 | 0.022 | 0.022 | 0 | 0.11 | 0.02 | 0.22 | 0.08 | 0.13 | 0.1 |
| 6 | 165609467 | rs9348003 | 1.071 | 0.494 | -2.246 | -1.041 | -0.021 | -0.057 | -0.057 | 0.013 | 0.09 | 0.01 | 0.14 | 0.23 | 0.09 | 0.06 |
| 6 | 165612858 | rs1764053 | -0.776 | 0.556 | -2.278 | -1.14 | 0.055 | 0.04 | 0.04 | 0.452 | 0.46 | 0.72 | 0.5 | 0.39 | 0.32 | 0.24 |

| | | | | | | | | | | | | | | | | |
|---|---|---|---|---|---|---|---|---|---|---|---|---|---|---|---|---|
| 6 | 165616420 | rs12526681 | 0.84 | 1.003 | -2.318 | -1.098 | -0.027 | -0.013 | -0.013 | 0.121 | 0.14 | 0.08 | 0.17 | 0.17 | 0.26 | 0.32 |
| 6 | 165621452 | rs1923608 | -0.033 | 0.779 | -1.669 | -1.204 | -0.001 | -0.01 | -0.01 | 0 | 0.19 | 0.16 | 0.35 | 0.33 | 0.19 | 0.16 |
| 6 | 165622183 | rs502201 | -1.507 | 0.535 | -1.65 | -1.329 | -0.061 | -0.06 | -0.06 | 0.045 | 0.61 | 0.9 | 0.68 | 0.58 | 0.56 | 0.56 |
| 6 | 165622459 | rs9654572 | 0.738 | 0.417 | -1.563 | -1.24 | -0.003 | 0.029 | 0.029 | 0 | 0.11 | 0.14 | 0.16 | 0.25 | 0.11 | 0.06 |
| 6 | 165623506 | rs580256 | 0.588 | 0.223 | -1.187 | -1.328 | -0.035 | -0.023 | -0.023 | 0 | 0.12 | 0.29 | 0.06 | 0.07 | 0.04 | 0.05 |
| 6 | 165625891 | rs1744506 | -1.648 | 0.169 | -1.703 | -1.495 | -0.031 | -0.125 | -0.125 | 1.231 | 0.64 | 0.94 | 0.58 | 0.5 | 0.67 | 0.57 |
| 6 | 165631791 | rs2197100 | 0.989 | 0.622 | -1.66 | -1.588 | -0.036 | -0.012 | -0.012 | 0.025 | 0.15 | 0.08 | 0.22 | 0.17 | 0.27 | 0.36 |
| 6 | 165633342 | rs475691 | -1.151 | 0.483 | -1.739 | -1.629 | -0.055 | -0.053 | -0.053 | 0.693 | 0.53 | 0.91 | 0.52 | 0.49 | 0.48 | 0.49 |
| 6 | 165633751 | rs1441890 | 0.929 | 0.25 | -1.749 | -1.62 | -0.03 | 0.009 | 0.009 | 0 | 0.1 | 0.01 | 0.21 | 0.08 | 0.12 | 0.12 |
| 6 | 165634127 | rs1319746 | 0.44 | -0.122 | -0.642 | -1.31 | -0.047 | 0.064 | 0.064 | 0 | 0.12 | 0.01 | 0.07 | 0 | 0.21 | 0.08 |
| 6 | 165638896 | rs7740603 | -0.514 | -0.124 | 0.026 | -1.199 | 0.009 | -0.036 | -0.036 | 0.15 | 0.29 | 0.43 | 0.28 | 0.17 | 0.31 | 0.38 |
| 6 | 165638939 | rs7740635 | 0.961 | -0.187 | 0.115 | -0.868 | -0.008 | -0.028 | -0.028 | 0.139 | 0.1 | 0.09 | 0.04 | 0 | 0.06 | 0.02 |
| 6 | 165640017 | rs16897566 | 0.469 | -0.125 | 0.113 | -0.724 | 0 | 0.006 | 0.006 | 0 | 0.11 | 0.14 | 0.16 | 0.25 | 0.1 | 0.06 |
| 6 | 165643282 | rs7772552 | -0.4 | -0.278 | 0.511 | -1.24 | 0.02 | -0.032 | -0.032 | 0 | 0.13 | 0.27 | 0.06 | 0.07 | 0.08 | 0.07 |
| 6 | 165645984 | rs7751240 | 0.113 | -0.424 | 0.611 | -1.363 | -0.045 | -0.096 | -0.096 | 0.358 | 0.15 | 0.38 | 0.07 | 0 | 0.06 | 0.02 |
| 6 | 165647395 | rs1764059 | -2.541 | -0.454 | 0.159 | -1.506 | -0.057 | 0.099 | 0.099 | 0 | 0.58 | 0.86 | 0.42 | 0.24 | 0.54 | 0.51 |
| 6 | 165647621 | rs16897582 | 1.321 | -0.33 | 0.524 | -1.262 | -0.009 | -0.033 | -0.033 | 0 | 0.11 | 0.14 | 0.05 | 0 | 0.07 | 0.02 |
| 6 | 165648938 | rs4709073 | 0.855 | -0.21 | 0.212 | -1.243 | -0.017 | -0.021 | -0.021 | 0.092 | 0.08 | 0.03 | 0.16 | 0.17 | 0.15 | 0.31 |
| 6 | 165657661 | rs1744493 | -1.75 | -0.127 | 0.19 | -1.254 | 0.016 | 0.097 | 0.097 | 0.123 | 0.43 | 0.78 | 0.32 | 0.25 | 0.35 | 0.41 |
| 6 | 165658548 | rs2119729 | 3.184 | -0.051 | -0.019 | -1.043 | -0.095 | -0.192 | -0.192 | 0.052 | 0.32 | 0.02 | 0.4 | 0.45 | 0.33 | 0.4 |
| 6 | 165658888 | rs1764043 | -2.405 | 0.612 | 0.954 | -1.024 | -0.053 | 0.11 | 0.11 | 0 | 0.57 | 0.88 | 0.42 | 0.24 | 0.55 | 0.5 |
| 6 | 165661646 | rs1157279 | -0.459 | 0.568 | 1.226 | -1.02 | -0.022 | -0.026 | -0.026 | 0 | 0.25 | 0.54 | 0.24 | 0.18 | 0.22 | 0.34 |
| 6 | 165662789 | rs11756728 | 1.017 | 0.687 | 1.134 | -0.978 | 0.002 | -0.004 | -0.004 | 0 | 0.07 | 0.04 | 0.03 | 0 | 0.05 | 0.01 |
| 6 | 165663792 | rs559156 | -0.643 | 0.793 | 1.194 | -0.948 | -0.003 | -0.051 | -0.051 | 0 | 0.27 | 0.29 | 0.12 | 0.07 | 0.2 | 0.14 |
| 6 | 165664853 | rs1998394 | -1.634 | 0.856 | 1.841 | -0.45 | 0.001 | -0.021 | -0.021 | 0 | 0.3 | 0.61 | 0.31 | 0.17 | 0.35 | 0.36 |
| 6 | 165665299 | rs4709075 | -1.464 | 0.659 | 1.965 | -0.243 | -0.015 | -0.018 | -0.018 | 0.296 | 0.31 | 0.58 | 0.26 | 0.21 | 0.26 | 0.4 |
| 6 | 165671448 | rs10946068 | -1.296 | 0.802 | 2.308 | -0.098 | 0.005 | 0.001 | 0.001 | 0 | 0.25 | 0.31 | 0.37 | 0.21 | 0.32 | 0.37 |
| 6 | 165673747 | rs4709927 | 0.628 | 0.838 | 2.501 | -0.039 | 0.033 | 0.047 | 0.047 | 0 | 0.87 | 0.75 | 0.83 | 0.77 | 0.87 | 0.93 |
| 6 | 165675433 | rs2146282 | -1.715 | 0.797 | 2.556 | -0.027 | -0.004 | -0.013 | -0.013 | 0.815 | 0.34 | 0.43 | 0.41 | 0.22 | 0.37 | 0.42 |
| 6 | 165678438 | rs4324759 | 2.237 | 0.799 | 2.052 | -0.027 | -0.018 | -0.012 | -0.012 | 0.106 | 0.53 | 0.3 | 0.42 | 0.55 | 0.5 | 0.51 |
| 6 | 165682170 | rs1105533 | -0.693 | 0.8 | 1.974 | -0.022 | -0.003 | 0.004 | 0.004 | 0 | 0.2 | 0.2 | 0.3 | 0.21 | 0.19 | 0.35 |
| 6 | 165683048 | rs6908035 | 0.975 | 0.792 | 1.856 | -0.029 | 0 | 0.039 | 0.039 | 0 | 0.07 | 0 | 0.04 | 0 | 0.05 | 0.02 |
| 6 | 165686789 | rs9356355 | 2.496 | 0.83 | 1.963 | -0.004 | -0.013 | -0.034 | -0.034 | 0 | 0.48 | 0.16 | 0.39 | 0.55 | 0.46 | 0.48 |
| 6 | 165688844 | rs9459344 | 1.044 | 0.624 | 1.999 | 0.141 | 0.005 | 0.007 | 0.007 | 0 | 0.77 | 0.48 | 0.76 | 0.76 | 0.81 | 0.88 |
| 6 | 165691210 | rs6909361 | -0.762 | 0.681 | 1.721 | 0.208 | -0.007 | 0.011 | 0.011 | 0 | 0.21 | 0.26 | 0.31 | 0.21 | 0.19 | 0.35 |
| 6 | 165691303 | rs6914041 | 0.968 | 0.685 | 1.679 | 0.203 | -0.003 | -0.014 | -0.014 | 0 | 0.73 | 0.5 | 0.75 | 0.76 | 0.78 | 0.85 |
| 6 | 165693624 | rs1413100 | 1.472 | 0.826 | 0.874 | 0.282 | 0.007 | -0.028 | -0.028 | 0 | 0.52 | 0.21 | 0.44 | 0.55 | 0.59 | 0.5 |
| 6 | 165706473 | rs1105228 | 1.115 | 0.44 | 1.367 | 0.018 | -0.016 | -0.031 | -0.031 | 0 | 0.65 | 0.5 | 0.62 | 0.81 | 0.69 | 0.56 |
| 6 | 165713961 | rs510579 | -1.543 | 0.397 | 1.635 | -0.039 | 0.019 | 0.068 | 0.068 | 0.008 | 0.4 | 0.82 | 0.44 | 0.21 | 0.38 | 0.46 |
| 6 | 165713998 | rs573814 | -1.188 | 0.489 | 1.424 | 0.028 | -0.009 | -0.033 | -0.033 | 0 | 0.36 | 0.47 | 0.4 | 0.2 | 0.37 | 0.45 |
| 6 | 165715673 | rs9459356 | -0.376 | 0.292 | 1.16 | 0.07 | -0.012 | -0.087 | -0.087 | 0.726 | 0.17 | 0.35 | 0.1 | 0.01 | 0.16 | 0.1 |
| 6 | 165716733 | rs484861 | -0.568 | 0.413 | 0.771 | 0.34 | -0.012 | 0.019 | 0.019 | 0 | 0.16 | 0.25 | 0.25 | 0.19 | 0.16 | 0.25 |
| 6 | 165722345 | rs646444 | -1.217 | 0.605 | 0.333 | 0.314 | -0.01 | 0.004 | 0.004 | 0 | 0.46 | 0.68 | 0.56 | 0.44 | 0.5 | 0.5 |
| 6 | 165723166 | rs575424 | -1.101 | 0.013 | 0.951 | 0.359 | -0.016 | 0.05 | 0.05 | 0 | 0.2 | 0.35 | 0.21 | 0.26 | 0.17 | 0.31 |
| 6 | 165724052 | rs504371 | -1.293 | -0.294 | 0.943 | 0.12 | -0.009 | 0.001 | 0.001 | 0 | 0.42 | 0.54 | 0.47 | 0.42 | 0.38 | 0.33 |
| 6 | 165732635 | rs533040 | -1.84 | -0.368 | 0.835 | -0.096 | 0.01 | 0.011 | 0.011 | 0 | 0.52 | 0.79 | 0.54 | 0.42 | 0.48 | 0.42 |
| 6 | 165744991 | rs9459370 | -0.415 | -0.395 | 1.111 | -0.158 | -0.008 | -0.04 | -0.04 | 0 | 0.14 | 0.2 | 0.08 | 0 | 0.09 | 0.05 |
| 6 | 165745732 | rs9457071 | -0.463 | -0.542 | 1.018 | -0.232 | 0.026 | 0.035 | 0.035 | 0 | 0.14 | 0.08 | 0.06 | 0 | 0.09 | 0.05 |
| 6 | 165746031 | rs9457072 | -0.11 | -0.469 | 1.053 | -0.195 | 0.018 | 0.025 | 0.025 | 0 | 0.13 | 0.08 | 0.06 | 0 | 0.09 | 0.05 |
| 6 | 165750845 | rs3799148 | 0.096 | -0.579 | 1.588 | -0.116 | -0.001 | 0.069 | 0.069 | 0 | 0.11 | 0.17 | 0.23 | 0.26 | 0.13 | 0.17 |
| 6 | 165750958 | rs1999548 | -0.223 | -0.591 | 1.557 | -0.131 | -0.002 | -0.027 | -0.027 | 0 | 0.13 | 0.18 | 0.08 | 0 | 0.09 | 0.05 |
| 6 | 165754461 | rs3778248 | 0.057 | -0.549 | 1.604 | -0.081 | -0.002 | 0.056 | 0.056 | 0 | 0.11 | 0.17 | 0.23 | 0.26 | 0.13 | 0.16 |
| 6 | 165754976 | rs525643 | -0.589 | -0.5 | 1.52 | -0.102 | -0.002 | 0.073 | 0.073 | 0 | 0.17 | 0.34 | 0.27 | 0.26 | 0.17 | 0.25 |
| 6 | 165759915 | rs484842 | -0.646 | -0.514 | 1.496 | -0.104 | -0.003 | 0.072 | 0.072 | 0 | 0.18 | 0.35 | 0.27 | 0.27 | 0.18 | 0.27 |
| 6 | 165763277 | rs9459375 | -0.161 | -0.459 | 1.501 | -0.142 | -0.002 | 0 | 0 | 0 | 0.18 | 0.16 | 0.2 | 0.16 | 0.22 | 0.14 |
| 6 | 165765865 | rs487489 | -1.569 | -0.526 | 1.293 | -0.13 | -0.001 | -0.008 | -0.008 | 0.312 | 0.37 | 0.67 | 0.48 | 0.43 | 0.39 | 0.41 |
| 6 | 165768220 | rs537436 | -1.656 | -0.557 | 1.27 | -0.146 | -0.001 | -0.01 | -0.01 | 0 | 0.38 | 0.69 | 0.48 | 0.43 | 0.4 | 0.41 |
| 6 | 165790374 | rs575669 | -2.033 | -0.593 | 1.243 | -0.181 | 0.019 | 0.02 | 0.02 | 0 | 0.5 | 0.77 | 0.55 | 0.43 | 0.49 | 0.45 |
| 6 | 165791239 | rs546311 | -0.534 | -0.407 | 1.173 | -0.138 | 0.023 | 0.069 | 0.069 | 0 | 0.18 | 0.35 | 0.27 | 0.26 | 0.2 | 0.27 |
| 6 | 165792989 | rs9459407 | -0.093 | -0.366 | 0.274 | -0.03 | 0.013 | 0.009 | 0.009 | 0 | 0.12 | 0.09 | 0.07 | 0 | 0.1 | 0.05 |
| 6 | 165798068 | rs572280 | -1.004 | -0.152 | 0.623 | 0.387 | -0.001 | -0.009 | -0.009 | 0.459 | 0.37 | 0.66 | 0.47 | 0.44 | 0.39 | 0.4 |
| 6 | 165798443 | rs16897780 | -1.227 | -0.118 | 0.755 | 1.004 | 0.013 | 0.227 | 0.227 | 0 | 0.07 | 0.18 | 0.18 | 0.3 | 0.08 | 0.23 |
| 6 | 165798851 | rs16897782 | -0.874 | -0.439 | 0.317 | 0.83 | -0.001 | 0.006 | 0.006 | 0 | 0.23 | 0.22 | 0.37 | 0.37 | 0.18 | 0.23 |
| 6 | 165798894 | rs220807 | -1.01 | -0.658 | -0.203 | 0.225 | -0.05 | -0.043 | -0.043 | 0 | 0.61 | 0.86 | 0.65 | 0.49 | 0.57 | 0.48 |
| 6 | 165798963 | rs9457084 | -0.504 | -1.115 | 0.38 | 0.143 | -0.02 | 0.029 | 0.029 | 0 | 0.31 | 0.59 | 0.42 | 0.41 | 0.21 | 0.35 |
| 6 | 165799038 | rs220806 | -0.017 | -0.363 | 0.256 | -0.627 | -0.028 | -0.051 | -0.051 | 0 | 0.36 | 0.47 | 0.28 | 0.16 | 0.26 | 0.29 |
| 6 | 165801094 | rs11969826 | 0.207 | -0.374 | 0.193 | -0.77 | -0.023 | -0.11 | -0.11 | 0 | 0.19 | 0.28 | 0.12 | 0 | 0.15 | 0.1 |
| 6 | 165801183 | rs17193 | -0.704 | 0.401 | -0.236 | -0.789 | -0.095 | -0.093 | -0.093 | 0 | 0.78 | 0.92 | 0.66 | 0.34 | 0.71 | 0.56 |
| 6 | 165801790 | rs1511746 | 0.389 | 0.333 | -0.095 | -0.819 | -0.031 | -0.131 | -0.131 | 0 | 0.2 | 0.35 | 0.13 | 0 | 0.15 | 0.21 |

| | | | | | | | | | | | | | | | | |
|---|---|---|---|---|---|---|---|---|---|---|---|---|---|---|---|---|
| 6 | 165802561 | rs220800 | -1.383 | 0.478 | -0.463 | -0.61 | -0.085 | -0.112 | -0.112 | 0.018 | 0.85 | 0.98 | 0.77 | 0.48 | 0.77 | 0.59 |
| 6 | 165808429 | rs3818177 | 0.577 | 0.265 | -0.471 | -0.745 | -0.038 | -0.022 | -0.022 | 0 | 0.3 | 0.13 | 0.4 | 0.25 | 0.41 | 0.18 |
| 6 | 165814173 | rs12528988 | 0.607 | 0.221 | -0.457 | -0.745 | -0.017 | -0.025 | -0.025 | 0.114 | 0.07 | 0.03 | 0.09 | 0.16 | 0.06 | 0.04 |
| 6 | 165815336 | rs220783 | 0.923 | -0.024 | -0.067 | -0.93 | -0.009 | -0.014 | -0.014 | 0 | 0.29 | 0.3 | 0.15 | 0.12 | 0.17 | 0.19 |
| 6 | 165816049 | rs573291 | 0.556 | 0.09 | 0.124 | -1.039 | -0.029 | -0.115 | -0.115 | 1.449 | 0.19 | 0.32 | 0.13 | 0 | 0.15 | 0.2 |
| 6 | 165817219 | rs541254 | 0.329 | 0.069 | 0.18 | -1.046 | -0.044 | -0.134 | -0.134 | 0 | 0.21 | 0.53 | 0.14 | 0 | 0.14 | 0.21 |
| 6 | 165819417 | rs538464 | 0.49 | 0.361 | 0.15 | -1.043 | -0.014 | -0.103 | -0.103 | 0 | 0.2 | 0.28 | 0.14 | 0 | 0.17 | 0.09 |
| 6 | 165820098 | rs220776 | -0.571 | 0.226 | 0.137 | -0.992 | -0.005 | -0.005 | -0.005 | 0 | 0.78 | 0.79 | 0.68 | 0.33 | 0.72 | 0.56 |
| 6 | 165821351 | rs220772 | 0.238 | 0.223 | 0.078 | -0.993 | 0.006 | -0.001 | -0.001 | 0 | 0.6 | 0.48 | 0.55 | 0.32 | 0.58 | 0.35 |
| 6 | 165821402 | rs220771 | 1.054 | -0.048 | 0.006 | -0.952 | 0.008 | 0.005 | 0.005 | 0 | 0.28 | 0.24 | 0.14 | 0.08 | 0.17 | 0.18 |
| 6 | 165823608 | rs220761 | -0.39 | 0.092 | -0.326 | -0.865 | 0.018 | 0.023 | 0.023 | 0 | 0.77 | 0.7 | 0.68 | 0.33 | 0.72 | 0.45 |
| 6 | 165827643 | rs220748 | 0.772 | -0.002 | -0.184 | -0.933 | -0.051 | -0.093 | -0.093 | 0 | 0.3 | 0.37 | 0.17 | 0.08 | 0.17 | 0.19 |
| 6 | 165831225 | rs525235 | 0.482 | 0.005 | -0.32 | -0.949 | 0.021 | 0.045 | 0.045 | 0 | 0.24 | 0.14 | 0.09 | 0 | 0.15 | 0.11 |
| 6 | 165831491 | rs578654 | 0.511 | -0.067 | 0.037 | -0.309 | -0.003 | 0.012 | 0.012 | 0 | 0.45 | 0.45 | 0.5 | 0.26 | 0.52 | 0.28 |
| 6 | 165832227 | rs220740 | -0.476 | -1.384 | 0.383 | -0.739 | -0.051 | -0.098 | -0.098 | 0 | 0.4 | 0.42 | 0.19 | 0.08 | 0.25 | 0.2 |
| 6 | 165832935 | rs3778249 | -0.145 | -0.98 | -0.018 | -0.249 | -0.036 | 0.008 | 0.008 | 0.153 | 0.14 | 0.32 | 0.12 | 0.15 | 0.11 | 0.14 |
| 6 | 165836746 | rs220813 | -0.506 | -0.61 | -0.486 | -0.455 | -0.039 | -0.057 | -0.057 | 0 | 0.29 | 0.32 | 0.14 | 0.1 | 0.15 | 0.28 |
| 6 | 165837769 | rs220810 | 0.359 | -0.363 | -1.163 | -0.312 | 0.034 | 0.103 | 0.103 | 0 | 0.17 | 0.02 | 0.08 | 0 | 0.1 | 0.07 |
| 6 | 165837800 | rs220809 | -0.005 | 1.235 | -0.518 | 0.795 | 0.046 | 0.062 | 0.062 | 0.125 | 0.77 | 0.66 | 0.67 | 0.47 | 0.68 | 0.56 |
| 6 | 165837992 | rs9459417 | -0.795 | 0.604 | -0.943 | 0.268 | 0.016 | 0.022 | 0.022 | 0 | 0.2 | 0.14 | 0.08 | 0.06 | 0.12 | 0.15 |
| 6 | 165844311 | rs470922 | 1.131 | 2.068 | -0.743 | 1.293 | 0.004 | 0.008 | 0.008 | 0 | 0.81 | 0.81 | 0.71 | 0.55 | 0.7 | 0.71 |
| 6 | 165845903 | rs9365885 | -1.185 | 2.444 | -0.602 | 1.733 | 0 | 0.021 | 0.021 | 0 | 0.12 | 0.14 | 0.19 | 0.38 | 0.19 | 0.18 |
| 6 | 165846042 | rs470278 | 0.308 | 3.082 | -1.206 | 1.585 | -0.055 | -0.037 | -0.037 | 0 | 0.78 | 0.91 | 0.65 | 0.78 | 0.66 | 0.73 |
| 6 | 165849238 | rs220821 | -1.03 | 2.215 | -0.558 | 1.164 | 0.002 | 0.022 | 0.022 | 0 | 0.32 | 0.56 | 0.44 | 0.44 | 0.32 | 0.3 |
| 6 | 165850480 | rs220817 | -0.186 | 2.203 | -0.338 | 1.197 | -0.015 | -0.017 | -0.017 | 0 | 0.2 | 0.09 | 0.31 | 0.21 | 0.22 | 0.21 |
| 6 | 165852979 | rs454165 | 0.347 | 2.016 | -0.406 | 1.055 | -0.008 | -0.007 | -0.007 | 3.194 | 0.84 | 0.84 | 0.77 | 0.62 | 0.73 | 0.72 |
| 6 | 165860246 | rs3778250 | 0.031 | 1.65 | -0.761 | 0.668 | -0.026 | -0.02 | -0.02 | 0.15 | 0.25 | 0.04 | 0.33 | 0.23 | 0.28 | 0.25 |
| 6 | 165861013 | rs1967290 | -0.276 | 1.722 | -0.131 | 0.677 | -0.008 | -0.016 | -0.016 | 0.868 | 0.3 | 0.39 | 0.36 | 0.23 | 0.29 | 0.23 |
| 6 | 165862711 | rs9457092 | -0.214 | 1.724 | -0.409 | 0.759 | -0.009 | -0.011 | -0.011 | 0 | 0.07 | 0.05 | 0.08 | 0.2 | 0.09 | 0.09 |
| 6 | 165862892 | rs16897883 | -0.307 | 1.718 | -0.412 | 0.756 | -0.009 | -0.013 | -0.013 | 0 | 0.08 | 0.05 | 0.08 | 0.2 | 0.09 | 0.17 |
| 6 | 165863036 | rs221749 | 1.509 | 1.198 | -0.95 | -0.078 | 0.015 | 0.031 | 0.031 | 0 | 0.31 | 0.21 | 0.26 | 0.2 | 0.24 | 0.34 |
| 6 | 165865657 | rs221748 | -0.069 | 1.184 | -0.881 | -0.073 | -0.03 | -0.013 | -0.013 | 0 | 0.68 | 0.77 | 0.7 | 0.64 | 0.61 | 0.66 |
| 6 | 165866460 | rs221746 | 0.362 | 1.14 | -0.894 | -0.076 | -0.041 | -0.023 | -0.023 | 0.023 | 0.63 | 0.72 | 0.63 | 0.43 | 0.53 | 0.57 |
| 6 | 165870785 | rs9364795 | -0.599 | 1.135 | -0.982 | -0.066 | -0.006 | -0.001 | -0.001 | 1.031 | 0.53 | 0.59 | 0.51 | 0.44 | 0.5 | 0.38 |
| 6 | 165871066 | rs7748041 | -0.001 | 1.126 | -0.987 | -0.112 | 0.006 | 0.015 | 0.015 | 0 | 0.46 | 0.54 | 0.43 | 0.22 | 0.41 | 0.29 |
| 6 | 165871819 | rs221743 | 0.895 | 0.498 | -1.061 | -0.248 | 0.026 | 0.043 | 0.043 | 0 | 0.32 | 0.17 | 0.25 | 0.2 | 0.23 | 0.35 |
| 6 | 165883486 | rs221735 | 0.216 | 0.784 | -0.577 | 0.049 | -0.008 | -0.005 | -0.005 | 0 | 0.58 | 0.4 | 0.43 | 0.43 | 0.47 | 0.51 |
| 6 | 165884927 | rs221726 | 0.833 | 0.001 | -0.254 | -0.643 | -0.005 | -0.065 | -0.065 | 0 | 0.81 | 0.75 | 0.9 | 1 | 0.83 | 0.91 |
| 6 | 165887531 | rs221724 | -0.439 | -0.514 | -0.033 | -0.781 | -0.003 | 0.011 | 0.011 | 0 | 0.56 | 0.45 | 0.61 | 0.21 | 0.6 | 0.47 |
| 6 | 165896944 | rs617739 | -0.901 | -1.421 | 0.587 | -0.115 | -0.035 | 0.025 | 0.025 | 0 | 0.26 | 0.41 | 0.26 | 0.28 | 0.22 | 0.25 |
| 6 | 165896984 | rs617817 | -0.901 | -1.421 | 0.587 | -0.115 | -0.035 | 0.025 | 0.025 | 0 | 0.26 | 0.41 | 0.26 | 0.28 | 0.22 | 0.25 |
| 6 | 165901124 | rs471544 | -1.292 | -1.506 | 0.498 | -0.215 | -0.024 | 0.006 | 0.006 | 0 | 0.3 | 0.4 | 0.29 | 0.28 | 0.24 | 0.28 |
| 6 | 165905549 | rs221730 | 0.857 | -1.744 | -0.086 | 0.115 | -0.143 | -0.062 | -0.062 | 0 | 0.35 | 0.03 | 0.48 | 0.43 | 0.45 | 0.3 |
| 6 | 165909462 | rs537557 | -2.151 | -2.372 | 1.696 | 0.844 | 0.054 | -0.002 | -0.002 | 2.791 | 0.34 | 0.8 | 0.31 | 0.28 | 0.24 | 0.28 |
| 6 | 165910706 | rs16897929 | -0.341 | -2.385 | 1.69 | 0.776 | 0.001 | -0.015 | -0.015 | 0 | 0.11 | 0.11 | 0.03 | 0 | 0.06 | 0.1 |
| 6 | 165911470 | rs494139 | -2.266 | -2.398 | 2.227 | 0.806 | -0.031 | 0.008 | 0.008 | 0 | 0.29 | 0.59 | 0.27 | 0.27 | 0.23 | 0.26 |
| 6 | 165912970 | rs676389 | -2.034 | -2.338 | 2.273 | 0.84 | 0.064 | 0.263 | 0.263 | 0.049 | 0.09 | 0.35 | 0.21 | 0.24 | 0.1 | 0.06 |
| 6 | 165918005 | rs7741165 | -1.121 | -2.33 | 2.162 | 0.827 | 0.045 | 0.247 | 0.247 | 0 | 0.05 | 0.19 | 0.16 | 0.24 | 0.05 | 0.07 |
| 6 | 165920969 | rs16897948 | -1.876 | -2.381 | 1.872 | 0.781 | -0.054 | 0.028 | 0.028 | 0 | 0.25 | 0.44 | 0.22 | 0.27 | 0.18 | 0.24 |
| 6 | 165924970 | rs611909 | 1.557 | -2.363 | 1.459 | 0.779 | -0.06 | -0.116 | -0.116 | 0.285 | 0.75 | 0.41 | 0.88 | 0.97 | 0.82 | 0.78 |
| 6 | 165930153 | rs686406 | 2.069 | -2.291 | 0.437 | 0.819 | 0.086 | -0.01 | -0.01 | 0.217 | 0.65 | 0.12 | 0.68 | 0.73 | 0.76 | 0.69 |
| 6 | 165932290 | rs569647 | 1.872 | -1.9 | 1.345 | 0.674 | -0.001 | 0 | 0 | 0 | 0.69 | 0.27 | 0.58 | 0.78 | 0.77 | 0.79 |
| 6 | 165935421 | rs519595 | 1.27 | -1.869 | 1.833 | 0.69 | -0.034 | 0.008 | 0.008 | 0 | 0.84 | 0.64 | 0.71 | 0.82 | 0.86 | 0.84 |
| 6 | 165935918 | rs515012 | 2.086 | -1.946 | 1.757 | 0.624 | -0.019 | -0.017 | -0.017 | 0.031 | 0.69 | 0.3 | 0.6 | 0.78 | 0.77 | 0.8 |
| 6 | 165943163 | rs576853 | 1.678 | -1.87 | 1.79 | 0.927 | -0.002 | -0.001 | -0.001 | 0 | 0.74 | 0.3 | 0.76 | 0.73 | 0.83 | 0.81 |
| 6 | 165945970 | rs6906719 | -1.665 | -1.86 | 2.027 | 0.966 | -0.052 | 0.057 | 0.057 | 0.096 | 0.18 | 0.53 | 0.19 | 0.25 | 0.09 | 0.13 |
| 6 | 165947458 | rs2031846 | -1.397 | -1.834 | 1.878 | 1.501 | -0.007 | 0.182 | 0.182 | 0 | 0.1 | 0.34 | 0.11 | 0.21 | 0.05 | 0.21 |
| 6 | 165948503 | rs484720 | 1.951 | -1.8 | 0.591 | 1.342 | 0.063 | 0.024 | 0.024 | 0.012 | 0.62 | 0.1 | 0.68 | 0.72 | 0.72 | 0.65 |
| 6 | 165948832 | rs481838 | 1.825 | -1.78 | 0.577 | 1.292 | 0.039 | -0.104 | -0.104 | 0 | 0.58 | 0.07 | 0.55 | 0.52 | 0.68 | 0.62 |
| 6 | 165953662 | rs9356369 | -1.437 | -1.652 | 1.537 | 1.367 | -0.022 | 0.319 | 0.319 | 0.157 | 0.11 | 0.43 | 0.12 | 0.33 | 0.06 | 0.11 |
| 6 | 165953796 | rs16897963 | -0.753 | -1.708 | 1.023 | 0.945 | 0 | -0.017 | -0.017 | 0 | 0.11 | 0.11 | 0.06 | 0 | 0.08 | 0.04 |
| 6 | 165955070 | rs1511744 | -1.672 | -2.162 | 1.418 | 1.1 | -0.081 | 0.103 | 0.103 | 0.009 | 0.23 | 0.47 | 0.19 | 0.38 | 0.09 | 0.25 |
| 6 | 165959283 | rs9347080 | -1.012 | -2.127 | 1.36 | 1.095 | -0.009 | 0.058 | 0.058 | 0 | 0.11 | 0.15 | 0.12 | 0.33 | 0.06 | 0.25 |
| 6 | 165959666 | rs7753004 | -0.79 | -2.295 | 1.385 | 0.819 | -0.004 | -0.047 | -0.047 | 0 | 0.19 | 0.28 | 0.1 | 0.03 | 0.16 | 0.1 |
| 6 | 165959846 | rs7772901 | -1.608 | -2.3 | 1.266 | 0.813 | -0.017 | 0.011 | 0.011 | 0 | 0.32 | 0.46 | 0.23 | 0.36 | 0.25 | 0.36 |
| 6 | 165961160 | rs6922908 | -0.769 | -2.31 | 1.407 | 0.537 | -0.047 | -0.042 | -0.042 | 1.11 | 0.13 | 0.25 | 0.07 | 0.05 | 0.05 | 0.03 |
| 6 | 165963574 | rs7761766 | -1.779 | -2.121 | 1.41 | 0.843 | -0.014 | 0.006 | 0.006 | 0 | 0.37 | 0.62 | 0.37 | 0.5 | 0.27 | 0.38 |
| 6 | 165965479 | rs634651 | -1.527 | -1.447 | 0.555 | 0.653 | -0.027 | -0.04 | -0.04 | 0.566 | 0.73 | 0.85 | 0.66 | 0.57 | 0.67 | 0.63 |
| 6 | 165969225 | rs6938249 | -1.354 | -1.628 | 1.948 | 0.88 | -0.032 | 0.143 | 0.143 | 0 | 0.11 | 0.23 | 0.11 | 0.34 | 0.03 | 0.25 |

| | | | | | | | | | | | | | | | | |
|---|---|---|---|---|---|---|---|---|---|---|---|---|---|---|---|---|
| 6 | 165969237 | rs1022179 | -2.227 | -1.626 | 1.348 | 0.856 | 0.074 | -0.048 | -0.048 | 1.808 | 0.37 | 0.77 | 0.39 | 0.46 | 0.2 | 0.32 |
| 6 | 165969801 | rs7760078 | -1.225 | -0.994 | 1.583 | 0.927 | -0.04 | -0.054 | -0.054 | 0.225 | 0.13 | 0.4 | 0.1 | 0.06 | 0.07 | 0.03 |
| 6 | 165992248 | rs943801 | 1.367 | -1.043 | 0.582 | 0.946 | 0.033 | 0.072 | 0.072 | 0 | 0.67 | 0.24 | 0.66 | 0.87 | 0.79 | 0.77 |
| 6 | 165995489 | rs16898017 | -0.668 | -1.041 | 1.151 | 0.909 | -0.028 | -0.059 | -0.059 | 0 | 0.13 | 0.29 | 0.06 | 0.01 | 0.06 | 0.12 |
| 6 | 166003186 | rs3008063 | 1.625 | -1.08 | 0.9 | 0.944 | 0.065 | -0.079 | -0.079 | 0.654 | 0.63 | 0.16 | 0.6 | 0.54 | 0.78 | 0.63 |
| 6 | 166007287 | rs2987298 | 1.09 | -0.945 | 0.84 | 1.035 | -0.035 | 0.119 | 0.119 | 0.033 | 0.75 | 0.45 | 0.66 | 0.55 | 0.84 | 0.74 |
| 6 | 166008387 | rs6917844 | -0.784 | -0.94 | 1.04 | 0.984 | -0.015 | -0.046 | -0.046 | 0.669 | 0.21 | 0.32 | 0.27 | 0.13 | 0.15 | 0.19 |
| 6 | 166011067 | rs2983505 | -0.75 | -0.8 | 0.437 | 0.62 | -0.023 | -0.09 | -0.09 | 0 | 0.2 | 0.36 | 0.12 | 0.05 | 0.15 | 0.19 |
| 6 | 166011637 | rs720701 | -1.024 | -0.389 | -0.609 | 0.882 | 0.065 | -0.022 | -0.022 | 0.02 | 0.44 | 0.77 | 0.46 | 0.51 | 0.31 | 0.49 |
| 6 | 166015231 | rs844157 | 0.778 | 0.738 | -0.143 | 1.048 | 0.013 | 0.027 | 0.027 | 0 | 0.68 | 0.55 | 0.4 | 0.4 | 0.59 | 0.62 |
| 6 | 166015431 | rs7762160 | 1.049 | 0.803 | -0.205 | 0.516 | 0.005 | 0.076 | 0.076 | 0 | 0.51 | 0.26 | 0.3 | 0.33 | 0.43 | 0.53 |
| 6 | 166016717 | rs12214904 | 0.356 | 0.818 | -0.341 | 0.43 | -0.016 | -0.002 | -0.002 | 0 | 0.13 | 0.02 | 0.07 | 0.07 | 0.13 | 0.09 |
| 6 | 166018290 | rs705789 | -1.143 | 0.505 | 0.551 | 0.304 | -0.047 | -0.061 | -0.061 | 0 | 0.32 | 0.53 | 0.17 | 0.14 | 0.24 | 0.19 |
| 6 | 166020095 | rs12198517 | 0.195 | 0.528 | 0.614 | 0.318 | 0.007 | 0.015 | 0.015 | 0.497 | 0.14 | 0.11 | 0.07 | 0.07 | 0.13 | 0.13 |
| 6 | 166027660 | rs3008013 | 0.389 | 0.743 | 1.027 | 0.372 | 0.024 | 0.04 | 0.04 | 0 | 0.81 | 0.78 | 0.48 | 0.46 | 0.67 | 0.67 |
| 6 | 166030770 | rs3008018 | -1.118 | 0.545 | 1.151 | 0.292 | -0.003 | -0.006 | -0.006 | 0 | 0.3 | 0.34 | 0.16 | 0.16 | 0.24 | 0.19 |
| 6 | 166031534 | rs12204986 | 0.381 | 0.549 | 1.129 | 0.328 | 0.006 | 0.011 | 0.011 | 0 | 0.13 | 0.05 | 0.07 | 0.07 | 0.13 | 0.09 |
| 6 | 166034053 | rs2983500 | -0.952 | 0.535 | 1.487 | 0.441 | -0.06 | -0.072 | -0.072 | 0.028 | 0.19 | 0.48 | 0.1 | 0.1 | 0.11 | 0.1 |
| 6 | 166038928 | rs9459462 | 0.541 | 1.287 | 0.911 | 0.842 | -0.007 | 0.002 | 0.002 | 0 | 0.84 | 0.88 | 0.5 | 0.49 | 0.69 | 0.66 |
| 6 | 166039571 | rs2983509 | 0.84 | 0.563 | 0.687 | 0.276 | 0.014 | 0.015 | 0.015 | 1.751 | 0.45 | 0.56 | 0.27 | 0.25 | 0.34 | 0.35 |
| 6 | 166042705 | rs9459464 | -0.539 | 0.947 | 0.369 | 0.337 | -0.004 | 0.045 | 0.045 | 0 | 0.39 | 0.25 | 0.68 | 0.71 | 0.58 | 0.39 |
| 6 | 166043862 | rs3008034 | 1.061 | 1.172 | 0.465 | 0.221 | 0.016 | 0.017 | 0.017 | 0.347 | 0.48 | 0.41 | 0.21 | 0.17 | 0.32 | 0.36 |
| 6 | 166048868 | rs9459465 | 0.607 | 1.416 | 0.446 | 0.193 | -0.018 | 0.065 | 0.065 | 0 | 0.23 | 0.1 | 0.12 | 0.02 | 0.24 | 0.12 |
| 6 | 166050841 | rs12529880 | 0.137 | 1.338 | 0.533 | 0.252 | -0.015 | 0.01 | 0.01 | 0 | 0.08 | 0.12 | 0.04 | 0.13 | 0.04 | 0.1 |
| 6 | 166051244 | rs2983515 | 0.183 | 1.459 | 0.413 | 0.285 | 0 | -0.006 | -0.006 | 0 | 0.86 | 0.84 | 0.89 | 0.88 | 0.87 | 0.72 |
| 6 | 166051508 | rs2983516 | 1.657 | 1.356 | 0.198 | 0.261 | 0.027 | 0.003 | 0.003 | 0.568 | 0.36 | 0.22 | 0.17 | 0.01 | 0.26 | 0.21 |
| 6 | 166054570 | rs9347083 | 0.158 | 2.151 | -0.06 | 1.375 | -0.041 | -0.015 | -0.015 | 0 | 0.07 | 0.02 | 0.46 | 0.38 | 0.25 | 0.12 |
| 6 | 166062379 | rs9365899 | -0.562 | 0.752 | 0.214 | 0.613 | -0.001 | -0.001 | -0.001 | 1.741 | 0.34 | 0.34 | 0.24 | 0.32 | 0.32 | 0.28 |
| 6 | 166063761 | rs2983526 | 0.356 | 0.896 | -0.132 | 0.5 | -0.019 | -0.019 | -0.019 | 0 | 0.76 | 0.77 | 0.5 | 0.44 | 0.61 | 0.71 |
| 6 | 166064152 | rs9365900 | -0.822 | -0.087 | -0.068 | 0.437 | -0.037 | 0.041 | 0.041 | 0 | 0.15 | 0.18 | 0.16 | 0.32 | 0.06 | 0.33 |
| 6 | 166065660 | rs10485103 | -0.085 | 0.221 | -0.518 | -0.302 | 0.003 | 0.002 | 0.002 | 0 | 0.1 | 0.08 | 0.09 | 0.08 | 0.04 | 0.29 |
| 6 | 166066233 | rs2983527 | 0.361 | 0.816 | -0.597 | -0.271 | 0.009 | -0.011 | -0.011 | 0.773 | 0.5 | 0.48 | 0.3 | 0.12 | 0.39 | 0.53 |
| 6 | 166069358 | rs875590 | -0.254 | 1.114 | -1.043 | 0.668 | -0.024 | -0.033 | -0.033 | 1.874 | 0.38 | 0.21 | 0.4 | 0.38 | 0.39 | 0.38 |
| 6 | 166069813 | rs2983530 | 0.884 | 0.996 | -0.144 | 1.422 | 0.004 | 0.093 | 0.093 | 0.26 | 0.76 | 0.44 | 0.72 | 0.52 | 0.79 | 0.58 |
| 6 | 166069843 | rs3008054 | 0.863 | 0.984 | -0.123 | 1.46 | 0 | 0.103 | 0.103 | 0.309 | 0.76 | 0.44 | 0.73 | 0.52 | 0.8 | 0.59 |
| 6 | 166070424 | rs911028 | 0.572 | 1.223 | 0.329 | 1.357 | 0.048 | 0.03 | 0.03 | 0.288 | 0.77 | 0.59 | 0.71 | 0.78 | 0.73 | 0.72 |
| 6 | 166077817 | rs3008061 | -1.752 | 0.571 | 0.856 | 1.426 | -0.042 | 0.118 | 0.118 | 0.92 | 0.17 | 0.42 | 0.16 | 0.34 | 0.12 | 0.29 |
| 6 | 166077866 | rs7738923 | -1.288 | 0.502 | 0.815 | 1.523 | -0.01 | 0.233 | 0.233 | 0 | 0.06 | 0.2 | 0.09 | 0.3 | 0.02 | 0.19 |
| 6 | 166088295 | rs9347084 | 0.094 | 0.53 | 0.616 | 0.62 | 0.005 | -0.076 | -0.076 | 0 | 0.42 | 0.5 | 0.36 | 0.19 | 0.42 | 0.46 |
| 6 | 166090378 | rs4709088 | 0.561 | 0.454 | 0.873 | 0.619 | -0.007 | 0.047 | 0.047 | 0.066 | 0.46 | 0.39 | 0.39 | 0.74 | 0.49 | 0.48 |
| 6 | 166092714 | rs9348032 | 1.031 | 0.357 | 0.831 | 0.696 | -0.016 | -0.003 | -0.003 | 0.105 | 0.88 | 0.85 | 0.75 | 0.91 | 0.9 | 0.93 |
| 6 | 166097078 | rs4235951 | 1.124 | 0.364 | 1 | 0.659 | -0.025 | -0.009 | -0.009 | 0 | 0.87 | 0.78 | 0.74 | 0.92 | 0.9 | 0.93 |
| 6 | 166099417 | rs6916862 | 1.374 | 0.314 | 1.41 | 0.621 | -0.019 | -0.008 | -0.008 | 0.72 | 0.87 | 0.82 | 0.75 | 0.9 | 0.89 | 0.89 |
| 6 | 166100083 | rs9365906 | -0.093 | 0.3 | 1.308 | 0.475 | -0.022 | 0.126 | 0.126 | 0.019 | 0.54 | 0.73 | 0.47 | 0.84 | 0.51 | 0.63 |
| 6 | 166100118 | rs9365907 | 0.215 | -0.001 | 1.155 | 0.764 | 0 | 0.05 | 0.05 | 0.167 | 0.46 | 0.29 | 0.35 | 0.34 | 0.43 | 0.45 |
| 6 | 166101951 | rs9689801 | 0.906 | -0.089 | 1.174 | 0.804 | -0.03 | -0.029 | -0.029 | 0 | 0.37 | 0.25 | 0.32 | 0.48 | 0.45 | 0.35 |
| 6 | 166103873 | rs9459474 | 1.86 | -0.465 | 1.408 | 0.977 | -0.076 | 0.085 | 0.085 | 0 | 0.76 | 0.49 | 0.61 | 0.58 | 0.86 | 0.73 |
| 6 | 166105413 | rs9356381 | -0.457 | -0.4 | 1.223 | 1.019 | -0.039 | 0.071 | 0.071 | 0.681 | 0.11 | 0.26 | 0.07 | 0.26 | 0.03 | 0.14 |
| 6 | 166105641 | rs4424058 | -1.258 | -0.4 | 1.222 | 0.998 | 0.038 | 0.04 | 0.04 | 0.07 | 0.06 | 0.28 | 0.08 | 0.1 | 0.02 | 0.15 |
| 6 | 166105884 | rs4532422 | 1.471 | -0.419 | 1.204 | 0.992 | -0.024 | 0.002 | 0.002 | 0.101 | 0.86 | 0.71 | 0.68 | 0.82 | 0.88 | 0.8 |
| 6 | 166106345 | rs9348034 | 0.299 | -0.196 | 0.73 | 1.073 | -0.013 | -0.026 | -0.026 | 0 | 0.92 | 0.99 | 0.76 | 0.92 | 0.9 | 0.95 |
| 6 | 166106444 | rs9365909 | -0.479 | -0.275 | 0.832 | 1.043 | -0.035 | 0.064 | 0.064 | 1.034 | 0.11 | 0.23 | 0.07 | 0.26 | 0.03 | 0.13 |
| 6 | 166108154 | rs9365910 | -0.209 | -0.245 | 0.598 | 1.059 | -0.017 | 0.035 | 0.035 | 0.949 | 0.1 | 0.15 | 0.07 | 0.26 | 0.03 | 0.13 |
| 6 | 166109230 | rs12191997 | 1.46 | 0.126 | 0.186 | 0.934 | -0.055 | 0.068 | 0.068 | 0 | 0.25 | 0.06 | 0.16 | 0.17 | 0.29 | 0.16 |
| 6 | 166109355 | rs12055769 | -0.078 | 0.312 | 0.469 | 1.185 | 0.001 | 0.052 | 0.052 | 0 | 0.11 | 0.1 | 0.14 | 0.36 | 0.15 | 0.19 |
| 6 | 166111816 | rs7760328 | -0.369 | 0.089 | 1.018 | 1.133 | -0.002 | -0.006 | -0.006 | 0.282 | 0.61 | 0.59 | 0.47 | 0.65 | 0.61 | 0.73 |
| 6 | 166114123 | rs9459484 | -1.332 | -0.053 | 0.787 | 1.135 | -0.003 | 0.029 | 0.029 | 0.08 | 0.06 | 0.24 | 0.04 | 0.06 | 0.02 | 0.04 |
| 6 | 166125295 | rs4463239 | 0.417 | -0.004 | 0.131 | 1.237 | 0.014 | 0.004 | 0.004 | 0.878 | 0.47 | 0.36 | 0.4 | 0.41 | 0.56 | 0.51 |
| 6 | 166128638 | rs9364813 | -0.727 | 0.124 | -0.845 | 1.128 | -0.009 | 0.007 | 0.007 | 0.143 | 0.13 | 0.16 | 0.07 | 0.25 | 0.06 | 0.22 |
| 6 | 166133803 | rs4235953 | -0.618 | 0.5 | -1.095 | 0.461 | -0.057 | 0.007 | 0.007 | 0 | 0.67 | 0.94 | 0.52 | 0.7 | 0.63 | 0.77 |
| 6 | 166140832 | rs4072988 | 0.505 | 0.702 | -0.522 | 0.109 | -0.008 | 0.108 | 0.108 | 2.258 | 0.33 | 0.12 | 0.22 | 0.09 | 0.34 | 0.24 |
| 6 | 166146245 | rs10455806 | 0.526 | 0.796 | -0.599 | 0.195 | -0.032 | 0.219 | 0.219 | 0.558 | 0.35 | 0.09 | 0.22 | 0.08 | 0.36 | 0.26 |
| 6 | 166146354 | rs1039001 | -1.078 | 0.388 | -0.131 | 0.337 | -0.051 | -0.006 | -0.006 | 0 | 0.17 | 0.24 | 0.08 | 0.17 | 0.1 | 0.34 |
| 6 | 166147109 | rs870448 | -0.051 | 0.816 | 0.161 | 0.623 | 0.021 | 0.066 | 0.066 | 0.649 | 0.52 | 0.34 | 0.31 | 0.25 | 0.47 | 0.6 |
| 6 | 166163634 | rs9356386 | -0.388 | 0.831 | 0.73 | 0.965 | 0.065 | 0.255 | 0.255 | 0.218 | 0.08 | 0.24 | 0.18 | 0.36 | 0.12 | 0.12 |
| 6 | 166164109 | rs11966353 | -1.309 | 0.755 | 0.713 | 0.501 | -0.04 | 0.001 | 0.001 | 0.119 | 0.13 | 0.39 | 0.08 | 0.12 | 0.06 | 0.24 |
| 6 | 166179399 | rs4709981 | 0.274 | 0.97 | 0.115 | 0.408 | -0.003 | 0.033 | 0.033 | 0.019 | 0.42 | 0.32 | 0.64 | 0.78 | 0.52 | 0.41 |
| 6 | 166181220 | rs2323029 | -0.618 | 1.204 | -0.884 | 0.355 | 0.021 | 0.127 | 0.127 | 3.036 | 0.59 | 0.8 | 0.74 | 0.89 | 0.59 | 0.66 |
| 6 | 166184203 | rs4709090 | -0.65 | 1.114 | -0.968 | 0.9 | -0.011 | -0.038 | -0.038 | 0.623 | 0.13 | 0.05 | 0.18 | 0.45 | 0.15 | 0.36 |

| | | | | | | | | | | | | | | | | |
|---|---|---|---|---|---|---|---|---|---|---|---|---|---|---|---|---|
| 6 | 166186011 | rs7748044 | -0.2 | 0.851 | -0.778 | -0.23 | 0.025 | 0.098 | 0.098 | 0.62 | 0.54 | 0.67 | 0.72 | 0.89 | 0.58 | 0.65 |
| 6 | 166196335 | rs4709984 | 1.033 | 1.14 | -0.455 | 0.849 | 0 | 0.003 | 0.003 | 0 | 0.84 | 0.8 | 0.85 | 0.75 | 0.84 | 0.69 |
| 6 | 166196458 | rs9348044 | 1.599 | 1.242 | 0.63 | 1.385 | 0.036 | 0.171 | 0.171 | 0 | 0.82 | 0.6 | 0.65 | 0.63 | 0.81 | 0.73 |
| 6 | 166196865 | rs4709985 | 0.18 | 1.118 | 0.451 | 0.144 | -0.007 | -0.082 | -0.082 | 0.018 | 0.63 | 0.5 | 0.73 | 0.86 | 0.62 | 0.7 |
| 6 | 166197239 | rs9364815 | 0.541 | 0.892 | 0.11 | 0.675 | -0.01 | -0.03 | -0.03 | 0 | 0.64 | 0.74 | 0.61 | 0.58 | 0.67 | 0.58 |
| 6 | 166197276 | rs12528870 | -1.342 | 1.056 | 0.914 | 0.795 | 0.07 | 0.156 | 0.156 | 0.106 | 0.08 | 0.36 | 0.08 | 0.21 | 0.05 | 0.14 |
| 6 | 166197889 | rs6928719 | 1.014 | 1.288 | 0.871 | 0.768 | -0.009 | 0.017 | 0.017 | 0.423 | 0.54 | 0.36 | 0.51 | 0.35 | 0.59 | 0.43 |
| 6 | 166198621 | rs6928737 | 0.749 | 1.292 | 0.393 | 0.694 | -0.011 | -0.018 | -0.018 | 0 | 0.6 | 0.73 | 0.59 | 0.57 | 0.64 | 0.58 |
| 6 | 166210622 | rs1494769 | 1.231 | 1.627 | 0.652 | 0.735 | 0.018 | 0.022 | 0.022 | 0.708 | 0.66 | 0.6 | 0.51 | 0.45 | 0.6 | 0.57 |
| 6 | 166211309 | rs6917116 | -1.144 | 1.718 | 0.388 | 0.612 | 0.005 | -0.024 | -0.024 | 1.832 | 0.22 | 0.27 | 0.13 | 0.11 | 0.24 | 0.21 |
| 6 | 166216338 | rs11755875 | -0.995 | 1.705 | 0.01 | 0.858 | -0.017 | -0.042 | -0.042 | 0 | 0.16 | 0.08 | 0.37 | 0.45 | 0.19 | 0.28 |
| 6 | 166224027 | rs13203335 | -1.536 | 1.548 | 0.073 | 0.686 | -0.009 | -0.029 | -0.029 | 0.63 | 0.21 | 0.12 | 0.39 | 0.43 | 0.24 | 0.32 |
| 6 | 166224203 | rs10455962 | -1.041 | 1.849 | -0.066 | 0.966 | -0.045 | -0.038 | -0.038 | 0 | 0.09 | 0.02 | 0.36 | 0.41 | 0.18 | 0.22 |
| 6 | 166227582 | rs9348048 | 0.079 | 1.686 | -0.385 | -0.26 | -0.026 | 0.048 | 0.048 | 0 | 0.24 | 0.11 | 0.18 | 0.02 | 0.3 | 0.14 |
| 6 | 166230553 | rs10455964 | 0.615 | 1.138 | -0.233 | -0.388 | 0.013 | 0.076 | 0.076 | 0.031 | 0.5 | 0.4 | 0.65 | 0.88 | 0.49 | 0.58 |
| 6 | 166231293 | rs12663089 | 0.047 | 1.186 | -0.421 | -0.283 | 0 | 0.007 | 0.007 | 0 | 0.21 | 0.21 | 0.33 | 0.57 | 0.17 | 0.41 |
| 6 | 166231573 | rs10806834 | -0.162 | 1.199 | -0.683 | -0.27 | 0.006 | -0.061 | -0.061 | 0 | 0.7 | 0.7 | 0.77 | 0.98 | 0.64 | 0.8 |
| 6 | 166231777 | rs4709988 | -0.686 | 0.828 | -1.388 | 0.338 | 0.026 | -0.026 | -0.026 | 0 | 0.61 | 0.85 | 0.68 | 0.63 | 0.63 | 0.66 |
| 6 | 166237461 | rs6920797 | 0.459 | 1.348 | -1.046 | -0.419 | -0.017 | 0.027 | 0.027 | 0.697 | 0.66 | 0.25 | 0.67 | 0.88 | 0.62 | 0.75 |
| 6 | 166244574 | rs6931483 | 0.126 | 0.945 | -1.366 | -1.669 | 0.008 | 0.038 | 0.038 | 0 | 0.29 | 0.2 | 0.2 | 0.03 | 0.29 | 0.08 |
| 6 | 166249068 | rs7741397 | 0.5 | 0.993 | -1.195 | -1.39 | 0.009 | -0.019 | -0.019 | 0.009 | 0.31 | 0.24 | 0.2 | 0.4 | 0.25 | 0.44 |
| 6 | 166249979 | rs12191225 | 0.416 | 1.243 | -0.893 | -1.592 | 0.002 | -0.013 | -0.013 | 0 | 0.37 | 0.31 | 0.28 | 0.07 | 0.4 | 0.25 |
| 6 | 166257570 | rs6915702 | -0.172 | 1.756 | -0.831 | -1.602 | 0.03 | 0.064 | 0.064 | 0 | 0.8 | 0.71 | 0.51 | 0.5 | 0.7 | 0.78 |
| 6 | 166260182 | rs4709989 | 0.073 | 1.472 | -0.62 | -1.566 | 0.04 | 0.12 | 0.12 | 0 | 0.15 | 0.24 | 0.45 | 0.5 | 0.25 | 0.21 |
| 6 | 166263488 | rs9348050 | 0.266 | 1.221 | -0.549 | -1.76 | -0.006 | -0.006 | -0.006 | 0 | 0.43 | 0.38 | 0.65 | 0.55 | 0.54 | 0.3 |
| 6 | 166266219 | rs13201944 | 0.812 | 0.906 | -0.657 | -1.744 | -0.003 | 0.026 | 0.026 | 0 | 0.07 | 0.03 | 0.02 | 0 | 0.04 | 0.01 |
| 6 | 166266322 | rs7751553 | 0.112 | 0.845 | -0.584 | -1.949 | 0.025 | 0.139 | 0.139 | 0 | 0.45 | 0.26 | 0.28 | 0.06 | 0.43 | 0.26 |
| 6 | 166275615 | rs2453392 | -0.878 | 0.889 | -0.521 | -1.921 | 0.003 | 0.021 | 0.021 | 0 | 0.66 | 0.78 | 0.76 | 0.97 | 0.63 | 0.88 |
| 6 | 166277064 | rs2498581 | 0.529 | 1.127 | -0.663 | -1.547 | -0.012 | -0.043 | -0.043 | 0.677 | 0.2 | 0.1 | 0.27 | 0.29 | 0.19 | 0.27 |
| 6 | 166287778 | rs2498575 | -1.859 | 1.091 | -0.597 | -1.59 | 0 | 0.109 | 0.109 | 0 | 0.73 | 0.92 | 0.84 | 0.97 | 0.73 | 0.92 |
| 6 | 166289033 | rs2280556 | 0.015 | 1.245 | -0.18 | -1.306 | 0.022 | 0.013 | 0.013 | 0 | 0.32 | 0.53 | 0.44 | 0.65 | 0.38 | 0.47 |
| 6 | 166289701 | rs2280559 | 0.672 | 1.099 | -0.043 | -1.304 | -0.001 | 0.063 | 0.063 | 0 | 0.24 | 0.33 | 0.17 | 0.42 | 0.23 | 0.41 |
| 6 | 166297510 | rs883299 | -0.161 | 0.821 | -0.234 | -1.535 | -0.002 | -0.08 | -0.08 | 1.821 | 0.46 | 0.48 | 0.69 | 0.91 | 0.5 | 0.69 |
| 6 | 166298612 | rs955075 | 0.764 | 0.949 | -0.596 | -0.907 | -0.023 | -0.048 | -0.048 | 3.628 | 0.18 | 0.06 | 0.29 | 0.28 | 0.2 | 0.26 |
| 6 | 166299347 | rs6932405 | -0.853 | 0.973 | 0.465 | -1.222 | 0.113 | 0.079 | 0.079 | 0 | 0.14 | 0.44 | 0.29 | 0.26 | 0.16 | 0.07 |
| 6 | 166299569 | rs4709991 | -0.15 | 0.777 | 0.21 | -1.452 | 0.032 | 0.094 | 0.094 | 0 | 0.54 | 0.68 | 0.73 | 0.92 | 0.55 | 0.69 |
| 6 | 166308198 | rs9364825 | -0.005 | 1.022 | 0.471 | -1.309 | 0.094 | 0.114 | 0.114 | 0 | 0.06 | 0.18 | 0.25 | 0.26 | 0.12 | 0.06 |
| 6 | 166308924 | rs10806835 | 0.327 | 0.863 | 0.534 | -1.477 | -0.001 | -0.014 | -0.014 | 3.265 | 0.47 | 0.48 | 0.48 | 0.66 | 0.42 | 0.6 |
| 6 | 166309288 | rs2461722 | -0.924 | 0.791 | 0.489 | -1.497 | 0.008 | -0.013 | -0.013 | 0 | 0.72 | 0.79 | 0.83 | 0.98 | 0.74 | 0.9 |
| 6 | 166323038 | rs2500482 | 0.178 | 0.83 | 0.698 | -1.284 | 0.015 | 0.036 | 0.036 | 0 | 0.63 | 0.77 | 0.8 | 0.96 | 0.65 | 0.73 |
| 6 | 166327142 | rs10946106 | 0.946 | 1.238 | 0.944 | -0.912 | -0.002 | -0.017 | -0.017 | 0 | 0.16 | 0.13 | 0.13 | 0.02 | 0.19 | 0.08 |
| 6 | 166327886 | rs9347099 | -0.316 | 1.348 | 1.266 | -0.686 | 0.1 | 0.146 | 0.146 | 0 | 0.05 | 0.2 | 0.24 | 0.28 | 0.1 | 0.06 |
| 6 | 166331460 | rs6899775 | -0.62 | 1.314 | 1.09 | -0.72 | 0.001 | -0.02 | -0.02 | 0 | 0.3 | 0.26 | 0.18 | 0.04 | 0.29 | 0.25 |
| 6 | 166333799 | rs9459531 | -1.328 | 1.38 | 1.097 | -0.748 | 0.008 | -0.024 | -0.024 | 0 | 0.4 | 0.49 | 0.46 | 0.32 | 0.47 | 0.35 |
| 6 | 166342650 | rs4472324 | -1.011 | 1.227 | 1.031 | -0.23 | 0.067 | 0.056 | 0.056 | 0 | 0.1 | 0.22 | 0.29 | 0.28 | 0.18 | 0.09 |
| 6 | 166346764 | rs660244 | -0.841 | -0.02 | 1.073 | -0.267 | -0.023 | -0.037 | -0.037 | 0 | 0.13 | 0.18 | 0.03 | 0.02 | 0.06 | 0.15 |
| 6 | 166354509 | rs167268 | -1.063 | 0.086 | 1.194 | -0.201 | -0.041 | 0.056 | 0.056 | 0.073 | 0.17 | 0.41 | 0.18 | 0.31 | 0.11 | 0.19 |
| 6 | 166355837 | rs300110 | -1.299 | 0.109 | 1.159 | -0.151 | -0.034 | 0.082 | 0.082 | 0.047 | 0.21 | 0.43 | 0.21 | 0.4 | 0.19 | 0.33 |
| 6 | 166362761 | rs6937549 | 0.662 | 0.248 | 0.556 | -0.487 | -0.011 | 0.01 | 0.01 | 0 | 0.17 | 0.09 | 0.12 | 0.01 | 0.19 | 0.1 |
| 6 | 166365759 | rs93622 | 1.045 | 0.287 | 0.208 | -0.227 | 0.011 | 0.015 | 0.015 | 0 | 0.55 | 0.17 | 0.61 | 0.57 | 0.55 | 0.54 |
| 6 | 166366474 | rs300117 | -0.183 | 0.249 | 0.447 | -0.245 | -0.01 | 0.014 | 0.014 | 0 | 0.3 | 0.48 | 0.3 | 0.37 | 0.3 | 0.29 |
| 6 | 166366573 | rs7751441 | -0.506 | 0.244 | 0.616 | -0.154 | 0.005 | 0.001 | 0.001 | 0.497 | 0.1 | 0.33 | 0.05 | 0.07 | 0.08 | 0.15 |
| 6 | 166367429 | rs213997 | 0.484 | 0.378 | 0.196 | -0.054 | -0.001 | 0 | 0 | 0 | 0.63 | 0.41 | 0.64 | 0.6 | 0.62 | 0.69 |
| 6 | 166368397 | rs3914144 | -0.585 | 0.2 | -0.169 | -0.314 | 0.002 | 0.023 | 0.023 | 0 | 0.1 | 0.03 | 0.04 | 0.04 | 0.08 | 0.03 |
| 6 | 166368450 | rs705791 | 1.258 | 0.033 | 0.167 | -1.091 | 0.008 | 0.018 | 0.018 | 0.245 | 0.56 | 0.36 | 0.76 | 0.91 | 0.6 | 0.68 |
| 6 | 166368956 | rs705792 | 0.719 | 0.347 | 0.683 | -1.028 | 0.005 | -0.012 | -0.012 | 0.556 | 0.85 | 0.72 | 0.89 | 0.91 | 0.83 | 0.81 |
| 6 | 166371492 | rs12203748 | -0.107 | 0.135 | 0.459 | -0.973 | -0.017 | 0.032 | 0.032 | 0 | 0.17 | 0.1 | 0.13 | 0.09 | 0.2 | 0.08 |
| 6 | 166374108 | rs697462 | 1.643 | 0.017 | 0.656 | -1.075 | 0.007 | 0.115 | 0.115 | 0.247 | 0.59 | 0.21 | 0.74 | 0.87 | 0.62 | 0.64 |
| 6 | 166376465 | rs6899687 | -0.419 | -0.099 | 0.507 | -1.179 | -0.011 | 0.064 | 0.064 | 0 | 0.2 | 0.11 | 0.13 | 0.03 | 0.2 | 0.08 |
| 6 | 166376825 | rs705795 | -0.488 | -0.042 | -0.156 | -1.571 | -0.001 | 0.02 | 0.02 | 0 | 0.76 | 0.87 | 0.86 | 0.81 | 0.73 | 0.82 |
| 6 | 166379282 | rs697467 | 1.098 | 0.248 | 0.546 | -1.41 | -0.001 | 0.038 | 0.038 | 0 | 0.68 | 0.72 | 0.72 | 0.89 | 0.66 | 0.69 |
| 6 | 166379896 | rs697471 | 1.528 | 0.272 | 0.81 | -1.481 | -0.001 | -0.002 | -0.002 | 0.337 | 0.59 | 0.38 | 0.65 | 0.85 | 0.56 | 0.65 |
| 6 | 166383562 | rs705802 | 1.322 | 0.704 | 1.058 | -1.359 | 0.026 | -0.099 | -0.099 | 0 | 0.69 | 0.45 | 0.79 | 0.93 | 0.64 | 0.81 |
| 6 | 166385941 | rs13437489 | -1.319 | 0.802 | 1.151 | -1.317 | 0.015 | -0.068 | -0.068 | 0.058 | 0.27 | 0.39 | 0.19 | 0.08 | 0.3 | 0.09 |
| 6 | 166387129 | rs705804 | 0.611 | 0.863 | 0.859 | -1.18 | -0.023 | 0.037 | 0.037 | 0 | 0.8 | 0.84 | 0.87 | 0.96 | 0.74 | 0.91 |
| 6 | 166396880 | rs697473 | 1.201 | 0.804 | 0.937 | -1.185 | 0.021 | -0.059 | -0.059 | 0 | 0.73 | 0.57 | 0.74 | 0.88 | 0.67 | 0.71 |
| 6 | 166403527 | rs705807 | 1.456 | 0.526 | 1.003 | -1.012 | 0.058 | 0.029 | 0.029 | 1.024 | 0.89 | 0.59 | 0.84 | 0.93 | 0.87 | 0.83 |
| 6 | 166408949 | rs300129 | 1.448 | 0.774 | 0.754 | -1.097 | 0.029 | -0.04 | -0.04 | 0 | 0.72 | 0.44 | 0.7 | 0.86 | 0.65 | 0.76 |
| 6 | 166413517 | rs12209946 | -0.016 | 0.606 | 0.065 | -1.09 | -0.009 | 0.026 | 0.026 | 0.198 | 0.07 | 0.01 | 0.03 | 0.02 | 0.05 | 0.08 |

| | | | | | | | | | | | | | | | | |
|---|---|---|---|---|---|---|---|---|---|---|---|---|---|---|---|---|
| 6 | 166420136 | rs300138 | -0.808 | 0.36 | 0.718 | -1.3 | -0.044 | -0.115 | -0.115 | 0 | 0.19 | 0.44 | 0.1 | 0.04 | 0.13 | 0.12 |
| 6 | 166421637 | rs697482 | 0.866 | 0.626 | 0.619 | -1.262 | 0.048 | 0.002 | 0.002 | 0 | 0.9 | 0.83 | 0.82 | 0.92 | 0.79 | 0.82 |
| 6 | 166422831 | rs300144 | -0.844 | 0.812 | 0.718 | -1.379 | -0.02 | -0.122 | -0.122 | 0.475 | 0.22 | 0.49 | 0.12 | 0.04 | 0.17 | 0.13 |
| 6 | 166436895 | rs9348064 | 2.093 | 0.208 | 0.08 | -1.432 | -0.091 | 0.075 | 0.075 | 6.278 | 0.46 | 0.08 | 0.47 | 0.67 | 0.48 | 0.55 |
| 6 | 166438811 | rs697487 | 0.2 | 0.241 | 0.264 | -1.436 | 0.094 | -0.014 | -0.014 | 0.747 | 0.93 | 0.83 | 0.92 | 0.97 | 0.85 | 0.97 |
| 6 | 166441028 | rs697491 | -1.489 | 0.133 | 0.32 | -1.495 | 0.026 | 0.031 | 0.031 | 6.888 | 0.44 | 0.61 | 0.35 | 0.25 | 0.33 | 0.29 |
| 6 | 166447606 | rs2323044 | 1.912 | 0.113 | 0.183 | -1.484 | -0.032 | 0.178 | 0.178 | 5.462 | 0.48 | 0.12 | 0.56 | 0.72 | 0.52 | 0.68 |
| 6 | 166455916 | rs777573 | -0.912 | -0.042 | 0.761 | -1.518 | -0.034 | -0.023 | -0.023 | 0.086 | 0.3 | 0.38 | 0.26 | 0.23 | 0.2 | 0.27 |
| 6 | 166465095 | rs420415 | 2.254 | 0.1 | 0.407 | -1.539 | -0.036 | 0.077 | 0.077 | 3.786 | 0.45 | 0.18 | 0.46 | 0.67 | 0.46 | 0.54 |
| 6 | 166468915 | rs300614 | -0.687 | 0.204 | 0.386 | -1.523 | 0.016 | -0.074 | -0.074 | 1.02 | 0.2 | 0.34 | 0.17 | 0.05 | 0.23 | 0.05 |
| 6 | 166470464 | rs4710003 | -1.128 | 0.163 | 0.504 | -1.5 | -0.004 | -0.032 | -0.032 | 0 | 0.41 | 0.41 | 0.4 | 0.23 | 0.42 | 0.21 |
| 6 | 166474887 | rs4458652 | 0.96 | 0.199 | 0.541 | -1.475 | -0.002 | -0.005 | -0.005 | 0 | 0.61 | 0.64 | 0.57 | 0.74 | 0.63 | 0.68 |
| 6 | 166475713 | rs4607409 | -0.071 | 0.124 | 0.435 | -1.524 | 0.009 | 0.005 | 0.005 | 0.961 | 0.82 | 0.88 | 0.77 | 0.8 | 0.83 | 0.85 |
| 6 | 166480503 | rs10946116 | 0.093 | 0.286 | 0.376 | -1.545 | 0.001 | 0.053 | 0.053 | 0 | 0.2 | 0.07 | 0.19 | 0.07 | 0.2 | 0.18 |
| 6 | 166482465 | rs1900055 | 0.793 | 0.267 | 0.437 | -1.562 | 0.025 | -0.033 | -0.033 | 0 | 0.75 | 0.62 | 0.68 | 0.77 | 0.67 | 0.82 |
| 6 | 166487606 | rs9356411 | 1.624 | 0.306 | 0.336 | -1.149 | 0.047 | 0.099 | 0.099 | 5.391 | 0.49 | 0.15 | 0.6 | 0.66 | 0.56 | 0.55 |
| 6 | 166492963 | rs6456037 | -0.754 | 0.121 | 0.4 | -1.02 | -0.023 | -0.011 | -0.011 | 0 | 0.3 | 0.39 | 0.29 | 0.26 | 0.24 | 0.28 |
| 6 | 166502703 | rs7757379 | -1.324 | 0.325 | 0.518 | -1.076 | -0.006 | -0.025 | -0.025 | 0.249 | 0.17 | 0.47 | 0.21 | 0.1 | 0.11 | 0.17 |
| 6 | 166504783 | rs2166251 | 1.648 | 0.569 | 0.388 | -1.16 | -0.012 | 0.032 | 0.032 | 3.135 | 0.7 | 0.2 | 0.67 | 0.85 | 0.66 | 0.65 |
| 6 | 166510397 | rs3127386 | 1.073 | 0.737 | 0.457 | -1.217 | 0.017 | -0.028 | -0.028 | 0 | 0.8 | 0.71 | 0.84 | 0.94 | 0.7 | 0.79 |
| 6 | 166510461 | rs9356421 | 2.04 | 0.535 | 0.148 | -1.511 | -0.012 | 0.207 | 0.207 | 2.316 | 0.51 | 0.11 | 0.47 | 0.82 | 0.47 | 0.56 |
| 6 | 166510520 | rs3099248 | 0.969 | 0.588 | 0.219 | -1.44 | 0.017 | -0.028 | -0.028 | 0.02 | 0.8 | 0.71 | 0.84 | 0.94 | 0.7 | 0.79 |
| 6 | 166510596 | rs3127389 | -0.771 | 0.528 | 0.32 | -1.302 | 0.046 | 0.03 | 0.03 | 4.203 | 0.11 | 0.5 | 0.18 | 0.09 | 0.07 | 0.14 |
| 6 | 166513487 | rs12196325 | 0.018 | 0.528 | 0.246 | -1.415 | -0.003 | -0.061 | -0.061 | 0 | 0.19 | 0.27 | 0.22 | 0.03 | 0.16 | 0.09 |
| 6 | 166517429 | rs1867596 | 1.609 | 0.441 | -0.799 | -0.942 | -0.062 | -0.054 | -0.054 | 0 | 0.26 | 0.05 | 0.3 | 0.3 | 0.29 | 0.4 |
| 6 | 166517762 | rs2028790 | 0.755 | 0.514 | -0.616 | -0.808 | -0.004 | -0.059 | -0.059 | 0.292 | 0.3 | 0.13 | 0.2 | 0.55 | 0.29 | 0.18 |
| 6 | 166517775 | rs9356424 | 0.888 | 0.178 | -0.739 | -0.788 | 0.04 | -0.114 | -0.114 | 0 | 0.29 | 0.06 | 0.18 | 0.55 | 0.22 | 0.17 |
| 6 | 166519460 | rs6914270 | -0.045 | 0.354 | -0.108 | -0.989 | 0.02 | 0.002 | 0.002 | 0 | 0.9 | 0.79 | 0.92 | 0.95 | 0.83 | 0.83 |
| 6 | 166520236 | rs2166248 | 0.764 | 0.199 | -0.175 | -1.221 | 0.005 | 0.04 | 0.04 | 0.258 | 0.57 | 0.33 | 0.64 | 0.88 | 0.57 | 0.53 |
| 6 | 166521387 | rs4504453 | 0.518 | 0.142 | -0.071 | -1.317 | 0.001 | -0.025 | -0.025 | 0 | 0.4 | 0.43 | 0.58 | 0.88 | 0.45 | 0.45 |
| 6 | 166523465 | rs9365951 | 0.473 | 0.12 | -0.085 | -0.768 | -0.013 | 0.166 | 0.166 | 0 | 0.11 | 0.21 | 0.23 | 0.55 | 0.1 | 0.12 |
| 6 | 166526466 | rs10946121 | 0.693 | 0.109 | -0.093 | -1.016 | -0.001 | -0.065 | -0.065 | 0 | 0.17 | 0.24 | 0.2 | 0.02 | 0.15 | 0.15 |
| 6 | 166526670 | rs10946122 | 0.768 | 0.102 | -0.136 | -1.018 | -0.001 | -0.034 | -0.034 | 0 | 0.17 | 0.19 | 0.2 | 0.02 | 0.14 | 0.12 |
| 6 | 166534145 | rs4709098 | -0.006 | -0.229 | -0.195 | -1.046 | 0.008 | 0.01 | 0.01 | 0.159 | 0.57 | 0.61 | 0.79 | 0.89 | 0.59 | 0.62 |
| 6 | 166540384 | rs3127428 | -1.283 | 0.025 | -0.359 | -1.002 | -0.094 | -0.05 | -0.05 | 2.199 | 0.68 | 0.94 | 0.62 | 0.68 | 0.6 | 0.63 |
| 6 | 166543262 | rs2166247 | 0.578 | 0.052 | -0.166 | -0.895 | 0 | -0.068 | -0.068 | 0.376 | 0.38 | 0.46 | 0.57 | 0.87 | 0.38 | 0.46 |
| 6 | 166545352 | rs9348077 | 0.349 | 0.051 | -0.044 | -0.71 | -0.007 | 0.341 | 0.341 | 0 | 0.11 | 0.34 | 0.12 | 0.51 | 0.1 | 0.1 |
| 6 | 166554170 | rs3099283 | -0.446 | -0.075 | -0.351 | -0.997 | 0.007 | 0.091 | 0.091 | 0 | 0.63 | 0.84 | 0.67 | 0.86 | 0.6 | 0.62 |
| 6 | 166556333 | rs9689603 | 0.506 | -0.12 | -0.231 | -1.067 | -0.006 | -0.044 | -0.044 | 0 | 0.42 | 0.52 | 0.45 | 0.84 | 0.38 | 0.46 |
| 6 | 166558164 | rs9459583 | 0.836 | -0.05 | -0.453 | -1.103 | -0.048 | 0.067 | 0.067 | 0 | 0.18 | 0.08 | 0.21 | 0.02 | 0.22 | 0.17 |
| 6 | 166561712 | rs10806845 | 0.698 | -0.077 | -0.485 | -1.019 | -0.006 | 0.051 | 0.051 | 1.507 | 0.43 | 0.16 | 0.52 | 0.35 | 0.5 | 0.5 |
| 6 | 166562346 | rs920962 | 0.808 | -0.1 | -0.451 | -1.006 | -0.02 | 0.018 | 0.018 | 0 | 0.18 | 0.11 | 0.21 | 0.03 | 0.22 | 0.17 |
| 6 | 166565375 | rs9348082 | -0.728 | -0.16 | -0.397 | -1.007 | 0 | -0.015 | -0.015 | 1.669 | 0.36 | 0.72 | 0.48 | 0.58 | 0.32 | 0.33 |
| 6 | 166566173 | rs10946129 | 1.818 | -0.161 | -0.431 | -1.033 | -0.07 | -0.067 | -0.067 | 0 | 0.25 | 0.05 | 0.32 | 0.32 | 0.28 | 0.32 |
| 6 | 166567345 | rs6914547 | -1.137 | -0.22 | -0.539 | -1.135 | -0.009 | -0.001 | -0.001 | 0 | 0.45 | 0.71 | 0.41 | 0.59 | 0.44 | 0.43 |
| 6 | 166567853 | rs9459589 | -0.216 | -0.196 | -0.495 | -1.152 | -0.038 | 0.056 | 0.056 | 0 | 0.28 | 0.14 | 0.27 | 0.07 | 0.34 | 0.29 |
| 6 | 166570112 | rs12207298 | 0.578 | -0.14 | -0.525 | -1.093 | -0.005 | 0.012 | 0.012 | 0 | 0.1 | 0.06 | 0.06 | 0.05 | 0.12 | 0.12 |
| 6 | 166570346 | rs9295323 | -0.489 | -0.172 | -0.498 | -1.068 | 0 | -0.005 | -0.005 | 0 | 0.35 | 0.66 | 0.34 | 0.53 | 0.32 | 0.3 |
| 6 | 166571443 | rs1056053 | -0.316 | -0.245 | -0.48 | -0.988 | 0.01 | 0.076 | 0.076 | 0 | 0.62 | 0.8 | 0.67 | 0.84 | 0.6 | 0.63 |
| 6 | 166571525 | rs3099270 | -1.089 | -0.259 | -0.486 | -0.997 | 0.012 | 0.025 | 0.025 | 0 | 0.72 | 0.81 | 0.73 | 0.89 | 0.71 | 0.75 |
| 6 | 166571579 | rs1056051 | -1.096 | -0.26 | -0.496 | -0.998 | 0.015 | 0.036 | 0.036 | 0 | 0.72 | 0.85 | 0.73 | 0.89 | 0.72 | 0.75 |
| 6 | 166572005 | rs3816300 | 0.636 | -0.283 | -0.149 | -0.898 | -0.014 | 0.194 | 0.194 | 0 | 0.11 | 0.22 | 0.12 | 0.52 | 0.1 | 0.11 |
| 6 | 166572045 | rs3127328 | -0.339 | -0.293 | -0.23 | -0.996 | -0.006 | -0.039 | -0.039 | 0 | 0.16 | 0.24 | 0.07 | 0.05 | 0.12 | 0.15 |
| 6 | 166573544 | rs2028791 | -0.384 | -0.284 | -0.245 | -0.987 | -0.006 | -0.024 | -0.024 | 0 | 0.16 | 0.27 | 0.2 | 0.09 | 0.12 | 0.17 |
| 6 | 166579270 | rs2305089 | 0.729 | -0.324 | -0.34 | -1.015 | 0.017 | 0.071 | 0.071 | 0.18 | 0.46 | 0.2 | 0.53 | 0.34 | 0.5 | 0.49 |
| 6 | 166581147 | rs3099266 | -0.351 | -0.239 | -0.458 | -1.03 | -0.001 | 0.053 | 0.053 | 0 | 0.63 | 0.84 | 0.65 | 0.79 | 0.58 | 0.62 |
| 6 | 166584582 | rs7773665 | 0.664 | -0.211 | -0.42 | -0.998 | -0.003 | -0.011 | -0.011 | 0 | 0.4 | 0.43 | 0.44 | 0.73 | 0.39 | 0.47 |
| 6 | 166584676 | rs7753771 | 0.476 | -0.245 | -0.369 | -0.841 | -0.019 | 0.29 | 0.29 | 0 | 0.11 | 0.34 | 0.1 | 0.43 | 0.09 | 0.1 |
| 6 | 166584876 | rs10946130 | 1.587 | -0.15 | -0.724 | -0.871 | -0.065 | -0.026 | -0.026 | 0 | 0.27 | 0.06 | 0.33 | 0.3 | 0.3 | 0.37 |
| 6 | 166585112 | rs2197450 | -0.289 | -0.099 | -0.68 | -0.898 | 0.003 | 0.052 | 0.052 | 0 | 0.62 | 0.8 | 0.64 | 0.78 | 0.58 | 0.62 |
| 6 | 166585603 | rs11962977 | 0.834 | 0.344 | -0.315 | -0.479 | -0.046 | 0.05 | 0.05 | 0.057 | 0.17 | 0.06 | 0.2 | 0.06 | 0.21 | 0.14 |
| 6 | 166586982 | rs9347107 | 0.382 | 0.263 | 0.049 | -0.186 | -0.02 | 0.179 | 0.179 | 0 | 0.1 | 0.22 | 0.08 | 0.43 | 0.09 | 0.1 |
| 6 | 166588221 | rs9348092 | -0.101 | 0.331 | -0.106 | -0.44 | 0.012 | 0.054 | 0.054 | 0 | 0.91 | 0.86 | 0.76 | 0.84 | 0.84 | 0.83 |
| 6 | 166592349 | rs3127348 | 1.03 | 0.495 | -0.573 | -1.124 | 0.016 | 0.003 | 0.003 | 0 | 0.44 | 0.37 | 0.23 | 0.05 | 0.36 | 0.19 |
| 6 | 166592755 | rs3099256 | 0.106 | 0.197 | -0.423 | -0.402 | 0.05 | 0.194 | 0.194 | 0 | 0.87 | 0.62 | 0.71 | 0.75 | 0.8 | 0.79 |
| 6 | 166592832 | rs12524688 | 1.184 | 0.433 | -1.223 | -0.185 | -0.025 | -0.032 | -0.032 | 0 | 0.13 | 0.05 | 0.23 | 0.29 | 0.17 | 0.18 |
| 6 | 166593891 | rs9348093 | 0.174 | 0.097 | -1.14 | -0.372 | 0.015 | -0.102 | -0.102 | 0 | 0.14 | 0.01 | 0.06 | 0.38 | 0.1 | 0.16 |
| 6 | 166603765 | rs10428822 | -0.313 | 0.781 | -0.483 | -0.434 | 0.001 | -0.024 | -0.024 | 0 | 0.13 | 0.23 | 0.09 | 0.06 | 0.15 | 0.16 |
| 6 | 166608241 | rs11759376 | -0.199 | 1.116 | -0.948 | -0.709 | -0.066 | 0.072 | 0.072 | 0 | 0.27 | 0.13 | 0.31 | 0.12 | 0.38 | 0.28 |

| | | | | | | | | | | | | | | | | |
|---|---|---|---|---|---|---|---|---|---|---|---|---|---|---|---|---|
| 6 | 166609388 | rs7750964 | 0.671 | 0.854 | -0.677 | -0.337 | 0.04 | 0.141 | 0.141 | 0 | 0.16 | 0.28 | 0.27 | 0.4 | 0.19 | 0.3 |
| 6 | 166610740 | rs4147236 | 1.248 | -0.001 | -1.104 | -0.733 | 0.106 | 0.133 | 0.133 | 0 | 0.35 | 0.14 | 0.27 | 0.1 | 0.23 | 0.21 |
| 6 | 166613351 | rs4710024 | 1.138 | 0.121 | -1.388 | -0.581 | 0.047 | 0.057 | 0.057 | 0 | 0.36 | 0.25 | 0.32 | 0.17 | 0.27 | 0.34 |
| 6 | 166615186 | rs4128605 | -0.544 | 0.826 | -1.591 | 0.088 | -0.01 | -0.004 | -0.004 | 0 | 0.42 | 0.31 | 0.41 | 0.42 | 0.47 | 0.48 |
| 6 | 166615985 | rs4709105 | -1.323 | 0.835 | -2.108 | 0.16 | -0.032 | -0.002 | -0.002 | 0 | 0.55 | 0.95 | 0.65 | 0.63 | 0.57 | 0.73 |
| 6 | 166617229 | rs13206417 | -0.518 | 0.915 | -1.088 | -0.122 | -0.006 | 0.005 | 0.005 | 0 | 0.12 | 0.18 | 0.2 | 0.12 | 0.09 | 0.16 |
| 6 | 166619821 | rs9459617 | -0.47 | 1.387 | -1.557 | -0.346 | -0.072 | 0.075 | 0.075 | 0 | 0.18 | 0.03 | 0.11 | 0.07 | 0.23 | 0.14 |
| 6 | 166623254 | rs12190692 | -0.196 | 1.763 | -1.673 | 0.063 | -0.108 | -0.071 | -0.071 | 0 | 0.18 | 0.02 | 0.19 | 0.23 | 0.3 | 0.37 |
| 6 | 166623917 | rs4591822 | -0.295 | 1.563 | -1.785 | -0.227 | -0.008 | 0.006 | 0.006 | 0 | 0.21 | 0.17 | 0.3 | 0.15 | 0.24 | 0.16 |
| 6 | 166624249 | rs4385290 | -1.156 | 1.295 | -0.831 | -0.307 | 0.001 | 0 | 0 | 0 | 0.39 | 0.37 | 0.5 | 0.38 | 0.54 | 0.52 |
| 6 | 166631191 | rs12526629 | -0.198 | 0.251 | -1.134 | -0.928 | -0.002 | -0.02 | -0.02 | 0 | 0.71 | 0.36 | 0.86 | 0.92 | 0.79 | 0.78 |
| 6 | 166634430 | rs7773598 | 0.001 | 0.633 | -0.796 | -0.71 | 0.011 | -0.005 | -0.005 | 0 | 0.8 | 0.76 | 0.69 | 0.86 | 0.75 | 0.85 |
| 6 | 166636119 | rs12663423 | 1.533 | 0.607 | -1.199 | -0.671 | 0.021 | -0.088 | -0.088 | 0.29 | 0.31 | 0.13 | 0.38 | 0.53 | 0.25 | 0.25 |
| 6 | 166640456 | rs1610310 | -0.458 | 0.431 | -0.536 | -1.246 | 0.004 | 0.078 | 0.078 | 0 | 0.48 | 0.59 | 0.3 | 0.22 | 0.47 | 0.53 |
| 6 | 166642850 | rs7753174 | 0.652 | 0.016 | -0.598 | -1.132 | 0 | -0.028 | -0.028 | 0 | 0.2 | 0.2 | 0.09 | 0.01 | 0.14 | 0.08 |
| 6 | 166651396 | rs10946137 | 0.467 | 0.517 | -0.889 | -0.93 | -0.098 | -0.033 | -0.033 | 0 | 0.23 | 0.06 | 0.21 | 0.22 | 0.36 | 0.4 |
| 6 | 166653608 | rs6456063 | -0.076 | 0.608 | -0.528 | -0.925 | 0.004 | -0.009 | -0.009 | 1.985 | 0.87 | 0.82 | 0.77 | 0.94 | 0.86 | 0.88 |
| 6 | 166654968 | rs9348102 | 1.564 | 0.244 | -0.834 | -0.786 | -0.002 | -0.096 | -0.096 | 0 | 0.31 | 0.08 | 0.4 | 0.56 | 0.25 | 0.23 |
| 6 | 166660733 | rs9356453 | -0.147 | 0.358 | -0.076 | -0.872 | -0.001 | -0.024 | -0.024 | 0 | 0.7 | 0.64 | 0.7 | 0.86 | 0.75 | 0.8 |
| 6 | 166662128 | rs1610315 | -0.085 | 0.33 | -0.082 | -0.871 | -0.003 | -0.023 | -0.023 | 0.033 | 0.69 | 0.63 | 0.71 | 0.86 | 0.75 | 0.77 |
| 6 | 166665154 | rs7740044 | -0.248 | 0.438 | 0.379 | -0.624 | 0.026 | -0.028 | -0.028 | 0.087 | 0.89 | 0.77 | 0.79 | 0.94 | 0.9 | 0.89 |
| 6 | 166667434 | rs9356458 | 0.146 | 0.481 | 0.094 | -0.763 | -0.001 | -0.014 | -0.014 | 0 | 0.42 | 0.43 | 0.27 | 0.31 | 0.43 | 0.53 |
| 6 | 166667861 | rs4709109 | 1.355 | 0.479 | -0.025 | -0.534 | -0.097 | -0.139 | -0.139 | 0.269 | 0.34 | 0.04 | 0.43 | 0.54 | 0.35 | 0.24 |
| 6 | 166669864 | rs9365985 | -0.428 | 0.423 | 0.307 | -0.552 | -0.008 | 0.006 | 0.006 | 0 | 0.56 | 0.57 | 0.49 | 0.41 | 0.57 | 0.7 |
| 6 | 166672508 | rs9356460 | 0.04 | 0.312 | 0.343 | -0.592 | 0 | -0.002 | -0.002 | 0 | 0.45 | 0.39 | 0.42 | 0.3 | 0.45 | 0.58 |
| 6 | 166673072 | rs4367352 | 0.429 | 0.446 | -0.16 | -0.475 | 0.003 | 0.068 | 0.068 | 0 | 0.44 | 0.21 | 0.5 | 0.71 | 0.45 | 0.37 |
| 6 | 166673508 | rs9365988 | 1.246 | 0.682 | -0.638 | -0.326 | -0.063 | -0.101 | -0.101 | 0 | 0.33 | 0.06 | 0.42 | 0.55 | 0.33 | 0.24 |
| 6 | 166674624 | rs6924775 | 0.453 | 0.917 | 0.093 | 0.206 | 0.012 | -0.004 | -0.004 | 0.704 | 0.92 | 0.87 | 0.93 | 0.94 | 0.91 | 0.94 |
| 6 | 166675798 | rs12207833 | -0.509 | 0.913 | 0.338 | 0.275 | 0.013 | -0.005 | -0.005 | 0 | 0.24 | 0.24 | 0.23 | 0.21 | 0.35 | 0.42 |
| 6 | 166676586 | rs7753126 | -0.936 | 0.198 | 0.258 | 0.326 | -0.022 | -0.006 | -0.006 | 0.377 | 0.18 | 0.71 | 0.14 | 0.17 | 0.17 | 0.21 |
| 6 | 166676758 | rs7774584 | -0.762 | 0.596 | -0.185 | 0.48 | -0.001 | 0 | 0 | 0 | 0.46 | 0.47 | 0.42 | 0.44 | 0.59 | 0.55 |
| 6 | 166677036 | rs6940834 | -0.528 | 1.002 | -0.703 | 0.752 | -0.016 | -0.037 | -0.037 | 0.818 | 0.4 | 0.25 | 0.54 | 0.55 | 0.44 | 0.33 |
| 6 | 166679074 | rs6910073 | -0.951 | 1.035 | -0.354 | 0.505 | 0.008 | 0 | 0 | 0 | 0.38 | 0.59 | 0.63 | 0.63 | 0.52 | 0.38 |
| 6 | 166684657 | rs6909266 | -0.887 | 1.037 | -0.742 | -0.168 | 0.019 | -0.02 | -0.02 | 0 | 0.23 | 0.33 | 0.31 | 0.22 | 0.31 | 0.18 |
| 6 | 166686851 | rs10946145 | -0.952 | 1.045 | -1.026 | -0.181 | 0 | -0.018 | -0.018 | 0.257 | 0.1 | 0.12 | 0.06 | 0 | 0.08 | 0.01 |
| 6 | 166689600 | rs9355566 | -0.681 | 1.256 | -1.355 | 0.104 | -0.041 | 0.024 | 0.024 | 0 | 0.31 | 0.16 | 0.34 | 0.3 | 0.41 | 0.19 |
| 6 | 166699558 | rs12206441 | -0.495 | 1.016 | -0.877 | -0.196 | -0.002 | -0.021 | -0.021 | 0 | 0.16 | 0.14 | 0.12 | 0 | 0.19 | 0.13 |
| 6 | 166699631 | rs13195100 | -0.792 | 0.647 | -0.346 | -0.173 | -0.012 | -0.002 | -0.002 | 0.237 | 0.05 | 0.15 | 0.04 | 0.03 | 0.01 | 0.01 |
| 6 | 166702780 | rs9365997 | -0.505 | 0.299 | -0.42 | 0.594 | 0.002 | 0.013 | 0.013 | 0 | 0.1 | 0.09 | 0.02 | 0.26 | 0.04 | 0.11 |
| 6 | 166706121 | rs6934656 | -0.593 | 0.694 | -0.283 | -0.338 | 0.016 | -0.038 | -0.038 | 0 | 0.72 | 0.67 | 0.56 | 0.91 | 0.61 | 0.76 |
| 6 | 166707250 | rs1033725 | 0.249 | -0.008 | -0.766 | -0.257 | 0.024 | -0.025 | -0.025 | 0 | 0.4 | 0.21 | 0.29 | 0.61 | 0.36 | 0.4 |
| 6 | 166710938 | rs6902135 | 0.226 | 0.241 | 0.021 | -0.377 | -0.002 | 0.002 | 0.002 | 0 | 0.42 | 0.35 | 0.39 | 0.6 | 0.44 | 0.4 |
| 6 | 166718436 | rs1998167 | -0.77 | 0.413 | 0.639 | -0.602 | 0.002 | -0.081 | -0.081 | 0 | 0.7 | 0.56 | 0.58 | 0.89 | 0.72 | 0.67 |
| 6 | 166718662 | rs1998168 | -0.953 | 0.291 | 0.867 | -0.665 | -0.015 | 0.066 | 0.066 | 0 | 0.73 | 0.81 | 0.59 | 0.92 | 0.72 | 0.77 |
| 6 | 166722701 | rs9459644 | -0.648 | 0.309 | 1.219 | -0.821 | 0.001 | 0.001 | 0.001 | 0.473 | 0.39 | 0.59 | 0.36 | 0.33 | 0.37 | 0.38 |
| 6 | 166724312 | rs10946155 | -0.389 | 0.323 | 1.222 | -0.826 | -0.004 | -0.01 | -0.01 | 0 | 0.37 | 0.48 | 0.35 | 0.3 | 0.36 | 0.28 |
| 6 | 166737980 | rs4710041 | 0.497 | 1.036 | 1.324 | -0.97 | 0.014 | -0.012 | -0.012 | 0.336 | 0.9 | 0.9 | 0.78 | 0.98 | 0.82 | 0.95 |
| 6 | 166739684 | rs10806847 | 0.598 | 1.032 | 1.821 | -0.976 | 0.026 | -0.015 | -0.015 | 0 | 0.88 | 0.83 | 0.76 | 0.95 | 0.77 | 0.85 |
| 6 | 166747572 | rs4144886 | 0.664 | 0.92 | 1.837 | -0.623 | -0.002 | -0.022 | -0.022 | 0.041 | 0.25 | 0.17 | 0.18 | 0.34 | 0.25 | 0.24 |
| 6 | 166751416 | rs9356478 | 0.598 | 0.989 | 1.638 | -0.621 | 0.039 | -0.023 | -0.023 | 0.526 | 0.87 | 0.78 | 0.75 | 0.96 | 0.75 | 0.85 |
| 6 | 166751690 | rs4709111 | 0 | 0.793 | 1.216 | -0.552 | 0.003 | 0 | 0 | 0 | 0.59 | 0.39 | 0.54 | 0.63 | 0.5 | 0.62 |
| 6 | 166761245 | rs9457165 | 1.03 | 0.588 | 0.472 | -0.603 | 0.015 | 0.038 | 0.038 | 0.41 | 0.21 | 0.13 | 0.31 | 0.08 | 0.13 | 0.21 |
| 6 | 166763633 | rs9885912 | 0.368 | 0.785 | 0.71 | -0.203 | 0 | 0.002 | 0.002 | 0 | 0.51 | 0.39 | 0.55 | 0.47 | 0.46 | 0.49 |
| 6 | 166769136 | rs9356480 | 0.113 | 0.53 | 0.268 | -0.311 | -0.004 | -0.059 | -0.059 | 0 | 0.31 | 0.17 | 0.18 | 0.51 | 0.32 | 0.45 |
| 6 | 166777855 | rs6456078 | 0.278 | 0.736 | 0.664 | -0.572 | -0.004 | -0.002 | -0.002 | 0 | 0.85 | 0.83 | 0.73 | 0.97 | 0.73 | 0.92 |
| 6 | 166778679 | rs3728 | 0.881 | 0.58 | 0.63 | -0.332 | -0.045 | -0.01 | -0.01 | 0 | 0.43 | 0.12 | 0.45 | 0.64 | 0.44 | 0.7 |
| 6 | 166782143 | rs6456079 | 0.956 | 0.878 | 1.937 | -0.27 | -0.006 | 0 | 0 | 0 | 0.84 | 0.83 | 0.73 | 0.97 | 0.72 | 0.92 |
| 6 | 166791235 | rs3734605 | 0.921 | 0.844 | 1.998 | -0.261 | 0.009 | -0.016 | -0.016 | 3.004 | 0.85 | 0.82 | 0.72 | 0.97 | 0.72 | 0.92 |
| 6 | 166792405 | rs2343591 | 1.619 | 0.8 | 2.311 | -0.28 | 0.04 | -0.104 | -0.104 | 0 | 0.78 | 0.43 | 0.69 | 0.96 | 0.72 | 0.92 |
| 6 | 166796433 | rs1546854 | 0.823 | 0.945 | 1.687 | -0.263 | 0.007 | -0.01 | -0.01 | 0.391 | 0.86 | 0.83 | 0.72 | 0.97 | 0.72 | 0.93 |
| 6 | 166799022 | rs3757195 | 0.997 | 0.957 | 1.597 | -0.257 | 0.048 | -0.045 | -0.045 | 0 | 0.85 | 0.74 | 0.71 | 0.97 | 0.72 | 0.93 |
| 6 | 166819774 | rs9355578 | 1.267 | 0.489 | 1.37 | 0.144 | 0.04 | -0.011 | -0.011 | 0 | 0.77 | 0.61 | 0.72 | 0.78 | 0.67 | 0.86 |
| 6 | 166826021 | rs2984 | 0.632 | 0.445 | 0.661 | 0.202 | 0.028 | 0.005 | 0.005 | 0 | 0.8 | 0.74 | 0.73 | 0.77 | 0.67 | 0.86 |
| 6 | 166826255 | rs909631 | 1.331 | 0.233 | 0.877 | 0.262 | 0.02 | 0.007 | 0.007 | 0 | 0.76 | 0.31 | 0.66 | 0.74 | 0.66 | 0.84 |
| 6 | 166826304 | rs760670 | 0.167 | 0.679 | -0.032 | 0.488 | 0.015 | 0.001 | 0.001 | 0.053 | 0.85 | 0.86 | 0.75 | 0.79 | 0.68 | 0.86 |
| 6 | 166830696 | rs6925364 | 0.387 | 0.623 | -0.057 | 0.35 | 0.043 | 0.021 | 0.021 | 0 | 0.8 | 0.74 | 0.7 | 0.75 | 0.65 | 0.85 |
| 6 | 166831694 | rs10946164 | 0.178 | 0.19 | -0.732 | -0.346 | -0.134 | 0.117 | 0.117 | 0 | 0.4 | 0.02 | 0.24 | 0.2 | 0.44 | 0.43 |
| 6 | 166838291 | rs16898917 | -1.353 | -0.025 | 1.037 | 0.428 | 0.007 | -0.002 | -0.002 | 0 | 0.07 | 0.42 | 0.03 | 0 | 0.01 | 0.01 |
| 6 | 166851320 | rs2049957 | -0.213 | 0.356 | 0.001 | 0.755 | -0.001 | -0.002 | -0.002 | 0 | 0.48 | 0.5 | 0.46 | 0.47 | 0.52 | 0.35 |

| | | | | | | | | | | | | | | | | |
|---|---|---|---|---|---|---|---|---|---|---|---|---|---|---|---|---|
| 6 | 166853513 | rs9355580 | -0.319 | 0.119 | -0.37 | 0.317 | -0.017 | 0.084 | 0.084 | 0 | 0.28 | 0.09 | 0.27 | 0.09 | 0.3 | 0.17 |
| 6 | 166855297 | rs1003642 | 0.439 | -0.121 | -0.217 | 1.049 | -0.004 | 0.037 | 0.037 | 0.344 | 0.92 | 0.91 | 0.86 | 0.82 | 0.91 | 0.81 |
| 6 | 166859080 | rs9295346 | -0.351 | -0.456 | 0.343 | 1.462 | 0.003 | 0.016 | 0.016 | 0 | 0.33 | 0.25 | 0.37 | 0.18 | 0.32 | 0.25 |
| 6 | 166859572 | rs6456083 | -0.592 | -1.39 | 0.322 | 1.445 | -0.098 | 0.016 | 0.016 | 1.666 | 0.17 | 0.61 | 0.08 | 0.15 | 0.04 | 0.13 |
| 6 | 166859725 | rs2246553 | 1.011 | -1.415 | 0.304 | 1.688 | -0.055 | 0.074 | 0.074 | 1.448 | 0.77 | 0.34 | 0.79 | 0.62 | 0.87 | 0.71 |
| 6 | 166861171 | rs2281052 | 0.275 | -0.936 | 0.395 | 1.917 | -0.007 | 0.03 | 0.03 | 0 | 0.65 | 0.78 | 0.6 | 0.78 | 0.6 | 0.67 |
| 6 | 166861958 | rs9457170 | -0.501 | -0.996 | 0.752 | 1.677 | 0.008 | 0.066 | 0.066 | 0.315 | 0.05 | 0.23 | 0.11 | 0.1 | 0.02 | 0.08 |
| 6 | 166865131 | rs2072828 | -0.428 | -0.742 | 0.515 | 1.665 | -0.001 | -0.027 | -0.027 | 0 | 0.35 | 0.25 | 0.43 | 0.41 | 0.39 | 0.37 |
| 6 | 166868072 | rs6912899 | -1.168 | -0.507 | 1.044 | 1.991 | 0.022 | 0.302 | 0.302 | 0.034 | 0.09 | 0.34 | 0.16 | 0.3 | 0.08 | 0.2 |
| 6 | 166873823 | rs12208871 | -0.096 | -0.247 | 0.244 | 1.127 | 0.004 | -0.019 | -0.019 | 0 | 0.09 | 0.09 | 0.18 | 0.03 | 0.14 | 0.1 |
| 6 | 166875819 | rs3817796 | -1.216 | -0.786 | 0.199 | 1.791 | 0.008 | -0.008 | -0.008 | 0 | 0.18 | 0.2 | 0.17 | 0.41 | 0.12 | 0.22 |
| 6 | 166876628 | rs3817792 | -1.262 | -0.889 | 0.418 | 1.864 | 0.002 | 0.009 | 0.009 | 0.773 | 0.17 | 0.19 | 0.16 | 0.39 | 0.12 | 0.27 |
| 6 | 166876708 | rs3817790 | -0.474 | -0.996 | 0.553 | 0.986 | -0.004 | -0.018 | -0.018 | 0 | 0.32 | 0.32 | 0.39 | 0.18 | 0.3 | 0.19 |
| 6 | 166877012 | rs1004700 | -1.585 | -1.311 | 0.457 | 0.919 | -0.025 | -0.021 | -0.021 | 0.003 | 0.29 | 0.39 | 0.36 | 0.23 | 0.22 | 0.19 |
| 6 | 166877118 | rs1004699 | -1.538 | -1.31 | 0.456 | 0.912 | -0.001 | -0.005 | -0.005 | 0.039 | 0.43 | 0.39 | 0.43 | 0.23 | 0.34 | 0.23 |
| 6 | 166877532 | rs2235287 | -0.929 | -1.326 | 0.578 | 0.827 | -0.015 | -0.026 | -0.026 | 0.006 | 0.21 | 0.4 | 0.34 | 0.18 | 0.19 | 0.14 |
| 6 | 166878094 | rs12202788 | -0.791 | -1.306 | 0.772 | 0.84 | -0.004 | -0.047 | -0.047 | 0.26 | 0.19 | 0.26 | 0.31 | 0.06 | 0.18 | 0.12 |
| 6 | 166878708 | rs9295348 | -1.546 | -1.441 | 1.002 | 0.852 | 0.001 | -0.002 | -0.002 | 0 | 0.33 | 0.32 | 0.46 | 0.45 | 0.3 | 0.38 |
| 6 | 166878799 | rs6905447 | -1.721 | -1.448 | 1.01 | 0.662 | 0.04 | 0.058 | 0.058 | 1.804 | 0.51 | 0.75 | 0.57 | 0.58 | 0.44 | 0.53 |
| 6 | 166882664 | rs16898963 | 0.476 | -1.357 | 1.055 | 0.462 | 0 | 0 | 0 | 0 | 0.15 | 0.13 | 0.11 | 0.13 | 0.14 | 0.14 |
| 6 | 166883981 | rs9459678 | -1.36 | -1.462 | 1.12 | 0.22 | 0.026 | 0.07 | 0.07 | 0 | 0.39 | 0.68 | 0.45 | 0.18 | 0.33 | 0.28 |
| 6 | 166884139 | rs6902946 | -1.048 | -1.845 | 0.858 | -0.448 | -0.011 | -0.06 | -0.06 | 1.517 | 0.43 | 0.49 | 0.45 | 0.15 | 0.37 | 0.31 |
| 6 | 166884211 | rs6926393 | -0.888 | -1.282 | 0.987 | -0.276 | -0.029 | 0.019 | 0.019 | 0 | 0.37 | 0.46 | 0.31 | 0.5 | 0.33 | 0.41 |
| 6 | 166884534 | rs9366015 | -1.854 | -0.309 | 1.029 | 0.079 | -0.006 | -0.018 | -0.018 | 0 | 0.31 | 0.3 | 0.37 | 0.43 | 0.33 | 0.38 |
| 6 | 166884765 | rs12199759 | -0.937 | -0.795 | 0.951 | -0.075 | -0.001 | -0.013 | -0.013 | 0.018 | 0.16 | 0.16 | 0.21 | 0.04 | 0.17 | 0.16 |
| 6 | 166884848 | rs2235283 | -1.618 | -1.101 | 0.891 | 0.242 | -0.007 | 0.003 | 0.003 | 2.981 | 0.38 | 0.43 | 0.48 | 0.59 | 0.33 | 0.47 |
| 6 | 166886340 | rs909635 | -1.776 | -0.922 | 0.225 | 0.175 | -0.15 | -0.01 | -0.01 | 0 | 0.63 | 0.97 | 0.58 | 0.65 | 0.48 | 0.59 |
| 6 | 166886680 | rs909634 | -0.543 | -0.702 | 1.631 | 0.971 | -0.037 | 0.038 | 0.038 | 0 | 0.31 | 0.43 | 0.21 | 0.49 | 0.21 | 0.28 |
| 6 | 166891620 | rs7742938 | -1.22 | -0.558 | 2.035 | 0.76 | -0.021 | 0.019 | 0.019 | 1.346 | 0.08 | 0.32 | 0.05 | 0.05 | 0.02 | 0 |
| 6 | 166892798 | rs6909920 | -1.505 | -0.417 | 2.341 | 0.709 | 0.006 | -0.048 | -0.048 | 0.123 | 0.17 | 0.58 | 0.22 | 0.09 | 0.14 | 0.17 |
| 6 | 166892866 | rs6926193 | -1.38 | -0.437 | 2.4 | 0.7 | 0.006 | -0.061 | -0.061 | 0.167 | 0.16 | 0.35 | 0.2 | 0.07 | 0.14 | 0.11 |
| 6 | 166892969 | rs6456086 | -1.747 | -0.761 | 2.331 | 0.749 | -0.019 | -0.034 | -0.034 | 0.039 | 0.18 | 0.64 | 0.21 | 0.09 | 0.12 | 0.17 |
| 6 | 166893239 | rs6926800 | -1.618 | -0.657 | 2.479 | 0.77 | -0.016 | -0.067 | -0.067 | 0.65 | 0.17 | 0.49 | 0.26 | 0.08 | 0.11 | 0.16 |
| 6 | 166893339 | rs4709117 | -0.675 | -0.388 | 2.198 | 1.082 | -0.005 | 0.004 | 0.004 | 0 | 0.33 | 0.33 | 0.28 | 0.53 | 0.26 | 0.39 |
| 6 | 166895934 | rs9347129 | -1.16 | -0.001 | 2.163 | 1.17 | 0.02 | -0.009 | -0.009 | 0 | 0.14 | 0.27 | 0.24 | 0.09 | 0.13 | 0.19 |
| 6 | 166898877 | rs9295351 | -1.291 | -0.005 | 1.869 | 1.168 | 0.014 | -0.014 | -0.014 | 0 | 0.14 | 0.3 | 0.24 | 0.09 | 0.13 | 0.19 |
| 6 | 166903347 | rs3778432 | -1.157 | -0.011 | 1.357 | 1.136 | 0.007 | -0.045 | -0.045 | 0 | 0.14 | 0.39 | 0.2 | 0.06 | 0.11 | 0.14 |
| 6 | 166904398 | rs2281045 | -0.606 | -0.133 | 0.485 | 1.322 | 0.007 | -0.112 | -0.112 | 0 | 0.29 | 0.08 | 0.31 | 0.52 | 0.28 | 0.37 |
| 6 | 166904927 | rs1569462 | -1.887 | -0.218 | 0.516 | 1.261 | 0.039 | 0.061 | 0.061 | 0.104 | 0.48 | 0.84 | 0.66 | 0.61 | 0.4 | 0.56 |
| 6 | 166905262 | rs3817781 | -0.387 | -0.096 | 0.339 | 1.352 | -0.01 | -0.136 | -0.136 | 0 | 0.28 | 0.05 | 0.43 | 0.5 | 0.28 | 0.37 |
| 6 | 166910553 | rs4710059 | -1.806 | -0.192 | 0.309 | 1.333 | -0.001 | 0.124 | 0.124 | 1 | 0.48 | 0.93 | 0.53 | 0.69 | 0.4 | 0.55 |
| 6 | 166911186 | rs6940371 | -1.598 | -0.208 | 0.341 | 1.357 | -0.023 | -0.016 | -0.016 | 0 | 0.44 | 0.91 | 0.51 | 0.59 | 0.38 | 0.54 |
| 6 | 166912157 | rs6923965 | -0.947 | -0.192 | 1.134 | 1.163 | 0.031 | -0.072 | -0.072 | 0 | 0.12 | 0.5 | 0.17 | 0.02 | 0.07 | 0.11 |
| 6 | 166913485 | rs9457173 | -0.662 | -0.325 | 0.584 | 1.444 | 0.01 | -0.045 | -0.045 | 0 | 0.28 | 0.08 | 0.32 | 0.61 | 0.27 | 0.36 |
| 6 | 166915130 | rs10946174 | -0.947 | -0.19 | 0.737 | 1.402 | 0.006 | -0.003 | -0.003 | 0 | 0.33 | 0.24 | 0.33 | 0.58 | 0.29 | 0.38 |
| 6 | 166915537 | rs2294323 | -1.336 | -0.222 | -0.165 | 1.954 | 0.006 | 0.002 | 0.002 | 0.793 | 0.28 | 0.7 | 0.18 | 0.27 | 0.12 | 0.16 |
| 6 | 166916557 | rs3778430 | 0.157 | -0.282 | -0.306 | 2.093 | -0.035 | -0.112 | -0.112 | 0 | 0.16 | 0.02 | 0.33 | 0.35 | 0.2 | 0.31 |
| 6 | 166917844 | rs2071941 | -1.336 | -0.329 | 0.404 | 1.635 | -0.036 | 0.003 | 0.003 | 0 | 0.25 | 0.56 | 0.17 | 0.27 | 0.14 | 0.16 |
| 6 | 166919755 | rs1894660 | -1.236 | 0.44 | 0.233 | 1.997 | 0.019 | 0.11 | 0.11 | 0 | 0.17 | 0.32 | 0.36 | 0.4 | 0.18 | 0.36 |
| 6 | 166922041 | rs411281 | 0.523 | 0.171 | -0.537 | 1.688 | 0.006 | -0.023 | -0.023 | 0 | 0.88 | 0.97 | 0.91 | 0.84 | 0.89 | 0.91 |
| 6 | 166922093 | rs411236 | -0.673 | -0.095 | -0.771 | 1.177 | 0.004 | 0.003 | 0.003 | 0 | 0.66 | 0.69 | 0.81 | 0.66 | 0.73 | 0.73 |
| 6 | 166922813 | rs3799670 | -0.152 | -0.36 | -1.164 | 1.398 | 0.042 | -0.064 | -0.064 | 0 | 0.15 | 0.03 | 0.09 | 0.18 | 0.09 | 0.11 |
| 6 | 166928580 | rs378373 | 0.263 | 0.184 | -1.933 | 1.143 | -0.107 | -0.072 | -0.072 | 1.288 | 0.2 | 0.03 | 0.3 | 0.23 | 0.34 | 0.32 |
| 6 | 166929956 | rs3799665 | 0.941 | -0.298 | -2.536 | 0.422 | -0.036 | -0.08 | -0.08 | 0.536 | 0.27 | 0.12 | 0.26 | 0.46 | 0.26 | 0.22 |
| 6 | 166930424 | rs9364852 | 0.457 | -0.901 | -1.522 | -0.121 | -0.001 | 0 | 0 | 0.344 | 0.46 | 0.54 | 0.37 | 0.57 | 0.4 | 0.35 |
| 6 | 166935195 | rs2187911 | -0.336 | -0.971 | -1.138 | -0.191 | -0.033 | -0.044 | -0.044 | 0 | 0.65 | 0.77 | 0.6 | 0.62 | 0.55 | 0.62 |
| 6 | 166936021 | rs743980 | 0.339 | -1.514 | -0.724 | -0.423 | -0.039 | -0.113 | -0.113 | 0 | 0.22 | 0.39 | 0.24 | 0.07 | 0.15 | 0.27 |
| 6 | 166936266 | rs3799658 | 1.093 | -1.502 | -0.824 | -0.387 | 0.012 | -0.045 | -0.045 | 0 | 0.14 | 0.04 | 0.09 | 0.15 | 0.11 | 0.06 |
| 6 | 166938431 | rs6939941 | -0.293 | -1.463 | -0.687 | -0.357 | -0.032 | -0.028 | -0.028 | 0 | 0.36 | 0.48 | 0.33 | 0.22 | 0.26 | 0.34 |
| 6 | 166940184 | rs6924732 | 0.23 | -1.234 | -0.707 | -0.43 | -0.02 | -0.061 | -0.061 | 0 | 0.22 | 0.29 | 0.25 | 0.07 | 0.16 | 0.28 |
| 6 | 166941578 | rs11969535 | 0.259 | -1.228 | -0.704 | -0.434 | -0.02 | -0.064 | -0.064 | 0 | 0.22 | 0.29 | 0.25 | 0.05 | 0.16 | 0.28 |
| 6 | 166943513 | rs11970312 | 0.289 | -1.229 | -0.727 | -0.441 | -0.021 | -0.059 | -0.059 | 0 | 0.21 | 0.28 | 0.25 | 0.05 | 0.15 | 0.27 |
| 6 | 166951139 | rs2187909 | 0.568 | -0.698 | -0.795 | -0.158 | -0.038 | -0.011 | -0.011 | 0 | 0.35 | 0.1 | 0.39 | 0.3 | 0.4 | 0.56 |
| 6 | 166952264 | rs1003857 | -0.034 | -0.659 | -0.292 | -0.306 | 0.035 | -0.1 | -0.1 | 0 | 0.81 | 0.58 | 0.85 | 0.94 | 0.79 | 0.87 |
| 6 | 166952981 | rs11755127 | 1.583 | -0.861 | -0.866 | -0.197 | -0.055 | -0.109 | -0.109 | 0 | 0.25 | 0.09 | 0.24 | 0.44 | 0.26 | 0.18 |
| 6 | 166954244 | rs3778409 | 1.112 | -0.779 | -0.772 | -0.197 | 0.003 | -0.047 | -0.047 | 0.338 | 0.14 | 0.03 | 0.08 | 0.15 | 0.11 | 0.06 |
| 6 | 166954433 | rs3799639 | 0.547 | -0.542 | -0.224 | -0.41 | 0.005 | -0.108 | -0.108 | 0 | 0.76 | 0.57 | 0.74 | 0.89 | 0.76 | 0.8 |
| 6 | 166955738 | rs763187 | -0.619 | -0.129 | -0.606 | -0.282 | 0 | -0.027 | -0.027 | 0.095 | 0.88 | 0.9 | 0.92 | 0.99 | 0.84 | 0.92 |
| 6 | 166956895 | rs6918886 | 0.175 | -0.422 | -0.592 | -0.211 | -0.005 | 0.015 | 0.015 | 0 | 0.39 | 0.28 | 0.52 | 0.36 | 0.43 | 0.6 |

| | | | | | | | | | | | | | | | | |
|---|---|---|---|---|---|---|---|---|---|---|---|---|---|---|---|---|
| 6 | 166963233 | rs6456093 | 0.243 | -0.434 | -0.622 | 0.039 | 0.01 | 0.011 | 0.011 | 0.572 | 0.43 | 0.65 | 0.39 | 0.58 | 0.35 | 0.26 |
| 6 | 166965035 | rs3778405 | -0.474 | -1.384 | -0.845 | 0.773 | 0.012 | -0.026 | -0.026 | 0.301 | 0.18 | 0.15 | 0.25 | 0.37 | 0.1 | 0.38 |
| 6 | 166966530 | rs9366027 | 0.912 | -1.331 | -0.786 | 0.716 | 0.019 | 0.006 | 0.006 | 0 | 0.11 | 0.15 | 0.06 | 0.13 | 0.1 | 0.15 |
| 6 | 166967389 | rs3799627 | 0.677 | -1.329 | -0.951 | 1.035 | 0.002 | -0.074 | -0.074 | 0 | 0.11 | 0.02 | 0.14 | 0.29 | 0.07 | 0.27 |
| 6 | 166969936 | rs971152 | -0.978 | -1.734 | -0.997 | 0.475 | 0.008 | 0.012 | 0.012 | 0.891 | 0.29 | 0.66 | 0.24 | 0.23 | 0.16 | 0.28 |
| 6 | 166971926 | rs9355584 | -0.88 | -2.019 | -1.651 | 0.448 | 0.041 | -0.051 | -0.051 | 0.124 | 0.44 | 0.77 | 0.36 | 0.43 | 0.28 | 0.36 |
| 6 | 166973105 | rs3823204 | 1.602 | -1.989 | -1.629 | 0.476 | -0.012 | -0.086 | -0.086 | 0 | 0.13 | 0.01 | 0.11 | 0.2 | 0.12 | 0.08 |
| 6 | 166974222 | rs3799620 | -1.291 | -2.106 | -1.713 | 0.338 | -0.005 | 0.103 | 0.103 | 0 | 0.56 | 0.79 | 0.51 | 0.66 | 0.35 | 0.63 |
| 6 | 166975390 | rs2032546 | 1.677 | -2.076 | -2.159 | -0.113 | -0.034 | -0.022 | -0.022 | 0 | 0.2 | 0.07 | 0.28 | 0.28 | 0.25 | 0.23 |
| 6 | 166976374 | rs3799619 | 1.055 | -2.074 | -1.48 | 0.324 | -0.015 | -0.08 | -0.08 | 2.101 | 0.13 | 0.02 | 0.14 | 0.28 | 0.1 | 0.29 |
| 6 | 166977609 | rs3823202 | 0.235 | -2.096 | -1.12 | 0.169 | -0.01 | 0.022 | 0.022 | 0.11 | 0.08 | 0.24 | 0.11 | 0.1 | 0.03 | 0.09 |
| 6 | 166978752 | rs3778403 | 0.327 | -2.164 | -1.239 | 0.123 | 0.013 | 0.019 | 0.019 | 0 | 0.44 | 0.3 | 0.51 | 0.58 | 0.4 | 0.36 |
| 6 | 166979046 | rs3778401 | -0.507 | -2.215 | -1.19 | 0.085 | 0.002 | -0.03 | -0.03 | 0.062 | 0.58 | 0.61 | 0.69 | 0.8 | 0.48 | 0.64 |
| 6 | 166979148 | rs3778400 | 1.444 | -2.141 | -1.472 | 0.131 | 0 | -0.068 | -0.068 | 0 | 0.11 | 0.02 | 0.14 | 0.22 | 0.07 | 0.28 |
| 6 | 166979176 | rs3778398 | -1.658 | -1.889 | -1.258 | 0.089 | -0.052 | 0.074 | 0.074 | 0 | 0.77 | 0.87 | 0.79 | 0.93 | 0.61 | 0.86 |
| 6 | 166981135 | rs3778394 | 0.649 | -2.02 | -1.236 | 0.256 | -0.003 | 0.004 | 0.004 | 0.014 | 0.23 | 0.26 | 0.23 | 0.31 | 0.14 | 0.13 |
| 6 | 166981608 | rs3799613 | 1.423 | -1.812 | -1.273 | 0.091 | -0.04 | -0.023 | -0.023 | 0 | 0.21 | 0.06 | 0.29 | 0.28 | 0.26 | 0.24 |
| 6 | 166982506 | rs3799611 | 0.91 | -1.93 | -1.265 | 0.098 | -0.031 | -0.049 | -0.049 | 0 | 0.24 | 0.1 | 0.38 | 0.37 | 0.27 | 0.3 |
| 6 | 166982766 | rs3799610 | -0.034 | -2.633 | -0.591 | 0.386 | 0.032 | 0.008 | 0.008 | 0 | 0.43 | 0.24 | 0.33 | 0.51 | 0.3 | 0.53 |
| 6 | 166983874 | rs7760101 | 0.768 | -2.324 | -0.514 | 0.361 | 0.006 | -0.018 | -0.018 | 0 | 0.06 | 0.23 | 0.03 | 0 | 0.01 | 0 |
| 6 | 166984094 | rs6456095 | -1.507 | -2.384 | -0.747 | 0.263 | -0.09 | 0.103 | 0.103 | 0 | 0.68 | 0.86 | 0.66 | 0.84 | 0.52 | 0.77 |
| 6 | 166984656 | rs12661820 | 0.144 | -2.287 | -0.702 | 0.399 | -0.014 | -0.01 | -0.01 | 0 | 0.41 | 0.43 | 0.43 | 0.58 | 0.29 | 0.52 |
| 6 | 166984801 | rs9459695 | 1.213 | -1.902 | -0.633 | 0.512 | -0.019 | -0.034 | -0.034 | 0 | 0.15 | 0.05 | 0.25 | 0.26 | 0.2 | 0.22 |
| 6 | 166986767 | rs11755586 | -0.561 | -1.86 | -0.421 | 0.58 | 0.019 | 0.026 | 0.026 | 0 | 0.52 | 0.43 | 0.47 | 0.65 | 0.34 | 0.61 |
| 6 | 166987743 | rs4709122 | -0.428 | -1.221 | -0.309 | 0.596 | -0.023 | -0.012 | -0.012 | 0 | 0.36 | 0.51 | 0.34 | 0.3 | 0.2 | 0.5 |
| 6 | 166996991 | rs960145 | -0.585 | -0.737 | -0.308 | 0.369 | -0.082 | 0.029 | 0.029 | 0.067 | 0.59 | 0.87 | 0.59 | 0.77 | 0.46 | 0.79 |
| 6 | 166998662 | rs4710072 | 0.487 | -0.728 | -0.184 | 0.462 | 0.014 | 0.006 | 0.006 | 0.493 | 0.15 | 0.17 | 0.14 | 0.16 | 0.17 | 0.19 |
| 6 | 167003381 | rs3778385 | 0.359 | -0.69 | -0.376 | 0.589 | -0.008 | 0.007 | 0.007 | 0 | 0.37 | 0.43 | 0.35 | 0.57 | 0.29 | 0.44 |
| 6 | 167012206 | rs3823196 | 0.637 | -0.234 | -1.173 | 0.886 | 0.014 | -0.077 | -0.077 | 0.607 | 0.13 | 0.02 | 0.09 | 0.23 | 0.09 | 0.05 |
| 6 | 167015116 | rs2032544 | 0.473 | -0.224 | -1.086 | 0.785 | 0.022 | -0.036 | -0.036 | 0 | 0.16 | 0.1 | 0.11 | 0.3 | 0.1 | 0.2 |
| 6 | 167016947 | rs3799585 | -0.89 | -0.083 | -1.093 | 0.544 | 0.002 | -0.031 | -0.031 | 0.057 | 0.75 | 0.83 | 0.71 | 0.98 | 0.58 | 0.85 |
| 6 | 167018721 | rs10946177 | 0.214 | -0.694 | -1.562 | 1.359 | 0.012 | -0.023 | -0.023 | 0 | 0.34 | 0.17 | 0.4 | 0.47 | 0.28 | 0.39 |
| 6 | 167020970 | rs9355586 | -0.627 | -0.361 | -0.617 | 1.732 | 0.021 | -0.031 | -0.031 | 0 | 0.16 | 0.1 | 0.11 | 0.28 | 0.1 | 0.2 |
| 6 | 167029135 | rs916390 | -1.177 | 0.15 | 0.153 | 1.4 | -0.006 | 0.021 | 0.021 | 0 | 0.34 | 0.53 | 0.32 | 0.41 | 0.31 | 0.37 |
| 6 | 167029183 | rs873070 | -0.647 | 0.146 | 0.118 | 1.38 | -0.015 | 0.01 | 0.01 | 0 | 0.13 | 0.08 | 0.11 | 0.15 | 0.18 | 0.08 |
| 6 | 167032325 | rs3778372 | -1.229 | 0.074 | 0.68 | 1.549 | 0.016 | 0.066 | 0.066 | 0 | 0.17 | 0.32 | 0.13 | 0.29 | 0.18 | 0.16 |
| 6 | 167033404 | rs3778369 | -0.961 | 0.085 | 0.684 | 1.549 | -0.006 | -0.004 | -0.004 | 0 | 0.15 | 0.12 | 0.11 | 0.19 | 0.18 | 0.15 |
| 6 | 167039406 | rs3799566 | -1.434 | 0.048 | 0.728 | 1.416 | -0.002 | -0.002 | -0.002 | 0 | 0.19 | 0.15 | 0.14 | 0.25 | 0.2 | 0.13 |
| 6 | 167043542 | rs4145432 | -0.634 | -0.031 | 0.812 | 1.632 | 0 | -0.001 | -0.001 | 0 | 0.11 | 0.09 | 0.16 | 0.18 | 0.11 | 0.17 |
| 6 | 167044976 | rs9364861 | -1.776 | -0.066 | 0.698 | 1.523 | -0.02 | -0.022 | -0.022 | 0 | 0.25 | 0.13 | 0.27 | 0.36 | 0.28 | 0.25 |
| 6 | 167045295 | rs9356498 | -0.661 | -0.11 | 0.664 | 1.297 | -0.049 | 0.004 | 0.004 | 0 | 0.14 | 0.02 | 0.11 | 0.18 | 0.18 | 0.09 |
| 6 | 167049121 | rs9459702 | -0.946 | -0.276 | 0.966 | 1.295 | -0.007 | 0.003 | 0.003 | 0 | 0.05 | 0.03 | 0.03 | 0.06 | 0.02 | 0.02 |
| 6 | 167054213 | rs9295357 | -2.452 | -0.139 | 1.284 | 1.445 | -0.013 | 0.032 | 0.032 | 0.187 | 0.34 | 0.45 | 0.32 | 0.44 | 0.28 | 0.37 |
| 6 | 167055626 | rs9366036 | -0.491 | -0.155 | 1.243 | 1.445 | -0.021 | -0.031 | -0.031 | 0 | 0.11 | 0.02 | 0.15 | 0.14 | 0.1 | 0.19 |
| 6 | 167058026 | rs9347140 | -1.914 | -0.21 | 1.327 | 1.429 | -0.021 | 0.03 | 0.03 | 0 | 0.19 | 0.3 | 0.21 | 0.2 | 0.13 | 0.21 |
| 6 | 167060732 | rs12333057 | -1.451 | -0.193 | 1.345 | 1.437 | -0.017 | 0.057 | 0.057 | 0 | 0.07 | 0.19 | 0.04 | 0.06 | 0.02 | 0.02 |
| 6 | 167064415 | rs10484525 | -2.443 | -0.189 | 1.199 | 1.441 | -0.006 | 0.015 | 0.015 | 0.274 | 0.33 | 0.38 | 0.31 | 0.43 | 0.28 | 0.37 |
| 6 | 167065170 | rs6933990 | -1.789 | -0.183 | 1.223 | 1.444 | -0.016 | 0.022 | 0.022 | 0.117 | 0.18 | 0.26 | 0.2 | 0.2 | 0.12 | 0.19 |
| 6 | 167065891 | rs6919949 | -0.844 | -0.055 | 1.062 | 1.385 | -0.003 | -0.006 | -0.006 | 0 | 0.15 | 0.11 | 0.11 | 0.23 | 0.16 | 0.18 |
| 6 | 167074367 | rs9457192 | -1.746 | -0.038 | 1.046 | 1.444 | 0 | 0.005 | 0.005 | 0.046 | 0.29 | 0.3 | 0.29 | 0.38 | 0.26 | 0.35 |
| 6 | 167081819 | rs6903559 | -1.066 | -0.263 | 1.056 | 0.372 | 0 | 0 | 0 | 0 | 0.42 | 0.45 | 0.24 | 0.32 | 0.33 | 0.41 |
| 6 | 167082896 | rs7752989 | -0.324 | -0.449 | 1.17 | 0.221 | 0.007 | 0.001 | 0.001 | 0 | 0.21 | 0.2 | 0.15 | 0.24 | 0.18 | 0.27 |
| 6 | 167083690 | rs6921577 | -2.076 | -0.382 | 1.506 | 0.209 | 0.001 | 0.003 | 0.003 | 0 | 0.52 | 0.6 | 0.37 | 0.38 | 0.44 | 0.48 |
| 6 | 167084126 | rs12198203 | -1.11 | -0.246 | 1.443 | 0.327 | 0.002 | 0.01 | 0.01 | 0 | 0.31 | 0.31 | 0.19 | 0.26 | 0.27 | 0.37 |
| 6 | 167085055 | rs4518470 | -1.19 | -0.236 | 1.667 | 0.373 | 0.005 | 0.036 | 0.036 | 0 | 0.14 | 0.33 | 0.15 | 0.12 | 0.11 | 0.08 |
| 6 | 167089020 | rs9348171 | -0.878 | -0.143 | 1.268 | 0.381 | 0.005 | 0.002 | 0.002 | 0.242 | 0.25 | 0.25 | 0.16 | 0.26 | 0.2 | 0.19 |
| 6 | 167091844 | rs2281144 | -1.292 | -0.09 | 1.61 | 0.553 | 0 | 0.043 | 0.043 | 0 | 0.17 | 0.5 | 0.17 | 0.13 | 0.12 | 0.18 |
| 6 | 167096277 | rs10484524 | -0.569 | -0.087 | 1.054 | 0.54 | -0.004 | 0.045 | 0.045 | 0.381 | 0.23 | 0.07 | 0.15 | 0.2 | 0.2 | 0.18 |
| 6 | 167096773 | rs9295361 | -1.284 | -0.065 | 1.354 | 0.412 | 0.01 | -0.009 | -0.009 | 0 | 0.23 | 0.54 | 0.11 | 0.14 | 0.2 | 0.27 |
| 6 | 167096845 | rs9283866 | -1.448 | -0.02 | 1.252 | 0.446 | 0.008 | 0.011 | 0.011 | 0.11 | 0.48 | 0.63 | 0.26 | 0.41 | 0.4 | 0.47 |
| 6 | 167100377 | rs926389 | -1.513 | 0.205 | 1.384 | 0.626 | 0.001 | -0.001 | -0.001 | 0 | 0.34 | 0.7 | 0.2 | 0.37 | 0.26 | 0.33 |
| 6 | 167101325 | rs9459715 | -0.117 | 0.287 | 1.505 | 0.587 | 0.019 | -0.04 | -0.04 | 0.237 | 0.17 | 0.08 | 0.08 | 0.35 | 0.09 | 0.17 |
| 6 | 167103095 | rs7770039 | -2.048 | 0.292 | 1.6 | 0.443 | 0.002 | -0.075 | -0.075 | 1.842 | 0.18 | 0.6 | 0.12 | 0.03 | 0.18 | 0.17 |
| 6 | 167107415 | rs6456101 | -1.41 | 0.359 | 1.838 | 0.515 | 0.029 | -0.035 | -0.035 | 0 | 0.08 | 0.31 | 0.04 | 0 | 0.08 | 0.03 |
| 6 | 167107444 | rs6456102 | -1.569 | 0.863 | 1.488 | 0.242 | -0.008 | -0.004 | -0.004 | 0 | 0.45 | 0.74 | 0.34 | 0.44 | 0.47 | 0.37 |
| 6 | 167108595 | rs909768 | 0.075 | 0.86 | 1.505 | 0.26 | 0.056 | -0.034 | -0.034 | 0.012 | 0.15 | 0.35 | 0.17 | 0.08 | 0.19 | 0.1 |
| 6 | 167108650 | rs12662678 | 0.211 | 0.915 | 0.697 | 0.313 | 0.019 | -0.08 | -0.08 | 6.244 | 0.17 | 0.03 | 0.06 | 0.34 | 0.09 | 0.17 |
| 6 | 167110134 | rs2345258 | -0.267 | 0.888 | 0.887 | 0.25 | 0.026 | -0.062 | -0.062 | 1.532 | 0.18 | 0.59 | 0.19 | 0.08 | 0.19 | 0.09 |
| 6 | 167110268 | rs12663470 | -0.493 | 0.493 | 0.717 | 0.654 | 0.007 | -0.024 | -0.024 | 0 | 0.23 | 0.16 | 0.12 | 0.37 | 0.21 | 0.25 |

| | | | | | | | | | | | | | | | | |
|---|---|---|---|---|---|---|---|---|---|---|---|---|---|---|---|---|
| 6 | 167113245 | rs16900973 | -1.311 | 0.792 | 1.112 | 0.338 | 0.009 | -0.025 | -0.025 | 1.185 | 0.06 | 0.13 | 0.03 | 0 | 0.08 | 0.03 |
| 6 | 167114049 | rs6911624 | -0.924 | 0.375 | 1.362 | 0.238 | 0 | -0.032 | -0.032 | 0 | 0.2 | 0.44 | 0.26 | 0.12 | 0.18 | 0.12 |
| 6 | 167116160 | rs3817828 | 1.738 | 0.908 | 1.729 | 0.354 | -0.008 | 0.02 | 0.02 | 1.225 | 0.69 | 0.22 | 0.66 | 0.79 | 0.68 | 0.83 |
| 6 | 167117615 | rs760837 | 0.645 | 1.022 | 1.807 | 0.321 | -0.002 | 0.001 | 0.001 | 0 | 0.92 | 0.95 | 0.93 | 0.96 | 0.88 | 0.97 |
| 6 | 167122054 | rs4710081 | 2.166 | 0.979 | 2.005 | 0.291 | -0.014 | 0.031 | 0.031 | 0.618 | 0.69 | 0.18 | 0.66 | 0.8 | 0.68 | 0.83 |
| 6 | 167129817 | rs7768395 | 1.168 | 1.032 | 2.16 | 0.314 | 0.02 | -0.04 | -0.04 | 0.003 | 0.89 | 0.73 | 0.9 | 0.96 | 0.87 | 0.96 |
| 6 | 167131911 | rs12660310 | -0.902 | 0.52 | 2.198 | 0.578 | -0.002 | -0.001 | -0.001 | 0 | 0.12 | 0.15 | 0.21 | 0.11 | 0.18 | 0.11 |
| 6 | 167139479 | rs880900 | -2.154 | 0.181 | 1.379 | 0.771 | 0.001 | -0.001 | -0.001 | 0.056 | 0.28 | 0.76 | 0.28 | 0.24 | 0.22 | 0.16 |
| 6 | 167147083 | rs16899333 | -1.273 | 0.145 | 1.207 | 0.685 | -0.007 | -0.017 | -0.017 | 1.117 | 0.18 | 0.61 | 0.23 | 0.12 | 0.17 | 0.1 |
| 6 | 167147741 | rs6456115 | -1.13 | 0.137 | 1.101 | 0.681 | 0.005 | -0.015 | -0.015 | 3.982 | 0.18 | 0.51 | 0.22 | 0.12 | 0.17 | 0.11 |
| 6 | 167150023 | rs6932950 | -1.995 | 0.099 | 1.017 | 0.692 | 0.017 | 0.019 | 0.019 | 1.013 | 0.26 | 0.78 | 0.28 | 0.16 | 0.2 | 0.13 |
| 6 | 167151463 | rs7770200 | -2 | 0.168 | 0.741 | 0.974 | -0.017 | 0.045 | 0.045 | 0.157 | 0.18 | 0.4 | 0.14 | 0.21 | 0.14 | 0.12 |
| 6 | 167152526 | rs6456121 | -1.845 | 0.384 | -0.432 | 0.921 | -0.004 | -0.065 | -0.065 | 3.977 | 0.34 | 0.88 | 0.36 | 0.32 | 0.29 | 0.22 |
| 6 | 167154016 | rs909765 | -0.58 | 0.413 | -0.403 | 0.914 | -0.005 | -0.003 | -0.003 | 0 | 0.07 | 0.05 | 0.05 | 0.08 | 0.1 | 0.05 |
| 6 | 167155002 | rs3734600 | -1.254 | 0.357 | -0.428 | 0.935 | 0.028 | -0.018 | -0.018 | 0 | 0.27 | 0.84 | 0.3 | 0.24 | 0.19 | 0.17 |
| 6 | 167155074 | rs1108536 | -0.658 | 0.734 | 0.048 | 0.843 | -0.004 | -0.003 | -0.003 | 0.061 | 0.06 | 0.04 | 0.05 | 0.08 | 0.09 | 0.03 |
| 6 | 167156675 | rs1973900 | -0.994 | 0.642 | 0.022 | 0.826 | -0.004 | -0.002 | -0.002 | 0 | 0.08 | 0.06 | 0.05 | 0.08 | 0.09 | 0.05 |
| 6 | 167159163 | rs7763415 | -1.089 | 0.433 | 0.819 | 1.056 | 0.011 | 0.073 | 0.073 | 0.697 | 0.13 | 0.44 | 0.21 | 0.19 | 0.09 | 0.14 |
| 6 | 167161523 | rs4710089 | -0.676 | 0.657 | 0.887 | 1.002 | -0.005 | -0.003 | -0.003 | 0 | 0.27 | 0.54 | 0.31 | 0.26 | 0.24 | 0.24 |
| 6 | 167161571 | rs1883361 | -0.311 | 0.644 | -0.613 | 0.693 | -0.05 | -0.055 | -0.055 | 2.297 | 0.64 | 0.9 | 0.61 | 0.68 | 0.58 | 0.68 |
| 6 | 167167501 | rs7740362 | 0.119 | 0.725 | -0.763 | 0.444 | -0.016 | -0.042 | -0.042 | 0 | 0.28 | 0.15 | 0.35 | 0.31 | 0.33 | 0.3 |
| 6 | 167169223 | rs1202674 | -0.031 | 0.628 | -0.505 | -0.066 | -0.029 | 0.089 | 0.089 | 0 | 0.72 | 0.9 | 0.83 | 1 | 0.68 | 0.88 |
| 6 | 167171618 | rs9355592 | -1.018 | 0.5 | 0.04 | 0.84 | -0.011 | 0.168 | 0.168 | 0.83 | 0.06 | 0.21 | 0.11 | 0.29 | 0.02 | 0.09 |
| 6 | 167172048 | rs656938 | 0.434 | -0.138 | 0.375 | 0.196 | -0.011 | -0.022 | -0.022 | 0 | 0.57 | 0.51 | 0.66 | 0.7 | 0.63 | 0.71 |
| 6 | 167175396 | rs505982 | -0.5 | -0.214 | 0.42 | 0.042 | 0.035 | 0.058 | 0.058 | 3.38 | 0.41 | 0.73 | 0.32 | 0.26 | 0.32 | 0.2 |
| 6 | 167175911 | rs9347154 | -0.66 | -0.196 | 0.237 | 0.123 | -0.018 | 0.099 | 0.099 | 0 | 0.06 | 0.14 | 0.11 | 0.25 | 0.02 | 0.08 |
| 6 | 167176934 | rs388372 | -0.772 | -0.028 | -0.616 | 0.15 | -0.04 | -0.08 | -0.08 | 0.242 | 0.68 | 0.88 | 0.72 | 0.48 | 0.58 | 0.67 |
| 6 | 167177037 | rs388033 | 1.278 | -0.03 | -0.817 | 0.081 | -0.062 | -0.023 | -0.023 | 2.508 | 0.2 | 0.02 | 0.17 | 0.14 | 0.18 | 0.2 |
| 6 | 167177422 | rs635808 | 0.589 | -0.267 | -0.011 | -0.107 | -0.01 | -0.149 | -0.149 | 0 | 0.26 | 0.44 | 0.15 | 0 | 0.23 | 0.1 |
| 6 | 167177733 | rs637191 | -1.294 | -0.295 | -0.208 | 0.854 | -0.034 | 0.067 | 0.067 | 0 | 0.14 | 0.34 | 0.22 | 0.28 | 0.09 | 0.18 |
| 6 | 167180528 | rs409161 | -0.069 | -0.085 | -1.5 | 0.871 | -0.016 | -0.096 | -0.096 | 0 | 0.3 | 0.06 | 0.31 | 0.61 | 0.27 | 0.38 |
| 6 | 167181925 | rs733294 | 0.723 | -0.422 | -0.763 | 0.282 | 0.004 | -0.022 | -0.022 | 0 | 0.24 | 0.26 | 0.19 | 0.12 | 0.27 | 0.1 |
| 6 | 167182277 | rs481830 | -0.434 | -0.447 | -0.437 | 0.068 | 0.005 | 0.006 | 0.006 | 0.161 | 0.5 | 0.57 | 0.53 | 0.7 | 0.39 | 0.57 |
| 6 | 167183294 | rs16899429 | 0.404 | -0.313 | 0.383 | 0.539 | -0.008 | 0.021 | 0.021 | 3.317 | 0.23 | 0.31 | 0.27 | 0.32 | 0.18 | 0.26 |
| 6 | 167184881 | rs2345067 | -1.26 | -0.924 | 0.439 | 0.291 | 0.055 | 0.172 | 0.172 | 0.12 | 0.52 | 0.73 | 0.56 | 0.84 | 0.41 | 0.59 |
| 6 | 167185674 | rs6456123 | -1.416 | -0.901 | 0.578 | 0.271 | -0.003 | -0.001 | -0.001 | 0.004 | 0.71 | 0.7 | 0.67 | 0.69 | 0.64 | 0.66 |
| 6 | 167186024 | rs17590608 | 1.704 | -0.751 | 0.522 | 0.405 | -0.023 | 0.052 | 0.052 | 0 | 0.21 | 0.1 | 0.13 | 0 | 0.24 | 0.09 |
| 6 | 167187771 | rs655734 | -1.751 | -0.842 | 1.091 | 0.475 | 0.038 | 0.242 | 0.242 | 3.167 | 0.07 | 0.48 | 0.08 | 0.15 | 0.01 | 0.01 |
| 6 | 167188350 | rs2072638 | 0.11 | -0.893 | 0.781 | 0.349 | -0.001 | 0.006 | 0.006 | 0 | 0.36 | 0.38 | 0.26 | 0.5 | 0.34 | 0.4 |
| 6 | 167188463 | rs2072640 | -0.488 | -1.008 | 0.772 | 0.384 | -0.001 | -0.059 | -0.059 | 0 | 0.12 | 0.03 | 0.12 | 0.5 | 0.09 | 0.31 |
| 6 | 167188492 | rs2072641 | 0.132 | -0.781 | 0.781 | 0.315 | -0.007 | 0.014 | 0.014 | 0 | 0.35 | 0.38 | 0.24 | 0.54 | 0.25 | 0.4 |
| 6 | 167189689 | rs533810 | -0.347 | -1.035 | 0.285 | -0.204 | 0.069 | -0.046 | -0.046 | 1.393 | 0.51 | 0.89 | 0.43 | 0.47 | 0.32 | 0.48 |
| 6 | 167190753 | rs6919216 | 1.586 | -0.931 | 0.794 | -0.305 | -0.015 | -0.106 | -0.106 | 0 | 0.22 | 0.31 | 0.13 | 0 | 0.15 | 0.09 |
| 6 | 167192368 | rs9457204 | 1.012 | -0.949 | 0.774 | -0.314 | -0.033 | -0.145 | -0.145 | 1.814 | 0.25 | 0.4 | 0.13 | 0 | 0.16 | 0.09 |
| 6 | 167196032 | rs845641 | 1.768 | -0.968 | 0.501 | -0.373 | -0.125 | 0.037 | 0.037 | 0 | 0.29 | 0.03 | 0.29 | 0.19 | 0.38 | 0.34 |
| 6 | 167199572 | rs6913892 | 1.514 | -0.804 | 0.888 | -0.593 | -0.002 | -0.046 | -0.046 | 0 | 0.21 | 0.19 | 0.12 | 0 | 0.15 | 0.09 |
| 6 | 167200813 | rs1927430 | 0.153 | -0.853 | 1.002 | -0.563 | -0.019 | -0.076 | -0.076 | 0.175 | 0.4 | 0.47 | 0.27 | 0.06 | 0.29 | 0.22 |
| 6 | 167209669 | rs10946189 | 0.282 | -0.816 | 0.882 | -0.543 | -0.001 | -0.024 | -0.024 | 0 | 0.38 | 0.34 | 0.27 | 0.06 | 0.29 | 0.22 |
| 6 | 167211587 | rs707768 | 1.386 | -1.034 | 0.833 | -0.309 | -0.132 | -0.126 | -0.126 | 1.616 | 0.35 | 0.08 | 0.49 | 0.54 | 0.49 | 0.51 |
| 6 | 167211690 | rs13437290 | 1.212 | -0.776 | 1.139 | -0.332 | -0.003 | -0.048 | -0.048 | 0 | 0.23 | 0.21 | 0.12 | 0 | 0.16 | 0.11 |
| 6 | 167215384 | rs7754623 | 0.622 | -0.859 | 1.342 | -0.213 | -0.055 | -0.156 | -0.156 | 0.296 | 0.27 | 0.4 | 0.12 | 0 | 0.12 | 0.11 |
| 6 | 167216861 | rs10946191 | 0.651 | -0.614 | 0.984 | -0.014 | -0.02 | 0.001 | 0.001 | 7.229 | 0.14 | 0.03 | 0.14 | 0.06 | 0.13 | 0.14 |
| 6 | 167217236 | rs707766 | 0.812 | -0.61 | 1.192 | -0.056 | -0.008 | -0.073 | -0.073 | 0 | 0.67 | 0.52 | 0.71 | 0.93 | 0.69 | 0.75 |
| 6 | 167218762 | rs2178806 | 0.715 | -0.493 | 1.254 | 0.232 | -0.04 | 0.061 | 0.061 | 0 | 0.26 | 0.41 | 0.2 | 0.37 | 0.15 | 0.23 |
| 6 | 167219157 | rs845665 | 1.362 | 0.081 | 2.705 | 1.409 | -0.005 | 0.048 | 0.048 | 0.015 | 0.7 | 0.74 | 0.72 | 0.91 | 0.69 | 0.75 |
| 6 | 167223099 | rs845670 | 0.991 | -0.023 | 3.301 | 1.991 | -0.01 | 0.031 | 0.031 | 0 | 0.86 | 0.87 | 0.87 | 0.98 | 0.82 | 0.88 |
| 6 | 167223262 | rs845671 | 2.665 | 0.011 | 3.475 | 2.129 | 0.002 | -0.006 | -0.006 | 0 | 0.65 | 0.26 | 0.54 | 0.64 | 0.65 | 0.62 |
| 6 | 167223842 | rs845674 | 2.875 | -0.002 | 3.491 | 2.136 | 0 | 0.002 | 0.002 | 0.204 | 0.65 | 0.34 | 0.55 | 0.64 | 0.64 | 0.63 |
| 6 | 167223962 | rs4710106 | -2.402 | 0.015 | 3.426 | 2.143 | 0.041 | 0.151 | 0.151 | 1.047 | 0.13 | 0.37 | 0.22 | 0.27 | 0.14 | 0.19 |
| 6 | 167229078 | rs1099650 | 1.982 | 0.09 | 3.365 | 2.129 | -0.012 | -0.067 | -0.067 | 0.21 | 0.76 | 0.49 | 0.75 | 0.91 | 0.78 | 0.81 |
| 6 | 167230097 | rs1309150 | 1.696 | 0.09 | 3.363 | 2.129 | -0.004 | -0.03 | -0.03 | 0 | 0.78 | 0.63 | 0.77 | 0.91 | 0.78 | 0.81 |
| 6 | 167232907 | rs6902718 | 2.296 | 0.112 | 3.321 | 2.254 | -0.003 | 0.025 | 0.025 | 0.292 | 0.73 | 0.32 | 0.65 | 0.54 | 0.74 | 0.71 |
| 6 | 167234990 | rs1099661 | 1.52 | 0.193 | 3.053 | 2.091 | 0.002 | 0.022 | 0.022 | 0 | 0.83 | 0.73 | 0.77 | 0.61 | 0.79 | 0.78 |
| 6 | 167237848 | rs1103031 | 1.493 | 0.15 | 2.893 | 2.006 | 0.004 | -0.1 | -0.1 | 0 | 0.82 | 0.59 | 0.85 | 0.98 | 0.82 | 0.88 |
| 6 | 167238612 | rs1202621 | 2.004 | 0.147 | 2.565 | 2.044 | 0.03 | 0.018 | 0.018 | 0.594 | 0.71 | 0.49 | 0.55 | 0.31 | 0.65 | 0.58 |
| 6 | 167245665 | rs9355598 | 1.545 | 0.156 | 1.438 | 1.691 | 0.001 | 0 | 0 | 0.044 | 0.75 | 0.7 | 0.66 | 0.71 | 0.68 | 0.69 |
| 6 | 167256652 | rs9459774 | -1.436 | -0.755 | 0.672 | 1.442 | 0.003 | 0.069 | 0.069 | 0 | 0.17 | 0.27 | 0.21 | 0.36 | 0.2 | 0.19 |
| 6 | 167267919 | rs2881194 | 1.556 | -1.339 | 0.206 | 0.773 | 0.054 | -0.037 | -0.037 | 0.392 | 0.75 | 0.19 | 0.8 | 0.66 | 0.92 | 0.76 |
| 6 | 167269979 | rs6918384 | -1.686 | -1.411 | 1.05 | 0.601 | 0.002 | 0.091 | 0.091 | 0 | 0.17 | 0.59 | 0.26 | 0.33 | 0.22 | 0.12 |

| | | | | | | | | | | | | | | | | |
|---|---|---|---|---|---|---|---|---|---|---|---|---|---|---|---|---|
| 6 | 167270648 | rs4710124 | 1.235 | -1.44 | 1.52 | 0.622 | -0.044 | -0.084 | -0.084 | 1.04 | 0.81 | 0.66 | 0.92 | 0.95 | 0.93 | 0.83 |
| 6 | 167271139 | rs9356529 | 2.394 | -1.58 | 1.193 | 0.531 | 0.048 | -0.053 | -0.053 | 0 | 0.62 | 0.13 | 0.66 | 0.63 | 0.71 | 0.69 |
| 6 | 167271460 | rs9348198 | 2.393 | -1.349 | 1.324 | 0.163 | 0.041 | 0.165 | 0.165 | 0.876 | 0.65 | 0.16 | 0.76 | 0.91 | 0.73 | 0.74 |
| 6 | 167272280 | rs1536645 | 1.62 | -1.333 | 1.647 | 0.206 | -0.022 | -0.028 | -0.028 | 0 | 0.77 | 0.3 | 0.88 | 0.91 | 0.93 | 0.79 |
| 6 | 167274198 | rs7747434 | 1.712 | -1.312 | 1.538 | 0.271 | 0.005 | -0.006 | -0.006 | 0 | 0.77 | 0.27 | 0.81 | 0.65 | 0.91 | 0.75 |
| 6 | 167279331 | rs12216339 | 1.79 | -1.391 | 2.363 | 0.092 | -0.003 | -0.001 | -0.001 | 0.018 | 0.63 | 0.37 | 0.63 | 0.64 | 0.68 | 0.69 |
| 6 | 167282950 | rs4710128 | 0.957 | -1.339 | 1.944 | -0.757 | 0.002 | 0.001 | 0.001 | 0 | 0.79 | 0.8 | 0.8 | 0.98 | 0.76 | 0.83 |
| 6 | 167287772 | rs13204302 | 1.344 | -1.304 | 1.559 | -0.856 | 0.021 | 0.052 | 0.052 | 0.131 | 0.63 | 0.32 | 0.66 | 0.83 | 0.66 | 0.73 |
| 6 | 167289844 | rs7766409 | 1.209 | -1.275 | 1.566 | -0.85 | 0.008 | -0.01 | -0.01 | 2.304 | 0.66 | 0.37 | 0.71 | 0.91 | 0.68 | 0.79 |
| 6 | 167298751 | rs9366069 | 1.163 | -1.095 | 1.263 | -0.752 | 0.019 | 0.054 | 0.054 | 0.017 | 0.65 | 0.32 | 0.67 | 0.82 | 0.66 | 0.74 |
| 6 | 167311818 | rs9459795 | -1.661 | -0.526 | 1.71 | 0.003 | 0.083 | 0.109 | 0.109 | 0.408 | 0.05 | 0.32 | 0.08 | 0.05 | 0.03 | 0.05 |
| 6 | 167321104 | rs13216122 | -1.69 | -0.883 | 0.962 | -0.048 | -0.015 | 0 | 0 | 0 | 0.14 | 0.2 | 0.07 | 0.15 | 0.08 | 0.12 |
| 6 | 167321693 | rs943688 | 0.798 | -0.879 | 0.689 | -0.842 | -0.001 | -0.039 | -0.039 | 1.852 | 0.84 | 0.84 | 0.92 | 0.99 | 0.88 | 0.86 |
| 6 | 167326008 | rs6933508 | 2.182 | -0.668 | 0.325 | -0.831 | 0.018 | 0.106 | 0.106 | 0.99 | 0.66 | 0.15 | 0.75 | 0.84 | 0.71 | 0.73 |
| 6 | 167327101 | rs6913013 | 1.57 | -1.308 | 0.656 | -0.851 | 0.01 | 0.01 | 0.01 | 0.066 | 0.77 | 0.26 | 0.89 | 0.94 | 0.92 | 0.86 |
| 6 | 167333973 | rs9355606 | 1.722 | -1.26 | 0.393 | -0.884 | 0.071 | 0.092 | 0.092 | 0.629 | 0.7 | 0.2 | 0.83 | 0.9 | 0.84 | 0.81 |
| 6 | 167334382 | rs10806874 | 1.591 | -1.289 | 0.502 | -0.865 | 0.051 | 0.059 | 0.059 | 0.613 | 0.71 | 0.25 | 0.84 | 0.9 | 0.84 | 0.81 |
| 6 | 167341774 | rs9355607 | 1.248 | -0.88 | 0.525 | -0.762 | 0.001 | -0.006 | -0.006 | 0 | 0.74 | 0.31 | 0.85 | 0.9 | 0.84 | 0.83 |
| 6 | 167344583 | rs13213697 | -1.309 | -0.86 | 0.54 | -1.119 | -0.016 | -0.096 | -0.096 | 0.061 | 0.28 | 0.36 | 0.15 | 0.01 | 0.19 | 0.14 |
| 6 | 167348416 | rs10946197 | -0.841 | -0.142 | -0.134 | -1.689 | -0.026 | 0.038 | 0.038 | 0 | 0.25 | 0.15 | 0.18 | 0 | 0.27 | 0.17 |
| 6 | 167351120 | rs2757048 | -1.195 | -0.627 | 0.272 | -0.51 | -0.05 | 0.224 | 0.224 | 0 | 0.19 | 0.52 | 0.25 | 0.49 | 0.13 | 0.39 |
| 6 | 167352104 | rs3777722 | -0.269 | -0.427 | -0.282 | -0.107 | 0.003 | -0.037 | -0.037 | 0 | 0.1 | 0.03 | 0.23 | 0.41 | 0.11 | 0.33 |
| 6 | 167358233 | rs1212550 | -0.88 | -0.482 | -0.138 | -0.488 | 0 | 0.038 | 0.038 | 0.248 | 0.43 | 0.66 | 0.3 | 0.27 | 0.36 | 0.27 |
| 6 | 167359558 | rs2769343 | -1.395 | -0.384 | 0.533 | -0.084 | -0.001 | 0.017 | 0.017 | 0.007 | 0.34 | 0.46 | 0.54 | 0.52 | 0.31 | 0.46 |
| 6 | 167360389 | rs2236313 | -1.802 | -0.742 | 0.776 | -0.262 | 0.006 | 0.025 | 0.025 | 0 | 0.49 | 0.63 | 0.59 | 0.64 | 0.44 | 0.66 |
| 6 | 167360724 | rs1079145 | -0.406 | -0.764 | 0.422 | -0.051 | 0.032 | -0.074 | -0.074 | 0.974 | 0.13 | 0.03 | 0.34 | 0.43 | 0.1 | 0.32 |
| 6 | 167363948 | rs3798307 | -0.841 | -0.541 | 0.586 | -0.127 | 0.03 | -0.075 | -0.075 | 0 | 0.3 | 0.14 | 0.44 | 0.43 | 0.22 | 0.4 |
| 6 | 167364112 | rs12197030 | -0.349 | -0.43 | 1.054 | -0.146 | 0.058 | 0.016 | 0.016 | 0 | 0.08 | 0.22 | 0.09 | 0.1 | 0.13 | 0.06 |
| 6 | 167373708 | rs9366076 | -0.707 | -0.496 | 0.882 | -0.022 | -0.009 | -0.083 | -0.083 | 0 | 0.29 | 0.1 | 0.42 | 0.42 | 0.26 | 0.41 |
| 6 | 167383075 | rs9355610 | -1.36 | -0.21 | 1.184 | 0.033 | 0 | -0.002 | -0.002 | 0 | 0.42 | 0.41 | 0.51 | 0.54 | 0.37 | 0.63 |
| 6 | 167397614 | rs435359 | 0.914 | -0.179 | 1.122 | 0.013 | 0.003 | -0.001 | -0.001 | 0 | 0.9 | 0.86 | 0.91 | 0.91 | 0.88 | 0.92 |
| 6 | 167398071 | rs394522 | 2.108 | -0.25 | 1.291 | -0.006 | -0.012 | 0.036 | 0.036 | 0 | 0.47 | 0.22 | 0.38 | 0.37 | 0.51 | 0.29 |
| 6 | 167403873 | rs933243 | -1.578 | -0.241 | 1.137 | 0.014 | -0.006 | 0.002 | 0.002 | 0 | 0.44 | 0.52 | 0.52 | 0.54 | 0.37 | 0.63 |
| 6 | 167407584 | rs4709142 | -0.366 | -0.203 | 1.191 | 0.052 | 0.003 | 0.021 | 0.021 | 0 | 0.06 | 0.14 | 0.05 | 0.12 | 0.05 | 0.07 |
| 6 | 167419105 | rs9459836 | 2.184 | 0.237 | 0.801 | -0.072 | -0.038 | 0.059 | 0.059 | 0 | 0.43 | 0.1 | 0.33 | 0.27 | 0.43 | 0.26 |
| 6 | 167435325 | rs7749278 | 1.844 | 0.222 | 0.998 | -0.053 | 0.006 | 0.002 | 0.002 | 0 | 0.47 | 0.36 | 0.36 | 0.45 | 0.45 | 0.33 |
| 6 | 167437988 | rs2301436 | 1.906 | 0.228 | 1.002 | -0.042 | 0.004 | 0.014 | 0.014 | 0 | 0.47 | 0.36 | 0.36 | 0.36 | 0.45 | 0.32 |
| 6 | 167455139 | rs162297 | -0.435 | 0.18 | 1.023 | 0.04 | -0.034 | -0.103 | -0.103 | 2.231 | 0.32 | 0.11 | 0.46 | 0.47 | 0.34 | 0.42 |
| 6 | 167455629 | rs162298 | -1.001 | 0.052 | 0.942 | 0.215 | 0.003 | 0.032 | 0.032 | 0 | 0.33 | 0.78 | 0.4 | 0.26 | 0.28 | 0.24 |
| 6 | 167461501 | rs12529876 | 1.47 | 0.406 | 0.978 | 0.263 | 0.013 | 0.011 | 0.011 | 0 | 0.46 | 0.36 | 0.35 | 0.36 | 0.4 | 0.31 |
| 6 | 167492552 | rs7757618 | 1.044 | 0.603 | 0.858 | 0.287 | 0.018 | -0.005 | -0.005 | 0.798 | 0.88 | 0.79 | 0.88 | 0.88 | 0.85 | 0.9 |
| 6 | 167500562 | rs204295 | -0.936 | 0.453 | 0.874 | 0.282 | -0.001 | -0.001 | -0.001 | 0 | 0.47 | 0.54 | 0.54 | 0.55 | 0.49 | 0.64 |
| 6 | 167502638 | rs1331301 | -0.029 | 0.49 | 0.812 | 0.269 | -0.073 | -0.074 | -0.074 | 0.238 | 0.32 | 0.14 | 0.46 | 0.42 | 0.41 | 0.38 |
| 6 | 167505791 | rs6902119 | 0.655 | -0.099 | 1.501 | 0.4 | 0.003 | 0.003 | 0.003 | 0 | 0.54 | 0.44 | 0.4 | 0.37 | 0.44 | 0.5 |
| 6 | 167507658 | rs150115 | -1.349 | -0.105 | 1.675 | 0.439 | 0.061 | 0.158 | 0.158 | 0 | 0.08 | 0.35 | 0.06 | 0.2 | 0.05 | 0.1 |
| 6 | 167507676 | rs86757 | -1.156 | -0.11 | 1.676 | 0.405 | 0.069 | 0.07 | 0.07 | 0 | 0.08 | 0.32 | 0.05 | 0.1 | 0.05 | 0.07 |
| 6 | 167509833 | rs150113 | -0.772 | -0.078 | 1.738 | 0.418 | 0.073 | 0.091 | 0.091 | 0 | 0.07 | 0.27 | 0.05 | 0.11 | 0.05 | 0.07 |
| 6 | 167513998 | rs4710181 | 1.236 | -0.141 | 1.131 | 0.513 | -0.023 | -0.007 | -0.007 | 0.438 | 0.46 | 0.09 | 0.38 | 0.41 | 0.42 | 0.42 |
| 6 | 167515233 | rs4710184 | 2.012 | -0.153 | 1.235 | 0.474 | -0.052 | -0.022 | -0.022 | 0 | 0.86 | 0.3 | 0.93 | 0.9 | 0.97 | 0.86 |
| 6 | 167516458 | rs975822 | -0.303 | -0.159 | 1.217 | 0.501 | -0.026 | -0.037 | -0.037 | 1.746 | 0.39 | 0.21 | 0.54 | 0.44 | 0.55 | 0.44 |
| 6 | 167521624 | rs6905911 | 1.252 | -0.082 | 1.092 | 0.717 | 0 | -0.055 | -0.055 | 0 | 0.46 | 0.1 | 0.37 | 0.46 | 0.4 | 0.42 |
| 6 | 167522300 | rs6456156 | 0.404 | -0.126 | 1.244 | 0.765 | -0.013 | -0.02 | -0.02 | 1.639 | 0.56 | 0.34 | 0.42 | 0.49 | 0.49 | 0.56 |
| 6 | 167522386 | rs4710185 | 1.496 | -0.29 | 0.898 | 0.772 | -0.03 | 0.031 | 0.031 | 0 | 0.41 | 0.03 | 0.32 | 0.25 | 0.35 | 0.31 |
| 6 | 167523395 | rs6907666 | -0.989 | -0.02 | 1.228 | 0.749 | 0 | -0.001 | -0.001 | 0 | 0.51 | 0.55 | 0.61 | 0.56 | 0.61 | 0.61 |
| 6 | 167524743 | rs1556413 | -0.052 | 0.059 | 1.107 | 0.786 | -0.053 | -0.041 | -0.041 | 0.821 | 0.37 | 0.2 | 0.52 | 0.43 | 0.48 | 0.41 |
| 6 | 167526506 | rs1015077 | -1.429 | 0.141 | 0.993 | 0.81 | -0.019 | 0.042 | 0.042 | 0.185 | 0.13 | 0.31 | 0.07 | 0.21 | 0.09 | 0.21 |
| 6 | 167527296 | rs6908364 | -0.174 | 0.234 | 1.099 | 0.89 | 0.024 | 0.013 | 0.013 | 0 | 0.37 | 0.43 | 0.53 | 0.39 | 0.48 | 0.4 |
| 6 | 167532793 | rs3093024 | -1.004 | 0.079 | 0.149 | 0.678 | 0.038 | 0.041 | 0.041 | 0.837 | 0.51 | 0.68 | 0.63 | 0.57 | 0.58 | 0.62 |
| 6 | 167535914 | rs3093021 | -0.426 | 0.135 | 0.06 | 0.648 | -0.044 | 0.021 | 0.021 | 2.495 | 0.11 | 0.36 | 0.04 | 0.1 | 0.05 | 0.14 |
| 6 | 167537594 | rs1855025 | -0.339 | 0.974 | -0.955 | 0.915 | -0.052 | -0.036 | -0.036 | 0 | 0.36 | 0.18 | 0.54 | 0.37 | 0.48 | 0.4 |
| 6 | 167544709 | rs3798315 | -0.267 | 1.17 | -0.878 | 0.907 | -0.032 | -0.029 | -0.029 | 0.542 | 0.1 | 0.04 | 0.11 | 0.14 | 0.13 | 0.13 |
| 6 | 167548281 | rs3093012 | -0.047 | 0.601 | -0.926 | 0.659 | 0.009 | 0.016 | 0.016 | 0.245 | 0.47 | 0.37 | 0.62 | 0.74 | 0.58 | 0.53 |
| 6 | 167548607 | rs3093010 | 0.426 | 0.664 | -1.589 | 0.596 | -0.011 | 0.011 | 0.011 | 0 | 0.32 | 0.12 | 0.46 | 0.29 | 0.34 | 0.27 |
| 6 | 167549477 | rs3093009 | -0.45 | 0.041 | -1.051 | 0.086 | -0.008 | -0.032 | -0.032 | 0 | 0.23 | 0.59 | 0.21 | 0.13 | 0.21 | 0.17 |
| 6 | 167549536 | rs3093008 | -0.069 | -0.175 | -0.312 | -0.092 | 0.034 | 0.025 | 0.025 | 2.251 | 0.06 | 0.36 | 0.04 | 0 | 0.01 | 0 |
| 6 | 167549775 | rs3093007 | -0.07 | -0.096 | -0.314 | -0.068 | 0.001 | -0.012 | -0.012 | 0 | 0.14 | 0.18 | 0.12 | 0.05 | 0.18 | 0.12 |
| 6 | 167550042 | rs2071171 | 0.224 | 0.394 | -0.707 | 0.02 | 0 | 0.002 | 0.002 | 0 | 0.61 | 0.41 | 0.71 | 0.54 | 0.62 | 0.6 |
| 6 | 167554185 | rs3093001 | 1.124 | 0.522 | -0.105 | 0.602 | 0.006 | 0.012 | 0.012 | 0.786 | 0.51 | 0.17 | 0.62 | 0.58 | 0.51 | 0.56 |
| 6 | 167554197 | rs3093000 | 0.024 | 0.502 | 0.156 | 0.605 | 0 | -0.001 | -0.001 | 0 | 0.69 | 0.68 | 0.79 | 0.63 | 0.69 | 0.68 |

| | | | | | | | | | | | | | | | | |
|---|---|---|---|---|---|---|---|---|---|---|---|---|---|---|---|---|
| 6 | 167555155 | rs4710188 | 0.74 | 0.537 | 0.113 | 0.585 | 0.012 | -0.017 | -0.017 | 0.178 | 0.61 | 0.3 | 0.71 | 0.54 | 0.65 | 0.62 |
| 6 | 167558739 | rs879840 | 0.609 | 0.592 | -0.05 | 0.602 | 0 | 0 | 0 | 0 | 0.62 | 0.62 | 0.74 | 0.54 | 0.61 | 0.58 |
| 6 | 167566686 | rs9459893 | -0.934 | 0.507 | -0.085 | 0.749 | 0.019 | 0.072 | 0.072 | 0 | 0.06 | 0.21 | 0.18 | 0.11 | 0.03 | 0.06 |
| 6 | 167571911 | rs2345968 | -0.191 | 0.431 | -0.186 | 0.709 | -0.006 | 0.024 | 0.024 | 0 | 0.07 | 0.15 | 0.18 | 0.11 | 0.04 | 0.06 |
| 6 | 167572542 | rs4710191 | 1.421 | 0.505 | -0.728 | 0.638 | -0.032 | -0.048 | -0.048 | 0.56 | 0.45 | 0.1 | 0.47 | 0.49 | 0.43 | 0.47 |
| 6 | 167576698 | rs1999375 | -1.055 | 0.412 | -0.251 | 0.811 | 0.002 | -0.027 | -0.027 | 0 | 0.4 | 0.69 | 0.42 | 0.5 | 0.39 | 0.43 |
| 6 | 167578405 | rs723558 | -0.461 | 0.715 | -0.521 | 1.004 | -0.026 | -0.042 | -0.042 | 0.003 | 0.28 | 0.15 | 0.2 | 0.38 | 0.35 | 0.37 |
| 6 | 167601309 | rs2345969 | -1.074 | 0.786 | 0.525 | 0.954 | 0.018 | 0.019 | 0.019 | 1.268 | 0.19 | 0.57 | 0.27 | 0.25 | 0.2 | 0.16 |
| 6 | 167611757 | rs9347182 | -1.077 | 0.681 | 0.165 | 1.167 | -0.009 | 0.003 | 0.003 | 0 | 0.38 | 0.33 | 0.43 | 0.62 | 0.43 | 0.51 |
| 6 | 167615010 | rs2180713 | -0.983 | 0.7 | 0.336 | 1.414 | -0.007 | -0.015 | -0.015 | 0 | 0.37 | 0.31 | 0.31 | 0.48 | 0.43 | 0.48 |
| 6 | 167616209 | rs10946220 | 0.092 | 0.825 | 0.239 | 1.071 | -0.009 | 0.013 | 0.013 | 1.358 | 0.11 | 0.09 | 0.06 | 0.09 | 0.12 | 0.08 |
| 6 | 167622828 | rs6904588 | 0.732 | 0.796 | 0.081 | 0.217 | -0.006 | 0.002 | 0.002 | 0 | 0.24 | 0.19 | 0.2 | 0 | 0.24 | 0.06 |
| 6 | 167624079 | rs9364896 | 1.023 | 0.623 | 0.288 | 0.497 | -0.018 | -0.023 | -0.023 | 0 | 0.57 | 0.25 | 0.56 | 0.56 | 0.53 | 0.54 |
| 6 | 167634544 | rs914331 | 0.971 | 0.709 | 0.478 | -0.058 | -0.007 | 0.011 | 0.011 | 0 | 0.6 | 0.64 | 0.7 | 0.71 | 0.53 | 0.57 |
| 6 | 167640643 | rs2094946 | 0.104 | 0.854 | 0.376 | -0.051 | 0.001 | -0.006 | -0.006 | 0 | 0.81 | 0.79 | 0.88 | 0.86 | 0.78 | 0.81 |
| 6 | 167644277 | rs7761658 | -0.198 | 0.601 | 0.683 | 0.234 | -0.002 | 0.001 | 0.001 | 0 | 0.21 | 0.19 | 0.18 | 0.15 | 0.26 | 0.24 |
| 6 | 167646739 | rs2018748 | 0.327 | 0.59 | 0.679 | 0.212 | 0.001 | -0.009 | -0.009 | 0 | 0.77 | 0.71 | 0.81 | 0.85 | 0.74 | 0.76 |
| 6 | 167651236 | rs6925016 | 0.042 | 0.302 | 0.218 | 0.095 | -0.022 | 0.058 | 0.058 | 0 | 0.4 | 0.3 | 0.32 | 0.17 | 0.45 | 0.3 |
| 6 | 167663244 | rs7749161 | -0.38 | 0.195 | -0.096 | 0.554 | -0.002 | -0.036 | -0.036 | 0 | 0.17 | 0.07 | 0.13 | 0.24 | 0.18 | 0.27 |
| 6 | 167663636 | rs7773576 | -0.242 | 0.132 | 0.394 | 0.17 | 0.006 | -0.095 | -0.095 | 0.108 | 0.79 | 0.53 | 0.86 | 0.9 | 0.84 | 0.85 |
| 6 | 167664918 | rs4709159 | 1.331 | 0.132 | -0.082 | 0.588 | -0.023 | -0.056 | -0.056 | 2.082 | 0.32 | 0.17 | 0.39 | 0.39 | 0.37 | 0.37 |
| 6 | 167665326 | rs6903113 | -0.326 | 0.151 | -0.06 | 0.725 | 0.001 | -0.035 | -0.035 | 0.552 | 0.18 | 0.07 | 0.13 | 0.25 | 0.18 | 0.28 |
| 6 | 167685688 | rs2092091 | -0.826 | 0.17 | 0.314 | 0.751 | -0.001 | -0.001 | -0.001 | 1.079 | 0.39 | 0.4 | 0.28 | 0.37 | 0.37 | 0.42 |
| 6 | 167687856 | rs1854852 | 0.988 | 0.209 | 0.063 | 0.716 | -0.009 | 0.033 | 0.033 | 0 | 0.26 | 0.11 | 0.23 | 0.17 | 0.28 | 0.15 |
| 6 | 167691225 | rs172283 | -1.618 | 0.156 | 0.144 | 0.77 | -0.034 | -0.084 | -0.084 | 0 | 0.69 | 0.88 | 0.63 | 0.62 | 0.67 | 0.63 |
| 6 | 167696948 | rs3010558 | -0.214 | 0.162 | 0.04 | 0.695 | -0.017 | -0.07 | -0.07 | 0.084 | 0.19 | 0.04 | 0.13 | 0.26 | 0.19 | 0.28 |
| 6 | 167698769 | rs206990 | 0.867 | 0.171 | 0.282 | 0.696 | -0.003 | 0.013 | 0.013 | 0 | 0.27 | 0.2 | 0.25 | 0.17 | 0.29 | 0.15 |
| 6 | 167699704 | rs2981956 | -0.74 | 0.193 | 0.276 | 0.704 | -0.004 | -0.007 | -0.007 | 0 | 0.4 | 0.48 | 0.29 | 0.39 | 0.37 | 0.42 |
| 6 | 167704244 | rs2989555 | -1.227 | 0.29 | 0.153 | 0.749 | 0.002 | 0.024 | 0.024 | 0.315 | 0.66 | 0.58 | 0.51 | 0.54 | 0.65 | 0.57 |
| 6 | 167704944 | rs2076005 | -1.274 | 0.284 | 0.149 | 0.746 | 0.002 | 0.026 | 0.026 | 0.433 | 0.66 | 0.58 | 0.51 | 0.54 | 0.65 | 0.57 |
| 6 | 167706981 | rs2981961 | -1.65 | 0.002 | 0.273 | 1.879 | 0.009 | 0.016 | 0.016 | 0.079 | 0.43 | 0.78 | 0.42 | 0.69 | 0.38 | 0.5 |
| 6 | 167707727 | rs2076006 | -0.225 | 0.348 | -0.284 | 1.691 | 0.011 | 0.066 | 0.066 | 1.033 | 0.2 | 0.38 | 0.17 | 0.36 | 0.18 | 0.17 |
| 6 | 167709501 | rs2076009 | -0.167 | 0.402 | -0.257 | 1.741 | -0.002 | 0.05 | 0.05 | 0.723 | 0.11 | 0.14 | 0.13 | 0.28 | 0.12 | 0.21 |
| 6 | 167709702 | rs2235197 | 0.073 | 0.368 | -0.419 | 1.644 | -0.007 | -0.033 | -0.033 | 0 | 0.1 | 0.04 | 0.1 | 0.14 | 0.12 | 0.17 |
| 6 | 167710324 | rs2981962 | -0.455 | 0.437 | -0.157 | 1.308 | 0.007 | -0.001 | -0.001 | 0 | 0.2 | 0.32 | 0.15 | 0.22 | 0.18 | 0.14 |
| 6 | 167710832 | rs3010544 | -0.421 | 0.403 | -0.261 | 1.328 | 0.005 | 0.001 | 0.001 | 0.715 | 0.19 | 0.3 | 0.15 | 0.22 | 0.18 | 0.14 |
| 6 | 167713357 | rs6923070 | -1.006 | 0.444 | -0.635 | 1.295 | -0.002 | 0.011 | 0.011 | 0 | 0.1 | 0.1 | 0.06 | 0.22 | 0.08 | 0.12 |
| 6 | 167715424 | rs13215796 | -0.653 | -0.074 | -0.609 | 1.119 | 0.008 | 0.002 | 0.002 | 0 | 0.34 | 0.29 | 0.31 | 0.29 | 0.28 | 0.39 |
| 6 | 167716043 | rs2981973 | 0.404 | 0.291 | -0.614 | 1.085 | -0.001 | 0 | 0 | 0 | 0.38 | 0.29 | 0.36 | 0.45 | 0.36 | 0.44 |
| 6 | 167718396 | rs13198547 | -0.044 | 0.141 | -0.606 | 0.569 | 0.016 | 0.02 | 0.02 | 0 | 0.33 | 0.25 | 0.21 | 0.22 | 0.28 | 0.33 |
| 6 | 167718445 | rs13198321 | -0.001 | 0.127 | -0.465 | 0.411 | 0.011 | 0.022 | 0.022 | 0 | 0.33 | 0.26 | 0.21 | 0.13 | 0.27 | 0.31 |
| 6 | 167721545 | rs9457296 | -0.469 | 0.144 | 0.493 | 0.363 | 0.006 | -0.032 | -0.032 | 0 | 0.06 | 0.1 | 0.04 | 0 | 0.06 | 0.03 |
| 6 | 167737138 | rs7754688 | -0.655 | -0.043 | 1.163 | 0.427 | 0.001 | -0.002 | -0.002 | 0 | 0.72 | 0.7 | 0.68 | 0.83 | 0.64 | 0.81 |
| 6 | 167737682 | rs3814080 | -0.175 | -0.115 | 1.23 | 0.418 | 0.031 | 0.037 | 0.037 | 0 | 0.42 | 0.68 | 0.41 | 0.31 | 0.36 | 0.43 |
| 6 | 167739769 | rs9456267 | -1.388 | -0.043 | 1.324 | 0.354 | 0.006 | -0.053 | -0.053 | 0.076 | 0.13 | 0.55 | 0.2 | 0.03 | 0.12 | 0.06 |
| 6 | 167740363 | rs4709169 | -0.651 | 0.024 | 0.934 | 0.335 | -0.113 | 0.01 | 0.01 | 0 | 0.73 | 0.96 | 0.7 | 0.83 | 0.64 | 0.81 |
| 6 | 167743729 | rs9347206 | 1.509 | 0.391 | 1.338 | 0.484 | 0.019 | -0.072 | -0.072 | 0.259 | 0.86 | 0.8 | 0.89 | 1 | 0.83 | 0.91 |
| 6 | 167750150 | rs2981977 | 0.811 | 0.578 | 0.965 | 0.409 | -0.003 | 0.004 | 0.004 | 0.231 | 0.66 | 0.42 | 0.56 | 0.77 | 0.6 | 0.76 |
| 6 | 167760702 | rs9457305 | -0.135 | 0.903 | 1.07 | 0.479 | -0.002 | 0.001 | 0.001 | 0 | 0.54 | 0.43 | 0.56 | 0.39 | 0.58 | 0.58 |
| 6 | 167768161 | rs9457309 | -0.711 | 0.766 | 1.535 | 0.645 | 0.078 | -0.028 | -0.028 | 2.997 | 0.08 | 0.3 | 0.07 | 0 | 0.1 | 0.05 |
| 6 | 167770736 | rs2981988 | 0.804 | 0.799 | 0.539 | 0.642 | 0.022 | 0.03 | 0.03 | 0 | 0.86 | 0.45 | 0.73 | 0.86 | 0.84 | 0.89 |
| 6 | 168348424 | rs669861 | -0.919 | 0.05 | 1.412 | 0.707 | -0.02 | 0.039 | 0.039 | 0 | 0.17 | 0.44 | 0.23 | 0.22 | 0.11 | 0.26 |
| 6 | 168349206 | rs12665666 | -0.278 | -0.222 | 0.677 | -0.062 | -0.007 | 0.008 | 0.008 | 0 | 0.12 | 0.25 | 0.22 | 0.2 | 0.1 | 0.25 |
| 6 | 168364813 | rs1418 | -0.603 | -0.138 | 0.146 | 0.103 | 0.004 | 0.01 | 0.01 | 0 | 0.43 | 0.42 | 0.56 | 0.8 | 0.41 | 0.64 |
| 6 | 168378848 | rs227249 | -0.469 | 0.436 | -0.241 | 0.167 | -0.009 | 0.045 | 0.045 | 0 | 0.44 | 0.33 | 0.54 | 0.76 | 0.55 | 0.54 |
| 6 | 168379782 | rs9346516 | 0.639 | 0.397 | 0.299 | 0.524 | -0.013 | -0.002 | -0.002 | 0.032 | 0.75 | 0.6 | 0.67 | 0.74 | 0.74 | 0.68 |
| 6 | 168385774 | rs9455927 | -1.093 | 0.594 | 0.29 | 1.339 | -0.015 | -0.027 | -0.027 | 0 | 0.16 | 0.09 | 0.32 | 0.38 | 0.26 | 0.38 |
| 6 | 168386575 | rs6900844 | 0.055 | 0.223 | -0.095 | 0.929 | 0.004 | -0.007 | -0.007 | 0.04 | 0.28 | 0.24 | 0.25 | 0.33 | 0.24 | 0.21 |
| 6 | 168388213 | rs2843011 | -0.023 | 0.171 | -0.575 | 0.421 | 0.004 | -0.038 | -0.038 | 0 | 0.8 | 0.93 | 0.82 | 0.74 | 0.8 | 0.76 |
| 6 | 168389324 | rs7741791 | 0.305 | 0.225 | 0.075 | -0.362 | -0.008 | -0.113 | -0.113 | 0.002 | 0.22 | 0.4 | 0.14 | 0.03 | 0.2 | 0.07 |
| 6 | 168389858 | rs4708617 | -0.207 | 0.135 | -0.052 | -0.033 | 0.011 | 0.054 | 0.054 | 0.435 | 0.11 | 0.2 | 0.09 | 0.22 | 0.09 | 0.14 |
| 6 | 168390030 | rs2516794 | 1.089 | -0.373 | -0.635 | -0.448 | -0.05 | -0.044 | -0.044 | 1.194 | 0.41 | 0.23 | 0.47 | 0.48 | 0.47 | 0.43 |
| 6 | 168390136 | rs2516795 | 0.28 | -0.178 | -0.6 | -0.15 | -0.014 | -0.049 | -0.049 | 0 | 0.77 | 0.91 | 0.77 | 0.6 | 0.76 | 0.77 |
| 6 | 168393776 | rs3734887 | -0.408 | -0.392 | -0.393 | -0.335 | -0.001 | -0.033 | -0.033 | 0 | 0.24 | 0.09 | 0.32 | 0.36 | 0.23 | 0.19 |
| 6 | 168394420 | rs2171982 | -1.053 | -0.604 | 0.029 | -0.615 | -0.009 | -0.05 | -0.05 | 0 | 0.81 | 0.77 | 0.89 | 0.97 | 0.85 | 0.92 |
| 6 | 168394717 | rs3752610 | 0.391 | -0.618 | -0.027 | -0.645 | 0.002 | 0.023 | 0.023 | 0 | 0.14 | 0.15 | 0.19 | 0.44 | 0.12 | 0.35 |
| 6 | 168394923 | rs3752611 | -0.306 | -0.758 | 0.334 | -0.758 | -0.017 | 0.049 | 0.049 | 0 | 0.22 | 0.28 | 0.21 | 0.44 | 0.14 | 0.39 |
| 6 | 168395957 | rs3778667 | -0.547 | -0.706 | 0.284 | -0.766 | 0.008 | -0.004 | -0.004 | 0.054 | 0.31 | 0.2 | 0.34 | 0.37 | 0.25 | 0.23 |
| 6 | 168399960 | rs9455929 | 1.647 | -0.531 | 0.332 | -0.782 | 0.022 | -0.072 | -0.072 | 0 | 0.29 | 0.39 | 0.31 | 0.06 | 0.34 | 0.12 |

| | | | | | | | | | | | | | | | |
|---|---|---|---|---|---|---|---|---|---|---|---|---|---|---|---|
| 6 | 168401923 | rs1812028 | -0.793 | -0.604 | 0.24 | -0.746 | 0.001 | -0.002 | -0.002 | 0 | 0.37 | 0.4 | 0.32 | 0.52 | 0.28 | 0.44 |
| 6 | 168404976 | rs2516808 | 0.405 | -0.5 | 0.464 | -0.745 | -0.002 | -0.018 | -0.018 | 0.839 | 0.57 | 0.48 | 0.73 | 0.78 | 0.57 | 0.64 |
| 6 | 168410351 | rs12194885 | 0.016 | 0.03 | 0.555 | -0.261 | 0.033 | 0.023 | 0.023 | 0.094 | 0.09 | 0.2 | 0.08 | 0.12 | 0.18 | 0.22 |
| 6 | 168410722 | rs2843003 | 1.334 | -0.136 | 0.615 | -0.585 | 0.027 | -0.088 | -0.088 | 1.149 | 0.28 | 0.39 | 0.28 | 0.07 | 0.34 | 0.14 |
| 6 | 168413195 | rs1858675 | -0.639 | -0.212 | -0.181 | -0.517 | -0.012 | -0.018 | -0.018 | 0 | 0.3 | 0.23 | 0.38 | 0.39 | 0.38 | 0.44 |
| 6 | 168413457 | rs227242 | 1.949 | -0.239 | -0.211 | -0.549 | -0.067 | 0.1 | 0.1 | 0.06 | 0.25 | 0.08 | 0.25 | 0.07 | 0.31 | 0.13 |
| 6 | 168415372 | rs9364385 | -0.771 | -0.184 | -0.127 | -0.421 | -0.005 | -0.009 | -0.009 | 0 | 0.31 | 0.26 | 0.38 | 0.39 | 0.38 | 0.44 |
| 6 | 168417597 | rs12198634 | 0.388 | -0.15 | -0.136 | -0.462 | 0.02 | 0.025 | 0.025 | 0 | 0.09 | 0.19 | 0.07 | 0.12 | 0.15 | 0.22 |
| 6 | 168417986 | rs3807061 | -0.853 | -0.177 | -0.194 | -0.345 | -0.006 | -0.008 | -0.008 | 0 | 0.31 | 0.27 | 0.39 | 0.42 | 0.41 | 0.44 |
| 6 | 168418669 | rs7744329 | -0.209 | -0.074 | -0.554 | -0.502 | -0.029 | -0.049 | -0.049 | 0 | 0.22 | 0.06 | 0.32 | 0.3 | 0.24 | 0.22 |
| 6 | 168420258 | rs227235 | 1.258 | -0.059 | -0.241 | -0.638 | -0.011 | 0.017 | 0.017 | 0 | 0.28 | 0.26 | 0.29 | 0.17 | 0.32 | 0.14 |
| 6 | 168427146 | rs9455933 | -1.078 | -0.309 | -0.339 | -0.682 | 0.001 | -0.037 | -0.037 | 0 | 0.47 | 0.54 | 0.5 | 0.89 | 0.45 | 0.77 |
| 6 | 168432093 | rs9364386 | -0.562 | 0.264 | -0.55 | 0.139 | -0.024 | -0.049 | -0.049 | 0 | 0.18 | 0.05 | 0.16 | 0.3 | 0.17 | 0.26 |
| 6 | 168432236 | rs9355156 | -0.562 | 0.264 | -0.55 | 0.139 | -0.024 | -0.049 | -0.049 | 1.865 | 0.18 | 0.05 | 0.16 | 0.3 | 0.17 | 0.26 |
| 6 | 168440640 | rs2073632 | 0.164 | 0.272 | -0.448 | 0.314 | -0.012 | -0.038 | -0.038 | 0 | 0.09 | 0.02 | 0.08 | 0.22 | 0.08 | 0.21 |
| 6 | 168442475 | rs4708629 | 0.655 | 0.384 | -0.29 | 0.192 | -0.001 | -0.009 | -0.009 | 0 | 0.56 | 0.52 | 0.57 | 0.59 | 0.58 | 0.6 |
| 6 | 168442765 | rs2073634 | -0.203 | 0.545 | -0.136 | 1.394 | -0.026 | -0.051 | -0.051 | 0.135 | 0.08 | 0.01 | 0.09 | 0.21 | 0.08 | 0.21 |
| 6 | 168450159 | rs225762 | -0.987 | 0.513 | -0.175 | 0.949 | -0.004 | 0.062 | 0.062 | 3.574 | 0.28 | 0.48 | 0.32 | 0.49 | 0.3 | 0.46 |
| 6 | 168455806 | rs4708631 | 0.591 | 0.569 | -0.575 | 0.613 | 0.001 | 0 | 0 | 0 | 0.86 | 0.84 | 0.82 | 0.85 | 0.89 | 0.93 |
| 6 | 168457279 | rs3734897 | 0.649 | 0.778 | -0.608 | 0.712 | -0.006 | -0.002 | -0.002 | 0 | 0.84 | 0.77 | 0.75 | 0.83 | 0.87 | 0.91 |
| 6 | 168457801 | rs3734898 | 0.575 | 0.777 | -0.617 | 0.709 | -0.002 | -0.001 | -0.001 | 0 | 0.86 | 0.82 | 0.82 | 0.85 | 0.89 | 0.93 |
| 6 | 168457966 | rs3734899 | 1.32 | 1.257 | -1.339 | 0.7 | -0.055 | -0.021 | -0.021 | 0.151 | 0.48 | 0.11 | 0.35 | 0.49 | 0.49 | 0.62 |
| 6 | 168458074 | rs3734900 | 1.205 | 1.42 | -1.204 | 0.785 | -0.016 | -0.045 | -0.045 | 0.178 | 0.53 | 0.15 | 0.43 | 0.53 | 0.55 | 0.71 |
| 6 | 168462626 | rs12197610 | 0.059 | 1.35 | -0.551 | 0.718 | 0.08 | -0.083 | -0.083 | 1.975 | 0.17 | 0.45 | 0.18 | 0.04 | 0.2 | 0.11 |
| 6 | 168463153 | rs3734901 | 0.352 | 1.262 | -0.837 | 0.882 | 0.006 | -0.038 | -0.038 | 0 | 0.07 | 0 | 0.02 | 0.16 | 0.06 | 0.04 |
| 6 | 168466504 | rs9295041 | 0.37 | 1.278 | -0.73 | 0.833 | 0 | -0.001 | -0.001 | 0 | 0.82 | 0.76 | 0.74 | 0.79 | 0.83 | 0.84 |
| 6 | 168468186 | rs6906979 | -0.175 | 1.284 | -0.788 | 0.82 | 0.006 | -0.001 | -0.001 | 0 | 0.86 | 0.88 | 0.82 | 0.84 | 0.89 | 0.93 |
| 6 | 168472299 | rs1490398 | 1.785 | 1.299 | -1 | 0.834 | -0.036 | 0.001 | 0.001 | 1.761 | 0.49 | 0.09 | 0.38 | 0.48 | 0.51 | 0.61 |
| 6 | 168474309 | rs1565444 | -0.381 | 0.954 | -0.827 | 0.786 | 0.004 | 0 | 0 | 0 | 0.85 | 0.84 | 0.81 | 0.84 | 0.89 | 0.92 |
| 6 | 168474949 | rs902391 | 0.886 | 0.944 | -0.796 | 0.786 | 0 | -0.001 | -0.001 | 0 | 0.76 | 0.74 | 0.72 | 0.71 | 0.8 | 0.79 |
| 6 | 168484085 | rs4708431 | -0.862 | 1.422 | -0.674 | 0.495 | 0.055 | -0.038 | -0.038 | 0 | 0.29 | 0.57 | 0.35 | 0.17 | 0.41 | 0.26 |
| 6 | 168486186 | rs4708639 | -0.288 | 1.283 | -0.813 | 0.589 | 0 | 0 | 0 | 0 | 0.86 | 0.82 | 0.84 | 0.86 | 0.89 | 0.93 |
| 6 | 168486306 | rs10455755 | 1.045 | 1.803 | -0.732 | 0.502 | 0 | 0.011 | 0.011 | 0.18 | 0.71 | 0.66 | 0.63 | 0.57 | 0.72 | 0.73 |
| 6 | 168488562 | rs9355052 | 1.615 | 1.418 | -1.375 | 0.359 | 0.011 | -0.052 | -0.052 | 0.672 | 0.43 | 0.09 | 0.29 | 0.5 | 0.36 | 0.57 |
| 6 | 168490016 | rs6907103 | 0.682 | 1.593 | -1.182 | 0.342 | -0.009 | 0.001 | 0.001 | 0 | 0.61 | 0.33 | 0.51 | 0.63 | 0.57 | 0.76 |
| 6 | 168490568 | rs9364220 | 1.438 | 1.662 | -1.468 | 0.22 | 0.06 | -0.044 | -0.044 | 0 | 0.46 | 0.15 | 0.38 | 0.54 | 0.37 | 0.57 |
| 6 | 168491848 | rs902385 | 0.516 | 1.476 | -1.943 | 0.022 | 0.003 | -0.004 | -0.004 | 0.487 | 0.59 | 0.63 | 0.71 | 0.6 | 0.64 | 0.7 |
| 6 | 168492160 | rs1873337 | 0.143 | 1.432 | -2.143 | -0.014 | 0.038 | -0.068 | -0.068 | 0.694 | 0.22 | 0.6 | 0.31 | 0.06 | 0.32 | 0.14 |
| 6 | 168492273 | rs6455482 | 0.104 | 0.993 | -2.529 | 0.386 | 0.002 | -0.024 | -0.024 | 0 | 0.77 | 0.85 | 0.81 | 0.71 | 0.78 | 0.84 |
| 6 | 168492963 | rs9346650 | 0.788 | 0.699 | -2.424 | 0.539 | 0.03 | -0.102 | -0.102 | 0 | 0.38 | 0.08 | 0.42 | 0.55 | 0.33 | 0.58 |
| 6 | 168499925 | rs9455952 | 0.171 | 0.846 | -1.665 | 0.822 | 0 | -0.043 | -0.043 | 1.884 | 0.84 | 0.94 | 0.9 | 0.8 | 0.86 | 0.92 |
| 6 | 168506075 | rs902382 | -0.293 | 1.407 | -1.509 | 0.506 | -0.025 | 0.023 | 0.023 | 0 | 0.1 | 0.04 | 0.15 | 0.03 | 0.16 | 0.1 |
| 6 | 168511016 | rs1473500 | -0.805 | 0.785 | -1.012 | 0.184 | -0.03 | -0.038 | -0.038 | 0 | 0.1 | 0.62 | 0.12 | 0.05 | 0.02 | 0.05 |
| 6 | 168511079 | rs12213265 | 0.447 | 0.37 | -1.802 | 0.732 | 0.004 | -0.124 | -0.124 | 0 | 0.19 | 0.02 | 0.26 | 0.37 | 0.15 | 0.32 |
| 6 | 168513790 | rs4708647 | -1.28 | 1.179 | -2.14 | 0.467 | 0.006 | 0.008 | 0.008 | 0 | 0.54 | 0.87 | 0.72 | 0.58 | 0.59 | 0.65 |
| 6 | 168515674 | rs4708435 | -1.19 | 1.161 | -2.099 | 0.457 | 0.026 | 0.027 | 0.027 | 0 | 0.52 | 0.81 | 0.71 | 0.58 | 0.59 | 0.66 |
| 6 | 168517579 | rs4708652 | -0.501 | 1.228 | -2.099 | 0.445 | 0.02 | -0.024 | -0.024 | 0.191 | 0.4 | 0.71 | 0.52 | 0.44 | 0.34 | 0.47 |
| 6 | 168519224 | rs9355166 | -0.194 | 1.665 | -1.618 | 0.062 | -0.003 | 0.001 | 0.001 | 0 | 0.13 | 0.11 | 0.19 | 0.13 | 0.25 | 0.18 |
| 6 | 168526240 | rs2171981 | -0.854 | 1.519 | -1.409 | 0.011 | 0.019 | -0.03 | -0.03 | 0 | 0.25 | 0.66 | 0.36 | 0.14 | 0.39 | 0.28 |
| 6 | 168526858 | rs4708660 | -0.75 | 0.885 | -1.649 | 0.612 | 0.01 | 0.009 | 0.009 | 0 | 0.52 | 0.84 | 0.71 | 0.46 | 0.57 | 0.68 |
| 6 | 168531451 | rs9455576 | -0.563 | 1.244 | -0.5 | 0.177 | 0.014 | -0.014 | -0.014 | 0 | 0.2 | 0.27 | 0.26 | 0.12 | 0.27 | 0.26 |
| 6 | 168533451 | rs1565441 | -0.31 | 1.008 | -0.542 | 0.155 | -0.003 | 0.002 | 0.002 | 0 | 0.42 | 0.6 | 0.48 | 0.37 | 0.45 | 0.36 |
| 6 | 168533648 | rs6937877 | 0.524 | 1.085 | -0.378 | 0.202 | -0.004 | -0.012 | -0.012 | 0 | 0.24 | 0.49 | 0.2 | 0.25 | 0.17 | 0.09 |
| 6 | 168533787 | rs746800 | -0.212 | 0.405 | -0.492 | 0.424 | -0.031 | -0.048 | -0.048 | 0 | 0.61 | 0.83 | 0.56 | 0.54 | 0.54 | 0.53 |
| 6 | 168535138 | rs6931528 | -0.14 | -0.12 | -0.351 | -0.014 | -0.007 | 0.015 | 0.015 | 0 | 0.85 | 0.95 | 0.9 | 0.88 | 0.82 | 0.87 |
| 6 | 168538682 | rs12215821 | 0.198 | 0.084 | -0.336 | -0.163 | -0.017 | 0.023 | 0.023 | 0 | 0.1 | 0.06 | 0.15 | 0.01 | 0.14 | 0.16 |
| 6 | 168542557 | rs1602283 | 0.27 | -0.509 | 0.247 | 0.214 | 0.002 | 0.002 | 0.002 | 0 | 0.84 | 0.83 | 0.87 | 0.85 | 0.82 | 0.86 |
| 6 | 168542900 | rs4708672 | 1.261 | -0.81 | 0.525 | 0.143 | 0.057 | 0.036 | 0.036 | 1.187 | 0.61 | 0.25 | 0.63 | 0.66 | 0.7 | 0.68 |
| 6 | 168543303 | rs6939959 | -0.172 | -0.588 | 0.357 | 0.218 | -0.047 | -0.021 | -0.021 | 0 | 0.19 | 0.33 | 0.13 | 0.14 | 0.11 | 0.12 |
| 6 | 168546852 | rs9355170 | 0.858 | -0.5 | 0.229 | 0.255 | -0.028 | -0.016 | -0.016 | 0.323 | 0.65 | 0.45 | 0.74 | 0.71 | 0.71 | 0.74 |
| 6 | 168548496 | rs980909 | -0.285 | -0.149 | -0.469 | 0.635 | -0.002 | -0.006 | -0.006 | 0 | 0.93 | 0.95 | 0.96 | 0.88 | 0.94 | 0.98 |
| 6 | 168548525 | rs6916885 | 0.017 | -0.093 | -0.346 | 0.598 | -0.012 | 0.005 | 0.005 | 0 | 0.76 | 0.7 | 0.82 | 0.73 | 0.83 | 0.85 |
| 6 | 168549899 | rs6933250 | -0.109 | 0.285 | -0.169 | 0.499 | -0.005 | -0.014 | -0.014 | 0 | 0.33 | 0.55 | 0.17 | 0.22 | 0.26 | 0.26 |
| 6 | 168549995 | rs6927982 | -0.245 | 0.342 | -0.813 | 0.385 | 0.022 | 0.146 | 0.146 | 0.12 | 0.43 | 0.85 | 0.26 | 0.22 | 0.37 | 0.38 |
| 6 | 168550184 | rs9364407 | -1.242 | 0.594 | 0.188 | 1.279 | 0.01 | 0.161 | 0.161 | 0 | 0.09 | 0.23 | 0.13 | 0.31 | 0.1 | 0.08 |
| 6 | 168552065 | rs12214678 | -0.453 | 1.013 | -0.299 | 0.218 | -0.009 | 0.008 | 0.008 | 0 | 0.17 | 0.1 | 0.15 | 0.01 | 0.2 | 0.14 |
| 6 | 168554222 | rs12199628 | -1.171 | 0.777 | 0.31 | 0.034 | 0.012 | -0.071 | -0.071 | 0 | 0.23 | 0.29 | 0.23 | 0.04 | 0.28 | 0.19 |
| 6 | 168554421 | rs10945476 | -1.183 | 0.772 | 0.34 | 0.031 | 0.013 | -0.075 | -0.075 | 0 | 0.24 | 0.29 | 0.28 | 0.05 | 0.28 | 0.19 |
| 6 | 168555227 | rs4708676 | -1.074 | 0.75 | 0.042 | 0.138 | 0.002 | 0.001 | 0.001 | 0.131 | 0.52 | 0.53 | 0.54 | 0.46 | 0.64 | 0.53 |

| | | | | | | | | | | | | | | | | |
|---|---|---|---|---|---|---|---|---|---|---|---|---|---|---|---|---|
| 6 | 168558846 | rs1809698 | 0.019 | 0.625 | 0.394 | 0.127 | 0.023 | -0.101 | -0.101 | 0 | 0.2 | 0.35 | 0.18 | 0.01 | 0.21 | 0.15 |
| 6 | 168559038 | rs1871431 | -0.952 | 0.547 | 0.615 | 0.108 | 0.022 | 0.029 | 0.029 | 0 | 0.49 | 0.7 | 0.49 | 0.44 | 0.58 | 0.49 |
| 6 | 168564920 | rs9346527 | -0.243 | 0.456 | 0.424 | 0.473 | -0.003 | -0.01 | -0.01 | 0 | 0.27 | 0.26 | 0.26 | 0.4 | 0.34 | 0.29 |
| 6 | 168571947 | rs9364414 | 0.116 | 0.485 | 0.074 | 0.552 | -0.054 | -0.075 | -0.075 | 0 | 0.22 | 0.12 | 0.21 | 0.39 | 0.32 | 0.25 |
| 6 | 168573159 | rs4708679 | -0.001 | 0.476 | 0.17 | 0.533 | -0.028 | -0.043 | -0.043 | 0 | 0.22 | 0.16 | 0.22 | 0.39 | 0.32 | 0.25 |
| 6 | 168576278 | rs6941655 | -0.117 | 0.099 | -0.009 | 0.274 | -0.01 | -0.046 | -0.046 | 0 | 0.15 | 0.38 | 0.08 | 0.04 | 0.09 | 0.11 |
| 6 | 168576745 | rs9364229 | -0.326 | 0.113 | -0.334 | 0.327 | 0.001 | 0.001 | 0.001 | 0 | 0.38 | 0.44 | 0.3 | 0.43 | 0.42 | 0.36 |
| 6 | 168577460 | rs13200002 | -1.132 | -0.296 | -0.835 | 1.077 | 0.004 | -0.013 | -0.013 | 0 | 0.19 | 0.16 | 0.12 | 0.3 | 0.16 | 0.37 |
| 6 | 168578718 | rs6455492 | -0.721 | -0.409 | -1.744 | 0.511 | -0.134 | 0.023 | 0.023 | 0.233 | 0.63 | 0.97 | 0.62 | 0.78 | 0.47 | 0.71 |
| 6 | 168579302 | rs7751205 | -0.283 | -0.871 | -1.496 | 0.279 | 0.009 | 0.012 | 0.012 | 0 | 0.14 | 0.14 | 0.05 | 0.05 | 0.08 | 0.18 |
| 6 | 168580694 | rs4708445 | -0.389 | -0.812 | -1.205 | 1.302 | 0.011 | 0.107 | 0.107 | 0.204 | 0.07 | 0.09 | 0.29 | 0.45 | 0.14 | 0.25 |
| 6 | 168580741 | rs11960954 | -0.821 | -1.857 | -0.682 | 0.888 | 0.039 | 0.021 | 0.021 | 0 | 0.4 | 0.7 | 0.19 | 0.3 | 0.19 | 0.37 |
| 6 | 168581362 | rs9455968 | -0.42 | -1.713 | 0.243 | 1.008 | -0.023 | 0.01 | 0.01 | 0 | 0.24 | 0.28 | 0.12 | 0.31 | 0.12 | 0.23 |
| 6 | 168582182 | rs9455971 | 0.352 | -1.704 | 0.133 | 0.863 | -0.001 | -0.013 | -0.013 | 0.766 | 0.06 | 0.05 | 0.02 | 0 | 0.04 | 0.02 |
| 6 | 168583006 | rs9455973 | -0.799 | -1.873 | 0.381 | 0.942 | -0.079 | 0.066 | 0.066 | 0 | 0.23 | 0.5 | 0.23 | 0.32 | 0.09 | 0.27 |
| 6 | 168589242 | rs9355178 | -1.68 | -1.762 | -0.017 | 0.965 | 0.115 | 0.02 | 0.02 | 0 | 0.49 | 0.8 | 0.44 | 0.57 | 0.3 | 0.43 |
| 6 | 168592134 | rs9283861 | -2.345 | -1.65 | -0.352 | 0.812 | -0.152 | 0.001 | 0.001 | 0.043 | 0.63 | 0.97 | 0.61 | 0.76 | 0.5 | 0.71 |
| 6 | 168593739 | rs12198918 | -0.68 | -1.8 | -0.131 | 0.694 | -0.001 | -0.001 | -0.001 | 0 | 0.23 | 0.23 | 0.2 | 0.18 | 0.26 | 0.37 |
| 6 | 168593956 | rs9346533 | -1.693 | -1.798 | -0.071 | 0.728 | 0.09 | 0.02 | 0.02 | 0 | 0.49 | 0.74 | 0.46 | 0.57 | 0.3 | 0.43 |
| 6 | 168595832 | rs12213783 | -1.144 | -1.981 | 0.294 | 0.995 | -0.003 | -0.003 | -0.003 | 0 | 0.26 | 0.31 | 0.33 | 0.25 | 0.28 | 0.38 |
| 6 | 168596251 | rs4708691 | -1.474 | -1.978 | 0.401 | 1.13 | 0.007 | -0.006 | -0.006 | 0 | 0.38 | 0.66 | 0.31 | 0.52 | 0.25 | 0.3 |
| 6 | 168597242 | rs6918784 | -1.385 | -2.01 | 0.758 | 1.02 | -0.017 | -0.011 | -0.011 | 0.009 | 0.36 | 0.59 | 0.21 | 0.26 | 0.23 | 0.29 |
| 6 | 168597347 | rs9455595 | -0.833 | -1.927 | 1.347 | 1.087 | 0.029 | 0.049 | 0.049 | 0 | 0.07 | 0.3 | 0.05 | 0.06 | 0.04 | 0.07 |
| 6 | 168599333 | rs2306286 | -0.442 | -1.917 | 1.05 | 1.057 | -0.021 | 0.068 | 0.068 | 0.332 | 0.13 | 0.2 | 0.25 | 0.32 | 0.07 | 0.13 |
| 6 | 168605875 | rs9455600 | -1.838 | -2.009 | 1.413 | 0.87 | -0.019 | -0.023 | -0.023 | 0.285 | 0.36 | 0.54 | 0.49 | 0.75 | 0.31 | 0.44 |
| 6 | 168609335 | rs9455983 | -1.537 | -2.027 | 1.38 | 0.884 | -0.001 | -0.003 | -0.003 | 0 | 0.4 | 0.56 | 0.5 | 0.74 | 0.33 | 0.47 |
| 6 | 168612631 | rs9455988 | -0.987 | -1.862 | 1.555 | 1.055 | 0 | 0 | 0 | 0 | 0.18 | 0.18 | 0.15 | 0.17 | 0.15 | 0.29 |
| 6 | 168617892 | rs9455992 | -1.295 | -1.943 | 1.676 | 1.079 | -0.035 | 0.006 | 0.006 | 0 | 0.25 | 0.49 | 0.2 | 0.26 | 0.19 | 0.21 |
| 6 | 168620659 | rs7744123 | -0.679 | -1.96 | 1.793 | 1.089 | -0.015 | -0.004 | -0.004 | 3.017 | 0.2 | 0.28 | 0.14 | 0.2 | 0.15 | 0.15 |
| 6 | 168620671 | rs7767297 | -0.613 | -1.954 | 1.597 | 1.092 | -0.028 | -0.006 | -0.006 | 0 | 0.2 | 0.35 | 0.15 | 0.21 | 0.15 | 0.15 |
| 6 | 168620970 | rs7760693 | -2.579 | -1.962 | 1.055 | 1.034 | -0.076 | 0.068 | 0.068 | 0 | 0.56 | 0.97 | 0.64 | 0.74 | 0.46 | 0.57 |
| 6 | 168621367 | rs9455995 | -1.218 | -1.85 | 1.23 | 0.928 | -0.026 | 0 | 0 | 0 | 0.25 | 0.56 | 0.2 | 0.27 | 0.19 | 0.21 |
| 6 | 168623762 | rs2170001 | 0.424 | -1.815 | 1.118 | 0.856 | 0 | -0.026 | -0.026 | 0.113 | 0.1 | 0.12 | 0.08 | 0 | 0.1 | 0.02 |
| 6 | 168624848 | rs4383786 | -1.547 | -1.797 | 1.164 | 0.993 | -0.001 | 0.002 | 0.002 | 0 | 0.31 | 0.61 | 0.31 | 0.43 | 0.29 | 0.37 |
| 6 | 168626744 | rs6913465 | -1.672 | -2.149 | 1.26 | 1.002 | -0.011 | -0.002 | -0.002 | 1.925 | 0.37 | 0.58 | 0.39 | 0.38 | 0.26 | 0.33 |
| 6 | 168627448 | rs4708701 | -1.894 | -2.264 | 1.326 | 1.004 | 0.058 | 0.016 | 0.016 | 0 | 0.47 | 0.71 | 0.54 | 0.57 | 0.36 | 0.4 |
| 6 | 168629381 | rs2291098 | -1.511 | -2.204 | 1.301 | 1.063 | -0.013 | 0.015 | 0.015 | 0.05 | 0.3 | 0.41 | 0.43 | 0.47 | 0.2 | 0.29 |
| 6 | 168630519 | rs9455613 | -0.248 | -2.135 | 1.166 | 1.068 | 0.009 | 0.087 | 0.087 | 0 | 0.05 | 0.19 | 0.09 | 0.16 | 0.03 | 0.11 |
| 6 | 168630983 | rs2127330 | -2.305 | -2.182 | 0.979 | 1.036 | 0.013 | 0.082 | 0.082 | 0 | 0.55 | 0.94 | 0.63 | 0.74 | 0.38 | 0.51 |
| 6 | 168632301 | rs2035273 | -0.214 | -2.099 | 1.177 | 1.135 | 0.014 | 0.098 | 0.098 | 0 | 0.05 | 0.18 | 0.1 | 0.16 | 0.03 | 0.11 |
| 6 | 168633971 | rs4708702 | -1.095 | -2.105 | 0.919 | 1.044 | 0.005 | -0.005 | -0.005 | 0 | 0.25 | 0.28 | 0.35 | 0.31 | 0.17 | 0.1 |
| 6 | 168651225 | rs4708705 | -0.772 | -2.029 | 1.241 | 1.109 | 0.009 | -0.011 | -0.011 | 0.023 | 0.31 | 0.25 | 0.45 | 0.58 | 0.27 | 0.38 |
| 6 | 168654978 | rs9355058 | -1.398 | -2.104 | 1.418 | 1.005 | 0.006 | 0.007 | 0.007 | 0.013 | 0.42 | 0.56 | 0.56 | 0.79 | 0.31 | 0.57 |
| 6 | 168664184 | rs9346673 | -0.774 | -1.689 | 1.214 | 1.105 | 0.004 | -0.009 | -0.009 | 1.458 | 0.32 | 0.29 | 0.45 | 0.58 | 0.29 | 0.39 |
| 6 | 168664479 | rs6911413 | -1.883 | -2.058 | 0.78 | 0.96 | 0.09 | 0.196 | 0.196 | 0 | 0.52 | 0.89 | 0.63 | 0.81 | 0.36 | 0.6 |
| 6 | 168665760 | rs1871428 | -1.916 | -2.122 | 0.719 | 0.95 | 0.044 | 0.182 | 0.182 | 0.035 | 0.52 | 0.96 | 0.64 | 0.81 | 0.36 | 0.6 |
| 6 | 168666768 | rs6924059 | -1.152 | -2.103 | 1.236 | 1.019 | -0.007 | 0.138 | 0.138 | 0 | 0.1 | 0.38 | 0.11 | 0.21 | 0.03 | 0.1 |
| 6 | 168670016 | rs9364421 | -1.684 | -2.057 | 0.622 | 0.739 | 0.09 | 0.146 | 0.146 | 0 | 0.51 | 0.8 | 0.6 | 0.78 | 0.36 | 0.57 |
| 6 | 168673955 | rs7773617 | -0.749 | -1.933 | 0.804 | 0.79 | -0.012 | 0.008 | 0.008 | 0 | 0.29 | 0.35 | 0.28 | 0.34 | 0.27 | 0.25 |
| 6 | 168685495 | rs10945492 | -0.421 | -1.951 | 0.174 | 0.242 | 0.015 | -0.002 | -0.002 | 0 | 0.36 | 0.2 | 0.31 | 0.39 | 0.35 | 0.31 |
| 6 | 168685559 | rs2054476 | -1.323 | -1.528 | -0.214 | 0.263 | -0.019 | 0.173 | 0.173 | 0 | 0.55 | 0.9 | 0.65 | 0.82 | 0.44 | 0.58 |
| 6 | 168686611 | rs10945493 | -0.896 | -0.66 | 0.664 | 1.03 | 0.003 | 0.12 | 0.12 | 0 | 0.05 | 0.08 | 0.28 | 0.35 | 0.08 | 0.14 |
| 6 | 168686848 | rs10945494 | -1.246 | -0.665 | 0.891 | 0.988 | 0.051 | 0.322 | 0.322 | 0 | 0.06 | 0.26 | 0.29 | 0.36 | 0.08 | 0.14 |
| 6 | 168689657 | rs6924136 | -0.635 | -0.696 | 0.481 | -0.249 | 0.033 | -0.007 | -0.007 | 0 | 0.1 | 0.24 | 0.07 | 0.06 | 0.13 | 0.05 |
| 6 | 168691883 | rs9455622 | 0.387 | -0.482 | -0.682 | -0.745 | -0.025 | 0.031 | 0.031 | 0 | 0.36 | 0.28 | 0.34 | 0.17 | 0.49 | 0.31 |
| 6 | 168693090 | rs1479516 | 0.38 | -0.514 | -0.686 | -0.741 | -0.02 | 0.024 | 0.024 | 0 | 0.36 | 0.29 | 0.37 | 0.17 | 0.49 | 0.33 |
| 6 | 168695624 | rs7738825 | -0.289 | -0.986 | -1.495 | -0.714 | 0.04 | -0.032 | -0.032 | 0 | 0.26 | 0.11 | 0.34 | 0.66 | 0.13 | 0.38 |
| 6 | 168699291 | rs2306285 | 0.541 | -0.496 | -1.686 | -1.062 | -0.163 | 0.037 | 0.037 | 0 | 0.29 | 0.04 | 0.39 | 0.2 | 0.49 | 0.3 |
| 6 | 168700846 | rs12202943 | -0.131 | -0.422 | -1.458 | -1.031 | -0.004 | 0.024 | 0.024 | 0 | 0.15 | 0.1 | 0.09 | 0.07 | 0.15 | 0.14 |
| 6 | 168703821 | rs7750101 | 0.056 | -1.038 | -0.729 | -1.173 | 0.002 | 0.001 | 0.001 | 0 | 0.47 | 0.56 | 0.47 | 0.26 | 0.59 | 0.47 |
| 6 | 168703878 | rs9456047 | -0.35 | -1.292 | -0.095 | -1.279 | -0.043 | -0.092 | -0.092 | 0 | 0.19 | 0.31 | 0.09 | 0.01 | 0.1 | 0.1 |
| 6 | 168705006 | rs7768092 | -0.698 | -1.207 | 0.079 | -1.296 | -0.06 | -0.138 | -0.138 | 0.02 | 0.22 | 0.43 | 0.09 | 0 | 0.1 | 0.1 |
| 6 | 168705822 | rs9346686 | -0.097 | -0.825 | -0.455 | -1.693 | 0.009 | 0.034 | 0.034 | 0 | 0.46 | 0.42 | 0.63 | 0.88 | 0.46 | 0.48 |
| 6 | 168705899 | rs13193695 | -0.451 | -1.306 | -0.788 | -0.532 | 0.053 | -0.025 | -0.025 | 0 | 0.27 | 0.11 | 0.46 | 0.3 | 0.19 | 0.25 |
| 6 | 168707435 | rs10945498 | -0.642 | -1.285 | 0.002 | -0.853 | -0.026 | -0.017 | -0.017 | 0 | 0.36 | 0.54 | 0.35 | 0.32 | 0.26 | 0.46 |
| 6 | 168708045 | rs12530360 | -0.109 | -1.173 | -0.223 | -0.865 | -0.007 | 0.002 | 0.002 | 0 | 0.14 | 0.09 | 0.11 | 0.12 | 0.14 | 0.17 |
| 6 | 168709385 | rs6925614 | -0.845 | -1.105 | -0.182 | -1.16 | 0 | 0 | 0 | 0 | 0.42 | 0.46 | 0.38 | 0.25 | 0.37 | 0.4 |
| 6 | 168711170 | rs1551988 | -0.33 | -1.062 | -0.368 | -1.333 | 0.01 | 0.01 | 0.01 | 0 | 0.32 | 0.33 | 0.27 | 0.13 | 0.22 | 0.31 |
| 6 | 168712643 | rs13207327 | -1.316 | -1.203 | -0.837 | -1.709 | -0.067 | 0.081 | 0.081 | 0 | 0.69 | 0.84 | 0.77 | 0.94 | 0.54 | 0.74 |

| | | | | | | | | | | | | | | | | |
|---|---|---|---|---|---|---|---|---|---|---|---|---|---|---|---|---|
| 6 | 168715141 | rs4549598 | -0.158 | -1.481 | -0.805 | -1.471 | 0.024 | 0.035 | 0.035 | 0 | 0.29 | 0.22 | 0.26 | 0.12 | 0.22 | 0.31 |
| 6 | 168719924 | rs4292501 | 0.415 | -1.043 | -1.34 | -1.449 | -0.012 | 0.251 | 0.251 | 0.589 | 0.49 | 0.13 | 0.65 | 0.82 | 0.55 | 0.53 |
| 6 | 168726618 | rs9346688 | 0.919 | -0.76 | -1.271 | -0.915 | 0.025 | 0.001 | 0.001 | 4.305 | 0.15 | 0.08 | 0.21 | 0.13 | 0.09 | 0.1 |
| 6 | 168731137 | rs9346539 | 0.403 | -0.846 | -0.944 | -1.133 | 0.008 | 0.022 | 0.022 | 0.147 | 0.62 | 0.33 | 0.6 | 0.87 | 0.66 | 0.8 |
| 6 | 168731524 | rs9346690 | 0.403 | -0.846 | -0.944 | -1.133 | 0.008 | 0.022 | 0.022 | 0.147 | 0.62 | 0.33 | 0.6 | 0.87 | 0.66 | 0.8 |
| 6 | 168738199 | rs4708456 | 0.411 | -0.815 | -0.726 | -1.113 | 0.026 | 0.048 | 0.048 | 2.524 | 0.45 | 0.33 | 0.53 | 0.69 | 0.64 | 0.56 |
| 6 | 168743094 | rs7744524 | 0.261 | -0.499 | -0.505 | -0.891 | 0.017 | -0.046 | -0.046 | 2.093 | 0.12 | 0.16 | 0.1 | 0 | 0.18 | 0.07 |
| 6 | 168743792 | rs4485988 | 0.019 | -0.73 | -0.207 | -0.762 | -0.018 | -0.058 | -0.058 | 5.48 | 0.9 | 0.73 | 0.93 | 1 | 0.94 | 0.97 |
| 6 | 168750052 | rs9364244 | 1.488 | -0.487 | -0.598 | -0.309 | -0.016 | 0.033 | 0.033 | 2.844 | 0.06 | 0.09 | 0.16 | 0.19 | 0.01 | 0.09 |
| 6 | 168756517 | rs9364245 | 0.908 | -0.243 | -0.881 | 0.64 | -0.003 | -0.048 | -0.048 | 2.161 | 0.11 | 0.03 | 0.12 | 0.33 | 0.1 | 0.09 |
| 6 | 168760569 | rs4708459 | -0.54 | -0.646 | -1.178 | 0.01 | 0.034 | 0.121 | 0.121 | 0.301 | 0.5 | 0.91 | 0.45 | 0.33 | 0.4 | 0.46 |
| 6 | 168774529 | rs1886399 | -0.863 | -0.087 | -0.878 | 0.301 | -0.001 | 0.002 | 0.002 | 0 | 0.75 | 0.8 | 0.81 | 0.77 | 0.72 | 0.78 |
| 6 | 168775942 | rs3892414 | -0.364 | -0.17 | -0.572 | 0.417 | -0.053 | 0.144 | 0.144 | 0 | 0.18 | 0.51 | 0.25 | 0.35 | 0.09 | 0.24 |
| 6 | 168786154 | rs11755288 | 0.388 | 0.266 | -0.958 | 0.429 | -0.092 | -0.08 | -0.08 | 0 | 0.32 | 0.14 | 0.45 | 0.42 | 0.46 | 0.33 |
| 6 | 168800562 | rs9346705 | 0.134 | 0.188 | -0.486 | 0.057 | 0.003 | 0.008 | 0.008 | 0.099 | 0.39 | 0.4 | 0.34 | 0.17 | 0.35 | 0.3 |
| 6 | 168801934 | rs4588663 | 0.685 | 0.293 | -0.69 | 0.317 | -0.005 | -0.009 | -0.009 | 0.046 | 0.26 | 0.24 | 0.42 | 0.59 | 0.29 | 0.38 |
| 6 | 168805887 | rs9364253 | -0.241 | -0.019 | -0.575 | 0.183 | 0.001 | 0.024 | 0.024 | 0 | 0.65 | 0.77 | 0.77 | 0.77 | 0.64 | 0.68 |
| 6 | 168808760 | rs2609299 | 0.015 | -0.399 | -0.154 | 0.443 | -0.009 | -0.015 | -0.015 | 10.079 | 0.3 | 0.31 | 0.24 | 0.23 | 0.27 | 0.25 |
| 6 | 168809019 | rs7747132 | -0.226 | -0.368 | 0.074 | 1.178 | -0.001 | 0.027 | 0.027 | 0 | 0.7 | 0.69 | 0.76 | 0.55 | 0.68 | 0.63 |
| 6 | 168814455 | rs2763264 | -0.226 | -0.245 | -0.098 | 0.984 | 0.01 | -0.004 | -0.004 | 0 | 0.48 | 0.64 | 0.41 | 0.48 | 0.43 | 0.53 |
| 6 | 168815037 | rs9364451 | -0.2 | -0.778 | -0.438 | 0.87 | -0.024 | 0.034 | 0.034 | 0 | 0.26 | 0.32 | 0.26 | 0.46 | 0.16 | 0.36 |
| 6 | 168829515 | rs2763269 | -0.761 | -0.575 | -0.911 | 0.417 | -0.011 | 0.001 | 0.001 | 0 | 0.71 | 0.83 | 0.79 | 0.69 | 0.68 | 0.72 |
| 6 | 168846625 | rs9346712 | 1 | -0.6 | -1.063 | 0.979 | 0.024 | -0.005 | -0.005 | 0 | 0.07 | 0.02 | 0.17 | 0.29 | 0.01 | 0.07 |
| 6 | 168847175 | rs2609334 | 1.097 | -0.375 | -1.301 | 0.411 | -0.095 | 0.118 | 0.118 | 0.599 | 0.18 | 0.02 | 0.12 | 0 | 0.26 | 0.16 |
| 6 | 168856841 | rs4708742 | 1.093 | -0.503 | -0.796 | 0.806 | -0.026 | -0.016 | -0.016 | 0 | 0.23 | 0.09 | 0.37 | 0.23 | 0.25 | 0.17 |
| 6 | 168874779 | rs9364457 | -0.886 | -0.67 | -0.052 | 1.161 | -0.051 | 0.164 | 0.164 | 0.01 | 0.19 | 0.55 | 0.25 | 0.45 | 0.12 | 0.24 |
| 6 | 168879958 | rs9364458 | -0.673 | -0.647 | 0.104 | 1.175 | -0.037 | 0.136 | 0.136 | 0 | 0.18 | 0.36 | 0.23 | 0.45 | 0.11 | 0.22 |
| 6 | 168889305 | rs1232314 | 0.917 | -0.344 | -0.462 | 0.924 | -0.057 | -0.076 | -0.076 | 0.041 | 0.17 | 0.02 | 0.14 | 0.35 | 0.21 | 0.19 |
| 6 | 168895058 | rs6938236 | -0.274 | -0.743 | 0.476 | 1.12 | -0.025 | 0.406 | 0.406 | 1.562 | 0.1 | 0.53 | 0.22 | 0.39 | 0.02 | 0.1 |
| 6 | 168898415 | rs2609326 | -0.073 | -0.894 | -0.567 | 0.014 | 0.013 | 0.016 | 0.016 | 0 | 0.52 | 0.44 | 0.51 | 0.78 | 0.62 | 0.45 |
| 6 | 168901450 | rs2609330 | -0.749 | -0.807 | -0.69 | -0.071 | -0.002 | 0.005 | 0.005 | 0 | 0.69 | 0.75 | 0.55 | 0.77 | 0.67 | 0.73 |
| 6 | 168902366 | rs2749246 | -1.059 | -0.861 | -0.727 | -0.027 | -0.006 | -0.004 | -0.004 | 0 | 0.75 | 0.81 | 0.6 | 0.77 | 0.73 | 0.81 |
| 6 | 168903232 | rs2609333 | -0.44 | -1.197 | -0.7 | -0.198 | 0.002 | 0.025 | 0.025 | 0 | 0.57 | 0.64 | 0.43 | 0.76 | 0.43 | 0.63 |
| 6 | 168905499 | rs2749250 | -0.377 | -1.021 | -0.623 | -0.125 | 0.015 | -0.001 | -0.001 | 0 | 0.52 | 0.62 | 0.27 | 0.56 | 0.42 | 0.61 |
| 6 | 168908013 | rs6901357 | -0.314 | -1.421 | -0.433 | -0.129 | -0.065 | -0.084 | -0.084 | 0.009 | 0.16 | 0.26 | 0.05 | 0.01 | 0.05 | 0.22 |
| 6 | 168911377 | rs2749258 | 0.385 | -1.021 | -1.624 | 0.153 | 0.125 | 0.003 | 0.003 | 1.557 | 0.52 | 0.12 | 0.4 | 0.44 | 0.69 | 0.38 |
| 6 | 168915232 | rs5016787 | -0.315 | -1.118 | -0.847 | 0.024 | 0.077 | 0.04 | 0.04 | 7.987 | 0.95 | 0.69 | 0.95 | 1 | 0.97 | 0.98 |
| 6 | 168915345 | rs2749266 | 0.973 | -0.45 | -1.364 | -0.466 | -0.117 | -0.144 | -0.144 | 0 | 0.23 | 0.02 | 0.21 | 0.43 | 0.31 | 0.25 |
| 6 | 168915546 | rs7740366 | 0.177 | -0.605 | -1.436 | -0.135 | -0.037 | 0.003 | 0.003 | 1.401 | 0.38 | 0.52 | 0.43 | 0.34 | 0.22 | 0.47 |
| 6 | 168917870 | rs2144745 | 1.153 | -0.824 | -1.662 | -0.104 | -0.002 | 0.003 | 0.003 | 0.874 | 0.18 | 0.24 | 0.28 | 0.22 | 0.15 | 0.18 |
| 6 | 168918704 | rs2144746 | 1.194 | -0.46 | -1.74 | -0.18 | -0.054 | -0.056 | -0.056 | 0 | 0.23 | 0.12 | 0.24 | 0.42 | 0.34 | 0.26 |
| 6 | 168918741 | rs2144747 | -0.052 | -0.69 | -1.697 | -0.161 | 0.129 | 0.045 | 0.045 | 2.859 | 0.93 | 0.62 | 0.92 | 0.96 | 0.92 | 0.92 |
| 6 | 168918946 | rs2180344 | 0.814 | -0.972 | -1.82 | -0.27 | 0.102 | -0.001 | -0.001 | 0.004 | 0.52 | 0.09 | 0.39 | 0.44 | 0.69 | 0.39 |
| 6 | 168924891 | rs12200380 | 0.911 | -0.905 | -1.521 | -0.816 | -0.007 | 0.005 | 0.005 | 0.17 | 0.19 | 0.27 | 0.29 | 0.21 | 0.14 | 0.21 |
| 6 | 168926062 | rs2763249 | -0.359 | -0.904 | -1.771 | -0.821 | -0.012 | -0.003 | -0.003 | 0 | 0.75 | 0.36 | 0.66 | 0.79 | 0.82 | 0.73 |
| 6 | 168926785 | rs6919839 | 1.085 | -0.524 | -1.806 | -0.597 | 0.001 | -0.001 | -0.001 | 0.691 | 0.17 | 0.17 | 0.26 | 0.19 | 0.14 | 0.18 |
| 6 | 168935525 | rs13205533 | -0.363 | -0.361 | -2.015 | -0.888 | 0.015 | 0.142 | 0.142 | 0.018 | 0.47 | 0.22 | 0.31 | 0.07 | 0.45 | 0.38 |
| 6 | 168941670 | rs17551813 | 0.188 | -0.382 | -1.609 | -0.227 | -0.007 | -0.007 | -0.007 | 0 | 0.1 | 0 | 0.1 | 0.06 | 0.06 | 0.21 |
| 6 | 168943376 | rs1402 | -0.268 | -0.385 | -1.501 | -0.23 | -0.018 | -0.019 | -0.019 | 0 | 0.15 | 0.18 | 0.13 | 0.08 | 0.08 | 0.22 |
| 6 | 168944598 | rs2180346 | -0.58 | 0.154 | -1.942 | 0.569 | 0.001 | -0.031 | -0.031 | 0 | 0.11 | 0.07 | 0.22 | 0.38 | 0.11 | 0.17 |
| 6 | 168948488 | rs4708477 | -0.275 | 0.309 | -2.117 | -0.695 | -0.05 | 0.184 | 0.184 | 0.043 | 0.67 | 0.93 | 0.78 | 1 | 0.61 | 0.84 |
| 6 | 168951165 | rs1884477 | -0.112 | 0.02 | -1.363 | 0.165 | 0.002 | 0.002 | 0.002 | 0.144 | 0.3 | 0.31 | 0.3 | 0.23 | 0.23 | 0.37 |
| 6 | 168960552 | rs10945513 | 0.104 | 0.254 | -1.505 | 0.354 | -0.001 | 0.049 | 0.049 | 0.899 | 0.45 | 0.27 | 0.52 | 0.35 | 0.44 | 0.3 |
| 6 | 168966538 | rs11969025 | 0.359 | 0.418 | -1.137 | 0.551 | 0.002 | 0.072 | 0.072 | 0.48 | 0.35 | 0.17 | 0.3 | 0.2 | 0.31 | 0.25 |
| 6 | 168967691 | rs2342630 | -0.349 | 0.286 | -1.207 | 1.048 | -0.052 | -0.01 | -0.01 | 0 | 0.51 | 0.87 | 0.45 | 0.52 | 0.43 | 0.51 |
| 6 | 168968434 | rs12195840 | -0.24 | 0.276 | -0.041 | 1.232 | 0.097 | 0.134 | 0.134 | 0.005 | 0.06 | 0.21 | 0.08 | 0.11 | 0.08 | 0.08 |
| 6 | 168970508 | rs6455530 | -0.418 | 0.226 | 0.262 | 1.218 | 0.015 | 0.202 | 0.202 | 0 | 0.1 | 0.44 | 0.07 | 0.2 | 0.05 | 0.17 |
| 6 | 168976562 | rs6934395 | -0.381 | 0.27 | 0.866 | 1.045 | 0.088 | 0.123 | 0.123 | 0.022 | 0.06 | 0.21 | 0.08 | 0.12 | 0.08 | 0.08 |
| 6 | 168980519 | rs4537111 | 0.25 | 0.224 | 0.849 | 0.998 | 0 | -0.002 | -0.002 | 0.199 | 0.42 | 0.42 | 0.37 | 0.32 | 0.39 | 0.33 |
| 6 | 168991469 | rs9455669 | -0.034 | 0.107 | 0.93 | 0.871 | -0.008 | -0.011 | -0.011 | 0.925 | 0.27 | 0.3 | 0.34 | 0.16 | 0.27 | 0.36 |
| 6 | 168991834 | rs2180342 | -0.296 | 0.435 | 0.815 | 0.866 | 0 | 0.002 | 0.002 | 0 | 0.81 | 0.81 | 0.68 | 0.62 | 0.79 | 0.82 |
| 6 | 168997347 | rs4708757 | 0.023 | 0.382 | 0.847 | 0.918 | 0.001 | -0.005 | -0.005 | 0 | 0.69 | 0.64 | 0.7 | 0.76 | 0.65 | 0.61 |
| 6 | 169002767 | rs2880405 | -0.58 | 0.334 | 0.751 | 0.908 | 0.008 | -0.003 | -0.003 | 0 | 0.46 | 0.57 | 0.58 | 0.51 | 0.45 | 0.51 |
| 6 | 169004019 | rs12530037 | 0.996 | 0.404 | 0.5 | 0.924 | -0.019 | 0.032 | 0.032 | 0 | 0.46 | 0.29 | 0.35 | 0.38 | 0.46 | 0.38 |
| 6 | 169005418 | rs9364471 | 0.41 | 0.325 | 0.136 | 0.926 | -0.002 | 0.004 | 0.004 | 0 | 0.13 | 0.12 | 0.14 | 0.35 | 0.16 | 0.19 |
| 6 | 169005721 | rs4708759 | 0.096 | -0.088 | 0.115 | 0.308 | 0.002 | 0.001 | 0.001 | 0.048 | 0.27 | 0.22 | 0.27 | 0.12 | 0.28 | 0.28 |
| 6 | 169006974 | rs9455673 | -0.405 | -0.46 | -0.002 | 0.405 | 0.06 | 0.048 | 0.048 | 0 | 0.36 | 0.25 | 0.33 | 0.23 | 0.22 | 0.37 |
| 6 | 169007180 | rs9456238 | 0.475 | -0.288 | -0.179 | 0.397 | 0.047 | 0.03 | 0.03 | 0.497 | 0.23 | 0.11 | 0.18 | 0.13 | 0.17 | 0.28 |
| 6 | 169008945 | rs3734421 | 0.243 | -0.229 | 0.209 | 0.437 | 0.003 | 0.003 | 0.003 | 0 | 0.28 | 0.23 | 0.14 | 0.13 | 0.2 | 0.28 |

| | | | | | | | | | | | | | | | | |
|---|---|---|---|---|---|---|---|---|---|---|---|---|---|---|---|---|
| 6 | 169009298 | rs6913757 | 0.125 | 0.184 | -0.197 | 0.567 | 0.002 | 0.006 | 0.006 | 0 | 0.65 | 0.37 | 0.49 | 0.58 | 0.63 | 0.63 |
| 6 | 169010437 | rs6932414 | 0.186 | 0.457 | 0.027 | 0.769 | 0.037 | 0.055 | 0.055 | 0 | 0.94 | 0.87 | 0.94 | 0.88 | 0.92 | 0.92 |
| 6 | 169014011 | rs9295077 | 0.862 | 0.37 | -0.021 | 0.72 | 0.01 | 0.01 | 0.01 | 0 | 0.27 | 0.2 | 0.14 | 0.13 | 0.2 | 0.28 |
| 6 | 169018179 | rs7761264 | 0.299 | 0.37 | 0.267 | 0.881 | -0.002 | -0.001 | -0.001 | 0 | 0.26 | 0.28 | 0.25 | 0.26 | 0.28 | 0.25 |
| 6 | 169021311 | rs2342639 | 0.444 | 0.382 | 0.388 | 0.75 | 0.038 | 0.057 | 0.057 | 0.023 | 0.94 | 0.83 | 0.93 | 0.88 | 0.92 | 0.92 |
| 6 | 169022053 | rs12197584 | 0.509 | 0.33 | 0.125 | 0.614 | 0.013 | 0.012 | 0.012 | 0 | 0.16 | 0.2 | 0.21 | 0.28 | 0.22 | 0.2 |
| 6 | 169027613 | rs17553738 | 0.595 | 0.468 | -0.089 | 0.333 | -0.009 | -0.065 | -0.065 | 0.391 | 0.19 | 0.25 | 0.2 | 0.01 | 0.13 | 0.14 |
| 6 | 169029167 | rs9455679 | 0.87 | 0.299 | -0.385 | 0.434 | 0.049 | 0.04 | 0.04 | 0.013 | 0.27 | 0.17 | 0.13 | 0.14 | 0.18 | 0.3 |
| 6 | 169033040 | rs2342645 | 0.251 | 0.564 | -0.254 | 0.503 | 0.016 | 0.003 | 0.003 | 0.597 | 0.43 | 0.3 | 0.35 | 0.43 | 0.42 | 0.49 |
| 6 | 169033082 | rs2342646 | -0.653 | 0.616 | -0.386 | 0.576 | -0.01 | 0 | 0 | 0.203 | 0.71 | 0.77 | 0.79 | 0.71 | 0.7 | 0.76 |
| 6 | 169039497 | rs10945529 | 0.433 | 0.115 | -1.409 | 0.318 | -0.002 | 0.058 | 0.058 | 1.657 | 0.3 | 0.03 | 0.14 | 0.15 | 0.24 | 0.36 |
| 6 | 169039594 | rs4708481 | -0.08 | 0.177 | -1.024 | 0.785 | 0.054 | 0.026 | 0.026 | 0.791 | 0.49 | 0.27 | 0.44 | 0.46 | 0.4 | 0.54 |
| 6 | 169047090 | rs6931082 | -0.212 | 0.477 | 0.053 | 1.197 | 0.027 | 0.065 | 0.065 | 1.599 | 0.12 | 0.3 | 0.06 | 0.19 | 0.12 | 0.12 |
| 6 | 169047112 | rs9295079 | 1.1 | 0.495 | -0.648 | 0.632 | 0.007 | 0.238 | 0.238 | 0.058 | 0.49 | 0.15 | 0.38 | 0.13 | 0.45 | 0.42 |
| 6 | 169047901 | rs4708483 | -0.156 | 0.3 | -0.281 | 1.196 | 0.003 | -0.006 | -0.006 | 0 | 0.76 | 0.8 | 0.84 | 0.5 | 0.82 | 0.74 |
| 6 | 169048303 | rs6937595 | 0.079 | 0.582 | -0.085 | 1.216 | 0.02 | 0.058 | 0.058 | 1.792 | 0.11 | 0.22 | 0.05 | 0.19 | 0.12 | 0.11 |
| 6 | 169048842 | rs9455680 | -0.653 | -0.188 | -0.212 | 1.133 | 0.009 | -0.002 | -0.002 | 0 | 0.18 | 0.12 | 0.24 | 0.17 | 0.14 | 0.25 |
| 6 | 169049030 | rs9455681 | 1.893 | -0.128 | -0.602 | 0.932 | -0.149 | 0.125 | 0.125 | 0.047 | 0.38 | 0.04 | 0.36 | 0.14 | 0.48 | 0.32 |
| 6 | 169049871 | rs1885633 | -0.709 | 0.302 | 0.631 | 1.186 | 0.044 | -0.062 | -0.062 | 0 | 0.13 | 0.23 | 0.17 | 0.01 | 0.22 | 0.11 |
| 6 | 169051020 | rs3818311 | -0.901 | 0.033 | 0.729 | 1.687 | 0.008 | 0.023 | 0.023 | 0 | 0.28 | 0.35 | 0.25 | 0.47 | 0.32 | 0.33 |
| 6 | 169058051 | rs4708373 | 0.18 | 0.059 | 0.992 | 1.65 | 0.06 | -0.04 | -0.04 | 0 | 0.08 | 0.16 | 0.13 | 0 | 0.16 | 0.05 |
| 6 | 169058581 | rs1124346 | -0.044 | 0.011 | 0.616 | 1.694 | -0.002 | 0.01 | 0.01 | 0 | 0.2 | 0.19 | 0.12 | 0.47 | 0.16 | 0.29 |
| 6 | 169074105 | rs196454 | -1.954 | -0.193 | 0.081 | 1.539 | -0.009 | 0.043 | 0.043 | 0.152 | 0.47 | 0.93 | 0.57 | 0.69 | 0.41 | 0.51 |
| 6 | 169075136 | rs1885634 | -0.754 | -0.189 | 0.486 | 1.634 | -0.023 | 0.126 | 0.126 | 2.928 | 0.11 | 0.5 | 0.09 | 0.19 | 0.04 | 0.12 |
| 6 | 169075499 | rs654391 | -0.52 | 0.008 | -0.025 | 1.592 | -0.046 | -0.114 | -0.114 | 0.865 | 0.26 | 0.11 | 0.43 | 0.49 | 0.3 | 0.3 |
| 6 | 169076700 | rs1322556 | -0.85 | -0.032 | -0.015 | 1.724 | -0.002 | -0.004 | -0.004 | 0 | 0.28 | 0.28 | 0.25 | 0.45 | 0.31 | 0.33 |
| 6 | 169078546 | rs601639 | -0.024 | -0.148 | -0.56 | 1.974 | 0.031 | -0.066 | -0.066 | 0 | 0.18 | 0.06 | 0.22 | 0.43 | 0.12 | 0.24 |
| 6 | 169086528 | rs16887281 | 0.441 | -0.302 | -0.649 | 2.003 | 0.04 | -0.105 | -0.105 | 0 | 0.17 | 0.02 | 0.18 | 0.44 | 0.11 | 0.26 |
| 6 | 169086561 | rs9346563 | -1.378 | 0.29 | -0.374 | 1.97 | -0.02 | 0.027 | 0.027 | 1.53 | 0.26 | 0.82 | 0.3 | 0.22 | 0.28 | 0.26 |
| 6 | 169094291 | rs647547 | -1.854 | 0.275 | -0.561 | 2.134 | -0.05 | -0.134 | -0.134 | 2.079 | 0.36 | 0.92 | 0.43 | 0.51 | 0.35 | 0.35 |
| 6 | 169118534 | rs196462 | -0.345 | 1.093 | 0.054 | 1.758 | 0 | -0.001 | -0.001 | 0.039 | 0.07 | 0.06 | 0.09 | 0.04 | 0.07 | 0.03 |
| 6 | 169121081 | rs9455695 | -0.093 | 1.203 | 0.409 | 1.554 | 0.022 | -0.032 | -0.032 | 0 | 0.09 | 0.15 | 0.13 | 0 | 0.13 | 0.04 |
| 6 | 169122136 | rs1322558 | 0.471 | 1.207 | 0.362 | 1.556 | -0.044 | 0.039 | 0.039 | 0 | 0.07 | 0.01 | 0.12 | 0 | 0.13 | 0.03 |
| 6 | 169127523 | rs1407515 | -0.222 | 1.181 | 0.69 | 1.536 | 0.015 | -0.027 | -0.027 | 0 | 0.09 | 0.13 | 0.13 | 0 | 0.13 | 0.04 |
| 6 | 169128134 | rs4708487 | -0.925 | 1.045 | 0.812 | 1.871 | 0 | 0.007 | 0.007 | 3.317 | 0.28 | 0.35 | 0.43 | 0.45 | 0.26 | 0.31 |
| 6 | 169128147 | rs16887303 | 0.229 | 1.006 | 0.804 | 1.883 | 0.006 | -0.008 | -0.008 | 4.197 | 0.18 | 0.18 | 0.29 | 0.45 | 0.13 | 0.28 |
| 6 | 169133034 | rs196489 | -1.334 | 0.98 | 1.293 | 1.683 | -0.002 | -0.002 | -0.002 | 0.039 | 0.34 | 0.36 | 0.44 | 0.69 | 0.27 | 0.4 |
| 6 | 169135214 | rs9346566 | -0.839 | 1.029 | 1.113 | 1.489 | 0.008 | 0.002 | 0.002 | 0 | 0.16 | 0.18 | 0.18 | 0.15 | 0.2 | 0.12 |
| 6 | 169139351 | rs196501 | 0.463 | 1.03 | 0.908 | 1.455 | -0.027 | -0.017 | -0.017 | 1.539 | 0.05 | 0.01 | 0.08 | 0.04 | 0.07 | 0.03 |
| 6 | 169140572 | rs196506 | -2.101 | 0.88 | 1.236 | 1.37 | -0.044 | 0.2 | 0.2 | 0.094 | 0.48 | 0.88 | 0.65 | 0.82 | 0.44 | 0.53 |
| 6 | 169153014 | rs1407514 | 0.007 | 0.915 | 1.86 | 1.637 | -0.016 | 0.047 | 0.047 | 2.337 | 0.2 | 0.24 | 0.29 | 0.47 | 0.14 | 0.28 |
| 6 | 169156639 | rs9364286 | -0.34 | 1.064 | 1.83 | 1.687 | 0.001 | 0.018 | 0.018 | 0.013 | 0.08 | 0.11 | 0.04 | 0.14 | 0.08 | 0.1 |
| 6 | 169158652 | rs911939 | 1.183 | 1.322 | 2.245 | 1.618 | 0.034 | 0 | 0 | 0 | 0.84 | 0.75 | 0.76 | 0.83 | 0.78 | 0.88 |
| 6 | 169161550 | rs9364289 | -1.308 | 1.148 | 2.253 | 1.684 | -0.007 | 0.005 | 0.005 | 0 | 0.1 | 0.18 | 0.11 | 0.16 | 0.09 | 0.15 |
| 6 | 169166139 | rs1575341 | -0.099 | 1.242 | 2.321 | 1.671 | -0.016 | -0.067 | -0.067 | 0.258 | 0.3 | 0.16 | 0.4 | 0.47 | 0.28 | 0.37 |
| 6 | 169168216 | rs9294911 | 1.998 | 1.326 | 2.692 | 1.629 | 0.011 | 0.118 | 0.118 | 0 | 0.47 | 0.27 | 0.33 | 0.18 | 0.49 | 0.4 |
| 6 | 169175473 | rs1040932 | -1.447 | 1.293 | 2.848 | 1.653 | -0.006 | -0.002 | -0.002 | 0 | 0.11 | 0.12 | 0.1 | 0.16 | 0.09 | 0.17 |
| 6 | 169182237 | rs927476 | 1.969 | 1.252 | 2.746 | 1.601 | -0.005 | 0.194 | 0.194 | 0.005 | 0.47 | 0.14 | 0.31 | 0.18 | 0.49 | 0.42 |
| 6 | 169191771 | rs6942031 | -1.156 | 1.286 | 2.891 | 1.69 | -0.005 | 0.001 | 0.001 | 0 | 0.42 | 0.48 | 0.52 | 0.63 | 0.37 | 0.52 |
| 6 | 169196780 | rs4585533 | -1.759 | 1.357 | 2.593 | 1.649 | 0.01 | 0.151 | 0.151 | 0.01 | 0.49 | 0.81 | 0.6 | 0.78 | 0.42 | 0.58 |
| 6 | 169199202 | rs6936903 | -1.418 | 1.479 | 2.761 | 1.65 | 0.005 | 0.019 | 0.019 | 0.191 | 0.46 | 0.62 | 0.58 | 0.68 | 0.41 | 0.57 |
| 6 | 169199944 | rs6899552 | -1.94 | 1.472 | 2.772 | 1.631 | 0.008 | 0.161 | 0.161 | 0 | 0.52 | 0.78 | 0.69 | 0.83 | 0.51 | 0.61 |
| 6 | 169200386 | rs9364294 | -1.438 | 1.53 | 2.858 | 1.651 | 0.098 | 0.083 | 0.083 | 0.219 | 0.06 | 0.3 | 0.12 | 0.14 | 0.09 | 0.09 |
| 6 | 169201047 | rs10806650 | -0.834 | 1.525 | 2.78 | 1.671 | 0.002 | 0.013 | 0.013 | 0 | 0.37 | 0.55 | 0.51 | 0.61 | 0.35 | 0.47 |
| 6 | 169202852 | rs9364299 | -0.678 | 1.529 | 2.636 | 1.673 | -0.002 | -0.005 | -0.005 | 0 | 0.35 | 0.33 | 0.49 | 0.61 | 0.35 | 0.47 |
| 6 | 169204833 | rs9294913 | -0.764 | 1.554 | 2.474 | 1.687 | 0.003 | 0.014 | 0.014 | 0 | 0.37 | 0.45 | 0.5 | 0.61 | 0.35 | 0.48 |
| 6 | 169211953 | rs4496770 | -1.466 | 1.438 | 1.954 | 1.681 | 0.017 | 0.09 | 0.09 | 0 | 0.16 | 0.3 | 0.18 | 0.27 | 0.16 | 0.13 |
| 6 | 169212299 | rs7454257 | 0.002 | 1.626 | 1.095 | 1.547 | -0.019 | -0.069 | -0.069 | 0.062 | 0.3 | 0.15 | 0.4 | 0.46 | 0.29 | 0.38 |
| 6 | 169220177 | rs4594925 | -0.267 | 1.704 | 1.042 | 1.587 | 0 | -0.028 | -0.028 | 0 | 0.33 | 0.27 | 0.41 | 0.46 | 0.29 | 0.38 |
| 6 | 169222409 | rs4708497 | -1.224 | 2.023 | 0.99 | 1.63 | -0.003 | 0.067 | 0.067 | 0 | 0.49 | 0.72 | 0.6 | 0.67 | 0.45 | 0.51 |
| 6 | 169228159 | rs4708499 | -1.039 | 2.047 | 0.482 | 1.62 | 0.117 | 0.137 | 0.137 | 0.416 | 0.13 | 0.38 | 0.26 | 0.29 | 0.23 | 0.14 |
| 6 | 169229013 | rs4708500 | -1.135 | 1.376 | -0.691 | 0.741 | 0.009 | 0.136 | 0.136 | 0.53 | 0.55 | 0.87 | 0.71 | 0.8 | 0.52 | 0.59 |
| 6 | 169237939 | rs6900380 | 0.971 | 2.177 | -0.102 | 1.521 | 0.003 | 0.005 | 0.005 | 0 | 0.78 | 0.82 | 0.75 | 0.62 | 0.68 | 0.66 |
| 6 | 169243539 | rs9364309 | -1.203 | 2.362 | -0.361 | 1.705 | -0.041 | -0.091 | -0.091 | 0 | 0.15 | 0.03 | 0.35 | 0.67 | 0.17 | 0.4 |
| 6 | 169249038 | rs9364180 | -1.989 | 1.711 | 0.31 | 1.435 | -0.014 | 0.028 | 0.028 | 0.449 | 0.23 | 0.29 | 0.4 | 0.68 | 0.18 | 0.43 |
| 6 | 169251249 | rs7453792 | 1.779 | 1.286 | 0.683 | 0.76 | 0.02 | 0.069 | 0.069 | 2.324 | 0.55 | 0.27 | 0.38 | 0.16 | 0.62 | 0.46 |
| 6 | 169252480 | rs9455734 | -0.706 | 1.29 | 0.698 | 0.888 | 0.004 | -0.049 | -0.049 | 0.029 | 0.09 | 0.2 | 0.07 | 0 | 0.06 | 0.06 |
| 6 | 169263416 | rs9364314 | -0.935 | 0.994 | 0.483 | 1.239 | 0.008 | -0.024 | -0.024 | 0 | 0.13 | 0.08 | 0.33 | 0.65 | 0.11 | 0.37 |
| 6 | 169264259 | rs9294923 | -0.905 | 0.994 | 0.483 | 1.239 | 0.008 | -0.023 | -0.023 | 0 | 0.13 | 0.08 | 0.33 | 0.65 | 0.11 | 0.37 |

| | | | | | | | | | | | | | | | | |
|---|---|---|---|---|---|---|---|---|---|---|---|---|---|---|---|---|
| 6 | 169270879 | rs13191839 | 0.057 | 0.836 | 0.876 | 1.062 | -0.002 | -0.034 | -0.034 | 0 | 0.15 | 0.22 | 0.09 | 0 | 0.12 | 0.05 |
| 6 | 169270957 | rs7756719 | 1.342 | 0.824 | 0.943 | 1.104 | 0 | -0.001 | -0.001 | 1.836 | 0.71 | 0.64 | 0.57 | 0.35 | 0.76 | 0.58 |
| 6 | 169274467 | rs6455435 | -0.191 | 0.885 | 1.39 | 1 | -0.024 | -0.099 | -0.099 | 0 | 0.16 | 0.37 | 0.11 | 0 | 0.13 | 0.05 |
| 6 | 169276167 | rs9294927 | -0.973 | 0.672 | 0.888 | 0.673 | -0.041 | -0.046 | -0.046 | 5.97 | 0.24 | 0.09 | 0.43 | 0.69 | 0.27 | 0.44 |
| 6 | 169277046 | rs9455754 | 1.206 | 0.681 | 0.896 | 0.71 | -0.003 | -0.014 | -0.014 | 0 | 0.59 | 0.48 | 0.45 | 0.31 | 0.61 | 0.51 |
| 6 | 169296305 | rs6920859 | -1.039 | 0.985 | 1.111 | 0.651 | 0.101 | -0.009 | -0.009 | 3.709 | 0.23 | 0.44 | 0.27 | 0.14 | 0.31 | 0.11 |
| 6 | 169296369 | rs6922341 | -1.066 | 1.083 | 1.168 | 0.712 | 0.222 | 0.132 | 0.132 | 0 | 0.12 | 0.43 | 0.22 | 0.14 | 0.21 | 0.09 |
| 6 | 169297420 | rs4708389 | -1.497 | 0.932 | 1.072 | 1.014 | 0.007 | 0.013 | 0.013 | 0.06 | 0.3 | 0.61 | 0.31 | 0.36 | 0.34 | 0.35 |
| 6 | 169306641 | rs4708542 | 0.021 | -0.381 | 0.651 | 0.595 | -0.001 | 0 | 0 | 0 | 0.7 | 0.69 | 0.64 | 0.59 | 0.73 | 0.83 |
| 6 | 169318610 | rs8180556 | 1.324 | -0.501 | 1.233 | 0.588 | 0.078 | 0.067 | 0.067 | 0.3 | 0.91 | 0.6 | 0.89 | 0.94 | 0.92 | 0.96 |
| 6 | 169320658 | rs9364328 | -0.389 | -0.683 | 1.131 | 0.563 | 0.024 | 0.047 | 0.047 | 0 | 0.24 | 0.13 | 0.14 | 0.12 | 0.21 | 0.13 |
| 6 | 169323526 | rs9294935 | 1.36 | -0.524 | 1.242 | 1.011 | 0.008 | 0.165 | 0.165 | 0.225 | 0.83 | 0.56 | 0.84 | 0.69 | 0.88 | 0.75 |
| 6 | 169327156 | rs6927830 | -1.334 | -0.645 | 0.822 | 0.843 | -0.016 | 0.018 | 0.018 | 0 | 0.38 | 0.43 | 0.24 | 0.46 | 0.3 | 0.35 |
| 6 | 169327747 | rs9346491 | -0.931 | -0.542 | 0.141 | 0.212 | 0.013 | 0.03 | 0.03 | 0 | 0.28 | 0.17 | 0.18 | 0.17 | 0.25 | 0.14 |
| 6 | 169334481 | rs9455504 | -0.303 | -0.548 | -0.021 | 0.261 | 0.044 | 0.011 | 0.011 | 0 | 0.22 | 0.08 | 0.1 | 0.15 | 0.17 | 0.11 |
| 6 | 169334898 | rs6455455 | 0.771 | -0.08 | 0.657 | 0.224 | 0.019 | -0.04 | -0.04 | 0 | 0.89 | 0.76 | 0.89 | 0.98 | 0.92 | 0.89 |
| 6 | 169338671 | rs4596451 | -0.31 | 0.024 | 0.251 | 0.162 | 0.004 | -0.008 | -0.008 | 1.488 | 0.34 | 0.35 | 0.39 | 0.28 | 0.45 | 0.19 |
| 6 | 169343146 | rs9346596 | -0.356 | -0.543 | 0.112 | 0.34 | 0.017 | -0.021 | -0.021 | 0 | 0.25 | 0.17 | 0.2 | 0.52 | 0.18 | 0.36 |
| 6 | 169347773 | rs9294937 | 0.592 | -0.529 | 0.332 | 0.665 | 0.006 | 0.003 | 0.003 | 0 | 0.49 | 0.42 | 0.44 | 0.42 | 0.45 | 0.52 |
| 6 | 169348035 | rs6905095 | -0.484 | -0.798 | 0.38 | 0.554 | -0.005 | -0.003 | -0.003 | 0.403 | 0.39 | 0.36 | 0.51 | 0.58 | 0.46 | 0.44 |
| 6 | 169352091 | rs2180332 | 0.786 | -0.704 | -0.264 | 0.599 | -0.091 | 0.001 | 0.001 | 0 | 0.21 | 0.11 | 0.34 | 0.27 | 0.37 | 0.17 |
| 6 | 169358695 | rs11969044 | 0.516 | -0.655 | -0.118 | 0.751 | -0.003 | 0.014 | 0.014 | 0 | 0.1 | 0.11 | 0.07 | 0.2 | 0.1 | 0.08 |
| 6 | 169362590 | rs3818701 | -0.519 | -0.094 | -0.504 | 0.675 | -0.086 | -0.076 | -0.076 | 0.551 | 0.34 | 0.16 | 0.46 | 0.56 | 0.42 | 0.4 |
| 6 | 169365642 | rs9455801 | -0.675 | -0.193 | -0.486 | 0.612 | -0.048 | -0.029 | -0.029 | 0.711 | 0.38 | 0.21 | 0.49 | 0.57 | 0.44 | 0.43 |
| 6 | 169368878 | rs13200536 | 0.569 | 0.029 | -0.535 | 0.731 | -0.1 | -0.008 | -0.008 | 0 | 0.2 | 0.08 | 0.31 | 0.26 | 0.33 | 0.12 |
| 6 | 169369886 | rs9455812 | -0.374 | -0.55 | 0.135 | 0.988 | -0.003 | 0.019 | 0.019 | 0 | 0.15 | 0.13 | 0.15 | 0.31 | 0.09 | 0.27 |
| 6 | 169371188 | rs7755281 | -0.611 | -0.244 | -0.024 | 0.426 | -0.033 | -0.072 | -0.072 | 0 | 0.12 | 0.26 | 0.04 | 0 | 0.08 | 0.03 |
| 6 | 169381348 | rs6931239 | -0.698 | 0.329 | -0.699 | 1.187 | -0.019 | -0.065 | -0.065 | 0 | 0.18 | 0.07 | 0.28 | 0.42 | 0.2 | 0.34 |
| 6 | 169382503 | rs9364337 | -0.966 | 0.062 | -0.21 | 1.044 | 0 | 0.035 | 0.035 | 0 | 0.08 | 0.15 | 0.05 | 0.08 | 0.06 | 0.06 |
| 6 | 169387386 | rs9294943 | -0.278 | -0.141 | -1.448 | 0.663 | 0.004 | 0.004 | 0.004 | 0.701 | 0.39 | 0.18 | 0.39 | 0.32 | 0.41 | 0.55 |
| 6 | 169393301 | rs1757974 | -0.157 | -0.339 | -1.031 | 0.206 | -0.025 | -0.062 | -0.062 | 0.666 | 0.62 | 0.44 | 0.72 | 0.87 | 0.72 | 0.67 |
| 6 | 169393766 | rs1757975 | -0.285 | -0.525 | -0.411 | 0.06 | -0.017 | -0.043 | -0.043 | 0 | 0.65 | 0.58 | 0.77 | 0.88 | 0.75 | 0.7 |
| 6 | 169394863 | rs9346498 | -0.283 | -0.629 | -0.31 | -0.054 | -0.028 | 0.055 | 0.055 | 0 | 0.43 | 0.7 | 0.55 | 0.76 | 0.49 | 0.42 |
| 6 | 169397298 | rs2016484 | -0.677 | -0.653 | -0.328 | -0.095 | 0.028 | 0.022 | 0.022 | 0.152 | 0.7 | 0.77 | 0.88 | 1 | 0.81 | 0.89 |
| 6 | 169398467 | rs7766847 | -0.015 | -0.605 | -0.272 | -0.04 | 0.005 | 0.002 | 0.002 | 0 | 0.38 | 0.54 | 0.43 | 0.63 | 0.42 | 0.22 |
| 6 | 169400285 | rs17181982 | 1.25 | -0.584 | -0.637 | -0.168 | -0.046 | -0.008 | -0.008 | 0 | 0.33 | 0.07 | 0.36 | 0.25 | 0.33 | 0.51 |
| 6 | 169402207 | rs9355031 | 0.096 | -0.465 | -0.406 | -0.217 | -0.001 | 0.02 | 0.02 | 0 | 0.1 | 0.24 | 0.06 | 0.08 | 0.08 | 0.06 |
| 6 | 169405314 | rs17182135 | 1.014 | -0.278 | -0.461 | -0.164 | -0.033 | -0.037 | -0.037 | 0 | 0.37 | 0.19 | 0.46 | 0.37 | 0.4 | 0.69 |
| 6 | 169405638 | rs9355119 | 0.207 | -0.168 | -0.494 | -0.189 | -0.004 | -0.004 | -0.004 | 0 | 0.47 | 0.4 | 0.52 | 0.45 | 0.48 | 0.75 |
| 6 | 169406072 | rs4708575 | 0.207 | -0.168 | -0.494 | -0.189 | -0.004 | -0.004 | -0.004 | 0 | 0.47 | 0.4 | 0.52 | 0.45 | 0.48 | 0.75 |
| 6 | 169410675 | rs9364195 | -0.773 | -0.272 | -0.688 | -0.304 | 0.037 | 0.066 | 0.066 | 0.434 | 0.72 | 0.83 | 0.89 | 1 | 0.81 | 0.9 |
| 6 | 169421120 | rs11752531 | 0.76 | -0.197 | -0.34 | -0.209 | 0.017 | 0.049 | 0.049 | 0 | 0.21 | 0.23 | 0.3 | 0.54 | 0.29 | 0.13 |
| 6 | 169422415 | rs11757721 | 0.76 | -0.004 | -0.34 | -0.209 | 0.013 | 0.049 | 0.049 | 0 | 0.21 | 0.23 | 0.3 | 0.54 | 0.29 | 0.13 |
| 6 | 169427629 | rs655307 | -0.684 | 0.022 | -0.392 | -0.165 | 0.038 | 0.064 | 0.064 | 0.443 | 0.71 | 0.82 | 0.89 | 1 | 0.81 | 0.91 |
| 6 | 169428312 | rs10945359 | 0.238 | 0.237 | -0.436 | 0.098 | -0.003 | -0.012 | -0.012 | 0.436 | 0.55 | 0.41 | 0.67 | 0.82 | 0.64 | 0.66 |
| 6 | 169428452 | rs866743 | 0.145 | 0.265 | -0.625 | 0.112 | 0.012 | 0.022 | 0.022 | 0.686 | 0.55 | 0.35 | 0.66 | 0.82 | 0.64 | 0.66 |
| 6 | 169429526 | rs12529420 | 0.629 | 0.254 | -1.344 | -0.042 | -0.056 | -0.005 | -0.005 | 0 | 0.38 | 0.07 | 0.37 | 0.27 | 0.37 | 0.56 |
| 6 | 169434121 | rs369417 | -0.994 | -0.003 | -1.227 | 0.047 | -0.107 | -0.051 | -0.051 | 0.395 | 0.73 | 0.97 | 0.83 | 0.83 | 0.66 | 0.64 |
| 6 | 169436715 | rs416598 | -0.122 | -0.215 | -0.452 | 0.383 | -0.005 | -0.001 | -0.001 | 0 | 0.88 | 0.88 | 0.95 | 0.91 | 0.89 | 0.9 |
| 6 | 169440779 | rs12110438 | 0.816 | -0.357 | -0.523 | 0.64 | -0.013 | -0.002 | -0.002 | 0 | 0.17 | 0.05 | 0.33 | 0.22 | 0.19 | 0.16 |
| 6 | 169440856 | rs7765333 | -0.435 | -0.486 | -0.423 | 0.504 | 0.009 | 0.022 | 0.022 | 0 | 0.55 | 0.76 | 0.73 | 0.75 | 0.57 | 0.54 |
| 6 | 169442109 | rs13203299 | 1.635 | -0.194 | -0.587 | 0.643 | -0.05 | -0.045 | -0.045 | 0.491 | 0.08 | 0.01 | 0.18 | 0.3 | 0.19 | 0.07 |
| 6 | 169444202 | rs17182988 | 1.205 | -0.661 | -0.214 | 0.2 | 0.01 | -0.013 | -0.013 | 0.062 | 0.18 | 0.22 | 0.06 | 0 | 0.08 | 0.07 |
| 6 | 169447055 | rs376307 | -0.776 | -0.424 | -0.209 | 0.439 | -0.001 | 0 | 0 | 0 | 0.89 | 0.91 | 0.95 | 0.92 | 0.89 | 0.91 |
| 6 | 169448440 | rs432401 | 0.179 | -0.39 | 0.046 | 0.535 | 0.006 | 0.028 | 0.028 | 0 | 0.84 | 0.77 | 0.83 | 0.76 | 0.82 | 0.71 |
| 6 | 169448796 | rs12193635 | 0.162 | -0.944 | -0.1 | 0.323 | 0.045 | 0.071 | 0.071 | 0.136 | 0.4 | 0.22 | 0.41 | 0.28 | 0.29 | 0.27 |
| 6 | 169449239 | rs12207730 | 1.258 | -1.105 | -0.068 | 0.235 | -0.005 | -0.035 | -0.035 | 0 | 0.18 | 0.18 | 0.06 | 0 | 0.07 | 0.07 |
| 6 | 169451803 | rs10945364 | 0.595 | -1.018 | -0.201 | 0.401 | 0.05 | 0.105 | 0.105 | 0.454 | 0.49 | 0.31 | 0.25 | 0.17 | 0.36 | 0.4 |
| 6 | 169454921 | rs380992 | -0.16 | -0.709 | -0.241 | 0.675 | 0.012 | 0.013 | 0.013 | 1.94 | 0.66 | 0.66 | 0.46 | 0.43 | 0.54 | 0.72 |
| 6 | 169458719 | rs9455530 | 1.199 | -0.477 | -0.871 | 0.883 | -0.077 | -0.043 | -0.043 | 0 | 0.3 | 0.03 | 0.18 | 0.17 | 0.29 | 0.34 |
| 6 | 169459130 | rs9294948 | 0.083 | -0.251 | 0.314 | 1.446 | -0.004 | 0.001 | 0.001 | 0 | 0.1 | 0.09 | 0.04 | 0.08 | 0.08 | 0.08 |
| 6 | 169462997 | rs369894 | 0.232 | -0.165 | 0.355 | 1.606 | 0.014 | 0.017 | 0.017 | 0 | 0.77 | 0.71 | 0.75 | 0.66 | 0.71 | 0.87 |
| 6 | 169463101 | rs9689200 | -1.403 | -0.027 | 0.433 | 1.984 | -0.026 | 0.116 | 0.116 | 0.181 | 0.23 | 0.45 | 0.28 | 0.47 | 0.17 | 0.43 |
| 6 | 169463997 | rs12526105 | 0.134 | 0.05 | -0.566 | 1.466 | 0.007 | -0.02 | -0.02 | 0 | 0.1 | 0.05 | 0.23 | 0.22 | 0.07 | 0.13 |
| 6 | 169464306 | rs446212 | 0.227 | 0.554 | -0.234 | 1.23 | -0.011 | -0.011 | -0.011 | 0 | 0.82 | 0.87 | 0.78 | 0.64 | 0.72 | 0.85 |
| 6 | 169467285 | rs9294949 | -0.913 | -0.463 | 0.17 | 0.529 | -0.07 | -0.035 | -0.035 | 0.011 | 0.2 | 0.37 | 0.08 | 0.1 | 0.11 | 0.17 |
| 6 | 169471078 | rs9346612 | 0.109 | -0.112 | -0.505 | 0.681 | 0.023 | -0.02 | -0.02 | 0 | 0.22 | 0.16 | 0.28 | 0.29 | 0.19 | 0.27 |
| 6 | 169471707 | rs4708582 | -0.286 | 0.537 | -0.43 | 0.181 | 0.001 | 0.001 | 0.001 | 0 | 0.53 | 0.49 | 0.49 | 0.47 | 0.49 | 0.49 |
| 6 | 169472759 | rs9355126 | 0.088 | 0.54 | -0.448 | 0.013 | -0.022 | -0.005 | -0.005 | 0 | 0.42 | 0.47 | 0.43 | 0.42 | 0.31 | 0.42 |

| | | | | | | | | | | | | | | | | |
|---|---|---|---|---|---|---|---|---|---|---|---|---|---|---|---|---|
| 6 | 169473174 | rs436474 | -1.117 | 1.231 | -0.531 | -0.063 | -0.02 | -0.033 | -0.033 | 0 | 0.79 | 0.84 | 0.65 | 0.58 | 0.71 | 0.78 |
| 6 | 169473816 | rs633668 | -1.241 | 1.19 | -0.551 | 0.137 | -0.093 | -0.117 | -0.117 | 0.884 | 0.8 | 0.96 | 0.66 | 0.65 | 0.71 | 0.79 |
| 6 | 169473856 | rs13201804 | 0.185 | 1.286 | -0.709 | -0.033 | -0.003 | -0.003 | -0.003 | 0 | 0.55 | 0.48 | 0.5 | 0.43 | 0.45 | 0.57 |
| 6 | 169474687 | rs10485332 | 1.018 | 1.352 | -0.781 | -0.018 | -0.027 | 0.065 | 0.065 | 2.532 | 0.13 | 0.04 | 0.07 | 0 | 0.14 | 0.15 |
| 6 | 169474908 | rs442432 | -1.19 | 1.422 | -0.682 | 0.093 | -0.073 | -0.096 | -0.096 | 0 | 0.79 | 0.93 | 0.65 | 0.65 | 0.7 | 0.79 |
| 6 | 169477665 | rs2183840 | 0.823 | 1.437 | -0.555 | 0.084 | -0.026 | 0.025 | 0.025 | 0 | 0.14 | 0.09 | 0.07 | 0.06 | 0.19 | 0.07 |
| 6 | 169485639 | rs12179836 | 0.009 | 1.199 | -0.461 | 0.079 | -0.049 | -0.058 | -0.058 | 0 | 0.37 | 0.27 | 0.44 | 0.49 | 0.49 | 0.29 |
| 6 | 169489561 | rs10945370 | 0.022 | 1.222 | -0.463 | 0.077 | -0.058 | -0.053 | -0.053 | 0 | 0.37 | 0.25 | 0.43 | 0.42 | 0.49 | 0.28 |
| 6 | 169491266 | rs2224644 | -0.085 | 1.149 | -0.159 | 0.04 | 0.005 | -0.005 | -0.005 | 0 | 0.91 | 0.88 | 0.95 | 0.94 | 0.93 | 0.86 |
| 6 | 169492964 | rs9455534 | -0.037 | 1.155 | -0.01 | 0.064 | -0.025 | -0.019 | -0.019 | 0 | 0.38 | 0.32 | 0.45 | 0.52 | 0.49 | 0.29 |
| 6 | 169495334 | rs9767300 | 0.482 | 0.782 | -0.001 | -0.057 | 0.02 | 0.036 | 0.036 | 0.013 | 0.06 | 0.17 | 0.04 | 0.07 | 0.05 | 0.12 |
| 6 | 169503245 | rs9406337 | 0.167 | 1.346 | 0.105 | 0.027 | -0.016 | -0.02 | -0.02 | 0 | 0.34 | 0.21 | 0.39 | 0.51 | 0.41 | 0.29 |
| 6 | 169506837 | rs833681 | -0.126 | 1.306 | 0.084 | -0.098 | -0.002 | -0.062 | -0.062 | 0 | 0.77 | 0.71 | 0.88 | 0.96 | 0.8 | 0.7 |
| 6 | 169508247 | rs9294955 | 0.439 | 1.465 | 0.044 | -0.1 | 0.002 | -0.049 | -0.049 | 0 | 0.63 | 0.5 | 0.8 | 0.83 | 0.64 | 0.62 |
| 6 | 169509545 | rs4708592 | 0.166 | 1.437 | 0.131 | -0.059 | -0.009 | 0.05 | 0.05 | 1.215 | 0.88 | 0.52 | 0.91 | 0.85 | 0.93 | 0.85 |
| 6 | 169514577 | rs1407553 | 0.228 | 1.464 | 0.048 | -0.149 | -0.009 | -0.002 | -0.002 | 0.466 | 0.56 | 0.63 | 0.54 | 0.48 | 0.52 | 0.57 |
| 6 | 169518030 | rs12525738 | 1.162 | 1.508 | -0.071 | -0.13 | -0.055 | 0.019 | 0.019 | 0 | 0.12 | 0.01 | 0.06 | 0.05 | 0.16 | 0.07 |
| 6 | 169519052 | rs2093524 | 0.174 | 1.504 | -0.003 | -0.129 | -0.001 | -0.001 | -0.001 | 0.579 | 0.56 | 0.5 | 0.54 | 0.58 | 0.55 | 0.5 |
| 6 | 169520560 | rs9355132 | 0.774 | 1.472 | -0.036 | -0.121 | 0.007 | 0 | 0 | 0 | 0.42 | 0.38 | 0.46 | 0.45 | 0.38 | 0.42 |
| 6 | 169532439 | rs7757988 | 0.752 | 1.487 | -0.132 | -0.156 | 0.007 | 0.006 | 0.006 | 0 | 0.42 | 0.41 | 0.46 | 0.43 | 0.38 | 0.42 |
| 6 | 169532735 | rs7762175 | 1.175 | 1.397 | -0.183 | -0.244 | -0.021 | -0.031 | -0.031 | 0 | 0.19 | 0.29 | 0.19 | 0.13 | 0.12 | 0.2 |
| 6 | 169539647 | rs13200360 | 0.542 | 1.405 | -0.261 | -0.084 | 0.003 | -0.001 | -0.001 | 0.002 | 0.44 | 0.59 | 0.53 | 0.48 | 0.38 | 0.44 |
| 6 | 169540055 | rs833680 | -0.352 | 1.189 | -0.325 | -0.246 | -0.001 | -0.011 | -0.011 | 0 | 0.79 | 0.81 | 0.88 | 0.86 | 0.8 | 0.71 |
| 6 | 169542869 | rs13217516 | 0.868 | 1.54 | -0.963 | 0.084 | -0.061 | -0.121 | -0.121 | 0 | 0.21 | 0.04 | 0.32 | 0.35 | 0.25 | 0.21 |
| 6 | 169546063 | rs9689078 | -0.349 | 1.816 | -0.911 | 0.004 | -0.012 | -0.006 | -0.006 | 0 | 0.44 | 0.31 | 0.46 | 0.51 | 0.5 | 0.44 |
| 6 | 169546486 | rs9766768 | -1.159 | 1.78 | -0.923 | -0.018 | -0.04 | -0.046 | -0.046 | 0.536 | 0.73 | 0.57 | 0.78 | 0.79 | 0.84 | 0.68 |
| 6 | 169548692 | rs9355134 | 0.191 | 2.57 | -0.498 | 0.598 | -0.029 | 0.004 | 0.004 | 0 | 0.14 | 0.07 | 0.07 | 0.1 | 0.17 | 0.08 |
| 6 | 169549501 | rs1885656 | -0.751 | 2.592 | -0.467 | 0.602 | 0.002 | -0.006 | -0.006 | 0 | 0.28 | 0.25 | 0.16 | 0.18 | 0.32 | 0.24 |
| 6 | 169554727 | rs9294963 | -1.15 | 2.466 | -0.009 | 1.14 | 0.022 | 0.024 | 0.024 | 0 | 0.36 | 0.42 | 0.42 | 0.46 | 0.44 | 0.29 |
| 6 | 169555451 | rs911970 | -0.859 | 1.848 | 0.47 | 1.438 | 0.023 | 0.1 | 0.1 | 0 | 0.22 | 0.35 | 0.35 | 0.36 | 0.27 | 0.21 |
| 6 | 169555623 | rs2343367 | -0.326 | 1.345 | -0.375 | 1.625 | -0.002 | -0.02 | -0.02 | 0 | 0.17 | 0.13 | 0.32 | 0.36 | 0.2 | 0.19 |
| 6 | 169556108 | rs2093525 | -0.461 | 1.362 | -0.433 | 1.633 | -0.001 | 0.036 | 0.036 | 0 | 0.18 | 0.21 | 0.32 | 0.35 | 0.21 | 0.2 |
| 6 | 169558178 | rs12210313 | -1.043 | 1.731 | -0.731 | 1.489 | -0.043 | -0.036 | -0.036 | 0.02 | 0.07 | 0.01 | 0.22 | 0.23 | 0.15 | 0.13 |
| 6 | 169558952 | rs2181284 | -0.813 | 1.238 | -0.297 | 1.026 | -0.01 | -0.029 | -0.029 | 0.711 | 0.05 | 0.1 | 0.02 | 0 | 0.04 | 0.02 |
| 6 | 169559115 | rs1985206 | -0.135 | 0.829 | -0.432 | 0.5 | 0.002 | 0.004 | 0.004 | 0 | 0.47 | 0.41 | 0.44 | 0.53 | 0.52 | 0.47 |
| 6 | 169559506 | rs9294965 | 0.117 | -0.066 | 0.139 | -0.127 | -0.011 | -0.063 | -0.063 | 0 | 0.11 | 0.24 | 0.08 | 0 | 0.08 | 0.02 |
| 6 | 169560597 | rs9455882 | -1.231 | -0.012 | 0.344 | 0.038 | 0 | 0 | 0 | 0 | 0.13 | 0.15 | 0.18 | 0.13 | 0.13 | 0.07 |
| 6 | 169561395 | rs9688959 | -0.477 | -0.629 | 0.343 | 0.055 | -0.056 | 0 | 0 | 0 | 0.68 | 0.58 | 0.66 | 0.68 | 0.82 | 0.6 |
| 6 | 169562308 | rs4708599 | 0.775 | -0.599 | -0.097 | -0.009 | 0.004 | -0.003 | -0.003 | 0 | 0.46 | 0.43 | 0.42 | 0.43 | 0.59 | 0.4 |
| 6 | 169563337 | rs6925188 | -0.62 | -0.639 | -0.434 | -0.174 | -0.004 | 0.012 | 0.012 | 0 | 0.16 | 0.11 | 0.2 | 0.13 | 0.17 | 0.09 |
| 6 | 169565035 | rs6910183 | -0.997 | -0.73 | -0.142 | -0.059 | -0.006 | -0.001 | -0.001 | 0 | 0.68 | 0.69 | 0.67 | 0.72 | 0.82 | 0.6 |
| 6 | 169571419 | rs9294970 | -0.934 | -0.791 | -0.157 | -0.1 | -0.002 | 0.021 | 0.021 | 0 | 0.26 | 0.12 | 0.25 | 0.25 | 0.25 | 0.22 |
| 6 | 169571693 | rs3763267 | 0.816 | -0.766 | -0.184 | -0.09 | -0.017 | -0.014 | -0.014 | 0 | 0.1 | 0 | 0.06 | 0.12 | 0.09 | 0.13 |
| 6 | 169577804 | rs1475139 | 0.995 | -0.927 | 0.002 | -0.113 | 0 | -0.001 | -0.001 | 0 | 0.4 | 0.37 | 0.41 | 0.42 | 0.57 | 0.39 |
| 6 | 169586374 | rs9689350 | -1.101 | -1.808 | 0.17 | -0.426 | -0.022 | -0.001 | -0.001 | 0.229 | 0.85 | 0.77 | 0.85 | 0.85 | 0.93 | 0.8 |
| 6 | 169589925 | rs1555523 | 1.138 | -1.76 | 0.158 | -0.445 | -0.009 | -0.003 | -0.003 | 0 | 0.16 | 0.18 | 0.18 | 0.12 | 0.1 | 0.18 |
| 6 | 169591548 | rs7743310 | -1.475 | -1.658 | 0.132 | -0.534 | 0.008 | 0.016 | 0.016 | 0 | 0.6 | 0.53 | 0.59 | 0.58 | 0.47 | 0.62 |
| 6 | 169591678 | rs7743597 | -0.984 | -1.619 | 0.088 | -0.514 | 0.022 | 0.013 | 0.013 | 0 | 0.42 | 0.32 | 0.41 | 0.45 | 0.36 | 0.44 |
| 6 | 169591793 | rs9505942 | -0.484 | -1.488 | 0.183 | -0.428 | 0 | 0 | 0 | 0 | 0.16 | 0.11 | 0.19 | 0.18 | 0.16 | 0.1 |
| 6 | 169593545 | rs6941695 | 0.942 | -1.443 | 0.042 | -0.482 | -0.008 | -0.003 | -0.003 | 0 | 0.16 | 0.18 | 0.18 | 0.13 | 0.1 | 0.2 |
| 6 | 169594139 | rs9406329 | 1.06 | -1.348 | 0.023 | -0.451 | 0 | 0 | 0 | 0.457 | 0.15 | 0.19 | 0.16 | 0.15 | 0.11 | 0.21 |
| 6 | 169595246 | rs6910371 | -0.128 | -1.198 | -0.103 | -0.499 | 0.006 | 0.004 | 0.004 | 0 | 0.25 | 0.2 | 0.23 | 0.17 | 0.18 | 0.23 |
| 6 | 169595259 | rs9406318 | -0.114 | -0.627 | -0.133 | -0.338 | -0.026 | -0.023 | -0.023 | 0 | 0.07 | 0.01 | 0.14 | 0.1 | 0.08 | 0.04 |
| 6 | 169598871 | rs12181211 | 0.614 | -0.255 | 0.084 | -1.004 | -0.008 | -0.084 | -0.084 | 2.179 | 0.21 | 0.3 | 0.15 | 0.02 | 0.19 | 0.03 |
| 6 | 169599832 | rs9505953 | 0.427 | -0.742 | 0.059 | -1.287 | 0.008 | 0.012 | 0.012 | 0 | 0.25 | 0.22 | 0.22 | 0.16 | 0.2 | 0.23 |
| 6 | 169600161 | rs7381269 | -1.199 | -0.735 | 0.147 | -1.412 | -0.014 | -0.016 | -0.016 | 0 | 0.78 | 0.82 | 0.72 | 0.47 | 0.66 | 0.64 |
| 6 | 169601313 | rs7775470 | 1.439 | -0.885 | 0.496 | -1.461 | -0.004 | -0.042 | -0.042 | 0 | 0.18 | 0.23 | 0.13 | 0.02 | 0.17 | 0.03 |
| 6 | 169606443 | rs9505933 | 0.107 | -1.169 | 0.426 | -1.567 | 0.021 | 0.017 | 0.017 | 0.45 | 0.43 | 0.42 | 0.5 | 0.36 | 0.32 | 0.44 |
| 6 | 169607718 | rs7764005 | -0.139 | -1.168 | 0.517 | -1.574 | -0.013 | 0.005 | 0.005 | 2.922 | 0.62 | 0.73 | 0.62 | 0.39 | 0.48 | 0.47 |
| 6 | 169609256 | rs12665286 | 1.376 | -1.122 | 0.227 | -1.377 | 0.014 | 0.009 | 0.009 | 0 | 0.17 | 0.13 | 0.27 | 0.13 | 0.11 | 0.21 |
| 6 | 169616423 | rs1132782 | 1.101 | -1.017 | 0.179 | -1.382 | -0.002 | 0.001 | 0.001 | 0 | 0.23 | 0.2 | 0.17 | 0.1 | 0.24 | 0.08 |
| 6 | 169617726 | rs8089 | 1.117 | -1.03 | 0.178 | -1.383 | -0.002 | 0 | 0 | 0 | 0.23 | 0.2 | 0.17 | 0.1 | 0.24 | 0.08 |
| 6 | 169617840 | rs12435 | 0.276 | -0.63 | 0.138 | -1.018 | -0.014 | -0.012 | -0.012 | 4.938 | 0.28 | 0.35 | 0.24 | 0.24 | 0.21 | 0.24 |
| 6 | 169620247 | rs9393165 | -0.86 | 0.288 | -0.207 | -0.345 | -0.041 | -0.043 | -0.043 | 0 | 0.8 | 0.89 | 0.72 | 0.47 | 0.65 | 0.64 |
| 6 | 169621136 | rs10945405 | -0.353 | 0.194 | -0.093 | -0.182 | -0.034 | -0.024 | -0.024 | 0 | 0.7 | 0.79 | 0.68 | 0.47 | 0.55 | 0.51 |
| 6 | 169625874 | rs9505888 | -0.049 | 0.372 | 0.112 | -0.343 | 0.015 | -0.014 | -0.014 | 0 | 0.65 | 0.54 | 0.58 | 0.32 | 0.58 | 0.49 |
| 6 | 169626065 | rs9505889 | -0.456 | -0.522 | -0.451 | -0.815 | -0.002 | 0.007 | 0.007 | 0 | 0.22 | 0.26 | 0.34 | 0.33 | 0.2 | 0.22 |
| 6 | 169630340 | rs6605521 | -0.553 | 0.347 | -1.16 | -0.351 | -0.072 | -0.085 | -0.085 | 0 | 0.69 | 0.87 | 0.66 | 0.5 | 0.56 | 0.59 |
| 6 | 169631907 | rs6940420 | -0.064 | -0.749 | -1.128 | -0.672 | -0.02 | 0.005 | 0.005 | 0 | 0.24 | 0.32 | 0.25 | 0.22 | 0.19 | 0.3 |

| | | | | | | | | | | | | | | | | |
|---|---|---|---|---|---|---|---|---|---|---|---|---|---|---|---|---|
| 6 | 169633185 | rs11756502 | -0.393 | -1.025 | -1.151 | -0.855 | -0.053 | 0.074 | 0.074 | 0 | 0.27 | 0.47 | 0.33 | 0.46 | 0.2 | 0.24 |
| 6 | 169637456 | rs9283850 | -0.442 | -0.821 | -1.888 | -1.383 | -0.03 | 0.188 | 0.188 | 0 | 0.54 | 0.75 | 0.66 | 0.9 | 0.52 | 0.49 |
| 6 | 169639901 | rs9379336 | 0.504 | -0.634 | -1.543 | -0.938 | -0.007 | -0.063 | -0.063 | 0.059 | 0.22 | 0.17 | 0.33 | 0.44 | 0.22 | 0.36 |
| 6 | 169640257 | rs4076564 | 0.434 | -0.517 | -1.149 | -0.618 | -0.008 | 0.18 | 0.18 | 0.024 | 0.08 | 0.39 | 0.12 | 0.19 | 0.04 | 0.04 |
| 6 | 169643685 | rs9379341 | 0.14 | -0.129 | -1.792 | -1.414 | 0.008 | 0.003 | 0.003 | 0.221 | 0.37 | 0.65 | 0.47 | 0.63 | 0.3 | 0.51 |
| 6 | 169646920 | rs9505896 | 0.216 | 0.025 | -1.881 | -1.563 | -0.007 | 0.067 | 0.067 | 0.043 | 0.27 | 0.1 | 0.17 | 0.02 | 0.26 | 0.19 |
| 6 | 169647022 | rs7382358 | 0.158 | -0.09 | -2.168 | -1.292 | 0.03 | 0.036 | 0.036 | 0.064 | 0.36 | 0.16 | 0.27 | 0.27 | 0.31 | 0.35 |
| 6 | 169648030 | rs11756439 | 0.158 | -0.09 | -2.183 | -1.292 | 0.031 | 0.036 | 0.036 | 0.064 | 0.36 | 0.16 | 0.27 | 0.27 | 0.31 | 0.35 |
| 6 | 169652568 | rs9717609 | 0.363 | 0.336 | -1.463 | -0.891 | -0.036 | 0.06 | 0.06 | 2.344 | 0.12 | 0.01 | 0.11 | 0.02 | 0.14 | 0.03 |
| 6 | 169656120 | rs6605524 | -0.286 | -0.071 | -1.298 | -1.125 | 0.021 | 0.199 | 0.199 | 4.784 | 0.51 | 0.74 | 0.63 | 0.87 | 0.47 | 0.67 |
| 6 | 169663638 | rs9717830 | -0.467 | -1.108 | -0.736 | -0.643 | 0.01 | 0.006 | 0.006 | 0.954 | 0.39 | 0.3 | 0.42 | 0.67 | 0.24 | 0.46 |
| 6 | 169667351 | rs9379335 | 0.191 | -0.978 | -0.799 | -0.149 | -0.016 | 0.014 | 0.014 | 0.084 | 0.07 | 0.05 | 0.14 | 0.15 | 0.01 | 0.04 |
| 6 | 169668998 | rs7382539 | -0.664 | -0.771 | -0.458 | -0.921 | 0.056 | 0.076 | 0.076 | 0 | 0.49 | 0.59 | 0.55 | 0.76 | 0.32 | 0.6 |
| 6 | 169672120 | rs9406324 | -1.088 | 0.049 | -0.089 | -0.623 | -0.006 | 0.016 | 0.016 | 0.011 | 0.09 | 0.2 | 0.13 | 0.08 | 0.08 | 0.11 |
| 6 | 169678116 | rs7761056 | -0.267 | 0.216 | -0.717 | -1.277 | 0.006 | 0.012 | 0.012 | 1.025 | 0.34 | 0.71 | 0.26 | 0.01 | 0.31 | 0.14 |
| 6 | 169697100 | rs9294983 | -0.079 | 0.406 | -0.682 | -1.032 | -0.006 | 0.064 | 0.064 | 0 | 0.54 | 0.67 | 0.6 | 0.8 | 0.48 | 0.62 |
| 6 | 169700931 | rs13215678 | -0.525 | 0.799 | -0.687 | -1.135 | -0.002 | -0.006 | -0.006 | 0.516 | 0.14 | 0.17 | 0.11 | 0.02 | 0.17 | 0.17 |
| 6 | 169713606 | rs11964794 | 0.379 | 0.532 | -0.765 | -1.631 | -0.05 | 0.066 | 0.066 | 0.844 | 0.36 | 0.2 | 0.25 | 0 | 0.44 | 0.18 |
| 6 | 169717377 | rs9379340 | -0.023 | -0.235 | -0.122 | -0.951 | 0 | -0.001 | -0.001 | 0 | 0.51 | 0.5 | 0.69 | 0.71 | 0.55 | 0.53 |
| 6 | 169717559 | rs9406330 | -0.329 | -0.601 | 0.114 | -1.146 | 0 | -0.004 | -0.004 | 0 | 0.58 | 0.58 | 0.71 | 0.71 | 0.57 | 0.56 |
| 6 | 169718094 | rs9406333 | -0.196 | -0.673 | 0.248 | -0.637 | -0.007 | 0.035 | 0.035 | 0 | 0.14 | 0.17 | 0.43 | 0.57 | 0.11 | 0.35 |
| 6 | 169718109 | rs9406320 | -0.29 | -0.582 | 0.322 | -0.598 | -0.001 | 0.001 | 0.001 | 0 | 0.58 | 0.57 | 0.71 | 0.63 | 0.58 | 0.55 |
| 6 | 169720683 | rs9688888 | -0.272 | -0.783 | -0.5 | -0.468 | 0.029 | -0.013 | -0.013 | 0 | 0.26 | 0.2 | 0.48 | 0.77 | 0.16 | 0.49 |
| 6 | 169723097 | rs12202886 | 0.762 | -0.358 | -0.476 | -0.976 | -0.053 | 0.093 | 0.093 | 0.119 | 0.38 | 0.18 | 0.26 | 0.03 | 0.45 | 0.17 |
| 6 | 169725428 | rs9283852 | -1.041 | -0.702 | 0.113 | -0.199 | -0.007 | 0.001 | 0.001 | 0 | 0.18 | 0.25 | 0.23 | 0.19 | 0.16 | 0.25 |
| 6 | 169726479 | rs9379343 | -0.721 | -0.942 | 0.003 | -0.846 | -0.001 | -0.019 | -0.019 | 0 | 0.37 | 0.41 | 0.6 | 0.88 | 0.36 | 0.65 |
| 6 | 169727601 | rs12190007 | 1.393 | -0.845 | -0.094 | -1.383 | -0.013 | -0.004 | -0.004 | 0 | 0.4 | 0.32 | 0.27 | 0.01 | 0.46 | 0.22 |
| 6 | 169733088 | rs7382645 | -0.742 | -1.094 | -0.108 | -0.495 | -0.008 | -0.002 | -0.002 | 0 | 0.38 | 0.59 | 0.37 | 0.38 | 0.27 | 0.55 |
| 6 | 169739409 | rs7750869 | -0.205 | -0.952 | -0.578 | -0.692 | -0.012 | 0.057 | 0.057 | 0 | 0.84 | 0.97 | 0.91 | 1 | 0.83 | 0.95 |
| 6 | 169742250 | rs7766683 | 0.097 | -1.351 | -0.15 | -0.991 | -0.001 | -0.074 | -0.074 | 0.954 | 0.12 | 0.48 | 0.06 | 0 | 0.05 | 0.05 |
| 6 | 169743497 | rs10455745 | 2.075 | -1.603 | -1.345 | -1.098 | -0.092 | 0.195 | 0.195 | 0 | 0.4 | 0.08 | 0.27 | 0.12 | 0.47 | 0.26 |
| 6 | 169756330 | rs1028295 | 0.449 | -1.603 | -1.352 | -0.836 | -0.026 | -0.047 | -0.047 | 0.174 | 0.1 | 0.02 | 0.27 | 0.59 | 0.12 | 0.23 |
| 6 | 169756921 | rs1028296 | 0.303 | -1.703 | -1.283 | -0.945 | 0.003 | 0.049 | 0.049 | 0.721 | 0.76 | 0.89 | 0.89 | 0.95 | 0.79 | 0.89 |
| 6 | 169763254 | rs7741251 | 0.478 | -1.708 | -1.261 | -0.947 | 0.005 | 0.029 | 0.029 | 0.685 | 0.75 | 0.86 | 0.88 | 0.95 | 0.79 | 0.9 |
| 6 | 169765845 | rs16887939 | 0.87 | -1.726 | -1.64 | -0.961 | -0.05 | -0.064 | -0.064 | 0 | 0.15 | 0.02 | 0.26 | 0.25 | 0.17 | 0.36 |
| 6 | 169765884 | rs9371116 | 0.45 | -1.643 | -1.58 | -0.857 | -0.026 | -0.05 | -0.05 | 0 | 0.1 | 0.02 | 0.27 | 0.59 | 0.12 | 0.22 |
| 6 | 169767163 | rs12216263 | 1.946 | -1.564 | -1.492 | -1.233 | -0.088 | 0.197 | 0.197 | 0 | 0.4 | 0.08 | 0.27 | 0.12 | 0.47 | 0.26 |
| 6 | 169776520 | rs4716370 | -0.435 | -1.262 | -1.322 | -0.916 | -0.008 | 0.031 | 0.031 | 0 | 0.85 | 0.96 | 0.92 | 0.95 | 0.84 | 0.96 |
| 6 | 169778429 | rs9283855 | 0.681 | -1.318 | -1.262 | -0.977 | 0.006 | 0.049 | 0.049 | 0.326 | 0.75 | 0.87 | 0.88 | 0.95 | 0.79 | 0.9 |
| 6 | 169779183 | rs9294994 | -0.317 | -1.238 | -1.264 | -0.891 | -0.001 | 0.027 | 0.027 | 0 | 0.84 | 0.93 | 0.91 | 0.95 | 0.84 | 0.95 |
| 6 | 169785373 | rs3006189 | -0.419 | -1.221 | -1.303 | -0.901 | -0.008 | 0.055 | 0.055 | 0 | 0.85 | 0.96 | 0.91 | 1 | 0.84 | 0.96 |
| 6 | 169789010 | rs9383524 | -0.594 | -1.106 | -1.525 | -0.997 | -0.039 | 0.034 | 0.034 | 0.439 | 0.28 | 0.11 | 0.55 | 0.84 | 0.29 | 0.6 |
| 6 | 169794808 | rs419944 | 0.648 | -0.883 | -1.251 | -0.651 | -0.01 | -0.014 | -0.014 | 0 | 0.08 | 0.01 | 0.02 | 0.04 | 0.05 | 0.06 |
| 6 | 169795607 | rs370834 | -0.316 | -0.931 | -1.329 | -0.798 | -0.035 | 0.095 | 0.095 | 0.032 | 0.85 | 0.99 | 0.92 | 1 | 0.84 | 0.96 |
| 6 | 169798329 | rs3006186 | 0.709 | -0.901 | -1.024 | -0.485 | -0.004 | 0.019 | 0.019 | 0.3 | 0.83 | 0.96 | 0.94 | 0.92 | 0.87 | 0.9 |
| 6 | 169798796 | rs9406339 | 0.393 | -0.838 | -1.402 | -0.133 | -0.034 | -0.104 | -0.104 | 0.702 | 0.15 | 0.03 | 0.27 | 0.52 | 0.15 | 0.22 |
| 6 | 169806814 | rs9478125 | -1.144 | -1.177 | 0.108 | -0.242 | -0.018 | -0.023 | -0.023 | 0.631 | 0.31 | 0.59 | 0.35 | 0.25 | 0.22 | 0.4 |
| 6 | 169810854 | rs16887985 | -1.515 | -1.34 | 0.113 | -0.344 | 0.035 | -0.014 | -0.014 | 0 | 0.37 | 0.75 | 0.38 | 0.38 | 0.25 | 0.45 |
| 6 | 169818001 | rs4469299 | 0.421 | -1.233 | 0.601 | -0.235 | 0.006 | -0.009 | -0.009 | 0 | 0.11 | 0.08 | 0.23 | 0.2 | 0.07 | 0.3 |
| 6 | 169857451 | rs1047910 | -1.359 | -1.246 | 0.819 | -0.332 | -0.009 | -0.015 | -0.015 | 0 | 0.3 | 0.61 | 0.35 | 0.24 | 0.22 | 0.42 |
| 6 | 169888764 | rs2865091 | -0.188 | -1.198 | 0.987 | -0.328 | 0 | -0.008 | -0.008 | 0.062 | 0.09 | 0.13 | 0.06 | 0.03 | 0.09 | 0.07 |
| 6 | 169898317 | rs4716360 | -1.54 | -1.262 | 1.041 | -0.336 | -0.007 | -0.013 | -0.013 | 0 | 0.37 | 0.59 | 0.36 | 0.29 | 0.27 | 0.47 |
| 6 | 169912559 | rs2997887 | 1.077 | -1.297 | 0.885 | -0.219 | 0.007 | 0.199 | 0.199 | 0 | 0.94 | 0.9 | 0.73 | 0.43 | 0.93 | 0.88 |
| 6 | 169914218 | rs9477987 | -1.789 | -1.319 | 0.937 | -0.271 | 0.035 | 0.057 | 0.057 | 0 | 0.4 | 0.81 | 0.39 | 0.29 | 0.27 | 0.47 |
| 6 | 169934212 | rs3006195 | 1.344 | -1.559 | 0.951 | -0.293 | -0.006 | -0.016 | -0.016 | 0 | 0.63 | 0.4 | 0.64 | 0.74 | 0.73 | 0.54 |
| 6 | 169958982 | rs3734905 | 0.523 | -1.52 | 0.776 | -0.222 | 0.009 | -0.02 | -0.02 | 0 | 0.11 | 0.06 | 0.21 | 0.21 | 0.07 | 0.3 |
| 6 | 169966644 | rs9396970 | -0.645 | -1.48 | 0.745 | -0.195 | -0.002 | 0.049 | 0.049 | 0 | 0.08 | 0.07 | 0.28 | 0.65 | 0.08 | 0.18 |
| 6 | 169967244 | rs3817865 | 1.323 | -1.587 | 0.658 | -0.332 | 0.047 | 0.034 | 0.034 | 0 | 0.58 | 0.18 | 0.61 | 0.67 | 0.73 | 0.52 |
| 6 | 169974663 | rs9295006 | 1.641 | -1.572 | 0.742 | -0.304 | 0.053 | 0.008 | 0.008 | 0 | 0.54 | 0.26 | 0.6 | 0.59 | 0.7 | 0.48 |
| 6 | 169977630 | rs9478060 | -1.195 | -1.363 | 1.243 | 0.207 | -0.01 | 0.027 | 0.027 | 0 | 0.29 | 0.43 | 0.45 | 0.34 | 0.23 | 0.42 |
| 6 | 169989238 | rs6932553 | -0.104 | -1.259 | 1.146 | -0.005 | 0.001 | -0.014 | -0.014 | 0.342 | 0.09 | 0.16 | 0.07 | 0.04 | 0.09 | 0.07 |
| 6 | 169994413 | rs2494673 | 0.679 | -1.293 | 1.15 | -0.014 | -0.019 | -0.012 | -0.012 | 0 | 0.86 | 0.74 | 0.93 | 0.9 | 0.91 | 0.89 |
| 6 | 169995365 | rs4716369 | -1.135 | -1.337 | 1.334 | -0.009 | -0.023 | -0.011 | -0.011 | 0 | 0.31 | 0.51 | 0.33 | 0.28 | 0.22 | 0.42 |
| 6 | 169998186 | rs2050062 | -0.167 | -0.895 | 1.485 | 0.578 | -0.003 | -0.026 | -0.026 | 0 | 0.92 | 0.9 | 0.97 | 0.98 | 0.95 | 0.95 |
| 6 | 170036309 | rs7775504 | -1.787 | -0.542 | 1.297 | 0.101 | -0.004 | -0.064 | -0.064 | 1.218 | 0.28 | 0.55 | 0.17 | 0.15 | 0.23 | 0.2 |
| 6 | 170042465 | rs7762296 | -2.094 | -0.543 | 1.209 | 0.058 | 0 | -0.016 | -0.016 | 0 | 0.33 | 0.61 | 0.49 | 0.74 | 0.31 | 0.34 |
| 6 | 170052049 | rs9478086 | -0.979 | -0.427 | 1.075 | 0.014 | 0.013 | -0.052 | -0.052 | 1.23 | 0.14 | 0.34 | 0.14 | 0.04 | 0.13 | 0.09 |
| 6 | 170058374 | rs3800544 | -0.223 | -0.421 | 0.973 | 0.051 | 0 | 0.001 | 0.001 | 0 | 0.08 | 0.1 | 0.05 | 0.09 | 0.07 | 0.07 |
| 6 | 170059627 | rs3823464 | -1.098 | -0.439 | 1.104 | 0.097 | 0.002 | 0.032 | 0.032 | 0.042 | 0.16 | 0.26 | 0.4 | 0.64 | 0.19 | 0.2 |

| | | | | | | | | | | | | | | | | |
|---|---|---|---|---|---|---|---|---|---|---|---|---|---|---|---|---|
| 6 | 170061501 | rs9396992 | -1.323 | -0.572 | 1.013 | 0.037 | 0 | 0.005 | 0.005 | 0 | 0.2 | 0.25 | 0.39 | 0.65 | 0.22 | 0.24 |
| 6 | 170068058 | rs4509126 | -1.166 | -0.648 | 1.046 | -0.274 | 0.001 | 0.057 | 0.057 | 0 | 0.15 | 0.27 | 0.18 | 0.29 | 0.12 | 0.24 |
| 6 | 170101841 | rs1137523 | -1.101 | -0.956 | 0.921 | -0.496 | -0.001 | 0.016 | 0.016 | 0 | 0.17 | 0.2 | 0.37 | 0.64 | 0.19 | 0.21 |
| 6 | 170107702 | rs1127489 | -1.089 | -0.668 | 0.89 | -0.439 | 0.008 | 0.058 | 0.058 | 0 | 0.15 | 0.28 | 0.18 | 0.3 | 0.12 | 0.25 |
| 6 | 170122234 | rs6931721 | -1.126 | -0.437 | 0.337 | -0.983 | -0.004 | 0.053 | 0.053 | 1.251 | 0.87 | 0.96 | 0.93 | 1 | 0.87 | 0.98 |
| 6 | 170138583 | rs7762260 | 0.086 | -0.394 | 0.426 | -0.922 | 0 | -0.002 | -0.002 | 0.736 | 0.09 | 0.13 | 0.09 | 0.05 | 0.11 | 0.09 |
| 6 | 170140184 | rs9383520 | -0.668 | -0.4 | 0.431 | -0.869 | -0.001 | 0.001 | 0.001 | 0 | 0.14 | 0.14 | 0.37 | 0.64 | 0.18 | 0.2 |
| 6 | 170141552 | rs7740218 | -0.866 | -0.262 | 0.54 | -0.871 | 0.017 | 0.071 | 0.071 | 0 | 0.16 | 0.33 | 0.2 | 0.29 | 0.13 | 0.24 |
| 6 | 170156683 | rs9478102 | 0.975 | -0.302 | 0.389 | -0.92 | 0.001 | 0.036 | 0.036 | 1.264 | 0.13 | 0.06 | 0.07 | 0.03 | 0.13 | 0.02 |
| 6 | 170162537 | rs11966349 | -0.477 | -0.175 | 0.52 | -0.792 | 0.018 | 0.091 | 0.091 | 0 | 0.11 | 0.24 | 0.17 | 0.27 | 0.1 | 0.19 |
| 6 | 170172316 | rs6937197 | 0.219 | -0.799 | 0.387 | -1.161 | -0.041 | -0.081 | -0.081 | 0 | 0.24 | 0.37 | 0.13 | 0.05 | 0.15 | 0.2 |
| 6 | 170174055 | rs4716397 | -1.233 | -0.801 | 0.39 | -1.216 | 0.019 | 0.039 | 0.039 | 0 | 0.25 | 0.44 | 0.3 | 0.33 | 0.24 | 0.29 |
| 6 | 170176647 | rs4716399 | -1.228 | -0.798 | 0.36 | -1.214 | 0.017 | 0.039 | 0.039 | 0 | 0.26 | 0.5 | 0.3 | 0.33 | 0.24 | 0.29 |
| 6 | 170184463 | rs9371127 | -0.407 | -0.718 | 0.51 | -1.048 | -0.037 | 0.016 | 0.016 | 0.096 | 0.28 | 0.39 | 0.4 | 0.65 | 0.22 | 0.31 |
| 6 | 170193933 | rs12214759 | -0.214 | -0.617 | -0.085 | -2.117 | 0.001 | -0.022 | -0.022 | 0.438 | 0.65 | 0.62 | 0.42 | 0.19 | 0.64 | 0.66 |
| 6 | 170198918 | rs6939878 | 0.169 | -0.943 | 0.568 | -1.258 | 0.003 | 0.006 | 0.006 | 0 | 0.08 | 0.14 | 0.06 | 0.1 | 0.08 | 0.07 |
| 6 | 170200081 | rs9478112 | 0.345 | -1.021 | 0.592 | -1.32 | -0.033 | -0.067 | -0.067 | 0 | 0.24 | 0.32 | 0.12 | 0.05 | 0.15 | 0.2 |
| 6 | 170202171 | rs3749889 | -0.046 | -0.92 | 0.45 | -1.243 | 0.003 | 0.008 | 0.008 | 0 | 0.09 | 0.17 | 0.07 | 0.1 | 0.08 | 0.07 |
| 6 | 170205573 | rs713367 | -0.046 | -0.906 | 0.45 | -1.243 | 0.004 | 0.008 | 0.008 | 0 | 0.09 | 0.17 | 0.07 | 0.1 | 0.08 | 0.07 |
| 6 | 170206725 | rs725290 | -0.088 | -0.943 | 0.445 | -1.275 | -0.027 | -0.071 | -0.071 | 0 | 0.28 | 0.35 | 0.14 | 0.06 | 0.18 | 0.25 |
| 6 | 170210754 | rs11965178 | -0.264 | -0.288 | 0.094 | -0.649 | 0.002 | 0.006 | 0.006 | 0 | 0.09 | 0.15 | 0.06 | 0.11 | 0.08 | 0.07 |
| 6 | 170212491 | rs16888431 | -0.644 | -0.368 | -0.028 | -0.729 | 0 | 0 | 0 | 2.192 | 0.18 | 0.2 | 0.14 | 0.15 | 0.18 | 0.16 |
| 6 | 170227259 | rs9371179 | -1.443 | -0.497 | 0.101 | -0.879 | -0.001 | -0.004 | -0.004 | 0.09 | 0.33 | 0.69 | 0.47 | 0.77 | 0.31 | 0.3 |
| 6 | 170228698 | rs7762762 | -0.473 | -0.385 | 0.073 | -0.834 | 0 | 0 | 0 | 0 | 0.18 | 0.21 | 0.15 | 0.15 | 0.19 | 0.16 |
| 6 | 170230182 | rs2864054 | -1.006 | -0.86 | -0.088 | -1.182 | -0.019 | 0.001 | 0.001 | 0 | 0.31 | 0.38 | 0.31 | 0.34 | 0.2 | 0.51 |
| 6 | 170232507 | rs4716407 | -1.612 | -0.761 | -1.007 | -1.635 | -0.129 | 0.244 | 0.244 | 0.111 | 0.62 | 0.93 | 0.74 | 0.99 | 0.51 | 0.67 |
| 6 | 170239266 | rs6908341 | -0.283 | -0.767 | -0.338 | -0.735 | 0 | 0 | 0 | 0 | 0.17 | 0.14 | 0.23 | 0.18 | 0.13 | 0.35 |
| 6 | 170247287 | rs12527469 | -0.182 | -0.818 | -0.279 | -0.64 | -0.006 | 0.001 | 0.001 | 0 | 0.13 | 0.12 | 0.22 | 0.17 | 0.11 | 0.34 |
| 6 | 170250768 | rs10455752 | 0.511 | -0.737 | -0.292 | -0.939 | 0.005 | -0.001 | -0.001 | 0 | 0.15 | 0.1 | 0.07 | 0 | 0.13 | 0.02 |
| 6 | 170255046 | rs4716412 | -1.063 | -1.391 | 0.076 | -0.762 | 0.012 | 0.008 | 0.008 | 0.602 | 0.38 | 0.6 | 0.33 | 0.32 | 0.3 | 0.37 |
| 6 | 170258821 | rs9383484 | -1.255 | -1.529 | -0.273 | -0.978 | 0.086 | 0.197 | 0.197 | 0.462 | 0.46 | 0.75 | 0.62 | 0.9 | 0.35 | 0.56 |
| 6 | 170261237 | rs714189 | -1.467 | -1.647 | 0.201 | -1.053 | 0.066 | 0.116 | 0.116 | 0.939 | 0.5 | 0.69 | 0.62 | 0.91 | 0.36 | 0.59 |
| 6 | 170267771 | rs7453586 | -1.39 | -1.735 | 0.153 | -0.997 | 0.083 | 0.171 | 0.171 | 1.39 | 0.51 | 0.74 | 0.63 | 0.9 | 0.37 | 0.59 |
| 6 | 170268791 | rs7749330 | -0.477 | -1.935 | 0.557 | -0.84 | -0.052 | 0.071 | 0.071 | 0.666 | 0.25 | 0.47 | 0.36 | 0.59 | 0.18 | 0.22 |
| 6 | 170269355 | rs7766161 | -0.878 | -1.803 | 0.341 | -1.263 | 0.005 | 0.003 | 0.003 | 2.609 | 0.25 | 0.22 | 0.4 | 0.76 | 0.2 | 0.41 |
| 6 | 170270028 | rs10806671 | -1.536 | -1.307 | 0.097 | -1.693 | 0.022 | 0.067 | 0.067 | 0.827 | 0.51 | 0.64 | 0.67 | 0.97 | 0.44 | 0.61 |
| 6 | 170272641 | rs7744688 | -0.901 | -1.404 | 0.116 | -1.654 | -0.015 | -0.041 | -0.041 | 0 | 0.33 | 0.52 | 0.44 | 0.79 | 0.26 | 0.41 |
| 6 | 170275554 | rs9637984 | 0.378 | -1.056 | -0.109 | -0.852 | -0.002 | 0.005 | 0.005 | 0 | 0.16 | 0.08 | 0.22 | 0.18 | 0.16 | 0.11 |
| 6 | 170279658 | rs4134409 | 0.378 | -1.056 | -0.109 | -0.852 | -0.002 | 0.005 | 0.005 | 0 | 0.16 | 0.08 | 0.22 | 0.18 | 0.16 | 0.11 |
| 6 | 170331956 | rs12183541 | -0.519 | -0.952 | 0.005 | -0.767 | 0.01 | 0.039 | 0.039 | 0 | 0.81 | 0.93 | 0.86 | 0.94 | 0.88 | 0.82 |
| 6 | 170332621 | rs6940159 | 0.31 | -0.823 | 0.153 | -0.663 | 0.007 | 0.024 | 0.024 | 0.042 | 0.49 | 0.58 | 0.43 | 0.15 | 0.59 | 0.31 |
| 6 | 170333717 | rs4710807 | 0.642 | -0.847 | 0.218 | -0.13 | 0.004 | -0.004 | -0.004 | 0.311 | 0.27 | 0.17 | 0.2 | 0.31 | 0.26 | 0.32 |
| 6 | 170334114 | rs2018894 | 0.091 | -0.582 | 0.102 | -0.261 | -0.001 | -0.032 | -0.032 | 1.375 | 0.58 | 0.57 | 0.72 | 0.9 | 0.66 | 0.71 |
| 6 | 170335567 | rs1008794 | 0.355 | -0.192 | 0.258 | -0.113 | 0 | -0.041 | -0.041 | 1.441 | 0.21 | 0.26 | 0.13 | 0.03 | 0.22 | 0.1 |
| 6 | 170337938 | rs11968737 | -0.277 | 0.541 | 0.803 | 0.595 | 0.079 | 0.002 | 0.002 | 0 | 0.05 | 0.23 | 0.06 | 0.03 | 0.11 | 0.06 |
| 6 | 170341197 | rs9459964 | 0.67 | 0.589 | 0.509 | 0.597 | -0.003 | 0.001 | 0.001 | 0 | 0.26 | 0.23 | 0.3 | 0.12 | 0.37 | 0.2 |
| 6 | 170342187 | rs6929872 | -0.51 | 0.307 | -0.337 | 0.196 | -0.002 | -0.001 | -0.001 | 0 | 0.56 | 0.54 | 0.55 | 0.45 | 0.74 | 0.6 |
| 6 | 170347127 | rs1466749 | -0.18 | 0.337 | -0.406 | 0.184 | -0.002 | -0.001 | -0.001 | 0 | 0.51 | 0.49 | 0.51 | 0.43 | 0.63 | 0.54 |
| 6 | 170351025 | rs6932737 | -0.769 | -0.003 | -0.729 | -0.099 | 0.021 | -0.009 | -0.009 | 1.914 | 0.6 | 0.66 | 0.64 | 0.52 | 0.77 | 0.67 |
| 6 | 170352345 | rs6456185 | -0.02 | 0.262 | -0.702 | 0.012 | 0.024 | 0.024 | 0.024 | 1.889 | 0.35 | 0.56 | 0.39 | 0.44 | 0.53 | 0.3 |
| 6 | 170353312 | rs1000104 | 0.127 | 0.016 | -0.336 | 0.497 | 0.014 | -0.002 | -0.002 | 0 | 0.43 | 0.38 | 0.43 | 0.49 | 0.26 | 0.4 |
| 6 | 170358566 | rs1532389 | 1.292 | 0.754 | 0.056 | 1.277 | 0.005 | 0.084 | 0.084 | 0 | 0.86 | 0.75 | 0.71 | 0.68 | 0.82 | 0.75 |
| 6 | 170384030 | rs9460225 | 1.15 | 0.581 | -0.12 | 1.203 | -0.006 | 0.042 | 0.042 | 0 | 0.86 | 0.79 | 0.76 | 0.69 | 0.86 | 0.71 |
| 6 | 170384298 | rs11753530 | 0.431 | 0.609 | -0.835 | -0.091 | -0.011 | 0.122 | 0.122 | 0 | 0.34 | 0.13 | 0.23 | 0.08 | 0.28 | 0.28 |
| 6 | 170401188 | rs9460269 | 0.321 | 1.21 | -0.862 | 0.658 | -0.088 | -0.03 | -0.03 | 0 | 0.18 | 0.03 | 0.28 | 0.17 | 0.33 | 0.13 |
| 6 | 170409966 | rs6456234 | 1.177 | 0.604 | 0.52 | 0.623 | 0.03 | -0.037 | -0.037 | 0 | 0.91 | 0.61 | 0.93 | 0.99 | 0.91 | 0.94 |
| 6 | 170412069 | rs883488 | 1.434 | 0.41 | 0.699 | 1.184 | -0.017 | 0.189 | 0.189 | 0.228 | 0.88 | 0.62 | 0.78 | 0.68 | 0.93 | 0.74 |
| 6 | 170414285 | rs9460045 | 1.52 | 0.344 | 0.539 | 1.51 | -0.022 | 0.118 | 0.118 | 0.295 | 0.88 | 0.72 | 0.78 | 0.42 | 0.92 | 0.71 |
| 6 | 170416717 | rs1109616 | -0.249 | 0.291 | -0.169 | 0.362 | -0.002 | 0.123 | 0.123 | 0.357 | 0.46 | 0.75 | 0.41 | 0.16 | 0.5 | 0.36 |
| 6 | 170422610 | rs9356682 | -0.536 | -0.109 | 0.26 | 0.751 | 0.002 | -0.007 | -0.007 | 0 | 0.18 | 0.15 | 0.26 | 0.35 | 0.13 | 0.34 |
| 6 | 170424092 | rs6936493 | -0.763 | -0.087 | 0.748 | 0.465 | 0.007 | 0.018 | 0.018 | 0.115 | 0.36 | 0.52 | 0.49 | 0.54 | 0.41 | 0.45 |
| 6 | 170425521 | rs10946292 | -1.007 | -0.103 | 1.035 | 0.534 | -0.029 | 0.111 | 0.111 | 0.015 | 0.22 | 0.5 | 0.29 | 0.37 | 0.14 | 0.35 |
| 6 | 170426027 | rs4710879 | 0.394 | -0.178 | 0.561 | 0.018 | 0.005 | 0.075 | 0.075 | 0 | 0.45 | 0.22 | 0.5 | 0.38 | 0.62 | 0.44 |
| 6 | 170427509 | rs13210275 | 0.394 | -0.178 | 0.565 | 0.015 | 0.005 | 0.077 | 0.077 | 0 | 0.45 | 0.22 | 0.51 | 0.38 | 0.62 | 0.44 |
| 6 | 170428032 | rs903630 | -1.814 | -0.263 | 0.858 | -0.115 | -0.012 | 0.01 | 0.01 | 0 | 0.74 | 0.65 | 0.87 | 0.74 | 0.87 | 0.85 |
| 6 | 170428556 | rs3012418 | -1.147 | -0.15 | 0.564 | -0.351 | -0.007 | -0.081 | -0.081 | 0.149 | 0.67 | 0.68 | 0.81 | 0.98 | 0.77 | 0.81 |
| 6 | 170431335 | rs3013251 | -0.262 | 0.008 | -0.123 | -0.174 | -0.008 | 0.031 | 0.031 | 0 | 0.19 | 0.29 | 0.29 | 0.6 | 0.15 | 0.38 |
| 6 | 170433036 | rs4710696 | 0.667 | 0.161 | -1.063 | -0.182 | -0.083 | 0.071 | 0.071 | 0 | 0.24 | 0.04 | 0.28 | 0.2 | 0.34 | 0.28 |
| 6 | 170438227 | rs9459963 | 1.29 | 0.252 | -0.993 | -0.192 | -0.049 | -0.037 | -0.037 | 0 | 0.14 | 0.03 | 0.2 | 0.17 | 0.27 | 0.1 |

| | | | | | | | | | | | | | | | | |
|---|---|---|---|---|---|---|---|---|---|---|---|---|---|---|---|---|
| 6 | 170439874 | rs4710698 | -0.043 | -0.676 | -0.788 | -0.154 | -0.02 | -0.045 | -0.045 | 0 | 0.45 | 0.12 | 0.58 | 0.51 | 0.63 | 0.43 |
| 6 | 170444518 | rs3012454 | -0.508 | -1.049 | -0.565 | -0.387 | 0.007 | 0.005 | 0.005 | 1.446 | 0.68 | 0.75 | 0.75 | 0.78 | 0.77 | 0.75 |
| 6 | 170444663 | rs4710755 | 1.014 | -1.016 | -0.888 | -0.494 | -0.01 | -0.01 | -0.01 | 0 | 0.18 | 0.11 | 0.21 | 0.3 | 0.21 | 0.16 |
| 6 | 170444698 | rs4710756 | 0.091 | -1.46 | -0.441 | -0.626 | 0.061 | 0.022 | 0.022 | 0.869 | 0.52 | 0.35 | 0.68 | 0.62 | 0.69 | 0.55 |
| 6 | 170444816 | rs12204719 | 1.6 | -1.301 | -0.44 | -0.845 | -0.026 | -0.028 | -0.028 | 0 | 0.19 | 0.11 | 0.26 | 0.26 | 0.31 | 0.18 |
| 6 | 170445169 | rs3013257 | 0.173 | -1.239 | -0.066 | -0.932 | -0.003 | -0.003 | -0.003 | 0 | 0.42 | 0.58 | 0.4 | 0.45 | 0.48 | 0.42 |
| 6 | 170445784 | rs2311179 | -0.081 | -1.383 | -0.231 | -1.06 | 0.015 | 0.026 | 0.026 | 0.29 | 0.57 | 0.39 | 0.68 | 0.76 | 0.7 | 0.58 |
| 6 | 170445870 | rs9366130 | 0.284 | -1.169 | 0.021 | -0.905 | -0.026 | -0.028 | -0.028 | 0.474 | 0.39 | 0.29 | 0.38 | 0.47 | 0.49 | 0.37 |
| 6 | 170445911 | rs4582363 | -0.189 | -1.119 | 0.183 | -0.891 | 0 | 0 | 0 | 0 | 0.47 | 0.51 | 0.51 | 0.51 | 0.55 | 0.57 |
| 6 | 170448999 | rs3012361 | -0.269 | -1.111 | 0.082 | -0.775 | 0.022 | 0.018 | 0.018 | 1.8 | 0.63 | 0.7 | 0.73 | 0.76 | 0.76 | 0.73 |
| 6 | 170449482 | rs2935065 | -2.196 | -1.243 | -0.062 | -0.888 | 0.006 | -0.012 | -0.012 | 0 | 0.82 | 0.88 | 0.85 | 0.78 | 0.9 | 0.91 |
| 6 | 170449680 | rs903625 | 0.625 | -1.168 | 0.055 | -0.893 | 0.026 | 0.062 | 0.062 | 0.013 | 0.32 | 0.44 | 0.31 | 0.43 | 0.39 | 0.45 |
| 6 | 170450340 | rs719960 | 0.214 | -1.156 | 0.055 | -0.865 | 0.008 | 0.009 | 0.009 | 0.064 | 0.42 | 0.48 | 0.38 | 0.43 | 0.49 | 0.5 |
| 6 | 170455060 | rs4710758 | 1.53 | -1.062 | 0.094 | -0.785 | -0.03 | -0.015 | -0.015 | 0.934 | 0.19 | 0.07 | 0.33 | 0.3 | 0.27 | 0.23 |
| 6 | 170466300 | rs2935100 | -1.322 | -1.212 | 0.284 | -0.719 | -0.007 | 0.001 | 0.001 | 3.36 | 0.76 | 0.72 | 0.83 | 0.76 | 0.89 | 0.86 |
| 6 | 170472702 | rs937327 | 0.043 | -1.157 | 0.25 | -0.748 | -0.005 | -0.006 | -0.006 | 0 | 0.38 | 0.65 | 0.34 | 0.48 | 0.41 | 0.47 |
| 6 | 170475212 | rs1021543 | -1.518 | -1.144 | 0.474 | -0.695 | -0.043 | -0.019 | -0.019 | 0.312 | 0.22 | 0.38 | 0.15 | 0.2 | 0.17 | 0.34 |
| 6 | 170488009 | rs2935105 | 0.085 | -0.843 | 0.222 | -0.572 | 0.014 | 0.021 | 0.021 | 0.101 | 0.35 | 0.53 | 0.34 | 0.46 | 0.41 | 0.48 |
| 6 | 170492449 | rs4540249 | -0.535 | -1.085 | 0.163 | -0.599 | -0.012 | -0.011 | -0.011 | 0 | 0.69 | 0.74 | 0.7 | 0.71 | 0.63 | 0.72 |
| 6 | 170496354 | rs2935090 | -0.098 | -1.223 | 0.069 | -0.641 | 0.003 | 0.001 | 0.001 | 4.204 | 0.72 | 0.74 | 0.77 | 0.79 | 0.84 | 0.83 |
| 6 | 170502363 | rs9459971 | -0.292 | -0.463 | -0.336 | -0.138 | 0.001 | 0.001 | 0.001 | 2.316 | 0.65 | 0.39 | 0.73 | 0.76 | 0.75 | 0.86 |
| 6 | 170516123 | rs9348260 | -0.039 | -0.213 | -0.624 | 0.122 | -0.013 | -0.021 | -0.021 | 0 | 0.58 | 0.44 | 0.66 | 0.76 | 0.69 | 0.74 |
| 6 | 170517351 | rs12196403 | -0.669 | 0.178 | -0.923 | 0.613 | 0.009 | 0.026 | 0.026 | 1.779 | 0.34 | 0.49 | 0.31 | 0.29 | 0.37 | 0.25 |
| 6 | 170526065 | rs9366181 | -0.851 | -0.112 | -1.59 | 0.326 | 0.024 | 0.038 | 0.038 | 0.649 | 0.27 | 0.82 | 0.18 | 0.14 | 0.17 | 0.08 |
| 6 | 170537154 | rs2983227 | 0.506 | 0.171 | -2.458 | 0.164 | -0.156 | 0.059 | 0.059 | 1.691 | 0.38 | 0.02 | 0.42 | 0.21 | 0.53 | 0.37 |
| 6 | 170538325 | rs745285 | -1.074 | -0.295 | -1.851 | 1.145 | 0.052 | -0.1 | -0.1 | 0 | 0.3 | 0.87 | 0.23 | 0.44 | 0.18 | 0.21 |
| 6 | 170543570 | rs4710771 | -0.495 | 0.004 | -1.387 | 0.643 | -0.001 | 0.002 | 0.002 | 0.006 | 0.12 | 0.13 | 0.1 | 0.09 | 0.12 | 0.03 |
| 6 | 170546108 | rs6456182 | -1.014 | -0.05 | -0.548 | 0.815 | -0.01 | 0.06 | 0.06 | 0 | 0.19 | 0.41 | 0.15 | 0.66 | 0.18 | 0.33 |
| 6 | 170546310 | rs6456183 | -1.308 | 0.153 | -0.65 | 0.044 | -0.002 | -0.028 | -0.028 | 0 | 0.29 | 0.56 | 0.24 | 0.79 | 0.3 | 0.38 |
| 6 | 170547690 | rs9460087 | -0.356 | -0.206 | -0.946 | -0.49 | -0.012 | -0.019 | -0.019 | 1.562 | 0.3 | 0.34 | 0.26 | 0.2 | 0.26 | 0.31 |
| 6 | 170548721 | rs9460089 | 0.025 | -0.264 | -1.204 | -0.461 | 0.027 | 0.072 | 0.072 | 0 | 0.21 | 0.07 | 0.22 | 0.17 | 0.18 | 0.22 |
| 6 | 170552613 | rs7763013 | -1.042 | 0.026 | -0.335 | 0.324 | 0.004 | 0.026 | 0.026 | 0.43 | 0.3 | 0.36 | 0.24 | 0.43 | 0.34 | 0.33 |
| 6 | 170554506 | rs6456189 | -0.706 | -0.204 | 0.464 | -0.292 | -0.009 | -0.102 | -0.102 | 0.105 | 0.23 | 0.4 | 0.16 | 0.07 | 0.2 | 0.15 |
| 6 | 170554601 | rs6456190 | -0.292 | -0.154 | 0.441 | -0.266 | 0 | -0.011 | -0.011 | 0 | 0.19 | 0.16 | 0.14 | 0.07 | 0.17 | 0.16 |
| 6 | 170558004 | rs3012428 | 1.083 | -0.125 | 0.411 | -0.166 | 0.002 | 0.003 | 0.003 | 0.615 | 0.51 | 0.47 | 0.55 | 0.61 | 0.58 | 0.62 |
| 6 | 170560679 | rs2144667 | 0.187 | -0.191 | 0.332 | -0.087 | 0.001 | -0.002 | -0.002 | 0 | 0.8 | 0.82 | 0.82 | 0.79 | 0.83 | 0.82 |
| 6 | 170563478 | rs7760268 | 0.294 | 0.272 | 0.114 | -0.733 | 0 | -0.015 | -0.015 | 0.668 | 0.22 | 0.25 | 0.22 | 0.11 | 0.26 | 0.1 |
| 6 | 170564749 | rs9348301 | 0.081 | 0.495 | 0.365 | -0.457 | 0.004 | 0.027 | 0.027 | 0.021 | 0.24 | 0.31 | 0.23 | 0.42 | 0.29 | 0.26 |
| 6 | 170566240 | rs7771267 | 0.371 | 0.487 | 0.408 | -0.943 | 0.006 | -0.045 | -0.045 | 0.749 | 0.23 | 0.31 | 0.22 | 0.11 | 0.26 | 0.1 |
| 6 | 170566631 | rs9348302 | -0.197 | 0.372 | 0.448 | -0.502 | 0.004 | 0.006 | 0.006 | 0 | 0.28 | 0.3 | 0.27 | 0.67 | 0.3 | 0.4 |
| 6 | 170567708 | rs3012372 | 0.283 | 0.279 | 0.199 | 0.382 | 0.002 | -0.002 | -0.002 | 0 | 0.78 | 0.81 | 0.81 | 0.79 | 0.83 | 0.81 |
| 6 | 170572741 | rs3012437 | 0.719 | 0.7 | 0.233 | -0.418 | 0.003 | -0.023 | -0.023 | 0 | 0.6 | 0.62 | 0.66 | 0.78 | 0.58 | 0.8 |
| 6 | 170579499 | rs9295393 | 0.372 | 0.782 | 0.61 | -0.93 | 0.004 | 0.003 | 0.003 | 0 | 0.67 | 0.76 | 0.66 | 0.68 | 0.69 | 0.74 |
| 6 | 170580521 | rs7756780 | 0.921 | 0.668 | 0.659 | -0.948 | 0.001 | 0.031 | 0.031 | 0 | 0.49 | 0.56 | 0.54 | 0.68 | 0.5 | 0.67 |
| 6 | 170585755 | rs9348305 | -0.169 | 0.525 | 0.285 | -1.081 | 0.008 | 0.054 | 0.054 | 0 | 0.22 | 0.12 | 0.18 | 0.04 | 0.2 | 0.24 |
| 6 | 170585917 | rs9366197 | -1.333 | 0.443 | 0.792 | -1.339 | 0.001 | -0.083 | -0.083 | 0 | 0.42 | 0.58 | 0.49 | 0.99 | 0.37 | 0.6 |
| 6 | 170589360 | rs9460103 | -0.988 | 1.057 | 0.346 | -1.018 | -0.001 | 0.002 | 0.002 | 0 | 0.38 | 0.41 | 0.36 | 0.28 | 0.4 | 0.37 |
| 6 | 170591923 | rs1033583 | -1.417 | 1.85 | 0.08 | -0.563 | -0.025 | -0.01 | -0.01 | 0.74 | 0.25 | 0.21 | 0.26 | 0.24 | 0.33 | 0.24 |
| 6 | 170620572 | rs2024694 | -1.489 | 1.454 | 0.899 | -0.656 | 0.031 | 0.004 | 0.004 | 2.424 | 0.06 | 0.25 | 0.04 | 0.02 | 0.06 | 0.06 |
| 6 | 170628231 | rs7774272 | -1.838 | 1.445 | 0.875 | -0.713 | 0.023 | -0.014 | -0.014 | 0 | 0.07 | 0.21 | 0.06 | 0.02 | 0.1 | 0.08 |
| 6 | 170641014 | rs9459991 | -2.202 | 1.323 | 1.014 | -0.68 | 0.01 | 0.256 | 0.256 | 2.176 | 0.1 | 0.3 | 0.12 | 0.59 | 0.13 | 0.25 |
| 7 | 123956 | rs3923903 | -0.124 | 0.201 | -1.425 | -0.153 | -0.016 | 0.086 | 0.086 | 0.022 | 0.47 | 0.27 | 0.58 | 0.68 | 0.48 | 0.4 |
| 7 | 127976 | rs4916960 | 0.352 | -0.742 | -0.484 | 0.493 | -0.007 | 0.011 | 0.011 | 0 | 0.71 | 0.58 | 0.72 | 0.65 | 0.76 | 0.66 |
| 7 | 753495 | rs7797121 | -0.769 | 0.084 | 0.203 | 0.689 | -0.019 | 0.04 | 0.04 | 0 | 0.17 | 0.22 | 0.21 | 0.27 | 0.09 | 0.28 |
| 7 | 795227 | rs7795258 | 0.412 | 0.986 | 0.038 | 0.937 | 0.003 | 0.028 | 0.028 | 0 | 0.84 | 0.77 | 0.78 | 0.61 | 0.82 | 0.86 |
| 7 | 825494 | rs9458 | -0.523 | 1.066 | 0.451 | 1.289 | 0.004 | 0.038 | 0.038 | 0.394 | 0.11 | 0.11 | 0.21 | 0.27 | 0.14 | 0.28 |
| 7 | 828493 | rs4397290 | -0.698 | 0.975 | 0.334 | 0.975 | 0.002 | 0.025 | 0.025 | 0 | 0.14 | 0.18 | 0.21 | 0.38 | 0.18 | 0.13 |
| 7 | 857352 | rs4275123 | -1.155 | 0.983 | 0.934 | 0.718 | 0.045 | 0.014 | 0.014 | 0 | 0.16 | 0.43 | 0.25 | 0.2 | 0.2 | 0.09 |
| 7 | 864389 | rs4719475 | -1.758 | 1.015 | 1.257 | 0.829 | 0.014 | -0.008 | -0.008 | 0 | 0.3 | 0.42 | 0.43 | 0.23 | 0.35 | 0.41 |
| 7 | 869147 | rs11769388 | -1.055 | 0.879 | 0.644 | 0.798 | -0.013 | -0.018 | -0.018 | 2.795 | 0.14 | 0.1 | 0.24 | 0.14 | 0.18 | 0.26 |
| 7 | 869660 | rs4481481 | -0.06 | 0.652 | 0.564 | 0.4 | 0.003 | 0.013 | 0.013 | 0 | 0.7 | 0.73 | 0.86 | 0.79 | 0.76 | 0.78 |
| 7 | 869846 | rs10246200 | 1.116 | 0.647 | 0.576 | 0.411 | -0.002 | 0.02 | 0.02 | 0 | 0.57 | 0.63 | 0.62 | 0.65 | 0.58 | 0.52 |
| 7 | 884665 | rs6960062 | -1.307 | 0.591 | 0.863 | 0.738 | 0.024 | 0.027 | 0.027 | 0 | 0.21 | 0.28 | 0.43 | 0.27 | 0.34 | 0.32 |
| 7 | 889358 | rs12669325 | 0.866 | 0.591 | 0.689 | 0.799 | -0.04 | -0.038 | -0.038 | 1.141 | 0.07 | 0.01 | 0.18 | 0.11 | 0.13 | 0.22 |
| 7 | 892902 | rs4076072 | -0.108 | 0.456 | 0.654 | 0.794 | -0.023 | -0.021 | -0.021 | 0 | 0.14 | 0.08 | 0.24 | 0.13 | 0.18 | 0.25 |
| 7 | 901574 | rs7809344 | -0.537 | 0.879 | 0.362 | 0.569 | -0.013 | -0.011 | -0.011 | 2.782 | 0.17 | 0.1 | 0.24 | 0.14 | 0.19 | 0.28 |
| 7 | 909350 | rs4544987 | -0.021 | 0.945 | 0.332 | 0.598 | 0.004 | 0.018 | 0.018 | 0 | 0.71 | 0.74 | 0.86 | 0.79 | 0.76 | 0.78 |
| 7 | 912755 | rs4722034 | 0.01 | 0.939 | 0.341 | 0.596 | 0.002 | 0.009 | 0.009 | 0 | 0.7 | 0.72 | 0.86 | 0.79 | 0.76 | 0.78 |
| 7 | 916678 | rs10950881 | -0.145 | 0.499 | -0.32 | 0.825 | -0.001 | 0.017 | 0.017 | 0 | 0.74 | 0.84 | 0.87 | 1 | 0.77 | 0.81 |

| | | | | | | | | | | | | | | | | |
|---|---|---|---|---|---|---|---|---|---|---|---|---|---|---|---|---|
| 7 | 925949 | rs10226357 | 1.249 | 0.802 | -0.622 | 1.082 | -0.002 | -0.016 | -0.016 | 0 | 0.55 | 0.48 | 0.6 | 0.69 | 0.58 | 0.54 |
| 7 | 930689 | rs13171 | -1.083 | 1.306 | -1.087 | 1.643 | -0.015 | -0.039 | -0.039 | 0 | 0.15 | 0.09 | 0.24 | 0.27 | 0.19 | 0.27 |
| 7 | 965643 | rs3757854 | -0.579 | 1.212 | -1.282 | 1.397 | -0.01 | 0.022 | 0.022 | 0 | 0.44 | 0.3 | 0.48 | 0.43 | 0.46 | 0.35 |
| 7 | 971124 | rs6956301 | 0.29 | 1.897 | -0.722 | 1.405 | 0.016 | 0.004 | 0.004 | 0.001 | 0.78 | 0.72 | 0.62 | 0.71 | 0.66 | 0.89 |
| 7 | 977616 | rs35585784 | -0.78 | 1.173 | -0.807 | 1.629 | -0.002 | -0.005 | -0.005 | 0 | 0.13 | 0.12 | 0.15 | 0.23 | 0.14 | 0.19 |
| 7 | 985840 | rs7811408 | -0.414 | 0.338 | -0.933 | 1.185 | -0.015 | -0.003 | -0.003 | 0.005 | 0.31 | 0.55 | 0.17 | 0.24 | 0.2 | 0.19 |
| 7 | 993173 | rs36015262 | -0.943 | 0.894 | -1.196 | 1.188 | -0.018 | -0.031 | -0.031 | 0.372 | 0.14 | 0.07 | 0.15 | 0.22 | 0.15 | 0.2 |
| 7 | 998802 | rs10951261 | 0.282 | 1.38 | -1.293 | 0.87 | -0.045 | -0.043 | -0.043 | 0 | 0.17 | 0.08 | 0.29 | 0.29 | 0.29 | 0.13 |
| 7 | 1003047 | rs7799459 | -0.824 | 0.898 | -0.676 | 0.6 | -0.027 | -0.037 | -0.037 | 0 | 0.11 | 0.19 | 0.15 | 0.02 | 0.06 | 0.03 |
| 7 | 1005950 | rs28448690 | -0.857 | 0.901 | -0.507 | 0.69 | -0.029 | -0.014 | -0.014 | 1.628 | 0.11 | 0.26 | 0.15 | 0.04 | 0.06 | 0.04 |
| 7 | 1009337 | rs10264122 | -1.715 | 1.025 | -0.92 | 0.596 | -0.027 | 0.03 | 0.03 | 0 | 0.56 | 0.92 | 0.67 | 0.61 | 0.59 | 0.47 |
| 7 | 1010494 | rs6942989 | -0.545 | 0.582 | -0.517 | 0.601 | -0.027 | -0.035 | -0.035 | 0 | 0.11 | 0.18 | 0.15 | 0.02 | 0.06 | 0.03 |
| 7 | 1011968 | rs6462277 | -0.146 | 0.679 | 0.378 | 0.588 | -0.031 | -0.106 | -0.106 | 0 | 0.16 | 0.41 | 0.07 | 0 | 0.08 | 0.13 |
| 7 | 1016948 | rs10272099 | -1.016 | 0.929 | 0.189 | 0.682 | 0 | -0.007 | -0.007 | 0 | 0.32 | 0.7 | 0.37 | 0.52 | 0.33 | 0.27 |
| 7 | 1019784 | rs10274964 | -1.16 | 0.919 | 0.847 | 0.652 | 0.024 | 0.006 | 0.006 | 0 | 0.35 | 0.49 | 0.45 | 0.59 | 0.44 | 0.4 |
| 7 | 1020272 | rs7807150 | -0.116 | 0.871 | 1.224 | 0.689 | -0.006 | -0.034 | -0.034 | 0.327 | 0.14 | 0.32 | 0.08 | 0.06 | 0.1 | 0.13 |
| 7 | 1022728 | rs12701220 | 0.87 | 1.019 | 0.834 | 0.903 | -0.027 | -0.048 | -0.048 | 0 | 0.13 | 0.07 | 0.25 | 0.3 | 0.22 | 0.13 |
| 7 | 1024581 | rs3824081 | -2.047 | 0.86 | 0.877 | 0.77 | -0.01 | 0.087 | 0.087 | 0.035 | 0.49 | 0.76 | 0.61 | 0.62 | 0.5 | 0.47 |
| 7 | 1031117 | rs11976017 | -1.829 | 0.892 | 1.039 | 0.177 | -0.006 | 0.009 | 0.009 | 3.006 | 0.34 | 0.61 | 0.35 | 0.31 | 0.27 | 0.34 |
| 7 | 1034574 | rs6462495 | -1.252 | 0.934 | 1.009 | 0.205 | -0.01 | -0.029 | -0.029 | 0 | 0.33 | 0.82 | 0.38 | 0.35 | 0.33 | 0.27 |
| 7 | 1034903 | rs1007766 | -2.099 | 0.936 | 0.968 | 0.198 | -0.078 | -0.067 | -0.067 | 0 | 0.48 | 0.91 | 0.61 | 0.44 | 0.48 | 0.46 |
| 7 | 1036799 | rs8591 | 1.021 | 1.036 | 1.032 | 0.242 | -0.019 | 0.007 | 0.007 | 0 | 0.13 | 0.07 | 0.24 | 0.14 | 0.21 | 0.11 |
| 7 | 1036851 | rs8145 | -0.715 | 1.028 | 1.165 | 0.236 | -0.002 | 0.012 | 0.012 | 0.593 | 0.18 | 0.1 | 0.24 | 0.09 | 0.17 | 0.21 |
| 7 | 1039457 | rs12701348 | 1.023 | 1.033 | 1.162 | 0.239 | -0.042 | -0.001 | -0.001 | 0.208 | 0.13 | 0.05 | 0.23 | 0.14 | 0.21 | 0.11 |
| 7 | 1044938 | rs9639696 | -1.74 | 0.999 | 1.578 | 0.246 | 0 | 0.022 | 0.022 | 0.597 | 0.33 | 0.51 | 0.35 | 0.3 | 0.28 | 0.34 |
| 7 | 1046637 | rs11977029 | -0.775 | 1.047 | 1.404 | 0.267 | -0.001 | 0.013 | 0.013 | 0.542 | 0.18 | 0.11 | 0.24 | 0.09 | 0.17 | 0.21 |
| 7 | 1051776 | rs10229964 | -1.566 | 1.044 | 1.623 | 0.183 | 0.025 | 0.042 | 0.042 | 0 | 0.34 | 0.47 | 0.55 | 0.42 | 0.42 | 0.33 |
| 7 | 1052353 | rs13224488 | 0.932 | 1.036 | 1.201 | 0.162 | -0.019 | 0.003 | 0.003 | 0 | 0.13 | 0.06 | 0.24 | 0.15 | 0.21 | 0.11 |
| 7 | 1053958 | rs1881116 | -0.597 | 1.025 | 1.438 | 0.104 | 0.008 | -0.057 | -0.057 | 2.55 | 0.26 | 0.48 | 0.32 | 0.16 | 0.28 | 0.23 |
| 7 | 1060288 | rs11763020 | 0.337 | 0.906 | 0.72 | -0.033 | 0.006 | 0.097 | 0.097 | 0 | 0.2 | 0.06 | 0.09 | 0 | 0.13 | 0.14 |
| 7 | 1081972 | rs10224368 | -0.895 | 0.89 | 1.032 | 0.143 | -0.001 | -0.001 | -0.001 | 0 | 0.37 | 0.34 | 0.37 | 0.28 | 0.33 | 0.3 |
| 7 | 1091644 | rs2363285 | -1.519 | 0.877 | 1.06 | 0.099 | -0.006 | -0.015 | -0.015 | 0 | 0.58 | 0.72 | 0.62 | 0.43 | 0.53 | 0.49 |
| 7 | 1094819 | rs4720335 | -0.401 | 0.897 | 0.927 | 0.05 | 0 | -0.001 | -0.001 | 0 | 0.11 | 0.1 | 0.13 | 0.05 | 0.06 | 0.05 |
| 7 | 1096766 | rs1881124 | 0.538 | 1.075 | 1.143 | -0.016 | 0 | -0.007 | -0.007 | 3.227 | 0.15 | 0.18 | 0.18 | 0.1 | 0.15 | 0.17 |
| 7 | 1096918 | rs1881125 | -1.512 | 0.933 | 1.014 | -0.497 | -0.011 | -0.018 | -0.018 | 0.356 | 0.61 | 0.67 | 0.57 | 0.36 | 0.53 | 0.48 |
| 7 | 1097952 | rs6953080 | -0.113 | 0.615 | 0.978 | -0.406 | -0.001 | -0.004 | -0.004 | 0 | 0.11 | 0.1 | 0.13 | 0.02 | 0.06 | 0.04 |
| 7 | 1102241 | rs12701823 | -0.192 | 0.677 | 1.132 | -0.457 | 0.011 | -0.009 | -0.009 | 0 | 0.22 | 0.29 | 0.29 | 0.17 | 0.26 | 0.29 |
| 7 | 1103697 | rs10280960 | -1.553 | 0.73 | 1.321 | -0.489 | 0.01 | 0.004 | 0.004 | 0 | 0.59 | 0.55 | 0.53 | 0.37 | 0.47 | 0.48 |
| 7 | 1105426 | rs12701834 | -0.198 | 0.71 | 1.141 | -0.521 | 0.018 | 0.039 | 0.039 | 0.348 | 0.39 | 0.28 | 0.26 | 0.1 | 0.28 | 0.31 |
| 7 | 1117436 | rs10274367 | -0.749 | 1.039 | 1.48 | -0.235 | 0.045 | -0.004 | -0.004 | 0.162 | 0.24 | 0.41 | 0.36 | 0.25 | 0.33 | 0.31 |
| 7 | 1121188 | rs12540123 | -0.781 | 1.043 | 1.584 | -0.129 | 0.027 | 0.038 | 0.038 | 0 | 0.24 | 0.39 | 0.34 | 0.36 | 0.34 | 0.31 |
| 7 | 1125893 | rs3808350 | -1.318 | 0.938 | 1.507 | -0.265 | -0.019 | -0.029 | -0.029 | 0 | 0.32 | 0.75 | 0.42 | 0.43 | 0.39 | 0.35 |
| 7 | 1126659 | rs3808351 | -0.919 | 0.762 | 1.619 | -0.356 | -0.005 | 0.012 | 0.012 | 0 | 0.28 | 0.27 | 0.31 | 0.18 | 0.3 | 0.33 |
| 7 | 1130584 | rs12702029 | -1.403 | 0.914 | 1.162 | -0.559 | -0.02 | -0.024 | -0.024 | 0 | 0.37 | 0.74 | 0.48 | 0.41 | 0.47 | 0.36 |
| 7 | 1131356 | rs3802141 | -1.816 | 0.77 | 1.213 | -0.526 | -0.021 | -0.057 | -0.057 | 0.44 | 0.66 | 0.82 | 0.65 | 0.44 | 0.6 | 0.59 |
| 7 | 1132972 | rs3808353 | 0.83 | 0.873 | 1.433 | -0.549 | -0.005 | 0.02 | 0.02 | 0 | 0.15 | 0.15 | 0.18 | 0.08 | 0.18 | 0.19 |
| 7 | 1133943 | rs10951737 | -1.374 | 0.887 | 1.539 | -0.465 | 0.049 | 0.016 | 0.016 | 0 | 0.14 | 0.5 | 0.17 | 0.09 | 0.14 | 0.15 |
| 7 | 1140418 | rs6463224 | 0.132 | 0.859 | 1.354 | -0.391 | 0.004 | 0.035 | 0.035 | 0 | 0.19 | 0.3 | 0.21 | 0.23 | 0.2 | 0.19 |
| 7 | 1141129 | rs13242034 | 0.723 | 0.951 | 1.39 | -0.388 | -0.001 | -0.001 | -0.001 | 0 | 0.16 | 0.21 | 0.21 | 0.15 | 0.2 | 0.19 |
| 7 | 1141245 | rs12702080 | -0.852 | 0.765 | 1.435 | -0.543 | -0.001 | -0.007 | -0.007 | 0.005 | 0.55 | 0.53 | 0.47 | 0.33 | 0.46 | 0.39 |
| 7 | 1141738 | rs10262070 | 0.348 | 0.73 | 1.132 | -0.521 | 0.051 | 0.128 | 0.128 | 0 | 0.34 | 0.16 | 0.21 | 0.03 | 0.21 | 0.19 |
| 7 | 1141925 | rs10262232 | 0.417 | 0.729 | 1.131 | -0.52 | 0.045 | 0.026 | 0.026 | 0 | 0.35 | 0.16 | 0.23 | 0.18 | 0.22 | 0.2 |
| 7 | 1142977 | rs10266519 | 1.058 | 0.618 | 0.994 | -0.612 | 0.036 | 0.137 | 0.137 | 0.155 | 0.27 | 0.05 | 0.18 | 0.03 | 0.14 | 0.16 |
| 7 | 1144527 | rs4236375 | -1.206 | 0.813 | 1.161 | -0.522 | -0.056 | -0.015 | -0.015 | 0.425 | 0.6 | 0.81 | 0.53 | 0.35 | 0.49 | 0.42 |
| 7 | 1151066 | rs6955290 | 0.455 | 0.805 | 1.269 | -0.534 | 0.017 | -0.051 | -0.051 | 0 | 0.18 | 0.34 | 0.25 | 0.14 | 0.24 | 0.19 |
| 7 | 1161646 | rs11766651 | -1.95 | 0.983 | 0.563 | -0.537 | -0.052 | -0.134 | -0.134 | 0 | 0.67 | 0.94 | 0.65 | 0.42 | 0.62 | 0.55 |
| 7 | 1162084 | rs7803208 | -0.247 | 0.934 | 0.671 | -0.553 | 0.018 | 0.074 | 0.074 | 0 | 0.23 | 0.49 | 0.29 | 0.33 | 0.26 | 0.23 |
| 7 | 1162222 | rs7793336 | -1.144 | 0.983 | 0.595 | -0.519 | -0.083 | -0.039 | -0.039 | 0 | 0.61 | 0.89 | 0.54 | 0.35 | 0.51 | 0.42 |
| 7 | 1169369 | rs6968630 | -1.17 | 0.974 | 0.57 | -0.455 | -0.086 | -0.051 | -0.051 | 0 | 0.61 | 0.89 | 0.54 | 0.37 | 0.51 | 0.42 |
| 7 | 1172588 | rs4305794 | -0.739 | 0.949 | 0.582 | -0.592 | 0.001 | -0.023 | -0.023 | 0 | 0.56 | 0.58 | 0.48 | 0.2 | 0.47 | 0.41 |
| 7 | 1172969 | rs13222294 | 0.628 | 1.081 | 0.253 | -0.43 | -0.001 | -0.01 | -0.01 | 0 | 0.17 | 0.14 | 0.19 | 0.08 | 0.2 | 0.19 |
| 7 | 1181139 | rs10278051 | -0.471 | 1.043 | -0.004 | -0.245 | -0.001 | 0 | 0 | 0 | 0.41 | 0.59 | 0.36 | 0.28 | 0.3 | 0.32 |
| 7 | 1183964 | rs28522260 | 0.647 | 0.637 | -0.003 | -0.264 | 0.032 | 0.129 | 0.129 | 0.258 | 0.27 | 0.07 | 0.18 | 0.03 | 0.15 | 0.16 |
| 7 | 1189178 | rs6943934 | -1.16 | 0.704 | 0.2 | -0.218 | 0.099 | 0.288 | 0.288 | 0.233 | 0.06 | 0.32 | 0.06 | 0.16 | 0.06 | 0.01 |
| 7 | 1192572 | rs1133122 | 0.67 | 0.677 | 0.163 | -0.294 | 0.031 | 0.128 | 0.128 | 0.309 | 0.27 | 0.07 | 0.18 | 0.03 | 0.15 | 0.16 |
| 7 | 1194274 | rs11763865 | 0.159 | 0.729 | 0.207 | -0.289 | -0.014 | -0.089 | -0.089 | 0 | 0.32 | 0.38 | 0.23 | 0.03 | 0.2 | 0.16 |
| 7 | 1198743 | rs4075937 | -0.678 | 0.519 | 0.101 | -0.225 | -0.001 | -0.019 | -0.019 | 0 | 0.59 | 0.46 | 0.57 | 0.25 | 0.57 | 0.53 |
| 7 | 1203957 | rs2960837 | -1.046 | 0.539 | 0.035 | -0.119 | 0.004 | -0.027 | -0.027 | 0 | 0.63 | 0.71 | 0.62 | 0.42 | 0.62 | 0.54 |
| 7 | 1213127 | rs1543985 | -1.554 | 0.56 | -0.308 | 0.199 | -0.039 | -0.092 | -0.092 | 0 | 0.68 | 0.92 | 0.65 | 0.47 | 0.62 | 0.6 |

| | | | | | | | | | | | | | | | | |
|---|---|---|---|---|---|---|---|---|---|---|---|---|---|---|---|---|
| 7 | 1213943 | rs10275401 | 0.768 | 0.496 | -0.355 | -0.224 | 0.033 | 0.09 | 0.09 | 0.495 | 0.26 | 0.1 | 0.18 | 0.03 | 0.14 | 0.15 |
| 7 | 1216013 | rs1880297 | -0.183 | 0.917 | -0.049 | -0.015 | 0.002 | -0.014 | -0.014 | 0 | 0.34 | 0.39 | 0.39 | 0.22 | 0.44 | 0.42 |
| 7 | 1216212 | rs1880296 | -0.09 | 1.225 | 0.028 | 0.472 | -0.002 | -0.034 | -0.034 | 0.308 | 0.57 | 0.52 | 0.57 | 0.24 | 0.57 | 0.57 |
| 7 | 1216438 | rs2949170 | -0.5 | 1.196 | -1.083 | 0.42 | -0.005 | -0.061 | -0.061 | 0 | 0.63 | 0.83 | 0.63 | 0.43 | 0.61 | 0.6 |
| 7 | 1221100 | rs17151792 | -0.751 | 2.08 | -0.4 | 1.199 | -0.013 | -0.011 | -0.011 | 0.147 | 0.14 | 0.15 | 0.27 | 0.28 | 0.26 | 0.16 |
| 7 | 1221289 | rs6961510 | 0.249 | 2.093 | -0.491 | 1.323 | 0.013 | 0.004 | 0.004 | 0 | 0.76 | 0.65 | 0.58 | 0.8 | 0.66 | 0.72 |
| 7 | 1224910 | rs12702457 | -0.564 | 1.976 | -0.148 | 0.808 | 0.042 | 0.005 | 0.005 | 0 | 0.18 | 0.26 | 0.28 | 0.21 | 0.28 | 0.17 |
| 7 | 1231678 | rs9640008 | 0.749 | 0.893 | -0.253 | 0.882 | -0.002 | -0.002 | -0.002 | 0 | 0.58 | 0.47 | 0.3 | 0.49 | 0.37 | 0.6 |
| 7 | 1243525 | rs7456553 | -0.303 | 0.222 | -0.69 | 0.678 | -0.024 | -0.023 | -0.023 | 0 | 0.39 | 0.18 | 0.35 | 0.43 | 0.39 | 0.28 |
| 7 | 1245798 | rs7782777 | -0.257 | 0.226 | -0.703 | 0.662 | -0.005 | -0.023 | -0.023 | 0 | 0.38 | 0.17 | 0.38 | 0.42 | 0.36 | 0.27 |
| 7 | 1246030 | rs11767578 | 0.388 | -0.101 | 0.434 | 0.303 | -0.011 | -0.004 | -0.004 | 0 | 0.92 | 0.54 | 0.94 | 0.99 | 0.99 | 0.98 |
| 7 | 1260636 | rs10081255 | 0.646 | -0.051 | -0.343 | 0.61 | 0.025 | -0.007 | -0.007 | 0.048 | 0.45 | 0.16 | 0.57 | 0.42 | 0.59 | 0.67 |
| 7 | 1274582 | rs4724799 | 0.864 | -1.086 | -0.546 | 0.663 | 0.106 | 0.014 | 0.014 | 1.969 | 0.59 | 0.1 | 0.67 | 0.32 | 0.78 | 0.69 |
| 7 | 1283597 | rs10235272 | 0.557 | -0.895 | -0.395 | 0.782 | 0.069 | -0.027 | -0.027 | 0 | 0.63 | 0.14 | 0.58 | 0.4 | 0.79 | 0.78 |
| 7 | 1288639 | rs13241427 | 0.688 | -1.486 | -0.006 | 0.685 | -0.037 | 0.003 | 0.003 | 0.563 | 0.65 | 0.56 | 0.62 | 0.67 | 0.8 | 0.83 |
| 7 | 1289279 | rs13242133 | 0.618 | -1.459 | 0.023 | 0.569 | -0.022 | -0.007 | -0.007 | 0 | 0.64 | 0.41 | 0.61 | 0.72 | 0.8 | 0.83 |
| 7 | 1303439 | rs1949783 | 1.051 | -0.643 | -0.748 | 0.86 | 0.075 | -0.002 | -0.002 | 0 | 0.57 | 0.1 | 0.54 | 0.36 | 0.72 | 0.74 |
| 7 | 1307284 | rs3750159 | -1.039 | -0.332 | -0.042 | 1.03 | 0.002 | 0.045 | 0.045 | 0.017 | 0.22 | 0.36 | 0.27 | 0.43 | 0.21 | 0.26 |
| 7 | 1316239 | rs2690262 | 0.62 | -0.531 | -0.424 | 0.367 | -0.003 | -0.004 | -0.004 | 0 | 0.55 | 0.44 | 0.49 | 0.66 | 0.67 | 0.66 |
| 7 | 1323967 | rs4724883 | -0.017 | -0.916 | -0.432 | 0.252 | -0.035 | -0.084 | -0.084 | 0 | 0.18 | 0.25 | 0.15 | 0 | 0.08 | 0.1 |
| 7 | 1324345 | rs10952002 | -0.8 | -0.041 | -1.463 | 0.61 | -0.065 | -0.003 | -0.003 | 0 | 0.54 | 0.91 | 0.52 | 0.47 | 0.47 | 0.61 |
| 7 | 1325122 | rs2690256 | 0.714 | -0.077 | -1.804 | 0.062 | -0.153 | 0.017 | 0.017 | 0 | 0.43 | 0.02 | 0.33 | 0.26 | 0.55 | 0.38 |
| 7 | 1328707 | rs11772032 | -0.282 | 0.212 | -1.271 | -0.18 | -0.02 | 0.012 | 0.012 | 0.23 | 0.22 | 0.12 | 0.19 | 0.22 | 0.21 | 0.3 |
| 7 | 1329569 | rs4598160 | 1.419 | -0.187 | -1.768 | -0.26 | -0.139 | -0.014 | -0.014 | 0 | 0.4 | 0.02 | 0.3 | 0.26 | 0.45 | 0.4 |
| 7 | 1331254 | rs11763092 | 0.13 | -0.507 | -0.189 | -0.421 | -0.028 | -0.122 | -0.122 | 0 | 0.26 | 0.36 | 0.25 | 0 | 0.23 | 0.11 |
| 7 | 1331920 | rs10240008 | 1.093 | -0.46 | 0.074 | -0.291 | 0.008 | 0.033 | 0.033 | 0 | 0.49 | 0.42 | 0.53 | 0.7 | 0.55 | 0.54 |
| 7 | 1333246 | rs1104950 | 1.558 | -0.216 | 0.091 | -0.227 | -0.031 | 0.042 | 0.042 | 0.687 | 0.42 | 0.22 | 0.31 | 0.26 | 0.48 | 0.37 |
| 7 | 1335231 | rs10952012 | 0.112 | -0.485 | 0.139 | -1.053 | 0.008 | -0.002 | -0.002 | 0 | 0.77 | 0.68 | 0.79 | 0.72 | 0.79 | 0.66 |
| 7 | 1335447 | rs12540710 | -0.289 | -0.711 | -0.518 | -1.198 | 0.015 | -0.004 | -0.004 | 0 | 0.26 | 0.21 | 0.24 | 0.01 | 0.23 | 0.17 |
| 7 | 1338394 | rs4724910 | -1.319 | 0.107 | -0.722 | -0.789 | 0.018 | 0.118 | 0.118 | 0.294 | 0.49 | 0.74 | 0.44 | 0.31 | 0.46 | 0.53 |
| 7 | 1342825 | rs13230410 | -0.966 | 0.693 | -0.737 | -0.14 | -0.008 | 0.021 | 0.021 | 0 | 0.27 | 0.18 | 0.22 | 0.26 | 0.25 | 0.35 |
| 7 | 1348538 | rs13241587 | -0.836 | 1.088 | -0.883 | -0.418 | -0.045 | 0.051 | 0.051 | 0 | 0.24 | 0.11 | 0.22 | 0.21 | 0.27 | 0.35 |
| 7 | 1360473 | rs7794099 | -1.57 | 0.948 | 0.204 | -0.254 | 0.063 | -0.012 | -0.012 | 0 | 0.06 | 0.33 | 0.05 | 0 | 0.05 | 0.01 |
| 7 | 1363454 | rs13221166 | -0.085 | 0.887 | -0.332 | -0.454 | 0.004 | 0.01 | 0.01 | 0 | 0.56 | 0.42 | 0.5 | 0.31 | 0.67 | 0.37 |
| 7 | 1380226 | rs1106296 | -0.334 | 0.823 | -0.321 | -0.036 | 0 | -0.001 | -0.001 | 0.243 | 0.39 | 0.58 | 0.39 | 0.48 | 0.35 | 0.29 |
| 7 | 1381028 | rs1880292 | 0.823 | 0.964 | -0.798 | -0.184 | 0.023 | -0.032 | -0.032 | 0.739 | 0.4 | 0.19 | 0.43 | 0.47 | 0.32 | 0.49 |
| 7 | 1382320 | rs7459420 | -0.462 | 0.383 | -0.275 | -0.39 | 0.011 | 0.036 | 0.036 | 0 | 0.39 | 0.23 | 0.36 | 0.31 | 0.37 | 0.32 |
| 7 | 1387185 | rs2361713 | 0.028 | -0.128 | -0.573 | -0.465 | 0.014 | -0.028 | -0.028 | 0 | 0.28 | 0.04 | 0.17 | 0.24 | 0.21 | 0.26 |
| 7 | 1389397 | rs7809940 | -0.971 | -0.174 | 0.364 | -0.542 | -0.015 | -0.012 | -0.012 | 0 | 0.4 | 0.49 | 0.37 | 0.29 | 0.32 | 0.39 |
| 7 | 1389594 | rs4442018 | -0.967 | -0.079 | 0.263 | -0.505 | -0.001 | -0.01 | -0.01 | 1.164 | 0.09 | 0.12 | 0.18 | 0.04 | 0.11 | 0.1 |
| 7 | 1410550 | rs3934914 | 0.928 | -0.257 | 0.434 | -0.593 | 0.023 | 0.035 | 0.035 | 0 | 0.58 | 0.31 | 0.51 | 0.72 | 0.62 | 0.63 |
| 7 | 1410607 | rs4288278 | -0.153 | -0.269 | 0.186 | -0.608 | -0.062 | -0.048 | -0.048 | 0.013 | 0.71 | 0.99 | 0.78 | 0.75 | 0.69 | 0.73 |
| 7 | 1420019 | rs10807960 | 0.808 | -0.038 | 1.147 | -0.476 | -0.005 | -0.011 | -0.011 | 0.059 | 0.66 | 0.6 | 0.55 | 0.72 | 0.64 | 0.65 |
| 7 | 1428476 | rs11514810 | -1.113 | -0.114 | 1.565 | -0.813 | 0.006 | -0.047 | -0.047 | 0 | 0.13 | 0.37 | 0.27 | 0.03 | 0.1 | 0.08 |
| 7 | 1438686 | rs10233864 | -1.279 | -0.296 | 1.514 | -0.799 | -0.019 | -0.04 | -0.04 | 0 | 0.08 | 0.3 | 0.05 | 0 | 0.02 | 0.08 |
| 7 | 1448188 | rs10267348 | -1.991 | -0.503 | 1.421 | -0.983 | -0.046 | -0.092 | -0.092 | 0 | 0.22 | 0.6 | 0.3 | 0.03 | 0.12 | 0.16 |
| 7 | 1458931 | rs10231967 | -2.238 | -0.649 | 1.273 | -1.127 | -0.045 | -0.109 | -0.109 | 0.05 | 0.26 | 0.57 | 0.31 | 0.03 | 0.14 | 0.17 |
| 7 | 1461066 | rs7807443 | -1.214 | -0.736 | 1.361 | -1.162 | -0.027 | -0.061 | -0.061 | 0 | 0.14 | 0.52 | 0.28 | 0.03 | 0.08 | 0.07 |
| 7 | 1462702 | rs6955047 | -1.54 | -0.656 | 1.49 | -1.12 | -0.049 | -0.101 | -0.101 | 0.544 | 0.2 | 0.42 | 0.28 | 0.03 | 0.11 | 0.12 |
| 7 | 1464603 | rs7800232 | -1.685 | -0.659 | 1.101 | -1.114 | -0.053 | -0.103 | -0.103 | 0 | 0.22 | 0.57 | 0.3 | 0.03 | 0.11 | 0.15 |
| 7 | 1473468 | rs12670417 | 0.575 | -0.249 | 0.222 | -0.817 | -0.043 | -0.052 | -0.052 | 0.14 | 0.23 | 0.08 | 0.19 | 0.34 | 0.28 | 0.46 |
| 7 | 1476596 | rs10262345 | -1.749 | -0.66 | 0.869 | -0.994 | -0.049 | -0.12 | -0.12 | 0 | 0.25 | 0.56 | 0.31 | 0.03 | 0.14 | 0.15 |
| 7 | 1477113 | rs10266642 | -1.752 | -0.657 | 0.871 | -0.993 | -0.049 | -0.12 | -0.12 | 0 | 0.25 | 0.56 | 0.31 | 0.03 | 0.14 | 0.15 |
| 7 | 1478972 | rs10253769 | -1.72 | -0.661 | 0.883 | -0.981 | -0.051 | -0.119 | -0.119 | 1.004 | 0.25 | 0.56 | 0.31 | 0.03 | 0.14 | 0.15 |
| 7 | 1493367 | rs12535457 | -1.174 | -0.395 | 1.407 | -0.731 | -0.026 | -0.054 | -0.054 | 0 | 0.14 | 0.56 | 0.28 | 0.03 | 0.08 | 0.07 |
| 7 | 1507011 | rs6962610 | 0.422 | 0.05 | 1.002 | -0.842 | 0.001 | 0.081 | 0.081 | 0 | 0.51 | 0.82 | 0.41 | 0.66 | 0.46 | 0.5 |
| 7 | 1508191 | rs11972070 | 0.582 | 0.004 | 1.609 | -1.018 | 0.003 | 0.018 | 0.018 | 0 | 0.48 | 0.45 | 0.38 | 0.71 | 0.45 | 0.47 |
| 7 | 1521875 | rs4560715 | 0.36 | 0.08 | 1.21 | -0.995 | -0.017 | 0.121 | 0.121 | 0 | 0.52 | 0.84 | 0.42 | 0.71 | 0.47 | 0.5 |
| 7 | 1529860 | rs6968011 | 0.545 | 0.073 | 1.42 | -0.999 | 0.011 | 0.092 | 0.092 | 0 | 0.5 | 0.72 | 0.41 | 0.71 | 0.46 | 0.47 |
| 7 | 1536717 | rs3735660 | 0.387 | 0.069 | 1.269 | -1.001 | -0.015 | 0.124 | 0.124 | 0 | 0.52 | 0.84 | 0.42 | 0.71 | 0.47 | 0.46 |
| 7 | 1542697 | rs3752714 | 0.449 | 0.087 | 1.285 | -0.99 | 0.004 | 0.109 | 0.109 | 0 | 0.51 | 0.77 | 0.42 | 0.71 | 0.46 | 0.44 |
| 7 | 1551716 | rs10282181 | 0.256 | 0.627 | 1.73 | -0.838 | 0 | -0.004 | -0.004 | 0 | 0.54 | 0.56 | 0.46 | 0.67 | 0.53 | 0.53 |
| 7 | 1552438 | rs6965558 | -0.129 | 0.441 | 1.645 | -0.978 | -0.039 | 0.001 | 0.001 | 0 | 0.63 | 0.77 | 0.54 | 0.66 | 0.56 | 0.51 |
| 7 | 1557967 | rs2117347 | 0.563 | 0.539 | 2.022 | -0.858 | 0 | -0.003 | -0.003 | 0 | 0.56 | 0.48 | 0.48 | 0.67 | 0.55 | 0.47 |
| 7 | 1561702 | rs11973950 | 0.477 | 0.488 | 2.023 | -0.85 | 0 | -0.012 | -0.012 | 0.158 | 0.57 | 0.51 | 0.49 | 0.67 | 0.57 | 0.49 |
| 7 | 1563025 | rs6463894 | 0.384 | 0.509 | 2.117 | -0.841 | -0.003 | 0.015 | 0.015 | 0 | 0.58 | 0.69 | 0.52 | 0.67 | 0.57 | 0.49 |
| 7 | 1563401 | rs6463900 | 0.384 | 0.509 | 2.117 | -0.841 | -0.003 | 0.015 | 0.015 | 0 | 0.58 | 0.69 | 0.52 | 0.67 | 0.57 | 0.49 |
| 7 | 1563732 | rs4725180 | 0.426 | 0.495 | 2.14 | -0.847 | 0 | -0.003 | -0.003 | 0 | 0.58 | 0.57 | 0.5 | 0.67 | 0.57 | 0.49 |
| 7 | 1565409 | rs6978445 | 0.213 | 0.583 | 2.023 | -0.847 | 0 | 0.03 | 0.03 | 0 | 0.6 | 0.8 | 0.7 | 0.7 | 0.63 | 0.53 |

| | | | | | | | | | | | | | | | | |
|---|---|---|---|---|---|---|---|---|---|---|---|---|---|---|---|---|
| 7 | 1569418 | rs3814477 | 0.33 | 0.606 | 2.084 | -0.843 | -0.003 | 0.01 | 0.01 | 0 | 0.59 | 0.66 | 0.51 | 0.66 | 0.57 | 0.55 |
| 7 | 1574403 | rs4720833 | 0.219 | 0.52 | 1.849 | -0.821 | 0.003 | 0.027 | 0.027 | 0 | 0.6 | 0.8 | 0.7 | 0.69 | 0.63 | 0.55 |
| 7 | 1580921 | rs3735656 | 0.289 | 0.514 | 1.978 | -0.824 | 0.019 | 0.042 | 0.042 | 0 | 0.53 | 0.71 | 0.53 | 0.66 | 0.57 | 0.54 |
| 7 | 1583288 | rs892523 | 0.304 | 0.438 | 1.941 | -0.864 | 0.013 | 0.021 | 0.021 | 0 | 0.53 | 0.6 | 0.67 | 0.69 | 0.62 | 0.51 |
| 7 | 1586866 | rs3824072 | 0.181 | 0.362 | 1.701 | -0.879 | 0.034 | 0.068 | 0.068 | 0.107 | 0.56 | 0.8 | 0.7 | 0.69 | 0.62 | 0.55 |
| 7 | 1590280 | rs2304359 | 0.102 | 0.546 | 1.648 | -0.81 | 0.002 | 0.003 | 0.003 | 0 | 0.54 | 0.56 | 0.65 | 0.69 | 0.63 | 0.54 |
| 7 | 1590443 | rs2304361 | 0.104 | 0.559 | 1.622 | -0.813 | 0.007 | 0.009 | 0.009 | 0 | 0.54 | 0.56 | 0.65 | 0.69 | 0.63 | 0.54 |
| 7 | 1596196 | rs7811444 | -2.061 | 1.472 | 2.245 | -0.505 | 0.046 | -0.051 | -0.051 | 0.043 | 0.2 | 0.52 | 0.35 | 0.11 | 0.24 | 0.16 |
| 7 | 1602673 | rs2033572 | 2.084 | 1.222 | 1.512 | -0.346 | -0.002 | -0.001 | -0.001 | 0.186 | 0.71 | 0.28 | 0.67 | 0.84 | 0.71 | 0.8 |
| 7 | 1604320 | rs12702933 | 1.757 | 1.308 | 1.652 | -0.373 | 0.126 | 0.025 | 0.025 | 0 | 0.87 | 0.57 | 0.82 | 0.88 | 0.82 | 0.84 |
| 7 | 1606615 | rs1865347 | 2.039 | 1.289 | 1.715 | -0.414 | 0.114 | 0.011 | 0.011 | 0 | 0.86 | 0.5 | 0.81 | 0.89 | 0.82 | 0.84 |
| 7 | 1606847 | rs10244477 | -0.686 | 1.431 | 0.429 | -0.343 | -0.018 | -0.081 | -0.081 | 2.127 | 0.17 | 0.27 | 0.13 | 0.01 | 0.09 | 0.02 |
| 7 | 1607259 | rs12664 | -0.69 | 1.43 | 0.429 | -0.344 | -0.018 | -0.081 | -0.081 | 2.09 | 0.17 | 0.27 | 0.13 | 0 | 0.09 | 0.02 |
| 7 | 1614261 | rs10274145 | 1.62 | 1.79 | 0.588 | 0.336 | -0.014 | -0.012 | -0.012 | 0 | 0.92 | 0.97 | 0.86 | 0.88 | 0.82 | 0.85 |
| 7 | 1620592 | rs4725234 | 1.514 | 1.551 | 0.441 | 0.509 | 0.074 | 0.047 | 0.047 | 0 | 0.9 | 0.69 | 0.83 | 0.88 | 0.85 | 0.87 |
| 7 | 1629188 | rs1275 | -0.909 | 1.57 | -0.627 | 0.901 | -0.027 | -0.069 | -0.069 | 0.532 | 0.28 | 0.14 | 0.44 | 0.53 | 0.4 | 0.4 |
| 7 | 1631580 | rs753112 | -0.574 | 1.659 | -0.607 | 0.969 | -0.019 | 0.022 | 0.022 | 0 | 0.19 | 0.11 | 0.31 | 0.14 | 0.27 | 0.21 |
| 7 | 1640842 | rs10257217 | -0.704 | 1.047 | -0.821 | 0.617 | 0.013 | -0.004 | -0.004 | 0 | 0.68 | 0.79 | 0.79 | 0.66 | 0.77 | 0.66 |
| 7 | 1655601 | rs10429166 | -0.998 | 1.056 | 0.494 | 0.335 | 0.014 | -0.035 | -0.035 | 0 | 0.39 | 0.51 | 0.47 | 0.2 | 0.53 | 0.41 |
| 7 | 1667380 | rs10247783 | 0.007 | 1.056 | 0.951 | 0.145 | -0.008 | -0.066 | -0.066 | 0 | 0.11 | 0.18 | 0.06 | 0 | 0.06 | 0.03 |
| 7 | 1677026 | rs9886201 | -1.662 | 0.941 | 1.537 | 0.726 | -0.016 | 0.046 | 0.046 | 0.161 | 0.24 | 0.57 | 0.18 | 0.42 | 0.18 | 0.2 |
| 7 | 1677399 | rs4464874 | -1.744 | 0.545 | 0.952 | 0.625 | -0.072 | -0.032 | -0.032 | 0 | 0.52 | 0.86 | 0.53 | 0.61 | 0.51 | 0.47 |
| 7 | 1677696 | rs4720882 | 0.115 | 0.49 | 0.935 | 0.607 | -0.087 | 0.022 | 0.022 | 0.459 | 0.24 | 0.01 | 0.31 | 0.19 | 0.32 | 0.27 |
| 7 | 1680165 | rs12703010 | -1.07 | 0.613 | 1.33 | 0.674 | 0.003 | -0.013 | -0.013 | 0 | 0.41 | 0.63 | 0.45 | 0.56 | 0.38 | 0.36 |
| 7 | 1685010 | rs7785046 | -1.581 | 0.543 | 0.75 | 0.707 | -0.069 | -0.029 | -0.029 | 0 | 0.51 | 0.86 | 0.52 | 0.61 | 0.51 | 0.48 |
| 7 | 1688369 | rs6464024 | -0.575 | 0.565 | 0.983 | 0.71 | 0 | -0.004 | -0.004 | 4.246 | 0.35 | 0.35 | 0.39 | 0.24 | 0.44 | 0.37 |
| 7 | 1689761 | rs10274735 | 0.333 | 0.541 | 0.239 | 0.778 | -0.001 | -0.065 | -0.065 | 0 | 0.11 | 0.17 | 0.05 | 0 | 0.07 | 0.02 |
| 7 | 1704673 | rs6952435 | -1.334 | 0.672 | -0.673 | 0.631 | -0.073 | -0.034 | -0.034 | 0 | 0.53 | 0.89 | 0.53 | 0.62 | 0.54 | 0.48 |
| 7 | 1714703 | rs10270726 | -0.953 | 0.591 | -0.321 | 0.706 | 0.005 | -0.049 | -0.049 | 0.004 | 0.4 | 0.77 | 0.41 | 0.56 | 0.35 | 0.36 |
| 7 | 1724090 | rs6950626 | -0.707 | 0.638 | -0.871 | -0.125 | 0.002 | 0.037 | 0.037 | 0 | 0.16 | 0.07 | 0.09 | 0.06 | 0.16 | 0.11 |
| 7 | 1733102 | rs4720922 | -0.19 | 0.341 | -0.983 | 0.101 | -0.088 | -0.024 | -0.024 | 3.246 | 0.23 | 0.01 | 0.28 | 0.16 | 0.27 | 0.28 |
| 7 | 1750265 | rs7780342 | -1.116 | 0.266 | -0.244 | 0.091 | -0.033 | -0.058 | -0.058 | 0 | 0.17 | 0.4 | 0.11 | 0.06 | 0.13 | 0.15 |
| 7 | 1750797 | rs4720940 | -0.907 | 0.689 | -1.083 | 0.176 | 0 | -0.011 | -0.011 | 0.036 | 0.17 | 0.22 | 0.29 | 0.06 | 0.23 | 0.18 |
| 7 | 1766670 | rs4720965 | -0.494 | 0.286 | -1.593 | 0.458 | -0.029 | -0.064 | -0.064 | 2.443 | 0.24 | 0.01 | 0.11 | 0.2 | 0.19 | 0.29 |
| 7 | 1778604 | rs12699191 | 0.09 | 0.706 | -0.896 | 0.365 | 0.001 | -0.009 | -0.009 | 0 | 0.08 | 0.08 | 0.04 | 0 | 0.05 | 0.02 |
| 7 | 1787236 | rs9177 | 0.045 | 1.02 | -0.856 | 0.761 | -0.004 | -0.004 | -0.004 | 0.05 | 0.41 | 0.39 | 0.63 | 0.5 | 0.45 | 0.4 |
| 7 | 1792688 | rs6949979 | -1.11 | 1.066 | -0.775 | 1.112 | -0.001 | 0.003 | 0.003 | 0.147 | 0.14 | 0.14 | 0.09 | 0.27 | 0.13 | 0.17 |
| 7 | 1792804 | rs4719278 | 0 | 1.569 | -0.181 | 0.739 | 0.029 | -0.038 | -0.038 | 0.399 | 0.21 | 0.4 | 0.24 | 0.18 | 0.27 | 0.15 |
| 7 | 1806723 | rs12538366 | -0.918 | 0.876 | -0.772 | 0.965 | -0.017 | -0.082 | -0.082 | 0 | 0.18 | 0.04 | 0.1 | 0.33 | 0.15 | 0.18 |
| 7 | 1809120 | rs10277267 | -1.526 | 1.107 | -0.339 | 0.697 | -0.003 | -0.01 | -0.01 | 0 | 0.26 | 0.26 | 0.37 | 0.34 | 0.24 | 0.25 |
| 7 | 1809618 | rs1919908 | -1.503 | 1.117 | -0.323 | 0.767 | -0.014 | -0.044 | -0.044 | 0 | 0.24 | 0.11 | 0.33 | 0.31 | 0.21 | 0.24 |
| 7 | 1813198 | rs13245980 | -1.729 | 0.776 | 0.652 | 0.486 | -0.01 | 0.019 | 0.019 | 0 | 0.3 | 0.59 | 0.38 | 0.36 | 0.24 | 0.29 |
| 7 | 1819139 | rs10227806 | -0.934 | 0.72 | 1.098 | 0.471 | 0.013 | -0.037 | -0.037 | 0.003 | 0.07 | 0.21 | 0.21 | 0.01 | 0.06 | 0.09 |
| 7 | 1821278 | rs1881858 | -1.667 | 0.964 | 1.087 | 0.48 | 0.003 | 0.037 | 0.037 | 0 | 0.29 | 0.53 | 0.39 | 0.37 | 0.26 | 0.29 |
| 7 | 1825304 | rs10950371 | -1.488 | 0.682 | 0.993 | 0.359 | -0.002 | 0.011 | 0.011 | 0 | 0.3 | 0.42 | 0.39 | 0.36 | 0.27 | 0.31 |
| 7 | 1827002 | rs6954989 | 1.012 | 0.435 | 0.331 | -0.122 | -0.013 | 0.04 | 0.04 | 0 | 0.34 | 0.24 | 0.27 | 0.16 | 0.38 | 0.21 |
| 7 | 1829899 | rs7789703 | -0.94 | 0.556 | 0.325 | -0.057 | -0.007 | 0.014 | 0.014 | 0 | 0.3 | 0.42 | 0.23 | 0.35 | 0.25 | 0.26 |
| 7 | 1837636 | rs2895215 | -1.069 | 0.495 | 0.06 | 0.017 | -0.006 | 0.005 | 0.005 | 0 | 0.32 | 0.59 | 0.4 | 0.36 | 0.25 | 0.32 |
| 7 | 1842407 | rs12699299 | 0.609 | 0.497 | 0.106 | -0.257 | 0 | 0.025 | 0.025 | 0 | 0.38 | 0.31 | 0.28 | 0.17 | 0.38 | 0.24 |
| 7 | 1849127 | rs4540313 | -0.008 | 0.448 | 0.248 | 0.214 | 0 | -0.002 | -0.002 | 0.055 | 0.49 | 0.46 | 0.46 | 0.28 | 0.42 | 0.31 |
| 7 | 1851461 | rs7807190 | -1.009 | 0.32 | 0.055 | 0.155 | -0.002 | -0.004 | -0.004 | 0.017 | 0.33 | 0.62 | 0.33 | 0.29 | 0.24 | 0.4 |
| 7 | 1866916 | rs13225346 | 0.502 | 0.197 | 0.282 | 0.402 | 0.01 | 0.015 | 0.015 | 0 | 0.4 | 0.42 | 0.37 | 0.45 | 0.43 | 0.26 |
| 7 | 1868092 | rs7799807 | -0.826 | 0.05 | 0.281 | 0.279 | -0.001 | -0.004 | -0.004 | 0.82 | 0.67 | 0.64 | 0.52 | 0.59 | 0.62 | 0.52 |
| 7 | 1876081 | rs4458759 | 0.537 | 0.194 | 0.362 | 0.313 | 0 | 0.002 | 0.002 | 0 | 0.41 | 0.52 | 0.29 | 0.43 | 0.41 | 0.19 |
| 7 | 1886937 | rs6954673 | 0.973 | 0.071 | 0.12 | 0.274 | -0.038 | -0.083 | -0.083 | 0.957 | 0.27 | 0.1 | 0.46 | 0.42 | 0.35 | 0.37 |
| 7 | 1887362 | rs6957894 | 0.696 | 0.016 | 0.418 | 0.25 | 0.001 | 0.008 | 0.008 | 0 | 0.29 | 0.37 | 0.48 | 0.42 | 0.36 | 0.45 |
| 7 | 1887523 | rs6978725 | -1.463 | -0.021 | 0.941 | 0.3 | 0.032 | 0.069 | 0.069 | 0 | 0.05 | 0.21 | 0.11 | 0.09 | 0.03 | 0.09 |
| 7 | 1889599 | rs10250550 | -1.188 | -0.017 | 0.685 | 0.215 | -0.002 | -0.002 | -0.002 | 0 | 0.63 | 0.68 | 0.74 | 0.65 | 0.6 | 0.83 |
| 7 | 1892309 | rs11764337 | -0.392 | -0.063 | 0.235 | 0.155 | 0.017 | 0.112 | 0.112 | 0 | 0.28 | 0.06 | 0.14 | 0.14 | 0.21 | 0.29 |
| 7 | 1912222 | rs4721135 | 0.794 | 0.07 | 0.314 | 0.271 | -0.007 | -0.023 | -0.023 | 0.968 | 0.3 | 0.29 | 0.47 | 0.44 | 0.4 | 0.37 |
| 7 | 1914679 | rs4719331 | -0.814 | -0.139 | 0.528 | 0.138 | -0.021 | -0.029 | -0.029 | 0.064 | 0.33 | 0.55 | 0.21 | 0.22 | 0.21 | 0.37 |
| 7 | 1920826 | rs10275045 | 0.874 | 0.007 | 0.307 | 0.269 | -0.059 | -0.071 | -0.071 | 0 | 0.29 | 0.14 | 0.48 | 0.44 | 0.42 | 0.42 |
| 7 | 1923385 | rs7783715 | -0.562 | -0.151 | 0.778 | 0.185 | 0.001 | 0.002 | 0.002 | 0.312 | 0.57 | 0.36 | 0.67 | 0.66 | 0.62 | 0.73 |
| 7 | 1934131 | rs6951956 | -0.965 | 0.013 | 1.094 | 0.011 | -0.012 | -0.061 | -0.061 | 0.017 | 0.33 | 0.42 | 0.28 | 0.14 | 0.23 | 0.43 |
| 7 | 1947895 | rs6970796 | -0.943 | 0.057 | 1.173 | 0.022 | 0.121 | 0.041 | 0.041 | 0 | 0.05 | 0.41 | 0.15 | 0.01 | 0.03 | 0.15 |
| 7 | 1962163 | rs10268609 | 0.182 | -0.009 | 0.677 | -0.029 | 0.027 | 0.067 | 0.067 | 0 | 0.73 | 0.87 | 0.86 | 0.87 | 0.81 | 0.73 |
| 7 | 1968245 | rs12699476 | 0.981 | 0.309 | 0.656 | -0.129 | -0.001 | -0.008 | -0.008 | 0.017 | 0.66 | 0.65 | 0.68 | 0.77 | 0.73 | 0.63 |
| 7 | 1968953 | rs12699477 | 1.166 | 0.45 | 0.317 | 0.057 | -0.076 | -0.169 | -0.169 | 0 | 0.26 | 0.03 | 0.42 | 0.45 | 0.37 | 0.36 |
| 7 | 1976556 | rs2280550 | 0.882 | 0.139 | 0.196 | 0.008 | -0.101 | -0.171 | -0.171 | 0.156 | 0.27 | 0.03 | 0.44 | 0.44 | 0.39 | 0.36 |

| | | | | | | | | | | | | | | | | |
|---|---|---|---|---|---|---|---|---|---|---|---|---|---|---|---|---|
| 7 | 1984122 | rs1554506 | 1.636 | 0.202 | 0.653 | 0.03 | 0.142 | 0.175 | 0.175 | 0 | 0.95 | 0.68 | 0.84 | 0.91 | 0.96 | 0.91 |
| 7 | 1987092 | rs10237340 | 2.126 | 0.182 | 0.661 | 0.02 | 0.103 | 0.14 | 0.14 | 0.184 | 0.94 | 0.64 | 0.83 | 0.9 | 0.96 | 0.91 |
| 7 | 1993899 | rs7804190 | -0.886 | 0.129 | 0.159 | -0.423 | 0.016 | 0.025 | 0.025 | 0.012 | 0.65 | 0.56 | 0.65 | 0.59 | 0.64 | 0.74 |
| 7 | 2004421 | rs12666575 | 1.286 | 0.262 | -0.078 | -0.438 | -0.086 | -0.173 | -0.173 | 5.545 | 0.27 | 0.03 | 0.44 | 0.45 | 0.39 | 0.33 |
| 7 | 2017445 | rs6461049 | -0.598 | 0.256 | 0.013 | -0.505 | 0.011 | 0.015 | 0.015 | 0 | 0.66 | 0.61 | 0.38 | 0.45 | 0.56 | 0.57 |
| 7 | 2018870 | rs3996329 | 0.185 | 0.097 | -0.308 | -0.781 | 0.009 | 0.068 | 0.068 | 0 | 0.24 | 0.06 | 0.11 | 0.12 | 0.19 | 0.28 |
| 7 | 2032803 | rs11764590 | -0.255 | 0.132 | -0.245 | -0.737 | 0.006 | 0.122 | 0.122 | 0 | 0.28 | 0.04 | 0.14 | 0.12 | 0.21 | 0.32 |
| 7 | 2039594 | rs10950503 | -0.436 | 0.113 | -0.135 | -0.434 | -0.008 | 0.051 | 0.051 | 0.41 | 0.56 | 0.25 | 0.63 | 0.72 | 0.6 | 0.65 |
| 7 | 2053747 | rs3996325 | -0.828 | 0.069 | -0.063 | -0.438 | -0.003 | -0.003 | -0.003 | 0 | 0.33 | 0.28 | 0.3 | 0.27 | 0.26 | 0.41 |
| 7 | 2057934 | rs883444 | 1.577 | 0.09 | 0.293 | -0.426 | 0.047 | 0.068 | 0.068 | 0.833 | 0.91 | 0.46 | 0.81 | 0.91 | 0.96 | 0.83 |
| 7 | 2068470 | rs4721321 | -1.023 | 0.02 | 0.181 | -0.416 | -0.007 | -0.008 | -0.008 | 0.436 | 0.36 | 0.56 | 0.33 | 0.28 | 0.26 | 0.47 |
| 7 | 2070517 | rs7778752 | -1.309 | 0.148 | 0.176 | -0.33 | -0.019 | 0.058 | 0.058 | 0 | 0.64 | 0.82 | 0.79 | 0.77 | 0.65 | 0.8 |
| 7 | 2094340 | rs6461108 | -1.628 | 0.304 | 0.334 | -0.363 | 0.109 | 0.061 | 0.061 | 0 | 0.09 | 0.47 | 0.19 | 0.09 | 0.09 | 0.17 |
| 7 | 2110346 | rs3823624 | -0.427 | 0.136 | -0.164 | -0.389 | 0.008 | 0.025 | 0.025 | 0 | 0.28 | 0.06 | 0.14 | 0.18 | 0.21 | 0.29 |
| 7 | 2111636 | rs10950536 | 0.413 | 0.598 | -0.197 | -1.032 | 0.002 | 0.03 | 0.03 | 0 | 0.48 | 0.55 | 0.7 | 0.79 | 0.52 | 0.63 |
| 7 | 2115163 | rs3729620 | 0.275 | 0.562 | -0.546 | -1.244 | -0.037 | 0.048 | 0.048 | 0.319 | 0.32 | 0.03 | 0.25 | 0.18 | 0.22 | 0.25 |
| 7 | 2119657 | rs7796547 | -1.053 | 0.686 | -0.291 | -1.276 | -0.007 | 0.013 | 0.013 | 0.861 | 0.71 | 0.79 | 0.84 | 0.77 | 0.66 | 0.79 |
| 7 | 2120378 | rs3800879 | -1.023 | 0.68 | -0.242 | -1.271 | 0.008 | -0.006 | -0.006 | 0.083 | 0.7 | 0.68 | 0.79 | 0.69 | 0.65 | 0.77 |
| 7 | 2120635 | rs3800881 | -0.756 | 0.701 | -0.231 | -1.275 | 0.017 | -0.01 | -0.01 | 0.49 | 0.68 | 0.65 | 0.75 | 0.68 | 0.62 | 0.68 |
| 7 | 2120758 | rs3800882 | 1.075 | 0.618 | -0.143 | -1.322 | -0.01 | 0.039 | 0.039 | 0 | 0.34 | 0.55 | 0.5 | 0.5 | 0.4 | 0.43 |
| 7 | 2121156 | rs1558461 | 2.545 | 0.583 | -0.478 | -1.281 | -0.047 | -0.014 | -0.014 | 0 | 0.14 | 0.01 | 0.13 | 0.08 | 0.11 | 0.09 |
| 7 | 2123963 | rs3800893 | 0.462 | 0.711 | -0.287 | -1.176 | -0.025 | 0.053 | 0.053 | 0.261 | 0.33 | 0.03 | 0.24 | 0.18 | 0.22 | 0.26 |
| 7 | 2125942 | rs3800895 | 0.034 | 0.715 | -0.223 | -1.181 | 0.001 | 0.037 | 0.037 | 0.026 | 0.37 | 0.22 | 0.3 | 0.17 | 0.27 | 0.36 |
| 7 | 2130014 | rs13242473 | 0.429 | 0.626 | -0.438 | -1.077 | -0.025 | 0.039 | 0.039 | 0.235 | 0.33 | 0.03 | 0.24 | 0.21 | 0.22 | 0.26 |
| 7 | 2135170 | rs3800897 | 1.629 | 0.143 | -0.3 | -0.947 | -0.033 | 0.054 | 0.054 | 0 | 0.11 | 0.13 | 0.07 | 0.36 | 0.03 | 0.08 |
| 7 | 2149967 | rs10224497 | 0.933 | 0.66 | -0.166 | -0.918 | -0.01 | -0.01 | -0.01 | 0 | 0.33 | 0.26 | 0.54 | 0.59 | 0.43 | 0.47 |
| 7 | 2159437 | rs3800908 | 0.585 | 0.566 | -0.209 | -0.941 | 0.001 | 0.001 | 0.001 | 0 | 0.37 | 0.54 | 0.57 | 0.59 | 0.43 | 0.53 |
| 7 | 2159817 | rs3778978 | 1.471 | 0.671 | -0.774 | -0.919 | -0.08 | -0.176 | -0.176 | 0.605 | 0.28 | 0.02 | 0.45 | 0.5 | 0.39 | 0.36 |
| 7 | 2167978 | rs3800918 | -1 | 0.738 | -0.616 | -1.109 | -0.012 | 0.026 | 0.026 | 1.007 | 0.73 | 0.8 | 0.84 | 0.78 | 0.72 | 0.81 |
| 7 | 2170759 | rs6461168 | -0.158 | 0.464 | -0.576 | -1.18 | 0.028 | 0.038 | 0.038 | 0 | 0.4 | 0.71 | 0.35 | 0.27 | 0.27 | 0.35 |
| 7 | 2172455 | rs3778991 | 1.406 | 0.73 | -0.691 | -0.803 | -0.089 | -0.179 | -0.179 | 0.147 | 0.28 | 0.02 | 0.44 | 0.49 | 0.37 | 0.36 |
| 7 | 2175855 | rs3778994 | 0.966 | 0.74 | -0.6 | -0.821 | -0.036 | -0.074 | -0.074 | 0 | 0.32 | 0.18 | 0.48 | 0.5 | 0.41 | 0.43 |
| 7 | 2175871 | rs3778995 | -0.545 | 0.56 | 0.196 | -0.798 | 0.006 | 0.051 | 0.051 | 0.56 | 0.1 | 0.25 | 0.15 | 0.2 | 0.09 | 0.16 |
| 7 | 2190452 | rs6461188 | 0.255 | 0.291 | 0.408 | -1.186 | -0.022 | -0.081 | -0.081 | 0 | 0.32 | 0.48 | 0.21 | 0.06 | 0.19 | 0.19 |
| 7 | 2212758 | rs6967286 | 0.642 | 0.652 | 0.481 | -0.94 | 0.037 | 0.085 | 0.085 | 0 | 0.23 | 0.39 | 0.44 | 0.54 | 0.37 | 0.32 |
| 7 | 2212923 | rs1637761 | 0.072 | 0.292 | 0.421 | -1.062 | -0.001 | 0.001 | 0.001 | 0.066 | 0.63 | 0.67 | 0.41 | 0.29 | 0.53 | 0.47 |
| 7 | 2220053 | rs1637759 | -0.378 | 0.163 | -0.025 | -1.278 | 0.001 | 0.02 | 0.02 | 0.354 | 0.59 | 0.88 | 0.7 | 0.61 | 0.61 | 0.65 |
| 7 | 2220092 | rs17790410 | 1.648 | 0.116 | -0.175 | -1.337 | -0.008 | 0.102 | 0.102 | 0 | 0.26 | 0.05 | 0.16 | 0.06 | 0.19 | 0.19 |
| 7 | 2226162 | rs10226475 | 0.905 | -0.041 | 0.059 | -1.299 | 0.006 | 0.018 | 0.018 | 2.098 | 0.26 | 0.23 | 0.45 | 0.55 | 0.38 | 0.38 |
| 7 | 2230225 | rs4719457 | 1.301 | 0.291 | -0.232 | -1.329 | -0.001 | -0.001 | -0.001 | 0 | 0.22 | 0.17 | 0.5 | 0.73 | 0.37 | 0.33 |
| 7 | 2231002 | rs17186614 | 1.395 | 0.281 | -0.936 | -1.341 | -0.04 | 0.047 | 0.047 | 1.518 | 0.27 | 0.04 | 0.15 | 0.19 | 0.29 | 0.18 |
| 7 | 2239569 | rs1637738 | -0.631 | 0.435 | -1.461 | -1.158 | 0.005 | 0.081 | 0.081 | 0.284 | 0.59 | 0.9 | 0.73 | 0.72 | 0.62 | 0.67 |
| 7 | 2240094 | rs11976342 | 1.126 | 0.759 | -1.5 | -0.792 | -0.065 | -0.102 | -0.102 | 0 | 0.2 | 0.04 | 0.42 | 0.62 | 0.36 | 0.3 |
| 7 | 2247730 | rs17187301 | 0.63 | 0.606 | -1.371 | -1.163 | -0.028 | 0.127 | 0.127 | 0 | 0.29 | 0.05 | 0.19 | 0.06 | 0.24 | 0.21 |
| 7 | 2248472 | rs1612548 | 0.586 | 0.596 | -0.791 | -1.221 | 0.003 | -0.071 | -0.071 | 0.012 | 0.3 | 0.56 | 0.26 | 0.1 | 0.23 | 0.32 |
| 7 | 2248959 | rs3800933 | 0.748 | 0.674 | -1.419 | -0.994 | -0.123 | 0.073 | 0.073 | 1.134 | 0.38 | 0.06 | 0.5 | 0.75 | 0.49 | 0.49 |
| 7 | 2261468 | rs6461233 | 0.276 | 0.627 | -0.592 | -0.594 | 0.021 | 0.047 | 0.047 | 0 | 0.2 | 0.26 | 0.39 | 0.51 | 0.34 | 0.37 |
| 7 | 2267122 | rs1639922 | 0.691 | 0.202 | -0.615 | -0.998 | -0.026 | -0.065 | -0.065 | 0 | 0.36 | 0.54 | 0.27 | 0.1 | 0.25 | 0.21 |
| 7 | 2268970 | rs1626670 | -0.991 | 0.309 | -0.754 | -1.016 | 0.013 | 0.022 | 0.022 | 0.432 | 0.61 | 0.84 | 0.71 | 0.62 | 0.63 | 0.62 |
| 7 | 2269947 | rs17791802 | 1.241 | 0.346 | -1.146 | -0.811 | -0.032 | 0.09 | 0.09 | 0 | 0.29 | 0.02 | 0.22 | 0.07 | 0.23 | 0.19 |
| 7 | 2272936 | rs3757440 | -0.055 | 0.733 | -0.617 | -0.437 | 0.036 | 0.075 | 0.075 | 0.053 | 0.22 | 0.32 | 0.4 | 0.51 | 0.35 | 0.37 |
| 7 | 2278226 | rs7799006 | 0.26 | 0.819 | -0.639 | -0.346 | 0.04 | 0.081 | 0.081 | 0 | 0.2 | 0.28 | 0.39 | 0.51 | 0.34 | 0.37 |
| 7 | 2285178 | rs33922585 | 0.077 | 0.596 | -1.114 | -0.259 | -0.1 | -0.136 | -0.136 | 0 | 0.21 | 0.04 | 0.38 | 0.54 | 0.37 | 0.31 |
| 7 | 2286347 | rs10232413 | -1.29 | 0.11 | -0.875 | -0.325 | -0.038 | 0.042 | 0.042 | 0 | 0.9 | 0.83 | 0.89 | 0.8 | 0.97 | 0.87 |
| 7 | 2287560 | rs4721505 | 0.26 | -0.091 | -0.735 | -0.784 | 0.014 | 0.018 | 0.018 | 0.068 | 0.52 | 0.48 | 0.61 | 0.73 | 0.71 | 0.57 |
| 7 | 2290522 | rs1799832 | 1.114 | -0.166 | -1.54 | -0.75 | 0.047 | 0.122 | 0.122 | 0.645 | 0.31 | 0.1 | 0.21 | 0.06 | 0.21 | 0.21 |
| 7 | 2293609 | rs4719471 | -1.313 | 0.109 | -1.368 | -0.498 | -0.031 | 0.031 | 0.031 | 0 | 0.9 | 0.86 | 0.89 | 0.8 | 0.97 | 0.87 |
| 7 | 2294123 | rs12534131 | 0.783 | 0.305 | -1.481 | -0.479 | -0.11 | -0.139 | -0.139 | 0.01 | 0.17 | 0.02 | 0.38 | 0.5 | 0.36 | 0.29 |
| 7 | 2298071 | rs2398669 | -1.291 | 0.237 | -1.41 | -0.474 | -0.023 | 0.014 | 0.014 | 0 | 0.91 | 0.87 | 0.89 | 0.81 | 0.97 | 0.88 |
| 7 | 2298227 | rs2398668 | 0.745 | 0.326 | -1.493 | -0.434 | -0.112 | -0.139 | -0.139 | 0 | 0.17 | 0.02 | 0.38 | 0.5 | 0.36 | 0.31 |
| 7 | 2301989 | rs4721524 | -1.189 | 0.132 | -1.251 | -0.545 | -0.033 | 0.036 | 0.036 | 0 | 0.9 | 0.85 | 0.89 | 0.8 | 0.97 | 0.87 |
| 7 | 2309355 | rs6947645 | 1.307 | 0.055 | -1.401 | -0.709 | -0.004 | -0.037 | -0.037 | 0 | 0.36 | 0.53 | 0.24 | 0.2 | 0.34 | 0.27 |
| 7 | 2310974 | rs12535174 | 0.374 | 0.071 | -1.593 | -0.7 | -0.102 | -0.126 | -0.126 | 0.114 | 0.22 | 0.06 | 0.4 | 0.54 | 0.39 | 0.34 |
| 7 | 2313607 | rs7456643 | 1.131 | 0.314 | -1.559 | -0.892 | -0.021 | 0.127 | 0.127 | 0 | 0.28 | 0.03 | 0.2 | 0.05 | 0.21 | 0.21 |
| 7 | 2315056 | rs4549678 | 0.366 | 0.581 | -0.962 | -0.416 | 0 | -0.02 | -0.02 | 0 | 0.4 | 0.55 | 0.28 | 0.2 | 0.39 | 0.36 |
| 7 | 2332968 | rs12699820 | -0.376 | 0.998 | -1.127 | 0.325 | -0.072 | -0.085 | -0.085 | 0 | 0.21 | 0.09 | 0.4 | 0.59 | 0.36 | 0.34 |
| 7 | 2346115 | rs10249052 | -0.736 | 0.419 | -0.294 | 0.378 | 0.008 | 0.028 | 0.028 | 0 | 0.3 | 0.37 | 0.47 | 0.61 | 0.33 | 0.42 |
| 7 | 2347753 | rs6963924 | 0.25 | -0.685 | -0.159 | -0.022 | 0.014 | 0.055 | 0.055 | 0.278 | 0.39 | 0.25 | 0.25 | 0.16 | 0.31 | 0.21 |
| 7 | 2349470 | rs10479970 | 0.007 | -0.927 | 0.194 | -0.037 | 0.025 | 0.059 | 0.059 | 0 | 0.42 | 0.69 | 0.36 | 0.24 | 0.31 | 0.26 |

| | | | | | | | | | | | | | | | | |
|---|---|---|---|---|---|---|---|---|---|---|---|---|---|---|---|---|
| 7 | 2351981 | rs4442022 | -0.177 | -0.741 | 0.287 | -0.07 | 0.039 | 0.077 | 0.077 | 0.567 | 0.44 | 0.69 | 0.36 | 0.24 | 0.31 | 0.27 |
| 7 | 2363129 | rs12670380 | 0.304 | -0.448 | 0.389 | -0.238 | -0.006 | -0.031 | -0.031 | 0 | 0.37 | 0.4 | 0.26 | 0.16 | 0.3 | 0.24 |
| 7 | 2370956 | rs12699839 | -0.428 | -0.419 | -0.302 | -0.077 | -0.003 | 0.014 | 0.014 | 0.972 | 0.48 | 0.34 | 0.57 | 0.6 | 0.51 | 0.6 |
| 7 | 2377332 | rs2895183 | -0.239 | -0.269 | -0.225 | -0.323 | -0.005 | 0.016 | 0.016 | 2.985 | 0.43 | 0.36 | 0.56 | 0.79 | 0.5 | 0.6 |
| 7 | 2382358 | rs12699855 | -0.239 | -0.232 | -0.208 | -0.404 | -0.005 | 0.007 | 0.007 | 2.768 | 0.43 | 0.36 | 0.56 | 0.67 | 0.5 | 0.6 |
| 7 | 2393947 | rs7801817 | -0.472 | -0.621 | 0.163 | -0.732 | -0.022 | -0.05 | -0.05 | 0.273 | 0.68 | 0.45 | 0.76 | 0.86 | 0.78 | 0.77 |
| 7 | 2395509 | rs3807430 | 0.942 | -0.37 | -0.428 | -0.593 | -0.077 | -0.013 | -0.013 | 1.695 | 0.41 | 0.14 | 0.59 | 0.57 | 0.51 | 0.36 |
| 7 | 2399179 | rs12533700 | 0.835 | -0.291 | -0.344 | -0.686 | -0.032 | 0.065 | 0.065 | 0 | 0.43 | 0.24 | 0.6 | 0.72 | 0.51 | 0.36 |
| 7 | 2405127 | rs7791679 | 0.151 | 0.323 | -0.592 | -0.464 | -0.001 | -0.012 | -0.012 | 0 | 0.76 | 0.81 | 0.85 | 0.88 | 0.74 | 0.59 |
| 7 | 2412032 | rs1010303 | 1.051 | 0.021 | -0.791 | -0.519 | -0.091 | 0.047 | 0.047 | 0.059 | 0.37 | 0.13 | 0.57 | 0.72 | 0.5 | 0.4 |
| 7 | 2414567 | rs1010302 | 0.47 | 0.179 | -0.907 | -0.54 | -0.026 | -0.053 | -0.053 | 1.48 | 0.37 | 0.61 | 0.27 | 0.14 | 0.22 | 0.19 |
| 7 | 2414825 | rs1548597 | 0.925 | 1.048 | -1.12 | -0.142 | -0.03 | 0.043 | 0.043 | 0.109 | 0.77 | 0.92 | 0.86 | 0.87 | 0.73 | 0.62 |
| 7 | 2424266 | rs4719524 | 1.018 | 0.988 | -1.243 | -0.178 | -0.035 | -0.008 | -0.008 | 0 | 0.76 | 0.96 | 0.84 | 0.74 | 0.73 | 0.61 |
| 7 | 2433729 | rs11974910 | -0.509 | 1.145 | -1.03 | -0.4 | 0.016 | -0.016 | -0.016 | 0 | 0.08 | 0.02 | 0.07 | 0.15 | 0.06 | 0.13 |
| 7 | 2442246 | rs12538383 | -0.506 | 1.105 | -0.634 | -0.454 | -0.012 | 0.021 | 0.021 | 0 | 0.14 | 0.29 | 0.17 | 0.17 | 0.13 | 0.15 |
| 7 | 2448493 | rs6952809 | 1.232 | 1.498 | -0.384 | -0.688 | 0.048 | -0.028 | -0.028 | 0 | 0.75 | 0.53 | 0.71 | 0.79 | 0.67 | 0.65 |
| 7 | 2449733 | rs10215904 | -0.311 | 1.409 | -0.791 | -0.633 | -0.012 | 0.013 | 0.013 | 0 | 0.14 | 0.29 | 0.18 | 0.15 | 0.13 | 0.14 |
| 7 | 2452661 | rs2270005 | -0.533 | 1.384 | -0.802 | -0.621 | -0.017 | 0.02 | 0.02 | 0 | 0.15 | 0.37 | 0.19 | 0.16 | 0.13 | 0.2 |
| 7 | 2457709 | rs6972852 | -0.143 | 1.567 | -1.297 | -0.675 | -0.014 | 0.012 | 0.012 | 0.888 | 0.89 | 0.95 | 0.9 | 0.94 | 0.81 | 0.79 |
| 7 | 2472455 | rs3735100 | -0.314 | 0.734 | -1.267 | -0.425 | 0.017 | -0.017 | -0.017 | 1.253 | 0.08 | 0.02 | 0.06 | 0.15 | 0.04 | 0.13 |
| 7 | 2479029 | rs2969078 | -1.015 | 1.074 | -1.129 | -0.726 | -0.001 | 0 | 0 | 0 | 0.21 | 0.26 | 0.3 | 0.19 | 0.25 | 0.18 |
| 7 | 2483135 | rs11772554 | -0.818 | 1.257 | -1.429 | -1.011 | -0.013 | 0.019 | 0.019 | 1.052 | 0.1 | 0.09 | 0.11 | 0.01 | 0.18 | 0.09 |
| 7 | 2486004 | rs2906175 | 0.373 | 0.884 | -1.938 | -1.724 | 0.053 | 0.099 | 0.099 | 0.378 | 0.53 | 0.68 | 0.45 | 0.77 | 0.34 | 0.47 |
| 7 | 2486810 | rs2906173 | -0.564 | 1.356 | -2.091 | -1.895 | -0.042 | 0.045 | 0.045 | 0.768 | 0.72 | 0.86 | 0.62 | 0.85 | 0.66 | 0.79 |
| 7 | 2498052 | rs886626 | -0.855 | 1.345 | -2.456 | -1.926 | -0.113 | 0.033 | 0.033 | 0.013 | 0.74 | 0.97 | 0.69 | 0.87 | 0.68 | 0.79 |
| 7 | 2499770 | rs11768384 | -0.624 | 0.831 | -2.651 | -2.268 | -0.043 | 0.108 | 0.108 | 0.16 | 0.3 | 0.13 | 0.22 | 0.06 | 0.4 | 0.35 |
| 7 | 2505567 | rs1362136 | 1.042 | 0.771 | -2.691 | -2.281 | 0.037 | 0.053 | 0.053 | 0.454 | 0.42 | 0.63 | 0.4 | 0.79 | 0.27 | 0.38 |
| 7 | 2508072 | rs2906166 | -1.148 | 0.862 | -2.711 | -2.249 | -0.113 | 0.033 | 0.033 | 0.083 | 0.74 | 0.96 | 0.69 | 0.87 | 0.68 | 0.79 |
| 7 | 2510094 | rs11764837 | 0.973 | 0.902 | -2.587 | -2.125 | -0.062 | 0.055 | 0.055 | 0.02 | 0.14 | 0.01 | 0.1 | 0.04 | 0.2 | 0.2 |
| 7 | 2513442 | rs2969071 | 0.291 | 0.363 | -2.208 | -1.79 | 0.049 | 0.07 | 0.07 | 0.456 | 0.43 | 0.64 | 0.41 | 0.79 | 0.29 | 0.38 |
| 7 | 2528599 | rs2906142 | 0.356 | 0.75 | -1.857 | -1.462 | 0.001 | -0.001 | -0.001 | 0.19 | 0.4 | 0.55 | 0.38 | 0.77 | 0.3 | 0.45 |
| 7 | 2533903 | rs4721731 | -0.56 | 0.991 | -1.858 | -0.727 | -0.042 | 0.025 | 0.025 | 0.177 | 0.07 | 0 | 0.15 | 0.01 | 0.15 | 0.08 |
| 7 | 2542296 | rs12700017 | 0.432 | 0.845 | -1.937 | -0.685 | -0.061 | -0.012 | -0.012 | 2.636 | 0.18 | 0.04 | 0.19 | 0.12 | 0.24 | 0.15 |
| 7 | 2552986 | rs286560 | 0.163 | 0.631 | -1.86 | -1.087 | -0.012 | 0.111 | 0.111 | 0.868 | 0.32 | 0.13 | 0.27 | 0.14 | 0.34 | 0.46 |
| 7 | 2556157 | rs1982157 | -0.886 | -0.259 | -1.019 | -0.459 | -0.019 | 0.009 | 0.009 | 0.03 | 0.63 | 0.63 | 0.59 | 0.61 | 0.71 | 0.71 |
| 7 | 2557266 | rs755179 | -0.306 | 0.433 | -0.428 | -1.162 | 0.013 | -0.068 | -0.068 | 0 | 0.37 | 0.48 | 0.36 | 0.17 | 0.46 | 0.44 |
| 7 | 2560777 | rs4632959 | -0.155 | 0.109 | -0.615 | -0.706 | -0.047 | 0.103 | 0.103 | 2.322 | 0.39 | 0.17 | 0.35 | 0.22 | 0.55 | 0.49 |
| 7 | 2580070 | rs1061545 | -2.007 | -0.233 | 0.477 | 0.188 | 0.006 | 0 | 0 | 0 | 0.37 | 0.73 | 0.41 | 0.37 | 0.35 | 0.34 |
| 7 | 2580275 | rs6461429 | -1.979 | -0.21 | 0.455 | 0.181 | 0.004 | 0 | 0 | 0 | 0.36 | 0.72 | 0.41 | 0.37 | 0.35 | 0.34 |
| 7 | 2587150 | rs6942871 | -0.769 | -0.236 | 0.659 | 0.162 | -0.025 | 0.19 | 0.19 | 0 | 0.07 | 0.27 | 0.07 | 0.28 | 0.02 | 0.13 |
| 7 | 2589613 | rs11766591 | -1.073 | -0.102 | 0.03 | 0.162 | -0.045 | 0.072 | 0.072 | 0 | 0.12 | 0.01 | 0.19 | 0.01 | 0.18 | 0.15 |
| 7 | 2590383 | rs4721781 | -1.689 | 0.002 | 0.084 | 0.244 | 0.009 | 0.002 | 0.002 | 0 | 0.37 | 0.74 | 0.42 | 0.37 | 0.36 | 0.34 |
| 7 | 2594338 | rs11773376 | -1.073 | 0.874 | 0.085 | 0.125 | -0.031 | 0.026 | 0.026 | 0 | 0.05 | 0.01 | 0.16 | 0.01 | 0.14 | 0.07 |
| 7 | 2600076 | rs4721814 | -0.749 | 0.636 | 0.11 | 0.223 | 0.012 | -0.038 | -0.038 | 3.532 | 0.16 | 0.45 | 0.2 | 0.09 | 0.13 | 0.04 |
| 7 | 2604478 | rs11773176 | -0.92 | 1.111 | -0.649 | 0.27 | -0.03 | 0.025 | 0.025 | 0 | 0.05 | 0.01 | 0.11 | 0.01 | 0.14 | 0.07 |
| 7 | 2605333 | rs1550202 | -1.084 | 0.092 | -1 | 0.255 | 0.032 | 0.032 | 0.032 | 0 | 0.45 | 0.81 | 0.45 | 0.39 | 0.39 | 0.45 |
| 7 | 2606934 | rs10488610 | -1.165 | -0.243 | -1.032 | 0.484 | 0.016 | -0.003 | -0.003 | 0 | 0.12 | 0.09 | 0.11 | 0.11 | 0.08 | 0.17 |
| 7 | 2607339 | rs2917756 | -1.869 | -0.058 | -0.325 | 0.684 | -0.003 | 0.011 | 0.011 | 4.25 | 0.27 | 0.65 | 0.33 | 0.34 | 0.31 | 0.39 |
| 7 | 2609309 | rs6959481 | -0.855 | -0.009 | 0.446 | 0.63 | 0.03 | -0.06 | -0.06 | 0.259 | 0.1 | 0.24 | 0.11 | 0 | 0.15 | 0.11 |
| 7 | 2609765 | rs1036504 | -0.484 | 0.028 | -0.371 | 0.264 | 0.008 | -0.011 | -0.011 | 0.112 | 0.57 | 0.66 | 0.61 | 0.51 | 0.61 | 0.61 |
| 7 | 2613424 | rs2917750 | -0.454 | 0.003 | 0.09 | 0.271 | -0.008 | 0.006 | 0.006 | 0 | 0.56 | 0.51 | 0.59 | 0.49 | 0.62 | 0.62 |
| 7 | 2622649 | rs2289165 | -1.5 | -0.014 | -0.074 | 0.307 | 0.037 | -0.098 | -0.098 | 0 | 0.66 | 0.9 | 0.72 | 0.49 | 0.76 | 0.73 |
| 7 | 2629020 | rs12700120 | 0.494 | 0.171 | 0.086 | 0.259 | -0.019 | 0.049 | 0.049 | 0 | 0.41 | 0.34 | 0.32 | 0.16 | 0.44 | 0.33 |
| 7 | 2632477 | rs2917730 | -1.445 | 0.051 | -0.1 | 0.407 | 0.037 | -0.085 | -0.085 | 0 | 0.66 | 0.91 | 0.72 | 0.46 | 0.76 | 0.73 |
| 7 | 2640141 | rs2969018 | -0.859 | 0.516 | 0.33 | 0.899 | 0.047 | 0.254 | 0.254 | 0.301 | 0.12 | 0.26 | 0.31 | 0.42 | 0.15 | 0.33 |
| 7 | 2640821 | rs2005394 | -1.361 | 0.964 | 0.194 | 0.428 | 0.002 | 0.002 | 0.002 | 0 | 0.38 | 0.56 | 0.56 | 0.47 | 0.45 | 0.5 |
| 7 | 2643352 | rs4721867 | 0.727 | 0.973 | -0.121 | 0.34 | -0.021 | 0.044 | 0.044 | 0 | 0.18 | 0.1 | 0.16 | 0.05 | 0.17 | 0.07 |
| 7 | 2643607 | rs4719601 | -1.698 | 0.945 | -0.154 | 0.313 | 0.069 | -0.003 | -0.003 | 0.229 | 0.6 | 0.79 | 0.71 | 0.59 | 0.71 | 0.76 |
| 7 | 2644519 | rs2293404 | -0.953 | 1.184 | -0.289 | 0.446 | 0.02 | 0.043 | 0.043 | 0 | 0.3 | 0.4 | 0.46 | 0.5 | 0.33 | 0.4 |
| 7 | 2644607 | rs2293405 | 0.713 | 1.194 | -0.187 | 0.446 | -0.012 | 0.031 | 0.031 | 0 | 0.18 | 0.1 | 0.15 | 0.05 | 0.17 | 0.07 |
| 7 | 2645552 | rs2293407 | -0.949 | 1.198 | -0.225 | 0.452 | 0.02 | 0.043 | 0.043 | 0 | 0.3 | 0.4 | 0.46 | 0.49 | 0.33 | 0.4 |
| 7 | 2647007 | rs17833498 | 0.698 | 1.272 | -0.665 | 0.478 | -0.02 | 0.042 | 0.042 | 0 | 0.18 | 0.09 | 0.15 | 0.05 | 0.17 | 0.07 |
| 7 | 2649673 | rs3735108 | -1.111 | 1.114 | -0.443 | 0.543 | -0.016 | -0.013 | -0.013 | 0.344 | 0.35 | 0.28 | 0.52 | 0.5 | 0.45 | 0.51 |
| 7 | 2649819 | rs3735110 | -0.95 | 1.181 | -0.841 | 0.574 | -0.014 | -0.012 | -0.012 | 0 | 0.35 | 0.3 | 0.53 | 0.5 | 0.44 | 0.51 |
| 7 | 2649932 | rs3735111 | -0.746 | 1.431 | -0.588 | 1.105 | 0.027 | 0.171 | 0.171 | 0.347 | 0.11 | 0.17 | 0.29 | 0.44 | 0.15 | 0.34 |
| 7 | 2652630 | rs1045753 | 0.338 | 1.127 | -1.077 | 0.533 | -0.078 | 0.07 | 0.07 | 0 | 0.19 | 0.01 | 0.15 | 0.09 | 0.24 | 0.24 |
| 7 | 2662818 | rs12537408 | -0.677 | 0.7 | -0.299 | 1.609 | 0.047 | 0.246 | 0.246 | 0 | 0.07 | 0.18 | 0.22 | 0.32 | 0.11 | 0.18 |
| 7 | 2664214 | rs17132513 | 0.321 | -0.944 | -1.675 | 0.975 | 0.121 | -0.043 | -0.043 | 0 | 0.31 | 0.07 | 0.18 | 0.33 | 0.18 | 0.28 |
| 7 | 2667885 | rs1568968 | -1.052 | -0.14 | -0.451 | 1.036 | 0.009 | 0.019 | 0.019 | 2.561 | 0.11 | 0.71 | 0.16 | 0.13 | 0.12 | 0.15 |

| | | | | | | | | | | | | | | | | |
|---|---|---|---|---|---|---|---|---|---|---|---|---|---|---|---|---|
| 7 | 2669587 | rs4721900 | -0.813 | 0.342 | -0.594 | 0.623 | -0.002 | -0.005 | -0.005 | 0.063 | 0.29 | 0.38 | 0.2 | 0.2 | 0.25 | 0.22 |
| 7 | 2672972 | rs12670408 | -1.513 | 0.12 | -0.822 | 0.699 | -0.022 | 0.023 | 0.023 | 0 | 0.5 | 0.84 | 0.5 | 0.5 | 0.51 | 0.54 |
| 7 | 2673288 | rs10254690 | -1.877 | 0.122 | -1.028 | 0.813 | -0.038 | 0.153 | 0.153 | 0 | 0.55 | 0.88 | 0.56 | 0.75 | 0.56 | 0.62 |
| 7 | 2675662 | rs1105421 | -1.784 | 0.058 | -1.153 | 0.814 | -0.067 | 0.169 | 0.169 | 0 | 0.55 | 0.9 | 0.57 | 0.78 | 0.56 | 0.62 |
| 7 | 2679031 | rs4721904 | -1.146 | -0.014 | -0.422 | 0.899 | -0.002 | 0.003 | 0.003 | 0 | 0.34 | 0.43 | 0.26 | 0.46 | 0.3 | 0.3 |
| 7 | 2679449 | rs11768734 | -0.429 | 0.446 | -0.125 | 0.779 | 0.006 | 0.005 | 0.005 | 0.653 | 0.16 | 0.33 | 0.14 | 0.19 | 0.17 | 0.14 |
| 7 | 2686059 | rs10950811 | -0.792 | 0.596 | -0.251 | 0.386 | 0.001 | 0.04 | 0.04 | 0 | 0.49 | 0.65 | 0.52 | 0.68 | 0.49 | 0.53 |
| 7 | 2689472 | rs3801077 | -0.88 | 0.675 | -0.447 | 0.413 | -0.014 | 0.091 | 0.091 | 0 | 0.53 | 0.78 | 0.59 | 0.68 | 0.49 | 0.57 |
| 7 | 2703464 | rs13188 | -0.892 | 0.269 | 0.33 | 0.566 | 0.007 | 0.011 | 0.011 | 0.025 | 0.21 | 0.16 | 0.16 | 0.08 | 0.14 | 0.14 |
| 7 | 2703652 | rs3801075 | -1.439 | 0.089 | 0.763 | 0.487 | -0.011 | -0.028 | -0.028 | 0 | 0.26 | 0.61 | 0.21 | 0.09 | 0.16 | 0.15 |
| 7 | 2706478 | rs7782028 | -0.826 | -0.17 | 0.756 | 0.626 | -0.022 | -0.025 | -0.025 | 1.148 | 0.2 | 0.26 | 0.16 | 0.1 | 0.12 | 0.09 |
| 7 | 2714357 | rs1724932 | 1.067 | 0.377 | 0.471 | 1.044 | -0.002 | -0.005 | -0.005 | 1.459 | 0.69 | 0.55 | 0.67 | 0.71 | 0.68 | 0.68 |
| 7 | 2715373 | rs1724933 | 1.421 | 0.304 | 0.071 | 1.003 | -0.002 | -0.002 | -0.002 | 0 | 0.66 | 0.36 | 0.66 | 0.7 | 0.68 | 0.69 |
| 7 | 2724098 | rs10264843 | -1.414 | 0.16 | -0.695 | 1.118 | 0.04 | 0 | 0 | 0 | 0.36 | 0.78 | 0.34 | 0.32 | 0.31 | 0.31 |
| 7 | 2727337 | rs7776527 | -0.636 | 0.9 | -0.842 | 1.373 | -0.044 | -0.091 | -0.091 | 0 | 0.19 | 0.04 | 0.21 | 0.31 | 0.27 | 0.19 |
| 7 | 2733676 | rs3996391 | -1.137 | 1.095 | -0.348 | 1.5 | 0.015 | 0.093 | 0.093 | 1.662 | 0.16 | 0.34 | 0.26 | 0.41 | 0.24 | 0.37 |
| 7 | 2734713 | rs10276467 | -0.343 | 1.082 | -0.625 | 1.639 | 0 | 0.085 | 0.085 | 0 | 0.07 | 0.1 | 0.12 | 0.41 | 0.1 | 0.23 |
| 7 | 2739017 | rs809547 | -0.921 | 0.955 | -1.554 | 0.266 | -0.008 | 0.134 | 0.134 | 0.094 | 0.61 | 0.93 | 0.65 | 0.87 | 0.73 | 0.73 |
| 7 | 2745124 | rs6461534 | 0.318 | 1.355 | -0.659 | 0.356 | 0.047 | -0.054 | -0.054 | 0 | 0.71 | 0.55 | 0.65 | 0.83 | 0.54 | 0.72 |
| 7 | 2748487 | rs13235873 | -1.065 | 1.087 | -0.426 | 0.998 | 0.008 | 0.093 | 0.093 | 2.061 | 0.21 | 0.32 | 0.19 | 0.57 | 0.29 | 0.38 |
| 7 | 2750047 | rs10229834 | -1.106 | 0.602 | -0.239 | 0.343 | -0.011 | -0.067 | -0.067 | 0 | 0.16 | 0.66 | 0.12 | 0.01 | 0.18 | 0.1 |
| 7 | 2752487 | rs7776970 | -0.087 | 0.172 | -0.646 | -0.492 | -0.005 | -0.052 | -0.052 | 0.986 | 0.37 | 0.56 | 0.27 | 0.09 | 0.34 | 0.26 |
| 7 | 2752932 | rs7778736 | 0.051 | 0.098 | -0.492 | -0.494 | -0.009 | -0.09 | -0.09 | 4.89 | 0.36 | 0.48 | 0.26 | 0.1 | 0.34 | 0.26 |
| 7 | 2753667 | rs11769575 | 0.884 | 0.092 | -0.885 | -0.587 | 0 | -0.002 | -0.002 | 1.287 | 0.15 | 0.12 | 0.14 | 0.12 | 0.14 | 0.18 |
| 7 | 2753725 | rs11976039 | 0.449 | 0.033 | -0.905 | -0.623 | -0.009 | -0.092 | -0.092 | 0 | 0.36 | 0.48 | 0.26 | 0.1 | 0.34 | 0.26 |
| 7 | 2753929 | rs10950842 | 0.359 | 0.04 | -1.336 | -1.113 | -0.007 | 0.013 | 0.013 | 0 | 0.58 | 0.6 | 0.49 | 0.1 | 0.47 | 0.42 |
| 7 | 2754625 | rs4719646 | -0.71 | 0.139 | -1.271 | -0.974 | -0.015 | -0.003 | -0.003 | 0 | 0.74 | 0.75 | 0.63 | 0.22 | 0.61 | 0.6 |
| 7 | 2757534 | rs6979597 | 0.663 | -0.322 | -1.361 | -1.144 | -0.011 | -0.126 | -0.126 | 1.778 | 0.33 | 0.47 | 0.25 | 0.07 | 0.29 | 0.26 |
| 7 | 2763102 | rs798544 | 1.392 | 0.36 | -1.401 | -0.302 | -0.019 | -0.012 | -0.012 | 0.087 | 0.16 | 0.14 | 0.27 | 0.21 | 0.29 | 0.15 |
| 7 | 2764665 | rs798539 | -1.553 | 0.439 | -1.51 | -0.224 | -0.049 | -0.033 | -0.033 | 0.804 | 0.91 | 0.99 | 0.88 | 0.41 | 0.87 | 0.74 |
| 7 | 2765723 | rs4722006 | 0.017 | 0.157 | -1.235 | -0.442 | 0.021 | 0.08 | 0.08 | 0 | 0.51 | 0.62 | 0.41 | 0.18 | 0.45 | 0.48 |
| 7 | 2768626 | rs12484 | 0.03 | 0.141 | -1.242 | -0.449 | 0.02 | 0.08 | 0.08 | 0 | 0.5 | 0.62 | 0.41 | 0.19 | 0.46 | 0.47 |
| 7 | 2768864 | rs798532 | -0.856 | 0.136 | -1.272 | -0.459 | 0.013 | -0.052 | -0.052 | 0 | 0.67 | 0.79 | 0.67 | 0.41 | 0.75 | 0.63 |
| 7 | 2768935 | rs7805092 | -0.489 | 0.114 | -1.26 | -0.354 | -0.028 | 0.126 | 0.126 | 0 | 0.54 | 0.83 | 0.42 | 0.19 | 0.47 | 0.48 |
| 7 | 2773605 | rs208358 | -0.962 | -0.138 | -1.053 | -0.591 | -0.011 | -0.085 | -0.085 | 0 | 0.7 | 0.6 | 0.76 | 0.95 | 0.75 | 0.74 |
| 7 | 2780863 | rs208354 | 1.876 | -0.028 | -1.336 | -0.412 | -0.043 | -0.02 | -0.02 | 0 | 0.08 | 0.01 | 0.11 | 0.58 | 0.12 | 0.25 |
| 7 | 2782152 | rs208353 | -1.54 | -0.175 | -0.721 | -0.506 | -0.002 | 0.035 | 0.035 | 0.121 | 0.93 | 0.99 | 0.95 | 1 | 0.91 | 0.92 |
| 7 | 2789704 | rs798503 | 0.35 | -0.243 | -0.644 | -0.297 | 0 | 0.01 | 0.01 | 0.061 | 0.41 | 0.41 | 0.48 | 0.32 | 0.41 | 0.26 |
| 7 | 2792524 | rs11761804 | 1.615 | -0.304 | -0.753 | -0.615 | 0.037 | 0.066 | 0.066 | 1.125 | 0.22 | 0.1 | 0.2 | 0 | 0.11 | 0.1 |
| 7 | 2797267 | rs798495 | 1.442 | -0.228 | -0.657 | -0.563 | -0.049 | -0.036 | -0.036 | 0 | 0.16 | 0.09 | 0.26 | 0.23 | 0.3 | 0.17 |
| 7 | 2799686 | rs208346 | 0.827 | -0.311 | -0.597 | -0.565 | -0.001 | 0.008 | 0.008 | 0 | 0.53 | 0.63 | 0.51 | 0.63 | 0.39 | 0.53 |
| 7 | 2803037 | rs798486 | 1.006 | -0.247 | -0.251 | -0.26 | -0.025 | -0.015 | -0.015 | 0 | 0.16 | 0.15 | 0.26 | 0.22 | 0.28 | 0.17 |
| 7 | 2811543 | rs1182206 | -1.089 | -0.29 | -0.239 | -0.281 | 0.004 | 0.006 | 0.006 | 0.411 | 0.61 | 0.53 | 0.64 | 0.36 | 0.61 | 0.47 |
| 7 | 2814734 | rs9332401 | 0.26 | -0.255 | -0.398 | -0.369 | 0 | -0.005 | -0.005 | 0 | 0.19 | 0.08 | 0.16 | 0.13 | 0.19 | 0.18 |
| 7 | 2825922 | rs2527684 | 0.256 | -0.318 | -0.83 | -0.443 | -0.01 | 0.043 | 0.043 | 0.437 | 0.38 | 0.3 | 0.47 | 0.23 | 0.4 | 0.25 |
| 7 | 2831083 | rs2533888 | -0.374 | -0.376 | -0.894 | -0.462 | -0.018 | 0.088 | 0.088 | 1.041 | 0.5 | 0.19 | 0.53 | 0.32 | 0.54 | 0.43 |
| 7 | 2836433 | rs757788 | 1.23 | -0.145 | -0.53 | -0.484 | 0.031 | 0.04 | 0.04 | 0.535 | 0.21 | 0.08 | 0.2 | 0 | 0.1 | 0.1 |
| 7 | 2860217 | rs2644295 | -0.091 | -0.08 | -0.495 | -0.486 | -0.003 | 0.018 | 0.018 | 0.055 | 0.39 | 0.33 | 0.48 | 0.23 | 0.4 | 0.25 |
| 7 | 2863574 | rs2107964 | 1.221 | -0.11 | -0.504 | -0.473 | -0.064 | -0.048 | -0.048 | 0.528 | 0.15 | 0.06 | 0.22 | 0.23 | 0.28 | 0.15 |
| 7 | 2869391 | rs1182190 | -0.3 | -0.121 | -0.477 | -0.552 | 0.001 | 0.016 | 0.016 | 0.004 | 0.41 | 0.41 | 0.49 | 0.2 | 0.41 | 0.27 |
| 7 | 2869985 | rs1182188 | 0.794 | 0.107 | -0.488 | -0.312 | -0.021 | -0.005 | -0.005 | 0 | 0.16 | 0.16 | 0.27 | 0.2 | 0.3 | 0.16 |
| 7 | 2873648 | rs1182179 | 0.885 | 0.136 | -0.553 | -0.299 | -0.029 | -0.009 | -0.009 | 0 | 0.16 | 0.12 | 0.26 | 0.2 | 0.3 | 0.16 |
| 7 | 2884283 | rs1636249 | 0.02 | 0.083 | -0.398 | -0.366 | 0.009 | 0.02 | 0.02 | 0 | 0.21 | 0.3 | 0.28 | 0.34 | 0.3 | 0.17 |
| 7 | 2894512 | rs11765386 | 1.145 | 0.001 | -0.503 | -0.69 | 0.012 | -0.015 | -0.015 | 0 | 0.22 | 0.19 | 0.19 | 0.01 | 0.09 | 0.09 |
| 7 | 2895627 | rs1713913 | 0.006 | 0.216 | -0.414 | -0.494 | 0.001 | 0.004 | 0.004 | 0 | 0.4 | 0.41 | 0.47 | 0.36 | 0.38 | 0.27 |
| 7 | 2904928 | rs2527521 | -0.442 | 0.586 | -0.245 | -0.82 | -0.002 | 0.007 | 0.007 | 0 | 0.49 | 0.45 | 0.58 | 0.44 | 0.48 | 0.31 |
| 7 | 2910399 | rs1859560 | -0.759 | 0.376 | -0.249 | -0.798 | 0.004 | 0.001 | 0.001 | 0 | 0.89 | 0.94 | 0.92 | 0.93 | 0.87 | 0.85 |
| 7 | 2911972 | rs1182144 | -0.602 | 0.685 | -0.266 | -0.806 | 0.001 | 0 | 0 | 0.603 | 0.53 | 0.52 | 0.61 | 0.45 | 0.53 | 0.34 |
| 7 | 2915378 | rs6977940 | 0.23 | -0.077 | 0.026 | -1.575 | 0.016 | 0.048 | 0.048 | 0 | 0.23 | 0.15 | 0.2 | 0 | 0.13 | 0.13 |
| 7 | 2918778 | rs6957938 | -0.882 | 0.803 | 1.121 | -0.59 | 0.004 | -0.043 | -0.043 | 0 | 0.12 | 0.14 | 0.1 | 0 | 0.12 | 0.05 |
| 7 | 2920067 | rs17132769 | -1.306 | 0.793 | 1.121 | -0.599 | 0.017 | -0.093 | -0.093 | 0 | 0.15 | 0.36 | 0.12 | 0 | 0.14 | 0.06 |
| 7 | 2921582 | rs2644270 | 0.814 | 0.671 | 0.563 | -0.405 | -0.005 | -0.005 | -0.005 | 2.421 | 0.86 | 0.82 | 0.89 | 0.92 | 0.85 | 0.79 |
| 7 | 2923584 | rs2644269 | 0.723 | 0.644 | 0.183 | -0.375 | 0.007 | 0.003 | 0.003 | 0 | 0.86 | 0.91 | 0.89 | 0.91 | 0.86 | 0.79 |
| 7 | 2926448 | rs11769099 | 2.097 | 0.623 | 0.388 | -0.5 | -0.01 | -0.066 | -0.066 | 0 | 0.7 | 0.53 | 0.75 | 0.9 | 0.72 | 0.7 |
| 7 | 2927572 | rs2644302 | 0.909 | 0.96 | 0.205 | -0.316 | -0.006 | -0.008 | -0.008 | 0 | 0.85 | 0.74 | 0.89 | 0.91 | 0.84 | 0.84 |
| 7 | 2929822 | rs1713924 | 1.798 | 1.066 | -0.598 | -0.132 | -0.002 | 0.008 | 0.008 | 0 | 0.77 | 0.8 | 0.81 | 0.88 | 0.74 | 0.78 |
| 7 | 2930621 | rs11766331 | -0.668 | 1.251 | -0.659 | -0.118 | -0.019 | 0.027 | 0.027 | 0 | 0.11 | 0.05 | 0.24 | 0.01 | 0.13 | 0.07 |
| 7 | 2930748 | rs1612849 | 0.76 | 0.815 | -0.508 | 0.229 | -0.001 | 0 | 0 | 0 | 0.87 | 0.87 | 0.89 | 0.9 | 0.84 | 0.85 |
| 7 | 2933195 | rs6948739 | -0.629 | 0.674 | -0.794 | -0.178 | -0.002 | 0.051 | 0.051 | 0 | 0.37 | 0.26 | 0.42 | 0.23 | 0.32 | 0.41 |

| | | | | | | | | | | | | | | | | |
|---|---|---|---|---|---|---|---|---|---|---|---|---|---|---|---|---|
| 7 | 2937450 | rs2527517 | -1.107 | 0.041 | -0.396 | 0.8 | -0.021 | 0.032 | 0.032 | 0 | 0.15 | 0.24 | 0.2 | 0.31 | 0.06 | 0.24 |
| 7 | 2938919 | rs2527516 | -1.244 | -0.007 | -0.015 | 0.756 | -0.04 | 0.068 | 0.068 | 0 | 0.16 | 0.33 | 0.22 | 0.32 | 0.07 | 0.27 |
| 7 | 2942273 | rs11982651 | -0.107 | 0.244 | -0.487 | 0.41 | -0.008 | 0.008 | 0.008 | 0.926 | 0.4 | 0.38 | 0.57 | 0.62 | 0.49 | 0.5 |
| 7 | 2943818 | rs7810813 | 0.084 | -0.007 | -0.932 | -0.064 | 0.006 | 0.01 | 0.01 | 0 | 0.19 | 0.12 | 0.16 | 0.2 | 0.15 | 0.27 |
| 7 | 2944263 | rs1621828 | -0.458 | -0.029 | -1.34 | -0.446 | 0.001 | 0.069 | 0.069 | 3.382 | 0.76 | 0.96 | 0.83 | 0.88 | 0.78 | 0.91 |
| 7 | 2944317 | rs3735118 | 0.468 | 0.128 | -1.009 | -0.294 | -0.043 | -0.035 | -0.035 | 0 | 0.1 | 0.02 | 0.1 | 0.15 | 0.11 | 0.25 |
| 7 | 2946461 | rs1124581 | -0.305 | 0.469 | -0.696 | -0.152 | 0.01 | 0.054 | 0.054 | 0.02 | 0.49 | 0.7 | 0.53 | 0.65 | 0.47 | 0.57 |
| 7 | 2951346 | rs2679258 | -0.494 | 0.306 | -0.761 | -0.166 | -0.03 | 0.168 | 0.168 | 1.158 | 0.55 | 0.85 | 0.66 | 0.82 | 0.5 | 0.75 |
| 7 | 2951824 | rs2301921 | -0.568 | 0.124 | -0.528 | 0.083 | -0.024 | 0.103 | 0.103 | 0 | 0.05 | 0.15 | 0.14 | 0.17 | 0.02 | 0.15 |
| 7 | 2955097 | rs2301920 | -0.248 | 0.293 | 0.302 | 0.164 | -0.008 | 0.051 | 0.051 | 0.776 | 0.23 | 0.38 | 0.38 | 0.6 | 0.22 | 0.41 |
| 7 | 2956085 | rs2679261 | -0.522 | 0.427 | 0.847 | -0.127 | 0 | -0.007 | -0.007 | 0.568 | 0.51 | 0.54 | 0.62 | 0.81 | 0.48 | 0.75 |
| 7 | 2956217 | rs6461700 | -0.956 | 0.789 | 1.169 | 0.172 | 0.017 | 0.071 | 0.071 | 0 | 0.05 | 0.19 | 0.18 | 0.13 | 0.06 | 0.15 |
| 7 | 2956267 | rs6461701 | -0.139 | 0.605 | 1.272 | 0.238 | -0.024 | 0.109 | 0.109 | 0.279 | 0.22 | 0.36 | 0.28 | 0.48 | 0.18 | 0.42 |
| 7 | 2956759 | rs3735120 | -0.828 | 0.629 | 1.127 | 0.154 | -0.001 | 0.122 | 0.122 | 0 | 0.05 | 0.2 | 0.15 | 0.16 | 0.03 | 0.15 |
| 7 | 2956918 | rs3735123 | -1.117 | 0.617 | 1.004 | 0.149 | 0.017 | 0.173 | 0.173 | 0 | 0.06 | 0.29 | 0.16 | 0.16 | 0.03 | 0.15 |
| 7 | 2959172 | rs3735126 | -0.948 | 0.619 | 0.978 | 0.052 | 0.024 | 0.169 | 0.169 | 0 | 0.06 | 0.29 | 0.15 | 0.15 | 0.03 | 0.15 |
| 7 | 2961409 | rs958331 | 0.291 | 0.809 | 0.808 | -0.022 | 0.002 | 0.053 | 0.053 | 1.205 | 0.19 | 0.28 | 0.26 | 0.48 | 0.18 | 0.29 |
| 7 | 2961414 | rs1878805 | 0.878 | 0.716 | 0.254 | -0.234 | -0.007 | 0.047 | 0.047 | 0.078 | 0.25 | 0.17 | 0.23 | 0.13 | 0.27 | 0.28 |
| 7 | 2961909 | rs1713911 | -1.974 | 0.807 | -0.107 | -0.544 | -0.06 | -0.023 | -0.023 | 0.267 | 0.83 | 0.97 | 0.86 | 0.84 | 0.75 | 0.83 |
| 7 | 2966138 | rs1182137 | 0.435 | 0.558 | 0.451 | -0.998 | 0.001 | 0.129 | 0.129 | 4.138 | 0.28 | 0.04 | 0.23 | 0.07 | 0.25 | 0.15 |
| 7 | 2966445 | rs1182136 | -1.319 | 0.645 | 0.705 | -1.009 | 0.006 | -0.009 | -0.009 | 0 | 0.64 | 0.6 | 0.69 | 0.37 | 0.62 | 0.55 |
| 7 | 2968361 | rs2527506 | 0.618 | 0.306 | 0.371 | -1.173 | 0.009 | 0.018 | 0.018 | 0 | 0.27 | 0.22 | 0.21 | 0.22 | 0.24 | 0.25 |
| 7 | 2969496 | rs4991791 | -1.347 | 0.48 | 0.891 | -0.912 | 0.038 | 0.087 | 0.087 | 0.029 | 0.06 | 0.23 | 0.25 | 0.15 | 0.06 | 0.15 |
| 7 | 2969680 | rs1621509 | -0.405 | 0.723 | 0.156 | -1.286 | -0.006 | 0.153 | 0.153 | 2.826 | 0.27 | 0.04 | 0.18 | 0.01 | 0.24 | 0.09 |
| 7 | 2972072 | rs1636167 | -0.689 | 0.965 | 0.416 | -0.996 | 0 | -0.142 | -0.142 | 0.026 | 0.31 | 0.5 | 0.35 | 0.05 | 0.3 | 0.17 |
| 7 | 2990009 | rs7806365 | -1.144 | 1.415 | 0.332 | -0.22 | 0.039 | -0.041 | -0.041 | 0 | 0.22 | 0.49 | 0.38 | 0.19 | 0.25 | 0.26 |
| 7 | 3008328 | rs6949660 | -0.965 | 1.74 | 0.55 | 0.021 | 0.016 | 0.008 | 0.008 | 0 | 0.17 | 0.28 | 0.11 | 0.15 | 0.18 | 0.2 |
| 7 | 3018962 | rs4719737 | -0.131 | 1.226 | -0.436 | -0.679 | 0 | 0.037 | 0.037 | 0 | 0.56 | 0.63 | 0.66 | 0.75 | 0.54 | 0.63 |
| 7 | 3020375 | rs1878808 | -0.065 | 1.073 | -0.663 | -0.849 | 0.039 | 0.132 | 0.132 | 0 | 0.49 | 0.25 | 0.63 | 0.74 | 0.55 | 0.63 |
| 7 | 3021239 | rs1878806 | -0.21 | 0.967 | -1.523 | -1.108 | 0.021 | 0.159 | 0.159 | 0.769 | 0.55 | 0.92 | 0.69 | 0.73 | 0.62 | 0.72 |
| 7 | 3031192 | rs1843933 | 0.096 | 0.666 | -1.245 | -1.654 | 0.014 | 0.062 | 0.062 | 0.027 | 0.32 | 0.23 | 0.27 | 0.17 | 0.27 | 0.19 |
| 7 | 3032228 | rs4722304 | -0.6 | 1.074 | -0.919 | -1.944 | -0.006 | 0.021 | 0.021 | 0.132 | 0.61 | 0.66 | 0.6 | 0.31 | 0.6 | 0.51 |
| 7 | 3032284 | rs6461749 | -0.021 | 1.149 | -0.684 | -1.827 | 0.01 | -0.063 | -0.063 | 0 | 0.29 | 0.43 | 0.33 | 0.14 | 0.33 | 0.31 |
| 7 | 3033865 | rs11769046 | -0.519 | 1.253 | -0.258 | -1.918 | 0.037 | -0.053 | -0.053 | 0 | 0.1 | 0.31 | 0.09 | 0.01 | 0.1 | 0.17 |
| 7 | 3037438 | rs2396971 | 0.189 | 0.94 | -0.933 | -2.185 | -0.02 | 0.159 | 0.159 | 0.497 | 0.38 | 0.12 | 0.38 | 0.08 | 0.39 | 0.27 |
| 7 | 3038397 | rs6953804 | 0.242 | 0.777 | -0.931 | -2.256 | -0.005 | 0.071 | 0.071 | 0 | 0.25 | 0.08 | 0.12 | 0.11 | 0.21 | 0.21 |
| 7 | 3043406 | rs13245486 | -0.768 | 0.6 | -0.308 | -2.127 | -0.001 | 0 | 0 | 0 | 0.66 | 0.69 | 0.63 | 0.32 | 0.67 | 0.59 |
| 7 | 3046735 | rs2029279 | 0.682 | 0.811 | -0.298 | -2.202 | -0.018 | 0.135 | 0.135 | 0.015 | 0.37 | 0.13 | 0.38 | 0.08 | 0.4 | 0.26 |
| 7 | 3047721 | rs11982868 | 0.514 | 0.785 | 0.13 | -2.203 | 0.01 | 0.028 | 0.028 | 0 | 0.21 | 0.43 | 0.21 | 0.24 | 0.19 | 0.28 |
| 7 | 3049009 | rs10236776 | 0.021 | 0.764 | 0.482 | -2.245 | -0.001 | -0.065 | -0.065 | 0 | 0.46 | 0.48 | 0.43 | 0.08 | 0.48 | 0.35 |
| 7 | 3051375 | rs6461781 | -0.169 | 1.011 | 0.079 | -1.768 | 0.007 | 0.015 | 0.015 | 0 | 0.25 | 0.39 | 0.24 | 0.29 | 0.26 | 0.36 |
| 7 | 3053209 | rs17237863 | 1.254 | 1.253 | -0.417 | -1.477 | -0.022 | 0.027 | 0.027 | 0 | 0.05 | 0.01 | 0.05 | 0.01 | 0.11 | 0.12 |
| 7 | 3054098 | rs12671372 | 0.504 | 0.97 | -0.418 | -1.599 | -0.007 | 0.012 | 0.012 | 0 | 0.24 | 0.2 | 0.32 | 0.09 | 0.3 | 0.22 |
| 7 | 3059377 | rs6947877 | -0.27 | 1.08 | -0.949 | -1.602 | 0.001 | -0.009 | -0.009 | 0 | 0.38 | 0.36 | 0.47 | 0.29 | 0.41 | 0.4 |
| 7 | 3060397 | rs10950998 | -0.813 | 1.098 | -1.538 | -1.562 | -0.002 | 0.011 | 0.011 | 0 | 0.48 | 0.52 | 0.54 | 0.29 | 0.53 | 0.49 |
| 7 | 3062728 | rs4722332 | -0.758 | 1.134 | -1.745 | -1.497 | 0 | 0.034 | 0.034 | 0 | 0.48 | 0.58 | 0.57 | 0.29 | 0.56 | 0.5 |
| 7 | 3063921 | rs1843939 | 0.344 | 1.364 | -2.14 | -1.322 | -0.007 | 0.056 | 0.056 | 0 | 0.24 | 0.09 | 0.14 | 0.11 | 0.22 | 0.22 |
| 7 | 3064776 | rs17238159 | -0.027 | 1.637 | -1.499 | -0.567 | -0.02 | -0.003 | -0.003 | 0.128 | 0.1 | 0 | 0.07 | 0.05 | 0.1 | 0.12 |
| 7 | 3067913 | rs17834873 | -0.681 | 1.835 | -0.98 | -0.327 | -0.016 | 0.038 | 0.038 | 0 | 0.07 | 0.01 | 0.05 | 0 | 0.1 | 0.09 |
| 7 | 3070171 | rs10280353 | -0.906 | 2.368 | -0.725 | -0.094 | 0.003 | -0.001 | -0.001 | 0 | 0.15 | 0.14 | 0.33 | 0.14 | 0.22 | 0.13 |
| 7 | 3071115 | rs10951005 | -0.805 | 2.173 | -0.666 | -0.303 | -0.004 | 0.004 | 0.004 | 0 | 0.22 | 0.18 | 0.35 | 0.08 | 0.25 | 0.15 |
| 7 | 3071195 | rs6461796 | -0.943 | 2.184 | -0.529 | -0.428 | -0.004 | 0.003 | 0.003 | 0 | 0.31 | 0.66 | 0.43 | 0.1 | 0.37 | 0.26 |
| 7 | 3072215 | rs6970813 | -0.142 | 1.693 | -0.712 | 0.083 | -0.014 | -0.012 | -0.012 | 0 | 0.9 | 0.96 | 0.87 | 0.85 | 0.86 | 0.87 |
| 7 | 3076053 | rs10951010 | 0.028 | 1.35 | -0.552 | 0.351 | 0 | 0.022 | 0.022 | 1.505 | 0.23 | 0.36 | 0.15 | 0.28 | 0.25 | 0.26 |
| 7 | 3076707 | rs4722353 | -0.444 | 1.227 | -0.854 | 0.303 | -0.035 | -0.025 | -0.025 | 0 | 0.9 | 0.99 | 0.86 | 0.85 | 0.86 | 0.89 |
| 7 | 3079604 | rs4610634 | -0.068 | 1.016 | -0.556 | -0.303 | 0.003 | -0.061 | -0.061 | 0 | 0.19 | 0.27 | 0.14 | 0.03 | 0.2 | 0.1 |
| 7 | 3081727 | rs6953573 | 0.43 | 0.702 | -0.765 | -0.397 | -0.017 | 0.055 | 0.055 | 0 | 0.72 | 0.84 | 0.73 | 0.87 | 0.67 | 0.77 |
| 7 | 3081758 | rs10951014 | 0.292 | 0.697 | -0.602 | -0.306 | -0.001 | 0 | 0 | 0 | 0.31 | 0.3 | 0.19 | 0.3 | 0.29 | 0.27 |
| 7 | 3081854 | rs10951015 | 0.093 | 0.039 | -1.029 | -0.503 | 0.01 | 0.111 | 0.111 | 0 | 0.48 | 0.33 | 0.63 | 0.77 | 0.55 | 0.53 |
| 7 | 3086236 | rs4722366 | 0.335 | -0.278 | -1.259 | 0.274 | 0.027 | -0.078 | -0.078 | 1.102 | 0.78 | 0.91 | 0.78 | 0.61 | 0.84 | 0.74 |
| 7 | 3087086 | rs9986734 | 0.461 | -0.35 | -1.279 | 0.242 | 0.031 | -0.079 | -0.079 | 0 | 0.77 | 0.91 | 0.78 | 0.61 | 0.84 | 0.73 |
| 7 | 3090282 | rs7805181 | 0.543 | -0.423 | -1.061 | 0.228 | 0.014 | -0.038 | -0.038 | 0 | 0.77 | 0.82 | 0.77 | 0.63 | 0.82 | 0.64 |
| 7 | 3093297 | rs11768102 | 0.297 | 0.434 | -0.702 | -0.371 | 0.039 | -0.065 | -0.065 | 0.102 | 0.14 | 0.21 | 0.2 | 0.02 | 0.21 | 0.11 |
| 7 | 3094100 | rs1976134 | -0.25 | 1.026 | -0.795 | 0.132 | -0.005 | -0.02 | -0.02 | 0.838 | 0.82 | 0.83 | 0.75 | 0.61 | 0.81 | 0.79 |
| 7 | 3096211 | rs6461814 | 0.336 | 0.348 | -0.814 | 0.089 | -0.003 | 0.008 | 0.008 | 0.051 | 0.54 | 0.42 | 0.46 | 0.59 | 0.52 | 0.54 |
| 7 | 3096236 | rs10951028 | 0.179 | 0.402 | -0.984 | 0.623 | 0.001 | -0.037 | -0.037 | 0 | 0.26 | 0.19 | 0.25 | 0.48 | 0.28 | 0.38 |
| 7 | 3101279 | rs2396964 | -0.897 | 0.455 | -0.648 | 0.674 | -0.003 | 0.016 | 0.016 | 0.578 | 0.24 | 0.29 | 0.25 | 0.34 | 0.23 | 0.29 |
| 7 | 3101370 | rs2396963 | -1.225 | 0.083 | -1.216 | 0.26 | 0.018 | -0.055 | -0.055 | 0 | 0.72 | 0.84 | 0.74 | 0.62 | 0.79 | 0.76 |
| 7 | 3102237 | rs7780988 | -0.248 | 0.142 | -0.936 | 0.156 | -0.015 | 0.029 | 0.029 | 0 | 0.23 | 0.3 | 0.24 | 0.35 | 0.19 | 0.2 |

| | | | | | | | | | | | | | | | | |
|---|---|---|---|---|---|---|---|---|---|---|---|---|---|---|---|---|
| 7 | 3107183 | rs4722382 | -1.119 | 0.524 | -1.009 | -0.01 | -0.003 | -0.066 | -0.066 | 0.613 | 0.65 | 0.82 | 0.66 | 0.57 | 0.68 | 0.76 |
| 7 | 3109177 | rs12531570 | -0.585 | 1.436 | -0.338 | 0.056 | 0.018 | -0.003 | -0.003 | 2.141 | 0.12 | 0.16 | 0.19 | 0.09 | 0.2 | 0.21 |
| 7 | 3113034 | rs1843938 | -0.866 | 1.053 | -0.026 | -0.05 | -0.009 | 0.015 | 0.015 | 0 | 0.39 | 0.41 | 0.4 | 0.55 | 0.37 | 0.31 |
| 7 | 3117874 | rs11772124 | -0.295 | 0.813 | -0.281 | -0.008 | 0.041 | -0.043 | -0.043 | 0 | 0.29 | 0.15 | 0.23 | 0.46 | 0.21 | 0.24 |
| 7 | 3119823 | rs13221367 | -1.856 | 1.084 | -0.282 | -0.113 | 0.013 | -0.017 | -0.017 | 0.58 | 0.59 | 0.77 | 0.53 | 0.61 | 0.61 | 0.67 |
| 7 | 3125220 | rs4722398 | 1.085 | 1.468 | -0.39 | -0.056 | -0.018 | 0.022 | 0.022 | 0 | 0.12 | 0.07 | 0.06 | 0.03 | 0.13 | 0.17 |
| 7 | 3125512 | rs7782384 | -0.833 | 1.16 | -0.451 | 0.004 | -0.012 | -0.047 | -0.047 | 0 | 0.29 | 0.18 | 0.35 | 0.55 | 0.35 | 0.3 |
| 7 | 3126241 | rs11767716 | -0.169 | 0.709 | -0.535 | 0.033 | 0.023 | -0.051 | -0.051 | 0 | 0.21 | 0.09 | 0.18 | 0.47 | 0.17 | 0.22 |
| 7 | 3128789 | rs4722404 | 1.583 | 0.996 | -0.187 | -0.088 | 0.003 | 0.027 | 0.027 | 0.019 | 0.55 | 0.23 | 0.55 | 0.36 | 0.48 | 0.41 |
| 7 | 3129705 | rs10951034 | 1.689 | 0.803 | -0.251 | -0.189 | 0.026 | 0.042 | 0.042 | 0 | 0.47 | 0.21 | 0.51 | 0.35 | 0.45 | 0.38 |
| 7 | 3131792 | rs12540595 | -1.193 | 1.047 | 0.207 | -0.234 | 0.001 | 0.016 | 0.016 | 0.142 | 0.36 | 0.58 | 0.42 | 0.43 | 0.32 | 0.36 |
| 7 | 3137057 | rs7788891 | -0.996 | 1.031 | 0.746 | -0.253 | -0.019 | 0.002 | 0.002 | 1.072 | 0.3 | 0.4 | 0.14 | 0.31 | 0.16 | 0.29 |
| 7 | 3137677 | rs1968924 | -1.736 | 1.118 | 0.668 | -0.315 | 0.002 | -0.001 | -0.001 | 0 | 0.48 | 0.74 | 0.44 | 0.47 | 0.46 | 0.55 |
| 7 | 3140464 | rs7809283 | 1.109 | 1.28 | 0.486 | -0.293 | -0.008 | 0.004 | 0.004 | 0 | 0.09 | 0.07 | 0.06 | 0.03 | 0.13 | 0.15 |
| 7 | 3145429 | rs4719797 | 0.176 | 1.227 | 0.586 | -0.311 | 0.001 | -0.019 | -0.019 | 0.23 | 0.95 | 0.91 | 0.96 | 1 | 0.93 | 0.86 |
| 7 | 3146495 | rs4719798 | 0.076 | 1.359 | 0.579 | -0.282 | -0.013 | -0.008 | -0.008 | 0 | 0.89 | 0.88 | 0.75 | 0.87 | 0.77 | 0.78 |
| 7 | 3146983 | rs6967398 | 0.989 | 1.338 | 0.9 | -0.315 | 0.074 | -0.048 | -0.048 | 0.56 | 0.1 | 0.28 | 0.08 | 0.03 | 0.13 | 0.15 |
| 7 | 3147020 | rs6943837 | -1.592 | 1.134 | 0.948 | -0.293 | 0.007 | -0.004 | -0.004 | 1.909 | 0.43 | 0.64 | 0.39 | 0.51 | 0.34 | 0.49 |
| 7 | 3148092 | rs1872881 | -1.454 | 0.6 | 0.884 | -0.111 | 0.002 | -0.008 | -0.008 | 1.824 | 0.35 | 0.31 | 0.4 | 0.61 | 0.33 | 0.36 |
| 7 | 3152962 | rs4722435 | 1.625 | 0.606 | 1.006 | -0.065 | -0.001 | -0.001 | -0.001 | 9.135 | 0.61 | 0.51 | 0.51 | 0.38 | 0.6 | 0.51 |
| 7 | 3153173 | rs9986945 | 0.584 | 0.49 | 0.683 | -0.059 | 0.014 | -0.043 | -0.043 | 0.21 | 0.13 | 0.05 | 0.18 | 0.28 | 0.1 | 0.15 |
| 7 | 3154810 | rs6958161 | -1.42 | 0.507 | 0.659 | -0.05 | 0.006 | -0.008 | -0.008 | 0 | 0.32 | 0.3 | 0.29 | 0.46 | 0.2 | 0.35 |
| 7 | 3155301 | rs10951052 | -1.038 | 0.656 | 0.681 | -0.042 | -0.003 | -0.004 | -0.004 | 0 | 0.26 | 0.29 | 0.36 | 0.45 | 0.3 | 0.36 |
| 7 | 3157702 | rs6955702 | -1.413 | 0.438 | 0.209 | -0.253 | 0.002 | -0.002 | -0.002 | 0.207 | 0.4 | 0.57 | 0.53 | 0.63 | 0.46 | 0.49 |
| 7 | 3157916 | rs10499319 | 2.422 | 0.7 | -0.242 | -0.363 | -0.009 | -0.02 | -0.02 | 0 | 0.37 | 0.13 | 0.37 | 0.36 | 0.34 | 0.26 |
| 7 | 3167399 | rs6461856 | 1.026 | 0.096 | 0.366 | -0.055 | -0.015 | 0 | 0 | 0.052 | 0.7 | 0.68 | 0.83 | 0.68 | 0.79 | 0.76 |
| 7 | 3167751 | rs6942973 | -0.641 | 0.116 | 0.592 | -0.025 | 0.105 | 0.037 | 0.037 | 0.094 | 0.06 | 0.36 | 0.05 | 0 | 0.05 | 0.14 |
| 7 | 3169494 | rs12674252 | 1.257 | -0.062 | 0.047 | -0.106 | 0.032 | 0.008 | 0.008 | 0.068 | 0.64 | 0.31 | 0.78 | 0.68 | 0.75 | 0.61 |
| 7 | 3175338 | rs936909 | 0.818 | 0.225 | 0.321 | -0.55 | -0.035 | -0.071 | -0.071 | 0.843 | 0.8 | 0.67 | 0.85 | 0.95 | 0.83 | 0.8 |
| 7 | 3176241 | rs2396955 | -0.102 | 1.091 | 0.326 | -0.234 | 0.012 | 0.02 | 0.02 | 0 | 0.08 | 0.11 | 0.3 | 0.26 | 0.23 | 0.12 |
| 7 | 3179991 | rs7779245 | 0.847 | 0.641 | 0.319 | -0.457 | -0.036 | -0.056 | -0.056 | 0 | 0.76 | 0.69 | 0.88 | 0.95 | 0.85 | 0.73 |
| 7 | 3180657 | rs6964682 | 0.9 | 0.578 | 0.385 | -0.483 | -0.076 | -0.106 | -0.106 | 0 | 0.74 | 0.47 | 0.86 | 0.95 | 0.85 | 0.72 |
| 7 | 3181785 | rs17175397 | -0.589 | 1.947 | -0.033 | -0.237 | -0.087 | -0.016 | -0.016 | 2.795 | 0.16 | 0.04 | 0.18 | 0.13 | 0.25 | 0.15 |
| 7 | 3185929 | rs17175551 | -1.529 | 1.698 | 0.216 | -0.536 | 0.002 | 0 | 0 | 0 | 0.22 | 0.23 | 0.33 | 0.26 | 0.32 | 0.25 |
| 7 | 3189341 | rs11771337 | 2.031 | 1.691 | 0.77 | -0.554 | 0.031 | -0.003 | -0.003 | 0 | 0.73 | 0.39 | 0.58 | 0.71 | 0.61 | 0.58 |
| 7 | 3190823 | rs4722477 | 2.038 | 1.661 | 0.903 | -0.547 | 0.062 | 0.008 | 0.008 | 0 | 0.75 | 0.52 | 0.59 | 0.71 | 0.61 | 0.66 |
| 7 | 3191990 | rs6943452 | -1.789 | 1.526 | 0.71 | -0.623 | 0.015 | -0.008 | -0.008 | 1.886 | 0.27 | 0.35 | 0.38 | 0.26 | 0.33 | 0.46 |
| 7 | 3194156 | rs7783979 | 0.439 | 1.189 | 0.491 | -0.549 | -0.001 | -0.023 | -0.023 | 0.016 | 0.92 | 0.74 | 0.92 | 0.97 | 0.89 | 0.9 |
| 7 | 3196435 | rs1074118 | -1.101 | 1.519 | -0.063 | -0.389 | -0.001 | 0 | 0 | 0 | 0.14 | 0.18 | 0.26 | 0.14 | 0.16 | 0.18 |
| 7 | 3197464 | rs6960859 | -0.451 | 1.315 | 0.096 | -0.236 | 0 | 0.051 | 0.051 | 0 | 0.22 | 0.34 | 0.28 | 0.37 | 0.18 | 0.33 |
| 7 | 3197867 | rs4634530 | -0.958 | 1.577 | -0.355 | -1.102 | -0.032 | 0.108 | 0.108 | 0 | 0.42 | 0.71 | 0.36 | 0.1 | 0.53 | 0.43 |
| 7 | 3201313 | rs12700653 | -0.568 | 1.76 | -0.144 | -1.165 | -0.002 | 0.002 | 0.002 | 0.222 | 0.4 | 0.52 | 0.42 | 0.48 | 0.36 | 0.64 |
| 7 | 3202157 | rs7801659 | -1.922 | 2.077 | -0.26 | -1.184 | -0.003 | -0.091 | -0.091 | 0 | 0.66 | 0.86 | 0.62 | 0.53 | 0.7 | 0.87 |
| 7 | 3204014 | rs6975435 | -0.192 | 2.428 | 0.131 | -1.223 | 0.019 | -0.02 | -0.02 | 0.142 | 0.9 | 0.83 | 0.93 | 0.96 | 0.83 | 0.86 |
| 7 | 3204177 | rs10755848 | 1.015 | 2.375 | 0.009 | -1.262 | 0.001 | -0.002 | -0.002 | 0.294 | 0.49 | 0.28 | 0.5 | 0.48 | 0.46 | 0.22 |
| 7 | 3208071 | rs10951095 | 1.009 | 2.594 | 0.565 | -1.053 | 0.082 | 0.004 | 0.004 | 0 | 0.84 | 0.49 | 0.8 | 0.85 | 0.74 | 0.75 |
| 7 | 3211607 | rs7777228 | 0.508 | 2.219 | 0.346 | -1.146 | 0.037 | -0.008 | -0.008 | 0 | 0.29 | 0.18 | 0.2 | 0.34 | 0.13 | 0.35 |
| 7 | 3212079 | rs10240660 | 1.693 | 2.495 | 0.352 | -1.217 | 0.02 | -0.042 | -0.042 | 0 | 0.66 | 0.46 | 0.59 | 0.79 | 0.42 | 0.51 |
| 7 | 3214048 | rs1525550 | 0.096 | 2.352 | 0.615 | -0.775 | 0.02 | -0.019 | -0.019 | 0 | 0.9 | 0.84 | 0.81 | 0.95 | 0.81 | 0.87 |
| 7 | 3216850 | rs6961724 | 0.754 | 2.045 | 0.64 | -0.769 | 0.029 | 0 | 0 | 0 | 0.84 | 0.66 | 0.72 | 0.85 | 0.81 | 0.74 |
| 7 | 3217887 | rs10951110 | 1.604 | 2.141 | 0.417 | -0.837 | -0.017 | -0.044 | -0.044 | 0 | 0.4 | 0.25 | 0.38 | 0.46 | 0.44 | 0.16 |
| 7 | 3218524 | rs7785633 | -0.177 | 2.034 | 0.336 | -0.862 | -0.008 | -0.004 | -0.004 | 0 | 0.39 | 0.12 | 0.29 | 0.38 | 0.34 | 0.51 |
| 7 | 3219163 | rs17133109 | 0.323 | 2.071 | 0.343 | -0.87 | -0.011 | 0.01 | 0.01 | 0 | 0.09 | 0.06 | 0.1 | 0.04 | 0.16 | 0.14 |
| 7 | 3220259 | rs10241890 | 1.606 | 2.129 | 0.354 | -0.889 | -0.005 | -0.014 | -0.014 | 0 | 0.4 | 0.36 | 0.38 | 0.47 | 0.44 | 0.16 |
| 7 | 3220416 | rs17799419 | 0.518 | 2.301 | 0.203 | -0.898 | -0.007 | -0.043 | -0.043 | 0 | 0.31 | 0.07 | 0.22 | 0.34 | 0.24 | 0.38 |
| 7 | 3220990 | rs4534049 | 0.897 | 1.512 | 0.334 | -0.824 | 0.005 | 0.002 | 0.002 | 0.046 | 0.5 | 0.53 | 0.5 | 0.5 | 0.6 | 0.28 |
| 7 | 3222960 | rs870226 | 0.408 | 1.238 | -0.248 | -0.702 | 0.024 | -0.061 | -0.061 | 0 | 0.31 | 0.06 | 0.21 | 0.34 | 0.21 | 0.37 |
| 7 | 3223846 | rs11768094 | 0.652 | 1.082 | 0.223 | -0.349 | 0.01 | -0.006 | -0.006 | 0 | 0.45 | 0.43 | 0.37 | 0.47 | 0.42 | 0.25 |
| 7 | 3224002 | rs4719830 | 0.711 | 0.647 | 0.215 | -0.184 | -0.016 | -0.013 | -0.013 | 3.052 | 0.42 | 0.32 | 0.43 | 0.52 | 0.51 | 0.33 |
| 7 | 3228573 | rs17133145 | 0.765 | 0.909 | 0.587 | -0.321 | -0.001 | 0.008 | 0.008 | 0 | 0.45 | 0.36 | 0.45 | 0.49 | 0.49 | 0.24 |
| 7 | 3229109 | rs4719836 | -0.219 | 0.972 | 0.773 | -0.37 | -0.01 | 0.046 | 0.046 | 0 | 0.6 | 0.82 | 0.71 | 0.66 | 0.66 | 0.47 |
| 7 | 3230292 | rs11771362 | 0.081 | 0.832 | 0.782 | -0.425 | -0.001 | -0.003 | -0.003 | 0 | 0.08 | 0.09 | 0.07 | 0 | 0.06 | 0.02 |
| 7 | 3230348 | rs6969544 | 0.94 | 0.772 | 1.207 | -0.401 | 0.005 | -0.001 | -0.001 | 0 | 0.42 | 0.58 | 0.4 | 0.49 | 0.44 | 0.25 |
| 7 | 3232216 | rs17133152 | 1.41 | 0.814 | 1.056 | -0.387 | -0.013 | -0.048 | -0.048 | 0 | 0.38 | 0.14 | 0.35 | 0.49 | 0.43 | 0.23 |
| 7 | 3232378 | rs10272892 | 0.492 | 0.872 | 1.209 | -0.396 | -0.001 | -0.004 | -0.004 | 0 | 0.48 | 0.49 | 0.51 | 0.53 | 0.55 | 0.37 |
| 7 | 3232907 | rs10270339 | -0.072 | 0.885 | 1.107 | -0.382 | 0.01 | -0.012 | -0.012 | 0 | 0.08 | 0.19 | 0.14 | 0.05 | 0.09 | 0.07 |
| 7 | 3233684 | rs10228720 | 0.096 | 0.889 | 0.952 | -0.38 | 0.005 | -0.011 | -0.011 | 3.427 | 0.08 | 0.15 | 0.14 | 0.05 | 0.09 | 0.07 |
| 7 | 3234248 | rs12673364 | 1.319 | 0.879 | 0.458 | -0.332 | 0.002 | -0.019 | -0.019 | 0.537 | 0.38 | 0.25 | 0.36 | 0.48 | 0.43 | 0.23 |
| 7 | 3239495 | rs12700693 | -1.329 | 1.052 | 0.178 | -0.406 | 0.032 | -0.08 | -0.08 | 0 | 0.22 | 0.39 | 0.29 | 0.08 | 0.26 | 0.22 |

| | | | | | | | | | | | | | | | | |
|---|---|---|---|---|---|---|---|---|---|---|---|---|---|---|---|---|
| 7 | 3241619 | rs10229652 | -0.426 | 0.51 | -0.8 | -0.064 | -0.014 | -0.025 | -0.025 | 0 | 0.66 | 0.95 | 0.77 | 0.66 | 0.75 | 0.6 |
| 7 | 3244898 | rs1404874 | 0.415 | 0.55 | -0.459 | 0.019 | 0.025 | 0.035 | 0.035 | 0 | 0.52 | 0.33 | 0.58 | 0.59 | 0.64 | 0.43 |
| 7 | 3246139 | rs17133183 | -1.081 | 0.608 | -0.229 | 0.047 | 0.023 | -0.011 | -0.011 | 0 | 0.14 | 0.31 | 0.19 | 0.13 | 0.13 | 0.12 |
| 7 | 3246357 | rs2334065 | 0.05 | 0.547 | -0.621 | 0.116 | -0.001 | -0.016 | -0.016 | 0 | 0.61 | 0.78 | 0.65 | 0.53 | 0.68 | 0.48 |
| 7 | 3248483 | rs4722574 | 0.016 | 0.135 | -0.871 | 0.03 | 0.005 | 0.014 | 0.014 | 0 | 0.62 | 0.87 | 0.73 | 0.62 | 0.72 | 0.49 |
| 7 | 3248674 | rs4719856 | 0.261 | 0.049 | -0.844 | -0.058 | 0.013 | 0.082 | 0.082 | 0 | 0.55 | 0.78 | 0.69 | 0.65 | 0.66 | 0.42 |
| 7 | 3251558 | rs4398802 | 0.823 | -0.071 | -0.886 | 0.463 | 0.008 | -0.014 | -0.014 | 0 | 0.34 | 0.18 | 0.22 | 0.36 | 0.37 | 0.19 |
| 7 | 3257986 | rs17133184 | -0.475 | 0.034 | -0.627 | 0.432 | -0.002 | 0.001 | 0.001 | 0 | 0.17 | 0.17 | 0.13 | 0.16 | 0.23 | 0.29 |
| 7 | 3258801 | rs6461923 | -0.715 | -0.014 | -0.148 | 0.478 | -0.003 | -0.007 | -0.007 | 0.013 | 0.69 | 0.69 | 0.56 | 0.54 | 0.68 | 0.66 |
| 7 | 3264548 | rs7795734 | -0.329 | 0.011 | -0.744 | 0.418 | 0 | 0.052 | 0.052 | 0 | 0.33 | 0.21 | 0.33 | 0.18 | 0.33 | 0.48 |
| 7 | 3264878 | rs4719863 | 0.537 | 0.121 | -0.488 | 0.785 | 0.002 | -0.023 | -0.023 | 0.22 | 0.32 | 0.17 | 0.21 | 0.37 | 0.35 | 0.17 |
| 7 | 3270895 | rs4719865 | 0.507 | 0.049 | -0.461 | 0.673 | 0.011 | 0.004 | 0.004 | 1.829 | 0.32 | 0.24 | 0.2 | 0.29 | 0.3 | 0.15 |
| 7 | 3271998 | rs11972265 | 0.222 | -0.023 | -0.331 | 0.718 | 0.008 | 0.011 | 0.011 | 3.608 | 0.34 | 0.28 | 0.28 | 0.76 | 0.3 | 0.2 |
| 7 | 3272526 | rs10951131 | 0.144 | 0.41 | -0.263 | 0.351 | 0.001 | 0.009 | 0.009 | 2.098 | 0.55 | 0.6 | 0.66 | 0.72 | 0.55 | 0.7 |
| 7 | 3276268 | rs940777 | -0.6 | 1.023 | -0.116 | 0.853 | -0.005 | -0.004 | -0.004 | 0 | 0.32 | 0.35 | 0.44 | 0.5 | 0.49 | 0.31 |
| 7 | 3284552 | rs16870824 | -0.047 | -0.362 | -0.074 | 0.366 | 0.01 | 0.009 | 0.009 | 0 | 0.34 | 0.29 | 0.16 | 0.31 | 0.19 | 0.38 |
| 7 | 3284594 | rs12700737 | -0.014 | -0.068 | 0.475 | 0.374 | 0.047 | 0.041 | 0.041 | 0 | 0.24 | 0.37 | 0.4 | 0.31 | 0.39 | 0.17 |
| 7 | 3285433 | rs11977745 | 0.526 | -0.103 | 0.441 | 0.372 | -0.004 | -0.008 | -0.008 | 0 | 0.31 | 0.29 | 0.13 | 0.18 | 0.15 | 0.35 |
| 7 | 3285554 | rs17133201 | 0.995 | -0.161 | 0.378 | 0.328 | 0.064 | 0.072 | 0.072 | 0 | 0.25 | 0.07 | 0.1 | 0.05 | 0.09 | 0.27 |
| 7 | 3286120 | rs16870829 | -0.096 | -0.018 | 0.323 | 0.315 | 0.022 | -0.042 | -0.042 | 0.133 | 0.08 | 0.25 | 0.07 | 0 | 0.09 | 0.02 |
| 7 | 3286230 | rs6947894 | -1.378 | -0.055 | -0.004 | 0.326 | -0.04 | -0.015 | -0.015 | 0 | 0.6 | 0.8 | 0.58 | 0.62 | 0.58 | 0.55 |
| 7 | 3286425 | rs17811536 | 0.806 | 0.083 | -0.144 | 0.634 | -0.027 | -0.03 | -0.03 | 0 | 0.16 | 0.15 | 0.33 | 0.31 | 0.3 | 0.15 |
| 7 | 3288471 | rs12700739 | -0.77 | 0.088 | -0.152 | 0.503 | 0 | 0.02 | 0.02 | 0 | 0.53 | 0.61 | 0.55 | 0.67 | 0.48 | 0.52 |
| 7 | 3295229 | rs6963845 | -0.18 | -0.568 | -0.513 | 0.438 | -0.004 | 0.039 | 0.039 | 0.024 | 0.06 | 0.15 | 0.07 | 0.18 | 0.05 | 0.08 |
| 7 | 3298532 | rs6946117 | -0.039 | -0.753 | -0.591 | 0.298 | 0.015 | 0.013 | 0.013 | 0 | 0.31 | 0.26 | 0.17 | 0.24 | 0.14 | 0.37 |
| 7 | 3307922 | rs12056190 | 0.207 | 0.389 | -0.894 | 0.494 | -0.047 | -0.035 | -0.035 | 0 | 0.64 | 0.75 | 0.63 | 0.45 | 0.51 | 0.67 |
| 7 | 3312870 | rs2049477 | -0.254 | 0.613 | -1.251 | 0.464 | -0.14 | -0.115 | -0.115 | 0 | 0.74 | 0.97 | 0.71 | 0.65 | 0.62 | 0.77 |
| 7 | 3313326 | rs17133206 | 0.897 | -0.204 | -0.761 | -0.211 | -0.007 | -0.038 | -0.038 | 0.351 | 0.25 | 0.27 | 0.15 | 0.05 | 0.14 | 0.27 |
| 7 | 3315572 | rs17133208 | -0.936 | 0.08 | -0.455 | -0.088 | -0.004 | 0.027 | 0.027 | 0 | 0.25 | 0.3 | 0.33 | 0.41 | 0.25 | 0.23 |
| 7 | 3318658 | rs10276303 | 1.292 | -0.037 | -0.367 | -0.352 | -0.017 | 0.058 | 0.058 | 0 | 0.28 | 0.15 | 0.28 | 0.11 | 0.31 | 0.39 |
| 7 | 3320172 | rs11773532 | 0.292 | -0.261 | 0.079 | -0.507 | -0.001 | 0 | 0 | 0 | 0.39 | 0.34 | 0.43 | 0.55 | 0.33 | 0.4 |
| 7 | 3320807 | rs10241023 | 1.446 | -0.315 | -0.08 | -0.502 | -0.04 | 0.106 | 0.106 | 0 | 0.27 | 0.11 | 0.23 | 0.07 | 0.31 | 0.38 |
| 7 | 3326129 | rs12700767 | -1.409 | -0.166 | -0.006 | -0.377 | -0.059 | -0.075 | -0.075 | 0 | 0.8 | 0.91 | 0.73 | 0.61 | 0.71 | 0.86 |
| 7 | 3332630 | rs11773762 | 0.427 | -0.301 | 0.281 | -0.428 | 0.017 | -0.029 | -0.029 | 0 | 0.37 | 0.25 | 0.41 | 0.51 | 0.32 | 0.39 |
| 7 | 3335041 | rs9986772 | -0.439 | -0.239 | 0.362 | -0.45 | 0 | -0.003 | -0.003 | 0 | 0.54 | 0.6 | 0.57 | 0.45 | 0.57 | 0.62 |
| 7 | 3344986 | rs1404872 | 0.331 | -0.374 | 0.499 | -0.523 | -0.039 | -0.099 | -0.099 | 0.084 | 0.15 | 0.38 | 0.07 | 0 | 0.08 | 0.08 |
| 7 | 3349625 | rs1880610 | -1.513 | -0.303 | 0.469 | -0.499 | -0.048 | -0.065 | -0.065 | 0.14 | 0.8 | 0.91 | 0.73 | 0.59 | 0.71 | 0.86 |
| 7 | 3351810 | rs7785994 | -0.705 | -0.364 | 0.462 | -0.545 | -0.046 | -0.051 | -0.051 | 0 | 0.62 | 0.81 | 0.51 | 0.55 | 0.52 | 0.48 |
| 7 | 3358580 | rs4316064 | -0.428 | -0.349 | 0.524 | -0.564 | -0.006 | -0.006 | -0.006 | 0 | 0.59 | 0.64 | 0.49 | 0.55 | 0.52 | 0.48 |
| 7 | 3359886 | rs9691979 | 0.34 | -0.33 | 0.57 | -0.391 | -0.03 | -0.062 | -0.062 | 0 | 0.39 | 0.49 | 0.21 | 0.16 | 0.23 | 0.34 |
| 7 | 3366165 | rs11979128 | -0.308 | -0.312 | 0.496 | -0.372 | 0.006 | 0.007 | 0.007 | 0 | 0.36 | 0.47 | 0.35 | 0.42 | 0.37 | 0.21 |
| 7 | 3368197 | rs4607498 | -0.791 | -0.336 | 0.422 | -0.352 | -0.079 | -0.079 | -0.079 | 1.702 | 0.62 | 0.86 | 0.5 | 0.55 | 0.51 | 0.48 |
| 7 | 3369615 | rs7457935 | 1.203 | -0.314 | 0.491 | -0.323 | -0.032 | -0.045 | -0.045 | 0 | 0.21 | 0.17 | 0.28 | 0.42 | 0.29 | 0.14 |
| 7 | 3371579 | rs11981207 | 0.893 | -0.309 | 0.443 | -0.321 | -0.022 | -0.056 | -0.056 | 0.041 | 0.12 | 0.22 | 0.05 | 0 | 0.07 | 0.07 |
| 7 | 3371961 | rs4416733 | -0.209 | -0.247 | 0.262 | -0.265 | 0.004 | 0.006 | 0.006 | 0 | 0.36 | 0.46 | 0.34 | 0.42 | 0.37 | 0.21 |
| 7 | 3380895 | rs4131987 | -0.633 | -0.167 | 0.134 | -0.462 | -0.057 | -0.05 | -0.05 | 0 | 0.62 | 0.81 | 0.48 | 0.53 | 0.5 | 0.46 |
| 7 | 3385384 | rs4544975 | -0.64 | -0.22 | 0.04 | -0.494 | -0.053 | -0.051 | -0.051 | 0.369 | 0.62 | 0.81 | 0.49 | 0.53 | 0.51 | 0.49 |
| 7 | 3390013 | rs7776559 | -0.109 | -0.391 | 0.19 | -0.503 | 0.008 | 0.011 | 0.011 | 0 | 0.35 | 0.47 | 0.34 | 0.39 | 0.37 | 0.21 |
| 7 | 3396065 | rs6947525 | -0.93 | -0.174 | -0.155 | -0.584 | -0.051 | -0.033 | -0.033 | 0.42 | 0.65 | 0.82 | 0.54 | 0.41 | 0.53 | 0.42 |
| 7 | 3400463 | rs4286835 | 1.289 | -0.198 | -0.109 | -0.551 | -0.046 | -0.062 | -0.062 | 1.115 | 0.2 | 0.12 | 0.28 | 0.4 | 0.3 | 0.14 |
| 7 | 3413738 | rs12673513 | -0.279 | -0.294 | -0.241 | -0.485 | -0.009 | -0.004 | -0.004 | 0 | 0.07 | 0.01 | 0.18 | 0.4 | 0.08 | 0.12 |
| 7 | 3413793 | rs12667184 | 0.184 | -0.445 | -0.224 | -0.663 | -0.016 | -0.007 | -0.007 | 0.51 | 0.42 | 0.23 | 0.55 | 0.6 | 0.58 | 0.66 |
| 7 | 3414624 | rs10258056 | 0.293 | -0.451 | -0.245 | -0.667 | -0.05 | -0.034 | -0.034 | 0 | 0.41 | 0.14 | 0.54 | 0.59 | 0.58 | 0.65 |
| 7 | 3420856 | rs939912 | -1.542 | -0.462 | -0.194 | -0.68 | -0.019 | -0.024 | -0.024 | 0 | 0.8 | 0.87 | 0.72 | 0.6 | 0.71 | 0.86 |
| 7 | 3420940 | rs4418248 | 1.766 | -0.634 | 0.032 | -0.8 | -0.04 | -0.077 | -0.077 | 0.139 | 0.23 | 0.24 | 0.11 | 0 | 0.07 | 0.12 |
| 7 | 3424878 | rs10278205 | 0.389 | -0.424 | -0.12 | -0.801 | -0.047 | -0.029 | -0.029 | 0.281 | 0.41 | 0.18 | 0.54 | 0.58 | 0.57 | 0.63 |
| 7 | 3431050 | rs6967281 | -1.48 | -0.318 | 0.077 | -0.733 | -0.04 | -0.053 | -0.053 | 0 | 0.8 | 0.86 | 0.73 | 0.59 | 0.71 | 0.87 |
| 7 | 3434275 | rs11971511 | 0.142 | -0.652 | 0.25 | -0.925 | 0.059 | 0.025 | 0.025 | 0.244 | 0.4 | 0.68 | 0.19 | 0.01 | 0.15 | 0.22 |
| 7 | 3445311 | rs6952184 | 0.093 | -0.582 | 0.38 | -0.871 | -0.037 | -0.104 | -0.104 | 0 | 0.15 | 0.42 | 0.07 | 0 | 0.08 | 0.1 |
| 7 | 3451138 | rs17133289 | 1.287 | -0.63 | 0.577 | -0.919 | -0.01 | -0.021 | -0.021 | 0 | 0.27 | 0.23 | 0.14 | 0.13 | 0.13 | 0.24 |
| 7 | 3451251 | rs6959573 | 0.104 | -0.948 | 0.428 | -1.099 | 0.066 | 0.07 | 0.07 | 0 | 0.38 | 0.74 | 0.19 | 0.13 | 0.18 | 0.33 |
| 7 | 3455249 | rs17133292 | 1.427 | -0.783 | 0.502 | -1.002 | 0.015 | 0.011 | 0.011 | 0 | 0.26 | 0.18 | 0.13 | 0.14 | 0.12 | 0.26 |
| 7 | 3459998 | rs17133303 | 0.247 | -0.713 | 0.489 | -0.958 | -0.026 | -0.067 | -0.067 | 0 | 0.13 | 0.23 | 0.06 | 0 | 0.07 | 0.07 |
| 7 | 3472629 | rs17133346 | 1.32 | -0.688 | 0.456 | -1.01 | 0.005 | -0.005 | -0.005 | 0 | 0.27 | 0.2 | 0.15 | 0.14 | 0.15 | 0.24 |
| 7 | 3472721 | rs17133347 | 1.08 | -0.441 | 0.196 | -0.743 | -0.031 | 0.009 | 0.009 | 0 | 0.07 | 0.01 | 0.25 | 0.38 | 0.12 | 0.1 |
| 7 | 3475716 | rs7802106 | -1.599 | -0.294 | 0.132 | -0.812 | -0.059 | -0.08 | -0.08 | 0 | 0.81 | 0.94 | 0.72 | 0.59 | 0.71 | 0.85 |
| 7 | 3476330 | rs10235617 | 1.945 | -0.348 | 0.034 | -1.002 | -0.075 | 0.129 | 0.129 | 0 | 0.25 | 0.03 | 0.21 | 0.06 | 0.3 | 0.33 |
| 7 | 3479432 | rs2204123 | -0.057 | -0.483 | 0.309 | -0.88 | 0.068 | 0.089 | 0.089 | 0.015 | 0.43 | 0.71 | 0.22 | 0.15 | 0.23 | 0.33 |
| 7 | 3481695 | rs10243467 | -0.199 | -0.43 | 0.541 | -0.809 | 0.001 | -0.002 | -0.002 | 1.531 | 0.51 | 0.36 | 0.55 | 0.46 | 0.55 | 0.59 |

| | | | | | | | | | | | | | | | | |
|---|---|---|---|---|---|---|---|---|---|---|---|---|---|---|---|---|
| 7 | 3482926 | rs6415253 | -1.407 | -0.335 | 0.392 | -0.768 | -0.029 | -0.041 | -0.041 | 0 | 0.8 | 0.88 | 0.72 | 0.59 | 0.71 | 0.85 |
| 7 | 3486358 | rs1357309 | -1.272 | -0.361 | 0.631 | -0.777 | 0.003 | 0.01 | 0.01 | 0 | 0.79 | 0.82 | 0.71 | 0.59 | 0.7 | 0.83 |
| 7 | 3488521 | rs17133370 | 1.512 | -0.477 | 0.742 | -0.871 | -0.01 | -0.032 | -0.032 | 0.779 | 0.26 | 0.25 | 0.15 | 0.1 | 0.14 | 0.24 |
| 7 | 3490865 | rs2178629 | -0.012 | 0.002 | 0.436 | -1.102 | -0.001 | -0.038 | -0.038 | 0 | 0.8 | 0.78 | 0.9 | 1 | 0.83 | 0.89 |
| 7 | 3494122 | rs2204122 | 0.206 | -0.164 | 0.343 | -1.064 | 0.07 | 0.076 | 0.076 | 0.166 | 0.41 | 0.72 | 0.21 | 0.11 | 0.2 | 0.32 |
| 7 | 3498739 | rs1851419 | -0.938 | 0.045 | -0.007 | -0.893 | -0.052 | -0.064 | -0.064 | 0 | 0.82 | 0.9 | 0.73 | 0.61 | 0.71 | 0.85 |
| 7 | 3504911 | rs17133377 | 0.628 | -0.099 | 0.093 | -0.97 | -0.01 | -0.036 | -0.036 | 0.08 | 0.12 | 0.18 | 0.05 | 0 | 0.07 | 0.07 |
| 7 | 3505231 | rs17133379 | 1.517 | -0.104 | -0.041 | -0.923 | 0.012 | 0.006 | 0.006 | 0.001 | 0.26 | 0.17 | 0.13 | 0.11 | 0.13 | 0.24 |
| 7 | 3510741 | rs1915981 | -0.759 | -0.02 | -0.263 | -0.839 | -0.069 | -0.081 | -0.081 | 1.6 | 0.8 | 0.9 | 0.71 | 0.59 | 0.67 | 0.84 |
| 7 | 3515437 | rs17133409 | 1.218 | -0.432 | 0.041 | -0.968 | -0.043 | -0.069 | -0.069 | 0.061 | 0.27 | 0.27 | 0.14 | 0.1 | 0.12 | 0.27 |
| 7 | 3521000 | rs10499335 | -0.284 | 0.365 | -0.281 | -0.919 | 0.022 | -0.022 | -0.022 | 0.018 | 0.55 | 0.74 | 0.52 | 0.55 | 0.42 | 0.47 |
| 7 | 3521573 | rs10242223 | -0.488 | 0.57 | -0.83 | -1.119 | -0.073 | -0.106 | -0.106 | 0 | 0.73 | 0.93 | 0.74 | 0.59 | 0.67 | 0.85 |
| 7 | 3526685 | rs1548210 | -0.229 | 0.677 | -0.438 | -0.97 | -0.004 | -0.026 | -0.026 | 0 | 0.39 | 0.55 | 0.36 | 0.27 | 0.41 | 0.49 |
| 7 | 3531146 | rs2141819 | 0.615 | 0.511 | -0.316 | -1.146 | -0.01 | -0.006 | -0.006 | 0 | 0.42 | 0.5 | 0.4 | 0.32 | 0.27 | 0.41 |
| 7 | 3534505 | rs7794763 | 0.078 | 1.183 | -0.302 | -1.162 | -0.02 | -0.002 | -0.002 | 0 | 0.65 | 0.69 | 0.66 | 0.38 | 0.56 | 0.81 |
| 7 | 3539520 | rs4719964 | 1.085 | 1.09 | -0.346 | -1.148 | -0.016 | 0.05 | 0.05 | 0 | 0.23 | 0.15 | 0.24 | 0.06 | 0.29 | 0.4 |
| 7 | 3544638 | rs882921 | 0.209 | 0.75 | 0.042 | -1.146 | -0.02 | -0.073 | -0.073 | 0 | 0.12 | 0.37 | 0.06 | 0 | 0.07 | 0.08 |
| 7 | 3544743 | rs1882426 | -1.497 | 0.876 | -0.511 | -1.175 | -0.033 | 0.035 | 0.035 | 0 | 0.87 | 0.99 | 0.94 | 0.98 | 0.89 | 0.95 |
| 7 | 3547206 | rs1522505 | 1.588 | 0.307 | -0.265 | -1.202 | 0.034 | -0.002 | -0.002 | 0 | 0.31 | 0.15 | 0.36 | 0.31 | 0.22 | 0.33 |
| 7 | 3556132 | rs1108879 | -0.144 | 0.531 | -0.282 | -1.223 | -0.042 | -0.01 | -0.01 | 0 | 0.67 | 0.76 | 0.66 | 0.38 | 0.57 | 0.81 |
| 7 | 3559826 | rs2705608 | -0.759 | 0.485 | -0.374 | -1.174 | -0.043 | -0.056 | -0.056 | 0 | 0.72 | 0.83 | 0.7 | 0.6 | 0.65 | 0.85 |
| 7 | 3560919 | rs2705609 | 1.519 | 0.377 | -0.607 | -1.266 | -0.048 | 0.073 | 0.073 | 0 | 0.2 | 0.05 | 0.22 | 0.06 | 0.27 | 0.34 |
| 7 | 3571289 | rs12111969 | 1.767 | 0.134 | -0.446 | -1.266 | 0.015 | -0.001 | -0.001 | 0 | 0.3 | 0.18 | 0.36 | 0.32 | 0.22 | 0.34 |
| 7 | 3571965 | rs11973763 | 0.669 | 0.158 | -0.251 | -1.29 | -0.02 | -0.054 | -0.054 | 1.81 | 0.11 | 0.24 | 0.04 | 0 | 0.06 | 0.06 |
| 7 | 3572028 | rs4722903 | 1.079 | 0.291 | -0.306 | -1.307 | -0.01 | 0.049 | 0.049 | 0 | 0.23 | 0.14 | 0.23 | 0.06 | 0.26 | 0.4 |
| 7 | 3581391 | rs2614950 | 0.753 | 0.24 | -0.201 | -1.375 | -0.014 | 0.027 | 0.027 | 0 | 0.25 | 0.18 | 0.28 | 0.06 | 0.32 | 0.42 |
| 7 | 3585560 | rs17133545 | 0.475 | 0.211 | -0.236 | -1.241 | -0.02 | -0.058 | -0.058 | 0 | 0.11 | 0.25 | 0.05 | 0 | 0.07 | 0.06 |
| 7 | 3598581 | rs10216009 | 1.576 | 0.113 | -0.413 | -1.192 | 0.01 | -0.001 | -0.001 | 0 | 0.31 | 0.2 | 0.36 | 0.34 | 0.21 | 0.33 |
| 7 | 3602348 | rs7811384 | -0.18 | 0.127 | -0.649 | -1.135 | -0.047 | -0.055 | -0.055 | 0 | 0.64 | 0.76 | 0.68 | 0.57 | 0.58 | 0.79 |
| 7 | 3609712 | rs2141823 | -0.24 | 0.108 | -0.593 | -1.038 | -0.045 | -0.033 | -0.033 | 0.074 | 0.64 | 0.81 | 0.67 | 0.59 | 0.55 | 0.83 |
| 7 | 3611608 | rs9791502 | 0.277 | -0.078 | -0.673 | -1.097 | 0.022 | -0.014 | -0.014 | 0 | 0.54 | 0.68 | 0.61 | 0.56 | 0.41 | 0.71 |
| 7 | 3614237 | rs16870889 | 0.041 | 0.151 | -0.572 | -0.675 | 0 | -0.004 | -0.004 | 0 | 0.06 | 0.05 | 0.03 | 0.18 | 0.07 | 0.04 |
| 7 | 3617414 | rs9654994 | 1.661 | -0.351 | -0.484 | -1.028 | -0.027 | 0.019 | 0.019 | 0 | 0.22 | 0.24 | 0.31 | 0.34 | 0.11 | 0.25 |
| 7 | 3618143 | rs10238338 | 2.398 | -0.432 | -0.975 | -1.133 | 0.045 | 0.084 | 0.084 | 0 | 0.17 | 0.05 | 0.13 | 0 | 0.07 | 0.09 |
| 7 | 3624238 | rs13230266 | 0.097 | 0.273 | -0.549 | -0.626 | -0.001 | -0.023 | -0.023 | 0.886 | 0.06 | 0.04 | 0.04 | 0.12 | 0.06 | 0.03 |
| 7 | 3627799 | rs2614967 | -0.97 | 0.195 | -0.616 | -0.708 | -0.042 | -0.053 | -0.053 | 0.011 | 0.72 | 0.84 | 0.72 | 0.52 | 0.59 | 0.83 |
| 7 | 3630272 | rs10951254 | 0.342 | -0.551 | -0.441 | -0.919 | -0.004 | 0.013 | 0.013 | 0 | 0.29 | 0.33 | 0.46 | 0.39 | 0.3 | 0.54 |
| 7 | 3631115 | rs10951255 | 1.012 | -0.363 | -0.705 | -0.892 | -0.044 | 0.118 | 0.118 | 0 | 0.23 | 0.04 | 0.26 | 0.05 | 0.26 | 0.34 |
| 7 | 3638547 | rs2341587 | -0.239 | -0.336 | -0.406 | -0.847 | 0.021 | -0.002 | -0.002 | 0 | 0.63 | 0.59 | 0.64 | 0.39 | 0.5 | 0.81 |
| 7 | 3639612 | rs13223065 | 0.832 | -0.305 | -0.798 | -0.801 | -0.002 | -0.003 | -0.003 | 0 | 0.23 | 0.2 | 0.26 | 0.05 | 0.25 | 0.36 |
| 7 | 3659530 | rs12535467 | 0.404 | -0.175 | -0.957 | -0.799 | -0.007 | -0.002 | -0.002 | 0 | 0.28 | 0.25 | 0.28 | 0.06 | 0.35 | 0.36 |
| 7 | 3676463 | rs10263338 | 0.651 | -0.458 | -1.058 | -0.767 | -0.005 | -0.001 | -0.001 | 0 | 0.39 | 0.62 | 0.4 | 0.36 | 0.24 | 0.48 |
| 7 | 3677429 | rs17133605 | -0.54 | -0.349 | -1.444 | -0.765 | -0.019 | -0.008 | -0.008 | 0 | 0.66 | 0.77 | 0.69 | 0.42 | 0.59 | 0.83 |
| 7 | 3682486 | rs12531560 | 0.858 | -0.1 | -1.55 | -0.713 | -0.088 | 0.121 | 0.121 | 0 | 0.25 | 0.04 | 0.27 | 0.06 | 0.34 | 0.32 |
| 7 | 3687542 | rs1025456 | -0.323 | -0.106 | -1.24 | -0.204 | -0.004 | 0.045 | 0.045 | 0 | 0.95 | 0.91 | 0.98 | 0.82 | 0.94 | 0.96 |
| 7 | 3696883 | rs2099301 | -0.323 | -0.111 | -1.24 | -0.193 | -0.005 | 0.035 | 0.035 | 0 | 0.95 | 0.91 | 0.98 | 0.83 | 0.94 | 0.97 |
| 7 | 3697421 | rs6462213 | -0.252 | -0.126 | -1.366 | -0.117 | -0.022 | -0.012 | -0.012 | 0 | 0.64 | 0.75 | 0.68 | 0.44 | 0.56 | 0.81 |
| 7 | 3705355 | rs882553 | 0.31 | -0.137 | -1.645 | -0.463 | -0.019 | 0.025 | 0.025 | 0 | 0.6 | 0.67 | 0.64 | 0.73 | 0.5 | 0.59 |
| 7 | 3710751 | rs752587 | -0.525 | -0.361 | -2.256 | -0.391 | -0.103 | -0.039 | -0.039 | 0.492 | 0.72 | 0.98 | 0.73 | 0.76 | 0.64 | 0.66 |
| 7 | 3713194 | rs13223746 | -0.392 | -0.455 | -2.136 | -0.412 | 0.055 | -0.027 | -0.027 | 0.282 | 0.59 | 0.26 | 0.56 | 0.48 | 0.73 | 0.52 |
| 7 | 3714732 | rs11764509 | 1.038 | -0.156 | -2.08 | -0.458 | -0.045 | -0.048 | -0.048 | 0 | 0.3 | 0.15 | 0.29 | 0.4 | 0.39 | 0.18 |
| 7 | 3718051 | rs7790719 | -0.168 | -1.164 | -2.044 | -0.214 | 0.068 | 0.078 | 0.078 | 0 | 0.63 | 0.86 | 0.68 | 0.81 | 0.74 | 0.7 |
| 7 | 3720117 | rs12536936 | 0.885 | -2.19 | -1.877 | -0.161 | 0.203 | -0.001 | -0.001 | 0 | 0.55 | 0.12 | 0.48 | 0.38 | 0.79 | 0.44 |
| 7 | 3727061 | rs10951278 | -0.331 | -2.192 | -1.534 | -0.172 | 0.05 | 0.028 | 0.028 | 0 | 0.68 | 0.77 | 0.73 | 0.78 | 0.87 | 0.74 |
| 7 | 3727993 | rs2217607 | 0.887 | -2.054 | -1.682 | -0.165 | 0.154 | 0.016 | 0.016 | 0 | 0.55 | 0.16 | 0.47 | 0.38 | 0.76 | 0.43 |
| 7 | 3728881 | rs2060185 | 0.852 | -2.077 | -1.641 | -0.13 | 0.16 | 0.011 | 0.011 | 0 | 0.56 | 0.16 | 0.47 | 0.38 | 0.77 | 0.44 |
| 7 | 3732253 | rs992141 | 0.299 | -1.907 | -1.237 | -0.73 | 0.057 | 0.036 | 0.036 | 0 | 0.66 | 0.74 | 0.73 | 0.78 | 0.86 | 0.74 |
| 7 | 3735435 | rs1529661 | 0.452 | -1.892 | -1.315 | -0.773 | 0.11 | 0.076 | 0.076 | 0.512 | 0.62 | 0.81 | 0.71 | 0.78 | 0.84 | 0.74 |
| 7 | 3736788 | rs1529657 | 0.205 | -1.879 | -1.498 | -0.751 | -0.002 | -0.036 | -0.036 | 0 | 0.94 | 1 | 0.93 | 0.83 | 0.94 | 0.94 |
| 7 | 3736942 | rs1529655 | 0.488 | -1.734 | -1.383 | -0.707 | 0.111 | 0.082 | 0.082 | 0.272 | 0.6 | 0.76 | 0.69 | 0.78 | 0.83 | 0.73 |
| 7 | 3743555 | rs10256504 | -0.277 | -1.438 | -1.004 | -0.855 | 0.053 | 0.053 | 0.053 | 0 | 0.24 | 0.19 | 0.18 | 0.07 | 0.08 | 0.17 |
| 7 | 3750555 | rs10248557 | 0.748 | -1.442 | -0.275 | -0.912 | -0.031 | -0.074 | -0.074 | 0 | 0.16 | 0.24 | 0.11 | 0 | 0.06 | 0.07 |
| 7 | 3751099 | rs10264235 | 0.551 | -1.499 | 0.07 | -0.936 | -0.05 | -0.106 | -0.106 | 2.15 | 0.16 | 0.34 | 0.13 | 0 | 0.06 | 0.09 |
| 7 | 3753766 | rs10261179 | 0.872 | -1.517 | 0.08 | -0.944 | 0.014 | 0 | 0 | 0 | 0.15 | 0.12 | 0.11 | 0 | 0.06 | 0.07 |
| 7 | 3753952 | rs6969732 | -1.157 | -1.725 | 0.785 | -0.939 | -0.003 | 0 | 0 | 0 | 0.3 | 0.69 | 0.26 | 0.15 | 0.14 | 0.21 |
| 7 | 3760988 | rs10268585 | -1.779 | -1.667 | -0.184 | -1.072 | -0.103 | -0.135 | -0.135 | 0 | 0.57 | 0.98 | 0.63 | 0.46 | 0.45 | 0.71 |
| 7 | 3763146 | rs6952117 | 0.91 | -1.625 | 0.474 | -1.067 | -0.045 | -0.095 | -0.095 | 0 | 0.17 | 0.29 | 0.13 | 0 | 0.06 | 0.09 |
| 7 | 3765004 | rs6953733 | 0.872 | -1.625 | 0.465 | -1.001 | -0.048 | -0.038 | -0.038 | 0 | 0.17 | 0.32 | 0.13 | 0.07 | 0.06 | 0.09 |
| 7 | 3766819 | rs2051918 | -1.3 | -1.54 | 0.685 | -0.737 | -0.05 | 0.131 | 0.131 | 0 | 0.16 | 0.4 | 0.17 | 0.32 | 0.09 | 0.17 |

| | | | | | | | | | | | | | | | | |
|---|---|---|---|---|---|---|---|---|---|---|---|---|---|---|---|---|
| 7 | 3792895 | rs6975070 | -1.079 | -1.094 | 0.353 | -1.673 | 0.033 | 0.046 | 0.046 | 0 | 0.43 | 0.69 | 0.36 | 0.18 | 0.27 | 0.3 |
| 7 | 3825425 | rs17324338 | 0.092 | -0.718 | 0.096 | -1.726 | -0.051 | 0.066 | 0.066 | 0 | 0.14 | 0.04 | 0.2 | 0.01 | 0.26 | 0.21 |
| 7 | 3836379 | rs11981385 | -0.591 | -1.476 | 0.979 | -1.667 | -0.054 | -0.052 | -0.052 | 2.242 | 0.33 | 0.54 | 0.27 | 0.15 | 0.14 | 0.22 |
| 7 | 3839029 | rs7787471 | -0.885 | -1.205 | 1.31 | -1.747 | 0.005 | 0.012 | 0.012 | 0.008 | 0.07 | 0.34 | 0.08 | 0.05 | 0.05 | 0.08 |
| 7 | 3848455 | rs13234640 | -0.75 | -1.19 | 1.388 | -1.743 | -0.012 | -0.006 | -0.006 | 0 | 0.07 | 0.26 | 0.08 | 0.05 | 0.06 | 0.05 |
| 7 | 3848651 | rs1548617 | -0.411 | -1.425 | 1.349 | -2.073 | 0.017 | 0.019 | 0.019 | 0 | 0.33 | 0.19 | 0.23 | 0.07 | 0.18 | 0.21 |
| 7 | 3849481 | rs6945778 | -1.081 | -1.457 | 1.69 | -2.086 | 0.043 | 0.06 | 0.06 | 0 | 0.42 | 0.62 | 0.32 | 0.12 | 0.24 | 0.26 |
| 7 | 3849844 | rs6945255 | -0.538 | -1.444 | 1.532 | -2.083 | 0.031 | 0.05 | 0.05 | 0 | 0.35 | 0.17 | 0.23 | 0.08 | 0.18 | 0.21 |
| 7 | 3849907 | rs6946432 | -0.539 | -1.437 | 1.552 | -2.081 | 0.023 | 0.03 | 0.03 | 0 | 0.35 | 0.2 | 0.23 | 0.08 | 0.18 | 0.21 |
| 7 | 3852261 | rs17324947 | -0.018 | -1.506 | 1.775 | -1.825 | 0.012 | 0.006 | 0.006 | 0 | 0.27 | 0.17 | 0.2 | 0.07 | 0.11 | 0.14 |
| 7 | 3858097 | rs13221624 | -0.746 | -1.503 | 1.74 | -1.825 | -0.029 | -0.051 | -0.051 | 0 | 0.35 | 0.54 | 0.29 | 0.12 | 0.17 | 0.26 |
| 7 | 3860836 | rs2341959 | -0.786 | -1.501 | 1.629 | -1.828 | -0.046 | -0.077 | -0.077 | 0.996 | 0.35 | 0.48 | 0.29 | 0.12 | 0.18 | 0.26 |
| 7 | 3863914 | rs1548614 | -0.221 | -1.484 | 1.419 | -1.815 | 0.051 | 0.069 | 0.069 | 0 | 0.29 | 0.12 | 0.21 | 0.07 | 0.12 | 0.18 |
| 7 | 3869969 | rs17259924 | -0.087 | -1.47 | 1.424 | -1.809 | 0.036 | 0.047 | 0.047 | 0 | 0.28 | 0.14 | 0.2 | 0.07 | 0.13 | 0.18 |
| 7 | 3876162 | rs6462361 | -0.559 | -1.433 | 1.382 | -1.793 | 0.043 | 0.054 | 0.054 | 0 | 0.34 | 0.19 | 0.26 | 0.12 | 0.18 | 0.26 |
| 7 | 3886254 | rs41871 | 0.477 | -0.77 | 0.922 | -1.924 | -0.052 | 0.047 | 0.047 | 0 | 0.13 | 0.04 | 0.2 | 0 | 0.27 | 0.21 |
| 7 | 3891289 | rs12532912 | -0.211 | -1.057 | 1.225 | -1.944 | -0.032 | -0.048 | -0.048 | 0 | 0.32 | 0.32 | 0.25 | 0.12 | 0.16 | 0.25 |
| 7 | 3900810 | rs10085568 | -0.78 | -1.204 | 0.951 | -2.042 | 0.076 | 0.097 | 0.097 | 0 | 0.4 | 0.74 | 0.32 | 0.12 | 0.2 | 0.31 |
| 7 | 3905689 | rs6959510 | -0.645 | -1.033 | 0.741 | -2.017 | -0.075 | 0.078 | 0.078 | 0 | 0.85 | 0.97 | 0.8 | 1 | 0.72 | 0.81 |
| 7 | 3908096 | rs17261199 | 1.369 | -1.205 | 0.668 | -2.121 | -0.012 | 0.244 | 0.244 | 0 | 0.46 | 0.1 | 0.47 | 0.87 | 0.51 | 0.48 |
| 7 | 3910657 | rs6961561 | -0.194 | -1.342 | 1.449 | -1.916 | -0.072 | -0.086 | -0.086 | 3.356 | 0.3 | 0.48 | 0.27 | 0.13 | 0.17 | 0.28 |
| 7 | 3910982 | rs12690804 | -0.095 | -1.309 | 1.246 | -1.901 | -0.015 | -0.02 | -0.02 | 0 | 0.28 | 0.3 | 0.24 | 0.13 | 0.16 | 0.27 |
| 7 | 3918251 | rs6958535 | -0.332 | -1.239 | 1.321 | -1.881 | -0.037 | -0.055 | -0.055 | 0 | 0.32 | 0.39 | 0.25 | 0.13 | 0.18 | 0.28 |
| 7 | 3925535 | rs7804166 | -0.485 | -1.125 | 1.105 | -1.916 | -0.023 | -0.044 | -0.044 | 0.455 | 0.34 | 0.34 | 0.27 | 0.13 | 0.2 | 0.28 |
| 7 | 3925820 | rs7800775 | -0.829 | -1.121 | 0.378 | -1.921 | 0.087 | 0.134 | 0.134 | 0.003 | 0.39 | 0.76 | 0.32 | 0.14 | 0.21 | 0.32 |
| 7 | 3935045 | rs2107969 | 1.702 | -1.003 | -0.383 | -1.35 | -0.165 | -0.132 | -0.132 | 0 | 0.37 | 0.03 | 0.29 | 0.57 | 0.44 | 0.21 |
| 7 | 3938385 | rs12532171 | -0.201 | -1.22 | 0.443 | -1.857 | 0.025 | 0.025 | 0.025 | 0 | 0.31 | 0.3 | 0.25 | 0.13 | 0.17 | 0.25 |
| 7 | 3955516 | rs4723269 | -0.679 | -0.745 | 0.593 | -1.877 | 0.026 | 0.03 | 0.03 | 0 | 0.39 | 0.65 | 0.38 | 0.2 | 0.27 | 0.33 |
| 7 | 3958048 | rs382301 | -0.626 | -0.357 | -0.01 | -2.147 | -0.086 | 0.109 | 0.109 | 0 | 0.8 | 0.96 | 0.79 | 0.98 | 0.68 | 0.72 |
| 7 | 3958295 | rs4723275 | -0.687 | -1.025 | 1.038 | -1.745 | 0.042 | 0.062 | 0.062 | 0 | 0.37 | 0.24 | 0.36 | 0.2 | 0.26 | 0.31 |
| 7 | 3967795 | rs17134054 | -0.889 | -1.164 | 1.567 | -1.626 | -0.005 | -0.006 | -0.006 | 0.018 | 0.39 | 0.55 | 0.37 | 0.22 | 0.29 | 0.3 |
| 7 | 3972185 | rs2044225 | -0.306 | -1.116 | 1.429 | -1.601 | 0.048 | 0.06 | 0.06 | 0 | 0.33 | 0.14 | 0.34 | 0.21 | 0.24 | 0.29 |
| 7 | 3978342 | rs4723284 | -0.975 | -1.158 | 1.764 | -1.615 | 0.062 | 0.071 | 0.071 | 0 | 0.39 | 0.75 | 0.41 | 0.22 | 0.28 | 0.37 |
| 7 | 3980563 | rs7801639 | -0.981 | -1.105 | 1.799 | -1.501 | 0.067 | 0.074 | 0.074 | 0 | 0.41 | 0.84 | 0.43 | 0.26 | 0.31 | 0.37 |
| 7 | 3982653 | rs11977815 | -0.773 | -1.218 | 1.782 | -1.687 | -0.025 | -0.034 | -0.034 | 0 | 0.37 | 0.51 | 0.39 | 0.21 | 0.29 | 0.35 |
| 7 | 3983402 | rs12536527 | 0.208 | -1.293 | 1.269 | -1.925 | 0.003 | 0.023 | 0.023 | 0 | 0.2 | 0.09 | 0.29 | 0.16 | 0.19 | 0.25 |
| 7 | 4003958 | rs4720128 | -1.158 | -0.863 | 0.514 | -1.788 | 0.051 | 0.093 | 0.093 | 0 | 0.39 | 0.75 | 0.4 | 0.19 | 0.28 | 0.37 |
| 7 | 4004884 | rs10277771 | -0.952 | -0.807 | 0.451 | -1.764 | -0.008 | -0.019 | -0.019 | 0 | 0.37 | 0.57 | 0.39 | 0.19 | 0.29 | 0.37 |
| 7 | 4022953 | rs7778932 | -1.264 | 0.102 | -0.324 | -1.646 | -0.117 | 0.041 | 0.041 | 0 | 0.67 | 0.96 | 0.59 | 0.24 | 0.56 | 0.83 |
| 7 | 4024024 | rs10215031 | -0.421 | 0.138 | 0.303 | -1.826 | -0.005 | 0.061 | 0.061 | 0.087 | 0.56 | 0.29 | 0.45 | 0.18 | 0.52 | 0.7 |
| 7 | 4027536 | rs6956208 | -1.027 | 0.037 | 1.329 | -1.492 | 0.009 | -0.021 | -0.021 | 0.427 | 0.21 | 0.3 | 0.27 | 0.16 | 0.21 | 0.28 |
| 7 | 4028352 | rs10270839 | -0.897 | -0.258 | 1.7 | -1.62 | -0.036 | -0.107 | -0.107 | 0 | 0.15 | 0.38 | 0.1 | 0 | 0.08 | 0.07 |
| 7 | 4031064 | rs17134117 | -0.6 | -0.19 | 1.639 | -1.614 | 0.001 | 0 | 0 | 0 | 0.08 | 0.12 | 0.13 | 0.11 | 0.07 | 0.15 |
| 7 | 4035307 | rs6967503 | -0.334 | -0.341 | 2.233 | -1.621 | -0.006 | -0.031 | -0.031 | 0 | 0.12 | 0.1 | 0.06 | 0 | 0.08 | 0.07 |
| 7 | 4038917 | rs6950410 | -1.773 | -0.265 | 2.22 | -1.677 | -0.009 | -0.093 | -0.093 | 0.944 | 0.29 | 0.39 | 0.17 | 0.06 | 0.24 | 0.28 |
| 7 | 4041311 | rs10241063 | -1.65 | -0.294 | 2.227 | -1.654 | -0.012 | -0.078 | -0.078 | 0 | 0.28 | 0.38 | 0.18 | 0.06 | 0.25 | 0.21 |
| 7 | 4042436 | rs6462522 | -1.557 | -0.382 | 2.151 | -1.659 | -0.013 | -0.076 | -0.076 | 0 | 0.28 | 0.34 | 0.17 | 0.06 | 0.24 | 0.2 |
| 7 | 4050276 | rs6942556 | -1.03 | -0.479 | 2.304 | -1.188 | -0.017 | 0.027 | 0.027 | 0 | 0.2 | 0.57 | 0.23 | 0.23 | 0.15 | 0.24 |
| 7 | 4054773 | rs7793935 | -0.986 | -0.407 | 2.286 | -1.297 | 0.013 | -0.016 | -0.016 | 0 | 0.09 | 0.15 | 0.14 | 0.06 | 0.09 | 0.17 |
| 7 | 4057967 | rs12154868 | 0.069 | -0.352 | 1.52 | -1.418 | -0.007 | 0.059 | 0.059 | 0 | 0.1 | 0 | 0.05 | 0 | 0.07 | 0.04 |
| 7 | 4062727 | rs10228937 | -2.277 | -0.352 | 1.607 | -1.344 | 0.021 | 0.027 | 0.027 | 0.1 | 0.4 | 0.8 | 0.43 | 0.33 | 0.32 | 0.46 |
| 7 | 4070095 | rs2342489 | -0.946 | -0.205 | 2.157 | -2.033 | -0.007 | -0.047 | -0.047 | 0 | 0.18 | 0.7 | 0.18 | 0.08 | 0.15 | 0.21 |
| 7 | 4075621 | rs7791029 | -1.177 | -0.245 | 2.53 | -2.05 | 0 | -0.007 | -0.007 | 0 | 0.19 | 0.77 | 0.18 | 0.08 | 0.15 | 0.21 |
| 7 | 4077620 | rs6462544 | -0.367 | -0.211 | 2.773 | -2.038 | 0.021 | -0.065 | -0.065 | 3.952 | 0.15 | 0.52 | 0.1 | 0.06 | 0.13 | 0.16 |
| 7 | 4080757 | rs10259751 | -1.204 | -0.21 | 2.77 | -2.037 | 0 | -0.008 | -0.008 | 0 | 0.19 | 0.77 | 0.18 | 0.07 | 0.15 | 0.21 |
| 7 | 4081194 | rs12155314 | -0.387 | -0.202 | 2.739 | -2.034 | 0.02 | -0.065 | -0.065 | 0 | 0.16 | 0.52 | 0.11 | 0.06 | 0.13 | 0.16 |
| 7 | 4085343 | rs4442012 | -0.885 | -0.204 | 2.635 | -2.036 | -0.005 | -0.055 | -0.055 | 1.009 | 0.18 | 0.69 | 0.17 | 0.06 | 0.15 | 0.17 |
| 7 | 4087468 | rs1004043 | -2.41 | -0.244 | 2.099 | -2.353 | 0.011 | 0.188 | 0.188 | 0 | 0.4 | 0.87 | 0.29 | 0.08 | 0.33 | 0.4 |
| 7 | 4088876 | rs17134285 | -0.365 | -0.159 | 2.146 | -2.241 | 0.005 | -0.009 | -0.009 | 0.308 | 0.14 | 0.14 | 0.13 | 0 | 0.1 | 0.03 |
| 7 | 4089691 | rs13226987 | -0.047 | -0.024 | 2.313 | -2.076 | -0.003 | -0.03 | -0.03 | 0.174 | 0.14 | 0.25 | 0.08 | 0.06 | 0.13 | 0.11 |
| 7 | 4105705 | rs7801843 | -0.445 | -0.031 | 2.325 | -2.076 | -0.007 | -0.055 | -0.055 | 0 | 0.16 | 0.58 | 0.1 | 0.07 | 0.13 | 0.11 |
| 7 | 4105936 | rs10231929 | -0.317 | -0.016 | 2.341 | -2.089 | -0.002 | -0.069 | -0.069 | 0.305 | 0.16 | 0.47 | 0.09 | 0.06 | 0.13 | 0.11 |
| 7 | 4106198 | rs6971090 | -0.322 | 0.007 | 1.561 | -2.075 | 0 | -0.059 | -0.059 | 0 | 0.16 | 0.48 | 0.1 | 0.08 | 0.13 | 0.15 |
| 7 | 4111112 | rs6462577 | -0.729 | 0.042 | 0.43 | -2.064 | -0.002 | 0.01 | 0.01 | 0.139 | 0.21 | 0.78 | 0.21 | 0.08 | 0.21 | 0.2 |
| 7 | 4115792 | rs7809492 | -1.349 | -0.068 | 0.267 | -2.212 | 0.018 | 0.103 | 0.103 | 0 | 0.4 | 0.75 | 0.28 | 0.08 | 0.32 | 0.41 |
| 7 | 4134107 | rs7784114 | -0.801 | 0.25 | 0.51 | -1.988 | -0.017 | -0.11 | -0.11 | 0 | 0.28 | 0.59 | 0.21 | 0.07 | 0.22 | 0.33 |
| 7 | 4135985 | rs7802118 | -0.596 | 0.214 | 0.725 | -2.035 | -0.014 | -0.101 | -0.101 | 0.288 | 0.26 | 0.44 | 0.2 | 0.07 | 0.21 | 0.32 |
| 7 | 4136969 | rs7790901 | -0.761 | 0.208 | 0.185 | -2.038 | -0.018 | -0.104 | -0.104 | 0 | 0.28 | 0.6 | 0.21 | 0.07 | 0.22 | 0.33 |
| 7 | 4139470 | rs10234430 | -0.762 | 0.142 | -0.442 | -2.579 | -0.004 | -0.106 | -0.106 | 0 | 0.41 | 0.57 | 0.27 | 0.08 | 0.35 | 0.37 |

| | | | | | | | | | | | | | | | | |
|---|---|---|---|---|---|---|---|---|---|---|---|---|---|---|---|---|
| 7 | 4139785 | rs882362 | -0.088 | 0.718 | -0.836 | -2.801 | 0 | -0.028 | -0.028 | 0 | 0.66 | 0.68 | 0.4 | 0.11 | 0.59 | 0.73 |
| 7 | 4141250 | rs7793159 | -0.406 | 0.025 | -1.016 | -2.495 | 0.007 | -0.008 | -0.008 | 0 | 0.45 | 0.54 | 0.25 | 0.13 | 0.34 | 0.63 |
| 7 | 4152089 | rs1062213 | -0.371 | 0.417 | -1.423 | -1.994 | -0.04 | 0.045 | 0.045 | 0 | 0.27 | 0.16 | 0.2 | 0.14 | 0.27 | 0.22 |
| 7 | 4161793 | rs8180800 | -1.066 | -0.256 | -1.961 | -2.337 | -0.028 | 0.18 | 0.18 | 0 | 0.62 | 0.87 | 0.68 | 0.9 | 0.63 | 0.7 |
| 7 | 4167274 | rs651659 | 0.238 | -0.776 | -1.089 | -2.352 | -0.024 | -0.066 | -0.066 | 0 | 0.19 | 0.24 | 0.09 | 0 | 0.13 | 0.16 |
| 7 | 4167991 | rs11763980 | 0.682 | -0.682 | -0.726 | -2.092 | -0.01 | 0.04 | 0.04 | 0 | 0.1 | 0.14 | 0.22 | 0.76 | 0.09 | 0.19 |
| 7 | 4171506 | rs632896 | 0.323 | -0.857 | -0.872 | -2.269 | 0.023 | 0.04 | 0.04 | 0 | 0.15 | 0.07 | 0.09 | 0.01 | 0.15 | 0.11 |
| 7 | 4172006 | rs13224870 | 1.676 | -0.971 | -0.736 | -2.327 | -0.052 | 0.181 | 0.181 | 0 | 0.37 | 0.11 | 0.31 | 0.1 | 0.39 | 0.39 |
| 7 | 4172492 | rs667645 | -0.854 | -1.015 | -0.523 | -2.375 | -0.014 | -0.011 | -0.011 | 4.182 | 0.3 | 0.54 | 0.43 | 0.75 | 0.31 | 0.25 |
| 7 | 4173613 | rs11766511 | 1.591 | -0.815 | -0.748 | -2.286 | -0.041 | 0.185 | 0.185 | 0 | 0.38 | 0.11 | 0.31 | 0.1 | 0.39 | 0.3 |
| 7 | 4175508 | rs1077473 | 0.634 | -0.729 | -0.298 | -2.218 | 0 | 0.118 | 0.118 | 0 | 0.54 | 0.71 | 0.65 | 0.88 | 0.56 | 0.62 |
| 7 | 4177275 | rs636392 | 0.254 | -0.344 | -0.405 | -2.156 | -0.002 | -0.024 | -0.024 | 1.12 | 0.16 | 0.13 | 0.16 | 0 | 0.2 | 0.1 |
| 7 | 4177712 | rs4236342 | 1.223 | -0.558 | -0.561 | -2.188 | -0.015 | 0.164 | 0.164 | 0 | 0.47 | 0.21 | 0.51 | 0.84 | 0.48 | 0.41 |
| 7 | 4182531 | rs1636412 | -0.478 | -0.464 | -0.532 | -2.028 | -0.006 | -0.04 | -0.04 | 0 | 0.2 | 0.23 | 0.16 | 0.05 | 0.18 | 0.23 |
| 7 | 4182850 | rs640945 | -0.117 | -1.38 | -1.082 | -2.267 | -0.024 | 0.108 | 0.108 | 0.054 | 0.64 | 0.92 | 0.78 | 0.84 | 0.72 | 0.67 |
| 7 | 4185685 | rs663486 | -0.327 | -1.585 | -1.214 | -2.095 | -0.017 | -0.05 | -0.05 | 0 | 0.72 | 0.96 | 0.77 | 0.73 | 0.79 | 0.69 |
| 7 | 4187971 | rs674329 | -0.053 | -1.365 | -0.99 | -2.092 | -0.045 | 0.047 | 0.047 | 0.137 | 0.66 | 0.96 | 0.78 | 0.78 | 0.7 | 0.67 |
| 7 | 4191811 | rs10951446 | 1.191 | -1.434 | -0.364 | -2.223 | 0.003 | -0.013 | -0.013 | 0.003 | 0.43 | 0.44 | 0.57 | 0.73 | 0.53 | 0.35 |
| 7 | 4192168 | rs663911 | -0.523 | -1.293 | -0.089 | -2.176 | -0.009 | -0.035 | -0.035 | 0 | 0.21 | 0.27 | 0.18 | 0.05 | 0.17 | 0.23 |
| 7 | 4195988 | rs619672 | -1.166 | -1.065 | 0.04 | -2.452 | -0.001 | -0.026 | -0.026 | 0 | 0.24 | 0.73 | 0.26 | 0 | 0.23 | 0.24 |
| 7 | 4203145 | rs634381 | -0.872 | -1.059 | 0.748 | -2.534 | 0 | -0.074 | -0.074 | 3.346 | 0.21 | 0.72 | 0.26 | 0 | 0.21 | 0.2 |
| 7 | 4206457 | rs935567 | 1.255 | -1.383 | 0.051 | -2.797 | -0.057 | 0.163 | 0.163 | 0.023 | 0.46 | 0.12 | 0.52 | 0.77 | 0.51 | 0.41 |
| 7 | 4208658 | rs666035 | 0.346 | -1.512 | 0.356 | -2.605 | -0.001 | -0.018 | -0.018 | 0 | 0.14 | 0.12 | 0.15 | 0 | 0.14 | 0.13 |
| 7 | 4211281 | rs623667 | -0.16 | -1.657 | 0.223 | -2.553 | 0.024 | 0.041 | 0.041 | 0.314 | 0.65 | 0.93 | 0.78 | 0.76 | 0.72 | 0.69 |
| 7 | 4211696 | rs631738 | 0.323 | -1.681 | 0.872 | -2.695 | -0.002 | -0.007 | -0.007 | 0 | 0.59 | 0.55 | 0.69 | 0.76 | 0.66 | 0.67 |
| 7 | 4212421 | rs9639711 | 1.273 | -1.729 | 0.487 | -2.704 | -0.013 | 0.06 | 0.06 | 0.016 | 0.43 | 0.27 | 0.53 | 0.77 | 0.51 | 0.46 |
| 7 | 4213571 | rs602780 | 0.504 | -1.592 | 0.068 | -2.63 | -0.001 | -0.014 | -0.014 | 0.018 | 0.13 | 0.12 | 0.15 | 0 | 0.13 | 0.13 |
| 7 | 4213975 | rs671694 | -0.02 | -2.024 | -0.147 | -2.513 | -0.021 | 0.027 | 0.027 | 0.368 | 0.66 | 0.98 | 0.78 | 0.76 | 0.73 | 0.68 |
| 7 | 4214725 | rs10235503 | 0.684 | -2.564 | 0.063 | -2.674 | 0.048 | 0.085 | 0.085 | 0 | 0.51 | 0.3 | 0.67 | 0.7 | 0.59 | 0.53 |
| 7 | 4216771 | rs665163 | 0.442 | -2.097 | 0.063 | -2.336 | -0.013 | -0.026 | -0.026 | 0.094 | 0.55 | 0.47 | 0.7 | 0.71 | 0.65 | 0.6 |
| 7 | 4222089 | rs10265031 | -0.428 | -2.835 | 0.043 | -2.345 | -0.054 | -0.075 | -0.075 | 0 | 0.18 | 0.29 | 0.05 | 0 | 0.06 | 0.05 |
| 7 | 4224124 | rs17134457 | 0.547 | -2.696 | 0.024 | -2.358 | -0.006 | -0.014 | -0.014 | 0.125 | 0.1 | 0.11 | 0.07 | 0.07 | 0.08 | 0.11 |
| 7 | 4224833 | rs17134460 | -0.645 | -2.872 | 0.144 | -2.433 | -0.038 | -0.055 | -0.055 | 0.231 | 0.17 | 0.29 | 0.1 | 0.07 | 0.09 | 0.19 |
| 7 | 4225870 | rs625537 | -1.311 | -2.333 | -0.535 | -2.603 | 0 | 0.03 | 0.03 | 0.038 | 0.66 | 0.74 | 0.35 | 0.26 | 0.38 | 0.51 |
| 7 | 4227904 | rs7786536 | -0.976 | -2.204 | -0.453 | -2.414 | 0.032 | 0.035 | 0.035 | 0.038 | 0.54 | 0.63 | 0.29 | 0.25 | 0.3 | 0.41 |
| 7 | 4228898 | rs16871145 | -0.926 | -1.617 | 0.063 | -2.077 | -0.097 | -0.155 | -0.155 | 0.051 | 0.24 | 0.43 | 0.08 | 0 | 0.08 | 0.16 |
| 7 | 4237382 | rs6978817 | -0.646 | 0.307 | 0.037 | -1.641 | 0.14 | -0.031 | -0.031 | 1.693 | 0.07 | 0.18 | 0.1 | 0.01 | 0.21 | 0.12 |
| 7 | 4237425 | rs17134485 | -0.107 | -0.608 | -0.24 | -1.978 | -0.038 | -0.022 | -0.022 | 0 | 0.39 | 0.54 | 0.27 | 0.29 | 0.27 | 0.37 |
| 7 | 4237801 | rs7810341 | -0.125 | -0.3 | -0.159 | -1.906 | 0.131 | -0.029 | -0.029 | 0 | 0.07 | 0.18 | 0.1 | 0.01 | 0.19 | 0.12 |
| 7 | 4240085 | rs4723454 | -0.239 | -0.477 | -0.263 | -2.154 | 0.003 | 0.003 | 0.003 | 0.612 | 0.5 | 0.54 | 0.27 | 0.28 | 0.32 | 0.41 |
| 7 | 4242376 | rs1806470 | 0.305 | -0.558 | -0.257 | -2.19 | -0.056 | -0.024 | -0.024 | 0 | 0.34 | 0.49 | 0.19 | 0.29 | 0.24 | 0.27 |
| 7 | 4243557 | rs649232 | -0.476 | -0.523 | -0.375 | -2.319 | 0.001 | 0.024 | 0.024 | 0 | 0.46 | 0.57 | 0.3 | 0.3 | 0.48 | 0.52 |
| 7 | 4243758 | rs17134567 | 0.056 | -0.646 | -0.247 | -2.114 | -0.036 | -0.021 | -0.021 | 0 | 0.37 | 0.52 | 0.21 | 0.29 | 0.27 | 0.37 |
| 7 | 4246412 | rs7812170 | -0.516 | -0.235 | -0.665 | -2.074 | 0.025 | -0.113 | -0.113 | 0 | 0.27 | 0.33 | 0.22 | 0.03 | 0.33 | 0.3 |
| 7 | 4251135 | rs10260900 | -1.056 | -0.191 | -1.44 | -1.63 | -0.167 | -0.07 | -0.07 | 0 | 0.65 | 0.98 | 0.43 | 0.37 | 0.58 | 0.65 |
| 7 | 4263659 | rs4723505 | -0.658 | -0.467 | -0.892 | -1.801 | 0.001 | -0.03 | -0.03 | 0 | 0.38 | 0.38 | 0.27 | 0.15 | 0.42 | 0.45 |
| 7 | 4264920 | rs13240660 | 0.277 | -0.137 | -1.082 | -1.849 | -0.026 | 0.023 | 0.023 | 0 | 0.1 | 0.07 | 0.09 | 0.01 | 0.25 | 0.09 |
| 7 | 4265655 | rs11761645 | 1.149 | -0.768 | -1.178 | -2.047 | 0.03 | 0.023 | 0.023 | 0 | 0.21 | 0.15 | 0.12 | 0.15 | 0.19 | 0.13 |
| 7 | 4266379 | rs610962 | -0.97 | -0.546 | -0.848 | -1.96 | -0.064 | 0.012 | 0.012 | 0 | 0.63 | 0.83 | 0.44 | 0.37 | 0.61 | 0.59 |
| 7 | 4268061 | rs10269222 | 0.626 | -0.774 | -0.65 | -2.002 | 0 | 0 | 0 | 0.55 | 0.11 | 0.12 | 0.06 | 0.12 | 0.05 | 0.13 |
| 7 | 4270682 | rs647910 | -1.037 | -0.422 | -0.532 | -2.033 | -0.026 | 0.009 | 0.009 | 0 | 0.64 | 0.78 | 0.44 | 0.37 | 0.62 | 0.59 |
| 7 | 4273289 | rs17134653 | -0.222 | -0.529 | -0.043 | -1.642 | -0.029 | -0.026 | -0.026 | 0 | 0.13 | 0.19 | 0.07 | 0.09 | 0.06 | 0.19 |
| 7 | 4279500 | rs10255750 | -0.122 | 0.04 | 0.009 | -1.956 | -0.02 | -0.033 | -0.033 | 17.728 | 0.28 | 0.44 | 0.18 | 0.19 | 0.26 | 0.15 |
| 7 | 4282613 | rs10265196 | -1.316 | 0.113 | -0.582 | -2.269 | -0.081 | 0.051 | 0.051 | 0 | 0.54 | 0.94 | 0.48 | 0.37 | 0.58 | 0.48 |
| 7 | 4283564 | rs17134661 | 0.408 | 0.166 | -0.005 | -2.066 | -0.009 | 0.037 | 0.037 | 0 | 0.07 | 0.17 | 0.05 | 0.15 | 0.04 | 0.09 |
| 7 | 4283769 | rs10479843 | -1.343 | 0.136 | 0.074 | -2.06 | 0.005 | -0.03 | -0.03 | 0 | 0.3 | 0.57 | 0.32 | 0.18 | 0.34 | 0.32 |
| 7 | 4286597 | rs929372 | -0.286 | 0.16 | 0 | -2.085 | 0.041 | -0.124 | -0.124 | 0 | 0.21 | 0.36 | 0.25 | 0.02 | 0.28 | 0.21 |
| 7 | 4293056 | rs12531029 | 0.308 | 0.371 | -0.26 | -2.106 | -0.017 | 0.018 | 0.018 | 0 | 0.16 | 0.08 | 0.22 | 0.04 | 0.27 | 0.09 |
| 7 | 4298625 | rs13228414 | 1.084 | 0.204 | -0.239 | -2.211 | 0.01 | 0.023 | 0.023 | 0 | 0.22 | 0.15 | 0.14 | 0.15 | 0.22 | 0.14 |
| 7 | 4301190 | rs13247068 | -0.031 | 0.431 | -0.171 | -2.091 | -0.012 | 0.001 | 0.001 | 0 | 0.18 | 0.11 | 0.26 | 0.03 | 0.29 | 0.22 |
| 7 | 4304727 | rs10270413 | 0.89 | 0.665 | 0.269 | -2.083 | 0.036 | -0.082 | -0.082 | 0 | 0.73 | 0.56 | 0.69 | 0.89 | 0.66 | 0.71 |
| 7 | 4305834 | rs1044701 | 0.87 | 0.564 | -0.455 | -2.114 | -0.02 | 0.05 | 0.05 | 0 | 0.43 | 0.09 | 0.51 | 0.64 | 0.4 | 0.51 |
| 7 | 4307664 | rs3801065 | -0.469 | 0.528 | 0.027 | -2.02 | -0.001 | 0.001 | 0.001 | 0 | 0.43 | 0.54 | 0.39 | 0.19 | 0.53 | 0.33 |
| 7 | 4308109 | rs1562538 | -0.537 | 0.52 | 0.023 | -2.025 | -0.002 | 0.016 | 0.016 | 0 | 0.45 | 0.57 | 0.39 | 0.19 | 0.53 | 0.33 |
| 7 | 4308220 | rs10227750 | -1.269 | 0.344 | 0.634 | -1.666 | 0.042 | 0.038 | 0.038 | 0.167 | 0.08 | 0.42 | 0.08 | 0.03 | 0.02 | 0.17 |
| 7 | 4310365 | rs6955448 | 0.095 | 0.69 | -0.017 | -0.917 | 0 | -0.002 | -0.002 | 0 | 0.31 | 0.33 | 0.28 | 0.34 | 0.36 | 0.28 |
| 7 | 4311316 | rs10237163 | -0.568 | 0.867 | -0.26 | -0.83 | -0.001 | 0.007 | 0.007 | 1.063 | 0.44 | 0.6 | 0.39 | 0.38 | 0.5 | 0.42 |
| 7 | 4321571 | rs10259929 | 0.298 | 1.021 | -0.74 | -0.054 | 0.001 | 0.003 | 0.003 | 0 | 0.21 | 0.15 | 0.41 | 0.51 | 0.19 | 0.27 |
| 7 | 4324730 | rs11762889 | -0.107 | 0.933 | -1.354 | -0.45 | 0.006 | -0.007 | -0.007 | 3.338 | 0.37 | 0.17 | 0.49 | 0.59 | 0.35 | 0.45 |

| | | | | | | | | | | | | | | | |
|---|---|---|---|---|---|---|---|---|---|---|---|---|---|---|---|
| 7 | 4324762 | rs4720226 | -0.125 | 1.229 | -1.143 | -0.543 | -0.01 | -0.004 | -0.004 | 0.148 | 0.17 | 0.18 | 0.28 | 0.21 | 0.34 | 0.15 |
| 7 | 4325870 | rs12701532 | 0.271 | 1 | -0.956 | -0.309 | 0.008 | 0.007 | 0.007 | 0 | 0.13 | 0.15 | 0.22 | 0.22 | 0.26 | 0.14 |
| 7 | 4327351 | rs4723588 | 0.645 | 1.016 | -1.082 | -0.655 | 0.006 | 0.017 | 0.017 | 0.783 | 0.18 | 0.15 | 0.11 | 0.01 | 0.16 | 0.16 |
| 7 | 4330892 | rs4720229 | -0.103 | 0.219 | -0.71 | -0.227 | -0.01 | 0.007 | 0.007 | 0 | 0.37 | 0.35 | 0.42 | 0.35 | 0.45 | 0.33 |
| 7 | 4332438 | rs4723604 | -0.547 | 0.003 | -0.745 | -0.042 | -0.001 | -0.024 | -0.024 | 0 | 0.66 | 0.66 | 0.8 | 0.88 | 0.71 | 0.69 |
| 7 | 4333275 | rs2189973 | 0.196 | 0.523 | -0.754 | 0.111 | 0.006 | -0.009 | -0.009 | 0 | 0.41 | 0.51 | 0.46 | 0.36 | 0.48 | 0.42 |
| 7 | 4333610 | rs10488357 | 0.196 | 0.523 | -0.872 | 0.111 | 0.003 | -0.005 | -0.005 | 0.099 | 0.41 | 0.47 | 0.46 | 0.36 | 0.48 | 0.42 |
| 7 | 4336477 | rs10232703 | -0.383 | 0.531 | -0.997 | 0.99 | -0.003 | 0.141 | 0.141 | 0 | 0.21 | 0.46 | 0.28 | 0.42 | 0.22 | 0.2 |
| 7 | 4337070 | rs11773770 | 0.056 | 0.622 | -2.245 | 0.825 | -0.011 | -0.069 | -0.069 | 1.454 | 0.17 | 0.05 | 0.29 | 0.63 | 0.22 | 0.2 |
| 7 | 4339768 | rs13241730 | -0.381 | 0.048 | -2.302 | 0.061 | 0.016 | 0.139 | 0.139 | 0 | 0.34 | 0.07 | 0.22 | 0.09 | 0.26 | 0.31 |
| 7 | 4349949 | rs2107834 | -0.737 | 0.893 | -1.998 | 0.776 | -0.053 | -0.048 | -0.048 | 0.044 | 0.13 | 0.02 | 0.3 | 0.26 | 0.21 | 0.18 |
| 7 | 4352417 | rs2097884 | 0.098 | -0.491 | -2.469 | 0.313 | 0.102 | 0.031 | 0.031 | 0 | 0.42 | 0.21 | 0.32 | 0.35 | 0.27 | 0.48 |
| 7 | 4356023 | rs12533379 | -0.593 | 0.042 | -2.319 | 0.17 | -0.038 | 0.02 | 0.02 | 0 | 0.14 | 0.03 | 0.24 | 0.1 | 0.19 | 0.21 |
| 7 | 4363764 | rs6462750 | 0.009 | -0.862 | -2.527 | -0.515 | 0.02 | 0.035 | 0.035 | 0 | 0.76 | 0.86 | 0.88 | 0.97 | 0.81 | 0.72 |
| 7 | 4366797 | rs6462752 | -0.413 | -1.931 | -3.166 | -0.692 | 0.025 | 0.009 | 0.009 | 0 | 0.39 | 0.73 | 0.48 | 0.38 | 0.32 | 0.36 |
| 7 | 4367700 | rs2158629 | 1.318 | -1.989 | -3.232 | -0.729 | -0.098 | -0.011 | -0.011 | 0 | 0.36 | 0.1 | 0.4 | 0.65 | 0.5 | 0.36 |
| 7 | 4370566 | rs4425656 | 0.138 | -2.269 | -3.237 | -1.149 | -0.028 | 0.02 | 0.02 | 0 | 0.17 | 0.02 | 0.25 | 0.11 | 0.19 | 0.21 |
| 7 | 4371785 | rs6462756 | -0.365 | -2.462 | -2.963 | -1.24 | 0.025 | 0.045 | 0.045 | 0 | 0.75 | 0.87 | 0.87 | 0.95 | 0.82 | 0.73 |
| 7 | 4372595 | rs12701554 | -0.121 | -2.506 | -2.243 | -1.125 | -0.014 | -0.051 | -0.051 | 0.099 | 0.73 | 0.68 | 0.84 | 0.89 | 0.82 | 0.75 |
| 7 | 4374804 | rs4723644 | 0.331 | -2.324 | -1.492 | -1.034 | 0.028 | 0.011 | 0.011 | 0.477 | 0.54 | 0.43 | 0.74 | 0.89 | 0.74 | 0.67 |
| 7 | 4379718 | rs10236427 | -0.389 | -1.819 | -1.437 | -0.011 | 0.001 | -0.05 | -0.05 | 0.523 | 0.27 | 0.13 | 0.39 | 0.39 | 0.24 | 0.28 |
| 7 | 4379761 | rs7794252 | -0.965 | -2.122 | -0.401 | 0.037 | -0.014 | 0.026 | 0.026 | 0 | 0.34 | 0.55 | 0.46 | 0.5 | 0.29 | 0.33 |
| 7 | 4381743 | rs17134702 | -0.387 | -1.951 | 0.262 | -0.033 | 0.002 | 0.064 | 0.064 | 2.831 | 0.16 | 0.28 | 0.24 | 0.31 | 0.14 | 0.22 |
| 7 | 4382159 | rs4723652 | -0.07 | -1.993 | 0.36 | 0.117 | -0.002 | -0.002 | -0.002 | 0.238 | 0.14 | 0.27 | 0.2 | 0.11 | 0.13 | 0.21 |
| 7 | 4384024 | rs7792601 | 0.013 | -1.302 | -0.518 | -0.597 | -0.008 | -0.08 | -0.08 | 0.561 | 0.72 | 0.67 | 0.83 | 0.96 | 0.78 | 0.8 |
| 7 | 4384233 | rs10268968 | 0.505 | -1.422 | -0.27 | -0.117 | -0.011 | -0.056 | -0.056 | 0 | 0.13 | 0.04 | 0.21 | 0.33 | 0.13 | 0.23 |
| 7 | 4386247 | rs6980275 | -0.071 | -1.601 | 0.011 | -0.486 | 0.037 | -0.01 | -0.01 | 0 | 0.07 | 0.29 | 0.06 | 0 | 0.06 | 0.12 |
| 7 | 4388142 | rs12534827 | 0.462 | -1.538 | -0.677 | -0.398 | 0 | -0.014 | -0.014 | 0 | 0.13 | 0.12 | 0.21 | 0.23 | 0.13 | 0.21 |
| 7 | 4389065 | rs10280936 | -1.6 | -1.674 | -1.025 | -0.582 | -0.096 | 0.013 | 0.013 | 0 | 0.57 | 0.92 | 0.53 | 0.33 | 0.46 | 0.56 |
| 7 | 4394389 | rs4723670 | 0.128 | -2.205 | -0.834 | -0.761 | 0.026 | 0.07 | 0.07 | 1.609 | 0.22 | 0.09 | 0.12 | 0.1 | 0.16 | 0.08 |
| 7 | 4398027 | rs4720253 | -0.231 | -2.219 | -0.86 | -0.331 | 0.006 | -0.093 | -0.093 | 0 | 0.22 | 0.11 | 0.3 | 0.34 | 0.17 | 0.3 |
| 7 | 4398350 | rs10240742 | -1.152 | -2.35 | -0.283 | -0.406 | -0.014 | 0.014 | 0.014 | 0 | 0.32 | 0.45 | 0.36 | 0.34 | 0.22 | 0.43 |
| 7 | 4399050 | rs2057891 | -2.064 | -2.31 | 0.024 | -0.426 | 0.001 | -0.044 | -0.044 | 0 | 0.59 | 0.8 | 0.51 | 0.44 | 0.4 | 0.51 |
| 7 | 4399743 | rs5021672 | -1.384 | -2.299 | 0.113 | -0.42 | 0.01 | -0.001 | -0.001 | 0 | 0.49 | 0.44 | 0.44 | 0.44 | 0.35 | 0.38 |
| 7 | 4401148 | rs10216277 | -0.245 | -2.139 | -0.244 | -0.374 | 0 | -0.043 | -0.043 | 0.575 | 0.09 | 0.34 | 0.06 | 0 | 0.05 | 0.13 |
| 7 | 4401550 | rs11760485 | -1.282 | -2.161 | -0.147 | -0.38 | 0.072 | -0.034 | -0.034 | 0 | 0.48 | 0.23 | 0.42 | 0.44 | 0.34 | 0.38 |
| 7 | 4402530 | rs6951614 | -1.848 | -2.028 | -0.153 | -0.257 | -0.004 | 0.002 | 0.002 | 0 | 0.57 | 0.59 | 0.49 | 0.51 | 0.4 | 0.52 |
| 7 | 4407330 | rs4236353 | -1.798 | -1.995 | -0.159 | -0.455 | -0.006 | 0.007 | 0.007 | 0 | 0.57 | 0.57 | 0.48 | 0.44 | 0.4 | 0.51 |
| 7 | 4408299 | rs11980802 | 0.282 | -2.369 | -0.124 | -0.482 | 0.021 | 0.052 | 0.052 | 0.779 | 0.23 | 0.11 | 0.12 | 0.09 | 0.18 | 0.07 |
| 7 | 4409754 | rs4723703 | -0.476 | -1.83 | 0.089 | -0.222 | -0.027 | -0.085 | -0.085 | 0 | 0.41 | 0.51 | 0.24 | 0.09 | 0.28 | 0.28 |
| 7 | 4411209 | rs10488360 | -0.866 | -1.834 | -0.058 | -0.183 | 0.096 | -0.021 | -0.021 | 3.541 | 0.48 | 0.32 | 0.42 | 0.43 | 0.3 | 0.36 |
| 7 | 4411496 | rs1557962 | 0.214 | -1.852 | -0.062 | -0.19 | 0.049 | 0.078 | 0.078 | 0.853 | 0.33 | 0.25 | 0.18 | 0.09 | 0.22 | 0.14 |
| 7 | 4411828 | rs7459304 | -1.062 | -1.418 | 0.434 | -0.129 | 0.026 | -0.017 | -0.017 | 0 | 0.56 | 0.71 | 0.5 | 0.43 | 0.37 | 0.49 |
| 7 | 4411996 | rs11763858 | -0.522 | -1.41 | 0.376 | -0.124 | 0.093 | -0.021 | -0.021 | 0 | 0.48 | 0.32 | 0.42 | 0.43 | 0.31 | 0.36 |
| 7 | 4413313 | rs2107842 | -0.658 | -0.643 | 0.251 | 0.328 | 0.019 | -0.003 | -0.003 | 0 | 0.51 | 0.62 | 0.49 | 0.43 | 0.37 | 0.49 |
| 7 | 4414803 | rs9639771 | -0.485 | -0.637 | 0.048 | 0.33 | 0.01 | -0.001 | -0.001 | 0 | 0.5 | 0.53 | 0.47 | 0.43 | 0.36 | 0.48 |
| 7 | 4418719 | rs314624 | -0.568 | -1.047 | -0.863 | 0.524 | 0.048 | -0.004 | -0.004 | 1.111 | 0.51 | 0.71 | 0.49 | 0.43 | 0.36 | 0.48 |
| 7 | 4419287 | rs314627 | -0.606 | -0.981 | -0.895 | 0.539 | 0.038 | -0.006 | -0.006 | 0 | 0.51 | 0.69 | 0.49 | 0.43 | 0.37 | 0.49 |
| 7 | 4424054 | rs314647 | -0.68 | -0.885 | -0.945 | 1.041 | 0.007 | -0.032 | -0.032 | 0.562 | 0.13 | 0.09 | 0.21 | 0.23 | 0.09 | 0.24 |
| 7 | 4425134 | rs314650 | -1.589 | -1.129 | -1.002 | 1.153 | 0.006 | -0.016 | -0.016 | 1.401 | 0.14 | 0.1 | 0.12 | 0.12 | 0.13 | 0.16 |
| 7 | 4434887 | rs6945672 | -1.384 | -2.422 | -1.354 | 0.993 | 0.152 | -0.05 | -0.05 | 0 | 0.43 | 0.8 | 0.43 | 0.5 | 0.31 | 0.51 |
| 7 | 4435534 | rs4723721 | -0.701 | -2.809 | -0.537 | 1.135 | -0.062 | -0.007 | -0.007 | 0 | 0.2 | 0.47 | 0.21 | 0.17 | 0.17 | 0.17 |
| 7 | 4435801 | rs11982232 | -1.107 | -1.896 | -0.44 | 1.178 | 0.004 | -0.007 | -0.007 | 0 | 0.21 | 0.2 | 0.21 | 0.33 | 0.14 | 0.34 |
| 7 | 4445873 | rs1447402 | -1.486 | -1.48 | 0.533 | 0.69 | 0.009 | -0.087 | -0.087 | 0.017 | 0.13 | 0.44 | 0.1 | 0 | 0.11 | 0.04 |
| 7 | 4452636 | rs10951550 | 0.818 | -1.375 | -0.549 | 0.634 | -0.024 | -0.017 | -0.017 | 0.657 | 0.42 | 0.24 | 0.45 | 0.53 | 0.47 | 0.35 |
| 7 | 4455097 | rs6958692 | -0.434 | -1.748 | -0.381 | 0.457 | -0.047 | 0.045 | 0.045 | 0.115 | 0.22 | 0.54 | 0.3 | 0.3 | 0.16 | 0.39 |
| 7 | 4457003 | rs917118 | -1.588 | -1.792 | -0.459 | 0.409 | 0.02 | 0.007 | 0.007 | 0.19 | 0.35 | 0.7 | 0.39 | 0.3 | 0.28 | 0.45 |
| 7 | 4457235 | rs12537127 | 1.606 | -2.028 | -0.549 | 0.627 | -0.111 | -0.078 | -0.078 | 0 | 0.35 | 0.11 | 0.41 | 0.45 | 0.41 | 0.31 |
| 7 | 4457707 | rs10261581 | -0.716 | -2.07 | -0.323 | 0.731 | -0.053 | 0.03 | 0.03 | 0.018 | 0.24 | 0.54 | 0.32 | 0.3 | 0.17 | 0.41 |
| 7 | 4463248 | rs7810278 | -1.376 | -2.065 | -0.779 | 0.713 | 0.072 | -0.047 | -0.047 | 0 | 0.35 | 0.83 | 0.4 | 0.38 | 0.3 | 0.44 |
| 7 | 4468250 | rs12674358 | 0.177 | -1.531 | -0.644 | 0.906 | 0 | 0.001 | 0.001 | 0 | 0.07 | 0.07 | 0.04 | 0.09 | 0.06 | 0.04 |
| 7 | 4469281 | rs17134816 | 1.248 | -1.181 | -0.771 | 0.796 | -0.077 | -0.07 | -0.07 | 0 | 0.33 | 0.14 | 0.39 | 0.45 | 0.38 | 0.28 |
| 7 | 4476950 | rs13247208 | 1.152 | -0.973 | -0.411 | 1.011 | -0.034 | -0.012 | -0.012 | 0.266 | 0.38 | 0.19 | 0.4 | 0.42 | 0.41 | 0.34 |
| 7 | 4477012 | rs314583 | 0.301 | -0.988 | -0.606 | 1.279 | 0.005 | -0.034 | -0.034 | 0 | 0.9 | 0.94 | 0.79 | 0.81 | 0.95 | 0.84 |
| 7 | 4477024 | rs314582 | 1.268 | -1.16 | -0.785 | 1.149 | -0.064 | 0.036 | 0.036 | 0 | 0.26 | 0.03 | 0.18 | 0.18 | 0.29 | 0.24 |
| 7 | 4478588 | rs7783362 | 0.089 | -1.194 | -0.351 | 1.107 | -0.008 | 0.008 | 0.008 | 0 | 0.59 | 0.76 | 0.55 | 0.64 | 0.58 | 0.58 |
| 7 | 4479140 | rs423854 | 1.273 | -1.135 | -0.487 | 1.165 | -0.074 | 0.006 | 0.006 | 0 | 0.26 | 0.01 | 0.18 | 0.18 | 0.28 | 0.25 |
| 7 | 4479253 | rs394647 | 1.255 | -1.136 | -0.486 | 1.161 | -0.064 | 0.019 | 0.019 | 0 | 0.26 | 0.03 | 0.18 | 0.18 | 0.3 | 0.25 |
| 7 | 4483857 | rs172707 | 1.725 | -1.659 | 0.356 | 1.139 | -0.009 | 0.035 | 0.035 | 0 | 0.71 | 0.47 | 0.63 | 0.61 | 0.73 | 0.6 |

| | | | | | | | | | | | | | | | | |
|---|---|---|---|---|---|---|---|---|---|---|---|---|---|---|---|---|
| 7 | 4487983 | rs314600 | -0.354 | -1.281 | 0.756 | 1.359 | 0.002 | -0.002 | -0.002 | 0 | 0.45 | 0.39 | 0.4 | 0.43 | 0.46 | 0.45 |
| 7 | 4490854 | rs314605 | 0.649 | -0.947 | 0.628 | 1.381 | -0.004 | 0.036 | 0.036 | 0 | 0.87 | 0.74 | 0.73 | 0.76 | 0.9 | 0.81 |
| 7 | 4495001 | rs6958864 | -0.731 | -0.968 | 0.734 | 1.463 | -0.01 | 0.067 | 0.067 | 0 | 0.12 | 0.22 | 0.09 | 0.18 | 0.1 | 0.08 |
| 7 | 4497688 | rs895710 | 1.228 | -1.017 | -0.133 | 1.309 | -0.037 | 0.019 | 0.019 | 0.039 | 0.37 | 0.1 | 0.28 | 0.31 | 0.37 | 0.3 |
| 7 | 4500723 | rs6952622 | -0.148 | -0.96 | -0.174 | 1.296 | -0.072 | 0.038 | 0.038 | 0 | 0.58 | 0.9 | 0.56 | 0.64 | 0.55 | 0.5 |
| 7 | 4501085 | rs10248664 | -1.089 | -0.982 | 0.298 | 1.319 | 0.018 | 0.08 | 0.08 | 0.01 | 0.15 | 0.28 | 0.16 | 0.3 | 0.24 | 0.24 |
| 7 | 4501781 | rs10951566 | 1.419 | -1.119 | -0.177 | 1.377 | -0.028 | 0.039 | 0.039 | 0.128 | 0.38 | 0.19 | 0.36 | 0.3 | 0.38 | 0.4 |
| 7 | 4502285 | rs17753780 | -0.779 | -1.059 | -0.237 | 1.41 | -0.041 | 0.062 | 0.062 | 0 | 0.65 | 0.91 | 0.61 | 0.77 | 0.66 | 0.69 |
| 7 | 4503061 | rs10951567 | 0.384 | -1.061 | 0.057 | 1.221 | -0.001 | -0.006 | -0.006 | 0 | 0.49 | 0.53 | 0.41 | 0.3 | 0.42 | 0.44 |
| 7 | 4503136 | rs10951568 | 1.021 | -1.017 | -0.824 | 1.211 | -0.003 | 0.029 | 0.029 | 0 | 0.41 | 0.33 | 0.38 | 0.29 | 0.38 | 0.4 |
| 7 | 4503153 | rs6964070 | -0.053 | -1.075 | -1.408 | 1.367 | -0.063 | -0.072 | -0.072 | 0 | 0.58 | 0.89 | 0.52 | 0.55 | 0.54 | 0.63 |
| 7 | 4508102 | rs1466066 | -1.093 | -0.602 | -1.213 | 1.334 | -0.048 | 0.051 | 0.051 | 1.101 | 0.63 | 0.76 | 0.61 | 0.76 | 0.57 | 0.62 |
| 7 | 4509683 | rs12155305 | -0.223 | 0.011 | -0.918 | 1.646 | 0 | 0.098 | 0.098 | 0 | 0.05 | 0.11 | 0.04 | 0.25 | 0.06 | 0.18 |
| 7 | 4513032 | rs12533818 | -0.332 | 0.133 | -1.404 | 0.868 | -0.038 | 0.009 | 0.009 | 0 | 0.34 | 0.11 | 0.44 | 0.3 | 0.34 | 0.34 |
| 7 | 4518110 | rs4720305 | 0.778 | -0.154 | -1.028 | 0.781 | 0.004 | 0.014 | 0.014 | 0 | 0.9 | 0.97 | 0.96 | 0.94 | 0.93 | 0.9 |
| 7 | 4525159 | rs6957311 | 0.696 | 0.071 | -1.285 | 0.885 | -0.039 | 0.03 | 0.03 | 0.576 | 0.64 | 0.83 | 0.6 | 0.71 | 0.61 | 0.45 |
| 7 | 4526692 | rs4723803 | -0.092 | 0.234 | -1.153 | 0.921 | -0.111 | 0.004 | 0.004 | 0.705 | 0.27 | 0.03 | 0.38 | 0.23 | 0.34 | 0.5 |
| 7 | 4530940 | rs11971741 | 1.091 | -0.336 | -0.221 | 0.575 | 0.004 | 0.001 | 0.001 | 0.019 | 0.52 | 0.39 | 0.53 | 0.48 | 0.57 | 0.37 |
| 7 | 4534797 | rs11771140 | 1.162 | -0.421 | -0.171 | 0.525 | 0.001 | 0 | 0 | 0 | 0.48 | 0.42 | 0.52 | 0.48 | 0.57 | 0.4 |
| 7 | 4539599 | rs7799050 | 0.898 | -0.542 | 0.084 | 0.51 | -0.008 | -0.009 | -0.009 | 0.581 | 0.55 | 0.38 | 0.5 | 0.5 | 0.53 | 0.36 |
| 7 | 4540687 | rs6975898 | 0.253 | -0.747 | 0.43 | 0.538 | 0.006 | -0.006 | -0.006 | 0.012 | 0.69 | 0.64 | 0.56 | 0.77 | 0.61 | 0.67 |
| 7 | 4544188 | rs7784269 | 1.348 | -1.072 | 0.628 | 0.452 | -0.004 | -0.012 | -0.012 | 0 | 0.55 | 0.28 | 0.49 | 0.52 | 0.58 | 0.62 |
| 7 | 4548541 | rs2197681 | 1.876 | -1.132 | 0.397 | 0.384 | -0.064 | -0.06 | -0.06 | 0 | 0.43 | 0.12 | 0.46 | 0.5 | 0.51 | 0.39 |
| 7 | 4552959 | rs1447398 | -0.196 | -0.543 | 0.69 | 0.575 | -0.027 | 0.061 | 0.061 | 0 | 0.75 | 0.82 | 0.63 | 0.87 | 0.71 | 0.77 |
| 7 | 4553216 | rs1447399 | -0.22 | -0.543 | 0.69 | 0.586 | -0.027 | 0.058 | 0.058 | 0 | 0.75 | 0.82 | 0.63 | 0.85 | 0.7 | 0.76 |
| 7 | 4553521 | rs10265380 | -1.434 | -0.602 | 0.801 | 0.617 | -0.001 | 0.1 | 0.1 | 0 | 0.18 | 0.31 | 0.12 | 0.33 | 0.14 | 0.17 |
| 7 | 4554285 | rs879946 | -0.272 | -0.454 | 0.586 | 0.43 | 0.05 | 0.046 | 0.046 | 0 | 0.08 | 0.21 | 0.08 | 0.1 | 0.11 | 0.11 |
| 7 | 4558774 | rs6977884 | -0.025 | -0.382 | -0.163 | 0.258 | -0.034 | 0.069 | 0.069 | 0 | 0.75 | 0.85 | 0.64 | 0.87 | 0.72 | 0.8 |
| 7 | 4560681 | rs10276022 | 0.981 | -0.292 | 0.067 | 0.413 | -0.002 | 0.002 | 0.002 | 0 | 0.55 | 0.32 | 0.5 | 0.58 | 0.57 | 0.64 |
| 7 | 4563881 | rs17134910 | -1.075 | -0.344 | 0.681 | -0.076 | -0.007 | -0.114 | -0.114 | 0 | 0.21 | 0.42 | 0.2 | 0.07 | 0.19 | 0.29 |
| 7 | 4565594 | rs17134911 | -0.724 | -0.83 | 0.415 | 0.35 | 0.025 | -0.032 | -0.032 | 0 | 0.09 | 0.03 | 0.03 | 0.24 | 0.02 | 0.05 |
| 7 | 4569239 | rs1117769 | -0.855 | -0.678 | 0.955 | -0.056 | -0.021 | -0.082 | -0.082 | 0 | 0.14 | 0.55 | 0.15 | 0.05 | 0.09 | 0.24 |
| 7 | 4580498 | rs10266190 | -0.921 | -0.986 | 0.514 | -0.016 | -0.02 | -0.031 | -0.031 | 1.044 | 0.37 | 0.46 | 0.53 | 0.23 | 0.3 | 0.37 |
| 7 | 4584559 | rs10232337 | -0.994 | -1.049 | 0.577 | -0.072 | 0.001 | 0 | 0 | 0 | 0.38 | 0.58 | 0.55 | 0.23 | 0.3 | 0.37 |
| 7 | 4589079 | rs4723872 | -0.681 | -1.171 | 0.294 | -0.062 | 0.019 | 0.03 | 0.03 | 0 | 0.34 | 0.13 | 0.5 | 0.25 | 0.3 | 0.39 |
| 7 | 4589812 | rs4343988 | 1.491 | -0.895 | -0.033 | 0.349 | -0.119 | -0.021 | -0.021 | 0 | 0.42 | 0.06 | 0.31 | 0.35 | 0.51 | 0.38 |
| 7 | 4591812 | rs4236360 | -0.602 | -0.781 | 0.371 | 0.536 | 0.004 | 0.11 | 0.11 | 0.321 | 0.82 | 0.6 | 0.84 | 0.6 | 0.82 | 0.77 |
| 7 | 4592492 | rs4596557 | 0.181 | -1.14 | 0.148 | 0.464 | -0.005 | 0.004 | 0.004 | 0 | 0.24 | 0.1 | 0.48 | 0.22 | 0.27 | 0.33 |
| 7 | 4593385 | rs7812186 | 0.17 | -0.996 | 0.091 | 0.508 | 0.012 | 0.084 | 0.084 | 0.07 | 0.75 | 0.43 | 0.8 | 0.6 | 0.76 | 0.73 |
| 7 | 4598404 | rs10238077 | -0.311 | -0.991 | 0.059 | 0.511 | 0.013 | 0.092 | 0.092 | 0.015 | 0.78 | 0.57 | 0.82 | 0.59 | 0.76 | 0.74 |
| 7 | 4600429 | rs4720331 | -0.944 | -0.83 | 0.147 | 0.55 | -0.015 | -0.021 | -0.021 | 0 | 0.38 | 0.51 | 0.53 | 0.25 | 0.29 | 0.38 |
| 7 | 4609838 | rs10241876 | -0.298 | -0.571 | 0.312 | 0.666 | 0.02 | 0.109 | 0.109 | 0.677 | 0.79 | 0.56 | 0.82 | 0.59 | 0.77 | 0.75 |
| 7 | 4611655 | rs4723895 | 0.003 | -0.441 | -0.085 | 0.724 | 0.003 | 0.067 | 0.067 | 0.855 | 0.76 | 0.39 | 0.8 | 0.59 | 0.77 | 0.75 |
| 7 | 4613341 | rs1880121 | 0.003 | -0.441 | -0.25 | 0.724 | 0.001 | 0.053 | 0.053 | 0.855 | 0.76 | 0.37 | 0.8 | 0.59 | 0.77 | 0.75 |
| 7 | 4644965 | rs2343561 | 1.285 | -0.458 | -0.837 | 0.611 | -0.078 | -0.001 | -0.001 | 1.63 | 0.4 | 0.09 | 0.3 | 0.35 | 0.49 | 0.43 |
| 7 | 4647262 | rs12701792 | 0.647 | 0.016 | -0.581 | 0.503 | -0.003 | -0.003 | -0.003 | 0.176 | 0.55 | 0.36 | 0.34 | 0.53 | 0.54 | 0.64 |
| 7 | 4651184 | rs1015123 | -0.439 | 0.074 | -0.88 | 0.187 | -0.091 | 0.006 | 0.006 | 0.081 | 0.2 | 0.01 | 0.46 | 0.15 | 0.25 | 0.13 |
| 7 | 4658125 | rs10262589 | -1.372 | -0.11 | -0.123 | 0.55 | -0.01 | -0.006 | -0.006 | 0 | 0.14 | 0.13 | 0.42 | 0.19 | 0.2 | 0.25 |
| 7 | 4659324 | rs7786722 | -1.607 | -0.412 | 0.32 | 0.48 | -0.005 | -0.012 | -0.012 | 2.135 | 0.27 | 0.51 | 0.47 | 0.24 | 0.23 | 0.33 |
| 7 | 4667201 | rs10259797 | -1.585 | -0.471 | 0.053 | 0.532 | -0.007 | -0.006 | -0.006 | 0 | 0.27 | 0.61 | 0.48 | 0.22 | 0.24 | 0.33 |
| 7 | 4686519 | rs11977885 | -1.312 | -0.555 | -0.434 | 0.953 | -0.005 | 0.055 | 0.055 | 0 | 0.23 | 0.4 | 0.29 | 0.32 | 0.2 | 0.34 |
| 7 | 4691867 | rs17135014 | -0.863 | -0.402 | -0.814 | 1.011 | -0.001 | -0.034 | -0.034 | 0 | 0.1 | 0.02 | 0.11 | 0.14 | 0.08 | 0.11 |
| 7 | 4708134 | rs17828856 | 0.035 | -0.673 | -0.878 | 0.872 | -0.008 | -0.028 | -0.028 | 0.289 | 0.13 | 0.04 | 0.15 | 0.15 | 0.12 | 0.26 |
| 7 | 4711169 | rs17135019 | -1.093 | -0.706 | -0.73 | 0.824 | -0.005 | 0.013 | 0.013 | 0 | 0.22 | 0.27 | 0.2 | 0.28 | 0.18 | 0.34 |
| 7 | 4718631 | rs1001396 | 0.342 | -0.99 | -1.587 | 0.879 | 0 | -0.073 | -0.073 | 0 | 0.12 | 0 | 0.19 | 0.33 | 0.11 | 0.16 |
| 7 | 4723534 | rs9791644 | -0.299 | -0.716 | -1.591 | -0.478 | -0.005 | 0.018 | 0.018 | 0 | 0.17 | 0.06 | 0.1 | 0.09 | 0.16 | 0.17 |
| 7 | 4727980 | rs6964158 | -0.512 | -0.514 | -1.47 | -0.689 | 0.004 | 0.025 | 0.025 | 0 | 0.38 | 0.8 | 0.32 | 0.37 | 0.34 | 0.34 |
| 7 | 4740621 | rs10259192 | -0.835 | -0.323 | -0.444 | -1.058 | -0.028 | -0.084 | -0.084 | 0.249 | 0.16 | 0.64 | 0.14 | 0 | 0.1 | 0.06 |
| 7 | 4749527 | rs10259435 | -0.65 | -0.359 | -0.4 | -1.094 | -0.019 | -0.057 | -0.057 | 0 | 0.13 | 0.22 | 0.06 | 0 | 0.07 | 0.03 |
| 7 | 4759542 | rs4523152 | -0.828 | 0.187 | -0.408 | -1.412 | 0.005 | -0.061 | -0.061 | 3.972 | 0.36 | 0.46 | 0.22 | 0.13 | 0.36 | 0.28 |
| 7 | 4760696 | rs4623321 | 0.895 | 0.343 | -1.071 | -1.496 | 0.013 | 0.026 | 0.026 | 0 | 0.47 | 0.36 | 0.51 | 0.75 | 0.4 | 0.51 |
| 7 | 4765018 | rs6973046 | 0.447 | -0.136 | -1.134 | -1.844 | -0.013 | 0.06 | 0.06 | 0 | 0.44 | 0.36 | 0.57 | 0.83 | 0.46 | 0.69 |
| 7 | 4768038 | rs7788807 | 0.093 | -0.062 | -1.343 | -1.925 | 0.01 | 0.001 | 0.001 | 1.101 | 0.14 | 0.16 | 0.06 | 0 | 0.08 | 0.02 |
| 7 | 4780514 | rs3087749 | 0.275 | 0.251 | -1.335 | -1.711 | -0.012 | 0.086 | 0.086 | 0.068 | 0.45 | 0.32 | 0.56 | 0.85 | 0.48 | 0.7 |
| 7 | 4784652 | rs7784748 | -0.052 | 0.533 | -1.033 | -1.351 | 0 | -0.033 | -0.033 | 0 | 0.64 | 0.39 | 0.7 | 0.86 | 0.6 | 0.76 |
| 7 | 4787092 | rs12701915 | -0.332 | 0.754 | -1.934 | -0.692 | -0.031 | -0.029 | -0.029 | 0 | 0.42 | 0.06 | 0.44 | 0.64 | 0.41 | 0.43 |
| 7 | 4787612 | rs9690101 | 0.173 | 0.759 | -2.463 | -0.641 | -0.111 | -0.035 | -0.035 | 0.02 | 0.65 | 0.94 | 0.55 | 0.65 | 0.57 | 0.62 |
| 7 | 4789525 | rs11763558 | -0.964 | -0.199 | -0.553 | -0.813 | -0.019 | -0.063 | -0.063 | 0.542 | 0.15 | 0.39 | 0.07 | 0.07 | 0.09 | 0.25 |
| 7 | 4799694 | rs6463034 | -1.546 | -0.235 | 0.27 | -0.93 | -0.019 | -0.092 | -0.092 | 0.063 | 0.19 | 0.63 | 0.1 | 0.07 | 0.12 | 0.27 |

| | | | | | | | | | | | | | | | | |
|---|---|---|---|---|---|---|---|---|---|---|---|---|---|---|---|---|
| 7 | 4805137 | rs11976296 | -1.867 | -0.702 | 0.797 | -0.931 | -0.005 | -0.054 | -0.054 | 0 | 0.1 | 0.54 | 0.06 | 0 | 0.05 | 0.04 |
| 7 | 4805376 | rs17135086 | -2.176 | -0.814 | 1.229 | -0.979 | -0.012 | -0.048 | -0.048 | 0 | 0.1 | 0.56 | 0.06 | 0 | 0.03 | 0.03 |
| 7 | 4809660 | rs17135096 | -2.328 | -0.776 | 1.266 | -1.032 | -0.036 | -0.069 | -0.069 | 0 | 0.19 | 0.72 | 0.11 | 0.05 | 0.08 | 0.24 |
| 7 | 4810801 | rs11628 | -2.407 | -0.785 | 0.982 | -1.036 | -0.011 | -0.029 | -0.029 | 0 | 0.19 | 0.79 | 0.11 | 0.05 | 0.09 | 0.24 |
| 7 | 4813383 | rs10807911 | 0.9 | -0.26 | 0.489 | -0.983 | -0.074 | -0.048 | -0.048 | 0 | 0.15 | 0.03 | 0.26 | 0.2 | 0.3 | 0.11 |
| 7 | 4814613 | rs17135102 | -1.716 | -1.569 | 1.672 | -0.96 | -0.015 | -0.056 | -0.056 | 0 | 0.14 | 0.24 | 0.07 | 0.05 | 0.08 | 0.24 |
| 7 | 4815603 | rs3750014 | -2.767 | -1.61 | 1.426 | -0.979 | -0.001 | -0.018 | -0.018 | 0 | 0.2 | 0.8 | 0.11 | 0.05 | 0.09 | 0.24 |
| 7 | 4817952 | rs17135113 | -1.185 | -1.521 | 1.393 | -0.936 | -0.004 | -0.026 | -0.026 | 0.362 | 0.08 | 0.11 | 0.05 | 0 | 0.05 | 0.04 |
| 7 | 4820581 | rs17135115 | -0.284 | -1.597 | 1.326 | -0.914 | 0 | -0.001 | -0.001 | 0 | 0.09 | 0.01 | 0.03 | 0.07 | 0.07 | 0.21 |
| 7 | 4823971 | rs17135121 | -1.451 | -1.584 | 1.612 | -1.034 | -0.032 | -0.083 | -0.083 | 0 | 0.16 | 0.61 | 0.1 | 0.05 | 0.1 | 0.23 |
| 7 | 4825686 | rs7801122 | -1.356 | -1.444 | 1.514 | -0.962 | -0.021 | -0.064 | -0.064 | 0 | 0.19 | 0.29 | 0.1 | 0.05 | 0.12 | 0.25 |
| 7 | 4828593 | rs3750012 | -1.402 | -1.428 | 0.782 | -0.955 | -0.036 | -0.09 | -0.09 | 2.401 | 0.21 | 0.33 | 0.1 | 0.05 | 0.12 | 0.26 |
| 7 | 4845701 | rs6963791 | -1.371 | -1.556 | -0.552 | -0.966 | 0.025 | 0.01 | 0.01 | 0 | 0.35 | 0.71 | 0.18 | 0.05 | 0.21 | 0.33 |
| 7 | 4869751 | rs9655042 | -0.803 | -1.44 | -0.729 | 0.143 | 0.003 | -0.001 | -0.001 | 2.162 | 0.36 | 0.59 | 0.19 | 0.39 | 0.23 | 0.5 |
| 7 | 4876057 | rs3763384 | -0.224 | -0.985 | -1.438 | 0.015 | -0.04 | -0.051 | -0.051 | 2.546 | 0.32 | 0.04 | 0.29 | 0.33 | 0.29 | 0.4 |
| 7 | 4877012 | rs3763381 | -1.218 | -0.92 | -0.8 | 0.073 | -0.009 | 0.002 | 0.002 | 0.996 | 0.55 | 0.77 | 0.45 | 0.5 | 0.46 | 0.65 |
| 7 | 4880954 | rs13243520 | -0.908 | -1.301 | -0.917 | 0.021 | 0.05 | 0.032 | 0.032 | 0.019 | 0.52 | 0.83 | 0.42 | 0.45 | 0.38 | 0.58 |
| 7 | 4881925 | rs6958275 | -1.14 | -1.465 | -0.947 | -0.044 | -0.028 | 0.038 | 0.038 | 0.052 | 0.54 | 0.88 | 0.45 | 0.4 | 0.45 | 0.58 |
| 7 | 4883121 | rs17135213 | 0.119 | -1.774 | -1.08 | -0.082 | -0.062 | 0.001 | 0.001 | 0 | 0.34 | 0.02 | 0.3 | 0.27 | 0.31 | 0.41 |
| 7 | 4885221 | rs6964780 | -0.923 | -1.871 | -0.353 | -0.042 | 0.052 | 0.07 | 0.07 | 0 | 0.51 | 0.74 | 0.43 | 0.41 | 0.41 | 0.55 |
| 7 | 4893679 | rs10235539 | -0.865 | -2.16 | -0.116 | -0.102 | 0.05 | 0.045 | 0.045 | 0 | 0.48 | 0.67 | 0.37 | 0.41 | 0.34 | 0.54 |
| 7 | 4897687 | rs11972290 | -1.046 | -1.98 | 1.012 | -0.097 | 0.055 | 0.039 | 0.039 | 0.035 | 0.08 | 0.3 | 0.07 | 0.11 | 0.08 | 0.14 |
| 7 | 4899368 | rs17135241 | -1.005 | -1.956 | 1.135 | -0.094 | 0.029 | 0.015 | 0.015 | 0 | 0.07 | 0.17 | 0.06 | 0.11 | 0.08 | 0.14 |
| 7 | 4900095 | rs1553960 | -0.97 | -2.079 | 1.26 | -0.105 | 0.022 | 0.02 | 0.02 | 0 | 0.48 | 0.55 | 0.36 | 0.41 | 0.34 | 0.54 |
| 7 | 4900341 | rs17135247 | 0.146 | -2.081 | 1.25 | -0.093 | 0.035 | -0.015 | -0.015 | 0 | 0.37 | 0.03 | 0.26 | 0.31 | 0.25 | 0.4 |
| 7 | 4901191 | rs3750010 | -1.478 | -2.007 | 1.624 | -0.079 | 0.056 | 0.027 | 0.027 | 0 | 0.08 | 0.31 | 0.08 | 0.11 | 0.09 | 0.14 |
| 7 | 4903574 | rs7794427 | -0.024 | -2.237 | 1.302 | -0.025 | 0.032 | 0.018 | 0.018 | 0 | 0.38 | 0.21 | 0.28 | 0.3 | 0.25 | 0.4 |
| 7 | 4906887 | rs10240422 | -0.332 | -2.664 | 0.866 | -0.034 | 0.079 | 0.047 | 0.047 | 0 | 0.4 | 0.15 | 0.37 | 0.33 | 0.26 | 0.43 |
| 7 | 4909983 | rs10266401 | -1.506 | -2.912 | 0.254 | 0.586 | 0.008 | 0.094 | 0.094 | 0 | 0.58 | 0.96 | 0.64 | 0.81 | 0.36 | 0.62 |
| 7 | 4914358 | rs11973654 | -0.942 | -2.578 | 0.738 | 0.342 | 0.12 | 0.082 | 0.082 | 0.175 | 0.49 | 0.84 | 0.45 | 0.4 | 0.3 | 0.51 |
| 7 | 4918897 | rs6944694 | -1.714 | -2.578 | 1.232 | 0.368 | -0.024 | -0.012 | -0.012 | 0 | 0.14 | 0.55 | 0.07 | 0.14 | 0.06 | 0.14 |
| 7 | 4919277 | rs4724137 | -0.365 | -2.45 | 0.991 | 0.395 | 0.02 | 0.011 | 0.011 | 0 | 0.39 | 0.4 | 0.4 | 0.35 | 0.27 | 0.46 |
| 7 | 4923925 | rs6463152 | -0.319 | -2.175 | 0.546 | 0.766 | 0.012 | -0.009 | -0.009 | 0 | 0.37 | 0.3 | 0.39 | 0.46 | 0.26 | 0.41 |
| 7 | 4924960 | rs7782302 | -0.665 | -1.59 | 0.317 | 0.669 | 0.018 | -0.001 | -0.001 | 0 | 0.57 | 0.65 | 0.47 | 0.46 | 0.33 | 0.45 |
| 7 | 4925012 | rs7782329 | -0.788 | -0.517 | 0.183 | 0.612 | 0.001 | -0.019 | -0.019 | 0 | 0.15 | 0.22 | 0.08 | 0.13 | 0.11 | 0.15 |
| 7 | 4929721 | rs393275 | -0.569 | 0.887 | -0.111 | 0.459 | 0.002 | -0.007 | -0.007 | 0 | 0.38 | 0.61 | 0.3 | 0.28 | 0.42 | 0.47 |
| 7 | 4931576 | rs13222836 | -0.904 | 0.935 | -0.835 | 0.969 | 0.066 | 0.003 | 0.003 | 0.356 | 0.48 | 0.27 | 0.55 | 0.58 | 0.56 | 0.65 |
| 7 | 4935672 | rs2344727 | -0.262 | 1.265 | -0.856 | 0.893 | -0.071 | 0.064 | 0.064 | 0.09 | 0.35 | 0.14 | 0.25 | 0.26 | 0.42 | 0.46 |
| 7 | 4937916 | rs10229311 | -0.529 | 0.983 | -0.157 | 1.3 | 0.085 | 0.213 | 0.213 | 0 | 0.13 | 0.48 | 0.34 | 0.32 | 0.15 | 0.14 |
| 7 | 4942184 | rs300532 | -0.469 | 0.996 | -1.35 | 0.63 | 0.008 | 0.02 | 0.02 | 2.335 | 0.31 | 0.22 | 0.16 | 0.26 | 0.25 | 0.43 |
| 7 | 4946175 | rs372454 | 0.864 | 0.894 | -2.174 | 0.262 | -0.066 | 0.232 | 0.232 | 1.161 | 0.46 | 0.04 | 0.32 | 0.17 | 0.48 | 0.38 |
| 7 | 4946226 | rs41399345 | -0.115 | 0.649 | -1.107 | 1.076 | 0.007 | -0.011 | -0.011 | 0 | 0.11 | 0.05 | 0.05 | 0.19 | 0.04 | 0.11 |
| 7 | 4959807 | rs932061 | 0.051 | 0.74 | -1.104 | 0.593 | 0.007 | 0.016 | 0.016 | 0 | 0.46 | 0.62 | 0.6 | 0.3 | 0.41 | 0.28 |
| 7 | 4962688 | rs12671040 | -0.243 | 1.02 | -1.516 | 0.567 | 0.01 | -0.011 | -0.011 | 0 | 0.63 | 0.56 | 0.45 | 0.67 | 0.45 | 0.53 |
| 7 | 4964002 | rs2089967 | -0.258 | 1.01 | -1.445 | 0.577 | 0.01 | -0.01 | -0.01 | 0 | 0.63 | 0.57 | 0.45 | 0.67 | 0.45 | 0.53 |
| 7 | 4967663 | rs13224720 | 0.914 | 1.889 | -1.862 | 0.538 | -0.075 | -0.075 | -0.075 | 0 | 0.12 | 0 | 0.12 | 0.26 | 0.19 | 0.28 |
| 7 | 4968125 | rs11761526 | 0.452 | 1.392 | -1.745 | 0.741 | 0.002 | -0.071 | -0.071 | 0 | 0.22 | 0.07 | 0.12 | 0.3 | 0.15 | 0.28 |
| 7 | 4968943 | rs2965040 | 0.067 | 0.68 | -1.757 | 0.016 | 0.023 | 0.111 | 0.111 | 0 | 0.5 | 0.72 | 0.48 | 0.7 | 0.56 | 0.6 |
| 7 | 4972973 | rs2965042 | -0.044 | 0.793 | -1.473 | -0.013 | -0.012 | -0.002 | -0.002 | 0 | 0.69 | 0.72 | 0.54 | 0.68 | 0.67 | 0.64 |
| 7 | 4974485 | rs4573156 | -0.66 | 0.62 | -1.275 | 0.003 | 0.008 | 0.03 | 0.03 | 0.107 | 0.45 | 0.71 | 0.58 | 0.65 | 0.38 | 0.48 |
| 7 | 4975010 | rs2137502 | 0.846 | 0.364 | -1.366 | 0.23 | -0.015 | 0.134 | 0.134 | 0 | 0.17 | 0.41 | 0.18 | 0.32 | 0.13 | 0.16 |
| 7 | 4975956 | rs10240589 | 0.813 | 0.667 | -1.821 | 0.429 | 0.014 | -0.058 | -0.058 | 0.495 | 0.2 | 0.07 | 0.12 | 0.32 | 0.14 | 0.28 |
| 7 | 4992952 | rs7800341 | 0.331 | 0.93 | -1.788 | 0.156 | -0.122 | -0.031 | -0.031 | 0.193 | 0.33 | 0.06 | 0.29 | 0.33 | 0.46 | 0.42 |
| 7 | 5003373 | rs907719 | 0.26 | 1.186 | -1.844 | 0.2 | -0.057 | 0.028 | 0.028 | 0 | 0.35 | 0.12 | 0.45 | 0.31 | 0.45 | 0.41 |
| 7 | 5010993 | rs2942562 | -0.12 | 0.6 | -2.366 | 0.341 | 0.011 | 0.08 | 0.08 | 0 | 0.52 | 0.13 | 0.59 | 0.71 | 0.59 | 0.72 |
| 7 | 5014218 | rs11981905 | 0.371 | -0.075 | -1.763 | -0.241 | 0.017 | 0.047 | 0.047 | 0.328 | 0.57 | 0.68 | 0.57 | 0.31 | 0.64 | 0.5 |
| 7 | 5022223 | rs6463205 | 0.803 | -0.26 | -1.712 | -0.398 | -0.009 | -0.034 | -0.034 | 2.296 | 0.13 | 0.2 | 0.1 | 0.01 | 0.1 | 0.08 |
| 7 | 5031771 | rs13235865 | 1.528 | -0.357 | -1.881 | -0.261 | -0.021 | -0.104 | -0.104 | 0 | 0.16 | 0.01 | 0.15 | 0.39 | 0.14 | 0.28 |
| 7 | 5036641 | rs9639892 | -0.389 | -0.358 | -1.042 | -0.429 | -0.002 | 0.019 | 0.019 | 0 | 0.48 | 0.46 | 0.43 | 0.64 | 0.44 | 0.51 |
| 7 | 5037483 | rs7826 | 1.424 | -0.133 | -1.642 | -0.71 | -0.001 | 0.089 | 0.089 | 0 | 0.27 | 0.12 | 0.21 | 0.06 | 0.32 | 0.2 |
| 7 | 5041931 | rs12671224 | 0.935 | -0.365 | -1.329 | -0.297 | -0.002 | -0.042 | -0.042 | 0.064 | 0.13 | 0.07 | 0.16 | 0.26 | 0.1 | 0.15 |
| 7 | 5045169 | rs4455734 | 0.872 | -0.48 | -1.328 | 0.091 | -0.002 | -0.028 | -0.028 | 0.204 | 0.1 | 0.06 | 0.05 | 0.16 | 0.08 | 0.14 |
| 7 | 5050816 | rs7786822 | -1.008 | -0.42 | -0.771 | -0.057 | 0.004 | 0.001 | 0.001 | 0.089 | 0.62 | 0.64 | 0.41 | 0.55 | 0.5 | 0.54 |
| 7 | 5052357 | rs6977895 | -0.846 | -0.477 | -0.55 | -0.165 | -0.002 | -0.001 | -0.001 | 0 | 0.59 | 0.45 | 0.4 | 0.42 | 0.5 | 0.49 |
| 7 | 5070502 | rs10258089 | 1.574 | -0.528 | -0.499 | -0.499 | 0.006 | -0.056 | -0.056 | 2.85 | 0.17 | 0.2 | 0.11 | 0 | 0.19 | 0.08 |
| 7 | 5100411 | rs10268122 | 0.371 | -1.197 | -0.284 | -0.834 | 0.004 | -0.012 | -0.012 | 0 | 0.18 | 0.2 | 0.09 | 0.01 | 0.11 | 0.07 |
| 7 | 5101752 | rs11979787 | 1.681 | -1.311 | -0.493 | -0.741 | 0.008 | 0.004 | 0.004 | 0 | 0.09 | 0.05 | 0.12 | 0.37 | 0.09 | 0.16 |
| 7 | 5104275 | rs12674330 | 1.461 | -1.232 | -0.472 | -0.757 | 0.007 | -0.073 | -0.073 | 0.482 | 0.16 | 0.04 | 0.15 | 0.37 | 0.13 | 0.23 |
| 7 | 5104359 | rs12668616 | 1.461 | -1.232 | -0.472 | -0.757 | 0.007 | -0.073 | -0.073 | 0 | 0.16 | 0.04 | 0.15 | 0.37 | 0.13 | 0.23 |

| | | | | | | | | | | | | | | | | |
|---|---|---|---|---|---|---|---|---|---|---|---|---|---|---|---|---|
| 7 | 5105882 | rs9690559 | 1.447 | -1.224 | -0.448 | -0.754 | 0.008 | -0.07 | -0.07 | 0.107 | 0.17 | 0.05 | 0.15 | 0.37 | 0.13 | 0.23 |
| 7 | 5106222 | rs7808531 | -0.217 | -1.07 | 0.053 | -0.979 | 0.001 | 0.001 | 0.001 | 0 | 0.46 | 0.47 | 0.33 | 0.54 | 0.41 | 0.48 |
| 7 | 5106686 | rs4585633 | -1.385 | -0.824 | -0.174 | -0.854 | -0.005 | -0.002 | -0.002 | 0 | 0.63 | 0.69 | 0.42 | 0.55 | 0.51 | 0.55 |
| 7 | 5107777 | rs7794683 | -1.561 | -0.799 | -0.561 | -0.848 | -0.065 | -0.037 | -0.037 | 0.164 | 0.64 | 0.89 | 0.43 | 0.55 | 0.51 | 0.55 |
| 7 | 5107872 | rs7778444 | -1.617 | -0.774 | -0.593 | -0.852 | -0.074 | -0.043 | -0.043 | 0 | 0.65 | 0.9 | 0.43 | 0.55 | 0.51 | 0.55 |
| 7 | 5108770 | rs1127434 | -1.588 | -0.734 | -0.576 | -0.835 | -0.067 | -0.04 | -0.04 | 0.013 | 0.65 | 0.9 | 0.43 | 0.56 | 0.52 | 0.55 |
| 7 | 5112554 | rs1130329 | -0.839 | -0.652 | -0.48 | -0.646 | 0.016 | 0.025 | 0.025 | 2.975 | 0.46 | 0.43 | 0.27 | 0.29 | 0.34 | 0.48 |
| 7 | 5120148 | rs6961543 | 0.116 | -0.185 | -0.44 | -0.305 | -0.022 | -0.076 | -0.076 | 0 | 0.26 | 0.41 | 0.16 | 0.03 | 0.17 | 0.18 |
| 7 | 5122645 | rs7795386 | 0.531 | 0.165 | -1.588 | -0.206 | -0.059 | 0.024 | 0.024 | 0 | 0.39 | 0.08 | 0.21 | 0.31 | 0.35 | 0.36 |
| 7 | 5124934 | rs6958550 | -0.304 | 1.702 | -0.576 | 0.762 | -0.009 | -0.004 | -0.004 | 0 | 0.64 | 0.72 | 0.4 | 0.59 | 0.51 | 0.53 |
| 7 | 5133936 | rs12667108 | 0.064 | 2.49 | -0.346 | 1.291 | -0.054 | -0.065 | -0.065 | 0 | 0.11 | 0.02 | 0.11 | 0.32 | 0.17 | 0.13 |
| 7 | 5144105 | rs4720492 | 1.139 | 2.374 | 0.156 | 1.107 | 0.046 | 0.065 | 0.065 | 0 | 0.7 | 0.52 | 0.48 | 0.6 | 0.64 | 0.57 |
| 7 | 5146319 | rs4724358 | 0.295 | 2.647 | 0.222 | 1.315 | 0.073 | 0.112 | 0.112 | 0 | 0.84 | 0.7 | 0.55 | 0.63 | 0.73 | 0.63 |
| 7 | 5150581 | rs7798496 | 0.822 | 2.019 | -0.036 | 0.488 | 0.01 | -0.034 | -0.034 | 0.852 | 0.61 | 0.5 | 0.38 | 0.26 | 0.49 | 0.47 |
| 7 | 5153283 | rs6463266 | 1.2 | 2.154 | 0.546 | 0.705 | 0.134 | 0.086 | 0.086 | 0 | 0.87 | 0.56 | 0.68 | 0.75 | 0.8 | 0.78 |
| 7 | 5167160 | rs9655048 | 0.497 | 1.323 | -0.783 | 0.275 | 0 | -0.001 | -0.001 | 0 | 0.4 | 0.61 | 0.38 | 0.57 | 0.44 | 0.3 |
| 7 | 5177152 | rs13227672 | -0.34 | 0.76 | -1.565 | 0.001 | 0.027 | 0.071 | 0.071 | 0 | 0.39 | 0.17 | 0.18 | 0.25 | 0.35 | 0.34 |
| 7 | 5198328 | rs10248323 | -0.337 | 0.235 | -1.781 | -0.494 | 0.04 | 0.165 | 0.165 | 0 | 0.29 | 0.07 | 0.13 | 0.01 | 0.21 | 0.22 |
| 7 | 5214016 | rs4724430 | -0.095 | 0.905 | -1.866 | 0.481 | -0.073 | -0.1 | -0.1 | 0 | 0.25 | 0.08 | 0.44 | 0.45 | 0.3 | 0.34 |
| 7 | 5215853 | rs4724432 | 0.341 | 0.213 | -0.855 | -0.162 | -0.039 | -0.086 | -0.086 | 0 | 0.83 | 0.64 | 0.89 | 0.95 | 0.88 | 0.82 |
| 7 | 5220675 | rs4352737 | -0.913 | -0.2 | -0.8 | -0.024 | -0.033 | 0.017 | 0.017 | 0 | 0.55 | 0.78 | 0.49 | 0.59 | 0.47 | 0.58 |
| 7 | 5222956 | rs7790378 | 0.53 | -0.122 | -0.353 | 0.154 | 0.016 | 0.021 | 0.021 | 0 | 0.52 | 0.66 | 0.44 | 0.62 | 0.64 | 0.52 |
| 7 | 5224407 | rs6955907 | -0.685 | 0.253 | -0.298 | 0.318 | 0.047 | 0.099 | 0.099 | 0.906 | 0.12 | 0.36 | 0.09 | 0.26 | 0.17 | 0.21 |
| 7 | 5240541 | rs4591933 | 0.303 | 0.044 | -0.587 | 0.125 | -0.001 | 0.01 | 0.01 | 0.017 | 0.63 | 0.71 | 0.47 | 0.72 | 0.66 | 0.59 |
| 7 | 5254575 | rs6956071 | 1.001 | -0.129 | -1.072 | -0.343 | -0.001 | 0.001 | 0.001 | 0 | 0.38 | 0.55 | 0.3 | 0.41 | 0.41 | 0.38 |
| 7 | 5255294 | rs4236378 | -0.553 | -0.181 | -1.21 | -0.461 | 0.012 | 0.033 | 0.033 | 0 | 0.79 | 0.86 | 0.86 | 0.96 | 0.83 | 0.89 |
| 7 | 5261560 | rs4724461 | 0.065 | -0.175 | -1.343 | -0.553 | 0.019 | 0.139 | 0.139 | 0 | 0.71 | 0.89 | 0.83 | 0.96 | 0.8 | 0.87 |
| 7 | 5267524 | rs4720539 | 0.472 | 0.285 | -1.041 | -0.133 | -0.002 | -0.001 | -0.001 | 0 | 0.59 | 0.57 | 0.42 | 0.41 | 0.57 | 0.46 |
| 7 | 5268516 | rs11761408 | -0.19 | 0.724 | -1.328 | 0.042 | 0.008 | 0.043 | 0.043 | 0 | 0.73 | 0.81 | 0.84 | 0.96 | 0.77 | 0.84 |
| 7 | 5272177 | rs12472 | 0.667 | 1.091 | -1.393 | 0.885 | -0.027 | -0.037 | -0.037 | 0 | 0.2 | 0.07 | 0.29 | 0.24 | 0.2 | 0.28 |
| 7 | 5272927 | rs6949942 | -0.613 | 1.165 | -1.354 | 0.869 | 0.011 | 0.044 | 0.044 | 0.191 | 0.11 | 0.17 | 0.17 | 0.3 | 0.15 | 0.17 |
| 7 | 5273464 | rs13244634 | 0.139 | 0.789 | -1.931 | 0.549 | -0.03 | -0.039 | -0.039 | 0.344 | 0.24 | 0.07 | 0.4 | 0.25 | 0.25 | 0.29 |
| 7 | 5278439 | rs7808703 | 0.108 | 0.243 | -1.777 | 0.115 | 0.022 | 0.016 | 0.016 | 0 | 0.84 | 0.95 | 0.95 | 0.88 | 0.9 | 0.77 |
| 7 | 5279887 | rs13247861 | -0.413 | 1.21 | -1.395 | 0.724 | -0.054 | -0.012 | -0.012 | 0 | 0.34 | 0.18 | 0.55 | 0.61 | 0.47 | 0.39 |
| 7 | 5303440 | rs6463343 | -0.129 | 0.967 | -1.056 | 0.81 | 0.002 | -0.001 | -0.001 | 0 | 0.42 | 0.33 | 0.68 | 0.54 | 0.57 | 0.43 |
| 7 | 5319387 | rs10279978 | 0.151 | 0.864 | -1.083 | 0.848 | 0.003 | 0.003 | 0.003 | 0.103 | 0.36 | 0.36 | 0.38 | 0.37 | 0.35 | 0.3 |
| 7 | 5324257 | rs10269209 | -0.434 | 1.693 | -1.077 | 1.094 | -0.011 | -0.006 | -0.006 | 0.181 | 0.2 | 0.17 | 0.17 | 0.21 | 0.28 | 0.21 |
| 7 | 5328429 | rs6958502 | -0.395 | 0.798 | -1.801 | 0.974 | 0.026 | -0.067 | -0.067 | 0 | 0.72 | 0.91 | 0.78 | 0.59 | 0.82 | 0.49 |
| 7 | 5334209 | rs6965716 | 0.653 | 1.511 | -0.645 | 1.133 | 0.043 | -0.001 | -0.001 | 1.22 | 0.69 | 0.45 | 0.48 | 0.73 | 0.55 | 0.79 |
| 7 | 5338475 | rs3889348 | 0.636 | 1.484 | -0.393 | 0.831 | -0.015 | 0.001 | 0.001 | 0 | 0.7 | 0.79 | 0.72 | 0.74 | 0.65 | 0.77 |
| 7 | 5338714 | rs6950111 | -0.806 | 1.539 | -0.304 | 0.286 | 0.022 | -0.062 | -0.062 | 0 | 0.28 | 0.52 | 0.28 | 0.12 | 0.31 | 0.15 |
| 7 | 5347749 | rs9639976 | -0.5 | 2.289 | -0.791 | 0.581 | -0.054 | -0.03 | -0.03 | 0 | 0.14 | 0.06 | 0.17 | 0.12 | 0.26 | 0.08 |
| 7 | 5359680 | rs6943227 | 0.445 | 1.716 | -0.497 | -0.15 | 0.011 | -0.018 | -0.018 | 0 | 0.78 | 0.71 | 0.6 | 0.83 | 0.76 | 0.71 |
| 7 | 5366935 | rs10278799 | 0.172 | 0.989 | -0.422 | -0.298 | -0.009 | 0 | 0 | 0 | 0.12 | 0.18 | 0.09 | 0.13 | 0.06 | 0.15 |
| 7 | 5373370 | rs4486099 | 0.498 | 0.929 | -0.257 | -0.433 | -0.029 | -0.043 | -0.043 | 0.34 | 0.73 | 0.56 | 0.79 | 0.86 | 0.83 | 0.71 |
| 7 | 5375542 | rs10234709 | 0.801 | 0.05 | -0.642 | -1.534 | 0 | -0.062 | -0.062 | 0 | 0.38 | 0.41 | 0.34 | 0.07 | 0.44 | 0.22 |
| 7 | 5376032 | rs10253708 | 0.782 | -0.066 | -1.058 | -1.223 | -0.002 | -0.085 | -0.085 | 0 | 0.26 | 0.28 | 0.21 | 0.04 | 0.25 | 0.25 |
| 7 | 5376689 | rs6978911 | -0.174 | -0.01 | -2.085 | -0.569 | -0.037 | -0.057 | -0.057 | 0 | 0.4 | 0.08 | 0.31 | 0.59 | 0.38 | 0.3 |
| 7 | 5394951 | rs4724592 | -1.149 | 0.393 | -1.742 | -0.524 | -0.053 | 0.069 | 0.069 | 0 | 0.8 | 0.98 | 0.79 | 0.95 | 0.77 | 0.86 |
| 7 | 5396955 | rs7805732 | -0.263 | 0.351 | -1.368 | -0.394 | -0.012 | 0.03 | 0.03 | 0 | 0.61 | 0.87 | 0.46 | 0.29 | 0.64 | 0.6 |
| 7 | 5402606 | rs4357206 | 0.371 | 0.3 | -1.031 | -0.392 | -0.006 | 0.008 | 0.008 | 0.006 | 0.33 | 0.3 | 0.35 | 0.29 | 0.35 | 0.35 |
| 7 | 5406479 | rs4445136 | 0.42 | 0.099 | -1.147 | -0.553 | -0.009 | -0.01 | -0.01 | 0 | 0.38 | 0.66 | 0.53 | 0.5 | 0.41 | 0.46 |
| 7 | 5406582 | rs4626500 | -0.152 | 0.019 | -1.728 | -0.171 | -0.056 | -0.157 | -0.157 | 0 | 0.71 | 0.95 | 0.66 | 0.53 | 0.67 | 0.73 |
| 7 | 5409059 | rs6945065 | -1.055 | -0.188 | -0.462 | -0.63 | -0.016 | -0.037 | -0.037 | 0.199 | 0.14 | 0.2 | 0.03 | 0.05 | 0.08 | 0.16 |
| 7 | 5416248 | rs7804232 | -0.869 | -0.174 | -0.587 | -0.485 | -0.02 | -0.022 | -0.022 | 0 | 0.14 | 0.24 | 0.12 | 0.07 | 0.07 | 0.04 |
| 7 | 5427128 | rs4724662 | -0.963 | -0.285 | -1.843 | -1.025 | 0.019 | 0.144 | 0.144 | 0 | 0.39 | 0.86 | 0.38 | 0.17 | 0.26 | 0.41 |
| 7 | 5432672 | rs7791057 | -0.635 | -0.324 | -1.582 | -1.588 | 0.001 | 0.055 | 0.055 | 0.239 | 0.56 | 0.68 | 0.65 | 0.86 | 0.58 | 0.57 |
| 7 | 5436368 | rs10236222 | -0.074 | 0.434 | -1.635 | -0.948 | -0.13 | 0.058 | 0.058 | 0.023 | 0.33 | 0.1 | 0.43 | 0.17 | 0.49 | 0.39 |
| 7 | 5452756 | rs4628168 | 0.459 | 0.334 | -1.108 | -0.954 | 0.013 | 0.033 | 0.033 | 0 | 0.64 | 0.28 | 0.53 | 0.85 | 0.66 | 0.61 |
| 7 | 5473107 | rs13228828 | 1.005 | 0.479 | -0.468 | -0.522 | -0.007 | -0.002 | -0.002 | 0 | 0.93 | 0.81 | 0.86 | 0.96 | 0.93 | 0.9 |
| 7 | 5477922 | rs4720630 | 1.305 | 1.581 | -0.506 | -0.978 | -0.009 | 0.008 | 0.008 | 0 | 0.82 | 0.83 | 0.77 | 0.95 | 0.73 | 0.85 |
| 7 | 5480451 | rs11771525 | 1.233 | 1.389 | -0.163 | -1.023 | 0 | -0.002 | -0.002 | 0 | 0.73 | 0.76 | 0.46 | 0.19 | 0.66 | 0.53 |
| 7 | 5481120 | rs11760396 | 1.172 | 0.746 | -0.189 | -1.324 | -0.001 | -0.012 | -0.012 | 0 | 0.52 | 0.57 | 0.33 | 0.17 | 0.5 | 0.46 |
| 7 | 5483442 | rs6975264 | 0.595 | 0.602 | -0.004 | -1.329 | -0.003 | -0.019 | -0.019 | 0 | 0.66 | 0.68 | 0.65 | 0.94 | 0.57 | 0.79 |
| 7 | 5485083 | rs4724679 | -1.12 | 0.125 | 0.133 | -0.198 | -0.018 | 0.247 | 0.247 | 0 | 0.06 | 0.18 | 0.06 | 0.59 | 0.02 | 0.19 |
| 7 | 5490606 | rs4577859 | -0.579 | 0.382 | -0.123 | -0.414 | 0 | 0.07 | 0.07 | 0.07 | 0.1 | 0.15 | 0.09 | 0.62 | 0.11 | 0.39 |
| 7 | 5494677 | rs6964614 | -0.268 | 0.203 | -0.189 | -1.31 | 0 | -0.005 | -0.005 | 0 | 0.9 | 0.91 | 0.76 | 0.95 | 0.92 | 0.85 |
| 7 | 5496257 | rs7799592 | 0.795 | -0.054 | -0.222 | -1.917 | 0 | -0.008 | -0.008 | 0.034 | 0.67 | 0.73 | 0.66 | 0.85 | 0.71 | 0.67 |
| 7 | 5502964 | rs4454205 | 1.976 | -1.08 | 0.514 | -1.976 | -0.025 | -0.092 | -0.092 | 0 | 0.67 | 0.45 | 0.63 | 0.9 | 0.73 | 0.58 |

| | | | | | | | | | | | | | | | | |
|---|---|---|---|---|---|---|---|---|---|---|---|---|---|---|---|---|
| 7 | 5506499 | rs6948909 | -0.979 | -1.292 | 0.837 | -1.816 | -0.01 | -0.074 | -0.074 | 0 | 0.18 | 0.3 | 0.21 | 0.05 | 0.14 | 0.29 |
| 7 | 5509098 | rs4724690 | 0.018 | -1.672 | 0.921 | -2.114 | -0.009 | -0.027 | -0.027 | 4.571 | 0.61 | 0.55 | 0.63 | 0.8 | 0.69 | 0.79 |
| 7 | 5510159 | rs10951953 | 1.292 | -1.652 | 1.195 | -2.154 | 0.005 | 0.05 | 0.05 | 0 | 0.43 | 0.34 | 0.43 | 0.75 | 0.54 | 0.47 |
| 7 | 5512271 | rs4527789 | -0.197 | -1.671 | 0.872 | -2.075 | 0.014 | 0.016 | 0.016 | 1.469 | 0.91 | 0.97 | 0.91 | 0.95 | 0.92 | 0.92 |
| 7 | 5516293 | rs12535397 | -0.554 | -1.635 | 1.109 | -2.016 | 0.007 | 0.061 | 0.061 | 0.02 | 0.47 | 0.32 | 0.5 | 0.23 | 0.44 | 0.51 |
| 7 | 5516940 | rs7808325 | 1.479 | -1.654 | 1.335 | -1.966 | 0.011 | -0.02 | -0.02 | 6.539 | 0.41 | 0.47 | 0.36 | 0.66 | 0.48 | 0.39 |
| 7 | 5519542 | rs4416742 | -0.694 | -1.744 | 1.223 | -2.047 | -0.003 | 0 | 0 | 0 | 0.94 | 0.92 | 0.9 | 0.94 | 0.95 | 0.93 |
| 7 | 5520663 | rs10281893 | -0.501 | -1.855 | 1.184 | -2.11 | 0.018 | 0.07 | 0.07 | 0.001 | 0.46 | 0.28 | 0.49 | 0.22 | 0.4 | 0.51 |
| 7 | 5521468 | rs10951956 | -0.245 | -2.386 | 1.257 | -2.029 | -0.002 | -0.02 | -0.02 | 0 | 0.15 | 0.13 | 0.18 | 0.04 | 0.13 | 0.28 |
| 7 | 5524805 | rs10216189 | 1.566 | -2.503 | 1.249 | -1.976 | 0 | -0.008 | -0.008 | 0 | 0.42 | 0.53 | 0.38 | 0.71 | 0.51 | 0.45 |
| 7 | 5527223 | rs4352753 | -0.644 | -2.492 | 1.243 | -1.971 | 0.023 | 0.081 | 0.081 | 0 | 0.44 | 0.21 | 0.46 | 0.23 | 0.4 | 0.47 |
| 7 | 5528375 | rs10807949 | -0.74 | -2.524 | 1.223 | -1.987 | 0.018 | 0.046 | 0.046 | 0 | 0.47 | 0.35 | 0.48 | 0.23 | 0.4 | 0.47 |
| 7 | 5528883 | rs4724701 | 1.576 | -2.438 | 1.258 | -1.928 | 0.003 | -0.02 | -0.02 | 0 | 0.41 | 0.43 | 0.38 | 0.7 | 0.51 | 0.44 |
| 7 | 5530954 | rs11975313 | -0.439 | -2.396 | 1.104 | -1.929 | -0.009 | -0.054 | -0.054 | 0.657 | 0.16 | 0.27 | 0.21 | 0.04 | 0.14 | 0.23 |
| 7 | 5534598 | rs4320462 | 1.454 | -2.546 | 1.554 | -1.599 | 0.037 | 0.051 | 0.051 | 0.352 | 0.49 | 0.32 | 0.54 | 0.73 | 0.61 | 0.61 |
| 7 | 5534924 | rs7808849 | -0.467 | -2.147 | 0.592 | -1.418 | 0.053 | 0.109 | 0.109 | 0.265 | 0.35 | 0.1 | 0.3 | 0.19 | 0.28 | 0.26 |
| 7 | 5542513 | rs6463489 | -0.203 | -2.102 | 0.542 | -1.372 | -0.007 | -0.017 | -0.017 | 0.224 | 0.94 | 0.96 | 0.9 | 0.93 | 0.93 | 0.92 |
| 7 | 5545427 | rs1860883 | -0.723 | -2.015 | 0.704 | -1.322 | 0.046 | 0.069 | 0.069 | 0.534 | 0.42 | 0.22 | 0.39 | 0.26 | 0.35 | 0.34 |
| 7 | 5562740 | rs852432 | -0.918 | -1.726 | 0.878 | -1.155 | 0.001 | -0.005 | -0.005 | 2.906 | 0.39 | 0.39 | 0.32 | 0.19 | 0.33 | 0.27 |
| 7 | 5571625 | rs2966449 | -1.143 | -1.679 | 0.511 | -1.142 | -0.018 | -0.035 | -0.035 | 0 | 0.44 | 0.47 | 0.35 | 0.22 | 0.36 | 0.3 |
| 7 | 5572132 | rs852392 | -0.315 | -1.005 | 0.159 | -0.77 | 0.01 | 0.02 | 0.02 | 0 | 0.25 | 0.16 | 0.2 | 0.08 | 0.18 | 0.2 |
| 7 | 5572226 | rs852391 | -0.35 | -0.981 | 0.103 | -0.764 | 0.004 | 0.004 | 0.004 | 0 | 0.25 | 0.2 | 0.2 | 0.08 | 0.18 | 0.2 |
| 7 | 5578489 | rs7781531 | -0.935 | -0.448 | -0.026 | -0.577 | 0.008 | -0.02 | -0.02 | 0 | 0.05 | 0.17 | 0.02 | 0 | 0.04 | 0.02 |
| 7 | 5581695 | rs6963206 | -0.687 | -0.547 | -0.371 | -0.298 | 0.004 | -0.002 | -0.002 | 2.746 | 0.22 | 0.19 | 0.17 | 0.16 | 0.19 | 0.27 |
| 7 | 5582362 | rs6967973 | -0.545 | -0.467 | -0.387 | -0.281 | 0.002 | -0.003 | -0.003 | 0.449 | 0.21 | 0.19 | 0.17 | 0.16 | 0.19 | 0.27 |
| 7 | 5584934 | rs10951959 | -0.32 | -0.529 | -0.716 | -0.827 | 0.022 | 0.061 | 0.061 | 0 | 0.2 | 0.1 | 0.15 | 0.02 | 0.13 | 0.14 |
| 7 | 5592436 | rs852540 | -1.052 | -0.092 | -0.545 | -0.434 | -0.015 | -0.002 | -0.002 | 0.303 | 0.06 | 0.04 | 0.08 | 0.05 | 0.09 | 0.16 |
| 7 | 5596849 | rs1474445 | -1.092 | -0.919 | -0.56 | -0.642 | 0.006 | 0.003 | 0.003 | 0.279 | 0.25 | 0.25 | 0.18 | 0.19 | 0.22 | 0.28 |
| 7 | 5605130 | rs12666245 | -1.197 | -1.017 | -0.401 | -0.703 | -0.002 | -0.001 | -0.001 | 0 | 0.24 | 0.28 | 0.17 | 0.19 | 0.22 | 0.3 |
| 7 | 5607884 | rs3857755 | -0.801 | -1.034 | -0.207 | -0.662 | -0.012 | 0 | 0 | 0 | 0.2 | 0.29 | 0.15 | 0.18 | 0.17 | 0.28 |
| 7 | 5608932 | rs852499 | -1.388 | -1.367 | -0.083 | -0.838 | 0.014 | 0.005 | 0.005 | 0.003 | 0.3 | 0.74 | 0.2 | 0.23 | 0.21 | 0.33 |
| 7 | 5618237 | rs2693946 | -1.071 | -0.893 | -0.299 | -1.305 | 0.006 | 0.015 | 0.015 | 0 | 0.49 | 0.6 | 0.36 | 0.11 | 0.45 | 0.39 |
| 7 | 5619078 | rs6946228 | -1.239 | -1.421 | -0.277 | -1.399 | 0.003 | 0.003 | 0.003 | 1.073 | 0.33 | 0.68 | 0.22 | 0.16 | 0.27 | 0.3 |
| 7 | 5626564 | rs852380 | -0.727 | -1.187 | -0.178 | -1.34 | 0.055 | 0.096 | 0.096 | 0 | 0.51 | 0.64 | 0.33 | 0.15 | 0.34 | 0.37 |
| 7 | 5639126 | rs852479 | -1.554 | -0.879 | 0.062 | -0.934 | -0.012 | 0.023 | 0.023 | 0 | 0.1 | 0.16 | 0.06 | 0.18 | 0.08 | 0.11 |
| 7 | 5642746 | rs1640234 | -0.316 | -0.047 | -0.311 | -1.198 | -0.031 | -0.004 | -0.004 | 0 | 0.07 | 0.01 | 0.04 | 0.03 | 0.09 | 0.05 |
| 7 | 5644693 | rs7782428 | 0.77 | 0.363 | -0.089 | -1.775 | 0.011 | 0.028 | 0.028 | 0.49 | 0.49 | 0.43 | 0.48 | 0.81 | 0.44 | 0.55 |
| 7 | 5660686 | rs852441 | -1.749 | 0.313 | 0.876 | -1.675 | -0.011 | 0.054 | 0.054 | 0 | 0.1 | 0.34 | 0.11 | 0.1 | 0.06 | 0.13 |
| 7 | 5675305 | rs852376 | -1.84 | 0.257 | 1.407 | -1.7 | -0.01 | 0.065 | 0.065 | 0 | 0.11 | 0.5 | 0.12 | 0.1 | 0.06 | 0.13 |
| 7 | 5686742 | rs7780518 | -1.045 | 0.773 | 0.879 | -1.976 | -0.008 | 0.004 | 0.004 | 0 | 0.12 | 0.1 | 0.1 | 0.08 | 0.18 | 0.2 |
| 7 | 5697579 | rs852517 | -0.007 | 0.881 | 0.299 | -1.292 | -0.005 | 0.051 | 0.051 | 2.077 | 0.62 | 0.89 | 0.77 | 0.73 | 0.62 | 0.72 |
| 7 | 5697802 | rs6953970 | 1.341 | 0.663 | 0.127 | -1.188 | -0.043 | -0.057 | -0.057 | 0.194 | 0.39 | 0.11 | 0.3 | 0.59 | 0.41 | 0.43 |
| 7 | 5698964 | rs13247447 | 1.333 | 0.665 | 0.134 | -1.195 | -0.04 | -0.057 | -0.057 | 0 | 0.39 | 0.11 | 0.3 | 0.59 | 0.41 | 0.44 |
| 7 | 5710427 | rs4415230 | 0.567 | 0.683 | 0.164 | -1.765 | -0.005 | -0.005 | -0.005 | 0 | 0.07 | 0.04 | 0.05 | 0.06 | 0.1 | 0.12 |
| 7 | 5712600 | rs3779092 | 0.977 | 0.825 | -0.044 | -1.972 | -0.065 | 0.136 | 0.136 | 0 | 0.45 | 0.11 | 0.35 | 0.76 | 0.5 | 0.52 |
| 7 | 5716463 | rs3807583 | -0.103 | 0.772 | 0.059 | -1.898 | -0.013 | 0.001 | 0.001 | 0 | 0.09 | 0.05 | 0.08 | 0.06 | 0.15 | 0.17 |
| 7 | 5719557 | rs852397 | -1.35 | 0.666 | 0.64 | -1.84 | -0.006 | 0.053 | 0.053 | 0 | 0.1 | 0.54 | 0.12 | 0.08 | 0.06 | 0.13 |
| 7 | 5719739 | rs852396 | -1.207 | 0.665 | 0.709 | -1.838 | 0.014 | 0.066 | 0.066 | 0 | 0.08 | 0.5 | 0.08 | 0.05 | 0.04 | 0.1 |
| 7 | 5721448 | rs852394 | 0.435 | 0.571 | 0.473 | -1.775 | 0.04 | 0.138 | 0.138 | 1.297 | 0.5 | 0.82 | 0.41 | 0.26 | 0.47 | 0.42 |
| 7 | 5726022 | rs4724712 | 0.884 | 0.499 | 0.846 | -1.477 | -0.017 | 0.188 | 0.188 | 0 | 0.48 | 0.21 | 0.38 | 0.89 | 0.54 | 0.57 |
| 7 | 5732222 | rs3801009 | 0.731 | 0.512 | 0.982 | -1.357 | -0.001 | 0.015 | 0.015 | 0 | 0.06 | 0.11 | 0.04 | 0.13 | 0.06 | 0.08 |
| 7 | 5755231 | rs2287592 | 1.189 | 0.534 | 0.942 | -1.444 | -0.007 | 0.052 | 0.052 | 0 | 0.45 | 0.35 | 0.36 | 0.75 | 0.49 | 0.44 |
| 7 | 5772259 | rs13230831 | 2.203 | 0.453 | 0.851 | -1.312 | -0.047 | -0.046 | -0.046 | 0 | 0.32 | 0.06 | 0.27 | 0.69 | 0.32 | 0.31 |
| 7 | 5785086 | rs4724718 | 1.57 | 0.035 | 1.401 | -1.422 | 0.005 | 0.07 | 0.07 | 0 | 0.9 | 0.51 | 0.88 | 0.92 | 0.94 | 0.86 |
| 7 | 5785725 | rs10951960 | 0.1 | 0.059 | 1.334 | -1.555 | -0.052 | 0.169 | 0.169 | 1.439 | 0.38 | 0.13 | 0.4 | 0 | 0.41 | 0.24 |
| 7 | 5785999 | rs17135724 | 0.476 | -0.126 | 1.52 | -1.309 | 0 | 0 | 0 | 0 | 0.06 | 0.05 | 0.04 | 0.13 | 0.07 | 0.09 |
| 7 | 5792745 | rs6463513 | -1.429 | -0.131 | 1.497 | -1.339 | 0.033 | 0.057 | 0.057 | 0.193 | 0.09 | 0.47 | 0.11 | 0.05 | 0.05 | 0.09 |
| 7 | 5814178 | rs6944358 | -0.383 | -0.106 | 0.714 | -1.391 | -0.029 | 0.125 | 0.125 | 1.235 | 0.44 | 0.26 | 0.51 | 0.03 | 0.45 | 0.3 |
| 7 | 5814689 | rs12538870 | 1.569 | -0.107 | 0.673 | -1.342 | -0.041 | 0.077 | 0.077 | 0.91 | 0.39 | 0.11 | 0.32 | 0.75 | 0.42 | 0.47 |
| 7 | 5820350 | rs2008415 | -0.851 | -0.197 | 0.71 | -1.372 | 0.017 | 0.065 | 0.065 | 2.246 | 0.5 | 0.38 | 0.56 | 0.17 | 0.53 | 0.38 |
| 7 | 5823221 | rs13241373 | 1.996 | -0.24 | 1.418 | 0.014 | -0.027 | 0.006 | 0.006 | 0 | 0.86 | 0.41 | 0.9 | 0.84 | 0.9 | 0.85 |
| 7 | 5834567 | rs2345662 | 1.281 | -0.243 | 0.783 | 0.082 | -0.053 | 0.026 | 0.026 | 0 | 0.84 | 0.47 | 0.79 | 0.88 | 0.92 | 0.76 |
| 7 | 5850642 | rs4724739 | 1.464 | -0.204 | -0.548 | 0.092 | 0.053 | 0.057 | 0.057 | 0 | 0.54 | 0.07 | 0.56 | 0.67 | 0.59 | 0.48 |
| 7 | 5853467 | rs10249342 | -0.55 | -0.237 | 0.109 | -0.072 | -0.001 | -0.005 | -0.005 | 0 | 0.14 | 0.18 | 0.1 | 0.03 | 0.15 | 0.11 |
| 7 | 5854003 | rs6969131 | 0.004 | -0.298 | 0.718 | -0.083 | -0.006 | -0.005 | -0.005 | 0 | 0.94 | 0.83 | 0.95 | 0.96 | 0.97 | 0.91 |
| 7 | 5859558 | rs28404743 | -0.481 | -0.28 | 0.443 | -0.085 | -0.003 | -0.008 | -0.008 | 0.842 | 0.14 | 0.19 | 0.11 | 0.03 | 0.15 | 0.11 |
| 7 | 5871225 | rs7785084 | 1.845 | -0.335 | 0.971 | -0.141 | -0.046 | -0.089 | -0.089 | 0 | 0.76 | 0.42 | 0.84 | 0.94 | 0.8 | 0.79 |
| 7 | 5872994 | rs7791213 | -0.682 | -0.179 | 0.384 | -0.075 | -0.011 | -0.039 | -0.039 | 4.206 | 0.15 | 0.27 | 0.11 | 0.03 | 0.16 | 0.1 |
| 7 | 5878648 | rs888586 | -0.874 | -0.233 | -0.033 | -0.098 | -0.011 | -0.034 | -0.034 | 0 | 0.18 | 0.29 | 0.1 | 0.03 | 0.17 | 0.11 |

| | | | | | | | | | | | | | | | | |
|---|---|---|---|---|---|---|---|---|---|---|---|---|---|---|---|---|
| 7 | 5883522 | rs308092 | -1.808 | -0.07 | -0.866 | 0.677 | 0.02 | -0.039 | -0.039 | 0 | 0.44 | 0.87 | 0.42 | 0.48 | 0.45 | 0.45 |
| 7 | 5886060 | rs308085 | -1.811 | -0.016 | -0.815 | 0.825 | 0.021 | -0.049 | -0.049 | 0 | 0.44 | 0.85 | 0.48 | 0.42 | 0.45 | 0.45 |
| 7 | 5888663 | rs308080 | -1.73 | -0.054 | -0.844 | 0.747 | 0.026 | 0.011 | 0.011 | 0 | 0.45 | 0.86 | 0.47 | 0.54 | 0.44 | 0.49 |
| 7 | 5890168 | rs308076 | -1.782 | 0.228 | -0.874 | 0.673 | 0.015 | -0.012 | -0.012 | 0.098 | 0.46 | 0.85 | 0.62 | 0.49 | 0.46 | 0.52 |
| 7 | 5891221 | rs308074 | -0.256 | 0.68 | -0.585 | 0.345 | -0.017 | 0.009 | 0.009 | 0 | 0.06 | 0.01 | 0.2 | 0.04 | 0.05 | 0.06 |
| 7 | 5908742 | rs6953261 | -0.986 | 0.485 | -0.306 | 0.282 | -0.002 | -0.014 | -0.014 | 0 | 0.25 | 0.67 | 0.23 | 0.12 | 0.27 | 0.19 |
| 7 | 5911291 | rs9648583 | -1.074 | 1.256 | -0.351 | 0.6 | -0.008 | 0.01 | 0.01 | 0 | 0.16 | 0.09 | 0.41 | 0.11 | 0.18 | 0.23 |
| 7 | 5925156 | rs2881189 | -0.052 | 1.32 | -0.623 | 0.837 | 0 | -0.003 | -0.003 | 0 | 0.21 | 0.15 | 0.11 | 0.47 | 0.16 | 0.28 |
| 7 | 5929666 | rs7789795 | -0.183 | 1.582 | -0.583 | 0.452 | -0.033 | 0.006 | 0.006 | 0 | 0.09 | 0.02 | 0.11 | 0.05 | 0.09 | 0.1 |
| 7 | 6025136 | rs7811924 | -0.186 | 1.33 | 0.454 | 0.554 | 0.026 | 0.041 | 0.041 | 5.828 | 0.08 | 0.29 | 0.05 | 0.09 | 0.06 | 0.09 |
| 7 | 6026775 | rs2228006 | -0.357 | 0.805 | -0.148 | 0.215 | 0.007 | 0.041 | 0.041 | 0 | 0.84 | 0.94 | 0.77 | 0.88 | 0.88 | 0.89 |
| 7 | 6035507 | rs2286680 | 0.451 | 0.216 | -0.178 | 0.655 | -0.006 | -0.033 | -0.033 | 0 | 0.17 | 0.25 | 0.11 | 0.07 | 0.14 | 0.15 |
| 7 | 6036245 | rs10235277 | 0.053 | 0.299 | -0.364 | 0.537 | 0.018 | -0.099 | -0.099 | 2.954 | 0.77 | 0.92 | 0.7 | 0.54 | 0.79 | 0.64 |
| 7 | 6036515 | rs12112229 | -0.119 | 1.199 | 0.356 | 0.001 | 0.044 | -0.065 | -0.065 | 0.27 | 0.22 | 0.58 | 0.24 | 0.09 | 0.25 | 0.19 |
| 7 | 6046783 | rs7797466 | -0.152 | 0.981 | 0.318 | 0.114 | -0.016 | -0.028 | -0.028 | 0.175 | 0.24 | 0.34 | 0.15 | 0.16 | 0.18 | 0.24 |
| 7 | 6048973 | rs1062372 | -0.152 | 0.983 | 0.324 | 0.111 | -0.009 | -0.031 | -0.031 | 0.046 | 0.24 | 0.35 | 0.16 | 0.16 | 0.19 | 0.24 |
| 7 | 6053264 | rs1860461 | -0.23 | 0.691 | -0.397 | -0.667 | 0.005 | 0.024 | 0.024 | 0 | 0.53 | 0.45 | 0.45 | 0.78 | 0.47 | 0.61 |
| 7 | 6053286 | rs1860460 | -0.285 | 0.795 | -0.287 | -0.628 | 0.048 | -0.087 | -0.087 | 0.355 | 0.75 | 0.51 | 0.62 | 0.86 | 0.72 | 0.76 |
| 7 | 6054987 | rs6979676 | 0.054 | 0.192 | -0.588 | -0.645 | -0.01 | -0.03 | -0.03 | 0 | 0.34 | 0.35 | 0.19 | 0.16 | 0.24 | 0.26 |
| 7 | 6062828 | rs7781199 | 0.788 | 0.229 | -0.642 | -0.559 | -0.003 | -0.014 | -0.014 | 0 | 0.18 | 0.19 | 0.12 | 0.06 | 0.15 | 0.17 |
| 7 | 6063283 | rs4560 | 1.539 | 0.324 | -1.05 | -0.444 | -0.033 | -0.055 | -0.055 | 0 | 0.4 | 0.16 | 0.4 | 0.37 | 0.46 | 0.33 |
| 7 | 6065746 | rs10249759 | -0.132 | 0.282 | -0.875 | -0.794 | -0.004 | -0.024 | -0.024 | 1.709 | 0.63 | 0.62 | 0.64 | 0.81 | 0.68 | 0.65 |
| 7 | 6066461 | rs2639 | -0.185 | 0.337 | -0.859 | -0.8 | 0.092 | 0.226 | 0.226 | 1.863 | 0.16 | 0.44 | 0.22 | 0.44 | 0.2 | 0.3 |
| 7 | 6066946 | rs10254125 | -0.038 | -0.061 | -1.025 | -1.052 | -0.001 | -0.014 | -0.014 | 0 | 0.6 | 0.62 | 0.64 | 0.81 | 0.68 | 0.65 |
| 7 | 6070563 | rs17136342 | 2.729 | -0.051 | -1.334 | -1.158 | -0.016 | 0.051 | 0.051 | 0 | 0.23 | 0.14 | 0.2 | 0.06 | 0.28 | 0.16 |
| 7 | 6071277 | rs4724770 | -1.094 | -0.015 | -1.319 | -1.092 | 0 | 0.069 | 0.069 | 0 | 0.79 | 0.93 | 0.77 | 0.88 | 0.83 | 0.82 |
| 7 | 6078468 | rs2302330 | 0.274 | 0.587 | -0.885 | -0.859 | 0.1 | 0.018 | 0.018 | 3.406 | 0.11 | 0.38 | 0.16 | 0.11 | 0.13 | 0.14 |
| 7 | 6079662 | rs17136358 | 1.908 | 0.404 | -0.988 | -0.829 | 0.001 | -0.001 | -0.001 | 0 | 0.06 | 0.04 | 0.03 | 0.09 | 0.05 | 0.09 |
| 7 | 6090727 | rs852154 | 0.598 | 0.201 | -0.908 | -0.909 | 0.013 | 0.008 | 0.008 | 0.043 | 0.49 | 0.35 | 0.43 | 0.36 | 0.47 | 0.34 |
| 7 | 6095157 | rs852151 | 1.512 | 0.424 | -0.875 | -0.333 | -0.032 | -0.101 | -0.101 | 0 | 0.15 | 0.02 | 0.2 | 0.3 | 0.18 | 0.17 |
| 7 | 6114558 | rs13230531 | 0.887 | 0.76 | -0.77 | -1.149 | -0.003 | 0.002 | 0.002 | 0 | 0.1 | 0.05 | 0.19 | 0.08 | 0.22 | 0.18 |
| 7 | 6117152 | rs591982 | -0.531 | 0.227 | -0.485 | -1.295 | 0.005 | 0.04 | 0.04 | 0 | 0.49 | 0.59 | 0.54 | 0.8 | 0.53 | 0.64 |
| 7 | 6174379 | rs9648587 | 0.68 | 0.837 | 0.687 | -0.683 | 0.006 | 0.013 | 0.013 | 0 | 0.12 | 0.06 | 0.22 | 0.14 | 0.12 | 0.16 |
| 7 | 6189780 | rs3764814 | 0.246 | 1.099 | 1.32 | -0.233 | -0.036 | 0.29 | 0.29 | 0.003 | 0.14 | 0.43 | 0.18 | 0.43 | 0.09 | 0.2 |
| 7 | 6201305 | rs6959740 | 0.832 | 1.424 | 1.649 | -1.074 | 0.118 | -0.069 | -0.069 | 0.052 | 0.12 | 0.37 | 0.15 | 0.01 | 0.15 | 0.06 |
| 7 | 6203231 | rs6953241 | -1.179 | 1.476 | 1.442 | -1.036 | 0.2 | 0.081 | 0.081 | 0 | 0.07 | 0.31 | 0.17 | 0.07 | 0.15 | 0.05 |
| 7 | 6204999 | rs2301910 | 0.358 | 1.786 | 0.474 | -1.552 | -0.043 | 0.095 | 0.095 | 0 | 0.71 | 0.94 | 0.7 | 0.85 | 0.71 | 0.7 |
| 7 | 6207142 | rs3793202 | -1.757 | 1.815 | 1.262 | -1.124 | 0.145 | 0.02 | 0.02 | 0.12 | 0.1 | 0.62 | 0.2 | 0.06 | 0.17 | 0.13 |
| 7 | 6208369 | rs1017035 | 0.204 | 1.556 | 0.724 | -1.182 | -0.13 | 0.191 | 0.191 | 0 | 0.73 | 0.99 | 0.8 | 1 | 0.7 | 0.79 |
| 7 | 6213396 | rs17136059 | -2.222 | 1.452 | 2.067 | 0.037 | 0.186 | 0.031 | 0.031 | 0.134 | 0.1 | 0.43 | 0.18 | 0.06 | 0.18 | 0.15 |
| 7 | 6223280 | rs10240988 | -1.542 | 1.433 | 2.163 | 0.204 | 0.264 | 0.088 | 0.088 | 0 | 0.07 | 0.43 | 0.2 | 0.06 | 0.19 | 0.14 |
| 7 | 6237303 | rs6974814 | 0.523 | -0.719 | 1.664 | 0.174 | -0.002 | -0.013 | -0.013 | 0.123 | 0.09 | 0.09 | 0.02 | 0 | 0.02 | 0.01 |
| 7 | 6261683 | rs7791225 | -1.065 | -0.713 | 1.695 | 0.123 | -0.016 | -0.049 | -0.049 | 2.493 | 0.18 | 0.74 | 0.14 | 0.03 | 0.09 | 0.11 |
| 7 | 6274585 | rs10262194 | -1.503 | -0.71 | 1.714 | 0.23 | -0.023 | 0.03 | 0.03 | 0.443 | 0.21 | 0.67 | 0.22 | 0.34 | 0.14 | 0.33 |
| 7 | 6291937 | rs10255662 | -1.451 | -0.505 | 1.399 | -0.877 | 0.003 | -0.102 | -0.102 | 0.53 | 0.22 | 0.34 | 0.17 | 0.04 | 0.17 | 0.2 |
| 7 | 6296843 | rs6463540 | -1.256 | -0.724 | 1.276 | -0.554 | -0.018 | -0.05 | -0.05 | 0 | 0.24 | 0.71 | 0.14 | 0.04 | 0.09 | 0.11 |
| 7 | 6316798 | rs7780269 | -1.97 | 0.042 | 1.231 | 0.201 | 0.05 | 0.052 | 0.052 | 0.236 | 0.06 | 0.4 | 0.04 | 0.03 | 0.03 | 0.04 |
| 7 | 6319654 | rs9986688 | -1.626 | 0.092 | 0.341 | 0.204 | -0.02 | -0.022 | -0.022 | 0 | 0.07 | 0.25 | 0.04 | 0.03 | 0.03 | 0.04 |
| 7 | 6334747 | rs11764399 | -1.014 | 0.616 | -0.085 | 0.26 | 0 | -0.001 | -0.001 | 0 | 0.09 | 0.08 | 0.1 | 0.05 | 0.13 | 0.06 |
| 7 | 6347851 | rs6966217 | -0.659 | 0.359 | -0.651 | 0.759 | -0.001 | -0.106 | -0.106 | 0.092 | 0.23 | 0.07 | 0.25 | 0.59 | 0.25 | 0.49 |
| 7 | 6348344 | rs7802183 | 0.217 | -0.107 | -1.122 | 0.053 | 0.01 | 0.031 | 0.031 | 0 | 0.13 | 0.06 | 0.12 | 0.02 | 0.1 | 0.13 |
| 7 | 6352029 | rs10235034 | -1.253 | -0.235 | -0.492 | -0.407 | 0 | -0.001 | -0.001 | 0 | 0.42 | 0.4 | 0.38 | 0.64 | 0.41 | 0.57 |
| 7 | 6352485 | rs17136142 | -0.003 | -0.06 | -0.654 | -0.283 | -0.03 | -0.127 | -0.127 | 1.272 | 0.22 | 0.06 | 0.26 | 0.59 | 0.27 | 0.49 |
| 7 | 6353371 | rs4724794 | 0.03 | -0.148 | -0.666 | -0.281 | -0.015 | -0.136 | -0.136 | 0 | 0.22 | 0.06 | 0.25 | 0.59 | 0.25 | 0.49 |
| 7 | 6362809 | rs6463547 | -1.017 | -0.261 | 0.128 | 0.417 | 0.008 | 0.032 | 0.032 | 1.064 | 0.21 | 0.41 | 0.22 | 0.27 | 0.23 | 0.4 |
| 7 | 6362978 | rs2113628 | -0.323 | -0.281 | 0.068 | 0.558 | 0.015 | 0.295 | 0.295 | 0 | 0.07 | 0.28 | 0.18 | 0.27 | 0.04 | 0.09 |
| 7 | 6370194 | rs6952125 | -1.146 | -0.083 | -0.933 | -0.193 | -0.023 | 0.223 | 0.223 | 0.019 | 0.54 | 0.9 | 0.66 | 0.88 | 0.46 | 0.69 |
| 7 | 6377836 | rs4724795 | -1.224 | 0.41 | -0.278 | 0.729 | 0.002 | 0.008 | 0.008 | 0 | 0.18 | 0.23 | 0.22 | 0.35 | 0.29 | 0.27 |
| 7 | 6383524 | rs7791772 | -1.681 | 0.244 | -0.068 | 0.826 | 0 | -0.006 | -0.006 | 0 | 0.33 | 0.38 | 0.39 | 0.81 | 0.36 | 0.56 |
| 7 | 6384000 | rs4724798 | -1.765 | 0.107 | 0.577 | 0.756 | -0.004 | -0.051 | -0.051 | 0 | 0.36 | 0.54 | 0.4 | 0.85 | 0.37 | 0.59 |
| 7 | 6402827 | rs2042524 | -1.487 | 0.05 | 0.472 | 0.332 | -0.016 | 0.009 | 0.009 | 0 | 0.4 | 0.75 | 0.5 | 0.37 | 0.43 | 0.35 |
| 7 | 6410528 | rs4724800 | -0.751 | 0.505 | 0.954 | 0.546 | 0.073 | 0.091 | 0.091 | 0 | 0.13 | 0.29 | 0.22 | 0.27 | 0.26 | 0.14 |
| 7 | 6418275 | rs7784465 | -0.2 | 0.081 | 0.965 | 0.29 | 0.001 | -0.028 | -0.028 | 0.015 | 0.14 | 0.18 | 0.2 | 0.02 | 0.13 | 0.04 |
| 7 | 6420955 | rs10499344 | -0.265 | 0.06 | 0.863 | 0.316 | 0.002 | -0.04 | -0.04 | 0 | 0.14 | 0.22 | 0.21 | 0.03 | 0.13 | 0.04 |
| 7 | 6422556 | rs10951982 | -0.532 | 0.203 | 0.696 | 0.492 | -0.037 | -0.043 | -0.043 | 0.071 | 0.11 | 0.04 | 0.18 | 0.27 | 0.24 | 0.14 |
| 7 | 6426954 | rs3729790 | -0.487 | 0.221 | 0.722 | 0.505 | -0.027 | -0.032 | -0.032 | 0.486 | 0.11 | 0.05 | 0.18 | 0.26 | 0.22 | 0.14 |
| 7 | 6442371 | rs9374 | -0.486 | 0.205 | 0.731 | 0.505 | -0.02 | -0.023 | -0.023 | 0 | 0.11 | 0.05 | 0.18 | 0.27 | 0.23 | 0.14 |
| 7 | 6443839 | rs4720672 | -0.314 | 0.271 | 0.726 | 0.616 | -0.01 | -0.014 | -0.014 | 0 | 0.1 | 0.05 | 0.16 | 0.16 | 0.16 | 0.14 |
| 7 | 6449229 | rs836560 | -1.145 | -0.225 | 0.979 | 0.57 | -0.024 | 0.244 | 0.244 | 0 | 0.12 | 0.27 | 0.18 | 0.52 | 0.05 | 0.34 |

| | | | | | | | | | | | | | | | | |
|---|---|---|---|---|---|---|---|---|---|---|---|---|---|---|---|---|
| 7 | 6449496 | rs2303361 | -0.262 | -0.121 | 0.882 | 0.558 | -0.014 | -0.015 | -0.015 | 0.315 | 0.11 | 0.05 | 0.18 | 0.27 | 0.21 | 0.14 |
| 7 | 6451481 | rs866279 | -0.902 | -0.232 | 0.987 | 0.48 | -0.025 | -0.063 | -0.063 | 0.018 | 0.18 | 0.43 | 0.25 | 0.07 | 0.13 | 0.12 |
| 7 | 6456976 | rs702486 | -2.081 | -0.695 | -0.293 | 0.112 | -0.004 | 0.174 | 0.174 | 0.008 | 0.46 | 0.84 | 0.56 | 0.82 | 0.43 | 0.57 |
| 7 | 6462646 | rs836518 | -1.547 | -0.656 | -0.233 | 0.189 | 0.025 | -0.022 | -0.022 | 0 | 0.35 | 0.75 | 0.38 | 0.56 | 0.21 | 0.43 |
| 7 | 6465598 | rs836524 | -0.489 | -0.656 | -0.137 | 0.178 | -0.014 | 0.028 | 0.028 | 0 | 0.05 | 0.19 | 0.05 | 0.05 | 0.02 | 0.11 |
| 7 | 6474073 | rs836534 | -1.748 | -0.421 | -0.972 | 0.272 | 0.005 | 0.136 | 0.136 | 0.001 | 0.46 | 0.79 | 0.56 | 0.81 | 0.42 | 0.57 |
| 7 | 6474158 | rs836535 | -0.4 | -0.433 | -0.988 | 0.249 | 0.008 | 0.006 | 0.006 | 1.753 | 0.28 | 0.74 | 0.26 | 0.07 | 0.17 | 0.17 |
| 7 | 6475831 | rs3828946 | -1.33 | -0.303 | -1.027 | 0.299 | -0.029 | -0.037 | -0.037 | 0 | 0.18 | 0.08 | 0.3 | 0.74 | 0.25 | 0.4 |
| 7 | 6477002 | rs836540 | -0.952 | -0.295 | -0.928 | 0.13 | 0.053 | 0.002 | 0.002 | 0.855 | 0.39 | 0.81 | 0.4 | 0.64 | 0.22 | 0.55 |
| 7 | 6487156 | rs3828944 | 0.57 | 0.486 | -0.539 | 0.887 | -0.022 | -0.028 | -0.028 | 0 | 0.89 | 0.95 | 0.82 | 0.7 | 0.79 | 0.86 |
| 7 | 6495017 | rs7792987 | 1.52 | -0.249 | -0.029 | 0.894 | -0.02 | 0.058 | 0.058 | 0.009 | 0.77 | 0.61 | 0.63 | 0.53 | 0.78 | 0.65 |
| 7 | 6507517 | rs11980237 | -0.432 | -0.443 | 0.621 | 1.181 | -0.011 | 0.205 | 0.205 | 0 | 0.08 | 0.32 | 0.02 | 0.25 | 0.02 | 0.14 |
| 7 | 6512050 | rs6959558 | 0.6 | -1.156 | 0.382 | 1.078 | 0.006 | 0.005 | 0.005 | 0.152 | 0.85 | 0.9 | 0.85 | 0.58 | 0.93 | 0.65 |
| 7 | 6516035 | rs12702514 | 1.749 | -1.004 | -0.54 | -0.162 | -0.049 | 0.114 | 0.114 | 0 | 0.37 | 0.22 | 0.33 | 0.07 | 0.46 | 0.17 |
| 7 | 6516437 | rs17198191 | 0.781 | -1.034 | -0.785 | -0.558 | -0.029 | 0.055 | 0.055 | 3.896 | 0.2 | 0.05 | 0.12 | 0 | 0.21 | 0.12 |
| 7 | 6517440 | rs13222291 | 2.062 | -0.886 | -0.763 | -0.426 | -0.067 | 0.151 | 0.151 | 0 | 0.37 | 0.19 | 0.33 | 0.07 | 0.46 | 0.17 |
| 7 | 6520676 | rs6964421 | 0.786 | -0.859 | -0.703 | -0.323 | -0.005 | 0.006 | 0.006 | 0.778 | 0.57 | 0.57 | 0.68 | 0.54 | 0.68 | 0.51 |
| 7 | 6535517 | rs1523609 | -0.289 | -0.559 | -1.092 | -0.005 | -0.013 | -0.045 | -0.045 | 9.586 | 0.9 | 0.98 | 0.85 | 0.54 | 0.94 | 0.65 |
| 7 | 6537655 | rs4574760 | 1.955 | -0.454 | -1.294 | -0.419 | -0.101 | 0.163 | 0.163 | 0 | 0.33 | 0.09 | 0.3 | 0.06 | 0.43 | 0.13 |
| 7 | 6544247 | rs12702519 | 0.455 | -0.549 | -0.62 | 0.122 | -0.012 | 0.08 | 0.08 | 0.237 | 0.85 | 0.81 | 0.79 | 0.46 | 0.92 | 0.59 |
| 7 | 6544691 | rs2881698 | -0.302 | -0.24 | -0.67 | -0.221 | 0.054 | 0.111 | 0.111 | 0 | 0.1 | 0.33 | 0.28 | 0.17 | 0.19 | 0.15 |
| 7 | 6550494 | rs12702520 | 0.69 | -0.431 | -1.81 | -0.774 | 0.006 | 0.004 | 0.004 | 0.111 | 0.56 | 0.51 | 0.66 | 0.23 | 0.72 | 0.39 |
| 7 | 6557135 | rs10447683 | 0.799 | 0.232 | -2.927 | -0.737 | -0.148 | 0.295 | 0.295 | 0 | 0.4 | 0.02 | 0.34 | 0.05 | 0.51 | 0.24 |
| 7 | 6567114 | rs1105664 | 0.544 | 0.286 | -2.034 | 0.814 | -0.037 | -0.087 | -0.087 | 0 | 0.88 | 0.98 | 0.84 | 0.42 | 0.89 | 0.64 |
| 7 | 6578581 | rs13225983 | 0.82 | 0.662 | -2.285 | 0.135 | -0.121 | 0.043 | 0.043 | 0.891 | 0.31 | 0.06 | 0.31 | 0.22 | 0.48 | 0.15 |
| 7 | 6587526 | rs952042 | 0.215 | -0.398 | -1.732 | -0.154 | -0.001 | -0.004 | -0.004 | 0 | 0.35 | 0.37 | 0.36 | 0.27 | 0.29 | 0.29 |
| 7 | 6591796 | rs11762346 | 0.168 | -0.411 | -1.719 | -0.173 | -0.001 | -0.002 | -0.002 | 0.173 | 0.34 | 0.34 | 0.36 | 0.27 | 0.28 | 0.29 |
| 7 | 6613604 | rs7800401 | 0.586 | -0.541 | -1.168 | -0.171 | 0.011 | 0.012 | 0.012 | 0 | 0.06 | 0.19 | 0.1 | 0.05 | 0.07 | 0.07 |
| 7 | 6617945 | rs7793367 | -0.424 | -0.906 | -0.943 | -0.478 | 0.02 | 0.043 | 0.043 | 0 | 0.46 | 0.65 | 0.44 | 0.24 | 0.33 | 0.32 |
| 7 | 6620757 | rs4605951 | -1.488 | -0.967 | -1.066 | -0.378 | 0.008 | -0.134 | -0.134 | 0 | 0.82 | 0.98 | 0.83 | 0.53 | 0.86 | 0.51 |
| 7 | 6621277 | rs11559146 | 0.779 | -1.275 | -0.609 | -0.608 | -0.011 | -0.042 | -0.042 | 0 | 0.32 | 0.44 | 0.37 | 0.24 | 0.28 | 0.28 |
| 7 | 6625847 | rs7794450 | -0.842 | -1.188 | -0.665 | -0.578 | 0.032 | 0.121 | 0.121 | 0 | 0.49 | 0.8 | 0.52 | 0.29 | 0.4 | 0.37 |
| 7 | 6652740 | rs6973331 | 1.678 | -1.016 | -0.505 | -0.696 | -0.015 | -0.054 | -0.054 | 0 | 0.11 | 0.2 | 0.06 | 0 | 0.04 | 0.04 |
| 7 | 6656897 | rs1806552 | -1.899 | -0.804 | -0.708 | -0.628 | -0.102 | 0.011 | 0.011 | 0.109 | 0.72 | 0.93 | 0.69 | 0.76 | 0.54 | 0.86 |
| 7 | 6659509 | rs2346263 | 0.678 | -1.008 | -0.1 | -0.709 | -0.001 | -0.001 | -0.001 | 0.043 | 0.32 | 0.34 | 0.37 | 0.31 | 0.28 | 0.27 |
| 7 | 6662749 | rs2462655 | -1.497 | -0.122 | -0.126 | -0.854 | -0.076 | 0.067 | 0.067 | 0.37 | 0.71 | 0.87 | 0.69 | 0.85 | 0.54 | 0.88 |
| 7 | 6663670 | rs940871 | -1.576 | -0.452 | -0.217 | -0.819 | -0.126 | 0.066 | 0.066 | 0.369 | 0.71 | 0.93 | 0.68 | 0.85 | 0.51 | 0.87 |
| 7 | 6668027 | rs2244572 | 0.078 | -1.201 | 0.543 | -0.821 | -0.021 | -0.036 | -0.036 | 0 | 0.44 | 0.5 | 0.51 | 0.81 | 0.32 | 0.74 |
| 7 | 6681006 | rs4720680 | -0.533 | -1.206 | 0.368 | -0.736 | -0.023 | -0.008 | -0.008 | 0.393 | 0.36 | 0.45 | 0.27 | 0.31 | 0.26 | 0.31 |
| 7 | 6681619 | rs940870 | 1.252 | -1.149 | 0.081 | -0.638 | 0.028 | -0.067 | -0.067 | 0 | 0.3 | 0.13 | 0.35 | 0.52 | 0.24 | 0.52 |
| 7 | 6683454 | rs2253869 | -0.928 | -0.594 | 0.196 | -0.617 | -0.025 | 0.028 | 0.028 | 0.158 | 0.68 | 0.71 | 0.62 | 0.84 | 0.51 | 0.84 |
| 7 | 6694718 | rs2464877 | -0.874 | -0.481 | 0.242 | -1.192 | -0.021 | 0.016 | 0.016 | 1.362 | 0.67 | 0.71 | 0.64 | 0.96 | 0.51 | 0.86 |
| 7 | 6704332 | rs2251188 | -1.551 | -0.215 | -0.224 | -1.098 | -0.178 | 0.103 | 0.103 | 0.888 | 0.77 | 0.98 | 0.71 | 0.96 | 0.56 | 0.9 |
| 7 | 6706631 | rs6964079 | -0.222 | -0.449 | 1.33 | -0.452 | -0.012 | 0.207 | 0.207 | 0.049 | 0.08 | 0.28 | 0.11 | 0.29 | 0.05 | 0.17 |
| 7 | 6721661 | rs10951991 | 0.155 | -0.308 | 1.271 | -0.44 | -0.016 | 0.17 | 0.17 | 0 | 0.07 | 0.24 | 0.11 | 0.29 | 0.05 | 0.16 |
| 7 | 6734103 | rs3801033 | -1.468 | -0.509 | 1.011 | -0.697 | -0.028 | -0.009 | -0.009 | 2.077 | 0.35 | 0.55 | 0.25 | 0.32 | 0.24 | 0.29 |
| 7 | 6735534 | rs3801035 | 0.427 | -0.531 | 0.758 | -0.788 | -0.008 | 0.079 | 0.079 | 0.213 | 0.14 | 0.02 | 0.06 | 0.02 | 0.12 | 0.09 |
| 7 | 6743097 | rs17789894 | 0.426 | -0.408 | 0.589 | -0.784 | -0.026 | 0.079 | 0.079 | 0.222 | 0.14 | 0.02 | 0.08 | 0.02 | 0.14 | 0.09 |
| 7 | 6744957 | rs7798471 | -1.629 | -0.441 | 0.706 | -0.765 | -0.011 | -0.006 | -0.006 | 0 | 0.35 | 0.58 | 0.27 | 0.31 | 0.27 | 0.29 |
| 7 | 6746738 | rs9639027 | -1.787 | -0.525 | 0.305 | -0.85 | 0.098 | 0.087 | 0.087 | 0 | 0.46 | 0.91 | 0.34 | 0.32 | 0.3 | 0.38 |
| 7 | 6758018 | rs6944332 | -0.642 | -0.744 | 0.578 | -0.749 | -0.06 | 0.079 | 0.079 | 0.062 | 0.18 | 0.64 | 0.19 | 0.31 | 0.08 | 0.25 |
| 7 | 6773892 | rs13224907 | -0.743 | -0.494 | -0.313 | -0.631 | -0.014 | 0.017 | 0.017 | 0 | 0.23 | 0.27 | 0.18 | 0.34 | 0.17 | 0.26 |
| 7 | 6779270 | rs7779296 | -0.851 | -0.038 | -0.446 | -0.341 | 0.112 | 0.056 | 0.056 | 0 | 0.47 | 0.88 | 0.33 | 0.35 | 0.29 | 0.39 |
| 7 | 6780231 | rs10215807 | -0.826 | 1.266 | 0.534 | 0.849 | -0.015 | 0.199 | 0.199 | 0 | 0.08 | 0.23 | 0.11 | 0.31 | 0.05 | 0.16 |
| 7 | 6870635 | rs10242703 | 0.158 | 1.908 | 0.045 | -0.137 | 0.003 | -0.001 | -0.001 | 0 | 0.56 | 0.47 | 0.54 | 0.48 | 0.45 | 0.7 |
| 7 | 6929662 | rs4236396 | -1.246 | 1.314 | 0.356 | -0.419 | 0 | -0.051 | -0.051 | 0 | 0.26 | 0.24 | 0.24 | 0 | 0.25 | 0.15 |
| 7 | 6937338 | rs6463598 | -1.509 | 1.201 | 0.743 | 0.058 | -0.003 | -0.025 | -0.025 | 0 | 0.3 | 0.44 | 0.33 | 0.2 | 0.27 | 0.36 |
| 7 | 6955934 | rs6950893 | -1.262 | 1.524 | 0.477 | -0.099 | 0.005 | -0.015 | -0.015 | 0.179 | 0.23 | 0.28 | 0.2 | 0.17 | 0.26 | 0.26 |
| 7 | 6985071 | rs9691350 | -1.618 | 1.286 | 0.868 | -0.274 | 0.053 | -0.063 | -0.063 | 3.954 | 0.11 | 0.55 | 0.17 | 0 | 0.08 | 0.11 |
| 7 | 6986621 | rs10253555 | -2.021 | 1.751 | 0.73 | -0.482 | 0.043 | -0.103 | -0.103 | 0 | 0.17 | 0.65 | 0.24 | 0 | 0.19 | 0.15 |
| 7 | 7040309 | rs4074750 | -0.769 | 0.538 | 0.111 | -0.742 | 0.001 | -0.005 | -0.005 | 1.459 | 0.4 | 0.62 | 0.34 | 0.88 | 0.31 | 0.4 |
| 7 | 7040500 | rs4074748 | -0.791 | 0.524 | 0.022 | -0.755 | 0.015 | 0.018 | 0.018 | 0 | 0.41 | 0.66 | 0.34 | 0.88 | 0.31 | 0.4 |
| 7 | 7044010 | rs6945669 | -0.799 | 0.546 | -1.149 | -0.942 | 0.05 | 0.219 | 0.219 | 0.869 | 0.46 | 0.97 | 0.49 | 0.89 | 0.38 | 0.38 |
| 7 | 7052237 | rs3924856 | -1.029 | 0.476 | -0.663 | -0.262 | 0.006 | -0.011 | -0.011 | 0.003 | 0.13 | 0.1 | 0.21 | 0.75 | 0.11 | 0.27 |
| 7 | 7052481 | rs4513887 | -1.566 | 0.538 | 0.12 | 0.961 | 0.029 | 0.278 | 0.278 | 0.799 | 0.08 | 0.25 | 0.09 | 0.44 | 0.06 | 0.18 |
| 7 | 7052722 | rs6979563 | 0.088 | 1.031 | -0.744 | -0.326 | -0.031 | 0.068 | 0.068 | 0.151 | 0.29 | 0.17 | 0.4 | 0.07 | 0.37 | 0.43 |
| 7 | 7053611 | rs10261824 | -0.875 | 0.661 | -1.103 | -0.16 | 0.035 | 0.102 | 0.102 | 0 | 0.51 | 0.89 | 0.39 | 0.67 | 0.45 | 0.55 |
| 7 | 7054045 | rs4724913 | 1.102 | 0.766 | -0.098 | -0.248 | -0.001 | -0.001 | -0.001 | 0.391 | 0.63 | 0.44 | 0.71 | 0.38 | 0.67 | 0.53 |
| 7 | 7054164 | rs12671158 | -0.327 | 0.722 | 0.015 | -0.089 | -0.003 | -0.005 | -0.005 | 0 | 0.13 | 0.09 | 0.12 | 0.5 | 0.14 | 0.22 |

| | | | | | | | | | | | | | | | | |
|---|---|---|---|---|---|---|---|---|---|---|---|---|---|---|---|---|
| 7 | 7059936 | rs4075591 | 1.519 | 0.495 | -0.213 | -0.181 | 0.025 | 0.059 | 0.059 | 0 | 0.6 | 0.12 | 0.59 | 0.72 | 0.69 | 0.67 |
| 7 | 7060257 | rs4407769 | 0.448 | 0.381 | 0.672 | -0.236 | 0.023 | 0 | 0 | 0 | 0.89 | 0.79 | 0.87 | 0.89 | 0.87 | 0.81 |
| 7 | 7071374 | rs7781831 | -0.156 | 0.412 | 0.635 | -0.21 | -0.032 | -0.018 | -0.018 | 0 | 0.92 | 0.98 | 0.89 | 0.89 | 0.87 | 0.83 |
| 7 | 7076459 | rs7780465 | -0.906 | 0.242 | 1.314 | -0.281 | -0.054 | -0.169 | -0.169 | 0.535 | 0.25 | 0.47 | 0.12 | 0.01 | 0.14 | 0.12 |
| 7 | 7087248 | rs7788270 | -0.936 | 0.372 | 2.041 | -0.071 | -0.03 | -0.002 | -0.002 | 0 | 0.2 | 0.5 | 0.25 | 0.17 | 0.14 | 0.13 |
| 7 | 7091625 | rs13236240 | 2.41 | 0.436 | 1.511 | -0.112 | -0.001 | 0.199 | 0.199 | 0 | 0.51 | 0.16 | 0.49 | 0.21 | 0.57 | 0.47 |
| 7 | 7093149 | rs6967320 | 1.44 | 0.492 | 1.629 | -0.051 | 0.005 | -0.002 | -0.002 | 0.124 | 0.71 | 0.64 | 0.62 | 0.71 | 0.67 | 0.66 |
| 7 | 7093312 | rs16876441 | 0.1 | 0.455 | 1.262 | 0.044 | -0.012 | -0.002 | -0.002 | 0 | 0.08 | 0.01 | 0.1 | 0.5 | 0.08 | 0.15 |
| 7 | 7097895 | rs12702572 | -0.046 | 0.424 | 1.653 | -0.531 | -0.005 | -0.076 | -0.076 | 0 | 0.16 | 0.25 | 0.12 | 0 | 0.14 | 0.1 |
| 7 | 7116397 | rs10271901 | -0.872 | 0.425 | 1.52 | -0.515 | -0.014 | -0.062 | -0.062 | 1.047 | 0.12 | 0.21 | 0.05 | 0.01 | 0.05 | 0.07 |
| 7 | 7117799 | rs2205542 | -0.499 | 0.53 | 1.755 | -0.205 | 0.003 | 0.142 | 0.142 | 0 | 0.1 | 0.21 | 0.12 | 0.54 | 0.12 | 0.16 |
| 7 | 7117866 | rs2205541 | -0.471 | 0.53 | 1.756 | -0.227 | 0.004 | 0.142 | 0.142 | 0 | 0.1 | 0.21 | 0.12 | 0.54 | 0.12 | 0.16 |
| 7 | 7119469 | rs2280918 | -0.297 | 0.461 | 1.346 | -0.27 | 0.004 | -0.001 | -0.001 | 1.494 | 0.2 | 0.06 | 0.19 | 0.18 | 0.15 | 0.13 |
| 7 | 7120438 | rs4720720 | 2.137 | 0.554 | 1.445 | -0.359 | 0.005 | 0.049 | 0.049 | 0 | 0.57 | 0.31 | 0.46 | 0.16 | 0.61 | 0.47 |
| 7 | 7120557 | rs4720721 | 2.118 | 0.553 | 1.459 | -0.359 | 0.004 | 0.036 | 0.036 | 0 | 0.57 | 0.32 | 0.51 | 0.16 | 0.61 | 0.47 |
| 7 | 7123008 | rs10256956 | 2.569 | 0.612 | 1.161 | -0.508 | -0.03 | 0.312 | 0.312 | 0.272 | 0.5 | 0.12 | 0.48 | 0.14 | 0.57 | 0.44 |
| 7 | 7129858 | rs10260787 | 2.563 | 0.504 | 1.169 | -0.499 | -0.022 | 0.303 | 0.303 | 0.326 | 0.51 | 0.12 | 0.48 | 0.14 | 0.57 | 0.44 |
| 7 | 7130532 | rs1319627 | -1.601 | 0.512 | 1.276 | -0.471 | -0.004 | -0.02 | -0.02 | 1.652 | 0.33 | 0.48 | 0.38 | 0.3 | 0.28 | 0.36 |
| 7 | 7130847 | rs1160673 | 0.687 | 0.502 | 0.988 | -0.352 | 0.001 | 0.149 | 0.149 | 0 | 0.89 | 0.73 | 0.87 | 0.44 | 0.88 | 0.84 |
| 7 | 7136077 | rs17163287 | 1.069 | 0.528 | -0.202 | -0.492 | -0.003 | 0.059 | 0.059 | 0 | 0.7 | 0.9 | 0.68 | 0.25 | 0.72 | 0.71 |
| 7 | 7137541 | rs6963476 | 2.153 | 0.539 | -0.586 | -0.525 | -0.012 | 0.29 | 0.29 | 0.211 | 0.51 | 0.13 | 0.47 | 0.14 | 0.57 | 0.43 |
| 7 | 7138467 | rs974583 | 0.874 | 0.901 | -0.724 | -0.364 | -0.034 | 0.026 | 0.026 | 0 | 0.72 | 0.96 | 0.7 | 0.26 | 0.74 | 0.72 |
| 7 | 7141137 | rs1347025 | -1.03 | 1.014 | 0.2 | 0.26 | 0.032 | -0.041 | -0.041 | 0 | 0.16 | 0.54 | 0.17 | 0.12 | 0.13 | 0.26 |
| 7 | 7141565 | rs4724942 | 0.777 | 1.488 | -0.484 | 0.516 | -0.044 | -0.053 | -0.053 | 0 | 0.91 | 0.99 | 0.88 | 0.43 | 0.88 | 0.84 |
| 7 | 7150307 | rs1432193 | 1.31 | 1.18 | -0.366 | -0.089 | 0.006 | -0.083 | -0.083 | 0 | 0.62 | 0.54 | 0.51 | 0.13 | 0.68 | 0.49 |
| 7 | 7163442 | rs7801523 | 0.888 | 1.187 | -0.925 | 0.68 | -0.042 | -0.084 | -0.084 | 0 | 0.88 | 0.99 | 0.76 | 0.45 | 0.86 | 0.77 |
| 7 | 7167595 | rs7457456 | -0.811 | 1.218 | -0.973 | -0.023 | -0.019 | 0.051 | 0.051 | 1.045 | 0.2 | 0.11 | 0.26 | 0.09 | 0.2 | 0.28 |
| 7 | 7168493 | rs2163640 | -1.359 | 0.72 | 0.197 | 1.026 | 0.008 | 0.309 | 0.309 | 1.369 | 0.13 | 0.4 | 0.16 | 0.35 | 0.1 | 0.17 |
| 7 | 7170481 | rs17308158 | 0.387 | 0.859 | -0.474 | -0.575 | 0 | 0.003 | 0.003 | 0.549 | 0.45 | 0.42 | 0.3 | 0.08 | 0.38 | 0.35 |
| 7 | 7170935 | rs17248900 | -0.463 | 0.553 | -0.913 | -1.134 | 0.017 | 0.046 | 0.046 | 0 | 0.28 | 0.21 | 0.19 | 0.04 | 0.25 | 0.31 |
| 7 | 7171376 | rs6970013 | -0.322 | 0.37 | -0.52 | -0.327 | -0.002 | -0.001 | -0.001 | 0 | 0.43 | 0.44 | 0.4 | 0.41 | 0.44 | 0.23 |
| 7 | 7173681 | rs2060163 | -0.664 | 0.497 | -0.567 | -0.379 | 0.003 | -0.011 | -0.011 | 0.023 | 0.27 | 0.39 | 0.5 | 0.71 | 0.28 | 0.35 |
| 7 | 7174964 | rs6463650 | -1.158 | 0.539 | -1.459 | -0.286 | -0.135 | 0.089 | 0.089 | 0.193 | 0.58 | 0.96 | 0.7 | 0.76 | 0.54 | 0.53 |
| 7 | 7178545 | rs1025444 | -0.524 | 0.564 | -1.06 | 0.579 | -0.009 | -0.029 | -0.029 | 0.126 | 0.09 | 0.06 | 0.35 | 0.39 | 0.1 | 0.24 |
| 7 | 7185752 | rs12531403 | 0.489 | 0.66 | -0.986 | 0.681 | -0.014 | 0.029 | 0.029 | 0 | 0.06 | 0.04 | 0.35 | 0.38 | 0.09 | 0.23 |
| 7 | 7186554 | rs724254 | -0.985 | 0.569 | -0.974 | 0.349 | -0.022 | 0.164 | 0.164 | 0 | 0.52 | 0.84 | 0.67 | 0.75 | 0.51 | 0.48 |
| 7 | 7187869 | rs1574687 | -0.444 | 0.808 | 0.005 | 0.521 | 0.002 | 0.044 | 0.044 | 0 | 0.19 | 0.23 | 0.14 | 0.37 | 0.19 | 0.15 |
| 7 | 7194759 | rs10244129 | 0.571 | 0.873 | 0.378 | 0.498 | 0.005 | 0.063 | 0.063 | 0 | 0.77 | 0.66 | 0.8 | 0.49 | 0.77 | 0.83 |
| 7 | 7196848 | rs2163638 | 0.799 | 0.826 | -0.059 | 0.112 | -0.017 | 0.033 | 0.033 | 0 | 0.17 | 0.13 | 0.14 | 0 | 0.21 | 0.06 |
| 7 | 7196878 | rs11773545 | 0.294 | 0.965 | -0.044 | 0.964 | 0 | 0.016 | 0.016 | 0 | 0.08 | 0.07 | 0.06 | 0.18 | 0.1 | 0.02 |
| 7 | 7205749 | rs4720723 | 0.116 | 1.348 | -0.066 | 1.145 | -0.002 | 0.017 | 0.017 | 0 | 0.13 | 0.11 | 0.34 | 0.39 | 0.19 | 0.34 |
| 7 | 7212769 | rs1594575 | -0.922 | 0.825 | 0.368 | 1.004 | -0.002 | 0.024 | 0.024 | 0.021 | 0.34 | 0.41 | 0.35 | 0.49 | 0.32 | 0.24 |
| 7 | 7215822 | rs11760845 | 0.183 | 0.728 | 0.558 | 0.953 | -0.013 | 0.004 | 0.004 | 0 | 0.14 | 0.2 | 0.09 | 0.16 | 0.09 | 0.14 |
| 7 | 7217796 | rs11762665 | -0.07 | 0.776 | 0.241 | 1.054 | -0.001 | 0.012 | 0.012 | 0 | 0.09 | 0.08 | 0.07 | 0.2 | 0.12 | 0.03 |
| 7 | 7227509 | rs12671825 | -0.508 | 0.975 | 0.324 | 1.063 | 0.013 | 0.141 | 0.141 | 0 | 0.2 | 0.31 | 0.39 | 0.42 | 0.24 | 0.37 |
| 7 | 7250753 | rs11765592 | 0.681 | 1.099 | 0.035 | 1.076 | -0.037 | -0.04 | -0.04 | 0 | 0.07 | 0.01 | 0.05 | 0.2 | 0.11 | 0.03 |
| 7 | 7252775 | rs13246962 | 1.114 | 1.113 | 0.206 | 0.448 | -0.004 | -0.001 | -0.001 | 0.894 | 0.17 | 0.12 | 0.15 | 0 | 0.21 | 0.07 |
| 7 | 7254279 | rs10260052 | 0.729 | 0.927 | 0.336 | 0.442 | -0.002 | 0 | 0 | 0 | 0.13 | 0.16 | 0.08 | 0.17 | 0.07 | 0.14 |
| 7 | 7255017 | rs10238682 | -1.212 | 0.851 | 0.734 | 0.357 | -0.023 | 0.129 | 0.129 | 0 | 0.26 | 0.53 | 0.27 | 0.52 | 0.2 | 0.18 |
| 7 | 7255479 | rs2108790 | -2.328 | 0.842 | 0.433 | 0.018 | -0.016 | 0.161 | 0.161 | 0.052 | 0.61 | 0.85 | 0.79 | 0.93 | 0.64 | 0.58 |
| 7 | 7258339 | rs6960962 | -0.213 | 1.04 | 0.793 | 0.347 | -0.009 | -0.026 | -0.026 | 0.022 | 0.28 | 0.5 | 0.27 | 0.17 | 0.26 | 0.11 |
| 7 | 7260653 | rs11772534 | -0.381 | 1.043 | 0.733 | 0.434 | -0.004 | 0.099 | 0.099 | 0 | 0.18 | 0.28 | 0.22 | 0.53 | 0.17 | 0.07 |
| 7 | 7263117 | rs10272173 | 1.143 | 1.213 | 0.667 | 0.352 | 0.012 | -0.068 | -0.068 | 2.937 | 0.17 | 0.24 | 0.16 | 0 | 0.22 | 0.08 |
| 7 | 7266939 | rs2159191 | -1.371 | 1.172 | 0.516 | 0.401 | -0.017 | -0.028 | -0.028 | 4.856 | 0.39 | 0.76 | 0.41 | 0.52 | 0.4 | 0.17 |
| 7 | 7267787 | rs6960832 | -1.662 | 1.083 | 0.411 | 0.336 | -0.007 | 0.112 | 0.112 | 0 | 0.48 | 0.83 | 0.45 | 0.7 | 0.45 | 0.29 |
| 7 | 7268431 | rs10259085 | -1.662 | 1.134 | 0.412 | 0.337 | -0.024 | 0.112 | 0.112 | 0 | 0.48 | 0.83 | 0.45 | 0.7 | 0.47 | 0.29 |
| 7 | 7268794 | rs12539501 | 0.962 | 1.396 | 0.192 | 0.32 | -0.029 | -0.032 | -0.032 | 0.294 | 0.11 | 0.04 | 0.34 | 0.24 | 0.2 | 0.33 |
| 7 | 7269171 | rs11772919 | 0.027 | 1.408 | 0.264 | 0.332 | 0 | 0.105 | 0.105 | 0 | 0.17 | 0.25 | 0.2 | 0.52 | 0.17 | 0.07 |
| 7 | 7271879 | rs11972596 | 0.993 | 1.501 | 0.126 | 0.367 | -0.042 | -0.046 | -0.046 | 0 | 0.11 | 0.05 | 0.34 | 0.23 | 0.2 | 0.33 |
| 7 | 7273559 | rs1008898 | -1.016 | 1.165 | 0.317 | 0.526 | 0.006 | 0.148 | 0.148 | 0 | 0.22 | 0.47 | 0.26 | 0.53 | 0.2 | 0.19 |
| 7 | 7276879 | rs13226799 | 1.313 | 1.214 | -0.567 | 0.074 | 0.008 | -0.007 | -0.007 | 0 | 0.11 | 0.11 | 0.06 | 0.18 | 0.06 | 0.13 |
| 7 | 7279010 | rs10279432 | -2.354 | 1.194 | -0.76 | -0.115 | -0.076 | 0.258 | 0.258 | 0.014 | 0.62 | 0.97 | 0.8 | 0.93 | 0.65 | 0.65 |
| 7 | 7281805 | rs13245879 | 1.301 | 1.149 | -0.684 | -0.031 | -0.001 | -0.009 | -0.009 | 0 | 0.11 | 0.11 | 0.08 | 0.18 | 0.09 | 0.14 |
| 7 | 7282436 | rs7794210 | -1.161 | 1.085 | -0.54 | 0 | -0.017 | -0.01 | -0.01 | 0.746 | 0.31 | 0.42 | 0.29 | 0.71 | 0.25 | 0.25 |
| 7 | 7283569 | rs1047763 | -0.45 | 1.247 | -0.729 | 0.121 | -0.001 | 0.099 | 0.099 | 0 | 0.19 | 0.29 | 0.23 | 0.53 | 0.18 | 0.09 |
| 7 | 7284546 | rs10251492 | -2.353 | 1.194 | -0.976 | -0.03 | -0.081 | 0.233 | 0.233 | 0.004 | 0.65 | 0.97 | 0.81 | 0.93 | 0.66 | 0.66 |
| 7 | 7286258 | rs5010184 | 0.248 | 1.036 | -0.672 | 0.213 | -0.001 | 0.065 | 0.065 | 0 | 0.16 | 0.19 | 0.2 | 0.53 | 0.16 | 0.07 |
| 7 | 7292458 | rs7778301 | -0.07 | 0.734 | -0.755 | 0.048 | -0.034 | 0.08 | 0.08 | 0 | 0.18 | 0.29 | 0.2 | 0.63 | 0.13 | 0.1 |
| 7 | 7292892 | rs2108780 | -1.546 | 1.224 | -1.705 | -0.53 | -0.119 | 0.272 | 0.272 | 0.093 | 0.61 | 0.97 | 0.78 | 0.93 | 0.62 | 0.64 |

| | | | | | | | | | | | | | | | | |
|---|---|---|---|---|---|---|---|---|---|---|---|---|---|---|---|---|
| 7 | 7298188 | rs17252812 | 0.805 | 1.804 | -1.022 | -0.021 | -0.032 | 0.06 | 0.06 | 0 | 0.15 | 0.06 | 0.14 | 0 | 0.2 | 0.08 |
| 7 | 7301540 | rs13237081 | 0.664 | 2.126 | -0.944 | 0.531 | -0.044 | -0.04 | -0.04 | 0 | 0.08 | 0.03 | 0.3 | 0.17 | 0.19 | 0.25 |
| 7 | 7302293 | rs1299548 | -0.074 | 1.354 | -1.136 | 0.194 | -0.025 | 0.053 | 0.053 | 0.041 | 0.6 | 0.72 | 0.39 | 0.24 | 0.47 | 0.38 |
| 7 | 7304142 | rs10280773 | -0.118 | 1.142 | -0.329 | 0.868 | 0.001 | 0.259 | 0.259 | 0 | 0.07 | 0.19 | 0.15 | 0.48 | 0.07 | 0.05 |
| 7 | 7307032 | rs4724965 | 0.042 | 1.505 | 0.134 | 0.449 | 0.07 | 0.047 | 0.047 | 0 | 0.12 | 0.26 | 0.29 | 0.15 | 0.19 | 0.25 |
| 7 | 7309519 | rs10277495 | -0.502 | 1.354 | 0.652 | 0.902 | 0.015 | 0.029 | 0.029 | 0 | 0.13 | 0.24 | 0.27 | 0.15 | 0.16 | 0.24 |
| 7 | 7309566 | rs6970785 | -0.829 | 1.095 | -0.035 | 0.226 | 0 | 0.001 | 0.001 | 0 | 0.4 | 0.65 | 0.57 | 0.6 | 0.42 | 0.38 |
| 7 | 7310631 | rs16877378 | 1.925 | 1.111 | -0.177 | 0.356 | -0.024 | -0.01 | -0.01 | 0 | 0.07 | 0.02 | 0.1 | 0.26 | 0.12 | 0.03 |
| 7 | 7311593 | rs1294656 | 0.416 | 1.066 | -0.412 | -0.056 | -0.013 | 0.018 | 0.018 | 0 | 0.38 | 0.23 | 0.3 | 0.28 | 0.39 | 0.25 |
| 7 | 7311749 | rs1922630 | 0.562 | 1.181 | -0.377 | 0.077 | 0 | 0.001 | 0.001 | 0 | 0.1 | 0.1 | 0.24 | 0.16 | 0.15 | 0.24 |
| 7 | 7313044 | rs1796120 | -1.89 | 0.804 | -0.439 | -0.266 | 0.007 | 0.024 | 0.024 | 1.456 | 0.75 | 0.87 | 0.85 | 0.88 | 0.79 | 0.6 |
| 7 | 7313151 | rs17165881 | 1.729 | 0.937 | -0.527 | 0.138 | -0.002 | 0.013 | 0.013 | 0 | 0.09 | 0.07 | 0.1 | 0.24 | 0.12 | 0.03 |
| 7 | 7313294 | rs6973768 | 2.29 | 0.813 | -0.602 | 0.221 | 0 | 0.01 | 0.01 | 0.006 | 0.07 | 0.07 | 0.05 | 0.19 | 0.04 | 0.02 |
| 7 | 7313303 | rs1796123 | 0.972 | 0.98 | -0.825 | -0.279 | -0.017 | 0.123 | 0.123 | 0 | 0.28 | 0.12 | 0.19 | 0.03 | 0.26 | 0.23 |
| 7 | 7313715 | rs1541457 | -0.166 | 1.271 | -0.281 | 0.18 | 0.028 | 0.033 | 0.033 | 0 | 0.15 | 0.18 | 0.33 | 0.26 | 0.24 | 0.25 |
| 7 | 7315250 | rs1638201 | 2.261 | 0.454 | -0.78 | 0.126 | -0.004 | -0.062 | -0.062 | 0 | 0.1 | 0.02 | 0.09 | 0.24 | 0.07 | 0.04 |
| 7 | 7315815 | rs1796119 | -0.06 | 0.373 | -0.662 | -0.242 | -0.068 | -0.028 | -0.028 | 0.024 | 0.35 | 0.16 | 0.49 | 0.52 | 0.49 | 0.37 |
| 7 | 7316391 | rs722037 | 0.248 | 0.576 | -0.839 | -0.107 | -0.105 | -0.083 | -0.083 | 0 | 0.27 | 0.09 | 0.45 | 0.52 | 0.39 | 0.32 |
| 7 | 7316977 | rs1638214 | 0.165 | 0.566 | -0.743 | -0.112 | -0.068 | -0.053 | -0.053 | 5.257 | 0.28 | 0.15 | 0.44 | 0.52 | 0.39 | 0.31 |
| 7 | 7317590 | rs2270080 | 1.162 | 0.473 | -0.543 | 0.133 | -0.002 | 0.009 | 0.009 | 3.046 | 0.08 | 0.11 | 0.17 | 0.26 | 0.09 | 0.03 |
| 7 | 7324800 | rs1638202 | -1.925 | 0.062 | -0.559 | -0.304 | 0.007 | 0.027 | 0.027 | 0.023 | 0.74 | 0.84 | 0.84 | 0.88 | 0.77 | 0.61 |
| 7 | 7326948 | rs6463666 | -0.477 | -0.008 | -0.384 | -0.152 | -0.003 | 0.001 | 0.001 | 2.941 | 0.7 | 0.72 | 0.78 | 0.67 | 0.76 | 0.9 |
| 7 | 7328775 | rs2141911 | -1.606 | -0.071 | -0.675 | -0.145 | -0.038 | -0.081 | -0.081 | 0 | 0.82 | 0.96 | 0.85 | 0.75 | 0.81 | 0.93 |
| 7 | 7330622 | rs7784370 | 0.562 | -0.042 | -0.152 | -0.284 | 0 | 0.001 | 0.001 | 0.094 | 0.44 | 0.53 | 0.39 | 0.38 | 0.43 | 0.48 |
| 7 | 7332321 | rs6463669 | -0.212 | -0.262 | -0.175 | -0.285 | 0.005 | 0.052 | 0.052 | 0.242 | 0.47 | 0.32 | 0.54 | 0.62 | 0.51 | 0.48 |
| 7 | 7332580 | rs10273515 | 1.591 | -0.118 | -0.164 | 0.027 | 0.001 | 0.018 | 0.018 | 4.708 | 0.07 | 0.11 | 0.13 | 0.22 | 0.08 | 0.02 |
| 7 | 7333294 | rs1317198 | -0.961 | -0.249 | -0.152 | -0.394 | -0.002 | 0 | 0 | 0.449 | 0.4 | 0.61 | 0.6 | 0.63 | 0.5 | 0.51 |
| 7 | 7335868 | rs17166197 | 1.023 | -0.524 | -0.451 | -0.116 | 0.026 | -0.001 | -0.001 | 0 | 0.15 | 0.07 | 0.1 | 0.14 | 0.08 | 0.17 |
| 7 | 7341918 | rs2348033 | 0.527 | -0.388 | -0.341 | -0.033 | 0.002 | -0.003 | -0.003 | 0 | 0.18 | 0.17 | 0.18 | 0.33 | 0.11 | 0.09 |
| 7 | 7342827 | rs2348034 | 0.12 | -0.076 | -0.383 | -0.117 | 0.03 | 0.026 | 0.026 | 0 | 0.3 | 0.42 | 0.46 | 0.31 | 0.42 | 0.42 |
| 7 | 7343396 | rs6973410 | -1.745 | -0.229 | -1.006 | -0.398 | -0.039 | 0.157 | 0.157 | 0.134 | 0.7 | 0.94 | 0.82 | 0.9 | 0.71 | 0.58 |
| 7 | 7343909 | rs1548774 | 0.519 | -0.158 | -0.683 | -0.403 | -0.032 | -0.13 | -0.13 | 0 | 0.25 | 0.48 | 0.18 | 0.03 | 0.24 | 0.22 |
| 7 | 7343980 | rs12673989 | 1.069 | 0.371 | -0.952 | -0.038 | -0.025 | -0.047 | -0.047 | 0 | 0.19 | 0.05 | 0.16 | 0.24 | 0.19 | 0.07 |
| 7 | 7344571 | rs7784265 | -0.827 | 0.301 | -1.012 | -0.515 | -0.154 | -0.07 | -0.07 | 0.275 | 0.66 | 0.93 | 0.54 | 0.65 | 0.56 | 0.38 |
| 7 | 7345002 | rs6463679 | -0.607 | -0.748 | -0.719 | -1.688 | 0.001 | 0.051 | 0.051 | 0.25 | 0.64 | 0.8 | 0.76 | 0.9 | 0.62 | 0.52 |
| 7 | 7345085 | rs7801475 | 0.531 | 0.13 | 0.152 | -0.857 | 0.008 | -0.057 | -0.057 | 0 | 0.1 | 0.35 | 0.08 | 0 | 0.12 | 0.05 |
| 7 | 7345882 | rs10265493 | 0.244 | 0.219 | -0.189 | -0.844 | -0.017 | -0.08 | -0.08 | 1.401 | 0.12 | 0.3 | 0.1 | 0 | 0.14 | 0.05 |
| 7 | 7347708 | rs6966989 | -0.104 | 0.856 | -0.795 | -0.253 | -0.002 | -0.003 | -0.003 | 0 | 0.26 | 0.24 | 0.4 | 0.27 | 0.36 | 0.29 |
| 7 | 7348198 | rs17166296 | 0.375 | 1.287 | -1.1 | 0.366 | 0.003 | 0.12 | 0.12 | 0.447 | 0.09 | 0.23 | 0.14 | 0.38 | 0.09 | 0.05 |
| 7 | 7348633 | rs10486156 | 0.438 | 1.429 | -1.312 | 0.349 | -0.022 | -0.021 | -0.021 | 0 | 0.15 | 0.07 | 0.33 | 0.27 | 0.29 | 0.25 |
| 7 | 7349495 | rs6956860 | 0.053 | 0.803 | -1.25 | -0.06 | -0.002 | 0.061 | 0.061 | 0.15 | 0.11 | 0.2 | 0.14 | 0.37 | 0.09 | 0.05 |
| 7 | 7350683 | rs10486157 | 0.1 | 0.583 | -1.802 | -1.348 | 0.023 | 0.11 | 0.11 | 0.016 | 0.37 | 0.17 | 0.25 | 0.19 | 0.28 | 0.37 |
| 7 | 7352663 | rs2348038 | 0.415 | 0.902 | -1.238 | -0.733 | -0.003 | 0.044 | 0.044 | 0 | 0.85 | 0.93 | 0.91 | 1 | 0.82 | 0.87 |
| 7 | 7355509 | rs10081184 | 0.801 | 1.4 | -0.557 | -0.48 | 0.063 | -0.071 | -0.071 | 0 | 0.75 | 0.6 | 0.74 | 0.91 | 0.69 | 0.64 |
| 7 | 7361523 | rs1882600 | 0.579 | -0.018 | -1.388 | -0.371 | 0.087 | 0.119 | 0.119 | 0 | 0.52 | 0.17 | 0.55 | 0.72 | 0.64 | 0.71 |
| 7 | 7365006 | rs4724972 | 0.423 | -0.532 | -1.117 | -0.591 | 0.002 | 0.03 | 0.03 | 0.553 | 0.79 | 0.89 | 0.83 | 0.89 | 0.8 | 0.92 |
| 7 | 7365426 | rs1294608 | 1.287 | -0.909 | -0.258 | -0.789 | -0.005 | -0.018 | -0.018 | 0 | 0.69 | 0.35 | 0.77 | 0.88 | 0.75 | 0.82 |
| 7 | 7367474 | rs7811194 | -0.918 | -0.922 | 1.079 | -0.741 | 0.048 | -0.003 | -0.003 | 0 | 0.08 | 0.5 | 0.05 | 0 | 0.05 | 0.02 |
| 7 | 7368604 | rs2077279 | -1.569 | -1.017 | 1.397 | -0.83 | -0.018 | 0.013 | 0.013 | 0.225 | 0.15 | 0.5 | 0.14 | 0.12 | 0.13 | 0.09 |
| 7 | 7389173 | rs2348039 | 1.944 | -1.022 | 2.079 | -0.899 | -0.015 | -0.06 | -0.06 | 0 | 0.73 | 0.42 | 0.77 | 0.87 | 0.76 | 0.82 |
| 7 | 7403978 | rs6974119 | -0.02 | -1.01 | 2.654 | -0.895 | -0.004 | -0.008 | -0.008 | 0 | 0.9 | 0.92 | 0.92 | 1 | 0.9 | 0.98 |
| 7 | 7405069 | rs9640038 | 2.281 | -1.109 | 2.714 | -0.952 | 0.014 | 0.039 | 0.039 | 2.103 | 0.7 | 0.19 | 0.74 | 0.87 | 0.73 | 0.81 |
| 7 | 7413816 | rs7808775 | 2.207 | -1.124 | 2.71 | -0.927 | 0.012 | 0.03 | 0.03 | 1.507 | 0.72 | 0.19 | 0.76 | 0.87 | 0.76 | 0.88 |
| 7 | 7415217 | rs1294651 | 1.656 | -1.137 | 2.859 | -0.54 | -0.002 | -0.001 | -0.001 | 0 | 0.78 | 0.72 | 0.79 | 0.8 | 0.79 | 0.88 |
| 7 | 7420613 | rs1294625 | 2.276 | -1.137 | 2.932 | -0.638 | -0.013 | -0.027 | -0.027 | 0 | 0.75 | 0.3 | 0.78 | 0.89 | 0.78 | 0.86 |
| 7 | 7424599 | rs1294632 | 0.523 | -1.093 | 2.828 | -0.633 | 0 | -0.045 | -0.045 | 1.034 | 0.91 | 0.82 | 0.92 | 1 | 0.92 | 0.96 |
| 7 | 7425664 | rs1294634 | 2.45 | -1.172 | 2.936 | 0.352 | -0.027 | 0.044 | 0.044 | 0.162 | 0.72 | 0.42 | 0.67 | 0.66 | 0.75 | 0.74 |
| 7 | 7426828 | rs17167202 | -1.058 | -1.176 | 2.967 | 0.356 | -0.029 | 0.026 | 0.026 | 0 | 0.12 | 0.35 | 0.11 | 0.12 | 0.09 | 0.07 |
| 7 | 7427535 | rs11526212 | 2.637 | -1.168 | 2.638 | 0.361 | -0.002 | 0.001 | 0.001 | 0.726 | 0.68 | 0.34 | 0.63 | 0.65 | 0.7 | 0.73 |
| 7 | 7435730 | rs10952057 | 1.211 | -1.315 | 2.452 | -0.077 | -0.004 | -0.066 | -0.066 | 0 | 0.85 | 0.78 | 0.91 | 1 | 0.9 | 0.98 |
| 7 | 7437126 | rs4724986 | 2.116 | -1.384 | 2.271 | 0.042 | 0.01 | 0.012 | 0.012 | 0.803 | 0.75 | 0.19 | 0.79 | 0.87 | 0.8 | 0.88 |
| 7 | 7438102 | rs17167230 | -0.751 | -1.338 | 2.126 | 0.253 | -0.008 | 0.003 | 0.003 | 0 | 0.1 | 0.13 | 0.09 | 0.13 | 0.09 | 0.08 |
| 7 | 7438529 | rs12702612 | -1.234 | -1.34 | 2.214 | 0.253 | -0.026 | 0.034 | 0.034 | 0 | 0.12 | 0.36 | 0.1 | 0.13 | 0.09 | 0.08 |
| 7 | 7441152 | rs10235505 | -0.485 | -2.278 | 0.918 | -0.144 | 0.133 | 0.106 | 0.106 | 3.762 | 0.49 | 0.78 | 0.3 | 0.28 | 0.26 | 0.33 |
| 7 | 7441978 | rs6979399 | -0.374 | -2.718 | 0.899 | -0.363 | 0.003 | 0.003 | 0.003 | 0 | 0.15 | 0.15 | 0.11 | 0.08 | 0.14 | 0.15 |
| 7 | 7442169 | rs6962404 | -0.396 | -2.72 | 0.885 | -0.364 | 0.006 | 0.007 | 0.007 | 0 | 0.15 | 0.12 | 0.11 | 0.08 | 0.14 | 0.15 |
| 7 | 7443807 | rs17167323 | 0.831 | -2.988 | 1.341 | -0.459 | -0.014 | -0.004 | -0.004 | 0 | 0.25 | 0.26 | 0.15 | 0.22 | 0.11 | 0.15 |
| 7 | 7447324 | rs1294597 | 0.993 | -2.994 | 1.427 | -0.462 | 0.007 | 0.004 | 0.004 | 4.439 | 0.23 | 0.21 | 0.15 | 0.22 | 0.11 | 0.14 |
| 7 | 7470558 | rs10234561 | 0.905 | -3.041 | 1.585 | -0.379 | -0.036 | -0.016 | -0.016 | 0 | 0.24 | 0.34 | 0.16 | 0.21 | 0.12 | 0.15 |

| | | | | | | | | | | | | | | | | |
|---|---|---|---|---|---|---|---|---|---|---|---|---|---|---|---|---|
| 7 | 7478148 | rs6946354 | -1.14 | -3.31 | 1.565 | -0.729 | -0.024 | -0.041 | -0.041 | 1.094 | 0.22 | 0.32 | 0.12 | 0.04 | 0.18 | 0.19 |
| 7 | 7480382 | rs16878052 | 0.885 | -3.205 | 1.671 | -0.61 | -0.04 | -0.04 | -0.04 | 0 | 0.24 | 0.36 | 0.16 | 0.17 | 0.12 | 0.16 |
| 7 | 7488601 | rs10255201 | 0.785 | -3.214 | 1.615 | -0.614 | -0.057 | -0.057 | -0.057 | 0 | 0.25 | 0.4 | 0.16 | 0.14 | 0.11 | 0.14 |
| 7 | 7490418 | rs10263223 | 0.157 | -3.306 | 1.484 | -0.655 | -0.075 | -0.076 | -0.076 | 0.039 | 0.32 | 0.49 | 0.18 | 0.17 | 0.12 | 0.17 |
| 7 | 7492402 | rs17167922 | -1.072 | -3.341 | 1.025 | -0.672 | 0.135 | 0.144 | 0.144 | 2.436 | 0.51 | 0.8 | 0.3 | 0.21 | 0.28 | 0.33 |
| 7 | 7492995 | rs17167926 | 0.255 | -3.18 | 0.804 | -0.611 | 0.004 | -0.001 | -0.001 | 0 | 0.08 | 0.06 | 0.07 | 0.04 | 0.09 | 0.05 |
| 7 | 7493244 | rs17459676 | 0.687 | -3.007 | 0.714 | -0.59 | -0.001 | 0.02 | 0.02 | 0 | 0.08 | 0.04 | 0.03 | 0 | 0.06 | 0.11 |
| 7 | 7497676 | rs12234520 | 1.665 | -2.287 | -0.312 | -0.083 | -0.055 | -0.109 | -0.109 | 0 | 0.17 | 0 | 0.14 | 0.36 | 0.18 | 0.32 |
| 7 | 7498405 | rs6963106 | 1.404 | -2.605 | 0.087 | -0.231 | 0.052 | 0.054 | 0.054 | 5.084 | 0.23 | 0.09 | 0.15 | 0.14 | 0.11 | 0.14 |
| 7 | 7517366 | rs6943692 | -0.989 | -1.627 | -0.365 | -0.145 | -0.029 | -0.072 | -0.072 | 0 | 0.62 | 0.89 | 0.44 | 0.48 | 0.43 | 0.5 |
| 7 | 7528385 | rs10250756 | 1.127 | -1.179 | 0.46 | -0.221 | 0.001 | 0.001 | 0.001 | 4.05 | 0.25 | 0.25 | 0.17 | 0.17 | 0.12 | 0.14 |
| 7 | 7531006 | rs6946913 | 0.839 | -1.036 | 0.436 | -0.238 | -0.075 | -0.073 | -0.073 | 0 | 0.27 | 0.54 | 0.17 | 0.17 | 0.12 | 0.17 |
| 7 | 7531819 | rs11772657 | 0.007 | -1.084 | 0.074 | -0.383 | 0.01 | 0.005 | 0.005 | 0.184 | 0.62 | 0.33 | 0.77 | 0.83 | 0.79 | 0.73 |
| 7 | 7533908 | rs972766 | 1.34 | -1.161 | -0.065 | -0.13 | 0.059 | 0.05 | 0.05 | 5.106 | 0.23 | 0.09 | 0.15 | 0.15 | 0.11 | 0.16 |
| 7 | 7534979 | rs13312659 | 0.18 | -0.96 | -0.006 | 0.122 | 0.013 | 0.009 | 0.009 | 1.063 | 0.59 | 0.31 | 0.66 | 0.62 | 0.7 | 0.66 |
| 7 | 7539233 | rs11971297 | 0.863 | -0.691 | -0.398 | 0.253 | -0.003 | -0.003 | -0.003 | 0 | 0.13 | 0.11 | 0.14 | 0.21 | 0.15 | 0.17 |
| 7 | 7553733 | rs7795005 | -1.192 | -0.153 | -1.048 | 0.236 | -0.068 | -0.108 | -0.108 | 0.019 | 0.7 | 0.93 | 0.69 | 0.55 | 0.66 | 0.58 |
| 7 | 7554505 | rs17168369 | 1.117 | -0.628 | -0.254 | 0.267 | 0.055 | 0.048 | 0.048 | 7.105 | 0.22 | 0.06 | 0.14 | 0.14 | 0.1 | 0.13 |
| 7 | 7555713 | rs7791808 | 0.41 | -0.634 | -0.14 | 0.264 | 0.004 | -0.001 | -0.001 | 0 | 0.9 | 0.92 | 0.94 | 0.95 | 0.91 | 0.94 |
| 7 | 7555825 | rs7778306 | 0.873 | -0.804 | 0.484 | 0.287 | 0.064 | 0.039 | 0.039 | 0.01 | 0.52 | 0.24 | 0.68 | 0.67 | 0.64 | 0.6 |
| 7 | 7559541 | rs17151557 | 0.104 | -0.75 | 1.213 | 0.266 | -0.01 | -0.008 | -0.008 | 0 | 0.35 | 0.41 | 0.23 | 0.28 | 0.26 | 0.34 |
| 7 | 7569303 | rs7782245 | 1.45 | -0.928 | 0.99 | 0.275 | 0.062 | 0.087 | 0.087 | 6.972 | 0.21 | 0.02 | 0.07 | 0.06 | 0.08 | 0.14 |
| 7 | 7570959 | rs17168516 | 0.034 | -0.844 | 1.426 | 0.325 | -0.009 | -0.005 | -0.005 | 0 | 0.35 | 0.6 | 0.23 | 0.28 | 0.23 | 0.32 |
| 7 | 7588857 | rs12532693 | 0.525 | -0.685 | 1.129 | 0.377 | 0.041 | 0.066 | 0.066 | 3.427 | 0.57 | 0.13 | 0.7 | 0.71 | 0.69 | 0.66 |
| 7 | 7590175 | rs4724999 | 0.881 | -0.648 | 1.176 | 0.407 | 0.076 | 0.054 | 0.054 | 1.025 | 0.5 | 0.16 | 0.66 | 0.66 | 0.63 | 0.61 |
| 7 | 7618720 | rs1544465 | 0.374 | -0.573 | 1.715 | 0.019 | -0.008 | -0.004 | -0.004 | 0 | 0.59 | 0.45 | 0.73 | 0.66 | 0.71 | 0.64 |
| 7 | 7645806 | rs2286209 | 0.54 | -0.558 | 1.625 | -0.005 | 0.012 | 0.008 | 0.008 | 1.979 | 0.58 | 0.4 | 0.74 | 0.67 | 0.71 | 0.69 |
| 7 | 7646687 | rs4222 | 0.484 | -0.323 | 1.756 | -0.009 | -0.038 | -0.022 | -0.022 | 0 | 0.66 | 0.56 | 0.8 | 0.74 | 0.77 | 0.67 |
| 7 | 7653628 | rs10250880 | 0.153 | -0.24 | 1.71 | -0.199 | -0.027 | -0.054 | -0.054 | 0 | 0.27 | 0.35 | 0.13 | 0.08 | 0.13 | 0.18 |
| 7 | 7655026 | rs11772656 | 0.078 | -0.26 | 1.54 | -0.185 | -0.056 | -0.116 | -0.116 | 0 | 0.69 | 0.45 | 0.85 | 0.93 | 0.84 | 0.81 |
| 7 | 7657304 | rs1557996 | 0.08 | -0.241 | 1.41 | -0.177 | -0.047 | -0.1 | -0.1 | 1.015 | 0.7 | 0.55 | 0.85 | 0.92 | 0.84 | 0.81 |
| 7 | 7657786 | rs11975945 | 0.541 | 0.164 | 0.916 | 0.218 | 0.006 | 0.038 | 0.038 | 0 | 0.1 | 0.08 | 0.12 | 0.28 | 0.15 | 0.17 |
| 7 | 7660540 | rs12702622 | 0.215 | 0.146 | 1.341 | -0.085 | 0.097 | 0.284 | 0.284 | 0 | 0.86 | 0.59 | 0.78 | 0.6 | 0.82 | 0.78 |
| 7 | 7663794 | rs886727 | 0.858 | -0.156 | 1.015 | -0.322 | 0.034 | -0.023 | -0.023 | 0 | 0.58 | 0.3 | 0.67 | 0.54 | 0.71 | 0.66 |
| 7 | 7664696 | rs10265974 | -0.388 | 0.101 | 0.365 | -2.002 | -0.021 | 0.031 | 0.031 | 0.572 | 0.65 | 0.79 | 0.47 | 0.16 | 0.54 | 0.37 |
| 7 | 7668666 | rs10254096 | -0.255 | -0.06 | 0.349 | -2.029 | -0.037 | 0.069 | 0.069 | 0.336 | 0.63 | 0.85 | 0.52 | 0.17 | 0.55 | 0.34 |
| 7 | 7669560 | rs7798204 | 0.233 | 0.027 | 0.254 | -2.001 | -0.016 | 0.044 | 0.044 | 0.277 | 0.36 | 0.19 | 0.48 | 0.83 | 0.43 | 0.66 |
| 7 | 7670797 | rs10280932 | 0.579 | 0.084 | 0.086 | -1.982 | 0.001 | -0.015 | -0.015 | 0 | 0.4 | 0.64 | 0.44 | 0.1 | 0.44 | 0.22 |
| 7 | 7672023 | rs6962587 | 0.966 | 0.204 | -0.82 | -1.886 | -0.027 | 0.047 | 0.047 | 1.036 | 0.34 | 0.27 | 0.41 | 0.1 | 0.43 | 0.21 |
| 7 | 7672891 | rs17466674 | -0.141 | 1.096 | -1.015 | -1.378 | -0.005 | 0.006 | 0.006 | 0 | 0.14 | 0.11 | 0.13 | 0.1 | 0.15 | 0.13 |
| 7 | 7677757 | rs6463709 | 0.469 | 1.056 | -1.37 | -1.091 | -0.055 | -0.045 | -0.045 | 0 | 0.23 | 0.02 | 0.36 | 0.76 | 0.3 | 0.56 |
| 7 | 7679885 | rs3735460 | 0.105 | 0.608 | -1.12 | -1.283 | -0.02 | 0.012 | 0.012 | 0 | 0.11 | 0.01 | 0.18 | 0.09 | 0.09 | 0.09 |
| 7 | 7680632 | rs10251718 | -1.048 | 0.317 | -0.194 | -1.409 | -0.019 | -0.052 | -0.052 | 0.91 | 0.1 | 0.35 | 0.06 | 0.01 | 0.03 | 0.07 |
| 7 | 7680697 | rs17169194 | 0.59 | 0.814 | -0.778 | -1.466 | -0.035 | 0.005 | 0.005 | 0 | 0.09 | 0.02 | 0.07 | 0.06 | 0.1 | 0.03 |
| 7 | 7681020 | rs7792009 | 0.817 | 0.843 | -1.224 | -1.787 | -0.046 | -0.009 | -0.009 | 0 | 0.23 | 0.03 | 0.3 | 0.76 | 0.31 | 0.55 |
| 7 | 7681131 | rs17169196 | -0.609 | 0.294 | -0.585 | -2.056 | -0.022 | -0.03 | -0.03 | 0.039 | 0.08 | 0.33 | 0.03 | 0.01 | 0.01 | 0.05 |
| 7 | 7681576 | rs3757527 | -0.435 | 0.445 | -1.44 | -2.783 | -0.013 | 0.19 | 0.19 | 1.035 | 0.6 | 0.13 | 0.54 | 0.87 | 0.64 | 0.75 |
| 7 | 7684270 | rs4720745 | 0.84 | 0.342 | -0.785 | -3.034 | 0.016 | 0.043 | 0.043 | 0 | 0.52 | 0.61 | 0.66 | 0.89 | 0.64 | 0.69 |
| 7 | 7687826 | rs1859604 | 0.604 | 0.321 | -0.579 | -3.028 | 0.016 | 0.132 | 0.132 | 0.307 | 0.54 | 0.73 | 0.67 | 0.89 | 0.64 | 0.69 |
| 7 | 7687840 | rs1859605 | 1.378 | 0.409 | -0.73 | -3.076 | -0.111 | 0.028 | 0.028 | 0 | 0.25 | 0.04 | 0.34 | 0.8 | 0.39 | 0.54 |
| 7 | 7688023 | rs10230651 | 1.34 | 0.388 | -0.691 | -3.046 | -0.014 | -0.056 | -0.056 | 0 | 0.28 | 0.56 | 0.32 | 0.08 | 0.25 | 0.15 |
| 7 | 7690424 | rs17545935 | 1.106 | 0.535 | -1.279 | -3.003 | -0.028 | 0.102 | 0.102 | 0.26 | 0.19 | 0.03 | 0.12 | 0 | 0.17 | 0.1 |
| 7 | 7691176 | rs12668835 | 1.172 | 0.793 | -1.119 | -2.813 | -0.115 | 0.026 | 0.026 | 0 | 0.25 | 0.04 | 0.34 | 0.8 | 0.39 | 0.54 |
| 7 | 7692483 | rs886724 | 1.086 | 0.637 | -0.666 | -2.842 | -0.029 | 0.027 | 0.027 | 0 | 0.26 | 0.19 | 0.35 | 0.81 | 0.39 | 0.54 |
| 7 | 7703374 | rs7781054 | 1.141 | 0.21 | 0.354 | -2.598 | 0 | -0.013 | -0.013 | 0 | 0.61 | 0.57 | 0.69 | 0.84 | 0.69 | 0.62 |
| 7 | 7703579 | rs1012997 | 0.149 | 0.175 | 0.348 | -2.643 | 0.022 | 0.017 | 0.017 | 0 | 0.89 | 0.72 | 0.84 | 0.85 | 0.87 | 0.76 |
| 7 | 7703960 | rs1557997 | 1.286 | 0.173 | 0.407 | -2.644 | -0.003 | -0.038 | -0.038 | 0 | 0.61 | 0.52 | 0.69 | 0.84 | 0.69 | 0.62 |
| 7 | 7705275 | rs4720750 | 0.803 | 0.154 | 0.341 | -2.645 | 0 | -0.005 | -0.005 | 0 | 0.8 | 0.74 | 0.8 | 0.79 | 0.79 | 0.71 |
| 7 | 7709154 | rs7791564 | 1.636 | 0.136 | 0.342 | -2.639 | 0.006 | 0.017 | 0.017 | 0 | 0.52 | 0.52 | 0.64 | 0.78 | 0.61 | 0.58 |
| 7 | 7710914 | rs13227585 | -0.012 | 0.115 | -0.061 | -2.69 | 0.007 | 0.071 | 0.071 | 0.664 | 0.25 | 0.12 | 0.14 | 0.01 | 0.19 | 0.13 |
| 7 | 7719987 | rs10264890 | 1.566 | 0.174 | 0.244 | -2.558 | 0.015 | 0.095 | 0.095 | 0 | 0.48 | 0.3 | 0.62 | 0.78 | 0.61 | 0.59 |
| 7 | 7749866 | rs1045580 | -0.735 | 0.459 | -0.546 | -2.436 | -0.006 | 0.185 | 0.185 | 0.159 | 0.34 | 0.12 | 0.21 | 0.07 | 0.29 | 0.22 |
| 7 | 7750990 | rs11773126 | -1.108 | 0.435 | -0.226 | -2.413 | 0.007 | 0.041 | 0.041 | 1.495 | 0.39 | 0.29 | 0.26 | 0.17 | 0.33 | 0.34 |
| 7 | 7752559 | rs13242539 | -0.713 | 0.708 | -0.371 | -2.439 | -0.021 | 0.007 | 0.007 | 0 | 0.15 | 0.07 | 0.17 | 0.13 | 0.19 | 0.21 |
| 7 | 7754563 | rs17136498 | 0.989 | 0.639 | -0.447 | -2.436 | -0.007 | 0.013 | 0.013 | 0 | 0.08 | 0.01 | 0.05 | 0.06 | 0.07 | 0.05 |
| 7 | 7755797 | rs2160138 | -1.486 | 0.596 | -0.271 | -2.487 | -0.002 | 0.086 | 0.086 | 0.625 | 0.48 | 0.31 | 0.38 | 0.2 | 0.43 | 0.42 |
| 7 | 7769363 | rs4720751 | -1.654 | 0.505 | 0.524 | -2.085 | 0.015 | -0.017 | -0.017 | 0 | 0.26 | 0.42 | 0.24 | 0.24 | 0.25 | 0.26 |
| 7 | 7769505 | rs11760820 | 0.539 | 0.496 | 0.726 | -2.104 | -0.002 | 0.002 | 0.002 | 0 | 0.08 | 0.16 | 0.06 | 0.08 | 0.07 | 0.03 |
| 7 | 7770111 | rs11978146 | -1.647 | 0.517 | 0.793 | -2.08 | 0.037 | -0.014 | -0.014 | 0 | 0.19 | 0.4 | 0.19 | 0.16 | 0.19 | 0.23 |

| | | | | | | | | | | | | | | | | |
|---|---|---|---|---|---|---|---|---|---|---|---|---|---|---|---|---|
| 7 | 7770203 | rs11971584 | -0.634 | 0.48 | 0.693 | -2.143 | -0.001 | -0.038 | -0.038 | 0 | 0.12 | 0.22 | 0.12 | 0.03 | 0.1 | 0.1 |
| 7 | 7771501 | rs7804204 | -1.36 | 0.265 | 0.777 | -2.268 | -0.015 | -0.09 | -0.09 | 0.675 | 0.16 | 0.41 | 0.13 | 0.04 | 0.1 | 0.14 |
| 7 | 7772839 | rs17136863 | 1.018 | -0.135 | 0.296 | -2.425 | 0.015 | 0.049 | 0.049 | 1.113 | 0.09 | 0.01 | 0.02 | 0 | 0.04 | 0.01 |
| 7 | 7773841 | rs17136898 | -1.252 | -0.119 | 0.384 | -2.417 | 0.015 | 0.035 | 0.035 | 0 | 0.19 | 0.16 | 0.11 | 0.1 | 0.14 | 0.16 |
| 7 | 7774684 | rs10952069 | -2.418 | -0.137 | 1.183 | -2.471 | -0.009 | -0.035 | -0.035 | 0.023 | 0.29 | 0.51 | 0.19 | 0.23 | 0.23 | 0.29 |
| 7 | 7777975 | rs17137001 | -0.376 | -0.15 | 1.258 | -2.47 | 0 | -0.014 | -0.014 | 0 | 0.1 | 0.13 | 0.06 | 0.02 | 0.07 | 0.12 |
| 7 | 7778310 | rs2241075 | -1.406 | -0.15 | 1.351 | -2.475 | 0.017 | 0.018 | 0.018 | 0 | 0.18 | 0.36 | 0.13 | 0.21 | 0.16 | 0.17 |
| 7 | 7779091 | rs13237260 | -2.426 | -0.136 | 1.316 | -2.468 | -0.008 | -0.038 | -0.038 | 0 | 0.28 | 0.48 | 0.18 | 0.23 | 0.23 | 0.29 |
| 7 | 7779324 | rs10486187 | -0.775 | -0.024 | 1.341 | -2.391 | 0.033 | 0.031 | 0.031 | 0 | 0.08 | 0.16 | 0.07 | 0.13 | 0.1 | 0.13 |
| 7 | 7782919 | rs2024372 | -2.182 | 0.045 | 0.556 | -3.023 | -0.003 | 0.012 | 0.012 | 0.509 | 0.58 | 0.57 | 0.37 | 0.24 | 0.49 | 0.45 |
| 7 | 7785047 | rs717297 | -1.188 | -0.212 | 0.993 | -2.903 | -0.017 | -0.145 | -0.145 | 0.043 | 0.23 | 0.44 | 0.15 | 0.01 | 0.19 | 0.07 |
| 7 | 7786415 | rs17137154 | 0.552 | 0.023 | 0.445 | -2.832 | -0.025 | -0.011 | -0.011 | 0 | 0.09 | 0 | 0.06 | 0.08 | 0.09 | 0.03 |
| 7 | 7787006 | rs11984316 | -1.312 | -0.162 | 1.076 | -2.869 | -0.003 | -0.083 | -0.083 | 0.315 | 0.14 | 0.49 | 0.1 | 0.03 | 0.1 | 0.11 |
| 7 | 7788142 | rs1541549 | -2.008 | -0.291 | 0.996 | -2.942 | 0.004 | -0.02 | -0.02 | 0 | 0.33 | 0.63 | 0.22 | 0.03 | 0.28 | 0.16 |
| 7 | 7788155 | rs1541550 | -0.471 | -0.097 | 1.069 | -2.778 | 0.027 | -0.033 | -0.033 | 0 | 0.1 | 0.35 | 0.07 | 0.03 | 0.07 | 0.08 |
| 7 | 7790168 | rs17137233 | -0.554 | 0.189 | 0.431 | -2.453 | -0.001 | -0.017 | -0.017 | 0 | 0.18 | 0.12 | 0.12 | 0 | 0.19 | 0.04 |
| 7 | 7792702 | rs17137257 | -0.686 | 0.369 | 0.438 | -2.042 | 0.003 | -0.004 | -0.004 | 0 | 0.13 | 0.18 | 0.11 | 0.16 | 0.14 | 0.14 |
| 7 | 7794484 | rs13230086 | -1.399 | 0.515 | 0.225 | -2.38 | -0.001 | -0.042 | -0.042 | 0 | 0.34 | 0.58 | 0.3 | 0.19 | 0.29 | 0.37 |
| 7 | 7795531 | rs13234855 | -1.217 | 0.405 | -0.218 | -2.087 | -0.021 | 0.048 | 0.048 | 0 | 0.38 | 0.27 | 0.32 | 0.22 | 0.43 | 0.29 |
| 7 | 7798217 | rs17151585 | 0.378 | 0.032 | 0.336 | -2.062 | -0.004 | -0.034 | -0.034 | 0 | 0.09 | 0.11 | 0.1 | 0 | 0.06 | 0.08 |
| 7 | 7798728 | rs6463715 | -0.591 | -0.416 | 0.622 | -2.366 | -0.022 | -0.059 | -0.059 | 0 | 0.12 | 0.16 | 0.05 | 0 | 0.07 | 0.08 |
| 7 | 7798734 | rs6463716 | -2.064 | -0.311 | 0.972 | -2.457 | -0.018 | -0.135 | -0.135 | 0 | 0.29 | 0.49 | 0.26 | 0.04 | 0.23 | 0.18 |
| 7 | 7800214 | rs11764499 | -0.689 | -0.867 | 0.085 | -2.727 | 0.019 | 0.093 | 0.093 | 0.773 | 0.26 | 0.08 | 0.24 | 0.09 | 0.16 | 0.19 |
| 7 | 7800451 | rs10258735 | -1.864 | -0.815 | 0.204 | -2.836 | -0.008 | -0.122 | -0.122 | 4.209 | 0.33 | 0.5 | 0.28 | 0.1 | 0.3 | 0.33 |
| 7 | 7800487 | rs11765896 | 0.307 | -0.958 | 0.122 | -2.682 | 0.007 | 0.061 | 0.061 | 0.685 | 0.17 | 0 | 0.06 | 0.08 | 0.09 | 0.11 |
| 7 | 7800687 | rs17137412 | -0.396 | -0.689 | 0.65 | -2.375 | 0.073 | 0.01 | 0.01 | 0.1 | 0.07 | 0.32 | 0.09 | 0.03 | 0.11 | 0.05 |
| 7 | 7801684 | rs17137415 | -1.676 | -0.729 | 0.347 | -2.447 | -0.007 | -0.115 | -0.115 | 0.007 | 0.34 | 0.56 | 0.31 | 0.06 | 0.33 | 0.26 |
| 7 | 7804403 | rs9692542 | -1.71 | -0.981 | 0.038 | -1.762 | 0.026 | 0.039 | 0.039 | 0 | 0.36 | 0.73 | 0.31 | 0.22 | 0.29 | 0.32 |
| 7 | 7804891 | rs17137468 | -1.106 | -1.226 | 0.462 | -1.582 | -0.023 | -0.035 | -0.035 | 0.132 | 0.14 | 0.3 | 0.09 | 0.09 | 0.08 | 0.07 |
| 7 | 7805234 | rs10085885 | -1.345 | -1.062 | -0.113 | -1.548 | -0.035 | -0.037 | -0.037 | 0 | 0.25 | 0.6 | 0.21 | 0.14 | 0.15 | 0.25 |
| 7 | 7806677 | rs11761203 | -0.235 | -0.56 | -0.986 | -1.207 | 0.01 | 0.057 | 0.057 | 1.026 | 0.18 | 0.02 | 0.06 | 0.1 | 0.11 | 0.18 |
| 7 | 7807571 | rs10085476 | -1.306 | -0.186 | -0.134 | -0.734 | -0.011 | -0.02 | -0.02 | 2.376 | 0.24 | 0.32 | 0.22 | 0.18 | 0.17 | 0.22 |
| 7 | 7808253 | rs10276762 | -1.111 | 0.371 | -0.46 | 0.661 | -0.004 | 0.072 | 0.072 | 0 | 0.27 | 0.56 | 0.28 | 0.59 | 0.23 | 0.48 |
| 7 | 7809142 | rs6955585 | -0.795 | 1.042 | -1.087 | 1.131 | -0.016 | -0.041 | -0.041 | 0 | 0.21 | 0.14 | 0.29 | 0.39 | 0.23 | 0.37 |
| 7 | 7810122 | rs7357092 | 0.074 | 0.575 | -1.498 | 1.032 | 0 | -0.019 | -0.019 | 0.01 | 0.78 | 0.91 | 0.89 | 0.82 | 0.83 | 0.82 |
| 7 | 7811042 | rs11768218 | -0.084 | 0.944 | -1.456 | 0.83 | -0.05 | 0.015 | 0.015 | 0 | 0.39 | 0.23 | 0.39 | 0.37 | 0.47 | 0.36 |
| 7 | 7813890 | rs11771147 | -0.109 | 0.737 | -0.302 | 0.77 | 0.007 | -0.004 | -0.004 | 0 | 0.41 | 0.42 | 0.41 | 0.4 | 0.47 | 0.37 |
| 7 | 7814659 | rs11764567 | -0.417 | 0.786 | -0.23 | 0.868 | -0.01 | 0.003 | 0.003 | 0.059 | 0.46 | 0.66 | 0.48 | 0.5 | 0.49 | 0.46 |
| 7 | 7818884 | rs11978293 | -0.163 | 0.767 | -0.208 | 0.852 | -0.008 | 0.017 | 0.017 | 0 | 0.43 | 0.36 | 0.45 | 0.44 | 0.52 | 0.4 |
| 7 | 7821594 | rs6944658 | -0.176 | 0.732 | -0.019 | 0.835 | 0 | 0.005 | 0.005 | 0 | 0.45 | 0.46 | 0.5 | 0.43 | 0.52 | 0.44 |
| 7 | 7822303 | rs10272400 | 0.41 | 0.853 | 0.082 | 1.002 | -0.011 | 0.004 | 0.004 | 0 | 0.4 | 0.55 | 0.42 | 0.55 | 0.34 | 0.44 |
| 7 | 7822787 | rs10275971 | 0.558 | 0.272 | -0.106 | 0.915 | 0.012 | -0.02 | -0.02 | 0 | 0.66 | 0.29 | 0.78 | 0.61 | 0.75 | 0.7 |
| 7 | 7827038 | rs6958354 | 0.444 | 0.024 | -0.505 | 0.006 | 0 | -0.002 | -0.002 | 0 | 0.53 | 0.51 | 0.61 | 0.74 | 0.53 | 0.63 |
| 7 | 7828731 | rs11983544 | 0.132 | -0.109 | -0.284 | -0.132 | -0.001 | 0.014 | 0.014 | 0 | 0.64 | 0.71 | 0.76 | 0.77 | 0.68 | 0.76 |
| 7 | 7829515 | rs4725029 | 0.089 | -0.545 | -0.432 | -0.336 | -0.003 | -0.018 | -0.018 | 0 | 0.87 | 0.85 | 0.91 | 0.99 | 0.92 | 0.94 |
| 7 | 7831739 | rs6966185 | 0.301 | -0.704 | -0.6 | -0.175 | 0.01 | 0.006 | 0.006 | 1.291 | 0.44 | 0.36 | 0.52 | 0.67 | 0.58 | 0.69 |
| 7 | 7833725 | rs6957398 | 1.022 | -1.163 | -0.369 | -0.211 | -0.047 | -0.014 | -0.014 | 0.059 | 0.23 | 0.32 | 0.21 | 0.15 | 0.15 | 0.1 |
| 7 | 7834469 | rs10278342 | 1.351 | -0.193 | -0.875 | 0.424 | -0.085 | -0.061 | -0.061 | 1.191 | 0.28 | 0.11 | 0.38 | 0.39 | 0.36 | 0.4 |
| 7 | 7838994 | rs6975739 | 0.089 | -0.231 | -0.242 | 0.711 | 0.012 | 0.094 | 0.094 | 0.015 | 0.09 | 0.29 | 0.06 | 0.15 | 0.08 | 0.09 |
| 7 | 7839445 | rs6960751 | 1.159 | -0.275 | -0.463 | 0.627 | -0.001 | 0.001 | 0.001 | 0.751 | 0.63 | 0.38 | 0.67 | 0.61 | 0.63 | 0.69 |
| 7 | 7840613 | rs10244530 | 0.139 | -0.259 | -0.537 | 0.443 | -0.011 | 0.007 | 0.007 | 0 | 0.16 | 0.3 | 0.1 | 0.16 | 0.1 | 0.15 |
| 7 | 7841423 | rs17137990 | 2.226 | -0.47 | -0.617 | 0.35 | 0.041 | -0.015 | -0.015 | 0.189 | 0.08 | 0 | 0.03 | 0.08 | 0.03 | 0.02 |
| 7 | 7841524 | rs6463722 | 0.332 | -0.148 | -0.759 | 0.258 | 0.003 | 0.005 | 0.005 | 0.023 | 0.46 | 0.58 | 0.37 | 0.26 | 0.43 | 0.23 |
| 7 | 7842281 | rs12671658 | 0.268 | -0.15 | -0.842 | 0.258 | 0.016 | 0.035 | 0.035 | 0.005 | 0.47 | 0.62 | 0.38 | 0.26 | 0.43 | 0.23 |
| 7 | 7843721 | rs6943366 | 1.478 | -0.171 | -0.842 | 0.3 | -0.012 | -0.009 | -0.009 | 0 | 0.27 | 0.28 | 0.23 | 0.19 | 0.2 | 0.1 |
| 7 | 7844955 | rs7793050 | 2.014 | 0.311 | -1.305 | 0.632 | -0.072 | -0.062 | -0.062 | 0 | 0.22 | 0.03 | 0.23 | 0.22 | 0.23 | 0.32 |
| 7 | 7845138 | rs4725030 | -0.253 | -0.298 | -0.828 | 0.826 | -0.008 | 0.013 | 0.013 | 0 | 0.18 | 0.22 | 0.18 | 0.21 | 0.14 | 0.14 |
| 7 | 7845273 | rs4725031 | 0.255 | -0.492 | -0.76 | 0.828 | -0.042 | 0.001 | 0.001 | 0.165 | 0.09 | 0.21 | 0.04 | 0.06 | 0.02 | 0.05 |
| 7 | 7845574 | rs6946178 | -1.688 | -0.284 | -0.878 | 0.962 | -0.013 | -0.01 | -0.01 | 0 | 0.94 | 0.98 | 0.89 | 0.77 | 0.88 | 0.82 |
| 7 | 7846426 | rs6969660 | 0.875 | -0.115 | -0.625 | 0.877 | 0 | 0.001 | 0.001 | 0.94 | 0.63 | 0.6 | 0.56 | 0.46 | 0.57 | 0.58 |
| 7 | 7846920 | rs7801961 | -0.512 | 0.097 | -0.9 | 0.657 | -0.036 | -0.027 | -0.027 | 0 | 0.87 | 0.96 | 0.83 | 0.77 | 0.81 | 0.76 |
| 7 | 7848087 | rs10237037 | 0.628 | -0.069 | -0.758 | 0.591 | -0.02 | -0.003 | -0.003 | 0 | 0.08 | 0.12 | 0.04 | 0.06 | 0.02 | 0.06 |
| 7 | 7848912 | rs10259199 | 0.918 | -0.187 | -0.591 | 0.419 | -0.017 | -0.041 | -0.041 | 0.306 | 0.18 | 0.27 | 0.11 | 0.07 | 0.15 | 0.06 |
| 7 | 7849547 | rs6971634 | 1.197 | -0.059 | -0.739 | 0.733 | 0.002 | 0.001 | 0.001 | 0 | 0.25 | 0.22 | 0.26 | 0.19 | 0.23 | 0.15 |
| 7 | 7851307 | rs10249817 | 0.971 | -0.012 | -0.674 | 0.694 | 0.004 | -0.015 | -0.015 | 0 | 0.45 | 0.51 | 0.34 | 0.31 | 0.47 | 0.48 |
| 7 | 7851688 | rs12702654 | -0.786 | 0.171 | -0.55 | 0.389 | -0.005 | 0.002 | 0.002 | 0 | 0.94 | 0.97 | 0.89 | 0.78 | 0.88 | 0.84 |
| 7 | 7854541 | rs4725035 | 0.148 | 0.145 | -0.609 | 0.387 | -0.004 | 0.01 | 0.01 | 0 | 0.62 | 0.71 | 0.57 | 0.64 | 0.6 | 0.64 |
| 7 | 7854806 | rs4725037 | -0.504 | -0.025 | -0.49 | 0.794 | -0.013 | -0.01 | -0.01 | 0 | 0.94 | 0.98 | 0.89 | 0.9 | 0.9 | 0.86 |
| 7 | 7856669 | rs6956438 | 0.612 | -0.167 | -0.296 | 0.798 | -0.007 | 0.068 | 0.068 | 0.048 | 0.87 | 0.75 | 0.81 | 0.7 | 0.87 | 0.79 |

| | | | | | | | | | | | | | | | | |
|---|---|---|---|---|---|---|---|---|---|---|---|---|---|---|---|---|
| 7 | 7856810 | rs6943254 | 0.203 | -0.209 | -0.562 | 0.503 | -0.021 | 0.004 | 0.004 | 0.227 | 0.62 | 0.49 | 0.56 | 0.56 | 0.67 | 0.64 |
| 7 | 7859202 | rs2194656 | 0.311 | 0.215 | -0.772 | 0.143 | -0.002 | 0.015 | 0.015 | 2.633 | 0.32 | 0.37 | 0.26 | 0.46 | 0.35 | 0.4 |
| 7 | 7859336 | rs2194658 | -0.432 | 0.576 | -1.154 | -0.055 | 0.002 | 0.015 | 0.015 | 0.044 | 0.65 | 0.58 | 0.57 | 0.6 | 0.63 | 0.62 |
| 7 | 7861786 | rs11762146 | 0.072 | 0.587 | -1.589 | -0.059 | -0.003 | -0.024 | -0.024 | 3.018 | 0.12 | 0.06 | 0.14 | 0.2 | 0.13 | 0.14 |
| 7 | 7863237 | rs2881964 | -1.091 | 0.685 | -1.523 | -0.039 | 0.012 | -0.049 | -0.049 | 0 | 0.75 | 0.86 | 0.72 | 0.62 | 0.76 | 0.77 |
| 7 | 7863320 | rs7804567 | 0.344 | 0.801 | -1.5 | -0.002 | -0.007 | -0.012 | -0.012 | 0 | 0.07 | 0.06 | 0.14 | 0.2 | 0.11 | 0.09 |
| 7 | 7864751 | rs12702661 | -1.256 | 0.909 | -1.458 | 0.21 | 0.007 | -0.015 | -0.015 | 1.637 | 0.77 | 0.84 | 0.75 | 0.78 | 0.76 | 0.8 |
| 7 | 7864890 | rs17494328 | 0.932 | 1.431 | -1.399 | 0.306 | -0.006 | -0.012 | -0.012 | 0 | 0.31 | 0.25 | 0.35 | 0.6 | 0.4 | 0.43 |
| 7 | 7866979 | rs12533670 | -0.154 | 1.34 | -1.104 | -0.373 | 0.01 | -0.098 | -0.098 | 0.128 | 0.17 | 0.32 | 0.26 | 0 | 0.14 | 0.15 |
| 7 | 7868613 | rs17494426 | 0.043 | 1.174 | -1.489 | 0.394 | -0.039 | 0.023 | 0.023 | 1.212 | 0.06 | 0.01 | 0.05 | 0.01 | 0.11 | 0.08 |
| 7 | 7868740 | rs35795180 | 0.123 | 0.233 | -1.21 | 0.651 | 0.008 | -0.042 | -0.042 | 0.804 | 0.08 | 0 | 0.01 | 0.16 | 0.05 | 0.07 |
| 7 | 7870047 | rs10952075 | 0.059 | 0.508 | -0.977 | 0.108 | 0 | -0.01 | -0.01 | 0.145 | 0.57 | 0.46 | 0.66 | 0.74 | 0.54 | 0.57 |
| 7 | 7870208 | rs11974817 | 0.076 | 0.12 | -0.663 | 0.665 | -0.004 | -0.003 | -0.003 | 0 | 0.58 | 0.61 | 0.55 | 0.52 | 0.52 | 0.57 |
| 7 | 7871121 | rs17560422 | 0.038 | -0.504 | -1.184 | 0.532 | 0 | -0.041 | -0.041 | 0.279 | 0.2 | 0.05 | 0.12 | 0.22 | 0.15 | 0.26 |
| 7 | 7871531 | rs17494629 | 0.706 | -0.338 | -1.768 | 0 | -0.05 | 0.115 | 0.115 | 0 | 0.18 | 0.01 | 0.1 | 0.01 | 0.22 | 0.13 |
| 7 | 7872122 | rs10233915 | -0.254 | -0.584 | -1.963 | -0.012 | -0.033 | -0.054 | -0.054 | 0.763 | 0.33 | 0.82 | 0.53 | 0.57 | 0.37 | 0.28 |
| 7 | 7873385 | rs10241412 | 0.011 | -0.741 | -0.747 | 0.076 | 0.02 | 0.028 | 0.028 | 0 | 0.3 | 0.53 | 0.51 | 0.58 | 0.37 | 0.3 |
| 7 | 7875445 | rs2042545 | -0.617 | -0.862 | -0.962 | 0.022 | -0.028 | 0.013 | 0.013 | 0 | 0.43 | 0.79 | 0.62 | 0.6 | 0.51 | 0.44 |
| 7 | 7876466 | rs10260454 | -1.782 | -0.757 | -1.438 | 0.029 | -0.084 | -0.007 | -0.007 | 0 | 0.72 | 0.98 | 0.73 | 0.77 | 0.7 | 0.7 |
| 7 | 7880597 | rs758994 | -0.396 | -0.056 | -0.818 | 0.368 | -0.019 | -0.024 | -0.024 | 0.283 | 0.78 | 0.84 | 0.65 | 0.63 | 0.7 | 0.75 |
| 7 | 7882474 | rs7795698 | 0.814 | -0.263 | -0.51 | -0.064 | 0 | -0.031 | -0.031 | 0 | 0.52 | 0.5 | 0.4 | 0.2 | 0.53 | 0.44 |
| 7 | 7883323 | rs7780564 | 0.521 | 0.366 | -0.281 | 0.114 | 0.025 | 0.004 | 0.004 | 0 | 0.63 | 0.53 | 0.44 | 0.4 | 0.55 | 0.61 |
| 7 | 7888710 | rs13241138 | -0.319 | 0.404 | -0.152 | -0.617 | 0.006 | -0.1 | -0.1 | 0 | 0.37 | 0.5 | 0.4 | 0.05 | 0.42 | 0.3 |
| 7 | 7897979 | rs4270868 | 1.403 | 0.081 | -0.897 | -0.574 | -0.064 | 0.065 | 0.065 | 0 | 0.32 | 0.03 | 0.27 | 0.18 | 0.32 | 0.31 |
| 7 | 7898658 | rs6974910 | -0.432 | -0.083 | -0.579 | -0.806 | -0.002 | -0.06 | -0.06 | 1.909 | 0.22 | 0.66 | 0.33 | 0.05 | 0.21 | 0.15 |
| 7 | 7901243 | rs6965873 | -0.235 | -0.02 | -0.59 | -0.972 | -0.002 | -0.065 | -0.065 | 0 | 0.29 | 0.62 | 0.33 | 0 | 0.26 | 0.17 |
| 7 | 7901510 | rs7790025 | -0.159 | 0.041 | -0.411 | -0.942 | -0.003 | -0.023 | -0.023 | 0 | 0.7 | 0.61 | 0.68 | 0.18 | 0.72 | 0.56 |
| 7 | 7901922 | rs6953475 | 0.162 | -0.014 | 0.154 | -0.95 | 0.005 | -0.034 | -0.034 | 0 | 0.08 | 0.21 | 0.22 | 0.01 | 0.06 | 0.05 |
| 7 | 7902369 | rs17562004 | 1.81 | 0.032 | -0.438 | -0.936 | -0.069 | 0.082 | 0.082 | 0 | 0.32 | 0.03 | 0.27 | 0.17 | 0.32 | 0.28 |
| 7 | 7902687 | rs7808919 | -0.132 | 0.057 | -0.41 | -0.928 | -0.003 | -0.022 | -0.022 | 0 | 0.7 | 0.6 | 0.67 | 0.17 | 0.72 | 0.55 |
| 7 | 7903077 | rs6463733 | -0.784 | -0.025 | -0.5 | -0.643 | 0.022 | 0.092 | 0.092 | 0 | 0.5 | 0.61 | 0.45 | 0.17 | 0.44 | 0.45 |
| 7 | 7905999 | rs6463736 | -0.82 | -0.009 | -0.494 | -0.589 | 0.046 | 0.079 | 0.079 | 0 | 0.83 | 0.64 | 0.74 | 0.33 | 0.76 | 0.78 |
| 7 | 7906204 | rs2109971 | -1.05 | -0.014 | -0.394 | -0.14 | 0.018 | 0.045 | 0.045 | 0 | 0.84 | 0.77 | 0.78 | 0.59 | 0.76 | 0.77 |
| 7 | 7906852 | rs6463740 | -1.118 | -0.024 | -0.302 | -0.224 | 0.004 | 0.028 | 0.028 | 0 | 0.53 | 0.75 | 0.52 | 0.39 | 0.47 | 0.5 |
| 7 | 7907142 | rs17562330 | 0.908 | 0.148 | -0.439 | -0.48 | -0.003 | -0.009 | -0.009 | 0 | 0.19 | 0.14 | 0.12 | 0.23 | 0.2 | 0.15 |
| 7 | 7908235 | rs6969177 | -0.666 | 0.057 | -0.546 | -0.459 | -0.042 | 0.13 | 0.13 | 0.174 | 0.44 | 0.85 | 0.44 | 0.23 | 0.42 | 0.31 |
| 7 | 7908970 | rs6954729 | -1.398 | 0.277 | -0.297 | -0.718 | -0.002 | -0.028 | -0.028 | 0.041 | 0.67 | 0.72 | 0.71 | 0.49 | 0.66 | 0.67 |
| 7 | 7911371 | rs732855 | -0.396 | 0.232 | -0.118 | -0.742 | 0.008 | -0.025 | -0.025 | 0 | 0.39 | 0.26 | 0.38 | 0.49 | 0.37 | 0.48 |
| 7 | 7913819 | rs10255964 | 1.451 | 0.31 | -0.16 | -0.628 | -0.001 | -0.024 | -0.024 | 0 | 0.31 | 0.21 | 0.28 | 0.5 | 0.31 | 0.35 |
| 7 | 7913945 | rs10241098 | 0.62 | 0.295 | 0.018 | -0.621 | -0.001 | -0.001 | -0.001 | 0 | 0.53 | 0.46 | 0.39 | 0.5 | 0.53 | 0.47 |
| 7 | 7916567 | rs3735459 | -0.64 | 0.425 | -0.235 | -0.173 | 0.005 | 0.032 | 0.032 | 0 | 0.86 | 0.85 | 0.79 | 0.68 | 0.81 | 0.81 |
| 7 | 7917360 | rs1558475 | 0.125 | 0.194 | -0.318 | -0.452 | -0.034 | -0.011 | -0.011 | 0.574 | 0.19 | 0.29 | 0.23 | 0.16 | 0.1 | 0.22 |
| 7 | 7918285 | rs17497014 | -0.672 | 0.37 | -0.749 | -0.715 | 0 | 0.168 | 0.168 | 0 | 0.48 | 0.27 | 0.39 | 0.13 | 0.46 | 0.45 |
| 7 | 7918642 | rs10605 | 0.485 | 0.483 | -1.089 | -0.643 | 0.024 | 0.022 | 0.022 | 0 | 0.54 | 0.73 | 0.6 | 0.64 | 0.4 | 0.55 |
| 7 | 7923522 | rs17563175 | 1.093 | 0.601 | -1.292 | -0.796 | -0.033 | 0.07 | 0.07 | 0 | 0.19 | 0.09 | 0.09 | 0 | 0.23 | 0.12 |
| 7 | 7924284 | rs11761799 | 0.818 | 0.426 | -0.443 | 0.147 | 0.035 | 0.064 | 0.064 | 0 | 0.72 | 0.58 | 0.67 | 0.62 | 0.63 | 0.71 |
| 7 | 7924648 | rs17563229 | -0.343 | 0.676 | -1.606 | 0.291 | 0.012 | 0.089 | 0.089 | 0.096 | 0.28 | 0.09 | 0.28 | 0.11 | 0.23 | 0.38 |
| 7 | 7924797 | rs7778134 | 0.324 | 1.104 | -0.847 | 1.115 | -0.011 | -0.021 | -0.021 | 0 | 0.83 | 0.87 | 0.76 | 0.62 | 0.77 | 0.81 |
| 7 | 7928202 | rs2348769 | -1.036 | 0.754 | -0.886 | 0.588 | -0.006 | -0.003 | -0.003 | 0 | 0.25 | 0.18 | 0.38 | 0.71 | 0.29 | 0.24 |
| 7 | 7929851 | rs10278724 | 0.16 | 0.754 | -0.965 | 0.019 | -0.001 | -0.017 | -0.017 | 0 | 0.1 | 0.13 | 0.07 | 0 | 0.13 | 0.13 |
| 7 | 7930948 | rs10237424 | -0.361 | -0.54 | -0.791 | -0.672 | -0.015 | -0.015 | -0.015 | 0 | 0.31 | 0.3 | 0.26 | 0.18 | 0.17 | 0.31 |
| 7 | 7934974 | rs17139306 | 0.61 | -0.852 | -1.277 | -0.746 | 0.053 | 0.008 | 0.008 | 2.083 | 0.14 | 0.01 | 0.16 | 0.12 | 0.07 | 0.26 |
| 7 | 7937614 | rs6463745 | -1.22 | -0.953 | -0.543 | -0.969 | -0.01 | -0.041 | -0.041 | 0 | 0.44 | 0.48 | 0.35 | 0.18 | 0.36 | 0.47 |
| 7 | 7938344 | rs7785709 | -1.287 | -1.318 | -0.893 | 0.114 | 0.001 | -0.001 | -0.001 | 0 | 0.46 | 0.47 | 0.5 | 0.51 | 0.35 | 0.47 |
| 7 | 7938596 | rs10486201 | -0.528 | -0.863 | -1.45 | 0.003 | -0.007 | 0.05 | 0.05 | 0 | 0.19 | 0.01 | 0.1 | 0.08 | 0.16 | 0.1 |
| 7 | 7942576 | rs4725044 | -0.792 | -1.273 | -0.699 | 0.174 | -0.031 | -0.006 | -0.006 | 0 | 0.24 | 0.66 | 0.23 | 0.26 | 0.1 | 0.43 |
| 7 | 7943482 | rs4720759 | 0.264 | -1.351 | -1.064 | 0.126 | 0.017 | 0.016 | 0.016 | 2.902 | 0.15 | 0 | 0.18 | 0.1 | 0.08 | 0.33 |
| 7 | 7946400 | rs4725047 | 0.14 | -1.114 | -1.811 | 0.109 | -0.013 | -0.079 | -0.079 | 0 | 0.68 | 0.9 | 0.58 | 0.44 | 0.64 | 0.59 |
| 7 | 7949059 | rs3919526 | -0.73 | -0.034 | -2.479 | 0.249 | -0.03 | -0.084 | -0.084 | 0 | 0.78 | 0.97 | 0.75 | 0.72 | 0.76 | 0.84 |
| 7 | 7949712 | rs10243568 | -0.535 | -0.222 | -2.475 | 0.034 | -0.019 | 0.092 | 0.092 | 3.31 | 0.37 | 0.82 | 0.36 | 0.23 | 0.35 | 0.42 |
| 7 | 7949823 | rs12702676 | 0.716 | -0.351 | -2.493 | 0.055 | -0.016 | -0.021 | -0.021 | 0 | 0.56 | 0.85 | 0.47 | 0.44 | 0.38 | 0.55 |
| 7 | 7949890 | rs12702677 | 1.88 | 0.136 | -2.361 | 0.025 | -0.053 | 0.01 | 0.01 | 1.912 | 0.31 | 0.04 | 0.22 | 0.21 | 0.28 | 0.22 |
| 7 | 7952201 | rs6463746 | 1.914 | -0.464 | -2.273 | 0.073 | -0.09 | 0.014 | 0.014 | 0.872 | 0.3 | 0.06 | 0.23 | 0.2 | 0.33 | 0.25 |
| 7 | 7953906 | rs6463747 | 1.863 | -0.378 | -2.343 | 0.066 | -0.109 | -0.009 | -0.009 | 0 | 0.31 | 0.05 | 0.22 | 0.22 | 0.33 | 0.24 |
| 7 | 7954298 | rs6950643 | 1.069 | -0.483 | -2.333 | 0.041 | -0.035 | 0.075 | 0.075 | 0.621 | 0.41 | 0.06 | 0.32 | 0.21 | 0.54 | 0.32 |
| 7 | 7957841 | rs6463749 | -0.669 | -0.286 | -1.295 | 0.475 | 0.005 | 0.007 | 0.007 | 6.921 | 0.86 | 0.74 | 0.85 | 0.83 | 0.83 | 0.92 |
| 7 | 7957904 | rs6463750 | -0.912 | -0.21 | -0.524 | 0.494 | 0.019 | 0.025 | 0.025 | 0 | 0.27 | 0.4 | 0.3 | 0.38 | 0.34 | 0.27 |
| 7 | 7967220 | rs2041566 | 0.712 | -0.492 | -0.738 | -0.057 | -0.004 | -0.006 | -0.006 | 3.581 | 0.23 | 0.21 | 0.22 | 0.1 | 0.15 | 0.37 |
| 7 | 7967424 | rs13226746 | 1.34 | -0.659 | -1.016 | 0.193 | -0.085 | -0.106 | -0.106 | 0 | 0.33 | 0.07 | 0.34 | 0.45 | 0.37 | 0.35 |

| | | | | | | | | | | | | | | | | |
|---|---|---|---|---|---|---|---|---|---|---|---|---|---|---|---|---|
| 7 | 7968608 | rs10237854 | -0.195 | -0.716 | -0.167 | 0.23 | 0.03 | -0.027 | -0.027 | 0 | 0.74 | 0.64 | 0.71 | 0.83 | 0.61 | 0.85 |
| 7 | 7969336 | rs10252010 | 0.56 | -0.903 | -0.107 | 0.169 | 0.022 | -0.023 | -0.023 | 0 | 0.18 | 0.16 | 0.15 | 0.27 | 0.11 | 0.15 |
| 7 | 7970560 | rs7796084 | 0.168 | -0.966 | 0.214 | 0.34 | -0.006 | -0.004 | -0.004 | 0 | 0.7 | 0.72 | 0.59 | 0.61 | 0.57 | 0.74 |
| 7 | 7971917 | rs12702682 | 2.125 | -1.013 | 0.117 | 0.308 | -0.05 | 0.001 | 0.001 | 0.694 | 0.27 | 0.11 | 0.2 | 0.25 | 0.31 | 0.22 |
| 7 | 7972285 | rs10251201 | 0.156 | -1.107 | 0.164 | 0.359 | -0.018 | 0.013 | 0.013 | 0.19 | 0.62 | 0.67 | 0.6 | 0.71 | 0.51 | 0.72 |
| 7 | 7972762 | rs12670023 | 1.469 | -1.11 | 0.165 | 0.326 | 0.032 | 0.026 | 0.026 | 4.358 | 0.2 | 0.1 | 0.21 | 0.12 | 0.14 | 0.36 |
| 7 | 7973654 | rs6966273 | 0.739 | -0.841 | 0.117 | 0.31 | 0.006 | -0.006 | -0.006 | 0 | 0.54 | 0.61 | 0.44 | 0.51 | 0.44 | 0.6 |
| 7 | 7974827 | rs11974862 | 0.388 | -1.209 | 0.167 | 0.347 | -0.093 | -0.013 | -0.013 | 0.218 | 0.28 | 0.51 | 0.26 | 0.27 | 0.15 | 0.38 |
| 7 | 7975662 | rs7782224 | 0.328 | -1.199 | 0.171 | 0.352 | -0.094 | -0.015 | -0.015 | 0 | 0.28 | 0.52 | 0.26 | 0.27 | 0.15 | 0.38 |
| 7 | 7978565 | rs4725051 | 0.821 | -0.936 | 0.248 | 0.037 | 0.004 | 0.022 | 0.022 | 1.319 | 0.51 | 0.58 | 0.44 | 0.36 | 0.46 | 0.58 |
| 7 | 7978892 | rs13235362 | 2.029 | -0.78 | -0.292 | 0.078 | -0.058 | -0.006 | -0.006 | 3.015 | 0.26 | 0.09 | 0.2 | 0.26 | 0.31 | 0.22 |
| 7 | 7980355 | rs6950165 | -1.038 | -0.747 | -0.243 | 0.067 | 0.003 | 0.042 | 0.042 | 0.355 | 0.61 | 0.77 | 0.69 | 0.74 | 0.45 | 0.73 |
| 7 | 7982218 | rs16872177 | 0.14 | -0.724 | -0.344 | 0.169 | 0.054 | -0.009 | -0.009 | 0.207 | 0.37 | 0.25 | 0.35 | 0.37 | 0.26 | 0.5 |
| 7 | 7987137 | rs17140599 | 0.083 | 0.249 | 0.315 | 0.755 | -0.006 | 0.034 | 0.034 | 0 | 0.08 | 0.08 | 0.14 | 0.2 | 0.05 | 0.12 |
| 7 | 7987940 | rs10255002 | 0.689 | 0.309 | 0.248 | 0.704 | 0.001 | 0.001 | 0.001 | 2.275 | 0.13 | 0.16 | 0.17 | 0.17 | 0.15 | 0.11 |
| 7 | 7988212 | rs6947392 | -0.05 | 0.436 | 0.059 | 0.829 | 0.037 | -0.05 | -0.05 | 0.132 | 0.88 | 0.84 | 0.89 | 1 | 0.78 | 0.96 |
| 7 | 7991437 | rs1990367 | 1.087 | 0.657 | -1.027 | 1.134 | -0.006 | -0.015 | -0.015 | 0 | 0.64 | 0.72 | 0.61 | 0.53 | 0.61 | 0.7 |
| 7 | 7996282 | rs37949 | 0.455 | 1.074 | -0.598 | 0.846 | 0.009 | 0.008 | 0.008 | 0 | 0.84 | 0.84 | 0.77 | 0.79 | 0.79 | 0.84 |
| 7 | 8000032 | rs37955 | -0.963 | 0.427 | -0.869 | 0.573 | 0.001 | 0.001 | 0.001 | 0 | 0.18 | 0.12 | 0.12 | 0.13 | 0.14 | 0.12 |
| 7 | 8000971 | rs37957 | 0.145 | 0.869 | -1.393 | 0.529 | -0.028 | 0.019 | 0.019 | 0 | 0.71 | 0.92 | 0.77 | 0.77 | 0.7 | 0.69 |
| 7 | 8002389 | rs37959 | 0.489 | 0.398 | -0.961 | 0.529 | -0.006 | -0.01 | -0.01 | 0.34 | 0.41 | 0.58 | 0.36 | 0.31 | 0.36 | 0.24 |
| 7 | 8005762 | rs6956133 | -0.516 | -0.491 | -0.525 | 0.919 | -0.053 | 0.023 | 0.023 | 0.635 | 0.13 | 0.21 | 0.13 | 0.19 | 0.05 | 0.12 |
| 7 | 8005920 | rs37967 | 0.626 | -0.361 | -0.27 | 0.999 | -0.001 | -0.027 | -0.027 | 0 | 0.45 | 0.51 | 0.38 | 0.21 | 0.43 | 0.28 |
| 7 | 8005973 | rs37968 | -1 | 0 | -0.539 | 1.478 | -0.043 | 0.005 | 0.005 | 0 | 0.72 | 0.89 | 0.77 | 0.75 | 0.69 | 0.68 |
| 7 | 8007509 | rs37972 | -0.443 | 0.091 | -0.539 | 1.606 | -0.069 | -0.043 | -0.043 | 0 | 0.65 | 0.82 | 0.57 | 0.56 | 0.56 | 0.49 |
| 7 | 8007876 | rs37973 | -0.529 | 0.089 | -0.502 | 1.603 | -0.058 | -0.051 | -0.051 | 0 | 0.65 | 0.8 | 0.56 | 0.52 | 0.56 | 0.49 |
| 7 | 8014957 | rs37976 | -0.411 | 0.742 | -0.597 | 1.553 | -0.054 | -0.054 | -0.054 | 0.982 | 0.64 | 0.8 | 0.54 | 0.53 | 0.54 | 0.5 |
| 7 | 8019934 | rs715377 | -0.859 | 0.524 | 0.089 | 1.452 | -0.018 | 0.035 | 0.035 | 0.231 | 0.12 | 0.34 | 0.09 | 0.17 | 0.1 | 0.09 |
| 7 | 8021570 | rs12537880 | 0.553 | 0.402 | -0.107 | 1.446 | -0.007 | -0.008 | -0.008 | 0 | 0.1 | 0.09 | 0.21 | 0.22 | 0.14 | 0.19 |
| 7 | 8027174 | rs37990 | -1.022 | 0.321 | -0.087 | 1.427 | -0.01 | 0.022 | 0.022 | 0 | 0.72 | 0.79 | 0.74 | 0.79 | 0.67 | 0.69 |
| 7 | 8042389 | rs38010 | 0.686 | 0.066 | 0.09 | 1.515 | 0.013 | 0.077 | 0.077 | 1.726 | 0.5 | 0.7 | 0.38 | 0.35 | 0.42 | 0.36 |
| 7 | 8044483 | rs2041362 | 1.128 | 0.088 | 0.05 | 1.527 | -0.002 | 0.002 | 0.002 | 0 | 0.08 | 0.1 | 0.21 | 0.16 | 0.13 | 0.17 |
| 7 | 8055664 | rs17141876 | 1.32 | 0.037 | 0.033 | 1.5 | 0.032 | -0.03 | -0.03 | 0 | 0.09 | 0.01 | 0.11 | 0.17 | 0.03 | 0.12 |
| 7 | 8059186 | rs7798367 | -1.152 | 0.124 | -0.023 | 1.444 | -0.031 | 0.02 | 0.02 | 0 | 0.72 | 0.9 | 0.76 | 0.78 | 0.68 | 0.69 |
| 7 | 8060056 | rs1476823 | -0.351 | 0.126 | 0.003 | 1.47 | -0.043 | -0.032 | -0.032 | 1.648 | 0.63 | 0.78 | 0.54 | 0.61 | 0.54 | 0.53 |
| 7 | 8062590 | rs10243846 | 1.336 | 0.046 | 0.159 | 1.438 | 0.004 | -0.022 | -0.022 | 0 | 0.42 | 0.42 | 0.34 | 0.25 | 0.4 | 0.31 |
| 7 | 8074608 | rs13245462 | -0.867 | 0.033 | 0.162 | 1.441 | -0.014 | 0.021 | 0.021 | 0.241 | 0.17 | 0.23 | 0.18 | 0.35 | 0.14 | 0.22 |
| 7 | 8080481 | rs7801924 | 0.203 | 0.044 | 0.151 | 1.455 | 0.005 | -0.027 | -0.027 | 0 | 0.58 | 0.7 | 0.58 | 0.52 | 0.54 | 0.54 |
| 7 | 8086639 | rs4725061 | 1.299 | 0.033 | 0.25 | 1.473 | 0.001 | -0.009 | -0.009 | 0 | 0.42 | 0.4 | 0.34 | 0.26 | 0.4 | 0.31 |
| 7 | 8088394 | rs10226865 | -0.39 | 0.038 | 0.181 | 1.479 | -0.05 | -0.048 | -0.048 | 0.047 | 0.63 | 0.79 | 0.55 | 0.61 | 0.55 | 0.51 |
| 7 | 8098140 | rs9648623 | 0.64 | 0.002 | 0.351 | 0.832 | 0.053 | -0.072 | -0.072 | 0.017 | 0.16 | 0.28 | 0.11 | 0.02 | 0.16 | 0.15 |
| 7 | 8101099 | rs11978165 | 0.095 | -0.661 | -0.026 | 0.723 | 0.03 | -0.004 | -0.004 | 0.637 | 0.56 | 0.67 | 0.59 | 0.43 | 0.58 | 0.47 |
| 7 | 8101765 | rs6463755 | -1.318 | -0.401 | 0.088 | 0.757 | -0.017 | -0.025 | -0.025 | 0 | 0.73 | 0.85 | 0.75 | 0.68 | 0.67 | 0.66 |
| 7 | 8112014 | rs2191318 | -0.334 | -0.197 | 0.097 | 0.739 | -0.017 | 0.02 | 0.02 | 0 | 0.62 | 0.68 | 0.67 | 0.71 | 0.55 | 0.61 |
| 7 | 8113312 | rs11976862 | 0.962 | -0.302 | 0.091 | 0.795 | 0 | -0.013 | -0.013 | 0 | 0.43 | 0.4 | 0.35 | 0.31 | 0.42 | 0.34 |
| 7 | 8113751 | rs4236406 | 1.186 | -0.375 | -0.045 | 0.5 | 0 | -0.005 | -0.005 | 0 | 0.43 | 0.4 | 0.33 | 0.33 | 0.38 | 0.28 |
| 7 | 8115794 | rs17142727 | -0.07 | -0.44 | -0.277 | 0.486 | -0.005 | -0.032 | -0.032 | 0 | 0.15 | 0.04 | 0.18 | 0.23 | 0.16 | 0.21 |
| 7 | 8115979 | rs10253260 | -1.077 | -0.389 | -0.259 | 0.497 | -0.024 | 0.032 | 0.032 | 0 | 0.72 | 0.88 | 0.75 | 0.76 | 0.66 | 0.67 |
| 7 | 8116932 | rs10952084 | 0.65 | -0.363 | -0.206 | 0.457 | 0.001 | 0.014 | 0.014 | 0 | 0.09 | 0.1 | 0.21 | 0.24 | 0.13 | 0.19 |
| 7 | 8116948 | rs6964558 | 0.898 | -0.347 | -0.197 | 0.363 | 0.004 | 0.01 | 0.01 | 0 | 0.09 | 0.1 | 0.21 | 0.17 | 0.13 | 0.17 |
| 7 | 8122313 | rs1468594 | -0.678 | -0.119 | -0.503 | -0.104 | 0.001 | 0.001 | 0.001 | 0 | 0.82 | 0.79 | 0.78 | 0.73 | 0.74 | 0.76 |
| 7 | 8123427 | rs10257285 | -0.346 | -0.012 | -0.378 | 0.063 | 0.002 | 0.013 | 0.013 | 0 | 0.68 | 0.71 | 0.73 | 0.73 | 0.66 | 0.69 |
| 7 | 8127710 | rs4725068 | 1.315 | 0.35 | -0.357 | 0.177 | -0.001 | -0.008 | -0.008 | 0 | 0.41 | 0.39 | 0.31 | 0.3 | 0.36 | 0.28 |
| 7 | 8129728 | rs6967565 | -1.486 | 0.653 | 0.001 | 0.175 | 0.006 | 0.025 | 0.025 | 0 | 0.26 | 0.33 | 0.4 | 0.4 | 0.29 | 0.37 |
| 7 | 8134294 | rs6952006 | 0.448 | 0.978 | 0.092 | -0.299 | 0.014 | 0.037 | 0.037 | 1.171 | 0.87 | 0.72 | 0.7 | 0.84 | 0.84 | 0.83 |
| 7 | 8142830 | rs7803536 | -0.974 | 0.759 | 0.505 | -0.705 | 0.027 | -0.112 | -0.112 | 0 | 0.21 | 0.31 | 0.14 | 0 | 0.22 | 0.11 |
| 7 | 8143709 | rs6977396 | -1.214 | 0.695 | 1.043 | -0.596 | 0.05 | 0.008 | 0.008 | 0 | 0.1 | 0.39 | 0.1 | 0.09 | 0.11 | 0.09 |
| 7 | 8145013 | rs9655062 | 0.02 | 0.798 | 0.921 | -0.555 | 0.059 | -0.029 | -0.029 | 0.806 | 0.06 | 0.14 | 0.07 | 0 | 0.09 | 0.05 |
| 7 | 8145201 | rs11976736 | 0.241 | 0.54 | 0.665 | -0.825 | 0.002 | 0.001 | 0.001 | 0 | 0.25 | 0.22 | 0.17 | 0.28 | 0.23 | 0.32 |
| 7 | 8145640 | rs6959703 | 0.756 | 0.004 | 0.467 | -0.935 | -0.024 | -0.021 | -0.021 | 0 | 0.74 | 0.59 | 0.76 | 0.85 | 0.82 | 0.84 |
| 7 | 8145733 | rs17401797 | 1.83 | 0.425 | 0.316 | -0.917 | -0.03 | -0.01 | -0.01 | 0 | 0.41 | 0.19 | 0.52 | 0.57 | 0.54 | 0.45 |
| 7 | 8147385 | rs2109386 | 0.351 | 1.117 | 0.731 | -0.016 | -0.022 | 0.136 | 0.136 | 0.888 | 0.91 | 0.66 | 0.83 | 0.85 | 0.97 | 0.91 |
| 7 | 8149596 | rs7777031 | 0.975 | 1.172 | 0.394 | -0.131 | 0.005 | 0.08 | 0.08 | 0 | 0.74 | 0.49 | 0.67 | 0.57 | 0.75 | 0.61 |
| 7 | 8150562 | rs12702696 | 2.162 | 1.256 | -0.766 | -0.099 | -0.053 | -0.031 | -0.031 | 0.002 | 0.47 | 0.05 | 0.49 | 0.57 | 0.51 | 0.5 |
| 7 | 8152340 | rs12702698 | 2.041 | 1.394 | -0.442 | -0.046 | -0.018 | 0.089 | 0.089 | 0 | 0.51 | 0.12 | 0.52 | 0.67 | 0.54 | 0.55 |
| 7 | 8154535 | rs3807853 | 0.296 | 1.63 | 0.137 | 0.385 | -0.005 | -0.005 | -0.005 | 0 | 0.11 | 0.07 | 0.08 | 0.18 | 0.14 | 0.15 |
| 7 | 8156375 | rs3807852 | 1.961 | 1.569 | 0.19 | 0.358 | -0.01 | -0.013 | -0.013 | 0 | 0.64 | 0.25 | 0.62 | 0.61 | 0.65 | 0.64 |
| 7 | 8159569 | rs17143355 | -0.486 | 1.753 | 0.531 | 0.242 | 0.042 | -0.04 | -0.04 | 0 | 0.07 | 0.14 | 0.06 | 0 | 0.11 | 0.06 |
| 7 | 8160668 | rs3807846 | -1.931 | 1.562 | 0.779 | 0.319 | 0 | -0.001 | -0.001 | 0 | 0.31 | 0.65 | 0.23 | 0.29 | 0.28 | 0.29 |

| | | | | | | | | | | | | | | | | |
|---|---|---|---|---|---|---|---|---|---|---|---|---|---|---|---|---|
| 7 | 8161527 | rs10268598 | -2.039 | 1.762 | 0.932 | 0.343 | 0.087 | 0.077 | 0.077 | 0 | 0.2 | 0.58 | 0.29 | 0.3 | 0.26 | 0.27 |
| 7 | 8161981 | rs6965329 | -2.05 | 1.605 | 0.869 | 0.395 | -0.01 | 0.003 | 0.003 | 0.224 | 0.4 | 0.6 | 0.39 | 0.33 | 0.42 | 0.4 |
| 7 | 8162084 | rs3823839 | -2.219 | 1.629 | 0.043 | 0.341 | -0.102 | -0.061 | -0.061 | 0 | 0.46 | 0.96 | 0.44 | 0.42 | 0.41 | 0.43 |
| 7 | 8165121 | rs10259684 | -2.041 | 1.378 | 0.587 | 0.209 | 0.003 | 0.013 | 0.013 | 0.027 | 0.4 | 0.75 | 0.42 | 0.38 | 0.38 | 0.45 |
| 7 | 8166023 | rs12532271 | -1.391 | 1.105 | 0.897 | 0.346 | -0.002 | 0.015 | 0.015 | 0.282 | 0.24 | 0.36 | 0.32 | 0.32 | 0.23 | 0.28 |
| 7 | 8168129 | rs6463769 | 1.734 | 1.854 | 1.146 | 0.138 | 0.036 | -0.079 | -0.079 | 0 | 0.79 | 0.68 | 0.84 | 0.94 | 0.75 | 0.73 |
| 7 | 8179655 | rs3807843 | -1.592 | 2.132 | 1.431 | 0.193 | 0.078 | -0.087 | -0.087 | 0.33 | 0.14 | 0.34 | 0.13 | 0.01 | 0.22 | 0.1 |
| 7 | 8179846 | rs887848 | 2.341 | 2.061 | 1.494 | 0.125 | 0.039 | -0.062 | -0.062 | 0 | 0.81 | 0.69 | 0.81 | 0.93 | 0.72 | 0.74 |
| 7 | 8184961 | rs2110332 | 2.999 | 1.766 | 1.8 | 0.193 | 0.126 | 0.013 | 0.013 | 0 | 0.93 | 0.65 | 0.93 | 1 | 0.92 | 0.96 |
| 7 | 8187094 | rs10156091 | 1.798 | 1.787 | 1.626 | 0.189 | 0.018 | -0.045 | -0.045 | 0.119 | 0.88 | 0.74 | 0.87 | 0.94 | 0.88 | 0.81 |
| 7 | 8189620 | rs3757524 | -1.848 | 1.533 | 2.11 | 0.962 | -0.017 | 0.133 | 0.133 | 5.194 | 0.15 | 0.54 | 0.25 | 0.24 | 0.09 | 0.18 |
| 7 | 8191966 | rs3807839 | 2.732 | 1.58 | 1.709 | 1.452 | 0.093 | 0.132 | 0.132 | 0 | 0.81 | 0.58 | 0.73 | 0.68 | 0.74 | 0.71 |
| 7 | 8194701 | rs13226206 | -0.534 | 1.407 | 1.15 | 1.169 | 0.004 | 0.002 | 0.002 | 0 | 0.07 | 0.06 | 0.07 | 0.04 | 0.16 | 0.07 |
| 7 | 8196112 | rs3757523 | 0.833 | 0.855 | 0.85 | 1.612 | -0.015 | -0.014 | -0.014 | 0 | 0.83 | 0.92 | 0.83 | 0.78 | 0.83 | 0.85 |
| 7 | 8199401 | rs2348895 | -2.141 | 1.083 | 1.214 | 1.417 | -0.019 | 0.137 | 0.137 | 0.165 | 0.11 | 0.4 | 0.09 | 0.17 | 0.08 | 0.1 |
| 7 | 8204811 | rs6463778 | 1.133 | 1.195 | 0.143 | 1.352 | -0.021 | -0.05 | -0.05 | 0 | 0.82 | 0.9 | 0.75 | 0.57 | 0.77 | 0.73 |
| 7 | 8214451 | rs2058518 | 1.401 | 1.302 | 0.726 | 0.377 | 0.032 | -0.053 | -0.053 | 0.739 | 0.75 | 0.6 | 0.83 | 0.9 | 0.7 | 0.83 |
| 7 | 8214574 | rs2058519 | 1.378 | 1.197 | 0.726 | 0.373 | 0.026 | -0.055 | -0.055 | 0.086 | 0.76 | 0.6 | 0.83 | 0.9 | 0.73 | 0.83 |
| 7 | 8214666 | rs7804613 | -1.073 | 1.126 | 0.705 | 0.117 | 0.006 | -0.087 | -0.087 | 0 | 0.21 | 0.34 | 0.15 | 0 | 0.23 | 0.11 |
| 7 | 8215208 | rs7809298 | -1.543 | 0.847 | 0.679 | 0.007 | -0.005 | -0.137 | -0.137 | 0 | 0.28 | 0.42 | 0.17 | 0 | 0.25 | 0.11 |
| 7 | 8215475 | rs3779370 | 2.204 | 0.696 | 1.131 | 0.506 | 0.049 | -0.041 | -0.041 | 0 | 0.78 | 0.46 | 0.8 | 0.88 | 0.73 | 0.86 |
| 7 | 8215543 | rs3779368 | -1.806 | 0.693 | 0.416 | 0.6 | 0.031 | -0.024 | -0.024 | 0.794 | 0.18 | 0.33 | 0.17 | 0.12 | 0.22 | 0.14 |
| 7 | 8217022 | rs881451 | 0.378 | 0.106 | -0.566 | 0.774 | 0.047 | 0.016 | 0.016 | 10.277 | 0.81 | 0.92 | 0.95 | 0.85 | 0.94 | 0.89 |
| 7 | 8220016 | rs12668294 | 1.587 | 0.548 | -1.23 | 0.959 | -0.043 | 0.055 | 0.055 | 5.663 | 0.5 | 0.03 | 0.63 | 0.64 | 0.6 | 0.71 |
| 7 | 8222595 | rs3823833 | 1.575 | 0.721 | -1.254 | 1.167 | -0.103 | -0.036 | -0.036 | 0 | 0.49 | 0.03 | 0.56 | 0.54 | 0.55 | 0.7 |
| 7 | 8225890 | rs10271463 | -1.37 | 0.456 | -0.883 | 1.034 | 0.063 | 0.11 | 0.11 | 4.396 | 0.46 | 0.79 | 0.31 | 0.26 | 0.32 | 0.21 |
| 7 | 8233642 | rs3807826 | -1.578 | 0.43 | -1.349 | 1.297 | 0.031 | -0.016 | -0.016 | 0 | 0.47 | 0.96 | 0.42 | 0.41 | 0.35 | 0.23 |
| 7 | 8233708 | rs3823832 | 0.96 | 0.316 | -1.257 | 1.247 | 0.017 | -0.009 | -0.009 | 0 | 0.78 | 0.17 | 0.79 | 0.76 | 0.87 | 0.9 |
| 7 | 8237811 | rs10952094 | 1.97 | 0.417 | -1.553 | 1.325 | -0.037 | -0.066 | -0.066 | 0 | 0.5 | 0.03 | 0.51 | 0.51 | 0.59 | 0.75 |
| 7 | 8239909 | rs17402321 | -0.401 | 0.402 | -1.493 | 1.098 | -0.03 | 0.019 | 0.019 | 7.451 | 0.21 | 0.07 | 0.2 | 0.17 | 0.21 | 0.13 |
| 7 | 8241483 | rs2348900 | 1.881 | 0.366 | -1.493 | 1.112 | -0.031 | 0.003 | 0.003 | 8.747 | 0.52 | 0.03 | 0.57 | 0.61 | 0.61 | 0.76 |
| 7 | 8241749 | rs3823822 | -0.707 | 0.88 | -0.946 | 1.163 | 0 | 0.043 | 0.043 | 8.279 | 0.2 | 0.06 | 0.15 | 0.12 | 0.14 | 0.07 |
| 7 | 8242599 | rs17402432 | -0.498 | 0.98 | -0.914 | 1.207 | -0.006 | 0.036 | 0.036 | 7.869 | 0.19 | 0.06 | 0.15 | 0.12 | 0.14 | 0.08 |
| 7 | 8243084 | rs10486212 | -0.336 | 0.232 | -0.55 | 2.023 | 0.03 | -0.037 | -0.037 | 0.016 | 0.18 | 0.1 | 0.11 | 0.38 | 0.07 | 0.07 |
| 7 | 8243602 | rs16872827 | -1.904 | 0.939 | 0.573 | 1.888 | -0.009 | 0.042 | 0.042 | 0 | 0.13 | 0.23 | 0.17 | 0.29 | 0.14 | 0.13 |
| 7 | 8244012 | rs10952097 | 1.426 | 0.808 | 1.21 | 1.381 | 0.032 | -0.012 | -0.012 | 1.669 | 0.87 | 0.39 | 0.9 | 0.89 | 0.89 | 0.91 |
| 7 | 8247347 | rs3807822 | -1.641 | 1.172 | 0.808 | 1.319 | -0.004 | -0.007 | -0.007 | 0 | 0.05 | 0.02 | 0.12 | 0.12 | 0.05 | 0.07 |
| 7 | 8249663 | rs1981715 | 2.233 | 1.197 | 1.498 | 1.338 | 0.045 | 0.13 | 0.13 | 0.018 | 0.87 | 0.64 | 0.8 | 0.67 | 0.87 | 0.9 |
| 7 | 8250553 | rs16870299 | -1.198 | 1.292 | 1.089 | 0.94 | -0.006 | -0.007 | -0.007 | 3.051 | 0.14 | 0.11 | 0.22 | 0.25 | 0.16 | 0.1 |
| 7 | 8253685 | rs3779363 | -2.21 | 1.113 | 1.366 | 0.589 | 0.002 | -0.005 | -0.005 | 0 | 0.32 | 0.64 | 0.37 | 0.35 | 0.35 | 0.22 |
| 7 | 8256455 | rs7806154 | -1.628 | 0.902 | 1.404 | 0.427 | 0 | 0.002 | 0.002 | 0.187 | 0.22 | 0.72 | 0.27 | 0.27 | 0.15 | 0.15 |
| 7 | 8257560 | rs11981246 | -1.078 | 1.093 | 1.282 | 0.368 | 0.006 | -0.081 | -0.081 | 6.303 | 0.15 | 0.29 | 0.14 | 0.01 | 0.18 | 0.1 |
| 7 | 8258516 | rs3823818 | -1.55 | 1.008 | 0.609 | 0.572 | -0.005 | 0.006 | 0.006 | 0 | 0.33 | 0.71 | 0.39 | 0.32 | 0.32 | 0.27 |
| 7 | 8264410 | rs7796406 | -1.85 | 0.134 | 1.171 | 0.367 | -0.004 | -0.021 | -0.021 | 0 | 0.07 | 0.34 | 0.03 | 0 | 0.01 | 0.02 |
| 7 | 8269678 | rs6977682 | -2.289 | -0.37 | 1.637 | 0.163 | -0.049 | -0.055 | -0.055 | 0 | 0.19 | 0.65 | 0.12 | 0.08 | 0.07 | 0.1 |
| 7 | 8274964 | rs3807816 | -0.978 | -0.524 | 0.256 | -0.64 | 0.043 | 0.169 | 0.169 | 0 | 0.49 | 0.69 | 0.44 | 0.12 | 0.44 | 0.27 |
| 7 | 8275059 | rs3807815 | -0.958 | 0.167 | 0.224 | -0.283 | 0 | 0.025 | 0.025 | 0 | 0.56 | 0.65 | 0.61 | 0.32 | 0.6 | 0.5 |
| 7 | 8276621 | rs3779360 | -0.377 | -0.507 | -0.325 | -0.701 | -0.01 | 0.045 | 0.045 | 0 | 0.58 | 0.74 | 0.51 | 0.15 | 0.49 | 0.33 |
| 7 | 8287792 | rs3807814 | -0.544 | 0.04 | -0.36 | 0.296 | -0.007 | 0.012 | 0.012 | 0 | 0.37 | 0.47 | 0.26 | 0.47 | 0.32 | 0.22 |
| 7 | 8294796 | rs3807809 | -1.098 | 0.072 | -0.541 | 0.647 | 0.009 | -0.008 | -0.008 | 0 | 0.45 | 0.86 | 0.36 | 0.59 | 0.39 | 0.25 |
| 7 | 8298573 | rs3779356 | -0.677 | -0.151 | -0.618 | 0.634 | 0.001 | -0.001 | -0.001 | 0.515 | 0.41 | 0.6 | 0.34 | 0.61 | 0.37 | 0.29 |
| 7 | 8299942 | rs3807807 | -0.742 | -0.103 | -0.354 | 1.217 | -0.025 | -0.024 | -0.024 | 0 | 0.6 | 0.88 | 0.63 | 0.62 | 0.61 | 0.42 |
| 7 | 8302284 | rs9472 | -0.332 | 0.092 | -0.26 | 1.372 | 0.008 | 0.008 | 0.008 | 0 | 0.52 | 0.71 | 0.55 | 0.56 | 0.56 | 0.39 |
| 7 | 8307886 | rs6965830 | -0.568 | -0.228 | -0.564 | 0.879 | -0.013 | 0.045 | 0.045 | 0 | 0.52 | 0.83 | 0.57 | 0.64 | 0.5 | 0.37 |
| 7 | 8308993 | rs1558557 | -0.076 | -0.093 | -0.833 | 0.934 | -0.001 | -0.001 | -0.001 | 0 | 0.43 | 0.42 | 0.49 | 0.55 | 0.49 | 0.36 |
| 7 | 8313971 | rs6463790 | -0.442 | 0.113 | -1.536 | 0.535 | -0.03 | -0.026 | -0.026 | 0 | 0.51 | 0.89 | 0.57 | 0.58 | 0.5 | 0.36 |
| 7 | 8318827 | rs2349178 | -0.417 | 0.314 | -1.268 | 0.396 | 0 | 0.001 | 0.001 | 0 | 0.49 | 0.48 | 0.5 | 0.45 | 0.52 | 0.38 |
| 7 | 8322859 | rs12333784 | -0.593 | 0.004 | -1.879 | 0.566 | 0.042 | -0.082 | -0.082 | 0 | 0.38 | 0.85 | 0.3 | 0.52 | 0.3 | 0.25 |
| 7 | 8326998 | rs6969069 | -1.076 | -0.213 | -1.898 | -0.01 | 0.03 | 0.064 | 0.064 | 0 | 0.22 | 0.07 | 0.19 | 0.1 | 0.15 | 0.15 |
| 7 | 8327061 | rs7801871 | -1.005 | 0.12 | -0.633 | 0.047 | -0.005 | -0.005 | -0.005 | 0 | 0.12 | 0.33 | 0.12 | 0.1 | 0.07 | 0.11 |
| 7 | 8328354 | rs2056449 | 0.039 | 0.203 | -1.546 | 0.075 | -0.011 | 0.015 | 0.015 | 0 | 0.63 | 0.89 | 0.74 | 0.66 | 0.68 | 0.53 |
| 7 | 8328880 | rs12670364 | -1.607 | 0.61 | -0.531 | 0.277 | 0.028 | 0.018 | 0.018 | 0 | 0.11 | 0.22 | 0.14 | 0.1 | 0.1 | 0.18 |
| 7 | 8329762 | rs7797010 | -1.461 | 0.498 | -0.563 | 0.307 | 0.002 | 0.02 | 0.02 | 0 | 0.09 | 0.16 | 0.12 | 0.1 | 0.08 | 0.16 |
| 7 | 8337432 | rs12666065 | -0.487 | 0.039 | -1.12 | 1.033 | 0.021 | -0.057 | -0.057 | 0 | 0.17 | 0.11 | 0.1 | 0.4 | 0.12 | 0.25 |
| 7 | 8345416 | rs720964 | 0.384 | -0.004 | -1.266 | 0.376 | 0.002 | -0.01 | -0.01 | 1.408 | 0.31 | 0.27 | 0.33 | 0.43 | 0.32 | 0.26 |
| 7 | 8355886 | rs12668479 | 1.13 | -0.016 | -1.453 | 0.461 | -0.003 | -0.057 | -0.057 | 0.212 | 0.1 | 0.02 | 0.06 | 0.35 | 0.08 | 0.16 |
| 7 | 8356809 | rs6968572 | -0.898 | -0.314 | -1.088 | 0.098 | -0.002 | -0.042 | -0.042 | 0 | 0.54 | 0.59 | 0.55 | 0.88 | 0.5 | 0.53 |
| 7 | 8357822 | rs2215890 | -0.991 | 1.106 | -1.256 | -0.107 | -0.043 | 0.039 | 0.039 | 0 | 0.9 | 0.94 | 0.75 | 0.99 | 0.82 | 0.72 |
| 7 | 8359623 | rs765370 | 0.236 | 0.796 | -1.138 | 0.052 | -0.012 | 0.027 | 0.027 | 0 | 0.37 | 0.5 | 0.45 | 0.52 | 0.38 | 0.28 |

| | | | | | | | | | | | | | | | | |
|---|---|---|---|---|---|---|---|---|---|---|---|---|---|---|---|---|
| 7 | 8362318 | rs10226403 | -0.347 | 0.96 | -1.388 | -0.64 | -0.031 | 0.027 | 0.027 | 0 | 0.9 | 0.95 | 0.77 | 0.99 | 0.82 | 0.73 |
| 7 | 8362456 | rs10239677 | -0.663 | -0.818 | -2.138 | -0.691 | 0.107 | -0.019 | -0.019 | 0.067 | 0.37 | 0.83 | 0.36 | 0.4 | 0.27 | 0.31 |
| 7 | 8363127 | rs13236357 | -0.701 | -0.46 | -2.085 | -1.426 | -0.081 | 0.174 | 0.174 | 0 | 0.55 | 0.87 | 0.59 | 0.86 | 0.53 | 0.63 |
| 7 | 8364216 | rs17147407 | 0.564 | -0.536 | -1.651 | -2.146 | 0 | -0.003 | -0.003 | 0 | 0.13 | 0.17 | 0.11 | 0.14 | 0.18 | 0.14 |
| 7 | 8368184 | rs11763789 | 0.089 | -0.6 | -1.723 | -2.383 | -0.007 | 0.033 | 0.033 | 0 | 0.16 | 0.11 | 0.12 | 0.08 | 0.23 | 0.16 |
| 7 | 8368772 | rs2024364 | 0.589 | -0.772 | -1.073 | -2.48 | -0.001 | 0.02 | 0.02 | 0 | 0.3 | 0.26 | 0.38 | 0.8 | 0.26 | 0.41 |
| 7 | 8375933 | rs12702719 | 0.812 | -0.834 | -0.64 | -2.419 | -0.001 | 0.001 | 0.001 | 0 | 0.27 | 0.23 | 0.36 | 0.74 | 0.24 | 0.37 |
| 7 | 8382310 | rs6463799 | -0.334 | -1.053 | -0.578 | -2.498 | -0.003 | 0.067 | 0.067 | 0 | 0.42 | 0.64 | 0.49 | 0.92 | 0.44 | 0.48 |
| 7 | 8382895 | rs6975631 | 1.09 | -0.93 | -0.364 | -2.651 | 0.003 | 0.077 | 0.077 | 2.007 | 0.08 | 0.35 | 0.05 | 0.12 | 0.03 | 0.07 |
| 7 | 8383945 | rs882168 | -1.244 | -1.141 | -0.836 | -2.936 | -0.037 | 0.108 | 0.108 | 0 | 0.59 | 0.81 | 0.57 | 0.99 | 0.56 | 0.62 |
| 7 | 8387803 | rs9691069 | 0.604 | -0.865 | -1.613 | -3.072 | 0.006 | 0.078 | 0.078 | 0.014 | 0.55 | 0.64 | 0.48 | 0.86 | 0.42 | 0.55 |
| 7 | 8394248 | rs717698 | 0.3 | -0.219 | -2.053 | -2.875 | -0.087 | 0.117 | 0.117 | 0 | 0.65 | 0.84 | 0.53 | 0.86 | 0.5 | 0.64 |
| 7 | 8396670 | rs974644 | 0.699 | -0.252 | -2.093 | -2.913 | -0.002 | 0.013 | 0.013 | 0 | 0.15 | 0.09 | 0.13 | 0.12 | 0.2 | 0.09 |
| 7 | 8396706 | rs974645 | -0.378 | -0.364 | -2.215 | -3.013 | -0.028 | 0.099 | 0.099 | 0.01 | 0.61 | 0.84 | 0.35 | 0.26 | 0.5 | 0.41 |
| 7 | 8396913 | rs10479878 | 1.388 | -0.483 | -2.07 | -3.006 | 0.005 | 0.007 | 0.007 | 0 | 0.47 | 0.48 | 0.43 | 0.83 | 0.35 | 0.48 |
| 7 | 8398372 | rs10242397 | 1.839 | -0.502 | -1.898 | -2.901 | -0.061 | -0.111 | -0.111 | 0 | 0.28 | 0.42 | 0.13 | 0.11 | 0.16 | 0.17 |
| 7 | 8404148 | rs10264110 | -0.102 | -0.367 | -2.056 | -2.757 | 0.022 | 0.032 | 0.032 | 0 | 0.17 | 0.17 | 0.09 | 0.03 | 0.14 | 0.16 |
| 7 | 8416721 | rs7779531 | -0.101 | -0.305 | -2.048 | -2.751 | -0.007 | -0.024 | -0.024 | 0.159 | 0.18 | 0.23 | 0.08 | 0.03 | 0.15 | 0.16 |
| 7 | 8416819 | rs10239820 | 0.887 | -0.34 | -2.204 | -2.884 | -0.014 | -0.047 | -0.047 | 0 | 0.42 | 0.46 | 0.24 | 0.19 | 0.35 | 0.26 |
| 7 | 8422236 | rs17148140 | 1.739 | -0.979 | -1.746 | -3.001 | -0.063 | -0.154 | -0.154 | 0 | 0.26 | 0.39 | 0.12 | 0.05 | 0.15 | 0.16 |
| 7 | 8423035 | rs12702726 | 0.158 | -0.255 | -2.132 | -2.641 | -0.013 | -0.019 | -0.019 | 0 | 0.12 | 0.2 | 0.06 | 0.03 | 0.09 | 0.1 |
| 7 | 8423108 | rs42953 | -1.385 | -0.111 | -2.166 | -2.653 | -0.028 | 0.077 | 0.077 | 0 | 0.38 | 0.32 | 0.44 | 0.1 | 0.45 | 0.43 |
| 7 | 8426538 | rs10259079 | -0.532 | 0.014 | -2.827 | -2.53 | 0.001 | -0.009 | -0.009 | 0 | 0.17 | 0.15 | 0.09 | 0.03 | 0.14 | 0.17 |
| 7 | 8426887 | rs6967204 | 1.32 | 0.037 | -2.91 | -2.563 | -0.027 | 0.082 | 0.082 | 0 | 0.61 | 0.7 | 0.56 | 0.9 | 0.53 | 0.58 |
| 7 | 8433402 | rs2214695 | -0.56 | -0.217 | -2.734 | -2.392 | -0.063 | 0.05 | 0.05 | 0 | 0.83 | 0.95 | 0.67 | 0.93 | 0.73 | 0.77 |
| 7 | 8437955 | rs4628179 | 0.933 | -0.339 | -2.585 | -2.193 | -0.118 | 0.073 | 0.073 | 0 | 0.32 | 0.07 | 0.38 | 0.79 | 0.4 | 0.41 |
| 7 | 8449905 | rs10447691 | -0.578 | -0.144 | -1.358 | -1.162 | 0.004 | 0.003 | 0.003 | 0 | 0.11 | 0.11 | 0.06 | 0.04 | 0.09 | 0.11 |
| 7 | 8453313 | rs1007096 | -1.775 | 0.009 | -1.394 | -1.139 | 0.006 | 0.017 | 0.017 | 0 | 0.32 | 0.34 | 0.21 | 0.11 | 0.31 | 0.35 |
| 7 | 8464643 | rs12702747 | -1.587 | -0.126 | -0.905 | -0.736 | -0.007 | -0.031 | -0.031 | 2.385 | 0.19 | 0.28 | 0.1 | 0.04 | 0.16 | 0.19 |
| 7 | 8472460 | rs12702749 | -0.712 | 0.136 | -1.137 | -0.275 | 0.011 | 0.016 | 0.016 | 0 | 0.08 | 0.02 | 0.04 | 0.04 | 0.08 | 0.12 |
| 7 | 8472781 | rs11768154 | -0.679 | 0.144 | -1.134 | -0.271 | 0.01 | 0.016 | 0.016 | 0 | 0.08 | 0.02 | 0.04 | 0.04 | 0.08 | 0.12 |
| 7 | 8477442 | rs4141173 | -0.406 | 0.493 | -1.718 | -0.489 | -0.054 | 0.048 | 0.048 | 0 | 0.28 | 0.07 | 0.32 | 0.75 | 0.32 | 0.54 |
| 7 | 8484510 | rs7806226 | 0.597 | 0.06 | -0.973 | -0.923 | -0.02 | -0.052 | -0.052 | 0 | 0.81 | 0.39 | 0.63 | 0.9 | 0.86 | 0.85 |
| 7 | 8484908 | rs7805772 | 1.173 | -0.008 | -0.244 | -1.282 | -0.001 | -0.039 | -0.039 | 0 | 0.78 | 0.74 | 0.64 | 0.9 | 0.8 | 0.82 |
| 7 | 8488691 | rs10270530 | -0.275 | -0.829 | 0.115 | -0.874 | 0.012 | 0.055 | 0.055 | 0 | 0.42 | 0.75 | 0.46 | 0.75 | 0.39 | 0.53 |
| 7 | 8489911 | rs6945018 | -1.107 | -0.775 | -0.057 | -0.795 | -0.127 | 0.217 | 0.217 | 0 | 0.59 | 0.92 | 0.53 | 0.87 | 0.48 | 0.67 |
| 7 | 8503543 | rs2040891 | -1.133 | -0.792 | -0.06 | -0.358 | -0.12 | 0.121 | 0.121 | 0 | 0.59 | 0.93 | 0.52 | 0.79 | 0.46 | 0.68 |
| 7 | 8505405 | rs7796918 | -1.029 | -0.915 | 0.407 | -0.334 | -0.068 | 0.042 | 0.042 | 0.993 | 0.15 | 0.53 | 0.07 | 0.18 | 0.03 | 0.13 |
| 7 | 8508026 | rs10282272 | -0.944 | -0.899 | -0.223 | -0.326 | -0.062 | 0.016 | 0.016 | 1.358 | 0.15 | 0.65 | 0.07 | 0.18 | 0.03 | 0.14 |
| 7 | 8522345 | rs10247842 | -1.003 | -0.605 | -1.123 | -0.277 | -0.071 | 0.103 | 0.103 | 0 | 0.58 | 0.86 | 0.51 | 0.76 | 0.45 | 0.68 |
| 7 | 8527231 | rs10243563 | -0.646 | -0.318 | -0.338 | 0.24 | -0.025 | -0.039 | -0.039 | 0 | 0.1 | 0.21 | 0.05 | 0.02 | 0.03 | 0.08 |
| 7 | 8528063 | rs10486228 | -0.154 | 0.262 | -0.878 | 0.592 | -0.042 | -0.081 | -0.081 | 0 | 0.14 | 0.03 | 0.3 | 0.6 | 0.18 | 0.26 |
| 7 | 8532504 | rs6959225 | -0.73 | -0.624 | -0.426 | 0.71 | -0.058 | 0.047 | 0.047 | 3.318 | 0.33 | 0.48 | 0.29 | 0.46 | 0.21 | 0.24 |
| 7 | 8532519 | rs6977578 | 0.008 | -0.425 | -1.277 | -0.455 | -0.004 | 0.007 | 0.007 | 0.021 | 0.59 | 0.44 | 0.52 | 0.87 | 0.48 | 0.71 |
| 7 | 8532733 | rs10486231 | -0.44 | 0.099 | -1.279 | -0.44 | 0.002 | -0.003 | -0.003 | 0.589 | 0.09 | 0.08 | 0.04 | 0 | 0.06 | 0.09 |
| 7 | 8532844 | rs6978212 | 1.008 | 0.819 | -1.242 | -0.504 | -0.006 | 0.025 | 0.025 | 0.004 | 0.59 | 0.62 | 0.51 | 0.76 | 0.48 | 0.59 |
| 7 | 8533591 | rs4475381 | -0.236 | 1.03 | -1.127 | -0.328 | 0 | 0.003 | 0.003 | 0 | 0.11 | 0.16 | 0.17 | 0.17 | 0.13 | 0.12 |
| 7 | 8535486 | rs1859512 | -0.041 | 1.403 | -1.165 | -0.434 | 0 | -0.028 | -0.028 | 0.123 | 0.1 | 0.12 | 0.05 | 0 | 0.08 | 0.08 |
| 7 | 8539146 | rs4725098 | -0.964 | 1.947 | -1.353 | -0.35 | -0.082 | 0.028 | 0.028 | 0.569 | 0.84 | 0.97 | 0.76 | 0.94 | 0.7 | 0.83 |
| 7 | 8540236 | rs757704 | 0.986 | 1.6 | -0.436 | -0.589 | 0.031 | -0.033 | -0.033 | 0 | 0.66 | 0.59 | 0.53 | 0.78 | 0.49 | 0.66 |
| 7 | 8540295 | rs757705 | 0.744 | 1.072 | -0.607 | -1.075 | -0.018 | 0.072 | 0.072 | 0.014 | 0.6 | 0.73 | 0.6 | 0.9 | 0.49 | 0.58 |
| 7 | 8540931 | rs4141172 | -0.988 | 1.001 | -0.164 | -0.096 | 0.012 | 0.045 | 0.045 | 0 | 0.23 | 0.38 | 0.25 | 0.39 | 0.28 | 0.26 |
| 7 | 8542553 | rs17149860 | -0.43 | 0.614 | 0.119 | 0.194 | -0.007 | -0.001 | -0.001 | 0 | 0.22 | 0.2 | 0.21 | 0.21 | 0.1 | 0.2 |
| 7 | 8543538 | rs12534716 | -0.346 | 0.604 | 0.135 | 0.189 | -0.004 | 0 | 0 | 0 | 0.22 | 0.19 | 0.21 | 0.21 | 0.1 | 0.2 |
| 7 | 8544878 | rs10233230 | -0.842 | 0.749 | 0.137 | 0.366 | 0.006 | 0.019 | 0.019 | 0 | 0.24 | 0.38 | 0.22 | 0.34 | 0.27 | 0.26 |
| 7 | 8546068 | rs2189903 | -0.567 | 0.76 | 0.11 | 0.399 | -0.003 | 0.021 | 0.021 | 0 | 0.09 | 0.11 | 0.17 | 0.22 | 0.08 | 0.13 |
| 7 | 8547007 | rs17150003 | -0.131 | 0.809 | 0.245 | 0.381 | -0.026 | -0.048 | -0.048 | 0.021 | 0.19 | 0.15 | 0.21 | 0.36 | 0.25 | 0.24 |
| 7 | 8548572 | rs4725100 | -1.442 | 0.98 | 0.081 | 0.528 | 0.016 | 0.007 | 0.007 | 0 | 0.81 | 0.82 | 0.71 | 0.69 | 0.64 | 0.65 |
| 7 | 8549058 | rs7801099 | 0.989 | 0.718 | 0.299 | 0.391 | -0.004 | -0.003 | -0.003 | 0 | 0.55 | 0.6 | 0.51 | 0.48 | 0.52 | 0.43 |
| 7 | 8550648 | rs6978582 | 2.421 | 0.385 | 0.304 | 0.253 | 0.02 | 0.068 | 0.068 | 1.544 | 0.36 | 0.2 | 0.28 | 0.13 | 0.3 | 0.2 |
| 7 | 8550956 | rs2349485 | 0.237 | 0.645 | 0.56 | 0.292 | 0.022 | -0.002 | -0.002 | 0.023 | 0.71 | 0.66 | 0.58 | 0.71 | 0.62 | 0.65 |
| 7 | 8551289 | rs1013868 | -0.7 | 0.623 | 0.374 | 0.437 | -0.001 | -0.001 | -0.001 | 0 | 0.77 | 0.79 | 0.61 | 0.71 | 0.66 | 0.69 |
| 7 | 8560202 | rs2057750 | 0.249 | 0.457 | 0.422 | 0.756 | -0.003 | 0.011 | 0.011 | 0.47 | 0.57 | 0.64 | 0.51 | 0.71 | 0.55 | 0.54 |
| 7 | 8560291 | rs10255873 | 1.492 | 0.446 | 0.616 | 0.862 | -0.005 | -0.001 | -0.001 | 0 | 0.39 | 0.4 | 0.3 | 0.37 | 0.31 | 0.3 |
| 7 | 8560567 | rs4472426 | -0.214 | 0.607 | 0.042 | 0.936 | -0.003 | -0.004 | -0.004 | 0 | 0.19 | 0.22 | 0.21 | 0.35 | 0.24 | 0.24 |
| 7 | 8560944 | rs4640975 | -0.179 | 0.609 | -0.043 | 0.936 | -0.007 | -0.015 | -0.015 | 0 | 0.18 | 0.2 | 0.21 | 0.34 | 0.24 | 0.24 |
| 7 | 8561945 | rs2040765 | 1.689 | 0.567 | 0.008 | 0.89 | 0.013 | 0 | 0 | 0.17 | 0.37 | 0.26 | 0.28 | 0.37 | 0.31 | 0.32 |
| 7 | 8562722 | rs17150410 | -0.519 | 0.73 | -0.421 | 1.024 | -0.001 | 0.003 | 0.003 | 0 | 0.19 | 0.23 | 0.21 | 0.34 | 0.24 | 0.24 |
| 7 | 8571615 | rs11764549 | 0.983 | 0.905 | -0.425 | 0.666 | 0.005 | 0.035 | 0.035 | 1.92 | 0.45 | 0.24 | 0.47 | 0.33 | 0.41 | 0.35 |

| | | | | | | | | | | | | | | | | |
|---|---|---|---|---|---|---|---|---|---|---|---|---|---|---|---|---|
| 7 | 8573545 | rs17150705 | 0.532 | 0.663 | -0.953 | 1.09 | -0.002 | -0.02 | -0.02 | 0.238 | 0.07 | 0.01 | 0.08 | 0.28 | 0.06 | 0.07 |
| 7 | 8575468 | rs17150761 | 1.343 | 0.631 | -0.936 | 0.986 | -0.036 | -0.05 | -0.05 | 0.008 | 0.38 | 0.03 | 0.41 | 0.33 | 0.32 | 0.34 |
| 7 | 8576305 | rs7809396 | -0.964 | 1.036 | -0.621 | 1.248 | -0.007 | -0.018 | -0.018 | 0 | 0.2 | 0.21 | 0.19 | 0.33 | 0.24 | 0.24 |
| 7 | 8579216 | rs6955389 | 0.411 | 0.473 | -0.54 | 0.959 | 0.051 | 0.004 | 0.004 | 0.866 | 0.49 | 0.36 | 0.34 | 0.47 | 0.36 | 0.39 |
| 7 | 8582096 | rs7799856 | 0.004 | 0.869 | 0.097 | 0.966 | -0.011 | 0.013 | 0.013 | 0.397 | 0.62 | 0.71 | 0.42 | 0.65 | 0.45 | 0.52 |
| 7 | 8584243 | rs10486239 | -0.414 | 0.869 | 0.553 | 0.658 | -0.003 | -0.068 | -0.068 | 1.232 | 0.12 | 0.21 | 0.06 | 0 | 0.08 | 0.05 |
| 7 | 8584590 | rs4720785 | 0.559 | 0.317 | 0.443 | 0.397 | 0.044 | -0.011 | -0.011 | 0 | 0.46 | 0.24 | 0.33 | 0.45 | 0.35 | 0.4 |
| 7 | 8585496 | rs1859274 | 0.37 | 0.31 | 0.528 | 0.396 | 0.024 | -0.002 | -0.002 | 0 | 0.47 | 0.35 | 0.35 | 0.45 | 0.36 | 0.44 |
| 7 | 8587179 | rs6969718 | 0.863 | 0.21 | 0.728 | 0.495 | -0.004 | 0.001 | 0.001 | 0 | 0.39 | 0.6 | 0.32 | 0.44 | 0.27 | 0.35 |
| 7 | 8588980 | rs7812117 | 0.649 | 0.567 | 0.651 | 0.429 | 0.071 | -0.014 | -0.014 | 0.236 | 0.49 | 0.23 | 0.43 | 0.47 | 0.41 | 0.36 |
| 7 | 8594363 | rs2189461 | -0.239 | 1.107 | 1.167 | 0.094 | 0 | 0.01 | 0.01 | 0.016 | 0.66 | 0.61 | 0.6 | 0.5 | 0.64 | 0.51 |
| 7 | 8595389 | rs6963644 | -0.821 | 1.356 | 1.464 | 0.236 | -0.005 | -0.09 | -0.09 | 0.425 | 0.14 | 0.35 | 0.06 | 0 | 0.08 | 0.05 |
| 7 | 8597116 | rs6952939 | 0.749 | 1.39 | 1.382 | 0.221 | -0.007 | -0.007 | -0.007 | 0.817 | 0.43 | 0.39 | 0.5 | 0.51 | 0.49 | 0.42 |
| 7 | 8606260 | rs10227474 | -1.337 | 1.394 | 2.24 | 0.124 | 0.008 | -0.116 | -0.116 | 0 | 0.15 | 0.44 | 0.08 | 0 | 0.1 | 0.06 |
| 7 | 8623087 | rs6970999 | -1.271 | 1.44 | 2.192 | 0.517 | 0.001 | 0.001 | 0.001 | 0 | 0.3 | 0.72 | 0.32 | 0.28 | 0.25 | 0.3 |
| 7 | 8628692 | rs10249831 | -0.355 | 1.445 | 2.388 | 0.483 | 0.042 | -0.126 | -0.126 | 0 | 0.19 | 0.43 | 0.24 | 0.01 | 0.23 | 0.1 |
| 7 | 8633143 | rs10226547 | -1.694 | 1.466 | 2.24 | 0.507 | 0.009 | 0.013 | 0.013 | 0 | 0.32 | 0.78 | 0.33 | 0.28 | 0.25 | 0.3 |
| 7 | 8635885 | rs4538793 | -1.606 | 1.734 | 1.842 | 0.747 | 0.008 | 0.136 | 0.136 | 0 | 0.11 | 0.29 | 0.1 | 0.27 | 0.07 | 0.2 |
| 7 | 8637302 | rs10231458 | -0.216 | 1.749 | 1.303 | 0.21 | 0.01 | -0.069 | -0.069 | 0.147 | 0.19 | 0.33 | 0.11 | 0 | 0.22 | 0.11 |
| 7 | 8637693 | rs10265132 | -0.813 | 1.465 | 1.183 | 0.401 | 0.039 | -0.023 | -0.023 | 0 | 0.26 | 0.55 | 0.25 | 0.22 | 0.31 | 0.24 |
| 7 | 8638143 | rs6974213 | 1.381 | 1.279 | 0.458 | 0.41 | -0.013 | -0.129 | -0.129 | 0.446 | 0.42 | 0.03 | 0.28 | 0.5 | 0.39 | 0.35 |
| 7 | 8639507 | rs10232665 | -0.248 | 1.329 | 1.299 | 0.114 | 0.034 | -0.112 | -0.112 | 0.038 | 0.21 | 0.4 | 0.14 | 0 | 0.26 | 0.11 |
| 7 | 8641237 | rs10278529 | -1.434 | 1.364 | 1.241 | 0.144 | 0.009 | -0.034 | -0.034 | 0 | 0.32 | 0.61 | 0.3 | 0.2 | 0.31 | 0.27 |
| 7 | 8667171 | rs4492284 | -0.59 | 1.226 | 0.758 | 0.356 | 0.001 | 0.001 | 0.001 | 0 | 0.07 | 0.06 | 0.15 | 0.2 | 0.04 | 0.16 |
| 7 | 8673137 | rs6945631 | -1.158 | 1.357 | 1.114 | 0.169 | 0.021 | -0.032 | -0.032 | 0.086 | 0.27 | 0.49 | 0.28 | 0.2 | 0.29 | 0.26 |
| 7 | 8673637 | rs10231165 | -0.287 | 1.348 | 1.181 | 0.163 | 0.047 | -0.11 | -0.11 | 0 | 0.2 | 0.38 | 0.14 | 0 | 0.27 | 0.1 |
| 7 | 8675522 | rs10276202 | -1.06 | 1.378 | 1.141 | 0.18 | 0.027 | -0.033 | -0.033 | 0.191 | 0.27 | 0.51 | 0.28 | 0.2 | 0.29 | 0.26 |
| 7 | 8683681 | rs338 | 0.046 | 1.437 | 1.058 | 0.133 | 0.056 | -0.117 | -0.117 | 0 | 0.8 | 0.6 | 0.86 | 1 | 0.73 | 0.9 |
| 7 | 8697034 | rs7802092 | 0.837 | 1.456 | 0.97 | 0.201 | 0.036 | -0.035 | -0.035 | 0 | 0.72 | 0.43 | 0.71 | 0.8 | 0.68 | 0.71 |
| 7 | 8698007 | rs10952119 | 0.915 | 1.142 | 0.75 | 0.263 | 0.007 | -0.021 | -0.021 | 0 | 0.7 | 0.42 | 0.7 | 0.74 | 0.7 | 0.72 |
| 7 | 8720903 | rs11769942 | -0.456 | 0.77 | -0.07 | 0.57 | -0.012 | -0.012 | -0.012 | 0 | 0.4 | 0.48 | 0.46 | 0.29 | 0.33 | 0.39 |
| 7 | 8721760 | rs2057789 | -1.121 | 1.577 | -0.108 | 0.254 | 0.034 | -0.032 | -0.032 | 0.044 | 0.3 | 0.43 | 0.35 | 0.22 | 0.34 | 0.4 |
| 7 | 8721852 | rs12702765 | -0.941 | 1.48 | -0.727 | 0.419 | -0.031 | -0.011 | -0.011 | 3.011 | 0.19 | 0.08 | 0.19 | 0.18 | 0.25 | 0.19 |
| 7 | 8729005 | rs2189629 | 0.284 | 1.334 | -0.5 | -0.513 | 0.002 | -0.082 | -0.082 | 0 | 0.63 | 0.44 | 0.69 | 0.91 | 0.64 | 0.59 |
| 7 | 8729206 | rs2189630 | 0.69 | 0.992 | -1.281 | -1.131 | 0.008 | -0.024 | -0.024 | 0 | 0.3 | 0.37 | 0.45 | 0.8 | 0.38 | 0.36 |
| 7 | 8730421 | rs12702766 | 0.59 | 0.922 | -1.438 | -1.054 | 0.01 | 0.127 | 0.127 | 0 | 0.3 | 0.07 | 0.13 | 0.1 | 0.24 | 0.18 |
| 7 | 8732406 | rs12702767 | 0.466 | 0.715 | -1.173 | -1.076 | 0.027 | 0.062 | 0.062 | 0 | 0.31 | 0.18 | 0.23 | 0.1 | 0.22 | 0.19 |
| 7 | 8732867 | rs17153994 | 0.57 | 0.89 | -0.966 | -0.309 | 0.019 | 0.022 | 0.022 | 0 | 0.26 | 0.28 | 0.34 | 0.47 | 0.45 | 0.34 |
| 7 | 8736171 | rs7780955 | 0.364 | 0.077 | -0.285 | -0.082 | -0.057 | 0.081 | 0.081 | 3.349 | 0.14 | 0.32 | 0.1 | 0.17 | 0.04 | 0.09 |
| 7 | 8736299 | rs2189631 | -0.464 | -0.115 | -0.406 | -0.231 | 0.001 | 0.066 | 0.066 | 0 | 0.7 | 0.77 | 0.78 | 0.92 | 0.74 | 0.68 |
| 7 | 8737023 | rs12536578 | -0.042 | -0.076 | 0.014 | 0.483 | -0.04 | 0.047 | 0.047 | 2.702 | 0.13 | 0.18 | 0.09 | 0.17 | 0.04 | 0.09 |
| 7 | 8737318 | rs7782383 | -0.417 | -0.059 | -0.128 | 0.499 | -0.074 | 0.212 | 0.212 | 2.89 | 0.15 | 0.43 | 0.2 | 0.35 | 0.06 | 0.14 |
| 7 | 8737398 | rs7786015 | -0.712 | -0.003 | -0.138 | 0.525 | -0.1 | 0.083 | 0.083 | 0.039 | 0.24 | 0.46 | 0.22 | 0.35 | 0.09 | 0.19 |
| 7 | 8739365 | rs11536481 | -0.572 | 0.122 | -0.452 | 0.41 | -0.02 | -0.006 | -0.006 | 0 | 0.25 | 0.24 | 0.15 | 0.15 | 0.11 | 0.15 |
| 7 | 8751432 | rs1017002 | 0.031 | 1.188 | -1.736 | 0.133 | -0.057 | 0.045 | 0.045 | 0 | 0.67 | 0.86 | 0.73 | 0.78 | 0.59 | 0.67 |
| 7 | 8751843 | rs1017003 | 0.355 | 0.194 | -1.012 | 0.895 | 0.071 | 0.049 | 0.049 | 0 | 0.46 | 0.71 | 0.45 | 0.64 | 0.34 | 0.48 |
| 7 | 8755731 | rs1017005 | 0.013 | -0.388 | -1.26 | 0.998 | 0.008 | -0.032 | -0.032 | 0 | 0.17 | 0.12 | 0.28 | 0.31 | 0.17 | 0.24 |
| 7 | 8757367 | rs4141002 | 0.303 | -0.457 | -1.32 | 0.91 | 0.001 | -0.008 | -0.008 | 0.282 | 0.13 | 0.04 | 0.18 | 0.13 | 0.11 | 0.19 |
| 7 | 8764246 | rs17154637 | -0.111 | -0.961 | -1.241 | 0.354 | 0.021 | 0.033 | 0.033 | 0.238 | 0.46 | 0.67 | 0.43 | 0.4 | 0.38 | 0.44 |
| 7 | 8765540 | rs7779855 | 0.238 | -0.993 | -1.296 | 0.238 | 0.003 | -0.031 | -0.031 | 0 | 0.15 | 0.19 | 0.22 | 0.02 | 0.15 | 0.2 |
| 7 | 8772150 | rs12234501 | 1.124 | -0.938 | -1.039 | 0.247 | -0.033 | 0.007 | 0.007 | 0 | 0.26 | 0.33 | 0.15 | 0.34 | 0.16 | 0.2 |
| 7 | 8780037 | rs6463839 | -0.08 | -0.261 | -1.663 | 0.401 | 0.09 | -0.024 | -0.024 | 0 | 0.48 | 0.84 | 0.49 | 0.5 | 0.34 | 0.43 |
| 7 | 8784440 | rs1557876 | -0.992 | 0.012 | -1.702 | 0.424 | -0.096 | 0.005 | 0.005 | 0 | 0.67 | 0.91 | 0.71 | 0.7 | 0.54 | 0.67 |
| 7 | 8785903 | rs17334835 | -0.719 | 0.224 | -1.785 | 0.097 | -0.058 | -0.044 | -0.044 | 0.297 | 0.12 | 0.01 | 0.24 | 0.14 | 0.16 | 0.24 |
| 7 | 8785927 | rs17154753 | 0.292 | -0.265 | -1.557 | 0.271 | -0.005 | -0.019 | -0.019 | 0.926 | 0.11 | 0.06 | 0.17 | 0.19 | 0.15 | 0.17 |
| 7 | 8786057 | rs886505 | -0.233 | -0.056 | -0.996 | -0.555 | 0 | -0.008 | -0.008 | 0 | 0.62 | 0.63 | 0.54 | 0.85 | 0.5 | 0.57 |
| 7 | 8788531 | rs7806959 | 0.08 | -0.189 | -1.312 | -0.524 | 0.039 | 0.153 | 0.153 | 0 | 0.51 | 0.77 | 0.49 | 0.82 | 0.4 | 0.42 |
| 7 | 8790170 | rs10240887 | -0.078 | 0.265 | -1.369 | -1.026 | 0.024 | 0.073 | 0.073 | 0 | 0.61 | 0.74 | 0.69 | 0.88 | 0.64 | 0.66 |
| 7 | 8790549 | rs10225984 | -0.826 | 0.22 | -0.597 | -0.752 | 0.002 | 0.002 | 0.002 | 0 | 0.3 | 0.34 | 0.36 | 0.38 | 0.27 | 0.3 |
| 7 | 8794418 | rs2189621 | -0.433 | 0.379 | -1.312 | -0.967 | -0.002 | 0.081 | 0.081 | 0 | 0.6 | 0.79 | 0.49 | 0.81 | 0.63 | 0.6 |
| 7 | 8795294 | rs6975676 | -1.182 | 0.412 | -1.454 | -1.032 | -0.014 | 0.087 | 0.087 | 0 | 0.72 | 0.89 | 0.53 | 0.94 | 0.69 | 0.78 |
| 7 | 8796967 | rs6965209 | 0.211 | 0.231 | -1.054 | -0.631 | 0.008 | 0.009 | 0.009 | 0 | 0.34 | 0.31 | 0.19 | 0.2 | 0.28 | 0.41 |
| 7 | 8797321 | rs6463841 | -0.427 | 0.691 | -0.987 | -0.045 | -0.001 | -0.001 | -0.001 | 0 | 0.48 | 0.51 | 0.4 | 0.49 | 0.54 | 0.46 |
| 7 | 8797698 | rs10248396 | 1.435 | 1.033 | -0.961 | -0.007 | -0.019 | -0.017 | -0.017 | 0 | 0.11 | 0.06 | 0.2 | 0.31 | 0.24 | 0.06 |
| 7 | 8797737 | rs9791719 | 0.21 | 0.621 | -0.955 | 0.171 | -0.012 | 0.008 | 0.008 | 0 | 0.4 | 0.44 | 0.26 | 0.49 | 0.34 | 0.52 |
| 7 | 8799415 | rs12702774 | -0.152 | 1.337 | -0.454 | 0.633 | 0.029 | 0.04 | 0.04 | 0 | 0.16 | 0.19 | 0.24 | 0.32 | 0.26 | 0.09 |
| 7 | 8800804 | rs6973037 | -0.663 | 0.012 | -0.13 | 0.505 | -0.02 | 0.089 | 0.089 | 0 | 0.18 | 0.36 | 0.15 | 0.34 | 0.16 | 0.3 |
| 7 | 8800995 | rs1468298 | 0.092 | 0.166 | -0.503 | 0.303 | 0.013 | -0.011 | -0.011 | 0.069 | 0.14 | 0.18 | 0.07 | 0.11 | 0.12 | 0.16 |
| 7 | 8801357 | rs6973715 | -2.519 | -0.15 | -1.341 | 0.154 | -0.119 | 0.095 | 0.095 | 0.006 | 0.73 | 0.98 | 0.66 | 0.94 | 0.68 | 0.81 |

| | | | | | | | | | | | | | | | | |
|---|---|---|---|---|---|---|---|---|---|---|---|---|---|---|---|---|
| 7 | 8802039 | rs10268580 | -0.872 | -0.969 | -0.813 | 0.414 | -0.036 | 0.073 | 0.073 | 0 | 0.23 | 0.51 | 0.18 | 0.35 | 0.2 | 0.32 |
| 7 | 8802582 | rs1468301 | -0.87 | -0.94 | -0.809 | 0.343 | 0.003 | 0.062 | 0.062 | 0 | 0.3 | 0.5 | 0.22 | 0.46 | 0.28 | 0.46 |
| 7 | 8806028 | rs12668192 | 0.138 | -1.14 | -0.932 | 0.174 | -0.004 | 0.015 | 0.015 | 0 | 0.11 | 0 | 0.07 | 0.06 | 0.09 | 0.16 |
| 7 | 8806637 | rs10246356 | 0.079 | -1.398 | -0.844 | 0.161 | 0.031 | -0.02 | -0.02 | 0 | 0.42 | 0.3 | 0.38 | 0.48 | 0.37 | 0.41 |
| 7 | 8808404 | rs7807056 | 0.334 | -1.402 | -0.873 | 0.144 | 0.041 | -0.06 | -0.06 | 0 | 0.38 | 0.18 | 0.36 | 0.47 | 0.36 | 0.4 |
| 7 | 8815515 | rs1075126 | -0.599 | -1.49 | -0.556 | 0.086 | -0.038 | -0.06 | -0.06 | 1.002 | 0.19 | 0.42 | 0.12 | 0.06 | 0.12 | 0.19 |
| 7 | 8816498 | rs6975224 | 0.013 | -1.474 | -0.451 | 0.121 | -0.003 | 0 | 0 | 0 | 0.42 | 0.41 | 0.4 | 0.48 | 0.36 | 0.42 |
| 7 | 8817762 | rs10280502 | 1.407 | -1.643 | -0.737 | 0.224 | 0.005 | -0.042 | -0.042 | 0 | 0.13 | 0.05 | 0.18 | 0.31 | 0.15 | 0.1 |
| 7 | 8821834 | rs10228121 | -1.054 | -1.749 | -0.596 | 0.141 | 0.005 | -0.002 | -0.002 | 1.07 | 0.54 | 0.41 | 0.47 | 0.53 | 0.44 | 0.56 |
| 7 | 8822108 | rs10228390 | -2.861 | -1.616 | -0.837 | 0.19 | -0.152 | -0.115 | -0.115 | 0 | 0.74 | 0.96 | 0.57 | 0.65 | 0.6 | 0.77 |
| 7 | 8822413 | rs12702779 | 1.059 | -1.305 | -0.281 | 0.628 | -0.003 | 0 | 0 | 0 | 0.1 | 0.12 | 0.04 | 0.11 | 0.1 | 0.17 |
| 7 | 8823954 | rs886577 | -1.226 | -1.354 | -0.243 | 0.621 | 0 | -0.001 | -0.001 | 0 | 0.54 | 0.5 | 0.46 | 0.59 | 0.49 | 0.6 |
| 7 | 8827959 | rs11972509 | 0.769 | -1.231 | -0.273 | 0.681 | -0.004 | -0.052 | -0.052 | 0 | 0.15 | 0.05 | 0.19 | 0.38 | 0.17 | 0.11 |
| 7 | 8830494 | rs11977634 | -1.174 | -1.228 | -0.018 | 0.599 | -0.001 | -0.002 | -0.002 | 2.891 | 0.54 | 0.46 | 0.46 | 0.6 | 0.48 | 0.59 |
| 7 | 8833563 | rs7791419 | -1.441 | -1.217 | 0.009 | 0.616 | 0.004 | 0 | 0 | 0.65 | 0.58 | 0.54 | 0.47 | 0.6 | 0.5 | 0.6 |
| 7 | 8837843 | rs6951762 | 0.039 | -1.067 | 0.054 | 0.686 | 0.044 | 0.022 | 0.022 | 0 | 0.14 | 0.35 | 0.1 | 0.18 | 0.15 | 0.2 |
| 7 | 8844197 | rs10272895 | -0.355 | -1.194 | -0.132 | 0.605 | 0.047 | -0.043 | -0.043 | 0 | 0.42 | 0.22 | 0.38 | 0.45 | 0.37 | 0.44 |
| 7 | 8844448 | rs12702782 | 0.274 | -1.152 | -0.069 | 0.18 | 0.03 | -0.003 | -0.003 | 0 | 0.13 | 0.31 | 0.07 | 0.17 | 0.13 | 0.18 |
| 7 | 8846057 | rs12536728 | 0.357 | -1.442 | -0.758 | 0.168 | 0.052 | 0.067 | 0.067 | 0 | 0.27 | 0.13 | 0.19 | 0.12 | 0.18 | 0.31 |
| 7 | 8849533 | rs12702783 | -0.115 | -0.598 | -0.548 | 0.5 | 0.001 | 0.004 | 0.004 | 0 | 0.07 | 0.03 | 0.05 | 0.05 | 0.07 | 0.12 |
| 7 | 8850785 | rs1034755 | -0.521 | -0.854 | -0.368 | 0.521 | 0.027 | 0.015 | 0.015 | 0.071 | 0.45 | 0.37 | 0.36 | 0.31 | 0.34 | 0.47 |
| 7 | 8851998 | rs6964741 | -0.339 | -0.186 | -0.166 | 0.322 | 0.046 | -0.054 | -0.054 | 0 | 0.1 | 0.21 | 0.29 | 0 | 0.13 | 0.06 |
| 7 | 8853900 | rs9785044 | -0.874 | -0.191 | -0.32 | 0.709 | -0.005 | -0.009 | -0.009 | 0 | 0.19 | 0.2 | 0.19 | 0.39 | 0.22 | 0.27 |
| 7 | 8854128 | rs17155853 | 0.264 | -0.281 | -0.28 | 0.958 | -0.003 | 0.028 | 0.028 | 0 | 0.08 | 0.06 | 0.05 | 0.17 | 0.11 | 0.13 |
| 7 | 8855423 | rs9785055 | 0.289 | -0.115 | -0.461 | 0.992 | -0.007 | 0 | 0 | 0 | 0.13 | 0.11 | 0.08 | 0.18 | 0.13 | 0.22 |
| 7 | 8856074 | rs10486259 | -0.235 | -0.127 | -0.445 | 1.006 | -0.007 | -0.016 | -0.016 | 0 | 0.16 | 0.15 | 0.19 | 0.39 | 0.18 | 0.27 |
| 7 | 8867550 | rs2349786 | 0.322 | -0.124 | -0.37 | 0.87 | 0.025 | -0.042 | -0.042 | 0 | 0.08 | 0.19 | 0.25 | 0 | 0.08 | 0.05 |
| 7 | 8872635 | rs10252753 | 1.292 | -0.461 | -0.85 | 0.871 | 0.013 | -0.07 | -0.07 | 0 | 0.18 | 0.02 | 0.09 | 0.34 | 0.14 | 0.31 |
| 7 | 8876292 | rs6946006 | 0.5 | -0.335 | -0.511 | 0.993 | 0.013 | -0.048 | -0.048 | 0 | 0.15 | 0.07 | 0.17 | 0.21 | 0.13 | 0.09 |
| 7 | 8877256 | rs978514 | -0.94 | -0.232 | -0.124 | 1.099 | 0.006 | 0.002 | 0.002 | 0 | 0.37 | 0.45 | 0.53 | 0.4 | 0.38 | 0.37 |
| 7 | 8889394 | rs1154814 | -1.1 | -0.053 | 0.118 | 1.089 | 0.015 | 0.039 | 0.039 | 0 | 0.5 | 0.71 | 0.6 | 0.57 | 0.48 | 0.5 |
| 7 | 8890284 | rs7788194 | -0.873 | 0.294 | 0.449 | 1.224 | 0.057 | 0.155 | 0.155 | 0 | 0.1 | 0.29 | 0.36 | 0.22 | 0.11 | 0.08 |
| 7 | 8894344 | rs10486264 | -0.504 | -0.342 | -0.422 | 0.737 | 0.002 | 0.001 | 0.001 | 0 | 0.29 | 0.31 | 0.22 | 0.2 | 0.23 | 0.31 |
| 7 | 8899901 | rs1859416 | 0.098 | -0.29 | -0.275 | 1.088 | -0.014 | 0.111 | 0.111 | 0.491 | 0.25 | 0.35 | 0.26 | 0.47 | 0.25 | 0.41 |
| 7 | 8900786 | rs6967307 | 0.008 | 0.298 | -0.838 | 1.039 | 0.016 | 0.004 | 0.004 | 0 | 0.47 | 0.6 | 0.4 | 0.46 | 0.48 | 0.47 |
| 7 | 8901276 | rs13239626 | 0.605 | -0.083 | -1.312 | 0.602 | -0.127 | 0.216 | 0.216 | 0 | 0.42 | 0.04 | 0.49 | 0.11 | 0.49 | 0.19 |
| 7 | 8910032 | rs17156629 | 0.315 | -0.199 | -1.132 | 0.774 | 0.011 | -0.016 | -0.016 | 0 | 0.14 | 0.08 | 0.1 | 0.22 | 0.12 | 0.27 |
| 7 | 8919987 | rs2214621 | -1.071 | -0.504 | -0.408 | 0.963 | -0.03 | -0.028 | -0.028 | 0 | 0.68 | 0.65 | 0.8 | 0.74 | 0.73 | 0.56 |
| 7 | 8920903 | rs17156731 | -0.215 | -0.326 | -0.216 | 1.332 | 0.006 | 0.137 | 0.137 | 0 | 0.09 | 0.17 | 0.17 | 0.28 | 0.07 | 0.08 |
| 7 | 8924442 | rs6945904 | -0.113 | -0.425 | -0.349 | 1.18 | 0.013 | 0.008 | 0.008 | 0.404 | 0.57 | 0.65 | 0.73 | 0.58 | 0.62 | 0.29 |
| 7 | 8930961 | rs2113849 | 0.027 | -0.514 | 0.086 | 1.118 | -0.032 | -0.017 | -0.017 | 0 | 0.57 | 0.5 | 0.65 | 0.3 | 0.62 | 0.28 |
| 7 | 8935483 | rs4725142 | 0.025 | -0.579 | 0.02 | 1.086 | -0.001 | 0.001 | 0.001 | 0.719 | 0.57 | 0.6 | 0.65 | 0.3 | 0.62 | 0.26 |
| 7 | 8935674 | rs4725143 | -1.267 | -0.554 | 0.046 | 1.211 | 0.005 | 0.029 | 0.029 | 0 | 0.23 | 0.43 | 0.33 | 0.63 | 0.22 | 0.35 |
| 7 | 8937076 | rs17156940 | 1.197 | -0.531 | -0.135 | 0.975 | 0.005 | -0.048 | -0.048 | 0 | 0.1 | 0.01 | 0.07 | 0.17 | 0.1 | 0.24 |
| 7 | 8938582 | rs6463853 | 0.653 | -0.724 | -0.19 | 0.765 | 0.036 | 0.216 | 0.216 | 0.02 | 0.48 | 0.16 | 0.52 | 0.12 | 0.55 | 0.26 |
| 7 | 8942119 | rs11770027 | -0.355 | -0.85 | -0.024 | 0.838 | -0.004 | 0 | 0 | 0.424 | 0.6 | 0.6 | 0.69 | 0.3 | 0.66 | 0.32 |
| 7 | 8944613 | rs10256000 | -1.052 | -0.51 | 0.196 | 1.024 | -0.037 | 0.031 | 0.031 | 0.043 | 0.68 | 0.63 | 0.75 | 0.47 | 0.76 | 0.58 |
| 7 | 8945202 | rs4725145 | 0.059 | -0.318 | 0.226 | 1.166 | 0.031 | 0.045 | 0.045 | 0 | 0.5 | 0.4 | 0.58 | 0.26 | 0.58 | 0.35 |
| 7 | 8945240 | rs10256828 | 0.103 | -0.258 | 0.152 | 0.982 | 0 | 0.004 | 0.004 | 0 | 0.2 | 0.27 | 0.19 | 0.34 | 0.2 | 0.24 |
| 7 | 8946172 | rs1017266 | -0.304 | -0.106 | 0.17 | 0.807 | 0.013 | 0.01 | 0.01 | 0 | 0.56 | 0.63 | 0.66 | 0.29 | 0.62 | 0.33 |
| 7 | 8947944 | rs12702794 | 0.305 | -0.042 | -0.021 | 0.842 | 0.005 | 0.096 | 0.096 | 0 | 0.45 | 0.26 | 0.51 | 0.13 | 0.53 | 0.29 |
| 7 | 8948182 | rs12702795 | 0.333 | -0.042 | -0.025 | 0.842 | 0.003 | 0.09 | 0.09 | 0 | 0.44 | 0.26 | 0.5 | 0.12 | 0.51 | 0.28 |
| 7 | 8950864 | rs12702797 | 0.383 | -0.155 | 0.119 | 0.66 | 0.003 | 0.09 | 0.09 | 0 | 0.44 | 0.26 | 0.52 | 0.13 | 0.53 | 0.28 |
| 7 | 8953858 | rs10486269 | 1.138 | 0.622 | -0.016 | 0.506 | -0.073 | 0.126 | 0.126 | 0 | 0.28 | 0.07 | 0.36 | 0.08 | 0.33 | 0.2 |
| 7 | 8957041 | rs10266202 | -0.581 | 0.512 | 0.393 | 0.84 | 0.014 | 0 | 0 | 0 | 0.56 | 0.65 | 0.63 | 0.55 | 0.58 | 0.51 |
| 7 | 8959256 | rs11561884 | 0.033 | 0.664 | 0.544 | 0.917 | 0.039 | 0.175 | 0.175 | 0 | 0.08 | 0.12 | 0.1 | 0.3 | 0.08 | 0.07 |
| 7 | 8959373 | rs10247578 | 0.558 | 0.65 | 0.564 | 0.907 | -0.002 | -0.001 | -0.001 | 0.016 | 0.36 | 0.36 | 0.44 | 0.21 | 0.4 | 0.28 |
| 7 | 8959527 | rs12154520 | 0.291 | 0.657 | 0.623 | 0.899 | 0.028 | 0.009 | 0.009 | 0 | 0.38 | 0.48 | 0.48 | 0.39 | 0.41 | 0.27 |
| 7 | 8960296 | rs2016550 | -0.512 | 0.671 | 0.673 | 0.966 | 0.023 | 0.046 | 0.046 | 0 | 0.1 | 0.19 | 0.14 | 0.18 | 0.1 | 0.05 |
| 7 | 8963814 | rs2113851 | -0.311 | 0.668 | 0.948 | 0.48 | 0.003 | 0.004 | 0.004 | 0 | 0.48 | 0.55 | 0.6 | 0.28 | 0.52 | 0.27 |
| 7 | 8965531 | rs17157269 | 0.258 | 0.565 | 0.84 | 0.407 | -0.001 | -0.016 | -0.016 | 1.009 | 0.09 | 0.09 | 0.07 | 0.03 | 0.13 | 0.18 |
| 7 | 8966195 | rs17157273 | 0.782 | 0.678 | 0.798 | 0.723 | -0.012 | 0.089 | 0.089 | 0 | 0.17 | 0.32 | 0.12 | 0.3 | 0.15 | 0.21 |
| 7 | 8968224 | rs7783918 | -0.503 | 0.672 | 0.393 | 0.582 | 0.024 | 0.073 | 0.073 | 0 | 0.55 | 0.72 | 0.69 | 0.72 | 0.58 | 0.47 |
| 7 | 8968422 | rs17336189 | -0.141 | 0.996 | 0.26 | 0.142 | 0.007 | -0.015 | -0.015 | 3.12 | 0.41 | 0.49 | 0.53 | 0.28 | 0.44 | 0.26 |
| 7 | 8968569 | rs13230035 | 0.029 | 0.98 | 0.088 | 0.151 | 0.015 | -0.022 | -0.022 | 0 | 0.38 | 0.56 | 0.48 | 0.27 | 0.39 | 0.22 |
| 7 | 8972810 | rs5007445 | 0.035 | 0.87 | -0.145 | -0.09 | 0.001 | -0.044 | -0.044 | 0 | 0.37 | 0.65 | 0.45 | 0.1 | 0.36 | 0.23 |
| 7 | 8973252 | rs7809435 | 0.433 | 0.969 | -0.055 | -0.032 | 0.005 | -0.067 | -0.067 | 2.229 | 0.31 | 0.62 | 0.38 | 0.1 | 0.31 | 0.2 |
| 7 | 8974051 | rs12216543 | 0.658 | 1.126 | -0.457 | 0.055 | -0.01 | 0.019 | 0.019 | 0.055 | 0.28 | 0.27 | 0.36 | 0.1 | 0.31 | 0.21 |
| 7 | 8978004 | rs12702801 | 0.972 | 0.128 | -0.302 | 0.204 | -0.072 | 0.059 | 0.059 | 0 | 0.34 | 0.19 | 0.45 | 0.21 | 0.45 | 0.23 |

| | | | | | | | | | | | | | | | | |
|---|---|---|---|---|---|---|---|---|---|---|---|---|---|---|---|---|
| 7 | 8982058 | rs11984397 | 0.522 | 0.124 | -0.22 | 0.375 | -0.038 | 0.036 | 0.036 | 0 | 0.41 | 0.26 | 0.52 | 0.32 | 0.49 | 0.25 |
| 7 | 8982208 | rs6959207 | -0.632 | -0.132 | -0.011 | 0.421 | 0.004 | -0.004 | -0.004 | 0.674 | 0.68 | 0.72 | 0.75 | 0.61 | 0.71 | 0.55 |
| 7 | 8982831 | rs6964000 | 0.07 | 0.132 | -0.088 | -0.106 | 0.01 | -0.026 | -0.026 | 0 | 0.05 | 0.14 | 0.05 | 0 | 0.06 | 0.03 |
| 7 | 8985391 | rs17157517 | 0.043 | 0.078 | -0.349 | -0.256 | 0.003 | 0.07 | 0.07 | 0 | 0.44 | 0.36 | 0.45 | 0.26 | 0.56 | 0.38 |
| 7 | 8985708 | rs12702805 | 0.092 | 0.089 | -0.428 | -0.254 | -0.005 | 0.084 | 0.084 | 0 | 0.44 | 0.29 | 0.44 | 0.26 | 0.56 | 0.38 |
| 7 | 8989369 | rs10277839 | 0.603 | 0.149 | -0.643 | -0.203 | -0.098 | 0.068 | 0.068 | 0 | 0.37 | 0.14 | 0.38 | 0.26 | 0.5 | 0.34 |
| 7 | 8991603 | rs10155933 | -0.569 | -0.704 | 0.058 | 0.12 | -0.015 | 0.049 | 0.049 | 0 | 0.68 | 0.57 | 0.68 | 0.5 | 0.71 | 0.68 |
| 7 | 8992884 | rs10250332 | 0.538 | -0.24 | -0.468 | -0.207 | -0.052 | 0.031 | 0.031 | 0 | 0.34 | 0.19 | 0.28 | 0.29 | 0.41 | 0.25 |
| 7 | 8995047 | rs17414932 | -0.161 | -0.251 | 0.022 | -0.093 | 0.005 | -0.004 | -0.004 | 0 | 0.6 | 0.54 | 0.58 | 0.61 | 0.52 | 0.69 |
| 7 | 8995694 | rs2192597 | 0.009 | -0.203 | -0.007 | -0.006 | 0.106 | 0.151 | 0.151 | 0 | 0.94 | 0.66 | 0.86 | 0.89 | 0.93 | 0.95 |
| 7 | 8997725 | rs4725148 | 0.436 | -0.41 | -0.374 | -0.085 | -0.024 | 0.014 | 0.014 | 0 | 0.38 | 0.41 | 0.21 | 0.44 | 0.31 | 0.4 |
| 7 | 9001625 | rs2024376 | 0.286 | -0.381 | -0.167 | -0.078 | -0.001 | -0.006 | -0.006 | 0 | 0.71 | 0.54 | 0.56 | 0.76 | 0.72 | 0.65 |
| 7 | 9003918 | rs12702819 | 0.472 | 0.033 | -1.535 | -0.318 | -0.197 | 0.022 | 0.022 | 0 | 0.32 | 0.03 | 0.28 | 0.23 | 0.45 | 0.16 |
| 7 | 9006608 | rs10486277 | 0.512 | 0.188 | -1.18 | -0.297 | 0 | -0.007 | -0.007 | 0.087 | 0.07 | 0.06 | 0.03 | 0.01 | 0.05 | 0.18 |
| 7 | 9006896 | rs6961566 | -0.448 | -0.737 | -0.799 | -0.466 | -0.001 | 0.018 | 0.018 | 0 | 0.7 | 0.68 | 0.78 | 0.47 | 0.7 | 0.61 |
| 7 | 9009137 | rs10234242 | 0.021 | -0.746 | -0.231 | -0.553 | -0.02 | -0.027 | -0.027 | 0.78 | 0.07 | 0.26 | 0.04 | 0 | 0 | 0.04 |
| 7 | 9010195 | rs6976963 | -0.949 | -0.78 | -0.232 | -0.581 | -0.048 | -0.088 | -0.088 | 0.098 | 0.14 | 0.34 | 0.07 | 0.01 | 0.06 | 0.23 |
| 7 | 9015025 | rs10237048 | -0.22 | -0.725 | -0.316 | -0.553 | -0.018 | -0.035 | -0.035 | 1.806 | 0.08 | 0.34 | 0.05 | 0 | 0.01 | 0.05 |
| 7 | 9015608 | rs4725150 | 0.124 | -0.416 | -0.397 | -0.31 | -0.004 | 0.024 | 0.024 | 0 | 0.64 | 0.58 | 0.75 | 0.47 | 0.65 | 0.42 |
| 7 | 9015651 | rs16875030 | 0.027 | -0.17 | -0.306 | 0.006 | -0.008 | -0.025 | -0.025 | 0 | 0.07 | 0.11 | 0.03 | 0.01 | 0.05 | 0.19 |
| 7 | 9016366 | rs11768503 | -0.446 | -0.112 | -0.584 | 0.101 | 0.001 | -0.007 | -0.007 | 0 | 0.14 | 0.11 | 0.4 | 0.24 | 0.17 | 0.19 |
| 7 | 9018291 | rs10249915 | 0.596 | 0.134 | -0.705 | -0.147 | -0.009 | -0.012 | -0.012 | 0 | 0.62 | 0.56 | 0.47 | 0.71 | 0.65 | 0.51 |
| 7 | 9019492 | rs6949865 | 0.932 | 1.256 | -0.851 | -0.406 | -0.015 | 0.031 | 0.031 | 0 | 0.79 | 0.85 | 0.68 | 0.95 | 0.75 | 0.75 |
| 7 | 9019746 | rs4720814 | 1.04 | 1.266 | -0.025 | -0.017 | 0.028 | 0.025 | 0.025 | 0.584 | 0.82 | 0.51 | 0.68 | 0.78 | 0.81 | 0.8 |
| 7 | 9019826 | rs4720815 | 0.376 | 1.272 | -0.574 | -0.065 | -0.019 | 0.023 | 0.023 | 0 | 0.9 | 0.98 | 0.82 | 0.89 | 0.88 | 0.86 |
| 7 | 9021418 | rs887937 | 1.007 | 1.27 | -0.571 | 0.091 | 0 | 0.014 | 0.014 | 0 | 0.76 | 0.84 | 0.73 | 0.8 | 0.77 | 0.77 |
| 7 | 9021912 | rs9986680 | 0.872 | 1.395 | -0.628 | 0.484 | 0.035 | 0.032 | 0.032 | 0 | 0.42 | 0.24 | 0.24 | 0.28 | 0.28 | 0.3 |
| 7 | 9022420 | rs887938 | 0.446 | 1.715 | -0.789 | 0.257 | 0.004 | 0.04 | 0.04 | 0 | 0.47 | 0.34 | 0.54 | 0.37 | 0.42 | 0.38 |
| 7 | 9022706 | rs10486281 | 0.367 | 1.808 | -0.955 | 0.343 | -0.015 | -0.014 | -0.014 | 3.38 | 0.17 | 0.12 | 0.4 | 0.34 | 0.29 | 0.24 |
| 7 | 9024381 | rs6962574 | 1.229 | 0.524 | -0.948 | -0.301 | 0.058 | 0.07 | 0.07 | 0.025 | 0.3 | 0.12 | 0.13 | 0.03 | 0.13 | 0.14 |
| 7 | 9024438 | rs10486282 | -0.575 | 1.689 | -0.546 | 0.566 | -0.082 | -0.067 | -0.067 | 0 | 0.29 | 0.1 | 0.39 | 0.59 | 0.42 | 0.37 |
| 7 | 9032295 | rs10255740 | -0.692 | 1.321 | -0.029 | 0.274 | 0.026 | -0.011 | -0.011 | 0 | 0.37 | 0.5 | 0.6 | 0.6 | 0.49 | 0.43 |
| 7 | 9035815 | rs10236407 | -1.4 | 0.647 | 1.006 | 0.364 | 0.019 | 0.15 | 0.15 | 0.367 | 0.09 | 0.44 | 0.11 | 0.17 | 0.03 | 0.11 |
| 7 | 9039807 | rs4720816 | -1.486 | 0.123 | 0.815 | 0.032 | -0.057 | 0.014 | 0.014 | 0 | 0.16 | 0.48 | 0.13 | 0.17 | 0.06 | 0.1 |
| 7 | 9041670 | rs17158323 | -0.264 | 0.762 | 0.856 | -0.015 | 0.007 | -0.002 | -0.002 | 0 | 0.14 | 0.21 | 0.19 | 0.15 | 0.25 | 0.26 |
| 7 | 9042560 | rs17158339 | -0.924 | 0.577 | 1.002 | 0.068 | 0.022 | 0.147 | 0.147 | 0.232 | 0.08 | 0.41 | 0.1 | 0.17 | 0.03 | 0.08 |
| 7 | 9047500 | rs12702828 | -1.202 | 0.589 | -0.028 | -0.417 | -0.038 | 0.044 | 0.044 | 0.039 | 0.41 | 0.87 | 0.45 | 0.38 | 0.48 | 0.41 |
| 7 | 9048143 | rs11983401 | -0.364 | 0.563 | 0.312 | -0.318 | -0.003 | 0.003 | 0.003 | 0 | 0.23 | 0.19 | 0.25 | 0.23 | 0.25 | 0.18 |
| 7 | 9049793 | rs13241564 | -0.63 | 0.561 | 0.442 | -0.337 | 0.001 | -0.003 | -0.003 | 0 | 0.24 | 0.27 | 0.25 | 0.23 | 0.25 | 0.19 |
| 7 | 9052374 | rs13235007 | 0.335 | 0.715 | -0.097 | -0.501 | -0.032 | 0.062 | 0.062 | 0 | 0.17 | 0.05 | 0.16 | 0.05 | 0.22 | 0.11 |
| 7 | 9058629 | rs10232114 | -0.645 | 0.395 | 0.011 | -0.458 | 0.001 | 0.005 | 0.005 | 0 | 0.42 | 0.54 | 0.33 | 0.19 | 0.4 | 0.33 |
| 7 | 9061607 | rs12533648 | 0.302 | 0.589 | 0.218 | -0.598 | 0.003 | -0.004 | -0.004 | 0 | 0.23 | 0.2 | 0.15 | 0.05 | 0.21 | 0.14 |
| 7 | 9062629 | rs12702834 | -0.705 | 1.014 | 0.353 | 0.246 | 0.003 | 0.004 | 0.004 | 0 | 0.34 | 0.37 | 0.46 | 0.4 | 0.38 | 0.3 |
| 7 | 9062830 | rs10259251 | 0.635 | 0.998 | 0.146 | 0.401 | 0.002 | 0.011 | 0.011 | 0.057 | 0.06 | 0.06 | 0.17 | 0.16 | 0.09 | 0.13 |
| 7 | 9064295 | rs7792941 | 0.202 | 0.509 | 0.276 | 0.229 | -0.012 | -0.006 | -0.006 | 0.087 | 0.19 | 0.24 | 0.14 | 0.14 | 0.15 | 0.19 |
| 7 | 9066820 | rs10952132 | -0.012 | 0.269 | 0.013 | 0.183 | -0.01 | -0.016 | -0.016 | 0 | 0.28 | 0.3 | 0.19 | 0.13 | 0.2 | 0.22 |
| 7 | 9067203 | rs17158465 | -0.004 | 0.512 | -0.279 | 0.401 | -0.002 | -0.002 | -0.002 | 0 | 0.08 | 0.12 | 0.18 | 0.15 | 0.11 | 0.13 |
| 7 | 9068028 | rs17158491 | -0.342 | 0.517 | -0.212 | 0.429 | 0.002 | 0.007 | 0.007 | 0 | 0.09 | 0.14 | 0.24 | 0.17 | 0.12 | 0.16 |
| 7 | 9068698 | rs17158514 | 1.505 | 0.969 | -0.527 | 0.168 | -0.008 | -0.08 | -0.08 | 0.13 | 0.27 | 0.08 | 0.26 | 0.54 | 0.24 | 0.33 |
| 7 | 9072210 | rs17416081 | 0.926 | 1.339 | -0.636 | -0.075 | -0.054 | 0.011 | 0.011 | 0 | 0.09 | 0.01 | 0.15 | 0.05 | 0.2 | 0.16 |
| 7 | 9072385 | rs1030008 | 0.526 | 0.64 | -0.114 | -0.155 | -0.014 | -0.068 | -0.068 | 0 | 0.27 | 0.34 | 0.23 | 0.09 | 0.24 | 0.18 |
| 7 | 9073500 | rs10272373 | 0.714 | 0.674 | -0.137 | -0.139 | -0.009 | -0.054 | -0.054 | 0 | 0.25 | 0.31 | 0.22 | 0.09 | 0.23 | 0.18 |
| 7 | 9078497 | rs10952134 | 0.183 | 0.359 | -0.021 | -0.067 | 0.03 | -0.021 | -0.021 | 1.352 | 0.76 | 0.59 | 0.63 | 0.81 | 0.65 | 0.72 |
| 7 | 9079397 | rs12702842 | 1.08 | 0.367 | -0.439 | -0.251 | -0.077 | 0.092 | 0.092 | 0 | 0.23 | 0.01 | 0.21 | 0.07 | 0.27 | 0.17 |
| 7 | 9081052 | rs11983119 | 0.013 | 0.299 | -0.179 | 0.616 | -0.005 | -0.005 | -0.005 | 0 | 0.25 | 0.27 | 0.18 | 0.16 | 0.2 | 0.16 |
| 7 | 9084859 | rs726369 | -0.319 | 0.248 | -0.02 | 0.653 | 0.05 | 0.292 | 0.292 | 1.002 | 0.06 | 0.39 | 0.08 | 0.24 | 0.03 | 0.06 |
| 7 | 9086743 | rs11561890 | 1.088 | 0.379 | -1.262 | 0.278 | -0.052 | 0.008 | 0.008 | 0.021 | 0.5 | 0.06 | 0.4 | 0.46 | 0.46 | 0.56 |
| 7 | 9087523 | rs6463871 | -0.413 | -0.129 | -0.686 | 0.068 | -0.005 | -0.009 | -0.009 | 0.305 | 0.24 | 0.24 | 0.16 | 0.13 | 0.18 | 0.17 |
| 7 | 9097809 | rs2107749 | 0.259 | -0.212 | -0.99 | -0.043 | 0.009 | 0.028 | 0.028 | 0 | 0.8 | 0.83 | 0.88 | 0.92 | 0.86 | 0.79 |
| 7 | 9097947 | rs13242973 | 1.369 | -0.435 | -1.364 | 0.184 | 0.025 | -0.007 | -0.007 | 0 | 0.55 | 0.2 | 0.65 | 0.54 | 0.64 | 0.65 |
| 7 | 9101172 | rs4725158 | -0.992 | -0.278 | -1.358 | 0.237 | -0.003 | 0.004 | 0.004 | 0 | 0.94 | 0.86 | 0.94 | 0.92 | 0.96 | 0.92 |
| 7 | 9102112 | rs1476620 | 0.549 | -0.449 | -0.882 | 0.702 | -0.009 | 0.012 | 0.012 | 0 | 0.68 | 0.4 | 0.7 | 0.55 | 0.75 | 0.66 |
| 7 | 9102563 | rs17158852 | 1.901 | -0.34 | -2.295 | 0.61 | -0.122 | -0.042 | -0.042 | 0 | 0.37 | 0.03 | 0.4 | 0.31 | 0.41 | 0.42 |
| 7 | 9104184 | rs2214675 | -0.854 | 0.725 | -1.89 | -0.104 | -0.015 | 0.001 | 0.001 | 0 | 0.91 | 0.95 | 0.86 | 0.93 | 0.87 | 0.87 |
| 7 | 9104637 | rs12539486 | 0.508 | 0.667 | -1.567 | -0.099 | -0.001 | 0.007 | 0.007 | 0 | 0.09 | 0.07 | 0.1 | 0.17 | 0.1 | 0.1 |
| 7 | 9105028 | rs13227829 | 0.486 | 0.7 | -1.654 | -0.156 | -0.041 | -0.016 | -0.016 | 0 | 0.09 | 0.02 | 0.12 | 0.1 | 0.18 | 0.05 |
| 7 | 9108862 | rs11765761 | 0.289 | 0.57 | -1.64 | -0.168 | -0.054 | 0.002 | 0.002 | 0 | 0.13 | 0.02 | 0.15 | 0.1 | 0.22 | 0.05 |
| 7 | 9108917 | rs10247883 | -0.674 | 0.415 | -1.579 | -0.235 | 0.001 | 0 | 0 | 0 | 0.3 | 0.28 | 0.23 | 0.26 | 0.24 | 0.36 |
| 7 | 9110978 | rs12702844 | -1.493 | 0.927 | -1.526 | 0.268 | -0.022 | -0.009 | -0.009 | 0 | 0.23 | 0.12 | 0.25 | 0.26 | 0.32 | 0.17 |

| | | | | | | | | | | | | | | | | |
|---|---|---|---|---|---|---|---|---|---|---|---|---|---|---|---|---|
| 7 | 9119996 | rs12702864 | -0.601 | 1.245 | -1.494 | -0.007 | -0.038 | 0.011 | 0.011 | 0 | 0.13 | 0.05 | 0.15 | 0.1 | 0.22 | 0.05 |
| 7 | 9122800 | rs7810082 | 0.566 | 0.569 | -1.218 | -0.276 | -0.001 | -0.014 | -0.014 | 0.263 | 0.7 | 0.66 | 0.78 | 0.89 | 0.77 | 0.87 |
| 7 | 9125322 | rs10255611 | -0.654 | 0.197 | -1.661 | 0.088 | 0.01 | -0.031 | -0.031 | 0 | 0.15 | 0.08 | 0.09 | 0.14 | 0.1 | 0.15 |
| 7 | 9125379 | rs6947677 | -0.031 | 0.157 | -2.425 | -0.401 | -0.02 | -0.009 | -0.009 | 0 | 0.55 | 0.79 | 0.53 | 0.57 | 0.54 | 0.62 |
| 7 | 9126305 | rs2158573 | 0.117 | 0.044 | -1.577 | -0.707 | -0.001 | -0.004 | -0.004 | 0 | 0.56 | 0.54 | 0.54 | 0.63 | 0.58 | 0.64 |
| 7 | 9127173 | rs7791718 | -0.608 | -0.014 | -1.74 | -0.838 | -0.003 | -0.003 | -0.003 | 0.015 | 0.7 | 0.59 | 0.6 | 0.69 | 0.71 | 0.74 |
| 7 | 9128070 | rs6963727 | -1.097 | -0.029 | -1.639 | -0.845 | -0.011 | -0.012 | -0.012 | 0 | 0.75 | 0.81 | 0.61 | 0.69 | 0.71 | 0.74 |
| 7 | 9128241 | rs6463875 | 0.096 | 0.177 | -1.965 | -1.023 | 0.001 | -0.004 | -0.004 | 0 | 0.59 | 0.54 | 0.46 | 0.63 | 0.58 | 0.66 |
| 7 | 9128802 | rs10227982 | 0.195 | 0.119 | -2.227 | -0.977 | -0.004 | 0.031 | 0.031 | 0 | 0.16 | 0.06 | 0.06 | 0.05 | 0.14 | 0.12 |
| 7 | 9131069 | rs2107753 | -0.927 | 0.079 | -2.338 | -0.993 | -0.033 | -0.022 | -0.022 | 2.656 | 0.71 | 0.87 | 0.61 | 0.68 | 0.69 | 0.73 |
| 7 | 9135219 | rs12154452 | -0.616 | -0.135 | -2.207 | -1.067 | -0.002 | 0.012 | 0.012 | 0 | 0.18 | 0.18 | 0.16 | 0.1 | 0.15 | 0.18 |
| 7 | 9136704 | rs2286110 | -0.775 | -0.268 | -2.455 | -1.155 | -0.065 | 0.073 | 0.073 | 0 | 0.25 | 0.04 | 0.19 | 0.11 | 0.25 | 0.17 |
| 7 | 9136820 | rs2286111 | -1.13 | -0.859 | -2.699 | -1.829 | -0.082 | 0.248 | 0.248 | 0 | 0.54 | 0.92 | 0.39 | 0.16 | 0.47 | 0.33 |
| 7 | 9137229 | rs6945878 | 1.88 | -0.828 | -1.853 | -1.697 | 0.038 | -0.038 | -0.038 | 0 | 0.11 | 0.26 | 0.12 | 0.03 | 0.15 | 0.05 |
| 7 | 9138344 | rs12702865 | 1.88 | -0.838 | -1.882 | -1.701 | 0.031 | -0.063 | -0.063 | 1.911 | 0.11 | 0.24 | 0.12 | 0.01 | 0.15 | 0.05 |
| 7 | 9138376 | rs11514814 | 0.721 | -0.851 | -2.001 | -1.75 | -0.023 | 0.044 | 0.044 | 0 | 0.13 | 0.05 | 0.14 | 0.05 | 0.13 | 0.09 |
| 7 | 9138892 | rs6956172 | -0.605 | -1.08 | -1.68 | -1.846 | 0 | -0.021 | -0.021 | 0 | 0.32 | 0.65 | 0.28 | 0.09 | 0.28 | 0.16 |
| 7 | 9144240 | rs12538490 | 0.286 | -1.034 | -1.602 | -1.776 | -0.02 | -0.079 | -0.079 | 0.765 | 0.19 | 0.62 | 0.15 | 0.03 | 0.14 | 0.07 |
| 7 | 9144405 | rs2189869 | -1.454 | -1.068 | -2.179 | -1.832 | -0.125 | -0.054 | -0.054 | 0.016 | 0.72 | 0.93 | 0.49 | 0.64 | 0.58 | 0.65 |
| 7 | 9144719 | rs17159419 | 1.138 | -0.943 | -1.432 | -2.097 | -0.008 | -0.012 | -0.012 | 0 | 0.08 | 0.37 | 0.02 | 0 | 0 | 0.02 |
| 7 | 9146717 | rs11978878 | -1.452 | -0.56 | -1.478 | -1.981 | 0 | -0.025 | -0.025 | 0 | 0.55 | 0.55 | 0.4 | 0.19 | 0.51 | 0.34 |
| 7 | 9147398 | rs10247350 | -0.63 | -0.674 | -1.049 | -1.594 | 0.003 | 0.018 | 0.018 | 0 | 0.24 | 0.17 | 0.13 | 0.1 | 0.21 | 0.17 |
| 7 | 9150520 | rs293162 | -0.563 | -0.591 | -1.197 | -1.333 | -0.006 | -0.034 | -0.034 | 0 | 0.32 | 0.36 | 0.18 | 0.08 | 0.24 | 0.13 |
| 7 | 9154242 | rs10273445 | -0.049 | -0.185 | -1.007 | -0.649 | 0.003 | -0.011 | -0.011 | 0 | 0.07 | 0.11 | 0.05 | 0.04 | 0.1 | 0.08 |
| 7 | 9154302 | rs172399 | -1.321 | -0.459 | -1.32 | -0.611 | 0.025 | 0.042 | 0.042 | 0 | 0.35 | 0.73 | 0.16 | 0.2 | 0.24 | 0.22 |
| 7 | 9154681 | rs6961225 | -0.883 | -0.154 | -1.108 | -0.539 | -0.003 | -0.007 | -0.007 | 0.014 | 0.07 | 0.08 | 0.05 | 0 | 0.06 | 0.05 |
| 7 | 9157560 | rs293169 | -1.467 | -0.969 | -1.315 | -0.259 | 0.063 | 0.019 | 0.019 | 0 | 0.31 | 0.79 | 0.14 | 0.27 | 0.14 | 0.21 |
| 7 | 9158650 | rs293173 | 0.251 | -0.854 | -0.969 | -0.106 | -0.008 | 0.071 | 0.071 | 0 | 0.09 | 0.16 | 0.06 | 0.19 | 0.06 | 0.04 |
| 7 | 9159119 | rs293177 | -1.642 | -0.57 | -1.772 | -0.24 | -0.134 | -0.12 | -0.12 | 0 | 0.61 | 0.98 | 0.54 | 0.47 | 0.51 | 0.5 |
| 7 | 9161198 | rs293184 | 0.295 | -0.764 | -0.762 | -0.103 | 0.004 | 0.101 | 0.101 | 0 | 0.09 | 0.21 | 0.06 | 0.19 | 0.06 | 0.05 |
| 7 | 9161403 | rs293185 | -0.993 | -0.791 | -0.751 | -0.158 | -0.04 | 0.014 | 0.014 | 0.154 | 0.22 | 0.35 | 0.1 | 0.27 | 0.13 | 0.2 |
| 7 | 9167743 | rs2107755 | 0.407 | -0.28 | -1.178 | -0.192 | -0.029 | -0.016 | -0.016 | 0 | 0.07 | 0.01 | 0.1 | 0.06 | 0.09 | 0.08 |
| 7 | 9168502 | rs293192 | -1.122 | -0.785 | -0.83 | -0.223 | -0.007 | 0.024 | 0.024 | 0.204 | 0.3 | 0.43 | 0.27 | 0.31 | 0.26 | 0.23 |
| 7 | 9169798 | rs188374 | -1.68 | -0.656 | -1.662 | -0.523 | -0.127 | -0.065 | -0.065 | 0.124 | 0.55 | 0.98 | 0.42 | 0.36 | 0.43 | 0.4 |
| 7 | 9170998 | rs293195 | -1.341 | -0.918 | -1.487 | -0.487 | 0.018 | 0.006 | 0.006 | 0 | 0.45 | 0.93 | 0.32 | 0.29 | 0.31 | 0.24 |
| 7 | 9175023 | rs4585636 | -1.285 | -0.812 | -0.591 | -0.696 | -0.024 | -0.041 | -0.041 | 0 | 0.15 | 0.42 | 0.18 | 0.07 | 0.11 | 0.12 |
| 7 | 9176026 | rs991970 | 0.028 | -1.233 | -1.326 | -0.523 | 0.025 | -0.049 | -0.049 | 0 | 0.22 | 0.11 | 0.11 | 0.26 | 0.15 | 0.1 |
| 7 | 9189486 | rs1371736 | -0.7 | -1.204 | -1.137 | -0.491 | -0.015 | 0.007 | 0.007 | 0 | 0.31 | 0.35 | 0.19 | 0.34 | 0.19 | 0.19 |
| 7 | 9189617 | rs293134 | -2.199 | -0.972 | -1.548 | -0.765 | -0.15 | -0.131 | -0.131 | 0 | 0.6 | 0.98 | 0.55 | 0.43 | 0.54 | 0.42 |
| 7 | 9193124 | rs1440253 | -1.419 | -1.359 | -0.858 | -0.424 | 0.004 | 0 | 0 | 0 | 0.39 | 0.36 | 0.36 | 0.34 | 0.27 | 0.23 |
| 7 | 9193295 | rs10245058 | -0.461 | -1.27 | -1.136 | -0.376 | -0.004 | 0.001 | 0.001 | 0 | 0.24 | 0.27 | 0.12 | 0.27 | 0.15 | 0.1 |
| 7 | 9197108 | rs7791753 | -1.022 | -0.788 | -1.619 | -0.102 | 0.073 | 0.02 | 0.02 | 1.122 | 0.44 | 0.86 | 0.36 | 0.34 | 0.28 | 0.23 |
| 7 | 9197737 | rs10243151 | -1.028 | -0.74 | -1.614 | -1.035 | 0.07 | 0.237 | 0.237 | 1.019 | 0.44 | 0.86 | 0.4 | 0.09 | 0.28 | 0.23 |
| 7 | 9199121 | rs6969212 | 0.558 | -0.686 | -0.721 | -0.71 | -0.032 | -0.062 | -0.062 | 0 | 0.72 | 0.56 | 0.83 | 0.9 | 0.81 | 0.87 |
| 7 | 9201869 | rs2165272 | -0.05 | 0.775 | -0.971 | -0.325 | 0.015 | 0.046 | 0.046 | 0.232 | 0.09 | 0.15 | 0.18 | 0.21 | 0.11 | 0.11 |
| 7 | 9214074 | rs9640103 | 0.689 | 0.708 | -1.134 | -0.233 | 0.035 | -0.036 | -0.036 | 0 | 0.36 | 0.24 | 0.32 | 0.51 | 0.24 | 0.43 |
| 7 | 9214578 | rs2107748 | 0.759 | 0.719 | -1.131 | -0.231 | 0.015 | 0.044 | 0.044 | 0.339 | 0.09 | 0.16 | 0.21 | 0.2 | 0.12 | 0.11 |
| 7 | 9215329 | rs2057847 | -0.057 | 0.712 | -1.116 | -0.265 | 0.007 | 0.01 | 0.01 | 0 | 0.46 | 0.4 | 0.53 | 0.71 | 0.36 | 0.54 |
| 7 | 9216593 | rs395158 | -0.921 | 0.673 | -0.517 | -0.439 | 0.02 | -0.077 | -0.077 | 0 | 0.84 | 0.76 | 0.81 | 1 | 0.82 | 0.9 |
| 7 | 9216829 | rs17160189 | 0.214 | 0.228 | -0.828 | -0.354 | 0.048 | -0.027 | -0.027 | 0 | 0.4 | 0.23 | 0.32 | 0.52 | 0.25 | 0.43 |
| 7 | 9218021 | rs293136 | -0.039 | 0.546 | -0.47 | 0.454 | -0.006 | 0.031 | 0.031 | 0.155 | 0.36 | 0.25 | 0.24 | 0.2 | 0.4 | 0.3 |
| 7 | 9223595 | rs7796895 | -0.053 | -0.028 | -0.501 | 0.485 | -0.001 | 0.008 | 0.008 | 0 | 0.32 | 0.26 | 0.2 | 0.29 | 0.34 | 0.21 |
| 7 | 9224481 | rs17160259 | -0.009 | -0.375 | -0.027 | 0.442 | -0.012 | -0.007 | -0.007 | 0 | 0.09 | 0.13 | 0.04 | 0.03 | 0.06 | 0.08 |
| 7 | 9229882 | rs17160303 | -1.527 | -0.766 | 0.58 | -0.091 | -0.037 | -0.118 | -0.118 | 0 | 0.24 | 0.42 | 0.12 | 0.03 | 0.14 | 0.07 |
| 7 | 9231945 | rs17792080 | 0.17 | -0.689 | 0.66 | -0.086 | -0.002 | -0.008 | -0.008 | 0.998 | 0.08 | 0.06 | 0.05 | 0 | 0.07 | 0.01 |
| 7 | 9233910 | rs1285453 | -0.386 | -0.944 | 0.279 | -0.011 | -0.004 | 0.012 | 0.012 | 0.022 | 0.55 | 0.64 | 0.7 | 0.74 | 0.53 | 0.6 |
| 7 | 9233987 | rs1285454 | 0.33 | -0.998 | 0.356 | -0.025 | 0 | 0.006 | 0.006 | 1.308 | 0.43 | 0.49 | 0.65 | 0.71 | 0.46 | 0.54 |
| 7 | 9234364 | rs17203026 | 0.61 | -0.864 | 0.34 | -0.048 | 0.008 | -0.043 | -0.043 | 0 | 0.09 | 0.13 | 0.06 | 0 | 0.07 | 0.01 |
| 7 | 9235662 | rs1529234 | -1.208 | -1.057 | 0.445 | 0.474 | -0.008 | 0.034 | 0.034 | 0.404 | 0.3 | 0.52 | 0.42 | 0.46 | 0.27 | 0.29 |
| 7 | 9244093 | rs1440238 | -1.552 | -1.029 | 0.544 | 0.506 | -0.073 | -0.021 | -0.021 | 0.102 | 0.66 | 0.92 | 0.62 | 0.7 | 0.59 | 0.6 |
| 7 | 9247266 | rs4720829 | 0.026 | -1.244 | 0.595 | 0.578 | -0.01 | 0.006 | 0.006 | 0 | 0.45 | 0.19 | 0.47 | 0.62 | 0.47 | 0.49 |
| 7 | 9248557 | rs1285376 | -1.587 | -1.268 | 0.779 | 0.589 | -0.006 | 0.004 | 0.004 | 0.006 | 0.65 | 0.67 | 0.6 | 0.71 | 0.59 | 0.6 |
| 7 | 9248649 | rs1285377 | -0.682 | -1.317 | 0.804 | 0.556 | -0.003 | -0.001 | -0.001 | 0.087 | 0.18 | 0.19 | 0.08 | 0.12 | 0.1 | 0.15 |
| 7 | 9251214 | rs1285432 | -0.596 | -1.301 | 0.794 | 0.453 | 0.006 | 0.021 | 0.021 | 1.134 | 0.49 | 0.54 | 0.33 | 0.33 | 0.42 | 0.37 |
| 7 | 9253413 | rs2141531 | 1.255 | -1.101 | 0.476 | 0.472 | -0.074 | -0.019 | -0.019 | 0.008 | 0.28 | 0.02 | 0.18 | 0.25 | 0.3 | 0.24 |
| 7 | 9255592 | rs4329191 | 0.683 | -0.977 | 0.699 | 0.62 | 0.032 | -0.059 | -0.059 | 0.458 | 0.1 | 0.27 | 0.07 | 0 | 0.07 | 0.01 |
| 7 | 9256611 | rs17160634 | 0.604 | -0.853 | 0.428 | 0.727 | -0.001 | 0.013 | 0.013 | 0 | 0.17 | 0.17 | 0.28 | 0.38 | 0.17 | 0.25 |
| 7 | 9257996 | rs1285415 | -1.111 | -0.419 | -1.186 | 0.635 | -0.048 | 0.006 | 0.006 | 0.385 | 0.66 | 0.88 | 0.62 | 0.71 | 0.59 | 0.59 |
| 7 | 9266388 | rs1285407 | -0.098 | -0.13 | -0.89 | 0.514 | -0.066 | 0.022 | 0.022 | 0.01 | 0.32 | 0.08 | 0.23 | 0.24 | 0.36 | 0.31 |

| | | | | | | | | | | | | | | | | |
|---|---|---|---|---|---|---|---|---|---|---|---|---|---|---|---|---|
| 7 | 9267929 | rs1285405 | -0.15 | -0.141 | -0.856 | 0.523 | -0.048 | 0.034 | 0.034 | 0 | 0.32 | 0.12 | 0.24 | 0.25 | 0.36 | 0.31 |
| 7 | 9283620 | rs7777070 | 1.131 | -0.705 | -0.506 | 1.152 | 0.032 | -0.025 | -0.025 | 0 | 0.63 | 0.19 | 0.47 | 0.44 | 0.7 | 0.78 |
| 7 | 9283806 | rs1155197 | -1.004 | -0.974 | 0.509 | 1.044 | -0.052 | 0.079 | 0.079 | 0 | 0.22 | 0.39 | 0.41 | 0.54 | 0.15 | 0.18 |
| 7 | 9284038 | rs2119223 | -0.929 | -1.038 | 0.566 | 1.658 | -0.05 | 0.08 | 0.08 | 0 | 0.19 | 0.34 | 0.28 | 0.34 | 0.13 | 0.15 |
| 7 | 9285814 | rs2119217 | -1.246 | -1.103 | 0.51 | 1.875 | -0.04 | 0.228 | 0.228 | 0.304 | 0.14 | 0.38 | 0.24 | 0.46 | 0.07 | 0.07 |
| 7 | 9285968 | rs2165270 | -0.103 | -1.026 | 1.133 | 1.769 | 0.042 | -0.023 | -0.023 | 0.063 | 0.93 | 0.77 | 0.95 | 0.99 | 0.93 | 0.97 |
| 7 | 9287376 | rs1529242 | -0.942 | -1.055 | 1.143 | 1.755 | 0 | -0.004 | -0.004 | 0.142 | 0.13 | 0.15 | 0.23 | 0.08 | 0.15 | 0.12 |
| 7 | 9298621 | rs2699120 | -0.748 | -1.026 | 1.106 | 1.799 | -0.034 | 0.143 | 0.143 | 0.041 | 0.12 | 0.33 | 0.11 | 0.27 | 0.06 | 0.05 |
| 7 | 9301471 | rs1285396 | 1.336 | -1.066 | 1.109 | 1.68 | 0.046 | 0.023 | 0.023 | 0 | 0.82 | 0.55 | 0.75 | 0.79 | 0.82 | 0.88 |
| 7 | 9301716 | rs12702875 | 2.082 | -1.086 | 0.514 | 1.456 | 0.017 | -0.068 | -0.068 | 0 | 0.68 | 0.16 | 0.64 | 0.52 | 0.76 | 0.83 |
| 7 | 9323220 | rs1919796 | -1.406 | -1.096 | 0.847 | 1.588 | -0.015 | 0.142 | 0.142 | 0.876 | 0.22 | 0.57 | 0.3 | 0.46 | 0.17 | 0.14 |
| 7 | 9325970 | rs6963754 | 1.672 | -1.131 | 0.411 | 1.581 | 0.01 | -0.04 | -0.04 | 0 | 0.69 | 0.19 | 0.65 | 0.54 | 0.77 | 0.84 |
| 7 | 9326836 | rs4613895 | 0.324 | -1.067 | 0.152 | 1.495 | 0.008 | -0.01 | -0.01 | 0 | 0.05 | 0.03 | 0.16 | 0.07 | 0.07 | 0.09 |
| 7 | 9334068 | rs9648119 | 1.882 | -1.266 | -0.003 | 1.541 | 0.037 | -0.043 | -0.043 | 0 | 0.62 | 0.11 | 0.47 | 0.44 | 0.7 | 0.72 |
| 7 | 9334567 | rs10228006 | 0.174 | -1.049 | -0.448 | 0.78 | 0.001 | 0.057 | 0.057 | 0.463 | 0.46 | 0.37 | 0.41 | 0.32 | 0.5 | 0.59 |
| 7 | 9340431 | rs1569245 | -0.426 | -0.81 | -0.205 | 0.996 | 0.001 | 0.008 | 0.008 | 0 | 0.52 | 0.55 | 0.62 | 0.7 | 0.44 | 0.46 |
| 7 | 9340931 | rs1358657 | 1.243 | -0.997 | -0.373 | 0.781 | -0.013 | 0.053 | 0.053 | 0 | 0.34 | 0.24 | 0.21 | 0.11 | 0.4 | 0.3 |
| 7 | 9341294 | rs10253855 | 0.312 | -1.107 | -0.238 | 0.802 | 0 | 0.003 | 0.003 | 0 | 0.43 | 0.41 | 0.51 | 0.61 | 0.39 | 0.37 |
| 7 | 9350385 | rs10272334 | 1.302 | -0.862 | -0.808 | 1.076 | -0.015 | 0.007 | 0.007 | 0 | 0.76 | 0.65 | 0.73 | 0.72 | 0.79 | 0.67 |
| 7 | 9351067 | rs9918506 | 0.468 | -0.749 | -0.743 | 0.981 | 0.002 | -0.035 | -0.035 | 0 | 0.87 | 0.77 | 0.83 | 0.95 | 0.86 | 0.89 |
| 7 | 9352249 | rs1919791 | 1.25 | -1.021 | -0.729 | 1.321 | -0.041 | 0.01 | 0.01 | 0.388 | 0.26 | 0.12 | 0.21 | 0.23 | 0.3 | 0.28 |
| 7 | 9353334 | rs6961903 | 0.92 | -1.213 | -0.433 | 1.43 | 0.002 | -0.035 | -0.035 | 0 | 0.87 | 0.78 | 0.83 | 0.95 | 0.86 | 0.89 |
| 7 | 9356570 | rs17161552 | 0.184 | -1.213 | -0.46 | 1.685 | 0.001 | -0.047 | -0.047 | 0 | 0.09 | 0 | 0.14 | 0.15 | 0.04 | 0.03 |
| 7 | 9360629 | rs6971374 | 1.323 | -0.873 | -0.004 | 1.323 | -0.002 | 0.001 | 0.001 | 0 | 0.85 | 0.74 | 0.83 | 0.85 | 0.86 | 0.82 |
| 7 | 9362843 | rs10952159 | 1.953 | -0.772 | -0.126 | 1.333 | -0.008 | 0.036 | 0.036 | 0 | 0.76 | 0.65 | 0.72 | 0.63 | 0.79 | 0.6 |
| 7 | 9367328 | rs6463905 | 0.326 | -0.657 | -0.147 | 1.309 | -0.014 | -0.015 | -0.015 | 2.735 | 0.92 | 0.97 | 0.94 | 0.86 | 0.91 | 0.89 |
| 7 | 9368357 | rs3935319 | 0.105 | -0.624 | -0.137 | 1.326 | -0.015 | -0.017 | -0.017 | 0 | 0.93 | 0.97 | 0.94 | 0.86 | 0.91 | 0.89 |
| 7 | 9377055 | rs10241261 | 1.501 | -0.61 | -0.107 | 1.319 | -0.012 | 0.033 | 0.033 | 0 | 0.76 | 0.65 | 0.71 | 0.62 | 0.79 | 0.59 |
| 7 | 9379478 | rs12672847 | -0.837 | -0.273 | -0.166 | 1.205 | -0.003 | 0.04 | 0.04 | 0 | 0.09 | 0.17 | 0.11 | 0.18 | 0.07 | 0.22 |
| 7 | 9385303 | rs4313076 | -0.103 | -0.398 | -0.543 | 0.9 | -0.025 | -0.004 | -0.004 | 0 | 0.69 | 0.55 | 0.68 | 0.71 | 0.73 | 0.78 |
| 7 | 9387898 | rs11773271 | 0.227 | 0.216 | -1.02 | 0.167 | -0.002 | -0.007 | -0.007 | 0 | 0.41 | 0.33 | 0.31 | 0.15 | 0.52 | 0.36 |
| 7 | 9388118 | rs17161591 | -0.964 | 0.492 | -1.862 | 0.811 | -0.04 | -0.151 | -0.151 | 0 | 0.25 | 0.04 | 0.42 | 0.44 | 0.26 | 0.18 |
| 7 | 9388999 | rs17161595 | 0.108 | 0.475 | -1.705 | 0.482 | -0.006 | -0.019 | -0.019 | 0 | 0.11 | 0.03 | 0.29 | 0.15 | 0.13 | 0.11 |
| 7 | 9391363 | rs12702896 | 0.741 | 0.386 | -1.434 | -0.265 | 0 | -0.006 | -0.006 | 0 | 0.72 | 0.73 | 0.65 | 0.84 | 0.73 | 0.74 |
| 7 | 9392500 | rs2350839 | 0.254 | 0.544 | -1.598 | -0.465 | -0.007 | 0.014 | 0.014 | 0 | 0.8 | 0.86 | 0.9 | 0.99 | 0.8 | 0.83 |
| 7 | 9400774 | rs11978280 | 0.529 | 0.674 | -1.935 | -0.611 | -0.001 | 0.012 | 0.012 | 0 | 0.16 | 0.07 | 0.12 | 0.12 | 0.12 | 0.28 |
| 7 | 9403589 | rs13244949 | 0.974 | 1.254 | -2.412 | -0.663 | -0.047 | -0.039 | -0.039 | 1.999 | 0.12 | 0.03 | 0.11 | 0.11 | 0.16 | 0.05 |
| 7 | 9404231 | rs10952163 | -1.407 | 1.457 | -2.453 | -0.824 | -0.014 | 0.027 | 0.027 | 0.005 | 0.86 | 0.92 | 0.94 | 0.98 | 0.86 | 0.83 |
| 7 | 9408960 | rs7780856 | 2.278 | 0.911 | -2.593 | -0.831 | -0.035 | -0.108 | -0.108 | 0.164 | 0.3 | 0.11 | 0.28 | 0.55 | 0.33 | 0.32 |
| 7 | 9412730 | rs12702898 | 0.076 | 0.486 | -1.461 | -1.017 | -0.003 | -0.023 | -0.023 | 0 | 0.67 | 0.39 | 0.6 | 0.84 | 0.71 | 0.7 |
| 7 | 9414589 | rs7787758 | 0.875 | 0.435 | -1.531 | -0.964 | 0.016 | 0.105 | 0.105 | 0 | 0.51 | 0.3 | 0.48 | 0.72 | 0.58 | 0.41 |
| 7 | 9434108 | rs2350842 | -0.734 | 0.246 | -1.028 | -0.697 | -0.004 | -0.034 | -0.034 | 0.39 | 0.24 | 0.26 | 0.19 | 0.08 | 0.2 | 0.33 |
| 7 | 9447340 | rs4509227 | 0.471 | -0.123 | -1.093 | -0.744 | 0.016 | 0.026 | 0.026 | 0 | 0.14 | 0.02 | 0.11 | 0.08 | 0.09 | 0.3 |
| 7 | 9448650 | rs10266127 | -1.721 | -0.458 | -1.183 | -0.706 | 0.076 | 0.028 | 0.028 | 0 | 0.41 | 0.78 | 0.52 | 0.32 | 0.32 | 0.47 |
| 7 | 9458495 | rs2350835 | -0.625 | -0.286 | -1.159 | -0.876 | -0.009 | -0.025 | -0.025 | 0 | 0.21 | 0.23 | 0.17 | 0.08 | 0.16 | 0.33 |
| 7 | 9474205 | rs11976821 | 0.746 | -0.005 | -1.112 | -0.749 | -0.001 | -0.011 | -0.011 | 3.73 | 0.05 | 0.05 | 0.04 | 0 | 0.06 | 0.03 |
| 7 | 9487409 | rs2713328 | -0.435 | -0.098 | -1.04 | -0.772 | 0.002 | -0.009 | -0.009 | 0 | 0.85 | 0.92 | 0.67 | 0.82 | 0.83 | 0.87 |
| 7 | 9490281 | rs16876033 | -0.34 | -0.114 | -0.357 | -0.754 | 0.021 | -0.029 | -0.029 | 0 | 0.08 | 0.27 | 0.06 | 0 | 0.06 | 0.03 |
| 7 | 9496975 | rs10235237 | 1.319 | 0.022 | -1.251 | -1.027 | 0.003 | -0.003 | -0.003 | 0.026 | 0.4 | 0.35 | 0.31 | 0.56 | 0.37 | 0.35 |
| 7 | 9514529 | rs7782202 | -0.827 | -0.727 | -0.848 | -1.095 | -0.08 | -0.052 | -0.052 | 0 | 0.23 | 0.56 | 0.16 | 0.14 | 0.11 | 0.35 |
| 7 | 9517438 | rs2251806 | 1.676 | -0.495 | -1.554 | -1.112 | 0.071 | 0.042 | 0.042 | 0.014 | 0.6 | 0.19 | 0.48 | 0.66 | 0.68 | 0.54 |
| 7 | 9518522 | rs2709097 | -0.3 | 0.108 | -1.305 | -0.928 | -0.008 | -0.014 | -0.014 | 0 | 0.86 | 0.94 | 0.67 | 0.8 | 0.82 | 0.89 |
| 7 | 9531079 | rs2713320 | 0.075 | -0.069 | -1.353 | -0.895 | 0.006 | -0.014 | -0.014 | 0 | 0.86 | 0.94 | 0.7 | 0.78 | 0.85 | 0.87 |
| 7 | 9532166 | rs2713319 | -1.229 | 0.523 | -1.306 | -1.529 | 0.017 | 0.084 | 0.084 | 0 | 0.49 | 0.68 | 0.41 | 0.2 | 0.41 | 0.42 |
| 7 | 9534063 | rs2350849 | 0.249 | 0.853 | -0.839 | -1.438 | -0.02 | 0.041 | 0.041 | 0 | 0.75 | 0.86 | 0.81 | 0.95 | 0.71 | 0.84 |
| 7 | 9538003 | rs17161702 | -0.561 | 0.736 | -0.19 | -0.644 | 0.006 | -0.038 | -0.038 | 0 | 0.08 | 0.13 | 0.06 | 0 | 0.09 | 0.07 |
| 7 | 9539279 | rs9918508 | -0.369 | 0.777 | -0.246 | -0.626 | 0.006 | -0.033 | -0.033 | 0.275 | 0.08 | 0.12 | 0.06 | 0 | 0.09 | 0.07 |
| 7 | 9541999 | rs2709064 | 0.406 | 0.736 | -0.212 | -0.766 | -0.001 | -0.012 | -0.012 | 0.183 | 0.58 | 0.41 | 0.46 | 0.84 | 0.6 | 0.52 |
| 7 | 9542315 | rs2713344 | 1.506 | 0.675 | -0.48 | -0.218 | 0.024 | -0.044 | -0.044 | 1.64 | 0.32 | 0.14 | 0.23 | 0.38 | 0.28 | 0.23 |
| 7 | 9550478 | rs12702917 | 0.702 | 0.452 | 0.147 | -0.397 | 0.015 | 0.085 | 0.085 | 0 | 0.48 | 0.31 | 0.39 | 0.75 | 0.47 | 0.43 |
| 7 | 9569435 | rs10245938 | 0.68 | 0.432 | 0.057 | -0.441 | 0.012 | 0.146 | 0.146 | 0 | 0.5 | 0.22 | 0.43 | 0.78 | 0.5 | 0.47 |
| 7 | 9571002 | rs6463923 | 0.526 | 0.398 | 0.501 | -0.674 | 0.06 | -0.121 | -0.121 | 0 | 0.73 | 0.56 | 0.63 | 0.95 | 0.65 | 0.75 |
| 7 | 9572098 | rs17161716 | 0.301 | 0.149 | 0.636 | -0.521 | -0.023 | -0.021 | -0.021 | 0 | 0.24 | 0.35 | 0.2 | 0.17 | 0.15 | 0.3 |
| 7 | 9579438 | rs2709020 | -0.672 | 0.191 | 0.546 | -0.573 | -0.005 | 0.02 | 0.02 | 0.016 | 0.42 | 0.65 | 0.54 | 0.22 | 0.44 | 0.52 |
| 7 | 9582913 | rs968682 | -0.872 | 0.246 | 0.202 | -0.372 | -0.049 | -0.096 | -0.096 | 0 | 0.8 | 0.97 | 0.79 | 0.61 | 0.73 | 0.75 |
| 7 | 9583655 | rs10267144 | 2.206 | 0.099 | 0.448 | -0.48 | 0.018 | -0.041 | -0.041 | 0 | 0.28 | 0.2 | 0.21 | 0.39 | 0.23 | 0.2 |
| 7 | 9584460 | rs2709014 | -0.085 | 0.202 | 0.587 | -0.496 | 0 | 0.005 | 0.005 | 0.033 | 0.74 | 0.74 | 0.75 | 0.56 | 0.68 | 0.67 |
| 7 | 9584964 | rs9648634 | 0.036 | 0.303 | 0.522 | -0.498 | 0.021 | 0.007 | 0.007 | 0 | 0.07 | 0.18 | 0.05 | 0.05 | 0.08 | 0.09 |
| 7 | 9586778 | rs10276221 | 0.637 | 0.242 | 0.72 | -0.547 | 0.005 | 0.002 | 0.002 | 1.683 | 0.6 | 0.47 | 0.45 | 0.56 | 0.48 | 0.54 |

| | | | | | | | | | | | | | | | | |
|---|---|---|---|---|---|---|---|---|---|---|---|---|---|---|---|---|
| 7 | 9586925 | rs2709012 | 1.334 | 0.259 | 0.726 | -0.551 | 0.007 | -0.027 | -0.027 | 0 | 0.12 | 0.12 | 0.29 | 0 | 0.21 | 0.13 |
| 7 | 9587536 | rs10216270 | 0.146 | 0.116 | 0.68 | -0.502 | -0.003 | -0.001 | -0.001 | 0.14 | 0.66 | 0.66 | 0.49 | 0.61 | 0.53 | 0.63 |
| 7 | 9591668 | rs1505335 | 1.15 | 0.148 | 0.713 | -0.401 | 0.004 | -0.003 | -0.003 | 0 | 0.37 | 0.37 | 0.25 | 0.39 | 0.3 | 0.24 |
| 7 | 9592236 | rs2350865 | 1.638 | 0.337 | 0.881 | -0.232 | -0.009 | 0.011 | 0.011 | 0.076 | 0.3 | 0.36 | 0.22 | 0.39 | 0.23 | 0.2 |
| 7 | 9593806 | rs6964683 | 1.192 | 0.422 | 0.632 | -0.212 | -0.019 | 0.023 | 0.023 | 0.548 | 0.36 | 0.55 | 0.26 | 0.44 | 0.29 | 0.29 |
| 7 | 9597813 | rs2709004 | 0.036 | 0.578 | 0.479 | -0.255 | 0.001 | 0.005 | 0.005 | 0 | 0.73 | 0.73 | 0.74 | 0.6 | 0.67 | 0.72 |
| 7 | 9599465 | rs17161756 | 1.151 | 0.429 | 0.205 | -0.378 | -0.01 | -0.021 | -0.021 | 0 | 0.21 | 0.01 | 0.19 | 0.18 | 0.17 | 0.28 |
| 7 | 9606410 | rs2709070 | -0.264 | 0.47 | 0.317 | -0.293 | -0.029 | -0.067 | -0.067 | 0 | 0.75 | 0.9 | 0.66 | 0.55 | 0.67 | 0.71 |
| 7 | 9607139 | rs6975367 | 0.108 | 0.465 | 0.696 | -0.354 | 0.045 | 0.032 | 0.032 | 0.065 | 0.07 | 0.29 | 0.04 | 0.04 | 0.05 | 0.09 |
| 7 | 9608572 | rs10225720 | 1.963 | 0.417 | 0.691 | -0.428 | 0.01 | -0.014 | -0.014 | 0.189 | 0.29 | 0.22 | 0.23 | 0.36 | 0.23 | 0.19 |
| 7 | 9614561 | rs2938841 | 0.062 | 0.506 | 0.58 | -0.461 | -0.038 | 0.11 | 0.11 | 1.065 | 0.38 | 0.17 | 0.46 | 0.18 | 0.39 | 0.43 |
| 7 | 9616980 | rs17161791 | 1.343 | 0.468 | 0.776 | -0.356 | -0.015 | 0.013 | 0.013 | 0.493 | 0.35 | 0.43 | 0.27 | 0.41 | 0.28 | 0.28 |
| 7 | 9618902 | rs9942646 | 0.592 | 0.516 | 0.142 | -0.239 | 0.01 | 0.041 | 0.041 | 0.022 | 0.27 | 0.09 | 0.22 | 0.18 | 0.23 | 0.33 |
| 7 | 9620224 | rs1566917 | 1.571 | 0.419 | 0.339 | -0.257 | -0.024 | 0.016 | 0.016 | 0.32 | 0.31 | 0.41 | 0.24 | 0.37 | 0.23 | 0.19 |
| 7 | 9622923 | rs10280615 | 1.575 | 0.511 | -0.355 | -0.216 | 0.009 | -0.01 | -0.01 | 0 | 0.3 | 0.24 | 0.23 | 0.37 | 0.23 | 0.19 |
| 7 | 9629549 | rs12666565 | 0.748 | 0.633 | -0.94 | 0.414 | -0.01 | -0.015 | -0.015 | 0 | 0.16 | 0.09 | 0.17 | 0.25 | 0.21 | 0.18 |
| 7 | 9633177 | rs2657237 | 0.265 | 0.59 | -0.722 | 0.072 | -0.004 | 0.017 | 0.017 | 0 | 0.53 | 0.78 | 0.56 | 0.4 | 0.52 | 0.43 |
| 7 | 9636392 | rs2709047 | 0.32 | 0.611 | -0.699 | 0.081 | 0.001 | 0.02 | 0.02 | 0 | 0.52 | 0.74 | 0.55 | 0.39 | 0.52 | 0.43 |
| 7 | 9642449 | rs2709050 | -0.986 | 0.62 | -0.582 | -0.301 | -0.009 | 0.107 | 0.107 | 0 | 0.53 | 0.82 | 0.62 | 0.29 | 0.52 | 0.53 |
| 7 | 9642956 | rs17161870 | 0.308 | 0.643 | -0.157 | -0.473 | -0.002 | -0.002 | -0.002 | 0 | 0.26 | 0.27 | 0.21 | 0.21 | 0.19 | 0.3 |
| 7 | 9643283 | rs1354606 | -0.366 | 0.692 | 0.155 | -0.358 | 0.051 | -0.063 | -0.063 | 0.001 | 0.22 | 0.55 | 0.39 | 0.08 | 0.29 | 0.2 |
| 7 | 9644596 | rs3857762 | -0.195 | 0.435 | 0.266 | -0.332 | -0.008 | -0.004 | -0.004 | 0 | 0.3 | 0.36 | 0.3 | 0.26 | 0.23 | 0.34 |
| 7 | 9649124 | rs6463932 | -0.296 | 0.449 | 0.229 | -0.442 | 0.019 | 0.011 | 0.011 | 0 | 0.37 | 0.72 | 0.34 | 0.3 | 0.27 | 0.39 |
| 7 | 9650438 | rs10268421 | -0.234 | 0.505 | 0.033 | -0.417 | -0.017 | -0.008 | -0.008 | 0.043 | 0.35 | 0.47 | 0.33 | 0.29 | 0.26 | 0.38 |
| 7 | 9652280 | rs1032873 | 0.523 | 0.81 | -0.863 | -0.08 | -0.038 | 0.088 | 0.088 | 0 | 0.47 | 0.13 | 0.39 | 0.7 | 0.52 | 0.47 |
| 7 | 9662574 | rs7790648 | 0.669 | 1.063 | -0.361 | 0.38 | 0.02 | 0.019 | 0.019 | 1.102 | 0.66 | 0.57 | 0.52 | 0.49 | 0.48 | 0.61 |
| 7 | 9667148 | rs10236139 | 0.275 | 1.558 | -0.114 | 0.608 | 0.008 | -0.001 | -0.001 | 0 | 0.87 | 0.86 | 0.87 | 0.89 | 0.83 | 0.89 |
| 7 | 9667319 | rs13242542 | 0.963 | 0.779 | -0.048 | 0.134 | -0.012 | -0.018 | -0.018 | 0.171 | 0.19 | 0.19 | 0.17 | 0.11 | 0.11 | 0.11 |
| 7 | 9667987 | rs2882606 | 0.247 | 1.089 | 0.09 | 0.264 | -0.004 | -0.001 | -0.001 | 0 | 0.19 | 0.2 | 0.36 | 0.4 | 0.31 | 0.27 |
| 7 | 9670151 | rs9648636 | 0.167 | 1.1 | 0.061 | 0.01 | 0.008 | -0.001 | -0.001 | 0.777 | 0.87 | 0.86 | 0.87 | 0.89 | 0.83 | 0.89 |
| 7 | 9680160 | rs6974358 | 0.858 | 1.209 | 0.32 | 0.013 | 0.075 | 0 | 0 | 0 | 0.87 | 0.67 | 0.87 | 0.92 | 0.84 | 0.91 |
| 7 | 9682864 | rs6463935 | 0.098 | 0.884 | -0.505 | -0.634 | -0.018 | 0.048 | 0.048 | 0.063 | 0.7 | 0.82 | 0.79 | 0.93 | 0.69 | 0.72 |
| 7 | 9682996 | rs10464249 | 1.112 | 0.85 | -0.018 | -0.404 | -0.002 | -0.002 | -0.002 | 0 | 0.18 | 0.17 | 0.26 | 0.18 | 0.16 | 0.26 |
| 7 | 9683817 | rs17807466 | 1.021 | 0.976 | 0.023 | -0.433 | 0.011 | -0.055 | -0.055 | 0 | 0.82 | 0.76 | 0.85 | 0.98 | 0.79 | 0.83 |
| 7 | 9687570 | rs1394778 | 0.209 | 1.136 | 0.014 | -0.43 | 0.02 | -0.035 | -0.035 | 0 | 0.9 | 0.83 | 0.89 | 0.99 | 0.87 | 0.93 |
| 7 | 9689736 | rs1505341 | 1.129 | 1.112 | 0.013 | -0.427 | 0.011 | -0.052 | -0.052 | 0 | 0.82 | 0.75 | 0.85 | 0.98 | 0.79 | 0.84 |
| 7 | 9693082 | rs4273764 | -0.48 | 0.895 | 0.122 | -0.37 | -0.004 | -0.057 | -0.057 | 0 | 0.12 | 0.25 | 0.07 | 0 | 0.11 | 0.1 |
| 7 | 9693670 | rs11974998 | -0.293 | 0.796 | 0.517 | -0.054 | -0.005 | -0.008 | -0.008 | 0 | 0.11 | 0.33 | 0.06 | 0.05 | 0.09 | 0.1 |
| 7 | 9694577 | rs7796062 | -0.531 | 1.106 | 0.27 | -0.133 | 0 | 0 | 0 | 0 | 0.9 | 0.93 | 0.89 | 0.98 | 0.85 | 0.94 |
| 7 | 9694836 | rs7796507 | -0.147 | 0.953 | 0.52 | -0.106 | -0.006 | -0.009 | -0.009 | 0.132 | 0.11 | 0.32 | 0.06 | 0.05 | 0.09 | 0.1 |
| 7 | 9700425 | rs12670039 | -0.93 | 0.487 | -0.407 | -0.057 | -0.045 | -0.007 | -0.007 | 0 | 0.63 | 0.89 | 0.73 | 0.71 | 0.62 | 0.75 |
| 7 | 9701185 | rs17161902 | 1.293 | 0.349 | -0.534 | 0.066 | 0.031 | 0.007 | 0.007 | 1.013 | 0.29 | 0.11 | 0.27 | 0.27 | 0.21 | 0.4 |
| 7 | 9704732 | rs11767294 | 0.63 | 0.456 | 0.032 | 0.061 | 0.025 | 0.062 | 0.062 | 0 | 0.19 | 0.33 | 0.39 | 0.43 | 0.3 | 0.25 |
| 7 | 9708222 | rs6954232 | 0.519 | -0.13 | 0.111 | -0.32 | 0.004 | 0.006 | 0.006 | 0 | 0.36 | 0.31 | 0.32 | 0.24 | 0.3 | 0.48 |
| 7 | 9709384 | rs13223076 | 2.864 | -0.162 | -0.152 | -0.42 | -0.032 | 0.012 | 0.012 | 0.056 | 0.11 | 0.01 | 0.07 | 0.05 | 0.11 | 0.14 |
| 7 | 9711631 | rs6973468 | 0.853 | -0.279 | -0.095 | -0.491 | -0.015 | -0.005 | -0.005 | 5.987 | 0.25 | 0.31 | 0.25 | 0.24 | 0.19 | 0.34 |
| 7 | 9711659 | rs10224816 | 2.941 | -0.283 | -0.256 | -0.511 | -0.016 | 0.03 | 0.03 | 0 | 0.11 | 0.02 | 0.08 | 0.04 | 0.11 | 0.14 |
| 7 | 9712126 | rs957270 | 1.999 | -0.398 | -0.459 | -0.479 | 0.008 | -0.013 | -0.013 | 0 | 0.29 | 0.04 | 0.16 | 0.23 | 0.22 | 0.19 |
| 7 | 9713606 | rs11764744 | -0.853 | 0.131 | 0.025 | -0.339 | 0.011 | -0.035 | -0.035 | 0 | 0.77 | 0.74 | 0.79 | 0.95 | 0.73 | 0.8 |
| 7 | 9714758 | rs11525821 | 1.207 | 0.063 | -0.288 | -0.324 | 0.001 | 0.018 | 0.018 | 0 | 0.17 | 0.18 | 0.36 | 0.48 | 0.28 | 0.27 |
| 7 | 9715652 | rs1400927 | 1.399 | -0.694 | -0.094 | -0.402 | 0.025 | 0.053 | 0.053 | 0.407 | 0.38 | 0.16 | 0.21 | 0.23 | 0.31 | 0.28 |
| 7 | 9716062 | rs7799978 | -0.667 | -0.621 | 0.028 | -0.402 | 0.072 | -0.076 | -0.076 | 0.002 | 0.85 | 0.57 | 0.85 | 0.98 | 0.81 | 0.93 |
| 7 | 9716162 | rs7787594 | 0.145 | -0.622 | -0.013 | -0.378 | 0.002 | -0.01 | -0.01 | 0 | 0.57 | 0.37 | 0.41 | 0.47 | 0.42 | 0.53 |
| 7 | 9722307 | rs6944539 | 0.235 | -0.409 | -0.278 | -0.223 | -0.005 | 0.011 | 0.011 | 0 | 0.45 | 0.36 | 0.64 | 0.71 | 0.51 | 0.66 |
| 7 | 9734323 | rs320794 | 0.03 | -0.585 | -0.627 | -0.291 | -0.006 | 0.067 | 0.067 | 0 | 0.52 | 0.27 | 0.34 | 0.27 | 0.5 | 0.39 |
| 7 | 9735866 | rs320797 | 0.884 | -0.593 | -0.589 | -0.52 | 0.014 | -0.036 | -0.036 | 0 | 0.14 | 0.21 | 0.12 | 0.02 | 0.18 | 0.11 |
| 7 | 9737511 | rs6976386 | 1.968 | -0.792 | -0.43 | -0.38 | -0.005 | 0.002 | 0.002 | 0 | 0.15 | 0.2 | 0.2 | 0.18 | 0.11 | 0.24 |
| 7 | 9743804 | rs535495 | 0.572 | -0.769 | 0.176 | -0.47 | 0.031 | -0.071 | -0.071 | 0.011 | 0.16 | 0.38 | 0.13 | 0.02 | 0.18 | 0.11 |
| 7 | 9744170 | rs17808709 | 0.621 | -0.985 | -0.527 | -0.708 | 0.001 | 0.003 | 0.003 | 0 | 0.34 | 0.05 | 0.31 | 0.24 | 0.26 | 0.22 |
| 7 | 9744591 | rs13222133 | 0.81 | -0.981 | -0.479 | -0.516 | -0.001 | -0.011 | -0.011 | 2.109 | 0.09 | 0.06 | 0.19 | 0.13 | 0.07 | 0.07 |
| 7 | 9744833 | rs667922 | 0.539 | -0.868 | -0.306 | -0.785 | -0.001 | -0.008 | -0.008 | 0 | 0.15 | 0.15 | 0.13 | 0.02 | 0.19 | 0.11 |
| 7 | 9745524 | rs681489 | 0.109 | -1.082 | -0.252 | -0.604 | 0.001 | 0.008 | 0.008 | 0 | 0.42 | 0.3 | 0.43 | 0.3 | 0.43 | 0.33 |
| 7 | 9745966 | rs7787122 | 0.972 | -1.058 | -0.624 | -0.507 | 0.008 | 0.007 | 0.007 | 0 | 0.25 | 0.26 | 0.14 | 0.18 | 0.18 | 0.34 |
| 7 | 9748950 | rs320778 | -0.536 | -0.77 | -0.704 | -0.722 | -0.018 | -0.027 | -0.027 | 0 | 0.72 | 0.8 | 0.59 | 0.46 | 0.64 | 0.66 |
| 7 | 9757755 | rs320788 | -0.561 | -0.846 | -0.787 | -0.747 | -0.031 | -0.04 | -0.04 | 0.064 | 0.72 | 0.79 | 0.58 | 0.51 | 0.64 | 0.67 |
| 7 | 9771599 | rs320818 | -0.924 | -0.661 | -1.106 | -1.025 | -0.111 | 0.107 | 0.107 | 0.878 | 0.77 | 0.96 | 0.83 | 0.95 | 0.68 | 0.79 |
| 7 | 9775747 | rs4725227 | -0.077 | -1.17 | -0.749 | -0.965 | 0.032 | -0.021 | -0.021 | 0 | 0.41 | 0.75 | 0.45 | 0.57 | 0.31 | 0.48 |
| 7 | 9780728 | rs13232914 | 0.763 | -1.267 | -0.459 | -0.964 | -0.043 | -0.042 | -0.042 | 0 | 0.3 | 0.55 | 0.15 | 0.18 | 0.2 | 0.36 |
| 7 | 9791575 | rs10263442 | -0.5 | -0.994 | -0.091 | -0.983 | 0 | 0.146 | 0.146 | 0 | 0.1 | 0.24 | 0.29 | 0.4 | 0.11 | 0.13 |

| | | | | | | | | | | | | | | | | |
|---|---|---|---|---|---|---|---|---|---|---|---|---|---|---|---|---|
| 7 | 9805492 | rs494811 | -1.182 | -0.805 | -0.471 | -1.1 | -0.109 | 0.097 | 0.097 | 0.145 | 0.78 | 0.95 | 0.83 | 0.94 | 0.69 | 0.78 |
| 7 | 9807051 | rs7799117 | 0.392 | -1.544 | 0.538 | -0.88 | -0.011 | -0.017 | -0.017 | 0 | 0.33 | 0.37 | 0.16 | 0.16 | 0.22 | 0.34 |
| 7 | 9808465 | rs1877884 | -0.427 | -1.314 | 0.247 | -0.727 | 0.002 | 0.105 | 0.105 | 0 | 0.07 | 0.13 | 0.27 | 0.38 | 0.09 | 0.12 |
| 7 | 9815039 | rs1319747 | -0.209 | -1.402 | 0.064 | -0.809 | -0.01 | 0.006 | 0.006 | 0 | 0.4 | 0.61 | 0.45 | 0.56 | 0.31 | 0.46 |
| 7 | 9819610 | rs1399490 | -0.176 | -0.922 | -0.026 | -0.325 | -0.001 | 0.01 | 0.01 | 0.218 | 0.16 | 0.22 | 0.32 | 0.4 | 0.15 | 0.21 |
| 7 | 9826825 | rs1399487 | -0.684 | -0.485 | -0.427 | -0.645 | -0.104 | 0.082 | 0.082 | 0 | 0.78 | 0.96 | 0.79 | 0.93 | 0.62 | 0.79 |
| 7 | 9830163 | rs6957346 | 1.391 | -0.75 | 0.109 | -0.725 | -0.016 | -0.037 | -0.037 | 1.608 | 0.33 | 0.4 | 0.18 | 0.19 | 0.23 | 0.34 |
| 7 | 9831245 | rs6943135 | 1.081 | -0.437 | -0.316 | -0.488 | 0.005 | 0.022 | 0.022 | 0 | 0.1 | 0.06 | 0.06 | 0.01 | 0.08 | 0.11 |
| 7 | 9837098 | rs12672973 | 0.917 | -0.374 | -0.322 | -0.342 | 0.003 | -0.031 | -0.031 | 0.551 | 0.16 | 0.12 | 0.22 | 0.32 | 0.16 | 0.21 |
| 7 | 9843763 | rs12702944 | 0.988 | -0.236 | -0.065 | -0.815 | 0.006 | 0.031 | 0.031 | 0 | 0.1 | 0.05 | 0.06 | 0.01 | 0.08 | 0.11 |
| 7 | 9846607 | rs13243340 | 1.322 | -0.012 | 0.527 | -0.765 | -0.014 | -0.033 | -0.033 | 0.028 | 0.29 | 0.42 | 0.17 | 0.2 | 0.22 | 0.36 |
| 7 | 9849527 | rs17162067 | 0.932 | -0.014 | 0.69 | -0.708 | -0.009 | -0.004 | -0.004 | 0.573 | 0.33 | 0.56 | 0.44 | 0.7 | 0.29 | 0.46 |
| 7 | 9851332 | rs7779079 | -0.16 | 0.018 | 0.639 | -0.835 | 0.005 | -0.029 | -0.029 | 0 | 0.72 | 0.74 | 0.75 | 0.93 | 0.59 | 0.78 |
| 7 | 9852667 | rs929544 | 0.542 | -0.087 | 0.624 | -0.834 | 0 | -0.001 | -0.001 | 0 | 0.23 | 0.2 | 0.49 | 0.74 | 0.25 | 0.3 |
| 7 | 9853606 | rs161613 | 0.253 | 0.034 | 0.559 | -0.928 | 0.019 | -0.04 | -0.04 | 0 | 0.25 | 0.26 | 0.22 | 0.07 | 0.34 | 0.22 |
| 7 | 9858697 | rs12702945 | 1.132 | -0.151 | 0.454 | -0.972 | 0.036 | 0.053 | 0.053 | 0.01 | 0.2 | 0.08 | 0.09 | 0.01 | 0.12 | 0.12 |
| 7 | 9862917 | rs11976597 | -0.24 | 0.283 | 0.54 | -0.866 | 0.001 | 0 | 0 | 0 | 0.72 | 0.72 | 0.58 | 0.67 | 0.57 | 0.75 |
| 7 | 9867749 | rs726262 | -1.001 | 0.226 | 0.45 | -0.869 | 0.023 | 0.031 | 0.031 | 0.29 | 0.78 | 0.89 | 0.89 | 0.99 | 0.82 | 0.88 |
| 7 | 9868553 | rs161089 | 0.361 | 0.218 | 0.54 | -0.861 | 0.016 | -0.006 | -0.006 | 0 | 0.56 | 0.65 | 0.58 | 0.5 | 0.6 | 0.62 |
| 7 | 9870197 | rs1709592 | -1.123 | 0.237 | 0.376 | -0.855 | 0.027 | 0.053 | 0.053 | 0.62 | 0.79 | 0.91 | 0.89 | 0.99 | 0.82 | 0.88 |
| 7 | 9870471 | rs1625105 | -0.831 | 0.383 | 0.4 | -0.636 | 0.003 | -0.008 | -0.008 | 0 | 0.5 | 0.53 | 0.73 | 0.81 | 0.6 | 0.53 |
| 7 | 9870851 | rs1639734 | -0.535 | 0.397 | 0.404 | -0.632 | 0.009 | 0.057 | 0.057 | 0.33 | 0.46 | 0.36 | 0.6 | 0.71 | 0.55 | 0.51 |
| 7 | 9873984 | rs320810 | 0.175 | 0.421 | 0.266 | -0.616 | -0.012 | -0.003 | -0.003 | 0 | 0.7 | 0.72 | 0.56 | 0.67 | 0.57 | 0.74 |
| 7 | 9875208 | rs1620397 | -0.554 | 0.488 | -0.078 | -0.72 | 0.004 | 0.107 | 0.107 | 0.331 | 0.45 | 0.21 | 0.59 | 0.71 | 0.55 | 0.51 |
| 7 | 9877720 | rs1639730 | 0.54 | 0.537 | -0.162 | -0.689 | -0.027 | -0.006 | -0.006 | 0.998 | 0.24 | 0.15 | 0.3 | 0.22 | 0.33 | 0.25 |
| 7 | 9878251 | rs12667860 | 0.101 | -0.335 | -0.006 | -0.155 | -0.001 | -0.011 | -0.011 | 0 | 0.22 | 0.16 | 0.3 | 0.49 | 0.22 | 0.26 |
| 7 | 9881778 | rs12056077 | 0.336 | -0.253 | -0.434 | -0.118 | -0.011 | -0.062 | -0.062 | 0.215 | 0.2 | 0.09 | 0.29 | 0.49 | 0.22 | 0.25 |
| 7 | 9882397 | rs970275 | 0.959 | -0.885 | 0.343 | 0.509 | 0.119 | 0.088 | 0.088 | 0.354 | 0.93 | 0.54 | 0.84 | 0.9 | 0.9 | 0.98 |
| 7 | 9889287 | rs1514879 | 0.485 | -0.903 | -0.179 | 0.303 | -0.016 | -0.02 | -0.02 | 0 | 0.7 | 0.44 | 0.45 | 0.28 | 0.7 | 0.68 |
| 7 | 9893744 | rs972124 | 0.227 | -0.151 | -0.886 | 0.117 | 0.025 | 0.031 | 0.031 | 0 | 0.56 | 0.61 | 0.65 | 0.78 | 0.65 | 0.64 |
| 7 | 9898941 | rs12702950 | -0.872 | 0.426 | -0.922 | 0.52 | -0.004 | -0.004 | -0.004 | 0 | 0.07 | 0.03 | 0.05 | 0.08 | 0.11 | 0.13 |
| 7 | 9899380 | rs7776903 | 0.526 | 0.25 | -0.194 | 0.75 | 0.006 | 0.064 | 0.064 | 0.165 | 0.82 | 0.61 | 0.68 | 0.69 | 0.82 | 0.62 |
| 7 | 9903357 | rs728260 | 0.178 | 0.049 | -1.075 | 0.004 | -0.025 | 0.13 | 0.13 | 0 | 0.43 | 0.19 | 0.39 | 0.21 | 0.53 | 0.4 |
| 7 | 9905464 | rs17162125 | 0.382 | -0.64 | -0.222 | 0.104 | -0.011 | 0.068 | 0.068 | 0 | 0.27 | 0.42 | 0.24 | 0.44 | 0.21 | 0.23 |
| 7 | 9906364 | rs2351368 | 0.714 | -0.846 | -0.417 | 0.032 | 0.006 | 0.065 | 0.065 | 0 | 0.06 | 0.03 | 0.11 | 0.39 | 0.05 | 0.09 |
| 7 | 9909504 | rs10254084 | -0.158 | -0.96 | -0.289 | -0.008 | -0.015 | 0.073 | 0.073 | 0 | 0.28 | 0.44 | 0.24 | 0.44 | 0.21 | 0.25 |
| 7 | 9917356 | rs10487881 | 0.434 | -1.273 | -0.506 | -0.154 | 0.013 | -0.042 | -0.042 | 0 | 0.24 | 0.1 | 0.23 | 0.31 | 0.25 | 0.31 |
| 7 | 9917576 | rs10265396 | -0.766 | -1.18 | -0.292 | -0.189 | 0.035 | 0.043 | 0.043 | 0 | 0.49 | 0.73 | 0.5 | 0.39 | 0.41 | 0.47 |
| 7 | 9922243 | rs2971100 | -0.632 | -1.197 | -0.183 | 0.158 | -0.007 | 0.143 | 0.143 | 0 | 0.94 | 0.9 | 0.89 | 0.64 | 0.95 | 0.92 |
| 7 | 9922676 | rs4725239 | 0.485 | -1.154 | -0.184 | 0.053 | 0.012 | -0.027 | -0.027 | 0 | 0.26 | 0.14 | 0.25 | 0.29 | 0.27 | 0.34 |
| 7 | 9923279 | rs17162177 | 0.321 | -1.292 | -0.159 | -0.223 | -0.026 | -0.103 | -0.103 | 0.004 | 0.24 | 0.57 | 0.26 | 0.09 | 0.14 | 0.11 |
| 7 | 9926085 | rs2915132 | 0.921 | -1.175 | -0.37 | -0.209 | -0.024 | 0.105 | 0.105 | 0 | 0.43 | 0.16 | 0.39 | 0.25 | 0.54 | 0.45 |
| 7 | 9939876 | rs2351378 | 0.532 | -1.394 | -0.748 | -0.508 | 0.02 | -0.043 | -0.043 | 0 | 0.65 | 0.22 | 0.59 | 0.54 | 0.76 | 0.81 |
| 7 | 9944175 | rs7797070 | 0.152 | -1.527 | -0.939 | -0.478 | 0.007 | 0.064 | 0.064 | 0 | 0.37 | 0.05 | 0.3 | 0.25 | 0.35 | 0.31 |
| 7 | 9944520 | rs2351379 | -0.179 | -1.358 | -0.686 | -0.459 | -0.006 | 0.034 | 0.034 | 0 | 0.72 | 0.34 | 0.64 | 0.54 | 0.79 | 0.82 |
| 7 | 9951880 | rs6958845 | -0.112 | -1.388 | -0.585 | -0.384 | -0.006 | 0.037 | 0.037 | 0.009 | 0.86 | 0.83 | 0.84 | 0.63 | 0.9 | 0.91 |
| 7 | 9952849 | rs2915154 | 0.45 | -1.464 | -0.661 | -0.443 | 0.039 | 0.025 | 0.025 | 0 | 0.54 | 0.68 | 0.57 | 0.39 | 0.61 | 0.61 |
| 7 | 9955785 | rs2351383 | 0.003 | -1.415 | -0.608 | -0.236 | -0.012 | 0.087 | 0.087 | 0 | 0.91 | 0.87 | 0.88 | 0.63 | 0.95 | 0.9 |
| 7 | 9956258 | rs10277580 | 1.172 | -1.359 | -0.047 | -0.292 | 0.012 | -0.059 | -0.059 | 0 | 0.18 | 0.34 | 0.12 | 0.09 | 0.13 | 0.1 |
| 7 | 9957558 | rs2915156 | 0.304 | -1.387 | -0.568 | -0.146 | -0.013 | 0.059 | 0.059 | 0.007 | 0.85 | 0.8 | 0.83 | 0.64 | 0.9 | 0.91 |
| 7 | 9957795 | rs2351384 | 0.349 | -1.186 | -0.911 | -0.423 | -0.008 | 0.013 | 0.013 | 0 | 0.58 | 0.59 | 0.53 | 0.36 | 0.53 | 0.56 |
| 7 | 9965912 | rs1557978 | -1.299 | -0.673 | -1.053 | -0.252 | -0.04 | -0.023 | -0.023 | 0 | 0.77 | 0.85 | 0.71 | 0.72 | 0.64 | 0.68 |
| 7 | 9968807 | rs9648642 | -0.81 | -0.736 | -0.89 | -0.15 | -0.027 | 0.041 | 0.041 | 0 | 0.63 | 0.73 | 0.62 | 0.79 | 0.57 | 0.76 |
| 7 | 9977206 | rs17162231 | 0.542 | -0.36 | -1.024 | 0.803 | -0.003 | 0.042 | 0.042 | 0.386 | 0.11 | 0.12 | 0.08 | 0.34 | 0.05 | 0.07 |
| 7 | 9978539 | rs1016019 | -0.192 | 0.547 | -1.032 | 0.231 | -0.009 | 0.007 | 0.007 | 0.017 | 0.72 | 0.79 | 0.63 | 0.8 | 0.66 | 0.66 |
| 7 | 9988147 | rs10255183 | -0.729 | -0.983 | -0.844 | 0.232 | 0.012 | 0.009 | 0.009 | 1.073 | 0.25 | 0.19 | 0.23 | 0.19 | 0.17 | 0.35 |
| 7 | 9990303 | rs1859588 | 1.289 | -1.42 | -0.476 | 0.284 | -0.007 | -0.002 | -0.002 | 0 | 0.79 | 0.78 | 0.78 | 0.77 | 0.82 | 0.66 |
| 7 | 9994397 | rs1859586 | 0.212 | -1.244 | 0.359 | 0.624 | 0.023 | 0.052 | 0.052 | 0 | 0.28 | 0.44 | 0.44 | 0.42 | 0.34 | 0.2 |
| 7 | 9998779 | rs6955008 | -0.065 | -1.384 | 0.924 | 0.578 | 0.008 | 0.043 | 0.043 | 0 | 0.31 | 0.57 | 0.45 | 0.47 | 0.35 | 0.26 |
| 7 | 10000210 | rs2906453 | 1.209 | -1.983 | 1.123 | 0.292 | 0 | -0.024 | -0.024 | 2.56 | 0.44 | 0.45 | 0.28 | 0.11 | 0.48 | 0.44 |
| 7 | 10001709 | rs995857 | 0.577 | -2.023 | 0.898 | 0.516 | 0.001 | 0 | 0 | 0 | 0.85 | 0.87 | 0.82 | 0.83 | 0.88 | 0.74 |
| 7 | 10003616 | rs17162294 | 0.507 | -1.962 | 0.825 | 0.734 | -0.003 | 0.016 | 0.016 | 0.786 | 0.12 | 0.13 | 0.11 | 0.29 | 0.05 | 0.07 |
| 7 | 10004014 | rs12531709 | 1.101 | -1.578 | 0.95 | 0.524 | -0.028 | 0.038 | 0.038 | 0 | 0.87 | 0.68 | 0.85 | 0.75 | 0.92 | 0.7 |
| 7 | 10006341 | rs2971030 | 0.839 | -1.585 | 0.768 | 0.429 | -0.003 | 0.003 | 0.003 | 0 | 0.56 | 0.5 | 0.42 | 0.47 | 0.57 | 0.72 |
| 7 | 10007255 | rs2971028 | 0.732 | -1.603 | 0.901 | 0.461 | 0 | 0 | 0 | 0 | 0.57 | 0.56 | 0.43 | 0.47 | 0.58 | 0.73 |
| 7 | 10042560 | rs6463960 | 0.479 | -1.696 | 0.933 | 0.393 | -0.012 | -0.042 | -0.042 | 0 | 0.61 | 0.77 | 0.44 | 0.48 | 0.57 | 0.72 |
| 7 | 10044734 | rs2906408 | 0.85 | -1.593 | 0.918 | 0.319 | -0.001 | 0.001 | 0.001 | 0 | 0.57 | 0.55 | 0.41 | 0.48 | 0.58 | 0.73 |
| 7 | 10054784 | rs2906405 | 0.822 | -1.685 | 1.068 | 0.399 | -0.002 | 0.008 | 0.008 | 0 | 0.58 | 0.51 | 0.42 | 0.48 | 0.57 | 0.72 |
| 7 | 10059674 | rs7803616 | -0.102 | -1.576 | 1.068 | 0.401 | 0.013 | -0.044 | -0.044 | 0.119 | 0.18 | 0.31 | 0.15 | 0.07 | 0.2 | 0.08 |

| | | | | | | | | | | | | | | | | |
|---|---|---|---|---|---|---|---|---|---|---|---|---|---|---|---|---|
| 7 | 10062341 | rs12536973 | 0.771 | -1.754 | 1.012 | 0.45 | 0.001 | -0.018 | -0.018 | 0 | 0.14 | 0.11 | 0.28 | 0.35 | 0.12 | 0.13 |
| 7 | 10064734 | rs2906403 | 0.591 | -1.761 | 1.034 | 0.45 | 0.01 | -0.039 | -0.039 | 0.789 | 0.74 | 0.8 | 0.56 | 0.55 | 0.77 | 0.81 |
| 7 | 10069869 | rs6957830 | 1.388 | -1.82 | 1.074 | 0.415 | -0.003 | 0.002 | 0.002 | 0 | 0.55 | 0.51 | 0.42 | 0.49 | 0.56 | 0.72 |
| 7 | 10071092 | rs4413687 | 0.352 | -1.752 | 1.112 | 0.329 | 0.007 | 0.018 | 0.018 | 0 | 0.87 | 0.92 | 0.84 | 0.9 | 0.88 | 0.93 |
| 7 | 10072261 | rs2107970 | -0.245 | -1.752 | 1.113 | 0.33 | 0.003 | 0.008 | 0.008 | 0 | 0.9 | 0.92 | 0.84 | 0.9 | 0.89 | 0.93 |
| 7 | 10073500 | rs2971117 | 1.149 | -1.758 | 1.158 | 0.327 | -0.004 | -0.044 | -0.044 | 0 | 0.69 | 0.62 | 0.69 | 0.83 | 0.68 | 0.85 |
| 7 | 10074972 | rs2190146 | -0.147 | -2.046 | 1.143 | 0.338 | 0.005 | 0.005 | 0.005 | 0 | 0.89 | 0.9 | 0.84 | 0.9 | 0.9 | 0.94 |
| 7 | 10077485 | rs17162474 | -0.125 | -2.029 | 1.165 | 0.343 | 0.007 | -0.037 | -0.037 | 0 | 0.19 | 0.29 | 0.15 | 0.08 | 0.2 | 0.08 |
| 7 | 10079669 | rs2971111 | 1.159 | -2.374 | 1.287 | 0.386 | 0.006 | 0.004 | 0.004 | 0 | 0.61 | 0.38 | 0.42 | 0.15 | 0.71 | 0.54 |
| 7 | 10081250 | rs12333521 | -1 | -2.148 | 1.709 | 0.868 | -0.068 | 0.155 | 0.155 | 0 | 0.19 | 0.51 | 0.16 | 0.41 | 0.1 | 0.32 |
| 7 | 10086833 | rs2906374 | 2.209 | -2.128 | 1.457 | 0.832 | 0.083 | 0.007 | 0.007 | 0 | 0.5 | 0.21 | 0.54 | 0.43 | 0.59 | 0.53 |
| 7 | 10089086 | rs10257866 | -0.927 | -1.637 | 1.977 | 1.183 | -0.005 | -0.021 | -0.021 | 0 | 0.07 | 0.27 | 0.02 | 0 | 0.02 | 0.05 |
| 7 | 10091865 | rs10241626 | -1.61 | -1.561 | 1.806 | 1.206 | -0.005 | 0.008 | 0.008 | 0 | 0.32 | 0.6 | 0.26 | 0.48 | 0.27 | 0.4 |
| 7 | 10093255 | rs11980221 | -0.299 | -1.614 | 1.851 | 1.232 | -0.043 | 0.157 | 0.157 | 0 | 0.13 | 0.26 | 0.12 | 0.41 | 0.07 | 0.28 |
| 7 | 10093480 | rs6973780 | -1.422 | -1.479 | 1.086 | 1.183 | -0.004 | 0.005 | 0.005 | 0 | 0.33 | 0.6 | 0.27 | 0.48 | 0.29 | 0.4 |
| 7 | 10096276 | rs41426 | -0.889 | -1.948 | 1.831 | 1.716 | -0.027 | 0.075 | 0.075 | 0 | 0.14 | 0.2 | 0.13 | 0.41 | 0.08 | 0.29 |
| 7 | 10106803 | rs10239257 | -0.028 | -1.982 | 1.74 | 1.886 | 0.022 | -0.03 | -0.03 | 0.009 | 0.9 | 0.96 | 0.91 | 0.76 | 0.95 | 0.95 |
| 7 | 10113156 | rs6972783 | 2.31 | -2.266 | 2.311 | 1.959 | -0.003 | -0.002 | -0.002 | 0 | 0.72 | 0.34 | 0.59 | 0.36 | 0.82 | 0.64 |
| 7 | 10116825 | rs6952221 | 1.232 | -2.238 | 3.025 | 2.094 | -0.061 | 0.159 | 0.159 | 0 | 0.83 | 0.63 | 0.65 | 0.47 | 0.91 | 0.83 |
| 7 | 10121066 | rs12702976 | 2.347 | -2.254 | 2.727 | 2.177 | -0.009 | 0 | 0 | 0 | 0.72 | 0.32 | 0.57 | 0.37 | 0.81 | 0.62 |
| 7 | 10121205 | rs988418 | 0.515 | -2.261 | 2.802 | 2.328 | -0.018 | 0.073 | 0.073 | 0 | 0.84 | 0.78 | 0.65 | 0.54 | 0.89 | 0.87 |
| 7 | 10122252 | rs7811665 | 2.285 | -2.24 | 2.386 | 2.309 | -0.009 | 0 | 0 | 0 | 0.72 | 0.32 | 0.57 | 0.37 | 0.81 | 0.62 |
| 7 | 10124331 | rs12702978 | 1.141 | -1.694 | 1.894 | 2.107 | -0.032 | 0.196 | 0.196 | 0 | 0.81 | 0.59 | 0.62 | 0.53 | 0.86 | 0.86 |
| 7 | 10128774 | rs2191852 | 1.052 | -1.65 | 1.135 | 1.246 | 0.011 | 0.001 | 0.001 | 0 | 0.8 | 0.85 | 0.83 | 0.81 | 0.83 | 0.86 |
| 7 | 10129376 | rs17162571 | -1.42 | -1.659 | 1.324 | 1.053 | -0.009 | 0.001 | 0.001 | 0.116 | 0.1 | 0.15 | 0.13 | 0.13 | 0.06 | 0.2 |
| 7 | 10130272 | rs728220 | -1.278 | -1.694 | 1.552 | 1.041 | -0.011 | 0.026 | 0.026 | 0 | 0.08 | 0.24 | 0.04 | 0.09 | 0.04 | 0.17 |
| 7 | 10130672 | rs10251294 | -2.382 | -1.807 | 2.1 | 1.017 | -0.054 | 0.019 | 0.019 | 2.796 | 0.15 | 0.55 | 0.16 | 0.16 | 0.06 | 0.24 |
| 7 | 10137638 | rs10271818 | -1.519 | -1.659 | 1.408 | 1.049 | -0.03 | 0.002 | 0.002 | 0 | 0.27 | 0.62 | 0.19 | 0.3 | 0.18 | 0.36 |
| 7 | 10140489 | rs10255407 | -1.413 | -1.624 | 1.878 | 1.042 | -0.044 | 0.013 | 0.013 | 0 | 0.26 | 0.48 | 0.18 | 0.3 | 0.18 | 0.37 |
| 7 | 10146653 | rs7785204 | -2.411 | -2.013 | 1.583 | 1.049 | 0.066 | -0.018 | -0.018 | 0 | 0.35 | 0.84 | 0.26 | 0.37 | 0.25 | 0.4 |
| 7 | 10148521 | rs17162597 | 0.083 | -2.03 | 1.536 | 1.101 | -0.018 | -0.036 | -0.036 | 0 | 0.13 | 0.01 | 0.12 | 0.17 | 0.11 | 0.1 |
| 7 | 10154686 | rs17162631 | -0.858 | -2.373 | 2.125 | 0.885 | -0.008 | -0.028 | -0.028 | 0 | 0.09 | 0.11 | 0.05 | 0 | 0.08 | 0.03 |
| 7 | 10155133 | rs6963093 | -1.821 | -2.373 | 2.113 | 0.887 | -0.033 | -0.001 | -0.001 | 0 | 0.13 | 0.31 | 0.08 | 0.08 | 0.1 | 0.06 |
| 7 | 10161358 | rs10254939 | -2.566 | -2.406 | 2.321 | 0.886 | -0.031 | -0.01 | -0.01 | 0.049 | 0.17 | 0.71 | 0.1 | 0.11 | 0.1 | 0.12 |
| 7 | 10162137 | rs2041510 | -0.818 | -2.364 | 2.227 | 0.856 | -0.012 | -0.034 | -0.034 | 2.275 | 0.09 | 0.11 | 0.04 | 0 | 0.08 | 0.03 |
| 7 | 10172649 | rs17162693 | -2.421 | -2.694 | 2.503 | 1.016 | -0.03 | 0.003 | 0.003 | 0 | 0.37 | 0.46 | 0.43 | 0.4 | 0.31 | 0.33 |
| 7 | 10177323 | rs4725259 | -2.279 | -2.587 | 2.466 | 1.034 | -0.027 | 0 | 0 | 0 | 0.2 | 0.7 | 0.14 | 0.2 | 0.15 | 0.27 |
| 7 | 10182235 | rs2191849 | -1.957 | -2.54 | 2.509 | 1.034 | -0.044 | 0.023 | 0.023 | 0 | 0.18 | 0.51 | 0.13 | 0.21 | 0.14 | 0.25 |
| 7 | 10185972 | rs13240249 | 2.141 | -2.124 | 1.695 | 0.73 | 0.027 | 0.001 | 0.001 | 0 | 0.6 | 0.26 | 0.54 | 0.59 | 0.68 | 0.58 |
| 7 | 10187111 | rs222238 | 1.749 | -2.083 | 1.752 | 0.7 | -0.069 | 0.002 | 0.002 | 0 | 0.79 | 0.5 | 0.82 | 0.8 | 0.85 | 0.71 |
| 7 | 10191885 | rs974343 | 0.894 | -1.509 | 0.578 | 0.635 | 0.042 | -0.001 | -0.001 | 0 | 0.29 | 0.39 | 0.43 | 0.23 | 0.4 | 0.34 |
| 7 | 10205398 | rs222271 | 0.55 | -2.291 | 0.167 | 0.975 | 0.024 | -0.023 | -0.023 | 0 | 0.79 | 0.88 | 0.79 | 0.7 | 0.83 | 0.7 |
| 7 | 10211607 | rs222289 | 0.776 | -2.522 | -0.405 | 0.567 | -0.005 | 0.003 | 0.003 | 0.219 | 0.24 | 0.2 | 0.12 | 0.27 | 0.23 | 0.14 |
| 7 | 10216075 | rs222307 | -0.409 | -2.644 | -0.445 | 0.523 | 0.002 | 0.013 | 0.013 | 0 | 0.49 | 0.46 | 0.35 | 0.48 | 0.44 | 0.37 |
| 7 | 10220549 | rs222314 | 0.78 | -2.484 | -0.395 | 0.324 | -0.007 | -0.001 | -0.001 | 0 | 0.22 | 0.15 | 0.1 | 0.27 | 0.21 | 0.14 |
| 7 | 10222003 | rs222315 | 0.125 | -2.873 | -0.641 | 0.225 | 0.018 | 0.044 | 0.044 | 0 | 0.75 | 0.91 | 0.77 | 0.8 | 0.74 | 0.53 |
| 7 | 10226430 | rs10236225 | 0.909 | -2.636 | 0.276 | 0.337 | -0.004 | -0.007 | -0.007 | 0 | 0.44 | 0.39 | 0.59 | 0.41 | 0.52 | 0.39 |
| 7 | 10233678 | rs17162780 | -0.512 | -2.604 | 0.218 | 0.247 | -0.011 | -0.006 | -0.006 | 0 | 0.09 | 0.16 | 0.06 | 0.11 | 0.07 | 0.27 |
| 7 | 10245417 | rs9692033 | -0.043 | -2.789 | 0.302 | 0.365 | -0.022 | 0.032 | 0.032 | 0 | 0.24 | 0.41 | 0.25 | 0.29 | 0.18 | 0.27 |
| 7 | 10246371 | rs2107809 | 0.132 | -2.775 | 0.375 | 0.374 | -0.009 | 0.011 | 0.011 | 0.541 | 0.61 | 0.49 | 0.74 | 0.43 | 0.69 | 0.6 |
| 7 | 10249494 | rs222331 | 1.305 | -2.748 | 0.032 | 0.512 | -0.006 | -0.008 | -0.008 | 0.511 | 0.19 | 0.14 | 0.08 | 0.3 | 0.16 | 0.13 |
| 7 | 10252414 | rs2189952 | -0.533 | -2.587 | 0.198 | 0.415 | -0.016 | 0.011 | 0.011 | 0 | 0.31 | 0.39 | 0.28 | 0.33 | 0.24 | 0.29 |
| 7 | 10260962 | rs6959382 | -0.204 | -2.366 | 0.431 | 0.432 | -0.001 | 0.007 | 0.007 | 0 | 0.62 | 0.63 | 0.68 | 0.46 | 0.65 | 0.58 |
| 7 | 10263631 | rs6950145 | 0.317 | -2.138 | 0.354 | 0.356 | 0 | 0.001 | 0.001 | 0 | 0.1 | 0.09 | 0.06 | 0.12 | 0.11 | 0.21 |
| 7 | 10265600 | rs7802767 | 1.842 | -2.127 | 0.372 | 0.298 | 0.012 | -0.021 | -0.021 | 0 | 0.29 | 0.39 | 0.42 | 0.13 | 0.39 | 0.26 |
| 7 | 10267153 | rs7792558 | 1.689 | -1.91 | 0.341 | 0.272 | -0.002 | -0.003 | -0.003 | 0.14 | 0.3 | 0.34 | 0.45 | 0.13 | 0.42 | 0.27 |
| 7 | 10268690 | rs1024521 | -0.855 | -2.011 | 0.101 | 0.309 | -0.013 | -0.004 | -0.004 | 0.522 | 0.18 | 0.22 | 0.1 | 0.13 | 0.17 | 0.24 |
| 7 | 10276673 | rs2017532 | -0.057 | -1.9 | 0.053 | 0.24 | 0.003 | -0.009 | -0.009 | 0 | 0.59 | 0.37 | 0.63 | 0.47 | 0.63 | 0.58 |
| 7 | 10278806 | rs10225194 | 2.031 | -1.602 | -0.237 | 0.138 | -0.038 | 0.048 | 0.048 | 0 | 0.27 | 0.19 | 0.37 | 0.13 | 0.37 | 0.27 |
| 7 | 10279486 | rs17162816 | 1.272 | -1.633 | -0.128 | 0.189 | 0.009 | -0.018 | -0.018 | 0.159 | 0.17 | 0.07 | 0.18 | 0.22 | 0.11 | 0.09 |
| 7 | 10287951 | rs6943027 | -0.003 | -1.753 | 0.396 | 0.199 | -0.038 | -0.009 | -0.009 | 0 | 0.13 | 0.39 | 0.05 | 0.06 | 0.09 | 0.11 |
| 7 | 10293921 | rs10265578 | -0.984 | -1.77 | 0.051 | 0.175 | -0.067 | 0.01 | 0.01 | 0 | 0.71 | 0.9 | 0.71 | 0.66 | 0.64 | 0.67 |
| 7 | 10295322 | rs12164076 | 2.671 | -1.819 | 0.027 | 0.142 | -0.084 | 0.058 | 0.058 | 0 | 0.23 | 0.08 | 0.34 | 0.13 | 0.33 | 0.25 |
| 7 | 10297082 | rs7808481 | 1.372 | -1.934 | 0.278 | 0.247 | -0.027 | 0.081 | 0.081 | 0.352 | 0.08 | 0.22 | 0.07 | 0.12 | 0.01 | 0.01 |
| 7 | 10298164 | rs2189945 | 0.101 | -1.923 | 0.275 | 0.236 | 0.009 | -0.024 | -0.024 | 1.513 | 0.27 | 0.17 | 0.25 | 0.35 | 0.23 | 0.31 |
| 7 | 10298689 | rs17162825 | 1.257 | -1.9 | 0.298 | 0.307 | -0.002 | -0.02 | -0.02 | 0.662 | 0.09 | 0.03 | 0.06 | 0.14 | 0.12 | 0.22 |
| 7 | 10299080 | rs17162827 | 1.529 | -1.878 | 0.536 | 0.276 | 0.025 | 0.083 | 0.083 | 0 | 0.06 | 0.33 | 0.02 | 0.06 | 0.04 | 0.09 |
| 7 | 10299150 | rs17162828 | -0.429 | -1.704 | 0.479 | 0.254 | 0.027 | -0.024 | -0.024 | 0.64 | 0.33 | 0.19 | 0.27 | 0.35 | 0.28 | 0.33 |
| 7 | 10299937 | rs10265451 | 1.676 | -1.681 | 0.536 | 0.273 | 0.001 | 0.048 | 0.048 | 0 | 0.06 | 0.21 | 0.01 | 0.06 | 0.04 | 0.09 |

| | | | | | | | | | | | | | | | | |
|---|---|---|---|---|---|---|---|---|---|---|---|---|---|---|---|---|
| 7 | 10301776 | rs10228140 | -1.381 | -1.673 | 0.269 | 0.239 | 0.107 | 0.045 | 0.045 | 0 | 0.47 | 0.73 | 0.36 | 0.53 | 0.32 | 0.42 |
| 7 | 10302022 | rs10952197 | 1.788 | -1.452 | 0.021 | 0.189 | -0.002 | 0 | 0 | 0.016 | 0.31 | 0.31 | 0.42 | 0.25 | 0.34 | 0.26 |
| 7 | 10303377 | rs766549 | -0.804 | -1.032 | -0.552 | -0.12 | -0.071 | 0.005 | 0.005 | 0 | 0.71 | 0.92 | 0.72 | 0.66 | 0.65 | 0.67 |
| 7 | 10318296 | rs2351841 | 0.387 | -1.017 | -0.368 | -0.348 | -0.022 | 0.07 | 0.07 | 0 | 0.45 | 0.75 | 0.47 | 0.28 | 0.5 | 0.59 |
| 7 | 10318768 | rs10224072 | -0.956 | -0.93 | -0.687 | -0.374 | -0.031 | -0.051 | -0.051 | 0.019 | 0.79 | 0.92 | 0.79 | 0.62 | 0.75 | 0.72 |
| 7 | 10319806 | rs16876522 | 0.974 | -0.875 | 0.117 | -0.243 | 0.002 | -0.021 | -0.021 | 1.635 | 0.08 | 0.32 | 0.02 | 0.01 | 0.04 | 0.09 |
| 7 | 10326657 | rs16876525 | 1.402 | -0.811 | -0.733 | -0.074 | 0.02 | -0.047 | -0.047 | 0.169 | 0.19 | 0.07 | 0.22 | 0.35 | 0.12 | 0.09 |
| 7 | 10327371 | rs10235788 | -1.089 | -0.607 | -1.045 | -0.23 | -0.061 | 0.04 | 0.04 | 0.468 | 0.63 | 0.86 | 0.56 | 0.75 | 0.48 | 0.47 |
| 7 | 10337280 | rs7791241 | 0.78 | -0.679 | -0.647 | -0.121 | -0.002 | -0.003 | -0.003 | 0 | 0.47 | 0.39 | 0.58 | 0.48 | 0.47 | 0.38 |
| 7 | 10337941 | rs2351852 | 1.083 | -0.897 | -0.601 | -0.16 | -0.015 | 0.028 | 0.028 | 0 | 0.22 | 0.25 | 0.22 | 0.35 | 0.12 | 0.12 |
| 7 | 10339651 | rs6977174 | -0.44 | -0.842 | -0.122 | -0.268 | 0.026 | 0.096 | 0.096 | 0 | 0.87 | 0.6 | 0.9 | 0.84 | 0.86 | 0.79 |
| 7 | 10342311 | rs12536534 | 1.283 | -0.766 | -0.88 | -0.513 | -0.064 | 0.086 | 0.086 | 0 | 0.33 | 0.23 | 0.43 | 0.12 | 0.42 | 0.29 |
| 7 | 10343038 | rs12539924 | 2.104 | -0.632 | -1.124 | -0.415 | -0.065 | 0.064 | 0.064 | 0.028 | 0.24 | 0.13 | 0.35 | 0.12 | 0.34 | 0.27 |
| 7 | 10347299 | rs10226514 | 0.087 | -0.499 | -1.33 | -0.276 | 0.014 | 0.007 | 0.007 | 0 | 0.34 | 0.2 | 0.26 | 0.38 | 0.33 | 0.39 |
| 7 | 10349047 | rs6463988 | -0.841 | -0.64 | -1.831 | -0.299 | -0.017 | -0.028 | -0.028 | 0.043 | 0.88 | 0.97 | 0.92 | 0.87 | 0.82 | 0.79 |
| 7 | 10352241 | rs10488128 | 0.154 | -0.777 | -1.596 | -0.201 | -0.006 | -0.021 | -0.021 | 0.556 | 0.33 | 0.06 | 0.25 | 0.38 | 0.32 | 0.39 |
| 7 | 10356078 | rs2116615 | -0.965 | -0.265 | -0.79 | -0.27 | 0.017 | -0.02 | -0.02 | 0 | 0.65 | 0.62 | 0.59 | 0.8 | 0.53 | 0.55 |
| 7 | 10356213 | rs2163683 | -0.128 | -0.586 | -1.301 | -0.056 | -0.002 | 0.011 | 0.011 | 0 | 0.33 | 0.18 | 0.26 | 0.3 | 0.32 | 0.4 |
| 7 | 10360116 | rs17162921 | 0.367 | -0.273 | -1.198 | 0.152 | -0.014 | -0.013 | -0.013 | 0.27 | 0.21 | 0.15 | 0.27 | 0.19 | 0.24 | 0.3 |
| 7 | 10360414 | rs4725269 | 0.159 | -0.537 | -0.965 | 0.154 | 0.003 | 0.01 | 0.01 | 0 | 0.47 | 0.27 | 0.44 | 0.65 | 0.47 | 0.52 |
| 7 | 10361188 | rs10488129 | 0.898 | -0.269 | -0.995 | -0.181 | -0.028 | 0.06 | 0.06 | 0 | 0.29 | 0.14 | 0.34 | 0.06 | 0.33 | 0.25 |
| 7 | 10366844 | rs4725270 | 0.124 | -0.106 | -0.26 | 0.285 | 0.021 | 0.052 | 0.052 | 0 | 0.27 | 0.54 | 0.36 | 0.4 | 0.33 | 0.34 |
| 7 | 10368887 | rs10259967 | 0.779 | -0.079 | -0.744 | 0.358 | -0.013 | -0.033 | -0.033 | 0 | 0.21 | 0.15 | 0.29 | 0.35 | 0.25 | 0.31 |
| 7 | 10383334 | rs13236695 | 1.106 | -0.081 | -0.991 | -0.006 | -0.108 | 0.089 | 0.089 | 0 | 0.26 | 0.02 | 0.31 | 0.06 | 0.31 | 0.24 |
| 7 | 10383515 | rs6463991 | -1.674 | -0.542 | -1.09 | -0.101 | -0.001 | 0.031 | 0.031 | 0 | 0.73 | 0.91 | 0.78 | 0.84 | 0.79 | 0.83 |
| 7 | 10383638 | rs17442517 | 1.114 | -0.469 | -0.601 | -0.182 | -0.012 | 0.019 | 0.019 | 0 | 0.28 | 0.21 | 0.32 | 0.06 | 0.32 | 0.24 |
| 7 | 10386633 | rs6973363 | -1.694 | -0.592 | -0.517 | -0.204 | -0.035 | 0.04 | 0.04 | 0 | 0.75 | 0.96 | 0.78 | 0.87 | 0.79 | 0.83 |
| 7 | 10390212 | rs1549863 | -0.34 | -0.764 | -0.053 | -0.155 | 0.039 | -0.004 | -0.004 | 0 | 0.58 | 0.75 | 0.72 | 0.41 | 0.69 | 0.55 |
| 7 | 10394558 | rs11765391 | 0.67 | -0.676 | -0.149 | -0.208 | 0.021 | 0.031 | 0.031 | 0 | 0.27 | 0.46 | 0.35 | 0.32 | 0.32 | 0.31 |
| 7 | 10395132 | rs1295349 | 1.429 | -0.697 | -0.149 | -0.223 | -0.053 | 0.097 | 0.097 | 0 | 0.27 | 0.11 | 0.31 | 0.06 | 0.31 | 0.23 |
| 7 | 10395970 | rs1589764 | -0.615 | -0.734 | -0.102 | -0.21 | 0.006 | 0.027 | 0.027 | 0 | 0.46 | 0.62 | 0.45 | 0.78 | 0.47 | 0.6 |
| 7 | 10396118 | rs1295348 | -1.538 | -0.778 | -0.027 | -0.115 | 0.003 | 0.013 | 0.013 | 0 | 0.73 | 0.82 | 0.77 | 0.84 | 0.78 | 0.83 |
| 7 | 10426456 | rs12532295 | 0.51 | -0.686 | -0.057 | 0.102 | -0.007 | 0.028 | 0.028 | 0 | 0.15 | 0.14 | 0.06 | 0.43 | 0.11 | 0.27 |
| 7 | 10439066 | rs1581512 | 0.484 | -0.661 | 0.061 | 0.111 | -0.016 | 0.049 | 0.049 | 0 | 0.15 | 0.16 | 0.06 | 0.42 | 0.1 | 0.28 |
| 7 | 10458031 | rs17162989 | -0.025 | -0.489 | -0.08 | -0.195 | -0.012 | 0.004 | 0.004 | 0 | 0.37 | 0.29 | 0.4 | 0.37 | 0.44 | 0.42 |
| 7 | 10458795 | rs2049387 | -0.92 | -0.472 | 0.131 | -0.231 | -0.019 | 0.031 | 0.031 | 0 | 0.3 | 0.4 | 0.18 | 0.46 | 0.29 | 0.4 |
| 7 | 10459280 | rs17239281 | 1.771 | -0.256 | -0.046 | -0.51 | 0.004 | 0.02 | 0.02 | 0.074 | 0.09 | 0.07 | 0.03 | 0.04 | 0.08 | 0.11 |
| 7 | 10460056 | rs1524242 | -1.912 | -0.12 | -0.129 | -0.675 | -0.033 | 0.039 | 0.039 | 0 | 0.82 | 0.9 | 0.72 | 0.95 | 0.76 | 0.88 |
| 7 | 10460155 | rs17817981 | 1.049 | -0.066 | -0.032 | -0.468 | -0.003 | 0.041 | 0.041 | 0 | 0.12 | 0.08 | 0.07 | 0.04 | 0.12 | 0.12 |
| 7 | 10460817 | rs2049388 | 0.199 | -0.073 | -0.006 | -0.466 | 0.001 | -0.044 | -0.044 | 0 | 0.15 | 0.22 | 0.12 | 0.04 | 0.19 | 0.13 |
| 7 | 10467703 | rs2882836 | -0.407 | -0.018 | -0.056 | -0.444 | -0.008 | 0.047 | 0.047 | 0 | 0.45 | 0.39 | 0.47 | 0.75 | 0.51 | 0.6 |
| 7 | 10468765 | rs6946433 | -0.977 | -0.076 | 0.105 | -0.558 | -0.002 | -0.01 | -0.01 | 0 | 0.66 | 0.7 | 0.76 | 0.8 | 0.73 | 0.74 |
| 7 | 10469534 | rs765805 | 1.572 | 0.002 | -0.316 | -0.498 | -0.002 | -0.003 | -0.003 | 0 | 0.19 | 0.16 | 0.27 | 0.05 | 0.22 | 0.14 |
| 7 | 10473103 | rs12056273 | 0.734 | -0.221 | -0.465 | -0.273 | 0.028 | -0.067 | -0.067 | 0.179 | 0.15 | 0.03 | 0.06 | 0.44 | 0.11 | 0.29 |
| 7 | 10474405 | rs10952211 | -0.272 | -0.042 | -0.332 | -0.341 | 0 | -0.001 | -0.001 | 0.041 | 0.41 | 0.43 | 0.43 | 0.48 | 0.47 | 0.44 |
| 7 | 10474663 | rs2177511 | 1.178 | -0.112 | -0.467 | -0.084 | -0.045 | -0.066 | -0.066 | 0 | 0.22 | 0.07 | 0.3 | 0.32 | 0.26 | 0.28 |
| 7 | 10478374 | rs1524240 | -1.14 | -0.583 | -0.072 | -0.018 | -0.049 | 0.059 | 0.059 | 0 | 0.3 | 0.49 | 0.13 | 0.46 | 0.22 | 0.38 |
| 7 | 10478657 | rs17163018 | 0.158 | -0.342 | -0.562 | 0.342 | 0.008 | 0.001 | 0.001 | 0 | 0.13 | 0.08 | 0.06 | 0.43 | 0.1 | 0.27 |
| 7 | 10481065 | rs7797318 | -0.023 | -0.141 | -0.513 | 0.041 | 0.004 | -0.002 | -0.002 | 0.046 | 0.34 | 0.42 | 0.42 | 0.33 | 0.39 | 0.34 |
| 7 | 10491845 | rs6464009 | -1.136 | -0.087 | -0.921 | -0.11 | -0.001 | -0.017 | -0.017 | 0.236 | 0.75 | 0.88 | 0.79 | 0.66 | 0.78 | 0.8 |
| 7 | 10492168 | rs3801369 | 0.278 | -0.11 | -1.309 | -0.034 | -0.04 | -0.086 | -0.086 | 0 | 0.42 | 0.09 | 0.31 | 0.5 | 0.42 | 0.62 |
| 7 | 10494149 | rs2352114 | -0.897 | 0.135 | -0.864 | 0.589 | -0.004 | -0.025 | -0.025 | 0.013 | 0.75 | 0.91 | 0.79 | 0.66 | 0.78 | 0.8 |
| 7 | 10494987 | rs7796037 | -0.897 | 0.135 | -0.864 | 0.589 | -0.004 | -0.025 | -0.025 | 0 | 0.75 | 0.91 | 0.79 | 0.66 | 0.78 | 0.8 |
| 7 | 10516229 | rs17151708 | -0.069 | 0.318 | -0.865 | 1.049 | -0.01 | -0.013 | -0.013 | 0.256 | 0.09 | 0.02 | 0.07 | 0.38 | 0.11 | 0.25 |
| 7 | 10522050 | rs2222519 | -0.49 | 0.287 | -1.121 | 0.319 | -0.018 | -0.022 | -0.022 | 0 | 0.53 | 0.86 | 0.43 | 0.49 | 0.53 | 0.67 |
| 7 | 10524197 | rs6958833 | 0.818 | 0.156 | -1.229 | -0.43 | -0.037 | 0.046 | 0.046 | 3.166 | 0.21 | 0.1 | 0.2 | 0.11 | 0.22 | 0.31 |
| 7 | 10525322 | rs2352127 | 0.785 | 0.158 | -1.172 | -0.429 | -0.031 | 0.043 | 0.043 | 3.163 | 0.22 | 0.15 | 0.2 | 0.11 | 0.22 | 0.31 |
| 7 | 10525609 | rs2352128 | -1.481 | -0.081 | -1.609 | -0.72 | 0.007 | -0.021 | -0.021 | 0 | 0.75 | 0.9 | 0.78 | 0.68 | 0.75 | 0.89 |
| 7 | 10525926 | rs4725290 | 0.182 | -0.216 | -0.682 | -0.804 | -0.003 | 0.01 | 0.01 | 0 | 0.24 | 0.23 | 0.28 | 0.5 | 0.25 | 0.36 |
| 7 | 10554750 | rs1013719 | 1.471 | -0.187 | -0.915 | -0.847 | -0.04 | 0.106 | 0.106 | 0 | 0.23 | 0.08 | 0.24 | 0.05 | 0.23 | 0.19 |
| 7 | 10555821 | rs12111656 | 1.19 | -0.189 | -0.736 | -0.831 | -0.028 | 0.037 | 0.037 | 3.792 | 0.21 | 0.13 | 0.2 | 0.11 | 0.22 | 0.31 |
| 7 | 10558779 | rs1990103 | 0.279 | -0.322 | -0.288 | -0.766 | 0.009 | 0.029 | 0.029 | 0.107 | 0.5 | 0.39 | 0.45 | 0.45 | 0.52 | 0.44 |
| 7 | 10576502 | rs17163180 | 0.103 | -0.275 | -0.188 | -0.778 | 0 | 0.001 | 0.001 | 0.002 | 0.52 | 0.44 | 0.46 | 0.45 | 0.53 | 0.45 |
| 7 | 10580190 | rs13232957 | 1.38 | -0.293 | -0.223 | -0.8 | 0.011 | -0.012 | -0.012 | 0.114 | 0.26 | 0.12 | 0.23 | 0.34 | 0.25 | 0.17 |
| 7 | 10581402 | rs12533967 | 1.471 | -0.194 | -0.133 | -0.823 | -0.017 | 0.063 | 0.063 | 0 | 0.22 | 0.12 | 0.25 | 0.05 | 0.22 | 0.19 |
| 7 | 10583506 | rs17163213 | 0.69 | -0.179 | -0.107 | -0.752 | 0.019 | -0.041 | -0.041 | 0 | 0.24 | 0.32 | 0.22 | 0.11 | 0.27 | 0.31 |
| 7 | 10587973 | rs10486086 | -0.147 | -0.444 | -0.124 | -0.624 | -0.038 | 0.098 | 0.098 | 0.856 | 0.78 | 0.46 | 0.74 | 0.49 | 0.83 | 0.71 |
| 7 | 10593658 | rs17163246 | 0.237 | -0.416 | -0.209 | -0.704 | 0.006 | 0.006 | 0.006 | 0.135 | 0.56 | 0.39 | 0.51 | 0.45 | 0.6 | 0.52 |
| 7 | 10601877 | rs1016892 | -0.363 | -0.503 | -0.23 | -0.457 | -0.023 | 0.056 | 0.056 | 0 | 0.78 | 0.69 | 0.75 | 0.49 | 0.81 | 0.71 |

| | | | | | | | | | | | | | | | | |
|---|---|---|---|---|---|---|---|---|---|---|---|---|---|---|---|---|
| 7 | 10605795 | rs12666585 | -0.064 | -0.126 | -0.824 | -0.866 | 0.057 | 0.137 | 0.137 | 0.321 | 0.49 | 0.31 | 0.51 | 0.15 | 0.57 | 0.51 |
| 7 | 10609052 | rs17583347 | 0.582 | -0.003 | -1.131 | -0.751 | -0.061 | 0.112 | 0.112 | 0 | 0.23 | 0.03 | 0.25 | 0.09 | 0.26 | 0.23 |
| 7 | 10609528 | rs6947198 | -0.104 | -0.231 | -0.825 | -0.282 | -0.013 | -0.014 | -0.014 | 0 | 0.65 | 0.71 | 0.68 | 0.46 | 0.56 | 0.48 |
| 7 | 10614934 | rs6970987 | 1.123 | -0.625 | -0.665 | -0.435 | -0.007 | 0.004 | 0.004 | 0.004 | 0.23 | 0.19 | 0.3 | 0.27 | 0.17 | 0.15 |
| 7 | 10621699 | rs1012998 | -0.498 | -0.603 | -0.729 | -0.431 | -0.02 | -0.033 | -0.033 | 0 | 0.66 | 0.75 | 0.68 | 0.48 | 0.56 | 0.5 |
| 7 | 10622226 | rs7458352 | 1.055 | -0.71 | -0.741 | -0.258 | 0.022 | -0.032 | -0.032 | 0.17 | 0.22 | 0.07 | 0.29 | 0.27 | 0.16 | 0.15 |
| 7 | 10638180 | rs6962211 | -0.852 | -1.047 | -0.502 | -0.411 | 0.01 | 0.003 | 0.003 | 0 | 0.73 | 0.79 | 0.69 | 0.76 | 0.83 | 0.82 |
| 7 | 10642676 | rs7797298 | 1.482 | -0.852 | -0.876 | -0.788 | -0.079 | 0.11 | 0.11 | 0 | 0.22 | 0.04 | 0.25 | 0.04 | 0.3 | 0.18 |
| 7 | 10643492 | rs4339546 | 0.852 | -0.504 | -1.197 | -0.503 | -0.056 | -0.085 | -0.085 | 0 | 0.12 | 0.01 | 0.21 | 0.44 | 0.24 | 0.28 |
| 7 | 10645892 | rs7804426 | -1.333 | -0.69 | -0.928 | -0.608 | 0.026 | 0.008 | 0.008 | 0.002 | 0.73 | 0.84 | 0.69 | 0.77 | 0.83 | 0.82 |
| 7 | 10646407 | rs7776782 | 1.263 | -0.959 | -0.36 | -0.761 | -0.009 | 0.001 | 0.001 | 0 | 0.15 | 0.17 | 0.08 | 0.16 | 0.08 | 0.17 |
| 7 | 10649207 | rs10952224 | -0.591 | -0.868 | -0.144 | -0.933 | -0.001 | 0.008 | 0.008 | 0 | 0.47 | 0.56 | 0.41 | 0.7 | 0.5 | 0.62 |
| 7 | 10660424 | rs757819 | -1.331 | -0.894 | 0.019 | -0.907 | -0.014 | -0.004 | -0.004 | 0.469 | 0.73 | 0.72 | 0.68 | 0.76 | 0.83 | 0.82 |
| 7 | 10675914 | rs17163330 | 0.755 | -0.899 | 0.237 | -0.865 | 0 | -0.03 | -0.03 | 0 | 0.22 | 0.29 | 0.12 | 0.1 | 0.21 | 0.16 |
| 7 | 10678301 | rs6962705 | -0.918 | -0.912 | 0.103 | -0.905 | -0.001 | 0.014 | 0.014 | 0 | 0.51 | 0.56 | 0.42 | 0.71 | 0.53 | 0.66 |
| 7 | 10679384 | rs10263926 | 0.005 | -0.933 | 0.127 | -0.92 | 0.001 | -0.02 | -0.02 | 0 | 0.28 | 0.19 | 0.29 | 0.61 | 0.31 | 0.47 |
| 7 | 10680663 | rs766653 | 0.705 | -0.906 | 0.122 | -0.921 | 0.001 | -0.05 | -0.05 | 0 | 0.23 | 0.33 | 0.12 | 0.09 | 0.21 | 0.16 |
| 7 | 10683531 | rs7790096 | -0.659 | -0.78 | -0.14 | -0.946 | -0.001 | 0.015 | 0.015 | 0 | 0.51 | 0.55 | 0.42 | 0.71 | 0.52 | 0.63 |
| 7 | 10687242 | rs10228538 | -0.96 | -0.711 | -0.569 | -0.934 | -0.019 | 0.092 | 0.092 | 0 | 0.54 | 0.74 | 0.44 | 0.71 | 0.56 | 0.64 |
| 7 | 10698134 | rs11973940 | 1.49 | -0.934 | -0.82 | -0.429 | -0.01 | -0.011 | -0.011 | 1.154 | 0.16 | 0.17 | 0.12 | 0.25 | 0.2 | 0.16 |
| 7 | 10698353 | rs9655075 | -0.02 | -0.642 | -0.343 | -0.331 | -0.013 | 0.112 | 0.112 | 0 | 0.27 | 0.44 | 0.29 | 0.5 | 0.28 | 0.46 |
| 7 | 10701741 | rs2352452 | 0.079 | -0.651 | -0.295 | -0.338 | -0.008 | 0.126 | 0.126 | 0 | 0.26 | 0.4 | 0.3 | 0.5 | 0.28 | 0.44 |
| 7 | 10705605 | rs17163393 | 1.292 | -0.906 | -0.699 | -0.624 | -0.001 | 0.001 | 0.001 | 0 | 0.26 | 0.27 | 0.15 | 0.25 | 0.24 | 0.17 |
| 7 | 10713431 | rs7793387 | 1.874 | -0.854 | -0.69 | -0.478 | 0.004 | -0.005 | -0.005 | 0 | 0.14 | 0.07 | 0.12 | 0.13 | 0.13 | 0.16 |
| 7 | 10717127 | rs17586854 | -0.883 | -0.966 | -0.863 | -0.497 | -0.016 | 0.033 | 0.033 | 0.024 | 0.59 | 0.69 | 0.48 | 0.75 | 0.58 | 0.63 |
| 7 | 10720002 | rs2040754 | -0.857 | -0.993 | -0.875 | -0.511 | -0.056 | 0.083 | 0.083 | 0 | 0.59 | 0.84 | 0.49 | 0.75 | 0.58 | 0.63 |
| 7 | 10720330 | rs763397 | -1.831 | -0.865 | -1.1 | -0.615 | 0.003 | -0.043 | -0.043 | 0 | 0.83 | 0.93 | 0.74 | 0.79 | 0.87 | 0.83 |
| 7 | 10721320 | rs10258294 | -0.398 | -0.963 | -1.074 | -0.622 | -0.018 | 0.027 | 0.027 | 0 | 0.35 | 0.48 | 0.34 | 0.5 | 0.34 | 0.48 |
| 7 | 10721997 | rs2189437 | 1.634 | -0.905 | -1.206 | -0.582 | -0.045 | -0.062 | -0.062 | 0.038 | 0.1 | 0.01 | 0.2 | 0.36 | 0.19 | 0.28 |
| 7 | 10732220 | rs11773736 | 0.591 | -0.796 | -1.024 | -0.719 | -0.001 | -0.002 | -0.002 | 0.033 | 0.33 | 0.4 | 0.19 | 0.25 | 0.3 | 0.18 |
| 7 | 10747684 | rs2285914 | 0.115 | -0.81 | -1.025 | -0.708 | 0.013 | 0.001 | 0.001 | 0 | 0.5 | 0.52 | 0.54 | 0.55 | 0.58 | 0.64 |
| 7 | 10749891 | rs17587569 | 2.272 | -0.519 | -1.148 | -0.624 | -0.086 | 0.103 | 0.103 | 0 | 0.19 | 0.03 | 0.23 | 0.05 | 0.29 | 0.19 |
| 7 | 10749914 | rs17528240 | 1.14 | -0.358 | -0.699 | -0.126 | -0.01 | 0.003 | 0.003 | 0 | 0.25 | 0.21 | 0.14 | 0.25 | 0.24 | 0.16 |
| 7 | 10754626 | rs2189532 | -1.255 | -0.298 | -0.89 | -0.215 | 0.006 | -0.036 | -0.036 | 0.775 | 0.83 | 0.92 | 0.74 | 0.79 | 0.87 | 0.83 |
| 7 | 10781516 | rs10950313 | 0.91 | 0.013 | -0.761 | -0.307 | -0.002 | -0.031 | -0.031 | 0 | 0.27 | 0.25 | 0.17 | 0.39 | 0.25 | 0.15 |
| 7 | 10794857 | rs13224993 | 1.535 | 0.134 | -0.859 | -0.777 | -0.015 | 0.018 | 0.018 | 0 | 0.22 | 0.15 | 0.26 | 0.07 | 0.28 | 0.22 |
| 7 | 10795065 | rs7784127 | -0.003 | -0.085 | -0.58 | -1.071 | -0.012 | 0.011 | 0.011 | 0 | 0.38 | 0.44 | 0.47 | 0.54 | 0.36 | 0.53 |
| 7 | 10796892 | rs2214543 | 1.704 | -0.033 | -0.569 | -1.064 | -0.033 | 0.06 | 0.06 | 0 | 0.22 | 0.13 | 0.26 | 0.07 | 0.28 | 0.22 |
| 7 | 10804070 | rs4720903 | -1.989 | -0.05 | -0.468 | -0.999 | 0.006 | -0.042 | -0.042 | 0 | 0.92 | 0.88 | 0.93 | 1 | 0.92 | 0.91 |
| 7 | 10812405 | rs10259958 | -0.137 | -0.123 | -0.461 | -0.998 | -0.013 | 0.01 | 0.01 | 0 | 0.38 | 0.42 | 0.48 | 0.54 | 0.36 | 0.52 |
| 7 | 10817087 | rs7384091 | 1.705 | -0.128 | -0.548 | -0.9 | -0.054 | 0.096 | 0.096 | 0 | 0.21 | 0.06 | 0.25 | 0.07 | 0.28 | 0.22 |
| 7 | 10819155 | rs1971643 | -0.39 | -0.377 | -0.676 | -0.805 | -0.027 | 0.145 | 0.145 | 0 | 0.57 | 0.84 | 0.48 | 0.76 | 0.41 | 0.44 |
| 7 | 10819327 | rs1971642 | 0.518 | -0.405 | -0.219 | -0.72 | -0.016 | 0.023 | 0.023 | 0 | 0.26 | 0.29 | 0.29 | 0.37 | 0.16 | 0.29 |
| 7 | 10836089 | rs12673360 | 1.103 | -0.379 | -0.22 | -0.766 | 0.003 | -0.02 | -0.02 | 0.011 | 0.29 | 0.28 | 0.16 | 0.38 | 0.26 | 0.18 |
| 7 | 10836487 | rs979058 | -0.299 | -0.414 | -0.033 | -0.867 | 0 | -0.011 | -0.011 | 0 | 0.57 | 0.59 | 0.49 | 0.76 | 0.46 | 0.47 |
| 7 | 10845291 | rs12671402 | 1.78 | -0.176 | -0.243 | -0.933 | -0.036 | 0.058 | 0.058 | 0 | 0.2 | 0.1 | 0.32 | 0.07 | 0.26 | 0.17 |
| 7 | 10854493 | rs10486102 | 1.594 | -0.169 | -0.159 | -0.912 | -0.001 | -0.02 | -0.02 | 0 | 0.22 | 0.21 | 0.26 | 0.07 | 0.27 | 0.19 |
| 7 | 10855494 | rs4720910 | 0.617 | -0.7 | -0.171 | -0.763 | 0.02 | 0.017 | 0.017 | 0 | 0.5 | 0.4 | 0.42 | 0.45 | 0.52 | 0.35 |
| 7 | 10856098 | rs2057771 | 0.77 | -0.603 | -0.869 | -0.65 | 0.008 | -0.013 | -0.013 | 0 | 0.24 | 0.2 | 0.29 | 0.37 | 0.17 | 0.28 |
| 7 | 10857770 | rs17163658 | 1.946 | -0.682 | -1.029 | -0.584 | -0.006 | -0.023 | -0.023 | 0 | 0.14 | 0 | 0.1 | 0.13 | 0.1 | 0.15 |
| 7 | 10863492 | rs12531454 | 0.334 | -0.443 | -0.783 | -0.745 | 0.007 | 0.002 | 0.002 | 0 | 0.55 | 0.68 | 0.48 | 0.46 | 0.56 | 0.37 |
| 7 | 10875225 | rs17533461 | 0.807 | 0.813 | -0.746 | -0.479 | 0.06 | 0.101 | 0.101 | 0 | 0.2 | 0.34 | 0.17 | 0.33 | 0.27 | 0.17 |
| 7 | 10875894 | rs4720913 | -0.457 | 0.059 | -0.418 | -0.731 | 0.001 | 0.002 | 0.002 | 0.278 | 0.62 | 0.63 | 0.6 | 0.5 | 0.46 | 0.48 |
| 7 | 10879745 | rs17163678 | 0.444 | 0.231 | -1.39 | -0.59 | 0.027 | -0.016 | -0.016 | 0 | 0.37 | 0.3 | 0.41 | 0.43 | 0.24 | 0.29 |
| 7 | 10881827 | rs6973809 | 0.608 | 0.89 | -1.297 | -0.759 | -0.019 | 0.086 | 0.086 | 0 | 0.19 | 0.06 | 0.22 | 0.07 | 0.2 | 0.18 |
| 7 | 10884712 | rs12699156 | 0.499 | 0.981 | -0.9 | -0.674 | -0.002 | 0.015 | 0.015 | 0 | 0.55 | 0.55 | 0.5 | 0.8 | 0.49 | 0.64 |
| 7 | 10888816 | rs4520062 | 0.904 | 1.223 | -1.367 | -0.856 | -0.054 | 0.099 | 0.099 | 0 | 0.2 | 0.02 | 0.21 | 0.04 | 0.22 | 0.18 |
| 7 | 10889618 | rs968793 | -0.143 | 1.044 | -0.787 | -0.607 | 0.024 | -0.005 | -0.005 | 0 | 0.8 | 0.72 | 0.71 | 0.84 | 0.72 | 0.83 |
| 7 | 10892304 | rs1468493 | -0.155 | 1.089 | -0.431 | -0.877 | 0.116 | -0.06 | -0.06 | 1.329 | 0.87 | 0.63 | 0.87 | 0.96 | 0.77 | 0.91 |
| 7 | 10896020 | rs7787498 | -0.618 | 0.512 | -1.444 | -0.977 | -0.113 | 0.067 | 0.067 | 0 | 0.69 | 0.96 | 0.67 | 0.83 | 0.6 | 0.66 |
| 7 | 10896850 | rs10950321 | 1.073 | -0.02 | -2.089 | -0.96 | -0.029 | -0.098 | -0.098 | 0 | 0.23 | 0.06 | 0.25 | 0.59 | 0.24 | 0.33 |
| 7 | 10897225 | rs12375209 | 1.206 | -0.03 | -1.294 | -1.017 | 0 | -0.008 | -0.008 | 0.018 | 0.26 | 0.22 | 0.22 | 0.11 | 0.21 | 0.19 |
| 7 | 10902381 | rs6952827 | -0.275 | -0.089 | -1.028 | -1.006 | 0.017 | 0.058 | 0.058 | 0 | 0.43 | 0.72 | 0.45 | 0.71 | 0.38 | 0.47 |
| 7 | 10904752 | rs10239619 | -0.459 | -0.355 | -1.328 | -1.257 | 0.03 | 0.125 | 0.125 | 0 | 0.51 | 0.74 | 0.57 | 0.71 | 0.43 | 0.51 |
| 7 | 10904755 | rs10269406 | 0.003 | -0.011 | -0.597 | -1.243 | 0.01 | -0.029 | -0.029 | 0 | 0.19 | 0.3 | 0.22 | 0.12 | 0.2 | 0.16 |
| 7 | 10910808 | rs17163715 | 1.414 | -0.139 | -0.746 | -1.307 | 0 | -0.002 | -0.002 | 0.237 | 0.2 | 0.19 | 0.18 | 0.13 | 0.16 | 0.15 |
| 7 | 10912869 | rs10273132 | 1.235 | -0.133 | -0.514 | -1.261 | 0.043 | -0.041 | -0.041 | 0 | 0.1 | 0.26 | 0.1 | 0.04 | 0.14 | 0.08 |
| 7 | 10912949 | rs6460763 | 0.572 | -0.552 | -0.789 | -1.406 | -0.004 | -0.004 | -0.004 | 0 | 0.41 | 0.46 | 0.34 | 0.69 | 0.29 | 0.48 |
| 7 | 10915448 | rs4720914 | 1.152 | -0.187 | -0.829 | -1.278 | 0.009 | -0.01 | -0.01 | 0 | 0.08 | 0.03 | 0.12 | 0.08 | 0.06 | 0.09 |

| | | | | | | | | | | | | | | | | |
|---|---|---|---|---|---|---|---|---|---|---|---|---|---|---|---|---|
| 7 | 10920358 | rs6966010 | 0.404 | -0.218 | -0.535 | -1.481 | 0.001 | -0.037 | -0.037 | 0 | 0.59 | 0.58 | 0.51 | 0.81 | 0.44 | 0.61 |
| 7 | 10928142 | rs7357330 | -1.285 | 0.137 | -0.791 | -1.342 | -0.044 | 0.049 | 0.049 | 0 | 0.78 | 0.9 | 0.73 | 0.94 | 0.64 | 0.79 |
| 7 | 10930439 | rs10215108 | -0.334 | 0.109 | -0.48 | -1.348 | 0.004 | -0.009 | -0.009 | 0 | 0.71 | 0.75 | 0.6 | 0.85 | 0.58 | 0.71 |
| 7 | 10938739 | rs10226904 | 0.16 | -0.435 | -0.598 | -1.653 | 0.009 | 0.011 | 0.011 | 0 | 0.5 | 0.6 | 0.43 | 0.77 | 0.43 | 0.57 |
| 7 | 10944986 | rs12699162 | 1.229 | -0.15 | -0.788 | -1.446 | 0.003 | -0.015 | -0.015 | 0 | 0.21 | 0.17 | 0.27 | 0.07 | 0.36 | 0.2 |
| 7 | 10945193 | rs16876733 | 1.423 | -0.285 | -0.815 | -1.216 | 0 | -0.014 | -0.014 | 0 | 0.07 | 0.03 | 0.12 | 0.09 | 0.06 | 0.1 |
| 7 | 10950136 | rs7777849 | 0.21 | -0.602 | -1.044 | -1.401 | -0.03 | 0.089 | 0.089 | 0.096 | 0.4 | 0.22 | 0.35 | 0.16 | 0.43 | 0.22 |
| 7 | 10952613 | rs12699164 | -0.823 | -0.476 | -1.018 | -0.356 | -0.003 | -0.02 | -0.02 | 0 | 0.45 | 0.88 | 0.6 | 0.57 | 0.55 | 0.5 |
| 7 | 10952823 | rs13229903 | -0.871 | -0.517 | -1.044 | -0.361 | 0.052 | -0.03 | -0.03 | 0.015 | 0.55 | 0.9 | 0.65 | 0.57 | 0.61 | 0.51 |
| 7 | 10954219 | rs12699165 | 0.931 | 0.266 | -0.528 | 0.144 | 0.015 | -0.006 | -0.006 | 0 | 0.2 | 0.19 | 0.27 | 0.13 | 0.34 | 0.18 |
| 7 | 10954938 | rs10950329 | 0.306 | 0.245 | 0.062 | 0.243 | 0.027 | 0.151 | 0.151 | 0 | 0.12 | 0.24 | 0.17 | 0.31 | 0.14 | 0.22 |
| 7 | 10956540 | rs12536452 | 1.134 | 0.426 | 0.337 | 0.422 | -0.05 | 0.018 | 0.018 | 0.052 | 0.18 | 0.08 | 0.26 | 0.13 | 0.33 | 0.18 |
| 7 | 10956980 | rs6970457 | 0.539 | 0.417 | 0.362 | 0.449 | 0.04 | 0.229 | 0.229 | 0 | 0.09 | 0.24 | 0.15 | 0.31 | 0.11 | 0.19 |
| 7 | 10957622 | rs10486106 | 0.936 | 0.463 | 0.016 | 0.602 | 0.014 | -0.002 | -0.002 | 0 | 0.18 | 0.17 | 0.27 | 0.15 | 0.33 | 0.15 |
| 7 | 10957672 | rs17163764 | 0.331 | 0.142 | 0.185 | 0.367 | -0.004 | -0.003 | -0.003 | 0.173 | 0.13 | 0.14 | 0.15 | 0.13 | 0.09 | 0.11 |
| 7 | 10973362 | rs7789764 | -0.531 | 0.382 | -0.092 | 0.595 | -0.097 | -0.036 | -0.036 | 2.228 | 0.74 | 0.91 | 0.55 | 0.69 | 0.57 | 0.71 |
| 7 | 10975011 | rs218979 | -0.81 | -0.093 | 0.146 | 0.425 | -0.012 | 0.006 | 0.006 | 0 | 0.24 | 0.64 | 0.17 | 0.26 | 0.16 | 0.21 |
| 7 | 10975177 | rs218978 | -0.577 | -0.061 | 0.22 | 0.205 | -0.032 | -0.058 | -0.058 | 2.351 | 0.22 | 0.49 | 0.16 | 0.13 | 0.16 | 0.2 |
| 7 | 10979396 | rs1616965 | -0.629 | 0.141 | -0.59 | 0.173 | -0.132 | -0.055 | -0.055 | 0 | 0.74 | 0.94 | 0.65 | 0.69 | 0.57 | 0.7 |
| 7 | 10986944 | rs218970 | 1.406 | 0.028 | -0.608 | -0.457 | -0.059 | -0.017 | -0.017 | 0 | 0.13 | 0.02 | 0.25 | 0.1 | 0.25 | 0.27 |
| 7 | 10987822 | rs218968 | 1.232 | -0.029 | -0.05 | -0.12 | 0.003 | 0.242 | 0.242 | 0.345 | 0.47 | 0.13 | 0.36 | 0.15 | 0.43 | 0.47 |
| 7 | 10991660 | rs1074416 | 0.603 | 0.429 | 0.742 | 0.091 | 0 | -0.001 | -0.001 | 0 | 0.6 | 0.57 | 0.58 | 0.38 | 0.58 | 0.49 |
| 7 | 11012074 | rs6945372 | -1.519 | 0.504 | 0.802 | 0.233 | -0.001 | 0.004 | 0.004 | 0.496 | 0.23 | 0.7 | 0.29 | 0.35 | 0.21 | 0.08 |
| 7 | 11015444 | rs7807585 | -0.441 | 0.574 | 0.763 | 0.033 | -0.024 | -0.036 | -0.036 | 0 | 0.75 | 0.83 | 0.57 | 0.5 | 0.66 | 0.55 |
| 7 | 11015472 | rs10464254 | 0.472 | -0.181 | 1.504 | 0.275 | 0.102 | 0.203 | 0.203 | 0 | 0.06 | 0.25 | 0.19 | 0.23 | 0.1 | 0.02 |
| 7 | 11033756 | rs6950505 | -1.192 | -0.561 | 1.458 | 0.132 | -0.031 | -0.004 | -0.004 | 1.004 | 0.17 | 0.55 | 0.11 | 0.11 | 0.11 | 0.06 |
| 7 | 11036554 | rs6956108 | 0.131 | -0.484 | 1.887 | 0.199 | 0.005 | 0.116 | 0.116 | 0.421 | 0.07 | 0.28 | 0.03 | 0.1 | 0.03 | 0.02 |
| 7 | 11075263 | rs2353340 | -0.334 | -0.239 | 1.805 | -0.08 | 0.002 | -0.001 | -0.001 | 0.234 | 0.3 | 0.29 | 0.55 | 0.75 | 0.34 | 0.44 |
| 7 | 11091200 | rs17163925 | 0.467 | -0.294 | 1.9 | -0.106 | 0.015 | -0.024 | -0.024 | 0 | 0.05 | 0.11 | 0.06 | 0 | 0.07 | 0.04 |
| 7 | 11093709 | rs3801441 | -0.263 | -0.303 | 2.071 | -0.158 | 0 | 0 | 0 | 0 | 0.29 | 0.28 | 0.55 | 0.75 | 0.33 | 0.42 |
| 7 | 11113019 | rs7782589 | -0.274 | -0.285 | 1.956 | -0.153 | 0.002 | 0 | 0 | 0.05 | 0.29 | 0.3 | 0.55 | 0.75 | 0.33 | 0.43 |
| 7 | 11118333 | rs6971841 | -0.429 | -0.169 | 1.834 | -0.091 | 0.004 | -0.056 | -0.056 | 0.006 | 0.1 | 0.23 | 0.08 | 0 | 0.09 | 0.05 |
| 7 | 11140383 | rs7790568 | 0.074 | -0.593 | 1.634 | -0.346 | -0.036 | -0.102 | -0.102 | 0.548 | 0.2 | 0.32 | 0.09 | 0 | 0.11 | 0.04 |
| 7 | 11170396 | rs7790383 | -0.96 | -0.08 | 1.901 | 0.156 | 0.012 | -0.056 | -0.056 | 0 | 0.35 | 0.45 | 0.65 | 0.85 | 0.41 | 0.46 |
| 7 | 11171906 | rs4719258 | -1.052 | 0.003 | 1.699 | 0.193 | 0.001 | -0.012 | -0.012 | 0 | 0.34 | 0.39 | 0.65 | 0.85 | 0.39 | 0.43 |
| 7 | 11191734 | rs10499381 | 0.156 | -0.163 | 1.534 | 0.494 | -0.008 | -0.038 | -0.038 | 0 | 0.21 | 0.12 | 0.45 | 0.51 | 0.25 | 0.38 |
| 7 | 11203816 | rs10245045 | -1.164 | -0.079 | 1.177 | 0.419 | -0.006 | 0.002 | 0.002 | 0.032 | 0.35 | 0.67 | 0.55 | 0.62 | 0.36 | 0.37 |
| 7 | 11210291 | rs12668838 | -0.178 | -0.096 | 0.741 | 0.411 | -0.035 | 0.036 | 0.036 | 0 | 0.1 | 0.24 | 0.09 | 0.21 | 0.04 | 0.03 |
| 7 | 11210931 | rs12154393 | -0.947 | 0.332 | 0.775 | 0.454 | 0.013 | 0.027 | 0.027 | 0 | 0.33 | 0.37 | 0.56 | 0.52 | 0.4 | 0.41 |
| 7 | 11215293 | rs6968332 | -0.711 | 0.241 | 0.621 | 0.481 | 0.007 | 0.025 | 0.025 | 0 | 0.3 | 0.3 | 0.54 | 0.51 | 0.35 | 0.4 |
| 7 | 11216782 | rs10234187 | -0.603 | -0.007 | 0.497 | 0.695 | 0.004 | 0.049 | 0.049 | 0 | 0.28 | 0.3 | 0.5 | 0.51 | 0.32 | 0.37 |
| 7 | 11220846 | rs4720945 | -0.608 | 0.057 | 0.388 | 0.268 | -0.003 | 0 | 0 | 0.284 | 0.29 | 0.42 | 0.57 | 0.72 | 0.31 | 0.38 |
| 7 | 11228260 | rs10229042 | -0.615 | -0.173 | 0.556 | -0.208 | -0.028 | -0.136 | -0.136 | 1.245 | 0.19 | 0.45 | 0.08 | 0 | 0.13 | 0.09 |
| 7 | 11231727 | rs10440995 | -0.187 | -0.377 | -0.888 | -0.037 | 0.035 | 0.045 | 0.045 | 2.956 | 0.49 | 0.38 | 0.75 | 0.65 | 0.72 | 0.73 |
| 7 | 11232243 | rs4236261 | 0.09 | 0.443 | -1.285 | 0.579 | -0.027 | -0.007 | -0.007 | 0.564 | 0.18 | 0.08 | 0.38 | 0.2 | 0.28 | 0.27 |
| 7 | 11232753 | rs4720947 | -0.158 | 0.38 | -1.398 | 0.564 | -0.004 | 0.001 | 0.001 | 0.72 | 0.21 | 0.19 | 0.4 | 0.21 | 0.29 | 0.27 |
| 7 | 11233115 | rs6960105 | -0.389 | 0.458 | -2.114 | 0.506 | -0.016 | 0.007 | 0.007 | 0.835 | 0.49 | 0.87 | 0.6 | 0.65 | 0.5 | 0.45 |
| 7 | 11239345 | rs10226582 | -0.305 | 0.716 | -2.07 | 0.327 | -0.035 | -0.077 | -0.077 | 0 | 0.36 | 0.15 | 0.46 | 0.51 | 0.45 | 0.49 |
| 7 | 11240247 | rs2001817 | -0.854 | 0.63 | -2.635 | -0.22 | 0.011 | -0.056 | -0.056 | 0.286 | 0.59 | 0.89 | 0.69 | 0.6 | 0.68 | 0.71 |
| 7 | 11246692 | rs17164013 | 0.784 | 0.968 | -2.725 | -0.428 | -0.082 | -0.107 | -0.107 | 0 | 0.17 | 0.04 | 0.31 | 0.53 | 0.35 | 0.39 |
| 7 | 11248377 | rs6949976 | -0.798 | 0.808 | -1.412 | -0.749 | 0 | 0 | 0 | 0 | 0.46 | 0.44 | 0.43 | 0.55 | 0.54 | 0.63 |
| 7 | 11250828 | rs7794421 | -0.827 | 0.4 | -0.702 | -1.273 | -0.008 | -0.068 | -0.068 | 0 | 0.29 | 0.34 | 0.19 | 0.09 | 0.25 | 0.25 |
| 7 | 11252396 | rs17164021 | -0.535 | 0.587 | -0.394 | -1.216 | 0.01 | -0.029 | -0.029 | 0 | 0.18 | 0.2 | 0.15 | 0.07 | 0.2 | 0.18 |
| 7 | 11253503 | rs11983466 | 1.282 | -0.15 | -0.325 | -1.889 | -0.004 | -0.05 | -0.05 | 0 | 0.18 | 0.24 | 0.1 | 0 | 0.14 | 0.03 |
| 7 | 11253656 | rs6959531 | -0.124 | -0.096 | -0.264 | -1.809 | 0.018 | -0.036 | -0.036 | 0.026 | 0.18 | 0.3 | 0.16 | 0.09 | 0.18 | 0.18 |
| 7 | 11255948 | rs2525750 | 1.995 | -0.338 | -0.79 | -1.958 | -0.065 | 0.141 | 0.141 | 0 | 0.23 | 0.01 | 0.1 | 0.04 | 0.22 | 0.15 |
| 7 | 11258798 | rs10232432 | 1.183 | -0.6 | -0.426 | -1.806 | -0.061 | -0.044 | -0.044 | 2.931 | 0.19 | 0.29 | 0.1 | 0.05 | 0.08 | 0.11 |
| 7 | 11259361 | rs2714402 | 0.049 | -0.476 | -0.371 | -1.599 | -0.001 | -0.041 | -0.041 | 0 | 0.59 | 0.47 | 0.36 | 0.16 | 0.54 | 0.36 |
| 7 | 11262573 | rs2253825 | 0.662 | -0.404 | -0.251 | -1.642 | -0.019 | 0.066 | 0.066 | 0 | 0.41 | 0.23 | 0.24 | 0.1 | 0.41 | 0.34 |
| 7 | 11266778 | rs2189545 | 1.133 | -0.392 | -0.294 | -1.6 | -0.011 | 0.084 | 0.084 | 0 | 0.34 | 0.19 | 0.19 | 0.14 | 0.32 | 0.19 |
| 7 | 11267218 | rs2525744 | 0.46 | -0.471 | -0.158 | -1.429 | -0.001 | 0 | 0 | 3.389 | 0.44 | 0.4 | 0.29 | 0.21 | 0.44 | 0.36 |
| 7 | 11270607 | rs17164067 | 1.397 | -0.296 | -0.473 | -1.339 | -0.024 | -0.057 | -0.057 | 0.429 | 0.16 | 0.24 | 0.09 | 0 | 0.1 | 0.02 |
| 7 | 11272237 | rs2714411 | -0.3 | -0.177 | -0.359 | -1.207 | -0.012 | 0.028 | 0.028 | 0 | 0.64 | 0.69 | 0.38 | 0.18 | 0.53 | 0.46 |
| 7 | 11275420 | rs9791443 | -0.544 | 0.625 | -0.452 | -1.291 | 0.005 | 0.043 | 0.043 | 0 | 0.59 | 0.71 | 0.7 | 0.81 | 0.49 | 0.67 |
| 7 | 11276492 | rs12055949 | 0.235 | 0.11 | -0.384 | -1.379 | -0.019 | 0.079 | 0.079 | 0 | 0.44 | 0.28 | 0.27 | 0.1 | 0.44 | 0.34 |
| 7 | 11279164 | rs17164101 | 0.772 | 0.003 | -0.592 | -0.517 | 0.004 | -0.006 | -0.006 | 0 | 0.18 | 0.21 | 0.3 | 0.37 | 0.12 | 0.15 |
| 7 | 11285157 | rs10950342 | 0.329 | 0.131 | -0.486 | -0.359 | 0 | -0.008 | -0.008 | 0 | 0.26 | 0.26 | 0.32 | 0.48 | 0.25 | 0.17 |
| 7 | 11288583 | rs4236262 | -1.225 | 0.528 | -0.767 | -0.597 | -0.08 | -0.096 | -0.096 | 0.155 | 0.83 | 0.97 | 0.62 | 0.55 | 0.67 | 0.58 |
| 7 | 11289766 | rs17164144 | 0.992 | 0.433 | -0.709 | -0.758 | 0.005 | -0.016 | -0.016 | 0 | 0.09 | 0.12 | 0.07 | 0.02 | 0.09 | 0.13 |

| | | | | | | | | | | | | | | | | |
|---|---|---|---|---|---|---|---|---|---|---|---|---|---|---|---|---|
| 7 | 11290187 | rs17539729 | 1.193 | -0.224 | -1.015 | -0.895 | 0.063 | 0.072 | 0.072 | 0 | 0.21 | 0.07 | 0.1 | 0.01 | 0.13 | 0.11 |
| 7 | 11291400 | rs4720952 | -0.222 | 0.112 | -0.765 | -0.714 | 0.06 | 0.059 | 0.059 | 0.121 | 0.73 | 0.65 | 0.54 | 0.55 | 0.57 | 0.44 |
| 7 | 11294041 | rs10486113 | 0.62 | 0.123 | -1.57 | -0.847 | -0.001 | 0.006 | 0.006 | 0 | 0.23 | 0.22 | 0.19 | 0.18 | 0.25 | 0.07 |
| 7 | 11295119 | rs7812248 | 0.466 | 0.697 | -1.368 | -0.578 | 0.045 | 0.084 | 0.084 | 0 | 0.42 | 0.31 | 0.26 | 0.14 | 0.34 | 0.16 |
| 7 | 11295501 | rs10950344 | -0.741 | 1.407 | -1.973 | 0.175 | -0.049 | -0.054 | -0.054 | 0 | 0.87 | 0.97 | 0.71 | 0.51 | 0.75 | 0.61 |
| 7 | 11297388 | rs17540362 | 0.262 | 0.031 | -1.503 | -0.52 | -0.007 | 0.104 | 0.104 | 0 | 0.23 | 0.08 | 0.11 | 0.07 | 0.2 | 0.17 |
| 7 | 11298629 | rs10281406 | -0.31 | -0.329 | -0.885 | 0.02 | -0.008 | -0.024 | -0.024 | 0 | 0.1 | 0.12 | 0.13 | 0.01 | 0.07 | 0.1 |
| 7 | 11298722 | rs10281444 | -0.846 | -1.146 | -1.124 | 0.383 | 0.053 | 0.006 | 0.006 | 0.452 | 0.41 | 0.78 | 0.48 | 0.5 | 0.27 | 0.38 |
| 7 | 11298815 | rs11766703 | -0.083 | -0.23 | -0.708 | 0.646 | -0.018 | -0.005 | -0.005 | 0 | 0.14 | 0.26 | 0.09 | 0.17 | 0.07 | 0.11 |
| 7 | 11300347 | rs6460798 | 0.283 | 1.067 | -1.222 | 0.069 | -0.026 | 0.029 | 0.029 | 0.017 | 0.28 | 0.26 | 0.25 | 0.21 | 0.36 | 0.22 |
| 7 | 11301682 | rs12532526 | 0.32 | 1.248 | -1.355 | 0.099 | -0.02 | 0.004 | 0.004 | 0 | 0.07 | 0.03 | 0.16 | 0.06 | 0.08 | 0.11 |
| 7 | 11301960 | rs7811390 | -0.029 | 0.73 | -1.845 | -0.076 | 0.023 | -0.015 | -0.015 | 0 | 0.83 | 0.94 | 0.83 | 0.8 | 0.87 | 0.66 |
| 7 | 11304883 | rs7810741 | -0.598 | 0.87 | -1.4 | 0.028 | 0.004 | -0.018 | -0.018 | 0 | 0.89 | 0.95 | 0.84 | 0.77 | 0.9 | 0.68 |
| 7 | 11305971 | rs6946234 | 0.391 | 0.552 | -0.767 | -0.761 | -0.013 | -0.009 | -0.009 | 0 | 0.37 | 0.57 | 0.31 | 0.29 | 0.3 | 0.26 |
| 7 | 11306897 | rs7784717 | -0.486 | 0.58 | -0.71 | -0.754 | -0.009 | -0.04 | -0.04 | 0.24 | 0.77 | 0.89 | 0.56 | 0.39 | 0.76 | 0.52 |
| 7 | 11313841 | rs17541282 | 0.903 | 0.377 | -0.631 | -1.089 | -0.003 | -0.032 | -0.032 | 0.954 | 0.13 | 0.11 | 0.08 | 0 | 0.1 | 0.03 |
| 7 | 11314860 | rs10254054 | 0.833 | 0.304 | -0.102 | -1.385 | 0.016 | -0.062 | -0.062 | 0 | 0.34 | 0.6 | 0.29 | 0.15 | 0.41 | 0.2 |
| 7 | 11315293 | rs17613557 | 0.911 | 0.328 | -0.276 | -1.377 | -0.002 | -0.033 | -0.033 | 2.442 | 0.13 | 0.12 | 0.09 | 0 | 0.11 | 0.03 |
| 7 | 11316190 | rs17164201 | 0.847 | 0.511 | -0.391 | -1.381 | -0.012 | 0.015 | 0.015 | 0 | 0.07 | 0.07 | 0.08 | 0.02 | 0.1 | 0.06 |
| 7 | 11316581 | rs11765838 | 1.874 | 0.534 | -0.43 | -1.36 | -0.096 | 0.151 | 0.151 | 0 | 0.23 | 0.02 | 0.12 | 0.01 | 0.28 | 0.12 |
| 7 | 11321893 | rs10253872 | -0.094 | 0.504 | -0.169 | -1.26 | 0.009 | -0.011 | -0.011 | 0 | 0.69 | 0.79 | 0.54 | 0.39 | 0.73 | 0.48 |
| 7 | 11326981 | rs9771478 | -0.201 | 0.142 | -0.155 | -0.997 | -0.01 | 0.089 | 0.089 | 0.893 | 0.11 | 0.13 | 0.17 | 0.36 | 0.08 | 0.12 |
| 7 | 11328797 | rs10486122 | 1.513 | 0.211 | 0.099 | -1.073 | -0.035 | 0.033 | 0.033 | 0 | 0.27 | 0.21 | 0.27 | 0.15 | 0.41 | 0.19 |
| 7 | 11329134 | rs2214547 | -0.282 | 0.134 | 0.115 | -1.007 | -0.004 | 0.056 | 0.056 | 1.061 | 0.11 | 0.11 | 0.17 | 0.36 | 0.08 | 0.12 |
| 7 | 11334427 | rs11769360 | 0.912 | 0.11 | 0.37 | -1.189 | -0.002 | -0.002 | -0.002 | 0.024 | 0.27 | 0.26 | 0.19 | 0.25 | 0.24 | 0.26 |
| 7 | 11335132 | rs6978700 | 1.478 | 0.275 | 0.081 | -1.091 | -0.026 | 0.024 | 0.024 | 0 | 0.26 | 0.24 | 0.27 | 0.15 | 0.4 | 0.19 |
| 7 | 11343074 | rs2189349 | 0.584 | -0.015 | 0.143 | -1.157 | -0.001 | -0.001 | -0.001 | 0 | 0.4 | 0.46 | 0.39 | 0.44 | 0.41 | 0.29 |
| 7 | 11349162 | rs13237362 | -0.336 | 0.248 | 0.394 | -1.025 | 0.008 | 0.066 | 0.066 | 0 | 0.48 | 0.67 | 0.42 | 0.38 | 0.46 | 0.37 |
| 7 | 11349410 | rs10260325 | 1.156 | 0.259 | 0.364 | -1.032 | -0.024 | -0.042 | -0.042 | 0 | 0.31 | 0.25 | 0.29 | 0.37 | 0.37 | 0.24 |
| 7 | 11349438 | rs4454185 | -0.539 | 0.249 | 0.509 | -1.028 | -0.011 | 0.007 | 0.007 | 0.025 | 0.77 | 0.7 | 0.64 | 0.68 | 0.82 | 0.55 |
| 7 | 11351104 | rs5021557 | 1.118 | 0.31 | 0.31 | -1.005 | -0.026 | -0.053 | -0.053 | 0.513 | 0.3 | 0.2 | 0.28 | 0.38 | 0.35 | 0.23 |
| 7 | 11354086 | rs11762253 | 0.108 | 0.142 | 0.238 | -1.061 | 0.046 | 0.1 | 0.1 | 0 | 0.43 | 0.18 | 0.26 | 0.21 | 0.34 | 0.31 |
| 7 | 11357010 | rs10252651 | -0.413 | 0.192 | 0.315 | -1.042 | 0.001 | 0.001 | 0.001 | 0.581 | 0.76 | 0.78 | 0.63 | 0.65 | 0.71 | 0.58 |
| 7 | 11358235 | rs718392 | -1.176 | 0.26 | 0.118 | -1.031 | -0.05 | -0.047 | -0.047 | 0 | 0.81 | 0.92 | 0.65 | 0.66 | 0.72 | 0.61 |
| 7 | 11358787 | rs17621503 | 0.658 | 0.141 | 0.168 | -1.089 | 0.006 | 0.007 | 0.007 | 0 | 0.31 | 0.23 | 0.16 | 0.18 | 0.21 | 0.24 |
| 7 | 11369405 | rs916906 | 0.264 | 0.394 | 0.057 | -1.004 | 0.002 | -0.006 | -0.006 | 1.15 | 0.63 | 0.6 | 0.53 | 0.64 | 0.59 | 0.53 |
| 7 | 11369482 | rs916908 | 0.128 | 0.433 | 0.058 | -1.009 | -0.004 | -0.013 | -0.013 | 0.002 | 0.64 | 0.62 | 0.6 | 0.74 | 0.67 | 0.54 |
| 7 | 11370712 | rs4719266 | -1.579 | 0.318 | 0.048 | -0.996 | -0.004 | -0.024 | -0.024 | 0 | 0.83 | 0.93 | 0.73 | 0.76 | 0.84 | 0.63 |
| 7 | 11370880 | rs4719267 | -1.536 | 0.356 | 0.058 | -0.979 | -0.001 | -0.019 | -0.019 | 0 | 0.83 | 0.91 | 0.73 | 0.76 | 0.84 | 0.63 |
| 7 | 11371531 | rs10233772 | 0.379 | 0.741 | 0.273 | -0.842 | 0.006 | 0.011 | 0.011 | 0 | 0.17 | 0.12 | 0.12 | 0.02 | 0.15 | 0.09 |
| 7 | 11371740 | rs11762496 | 0.409 | 0.66 | 0.294 | -0.828 | 0.01 | 0.031 | 0.031 | 0 | 0.35 | 0.44 | 0.37 | 0.46 | 0.38 | 0.31 |
| 7 | 11381458 | rs16876820 | -0.395 | 0.725 | 0.364 | -1.238 | 0 | -0.097 | -0.097 | 4.043 | 0.19 | 0.32 | 0.16 | 0.02 | 0.18 | 0.1 |
| 7 | 11383722 | rs10261632 | 0.482 | 0.507 | 0.031 | -1.142 | -0.002 | -0.001 | -0.001 | 0.008 | 0.36 | 0.39 | 0.4 | 0.64 | 0.39 | 0.37 |
| 7 | 11384060 | rs10262124 | 0.817 | 0.52 | 0.116 | -0.48 | 0.008 | 0.01 | 0.01 | 0.333 | 0.65 | 0.87 | 0.63 | 0.71 | 0.73 | 0.56 |
| 7 | 11384111 | rs10262369 | -0.928 | 0.274 | 0.81 | 0.098 | 0.074 | 0.002 | 0.002 | 0 | 0.11 | 0.39 | 0.15 | 0.12 | 0.16 | 0.08 |
| 7 | 11384188 | rs6976538 | -0.312 | 0.584 | -0.588 | 0.273 | 0.029 | -0.049 | -0.049 | 0.048 | 0.81 | 0.92 | 0.74 | 0.71 | 0.88 | 0.61 |
| 7 | 11386860 | rs7796596 | 0.744 | 0.706 | -0.343 | 0.398 | -0.005 | 0 | 0 | 0 | 0.64 | 0.52 | 0.55 | 0.59 | 0.67 | 0.52 |
| 7 | 11387291 | rs7797181 | 0.153 | 0.28 | -1.456 | 0.586 | 0.014 | -0.037 | -0.037 | 0 | 0.74 | 0.9 | 0.69 | 0.7 | 0.81 | 0.6 |
| 7 | 11393134 | rs6958567 | -0.434 | 0.287 | -0.79 | 0.491 | -0.018 | -0.011 | -0.011 | 0 | 0.33 | 0.55 | 0.35 | 0.27 | 0.27 | 0.2 |
| 7 | 11393694 | rs1076224 | -0.911 | 1.163 | -0.85 | 0.08 | -0.016 | 0.033 | 0.033 | 0.205 | 0.33 | 0.76 | 0.39 | 0.15 | 0.38 | 0.17 |
| 7 | 11394610 | rs739773 | -0.282 | 1.335 | 0.305 | -0.156 | 0.063 | -0.099 | -0.099 | 0.007 | 0.15 | 0.33 | 0.11 | 0 | 0.2 | 0.07 |
| 7 | 11394751 | rs739774 | -1.72 | 1.418 | 0.337 | 0.051 | 0.006 | -0.018 | -0.018 | 0 | 0.3 | 0.62 | 0.3 | 0.17 | 0.38 | 0.13 |
| 7 | 11395720 | rs7797993 | -2.089 | 1.167 | 0.179 | 0.1 | -0.016 | 0.042 | 0.042 | 1.076 | 0.36 | 0.77 | 0.45 | 0.3 | 0.39 | 0.19 |
| 7 | 11404383 | rs17623238 | 1.324 | 0.59 | -0.663 | 0.162 | -0.033 | 0.09 | 0.09 | 0 | 0.4 | 0.14 | 0.28 | 0.25 | 0.4 | 0.36 |
| 7 | 11412531 | rs7777246 | -1.101 | 0.672 | 0.274 | 0.109 | 0.024 | -0.036 | -0.036 | 0 | 0.26 | 0.37 | 0.26 | 0.16 | 0.35 | 0.11 |
| 7 | 11418670 | rs2189352 | -0.776 | 0.694 | 0.394 | 0.009 | 0.038 | 0.007 | 0.007 | 1.606 | 0.2 | 0.59 | 0.33 | 0.23 | 0.24 | 0.11 |
| 7 | 11418931 | rs12670599 | -1.837 | 0.662 | 0.698 | 0.008 | 0.015 | -0.018 | -0.018 | 0.661 | 0.29 | 0.61 | 0.4 | 0.25 | 0.36 | 0.13 |
| 7 | 11420981 | rs7810604 | -1.832 | 0.613 | 0.701 | -0.002 | 0.019 | -0.018 | -0.018 | 0 | 0.29 | 0.59 | 0.41 | 0.25 | 0.37 | 0.13 |
| 7 | 11446902 | rs6956916 | -1.592 | 0.688 | 0.806 | 0.044 | 0.027 | -0.019 | -0.019 | 0 | 0.28 | 0.52 | 0.41 | 0.25 | 0.36 | 0.15 |
| 7 | 11450446 | rs17164548 | -0.103 | 0.647 | 0.771 | 0.053 | -0.001 | -0.011 | -0.011 | 0.381 | 0.07 | 0.09 | 0.08 | 0.03 | 0.1 | 0.03 |
| 7 | 11473561 | rs7782229 | -0.329 | 0.588 | 0.83 | 0.333 | 0.003 | -0.007 | -0.007 | 0 | 0.66 | 0.71 | 0.66 | 0.57 | 0.71 | 0.46 |
| 7 | 11485527 | rs10262052 | 0.414 | 0.802 | 0.817 | 0.325 | -0.005 | -0.003 | -0.003 | 0 | 0.57 | 0.64 | 0.53 | 0.54 | 0.52 | 0.38 |
| 7 | 11490005 | rs40244 | 0.512 | 1.249 | 0.901 | 0.273 | 0.002 | -0.003 | -0.003 | 0 | 0.8 | 0.78 | 0.78 | 0.86 | 0.71 | 0.88 |
| 7 | 11493935 | rs2107140 | 1.325 | 1.207 | 0.863 | 0.271 | -0.013 | -0.006 | -0.006 | 0 | 0.42 | 0.21 | 0.41 | 0.43 | 0.42 | 0.35 |
| 7 | 11494566 | rs39182 | 0.354 | 1.211 | 0.683 | 0.277 | 0.001 | -0.003 | -0.003 | 0 | 0.33 | 0.31 | 0.35 | 0.4 | 0.28 | 0.48 |
| 7 | 11496226 | rs17164591 | 1.45 | 1.566 | 0.622 | 0.307 | -0.037 | -0.018 | -0.018 | 0 | 0.4 | 0.09 | 0.24 | 0.38 | 0.38 | 0.34 |
| 7 | 11497040 | rs39183 | 0.563 | 1.316 | 0.634 | 0.284 | -0.005 | 0.008 | 0.008 | 0 | 0.31 | 0.31 | 0.3 | 0.38 | 0.23 | 0.44 |
| 7 | 11498778 | rs7792678 | -0.183 | 1.364 | 0.83 | 0.247 | -0.005 | 0.007 | 0.007 | 2.61 | 0.16 | 0.48 | 0.14 | 0.13 | 0.12 | 0.04 |
| 7 | 11501528 | rs1014184 | -0.109 | 1.78 | 0.382 | 0.083 | -0.001 | -0.017 | -0.017 | 1.657 | 0.08 | 0.07 | 0.08 | 0.16 | 0.1 | 0.11 |

| | | | | | | | | | | | | | | | | |
|---|---|---|---|---|---|---|---|---|---|---|---|---|---|---|---|---|
| 7 | 11503182 | rs12699192 | 0.929 | 1.674 | 0.197 | -0.078 | -0.11 | 0.146 | 0.146 | 0 | 0.42 | 0.07 | 0.28 | 0.26 | 0.46 | 0.34 |
| 7 | 11503363 | rs39186 | -0.359 | 1.714 | 0.347 | -0.108 | 0 | -0.008 | -0.008 | 0 | 0.45 | 0.53 | 0.61 | 0.62 | 0.47 | 0.62 |
| 7 | 11503849 | rs12538725 | 1.726 | 1.875 | 0.269 | -0.139 | -0.03 | 0.08 | 0.08 | 0 | 0.34 | 0.05 | 0.18 | 0.23 | 0.3 | 0.28 |
| 7 | 11504580 | rs39190 | 0.44 | 1.713 | 0.578 | 0.188 | -0.001 | -0.001 | -0.001 | 0 | 0.35 | 0.35 | 0.51 | 0.45 | 0.37 | 0.52 |
| 7 | 11505200 | rs11979605 | 0.545 | 1.219 | 0.974 | 0.162 | -0.018 | 0.012 | 0.012 | 0.094 | 0.1 | 0.18 | 0.1 | 0.11 | 0.07 | 0.03 |
| 7 | 11505293 | rs11983224 | -0.575 | 1.264 | 0.839 | 0.147 | -0.03 | 0.01 | 0.01 | 0.501 | 0.15 | 0.45 | 0.15 | 0.13 | 0.12 | 0.04 |
| 7 | 11506358 | rs39757 | -0.2 | 1.152 | 0.359 | -0.018 | -0.005 | -0.006 | -0.006 | 0 | 0.16 | 0.23 | 0.09 | 0.1 | 0.14 | 0.07 |
| 7 | 11507986 | rs17555491 | 0.782 | 1.07 | -0.296 | -0.059 | 0.007 | 0.162 | 0.162 | 0 | 0.49 | 0.21 | 0.25 | 0.2 | 0.42 | 0.46 |
| 7 | 11510053 | rs10252863 | 0.223 | 0.909 | 0.043 | 0.462 | 0.003 | 0.011 | 0.011 | 0 | 0.24 | 0.27 | 0.54 | 0.39 | 0.35 | 0.38 |
| 7 | 11512299 | rs1029466 | 0.166 | 0.935 | 0.377 | 0.323 | 0.016 | -0.025 | -0.025 | 3.021 | 0.79 | 0.68 | 0.84 | 0.9 | 0.76 | 0.89 |
| 7 | 11514865 | rs17164640 | -0.942 | 1.001 | 0.381 | 0.136 | 0.013 | -0.03 | -0.03 | 0 | 0.16 | 0.36 | 0.15 | 0.13 | 0.13 | 0.14 |
| 7 | 11518918 | rs4720989 | -0.082 | 0.833 | 0.003 | 0.274 | -0.002 | -0.012 | -0.012 | 0 | 0.74 | 0.77 | 0.81 | 0.69 | 0.74 | 0.84 |
| 7 | 11521542 | rs3735502 | 1.676 | 0.526 | -0.529 | -0.123 | -0.019 | 0.03 | 0.03 | 0 | 0.27 | 0.12 | 0.18 | 0.18 | 0.25 | 0.21 |
| 7 | 11522119 | rs2107141 | -0.67 | 0.704 | -0.383 | -0.145 | -0.002 | -0.003 | -0.003 | 0.187 | 0.73 | 0.73 | 0.78 | 0.69 | 0.71 | 0.83 |
| 7 | 11522155 | rs4720990 | 0.362 | 0.6 | -0.171 | -0.208 | 0 | -0.005 | -0.005 | 0 | 0.35 | 0.32 | 0.52 | 0.39 | 0.39 | 0.5 |
| 7 | 11522422 | rs17164653 | 0.827 | 0.962 | -0.012 | -0.027 | -0.019 | -0.015 | -0.015 | 0 | 0.35 | 0.42 | 0.17 | 0.3 | 0.27 | 0.31 |
| 7 | 11523432 | rs6971882 | 0.015 | 0.809 | -0.189 | -0.17 | -0.008 | 0.011 | 0.011 | 0 | 0.43 | 0.32 | 0.61 | 0.39 | 0.45 | 0.54 |
| 7 | 11524390 | rs10486134 | 2.187 | 0.844 | -0.134 | -0.092 | 0.015 | 0.016 | 0.016 | 0 | 0.22 | 0.1 | 0.09 | 0.15 | 0.17 | 0.17 |
| 7 | 11525339 | rs13240495 | 0.049 | 1.148 | -0.114 | 0.047 | 0 | 0.005 | 0.005 | 0 | 0.4 | 0.3 | 0.54 | 0.38 | 0.42 | 0.51 |
| 7 | 11528500 | rs10486135 | 0.458 | 1.036 | -0.03 | -0.03 | 0.002 | 0.002 | 0.002 | 0 | 0.51 | 0.48 | 0.33 | 0.31 | 0.39 | 0.47 |
| 7 | 11529374 | rs10232236 | -1.006 | 1.12 | 0.008 | 0.019 | -0.005 | -0.008 | -0.008 | 0 | 0.8 | 0.79 | 0.83 | 0.66 | 0.75 | 0.86 |
| 7 | 11532297 | rs10950351 | 0.455 | 0.788 | 0.027 | -0.011 | 0 | -0.002 | -0.002 | 0.896 | 0.5 | 0.48 | 0.32 | 0.31 | 0.38 | 0.45 |
| 7 | 11534319 | rs2189311 | 0.468 | 0.229 | -0.165 | 0.134 | -0.001 | -0.001 | -0.001 | 0 | 0.67 | 0.67 | 0.77 | 0.63 | 0.68 | 0.72 |
| 7 | 11535903 | rs10281072 | 0.224 | -0.458 | -0.358 | -0.038 | -0.029 | -0.055 | -0.055 | 0 | 0.15 | 0.2 | 0.06 | 0.04 | 0.06 | 0.15 |
| 7 | 11544610 | rs6943841 | 0.647 | 0.274 | -0.754 | 0.366 | 0.04 | 0.046 | 0.046 | 0 | 0.45 | 0.31 | 0.19 | 0.3 | 0.31 | 0.45 |
| 7 | 11545990 | rs11764433 | -0.498 | 0.913 | -0.737 | 0.873 | 0.001 | 0.02 | 0.02 | 0 | 0.22 | 0.31 | 0.2 | 0.35 | 0.24 | 0.15 |
| 7 | 11548082 | rs10250329 | 0.467 | 1.213 | -1.411 | 0.774 | -0.049 | -0.015 | -0.015 | 5.465 | 0.67 | 0.82 | 0.62 | 0.65 | 0.46 | 0.66 |
| 7 | 11548636 | rs6460818 | -0.058 | 0.58 | -1.275 | 0.621 | -0.002 | 0.019 | 0.019 | 0 | 0.51 | 0.57 | 0.44 | 0.38 | 0.48 | 0.42 |
| 7 | 11554096 | rs6972615 | -0.028 | 0.057 | -1.48 | 0.407 | 0.022 | 0.014 | 0.014 | 0 | 0.32 | 0.18 | 0.21 | 0.24 | 0.24 | 0.19 |
| 7 | 11555918 | rs10486138 | 0.769 | 0.028 | -0.767 | 0.455 | 0.009 | 0.027 | 0.027 | 0 | 0.09 | 0.22 | 0.11 | 0.11 | 0.09 | 0.07 |
| 7 | 11557685 | rs6460819 | -0.431 | 0.33 | -0.77 | 1.007 | 0 | 0.036 | 0.036 | 0 | 0.9 | 0.9 | 0.7 | 0.58 | 0.88 | 0.77 |
| 7 | 11559050 | rs2189320 | 0.164 | 0.178 | -0.72 | 0.527 | -0.017 | -0.021 | -0.021 | 0 | 0.18 | 0.29 | 0.15 | 0.11 | 0.13 | 0.09 |
| 7 | 11560362 | rs1023525 | 1.084 | 0.184 | -0.756 | 0.522 | -0.002 | -0.016 | -0.016 | 0.063 | 0.23 | 0.12 | 0.18 | 0.23 | 0.2 | 0.17 |
| 7 | 11560777 | rs10215244 | -0.114 | 0.185 | -0.38 | 0.427 | -0.017 | -0.021 | -0.021 | 0.336 | 0.18 | 0.29 | 0.15 | 0.11 | 0.13 | 0.09 |
| 7 | 11563097 | rs11982706 | -0.74 | 0.247 | -0.286 | 0.315 | -0.003 | -0.062 | -0.062 | 1.422 | 0.24 | 0.44 | 0.2 | 0.11 | 0.21 | 0.17 |
| 7 | 11568530 | rs12673692 | 0.661 | 0.238 | -0.369 | 0.592 | -0.003 | -0.002 | -0.002 | 0 | 0.37 | 0.41 | 0.24 | 0.36 | 0.29 | 0.38 |
| 7 | 11568650 | rs2354950 | 0.064 | 0.313 | -0.428 | 0.443 | -0.009 | -0.021 | -0.021 | 0 | 0.36 | 0.27 | 0.31 | 0.46 | 0.43 | 0.44 |
| 7 | 11569463 | rs12699202 | 0.364 | 0.21 | -0.557 | 0.686 | -0.103 | -0.058 | -0.058 | 2.403 | 0.3 | 0.04 | 0.2 | 0.25 | 0.37 | 0.3 |
| 7 | 11569734 | rs6963353 | 0.262 | 0.253 | -0.442 | 0.629 | 0.015 | -0.043 | -0.043 | 0 | 0.66 | 0.56 | 0.51 | 0.8 | 0.64 | 0.75 |
| 7 | 11569853 | rs12674351 | 1.695 | 0.241 | -0.424 | 0.641 | -0.006 | -0.001 | -0.001 | 0 | 0.32 | 0.35 | 0.2 | 0.35 | 0.23 | 0.32 |
| 7 | 11569988 | rs11766825 | -0.41 | 0.219 | -0.381 | 0.609 | 0.103 | 0.019 | 0.019 | 0 | 0.09 | 0.37 | 0.09 | 0.03 | 0.1 | 0.07 |
| 7 | 11570049 | rs6964213 | 1.337 | 0.168 | -0.371 | 0.629 | -0.034 | 0.039 | 0.039 | 0 | 0.36 | 0.51 | 0.31 | 0.54 | 0.27 | 0.45 |
| 7 | 11571680 | rs2023864 | -0.768 | 0.278 | -0.443 | 0.608 | -0.059 | 0.038 | 0.038 | 0.909 | 0.75 | 0.93 | 0.58 | 0.82 | 0.74 | 0.83 |
| 7 | 11574741 | rs1548504 | 1.529 | 0.344 | -0.377 | 0.532 | -0.005 | -0.001 | -0.001 | 0 | 0.33 | 0.38 | 0.21 | 0.35 | 0.24 | 0.32 |
| 7 | 11577625 | rs53 | 1.118 | 0.467 | -0.558 | 0.548 | -0.069 | -0.004 | -0.004 | 0 | 0.2 | 0.01 | 0.38 | 0.18 | 0.24 | 0.14 |
| 7 | 11578234 | rs6979125 | 1.729 | 0.408 | -0.421 | 0.63 | 0.001 | 0.001 | 0.001 | 0 | 0.33 | 0.32 | 0.17 | 0.32 | 0.22 | 0.32 |
| 7 | 11578386 | rs51 | -0.144 | 0.456 | -0.293 | 0.662 | 0.001 | 0.002 | 0.002 | 0 | 0.66 | 0.64 | 0.76 | 0.7 | 0.61 | 0.68 |
| 7 | 11579793 | rs10242744 | -1.336 | 0.477 | -0.176 | 0.52 | 0 | 0.065 | 0.065 | 0 | 0.18 | 0.52 | 0.21 | 0.22 | 0.16 | 0.22 |
| 7 | 11581003 | rs2072139 | 1.427 | 1.134 | -0.261 | 0.43 | -0.001 | -0.001 | -0.001 | 1.075 | 0.13 | 0.14 | 0.17 | 0.18 | 0.17 | 0.16 |
| 7 | 11581121 | rs47 | -0.753 | 1.083 | -0.322 | 0.49 | -0.045 | -0.107 | -0.107 | 0 | 0.79 | 0.99 | 0.87 | 0.76 | 0.77 | 0.72 |
| 7 | 11581134 | rs2074603 | 0.917 | 1.071 | -0.301 | 0.475 | -0.001 | 0 | 0 | 0 | 0.47 | 0.49 | 0.38 | 0.54 | 0.42 | 0.49 |
| 7 | 11581381 | rs10250564 | -0.279 | 1.07 | -0.335 | 0.247 | 0.047 | 0.022 | 0.022 | 0 | 0.23 | 0.52 | 0.26 | 0.22 | 0.27 | 0.23 |
| 7 | 11582043 | rs6962022 | 1.954 | 1.043 | -0.348 | 0.321 | 0.003 | 0.003 | 0.003 | 0 | 0.32 | 0.3 | 0.21 | 0.34 | 0.24 | 0.32 |
| 7 | 11583169 | rs12699208 | 1.486 | 1.142 | -0.375 | 0.394 | -0.01 | -0.012 | -0.012 | 0.013 | 0.13 | 0.09 | 0.16 | 0.18 | 0.17 | 0.16 |
| 7 | 11584063 | rs2354963 | -1.664 | 1.192 | -0.415 | 0.423 | -0.06 | -0.108 | -0.108 | 1.025 | 0.85 | 0.99 | 0.89 | 0.75 | 0.79 | 0.74 |
| 7 | 11585820 | rs6974688 | -1.659 | 1.205 | -0.409 | 0.399 | -0.06 | -0.109 | -0.109 | 0 | 0.85 | 0.98 | 0.89 | 0.75 | 0.79 | 0.73 |
| 7 | 11586291 | rs12699211 | -0.688 | 1.202 | -0.279 | 0.348 | 0.018 | -0.013 | -0.013 | 0 | 0.28 | 0.51 | 0.29 | 0.22 | 0.31 | 0.25 |
| 7 | 11588284 | rs40 | -0.076 | 1.162 | -0.287 | 0.31 | -0.024 | 0.009 | 0.009 | 0 | 0.37 | 0.24 | 0.57 | 0.37 | 0.43 | 0.31 |
| 7 | 11592760 | rs12699224 | 1.375 | 1.125 | -0.203 | 0.337 | 0.006 | 0.008 | 0.008 | 0 | 0.14 | 0.17 | 0.17 | 0.19 | 0.18 | 0.16 |
| 7 | 11593797 | rs7792687 | 1.421 | 1.128 | -0.211 | 0.338 | 0.003 | 0.004 | 0.004 | 0 | 0.14 | 0.15 | 0.17 | 0.19 | 0.18 | 0.15 |
| 7 | 11595832 | rs10280808 | -0.203 | 0.89 | -0.19 | 0.241 | -0.049 | -0.018 | -0.018 | 0 | 0.12 | 0.35 | 0.06 | 0.03 | 0.05 | 0.04 |
| 7 | 11597156 | rs19 | 0.864 | 0.926 | -0.227 | 0.236 | -0.005 | 0.007 | 0.007 | 0 | 0.54 | 0.41 | 0.61 | 0.53 | 0.49 | 0.48 |
| 7 | 11597387 | rs10229585 | -0.214 | 0.866 | -0.188 | 0.226 | -0.049 | -0.017 | -0.017 | 1.088 | 0.12 | 0.36 | 0.06 | 0.03 | 0.05 | 0.04 |
| 7 | 11600040 | rs2107479 | 0.753 | 0.939 | -0.278 | 0.22 | -0.014 | 0.015 | 0.015 | 0.772 | 0.21 | 0.12 | 0.4 | 0.19 | 0.25 | 0.15 |
| 7 | 11600616 | rs1034740 | 1.877 | 0.877 | -0.285 | 0.225 | 0.007 | 0.006 | 0.006 | 0 | 0.33 | 0.3 | 0.22 | 0.34 | 0.25 | 0.34 |
| 7 | 11602761 | rs6979760 | 0.655 | 0.631 | -0.011 | 0.552 | -0.001 | -0.004 | -0.004 | 0 | 0.6 | 0.53 | 0.67 | 0.53 | 0.57 | 0.54 |
| 7 | 11602899 | rs16 | 0.655 | 0.631 | -0.011 | 0.545 | -0.001 | -0.001 | -0.001 | 0 | 0.6 | 0.53 | 0.67 | 0.54 | 0.57 | 0.54 |
| 7 | 11603287 | rs6980325 | -0.449 | 0.607 | 0.009 | 0.594 | -0.021 | 0.01 | 0.01 | 0 | 0.25 | 0.44 | 0.22 | 0.23 | 0.22 | 0.19 |
| 7 | 11604426 | rs7805116 | 0.33 | 0.555 | -0.041 | 0.6 | 0.001 | -0.001 | -0.001 | 0 | 0.19 | 0.14 | 0.18 | 0.2 | 0.19 | 0.19 |

| | | | | | | | | | | | | | | | | |
|---|---|---|---|---|---|---|---|---|---|---|---|---|---|---|---|---|
| 7 | 11605462 | rs10234091 | 0.523 | 0.659 | 0.018 | 0.572 | -0.006 | -0.004 | -0.004 | 0 | 0.59 | 0.61 | 0.47 | 0.54 | 0.52 | 0.6 |
| 7 | 11607706 | rs70 | -1.201 | 0.711 | -0.105 | 0.618 | -0.02 | -0.052 | -0.052 | 0.617 | 0.84 | 0.93 | 0.89 | 0.77 | 0.8 | 0.76 |
| 7 | 11610954 | rs6949833 | 0.962 | 0.825 | 0.018 | 0.777 | 0.004 | 0.006 | 0.006 | 0 | 0.44 | 0.56 | 0.4 | 0.38 | 0.35 | 0.45 |
| 7 | 11611226 | rs12699239 | 0.039 | 1.01 | -0.023 | 0.792 | -0.019 | -0.027 | -0.027 | 0.009 | 0.66 | 0.81 | 0.63 | 0.64 | 0.61 | 0.64 |
| 7 | 11611400 | rs78 | 1.659 | 1.293 | 0.093 | 0.712 | -0.012 | -0.004 | -0.004 | 0 | 0.12 | 0.07 | 0.24 | 0.18 | 0.17 | 0.08 |
| 7 | 11611590 | rs12699240 | 0.214 | 1.242 | 0.139 | 0.723 | -0.038 | -0.039 | -0.039 | 1.345 | 0.19 | 0.07 | 0.22 | 0.26 | 0.25 | 0.2 |
| 7 | 11613179 | rs11769542 | -0.008 | 1.154 | 0.145 | 0.722 | -0.002 | -0.007 | -0.007 | 0.021 | 0.62 | 0.7 | 0.61 | 0.62 | 0.6 | 0.61 |
| 7 | 11614579 | rs2040816 | 0.797 | 0.836 | 0.206 | 0.856 | 0.001 | 0.004 | 0.004 | 0 | 0.14 | 0.18 | 0.18 | 0.21 | 0.17 | 0.16 |
| 7 | 11615654 | rs12699246 | 0.264 | 0.855 | 0.229 | 0.836 | 0 | 0.002 | 0.002 | 0 | 0.18 | 0.2 | 0.18 | 0.24 | 0.2 | 0.16 |
| 7 | 11616177 | rs10486153 | 0.427 | 0.824 | 0.202 | 0.828 | -0.002 | -0.009 | -0.009 | 0 | 0.17 | 0.15 | 0.18 | 0.23 | 0.19 | 0.16 |
| 7 | 11616500 | rs17164813 | 0.927 | 0.862 | 0.006 | 0.698 | 0.02 | 0.03 | 0.03 | 0 | 0.4 | 0.33 | 0.24 | 0.33 | 0.31 | 0.4 |
| 7 | 11619123 | rs17164819 | 1.396 | 1.017 | 0.056 | 0.711 | 0.002 | 0.004 | 0.004 | 0 | 0.35 | 0.34 | 0.24 | 0.34 | 0.29 | 0.38 |
| 7 | 11621813 | rs1859364 | 0.547 | 1.179 | -0.338 | 1.103 | 0.001 | 0.003 | 0.003 | 0 | 0.58 | 0.59 | 0.68 | 0.56 | 0.57 | 0.53 |
| 7 | 11622592 | rs10950355 | -0.254 | 0.563 | -0.035 | 1.057 | 0.004 | 0.027 | 0.027 | 0.469 | 0.76 | 0.68 | 0.82 | 0.77 | 0.74 | 0.7 |
| 7 | 11622892 | rs2189638 | -0.061 | 0.224 | 0.131 | 0.585 | -0.009 | -0.02 | -0.02 | 3.732 | 0.42 | 0.49 | 0.26 | 0.38 | 0.35 | 0.43 |
| 7 | 11627116 | rs17164850 | -1.263 | 0.43 | 0.413 | 0.932 | -0.011 | 0.022 | 0.022 | 0 | 0.07 | 0.26 | 0.03 | 0.04 | 0.05 | 0.03 |
| 7 | 11630298 | rs2074602 | -0.668 | 0.997 | -0.337 | 0.929 | -0.047 | -0.015 | -0.015 | 0 | 0.12 | 0.03 | 0.16 | 0.08 | 0.2 | 0.16 |
| 7 | 11642396 | rs2681049 | -1.786 | 1.207 | 0.648 | 1.335 | 0.05 | 0.051 | 0.051 | 0 | 0.23 | 0.4 | 0.31 | 0.31 | 0.34 | 0.35 |
| 7 | 11642451 | rs16869985 | -1.2 | 1.193 | 0.762 | 1.327 | 0.045 | 0.135 | 0.135 | 0 | 0.08 | 0.24 | 0.13 | 0.24 | 0.09 | 0.16 |
| 7 | 11647158 | rs2681054 | -2.004 | 1.369 | 0.62 | 1.207 | 0 | -0.001 | -0.001 | 0 | 0.24 | 0.23 | 0.33 | 0.31 | 0.35 | 0.4 |
| 7 | 11649165 | rs2681051 | -0.799 | 0.793 | 0.452 | 1.265 | -0.071 | -0.005 | -0.005 | 0 | 0.15 | 0.03 | 0.2 | 0.09 | 0.26 | 0.24 |
| 7 | 11649473 | rs2681050 | -1.751 | -0.048 | 0.595 | 1.451 | 0.024 | -0.059 | -0.059 | 0 | 0.36 | 0.8 | 0.33 | 0.56 | 0.36 | 0.57 |
| 7 | 11665731 | rs3846988 | -1.479 | -0.687 | 1.195 | 1.219 | -0.077 | 0.032 | 0.032 | 0 | 0.21 | 0.52 | 0.1 | 0.31 | 0.1 | 0.21 |
| 7 | 11669209 | rs17164930 | -1.201 | -0.458 | 0.169 | 0.653 | -0.035 | -0.037 | -0.037 | 0 | 0.18 | 0.18 | 0.06 | 0.03 | 0.08 | 0.12 |
| 7 | 11676377 | rs2074598 | -0.909 | 0.142 | -0.543 | 0.89 | 0.003 | 0.001 | 0.001 | 0 | 0.29 | 0.16 | 0.33 | 0.28 | 0.25 | 0.35 |
| 7 | 11676694 | rs2074597 | -0.931 | 0.197 | -0.012 | 1.498 | -0.006 | 0.176 | 0.176 | 0 | 0.21 | 0.5 | 0.26 | 0.45 | 0.2 | 0.23 |
| 7 | 11677088 | rs17576766 | -0.475 | 0.553 | -0.75 | 1.339 | -0.008 | -0.019 | -0.019 | 0 | 0.2 | 0.1 | 0.29 | 0.25 | 0.26 | 0.17 |
| 7 | 11677126 | rs12154996 | -0.042 | 1.074 | -1.634 | 0.391 | -0.04 | 0.065 | 0.065 | 0.226 | 0.24 | 0.08 | 0.18 | 0.08 | 0.29 | 0.21 |
| 7 | 11677840 | rs2526099 | -0.714 | 1.985 | -2.01 | 0.469 | -0.102 | 0.074 | 0.074 | 0.044 | 0.33 | 0.12 | 0.36 | 0.2 | 0.43 | 0.26 |
| 7 | 11678246 | rs2526098 | -0.64 | 1.534 | -2.267 | 0.557 | -0.001 | -0.063 | -0.063 | 0 | 0.75 | 0.91 | 0.68 | 0.68 | 0.72 | 0.76 |
| 7 | 11681776 | rs6460831 | 1.105 | 0.956 | -1.32 | 0.235 | -0.006 | 0.002 | 0.002 | 0.283 | 0.46 | 0.43 | 0.31 | 0.52 | 0.35 | 0.6 |
| 7 | 11682138 | rs1725449 | -0.083 | 1.443 | -1.119 | 0.459 | 0.01 | 0.002 | 0.002 | 0 | 0.08 | 0.1 | 0.23 | 0.16 | 0.18 | 0.05 |
| 7 | 11684576 | rs12699252 | 0.844 | 1.33 | -1.139 | 0.3 | 0 | 0 | 0 | 0 | 0.54 | 0.47 | 0.38 | 0.53 | 0.46 | 0.63 |
| 7 | 11703899 | rs2077983 | -0.581 | 1.425 | -1.238 | 0.288 | -0.007 | 0.008 | 0.008 | 0 | 0.14 | 0.1 | 0.12 | 0.13 | 0.17 | 0.15 |
| 7 | 11711073 | rs2049535 | 0.679 | 1.124 | -0.468 | 0.537 | -0.001 | -0.001 | -0.001 | 0 | 0.58 | 0.42 | 0.46 | 0.54 | 0.58 | 0.68 |
| 7 | 11725406 | rs1589944 | -0.385 | 0.888 | -0.367 | 0.612 | -0.001 | -0.018 | -0.018 | 0 | 0.35 | 0.27 | 0.38 | 0.41 | 0.39 | 0.37 |
| 7 | 11730333 | rs1568974 | -0.047 | 0.618 | 0.068 | 0.062 | 0.029 | -0.037 | -0.037 | 1.554 | 0.23 | 0.33 | 0.34 | 0.15 | 0.28 | 0.1 |
| 7 | 11733150 | rs17165047 | 1.638 | 0.675 | 0.095 | 0.038 | -0.007 | -0.005 | -0.005 | 0.388 | 0.15 | 0.13 | 0.11 | 0.14 | 0.11 | 0.07 |
| 7 | 11737891 | rs1405321 | 0.528 | 0.784 | 0.118 | 0.045 | 0.063 | -0.022 | -0.022 | 0.265 | 0.06 | 0.1 | 0.21 | 0.01 | 0.17 | 0.04 |
| 7 | 11740020 | rs2883585 | -0.609 | 0.773 | 0.131 | -0.012 | 0.032 | -0.04 | -0.04 | 1.066 | 0.29 | 0.35 | 0.39 | 0.17 | 0.39 | 0.22 |
| 7 | 11741736 | rs1526554 | -0.488 | 0.582 | 0.035 | 0.048 | 0.023 | -0.1 | -0.1 | 0 | 0.67 | 0.91 | 0.72 | 0.59 | 0.69 | 0.71 |
| 7 | 11742205 | rs10257747 | -0.967 | 0.453 | 0.286 | 0.147 | 0.012 | -0.009 | -0.009 | 2.807 | 0.57 | 0.64 | 0.67 | 0.46 | 0.67 | 0.43 |
| 7 | 11745609 | rs10950364 | 0.36 | 0.497 | 0 | 0.175 | 0.003 | 0.006 | 0.006 | 0 | 0.49 | 0.59 | 0.39 | 0.43 | 0.42 | 0.65 |
| 7 | 11746175 | rs17245738 | -0.634 | 0.57 | -0.147 | 0.108 | -0.014 | 0.006 | 0.006 | 0 | 0.15 | 0.13 | 0.28 | 0.03 | 0.28 | 0.2 |
| 7 | 11749647 | rs10225038 | -0.222 | 0.505 | -0.193 | 0.147 | -0.003 | -0.012 | -0.012 | 0 | 0.45 | 0.32 | 0.58 | 0.46 | 0.55 | 0.28 |
| 7 | 11753029 | rs1526549 | -0.155 | 0.757 | 0.326 | 0.919 | 0.031 | -0.106 | -0.106 | 0.532 | 0.69 | 0.91 | 0.73 | 0.56 | 0.71 | 0.71 |
| 7 | 11753282 | rs2355068 | 0.281 | 0.821 | 0.349 | 0.938 | -0.01 | -0.1 | -0.1 | 0.256 | 0.63 | 0.9 | 0.67 | 0.54 | 0.6 | 0.59 |
| 7 | 11756776 | rs1526543 | -0.099 | 0.85 | 0.361 | 0.847 | 0.016 | -0.006 | -0.006 | 0.111 | 0.06 | 0.08 | 0.21 | 0.01 | 0.17 | 0.04 |
| 7 | 11759464 | rs12699254 | 0.902 | 0.72 | 0.485 | 1.039 | 0 | -0.002 | -0.002 | 0.106 | 0.23 | 0.18 | 0.27 | 0.3 | 0.27 | 0.19 |
| 7 | 11759701 | rs17165078 | 1.134 | 0.642 | 0.633 | 1.069 | 0.034 | 0.02 | 0.02 | 0.162 | 0.48 | 0.32 | 0.36 | 0.5 | 0.36 | 0.5 |
| 7 | 11763981 | rs1526561 | 1.376 | 0.514 | 0.407 | 1.074 | 0.013 | 0.023 | 0.023 | 0 | 0.46 | 0.37 | 0.33 | 0.39 | 0.33 | 0.49 |
| 7 | 11765421 | rs10231490 | 0.757 | -0.359 | 0.564 | 1.271 | -0.005 | 0.026 | 0.026 | 0 | 0.25 | 0.31 | 0.29 | 0.3 | 0.27 | 0.16 |
| 7 | 11766879 | rs9691790 | 0.147 | -0.261 | 0.16 | 0.996 | -0.043 | -0.048 | -0.048 | 0 | 0.82 | 0.99 | 0.89 | 0.83 | 0.8 | 0.71 |
| 7 | 11767633 | rs13307585 | 1.915 | -0.462 | -0.034 | 0.985 | -0.132 | 0.121 | 0.121 | 0 | 0.37 | 0.03 | 0.42 | 0.24 | 0.39 | 0.28 |
| 7 | 11769311 | rs1526558 | 0.048 | -0.358 | 0.265 | 0.998 | 0.003 | -0.009 | -0.009 | 0.246 | 0.36 | 0.64 | 0.38 | 0.46 | 0.36 | 0.41 |
| 7 | 11771288 | rs10499409 | 1.075 | -0.07 | 0.452 | 0.887 | -0.012 | 0.036 | 0.036 | 0 | 0.45 | 0.33 | 0.51 | 0.37 | 0.45 | 0.3 |
| 7 | 11772735 | rs4721007 | -0.098 | -0.147 | 0.466 | 1.417 | 0.008 | -0.025 | -0.025 | 0 | 0.7 | 0.76 | 0.71 | 0.6 | 0.71 | 0.76 |
| 7 | 11774241 | rs1526538 | 0.792 | -0.051 | 0.087 | 1.183 | 0.034 | 0.04 | 0.04 | 0 | 0.57 | 0.16 | 0.58 | 0.42 | 0.59 | 0.5 |
| 7 | 11774458 | rs6956994 | 0.685 | -0.042 | 0.15 | 1.166 | 0.024 | 0.03 | 0.03 | 0 | 0.58 | 0.28 | 0.57 | 0.42 | 0.59 | 0.48 |
| 7 | 11779718 | rs1526520 | -0.829 | -0.213 | -0.475 | 1.618 | -0.026 | -0.105 | -0.105 | 0 | 0.42 | 0.95 | 0.49 | 0.54 | 0.42 | 0.47 |
| 7 | 11783957 | rs1467344 | -0.18 | -0.156 | 0.359 | 1.369 | -0.004 | 0.04 | 0.04 | 0.671 | 0.13 | 0.63 | 0.13 | 0.16 | 0.11 | 0.26 |
| 7 | 11794239 | rs6460840 | -0.599 | -0.356 | 0.03 | 1.545 | -0.037 | -0.096 | -0.096 | 0 | 0.76 | 0.96 | 0.74 | 0.71 | 0.74 | 0.76 |
| 7 | 11796710 | rs7795351 | 0.3 | -0.273 | 0.271 | 1.294 | -0.007 | -0.061 | -0.061 | 0.762 | 0.63 | 0.86 | 0.64 | 0.63 | 0.63 | 0.61 |
| 7 | 11803023 | rs1405315 | 1.127 | -0.267 | 0.559 | 1.169 | -0.001 | 0.059 | 0.059 | 0 | 0.48 | 0.39 | 0.45 | 0.3 | 0.46 | 0.33 |
| 7 | 11808215 | rs34067420 | 1.545 | 0.971 | 0.639 | 1.451 | 0.027 | 0.042 | 0.042 | 0.269 | 0.49 | 0.44 | 0.27 | 0.25 | 0.34 | 0.25 |
| 7 | 11809623 | rs1526518 | 0.9 | 0.936 | 0.836 | 1.479 | 0.036 | 0.046 | 0.046 | 0 | 0.11 | 0.29 | 0.14 | 0.15 | 0.14 | 0.26 |
| 7 | 11817290 | rs7783055 | -1.117 | 0.96 | 0.832 | 1.538 | 0.005 | -0.017 | -0.017 | 0 | 0.42 | 0.49 | 0.48 | 0.62 | 0.48 | 0.6 |
| 7 | 11819562 | rs10267051 | 0.171 | 1.063 | 0.694 | 1.689 | 0 | -0.028 | -0.028 | 0.045 | 0.24 | 0.15 | 0.27 | 0.37 | 0.26 | 0.24 |
| 7 | 11821125 | rs10242204 | -0.008 | 1.341 | 0.608 | 1.732 | -0.053 | -0.043 | -0.043 | 0 | 0.92 | 0.99 | 0.75 | 0.87 | 0.8 | 0.89 |

| | | | | | | | | | | | | | | | | |
|---|---|---|---|---|---|---|---|---|---|---|---|---|---|---|---|---|
| 7 | 11821993 | rs13437787 | -0.614 | 1.225 | 0.662 | 1.731 | -0.004 | -0.06 | -0.06 | 0.002 | 0.31 | 0.19 | 0.33 | 0.42 | 0.35 | 0.39 |
| 7 | 11822142 | rs12673487 | 0.723 | 1.366 | 0.866 | 1.78 | 0.033 | 0.038 | 0.038 | 0 | 0.12 | 0.29 | 0.14 | 0.17 | 0.15 | 0.27 |
| 7 | 11822611 | rs7793467 | -1.144 | 1.353 | 0.661 | 1.794 | -0.001 | 0.02 | 0.02 | 0.005 | 0.41 | 0.69 | 0.47 | 0.64 | 0.46 | 0.52 |
| 7 | 11822779 | rs6460841 | 0.882 | 1.541 | 0.016 | 1.948 | -0.031 | -0.015 | -0.015 | 0 | 0.66 | 0.74 | 0.47 | 0.47 | 0.51 | 0.62 |
| 7 | 11825210 | rs4721012 | -0.598 | 1.625 | -0.295 | 1.96 | -0.027 | -0.085 | -0.085 | 0.333 | 0.29 | 0.12 | 0.33 | 0.45 | 0.35 | 0.38 |
| 7 | 11825839 | rs7782151 | 0.884 | 1.615 | -0.205 | 1.905 | -0.015 | -0.013 | -0.013 | 0.219 | 0.88 | 0.91 | 0.65 | 0.71 | 0.76 | 0.87 |
| 7 | 11829124 | rs2355088 | -0.386 | 1.493 | -0.137 | 1.746 | 0.001 | -0.024 | -0.024 | 0 | 0.28 | 0.24 | 0.33 | 0.47 | 0.32 | 0.27 |
| 7 | 11834671 | rs6967913 | 0.841 | 2.66 | -0.066 | 2.009 | 0.002 | -0.003 | -0.003 | 2.01 | 0.9 | 0.92 | 0.72 | 0.87 | 0.78 | 0.89 |
| 7 | 11834856 | rs12531067 | 0.594 | 2.442 | -0.188 | 2.283 | -0.075 | -0.099 | -0.099 | 0 | 0.15 | 0.03 | 0.25 | 0.32 | 0.26 | 0.21 |
| 7 | 11845867 | rs17165231 | -0.596 | 2.295 | 0.053 | 1.736 | -0.004 | 0 | 0 | 0.732 | 0.08 | 0.09 | 0.06 | 0.06 | 0.08 | 0.12 |
| 7 | 11846108 | rs1815109 | 1.061 | 2.649 | 0.154 | 1.564 | 0.096 | 0.062 | 0.062 | 0 | 0.78 | 0.48 | 0.59 | 0.71 | 0.67 | 0.59 |
| 7 | 11846574 | rs7798977 | 0.643 | 2.865 | -0.438 | 1.69 | 0.009 | -0.002 | -0.002 | 0 | 0.93 | 0.95 | 0.74 | 0.84 | 0.81 | 0.89 |
| 7 | 11847740 | rs11768257 | 0.677 | 2.801 | -0.384 | 1.701 | 0.038 | 0.024 | 0.024 | 0 | 0.93 | 0.89 | 0.78 | 0.84 | 0.82 | 0.9 |
| 7 | 11852635 | rs9639102 | -0.497 | 1.782 | -0.42 | 1.523 | -0.002 | -0.002 | -0.002 | 0 | 0.25 | 0.27 | 0.34 | 0.32 | 0.35 | 0.27 |
| 7 | 11853708 | rs2883595 | 0.534 | 1.594 | -0.036 | 1.305 | 0.022 | 0.016 | 0.016 | 0 | 0.82 | 0.66 | 0.83 | 0.83 | 0.81 | 0.6 |
| 7 | 11855636 | rs6947930 | -0.587 | 1.441 | 0.032 | 1.665 | 0.003 | 0.032 | 0.032 | 0 | 0.32 | 0.42 | 0.45 | 0.48 | 0.31 | 0.24 |
| 7 | 11855947 | rs10270255 | -0.551 | 1.444 | 0.161 | 1.775 | 0.001 | 0.013 | 0.013 | 0 | 0.32 | 0.36 | 0.45 | 0.52 | 0.31 | 0.24 |
| 7 | 11856057 | rs12699276 | 0.077 | 1.242 | -0.799 | 1.228 | 0.003 | 0.006 | 0.006 | 0 | 0.39 | 0.63 | 0.36 | 0.32 | 0.39 | 0.53 |
| 7 | 11856806 | rs6460848 | -0.962 | 1.075 | -0.196 | 1.278 | 0 | 0.011 | 0.011 | 0 | 0.64 | 0.72 | 0.76 | 0.65 | 0.64 | 0.67 |
| 7 | 11857504 | rs4463318 | 0.51 | 1.004 | -0.18 | 1.129 | -0.025 | 0.084 | 0.084 | 0 | 0.29 | 0.07 | 0.27 | 0.1 | 0.3 | 0.42 |
| 7 | 11860433 | rs11972583 | -0.072 | 0.833 | 0.349 | 1.201 | -0.003 | 0.035 | 0.035 | 0.057 | 0.34 | 0.52 | 0.46 | 0.53 | 0.31 | 0.28 |
| 7 | 11860937 | rs7807756 | -0.683 | 0.705 | 0.987 | 1.159 | -0.005 | -0.007 | -0.007 | 0 | 0.61 | 0.5 | 0.74 | 0.63 | 0.64 | 0.68 |
| 7 | 11884682 | rs6942789 | -1.084 | 0.718 | 0.91 | 1.056 | -0.008 | 0.034 | 0.034 | 0 | 0.66 | 0.76 | 0.77 | 0.81 | 0.66 | 0.67 |
| 7 | 11885016 | rs6943249 | -1.602 | 0.693 | 1.177 | 1.022 | -0.003 | 0.055 | 0.055 | 0 | 0.7 | 0.83 | 0.81 | 0.87 | 0.7 | 0.78 |
| 7 | 11892575 | rs10950369 | -1.31 | 0.681 | 1.276 | 1.037 | -0.024 | 0.062 | 0.062 | 0 | 0.67 | 0.78 | 0.78 | 0.83 | 0.67 | 0.68 |
| 7 | 11893555 | rs10259581 | -1.594 | 0.684 | 1.293 | 1.04 | -0.002 | -0.043 | -0.043 | 1.901 | 0.69 | 0.63 | 0.78 | 0.89 | 0.71 | 0.79 |
| 7 | 11895273 | rs7458129 | -0.804 | 0.661 | 1.295 | 1.027 | -0.001 | 0.018 | 0.018 | 0 | 0.63 | 0.33 | 0.73 | 0.83 | 0.65 | 0.68 |
| 7 | 11904138 | rs7810903 | 0.963 | 0.807 | 0.675 | 1.046 | -0.082 | 0.034 | 0.034 | 0 | 0.25 | 0.02 | 0.26 | 0.1 | 0.3 | 0.4 |
| 7 | 11911677 | rs4307228 | -1.301 | 0.327 | 0.831 | 0.888 | -0.012 | 0.022 | 0.022 | 5.727 | 0.23 | 0.44 | 0.13 | 0.3 | 0.18 | 0.2 |
| 7 | 11911725 | rs6966165 | -0.076 | 0.235 | 0.79 | 0.865 | -0.001 | 0.009 | 0.009 | 0 | 0.48 | 0.54 | 0.69 | 0.58 | 0.53 | 0.59 |
| 7 | 11913955 | rs10215881 | -0.282 | 0.28 | 0.721 | 0.885 | 0.001 | 0.03 | 0.03 | 0 | 0.5 | 0.61 | 0.7 | 0.58 | 0.53 | 0.59 |
| 7 | 11915630 | rs13244014 | -1.017 | 0.389 | 0.922 | 0.977 | -0.004 | 0.018 | 0.018 | 6.64 | 0.21 | 0.36 | 0.1 | 0.3 | 0.18 | 0.2 |
| 7 | 11916354 | rs10253052 | 0.43 | 0.682 | 0.781 | 1.155 | -0.001 | 0.017 | 0.017 | 3.201 | 0.06 | 0.06 | 0.04 | 0.22 | 0.06 | 0.07 |
| 7 | 11925539 | rs2355129 | -0.929 | 0.642 | 1.355 | 1.176 | 0.004 | -0.006 | -0.006 | 0.017 | 0.63 | 0.47 | 0.74 | 0.6 | 0.64 | 0.63 |
| 7 | 11926347 | rs9655091 | -1.566 | 0.712 | 1.341 | 1.123 | -0.01 | -0.113 | -0.113 | 0 | 0.69 | 0.6 | 0.78 | 0.94 | 0.73 | 0.8 |
| 7 | 11930135 | rs1815106 | -1.662 | 0.72 | 1.416 | 1.135 | -0.003 | -0.051 | -0.051 | 0.661 | 0.69 | 0.72 | 0.79 | 0.94 | 0.73 | 0.81 |
| 7 | 11931567 | rs2355145 | 0.328 | 0.533 | 1.365 | 1.348 | -0.025 | 0.039 | 0.039 | 0 | 0.41 | 0.2 | 0.41 | 0.27 | 0.45 | 0.5 |
| 7 | 11932740 | rs2355147 | -1.075 | 0.561 | 1.333 | 1.367 | 0.002 | -0.004 | -0.004 | 0 | 0.62 | 0.44 | 0.74 | 0.6 | 0.64 | 0.62 |
| 7 | 11932891 | rs1961842 | -1.204 | 0.624 | 1.093 | 1.489 | 0.051 | 0.16 | 0.16 | 0 | 0.06 | 0.16 | 0.04 | 0.34 | 0.09 | 0.18 |
| 7 | 11933414 | rs10242930 | -0.284 | 0.542 | 0.854 | 1.407 | 0.011 | -0.035 | -0.035 | 0.822 | 0.2 | 0.13 | 0.31 | 0.32 | 0.18 | 0.12 |
| 7 | 11938060 | rs17165338 | -0.264 | 0.601 | 0.746 | 1.693 | -0.035 | -0.046 | -0.046 | 0 | 0.08 | 0.01 | 0.16 | 0.21 | 0.09 | 0.14 |
| 7 | 11938607 | rs2355757 | -1.45 | 0.501 | 1.256 | 1.671 | -0.006 | -0.016 | -0.016 | 0 | 0.66 | 0.61 | 0.77 | 0.67 | 0.69 | 0.72 |
| 7 | 11941048 | rs10250872 | -0.64 | 0.615 | 1.223 | 1.659 | 0.005 | 0.005 | 0.005 | 0 | 0.54 | 0.69 | 0.61 | 0.49 | 0.55 | 0.63 |
| 7 | 11950061 | rs6943159 | -1.105 | 0.675 | 1.165 | 1.577 | -0.005 | -0.011 | -0.011 | 0 | 0.4 | 0.55 | 0.42 | 0.73 | 0.36 | 0.39 |
| 7 | 11952193 | rs12531867 | 0.275 | 0.696 | 0.665 | 1.61 | 0.021 | -0.013 | -0.013 | 0 | 0.25 | 0.12 | 0.2 | 0.25 | 0.22 | 0.18 |
| 7 | 11954107 | rs17165392 | -0.216 | 0.779 | 0.68 | 1.625 | -0.021 | -0.026 | -0.026 | 0 | 0.08 | 0.04 | 0.16 | 0.21 | 0.09 | 0.11 |
| 7 | 11954575 | rs10234648 | 0.256 | 0.664 | 0.633 | 1.474 | 0.021 | -0.009 | -0.009 | 0 | 0.25 | 0.13 | 0.21 | 0.25 | 0.21 | 0.19 |
| 7 | 11958023 | rs1880312 | -1.924 | 0.628 | 0.631 | 1.38 | -0.001 | 0.06 | 0.06 | 0 | 0.71 | 0.84 | 0.82 | 0.95 | 0.73 | 0.8 |
| 7 | 11960047 | rs7801347 | 0.143 | 0.78 | 0.445 | 1.544 | 0.022 | -0.014 | -0.014 | 0.393 | 0.25 | 0.12 | 0.2 | 0.25 | 0.21 | 0.18 |
| 7 | 11965352 | rs10228842 | -1.12 | 0.929 | 0.532 | 1.719 | 0.014 | 0.106 | 0.106 | 0.001 | 0.12 | 0.19 | 0.19 | 0.48 | 0.14 | 0.18 |
| 7 | 11975557 | rs12699288 | 0.153 | 0.737 | 0.535 | 1.408 | -0.029 | 0.012 | 0.012 | 0 | 0.34 | 0.18 | 0.39 | 0.25 | 0.42 | 0.45 |
| 7 | 11982513 | rs11983339 | -0.583 | 0.611 | 0.782 | 1.514 | -0.012 | 0.023 | 0.023 | 0 | 0.34 | 0.45 | 0.39 | 0.48 | 0.27 | 0.31 |
| 7 | 11993510 | rs11975215 | -0.496 | 0.631 | 0.759 | 1.518 | 0.007 | -0.02 | -0.02 | 0 | 0.34 | 0.32 | 0.39 | 0.48 | 0.28 | 0.31 |
| 7 | 12000154 | rs2966628 | -1.147 | 0.478 | 0.821 | 1.396 | -0.007 | -0.04 | -0.04 | 0 | 0.67 | 0.56 | 0.79 | 0.73 | 0.71 | 0.75 |
| 7 | 12012769 | rs2192739 | -0.545 | 0.456 | 0.941 | 1.678 | -0.011 | 0.041 | 0.041 | 0 | 0.05 | 0.09 | 0.03 | 0.22 | 0.03 | 0.09 |
| 7 | 12013256 | rs10228515 | -0.865 | 0.658 | 1.164 | 1.742 | -0.004 | 0.003 | 0.003 | 0 | 0.43 | 0.7 | 0.44 | 0.59 | 0.42 | 0.61 |
| 7 | 12015739 | rs7809831 | 0.005 | 0.864 | 1.464 | 1.685 | -0.004 | 0.017 | 0.017 | 0 | 0.32 | 0.25 | 0.37 | 0.21 | 0.34 | 0.4 |
| 7 | 12017274 | rs13312476 | -0.371 | 0.859 | 1.139 | 1.683 | -0.002 | -0.019 | -0.019 | 0 | 0.35 | 0.55 | 0.39 | 0.21 | 0.34 | 0.41 |
| 7 | 12018593 | rs4412264 | -0.515 | 0.916 | 0.997 | 1.658 | -0.001 | -0.001 | -0.001 | 0 | 0.38 | 0.6 | 0.41 | 0.36 | 0.39 | 0.51 |
| 7 | 12025058 | rs7810106 | -2.099 | 0.898 | 1.434 | 1.451 | 0.084 | 0.177 | 0.177 | 0.358 | 0.07 | 0.58 | 0.07 | 0.15 | 0.06 | 0.11 |
| 7 | 12029512 | rs6965929 | -1.841 | 0.941 | 1.364 | 1.471 | 0.146 | 0.262 | 0.262 | 0.063 | 0.07 | 0.48 | 0.06 | 0.15 | 0.06 | 0.11 |
| 7 | 12032301 | rs6960153 | -0.717 | 0.853 | 0.88 | 1.431 | -0.001 | -0.001 | -0.001 | 0.44 | 0.38 | 0.41 | 0.39 | 0.54 | 0.39 | 0.59 |
| 7 | 12032343 | rs6964035 | -0.031 | 0.93 | 0.707 | 1.572 | -0.019 | -0.046 | -0.046 | 0 | 0.29 | 0.12 | 0.31 | 0.34 | 0.29 | 0.43 |
| 7 | 12032707 | rs10237588 | -0.621 | 0.938 | 0.999 | 1.312 | 0.002 | 0.056 | 0.056 | 0 | 0.26 | 0.37 | 0.29 | 0.53 | 0.22 | 0.55 |
| 7 | 12033933 | rs2192735 | 0.493 | 1.001 | 0.583 | 1.276 | 0.007 | -0.037 | -0.037 | 0 | 0.18 | 0.09 | 0.23 | 0.33 | 0.13 | 0.38 |
| 7 | 12042417 | rs12666416 | -1.214 | 1.085 | 0.719 | 0.884 | -0.004 | 0.035 | 0.035 | 0 | 0.48 | 0.38 | 0.52 | 0.69 | 0.51 | 0.63 |
| 7 | 12046503 | rs719680 | -1.509 | 1.019 | 0.4 | 0.736 | -0.004 | 0.047 | 0.047 | 0 | 0.53 | 0.72 | 0.56 | 0.7 | 0.52 | 0.65 |
| 7 | 12049086 | rs2215987 | 0.814 | 0.844 | -0.543 | 0.464 | -0.002 | -0.041 | -0.041 | 0 | 0.18 | 0.06 | 0.25 | 0.24 | 0.18 | 0.16 |
| 7 | 12049120 | rs2215986 | -0.477 | 0.817 | -0.589 | 0.491 | -0.008 | -0.073 | -0.073 | 3.621 | 0.4 | 0.19 | 0.46 | 0.5 | 0.41 | 0.46 |

| | | | | | | | | | | | | | | | | |
|---|---|---|---|---|---|---|---|---|---|---|---|---|---|---|---|---|
| 7 | 12050855 | rs2215985 | -0.352 | 0.801 | -0.589 | 0.491 | -0.012 | -0.072 | -0.072 | 0 | 0.4 | 0.19 | 0.46 | 0.5 | 0.42 | 0.46 |
| 7 | 12052718 | rs12673618 | 0.041 | 0.633 | -0.389 | 0.584 | 0.005 | -0.016 | -0.016 | 0.039 | 0.29 | 0.31 | 0.38 | 0.51 | 0.25 | 0.43 |
| 7 | 12053230 | rs10248034 | 0.334 | 0.661 | -0.513 | 0.504 | 0.011 | -0.006 | -0.006 | 0 | 0.19 | 0.14 | 0.26 | 0.19 | 0.15 | 0.16 |
| 7 | 12058208 | rs6956188 | -2.104 | 0.939 | -0.687 | 0.096 | -0.038 | 0.111 | 0.111 | 0.373 | 0.69 | 0.89 | 0.82 | 0.94 | 0.67 | 0.87 |
| 7 | 12061045 | rs2058644 | -1.129 | 0.961 | -0.471 | 0.493 | 0.001 | 0.024 | 0.024 | 0.027 | 0.42 | 0.7 | 0.46 | 0.32 | 0.42 | 0.42 |
| 7 | 12064193 | rs6950290 | -0.6 | 0.947 | 0.191 | 0.435 | 0.008 | -0.003 | -0.003 | 0 | 0.2 | 0.25 | 0.2 | 0.17 | 0.29 | 0.17 |
| 7 | 12066984 | rs2966608 | -1.228 | 0.918 | 0.214 | 0.459 | 0 | 0.009 | 0.009 | 0.026 | 0.42 | 0.68 | 0.46 | 0.36 | 0.42 | 0.43 |
| 7 | 12068095 | rs6943558 | 0.067 | 0.917 | 0.167 | 0.414 | -0.011 | 0.005 | 0.005 | 0 | 0.15 | 0.14 | 0.15 | 0.09 | 0.26 | 0.12 |
| 7 | 12070338 | rs4236266 | -1.085 | 0.911 | -0.05 | 0.467 | -0.007 | -0.014 | -0.014 | 0.011 | 0.42 | 0.74 | 0.47 | 0.4 | 0.44 | 0.43 |
| 7 | 12071117 | rs6979677 | -1.101 | 0.9 | -0.063 | 0.463 | -0.006 | -0.013 | -0.013 | 0 | 0.42 | 0.74 | 0.47 | 0.4 | 0.44 | 0.43 |
| 7 | 12071741 | rs7780167 | -1.176 | 0.908 | -0.067 | 0.471 | -0.007 | -0.012 | -0.012 | 0.584 | 0.43 | 0.76 | 0.48 | 0.4 | 0.45 | 0.43 |
| 7 | 12072864 | rs2966604 | -1.985 | 1.024 | -0.257 | 0.305 | -0.02 | 0.072 | 0.072 | 0.304 | 0.69 | 0.85 | 0.82 | 0.94 | 0.67 | 0.87 |
| 7 | 12075590 | rs1861276 | -0.859 | 1.085 | 0.195 | 0.681 | -0.007 | 0.019 | 0.019 | 0.289 | 0.31 | 0.58 | 0.37 | 0.35 | 0.27 | 0.42 |
| 7 | 12077909 | rs2215983 | -0.481 | 1.505 | 0.003 | 0.452 | 0.014 | 0.01 | 0.01 | 0 | 0.13 | 0.23 | 0.1 | 0.12 | 0.14 | 0.13 |
| 7 | 12078163 | rs976852 | -0.22 | 1.257 | -0.467 | 0.463 | -0.044 | -0.124 | -0.124 | 0 | 0.38 | 0.07 | 0.46 | 0.5 | 0.4 | 0.48 |
| 7 | 12084381 | rs1468786 | -1.114 | 1.162 | -0.236 | -0.034 | 0.007 | 0.017 | 0.017 | 2.033 | 0.53 | 0.69 | 0.57 | 0.39 | 0.58 | 0.51 |
| 7 | 12090999 | rs714161 | -2.43 | 1.558 | -0.204 | 0.115 | 0.01 | 0.08 | 0.08 | 2.998 | 0.73 | 0.9 | 0.82 | 0.91 | 0.77 | 0.88 |
| 7 | 12091807 | rs2215994 | -0.763 | 1.574 | 0.63 | 0.084 | 0.055 | 0.052 | 0.052 | 0 | 0.06 | 0.31 | 0.06 | 0.03 | 0.05 | 0.06 |
| 7 | 12091894 | rs17647913 | 1.369 | 1.595 | 0.478 | -0.001 | -0.006 | 0.011 | 0.011 | 0 | 0.17 | 0.08 | 0.22 | 0.12 | 0.15 | 0.18 |
| 7 | 12094966 | rs1035147 | 0.808 | 1.774 | 0.33 | -0.028 | -0.043 | -0.022 | -0.022 | 0 | 0.17 | 0.03 | 0.15 | 0.12 | 0.18 | 0.11 |
| 7 | 12096768 | rs10247762 | -0.117 | 1.746 | 0.606 | 0.001 | 0.005 | -0.001 | -0.001 | 0 | 0.11 | 0.14 | 0.15 | 0.13 | 0.21 | 0.17 |
| 7 | 12097701 | rs10251790 | 1.371 | 1.596 | 0.634 | 0.047 | -0.01 | -0.004 | -0.004 | 0.045 | 0.06 | 0.05 | 0.09 | 0.1 | 0.13 | 0.11 |
| 7 | 12103271 | rs7785860 | -0.007 | 1.582 | 0.712 | 0.039 | 0.025 | 0.014 | 0.014 | 0 | 0.09 | 0.2 | 0.13 | 0.13 | 0.18 | 0.16 |
| 7 | 12106056 | rs7801745 | -1.173 | 1.668 | 0.595 | -0.033 | 0.011 | -0.013 | -0.013 | 0 | 0.34 | 0.54 | 0.36 | 0.27 | 0.44 | 0.34 |
| 7 | 12108746 | rs6460873 | -1.476 | 1.687 | 0.556 | 0.326 | 0.036 | 0.005 | 0.005 | 0.722 | 0.53 | 0.65 | 0.61 | 0.51 | 0.65 | 0.52 |
| 7 | 12113859 | rs2160222 | -1.46 | 1.697 | 0.547 | 0.331 | 0.032 | 0.005 | 0.005 | 0.571 | 0.52 | 0.62 | 0.61 | 0.51 | 0.65 | 0.53 |
| 7 | 12115724 | rs6460878 | -1.16 | 1.717 | 0.529 | 0.373 | 0.012 | 0.017 | 0.017 | 0.209 | 0.34 | 0.53 | 0.36 | 0.4 | 0.44 | 0.34 |
| 7 | 12115931 | rs11976213 | -0.793 | 1.678 | 0.554 | 0.384 | 0.003 | 0.054 | 0.054 | 0 | 0.29 | 0.51 | 0.28 | 0.39 | 0.32 | 0.32 |
| 7 | 12119196 | rs12536809 | 0.322 | 1.728 | 0.38 | 0.339 | -0.012 | -0.024 | -0.024 | 0 | 0.6 | 0.78 | 0.6 | 0.4 | 0.62 | 0.43 |
| 7 | 12119520 | rs17648860 | 2.515 | 1.657 | 0.394 | 0.296 | 0.015 | 0.087 | 0.087 | 1.955 | 0.27 | 0.12 | 0.12 | 0.1 | 0.24 | 0.11 |
| 7 | 12121796 | rs1990602 | -1.525 | 1.654 | 0.379 | 0.372 | 0 | -0.012 | -0.012 | 0 | 0.81 | 0.87 | 0.77 | 0.61 | 0.84 | 0.66 |
| 7 | 12121945 | rs7783906 | 0.453 | 1.482 | 0.273 | 0.28 | -0.021 | -0.018 | -0.018 | 0.525 | 0.2 | 0.09 | 0.17 | 0.21 | 0.2 | 0.23 |
| 7 | 12123385 | rs2192745 | 0.379 | 1.506 | 0.609 | 0.277 | 0 | 0 | 0 | 0 | 0.54 | 0.59 | 0.4 | 0.46 | 0.5 | 0.4 |
| 7 | 12123472 | rs2192744 | -0.261 | 1.569 | 0.541 | 0.315 | 0.003 | 0.013 | 0.013 | 0 | 0.72 | 0.69 | 0.64 | 0.57 | 0.67 | 0.58 |
| 7 | 12126942 | rs7782424 | -0.894 | 1.594 | 0.594 | 0.335 | 0.015 | 0.021 | 0.021 | 0.039 | 0.32 | 0.48 | 0.38 | 0.36 | 0.39 | 0.31 |
| 7 | 12126980 | rs4721027 | 1.314 | 1.681 | 0.458 | 0.305 | -0.006 | 0.003 | 0.003 | 0 | 0.18 | 0.14 | 0.26 | 0.12 | 0.2 | 0.18 |
| 7 | 12127178 | rs7801957 | -0.364 | 1.557 | 0.523 | 0.499 | 0.058 | 0.075 | 0.075 | 0 | 0.11 | 0.34 | 0.2 | 0.15 | 0.19 | 0.09 |
| 7 | 12128426 | rs6958807 | -0.258 | 1.469 | -0.007 | 0.47 | -0.046 | 0.01 | 0.01 | 0 | 0.25 | 0.11 | 0.25 | 0.21 | 0.33 | 0.25 |
| 7 | 12130831 | rs2215992 | -1.101 | 1.215 | 0.242 | 0.625 | -0.006 | 0.012 | 0.012 | 0 | 0.8 | 0.82 | 0.77 | 0.63 | 0.87 | 0.74 |
| 7 | 12130843 | rs12699307 | 0.499 | 1.356 | 0.118 | 0.408 | -0.013 | 0.009 | 0.009 | 0 | 0.23 | 0.19 | 0.15 | 0.16 | 0.26 | 0.28 |
| 7 | 12133937 | rs12533558 | 1.242 | 0.861 | 0.35 | 0.53 | -0.027 | -0.014 | -0.014 | 0 | 0.32 | 0.46 | 0.23 | 0.25 | 0.27 | 0.22 |
| 7 | 12136802 | rs10807750 | -0.591 | 1.034 | 0.37 | 0.805 | 0.005 | 0.111 | 0.111 | 0 | 0.78 | 0.59 | 0.71 | 0.61 | 0.81 | 0.72 |
| 7 | 12142277 | rs6963473 | 0.486 | 1.038 | 0.389 | 0.644 | 0.005 | -0.002 | -0.002 | 0 | 0.36 | 0.65 | 0.44 | 0.4 | 0.34 | 0.34 |
| 7 | 12142345 | rs6949487 | 0.476 | 1.036 | 0.388 | 0.643 | 0.005 | -0.002 | -0.002 | 0 | 0.36 | 0.65 | 0.44 | 0.4 | 0.34 | 0.34 |
| 7 | 12145295 | rs2110713 | 1.045 | 0.905 | 0.44 | 0.585 | -0.022 | -0.019 | -0.019 | 0 | 0.28 | 0.56 | 0.32 | 0.23 | 0.2 | 0.19 |
| 7 | 12148307 | rs17165577 | 0.897 | 1.127 | -0.219 | 0.445 | -0.037 | 0.03 | 0.03 | 0 | 0.14 | 0.01 | 0.12 | 0.07 | 0.18 | 0.1 |
| 7 | 12157201 | rs11771114 | -0.106 | 1.122 | 0.392 | 0.505 | -0.001 | 0.008 | 0.008 | 0 | 0.51 | 0.54 | 0.49 | 0.35 | 0.53 | 0.4 |
| 7 | 12164100 | rs1003433 | -1.215 | 1.09 | 0.032 | 0.566 | -0.095 | -0.152 | -0.152 | 0 | 0.64 | 0.97 | 0.69 | 0.54 | 0.67 | 0.57 |
| 7 | 12166753 | rs4719294 | -0.248 | 0.935 | 0.388 | 0.458 | -0.004 | 0.029 | 0.029 | 0 | 0.53 | 0.63 | 0.5 | 0.36 | 0.53 | 0.41 |
| 7 | 12166784 | rs4721036 | -0.138 | 0.942 | 0.024 | 0.353 | -0.043 | 0.026 | 0.026 | 0.015 | 0.25 | 0.13 | 0.23 | 0.16 | 0.34 | 0.21 |
| 7 | 12170496 | rs4721042 | -0.372 | 1.14 | 0.231 | 0.536 | -0.015 | -0.007 | -0.007 | 0 | 0.56 | 0.68 | 0.51 | 0.5 | 0.54 | 0.55 |
| 7 | 12172083 | rs11770347 | 0.034 | 1.045 | -0.128 | 0.254 | 0.018 | 0.036 | 0.036 | 0 | 0.48 | 0.64 | 0.45 | 0.36 | 0.43 | 0.4 |
| 7 | 12173903 | rs2058641 | -0.772 | 0.884 | -0.039 | 0.281 | -0.016 | 0.014 | 0.014 | 0.51 | 0.38 | 0.65 | 0.52 | 0.3 | 0.51 | 0.34 |
| 7 | 12175112 | rs17652095 | 0.611 | 1.143 | -0.012 | 0.21 | -0.024 | -0.009 | -0.009 | 0.236 | 0.11 | 0.08 | 0.13 | 0.11 | 0.18 | 0.14 |
| 7 | 12175556 | rs6955418 | -1.045 | 1.1 | 0.004 | 0.228 | -0.017 | 0.016 | 0.016 | 0.51 | 0.38 | 0.65 | 0.52 | 0.3 | 0.51 | 0.34 |
| 7 | 12178420 | rs7786633 | -0.129 | 0.705 | 0.558 | 0.696 | 0.03 | 0.253 | 0.253 | 0 | 0.05 | 0.3 | 0.04 | 0.18 | 0.01 | 0.2 |
| 7 | 12180568 | rs17165656 | 1.408 | 0.672 | -0.12 | 0.81 | -0.047 | -0.048 | -0.048 | 0.024 | 0.21 | 0.02 | 0.22 | 0.2 | 0.2 | 0.22 |
| 7 | 12185607 | rs2111480 | -0.949 | 0.341 | -0.41 | 0.429 | -0.091 | -0.147 | -0.147 | 0.451 | 0.58 | 0.97 | 0.62 | 0.49 | 0.59 | 0.56 |
| 7 | 12188369 | rs11983376 | 1.017 | 0.65 | -0.215 | 0.046 | -0.004 | -0.001 | -0.001 | 0 | 0.12 | 0.14 | 0.14 | 0.1 | 0.18 | 0.13 |
| 7 | 12189920 | rs10224075 | 0.872 | 0.523 | -0.101 | -0.051 | -0.044 | 0.083 | 0.083 | 0 | 0.22 | 0.13 | 0.19 | 0.06 | 0.31 | 0.17 |
| 7 | 12190097 | rs4721049 | -0.362 | 0.495 | 0.278 | 0.047 | 0.004 | 0.006 | 0.006 | 0.047 | 0.5 | 0.62 | 0.41 | 0.17 | 0.58 | 0.42 |
| 7 | 12190364 | rs4721050 | -0.329 | 0.533 | 0.008 | 0.013 | 0.02 | 0.116 | 0.116 | 0.303 | 0.48 | 0.38 | 0.39 | 0.16 | 0.58 | 0.33 |
| 7 | 12192744 | rs7795554 | 0.401 | 0.526 | 0.088 | -0.02 | -0.003 | 0.003 | 0.003 | 0.482 | 0.38 | 0.33 | 0.26 | 0.06 | 0.39 | 0.19 |
| 7 | 12195083 | rs6963608 | -0.321 | 0.529 | 0.139 | 0.088 | 0.013 | 0.058 | 0.058 | 0.145 | 0.49 | 0.42 | 0.39 | 0.16 | 0.58 | 0.33 |
| 7 | 12196391 | rs12536822 | 0.43 | 0.497 | -0.018 | 0.213 | -0.006 | -0.013 | -0.013 | 0.763 | 0.3 | 0.48 | 0.39 | 0.25 | 0.27 | 0.28 |
| 7 | 12200060 | rs6952272 | 1.617 | 0.652 | -0.94 | 0.183 | -0.018 | -0.05 | -0.05 | 0 | 0.2 | 0.02 | 0.21 | 0.19 | 0.18 | 0.21 |
| 7 | 12200775 | rs4281019 | -0.433 | 0.811 | -0.593 | -0.154 | 0.019 | 0.163 | 0.163 | 0.157 | 0.52 | 0.82 | 0.41 | 0.19 | 0.57 | 0.42 |
| 7 | 12201160 | rs10247990 | 0.866 | 0.888 | -0.705 | -0.16 | -0.047 | 0.08 | 0.08 | 0.007 | 0.21 | 0.12 | 0.17 | 0.06 | 0.29 | 0.18 |
| 7 | 12212027 | rs1369491 | 0.418 | 0.713 | -0.324 | -0.03 | -0.005 | -0.034 | -0.034 | 0.644 | 0.31 | 0.59 | 0.23 | 0.13 | 0.27 | 0.24 |

| | | | | | | | | | | | | | | | | |
|---|---|---|---|---|---|---|---|---|---|---|---|---|---|---|---|---|
| 7 | 12213484 | rs1435536 | -1.253 | 0.592 | -0.659 | 0.304 | -0.034 | -0.079 | -0.079 | 0 | 0.77 | 0.97 | 0.82 | 0.67 | 0.78 | 0.76 |
| 7 | 12217878 | rs7780991 | 1.776 | 0.499 | -0.342 | 0.103 | -0.023 | -0.067 | -0.067 | 0.038 | 0.17 | 0.27 | 0.08 | 0.02 | 0.09 | 0.03 |
| 7 | 12218360 | rs6969617 | -1.195 | 0.718 | -0.819 | 0.248 | -0.026 | -0.058 | -0.058 | 0.497 | 0.77 | 0.89 | 0.8 | 0.63 | 0.76 | 0.75 |
| 7 | 12226725 | rs1020007 | 1.715 | 0.295 | -0.464 | 0.286 | -0.013 | -0.046 | -0.046 | 0.324 | 0.17 | 0.28 | 0.08 | 0.02 | 0.09 | 0.11 |
| 7 | 12227023 | rs17149891 | 0.672 | 0.503 | -1.048 | 0.469 | -0.005 | -0.106 | -0.106 | 0 | 0.25 | 0.07 | 0.41 | 0.47 | 0.23 | 0.34 |
| 7 | 12227892 | rs1468915 | 0.791 | 0.438 | -1.078 | 0.546 | -0.01 | -0.079 | -0.079 | 0 | 0.24 | 0.05 | 0.26 | 0.35 | 0.23 | 0.33 |
| 7 | 12240236 | rs17595402 | 1.405 | 0.652 | -0.994 | 0.444 | -0.015 | -0.023 | -0.023 | 0 | 0.06 | 0.03 | 0.06 | 0.1 | 0.07 | 0.11 |
| 7 | 12243680 | rs10258615 | 0.822 | 0.885 | -1.332 | -0.112 | -0.029 | 0.067 | 0.067 | 0 | 0.19 | 0.11 | 0.15 | 0.05 | 0.25 | 0.12 |
| 7 | 12249893 | rs1435528 | -1.15 | 0.797 | -1.689 | 0.164 | -0.064 | -0.046 | -0.046 | 0.043 | 0.64 | 0.98 | 0.77 | 0.69 | 0.71 | 0.75 |
| 7 | 12257728 | rs6460903 | -0.559 | 0.812 | -1.552 | 0.108 | -0.059 | 0.013 | 0.013 | 0 | 0.61 | 0.81 | 0.66 | 0.65 | 0.54 | 0.66 |
| 7 | 12260546 | rs12666249 | 0.421 | 0.716 | -0.969 | 0.201 | 0.031 | -0.033 | -0.033 | 0 | 0.43 | 0.24 | 0.47 | 0.53 | 0.3 | 0.43 |
| 7 | 12265988 | rs6966915 | -0.356 | 0.733 | -0.64 | 0.169 | -0.003 | 0.043 | 0.043 | 0 | 0.54 | 0.67 | 0.57 | 0.65 | 0.4 | 0.64 |
| 7 | 12270900 | rs16877361 | 1.018 | 0.735 | -0.06 | 0.181 | 0.028 | 0.055 | 0.055 | 0 | 0.1 | 0.27 | 0.08 | 0.12 | 0.1 | 0.2 |
| 7 | 12270967 | rs2302632 | 0.383 | 0.692 | -0.032 | 0.313 | -0.011 | -0.01 | -0.01 | 0 | 0.43 | 0.44 | 0.25 | 0.34 | 0.3 | 0.42 |
| 7 | 12271666 | rs1042946 | 2.808 | 0.722 | -0.092 | 0.211 | -0.008 | 0.088 | 0.088 | 1.648 | 0.14 | 0.01 | 0.05 | 0 | 0.08 | 0.02 |
| 7 | 12273152 | rs14978 | 0.262 | 0.716 | -0.093 | 0.224 | -0.006 | 0.003 | 0.003 | 1.137 | 0.44 | 0.41 | 0.49 | 0.53 | 0.3 | 0.44 |
| 7 | 12274220 | rs10488193 | 1.567 | 0.816 | -0.523 | 0.257 | -0.025 | -0.025 | -0.025 | 1.533 | 0.08 | 0.02 | 0.06 | 0.09 | 0.11 | 0.18 |
| 7 | 12275818 | rs1060700 | -0.278 | 0.843 | -0.464 | 0.213 | -0.006 | 0.054 | 0.054 | 0 | 0.54 | 0.72 | 0.57 | 0.65 | 0.41 | 0.64 |
| 7 | 12277081 | rs10488192 | 1.496 | 0.75 | -0.641 | 0.444 | -0.038 | -0.046 | -0.046 | 2.932 | 0.21 | 0.01 | 0.15 | 0.19 | 0.19 | 0.21 |
| 7 | 12286409 | rs6945902 | 0.159 | 0.581 | -0.293 | 0.545 | 0.004 | -0.008 | -0.008 | 0.105 | 0.32 | 0.67 | 0.42 | 0.43 | 0.22 | 0.42 |
| 7 | 12286682 | rs10225421 | -0.117 | 0.152 | -0.031 | 0.161 | -0.005 | 0.032 | 0.032 | 0 | 0.46 | 0.38 | 0.4 | 0.26 | 0.57 | 0.35 |
| 7 | 12298311 | rs10234185 | -1.271 | -0.022 | -0.505 | 0.085 | -0.087 | -0.132 | -0.132 | 0.135 | 0.7 | 0.96 | 0.74 | 0.59 | 0.67 | 0.57 |
| 7 | 12299586 | rs6949562 | -0.284 | -0.166 | 0.074 | -0.163 | 0.007 | 0.012 | 0.012 | 0.413 | 0.28 | 0.7 | 0.21 | 0.16 | 0.25 | 0.23 |
| 7 | 12300751 | rs4721066 | 1.44 | 0.074 | -0.254 | -0.408 | -0.055 | 0.097 | 0.097 | 0 | 0.16 | 0.02 | 0.1 | 0 | 0.2 | 0.1 |
| 7 | 12304073 | rs2216050 | -0.946 | -0.005 | 0.36 | -0.078 | -0.018 | -0.009 | -0.009 | 0 | 0.17 | 0.66 | 0.16 | 0.13 | 0.16 | 0.19 |
| 7 | 12308175 | rs7808979 | -0.967 | -0.121 | 0.184 | -0.152 | -0.001 | -0.001 | -0.001 | 0.435 | 0.22 | 0.74 | 0.21 | 0.17 | 0.24 | 0.23 |
| 7 | 12320341 | rs4496864 | -0.364 | -0.55 | 0.715 | 0.301 | 0.013 | 0.231 | 0.231 | 1.474 | 0.08 | 0.53 | 0.05 | 0.18 | 0.02 | 0.19 |
| 7 | 12322548 | rs10272812 | 0.264 | -0.568 | 1.115 | 0.313 | 0.01 | 0.13 | 0.13 | 0.758 | 0.07 | 0.32 | 0.04 | 0.14 | 0.02 | 0.16 |
| 7 | 12323757 | rs10247702 | -0.159 | -0.568 | 0.995 | 0.301 | 0.025 | 0.277 | 0.277 | 1.057 | 0.07 | 0.41 | 0.04 | 0.19 | 0.02 | 0.19 |
| 7 | 12335168 | rs2160271 | -0.936 | -0.618 | 0.88 | 0.282 | -0.04 | 0.117 | 0.117 | 3.21 | 0.1 | 0.66 | 0.06 | 0.19 | 0.03 | 0.2 |
| 7 | 12341939 | rs1861311 | 0.009 | -0.817 | 0.269 | 0.082 | 0.012 | 0.023 | 0.023 | 3.228 | 0.45 | 0.28 | 0.54 | 0.44 | 0.4 | 0.33 |
| 7 | 12342641 | rs1079506 | 0.612 | -0.812 | 0.139 | 0.171 | 0.005 | 0.125 | 0.125 | 0.885 | 0.06 | 0.28 | 0.04 | 0.14 | 0.02 | 0.16 |
| 7 | 12347689 | rs7795828 | 0.333 | -0.89 | -0.901 | 0.014 | -0.026 | -0.061 | -0.061 | 0 | 0.36 | 0.09 | 0.45 | 0.43 | 0.32 | 0.31 |
| 7 | 12357429 | rs12668545 | 0.771 | -1.056 | -1.028 | 0.077 | 0.017 | 0.014 | 0.014 | 0 | 0.26 | 0.12 | 0.39 | 0.27 | 0.22 | 0.18 |
| 7 | 12360805 | rs2356167 | -0.201 | -0.964 | -0.999 | -0.009 | -0.036 | -0.021 | -0.021 | 0.094 | 0.43 | 0.1 | 0.52 | 0.43 | 0.39 | 0.31 |
| 7 | 12373082 | rs6460929 | -1.651 | -0.78 | -1.096 | -0.132 | -0.115 | -0.099 | -0.099 | 0.021 | 0.62 | 0.94 | 0.7 | 0.62 | 0.59 | 0.56 |
| 7 | 12376811 | rs2192828 | -1.651 | -0.791 | -1.096 | -0.132 | -0.119 | -0.099 | -0.099 | 0 | 0.62 | 0.94 | 0.7 | 0.62 | 0.59 | 0.56 |
| 7 | 12385391 | rs10262667 | -0.411 | -0.764 | -0.562 | -0.075 | 0.008 | 0 | 0 | 0.445 | 0.16 | 0.56 | 0.16 | 0.14 | 0.19 | 0.2 |
| 7 | 12391319 | rs6967385 | -1.053 | -0.946 | -1.074 | 0.005 | -0.036 | -0.067 | -0.067 | 0.374 | 0.54 | 0.93 | 0.59 | 0.59 | 0.41 | 0.5 |
| 7 | 12391862 | rs7804218 | 0.28 | -0.791 | -0.469 | 0.241 | -0.006 | 0.014 | 0.014 | 7.816 | 0.13 | 0.15 | 0.09 | 0.19 | 0.1 | 0.14 |
| 7 | 12398651 | rs10251962 | 0.355 | -0.812 | -0.177 | 0.264 | -0.011 | 0.07 | 0.07 | 0.387 | 0.06 | 0.18 | 0.03 | 0.11 | 0.02 | 0.18 |
| 7 | 12400837 | rs2053380 | -0.349 | -0.884 | -0.951 | 0.107 | -0.049 | 0.013 | 0.013 | 0 | 0.7 | 0.9 | 0.55 | 0.72 | 0.69 | 0.84 |
| 7 | 12402989 | rs17165896 | 0.17 | -0.854 | -0.866 | 0.298 | 0.014 | -0.075 | -0.075 | 0.02 | 0.16 | 0.04 | 0.09 | 0.31 | 0.12 | 0.26 |
| 7 | 12405098 | rs2033871 | 1.631 | -0.691 | -0.695 | 0.098 | 0 | -0.014 | -0.014 | 0 | 0.07 | 0.08 | 0.1 | 0 | 0.14 | 0.03 |
| 7 | 12406826 | rs1371608 | 0.83 | -1.195 | -0.322 | -0.095 | -0.008 | -0.004 | -0.004 | 0 | 0.43 | 0.57 | 0.36 | 0.51 | 0.45 | 0.65 |
| 7 | 12409263 | rs17165906 | 0.866 | -1.18 | -0.211 | -0.059 | -0.006 | 0.015 | 0.015 | 8.748 | 0.13 | 0.17 | 0.08 | 0.2 | 0.1 | 0.19 |
| 7 | 12409327 | rs17165910 | 0.797 | -1.182 | -0.212 | -0.06 | -0.006 | 0.013 | 0.013 | 8.502 | 0.14 | 0.18 | 0.08 | 0.2 | 0.1 | 0.19 |
| 7 | 12409522 | rs3823844 | 0.633 | -1.196 | -0.211 | -0.066 | -0.013 | 0.027 | 0.027 | 7.81 | 0.14 | 0.23 | 0.09 | 0.2 | 0.1 | 0.21 |
| 7 | 12410112 | rs963323 | -0.13 | -1.506 | 0.156 | -0.316 | -0.012 | -0.021 | -0.021 | 0 | 0.76 | 0.7 | 0.85 | 0.82 | 0.82 | 0.76 |
| 7 | 12410811 | rs7786075 | -0.31 | -1.496 | 0.287 | -0.311 | -0.002 | -0.004 | -0.004 | 0.338 | 0.77 | 0.81 | 0.86 | 0.82 | 0.82 | 0.76 |
| 7 | 12415923 | rs17165942 | -0.078 | -1.104 | 0.654 | -0.083 | -0.002 | 0.006 | 0.006 | 0.747 | 0.2 | 0.3 | 0.18 | 0.21 | 0.24 | 0.24 |
| 7 | 12416367 | rs1440024 | -0.303 | -1.112 | 0.637 | -0.087 | -0.004 | 0.009 | 0.009 | 3.074 | 0.21 | 0.39 | 0.19 | 0.21 | 0.24 | 0.24 |
| 7 | 12418144 | rs4721095 | 0.998 | -1.092 | 0.54 | -0.06 | 0.007 | -0.026 | -0.026 | 0 | 0.13 | 0.07 | 0.08 | 0.2 | 0.1 | 0.21 |
| 7 | 12418602 | rs10270435 | 0.3 | -1.154 | 0.639 | -0.089 | -0.02 | 0.066 | 0.066 | 0.277 | 0.07 | 0.24 | 0.03 | 0.12 | 0.02 | 0.19 |
| 7 | 12418964 | rs10270931 | 0.617 | -1.16 | 0.506 | -0.2 | 0.017 | 0.039 | 0.039 | 0 | 0.48 | 0.34 | 0.43 | 0.59 | 0.55 | 0.56 |
| 7 | 12419108 | rs6967241 | 0.18 | -1.1 | 0.199 | -0.125 | -0.003 | -0.003 | -0.003 | 0 | 0.55 | 0.57 | 0.53 | 0.59 | 0.68 | 0.66 |
| 7 | 12420447 | rs6959370 | 0.155 | -1.051 | -0.031 | -0.073 | 0.009 | 0.011 | 0.011 | 0 | 0.55 | 0.48 | 0.67 | 0.63 | 0.69 | 0.67 |
| 7 | 12422721 | rs12699367 | 1.295 | -0.653 | -0.554 | 0.063 | -0.125 | -0.103 | -0.103 | 0.327 | 0.34 | 0.07 | 0.33 | 0.41 | 0.44 | 0.42 |
| 7 | 12428608 | rs6954047 | 0.108 | -0.814 | -0.051 | -0.192 | -0.018 | -0.006 | -0.006 | 0.764 | 0.38 | 0.52 | 0.5 | 0.41 | 0.35 | 0.38 |
| 7 | 12429488 | rs848013 | -0.646 | -0.848 | 0.153 | 0.062 | 0.012 | 0.008 | 0.008 | 0 | 0.48 | 0.66 | 0.57 | 0.49 | 0.44 | 0.42 |
| 7 | 12430812 | rs848011 | -0.663 | -0.83 | 0.125 | 0.058 | 0.014 | 0.011 | 0.011 | 0 | 0.48 | 0.7 | 0.58 | 0.49 | 0.45 | 0.42 |
| 7 | 12433195 | rs12669558 | 0.81 | -1.015 | 0.022 | 0.048 | 0.007 | -0.004 | -0.004 | 0 | 0.3 | 0.3 | 0.4 | 0.41 | 0.2 | 0.35 |
| 7 | 12433347 | rs17165995 | 0.218 | -0.959 | -0.032 | 0.07 | -0.023 | -0.016 | -0.016 | 0 | 0.17 | 0.4 | 0.12 | 0.13 | 0.11 | 0.15 |
| 7 | 12443795 | rs6968277 | 0.528 | -0.91 | -0.27 | 0.138 | 0.001 | 0.001 | 0.001 | 0.793 | 0.12 | 0.26 | 0.13 | 0.11 | 0.16 | 0.2 |
| 7 | 12444157 | rs9639128 | 1.317 | -0.948 | -0.637 | 0.141 | 0.024 | -0.009 | -0.009 | 0.16 | 0.25 | 0.07 | 0.37 | 0.29 | 0.18 | 0.18 |
| 7 | 12462428 | rs17444259 | 0.108 | -0.883 | -0.27 | 0.141 | 0 | 0.001 | 0.001 | 0 | 0.13 | 0.13 | 0.13 | 0.12 | 0.16 | 0.25 |
| 7 | 12463056 | rs1371607 | -0.61 | -0.958 | -0.433 | 0.138 | -0.006 | -0.013 | -0.013 | 0 | 0.24 | 0.1 | 0.21 | 0.25 | 0.27 | 0.36 |
| 7 | 12463109 | rs847901 | -1.943 | -0.999 | -0.443 | 0.079 | 0.003 | 0.019 | 0.019 | 1.26 | 0.39 | 0.82 | 0.31 | 0.33 | 0.37 | 0.41 |
| 7 | 12473521 | rs920428 | -0.751 | -0.955 | -0.339 | 0.117 | 0.003 | 0 | 0 | 0 | 0.25 | 0.23 | 0.22 | 0.25 | 0.27 | 0.35 |

| | | | | | | | | | | | | | | | | |
|---|---|---|---|---|---|---|---|---|---|---|---|---|---|---|---|---|
| 7 | 12476691 | rs17166028 | 1.561 | -0.909 | -0.436 | 0.135 | -0.026 | -0.038 | -0.038 | 0 | 0.11 | 0 | 0.08 | 0.13 | 0.11 | 0.12 |
| 7 | 12477478 | rs702474 | -2.268 | -0.505 | -0.986 | -0.343 | -0.027 | 0.075 | 0.075 | 0 | 0.86 | 0.99 | 0.9 | 0.97 | 0.86 | 0.82 |
| 7 | 12479584 | rs847976 | 1.512 | -0.34 | -0.254 | -0.177 | 0.023 | -0.001 | -0.001 | 0 | 0.1 | 0.18 | 0.06 | 0.08 | 0.1 | 0.05 |
| 7 | 12481461 | rs17542243 | -0.438 | -0.409 | -0.247 | -0.159 | 0.001 | -0.001 | -0.001 | 0 | 0.25 | 0.25 | 0.23 | 0.26 | 0.27 | 0.35 |
| 7 | 12482879 | rs1371601 | -0.391 | 0.099 | -0.375 | -0.285 | -0.013 | -0.019 | -0.019 | 0 | 0.56 | 0.32 | 0.65 | 0.55 | 0.59 | 0.56 |
| 7 | 12484517 | rs1440019 | 0.755 | 0.061 | -0.394 | -0.308 | -0.004 | 0.011 | 0.011 | 0.391 | 0.16 | 0.22 | 0.12 | 0.26 | 0.13 | 0.28 |
| 7 | 12484649 | rs1440018 | -0.836 | -0.003 | -0.549 | -0.348 | -0.007 | -0.009 | -0.009 | 0.052 | 0.6 | 0.67 | 0.67 | 0.55 | 0.59 | 0.56 |
| 7 | 12485026 | rs12699372 | 0.757 | 0.03 | -0.737 | -0.344 | -0.059 | 0.095 | 0.095 | 0 | 0.4 | 0.1 | 0.52 | 0.29 | 0.46 | 0.28 |
| 7 | 12487380 | rs1595804 | 0.198 | -0.012 | -0.685 | -0.281 | -0.057 | 0.044 | 0.044 | 0.586 | 0.51 | 0.1 | 0.6 | 0.44 | 0.57 | 0.39 |
| 7 | 12491847 | rs1100135 | -1.454 | 0.123 | -0.629 | -0.252 | -0.05 | -0.115 | -0.115 | 0.112 | 0.69 | 0.95 | 0.74 | 0.63 | 0.68 | 0.6 |
| 7 | 12498138 | rs12669338 | 1.526 | 0.17 | -0.578 | -0.241 | -0.04 | 0.025 | 0.025 | 0.016 | 0.31 | 0.08 | 0.43 | 0.3 | 0.32 | 0.21 |
| 7 | 12498511 | rs12667501 | -0.322 | 0.134 | -0.317 | -0.233 | -0.003 | -0.004 | -0.004 | 0 | 0.57 | 0.43 | 0.66 | 0.55 | 0.59 | 0.55 |
| 7 | 12505715 | rs848022 | -1.379 | 0.251 | -0.762 | -0.106 | 0.018 | 0.005 | 0.005 | 0.003 | 0.76 | 0.87 | 0.85 | 0.78 | 0.8 | 0.75 |
| 7 | 12507237 | rs2356486 | -0.574 | 0.769 | -0.607 | 0.062 | -0.002 | 0.001 | 0.001 | 0 | 0.42 | 0.62 | 0.67 | 0.61 | 0.48 | 0.55 |
| 7 | 12507250 | rs2883954 | -0.574 | 0.769 | -0.607 | 0.062 | -0.002 | 0.001 | 0.001 | 0 | 0.42 | 0.62 | 0.67 | 0.61 | 0.48 | 0.55 |
| 7 | 12511312 | rs11275 | -0.137 | 0.795 | -0.348 | 0.184 | 0.003 | 0.003 | 0.003 | 0 | 0.24 | 0.41 | 0.17 | 0.22 | 0.21 | 0.18 |
| 7 | 12519626 | rs10488154 | 0.747 | 1.222 | -0.407 | 0.212 | -0.007 | -0.008 | -0.008 | 0 | 0.19 | 0.16 | 0.46 | 0.48 | 0.21 | 0.42 |
| 7 | 12521871 | rs17544637 | 2.018 | 1.217 | -0.467 | 0.027 | 0.008 | -0.03 | -0.03 | 0 | 0.08 | 0.1 | 0.08 | 0 | 0.13 | 0.02 |
| 7 | 12522236 | rs6460955 | -0.781 | 0.848 | -0.672 | -0.204 | 0.007 | 0 | 0 | 0.014 | 0.68 | 0.81 | 0.8 | 0.69 | 0.71 | 0.71 |
| 7 | 12531907 | rs17166104 | 1.725 | 0.902 | -0.587 | -0.461 | 0.011 | 0.043 | 0.043 | 0.74 | 0.27 | 0.18 | 0.14 | 0.08 | 0.25 | 0.15 |
| 7 | 12533065 | rs847919 | -2.093 | 0.81 | -0.795 | -0.269 | -0.01 | -0.033 | -0.033 | 3.955 | 0.78 | 0.97 | 0.86 | 0.78 | 0.8 | 0.77 |
| 7 | 12533124 | rs847918 | 0.193 | 0.962 | -0.523 | -0.208 | 0.018 | 0.047 | 0.047 | 0 | 0.28 | 0.32 | 0.52 | 0.48 | 0.29 | 0.44 |
| 7 | 12534936 | rs847914 | -0.123 | 0.886 | -0.696 | -0.169 | 0 | -0.001 | -0.001 | 0.026 | 0.55 | 0.5 | 0.66 | 0.55 | 0.54 | 0.58 |
| 7 | 12543120 | rs13246109 | -1.696 | 0.958 | -0.572 | -0.192 | 0.082 | -0.031 | -0.031 | 0 | 0.91 | 0.68 | 0.94 | 1 | 0.92 | 0.98 |
| 7 | 12543665 | rs11977987 | 1.312 | 0.996 | -0.631 | -0.192 | -0.001 | 0.05 | 0.05 | 0 | 0.14 | 0.3 | 0.11 | 0.23 | 0.13 | 0.16 |
| 7 | 12543794 | rs17166119 | 0.793 | 1.043 | -1.145 | -0.267 | -0.027 | -0.072 | -0.072 | 0 | 0.22 | 0.11 | 0.49 | 0.48 | 0.27 | 0.43 |
| 7 | 12544057 | rs4721111 | -2.813 | 1.055 | -1.098 | -0.289 | -0.034 | 0.035 | 0.035 | 0 | 0.93 | 1 | 0.96 | 1 | 0.92 | 0.98 |
| 7 | 12547738 | rs11983536 | 1.877 | 0.862 | -1.046 | -0.243 | 0.002 | 0.109 | 0.109 | 0.714 | 0.24 | 0.07 | 0.1 | 0.08 | 0.19 | 0.14 |
| 7 | 12547917 | rs10249671 | -0.042 | 0.981 | -0.617 | 0.143 | 0.011 | 0.011 | 0.011 | 0.046 | 0.22 | 0.44 | 0.2 | 0.23 | 0.26 | 0.18 |
| 7 | 12553243 | rs12531198 | 0.181 | 1.203 | -0.869 | 0.32 | -0.002 | -0.013 | -0.013 | 0.803 | 0.22 | 0.16 | 0.13 | 0.3 | 0.18 | 0.36 |
| 7 | 12562063 | rs7801809 | 0.262 | 0.723 | -0.762 | -0.151 | -0.002 | -0.002 | -0.002 | 0 | 0.53 | 0.54 | 0.61 | 0.54 | 0.46 | 0.48 |
| 7 | 12562500 | rs1992309 | 2.077 | 0.745 | -1.049 | -0.17 | 0.003 | -0.036 | -0.036 | 0.629 | 0.07 | 0.11 | 0.03 | 0 | 0.05 | 0.02 |
| 7 | 12568567 | rs4291161 | -0.788 | 0.945 | -1.486 | -0.133 | -0.048 | -0.084 | -0.084 | 3.988 | 0.7 | 0.86 | 0.74 | 0.58 | 0.68 | 0.54 |
| 7 | 12570377 | rs4365982 | -0.236 | 0.675 | -1.216 | -0.024 | -0.047 | -0.022 | -0.022 | 0 | 0.61 | 0.77 | 0.65 | 0.58 | 0.52 | 0.51 |
| 7 | 12570647 | rs11771290 | 1.385 | 0.44 | -1.482 | 0.066 | -0.02 | -0.058 | -0.058 | 0 | 0.26 | 0.03 | 0.17 | 0.25 | 0.24 | 0.22 |
| 7 | 12572742 | rs10250755 | 0.555 | 0.54 | -0.88 | 0.204 | -0.003 | -0.02 | -0.02 | 0 | 0.42 | 0.26 | 0.46 | 0.44 | 0.46 | 0.37 |
| 7 | 12580007 | rs2159043 | 1.144 | 0.666 | -0.84 | 0.32 | 0.003 | 0.028 | 0.028 | 0 | 0.14 | 0.16 | 0.26 | 0.3 | 0.2 | 0.09 |
| 7 | 12581042 | rs10262356 | -1.253 | 0.579 | -0.599 | 0.15 | 0 | 0.005 | 0.005 | 0 | 0.77 | 0.8 | 0.71 | 0.62 | 0.72 | 0.61 |
| 7 | 12583080 | rs2108554 | 1.626 | 0.419 | -0.501 | -0.217 | -0.009 | 0.014 | 0.014 | 2.434 | 0.2 | 0.15 | 0.19 | 0.08 | 0.19 | 0.14 |
| 7 | 12587422 | rs10268418 | -0.252 | -0.224 | 0.321 | 0.237 | -0.008 | 0.012 | 0.012 | 1.174 | 0.66 | 0.51 | 0.67 | 0.61 | 0.66 | 0.57 |
| 7 | 12593214 | rs1034854 | 0.589 | 0.15 | 0.489 | 0.329 | -0.005 | 0.022 | 0.022 | 0.313 | 0.62 | 0.46 | 0.63 | 0.53 | 0.61 | 0.44 |
| 7 | 12596535 | rs11760555 | 0.535 | 0.155 | 0.309 | 0.335 | 0 | -0.002 | -0.002 | 0 | 0.62 | 0.62 | 0.64 | 0.53 | 0.6 | 0.43 |
| 7 | 12596616 | rs11764174 | 0.535 | 0.155 | 0.309 | 0.335 | 0 | -0.002 | -0.002 | 0.172 | 0.62 | 0.63 | 0.64 | 0.53 | 0.6 | 0.44 |
| 7 | 12604371 | rs6954111 | 0.471 | 0.151 | 0.471 | -0.421 | 0 | -0.016 | -0.016 | 0 | 0.52 | 0.52 | 0.41 | 0.24 | 0.46 | 0.37 |
| 7 | 12609679 | rs3735222 | 1.067 | 0.148 | 0.504 | -0.432 | -0.001 | -0.015 | -0.015 | 0 | 0.48 | 0.5 | 0.38 | 0.24 | 0.4 | 0.34 |
| 7 | 12610903 | rs2240573 | 1.148 | 0.015 | 0.491 | -0.293 | 0.002 | -0.003 | -0.003 | 1.54 | 0.1 | 0.07 | 0.03 | 0.16 | 0.05 | 0.17 |
| 7 | 12611179 | rs12699405 | 2.497 | 0.193 | 0.439 | -0.409 | -0.008 | 0.003 | 0.003 | 0 | 0.06 | 0.02 | 0.05 | 0 | 0.08 | 0.02 |
| 7 | 12612681 | rs7811582 | 0.471 | 0.07 | 0.368 | -0.395 | -0.002 | 0.022 | 0.022 | 1.355 | 0.53 | 0.6 | 0.42 | 0.25 | 0.46 | 0.38 |
| 7 | 12613059 | rs4458754 | 1.721 | -0.058 | 0.347 | -0.466 | -0.034 | 0.156 | 0.156 | 1.48 | 0.43 | 0.15 | 0.35 | 0.21 | 0.38 | 0.26 |
| 7 | 12614638 | rs2215063 | -0.263 | 0.045 | 0.321 | -0.398 | -0.013 | -0.003 | -0.003 | 0 | 0.63 | 0.71 | 0.46 | 0.4 | 0.52 | 0.54 |
| 7 | 12615772 | rs1990225 | 0.313 | 0.087 | 0.299 | -0.377 | -0.001 | 0 | 0 | 0 | 0.57 | 0.58 | 0.42 | 0.4 | 0.46 | 0.51 |
| 7 | 12618764 | rs17166188 | -1.022 | 0.503 | 0.266 | -0.8 | 0.03 | -0.017 | -0.017 | 0 | 0.1 | 0.41 | 0.07 | 0.03 | 0.08 | 0.11 |
| 7 | 12620804 | rs17283717 | 1.109 | 0.629 | -0.814 | -0.766 | -0.021 | 0.114 | 0.114 | 3.478 | 0.4 | 0.18 | 0.34 | 0.21 | 0.36 | 0.23 |
| 7 | 12631920 | rs6967031 | -0.472 | 0.169 | -0.699 | -0.685 | 0.002 | 0.097 | 0.097 | 2.778 | 0.45 | 0.28 | 0.56 | 0.79 | 0.52 | 0.66 |
| 7 | 12632085 | rs11764374 | 0.023 | -0.306 | -0.859 | -0.509 | -0.001 | 0.036 | 0.036 | 0.637 | 0.34 | 0.16 | 0.47 | 0.68 | 0.4 | 0.59 |
| 7 | 12643396 | rs849776 | 0.428 | -0.183 | 0.377 | -0.386 | 0 | -0.098 | -0.098 | 0 | 0.82 | 0.63 | 0.89 | 1 | 0.83 | 0.94 |
| 7 | 12644334 | rs849778 | -0.207 | -0.237 | 0.499 | -0.361 | -0.013 | -0.068 | -0.068 | 0.304 | 0.86 | 0.73 | 0.92 | 1 | 0.9 | 0.98 |
| 7 | 12645380 | rs10268615 | 1.108 | -0.364 | 0.025 | 0.018 | 0.003 | -0.01 | -0.01 | 0.103 | 0.28 | 0.25 | 0.15 | 0.38 | 0.23 | 0.39 |
| 7 | 12645621 | rs849780 | -1.837 | -0.391 | -0.425 | -0.06 | -0.004 | 0.004 | 0.004 | 0 | 0.93 | 0.96 | 0.95 | 1 | 0.92 | 0.99 |
| 7 | 12647933 | rs6978454 | -0.01 | -0.436 | -0.362 | -0.124 | -0.002 | -0.002 | -0.002 | 0 | 0.44 | 0.32 | 0.66 | 0.52 | 0.47 | 0.43 |
| 7 | 12654570 | rs991317 | 0.106 | -0.154 | -0.106 | 0.015 | -0.008 | -0.007 | -0.007 | 0 | 0.43 | 0.28 | 0.66 | 0.52 | 0.47 | 0.41 |
| 7 | 12654696 | rs2691819 | 0.995 | -0.144 | -0.124 | 0.027 | -0.001 | 0.012 | 0.012 | 0 | 0.32 | 0.48 | 0.2 | 0.39 | 0.31 | 0.42 |
| 7 | 12656415 | rs849798 | 1.541 | -0.122 | -0.49 | 0.06 | 0.003 | 0.024 | 0.024 | 0 | 0.14 | 0.06 | 0.07 | 0.09 | 0.13 | 0.14 |
| 7 | 12665739 | rs886890 | 0.045 | -0.117 | -0.53 | 0.068 | -0.003 | 0 | 0 | 0.09 | 0.45 | 0.36 | 0.67 | 0.52 | 0.49 | 0.42 |
| 7 | 12666907 | rs849758 | -0.284 | 0.109 | -0.602 | 0.086 | 0 | -0.003 | -0.003 | 0 | 0.58 | 0.4 | 0.73 | 0.61 | 0.59 | 0.55 |
| 7 | 12669016 | rs2529785 | -0.148 | -0.245 | -0.971 | -0.043 | -0.002 | -0.002 | -0.002 | 0.652 | 0.72 | 0.76 | 0.86 | 0.74 | 0.77 | 0.77 |
| 7 | 12669251 | rs731257 | 0.537 | 0.071 | -2.186 | 0.064 | -0.033 | -0.078 | -0.078 | 0 | 0.12 | 0.02 | 0.39 | 0.35 | 0.15 | 0.15 |
| 7 | 12670079 | rs11980671 | 1.195 | 0.37 | -2.053 | -0.227 | -0.021 | 0.023 | 0.023 | 0 | 0.22 | 0.1 | 0.22 | 0.01 | 0.29 | 0.16 |
| 7 | 12677573 | rs849763 | 0.514 | 0.255 | -1.924 | 0.072 | 0 | -0.005 | -0.005 | 0.068 | 0.64 | 0.32 | 0.44 | 0.56 | 0.65 | 0.68 |

| | | | | | | | | | | | | | | | | |
|---|---|---|---|---|---|---|---|---|---|---|---|---|---|---|---|---|
| 7 | 12680454 | rs2074866 | 0.839 | 0.219 | -1.945 | 0.115 | 0.025 | -0.063 | -0.063 | 0 | 0.33 | 0.12 | 0.17 | 0.41 | 0.25 | 0.45 |
| 7 | 12682919 | rs2691815 | 0.079 | 0.791 | -1.518 | 0.082 | -0.027 | 0.048 | 0.048 | 0 | 0.37 | 0.22 | 0.31 | 0.2 | 0.45 | 0.28 |
| 7 | 12685530 | rs2691808 | 0.257 | 0.567 | -2.406 | 0.433 | -0.004 | -0.004 | -0.004 | 0 | 0.39 | 0.09 | 0.21 | 0.34 | 0.34 | 0.46 |
| 7 | 12686194 | rs6972755 | -0.099 | 1.054 | -1.912 | 0.308 | -0.025 | 0.022 | 0.022 | 0 | 0.34 | 0.77 | 0.53 | 0.23 | 0.41 | 0.24 |
| 7 | 12689475 | rs1180398 | -0.215 | 0.611 | -2.092 | 0.631 | -0.004 | -0.141 | -0.141 | 0 | 0.33 | 0.04 | 0.27 | 0.49 | 0.29 | 0.35 |
| 7 | 12690418 | rs1017405 | -0.572 | 1.127 | -0.707 | -0.083 | 0.061 | -0.058 | -0.058 | 0.577 | 0.15 | 0.48 | 0.3 | 0.06 | 0.17 | 0.05 |
| 7 | 12691988 | rs10225719 | -0.465 | 1.269 | -0.51 | -0.314 | 0 | -0.002 | -0.002 | 0 | 0.41 | 0.59 | 0.46 | 0.32 | 0.46 | 0.36 |
| 7 | 12693140 | rs2033604 | -0.706 | 1.966 | -0.526 | -0.149 | 0.027 | 0.011 | 0.011 | 0.028 | 0.46 | 0.66 | 0.67 | 0.44 | 0.67 | 0.53 |
| 7 | 12693392 | rs10488225 | -1.094 | 2.22 | -0.646 | -0.043 | -0.005 | -0.004 | -0.004 | 0 | 0.06 | 0.03 | 0.2 | 0.09 | 0.2 | 0.17 |
| 7 | 12695507 | rs1017404 | -0.356 | 2.422 | -0.22 | -0.22 | 0.023 | -0.021 | -0.021 | 1.208 | 0.16 | 0.23 | 0.26 | 0.15 | 0.24 | 0.18 |
| 7 | 12697797 | rs2883999 | 0.083 | 1.791 | -0.79 | -0.449 | -0.027 | -0.042 | -0.042 | 0 | 0.32 | 0.11 | 0.35 | 0.38 | 0.36 | 0.33 |
| 7 | 12705331 | rs10247152 | -0.624 | 1.762 | -0.943 | -0.475 | -0.016 | 0.019 | 0.019 | 0.555 | 0.14 | 0.11 | 0.14 | 0.04 | 0.22 | 0.21 |
| 7 | 12705376 | rs11979770 | -1.398 | 1.5 | -1.145 | -0.527 | 0.021 | -0.138 | -0.138 | 3.117 | 0.65 | 0.95 | 0.63 | 0.51 | 0.75 | 0.71 |
| 7 | 12705413 | rs10246960 | -0.637 | 2.041 | -0.063 | -0.485 | 0.043 | -0.018 | -0.018 | 0 | 0.1 | 0.25 | 0.12 | 0.04 | 0.16 | 0.2 |
| 7 | 12708420 | rs4445118 | -1.075 | 2.08 | 0.301 | -0.569 | 0.043 | -0.027 | -0.027 | 0 | 0.3 | 0.44 | 0.39 | 0.23 | 0.52 | 0.45 |
| 7 | 12708998 | rs4721127 | -0.708 | 1.154 | 0.54 | -0.515 | -0.007 | 0.123 | 0.123 | 0.1 | 0.79 | 0.58 | 0.63 | 0.42 | 0.81 | 0.72 |
| 7 | 12712299 | rs6952054 | -0.603 | 0.477 | -0.208 | -1.072 | 0.07 | 0.006 | 0.006 | 0.232 | 0.58 | 0.78 | 0.5 | 0.4 | 0.69 | 0.68 |
| 7 | 12713070 | rs10488226 | 0.997 | 0.633 | -0.684 | -1.341 | -0.081 | 0.058 | 0.058 | 2.38 | 0.25 | 0.08 | 0.24 | 0.16 | 0.38 | 0.28 |
| 7 | 12713516 | rs12540556 | 0.728 | 1.097 | 0.271 | -0.996 | 0.013 | -0.001 | -0.001 | 0.256 | 0.07 | 0.11 | 0.04 | 0.04 | 0.05 | 0.21 |
| 7 | 12718149 | rs17615676 | 0.415 | 1.09 | 0.243 | -1.08 | -0.009 | 0.005 | 0.005 | 0 | 0.18 | 0.13 | 0.12 | 0.02 | 0.22 | 0.18 |
| 7 | 12721956 | rs6460972 | -1.252 | 0.849 | -0.102 | -1.117 | -0.04 | -0.134 | -0.134 | 0.862 | 0.77 | 0.98 | 0.58 | 0.4 | 0.78 | 0.7 |
| 7 | 12725678 | rs3757443 | -0.855 | 1.29 | 1.413 | -0.558 | 0.009 | -0.012 | -0.012 | 0 | 0.1 | 0.28 | 0.05 | 0.03 | 0.05 | 0.21 |
| 7 | 12728977 | rs2241748 | -1.276 | 1.318 | 1.426 | -0.497 | 0.016 | 0.106 | 0.106 | 0.131 | 0.13 | 0.47 | 0.11 | 0.19 | 0.09 | 0.22 |
| 7 | 12729269 | rs1058268 | -0.953 | 1.473 | 1.179 | -0.469 | 0.054 | 0.115 | 0.115 | 0 | 0.08 | 0.3 | 0.14 | 0.15 | 0.11 | 0.03 |
| 7 | 12732118 | rs7797379 | -1.499 | 1.155 | 0.218 | -0.507 | 0.002 | 0.052 | 0.052 | 0.408 | 0.35 | 0.74 | 0.26 | 0.21 | 0.31 | 0.41 |
| 7 | 12735954 | rs12534728 | -1.366 | 1.267 | 0.259 | -0.476 | -0.012 | 0.038 | 0.038 | 1.185 | 0.34 | 0.72 | 0.25 | 0.2 | 0.35 | 0.4 |
| 7 | 12743015 | rs7789919 | -1.052 | 1.358 | 0.595 | -0.082 | 0.011 | -0.002 | -0.002 | 0 | 0.39 | 0.48 | 0.47 | 0.39 | 0.47 | 0.42 |
| 7 | 12747443 | rs10950409 | -1.418 | 1.546 | 1.247 | -0.115 | 0.018 | 0.076 | 0.076 | 1.722 | 0.19 | 0.54 | 0.32 | 0.32 | 0.2 | 0.2 |
| 7 | 12754269 | rs17616207 | -0.144 | 1.659 | 0.968 | -0.107 | -0.002 | 0.001 | 0.001 | 0 | 0.05 | 0.07 | 0.22 | 0.06 | 0.09 | 0.07 |
| 7 | 12756145 | rs6970019 | -0.273 | 1.58 | 1.334 | -0.06 | 0.004 | -0.031 | -0.031 | 0 | 0.11 | 0.24 | 0.06 | 0.05 | 0.1 | 0.13 |
| 7 | 12756756 | rs4410798 | -1.131 | 1.695 | 1.68 | -0.049 | 0.032 | 0.11 | 0.11 | 0.967 | 0.16 | 0.42 | 0.13 | 0.28 | 0.14 | 0.15 |
| 7 | 12758408 | rs11981235 | -1.098 | 1.704 | 1.527 | -0.045 | 0.038 | 0.12 | 0.12 | 1.178 | 0.15 | 0.43 | 0.14 | 0.28 | 0.14 | 0.15 |
| 7 | 12761041 | rs7810893 | -1.565 | 2.03 | 1.115 | -0.615 | 0.012 | -0.032 | -0.032 | 0 | 0.28 | 0.44 | 0.4 | 0.14 | 0.38 | 0.31 |
| 7 | 12762235 | rs17166325 | -1.877 | 1.95 | 0.937 | -0.557 | 0.002 | -0.002 | -0.002 | 0.241 | 0.47 | 0.64 | 0.37 | 0.19 | 0.55 | 0.59 |
| 7 | 12765509 | rs6956267 | -2.401 | 1.888 | 0.432 | -0.577 | 0.044 | 0.11 | 0.11 | 0 | 0.56 | 0.82 | 0.45 | 0.23 | 0.66 | 0.67 |
| 7 | 12777308 | rs1524937 | -0.585 | 2.177 | 0.239 | -0.43 | 0 | -0.003 | -0.003 | 0 | 0.25 | 0.21 | 0.16 | 0.08 | 0.25 | 0.33 |
| 7 | 12781543 | rs17616845 | -0.371 | 2.178 | 0.182 | -0.33 | -0.052 | 0.033 | 0.033 | 2.754 | 0.15 | 0.06 | 0.16 | 0.09 | 0.23 | 0.25 |
| 7 | 12785780 | rs17616899 | -1.043 | 2.292 | 0.415 | -0.237 | -0.007 | 0.047 | 0.047 | 0.237 | 0.32 | 0.22 | 0.21 | 0.14 | 0.36 | 0.41 |
| 7 | 12793361 | rs10260162 | -0.943 | 2.043 | 1.056 | 0.322 | 0.104 | 0.038 | 0.038 | 2.987 | 0.12 | 0.23 | 0.19 | 0.18 | 0.26 | 0.2 |
| 7 | 12801051 | rs17617404 | -0.198 | 2.126 | 1.103 | 0.351 | 0.05 | 0.017 | 0.017 | 2.717 | 0.1 | 0.12 | 0.17 | 0.19 | 0.24 | 0.2 |
| 7 | 12801837 | rs849742 | -1.329 | 1.756 | 1.049 | 0.993 | 0.003 | -0.006 | -0.006 | 0 | 0.3 | 0.27 | 0.4 | 0.4 | 0.27 | 0.43 |
| 7 | 12804200 | rs702478 | -0.788 | 1.772 | 1.042 | 1.239 | -0.005 | 0.015 | 0.015 | 0 | 0.15 | 0.19 | 0.25 | 0.27 | 0.13 | 0.21 |
| 7 | 12804435 | rs702479 | -1.795 | 1.733 | 1.335 | 1.348 | -0.01 | 0.021 | 0.021 | 0.076 | 0.32 | 0.39 | 0.42 | 0.44 | 0.29 | 0.46 |
| 7 | 12805598 | rs849745 | 0.297 | 1.739 | 0.956 | 1.297 | -0.004 | -0.008 | -0.008 | 0 | 0.17 | 0.08 | 0.17 | 0.13 | 0.19 | 0.24 |
| 7 | 12808491 | rs12666603 | 0.454 | 1.714 | 0.965 | 1.297 | 0 | -0.011 | -0.011 | 0.047 | 0.16 | 0.08 | 0.16 | 0.13 | 0.17 | 0.24 |
| 7 | 12813460 | rs10215532 | -2.315 | 1.783 | 0.974 | 1.147 | 0.046 | 0.081 | 0.081 | 1.449 | 0.5 | 0.75 | 0.67 | 0.63 | 0.6 | 0.66 |
| 7 | 12817829 | rs4719334 | -1.538 | 2.126 | 1.111 | 1.207 | 0.095 | 0.027 | 0.027 | 1.411 | 0.17 | 0.29 | 0.23 | 0.23 | 0.3 | 0.21 |
| 7 | 12819414 | rs10281877 | -2.215 | 1.875 | 1.03 | 1.175 | 0.055 | 0.07 | 0.07 | 0.231 | 0.3 | 0.45 | 0.31 | 0.49 | 0.42 | 0.38 |
| 7 | 12820470 | rs10227287 | -1.354 | 1.871 | 0.661 | 1.096 | 0.105 | 0.045 | 0.045 | 0.345 | 0.14 | 0.29 | 0.22 | 0.18 | 0.27 | 0.17 |
| 7 | 12820572 | rs2356691 | -1.739 | 1.023 | 0.544 | 1.018 | 0.009 | 0.106 | 0.106 | 0.053 | 0.26 | 0.48 | 0.3 | 0.46 | 0.31 | 0.35 |
| 7 | 12821230 | rs12666534 | 0.983 | 0.492 | 0.182 | 0.279 | -0.001 | 0.061 | 0.061 | 0.474 | 0.51 | 0.36 | 0.49 | 0.35 | 0.51 | 0.31 |
| 7 | 12821521 | rs6974659 | 0.412 | 0.165 | 0.776 | 0.098 | 0.002 | 0.007 | 0.007 | 0.443 | 0.73 | 0.58 | 0.64 | 0.4 | 0.74 | 0.56 |
| 7 | 12822398 | rs16877556 | 0.209 | -0.23 | 0.447 | -0.371 | -0.001 | -0.007 | -0.007 | 0 | 0.53 | 0.49 | 0.52 | 0.4 | 0.55 | 0.55 |
| 7 | 12839631 | rs1990206 | -0.194 | -0.726 | 0.545 | -0.049 | 0.024 | 0.005 | 0.005 | 0.836 | 0.45 | 0.68 | 0.39 | 0.38 | 0.38 | 0.33 |
| 7 | 12840583 | rs13233019 | 1.936 | -1.001 | 0.463 | -0.119 | -0.044 | 0.122 | 0.122 | 1.685 | 0.19 | 0.01 | 0.14 | 0 | 0.19 | 0.02 |
| 7 | 12841226 | rs7797619 | -1.269 | -0.805 | 0.537 | 0.014 | -0.017 | -0.032 | -0.032 | 0.593 | 0.84 | 0.94 | 0.82 | 0.77 | 0.83 | 0.79 |
| 7 | 12844796 | rs7790473 | -0.416 | -0.85 | 0.947 | -0.138 | -0.023 | 0.057 | 0.057 | 0 | 0.25 | 0.42 | 0.26 | 0.37 | 0.22 | 0.33 |
| 7 | 12846898 | rs10240278 | -0.233 | -0.813 | 0.957 | -0.178 | 0.031 | 0.006 | 0.006 | 0.627 | 0.47 | 0.7 | 0.43 | 0.4 | 0.41 | 0.36 |
| 7 | 12848814 | rs10278168 | -0.055 | -0.913 | 0.99 | -0.092 | -0.008 | -0.005 | -0.005 | 0 | 0.45 | 0.55 | 0.39 | 0.35 | 0.4 | 0.34 |
| 7 | 12856667 | rs10230511 | -0.706 | -0.871 | 1.071 | -0.389 | 0.004 | 0.113 | 0.113 | 0.669 | 0.15 | 0.22 | 0.19 | 0.41 | 0.14 | 0.27 |
| 7 | 12856719 | rs17680996 | 0.609 | -0.931 | 1.154 | -0.446 | 0.014 | 0.024 | 0.024 | 0 | 0.15 | 0.15 | 0.1 | 0.06 | 0.15 | 0.16 |
| 7 | 12859244 | rs4141175 | -0.088 | -0.958 | 1.21 | -0.437 | -0.006 | -0.013 | -0.013 | 0.003 | 0.56 | 0.55 | 0.66 | 0.71 | 0.63 | 0.75 |
| 7 | 12860493 | rs4530933 | -0.701 | -0.91 | 1.289 | -0.362 | 0.035 | 0.256 | 0.256 | 0.54 | 0.11 | 0.31 | 0.19 | 0.34 | 0.11 | 0.21 |
| 7 | 12866971 | rs6960791 | 2.299 | -0.907 | 1.277 | -0.396 | -0.02 | -0.014 | -0.014 | 0 | 0.06 | 0.17 | 0.02 | 0.05 | 0.02 | 0.01 |
| 7 | 12868301 | rs13222475 | 1.176 | -0.934 | 1.292 | -0.454 | -0.015 | -0.081 | -0.081 | 0.501 | 0.27 | 0.34 | 0.18 | 0.05 | 0.21 | 0.03 |
| 7 | 12870484 | rs10236747 | -1.07 | -0.992 | 1.23 | -0.43 | -0.005 | -0.023 | -0.023 | 5.199 | 0.83 | 0.93 | 0.82 | 0.78 | 0.84 | 0.81 |
| 7 | 12872311 | rs6460980 | -0.854 | -1.111 | 1.297 | -0.447 | -0.002 | 0.038 | 0.038 | 0 | 0.26 | 0.32 | 0.28 | 0.42 | 0.27 | 0.43 |
| 7 | 12875858 | rs12699421 | 1.902 | -1.058 | 1.209 | -0.461 | 0.002 | -0.016 | -0.016 | 0 | 0.21 | 0.17 | 0.15 | 0.02 | 0.19 | 0.03 |
| 7 | 12877446 | rs11771731 | 1.844 | -1.086 | 1.09 | -0.459 | -0.035 | 0.004 | 0.004 | 0 | 0.28 | 0.16 | 0.35 | 0.29 | 0.35 | 0.3 |

| | | | | | | | | | | | | | | | | |
|---|---|---|---|---|---|---|---|---|---|---|---|---|---|---|---|---|
| 7 | 12886456 | rs2109293 | 1.404 | -1.226 | 1.195 | -0.434 | 0.003 | 0.013 | 0.013 | 0 | 0.1 | 0.06 | 0.09 | 0.01 | 0.13 | 0.14 |
| 7 | 12896263 | rs6956076 | -1.076 | -1.024 | 1.304 | -0.275 | -0.009 | 0.015 | 0.015 | 0 | 0.34 | 0.65 | 0.32 | 0.46 | 0.29 | 0.45 |
| 7 | 12900211 | rs10233257 | 1.783 | -0.975 | 1.245 | -0.42 | 0.005 | 0.008 | 0.008 | 1.513 | 0.2 | 0.12 | 0.14 | 0.02 | 0.19 | 0.03 |
| 7 | 12906866 | rs6959108 | 1.156 | -0.988 | 1.248 | -0.392 | 0.004 | 0.018 | 0.018 | 0 | 0.15 | 0.1 | 0.11 | 0.08 | 0.16 | 0.22 |
| 7 | 12907178 | rs6979870 | 1.041 | -0.775 | 1.171 | -0.272 | 0.001 | -0.003 | -0.003 | 0 | 0.18 | 0.16 | 0.13 | 0.07 | 0.19 | 0.24 |
| 7 | 12908084 | rs12665879 | 0.033 | -0.793 | 1.234 | -0.178 | -0.002 | 0.025 | 0.025 | 0 | 0.26 | 0.31 | 0.29 | 0.42 | 0.29 | 0.44 |
| 7 | 12908159 | rs10224102 | -0.292 | -0.796 | 1.221 | -0.196 | 0.006 | -0.004 | -0.004 | 0.104 | 0.48 | 0.61 | 0.44 | 0.43 | 0.48 | 0.46 |
| 7 | 12911330 | rs7781158 | 1.814 | -0.782 | 1.201 | -0.191 | -0.027 | -0.007 | -0.007 | 0 | 0.06 | 0.16 | 0.02 | 0.05 | 0.02 | 0.01 |
| 7 | 12912107 | rs758676 | 0.182 | -0.761 | 1.103 | -0.17 | -0.001 | -0.017 | -0.017 | 0 | 0.25 | 0.25 | 0.18 | 0.15 | 0.28 | 0.35 |
| 7 | 12917100 | rs7780771 | -0.704 | -0.759 | 1.055 | -0.135 | 0.006 | -0.012 | -0.012 | 0 | 0.54 | 0.77 | 0.47 | 0.48 | 0.49 | 0.49 |
| 7 | 12919380 | rs10280534 | 0.116 | -0.79 | 0.968 | -0.034 | -0.023 | 0.412 | 0.412 | 0 | 0.09 | 0.5 | 0.15 | 0.32 | 0.03 | 0.12 |
| 7 | 12925433 | rs12699429 | 0.014 | -0.712 | 0.751 | -0.355 | 0.006 | 0.056 | 0.056 | 0 | 0.46 | 0.35 | 0.33 | 0.15 | 0.47 | 0.37 |
| 7 | 12927594 | rs10950424 | 0.858 | -0.483 | 0.853 | -0.282 | 0.005 | 0.008 | 0.008 | 0 | 0.51 | 0.54 | 0.63 | 0.6 | 0.55 | 0.43 |
| 7 | 12927606 | rs12532245 | 1.428 | -0.494 | 0.908 | -0.227 | -0.007 | 0.044 | 0.044 | 0.826 | 0.22 | 0.39 | 0.26 | 0.29 | 0.2 | 0.09 |
| 7 | 12930953 | rs10488237 | 0.76 | -0.412 | 0.638 | -0.45 | 0.01 | 0 | 0 | 0 | 0.36 | 0.35 | 0.44 | 0.38 | 0.43 | 0.45 |
| 7 | 12933049 | rs10950426 | 1.426 | -0.369 | 0.457 | -0.443 | -0.061 | -0.008 | -0.008 | 0 | 0.29 | 0.12 | 0.36 | 0.31 | 0.35 | 0.34 |
| 7 | 12934357 | rs4719341 | 0.421 | -0.302 | 0.529 | -0.393 | 0.018 | 0.006 | 0.006 | 0.037 | 0.07 | 0.11 | 0.06 | 0.07 | 0.08 | 0.11 |
| 7 | 12934776 | rs10256019 | -1.129 | -0.431 | 0.44 | -0.392 | -0.008 | -0.007 | -0.007 | 0 | 0.85 | 0.95 | 0.92 | 0.84 | 0.85 | 0.83 |
| 7 | 12937656 | rs10252824 | 0.97 | -0.426 | 0.5 | -0.384 | -0.007 | -0.029 | -0.029 | 0.228 | 0.21 | 0.26 | 0.14 | 0.08 | 0.2 | 0.26 |
| 7 | 12941929 | rs10488241 | -0.245 | 0.232 | 0.466 | -0.077 | 0.109 | 0.08 | 0.08 | 0.214 | 0.08 | 0.37 | 0.08 | 0.07 | 0.08 | 0.11 |
| 7 | 12948857 | rs6957008 | -0.198 | 0.292 | 0.338 | -0.05 | 0.108 | 0.08 | 0.08 | 0.034 | 0.08 | 0.35 | 0.08 | 0.07 | 0.08 | 0.1 |
| 7 | 12951820 | rs17166509 | 0.48 | 0.288 | 0.066 | -0.877 | 0.108 | 0.103 | 0.103 | 0 | 0.06 | 0.31 | 0.08 | 0.07 | 0.06 | 0.09 |
| 7 | 12953229 | rs11514702 | 1.08 | 0.191 | 0.215 | -0.733 | 0 | -0.075 | -0.075 | 1.82 | 0.25 | 0.34 | 0.21 | 0.07 | 0.24 | 0.11 |
| 7 | 12953982 | rs1008368 | 0.923 | 0.172 | 0.201 | -0.753 | 0.01 | -0.002 | -0.002 | 0.078 | 0.29 | 0.3 | 0.38 | 0.31 | 0.36 | 0.33 |
| 7 | 12954300 | rs1548741 | 0.288 | 0.186 | 0.21 | -0.743 | -0.002 | -0.002 | -0.002 | 0.699 | 0.53 | 0.47 | 0.58 | 0.38 | 0.6 | 0.44 |
| 7 | 12955438 | rs1465277 | 0.276 | 0.206 | -0.011 | -0.734 | 0.007 | 0.007 | 0.007 | 0 | 0.54 | 0.5 | 0.58 | 0.39 | 0.6 | 0.44 |
| 7 | 12955892 | rs9632555 | 0.716 | 0.227 | -0.333 | -0.733 | 0.009 | 0.01 | 0.01 | 0 | 0.27 | 0.29 | 0.12 | 0.14 | 0.15 | 0.3 |
| 7 | 12956462 | rs9969214 | 0.849 | 0.372 | -0.937 | -0.574 | -0.035 | 0.001 | 0.001 | 0 | 0.29 | 0.17 | 0.37 | 0.32 | 0.35 | 0.33 |
| 7 | 12958992 | rs1160324 | -0.122 | -0.142 | -0.688 | 0.201 | -0.002 | 0.002 | 0.002 | 0.135 | 0.83 | 0.82 | 0.74 | 0.53 | 0.82 | 0.76 |
| 7 | 12959896 | rs4721160 | 0.118 | 0.347 | -0.876 | 0.112 | -0.009 | -0.008 | -0.008 | 0 | 0.54 | 0.43 | 0.58 | 0.38 | 0.6 | 0.44 |
| 7 | 12963669 | rs34080 | -0.488 | 1.068 | -1.21 | 0.436 | -0.015 | -0.015 | -0.015 | 0.998 | 0.26 | 0.19 | 0.33 | 0.31 | 0.25 | 0.33 |
| 7 | 12967449 | rs10252492 | -0.458 | 0.841 | -1.819 | -0.606 | -0.007 | 0.084 | 0.084 | 0 | 0.28 | 0.16 | 0.12 | 0.05 | 0.3 | 0.3 |
| 7 | 12972070 | rs34068 | 0.057 | 0.864 | -1.718 | 0.627 | 0.001 | 0.111 | 0.111 | 0 | 0.93 | 0.9 | 0.75 | 0.63 | 0.92 | 0.81 |
| 7 | 12972239 | rs34067 | 1.011 | 0.666 | -1.443 | 0.239 | -0.001 | 0.083 | 0.083 | 0 | 0.86 | 0.8 | 0.69 | 0.63 | 0.86 | 0.73 |
| 7 | 12972685 | rs17166539 | 0.056 | 0.541 | -0.38 | 0.175 | 0.006 | -0.031 | -0.031 | 0 | 0.07 | 0.16 | 0.06 | 0 | 0.06 | 0.08 |
| 7 | 12982479 | rs246828 | -0.529 | 0.677 | -0.436 | 0.302 | 0.004 | 0.133 | 0.133 | 0 | 0.95 | 0.93 | 0.76 | 0.64 | 0.93 | 0.88 |
| 7 | 12990038 | rs4721166 | -1.444 | 0.238 | -0.053 | 0.006 | 0.006 | -0.029 | -0.029 | 0 | 0.49 | 0.87 | 0.5 | 0.46 | 0.46 | 0.56 |
| 7 | 12996680 | rs4721168 | -2.019 | 0.128 | 0.627 | 0.043 | 0.002 | 0.247 | 0.247 | 0.178 | 0.17 | 0.52 | 0.31 | 0.41 | 0.11 | 0.22 |
| 7 | 12997006 | rs10278270 | -1.182 | 0.108 | 1.237 | -0.218 | 0.017 | -0.068 | -0.068 | 0 | 0.11 | 0.27 | 0.08 | 0 | 0.09 | 0.12 |
| 7 | 12997766 | rs6460991 | -1.784 | -0.085 | 0.656 | 0.213 | -0.018 | 0.02 | 0.02 | 0.645 | 0.49 | 0.87 | 0.49 | 0.55 | 0.48 | 0.58 |
| 7 | 12997879 | rs6964353 | 0.179 | -0.141 | 0.674 | 0.169 | -0.011 | 0.111 | 0.111 | 0 | 0.27 | 0.09 | 0.11 | 0.05 | 0.28 | 0.29 |
| 7 | 13006567 | rs4719343 | -0.139 | 0.086 | 0.699 | 0.323 | -0.001 | 0.101 | 0.101 | 0 | 0.07 | 0.07 | 0.25 | 0.45 | 0.09 | 0.14 |
| 7 | 13008008 | rs4719344 | -2.485 | -0.095 | 1.235 | 0.163 | -0.003 | 0.078 | 0.078 | 0.361 | 0.24 | 0.58 | 0.37 | 0.49 | 0.2 | 0.26 |
| 7 | 13008198 | rs10267804 | -1.206 | -0.093 | 1.42 | 0.152 | -0.005 | -0.069 | -0.069 | 0 | 0.12 | 0.23 | 0.08 | 0 | 0.08 | 0.09 |
| 7 | 13010195 | rs7789443 | -1.749 | -0.386 | 1.106 | 0.14 | -0.022 | 0.048 | 0.048 | 0.698 | 0.49 | 0.87 | 0.51 | 0.57 | 0.49 | 0.6 |
| 7 | 13018255 | rs10228675 | -1.705 | -0.567 | 0.832 | 0.3 | -0.104 | 0.09 | 0.09 | 1.168 | 0.51 | 0.97 | 0.59 | 0.71 | 0.5 | 0.65 |
| 7 | 13020308 | rs7794754 | -1.791 | -0.563 | 0.829 | 0.28 | -0.085 | 0.055 | 0.055 | 1.352 | 0.5 | 0.97 | 0.59 | 0.68 | 0.49 | 0.62 |
| 7 | 13021705 | rs10258251 | -1.221 | -0.377 | 1.249 | 0.277 | 0.037 | 0.281 | 0.281 | 0 | 0.12 | 0.32 | 0.3 | 0.51 | 0.12 | 0.2 |
| 7 | 13027695 | rs4719345 | -0.968 | 0.015 | 0.744 | 0.028 | 0.01 | 0.145 | 0.145 | 0 | 0.11 | 0.21 | 0.29 | 0.51 | 0.12 | 0.23 |
| 7 | 13031134 | rs6460994 | -0.493 | 0.03 | 0.839 | -0.445 | -0.002 | -0.065 | -0.065 | 0.016 | 0.35 | 0.43 | 0.21 | 0.06 | 0.36 | 0.42 |
| 7 | 13033428 | rs17166577 | -1.287 | 0.444 | 1.186 | 0.017 | 0.057 | -0.014 | -0.014 | 0 | 0.09 | 0.34 | 0.08 | 0.02 | 0.12 | 0.08 |
| 7 | 13048002 | rs1861238 | -1.572 | 0.302 | 0.702 | 0.205 | -0.003 | -0.015 | -0.015 | 0 | 0.3 | 0.73 | 0.4 | 0.51 | 0.27 | 0.31 |
| 7 | 13052836 | rs763534 | -1.206 | 0.346 | 1.484 | 0.284 | 0.016 | 0.202 | 0.202 | 0 | 0.2 | 0.47 | 0.34 | 0.41 | 0.17 | 0.26 |
| 7 | 13071287 | rs12537712 | -1.1 | 0.353 | 1.87 | 0.316 | 0.034 | 0.133 | 0.133 | 0 | 0.18 | 0.35 | 0.34 | 0.42 | 0.2 | 0.27 |
| 7 | 13074317 | rs17166626 | -0.773 | 0.291 | 2.179 | 0.295 | 0.021 | 0.111 | 0.111 | 0 | 0.15 | 0.28 | 0.32 | 0.43 | 0.17 | 0.25 |
| 7 | 13077065 | rs7778616 | -2.652 | 0.355 | 2.152 | 0.273 | -0.003 | -0.004 | -0.004 | 0 | 0.34 | 0.69 | 0.43 | 0.51 | 0.35 | 0.33 |
| 7 | 13083793 | rs17630382 | -0.176 | 0.306 | 2.126 | 0.11 | -0.011 | 0.021 | 0.021 | 0.109 | 0.12 | 0.05 | 0.07 | 0.09 | 0.15 | 0.07 |
| 7 | 13086613 | rs10227653 | -1.011 | 0.185 | 2.221 | 0.089 | -0.038 | -0.047 | -0.047 | 0.046 | 0.16 | 0.32 | 0.08 | 0.09 | 0.09 | 0.08 |
| 7 | 13089147 | rs13232408 | 0.316 | -0.207 | 1.185 | -0.315 | 0.031 | 0.056 | 0.056 | 0.37 | 0.24 | 0.09 | 0.16 | 0.08 | 0.21 | 0.29 |
| 7 | 13097797 | rs6962531 | -0.861 | -0.146 | 1.463 | -0.554 | -0.006 | -0.071 | -0.071 | 0 | 0.19 | 0.38 | 0.11 | 0.09 | 0.17 | 0.11 |
| 7 | 13103636 | rs2160193 | 0.533 | -0.119 | 1.556 | -0.312 | -0.001 | 0.034 | 0.034 | 0 | 0.14 | 0.18 | 0.31 | 0.38 | 0.16 | 0.22 |
| 7 | 13104434 | rs2058613 | -2.154 | -0.161 | 1.54 | -0.294 | 0.001 | -0.007 | -0.007 | 0 | 0.33 | 0.7 | 0.42 | 0.48 | 0.28 | 0.31 |
| 7 | 13105009 | rs7778887 | -0.019 | -0.199 | 1.415 | -0.299 | 0.041 | 0.233 | 0.233 | 0 | 0.16 | 0.43 | 0.33 | 0.38 | 0.17 | 0.23 |
| 7 | 13110857 | rs17166712 | 0.127 | -0.204 | 1.215 | -0.294 | 0.044 | 0.237 | 0.237 | 0 | 0.16 | 0.43 | 0.33 | 0.38 | 0.16 | 0.22 |
| 7 | 13116278 | rs6958548 | -1.186 | -0.295 | 0.969 | -0.383 | 0.001 | -0.002 | -0.002 | 0 | 0.43 | 0.61 | 0.47 | 0.44 | 0.4 | 0.53 |
| 7 | 13117948 | rs7782472 | -0.687 | -0.159 | 0.526 | -0.406 | -0.006 | 0.018 | 0.018 | 0.051 | 0.88 | 0.93 | 0.93 | 0.91 | 0.86 | 0.91 |
| 7 | 13124156 | rs2041231 | 0.547 | -0.279 | 0.527 | -0.517 | 0.022 | 0.044 | 0.044 | 0.155 | 0.24 | 0.14 | 0.12 | 0.06 | 0.21 | 0.28 |
| 7 | 13131480 | rs7808749 | 1.558 | -0.256 | 0.878 | -0.409 | -0.004 | 0.007 | 0.007 | 0 | 0.46 | 0.34 | 0.46 | 0.48 | 0.47 | 0.4 |
| 7 | 13152162 | rs1476772 | 1.838 | -0.767 | 0.839 | -0.454 | -0.089 | -0.047 | -0.047 | 0.437 | 0.41 | 0.09 | 0.43 | 0.48 | 0.45 | 0.39 |

| | | | | | | | | | | | | | | | | |
|---|---|---|---|---|---|---|---|---|---|---|---|---|---|---|---|---|
| 7 | 13176846 | rs11984152 | -1.367 | -0.573 | 1.225 | -0.252 | 0.023 | -0.027 | -0.027 | 0.089 | 0.07 | 0.21 | 0.04 | 0 | 0.06 | 0.03 |
| 7 | 13184540 | rs12699451 | 1.732 | -0.696 | 0.907 | -0.179 | 0 | -0.003 | -0.003 | 0 | 0.64 | 0.34 | 0.55 | 0.5 | 0.66 | 0.65 |
| 7 | 13185882 | rs2191041 | 0.225 | -0.508 | 0.62 | -0.133 | 0 | -0.002 | -0.002 | 0 | 0.85 | 0.87 | 0.91 | 0.88 | 0.82 | 0.88 |
| 7 | 13188639 | rs10234652 | 0.006 | -0.472 | 0.424 | -0.193 | -0.005 | 0.025 | 0.025 | 0 | 0.86 | 0.93 | 0.92 | 0.94 | 0.85 | 0.89 |
| 7 | 13190921 | rs7784835 | -1.001 | -0.652 | 0.685 | -0.255 | 0.016 | -0.021 | -0.021 | 0.134 | 0.4 | 0.73 | 0.43 | 0.44 | 0.34 | 0.47 |
| 7 | 13190950 | rs7804111 | -0.329 | -0.65 | 0.826 | -0.254 | -0.001 | -0.003 | -0.003 | 0 | 0.33 | 0.39 | 0.39 | 0.44 | 0.34 | 0.47 |
| 7 | 13193611 | rs12666139 | -0.005 | -0.648 | 0.895 | -0.261 | 0.001 | -0.026 | -0.026 | 0 | 0.31 | 0.3 | 0.37 | 0.44 | 0.29 | 0.44 |
| 7 | 13194971 | rs718814 | -0.345 | -0.643 | 0.901 | -0.256 | -0.001 | 0.018 | 0.018 | 1.363 | 0.34 | 0.45 | 0.39 | 0.44 | 0.34 | 0.45 |
| 7 | 13198073 | rs10279792 | 0.592 | -0.583 | 0.685 | -0.004 | 0.029 | 0.003 | 0.003 | 0 | 0.89 | 0.67 | 0.91 | 0.95 | 0.88 | 0.93 |
| 7 | 13200007 | rs1860269 | -0.788 | -0.661 | 0.496 | -0.028 | -0.013 | 0.016 | 0.016 | 3.34 | 0.38 | 0.57 | 0.4 | 0.46 | 0.31 | 0.47 |
| 7 | 13211495 | rs2191038 | -1.072 | -0.491 | 0.651 | 0.157 | -0.011 | 0.03 | 0.03 | 0.693 | 0.34 | 0.59 | 0.41 | 0.46 | 0.28 | 0.49 |
| 7 | 13214447 | rs2058156 | -0.676 | -0.417 | 0.019 | 0.139 | -0.017 | -0.005 | -0.005 | 0 | 0.93 | 0.97 | 0.94 | 0.95 | 0.88 | 0.95 |
| 7 | 13218051 | rs11760715 | 1.856 | -0.293 | -0.081 | 0.219 | -0.046 | 0.078 | 0.078 | 1.268 | 0.49 | 0.11 | 0.37 | 0.39 | 0.47 | 0.37 |
| 7 | 13226751 | rs4719355 | 0.709 | -0.363 | 0.512 | 0.21 | 0.015 | -0.04 | -0.04 | 0 | 0.85 | 0.48 | 0.88 | 0.95 | 0.86 | 0.93 |
| 7 | 13227326 | rs12666192 | 1.031 | -0.303 | -0.223 | 0.25 | -0.007 | 0.051 | 0.051 | 0.467 | 0.14 | 0.05 | 0.1 | 0.06 | 0.15 | 0.2 |
| 7 | 13227685 | rs12666706 | -0.244 | -0.255 | -0.172 | 0.316 | -0.008 | -0.059 | -0.059 | 0 | 0.26 | 0.17 | 0.36 | 0.46 | 0.26 | 0.43 |
| 7 | 13231065 | rs12672883 | -0.23 | -0.172 | -0.194 | 0.395 | -0.012 | -0.059 | -0.059 | 0 | 0.25 | 0.17 | 0.35 | 0.5 | 0.26 | 0.44 |
| 7 | 13237848 | rs194025 | 1.249 | -0.43 | -0.487 | 0.338 | 0.02 | 0.029 | 0.029 | 1.75 | 0.53 | 0.21 | 0.51 | 0.49 | 0.59 | 0.48 |
| 7 | 13238496 | rs16877717 | 0.263 | -0.216 | -0.441 | 0.696 | 0.018 | 0.126 | 0.126 | 0 | 0.05 | 0.13 | 0.18 | 0.22 | 0.07 | 0.1 |
| 7 | 13248712 | rs4721192 | -0.604 | -0.448 | -0.437 | 0.406 | 0.001 | 0.012 | 0.012 | 0.24 | 0.36 | 0.67 | 0.3 | 0.26 | 0.37 | 0.4 |
| 7 | 13249235 | rs10243842 | -1.495 | -0.447 | -1.288 | 0.018 | -0.049 | 0.022 | 0.022 | 0.099 | 0.6 | 0.88 | 0.6 | 0.57 | 0.57 | 0.64 |
| 7 | 13253069 | rs6973933 | -0.919 | -0.178 | -1.224 | 0.016 | -0.01 | 0.066 | 0.066 | 1.203 | 0.53 | 0.87 | 0.57 | 0.57 | 0.55 | 0.58 |
| 7 | 13255407 | rs6942409 | -1.432 | -0.25 | -1.107 | -0.052 | -0.05 | 0.056 | 0.056 | 0.249 | 0.6 | 0.85 | 0.6 | 0.67 | 0.58 | 0.65 |
| 7 | 13262586 | rs11766252 | -0.595 | -0.857 | -0.056 | -0.308 | -0.036 | 0.125 | 0.125 | 0.156 | 0.16 | 0.45 | 0.1 | 0.3 | 0.09 | 0.13 |
| 7 | 13263062 | rs10279810 | -1.026 | -0.945 | 0.142 | -0.531 | -0.004 | -0.118 | -0.118 | 0.248 | 0.19 | 0.39 | 0.13 | 0.02 | 0.21 | 0.11 |
| 7 | 13263382 | rs17166905 | 0.403 | -0.929 | 0.063 | -0.453 | -0.008 | 0.123 | 0.123 | 0 | 0.05 | 0.11 | 0.04 | 0.29 | 0.02 | 0.1 |
| 7 | 13265700 | rs7781725 | -1.926 | -1.042 | 0.116 | -0.637 | -0.012 | -0.01 | -0.01 | 0.047 | 0.33 | 0.56 | 0.25 | 0.31 | 0.31 | 0.29 |
| 7 | 13267944 | rs10215522 | 0.051 | -1.046 | 0.46 | -0.586 | -0.031 | 0.107 | 0.107 | 0.161 | 0.12 | 0.25 | 0.05 | 0.3 | 0.04 | 0.12 |
| 7 | 13268915 | rs7780368 | 1.259 | -1.165 | 0.527 | -0.743 | -0.003 | -0.024 | -0.024 | 0 | 0.15 | 0.2 | 0.09 | 0.08 | 0.13 | 0.18 |
| 7 | 13269300 | rs10253676 | -0.048 | -0.84 | 0.648 | -0.293 | -0.027 | 0.102 | 0.102 | 0 | 0.11 | 0.24 | 0.05 | 0.3 | 0.04 | 0.12 |
| 7 | 13270959 | rs17166908 | -1.676 | -0.803 | 0.51 | -0.327 | -0.001 | -0.001 | -0.001 | 0.011 | 0.3 | 0.31 | 0.26 | 0.31 | 0.36 | 0.33 |
| 7 | 13272254 | rs10281788 | -1.56 | -0.626 | 0.135 | -0.081 | -0.012 | -0.009 | -0.009 | 0.101 | 0.33 | 0.57 | 0.26 | 0.31 | 0.31 | 0.29 |
| 7 | 13274262 | rs2884164 | -0.582 | -1.25 | -0.055 | 0.009 | -0.048 | 0.025 | 0.025 | 0 | 0.25 | 0.37 | 0.19 | 0.34 | 0.15 | 0.26 |
| 7 | 13274561 | rs4329178 | 0.24 | -0.143 | -0.541 | 0.655 | -0.002 | -0.001 | -0.001 | 0 | 0.78 | 0.82 | 0.59 | 0.49 | 0.75 | 0.78 |
| 7 | 13274616 | rs4329179 | -1.59 | -0.081 | 0.1 | 0.823 | -0.005 | 0.074 | 0.074 | 0.062 | 0.17 | 0.68 | 0.34 | 0.42 | 0.15 | 0.13 |
| 7 | 13274710 | rs11767642 | -0.83 | -0.038 | -0.569 | 0.322 | 0.007 | 0.014 | 0.014 | 0.224 | 0.37 | 0.71 | 0.35 | 0.25 | 0.34 | 0.43 |
| 7 | 13275578 | rs17166921 | -0.126 | -0.153 | -0.136 | 0.394 | -0.004 | -0.004 | -0.004 | 4.593 | 0.17 | 0.19 | 0.15 | 0.16 | 0.15 | 0.15 |
| 7 | 13276093 | rs13227738 | 0.179 | -0.193 | -0.141 | 0.316 | 0.006 | -0.054 | -0.054 | 0 | 0.19 | 0.42 | 0.13 | 0.11 | 0.16 | 0.22 |
| 7 | 13276128 | rs7806310 | -2.03 | 0.29 | -0.374 | 0.552 | -0.007 | -0.027 | -0.027 | 0.034 | 0.51 | 0.88 | 0.46 | 0.53 | 0.5 | 0.53 |
| 7 | 13276497 | rs12699461 | -1.029 | -0.148 | -0.803 | 0.527 | 0 | -0.001 | -0.001 | 0 | 0.45 | 0.48 | 0.4 | 0.53 | 0.45 | 0.53 |
| 7 | 13276543 | rs4721197 | -0.233 | 0.044 | -0.779 | 0.521 | 0.027 | -0.009 | -0.009 | 0 | 0.33 | 0.21 | 0.26 | 0.34 | 0.25 | 0.31 |
| 7 | 13277948 | rs7780526 | -2.326 | 0.303 | -0.79 | 0.549 | -0.105 | 0.021 | 0.021 | 0.505 | 0.66 | 0.98 | 0.58 | 0.71 | 0.61 | 0.7 |
| 7 | 13282527 | rs10950444 | -1.18 | 0.396 | -0.576 | 0.391 | 0.002 | -0.023 | -0.023 | 0 | 0.55 | 0.53 | 0.51 | 0.72 | 0.53 | 0.66 |
| 7 | 13283419 | rs10234195 | -1.118 | 0.989 | 0.115 | 0.616 | -0.006 | -0.066 | -0.066 | 0 | 0.11 | 0.59 | 0.09 | 0 | 0.06 | 0.02 |
| 7 | 13285891 | rs10279866 | -0.086 | 1.03 | 0.047 | 0.708 | -0.006 | -0.009 | -0.009 | 0 | 0.17 | 0.24 | 0.15 | 0.17 | 0.15 | 0.16 |
| 7 | 13287995 | rs7787319 | -1.225 | 1.249 | 0 | 0.92 | 0.172 | 0.298 | 0.298 | 0 | 0.1 | 0.46 | 0.2 | 0.25 | 0.14 | 0.13 |
| 7 | 13289250 | rs10249615 | -2.101 | 1.244 | -0.07 | 0.89 | 0.042 | 0.068 | 0.068 | 0 | 0.19 | 0.61 | 0.24 | 0.3 | 0.23 | 0.18 |
| 7 | 13290030 | rs4719358 | 0.417 | 1.089 | -0.297 | 0.797 | 0 | -0.008 | -0.008 | 0 | 0.15 | 0.08 | 0.14 | 0.18 | 0.14 | 0.16 |
| 7 | 13293042 | rs10265874 | -0.715 | 0.672 | -0.311 | 0.626 | 0.002 | 0.014 | 0.014 | 0 | 0.42 | 0.63 | 0.34 | 0.29 | 0.41 | 0.44 |
| 7 | 13293713 | rs10254246 | 0.613 | 0.78 | -0.225 | 0.597 | 0.03 | -0.056 | -0.056 | 0 | 0.2 | 0.31 | 0.2 | 0.09 | 0.26 | 0.2 |
| 7 | 13293842 | rs17704933 | 0.686 | 0.889 | -0.364 | 0.803 | -0.027 | -0.006 | -0.006 | 0 | 0.05 | 0.01 | 0.15 | 0.19 | 0.13 | 0.13 |
| 7 | 13295482 | rs759181 | -1.684 | 0.618 | -0.353 | 0.656 | 0.001 | 0.003 | 0.003 | 0 | 0.44 | 0.57 | 0.36 | 0.6 | 0.38 | 0.49 |
| 7 | 13297072 | rs7787128 | 0.351 | 0.562 | -0.272 | 0.647 | -0.017 | 0.005 | 0.005 | 0 | 0.22 | 0.28 | 0.13 | 0.18 | 0.15 | 0.22 |
| 7 | 13300179 | rs13239242 | 0.465 | 1.142 | 0 | 0.56 | 0.034 | -0.061 | -0.061 | 0 | 0.2 | 0.33 | 0.21 | 0.1 | 0.26 | 0.21 |
| 7 | 13302054 | rs17166951 | 0.255 | 1.064 | -0.333 | 0.68 | 0 | -0.004 | -0.004 | 0 | 0.09 | 0.09 | 0.04 | 0.22 | 0.07 | 0.13 |
| 7 | 13303290 | rs1035084 | -1.872 | 0.957 | -0.483 | 0.631 | 0.049 | -0.019 | -0.019 | 0 | 0.5 | 0.87 | 0.49 | 0.52 | 0.56 | 0.56 |
| 7 | 13305542 | rs7780448 | -0.266 | 1.012 | -0.132 | 0.371 | 0.039 | -0.092 | -0.092 | 0 | 0.27 | 0.47 | 0.25 | 0.12 | 0.32 | 0.28 |
| 7 | 13306966 | rs11975497 | -1.589 | 0.998 | -0.24 | 0.423 | -0.005 | 0.035 | 0.035 | 0.004 | 0.43 | 0.65 | 0.31 | 0.33 | 0.42 | 0.44 |
| 7 | 13315977 | rs6954677 | -1.24 | 0.848 | -0.206 | 0.38 | -0.002 | 0.028 | 0.028 | 0 | 0.38 | 0.7 | 0.3 | 0.32 | 0.36 | 0.34 |
| 7 | 13316251 | rs6461019 | -0.256 | 0.837 | -0.183 | 0.37 | -0.001 | 0.003 | 0.003 | 0 | 0.12 | 0.14 | 0.06 | 0.22 | 0.1 | 0.15 |
| 7 | 13316690 | rs12374759 | -0.133 | 0.859 | -0.13 | 0.33 | 0.015 | -0.095 | -0.095 | 0.451 | 0.25 | 0.55 | 0.24 | 0.09 | 0.27 | 0.19 |
| 7 | 13319066 | rs7803179 | -1.165 | 0.924 | -0.024 | 0.53 | -0.002 | 0 | 0 | 0 | 0.37 | 0.34 | 0.41 | 0.27 | 0.49 | 0.43 |
| 7 | 13319336 | rs7788518 | -1.92 | 0.948 | -0.094 | 0.579 | 0.035 | -0.005 | -0.005 | 1.932 | 0.48 | 0.71 | 0.47 | 0.49 | 0.56 | 0.57 |
| 7 | 13321206 | rs12699473 | 0.506 | 1.248 | -0.245 | 0.494 | -0.008 | 0.004 | 0.004 | 0 | 0.19 | 0.16 | 0.2 | 0.09 | 0.26 | 0.19 |
| 7 | 13325060 | rs10488260 | -0.102 | 1.308 | -0.713 | 0.715 | -0.058 | -0.06 | -0.06 | 0.108 | 0.11 | 0.02 | 0.17 | 0.19 | 0.2 | 0.24 |
| 7 | 13333678 | rs13234975 | 0.279 | 1.531 | -0.497 | 0.402 | 0.1 | -0.074 | -0.074 | 0.448 | 0.18 | 0.48 | 0.25 | 0.1 | 0.29 | 0.16 |
| 7 | 13333829 | rs12154889 | -1.589 | 0.858 | -1.222 | 0.701 | 0.059 | 0.012 | 0.012 | 0.957 | 0.49 | 0.86 | 0.67 | 0.55 | 0.58 | 0.56 |
| 7 | 13336448 | rs2192325 | -2.03 | 0.628 | -1.637 | 0.469 | -0.036 | 0.021 | 0.021 | 0 | 0.68 | 0.99 | 0.86 | 0.71 | 0.72 | 0.71 |
| 7 | 13348832 | rs17707307 | -0.805 | 1.355 | -0.792 | -0.112 | -0.036 | -0.042 | -0.042 | 0 | 0.16 | 0.12 | 0.37 | 0.24 | 0.23 | 0.26 |

| | | | | | | | | | | | | | | | | |
|---|---|---|---|---|---|---|---|---|---|---|---|---|---|---|---|---|
| 7 | 13358803 | rs4721213 | 0.057 | 0.865 | -0.557 | -0.341 | 0.004 | 0.015 | 0.015 | 0 | 0.21 | 0.13 | 0.19 | 0.09 | 0.19 | 0.2 |
| 7 | 13373793 | rs987101 | -1.848 | 0.622 | -0.913 | -0.449 | 0.021 | -0.056 | -0.056 | 0 | 0.69 | 0.89 | 0.77 | 0.36 | 0.73 | 0.64 |
| 7 | 13395594 | rs2116629 | 1.021 | 1.088 | -0.888 | -0.686 | -0.037 | 0.066 | 0.066 | 0.477 | 0.24 | 0.12 | 0.22 | 0 | 0.28 | 0.07 |
| 7 | 13412439 | rs1594609 | 0.311 | 0.92 | -0.749 | -0.477 | -0.003 | 0.002 | 0.002 | 0 | 0.11 | 0.16 | 0.25 | 0.12 | 0.08 | 0.24 |
| 7 | 13415675 | rs4283932 | -0.127 | 1.031 | -0.634 | -0.167 | -0.005 | 0.017 | 0.017 | 0.617 | 0.45 | 0.62 | 0.56 | 0.21 | 0.46 | 0.39 |
| 7 | 13429017 | rs2568602 | -0.969 | 0.945 | -0.785 | 0.375 | 0.014 | -0.038 | -0.038 | 1.112 | 0.69 | 0.88 | 0.77 | 0.35 | 0.73 | 0.64 |
| 7 | 13433240 | rs2436219 | -0.135 | 0.993 | -0.512 | -0.014 | -0.002 | -0.012 | -0.012 | 0 | 0.57 | 0.54 | 0.66 | 0.23 | 0.59 | 0.49 |
| 7 | 13439084 | rs720543 | -0.361 | 0.724 | -0.105 | -0.657 | 0.003 | -0.046 | -0.046 | 1.084 | 0.26 | 0.51 | 0.3 | 0.13 | 0.25 | 0.24 |
| 7 | 13439978 | rs2568609 | -0.761 | 1.079 | -0.112 | -0.741 | 0.017 | -0.027 | -0.027 | 0.83 | 0.29 | 0.55 | 0.36 | 0.19 | 0.32 | 0.31 |
| 7 | 13441668 | rs17714678 | 0.758 | 1.195 | -0.494 | -0.861 | 0.007 | -0.014 | -0.014 | 0 | 0.28 | 0.31 | 0.38 | 0.12 | 0.31 | 0.16 |
| 7 | 13441694 | rs10262962 | -0.83 | 0.967 | -0.148 | -0.891 | 0.037 | 0.015 | 0.015 | 0.411 | 0.09 | 0.29 | 0.1 | 0.07 | 0.07 | 0.1 |
| 7 | 13443147 | rs2568610 | 0.37 | 0.961 | -0.312 | -0.889 | 0 | -0.001 | -0.001 | 0.168 | 0.21 | 0.26 | 0.26 | 0.13 | 0.25 | 0.22 |
| 7 | 13444996 | rs2568612 | -1.988 | 0.553 | -0.798 | -1.173 | -0.021 | -0.143 | -0.143 | 2.401 | 0.71 | 0.99 | 0.78 | 0.33 | 0.74 | 0.61 |
| 7 | 13445626 | rs11971750 | 0.351 | 0.68 | -0.566 | -1.188 | 0.001 | -0.006 | -0.006 | 0 | 0.34 | 0.34 | 0.4 | 0.14 | 0.4 | 0.22 |
| 7 | 13445846 | rs2723418 | 0.053 | 0.519 | -0.177 | -1.084 | 0.005 | -0.019 | -0.019 | 0 | 0.23 | 0.34 | 0.27 | 0.12 | 0.25 | 0.22 |
| 7 | 13445967 | rs2723419 | 0.053 | 0.519 | -0.177 | -1.084 | 0.005 | -0.019 | -0.019 | 0 | 0.23 | 0.34 | 0.27 | 0.12 | 0.25 | 0.22 |
| 7 | 13449643 | rs17715032 | 0.654 | 0.695 | -0.451 | -1.099 | -0.028 | 0.022 | 0.022 | 0 | 0.07 | 0.01 | 0.03 | 0.02 | 0.09 | 0.07 |
| 7 | 13450842 | rs6959532 | -0.115 | 0.5 | -0.307 | -1.278 | -0.004 | -0.041 | -0.041 | 0 | 0.08 | 0.17 | 0.05 | 0.01 | 0.05 | 0.02 |
| 7 | 13453726 | rs1432511 | -0.989 | 0.378 | -0.074 | -1.351 | -0.005 | -0.03 | -0.03 | 0 | 0.34 | 0.52 | 0.31 | 0.19 | 0.31 | 0.31 |
| 7 | 13455926 | rs2568617 | 0.936 | 0.438 | -0.046 | -1.339 | 0.001 | 0.005 | 0.005 | 0 | 0.18 | 0.17 | 0.18 | 0.12 | 0.18 | 0.19 |
| 7 | 13457236 | rs17658851 | 1.373 | 0.533 | -0.017 | -1.608 | 0.016 | -0.045 | -0.045 | 1.887 | 0.22 | 0.28 | 0.21 | 0.06 | 0.28 | 0.11 |
| 7 | 13459064 | rs17658983 | 0.176 | 0.372 | 0.03 | -1.583 | 0 | 0.007 | 0.007 | 0 | 0.23 | 0.22 | 0.22 | 0.13 | 0.25 | 0.22 |
| 7 | 13460486 | rs10479915 | -0.22 | 0.311 | -0.064 | -1.541 | 0.012 | -0.014 | -0.014 | 0.994 | 0.38 | 0.48 | 0.47 | 0.2 | 0.47 | 0.36 |
| 7 | 13462040 | rs1432514 | -1.823 | 0.316 | -0.396 | -1.434 | 0.013 | -0.093 | -0.093 | 0 | 0.65 | 0.94 | 0.72 | 0.32 | 0.71 | 0.54 |
| 7 | 13465676 | rs2568624 | 0.613 | 0.337 | -0.191 | -1.292 | 0.003 | 0.007 | 0.007 | 0 | 0.2 | 0.19 | 0.19 | 0.12 | 0.19 | 0.19 |
| 7 | 13466283 | rs1583258 | 0.257 | 0.363 | -0.152 | -1.27 | 0 | 0.003 | 0.003 | 0 | 0.22 | 0.24 | 0.21 | 0.14 | 0.22 | 0.21 |
| 7 | 13469125 | rs1594613 | -0.479 | 0.259 | 0.008 | -1.405 | 0.002 | -0.008 | -0.008 | 0 | 0.4 | 0.42 | 0.5 | 0.21 | 0.51 | 0.35 |
| 7 | 13469186 | rs1991762 | 0.913 | 0.396 | -0.099 | -1.432 | -0.05 | 0.076 | 0.076 | 0 | 0.25 | 0.1 | 0.21 | 0.06 | 0.32 | 0.13 |
| 7 | 13469277 | rs1991763 | -1.538 | 0.284 | -0.024 | -1.41 | 0.04 | -0.018 | -0.018 | 0 | 0.62 | 0.79 | 0.72 | 0.33 | 0.72 | 0.54 |
| 7 | 13473994 | rs2568630 | -0.026 | 0.384 | 0.072 | -1.436 | -0.01 | 0.027 | 0.027 | 0 | 0.25 | 0.57 | 0.38 | 0.23 | 0.24 | 0.28 |
| 7 | 13474180 | rs13237947 | 0.917 | 0.387 | -0.052 | -1.474 | -0.057 | 0.094 | 0.094 | 0 | 0.25 | 0.07 | 0.2 | 0.07 | 0.3 | 0.13 |
| 7 | 13476506 | rs6957197 | -1.114 | -0.328 | -0.235 | -1.343 | 0 | 0.006 | 0.006 | 0.133 | 0.54 | 0.56 | 0.55 | 0.32 | 0.52 | 0.42 |
| 7 | 13477214 | rs7791472 | 1.254 | -0.227 | -0.261 | -1.295 | -0.034 | 0.035 | 0.035 | 0 | 0.25 | 0.12 | 0.2 | 0.18 | 0.3 | 0.14 |
| 7 | 13478678 | rs10807764 | -0.573 | -0.304 | -0.236 | -1.288 | 0.001 | 0.002 | 0.002 | 0 | 0.22 | 0.18 | 0.3 | 0.14 | 0.21 | 0.24 |
| 7 | 13479783 | rs2568632 | 0.814 | -0.155 | -0.176 | -1.238 | 0 | -0.02 | -0.02 | 0 | 0.21 | 0.41 | 0.21 | 0.12 | 0.23 | 0.16 |
| 7 | 13480718 | rs2568633 | -0.21 | 0.044 | -0.053 | -1.143 | -0.007 | -0.009 | -0.009 | 0 | 0.56 | 0.68 | 0.61 | 0.39 | 0.54 | 0.44 |
| 7 | 13481626 | rs1432486 | -0.917 | 0.093 | -0.078 | -0.882 | 0.017 | 0.025 | 0.025 | 0 | 0.51 | 0.69 | 0.57 | 0.36 | 0.44 | 0.49 |
| 7 | 13482749 | rs10243359 | 0.478 | 0.203 | -0.027 | -0.867 | 0.006 | -0.01 | -0.01 | 0.02 | 0.06 | 0.13 | 0.05 | 0.04 | 0.06 | 0.06 |
| 7 | 13483020 | rs7785549 | 1.097 | 0.405 | 0.007 | -0.873 | -0.026 | 0.025 | 0.025 | 0 | 0.23 | 0.16 | 0.2 | 0.15 | 0.34 | 0.12 |
| 7 | 13485785 | rs10233716 | -0.154 | 0.202 | 0.165 | -0.779 | 0.016 | 0.083 | 0.083 | 0 | 0.05 | 0.13 | 0.08 | 0.11 | 0.06 | 0.07 |
| 7 | 13492428 | rs878644 | -0.903 | 0.036 | 0.15 | -0.921 | 0.004 | 0.012 | 0.012 | 0 | 0.47 | 0.6 | 0.42 | 0.36 | 0.53 | 0.36 |
| 7 | 13492885 | rs2723437 | 1.153 | -0.074 | 0.165 | -0.975 | 0.004 | 0.016 | 0.016 | 0 | 0.26 | 0.22 | 0.37 | 0.15 | 0.25 | 0.24 |
| 7 | 13493700 | rs10155975 | -1.274 | -0.013 | 0.435 | -0.983 | 0.001 | 0.029 | 0.029 | 0 | 0.24 | 0.49 | 0.23 | 0.21 | 0.22 | 0.25 |
| 7 | 13493720 | rs6962487 | -0.755 | 0.001 | 0.16 | -0.979 | -0.001 | 0.005 | 0.005 | 0 | 0.45 | 0.58 | 0.42 | 0.36 | 0.52 | 0.36 |
| 7 | 13497326 | rs10242586 | -0.54 | 0.086 | 0.016 | -0.853 | 0.009 | 0.006 | 0.006 | 0 | 0.16 | 0.21 | 0.16 | 0.14 | 0.21 | 0.19 |
| 7 | 13498299 | rs17167171 | 0.47 | 0.012 | -0.118 | -0.774 | -0.016 | 0.012 | 0.012 | 0.811 | 0.08 | 0.16 | 0.07 | 0.08 | 0.03 | 0.06 |
| 7 | 13500618 | rs2568574 | -1.962 | -0.011 | -0.646 | -0.817 | -0.024 | -0.162 | -0.162 | 0 | 0.76 | 0.96 | 0.81 | 0.5 | 0.79 | 0.6 |
| 7 | 13502530 | rs10266590 | 1.13 | 0.358 | -0.097 | -0.987 | -0.022 | 0.028 | 0.028 | 0 | 0.22 | 0.14 | 0.18 | 0.15 | 0.29 | 0.11 |
| 7 | 13507623 | rs1432496 | 0.517 | 0.279 | 0.218 | -1.005 | 0.007 | -0.049 | -0.049 | 0 | 0.3 | 0.4 | 0.26 | 0.13 | 0.35 | 0.17 |
| 7 | 13509696 | rs17167190 | 0.486 | 0.1 | 0.379 | -0.974 | -0.014 | 0.006 | 0.006 | 0.012 | 0.07 | 0.13 | 0.07 | 0.08 | 0.03 | 0.06 |
| 7 | 13511472 | rs2723457 | -0.758 | -0.215 | -0.041 | -0.962 | 0.034 | -0.014 | -0.014 | 1.262 | 0.65 | 0.76 | 0.71 | 0.36 | 0.77 | 0.61 |
| 7 | 13511606 | rs2723456 | 1.039 | -0.116 | -0.054 | -0.928 | -0.003 | 0.031 | 0.031 | 0 | 0.24 | 0.15 | 0.3 | 0.1 | 0.24 | 0.31 |
| 7 | 13511976 | rs2723454 | -0.146 | -0.086 | 0.062 | -1.009 | 0.004 | -0.033 | -0.033 | 0 | 0.62 | 0.57 | 0.67 | 0.79 | 0.58 | 0.74 |
| 7 | 13513686 | rs7781734 | 0.467 | 0.687 | 0.014 | -0.978 | 0.008 | -0.027 | -0.027 | 0 | 0.08 | 0.15 | 0.04 | 0.01 | 0.09 | 0.07 |
| 7 | 13516664 | rs2357504 | -1.265 | 0.581 | -0.055 | -0.999 | 0.04 | -0.012 | -0.012 | 0.078 | 0.65 | 0.76 | 0.71 | 0.36 | 0.78 | 0.61 |
| 7 | 13518160 | rs2723438 | 0.648 | 0.722 | -0.366 | -1.096 | 0.001 | 0.003 | 0.003 | 0 | 0.28 | 0.25 | 0.31 | 0.08 | 0.24 | 0.32 |
| 7 | 13518945 | rs2163689 | 0.768 | 0.772 | -0.957 | -1.145 | -0.041 | 0.049 | 0.049 | 0 | 0.28 | 0.16 | 0.26 | 0.14 | 0.41 | 0.2 |
| 7 | 13520443 | rs1347075 | -0.067 | 1.122 | -0.528 | -0.782 | -0.005 | -0.007 | -0.007 | 0.203 | 0.69 | 0.57 | 0.59 | 0.79 | 0.72 | 0.65 |
| 7 | 13522169 | rs2568584 | 0.392 | 0.995 | -0.465 | -0.618 | 0.006 | -0.019 | -0.019 | 0 | 0.6 | 0.52 | 0.55 | 0.77 | 0.59 | 0.55 |
| 7 | 13527659 | rs6461030 | 0.364 | 0.958 | -0.431 | -0.572 | 0.004 | -0.016 | -0.016 | 0 | 0.61 | 0.51 | 0.56 | 0.76 | 0.62 | 0.55 |
| 7 | 13529096 | rs10807765 | -0.787 | 1.238 | -0.513 | -1.21 | -0.002 | -0.005 | -0.005 | 0 | 0.45 | 0.54 | 0.45 | 0.36 | 0.54 | 0.31 |
| 7 | 13529463 | rs11973042 | 0.041 | 1.562 | -0.31 | -1.031 | 0.028 | 0.048 | 0.048 | 0 | 0.19 | 0.35 | 0.2 | 0.29 | 0.16 | 0.19 |
| 7 | 13543729 | rs13307587 | 0.51 | 2.004 | -0.725 | -1.522 | -0.057 | 0.09 | 0.09 | 0 | 0.21 | 0.06 | 0.21 | 0 | 0.34 | 0.08 |
| 7 | 13546297 | rs2692477 | -0.134 | 1.651 | -0.644 | -1.935 | -0.009 | -0.074 | -0.074 | 0 | 0.19 | 0.29 | 0.11 | 0.03 | 0.17 | 0.19 |
| 7 | 13548945 | rs2462613 | -0.542 | 1.802 | -0.33 | -1.947 | -0.008 | -0.013 | -0.013 | 0.467 | 0.4 | 0.35 | 0.33 | 0.03 | 0.52 | 0.25 |
| 7 | 13551373 | rs2723398 | -0.348 | 1.786 | -0.375 | -1.956 | -0.023 | 0.06 | 0.06 | 0 | 0.38 | 0.25 | 0.32 | 0.03 | 0.5 | 0.24 |
| 7 | 13557623 | rs2692464 | -0.55 | 1.766 | -0.372 | -1.939 | -0.008 | 0.003 | 0.003 | 0 | 0.39 | 0.33 | 0.31 | 0.03 | 0.49 | 0.24 |
| 7 | 13567908 | rs2692472 | 0.346 | 1.706 | -0.349 | -1.829 | -0.005 | -0.049 | -0.049 | 0 | 0.17 | 0.26 | 0.1 | 0.03 | 0.15 | 0.17 |
| 7 | 13569530 | rs17167207 | -0.637 | 1.766 | -0.812 | -1.773 | 0.01 | -0.025 | -0.025 | 0.018 | 0.1 | 0.18 | 0.14 | 0.08 | 0.1 | 0.11 |

| | | | | | | | | | | | | | | | | |
|---|---|---|---|---|---|---|---|---|---|---|---|---|---|---|---|---|
| 7 | 13571660 | rs12699513 | -0.238 | 1.806 | -0.983 | -1.908 | -0.008 | 0.02 | 0.02 | 0 | 0.32 | 0.25 | 0.32 | 0.08 | 0.41 | 0.16 |
| 7 | 13576298 | rs6951342 | 0.267 | 1.887 | -1.076 | -1.851 | -0.015 | 0.051 | 0.051 | 0 | 0.27 | 0.15 | 0.21 | 0 | 0.35 | 0.08 |
| 7 | 13577457 | rs6978901 | 0.761 | 1.874 | -1.224 | -1.81 | -0.077 | 0.153 | 0.153 | 0 | 0.23 | 0.01 | 0.18 | 0 | 0.31 | 0.06 |
| 7 | 13577494 | rs7810495 | 1.727 | 2.098 | -1.305 | -1.879 | 0.029 | 0.043 | 0.043 | 0 | 0.38 | 0.26 | 0.41 | 0.7 | 0.28 | 0.51 |
| 7 | 13579371 | rs9638696 | 1.105 | 2.128 | -1.266 | -1.841 | -0.023 | 0.056 | 0.056 | 0.346 | 0.6 | 0.28 | 0.6 | 0.7 | 0.59 | 0.58 |
| 7 | 13591197 | rs1880631 | 1.989 | 2.08 | -1.403 | -1.811 | 0.048 | 0.045 | 0.045 | 0 | 0.35 | 0.15 | 0.38 | 0.69 | 0.22 | 0.5 |
| 7 | 13594968 | rs10232684 | 0.422 | 2.668 | -1.313 | -1.538 | -0.024 | 0.089 | 0.089 | 0 | 0.24 | 0.09 | 0.21 | 0 | 0.33 | 0.06 |
| 7 | 13599216 | rs2222618 | -0.749 | 2.872 | -0.717 | -0.706 | 0.03 | -0.015 | -0.015 | 0 | 0.08 | 0.16 | 0.15 | 0.08 | 0.14 | 0.16 |
| 7 | 13601058 | rs17280514 | -0.502 | 2.426 | -1.195 | -0.455 | 0.008 | 0.01 | 0.01 | 0.016 | 0.09 | 0.11 | 0.13 | 0.17 | 0.12 | 0.07 |
| 7 | 13632228 | rs12668467 | 0.678 | 1.074 | -1.291 | -0.681 | 0.038 | 0.077 | 0.077 | 0 | 0.38 | 0.1 | 0.38 | 0.7 | 0.29 | 0.45 |
| 7 | 13634481 | rs6461037 | -1.185 | 1.415 | -0.147 | -0.59 | 0.023 | -0.004 | -0.004 | 0 | 0.24 | 0.48 | 0.22 | 0.23 | 0.26 | 0.24 |
| 7 | 13635158 | rs962177 | -1.102 | 1.388 | 0.014 | -0.592 | 0.02 | -0.007 | -0.007 | 0.06 | 0.23 | 0.4 | 0.2 | 0.22 | 0.25 | 0.24 |
| 7 | 13635955 | rs13246040 | 0.557 | 1.887 | -1.077 | -0.718 | -0.049 | 0.014 | 0.014 | 0 | 0.19 | 0.06 | 0.27 | 0.11 | 0.32 | 0.19 |
| 7 | 13638892 | rs6977948 | 0.525 | 1.739 | -1.128 | -0.926 | -0.017 | 0.038 | 0.038 | 0 | 0.16 | 0.07 | 0.1 | 0.02 | 0.15 | 0.13 |
| 7 | 13641674 | rs10230686 | -0.249 | 1.522 | -1.036 | -0.756 | -0.003 | 0.008 | 0.008 | 0 | 0.14 | 0.1 | 0.1 | 0.06 | 0.14 | 0.19 |
| 7 | 13643530 | rs6946350 | 1.02 | 1.285 | -1.093 | -0.555 | 0.062 | -0.007 | -0.007 | 0.336 | 0.4 | 0.17 | 0.39 | 0.65 | 0.28 | 0.45 |
| 7 | 13654199 | rs17581161 | -0.305 | 2.468 | -0.904 | -0.642 | -0.011 | 0.001 | 0.001 | 0 | 0.19 | 0.14 | 0.23 | 0.11 | 0.27 | 0.31 |
| 7 | 13654292 | rs17647702 | 0.571 | 2.555 | -0.896 | -0.599 | 0.006 | 0.008 | 0.008 | 0.003 | 0.06 | 0.06 | 0.14 | 0.1 | 0.14 | 0.18 |
| 7 | 13655987 | rs12699541 | -1.086 | 2.11 | -0.499 | -0.918 | 0.002 | -0.001 | -0.001 | 0 | 0.51 | 0.56 | 0.49 | 0.17 | 0.63 | 0.52 |
| 7 | 13656465 | rs7792011 | 0.521 | 2.107 | -0.562 | -0.944 | -0.004 | -0.008 | -0.008 | 0 | 0.07 | 0.16 | 0.05 | 0.05 | 0.04 | 0.08 |
| 7 | 13656560 | rs1581568 | 0.796 | 1.938 | -0.776 | -0.773 | -0.018 | 0.1 | 0.1 | 0 | 0.61 | 0.16 | 0.64 | 0.84 | 0.56 | 0.63 |
| 7 | 13656650 | rs1581567 | 0.258 | 1.781 | -0.683 | -0.661 | -0.001 | -0.002 | -0.002 | 0.249 | 0.68 | 0.32 | 0.68 | 0.89 | 0.59 | 0.71 |
| 7 | 13662014 | rs10807768 | -0.2 | 1.293 | -0.363 | -0.311 | -0.001 | -0.008 | -0.008 | 0 | 0.48 | 0.5 | 0.44 | 0.17 | 0.55 | 0.52 |
| 7 | 13674145 | rs10239445 | -0.089 | 1.036 | -0.933 | -0.26 | 0.008 | 0.071 | 0.071 | 0 | 0.52 | 0.68 | 0.41 | 0.28 | 0.55 | 0.51 |
| 7 | 13674179 | rs10269275 | -0.231 | 0.863 | -0.848 | -0.176 | 0.014 | 0.056 | 0.056 | 0 | 0.54 | 0.68 | 0.46 | 0.28 | 0.61 | 0.52 |
| 7 | 13679931 | rs11981893 | -0.698 | 0.647 | -0.377 | -0.293 | -0.012 | -0.062 | -0.062 | 0.383 | 0.27 | 0.56 | 0.2 | 0.13 | 0.28 | 0.22 |
| 7 | 13680095 | rs10499431 | -0.196 | 0.533 | -0.208 | -0.41 | -0.009 | 0.002 | 0.002 | 0 | 0.14 | 0.1 | 0.14 | 0.1 | 0.2 | 0.12 |
| 7 | 13682477 | rs10499436 | -0.023 | 1.219 | -1.122 | -0.275 | -0.071 | -0.06 | -0.06 | 0.005 | 0.15 | 0.01 | 0.13 | 0.19 | 0.18 | 0.29 |
| 7 | 13683080 | rs1525683 | -1.178 | 0.682 | -0.782 | -0.668 | -0.007 | -0.033 | -0.033 | 0 | 0.31 | 0.64 | 0.23 | 0.14 | 0.3 | 0.23 |
| 7 | 13685734 | rs891863 | -1.635 | 0.726 | -0.763 | -0.621 | -0.01 | 0.017 | 0.017 | 0 | 0.44 | 0.69 | 0.36 | 0.33 | 0.49 | 0.52 |
| 7 | 13692843 | rs7786891 | -0.33 | 0.552 | -0.1 | -0.532 | 0.005 | -0.021 | -0.021 | 0.147 | 0.2 | 0.39 | 0.18 | 0.11 | 0.22 | 0.14 |
| 7 | 13693060 | rs17167319 | 1.701 | 0.476 | -0.789 | -1.106 | 0.009 | -0.012 | -0.012 | 1.961 | 0.37 | 0.23 | 0.5 | 0.61 | 0.35 | 0.38 |
| 7 | 13696843 | rs17590581 | 1.449 | 0.07 | -1.553 | -1.838 | -0.045 | 0.109 | 0.109 | 0 | 0.17 | 0.01 | 0.17 | 0 | 0.21 | 0.07 |
| 7 | 13697970 | rs10243935 | -0.749 | 0.465 | -1.273 | -1.561 | -0.006 | 0.018 | 0.018 | 0 | 0.45 | 0.67 | 0.37 | 0.34 | 0.51 | 0.53 |
| 7 | 13701869 | rs10260350 | -0.742 | 0.447 | -1.131 | -1.474 | 0 | 0.011 | 0.011 | 0 | 0.46 | 0.64 | 0.37 | 0.34 | 0.52 | 0.53 |
| 7 | 13705124 | rs3864419 | 1.128 | 0.448 | -1.178 | -1.348 | -0.026 | 0 | 0 | 0 | 0.15 | 0.04 | 0.14 | 0.1 | 0.2 | 0.12 |
| 7 | 13722884 | rs12672414 | 1.153 | 0.431 | -1.148 | -0.643 | -0.009 | -0.082 | -0.082 | 0 | 0.34 | 0.12 | 0.44 | 0.48 | 0.35 | 0.42 |
| 7 | 13725443 | rs9655107 | -0.256 | 0.781 | -0.488 | -0.662 | 0.006 | 0 | 0 | 0 | 0.31 | 0.42 | 0.35 | 0.34 | 0.37 | 0.29 |
| 7 | 13728286 | rs1942144 | 0.792 | 0.754 | -0.431 | -0.621 | 0.009 | 0.028 | 0.028 | 0 | 0.11 | 0.21 | 0.09 | 0.23 | 0.14 | 0.16 |
| 7 | 13728505 | rs6461043 | 0.899 | 0.635 | -0.42 | -0.964 | 0.024 | 0.069 | 0.069 | 0 | 0.2 | 0.06 | 0.11 | 0.04 | 0.14 | 0.13 |
| 7 | 13729014 | rs7783357 | -0.011 | 0.743 | -0.364 | -0.964 | 0.001 | 0.1 | 0.1 | 0 | 0.37 | 0.13 | 0.34 | 0.15 | 0.34 | 0.25 |
| 7 | 13731213 | rs1893403 | -0.831 | 0.888 | -0.005 | -0.699 | -0.001 | 0.002 | 0.002 | 0 | 0.54 | 0.6 | 0.47 | 0.26 | 0.52 | 0.41 |
| 7 | 13731897 | rs17167374 | 0.609 | 0.903 | -0.022 | -0.681 | 0.007 | -0.015 | -0.015 | 0 | 0.2 | 0.29 | 0.26 | 0.11 | 0.22 | 0.12 |
| 7 | 13734285 | rs2848839 | -0.135 | 0.734 | -0.326 | -0.583 | -0.013 | -0.045 | -0.045 | 0 | 0.37 | 0.48 | 0.23 | 0.16 | 0.31 | 0.29 |
| 7 | 13741691 | rs10257635 | 0.494 | 0.917 | -0.588 | 0.139 | -0.027 | -0.032 | -0.032 | 1.973 | 0.18 | 0.05 | 0.14 | 0.14 | 0.19 | 0.15 |
| 7 | 13748392 | rs2848836 | -0.362 | 0.922 | 0 | 0.805 | -0.008 | -0.071 | -0.071 | 0.162 | 0.29 | 0.18 | 0.23 | 0.54 | 0.28 | 0.37 |
| 7 | 13753853 | rs2568639 | -0.543 | 1.236 | -0.418 | 0.839 | -0.046 | -0.042 | -0.042 | 4.096 | 0.65 | 0.75 | 0.44 | 0.52 | 0.57 | 0.55 |
| 7 | 13755474 | rs10486076 | 0.635 | 1.058 | -0.357 | 0.768 | -0.005 | 0.003 | 0.003 | 0 | 0.27 | 0.24 | 0.21 | 0.29 | 0.24 | 0.24 |
| 7 | 13756470 | rs9639165 | 0.202 | 1.13 | -0.344 | 0.838 | 0.002 | 0.014 | 0.014 | 0 | 0.31 | 0.34 | 0.28 | 0.64 | 0.31 | 0.46 |
| 7 | 13760017 | rs2528494 | 1.125 | 1.065 | -0.017 | 0.75 | 0.002 | 0.002 | 0.002 | 0.175 | 0.26 | 0.25 | 0.18 | 0.17 | 0.21 | 0.15 |
| 7 | 13761404 | rs2568654 | -0.412 | 1.039 | 0.002 | 0.695 | -0.013 | 0.038 | 0.038 | 0 | 0.57 | 0.64 | 0.47 | 0.85 | 0.52 | 0.62 |
| 7 | 13765043 | rs2568644 | -0.448 | 1.238 | -0.078 | 0.775 | -0.002 | 0 | 0 | 1.597 | 0.43 | 0.6 | 0.29 | 0.6 | 0.39 | 0.53 |
| 7 | 13765237 | rs2568643 | 0.02 | 0.934 | -0.187 | 0.668 | 0.007 | 0.007 | 0.007 | 0.032 | 0.37 | 0.36 | 0.22 | 0.24 | 0.31 | 0.31 |
| 7 | 13769291 | rs2528492 | 0.229 | 1.246 | 0.177 | 0.486 | 0.007 | -0.04 | -0.04 | 0 | 0.57 | 0.45 | 0.45 | 0.83 | 0.52 | 0.61 |
| 7 | 13770431 | rs6964915 | -0.048 | 0.951 | 0.062 | 0.498 | 0.014 | -0.048 | -0.048 | 0.076 | 0.15 | 0.51 | 0.07 | 0.05 | 0.12 | 0.16 |
| 7 | 13770745 | rs6965391 | 0.193 | 0.94 | 0.067 | 0.499 | -0.005 | -0.039 | -0.039 | 0 | 0.14 | 0.44 | 0.06 | 0.05 | 0.1 | 0.15 |
| 7 | 13773054 | rs2156013 | 0.552 | 1.089 | -0.035 | 0.644 | 0.031 | 0.002 | 0.002 | 1.196 | 0.49 | 0.32 | 0.36 | 0.46 | 0.41 | 0.38 |
| 7 | 13773221 | rs2848845 | 0.167 | 0.73 | -0.181 | 0.554 | -0.002 | 0.006 | 0.006 | 0.109 | 0.54 | 0.39 | 0.43 | 0.81 | 0.49 | 0.6 |
| 7 | 13777635 | rs2848837 | -0.639 | 1.241 | -0.957 | 0.995 | -0.063 | 0.072 | 0.072 | 0.142 | 0.69 | 0.89 | 0.5 | 0.86 | 0.59 | 0.76 |
| 7 | 13777723 | rs6974412 | -0.673 | 0.911 | -0.353 | 1.731 | 0.03 | 0.173 | 0.173 | 0 | 0.05 | 0.16 | 0.07 | 0.4 | 0.07 | 0.23 |
| 7 | 13779769 | rs17167480 | -0.876 | 0.798 | -0.425 | 1.586 | 0.011 | 0.119 | 0.119 | 0 | 0.05 | 0.14 | 0.09 | 0.4 | 0.06 | 0.22 |
| 7 | 13780484 | rs4719379 | 0.262 | -0.666 | -0.696 | 0.231 | 0.065 | 0.075 | 0.075 | 0.06 | 0.52 | 0.36 | 0.67 | 0.83 | 0.72 | 0.71 |
| 7 | 13780997 | rs12699544 | -0.143 | -0.997 | -0.671 | 0.026 | -0.002 | 0 | 0 | 0 | 0.81 | 0.77 | 0.84 | 0.85 | 0.82 | 0.77 |
| 7 | 13783853 | rs2190320 | 0.215 | -1.521 | -1.07 | -0.122 | 0.009 | 0.025 | 0.025 | 0 | 0.82 | 0.91 | 0.7 | 0.95 | 0.85 | 0.89 |
| 7 | 13786362 | rs11979631 | 1.02 | -1.52 | -0.803 | 0.114 | -0.011 | -0.001 | -0.001 | 0 | 0.47 | 0.44 | 0.49 | 0.63 | 0.68 | 0.57 |
| 7 | 13786456 | rs7456569 | 1.243 | -0.136 | -1.204 | -0.196 | 0.022 | -0.062 | -0.062 | 0 | 0.22 | 0.28 | 0.18 | 0.03 | 0.31 | 0.17 |
| 7 | 13792450 | rs6949771 | -0.031 | -0.876 | -1.162 | 0.581 | -0.008 | 0.009 | 0.009 | 0 | 0.9 | 0.96 | 0.73 | 0.96 | 0.88 | 0.91 |
| 7 | 13793914 | rs12673493 | 0.266 | -1.191 | -1.061 | 0.666 | 0.034 | -0.031 | -0.031 | 0 | 0.2 | 0.03 | 0.12 | 0.16 | 0.08 | 0.14 |
| 7 | 13794203 | rs6956462 | -0.391 | -1.006 | -0.566 | 0.679 | -0.04 | 0.012 | 0.012 | 0 | 0.35 | 0.47 | 0.23 | 0.35 | 0.17 | 0.33 |

| | | | | | | | | | | | | | | | |
|---|---|---|---|---|---|---|---|---|---|---|---|---|---|---|---|
| 7 | 13801382 | rs13224659 | -0.223 | 0.175 | -0.501 | 0.369 | 0.013 | -0.026 | -0.026 | 0 | 0.06 | 0.08 | 0.08 | 0 | 0.11 | 0.03 |
| 7 | 13801764 | rs11768005 | -0.433 | -0.038 | -0.853 | 1.186 | -0.009 | -0.035 | -0.035 | 0 | 0.16 | 0.09 | 0.39 | 0.43 | 0.23 | 0.17 |
| 7 | 13801825 | rs6461048 | -0.362 | 0.292 | -1.051 | 0.609 | -0.023 | 0.009 | 0.009 | 0.364 | 0.65 | 0.68 | 0.64 | 0.74 | 0.59 | 0.59 |
| 7 | 13804535 | rs12112961 | -0.339 | -0.133 | -0.733 | -0.078 | 0 | 0.006 | 0.006 | 0 | 0.41 | 0.41 | 0.52 | 0.54 | 0.44 | 0.43 |
| 7 | 13815500 | rs17732862 | 0.61 | -0.035 | -0.738 | -0.531 | -0.01 | 0.05 | 0.05 | 0 | 0.22 | 0.1 | 0.11 | 0.01 | 0.21 | 0.2 |
| 7 | 13816640 | rs2190317 | -1.132 | -0.088 | -0.886 | -0.414 | 0.038 | -0.031 | -0.031 | 0.168 | 0.83 | 1 | 0.95 | 0.87 | 0.91 | 0.81 |
| 7 | 13822381 | rs11980595 | 0.196 | 0.752 | -0.223 | -0.504 | -0.001 | -0.05 | -0.05 | 0.196 | 0.23 | 0.23 | 0.12 | 0.03 | 0.21 | 0.23 |
| 7 | 13828099 | rs7794026 | -0.473 | 1.041 | -0.027 | -0.256 | 0 | 0.011 | 0.011 | 0.054 | 0.42 | 0.65 | 0.51 | 0.23 | 0.51 | 0.34 |
| 7 | 13831393 | rs12699558 | 0.361 | 1.283 | 0.07 | -0.426 | -0.015 | 0.033 | 0.033 | 0 | 0.25 | 0.15 | 0.18 | 0.01 | 0.28 | 0.2 |
| 7 | 13833814 | rs10486070 | -0.251 | 1.224 | 0.363 | -0.453 | 0.011 | -0.125 | -0.125 | 3.196 | 0.3 | 0.48 | 0.22 | 0.07 | 0.29 | 0.21 |
| 7 | 13836003 | rs2051930 | 0.27 | 1.395 | -0.309 | -0.423 | -0.013 | 0.022 | 0.022 | 0 | 0.26 | 0.18 | 0.19 | 0.01 | 0.28 | 0.2 |
| 7 | 13841798 | rs10256467 | 0.145 | 1.327 | -0.259 | -0.409 | -0.004 | -0.023 | -0.023 | 0 | 0.27 | 0.24 | 0.2 | 0.01 | 0.3 | 0.2 |
| 7 | 13857235 | rs9792016 | 1.05 | 0.452 | -0.723 | 0.167 | 0.032 | -0.059 | -0.059 | 0 | 0.14 | 0 | 0.03 | 0.19 | 0.08 | 0.19 |
| 7 | 13858654 | rs6965481 | 0.209 | 0.799 | -0.72 | 0.521 | -0.001 | -0.001 | -0.001 | 1.238 | 0.13 | 0.13 | 0.19 | 0.18 | 0.16 | 0.17 |
| 7 | 13863636 | rs7806427 | 0.3 | 0.672 | -0.614 | 0.235 | 0.018 | 0.004 | 0.004 | 0.109 | 0.58 | 0.66 | 0.63 | 0.67 | 0.65 | 0.56 |
| 7 | 13864009 | rs12673394 | 0.853 | 0.673 | -0.381 | 0.215 | -0.001 | -0.001 | -0.001 | 0 | 0.46 | 0.5 | 0.34 | 0.47 | 0.45 | 0.49 |
| 7 | 13868825 | rs4560697 | -0.915 | 0.599 | -0.405 | 0.174 | 0.011 | 0.005 | 0.005 | 0 | 0.76 | 0.83 | 0.83 | 0.79 | 0.82 | 0.77 |
| 7 | 13871533 | rs7784117 | -1.01 | 0.72 | -0.565 | 0.036 | 0.013 | 0.013 | 0.013 | 0 | 0.76 | 0.84 | 0.84 | 0.83 | 0.83 | 0.79 |
| 7 | 13872906 | rs10950492 | 0.728 | 0.768 | -0.61 | 0.034 | 0 | 0 | 0 | 0 | 0.48 | 0.51 | 0.36 | 0.48 | 0.47 | 0.49 |
| 7 | 13875291 | rs10486069 | 0.603 | 1.009 | -0.658 | -0.06 | -0.063 | -0.071 | -0.071 | 0 | 0.16 | 0.02 | 0.19 | 0.17 | 0.23 | 0.2 |
| 7 | 13876094 | rs10281229 | -0.468 | 0.727 | -0.53 | -0.056 | 0.001 | 0.002 | 0.002 | 0 | 0.29 | 0.34 | 0.49 | 0.36 | 0.36 | 0.3 |
| 7 | 13876668 | rs7800028 | -0.111 | 0.758 | -0.537 | -0.044 | -0.02 | -0.024 | -0.024 | 0.733 | 0.26 | 0.2 | 0.47 | 0.32 | 0.36 | 0.29 |
| 7 | 13877930 | rs16870032 | 1.348 | 0.79 | -0.85 | -0.258 | 0.01 | -0.02 | -0.02 | 0.359 | 0.28 | 0.2 | 0.19 | 0.46 | 0.22 | 0.31 |
| 7 | 13877971 | rs10230244 | -0.52 | 1.044 | -0.935 | -0.283 | 0 | 0 | 0 | 0 | 0.46 | 0.57 | 0.63 | 0.32 | 0.59 | 0.45 |
| 7 | 13879464 | rs9986861 | -0.654 | 0.844 | -1.236 | -0.583 | 0.013 | 0.019 | 0.019 | 0 | 0.77 | 0.88 | 0.84 | 0.85 | 0.82 | 0.78 |
| 7 | 13880786 | rs10950493 | 0.008 | 1.477 | -0.906 | -0.235 | 0.018 | -0.056 | -0.056 | 0 | 0.18 | 0.25 | 0.16 | 0.02 | 0.23 | 0.16 |
| 7 | 13882718 | rs11766393 | -0.723 | 0.831 | -1.214 | 0.428 | -0.027 | -0.04 | -0.04 | 0 | 0.14 | 0.01 | 0.07 | 0.12 | 0.14 | 0.13 |
| 7 | 13883535 | rs2108150 | -1.154 | 0.82 | -0.584 | 0.494 | 0.003 | 0.015 | 0.015 | 0.611 | 0.22 | 0.7 | 0.23 | 0.24 | 0.23 | 0.25 |
| 7 | 13883729 | rs7799534 | -0.144 | 0.646 | -1.04 | 0.269 | 0.008 | 0.038 | 0.038 | 0 | 0.54 | 0.38 | 0.4 | 0.33 | 0.56 | 0.61 |
| 7 | 13886654 | rs7787720 | -0.3 | 0.258 | -0.904 | 0.307 | -0.003 | -0.001 | -0.001 | 0 | 0.39 | 0.38 | 0.39 | 0.4 | 0.47 | 0.52 |
| 7 | 13887745 | rs17167574 | 0.881 | -0.408 | -1.723 | 0.129 | 0.036 | -0.131 | -0.131 | 0.119 | 0.2 | 0.01 | 0.11 | 0.37 | 0.1 | 0.21 |
| 7 | 13889107 | rs12530627 | -0.444 | -0.276 | -0.791 | -0.254 | 0.017 | -0.065 | -0.065 | 0 | 0.63 | 0.61 | 0.57 | 0.89 | 0.57 | 0.76 |
| 7 | 13894017 | rs17677972 | 0.436 | -0.161 | -0.493 | 0.162 | 0.001 | 0.001 | 0.001 | 0.623 | 0.1 | 0.08 | 0.07 | 0.02 | 0.09 | 0.03 |
| 7 | 13899446 | rs2190324 | -0.861 | -0.381 | -0.089 | -0.016 | -0.037 | -0.07 | -0.07 | 0 | 0.18 | 0.45 | 0.18 | 0.03 | 0.14 | 0.06 |
| 7 | 13900731 | rs7795991 | -0.772 | -0.364 | 0.084 | -0.106 | 0.002 | -0.001 | -0.001 | 0 | 0.43 | 0.55 | 0.5 | 0.4 | 0.52 | 0.57 |
| 7 | 13900784 | rs252007 | 0.295 | -0.143 | -0.752 | -0.514 | -0.038 | 0.084 | 0.084 | 0 | 0.86 | 0.97 | 0.86 | 1 | 0.85 | 0.88 |
| 7 | 13902115 | rs17150029 | -0.591 | -0.017 | -0.217 | -0.351 | -0.001 | -0.002 | -0.002 | 1.731 | 0.22 | 0.24 | 0.21 | 0.14 | 0.21 | 0.16 |
| 7 | 13903105 | rs252001 | -0.533 | 0.068 | -0.455 | -0.363 | -0.039 | 0.061 | 0.061 | 0 | 0.9 | 0.99 | 0.88 | 1 | 0.88 | 0.88 |
| 7 | 13903309 | rs252000 | 0.8 | -0.004 | -0.166 | -0.181 | 0.038 | 0.01 | 0.01 | 0 | 0.44 | 0.21 | 0.35 | 0.37 | 0.35 | 0.28 |
| 7 | 13907827 | rs251994 | 1.12 | -0.587 | -0.753 | -0.288 | -0.047 | -0.056 | -0.056 | 0.067 | 0.37 | 0.07 | 0.4 | 0.33 | 0.35 | 0.28 |
| 7 | 13908250 | rs251993 | 0.3 | -0.756 | -0.182 | -0.414 | -0.062 | 0.003 | 0.003 | 1.058 | 0.73 | 0.57 | 0.84 | 0.68 | 0.88 | 0.87 |
| 7 | 13910748 | rs6962343 | 0.748 | -0.213 | -0.49 | -0.272 | -0.104 | -0.078 | -0.078 | 2.129 | 0.25 | 0.05 | 0.34 | 0.36 | 0.42 | 0.54 |
| 7 | 13911645 | rs722661 | -0.737 | -0.354 | -0.212 | -0.289 | 0.002 | -0.01 | -0.01 | 0 | 0.44 | 0.31 | 0.52 | 0.54 | 0.6 | 0.63 |
| 7 | 13912417 | rs12531613 | 0.479 | -0.071 | -0.51 | -0.364 | -0.026 | 0.039 | 0.039 | 0 | 0.07 | 0.01 | 0.05 | 0 | 0.1 | 0.05 |
| 7 | 13915696 | rs251989 | -0.116 | 0.173 | -0.615 | -0.178 | -0.017 | 0.023 | 0.023 | 0 | 0.68 | 0.79 | 0.66 | 0.77 | 0.62 | 0.59 |
| 7 | 13923962 | rs10486067 | 0.901 | -0.34 | -0.631 | -0.137 | 0.013 | 0.029 | 0.029 | 1.703 | 0.13 | 0.05 | 0.07 | 0.07 | 0.1 | 0.05 |
| 7 | 13931182 | rs9785000 | -1.718 | -0.352 | -0.41 | -0.001 | 0.008 | -0.036 | -0.036 | 0 | 0.71 | 0.92 | 0.75 | 0.72 | 0.76 | 0.81 |
| 7 | 13934688 | rs3735344 | 1.026 | -0.606 | 0.07 | -0.093 | 0.018 | 0.002 | 0.002 | 0 | 0.26 | 0.37 | 0.37 | 0.25 | 0.4 | 0.38 |
| 7 | 13935307 | rs3823702 | 2.056 | -0.626 | -0.365 | -0.187 | 0.073 | -0.109 | -0.109 | 0.553 | 0.25 | 0.01 | 0.09 | 0.28 | 0.11 | 0.17 |
| 7 | 13938185 | rs10268584 | 0.001 | -0.605 | 0.45 | -0.138 | 0.007 | -0.014 | -0.014 | 0.221 | 0.38 | 0.49 | 0.44 | 0.26 | 0.42 | 0.51 |
| 7 | 13942714 | rs6461056 | 1.179 | -0.319 | -0.079 | 0.03 | -0.015 | -0.002 | -0.002 | 2.99 | 0.23 | 0.24 | 0.29 | 0.24 | 0.35 | 0.37 |
| 7 | 13945675 | rs1859508 | -0.89 | -0.511 | -0.307 | 0.059 | 0.036 | 0.006 | 0.006 | 0.864 | 0.57 | 0.86 | 0.66 | 0.66 | 0.65 | 0.76 |
| 7 | 13947289 | rs10950497 | -1.2 | -1.05 | 0.285 | 0.237 | -0.016 | 0.014 | 0.014 | 0 | 0.31 | 0.4 | 0.33 | 0.44 | 0.27 | 0.36 |
| 7 | 13948517 | rs2237292 | -1.117 | -0.879 | 0.212 | 0.307 | -0.007 | -0.013 | -0.013 | 0 | 0.28 | 0.34 | 0.31 | 0.29 | 0.25 | 0.33 |
| 7 | 13955324 | rs3801091 | 0.217 | -1.609 | -0.264 | -0.071 | 0.014 | 0.034 | 0.034 | 2.252 | 0.13 | 0.04 | 0.07 | 0.06 | 0.1 | 0.06 |
| 7 | 13961306 | rs17167656 | -1.137 | -1.133 | 0.27 | 0.294 | -0.063 | -0.018 | -0.018 | 0 | 0.19 | 0.32 | 0.14 | 0.15 | 0.08 | 0.11 |
| 7 | 13967377 | rs10085783 | -0.957 | -0.595 | 0.023 | 0.288 | -0.024 | -0.051 | -0.051 | 0.476 | 0.09 | 0.22 | 0.1 | 0 | 0.03 | 0.03 |
| 7 | 13969085 | rs10242994 | -0.35 | -0.89 | -0.598 | 0.869 | -0.01 | 0.018 | 0.018 | 0 | 0.17 | 0.21 | 0.12 | 0.26 | 0.13 | 0.18 |
| 7 | 13974461 | rs12699566 | -0.291 | -0.172 | -1.248 | 0.937 | -0.029 | -0.083 | -0.083 | 0.581 | 0.25 | 0.09 | 0.29 | 0.45 | 0.28 | 0.26 |
| 7 | 13975769 | rs17167675 | 0.246 | -0.247 | -1.242 | 0.929 | -0.001 | -0.04 | -0.04 | 0.742 | 0.34 | 0.16 | 0.35 | 0.42 | 0.35 | 0.36 |
| 7 | 13978809 | rs9639168 | 0.242 | -0.278 | -1.233 | 0.929 | 0.003 | -0.035 | -0.035 | 0 | 0.34 | 0.18 | 0.35 | 0.42 | 0.34 | 0.36 |
| 7 | 13982420 | rs28615403 | 0.148 | -0.615 | -1.15 | 0.668 | 0.005 | -0.019 | -0.019 | 0.002 | 0.34 | 0.29 | 0.36 | 0.52 | 0.35 | 0.35 |
| 7 | 13990992 | rs2282872 | 0.302 | -0.262 | -0.637 | 0.479 | -0.005 | -0.001 | -0.001 | 0 | 0.36 | 0.37 | 0.52 | 0.34 | 0.42 | 0.4 |
| 7 | 13991221 | rs7790331 | -0.034 | -0.167 | -0.101 | 0.372 | 0.007 | -0.007 | -0.007 | 0 | 0.41 | 0.53 | 0.56 | 0.35 | 0.49 | 0.54 |
| 7 | 13992460 | rs6946355 | -0.455 | -0.627 | -0.513 | 0.303 | -0.013 | 0.007 | 0.007 | 0.016 | 0.49 | 0.8 | 0.61 | 0.43 | 0.52 | 0.48 |
| 7 | 13995209 | rs7800918 | -1.519 | -0.538 | 0.405 | 0.298 | -0.004 | -0.022 | -0.022 | 0 | 0.17 | 0.53 | 0.11 | 0.09 | 0.15 | 0.23 |
| 7 | 14002043 | rs10226058 | -1.382 | -0.649 | 0.888 | 0.496 | 0.017 | 0.009 | 0.009 | 0 | 0.12 | 0.36 | 0.08 | 0.07 | 0.09 | 0.25 |
| 7 | 14004472 | rs17167697 | -1.579 | -0.752 | 1.037 | 0.897 | 0.015 | 0.185 | 0.185 | 0 | 0.13 | 0.43 | 0.09 | 0.2 | 0.1 | 0.26 |
| 7 | 14004670 | rs757700 | 0.859 | -1.294 | 0.214 | 0.626 | 0.004 | 0 | 0 | 0.012 | 0.56 | 0.45 | 0.69 | 0.53 | 0.62 | 0.39 |

| | | | | | | | | | | | | | | | | |
|---|---|---|---|---|---|---|---|---|---|---|---|---|---|---|---|---|
| 7 | 14006895 | rs10278571 | -1.32 | -1.12 | 0.491 | 0.724 | 0.015 | 0.135 | 0.135 | 0 | 0.13 | 0.45 | 0.09 | 0.15 | 0.09 | 0.21 |
| 7 | 14013029 | rs17167711 | -1.039 | -0.955 | 0.066 | 0.519 | 0.043 | -0.008 | -0.008 | 0.473 | 0.06 | 0.23 | 0.06 | 0.01 | 0.05 | 0.19 |
| 7 | 14013219 | rs1476622 | -0.164 | -1.149 | -0.649 | 0.571 | -0.044 | -0.019 | -0.019 | 0.038 | 0.72 | 0.98 | 0.78 | 0.67 | 0.71 | 0.63 |
| 7 | 14020304 | rs17687519 | 0.22 | -1.711 | -0.825 | -0.243 | 0.019 | 0.07 | 0.07 | 0.852 | 0.25 | 0.1 | 0.13 | 0.01 | 0.17 | 0.24 |
| 7 | 14021095 | rs4721290 | 0.532 | -1.248 | -0.408 | 0.342 | 0.001 | 0 | 0 | 0 | 0.43 | 0.49 | 0.61 | 0.44 | 0.51 | 0.37 |
| 7 | 14026802 | rs3823706 | 0.904 | -1.656 | -0.792 | 0.367 | -0.027 | 0.04 | 0.04 | 0.172 | 0.13 | 0.24 | 0.16 | 0.2 | 0.08 | 0.09 |
| 7 | 14027557 | rs7789916 | -0.399 | -1.429 | -1.365 | 0.233 | -0.053 | -0.086 | -0.086 | 0.008 | 0.7 | 0.97 | 0.77 | 0.59 | 0.71 | 0.61 |
| 7 | 14027991 | rs6977237 | 0.391 | -1.106 | -1.077 | 0.376 | -0.007 | 0.036 | 0.036 | 0 | 0.46 | 0.68 | 0.63 | 0.59 | 0.54 | 0.38 |
| 7 | 14029739 | rs2073533 | 0.359 | -0.856 | -0.663 | 0.959 | -0.003 | -0.016 | -0.016 | 0.19 | 0.68 | 0.73 | 0.71 | 0.51 | 0.68 | 0.54 |
| 7 | 14040129 | rs1990077 | -0.628 | -1.38 | -0.762 | 0.795 | 0.054 | 0.039 | 0.039 | 0.056 | 0.47 | 0.77 | 0.5 | 0.62 | 0.4 | 0.49 |
| 7 | 14048098 | rs11978520 | 0.284 | -2.236 | 0.419 | 0.663 | -0.026 | 0.014 | 0.014 | 0 | 0.09 | 0.21 | 0.06 | 0.08 | 0.05 | 0.08 |
| 7 | 14059902 | rs7800203 | -0.226 | -2.315 | 1.47 | 0.784 | -0.024 | 0.098 | 0.098 | 0 | 0.1 | 0.45 | 0.08 | 0.12 | 0.04 | 0.11 |
| 7 | 14060101 | rs9784978 | 0.325 | -2.713 | 1.114 | 0.779 | 0.02 | -0.028 | -0.028 | 0 | 0.1 | 0.05 | 0.08 | 0.17 | 0.06 | 0.09 |
| 7 | 14061054 | rs7788676 | -0.25 | -2.579 | 1.472 | 0.357 | -0.038 | -0.077 | -0.077 | 0 | 0.24 | 0.38 | 0.14 | 0.08 | 0.15 | 0.11 |
| 7 | 14066134 | rs2190066 | -0.831 | -2.619 | 1.315 | 0.672 | -0.005 | 0 | 0 | 0 | 0.29 | 0.66 | 0.29 | 0.25 | 0.19 | 0.16 |
| 7 | 14075795 | rs11983337 | 0.06 | -2.442 | 1.61 | 0.643 | -0.028 | 0.012 | 0.012 | 0.066 | 0.21 | 0.38 | 0.22 | 0.16 | 0.15 | 0.1 |
| 7 | 14090857 | rs10259703 | -0.308 | -2.544 | 1.534 | 0.72 | -0.033 | 0.126 | 0.126 | 0.333 | 0.11 | 0.57 | 0.21 | 0.22 | 0.07 | 0.08 |
| 7 | 14091524 | rs12532319 | -0.298 | -2.554 | 1.592 | 0.636 | -0.027 | 0.011 | 0.011 | 0.238 | 0.11 | 0.53 | 0.1 | 0.09 | 0.06 | 0.08 |
| 7 | 14092993 | rs10238440 | -1.004 | -2.614 | 1.206 | 0.669 | 0.034 | -0.006 | -0.006 | 0 | 0.29 | 0.74 | 0.29 | 0.25 | 0.18 | 0.18 |
| 7 | 14102031 | rs6965533 | -0.483 | -2.497 | 1.602 | 0.573 | -0.035 | 0.025 | 0.025 | 1.501 | 0.12 | 0.59 | 0.1 | 0.12 | 0.06 | 0.12 |
| 7 | 14105064 | rs7797625 | -0.207 | -2.662 | 0.636 | 0.423 | 0.026 | 0.012 | 0.012 | 0.937 | 0.24 | 0.16 | 0.28 | 0.13 | 0.15 | 0.13 |
| 7 | 14110020 | rs2158664 | -0.109 | -2.661 | 0.417 | 0.359 | 0.032 | 0.018 | 0.018 | 0.901 | 0.23 | 0.1 | 0.28 | 0.13 | 0.15 | 0.11 |
| 7 | 14111193 | rs10278657 | -0.757 | -2.576 | 0.428 | 0.49 | 0.067 | 0.014 | 0.014 | 1.006 | 0.36 | 0.75 | 0.38 | 0.26 | 0.21 | 0.27 |
| 7 | 14113503 | rs2884306 | -1.468 | -2.767 | -0.45 | 0.251 | -0.005 | -0.086 | -0.086 | 0.159 | 0.61 | 0.91 | 0.54 | 0.49 | 0.38 | 0.55 |
| 7 | 14114798 | rs969419 | -0.891 | -2.976 | -0.053 | 0.14 | -0.001 | -0.001 | -0.001 | 0 | 0.31 | 0.64 | 0.32 | 0.41 | 0.21 | 0.39 |
| 7 | 14115074 | rs6972164 | -0.96 | -3.152 | -0.221 | 0.114 | 0.182 | 0.045 | 0.045 | 0 | 0.52 | 0.82 | 0.35 | 0.35 | 0.26 | 0.38 |
| 7 | 14117692 | rs2158662 | -1.662 | -2.814 | -0.438 | 0.194 | -0.041 | -0.108 | -0.108 | 0 | 0.62 | 0.91 | 0.54 | 0.48 | 0.41 | 0.59 |
| 7 | 14121020 | rs17167836 | -0.704 | -2.951 | -0.065 | 0.011 | 0.038 | 0.041 | 0.041 | 0 | 0.51 | 0.64 | 0.44 | 0.26 | 0.34 | 0.43 |
| 7 | 14124065 | rs1029544 | 0.206 | -2.947 | -0.16 | 0.056 | -0.018 | -0.014 | -0.014 | 0 | 0.33 | 0.36 | 0.23 | 0.12 | 0.18 | 0.22 |
| 7 | 14133305 | rs10267453 | -0.2 | -2.408 | -0.311 | 0.239 | 0.002 | 0.002 | 0.002 | 0 | 0.27 | 0.28 | 0.33 | 0.18 | 0.21 | 0.33 |
| 7 | 14133846 | rs6461068 | -0.138 | -1.298 | -0.556 | 0.807 | -0.001 | 0.179 | 0.179 | 0 | 0.08 | 0.28 | 0.08 | 0.25 | 0.05 | 0.15 |
| 7 | 14136740 | rs10085723 | -0.371 | -0.93 | -1.698 | 0.447 | -0.027 | -0.093 | -0.093 | 0 | 0.61 | 0.92 | 0.53 | 0.49 | 0.38 | 0.52 |
| 7 | 14139711 | rs2107888 | 0.209 | -0.206 | -1.431 | 0.523 | -0.047 | -0.073 | -0.073 | 0 | 0.38 | 0.14 | 0.42 | 0.54 | 0.53 | 0.47 |
| 7 | 14143121 | rs10243424 | -0.792 | 0.079 | -0.652 | 0.236 | -0.003 | -0.002 | -0.002 | 0 | 0.26 | 0.29 | 0.18 | 0.19 | 0.23 | 0.4 |
| 7 | 14143806 | rs2357852 | -0.835 | -0.006 | -0.626 | 0.612 | -0.005 | 0.007 | 0.007 | 0.164 | 0.26 | 0.32 | 0.19 | 0.3 | 0.22 | 0.4 |
| 7 | 14144663 | rs12533882 | -0.764 | -0.001 | -0.643 | 0.561 | -0.003 | -0.003 | -0.003 | 0.373 | 0.25 | 0.29 | 0.17 | 0.18 | 0.22 | 0.4 |
| 7 | 14146320 | rs7790836 | -1.477 | 0.531 | -1.046 | 0.325 | -0.08 | -0.071 | -0.071 | 0.32 | 0.65 | 0.87 | 0.47 | 0.54 | 0.47 | 0.67 |
| 7 | 14149236 | rs10231142 | -1.258 | 0.44 | -0.922 | 0.409 | 0.025 | -0.062 | -0.062 | 0 | 0.58 | 0.86 | 0.39 | 0.51 | 0.35 | 0.65 |
| 7 | 14151155 | rs6946604 | -0.974 | 0.054 | 0.07 | 0.225 | -0.039 | -0.001 | -0.001 | 0 | 0.32 | 0.45 | 0.21 | 0.28 | 0.22 | 0.32 |
| 7 | 14151697 | rs9638710 | -1.354 | 0.307 | -0.011 | 0.068 | 0.004 | 0.008 | 0.008 | 0.076 | 0.44 | 0.62 | 0.25 | 0.28 | 0.36 | 0.42 |
| 7 | 14156986 | rs7783337 | -1.028 | -0.949 | 0.29 | -0.184 | -0.013 | -0.002 | -0.002 | 0 | 0.21 | 0.24 | 0.19 | 0.24 | 0.15 | 0.18 |
| 7 | 14164274 | rs17167896 | -0.269 | -1.268 | -0.115 | -0.127 | -0.004 | 0.003 | 0.003 | 0 | 0.17 | 0.16 | 0.1 | 0.07 | 0.18 | 0.15 |
| 7 | 14170925 | rs17167907 | -0.569 | -1.118 | -0.38 | -0.022 | 0.004 | 0 | 0 | 0 | 0.15 | 0.18 | 0.12 | 0.16 | 0.18 | 0.2 |
| 7 | 14171253 | rs7792557 | -0.265 | -1.094 | -0.755 | 0.235 | -0.009 | 0.005 | 0.005 | 2.004 | 0.18 | 0.23 | 0.22 | 0.21 | 0.12 | 0.26 |
| 7 | 14172654 | rs726297 | -1.324 | -1.366 | -1.302 | 0.139 | 0.046 | -0.022 | -0.022 | 0 | 0.43 | 0.88 | 0.39 | 0.39 | 0.32 | 0.48 |
| 7 | 14173733 | rs7789750 | -1.274 | -0.648 | -0.532 | 0.017 | 0.031 | 0.034 | 0.034 | 0.619 | 0.34 | 0.76 | 0.2 | 0.22 | 0.24 | 0.23 |
| 7 | 14178260 | rs17167920 | -1.107 | 0.207 | -0.001 | 0.558 | -0.02 | 0.001 | 0.001 | 0.196 | 0.19 | 0.57 | 0.14 | 0.2 | 0.13 | 0.18 |
| 7 | 14180681 | rs10238563 | -1.06 | 0.858 | -0.898 | 0.609 | -0.009 | 0.005 | 0.005 | 0.812 | 0.28 | 0.84 | 0.32 | 0.26 | 0.31 | 0.22 |
| 7 | 14181736 | rs10950509 | 0.377 | 1.178 | -1.082 | 0.662 | 0.007 | 0.027 | 0.027 | 2.675 | 0.51 | 0.34 | 0.51 | 0.42 | 0.46 | 0.42 |
| 7 | 14182592 | rs714179 | 0.753 | 0.948 | -1.927 | 0.593 | 0.003 | 0.058 | 0.058 | 1.795 | 0.36 | 0.03 | 0.16 | 0.23 | 0.27 | 0.32 |
| 7 | 14185876 | rs979868 | -0.311 | 1.432 | -1.776 | 0.432 | -0.054 | 0.052 | 0.052 | 0.681 | 0.89 | 0.99 | 0.92 | 0.99 | 0.84 | 0.89 |
| 7 | 14187523 | rs3735345 | 0.216 | 0.727 | -1.714 | 0.533 | 0.019 | -0.003 | -0.003 | 0 | 0.13 | 0.06 | 0.11 | 0.11 | 0.07 | 0.05 |
| 7 | 14187830 | rs4719392 | 1.03 | 1.231 | -1.89 | 0.726 | 0.009 | 0.033 | 0.033 | 2.011 | 0.43 | 0.09 | 0.35 | 0.33 | 0.35 | 0.43 |
| 7 | 14187931 | rs17167930 | -0.451 | 0.933 | -0.859 | 1.083 | -0.03 | 0.06 | 0.06 | 0.043 | 0.17 | 0.34 | 0.2 | 0.24 | 0.13 | 0.11 |
| 7 | 14192611 | rs7777178 | 0.676 | 1.426 | -0.75 | 1.13 | 0.01 | 0.041 | 0.041 | 0.543 | 0.5 | 0.24 | 0.41 | 0.44 | 0.43 | 0.53 |
| 7 | 14196202 | rs10273051 | 0.358 | 1.388 | -0.825 | 1.221 | -0.013 | 0.001 | 0.001 | 0 | 0.11 | 0.14 | 0.15 | 0.12 | 0.05 | 0.02 |
| 7 | 14198112 | rs6962724 | 0.337 | 1.345 | -0.914 | 1.157 | -0.043 | -0.033 | -0.033 | 0.418 | 0.59 | 0.2 | 0.51 | 0.54 | 0.47 | 0.56 |
| 7 | 14198643 | rs12538534 | 0.673 | 1.424 | -0.991 | 0.97 | 0.007 | 0.06 | 0.06 | 0.866 | 0.47 | 0.1 | 0.35 | 0.34 | 0.37 | 0.48 |
| 7 | 14202305 | rs12699571 | -0.017 | 1.678 | -0.567 | 0.905 | 0.006 | 0.019 | 0.019 | 0 | 0.37 | 0.49 | 0.46 | 0.53 | 0.41 | 0.39 |
| 7 | 14207966 | rs969013 | 0.092 | 1.31 | -0.982 | 0.676 | -0.045 | 0.054 | 0.054 | 0.029 | 0.9 | 0.99 | 0.91 | 1 | 0.84 | 0.91 |
| 7 | 14209139 | rs7780524 | -0.223 | 1.023 | -1.131 | 0.734 | 0.011 | 0.076 | 0.076 | 0.059 | 0.48 | 0.73 | 0.68 | 0.63 | 0.48 | 0.46 |
| 7 | 14209220 | rs7799696 | 0.418 | 0.359 | -0.688 | 0.762 | -0.012 | -0.001 | -0.001 | 0 | 0.67 | 0.31 | 0.67 | 0.65 | 0.59 | 0.65 |
| 7 | 14210392 | rs992586 | -0.477 | 0.43 | -0.614 | 0.937 | -0.03 | -0.009 | -0.009 | 0.273 | 0.08 | 0.01 | 0.2 | 0.04 | 0.13 | 0.08 |
| 7 | 14212632 | rs7800027 | -0.823 | 0.246 | -0.331 | 1.25 | -0.068 | -0.097 | -0.097 | 1.463 | 0.65 | 0.96 | 0.7 | 0.63 | 0.68 | 0.74 |
| 7 | 14214561 | rs6969690 | -1.355 | 0.178 | 1.097 | 1.897 | 0.017 | 0.061 | 0.061 | 0 | 0.2 | 0.65 | 0.25 | 0.33 | 0.24 | 0.14 |
| 7 | 14214626 | rs6949550 | -2.277 | 0.154 | 1.622 | 1.928 | -0.001 | 0.002 | 0.002 | 0 | 0.28 | 0.66 | 0.5 | 0.46 | 0.34 | 0.26 |
| 7 | 14216609 | rs2286248 | -2.268 | 0.068 | 1.563 | 1.77 | -0.008 | 0.014 | 0.014 | 0.008 | 0.15 | 0.3 | 0.32 | 0.14 | 0.15 | 0.16 |
| 7 | 14217673 | rs6966154 | -0.931 | 0.291 | 1.818 | 1.907 | 0.039 | 0.098 | 0.098 | 0 | 0.16 | 0.54 | 0.2 | 0.27 | 0.21 | 0.13 |
| 7 | 14219213 | rs4719393 | -2.337 | 0.299 | 1.205 | 1.861 | -0.005 | -0.053 | -0.053 | 0 | 0.28 | 0.77 | 0.49 | 0.42 | 0.33 | 0.28 |

| | | | | | | | | | | | | | | | | |
|---|---|---|---|---|---|---|---|---|---|---|---|---|---|---|---|---|
| 7 | 14222693 | rs17695940 | -0.277 | 0.43 | 1.156 | 1.321 | -0.013 | 0 | 0 | 0 | 0.05 | 0.03 | 0.2 | 0.05 | 0.1 | 0.07 |
| 7 | 14226741 | rs10240586 | -2.13 | 0.347 | 1.552 | 1.315 | 0.004 | -0.013 | -0.013 | 0 | 0.25 | 0.69 | 0.4 | 0.16 | 0.31 | 0.25 |
| 7 | 14228703 | rs2068399 | -1.645 | 0.396 | 1.609 | 1.355 | -0.005 | -0.008 | -0.008 | 0 | 0.19 | 0.59 | 0.2 | 0.13 | 0.2 | 0.16 |
| 7 | 14229044 | rs2079459 | -0.693 | 0.609 | 1.047 | 1.124 | 0.022 | -0.04 | -0.04 | 0 | 0.13 | 0.25 | 0.11 | 0.04 | 0.19 | 0.11 |
| 7 | 14233250 | rs6461070 | -1.955 | 0.344 | 0.19 | 1.407 | 0.003 | -0.059 | -0.059 | 0 | 0.39 | 0.83 | 0.53 | 0.49 | 0.39 | 0.32 |
| 7 | 14234531 | rs10486052 | 0.056 | 0.2 | 0.331 | 1.676 | -0.029 | -0.022 | -0.022 | 0 | 0.06 | 0.01 | 0.2 | 0.09 | 0.1 | 0.06 |
| 7 | 14235801 | rs6967846 | -2.079 | -0.065 | 1.135 | 1.844 | -0.035 | 0.1 | 0.1 | 0 | 0.24 | 0.37 | 0.37 | 0.46 | 0.17 | 0.19 |
| 7 | 14239703 | rs11982041 | -2.474 | -0.037 | 1.371 | 1.846 | -0.008 | 0.028 | 0.028 | 0 | 0.31 | 0.59 | 0.46 | 0.54 | 0.31 | 0.25 |
| 7 | 14240130 | rs6971997 | -1.387 | 0.179 | 0.735 | 1.986 | 0.001 | 0.016 | 0.016 | 0 | 0.17 | 0.2 | 0.19 | 0.36 | 0.21 | 0.13 |
| 7 | 14241915 | rs17167959 | -0.81 | -0.082 | 0.413 | 1.975 | 0.003 | -0.04 | -0.04 | 0 | 0.1 | 0.02 | 0.1 | 0.3 | 0.07 | 0.08 |
| 7 | 14243631 | rs10486049 | -0.791 | -0.345 | 0.413 | 1.764 | 0.024 | -0.044 | -0.044 | 0 | 0.1 | 0.02 | 0.1 | 0.26 | 0.05 | 0.08 |
| 7 | 14244402 | rs10486048 | -0.186 | -0.027 | 0.455 | 1.881 | -0.029 | -0.022 | -0.022 | 0.064 | 0.06 | 0.01 | 0.2 | 0.08 | 0.1 | 0.06 |
| 7 | 14245377 | rs10274998 | -1.417 | -0.027 | 0.612 | 1.728 | -0.033 | 0.073 | 0.073 | 0 | 0.24 | 0.6 | 0.4 | 0.45 | 0.17 | 0.18 |
| 7 | 14254864 | rs17708126 | 0.154 | 0.833 | -1.021 | 0.013 | -0.011 | 0.101 | 0.101 | 2.317 | 0.17 | 0.01 | 0.09 | 0.03 | 0.16 | 0.1 |
| 7 | 14261762 | rs11772529 | -1.043 | 0.297 | 0.308 | 1.278 | 0.023 | 0.237 | 0.237 | 0 | 0.13 | 0.37 | 0.34 | 0.35 | 0.12 | 0.07 |
| 7 | 14265179 | rs7778972 | -0.772 | 0.167 | 0.426 | 1.956 | -0.02 | 0.141 | 0.141 | 0 | 0.05 | 0.13 | 0.09 | 0.26 | 0.02 | 0.04 |
| 7 | 14269425 | rs12539033 | 0.362 | 0.374 | 0.172 | 0.945 | -0.002 | -0.003 | -0.003 | 0 | 0.16 | 0.14 | 0.11 | 0.04 | 0.2 | 0.2 |
| 7 | 14270475 | rs985105 | 0.387 | -0.057 | 0.07 | 1.071 | 0.02 | 0.009 | 0.009 | 0 | 0.31 | 0.18 | 0.21 | 0.26 | 0.26 | 0.24 |
| 7 | 14272999 | rs7783422 | 0.208 | -0.145 | 0.702 | 1.093 | -0.017 | -0.006 | -0.006 | 0 | 0.32 | 0.36 | 0.22 | 0.28 | 0.25 | 0.25 |
| 7 | 14273653 | rs976760 | -0.517 | 0.389 | 0.587 | 0.88 | 0 | -0.001 | -0.001 | 0.015 | 0.67 | 0.65 | 0.59 | 0.41 | 0.74 | 0.58 |
| 7 | 14274343 | rs10241087 | -0.212 | 0.678 | 0.868 | 0.791 | 0.029 | 0.007 | 0.007 | 0.156 | 0.15 | 0.55 | 0.19 | 0.11 | 0.16 | 0.15 |
| 7 | 14276567 | rs1859732 | 0.408 | 1.099 | 0.643 | 0.588 | 0.002 | 0.005 | 0.005 | 0.041 | 0.39 | 0.32 | 0.35 | 0.37 | 0.41 | 0.34 |
| 7 | 14276766 | rs17708877 | 0.167 | 1.123 | 0.311 | 0.909 | -0.027 | 0.034 | 0.034 | 0 | 0.09 | 0.04 | 0.09 | 0 | 0.15 | 0.07 |
| 7 | 14278264 | rs7799548 | -0.206 | 0.816 | 0.32 | 1.102 | 0.008 | 0 | 0 | 0 | 0.16 | 0.24 | 0.3 | 0.17 | 0.19 | 0.07 |
| 7 | 14279074 | rs10229230 | -0.033 | 0.823 | -0.013 | 1.087 | -0.078 | -0.14 | -0.14 | 0.046 | 0.61 | 0.91 | 0.69 | 0.54 | 0.61 | 0.44 |
| 7 | 14280528 | rs10252549 | 0.896 | 0.956 | 0.209 | 0.964 | 0.006 | 0.039 | 0.039 | 0 | 0.45 | 0.71 | 0.4 | 0.37 | 0.46 | 0.39 |
| 7 | 14280548 | rs6943375 | 0.494 | 0.474 | 0.161 | 1.097 | -0.009 | 0.097 | 0.097 | 0 | 0.51 | 0.84 | 0.47 | 0.34 | 0.56 | 0.42 |
| 7 | 14283147 | rs10950513 | 0.879 | 0.567 | 0.473 | 1.092 | -0.002 | 0.034 | 0.034 | 0 | 0.46 | 0.67 | 0.41 | 0.33 | 0.48 | 0.38 |
| 7 | 14283396 | rs12673837 | 0.12 | 0.311 | -0.033 | 1.111 | -0.015 | -0.028 | -0.028 | 0 | 0.26 | 0.16 | 0.23 | 0.38 | 0.29 | 0.35 |
| 7 | 14283887 | rs6960036 | 0.288 | -0.133 | 0.129 | 0.864 | -0.038 | -0.024 | -0.024 | 0.1 | 0.75 | 0.62 | 0.83 | 0.81 | 0.86 | 0.78 |
| 7 | 14283976 | rs38291 | -0.129 | 0.875 | -0.194 | 1.396 | -0.066 | -0.074 | -0.074 | 0 | 0.78 | 0.97 | 0.82 | 0.73 | 0.77 | 0.67 |
| 7 | 14286113 | rs17168019 | 0.442 | -0.069 | 0.402 | 0.965 | -0.005 | 0.056 | 0.056 | 0 | 0.09 | 0.21 | 0.06 | 0.18 | 0.05 | 0.02 |
| 7 | 14288573 | rs12537544 | 0.304 | -0.127 | 0.201 | 1.064 | -0.005 | -0.034 | -0.034 | 0 | 0.17 | 0.04 | 0.13 | 0.21 | 0.14 | 0.19 |
| 7 | 14288622 | rs10950514 | -0.34 | -0.061 | 0.313 | 0.931 | -0.007 | 0.016 | 0.016 | 0 | 0.26 | 0.3 | 0.2 | 0.36 | 0.19 | 0.2 |
| 7 | 14292511 | rs10235178 | -0.986 | 0.191 | 0.076 | 0.769 | 0.015 | 0.014 | 0.014 | 0 | 0.34 | 0.47 | 0.31 | 0.39 | 0.37 | 0.24 |
| 7 | 14294598 | rs1011539 | 0.221 | 0.387 | -0.03 | 0.407 | 0.014 | -0.032 | -0.032 | 1.981 | 0.09 | 0.13 | 0.1 | 0 | 0.17 | 0.04 |
| 7 | 14296948 | rs6955524 | 1.271 | -0.144 | -0.349 | -0.32 | -0.024 | 0.018 | 0.018 | 0 | 0.32 | 0.23 | 0.31 | 0.01 | 0.41 | 0.29 |
| 7 | 14302653 | rs38279 | -0.466 | -1.534 | 0.211 | 0.581 | -0.013 | 0.073 | 0.073 | 0.047 | 0.29 | 0.42 | 0.24 | 0.54 | 0.23 | 0.22 |
| 7 | 14303025 | rs38278 | -1.457 | -1.6 | -0.203 | 0.352 | 0.037 | 0.205 | 0.205 | 0 | 0.54 | 0.82 | 0.59 | 0.99 | 0.39 | 0.63 |
| 7 | 14308129 | rs40450 | -1.473 | -1.709 | -0.21 | 0.35 | 0.042 | 0.198 | 0.198 | 0 | 0.54 | 0.81 | 0.59 | 0.99 | 0.38 | 0.62 |
| 7 | 14308206 | rs17168036 | -0.722 | -1.062 | 0.364 | 1.578 | -0.008 | 0.147 | 0.147 | 0 | 0.12 | 0.29 | 0.26 | 0.33 | 0.08 | 0.24 |
| 7 | 14308865 | rs38262 | -1.182 | -1.087 | 0.263 | 1.696 | -0.009 | 0.052 | 0.052 | 0 | 0.21 | 0.25 | 0.31 | 0.49 | 0.15 | 0.26 |
| 7 | 14310025 | rs38259 | 0.322 | -0.573 | -0.123 | 1.436 | 0.001 | -0.041 | -0.041 | 0 | 0.1 | 0 | 0.05 | 0.17 | 0.06 | 0.02 |
| 7 | 14310702 | rs6955426 | -0.481 | -0.258 | 0.391 | 1.366 | -0.021 | 0.106 | 0.106 | 0 | 0.16 | 0.33 | 0.18 | 0.3 | 0.11 | 0.18 |
| 7 | 14317221 | rs7793052 | 0.627 | -0.401 | 0.209 | 0.963 | -0.03 | 0.033 | 0.033 | 0.861 | 0.77 | 0.57 | 0.86 | 0.69 | 0.84 | 0.85 |
| 7 | 14329729 | rs4463321 | -0.106 | -0.495 | -0.169 | 0.85 | -0.028 | -0.026 | -0.026 | 0.016 | 0.83 | 0.99 | 0.87 | 0.81 | 0.84 | 0.93 |
| 7 | 14332000 | rs17711148 | 0.406 | 0.089 | -0.608 | 0.672 | -0.057 | 0.025 | 0.025 | 0 | 0.17 | 0.04 | 0.25 | 0.13 | 0.26 | 0.14 |
| 7 | 14339146 | rs969690 | 1.806 | 0.108 | -0.811 | 0.718 | -0.029 | -0.021 | -0.021 | 0.128 | 0.49 | 0.14 | 0.5 | 0.44 | 0.49 | 0.6 |
| 7 | 14349699 | rs10231874 | 1.232 | 0.151 | -0.959 | 0.742 | -0.005 | 0.002 | 0.002 | 1.513 | 0.68 | 0.41 | 0.77 | 0.59 | 0.75 | 0.74 |
| 7 | 14350339 | rs6980003 | -0.766 | 0.128 | -0.853 | 0.815 | -0.001 | 0.055 | 0.055 | 0.834 | 0.14 | 0.57 | 0.1 | 0.22 | 0.09 | 0.19 |
| 7 | 14351735 | rs6947472 | 1.5 | 0.125 | -1.221 | 0.804 | -0.034 | -0.034 | -0.034 | 0 | 0.48 | 0.14 | 0.49 | 0.43 | 0.49 | 0.59 |
| 7 | 14359852 | rs6944473 | -0.886 | 0.024 | -0.614 | 0.982 | 0.009 | 0.083 | 0.083 | 0.444 | 0.09 | 0.47 | 0.06 | 0.08 | 0.05 | 0.13 |
| 7 | 14360009 | rs10215596 | 0.054 | 0.489 | -1.834 | 0.948 | -0.017 | -0.018 | -0.018 | 0.09 | 0.08 | 0.05 | 0.06 | 0.14 | 0.08 | 0.11 |
| 7 | 14360606 | rs6415237 | -0.073 | 0.399 | -1.834 | 0.748 | -0.026 | -0.037 | -0.037 | 0 | 0.83 | 0.99 | 0.88 | 0.8 | 0.84 | 0.94 |
| 7 | 14364565 | rs1012221 | 0.456 | 0.354 | -1.889 | 0.737 | -0.029 | -0.029 | -0.029 | 0.196 | 0.79 | 0.99 | 0.86 | 0.8 | 0.8 | 0.89 |
| 7 | 14371050 | rs17712036 | -0.03 | 0.616 | -2.225 | 0.358 | -0.044 | 0.086 | 0.086 | 0.139 | 0.21 | 0.07 | 0.17 | 0 | 0.3 | 0.19 |
| 7 | 14371506 | rs917324 | -0.389 | 0.555 | -2.421 | 0.412 | -0.038 | -0.044 | -0.044 | 0.105 | 0.84 | 0.99 | 0.87 | 0.81 | 0.83 | 0.94 |
| 7 | 14378748 | rs2190671 | 0.306 | 0.892 | -2.03 | 0.278 | -0.049 | 0.076 | 0.076 | 0 | 0.16 | 0.04 | 0.12 | 0 | 0.22 | 0.13 |
| 7 | 14379251 | rs2108510 | -0.524 | 0.519 | -1.483 | 1.169 | 0.024 | -0.032 | -0.032 | 0.161 | 0.08 | 0.02 | 0.14 | 0.3 | 0.04 | 0.16 |
| 7 | 14379559 | rs13242229 | -0.36 | 1.048 | -1.415 | 0.656 | -0.025 | -0.051 | -0.051 | 0.916 | 0.25 | 0.1 | 0.24 | 0.38 | 0.28 | 0.4 |
| 7 | 14390063 | rs17168087 | 0.769 | 1.111 | -0.399 | 0.222 | 0.023 | 0.047 | 0.047 | 0 | 0.76 | 0.56 | 0.66 | 0.35 | 0.74 | 0.81 |
| 7 | 14391426 | rs6971911 | -0.125 | 1.07 | -0.553 | -0.494 | -0.007 | -0.005 | -0.005 | 0.157 | 0.38 | 0.5 | 0.46 | 0.34 | 0.34 | 0.48 |
| 7 | 14398086 | rs6977358 | -0.723 | 1.01 | -0.636 | -0.558 | -0.066 | -0.1 | -0.1 | 0 | 0.84 | 0.99 | 0.71 | 0.39 | 0.77 | 0.85 |
| 7 | 14401235 | rs10232313 | -1.232 | 0.57 | 0.365 | -0.064 | 0.041 | 0.034 | 0.034 | 0 | 0.06 | 0.26 | 0.04 | 0.05 | 0.04 | 0.12 |
| 7 | 14404866 | rs4721330 | -0.429 | 0.663 | 0.448 | -0.138 | 0 | -0.01 | -0.01 | 0 | 0.37 | 0.43 | 0.45 | 0.3 | 0.35 | 0.48 |
| 7 | 14410451 | rs2116532 | 0.971 | 0.634 | 0.269 | -0.136 | -0.006 | 0.017 | 0.017 | 0.22 | 0.56 | 0.3 | 0.51 | 0.61 | 0.61 | 0.39 |
| 7 | 14415253 | rs6947526 | 0.548 | 0.558 | 0.185 | -0.135 | -0.001 | -0.003 | -0.003 | 0 | 0.6 | 0.44 | 0.54 | 0.64 | 0.65 | 0.53 |
| 7 | 14415780 | rs12699609 | 0.909 | 0.674 | -0.482 | -0.095 | -0.014 | 0.036 | 0.036 | 0.729 | 0.52 | 0.13 | 0.49 | 0.6 | 0.6 | 0.4 |
| 7 | 14415835 | rs17775555 | 0.682 | 0.829 | -0.662 | -0.122 | -0.032 | 0.069 | 0.069 | 0.266 | 0.16 | 0.02 | 0.1 | 0 | 0.16 | 0.09 |

| | | | | | | | | | | | | | | | | |
|---|---|---|---|---|---|---|---|---|---|---|---|---|---|---|---|---|
| 7 | 14423100 | rs7785682 | 0.142 | 0.984 | -0.318 | 0.434 | 0.028 | 0.066 | 0.066 | 0 | 0.88 | 0.75 | 0.69 | 0.75 | 0.85 | 0.76 |
| 7 | 14424939 | rs10245285 | 0.227 | 0.842 | -1.165 | 0.37 | 0.011 | 0.049 | 0.049 | 0.028 | 0.5 | 0.13 | 0.44 | 0.56 | 0.6 | 0.39 |
| 7 | 14436385 | rs10215028 | -1.571 | 1.021 | 1.64 | 0.87 | 0.06 | 0.133 | 0.133 | 1.738 | 0.1 | 0.34 | 0.14 | 0.23 | 0.08 | 0.23 |
| 7 | 14437333 | rs12699614 | -1.719 | 0.836 | 1.48 | 0.808 | 0.001 | 0.014 | 0.014 | 0 | 0.09 | 0.11 | 0.12 | 0.23 | 0.08 | 0.25 |
| 7 | 14453061 | rs891808 | 2.189 | 1.294 | 1.786 | 0.912 | 0.054 | 0.106 | 0.106 | 0.015 | 0.86 | 0.63 | 0.66 | 0.66 | 0.85 | 0.71 |
| 7 | 14453857 | rs11769240 | -0.193 | 2.152 | 1.289 | 0.562 | -0.002 | -0.001 | -0.001 | 0 | 0.15 | 0.19 | 0.11 | 0.06 | 0.21 | 0.18 |
| 7 | 14454355 | rs11761424 | -1.599 | 2.467 | 0.953 | 1.146 | -0.015 | -0.023 | -0.023 | 0.148 | 0.25 | 0.18 | 0.38 | 0.47 | 0.34 | 0.42 |
| 7 | 14464118 | rs1431534 | -1.612 | 2.31 | 0.779 | 1.128 | -0.031 | -0.082 | -0.082 | 0 | 0.2 | 0.08 | 0.34 | 0.48 | 0.26 | 0.34 |
| 7 | 14465236 | rs7811768 | 0.277 | 2.337 | 0.802 | 1.13 | -0.002 | -0.003 | -0.003 | 0 | 0.15 | 0.19 | 0.11 | 0.05 | 0.21 | 0.18 |
| 7 | 14465892 | rs2043221 | -1.31 | 1.986 | 0.673 | 1.208 | -0.017 | -0.075 | -0.075 | 0 | 0.15 | 0.05 | 0.3 | 0.48 | 0.18 | 0.3 |
| 7 | 14470797 | rs1367775 | -1.135 | 1.579 | 0.764 | 1.286 | -0.002 | 0.028 | 0.028 | 0 | 0.11 | 0.1 | 0.19 | 0.29 | 0.11 | 0.09 |
| 7 | 14471998 | rs196755 | 0.895 | 0.945 | 0.026 | 0.632 | -0.031 | 0.131 | 0.131 | 0 | 0.21 | 0.02 | 0.13 | 0 | 0.2 | 0.16 |
| 7 | 14476181 | rs16878172 | -0.229 | 1.395 | 0.847 | 1.449 | 0.004 | 0.007 | 0.007 | 0 | 0.08 | 0.04 | 0.18 | 0.29 | 0.08 | 0.07 |
| 7 | 14478991 | rs12531006 | -1.088 | 1.3 | 1.443 | 1.64 | -0.002 | 0.013 | 0.013 | 0 | 0.11 | 0.1 | 0.07 | 0.24 | 0.11 | 0.08 |
| 7 | 14485818 | rs9690425 | -0.581 | 1.443 | 1.518 | 1.695 | -0.018 | -0.006 | -0.006 | 1.33 | 0.25 | 0.24 | 0.39 | 0.29 | 0.36 | 0.43 |
| 7 | 14486957 | rs10253325 | -2.182 | 1.342 | 1.611 | 1.694 | -0.031 | 0.098 | 0.098 | 0 | 0.17 | 0.34 | 0.25 | 0.36 | 0.12 | 0.13 |
| 7 | 14490193 | rs17651211 | -1.204 | 1.389 | 1.741 | 1.712 | -0.01 | 0.072 | 0.072 | 0 | 0.12 | 0.24 | 0.08 | 0.24 | 0.09 | 0.08 |
| 7 | 14494842 | rs1431520 | -0.72 | 1.465 | 1.353 | 1.761 | 0.037 | 0.028 | 0.028 | 0 | 0.27 | 0.38 | 0.56 | 0.39 | 0.36 | 0.47 |
| 7 | 14504188 | rs17168203 | -0.62 | 1.463 | 1.24 | 1.791 | -0.001 | 0.015 | 0.015 | 0 | 0.09 | 0.08 | 0.06 | 0.24 | 0.09 | 0.09 |
| 7 | 14505694 | rs10499441 | -0.717 | 1.477 | 1.225 | 1.409 | -0.017 | -0.012 | -0.012 | 0 | 0.27 | 0.26 | 0.55 | 0.39 | 0.36 | 0.47 |
| 7 | 14514544 | rs1978671 | -0.473 | 1.471 | 1.125 | 1.404 | -0.07 | -0.026 | -0.026 | 1.723 | 0.25 | 0.15 | 0.39 | 0.29 | 0.36 | 0.44 |
| 7 | 14526146 | rs12699622 | -1.542 | 1.447 | 1.165 | 1.678 | -0.037 | 0.046 | 0.046 | 0 | 0.35 | 0.24 | 0.44 | 0.73 | 0.44 | 0.52 |
| 7 | 14538449 | rs10264038 | -2.25 | 1.468 | 1.654 | 1.88 | -0.021 | 0.221 | 0.221 | 0 | 0.16 | 0.42 | 0.14 | 0.55 | 0.12 | 0.13 |
| 7 | 14549462 | rs9639203 | -1.285 | 1.722 | 1.63 | 1.801 | -0.044 | 0.037 | 0.037 | 0.008 | 0.33 | 0.24 | 0.57 | 0.72 | 0.44 | 0.51 |
| 7 | 14554546 | rs12531250 | -1.76 | 1.714 | 1.712 | 1.85 | -0.016 | 0.144 | 0.144 | 0 | 0.14 | 0.22 | 0.09 | 0.45 | 0.11 | 0.11 |
| 7 | 14554729 | rs12699624 | -0.623 | 1.807 | 1.589 | 1.787 | -0.057 | -0.013 | -0.013 | 0 | 0.25 | 0.18 | 0.49 | 0.27 | 0.35 | 0.42 |
| 7 | 14558320 | rs10270441 | -0.916 | 1.846 | 1.784 | 1.781 | 0.044 | 0.006 | 0.006 | 0 | 0.27 | 0.4 | 0.54 | 0.3 | 0.38 | 0.45 |
| 7 | 14562924 | rs12699628 | -0.917 | 1.793 | 1.499 | 1.78 | -0.025 | 0 | 0 | 0.758 | 0.27 | 0.26 | 0.52 | 0.28 | 0.36 | 0.45 |
| 7 | 14563045 | rs1367781 | -1.153 | 1.767 | 1.51 | 1.828 | 0.026 | 0.012 | 0.012 | 0.075 | 0.29 | 0.4 | 0.57 | 0.38 | 0.36 | 0.47 |
| 7 | 14564068 | rs4719409 | -0.678 | 1.826 | 1.153 | 1.715 | -0.061 | -0.016 | -0.016 | 0.929 | 0.26 | 0.17 | 0.51 | 0.28 | 0.36 | 0.44 |
| 7 | 14569676 | rs1991687 | -2.072 | 1.444 | 1.318 | 1.988 | -0.025 | 0.222 | 0.222 | 0 | 0.14 | 0.39 | 0.15 | 0.55 | 0.09 | 0.14 |
| 7 | 14591352 | rs12699645 | -0.901 | 1.685 | 0.832 | 1.984 | -0.059 | -0.051 | -0.051 | 0 | 0.32 | 0.2 | 0.57 | 0.49 | 0.46 | 0.52 |
| 7 | 14591967 | rs10257158 | -0.885 | 1.426 | 1.24 | 1.988 | -0.016 | 0.242 | 0.242 | 0 | 0.09 | 0.25 | 0.11 | 0.33 | 0.06 | 0.06 |
| 7 | 14592662 | rs6956840 | -1.636 | 1.373 | 0.944 | 1.662 | 0.007 | 0.045 | 0.045 | 0 | 0.46 | 0.64 | 0.69 | 0.84 | 0.54 | 0.6 |
| 7 | 14598424 | rs4719411 | -1.336 | 1.659 | 0.775 | 1.729 | 0.005 | -0.013 | -0.013 | 0 | 0.42 | 0.51 | 0.59 | 0.58 | 0.52 | 0.59 |
| 7 | 14614930 | rs13438451 | -1.561 | 1.007 | 0.867 | 2.305 | -0.046 | 0.177 | 0.177 | 0 | 0.16 | 0.4 | 0.21 | 0.65 | 0.13 | 0.13 |
| 7 | 14618505 | rs12674445 | -1.289 | 1.536 | -0.023 | 2.275 | -0.039 | -0.078 | -0.078 | 0 | 0.11 | 0.01 | 0.19 | 0.34 | 0.14 | 0.15 |
| 7 | 14619026 | rs17168277 | -1.806 | 1.573 | 0.083 | 2.091 | -0.007 | -0.028 | -0.028 | 0.015 | 0.16 | 0.11 | 0.23 | 0.4 | 0.18 | 0.18 |
| 7 | 14622333 | rs17595206 | -1.518 | 2.143 | -0.093 | 1.099 | -0.035 | -0.012 | -0.012 | 0.364 | 0.3 | 0.21 | 0.37 | 0.68 | 0.38 | 0.46 |
| 7 | 14636250 | rs1357869 | 1.858 | 2.602 | 0.657 | 0.81 | 0.011 | -0.012 | -0.012 | 0 | 0.62 | 0.51 | 0.52 | 0.31 | 0.5 | 0.43 |
| 7 | 14639749 | rs6944740 | 1.462 | 2.712 | 0.874 | 0.83 | -0.008 | 0.002 | 0.002 | 1.321 | 0.64 | 0.63 | 0.55 | 0.32 | 0.52 | 0.46 |
| 7 | 14642523 | rs10236011 | 1.253 | 2.426 | 0.873 | 0.825 | 0.001 | 0 | 0 | 0.051 | 0.69 | 0.64 | 0.79 | 0.31 | 0.67 | 0.52 |
| 7 | 14644667 | rs17150062 | -1.487 | 2.207 | 1.273 | 0.752 | 0.019 | 0.014 | 0.014 | 0.021 | 0.3 | 0.51 | 0.21 | 0.7 | 0.32 | 0.44 |
| 7 | 14653054 | rs1525088 | -1.256 | 2.234 | 1.026 | 0.75 | 0.001 | 0.001 | 0.001 | 0.007 | 0.27 | 0.28 | 0.19 | 0.7 | 0.3 | 0.42 |
| 7 | 14670478 | rs10271056 | -1.759 | 2.341 | 1.233 | 0.893 | 0.01 | 0.143 | 0.143 | 0.049 | 0.2 | 0.44 | 0.15 | 0.58 | 0.21 | 0.29 |
| 7 | 14671325 | rs12672921 | -1.764 | 2.323 | 1.239 | 0.892 | -0.002 | 0.117 | 0.117 | 0.01 | 0.2 | 0.4 | 0.13 | 0.58 | 0.2 | 0.29 |
| 7 | 14673868 | rs10254283 | -1.852 | 2.317 | 1.18 | 0.889 | -0.002 | 0.148 | 0.148 | 0.055 | 0.2 | 0.45 | 0.14 | 0.58 | 0.2 | 0.29 |
| 7 | 14675706 | rs17168320 | -1.151 | 1.966 | 0.367 | 0.983 | 0.033 | -0.084 | -0.084 | 0.041 | 0.13 | 0.01 | 0.06 | 0.49 | 0.09 | 0.15 |
| 7 | 14684726 | rs1404621 | 0.992 | 1.991 | 0.417 | 0.164 | -0.026 | -0.01 | -0.01 | 0 | 0.8 | 0.83 | 0.86 | 0.79 | 0.76 | 0.63 |
| 7 | 14697617 | rs10950533 | 1.007 | 1.962 | 0.372 | 0.167 | -0.039 | -0.017 | -0.017 | 0 | 0.8 | 0.87 | 0.87 | 0.78 | 0.76 | 0.62 |
| 7 | 14698853 | rs2049449 | -1.236 | 1.61 | 0.989 | -0.208 | -0.056 | 0.048 | 0.048 | 0 | 0.21 | 0.47 | 0.39 | 0.74 | 0.16 | 0.25 |
| 7 | 14702076 | rs1357858 | 0.659 | 1.792 | 0.936 | -0.594 | -0.014 | 0.02 | 0.02 | 0 | 0.82 | 0.88 | 0.89 | 0.89 | 0.84 | 0.75 |
| 7 | 14703078 | rs10279786 | 0.785 | 1.787 | 1.163 | -0.596 | 0.013 | -0.028 | -0.028 | 0 | 0.81 | 0.66 | 0.86 | 0.89 | 0.84 | 0.76 |
| 7 | 14703526 | rs11767076 | -1.064 | 1.237 | 0.85 | -0.409 | -0.001 | -0.001 | -0.001 | 0 | 0.22 | 0.28 | 0.39 | 0.83 | 0.24 | 0.39 |
| 7 | 14716865 | rs1525079 | 0.958 | 0.545 | 0.766 | -1.301 | 0 | -0.041 | -0.041 | 0.274 | 0.43 | 0.39 | 0.37 | 0.08 | 0.38 | 0.3 |
| 7 | 14717872 | rs6956435 | -0.612 | 0.26 | 0.616 | -1.505 | 0.019 | 0.178 | 0.178 | 0 | 0.51 | 0.77 | 0.65 | 0.92 | 0.46 | 0.66 |
| 7 | 14719872 | rs1357861 | -0.601 | 0.229 | 1.085 | -1.354 | 0.004 | 0.021 | 0.021 | 0 | 0.48 | 0.47 | 0.53 | 0.82 | 0.47 | 0.64 |
| 7 | 14726723 | rs1404613 | -0.45 | 0.23 | 0.974 | -1.391 | 0.006 | 0.069 | 0.069 | 0 | 0.47 | 0.67 | 0.63 | 0.86 | 0.45 | 0.62 |
| 7 | 14731648 | rs17668719 | 1.571 | 0.435 | 0.192 | -1.726 | -0.013 | 0.152 | 0.152 | 0.115 | 0.33 | 0.1 | 0.21 | 0.01 | 0.34 | 0.2 |
| 7 | 14737999 | rs1568846 | -0.306 | 0.061 | 0.47 | -1.454 | -0.036 | 0.199 | 0.199 | 0 | 0.42 | 0.15 | 0.59 | 0.88 | 0.44 | 0.61 |
| 7 | 14751257 | rs10259838 | -0.73 | 0.086 | 1.007 | -1.483 | -0.001 | -0.001 | -0.001 | 0 | 0.47 | 0.53 | 0.65 | 0.88 | 0.48 | 0.66 |
| 7 | 14758273 | rs1997040 | -0.356 | 0.199 | -0.255 | -1.326 | -0.04 | 0.176 | 0.176 | 0 | 0.4 | 0.15 | 0.59 | 0.88 | 0.44 | 0.61 |
| 7 | 14758773 | rs12699658 | -0.346 | 0.295 | -0.196 | -1.283 | -0.019 | 0.16 | 0.16 | 0 | 0.41 | 0.19 | 0.59 | 0.88 | 0.44 | 0.62 |
| 7 | 14778015 | rs1525095 | 1.267 | 0.445 | 0.586 | 0.076 | 0.011 | 0.099 | 0.099 | 0.053 | 0.82 | 0.36 | 0.68 | 0.65 | 0.86 | 0.78 |
| 7 | 14789057 | rs1880553 | -1.355 | 0.233 | 1.308 | 0.322 | 0.036 | 0.178 | 0.178 | 0 | 0.16 | 0.49 | 0.3 | 0.36 | 0.13 | 0.23 |
| 7 | 14801502 | rs6971441 | -2.404 | 0.228 | 2.348 | 0.521 | 0.076 | 0.285 | 0.285 | 0.188 | 0.12 | 0.48 | 0.36 | 0.31 | 0.1 | 0.17 |
| 7 | 14806558 | rs2358067 | -1.702 | 0.219 | 2.389 | 0.51 | 0.017 | 0.097 | 0.097 | 0 | 0.15 | 0.27 | 0.28 | 0.36 | 0.13 | 0.23 |
| 7 | 14809355 | rs12333979 | -0.823 | 0.196 | 2.2 | 0.471 | -0.015 | 0.01 | 0.01 | 0 | 0.11 | 0.28 | 0.1 | 0.12 | 0.07 | 0.13 |
| 7 | 14811545 | rs12333881 | -0.794 | 0.196 | 2.234 | 0.471 | -0.013 | 0.012 | 0.012 | 0 | 0.11 | 0.28 | 0.1 | 0.12 | 0.07 | 0.13 |

| | | | | | | | | | | | | | | | | |
|---|---|---|---|---|---|---|---|---|---|---|---|---|---|---|---|---|
| 7 | 14818008 | rs11982632 | -1.366 | 0.387 | 1.961 | 0.452 | -0.003 | 0.01 | 0.01 | 0 | 0.13 | 0.46 | 0.11 | 0.11 | 0.09 | 0.13 |
| 7 | 14818061 | rs7788183 | -0.291 | 1.214 | 1.146 | 0.636 | -0.098 | -0.108 | -0.108 | 0.043 | 0.82 | 0.97 | 0.78 | 0.6 | 0.73 | 0.73 |
| 7 | 14818166 | rs2358068 | -1.746 | 1.04 | 1.513 | 0.738 | 0.114 | 0.348 | 0.348 | 0.086 | 0.11 | 0.39 | 0.35 | 0.32 | 0.1 | 0.16 |
| 7 | 14832091 | rs7796443 | -1.809 | 0.384 | 0.842 | 0.356 | -0.001 | 0.001 | 0.001 | 0 | 0.28 | 0.65 | 0.35 | 0.36 | 0.24 | 0.33 |
| 7 | 14840654 | rs6944904 | -1.075 | 0.232 | 0.536 | 0.252 | -0.023 | 0.019 | 0.019 | 0.007 | 0.17 | 0.68 | 0.26 | 0.18 | 0.11 | 0.19 |
| 7 | 14842034 | rs7784714 | -1.686 | 0.248 | 0.479 | 0.236 | -0.008 | -0.004 | -0.004 | 0 | 0.27 | 0.69 | 0.28 | 0.18 | 0.2 | 0.28 |
| 7 | 14850821 | rs10487780 | 1.702 | -0.409 | -0.86 | -0.377 | -0.097 | 0.032 | 0.032 | 1.342 | 0.23 | 0.03 | 0.2 | 0.13 | 0.34 | 0.19 |
| 7 | 14851617 | rs10277367 | -1.41 | -0.502 | -0.848 | -0.32 | 0.007 | -0.018 | -0.018 | 0 | 0.34 | 0.69 | 0.48 | 0.43 | 0.27 | 0.4 |
| 7 | 14852294 | rs766366 | -0.704 | -0.647 | -1.034 | -0.308 | -0.004 | 0.011 | 0.011 | 0 | 0.56 | 0.84 | 0.69 | 0.56 | 0.61 | 0.59 |
| 7 | 14856115 | rs7784940 | 0.512 | -1.511 | -0.946 | -0.639 | 0.068 | 0.088 | 0.088 | 0 | 0.32 | 0.2 | 0.12 | 0.07 | 0.2 | 0.25 |
| 7 | 14856700 | rs10260512 | 0.738 | -1.061 | -0.619 | -0.394 | -0.006 | 0.001 | 0.001 | 0 | 0.54 | 0.37 | 0.51 | 0.4 | 0.54 | 0.42 |
| 7 | 14856726 | rs16878281 | 0.294 | -0.745 | -0.666 | -0.305 | -0.014 | -0.002 | -0.002 | 0 | 0.11 | 0.07 | 0.09 | 0.12 | 0.13 | 0.11 |
| 7 | 14861101 | rs2041407 | 1.429 | -1.437 | -0.228 | -0.117 | -0.071 | 0.08 | 0.08 | 0 | 0.31 | 0.14 | 0.3 | 0.13 | 0.49 | 0.18 |
| 7 | 14866959 | rs17368281 | 1.617 | -1.39 | -0.411 | -0.094 | -0.18 | 0.122 | 0.122 | 0 | 0.3 | 0.03 | 0.27 | 0.09 | 0.48 | 0.18 |
| 7 | 14868418 | rs11978840 | 1.413 | -1.771 | 0.085 | -0.107 | 0.005 | 0.004 | 0.004 | 0 | 0.26 | 0.29 | 0.15 | 0.19 | 0.16 | 0.22 |
| 7 | 14868813 | rs10085416 | 1.024 | -1.817 | 0.664 | -0.02 | 0.005 | 0 | 0 | 0 | 0.38 | 0.41 | 0.32 | 0.37 | 0.51 | 0.31 |
| 7 | 14871780 | rs6461139 | -1.779 | -1.285 | 0.659 | 0.018 | -0.054 | 0.06 | 0.06 | 0 | 0.9 | 1 | 0.9 | 1 | 0.85 | 0.97 |
| 7 | 14871856 | rs6461140 | -0.759 | -1.522 | 0.698 | 0.027 | 0.035 | 0.016 | 0.016 | 0 | 0.49 | 0.61 | 0.56 | 0.63 | 0.33 | 0.66 |
| 7 | 14874381 | rs2041409 | -0.608 | -1.528 | 0.895 | 0.031 | -0.001 | 0 | 0 | 0 | 0.48 | 0.51 | 0.55 | 0.6 | 0.32 | 0.66 |
| 7 | 14874799 | rs17168465 | 0.28 | -1.459 | 0.832 | -0.019 | 0.001 | 0.004 | 0.004 | 0 | 0.12 | 0.07 | 0.24 | 0.43 | 0.09 | 0.35 |
| 7 | 14876917 | rs10241170 | 0.316 | -1.434 | 1.063 | 0.067 | 0.019 | 0.261 | 0.261 | 0 | 0.07 | 0.26 | 0.03 | 0.28 | 0.03 | 0.11 |
| 7 | 14880482 | rs2286767 | 1.001 | -1.54 | 0.794 | -0.259 | -0.015 | -0.01 | -0.01 | 0 | 0.28 | 0.33 | 0.16 | 0.21 | 0.16 | 0.28 |
| 7 | 14884246 | rs4721380 | 1.713 | -1.223 | 0.491 | -0.277 | -0.182 | 0.175 | 0.175 | 1.255 | 0.34 | 0.03 | 0.27 | 0.09 | 0.48 | 0.18 |
| 7 | 14888418 | rs1117750 | 0.306 | -1.14 | 0.886 | 0.062 | 0.04 | 0.322 | 0.322 | 0 | 0.07 | 0.3 | 0.03 | 0.29 | 0.03 | 0.11 |
| 7 | 14891202 | rs975689 | 1.703 | -1.045 | 0.678 | -0.024 | -0.125 | 0.167 | 0.167 | 0.669 | 0.34 | 0.09 | 0.26 | 0.08 | 0.48 | 0.18 |
| 7 | 14898152 | rs994369 | -0.391 | -1.181 | 0.9 | -0.058 | -0.01 | -0.005 | -0.005 | 0.006 | 0.47 | 0.52 | 0.57 | 0.64 | 0.32 | 0.65 |
| 7 | 14898282 | rs17168486 | 0.206 | -1.311 | 0.68 | -0.007 | -0.004 | -0.04 | -0.04 | 0.142 | 0.16 | 0.08 | 0.39 | 0.46 | 0.17 | 0.36 |
| 7 | 14899267 | rs17310345 | 0.858 | -1.323 | 0.724 | -0.105 | 0.021 | 0.015 | 0.015 | 0 | 0.29 | 0.24 | 0.17 | 0.2 | 0.16 | 0.29 |
| 7 | 14901576 | rs4634526 | 1.62 | -1.1 | 0.308 | -0.304 | -0.103 | 0.16 | 0.16 | 0.092 | 0.34 | 0.12 | 0.26 | 0.07 | 0.48 | 0.18 |
| 7 | 14905838 | rs7788248 | 0.667 | -1.385 | 0.634 | -0.132 | -0.007 | 0 | 0 | 0 | 0.3 | 0.28 | 0.1 | 0.29 | 0.13 | 0.25 |
| 7 | 14909563 | rs4721385 | 0.502 | -1.159 | 1.122 | -0.084 | 0.018 | 0.064 | 0.064 | 0 | 0.79 | 0.61 | 0.83 | 0.71 | 0.8 | 0.84 |
| 7 | 14912174 | rs10270993 | 0.484 | -1.302 | 0.991 | 0.016 | 0.007 | 0.183 | 0.183 | 0 | 0.05 | 0.15 | 0.02 | 0.28 | 0.03 | 0.12 |
| 7 | 14915289 | rs6975734 | 0 | -1.366 | 1.056 | 0.018 | 0.007 | 0.211 | 0.211 | 0 | 0.93 | 0.78 | 0.97 | 0.7 | 0.97 | 0.88 |
| 7 | 14918063 | rs2358073 | -0.754 | -1.55 | 1.033 | -0.222 | 0.021 | 0.017 | 0.017 | 0 | 0.52 | 0.62 | 0.55 | 0.92 | 0.35 | 0.77 |
| 7 | 14922007 | rs10276674 | -0.489 | -1.488 | 0.885 | 0.111 | -0.007 | 0.069 | 0.069 | 0 | 0.2 | 0.25 | 0.4 | 0.45 | 0.18 | 0.48 |
| 7 | 14926914 | rs6972713 | 1.518 | -1.396 | 1.024 | -0.446 | -0.001 | 0.029 | 0.029 | 0.282 | 0.6 | 0.36 | 0.43 | 0.26 | 0.63 | 0.38 |
| 7 | 14927543 | rs217582 | 0.583 | -1.224 | 0.981 | -0.531 | 0.014 | -0.066 | -0.066 | 0.314 | 0.14 | 0.19 | 0.14 | 0 | 0.16 | 0.04 |
| 7 | 14928665 | rs170094 | 0.623 | -1.409 | 0.629 | -0.559 | 0.041 | 0.072 | 0.072 | 0.011 | 0.5 | 0.57 | 0.44 | 0.11 | 0.64 | 0.22 |
| 7 | 14931048 | rs2058281 | -0.839 | -1.375 | 0.401 | -0.637 | 0.012 | -0.009 | -0.009 | 0 | 0.5 | 0.48 | 0.56 | 0.92 | 0.36 | 0.77 |
| 7 | 14931662 | rs6961664 | 0.674 | -1.762 | 0.189 | -0.486 | -0.001 | 0.003 | 0.003 | 0 | 0.27 | 0.26 | 0.14 | 0.14 | 0.12 | 0.19 |
| 7 | 14933955 | rs2109396 | 0.191 | -1.644 | -0.372 | -0.805 | -0.028 | -0.129 | -0.129 | 0 | 0.62 | 0.5 | 0.8 | 0.94 | 0.73 | 0.73 |
| 7 | 14934554 | rs6968886 | 0.792 | -1.361 | -1.08 | -0.421 | 0.051 | 0.04 | 0.04 | 0.485 | 0.49 | 0.24 | 0.54 | 0.6 | 0.59 | 0.51 |
| 7 | 14939924 | rs217598 | -0.008 | -1.347 | -1.107 | -0.369 | 0.017 | -0.012 | -0.012 | 0.072 | 0.65 | 0.7 | 0.82 | 0.58 | 0.75 | 0.56 |
| 7 | 14941556 | rs4721386 | 1.346 | -0.991 | -1.869 | -0.459 | 0.026 | -0.02 | -0.02 | 0.284 | 0.57 | 0.16 | 0.55 | 0.56 | 0.62 | 0.6 |
| 7 | 14941720 | rs4721388 | 0.712 | -0.471 | -1.384 | -0.134 | 0.003 | -0.005 | -0.005 | 0 | 0.66 | 0.34 | 0.7 | 0.6 | 0.71 | 0.55 |
| 7 | 14943872 | rs7811739 | 0.761 | -0.395 | -1.959 | -0.972 | -0.059 | 0.156 | 0.156 | 1.787 | 0.24 | 0.01 | 0.13 | 0.01 | 0.25 | 0.07 |
| 7 | 14950641 | rs6950662 | -0.492 | -2.046 | -0.435 | -0.472 | -0.096 | -0.007 | -0.007 | 1.532 | 0.27 | 0.47 | 0.16 | 0.8 | 0.1 | 0.49 |
| 7 | 14951097 | rs36845 | -1.838 | -2.182 | -0.358 | -0.76 | 0.016 | 0.179 | 0.179 | 0 | 0.56 | 0.76 | 0.5 | 0.92 | 0.41 | 0.74 |
| 7 | 14957222 | rs10260034 | -0.286 | -2.284 | 0.212 | -0.62 | -0.087 | 0.001 | 0.001 | 0 | 0.27 | 0.37 | 0.16 | 0.79 | 0.1 | 0.48 |
| 7 | 14960988 | rs10276360 | -1.187 | -2.324 | 0.427 | -0.718 | 0 | -0.021 | -0.021 | 0.723 | 0.43 | 0.54 | 0.35 | 0.92 | 0.23 | 0.67 |
| 7 | 14961214 | rs10276618 | -0.284 | -2.344 | 0.612 | -0.704 | -0.04 | -0.005 | -0.005 | 0 | 0.25 | 0.27 | 0.15 | 0.83 | 0.1 | 0.48 |
| 7 | 14961556 | rs7794079 | 0.27 | -2.017 | 0.299 | -0.538 | 0.008 | 0.01 | 0.01 | 0 | 0.16 | 0.12 | 0.19 | 0.09 | 0.13 | 0.19 |
| 7 | 14972555 | rs36866 | 0.262 | -1.883 | 0.278 | -0.546 | -0.028 | 0.03 | 0.03 | 0 | 0.11 | 0.06 | 0.14 | 0 | 0.18 | 0.06 |
| 7 | 14972884 | rs7787411 | -0.393 | -2.24 | 0.524 | -0.586 | -0.087 | -0.014 | -0.014 | 0.187 | 0.27 | 0.38 | 0.16 | 0.82 | 0.1 | 0.47 |
| 7 | 14976097 | rs36873 | -1.621 | -2.099 | 0.604 | -0.003 | 0.027 | 0.084 | 0.084 | 0 | 0.55 | 0.76 | 0.49 | 0.78 | 0.39 | 0.73 |
| 7 | 14988867 | rs40628 | -1.723 | -2.392 | 0.636 | -0.649 | 0.021 | 0.196 | 0.196 | 0.054 | 0.57 | 0.86 | 0.51 | 0.89 | 0.4 | 0.74 |
| 7 | 14991989 | rs36893 | -0.843 | -2.6 | 0.769 | -0.637 | -0.005 | -0.044 | -0.044 | 0 | 0.31 | 0.42 | 0.34 | 0.1 | 0.3 | 0.26 |
| 7 | 14995522 | rs36897 | -1.892 | -2.576 | 0.419 | -0.631 | 0.025 | 0.209 | 0.209 | 0 | 0.57 | 0.84 | 0.51 | 0.91 | 0.39 | 0.74 |
| 7 | 14996843 | rs7811800 | -0.592 | -2.803 | 0.622 | -0.466 | -0.078 | -0.001 | -0.001 | 0.332 | 0.25 | 0.34 | 0.17 | 0.81 | 0.1 | 0.47 |
| 7 | 14998466 | rs9648193 | -0.513 | -2.624 | 0.75 | 0.012 | 0.029 | -0.037 | -0.037 | 0.586 | 0.19 | 0.08 | 0.13 | 0.61 | 0.09 | 0.39 |
| 7 | 15000359 | rs36911 | -0.979 | -2.063 | 0.799 | -0.067 | 0.008 | -0.063 | -0.063 | 0 | 0.31 | 0.57 | 0.36 | 0.11 | 0.31 | 0.29 |
| 7 | 15000858 | rs36912 | -0.988 | -2.063 | 0.609 | -0.07 | 0.006 | -0.05 | -0.05 | 1.747 | 0.32 | 0.59 | 0.36 | 0.11 | 0.31 | 0.29 |
| 7 | 15002829 | rs36916 | -1.436 | -1.673 | -0.023 | 0.127 | 0.047 | 0.186 | 0.186 | 0.03 | 0.51 | 0.78 | 0.5 | 0.83 | 0.38 | 0.73 |
| 7 | 15005958 | rs17168560 | -0.378 | -1.44 | -0.837 | 0.615 | 0.029 | -0.043 | -0.043 | 0 | 0.1 | 0.01 | 0.1 | 0.58 | 0.06 | 0.33 |
| 7 | 15010824 | rs17168564 | -0.166 | -1.082 | -0.422 | 0.323 | -0.004 | 0.003 | 0.003 | 0 | 0.16 | 0.16 | 0.19 | 0.18 | 0.12 | 0.24 |
| 7 | 15012831 | rs7805162 | -0.719 | -1.136 | -0.506 | 0.21 | 0.006 | -0.001 | -0.001 | 0.084 | 0.37 | 0.67 | 0.31 | 0.66 | 0.29 | 0.44 |
| 7 | 15013768 | rs10950544 | -0.942 | -0.462 | -0.588 | 0.667 | 0.018 | 0.002 | 0.002 | 0.067 | 0.34 | 0.74 | 0.34 | 0.34 | 0.24 | 0.31 |
| 7 | 15021223 | rs7800343 | 0.338 | 0.909 | -1.052 | 1.283 | -0.123 | -0.165 | -0.165 | 0 | 0.75 | 0.97 | 0.76 | 0.5 | 0.66 | 0.64 |
| 7 | 15025332 | rs917037 | -0.757 | 0.496 | -0.651 | 1.216 | -0.003 | 0.018 | 0.018 | 0 | 0.12 | 0.15 | 0.13 | 0.16 | 0.09 | 0.09 |

| | | | | | | | | | | | | | | | | |
|---|---|---|---|---|---|---|---|---|---|---|---|---|---|---|---|---|
| 7 | 15026787 | rs12670968 | -0.301 | 0.758 | -0.851 | 1.35 | -0.092 | -0.187 | -0.187 | 0.135 | 0.27 | 0.03 | 0.22 | 0.47 | 0.32 | 0.38 |
| 7 | 15036251 | rs6953636 | -0.625 | 0.5 | 0.05 | 0.606 | 0.014 | -0.033 | -0.033 | 0 | 0.13 | 0.25 | 0.07 | 0.05 | 0.12 | 0.1 |
| 7 | 15042907 | rs10487790 | -1.138 | 0.212 | 0.402 | 0.628 | -0.002 | -0.021 | -0.021 | 0.374 | 0.18 | 0.37 | 0.08 | 0.12 | 0.14 | 0.15 |
| 7 | 15042959 | rs10244143 | 1.476 | 0.243 | 0.418 | 0.944 | 0.006 | 0.02 | 0.02 | 0 | 0.5 | 0.44 | 0.56 | 0.21 | 0.49 | 0.44 |
| 7 | 15045529 | rs9639217 | -0.527 | 0.253 | 0.343 | 1.133 | -0.011 | 0.005 | 0.005 | 0.181 | 0.36 | 0.27 | 0.38 | 0.68 | 0.4 | 0.45 |
| 7 | 15048069 | rs10243242 | -0.797 | 0 | 0.737 | 1.286 | -0.011 | 0.035 | 0.035 | 0 | 0.08 | 0.17 | 0.05 | 0.18 | 0.05 | 0.02 |
| 7 | 15048611 | rs6976086 | 1.241 | -0.203 | 1.191 | 1.286 | -0.002 | -0.007 | -0.007 | 1.256 | 0.46 | 0.43 | 0.53 | 0.26 | 0.47 | 0.4 |
| 7 | 15048814 | rs6976381 | -0.35 | -0.272 | 1.242 | 1.339 | -0.086 | -0.098 | -0.098 | 0 | 0.4 | 0.1 | 0.36 | 0.49 | 0.48 | 0.52 |
| 7 | 15051233 | rs10950547 | 1.571 | -0.41 | 1.424 | 1.318 | -0.042 | 0.075 | 0.075 | 0.047 | 0.86 | 0.6 | 0.9 | 0.75 | 0.94 | 0.92 |
| 7 | 15051754 | rs6972083 | 1.536 | -0.406 | 1.416 | 1.32 | -0.04 | 0.061 | 0.061 | 0.043 | 0.87 | 0.63 | 0.91 | 0.75 | 0.94 | 0.93 |
| 7 | 15053589 | rs10267273 | 1.288 | -0.376 | 1.583 | 1.301 | 0 | 0.001 | 0.001 | 0.02 | 0.46 | 0.42 | 0.54 | 0.27 | 0.47 | 0.42 |
| 7 | 15054232 | rs10238625 | 1.252 | -0.371 | 1.566 | 1.303 | 0.001 | 0.013 | 0.013 | 0.009 | 0.46 | 0.52 | 0.54 | 0.27 | 0.47 | 0.42 |
| 7 | 15054370 | rs10264723 | 0.704 | -0.363 | 1.531 | 1.352 | -0.001 | 0.013 | 0.013 | 0.021 | 0.53 | 0.59 | 0.57 | 0.37 | 0.49 | 0.43 |
| 7 | 15059272 | rs11514706 | -0.224 | -0.35 | 1.335 | 1.328 | -0.103 | -0.104 | -0.104 | 0 | 0.39 | 0.08 | 0.36 | 0.48 | 0.47 | 0.53 |
| 7 | 15059906 | rs4721400 | 1.046 | -0.085 | 1.394 | 1.094 | 0.005 | 0.027 | 0.027 | 0.04 | 0.46 | 0.39 | 0.53 | 0.26 | 0.47 | 0.39 |
| 7 | 15063085 | rs7798129 | -0.747 | -0.177 | 1.143 | 1.106 | -0.027 | 0.03 | 0.03 | 0 | 0.06 | 0.26 | 0.05 | 0.08 | 0.01 | 0.05 |
| 7 | 15063833 | rs10244051 | -0.924 | 0.07 | 0.76 | 0.948 | 0.002 | -0.011 | -0.011 | 1.397 | 0.57 | 0.56 | 0.46 | 0.69 | 0.51 | 0.64 |
| 7 | 15070803 | rs17168608 | -1.735 | -0.334 | 1.012 | 1.028 | -0.02 | 0.014 | 0.014 | 0 | 0.14 | 0.38 | 0.17 | 0.18 | 0.11 | 0.08 |
| 7 | 15072029 | rs17377955 | -0.803 | -0.411 | 1.33 | 1.098 | 0.001 | 0.005 | 0.005 | 0 | 0.17 | 0.23 | 0.13 | 0.16 | 0.15 | 0.17 |
| 7 | 15073547 | rs720872 | -1.018 | -0.363 | 1.461 | 1.005 | 0.02 | -0.015 | -0.015 | 0 | 0.05 | 0.21 | 0.03 | 0.01 | 0.05 | 0.03 |
| 7 | 15074567 | rs7804568 | -1.862 | -0.372 | 1.399 | 1.046 | -0.016 | -0.022 | -0.022 | 0 | 0.13 | 0.35 | 0.07 | 0.08 | 0.08 | 0.07 |
| 7 | 15078991 | rs218066 | 1.107 | -0.517 | 0.627 | 0.808 | -0.005 | 0.092 | 0.092 | 3.374 | 0.47 | 0.25 | 0.57 | 0.3 | 0.54 | 0.39 |
| 7 | 15082439 | rs218077 | 1.076 | -0.564 | 1.111 | 0.678 | -0.001 | 0.054 | 0.054 | 2.714 | 0.47 | 0.37 | 0.52 | 0.2 | 0.54 | 0.35 |
| 7 | 15082534 | rs218078 | 0.726 | -0.568 | 1.293 | 0.679 | 0.012 | 0.01 | 0.01 | 0 | 0.94 | 0.8 | 0.9 | 0.94 | 0.93 | 0.96 |
| 7 | 15082905 | rs2041371 | -0.99 | -0.598 | 1.274 | 0.638 | -0.001 | -0.018 | -0.018 | 5.577 | 0.51 | 0.49 | 0.5 | 0.81 | 0.47 | 0.64 |
| 7 | 15090452 | rs6953736 | -0.085 | -0.598 | 1.257 | 1.013 | -0.005 | 0.013 | 0.013 | 1.423 | 0.34 | 0.31 | 0.34 | 0.49 | 0.32 | 0.52 |
| 7 | 15098004 | rs2159808 | -0.034 | -0.601 | 1.156 | 1.236 | 0.011 | -0.03 | -0.03 | 0 | 0.34 | 0.21 | 0.3 | 0.46 | 0.28 | 0.53 |
| 7 | 15098712 | rs7801401 | -2.191 | -1.026 | 2.059 | 1.943 | -0.009 | 0.019 | 0.019 | 0.066 | 0.18 | 0.24 | 0.14 | 0.3 | 0.1 | 0.15 |
| 7 | 15100749 | rs17348054 | -2.104 | -0.851 | 2.004 | 1.816 | 0.057 | 0.229 | 0.229 | 2.642 | 0.09 | 0.51 | 0.24 | 0.24 | 0.09 | 0.09 |
| 7 | 15105524 | rs10272051 | -1.502 | -0.855 | 1.908 | 1.745 | -0.006 | -0.004 | -0.004 | 0.039 | 0.14 | 0.16 | 0.06 | 0.09 | 0.08 | 0.05 |
| 7 | 15108184 | rs12386767 | -0.827 | -0.861 | 1.831 | 1.749 | 0.029 | 0.014 | 0.014 | 0.36 | 0.11 | 0.07 | 0.02 | 0.08 | 0.02 | 0.02 |
| 7 | 15108838 | rs8180732 | -2.385 | -0.821 | 2.286 | 1.831 | 0.065 | 0.3 | 0.3 | 2 | 0.11 | 0.44 | 0.28 | 0.43 | 0.15 | 0.21 |
| 7 | 15111567 | rs6945149 | -2.691 | -0.832 | 1.758 | 1.781 | -0.007 | 0.037 | 0.037 | 0 | 0.25 | 0.67 | 0.35 | 0.48 | 0.23 | 0.24 |
| 7 | 15114140 | rs10262813 | -2.838 | -0.841 | 1.862 | 1.85 | -0.007 | 0.038 | 0.038 | 0 | 0.27 | 0.44 | 0.39 | 0.54 | 0.25 | 0.27 |
| 7 | 15114926 | rs6971261 | -1.63 | -0.541 | 1.818 | 2.03 | 0.009 | -0.019 | -0.019 | 0 | 0.15 | 0.17 | 0.23 | 0.27 | 0.11 | 0.07 |
| 7 | 15121951 | rs4598156 | -0.708 | -0.514 | 2.108 | 1.905 | 0.122 | 0.068 | 0.068 | 0.104 | 0.05 | 0.36 | 0.05 | 0.01 | 0.05 | 0.03 |
| 7 | 15124846 | rs917796 | -1.56 | -0.558 | 1.484 | 2.08 | -0.003 | 0.005 | 0.005 | 0 | 0.16 | 0.23 | 0.24 | 0.26 | 0.11 | 0.07 |
| 7 | 15125363 | rs12112796 | -0.825 | -0.571 | 1.551 | 1.857 | 0.122 | 0.067 | 0.067 | 0.198 | 0.05 | 0.38 | 0.05 | 0.01 | 0.05 | 0.03 |
| 7 | 15126153 | rs6964377 | -2.27 | -0.558 | 0.724 | 1.899 | -0.004 | 0.06 | 0.06 | 0 | 0.24 | 0.66 | 0.33 | 0.51 | 0.22 | 0.19 |
| 7 | 15131687 | rs7791047 | -2.001 | -0.204 | 0.178 | 1.826 | 0.003 | 0.033 | 0.033 | 0 | 0.24 | 0.64 | 0.4 | 0.49 | 0.27 | 0.21 |
| 7 | 15140690 | rs9648196 | -1.874 | -1.021 | -0.866 | 1.689 | 0.058 | -0.126 | -0.126 | 0.4 | 0.28 | 0.91 | 0.23 | 0.5 | 0.2 | 0.2 |
| 7 | 15145681 | rs7808380 | -0.624 | -1.075 | -1.149 | 0.892 | 0.034 | 0.047 | 0.047 | 0 | 0.52 | 0.77 | 0.58 | 0.34 | 0.41 | 0.59 |
| 7 | 15146113 | rs763868 | -0.502 | -1.265 | -1.3 | 0.753 | 0.051 | 0.051 | 0.051 | 0 | 0.48 | 0.74 | 0.56 | 0.33 | 0.38 | 0.54 |
| 7 | 15148454 | rs2191994 | -1.096 | -1.745 | -0.725 | 0.482 | -0.038 | -0.096 | -0.096 | 0 | 0.16 | 0.59 | 0.09 | 0.01 | 0.09 | 0.14 |
| 7 | 15157688 | rs2041556 | 1.494 | -1.676 | -0.749 | 0.188 | -0.068 | -0.009 | -0.009 | 0 | 0.37 | 0.15 | 0.29 | 0.68 | 0.43 | 0.34 |
| 7 | 15159059 | rs1547962 | -0.509 | -1.837 | -0.028 | 0.053 | 0.002 | 0.003 | 0.003 | 0 | 0.49 | 0.45 | 0.58 | 0.3 | 0.45 | 0.53 |
| 7 | 15163610 | rs17387298 | 1.698 | -1.791 | -0.104 | 0.136 | -0.161 | -0.087 | -0.087 | 0 | 0.36 | 0.06 | 0.3 | 0.66 | 0.43 | 0.34 |
| 7 | 15169269 | rs2080041 | 1.815 | -1.898 | -0.237 | 0.515 | -0.167 | -0.166 | -0.166 | 0 | 0.33 | 0.06 | 0.26 | 0.55 | 0.42 | 0.31 |
| 7 | 15172463 | rs764702 | 1.696 | -2.116 | -0.293 | 0.127 | -0.063 | -0.018 | -0.018 | 0 | 0.33 | 0.16 | 0.34 | 0.68 | 0.43 | 0.31 |
| 7 | 15174347 | rs2961246 | 1.48 | -2.177 | -0.097 | 0.049 | -0.026 | -0.109 | -0.109 | 0 | 0.82 | 0.54 | 0.87 | 0.98 | 0.88 | 0.84 |
| 7 | 15177384 | rs9639223 | 0.574 | -2.027 | 0.19 | 0.269 | 0 | -0.009 | -0.009 | 0 | 0.45 | 0.54 | 0.39 | 0.66 | 0.48 | 0.39 |
| 7 | 15178883 | rs9639226 | 0.525 | -2.088 | 0.2 | 0.253 | -0.002 | 0.012 | 0.012 | 0.283 | 0.44 | 0.44 | 0.4 | 0.66 | 0.49 | 0.39 |
| 7 | 15181337 | rs2961256 | 0.924 | -2.035 | 0.455 | 0.262 | 0.002 | -0.033 | -0.033 | 0 | 0.91 | 0.74 | 0.77 | 0.99 | 0.95 | 0.94 |
| 7 | 15191273 | rs2908320 | 0.671 | -2.606 | -0.049 | -0.193 | 0.006 | 0.004 | 0.004 | 0 | 0.6 | 0.37 | 0.5 | 0.71 | 0.7 | 0.48 |
| 7 | 15196418 | rs12699686 | 0.497 | -2.759 | -0.016 | -0.334 | -0.012 | -0.095 | -0.095 | 0 | 0.74 | 0.61 | 0.65 | 0.98 | 0.77 | 0.83 |
| 7 | 15198589 | rs6966824 | 1.169 | -2.602 | -0.177 | -0.23 | -0.002 | 0.004 | 0.004 | 1.637 | 0.43 | 0.4 | 0.39 | 0.7 | 0.49 | 0.39 |
| 7 | 15199905 | rs12699688 | 2.642 | -2.422 | -0.889 | 0.03 | -0.114 | -0.096 | -0.096 | 0 | 0.23 | 0.04 | 0.26 | 0.33 | 0.34 | 0.2 |
| 7 | 15205107 | rs2908333 | 0.627 | -2.498 | -0.457 | 0.035 | 0.011 | 0.009 | 0.009 | 0 | 0.57 | 0.4 | 0.48 | 0.71 | 0.66 | 0.46 |
| 7 | 15206460 | rs2908336 | 0.135 | -2.479 | -0.609 | -0.002 | 0.009 | 0.006 | 0.006 | 0 | 0.62 | 0.65 | 0.53 | 0.72 | 0.7 | 0.5 |
| 7 | 15207942 | rs2908337 | 0.675 | -1.688 | -1.148 | -0.361 | -0.041 | 0.099 | 0.099 | 0.279 | 0.16 | 0.03 | 0.1 | 0 | 0.19 | 0.08 |
| 7 | 15213969 | rs11768872 | 1.275 | -1.515 | -1.12 | -0.158 | -0.086 | 0.085 | 0.085 | 0.125 | 0.38 | 0.12 | 0.33 | 0.29 | 0.48 | 0.31 |
| 7 | 15215277 | rs10243870 | 0.503 | -1.601 | -0.884 | 0.125 | 0.001 | 0.111 | 0.111 | 0 | 0.14 | 0.18 | 0.2 | 0.49 | 0.14 | 0.19 |
| 7 | 15221999 | rs4489212 | 1.258 | -1.412 | -1.349 | -0.213 | -0.089 | 0.077 | 0.077 | 1.208 | 0.38 | 0.12 | 0.31 | 0.3 | 0.48 | 0.31 |
| 7 | 15225539 | rs6962150 | -0.43 | -1.56 | -0.606 | -0.464 | -0.032 | -0.048 | -0.048 | 0.009 | 0.25 | 0.32 | 0.28 | 0.13 | 0.16 | 0.35 |
| 7 | 15230731 | rs7784727 | -0.587 | -1.175 | 0.1 | 0.394 | 0.015 | 0.102 | 0.102 | 0.149 | 0.17 | 0.43 | 0.16 | 0.22 | 0.17 | 0.13 |
| 7 | 15232798 | rs10237214 | -0.468 | -1.463 | -0.044 | 0.309 | -0.016 | -0.002 | -0.002 | 0.015 | 0.17 | 0.19 | 0.29 | 0.14 | 0.1 | 0.3 |
| 7 | 15233341 | rs10225336 | -1.257 | -1.68 | -0.267 | 0.012 | 0.036 | 0.043 | 0.043 | 0 | 0.45 | 0.76 | 0.5 | 0.3 | 0.41 | 0.48 |
| 7 | 15235144 | rs6944905 | 0.695 | -1.578 | -0.837 | 0.555 | -0.033 | -0.084 | -0.084 | 0 | 0.13 | 0.05 | 0.18 | 0.36 | 0.14 | 0.2 |
| 7 | 15236377 | rs4721416 | -1.47 | -1.606 | -0.025 | 0.233 | 0 | 0.003 | 0.003 | 0 | 0.53 | 0.57 | 0.59 | 0.63 | 0.48 | 0.61 |

| | | | | | | | | | | | | | | | | |
|---|---|---|---|---|---|---|---|---|---|---|---|---|---|---|---|---|
| 7 | 15241437 | rs7777171 | -1.183 | -1.563 | -0.541 | 0.293 | 0.031 | -0.022 | -0.022 | 0.377 | 0.39 | 0.71 | 0.52 | 0.51 | 0.29 | 0.56 |
| 7 | 15249864 | rs4551238 | -1.904 | -1.006 | -1.003 | 0.127 | -0.091 | 0.021 | 0.021 | 0 | 0.62 | 0.9 | 0.7 | 0.64 | 0.55 | 0.69 |
| 7 | 15251070 | rs4636099 | 1.207 | -1.06 | -0.528 | 0.037 | 0.012 | -0.003 | -0.003 | 0.603 | 0.14 | 0.06 | 0.07 | 0.11 | 0.11 | 0.09 |
| 7 | 15255144 | rs10081260 | 0.267 | -1.093 | 0.009 | -0.168 | 0.019 | -0.032 | -0.032 | 0 | 0.09 | 0.16 | 0.07 | 0.01 | 0.1 | 0.1 |
| 7 | 15258456 | rs6977610 | -0.696 | -1.725 | 0.175 | -0.519 | -0.04 | -0.119 | -0.119 | 0 | 0.25 | 0.41 | 0.3 | 0.05 | 0.16 | 0.32 |
| 7 | 15262599 | rs12674087 | 1.583 | -1.646 | -0.114 | -0.341 | 0.014 | -0.042 | -0.042 | 0 | 0.15 | 0.07 | 0.09 | 0.18 | 0.14 | 0.11 |
| 7 | 15264585 | rs7777548 | 1.018 | -1.702 | -0.084 | -0.074 | 0.004 | 0.002 | 0.002 | 0 | 0.63 | 0.69 | 0.63 | 0.67 | 0.69 | 0.5 |
| 7 | 15270751 | rs10248417 | -0.168 | -1.382 | 0.13 | -0.011 | -0.01 | -0.018 | -0.018 | 0 | 0.8 | 0.87 | 0.86 | 0.73 | 0.73 | 0.77 |
| 7 | 15272715 | rs6461161 | 0.629 | -1.473 | 0.218 | -0.041 | -0.002 | 0.011 | 0.011 | 0 | 0.63 | 0.45 | 0.75 | 0.54 | 0.59 | 0.66 |
| 7 | 15279488 | rs10263318 | -0.482 | -1.419 | -0.279 | -0.133 | -0.022 | -0.076 | -0.076 | 0 | 0.25 | 0.32 | 0.29 | 0.05 | 0.16 | 0.31 |
| 7 | 15282064 | rs9784982 | -1.476 | -1.243 | -1.158 | 0.058 | -0.039 | -0.075 | -0.075 | 0.908 | 0.89 | 0.99 | 0.92 | 0.73 | 0.86 | 0.85 |
| 7 | 15284032 | rs7812160 | 0.468 | -1.377 | -1.175 | -0.037 | -0.018 | -0.077 | -0.077 | 0 | 0.66 | 0.81 | 0.78 | 0.55 | 0.59 | 0.65 |
| 7 | 15318473 | rs10258070 | -1.288 | -1.745 | -1.351 | -0.229 | 0.111 | 0.1 | 0.1 | 0 | 0.45 | 0.87 | 0.4 | 0.25 | 0.27 | 0.43 |
| 7 | 15318794 | rs12333630 | -0.882 | -1.611 | -1.219 | -0.199 | 0.021 | 0.029 | 0.029 | 0 | 0.27 | 0.77 | 0.33 | 0.05 | 0.17 | 0.3 |
| 7 | 15331080 | rs12234378 | 0.219 | -1.377 | -1.102 | -0.036 | 0.012 | 0.002 | 0.002 | 0 | 0.25 | 0.17 | 0.13 | 0.2 | 0.22 | 0.22 |
| 7 | 15340329 | rs4377857 | -0.022 | -1.273 | -1.054 | 0.068 | -0.025 | -0.043 | -0.043 | 0 | 0.8 | 0.89 | 0.86 | 0.72 | 0.75 | 0.72 |
| 7 | 15351277 | rs4623313 | -0.354 | -1.224 | -0.972 | 0.069 | 0.002 | -0.01 | -0.01 | 0 | 0.85 | 0.89 | 0.92 | 0.73 | 0.86 | 0.8 |
| 7 | 15353729 | rs10279259 | -1.102 | -1.485 | -1.009 | -0.248 | 0.005 | -0.007 | -0.007 | 0 | 0.26 | 0.69 | 0.32 | 0.07 | 0.14 | 0.32 |
| 7 | 15357403 | rs4255036 | 2.167 | -1.215 | -1.09 | -0.177 | -0.114 | -0.09 | -0.09 | 0 | 0.37 | 0.08 | 0.48 | 0.5 | 0.48 | 0.34 |
| 7 | 15359354 | rs10266208 | 1.093 | -1.123 | -0.972 | 0.021 | 0.023 | -0.007 | -0.007 | 0 | 0.51 | 0.79 | 0.59 | 0.5 | 0.54 | 0.44 |
| 7 | 15365939 | rs7781413 | -0.517 | -0.659 | -0.506 | 0.179 | -0.028 | 0.016 | 0.016 | 0 | 0.37 | 0.48 | 0.34 | 0.51 | 0.24 | 0.4 |
| 7 | 15375586 | rs4074352 | -1.281 | -0.167 | -0.968 | 0.185 | 0.014 | -0.004 | -0.004 | 0.765 | 0.53 | 0.8 | 0.5 | 0.53 | 0.45 | 0.5 |
| 7 | 15377889 | rs4467844 | 1.692 | -0.129 | -0.574 | 0.438 | -0.004 | 0.085 | 0.085 | 0 | 0.16 | 0.21 | 0.1 | 0.45 | 0.12 | 0.18 |
| 7 | 15397449 | rs6957765 | 1.101 | -0.145 | -0.586 | 0.495 | -0.007 | 0.014 | 0.014 | 0 | 0.2 | 0.25 | 0.12 | 0.24 | 0.16 | 0.19 |
| 7 | 15399672 | rs10227084 | -0.589 | -0.282 | -0.416 | 0.207 | -0.009 | 0.012 | 0.012 | 0 | 0.41 | 0.5 | 0.42 | 0.49 | 0.4 | 0.43 |
| 7 | 15399963 | rs4599703 | 1.102 | -0.489 | -1.009 | 0.418 | -0.005 | 0.044 | 0.044 | 0 | 0.2 | 0.24 | 0.12 | 0.4 | 0.17 | 0.19 |
| 7 | 15404211 | rs7786312 | -0.322 | 0.771 | -1.608 | 0.611 | -0.005 | -0.011 | -0.011 | 0 | 0.51 | 0.78 | 0.51 | 0.5 | 0.52 | 0.51 |
| 7 | 15413574 | rs4472413 | 0.351 | 0.816 | -1.4 | 0.316 | 0 | 0.03 | 0.03 | 0 | 0.42 | 0.32 | 0.45 | 0.69 | 0.43 | 0.41 |
| 7 | 15421880 | rs10269850 | -1.003 | 0.856 | -1.94 | 0.27 | -0.031 | 0.002 | 0.002 | 1.716 | 0.39 | 0.87 | 0.31 | 0.32 | 0.37 | 0.4 |
| 7 | 15424383 | rs4484577 | 0.398 | 0.584 | -1.761 | 0.447 | 0 | -0.002 | -0.002 | 0.145 | 0.78 | 0.81 | 0.59 | 0.35 | 0.8 | 0.64 |
| 7 | 15428399 | rs4286841 | -0.656 | 1.21 | -2.105 | 0.515 | -0.078 | -0.079 | -0.079 | 0 | 0.87 | 0.99 | 0.76 | 0.72 | 0.77 | 0.81 |
| 7 | 15434065 | rs10239038 | 0.904 | 0.429 | -1.529 | -0.345 | -0.001 | 0 | 0 | 0.357 | 0.62 | 0.63 | 0.41 | 0.33 | 0.51 | 0.5 |
| 7 | 15434633 | rs6960653 | 1.433 | 0.103 | -1.508 | -0.524 | -0.015 | -0.016 | -0.016 | 0 | 0.43 | 0.47 | 0.28 | 0.26 | 0.34 | 0.27 |
| 7 | 15437962 | rs13242762 | 2.291 | -0.234 | -2.19 | -0.586 | -0.036 | 0.016 | 0.016 | 1.141 | 0.33 | 0.08 | 0.23 | 0.25 | 0.34 | 0.26 |
| 7 | 15440108 | rs4721433 | 0.73 | -0.362 | -2.053 | -0.695 | -0.002 | 0.06 | 0.06 | 0 | 0.16 | 0.02 | 0.07 | 0.07 | 0.14 | 0.2 |
| 7 | 15455333 | rs10263571 | 0.802 | -0.987 | -1.881 | -0.664 | 0.021 | 0.044 | 0.044 | 0 | 0.15 | 0.05 | 0.07 | 0.07 | 0.12 | 0.15 |
| 7 | 15456251 | rs4472405 | -0.52 | -0.443 | -1.888 | -0.634 | -0.043 | -0.113 | -0.113 | 0 | 0.8 | 0.96 | 0.59 | 0.58 | 0.77 | 0.69 |
| 7 | 15464307 | rs4431490 | 1.386 | -0.045 | -1.47 | -1.318 | -0.066 | 0.004 | 0.004 | 1.091 | 0.34 | 0.08 | 0.25 | 0.28 | 0.38 | 0.33 |
| 7 | 15468199 | rs11765688 | 0.458 | 0.109 | -1.285 | -1.321 | -0.002 | -0.013 | -0.013 | 0.176 | 0.13 | 0.14 | 0.13 | 0 | 0.22 | 0.09 |
| 7 | 15473799 | rs4615456 | 0.604 | -0.456 | -0.89 | -1.173 | 0 | 0.016 | 0.016 | 0.046 | 0.43 | 0.33 | 0.29 | 0.28 | 0.42 | 0.36 |
| 7 | 15478152 | rs6944475 | 0.914 | -0.532 | -0.714 | -1.009 | -0.001 | 0.001 | 0.001 | 0 | 0.37 | 0.33 | 0.28 | 0.27 | 0.38 | 0.31 |
| 7 | 15479473 | rs7787350 | 0.535 | -1.824 | -0.213 | -1.275 | 0.07 | 0.036 | 0.036 | 0 | 0.51 | 0.37 | 0.43 | 0.35 | 0.63 | 0.4 |
| 7 | 15484101 | rs10278280 | 0.922 | -1.8 | 0.293 | -1.383 | -0.004 | 0.006 | 0.006 | 0 | 0.42 | 0.4 | 0.38 | 0.33 | 0.46 | 0.32 |
| 7 | 15486094 | rs10215514 | 0.146 | -1.952 | 0.392 | -1.342 | 0.016 | -0.025 | -0.025 | 0.82 | 0.81 | 0.88 | 0.6 | 0.38 | 0.83 | 0.66 |
| 7 | 15486895 | rs6461176 | 1.089 | -2.018 | 0.775 | -1.363 | 0.012 | 0.04 | 0.04 | 0 | 0.71 | 0.57 | 0.51 | 0.38 | 0.65 | 0.62 |
| 7 | 15495150 | rs4628172 | 0.095 | -1.812 | 0.427 | -1.301 | 0.016 | -0.038 | -0.038 | 0.219 | 0.81 | 0.89 | 0.6 | 0.38 | 0.82 | 0.66 |
| 7 | 15495188 | rs11773289 | 0.671 | -1.712 | 0.784 | -1.321 | 0.04 | -0.041 | -0.041 | 0 | 0.09 | 0.13 | 0.08 | 0 | 0.19 | 0.05 |
| 7 | 15503313 | rs17168763 | -0.026 | -1.954 | 0.422 | -1.25 | 0.001 | 0.011 | 0.011 | 0 | 0.15 | 0.13 | 0.39 | 0.62 | 0.15 | 0.29 |
| 7 | 15507516 | rs1529893 | -0.346 | -2.187 | 0.814 | -1.216 | 0.035 | 0.034 | 0.034 | 0 | 0.75 | 0.85 | 0.88 | 0.96 | 0.83 | 0.71 |
| 7 | 15509401 | rs17168775 | -0.413 | -2.239 | 0.748 | -1.156 | 0.013 | -0.011 | -0.011 | 1.826 | 0.13 | 0.08 | 0.32 | 0.62 | 0.12 | 0.29 |
| 7 | 15509538 | rs12530666 | 1.481 | -2.349 | 1 | -1.328 | 0.021 | 0.031 | 0.031 | 0 | 0.47 | 0.28 | 0.46 | 0.33 | 0.53 | 0.4 |
| 7 | 15512916 | rs10950562 | 1.596 | -2.24 | 1.065 | -1.29 | -0.009 | 0.012 | 0.012 | 0 | 0.42 | 0.28 | 0.45 | 0.33 | 0.5 | 0.36 |
| 7 | 15521358 | rs6461187 | -1.596 | -2.145 | 0.926 | -1.253 | 0 | 0.014 | 0.014 | 0.093 | 0.32 | 0.58 | 0.43 | 0.62 | 0.34 | 0.35 |
| 7 | 15534475 | rs17150113 | 1.108 | -1.92 | 0.929 | -1.866 | 0 | -0.046 | -0.046 | 0 | 0.6 | 0.37 | 0.77 | 0.94 | 0.64 | 0.67 |
| 7 | 15534763 | rs1107506 | 0.278 | -1.918 | 0.817 | -1.869 | -0.073 | -0.124 | -0.124 | 0 | 0.7 | 0.49 | 0.84 | 0.94 | 0.83 | 0.73 |
| 7 | 15547950 | rs1992025 | 0.263 | -2.658 | 0.441 | -1.371 | 0.021 | -0.009 | -0.009 | 0.043 | 0.69 | 0.73 | 0.86 | 0.96 | 0.79 | 0.71 |
| 7 | 15549468 | rs12699727 | 0.41 | -2.722 | 0.453 | -1.354 | -0.018 | -0.074 | -0.074 | 0 | 0.67 | 0.59 | 0.85 | 0.95 | 0.8 | 0.71 |
| 7 | 15582208 | rs2196395 | 1.028 | -2.488 | -0.271 | -1.088 | 0.181 | 0.243 | 0.243 | 0 | 0.5 | 0.17 | 0.69 | 0.83 | 0.68 | 0.51 |
| 7 | 15593768 | rs916943 | -1.5 | -1.382 | -0.306 | -0.602 | 0.009 | 0.034 | 0.034 | 0 | 0.12 | 0.06 | 0.05 | 0 | 0.09 | 0.09 |
| 7 | 15593919 | rs17329829 | -1.438 | -1.346 | -0.008 | -0.028 | -0.008 | 0.037 | 0.037 | 0 | 0.08 | 0.14 | 0.06 | 0.12 | 0.05 | 0.04 |
| 7 | 15598156 | rs978897 | -1.225 | -1.392 | 0.239 | -0.627 | -0.025 | -0.101 | -0.101 | 0.013 | 0.2 | 0.59 | 0.1 | 0 | 0.14 | 0.13 |
| 7 | 15603853 | rs17389549 | -0.21 | -1.496 | -0.618 | -0.275 | 0.03 | 0.054 | 0.054 | 0 | 0.22 | 0.07 | 0.15 | 0.16 | 0.16 | 0.28 |
| 7 | 15607046 | rs7782511 | -1.209 | -1.723 | 0.204 | 0.66 | -0.072 | 0.088 | 0.088 | 0 | 0.27 | 0.44 | 0.36 | 0.44 | 0.18 | 0.21 |
| 7 | 15607231 | rs7801643 | -1.442 | -1.085 | 0.523 | 0.932 | 0.019 | -0.02 | -0.02 | 0 | 0.31 | 0.71 | 0.48 | 0.63 | 0.29 | 0.23 |
| 7 | 15607614 | rs7786817 | -0.716 | -1.312 | 1.497 | 0.667 | -0.025 | 0.188 | 0.188 | 0 | 0.13 | 0.23 | 0.18 | 0.43 | 0.1 | 0.11 |
| 7 | 15610187 | rs13235486 | -0.848 | -1.383 | 1.989 | 0.55 | -0.032 | -0.071 | -0.071 | 0 | 0.14 | 0.28 | 0.19 | 0 | 0.08 | 0.1 |
| 7 | 15613379 | rs6946895 | -1.126 | -1.349 | 1.855 | 0.564 | 0.03 | -0.021 | -0.021 | 0 | 0.06 | 0.13 | 0.05 | 0.02 | 0.12 | 0.02 |
| 7 | 15617256 | rs10246891 | -2.045 | -1.597 | 1.973 | 0.61 | -0.022 | 0.017 | 0.017 | 0 | 0.27 | 0.68 | 0.38 | 0.44 | 0.16 | 0.21 |
| 7 | 15620083 | rs17168843 | -0.415 | -1.523 | 1.661 | 0.878 | 0.01 | -0.017 | -0.017 | 0.031 | 0.1 | 0.03 | 0.05 | 0.1 | 0.09 | 0.09 |

| | | | | | | | | | | | | | | | | |
|---|---|---|---|---|---|---|---|---|---|---|---|---|---|---|---|---|
| 7 | 15621650 | rs6944184 | -2.346 | -1.664 | 2.181 | 0.787 | -0.036 | -0.068 | -0.068 | 0 | 0.26 | 0.45 | 0.14 | 0.13 | 0.24 | 0.2 |
| 7 | 15630207 | rs10950569 | -1.122 | -2.208 | 1.049 | -0.136 | 0.056 | 0.104 | 0.104 | 0 | 0.67 | 0.84 | 0.81 | 0.89 | 0.79 | 0.59 |
| 7 | 15634235 | rs17168857 | 0.657 | -2.168 | 0.949 | -0.097 | 0.002 | -0.025 | -0.025 | 0 | 0.09 | 0.01 | 0.04 | 0.1 | 0.09 | 0.09 |
| 7 | 15641042 | rs17168881 | -1.339 | -2.202 | 1.387 | 0.173 | -0.026 | 0.194 | 0.194 | 0.071 | 0.18 | 0.46 | 0.22 | 0.39 | 0.16 | 0.14 |
| 7 | 15643347 | rs6946852 | 0.303 | -2.28 | 1.437 | 0.144 | 0.01 | -0.001 | -0.001 | 0.028 | 0.49 | 0.47 | 0.6 | 0.5 | 0.63 | 0.44 |
| 7 | 15644223 | rs7458065 | -1.152 | -2.181 | 1.101 | 0.029 | 0.056 | 0.145 | 0.145 | 0 | 0.68 | 0.94 | 0.82 | 0.89 | 0.79 | 0.58 |
| 7 | 15647727 | rs17168895 | -0.405 | -2.08 | 1.343 | 0.09 | -0.002 | -0.059 | -0.059 | 0 | 0.12 | 0.26 | 0.08 | 0.03 | 0.13 | 0.1 |
| 7 | 15648861 | rs6946099 | 1.342 | -1.819 | 1.348 | 0.142 | -0.055 | -0.034 | -0.034 | 0 | 0.34 | 0.16 | 0.51 | 0.4 | 0.51 | 0.35 |
| 7 | 15651158 | rs10777 | -1.015 | -1.797 | 1.434 | 0.183 | 0.045 | 0.099 | 0.099 | 0 | 0.66 | 0.81 | 0.81 | 0.89 | 0.77 | 0.58 |
| 7 | 15652477 | rs2237494 | 0.728 | -1.674 | 1.396 | 0.295 | 0.014 | 0.002 | 0.002 | 0 | 0.43 | 0.24 | 0.56 | 0.42 | 0.57 | 0.44 |
| 7 | 15656707 | rs7793334 | 0.326 | -1.639 | 1.239 | 0.059 | 0.004 | -0.003 | -0.003 | 0 | 0.48 | 0.47 | 0.73 | 0.79 | 0.65 | 0.48 |
| 7 | 15657307 | rs10266294 | 0.364 | -1.638 | 1.24 | 0.059 | 0.003 | -0.005 | -0.005 | 0.551 | 0.48 | 0.46 | 0.73 | 0.79 | 0.65 | 0.48 |
| 7 | 15658957 | rs10486768 | 0.305 | -1.605 | 0.663 | 0.115 | -0.004 | -0.011 | -0.011 | 0 | 0.11 | 0.01 | 0.05 | 0.09 | 0.1 | 0.09 |
| 7 | 15668526 | rs1548691 | 0.265 | -1.68 | 0.752 | 0.07 | 0.053 | 0.121 | 0.121 | 0 | 0.51 | 0.67 | 0.73 | 0.79 | 0.63 | 0.47 |
| 7 | 15671471 | rs16878569 | -0.109 | -1.316 | 0.404 | 0.356 | 0.009 | -0.003 | -0.003 | 0.004 | 0.11 | 0.06 | 0.05 | 0.09 | 0.1 | 0.09 |
| 7 | 15674830 | rs10281319 | -0.252 | -1.604 | 0.645 | 0.244 | 0.007 | -0.002 | -0.002 | 0 | 0.54 | 0.63 | 0.61 | 0.45 | 0.63 | 0.45 |
| 7 | 15680072 | rs13438001 | -0.861 | -1.8 | 0.946 | 0.215 | -0.047 | -0.02 | -0.02 | 0 | 0.2 | 0.48 | 0.09 | 0.15 | 0.13 | 0.16 |
| 7 | 15692093 | rs17336581 | 1.397 | -0.374 | -0.133 | -0.069 | -0.118 | -0.104 | -0.104 | 0.033 | 0.22 | 0.01 | 0.29 | 0.33 | 0.32 | 0.17 |
| 7 | 15692150 | rs10486771 | -0.211 | -0.921 | -0.094 | -0.136 | 0.032 | -0.001 | -0.001 | 0 | 0.19 | 0.05 | 0.15 | 0.17 | 0.13 | 0.16 |
| 7 | 15696151 | rs12056299 | -1.834 | -1.211 | -0.371 | -0.332 | 0.009 | 0.106 | 0.106 | 0 | 0.71 | 0.93 | 0.82 | 0.89 | 0.79 | 0.63 |
| 7 | 15698178 | rs7787043 | 0.001 | -1.262 | -0.173 | -0.275 | 0.024 | 0.042 | 0.042 | 0 | 0.44 | 0.35 | 0.63 | 0.7 | 0.65 | 0.47 |
| 7 | 15699036 | rs10278557 | 1.334 | -0.672 | -0.551 | -0.215 | -0.035 | -0.023 | -0.023 | 0 | 0.26 | 0.13 | 0.27 | 0.31 | 0.26 | 0.24 |
| 7 | 15699816 | rs10282109 | -1.588 | -0.411 | -0.018 | -0.034 | 0.007 | 0.014 | 0.014 | 0 | 0.69 | 0.78 | 0.81 | 0.86 | 0.78 | 0.61 |
| 7 | 15702723 | rs6461201 | 0.665 | 0.225 | -0.222 | 0.408 | -0.043 | -0.043 | -0.043 | 0 | 0.12 | 0.04 | 0.29 | 0.37 | 0.28 | 0.2 |
| 7 | 15713934 | rs6461202 | -1.146 | -0.29 | 0.176 | 0.146 | -0.011 | -0.004 | -0.004 | 3.852 | 0.44 | 0.62 | 0.48 | 0.5 | 0.54 | 0.4 |
| 7 | 15716973 | rs6970462 | 1.06 | 0.189 | -0.536 | 0.031 | -0.055 | 0.049 | 0.049 | 0 | 0.11 | 0.01 | 0.23 | 0.03 | 0.23 | 0.17 |
| 7 | 15721178 | rs3801427 | -1.668 | -0.465 | -0.509 | -0.099 | 0.043 | 0.072 | 0.072 | 0 | 0.66 | 0.86 | 0.73 | 0.85 | 0.77 | 0.59 |
| 7 | 15721466 | rs6959056 | -0.41 | -0.396 | -0.431 | 0.036 | -0.066 | -0.07 | -0.07 | 0.508 | 0.37 | 0.11 | 0.42 | 0.48 | 0.48 | 0.37 |
| 7 | 15722836 | rs3801430 | 0.754 | -0.257 | 0.279 | 0.238 | 0.022 | 0.104 | 0.104 | 0 | 0.21 | 0.4 | 0.26 | 0.35 | 0.21 | 0.16 |
| 7 | 15732542 | rs10486774 | 0.294 | -0.79 | -0.236 | 0.092 | 0.014 | 0.011 | 0.011 | 0 | 0.18 | 0.09 | 0.08 | 0.16 | 0.14 | 0.13 |
| 7 | 15734715 | rs726395 | 1.2 | -0.679 | -0.145 | -0.054 | -0.004 | -0.001 | -0.001 | 0.807 | 0.54 | 0.41 | 0.58 | 0.49 | 0.52 | 0.61 |
| 7 | 15734821 | rs12056115 | 0.55 | -0.478 | -0.277 | 0.057 | -0.063 | -0.058 | -0.058 | 0.717 | 0.2 | 0.06 | 0.3 | 0.35 | 0.32 | 0.24 |
| 7 | 15735165 | rs11972297 | 1.404 | -0.553 | -0.3 | 0.099 | -0.053 | -0.083 | -0.083 | 0 | 0.17 | 0.05 | 0.3 | 0.35 | 0.28 | 0.15 |
| 7 | 15735559 | rs17168992 | 0.94 | -0.521 | -0.115 | 0.124 | 0.016 | 0.052 | 0.052 | 0 | 0.19 | 0.28 | 0.33 | 0.35 | 0.28 | 0.15 |
| 7 | 15743093 | rs11773055 | -0.427 | -0.941 | -0.076 | 0.194 | 0.001 | -0.005 | -0.005 | 0 | 0.39 | 0.64 | 0.58 | 0.56 | 0.61 | 0.36 |
| 7 | 15745324 | rs6942413 | -1.488 | -1.299 | -0.705 | 0.039 | -0.012 | 0.026 | 0.026 | 0 | 0.54 | 0.69 | 0.52 | 0.67 | 0.51 | 0.44 |
| 7 | 15746973 | rs11771752 | 0.022 | -0.558 | -0.198 | 0.377 | -0.019 | -0.016 | -0.016 | 1.681 | 0.25 | 0.19 | 0.32 | 0.38 | 0.33 | 0.25 |
| 7 | 15747273 | rs17169004 | -0.039 | -0.497 | 0.069 | 0.614 | 0.028 | -0.024 | -0.024 | 0 | 0.21 | 0.12 | 0.15 | 0.3 | 0.14 | 0.17 |
| 7 | 15752122 | rs17169008 | 0.584 | 0.418 | 0.407 | 0.901 | 0.039 | 0.053 | 0.053 | 0 | 0.17 | 0.3 | 0.27 | 0.22 | 0.28 | 0.12 |
| 7 | 15758208 | rs632846 | -0.084 | 0.205 | -0.193 | 0.667 | -0.032 | -0.015 | -0.015 | 0 | 0.3 | 0.13 | 0.43 | 0.35 | 0.4 | 0.26 |
| 7 | 15758899 | rs10226819 | -0.735 | -0.334 | -0.161 | 0.348 | -0.017 | -0.083 | -0.083 | 0 | 0.35 | 0.91 | 0.44 | 0.51 | 0.35 | 0.26 |
| 7 | 15760654 | rs10264045 | -0.663 | -0.438 | 0.674 | 0.606 | 0.02 | 0.09 | 0.09 | 1.307 | 0.11 | 0.56 | 0.08 | 0.13 | 0.05 | 0.07 |
| 7 | 15760677 | rs531653 | -1.186 | -0.544 | 0.694 | 0.705 | 0.081 | 0.037 | 0.037 | 0 | 0.54 | 0.88 | 0.75 | 0.62 | 0.72 | 0.43 |
| 7 | 15761479 | rs10254339 | -0.746 | -0.37 | 1.137 | 0.698 | 0.029 | 0.113 | 0.113 | 1.016 | 0.1 | 0.56 | 0.08 | 0.13 | 0.05 | 0.07 |
| 7 | 15761537 | rs10268121 | -0.919 | -0.368 | 1.132 | 0.699 | 0.012 | 0.086 | 0.086 | 1.383 | 0.11 | 0.59 | 0.08 | 0.13 | 0.05 | 0.07 |
| 7 | 15762763 | rs10272623 | -0.726 | -0.431 | 1.129 | 0.589 | 0.006 | 0.034 | 0.034 | 0.466 | 0.16 | 0.46 | 0.09 | 0.14 | 0.1 | 0.15 |
| 7 | 15771599 | rs17340395 | 1.194 | 0.04 | 0.682 | 0.908 | -0.001 | -0.001 | -0.001 | 0.192 | 0.15 | 0.19 | 0.24 | 0.18 | 0.25 | 0.1 |
| 7 | 15775095 | rs10249588 | -0.917 | -0.098 | 0.783 | 1.062 | -0.039 | 0.118 | 0.118 | 0 | 0.17 | 0.37 | 0.17 | 0.32 | 0.08 | 0.16 |
| 7 | 15775401 | rs10282201 | 0.563 | 0.207 | -0.224 | 1.005 | 0 | -0.027 | -0.027 | 0 | 0.09 | 0.05 | 0.13 | 0.21 | 0.07 | 0.11 |
| 7 | 15775437 | rs37409 | 0.023 | -0.021 | -0.24 | 0.72 | -0.051 | -0.027 | -0.027 | 0 | 0.29 | 0.1 | 0.42 | 0.36 | 0.41 | 0.26 |
| 7 | 15776389 | rs12670755 | -0.544 | 0.303 | 0.691 | 0.961 | 0.083 | 0.03 | 0.03 | 0 | 0.27 | 0.44 | 0.47 | 0.62 | 0.49 | 0.24 |
| 7 | 15782093 | rs37414 | 1.165 | -1.354 | 0.767 | 0.911 | -0.038 | 0.01 | 0.01 | 0.232 | 0.78 | 0.3 | 0.88 | 0.7 | 0.93 | 0.76 |
| 7 | 15788241 | rs37423 | 1.38 | -1.419 | 0.653 | 0.903 | -0.014 | -0.003 | -0.003 | 0 | 0.76 | 0.23 | 0.89 | 0.69 | 0.92 | 0.74 |
| 7 | 15794307 | rs6461207 | -1.55 | -0.492 | 1.145 | 1.085 | 0.008 | 0.166 | 0.166 | 1.495 | 0.07 | 0.55 | 0.06 | 0.11 | 0.01 | 0.01 |
| 7 | 15805403 | rs17169067 | 0.117 | -0.395 | 0.737 | 1.007 | -0.04 | -0.077 | -0.077 | 0.567 | 0.13 | 0.01 | 0.15 | 0.28 | 0.18 | 0.18 |
| 7 | 15806208 | rs10248805 | -0.976 | -0.3 | 1.365 | 0.395 | 0.028 | 0.055 | 0.055 | 0 | 0.19 | 0.56 | 0.32 | 0.23 | 0.22 | 0.11 |
| 7 | 15807092 | rs1942653 | -1.785 | -1.1 | 2.05 | 0.963 | 0.038 | 0.374 | 0.374 | 0.742 | 0.06 | 0.46 | 0.05 | 0.22 | 0.01 | 0.04 |
| 7 | 15815683 | rs13234197 | -1.05 | -0.775 | 0.93 | 0.407 | -0.018 | -0.071 | -0.071 | 0 | 0.13 | 0.26 | 0.04 | 0 | 0.08 | 0.14 |
| 7 | 15833180 | rs12113594 | -1.053 | -0.824 | 0.571 | 0.847 | -0.026 | 0.067 | 0.067 | 0 | 0.13 | 0.28 | 0.08 | 0.24 | 0.07 | 0.26 |
| 7 | 15837740 | rs12533562 | -1.733 | -1.119 | 0.748 | 0.748 | 0.01 | -0.004 | -0.004 | 0.112 | 0.33 | 0.71 | 0.28 | 0.33 | 0.24 | 0.35 |
| 7 | 15839429 | rs10278213 | -1.81 | -1.043 | 0.681 | 0.724 | 0.018 | -0.004 | -0.004 | 0 | 0.36 | 0.71 | 0.29 | 0.34 | 0.27 | 0.36 |
| 7 | 15852195 | rs10950582 | -0.161 | -1.016 | 1.07 | 0.68 | 0.036 | -0.003 | -0.003 | 0 | 0.1 | 0.19 | 0.16 | 0.07 | 0.15 | 0.05 |
| 7 | 15859501 | rs12531519 | -1.528 | -1.122 | 1.055 | 0.69 | -0.001 | 0.073 | 0.073 | 0.188 | 0.2 | 0.57 | 0.22 | 0.3 | 0.17 | 0.14 |
| 7 | 15868471 | rs11768573 | -0.045 | -0.952 | 0.656 | 1.145 | 0.031 | 0.044 | 0.044 | 0 | 0.09 | 0.15 | 0.2 | 0.14 | 0.15 | 0.06 |
| 7 | 15872713 | rs6974999 | -1.999 | -1.127 | 0.536 | 1.019 | 0.037 | -0.039 | -0.039 | 0.018 | 0.37 | 0.79 | 0.32 | 0.49 | 0.26 | 0.39 |
| 7 | 15874219 | rs10486781 | 0.166 | -0.786 | 0.036 | 0.152 | 0.026 | -0.001 | -0.001 | 0 | 0.08 | 0.12 | 0.16 | 0.05 | 0.15 | 0.04 |
| 7 | 15877633 | rs13227386 | -1.287 | -1.545 | -0.137 | 0.191 | -0.047 | -0.034 | -0.034 | 0.581 | 0.23 | 0.28 | 0.07 | 0.11 | 0.1 | 0.18 |
| 7 | 15881354 | rs1942638 | -0.085 | -1.301 | -1.082 | 0.167 | 0.108 | 0.071 | 0.071 | 0 | 0.48 | 0.73 | 0.38 | 0.63 | 0.31 | 0.37 |
| 7 | 15881632 | rs1942637 | -0.351 | -1.995 | -0.668 | -0.26 | -0.015 | -0.007 | -0.007 | 0 | 0.45 | 0.52 | 0.54 | 0.66 | 0.25 | 0.62 |

| | | | | | | | | | | | | | | | | |
|---|---|---|---|---|---|---|---|---|---|---|---|---|---|---|---|---|
| 7 | 15881942 | rs1942636 | 0.423 | 0.125 | -0.717 | 0.025 | 0.052 | 0.017 | 0.017 | 0 | 0.09 | 0.21 | 0.22 | 0.18 | 0.23 | 0.06 |
| 7 | 15883552 | rs10226196 | -0.703 | -0.216 | -1 | -0.076 | -0.026 | -0.001 | -0.001 | 0.161 | 0.2 | 0.08 | 0.43 | 0.23 | 0.28 | 0.42 |
| 7 | 15883805 | rs17169105 | -0.317 | -0.466 | -0.093 | 0.836 | 0.002 | 0.164 | 0.164 | 0 | 0.09 | 0.26 | 0.04 | 0.26 | 0.04 | 0.12 |
| 7 | 15884675 | rs10224052 | 0.747 | -0.92 | -0.371 | 0.769 | 0.018 | 0.017 | 0.017 | 0 | 0.31 | 0.23 | 0.21 | 0.16 | 0.23 | 0.16 |
| 7 | 15889830 | rs10258707 | 0.218 | -1.096 | -0.265 | 0.737 | -0.01 | -0.023 | -0.023 | 0 | 0.4 | 0.41 | 0.27 | 0.15 | 0.27 | 0.23 |
| 7 | 15891058 | rs38168 | 0.071 | -0.665 | -0.289 | 0.986 | 0.001 | 0.02 | 0.02 | 0 | 0.51 | 0.58 | 0.46 | 0.59 | 0.48 | 0.67 |
| 7 | 15891270 | rs17169111 | -0.046 | -0.801 | -0.142 | 1.22 | 0.003 | 0.008 | 0.008 | 0 | 0.08 | 0.07 | 0.04 | 0.27 | 0.06 | 0.13 |
| 7 | 15891740 | rs7797334 | 0.679 | -0.838 | -0.057 | 1.243 | -0.01 | 0.049 | 0.049 | 0 | 0.06 | 0.1 | 0.02 | 0.25 | 0.03 | 0.09 |
| 7 | 15892861 | rs1942634 | 1.031 | -0.79 | -0.129 | 1.054 | 0.023 | 0.114 | 0.114 | 0.109 | 0.3 | 0.07 | 0.2 | 0.09 | 0.22 | 0.15 |
| 7 | 15893300 | rs38169 | -0.761 | -0.851 | 0.255 | 1.06 | 0.017 | 0.002 | 0.002 | 0 | 0.84 | 0.78 | 0.66 | 0.76 | 0.74 | 0.6 |
| 7 | 15893530 | rs38171 | 0.187 | -0.871 | 0.327 | 1.034 | 0.001 | -0.006 | -0.006 | 0 | 0.44 | 0.44 | 0.38 | 0.63 | 0.46 | 0.33 |
| 7 | 15895516 | rs38173 | 0.581 | -0.752 | 0.421 | 1.056 | 0.028 | 0.109 | 0.109 | 0 | 0.75 | 0.53 | 0.62 | 0.5 | 0.7 | 0.51 |
| 7 | 15897219 | rs38175 | 0.582 | -0.741 | 0.42 | 1.009 | 0.019 | 0.082 | 0.082 | 0 | 0.74 | 0.47 | 0.62 | 0.49 | 0.69 | 0.5 |
| 7 | 15899838 | rs6959214 | 1.434 | -1.103 | -0.151 | 0.877 | 0.042 | 0.134 | 0.134 | 0.042 | 0.28 | 0.08 | 0.14 | 0.01 | 0.19 | 0.13 |
| 7 | 15901834 | rs28273 | 0.553 | 0.136 | -0.233 | 0.751 | 0.002 | 0.005 | 0.005 | 0 | 0.66 | 0.6 | 0.58 | 0.49 | 0.61 | 0.48 |
| 7 | 15904870 | rs38194 | 0.269 | 0.181 | -0.617 | 0.814 | -0.009 | -0.018 | -0.018 | 0 | 0.68 | 0.73 | 0.6 | 0.51 | 0.62 | 0.51 |
| 7 | 15906295 | rs1942645 | 0.965 | 0.554 | -0.718 | 0.892 | -0.007 | 0.06 | 0.06 | 0 | 0.57 | 0.65 | 0.43 | 0.25 | 0.44 | 0.43 |
| 7 | 15913289 | rs38204 | -0.427 | 0.564 | -0.743 | 0.705 | -0.022 | 0.02 | 0.02 | 0 | 0.75 | 0.82 | 0.62 | 0.74 | 0.66 | 0.58 |
| 7 | 15917071 | rs38212 | 0.113 | 0.281 | -0.682 | 1 | -0.012 | -0.057 | -0.057 | 0.248 | 0.16 | 0.08 | 0.21 | 0.37 | 0.17 | 0.08 |
| 7 | 15927187 | rs38224 | 0.307 | 0.652 | -0.556 | 0.915 | -0.014 | -0.013 | -0.013 | 0 | 0.68 | 0.72 | 0.6 | 0.53 | 0.61 | 0.46 |
| 7 | 15928175 | rs9655121 | 1.09 | 0.628 | -0.599 | 0.857 | 0.006 | 0.093 | 0.093 | 0 | 0.57 | 0.7 | 0.39 | 0.16 | 0.44 | 0.39 |
| 7 | 15928418 | rs38227 | -0.475 | 0.694 | -0.6 | 1.193 | -0.023 | -0.06 | -0.06 | 0 | 0.2 | 0.14 | 0.18 | 0.51 | 0.22 | 0.2 |
| 7 | 15929081 | rs38230 | -0.475 | 0.694 | -0.6 | 1.195 | -0.023 | -0.06 | -0.06 | 0 | 0.2 | 0.13 | 0.18 | 0.51 | 0.22 | 0.2 |
| 7 | 15932674 | rs38237 | -0.009 | 0.403 | -0.436 | 0.861 | -0.006 | -0.032 | -0.032 | 0 | 0.31 | 0.23 | 0.35 | 0.47 | 0.31 | 0.3 |
| 7 | 15932916 | rs38238 | -0.522 | 0.623 | -0.443 | 0.887 | -0.013 | 0.007 | 0.007 | 0 | 0.77 | 0.8 | 0.62 | 0.74 | 0.66 | 0.61 |
| 7 | 15941283 | rs38246 | -1.084 | 0.767 | -0.506 | 0.839 | -0.015 | 0.011 | 0.011 | 0 | 0.83 | 0.84 | 0.78 | 0.9 | 0.72 | 0.91 |
| 7 | 15942119 | rs38247 | 0.312 | 0.531 | -0.492 | 1.083 | -0.007 | -0.043 | -0.043 | 0 | 0.16 | 0.09 | 0.17 | 0.4 | 0.18 | 0.08 |
| 7 | 15948435 | rs38255 | -0.224 | 0.406 | -0.474 | 0.992 | -0.002 | -0.044 | -0.044 | 0.083 | 0.21 | 0.12 | 0.23 | 0.6 | 0.21 | 0.16 |
| 7 | 15956470 | rs2215169 | 0.005 | 0.515 | -0.297 | 0.894 | -0.005 | -0.008 | -0.008 | 0 | 0.73 | 0.73 | 0.6 | 0.51 | 0.64 | 0.54 |
| 7 | 15959199 | rs6965426 | -0.747 | 0.534 | -0.362 | 0.88 | -0.024 | 0.006 | 0.006 | 0 | 0.79 | 0.84 | 0.63 | 0.74 | 0.67 | 0.6 |
| 7 | 15959268 | rs16878646 | 0.7 | 0.43 | -0.426 | 1.046 | -0.049 | -0.105 | -0.105 | 0.339 | 0.15 | 0.02 | 0.15 | 0.31 | 0.17 | 0.11 |
| 7 | 15964744 | rs1072964 | 2.046 | 0.01 | 0.258 | 0.77 | -0.022 | 0.018 | 0.018 | 0 | 0.07 | 0.16 | 0.1 | 0.09 | 0.03 | 0.04 |
| 7 | 15973525 | rs1406100 | 1.344 | 0.115 | 0.126 | 0.873 | -0.006 | 0.061 | 0.061 | 0 | 0.06 | 0.09 | 0.03 | 0.23 | 0.04 | 0.07 |
| 7 | 15988412 | rs12674430 | 0.097 | -0.016 | -0.033 | 0.619 | -0.015 | -0.08 | -0.08 | 0 | 0.32 | 0.12 | 0.3 | 0.4 | 0.3 | 0.3 |
| 7 | 15992365 | rs800718 | -2.491 | 0.102 | 0.028 | 0.458 | -0.062 | 0.029 | 0.029 | 0.014 | 0.89 | 0.98 | 0.81 | 0.89 | 0.75 | 0.93 |
| 7 | 15993845 | rs17169172 | 1.793 | -0.236 | 0.672 | 0.435 | -0.002 | 0.002 | 0.002 | 0 | 0.09 | 0.1 | 0.17 | 0.16 | 0.07 | 0.34 |
| 7 | 15994459 | rs10241996 | -0.07 | -0.249 | 0.944 | 0.378 | -0.004 | -0.036 | -0.036 | 0 | 0.36 | 0.21 | 0.32 | 0.41 | 0.32 | 0.34 |
| 7 | 15994571 | rs10255820 | -0.566 | -0.273 | 0.978 | 0.098 | 0.006 | 0.028 | 0.028 | 0.242 | 0.42 | 0.33 | 0.34 | 0.64 | 0.35 | 0.4 |
| 7 | 15998044 | rs1997264 | -1.202 | -0.273 | 0.935 | 0.078 | 0.004 | 0.029 | 0.029 | 0.327 | 0.52 | 0.6 | 0.52 | 0.78 | 0.43 | 0.75 |
| 7 | 15998111 | rs1997263 | -0.905 | -0.252 | 0.69 | 0.086 | 0.009 | 0.069 | 0.069 | 0.006 | 0.49 | 0.38 | 0.51 | 0.78 | 0.42 | 0.75 |
| 7 | 16007799 | rs12670350 | 0.68 | 0.011 | 0.589 | 0.554 | -0.017 | 0.09 | 0.09 | 0 | 0.08 | 0.13 | 0.03 | 0.27 | 0.04 | 0.07 |
| 7 | 16016504 | rs1918857 | -1.303 | 0.101 | 0.601 | 0.294 | 0.026 | -0.013 | -0.013 | 3.429 | 0.8 | 0.7 | 0.69 | 0.78 | 0.71 | 0.89 |
| 7 | 16017501 | rs17457220 | 2.059 | -0.476 | -0.416 | -0.08 | 0.006 | 0.068 | 0.068 | 0 | 0.31 | 0.17 | 0.2 | 0.01 | 0.3 | 0.21 |
| 7 | 16025206 | rs11769915 | 1.315 | -0.323 | -0.734 | 0.096 | -0.013 | -0.044 | -0.044 | 0 | 0.15 | 0.06 | 0.16 | 0.27 | 0.17 | 0.09 |
| 7 | 16025340 | rs13234986 | 0.726 | -0.606 | -0.431 | -0.035 | -0.011 | -0.032 | -0.032 | 0 | 0.28 | 0.15 | 0.29 | 0.36 | 0.3 | 0.27 |
| 7 | 16027001 | rs6952901 | -0.085 | -0.66 | -0.176 | 0.006 | -0.006 | 0.029 | 0.029 | 0 | 0.2 | 0.25 | 0.21 | 0.43 | 0.14 | 0.45 |
| 7 | 16027025 | rs6953206 | -0.097 | -0.729 | -0.173 | 0.007 | -0.011 | 0.032 | 0.032 | 0 | 0.2 | 0.25 | 0.2 | 0.42 | 0.13 | 0.45 |
| 7 | 16027189 | rs6953523 | 0.01 | -0.635 | -0.092 | -0.102 | 0.001 | 0.031 | 0.031 | 0.305 | 0.7 | 0.42 | 0.66 | 0.55 | 0.69 | 0.82 |
| 7 | 16027998 | rs6461213 | -1.016 | -0.402 | 0.052 | -0.09 | 0.037 | -0.035 | -0.035 | 0 | 0.78 | 0.48 | 0.69 | 0.8 | 0.73 | 0.9 |
| 7 | 16036970 | rs17169198 | -0.148 | -0.403 | -0.405 | -0.035 | -0.01 | -0.032 | -0.032 | 0 | 0.39 | 0.22 | 0.47 | 0.49 | 0.39 | 0.63 |
| 7 | 16038502 | rs10257485 | 0.866 | -0.6 | -0.413 | -0.077 | 0.033 | 0.025 | 0.025 | 0 | 0.4 | 0.2 | 0.22 | 0.27 | 0.34 | 0.28 |
| 7 | 16043897 | rs10263175 | -0.604 | -1.149 | 0.363 | -0.03 | -0.072 | 0.144 | 0.144 | 0.796 | 0.15 | 0.56 | 0.06 | 0.31 | 0.05 | 0.09 |
| 7 | 16045110 | rs17169204 | 2.783 | -1.646 | -0.123 | -0.555 | -0.006 | 0.028 | 0.028 | 0.195 | 0.1 | 0.05 | 0.08 | 0 | 0.11 | 0.07 |
| 7 | 16045952 | rs800709 | -2.854 | -1.492 | -0.207 | -0.728 | -0.01 | 0.008 | 0.008 | 0 | 0.88 | 0.9 | 0.81 | 0.91 | 0.76 | 0.94 |
| 7 | 16054913 | rs2389577 | -1.962 | -1.836 | -0.557 | -0.838 | 0.015 | 0.021 | 0.021 | 0 | 0.77 | 0.86 | 0.81 | 0.85 | 0.82 | 0.83 |
| 7 | 16058837 | rs13247199 | -0.129 | -1.936 | -0.297 | -0.698 | -0.015 | 0.139 | 0.139 | 0 | 0.44 | 0.17 | 0.51 | 0.76 | 0.46 | 0.66 |
| 7 | 16059180 | rs2030972 | 0.888 | -2.192 | -0.503 | -0.753 | -0.088 | -0.112 | -0.112 | 0 | 0.32 | 0.07 | 0.46 | 0.45 | 0.41 | 0.58 |
| 7 | 16060201 | rs11761043 | -0.773 | -1.695 | 0.412 | -1.276 | 0.002 | -0.02 | -0.02 | 0 | 0.59 | 0.42 | 0.6 | 0.76 | 0.56 | 0.73 |
| 7 | 16060334 | rs10950597 | -0.403 | -1.601 | 0.452 | -1.17 | 0.009 | 0.055 | 0.055 | 0 | 0.49 | 0.36 | 0.51 | 0.75 | 0.46 | 0.67 |
| 7 | 16062239 | rs1528144 | 2.171 | -1.608 | -0.092 | -1.119 | 0.003 | 0.001 | 0.001 | 0 | 0.09 | 0.03 | 0.17 | 0.1 | 0.08 | 0.35 |
| 7 | 16062958 | rs1528146 | 0.203 | -1.493 | 0.293 | -1.199 | 0.008 | 0.035 | 0.035 | 0 | 0.5 | 0.46 | 0.47 | 0.24 | 0.54 | 0.33 |
| 7 | 16076502 | rs6461217 | -0.296 | -1.148 | -0.183 | -0.97 | -0.001 | -0.016 | -0.016 | 0 | 0.53 | 0.47 | 0.62 | 0.84 | 0.52 | 0.72 |
| 7 | 16085019 | rs13231687 | -0.167 | -1.216 | -0.375 | -0.947 | 0.005 | 0.097 | 0.097 | 0 | 0.52 | 0.36 | 0.61 | 0.84 | 0.53 | 0.74 |
| 7 | 16089752 | rs9648211 | -0.045 | -1.14 | -0.818 | -0.638 | -0.048 | 0.083 | 0.083 | 0.039 | 0.4 | 0.13 | 0.47 | 0.72 | 0.42 | 0.66 |
| 7 | 16100729 | rs1881241 | -0.525 | -0.861 | 0.124 | -0.106 | -0.022 | 0.055 | 0.055 | 0.022 | 0.81 | 0.51 | 0.8 | 0.74 | 0.85 | 0.83 |
| 7 | 16103718 | rs17284526 | 1.437 | -0.406 | -0.731 | -0.053 | -0.074 | 0.019 | 0.019 | 0.049 | 0.2 | 0.01 | 0.24 | 0.14 | 0.23 | 0.46 |
| 7 | 16104086 | rs6959719 | 0.059 | -0.58 | -0.746 | 0.349 | -0.063 | -0.008 | -0.008 | 0.069 | 0.4 | 0.15 | 0.44 | 0.34 | 0.46 | 0.62 |
| 7 | 16107358 | rs1528149 | 1.549 | -0.707 | -0.963 | 0.233 | -0.088 | 0.24 | 0.24 | 1.384 | 0.35 | 0.04 | 0.35 | 0.07 | 0.38 | 0.15 |
| 7 | 16115114 | rs10244722 | 2.092 | -0.595 | -0.596 | 0.384 | 0.002 | -0.004 | -0.004 | 0 | 0.07 | 0.06 | 0.1 | 0.12 | 0.06 | 0.06 |

| | | | | | | | | | | | | | | | | |
|---|---|---|---|---|---|---|---|---|---|---|---|---|---|---|---|---|
| 7 | 16119502 | rs12537658 | -0.048 | -0.569 | -0.44 | 0.23 | 0.001 | -0.003 | -0.003 | 0 | 0.39 | 0.38 | 0.42 | 0.65 | 0.43 | 0.66 |
| 7 | 16124957 | rs11773181 | 0.121 | -1.179 | -1.138 | 0.323 | 0.051 | -0.017 | -0.017 | 0 | 0.71 | 0.11 | 0.75 | 0.71 | 0.81 | 0.8 |
| 7 | 16127297 | rs12539174 | 0.575 | -1.054 | -1.19 | 0.538 | -0.098 | -0.039 | -0.039 | 0 | 0.32 | 0.07 | 0.38 | 0.31 | 0.41 | 0.6 |
| 7 | 16133713 | rs17169263 | 0.37 | -1.186 | -1.3 | 0.486 | 0.088 | -0.097 | -0.097 | 0.044 | 0.68 | 0.1 | 0.76 | 0.41 | 0.83 | 0.74 |
| 7 | 16140077 | rs3857708 | 2.418 | -1.198 | -1.323 | 0.292 | -0.115 | 0.145 | 0.145 | 3.589 | 0.26 | 0.02 | 0.3 | 0.07 | 0.33 | 0.1 |
| 7 | 16144157 | rs10950603 | 1.182 | -0.716 | -0.719 | 0.925 | -0.072 | -0.073 | -0.073 | 0.055 | 0.13 | 0.03 | 0.16 | 0.18 | 0.2 | 0.14 |
| 7 | 16149832 | rs11773405 | 1.461 | -0.677 | -0.703 | 0.541 | -0.057 | 0.003 | 0.003 | 0.767 | 0.2 | 0.02 | 0.23 | 0.16 | 0.22 | 0.46 |
| 7 | 16159827 | rs11760717 | 0.478 | -0.877 | -0.149 | 0.575 | -0.045 | 0.014 | 0.014 | 0.083 | 0.36 | 0.19 | 0.4 | 0.63 | 0.42 | 0.64 |
| 7 | 16163218 | rs4721475 | -0.319 | -0.985 | 0.445 | 0.529 | 0 | -0.024 | -0.024 | 0.019 | 0.45 | 0.47 | 0.52 | 0.77 | 0.47 | 0.72 |
| 7 | 16168502 | rs10269890 | 1.015 | -1.004 | 0.763 | 0.55 | -0.025 | 0.019 | 0.019 | 0 | 0.1 | 0.22 | 0.11 | 0.12 | 0.05 | 0.06 |
| 7 | 16168708 | rs6968537 | -0.421 | -0.959 | 0.651 | 0.51 | -0.001 | -0.018 | -0.018 | 0 | 0.46 | 0.53 | 0.53 | 0.75 | 0.47 | 0.72 |
| 7 | 16209097 | rs17169296 | 1.842 | -0.94 | 0.36 | 0.721 | 0.009 | -0.012 | -0.012 | 0 | 0.08 | 0.06 | 0.1 | 0.13 | 0.06 | 0.09 |
| 7 | 16209427 | rs1527206 | -0.367 | -0.919 | 0.447 | 0.634 | -0.001 | 0 | 0 | 0 | 0.45 | 0.46 | 0.51 | 0.75 | 0.46 | 0.72 |
| 7 | 16211553 | rs1012401 | -1.164 | -1.098 | 0.643 | 0.482 | -0.028 | -0.01 | -0.01 | 0 | 0.83 | 0.59 | 0.9 | 0.84 | 0.9 | 0.87 |
| 7 | 16211816 | rs1012402 | -1.382 | -1.11 | 0.828 | 0.476 | -0.021 | -0.007 | -0.007 | 0 | 0.84 | 0.66 | 0.91 | 0.84 | 0.9 | 0.87 |
| 7 | 16213150 | rs1527199 | 0.063 | -1.013 | 0.389 | 0.688 | -0.006 | -0.003 | -0.003 | 0 | 0.42 | 0.38 | 0.49 | 0.42 | 0.48 | 0.65 |
| 7 | 16218633 | rs17367660 | 1.335 | -1.107 | 0.131 | 0.411 | -0.084 | 0.194 | 0.194 | 1.038 | 0.35 | 0.08 | 0.38 | 0.06 | 0.43 | 0.15 |
| 7 | 16225174 | rs7809306 | 1.107 | -0.94 | 0.096 | 0.633 | -0.022 | -0.112 | -0.112 | 0.381 | 0.27 | 0.1 | 0.28 | 0.51 | 0.3 | 0.52 |
| 7 | 16229021 | rs10487915 | 1.446 | -0.985 | 0.093 | 0.52 | -0.059 | 0.145 | 0.145 | 2.132 | 0.35 | 0.13 | 0.38 | 0.09 | 0.43 | 0.15 |
| 7 | 16233634 | rs12668400 | 0.897 | -0.922 | 0.216 | 0.629 | 0.001 | -0.02 | -0.02 | 0 | 0.29 | 0.25 | 0.28 | 0.51 | 0.3 | 0.52 |
| 7 | 16238691 | rs6461229 | 0.971 | -0.91 | 0.196 | 0.687 | 0 | -0.032 | -0.032 | 0 | 0.28 | 0.23 | 0.28 | 0.52 | 0.29 | 0.52 |
| 7 | 16242073 | rs1527226 | 0.101 | -0.934 | 0.512 | 0.711 | -0.007 | 0.004 | 0.004 | 0 | 0.73 | 0.6 | 0.78 | 0.74 | 0.77 | 0.76 |
| 7 | 16245357 | rs1918259 | 1.222 | -0.972 | 0.514 | 0.748 | -0.011 | 0.002 | 0.002 | 1.927 | 0.36 | 0.3 | 0.39 | 0.06 | 0.42 | 0.15 |
| 7 | 16245752 | rs1527219 | 0.82 | -0.985 | 0.598 | 0.768 | 0.004 | 0.002 | 0.002 | 3.684 | 0.4 | 0.48 | 0.42 | 0.38 | 0.43 | 0.22 |
| 7 | 16251425 | rs1527222 | 1.124 | -0.705 | 0.428 | 0.905 | -0.01 | 0.004 | 0.004 | 0 | 0.09 | 0.11 | 0.11 | 0.13 | 0.06 | 0.1 |
| 7 | 16262371 | rs12216621 | 1.433 | -0.624 | 0.121 | 1.201 | -0.052 | 0.02 | 0.02 | 0.68 | 0.23 | 0.02 | 0.24 | 0.21 | 0.26 | 0.45 |
| 7 | 16270332 | rs6956986 | 1.18 | -0.746 | 0.328 | 1.144 | -0.005 | 0.009 | 0.009 | 4.726 | 0.38 | 0.37 | 0.45 | 0.18 | 0.44 | 0.16 |
| 7 | 16271635 | rs1918265 | 1.007 | -0.769 | 0.019 | 1.24 | -0.003 | -0.052 | -0.052 | 0 | 0.26 | 0.13 | 0.26 | 0.42 | 0.27 | 0.52 |
| 7 | 16274120 | rs6974124 | 1.228 | -0.648 | 0.114 | 1.139 | -0.042 | 0.076 | 0.076 | 4.105 | 0.37 | 0.17 | 0.43 | 0.18 | 0.44 | 0.16 |
| 7 | 16284326 | rs10486791 | -0.362 | -0.666 | 0.594 | 1.269 | 0.009 | 0.033 | 0.033 | 0 | 0.19 | 0.32 | 0.19 | 0.25 | 0.19 | 0.2 |
| 7 | 16284902 | rs12532139 | 0.862 | -0.609 | 0.556 | 1.419 | -0.037 | -0.043 | -0.043 | 2.754 | 0.09 | 0.03 | 0.12 | 0.2 | 0.14 | 0.11 |
| 7 | 16297326 | rs9918580 | 0.204 | -0.758 | 0.554 | 1.196 | 0.018 | -0.011 | -0.011 | 0 | 0.55 | 0.35 | 0.62 | 0.44 | 0.62 | 0.36 |
| 7 | 16297549 | rs1295154 | -1.717 | -0.783 | 0.275 | 1.237 | -0.052 | 0.025 | 0.025 | 0.895 | 0.77 | 0.98 | 0.76 | 0.8 | 0.73 | 0.55 |
| 7 | 16301026 | rs17459061 | 0.426 | -0.652 | 0.114 | 1.115 | -0.064 | -0.025 | -0.025 | 1.149 | 0.14 | 0.03 | 0.15 | 0.11 | 0.2 | 0.14 |
| 7 | 16301850 | rs17459166 | 1.017 | -0.733 | 0.479 | 0.944 | 0 | 0.004 | 0.004 | 2.946 | 0.41 | 0.39 | 0.49 | 0.33 | 0.42 | 0.23 |
| 7 | 16304817 | rs1295143 | 0.466 | -0.865 | 0.708 | 1.14 | -0.038 | -0.024 | -0.024 | 0.024 | 0.18 | 0.42 | 0.1 | 0.15 | 0.1 | 0.13 |
| 7 | 16306703 | rs6954901 | 0.79 | -0.804 | 0.652 | 1.111 | -0.008 | 0.03 | 0.03 | 1.719 | 0.46 | 0.33 | 0.55 | 0.33 | 0.48 | 0.25 |
| 7 | 16310888 | rs10247201 | 0.907 | -0.833 | 0.527 | 1.1 | -0.025 | 0.015 | 0.015 | 0 | 0.1 | 0.21 | 0.11 | 0.13 | 0.05 | 0.1 |
| 7 | 16311996 | rs10280613 | 1.06 | -0.845 | 0.357 | 1.095 | -0.017 | 0.005 | 0.005 | 0 | 0.1 | 0.13 | 0.1 | 0.14 | 0.05 | 0.1 |
| 7 | 16312057 | rs11766565 | 0.542 | -0.846 | 0.057 | 1.198 | 0.024 | 0.029 | 0.029 | 4.035 | 0.48 | 0.35 | 0.55 | 0.52 | 0.57 | 0.31 |
| 7 | 16312781 | rs1295133 | 0.671 | -0.528 | -0.407 | 1.154 | -0.016 | -0.012 | -0.012 | 0 | 0.17 | 0.24 | 0.09 | 0.14 | 0.1 | 0.12 |
| 7 | 16313122 | rs1295132 | -0.142 | 0.063 | -1.012 | 1.314 | -0.078 | -0.086 | -0.086 | 0.942 | 0.65 | 0.91 | 0.65 | 0.46 | 0.58 | 0.37 |
| 7 | 16319042 | rs12531451 | -0.498 | 0.097 | -0.877 | 1.237 | -0.069 | -0.1 | -0.1 | 0 | 0.19 | 0.06 | 0.21 | 0.38 | 0.27 | 0.21 |
| 7 | 16321111 | rs10282434 | -0.485 | -0.33 | -0.585 | 0.761 | -0.025 | -0.029 | -0.029 | 0.796 | 0.62 | 0.55 | 0.73 | 0.77 | 0.72 | 0.45 |
| 7 | 16322506 | rs12666960 | 1.066 | -0.186 | -0.701 | 0.833 | -0.018 | -0.046 | -0.046 | 0 | 0.09 | 0.02 | 0.07 | 0.25 | 0.12 | 0.09 |
| 7 | 16323312 | rs12539671 | -0.223 | -0.204 | -0.682 | 0.812 | -0.069 | -0.063 | -0.063 | 0 | 0.19 | 0.05 | 0.21 | 0.29 | 0.27 | 0.2 |
| 7 | 16326646 | rs35681285 | 1.438 | -0.441 | -0.35 | 0.686 | -0.023 | -0.042 | -0.042 | 0 | 0.1 | 0.2 | 0.06 | 0.02 | 0.04 | 0.04 |
| 7 | 16342982 | rs2389594 | -1.126 | -0.382 | -0.678 | 0.77 | 0.018 | 0.016 | 0.016 | 0 | 0.72 | 0.8 | 0.79 | 0.79 | 0.76 | 0.49 |
| 7 | 16346213 | rs6975353 | -1.343 | -0.385 | -1.066 | 0.769 | -0.009 | -0.012 | -0.012 | 0.16 | 0.74 | 0.99 | 0.82 | 0.8 | 0.77 | 0.51 |
| 7 | 16348830 | rs17169417 | 0.7 | -0.164 | -0.6 | 0.877 | -0.011 | -0.034 | -0.034 | 0.195 | 0.11 | 0.04 | 0.11 | 0.28 | 0.12 | 0.14 |
| 7 | 16351845 | rs2189733 | 2.018 | -0.102 | -1.016 | 0.541 | -0.081 | 0.045 | 0.045 | 0.126 | 0.21 | 0.04 | 0.31 | 0.1 | 0.31 | 0.1 |
| 7 | 16357305 | rs10950606 | 1.315 | -0.008 | -1.02 | 0.456 | -0.033 | 0.03 | 0.03 | 0 | 0.08 | 0.03 | 0.1 | 0 | 0.15 | 0.06 |
| 7 | 16358315 | rs984810 | 0.758 | -0.924 | -0.409 | 0.659 | 0.001 | -0.002 | -0.002 | 0 | 0.69 | 0.31 | 0.66 | 0.55 | 0.8 | 0.74 |
| 7 | 16360570 | rs6968164 | 0.258 | -0.875 | -0.193 | 0.738 | 0 | 0.034 | 0.034 | 0 | 0.48 | 0.34 | 0.35 | 0.37 | 0.49 | 0.65 |
| 7 | 16366339 | rs2158485 | -1.296 | -0.923 | 0.21 | 0.776 | -0.01 | 0.018 | 0.018 | 0 | 0.93 | 0.84 | 0.86 | 0.85 | 0.96 | 0.93 |
| 7 | 16366674 | rs11761604 | 0.618 | -1.048 | 0.157 | 0.698 | -0.028 | 0.034 | 0.034 | 0 | 0.24 | 0.13 | 0.24 | 0.15 | 0.29 | 0.18 |
| 7 | 16367282 | rs6973038 | 0.192 | -1.036 | 0.152 | 0.727 | 0 | 0.033 | 0.033 | 0 | 0.48 | 0.34 | 0.35 | 0.36 | 0.49 | 0.64 |
| 7 | 16370188 | rs7784596 | -0.26 | -1.07 | 0.381 | 0.706 | 0 | -0.001 | -0.001 | 0 | 0.59 | 0.55 | 0.41 | 0.38 | 0.53 | 0.68 |
| 7 | 16375281 | rs6949613 | -1.263 | -1.031 | 0.411 | 0.686 | -0.01 | 0.015 | 0.015 | 0 | 0.92 | 0.84 | 0.86 | 0.85 | 0.96 | 0.93 |
| 7 | 16377743 | rs1921846 | -0.046 | -1.073 | 0.371 | 0.641 | 0.002 | -0.063 | -0.063 | 0 | 0.36 | 0.48 | 0.3 | 0.17 | 0.33 | 0.22 |
| 7 | 16381366 | rs2526604 | -0.792 | -1.078 | 0.347 | 0.717 | -0.004 | 0.002 | 0.002 | 0 | 0.46 | 0.55 | 0.42 | 0.45 | 0.45 | 0.36 |
| 7 | 16383169 | rs2526600 | 1.07 | -1.006 | 0.346 | 0.752 | 0.006 | -0.019 | -0.019 | 0 | 0.31 | 0.42 | 0.47 | 0.24 | 0.35 | 0.17 |
| 7 | 16383860 | rs4537216 | 1.636 | -0.973 | 0.384 | 0.629 | 0 | -0.007 | -0.007 | 0 | 0.25 | 0.3 | 0.34 | 0.09 | 0.31 | 0.1 |
| 7 | 16384664 | rs2107587 | -0.521 | -0.992 | 0.24 | 0.726 | 0 | 0.002 | 0.002 | 0 | 0.62 | 0.38 | 0.53 | 0.75 | 0.63 | 0.8 |
| 7 | 16386494 | rs4146016 | -0.043 | -1.012 | 0.239 | 0.608 | -0.003 | -0.001 | -0.001 | 0 | 0.54 | 0.36 | 0.48 | 0.51 | 0.53 | 0.71 |
| 7 | 16388295 | rs12666868 | 1.14 | -1.001 | 0.197 | 0.66 | -0.011 | -0.019 | -0.019 | 0 | 0.11 | 0.04 | 0.12 | 0.38 | 0.12 | 0.15 |
| 7 | 16388624 | rs4615457 | 1.141 | -1.003 | 0.196 | 0.659 | -0.011 | -0.019 | -0.019 | 0 | 0.11 | 0.04 | 0.12 | 0.38 | 0.12 | 0.15 |
| 7 | 16388904 | rs7810150 | -0.099 | -1.014 | 0.343 | 0.636 | -0.003 | 0 | 0 | 0 | 0.54 | 0.39 | 0.48 | 0.51 | 0.53 | 0.71 |
| 7 | 16389412 | rs4565368 | 0.883 | -0.882 | 0.07 | 0.707 | 0.013 | -0.005 | -0.005 | 0 | 0.29 | 0.23 | 0.24 | 0.36 | 0.25 | 0.53 |

| | | | | | | | | | | | | | | | | |
|---|---|---|---|---|---|---|---|---|---|---|---|---|---|---|---|---|
| 7 | 16399003 | rs2222862 | 0.485 | -0.792 | 0.089 | 0.751 | -0.004 | -0.002 | -0.002 | 2.244 | 0.35 | 0.36 | 0.46 | 0.38 | 0.44 | 0.24 |
| 7 | 16399822 | rs10950616 | 0.391 | -1.306 | 0.177 | 1.181 | -0.011 | -0.019 | -0.019 | 0 | 0.11 | 0.04 | 0.12 | 0.38 | 0.12 | 0.14 |
| 7 | 16401190 | rs7789712 | -0.677 | -1.296 | 0.497 | 0.605 | -0.027 | -0.041 | -0.041 | 0 | 0.89 | 0.82 | 0.94 | 0.98 | 0.96 | 0.96 |
| 7 | 16406272 | rs12699786 | 0.273 | -0.968 | 0.352 | 1.012 | 0.013 | -0.001 | -0.001 | 0.139 | 0.23 | 0.17 | 0.11 | 0.22 | 0.2 | 0.46 |
| 7 | 16408071 | rs12671637 | -0.63 | -0.379 | 0.966 | 1.197 | 0.003 | 0.011 | 0.011 | 0 | 0.16 | 0.12 | 0.07 | 0.08 | 0.11 | 0.26 |
| 7 | 16431594 | rs11972185 | -1.128 | -0.679 | 1.612 | 1.452 | -0.048 | 0.038 | 0.038 | 0.748 | 0.19 | 0.58 | 0.18 | 0.3 | 0.07 | 0.17 |
| 7 | 16436708 | rs17169453 | -0.249 | -0.702 | 1.468 | 1.42 | -0.014 | 0.012 | 0.012 | 0 | 0.16 | 0.04 | 0.07 | 0.08 | 0.11 | 0.26 |
| 7 | 16437070 | rs4351323 | -0.713 | -0.74 | 1.935 | 1.429 | -0.045 | 0.107 | 0.107 | 0.16 | 0.17 | 0.4 | 0.17 | 0.3 | 0.07 | 0.17 |
| 7 | 16437330 | rs17617900 | -1.258 | -0.177 | 1.821 | 1.579 | -0.008 | -0.006 | -0.006 | 0.168 | 0.26 | 0.22 | 0.32 | 0.26 | 0.31 | 0.38 |
| 7 | 16437810 | rs6976548 | -2.174 | -0.175 | 1.702 | 1.579 | -0.004 | 0.004 | 0.004 | 0 | 0.43 | 0.69 | 0.5 | 0.56 | 0.38 | 0.55 |
| 7 | 16438182 | rs6461247 | 0.532 | -0.149 | 1.708 | 1.583 | -0.011 | -0.009 | -0.009 | 0 | 0.08 | 0.04 | 0.25 | 0.17 | 0.2 | 0.11 |
| 7 | 16438357 | rs6461248 | -2.061 | -0.304 | 1.588 | 1.497 | -0.009 | 0.002 | 0.002 | 0 | 0.43 | 0.75 | 0.5 | 0.56 | 0.38 | 0.55 |
| 7 | 16439560 | rs11974818 | -0.219 | -0.292 | 1.554 | 1.471 | 0.003 | 0.014 | 0.014 | 0.017 | 0.16 | 0.1 | 0.07 | 0.08 | 0.11 | 0.26 |
| 7 | 16440023 | rs6949292 | -1.499 | -0.19 | 1.444 | 1.464 | -0.005 | 0.001 | 0.001 | 0 | 0.42 | 0.69 | 0.5 | 0.56 | 0.38 | 0.55 |
| 7 | 16469491 | rs10240752 | -1.596 | -0.093 | 0.528 | 1.097 | -0.015 | -0.033 | -0.033 | 0 | 0.44 | 0.84 | 0.51 | 0.45 | 0.38 | 0.54 |
| 7 | 16471712 | rs10249569 | -1.207 | -0.807 | 0.926 | 0.881 | -0.026 | -0.004 | -0.004 | 0 | 0.31 | 0.6 | 0.13 | 0.25 | 0.14 | 0.38 |
| 7 | 16484541 | rs1921835 | 0.576 | 0.058 | -0.302 | 0.644 | 0.001 | 0.038 | 0.038 | 0 | 0.9 | 0.96 | 0.95 | 1 | 0.94 | 0.91 |
| 7 | 16490580 | rs10240946 | -0.566 | -0.301 | 0.236 | 0.762 | -0.019 | 0.02 | 0.02 | 0 | 0.08 | 0.16 | 0.03 | 0.11 | 0.02 | 0.08 |
| 7 | 16494407 | rs4721503 | 1.669 | -0.905 | 0.198 | 0.469 | -0.01 | -0.013 | -0.013 | 0 | 0.8 | 0.77 | 0.93 | 0.9 | 0.91 | 0.84 |
| 7 | 16500142 | rs10270805 | 0.678 | -0.9 | 0.147 | 0.455 | 0.029 | 0.063 | 0.063 | 0.472 | 0.85 | 0.94 | 0.94 | 1 | 0.93 | 0.87 |
| 7 | 16500884 | rs12699798 | 1.97 | -0.68 | 0.486 | 0.879 | 0.01 | 0.019 | 0.019 | 0 | 0.55 | 0.35 | 0.56 | 0.43 | 0.6 | 0.42 |
| 7 | 16503724 | rs17619858 | -0.445 | -0.687 | 0.225 | 1.508 | -0.032 | -0.027 | -0.027 | 0.263 | 0.15 | 0.07 | 0.27 | 0.18 | 0.25 | 0.26 |
| 7 | 16507247 | rs2528477 | -0.38 | -0.734 | 0.208 | 1.442 | 0.039 | 0.09 | 0.09 | 0.398 | 0.17 | 0.02 | 0.05 | 0.02 | 0.07 | 0.18 |
| 7 | 16509086 | rs1981596 | 0.185 | -0.687 | 0.789 | 1.59 | -0.024 | 0.161 | 0.161 | 0.106 | 0.1 | 0.19 | 0.07 | 0.29 | 0.05 | 0.16 |
| 7 | 16514246 | rs2528474 | -1.734 | -0.605 | 0.44 | 1.643 | -0.019 | -0.006 | -0.006 | 0.152 | 0.31 | 0.15 | 0.34 | 0.28 | 0.34 | 0.42 |
| 7 | 16518195 | rs2528472 | -2.496 | -0.666 | 0.724 | 1.572 | 0.014 | 0.03 | 0.03 | 0 | 0.44 | 0.77 | 0.44 | 0.52 | 0.39 | 0.57 |
| 7 | 16518247 | rs2528471 | -1.664 | -0.361 | 0.428 | 1.401 | -0.045 | -0.006 | -0.006 | 0 | 0.29 | 0.09 | 0.33 | 0.22 | 0.34 | 0.41 |
| 7 | 16519510 | rs2528468 | -0.251 | -0.409 | 0.371 | 1.301 | 0.024 | 0.077 | 0.077 | 0.27 | 0.16 | 0.01 | 0.04 | 0.02 | 0.07 | 0.18 |
| 7 | 16519589 | rs2528467 | -1.161 | -0.326 | 0.401 | 1.374 | -0.046 | -0.001 | -0.001 | 0 | 0.26 | 0.08 | 0.29 | 0.18 | 0.28 | 0.35 |
| 7 | 16520193 | rs2704668 | -1.757 | -0.528 | 0.97 | 1.615 | -0.001 | 0.002 | 0.002 | 0 | 0.38 | 0.35 | 0.43 | 0.48 | 0.37 | 0.55 |
| 7 | 16520887 | rs2528465 | -1.458 | 0.18 | 0.896 | 1.653 | -0.011 | -0.002 | -0.002 | 0 | 0.4 | 0.31 | 0.45 | 0.32 | 0.41 | 0.58 |
| 7 | 16521647 | rs11770436 | 1.923 | 0.045 | 0.245 | 1.155 | 0.02 | -0.011 | -0.011 | 3.96 | 0.41 | 0.22 | 0.26 | 0.48 | 0.38 | 0.28 |
| 7 | 16526893 | rs2704674 | 0.993 | 0.089 | 0.234 | 1.096 | 0.03 | 0.031 | 0.031 | 0.262 | 0.64 | 0.53 | 0.5 | 0.58 | 0.51 | 0.47 |
| 7 | 16529037 | rs6461258 | 1.616 | -0.082 | 0.154 | 1.022 | 0.005 | 0 | 0 | 2.175 | 0.49 | 0.54 | 0.45 | 0.58 | 0.44 | 0.34 |
| 7 | 16531768 | rs2158794 | -0.451 | 0.058 | 0.187 | 1.109 | 0.004 | 0.013 | 0.013 | 0 | 0.91 | 0.87 | 0.94 | 0.86 | 0.88 | 0.93 |
| 7 | 16531836 | rs2158793 | 1.45 | 0.238 | 0.147 | 1.253 | 0.001 | -0.001 | -0.001 | 0.278 | 0.59 | 0.57 | 0.65 | 0.64 | 0.52 | 0.49 |
| 7 | 16532088 | rs1990130 | 1.081 | 0.288 | -0.111 | 1.161 | 0.001 | -0.02 | -0.02 | 0 | 0.09 | 0.03 | 0.2 | 0.07 | 0.09 | 0.15 |
| 7 | 16534157 | rs11980729 | -0.903 | 0.367 | 0.012 | 1.084 | -0.006 | 0.004 | 0.004 | 0 | 0.32 | 0.3 | 0.28 | 0.22 | 0.36 | 0.45 |
| 7 | 16538712 | rs7794899 | -0.221 | 0.304 | -0.107 | 1.051 | 0.004 | 0.117 | 0.117 | 0 | 0.79 | 0.98 | 0.9 | 1 | 0.83 | 0.83 |
| 7 | 16541388 | rs917221 | 1.132 | 0.192 | 0.499 | 1.168 | -0.013 | 0.054 | 0.054 | 0 | 0.56 | 0.32 | 0.63 | 0.79 | 0.53 | 0.49 |
| 7 | 16542364 | rs9639262 | -1.249 | 0.388 | 0.843 | 1.473 | 0.085 | 0.492 | 0.492 | 0.855 | 0.07 | 0.42 | 0.05 | 0.36 | 0.05 | 0.08 |
| 7 | 16544872 | rs886757 | 0.666 | -0.095 | -0.502 | 0.01 | -0.028 | 0.059 | 0.059 | 0 | 0.3 | 0.16 | 0.4 | 0.22 | 0.36 | 0.21 |
| 7 | 16550888 | rs17169520 | 0.319 | 0.204 | -0.29 | -0.194 | 0.024 | -0.024 | -0.024 | 0.187 | 0.25 | 0.39 | 0.37 | 0.24 | 0.29 | 0.23 |
| 7 | 16552572 | rs10242167 | -0.103 | -0.109 | -1.377 | -0.28 | -0.024 | -0.003 | -0.003 | 0 | 0.73 | 0.96 | 0.7 | 0.79 | 0.75 | 0.71 |
| 7 | 16555274 | rs7804057 | 1.345 | -0.094 | -0.988 | 0.04 | -0.021 | -0.062 | -0.062 | 0 | 0.28 | 0.15 | 0.22 | 0.41 | 0.31 | 0.39 |
| 7 | 16557858 | rs4721511 | 0.897 | -0.235 | -1.134 | -0.11 | 0.008 | -0.042 | -0.042 | 0.007 | 0.12 | 0.03 | 0.16 | 0.2 | 0.1 | 0.17 |
| 7 | 16560564 | rs7797565 | 0.128 | -0.396 | -0.88 | -0.358 | 0.019 | 0.05 | 0.05 | 0 | 0.65 | 0.83 | 0.66 | 0.8 | 0.71 | 0.63 |
| 7 | 16561687 | rs2190310 | -0.029 | -0.522 | -0.701 | -0.533 | -0.023 | 0.116 | 0.116 | 1.258 | 0.54 | 0.75 | 0.47 | 0.75 | 0.52 | 0.56 |
| 7 | 16564431 | rs11765776 | 1.073 | -0.236 | -0.674 | -0.347 | -0.018 | 0.006 | 0.006 | 0 | 0.22 | 0.16 | 0.34 | 0.25 | 0.29 | 0.25 |
| 7 | 16566391 | rs6461263 | -0.359 | -0.437 | 0.121 | -0.474 | -0.016 | -0.111 | -0.111 | 0 | 0.7 | 0.64 | 0.78 | 1 | 0.78 | 0.74 |
| 7 | 16567628 | rs6974206 | 0.258 | -0.291 | 0.059 | -0.207 | -0.005 | -0.009 | -0.009 | 0.152 | 0.33 | 0.3 | 0.48 | 0.45 | 0.36 | 0.38 |
| 7 | 16572051 | rs1404171 | -0.398 | -0.129 | -0.078 | -0.155 | -0.003 | -0.002 | -0.002 | 0 | 0.43 | 0.41 | 0.54 | 0.6 | 0.48 | 0.4 |
| 7 | 16573508 | rs886752 | -0.354 | -0.024 | -0.284 | 0.152 | 0.005 | 0.037 | 0.037 | 0.993 | 0.52 | 0.39 | 0.58 | 0.86 | 0.56 | 0.56 |
| 7 | 16573947 | rs17169536 | 1.846 | -0.057 | -0.917 | 0.201 | -0.053 | -0.099 | -0.099 | 0 | 0.25 | 0.05 | 0.22 | 0.4 | 0.3 | 0.33 |
| 7 | 16576251 | rs12671061 | 1.71 | 0.042 | -0.832 | 0.323 | 0.01 | -0.037 | -0.037 | 0 | 0.11 | 0.03 | 0.17 | 0.2 | 0.08 | 0.16 |
| 7 | 16576978 | rs12669350 | 0.147 | 0.052 | -0.849 | 0.315 | -0.004 | -0.079 | -0.079 | 0 | 0.19 | 0.07 | 0.2 | 0.35 | 0.2 | 0.18 |
| 7 | 16590306 | rs6942710 | 0.277 | 0.413 | -0.284 | 0.401 | 0 | 0 | 0 | 0 | 0.37 | 0.4 | 0.3 | 0.55 | 0.42 | 0.35 |
| 7 | 16590696 | rs10224153 | 0.824 | 0.234 | -0.524 | 0.367 | -0.002 | -0.009 | -0.009 | 0 | 0.09 | 0.1 | 0.06 | 0.15 | 0.12 | 0.03 |
| 7 | 16592915 | rs518011 | -0.947 | 0.637 | -0.68 | 0.702 | -0.02 | -0.033 | -0.033 | 0.267 | 0.72 | 0.83 | 0.65 | 0.54 | 0.64 | 0.65 |
| 7 | 16592993 | rs477644 | -0.435 | 0.093 | -0.683 | 0.507 | 0.041 | 0.04 | 0.04 | 0 | 0.5 | 0.67 | 0.41 | 0.35 | 0.36 | 0.41 |
| 7 | 16594454 | rs17169541 | 0.756 | 0.249 | -0.921 | 1.008 | -0.013 | -0.066 | -0.066 | 0 | 0.12 | 0.04 | 0.29 | 0.35 | 0.12 | 0.19 |
| 7 | 16596858 | rs10256540 | 0.12 | 0.18 | -0.466 | 0.854 | 0 | 0.042 | 0.042 | 0 | 0.06 | 0.13 | 0.06 | 0.14 | 0.08 | 0.03 |
| 7 | 16599762 | rs7780726 | 0.401 | -0.258 | -0.186 | -0.073 | -0.03 | -0.143 | -0.143 | 0.543 | 0.27 | 0.36 | 0.13 | 0 | 0.19 | 0.27 |
| 7 | 16602243 | rs7810696 | -0.307 | 0.277 | -0.659 | 0.148 | 0.014 | 0.074 | 0.074 | 0 | 0.5 | 0.57 | 0.32 | 0.2 | 0.46 | 0.53 |
| 7 | 16603225 | rs2389674 | -0.228 | -0.124 | -0.671 | 0.306 | -0.006 | -0.003 | -0.003 | 0 | 0.57 | 0.57 | 0.58 | 0.55 | 0.49 | 0.73 |
| 7 | 16605357 | rs706021 | -0.161 | -0.172 | -0.552 | 0.372 | -0.018 | -0.018 | -0.018 | 0.044 | 0.6 | 0.36 | 0.51 | 0.52 | 0.48 | 0.68 |
| 7 | 16607871 | rs520485 | -0.544 | -0.27 | -1.07 | 0.274 | -0.129 | 0.081 | 0.081 | 0 | 0.66 | 0.91 | 0.59 | 0.83 | 0.54 | 0.67 |
| 7 | 16615490 | rs569022 | -0.137 | -0.712 | -0.361 | 0.377 | 0.04 | 0.044 | 0.044 | 0.042 | 0.42 | 0.33 | 0.2 | 0.21 | 0.27 | 0.46 |
| 7 | 16620928 | rs562687 | -0.579 | -0.607 | -0.299 | 0.275 | 0.068 | 0.01 | 0.01 | 0 | 0.49 | 0.69 | 0.28 | 0.51 | 0.34 | 0.49 |

| | | | | | | | | | | | | | | | | |
|---|---|---|---|---|---|---|---|---|---|---|---|---|---|---|---|---|
| 7 | 16622785 | rs541415 | -0.393 | -0.873 | -0.347 | 0.342 | -0.062 | 0.027 | 0.027 | 0 | 0.34 | 0.49 | 0.17 | 0.39 | 0.18 | 0.38 |
| 7 | 16627015 | rs6942828 | 0.624 | 0.606 | -0.587 | -0.07 | -0.003 | 0.005 | 0.005 | 0 | 0.79 | 0.81 | 0.63 | 0.81 | 0.76 | 0.81 |
| 7 | 16639379 | rs4721517 | 0.008 | 0.548 | -0.514 | -0.178 | -0.001 | 0 | 0 | 0 | 0.14 | 0.13 | 0.35 | 0.1 | 0.17 | 0.13 |
| 7 | 16640300 | rs1044983 | 0.721 | 1.104 | -0.252 | -0.018 | 0.05 | 0.193 | 0.193 | 0 | 0.18 | 0.36 | 0.22 | 0.48 | 0.22 | 0.32 |
| 7 | 16640358 | rs543387 | -0.771 | 1.191 | -0.732 | -0.29 | 0.016 | 0.001 | 0.001 | 0.175 | 0.86 | 0.84 | 0.84 | 0.84 | 0.72 | 0.91 |
| 7 | 16641749 | rs523615 | 0.266 | 1.007 | -0.692 | -0.211 | 0.017 | 0.017 | 0.017 | 0.685 | 0.34 | 0.49 | 0.57 | 0.58 | 0.38 | 0.46 |
| 7 | 16647832 | rs13244745 | -0.236 | 0.979 | -0.895 | 0.37 | -0.002 | 0.001 | 0.001 | 0.055 | 0.18 | 0.2 | 0.15 | 0.25 | 0.14 | 0.12 |
| 7 | 16652523 | rs2177539 | -0.134 | 1.227 | -1.315 | 0.326 | -0.01 | -0.001 | -0.001 | 0.049 | 0.16 | 0.13 | 0.16 | 0.17 | 0.26 | 0.13 |
| 7 | 16653608 | rs11765150 | -0.422 | 0.086 | -1.237 | 0.28 | 0.002 | -0.015 | -0.015 | 0.596 | 0.17 | 0.09 | 0.14 | 0.24 | 0.14 | 0.1 |
| 7 | 16654334 | rs10272016 | -0.716 | -0.218 | -1.267 | -0.291 | 0.018 | 0.035 | 0.035 | 0.085 | 0.2 | 0.11 | 0.08 | 0.07 | 0.12 | 0.1 |
| 7 | 16655632 | rs9638734 | -0.462 | -0.214 | -1.609 | -0.211 | 0.025 | 0.052 | 0.052 | 0 | 0.26 | 0.04 | 0.15 | 0.13 | 0.2 | 0.11 |
| 7 | 16656292 | rs10261839 | 0.06 | -0.033 | -1.099 | 0.015 | 0.002 | 0.002 | 0.002 | 0 | 0.49 | 0.53 | 0.66 | 0.47 | 0.48 | 0.59 |
| 7 | 16656546 | rs10264878 | -1.461 | -0.549 | -0.849 | 0.136 | 0.004 | -0.01 | -0.01 | 0 | 0.76 | 0.76 | 0.86 | 0.61 | 0.81 | 0.67 |
| 7 | 16662008 | rs1404398 | -1.428 | -0.728 | -0.917 | 0.033 | 0.011 | -0.033 | -0.033 | 0 | 0.76 | 0.81 | 0.87 | 0.59 | 0.82 | 0.67 |
| 7 | 16663446 | rs2286223 | 0.093 | -0.647 | -0.772 | 0.024 | 0.011 | 0.023 | 0.023 | 0 | 0.21 | 0.1 | 0.21 | 0.14 | 0.22 | 0.07 |
| 7 | 16666301 | rs1404396 | 0.881 | -0.128 | -0.891 | 0.18 | -0.061 | 0.002 | 0.002 | 1.409 | 0.24 | 0.12 | 0.3 | 0.24 | 0.39 | 0.36 |
| 7 | 16666454 | rs3815236 | 1.065 | -0.27 | -0.868 | 0.187 | -0.006 | 0.019 | 0.019 | 0 | 0.21 | 0.13 | 0.29 | 0.11 | 0.2 | 0.23 |
| 7 | 16667920 | rs10486794 | 0.209 | -0.222 | -0.938 | 0.415 | -0.085 | 0.013 | 0.013 | 1.082 | 0.28 | 0.07 | 0.3 | 0.24 | 0.4 | 0.36 |
| 7 | 16670471 | rs11981854 | 0.261 | 0.048 | -0.066 | 0.638 | 0.004 | 0.107 | 0.107 | 1.868 | 0.07 | 0.29 | 0.06 | 0.11 | 0.02 | 0.02 |
| 7 | 16674634 | rs12670185 | 0.546 | 0.179 | -0.766 | 0.697 | -0.013 | 0.069 | 0.069 | 1.294 | 0.07 | 0.2 | 0.05 | 0.11 | 0.02 | 0.02 |
| 7 | 16676955 | rs4719474 | 0.236 | 0.616 | -1.735 | 0.556 | 0.02 | 0.019 | 0.019 | 0.136 | 0.46 | 0.64 | 0.55 | 0.36 | 0.39 | 0.3 |
| 7 | 16681028 | rs6976832 | 0.104 | 0.313 | -1.847 | 0.751 | 0.018 | -0.044 | -0.044 | 0 | 0.7 | 0.83 | 0.86 | 0.62 | 0.77 | 0.66 |
| 7 | 16686940 | rs706024 | -0.066 | 0.701 | -2.399 | 0.39 | -0.076 | 0.078 | 0.078 | 0 | 0.62 | 0.92 | 0.66 | 0.7 | 0.59 | 0.58 |
| 7 | 16691244 | rs4721525 | 0.077 | 0.612 | -2.832 | 0.708 | -0.026 | -0.039 | -0.039 | 0 | 0.18 | 0.02 | 0.18 | 0.2 | 0.16 | 0.17 |
| 7 | 16700110 | rs4721526 | -0.63 | 0.127 | -2.201 | 0.663 | 0.001 | 0.013 | 0.013 | 0 | 0.25 | 0.13 | 0.22 | 0.19 | 0.21 | 0.27 |
| 7 | 16700447 | rs11771285 | 0.579 | 0.122 | -1.771 | 0.703 | -0.015 | -0.012 | -0.012 | 0 | 0.24 | 0.18 | 0.31 | 0.25 | 0.3 | 0.21 |
| 7 | 16700798 | rs696279 | -0.865 | 0.047 | -1.885 | 0.613 | -0.012 | -0.036 | -0.036 | 0 | 0.54 | 0.76 | 0.55 | 0.44 | 0.51 | 0.5 |
| 7 | 16702867 | rs17625938 | 0.538 | 0.474 | -1.536 | 0.586 | -0.02 | -0.017 | -0.017 | 3.565 | 0.22 | 0.15 | 0.32 | 0.25 | 0.3 | 0.2 |
| 7 | 16703752 | rs10499471 | 0.168 | 0.662 | -0.277 | 0.536 | 0.032 | 0.011 | 0.011 | 0 | 0.25 | 0.35 | 0.34 | 0.25 | 0.29 | 0.21 |
| 7 | 16704191 | rs1841217 | -0.074 | 0.303 | -0.204 | 0.354 | -0.003 | 0.038 | 0.038 | 0 | 0.17 | 0.06 | 0.17 | 0.04 | 0.15 | 0.19 |
| 7 | 16704212 | rs10499472 | -0.018 | 0.293 | 0.942 | 0.694 | -0.027 | 0.031 | 0.031 | 0 | 0.33 | 0.52 | 0.23 | 0.49 | 0.26 | 0.31 |
| 7 | 16707747 | rs3779048 | 0.358 | 0.393 | 0.495 | 0.528 | -0.036 | -0.084 | -0.084 | 0 | 0.28 | 0.04 | 0.18 | 0.38 | 0.27 | 0.34 |
| 7 | 16708028 | rs17169573 | -1.339 | 0.339 | 1.21 | 0.377 | -0.011 | -0.098 | -0.098 | 0 | 0.14 | 0.58 | 0.1 | 0 | 0.08 | 0.07 |
| 7 | 16710153 | rs11973867 | -1.338 | 0.515 | 0.881 | 0.495 | 0.108 | 0.013 | 0.013 | 0 | 0.06 | 0.29 | 0.05 | 0 | 0.06 | 0.01 |
| 7 | 16710566 | rs11763990 | 0.296 | 0.378 | -0.15 | 0.698 | -0.039 | -0.086 | -0.086 | 0.076 | 0.29 | 0.04 | 0.18 | 0.39 | 0.27 | 0.34 |
| 7 | 16719073 | rs10238254 | 0.228 | 0.209 | -0.031 | 0.715 | 0.052 | -0.053 | -0.053 | 0 | 0.29 | 0.11 | 0.13 | 0.38 | 0.2 | 0.25 |
| 7 | 16724005 | rs10243614 | 0.243 | 0.188 | 0.209 | 0.706 | 0.027 | -0.018 | -0.018 | 0 | 0.3 | 0.19 | 0.13 | 0.38 | 0.2 | 0.27 |
| 7 | 16730542 | rs17169592 | 0.106 | 0.632 | 0.402 | 0.73 | 0.049 | -0.016 | -0.016 | 0 | 0.06 | 0.08 | 0.07 | 0.03 | 0.11 | 0.15 |
| 7 | 16731253 | rs696282 | -0.331 | 0.518 | 0.436 | 0.684 | -0.01 | -0.001 | -0.001 | 0.01 | 0.41 | 0.51 | 0.28 | 0.39 | 0.34 | 0.42 |
| 7 | 16734307 | rs6977880 | -1.096 | 0.383 | 1.079 | 0.759 | 0.015 | 0.185 | 0.185 | 0 | 0.09 | 0.46 | 0.08 | 0.18 | 0.05 | 0.07 |
| 7 | 16735586 | rs706041 | 0.16 | 1.473 | -0.755 | 0.095 | -0.026 | 0.058 | 0.058 | 0 | 0.7 | 0.81 | 0.69 | 0.83 | 0.65 | 0.83 |
| 7 | 16735782 | rs706042 | -1.572 | 1.823 | -0.101 | 0.28 | 0.031 | -0.003 | -0.003 | 0.008 | 0.13 | 0.28 | 0.15 | 0.13 | 0.16 | 0.15 |
| 7 | 16738006 | rs12539223 | -0.586 | 2.074 | -0.792 | 0.527 | -0.071 | -0.034 | -0.034 | 0 | 0.22 | 0.08 | 0.31 | 0.25 | 0.35 | 0.31 |
| 7 | 16740101 | rs818323 | -1.04 | 1.317 | -1.581 | 0.534 | -0.051 | -0.108 | -0.108 | 0 | 0.65 | 0.93 | 0.67 | 0.46 | 0.61 | 0.61 |
| 7 | 16740554 | rs1459558 | -0.18 | 1.213 | -0.954 | 0.279 | -0.101 | -0.051 | -0.051 | 0 | 0.26 | 0.07 | 0.34 | 0.27 | 0.42 | 0.32 |
| 7 | 16745589 | rs1149510 | -1.175 | 0.267 | -0.568 | -0.019 | 0.006 | 0.092 | 0.092 | 0 | 0.5 | 0.82 | 0.51 | 0.28 | 0.54 | 0.48 |
| 7 | 16747549 | rs17627275 | -0.317 | 0.342 | -0.181 | 0.068 | -0.055 | -0.035 | -0.035 | 0 | 0.25 | 0.12 | 0.33 | 0.28 | 0.42 | 0.36 |
| 7 | 16748217 | rs1674815 | -0.579 | 0.26 | 0.308 | -0.173 | 0.014 | 0.001 | 0.001 | 0 | 0.27 | 0.63 | 0.32 | 0.24 | 0.34 | 0.35 |
| 7 | 16754957 | rs706045 | -1.221 | 0.207 | 0.364 | -0.265 | -0.074 | 0.121 | 0.121 | 0 | 0.45 | 0.89 | 0.49 | 0.23 | 0.5 | 0.42 |
| 7 | 16755448 | rs818489 | -1.229 | 0.207 | 0.364 | -0.265 | -0.073 | 0.122 | 0.122 | 0 | 0.45 | 0.89 | 0.49 | 0.23 | 0.5 | 0.42 |
| 7 | 16761612 | rs1144628 | 0.027 | 0.34 | 0.646 | -0.23 | -0.018 | -0.121 | -0.121 | 0 | 0.22 | 0.37 | 0.18 | 0 | 0.18 | 0.08 |
| 7 | 16761748 | rs17684089 | 0.178 | 0.332 | 0.086 | -0.056 | -0.056 | -0.032 | -0.032 | 0 | 0.19 | 0.07 | 0.27 | 0.22 | 0.32 | 0.34 |
| 7 | 16761784 | rs11770872 | 0.723 | -0.109 | 0.055 | 0.366 | 0.029 | -0.063 | -0.063 | 0 | 0.11 | 0.01 | 0.14 | 0.24 | 0.05 | 0.18 |
| 7 | 16764156 | rs697521 | 0.729 | 0.056 | 0.414 | -0.002 | -0.016 | -0.046 | -0.046 | 0 | 0.15 | 0.17 | 0.1 | 0 | 0.08 | 0.07 |
| 7 | 16765190 | rs7787867 | -0.168 | 0.708 | 0.655 | 0.156 | 0.028 | 0.005 | 0.005 | 0 | 0.22 | 0.27 | 0.28 | 0.22 | 0.34 | 0.32 |
| 7 | 16765898 | rs697523 | -0.141 | 0.182 | 0.867 | -0.097 | -0.053 | -0.111 | -0.111 | 0.658 | 0.2 | 0.34 | 0.11 | 0 | 0.1 | 0.07 |
| 7 | 16766403 | rs2389676 | -1.277 | 0.876 | 1.074 | 0.117 | 0.064 | 0.051 | 0.051 | 0 | 0.18 | 0.34 | 0.32 | 0.25 | 0.26 | 0.26 |
| 7 | 16767532 | rs697526 | -2.051 | 0.459 | 0.346 | 0.366 | -0.1 | -0.076 | -0.076 | 0 | 0.6 | 0.92 | 0.66 | 0.54 | 0.57 | 0.59 |
| 7 | 16771045 | rs11531480 | -1.101 | 0.528 | 0.888 | 0.168 | 0.06 | 0.057 | 0.057 | 0 | 0.18 | 0.34 | 0.3 | 0.25 | 0.25 | 0.25 |
| 7 | 16776257 | rs12672531 | -0.497 | 0.781 | 0.545 | 0.267 | -0.042 | -0.035 | -0.035 | 0 | 0.2 | 0.03 | 0.19 | 0.21 | 0.18 | 0.25 |
| 7 | 16776509 | rs1168059 | -1.661 | 1.015 | 1.092 | 0.409 | 0.012 | 0.018 | 0.018 | 0 | 0.34 | 0.41 | 0.4 | 0.4 | 0.38 | 0.43 |
| 7 | 16777700 | rs6956775 | 0.528 | 1.515 | 1.447 | 0.619 | 0.054 | 0.079 | 0.079 | 0 | 0.07 | 0.14 | 0.19 | 0.2 | 0.16 | 0.12 |
| 7 | 16780009 | rs7800565 | 0.377 | 1.455 | 1.048 | 0.532 | 0.002 | 0.033 | 0.033 | 0 | 0.2 | 0.09 | 0.13 | 0.08 | 0.19 | 0.16 |
| 7 | 16782963 | rs1723808 | -1.993 | 1.692 | 1.288 | 0.782 | 0.068 | 0.066 | 0.066 | 0.027 | 0.14 | 0.62 | 0.26 | 0.22 | 0.22 | 0.14 |
| 7 | 16784421 | rs1723803 | -1.479 | 1.864 | 1.348 | 0.948 | 0.245 | 0.31 | 0.31 | 0.109 | 0.07 | 0.46 | 0.2 | 0.2 | 0.15 | 0.13 |
| 7 | 16785522 | rs1674801 | -1.651 | 2.593 | 0.999 | 0.864 | 0.141 | 0.133 | 0.133 | 0 | 0.21 | 0.49 | 0.41 | 0.38 | 0.37 | 0.32 |
| 7 | 16785553 | rs1674802 | -1.025 | 2.051 | 0.608 | 0.807 | 0.004 | -0.024 | -0.024 | 0.015 | 0.43 | 0.76 | 0.57 | 0.47 | 0.56 | 0.47 |
| 7 | 16788739 | rs2693095 | -1.248 | 2.365 | 1.006 | 0.966 | 0.239 | 0.229 | 0.229 | 0.012 | 0.06 | 0.35 | 0.2 | 0.19 | 0.17 | 0.11 |
| 7 | 16789764 | rs706053 | -1.526 | 2.312 | 1.056 | 1.083 | 0.202 | 0.215 | 0.215 | 0.013 | 0.07 | 0.33 | 0.23 | 0.2 | 0.17 | 0.11 |

| | | | | | | | | | | | | | | | | |
|---|---|---|---|---|---|---|---|---|---|---|---|---|---|---|---|---|
| 7 | 16795114 | rs818803 | -0.089 | 1.847 | 0.689 | 0.196 | 0.001 | -0.013 | -0.013 | 0.022 | 0.39 | 0.6 | 0.39 | 0.19 | 0.42 | 0.18 |
| 7 | 16795564 | rs818805 | -1.104 | 1.882 | 0.887 | 0.074 | 0.048 | -0.024 | -0.024 | 0 | 0.09 | 0.38 | 0.07 | 0 | 0.06 | 0.01 |
| 7 | 16796532 | rs818806 | -0.744 | 2.007 | 0.041 | -0.487 | -0.048 | 0.057 | 0.057 | 0 | 0.89 | 0.98 | 0.79 | 1 | 0.82 | 0.88 |
| 7 | 16802589 | rs3807490 | 0.314 | 1.907 | 0.602 | -0.343 | 0.005 | -0.024 | -0.024 | 0.77 | 0.38 | 0.46 | 0.38 | 0.2 | 0.41 | 0.18 |
| 7 | 16803990 | rs706057 | -0.624 | 1.437 | -0.048 | -0.27 | -0.021 | -0.074 | -0.074 | 0 | 0.6 | 0.88 | 0.6 | 0.43 | 0.62 | 0.48 |
| 7 | 16807354 | rs12530923 | 1.38 | 1.273 | 0.082 | -0.292 | -0.036 | 0.034 | 0.034 | 0 | 0.31 | 0.15 | 0.33 | 0.2 | 0.37 | 0.17 |
| 7 | 16811167 | rs706065 | 0.229 | 1.211 | 0.191 | -0.25 | -0.001 | 0.009 | 0.009 | 0 | 0.47 | 0.41 | 0.51 | 0.4 | 0.53 | 0.28 |
| 7 | 16811808 | rs11982151 | 0.21 | 1.707 | 0.253 | 0.121 | 0.001 | 0.001 | 0.001 | 0 | 0.15 | 0.19 | 0.17 | 0.19 | 0.16 | 0.11 |
| 7 | 16814399 | rs697528 | -1.258 | 1.678 | -0.24 | -0.008 | -0.023 | 0.021 | 0.021 | 0 | 0.94 | 0.98 | 0.82 | 1 | 0.87 | 0.95 |
| 7 | 16815345 | rs747347 | 0.555 | 0.917 | -0.068 | -0.018 | 0.001 | 0.003 | 0.003 | 0 | 0.46 | 0.52 | 0.36 | 0.23 | 0.41 | 0.27 |
| 7 | 16815542 | rs3816599 | 0.301 | 0.941 | -0.082 | 0.001 | -0.008 | -0.005 | -0.005 | 0 | 0.16 | 0.22 | 0.17 | 0.18 | 0.14 | 0.11 |
| 7 | 16816000 | rs2290837 | 0.443 | 1.153 | -0.573 | -0.084 | -0.006 | -0.035 | -0.035 | 0 | 0.42 | 0.48 | 0.35 | 0.2 | 0.39 | 0.23 |
| 7 | 16818799 | rs6461276 | 0.483 | 1.699 | -1.647 | 0.194 | 0.001 | 0.06 | 0.06 | 0 | 0.42 | 0.26 | 0.34 | 0.24 | 0.4 | 0.27 |
| 7 | 16821216 | rs3807495 | 0.637 | 1.661 | -1.63 | -0.006 | -0.013 | 0.092 | 0.092 | 0.055 | 0.37 | 0.23 | 0.32 | 0.17 | 0.39 | 0.19 |
| 7 | 16821355 | rs3807496 | 0.533 | 1.62 | -1.632 | 0.795 | -0.015 | -0.07 | -0.07 | 0 | 0.62 | 0.9 | 0.55 | 0.61 | 0.61 | 0.52 |
| 7 | 16821637 | rs2389700 | 0.862 | 1.749 | -0.809 | 0.671 | -0.01 | 0.012 | 0.012 | 0 | 0.9 | 0.91 | 0.92 | 0.97 | 0.86 | 0.88 |
| 7 | 16823169 | rs3807498 | -1.435 | 1.682 | -0.297 | 1.104 | 0.006 | 0.037 | 0.037 | 0 | 0.06 | 0.13 | 0.11 | 0.18 | 0.07 | 0.15 |
| 7 | 16823978 | rs6616 | 2.32 | 1.964 | 0.361 | 1.255 | 0.022 | 0.018 | 0.018 | 0 | 0.81 | 0.35 | 0.77 | 0.71 | 0.73 | 0.67 |
| 7 | 16824662 | rs1037791 | 2.357 | 2.137 | 1.333 | 1.279 | 0.137 | 0.128 | 0.128 | 0 | 0.86 | 0.63 | 0.77 | 0.7 | 0.75 | 0.73 |
| 7 | 16825385 | rs2125711 | 2.125 | 2.113 | 1.473 | 1.279 | 0.112 | 0.052 | 0.052 | 0 | 0.87 | 0.56 | 0.85 | 0.87 | 0.79 | 0.82 |
| 7 | 16825517 | rs17629192 | -0.36 | 1.983 | 0.806 | 1.165 | -0.024 | 0.02 | 0.02 | 0 | 0.17 | 0.07 | 0.2 | 0.11 | 0.2 | 0.15 |
| 7 | 16825580 | rs4629751 | 2.771 | 2.07 | 1.102 | 2.001 | 0.005 | 0.021 | 0.021 | 0.311 | 0.76 | 0.29 | 0.75 | 0.58 | 0.72 | 0.72 |
| 7 | 16826463 | rs10247482 | 0.181 | 2.197 | 0.765 | 1.267 | 0.021 | -0.005 | -0.005 | 0.05 | 0.94 | 0.92 | 0.9 | 0.97 | 0.84 | 0.95 |
| 7 | 16830910 | rs940827 | 2.18 | 2.148 | 0.938 | 1.308 | 0.175 | 0.093 | 0.093 | 0 | 0.86 | 0.52 | 0.74 | 0.76 | 0.73 | 0.72 |
| 7 | 16831537 | rs15504 | -1.432 | 1.499 | 0.615 | 0.993 | 0.066 | 0.145 | 0.145 | 0.045 | 0.08 | 0.47 | 0.07 | 0.09 | 0.04 | 0.13 |
| 7 | 16837404 | rs819010 | 1.207 | 1.989 | -0.408 | 0.618 | -0.037 | 0.027 | 0.027 | 0.003 | 0.69 | 0.76 | 0.64 | 0.71 | 0.56 | 0.7 |
| 7 | 16837618 | rs2280654 | -0.939 | 2.034 | -0.493 | 0.668 | -0.012 | -0.006 | -0.006 | 0.021 | 0.15 | 0.16 | 0.22 | 0.28 | 0.22 | 0.35 |
| 7 | 16839835 | rs11773294 | -0.822 | 1.11 | -0.37 | 0.498 | 0.013 | 0.048 | 0.048 | 0.091 | 0.15 | 0.06 | 0.06 | 0 | 0.11 | 0.08 |
| 7 | 16843465 | rs706071 | 1.336 | 1.419 | -0.383 | 0.753 | -0.038 | 0.02 | 0.02 | 0.166 | 0.8 | 0.89 | 0.84 | 0.9 | 0.72 | 0.78 |
| 7 | 16844064 | rs17150186 | -0.236 | 1.253 | -0.587 | 0.452 | 0 | 0.01 | 0.01 | 0 | 0.2 | 0.13 | 0.18 | 0.21 | 0.19 | 0.27 |
| 7 | 16846146 | rs1459564 | 2.406 | 0.275 | -1.114 | -0.185 | -0.047 | 0.113 | 0.113 | 0.095 | 0.32 | 0.08 | 0.4 | 0.17 | 0.37 | 0.21 |
| 7 | 16846354 | rs706075 | -0.436 | 0.115 | -0.875 | 0.214 | 0 | -0.002 | -0.002 | 0 | 0.34 | 0.64 | 0.23 | 0.39 | 0.22 | 0.3 |
| 7 | 16846624 | rs706076 | 0.153 | -0.036 | -0.378 | 0.136 | -0.03 | 0.048 | 0.048 | 0 | 0.27 | 0.35 | 0.19 | 0.4 | 0.19 | 0.3 |
| 7 | 16847745 | rs706078 | -1.984 | 0.047 | -0.422 | 0.133 | -0.041 | -0.026 | -0.026 | 0 | 0.61 | 0.77 | 0.43 | 0.6 | 0.46 | 0.64 |
| 7 | 16848712 | rs6960534 | 0.377 | 0.022 | -0.378 | 0.115 | -0.001 | -0.056 | -0.056 | 0 | 0.1 | 0.27 | 0.04 | 0 | 0.05 | 0.06 |
| 7 | 16848890 | rs1563780 | 0.905 | 0.048 | -0.399 | 0.135 | 0.012 | -0.009 | -0.009 | 0.545 | 0.64 | 0.59 | 0.68 | 0.7 | 0.58 | 0.53 |
| 7 | 16849012 | rs1563781 | 2.5 | 0.037 | -0.508 | 0.116 | -0.019 | 0.01 | 0.01 | 0.067 | 0.33 | 0.18 | 0.46 | 0.3 | 0.36 | 0.22 |
| 7 | 16851612 | rs10085446 | -0.925 | 0.04 | -0.402 | 0.104 | 0 | -0.002 | -0.002 | 2.663 | 0.33 | 0.34 | 0.24 | 0.2 | 0.27 | 0.35 |
| 7 | 16855007 | rs10224320 | -0.851 | 0.035 | -0.471 | 0.102 | 0.001 | 0.001 | 0.001 | 0 | 0.32 | 0.33 | 0.24 | 0.2 | 0.27 | 0.35 |
| 7 | 16856549 | rs7807123 | 1.345 | 0.047 | -0.645 | 0.195 | 0.01 | 0.012 | 0.012 | 0 | 0.53 | 0.33 | 0.64 | 0.49 | 0.56 | 0.49 |
| 7 | 16856582 | rs7807024 | -0.281 | 0.257 | -0.636 | 0.167 | -0.006 | -0.01 | -0.01 | 0 | 0.94 | 0.95 | 0.9 | 0.89 | 0.83 | 0.86 |
| 7 | 16861849 | rs10253216 | 0.037 | 0.107 | -0.614 | 0.151 | -0.026 | 0.033 | 0.033 | 0 | 0.26 | 0.6 | 0.26 | 0.52 | 0.16 | 0.28 |
| 7 | 16862235 | rs7797977 | 0.636 | 0.145 | -0.632 | 0.129 | -0.001 | 0.006 | 0.006 | 0 | 0.67 | 0.32 | 0.63 | 0.37 | 0.66 | 0.56 |
| 7 | 16863727 | rs11768848 | 0.541 | 0.153 | -0.598 | 0.133 | -0.001 | 0.005 | 0.005 | 0 | 0.68 | 0.33 | 0.63 | 0.37 | 0.67 | 0.58 |
| 7 | 16875057 | rs7809532 | 0.504 | -0.117 | -0.759 | 0.031 | 0.006 | 0 | 0 | 0 | 0.72 | 0.69 | 0.68 | 0.7 | 0.61 | 0.58 |
| 7 | 16878113 | rs10232481 | -0.103 | -0.323 | -0.354 | 0.317 | -0.031 | 0.041 | 0.041 | 0 | 0.25 | 0.59 | 0.26 | 0.52 | 0.16 | 0.29 |
| 7 | 16882053 | rs1404966 | -0.662 | -0.297 | -0.536 | 0.325 | 0 | -0.003 | -0.003 | 0.005 | 0.3 | 0.67 | 0.27 | 0.53 | 0.16 | 0.28 |
| 7 | 16884241 | rs17136737 | -0.563 | -0.149 | -0.283 | 0.727 | -0.049 | 0.104 | 0.104 | 0 | 0.27 | 0.44 | 0.26 | 0.49 | 0.17 | 0.3 |
| 7 | 16886324 | rs12666607 | -0.266 | -0.125 | -0.375 | 0.392 | -0.002 | 0.01 | 0.01 | 0.16 | 0.22 | 0.18 | 0.17 | 0.19 | 0.22 | 0.36 |
| 7 | 16886936 | rs1525742 | 0.248 | 0.085 | -0.476 | 0.265 | 0.05 | -0.028 | -0.028 | 0 | 0.82 | 0.66 | 0.81 | 0.91 | 0.73 | 0.81 |
| 7 | 16886964 | rs1525741 | 1.228 | -0.017 | -0.479 | 0.27 | 0.006 | -0.018 | -0.018 | 0 | 0.59 | 0.46 | 0.64 | 0.72 | 0.51 | 0.46 |
| 7 | 16889926 | rs1525738 | -1.227 | -0.19 | -0.965 | 0.03 | 0.003 | 0.084 | 0.084 | 0 | 0.51 | 0.66 | 0.52 | 0.81 | 0.53 | 0.75 |
| 7 | 16890368 | rs6978138 | 1.068 | -0.125 | -0.682 | 0.231 | 0.004 | 0.003 | 0.003 | 0 | 0.61 | 0.4 | 0.69 | 0.66 | 0.67 | 0.58 |
| 7 | 16890982 | rs12534941 | -0.263 | -0.101 | -0.822 | 0.208 | 0.029 | 0.086 | 0.086 | 0 | 0.28 | 0.09 | 0.16 | 0.09 | 0.23 | 0.21 |
| 7 | 16895475 | rs10950635 | -0.75 | 0.58 | 0.029 | 0.813 | -0.024 | -0.008 | -0.008 | 0 | 0.22 | 0.17 | 0.24 | 0.29 | 0.33 | 0.41 |
| 7 | 16899197 | rs11798 | 0.745 | 0.649 | -0.173 | 0.866 | 0.031 | -0.005 | -0.005 | 0 | 0.74 | 0.46 | 0.72 | 0.74 | 0.66 | 0.59 |
| 7 | 16899376 | rs4719484 | 0.753 | 0.651 | -0.173 | 0.837 | 0.03 | 0.001 | 0.001 | 0.1 | 0.74 | 0.45 | 0.72 | 0.72 | 0.66 | 0.59 |
| 7 | 16903770 | rs10280750 | -0.561 | -0.312 | -0.017 | 1.198 | -0.04 | 0.066 | 0.066 | 0 | 0.18 | 0.32 | 0.15 | 0.3 | 0.08 | 0.18 |
| 7 | 16904410 | rs11975058 | -0.278 | 0.093 | 0.126 | 0.985 | 0.005 | 0.005 | 0.005 | 0 | 0.25 | 0.21 | 0.22 | 0.34 | 0.28 | 0.33 |
| 7 | 16918471 | rs11496005 | 1.092 | 0.472 | -0.738 | 0.544 | -0.018 | 0.007 | 0.007 | 0.158 | 0.25 | 0.15 | 0.32 | 0.2 | 0.3 | 0.2 |
| 7 | 16919172 | rs10249546 | 0.362 | -0.22 | -1.131 | 0.262 | 0.007 | 0.012 | 0.012 | 0 | 0.23 | 0.18 | 0.24 | 0.21 | 0.23 | 0.27 |
| 7 | 16919476 | rs4370433 | 0.2 | 0.156 | -0.815 | 0.338 | -0.001 | 0.001 | 0.001 | 0.234 | 0.55 | 0.51 | 0.62 | 0.58 | 0.49 | 0.51 |
| 7 | 16922366 | rs10950638 | 0.905 | 0.202 | -1.372 | 0.266 | -0.062 | 0.006 | 0.006 | 0 | 0.21 | 0.03 | 0.27 | 0.14 | 0.23 | 0.2 |
| 7 | 16925518 | rs10950639 | 0.499 | -0.753 | -1.132 | -0.046 | 0.007 | 0.009 | 0.009 | 0 | 0.28 | 0.25 | 0.24 | 0.14 | 0.23 | 0.2 |
| 7 | 16925645 | rs17136772 | 0.304 | -1.31 | -0.483 | 0.217 | -0.04 | 0.049 | 0.049 | 0 | 0.16 | 0.3 | 0.05 | 0.26 | 0.06 | 0.12 |
| 7 | 16925781 | rs6968539 | 0.604 | -0.75 | -0.09 | 0.259 | 0.025 | 0.119 | 0.119 | 0 | 0.87 | 0.61 | 0.84 | 0.72 | 0.88 | 0.7 |
| 7 | 16926273 | rs6969383 | -0.15 | -0.645 | -0.508 | 0.095 | 0.004 | 0.003 | 0.003 | 0 | 0.61 | 0.41 | 0.57 | 0.57 | 0.66 | 0.5 |
| 7 | 16928306 | rs2190271 | 1.063 | -0.222 | -0.697 | 0.148 | -0.113 | -0.044 | -0.044 | 1.103 | 0.17 | 0.01 | 0.26 | 0.17 | 0.32 | 0.18 |
| 7 | 16931466 | rs6962438 | 1.015 | -0.657 | -0.128 | 0.085 | -0.015 | 0.03 | 0.03 | 0.062 | 0.26 | 0.13 | 0.3 | 0.14 | 0.26 | 0.21 |

| | | | | | | | | | | | | | | | | |
|---|---|---|---|---|---|---|---|---|---|---|---|---|---|---|---|---|
| 7 | 16932506 | rs7800062 | 0.049 | -0.688 | 0.263 | 0.117 | 0.062 | 0.305 | 0.305 | 0 | 0.93 | 0.58 | 0.93 | 0.71 | 0.99 | 0.75 |
| 7 | 16932544 | rs7800176 | 0.897 | -0.675 | 0.054 | 0.118 | -0.013 | 0.036 | 0.036 | 0.004 | 0.27 | 0.13 | 0.3 | 0.14 | 0.26 | 0.22 |
| 7 | 16932926 | rs4582450 | 0.669 | -0.447 | 0.617 | 0.55 | 0.04 | 0.229 | 0.229 | 0 | 0.92 | 0.69 | 0.91 | 0.71 | 0.96 | 0.74 |
| 7 | 16946486 | rs4559136 | 0.607 | -0.341 | 0.439 | 0.674 | 0.06 | 0.207 | 0.207 | 0 | 0.92 | 0.62 | 0.9 | 0.77 | 0.94 | 0.89 |
| 7 | 16947921 | rs847375 | 0.655 | -0.769 | -0.402 | 0.192 | -0.004 | 0.051 | 0.051 | 0 | 0.31 | 0.15 | 0.39 | 0.22 | 0.29 | 0.36 |
| 7 | 16955730 | rs4563783 | 0.095 | -1.268 | -0.474 | 0.126 | 0.03 | -0.002 | -0.002 | 0.057 | 0.64 | 0.3 | 0.74 | 0.59 | 0.76 | 0.62 |
| 7 | 16957686 | rs12532502 | 1.197 | -1.321 | -0.165 | 0.11 | 0.009 | 0.006 | 0.006 | 0 | 0.21 | 0.18 | 0.13 | 0.14 | 0.16 | 0.16 |
| 7 | 16959444 | rs4320447 | 0.352 | -1.191 | 0.035 | 0.21 | 0.013 | 0.149 | 0.149 | 0.015 | 0.9 | 0.56 | 0.9 | 0.77 | 0.94 | 0.87 |
| 7 | 16959685 | rs4534036 | 1.334 | -1.211 | -0.014 | 0.181 | -0.027 | 0.078 | 0.078 | 0 | 0.86 | 0.44 | 0.86 | 0.73 | 0.91 | 0.79 |
| 7 | 16963044 | rs17136841 | -0.415 | -1.22 | 0.562 | 0.204 | -0.004 | 0.153 | 0.153 | 0 | 0.07 | 0.21 | 0.06 | 0.27 | 0.02 | 0.18 |
| 7 | 16964510 | rs13237319 | 1.242 | -1.404 | 0.268 | 0.109 | 0.021 | 0.016 | 0.016 | 0 | 0.21 | 0.06 | 0.15 | 0.15 | 0.18 | 0.18 |
| 7 | 16965115 | rs847408 | -0.241 | -1.542 | 0.356 | 0.08 | -0.035 | 0.008 | 0.008 | 0.081 | 0.33 | 0.56 | 0.25 | 0.43 | 0.23 | 0.43 |
| 7 | 16966355 | rs847403 | -0.236 | -1.549 | 0.355 | 0.082 | -0.039 | 0.009 | 0.009 | 0.1 | 0.33 | 0.55 | 0.25 | 0.43 | 0.23 | 0.43 |
| 7 | 16968409 | rs4333473 | -1.098 | -2.055 | -0.331 | 0.365 | 0.038 | -0.056 | -0.056 | 0 | 0.85 | 0.95 | 0.96 | 0.78 | 0.94 | 0.84 |
| 7 | 16968509 | rs847396 | 1.147 | -1.929 | -0.237 | 0.324 | 0.032 | 0.03 | 0.03 | 0 | 0.24 | 0.1 | 0.17 | 0.16 | 0.21 | 0.25 |
| 7 | 16970321 | rs847393 | 1.326 | -1.957 | -0.228 | 0.327 | 0.035 | 0.024 | 0.024 | 0 | 0.23 | 0.11 | 0.14 | 0.16 | 0.19 | 0.23 |
| 7 | 16973077 | rs10276006 | 0.151 | -1.81 | 0.027 | 0.465 | 0.022 | 0.177 | 0.177 | 0 | 0.06 | 0.29 | 0.06 | 0.22 | 0.02 | 0.11 |
| 7 | 16975024 | rs2108091 | -0.603 | -1.11 | -0.079 | 0.555 | 0.047 | 0.204 | 0.204 | 0 | 0.08 | 0.35 | 0.11 | 0.33 | 0.06 | 0.21 |
| 7 | 16975053 | rs847386 | -0.047 | -1.152 | -0.442 | 0.372 | -0.077 | -0.089 | -0.089 | 0 | 0.63 | 0.84 | 0.68 | 0.57 | 0.57 | 0.68 |
| 7 | 16977505 | rs1990114 | -1.139 | -1.027 | -0.264 | 0.427 | 0.048 | 0.211 | 0.211 | 0 | 0.11 | 0.46 | 0.16 | 0.33 | 0.09 | 0.24 |
| 7 | 16979542 | rs10487585 | -0.686 | -0.986 | -0.491 | 0.43 | 0.065 | 0.231 | 0.231 | 0 | 0.09 | 0.39 | 0.11 | 0.33 | 0.07 | 0.22 |
| 7 | 16984280 | rs1636744 | 0.755 | -0.982 | -1.564 | 0.094 | -0.094 | -0.044 | -0.044 | 0 | 0.22 | 0.09 | 0.34 | 0.25 | 0.41 | 0.26 |
| 7 | 16988435 | rs4543439 | -1.202 | -1.332 | -1.789 | 0.235 | 0.048 | -0.095 | -0.095 | 0 | 0.85 | 0.96 | 0.93 | 0.55 | 0.94 | 0.83 |
| 7 | 16989462 | rs1403804 | 1.388 | -1.22 | -1.823 | -0.011 | 0.021 | 0.052 | 0.052 | 0 | 0.24 | 0.11 | 0.17 | 0.11 | 0.21 | 0.28 |
| 7 | 16994809 | rs13237658 | 1.661 | -0.829 | -1.716 | 0.007 | -0.053 | 0.028 | 0.028 | 0 | 0.24 | 0.02 | 0.35 | 0.14 | 0.24 | 0.2 |
| 7 | 17004301 | rs1721019 | 0.751 | -0.338 | -1.222 | 0.252 | 0.058 | 0.024 | 0.024 | 0 | 0.21 | 0.28 | 0.3 | 0.27 | 0.41 | 0.26 |
| 7 | 17004387 | rs6461292 | -0.207 | -0.972 | -1.841 | 0.296 | 0.077 | -0.066 | -0.066 | 0.041 | 0.6 | 0.86 | 0.76 | 0.45 | 0.72 | 0.62 |
| 7 | 17008167 | rs1622628 | 0.16 | -0.753 | -1.981 | 0.176 | 0.02 | 0.053 | 0.053 | 0 | 0.52 | 0.84 | 0.56 | 0.32 | 0.61 | 0.48 |
| 7 | 17009612 | rs1636731 | -0.428 | -0.884 | -1.79 | 0.12 | 0.041 | 0.065 | 0.065 | 0 | 0.5 | 0.29 | 0.46 | 0.68 | 0.4 | 0.53 |
| 7 | 17015695 | rs1523635 | 0.228 | -0.722 | -2.056 | 0.152 | -0.039 | -0.074 | -0.074 | 0 | 0.26 | 0.1 | 0.36 | 0.65 | 0.28 | 0.37 |
| 7 | 17018432 | rs847440 | -0.855 | -0.512 | -1.372 | -0.152 | 0.003 | 0.026 | 0.026 | 0 | 0.54 | 0.63 | 0.62 | 0.79 | 0.53 | 0.5 |
| 7 | 17025992 | rs763783 | 0.163 | -1.171 | -1.417 | 1.044 | 0.015 | -0.021 | -0.021 | 0 | 0.11 | 0.06 | 0.07 | 0.33 | 0.04 | 0.11 |
| 7 | 17028225 | rs763522 | 1.5 | -1.376 | -1.537 | 0.955 | 0.006 | -0.003 | -0.003 | 0 | 0.11 | 0.03 | 0.05 | 0.09 | 0.07 | 0.1 |
| 7 | 17028698 | rs4529355 | -1.294 | -1.911 | -1.178 | 0.487 | 0.041 | 0.056 | 0.056 | 0.566 | 0.52 | 0.71 | 0.38 | 0.77 | 0.35 | 0.46 |
| 7 | 17032172 | rs17137033 | 0.573 | -2.064 | -0.998 | 0.467 | 0.008 | -0.009 | -0.009 | 0 | 0.11 | 0.08 | 0.03 | 0.29 | 0.06 | 0.13 |
| 7 | 17033088 | rs9638738 | -1.404 | -1.985 | -1.029 | 0.381 | 0.042 | 0.072 | 0.072 | 0.696 | 0.53 | 0.74 | 0.39 | 0.78 | 0.37 | 0.46 |
| 7 | 17039332 | rs847423 | 0.371 | -1.579 | -0.746 | 0.43 | 0.033 | -0.06 | -0.06 | 0.072 | 0.17 | 0.07 | 0.11 | 0.35 | 0.1 | 0.12 |
| 7 | 17040226 | rs7808216 | 0.179 | -1.585 | -0.433 | 0.372 | 0 | 0 | 0 | 0 | 0.24 | 0.26 | 0.27 | 0.39 | 0.23 | 0.19 |
| 7 | 17048140 | rs7786255 | 1.506 | -1.363 | -0.617 | 0.28 | 0.005 | 0.001 | 0.001 | 0 | 0.11 | 0.04 | 0.06 | 0.08 | 0.07 | 0.1 |
| 7 | 17050971 | rs4436007 | -0.641 | -1.338 | -0.438 | 0.116 | 0.045 | 0.011 | 0.011 | 0 | 0.93 | 0.74 | 0.95 | 0.98 | 0.93 | 0.89 |
| 7 | 17052786 | rs2158767 | 1.049 | -0.909 | -1.235 | -0.036 | -0.104 | 0.023 | 0.023 | 0 | 0.19 | 0.04 | 0.44 | 0.13 | 0.32 | 0.28 |
| 7 | 17054346 | rs3944100 | 0.048 | -0.893 | -1.266 | -0.068 | -0.102 | 0.047 | 0.047 | 0.012 | 0.33 | 0.08 | 0.48 | 0.21 | 0.36 | 0.38 |
| 7 | 17059433 | rs12670784 | 0.141 | -1.333 | -1.224 | -0.039 | -0.118 | 0.044 | 0.044 | 0.026 | 0.32 | 0.08 | 0.49 | 0.21 | 0.38 | 0.37 |
| 7 | 17067445 | rs2163069 | -0.998 | -1.648 | -0.907 | -0.041 | 0.001 | 0 | 0 | 0 | 0.6 | 0.68 | 0.78 | 0.62 | 0.62 | 0.58 |
| 7 | 17069968 | rs1523632 | 0.003 | -1.511 | -1.32 | 0.044 | -0.03 | -0.057 | -0.057 | 0.365 | 0.4 | 0.19 | 0.7 | 0.53 | 0.54 | 0.47 |
| 7 | 17079154 | rs11772758 | 0.144 | -1.353 | -1.128 | -0.105 | 0.014 | 0.034 | 0.034 | 0 | 0.25 | 0.16 | 0.2 | 0.12 | 0.2 | 0.21 |
| 7 | 17081157 | rs518136 | -0.583 | -1.313 | -0.695 | 0.079 | 0.024 | 0.026 | 0.026 | 0.186 | 0.44 | 0.63 | 0.33 | 0.26 | 0.35 | 0.38 |
| 7 | 17081981 | rs17345341 | 2.232 | -0.515 | -0.911 | 0.239 | -0.036 | 0.018 | 0.018 | 0 | 0.09 | 0.01 | 0.14 | 0.03 | 0.15 | 0.07 |
| 7 | 17082894 | rs4721562 | 0.984 | -0.122 | -0.638 | 0.497 | -0.114 | 0.005 | 0.005 | 0 | 0.16 | 0.03 | 0.41 | 0.11 | 0.31 | 0.27 |
| 7 | 17085321 | rs6961860 | -0.235 | -0.785 | 0.062 | 0.619 | 0.073 | 0.056 | 0.056 | 0 | 0.28 | 0.52 | 0.44 | 0.44 | 0.36 | 0.4 |
| 7 | 17095077 | rs546484 | 0.613 | -1.01 | 0.168 | 0.512 | -0.001 | -0.014 | -0.014 | 0 | 0.55 | 0.38 | 0.35 | 0.21 | 0.46 | 0.42 |
| 7 | 17095636 | rs11765040 | 1.399 | -1.034 | 0.14 | 0.638 | -0.076 | -0.062 | -0.062 | 0.468 | 0.17 | 0.07 | 0.33 | 0.36 | 0.32 | 0.27 |
| 7 | 17096005 | rs13227633 | 0.626 | -1.08 | 0.309 | 0.631 | 0 | 0 | 0 | 0 | 0.56 | 0.38 | 0.35 | 0.21 | 0.47 | 0.42 |
| 7 | 17098097 | rs530947 | -0.839 | -1.531 | -0.14 | 0.542 | 0.019 | 0.003 | 0.003 | 0 | 0.61 | 0.92 | 0.79 | 0.71 | 0.67 | 0.67 |
| 7 | 17099057 | rs10228079 | 0.182 | -1.434 | 0.137 | 0.605 | -0.007 | 0.092 | 0.092 | 0 | 0.13 | 0.29 | 0.23 | 0.28 | 0.12 | 0.19 |
| 7 | 17099149 | rs10257625 | 0.898 | -1.432 | 0.108 | 0.613 | -0.001 | 0.024 | 0.024 | 0 | 0.1 | 0.12 | 0.21 | 0.23 | 0.12 | 0.16 |
| 7 | 17103344 | rs6974162 | 0.112 | -1.44 | -0.265 | 0.587 | -0.01 | 0.092 | 0.092 | 0 | 0.13 | 0.3 | 0.23 | 0.28 | 0.12 | 0.19 |
| 7 | 17104923 | rs540509 | 1.563 | -1.482 | -0.824 | 0.428 | 0.011 | 0.045 | 0.045 | 0 | 0.26 | 0.11 | 0.14 | 0.13 | 0.23 | 0.23 |
| 7 | 17105122 | rs672365 | -2.044 | -1.469 | -0.815 | 0.344 | 0.007 | -0.017 | -0.017 | 0 | 0.87 | 0.93 | 0.92 | 0.84 | 0.9 | 0.9 |
| 7 | 17109277 | rs13224356 | 2.061 | -1.533 | -0.91 | 0.169 | -0.019 | 0.099 | 0.099 | 0.019 | 0.19 | 0.02 | 0.16 | 0.02 | 0.18 | 0.11 |
| 7 | 17110253 | rs819367 | -1.293 | -0.984 | -0.517 | 0.572 | 0.01 | 0.036 | 0.036 | 0 | 0.73 | 0.85 | 0.85 | 0.89 | 0.76 | 0.76 |
| 7 | 17110307 | rs4265079 | 1.439 | -0.868 | -0.656 | 0.758 | -0.04 | -0.028 | -0.028 | 0 | 0.17 | 0.14 | 0.34 | 0.34 | 0.32 | 0.25 |
| 7 | 17111402 | rs11981130 | 1.575 | -0.762 | -0.862 | 0.534 | 0.015 | 0.03 | 0.03 | 0 | 0.28 | 0.18 | 0.19 | 0.07 | 0.22 | 0.21 |
| 7 | 17112008 | rs819365 | 0.158 | -0.742 | -0.953 | 0.64 | 0.027 | 0.016 | 0.016 | 0 | 0.4 | 0.13 | 0.23 | 0.27 | 0.34 | 0.35 |
| 7 | 17116201 | rs10499481 | 0.634 | -0.264 | -1.039 | 0.781 | -0.011 | -0.057 | -0.057 | 0 | 0.1 | 0.02 | 0.2 | 0.25 | 0.11 | 0.17 |
| 7 | 17117736 | rs17137193 | 1.487 | -0.427 | -0.787 | 0.412 | 0.007 | -0.01 | -0.01 | 0 | 0.13 | 0.07 | 0.08 | 0.15 | 0.1 | 0.1 |
| 7 | 17126744 | rs6970379 | -0.582 | -0.299 | -0.645 | 0.142 | 0.009 | 0.016 | 0.016 | 0 | 0.73 | 0.87 | 0.85 | 0.86 | 0.76 | 0.76 |
| 7 | 17127579 | rs10247820 | 0.309 | -0.614 | -0.528 | -0.006 | 0.014 | 0.021 | 0.021 | 0 | 0.3 | 0.2 | 0.2 | 0.08 | 0.23 | 0.22 |
| 7 | 17129802 | rs7801215 | 0.035 | 0.233 | -0.083 | 0.809 | -0.016 | -0.038 | -0.038 | 0 | 0.78 | 0.74 | 0.88 | 0.93 | 0.82 | 0.8 |

| | | | | | | | | | | | | | | | | |
|---|---|---|---|---|---|---|---|---|---|---|---|---|---|---|---|---|
| 7 | 17132458 | rs6946140 | -1.022 | 1.055 | -0.316 | 1.667 | 0.003 | 0.026 | 0.026 | 0 | 0.1 | 0.11 | 0.12 | 0.31 | 0.11 | 0.16 |
| 7 | 17133514 | rs7801891 | -1.457 | 0.772 | -0.143 | 1.395 | -0.004 | 0.02 | 0.02 | 0 | 0.06 | 0.12 | 0.05 | 0.08 | 0.06 | 0.08 |
| 7 | 17144007 | rs10233878 | -0.507 | 0.646 | 0.092 | 1.636 | 0.036 | 0.037 | 0.037 | 0 | 0.53 | 0.74 | 0.73 | 0.59 | 0.63 | 0.51 |
| 7 | 17163262 | rs10085375 | -0.443 | 0.806 | 0.432 | 1.46 | 0.033 | 0.009 | 0.009 | 0 | 0.51 | 0.62 | 0.66 | 0.46 | 0.63 | 0.47 |
| 7 | 17169922 | rs6973953 | 0.444 | 1.261 | 0.076 | 1.398 | -0.005 | -0.01 | -0.01 | 1.59 | 0.18 | 0.12 | 0.31 | 0.18 | 0.21 | 0.23 |
| 7 | 17170680 | rs1997091 | -0.554 | 1.611 | 0.187 | 1.298 | -0.001 | -0.001 | -0.001 | 0 | 0.3 | 0.31 | 0.42 | 0.38 | 0.4 | 0.36 |
| 7 | 17172181 | rs4721576 | -0.665 | 1.631 | 0.339 | 0.976 | 0.01 | 0.046 | 0.046 | 0.662 | 0.19 | 0.42 | 0.23 | 0.26 | 0.23 | 0.12 |
| 7 | 17173879 | rs7341466 | -0.395 | 1.565 | -0.018 | 1.021 | -0.003 | -0.005 | -0.005 | 0 | 0.05 | 0.05 | 0.06 | 0.19 | 0.08 | 0.06 |
| 7 | 17176784 | rs1404962 | -1.113 | 1.188 | -0.334 | 0.929 | -0.002 | 0.016 | 0.016 | 0.223 | 0.41 | 0.53 | 0.36 | 0.46 | 0.38 | 0.28 |
| 7 | 17183451 | rs17137264 | -0.792 | 1.23 | -0.363 | 0.84 | -0.02 | -0.023 | -0.023 | 0 | 0.06 | 0.04 | 0.09 | 0.15 | 0.11 | 0.04 |
| 7 | 17183694 | rs11763349 | 0.657 | 1.164 | -0.421 | 0.802 | -0.07 | 0.014 | 0.014 | 2.472 | 0.13 | 0.01 | 0.2 | 0.07 | 0.19 | 0.08 |
| 7 | 17184107 | rs6944767 | 0.768 | 0.857 | -0.216 | 0.475 | -0.019 | -0.057 | -0.057 | 0 | 0.13 | 0.16 | 0.04 | 0 | 0.07 | 0.11 |
| 7 | 17186779 | rs6461299 | 0.76 | 0.94 | -0.439 | 0.392 | 0.028 | 0.001 | 0.001 | 0 | 0.51 | 0.37 | 0.31 | 0.49 | 0.42 | 0.35 |
| 7 | 17192272 | rs11766104 | 0.622 | 0.804 | -0.171 | 0.434 | 0.003 | -0.038 | -0.038 | 0.363 | 0.13 | 0.19 | 0.07 | 0.01 | 0.17 | 0.06 |
| 7 | 17196492 | rs11772495 | 0.313 | 0.342 | -0.004 | 0.298 | -0.006 | -0.016 | -0.016 | 0 | 0.64 | 0.43 | 0.58 | 0.27 | 0.68 | 0.48 |
| 7 | 17196857 | rs17137304 | 0.477 | 0.272 | -0.103 | 0.648 | 0.021 | -0.076 | -0.076 | 0 | 0.19 | 0.04 | 0.06 | 0.3 | 0.14 | 0.21 |
| 7 | 17197571 | rs1525735 | 0.788 | 0.413 | -0.219 | 0.32 | 0.015 | 0.067 | 0.067 | 0 | 0.5 | 0.39 | 0.46 | 0.14 | 0.53 | 0.41 |
| 7 | 17198181 | rs6461304 | 0.897 | 0.372 | -0.163 | 0.294 | -0.006 | 0.131 | 0.131 | 0 | 0.45 | 0.28 | 0.44 | 0.14 | 0.52 | 0.4 |
| 7 | 17200506 | rs11773165 | -0.228 | 0.429 | -0.211 | 0.368 | -0.002 | -0.01 | -0.01 | 0 | 0.19 | 0.23 | 0.15 | 0.19 | 0.17 | 0.07 |
| 7 | 17204692 | rs17150193 | 0.657 | 0.476 | -0.39 | 0.377 | -0.03 | 0.008 | 0.008 | 0 | 0.14 | 0.05 | 0.12 | 0.16 | 0.15 | 0.06 |
| 7 | 17207578 | rs11773260 | 0.885 | 0.358 | -0.195 | 0.486 | 0.001 | 0.02 | 0.02 | 0 | 0.46 | 0.41 | 0.44 | 0.36 | 0.51 | 0.43 |
| 7 | 17207733 | rs10499489 | 1.435 | 0.851 | -0.652 | 0.007 | -0.059 | 0.034 | 0.034 | 1.033 | 0.09 | 0.01 | 0.11 | 0.02 | 0.2 | 0.08 |
| 7 | 17212073 | rs11770386 | 0.15 | 0.537 | -0.297 | 0.093 | -0.006 | -0.006 | -0.006 | 0 | 0.18 | 0.22 | 0.11 | 0.18 | 0.14 | 0.06 |
| 7 | 17214399 | rs17644896 | 1.625 | 0.565 | -0.502 | -0.072 | -0.003 | 0.025 | 0.025 | 0 | 0.31 | 0.23 | 0.28 | 0.1 | 0.31 | 0.31 |
| 7 | 17221063 | rs1464906 | 0.168 | -0.209 | -0.345 | 0.281 | 0.01 | 0.005 | 0.005 | 0.242 | 0.6 | 0.69 | 0.54 | 0.65 | 0.64 | 0.48 |
| 7 | 17221154 | rs12670338 | -0.591 | 0.048 | 0.399 | 0.959 | -0.033 | 0.076 | 0.076 | 0.88 | 0.21 | 0.42 | 0.13 | 0.38 | 0.14 | 0.11 |
| 7 | 17235800 | rs6950788 | 0.834 | 0.321 | 0.529 | 0.278 | -0.035 | -0.112 | -0.112 | 0 | 0.75 | 0.47 | 0.87 | 0.97 | 0.79 | 0.93 |
| 7 | 17238185 | rs10499496 | -0.635 | 0.316 | 0.832 | 0.314 | 0.033 | 0.001 | 0.001 | 0 | 0.06 | 0.24 | 0.04 | 0 | 0.05 | 0.02 |
| 7 | 17244315 | rs4073703 | 0.372 | 0.458 | 0.444 | 0.414 | -0.06 | -0.053 | -0.053 | 0 | 0.16 | 0.02 | 0.12 | 0.25 | 0.19 | 0.13 |
| 7 | 17245862 | rs4145933 | 1.315 | 0.027 | 1.197 | 0.459 | 0.055 | 0.012 | 0.012 | 0.346 | 0.93 | 0.66 | 0.96 | 1 | 0.94 | 0.98 |
| 7 | 17253922 | rs4552804 | 0.344 | 0.084 | 0.701 | 0.471 | -0.034 | -0.096 | -0.096 | 3.048 | 0.8 | 0.66 | 0.91 | 0.98 | 0.85 | 0.95 |
| 7 | 17254446 | rs4524681 | 1.543 | 0.082 | 0.67 | 0.575 | 0.016 | 0.008 | 0.008 | 0 | 0.6 | 0.32 | 0.49 | 0.65 | 0.6 | 0.64 |
| 7 | 17262556 | rs4391312 | 0.297 | 0.077 | 0.382 | 0.448 | -0.035 | -0.098 | -0.098 | 0.93 | 0.79 | 0.66 | 0.92 | 0.98 | 0.85 | 0.95 |
| 7 | 17273749 | rs1859022 | 0.994 | 1.014 | 0.118 | 1.262 | 0.013 | 0.046 | 0.046 | 1.709 | 0.78 | 0.71 | 0.56 | 0.66 | 0.71 | 0.63 |
| 7 | 17277084 | rs1476079 | -0.85 | 1.04 | -0.364 | 1.395 | -0.001 | 0.007 | 0.007 | 0.423 | 0.19 | 0.19 | 0.41 | 0.36 | 0.23 | 0.31 |
| 7 | 17279294 | rs1476081 | -1.081 | 0.875 | -0.104 | 1.326 | 0.006 | 0.034 | 0.034 | 0 | 0.26 | 0.38 | 0.42 | 0.38 | 0.3 | 0.31 |
| 7 | 17280135 | rs10279112 | -0.086 | 1.526 | -1.081 | 1.557 | -0.088 | -0.101 | -0.101 | 0.626 | 0.83 | 0.98 | 0.74 | 0.69 | 0.77 | 0.82 |
| 7 | 17287106 | rs6968554 | 1.114 | 1.167 | -1.047 | 1.316 | 0.004 | 0.013 | 0.013 | 0.432 | 0.56 | 0.73 | 0.4 | 0.4 | 0.62 | 0.47 |
| 7 | 17296072 | rs17137472 | 1.708 | 0.471 | -0.712 | 0.758 | 0.008 | -0.002 | -0.002 | 0 | 0.39 | 0.41 | 0.3 | 0.38 | 0.49 | 0.38 |
| 7 | 17303778 | rs10275488 | 0.769 | 0.387 | -0.412 | 0.913 | 0 | 0.006 | 0.006 | 0 | 0.13 | 0.14 | 0.34 | 0.26 | 0.12 | 0.2 |
| 7 | 17318312 | rs13236243 | 1.096 | 0.034 | -0.491 | 0.887 | -0.004 | 0.012 | 0.012 | 0 | 0.68 | 0.55 | 0.5 | 0.64 | 0.73 | 0.62 |
| 7 | 17322683 | rs4719495 | -0.535 | 0.051 | -0.383 | 0.925 | 0.009 | 0.049 | 0.049 | 0 | 0.83 | 0.77 | 0.76 | 0.68 | 0.82 | 0.78 |
| 7 | 17326058 | rs2301677 | -1.232 | 0.024 | 0.265 | 0.779 | -0.018 | -0.012 | -0.012 | 0 | 0.27 | 0.55 | 0.43 | 0.27 | 0.22 | 0.25 |
| 7 | 17329402 | rs765623 | -0.159 | 0.091 | 0.047 | 0.84 | -0.028 | -0.058 | -0.058 | 1.081 | 0.81 | 0.92 | 0.74 | 0.67 | 0.78 | 0.72 |
| 7 | 17332989 | rs10250822 | -0.175 | 0.149 | 0.066 | 0.866 | -0.023 | -0.054 | -0.054 | 0 | 0.81 | 0.91 | 0.75 | 0.64 | 0.78 | 0.72 |
| 7 | 17334899 | rs4719497 | 1.696 | 0.299 | -0.038 | 0.741 | -0.042 | 0.046 | 0.046 | 0 | 0.08 | 0.01 | 0.08 | 0 | 0.12 | 0.04 |
| 7 | 17336035 | rs3757824 | -0.36 | 0.141 | 0.128 | 0.756 | -0.003 | -0.004 | -0.004 | 0 | 0.18 | 0.15 | 0.39 | 0.27 | 0.21 | 0.28 |
| 7 | 17341796 | rs12532771 | 1.542 | 0.049 | 0.442 | 0.53 | -0.002 | 0.002 | 0.002 | 0 | 0.61 | 0.45 | 0.36 | 0.41 | 0.65 | 0.54 |
| 7 | 17345614 | rs2282885 | 1.922 | 0.161 | -0.718 | 0.054 | -0.118 | 0.021 | 0.021 | 0.749 | 0.31 | 0.02 | 0.21 | 0.2 | 0.4 | 0.21 |
| 7 | 17348119 | rs2237298 | 0.774 | 0.096 | -0.151 | 0.242 | 0.002 | 0.008 | 0.008 | 0 | 0.63 | 0.52 | 0.36 | 0.4 | 0.65 | 0.55 |
| 7 | 17349626 | rs17779352 | 2.663 | -0.118 | -0.495 | -0.065 | -0.005 | 0.02 | 0.02 | 0.187 | 0.07 | 0.02 | 0.03 | 0.04 | 0.07 | 0.1 |
| 7 | 17357873 | rs1476080 | 0.611 | -0.119 | -0.251 | 0.074 | 0 | 0.001 | 0.001 | 0 | 0.63 | 0.57 | 0.38 | 0.39 | 0.66 | 0.59 |
| 7 | 17359594 | rs2237297 | 1.272 | -0.05 | -0.309 | 0.539 | 0.003 | 0.128 | 0.128 | 0 | 0.09 | 0.15 | 0.14 | 0.36 | 0.1 | 0.16 |
| 7 | 17360521 | rs17137566 | 0.511 | -0.04 | -0.366 | 0.428 | 0.003 | 0.05 | 0.05 | 0 | 0.14 | 0.15 | 0.17 | 0.38 | 0.18 | 0.17 |
| 7 | 17368085 | rs6960165 | -1.189 | -0.065 | -0.584 | 0.332 | -0.067 | -0.046 | -0.046 | 1.211 | 0.83 | 0.97 | 0.75 | 0.8 | 0.77 | 0.85 |
| 7 | 17368420 | rs2158041 | -1.282 | -0.098 | -0.579 | 0.239 | -0.065 | -0.064 | -0.064 | 0 | 0.84 | 0.97 | 0.75 | 0.76 | 0.78 | 0.81 |
| 7 | 17371363 | rs7811989 | -0.701 | -0.105 | -0.395 | 0.243 | -0.017 | -0.009 | -0.009 | 2.325 | 0.79 | 0.89 | 0.53 | 0.76 | 0.76 | 0.7 |
| 7 | 17379110 | rs2066853 | 0.052 | -0.214 | -0.183 | 0.286 | -0.046 | 0.159 | 0.159 | 0.203 | 0.18 | 0.46 | 0.17 | 0.37 | 0.11 | 0.15 |
| 7 | 17380662 | rs2040623 | 0.095 | -0.147 | -0.29 | 0.3 | -0.001 | 0.011 | 0.011 | 0 | 0.18 | 0.16 | 0.38 | 0.4 | 0.21 | 0.29 |
| 7 | 17381740 | rs2106728 | 1.783 | -0.397 | -0.645 | 0.018 | -0.104 | 0.027 | 0.027 | 0.636 | 0.29 | 0.03 | 0.21 | 0.19 | 0.4 | 0.21 |
| 7 | 17399858 | rs10950657 | 1.672 | -0.376 | -0.083 | 0.255 | 0 | 0.043 | 0.043 | 0 | 0.53 | 0.27 | 0.3 | 0.35 | 0.55 | 0.46 |
| 7 | 17403696 | rs10274243 | -0.024 | -0.4 | -0.058 | 0.448 | -0.032 | 0.11 | 0.11 | 0 | 0.16 | 0.31 | 0.15 | 0.38 | 0.1 | 0.16 |
| 7 | 17405718 | rs7780687 | -0.518 | -0.471 | -0.232 | 0.473 | -0.027 | 0.071 | 0.071 | 0 | 0.22 | 0.4 | 0.2 | 0.39 | 0.19 | 0.19 |
| 7 | 17407061 | rs2053998 | 0.296 | -0.049 | -0.78 | 0.764 | -0.001 | 0.005 | 0.005 | 0 | 0.18 | 0.16 | 0.39 | 0.38 | 0.21 | 0.26 |
| 7 | 17425895 | rs12667669 | 0.651 | 0.007 | -0.95 | 0.792 | -0.059 | -0.102 | -0.102 | 0.875 | 0.15 | 0.01 | 0.38 | 0.37 | 0.21 | 0.24 |
| 7 | 17432838 | rs9639288 | 0.47 | 0.006 | -0.697 | 0.776 | 0 | 0.012 | 0.012 | 0 | 0.17 | 0.16 | 0.38 | 0.37 | 0.21 | 0.25 |
| 7 | 17436278 | rs6947693 | -0.456 | -0.174 | -1.312 | 0.568 | -0.044 | -0.038 | -0.038 | 0 | 0.78 | 0.93 | 0.75 | 0.76 | 0.76 | 0.76 |
| 7 | 17439609 | rs6461314 | 2.41 | 0.031 | -1.372 | 0.61 | -0.008 | -0.004 | -0.004 | 1.777 | 0.06 | 0.03 | 0.07 | 0.14 | 0.09 | 0.11 |
| 7 | 17442679 | rs1077773 | 1.193 | -0.483 | -1.176 | 0.256 | 0.005 | 0.005 | 0.005 | 0 | 0.52 | 0.35 | 0.58 | 0.58 | 0.57 | 0.55 |

| | | | | | | | | | | | | | | | | |
|---|---|---|---|---|---|---|---|---|---|---|---|---|---|---|---|---|
| 7 | 17448444 | rs6461315 | -0.452 | -0.452 | -0.926 | -0.005 | -0.021 | -0.01 | -0.01 | 0.014 | 0.76 | 0.85 | 0.72 | 0.76 | 0.72 | 0.75 |
| 7 | 17450853 | rs7794392 | 2.168 | -0.523 | -1.038 | -0.024 | -0.023 | 0.051 | 0.051 | 0 | 0.33 | 0.15 | 0.2 | 0.2 | 0.37 | 0.3 |
| 7 | 17451588 | rs1111547 | 0.523 | -0.389 | -0.406 | 0.111 | -0.015 | -0.017 | -0.017 | 0.248 | 0.69 | 0.44 | 0.7 | 0.77 | 0.75 | 0.77 |
| 7 | 17453479 | rs6977113 | -0.416 | -0.558 | -0.024 | 0.172 | -0.031 | -0.054 | -0.054 | 0 | 0.13 | 0.38 | 0.06 | 0.04 | 0.04 | 0.08 |
| 7 | 17462221 | rs4721602 | -0.465 | -0.412 | -0.278 | 0.295 | -0.014 | -0.026 | -0.026 | 0 | 0.27 | 0.53 | 0.26 | 0.19 | 0.2 | 0.19 |
| 7 | 17464410 | rs1078032 | -0.502 | -0.134 | -0.584 | 0.285 | 0.008 | 0 | 0 | 0.345 | 0.4 | 0.53 | 0.41 | 0.37 | 0.45 | 0.28 |
| 7 | 17464427 | rs1078031 | -0.209 | -0.082 | -0.555 | 0.314 | 0.015 | 0.004 | 0.004 | 0.059 | 0.38 | 0.52 | 0.4 | 0.37 | 0.45 | 0.25 |
| 7 | 17467225 | rs4721604 | -0.148 | -0.115 | -0.468 | 0.381 | -0.004 | 0 | 0 | 0.326 | 0.37 | 0.63 | 0.41 | 0.34 | 0.44 | 0.3 |
| 7 | 17472677 | rs12699853 | -0.127 | -0.847 | -0.079 | 0.231 | 0.005 | 0.001 | 0.001 | 3.58 | 0.39 | 0.43 | 0.41 | 0.37 | 0.46 | 0.34 |
| 7 | 17475607 | rs2034684 | 1.46 | -0.721 | 0.207 | 0.31 | 0.019 | 0.015 | 0.015 | 3.6 | 0.18 | 0.3 | 0.19 | 0.19 | 0.26 | 0.07 |
| 7 | 17481311 | rs10499499 | 1.011 | -0.742 | -0.323 | 0.521 | -0.071 | -0.017 | -0.017 | 0.02 | 0.17 | 0.01 | 0.23 | 0.14 | 0.21 | 0.1 |
| 7 | 17490691 | rs655478 | -0.037 | -1.042 | -0.215 | 0.507 | 0.002 | -0.004 | -0.004 | 0 | 0.71 | 0.8 | 0.72 | 0.72 | 0.73 | 0.65 |
| 7 | 17492960 | rs1545534 | -0.8 | -1.091 | -0.026 | 0.606 | 0.005 | 0.008 | 0.008 | 0.259 | 0.52 | 0.59 | 0.49 | 0.54 | 0.58 | 0.47 |
| 7 | 17493007 | rs1545535 | 0.653 | -0.951 | -0.382 | 0.713 | -0.017 | -0.018 | -0.018 | 0 | 0.23 | 0.19 | 0.29 | 0.27 | 0.3 | 0.31 |
| 7 | 17495669 | rs1868361 | 1.731 | -1.057 | -0.68 | 0.584 | -0.011 | -0.029 | -0.029 | 0 | 0.13 | 0.07 | 0.33 | 0.33 | 0.16 | 0.24 |
| 7 | 17497568 | rs497266 | -0.104 | -1.197 | -0.487 | 0.549 | -0.008 | -0.007 | -0.007 | 0 | 0.65 | 0.43 | 0.75 | 0.73 | 0.69 | 0.71 |
| 7 | 17499411 | rs555952 | -0.469 | -1.326 | -0.367 | 0.525 | -0.044 | -0.002 | -0.002 | 0 | 0.84 | 0.56 | 0.9 | 0.89 | 0.93 | 0.77 |
| 7 | 17499822 | rs559613 | 1.47 | -1.323 | -0.501 | 0.5 | 0 | 0 | 0 | 0.553 | 0.32 | 0.33 | 0.16 | 0.21 | 0.3 | 0.33 |
| 7 | 17500595 | rs603819 | 0.131 | -1.105 | -0.213 | 0.539 | -0.008 | -0.006 | -0.006 | 0 | 0.68 | 0.53 | 0.67 | 0.72 | 0.72 | 0.67 |
| 7 | 17501082 | rs606056 | 0.959 | -0.982 | -0.131 | 0.582 | -0.001 | -0.001 | -0.001 | 0 | 0.46 | 0.51 | 0.5 | 0.5 | 0.45 | 0.57 |
| 7 | 17504506 | rs662356 | 0.592 | -0.936 | -0.204 | 0.559 | 0.001 | -0.001 | -0.001 | 0 | 0.49 | 0.43 | 0.54 | 0.54 | 0.48 | 0.6 |
| 7 | 17504646 | rs13244652 | 0.284 | -0.866 | -0.498 | 0.613 | -0.075 | -0.035 | -0.035 | 0.773 | 0.38 | 0.12 | 0.39 | 0.39 | 0.47 | 0.2 |
| 7 | 17505619 | rs677641 | 0.565 | -0.718 | -0.299 | 0.63 | 0.001 | 0.001 | 0.001 | 3.825 | 0.48 | 0.55 | 0.54 | 0.48 | 0.46 | 0.59 |
| 7 | 17506531 | rs1059788 | 1.323 | -0.647 | -0.431 | 0.639 | -0.018 | -0.012 | -0.012 | 0.603 | 0.21 | 0.11 | 0.16 | 0.22 | 0.26 | 0.1 |
| 7 | 17506813 | rs1059791 | 1.339 | -0.646 | -0.436 | 0.64 | -0.019 | -0.013 | -0.013 | 0 | 0.21 | 0.09 | 0.16 | 0.22 | 0.26 | 0.1 |
| 7 | 17509011 | rs536532 | -0.271 | -0.678 | -0.331 | 0.572 | 0 | 0 | 0 | 0 | 0.69 | 0.7 | 0.68 | 0.7 | 0.71 | 0.68 |
| 7 | 17509089 | rs632859 | 0.794 | -0.657 | -0.348 | 0.591 | 0 | 0 | 0 | 0 | 0.46 | 0.48 | 0.47 | 0.46 | 0.45 | 0.57 |
| 7 | 17510224 | rs7800225 | 0.94 | -0.519 | -0.485 | 0.594 | -0.02 | -0.014 | -0.014 | 0 | 0.23 | 0.13 | 0.2 | 0.24 | 0.28 | 0.11 |
| 7 | 17511491 | rs512302 | 0.258 | -0.725 | -0.564 | 0.485 | 0.005 | -0.004 | -0.004 | 0.659 | 0.63 | 0.63 | 0.46 | 0.59 | 0.67 | 0.57 |
| 7 | 17513099 | rs17137735 | 1.265 | -0.273 | -1.098 | 0.358 | -0.034 | 0.001 | 0.001 | 0 | 0.17 | 0.04 | 0.39 | 0.14 | 0.2 | 0.14 |
| 7 | 17514336 | rs524715 | 0.521 | -0.523 | -1.015 | 0.314 | -0.023 | -0.008 | -0.008 | 0.049 | 0.42 | 0.28 | 0.36 | 0.46 | 0.47 | 0.54 |
| 7 | 17514416 | rs10269035 | 1.831 | -0.67 | -1.373 | 0.248 | -0.004 | -0.01 | -0.01 | 0 | 0.17 | 0.16 | 0.13 | 0.24 | 0.22 | 0.09 |
| 7 | 17514653 | rs611935 | -1.65 | -0.649 | -1.58 | -0.139 | 0.023 | 0.008 | 0.008 | 1.463 | 0.88 | 0.94 | 0.96 | 0.92 | 0.94 | 0.8 |
| 7 | 17515400 | rs12670838 | -0.538 | -0.638 | -1.459 | -0.205 | 0.01 | 0.004 | 0.004 | 1.253 | 0.81 | 0.81 | 0.9 | 0.83 | 0.92 | 0.76 |
| 7 | 17515440 | rs17704807 | -0.105 | -0.043 | -1.78 | -0.044 | -0.064 | 0.101 | 0.101 | 0.189 | 0.38 | 0.12 | 0.53 | 0.21 | 0.43 | 0.4 |
| 7 | 17518574 | rs4721619 | -0.341 | 0.458 | -1.326 | 0.467 | -0.022 | 0.016 | 0.016 | 0 | 0.29 | 0.17 | 0.49 | 0.24 | 0.3 | 0.39 |
| 7 | 17520523 | rs1979098 | -0.131 | 0.421 | -1.206 | 0.525 | -0.006 | 0.007 | 0.007 | 0 | 0.28 | 0.23 | 0.47 | 0.25 | 0.27 | 0.36 |
| 7 | 17521073 | rs6960468 | -0.284 | 0.397 | -0.722 | 0.688 | -0.007 | 0.004 | 0.004 | 0 | 0.27 | 0.5 | 0.46 | 0.29 | 0.24 | 0.31 |
| 7 | 17521686 | rs1448666 | -0.932 | 0.591 | -1.571 | 0.608 | 0.005 | 0.147 | 0.147 | 0.743 | 0.55 | 0.3 | 0.7 | 0.78 | 0.62 | 0.57 |
| 7 | 17522125 | rs1448670 | -0.219 | 0.262 | -1.111 | 1.022 | 0.005 | -0.011 | -0.011 | 0 | 0.23 | 0.15 | 0.4 | 0.24 | 0.2 | 0.29 |
| 7 | 17522503 | rs17137763 | -0.595 | 0.4 | -0.445 | 1.41 | 0.003 | 0.142 | 0.142 | 0.045 | 0.13 | 0.29 | 0.14 | 0.31 | 0.12 | 0.18 |
| 7 | 17524639 | rs16871596 | -0.345 | 0.587 | -1.138 | 0.894 | 0.006 | -0.007 | -0.007 | 1.081 | 0.45 | 0.56 | 0.27 | 0.48 | 0.38 | 0.23 |
| 7 | 17527048 | rs13247700 | -0.222 | 0.475 | -1.832 | 0.62 | -0.037 | -0.037 | -0.037 | 0 | 0.17 | 0.03 | 0.24 | 0.18 | 0.15 | 0.41 |
| 7 | 17528021 | rs516794 | -0.5 | 0.318 | -2.554 | 0.48 | -0.089 | -0.049 | -0.049 | 0.154 | 0.62 | 0.86 | 0.69 | 0.57 | 0.51 | 0.65 |
| 7 | 17528130 | rs517723 | 1.378 | 0.035 | -2.572 | 0.439 | 0.025 | -0.008 | -0.008 | 0 | 0.17 | 0.06 | 0.17 | 0.2 | 0.12 | 0.09 |
| 7 | 17532188 | rs13239692 | -0.476 | 0.489 | -1.449 | 0.158 | -0.021 | 0.019 | 0.019 | 0 | 0.33 | 0.49 | 0.34 | 0.38 | 0.25 | 0.55 |
| 7 | 17533687 | rs12699858 | -0.998 | 1.131 | -1.07 | -0.067 | -0.002 | 0.001 | 0.001 | 0.093 | 0.28 | 0.32 | 0.41 | 0.18 | 0.32 | 0.46 |
| 7 | 17540774 | rs10259109 | -0.905 | 1.392 | -1.121 | -0.208 | 0.013 | -0.029 | -0.029 | 0 | 0.27 | 0.35 | 0.41 | 0.12 | 0.3 | 0.43 |
| 7 | 17544284 | rs1349137 | -0.461 | 1.804 | -1.729 | 0.192 | -0.041 | 0.011 | 0.011 | 0.002 | 0.16 | 0.1 | 0.33 | 0.12 | 0.22 | 0.39 |
| 7 | 17551902 | rs17706320 | -0.698 | 1.924 | -2.039 | 0.056 | -0.072 | 0.02 | 0.02 | 0 | 0.23 | 0.06 | 0.29 | 0.18 | 0.33 | 0.34 |
| 7 | 17552884 | rs11761325 | 0.168 | 1.985 | -2.076 | 0.041 | -0.032 | 0.019 | 0.019 | 0 | 0.13 | 0.05 | 0.09 | 0.08 | 0.21 | 0.1 |
| 7 | 17554444 | rs7785619 | 0.463 | 1.994 | -2.285 | 0.576 | -0.002 | -0.021 | -0.021 | 0.149 | 0.09 | 0.05 | 0.11 | 0.23 | 0.08 | 0.14 |
| 7 | 17559127 | rs12699863 | 0.794 | 2.003 | -2.302 | 0.567 | -0.039 | -0.019 | -0.019 | 0.403 | 0.09 | 0 | 0.2 | 0.1 | 0.12 | 0.23 |
| 7 | 17561583 | rs10499504 | 0.52 | 2.165 | -2.319 | 0.548 | -0.057 | -0.011 | -0.011 | 0 | 0.1 | 0.01 | 0.23 | 0.1 | 0.18 | 0.24 |
| 7 | 17561865 | rs6461323 | -1.17 | 2.126 | -1.79 | -0.059 | -0.002 | 0 | 0 | 0 | 0.35 | 0.62 | 0.54 | 0.34 | 0.43 | 0.41 |
| 7 | 17563282 | rs11772776 | 0.686 | 1.675 | -2.003 | 0.09 | -0.011 | 0.012 | 0.012 | 0 | 0.12 | 0.05 | 0.06 | 0.08 | 0.15 | 0.1 |
| 7 | 17565087 | rs17137807 | -0.598 | 1.749 | -2.001 | 0.124 | -0.055 | -0.072 | -0.072 | 0 | 0.2 | 0.05 | 0.34 | 0.33 | 0.25 | 0.37 |
| 7 | 17566157 | rs4721626 | 0.872 | 1.304 | -1.783 | 0.211 | -0.001 | 0.001 | 0.001 | 0.019 | 0.65 | 0.63 | 0.64 | 0.64 | 0.61 | 0.55 |
| 7 | 17566398 | rs1375238 | 1.179 | 0.519 | -2.207 | -0.284 | -0.003 | 0.005 | 0.005 | 2.118 | 0.37 | 0.53 | 0.44 | 0.39 | 0.37 | 0.39 |
| 7 | 17566650 | rs1031274 | -0.568 | 0.277 | -1.191 | 0.407 | -0.006 | 0.016 | 0.016 | 0.092 | 0.9 | 0.84 | 0.95 | 0.83 | 0.94 | 0.88 |
| 7 | 17566664 | rs1031275 | 0.222 | 0.326 | -1.246 | 0.444 | -0.011 | -0.002 | -0.002 | 0.316 | 0.74 | 0.61 | 0.87 | 0.75 | 0.79 | 0.8 |
| 7 | 17566678 | rs1031276 | 0.701 | 0.534 | -1.206 | 0.316 | -0.005 | 0.013 | 0.013 | 0.005 | 0.65 | 0.58 | 0.77 | 0.51 | 0.71 | 0.67 |
| 7 | 17571110 | rs10229888 | 0.127 | 0.706 | -1.026 | 0.093 | 0.018 | 0.049 | 0.049 | 0 | 0.48 | 0.67 | 0.6 | 0.65 | 0.55 | 0.62 |
| 7 | 17572925 | rs974821 | 0.326 | 0.471 | -1.086 | 0.217 | 0 | -0.003 | -0.003 | 0 | 0.92 | 0.91 | 0.93 | 0.81 | 0.93 | 0.85 |
| 7 | 17574471 | rs10268888 | 0.883 | 0.591 | -0.804 | 0.296 | 0.002 | 0.017 | 0.017 | 0 | 0.1 | 0.17 | 0.15 | 0.17 | 0.1 | 0.07 |
| 7 | 17574672 | rs11760243 | 1.246 | 0.41 | -1.382 | -0.192 | 0.007 | 0.171 | 0.171 | 1.383 | 0.25 | 0.01 | 0.11 | 0 | 0.17 | 0.04 |
| 7 | 17575717 | rs4721629 | 0.267 | 0.417 | -0.948 | 0.014 | 0.002 | -0.003 | -0.003 | 0 | 0.88 | 0.9 | 0.92 | 0.84 | 0.86 | 0.83 |
| 7 | 17576693 | rs1448655 | 0.369 | 0.396 | -0.815 | -0.024 | -0.003 | -0.002 | -0.002 | 0 | 0.87 | 0.84 | 0.91 | 0.86 | 0.86 | 0.83 |
| 7 | 17582606 | rs12699868 | 1.22 | 0.683 | -0.567 | 0.169 | -0.005 | -0.012 | -0.012 | 0 | 0.3 | 0.25 | 0.35 | 0.46 | 0.33 | 0.48 |

| | | | | | | | | | | | | | | | | |
|---|---|---|---|---|---|---|---|---|---|---|---|---|---|---|---|---|
| 7 | 17583429 | rs10247400 | 0.891 | 0.338 | -0.323 | -0.242 | 0.012 | 0.004 | 0.004 | 2.851 | 0.68 | 0.6 | 0.64 | 0.68 | 0.61 | 0.59 |
| 7 | 17585609 | rs968170 | 0.632 | 0.37 | -0.116 | -0.067 | -0.006 | 0.02 | 0.02 | 0.204 | 0.89 | 0.8 | 0.92 | 0.82 | 0.87 | 0.77 |
| 7 | 17588072 | rs10242846 | 0.171 | 0.656 | -0.503 | -0.142 | -0.002 | 0.018 | 0.018 | 0 | 0.47 | 0.41 | 0.54 | 0.71 | 0.52 | 0.52 |
| 7 | 17593328 | rs4721630 | 0.93 | 0.617 | -0.159 | -0.006 | 0.004 | -0.039 | -0.039 | 0 | 0.89 | 0.76 | 0.93 | 0.94 | 0.89 | 0.88 |
| 7 | 17594314 | rs6963617 | -0.655 | 0.302 | -1.033 | -0.461 | 0.018 | 0.1 | 0.1 | 0 | 0.32 | 0.17 | 0.29 | 0.09 | 0.29 | 0.23 |
| 7 | 17599438 | rs13221764 | 0.867 | 0.618 | -0.557 | 0.176 | -0.005 | 0.011 | 0.011 | 0 | 0.55 | 0.34 | 0.67 | 0.38 | 0.54 | 0.51 |
| 7 | 17604524 | rs7807962 | -0.662 | 0.863 | 0.345 | 0.754 | 0.056 | 0.107 | 0.107 | 0.016 | 0.09 | 0.29 | 0.12 | 0.16 | 0.09 | 0.09 |
| 7 | 17605070 | rs1448673 | 0.27 | 0.837 | 0.412 | 0.732 | -0.003 | -0.014 | -0.014 | 0 | 0.71 | 0.72 | 0.79 | 0.56 | 0.67 | 0.62 |
| 7 | 17612033 | rs6461326 | 0.696 | 0.638 | 0.505 | 0.691 | -0.002 | -0.006 | -0.006 | 0.297 | 0.64 | 0.67 | 0.78 | 0.54 | 0.62 | 0.59 |
| 7 | 17614091 | rs10270855 | -1.759 | 0.829 | 0.616 | 0.934 | 0.026 | 0.056 | 0.056 | 0.282 | 0.2 | 0.64 | 0.22 | 0.33 | 0.24 | 0.26 |
| 7 | 17621253 | rs1448672 | 1.842 | 0.672 | -0.124 | 1.133 | -0.122 | -0.08 | -0.08 | 0 | 0.48 | 0.03 | 0.62 | 0.43 | 0.54 | 0.43 |
| 7 | 17623225 | rs12699878 | 1.453 | 0.638 | 0.834 | 1.229 | 0.017 | 0.004 | 0.004 | 0 | 0.58 | 0.35 | 0.68 | 0.56 | 0.68 | 0.6 |
| 7 | 17625437 | rs1562630 | 1.298 | 0.615 | 1.279 | 1.232 | -0.012 | -0.003 | -0.003 | 0.272 | 0.59 | 0.52 | 0.7 | 0.57 | 0.68 | 0.6 |
| 7 | 17626916 | rs16871695 | -0.342 | 0.617 | 1.391 | 1.298 | 0.056 | 0.049 | 0.049 | 0 | 0.08 | 0.22 | 0.06 | 0.13 | 0.15 | 0.18 |
| 7 | 17626941 | rs6944072 | 1.069 | 0.544 | 1.445 | 1.233 | -0.011 | 0.002 | 0.002 | 0 | 0.63 | 0.42 | 0.71 | 0.57 | 0.72 | 0.64 |
| 7 | 17629311 | rs17137913 | -0.412 | 0.568 | 1.663 | 1.334 | 0.016 | 0.08 | 0.08 | 0.014 | 0.11 | 0.22 | 0.16 | 0.23 | 0.11 | 0.1 |
| 7 | 17629654 | rs17714316 | 0.346 | 0.306 | 1.24 | 1.021 | 0.01 | 0.033 | 0.033 | 0 | 0.14 | 0.08 | 0.08 | 0.02 | 0.09 | 0.13 |
| 7 | 17632351 | rs13228576 | 1.271 | 0.583 | 1.321 | 1.295 | 0.021 | 0.003 | 0.003 | 0.024 | 0.63 | 0.31 | 0.7 | 0.59 | 0.72 | 0.64 |
| 7 | 17640215 | rs17137965 | -0.586 | 0.68 | 1.486 | 1.07 | 0.079 | 0.078 | 0.078 | 0 | 0.09 | 0.25 | 0.07 | 0.14 | 0.15 | 0.18 |
| 7 | 17641577 | rs1448664 | -1.042 | 0.646 | 1.543 | 1.095 | 0.049 | 0.068 | 0.068 | 0 | 0.1 | 0.29 | 0.07 | 0.15 | 0.14 | 0.17 |
| 7 | 17644213 | rs2892855 | -2.133 | 0.766 | 1.314 | 1.786 | -0.127 | 0.057 | 0.057 | 0 | 0.51 | 0.97 | 0.38 | 0.69 | 0.47 | 0.6 |
| 7 | 17644647 | rs10950666 | -1.188 | 0.842 | 1.474 | 1.745 | -0.003 | 0.031 | 0.031 | 0 | 0.35 | 0.52 | 0.3 | 0.53 | 0.35 | 0.39 |
| 7 | 17651185 | rs11486792 | -1.23 | 0.901 | 1.478 | 1.746 | -0.001 | 0.022 | 0.022 | 0 | 0.36 | 0.53 | 0.29 | 0.48 | 0.35 | 0.39 |
| 7 | 17658420 | rs7803926 | -2.348 | 0.844 | 1.227 | 1.836 | -0.118 | 0.096 | 0.096 | 0 | 0.52 | 0.97 | 0.39 | 0.7 | 0.48 | 0.58 |
| 7 | 17661266 | rs2389827 | -0.579 | 0.848 | 1.468 | 1.852 | 0.007 | 0.113 | 0.113 | 0 | 0.09 | 0.22 | 0.06 | 0.23 | 0.07 | 0.15 |
| 7 | 17662408 | rs10246213 | -1.478 | 0.849 | 1.539 | 1.8 | 0.002 | 0.008 | 0.008 | 0 | 0.4 | 0.44 | 0.29 | 0.46 | 0.38 | 0.42 |
| 7 | 17662517 | rs7788582 | -1.165 | 0.858 | 1.488 | 1.795 | 0.002 | 0.03 | 0.03 | 0 | 0.35 | 0.5 | 0.3 | 0.46 | 0.36 | 0.39 |
| 7 | 17663745 | rs17138004 | -0.835 | 0.801 | 1.561 | 1.666 | 0.001 | -0.034 | -0.034 | 0 | 0.12 | 0.17 | 0.05 | 0 | 0.11 | 0.05 |
| 7 | 17667816 | rs2192073 | -1.8 | 0.791 | 1.603 | 1.703 | 0.002 | 0.037 | 0.037 | 0 | 0.39 | 0.75 | 0.32 | 0.69 | 0.35 | 0.52 |
| 7 | 17668920 | rs10249622 | -1.824 | 0.792 | 1.67 | 1.79 | -0.001 | -0.001 | -0.001 | 0 | 0.43 | 0.53 | 0.32 | 0.66 | 0.4 | 0.56 |
| 7 | 17668993 | rs10265182 | -0.202 | 0.78 | 1.498 | 1.641 | 0.003 | -0.029 | -0.029 | 0.421 | 0.06 | 0.14 | 0.04 | 0 | 0.06 | 0.02 |
| 7 | 17675093 | rs7796629 | -1.034 | 0.777 | 1.589 | 1.709 | 0.008 | 0.043 | 0.043 | 0.097 | 0.15 | 0.28 | 0.09 | 0.23 | 0.12 | 0.19 |
| 7 | 17676305 | rs6969402 | -1.862 | 0.775 | 1.527 | 1.706 | 0.012 | 0.028 | 0.028 | 0.252 | 0.21 | 0.47 | 0.13 | 0.23 | 0.19 | 0.21 |
| 7 | 17677721 | rs6961577 | -2.364 | 0.878 | 1.06 | 1.701 | -0.112 | 0.083 | 0.083 | 0 | 0.51 | 0.97 | 0.39 | 0.69 | 0.47 | 0.58 |
| 7 | 17681020 | rs6958106 | -1.548 | 0.831 | 1.399 | 1.788 | 0.004 | 0.054 | 0.054 | 0.021 | 0.14 | 0.3 | 0.09 | 0.23 | 0.14 | 0.17 |
| 7 | 17686090 | rs723724 | -1.598 | 0.831 | 1.435 | 1.788 | 0.007 | 0.069 | 0.069 | 0.085 | 0.14 | 0.35 | 0.1 | 0.23 | 0.14 | 0.17 |
| 7 | 17687722 | rs10225730 | -0.254 | 0.82 | 1.444 | 1.742 | -0.002 | -0.03 | -0.03 | 0 | 0.06 | 0.18 | 0.02 | 0 | 0.04 | 0.02 |
| 7 | 17690211 | rs765237 | -1.175 | 0.923 | 1.69 | 1.727 | -0.001 | 0.044 | 0.044 | 0 | 0.3 | 0.4 | 0.28 | 0.46 | 0.28 | 0.34 |
| 7 | 17690608 | rs2159853 | -2.021 | 0.978 | 1.364 | 1.725 | 0.004 | 0.012 | 0.012 | 0.325 | 0.19 | 0.3 | 0.11 | 0.24 | 0.22 | 0.24 |
| 7 | 17691608 | rs12699882 | 1.811 | 0.223 | -0.402 | 0.441 | -0.12 | 0.067 | 0.067 | 0 | 0.21 | 0.02 | 0.46 | 0.08 | 0.32 | 0.28 |
| 7 | 17698677 | rs4721639 | -0.63 | 0.322 | 0.279 | 0.747 | 0.002 | 0.007 | 0.007 | 0 | 0.29 | 0.29 | 0.27 | 0.63 | 0.29 | 0.37 |
| 7 | 17698736 | rs4719515 | -0.084 | 0.404 | -0.028 | 1.165 | -0.028 | -0.04 | -0.04 | 0 | 0.07 | 0.01 | 0.04 | 0.15 | 0.12 | 0.11 |
| 7 | 17699814 | rs10950670 | -1.923 | 0.047 | -0.157 | 0.707 | -0.041 | 0.087 | 0.087 | 0 | 0.55 | 0.88 | 0.4 | 0.69 | 0.51 | 0.62 |
| 7 | 17702281 | rs2389829 | -0.265 | 0.124 | 0.546 | 0.642 | 0.007 | -0.024 | -0.024 | 0 | 0.14 | 0.3 | 0.05 | 0.06 | 0.09 | 0.17 |
| 7 | 17703416 | rs7781652 | -2.003 | 0.249 | 0.561 | 0.596 | -0.003 | 0.003 | 0.003 | 0.4 | 0.3 | 0.74 | 0.17 | 0.22 | 0.29 | 0.29 |
| 7 | 17705199 | rs10260737 | -1.636 | 0.283 | 0.789 | 0.559 | 0.051 | -0.042 | -0.042 | 0 | 0.1 | 0.46 | 0.07 | 0 | 0.07 | 0.02 |
| 7 | 17711135 | rs4721643 | 0.03 | -0.116 | -0.041 | 0.551 | -0.007 | -0.006 | -0.006 | 0 | 0.56 | 0.45 | 0.73 | 0.74 | 0.63 | 0.65 |
| 7 | 17712278 | rs6973747 | 0.135 | -0.023 | -0.021 | 0.408 | 0.009 | 0.017 | 0.017 | 0 | 0.49 | 0.56 | 0.64 | 0.32 | 0.58 | 0.5 |
| 7 | 17712428 | rs13230396 | 1.502 | 0.864 | -0.594 | 0.024 | 0.004 | 0.008 | 0.008 | 0 | 0.27 | 0.17 | 0.43 | 0.27 | 0.23 | 0.3 |
| 7 | 17712942 | rs6461336 | -0.79 | 0.984 | -0.48 | -0.038 | -0.017 | 0.025 | 0.025 | 0 | 0.61 | 0.65 | 0.44 | 0.69 | 0.58 | 0.58 |
| 7 | 17713450 | rs10253857 | -1.07 | 0.276 | -0.261 | 0.247 | -0.02 | 0.006 | 0.006 | 0 | 0.69 | 0.67 | 0.84 | 0.63 | 0.76 | 0.78 |
| 7 | 17715616 | rs2723543 | -0.686 | 0.396 | -0.326 | 0.418 | -0.048 | 0.035 | 0.035 | 0 | 0.65 | 0.58 | 0.81 | 0.53 | 0.72 | 0.69 |
| 7 | 17718157 | rs2723548 | 0.441 | 0.671 | -0.433 | 0.316 | 0.001 | 0.026 | 0.026 | 0 | 0.48 | 0.45 | 0.57 | 0.32 | 0.42 | 0.49 |
| 7 | 17719652 | rs956758 | 1.264 | 0.45 | -0.485 | 0.261 | -0.051 | -0.001 | -0.001 | 0 | 0.39 | 0.09 | 0.5 | 0.32 | 0.35 | 0.43 |
| 7 | 17719956 | rs2075083 | 0.574 | 0.477 | -0.514 | 0.21 | 0.003 | -0.029 | -0.029 | 0 | 0.45 | 0.17 | 0.63 | 0.47 | 0.48 | 0.51 |
| 7 | 17720158 | rs2075084 | -0.733 | 0.543 | -0.23 | 0.618 | -0.042 | 0.011 | 0.011 | 0 | 0.67 | 0.61 | 0.83 | 0.61 | 0.75 | 0.77 |
| 7 | 17721858 | rs6960473 | -0.756 | 2.065 | 0.175 | 1.08 | 0.009 | 0.009 | 0.009 | 0 | 0.09 | 0.1 | 0.13 | 0.15 | 0.13 | 0.09 |
| 7 | 17722076 | rs2723549 | -0.754 | 1.456 | 0.076 | 0.837 | 0.009 | 0.036 | 0.036 | 0.017 | 0.07 | 0.19 | 0.06 | 0.1 | 0.09 | 0.14 |
| 7 | 17723110 | rs4721648 | 0.765 | 1.313 | -0.176 | 0.788 | -0.018 | 0.031 | 0.031 | 0 | 0.68 | 0.55 | 0.69 | 0.4 | 0.67 | 0.6 |
| 7 | 17723272 | rs13222101 | 0.221 | 2.021 | 0.033 | 0.93 | -0.006 | 0.005 | 0.005 | 0 | 0.09 | 0.08 | 0.11 | 0.04 | 0.17 | 0.11 |
| 7 | 17723962 | rs1558486 | 0.307 | 1.773 | 0.334 | 1.259 | 0.014 | 0.146 | 0.146 | 0 | 0.83 | 0.43 | 0.83 | 0.6 | 0.8 | 0.85 |
| 7 | 17725680 | rs11765014 | 1.214 | 1.999 | 0.083 | 1.157 | 0.009 | 0.062 | 0.062 | 0 | 0.72 | 0.38 | 0.78 | 0.5 | 0.7 | 0.71 |
| 7 | 17726121 | rs13240172 | -0.115 | 1.968 | 0.114 | 1.187 | 0.034 | 0.167 | 0.167 | 0 | 0.85 | 0.64 | 0.85 | 0.6 | 0.8 | 0.86 |
| 7 | 17727154 | rs6959390 | 1.056 | 1.92 | -0.558 | 0.881 | 0.009 | -0.001 | -0.001 | 0.084 | 0.64 | 0.39 | 0.68 | 0.33 | 0.54 | 0.56 |
| 7 | 17727264 | rs6959560 | 0.78 | 1.932 | -0.56 | 0.863 | 0.027 | 0.036 | 0.036 | 0 | 0.69 | 0.41 | 0.75 | 0.4 | 0.64 | 0.69 |
| 7 | 17727916 | rs2723557 | -1.226 | 0.798 | -1.669 | -0.044 | 0.115 | 0.186 | 0.186 | 0 | 0.63 | 0.91 | 0.63 | 0.93 | 0.74 | 0.75 |
| 7 | 17728255 | rs6964887 | -0.781 | 1.698 | -1.229 | 0.649 | 0.027 | 0.018 | 0.018 | 0 | 0.47 | 0.72 | 0.39 | 0.55 | 0.57 | 0.48 |
| 7 | 17730469 | rs2723559 | -1.221 | 1.593 | -1.539 | 0.682 | 0.124 | 0.019 | 0.019 | 0.153 | 0.56 | 0.89 | 0.54 | 0.63 | 0.68 | 0.64 |
| 7 | 17731550 | rs2058452 | -0.846 | 2.322 | -1.509 | 0.612 | -0.047 | 0.022 | 0.022 | 0 | 0.27 | 0.22 | 0.29 | 0.2 | 0.37 | 0.33 |

| | | | | | | | | | | | | | | | | |
|---|---|---|---|---|---|---|---|---|---|---|---|---|---|---|---|---|
| 7 | 17732490 | rs2537609 | -1.443 | 1.989 | -1.439 | 1.025 | 0.126 | -0.072 | -0.072 | 1.081 | 0.55 | 0.87 | 0.53 | 0.49 | 0.67 | 0.58 |
| 7 | 17738031 | rs10488472 | 0.68 | 2.205 | -1.361 | 1.065 | -0.041 | -0.005 | -0.005 | 0 | 0.08 | 0.01 | 0.13 | 0.08 | 0.11 | 0.16 |
| 7 | 17739875 | rs2389833 | 0.047 | 2.203 | -1.397 | 1.029 | -0.018 | -0.018 | -0.018 | 0 | 0.1 | 0.09 | 0.16 | 0.22 | 0.13 | 0.22 |
| 7 | 17743719 | rs4486086 | -0.909 | 2.482 | -1.325 | 0.963 | -0.009 | -0.02 | -0.02 | 3.682 | 0.42 | 0.68 | 0.41 | 0.51 | 0.47 | 0.49 |
| 7 | 17746628 | rs6953198 | -1.583 | 2.43 | -1.473 | 0.944 | 0.113 | 0.044 | 0.044 | 0.137 | 0.54 | 0.76 | 0.54 | 0.63 | 0.68 | 0.64 |
| 7 | 17746655 | rs6970817 | -0.966 | 2.451 | -1.382 | 0.829 | 0.007 | -0.006 | -0.006 | 0 | 0.45 | 0.75 | 0.39 | 0.41 | 0.55 | 0.42 |
| 7 | 17749477 | rs10256609 | 0.275 | 2.455 | -1.318 | 0.828 | -0.032 | -0.004 | -0.004 | 0 | 0.13 | 0.07 | 0.13 | 0.12 | 0.22 | 0.15 |
| 7 | 17750811 | rs2041589 | 0.275 | 2.455 | -1.318 | 0.828 | -0.032 | -0.004 | -0.004 | 2.278 | 0.13 | 0.07 | 0.13 | 0.12 | 0.22 | 0.15 |
| 7 | 17755584 | rs7786114 | -0.361 | 2.383 | -1.308 | 0.805 | -0.007 | -0.001 | -0.001 | 0 | 0.33 | 0.68 | 0.26 | 0.29 | 0.33 | 0.26 |
| 7 | 17760799 | rs10265355 | -0.743 | 2.41 | -1.265 | 0.81 | -0.008 | 0.004 | 0.004 | 0.131 | 0.41 | 0.61 | 0.4 | 0.37 | 0.46 | 0.42 |
| 7 | 17764375 | rs2723580 | -1.788 | 1.112 | -1.868 | 0.364 | 0.094 | 0.064 | 0.064 | 0 | 0.66 | 0.78 | 0.58 | 0.81 | 0.81 | 0.63 |
| 7 | 17766429 | rs17138233 | 1.087 | 2.026 | -1.735 | 0.582 | -0.036 | -0.036 | -0.036 | 0.448 | 0.06 | 0 | 0.04 | 0.12 | 0.12 | 0.1 |
| 7 | 17768277 | rs13226195 | -0.301 | 2.31 | -1.509 | 0.513 | -0.028 | 0.003 | 0.003 | 0 | 0.15 | 0.11 | 0.19 | 0.12 | 0.29 | 0.17 |
| 7 | 17770454 | rs2110014 | 0.121 | 1.771 | -1.641 | 0.767 | -0.034 | -0.122 | -0.122 | 0 | 0.17 | 0.01 | 0.15 | 0.41 | 0.17 | 0.28 |
| 7 | 17780891 | rs3919528 | -0.39 | 2.375 | -1.711 | 0.426 | -0.059 | 0.032 | 0.032 | 0 | 0.21 | 0.17 | 0.23 | 0.11 | 0.38 | 0.2 |
| 7 | 17781415 | rs2723530 | -0.766 | 1.86 | -1.96 | 0.151 | -0.002 | 0.087 | 0.087 | 0.036 | 0.42 | 0.71 | 0.37 | 0.82 | 0.42 | 0.54 |
| 7 | 17781483 | rs2537575 | 0.62 | 1.758 | -0.844 | 0.341 | 0.026 | 0.073 | 0.073 | 0.154 | 0.21 | 0.37 | 0.19 | 0.31 | 0.24 | 0.18 |
| 7 | 17782609 | rs2098223 | -0.135 | 1.759 | -1.191 | 0.34 | -0.055 | 0.03 | 0.03 | 0 | 0.21 | 0.16 | 0.23 | 0.11 | 0.38 | 0.2 |
| 7 | 17783473 | rs12536448 | 0.992 | 1.948 | -1.124 | 0.345 | -0.025 | -0.009 | -0.009 | 0 | 0.11 | 0.06 | 0.14 | 0.14 | 0.2 | 0.14 |
| 7 | 17784344 | rs2723539 | -1.443 | 1.127 | -1.161 | 0.399 | 0.136 | 0.03 | 0.03 | 0.037 | 0.52 | 0.93 | 0.51 | 0.62 | 0.66 | 0.57 |
| 7 | 17786633 | rs2723535 | -2.119 | 1.069 | -1.178 | 0.324 | 0.139 | 0.197 | 0.197 | 0 | 0.63 | 0.92 | 0.59 | 0.93 | 0.79 | 0.73 |
| 7 | 17794682 | rs1524780 | -0.96 | 1.254 | -1.101 | 0.43 | -0.014 | -0.054 | -0.054 | 0.069 | 0.4 | 0.2 | 0.47 | 0.5 | 0.55 | 0.51 |
| 7 | 17796008 | rs16871818 | 0.212 | 1.428 | -1.287 | 0.445 | -0.018 | -0.09 | -0.09 | 0.099 | 0.2 | 0.03 | 0.15 | 0.38 | 0.18 | 0.32 |
| 7 | 17798274 | rs2140711 | 0.321 | 1.412 | -1.292 | 0.379 | -0.024 | -0.081 | -0.081 | 0 | 0.18 | 0.03 | 0.15 | 0.36 | 0.18 | 0.31 |
| 7 | 17798958 | rs2389852 | -0.644 | 0.826 | -0.873 | 0.118 | 0.008 | -0.003 | -0.003 | 0 | 0.42 | 0.37 | 0.39 | 0.61 | 0.58 | 0.54 |
| 7 | 17801437 | rs2537583 | -0.048 | 0.834 | -0.701 | 0.041 | 0.027 | 0.114 | 0.114 | 0.813 | 0.2 | 0.5 | 0.2 | 0.31 | 0.21 | 0.18 |
| 7 | 17807563 | rs1830005 | -0.738 | 1.003 | -0.915 | 0.066 | 0.002 | 0.002 | 0.002 | 0 | 0.43 | 0.43 | 0.41 | 0.67 | 0.6 | 0.56 |
| 7 | 17808141 | rs10271676 | -0.629 | 1.153 | -0.856 | 0.138 | -0.001 | 0 | 0 | 0 | 0.4 | 0.42 | 0.39 | 0.62 | 0.56 | 0.54 |
| 7 | 17809788 | rs17138250 | -0.009 | 1.297 | -0.501 | 0.733 | 0 | 0.024 | 0.024 | 0 | 0.15 | 0.14 | 0.15 | 0.3 | 0.18 | 0.17 |
| 7 | 17811797 | rs2254737 | -0.147 | 0.382 | -1.224 | -0.08 | 0.031 | 0.094 | 0.094 | 0 | 0.55 | 0.66 | 0.5 | 0.81 | 0.42 | 0.56 |
| 7 | 17813444 | rs10488475 | 1.464 | 0.664 | -1.518 | -0.467 | -0.02 | 0.051 | 0.051 | 0 | 0.12 | 0.02 | 0.08 | 0 | 0.1 | 0.15 |
| 7 | 17813677 | rs2723520 | 0.414 | 1.83 | -1.788 | -0.676 | -0.016 | 0.065 | 0.065 | 0 | 0.57 | 0.65 | 0.72 | 0.82 | 0.54 | 0.63 |
| 7 | 17815402 | rs2723512 | 1.508 | -0.468 | -1.536 | -0.351 | -0.008 | 0.001 | 0.001 | 0.223 | 0.27 | 0.27 | 0.2 | 0.63 | 0.14 | 0.32 |
| 7 | 17822386 | rs2691619 | -1.898 | -0.078 | -2.035 | -0.232 | -0.027 | 0.03 | 0.03 | 0.418 | 0.82 | 0.99 | 0.92 | 0.88 | 0.83 | 0.84 |
| 7 | 17826628 | rs1852011 | -0.555 | -1.177 | -1.828 | -0.145 | -0.041 | 0.028 | 0.028 | 0 | 0.66 | 0.69 | 0.57 | 0.75 | 0.53 | 0.71 |
| 7 | 17831806 | rs2691583 | -0.577 | -1.212 | -1.856 | -0.165 | -0.04 | 0.027 | 0.027 | 0 | 0.65 | 0.7 | 0.56 | 0.74 | 0.52 | 0.71 |
| 7 | 17872093 | rs1524776 | -0.709 | -1.266 | -1.806 | 0.134 | -0.102 | -0.02 | -0.02 | 0 | 0.66 | 0.85 | 0.58 | 0.64 | 0.52 | 0.7 |
| 7 | 17874271 | rs4721660 | -2.345 | -1.399 | -1.938 | 0.034 | 0.008 | -0.047 | -0.047 | 0.109 | 0.87 | 0.99 | 0.96 | 0.78 | 0.93 | 0.87 |
| 7 | 17875431 | rs2691601 | -0.563 | -1.489 | -1.805 | 0.022 | -0.102 | 0.017 | 0.017 | 0 | 0.66 | 0.86 | 0.58 | 0.71 | 0.52 | 0.71 |
| 7 | 17908546 | rs12699916 | 1.599 | -1.41 | -1.767 | 0.023 | -0.133 | -0.03 | -0.03 | 0 | 0.19 | 0.02 | 0.36 | 0.15 | 0.39 | 0.16 |
| 7 | 17911038 | rs10282707 | -0.251 | -2.074 | -1.383 | 0.757 | 0.076 | -0.007 | -0.007 | 0 | 0.52 | 0.79 | 0.48 | 0.5 | 0.41 | 0.46 |
| 7 | 17911752 | rs1917368 | -0.293 | -2.171 | -1.396 | 0.733 | 0.081 | -0.008 | -0.008 | 0 | 0.52 | 0.84 | 0.49 | 0.43 | 0.4 | 0.46 |
| 7 | 17912336 | rs4721664 | 1.183 | -2.141 | -1.161 | 0.173 | 0.017 | -0.024 | -0.024 | 0 | 0.08 | 0.17 | 0.09 | 0.02 | 0.11 | 0.02 |
| 7 | 17927789 | rs2714878 | -0.499 | -2.269 | -1.195 | 0.186 | 0.003 | 0.009 | 0.009 | 0 | 0.56 | 0.53 | 0.47 | 0.71 | 0.41 | 0.69 |
| 7 | 17929655 | rs4291154 | 0.208 | -2.793 | -1.139 | -0.093 | 0.035 | -0.02 | -0.02 | 0 | 0.38 | 0.73 | 0.31 | 0.57 | 0.28 | 0.44 |
| 7 | 17948140 | rs2714862 | 0.551 | -2.539 | -1.422 | -0.009 | -0.04 | 0.014 | 0.014 | 0 | 0.38 | 0.43 | 0.3 | 0.52 | 0.25 | 0.41 |
| 7 | 17959134 | rs2714858 | -0.058 | -2.204 | -1.459 | 0.306 | 0 | 0 | 0 | 0.008 | 0.42 | 0.52 | 0.45 | 0.57 | 0.35 | 0.62 |
| 7 | 17965838 | rs17138455 | 0.508 | -2.271 | -1.686 | 0.433 | 0.087 | -0.006 | -0.006 | 0.071 | 0.27 | 0.06 | 0.21 | 0.27 | 0.15 | 0.2 |
| 7 | 17968205 | rs10085395 | 1.147 | -1.42 | -1.289 | 0.361 | 0.012 | 0.013 | 0.013 | 0 | 0.23 | 0.18 | 0.12 | 0.07 | 0.17 | 0.08 |
| 7 | 17969900 | rs12699918 | 0.144 | 0.061 | -1.374 | 0.783 | -0.119 | -0.021 | -0.021 | 0 | 0.33 | 0.1 | 0.42 | 0.36 | 0.48 | 0.29 |
| 7 | 17978129 | rs6973480 | 0.015 | 0.112 | -1.317 | 0.776 | -0.037 | 0.01 | 0.01 | 0 | 0.36 | 0.31 | 0.57 | 0.35 | 0.5 | 0.3 |
| 7 | 17983297 | rs4543441 | 0.018 | 0.599 | -1.121 | 0.998 | 0.005 | -0.001 | -0.001 | 1.559 | 0.3 | 0.34 | 0.53 | 0.33 | 0.49 | 0.25 |
| 7 | 17992014 | rs4466298 | -0.376 | 0.391 | -1.244 | 1.415 | -0.001 | -0.012 | -0.012 | 0 | 0.37 | 0.31 | 0.58 | 0.59 | 0.57 | 0.34 |
| 7 | 17994494 | rs12699921 | -0.005 | 0.694 | -1.184 | 1.124 | -0.086 | -0.014 | -0.014 | 0.073 | 0.29 | 0.16 | 0.51 | 0.33 | 0.49 | 0.22 |
| 7 | 18002126 | rs13224298 | 0.027 | 0.66 | -1.104 | 1.122 | -0.061 | -0.008 | -0.008 | 0 | 0.28 | 0.18 | 0.52 | 0.34 | 0.51 | 0.22 |
| 7 | 18003670 | rs11766489 | -0.122 | 0.566 | -0.987 | 1.076 | 0.019 | -0.003 | -0.003 | 0 | 0.31 | 0.39 | 0.54 | 0.33 | 0.52 | 0.24 |
| 7 | 18005237 | rs9648238 | 0.097 | 0.597 | -1.238 | 1.09 | -0.116 | -0.028 | -0.028 | 0 | 0.28 | 0.09 | 0.51 | 0.33 | 0.52 | 0.22 |
| 7 | 18017943 | rs12669067 | -0.928 | 0.037 | -1.583 | 1.156 | 0.014 | -0.006 | -0.006 | 0 | 0.45 | 0.39 | 0.6 | 0.53 | 0.62 | 0.35 |
| 7 | 18022242 | rs11535160 | 0.618 | -0.104 | -1.644 | 0.942 | -0.09 | 0.019 | 0.019 | 0.679 | 0.26 | 0.1 | 0.49 | 0.25 | 0.53 | 0.19 |
| 7 | 18022771 | rs10499509 | -0.889 | -0.115 | -1.658 | 1.007 | 0.028 | 0.003 | 0.003 | 0 | 0.44 | 0.37 | 0.62 | 0.47 | 0.65 | 0.33 |
| 7 | 18023617 | rs12530608 | 0.146 | -0.111 | -1.766 | 0.993 | 0.011 | -0.027 | -0.027 | 0 | 0.16 | 0.12 | 0.11 | 0.26 | 0.13 | 0.12 |
| 7 | 18025423 | rs6947751 | -0.011 | 0.211 | -1.427 | 0.501 | -0.055 | 0.041 | 0.041 | 0.482 | 0.29 | 0.2 | 0.5 | 0.21 | 0.52 | 0.19 |
| 7 | 18025440 | rs6947763 | -0.128 | 0.215 | -1.427 | 0.503 | -0.03 | 0.026 | 0.026 | 0.171 | 0.3 | 0.28 | 0.51 | 0.21 | 0.52 | 0.19 |
| 7 | 18034431 | rs12699936 | -0.307 | -0.1 | -1.539 | 0.334 | -0.038 | 0.053 | 0.053 | 0 | 0.37 | 0.2 | 0.55 | 0.27 | 0.55 | 0.32 |
| 7 | 18035383 | rs12155344 | -0.291 | -0.078 | -1.546 | 0.334 | -0.055 | 0.06 | 0.06 | 0.183 | 0.37 | 0.16 | 0.54 | 0.27 | 0.55 | 0.31 |
| 7 | 18036309 | rs12155045 | -0.113 | -0.095 | -1.559 | 0.326 | -0.082 | 0.051 | 0.051 | 0.519 | 0.35 | 0.11 | 0.54 | 0.27 | 0.55 | 0.3 |
| 7 | 18036784 | rs12699939 | -0.293 | -0.084 | -1.554 | 0.331 | -0.055 | 0.06 | 0.06 | 0.258 | 0.37 | 0.16 | 0.54 | 0.28 | 0.55 | 0.3 |
| 7 | 18039252 | rs4721679 | -0.274 | -0.114 | -1.52 | 0.318 | -0.04 | 0.054 | 0.054 | 0.271 | 0.37 | 0.2 | 0.54 | 0.28 | 0.55 | 0.28 |
| 7 | 18040264 | rs6957876 | -0.326 | -0.15 | -1.341 | 0.287 | -0.017 | 0.033 | 0.033 | 0.176 | 0.37 | 0.31 | 0.56 | 0.28 | 0.55 | 0.28 |

| | | | | | | | | | | | | | | | | |
|---|---|---|---|---|---|---|---|---|---|---|---|---|---|---|---|---|
| 7 | 18041085 | rs4568518 | -0.426 | -0.182 | -1.021 | 0.258 | -0.018 | 0.038 | 0.038 | 0.153 | 0.39 | 0.29 | 0.56 | 0.28 | 0.56 | 0.31 |
| 7 | 18044262 | rs12699949 | -0.663 | -0.485 | -0.898 | 0.286 | -0.006 | -0.001 | -0.001 | 0.808 | 0.58 | 0.53 | 0.69 | 0.61 | 0.67 | 0.57 |
| 7 | 18045091 | rs6461367 | -0.43 | -0.447 | -1.16 | 0.303 | 0.021 | 0.006 | 0.006 | 0 | 0.56 | 0.33 | 0.66 | 0.61 | 0.66 | 0.55 |
| 7 | 18055120 | rs13232596 | 0.304 | 0.772 | -1.356 | 0.241 | -0.047 | -0.019 | -0.019 | 0.278 | 0.07 | 0.01 | 0.09 | 0.06 | 0.16 | 0.06 |
| 7 | 18058318 | rs10950697 | -0.36 | -0.313 | -1.199 | 0.19 | 0.042 | 0.006 | 0.006 | 0.106 | 0.54 | 0.19 | 0.65 | 0.61 | 0.66 | 0.57 |
| 7 | 18062610 | rs7781290 | -0.475 | -0.257 | -1.174 | 0.201 | 0.007 | 0.001 | 0.001 | 0 | 0.57 | 0.43 | 0.68 | 0.61 | 0.67 | 0.58 |
| 7 | 18062796 | rs7781217 | -0.427 | -0.154 | -1.631 | 0.247 | 0.039 | 0.001 | 0.001 | 0.198 | 0.55 | 0.19 | 0.64 | 0.61 | 0.66 | 0.57 |
| 7 | 18066569 | rs3800962 | 1.128 | 0.136 | -1.283 | 0.231 | -0.035 | 0.125 | 0.125 | 2.65 | 0.38 | 0.2 | 0.4 | 0.15 | 0.42 | 0.22 |
| 7 | 18069638 | rs6976627 | -0.356 | 0.246 | -0.514 | 0.378 | 0.017 | -0.071 | -0.071 | 0 | 0.86 | 0.76 | 0.89 | 1 | 0.83 | 0.93 |
| 7 | 18083032 | rs4416730 | 0.132 | -0.113 | -1.128 | 1.079 | -0.007 | -0.08 | -0.08 | 0.123 | 0.14 | 0.03 | 0.14 | 0.41 | 0.12 | 0.22 |
| 7 | 18085049 | rs17138549 | 0.566 | -0.204 | -1.162 | 1.121 | 0.009 | -0.009 | -0.009 | 0 | 0.07 | 0.04 | 0.09 | 0.2 | 0.04 | 0.14 |
| 7 | 18085785 | rs17138554 | 0.176 | -0.326 | -0.992 | 0.794 | -0.004 | -0.001 | -0.001 | 0.033 | 0.13 | 0.14 | 0.1 | 0.1 | 0.1 | 0.18 |
| 7 | 18088558 | rs6963953 | -0.652 | -0.245 | -0.652 | 1.127 | -0.02 | 0.052 | 0.052 | 0 | 0.22 | 0.37 | 0.24 | 0.61 | 0.17 | 0.36 |
| 7 | 18090280 | rs10257697 | 0.187 | 0.26 | -1.313 | 0.801 | 0.008 | 0.028 | 0.028 | 0 | 0.55 | 0.28 | 0.59 | 0.66 | 0.59 | 0.49 |
| 7 | 18093358 | rs7804628 | 0.587 | 0.317 | -1.787 | 0.461 | -0.105 | 0.097 | 0.097 | 0.907 | 0.41 | 0.08 | 0.44 | 0.26 | 0.48 | 0.26 |
| 7 | 18094115 | rs4449697 | 0.228 | -0.513 | -1.868 | 0.296 | 0.059 | 0.055 | 0.055 | 0.058 | 0.34 | 0.1 | 0.2 | 0.2 | 0.29 | 0.24 |
| 7 | 18095821 | rs10275615 | -0.13 | -0.784 | -0.977 | 0.754 | -0.048 | 0.024 | 0.024 | 0.033 | 0.26 | 0.4 | 0.26 | 0.32 | 0.18 | 0.38 |
| 7 | 18101378 | rs6948797 | 1.233 | -0.439 | -0.76 | 0.561 | -0.007 | -0.031 | -0.031 | 0.41 | 0.09 | 0.13 | 0.12 | 0.02 | 0.06 | 0.12 |
| 7 | 18104972 | rs6966799 | -0.357 | -0.04 | -0.513 | 0.441 | 0.011 | -0.015 | -0.015 | 0 | 0.38 | 0.58 | 0.49 | 0.24 | 0.39 | 0.42 |
| 7 | 18110113 | rs7808162 | 1.091 | 0.149 | -0.6 | 0.684 | -0.007 | -0.032 | -0.032 | 0 | 0.09 | 0.14 | 0.12 | 0.02 | 0.06 | 0.12 |
| 7 | 18113008 | rs2731568 | -0.107 | 0.317 | -1.054 | 1.208 | -0.072 | -0.107 | -0.107 | 1.878 | 0.21 | 0.02 | 0.18 | 0.33 | 0.27 | 0.21 |
| 7 | 18122596 | rs1540880 | -0.8 | 0.216 | -0.042 | 1.178 | 0.011 | 0.1 | 0.1 | 0 | 0.06 | 0.28 | 0.07 | 0.1 | 0.02 | 0.12 |
| 7 | 18128248 | rs2049726 | -0.344 | 0.361 | -0.721 | 1.144 | 0.011 | -0.024 | -0.024 | 0 | 0.55 | 0.78 | 0.69 | 0.53 | 0.54 | 0.78 |
| 7 | 18150005 | rs1540882 | 0.461 | 0.179 | -0.764 | 1.413 | 0.051 | -0.008 | -0.008 | 0 | 0.57 | 0.21 | 0.67 | 0.56 | 0.7 | 0.52 |
| 7 | 18150431 | rs2731564 | 0.286 | -0.288 | -0.014 | 1.358 | -0.027 | -0.007 | -0.007 | 0 | 0.83 | 0.59 | 0.85 | 0.85 | 0.88 | 0.85 |
| 7 | 18158464 | rs4721694 | 0.323 | -0.323 | -0.021 | 1.296 | 0.006 | -0.003 | -0.003 | 0.074 | 0.26 | 0.21 | 0.17 | 0.29 | 0.18 | 0.37 |
| 7 | 18161620 | rs2697908 | 0.907 | -0.395 | -0.469 | 1.423 | 0.062 | -0.028 | -0.028 | 0 | 0.55 | 0.22 | 0.67 | 0.54 | 0.7 | 0.43 |
| 7 | 18163689 | rs1569174 | 0.599 | -0.431 | -0.703 | 1.4 | -0.013 | -0.055 | -0.055 | 0 | 0.23 | 0.04 | 0.14 | 0.27 | 0.17 | 0.37 |
| 7 | 18164980 | rs12535940 | 0.629 | -0.484 | -0.671 | 1.291 | -0.006 | -0.038 | -0.038 | 1.276 | 0.23 | 0.05 | 0.14 | 0.27 | 0.17 | 0.37 |
| 7 | 18171029 | rs1178370 | -1.01 | -0.483 | -0.507 | 1.354 | 0.074 | 0.018 | 0.018 | 1.086 | 0.45 | 0.79 | 0.3 | 0.39 | 0.28 | 0.55 |
| 7 | 18172374 | rs4490725 | -0.518 | -0.284 | -0.543 | 1.445 | -0.015 | -0.031 | -0.031 | 0.894 | 0.06 | 0.01 | 0.04 | 0.17 | 0.06 | 0.13 |
| 7 | 18186306 | rs12699970 | 0.407 | -0.164 | -0.456 | 1.352 | 0.008 | 0.015 | 0.015 | 1.422 | 0.25 | 0.14 | 0.15 | 0.22 | 0.18 | 0.36 |
| 7 | 18189670 | rs1178356 | -1.641 | 0.358 | -1.305 | 1.307 | -0.013 | -0.027 | -0.027 | 0.454 | 0.69 | 0.9 | 0.52 | 0.67 | 0.69 | 0.76 |
| 7 | 18193589 | rs12171593 | -0.049 | -0.131 | -0.19 | 1.563 | -0.056 | 0.056 | 0.056 | 0 | 0.13 | 0.34 | 0.1 | 0.14 | 0.05 | 0.12 |
| 7 | 18200205 | rs7788797 | 1.051 | -0.092 | -0.293 | 1.617 | 0.01 | -0.018 | -0.018 | 1.503 | 0.25 | 0.13 | 0.16 | 0.3 | 0.19 | 0.36 |
| 7 | 18215216 | rs1961051 | 1.06 | -0.075 | -0.306 | 1.605 | -0.02 | 0.001 | 0.001 | 1.336 | 0.24 | 0.04 | 0.14 | 0.22 | 0.19 | 0.35 |
| 7 | 18216046 | rs17138718 | 1.052 | -0.076 | -0.302 | 1.614 | 0.009 | 0.013 | 0.013 | 1.875 | 0.25 | 0.12 | 0.15 | 0.24 | 0.19 | 0.35 |
| 7 | 18218278 | rs1178344 | -0.888 | -0.21 | -0.099 | 1.574 | 0.059 | 0.002 | 0.002 | 0.512 | 0.44 | 0.75 | 0.29 | 0.41 | 0.28 | 0.5 |
| 7 | 18221192 | rs1406361 | -0.252 | 0.503 | 0.14 | 1.396 | 0.047 | -0.019 | -0.019 | 0.014 | 0.14 | 0.36 | 0.15 | 0.1 | 0.21 | 0.13 |
| 7 | 18221376 | rs16871944 | 1.071 | 0.574 | -0.214 | 1.407 | 0.011 | -0.006 | -0.006 | 1.408 | 0.25 | 0.12 | 0.14 | 0.28 | 0.19 | 0.35 |
| 7 | 18221400 | rs1178339 | -1.227 | 0.375 | -0.364 | 1.223 | 0.001 | 0.015 | 0.015 | 0 | 0.22 | 0.56 | 0.2 | 0.24 | 0.24 | 0.21 |
| 7 | 18223455 | rs2058074 | 0.795 | 0.178 | -0.563 | 0.914 | 0.008 | -0.03 | -0.03 | 0 | 0.26 | 0.07 | 0.15 | 0.31 | 0.19 | 0.36 |
| 7 | 18224597 | rs17138740 | 0.776 | 0.211 | -0.527 | 0.815 | 0.013 | -0.014 | -0.014 | 0 | 0.26 | 0.16 | 0.16 | 0.32 | 0.19 | 0.36 |
| 7 | 18233458 | rs1178328 | -0.601 | 0.354 | -0.444 | 0.854 | 0.029 | -0.009 | -0.009 | 0 | 0.44 | 0.69 | 0.29 | 0.5 | 0.36 | 0.59 |
| 7 | 18236056 | rs4721701 | -0.355 | 0.174 | -0.002 | 0.765 | -0.008 | 0.003 | 0.003 | 0.04 | 0.42 | 0.49 | 0.29 | 0.49 | 0.36 | 0.58 |
| 7 | 18258567 | rs1178375 | -0.41 | 0.118 | 0.354 | 0.754 | 0.022 | 0.015 | 0.015 | 0 | 0.06 | 0.22 | 0.03 | 0.02 | 0.03 | 0.03 |
| 7 | 18266604 | rs2567209 | 0.004 | 0.435 | -0.032 | 0.726 | 0.004 | 0.006 | 0.006 | 0 | 0.75 | 0.68 | 0.8 | 0.77 | 0.7 | 0.86 |
| 7 | 18270430 | rs1007192 | 0.269 | 0.218 | -0.355 | 0.568 | 0.003 | 0.002 | 0.002 | 0.136 | 0.7 | 0.52 | 0.79 | 0.77 | 0.69 | 0.84 |
| 7 | 18281894 | rs6461374 | -0.515 | 0.376 | -0.548 | 0.662 | 0.005 | 0.006 | 0.006 | 0 | 0.78 | 0.75 | 0.82 | 0.77 | 0.72 | 0.87 |
| 7 | 18288593 | rs917325 | 0.823 | 0.343 | -1.065 | 0.753 | -0.103 | -0.06 | -0.06 | 0 | 0.39 | 0.1 | 0.59 | 0.47 | 0.49 | 0.44 |
| 7 | 18301455 | rs17346782 | 1.779 | 0.784 | -1.363 | 0.526 | -0.066 | -0.024 | -0.024 | 0 | 0.13 | 0.01 | 0.25 | 0.14 | 0.17 | 0.14 |
| 7 | 18310791 | rs302178 | 0.781 | 0.327 | -1.068 | 0.69 | -0.069 | -0.057 | -0.057 | 0.128 | 0.38 | 0.14 | 0.59 | 0.49 | 0.47 | 0.44 |
| 7 | 18314737 | rs6950126 | 0.555 | 0.188 | -0.859 | 0.635 | -0.029 | -0.024 | -0.024 | 0.544 | 0.35 | 0.44 | 0.2 | 0.28 | 0.22 | 0.4 |
| 7 | 18316305 | rs6956728 | 0.891 | 0.328 | -1.101 | 0.721 | -0.096 | -0.082 | -0.082 | 0.275 | 0.37 | 0.12 | 0.58 | 0.49 | 0.47 | 0.44 |
| 7 | 18326622 | rs17138914 | 1.363 | -0.05 | -0.714 | 1.187 | 0.037 | -0.064 | -0.064 | 0.893 | 0.1 | 0 | 0.04 | 0.17 | 0.04 | 0.08 |
| 7 | 18328568 | rs625733 | 0.279 | -0.423 | -0.032 | 0.928 | -0.041 | -0.054 | -0.054 | 2.409 | 0.1 | 0.33 | 0.05 | 0 | 0.03 | 0.03 |
| 7 | 18328833 | rs11769668 | 0.284 | -0.132 | -0.035 | 0.918 | -0.005 | -0.021 | -0.021 | 0 | 0.21 | 0.16 | 0.21 | 0.23 | 0.24 | 0.14 |
| 7 | 18329050 | rs623532 | -1.045 | -0.088 | -0.054 | 0.926 | 0.001 | 0 | 0 | 0.053 | 0.37 | 0.63 | 0.28 | 0.25 | 0.3 | 0.2 |
| 7 | 18330980 | rs1015088 | -0.917 | -0.07 | -0.062 | 0.933 | 0.015 | 0.005 | 0.005 | 0.281 | 0.36 | 0.68 | 0.29 | 0.24 | 0.29 | 0.19 |
| 7 | 18333696 | rs2960783 | 1.919 | -0.232 | -0.274 | 0.775 | -0.044 | -0.07 | -0.07 | 0.038 | 0.14 | 0.01 | 0.11 | 0.26 | 0.15 | 0.25 |
| 7 | 18336292 | rs2915399 | -1.643 | -0.237 | -0.389 | 0.844 | -0.038 | 0.077 | 0.077 | 0 | 0.59 | 0.83 | 0.47 | 0.72 | 0.55 | 0.66 |
| 7 | 18338350 | rs582937 | 1.406 | 0.152 | -0.201 | 0.904 | 0.011 | 0.035 | 0.035 | 0.545 | 0.16 | 0.29 | 0.13 | 0.26 | 0.17 | 0.25 |
| 7 | 18343723 | rs614304 | 1.887 | 0.142 | -0.233 | 0.927 | -0.01 | -0.024 | -0.024 | 0.094 | 0.14 | 0.08 | 0.12 | 0.26 | 0.15 | 0.25 |
| 7 | 18344095 | rs7806296 | -0.905 | 0.119 | -0.303 | 0.935 | 0 | -0.001 | -0.001 | 0 | 0.4 | 0.59 | 0.36 | 0.46 | 0.38 | 0.36 |
| 7 | 18347657 | rs636246 | 1.723 | 0.25 | -0.379 | 0.982 | 0.003 | 0.013 | 0.013 | 1.217 | 0.14 | 0.2 | 0.12 | 0.26 | 0.15 | 0.25 |
| 7 | 18364750 | rs604804 | 1.746 | 0.3 | -0.336 | 1.148 | 0.005 | 0.014 | 0.014 | 0.936 | 0.14 | 0.2 | 0.12 | 0.26 | 0.16 | 0.24 |
| 7 | 18370302 | rs4721708 | 0.185 | 0.22 | -0.253 | 1.224 | -0.004 | 0.019 | 0.019 | 0 | 0.69 | 0.4 | 0.73 | 0.55 | 0.71 | 0.54 |
| 7 | 18382590 | rs1318776 | 1.154 | 0.209 | -0.415 | 1.028 | -0.023 | 0.11 | 0.11 | 0 | 0.46 | 0.2 | 0.52 | 0.29 | 0.46 | 0.41 |
| 7 | 18383716 | rs17347075 | 1.372 | 0.219 | -0.667 | 1.018 | -0.063 | 0.105 | 0.105 | 0 | 0.42 | 0.11 | 0.5 | 0.27 | 0.44 | 0.38 |

| | | | | | | | | | | | | | | | | |
|---|---|---|---|---|---|---|---|---|---|---|---|---|---|---|---|---|
| 7 | 18384157 | rs17347096 | 1.247 | 0.255 | -0.681 | 1.028 | -0.069 | 0.105 | 0.105 | 0 | 0.43 | 0.09 | 0.5 | 0.27 | 0.46 | 0.39 |
| 7 | 18388143 | rs17419024 | 0.949 | 0.092 | -0.835 | 1.081 | -0.013 | -0.034 | -0.034 | 0 | 0.51 | 0.14 | 0.61 | 0.5 | 0.55 | 0.57 |
| 7 | 18391557 | rs10486304 | -0.255 | 0.549 | -0.474 | 1.171 | -0.001 | 0.005 | 0.005 | 0 | 0.24 | 0.26 | 0.21 | 0.23 | 0.23 | 0.14 |
| 7 | 18392320 | rs2389941 | 1.229 | 1.286 | -0.671 | 1.21 | -0.003 | -0.009 | -0.009 | 0 | 0.12 | 0.08 | 0.09 | 0.18 | 0.11 | 0.19 |
| 7 | 18392543 | rs655828 | 0.826 | 1.298 | -0.624 | 1.106 | -0.006 | -0.017 | -0.017 | 0 | 0.15 | 0.1 | 0.23 | 0.29 | 0.17 | 0.26 |
| 7 | 18394098 | rs17347159 | 0.428 | 1.972 | -0.29 | 0.739 | -0.051 | 0.048 | 0.048 | 0.039 | 0.12 | 0.02 | 0.13 | 0.03 | 0.19 | 0.14 |
| 7 | 18399925 | rs1367697 | -0.02 | 1.645 | 0.095 | 0.693 | -0.011 | -0.042 | -0.042 | 0.497 | 0.92 | 0.88 | 0.96 | 1 | 0.94 | 0.93 |
| 7 | 18402530 | rs10255111 | 0.182 | 2.151 | 0.186 | 0.863 | 0.033 | -0.004 | -0.004 | 0.017 | 0.18 | 0.35 | 0.33 | 0.16 | 0.22 | 0.16 |
| 7 | 18402946 | rs17419156 | 0.992 | 1.461 | -0.667 | 1.095 | -0.013 | -0.006 | -0.006 | 0 | 0.54 | 0.12 | 0.45 | 0.58 | 0.49 | 0.5 |
| 7 | 18403678 | rs7788359 | -0.181 | 1.318 | 0.212 | 1.052 | -0.002 | -0.036 | -0.036 | 0 | 0.07 | 0.12 | 0.04 | 0 | 0.06 | 0.07 |
| 7 | 18404917 | rs1365893 | -0.977 | 1.913 | 0.791 | 1.709 | 0.027 | 0.071 | 0.071 | 0 | 0.22 | 0.43 | 0.29 | 0.42 | 0.25 | 0.29 |
| 7 | 18411672 | rs10236525 | 0.469 | 1.972 | 0.939 | 1.812 | 0.031 | 0.015 | 0.015 | 0 | 0.12 | 0.16 | 0.25 | 0.21 | 0.23 | 0.17 |
| 7 | 18420005 | rs17139138 | -0.977 | 1.901 | 0.973 | 1.824 | -0.024 | 0.076 | 0.076 | 0 | 0.09 | 0.26 | 0.06 | 0.22 | 0.04 | 0.13 |
| 7 | 18422684 | rs7786745 | 0.626 | 2 | 0.89 | 1.784 | 0.008 | 0.001 | 0.001 | 0 | 0.12 | 0.14 | 0.26 | 0.22 | 0.23 | 0.17 |
| 7 | 18436019 | rs10225649 | -1.2 | 1.886 | 0.963 | 1.858 | 0.044 | 0.109 | 0.109 | 0 | 0.23 | 0.45 | 0.33 | 0.45 | 0.27 | 0.31 |
| 7 | 18450104 | rs2162563 | 0.097 | 1.768 | 1.05 | 1.582 | 0.033 | 0.008 | 0.008 | 0 | 0.83 | 0.75 | 0.72 | 0.8 | 0.75 | 0.81 |
| 7 | 18455288 | rs13228023 | 1.364 | 1.891 | 1.141 | 1.665 | -0.005 | -0.002 | -0.002 | 0 | 0.66 | 0.4 | 0.6 | 0.34 | 0.62 | 0.6 |
| 7 | 18504411 | rs213272 | -0.94 | 1.932 | 1.382 | 1.707 | 0.019 | 0.02 | 0.02 | 0 | 0.3 | 0.6 | 0.37 | 0.6 | 0.33 | 0.3 |
| 7 | 18504508 | rs213273 | -0.988 | 1.752 | 0.957 | 1.521 | -0.01 | 0.032 | 0.032 | 0 | 0.94 | 0.99 | 0.97 | 1 | 0.94 | 0.93 |
| 7 | 18517279 | rs17347800 | 0.991 | 1.878 | 0.639 | 1.585 | -0.012 | -0.018 | -0.018 | 0 | 0.06 | 0.02 | 0.04 | 0.1 | 0.08 | 0.16 |
| 7 | 18522740 | rs10243618 | -0.352 | 1.045 | 0.061 | 0.878 | -0.035 | 0.012 | 0.012 | 0.112 | 0.33 | 0.53 | 0.22 | 0.36 | 0.26 | 0.34 |
| 7 | 18548419 | rs17432448 | 1.105 | 0.75 | -0.557 | 0.237 | 0.02 | 0.135 | 0.135 | 0 | 0.41 | 0.13 | 0.47 | 0.16 | 0.37 | 0.34 |
| 7 | 18579254 | rs2704289 | 0.787 | 1.372 | -0.61 | 0.659 | -0.085 | -0.044 | -0.044 | 0 | 0.19 | 0.01 | 0.2 | 0.23 | 0.27 | 0.13 |
| 7 | 18590659 | rs11543651 | 0.198 | -0.115 | -0.395 | -0.35 | -0.002 | -0.042 | -0.042 | 0.75 | 0.31 | 0.32 | 0.42 | 0.1 | 0.3 | 0.28 |
| 7 | 18601365 | rs801763 | -0.163 | -1.371 | -0.977 | -0.747 | 0.06 | 0.062 | 0.062 | 0.089 | 0.34 | 0.79 | 0.24 | 0.08 | 0.13 | 0.26 |
| 7 | 18604337 | rs17150238 | -0.389 | -0.792 | -0.758 | -0.825 | 0.001 | -0.026 | -0.026 | 0 | 0.23 | 0.33 | 0.36 | 0.07 | 0.22 | 0.24 |
| 7 | 18614519 | rs2520456 | 1.598 | -0.36 | -1.595 | -0.422 | -0.116 | -0.077 | -0.077 | 3.087 | 0.24 | 0.01 | 0.22 | 0.3 | 0.32 | 0.26 |
| 7 | 18616914 | rs3807917 | 0.841 | -0.367 | -1.59 | -0.192 | -0.11 | -0.108 | -0.108 | 0 | 0.31 | 0.13 | 0.41 | 0.46 | 0.4 | 0.36 |
| 7 | 18619934 | rs1615309 | -0.029 | -0.588 | -1.386 | -0.498 | 0.011 | -0.029 | -0.029 | 0 | 0.56 | 0.76 | 0.48 | 0.51 | 0.46 | 0.49 |
| 7 | 18621630 | rs1726597 | -0.277 | -0.985 | -1.665 | -0.635 | 0.019 | 0.031 | 0.031 | 0 | 0.32 | 0.71 | 0.36 | 0.16 | 0.21 | 0.27 |
| 7 | 18621925 | rs13234273 | 1.798 | -0.958 | -1.671 | -0.714 | 0.007 | 0.026 | 0.026 | 0 | 0.13 | 0.06 | 0.11 | 0.04 | 0.09 | 0.15 |
| 7 | 18622929 | rs2520458 | 1.31 | -0.729 | -1.388 | -0.423 | -0.078 | -0.097 | -0.097 | 0.188 | 0.3 | 0.09 | 0.28 | 0.53 | 0.38 | 0.38 |
| 7 | 18624800 | rs2073974 | 0.22 | -0.76 | -0.688 | -0.4 | -0.001 | 0.001 | 0.001 | 0.098 | 0.4 | 0.4 | 0.57 | 0.63 | 0.52 | 0.53 |
| 7 | 18626623 | rs2695029 | -0.35 | -0.743 | -0.915 | -0.4 | 0.009 | 0.032 | 0.032 | 0 | 0.57 | 0.62 | 0.58 | 0.66 | 0.54 | 0.62 |
| 7 | 18628779 | rs1726614 | -1.182 | -0.667 | -0.869 | -0.46 | -0.001 | -0.044 | -0.044 | 0 | 0.91 | 0.81 | 0.95 | 1 | 0.92 | 0.92 |
| 7 | 18629590 | rs3807907 | 0.667 | -0.695 | -0.9 | -0.227 | -0.082 | -0.059 | -0.059 | 0 | 0.36 | 0.12 | 0.47 | 0.52 | 0.47 | 0.44 |
| 7 | 18630208 | rs1726610 | -0.795 | -0.261 | -0.737 | -0.824 | -0.001 | -0.006 | -0.006 | 0 | 0.69 | 0.63 | 0.66 | 0.87 | 0.63 | 0.78 |
| 7 | 18630523 | rs2106506 | 0.465 | -0.198 | -0.729 | -0.703 | 0.013 | 0.013 | 0.013 | 0 | 0.25 | 0.18 | 0.24 | 0.04 | 0.15 | 0.19 |
| 7 | 18631680 | rs2695027 | 0.97 | 0.236 | -1.293 | -0.249 | -0.044 | 0.023 | 0.023 | 0 | 0.36 | 0.13 | 0.3 | 0.71 | 0.38 | 0.52 |
| 7 | 18633807 | rs3213615 | -1.068 | 0.956 | -0.792 | -0.025 | -0.016 | -0.011 | -0.011 | 0 | 0.18 | 0.1 | 0.34 | 0.31 | 0.31 | 0.31 |
| 7 | 18633901 | rs3213616 | -0.057 | -0.15 | -0.339 | -0.155 | 0.001 | -0.002 | -0.002 | 0.228 | 0.74 | 0.25 | 0.55 | 0.58 | 0.81 | 0.77 |
| 7 | 18635277 | rs3814993 | 0.386 | -0.21 | -0.122 | -0.459 | -0.001 | -0.053 | -0.053 | 0 | 0.27 | 0.37 | 0.25 | 0.09 | 0.28 | 0.21 |
| 7 | 18635619 | rs2695026 | 0.5 | -0.57 | -0.215 | -0.332 | 0.021 | -0.03 | -0.03 | 0 | 0.68 | 0.22 | 0.53 | 0.5 | 0.79 | 0.75 |
| 7 | 18638652 | rs801534 | -0.624 | -0.256 | -0.132 | -0.291 | 0.004 | 0.004 | 0.004 | 3.782 | 0.92 | 0.87 | 0.84 | 0.87 | 0.9 | 0.97 |
| 7 | 18644442 | rs801511 | -1.097 | -0.252 | -0.131 | -0.274 | -0.002 | -0.002 | -0.002 | 0 | 0.93 | 0.94 | 0.84 | 0.86 | 0.89 | 0.95 |
| 7 | 18647887 | rs2254993 | -0.465 | -0.499 | -0.112 | -0.278 | 0.003 | 0.007 | 0.007 | 3.486 | 0.92 | 0.85 | 0.85 | 0.87 | 0.9 | 0.97 |
| 7 | 18648426 | rs1726599 | 0.662 | -0.842 | 0.091 | -0.4 | -0.054 | 0.052 | 0.052 | 0.123 | 0.21 | 0.55 | 0.27 | 0.29 | 0.1 | 0.16 |
| 7 | 18650086 | rs1405618 | -0.517 | -0.712 | -0.375 | -0.55 | -0.03 | 0.04 | 0.04 | 0 | 0.73 | 0.92 | 0.7 | 0.82 | 0.68 | 0.79 |
| 7 | 18659363 | rs2243880 | -0.321 | -0.679 | -0.017 | -0.576 | -0.017 | 0.087 | 0.087 | 2.773 | 0.77 | 0.33 | 0.6 | 0.51 | 0.83 | 0.85 |
| 7 | 18661409 | rs1554774 | 0.784 | -0.387 | -0.293 | -0.712 | 0.017 | -0.027 | -0.027 | 0.06 | 0.62 | 0.2 | 0.5 | 0.44 | 0.72 | 0.72 |
| 7 | 18664580 | rs3801986 | 1.016 | -0.026 | 0.529 | -0.49 | -0.003 | 0.007 | 0.007 | 0.104 | 0.49 | 0.36 | 0.5 | 0.52 | 0.52 | 0.62 |
| 7 | 18666217 | rs6963748 | -0.261 | 0.474 | 0.871 | 0.041 | -0.013 | 0.038 | 0.038 | 0 | 0.25 | 0.12 | 0.2 | 0.03 | 0.28 | 0.14 |
| 7 | 18666597 | rs17432977 | 0.694 | 0.537 | 0.869 | 0.079 | -0.003 | -0.007 | -0.007 | 0.878 | 0.19 | 0.16 | 0.17 | 0.01 | 0.22 | 0.1 |
| 7 | 18666901 | rs2188320 | -0.882 | 0.459 | 1.096 | 0.202 | 0.003 | 0 | 0 | 0 | 0.34 | 0.44 | 0.43 | 0.36 | 0.36 | 0.34 |
| 7 | 18672384 | rs801540 | 0.522 | 0.535 | 0.516 | 0.243 | -0.017 | 0.029 | 0.029 | 0 | 0.64 | 0.77 | 0.64 | 0.77 | 0.59 | 0.74 |
| 7 | 18674040 | rs2285430 | 1.172 | 0.291 | 0.144 | 0.419 | -0.005 | -0.019 | -0.019 | 2.648 | 0.55 | 0.22 | 0.38 | 0.45 | 0.56 | 0.62 |
| 7 | 18679534 | rs2519890 | 0.895 | 0.271 | 0.611 | 0.798 | 0.011 | 0.012 | 0.012 | 1.701 | 0.88 | 0.84 | 0.83 | 0.85 | 0.87 | 0.91 |
| 7 | 18683672 | rs3801983 | -1.693 | -0.709 | 0.712 | 0.987 | 0.001 | -0.003 | -0.003 | 0 | 0.23 | 0.18 | 0.36 | 0.5 | 0.19 | 0.35 |
| 7 | 18689386 | rs6958814 | -0.071 | -0.82 | 1.331 | 1.025 | 0 | -0.001 | -0.001 | 0 | 0.11 | 0.1 | 0.13 | 0.13 | 0.07 | 0.1 |
| 7 | 18691879 | rs2285438 | -0.561 | -0.835 | 1.319 | 1.071 | -0.002 | -0.001 | -0.001 | 0 | 0.05 | 0.05 | 0.06 | 0.1 | 0.02 | 0.06 |
| 7 | 18698661 | rs2285440 | -2.228 | -0.96 | 2.162 | 1.093 | 0.019 | 0.229 | 0.229 | 0.106 | 0.08 | 0.27 | 0.26 | 0.39 | 0.06 | 0.22 |
| 7 | 18699475 | rs2269754 | -3.071 | -1.058 | 2.581 | 0.952 | 0.014 | -0.035 | -0.035 | 0 | 0.3 | 0.8 | 0.45 | 0.53 | 0.2 | 0.33 |
| 7 | 18703062 | rs801521 | -1.641 | -1.128 | 3.128 | 0.967 | -0.038 | 0.007 | 0.007 | 0.558 | 0.19 | 0.54 | 0.17 | 0.17 | 0.09 | 0.12 |
| 7 | 18704889 | rs2285443 | -2.432 | -1.079 | 3.183 | 0.968 | -0.029 | 0.079 | 0.079 | 0.168 | 0.24 | 0.61 | 0.36 | 0.47 | 0.15 | 0.25 |
| 7 | 18706703 | rs17348528 | -0.512 | -1.064 | 3.201 | 0.99 | 0 | -0.003 | -0.003 | 0 | 0.13 | 0.13 | 0.16 | 0.17 | 0.1 | 0.1 |
| 7 | 18706918 | rs801524 | -2.843 | -1.078 | 3.295 | 1.042 | 0.021 | -0.062 | -0.062 | 0.633 | 0.27 | 0.82 | 0.41 | 0.51 | 0.18 | 0.27 |
| 7 | 18707842 | rs706086 | -1.922 | -1.088 | 3.365 | 1.004 | -0.048 | -0.001 | -0.001 | 0 | 0.2 | 0.5 | 0.18 | 0.17 | 0.09 | 0.1 |
| 7 | 18712720 | rs17139560 | -0.738 | -1.069 | 3.341 | 1.009 | -0.025 | 0.019 | 0.019 | 1.09 | 0.14 | 0.23 | 0.14 | 0.16 | 0.07 | 0.09 |
| 7 | 18713515 | rs706087 | -1.563 | -1.757 | 2.958 | 0.728 | -0.033 | 0.064 | 0.064 | 0.623 | 0.3 | 0.45 | 0.42 | 0.51 | 0.18 | 0.32 |

| | | | | | | | | | | | | | | | | |
|---|---|---|---|---|---|---|---|---|---|---|---|---|---|---|---|---|
| 7 | 18714254 | rs17139571 | -2.057 | -1.527 | 2.917 | 0.827 | 0.062 | 0.337 | 0.337 | 0.102 | 0.06 | 0.3 | 0.25 | 0.36 | 0.06 | 0.22 |
| 7 | 18718573 | rs4484585 | -2.256 | -1.51 | 2.951 | 0.802 | 0.084 | 0.341 | 0.341 | 0 | 0.06 | 0.39 | 0.25 | 0.34 | 0.06 | 0.19 |
| 7 | 18720135 | rs10257046 | -0.603 | -1.578 | 3.003 | 0.74 | -0.023 | 0.055 | 0.055 | 0.85 | 0.14 | 0.33 | 0.15 | 0.17 | 0.07 | 0.09 |
| 7 | 18722234 | rs6962675 | -1.918 | -1.666 | 2.853 | 0.675 | -0.021 | -0.045 | -0.045 | 0 | 0.08 | 0.34 | 0.04 | 0 | 0.03 | 0.01 |
| 7 | 18724179 | rs2022090 | -1.749 | -1.612 | 2.851 | 0.712 | -0.048 | -0.017 | -0.017 | 0 | 0.2 | 0.5 | 0.19 | 0.16 | 0.11 | 0.11 |
| 7 | 18724512 | rs17139630 | -2.357 | -1.436 | 2.775 | 0.856 | 0.091 | 0.366 | 0.366 | 0.256 | 0.07 | 0.39 | 0.26 | 0.36 | 0.06 | 0.24 |
| 7 | 18728485 | rs6950376 | -0.031 | -1.436 | 2.211 | 0.822 | 0.001 | -0.008 | -0.008 | 0.266 | 0.12 | 0.09 | 0.15 | 0.16 | 0.09 | 0.1 |
| 7 | 18729827 | rs10276613 | -1.894 | -1.53 | 2.231 | 0.827 | -0.014 | 0.061 | 0.061 | 1.926 | 0.22 | 0.59 | 0.41 | 0.53 | 0.14 | 0.36 |
| 7 | 18734065 | rs12669789 | 0.125 | -1.681 | 1.985 | 0.792 | -0.019 | 0.026 | 0.026 | 0 | 0.15 | 0.16 | 0.2 | 0.31 | 0.07 | 0.18 |
| 7 | 18734921 | rs1178098 | -1.56 | -2.134 | 1.428 | 0.142 | 0.009 | 0.001 | 0.001 | 0 | 0.45 | 0.39 | 0.44 | 0.48 | 0.28 | 0.41 |
| 7 | 18738911 | rs212666 | -1.739 | -1.948 | 1.251 | 0.255 | 0.038 | -0.048 | -0.048 | 0.31 | 0.4 | 0.8 | 0.5 | 0.53 | 0.27 | 0.41 |
| 7 | 18739670 | rs10236906 | -0.107 | -1.873 | 1.315 | 0.331 | -0.018 | 0.016 | 0.016 | 0.075 | 0.15 | 0.22 | 0.18 | 0.21 | 0.09 | 0.15 |
| 7 | 18740164 | rs4721721 | -0.201 | -1.825 | 1.303 | 0.341 | -0.006 | 0.016 | 0.016 | 0.063 | 0.16 | 0.23 | 0.19 | 0.21 | 0.11 | 0.15 |
| 7 | 18741367 | rs1178103 | -0.058 | -1.871 | 0.478 | 0.157 | -0.001 | -0.002 | -0.002 | 0.107 | 0.19 | 0.13 | 0.25 | 0.29 | 0.18 | 0.23 |
| 7 | 18750392 | rs1178117 | -1.657 | -2.105 | 0.064 | 0.357 | 0.045 | -0.042 | -0.042 | 0 | 0.4 | 0.79 | 0.4 | 0.49 | 0.27 | 0.43 |
| 7 | 18751998 | rs1178118 | -0.616 | -1.614 | 0.038 | 0.065 | -0.003 | 0.005 | 0.005 | 0.096 | 0.22 | 0.25 | 0.25 | 0.32 | 0.18 | 0.28 |
| 7 | 18753320 | rs1178119 | -0.463 | -1.282 | -0.807 | 0.068 | -0.007 | -0.024 | -0.024 | 0 | 0.28 | 0.03 | 0.25 | 0.29 | 0.21 | 0.25 |
| 7 | 18763117 | rs212664 | -0.259 | -0.8 | -1.004 | -0.677 | -0.026 | 0.186 | 0.186 | 0 | 0.66 | 0.95 | 0.8 | 0.96 | 0.72 | 0.74 |
| 7 | 18767343 | rs1178127 | -0.709 | -0.702 | -0.219 | 0.31 | 0.007 | -0.045 | -0.045 | 0 | 0.31 | 0.21 | 0.3 | 0.52 | 0.26 | 0.39 |
| 7 | 18768334 | rs1178129 | -0.272 | -0.858 | 0.116 | 0.474 | -0.013 | -0.103 | -0.103 | 5.205 | 0.22 | 0.07 | 0.17 | 0.41 | 0.21 | 0.24 |
| 7 | 18768788 | rs1178130 | -0.896 | -0.963 | 1.193 | 0.405 | -0.01 | 0.043 | 0.043 | 0 | 0.27 | 0.55 | 0.3 | 0.51 | 0.23 | 0.34 |
| 7 | 18778113 | rs1178148 | -0.327 | -0.932 | 1.189 | 0.538 | -0.004 | -0.089 | -0.089 | 0 | 0.23 | 0.08 | 0.17 | 0.55 | 0.21 | 0.24 |
| 7 | 18783170 | rs1178156 | -0.611 | -0.998 | 1.781 | 0.5 | -0.008 | 0.046 | 0.046 | 0 | 0.25 | 0.39 | 0.29 | 0.64 | 0.22 | 0.31 |
| 7 | 18786967 | rs1178162 | -0.79 | -0.983 | 1.914 | 0.496 | -0.014 | 0.044 | 0.044 | 0 | 0.26 | 0.44 | 0.3 | 0.64 | 0.22 | 0.3 |
| 7 | 18787563 | rs1178163 | -0.018 | -0.983 | 2.008 | 0.536 | 0.005 | -0.039 | -0.039 | 1.416 | 0.2 | 0.13 | 0.16 | 0.4 | 0.18 | 0.14 |
| 7 | 18788589 | rs1178164 | -1.24 | -1.027 | 2.129 | 0.497 | -0.013 | 0.016 | 0.016 | 0 | 0.28 | 0.59 | 0.33 | 0.64 | 0.23 | 0.37 |
| 7 | 18794541 | rs1178172 | -1.083 | -1.015 | 2.183 | 0.503 | -0.014 | 0.029 | 0.029 | 0 | 0.27 | 0.53 | 0.32 | 0.64 | 0.23 | 0.34 |
| 7 | 18795250 | rs17139817 | -1.374 | -0.956 | 1.83 | 0.549 | 0 | 0.085 | 0.085 | 0 | 0.06 | 0.17 | 0.14 | 0.25 | 0.04 | 0.17 |
| 7 | 18799718 | rs1178178 | -0.005 | -0.929 | 1.47 | 0.446 | -0.011 | -0.104 | -0.104 | 0 | 0.19 | 0.05 | 0.16 | 0.4 | 0.18 | 0.18 |
| 7 | 18805880 | rs12056043 | -2.372 | -0.848 | 1.821 | 0.61 | 0.054 | 0.22 | 0.22 | 0.157 | 0.08 | 0.43 | 0.24 | 0.24 | 0.05 | 0.19 |
| 7 | 18806051 | rs6959028 | 0.264 | -0.809 | 1.068 | 0.272 | -0.057 | -0.06 | -0.06 | 0 | 0.6 | 0.91 | 0.68 | 0.59 | 0.58 | 0.62 |
| 7 | 18808396 | rs13241957 | 0.764 | -0.775 | 1.279 | -0.182 | 0 | 0.004 | 0.004 | 0 | 0.5 | 0.56 | 0.56 | 0.44 | 0.53 | 0.56 |
| 7 | 18814931 | rs10234685 | -0.927 | -0.516 | 1.503 | -0.176 | 0.019 | 0.002 | 0.002 | 0 | 0.07 | 0.18 | 0.1 | 0.04 | 0.06 | 0.16 |
| 7 | 18819266 | rs2588599 | 0.317 | -0.6 | 1.645 | -0.075 | -0.009 | 0.044 | 0.044 | 0.001 | 0.8 | 0.73 | 0.69 | 0.4 | 0.83 | 0.82 |
| 7 | 18820271 | rs2588604 | 0.341 | -0.606 | 1.691 | -0.078 | -0.015 | 0.075 | 0.075 | 0.005 | 0.8 | 0.66 | 0.69 | 0.4 | 0.82 | 0.81 |
| 7 | 18823893 | rs12699981 | 0.23 | -0.629 | 1.627 | -0.063 | -0.011 | 0.049 | 0.049 | 0.011 | 0.81 | 0.73 | 0.7 | 0.41 | 0.84 | 0.83 |
| 7 | 18834375 | rs6972422 | 0.331 | -0.637 | 1.007 | -0.485 | -0.056 | 0.118 | 0.118 | 0 | 0.22 | 0.04 | 0.16 | 0.02 | 0.26 | 0.2 |
| 7 | 18835449 | rs10248070 | -0.171 | -0.559 | 1.331 | -0.399 | 0.019 | 0.018 | 0.018 | 0 | 0.06 | 0.14 | 0.09 | 0.06 | 0.05 | 0.12 |
| 7 | 18836667 | rs12540872 | 1.349 | -0.54 | 1.478 | -0.339 | -0.027 | 0.077 | 0.077 | 2.023 | 0.49 | 0.16 | 0.4 | 0.39 | 0.52 | 0.5 |
| 7 | 18836701 | rs6963568 | 0.911 | -0.35 | 1.643 | -0.171 | -0.003 | 0.022 | 0.022 | 0.714 | 0.73 | 0.37 | 0.58 | 0.42 | 0.77 | 0.71 |
| 7 | 18838616 | rs12531908 | 0.844 | -0.117 | 1.526 | -0.292 | -0.002 | 0.026 | 0.026 | 0.361 | 0.71 | 0.54 | 0.57 | 0.37 | 0.73 | 0.73 |
| 7 | 18843093 | rs2440598 | 0.357 | -0.345 | 1.263 | -0.569 | 0.006 | -0.075 | -0.075 | 0.042 | 0.21 | 0.39 | 0.17 | 0.04 | 0.23 | 0.2 |
| 7 | 18845072 | rs10215908 | 0.028 | -0.369 | 0.967 | -0.482 | 0.031 | 0.029 | 0.029 | 0 | 0.05 | 0.13 | 0.09 | 0.06 | 0.06 | 0.14 |
| 7 | 18847958 | rs2520346 | -0.526 | -0.493 | 0.583 | -0.463 | 0.01 | -0.11 | -0.11 | 0 | 0.82 | 0.98 | 0.72 | 0.4 | 0.85 | 0.92 |
| 7 | 18848387 | rs7777331 | -1.151 | -0.059 | 1.356 | -0.732 | 0.005 | 0.002 | 0.002 | 0 | 0.08 | 0.13 | 0.11 | 0.05 | 0.08 | 0.21 |
| 7 | 18854050 | rs2520354 | -1.36 | 0.185 | 0.998 | -0.832 | -0.019 | 0.096 | 0.096 | 4.081 | 0.33 | 0.88 | 0.39 | 0.09 | 0.34 | 0.41 |
| 7 | 18854401 | rs12699985 | -0.613 | 0.228 | 0.883 | -0.858 | 0.003 | -0.055 | -0.055 | 0 | 0.24 | 0.37 | 0.28 | 0.04 | 0.28 | 0.24 |
| 7 | 18860014 | rs13241157 | -0.524 | 0.484 | 0.496 | -0.711 | 0.035 | -0.133 | -0.133 | 11.897 | 0.24 | 0.56 | 0.3 | 0.04 | 0.3 | 0.21 |
| 7 | 18861140 | rs2520361 | -0.827 | -0.3 | -0.095 | -0.158 | -0.019 | -0.025 | -0.025 | 0 | 0.28 | 0.57 | 0.24 | 0.22 | 0.17 | 0.19 |
| 7 | 18863899 | rs2520362 | -0.223 | -0.222 | -0.407 | 0.042 | -0.003 | 0.003 | 0.003 | 0 | 0.55 | 0.45 | 0.49 | 0.45 | 0.51 | 0.75 |
| 7 | 18864040 | rs7808313 | -0.504 | -0.474 | -0.006 | -0.1 | -0.005 | -0.054 | -0.054 | 0.137 | 0.12 | 0.19 | 0.05 | 0 | 0.09 | 0.05 |
| 7 | 18866190 | rs3852253 | -1.08 | -1.462 | 0.072 | 0.155 | -0.035 | -0.005 | -0.005 | 2.294 | 0.19 | 0.69 | 0.08 | 0.17 | 0.06 | 0.02 |
| 7 | 18866447 | rs3852255 | -0.82 | -1.44 | 1.128 | 0.166 | -0.076 | -0.003 | -0.003 | 0 | 0.17 | 0.5 | 0.06 | 0.15 | 0.06 | 0.02 |
| 7 | 18868031 | rs17349860 | 0.18 | -2.127 | 0.373 | 0.168 | 0.031 | 0.022 | 0.022 | 0 | 0.38 | 0.24 | 0.35 | 0.33 | 0.34 | 0.54 |
| 7 | 18874385 | rs2016515 | 0.694 | -2.465 | 0.703 | 0.014 | 0.012 | -0.018 | -0.018 | 0 | 0.3 | 0.12 | 0.32 | 0.34 | 0.3 | 0.51 |
| 7 | 18877874 | rs2073963 | -0.457 | -2.08 | 1.032 | -0.102 | 0.001 | -0.001 | -0.001 | 0.307 | 0.45 | 0.52 | 0.39 | 0.49 | 0.39 | 0.59 |
| 7 | 18882203 | rs12056282 | -1.216 | -1.786 | 1.077 | -0.064 | 0.021 | 0.12 | 0.12 | 1.158 | 0.1 | 0.39 | 0.05 | 0.15 | 0.04 | 0.03 |
| 7 | 18886177 | rs17140133 | -2.013 | -1.805 | 1.107 | -0.062 | 0.03 | 0.118 | 0.118 | 0.312 | 0.07 | 0.3 | 0.04 | 0.14 | 0.03 | 0.02 |
| 7 | 18890000 | rs756853 | 0.457 | -2.143 | 0.757 | -0.052 | -0.005 | 0 | 0 | 0.302 | 0.53 | 0.51 | 0.55 | 0.47 | 0.59 | 0.36 |
| 7 | 18894855 | rs13239638 | 0.951 | -2.193 | 0.614 | 0.133 | -0.042 | -0.003 | -0.003 | 0 | 0.24 | 0.07 | 0.24 | 0.21 | 0.28 | 0.39 |
| 7 | 18894897 | rs10230371 | -1.834 | -2.841 | 0.349 | 0.251 | -0.124 | -0.086 | -0.086 | 0 | 0.64 | 0.95 | 0.5 | 0.68 | 0.45 | 0.71 |
| 7 | 18895297 | rs12699994 | -0.197 | -3.083 | 0.56 | -0.336 | 0.023 | 0.023 | 0.023 | 0.031 | 0.4 | 0.24 | 0.39 | 0.32 | 0.36 | 0.57 |
| 7 | 18910232 | rs6461390 | -1.374 | -3.051 | 0.339 | -0.283 | 0.015 | -0.019 | -0.019 | 0.014 | 0.6 | 0.66 | 0.45 | 0.48 | 0.39 | 0.58 |
| 7 | 18913380 | rs7794241 | -0.519 | -3.039 | 0.401 | -0.427 | 0.018 | 0.015 | 0.015 | 0 | 0.48 | 0.42 | 0.41 | 0.33 | 0.36 | 0.56 |
| 7 | 18915133 | rs6959501 | -0.911 | -2.941 | -0.121 | -0.49 | 0.016 | 0.02 | 0.02 | 0 | 0.14 | 0.1 | 0.08 | 0.04 | 0.1 | 0.16 |
| 7 | 18924850 | rs10252945 | 0.216 | -2.505 | -0.348 | -0.327 | 0.005 | 0.032 | 0.032 | 0 | 0.29 | 0.2 | 0.25 | 0.17 | 0.25 | 0.35 |
| 7 | 18928240 | rs17350341 | 0.684 | -2.495 | -0.528 | -0.315 | -0.019 | 0.043 | 0.043 | 0.008 | 0.25 | 0.08 | 0.22 | 0.15 | 0.25 | 0.34 |
| 7 | 18928503 | rs17350355 | 0.728 | -2.619 | -0.399 | -0.292 | 0.073 | 0.066 | 0.066 | 0.006 | 0.48 | 0.27 | 0.61 | 0.64 | 0.62 | 0.5 |
| 7 | 18931219 | rs10278449 | 0.136 | -1.881 | -0.773 | -0.637 | -0.064 | -0.051 | -0.051 | 0 | 0.72 | 0.48 | 0.75 | 0.85 | 0.85 | 0.82 |

| | | | | | | | | | | | | | | | | |
|---|---|---|---|---|---|---|---|---|---|---|---|---|---|---|---|---|
| 7 | 18931305 | rs2190278 | -0.188 | -0.4 | -1.111 | -0.209 | -0.012 | 0.03 | 0.03 | 0.175 | 0.25 | 0.11 | 0.22 | 0.17 | 0.25 | 0.35 |
| 7 | 18934838 | rs10272709 | -0.96 | -0.262 | -0.399 | -0.119 | -0.012 | -0.052 | -0.052 | 0.142 | 0.18 | 0.4 | 0.21 | 0.07 | 0.13 | 0.14 |
| 7 | 18935310 | rs11764843 | 0.423 | -0.089 | -1.849 | -0.253 | 0.026 | 0.123 | 0.123 | 0.901 | 0.55 | 0.11 | 0.46 | 0.22 | 0.64 | 0.67 |
| 7 | 18938081 | rs13438248 | -1.547 | 0.391 | -0.65 | -0.387 | 0.024 | -0.086 | -0.086 | 0 | 0.14 | 0.33 | 0.12 | 0 | 0.15 | 0.03 |
| 7 | 18938376 | rs6954476 | -1.47 | -0.099 | -0.386 | -0.436 | -0.047 | -0.066 | -0.066 | 0.181 | 0.11 | 0.43 | 0.07 | 0 | 0.03 | 0.02 |
| 7 | 18938958 | rs2301949 | -0.817 | 1.245 | -1.196 | 0.593 | -0.03 | -0.035 | -0.035 | 0 | 0.16 | 0.07 | 0.25 | 0.42 | 0.26 | 0.16 |
| 7 | 18943707 | rs17140251 | 0.018 | 0.748 | -1.499 | 0.503 | 0.002 | -0.032 | -0.032 | 0 | 0.07 | 0 | 0.09 | 0.12 | 0.06 | 0.3 |
| 7 | 18947955 | rs1117533 | -0.619 | 0.595 | -2.335 | 0.03 | -0.077 | -0.094 | -0.094 | 0 | 0.35 | 0.08 | 0.35 | 0.4 | 0.36 | 0.28 |
| 7 | 18952433 | rs13232564 | 0.271 | 0.881 | -2.419 | 0.156 | -0.053 | -0.082 | -0.082 | 1.397 | 0.12 | 0.03 | 0.32 | 0.34 | 0.17 | 0.13 |
| 7 | 18952826 | rs2190274 | -0.543 | 0.868 | -2.516 | 0.076 | -0.003 | 0.003 | 0.003 | 0 | 0.89 | 0.88 | 0.92 | 0.91 | 0.88 | 0.91 |
| 7 | 18953143 | rs4379363 | -0.339 | 1.316 | -2.601 | -0.318 | -0.021 | 0.03 | 0.03 | 0 | 0.18 | 0.07 | 0.17 | 0.13 | 0.21 | 0.37 |
| 7 | 18957681 | rs13231656 | -0.323 | 0.376 | -2.617 | -0.09 | -0.009 | -0.101 | -0.101 | 0 | 0.22 | 0.09 | 0.26 | 0.38 | 0.2 | 0.14 |
| 7 | 18958533 | rs6967021 | 1.811 | 0.287 | -2.413 | 0.162 | -0.002 | 0.08 | 0.08 | 0 | 0.51 | 0.64 | 0.37 | 0.28 | 0.49 | 0.38 |
| 7 | 18959138 | rs2214850 | -0.043 | 0.47 | -2.026 | 0.345 | 0 | 0.037 | 0.037 | 0 | 0.86 | 0.71 | 0.76 | 0.77 | 0.87 | 0.83 |
| 7 | 18959381 | rs6972136 | -0.647 | 0.594 | -2.17 | 0.363 | -0.042 | -0.088 | -0.088 | 0.024 | 0.25 | 0.12 | 0.31 | 0.38 | 0.29 | 0.15 |
| 7 | 18963677 | rs6973918 | -0.714 | 0.197 | -2.061 | 0.344 | -0.002 | 0.003 | 0.003 | 0 | 0.89 | 0.88 | 0.94 | 0.92 | 0.88 | 0.9 |
| 7 | 18964076 | rs12700014 | 1.659 | 0.037 | -1.727 | 0.269 | 0.008 | -0.021 | -0.021 | 0 | 0.4 | 0.41 | 0.34 | 0.26 | 0.5 | 0.27 |
| 7 | 18965590 | rs10246722 | -1.035 | 0.573 | -1.72 | 0.981 | -0.017 | -0.081 | -0.081 | 0.094 | 0.23 | 0.1 | 0.27 | 0.37 | 0.21 | 0.15 |
| 7 | 18965622 | rs7789752 | 0.429 | 0.416 | -1.623 | 1.181 | 0.023 | 0.029 | 0.029 | 0 | 0.78 | 0.89 | 0.91 | 0.87 | 0.87 | 0.88 |
| 7 | 18968003 | rs10226243 | -0.475 | 1.166 | -1.064 | 1.681 | -0.051 | -0.085 | -0.085 | 0 | 0.13 | 0.05 | 0.32 | 0.34 | 0.18 | 0.13 |
| 7 | 18974723 | rs10248565 | -1.209 | 0.815 | -0.34 | 0.56 | 0.01 | -0.051 | -0.051 | 0 | 0.1 | 0.22 | 0.1 | 0.01 | 0.12 | 0.04 |
| 7 | 18975056 | rs2717356 | -1.333 | -0.086 | -0.116 | 0.261 | -0.015 | -0.022 | -0.022 | 1.451 | 0.23 | 0.66 | 0.24 | 0.13 | 0.18 | 0.19 |
| 7 | 18975189 | rs2240419 | -0.278 | 0.096 | -1.264 | 0.195 | -0.037 | -0.014 | -0.014 | 0 | 0.77 | 0.94 | 0.77 | 0.75 | 0.76 | 0.85 |
| 7 | 18977047 | rs17140333 | 0.43 | -0.447 | -1.304 | 0.739 | 0.009 | -0.017 | -0.017 | 0 | 0.13 | 0.09 | 0.18 | 0.35 | 0.1 | 0.22 |
| 7 | 18977511 | rs17140345 | -0.121 | 0.516 | -1.458 | 0.652 | -0.013 | -0.04 | -0.04 | 0 | 0.17 | 0.09 | 0.33 | 0.38 | 0.18 | 0.26 |
| 7 | 18977653 | rs1034805 | 1.582 | 0.617 | -2.116 | 0.301 | -0.067 | -0.044 | -0.044 | 0 | 0.26 | 0.09 | 0.23 | 0.26 | 0.34 | 0.31 |
| 7 | 18979366 | rs11767043 | 0.397 | -0.168 | -1.74 | -0.073 | 0.001 | -0.022 | -0.022 | 0 | 0.21 | 0.19 | 0.11 | 0.01 | 0.18 | 0.23 |
| 7 | 18979516 | rs10223990 | 1.826 | -0.088 | -1.93 | -0.005 | -0.109 | -0.036 | -0.036 | 0 | 0.26 | 0.11 | 0.27 | 0.26 | 0.41 | 0.3 |
| 7 | 18980267 | rs2853552 | -0.717 | -0.218 | -0.962 | 0.069 | 0.003 | 0.008 | 0.008 | 4.993 | 0.68 | 0.77 | 0.71 | 0.73 | 0.71 | 0.82 |
| 7 | 18986970 | rs2528406 | 0.252 | -0.912 | -0.148 | -0.012 | -0.018 | -0.062 | -0.062 | 0.248 | 0.22 | 0.28 | 0.09 | 0.02 | 0.15 | 0.18 |
| 7 | 18989232 | rs2051920 | -0.345 | -0.711 | -0.369 | 0.504 | 0.002 | -0.014 | -0.014 | 0 | 0.76 | 0.81 | 0.63 | 0.61 | 0.76 | 0.74 |
| 7 | 18991904 | rs10282136 | 1.693 | -0.536 | -0.355 | 0.293 | -0.1 | 0.026 | 0.026 | 0 | 0.31 | 0.13 | 0.28 | 0.25 | 0.46 | 0.3 |
| 7 | 18993249 | rs11505418 | 1.535 | -0.547 | -0.277 | 0.309 | -0.065 | 0.021 | 0.021 | 0 | 0.32 | 0.2 | 0.29 | 0.27 | 0.46 | 0.32 |
| 7 | 19001691 | rs6461396 | -0.327 | -0.302 | -0.204 | 0.508 | 0 | -0.002 | -0.002 | 0 | 0.45 | 0.46 | 0.38 | 0.47 | 0.41 | 0.56 |
| 7 | 19003032 | rs7808621 | -0.937 | -0.493 | -0.261 | 0.563 | 0 | -0.002 | -0.002 | 0 | 0.79 | 0.82 | 0.66 | 0.72 | 0.81 | 0.85 |
| 7 | 19007884 | rs17140423 | -0.432 | -0.576 | -0.461 | 0.592 | 0.013 | 0.003 | 0.003 | 0.236 | 0.44 | 0.43 | 0.48 | 0.58 | 0.32 | 0.52 |
| 7 | 19013256 | rs4141042 | 0.918 | -0.552 | -0.439 | 0.631 | 0 | 0.011 | 0.011 | 0 | 0.16 | 0.16 | 0.21 | 0.4 | 0.15 | 0.31 |
| 7 | 19014843 | rs2526640 | -1.114 | -0.545 | -0.444 | 0.503 | -0.012 | 0.021 | 0.021 | 0.213 | 0.8 | 0.91 | 0.85 | 0.86 | 0.8 | 0.87 |
| 7 | 19016165 | rs2526639 | -0.193 | -0.964 | -0.509 | 0.472 | 0.018 | 0.002 | 0.002 | 4.524 | 0.69 | 0.82 | 0.76 | 0.7 | 0.76 | 0.73 |
| 7 | 19016871 | rs12533452 | 1.086 | -0.687 | -0.464 | 0.336 | -0.004 | -0.005 | -0.005 | 0 | 0.16 | 0.16 | 0.2 | 0.31 | 0.18 | 0.24 |
| 7 | 19017705 | rs4413678 | 1.802 | -0.702 | -0.703 | 0.104 | 0.008 | -0.002 | -0.002 | 0 | 0.11 | 0.06 | 0.03 | 0 | 0.05 | 0.01 |
| 7 | 19018009 | rs2240279 | -0.588 | -0.323 | -0.26 | 0.303 | -0.009 | -0.012 | -0.012 | 0 | 0.43 | 0.52 | 0.35 | 0.33 | 0.36 | 0.46 |
| 7 | 19019671 | rs10268180 | 1.834 | -0.208 | -0.713 | -0.491 | -0.028 | 0.024 | 0.024 | 0 | 0.34 | 0.19 | 0.27 | 0.28 | 0.39 | 0.35 |
| 7 | 19019880 | rs2717351 | -0.099 | -0.231 | -0.3 | 0.045 | 0.019 | 0.115 | 0.115 | 0 | 0.2 | 0.39 | 0.46 | 0.55 | 0.23 | 0.34 |
| 7 | 19021592 | rs2158495 | -0.1 | -0.782 | -0.819 | -0.311 | 0.006 | -0.002 | -0.002 | 0 | 0.22 | 0.2 | 0.11 | 0.02 | 0.17 | 0.19 |
| 7 | 19030853 | rs2717344 | 0.201 | -0.458 | -0.877 | 0.115 | 0.001 | 0 | 0 | 0 | 0.69 | 0.77 | 0.46 | 0.29 | 0.67 | 0.47 |
| 7 | 19034191 | rs7788833 | -1.263 | -1.77 | -0.459 | 0.266 | -0.003 | 0.002 | 0.002 | 0.336 | 0.28 | 0.29 | 0.35 | 0.39 | 0.22 | 0.35 |
| 7 | 19034579 | rs10227612 | -1.654 | -2.189 | -0.239 | -0.009 | -0.018 | -0.001 | -0.001 | 0.005 | 0.38 | 0.46 | 0.44 | 0.4 | 0.29 | 0.42 |
| 7 | 19035920 | rs2074633 | -1.341 | -2.296 | -0.085 | 0.136 | 0.006 | -0.002 | -0.002 | 0 | 0.3 | 0.27 | 0.25 | 0.36 | 0.22 | 0.35 |
| 7 | 19036578 | rs2526630 | 1.409 | -2.078 | -0.044 | -0.013 | -0.027 | 0.02 | 0.02 | 0.466 | 0.45 | 0.27 | 0.42 | 0.38 | 0.51 | 0.28 |
| 7 | 19036738 | rs7812296 | -1.607 | -2.138 | 0.041 | -0.005 | -0.04 | 0.018 | 0.018 | 0.733 | 0.56 | 0.8 | 0.58 | 0.62 | 0.49 | 0.72 |
| 7 | 19036775 | rs2023938 | 0.213 | -2.217 | 0.078 | -0.055 | -0.008 | -0.06 | -0.06 | 0 | 0.14 | 0.22 | 0.08 | 0 | 0.1 | 0.1 |
| 7 | 19036909 | rs7776735 | -0.746 | -2.188 | 0.079 | -0.025 | -0.003 | -0.003 | -0.003 | 0.016 | 0.41 | 0.53 | 0.5 | 0.62 | 0.39 | 0.62 |
| 7 | 19038646 | rs2158493 | -0.974 | -2.367 | -0.155 | -0.228 | -0.001 | -0.003 | -0.003 | 1.342 | 0.49 | 0.46 | 0.51 | 0.63 | 0.44 | 0.63 |
| 7 | 19038678 | rs2526629 | -1.419 | -2.351 | -0.087 | -0.224 | -0.034 | 0.028 | 0.028 | 0.828 | 0.55 | 0.76 | 0.58 | 0.62 | 0.49 | 0.72 |
| 7 | 19040331 | rs7792656 | -0.308 | -2.425 | -0.289 | -0.261 | -0.014 | -0.075 | -0.075 | 0 | 0.18 | 0.25 | 0.09 | 0 | 0.11 | 0.11 |
| 7 | 19042101 | rs10241753 | -0.263 | -2.48 | -0.206 | -0.11 | -0.004 | -0.01 | -0.01 | 0 | 0.36 | 0.26 | 0.48 | 0.61 | 0.38 | 0.6 |
| 7 | 19043360 | rs2526625 | -0.613 | -2.472 | -0.197 | -0.033 | 0.001 | 0.001 | 0.001 | 0 | 0.75 | 0.71 | 0.68 | 0.77 | 0.73 | 0.64 |
| 7 | 19048472 | rs12669496 | 0.562 | -2.106 | -0.263 | 0.307 | -0.002 | 0.006 | 0.006 | 0 | 0.34 | 0.23 | 0.34 | 0.23 | 0.32 | 0.37 |
| 7 | 19049162 | rs1548577 | -1.038 | -2.083 | -0.673 | 0.33 | -0.123 | -0.06 | -0.06 | 0 | 0.59 | 0.9 | 0.59 | 0.6 | 0.48 | 0.7 |
| 7 | 19049388 | rs2107595 | -0.709 | -2.029 | -0.601 | 0.467 | 0.002 | -0.003 | -0.003 | 0 | 0.22 | 0.21 | 0.22 | 0.36 | 0.17 | 0.31 |
| 7 | 19055280 | rs4721740 | -0.087 | -1.869 | -0.661 | 0.444 | 0.002 | 0.001 | 0.001 | 0.172 | 0.44 | 0.35 | 0.41 | 0.58 | 0.41 | 0.61 |
| 7 | 19055703 | rs10255384 | 0.345 | -1.732 | -0.554 | 0.487 | -0.003 | -0.068 | -0.068 | 0 | 0.13 | 0.23 | 0.07 | 0 | 0.09 | 0.1 |
| 7 | 19055797 | rs12530920 | 0.619 | -1.71 | -0.562 | 0.521 | -0.01 | -0.069 | -0.069 | 0 | 0.1 | 0 | 0.14 | 0.39 | 0.1 | 0.24 |
| 7 | 19059427 | rs7783974 | 0.045 | -1.331 | -0.578 | 0.027 | -0.005 | -0.068 | -0.068 | 0.94 | 0.13 | 0.22 | 0.07 | 0 | 0.09 | 0.11 |
| 7 | 19060012 | rs2906044 | -1.248 | -1.241 | -1.533 | 0.059 | -0.121 | -0.091 | -0.091 | 0.213 | 0.67 | 0.95 | 0.64 | 0.63 | 0.57 | 0.74 |
| 7 | 19060121 | rs12533919 | -0.333 | -0.78 | -1.099 | 0.499 | 0.005 | 0.029 | 0.029 | 0 | 0.51 | 0.32 | 0.45 | 0.62 | 0.48 | 0.63 |
| 7 | 19064020 | rs2526614 | -0.607 | -1.112 | -1.369 | 0.567 | -0.124 | -0.084 | -0.084 | 0 | 0.58 | 0.96 | 0.48 | 0.56 | 0.42 | 0.63 |
| 7 | 19070165 | rs1548576 | -1.276 | -1.493 | 0.701 | 0.112 | -0.04 | -0.056 | -0.056 | 0.485 | 0.12 | 0.77 | 0.07 | 0.01 | 0.04 | 0.06 |

| | | | | | | | | | | | | | | | | |
|---|---|---|---|---|---|---|---|---|---|---|---|---|---|---|---|---|
| 7 | 19070930 | rs2526644 | -0.552 | -1.911 | 0.444 | 0.149 | 0.044 | -0.006 | -0.006 | 0 | 0.15 | 0.05 | 0.08 | 0.14 | 0.11 | 0.3 |
| 7 | 19095905 | rs2717364 | 0.843 | -1.19 | 1.288 | 0.166 | 0.008 | -0.019 | -0.019 | 0 | 0.92 | 0.53 | 0.96 | 0.98 | 0.96 | 0.95 |
| 7 | 19100076 | rs2521735 | 1.334 | -1.194 | 1.451 | 0.164 | -0.049 | -0.069 | -0.069 | 0.017 | 0.89 | 0.28 | 0.94 | 0.98 | 0.96 | 0.95 |
| 7 | 19102313 | rs2717336 | -0.04 | -1.193 | 1.359 | 0.165 | -0.02 | -0.024 | -0.024 | 0 | 0.95 | 0.91 | 0.99 | 1 | 0.98 | 0.95 |
| 7 | 19104765 | rs2521736 | -0.04 | -1.191 | 0.909 | 0.166 | -0.02 | -0.024 | -0.024 | 0 | 0.95 | 0.92 | 0.99 | 1 | 0.98 | 0.95 |
| 7 | 19107986 | rs10253091 | -1.14 | -1.185 | 1.017 | 0.171 | 0.02 | 0.007 | 0.007 | 10.342 | 0.06 | 0.64 | 0.06 | 0.01 | 0.02 | 0 |
| 7 | 19128201 | rs2892913 | -1.69 | -1.203 | 0.638 | 0.142 | -0.029 | -0.044 | -0.044 | 0 | 0.12 | 0.79 | 0.07 | 0.02 | 0.04 | 0.06 |
| 7 | 19131207 | rs17435999 | -0.353 | -0.46 | -0.56 | 0.492 | 0 | 0.001 | 0.001 | 0.127 | 0.25 | 0.19 | 0.12 | 0.11 | 0.19 | 0.1 |
| 7 | 19133072 | rs10225279 | -0.835 | -0.398 | -1.525 | 0.658 | -0.091 | 0.036 | 0.036 | 0 | 0.58 | 0.88 | 0.61 | 0.64 | 0.47 | 0.66 |
| 7 | 19133464 | rs10228406 | 0.171 | -0.674 | -0.926 | 0.5 | -0.016 | -0.101 | -0.101 | 0 | 0.26 | 0.44 | 0.15 | 0.07 | 0.19 | 0.1 |
| 7 | 19134995 | rs7801041 | 0.098 | -0.642 | -0.99 | 0.521 | -0.014 | -0.095 | -0.095 | 0 | 0.27 | 0.46 | 0.15 | 0.07 | 0.19 | 0.1 |
| 7 | 19135210 | rs2390045 | -0.116 | -0.588 | -1.328 | 0.943 | -0.003 | -0.002 | -0.002 | 0 | 0.31 | 0.35 | 0.39 | 0.25 | 0.27 | 0.5 |
| 7 | 19137857 | rs6964466 | -0.53 | 0.028 | -1.861 | 0.917 | -0.068 | 0.04 | 0.04 | 0.045 | 0.54 | 0.92 | 0.58 | 0.63 | 0.42 | 0.64 |
| 7 | 19148847 | rs985246 | -0.823 | -1.07 | -0.685 | 1.122 | -0.042 | -0.014 | -0.014 | 0 | 0.37 | 0.45 | 0.5 | 0.69 | 0.21 | 0.52 |
| 7 | 19153843 | rs4721746 | -0.63 | -1.292 | -0.194 | 1.461 | -0.015 | 0.009 | 0.009 | 0.107 | 0.08 | 0.14 | 0.04 | 0.11 | 0.03 | 0.1 |
| 7 | 19160897 | rs10275272 | -2.012 | -2.101 | 0.563 | 1.887 | -0.046 | 0.029 | 0.029 | 0.255 | 0.22 | 0.43 | 0.17 | 0.27 | 0.12 | 0.33 |
| 7 | 19161218 | rs2189000 | -1.902 | -2.107 | 0.645 | 1.885 | -0.04 | 0.034 | 0.034 | 0.01 | 0.21 | 0.41 | 0.17 | 0.27 | 0.12 | 0.33 |
| 7 | 19163912 | rs17140698 | -0.263 | -2.122 | 0.834 | 1.886 | -0.002 | 0.013 | 0.013 | 0 | 0.1 | 0.11 | 0.1 | 0.16 | 0.09 | 0.22 |
| 7 | 19165264 | rs994823 | -0.526 | -2.15 | 0.897 | 1.873 | -0.001 | 0.042 | 0.042 | 0.193 | 0.1 | 0.15 | 0.12 | 0.26 | 0.09 | 0.21 |
| 7 | 19166673 | rs4721748 | -0.837 | -2.15 | 0.855 | 1.84 | -0.01 | -0.024 | -0.024 | 0.221 | 0.08 | 0.07 | 0.03 | 0.01 | 0.03 | 0.1 |
| 7 | 19172600 | rs1157350 | -2.45 | -2.478 | 1.472 | 2.739 | 0.063 | 0.065 | 0.065 | 0 | 0.45 | 0.71 | 0.58 | 0.7 | 0.29 | 0.54 |
| 7 | 19178075 | rs11975410 | -2.292 | -2.225 | 1.332 | 2.598 | -0.042 | 0.034 | 0.034 | 0.298 | 0.22 | 0.44 | 0.17 | 0.29 | 0.13 | 0.32 |
| 7 | 19186465 | rs10257807 | -0.462 | -2.21 | 1.193 | 2.66 | 0 | 0.001 | 0.001 | 0.934 | 0.09 | 0.07 | 0.08 | 0.17 | 0.06 | 0.19 |
| 7 | 19186799 | rs10228726 | -2.82 | -2.198 | 1.433 | 2.613 | -0.005 | 0.021 | 0.021 | 0 | 0.24 | 0.4 | 0.22 | 0.3 | 0.2 | 0.37 |
| 7 | 19191322 | rs10275878 | -2.146 | -1.917 | 1.188 | 2.678 | -0.001 | -0.004 | -0.004 | 0 | 0.3 | 0.33 | 0.22 | 0.21 | 0.26 | 0.21 |
| 7 | 19193072 | rs17351688 | -2.196 | -1.911 | 1.194 | 2.676 | -0.001 | -0.007 | -0.007 | 0.038 | 0.3 | 0.36 | 0.23 | 0.21 | 0.27 | 0.21 |
| 7 | 19194166 | rs17140750 | -0.173 | -2.051 | 1.117 | 3.178 | 0.026 | -0.016 | -0.016 | 0 | 0.08 | 0.03 | 0.06 | 0.3 | 0.03 | 0.24 |
| 7 | 19195769 | rs2390047 | -2.881 | -2.167 | 1.075 | 3.065 | -0.018 | -0.006 | -0.006 | 0 | 0.4 | 0.51 | 0.3 | 0.38 | 0.3 | 0.42 |
| 7 | 19198132 | rs7780145 | -2.729 | -2.245 | 1.077 | 3.018 | -0.024 | 0.036 | 0.036 | 0 | 0.36 | 0.52 | 0.28 | 0.51 | 0.28 | 0.4 |
| 7 | 19200324 | rs13242837 | -2.248 | -2.24 | 1.067 | 2.657 | 0.001 | 0.016 | 0.016 | 0.034 | 0.31 | 0.26 | 0.2 | 0.09 | 0.28 | 0.21 |
| 7 | 19206436 | rs10486336 | -2.831 | -2.331 | 1.002 | 2.858 | -0.01 | 0.058 | 0.058 | 0 | 0.3 | 0.51 | 0.27 | 0.44 | 0.27 | 0.37 |
| 7 | 19207222 | rs10486337 | -0.578 | -2.271 | 0.903 | 2.824 | -0.008 | -0.014 | -0.014 | 0 | 0.06 | 0.08 | 0.04 | 0.01 | 0.05 | 0.07 |
| 7 | 19209937 | rs12666922 | -2.201 | -2.376 | 0.867 | 2.803 | 0.004 | -0.004 | -0.004 | 0 | 0.27 | 0.2 | 0.2 | 0.27 | 0.22 | 0.31 |
| 7 | 19210250 | rs7806506 | -1.477 | -2.27 | 0.727 | 2.53 | -0.034 | 0.048 | 0.048 | 0 | 0.2 | 0.06 | 0.13 | 0.08 | 0.19 | 0.1 |
| 7 | 19211709 | rs7798717 | -1.855 | -2.176 | 0.861 | 2.505 | 0.003 | 0.003 | 0.003 | 0 | 0.12 | 0.12 | 0.06 | 0.05 | 0.08 | 0.12 |
| 7 | 19213238 | rs17140802 | -1.838 | -2.165 | 0.868 | 2.403 | -0.026 | 0.003 | 0.003 | 0 | 0.12 | 0.26 | 0.09 | 0.12 | 0.07 | 0.12 |
| 7 | 19215459 | rs10248721 | -2.166 | -2.224 | 0.762 | 2.693 | 0.009 | -0.019 | -0.019 | 0 | 0.29 | 0.22 | 0.3 | 0.4 | 0.23 | 0.39 |
| 7 | 19241659 | rs7779453 | -0.457 | -1.886 | 0.175 | 1.746 | -0.072 | 0.059 | 0.059 | 1.36 | 0.39 | 0.11 | 0.37 | 0.22 | 0.46 | 0.42 |
| 7 | 19244484 | rs17140852 | 0.254 | -1.756 | 0.675 | 1.731 | -0.023 | -0.004 | -0.004 | 0 | 0.1 | 0.17 | 0.02 | 0.05 | 0.03 | 0.04 |
| 7 | 19249169 | rs11535164 | -0.373 | -1.807 | 0.822 | 1.837 | -0.063 | 0.061 | 0.061 | 0.284 | 0.12 | 0.43 | 0.06 | 0.15 | 0.04 | 0.04 |
| 7 | 19250387 | rs10253644 | -0.695 | -1.816 | 0.891 | 1.822 | -0.077 | 0.052 | 0.052 | 0.645 | 0.14 | 0.48 | 0.07 | 0.15 | 0.04 | 0.04 |
| 7 | 19250811 | rs3922356 | -1.907 | -1.79 | 0.246 | 1.534 | -0.018 | 0.048 | 0.048 | 0 | 0.64 | 0.83 | 0.66 | 0.82 | 0.6 | 0.81 |
| 7 | 19252156 | rs4141213 | -0.437 | -1.934 | 0.129 | 1.942 | -0.001 | -0.031 | -0.031 | 0 | 0.09 | 0.03 | 0.18 | 0.27 | 0.12 | 0.24 |
| 7 | 19254640 | rs10264031 | -1.179 | -2.002 | 0.322 | 1.442 | 0.05 | 0.014 | 0.014 | 3.409 | 0.5 | 0.74 | 0.42 | 0.52 | 0.37 | 0.5 |
| 7 | 19254715 | rs10248905 | -0.451 | -1.37 | 0.442 | 1.565 | 0.001 | 0.032 | 0.032 | 0.161 | 0.36 | 0.45 | 0.38 | 0.48 | 0.32 | 0.45 |
| 7 | 19256283 | rs10253752 | -0.281 | -1.372 | -0.099 | 1.539 | -0.022 | 0.019 | 0.019 | 0 | 0.07 | 0.08 | 0.06 | 0.19 | 0.01 | 0.1 |
| 7 | 19261248 | rs7787452 | -0.097 | -1.189 | -0.194 | 1.334 | -0.018 | 0.018 | 0.018 | 1.624 | 0.3 | 0.16 | 0.28 | 0.19 | 0.31 | 0.34 |
| 7 | 19274183 | rs975510 | -0.746 | -1.196 | -0.042 | 1.292 | -0.005 | 0.016 | 0.016 | 0.939 | 0.4 | 0.43 | 0.39 | 0.48 | 0.35 | 0.45 |
| 7 | 19291993 | rs7792037 | -1.946 | -1.61 | 0.483 | 1.771 | -0.046 | 0.119 | 0.119 | 0 | 0.14 | 0.65 | 0.15 | 0.32 | 0.06 | 0.18 |
| 7 | 19293207 | rs4499987 | -0.805 | -1.6 | -0.265 | 1.805 | -0.024 | -0.029 | -0.029 | 0 | 0.2 | 0.04 | 0.32 | 0.28 | 0.21 | 0.24 |
| 7 | 19302302 | rs11974404 | -0.35 | -1.522 | 0.747 | 1.898 | 0.089 | 0.085 | 0.085 | 0.84 | 0.05 | 0.34 | 0.06 | 0.03 | 0.04 | 0.07 |
| 7 | 19305590 | rs7802361 | -1.666 | -0.949 | 0.347 | 1.745 | 0.001 | -0.051 | -0.051 | 1.93 | 0.34 | 0.78 | 0.31 | 0.46 | 0.33 | 0.51 |
| 7 | 19322409 | rs17141042 | -0.076 | -0.989 | 0.495 | 1.931 | -0.037 | 0.053 | 0.053 | 0 | 0.07 | 0.16 | 0.07 | 0.18 | 0.01 | 0.11 |
| 7 | 19324368 | rs2191893 | -0.563 | -0.977 | 0.497 | 1.952 | 0.068 | 0.192 | 0.192 | 2.172 | 0.06 | 0.35 | 0.08 | 0.14 | 0.05 | 0.08 |
| 7 | 19326733 | rs929554 | 0.253 | -0.987 | 0.062 | 1.877 | -0.031 | 0.006 | 0.006 | 0 | 0.2 | 0.08 | 0.15 | 0.14 | 0.28 | 0.33 |
| 7 | 19327430 | rs10486343 | 0.649 | -1.003 | -0.148 | 1.971 | -0.008 | -0.036 | -0.036 | 0 | 0.11 | 0 | 0.07 | 0.15 | 0.09 | 0.13 |
| 7 | 19330700 | rs985545 | 2.563 | -0.974 | -0.189 | 1.871 | -0.083 | 0.042 | 0.042 | 0 | 0.44 | 0.1 | 0.36 | 0.27 | 0.46 | 0.24 |
| 7 | 19333027 | rs10281369 | 2.588 | -0.971 | -0.17 | 1.821 | -0.084 | 0.06 | 0.06 | 0 | 0.44 | 0.1 | 0.36 | 0.25 | 0.46 | 0.23 |
| 7 | 19338209 | rs11763201 | -1.05 | -0.923 | -0.08 | 1.894 | -0.022 | -0.039 | -0.039 | 0.844 | 0.19 | 0.04 | 0.32 | 0.3 | 0.21 | 0.26 |
| 7 | 19340145 | rs9638759 | -1.832 | -1.003 | -0.147 | 1.931 | -0.01 | -0.057 | -0.057 | 0 | 0.15 | 0.06 | 0.3 | 0.41 | 0.18 | 0.24 |
| 7 | 19340930 | rs9639327 | -2.533 | -1.005 | 0.258 | 1.942 | -0.024 | 0.082 | 0.082 | 0.001 | 0.25 | 0.45 | 0.37 | 0.44 | 0.23 | 0.31 |
| 7 | 19346344 | rs2052149 | 2.747 | -1.16 | -0.253 | 1.847 | -0.04 | 0.102 | 0.102 | 0 | 0.45 | 0.19 | 0.39 | 0.23 | 0.49 | 0.23 |
| 7 | 19346506 | rs6980408 | -1.236 | -1.121 | -0.116 | 1.902 | -0.004 | 0.166 | 0.166 | 0 | 0.12 | 0.51 | 0.15 | 0.28 | 0.08 | 0.17 |
| 7 | 19347037 | rs17437296 | -2.325 | -1.126 | -0.11 | 1.904 | 0.002 | 0.048 | 0.048 | 0 | 0.5 | 0.63 | 0.58 | 0.66 | 0.51 | 0.77 |
| 7 | 19349040 | rs17141117 | -1.856 | -0.886 | 0.165 | 2.091 | -0.016 | 0.109 | 0.109 | 0 | 0.14 | 0.51 | 0.17 | 0.28 | 0.13 | 0.19 |
| 7 | 19349731 | rs17141120 | -1.034 | -0.578 | -0.272 | 1.864 | -0.011 | -0.031 | -0.031 | 0 | 0.16 | 0.04 | 0.29 | 0.25 | 0.14 | 0.26 |
| 7 | 19353971 | rs1990451 | 1.607 | -0.71 | -0.018 | 1.592 | 0.003 | 0.005 | 0.005 | 0 | 0.61 | 0.38 | 0.58 | 0.38 | 0.65 | 0.27 |
| 7 | 19355187 | rs13243246 | 2.331 | -0.689 | -0.062 | 1.555 | -0.008 | 0.092 | 0.092 | 0 | 0.48 | 0.18 | 0.41 | 0.31 | 0.55 | 0.22 |
| 7 | 19359782 | rs2390081 | -0.997 | -0.695 | 0.025 | 1.461 | 0.01 | -0.016 | -0.016 | 0 | 0.08 | 0.19 | 0.16 | 0.07 | 0.09 | 0.05 |

| | | | | | | | | | | | | | | | | |
|---|---|---|---|---|---|---|---|---|---|---|---|---|---|---|---|---|
| 7 | 19363844 | rs1465297 | 2.44 | -0.645 | -0.059 | 1.581 | -0.021 | 0.165 | 0.165 | 0 | 0.47 | 0.16 | 0.4 | 0.2 | 0.54 | 0.22 |
| 7 | 19369093 | rs10254503 | -1.025 | -0.589 | 0.049 | 2.007 | -0.001 | 0.013 | 0.013 | 0 | 0.26 | 0.3 | 0.28 | 0.46 | 0.24 | 0.55 |
| 7 | 19369899 | rs17141144 | -0.214 | -0.598 | 0.065 | 1.944 | 0.013 | 0.011 | 0.011 | 0 | 0.05 | 0.19 | 0.16 | 0.05 | 0.06 | 0.05 |
| 7 | 19370674 | rs9655164 | -0.381 | -0.619 | 0.046 | 1.984 | -0.01 | -0.07 | -0.07 | 0 | 0.21 | 0.1 | 0.13 | 0.4 | 0.19 | 0.52 |
| 7 | 19370787 | rs10262398 | 1.128 | -0.68 | 0.733 | 2.033 | 0.006 | 0.048 | 0.048 | 0 | 0.84 | 0.75 | 0.75 | 0.71 | 0.82 | 0.85 |
| 7 | 19371245 | rs9654950 | -1.152 | -0.636 | -0.049 | 1.993 | 0 | 0.003 | 0.003 | 0.03 | 0.27 | 0.3 | 0.31 | 0.37 | 0.27 | 0.57 |
| 7 | 19371790 | rs7796931 | 2.07 | -0.646 | -0.043 | 1.975 | 0 | 0.009 | 0.009 | 0 | 0.6 | 0.4 | 0.59 | 0.27 | 0.62 | 0.28 |
| 7 | 19373677 | rs4721754 | 1.714 | -0.62 | 0.029 | 2.018 | -0.006 | 0.103 | 0.103 | 0.055 | 0.79 | 0.42 | 0.72 | 0.59 | 0.81 | 0.8 |
| 7 | 19374370 | rs978068 | -1.555 | -0.679 | 0.285 | 1.999 | 0.009 | 0.083 | 0.083 | 0.44 | 0.07 | 0.29 | 0.16 | 0.16 | 0.06 | 0.05 |
| 7 | 19378959 | rs10155945 | -0.775 | -0.604 | -0.159 | 1.896 | -0.057 | -0.097 | -0.097 | 0 | 0.25 | 0.01 | 0.16 | 0.31 | 0.21 | 0.52 |
| 7 | 19389750 | rs6945594 | -1.225 | 0.179 | 0.683 | 2.178 | 0.002 | 0.006 | 0.006 | 0 | 0.26 | 0.26 | 0.31 | 0.4 | 0.31 | 0.56 |
| 7 | 19393520 | rs6966644 | 1.635 | 0.173 | 1.022 | 2.206 | 0.036 | 0.152 | 0.152 | 0 | 0.84 | 0.52 | 0.7 | 0.7 | 0.86 | 0.79 |
| 7 | 19396641 | rs7788306 | 1.504 | 0.241 | 1.054 | 2.199 | 0.025 | 0.158 | 0.158 | 1.204 | 0.85 | 0.55 | 0.75 | 0.69 | 0.87 | 0.81 |
| 7 | 19397441 | rs10249433 | -2.108 | 0.082 | 1.131 | 2.02 | -0.012 | -0.048 | -0.048 | 0 | 0.09 | 0.23 | 0.04 | 0 | 0.06 | 0.03 |
| 7 | 19399831 | rs9639329 | -1.146 | -0.091 | 0.766 | 1.88 | -0.014 | -0.025 | -0.025 | 0 | 0.31 | 0.17 | 0.38 | 0.39 | 0.34 | 0.54 |
| 7 | 19411289 | rs727919 | 1.56 | 0.042 | 0.586 | 1.716 | 0.011 | 0.096 | 0.096 | 0.217 | 0.78 | 0.54 | 0.7 | 0.65 | 0.81 | 0.8 |
| 7 | 19413024 | rs960046 | 3.536 | 0.179 | 0.45 | 1.454 | -0.068 | 0.107 | 0.107 | 0 | 0.39 | 0.09 | 0.28 | 0.18 | 0.42 | 0.22 |
| 7 | 19413271 | rs960044 | 2.861 | 0.177 | 0.5 | 1.467 | -0.011 | 0.09 | 0.09 | 0 | 0.45 | 0.21 | 0.32 | 0.26 | 0.48 | 0.25 |
| 7 | 19415207 | rs10950719 | 2.304 | 0.499 | 0.626 | 1.696 | -0.003 | 0.003 | 0.003 | 0 | 0.58 | 0.43 | 0.49 | 0.34 | 0.57 | 0.3 |
| 7 | 19418564 | rs2215621 | -1.549 | 0.236 | 0.856 | 2.084 | -0.016 | -0.056 | -0.056 | 0.66 | 0.28 | 0.14 | 0.35 | 0.47 | 0.31 | 0.57 |
| 7 | 19421606 | rs17437711 | -1.248 | 0.285 | 0.919 | 2.031 | -0.002 | -0.025 | -0.025 | 0 | 0.16 | 0.17 | 0.19 | 0.06 | 0.14 | 0.07 |
| 7 | 19423063 | rs10447604 | 1.374 | 0.302 | 1.593 | 2.2 | 0.032 | 0.129 | 0.129 | 6.908 | 0.9 | 0.56 | 0.8 | 0.88 | 0.94 | 0.93 |
| 7 | 19423672 | rs981704 | 2.137 | 0.283 | 1.633 | 2.205 | -0.019 | 0.075 | 0.075 | 10.802 | 0.87 | 0.4 | 0.78 | 0.82 | 0.94 | 0.93 |
| 7 | 19437424 | rs17437739 | 2.808 | 0.107 | 1.529 | 2.232 | 0.002 | 0.037 | 0.037 | 0 | 0.55 | 0.33 | 0.41 | 0.34 | 0.57 | 0.33 |
| 7 | 19437987 | rs6461427 | 2.474 | 0.41 | 1.884 | 2.451 | -0.029 | 0.049 | 0.049 | 10.37 | 0.87 | 0.35 | 0.78 | 0.82 | 0.94 | 0.93 |
| 7 | 19439393 | rs10227301 | 1.705 | 0.438 | 1.813 | 2.471 | 0.02 | 0.14 | 0.14 | 7.212 | 0.89 | 0.5 | 0.79 | 0.88 | 0.95 | 0.93 |
| 7 | 19447111 | rs4721765 | 2.388 | 0.637 | 1.769 | 2.36 | 0.002 | 0.156 | 0.156 | 4.644 | 0.84 | 0.49 | 0.69 | 0.72 | 0.88 | 0.9 |
| 7 | 19448460 | rs10242998 | 3.541 | 0.63 | 1.771 | 2.291 | -0.005 | 0.031 | 0.031 | 3.325 | 0.75 | 0.36 | 0.64 | 0.66 | 0.78 | 0.84 |
| 7 | 19448729 | rs10243311 | -1.011 | 0.746 | 1.731 | 2.286 | -0.014 | -0.025 | -0.025 | 0 | 0.24 | 0.16 | 0.34 | 0.48 | 0.31 | 0.57 |
| 7 | 19448849 | rs7341436 | -1.117 | 0.766 | 1.694 | 2.21 | 0.005 | -0.025 | -0.025 | 1.677 | 0.07 | 0.1 | 0.05 | 0 | 0.09 | 0.05 |
| 7 | 19451194 | rs10252231 | 1.79 | 0.718 | 1.651 | 2.317 | 0.002 | 0.154 | 0.154 | 10.093 | 0.89 | 0.58 | 0.77 | 0.83 | 0.95 | 0.93 |
| 7 | 19454846 | rs1024768 | -2.627 | 0.854 | 1.464 | 2.299 | 0.004 | -0.003 | -0.003 | 0 | 0.4 | 0.45 | 0.47 | 0.7 | 0.48 | 0.66 |
| 7 | 19462542 | rs10279302 | -1.326 | 0.782 | 1.31 | 2 | -0.02 | -0.042 | -0.042 | 0 | 0.24 | 0.16 | 0.33 | 0.46 | 0.31 | 0.57 |
| 7 | 19464027 | rs2192473 | 1.601 | 0.725 | 1.764 | 2.142 | 0.001 | 0.047 | 0.047 | 0 | 0.78 | 0.72 | 0.72 | 0.54 | 0.79 | 0.52 |
| 7 | 19464080 | rs6966182 | 1.764 | 0.642 | 2.087 | 2.012 | 0.006 | 0.097 | 0.097 | 0 | 0.72 | 0.53 | 0.67 | 0.5 | 0.71 | 0.4 |
| 7 | 19468258 | rs10279064 | -1.852 | 0.385 | 1.778 | 1.916 | 0.028 | 0.076 | 0.076 | 2.02 | 0.15 | 0.36 | 0.25 | 0.17 | 0.14 | 0.17 |
| 7 | 19471964 | rs6942577 | 1.691 | 0.439 | 1.727 | 1.813 | -0.009 | -0.005 | -0.005 | 0 | 0.93 | 0.82 | 0.84 | 0.93 | 0.94 | 0.92 |
| 7 | 19476939 | rs7811056 | 2.121 | 0.568 | 1.513 | 1.882 | -0.026 | 0.041 | 0.041 | 0 | 0.54 | 0.27 | 0.39 | 0.35 | 0.53 | 0.25 |
| 7 | 19477272 | rs6979828 | -0.844 | 0.538 | 1.64 | 1.901 | 0.011 | 0.02 | 0.02 | 0 | 0.28 | 0.37 | 0.19 | 0.38 | 0.27 | 0.5 |
| 7 | 19477609 | rs6461430 | -1.28 | 0.451 | 0.757 | 1.35 | -0.014 | -0.004 | -0.004 | 0 | 0.4 | 0.62 | 0.45 | 0.58 | 0.4 | 0.67 |
| 7 | 19478176 | rs10268699 | 1.502 | 0.114 | 1.279 | 1.102 | -0.002 | 0.001 | 0.001 | 0 | 0.95 | 0.9 | 0.84 | 0.93 | 0.94 | 0.92 |
| 7 | 19481290 | rs17141315 | -2.157 | 0.111 | 1.389 | 1.14 | -0.001 | 0.008 | 0.008 | 0 | 0.16 | 0.3 | 0.31 | 0.21 | 0.13 | 0.17 |
| 7 | 19481547 | rs17141316 | -2.042 | 0.045 | 1.38 | 1.049 | 0 | -0.04 | -0.04 | 0 | 0.15 | 0.26 | 0.21 | 0.09 | 0.12 | 0.14 |
| 7 | 19481770 | rs17141317 | -0.802 | -0.384 | 1.616 | 1.24 | -0.003 | 0.039 | 0.039 | 0 | 0.09 | 0.14 | 0.07 | 0.13 | 0.07 | 0.06 |
| 7 | 19481867 | rs17141318 | -2.026 | -0.53 | 2.043 | 1.177 | -0.007 | 0.01 | 0.01 | 0.137 | 0.14 | 0.35 | 0.22 | 0.18 | 0.1 | 0.14 |
| 7 | 19482653 | rs10229992 | -2.632 | -0.741 | 1.718 | 1.095 | -0.018 | 0.003 | 0.003 | 0 | 0.31 | 0.58 | 0.4 | 0.35 | 0.2 | 0.26 |
| 7 | 19496496 | rs2192472 | 1.631 | -0.842 | 1.787 | 1.062 | -0.036 | -0.006 | -0.006 | 0 | 0.86 | 0.6 | 0.74 | 0.85 | 0.92 | 0.83 |
| 7 | 19498089 | rs7792289 | -0.822 | -0.715 | 1.621 | 1.18 | -0.001 | 0.01 | 0.01 | 0 | 0.1 | 0.09 | 0.07 | 0.15 | 0.09 | 0.05 |
| 7 | 19509870 | rs2110470 | 2.553 | -0.907 | 1.917 | 1.112 | -0.034 | 0.004 | 0.004 | 0.03 | 0.72 | 0.37 | 0.66 | 0.7 | 0.82 | 0.78 |
| 7 | 19518160 | rs1548825 | 2.306 | -0.744 | 1.882 | 1.237 | -0.039 | 0.057 | 0.057 | 0.156 | 0.77 | 0.49 | 0.68 | 0.69 | 0.84 | 0.78 |
| 7 | 19522934 | rs10274448 | 1.017 | -0.643 | 1.8 | 1.009 | -0.002 | -0.004 | -0.004 | 0 | 0.93 | 0.93 | 0.86 | 0.91 | 0.93 | 0.9 |
| 7 | 19527095 | rs4719561 | 2.214 | -0.664 | 1.895 | 1.004 | -0.023 | -0.002 | -0.002 | 0.576 | 0.85 | 0.55 | 0.72 | 0.84 | 0.9 | 0.84 |
| 7 | 19528428 | rs2390117 | 1.129 | -0.637 | 1.805 | 0.988 | -0.001 | 0.01 | 0.01 | 0 | 0.92 | 0.71 | 0.86 | 0.92 | 0.96 | 0.9 |
| 7 | 19530893 | rs12700054 | 1.864 | -0.643 | 1.699 | 0.974 | -0.01 | -0.033 | -0.033 | 0 | 0.88 | 0.59 | 0.84 | 0.92 | 0.93 | 0.86 |
| 7 | 19531380 | rs17141444 | -2.283 | -0.685 | 1.696 | 0.98 | -0.033 | -0.005 | -0.005 | 0.537 | 0.13 | 0.39 | 0.26 | 0.14 | 0.07 | 0.13 |
| 7 | 19541920 | rs7810842 | -2.216 | -0.678 | 1.7 | 0.983 | -0.029 | -0.001 | -0.001 | 0 | 0.13 | 0.39 | 0.26 | 0.14 | 0.07 | 0.13 |
| 7 | 19544807 | rs6966240 | -2.255 | -0.673 | 1.776 | 1.006 | -0.023 | 0.007 | 0.007 | 1.023 | 0.13 | 0.41 | 0.26 | 0.14 | 0.07 | 0.13 |
| 7 | 19546965 | rs17141494 | -0.478 | -0.558 | 1.866 | 0.998 | 0.008 | 0.015 | 0.015 | 0.326 | 0.06 | 0.22 | 0.13 | 0.06 | 0.02 | 0.07 |
| 7 | 19547138 | rs10234065 | -2.345 | -0.821 | 1.826 | 1.14 | -0.041 | 0.014 | 0.014 | 0 | 0.25 | 0.59 | 0.34 | 0.28 | 0.16 | 0.19 |
| 7 | 19547949 | rs6944632 | 1.793 | -0.859 | 0.911 | 0.901 | 0.011 | 0.026 | 0.026 | 0.046 | 0.57 | 0.34 | 0.46 | 0.26 | 0.63 | 0.33 |
| 7 | 19547969 | rs16872296 | -1.269 | -1.271 | 1.305 | 0.978 | 0.001 | -0.021 | -0.021 | 0.32 | 0.1 | 0.37 | 0.15 | 0.06 | 0.05 | 0.09 |
| 7 | 19548842 | rs4721778 | 1.509 | -2.119 | 0.219 | 0.322 | 0.063 | 0.179 | 0.179 | 0 | 0.48 | 0.1 | 0.47 | 0.18 | 0.66 | 0.28 |
| 7 | 19549650 | rs10245919 | 0.768 | -2.166 | 0.938 | 0.322 | 0.012 | 0.018 | 0.018 | 0 | 0.59 | 0.63 | 0.62 | 0.25 | 0.69 | 0.36 |
| 7 | 19558261 | rs1476944 | 1.601 | -2.145 | 0.677 | 0.32 | 0.074 | 0.185 | 0.185 | 0 | 0.51 | 0.1 | 0.48 | 0.18 | 0.66 | 0.28 |
| 7 | 19563146 | rs10235171 | -0.426 | -2.11 | 0.792 | 0.759 | 0.002 | -0.002 | -0.002 | 0 | 0.85 | 0.89 | 0.83 | 0.5 | 0.87 | 0.5 |
| 7 | 19566286 | rs6461441 | -0.078 | -2.145 | 0.763 | 0.814 | 0.002 | -0.027 | -0.027 | 0 | 0.83 | 0.88 | 0.8 | 0.45 | 0.85 | 0.49 |
| 7 | 19573641 | rs17141561 | 0.214 | -2.061 | 0.724 | 1.227 | -0.007 | -0.04 | -0.04 | 0 | 0.15 | 0.07 | 0.11 | 0.48 | 0.15 | 0.5 |
| 7 | 19576001 | rs6971655 | 0.957 | -2.191 | 0.891 | 1.038 | 0.012 | 0.007 | 0.007 | 1.981 | 0.71 | 0.28 | 0.77 | 0.75 | 0.84 | 0.8 |
| 7 | 19578620 | rs12112355 | 0.04 | -2.16 | 1.52 | 1.063 | 0.002 | 0.049 | 0.049 | 0.252 | 0.06 | 0.27 | 0.13 | 0.07 | 0.01 | 0.06 |

| | | | | | | | | | | | | | | | | |
|---|---|---|---|---|---|---|---|---|---|---|---|---|---|---|---|---|
| 7 | 19578876 | rs2024353 | -0.855 | -2.296 | 1.633 | 1.164 | 0.012 | -0.01 | -0.01 | 0 | 0.31 | 0.18 | 0.25 | 0.69 | 0.27 | 0.61 |
| 7 | 19581830 | rs2192480 | 0.387 | -2.362 | 1.77 | 1.135 | 0.001 | -0.001 | -0.001 | 0.357 | 0.16 | 0.17 | 0.07 | 0.17 | 0.12 | 0.11 |
| 7 | 19587538 | rs10441042 | -0.176 | -2.321 | 1.824 | 0.93 | 0.003 | 0.031 | 0.031 | 0.349 | 0.07 | 0.34 | 0.05 | 0.06 | 0.01 | 0.06 |
| 7 | 19592673 | rs17141582 | -0.46 | -2.321 | 1.864 | 0.93 | -0.009 | 0.019 | 0.019 | 0.961 | 0.08 | 0.35 | 0.05 | 0.06 | 0.01 | 0.06 |
| 7 | 19599867 | rs17354370 | -0.836 | -2.186 | 1.63 | 0.976 | 0.003 | -0.005 | -0.005 | 0 | 0.31 | 0.19 | 0.36 | 0.71 | 0.28 | 0.61 |
| 7 | 19611149 | rs714392 | 1.541 | -2.229 | 1.7 | 0.972 | -0.008 | -0.018 | -0.018 | 0 | 0.59 | 0.54 | 0.52 | 0.21 | 0.66 | 0.29 |
| 7 | 19611630 | rs11486800 | 0.042 | -2.136 | 1.637 | 1.093 | -0.001 | 0.054 | 0.054 | 0 | 0.16 | 0.19 | 0.3 | 0.47 | 0.15 | 0.49 |
| 7 | 19617468 | rs9655166 | -0.519 | -2.13 | 1.811 | 0.833 | 0.031 | 0.058 | 0.058 | 0 | 0.05 | 0.3 | 0.1 | 0.05 | 0.02 | 0.02 |
| 7 | 19626493 | rs6962668 | -0.645 | -2.139 | 1.874 | 0.809 | 0.024 | -0.019 | -0.019 | 0 | 0.07 | 0.3 | 0.04 | 0 | 0.04 | 0.04 |
| 7 | 19627850 | rs12113592 | -0.826 | -2.221 | 1.193 | 0.954 | -0.012 | 0.051 | 0.051 | 0 | 0.22 | 0.24 | 0.24 | 0.56 | 0.17 | 0.53 |
| 7 | 19629693 | rs7810672 | -0.299 | -2.361 | 1.308 | 0.907 | -0.003 | 0.045 | 0.045 | 0.351 | 0.07 | 0.3 | 0.13 | 0.07 | 0.01 | 0.06 |
| 7 | 19632320 | rs4721805 | 0.355 | -2.412 | 1.035 | 0.904 | -0.021 | 0.03 | 0.03 | 0.572 | 0.17 | 0.33 | 0.17 | 0.19 | 0.13 | 0.11 |
| 7 | 19642100 | rs2390151 | -0.722 | -2.356 | 0.453 | 0.768 | -0.039 | -0.008 | -0.008 | 0 | 0.25 | 0.49 | 0.19 | 0.21 | 0.19 | 0.16 |
| 7 | 19644307 | rs12668640 | -0.713 | -2.211 | 0.051 | 0.911 | -0.008 | 0.036 | 0.036 | 0 | 0.21 | 0.22 | 0.31 | 0.59 | 0.16 | 0.5 |
| 7 | 19649160 | rs11978390 | -0.888 | -1.9 | -0.035 | 0.552 | -0.026 | 0.001 | 0.001 | 0 | 0.09 | 0.15 | 0.19 | 0.12 | 0.03 | 0.09 |
| 7 | 19650053 | rs10245106 | -0.988 | -2.542 | -0.341 | 0.198 | 0.127 | 0.15 | 0.15 | 0 | 0.5 | 0.79 | 0.49 | 0.81 | 0.31 | 0.67 |
| 7 | 19653149 | rs12700063 | 0.716 | -2.282 | -0.255 | 0.237 | 0.017 | 0.016 | 0.016 | 0 | 0.56 | 0.38 | 0.61 | 0.26 | 0.68 | 0.41 |
| 7 | 19654201 | rs4307235 | 1.047 | -2.152 | -0.845 | 0.151 | 0.055 | 0.122 | 0.122 | 0 | 0.46 | 0.16 | 0.48 | 0.19 | 0.64 | 0.29 |
| 7 | 19658389 | rs2892937 | -0.69 | -2.096 | -0.679 | 0.384 | -0.003 | -0.008 | -0.008 | 0 | 0.06 | 0.03 | 0.02 | 0.13 | 0.06 | 0.09 |
| 7 | 19660132 | rs13231224 | 0.605 | -2.739 | -1.019 | -0.152 | -0.011 | -0.007 | -0.007 | 0 | 0.63 | 0.39 | 0.58 | 0.28 | 0.69 | 0.62 |
| 7 | 19661286 | rs13235383 | 1.578 | -3.13 | -1.166 | -0.375 | 0.058 | 0.121 | 0.121 | 0 | 0.47 | 0.25 | 0.42 | 0.09 | 0.6 | 0.45 |
| 7 | 19661865 | rs17141740 | 0.69 | -3.08 | -1.161 | -0.407 | 0.025 | 0.032 | 0.032 | 0 | 0.61 | 0.29 | 0.49 | 0.2 | 0.68 | 0.58 |
| 7 | 19674247 | rs10257645 | -0.27 | -2.881 | -1.379 | -0.541 | -0.02 | -0.031 | -0.031 | 0 | 0.3 | 0.59 | 0.24 | 0.16 | 0.22 | 0.06 |
| 7 | 19682148 | rs10282743 | -2.118 | -2.434 | -1.341 | -0.407 | -0.035 | 0.091 | 0.091 | 0 | 0.63 | 0.79 | 0.62 | 0.94 | 0.51 | 0.58 |
| 7 | 19687779 | rs1476882 | -0.523 | -2.59 | -1.188 | 0.035 | 0.011 | -0.002 | -0.002 | 0 | 0.15 | 0.11 | 0.15 | 0.19 | 0.1 | 0.18 |
| 7 | 19691229 | rs6461452 | -0.731 | -2.564 | -1.142 | 0.075 | 0.013 | 0.012 | 0.012 | 0 | 0.38 | 0.66 | 0.41 | 0.75 | 0.28 | 0.37 |
| 7 | 19711762 | rs6974308 | -0.729 | -2.386 | -1.615 | 0.153 | 0.043 | 0.043 | 0.043 | 0.625 | 0.39 | 0.71 | 0.28 | 0.17 | 0.18 | 0.17 |
| 7 | 19716163 | rs12536095 | -1.819 | -0.689 | -1.003 | 1.377 | -0.002 | -0.001 | -0.001 | 0 | 0.16 | 0.17 | 0.31 | 0.75 | 0.22 | 0.4 |
| 7 | 19719456 | rs11771348 | -0.539 | -0.717 | -0.802 | 1.559 | -0.032 | -0.043 | -0.043 | 0 | 0.09 | 0.01 | 0.2 | 0.57 | 0.13 | 0.27 |
| 7 | 19730538 | rs12667746 | -1.664 | -0.86 | -0.745 | 1.33 | 0.002 | -0.003 | -0.003 | 0 | 0.2 | 0.2 | 0.34 | 0.61 | 0.16 | 0.35 |
| 7 | 19747283 | rs3948699 | -1.148 | -0.557 | -1.069 | 1.526 | -0.023 | -0.1 | -0.1 | 6.4 | 0.15 | 0.04 | 0.2 | 0.6 | 0.12 | 0.29 |
| 7 | 19751749 | rs17141813 | -0.86 | -0.405 | -1.402 | 1.239 | -0.065 | -0.103 | -0.103 | 0 | 0.14 | 0.02 | 0.17 | 0.58 | 0.16 | 0.24 |
| 7 | 19753807 | rs11972191 | -1.51 | -0.605 | -0.904 | 1.215 | 0 | 0.007 | 0.007 | 0.285 | 0.17 | 0.23 | 0.21 | 0.58 | 0.12 | 0.28 |
| 7 | 19755224 | rs10486362 | -0.028 | -0.605 | -0.923 | 1.233 | -0.032 | 0.025 | 0.025 | 0 | 0.06 | 0.01 | 0.15 | 0.57 | 0.08 | 0.22 |
| 7 | 19757731 | rs10239591 | -2.186 | -1.655 | -1.8 | 0.402 | -0.04 | 0.309 | 0.309 | 0.235 | 0.54 | 0.94 | 0.55 | 0.95 | 0.39 | 0.61 |
| 7 | 19758362 | rs7802120 | -1.653 | -1.133 | -1.532 | 0.657 | -0.012 | -0.058 | -0.058 | 0 | 0.25 | 0.16 | 0.25 | 0.59 | 0.21 | 0.31 |
| 7 | 19760190 | rs2285925 | -0.867 | -1.268 | -1.56 | 0.944 | -0.005 | -0.023 | -0.023 | 0 | 0.16 | 0.15 | 0.18 | 0.55 | 0.16 | 0.25 |
| 7 | 19760753 | rs2285928 | -1.381 | -1.266 | -1.561 | 0.777 | -0.047 | -0.133 | -0.133 | 0.972 | 0.23 | 0.03 | 0.23 | 0.59 | 0.2 | 0.31 |
| 7 | 19763478 | rs10435067 | -0.733 | -1.141 | -1.448 | 0.878 | -0.009 | -0.026 | -0.026 | 0 | 0.15 | 0.14 | 0.18 | 0.59 | 0.17 | 0.25 |
| 7 | 19767162 | rs10240229 | -0.646 | -1.294 | 0.076 | 1.302 | -0.018 | 0.045 | 0.045 | 0 | 0.18 | 0.33 | 0.18 | 0.28 | 0.15 | 0.14 |
| 7 | 19771693 | rs4721822 | -0.579 | -1.101 | -0.08 | 1.639 | -0.007 | 0.062 | 0.062 | 0 | 0.08 | 0.1 | 0.16 | 0.55 | 0.1 | 0.23 |
| 7 | 19783977 | rs2158379 | -1.678 | -1.406 | 0.738 | 1.858 | -0.007 | 0.171 | 0.171 | 0 | 0.19 | 0.51 | 0.31 | 0.43 | 0.13 | 0.26 |
| 7 | 19788403 | rs12666488 | -0.74 | -1.619 | 1.086 | 1.905 | 0.024 | 0.212 | 0.212 | 0 | 0.09 | 0.22 | 0.24 | 0.41 | 0.1 | 0.22 |
| 7 | 19790755 | rs17141889 | -0.822 | -1.764 | 1.274 | 1.76 | -0.005 | 0.198 | 0.198 | 0 | 0.19 | 0.34 | 0.18 | 0.56 | 0.18 | 0.18 |
| 7 | 19791171 | rs10252597 | -0.624 | -1.615 | 1.375 | 1.844 | 0.008 | 0.127 | 0.127 | 0 | 0.09 | 0.14 | 0.24 | 0.41 | 0.1 | 0.22 |
| 7 | 19798366 | rs7790076 | -0.602 | -1.685 | 1.88 | 1.864 | 0.005 | 0.053 | 0.053 | 0 | 0.17 | 0.25 | 0.18 | 0.29 | 0.19 | 0.16 |
| 7 | 19798833 | rs7791038 | -1.999 | -1.609 | 1.702 | 1.938 | -0.028 | 0.105 | 0.105 | 0 | 0.22 | 0.52 | 0.33 | 0.6 | 0.13 | 0.3 |
| 7 | 19801016 | rs17141931 | -0.682 | -1.565 | 1.309 | 1.848 | -0.031 | 0.011 | 0.011 | 0.408 | 0.12 | 0.26 | 0.08 | 0.17 | 0.04 | 0.09 |
| 7 | 19803925 | rs714238 | -1.638 | -1.559 | 1.496 | 1.893 | -0.014 | 0.092 | 0.092 | 0 | 0.2 | 0.36 | 0.33 | 0.46 | 0.13 | 0.28 |
| 7 | 19805480 | rs6977660 | -1.835 | -1.572 | 1.699 | 1.926 | -0.025 | 0.105 | 0.105 | 0 | 0.21 | 0.48 | 0.34 | 0.58 | 0.14 | 0.32 |
| 7 | 19810957 | rs3922342 | -0.956 | -1.59 | 1.743 | 1.759 | 0.006 | 0.138 | 0.138 | 0 | 0.2 | 0.39 | 0.17 | 0.41 | 0.21 | 0.2 |
| 7 | 19813466 | rs12672257 | -0.816 | -1.383 | 1.74 | 2.209 | 0.011 | 0.175 | 0.175 | 0 | 0.1 | 0.21 | 0.23 | 0.44 | 0.11 | 0.22 |
| 7 | 19816227 | rs10224938 | -0.684 | -1.375 | 1.72 | 2.213 | 0.012 | 0.188 | 0.188 | 0 | 0.1 | 0.22 | 0.23 | 0.45 | 0.11 | 0.22 |
| 7 | 19816825 | rs12531373 | -2.054 | -1.413 | 1.708 | 2.052 | -0.023 | 0.101 | 0.101 | 0 | 0.22 | 0.46 | 0.33 | 0.6 | 0.15 | 0.34 |
| 7 | 19843797 | rs10267417 | -0.667 | -1.354 | 1.503 | 2.336 | 0.01 | 0.115 | 0.115 | 0 | 0.1 | 0.21 | 0.21 | 0.34 | 0.11 | 0.22 |
| 7 | 19849629 | rs7781975 | -1.821 | -1.316 | 1.371 | 2.164 | -0.023 | 0.105 | 0.105 | 0 | 0.22 | 0.55 | 0.34 | 0.6 | 0.15 | 0.35 |
| 7 | 19854686 | rs16872385 | -1.741 | -1.347 | 1.281 | 1.925 | -0.028 | 0.057 | 0.057 | 0 | 0.22 | 0.42 | 0.11 | 0.37 | 0.13 | 0.29 |
| 7 | 19856478 | rs10277445 | -1.878 | -1.518 | 1.41 | 2.041 | -0.023 | 0.095 | 0.095 | 0 | 0.26 | 0.52 | 0.32 | 0.55 | 0.2 | 0.36 |
| 7 | 19861549 | rs6946403 | -1.007 | -1.469 | 1.396 | 2.086 | -0.007 | 0.128 | 0.128 | 0 | 0.15 | 0.31 | 0.26 | 0.38 | 0.12 | 0.25 |
| 7 | 19868190 | rs7784571 | -1.377 | -1.495 | 1.345 | 1.957 | -0.009 | 0.001 | 0.001 | 0 | 0.15 | 0.24 | 0.09 | 0.17 | 0.11 | 0.1 |
| 7 | 19874391 | rs17453222 | -0.36 | -1.457 | 1.213 | 1.729 | 0.008 | -0.015 | -0.015 | 0 | 0.09 | 0.2 | 0.08 | 0.08 | 0.11 | 0.06 |
| 7 | 19876806 | rs13239454 | -0.238 | -1.456 | 1.206 | 1.729 | 0.004 | -0.012 | -0.012 | 0 | 0.08 | 0.15 | 0.07 | 0.07 | 0.11 | 0.06 |
| 7 | 19894101 | rs13243056 | -0.774 | -1.49 | 1.288 | 1.822 | 0.001 | 0.16 | 0.16 | 0 | 0.14 | 0.31 | 0.26 | 0.39 | 0.12 | 0.26 |
| 7 | 19900526 | rs756479 | -2.203 | -1.517 | 1.295 | 1.803 | -0.02 | 0.066 | 0.066 | 0 | 0.28 | 0.57 | 0.35 | 0.47 | 0.23 | 0.32 |
| 7 | 19901724 | rs4548069 | -2.177 | -2.059 | 0.543 | 1.289 | 0.014 | 0.28 | 0.28 | 0.081 | 0.52 | 0.94 | 0.47 | 0.91 | 0.36 | 0.66 |
| 7 | 19911915 | rs10228558 | -1.062 | -1.31 | 1.368 | 1.855 | 0.004 | 0.205 | 0.205 | 0 | 0.14 | 0.4 | 0.26 | 0.39 | 0.12 | 0.25 |
| 7 | 19916190 | rs10253272 | -1.086 | -1.409 | 1.37 | 1.79 | -0.017 | 0.21 | 0.21 | 0 | 0.16 | 0.43 | 0.26 | 0.38 | 0.11 | 0.28 |
| 7 | 19939084 | rs6954874 | -2.701 | -1.503 | 1.405 | 1.8 | -0.016 | -0.024 | -0.024 | 0 | 0.32 | 0.61 | 0.36 | 0.77 | 0.25 | 0.54 |
| 7 | 19939331 | rs13228500 | -0.938 | -1.576 | 1.415 | 1.78 | -0.01 | 0.168 | 0.168 | 0 | 0.15 | 0.32 | 0.25 | 0.38 | 0.11 | 0.29 |

| | | | | | | | | | | | | | | | | |
|---|---|---|---|---|---|---|---|---|---|---|---|---|---|---|---|---|
| 7 | 19953422 | rs6968110 | -0.26 | -1.555 | 1.352 | 1.746 | 0 | -0.002 | -0.002 | 0 | 0.08 | 0.08 | 0.07 | 0.09 | 0.1 | 0.05 |
| 7 | 19959535 | rs10486375 | -0.964 | -1.576 | 1.277 | 1.762 | -0.028 | 0.081 | 0.081 | 0 | 0.08 | 0.16 | 0.04 | 0.28 | 0.03 | 0.18 |
| 7 | 19959826 | rs10259096 | -1.943 | -1.607 | 1.316 | 1.818 | -0.051 | 0.052 | 0.052 | 0 | 0.23 | 0.46 | 0.29 | 0.63 | 0.14 | 0.46 |
| 7 | 19959995 | rs13240921 | -0.629 | -1.58 | 1.323 | 1.703 | 0.007 | -0.033 | -0.033 | 1.048 | 0.1 | 0.2 | 0.07 | 0.02 | 0.1 | 0.05 |
| 7 | 19962374 | rs13237702 | -1.862 | -1.575 | 1.112 | 1.726 | -0.028 | 0.077 | 0.077 | 0 | 0.18 | 0.41 | 0.12 | 0.31 | 0.13 | 0.2 |
| 7 | 19970430 | rs2390203 | -1.774 | -1.291 | 0.669 | 2 | -0.037 | 0.029 | 0.029 | 0 | 0.23 | 0.41 | 0.29 | 0.66 | 0.15 | 0.48 |
| 7 | 19972439 | rs10261330 | -1.453 | -1.177 | 0.501 | 1.481 | -0.033 | 0.074 | 0.074 | 0 | 0.21 | 0.35 | 0.26 | 0.37 | 0.13 | 0.29 |
| 7 | 19979773 | rs1558043 | -1.361 | -0.864 | 0.484 | 1.367 | -0.011 | 0.082 | 0.082 | 0 | 0.17 | 0.31 | 0.26 | 0.36 | 0.12 | 0.35 |
| 7 | 19981460 | rs7790654 | -1.296 | -1.716 | 0.179 | 0.534 | 0 | 0 | 0 | 0 | 0.4 | 0.43 | 0.34 | 0.33 | 0.25 | 0.49 |
| 7 | 19983182 | rs13247232 | -0.887 | -0.929 | 0.136 | 0.409 | 0.002 | -0.001 | -0.001 | 0 | 0.36 | 0.39 | 0.42 | 0.33 | 0.3 | 0.52 |
| 7 | 19986136 | rs10486378 | -1.08 | -0.932 | 0.184 | 0.408 | 0.001 | 0 | 0 | 0.408 | 0.38 | 0.42 | 0.42 | 0.34 | 0.31 | 0.53 |
| 7 | 19997260 | rs10950785 | -0.636 | -1.069 | 0.118 | 0.339 | -0.011 | 0.013 | 0.013 | 0 | 0.25 | 0.35 | 0.32 | 0.32 | 0.21 | 0.45 |
| 7 | 19999873 | rs2390205 | -1.404 | -1 | 0.465 | 0.712 | -0.004 | 0.009 | 0.009 | 0.003 | 0.35 | 0.55 | 0.47 | 0.58 | 0.34 | 0.58 |
| 7 | 20001275 | rs13221825 | -1.284 | -1 | 0.423 | 0.711 | -0.016 | 0.009 | 0.009 | 0.115 | 0.34 | 0.56 | 0.4 | 0.6 | 0.29 | 0.58 |
| 7 | 20003094 | rs17142269 | -0.407 | -0.991 | 0.502 | 0.813 | -0.003 | 0.1 | 0.1 | 0.723 | 0.1 | 0.24 | 0.07 | 0.26 | 0.08 | 0.16 |
| 7 | 20005985 | rs6957312 | -1.132 | -0.77 | 0.533 | 0.591 | 0 | 0 | 0 | 0 | 0.38 | 0.4 | 0.47 | 0.39 | 0.38 | 0.49 |
| 7 | 20006793 | rs6980441 | -0.469 | -0.732 | 0.544 | 0.552 | -0.001 | 0.009 | 0.009 | 0 | 0.28 | 0.36 | 0.4 | 0.34 | 0.27 | 0.45 |
| 7 | 20010422 | rs11980927 | -1.728 | -0.205 | 0.285 | 0.588 | -0.01 | 0.023 | 0.023 | 0 | 0.6 | 0.68 | 0.58 | 0.74 | 0.5 | 0.77 |
| 7 | 20010778 | rs10950788 | -0.539 | -0.139 | 0.039 | 0.599 | -0.001 | -0.001 | -0.001 | 0 | 0.4 | 0.45 | 0.42 | 0.35 | 0.33 | 0.49 |
| 7 | 20018112 | rs6973418 | 1.015 | -0.229 | -0.371 | 0.787 | -0.001 | 0.012 | 0.012 | 0 | 0.79 | 0.78 | 0.84 | 0.61 | 0.82 | 0.65 |
| 7 | 20022789 | rs10280414 | 0.446 | 0.257 | -0.832 | 0.077 | -0.002 | 0.025 | 0.025 | 0 | 0.34 | 0.72 | 0.46 | 0.2 | 0.36 | 0.23 |
| 7 | 20025660 | rs2214920 | -1.042 | -0.248 | -1.325 | 0.144 | -0.106 | -0.004 | -0.004 | 0 | 0.67 | 0.88 | 0.46 | 0.66 | 0.53 | 0.72 |
| 7 | 20026035 | rs4721854 | -0.164 | -1.03 | -0.964 | -0.25 | 0.023 | 0.012 | 0.012 | 0 | 0.44 | 0.6 | 0.29 | 0.63 | 0.26 | 0.64 |
| 7 | 20030908 | rs17142283 | -0.748 | -1.275 | -0.209 | 0.523 | 0.019 | 0.23 | 0.23 | 0 | 0.11 | 0.42 | 0.11 | 0.3 | 0.08 | 0.25 |
| 7 | 20032182 | rs2190414 | -1.777 | -1.504 | -0.491 | 0.396 | -0.025 | 0.029 | 0.029 | 0.005 | 0.19 | 0.69 | 0.16 | 0.32 | 0.14 | 0.29 |
| 7 | 20032762 | rs11978406 | -1.837 | -1.943 | -1.15 | -0.126 | -0.014 | 0.106 | 0.106 | 0.274 | 0.54 | 0.88 | 0.4 | 0.7 | 0.47 | 0.68 |
| 7 | 20036179 | rs4530935 | -1.169 | -2.005 | -1.129 | -0.158 | 0.025 | 0.101 | 0.101 | 0.158 | 0.52 | 0.83 | 0.4 | 0.68 | 0.44 | 0.65 |
| 7 | 20039263 | rs10244904 | 0.904 | -2.26 | -0.961 | -0.272 | -0.019 | 0.036 | 0.036 | 0 | 0.14 | 0.04 | 0.12 | 0.04 | 0.2 | 0.06 |
| 7 | 20043654 | rs38157 | -1.917 | -2.508 | -1.107 | -0.257 | 0.016 | 0.18 | 0.18 | 0.122 | 0.54 | 0.88 | 0.4 | 0.78 | 0.44 | 0.68 |
| 7 | 20044127 | rs38155 | -2.3 | -1.856 | -0.101 | 0.358 | 0.012 | -0.022 | -0.022 | 0 | 0.38 | 0.79 | 0.29 | 0.47 | 0.37 | 0.35 |
| 7 | 20044438 | rs2158860 | -1.894 | -1.878 | 0.255 | 0.358 | 0.006 | -0.006 | -0.006 | 0 | 0.18 | 0.2 | 0.12 | 0.43 | 0.15 | 0.31 |
| 7 | 20047011 | rs11971303 | -1.268 | -1.883 | 0.252 | 0.395 | -0.004 | 0.072 | 0.072 | 0 | 0.1 | 0.17 | 0.1 | 0.43 | 0.09 | 0.24 |
| 7 | 20049554 | rs11980953 | -1.066 | -1.931 | 0.482 | 0.385 | -0.003 | 0.069 | 0.069 | 0 | 0.09 | 0.14 | 0.09 | 0.42 | 0.08 | 0.24 |
| 7 | 20050490 | rs4721857 | 0.415 | -1.867 | 0.361 | 0.375 | -0.025 | 0.035 | 0.035 | 0 | 0.15 | 0.06 | 0.12 | 0.04 | 0.2 | 0.05 |
| 7 | 20052543 | rs38147 | -2.036 | -1.838 | 0.289 | 0.339 | 0.01 | -0.014 | -0.014 | 0.095 | 0.37 | 0.76 | 0.27 | 0.46 | 0.35 | 0.36 |
| 7 | 20066243 | rs6957367 | -0.573 | -1.562 | 0.246 | 0.591 | 0.021 | 0.024 | 0.024 | 0.325 | 0.12 | 0.03 | 0.04 | 0 | 0.07 | 0.09 |
| 7 | 20066718 | rs2190416 | -0.563 | -1.56 | 0.248 | 0.799 | -0.002 | 0.121 | 0.121 | 0 | 0.07 | 0.13 | 0.08 | 0.47 | 0.09 | 0.24 |
| 7 | 20069620 | rs10263340 | -1.48 | -1.555 | 0.098 | 0.63 | -0.004 | 0 | 0 | 0 | 0.47 | 0.55 | 0.3 | 0.5 | 0.39 | 0.48 |
| 7 | 20071322 | rs10268448 | -1.441 | -1.453 | 0.025 | 0.66 | -0.011 | 0.003 | 0.003 | 0 | 0.45 | 0.5 | 0.29 | 0.5 | 0.37 | 0.46 |
| 7 | 20072606 | rs17362999 | -0.097 | -1.061 | -0.826 | 0.094 | -0.047 | 0.152 | 0.152 | 0.476 | 0.24 | 0.03 | 0.14 | 0.02 | 0.23 | 0.12 |
| 7 | 20082960 | rs4721861 | 1.235 | -1.319 | 0.088 | 1.288 | -0.036 | 0.069 | 0.069 | 2.752 | 0.76 | 0.42 | 0.76 | 0.45 | 0.83 | 0.52 |
| 7 | 20083533 | rs1035087 | 0.789 | -1.209 | 0.117 | 1.586 | -0.001 | 0.15 | 0.15 | 0.387 | 0.93 | 0.82 | 0.89 | 0.66 | 0.93 | 0.8 |
| 7 | 20085640 | rs1019249 | 1.611 | -1.497 | 0.095 | 1.183 | 0.015 | -0.002 | -0.002 | 0 | 0.62 | 0.34 | 0.6 | 0.45 | 0.7 | 0.44 |
| 7 | 20085995 | rs1019248 | 0.332 | -1.577 | 0.095 | 1.287 | 0 | 0.016 | 0.016 | 0 | 0.87 | 0.85 | 0.84 | 0.59 | 0.88 | 0.75 |
| 7 | 20088988 | rs10240011 | 1.95 | -1.55 | -0.383 | 0.983 | -0.049 | 0.135 | 0.135 | 0 | 0.42 | 0.16 | 0.42 | 0.21 | 0.53 | 0.27 |
| 7 | 20093641 | rs4721864 | 1.654 | -1.355 | -0.154 | 0.91 | 0.024 | 0.006 | 0.006 | 0.614 | 0.61 | 0.31 | 0.58 | 0.43 | 0.68 | 0.44 |
| 7 | 20104224 | rs13231772 | -0.179 | -1.322 | 0.094 | 0.846 | -0.003 | 0.001 | 0.001 | 0.074 | 0.13 | 0.12 | 0.15 | 0.14 | 0.07 | 0.21 |
| 7 | 20114589 | rs6954985 | 1.07 | -1.271 | -0.004 | 0.909 | 0.008 | -0.012 | -0.012 | 0 | 0.71 | 0.72 | 0.77 | 0.58 | 0.82 | 0.63 |
| 7 | 20116799 | rs1012811 | -0.179 | -1.199 | 0.015 | 0.361 | -0.003 | -0.057 | -0.057 | 0 | 0.25 | 0.25 | 0.13 | 0.06 | 0.25 | 0.12 |
| 7 | 20117042 | rs1012813 | 1.907 | -1.289 | -0.193 | 0.402 | -0.067 | -0.004 | -0.004 | 0 | 0.41 | 0.16 | 0.52 | 0.41 | 0.5 | 0.38 |
| 7 | 20117668 | rs17142393 | -0.492 | -1.624 | -0.01 | 0.507 | 0.022 | -0.031 | -0.031 | 0 | 0.22 | 0.14 | 0.13 | 0.37 | 0.15 | 0.36 |
| 7 | 20118352 | rs4721875 | 0.118 | -1.296 | 0.131 | 0.145 | -0.006 | -0.085 | -0.085 | 0 | 0.75 | 0.64 | 0.83 | 0.93 | 0.77 | 0.92 |
| 7 | 20118821 | rs2098283 | 0.402 | -1.265 | -0.062 | 0.158 | -0.016 | -0.116 | -0.116 | 0.54 | 0.72 | 0.51 | 0.81 | 0.94 | 0.77 | 0.91 |
| 7 | 20120813 | rs4721877 | -0.592 | -0.893 | -0.634 | 0.126 | 0.002 | -0.026 | -0.026 | 0 | 0.43 | 0.34 | 0.23 | 0.08 | 0.35 | 0.22 |
| 7 | 20123880 | rs11760416 | 1.132 | -0.628 | -1.017 | 0.784 | 0.047 | -0.03 | -0.03 | 0 | 0.56 | 0.18 | 0.53 | 0.49 | 0.64 | 0.44 |
| 7 | 20123914 | rs11970927 | -0.517 | -0.124 | -0.454 | 1.377 | 0 | 0.059 | 0.059 | 0 | 0.13 | 0.16 | 0.13 | 0.39 | 0.12 | 0.29 |
| 7 | 20124384 | rs726916 | 0.441 | 0.368 | -0.582 | 0.645 | 0.001 | -0.01 | -0.01 | 2.63 | 0.35 | 0.32 | 0.39 | 0.23 | 0.36 | 0.34 |
| 7 | 20124541 | rs7811906 | 0.64 | 0.457 | -0.375 | 1.384 | -0.001 | 0.01 | 0.01 | 0 | 0.74 | 0.4 | 0.63 | 0.64 | 0.77 | 0.71 |
| 7 | 20125192 | rs10281498 | -1.322 | 0.368 | -0.375 | 1.96 | -0.006 | 0.051 | 0.051 | 0.593 | 0.13 | 0.12 | 0.29 | 0.47 | 0.14 | 0.29 |
| 7 | 20125586 | rs7780826 | 0.946 | 0.708 | -0.528 | 1.211 | -0.024 | 0.03 | 0.03 | 0 | 0.58 | 0.32 | 0.42 | 0.3 | 0.46 | 0.45 |
| 7 | 20127561 | rs4721879 | 1.701 | 0.157 | -0.952 | 0.829 | -0.025 | 0.107 | 0.107 | 1.121 | 0.35 | 0.16 | 0.21 | 0.13 | 0.36 | 0.33 |
| 7 | 20139644 | rs2892970 | 0.129 | 0.332 | -1.103 | 1.385 | 0.039 | -0.018 | -0.018 | 0 | 0.25 | 0.1 | 0.23 | 0.24 | 0.12 | 0.23 |
| 7 | 20140288 | rs6957548 | 1.823 | 0.297 | -0.553 | 1.356 | -0.026 | 0.121 | 0.121 | 0.995 | 0.35 | 0.16 | 0.21 | 0.12 | 0.36 | 0.33 |
| 7 | 20142868 | rs12536790 | -0.026 | 0.423 | 0.495 | 1.288 | -0.003 | -0.005 | -0.005 | 0 | 0.26 | 0.2 | 0.4 | 0.47 | 0.39 | 0.28 |
| 7 | 20143669 | rs2390252 | 1.054 | 0.399 | 0.575 | 1.229 | -0.003 | -0.001 | -0.001 | 0.809 | 0.62 | 0.42 | 0.45 | 0.36 | 0.48 | 0.56 |
| 7 | 20144829 | rs11979495 | -0.087 | 0.4 | 0.31 | 1.386 | -0.003 | -0.004 | -0.004 | 0.141 | 0.26 | 0.21 | 0.33 | 0.44 | 0.39 | 0.28 |
| 7 | 20147715 | rs2110328 | -1.217 | 0.408 | 0.308 | 1.246 | 0 | 0.007 | 0.007 | 0.994 | 0.53 | 0.4 | 0.64 | 0.73 | 0.51 | 0.5 |
| 7 | 20149537 | rs17142427 | -0.151 | 0.148 | -0.006 | 1.121 | 0.033 | -0.023 | -0.023 | 0 | 0.24 | 0.1 | 0.29 | 0.24 | 0.12 | 0.23 |
| 7 | 20152801 | rs2390254 | 0.534 | 0.696 | 0.158 | 0.851 | 0.046 | -0.039 | -0.039 | 0.008 | 0.92 | 0.83 | 0.92 | 0.99 | 0.9 | 0.96 |

| | | | | | | | | | | | | | | | | |
|---|---|---|---|---|---|---|---|---|---|---|---|---|---|---|---|---|
| 7 | 20154816 | rs10235691 | 1.135 | 0.299 | -0.087 | 1.194 | 0.002 | -0.002 | -0.002 | 0.014 | 0.62 | 0.37 | 0.47 | 0.56 | 0.74 | 0.68 |
| 7 | 20156313 | rs10226814 | -1.668 | 0.311 | -0.155 | 0.543 | -0.011 | 0.067 | 0.067 | 0.021 | 0.6 | 0.71 | 0.63 | 0.8 | 0.6 | 0.68 |
| 7 | 20156919 | rs4719603 | -0.635 | 0.04 | -0.242 | 0.969 | 0.011 | -0.029 | -0.029 | 0 | 0.26 | 0.16 | 0.24 | 0.47 | 0.21 | 0.41 |
| 7 | 20157230 | rs10244572 | 0.114 | 0.009 | 0.017 | 0.32 | -0.017 | 0.033 | 0.033 | 0 | 0.16 | 0.21 | 0.11 | 0.27 | 0.13 | 0.23 |
| 7 | 20170125 | rs11765671 | 0.163 | 0.062 | 0.05 | 0.295 | -0.01 | 0.009 | 0.009 | 0 | 0.19 | 0.15 | 0.15 | 0.28 | 0.13 | 0.27 |
| 7 | 20177882 | rs2390256 | -1.268 | 0.167 | 0.031 | 0.056 | 0.021 | 0.013 | 0.013 | 0 | 0.35 | 0.7 | 0.3 | 0.28 | 0.31 | 0.31 |
| 7 | 20182020 | rs10231585 | -1.756 | 0.333 | 0.027 | -0.531 | 0.025 | 0.191 | 0.191 | 0 | 0.48 | 0.75 | 0.52 | 0.81 | 0.45 | 0.62 |
| 7 | 20182336 | rs10275612 | -2.537 | 0.359 | -0.176 | -0.657 | -0.066 | 0.126 | 0.126 | 0.131 | 0.63 | 0.85 | 0.69 | 0.85 | 0.59 | 0.71 |
| 7 | 20182948 | rs6980121 | -1.376 | 0.32 | 0.41 | -0.672 | -0.001 | -0.014 | -0.014 | 0 | 0.44 | 0.54 | 0.39 | 0.79 | 0.41 | 0.62 |
| 7 | 20183621 | rs6946746 | 0.469 | 0.363 | 0.516 | -0.564 | -0.001 | -0.007 | -0.007 | 0.809 | 0.16 | 0.18 | 0.19 | 0.06 | 0.14 | 0.09 |
| 7 | 20184421 | rs4721885 | -1.075 | 0.405 | 0.721 | -0.548 | -0.018 | -0.035 | -0.035 | 1.094 | 0.38 | 0.41 | 0.44 | 0.79 | 0.29 | 0.55 |
| 7 | 20185196 | rs10236093 | -0.629 | 0.512 | 0.583 | -0.678 | 0.059 | -0.04 | -0.04 | 0.02 | 0.07 | 0.21 | 0.06 | 0 | 0.16 | 0.07 |
| 7 | 20198440 | rs975263 | -0.477 | 0.362 | 0.633 | -0.732 | -0.018 | -0.019 | -0.019 | 0 | 0.29 | 0.28 | 0.33 | 0.78 | 0.24 | 0.51 |
| 7 | 20204135 | rs1990172 | -0.779 | 0.34 | 0.494 | -0.934 | 0.024 | 0.047 | 0.047 | 0.012 | 0.33 | 0.16 | 0.34 | 0.82 | 0.27 | 0.53 |
| 7 | 20209279 | rs3094999 | -1.943 | 0.644 | 0.095 | -0.829 | -0.01 | 0.05 | 0.05 | 1.19 | 0.55 | 0.58 | 0.57 | 0.85 | 0.54 | 0.67 |
| 7 | 20211985 | rs17142532 | 0.49 | 0.704 | -0.047 | -0.476 | -0.016 | 0.003 | 0.003 | 0 | 0.09 | 0.03 | 0.21 | 0.6 | 0.12 | 0.25 |
| 7 | 20212351 | rs10085691 | -0.548 | 0.771 | 0.021 | -1.562 | 0.027 | -0.032 | -0.032 | 0.095 | 0.07 | 0.14 | 0.06 | 0 | 0.16 | 0.07 |
| 7 | 20217706 | rs2390285 | -1.345 | 0.443 | -0.171 | -1.668 | 0.071 | 0.164 | 0.164 | 3.673 | 0.49 | 0.69 | 0.55 | 0.88 | 0.38 | 0.59 |
| 7 | 20222287 | rs3094993 | -0.292 | 0.392 | -0.075 | -1.694 | -0.038 | -0.058 | -0.058 | 0 | 0.33 | 0.46 | 0.34 | 0.81 | 0.25 | 0.5 |
| 7 | 20223145 | rs3114470 | -0.302 | 0.389 | -0.076 | -1.696 | -0.037 | -0.057 | -0.057 | 0 | 0.33 | 0.47 | 0.34 | 0.82 | 0.24 | 0.5 |
| 7 | 20225747 | rs17142558 | -0.114 | 0.42 | -0.052 | -1.698 | -0.011 | -0.067 | -0.067 | 2.877 | 0.17 | 0.29 | 0.19 | 0.04 | 0.14 | 0.09 |
| 7 | 20228202 | rs10277079 | -0.041 | 0.444 | -0.063 | -1.69 | -0.008 | -0.049 | -0.049 | 0.038 | 0.17 | 0.23 | 0.19 | 0.04 | 0.14 | 0.09 |
| 7 | 20233664 | rs3095012 | -0.278 | 0.389 | -0.049 | -1.658 | -0.039 | -0.056 | -0.056 | 0.657 | 0.33 | 0.45 | 0.34 | 0.81 | 0.25 | 0.5 |
| 7 | 20235436 | rs3095213 | -1.908 | 0.508 | 0.036 | -1.565 | -0.059 | 0.191 | 0.191 | 0.513 | 0.63 | 0.85 | 0.72 | 1 | 0.59 | 0.78 |
| 7 | 20236136 | rs3095212 | -0.173 | 0.864 | 0.249 | -1.243 | -0.007 | -0.059 | -0.059 | 0 | 0.36 | 0.45 | 0.39 | 0.82 | 0.34 | 0.52 |
| 7 | 20240680 | rs10249826 | 0.333 | 0.921 | 0.099 | -1.234 | 0.001 | 0.04 | 0.04 | 0 | 0.11 | 0.03 | 0.16 | 0.05 | 0.11 | 0.09 |
| 7 | 20251640 | rs3095009 | -0.556 | 1.121 | 0.505 | -1.045 | 0.002 | 0.026 | 0.026 | 0 | 0.47 | 0.54 | 0.56 | 0.87 | 0.46 | 0.62 |
| 7 | 20252033 | rs2190434 | -0.865 | 1.02 | 0.414 | -0.987 | 0.008 | 0.172 | 0.172 | 0 | 0.49 | 0.72 | 0.59 | 0.87 | 0.56 | 0.66 |
| 7 | 20255705 | rs3095008 | -0.502 | 0.863 | 0.427 | -1.08 | -0.001 | -0.043 | -0.043 | 0 | 0.44 | 0.48 | 0.62 | 0.89 | 0.43 | 0.6 |
| 7 | 20256825 | rs3095006 | -1.191 | 0.355 | -0.04 | -1.405 | 0.01 | 0.019 | 0.019 | 0 | 0.63 | 0.67 | 0.78 | 0.9 | 0.7 | 0.67 |
| 7 | 20258779 | rs10081272 | 0.483 | 0.834 | -0.591 | -1.177 | -0.012 | 0.013 | 0.013 | 0 | 0.17 | 0.11 | 0.11 | 0 | 0.23 | 0.07 |
| 7 | 20266666 | rs1029600 | -0.755 | 0.914 | -0.125 | -0.505 | -0.001 | -0.03 | -0.03 | 0 | 0.44 | 0.38 | 0.64 | 0.9 | 0.47 | 0.6 |
| 7 | 20280329 | rs2041017 | -1.431 | 0.266 | 0.497 | 0.767 | 0.011 | 0.148 | 0.148 | 1.441 | 0.13 | 0.51 | 0.09 | 0.31 | 0.1 | 0.25 |
| 7 | 20281553 | rs2041015 | -1.697 | 0.262 | -0.091 | 1.154 | -0.001 | 0 | 0 | 0 | 0.38 | 0.64 | 0.39 | 0.65 | 0.33 | 0.51 |
| 7 | 20282736 | rs11983202 | -1.118 | 0.133 | 0.272 | 1.186 | -0.001 | 0.027 | 0.027 | 0 | 0.26 | 0.42 | 0.24 | 0.31 | 0.26 | 0.32 |
| 7 | 20291878 | rs10486388 | -1.655 | 0.225 | 0.406 | 0.997 | 0.011 | 0.023 | 0.023 | 0.017 | 0.45 | 0.65 | 0.44 | 0.79 | 0.37 | 0.54 |
| 7 | 20292265 | rs4721892 | -1.696 | 0.258 | 0.411 | 0.937 | 0.018 | 0.056 | 0.056 | 0.017 | 0.47 | 0.71 | 0.46 | 0.8 | 0.41 | 0.58 |
| 7 | 20294461 | rs17142623 | 0.172 | 0.133 | 0.224 | 1.348 | -0.021 | -0.071 | -0.071 | 0.131 | 0.12 | 0 | 0.14 | 0.33 | 0.09 | 0.26 |
| 7 | 20297766 | rs17142627 | 0.492 | 0.104 | 0.586 | 0.827 | -0.007 | -0.006 | -0.006 | 0 | 0.08 | 0.1 | 0.06 | 0.05 | 0.07 | 0.07 |
| 7 | 20297926 | rs11763270 | -1.382 | 0.067 | 1.039 | 0.783 | 0.001 | 0.001 | 0.001 | 0.006 | 0.36 | 0.4 | 0.36 | 0.75 | 0.29 | 0.44 |
| 7 | 20303264 | rs6461483 | -1.792 | 0.129 | 1.029 | 0.723 | 0.003 | 0.004 | 0.004 | 0 | 0.41 | 0.64 | 0.4 | 0.79 | 0.31 | 0.53 |
| 7 | 20309378 | rs2892978 | -1.752 | 0.122 | 1.052 | 0.719 | -0.003 | -0.005 | -0.005 | 0 | 0.4 | 0.62 | 0.39 | 0.79 | 0.31 | 0.53 |
| 7 | 20315214 | rs2390288 | -1.866 | 0.118 | 0.983 | 0.718 | 0.008 | 0.017 | 0.017 | 0 | 0.42 | 0.69 | 0.46 | 0.79 | 0.35 | 0.56 |
| 7 | 20319981 | rs2529004 | -1.712 | 0.794 | 0.779 | 0.738 | 0 | -0.005 | -0.005 | 0 | 0.4 | 0.63 | 0.5 | 0.79 | 0.44 | 0.57 |
| 7 | 20326797 | rs17142673 | -0.96 | 0.748 | 0.915 | 0.767 | 0.004 | -0.006 | -0.006 | 2.84 | 0.27 | 0.2 | 0.33 | 0.73 | 0.21 | 0.42 |
| 7 | 20327258 | rs17142674 | 0.519 | 0.845 | 0.735 | 0.706 | 0.001 | -0.001 | -0.001 | 0.002 | 0.07 | 0.08 | 0.07 | 0.03 | 0.08 | 0.07 |
| 7 | 20331713 | rs3095208 | -0.84 | 1.012 | 0.812 | 0.687 | 0.08 | -0.062 | -0.062 | 0 | 0.12 | 0.32 | 0.15 | 0.03 | 0.24 | 0.11 |
| 7 | 20337516 | rs3095203 | -1.66 | 0.968 | 0.465 | 0.723 | 0 | 0.001 | 0.001 | 0 | 0.4 | 0.65 | 0.48 | 0.79 | 0.39 | 0.53 |
| 7 | 20338424 | rs17142692 | 0.187 | 0.641 | -0.292 | 0.634 | 0.004 | 0.011 | 0.011 | 0 | 0.07 | 0.04 | 0.06 | 0.03 | 0.08 | 0.07 |
| 7 | 20344366 | rs9791961 | -0.699 | 0.381 | -0.542 | 0.741 | -0.022 | 0.027 | 0.027 | 0 | 0.34 | 0.41 | 0.44 | 0.43 | 0.32 | 0.28 |
| 7 | 20345864 | rs3114429 | -1.112 | 0.284 | -1.118 | 0.264 | -0.001 | -0.001 | -0.001 | 3.259 | 0.47 | 0.56 | 0.46 | 0.56 | 0.49 | 0.59 |
| 7 | 20346445 | rs3095224 | -0.643 | 0.161 | -1.106 | 0.273 | -0.014 | 0.026 | 0.026 | 0.031 | 0.35 | 0.52 | 0.37 | 0.54 | 0.32 | 0.52 |
| 7 | 20347570 | rs3114431 | -1.047 | 0.262 | -1.068 | 0.316 | 0.002 | 0.002 | 0.002 | 0.003 | 0.48 | 0.52 | 0.43 | 0.55 | 0.44 | 0.57 |
| 7 | 20348727 | rs1558310 | 0.811 | 0.3 | -1.275 | 0.847 | 0.012 | -0.082 | -0.082 | 0 | 0.13 | 0.03 | 0.21 | 0.39 | 0.1 | 0.2 |
| 7 | 20351817 | rs990528 | 0.362 | 0.684 | -1.025 | 0.812 | -0.023 | -0.014 | -0.014 | 0 | 0.14 | 0.04 | 0.07 | 0.14 | 0.15 | 0.29 |
| 7 | 20354037 | rs975130 | -0.064 | 0.557 | -0.366 | 1.266 | 0.01 | 0.248 | 0.248 | 0.206 | 0.15 | 0.44 | 0.25 | 0.37 | 0.11 | 0.14 |
| 7 | 20356506 | rs206182 | 1.272 | 0.612 | -0.412 | 1.202 | -0.006 | 0 | 0 | 0 | 0.58 | 0.39 | 0.49 | 0.44 | 0.55 | 0.62 |
| 7 | 20357385 | rs206184 | 0.873 | 0.632 | -0.366 | 1.242 | -0.006 | 0.004 | 0.004 | 0 | 0.61 | 0.53 | 0.57 | 0.49 | 0.66 | 0.65 |
| 7 | 20358290 | rs206187 | -0.275 | 0.373 | -0.583 | 1.275 | -0.015 | -0.017 | -0.017 | 1.732 | 0.91 | 0.99 | 0.86 | 0.86 | 0.89 | 0.88 |
| 7 | 20360101 | rs206190 | 1.407 | 0.397 | -0.545 | 1.263 | -0.005 | -0.001 | -0.001 | 0.541 | 0.58 | 0.4 | 0.48 | 0.44 | 0.56 | 0.61 |
| 7 | 20360930 | rs206192 | 0.871 | -0.56 | -0.74 | 1.11 | -0.003 | 0.001 | 0.001 | 0 | 0.63 | 0.61 | 0.57 | 0.49 | 0.69 | 0.66 |
| 7 | 20361053 | rs206193 | 0.12 | 0.043 | -1.271 | 1.17 | -0.022 | -0.039 | -0.039 | 0 | 0.91 | 0.98 | 0.86 | 0.82 | 0.89 | 0.87 |
| 7 | 20366528 | rs6974002 | -0.506 | -0.381 | -0.943 | 1.011 | -0.013 | 0.007 | 0.007 | 0.398 | 0.3 | 0.36 | 0.28 | 0.37 | 0.22 | 0.23 |
| 7 | 20371758 | rs206198 | -0.216 | 0.024 | -2.192 | 0.305 | -0.002 | -0.003 | -0.003 | 0 | 0.44 | 0.78 | 0.63 | 0.62 | 0.39 | 0.7 |
| 7 | 20373272 | rs206202 | -0.729 | -0.292 | -2.755 | -0.298 | 0.016 | 0.064 | 0.064 | 0 | 0.83 | 0.96 | 0.87 | 0.99 | 0.86 | 0.9 |
| 7 | 20374252 | rs206203 | 0.168 | -0.19 | -2.952 | -0.383 | 0.014 | -0.071 | -0.071 | 0 | 0.79 | 0.91 | 0.82 | 0.66 | 0.82 | 0.87 |
| 7 | 20376018 | rs10486391 | -0.064 | -0.284 | -2.746 | -0.595 | -0.016 | -0.015 | -0.015 | 0 | 0.38 | 0.25 | 0.23 | 0.35 | 0.46 | 0.19 |
| 7 | 20376639 | rs206204 | -1.47 | -0.285 | -2.727 | -0.56 | 0.023 | 0.061 | 0.061 | 0 | 0.88 | 0.97 | 0.9 | 0.99 | 0.92 | 0.95 |
| 7 | 20383571 | rs3807934 | 0.71 | 0.003 | -2.705 | -0.555 | -0.009 | 0.021 | 0.021 | 0 | 0.47 | 0.68 | 0.65 | 0.63 | 0.44 | 0.76 |

| | | | | | | | | | | | | | | | | |
|---|---|---|---|---|---|---|---|---|---|---|---|---|---|---|---|---|
| 7 | 20389202 | rs17150347 | 1.152 | 0.139 | -2.814 | -0.491 | -0.012 | -0.002 | -0.002 | 1.154 | 0.19 | 0.11 | 0.24 | 0.22 | 0.19 | 0.35 |
| 7 | 20392652 | rs3807936 | 0.869 | 0.272 | -2.768 | -0.728 | 0.002 | 0.113 | 0.113 | 0 | 0.08 | 0.1 | 0.08 | 0.37 | 0.09 | 0.08 |
| 7 | 20393056 | rs13233253 | 1.222 | -0.02 | -2.975 | -1.027 | -0.075 | 0.141 | 0.141 | 0 | 0.23 | 0.06 | 0.13 | 0 | 0.35 | 0.08 |
| 7 | 20395067 | rs3757724 | 1.081 | 0.225 | -2.65 | -0.767 | 0.012 | 0.026 | 0.026 | 0.094 | 0.05 | 0.06 | 0.14 | 0.16 | 0.06 | 0.2 |
| 7 | 20395186 | rs9654959 | 1.997 | 0.249 | -2.781 | -0.982 | -0.044 | 0.081 | 0.081 | 0 | 0.13 | 0 | 0.06 | 0 | 0.11 | 0.04 |
| 7 | 20397974 | rs10265149 | 0.821 | 0.316 | -2.157 | -0.348 | 0.001 | 0.027 | 0.027 | 0.033 | 0.51 | 0.79 | 0.69 | 0.64 | 0.48 | 0.76 |
| 7 | 20408459 | rs17365098 | -0.027 | 1.262 | -1.444 | 0.084 | -0.011 | 0.093 | 0.093 | 0 | 0.3 | 0.19 | 0.16 | 0 | 0.35 | 0.09 |
| 7 | 20413590 | rs1023522 | 1.031 | 1.231 | -1.417 | 0.156 | -0.017 | -0.011 | -0.011 | 0.1 | 0.55 | 0.68 | 0.66 | 0.55 | 0.47 | 0.77 |
| 7 | 20418678 | rs3735619 | -0.318 | 1.345 | -1.223 | 0.198 | 0.001 | 0.086 | 0.086 | 0 | 0.08 | 0.1 | 0.07 | 0.41 | 0.1 | 0.11 |
| 7 | 20428173 | rs6954259 | -0.412 | 1.193 | -1.185 | 0.235 | -0.01 | 0.148 | 0.148 | 0 | 0.09 | 0.19 | 0.06 | 0.37 | 0.07 | 0.09 |
| 7 | 20429735 | rs2189300 | 1.288 | 1.243 | -1.17 | 0.18 | 0.005 | 0.092 | 0.092 | 0.083 | 0.86 | 0.72 | 0.83 | 0.62 | 0.85 | 0.86 |
| 7 | 20433107 | rs1014038 | 0.387 | 1.268 | -1.168 | -0.018 | 0.006 | -0.018 | -0.018 | 0 | 0.95 | 0.91 | 0.89 | 0.99 | 0.92 | 0.95 |
| 7 | 20437293 | rs717861 | 0.43 | 1.233 | -1.282 | -0.172 | -0.038 | 0.168 | 0.168 | 0 | 0.65 | 0.87 | 0.75 | 0.99 | 0.57 | 0.85 |
| 7 | 20437823 | rs10263803 | 0.553 | 1.236 | -1.281 | -0.17 | -0.04 | 0.164 | 0.164 | 0 | 0.65 | 0.88 | 0.76 | 0.99 | 0.57 | 0.85 |
| 7 | 20440465 | rs3807950 | 0.788 | 1.31 | -1.264 | -0.03 | -0.013 | -0.007 | -0.007 | 0 | 0.88 | 0.93 | 0.83 | 0.57 | 0.83 | 0.84 |
| 7 | 20441065 | rs13242474 | 0.213 | 1.208 | -1.113 | -0.149 | -0.009 | 0.039 | 0.039 | 0 | 0.23 | 0.15 | 0.12 | 0 | 0.27 | 0.08 |
| 7 | 20442783 | rs2158252 | 1.787 | 1.312 | -1.032 | -0.07 | 0.012 | 0.004 | 0.004 | 0 | 0.49 | 0.57 | 0.53 | 0.41 | 0.41 | 0.57 |
| 7 | 20442796 | rs3823974 | -1.345 | 0.891 | -1 | 0.029 | 0.002 | -0.017 | -0.017 | 0.054 | 0.4 | 0.18 | 0.22 | 0.41 | 0.43 | 0.19 |
| 7 | 20443021 | rs3807955 | -0.023 | 0.771 | -0.863 | 0.192 | -0.003 | 0.083 | 0.083 | 0 | 0.08 | 0.09 | 0.06 | 0.4 | 0.07 | 0.09 |
| 7 | 20444532 | rs2074424 | 1.766 | 0.842 | -0.548 | -0.875 | 0.019 | 0.01 | 0.01 | 0.763 | 0.87 | 0.77 | 0.76 | 0.83 | 0.86 | 0.74 |
| 7 | 20451427 | rs7782524 | 0.997 | 0.507 | -0.663 | -0.982 | 0.003 | 0.066 | 0.066 | 0 | 0.64 | 0.74 | 0.78 | 0.99 | 0.62 | 0.85 |
| 7 | 20453875 | rs4719613 | 1.175 | 0.58 | -0.659 | -0.95 | 0.009 | 0.03 | 0.03 | 0 | 0.63 | 0.72 | 0.77 | 0.99 | 0.63 | 0.85 |
| 7 | 20461417 | rs12534544 | 1.583 | 0.713 | -0.568 | -0.871 | 0 | -0.003 | -0.003 | 0 | 0.58 | 0.59 | 0.63 | 0.82 | 0.55 | 0.66 |
| 7 | 20464105 | rs2107128 | 1.651 | 0.914 | -0.384 | -0.701 | -0.001 | -0.009 | -0.009 | 0.161 | 0.8 | 0.63 | 0.73 | 0.81 | 0.79 | 0.71 |
| 7 | 20468022 | rs11976987 | -1.226 | 1.092 | -0.297 | -0.459 | 0.028 | 0.101 | 0.101 | 0.047 | 0.08 | 0.22 | 0.18 | 0.23 | 0.1 | 0.21 |
| 7 | 20468718 | rs11980974 | -1.064 | 1.102 | -0.344 | -0.475 | 0.03 | 0.107 | 0.107 | 0.06 | 0.08 | 0.21 | 0.18 | 0.25 | 0.1 | 0.21 |
| 7 | 20469685 | rs7803406 | -0.656 | 0.946 | -0.443 | -0.469 | -0.003 | 0.063 | 0.063 | 0 | 0.07 | 0.13 | 0.22 | 0.24 | 0.07 | 0.19 |
| 7 | 20470275 | rs4719617 | 1.398 | 0.913 | -0.602 | -0.576 | 0 | 0.012 | 0.012 | 0 | 0.56 | 0.39 | 0.58 | 0.73 | 0.53 | 0.65 |
| 7 | 20476194 | rs11979620 | -0.914 | 1.413 | -1.003 | 0.016 | 0.036 | 0.125 | 0.125 | 0 | 0.1 | 0.27 | 0.21 | 0.25 | 0.11 | 0.2 |
| 7 | 20478448 | rs17456488 | -0.508 | 1.351 | -1.832 | -0.551 | -0.029 | 0.062 | 0.062 | 0 | 0.23 | 0.13 | 0.13 | 0 | 0.31 | 0.08 |
| 7 | 20489873 | rs7796492 | -0.212 | 1.148 | -2.228 | -0.181 | 0.024 | 0.067 | 0.067 | 0.675 | 0.49 | 0.21 | 0.46 | 0.6 | 0.47 | 0.38 |
| 7 | 20490448 | rs17142863 | -0.456 | 1.367 | -1.426 | -0.418 | 0.014 | -0.033 | -0.033 | 2.031 | 0.09 | 0.19 | 0.06 | 0.01 | 0.1 | 0.18 |
| 7 | 20490812 | rs11971617 | -0.431 | 1.296 | -1.146 | -0.436 | 0.016 | -0.033 | -0.033 | 0 | 0.09 | 0.22 | 0.07 | 0.01 | 0.1 | 0.18 |
| 7 | 20494612 | rs6961781 | -0.41 | 1.468 | -2.048 | -0.8 | 0.009 | 0.049 | 0.049 | 0 | 0.52 | 0.4 | 0.38 | 0.76 | 0.48 | 0.65 |
| 7 | 20497113 | rs2188869 | -0.374 | 1.373 | -2.084 | -0.804 | 0.026 | 0.066 | 0.066 | 0 | 0.52 | 0.38 | 0.36 | 0.76 | 0.44 | 0.65 |
| 7 | 20497135 | rs2188870 | 0.203 | 1.16 | -2.402 | -0.734 | 0.009 | 0.036 | 0.036 | 0.476 | 0.41 | 0.03 | 0.3 | 0.75 | 0.34 | 0.47 |
| 7 | 20499625 | rs4141372 | -0.053 | 1.301 | -2.47 | -0.689 | -0.025 | -0.085 | -0.085 | 0.107 | 0.44 | 0.05 | 0.3 | 0.56 | 0.4 | 0.5 |
| 7 | 20503965 | rs1859044 | -1.098 | 1.783 | -1.767 | -0.826 | 0.006 | 0 | 0 | 0 | 0.18 | 0.27 | 0.23 | 0.22 | 0.23 | 0.22 |
| 7 | 20512082 | rs7793958 | -0.886 | 0.991 | -2.479 | -1.746 | 0.031 | 0.107 | 0.107 | 0.201 | 0.78 | 0.95 | 0.87 | 0.99 | 0.87 | 0.8 |
| 7 | 20512666 | rs2188872 | -0.894 | 0.975 | -2.486 | -1.753 | 0.031 | 0.107 | 0.107 | 0.174 | 0.78 | 0.95 | 0.87 | 0.99 | 0.88 | 0.8 |
| 7 | 20517352 | rs2893003 | -0.08 | 0.967 | -1.592 | -2.144 | -0.001 | -0.026 | -0.026 | 0 | 0.45 | 0.51 | 0.35 | 0.3 | 0.42 | 0.39 |
| 7 | 20518908 | rs17456802 | 1.354 | 0.785 | -1.398 | -2.268 | 0.005 | -0.003 | -0.003 | 0.022 | 0.27 | 0.17 | 0.12 | 0.02 | 0.19 | 0.16 |
| 7 | 20519106 | rs10273151 | 0.71 | 0.803 | -1.443 | -2.257 | -0.016 | 0.066 | 0.066 | 0 | 0.12 | 0 | 0.06 | 0.01 | 0.1 | 0.13 |
| 7 | 20523492 | rs17142970 | 0.994 | 0.668 | -1.142 | -2.342 | 0.002 | -0.042 | -0.042 | 0.063 | 0.31 | 0.27 | 0.12 | 0.03 | 0.19 | 0.19 |
| 7 | 20527424 | rs7784579 | -0.335 | 1.005 | -0.62 | -2.187 | 0 | 0.002 | 0.002 | 0 | 0.39 | 0.39 | 0.48 | 0.76 | 0.46 | 0.41 |
| 7 | 20527578 | rs10272302 | -0.365 | 1.076 | -0.412 | -2.122 | -0.001 | -0.011 | -0.011 | 0 | 0.49 | 0.52 | 0.35 | 0.32 | 0.42 | 0.43 |
| 7 | 20530767 | rs10226404 | -0.362 | 1.137 | -0.494 | -2.04 | 0.004 | -0.003 | -0.003 | 0 | 0.19 | 0.28 | 0.22 | 0.22 | 0.24 | 0.19 |
| 7 | 20535864 | rs12535843 | -0.403 | 1.18 | -0.629 | -1.838 | -0.015 | -0.012 | -0.012 | 0.963 | 0.19 | 0.12 | 0.22 | 0.26 | 0.25 | 0.2 |
| 7 | 20550979 | rs1012869 | 1.111 | 1.103 | -0.469 | -1.903 | 0.006 | 0 | 0 | 0 | 0.29 | 0.22 | 0.13 | 0.1 | 0.17 | 0.17 |
| 7 | 20552668 | rs10273566 | -0.257 | 1.172 | -0.676 | -1.798 | -0.029 | -0.027 | -0.027 | 0 | 0.19 | 0.09 | 0.22 | 0.26 | 0.25 | 0.19 |
| 7 | 20559001 | rs10233470 | -0.288 | 1.183 | -0.72 | -1.8 | -0.027 | 0.023 | 0.023 | 0 | 0.19 | 0.08 | 0.22 | 0.09 | 0.25 | 0.17 |
| 7 | 20560996 | rs6461503 | -0.553 | 1.801 | -0.349 | -1.965 | 0.008 | 0.062 | 0.062 | 0 | 0.55 | 0.76 | 0.41 | 0.37 | 0.5 | 0.48 |
| 7 | 20572445 | rs6461504 | 1.129 | 1.461 | -0.493 | -1.951 | -0.007 | -0.01 | -0.01 | 0 | 0.29 | 0.33 | 0.13 | 0.27 | 0.17 | 0.18 |
| 7 | 20572700 | rs916790 | -0.257 | 1.536 | -0.592 | -1.923 | 0.001 | 0.003 | 0.003 | 0 | 0.5 | 0.52 | 0.34 | 0.36 | 0.42 | 0.45 |
| 7 | 20576180 | rs10260395 | -0.355 | 1.839 | -0.537 | -1.673 | -0.023 | 0.023 | 0.023 | 0 | 0.19 | 0.1 | 0.22 | 0.09 | 0.25 | 0.18 |
| 7 | 20583122 | rs1989886 | 0.048 | 1.83 | -0.256 | -1.504 | 0.045 | 0.018 | 0.018 | 0 | 0.26 | 0.38 | 0.51 | 0.63 | 0.45 | 0.36 |
| 7 | 20583673 | rs10234192 | 1.228 | 1.139 | -0.839 | -1.228 | 0.005 | -0.104 | -0.104 | 0 | 0.16 | 0.03 | 0.25 | 0.5 | 0.21 | 0.21 |
| 7 | 20592679 | rs6977876 | 0.168 | 1.44 | -0.478 | -1.816 | 0.018 | -0.126 | -0.126 | 0.215 | 0.39 | 0.43 | 0.32 | 0.04 | 0.39 | 0.32 |
| 7 | 20596439 | rs999601 | 0.186 | 1.223 | -0.726 | -0.959 | -0.003 | -0.001 | -0.001 | 0.008 | 0.46 | 0.56 | 0.45 | 0.49 | 0.46 | 0.36 |
| 7 | 20596929 | rs17143056 | 0.041 | 1.947 | -0.518 | -0.763 | 0.013 | 0.003 | 0.003 | 0 | 0.07 | 0.06 | 0.17 | 0 | 0.15 | 0.06 |
| 7 | 20601159 | rs2188864 | 0.038 | 1.47 | -0.206 | -0.852 | -0.018 | -0.009 | -0.009 | 0 | 0.3 | 0.36 | 0.41 | 0.73 | 0.29 | 0.43 |
| 7 | 20604398 | rs2188865 | -0.945 | 1.694 | -0.422 | -1.05 | -0.026 | 0.05 | 0.05 | 0 | 0.65 | 0.86 | 0.7 | 0.73 | 0.62 | 0.74 |
| 7 | 20613366 | rs9885995 | -0.051 | 1.673 | -0.066 | -1.053 | 0.034 | -0.008 | -0.008 | 0 | 0.56 | 0.71 | 0.45 | 0.48 | 0.43 | 0.54 |
| 7 | 20618211 | rs17143104 | 0.894 | 1.623 | 0.026 | -1.029 | -0.039 | -0.135 | -0.135 | 0 | 0.28 | 0.42 | 0.14 | 0 | 0.18 | 0.19 |
| 7 | 20618597 | rs17143109 | 0.755 | 1.546 | -0.08 | -1.029 | 0.035 | -0.068 | -0.068 | 0 | 0.26 | 0.15 | 0.3 | 0.48 | 0.23 | 0.34 |
| 7 | 20625540 | rs10216228 | -0.994 | 1.824 | -0.113 | -1.05 | 0.032 | -0.001 | -0.001 | 0 | 0.95 | 0.95 | 0.85 | 1 | 0.87 | 0.94 |
| 7 | 20630890 | rs1858947 | -0.9 | 1.888 | -0.085 | -1.067 | -0.026 | 0.058 | 0.058 | 0 | 0.65 | 0.86 | 0.8 | 0.76 | 0.61 | 0.73 |
| 7 | 20633736 | rs10950819 | 0.273 | 1.891 | -0.065 | -1.049 | -0.02 | -0.028 | -0.028 | 0 | 0.31 | 0.21 | 0.54 | 0.5 | 0.36 | 0.46 |
| 7 | 20635309 | rs4721922 | 0.79 | 1.642 | -0.372 | -1.271 | -0.021 | -0.122 | -0.122 | 0 | 0.34 | 0.43 | 0.16 | 0 | 0.23 | 0.22 |

| | | | | | | | | | | | | | | | | |
|---|---|---|---|---|---|---|---|---|---|---|---|---|---|---|---|---|
| 7 | 20636479 | rs1858941 | -0.737 | 1.545 | -0.304 | -1.177 | -0.003 | -0.006 | -0.006 | 0 | 0.65 | 0.7 | 0.7 | 0.5 | 0.58 | 0.69 |
| 7 | 20636508 | rs1858942 | 0.006 | 0.812 | -0.588 | -0.975 | 0.017 | 0.038 | 0.038 | 0 | 0.54 | 0.64 | 0.63 | 0.76 | 0.45 | 0.67 |
| 7 | 20637310 | rs11971866 | 0.327 | 0.775 | -1.102 | -0.477 | 0.036 | -0.066 | -0.066 | 0.049 | 0.3 | 0.13 | 0.4 | 0.49 | 0.25 | 0.38 |
| 7 | 20638231 | rs10488581 | -0.794 | 1.67 | -0.495 | 0.119 | 0.014 | 0.058 | 0.058 | 0 | 0.08 | 0.15 | 0.05 | 0.3 | 0.15 | 0.12 |
| 7 | 20639237 | rs10236605 | 0.605 | 1.514 | -0.728 | 0.109 | -0.005 | 0.002 | 0.002 | 0 | 0.57 | 0.43 | 0.48 | 0.56 | 0.48 | 0.66 |
| 7 | 20639491 | rs6947113 | 0.014 | 1.892 | -0.612 | -0.15 | 0.031 | -0.036 | -0.036 | 0 | 0.31 | 0.36 | 0.3 | 0.15 | 0.42 | 0.14 |
| 7 | 20640509 | rs1858943 | -0.337 | 1.674 | -1.385 | -0.063 | -0.045 | 0.018 | 0.018 | 0.596 | 0.18 | 0.07 | 0.12 | 0.16 | 0.23 | 0.25 |
| 7 | 20640630 | rs7805271 | 0.088 | 1.469 | -1.231 | -0.009 | -0.005 | -0.003 | -0.003 | 0.02 | 0.13 | 0.19 | 0.12 | 0.1 | 0.08 | 0.06 |
| 7 | 20640689 | rs1858944 | -0.165 | 1.035 | -2.252 | -0.308 | -0.079 | 0.123 | 0.123 | 0 | 0.56 | 0.95 | 0.6 | 0.74 | 0.52 | 0.53 |
| 7 | 20641104 | rs7806034 | 0.966 | 0.684 | -1.366 | 0.125 | -0.012 | -0.01 | -0.01 | 0 | 0.27 | 0.36 | 0.17 | 0.18 | 0.24 | 0.14 |
| 7 | 20641893 | rs17215405 | -0.597 | 1.435 | -1.307 | -0.001 | 0.008 | 0.007 | 0.007 | 0.284 | 0.36 | 0.58 | 0.46 | 0.55 | 0.41 | 0.41 |
| 7 | 20642445 | rs17143177 | -0.187 | 1.5 | -0.375 | -0.75 | 0.074 | 0.024 | 0.024 | 0.06 | 0.13 | 0.31 | 0.19 | 0.14 | 0.16 | 0.11 |
| 7 | 20642697 | rs4719621 | -0.611 | 0.643 | -0.979 | -1.206 | 0.035 | 0.103 | 0.103 | 0 | 0.69 | 0.89 | 0.85 | 0.87 | 0.73 | 0.61 |
| 7 | 20647465 | rs6461509 | -1.385 | 0.543 | -0.981 | -1.187 | -0.018 | 0.069 | 0.069 | 0.197 | 0.87 | 0.99 | 0.92 | 0.99 | 0.85 | 0.89 |
| 7 | 20652652 | rs4457229 | -0.262 | 0.832 | -0.279 | -0.861 | 0.002 | 0.141 | 0.141 | 0.293 | 0.24 | 0.5 | 0.36 | 0.5 | 0.26 | 0.35 |
| 7 | 20665487 | rs11773775 | 1.776 | 0.677 | -0.135 | -0.825 | -0.002 | -0.01 | -0.01 | 0 | 0.06 | 0.11 | 0.04 | 0.04 | 0.06 | 0.13 |
| 7 | 20666750 | rs7802291 | 1.117 | 0.691 | -0.183 | -0.995 | -0.005 | 0.018 | 0.018 | 0.399 | 0.36 | 0.27 | 0.32 | 0.23 | 0.35 | 0.17 |
| 7 | 20669629 | rs1476485 | -0.518 | 0.556 | 0.166 | -0.925 | 0.015 | 0.025 | 0.025 | 0.022 | 0.16 | 0.41 | 0.24 | 0.19 | 0.16 | 0.27 |
| 7 | 20678713 | rs6461510 | -0.001 | 0.559 | 0.278 | -0.919 | 0.042 | 0.068 | 0.068 | 0 | 0.1 | 0.36 | 0.2 | 0.15 | 0.1 | 0.14 |
| 7 | 20679286 | rs2188594 | -0.766 | 0.534 | 0.305 | -0.921 | 0.006 | 0.018 | 0.018 | 0.191 | 0.17 | 0.53 | 0.24 | 0.2 | 0.16 | 0.27 |
| 7 | 20680353 | rs11773053 | 2.007 | 0.539 | 0.26 | -0.916 | -0.009 | -0.001 | -0.001 | 0 | 0.05 | 0 | 0.03 | 0.05 | 0.06 | 0.13 |
| 7 | 20680449 | rs2106563 | -1.231 | 0.517 | 0.079 | -1.176 | 0.011 | 0.123 | 0.123 | 0 | 0.66 | 0.89 | 0.82 | 0.87 | 0.7 | 0.6 |
| 7 | 20680551 | rs12700225 | 0.43 | 0.47 | 0.561 | -0.633 | 0.017 | 0.031 | 0.031 | 0 | 0.08 | 0.12 | 0.14 | 0.15 | 0.1 | 0.09 |
| 7 | 20684296 | rs6957205 | -0.996 | 0.501 | 0.469 | -0.707 | 0.015 | -0.047 | -0.047 | 0.283 | 0.22 | 0.54 | 0.27 | 0.15 | 0.24 | 0.17 |
| 7 | 20685508 | rs11983326 | -1.23 | 0.574 | 0.507 | -1.701 | 0.014 | -0.055 | -0.055 | 4.656 | 0.26 | 0.47 | 0.31 | 0.19 | 0.3 | 0.3 |
| 7 | 20687181 | rs2893006 | -0.674 | 0.502 | 0.54 | -1.711 | 0.002 | -0.107 | -0.107 | 0 | 0.2 | 0.4 | 0.17 | 0.04 | 0.21 | 0.21 |
| 7 | 20688135 | rs6950237 | -1.326 | 0.496 | 0.557 | -1.709 | 0.009 | -0.054 | -0.054 | 0 | 0.27 | 0.52 | 0.3 | 0.19 | 0.3 | 0.3 |
| 7 | 20689065 | rs6955287 | -1.367 | 0.505 | 0.581 | -1.71 | 0.006 | -0.047 | -0.047 | 0 | 0.27 | 0.53 | 0.3 | 0.2 | 0.3 | 0.3 |
| 7 | 20690173 | rs10950826 | -0.959 | 0.505 | 0.571 | -1.705 | 0.004 | 0.022 | 0.022 | 0.071 | 0.16 | 0.5 | 0.24 | 0.2 | 0.16 | 0.27 |
| 7 | 20692413 | rs2285557 | 0.053 | 0.531 | 0.376 | -1.701 | -0.046 | 0.078 | 0.078 | 0 | 0.16 | 0.03 | 0.09 | 0.04 | 0.21 | 0.16 |
| 7 | 20693159 | rs2285560 | 0.664 | 0.502 | 0.477 | -1.652 | 0.013 | 0.031 | 0.031 | 0 | 0.07 | 0.11 | 0.15 | 0.16 | 0.09 | 0.08 |
| 7 | 20693542 | rs1108864 | -1.586 | 0.327 | 0.222 | -1.916 | -0.005 | 0.099 | 0.099 | 2.18 | 0.78 | 0.99 | 0.89 | 0.96 | 0.8 | 0.94 |
| 7 | 20694735 | rs6958596 | 0.228 | 0.748 | 1.049 | -1.625 | 0.003 | 0.012 | 0.012 | 0.084 | 0.13 | 0.19 | 0.18 | 0.2 | 0.16 | 0.22 |
| 7 | 20698270 | rs2301641 | -0.981 | 0.724 | 0.931 | -1.623 | 0.013 | -0.037 | -0.037 | 0 | 0.29 | 0.58 | 0.31 | 0.2 | 0.33 | 0.31 |
| 7 | 20700361 | rs6952128 | -1.006 | 0.712 | 0.734 | -1.646 | 0 | -0.002 | -0.002 | 0 | 0.3 | 0.66 | 0.31 | 0.24 | 0.33 | 0.31 |
| 7 | 20703948 | rs1011559 | 0.327 | 0.63 | 0.701 | -1.66 | 0.006 | -0.058 | -0.058 | 0 | 0.15 | 0.31 | 0.12 | 0.02 | 0.15 | 0.13 |
| 7 | 20704137 | rs916739 | -0.044 | 0.231 | 0.589 | -1.551 | 0.037 | 0.05 | 0.05 | 0 | 0.15 | 0.33 | 0.19 | 0.22 | 0.17 | 0.18 |
| 7 | 20712495 | rs13240297 | 1.05 | 0.302 | -0.359 | -1.766 | -0.036 | 0.182 | 0.182 | 0 | 0.33 | 0.07 | 0.23 | 0.04 | 0.38 | 0.42 |
| 7 | 20721084 | rs11764760 | 0.051 | -0.31 | 0.05 | -0.83 | 0.02 | 0.033 | 0.033 | 0 | 0.13 | 0.17 | 0.18 | 0.23 | 0.18 | 0.18 |
| 7 | 20726621 | rs17143258 | -0.974 | -1.281 | -0.061 | -1.383 | -0.058 | -0.131 | -0.131 | 0.229 | 0.37 | 0.49 | 0.16 | 0.05 | 0.17 | 0.21 |
| 7 | 20726832 | rs10230205 | -0.93 | -1.229 | -0.223 | -1.36 | -0.03 | -0.084 | -0.084 | 1.52 | 0.36 | 0.44 | 0.15 | 0.05 | 0.17 | 0.22 |
| 7 | 20727011 | rs17143260 | -0.955 | -0.339 | 0.623 | -0.53 | -0.048 | -0.083 | -0.083 | 0.06 | 0.16 | 0.4 | 0.09 | 0.03 | 0.07 | 0.18 |
| 7 | 20731918 | rs2158857 | 0.065 | 0.15 | 0.273 | -0.282 | -0.008 | -0.019 | -0.019 | 0 | 0.8 | 0.78 | 0.92 | 0.91 | 0.84 | 0.92 |
| 7 | 20735885 | rs2158856 | 0.483 | 0.176 | 0.745 | -0.09 | -0.02 | -0.017 | -0.017 | 0.307 | 0.77 | 0.59 | 0.89 | 0.86 | 0.83 | 0.91 |
| 7 | 20736577 | rs17143272 | -0.62 | 0.719 | 0.739 | 0.305 | 0.029 | 0.163 | 0.163 | 0 | 0.15 | 0.25 | 0.17 | 0.55 | 0.17 | 0.22 |
| 7 | 20737300 | rs2024045 | -0.042 | 0.509 | 0.673 | -0.42 | 0.009 | 0.013 | 0.013 | 1.333 | 0.8 | 0.89 | 0.93 | 0.87 | 0.83 | 0.92 |
| 7 | 20739632 | rs11762180 | -0.356 | 1.059 | 0.24 | -0.691 | 0 | -0.011 | -0.011 | 0 | 0.16 | 0.14 | 0.24 | 0.02 | 0.24 | 0.16 |
| 7 | 20739839 | rs13229035 | -0.379 | 1.059 | 0.103 | -0.691 | -0.046 | 0.064 | 0.064 | 0.234 | 0.16 | 0.03 | 0.21 | 0.02 | 0.24 | 0.16 |
| 7 | 20740648 | rs17218211 | -0.276 | 1.072 | 0.12 | -0.685 | -0.007 | 0.004 | 0.004 | 0.054 | 0.15 | 0.12 | 0.24 | 0.02 | 0.24 | 0.16 |
| 7 | 20740718 | rs17218253 | -0.544 | 1.101 | 0.247 | -0.685 | 0.02 | -0.055 | -0.055 | 0 | 0.18 | 0.21 | 0.27 | 0.02 | 0.27 | 0.17 |
| 7 | 20741167 | rs4142216 | 1.346 | 1.051 | 0.312 | -0.76 | -0.001 | 0.009 | 0.009 | 0 | 0.58 | 0.42 | 0.65 | 0.84 | 0.54 | 0.74 |
| 7 | 20741668 | rs10488578 | -0.936 | 1.506 | -0.149 | -0.534 | 0.012 | -0.044 | -0.044 | 0 | 0.18 | 0.19 | 0.26 | 0.02 | 0.25 | 0.16 |
| 7 | 20744543 | rs2158855 | -1.148 | 1.357 | -0.515 | -0.509 | 0.005 | -0.04 | -0.04 | 0 | 0.21 | 0.2 | 0.27 | 0.02 | 0.28 | 0.17 |
| 7 | 20744563 | rs2158854 | -1.064 | 1.278 | -0.605 | -0.492 | 0.003 | -0.032 | -0.032 | 0.075 | 0.2 | 0.18 | 0.26 | 0.02 | 0.27 | 0.17 |
| 7 | 20754102 | rs7781446 | 0.799 | 1.428 | -1.314 | -0.442 | -0.076 | 0.085 | 0.085 | 0.283 | 0.77 | 0.91 | 0.79 | 0.95 | 0.62 | 0.71 |
| 7 | 20754482 | rs6461511 | 0.784 | 1.434 | -1.32 | -0.439 | -0.08 | 0.085 | 0.085 | 0.341 | 0.77 | 0.91 | 0.79 | 0.95 | 0.62 | 0.71 |
| 7 | 20754975 | rs6962189 | 0.92 | 1.255 | -1.502 | -0.518 | -0.116 | 0.099 | 0.099 | 0.854 | 0.76 | 0.97 | 0.78 | 0.95 | 0.62 | 0.68 |
| 7 | 20761221 | rs34972642 | -0.319 | 0.627 | -1.266 | 0.037 | 0.002 | -0.003 | -0.003 | 0 | 0.07 | 0.06 | 0.03 | 0.14 | 0.06 | 0.05 |
| 7 | 20761738 | rs10266338 | -1.268 | -0.042 | -0.415 | -0.17 | -0.037 | 0.018 | 0.018 | 0 | 0.13 | 0.22 | 0.05 | 0.13 | 0.08 | 0.05 |
| 7 | 20761824 | rs1023541 | 0.518 | 0.311 | -0.433 | -0.485 | -0.026 | 0.024 | 0.024 | 0 | 0.8 | 0.88 | 0.81 | 0.9 | 0.68 | 0.72 |
| 7 | 20763115 | rs10950830 | -0.292 | -0.178 | -0.35 | -0.264 | -0.03 | 0.043 | 0.043 | 0.107 | 0.29 | 0.5 | 0.21 | 0.33 | 0.22 | 0.22 |
| 7 | 20763907 | rs6461513 | 0.064 | 0.017 | -0.663 | -0.562 | -0.026 | 0.029 | 0.029 | 0 | 0.79 | 0.88 | 0.8 | 0.92 | 0.67 | 0.71 |
| 7 | 20764374 | rs10950831 | -0.456 | -0.253 | -1.148 | -0.348 | 0.054 | 0.067 | 0.067 | 0 | 0.44 | 0.7 | 0.6 | 0.72 | 0.31 | 0.37 |
| 7 | 20765631 | rs4721939 | -0.105 | 0.168 | -0.974 | -0.275 | -0.025 | 0.024 | 0.024 | 0 | 0.8 | 0.88 | 0.8 | 0.92 | 0.67 | 0.71 |
| 7 | 20767815 | rs4721940 | 0.953 | -0.295 | -0.882 | -0.219 | 0.006 | 0.08 | 0.08 | 0 | 0.42 | 0.18 | 0.19 | 0.25 | 0.38 | 0.36 |
| 7 | 20767954 | rs17143304 | -0.317 | -0.644 | 0.135 | -0.331 | -0.008 | -0.015 | -0.015 | 0.751 | 0.06 | 0.25 | 0.02 | 0 | 0.01 | 0.03 |
| 7 | 20768013 | rs10254317 | -0.656 | -0.108 | -0.162 | -0.502 | -0.013 | 0.037 | 0.037 | 0.001 | 0.41 | 0.79 | 0.67 | 0.69 | 0.43 | 0.36 |
| 7 | 20768899 | rs4721941 | -0.416 | -0.215 | -0.032 | -0.521 | 0.012 | 0.007 | 0.007 | 0.006 | 0.34 | 0.74 | 0.61 | 0.69 | 0.29 | 0.34 |
| 7 | 20769733 | rs11972818 | -0.502 | -0.119 | 0.267 | -0.445 | 0.007 | 0.029 | 0.029 | 0 | 0.07 | 0.31 | 0.03 | 0.04 | 0.01 | 0.03 |

| | | | | | | | | | | | | | | | | |
|---|---|---|---|---|---|---|---|---|---|---|---|---|---|---|---|---|
| 7 | 20774091 | rs6959944 | -0.056 | 0.259 | -0.255 | -0.386 | -0.009 | 0 | 0 | 1.407 | 0.08 | 0.07 | 0.09 | 0 | 0.15 | 0.02 |
| 7 | 20777041 | rs10269525 | -0.288 | 0.041 | -0.412 | -0.46 | -0.036 | 0.035 | 0.035 | 0 | 0.84 | 0.99 | 0.89 | 0.95 | 0.82 | 0.72 |
| 7 | 20777250 | rs9638774 | 0.663 | -0.495 | -0.76 | -0.869 | 0.02 | 0.079 | 0.079 | 0 | 0.68 | 0.91 | 0.83 | 0.86 | 0.71 | 0.58 |
| 7 | 20777911 | rs10244045 | -0.771 | -0.389 | -0.25 | -0.629 | -0.001 | 0 | 0 | 0 | 0.87 | 0.87 | 0.89 | 0.88 | 0.83 | 0.72 |
| 7 | 20778646 | rs6461515 | 0.295 | -0.512 | 0.507 | -0.659 | -0.013 | -0.018 | -0.018 | 0.226 | 0.81 | 0.64 | 0.86 | 0.82 | 0.82 | 0.71 |
| 7 | 20778773 | rs6461516 | -0.635 | -0.418 | 0.478 | -0.625 | 0.005 | -0.003 | -0.003 | 0 | 0.86 | 0.75 | 0.89 | 0.85 | 0.83 | 0.72 |
| 7 | 20782094 | rs10231520 | -0.385 | -0.384 | -0.474 | -0.252 | -0.041 | -0.028 | -0.028 | 4.498 | 0.35 | 0.14 | 0.26 | 0.29 | 0.36 | 0.29 |
| 7 | 20782474 | rs12669866 | -0.655 | -0.709 | -0.198 | -0.236 | -0.001 | -0.002 | -0.002 | 0 | 0.53 | 0.44 | 0.72 | 0.63 | 0.53 | 0.48 |
| 7 | 20784169 | rs6967583 | -0.611 | -0.419 | -0.251 | -0.028 | 0 | 0 | 0 | 0 | 0.84 | 0.82 | 0.88 | 0.85 | 0.84 | 0.72 |
| 7 | 20789195 | rs17220780 | -0.34 | -0.054 | -0.195 | 0.413 | 0.014 | 0.049 | 0.049 | 0 | 0.14 | 0.21 | 0.38 | 0.36 | 0.18 | 0.11 |
| 7 | 20790767 | rs4721944 | 1.529 | -0.092 | -0.86 | -0.07 | -0.078 | 0.024 | 0.024 | 0 | 0.3 | 0.04 | 0.21 | 0.19 | 0.35 | 0.28 |
| 7 | 20791190 | rs4721946 | 1.24 | -0.208 | -0.098 | -0.143 | 0.002 | -0.04 | -0.04 | 0 | 0.35 | 0.46 | 0.24 | 0.23 | 0.36 | 0.28 |
| 7 | 20794396 | rs2709747 | 1.127 | -0.294 | 0.045 | -0.128 | 0 | -0.022 | -0.022 | 0 | 0.37 | 0.56 | 0.25 | 0.22 | 0.36 | 0.28 |
| 7 | 20797030 | rs2528803 | 1.128 | -0.292 | -0.013 | -0.127 | 0.002 | -0.035 | -0.035 | 0 | 0.37 | 0.53 | 0.24 | 0.22 | 0.36 | 0.28 |
| 7 | 20815352 | rs1029595 | -1.163 | -0.363 | 0.039 | -0.124 | 0.002 | -0.035 | -0.035 | 0 | 0.63 | 0.51 | 0.76 | 0.78 | 0.63 | 0.73 |
| 7 | 20821273 | rs2286260 | 0.005 | -0.556 | -1.337 | 0.049 | -0.028 | -0.101 | -0.101 | 1.049 | 0.32 | 0.04 | 0.36 | 0.35 | 0.28 | 0.34 |
| 7 | 20831800 | rs16872619 | 0.516 | -0.718 | -1.364 | 0.288 | 0.018 | 0.005 | 0.005 | 0 | 0.2 | 0.04 | 0.22 | 0.16 | 0.16 | 0.18 |
| 7 | 20838951 | rs2529753 | 0.323 | -0.441 | -1.455 | 0.208 | -0.069 | -0.031 | -0.031 | 0 | 0.71 | 0.91 | 0.67 | 0.69 | 0.63 | 0.55 |
| 7 | 20844094 | rs2108520 | 0.791 | -0.322 | -0.997 | -0.041 | -0.017 | 0.019 | 0.019 | 0 | 0.57 | 0.77 | 0.48 | 0.36 | 0.55 | 0.49 |
| 7 | 20863175 | rs17143377 | 1.29 | -0.586 | -0.648 | 0 | 0.016 | 0.016 | 0.016 | 0 | 0.17 | 0.05 | 0.2 | 0.13 | 0.14 | 0.17 |
| 7 | 20867808 | rs2709722 | 0.757 | -0.676 | -0.758 | -0.137 | 0.001 | 0.036 | 0.036 | 0 | 0.52 | 0.75 | 0.44 | 0.38 | 0.47 | 0.42 |
| 7 | 20868521 | rs10256428 | 2.074 | -0.504 | -0.103 | -0.008 | -0.001 | 0 | 0 | 0.89 | 0.29 | 0.28 | 0.21 | 0.22 | 0.33 | 0.24 |
| 7 | 20878995 | rs2529763 | 1.24 | -0.44 | -0.241 | -0.023 | 0.001 | 0.008 | 0.008 | 0.005 | 0.4 | 0.67 | 0.36 | 0.27 | 0.39 | 0.25 |
| 7 | 20896062 | rs17143423 | 0.893 | -0.17 | -0.267 | 0.327 | -0.043 | -0.055 | -0.055 | 0 | 0.16 | 0.02 | 0.17 | 0.17 | 0.15 | 0.3 |
| 7 | 20903218 | rs17143431 | 2.223 | -0.589 | -0.707 | 0.032 | 0.009 | 0.044 | 0.044 | 0 | 0.17 | 0 | 0.06 | 0.07 | 0.11 | 0.15 |
| 7 | 20904270 | rs204587 | 1.607 | -0.743 | -0.741 | 0.107 | -0.02 | 0.042 | 0.042 | 0.036 | 0.5 | 0.22 | 0.46 | 0.36 | 0.55 | 0.42 |
| 7 | 20907104 | rs204590 | -0.669 | -0.557 | -0.755 | 0.349 | -0.021 | -0.017 | -0.017 | 0 | 0.9 | 0.96 | 0.78 | 0.77 | 0.82 | 0.85 |
| 7 | 20911982 | rs204596 | -0.022 | -0.564 | -0.495 | 0.365 | 0.069 | 0.036 | 0.036 | 0.387 | 0.92 | 0.79 | 0.79 | 0.84 | 0.84 | 0.86 |
| 7 | 20912112 | rs2529768 | -0.624 | -0.6 | -0.629 | 0.437 | -0.009 | 0.003 | 0.003 | 0 | 0.4 | 0.37 | 0.31 | 0.44 | 0.28 | 0.45 |
| 7 | 20921103 | rs17143451 | 1.548 | -0.552 | -0.718 | 0.503 | 0.028 | 0.022 | 0.022 | 0 | 0.22 | 0.1 | 0.15 | 0.13 | 0.15 | 0.19 |
| 7 | 20930177 | rs6947953 | 1.488 | -0.236 | -0.545 | 0.65 | -0.014 | 0.007 | 0.007 | 0 | 0.18 | 0.21 | 0.13 | 0.19 | 0.12 | 0.12 |
| 7 | 20930292 | rs6948125 | 0.04 | -0.157 | -0.615 | 0.694 | 0.001 | 0.005 | 0.005 | 0 | 0.29 | 0.26 | 0.34 | 0.42 | 0.3 | 0.29 |
| 7 | 20935024 | rs17143466 | 0.838 | 0.288 | -0.759 | 0.67 | -0.002 | 0.006 | 0.006 | 0 | 0.05 | 0.14 | 0.09 | 0.04 | 0.04 | 0.04 |
| 7 | 20940950 | rs1155096 | -0.751 | 0.304 | -0.997 | 0.652 | -0.031 | -0.04 | -0.04 | 0 | 0.61 | 0.78 | 0.43 | 0.56 | 0.41 | 0.6 |
| 7 | 20941262 | rs4140783 | -2.195 | 0.346 | -1.19 | 0.514 | -0.12 | -0.049 | -0.049 | 0 | 0.72 | 0.98 | 0.69 | 0.77 | 0.6 | 0.78 |
| 7 | 20944522 | rs6461521 | -1.058 | 0.144 | -0.593 | 0.149 | -0.016 | -0.01 | -0.01 | 0.24 | 0.62 | 0.66 | 0.56 | 0.59 | 0.48 | 0.62 |
| 7 | 20949268 | rs6955523 | 1.549 | -0.366 | -0.849 | 0.091 | 0.027 | 0.009 | 0.009 | 0 | 0.21 | 0.14 | 0.07 | 0.19 | 0.11 | 0.17 |
| 7 | 20950496 | rs2160093 | 0.129 | -0.079 | -0.781 | 0.156 | -0.003 | 0.002 | 0.002 | 0 | 0.24 | 0.27 | 0.21 | 0.25 | 0.23 | 0.15 |
| 7 | 20955370 | rs2709733 | -0.873 | -0.006 | -1.142 | 0.178 | -0.026 | -0.043 | -0.043 | 2.232 | 0.59 | 0.73 | 0.4 | 0.55 | 0.45 | 0.59 |
| 7 | 20960893 | rs17601859 | 1.288 | 0.351 | -1.591 | -0.234 | -0.023 | -0.045 | -0.045 | 0 | 0.08 | 0 | 0.09 | 0.11 | 0.12 | 0.05 |
| 7 | 20960966 | rs1008132 | -1.85 | 0.359 | -1.742 | -0.209 | -0.131 | -0.155 | -0.155 | 0 | 0.71 | 0.97 | 0.69 | 0.59 | 0.63 | 0.65 |
| 7 | 20961711 | rs10248211 | -0.449 | 0.204 | -1.103 | -0.262 | -0.004 | 0.009 | 0.009 | 0 | 0.32 | 0.2 | 0.31 | 0.26 | 0.38 | 0.18 |
| 7 | 20967833 | rs6962272 | 1.008 | 0.389 | -0.873 | -0.243 | 0.001 | 0.001 | 0.001 | 0 | 0.16 | 0.13 | 0.11 | 0.12 | 0.11 | 0.08 |
| 7 | 20970137 | rs6973043 | -0.109 | 0.502 | -0.717 | -0.219 | 0.018 | 0.029 | 0.029 | 0 | 0.47 | 0.24 | 0.36 | 0.39 | 0.46 | 0.31 |
| 7 | 20972111 | rs6945857 | -0.228 | 0.47 | -0.5 | -0.229 | 0.036 | 0.012 | 0.012 | 0 | 0.5 | 0.35 | 0.5 | 0.47 | 0.41 | 0.33 |
| 7 | 20978000 | rs2709725 | -0.695 | 0.539 | -0.543 | -0.243 | 0.009 | 0.003 | 0.003 | 0 | 0.53 | 0.56 | 0.52 | 0.44 | 0.42 | 0.34 |
| 7 | 20990451 | rs4721959 | 0.057 | 0.461 | -0.922 | -0.19 | 0.001 | 0 | 0 | 0 | 0.55 | 0.65 | 0.67 | 0.59 | 0.6 | 0.7 |
| 7 | 20993726 | rs2390406 | -0.18 | 0.151 | -1.027 | -0.406 | 0.009 | 0.021 | 0.021 | 0 | 0.43 | 0.41 | 0.25 | 0.27 | 0.35 | 0.19 |
| 7 | 20994491 | rs12155172 | -1.711 | 0.693 | -0.938 | -0.094 | -0.004 | -0.046 | -0.046 | 0 | 0.79 | 0.91 | 0.87 | 0.72 | 0.8 | 0.8 |
| 7 | 20998686 | rs6966904 | 1.36 | 0.794 | -1.221 | -0.279 | -0.06 | 0.034 | 0.034 | 0.011 | 0.3 | 0.05 | 0.24 | 0.24 | 0.31 | 0.36 |
| 7 | 21000345 | rs2390409 | -1.093 | 0.825 | -0.291 | 0.023 | -0.009 | -0.009 | -0.009 | 0.004 | 0.39 | 0.75 | 0.44 | 0.4 | 0.35 | 0.34 |
| 7 | 21015297 | rs12700247 | 0.809 | 0.823 | -0.454 | -0.059 | -0.047 | -0.026 | -0.026 | 0 | 0.08 | 0.01 | 0.23 | 0.09 | 0.13 | 0.09 |
| 7 | 21018240 | rs10270424 | -0.718 | 0.776 | 0.229 | -0.099 | 0.129 | 0.033 | 0.033 | 1.47 | 0.06 | 0.41 | 0.07 | 0 | 0.08 | 0.02 |
| 7 | 21019203 | rs17143600 | 1.067 | 0.701 | 0.218 | -0.107 | 0.012 | 0.005 | 0.005 | 0 | 0.17 | 0.11 | 0.07 | 0.14 | 0.11 | 0.14 |
| 7 | 21021498 | rs12111744 | 0.104 | 0.63 | 0.267 | -0.103 | -0.017 | -0.051 | -0.051 | 3.221 | 0.23 | 0.43 | 0.12 | 0.16 | 0.18 | 0.15 |
| 7 | 21023562 | rs17732809 | 0.934 | 0.71 | 0.154 | -0.119 | -0.014 | -0.006 | -0.006 | 0 | 0.08 | 0.03 | 0.23 | 0.09 | 0.12 | 0.08 |
| 7 | 21027080 | rs2390412 | 1.316 | 0.628 | 0.019 | -0.162 | -0.017 | 0.053 | 0.053 | 0 | 0.4 | 0.16 | 0.43 | 0.35 | 0.45 | 0.45 |
| 7 | 21030508 | rs17676368 | 1.072 | 0.485 | -0.029 | -0.225 | -0.017 | 0.099 | 0.099 | 0.132 | 0.47 | 0.17 | 0.48 | 0.35 | 0.52 | 0.49 |
| 7 | 21032355 | rs4598145 | -0.473 | 0.792 | 0.411 | -0.033 | 0.09 | -0.006 | -0.006 | 0 | 0.14 | 0.36 | 0.29 | 0.09 | 0.21 | 0.11 |
| 7 | 21033870 | rs10156011 | 1.062 | 0.871 | -0.414 | 0.007 | -0.022 | 0.049 | 0.049 | 0.036 | 0.4 | 0.16 | 0.43 | 0.35 | 0.45 | 0.45 |
| 7 | 21037046 | rs3810892 | 0.636 | 0.667 | -0.055 | 0.348 | 0.011 | 0.008 | 0.008 | 0 | 0.17 | 0.11 | 0.07 | 0.13 | 0.11 | 0.14 |
| 7 | 21047023 | rs10265596 | -1.281 | 0.79 | 0.085 | 0.462 | 0.08 | 0.003 | 0.003 | 0.045 | 0.05 | 0.24 | 0.05 | 0 | 0.08 | 0.02 |
| 7 | 21053970 | rs4721961 | 0.498 | 0.875 | -0.147 | 0.455 | -0.039 | 0.036 | 0.036 | 0 | 0.4 | 0.21 | 0.47 | 0.35 | 0.46 | 0.45 |
| 7 | 21059378 | rs9690626 | 0.478 | 1.019 | -0.243 | 0.57 | -0.008 | 0 | 0 | 0 | 0.23 | 0.08 | 0.27 | 0.23 | 0.2 | 0.23 |
| 7 | 21060304 | rs6946302 | -0.693 | 0.997 | -0.048 | 0.733 | -0.01 | 0.003 | 0.003 | 0 | 0.69 | 0.59 | 0.81 | 0.72 | 0.69 | 0.78 |
| 7 | 21064608 | rs6943750 | -0.001 | 1.116 | -0.723 | 0.826 | 0.047 | 0.074 | 0.074 | 0.018 | 0.07 | 0.24 | 0.1 | 0.12 | 0.1 | 0.11 |
| 7 | 21064647 | rs6963949 | -0.382 | 0.936 | -0.758 | 0.549 | -0.014 | -0.001 | -0.001 | 0 | 0.67 | 0.5 | 0.79 | 0.72 | 0.68 | 0.77 |
| 7 | 21066684 | rs6954678 | -0.12 | 1.287 | -1.148 | 0.363 | -0.016 | 0.017 | 0.017 | 0 | 0.37 | 0.19 | 0.43 | 0.36 | 0.38 | 0.42 |
| 7 | 21068214 | rs6461529 | 0.839 | 1.574 | -0.263 | 0.229 | 0.097 | -0.013 | -0.013 | 0 | 0.94 | 0.78 | 0.93 | 1 | 0.85 | 0.93 |

| | | | | | | | | | | | | | | | | |
|---|---|---|---|---|---|---|---|---|---|---|---|---|---|---|---|---|
| 7 | 21068570 | rs17676896 | -0.846 | 1.206 | -0.87 | 0.239 | -0.002 | 0.01 | 0.01 | 0 | 0.08 | 0.02 | 0.09 | 0.06 | 0.09 | 0.15 |
| 7 | 21072640 | rs34494536 | -1.015 | 0.947 | -1.132 | 0.212 | 0.009 | 0.007 | 0.007 | 0.02 | 0.18 | 0.09 | 0.17 | 0.09 | 0.17 | 0.14 |
| 7 | 21072954 | rs10270923 | 0.192 | 0.974 | -1.333 | -0.315 | 0 | 0.01 | 0.01 | 0 | 0.21 | 0.18 | 0.24 | 0.18 | 0.17 | 0.22 |
| 7 | 21080407 | rs6963278 | -0.242 | 0.729 | -1.292 | 0.077 | 0.01 | -0.019 | -0.019 | 0.78 | 0.44 | 0.26 | 0.59 | 0.49 | 0.38 | 0.5 |
| 7 | 21084329 | rs7795600 | -0.451 | 0.541 | -1.965 | -0.316 | 0 | -0.002 | -0.002 | 0 | 0.66 | 0.34 | 0.68 | 0.79 | 0.59 | 0.67 |
| 7 | 21085264 | rs13243044 | 1.677 | 0.786 | -1.049 | -0.439 | -0.001 | -0.015 | -0.015 | 0 | 0.19 | 0.25 | 0.11 | 0.13 | 0.17 | 0.2 |
| 7 | 21086373 | rs17734042 | 1.36 | 0.88 | -1.22 | -0.473 | -0.024 | 0.043 | 0.043 | 0 | 0.22 | 0.11 | 0.26 | 0.18 | 0.22 | 0.24 |
| 7 | 21087077 | rs7800398 | 0.308 | 0.983 | -1.093 | -0.325 | 0.003 | -0.002 | -0.002 | 0.282 | 0.45 | 0.49 | 0.5 | 0.42 | 0.51 | 0.47 |
| 7 | 21088583 | rs1019233 | 1.447 | 0.784 | -1.129 | -0.194 | -0.021 | -0.089 | -0.089 | 0 | 0.21 | 0.03 | 0.33 | 0.29 | 0.21 | 0.27 |
| 7 | 21090300 | rs10239242 | -1.594 | 0.825 | -0.403 | 0.093 | 0.002 | 0 | 0 | 0 | 0.86 | 0.85 | 0.96 | 0.86 | 0.91 | 0.88 |
| 7 | 21092593 | rs4719636 | -1.442 | 0.827 | -0.042 | -0.042 | -0.012 | -0.001 | -0.001 | 0.045 | 0.85 | 0.77 | 0.95 | 0.86 | 0.91 | 0.88 |
| 7 | 21093515 | rs4719637 | 0.332 | 0.832 | -0.086 | -0.073 | -0.006 | 0.023 | 0.023 | 0 | 0.44 | 0.41 | 0.36 | 0.29 | 0.51 | 0.44 |
| 7 | 21094423 | rs17734397 | 0.032 | 0.811 | -0.213 | -0.028 | 0.007 | 0.011 | 0.011 | 0 | 0.3 | 0.21 | 0.2 | 0.22 | 0.26 | 0.24 |
| 7 | 21094589 | rs917815 | -0.499 | 1.056 | 0.154 | 0.005 | -0.001 | -0.013 | -0.013 | 0.017 | 0.57 | 0.55 | 0.69 | 0.59 | 0.62 | 0.53 |
| 7 | 21094928 | rs917813 | -0.832 | 1.056 | 0.412 | -0.099 | 0.006 | 0.006 | 0.006 | 1.055 | 0.75 | 0.38 | 0.79 | 0.69 | 0.7 | 0.77 |
| 7 | 21096914 | rs10275129 | 0.533 | 1.143 | 0.399 | -0.025 | 0.018 | 0.025 | 0.025 | 0 | 0.77 | 0.5 | 0.75 | 0.76 | 0.68 | 0.75 |
| 7 | 21099912 | rs1981703 | 0.597 | 1.074 | 0.427 | -0.046 | 0.001 | 0 | 0 | 0 | 0.65 | 0.49 | 0.72 | 0.69 | 0.6 | 0.71 |
| 7 | 21103691 | rs2159851 | -0.381 | 1.09 | 0.229 | -0.101 | -0.004 | 0.039 | 0.039 | 0 | 0.3 | 0.08 | 0.2 | 0.16 | 0.27 | 0.27 |
| 7 | 21108002 | rs4719639 | 0.231 | 1.135 | 0.459 | -0.079 | 0.043 | -0.032 | -0.032 | 0 | 0.79 | 0.59 | 0.76 | 0.9 | 0.68 | 0.77 |
| 7 | 21109645 | rs12690773 | -0.17 | 1.152 | 0.327 | -0.064 | 0.021 | -0.018 | -0.018 | 0 | 0.2 | 0.28 | 0.23 | 0.1 | 0.32 | 0.23 |
| 7 | 21111223 | rs13228677 | 0.275 | 1.142 | 0.296 | -0.064 | -0.001 | 0.006 | 0.006 | 0 | 0.17 | 0.08 | 0.16 | 0.1 | 0.18 | 0.22 |
| 7 | 21111643 | rs1860875 | 0.781 | 0.801 | -0.04 | -0.258 | 0.019 | -0.007 | -0.007 | 0 | 0.44 | 0.35 | 0.56 | 0.56 | 0.39 | 0.47 |
| 7 | 21112893 | rs1558481 | -0.982 | 1.065 | 0.161 | -0.068 | 0.003 | -0.014 | -0.014 | 0 | 0.78 | 0.76 | 0.83 | 0.86 | 0.79 | 0.77 |
| 7 | 21113590 | rs17143815 | 1.279 | 0.915 | -0.19 | 0.062 | -0.021 | -0.119 | -0.119 | 0 | 0.2 | 0.04 | 0.44 | 0.44 | 0.21 | 0.27 |
| 7 | 21117480 | rs4719642 | 0.425 | 0.534 | -0.227 | 0.234 | 0.003 | -0.001 | -0.001 | 0 | 0.6 | 0.57 | 0.67 | 0.62 | 0.61 | 0.53 |
| 7 | 21135638 | rs17735470 | 0.189 | 0.186 | -0.888 | 0.375 | 0.006 | -0.021 | -0.021 | 0.007 | 0.15 | 0.02 | 0.14 | 0.12 | 0.13 | 0.17 |
| 7 | 21136170 | rs17678809 | 0.355 | 0.08 | -1.286 | -0.09 | 0.012 | 0.12 | 0.12 | 0.019 | 0.35 | 0.12 | 0.18 | 0.16 | 0.27 | 0.26 |
| 7 | 21138431 | rs1860874 | -1.805 | 0.54 | -1.347 | 0.362 | -0.025 | 0.003 | 0.003 | 0 | 0.85 | 0.89 | 0.63 | 0.93 | 0.65 | 0.91 |
| 7 | 21140343 | rs971992 | -0.548 | 0.105 | -1.042 | 0.435 | 0.016 | 0.003 | 0.003 | 0 | 0.49 | 0.44 | 0.3 | 0.44 | 0.34 | 0.47 |
| 7 | 21142122 | rs6962818 | 0.369 | 0.214 | -0.862 | 0.338 | -0.026 | -0.014 | -0.014 | 0 | 0.36 | 0.46 | 0.17 | 0.33 | 0.22 | 0.31 |
| 7 | 21143998 | rs10950838 | 0.739 | 0.824 | -1.211 | 0.393 | -0.143 | -0.004 | -0.004 | 0 | 0.2 | 0.02 | 0.41 | 0.08 | 0.43 | 0.12 |
| 7 | 21147678 | rs11974269 | 0.313 | 0.063 | -0.82 | 0.29 | -0.006 | -0.014 | -0.014 | 0.963 | 0.22 | 0.28 | 0.15 | 0.15 | 0.17 | 0.25 |
| 7 | 21148308 | rs17143871 | -0.337 | -0.145 | -0.767 | 0.668 | -0.01 | 0.017 | 0.017 | 0.625 | 0.26 | 0.55 | 0.27 | 0.3 | 0.21 | 0.32 |
| 7 | 21150461 | rs12700255 | 1.132 | -0.136 | -0.848 | 0.738 | -0.011 | 0.001 | 0.001 | 0 | 0.15 | 0.07 | 0.16 | 0.1 | 0.16 | 0.17 |
| 7 | 21150628 | rs17143873 | 2.447 | -0.246 | -1.297 | 0.836 | 0.01 | 0.002 | 0.002 | 1.763 | 0.14 | 0.03 | 0.04 | 0.08 | 0.09 | 0.03 |
| 7 | 21151172 | rs17679456 | 2.576 | -0.216 | -1.294 | 0.849 | 0.004 | -0.004 | -0.004 | 0 | 0.14 | 0.01 | 0.04 | 0.08 | 0.08 | 0.03 |
| 7 | 21154627 | rs10231764 | 0.594 | 0.334 | -0.661 | 1.146 | -0.013 | -0.012 | -0.012 | 0 | 0.39 | 0.31 | 0.56 | 0.5 | 0.54 | 0.4 |
| 7 | 21156235 | rs2390474 | 0.614 | 0.329 | -0.647 | 1.368 | -0.015 | 0.005 | 0.005 | 0 | 0.39 | 0.3 | 0.55 | 0.32 | 0.53 | 0.39 |
| 7 | 21158948 | rs12532292 | 1.9 | 0.552 | -0.616 | 1.332 | -0.04 | 0.004 | 0.004 | 0.404 | 0.16 | 0.09 | 0.36 | 0.08 | 0.36 | 0.09 |
| 7 | 21159575 | rs9655208 | 1.829 | 0.387 | -0.752 | 1.424 | -0.002 | -0.011 | -0.011 | 1.968 | 0.2 | 0.21 | 0.11 | 0.09 | 0.16 | 0.21 |
| 7 | 21160678 | rs9655209 | 1.511 | 0.459 | -0.722 | 1.404 | 0.001 | 0 | 0 | 0.07 | 0.23 | 0.22 | 0.2 | 0.26 | 0.18 | 0.28 |
| 7 | 21161819 | rs17679859 | 1.828 | 0.603 | -0.659 | 1.343 | -0.001 | 0.001 | 0.001 | 0.404 | 0.16 | 0.12 | 0.36 | 0.08 | 0.35 | 0.09 |
| 7 | 21163036 | rs10250250 | -0.382 | 0.425 | -0.564 | 1.29 | 0.009 | 0.024 | 0.024 | 0.592 | 0.47 | 0.64 | 0.41 | 0.71 | 0.37 | 0.61 |
| 7 | 21163355 | rs10950840 | -0.757 | 0.026 | -0.967 | 1.146 | 0.038 | 0.062 | 0.062 | 0 | 0.63 | 0.79 | 0.77 | 0.78 | 0.72 | 0.7 |
| 7 | 21163915 | rs17143888 | 1.407 | 0.025 | -1.011 | 1.212 | 0.006 | 0.004 | 0.004 | 0.705 | 0.22 | 0.19 | 0.2 | 0.24 | 0.18 | 0.28 |
| 7 | 21164766 | rs4721981 | -0.429 | 0.507 | -0.753 | 1.426 | -0.001 | -0.001 | -0.001 | 0 | 0.41 | 0.58 | 0.57 | 0.55 | 0.54 | 0.42 |
| 7 | 21168014 | rs17143912 | -0.409 | 0.215 | -0.835 | 0.777 | 0 | 0.001 | 0.001 | 0 | 0.08 | 0.08 | 0.08 | 0.15 | 0.07 | 0.17 |
| 7 | 21170601 | rs11772446 | 0.394 | 0.007 | -1.048 | 0.336 | 0.012 | 0.026 | 0.026 | 0.026 | 0.14 | 0.06 | 0.06 | 0.05 | 0.11 | 0.18 |
| 7 | 21171130 | rs10488365 | -0.208 | 0.889 | -1.527 | 0.522 | -0.106 | -0.065 | -0.065 | 0.657 | 0.21 | 0.05 | 0.26 | 0.33 | 0.28 | 0.34 |
| 7 | 21178262 | rs12538434 | -0.3 | 0.744 | -1.148 | 0.243 | -0.001 | 0 | 0 | 0.594 | 0.19 | 0.16 | 0.19 | 0.14 | 0.17 | 0.33 |
| 7 | 21179074 | rs6951681 | 0.346 | 1.257 | -0.305 | 0.618 | 0.12 | 0.169 | 0.169 | 0 | 0.06 | 0.28 | 0.1 | 0.18 | 0.09 | 0.04 |
| 7 | 21179706 | rs13229308 | -0.879 | 0.296 | -0.493 | 0.365 | 0.007 | 0.003 | 0.003 | 1.241 | 0.7 | 0.8 | 0.9 | 0.76 | 0.81 | 0.75 |
| 7 | 21181503 | rs12700256 | 1.547 | 0.182 | -0.497 | 0.26 | 0.009 | 0.01 | 0.01 | 0.809 | 0.18 | 0.1 | 0.06 | 0.13 | 0.11 | 0.21 |
| 7 | 21183179 | rs17143933 | 0.543 | -0.031 | -0.623 | -0.108 | -0.003 | -0.068 | -0.068 | 0 | 0.21 | 0.3 | 0.12 | 0 | 0.19 | 0.09 |
| 7 | 21185178 | rs10246207 | -0.635 | -0.04 | -0.581 | -0.007 | -0.001 | 0 | 0 | 0 | 0.39 | 0.41 | 0.43 | 0.36 | 0.39 | 0.55 |
| 7 | 21185221 | rs13241724 | 1.838 | -0.054 | -0.627 | -0.048 | 0.009 | 0.009 | 0.009 | 0.953 | 0.18 | 0.1 | 0.05 | 0.12 | 0.09 | 0.21 |
| 7 | 21192264 | rs4473915 | 0.767 | 0.152 | -0.388 | 0.107 | 0.102 | 0.218 | 0.218 | 0 | 0.07 | 0.24 | 0.2 | 0.38 | 0.11 | 0.17 |
| 7 | 21193137 | rs2215676 | -0.747 | 0.088 | -0.352 | 0.027 | 0 | -0.043 | -0.043 | 0.047 | 0.4 | 0.49 | 0.32 | 0.14 | 0.37 | 0.43 |
| 7 | 21193341 | rs2110007 | -2.6 | -0.051 | -0.439 | 0.009 | -0.011 | 0.003 | 0.003 | 0 | 0.88 | 0.88 | 0.96 | 0.88 | 0.92 | 0.96 |
| 7 | 21195371 | rs12700260 | 2.271 | 0.365 | -0.403 | 0.116 | -0.037 | -0.053 | -0.053 | 0 | 0.23 | 0.07 | 0.38 | 0.23 | 0.33 | 0.16 |
| 7 | 21197888 | rs4721986 | -1.38 | 0.748 | -0.1 | 0.172 | 0.005 | 0.008 | 0.008 | 0.083 | 0.46 | 0.72 | 0.52 | 0.53 | 0.48 | 0.6 |
| 7 | 21202731 | rs10269330 | -1.097 | 0.395 | -0.532 | 0.079 | -0.001 | 0.001 | 0.001 | 0 | 0.39 | 0.43 | 0.42 | 0.38 | 0.39 | 0.51 |
| 7 | 21202828 | rs7781186 | 0.148 | 0.475 | -0.779 | -0.106 | 0.001 | -0.027 | -0.027 | 0 | 0.2 | 0.23 | 0.12 | 0 | 0.19 | 0.09 |
| 7 | 21211121 | rs4470916 | 0.367 | 0.554 | -0.702 | 0.258 | 0.001 | -0.012 | -0.012 | 0 | 0.23 | 0.3 | 0.22 | 0.12 | 0.22 | 0.2 |
| 7 | 21211412 | rs4236302 | -0.758 | 1.134 | -0.572 | 0.605 | 0.022 | 0.012 | 0.012 | 0 | 0.26 | 0.4 | 0.3 | 0.37 | 0.26 | 0.38 |
| 7 | 21212389 | rs10224542 | 1.646 | 0.935 | -0.715 | 0.75 | -0.073 | -0.073 | -0.073 | 0 | 0.22 | 0.03 | 0.37 | 0.23 | 0.33 | 0.15 |
| 7 | 21215706 | rs7803836 | 0.252 | 0.755 | -0.158 | 0.765 | 0.021 | 0.123 | 0.123 | 0 | 0.07 | 0.29 | 0.16 | 0.22 | 0.04 | 0.2 |
| 7 | 21222278 | rs6966877 | 0.88 | 1 | -0.436 | 0.711 | -0.011 | -0.012 | -0.012 | 0 | 0.09 | 0.06 | 0.18 | 0.2 | 0.12 | 0.15 |
| 7 | 21222790 | rs6971109 | -1.114 | 1.008 | -0.2 | 0.548 | 0.021 | -0.009 | -0.009 | 0 | 0.24 | 0.41 | 0.18 | 0.21 | 0.24 | 0.23 |

| | | | | | | | | | | | | | | | | |
|---|---|---|---|---|---|---|---|---|---|---|---|---|---|---|---|---|
| 7 | 21225088 | rs10479785 | 0.439 | 0.713 | -0.3 | 0.663 | -0.016 | -0.013 | -0.013 | 1.845 | 0.38 | 0.27 | 0.58 | 0.58 | 0.46 | 0.46 |
| 7 | 21225260 | rs10479786 | 0.434 | 0.668 | -0.274 | 0.432 | -0.046 | -0.078 | -0.078 | 0.439 | 0.39 | 0.16 | 0.62 | 0.47 | 0.56 | 0.36 |
| 7 | 21228873 | rs10247670 | 0.22 | 0.681 | -0.186 | 0.452 | -0.008 | 0.017 | 0.017 | 0 | 0.4 | 0.3 | 0.64 | 0.62 | 0.57 | 0.42 |
| 7 | 21233327 | rs17144012 | -0.127 | 0.753 | -0.106 | 0.658 | 0.015 | 0.009 | 0.009 | 0 | 0.14 | 0.25 | 0.11 | 0.17 | 0.13 | 0.19 |
| 7 | 21234668 | rs12537582 | 0.569 | 0.328 | -0.634 | 0.71 | 0.004 | 0.004 | 0.004 | 0.898 | 0.18 | 0.13 | 0.07 | 0.17 | 0.12 | 0.21 |
| 7 | 21237935 | rs10247735 | 0.433 | 0.545 | -0.401 | 0.421 | 0 | -0.015 | -0.015 | 0.014 | 0.58 | 0.68 | 0.77 | 0.76 | 0.63 | 0.68 |
| 7 | 21238247 | rs6977771 | -0.858 | 0.315 | -0.785 | 0.316 | 0.022 | 0.051 | 0.051 | 0 | 0.48 | 0.61 | 0.42 | 0.37 | 0.53 | 0.5 |
| 7 | 21243188 | rs10499527 | 1.105 | 0.584 | -1.191 | 0.707 | 0.024 | 0.052 | 0.052 | 0.843 | 0.45 | 0.11 | 0.45 | 0.26 | 0.45 | 0.28 |
| 7 | 21245736 | rs10228903 | -0.342 | 0.734 | -0.631 | 1.258 | 0.004 | 0.045 | 0.045 | 0.825 | 0.16 | 0.21 | 0.29 | 0.43 | 0.17 | 0.32 |
| 7 | 21252028 | rs10240554 | -0.94 | 0.872 | -0.635 | 1.063 | -0.017 | 0 | 0 | 0 | 0.38 | 0.26 | 0.29 | 0.37 | 0.39 | 0.35 |
| 7 | 21257800 | rs4719652 | -0.505 | 1.131 | -0.214 | 1.161 | -0.007 | -0.003 | -0.003 | 0 | 0.29 | 0.22 | 0.26 | 0.3 | 0.32 | 0.26 |
| 7 | 21258856 | rs13309919 | 0.727 | 1.087 | -0.233 | 1.197 | 0.009 | 0.024 | 0.024 | 0.225 | 0.07 | 0.1 | 0.04 | 0.15 | 0.08 | 0.15 |
| 7 | 21259055 | rs6957594 | 2.055 | 0.614 | -0.637 | 0.914 | 0.015 | 0.058 | 0.058 | 1.183 | 0.38 | 0.11 | 0.4 | 0.24 | 0.38 | 0.21 |
| 7 | 21259339 | rs6975592 | -0.985 | 0.751 | 0.386 | 1.095 | 0.012 | -0.021 | -0.021 | 0.422 | 0.34 | 0.45 | 0.3 | 0.27 | 0.34 | 0.28 |
| 7 | 21260371 | rs4721994 | -0.28 | 0.659 | 0.271 | 1.1 | -0.002 | 0.006 | 0.006 | 0 | 0.25 | 0.18 | 0.17 | 0.18 | 0.26 | 0.16 |
| 7 | 21261834 | rs7786269 | 0.246 | 0.55 | 0.675 | 1.706 | 0.008 | 0.112 | 0.112 | 0.018 | 0.07 | 0.23 | 0.07 | 0.21 | 0.05 | 0.11 |
| 7 | 21262238 | rs4721998 | 0.437 | 0.571 | 0.694 | 1.659 | 0.015 | 0.238 | 0.238 | 1.095 | 0.9 | 0.67 | 0.86 | 0.56 | 0.93 | 0.69 |
| 7 | 21263531 | rs10499529 | 0.532 | 0.518 | 0.164 | 1.375 | -0.011 | 0.059 | 0.059 | 0 | 0.07 | 0.12 | 0.11 | 0.19 | 0.06 | 0.06 |
| 7 | 21265984 | rs7779989 | -0.407 | 0.421 | -0.079 | 1.153 | -0.008 | -0.003 | -0.003 | 0.745 | 0.34 | 0.32 | 0.22 | 0.27 | 0.24 | 0.25 |
| 7 | 21266123 | rs7780161 | -0.41 | 0.795 | -0.081 | 1.032 | 0.005 | 0.012 | 0.012 | 0.381 | 0.34 | 0.26 | 0.25 | 0.2 | 0.29 | 0.2 |
| 7 | 21266973 | rs10258822 | -0.171 | 0.274 | -0.048 | 1.052 | 0.008 | 0.008 | 0.008 | 0.687 | 0.32 | 0.25 | 0.21 | 0.19 | 0.24 | 0.19 |
| 7 | 21267438 | rs10246767 | -1.002 | 0.286 | -0.177 | 1.041 | 0.021 | 0.012 | 0.012 | 0.827 | 0.43 | 0.59 | 0.35 | 0.72 | 0.31 | 0.59 |
| 7 | 21276334 | rs6972011 | 0.216 | 0.823 | -0.164 | 1.012 | 0.035 | 0.001 | 0.001 | 0 | 0.77 | 0.65 | 0.82 | 0.81 | 0.73 | 0.78 |
| 7 | 21278363 | rs10240952 | 0.454 | 0.546 | -0.007 | 1.317 | -0.036 | 0.163 | 0.163 | 0.632 | 0.78 | 0.44 | 0.8 | 0.41 | 0.85 | 0.53 |
| 7 | 21280322 | rs2390498 | 0.557 | 0.694 | -0.509 | 0.765 | 0.004 | -0.027 | -0.027 | 0 | 0.67 | 0.57 | 0.79 | 0.8 | 0.68 | 0.75 |
| 7 | 21281641 | rs17688347 | 1.775 | 0.432 | -1.109 | 0.581 | -0.062 | 0.138 | 0.138 | 0.006 | 0.36 | 0.08 | 0.47 | 0.1 | 0.43 | 0.19 |
| 7 | 21284543 | rs10085845 | -1.324 | 0.268 | -0.321 | 0.917 | -0.017 | -0.002 | -0.002 | 0.926 | 0.89 | 0.84 | 0.95 | 0.92 | 0.93 | 0.93 |
| 7 | 21284666 | rs10807806 | -1.297 | 0.263 | -0.191 | 0.821 | -0.022 | -0.009 | -0.009 | 0 | 0.89 | 0.75 | 0.93 | 0.93 | 0.92 | 0.93 |
| 7 | 21285304 | rs4609107 | -0.701 | 0.172 | -0.503 | 0.98 | 0.013 | -0.056 | -0.056 | 0 | 0.29 | 0.13 | 0.28 | 0.5 | 0.26 | 0.4 |
| 7 | 21285699 | rs12673777 | -0.4 | 0.028 | -0.556 | 0.718 | -0.004 | 0.005 | 0.005 | 0 | 0.39 | 0.49 | 0.3 | 0.49 | 0.36 | 0.48 |
| 7 | 21286643 | rs4548059 | 2.3 | 1.172 | 0.123 | 1.091 | 0.018 | 0.019 | 0.019 | 2.111 | 0.38 | 0.24 | 0.45 | 0.27 | 0.37 | 0.26 |
| 7 | 21287199 | rs4418246 | 1.087 | 1.366 | 0.213 | 1.197 | 0.02 | 0.004 | 0.004 | 0 | 0.63 | 0.5 | 0.64 | 0.56 | 0.57 | 0.47 |
| 7 | 21287458 | rs4719656 | 1.432 | 1.143 | 0.352 | 1.141 | 0 | -0.001 | -0.001 | 0.344 | 0.54 | 0.43 | 0.61 | 0.55 | 0.53 | 0.55 |
| 7 | 21288213 | rs6970201 | -0.146 | 1.118 | 0.251 | 1.131 | 0.002 | 0.002 | 0.002 | 0 | 0.26 | 0.22 | 0.14 | 0.21 | 0.22 | 0.26 |
| 7 | 21289221 | rs12700267 | -1.134 | 1.202 | 0.055 | 0.914 | 0.006 | -0.003 | -0.003 | 0 | 0.37 | 0.32 | 0.29 | 0.4 | 0.32 | 0.39 |
| 7 | 21291229 | rs2700298 | -1.447 | 1.476 | 0.213 | 1.055 | -0.029 | -0.022 | -0.022 | 0 | 0.87 | 0.96 | 0.92 | 0.87 | 0.86 | 0.83 |
| 7 | 21291878 | rs2248634 | -0.395 | 1.454 | 0.267 | 1.041 | 0.033 | -0.001 | -0.001 | 0 | 0.83 | 0.68 | 0.86 | 0.84 | 0.81 | 0.78 |
| 7 | 21292628 | rs2700299 | 0.083 | 1.551 | -0.076 | 0.897 | 0.048 | -0.024 | -0.024 | 0 | 0.06 | 0.12 | 0.09 | 0 | 0.13 | 0.05 |
| 7 | 21294396 | rs2529142 | 0.709 | 1.43 | -0.041 | 0.958 | 0.001 | -0.019 | -0.019 | 0 | 0.72 | 0.79 | 0.77 | 0.67 | 0.75 | 0.71 |
| 7 | 21298242 | rs10277012 | -0.283 | 1.686 | -0.035 | 1.084 | 0.003 | 0 | 0 | 1.256 | 0.82 | 0.8 | 0.85 | 0.81 | 0.77 | 0.8 |
| 7 | 21302041 | rs1997391 | 1.276 | 1.651 | -0.177 | 1.183 | -0.032 | -0.03 | -0.03 | 0 | 0.67 | 0.76 | 0.68 | 0.42 | 0.64 | 0.52 |
| 7 | 21309288 | rs6461547 | 0.189 | 1.28 | -0.075 | 1.135 | 0.001 | 0.001 | 0.001 | 0 | 0.72 | 0.64 | 0.79 | 0.65 | 0.71 | 0.64 |
| 7 | 21312691 | rs6943932 | 2.022 | 1.349 | -0.127 | 1.09 | 0.018 | 0.023 | 0.023 | 0 | 0.4 | 0.25 | 0.45 | 0.27 | 0.41 | 0.3 |
| 7 | 21313936 | rs16872731 | 0.244 | 1.417 | -0.14 | 1.16 | -0.013 | -0.026 | -0.026 | 0 | 0.14 | 0.04 | 0.15 | 0.16 | 0.12 | 0.11 |
| 7 | 21315247 | rs9639371 | 0.432 | 1.468 | -0.077 | 0.971 | -0.002 | 0 | 0 | 0.885 | 0.16 | 0.15 | 0.13 | 0.42 | 0.17 | 0.33 |
| 7 | 21316043 | rs757968 | -0.338 | 1.472 | -0.136 | 0.946 | 0.035 | -0.023 | -0.023 | 0 | 0.77 | 0.93 | 0.83 | 0.72 | 0.81 | 0.72 |
| 7 | 21316604 | rs917269 | 0.242 | 1.549 | -0.099 | 0.934 | 0.006 | 0.007 | 0.007 | 0 | 0.67 | 0.89 | 0.82 | 0.7 | 0.7 | 0.68 |
| 7 | 21320232 | rs2097956 | 1.262 | 1.417 | 0.007 | 0.941 | 0 | 0 | 0 | 0 | 0.56 | 0.44 | 0.67 | 0.44 | 0.52 | 0.43 |
| 7 | 21323235 | rs6970402 | 1.977 | 1.338 | -0.056 | 0.87 | 0.008 | 0.018 | 0.018 | 0.074 | 0.4 | 0.32 | 0.47 | 0.16 | 0.4 | 0.31 |
| 7 | 21326835 | rs1637303 | -0.638 | 1.105 | -0.384 | 0.917 | 0.028 | 0.08 | 0.08 | 0 | 0.8 | 0.92 | 0.91 | 0.97 | 0.84 | 0.89 |
| 7 | 21327064 | rs2700281 | -0.218 | 1.109 | -0.389 | 1.001 | 0.03 | -0.021 | -0.021 | 0 | 0.77 | 0.91 | 0.88 | 0.73 | 0.81 | 0.73 |
| 7 | 21338231 | rs6976044 | -0.964 | 1.198 | -0.134 | 1.087 | -0.004 | 0.06 | 0.06 | 0 | 0.18 | 0.23 | 0.23 | 0.59 | 0.14 | 0.38 |
| 7 | 21345198 | rs2041023 | 1.008 | 1.421 | -0.421 | 1.158 | -0.033 | -0.013 | -0.013 | 0 | 0.05 | 0.01 | 0.17 | 0.25 | 0.1 | 0.06 |
| 7 | 21346861 | rs2529166 | -0.614 | 1.58 | -0.15 | 1.004 | 0.083 | 0.028 | 0.028 | 0 | 0.23 | 0.4 | 0.22 | 0.3 | 0.32 | 0.32 |
| 7 | 21350117 | rs2700289 | 0.567 | 1.517 | -0.304 | 0.94 | 0.017 | 0.031 | 0.031 | 0 | 0.65 | 0.77 | 0.72 | 0.74 | 0.72 | 0.79 |
| 7 | 21350308 | rs1637308 | 1.077 | 1.691 | -0.258 | 0.961 | 0.005 | -0.007 | -0.007 | 0 | 0.51 | 0.74 | 0.62 | 0.53 | 0.56 | 0.61 |
| 7 | 21351951 | rs6461551 | 1.013 | 1.361 | 0.253 | 0.859 | 0 | -0.007 | -0.007 | 0 | 0.55 | 0.53 | 0.68 | 0.74 | 0.53 | 0.65 |
| 7 | 21355088 | rs2529168 | 0.388 | 1.295 | -0.394 | 0.691 | -0.006 | -0.012 | -0.012 | 0.036 | 0.13 | 0.13 | 0.09 | 0 | 0.15 | 0.12 |
| 7 | 21359798 | rs2700295 | -0.4 | 1.235 | -0.346 | 0.71 | 0.097 | -0.082 | -0.082 | 0 | 0.17 | 0.44 | 0.15 | 0.1 | 0.2 | 0.23 |
| 7 | 21362447 | rs2041026 | 0.729 | 1.028 | -0.343 | 0.994 | -0.001 | 0.009 | 0.009 | 0 | 0.6 | 0.4 | 0.72 | 0.61 | 0.64 | 0.48 |
| 7 | 21372135 | rs12667306 | -1.039 | 0.841 | -0.187 | 1.85 | 0.105 | 0.318 | 0.318 | 0.795 | 0.07 | 0.37 | 0.15 | 0.28 | 0.06 | 0.26 |
| 7 | 21372342 | rs13229993 | 0.669 | 0.815 | -0.291 | 1.933 | -0.031 | -0.062 | -0.062 | 0.405 | 0.1 | 0.01 | 0.07 | 0.2 | 0.12 | 0.11 |
| 7 | 21373772 | rs2057999 | 1.247 | 0.744 | 0.153 | 1.577 | -0.01 | 0.061 | 0.061 | 0.478 | 0.81 | 0.48 | 0.78 | 0.69 | 0.85 | 0.67 |
| 7 | 21378687 | rs6461554 | -1.09 | 0.777 | 0.059 | 1.599 | 0.107 | 0.333 | 0.333 | 0.488 | 0.07 | 0.38 | 0.15 | 0.28 | 0.06 | 0.26 |
| 7 | 21378805 | rs6461555 | 0.092 | 0.631 | -1.171 | 0.957 | -0.005 | -0.018 | -0.018 | 0 | 0.88 | 0.86 | 0.93 | 0.97 | 0.91 | 0.93 |
| 7 | 21392007 | rs17144321 | -0.779 | 1.099 | -1.482 | 1.66 | -0.027 | -0.078 | -0.078 | 0 | 0.16 | 0.09 | 0.2 | 0.42 | 0.16 | 0.38 |
| 7 | 21396863 | rs10241715 | 0.087 | 0.786 | -1.814 | 1.506 | -0.043 | -0.081 | -0.081 | 0 | 0.84 | 0.99 | 0.89 | 0.79 | 0.83 | 0.83 |
| 7 | 21397405 | rs6974377 | -0.446 | 0.579 | -1.788 | 1.104 | 0.001 | 0.018 | 0.018 | 0 | 0.34 | 0.24 | 0.26 | 0.29 | 0.31 | 0.39 |
| 7 | 21397448 | rs6974533 | -0.506 | 0.504 | -1.703 | 1.069 | 0.002 | 0.014 | 0.014 | 0 | 0.34 | 0.26 | 0.27 | 0.29 | 0.31 | 0.4 |

| | | | | | | | | | | | | | | | | |
|---|---|---|---|---|---|---|---|---|---|---|---|---|---|---|---|---|
| 7 | 21398774 | rs7793253 | -0.225 | 0.107 | -1.453 | 0.76 | -0.016 | 0.03 | 0.03 | 0 | 0.27 | 0.6 | 0.38 | 0.44 | 0.19 | 0.41 |
| 7 | 21398857 | rs4722016 | -0.982 | 0.641 | -1.635 | 0.825 | -0.041 | -0.014 | -0.014 | 0 | 0.93 | 0.98 | 0.8 | 0.64 | 0.82 | 0.91 |
| 7 | 21399377 | rs10253780 | 1.591 | 0.415 | -1.663 | 0.525 | -0.035 | 0.171 | 0.171 | 0 | 0.34 | 0.12 | 0.24 | 0.02 | 0.34 | 0.16 |
| 7 | 21403759 | rs10486426 | 0.196 | 0.206 | -0.476 | 1.142 | 0.005 | 0.161 | 0.161 | 0 | 0.08 | 0.26 | 0.11 | 0.26 | 0.07 | 0.21 |
| 7 | 21406569 | rs7781656 | -0.965 | -0.036 | -0.65 | 0.953 | 0.012 | 0.009 | 0.009 | 0 | 0.51 | 0.66 | 0.51 | 0.58 | 0.43 | 0.71 |
| 7 | 21410237 | rs6965123 | 0.608 | 0.085 | -0.823 | 0.47 | 0 | 0 | 0 | 5.366 | 0.17 | 0.12 | 0.24 | 0.15 | 0.13 | 0.23 |
| 7 | 21413179 | rs7777432 | -0.724 | 0.099 | -0.733 | 0.424 | 0.01 | 0.044 | 0.044 | 0 | 0.39 | 0.25 | 0.38 | 0.2 | 0.34 | 0.45 |
| 7 | 21414350 | rs6962097 | 0.168 | -0.08 | -0.438 | 0.949 | -0.001 | 0 | 0 | 0 | 0.17 | 0.14 | 0.25 | 0.18 | 0.14 | 0.24 |
| 7 | 21421615 | rs12540638 | 0.479 | 0.513 | -0.556 | 1.402 | -0.033 | 0.005 | 0.005 | 0 | 0.07 | 0.03 | 0.1 | 0.47 | 0.15 | 0.17 |
| 7 | 21422747 | rs4722019 | 0.313 | -0.108 | -0.724 | 0.419 | 0.021 | -0.004 | -0.004 | 0 | 0.46 | 0.36 | 0.48 | 0.58 | 0.56 | 0.4 |
| 7 | 21424227 | rs7805954 | -0.993 | -0.657 | -0.448 | 0.064 | -0.007 | -0.031 | -0.031 | 0.011 | 0.76 | 0.78 | 0.86 | 0.92 | 0.82 | 0.75 |
| 7 | 21424543 | rs4722020 | -0.603 | -0.753 | -0.49 | 0.093 | 0.007 | 0.006 | 0.006 | 0 | 0.71 | 0.78 | 0.78 | 0.86 | 0.79 | 0.69 |
| 7 | 21426445 | rs11973690 | 0.056 | -0.712 | -0.648 | -0.048 | -0.012 | 0.032 | 0.032 | 0 | 0.25 | 0.39 | 0.31 | 0.28 | 0.22 | 0.26 |
| 7 | 21431809 | rs4389830 | -0.639 | -0.695 | -0.511 | 0.048 | -0.006 | -0.011 | -0.011 | 0 | 0.7 | 0.76 | 0.78 | 0.86 | 0.77 | 0.68 |
| 7 | 21433440 | rs17144391 | 0.081 | -0.694 | -0.321 | 0.035 | -0.025 | 0.051 | 0.051 | 0 | 0.23 | 0.47 | 0.29 | 0.28 | 0.16 | 0.25 |
| 7 | 21443004 | rs12672035 | 1.043 | -0.32 | -0.683 | 0.28 | -0.045 | -0.033 | -0.033 | 0 | 0.09 | 0.02 | 0.1 | 0.48 | 0.15 | 0.19 |
| 7 | 21443716 | rs6947410 | 1.089 | -0.414 | -0.639 | -0.082 | -0.093 | 0.161 | 0.161 | 0 | 0.33 | 0.15 | 0.31 | 0.04 | 0.41 | 0.18 |
| 7 | 21446473 | rs13226735 | 0.507 | -0.373 | -0.163 | 0.46 | -0.01 | 0.03 | 0.03 | 0 | 0.16 | 0.32 | 0.11 | 0.23 | 0.13 | 0.19 |
| 7 | 21449367 | rs12668671 | -0.169 | -0.334 | -0.632 | 0.375 | 0.027 | 0.024 | 0.024 | 0 | 0.52 | 0.43 | 0.58 | 0.66 | 0.63 | 0.5 |
| 7 | 21449418 | rs12666618 | 0.417 | 0.248 | -1.261 | 0.884 | -0.032 | -0.018 | -0.018 | 0 | 0.09 | 0.04 | 0.1 | 0.5 | 0.15 | 0.2 |
| 7 | 21450943 | rs253427 | 0.877 | -0.254 | -1.242 | 0.571 | 0.041 | 0.003 | 0.003 | 0 | 0.31 | 0.14 | 0.33 | 0.25 | 0.24 | 0.32 |
| 7 | 21451262 | rs253429 | 0.788 | -0.304 | -1.639 | 0.179 | 0.002 | -0.01 | -0.01 | 0 | 0.6 | 0.21 | 0.66 | 0.36 | 0.6 | 0.5 |
| 7 | 21456603 | rs39293 | 1.104 | -0.389 | -1.538 | 0.316 | 0.04 | -0.022 | -0.022 | 0.051 | 0.31 | 0.14 | 0.38 | 0.33 | 0.24 | 0.33 |
| 7 | 21460818 | rs39298 | -0.617 | -0.238 | -1.834 | 0.113 | -0.051 | 0.047 | 0.047 | 0 | 0.81 | 0.99 | 0.9 | 0.91 | 0.78 | 0.76 |
| 7 | 21462828 | rs10261327 | -0.016 | -0.122 | -0.948 | 0.302 | -0.001 | -0.007 | -0.007 | 0 | 0.42 | 0.57 | 0.43 | 0.53 | 0.52 | 0.37 |
| 7 | 21466395 | rs9986768 | 0.694 | -0.129 | -1.173 | -0.202 | -0.105 | 0.145 | 0.145 | 0.407 | 0.29 | 0.07 | 0.29 | 0.03 | 0.39 | 0.17 |
| 7 | 21470513 | rs39304 | 0.924 | -0.723 | -0.931 | 0.253 | 0.036 | 0.028 | 0.028 | 0 | 0.37 | 0.23 | 0.2 | 0.3 | 0.24 | 0.36 |
| 7 | 21474580 | rs7795411 | -1.03 | -0.623 | -0.188 | 0.369 | -0.006 | 0.219 | 0.219 | 0 | 0.09 | 0.25 | 0.33 | 0.63 | 0.04 | 0.21 |
| 7 | 21476132 | rs2282888 | -0.583 | -0.448 | -0.401 | 0.016 | -0.058 | 0.083 | 0.083 | 0.128 | 0.75 | 0.92 | 0.73 | 0.89 | 0.63 | 0.73 |
| 7 | 21476188 | rs2282889 | 0.854 | -0.377 | -0.131 | -0.056 | 0 | -0.028 | -0.028 | 0.185 | 0.63 | 0.53 | 0.39 | 0.25 | 0.59 | 0.49 |
| 7 | 21483605 | rs2237303 | -0.341 | -0.247 | -0.027 | 0.692 | -0.025 | -0.038 | -0.038 | 0 | 0.73 | 0.83 | 0.72 | 0.61 | 0.66 | 0.66 |
| 7 | 21485102 | rs2107448 | 0.918 | -0.306 | 0.078 | 0.509 | 0 | -0.037 | -0.037 | 0 | 0.62 | 0.48 | 0.38 | 0.22 | 0.59 | 0.47 |
| 7 | 21485397 | rs2390536 | 1.187 | -0.285 | -0.212 | 0.351 | -0.105 | 0.121 | 0.121 | 0 | 0.23 | 0.04 | 0.19 | 0.02 | 0.35 | 0.12 |
| 7 | 21488929 | rs2107449 | 0.634 | -0.126 | -0.202 | 0.349 | -0.085 | 0.119 | 0.119 | 0 | 0.27 | 0.08 | 0.19 | 0.04 | 0.37 | 0.16 |
| 7 | 21491021 | rs2158414 | -0.748 | 0.008 | -0.2 | 0.424 | -0.056 | 0.081 | 0.081 | 0 | 0.76 | 0.94 | 0.75 | 0.89 | 0.68 | 0.75 |
| 7 | 21496465 | rs6461565 | -1.47 | -0.368 | -0.284 | 0.39 | -0.101 | 0.065 | 0.065 | 0 | 0.8 | 0.99 | 0.76 | 0.92 | 0.7 | 0.78 |
| 7 | 21496496 | rs6461566 | -0.938 | 0.213 | -0.138 | 0.469 | 0.036 | 0.013 | 0.013 | 0 | 0.26 | 0.4 | 0.48 | 0.67 | 0.36 | 0.35 |
| 7 | 21498702 | rs6958382 | -0.49 | -0.169 | -0.104 | 0.593 | 0 | 0 | 0 | 0.017 | 0.06 | 0.07 | 0.02 | 0.01 | 0.06 | 0.07 |
| 7 | 21502073 | rs2282893 | 1.491 | -0.315 | -0.01 | 0.523 | 0.042 | 0.045 | 0.045 | 0 | 0.37 | 0.21 | 0.17 | 0.21 | 0.22 | 0.35 |
| 7 | 21509745 | rs17144483 | -1.919 | -0.156 | 0.187 | 1.068 | -0.008 | 0.087 | 0.087 | 0.144 | 0.09 | 0.19 | 0.09 | 0.37 | 0.05 | 0.16 |
| 7 | 21534152 | rs7811417 | -0.616 | -0.044 | -0.065 | 0.806 | -0.019 | -0.045 | -0.045 | 2.86 | 0.72 | 0.78 | 0.67 | 0.56 | 0.66 | 0.66 |
| 7 | 21535776 | rs6461569 | 0.398 | -0.036 | 0.069 | 0.671 | 0.028 | 0.02 | 0.02 | 0.437 | 0.51 | 0.55 | 0.53 | 0.6 | 0.33 | 0.58 |
| 7 | 21535971 | rs7811539 | 0.618 | -0.1 | 0.075 | 0.659 | 0.041 | 0.051 | 0.051 | 0.243 | 0.48 | 0.59 | 0.52 | 0.85 | 0.28 | 0.59 |
| 7 | 21540972 | rs11508471 | 1.472 | -0.094 | 0.028 | 0.615 | 0.047 | 0.052 | 0.052 | 0 | 0.39 | 0.24 | 0.18 | 0.2 | 0.24 | 0.36 |
| 7 | 21542891 | rs2390538 | 0.234 | 0.122 | 0.004 | 0.585 | -0.051 | 0.042 | 0.042 | 0 | 0.24 | 0.14 | 0.27 | 0.12 | 0.37 | 0.16 |
| 7 | 21548379 | rs17144546 | 0.544 | -0.393 | -0.085 | 0.443 | -0.025 | 0.057 | 0.057 | 0.521 | 0.08 | 0.17 | 0.06 | 0.19 | 0.03 | 0.2 |
| 7 | 21549891 | rs1018954 | -0.707 | 0.001 | -0.434 | 0.198 | -0.002 | -0.024 | -0.024 | 0.016 | 0.32 | 0.34 | 0.24 | 0.05 | 0.43 | 0.19 |
| 7 | 21551070 | rs3735440 | -1.291 | -0.017 | -0.85 | 0.246 | -0.047 | 0.072 | 0.072 | 0.314 | 0.76 | 0.93 | 0.74 | 0.89 | 0.67 | 0.76 |
| 7 | 21554464 | rs9648275 | -0.18 | 0.112 | -0.795 | 0.086 | -0.072 | 0.097 | 0.097 | 0 | 0.27 | 0.14 | 0.21 | 0.05 | 0.39 | 0.15 |
| 7 | 21556942 | rs10249014 | -0.368 | -0.099 | -0.721 | 0.144 | -0.059 | 0.098 | 0.098 | 0 | 0.29 | 0.17 | 0.21 | 0.05 | 0.39 | 0.15 |
| 7 | 21557854 | rs2023910 | -0.146 | -0.629 | -0.547 | 0.849 | -0.015 | -0.032 | -0.032 | 0.165 | 0.21 | 0.13 | 0.43 | 0.36 | 0.24 | 0.24 |
| 7 | 21558552 | rs2023911 | 0.451 | -0.765 | -0.432 | 0.594 | -0.013 | 0.003 | 0.003 | 0 | 0.43 | 0.47 | 0.5 | 0.52 | 0.27 | 0.49 |
| 7 | 21564835 | rs1473119 | -0.985 | -0.483 | -0.716 | -0.153 | -0.064 | 0.08 | 0.08 | 0.011 | 0.66 | 0.83 | 0.66 | 0.85 | 0.52 | 0.73 |
| 7 | 21569089 | rs10277163 | -0.148 | -0.298 | -0.421 | -0.107 | 0.002 | -0.021 | -0.021 | 1.708 | 0.2 | 0.23 | 0.16 | 0.06 | 0.25 | 0.1 |
| 7 | 21580913 | rs9639380 | -0.109 | -0.326 | -0.493 | 0.412 | -0.004 | 0.302 | 0.302 | 0 | 0.12 | 0.38 | 0.24 | 0.59 | 0.07 | 0.24 |
| 7 | 21581444 | rs6959819 | -1.072 | -0.485 | -0.892 | -0.032 | -0.05 | 0.087 | 0.087 | 0 | 0.64 | 0.8 | 0.6 | 0.86 | 0.52 | 0.67 |
| 7 | 21581561 | rs976516 | -0.303 | -0.294 | -1.265 | -0.227 | 0.003 | 0.014 | 0.014 | 0 | 0.52 | 0.43 | 0.36 | 0.28 | 0.46 | 0.45 |
| 7 | 21582516 | rs11760336 | 0.679 | -0.529 | -2.114 | -0.599 | 0.033 | 0.073 | 0.073 | 0 | 0.27 | 0.11 | 0.11 | 0.04 | 0.21 | 0.17 |
| 7 | 21582917 | rs2285942 | 0.746 | 0.405 | -1.668 | -0.12 | -0.031 | -0.001 | -0.001 | 0 | 0.07 | 0.01 | 0.07 | 0.05 | 0.14 | 0.05 |
| 7 | 21586595 | rs4722037 | -0.383 | -0.278 | -2.412 | -0.857 | -0.11 | 0.305 | 0.305 | 0 | 0.57 | 0.94 | 0.64 | 0.96 | 0.42 | 0.61 |
| 7 | 21589243 | rs4639414 | -0.14 | -0.152 | -1.832 | -0.423 | -0.014 | 0.011 | 0.011 | 0 | 0.13 | 0.07 | 0.1 | 0.09 | 0.2 | 0.16 |
| 7 | 21599233 | rs10950854 | -0.795 | -0.891 | -1.291 | -0.22 | 0.004 | 0.008 | 0.008 | 0 | 0.47 | 0.56 | 0.4 | 0.72 | 0.43 | 0.56 |
| 7 | 21603534 | rs10950855 | -0.027 | -1.26 | -0.76 | 0.666 | -0.037 | 0.15 | 0.15 | 0.181 | 0.1 | 0.3 | 0.29 | 0.34 | 0.03 | 0.15 |
| 7 | 21606129 | rs6977909 | 0.645 | -1.389 | -0.646 | 0.871 | 0.005 | 0 | 0 | 0 | 0.23 | 0.16 | 0.13 | 0.18 | 0.22 | 0.17 |
| 7 | 21606299 | rs6978629 | 0.048 | -1.341 | -0.625 | 0.902 | -0.036 | 0.111 | 0.111 | 0.237 | 0.1 | 0.29 | 0.13 | 0.29 | 0.03 | 0.12 |
| 7 | 21610296 | rs12669957 | -0.604 | -1.43 | -0.62 | 0.862 | -0.049 | 0.12 | 0.12 | 0 | 0.12 | 0.36 | 0.14 | 0.29 | 0.03 | 0.11 |
| 7 | 21612565 | rs6976617 | 0.607 | -1.342 | -0.573 | 0.458 | 0.008 | 0 | 0 | 0 | 0.23 | 0.14 | 0.13 | 0.18 | 0.22 | 0.17 |
| 7 | 21612705 | rs12673820 | -1.113 | -1.338 | -0.445 | 0.437 | 0.005 | 0.012 | 0.012 | 0 | 0.47 | 0.58 | 0.41 | 0.73 | 0.44 | 0.52 |
| 7 | 21615629 | rs12670286 | 0.429 | -1.389 | -0.402 | 0.671 | -0.035 | -0.084 | -0.084 | 0 | 0.13 | 0.02 | 0.17 | 0.48 | 0.17 | 0.18 |

| | | | | | | | | | | | | | | | | |
|---|---|---|---|---|---|---|---|---|---|---|---|---|---|---|---|---|
| 7 | 21619511 | rs12666258 | -0.108 | -1.452 | -0.355 | 0.375 | -0.005 | 0.002 | 0.002 | 0 | 0.22 | 0.26 | 0.13 | 0.2 | 0.23 | 0.19 |
| 7 | 21626856 | rs11765753 | 0.656 | -1.478 | -0.368 | 0.393 | -0.009 | 0.01 | 0.01 | 0 | 0.17 | 0.23 | 0.11 | 0.19 | 0.18 | 0.12 |
| 7 | 21628197 | rs12670130 | 0.834 | -1.457 | -0.541 | 0.402 | 0.004 | -0.007 | -0.007 | 0 | 0.17 | 0.12 | 0.11 | 0.19 | 0.18 | 0.12 |
| 7 | 21632107 | rs6947026 | 1.181 | -1.707 | -0.762 | -0.369 | -0.059 | 0.09 | 0.09 | 0 | 0.31 | 0.17 | 0.29 | 0.05 | 0.42 | 0.25 |
| 7 | 21639572 | rs17144747 | -0.173 | -1.407 | 0.005 | -0.252 | -0.015 | -0.026 | -0.026 | 0.195 | 0.08 | 0.17 | 0.04 | 0.02 | 0.04 | 0.08 |
| 7 | 21640361 | rs10269582 | 1.039 | -1.709 | -0.177 | -0.507 | -0.068 | 0.137 | 0.137 | 0 | 0.36 | 0.15 | 0.31 | 0.02 | 0.43 | 0.25 |
| 7 | 21640405 | rs10224537 | -1.64 | -1.722 | -0.052 | -0.388 | -0.001 | -0.001 | -0.001 | 0 | 0.74 | 0.76 | 0.7 | 0.81 | 0.82 | 0.72 |
| 7 | 21641218 | rs3827657 | -1.431 | -1.595 | -0.049 | -0.423 | 0 | 0 | 0 | 0.324 | 0.72 | 0.75 | 0.69 | 0.78 | 0.81 | 0.71 |
| 7 | 21641629 | rs7803642 | -0.773 | -1.045 | 0.404 | 0.383 | -0.001 | -0.003 | -0.003 | 0.631 | 0.29 | 0.17 | 0.35 | 0.75 | 0.36 | 0.4 |
| 7 | 21647686 | rs17144779 | -0.287 | -1.114 | 0.422 | 0.403 | -0.008 | -0.005 | -0.005 | 0 | 0.08 | 0.15 | 0.09 | 0.06 | 0.05 | 0.16 |
| 7 | 21648554 | rs9639382 | -0.792 | -1.134 | 0.108 | 0.524 | -0.025 | 0.03 | 0.03 | 0 | 0.23 | 0.46 | 0.22 | 0.25 | 0.22 | 0.28 |
| 7 | 21654602 | rs7779983 | -1.36 | -0.324 | -0.221 | 0.381 | -0.006 | 0.018 | 0.018 | 0.015 | 0.45 | 0.6 | 0.49 | 0.77 | 0.56 | 0.5 |
| 7 | 21657185 | rs8180768 | 0.239 | -0.876 | -0.271 | 0.785 | -0.002 | -0.019 | -0.019 | 0 | 0.13 | 0.09 | 0.11 | 0.18 | 0.18 | 0.13 |
| 7 | 21680313 | rs2222521 | -0.881 | -1.594 | 0.203 | 0.828 | -0.01 | 0.024 | 0.024 | 0.212 | 0.1 | 0.11 | 0.29 | 0.18 | 0.04 | 0.07 |
| 7 | 21680566 | rs10950865 | -0.854 | -1.952 | 0.174 | 0.367 | -0.016 | -0.027 | -0.027 | 0.446 | 0.65 | 0.79 | 0.77 | 0.43 | 0.64 | 0.62 |
| 7 | 21680751 | rs10274031 | -1.137 | -2.428 | 0.536 | 0.322 | -0.011 | -0.001 | -0.001 | 0 | 0.33 | 0.63 | 0.42 | 0.4 | 0.13 | 0.33 |
| 7 | 21680935 | rs10274412 | 0.457 | -2.416 | 0.994 | 0.273 | -0.029 | -0.01 | -0.01 | 0 | 0.18 | 0.21 | 0.07 | 0.17 | 0.08 | 0.17 |
| 7 | 21681379 | rs1568777 | -0.81 | -2.259 | 1.126 | 0.341 | 0.015 | 0.067 | 0.067 | 0.252 | 0.05 | 0.29 | 0.05 | 0.06 | 0.01 | 0.15 |
| 7 | 21681850 | rs17747517 | 0.335 | -2.264 | 0.699 | 0.326 | 0.052 | 0.033 | 0.033 | 0.673 | 0.21 | 0.1 | 0.07 | 0.17 | 0.1 | 0.2 |
| 7 | 21682636 | rs7796497 | -1.363 | -2.248 | 0.709 | 0.221 | 0.014 | 0 | 0 | 0 | 0.39 | 0.66 | 0.44 | 0.39 | 0.19 | 0.4 |
| 7 | 21684027 | rs13224416 | 1.28 | -2.307 | 0.355 | 0.479 | -0.1 | 0.009 | 0.009 | 0 | 0.31 | 0.11 | 0.39 | 0.22 | 0.52 | 0.39 |
| 7 | 21685178 | rs2893046 | -0.657 | -2.347 | 0.485 | 0.499 | -0.028 | 0.083 | 0.083 | 0 | 0.1 | 0.17 | 0.32 | 0.24 | 0.02 | 0.15 |
| 7 | 21690187 | rs10260278 | 0.393 | -2.279 | 0.563 | 0.467 | 0.001 | 0.001 | 0.001 | 0 | 0.43 | 0.44 | 0.71 | 0.47 | 0.54 | 0.55 |
| 7 | 21693607 | rs9639385 | -0.79 | -1.883 | 0.309 | 0.599 | -0.006 | 0.014 | 0.014 | 0 | 0.66 | 0.5 | 0.77 | 0.6 | 0.7 | 0.79 |
| 7 | 21693716 | rs7790091 | -0.075 | -1.84 | 0.334 | 0.602 | 0.008 | -0.006 | -0.006 | 0 | 0.58 | 0.32 | 0.73 | 0.54 | 0.64 | 0.63 |
| 7 | 21697133 | rs10238210 | 0.337 | -1.241 | -0.106 | 0.099 | 0.014 | -0.027 | -0.027 | 2.113 | 0.05 | 0.17 | 0.02 | 0 | 0.06 | 0.07 |
| 7 | 21698078 | rs10480009 | 0.337 | -1.241 | -0.114 | 0.099 | 0.028 | -0.02 | -0.02 | 2.576 | 0.05 | 0.21 | 0.02 | 0 | 0.06 | 0.07 |
| 7 | 21698100 | rs10499538 | 0.424 | -1.28 | -0.439 | 0.074 | 0.039 | 0.045 | 0.045 | 1.876 | 0.19 | 0.07 | 0.07 | 0 | 0.1 | 0.11 |
| 7 | 21704005 | rs6461593 | -0.534 | -1.206 | -0.443 | 0.108 | -0.041 | -0.133 | -0.133 | 0 | 0.28 | 0.41 | 0.11 | 0 | 0.17 | 0.18 |
| 7 | 21706255 | rs6975512 | -0.138 | 0.049 | -0.711 | 1.287 | -0.003 | 0 | 0 | 0 | 0.46 | 0.49 | 0.71 | 0.46 | 0.59 | 0.6 |
| 7 | 21716567 | rs2965387 | 0.104 | 0.539 | -1.205 | 1.108 | -0.003 | 0.004 | 0.004 | 0 | 0.58 | 0.74 | 0.42 | 0.42 | 0.57 | 0.57 |
| 7 | 21719768 | rs2965391 | 0.323 | 0.269 | -0.673 | 0.714 | 0.006 | -0.04 | -0.04 | 0 | 0.42 | 0.5 | 0.3 | 0.2 | 0.43 | 0.45 |
| 7 | 21728627 | rs2033884 | -0.77 | 0.616 | -0.584 | 1.586 | 0.005 | 0.034 | 0.034 | 0 | 0.26 | 0.32 | 0.25 | 0.48 | 0.29 | 0.29 |
| 7 | 21730647 | rs7790948 | -0.844 | 0.483 | -0.245 | 1.81 | -0.005 | 0.098 | 0.098 | 0.021 | 0.2 | 0.33 | 0.31 | 0.42 | 0.16 | 0.18 |
| 7 | 21741813 | rs418392 | 0.125 | 0.589 | -0.987 | 1.558 | -0.032 | -0.009 | -0.009 | 0 | 0.86 | 0.99 | 0.86 | 0.89 | 0.83 | 0.81 |
| 7 | 21747684 | rs17145085 | -0.502 | 0.701 | -0.567 | 2.945 | -0.031 | -0.032 | -0.032 | 0 | 0.06 | 0.01 | 0.09 | 0.23 | 0.09 | 0.12 |
| 7 | 21748121 | rs6972964 | 0.005 | -0.018 | -0.084 | 2.323 | -0.027 | 0.04 | 0.04 | 0 | 0.15 | 0.44 | 0.14 | 0.19 | 0.07 | 0.32 |
| 7 | 21749104 | rs12667433 | -0.792 | 0.004 | -0.648 | 1.932 | -0.018 | -0.036 | -0.036 | 0 | 0.1 | 0.03 | 0.11 | 0.21 | 0.11 | 0.12 |
| 7 | 21752786 | rs2906663 | -1.324 | -0.418 | -0.858 | 1.712 | -0.004 | -0.03 | -0.03 | 0 | 0.16 | 0.03 | 0.14 | 0.21 | 0.14 | 0.16 |
| 7 | 21754231 | rs764089 | -0.173 | -0.371 | -0.751 | 1.324 | -0.017 | 0.032 | 0.032 | 0 | 0.34 | 0.56 | 0.46 | 0.48 | 0.29 | 0.47 |
| 7 | 21754897 | rs2965406 | -0.818 | -0.36 | -0.711 | 1.098 | 0.003 | 0.005 | 0.005 | 0 | 0.46 | 0.56 | 0.59 | 0.69 | 0.42 | 0.62 |
| 7 | 21755427 | rs2965407 | -0.293 | -0.405 | -1.196 | 0.962 | 0.007 | -0.012 | -0.012 | 0 | 0.31 | 0.23 | 0.32 | 0.42 | 0.27 | 0.45 |
| 7 | 21757690 | rs2240471 | -0.052 | -0.124 | -1.354 | 0.412 | -0.043 | 0.105 | 0.105 | 0 | 0.36 | 0.18 | 0.31 | 0.18 | 0.43 | 0.42 |
| 7 | 21761038 | rs6976078 | -0.656 | 0.024 | -1.662 | 0.199 | 0.009 | 0.243 | 0.243 | 0 | 0.51 | 0.9 | 0.62 | 0.81 | 0.43 | 0.63 |
| 7 | 21761612 | rs12540711 | -0.822 | -0.331 | -1.765 | 0.148 | -0.126 | 0.207 | 0.207 | 0 | 0.58 | 0.95 | 0.68 | 0.87 | 0.55 | 0.67 |
| 7 | 21763145 | rs1029597 | -1.145 | -0.617 | -2.152 | -0.241 | 0.008 | 0.031 | 0.031 | 0.009 | 0.81 | 0.96 | 0.86 | 0.86 | 0.86 | 0.81 |
| 7 | 21763249 | rs2158870 | 0.136 | -0.654 | -1.162 | 0.258 | 0.028 | 0.016 | 0.016 | 0.062 | 0.26 | 0.34 | 0.2 | 0.28 | 0.33 | 0.22 |
| 7 | 21765452 | rs12536928 | 1.21 | -1.153 | -0.927 | 0.187 | -0.039 | 0.073 | 0.073 | 1.517 | 0.39 | 0.26 | 0.44 | 0.2 | 0.5 | 0.48 |
| 7 | 21765497 | rs11768670 | 1.191 | -1.15 | -0.573 | 0.18 | -0.019 | 0.024 | 0.024 | 0 | 0.19 | 0.12 | 0.15 | 0.06 | 0.26 | 0.08 |
| 7 | 21765909 | rs2074764 | 0.05 | -1.26 | -0.102 | 0.267 | -0.003 | 0 | 0 | 0 | 0.58 | 0.64 | 0.68 | 0.57 | 0.58 | 0.73 |
| 7 | 21766037 | rs16872875 | -0.174 | -1.3 | -0.007 | 0.309 | -0.065 | 0.144 | 0.144 | 0.395 | 0.17 | 0.35 | 0.22 | 0.38 | 0.07 | 0.21 |
| 7 | 21766996 | rs2906667 | 0.297 | -1.089 | -0.105 | 0.446 | -0.001 | -0.002 | -0.002 | 0 | 0.55 | 0.58 | 0.56 | 0.5 | 0.56 | 0.61 |
| 7 | 21768231 | rs2965375 | 0.173 | -0.971 | 0.072 | 0.543 | 0.015 | 0.049 | 0.049 | 0 | 0.68 | 0.56 | 0.75 | 0.43 | 0.68 | 0.71 |
| 7 | 21771290 | rs9639391 | 0.414 | -0.968 | -0.121 | 0.633 | 0 | 0.007 | 0.007 | 0 | 0.23 | 0.21 | 0.16 | 0.43 | 0.26 | 0.11 |
| 7 | 21773996 | rs6979397 | 1.09 | -1.475 | -0.316 | 0.396 | 0.061 | 0.067 | 0.067 | 0 | 0.57 | 0.26 | 0.57 | 0.29 | 0.75 | 0.58 |
| 7 | 21775387 | rs2072220 | 0.26 | -1.371 | 0.328 | 0.978 | 0.027 | 0.308 | 0.308 | 0 | 0.06 | 0.28 | 0.21 | 0.21 | 0.03 | 0.08 |
| 7 | 21777119 | rs2906681 | 0.781 | -1.451 | 0.186 | 0.864 | -0.001 | -0.001 | -0.001 | 0.131 | 0.54 | 0.47 | 0.67 | 0.38 | 0.58 | 0.64 |
| 7 | 21777455 | rs17145154 | 1.425 | -2.002 | 0.034 | 0.964 | 0.008 | -0.004 | -0.004 | 0 | 0.09 | 0.07 | 0.04 | 0.26 | 0.04 | 0.03 |
| 7 | 21778200 | rs2965365 | 0.803 | -1.9 | 0.155 | 0.925 | 0.017 | 0.027 | 0.027 | 0 | 0.58 | 0.6 | 0.57 | 0.24 | 0.7 | 0.56 |
| 7 | 21778449 | rs1109806 | 0.768 | -1.868 | 0.208 | 0.965 | 0.015 | 0.023 | 0.023 | 0 | 0.59 | 0.59 | 0.57 | 0.24 | 0.7 | 0.58 |
| 7 | 21779104 | rs933353 | -1.122 | -2.13 | 0.318 | 0.837 | 0.048 | 0.065 | 0.065 | 0.054 | 0.41 | 0.74 | 0.4 | 0.8 | 0.27 | 0.28 |
| 7 | 21779278 | rs9639393 | 1.674 | -2.134 | 0.342 | 0.789 | 0.08 | 0.18 | 0.18 | 0.108 | 0.5 | 0.22 | 0.52 | 0.08 | 0.65 | 0.49 |
| 7 | 21779331 | rs933354 | 0.266 | -2.107 | 0.442 | 1.085 | 0.001 | -0.003 | -0.003 | 0 | 0.22 | 0.24 | 0.29 | 0.26 | 0.19 | 0.15 |
| 7 | 21779665 | rs9639395 | 0.428 | -2.091 | 0.463 | 1.092 | 0 | -0.001 | -0.001 | 0 | 0.69 | 0.64 | 0.62 | 0.6 | 0.75 | 0.62 |
| 7 | 21780460 | rs2906690 | 0.618 | -2.06 | 0.593 | 1.108 | 0.042 | 0.122 | 0.122 | 0 | 0.05 | 0.28 | 0.03 | 0.09 | 0.03 | 0.05 |
| 7 | 21780930 | rs9639396 | 1.595 | -2.402 | 0.294 | 1.021 | 0.053 | 0.047 | 0.047 | 0.498 | 0.55 | 0.25 | 0.54 | 0.32 | 0.67 | 0.54 |
| 7 | 21781807 | rs1029598 | 0.138 | -2.067 | 0.401 | 1.158 | -0.003 | 0.008 | 0.008 | 0 | 0.22 | 0.29 | 0.3 | 0.27 | 0.18 | 0.16 |
| 7 | 21787705 | rs2965355 | -0.67 | -1.684 | 0.551 | 0.781 | 0 | 0 | 0 | 0 | 0.91 | 0.92 | 0.92 | 0.88 | 0.94 | 0.78 |
| 7 | 21789026 | rs4449698 | 1.397 | -1.859 | 0.453 | 0.617 | -0.014 | -0.048 | -0.048 | 0 | 0.55 | 0.48 | 0.55 | 0.14 | 0.68 | 0.54 |

| | | | | | | | | | | | | | | | | |
|---|---|---|---|---|---|---|---|---|---|---|---|---|---|---|---|---|
| 7 | 21798062 | rs6461598 | -0.834 | -1.61 | 0.192 | 0.671 | 0.001 | 0.007 | 0.007 | 0 | 0.9 | 0.96 | 0.91 | 0.89 | 0.93 | 0.77 |
| 7 | 21799157 | rs17277521 | 0.685 | -1.718 | 0.361 | 0.813 | -0.003 | 0.001 | 0.001 | 0 | 0.18 | 0.29 | 0.1 | 0.25 | 0.18 | 0.09 |
| 7 | 21799575 | rs10485976 | 0.885 | -1.74 | -0.156 | 1.058 | -0.011 | -0.071 | -0.071 | 0.197 | 0.12 | 0.02 | 0.05 | 0.26 | 0.08 | 0.04 |
| 7 | 21800369 | rs10281087 | 1.257 | -1.513 | -0.241 | 0.524 | -0.005 | -0.003 | -0.003 | 0 | 0.34 | 0.3 | 0.5 | 0.31 | 0.48 | 0.37 |
| 7 | 21801878 | rs10256021 | 0.051 | -1.485 | -0.506 | 0.557 | -0.004 | -0.037 | -0.037 | 0.282 | 0.41 | 0.23 | 0.5 | 0.44 | 0.33 | 0.35 |
| 7 | 21804225 | rs12700303 | 0.984 | -1.953 | -0.482 | 0.217 | 0.083 | 0.134 | 0.134 | 0.138 | 0.53 | 0.25 | 0.53 | 0.12 | 0.7 | 0.53 |
| 7 | 21804404 | rs4719674 | 1.288 | -1.538 | -0.278 | 0.392 | -0.022 | 0.034 | 0.034 | 0 | 0.21 | 0.14 | 0.25 | 0.05 | 0.22 | 0.21 |
| 7 | 21806539 | rs4722055 | -1.101 | -1.693 | -0.499 | 0.542 | -0.038 | 0.063 | 0.063 | 0.008 | 0.55 | 0.89 | 0.68 | 0.69 | 0.46 | 0.61 |
| 7 | 21807071 | rs17145316 | 0.91 | -1.806 | -0.129 | 0.955 | 0.001 | 0.016 | 0.016 | 0 | 0.09 | 0.04 | 0.09 | 0.08 | 0.06 | 0.21 |
| 7 | 21807211 | rs6978810 | 0.479 | -1.797 | 0.343 | 0.969 | -0.028 | 0.092 | 0.092 | 0 | 0.19 | 0.32 | 0.29 | 0.52 | 0.16 | 0.18 |
| 7 | 21810164 | rs11983051 | 0.352 | -1.817 | 0.371 | 0.931 | -0.004 | -0.014 | -0.014 | 0 | 0.11 | 0.27 | 0.11 | 0.09 | 0.06 | 0.24 |
| 7 | 21810234 | rs6977017 | -0.95 | -1.872 | -0.188 | 0.95 | 0.042 | -0.015 | -0.015 | 0 | 0.33 | 0.78 | 0.42 | 0.65 | 0.21 | 0.42 |
| 7 | 21812455 | rs17145347 | -1.267 | -1.457 | 0.194 | 1.056 | -0.02 | 0 | 0 | 0 | 0.29 | 0.55 | 0.25 | 0.29 | 0.23 | 0.38 |
| 7 | 21814769 | rs3926905 | -1.382 | -1.504 | 0.197 | 1.34 | -0.024 | 0.086 | 0.086 | 0.126 | 0.29 | 0.51 | 0.44 | 0.44 | 0.24 | 0.45 |
| 7 | 21814839 | rs17145367 | -0.299 | -1.417 | 0.033 | 1.258 | -0.009 | 0.034 | 0.034 | 0 | 0.08 | 0.09 | 0.08 | 0.13 | 0.05 | 0.12 |
| 7 | 21816750 | rs2189020 | 0.978 | -1.388 | -0.39 | 0.769 | -0.03 | 0.048 | 0.048 | 0.005 | 0.37 | 0.17 | 0.28 | 0.27 | 0.46 | 0.34 |
| 7 | 21817048 | rs2214324 | -0.103 | -1.549 | -0.138 | 0.836 | 0.005 | 0.019 | 0.019 | 0 | 0.49 | 0.55 | 0.39 | 0.61 | 0.55 | 0.43 |
| 7 | 21818323 | rs1557778 | 0.241 | -1.386 | -0.153 | 0.732 | -0.009 | -0.047 | -0.047 | 0 | 0.76 | 0.59 | 0.83 | 0.84 | 0.78 | 0.7 |
| 7 | 21818850 | rs2074330 | 0.409 | -1.352 | -0.207 | 0.823 | -0.008 | 0.066 | 0.066 | 0 | 0.09 | 0.11 | 0.09 | 0.34 | 0.08 | 0.1 |
| 7 | 21822910 | rs17358098 | 1.734 | -1.387 | -0.873 | 0.299 | -0.086 | 0.121 | 0.121 | 0.434 | 0.25 | 0.06 | 0.25 | 0.02 | 0.42 | 0.26 |
| 7 | 21824122 | rs6965750 | 1.107 | -1.662 | -0.671 | 0.337 | -0.001 | -0.029 | -0.029 | 0 | 0.12 | 0.23 | 0.07 | 0.06 | 0.14 | 0.1 |
| 7 | 21824682 | rs1859103 | 0.988 | -1.704 | -0.667 | 0.468 | -0.003 | 0.028 | 0.028 | 0 | 0.13 | 0.27 | 0.09 | 0.15 | 0.14 | 0.12 |
| 7 | 21825012 | rs961526 | -0.93 | -1.664 | -0.99 | 0.394 | -0.006 | 0.016 | 0.016 | 0 | 0.9 | 0.94 | 0.93 | 0.92 | 0.92 | 0.79 |
| 7 | 21826035 | rs11495974 | 1.638 | -1.453 | -1.07 | 0.314 | -0.11 | 0.145 | 0.145 | 0.907 | 0.26 | 0.06 | 0.28 | 0.02 | 0.45 | 0.26 |
| 7 | 21828407 | rs2074328 | 1.807 | -1.663 | -1.02 | 0.545 | 0.032 | -0.036 | -0.036 | 0 | 0.11 | 0.03 | 0.04 | 0.24 | 0.04 | 0.08 |
| 7 | 21830640 | rs10276109 | 0.809 | -1.624 | -0.818 | 0.527 | -0.003 | 0.006 | 0.006 | 0.126 | 0.37 | 0.36 | 0.45 | 0.48 | 0.29 | 0.33 |
| 7 | 21833447 | rs2893049 | -0.869 | -1.639 | -0.846 | -0.053 | -0.103 | 0.209 | 0.209 | 0.184 | 0.63 | 0.9 | 0.66 | 0.94 | 0.46 | 0.53 |
| 7 | 21835071 | rs10950875 | 0.335 | -1.642 | -0.778 | 0.037 | 0.01 | 0.018 | 0.018 | 0.47 | 0.46 | 0.35 | 0.48 | 0.73 | 0.32 | 0.39 |
| 7 | 21836072 | rs10230246 | 0.166 | -1.631 | -0.767 | 0.029 | 0.016 | 0.028 | 0.028 | 0.261 | 0.47 | 0.36 | 0.5 | 0.78 | 0.32 | 0.43 |
| 7 | 21839896 | rs6956349 | 1.72 | -1.526 | -0.853 | -0.134 | -0.13 | 0.163 | 0.163 | 0.431 | 0.26 | 0.04 | 0.27 | 0.01 | 0.45 | 0.26 |
| 7 | 21843579 | rs10233735 | 0.048 | -1.697 | -0.715 | 0.313 | -0.071 | 0.139 | 0.139 | 0 | 0.22 | 0.35 | 0.14 | 0.52 | 0.12 | 0.14 |
| 7 | 21845794 | rs7811898 | 0.453 | -0.937 | -0.251 | -0.418 | -0.008 | 0 | 0 | 0 | 0.17 | 0.24 | 0.14 | 0.16 | 0.14 | 0.14 |
| 7 | 21846551 | rs7780746 | 0.404 | -0.896 | -0.322 | -0.528 | 0.046 | 0.101 | 0.101 | 0.626 | 0.5 | 0.66 | 0.52 | 0.76 | 0.34 | 0.43 |
| 7 | 21852536 | rs2014193 | 0.664 | -0.806 | -0.196 | -0.844 | -0.003 | -0.028 | -0.028 | 0 | 0.47 | 0.45 | 0.5 | 0.81 | 0.34 | 0.44 |
| 7 | 21852615 | rs2014196 | -0.747 | -0.805 | -0.201 | -0.852 | -0.122 | 0.148 | 0.148 | 0.516 | 0.67 | 0.9 | 0.66 | 0.88 | 0.48 | 0.54 |
| 7 | 21852852 | rs12700308 | 0.572 | -0.585 | -0.18 | -0.722 | -0.004 | -0.03 | -0.03 | 0 | 0.47 | 0.45 | 0.5 | 0.81 | 0.34 | 0.43 |
| 7 | 21856151 | rs2214326 | 0.624 | -0.564 | -0.302 | -0.705 | 0.004 | -0.002 | -0.002 | 0 | 0.46 | 0.38 | 0.49 | 0.81 | 0.34 | 0.43 |
| 7 | 21858215 | rs9691045 | 0.185 | -0.305 | -0.22 | -0.152 | -0.01 | -0.038 | -0.038 | 4.855 | 0.16 | 0.25 | 0.14 | 0.07 | 0.14 | 0.11 |
| 7 | 21860316 | rs17145586 | 0.254 | -0.238 | -0.424 | -0.108 | -0.007 | -0.018 | -0.018 | 0.057 | 0.16 | 0.22 | 0.14 | 0.09 | 0.14 | 0.11 |
| 7 | 21861028 | rs2390592 | 0.105 | 0.057 | -0.139 | 0.354 | -0.001 | -0.002 | -0.002 | 1.085 | 0.91 | 0.83 | 0.92 | 0.88 | 0.92 | 0.79 |
| 7 | 21862963 | rs7781044 | 0.097 | 0.089 | -0.11 | 0.675 | -0.007 | -0.004 | -0.004 | 0 | 0.35 | 0.45 | 0.44 | 0.61 | 0.27 | 0.34 |
| 7 | 21863871 | rs10229866 | -0.624 | 0.433 | 0.192 | 0.811 | 0.019 | -0.003 | -0.003 | 0.169 | 0.18 | 0.32 | 0.13 | 0.13 | 0.2 | 0.1 |
| 7 | 21868744 | rs10485983 | -0.446 | 0.362 | 0.312 | 1.373 | 0.012 | 0.041 | 0.041 | 0 | 0.12 | 0.18 | 0.11 | 0.3 | 0.18 | 0.16 |
| 7 | 21869525 | rs7807695 | -1.313 | -0.578 | 0.615 | 0.82 | 0.007 | -0.045 | -0.045 | 0.433 | 0.09 | 0.28 | 0.02 | 0 | 0.05 | 0.05 |
| 7 | 21880186 | rs11763291 | 0.669 | -1.354 | -0.249 | 0.77 | 0.003 | 0.046 | 0.046 | 0.48 | 0.41 | 0.07 | 0.43 | 0.28 | 0.63 | 0.38 |
| 7 | 21881548 | rs10274154 | -0.44 | -1.642 | 0.078 | 1.166 | -0.008 | 0.168 | 0.168 | 0.088 | 0.11 | 0.44 | 0.04 | 0.25 | 0.05 | 0.13 |
| 7 | 21884455 | rs12700309 | 0.585 | -2.356 | -0.322 | 1.428 | 0.134 | -0.049 | -0.049 | 0 | 0.55 | 0.15 | 0.54 | 0.5 | 0.72 | 0.44 |
| 7 | 21888238 | rs10277904 | -1.242 | -2.667 | -0.06 | 1.19 | 0.02 | 0.014 | 0.014 | 0.078 | 0.76 | 0.85 | 0.86 | 0.95 | 0.86 | 0.71 |
| 7 | 21891451 | rs12700311 | -0.488 | -2.602 | 0.084 | 1.421 | 0.002 | 0 | 0 | 0.038 | 0.68 | 0.7 | 0.6 | 0.72 | 0.77 | 0.56 |
| 7 | 21892416 | rs2074326 | 1.283 | -2.762 | 0.153 | 1.401 | 0.091 | 0.103 | 0.103 | 0 | 0.46 | 0.26 | 0.47 | 0.25 | 0.67 | 0.37 |
| 7 | 21894596 | rs10254976 | -0.501 | -2.754 | 0.216 | 1.586 | 0.016 | -0.015 | -0.015 | 0 | 0.68 | 0.76 | 0.57 | 0.56 | 0.76 | 0.54 |
| 7 | 21895887 | rs16873036 | -0.852 | -2.78 | 0.255 | 1.598 | -0.058 | 0.124 | 0.124 | 0.536 | 0.17 | 0.39 | 0.09 | 0.32 | 0.07 | 0.05 |
| 7 | 21895985 | rs4719678 | 1.303 | -2.468 | -0.053 | 1.488 | 0.055 | 0.096 | 0.096 | 0.492 | 0.42 | 0.15 | 0.45 | 0.27 | 0.65 | 0.37 |
| 7 | 21897714 | rs2040664 | 0.406 | -2.454 | -0.016 | 1.507 | 0.07 | 0.009 | 0.009 | 0 | 0.53 | 0.71 | 0.49 | 0.47 | 0.7 | 0.5 |
| 7 | 21899373 | rs11765332 | 1.021 | -2.267 | -0.595 | 1.137 | 0.115 | 0.13 | 0.13 | 0.92 | 0.45 | 0.15 | 0.46 | 0.24 | 0.68 | 0.37 |
| 7 | 21900188 | rs10950880 | 0.09 | -2.394 | -0.389 | 1.119 | 0.016 | 0.013 | 0.013 | 1.076 | 0.61 | 0.7 | 0.8 | 0.73 | 0.82 | 0.68 |
| 7 | 21900924 | rs4722066 | 0.767 | -1.687 | -0.93 | 1.179 | 0.031 | 0.088 | 0.088 | 0 | 0.45 | 0.06 | 0.49 | 0.26 | 0.66 | 0.37 |
| 7 | 21901038 | rs17145715 | -1.155 | -1.866 | -0.394 | 1.409 | -0.008 | 0.044 | 0.044 | 0.051 | 0.17 | 0.3 | 0.12 | 0.24 | 0.11 | 0.1 |
| 7 | 21901540 | rs17145720 | -0.519 | -1.883 | -0.272 | 1.258 | 0.021 | -0.006 | -0.006 | 0.238 | 0.06 | 0.26 | 0.04 | 0 | 0.02 | 0.03 |
| 7 | 21901566 | rs4722067 | -0.641 | -2.09 | -0.2 | 1.399 | -0.048 | 0.059 | 0.059 | 0 | 0.72 | 0.58 | 0.65 | 0.54 | 0.82 | 0.61 |
| 7 | 21906980 | rs17145746 | -0.381 | -1.783 | -0.538 | 0.899 | -0.029 | -0.028 | -0.028 | 0.661 | 0.12 | 0.26 | 0.05 | 0.06 | 0.05 | 0.13 |
| 7 | 21908964 | rs10485987 | 0.535 | -2.241 | -1.179 | 0.971 | 0.16 | 0.022 | 0.022 | 0.079 | 0.54 | 0.12 | 0.55 | 0.34 | 0.75 | 0.43 |
| 7 | 21909081 | rs886313 | 1.313 | -1.09 | -1.756 | 0.663 | -0.173 | 0.119 | 0.119 | 0.49 | 0.37 | 0.03 | 0.35 | 0.16 | 0.46 | 0.25 |
| 7 | 21909867 | rs12539597 | -0.944 | -1.257 | -1.488 | 0.566 | 0.039 | -0.035 | -0.035 | 0.322 | 0.71 | 0.84 | 0.63 | 0.38 | 0.8 | 0.57 |
| 7 | 21910572 | rs11766053 | 1.311 | -1.089 | -1.626 | 0.502 | -0.165 | 0.121 | 0.121 | 0.452 | 0.37 | 0.04 | 0.35 | 0.16 | 0.45 | 0.24 |
| 7 | 21912150 | rs10254159 | -1.195 | -1.227 | -1.458 | 0.729 | 0.031 | -0.044 | -0.044 | 0 | 0.73 | 0.84 | 0.64 | 0.43 | 0.84 | 0.58 |
| 7 | 21912204 | rs10254192 | -1.328 | -1.181 | -1.439 | 0.834 | 0.023 | -0.042 | -0.042 | 0.313 | 0.73 | 0.85 | 0.64 | 0.45 | 0.84 | 0.61 |
| 7 | 21912476 | rs11761264 | 1.351 | -1.04 | -1.686 | 0.516 | -0.167 | 0.091 | 0.091 | 0.307 | 0.36 | 0.03 | 0.34 | 0.17 | 0.44 | 0.24 |
| 7 | 21915464 | rs2390595 | -1.268 | -0.905 | -1.021 | 0.455 | 0.001 | -0.006 | -0.006 | 1.182 | 0.3 | 0.69 | 0.3 | 0.17 | 0.38 | 0.28 |

| | | | | | | | | | | | | | | | | |
|---|---|---|---|---|---|---|---|---|---|---|---|---|---|---|---|---|
| 7 | 21919145 | rs10261104 | 0.32 | -0.743 | -1.202 | 0.222 | -0.014 | 0.024 | 0.024 | 0.109 | 0.1 | 0.04 | 0.08 | 0.05 | 0.1 | 0.07 |
| 7 | 21920046 | rs11763022 | 1.65 | -0.4 | -1.394 | 0.208 | -0.11 | 0.07 | 0.07 | 0.516 | 0.32 | 0.03 | 0.3 | 0.17 | 0.38 | 0.21 |
| 7 | 21920165 | rs11763025 | 1.58 | -0.411 | -1.396 | 0.206 | -0.112 | 0.074 | 0.074 | 0.477 | 0.32 | 0.03 | 0.3 | 0.17 | 0.38 | 0.21 |
| 7 | 21922555 | rs7350056 | -0.28 | -0.474 | -1.105 | 0.265 | -0.021 | 0.023 | 0.023 | 0 | 0.2 | 0.16 | 0.19 | 0.11 | 0.3 | 0.25 |
| 7 | 21922566 | rs7349999 | -0.22 | -0.447 | -0.893 | 0.304 | 0.004 | -0.003 | -0.003 | 0 | 0.17 | 0.23 | 0.2 | 0.12 | 0.26 | 0.18 |
| 7 | 21928527 | rs6461614 | -0.473 | -0.427 | -0.872 | 0.684 | 0 | 0 | 0 | 0 | 0.5 | 0.52 | 0.38 | 0.55 | 0.54 | 0.57 |
| 7 | 21929575 | rs4392794 | 0.088 | -0.454 | -0.809 | 0.697 | -0.013 | 0.135 | 0.135 | 0.364 | 0.83 | 0.59 | 0.67 | 0.72 | 0.89 | 0.8 |
| 7 | 21929756 | rs6949101 | 0.579 | -0.299 | -0.896 | 0.712 | -0.041 | 0.003 | 0.003 | 0 | 0.13 | 0.1 | 0.19 | 0.11 | 0.28 | 0.16 |
| 7 | 21932044 | rs12537531 | 0.609 | -0.426 | -0.849 | 0.693 | -0.016 | -0.022 | -0.022 | 0 | 0.34 | 0.25 | 0.3 | 0.47 | 0.42 | 0.41 |
| 7 | 21932317 | rs9639401 | 0.837 | -0.585 | -0.808 | 0.677 | 0.009 | -0.005 | -0.005 | 1.521 | 0.31 | 0.26 | 0.15 | 0.4 | 0.21 | 0.37 |
| 7 | 21932876 | rs12533084 | 2.157 | -0.576 | -0.985 | 0.725 | 0.03 | -0.071 | -0.071 | 0 | 0.2 | 0.04 | 0.11 | 0.35 | 0.14 | 0.25 |
| 7 | 21933483 | rs9639403 | 0.761 | -0.163 | -0.702 | 0.307 | -0.001 | -0.012 | -0.012 | 0 | 0.09 | 0.1 | 0.03 | 0.03 | 0.08 | 0.1 |
| 7 | 21935986 | rs1139225 | -0.924 | -0.05 | -0.477 | 0.318 | 0.022 | -0.004 | -0.004 | 0 | 0.07 | 0.22 | 0.06 | 0.03 | 0.07 | 0.07 |
| 7 | 21937988 | rs4636103 | 0.914 | -0.232 | -1.068 | 0.111 | -0.025 | 0.115 | 0.115 | 0.446 | 0.39 | 0.19 | 0.32 | 0.19 | 0.4 | 0.3 |
| 7 | 21939382 | rs35469450 | 0.608 | -0.053 | -1.051 | 0.231 | -0.054 | -0.001 | -0.001 | 0 | 0.13 | 0.06 | 0.18 | 0.1 | 0.29 | 0.16 |
| 7 | 21940960 | rs7971 | 1.025 | -0.171 | -1.019 | 0.187 | -0.049 | 0.111 | 0.111 | 0.359 | 0.35 | 0.09 | 0.3 | 0.17 | 0.36 | 0.23 |
| 7 | 21941669 | rs1128226 | 0.641 | -0.652 | -1.165 | -0.274 | -0.012 | 0.027 | 0.027 | 0 | 0.33 | 0.27 | 0.31 | 0.18 | 0.39 | 0.24 |
| 7 | 21945563 | rs12540021 | 1.176 | -0.031 | -1.438 | -0.013 | -0.023 | 0.078 | 0.078 | 0.251 | 0.31 | 0.08 | 0.25 | 0.15 | 0.28 | 0.22 |
| 7 | 21945687 | rs6461617 | 0.159 | -0.126 | -1.284 | -0.151 | -0.017 | 0.112 | 0.112 | 0 | 0.46 | 0.23 | 0.37 | 0.21 | 0.43 | 0.35 |
| 7 | 21945840 | rs4445125 | -1.373 | 0.199 | -1.055 | 0.027 | -0.016 | 0.012 | 0.012 | 0.262 | 0.35 | 0.3 | 0.31 | 0.24 | 0.49 | 0.4 |
| 7 | 21947138 | rs10238945 | -0.002 | 0.022 | -1.272 | -0.013 | -0.013 | 0.034 | 0.034 | 0.293 | 0.13 | 0.06 | 0.1 | 0.05 | 0.13 | 0.12 |
| 7 | 21949303 | rs12234262 | 0.096 | -0.353 | -1.378 | 0.279 | 0.032 | 0.042 | 0.042 | 0.778 | 0.52 | 0.26 | 0.39 | 0.46 | 0.6 | 0.6 |
| 7 | 21949886 | rs7811563 | 0.064 | -0.65 | -1.398 | 0.253 | 0.119 | 0.08 | 0.08 | 0.277 | 0.48 | 0.84 | 0.55 | 0.68 | 0.32 | 0.52 |
| 7 | 21951033 | rs4346898 | 1.029 | -0.335 | -1.439 | 0.064 | -0.116 | 0.048 | 0.048 | 0 | 0.33 | 0.07 | 0.26 | 0.21 | 0.41 | 0.29 |
| 7 | 21951105 | rs11978167 | 0.182 | -0.735 | -0.492 | 0.088 | 0.031 | -0.059 | -0.059 | 0 | 0.18 | 0.32 | 0.14 | 0.07 | 0.24 | 0.18 |
| 7 | 21952954 | rs6958427 | -0.647 | -0.858 | -0.867 | 0.338 | 0.005 | 0.001 | 0.001 | 0 | 0.91 | 0.92 | 0.97 | 0.95 | 0.95 | 0.91 |
| 7 | 21955785 | rs12700327 | 0.2 | -0.927 | -0.865 | 0.602 | 0.013 | 0.011 | 0.011 | 0 | 0.45 | 0.2 | 0.54 | 0.67 | 0.41 | 0.55 |
| 7 | 21956405 | rs1128250 | 1.366 | -1.039 | -0.813 | 0.697 | 0.066 | -0.1 | -0.1 | 0.501 | 0.28 | 0.11 | 0.42 | 0.58 | 0.18 | 0.36 |
| 7 | 21964362 | rs1406705 | 0.227 | -0.703 | -0.45 | 0.541 | -0.002 | -0.002 | -0.002 | 0 | 0.11 | 0.13 | 0.05 | 0.06 | 0.07 | 0.11 |
| 7 | 21979144 | rs7807075 | -0.267 | -0.728 | -0.42 | 0.65 | -0.004 | -0.001 | -0.001 | 0.546 | 0.54 | 0.44 | 0.69 | 0.49 | 0.59 | 0.56 |
| 7 | 21980773 | rs6946675 | 1.798 | -0.715 | -0.746 | 0.702 | 0.029 | -0.022 | -0.022 | 0 | 0.24 | 0.14 | 0.12 | 0.36 | 0.15 | 0.24 |
| 7 | 21981938 | rs12700337 | -1.825 | -0.792 | -0.781 | 0.72 | 0.012 | -0.015 | -0.015 | 0 | 0.75 | 0.78 | 0.88 | 0.54 | 0.86 | 0.76 |
| 7 | 21982300 | rs13226917 | 0.594 | -0.721 | -0.8 | 0.638 | -0.061 | 0.059 | 0.059 | 0.62 | 0.17 | 0.06 | 0.15 | 0.06 | 0.26 | 0.19 |
| 7 | 21982448 | rs10262937 | -1.76 | -0.813 | -0.772 | 0.742 | 0.012 | -0.014 | -0.014 | 0 | 0.76 | 0.8 | 0.89 | 0.56 | 0.87 | 0.76 |
| 7 | 21987884 | rs1534771 | 0.486 | -0.429 | -1.03 | 0.636 | -0.068 | 0.066 | 0.066 | 0 | 0.39 | 0.16 | 0.34 | 0.18 | 0.46 | 0.32 |
| 7 | 21990232 | rs1637075 | -0.028 | -0.198 | -0.648 | 0.8 | 0.004 | 0.002 | 0.002 | 0 | 0.5 | 0.48 | 0.31 | 0.5 | 0.46 | 0.49 |
| 7 | 21993660 | rs1316623 | -0.606 | 0.321 | -1.497 | 0.907 | -0.047 | -0.036 | -0.036 | 1.629 | 0.22 | 0.1 | 0.22 | 0.25 | 0.3 | 0.18 |
| 7 | 21993767 | rs2893061 | -0.044 | -0.412 | -1.4 | 0.853 | 0.013 | -0.027 | -0.027 | 0 | 0.06 | 0.03 | 0.02 | 0.08 | 0.04 | 0.03 |
| 7 | 21994453 | rs2699460 | 0.553 | -0.966 | -1.308 | 0.795 | 0.017 | -0.016 | -0.016 | 0 | 0.35 | 0.64 | 0.49 | 0.61 | 0.25 | 0.36 |
| 7 | 21995315 | rs4722071 | 0.104 | -0.111 | -1.633 | 0.232 | -0.045 | 0.04 | 0.04 | 0 | 0.23 | 0.13 | 0.21 | 0.08 | 0.36 | 0.12 |
| 7 | 22000425 | rs6461625 | -0.074 | 0.016 | -1.577 | 0.518 | -0.007 | -0.01 | -0.01 | 0 | 0.55 | 0.63 | 0.52 | 0.44 | 0.59 | 0.39 |
| 7 | 22001646 | rs1174939 | -0.745 | 0.09 | -1.457 | 0.104 | -0.002 | 0.022 | 0.022 | 0.233 | 0.35 | 0.7 | 0.29 | 0.08 | 0.39 | 0.16 |
| 7 | 22007118 | rs1174943 | -0.576 | 0.174 | -1.593 | 0.603 | -0.019 | 0.002 | 0.002 | 0 | 0.83 | 0.97 | 0.89 | 0.88 | 0.84 | 0.77 |
| 7 | 22010179 | rs4439015 | 0.124 | 0.276 | -1.718 | 1.023 | -0.006 | -0.06 | -0.06 | 0 | 0.11 | 0.03 | 0.18 | 0.24 | 0.12 | 0.11 |
| 7 | 22011292 | rs4722077 | 1.186 | 0.017 | -1.7 | 0.712 | -0.039 | 0.067 | 0.067 | 0.227 | 0.42 | 0.23 | 0.29 | 0.29 | 0.51 | 0.31 |
| 7 | 22018421 | rs11761561 | 1.154 | 0.193 | -1.562 | 0.829 | 0.013 | 0.092 | 0.092 | 0.741 | 0.5 | 0.17 | 0.35 | 0.32 | 0.57 | 0.5 |
| 7 | 22020039 | rs2527698 | 0.398 | 0.074 | -0.665 | 1.019 | -0.001 | -0.001 | -0.001 | 0 | 0.65 | 0.71 | 0.61 | 0.6 | 0.68 | 0.63 |
| 7 | 22020691 | rs7778988 | 1.236 | 0.168 | -0.459 | 0.797 | 0.012 | 0.018 | 0.018 | 0.662 | 0.57 | 0.3 | 0.54 | 0.6 | 0.62 | 0.46 |
| 7 | 22021924 | rs1177792 | -1.222 | 0.112 | -0.271 | 0.949 | 0 | -0.004 | -0.004 | 0 | 0.23 | 0.31 | 0.27 | 0.23 | 0.24 | 0.2 |
| 7 | 22024040 | rs1175000 | 1.482 | 0.217 | -0.417 | 0.962 | -0.013 | 0.004 | 0.004 | 1.313 | 0.5 | 0.22 | 0.32 | 0.47 | 0.47 | 0.5 |
| 7 | 22025485 | rs1174998 | -0.463 | 0.201 | -0.356 | 0.943 | 0.002 | -0.026 | -0.026 | 0.039 | 0.06 | 0.18 | 0.06 | 0.01 | 0.07 | 0.2 |
| 7 | 22025917 | rs1174995 | -0.168 | 0.252 | -0.227 | 0.988 | -0.004 | 0.011 | 0.011 | 0 | 0.73 | 0.75 | 0.7 | 0.61 | 0.79 | 0.71 |
| 7 | 22026842 | rs1174992 | 0.077 | 0.158 | -0.09 | 0.71 | -0.001 | -0.002 | -0.002 | 0 | 0.71 | 0.69 | 0.7 | 0.66 | 0.76 | 0.71 |
| 7 | 22027066 | rs1174991 | -0.413 | 0.154 | -0.103 | 0.71 | -0.001 | 0 | 0 | 0 | 0.75 | 0.71 | 0.75 | 0.75 | 0.79 | 0.76 |
| 7 | 22027212 | rs7783529 | 2.462 | 0.107 | -0.156 | 0.757 | -0.088 | -0.092 | -0.092 | 0 | 0.38 | 0.03 | 0.23 | 0.38 | 0.39 | 0.26 |
| 7 | 22027905 | rs12700339 | 1.024 | 0.293 | 0.071 | 0.237 | -0.005 | 0.059 | 0.059 | 0.377 | 0.58 | 0.21 | 0.57 | 0.66 | 0.61 | 0.48 |
| 7 | 22028717 | rs11976853 | 0.657 | 0.376 | 0.118 | 0.314 | 0.008 | -0.028 | -0.028 | 0.001 | 0.05 | 0.14 | 0.06 | 0 | 0.08 | 0.05 |
| 7 | 22029482 | rs2107973 | 0.978 | 0.467 | -0.026 | 0.465 | 0.002 | -0.009 | -0.009 | 0.01 | 0.54 | 0.57 | 0.47 | 0.61 | 0.54 | 0.56 |
| 7 | 22033912 | rs1730906 | 0.175 | 0.459 | -0.309 | 0.493 | -0.001 | 0.001 | 0.001 | 0.001 | 0.67 | 0.75 | 0.65 | 0.66 | 0.69 | 0.52 |
| 7 | 22037376 | rs1174954 | -0.945 | 0.471 | -0.313 | 0.538 | 0.006 | -0.039 | -0.039 | 0 | 0.18 | 0.31 | 0.24 | 0.06 | 0.22 | 0.15 |
| 7 | 22042280 | rs1636296 | -0.865 | 0.08 | -0.83 | 1.037 | 0.015 | -0.022 | -0.022 | 0 | 0.46 | 0.7 | 0.54 | 0.53 | 0.44 | 0.45 |
| 7 | 22042482 | rs1174957 | -0.203 | -0.067 | -0.324 | 1.104 | -0.006 | -0.001 | -0.001 | 0 | 0.19 | 0.27 | 0.23 | 0.16 | 0.19 | 0.19 |
| 7 | 22042832 | rs1174959 | -0.374 | -0.162 | -0.329 | 0.922 | -0.022 | 0.069 | 0.069 | 0 | 0.79 | 0.93 | 0.87 | 0.9 | 0.78 | 0.87 |
| 7 | 22043000 | rs10242914 | -0.501 | -0.22 | -0.397 | 0.939 | 0.009 | 0.036 | 0.036 | 0 | 0.07 | 0.23 | 0.15 | 0.12 | 0.06 | 0.16 |
| 7 | 22044116 | rs1174961 | 0.282 | -0.137 | -1.083 | 0.9 | -0.024 | 0.07 | 0.07 | 0 | 0.21 | 0.07 | 0.13 | 0.09 | 0.22 | 0.13 |
| 7 | 22045194 | rs1174963 | 0.229 | -0.303 | -0.987 | 1.134 | -0.002 | -0.024 | -0.024 | 0 | 0.21 | 0.1 | 0.23 | 0.28 | 0.19 | 0.26 |
| 7 | 22045272 | rs1174964 | 0.2 | -0.306 | -0.967 | 1.132 | -0.002 | -0.02 | -0.02 | 2.932 | 0.21 | 0.11 | 0.22 | 0.28 | 0.19 | 0.26 |
| 7 | 22045395 | rs12056027 | 1.639 | -0.384 | -1.148 | 1.139 | -0.021 | 0.098 | 0.098 | 2.046 | 0.5 | 0.06 | 0.37 | 0.3 | 0.55 | 0.47 |
| 7 | 22045685 | rs1730886 | 1.158 | -0.25 | -0.76 | 1.181 | 0.003 | -0.005 | -0.005 | 0 | 0.6 | 0.63 | 0.55 | 0.45 | 0.62 | 0.56 |

| | | | | | | | | | | | | | | | | |
|---|---|---|---|---|---|---|---|---|---|---|---|---|---|---|---|---|
| 7 | 22046015 | rs1174965 | 0.026 | 0.232 | -1.208 | 0.525 | -0.013 | 0.064 | 0.064 | 0 | 0.8 | 0.94 | 0.86 | 0.9 | 0.81 | 0.88 |
| 7 | 22046923 | rs6961921 | -0.393 | -0.393 | -0.824 | 0.744 | -0.006 | -0.001 | -0.001 | 0 | 0.33 | 0.34 | 0.32 | 0.25 | 0.27 | 0.3 |
| 7 | 22047162 | rs7795246 | 0.04 | 0.127 | -1.203 | 0.369 | -0.037 | 0.132 | 0.132 | 0 | 0.43 | 0.22 | 0.33 | 0.17 | 0.54 | 0.37 |
| 7 | 22047694 | rs7782821 | -0.6 | 0.027 | -0.392 | 1.285 | 0.006 | 0.134 | 0.134 | 0 | 0.06 | 0.2 | 0.08 | 0.23 | 0.05 | 0.22 |
| 7 | 22050169 | rs2699463 | 0.295 | -0.786 | -0.492 | 0.606 | -0.052 | -0.02 | -0.02 | 0 | 0.69 | 0.59 | 0.81 | 0.78 | 0.86 | 0.78 |
| 7 | 22051466 | rs6970233 | -0.468 | 0.038 | -1.146 | 0.611 | -0.023 | -0.03 | -0.03 | 0 | 0.32 | 0.76 | 0.3 | 0.38 | 0.37 | 0.29 |
| 7 | 22053732 | rs1637053 | -0.922 | 0.104 | -1.678 | 0.188 | -0.112 | -0.044 | -0.044 | 0 | 0.59 | 0.92 | 0.6 | 0.65 | 0.53 | 0.64 |
| 7 | 22057577 | rs1730913 | 0.125 | 0.412 | -0.131 | -0.174 | 0.046 | -0.022 | -0.022 | 0 | 0.77 | 0.54 | 0.66 | 0.84 | 0.69 | 0.87 |
| 7 | 22059791 | rs6947288 | 0.314 | 0.369 | 0.148 | -0.851 | 0.001 | -0.013 | -0.013 | 0 | 0.62 | 0.58 | 0.45 | 0.73 | 0.64 | 0.6 |
| 7 | 22072762 | rs1174981 | -0.174 | 0.273 | 0.25 | -1.07 | -0.01 | -0.009 | -0.009 | 0.939 | 0.22 | 0.35 | 0.26 | 0.24 | 0.16 | 0.3 |
| 7 | 22073513 | rs3857714 | -0.139 | 0.435 | 0.038 | -1.241 | -0.02 | 0.038 | 0.038 | 0 | 0.07 | 0.01 | 0.04 | 0 | 0.08 | 0.03 |
| 7 | 22074023 | rs1636319 | -1.461 | 0.197 | 0.474 | -1.476 | 0.003 | 0.051 | 0.051 | 0 | 0.52 | 0.64 | 0.49 | 0.27 | 0.43 | 0.46 |
| 7 | 22078387 | rs1174950 | -0.377 | 0.151 | 0.813 | -1.48 | -0.029 | -0.031 | -0.031 | 0 | 0.26 | 0.43 | 0.27 | 0.24 | 0.17 | 0.31 |
| 7 | 22079598 | rs1174952 | 0.839 | 0.111 | 0.73 | -1.489 | -0.004 | 0.01 | 0.01 | 0 | 0.17 | 0.17 | 0.17 | 0.03 | 0.18 | 0.12 |
| 7 | 22081348 | rs1174953 | -0.336 | 0.197 | 0.808 | -1.502 | -0.015 | -0.023 | -0.023 | 4.403 | 0.26 | 0.37 | 0.29 | 0.24 | 0.18 | 0.32 |
| 7 | 22082155 | rs1636327 | -1.212 | 0.255 | 0.676 | -1.501 | -0.024 | 0.056 | 0.056 | 0 | 0.84 | 0.93 | 0.83 | 0.97 | 0.81 | 0.88 |
| 7 | 22083233 | rs1619250 | 0.596 | 0.209 | 0.784 | -1.492 | -0.004 | -0.104 | -0.104 | 0 | 0.71 | 0.56 | 0.76 | 0.97 | 0.73 | 0.85 |
| 7 | 22086004 | rs952654 | 1.147 | 0.168 | 0.562 | -1.425 | 0 | 0.006 | 0.006 | 0 | 0.49 | 0.43 | 0.51 | 0.74 | 0.56 | 0.55 |
| 7 | 22088748 | rs1730932 | -0.579 | 0.346 | 0.531 | -1.287 | -0.007 | -0.034 | -0.034 | 0.161 | 0.27 | 0.45 | 0.29 | 0.22 | 0.2 | 0.32 |
| 7 | 22090928 | rs1637081 | -1.008 | 0.374 | 0.233 | -1.341 | 0 | 0 | 0 | 0 | 0.32 | 0.3 | 0.31 | 0.23 | 0.26 | 0.33 |
| 7 | 22092706 | rs10499562 | 0.977 | 0.471 | 0.2 | -1.473 | -0.001 | -0.005 | -0.005 | 0 | 0.53 | 0.52 | 0.51 | 0.76 | 0.55 | 0.56 |
| 7 | 22095150 | rs2528907 | -0.113 | 0.535 | -0.421 | -1.144 | -0.038 | 0.061 | 0.061 | 0 | 0.81 | 0.98 | 0.88 | 0.91 | 0.81 | 0.72 |
| 7 | 22096915 | rs10260796 | -1.024 | 0.145 | 0.205 | -1.12 | -0.015 | -0.056 | -0.056 | 0 | 0.1 | 0.28 | 0.05 | 0 | 0.06 | 0.03 |
| 7 | 22097449 | rs2699453 | -0.239 | 0.489 | -0.364 | -1.155 | -0.014 | 0.056 | 0.056 | 2.067 | 0.83 | 0.97 | 0.91 | 0.91 | 0.86 | 0.73 |
| 7 | 22097476 | rs1637078 | -0.354 | -0.068 | -0.101 | -1.219 | -0.023 | 0.003 | 0.003 | 0 | 0.14 | 0.28 | 0.19 | 0.16 | 0.06 | 0.12 |
| 7 | 22098259 | rs1833111 | -0.106 | 0.218 | -1.224 | -1.803 | -0.094 | 0.188 | 0.188 | 1.02 | 0.32 | 0.07 | 0.25 | 0.02 | 0.37 | 0.24 |
| 7 | 22100234 | rs1730900 | -1.19 | 0.546 | -0.737 | -1.036 | 0.004 | 0.008 | 0.008 | 0 | 0.78 | 0.78 | 0.64 | 0.71 | 0.72 | 0.82 |
| 7 | 22101207 | rs10234533 | -0.686 | 0.195 | -1.111 | -0.931 | -0.025 | 0.052 | 0.052 | 0 | 0.45 | 0.27 | 0.4 | 0.61 | 0.45 | 0.6 |
| 7 | 22104664 | rs4722082 | -0.158 | 0.308 | -0.178 | -0.287 | 0.064 | 0.165 | 0.165 | 0.096 | 0.16 | 0.38 | 0.24 | 0.55 | 0.24 | 0.22 |
| 7 | 22105864 | rs13223060 | 0.435 | -0.856 | -0.623 | -1.064 | 0.017 | 0.012 | 0.012 | 1.469 | 0.31 | 0.2 | 0.1 | 0.07 | 0.21 | 0.22 |
| 7 | 22107275 | rs12671298 | 0.14 | -1.12 | -0.289 | -0.439 | -0.018 | -0.006 | -0.006 | 0.583 | 0.18 | 0.29 | 0.13 | 0.18 | 0.09 | 0.2 |
| 7 | 22107371 | rs12671730 | 0.195 | -1.108 | -0.249 | -0.434 | -0.017 | -0.005 | -0.005 | 0.252 | 0.17 | 0.27 | 0.13 | 0.18 | 0.09 | 0.2 |
| 7 | 22108059 | rs10234682 | -1.75 | -1.203 | -1.039 | -0.908 | -0.187 | 0.159 | 0.159 | 0.153 | 0.72 | 0.97 | 0.63 | 0.94 | 0.55 | 0.65 |
| 7 | 22109436 | rs7784234 | 0.163 | -1.239 | -0.437 | -0.522 | -0.017 | -0.009 | -0.009 | 0 | 0.12 | 0.22 | 0.17 | 0.09 | 0.09 | 0.06 |
| 7 | 22109459 | rs10499559 | 1.819 | -1.09 | -0.323 | 0.08 | -0.004 | 0.05 | 0.05 | 0 | 0.08 | 0.05 | 0.07 | 0.33 | 0.15 | 0.1 |
| 7 | 22110092 | rs2214937 | 0.81 | -1.386 | -0.088 | -0.127 | 0.007 | -0.007 | -0.007 | 0.024 | 0.33 | 0.27 | 0.1 | 0.07 | 0.22 | 0.24 |
| 7 | 22112822 | rs17146140 | 1.308 | -0.852 | -0.014 | 0.286 | 0.023 | 0.119 | 0.119 | 0 | 0.09 | 0.13 | 0.08 | 0.33 | 0.15 | 0.1 |
| 7 | 22113119 | rs17146144 | 0.607 | -0.89 | -0.061 | 0.339 | 0.011 | 0.066 | 0.066 | 0 | 0.11 | 0.14 | 0.14 | 0.57 | 0.18 | 0.17 |
| 7 | 22113206 | rs6949977 | -0.095 | -0.408 | -0.169 | -0.029 | -0.003 | 0.001 | 0.001 | 0 | 0.29 | 0.36 | 0.44 | 0.29 | 0.26 | 0.26 |
| 7 | 22115454 | rs12539524 | 1.33 | -0.079 | -0.544 | 0.111 | 0.006 | 0.07 | 0.07 | 0 | 0.08 | 0.07 | 0.08 | 0.34 | 0.15 | 0.1 |
| 7 | 22116881 | rs7793513 | -0.157 | -0.9 | -0.562 | -0.491 | 0.001 | 0.001 | 0.001 | 0 | 0.46 | 0.54 | 0.59 | 0.55 | 0.49 | 0.37 |
| 7 | 22118542 | rs4722086 | -0.799 | -0.947 | -0.566 | -0.605 | 0.016 | 0.024 | 0.024 | 0 | 0.54 | 0.66 | 0.68 | 0.88 | 0.64 | 0.48 |
| 7 | 22121378 | rs12154589 | 1.062 | -0.946 | -0.636 | -0.645 | -0.005 | 0.013 | 0.013 | 0 | 0.18 | 0.15 | 0.22 | 0.1 | 0.24 | 0.11 |
| 7 | 22122151 | rs10271131 | 0.901 | -1.408 | -0.678 | -0.855 | -0.09 | -0.086 | -0.086 | 0 | 0.23 | 0.35 | 0.04 | 0.05 | 0.07 | 0.19 |
| 7 | 22122507 | rs7787246 | -2.057 | -1.199 | -0.76 | -0.825 | -0.102 | 0.118 | 0.118 | 0.504 | 0.77 | 0.98 | 0.72 | 0.93 | 0.68 | 0.69 |
| 7 | 22122563 | rs7787802 | -0.464 | -1.065 | -0.664 | -0.676 | 0.001 | 0.002 | 0.002 | 0 | 0.5 | 0.52 | 0.57 | 0.56 | 0.5 | 0.5 |
| 7 | 22127043 | rs4577865 | 0.536 | -1.209 | -0.982 | -0.675 | 0.001 | 0.007 | 0.007 | 0 | 0.27 | 0.27 | 0.22 | 0.21 | 0.23 | 0.33 |
| 7 | 22128464 | rs4722088 | -0.239 | -1.287 | -0.955 | -0.606 | 0.023 | -0.01 | -0.01 | 0 | 0.44 | 0.62 | 0.26 | 0.44 | 0.4 | 0.23 |
| 7 | 22128523 | rs4722089 | -0.797 | -1.26 | -1.081 | -0.635 | 0.045 | 0.114 | 0.114 | 0.481 | 0.49 | 0.7 | 0.42 | 0.7 | 0.45 | 0.28 |
| 7 | 22129599 | rs10267816 | 0.892 | -0.898 | -1.213 | -0.369 | -0.002 | 0.028 | 0.028 | 0 | 0.05 | 0.03 | 0.16 | 0.27 | 0.04 | 0.07 |
| 7 | 22131326 | rs1815976 | 1.194 | -0.799 | -1.181 | -0.66 | -0.028 | 0.016 | 0.016 | 0.267 | 0.21 | 0.06 | 0.19 | 0.18 | 0.2 | 0.19 |
| 7 | 22131772 | rs4722093 | 1.042 | -0.369 | -0.634 | -0.643 | 0.002 | -0.037 | -0.037 | 0 | 0.17 | 0.27 | 0.13 | 0.08 | 0.22 | 0.07 |
| 7 | 22133391 | rs1419761 | 1.375 | -0.325 | -0.591 | -0.602 | 0.003 | -0.027 | -0.027 | 0 | 0.15 | 0.22 | 0.12 | 0.08 | 0.21 | 0.07 |
| 7 | 22133922 | rs10269328 | -1.531 | -0.324 | -0.783 | -0.584 | 0 | 0.007 | 0.007 | 0 | 0.85 | 0.88 | 0.88 | 0.91 | 0.79 | 0.93 |
| 7 | 22134194 | rs4722096 | 0.444 | -0.506 | -1.08 | -0.744 | -0.013 | 0.148 | 0.148 | 0 | 0.41 | 0.21 | 0.41 | 0.11 | 0.39 | 0.55 |
| 7 | 22138141 | rs10264093 | 0.779 | 0.032 | -0.704 | 0.039 | 0.01 | 0.062 | 0.062 | 0 | 0.09 | 0.11 | 0.08 | 0.33 | 0.15 | 0.1 |
| 7 | 22140620 | rs1859800 | -0.682 | -0.044 | -0.871 | -0.357 | -0.023 | 0.083 | 0.083 | 0 | 0.6 | 0.76 | 0.6 | 0.85 | 0.49 | 0.61 |
| 7 | 22141711 | rs3915972 | -1.823 | 0.322 | -1.076 | -0.431 | -0.045 | 0.101 | 0.101 | 0.226 | 0.79 | 0.98 | 0.75 | 0.94 | 0.76 | 0.73 |
| 7 | 22142227 | rs13233663 | 1.317 | 0.35 | -0.91 | -0.38 | -0.003 | -0.013 | -0.013 | 0 | 0.17 | 0.18 | 0.13 | 0.08 | 0.2 | 0.1 |
| 7 | 22142742 | rs10225710 | 0.032 | 0.446 | -0.698 | -0.186 | 0 | 0 | 0 | 0 | 0.48 | 0.52 | 0.44 | 0.51 | 0.4 | 0.43 |
| 7 | 22145084 | rs16873129 | 0.292 | 0.668 | -0.76 | 0.299 | -0.01 | -0.012 | -0.012 | 0.342 | 0.07 | 0.06 | 0.03 | 0.16 | 0.08 | 0.18 |
| 7 | 22145115 | rs1054766 | -0.296 | 0.616 | -0.628 | 0.152 | 0.001 | 0.044 | 0.044 | 0.129 | 0.3 | 0.4 | 0.33 | 0.44 | 0.36 | 0.28 |
| 7 | 22146840 | rs7793103 | 0.303 | 0.689 | -0.61 | -0.342 | -0.011 | 0.028 | 0.028 | 2.316 | 0.25 | 0.36 | 0.16 | 0.59 | 0.2 | 0.27 |
| 7 | 22147562 | rs7798144 | -0.73 | 0.732 | -1.007 | -0.685 | -0.003 | -0.054 | -0.054 | 0 | 0.6 | 0.56 | 0.51 | 0.85 | 0.53 | 0.65 |
| 7 | 22147735 | rs10950889 | -0.33 | 0.951 | -0.386 | -0.543 | 0.014 | -0.021 | -0.021 | 0 | 0.37 | 0.48 | 0.37 | 0.26 | 0.33 | 0.39 |
| 7 | 22148016 | rs11760455 | -0.399 | 0.977 | -0.562 | -0.533 | 0.003 | -0.006 | -0.006 | 1.532 | 0.38 | 0.56 | 0.38 | 0.26 | 0.33 | 0.39 |
| 7 | 22148704 | rs3829757 | 0.089 | 0.946 | -0.849 | -0.528 | 0.004 | -0.016 | -0.016 | 0 | 0.26 | 0.28 | 0.32 | 0.11 | 0.25 | 0.18 |
| 7 | 22150888 | rs4722097 | 0.223 | 1.227 | -1.505 | -0.37 | -0.015 | 0.091 | 0.091 | 0.027 | 0.29 | 0.14 | 0.4 | 0.06 | 0.34 | 0.33 |
| 7 | 22151968 | rs4598147 | 0.252 | 1.514 | -1.061 | 0.676 | -0.013 | -0.016 | -0.016 | 2.144 | 0.66 | 0.71 | 0.71 | 0.37 | 0.57 | 0.71 |

| | | | | | | | | | | | | | | | | |
|---|---|---|---|---|---|---|---|---|---|---|---|---|---|---|---|---|
| 7 | 22154454 | rs13227860 | -0.416 | 1.861 | -0.834 | 0.841 | 0.005 | 0 | 0 | 0.653 | 0.14 | 0.17 | 0.25 | 0.22 | 0.3 | 0.07 |
| 7 | 22155376 | rs17146234 | -0.246 | 0.913 | -0.924 | 0.62 | 0.009 | 0.005 | 0.005 | 0.143 | 0.37 | 0.54 | 0.27 | 0.36 | 0.31 | 0.41 |
| 7 | 22157423 | rs1859801 | -0.414 | 1.168 | -0.996 | 0.667 | -0.003 | -0.002 | -0.002 | 0 | 0.44 | 0.57 | 0.49 | 0.51 | 0.55 | 0.35 |
| 7 | 22157782 | rs17707499 | -0.167 | 1.384 | -1.86 | 0.335 | -0.072 | 0.038 | 0.038 | 3.702 | 0.37 | 0.09 | 0.35 | 0.28 | 0.48 | 0.34 |
| 7 | 22158386 | rs7301 | -0.834 | 0.601 | -0.764 | 0.606 | -0.043 | 0.003 | 0.003 | 0 | 0.27 | 0.49 | 0.19 | 0.29 | 0.2 | 0.26 |
| 7 | 22158977 | rs3779069 | 0.189 | 0.971 | -2.2 | 0.531 | -0.018 | 0.057 | 0.057 | 1.123 | 0.52 | 0.08 | 0.43 | 0.63 | 0.47 | 0.59 |
| 7 | 22159182 | rs3779067 | 0.101 | 0.829 | -1.634 | 0.805 | 0.02 | 0.013 | 0.013 | 0.096 | 0.54 | 0.24 | 0.43 | 0.58 | 0.42 | 0.58 |
| 7 | 22162492 | rs10807809 | -0.085 | 0.766 | -0.72 | 0.733 | 0.009 | -0.013 | -0.013 | 0 | 0.58 | 0.45 | 0.5 | 0.7 | 0.51 | 0.76 |
| 7 | 22163039 | rs4722100 | -0.66 | 1.15 | -0.269 | 0.449 | 0.012 | -0.081 | -0.081 | 0 | 0.68 | 0.58 | 0.62 | 0.92 | 0.63 | 0.85 |
| 7 | 22163907 | rs10950892 | -0.374 | 0.516 | 0.065 | 1.161 | -0.003 | 0.055 | 0.055 | 0 | 0.09 | 0.15 | 0.18 | 0.28 | 0.11 | 0.25 |
| 7 | 22164266 | rs1859802 | 0.859 | 0.796 | 0.36 | 0.675 | -0.005 | 0 | 0 | 0 | 0.42 | 0.38 | 0.46 | 0.58 | 0.42 | 0.52 |
| 7 | 22164444 | rs1859803 | -0.791 | 0.53 | 0.995 | 0.638 | -0.014 | -0.065 | -0.065 | 0.433 | 0.67 | 0.55 | 0.66 | 0.89 | 0.7 | 0.83 |
| 7 | 22165180 | rs2302000 | -0.044 | 0.517 | 0.394 | 1.009 | -0.005 | 0.008 | 0.008 | 0.118 | 0.08 | 0.03 | 0.06 | 0.32 | 0.09 | 0.26 |
| 7 | 22165421 | rs17146285 | 1.682 | 0.355 | 0.414 | 0.635 | 0.004 | 0.03 | 0.03 | 0 | 0.23 | 0.16 | 0.29 | 0.11 | 0.22 | 0.14 |
| 7 | 22169078 | rs1859804 | 1.651 | 0.433 | 0.759 | 0.784 | -0.005 | 0.022 | 0.022 | 0.048 | 0.76 | 0.36 | 0.78 | 0.65 | 0.8 | 0.62 |
| 7 | 22171582 | rs6461638 | -0.319 | 0.401 | 0.833 | 0.65 | 0.003 | -0.018 | -0.018 | 0 | 0.58 | 0.38 | 0.53 | 0.9 | 0.6 | 0.75 |
| 7 | 22171815 | rs3807535 | 0.574 | 0.454 | 0.419 | 0.763 | -0.059 | -0.059 | -0.059 | 0 | 0.44 | 0.1 | 0.43 | 0.57 | 0.51 | 0.48 |
| 7 | 22173634 | rs3823665 | -0.889 | 1.124 | 0.777 | 1.149 | -0.001 | -0.015 | -0.015 | 0 | 0.14 | 0.1 | 0.09 | 0.36 | 0.14 | 0.28 |
| 7 | 22175472 | rs3901281 | 0.085 | 1.248 | 0.532 | 1.014 | -0.041 | -0.016 | -0.016 | 0 | 0.27 | 0.14 | 0.31 | 0.38 | 0.39 | 0.32 |
| 7 | 22177012 | rs4722101 | -1.156 | 1.173 | 0.584 | 0.871 | -0.011 | 0.038 | 0.038 | 2.133 | 0.43 | 0.69 | 0.44 | 0.41 | 0.52 | 0.5 |
| 7 | 22184018 | rs1859805 | -0.409 | 1.301 | 1.014 | 0.95 | 0.032 | -0.017 | -0.017 | 0.334 | 0.35 | 0.39 | 0.4 | 0.37 | 0.47 | 0.51 |
| 7 | 22184259 | rs883073 | 0.935 | 1.387 | 1.075 | 0.925 | 0.022 | -0.016 | -0.016 | 0 | 0.08 | 0.13 | 0.11 | 0.08 | 0.11 | 0.14 |
| 7 | 22187744 | rs3807531 | 0.422 | 1.371 | 1.037 | 0.957 | -0.024 | -0.003 | -0.003 | 0.194 | 0.26 | 0.14 | 0.27 | 0.31 | 0.34 | 0.33 |
| 7 | 22191936 | rs10276439 | -0.203 | 0.917 | 1.349 | 0.857 | 0.089 | -0.028 | -0.028 | 0.193 | 0.11 | 0.4 | 0.13 | 0.09 | 0.12 | 0.14 |
| 7 | 22196981 | rs1419756 | 0.921 | 0.756 | 1.065 | 1.017 | -0.019 | 0.08 | 0.08 | 0.726 | 0.77 | 0.67 | 0.85 | 0.6 | 0.82 | 0.71 |
| 7 | 22197725 | rs6959894 | 0.739 | 0.721 | 0.444 | 0.982 | 0.022 | -0.014 | -0.014 | 0 | 0.08 | 0.13 | 0.12 | 0.09 | 0.12 | 0.16 |
| 7 | 22202117 | rs12592 | -1.305 | 0.693 | 0.412 | 1.116 | 0.031 | 0.054 | 0.054 | 1.594 | 0.51 | 0.68 | 0.44 | 0.61 | 0.56 | 0.61 |
| 7 | 22202485 | rs1981601 | -1.351 | 0.703 | 0.293 | 1.048 | 0.029 | 0.087 | 0.087 | 1.768 | 0.51 | 0.7 | 0.44 | 0.66 | 0.56 | 0.62 |
| 7 | 22214726 | rs17146355 | 0.435 | 0.767 | -0.345 | 1.31 | 0.007 | -0.028 | -0.028 | 0 | 0.1 | 0.02 | 0.09 | 0.29 | 0.06 | 0.16 |
| 7 | 22216054 | rs7786882 | -0.432 | 0.93 | -0.197 | 0.731 | -0.05 | 0.026 | 0.026 | 1.61 | 0.35 | 0.22 | 0.31 | 0.32 | 0.48 | 0.4 |
| 7 | 22230393 | rs4722108 | -0.234 | 0.909 | -0.384 | 0.6 | -0.012 | 0.034 | 0.034 | 0 | 0.91 | 0.97 | 0.96 | 0.99 | 0.89 | 0.93 |
| 7 | 22233813 | rs3813383 | -0.618 | 0.755 | -0.027 | 0.538 | -0.01 | -0.004 | -0.004 | 0 | 0.16 | 0.21 | 0.13 | 0.14 | 0.13 | 0.24 |
| 7 | 22235825 | rs17146377 | 0.138 | 0.716 | -0.515 | 0.834 | 0.01 | -0.007 | -0.007 | 0 | 0.12 | 0.1 | 0.1 | 0.38 | 0.07 | 0.26 |
| 7 | 22237786 | rs13229289 | 0.3 | 1.062 | -0.638 | 0.579 | -0.025 | 0.019 | 0.019 | 0 | 0.19 | 0.08 | 0.18 | 0.13 | 0.34 | 0.18 |
| 7 | 22242535 | rs4719687 | -0.727 | 0.424 | -0.633 | 0.538 | -0.001 | -0.001 | -0.001 | 2.215 | 0.43 | 0.54 | 0.41 | 0.59 | 0.54 | 0.53 |
| 7 | 22249369 | rs1476686 | -1.466 | 0.166 | -1.505 | 0.634 | 0.062 | 0.068 | 0.068 | 1.851 | 0.59 | 0.88 | 0.57 | 0.76 | 0.68 | 0.76 |
| 7 | 22252165 | rs16873181 | 0.223 | 0.639 | -0.788 | 0.915 | -0.013 | 0.011 | 0.011 | 0.162 | 0.07 | 0.09 | 0.08 | 0.13 | 0.02 | 0.17 |
| 7 | 22254164 | rs16873182 | -0.697 | 1.611 | -0.977 | 0.728 | -0.025 | -0.006 | -0.006 | 0 | 0.15 | 0.09 | 0.12 | 0.16 | 0.21 | 0.2 |
| 7 | 22254262 | rs10278217 | -1.424 | 0.858 | -1.525 | 0.391 | 0.005 | 0.048 | 0.048 | 0.039 | 0.69 | 0.91 | 0.82 | 0.85 | 0.72 | 0.83 |
| 7 | 22256988 | rs1419759 | -1.515 | 0.668 | -0.828 | 0.354 | 0.007 | 0.042 | 0.042 | 0 | 0.64 | 0.69 | 0.65 | 0.85 | 0.69 | 0.83 |
| 7 | 22258795 | rs12534599 | 1.377 | 0.652 | -0.931 | 0.386 | 0.001 | -0.011 | -0.011 | 0 | 0.13 | 0.13 | 0.23 | 0.17 | 0.15 | 0.13 |
| 7 | 22268760 | rs17711814 | 0.071 | 0.78 | -0.769 | 0.946 | -0.026 | 0.01 | 0.01 | 0.168 | 0.18 | 0.08 | 0.17 | 0.15 | 0.23 | 0.26 |
| 7 | 22269345 | rs10499542 | -1.241 | 0.689 | -0.244 | 0.871 | 0.007 | 0 | 0 | 0 | 0.39 | 0.37 | 0.34 | 0.66 | 0.4 | 0.61 |
| 7 | 22271932 | rs2057987 | -1.664 | 0.666 | -0.178 | 0.744 | 0.006 | 0.065 | 0.065 | 0.012 | 0.53 | 0.58 | 0.57 | 0.83 | 0.55 | 0.74 |
| 7 | 22277446 | rs1419765 | -0.23 | 0.342 | 0.557 | 0.832 | 0.007 | -0.07 | -0.07 | 0 | 0.11 | 0.32 | 0.08 | 0 | 0.12 | 0.06 |
| 7 | 22279116 | rs11761102 | -0.233 | 0.358 | 0.427 | 0.836 | 0.001 | -0.045 | -0.045 | 0.002 | 0.11 | 0.16 | 0.08 | 0 | 0.13 | 0.11 |
| 7 | 22281041 | rs7795150 | -0.6 | 0.333 | 0.183 | 1.022 | -0.012 | -0.005 | -0.005 | 0.271 | 0.27 | 0.56 | 0.34 | 0.2 | 0.29 | 0.25 |
| 7 | 22283925 | rs4722116 | -1.754 | 0.29 | -0.177 | 0.553 | 0.029 | 0.109 | 0.109 | 0.183 | 0.51 | 0.75 | 0.5 | 0.7 | 0.46 | 0.55 |
| 7 | 22288276 | rs12700364 | 0.37 | 0.231 | -0.377 | 0.528 | -0.002 | 0.007 | 0.007 | 0 | 0.09 | 0.07 | 0.06 | 0 | 0.12 | 0.02 |
| 7 | 22289364 | rs10250461 | -0.29 | -0.138 | -0.181 | 0.874 | 0.02 | 0.01 | 0.01 | 0 | 0.18 | 0.07 | 0.1 | 0.17 | 0.11 | 0.14 |
| 7 | 22295360 | rs2214935 | -1.542 | -0.149 | 0.255 | 0.848 | 0.005 | 0.008 | 0.008 | 0.304 | 0.44 | 0.56 | 0.43 | 0.71 | 0.36 | 0.59 |
| 7 | 22304800 | rs4722120 | -0.549 | 0.062 | 0.156 | 1.241 | 0.01 | -0.004 | -0.004 | 0 | 0.19 | 0.11 | 0.11 | 0.27 | 0.14 | 0.23 |
| 7 | 22309509 | rs2528913 | 0.987 | 0.083 | 0.681 | 1.328 | 0.02 | 0.056 | 0.056 | 1.817 | 0.15 | 0.38 | 0.25 | 0.16 | 0.17 | 0.1 |
| 7 | 22311651 | rs17712784 | 0.535 | 0.233 | 0.491 | 1.07 | -0.01 | 0.009 | 0.009 | 0 | 0.09 | 0.05 | 0.17 | 0.01 | 0.13 | 0.09 |
| 7 | 22312065 | rs17146488 | -0.545 | 0.162 | 0.694 | 1.265 | 0.01 | 0.011 | 0.011 | 0 | 0.19 | 0.11 | 0.11 | 0.16 | 0.14 | 0.22 |
| 7 | 22316308 | rs1636899 | -1.678 | 0.131 | 0.712 | 1.413 | -0.004 | -0.011 | -0.011 | 0 | 0.39 | 0.53 | 0.44 | 0.67 | 0.37 | 0.62 |
| 7 | 22317659 | rs17146521 | -0.695 | 0.156 | 0.55 | 1.504 | -0.008 | 0.007 | 0.007 | 0 | 0.08 | 0.08 | 0.1 | 0.33 | 0.11 | 0.23 |
| 7 | 22322898 | rs1636892 | -1.255 | -0.218 | 0.909 | 1.514 | -0.007 | 0 | 0 | 0.525 | 0.36 | 0.59 | 0.37 | 0.33 | 0.32 | 0.32 |
| 7 | 22330700 | rs2302001 | -1.243 | -0.431 | 0.712 | 1.513 | 0.001 | -0.002 | -0.002 | 0 | 0.24 | 0.22 | 0.19 | 0.51 | 0.2 | 0.43 |
| 7 | 22339821 | rs2528645 | -1.728 | -0.396 | 0.708 | 1.375 | 0.011 | 0.029 | 0.029 | 1.556 | 0.45 | 0.74 | 0.49 | 0.66 | 0.43 | 0.62 |
| 7 | 22340865 | rs2686528 | 0.749 | -0.142 | 0.946 | 1.554 | 0.027 | 0.03 | 0.03 | 1.387 | 0.16 | 0.35 | 0.27 | 0.17 | 0.21 | 0.1 |
| 7 | 22341983 | rs1636897 | -1.353 | -0.212 | 0.745 | 1.572 | 0 | 0 | 0 | 0 | 0.34 | 0.66 | 0.39 | 0.33 | 0.36 | 0.35 |
| 7 | 22342416 | rs10950896 | -0.958 | -0.139 | -0.093 | 1.639 | -0.003 | 0.02 | 0.02 | 0 | 0.16 | 0.15 | 0.15 | 0.41 | 0.1 | 0.26 |
| 7 | 22344911 | rs1419767 | -0.837 | -0.142 | -0.193 | 1.308 | -0.001 | 0.001 | 0.001 | 2.774 | 0.29 | 0.32 | 0.33 | 0.24 | 0.29 | 0.29 |
| 7 | 22346329 | rs4719691 | -1.936 | -0.085 | -0.6 | 1.231 | 0.003 | -0.017 | -0.017 | 0.56 | 0.26 | 0.24 | 0.28 | 0.66 | 0.22 | 0.56 |
| 7 | 22347601 | rs13233212 | -0.372 | -0.153 | -0.59 | 1.39 | -0.007 | 0.081 | 0.081 | 0 | 0.12 | 0.16 | 0.15 | 0.43 | 0.07 | 0.27 |
| 7 | 22347861 | rs10499552 | -0.007 | -0.147 | -0.641 | 1.334 | 0.002 | -0.033 | -0.033 | 0 | 0.08 | 0.02 | 0.08 | 0.15 | 0.05 | 0.19 |
| 7 | 22352740 | rs4722126 | -1.477 | -0.144 | -0.599 | 1.317 | -0.001 | 0.015 | 0.015 | 0 | 0.19 | 0.2 | 0.22 | 0.56 | 0.13 | 0.42 |
| 7 | 22354617 | rs1636898 | -1.626 | -0.172 | -0.388 | 1.102 | 0 | -0.026 | -0.026 | 0.152 | 0.43 | 0.61 | 0.56 | 0.84 | 0.45 | 0.68 |

| | | | | | | | | | | | | | | | | |
|---|---|---|---|---|---|---|---|---|---|---|---|---|---|---|---|---|
| 7 | 22358205 | rs12333620 | 1.059 | 0.042 | -1.36 | 1.059 | 0.002 | 0.004 | 0.004 | 0 | 0.16 | 0.18 | 0.26 | 0.2 | 0.22 | 0.12 |
| 7 | 22358234 | rs12333614 | -1.323 | -0.338 | -1.825 | 0.764 | -0.001 | 0.026 | 0.026 | 0 | 0.51 | 0.36 | 0.51 | 0.56 | 0.5 | 0.56 |
| 7 | 22360535 | rs6947075 | -2.696 | -0.065 | -2.016 | 0.488 | -0.03 | 0.074 | 0.074 | 0.455 | 0.74 | 0.9 | 0.82 | 1 | 0.7 | 0.9 |
| 7 | 22365688 | rs6461643 | 1.106 | 0.216 | -1.785 | 0.953 | 0.002 | 0.004 | 0.004 | 0 | 0.15 | 0.18 | 0.26 | 0.2 | 0.22 | 0.12 |
| 7 | 22368678 | rs1522280 | -0.307 | 0.187 | -1.808 | 0.979 | -0.004 | -0.02 | -0.02 | 1.034 | 0.24 | 0.21 | 0.33 | 0.37 | 0.27 | 0.31 |
| 7 | 22374924 | rs1568571 | 1.035 | 0.261 | -1.765 | 0.852 | -0.008 | -0.016 | -0.016 | 0 | 0.16 | 0.12 | 0.25 | 0.21 | 0.22 | 0.14 |
| 7 | 22378982 | rs10226205 | -1.387 | 0.216 | -1.834 | 0.764 | -0.004 | -0.006 | -0.006 | 0 | 0.43 | 0.41 | 0.57 | 0.52 | 0.48 | 0.51 |
| 7 | 22380050 | rs17776545 | -0.929 | 0.241 | -2.163 | 0.45 | -0.017 | -0.016 | -0.016 | 1.126 | 0.16 | 0.11 | 0.25 | 0.24 | 0.21 | 0.3 |
| 7 | 22383866 | rs9639426 | 0.068 | 0.372 | -2.084 | 0.493 | -0.021 | -0.043 | -0.043 | 2.623 | 0.07 | 0.01 | 0.19 | 0.15 | 0.07 | 0.19 |
| 7 | 22385417 | rs4337997 | 0.986 | 0.501 | -2.105 | 0.554 | -0.018 | -0.019 | -0.019 | 0 | 0.16 | 0.07 | 0.13 | 0.17 | 0.19 | 0.14 |
| 7 | 22386924 | rs2041012 | -1.815 | 0.148 | -1.84 | 0.587 | -0.025 | -0.023 | -0.023 | 0 | 0.68 | 0.85 | 0.72 | 0.67 | 0.64 | 0.71 |
| 7 | 22389875 | rs2108288 | -1.153 | 0.23 | -1.64 | 0.98 | 0.004 | 0.004 | 0.004 | 0 | 0.19 | 0.18 | 0.23 | 0.17 | 0.18 | 0.17 |
| 7 | 22391090 | rs7798524 | -0.182 | 0.363 | -1.478 | 1.229 | 0.005 | -0.009 | -0.009 | 0.477 | 0.12 | 0.11 | 0.19 | 0.16 | 0.12 | 0.15 |
| 7 | 22391763 | rs17720371 | -0.383 | 0.387 | -1.702 | 1.139 | -0.021 | -0.002 | -0.002 | 1.787 | 0.38 | 0.26 | 0.43 | 0.38 | 0.42 | 0.42 |
| 7 | 22392134 | rs12539362 | -0.529 | 0.571 | -1.362 | 0.789 | 0.005 | 0.028 | 0.028 | 0 | 0.5 | 0.32 | 0.62 | 0.56 | 0.53 | 0.57 |
| 7 | 22405350 | rs10950899 | -0.805 | 1.274 | -1.03 | 0.796 | -0.029 | -0.031 | -0.031 | 0 | 0.05 | 0.02 | 0.22 | 0.18 | 0.1 | 0.19 |
| 7 | 22409383 | rs7779378 | -0.768 | 1.037 | -1.031 | 0.312 | -0.004 | 0.019 | 0.019 | 0 | 0.21 | 0.13 | 0.25 | 0.08 | 0.18 | 0.13 |
| 7 | 22410250 | rs6946978 | 0.913 | 0.855 | -0.122 | 0.224 | 0.002 | -0.048 | -0.048 | 0 | 0.77 | 0.64 | 0.73 | 0.92 | 0.82 | 0.87 |
| 7 | 22412741 | rs7794890 | 1.101 | 0.816 | -0.118 | 0.19 | -0.013 | 0.007 | 0.007 | 0.011 | 0.92 | 0.73 | 0.84 | 0.97 | 0.97 | 0.93 |
| 7 | 22414646 | rs6943931 | -1.421 | 0.745 | 0.149 | 0.351 | -0.003 | 0 | 0 | 0.272 | 0.29 | 0.25 | 0.33 | 0.36 | 0.34 | 0.39 |
| 7 | 22416306 | rs10228562 | 0.917 | 0.68 | 0.289 | 0.459 | -0.013 | 0.007 | 0.007 | 0.003 | 0.92 | 0.73 | 0.83 | 0.97 | 0.97 | 0.93 |
| 7 | 22422608 | rs2686485 | 1.312 | 0.651 | 0.422 | 0.446 | 0.025 | 0.057 | 0.057 | 0.116 | 0.93 | 0.64 | 0.83 | 0.97 | 0.97 | 0.95 |
| 7 | 22422917 | rs17146761 | -1.375 | 0.664 | 0.365 | 0.561 | 0.001 | -0.007 | -0.007 | 0.17 | 0.07 | 0.04 | 0.08 | 0.19 | 0.06 | 0.16 |
| 7 | 22431079 | rs4722136 | -0.907 | 0.676 | 0.372 | 0.558 | 0.005 | 0 | 0 | 0 | 0.07 | 0.04 | 0.14 | 0.03 | 0.03 | 0.07 |
| 7 | 22432872 | rs2528804 | 2.012 | 0.664 | 0.461 | 0.559 | -0.009 | 0.048 | 0.048 | 0.403 | 0.87 | 0.63 | 0.75 | 0.76 | 0.91 | 0.79 |
| 7 | 22433951 | rs2686498 | 0.301 | 0.576 | -0.167 | -0.083 | 0.002 | -0.081 | -0.081 | 0 | 0.72 | 0.57 | 0.69 | 0.89 | 0.68 | 0.79 |
| 7 | 22434643 | rs3094969 | 1.3 | 0.496 | -0.085 | 0.134 | 0 | 0.004 | 0.004 | 0 | 0.48 | 0.54 | 0.38 | 0.58 | 0.46 | 0.52 |
| 7 | 22437867 | rs3846995 | -1.295 | 0.508 | 0.183 | 0.294 | -0.001 | 0.002 | 0.002 | 0 | 0.53 | 0.47 | 0.58 | 0.43 | 0.5 | 0.51 |
| 7 | 22440260 | rs2686468 | 1.276 | 0.537 | 0.183 | 0.301 | -0.002 | 0.003 | 0.003 | 0 | 0.47 | 0.48 | 0.41 | 0.57 | 0.5 | 0.49 |
| 7 | 22446210 | rs2686507 | 1.353 | 0.526 | 0.178 | 0.297 | -0.001 | 0.001 | 0.001 | 0 | 0.46 | 0.47 | 0.4 | 0.56 | 0.5 | 0.49 |
| 7 | 22452083 | rs10258484 | -0.292 | 0.531 | 0.133 | 0.627 | -0.007 | 0.032 | 0.032 | 0.866 | 0.24 | 0.11 | 0.39 | 0.13 | 0.28 | 0.19 |
| 7 | 22455865 | rs2686518 | 1.023 | 0.447 | 0.245 | 0.624 | -0.009 | 0.063 | 0.063 | 0 | 0.55 | 0.74 | 0.59 | 0.77 | 0.54 | 0.67 |
| 7 | 22455947 | rs2686519 | 1.298 | 0.464 | 0.232 | 0.63 | 0.004 | 0.04 | 0.04 | 0 | 0.51 | 0.65 | 0.45 | 0.76 | 0.53 | 0.63 |
| 7 | 22456404 | rs2528824 | 1.298 | 0.464 | 0.232 | 0.63 | 0.004 | 0.04 | 0.04 | 0 | 0.51 | 0.65 | 0.45 | 0.76 | 0.53 | 0.63 |
| 7 | 22459324 | rs2177142 | 1.687 | 0.544 | 0.231 | 0.631 | -0.001 | 0 | 0 | 4.654 | 0.46 | 0.47 | 0.4 | 0.58 | 0.47 | 0.49 |
| 7 | 22471195 | rs740294 | 1.101 | 0.533 | 0.227 | 0.603 | -0.02 | -0.023 | -0.023 | 8.382 | 0.65 | 0.49 | 0.67 | 0.7 | 0.77 | 0.64 |
| 7 | 22473556 | rs10275929 | 0.387 | 0.526 | 0.122 | 0.597 | -0.051 | 0.041 | 0.041 | 0.692 | 0.21 | 0.05 | 0.26 | 0.11 | 0.29 | 0.15 |
| 7 | 22474012 | rs3813379 | -0.983 | 0.513 | 0.16 | 0.581 | -0.011 | 0.03 | 0.03 | 0.047 | 0.31 | 0.16 | 0.45 | 0.31 | 0.35 | 0.43 |
| 7 | 22474921 | rs10250942 | -0.158 | 0.505 | 0.148 | 0.577 | -0.053 | 0.002 | 0.002 | 0.363 | 0.24 | 0.07 | 0.3 | 0.29 | 0.33 | 0.31 |
| 7 | 22476834 | rs10237119 | 0.198 | 0.516 | 0.185 | 0.582 | -0.008 | 0.015 | 0.015 | 0.488 | 0.22 | 0.19 | 0.27 | 0.08 | 0.29 | 0.15 |
| 7 | 22477507 | rs7458118 | -1.213 | 0.509 | 0.221 | 0.579 | 0.001 | 0.013 | 0.013 | 0 | 0.33 | 0.3 | 0.46 | 0.28 | 0.36 | 0.43 |
| 7 | 22478797 | rs10485990 | -0.816 | 0.447 | 0.103 | 0.63 | -0.006 | 0.004 | 0.004 | 0 | 0.08 | 0.09 | 0.16 | 0.2 | 0.04 | 0.19 |
| 7 | 22479118 | rs2528845 | -0.321 | 0.614 | 0.051 | 0.647 | -0.001 | -0.006 | -0.006 | 0 | 0.78 | 0.81 | 0.86 | 0.89 | 0.83 | 0.91 |
| 7 | 22482415 | rs2686527 | 1.507 | 0.636 | 0.039 | 0.624 | -0.001 | 0.009 | 0.009 | 0.117 | 0.37 | 0.38 | 0.33 | 0.31 | 0.38 | 0.35 |
| 7 | 22484487 | rs3094974 | 0.256 | 0.681 | -0.475 | 0.663 | -0.009 | 0.008 | 0.008 | 0.117 | 0.71 | 0.8 | 0.55 | 0.69 | 0.65 | 0.66 |
| 7 | 22492390 | rs3114717 | 0.432 | 0.537 | -0.261 | 0.698 | -0.001 | -0.006 | -0.006 | 0 | 0.42 | 0.52 | 0.38 | 0.48 | 0.42 | 0.51 |
| 7 | 22512880 | rs10271156 | -0.363 | -0.398 | -0.404 | 0.847 | -0.007 | 0.002 | 0.002 | 0.108 | 0.38 | 0.53 | 0.36 | 0.45 | 0.23 | 0.3 |
| 7 | 22513195 | rs4722143 | -0.764 | 0.291 | -0.945 | 0.787 | 0.018 | 0.018 | 0.018 | 0 | 0.51 | 0.58 | 0.54 | 0.58 | 0.4 | 0.44 |
| 7 | 22513664 | rs17146953 | 0.553 | -0.244 | -0.687 | 0.744 | 0.03 | -0.031 | -0.031 | 1.467 | 0.27 | 0.16 | 0.16 | 0.4 | 0.16 | 0.24 |
| 7 | 22517358 | rs12540820 | -0.11 | -0.247 | -0.396 | 0.719 | 0.001 | 0.003 | 0.003 | 0 | 0.1 | 0.1 | 0.18 | 0.05 | 0.08 | 0.06 |
| 7 | 22517858 | rs17146956 | 0.532 | -0.261 | -0.252 | 0.734 | 0.014 | -0.015 | -0.015 | 0 | 0.27 | 0.19 | 0.15 | 0.4 | 0.15 | 0.24 |
| 7 | 22517994 | rs17298707 | -0.143 | -0.063 | -0.341 | 0.67 | -0.032 | -0.025 | -0.025 | 0 | 0.08 | 0.02 | 0.15 | 0.12 | 0.13 | 0.11 |
| 7 | 22521199 | rs10239540 | -0.013 | 0.161 | -0.343 | 0.615 | -0.021 | 0.026 | 0.026 | 0 | 0.07 | 0.04 | 0.04 | 0 | 0.09 | 0.04 |
| 7 | 22524201 | rs10252093 | -0.644 | 0.052 | -0.164 | 0.687 | 0.02 | -0.011 | -0.011 | 0.017 | 0.42 | 0.34 | 0.37 | 0.47 | 0.28 | 0.34 |
| 7 | 22525497 | rs4719696 | -1.134 | 0.016 | -0.134 | 0.639 | 0.013 | 0.016 | 0.016 | 0 | 0.49 | 0.56 | 0.54 | 0.58 | 0.4 | 0.43 |
| 7 | 22527691 | rs11761760 | 0.059 | 0.469 | 0.031 | 0.876 | 0.02 | -0.009 | -0.009 | 0 | 0.55 | 0.64 | 0.42 | 0.51 | 0.41 | 0.53 |
| 7 | 22528568 | rs17146982 | 0.846 | 0.417 | 0.184 | 0.846 | 0.021 | 0.002 | 0.002 | 0 | 0.43 | 0.37 | 0.21 | 0.46 | 0.32 | 0.47 |
| 7 | 22528662 | rs10234642 | 0.039 | 0.538 | 0.272 | 0.797 | 0.003 | -0.03 | -0.03 | 0 | 0.07 | 0.15 | 0.05 | 0 | 0.07 | 0.03 |
| 7 | 22534805 | rs6972788 | -0.324 | 0.14 | -0.063 | 0.727 | -0.026 | 0.005 | 0.005 | 7.475 | 0.58 | 0.74 | 0.6 | 0.61 | 0.54 | 0.61 |
| 7 | 22542093 | rs3857717 | 1.652 | 0.252 | 0.253 | 0.813 | 0.01 | -0.011 | -0.011 | 0.799 | 0.77 | 0.63 | 0.61 | 0.82 | 0.76 | 0.79 |
| 7 | 22543081 | rs12666499 | 0.84 | 0.426 | 0.207 | 0.702 | 0.018 | -0.02 | -0.02 | 0.011 | 0.83 | 0.78 | 0.76 | 0.92 | 0.77 | 0.83 |
| 7 | 22556193 | rs34374757 | 0.831 | -0.464 | -0.235 | 0.74 | 0.002 | -0.001 | -0.001 | 0 | 0.3 | 0.29 | 0.17 | 0.28 | 0.24 | 0.23 |
| 7 | 22580985 | rs886159 | -0.748 | -0.468 | 0.001 | 0.659 | 0.008 | 0.01 | 0.01 | 0.972 | 0.3 | 0.42 | 0.33 | 0.4 | 0.34 | 0.3 |
| 7 | 22581131 | rs13247988 | 1.582 | -0.91 | -0.204 | 0.343 | 0.028 | 0.024 | 0.024 | 13.064 | 0.33 | 0.25 | 0.18 | 0.2 | 0.21 | 0.24 |
| 7 | 22583326 | rs10261484 | -1.041 | -0.694 | 0.246 | 0.575 | -0.022 | -0.01 | -0.01 | 0 | 0.15 | 0.32 | 0.21 | 0.14 | 0.13 | 0.11 |
| 7 | 22584183 | rs4722158 | 0.752 | -0.349 | -0.141 | 0.604 | -0.006 | 0.018 | 0.018 | 0 | 0.06 | 0.02 | 0.14 | 0 | 0.08 | 0.04 |
| 7 | 22598742 | rs9655226 | 0.423 | 0.772 | 0.254 | 1.15 | 0.066 | 0.071 | 0.071 | 0 | 0.71 | 0.56 | 0.52 | 0.58 | 0.57 | 0.58 |
| 7 | 22601127 | rs17366136 | 1.247 | 0.421 | -0.332 | 0.992 | 0.072 | 0.105 | 0.105 | 4.7 | 0.49 | 0.3 | 0.35 | 0.24 | 0.35 | 0.32 |
| 7 | 22602932 | rs1122913 | -1.28 | 0.853 | -0.122 | 1.195 | 0.014 | 0.003 | 0.003 | 1.941 | 0.42 | 0.64 | 0.61 | 0.64 | 0.61 | 0.53 |

| | | | | | | | | | | | | | | | | |
|---|---|---|---|---|---|---|---|---|---|---|---|---|---|---|---|---|
| 7 | 22604319 | rs10275628 | 0.12 | 1.17 | -0.508 | 1.168 | -0.025 | -0.037 | -0.037 | 0 | 0.16 | 0.08 | 0.18 | 0.34 | 0.21 | 0.24 |
| 7 | 22612164 | rs2023718 | 0.42 | 1.79 | -0.319 | 1.046 | 0.01 | -0.001 | -0.001 | 0.44 | 0.13 | 0.16 | 0.25 | 0.1 | 0.29 | 0.23 |
| 7 | 22612299 | rs724080 | 0.407 | 1.145 | 0.213 | 1.342 | -0.022 | 0.072 | 0.072 | 0 | 0.87 | 0.78 | 0.76 | 0.75 | 0.91 | 0.81 |
| 7 | 22617721 | rs2905345 | 0.485 | 1.618 | -0.028 | 0.92 | -0.009 | -0.006 | -0.006 | 0 | 0.15 | 0.11 | 0.18 | 0.27 | 0.22 | 0.22 |
| 7 | 22618248 | rs2905346 | -1.387 | 1.477 | 0.151 | 0.814 | 0.001 | 0 | 0 | 0.404 | 0.33 | 0.32 | 0.44 | 0.39 | 0.51 | 0.46 |
| 7 | 22618338 | rs10156122 | 0.091 | 1.194 | 0.086 | 0.179 | 0.013 | -0.001 | -0.001 | 0 | 0.15 | 0.16 | 0.28 | 0.1 | 0.35 | 0.25 |
| 7 | 22634005 | rs756879 | -0.785 | -0.386 | 0.439 | -0.579 | -0.012 | -0.035 | -0.035 | 0.042 | 0.43 | 0.47 | 0.6 | 0.1 | 0.67 | 0.44 |
| 7 | 22643753 | rs4310110 | 0.77 | 0.335 | 0.344 | -0.55 | -0.027 | 0.025 | 0.025 | 0 | 0.06 | 0.02 | 0.06 | 0 | 0.16 | 0.07 |
| 7 | 22644935 | rs4265093 | 0.333 | -1.087 | 0.953 | -0.648 | -0.003 | 0.015 | 0.015 | 0 | 0.1 | 0.19 | 0.17 | 0.08 | 0.09 | 0.07 |
| 7 | 22648542 | rs2905309 | -1.283 | -1.325 | 0.846 | -0.753 | 0.087 | 0.13 | 0.13 | 0 | 0.56 | 0.76 | 0.6 | 0.12 | 0.72 | 0.44 |
| 7 | 22654968 | rs2961285 | -0.288 | -1.44 | 1.132 | -0.717 | 0.06 | 0.101 | 0.101 | 0 | 0.43 | 0.27 | 0.61 | 0.13 | 0.68 | 0.39 |
| 7 | 22659080 | rs2961286 | -1.069 | -1.424 | 1.096 | -0.864 | 0.092 | 0.083 | 0.083 | 0.003 | 0.55 | 0.74 | 0.74 | 0.17 | 0.72 | 0.46 |
| 7 | 22663453 | rs2961287 | -1.274 | -1.418 | 1.062 | -0.875 | 0.089 | 0.085 | 0.085 | 0 | 0.56 | 0.76 | 0.74 | 0.17 | 0.72 | 0.45 |
| 7 | 22669357 | rs7797706 | -0.181 | -1.41 | 1.388 | -0.782 | -0.001 | 0.021 | 0.021 | 0 | 0.12 | 0.32 | 0.17 | 0.08 | 0.09 | 0.07 |
| 7 | 22683622 | rs7793163 | 0.289 | -1.391 | 1.319 | -0.779 | -0.016 | 0.022 | 0.022 | 0 | 0.11 | 0.26 | 0.14 | 0.08 | 0.07 | 0.06 |
| 7 | 22687910 | rs2905316 | -0.413 | -2.462 | 0.581 | -1.382 | 0.038 | 0.041 | 0.041 | 0 | 0.63 | 0.32 | 0.69 | 0.1 | 0.81 | 0.54 |
| 7 | 22695031 | rs2961294 | 0.172 | -2.238 | 0.592 | -1.425 | 0.019 | 0.11 | 0.11 | 0 | 0.42 | 0.26 | 0.54 | 0.08 | 0.61 | 0.38 |
| 7 | 22698466 | rs1006509 | 0.175 | -2.229 | 0.565 | -1.421 | 0.022 | 0.123 | 0.123 | 0 | 0.42 | 0.25 | 0.53 | 0.08 | 0.61 | 0.38 |
| 7 | 22701167 | rs1989838 | -0.698 | -2.218 | 0.567 | -1.427 | 0.066 | 0.072 | 0.072 | 0 | 0.55 | 0.73 | 0.72 | 0.17 | 0.7 | 0.44 |
| 7 | 22704587 | rs2905321 | 0.25 | -1.939 | 0.341 | -1.381 | 0.009 | 0.078 | 0.078 | 0 | 0.41 | 0.28 | 0.39 | 0.07 | 0.6 | 0.35 |
| 7 | 22705474 | rs6962836 | 0.907 | -2.004 | 0.225 | -1.367 | -0.077 | -0.075 | -0.075 | 0.08 | 0.31 | 0.45 | 0.23 | 0.09 | 0.15 | 0.1 |
| 7 | 22706707 | rs2157958 | -0.318 | -1.426 | -0.197 | -1.703 | 0.009 | 0.009 | 0.009 | 0 | 0.45 | 0.38 | 0.59 | 0.09 | 0.64 | 0.37 |
| 7 | 22709003 | rs2905324 | 0.527 | -1.022 | 0.079 | -1.764 | 0.049 | 0.092 | 0.092 | 0 | 0.54 | 0.35 | 0.4 | 0.09 | 0.68 | 0.41 |
| 7 | 22712877 | rs2905325 | -0.815 | -1.32 | 0.142 | -1.408 | -0.019 | 0.068 | 0.068 | 0 | 0.89 | 0.83 | 0.69 | 0.3 | 0.96 | 0.65 |
| 7 | 22716524 | rs4719709 | -0.292 | -1.047 | 0.089 | -1.592 | -0.009 | 0.052 | 0.052 | 0 | 0.36 | 0.16 | 0.28 | 0.16 | 0.35 | 0.25 |
| 7 | 22731199 | rs1476483 | 0.559 | -1.53 | 0.024 | -1.634 | -0.01 | 0.021 | 0.021 | 0 | 0.17 | 0.1 | 0.13 | 0.03 | 0.2 | 0.13 |
| 7 | 22736446 | rs7802442 | -0.959 | -1.565 | 0.293 | -1.629 | 0.041 | 0.064 | 0.064 | 0 | 0.47 | 0.62 | 0.59 | 0.88 | 0.3 | 0.58 |
| 7 | 22740312 | rs4722167 | -0.357 | -1.304 | -0.198 | -1.498 | 0.015 | 0.027 | 0.027 | 0 | 0.34 | 0.19 | 0.25 | 0.16 | 0.25 | 0.23 |
| 7 | 22749157 | rs6949149 | 0.799 | -1.148 | -0.069 | -1.116 | -0.004 | 0.063 | 0.063 | 0 | 0.11 | 0.12 | 0.3 | 0.65 | 0.06 | 0.36 |
| 7 | 22752426 | rs6969502 | 0.634 | -1.552 | -0.277 | -1.689 | -0.021 | 0 | 0 | 0 | 0.42 | 0.57 | 0.42 | 0.66 | 0.2 | 0.44 |
| 7 | 22752873 | rs6969927 | 0.031 | -0.604 | -0.647 | -1.411 | 0.004 | -0.025 | -0.025 | 1.062 | 0.26 | 0.16 | 0.25 | 0.3 | 0.26 | 0.26 |
| 7 | 22757089 | rs6461662 | -0.207 | -0.396 | -0.853 | -1.998 | -0.099 | -0.017 | -0.017 | 3.063 | 0.72 | 0.87 | 0.51 | 0.21 | 0.53 | 0.43 |
| 7 | 22758082 | rs7801617 | 1.355 | -1.179 | -0.271 | -2.445 | -0.108 | -0.172 | -0.172 | 0.28 | 0.33 | 0.47 | 0.16 | 0 | 0.12 | 0.14 |
| 7 | 22758562 | rs7805828 | -0.325 | -1.074 | -0.279 | -2.368 | 0 | 0.001 | 0.001 | 0 | 0.4 | 0.38 | 0.36 | 0.22 | 0.41 | 0.3 |
| 7 | 22760488 | rs10499563 | 0.359 | -1.104 | -0.266 | -2.369 | -0.005 | -0.008 | -0.008 | 0.6 | 0.24 | 0.23 | 0.16 | 0.16 | 0.23 | 0.21 |
| 7 | 22761202 | rs2056576 | 0.511 | -1.151 | -0.217 | -2.619 | 0.032 | 0.041 | 0.041 | 1.077 | 0.46 | 0.41 | 0.29 | 0.06 | 0.33 | 0.19 |
| 7 | 22765232 | rs2069824 | 1.568 | -1.474 | -0.28 | -2.511 | 0.094 | 0.089 | 0.089 | 0.09 | 0.27 | 0.14 | 0.06 | 0 | 0.1 | 0.09 |
| 7 | 22768027 | rs2069837 | 1.114 | -1.843 | -0.327 | -2.428 | 0.018 | 0.003 | 0.003 | 0 | 0.18 | 0.12 | 0.08 | 0.14 | 0.09 | 0.16 |
| 7 | 22768124 | rs1474347 | -1.159 | -1.275 | -0.185 | -2.288 | -0.034 | 0.013 | 0.013 | 0 | 0.82 | 0.89 | 0.8 | 0.99 | 0.59 | 0.86 |
| 7 | 22768707 | rs1554606 | -0.823 | -1.352 | 0.082 | -2.317 | 0.07 | -0.093 | -0.093 | 0 | 0.78 | 0.69 | 0.71 | 0.99 | 0.57 | 0.8 |
| 7 | 22774231 | rs10242595 | 0.009 | -1.652 | -0.408 | -2.255 | -0.013 | -0.069 | -0.069 | 0.175 | 0.65 | 0.54 | 0.52 | 0.93 | 0.32 | 0.65 |
| 7 | 22785086 | rs1524105 | 0.107 | -1.611 | -0.445 | -2.249 | -0.014 | -0.109 | -0.109 | 0 | 0.65 | 0.57 | 0.6 | 0.94 | 0.33 | 0.66 |
| 7 | 22795153 | rs1524104 | 0.66 | -1.84 | -0.616 | -2.128 | 0.004 | 0.004 | 0.004 | 0 | 0.26 | 0.25 | 0.14 | 0.12 | 0.15 | 0.17 |
| 7 | 22798080 | rs7808122 | -0.802 | -1.021 | -1.009 | -2.434 | 0.048 | -0.08 | -0.08 | 0 | 0.78 | 0.74 | 0.74 | 0.99 | 0.57 | 0.81 |
| 7 | 22800653 | rs10229203 | 0.743 | -0.743 | -1.591 | -1.623 | 0.03 | 0.008 | 0.008 | 0.322 | 0.25 | 0.18 | 0.07 | 0 | 0.1 | 0.09 |
| 7 | 22805608 | rs7793526 | -0.382 | -0.704 | -1.593 | -1.59 | 0.022 | 0.074 | 0.074 | 0.096 | 0.4 | 0.27 | 0.26 | 0.08 | 0.37 | 0.23 |
| 7 | 22806718 | rs13236645 | -0.464 | 0.832 | -1.302 | -0.764 | -0.036 | 0.02 | 0.02 | 0 | 0.14 | 0.06 | 0.15 | 0.05 | 0.27 | 0.14 |
| 7 | 22811994 | rs17147267 | -0.173 | -0.796 | -1.328 | -1.057 | 0.044 | 0.046 | 0.046 | 0 | 0.31 | 0.17 | 0.14 | 0.04 | 0.19 | 0.14 |
| 7 | 22812578 | rs10265548 | -0.771 | 0.634 | -0.983 | -0.24 | 0.017 | 0.016 | 0.016 | 0 | 0.17 | 0.24 | 0.23 | 0.24 | 0.27 | 0.29 |
| 7 | 22812592 | rs1524102 | 0.28 | 0.474 | -1.326 | -1.206 | 0.014 | 0.084 | 0.084 | 0.094 | 0.38 | 0.24 | 0.46 | 0.11 | 0.34 | 0.28 |
| 7 | 22813879 | rs10270171 | 0.476 | 0.166 | -1.079 | -1.226 | -0.001 | 0.001 | 0.001 | 1.096 | 0.2 | 0.22 | 0.28 | 0.79 | 0.22 | 0.31 |
| 7 | 22814396 | rs12112489 | 0.385 | 0.134 | -1.241 | -1.678 | -0.011 | 0.076 | 0.076 | 0 | 0.45 | 0.34 | 0.46 | 0.17 | 0.49 | 0.43 |
| 7 | 22816495 | rs10260111 | -0.909 | 0.066 | -1.297 | -1.722 | -0.017 | -0.141 | -0.141 | 2.014 | 0.66 | 0.58 | 0.74 | 0.96 | 0.7 | 0.78 |
| 7 | 22816903 | rs2103109 | 0.371 | -0.06 | -1.325 | -1.765 | -0.052 | 0.133 | 0.133 | 0 | 0.43 | 0.17 | 0.45 | 0.16 | 0.49 | 0.43 |
| 7 | 22817187 | rs2091114 | -0.691 | -0.485 | -1.705 | -2.119 | -0.017 | 0.191 | 0.191 | 2.827 | 0.5 | 0.85 | 0.49 | 0.81 | 0.51 | 0.46 |
| 7 | 22821019 | rs13243243 | 1.736 | -0.245 | -1.644 | -1.98 | -0.031 | 0.023 | 0.023 | 0 | 0.1 | 0.04 | 0.11 | 0 | 0.19 | 0.05 |
| 7 | 22823906 | rs1524100 | -0.582 | -0.595 | -0.994 | -1.641 | 0.012 | 0.078 | 0.078 | 0.04 | 0.67 | 0.84 | 0.76 | 0.96 | 0.7 | 0.77 |
| 7 | 22826360 | rs7781074 | 1.176 | -0.324 | -0.869 | -1.303 | -0.019 | -0.093 | -0.093 | 0 | 0.18 | 0.04 | 0.2 | 0.59 | 0.2 | 0.27 |
| 7 | 22830007 | rs7810218 | 0.153 | -0.272 | -0.638 | -1.297 | -0.039 | -0.036 | -0.036 | 1.437 | 0.3 | 0.2 | 0.37 | 0.62 | 0.41 | 0.36 |
| 7 | 22831386 | rs7779573 | -1.184 | -0.352 | -0.546 | -1.442 | 0.017 | 0.037 | 0.037 | 0 | 0.54 | 0.59 | 0.53 | 0.86 | 0.59 | 0.6 |
| 7 | 22831954 | rs10242247 | 0.204 | -0.07 | -0.397 | -1.296 | -0.02 | -0.005 | -0.005 | 0 | 0.23 | 0.39 | 0.16 | 0.22 | 0.18 | 0.24 |
| 7 | 22842319 | rs10262258 | -0.267 | -0.284 | -0.486 | -1.255 | -0.003 | 0.005 | 0.005 | 0 | 0.37 | 0.63 | 0.27 | 0.23 | 0.38 | 0.28 |
| 7 | 22850333 | rs1029741 | -1.003 | -0.422 | -0.795 | -1.334 | 0.029 | 0.139 | 0.139 | 0 | 0.56 | 0.84 | 0.52 | 0.83 | 0.6 | 0.61 |
| 7 | 22861978 | rs4354222 | -0.992 | -0.421 | -0.797 | -1.334 | 0.029 | 0.148 | 0.148 | 0 | 0.55 | 0.86 | 0.52 | 0.85 | 0.6 | 0.61 |
| 7 | 22864724 | rs10225081 | 1.65 | -0.241 | -0.857 | -0.09 | -0.03 | -0.022 | -0.022 | 0.008 | 0.13 | 0.03 | 0.08 | 0.14 | 0.14 | 0.15 |
| 7 | 22870016 | rs10224419 | 0.959 | -0.251 | -0.567 | -0.152 | -0.025 | 0.014 | 0.014 | 0 | 0.24 | 0.14 | 0.16 | 0.19 | 0.27 | 0.15 |
| 7 | 22870074 | rs10224244 | 0.959 | -0.251 | -0.572 | -0.152 | -0.026 | 0.015 | 0.015 | 0 | 0.24 | 0.12 | 0.16 | 0.19 | 0.27 | 0.15 |
| 7 | 22877782 | rs17147339 | 1.206 | -0.264 | -0.106 | -0.097 | 0.039 | 0.405 | 0.405 | 0 | 0.06 | 0.24 | 0.14 | 0.45 | 0.09 | 0.12 |
| 7 | 22878983 | rs11767419 | 0.956 | -0.202 | -0.331 | -0.168 | -0.025 | 0.011 | 0.011 | 0 | 0.23 | 0.13 | 0.16 | 0.2 | 0.27 | 0.16 |

| | | | | | | | | | | | | | | | | |
|---|---|---|---|---|---|---|---|---|---|---|---|---|---|---|---|---|
| 7 | 22880856 | rs17147344 | -0.141 | -0.291 | -0.111 | -0.176 | 0.002 | -0.018 | -0.018 | 0 | 0.25 | 0.37 | 0.21 | 0.18 | 0.26 | 0.33 |
| 7 | 22882483 | rs10226018 | 0.251 | 0.383 | -0.395 | -0.203 | -0.052 | 0.026 | 0.026 | 0 | 0.37 | 0.18 | 0.24 | 0.3 | 0.41 | 0.32 |
| 7 | 22882621 | rs4329182 | 0.51 | 0.232 | -0.383 | -0.178 | 0.006 | 0.093 | 0.093 | 0.261 | 0.21 | 0.32 | 0.23 | 0.55 | 0.23 | 0.27 |
| 7 | 22882827 | rs4644138 | -1.603 | 0.224 | -0.754 | -0.393 | 0.047 | 0.208 | 0.208 | 0 | 0.58 | 0.85 | 0.6 | 0.98 | 0.66 | 0.74 |
| 7 | 22884898 | rs10264536 | -0.52 | 0.368 | -0.622 | -0.031 | 0.026 | 0.009 | 0.009 | 0 | 0.35 | 0.44 | 0.31 | 0.36 | 0.41 | 0.39 |
| 7 | 22894487 | rs717624 | -0.617 | 0.418 | -0.759 | 0.161 | -0.039 | 0.066 | 0.066 | 0.087 | 0.64 | 0.84 | 0.74 | 0.77 | 0.6 | 0.77 |
| 7 | 22894777 | rs3757446 | 1.625 | 0.385 | -0.806 | 0.256 | 0.002 | -0.004 | -0.004 | 0 | 0.12 | 0.07 | 0.06 | 0.13 | 0.1 | 0.15 |
| 7 | 22902620 | rs6461674 | -0.312 | 0.398 | -0.363 | 0.438 | 0.026 | 0.001 | 0.001 | 0 | 0.5 | 0.71 | 0.59 | 0.51 | 0.46 | 0.53 |
| 7 | 22905282 | rs17147383 | 0.827 | 0.221 | -0.377 | 0.58 | -0.026 | -0.066 | -0.066 | 1.994 | 0.33 | 0.1 | 0.43 | 0.43 | 0.36 | 0.35 |
| 7 | 22907231 | rs17147395 | 0.091 | 0.474 | -0.948 | 0.027 | 0.016 | 0.021 | 0.021 | 0.976 | 0.43 | 0.79 | 0.63 | 0.62 | 0.42 | 0.54 |
| 7 | 22914712 | rs2037607 | -0.231 | 0.599 | -0.755 | 0.331 | 0.002 | 0.007 | 0.007 | 0 | 0.56 | 0.63 | 0.71 | 0.62 | 0.6 | 0.67 |
| 7 | 22918729 | rs4722195 | 0.02 | 0.935 | -0.668 | 0.436 | -0.001 | 0.007 | 0.007 | 0.155 | 0.47 | 0.58 | 0.68 | 0.6 | 0.54 | 0.58 |
| 7 | 22923091 | rs4722196 | -0.776 | 0.988 | -0.575 | 0.468 | 0.028 | 0.011 | 0.011 | 0.193 | 0.48 | 0.41 | 0.6 | 0.52 | 0.58 | 0.58 |
| 7 | 22924527 | rs6461676 | 0.533 | 1.783 | -1.036 | 0.595 | -0.015 | -0.04 | -0.04 | 0 | 0.23 | 0.21 | 0.26 | 0.39 | 0.26 | 0.33 |
| 7 | 22932736 | rs1001026 | -0.58 | 1.553 | -0.719 | 0.278 | -0.003 | -0.006 | -0.006 | 0 | 0.35 | 0.36 | 0.39 | 0.09 | 0.39 | 0.27 |
| 7 | 22933477 | rs4722198 | -1.313 | 1.446 | -0.733 | 0.396 | -0.024 | 0.002 | 0.002 | 1.135 | 0.58 | 0.58 | 0.65 | 0.52 | 0.68 | 0.6 |
| 7 | 22942737 | rs11762836 | 1.148 | 1.598 | -0.936 | 0.239 | -0.013 | 0.031 | 0.031 | 0 | 0.06 | 0.01 | 0.03 | 0 | 0.07 | 0.05 |
| 7 | 22951843 | rs4719720 | 0.675 | 1.66 | -0.788 | 0.424 | 0.006 | 0.033 | 0.033 | 0 | 0.23 | 0.3 | 0.27 | 0.48 | 0.26 | 0.32 |
| 7 | 22959577 | rs11772647 | 1.23 | 1.632 | -0.799 | 0.416 | -0.013 | 0.03 | 0.03 | 0 | 0.06 | 0.01 | 0.03 | 0 | 0.07 | 0.05 |
| 7 | 22959756 | rs10277454 | -1.516 | 1.442 | -0.785 | 0.354 | -0.021 | -0.006 | -0.006 | 0 | 0.6 | 0.61 | 0.66 | 0.59 | 0.68 | 0.6 |
| 7 | 22975004 | rs3891325 | -1.093 | 1.611 | -0.761 | 0.467 | -0.018 | 0.029 | 0.029 | 0 | 0.39 | 0.35 | 0.42 | 0.18 | 0.46 | 0.39 |
| 7 | 22975659 | rs10247567 | -0.032 | 1.644 | -0.749 | 0.382 | -0.008 | -0.011 | -0.011 | 0 | 0.37 | 0.38 | 0.35 | 0.51 | 0.41 | 0.46 |
| 7 | 22980638 | rs7788668 | -0.362 | 1.707 | -0.792 | 0.372 | -0.025 | 0.029 | 0.029 | 0 | 0.25 | 0.18 | 0.31 | 0.1 | 0.31 | 0.25 |
| 7 | 22983890 | rs1583511 | -1.769 | 2.037 | -0.806 | 0.482 | -0.029 | -0.007 | -0.007 | 0.011 | 0.62 | 0.63 | 0.68 | 0.63 | 0.72 | 0.71 |
| 7 | 22984824 | rs2286494 | -1.708 | 1.964 | -0.888 | 0.483 | -0.045 | -0.012 | -0.012 | 0 | 0.62 | 0.57 | 0.67 | 0.63 | 0.72 | 0.71 |
| 7 | 22986107 | rs7779074 | 1.737 | 2.003 | -0.943 | 0.593 | -0.035 | -0.041 | -0.041 | 0 | 0.16 | 0.01 | 0.07 | 0.15 | 0.15 | 0.19 |
| 7 | 22988381 | rs10224604 | -0.844 | 2.094 | -0.91 | 0.624 | -0.146 | -0.082 | -0.082 | 2.856 | 0.28 | 0.08 | 0.41 | 0.33 | 0.41 | 0.35 |
| 7 | 22992629 | rs2058159 | 2.304 | 2.403 | -0.969 | 0.569 | -0.025 | 0.036 | 0.036 | 0.033 | 0.07 | 0 | 0.15 | 0 | 0.09 | 0.12 |
| 7 | 23002354 | rs6974381 | 1.644 | 2.095 | -0.902 | 0.59 | 0.001 | 0 | 0 | 0 | 0.17 | 0.17 | 0.09 | 0.15 | 0.15 | 0.19 |
| 7 | 23010063 | rs6461679 | -0.737 | 1.231 | -0.721 | 0.226 | 0.049 | 0.038 | 0.038 | 1.043 | 0.49 | 0.39 | 0.59 | 0.58 | 0.61 | 0.55 |
| 7 | 23010545 | rs10240943 | 1.348 | 1.257 | -0.723 | 0.258 | -0.016 | -0.033 | -0.033 | 0 | 0.22 | 0.18 | 0.19 | 0.37 | 0.25 | 0.29 |
| 7 | 23020672 | rs13240517 | 0.99 | 1.029 | -0.596 | 0.649 | -0.005 | -0.014 | -0.014 | 0 | 0.33 | 0.34 | 0.33 | 0.47 | 0.36 | 0.35 |
| 7 | 23024795 | rs17147527 | -0.873 | 1.157 | -0.809 | 0.681 | -0.112 | 0.026 | 0.026 | 0 | 0.23 | 0.05 | 0.3 | 0.12 | 0.31 | 0.25 |
| 7 | 23028041 | rs2286705 | -0.749 | 1.146 | -0.717 | 0.599 | -0.018 | -0.003 | -0.003 | 0 | 0.58 | 0.61 | 0.65 | 0.57 | 0.68 | 0.59 |
| 7 | 23028610 | rs2286707 | 0.058 | 1.122 | -0.775 | 0.601 | -0.1 | -0.011 | -0.011 | 0 | 0.17 | 0.01 | 0.26 | 0.1 | 0.25 | 0.2 |
| 7 | 23054378 | rs4719726 | -0.705 | 1.064 | -1.181 | 0.576 | -0.029 | -0.004 | -0.004 | 4.213 | 0.58 | 0.57 | 0.66 | 0.57 | 0.68 | 0.59 |
| 7 | 23067144 | rs17370204 | -0.433 | 1.605 | -1.556 | 0.806 | -0.024 | 0.066 | 0.066 | 0 | 0.13 | 0.04 | 0.06 | 0.01 | 0.12 | 0.08 |
| 7 | 23067738 | rs10488076 | -0.392 | 1.619 | -1.216 | 1.234 | -0.071 | -0.063 | -0.063 | 1.296 | 0.12 | 0.01 | 0.3 | 0.2 | 0.21 | 0.21 |
| 7 | 23095077 | rs764009 | 0.217 | 0.857 | -1.401 | 0.886 | -0.017 | 0.008 | 0.008 | 0 | 0.31 | 0.33 | 0.26 | 0.42 | 0.19 | 0.37 |
| 7 | 23100220 | rs10950938 | -0.934 | 0.818 | -1.399 | 0.914 | 0.005 | 0.002 | 0.002 | 0 | 0.49 | 0.4 | 0.53 | 0.53 | 0.43 | 0.58 |
| 7 | 23105157 | rs17147607 | -0.036 | 0.567 | -1.275 | 0.932 | -0.007 | 0.002 | 0.002 | 0 | 0.33 | 0.34 | 0.3 | 0.39 | 0.24 | 0.36 |
| 7 | 23106669 | rs4722224 | -0.319 | 0.733 | -1.066 | 0.903 | 0.008 | -0.003 | -0.003 | 0.349 | 0.48 | 0.38 | 0.52 | 0.49 | 0.44 | 0.58 |
| 7 | 23109033 | rs11532668 | -0.767 | 0.782 | -1.204 | 1.299 | 0.003 | -0.037 | -0.037 | 0 | 0.2 | 0.13 | 0.42 | 0.55 | 0.21 | 0.45 |
| 7 | 23116580 | rs6461687 | -0.149 | 0.879 | -1.562 | 0.487 | 0.012 | 0.037 | 0.037 | 0 | 0.42 | 0.8 | 0.39 | 0.35 | 0.39 | 0.37 |
| 7 | 23140624 | rs12700412 | 0.257 | 1.135 | -0.803 | 0.409 | -0.044 | 0.015 | 0.015 | 0 | 0.07 | 0 | 0.16 | 0.03 | 0.16 | 0.25 |
| 7 | 23143049 | rs7456915 | -0.074 | 0.399 | -0.782 | 0.211 | 0.001 | 0.011 | 0.011 | 0.517 | 0.36 | 0.77 | 0.32 | 0.34 | 0.34 | 0.26 |
| 7 | 23143851 | rs13438179 | -0.074 | 0.399 | -0.782 | 0.211 | 0.001 | 0.011 | 0.011 | 0.517 | 0.36 | 0.77 | 0.32 | 0.34 | 0.34 | 0.26 |
| 7 | 23144947 | rs10256524 | -0.064 | 0.4 | -0.784 | 0.212 | 0 | 0.007 | 0.007 | 0.339 | 0.36 | 0.71 | 0.32 | 0.34 | 0.34 | 0.26 |
| 7 | 23145295 | rs10256996 | -0.048 | 0.897 | -0.484 | 0.239 | 0.021 | -0.066 | -0.066 | 0 | 0.34 | 0.47 | 0.32 | 0.17 | 0.41 | 0.24 |
| 7 | 23145952 | rs11772019 | 0.139 | 0.922 | -0.704 | 0.042 | -0.044 | 0.052 | 0.052 | 0 | 0.09 | 0.01 | 0.09 | 0 | 0.14 | 0.04 |
| 7 | 23156079 | rs1558313 | -0.766 | 0.692 | -0.626 | 0.059 | 0 | 0.014 | 0.014 | 0.79 | 0.37 | 0.8 | 0.32 | 0.36 | 0.35 | 0.26 |
| 7 | 23164701 | rs15775 | -0.762 | 0.638 | -0.627 | 0.056 | 0.004 | 0.011 | 0.011 | 0.138 | 0.37 | 0.71 | 0.32 | 0.35 | 0.34 | 0.26 |
| 7 | 23175946 | rs6953996 | -0.294 | 0.738 | -0.661 | -0.094 | 0 | -0.061 | -0.061 | 0 | 0.33 | 0.46 | 0.29 | 0.18 | 0.33 | 0.23 |
| 7 | 23187179 | rs10247268 | -0.216 | 0.704 | -0.945 | -0.13 | 0.004 | -0.047 | -0.047 | 0 | 0.32 | 0.49 | 0.28 | 0.18 | 0.33 | 0.23 |
| 7 | 23194605 | rs955187 | 1.184 | 0.631 | -1.102 | -0.129 | 0.004 | 0.064 | 0.064 | 0.136 | 0.57 | 0.09 | 0.62 | 0.67 | 0.62 | 0.67 |
| 7 | 23214696 | rs706013 | -0.753 | 0.492 | -0.798 | -0.068 | -0.03 | 0.073 | 0.073 | 0.031 | 0.1 | 0.4 | 0.08 | 0.15 | 0.04 | 0.1 |
| 7 | 23220235 | rs10254544 | -0.29 | 0.449 | -0.768 | -0.305 | 0.004 | -0.06 | -0.06 | 0 | 0.32 | 0.47 | 0.28 | 0.18 | 0.32 | 0.23 |
| 7 | 23220450 | rs10242104 | -0.22 | 0.449 | -0.52 | -0.304 | 0.007 | -0.061 | -0.061 | 0 | 0.31 | 0.43 | 0.28 | 0.18 | 0.32 | 0.22 |
| 7 | 23220639 | rs858249 | -0.641 | 0.284 | -0.837 | -0.211 | -0.024 | -0.034 | -0.034 | 0 | 0.09 | 0.22 | 0.08 | 0.01 | 0.05 | 0.1 |
| 7 | 23228400 | rs1637224 | -0.443 | 0.286 | -0.831 | -0.228 | -0.008 | -0.018 | -0.018 | 0 | 0.08 | 0.14 | 0.08 | 0.01 | 0.05 | 0.1 |
| 7 | 23235483 | rs1728327 | -0.509 | 0.352 | -0.707 | -0.103 | -0.008 | -0.018 | -0.018 | 0 | 0.08 | 0.14 | 0.08 | 0.01 | 0.05 | 0.1 |
| 7 | 23239133 | rs858238 | -0.822 | 0.352 | -0.706 | -0.103 | -0.024 | -0.037 | -0.037 | 0 | 0.09 | 0.21 | 0.08 | 0.01 | 0.05 | 0.1 |
| 7 | 23247361 | rs858291 | -0.545 | 0.375 | -0.792 | -0.093 | -0.009 | -0.019 | -0.019 | 0 | 0.08 | 0.14 | 0.08 | 0.01 | 0.05 | 0.1 |
| 7 | 23260251 | rs1637193 | -1.105 | 0.815 | -1.159 | 0.014 | -0.019 | 0.081 | 0.081 | 0.318 | 0.44 | 0.96 | 0.4 | 0.3 | 0.39 | 0.39 |
| 7 | 23274764 | rs10268610 | -0.088 | 0.672 | -0.309 | 0.05 | 0.011 | -0.052 | -0.052 | 0.359 | 0.32 | 0.44 | 0.29 | 0.22 | 0.33 | 0.29 |
| 7 | 23286315 | rs858239 | -1.236 | 0.645 | -0.235 | 0.062 | -0.018 | 0.124 | 0.124 | 0.331 | 0.43 | 0.96 | 0.4 | 0.26 | 0.38 | 0.38 |
| 7 | 23292808 | rs1881203 | -0.303 | 0.725 | -0.179 | 0.084 | 0.012 | -0.077 | -0.077 | 0 | 0.32 | 0.44 | 0.31 | 0.15 | 0.34 | 0.29 |
| 7 | 23293098 | rs199348 | -0.811 | 0.718 | -0.092 | 0.08 | -0.002 | -0.044 | -0.044 | 0 | 0.39 | 0.53 | 0.36 | 0.13 | 0.38 | 0.37 |
| 7 | 23294144 | rs858275 | -0.897 | 0.717 | -0.031 | 0.081 | 0.007 | 0.056 | 0.056 | 0 | 0.41 | 0.72 | 0.38 | 0.28 | 0.38 | 0.38 |

| | | | | | | | | | | | | | | | | |
|---|---|---|---|---|---|---|---|---|---|---|---|---|---|---|---|---|
| 7 | 23296533 | rs199355 | -0.809 | 0.721 | -0.044 | 0.062 | -0.002 | -0.043 | -0.043 | 0 | 0.39 | 0.54 | 0.36 | 0.13 | 0.38 | 0.37 |
| 7 | 23296572 | rs199354 | 0.694 | 0.709 | -0.081 | 0.062 | 0.004 | 0.001 | 0.001 | 0 | 0.06 | 0.06 | 0.05 | 0.01 | 0.03 | 0.09 |
| 7 | 23298415 | rs10085838 | -0.11 | 0.75 | 0.212 | 0.036 | 0.013 | -0.073 | -0.073 | 0 | 0.31 | 0.4 | 0.31 | 0.12 | 0.34 | 0.28 |
| 7 | 23300049 | rs199351 | -1.176 | 0.687 | 0.024 | 0.035 | -0.023 | 0.049 | 0.049 | 0.437 | 0.43 | 0.96 | 0.39 | 0.35 | 0.38 | 0.39 |
| 7 | 23306500 | rs156428 | 0.605 | 0.741 | 0.091 | 0.028 | -0.015 | -0.021 | -0.021 | 0 | 0.06 | 0.08 | 0.05 | 0.01 | 0.03 | 0.09 |
| 7 | 23314547 | rs5850 | -0.308 | -0.008 | -0.203 | -0.327 | -0.001 | -0.054 | -0.054 | 0 | 0.36 | 0.41 | 0.3 | 0.12 | 0.34 | 0.28 |
| 7 | 23317600 | rs2268745 | -0.367 | -0.013 | -0.196 | -0.318 | -0.001 | -0.049 | -0.049 | 0 | 0.37 | 0.44 | 0.31 | 0.19 | 0.34 | 0.28 |
| 7 | 23330347 | rs886791 | -1.608 | 0.416 | -0.873 | 0.204 | -0.085 | -0.024 | -0.024 | 0 | 0.61 | 0.99 | 0.82 | 0.64 | 0.64 | 0.69 |
| 7 | 23337615 | rs199656 | -0.063 | 0.943 | -0.544 | 0.005 | 0.005 | -0.042 | -0.042 | 0.042 | 0.64 | 0.58 | 0.68 | 0.83 | 0.61 | 0.69 |
| 7 | 23339460 | rs149821 | 0.37 | 1.735 | -0.646 | 0.372 | 0.001 | 0.001 | 0.001 | 0 | 0.58 | 0.57 | 0.5 | 0.58 | 0.56 | 0.58 |
| 7 | 23340618 | rs156404 | 0.684 | 1.942 | -0.497 | -1.262 | -0.016 | -0.013 | -0.013 | 0 | 0.06 | 0.08 | 0.05 | 0.01 | 0.01 | 0.09 |
| 7 | 23348396 | rs156409 | 0.772 | 1.949 | -0.494 | -1.258 | -0.016 | -0.014 | -0.014 | 0 | 0.06 | 0.08 | 0.05 | 0.01 | 0.01 | 0.09 |
| 7 | 23353310 | rs156419 | 0.082 | 1.524 | -0.372 | -1.12 | -0.026 | -0.023 | -0.023 | 0 | 0.09 | 0.14 | 0.06 | 0.01 | 0.01 | 0.1 |
| 7 | 23354413 | rs156418 | -0.773 | 1.486 | -0.348 | -1.097 | -0.045 | 0.056 | 0.056 | 0 | 0.1 | 0.3 | 0.07 | 0.17 | 0.02 | 0.1 |
| 7 | 23355241 | rs17740440 | 1.526 | 1.564 | -0.437 | -1.234 | -0.034 | -0.006 | -0.006 | 0.263 | 0.05 | 0 | 0.15 | 0.03 | 0.14 | 0.18 |
| 7 | 23374134 | rs190647 | 0.482 | 0.677 | -0.219 | -1.115 | -0.023 | -0.021 | -0.021 | 5.634 | 0.07 | 0.12 | 0.06 | 0.01 | 0.01 | 0.09 |
| 7 | 23380185 | rs1964536 | -0.441 | 0.665 | -0.527 | -1.392 | 0.021 | -0.057 | -0.057 | 0 | 0.37 | 0.45 | 0.33 | 0.2 | 0.44 | 0.34 |
| 7 | 23389073 | rs7796541 | -0.309 | 0.053 | -0.241 | -1.253 | 0.005 | -0.06 | -0.06 | 0.285 | 0.34 | 0.42 | 0.3 | 0.11 | 0.37 | 0.3 |
| 7 | 23389751 | rs3807459 | -0.65 | -0.04 | -0.342 | -1.481 | 0.004 | -0.021 | -0.021 | 0 | 0.39 | 0.43 | 0.33 | 0.27 | 0.42 | 0.33 |
| 7 | 23394647 | rs274061 | -0.925 | -0.538 | -0.55 | -1.493 | -0.063 | -0.057 | -0.057 | 1.328 | 0.12 | 0.33 | 0.07 | 0.01 | 0.01 | 0.09 |
| 7 | 23398061 | rs274064 | -0.652 | -0.369 | -0.533 | -1.417 | -0.046 | -0.039 | -0.039 | 0.012 | 0.09 | 0.3 | 0.07 | 0.01 | 0.01 | 0.09 |
| 7 | 23401955 | rs433395 | -1.007 | -0.118 | -1.04 | -1.395 | 0.018 | 0.191 | 0.191 | 0.41 | 0.48 | 0.93 | 0.39 | 0.23 | 0.4 | 0.4 |
| 7 | 23415317 | rs6461708 | -0.972 | -0.846 | 0.01 | -0.946 | -0.01 | -0.002 | -0.002 | 0 | 0.22 | 0.68 | 0.18 | 0.19 | 0.12 | 0.25 |
| 7 | 23419593 | rs439590 | -1.112 | -1.096 | -0.092 | -1.017 | 0.028 | 0.004 | 0.004 | 2.163 | 0.3 | 0.75 | 0.17 | 0.24 | 0.16 | 0.29 |
| 7 | 23420767 | rs375092 | -1.221 | -1.014 | -0.224 | -1.213 | 0.07 | 0.029 | 0.029 | 3.32 | 0.31 | 0.84 | 0.23 | 0.2 | 0.15 | 0.3 |
| 7 | 23426058 | rs274029 | -0.446 | -0.794 | 0.521 | -0.939 | -0.08 | 0.046 | 0.046 | 0.136 | 0.15 | 0.38 | 0.05 | 0.13 | 0.03 | 0.1 |
| 7 | 23434735 | rs11762251 | -0.123 | 0.179 | -0.314 | -0.94 | -0.017 | 0.039 | 0.039 | 2.237 | 0.19 | 0.04 | 0.17 | 0.09 | 0.19 | 0.18 |
| 7 | 23436278 | rs274017 | -0.296 | 0.406 | -0.051 | -1.298 | 0.004 | -0.037 | -0.037 | 0 | 0.35 | 0.54 | 0.34 | 0.19 | 0.33 | 0.4 |
| 7 | 23436427 | rs12700428 | 0.443 | 0.685 | 0.163 | -1.188 | -0.01 | 0.009 | 0.009 | 0.012 | 0.21 | 0.16 | 0.27 | 0.11 | 0.29 | 0.34 |
| 7 | 23450198 | rs274035 | -1.237 | 0.623 | 0.114 | -1.408 | 0.013 | 0.034 | 0.034 | 0.486 | 0.58 | 0.7 | 0.57 | 0.28 | 0.6 | 0.57 |
| 7 | 23456221 | rs6651066 | 0.332 | 1.067 | -0.763 | -1.103 | 0.013 | -0.002 | -0.002 | 0 | 0.22 | 0.23 | 0.3 | 0.2 | 0.3 | 0.35 |
| 7 | 23464856 | rs12534523 | -0.617 | 0.453 | -1.027 | -1.024 | 0.011 | 0.063 | 0.063 | 3.161 | 0.3 | 0.14 | 0.21 | 0.1 | 0.28 | 0.2 |
| 7 | 23472061 | rs12700439 | -0.46 | 1.02 | -0.674 | -1.388 | 0.007 | -0.088 | -0.088 | 0 | 0.4 | 0.43 | 0.29 | 0.13 | 0.38 | 0.53 |
| 7 | 23473292 | rs6971100 | -0.758 | 1.483 | -0.359 | -1.528 | 0.011 | 0.076 | 0.076 | 0.134 | 0.47 | 0.63 | 0.39 | 0.16 | 0.49 | 0.53 |
| 7 | 23483815 | rs12540730 | -0.147 | 1.933 | -0.282 | -1.892 | -0.005 | 0.034 | 0.034 | 0 | 0.24 | 0.11 | 0.13 | 0.01 | 0.24 | 0.32 |
| 7 | 23495961 | rs4304239 | -1.273 | 1.994 | 0.18 | -1.975 | 0 | -0.097 | -0.097 | 0 | 0.38 | 0.55 | 0.27 | 0.05 | 0.34 | 0.38 |
| 7 | 23498358 | rs11770192 | -1.006 | 1.965 | 0.559 | -1.987 | 0.002 | -0.076 | -0.076 | 0 | 0.35 | 0.38 | 0.25 | 0.05 | 0.34 | 0.38 |
| 7 | 23502518 | rs12534005 | 0.685 | 1.869 | 0.657 | -1.966 | 0.054 | -0.013 | -0.013 | 0 | 0.08 | 0.21 | 0.11 | 0.04 | 0.12 | 0.02 |
| 7 | 23514868 | rs12700443 | -1.368 | 2.288 | 0.679 | -2.059 | 0.006 | -0.105 | -0.105 | 3.011 | 0.38 | 0.47 | 0.27 | 0.04 | 0.37 | 0.38 |
| 7 | 23519260 | rs10950949 | -1.492 | 2.215 | 0.637 | -2.086 | 0.003 | -0.088 | -0.088 | 0.044 | 0.39 | 0.55 | 0.27 | 0.04 | 0.37 | 0.38 |
| 7 | 23520963 | rs7779665 | -1.478 | 2.199 | 0.629 | -2.064 | 0.003 | -0.085 | -0.085 | 0 | 0.39 | 0.55 | 0.27 | 0.05 | 0.37 | 0.38 |
| 7 | 23523837 | rs10950951 | 0.024 | 2.196 | 0.479 | -2.154 | -0.007 | 0.011 | 0.011 | 0 | 0.2 | 0.11 | 0.12 | 0.01 | 0.21 | 0.3 |
| 7 | 23528927 | rs7776572 | -0.04 | 1.316 | -0.539 | -1.381 | -0.03 | -0.062 | -0.062 | 0.033 | 0.66 | 0.89 | 0.67 | 0.57 | 0.61 | 0.54 |
| 7 | 23539546 | rs2170681 | 1.186 | 0.179 | -0.901 | -1.781 | 0.025 | -0.033 | -0.033 | 0.239 | 0.15 | 0.12 | 0.24 | 0.34 | 0.11 | 0.12 |
| 7 | 23555814 | rs10259241 | 2.038 | 0.194 | -0.873 | -1.791 | 0.009 | -0.006 | -0.006 | 0 | 0.07 | 0.1 | 0.11 | 0.03 | 0.12 | 0.02 |
| 7 | 23564767 | rs17150424 | 1.342 | 0.077 | -0.892 | -1.768 | 0.03 | -0.043 | -0.043 | 0.84 | 0.14 | 0.09 | 0.23 | 0.34 | 0.12 | 0.12 |
| 7 | 23564901 | rs13240694 | 0.566 | 0.091 | -0.758 | -1.844 | 0 | -0.048 | -0.048 | 0.066 | 0.43 | 0.51 | 0.32 | 0.17 | 0.38 | 0.4 |
| 7 | 23568653 | rs4722238 | 0.595 | 0.109 | -0.667 | -1.837 | 0 | -0.043 | -0.043 | 2.166 | 0.43 | 0.46 | 0.32 | 0.18 | 0.38 | 0.4 |
| 7 | 23579399 | rs10249385 | 2.005 | 0.377 | -0.594 | -1.643 | 0.009 | -0.006 | -0.006 | 0 | 0.07 | 0.1 | 0.1 | 0.04 | 0.12 | 0.02 |
| 7 | 23586645 | rs13236851 | 2.033 | 0.657 | -0.797 | -1.578 | -0.044 | -0.015 | -0.015 | 1.218 | 0.07 | 0 | 0.11 | 0.11 | 0.15 | 0.17 |
| 7 | 23588135 | rs13241000 | -0.973 | 0.175 | -0.445 | -1.541 | 0.003 | 0.006 | 0.006 | 0 | 0.48 | 0.58 | 0.48 | 0.4 | 0.49 | 0.43 |
| 7 | 23589042 | rs11773115 | 0.825 | 0.204 | -0.708 | -1.662 | -0.011 | -0.003 | -0.003 | 1.541 | 0.19 | 0.1 | 0.13 | 0.01 | 0.26 | 0.29 |
| 7 | 23589186 | rs11773147 | 0.84 | 0.206 | -0.807 | -1.661 | -0.033 | 0.049 | 0.049 | 0 | 0.19 | 0.06 | 0.12 | 0.01 | 0.26 | 0.29 |
| 7 | 23597913 | rs12540601 | -0.093 | 0.223 | -0.725 | -1.108 | -0.012 | 0.043 | 0.043 | 0 | 0.36 | 0.17 | 0.35 | 0.25 | 0.39 | 0.4 |
| 7 | 23601224 | rs227940 | 0.845 | 0.243 | -0.455 | -1.147 | 0.001 | 0.018 | 0.018 | 0 | 0.52 | 0.39 | 0.55 | 0.7 | 0.52 | 0.59 |
| 7 | 23603318 | rs227944 | 1.664 | -0.237 | -0.893 | -0.885 | -0.015 | -0.138 | -0.138 | 6.926 | 0.18 | 0 | 0.19 | 0.42 | 0.18 | 0.23 |
| 7 | 23603597 | rs10246526 | 1.036 | -0.18 | -0.294 | -1.111 | 0.063 | -0.004 | -0.004 | 0.169 | 0.09 | 0.3 | 0.1 | 0.05 | 0.09 | 0.02 |
| 7 | 23607537 | rs13230456 | -0.108 | -0.698 | -0.871 | -1.32 | -0.065 | 0.172 | 0.172 | 0 | 0.3 | 0.1 | 0.21 | 0.02 | 0.34 | 0.32 |
| 7 | 23610629 | rs227951 | 0.82 | -1.071 | -0.39 | -1.3 | 0.03 | 0.104 | 0.104 | 0 | 0.5 | 0.24 | 0.5 | 0.73 | 0.51 | 0.54 |
| 7 | 23611517 | rs227952 | -0.578 | -0.865 | -0.237 | -0.864 | -0.016 | 0.014 | 0.014 | 0 | 0.86 | 0.73 | 0.81 | 0.77 | 0.87 | 0.91 |
| 7 | 23618026 | rs798645 | 0.37 | -0.823 | -0.203 | -1.054 | 0.001 | -0.002 | -0.002 | 0 | 0.73 | 0.64 | 0.61 | 0.75 | 0.65 | 0.79 |
| 7 | 23619734 | rs798646 | 1.009 | -0.957 | -0.369 | -1.238 | 0.009 | 0.004 | 0.004 | 0 | 0.42 | 0.42 | 0.43 | 0.57 | 0.45 | 0.52 |
| 7 | 23628533 | rs11980035 | 0.458 | -0.398 | -0.56 | -0.909 | -0.036 | 0.043 | 0.043 | 0 | 0.16 | 0.08 | 0.17 | 0.03 | 0.23 | 0.14 |
| 7 | 23630468 | rs116981 | -1.021 | -0.561 | -0.638 | -0.261 | 0.019 | -0.044 | -0.044 | 0 | 0.89 | 0.95 | 0.81 | 0.79 | 0.87 | 0.94 |
| 7 | 23637416 | rs227937 | 1.09 | -0.841 | -0.107 | -0.393 | 0.002 | 0.001 | 0.001 | 0 | 0.51 | 0.41 | 0.5 | 0.59 | 0.52 | 0.65 |
| 7 | 23640174 | rs17779181 | 1.49 | -0.8 | -0.376 | -0.385 | 0.005 | -0.028 | -0.028 | 0 | 0.17 | 0.02 | 0.1 | 0.16 | 0.12 | 0.14 |
| 7 | 23646317 | rs227917 | -0.333 | -0.886 | -0.254 | -0.284 | 0.043 | -0.069 | -0.069 | 0.379 | 0.72 | 0.88 | 0.7 | 0.62 | 0.75 | 0.8 |
| 7 | 23647378 | rs227919 | 1.084 | -1.011 | -0.079 | -0.426 | 0.001 | -0.006 | -0.006 | 0 | 0.66 | 0.47 | 0.6 | 0.71 | 0.61 | 0.76 |
| 7 | 23648714 | rs12700451 | 0.931 | -0.266 | 0.13 | -0.277 | -0.029 | 0.027 | 0.027 | 0 | 0.15 | 0.09 | 0.16 | 0.04 | 0.23 | 0.12 |

| | | | | | | | | | | | | | | | | |
|---|---|---|---|---|---|---|---|---|---|---|---|---|---|---|---|---|
| 7 | 23652561 | rs13227748 | 0.895 | -0.275 | 0.176 | -0.281 | -0.01 | 0.006 | 0.006 | 0 | 0.15 | 0.11 | 0.17 | 0.04 | 0.23 | 0.12 |
| 7 | 23667847 | rs13230595 | 0.915 | -0.262 | 0.128 | -0.275 | -0.021 | 0.019 | 0.019 | 0 | 0.15 | 0.1 | 0.16 | 0.03 | 0.23 | 0.12 |
| 7 | 23675080 | rs1010499 | -0.438 | -0.414 | 0.06 | -0.25 | 0.044 | -0.069 | -0.069 | 0 | 0.72 | 0.88 | 0.7 | 0.63 | 0.75 | 0.8 |
| 7 | 23684965 | rs10253843 | -0.481 | -0.404 | 0.021 | -0.252 | 0.044 | -0.093 | -0.093 | 0 | 0.72 | 0.92 | 0.71 | 0.63 | 0.74 | 0.78 |
| 7 | 23690640 | rs4722242 | 0.78 | -0.304 | 0.28 | -0.225 | 0 | -0.001 | -0.001 | 0 | 0.52 | 0.42 | 0.5 | 0.55 | 0.51 | 0.65 |
| 7 | 23710459 | rs1978252 | 0.707 | 0.547 | -0.066 | -0.306 | 0.005 | -0.014 | -0.014 | 0.35 | 0.12 | 0.13 | 0.14 | 0.04 | 0.15 | 0.07 |
| 7 | 23711767 | rs10248887 | -0.855 | -0.016 | 0.105 | -0.706 | -0.005 | -0.046 | -0.046 | 0 | 0.39 | 0.49 | 0.32 | 0.16 | 0.35 | 0.24 |
| 7 | 23720488 | rs12700456 | 1.64 | -0.312 | 0.016 | -0.574 | -0.021 | -0.008 | -0.008 | 0 | 0.46 | 0.25 | 0.5 | 0.61 | 0.49 | 0.67 |
| 7 | 23720720 | rs12700457 | 1.151 | -0.571 | 0.458 | -0.64 | 0.01 | -0.017 | -0.017 | 0 | 0.64 | 0.51 | 0.59 | 0.75 | 0.57 | 0.78 |
| 7 | 23720985 | rs4722244 | 1.865 | -0.547 | 0.374 | -0.628 | -0.021 | -0.005 | -0.005 | 0 | 0.46 | 0.25 | 0.5 | 0.61 | 0.48 | 0.67 |
| 7 | 23721573 | rs4722245 | 1.67 | -0.54 | 0.356 | -0.626 | -0.004 | 0.002 | 0.002 | 0.344 | 0.48 | 0.39 | 0.51 | 0.61 | 0.48 | 0.67 |
| 7 | 23727104 | rs11763528 | 1.393 | -0.571 | 0.221 | -0.592 | 0.021 | -0.027 | -0.027 | 0 | 0.15 | 0.02 | 0.07 | 0.14 | 0.09 | 0.11 |
| 7 | 23730983 | rs7782183 | 0.779 | -0.465 | 0.484 | -0.65 | 0.007 | -0.05 | -0.05 | 0 | 0.09 | 0.22 | 0.06 | 0 | 0.07 | 0.03 |
| 7 | 23736510 | rs1558308 | -0.982 | -0.482 | 0.312 | -0.655 | -0.024 | -0.058 | -0.058 | 0.661 | 0.87 | 0.94 | 0.8 | 0.78 | 0.79 | 0.87 |
| 7 | 23737795 | rs5029444 | 1.821 | -0.475 | 0.25 | -0.818 | 0.02 | 0.004 | 0.004 | 1.957 | 0.47 | 0.23 | 0.5 | 0.62 | 0.52 | 0.66 |
| 7 | 23739795 | rs6968174 | 1.256 | -0.511 | 0.282 | -0.841 | -0.018 | 0.028 | 0.028 | 0 | 0.14 | 0.1 | 0.14 | 0.03 | 0.17 | 0.07 |
| 7 | 23744549 | rs6974500 | 1.258 | -0.551 | 0.314 | -0.853 | 0.015 | -0.004 | -0.004 | 0 | 0.6 | 0.33 | 0.63 | 0.65 | 0.64 | 0.69 |
| 7 | 23745716 | rs6962385 | 1.218 | -0.456 | 0.248 | -0.83 | -0.018 | 0.028 | 0.028 | 0 | 0.14 | 0.1 | 0.14 | 0.03 | 0.16 | 0.07 |
| 7 | 23750270 | rs950934 | 1.071 | -0.563 | 0.404 | -0.849 | 0.035 | -0.021 | -0.021 | 0 | 0.06 | 0.21 | 0.05 | 0 | 0.06 | 0.03 |
| 7 | 23751403 | rs4722256 | 2.495 | 0.032 | -0.323 | -0.86 | 0.013 | -0.038 | -0.038 | 0.105 | 0.34 | 0.14 | 0.18 | 0.39 | 0.31 | 0.34 |
| 7 | 23751736 | rs7779633 | -0.642 | 0.044 | -0.138 | -0.886 | -0.019 | 0.014 | 0.014 | 0 | 0.37 | 0.5 | 0.35 | 0.36 | 0.35 | 0.31 |
| 7 | 23761242 | rs9654971 | 1.56 | 0.182 | -0.356 | -0.913 | -0.032 | 0.05 | 0.05 | 0.037 | 0.14 | 0.06 | 0.12 | 0.03 | 0.18 | 0.06 |
| 7 | 23763280 | rs10950957 | -0.636 | 0.019 | -0.218 | -0.924 | -0.039 | 0.093 | 0.093 | 0.039 | 0.83 | 0.55 | 0.78 | 0.79 | 0.84 | 0.9 |
| 7 | 23767480 | rs10231712 | -1.218 | 0.046 | -0.148 | -0.923 | 0.013 | -0.025 | -0.025 | 0 | 0.92 | 0.7 | 0.96 | 1 | 0.98 | 0.98 |
| 7 | 23768907 | rs12176610 | 2.861 | -0.19 | -0.526 | -1.031 | -0.038 | -0.132 | -0.132 | 2.143 | 0.3 | 0.02 | 0.16 | 0.4 | 0.29 | 0.33 |
| 7 | 23771484 | rs6958062 | 1.684 | 0.032 | -0.457 | -0.926 | -0.041 | 0.054 | 0.054 | 0.037 | 0.14 | 0.04 | 0.12 | 0.03 | 0.18 | 0.06 |
| 7 | 23772368 | rs2112035 | 1.915 | -0.014 | -0.348 | -0.928 | -0.015 | 0.003 | 0.003 | 0 | 0.4 | 0.41 | 0.19 | 0.39 | 0.32 | 0.35 |
| 7 | 23777938 | rs13246754 | -0.053 | 0.114 | -0.345 | -0.948 | -0.001 | -0.001 | -0.001 | 0 | 0.29 | 0.28 | 0.42 | 0.36 | 0.25 | 0.42 |
| 7 | 23794029 | rs4722266 | 1.373 | 0.342 | -0.32 | -0.925 | 0.023 | 0.004 | 0.004 | 0 | 0.12 | 0.2 | 0.31 | 0.21 | 0.13 | 0.28 |
| 7 | 23798717 | rs12670018 | 0.643 | 0.344 | -0.419 | -0.972 | 0.029 | -0.005 | -0.005 | 0.351 | 0.15 | 0.22 | 0.34 | 0.21 | 0.21 | 0.35 |
| 7 | 23802003 | rs10278268 | -0.277 | 0.309 | -0.443 | -1.029 | -0.003 | 0.04 | 0.04 | 0 | 0.69 | 0.79 | 0.76 | 0.82 | 0.73 | 0.81 |
| 7 | 23811800 | rs10247878 | 1.53 | 0.586 | -0.58 | -1.111 | -0.033 | 0.03 | 0.03 | 0 | 0.1 | 0.04 | 0.09 | 0.05 | 0.16 | 0.05 |
| 7 | 23821784 | rs6461736 | -0.866 | 0.61 | -0.52 | -1.086 | -0.002 | 0.02 | 0.02 | 0 | 0.4 | 0.33 | 0.44 | 0.28 | 0.36 | 0.41 |
| 7 | 23823355 | rs6461737 | -0.375 | 0.598 | -0.518 | -1.085 | 0.005 | 0.01 | 0.01 | 0 | 0.37 | 0.33 | 0.42 | 0.28 | 0.3 | 0.35 |
| 7 | 23823621 | rs12700465 | 0.578 | 0.655 | -0.344 | -1.116 | 0.042 | -0.052 | -0.052 | 0 | 0.17 | 0.25 | 0.18 | 0.11 | 0.19 | 0.16 |
| 7 | 23828088 | rs7782389 | -0.915 | 0.636 | -0.534 | -1.107 | -0.015 | 0.026 | 0.026 | 0 | 0.43 | 0.33 | 0.54 | 0.39 | 0.43 | 0.47 |
| 7 | 23830990 | rs10253205 | -0.562 | 0.52 | -0.486 | -1.08 | -0.006 | 0.008 | 0.008 | 0 | 0.25 | 0.48 | 0.29 | 0.22 | 0.27 | 0.21 |
| 7 | 23832103 | rs12666547 | 1.945 | 0.491 | -0.428 | -1.082 | 0.04 | 0.042 | 0.042 | 0 | 0.42 | 0.21 | 0.34 | 0.61 | 0.44 | 0.41 |
| 7 | 23833243 | rs7791283 | 0.429 | 0.517 | -0.422 | -1.084 | 0.039 | -0.004 | -0.004 | 0 | 0.15 | 0.22 | 0.34 | 0.21 | 0.22 | 0.34 |
| 7 | 23836281 | rs4418236 | 2.147 | 0.513 | -0.559 | -1.053 | -0.004 | 0.019 | 0.019 | 0 | 0.09 | 0.1 | 0.17 | 0.21 | 0.13 | 0.07 |
| 7 | 23840813 | rs12536714 | -0.856 | 0.517 | -0.601 | -1.055 | 0.011 | 0.038 | 0.038 | 0 | 0.46 | 0.33 | 0.56 | 0.39 | 0.49 | 0.53 |
| 7 | 23844242 | rs12700481 | 2.356 | 0.562 | -0.569 | -1.028 | -0.042 | 0 | 0 | 0 | 0.07 | 0 | 0.13 | 0.1 | 0.13 | 0.12 |
| 7 | 23845554 | rs10279416 | -0.257 | 0.238 | -0.456 | -0.974 | -0.036 | 0.046 | 0.046 | 0 | 0.21 | 0.47 | 0.25 | 0.22 | 0.2 | 0.14 |
| 7 | 23852833 | rs6949764 | 0.771 | 0.256 | -0.602 | -0.947 | 0.024 | -0.06 | -0.06 | 0 | 0.12 | 0.47 | 0.09 | 0.01 | 0.08 | 0.07 |
| 7 | 23856954 | rs2188851 | 0.45 | 0.176 | -0.868 | -0.864 | 0.023 | -0.018 | -0.018 | 0.061 | 0.18 | 0.13 | 0.2 | 0.21 | 0.16 | 0.09 |
| 7 | 23871591 | rs6461743 | -0.893 | 0.495 | -0.913 | -0.74 | 0.01 | 0.038 | 0.038 | 0 | 0.46 | 0.33 | 0.56 | 0.39 | 0.48 | 0.53 |
| 7 | 23873892 | rs6967709 | 0.458 | -0.43 | -0.899 | -0.542 | -0.043 | 0.015 | 0.015 | 0 | 0.15 | 0.05 | 0.24 | 0.11 | 0.22 | 0.13 |
| 7 | 23883181 | rs4719739 | 0.797 | -0.307 | -0.857 | -0.54 | -0.013 | -0.045 | -0.045 | 0 | 0.12 | 0.07 | 0.29 | 0.34 | 0.18 | 0.15 |
| 7 | 23892995 | rs12667136 | 0.797 | -0.302 | -0.857 | -0.54 | -0.005 | -0.045 | -0.045 | 0 | 0.12 | 0.07 | 0.29 | 0.33 | 0.17 | 0.14 |
| 7 | 23901883 | rs10276693 | 0.483 | -0.219 | -0.977 | -0.65 | -0.055 | 0.047 | 0.047 | 0 | 0.14 | 0.04 | 0.13 | 0.06 | 0.22 | 0.12 |
| 7 | 23903956 | rs10255647 | -0.825 | -0.478 | -0.463 | -0.781 | -0.008 | 0.054 | 0.054 | 0 | 0.25 | 0.4 | 0.36 | 0.33 | 0.25 | 0.18 |
| 7 | 23910147 | rs8180703 | -0.791 | -0.487 | -0.336 | -0.779 | -0.015 | 0.056 | 0.056 | 0 | 0.25 | 0.42 | 0.35 | 0.33 | 0.24 | 0.17 |
| 7 | 23911701 | rs2390847 | -1.807 | -0.763 | -0.471 | -1.021 | -0.01 | 0.005 | 0.005 | 0.077 | 0.56 | 0.65 | 0.59 | 0.39 | 0.5 | 0.35 |
| 7 | 23914001 | rs12530774 | 0.631 | -0.69 | -0.683 | -1.045 | 0.011 | 0.09 | 0.09 | 0.797 | 0.2 | 0.02 | 0.14 | 0.05 | 0.11 | 0.09 |
| 7 | 23914014 | rs17281622 | 0.016 | -0.846 | -0.605 | -0.926 | 0.001 | 0.044 | 0.044 | 0.003 | 0.31 | 0.25 | 0.24 | 0.06 | 0.26 | 0.17 |
| 7 | 23920231 | rs6970935 | -0.742 | -0.855 | -0.528 | -0.864 | -0.017 | 0.068 | 0.068 | 0 | 0.25 | 0.49 | 0.36 | 0.33 | 0.24 | 0.17 |
| 7 | 23920558 | rs6957300 | -0.117 | -0.694 | -1.026 | -0.794 | -0.005 | 0.121 | 0.121 | 0.03 | 0.3 | 0.11 | 0.22 | 0.06 | 0.25 | 0.17 |
| 7 | 23924545 | rs17282271 | -0.098 | -0.663 | -1.019 | -0.747 | -0.008 | 0.072 | 0.072 | 0.024 | 0.3 | 0.1 | 0.22 | 0.14 | 0.25 | 0.17 |
| 7 | 23925448 | rs4719742 | 0.476 | 0.08 | -0.481 | -0.65 | 0 | 0.011 | 0.011 | 0 | 0.13 | 0.14 | 0.29 | 0.25 | 0.18 | 0.13 |
| 7 | 23926738 | rs17148479 | -0.292 | -0.226 | -0.597 | -1.014 | -0.024 | -0.075 | -0.075 | 0 | 0.15 | 0.21 | 0.1 | 0 | 0.16 | 0.04 |
| 7 | 23928092 | rs12700497 | 0.59 | 0.447 | -1.054 | -0.562 | -0.049 | 0.063 | 0.063 | 0 | 0.13 | 0.04 | 0.11 | 0 | 0.2 | 0.16 |
| 7 | 23935130 | rs2390850 | 0.436 | 0.277 | -0.871 | -0.034 | -0.005 | 0.017 | 0.017 | 0 | 0.07 | 0.06 | 0.2 | 0.25 | 0.08 | 0.11 |
| 7 | 23937198 | rs10225926 | 0.491 | -0.239 | -0.758 | -0.6 | 0.01 | -0.033 | -0.033 | 2.525 | 0.65 | 0.49 | 0.71 | 0.78 | 0.6 | 0.6 |
| 7 | 23938560 | rs1548465 | -0.481 | -0.191 | -1.261 | -0.65 | -0.029 | 0.038 | 0.038 | 0.117 | 0.79 | 0.9 | 0.83 | 0.88 | 0.71 | 0.74 |
| 7 | 23941098 | rs6950415 | -0.231 | -0.84 | -1.489 | -1.253 | -0.007 | -0.012 | -0.012 | 0 | 0.18 | 0.76 | 0.19 | 0.1 | 0.14 | 0.16 |
| 7 | 23946488 | rs4722292 | 0.649 | -0.779 | -1.442 | -1.239 | 0 | -0.016 | -0.016 | 2.812 | 0.47 | 0.5 | 0.64 | 0.87 | 0.4 | 0.58 |
| 7 | 23955191 | rs2059447 | 0.115 | -0.62 | -1.629 | -1.347 | 0.022 | -0.096 | -0.096 | 0 | 0.68 | 0.54 | 0.73 | 0.93 | 0.62 | 0.72 |
| 7 | 23955245 | rs2059448 | -1.329 | -0.531 | -1.702 | -1.311 | -0.051 | 0.07 | 0.07 | 0.109 | 0.82 | 0.94 | 0.85 | 1 | 0.72 | 0.83 |
| 7 | 23956316 | rs12530590 | 0.601 | -0.92 | -1.594 | -0.987 | -0.059 | -0.027 | -0.027 | 0.29 | 0.13 | 0.38 | 0.07 | 0.08 | 0.04 | 0.09 |

| | | | | | | | | | | | | | | | | |
|---|---|---|---|---|---|---|---|---|---|---|---|---|---|---|---|---|
| 7 | 23959375 | rs2390851 | -0.569 | -0.285 | -1.495 | -0.724 | 0.022 | -0.088 | -0.088 | 0.897 | 0.69 | 0.58 | 0.74 | 0.94 | 0.62 | 0.71 |
| 7 | 23962975 | rs6973321 | -0.196 | -1.108 | -0.986 | -0.12 | -0.051 | -0.069 | -0.069 | 0 | 0.32 | 0.42 | 0.17 | 0.11 | 0.25 | 0.2 |
| 7 | 23965357 | rs17148539 | -0.549 | 0.222 | -1.344 | 0.917 | -0.016 | -0.055 | -0.055 | 0 | 0.14 | 0.06 | 0.26 | 0.4 | 0.13 | 0.22 |
| 7 | 23970771 | rs11975049 | -0.81 | 0.041 | -1.191 | 0.304 | 0.003 | 0.006 | 0.006 | 0 | 0.17 | 0.16 | 0.11 | 0.1 | 0.15 | 0.21 |
| 7 | 23980327 | rs11982318 | -0.828 | -0.139 | -0.745 | 0.57 | -0.021 | 0.03 | 0.03 | 0 | 0.12 | 0.26 | 0.1 | 0.13 | 0.1 | 0.13 |
| 7 | 23980973 | rs10261564 | 1.083 | -0.27 | -1.296 | 0.435 | -0.025 | 0.012 | 0.012 | 0 | 0.4 | 0.19 | 0.25 | 0.34 | 0.36 | 0.36 |
| 7 | 23981394 | rs12537767 | -0.001 | -0.224 | -1.256 | 0.485 | -0.02 | 0.025 | 0.025 | 0 | 0.12 | 0.25 | 0.1 | 0.13 | 0.1 | 0.13 |
| 7 | 23999091 | rs12669666 | 1.548 | -0.418 | -1.619 | 0.079 | -0.021 | -0.035 | -0.035 | 0 | 0.25 | 0.13 | 0.17 | 0.34 | 0.25 | 0.25 |
| 7 | 24005226 | rs1549395 | -0.972 | -0.772 | -1.927 | 0.185 | -0.128 | 0.026 | 0.026 | 0 | 0.73 | 0.98 | 0.65 | 0.8 | 0.6 | 0.81 |
| 7 | 24005443 | rs17148598 | 0.995 | -0.987 | -1.887 | 0.182 | -0.008 | -0.005 | -0.005 | 0 | 0.11 | 0.13 | 0.07 | 0.06 | 0.09 | 0.06 |
| 7 | 24005833 | rs1549398 | 0.024 | -0.728 | -1.423 | 0.207 | -0.004 | 0.005 | 0.005 | 0 | 0.61 | 0.68 | 0.56 | 0.67 | 0.52 | 0.69 |
| 7 | 24007152 | rs13221572 | 1.425 | -0.868 | -0.975 | 0.117 | -0.005 | -0.018 | -0.018 | 0 | 0.08 | 0.08 | 0.02 | 0 | 0.05 | 0.08 |
| 7 | 24007472 | rs7802177 | -0.317 | -0.821 | -1.092 | 0.007 | -0.009 | 0.016 | 0.016 | 0 | 0.67 | 0.73 | 0.61 | 0.73 | 0.63 | 0.75 |
| 7 | 24008462 | rs12532005 | -0.031 | -0.84 | -1.091 | -0.002 | 0.009 | 0.025 | 0.025 | 0 | 0.18 | 0.13 | 0.08 | 0.04 | 0.19 | 0.22 |
| 7 | 24013733 | rs7797227 | 1.012 | -0.872 | -0.549 | 0.092 | 0.028 | 0.307 | 0.307 | 0.13 | 0.05 | 0.25 | 0.23 | 0.29 | 0.02 | 0.11 |
| 7 | 24021963 | rs4722312 | 1.364 | -0.851 | -0.628 | 0.003 | -0.03 | 0.01 | 0.01 | 0 | 0.4 | 0.18 | 0.26 | 0.34 | 0.38 | 0.35 |
| 7 | 24035127 | rs10447620 | -0.743 | -0.842 | -0.549 | 0.03 | -0.026 | 0.047 | 0.047 | 0 | 0.26 | 0.39 | 0.33 | 0.39 | 0.24 | 0.42 |
| 7 | 24045162 | rs12531312 | 0.511 | -0.652 | -0.363 | 0.042 | -0.04 | -0.033 | -0.033 | 0 | 0.15 | 0.31 | 0.07 | 0.07 | 0.07 | 0.14 |
| 7 | 24048364 | rs7791476 | -0.337 | 0.04 | -0.317 | 0.282 | -0.028 | 0.015 | 0.015 | 0 | 0.75 | 0.84 | 0.65 | 0.79 | 0.66 | 0.83 |
| 7 | 24051036 | rs6461774 | -0.373 | 0.063 | -0.306 | 0.293 | -0.028 | 0.014 | 0.014 | 0 | 0.75 | 0.84 | 0.65 | 0.79 | 0.66 | 0.83 |
| 7 | 24052443 | rs7798628 | -0.492 | 0.076 | -0.269 | 0.3 | -0.049 | 0.024 | 0.024 | 0 | 0.77 | 0.92 | 0.68 | 0.82 | 0.67 | 0.84 |
| 7 | 24055011 | rs2216974 | 0.473 | 0.029 | -0.248 | 0.285 | -0.01 | 0.006 | 0.006 | 0 | 0.69 | 0.72 | 0.63 | 0.73 | 0.6 | 0.63 |
| 7 | 24058149 | rs6461777 | -0.303 | 0.093 | -0.195 | 0.384 | -0.043 | 0.043 | 0.043 | 0 | 0.76 | 0.91 | 0.68 | 0.83 | 0.67 | 0.84 |
| 7 | 24061437 | rs6947712 | -0.27 | 0.022 | -0.008 | 0.518 | -0.024 | 0.016 | 0.016 | 0 | 0.76 | 0.87 | 0.67 | 0.81 | 0.67 | 0.84 |
| 7 | 24062080 | rs10241053 | 0.064 | 0.111 | -0.194 | 0.372 | 0.019 | 0.067 | 0.067 | 0 | 0.7 | 0.52 | 0.43 | 0.52 | 0.65 | 0.75 |
| 7 | 24065914 | rs2390863 | 0.68 | -0.004 | -0.382 | 0.19 | -0.001 | -0.001 | -0.001 | 0 | 0.47 | 0.49 | 0.32 | 0.51 | 0.47 | 0.55 |
| 7 | 24069951 | rs719158 | 0.541 | 0.007 | -0.404 | 0.184 | -0.001 | 0 | 0 | 0 | 0.49 | 0.49 | 0.32 | 0.53 | 0.48 | 0.55 |
| 7 | 24075536 | rs17357907 | -0.189 | 0.012 | -0.559 | -0.021 | 0.007 | 0.06 | 0.06 | 0 | 0.29 | 0.2 | 0.16 | 0.07 | 0.3 | 0.34 |
| 7 | 24075809 | rs2042500 | -0.457 | -0.003 | -0.373 | 0.079 | 0.01 | 0.055 | 0.055 | 0 | 0.74 | 0.67 | 0.47 | 0.55 | 0.73 | 0.77 |
| 7 | 24076656 | rs10272622 | 0.273 | 0.011 | -0.501 | -0.014 | 0.015 | 0.056 | 0.056 | 0 | 0.27 | 0.17 | 0.15 | 0.03 | 0.26 | 0.23 |
| 7 | 24078362 | rs4722322 | 0.8 | 0.133 | -0.215 | 0.09 | -0.008 | -0.016 | -0.016 | 0 | 0.44 | 0.33 | 0.3 | 0.53 | 0.44 | 0.5 |
| 7 | 24079960 | rs11767698 | 1.131 | 0.425 | -0.134 | 0.247 | 0.001 | 0.002 | 0.002 | 0 | 0.1 | 0.07 | 0.23 | 0.18 | 0.15 | 0.12 |
| 7 | 24082848 | rs6946475 | 0.994 | 0.296 | 0.108 | 0.07 | -0.009 | -0.007 | -0.007 | 8.123 | 0.49 | 0.23 | 0.33 | 0.49 | 0.47 | 0.51 |
| 7 | 24091130 | rs272693 | -0.658 | 0.218 | 0.115 | 0.038 | -0.005 | -0.003 | -0.003 | 0 | 0.84 | 0.89 | 0.72 | 0.79 | 0.78 | 0.83 |
| 7 | 24091230 | rs1996616 | 1.16 | 0.189 | 0.044 | 0.057 | -0.012 | 0.004 | 0.004 | 0 | 0.48 | 0.24 | 0.33 | 0.47 | 0.47 | 0.48 |
| 7 | 24092278 | rs272696 | -0.511 | -0.04 | 0.132 | -0.06 | -0.015 | -0.002 | -0.002 | 0 | 0.34 | 0.57 | 0.39 | 0.32 | 0.31 | 0.35 |
| 7 | 24097729 | rs104987 | 0.198 | -0.002 | -0.449 | -0.054 | 0.013 | 0.026 | 0.026 | 0 | 0.65 | 0.84 | 0.6 | 0.72 | 0.65 | 0.69 |
| 7 | 24107060 | rs17660308 | 0.402 | 0.378 | -0.662 | -0.644 | -0.011 | 0.003 | 0.003 | 0 | 0.2 | 0.16 | 0.15 | 0 | 0.24 | 0.07 |
| 7 | 24107213 | rs272673 | -0.025 | -0.073 | -0.372 | -0.189 | -0.007 | 0.075 | 0.075 | 0 | 0.08 | 0.12 | 0.24 | 0.25 | 0.07 | 0.14 |
| 7 | 24113564 | rs2113502 | 0.934 | -0.355 | -0.371 | 0.312 | 0.014 | -0.001 | -0.001 | 0 | 0.31 | 0.3 | 0.18 | 0.29 | 0.21 | 0.36 |
| 7 | 24119660 | rs452276 | -0.764 | 0.017 | -0.13 | 0.292 | 0.037 | -0.003 | -0.003 | 0 | 0.83 | 0.82 | 0.71 | 0.81 | 0.77 | 0.83 |
| 7 | 24129888 | rs272669 | -0.111 | -0.255 | -0.079 | 0.097 | 0.004 | 0 | 0 | 0 | 0.48 | 0.43 | 0.71 | 0.51 | 0.64 | 0.49 |
| 7 | 24132490 | rs175374 | 0.824 | -0.508 | -0.128 | 0.213 | -0.049 | -0.032 | -0.032 | 1.594 | 0.34 | 0.12 | 0.44 | 0.34 | 0.45 | 0.24 |
| 7 | 24133719 | rs1908272 | -0.046 | -0.568 | -0.058 | 0.192 | 0.041 | 0.001 | 0.001 | 0.184 | 0.49 | 0.67 | 0.26 | 0.53 | 0.33 | 0.58 |
| 7 | 24137506 | rs13237798 | 1.184 | -0.725 | -0.154 | 0.215 | 0.015 | -0.001 | -0.001 | 0.019 | 0.31 | 0.27 | 0.17 | 0.31 | 0.2 | 0.35 |
| 7 | 24142928 | rs759869 | -0.809 | -0.543 | -0.682 | 0.193 | 0.054 | -0.021 | -0.021 | 0 | 0.8 | 0.69 | 0.69 | 0.83 | 0.74 | 0.73 |
| 7 | 24142977 | rs759870 | 0.066 | -0.768 | -0.603 | 0.32 | 0.018 | -0.002 | -0.002 | 0 | 0.46 | 0.61 | 0.25 | 0.49 | 0.29 | 0.49 |
| 7 | 24143458 | rs10251576 | 0.872 | 0.23 | -1.037 | 0.613 | 0.016 | -0.041 | -0.041 | 0 | 0.12 | 0.02 | 0.05 | 0.19 | 0.08 | 0.09 |
| 7 | 24145105 | rs272659 | 0.217 | 0.257 | -1.099 | 0.582 | -0.005 | 0.009 | 0.009 | 0.908 | 0.54 | 0.37 | 0.54 | 0.56 | 0.57 | 0.54 |
| 7 | 24148444 | rs17287854 | 0.485 | 0.501 | -1.565 | 0.623 | -0.019 | -0.016 | -0.016 | 0 | 0.09 | 0.01 | 0.1 | 0.1 | 0.12 | 0.05 |
| 7 | 24149942 | rs272711 | 0.436 | 0.056 | -1.34 | 0.613 | 0 | 0 | 0 | 2.462 | 0.46 | 0.53 | 0.52 | 0.47 | 0.55 | 0.41 |
| 7 | 24151175 | rs17288003 | -0.292 | -0.146 | -1.972 | 0.197 | 0.017 | 0.067 | 0.067 | 0.158 | 0.22 | 0.01 | 0.09 | 0.08 | 0.15 | 0.12 |
| 7 | 24152297 | rs1389653 | 0.123 | 0.144 | -1.401 | 0.793 | -0.007 | 0.001 | 0.001 | 0 | 0.46 | 0.71 | 0.49 | 0.41 | 0.53 | 0.35 |
| 7 | 24152502 | rs9638813 | -0.323 | 0.015 | -1.494 | 1.251 | 0.003 | -0.04 | -0.04 | 0 | 0.13 | 0.08 | 0.34 | 0.23 | 0.12 | 0.27 |
| 7 | 24152917 | rs272706 | -0.266 | 0.104 | -0.849 | 1.224 | 0 | -0.001 | -0.001 | 0 | 0.49 | 0.47 | 0.34 | 0.53 | 0.5 | 0.42 |
| 7 | 24166303 | rs172063 | -0.788 | 0.182 | -1.067 | 1.543 | -0.003 | -0.014 | -0.014 | 0 | 0.32 | 0.44 | 0.48 | 0.69 | 0.3 | 0.49 |
| 7 | 24168274 | rs272663 | 0.083 | 0.297 | -0.815 | 1.79 | -0.003 | 0.057 | 0.057 | 0 | 0.22 | 0.29 | 0.17 | 0.44 | 0.17 | 0.27 |
| 7 | 24171927 | rs6952104 | -0.804 | 0.257 | -0.634 | 1.51 | -0.018 | -0.012 | -0.012 | 0 | 0.52 | 0.76 | 0.38 | 0.5 | 0.53 | 0.41 |
| 7 | 24172966 | rs1978145 | -0.879 | 0.224 | -0.646 | 1.354 | -0.026 | -0.016 | -0.016 | 0 | 0.53 | 0.84 | 0.39 | 0.56 | 0.54 | 0.42 |
| 7 | 24177538 | rs198729 | 0.968 | 0.34 | -0.051 | 0.773 | 0.001 | 0.007 | 0.007 | 0 | 0.41 | 0.44 | 0.37 | 0.23 | 0.36 | 0.37 |
| 7 | 24181478 | rs6461782 | 1.417 | 0.446 | -0.285 | 0.661 | -0.042 | 0.026 | 0.026 | 0.035 | 0.16 | 0.04 | 0.1 | 0.07 | 0.22 | 0.05 |
| 7 | 24183209 | rs198700 | 0.644 | 0.368 | 0.318 | 0.659 | 0.001 | 0.02 | 0.02 | 0 | 0.58 | 0.72 | 0.5 | 0.3 | 0.58 | 0.42 |
| 7 | 24187619 | rs2905863 | -1.481 | 0.416 | 0.192 | 0.735 | -0.003 | 0.052 | 0.052 | 0 | 0.6 | 0.77 | 0.62 | 0.72 | 0.64 | 0.63 |
| 7 | 24191762 | rs198311 | -1.326 | 0.453 | 0.3 | 0.635 | -0.029 | 0.088 | 0.088 | 1.382 | 0.53 | 0.84 | 0.59 | 0.63 | 0.55 | 0.59 |
| 7 | 24192082 | rs198309 | -0.766 | 0.154 | 0.669 | 0.754 | -0.006 | 0.004 | 0.004 | 0 | 0.34 | 0.57 | 0.44 | 0.54 | 0.3 | 0.53 |
| 7 | 24195267 | rs198301 | -1.072 | 0.218 | 0.695 | 0.655 | 0.024 | 0.028 | 0.028 | 0.148 | 0.46 | 0.61 | 0.5 | 0.61 | 0.39 | 0.56 |
| 7 | 24196183 | rs198300 | -1.296 | 0.163 | 0.596 | 0.652 | 0.038 | 0.073 | 0.073 | 0.312 | 0.48 | 0.71 | 0.53 | 0.64 | 0.44 | 0.57 |
| 7 | 24200034 | rs17290428 | 1.348 | 0.077 | 0.541 | 0.667 | 0.005 | 0.003 | 0.003 | 0 | 0.06 | 0.01 | 0.02 | 0.05 | 0.06 | 0.05 |
| 7 | 24200098 | rs156282 | -1.242 | 0.084 | 0.592 | 0.664 | 0.043 | 0.063 | 0.063 | 0.233 | 0.47 | 0.69 | 0.51 | 0.63 | 0.4 | 0.57 |

| | | | | | | | | | | | | | | | | |
|---|---|---|---|---|---|---|---|---|---|---|---|---|---|---|---|---|
| 7 | 24207028 | rs2189494 | 1.7 | 0.077 | 0.41 | 0.667 | -0.019 | -0.009 | -0.009 | 0.137 | 0.11 | 0.01 | 0.05 | 0.08 | 0.1 | 0.04 |
| 7 | 24208722 | rs2097791 | 0.083 | 0.06 | 0.675 | 0.69 | 0.001 | -0.004 | -0.004 | 0.253 | 0.26 | 0.21 | 0.38 | 0.48 | 0.24 | 0.45 |
| 7 | 24216810 | rs156310 | -1.253 | 0.062 | 0.415 | 0.65 | 0.048 | 0.064 | 0.064 | 0.204 | 0.47 | 0.74 | 0.51 | 0.61 | 0.41 | 0.57 |
| 7 | 24219642 | rs17369508 | 1.321 | 0.078 | 0.325 | 0.692 | 0.004 | -0.009 | -0.009 | 0.423 | 0.06 | 0.01 | 0.02 | 0.09 | 0.06 | 0.06 |
| 7 | 24219795 | rs17369536 | 0.154 | 0.084 | 0.312 | 0.675 | -0.003 | 0.007 | 0.007 | 0 | 0.19 | 0.04 | 0.09 | 0.16 | 0.2 | 0.1 |
| 7 | 24232285 | rs756500 | -0.124 | 0.018 | 0.564 | 0.718 | -0.016 | 0.071 | 0.071 | 0.047 | 0.27 | 0.52 | 0.41 | 0.48 | 0.24 | 0.45 |
| 7 | 24234644 | rs17369837 | 0.183 | 0.028 | 0.452 | 0.647 | 0.004 | 0.01 | 0.01 | 0 | 0.19 | 0.07 | 0.1 | 0.16 | 0.2 | 0.1 |
| 7 | 24237549 | rs156289 | -0.67 | 0.062 | 0.584 | 0.69 | -0.011 | 0.011 | 0.011 | 0 | 0.33 | 0.55 | 0.43 | 0.57 | 0.3 | 0.51 |
| 7 | 24239113 | rs156284 | -0.751 | 0.063 | 0.388 | 0.691 | 0.007 | -0.006 | -0.006 | 0 | 0.33 | 0.66 | 0.44 | 0.57 | 0.3 | 0.51 |
| 7 | 24244012 | rs13245518 | -1.287 | 0.076 | 0.196 | 0.647 | 0.048 | 0.097 | 0.097 | 0.335 | 0.48 | 0.8 | 0.53 | 0.65 | 0.44 | 0.57 |
| 7 | 24244627 | rs10254428 | -1.265 | -0.037 | 0.005 | 0.449 | 0 | 0.042 | 0.042 | 2.714 | 0.57 | 0.71 | 0.61 | 0.77 | 0.6 | 0.66 |
| 7 | 24245778 | rs1494193 | -0.007 | -0.29 | 0.523 | 0.878 | 0.012 | -0.054 | -0.054 | 0 | 0.23 | 0.13 | 0.36 | 0.49 | 0.21 | 0.44 |
| 7 | 24249469 | rs198275 | -0.219 | -0.243 | 0.449 | 0.737 | -0.02 | 0.044 | 0.044 | 0.021 | 0.25 | 0.43 | 0.39 | 0.56 | 0.21 | 0.45 |
| 7 | 24259922 | rs3025153 | 0.755 | -0.007 | 0.126 | 0.765 | 0.004 | 0.002 | 0.002 | 0 | 0.06 | 0.01 | 0.02 | 0.05 | 0.07 | 0.06 |
| 7 | 24264513 | rs16213 | -0.601 | 0.073 | -0.062 | 0.677 | 0.01 | 0.016 | 0.016 | 0.408 | 0.43 | 0.73 | 0.51 | 0.61 | 0.4 | 0.5 |
| 7 | 24266046 | rs12055984 | -0.312 | 0.03 | 0.622 | 0.715 | 0.011 | 0.048 | 0.048 | 0 | 0.09 | 0.27 | 0.08 | 0.15 | 0.09 | 0.08 |
| 7 | 24266104 | rs10268827 | 0.705 | 0.396 | 0.079 | 0.313 | -0.001 | 0.007 | 0.007 | 0 | 0.23 | 0.19 | 0.19 | 0.15 | 0.22 | 0.2 |
| 7 | 24267358 | rs16210 | 0.784 | 0.227 | 0.199 | 0.628 | -0.009 | -0.015 | -0.015 | 0.896 | 0.09 | 0.04 | 0.13 | 0.16 | 0.12 | 0.05 |
| 7 | 24269205 | rs16207 | 0.996 | 0.208 | 0.653 | 0.533 | 0.023 | 0.068 | 0.068 | 0.076 | 0.77 | 0.45 | 0.72 | 0.66 | 0.71 | 0.81 |
| 7 | 24269361 | rs16206 | 0.594 | -0.15 | -0.001 | 0.433 | 0.009 | -0.014 | -0.014 | 0 | 0.22 | 0.17 | 0.28 | 0.46 | 0.15 | 0.42 |
| 7 | 24269503 | rs16205 | -0.221 | 0.287 | 0.051 | 0.306 | 0.019 | 0.031 | 0.031 | 0.287 | 0.87 | 0.78 | 0.81 | 0.81 | 0.81 | 0.89 |
| 7 | 24269523 | rs16204 | -0.221 | 0.287 | 0.051 | 0.306 | 0.019 | 0.031 | 0.031 | 0.287 | 0.87 | 0.78 | 0.81 | 0.81 | 0.81 | 0.89 |
| 7 | 24273131 | rs16484 | 1.262 | 0.241 | -0.097 | 0.29 | 0 | -0.005 | -0.005 | 0 | 0.06 | 0.1 | 0.05 | 0.04 | 0.09 | 0.06 |
| 7 | 24274753 | rs12666025 | -0.283 | 0.28 | -0.162 | 0.344 | 0.015 | 0.027 | 0.027 | 0.306 | 0.87 | 0.79 | 0.82 | 0.82 | 0.81 | 0.89 |
| 7 | 24280534 | rs16483 | 0.812 | 0.068 | -0.506 | 0.121 | -0.012 | 0.034 | 0.034 | 0 | 0.1 | 0.02 | 0.04 | 0.04 | 0.1 | 0.1 |
| 7 | 24282334 | rs6461787 | -1.036 | -0.214 | -0.227 | 0.022 | 0.018 | 0.031 | 0.031 | 0 | 0.45 | 0.62 | 0.54 | 0.8 | 0.4 | 0.61 |
| 7 | 24284281 | rs2006644 | -0.413 | -0.5 | 0.004 | 0.268 | -0.044 | 0.035 | 0.035 | 0 | 0.32 | 0.49 | 0.4 | 0.59 | 0.23 | 0.49 |
| 7 | 24284325 | rs2006643 | -0.783 | -0.609 | -0.09 | 0.187 | -0.037 | 0.014 | 0.014 | 0 | 0.37 | 0.57 | 0.41 | 0.59 | 0.27 | 0.58 |
| 7 | 24294750 | rs16180 | -1.298 | -0.384 | -0.204 | 0.18 | 0.021 | 0.025 | 0.025 | 0 | 0.47 | 0.63 | 0.51 | 0.73 | 0.38 | 0.61 |
| 7 | 24294993 | rs16179 | 0.426 | -0.78 | 0.081 | 0.62 | -0.025 | 0.183 | 0.183 | 0 | 0.1 | 0.27 | 0.28 | 0.4 | 0.06 | 0.2 |
| 7 | 24295850 | rs16177 | -0.691 | -0.797 | 0.127 | 0.604 | -0.041 | 0.137 | 0.137 | 0 | 0.17 | 0.41 | 0.31 | 0.41 | 0.11 | 0.29 |
| 7 | 24302220 | rs16166 | -1.366 | -0.841 | 0.058 | 0.549 | 0.019 | 0.016 | 0.016 | 0 | 0.46 | 0.63 | 0.48 | 0.73 | 0.35 | 0.6 |
| 7 | 24303807 | rs16162 | -0.609 | -0.884 | 0.157 | 0.585 | -0.046 | 0.143 | 0.143 | 0.028 | 0.16 | 0.35 | 0.3 | 0.42 | 0.09 | 0.28 |
| 7 | 24305843 | rs7804106 | 0.63 | -0.762 | 0.143 | 0.556 | 0.001 | 0.001 | 0.001 | 0 | 0.64 | 0.65 | 0.78 | 0.67 | 0.66 | 0.57 |
| 7 | 24306749 | rs6960016 | 1.828 | -0.248 | -0.329 | -0.015 | 0.002 | 0.001 | 0.001 | 0 | 0.28 | 0.27 | 0.32 | 0.27 | 0.3 | 0.27 |
| 7 | 24307220 | rs721262 | 0.888 | -0.253 | -0.322 | 0.249 | -0.001 | 0.022 | 0.022 | 0.673 | 0.48 | 0.36 | 0.67 | 0.64 | 0.51 | 0.5 |
| 7 | 24311728 | rs13223753 | 2.358 | -0.196 | -0.37 | 0.257 | -0.012 | 0.002 | 0.002 | 0 | 0.13 | 0.1 | 0.1 | 0 | 0.15 | 0.07 |
| 7 | 24312294 | rs10241609 | 1.969 | -0.253 | -0.4 | 0.32 | -0.007 | -0.014 | -0.014 | 0 | 0.08 | 0.02 | 0.07 | 0.16 | 0.1 | 0.04 |
| 7 | 24322062 | rs16479 | 1.262 | -0.237 | -0.351 | 0.296 | -0.003 | -0.008 | -0.008 | 0.111 | 0.05 | 0.07 | 0.02 | 0.02 | 0.04 | 0.09 |
| 7 | 24322325 | rs16149 | -0.022 | -0.343 | -0.257 | 0.285 | -0.006 | -0.002 | -0.002 | 0.199 | 0.3 | 0.36 | 0.19 | 0.32 | 0.26 | 0.33 |
| 7 | 24323410 | rs16147 | 0.83 | -0.261 | -0.323 | 0.287 | -0.001 | 0.02 | 0.02 | 0.645 | 0.48 | 0.36 | 0.67 | 0.64 | 0.51 | 0.5 |
| 7 | 24324759 | rs16141 | 0.956 | -0.081 | -0.379 | 0.2 | -0.002 | 0.017 | 0.017 | 1.126 | 0.47 | 0.37 | 0.64 | 0.66 | 0.49 | 0.49 |
| 7 | 24327526 | rs3779477 | 1.26 | -0.766 | -0.311 | 0.335 | -0.006 | 0.011 | 0.011 | 0.043 | 0.42 | 0.34 | 0.4 | 0.24 | 0.46 | 0.31 |
| 7 | 24329837 | rs16131 | 2.102 | -0.7 | -0.317 | 0.226 | -0.016 | 0.021 | 0.021 | 0 | 0.13 | 0.07 | 0.1 | 0.05 | 0.15 | 0.07 |
| 7 | 24330765 | rs16129 | 0.821 | -0.801 | -0.281 | 0.284 | -0.001 | 0.007 | 0.007 | 0.746 | 0.48 | 0.43 | 0.67 | 0.65 | 0.51 | 0.5 |
| 7 | 24332135 | rs16123 | 1.549 | -0.933 | -0.429 | 0.761 | -0.005 | 0.071 | 0.071 | 0 | 0.05 | 0.07 | 0.24 | 0.28 | 0.04 | 0.18 |
| 7 | 24334069 | rs16472 | -0.446 | -0.79 | -0.575 | 0.09 | 0 | -0.006 | -0.006 | 2.768 | 0.94 | 0.92 | 0.97 | 0.98 | 0.92 | 0.92 |
| 7 | 24335527 | rs17374047 | 0.171 | -0.702 | -1.323 | 0.194 | 0.039 | 0.031 | 0.031 | 0 | 0.22 | 0.08 | 0.1 | 0.17 | 0.16 | 0.31 |
| 7 | 24341544 | rs10269857 | 0.777 | -0.615 | -1.3 | 0.17 | 0.025 | 0.001 | 0.001 | 0.01 | 0.61 | 0.68 | 0.57 | 0.32 | 0.68 | 0.42 |
| 7 | 24343178 | rs16466 | 1.633 | -0.728 | -2.113 | -0.351 | -0.025 | 0.001 | 0.001 | 0.161 | 0.16 | 0.01 | 0.12 | 0.12 | 0.17 | 0.1 |
| 7 | 24344323 | rs16101 | 0.368 | -0.795 | -1.658 | -0.096 | 0.001 | -0.004 | -0.004 | 0 | 0.19 | 0.03 | 0.12 | 0.21 | 0.17 | 0.27 |
| 7 | 24345005 | rs16098 | -0.106 | -0.656 | -1.54 | 0.007 | 0.001 | -0.003 | -0.003 | 0 | 0.25 | 0.05 | 0.16 | 0.24 | 0.21 | 0.32 |
| 7 | 24347891 | rs16089 | -0.494 | -0.642 | -1.403 | 0.04 | 0.006 | 0.033 | 0.033 | 0.574 | 0.3 | 0.12 | 0.17 | 0.25 | 0.24 | 0.4 |
| 7 | 24348217 | rs16087 | -0.848 | -0.563 | -1.336 | -0.166 | 0.018 | 0.121 | 0.121 | 0.595 | 0.48 | 0.23 | 0.29 | 0.31 | 0.43 | 0.47 |
| 7 | 24349013 | rs16085 | -0.938 | -0.566 | -1.194 | -0.154 | 0.019 | 0.068 | 0.068 | 0.542 | 0.49 | 0.33 | 0.29 | 0.32 | 0.43 | 0.48 |
| 7 | 24350504 | rs16082 | 0.391 | -0.002 | -0.655 | -0.12 | -0.005 | -0.053 | -0.053 | 0 | 0.2 | 0.28 | 0.13 | 0.08 | 0.19 | 0.08 |
| 7 | 24353182 | rs16079 | -0.516 | 0.02 | -0.652 | 0.244 | -0.001 | -0.001 | -0.001 | 0 | 0.47 | 0.52 | 0.56 | 0.62 | 0.4 | 0.46 |
| 7 | 24355042 | rs10487603 | 0.169 | 0.175 | -0.645 | 0.332 | 0 | -0.005 | -0.005 | 0 | 0.13 | 0.2 | 0.13 | 0.09 | 0.13 | 0.04 |
| 7 | 24356504 | rs1012768 | 0.92 | 0.165 | -0.811 | 0.372 | -0.01 | 0.038 | 0.038 | 0 | 0.43 | 0.31 | 0.38 | 0.27 | 0.5 | 0.41 |
| 7 | 24358103 | rs17316431 | 0.973 | 0.811 | -1.138 | 0.124 | -0.084 | -0.027 | -0.027 | 0 | 0.14 | 0.01 | 0.17 | 0.1 | 0.2 | 0.19 |
| 7 | 24358497 | rs6461794 | -0.234 | -0.381 | -0.573 | 0.295 | 0.005 | 0.009 | 0.009 | 0 | 0.34 | 0.34 | 0.29 | 0.23 | 0.3 | 0.17 |
| 7 | 24364791 | rs10233456 | -1.26 | -0.429 | -0.578 | 0.404 | 0.002 | 0.027 | 0.027 | 0 | 0.92 | 0.89 | 0.95 | 0.83 | 0.94 | 0.86 |
| 7 | 24368686 | rs10487605 | 0.979 | -0.289 | -0.677 | 0.377 | 0.017 | 0.032 | 0.032 | 0 | 0.21 | 0.1 | 0.13 | 0.11 | 0.2 | 0.11 |
| 7 | 24370883 | rs2521624 | -0.821 | -0.33 | -0.751 | 0.377 | -0.007 | -0.005 | -0.005 | 0 | 0.43 | 0.49 | 0.51 | 0.63 | 0.38 | 0.43 |
| 7 | 24376523 | rs2521630 | -0.29 | -0.31 | -0.911 | 0.368 | 0.006 | 0.004 | 0.004 | 0 | 0.34 | 0.31 | 0.25 | 0.29 | 0.3 | 0.22 |
| 7 | 24378040 | rs2521634 | 0.455 | -0.233 | -0.854 | 0.339 | -0.018 | -0.01 | -0.01 | 0 | 0.24 | 0.36 | 0.17 | 0.19 | 0.24 | 0.16 |
| 7 | 24380385 | rs2245657 | -1.132 | -0.121 | -0.954 | 0.385 | 0.006 | 0.013 | 0.013 | 0 | 0.89 | 0.93 | 0.95 | 0.93 | 0.91 | 0.86 |
| 7 | 24381817 | rs2722401 | -1.229 | 0.227 | -0.528 | 0.713 | -0.001 | 0.155 | 0.155 | 0 | 0.24 | 0.49 | 0.4 | 0.53 | 0.21 | 0.32 |
| 7 | 24381949 | rs10260381 | 0.691 | 0.282 | -0.741 | 0.759 | -0.029 | -0.022 | -0.022 | 6.688 | 0.1 | 0.01 | 0.13 | 0.13 | 0.1 | 0.07 |

| | | | | | | | | | | | | | | | | |
|---|---|---|---|---|---|---|---|---|---|---|---|---|---|---|---|---|
| 7 | 24382726 | rs2722403 | -0.767 | 0.302 | -0.708 | 0.606 | -0.002 | 0.093 | 0.093 | 0 | 0.21 | 0.29 | 0.37 | 0.53 | 0.18 | 0.31 |
| 7 | 24382901 | rs2722404 | -0.894 | 0.522 | -0.715 | 0.612 | 0.013 | 0.022 | 0.022 | 0 | 0.49 | 0.61 | 0.55 | 0.65 | 0.46 | 0.44 |
| 7 | 24383941 | rs2390970 | -0.85 | 0.781 | -0.359 | 0.921 | 0.011 | 0.153 | 0.153 | 0.011 | 0.12 | 0.32 | 0.25 | 0.31 | 0.11 | 0.2 |
| 7 | 24390824 | rs6956026 | 0.075 | 1.221 | -1.541 | -0.146 | -0.087 | 0.112 | 0.112 | 0 | 0.67 | 0.9 | 0.69 | 0.84 | 0.56 | 0.79 |
| 7 | 24395900 | rs10238015 | 0.268 | 1.087 | -0.564 | -0.286 | -0.002 | 0.001 | 0.001 | 0.575 | 0.54 | 0.55 | 0.59 | 0.74 | 0.48 | 0.67 |
| 7 | 24402318 | rs6948996 | 0.275 | 0.925 | 0.366 | -0.189 | -0.002 | 0.011 | 0.011 | 0.548 | 0.54 | 0.55 | 0.55 | 0.69 | 0.48 | 0.66 |
| 7 | 24405250 | rs10253625 | 0.281 | 0.914 | 0.472 | -0.197 | -0.001 | -0.001 | -0.001 | 0 | 0.54 | 0.54 | 0.55 | 0.69 | 0.48 | 0.66 |
| 7 | 24405358 | rs10253966 | 0.251 | 0.901 | 0.741 | -0.203 | -0.004 | 0.019 | 0.019 | 0.615 | 0.54 | 0.59 | 0.55 | 0.69 | 0.48 | 0.66 |
| 7 | 24405877 | rs2390975 | 0.62 | 0.567 | 0.442 | -0.714 | -0.002 | 0.072 | 0.072 | 1.67 | 0.4 | 0.19 | 0.45 | 0.69 | 0.36 | 0.64 |
| 7 | 24406877 | rs17149360 | 0.419 | 0.957 | 0.42 | -0.993 | -0.004 | -0.014 | -0.014 | 0 | 0.36 | 0.37 | 0.32 | 0.23 | 0.3 | 0.18 |
| 7 | 24408359 | rs11970778 | 0.478 | 1.004 | 0.121 | -1.191 | -0.001 | -0.039 | -0.039 | 0 | 0.15 | 0.18 | 0.1 | 0 | 0.15 | 0.04 |
| 7 | 24411737 | rs34173602 | 0.512 | 1.377 | 0.377 | -1.209 | -0.008 | -0.027 | -0.027 | 0 | 0.37 | 0.4 | 0.31 | 0.24 | 0.32 | 0.2 |
| 7 | 24415851 | rs6950687 | 0.513 | 1.474 | 0.627 | -1.186 | -0.01 | -0.031 | -0.031 | 0.239 | 0.37 | 0.41 | 0.31 | 0.24 | 0.32 | 0.2 |
| 7 | 24417595 | rs7803032 | -0.349 | 1.211 | 0.081 | -1.202 | -0.062 | 0.107 | 0.107 | 0 | 0.6 | 0.8 | 0.47 | 0.25 | 0.56 | 0.33 |
| 7 | 24427853 | rs2521643 | -0.263 | 1.029 | 0.026 | -0.355 | -0.021 | 0.001 | 0.001 | 0.142 | 0.66 | 0.7 | 0.58 | 0.36 | 0.59 | 0.48 |
| 7 | 24428993 | rs4722352 | 0.836 | 0.534 | 0.044 | 0.411 | -0.019 | 0.052 | 0.052 | 0 | 0.08 | 0.15 | 0.13 | 0.24 | 0.05 | 0.13 |
| 7 | 24431105 | rs2722394 | 1.097 | 0.275 | -0.136 | 0.168 | 0.004 | 0.003 | 0.003 | 1.458 | 0.41 | 0.38 | 0.34 | 0.25 | 0.33 | 0.28 |
| 7 | 24431549 | rs10487611 | 0.275 | 0.794 | -0.73 | 0.383 | -0.031 | -0.057 | -0.057 | 0 | 0.24 | 0.1 | 0.24 | 0.34 | 0.26 | 0.37 |
| 7 | 24433001 | rs2521646 | 0.095 | 0.574 | -0.843 | 0.204 | -0.061 | -0.065 | -0.065 | 0 | 0.7 | 0.85 | 0.74 | 0.6 | 0.62 | 0.63 |
| 7 | 24434310 | rs11765036 | 0.309 | 0.637 | -1.031 | 0.333 | -0.073 | -0.018 | -0.018 | 0 | 0.15 | 0.01 | 0.15 | 0.12 | 0.2 | 0.18 |
| 7 | 24442486 | rs17149415 | -0.204 | -0.465 | -0.496 | -0.102 | -0.029 | -0.067 | -0.067 | 0 | 0.27 | 0.44 | 0.15 | 0.12 | 0.2 | 0.18 |
| 7 | 24442911 | rs4722357 | -1.693 | -0.462 | -0.424 | -0.049 | 0.008 | 0.013 | 0.013 | 1.192 | 0.67 | 0.73 | 0.69 | 0.8 | 0.7 | 0.84 |
| 7 | 24444385 | rs2106596 | -0.185 | -0.425 | -0.428 | 0.296 | -0.009 | -0.018 | -0.018 | 1.003 | 0.33 | 0.4 | 0.4 | 0.24 | 0.29 | 0.33 |
| 7 | 24444777 | rs4956 | -1.491 | -0.532 | -0.725 | 0.314 | 0.018 | 0.019 | 0.019 | 1.188 | 0.64 | 0.78 | 0.68 | 0.68 | 0.68 | 0.74 |
| 7 | 24445246 | rs92 | -0.401 | 0.103 | -0.642 | 0.611 | -0.078 | -0.052 | -0.052 | 0 | 0.33 | 0.19 | 0.29 | 0.57 | 0.43 | 0.53 |
| 7 | 24448899 | rs6956326 | -0.554 | 0.628 | -0.409 | 0.574 | 0.041 | 0.043 | 0.043 | 0 | 0.33 | 0.44 | 0.36 | 0.49 | 0.41 | 0.47 |
| 7 | 24454198 | rs9785009 | -1.112 | 0.288 | -0.647 | 0.319 | 0.064 | 0.054 | 0.054 | 0.459 | 0.52 | 0.74 | 0.48 | 0.63 | 0.6 | 0.63 |
| 7 | 24456585 | rs17149463 | -0.068 | 1.057 | -0.7 | 0.406 | -0.036 | -0.043 | -0.043 | 0 | 0.26 | 0.1 | 0.17 | 0.36 | 0.32 | 0.31 |
| 7 | 24462817 | rs9638820 | -0.937 | 0.549 | -0.389 | -0.398 | -0.001 | -0.05 | -0.05 | 0 | 0.45 | 0.47 | 0.56 | 0.88 | 0.52 | 0.75 |
| 7 | 24464343 | rs10228771 | -0.407 | 0.377 | -0.53 | -0.169 | -0.002 | -0.008 | -0.008 | 0 | 0.45 | 0.51 | 0.45 | 0.32 | 0.54 | 0.33 |
| 7 | 24465784 | rs6942803 | 0.474 | -0.26 | -0.524 | -0.726 | -0.003 | 0.045 | 0.045 | 0.364 | 0.55 | 0.65 | 0.36 | 0.24 | 0.57 | 0.38 |
| 7 | 24468285 | rs7796645 | 1.423 | -0.234 | -0.563 | -0.989 | 0.001 | -0.012 | -0.012 | 0 | 0.26 | 0.3 | 0.2 | 0.06 | 0.36 | 0.2 |
| 7 | 24468539 | rs12533743 | 1.414 | -0.189 | -0.545 | -0.976 | 0.007 | -0.018 | -0.018 | 0 | 0.24 | 0.3 | 0.2 | 0.06 | 0.35 | 0.2 |
| 7 | 24480784 | rs17327167 | 0.755 | 0.061 | -0.398 | -0.604 | -0.008 | 0.003 | 0.003 | 0.016 | 0.07 | 0.06 | 0.2 | 0 | 0.14 | 0.07 |
| 7 | 24488820 | rs10281745 | -0.736 | -0.546 | -0.426 | -0.529 | 0.009 | 0.011 | 0.011 | 1.513 | 0.42 | 0.67 | 0.4 | 0.36 | 0.37 | 0.39 |
| 7 | 24490073 | rs7782559 | 0.935 | -0.838 | -0.782 | -0.429 | 0.032 | -0.028 | -0.028 | 2.872 | 0.13 | 0.06 | 0.04 | 0.19 | 0.07 | 0.13 |
| 7 | 24491000 | rs17149501 | 1.02 | -0.504 | -0.55 | -0.667 | 0.003 | 0.007 | 0.007 | 14.852 | 0.17 | 0.18 | 0.11 | 0.06 | 0.14 | 0.11 |
| 7 | 24493240 | rs10224443 | -0.184 | -0.745 | -0.682 | -0.692 | 0.005 | 0.02 | 0.02 | 0 | 0.77 | 0.83 | 0.84 | 0.89 | 0.84 | 0.86 |
| 7 | 24504934 | rs17149516 | 0.726 | -0.732 | -0.251 | -0.541 | -0.006 | -0.028 | -0.028 | 0 | 0.06 | 0.16 | 0.02 | 0 | 0.03 | 0 |
| 7 | 24505207 | rs17149519 | 0.681 | -1.079 | -0.641 | -0.449 | 0.048 | -0.023 | -0.023 | 0 | 0.14 | 0.02 | 0.04 | 0.19 | 0.07 | 0.18 |
| 7 | 24513648 | rs7789043 | 0.646 | -1.101 | -0.408 | -0.468 | 0.015 | 0.001 | 0.001 | 0 | 0.14 | 0.09 | 0.05 | 0.18 | 0.07 | 0.17 |
| 7 | 24514729 | rs6961237 | -0.928 | -1.028 | -0.457 | 0.114 | 0.002 | 0.047 | 0.047 | 0 | 0.93 | 0.92 | 0.94 | 0.77 | 0.91 | 0.89 |
| 7 | 24515976 | rs739539 | 2.091 | -0.942 | -0.716 | -1.151 | -0.002 | 0.01 | 0.01 | 7.568 | 0.11 | 0.05 | 0.07 | 0 | 0.11 | 0.04 |
| 7 | 24516352 | rs17149538 | 0.269 | -1.093 | -0.424 | -0.921 | -0.029 | 0.006 | 0.006 | 0 | 0.17 | 0.24 | 0.08 | 0.25 | 0.1 | 0.19 |
| 7 | 24516645 | rs6943268 | 0.758 | -0.145 | -0.041 | -0.75 | -0.001 | -0.01 | -0.01 | 0 | 0.53 | 0.55 | 0.36 | 0.31 | 0.59 | 0.41 |
| 7 | 24517195 | rs719082 | 0.38 | -0.273 | 0.234 | -0.519 | -0.013 | -0.014 | -0.014 | 0 | 0.63 | 0.47 | 0.55 | 0.31 | 0.75 | 0.47 |
| 7 | 24521566 | rs11767990 | 0.322 | -0.199 | -0.442 | -0.524 | -0.004 | -0.004 | -0.004 | 0 | 0.64 | 0.65 | 0.57 | 0.3 | 0.75 | 0.46 |
| 7 | 24530016 | rs976681 | -0.545 | -0.168 | -0.18 | -0.335 | 0.008 | 0.008 | 0.008 | 0.926 | 0.26 | 0.41 | 0.36 | 0.65 | 0.25 | 0.47 |
| 7 | 24534070 | rs2521230 | -1.222 | -0.471 | -0.563 | -1.093 | -0.029 | 0.089 | 0.089 | 0.041 | 0.62 | 0.78 | 0.7 | 0.89 | 0.51 | 0.77 |
| 7 | 24540300 | rs17387919 | 0.102 | -1.069 | -0.44 | -1.154 | 0.01 | 0.009 | 0.009 | 0.237 | 0.19 | 0.1 | 0.22 | 0.2 | 0.14 | 0.22 |
| 7 | 24540490 | rs4236315 | -1.123 | -1.433 | -0.153 | -1.127 | 0.005 | 0.03 | 0.03 | 0 | 0.51 | 0.56 | 0.61 | 0.78 | 0.43 | 0.73 |
| 7 | 24541271 | rs4719771 | 1.086 | -1.151 | -0.555 | -0.999 | 0 | 0.056 | 0.056 | 0.642 | 0.15 | 0.17 | 0.15 | 0.51 | 0.13 | 0.38 |
| 7 | 24541524 | rs2521222 | -2.116 | -1.332 | -0.998 | -1.507 | -0.169 | 0.255 | 0.255 | 0 | 0.65 | 0.96 | 0.7 | 1 | 0.5 | 0.85 |
| 7 | 24547294 | rs11524615 | 0.53 | -1.176 | -0.439 | -0.729 | -0.002 | -0.039 | -0.039 | 0 | 0.13 | 0.04 | 0.23 | 0.27 | 0.11 | 0.3 |
| 7 | 24548327 | rs11771214 | 0.851 | -1.202 | -0.167 | -0.546 | -0.005 | 0.075 | 0.075 | 1.419 | 0.13 | 0.15 | 0.16 | 0.53 | 0.09 | 0.36 |
| 7 | 24549526 | rs11976703 | -0.273 | -1.483 | 0.115 | -0.719 | -0.042 | -0.014 | -0.014 | 0 | 0.15 | 0.33 | 0.09 | 0.12 | 0.07 | 0.04 |
| 7 | 24553708 | rs4722370 | 0.783 | -0.889 | -0.017 | -0.375 | -0.005 | 0.08 | 0.08 | 1.335 | 0.13 | 0.16 | 0.16 | 0.53 | 0.09 | 0.33 |
| 7 | 24556824 | rs4722372 | -0.607 | -1.004 | 0.065 | -0.487 | 0.006 | 0 | 0 | 0.412 | 0.31 | 0.24 | 0.39 | 0.68 | 0.24 | 0.57 |
| 7 | 24560690 | rs2711106 | -0.375 | -1.112 | 0.123 | -0.808 | -0.004 | -0.004 | -0.004 | 0 | 0.11 | 0.19 | 0.09 | 0.09 | 0.11 | 0.12 |
| 7 | 24561060 | rs2190436 | -1.353 | -1.154 | 0.371 | -0.828 | -0.001 | 0.007 | 0.007 | 0 | 0.17 | 0.3 | 0.2 | 0.18 | 0.19 | 0.15 |
| 7 | 24562158 | rs4719773 | 0.448 | -1.235 | 0.105 | -0.807 | -0.026 | -0.002 | -0.002 | 0 | 0.18 | 0.03 | 0.22 | 0.17 | 0.14 | 0.24 |
| 7 | 24569427 | rs2711098 | 0.236 | -0.99 | 0.546 | -0.433 | -0.007 | -0.041 | -0.041 | 5.885 | 0.08 | 0.14 | 0.04 | 0 | 0.06 | 0.05 |
| 7 | 24569933 | rs2711097 | -1.047 | -1.012 | 0.566 | -0.42 | -0.005 | 0.01 | 0.01 | 0 | 0.24 | 0.29 | 0.25 | 0.63 | 0.2 | 0.49 |
| 7 | 24578963 | rs2711095 | -1.317 | -0.605 | 0.195 | -0.635 | 0.008 | 0.023 | 0.023 | 0 | 0.17 | 0.39 | 0.2 | 0.18 | 0.19 | 0.15 |
| 7 | 24587380 | rs11767874 | -0.706 | -0.78 | -0.99 | -1.384 | 0.081 | 0.192 | 0.192 | 0 | 0.47 | 0.74 | 0.51 | 0.94 | 0.3 | 0.65 |
| 7 | 24600679 | rs2711125 | -0.53 | -0.391 | -0.86 | -1.21 | 0 | -0.009 | -0.009 | 0.172 | 0.13 | 0.15 | 0.17 | 0.06 | 0.16 | 0.15 |
| 7 | 24606157 | rs2529044 | -0.771 | -0.453 | -0.224 | -1.239 | 0.021 | -0.056 | -0.056 | 0.714 | 0.14 | 0.28 | 0.18 | 0.06 | 0.17 | 0.15 |
| 7 | 24625582 | rs11761560 | 0.038 | -0.994 | 0.296 | -0.902 | -0.019 | 0.082 | 0.082 | 0.801 | 0.18 | 0.41 | 0.19 | 0.67 | 0.11 | 0.36 |
| 7 | 24631245 | rs4722378 | 0.894 | -1.06 | 0.816 | -0.037 | 0 | 0.085 | 0.085 | 1.864 | 0.12 | 0.13 | 0.11 | 0.46 | 0.11 | 0.33 |

| | | | | | | | | | | | | | | | | |
|---|---|---|---|---|---|---|---|---|---|---|---|---|---|---|---|---|
| 7 | 24632111 | rs2711134 | 0.483 | -1.098 | 0.801 | -0.092 | -0.008 | -0.032 | -0.032 | 0 | 0.08 | 0.15 | 0.05 | 0 | 0.06 | 0.1 |
| 7 | 24637056 | rs12700538 | 0.857 | -0.963 | 0.791 | 0.028 | 0 | 0.061 | 0.061 | 1.864 | 0.12 | 0.12 | 0.11 | 0.46 | 0.11 | 0.33 |
| 7 | 24680053 | rs2711132 | -2.012 | -0.643 | -0.505 | -0.867 | -0.033 | 0.086 | 0.086 | 0.78 | 0.86 | 0.99 | 0.92 | 1 | 0.88 | 0.97 |
| 7 | 24703535 | rs11982399 | 0.287 | -0.526 | 0.179 | -0.308 | 0.018 | 0.231 | 0.231 | 0.021 | 0.06 | 0.3 | 0.09 | 0.21 | 0 | 0.02 |
| 7 | 24713291 | rs12700541 | 0.653 | 0.519 | 0.019 | -0.139 | 0.035 | -0.046 | -0.046 | 0 | 0.1 | 0.14 | 0.1 | 0 | 0.19 | 0.02 |
| 7 | 24724066 | rs11769204 | 1.112 | -0.099 | 0.017 | 0.104 | -0.003 | -0.007 | -0.007 | 0.195 | 0.12 | 0.04 | 0.1 | 0.49 | 0.11 | 0.33 |
| 7 | 24738118 | rs6957782 | 0.929 | -0.091 | 0.121 | 0.045 | 0.003 | -0.035 | -0.035 | 0 | 0.06 | 0.14 | 0.04 | 0 | 0.07 | 0.02 |
| 7 | 24738372 | rs17149888 | 0.326 | -0.221 | 0.356 | 0.096 | -0.006 | 0.168 | 0.168 | 0.011 | 0.06 | 0.26 | 0.08 | 0.2 | 0 | 0.02 |
| 7 | 24742436 | rs17149912 | -0.126 | -0.872 | -0.102 | 0.016 | 0.05 | 0 | 0 | 0 | 0.25 | 0.14 | 0.26 | 0.28 | 0.15 | 0.22 |
| 7 | 24742553 | rs2240005 | -1.749 | -0.99 | 0.254 | -0.154 | 0.024 | 0.013 | 0.013 | 1.004 | 0.24 | 0.52 | 0.29 | 0.27 | 0.25 | 0.23 |
| 7 | 24742585 | rs2074142 | -1.219 | -2.3 | 0.295 | 0.601 | -0.014 | -0.008 | -0.008 | 1.254 | 0.41 | 0.56 | 0.5 | 0.71 | 0.25 | 0.67 |
| 7 | 24746022 | rs2269812 | 0 | -1.549 | -0.097 | 0.516 | 0.05 | 0 | 0 | 0 | 0.23 | 0.08 | 0.26 | 0.2 | 0.14 | 0.2 |
| 7 | 24748565 | rs2721810 | -2.621 | -1.118 | -0.051 | 0.458 | -0.113 | 0.303 | 0.303 | 0 | 0.59 | 0.94 | 0.7 | 0.98 | 0.49 | 0.76 |
| 7 | 24749223 | rs2521755 | -2.545 | -1.082 | 0.06 | 0.472 | -0.053 | 0.213 | 0.213 | 0 | 0.58 | 0.86 | 0.69 | 0.98 | 0.49 | 0.76 |
| 7 | 24756951 | rs12540919 | 0.244 | -1.078 | 0.185 | 1.057 | 0.012 | -0.01 | -0.01 | 0.332 | 0.14 | 0.04 | 0.08 | 0.13 | 0.1 | 0.11 |
| 7 | 24757202 | rs2237306 | -0.935 | -1.117 | 0.552 | 1.276 | 0.021 | 0.169 | 0.169 | 0.583 | 0.14 | 0.5 | 0.27 | 0.35 | 0.13 | 0.21 |
| 7 | 24758161 | rs874540 | -1.936 | -1.124 | 0.218 | 1.147 | 0.084 | 0.084 | 0.084 | 0.174 | 0.5 | 0.83 | 0.62 | 0.67 | 0.36 | 0.51 |
| 7 | 24758753 | rs754555 | -0.266 | -1.088 | 0.291 | 0.675 | -0.001 | 0.023 | 0.023 | 0 | 0.19 | 0.2 | 0.14 | 0.44 | 0.15 | 0.35 |
| 7 | 24758795 | rs876305 | -2.264 | -1.091 | 0.289 | 0.669 | 0.009 | 0.249 | 0.249 | 0.393 | 0.54 | 0.83 | 0.66 | 0.98 | 0.41 | 0.75 |
| 7 | 24758818 | rs754554 | -0.324 | -1.015 | 0.292 | 0.75 | -0.001 | 0.023 | 0.023 | 0 | 0.19 | 0.2 | 0.14 | 0.44 | 0.15 | 0.35 |
| 7 | 24761850 | rs2237315 | -1.503 | -0.986 | 0.295 | 0.834 | 0.01 | -0.012 | -0.012 | 0 | 0.39 | 0.64 | 0.55 | 0.54 | 0.33 | 0.41 |
| 7 | 24769089 | rs1001464 | -0.624 | -0.701 | 0.309 | 0.845 | -0.045 | 0.03 | 0.03 | 0.723 | 0.19 | 0.42 | 0.24 | 0.73 | 0.09 | 0.38 |
| 7 | 24769400 | rs2721798 | -1.015 | -0.395 | -0.21 | 0.953 | 0.012 | 0.009 | 0.009 | 0 | 0.13 | 0.22 | 0.18 | 0.15 | 0.17 | 0.18 |
| 7 | 24772686 | rs2521766 | 0.014 | -0.311 | -0.327 | 0.429 | 0.004 | -0.033 | -0.033 | 0.714 | 0.08 | 0.15 | 0.06 | 0 | 0.07 | 0.1 |
| 7 | 24781080 | rs11509139 | 0.347 | -0.339 | -0.836 | 0.517 | -0.002 | 0.001 | 0.001 | 0 | 0.06 | 0.06 | 0.13 | 0.08 | 0.11 | 0.08 |
| 7 | 24781792 | rs2237324 | -1.315 | -0.519 | -0.835 | 0.389 | 0.007 | 0.013 | 0.013 | 0.059 | 0.33 | 0.69 | 0.34 | 0.73 | 0.25 | 0.52 |
| 7 | 24784057 | rs2721801 | -0.203 | -0.588 | -0.69 | 0.4 | -0.029 | -0.045 | -0.045 | 0.066 | 0.16 | 0.59 | 0.12 | 0.09 | 0.1 | 0.13 |
| 7 | 24784403 | rs2521768 | -1.735 | -0.456 | -1.63 | 0.476 | -0.084 | 0.139 | 0.139 | 0 | 0.57 | 0.93 | 0.62 | 0.77 | 0.45 | 0.72 |
| 7 | 24788209 | rs4722385 | -0.013 | -0.177 | -0.717 | 0.918 | 0.012 | -0.011 | -0.011 | 1.702 | 0.16 | 0.08 | 0.24 | 0.21 | 0.11 | 0.17 |
| 7 | 24803592 | rs4722388 | 0.13 | -0.053 | -0.769 | 0.941 | 0.004 | -0.011 | -0.011 | 0 | 0.14 | 0.09 | 0.24 | 0.21 | 0.11 | 0.17 |
| 7 | 24804756 | rs9648009 | 0.093 | -0.018 | -0.812 | 0.939 | 0.004 | -0.018 | -0.018 | 0 | 0.15 | 0.07 | 0.24 | 0.21 | 0.11 | 0.17 |
| 7 | 24805050 | rs6976667 | -1.368 | -0.235 | -0.696 | 0.906 | 0.007 | 0.014 | 0.014 | 1.798 | 0.41 | 0.32 | 0.51 | 0.69 | 0.36 | 0.61 |
| 7 | 24808368 | rs17150118 | 0.015 | -0.159 | -1.122 | 1.18 | 0.016 | -0.09 | -0.09 | 1.037 | 0.17 | 0.04 | 0.12 | 0.4 | 0.13 | 0.32 |
| 7 | 24818656 | rs1468185 | -1.068 | 0.041 | -0.87 | 0.938 | -0.001 | 0.003 | 0.003 | 4.229 | 0.28 | 0.25 | 0.21 | 0.41 | 0.25 | 0.38 |
| 7 | 24826613 | rs17150155 | -0.548 | 0.101 | -1.47 | 0.698 | 0.015 | -0.066 | -0.066 | 0 | 0.3 | 0.08 | 0.36 | 0.6 | 0.24 | 0.46 |
| 7 | 24828055 | rs2391000 | 0.525 | 0.489 | -1.149 | 0.338 | -0.027 | -0.011 | -0.011 | 0 | 0.06 | 0.02 | 0.13 | 0.07 | 0.11 | 0.1 |
| 7 | 24837746 | rs10486432 | 0.166 | 0.215 | -1.266 | 0.213 | -0.026 | -0.005 | -0.005 | 0.018 | 0.09 | 0.01 | 0.14 | 0.06 | 0.1 | 0.13 |
| 7 | 24843621 | rs10227466 | -1.79 | -0.218 | -1.561 | -0.377 | -0.067 | 0.189 | 0.189 | 0 | 0.6 | 0.84 | 0.72 | 0.98 | 0.5 | 0.77 |
| 7 | 24843810 | rs2285651 | 0.232 | -0.695 | -1.571 | 0.121 | 0.023 | -0.053 | -0.053 | 0.875 | 0.18 | 0.1 | 0.13 | 0.5 | 0.11 | 0.33 |
| 7 | 24848829 | rs1012775 | -2.042 | 0.554 | -1.927 | -0.597 | -0.078 | 0.238 | 0.238 | 0 | 0.65 | 0.94 | 0.8 | 0.98 | 0.63 | 0.79 |
| 7 | 24850995 | rs9648315 | 0.837 | 0.693 | -1.927 | -0.563 | -0.061 | 0.033 | 0.033 | 0 | 0.13 | 0.02 | 0.2 | 0.05 | 0.23 | 0.14 |
| 7 | 24853496 | rs739627 | -1.153 | 0.176 | -1.667 | -0.424 | 0.043 | 0.303 | 0.303 | 0 | 0.5 | 0.83 | 0.58 | 0.93 | 0.39 | 0.64 |
| 7 | 24856032 | rs10215692 | 0.503 | 0.008 | -1.525 | 0.068 | -0.017 | -0.065 | -0.065 | 0 | 0.13 | 0.01 | 0.22 | 0.24 | 0.12 | 0.21 |
| 7 | 24857983 | rs2106943 | -0.622 | 0.017 | -1.291 | -0.866 | 0.001 | -0.045 | -0.045 | 0.486 | 0.46 | 0.61 | 0.4 | 0.05 | 0.56 | 0.25 |
| 7 | 24863109 | rs6957090 | -1.559 | -0.088 | -1.622 | -0.629 | 0.004 | 0.102 | 0.102 | 0 | 0.65 | 0.83 | 0.77 | 0.83 | 0.66 | 0.74 |
| 7 | 24863792 | rs17150239 | 0.715 | 0.088 | -1.412 | -0.139 | 0.025 | -0.07 | -0.07 | 0 | 0.16 | 0.07 | 0.09 | 0.52 | 0.11 | 0.4 |
| 7 | 24866690 | rs7802069 | -1.233 | 0.534 | -0.481 | -0.206 | -0.006 | -0.004 | -0.004 | 0.374 | 0.46 | 0.48 | 0.66 | 0.34 | 0.63 | 0.39 |
| 7 | 24868863 | rs17291810 | 0.547 | 1.536 | -1.012 | -0.065 | -0.052 | 0.068 | 0.068 | 2.002 | 0.21 | 0.09 | 0.25 | 0.05 | 0.29 | 0.13 |
| 7 | 24870722 | rs13225320 | 0.633 | 1.648 | -1.161 | 0.029 | -0.005 | -0.017 | -0.017 | 0 | 0.08 | 0.02 | 0.29 | 0.26 | 0.09 | 0.15 |
| 7 | 24874459 | rs2285688 | 0.613 | 1.57 | -1.224 | 0.029 | -0.093 | 0.08 | 0.08 | 1.899 | 0.2 | 0.02 | 0.25 | 0.06 | 0.29 | 0.13 |
| 7 | 24874499 | rs2285689 | -0.701 | 1.076 | -0.4 | 0.692 | 0.01 | 0.009 | 0.009 | 0 | 0.17 | 0.14 | 0.36 | 0.26 | 0.24 | 0.19 |
| 7 | 24886767 | rs16873714 | -0.626 | 0.47 | -0.108 | 1.313 | 0 | 0.066 | 0.066 | 0 | 0.13 | 0.22 | 0.25 | 0.34 | 0.12 | 0.27 |
| 7 | 24890808 | rs10238784 | -0.871 | -0.37 | 0.326 | 1.454 | -0.024 | 0.062 | 0.062 | 0.102 | 0.21 | 0.25 | 0.23 | 0.46 | 0.13 | 0.38 |
| 7 | 24891093 | rs17150305 | -1.684 | -0.176 | 0.607 | 1.191 | -0.012 | 0.081 | 0.081 | 0.084 | 0.3 | 0.44 | 0.29 | 0.46 | 0.26 | 0.4 |
| 7 | 24892514 | rs6956360 | -0.39 | -0.057 | 0.038 | 0.635 | -0.005 | -0.011 | -0.011 | 0 | 0.34 | 0.34 | 0.46 | 0.5 | 0.36 | 0.38 |
| 7 | 24893147 | rs4470905 | -1.681 | -0.451 | -0.167 | 0.808 | 0.018 | 0.034 | 0.034 | 0 | 0.53 | 0.66 | 0.7 | 0.75 | 0.59 | 0.62 |
| 7 | 24894255 | rs17293003 | 0.626 | -0.493 | -0.028 | 0.741 | -0.008 | 0.017 | 0.017 | 3.888 | 0.24 | 0.19 | 0.21 | 0.15 | 0.26 | 0.12 |
| 7 | 24895243 | rs739709 | 0.239 | -0.534 | -0.149 | 0.889 | -0.051 | 0.014 | 0.014 | 0 | 0.14 | 0.26 | 0.08 | 0.14 | 0.05 | 0.15 |
| 7 | 24895575 | rs2007271 | -1.426 | -0.43 | -0.389 | 0.867 | 0.015 | 0.026 | 0.026 | 0 | 0.51 | 0.59 | 0.69 | 0.85 | 0.59 | 0.68 |
| 7 | 24896313 | rs10259998 | -1.205 | -0.308 | 0.091 | 1.338 | 0.006 | -0.035 | -0.035 | 2.018 | 0.25 | 0.21 | 0.43 | 0.57 | 0.22 | 0.46 |
| 7 | 24897854 | rs6955303 | -0.989 | -0.373 | 0.036 | 1.154 | -0.009 | -0.028 | -0.028 | 0 | 0.21 | 0.36 | 0.12 | 0.14 | 0.18 | 0.17 |
| 7 | 24898123 | rs17150327 | 0.263 | -0.118 | -0.483 | 1.345 | -0.02 | -0.049 | -0.049 | 0.117 | 0.08 | 0 | 0.16 | 0.23 | 0.09 | 0.18 |
| 7 | 24902762 | rs2285690 | -1.535 | -0.253 | -0.187 | 0.693 | 0.011 | 0.012 | 0.012 | 0.182 | 0.37 | 0.66 | 0.45 | 0.76 | 0.29 | 0.63 |
| 7 | 24908697 | rs17228163 | 0.596 | 0.119 | -0.273 | 0.383 | -0.004 | 0.008 | 0.008 | 0 | 0.23 | 0.17 | 0.25 | 0.15 | 0.26 | 0.2 |
| 7 | 24911061 | rs6952356 | -1.457 | -0.012 | 0.02 | 0.024 | -0.001 | -0.017 | -0.017 | 0 | 0.41 | 0.6 | 0.44 | 0.83 | 0.37 | 0.64 |
| 7 | 24911357 | rs6952815 | 0.562 | 0.05 | 0.345 | 0.3 | 0.033 | 0.027 | 0.027 | 0.114 | 0.07 | 0.17 | 0.06 | 0.08 | 0.09 | 0.16 |
| 7 | 24917974 | rs6979475 | 0.94 | 0.056 | 0.234 | 0.321 | 0.002 | -0.01 | -0.01 | 0.019 | 0.23 | 0.19 | 0.25 | 0.15 | 0.26 | 0.2 |
| 7 | 24918330 | rs879496 | 0.258 | -0.203 | 0.456 | 1.012 | -0.001 | 0.001 | 0.001 | 0 | 0.1 | 0.05 | 0.2 | 0.53 | 0.1 | 0.33 |
| 7 | 24921240 | rs8180767 | -2.207 | -0.209 | 0.546 | 0.725 | 0.01 | 0.051 | 0.051 | 0.08 | 0.42 | 0.73 | 0.45 | 0.81 | 0.36 | 0.64 |

| | | | | | | | | | | | | | | | | |
|---|---|---|---|---|---|---|---|---|---|---|---|---|---|---|---|---|
| 7 | 24923761 | rs17150421 | -0.634 | -0.479 | 0.538 | 0.867 | -0.004 | 0.003 | 0.003 | 0 | 0.17 | 0.24 | 0.17 | 0.18 | 0.16 | 0.21 |
| 7 | 24924015 | rs4141251 | -1.808 | -0.482 | 0.54 | 0.759 | -0.019 | -0.019 | -0.019 | 0 | 0.35 | 0.62 | 0.42 | 0.73 | 0.29 | 0.56 |
| 7 | 24925834 | rs6946141 | -1.664 | -0.139 | -0.005 | 0.528 | 0.007 | 0.142 | 0.142 | 0 | 0.57 | 0.76 | 0.66 | 0.92 | 0.58 | 0.76 |
| 7 | 24932346 | rs10486438 | 0.716 | 0.145 | 0.085 | 1.436 | -0.003 | 0.029 | 0.029 | 0 | 0.06 | 0.05 | 0.12 | 0.25 | 0.08 | 0.15 |
| 7 | 24933691 | rs10486439 | 0.522 | -0.015 | 0.008 | 1.462 | 0.025 | -0.022 | -0.022 | 0 | 0.06 | 0.02 | 0.09 | 0.11 | 0.04 | 0.06 |
| 7 | 24935581 | rs144 | 0.149 | 0.25 | 0.435 | 1.549 | -0.003 | -0.007 | -0.007 | 0.11 | 0.24 | 0.21 | 0.32 | 0.4 | 0.28 | 0.35 |
| 7 | 24936098 | rs136 | 0.161 | 0.252 | 0.436 | 1.547 | -0.003 | -0.007 | -0.007 | 0 | 0.24 | 0.21 | 0.32 | 0.41 | 0.28 | 0.35 |
| 7 | 24948424 | rs12216670 | 0.812 | 0.493 | 0.71 | 1.517 | 0.008 | 0.008 | 0.008 | 0 | 0.05 | 0.06 | 0.05 | 0.07 | 0.08 | 0.14 |
| 7 | 24949338 | rs143 | -0.958 | 0.402 | 0.85 | 1.427 | 0.008 | -0.007 | -0.007 | 0 | 0.36 | 0.36 | 0.49 | 0.73 | 0.44 | 0.64 |
| 7 | 24953647 | rs10215435 | 0.782 | 0.577 | 0.568 | 1.587 | 0.002 | 0.047 | 0.047 | 0 | 0.06 | 0.08 | 0.12 | 0.25 | 0.08 | 0.15 |
| 7 | 24961093 | rs7778780 | -1.7 | 0.654 | 0.886 | 1.466 | 0.071 | 0.171 | 0.171 | 0 | 0.2 | 0.47 | 0.24 | 0.4 | 0.25 | 0.34 |
| 7 | 24962199 | rs763852 | -0.738 | 0.611 | 0.734 | 1.254 | 0.034 | 0.191 | 0.191 | 0 | 0.16 | 0.36 | 0.28 | 0.49 | 0.19 | 0.31 |
| 7 | 24962751 | rs10486440 | -0.719 | 0.574 | 0.436 | 1.248 | 0.011 | 0.1 | 0.1 | 0 | 0.17 | 0.22 | 0.27 | 0.5 | 0.21 | 0.33 |
| 7 | 24964056 | rs117 | 0.022 | 0.46 | 0.002 | 1.031 | -0.005 | -0.005 | -0.005 | 0 | 0.25 | 0.18 | 0.32 | 0.36 | 0.28 | 0.36 |
| 7 | 24969081 | rs6461830 | -0.583 | 0.184 | 0.17 | 1.084 | -0.001 | 0.021 | 0.021 | 0 | 0.12 | 0.23 | 0.12 | 0.2 | 0.1 | 0.22 |
| 7 | 24970548 | rs2391007 | 0.715 | 0.308 | 0.066 | 1.126 | -0.001 | 0 | 0 | 0.018 | 0.64 | 0.66 | 0.62 | 0.56 | 0.63 | 0.62 |
| 7 | 24970805 | rs6951637 | -0.155 | 0.345 | -0.404 | 0.51 | -0.002 | 0.003 | 0.003 | 0 | 0.11 | 0.13 | 0.07 | 0.12 | 0.11 | 0.19 |
| 7 | 24971074 | rs4332039 | 1.759 | 0.321 | -0.991 | -0.172 | -0.016 | 0.101 | 0.101 | 0 | 0.29 | 0.11 | 0.17 | 0.05 | 0.29 | 0.23 |
| 7 | 24971213 | rs2391009 | -0.304 | 0.325 | -0.663 | 0.033 | -0.003 | 0.001 | 0.001 | 0 | 0.77 | 0.78 | 0.75 | 0.84 | 0.7 | 0.86 |
| 7 | 24972658 | rs10265239 | 0.278 | 0.1 | -0.585 | 0.057 | -0.002 | -0.034 | -0.034 | 0.052 | 0.2 | 0.21 | 0.18 | 0.03 | 0.17 | 0.12 |
| 7 | 24974501 | rs6952796 | 0.079 | 0.013 | -0.976 | 0.713 | -0.002 | 0.004 | 0.004 | 0 | 0.05 | 0.04 | 0.02 | 0.07 | 0.02 | 0.07 |
| 7 | 24979355 | rs757139 | 0.235 | 0.012 | -1.041 | 0.565 | 0.018 | 0.032 | 0.032 | 0 | 0.51 | 0.29 | 0.41 | 0.55 | 0.44 | 0.64 |
| 7 | 24979715 | rs10242782 | 0.509 | 0.085 | -0.908 | 0.198 | -0.007 | 0.119 | 0.119 | 0 | 0.42 | 0.24 | 0.22 | 0.07 | 0.41 | 0.24 |
| 7 | 24980171 | rs11771668 | 0.619 | -0.085 | -1.116 | 0.532 | -0.006 | -0.073 | -0.073 | 0.022 | 0.39 | 0.14 | 0.31 | 0.39 | 0.38 | 0.47 |
| 7 | 24982529 | rs4236316 | -0.077 | 0.074 | -0.566 | -0.091 | 0.004 | 0.006 | 0.006 | 0 | 0.27 | 0.24 | 0.3 | 0.07 | 0.21 | 0.13 |
| 7 | 24982863 | rs2710993 | -0.862 | 0.296 | -0.825 | -0.183 | -0.063 | 0.038 | 0.038 | 0 | 0.83 | 0.95 | 0.85 | 0.88 | 0.7 | 0.91 |
| 7 | 24984666 | rs2710995 | 0.618 | 0.134 | -0.06 | -0.15 | -0.001 | 0.009 | 0.009 | 0 | 0.53 | 0.55 | 0.52 | 0.82 | 0.49 | 0.77 |
| 7 | 24985941 | rs12533171 | 1.973 | -0.091 | -0.559 | -0.322 | -0.05 | 0.14 | 0.14 | 0 | 0.3 | 0.1 | 0.21 | 0.05 | 0.34 | 0.12 |
| 7 | 24988434 | rs7803029 | 1.298 | -0.066 | -0.14 | 0.138 | -0.041 | 0.035 | 0.035 | 0.147 | 0.38 | 0.22 | 0.39 | 0.26 | 0.46 | 0.29 |
| 7 | 24993561 | rs2710998 | -0.328 | -0.435 | 0.523 | 0.211 | -0.044 | 0.067 | 0.067 | 0 | 0.23 | 0.32 | 0.31 | 0.56 | 0.12 | 0.5 |
| 7 | 25007109 | rs2522240 | 0.527 | -0.258 | 0.56 | 0.186 | 0.002 | 0.002 | 0.002 | 0 | 0.53 | 0.44 | 0.51 | 0.57 | 0.46 | 0.65 |
| 7 | 25012699 | rs149 | -0.374 | -0.218 | 0.649 | 0.035 | -0.01 | -0.01 | -0.01 | 0 | 0.33 | 0.39 | 0.32 | 0.27 | 0.27 | 0.26 |
| 7 | 25013982 | rs10486445 | 1.941 | -0.061 | 0.445 | -0.134 | -0.066 | 0.18 | 0.18 | 0 | 0.34 | 0.15 | 0.23 | 0.01 | 0.39 | 0.14 |
| 7 | 25018426 | rs11973976 | 0.608 | -0.207 | 0.609 | -0.163 | -0.006 | -0.017 | -0.017 | 0 | 0.23 | 0.24 | 0.21 | 0.06 | 0.13 | 0.13 |
| 7 | 25021625 | rs146 | 0.985 | -0.113 | 0.389 | -0.034 | -0.003 | 0.002 | 0.002 | 0 | 0.08 | 0.05 | 0.11 | 0.19 | 0.14 | 0.12 |
| 7 | 25023441 | rs179 | 0.262 | -0.315 | 0.613 | 0.35 | -0.018 | -0.028 | -0.028 | 0.008 | 0.69 | 0.77 | 0.63 | 0.51 | 0.64 | 0.47 |
| 7 | 25023700 | rs4719787 | 0.804 | -0.36 | 0.994 | 0.31 | 0.005 | -0.015 | -0.015 | 0 | 0.63 | 0.47 | 0.52 | 0.28 | 0.61 | 0.37 |
| 7 | 25031419 | rs2711005 | 0.254 | -0.361 | 0.868 | 0.461 | 0.012 | -0.017 | -0.017 | 0 | 0.19 | 0.13 | 0.16 | 0.31 | 0.09 | 0.47 |
| 7 | 25031874 | rs17232369 | 1.602 | -0.535 | 0.619 | 0.298 | -0.011 | 0.024 | 0.024 | 0 | 0.15 | 0.01 | 0.17 | 0.06 | 0.11 | 0.12 |
| 7 | 25033791 | rs175 | -1.424 | -0.251 | 0.915 | 0.436 | 0.003 | 0.012 | 0.012 | 0 | 0.54 | 0.63 | 0.53 | 0.56 | 0.41 | 0.74 |
| 7 | 25034131 | rs173 | 0.639 | -0.248 | 0.583 | 0.436 | -0.011 | -0.012 | -0.012 | 0 | 0.09 | 0.05 | 0.08 | 0.17 | 0.13 | 0.13 |
| 7 | 25043942 | rs17298333 | 2.415 | -0.19 | 0.628 | 0.388 | -0.038 | 0.13 | 0.13 | 0 | 0.3 | 0.1 | 0.21 | 0.01 | 0.34 | 0.11 |
| 7 | 25045363 | rs2522226 | 0.621 | -0.304 | 0.732 | 0.432 | -0.003 | 0.013 | 0.013 | 0 | 0.17 | 0.17 | 0.21 | 0.32 | 0.09 | 0.43 |
| 7 | 25046341 | rs169 | -1.927 | -0.262 | 0.724 | 0.531 | -0.034 | -0.027 | -0.027 | 0 | 0.89 | 0.95 | 0.86 | 0.83 | 0.78 | 0.94 |
| 7 | 25047483 | rs17150754 | 0.911 | -0.268 | 0.821 | 0.529 | 0.026 | 0.036 | 0.036 | 0 | 0.08 | 0.13 | 0.09 | 0.18 | 0.14 | 0.13 |
| 7 | 25052796 | rs2522251 | 1.14 | -0.258 | 0.675 | 0.471 | -0.006 | 0.054 | 0.054 | 0 | 0.51 | 0.28 | 0.43 | 0.32 | 0.48 | 0.56 |
| 7 | 25055740 | rs2040672 | -1.7 | -0.234 | 0.466 | 0.297 | -0.054 | -0.045 | -0.045 | 0 | 0.9 | 0.98 | 0.87 | 0.82 | 0.78 | 0.94 |
| 7 | 25061014 | rs4722416 | 0.714 | -0.452 | 0.768 | 0.211 | -0.032 | -0.054 | -0.054 | 0 | 0.23 | 0.32 | 0.21 | 0.08 | 0.14 | 0.14 |
| 7 | 25067042 | rs4722417 | -0.038 | -0.203 | 0.743 | 0.503 | 0.003 | -0.017 | -0.017 | 0 | 0.74 | 0.8 | 0.71 | 0.52 | 0.78 | 0.41 |
| 7 | 25068223 | rs17150790 | 0.399 | -0.207 | 0.881 | 0.56 | 0.015 | 0.088 | 0.088 | 0 | 0.05 | 0.27 | 0.06 | 0.12 | 0.03 | 0.09 |
| 7 | 25068398 | rs2391081 | 2.251 | -0.108 | -0.045 | 0.419 | -0.046 | 0.092 | 0.092 | 0 | 0.3 | 0.1 | 0.26 | 0.14 | 0.34 | 0.1 |
| 7 | 25068898 | rs2001434 | -0.903 | -0.226 | -0.165 | 0.819 | -0.001 | -0.001 | -0.001 | 0 | 0.84 | 0.87 | 0.82 | 0.77 | 0.83 | 0.71 |
| 7 | 25069340 | rs4320455 | -0.271 | 0.123 | -0.522 | 0.058 | 0.039 | 0.016 | 0.016 | 0 | 0.2 | 0.32 | 0.23 | 0.31 | 0.31 | 0.14 |
| 7 | 25070863 | rs17725486 | 1.589 | 0.4 | -0.515 | 0.23 | -0.014 | 0.056 | 0.056 | 0 | 0.3 | 0.16 | 0.26 | 0.15 | 0.34 | 0.1 |
| 7 | 25071824 | rs10275560 | -0.239 | 0.067 | -0.439 | 0.314 | -0.006 | 0.054 | 0.054 | 0 | 0.7 | 0.52 | 0.66 | 0.54 | 0.76 | 0.36 |
| 7 | 25073439 | rs10085787 | -0.19 | 1.395 | -0.995 | 0.109 | 0.004 | 0.05 | 0.05 | 0 | 0.43 | 0.34 | 0.43 | 0.29 | 0.6 | 0.15 |
| 7 | 25074411 | rs7803101 | -0.522 | 1.938 | -0.838 | 0.692 | -0.076 | -0.051 | -0.051 | 0 | 0.12 | 0.01 | 0.15 | 0.16 | 0.24 | 0.04 |
| 7 | 25078543 | rs10282247 | -0.933 | 1.236 | -0.151 | 0.338 | -0.001 | -0.017 | -0.017 | 0 | 0.19 | 0.27 | 0.14 | 0.11 | 0.19 | 0.15 |
| 7 | 25088336 | rs2526401 | -0.949 | 0.97 | -0.435 | -0.077 | 0.031 | 0.125 | 0.125 | 0 | 0.48 | 0.71 | 0.59 | 0.79 | 0.39 | 0.73 |
| 7 | 25089059 | rs4236318 | 0.005 | 0.79 | -0.1 | 0.439 | 0.006 | -0.031 | -0.031 | 0 | 0.25 | 0.2 | 0.26 | 0.39 | 0.2 | 0.39 |
| 7 | 25094095 | rs255632 | -0.45 | 0.638 | 0.6 | 0.757 | -0.001 | 0.049 | 0.049 | 0.184 | 0.2 | 0.28 | 0.23 | 0.31 | 0.19 | 0.3 |
| 7 | 25096293 | rs39412 | -1.019 | 1.088 | -0.02 | 0.127 | 0.009 | 0.092 | 0.092 | 0 | 0.52 | 0.64 | 0.49 | 0.76 | 0.43 | 0.71 |
| 7 | 25099673 | rs39415 | 0.234 | 1.138 | -0.036 | 0.173 | 0.02 | -0.002 | -0.002 | 0.13 | 0.07 | 0.12 | 0.15 | 0.06 | 0.1 | 0.06 |
| 7 | 25100755 | rs39416 | -0.984 | 1.403 | -0.263 | 0.107 | 0.001 | 0.011 | 0.011 | 1.405 | 0.27 | 0.69 | 0.4 | 0.39 | 0.29 | 0.41 |
| 7 | 25101310 | rs4719793 | 0.181 | 1.014 | -0.166 | 0.278 | 0.003 | -0.008 | -0.008 | 0.059 | 0.28 | 0.24 | 0.16 | 0.41 | 0.22 | 0.34 |
| 7 | 25105848 | rs7789951 | 0.026 | 0.445 | -0.341 | 0.307 | -0.022 | 0.021 | 0.021 | 0 | 0.16 | 0.23 | 0.16 | 0.24 | 0.1 | 0.27 |
| 7 | 25109146 | rs39431 | -0.176 | 1.463 | -1.519 | -0.235 | -0.044 | 0.092 | 0.092 | 0 | 0.58 | 0.76 | 0.56 | 0.73 | 0.5 | 0.72 |
| 7 | 25115972 | rs2711014 | 0.596 | 1.743 | -1.744 | -0.475 | -0.067 | 0.095 | 0.095 | 0 | 0.73 | 0.89 | 0.78 | 0.93 | 0.63 | 0.72 |
| 7 | 25116332 | rs39436 | 0.34 | 1.324 | -1.983 | -0.82 | -0.029 | 0.048 | 0.048 | 0 | 0.59 | 0.29 | 0.51 | 0.29 | 0.52 | 0.55 |

| | | | | | | | | | | | | | | | | |
|---|---|---|---|---|---|---|---|---|---|---|---|---|---|---|---|---|
| 7 | 25119752 | rs39438 | 0.78 | 1.036 | -1.19 | 0.47 | 0.005 | 0.095 | 0.095 | 0 | 0.86 | 0.72 | 0.87 | 0.59 | 0.85 | 0.77 |
| 7 | 25128788 | rs39446 | 0.528 | 1.133 | -1.187 | 0.753 | 0.005 | 0.015 | 0.015 | 0 | 0.9 | 0.87 | 0.88 | 0.81 | 0.86 | 0.8 |
| 7 | 25132183 | rs10268780 | 0.141 | 0.3 | -1.007 | 0.039 | -0.003 | -0.06 | -0.06 | 0 | 0.17 | 0.33 | 0.13 | 0.11 | 0.14 | 0.1 |
| 7 | 25132379 | rs10239907 | -0.542 | -0.202 | -1.411 | 0.195 | 0.017 | -0.017 | -0.017 | 0 | 0.28 | 0.15 | 0.3 | 0.32 | 0.22 | 0.21 |
| 7 | 25132897 | rs12673033 | 0.358 | -0.743 | -1.283 | 0.044 | 0.031 | -0.026 | -0.026 | 0.34 | 0.17 | 0.08 | 0.19 | 0.21 | 0.09 | 0.11 |
| 7 | 25133849 | rs39453 | -0.549 | -0.752 | -1.323 | -0.123 | 0.021 | 0.04 | 0.04 | 0 | 0.4 | 0.2 | 0.37 | 0.28 | 0.37 | 0.23 |
| 7 | 25135566 | rs7800235 | -0.645 | -0.487 | -0.992 | 0.121 | 0.042 | 0.029 | 0.029 | 0 | 0.19 | 0.53 | 0.17 | 0.23 | 0.23 | 0.13 |
| 7 | 25138770 | rs17150913 | -0.346 | -0.511 | -0.434 | 0.119 | 0.048 | 0.049 | 0.049 | 0 | 0.16 | 0.36 | 0.15 | 0.27 | 0.22 | 0.11 |
| 7 | 25155861 | rs10951046 | -0.734 | -0.551 | -0.127 | 0.094 | 0.023 | 0.023 | 0.023 | 0 | 0.18 | 0.67 | 0.18 | 0.27 | 0.23 | 0.13 |
| 7 | 25157307 | rs12700583 | 2.575 | -0.793 | -0.622 | 0.016 | -0.107 | 0.146 | 0.146 | 0 | 0.37 | 0.04 | 0.33 | 0.16 | 0.38 | 0.43 |
| 7 | 25158246 | rs4719799 | -0.371 | -0.829 | -0.568 | 0.174 | 0.012 | -0.006 | -0.006 | 0 | 0.69 | 0.82 | 0.74 | 0.7 | 0.71 | 0.78 |
| 7 | 25159184 | rs12700584 | -0.497 | -0.859 | -0.785 | 0.16 | 0.015 | -0.026 | -0.026 | 0.022 | 0.7 | 0.9 | 0.75 | 0.7 | 0.71 | 0.78 |
| 7 | 25174332 | rs2717885 | -0.407 | -0.607 | -0.587 | 0.247 | 0.06 | 0.061 | 0.061 | 0 | 0.17 | 0.53 | 0.17 | 0.27 | 0.23 | 0.12 |
| 7 | 25174697 | rs2717886 | -0.407 | -0.607 | -0.587 | 0.247 | 0.06 | 0.061 | 0.061 | 0 | 0.17 | 0.53 | 0.17 | 0.27 | 0.23 | 0.12 |
| 7 | 25181912 | rs2285738 | 1.275 | -0.636 | -0.637 | 0.254 | 0.006 | 0.036 | 0.036 | 0 | 0.7 | 0.39 | 0.61 | 0.47 | 0.68 | 0.65 |
| 7 | 25182405 | rs11984293 | 0.271 | -0.577 | -0.56 | 0.166 | 0.083 | 0.067 | 0.067 | 0 | 0.13 | 0.42 | 0.12 | 0.16 | 0.17 | 0.09 |
| 7 | 25190106 | rs7806201 | 1.038 | -0.728 | -0.503 | 0.178 | 0.003 | 0.01 | 0.01 | 0 | 0.68 | 0.37 | 0.59 | 0.42 | 0.66 | 0.62 |
| 7 | 25194413 | rs2717900 | 0.62 | -1.05 | 0.056 | 0.326 | 0.002 | 0.002 | 0.002 | 0 | 0.55 | 0.59 | 0.5 | 0.42 | 0.61 | 0.55 |
| 7 | 25197078 | rs2717896 | -1.062 | -1.03 | 0.055 | 0.314 | 0.019 | 0.147 | 0.147 | 0.143 | 0.13 | 0.54 | 0.11 | 0.28 | 0.12 | 0.12 |
| 7 | 25202369 | rs1034691 | 1.782 | -0.925 | -0.504 | 0.241 | -0.112 | 0.226 | 0.226 | 0 | 0.42 | 0.05 | 0.39 | 0.14 | 0.49 | 0.43 |
| 7 | 25207974 | rs2717858 | -1.794 | -0.73 | 0.875 | 0.908 | -0.004 | 0.06 | 0.06 | 0.543 | 0.13 | 0.7 | 0.13 | 0.26 | 0.12 | 0.12 |
| 7 | 25207994 | rs2717857 | -1.157 | -0.58 | 0.299 | 1.01 | 0.023 | 0.129 | 0.129 | 0 | 0.11 | 0.34 | 0.09 | 0.26 | 0.12 | 0.1 |
| 7 | 25228115 | rs17151061 | -0.648 | -1.04 | -0.262 | 0.415 | 0.002 | -0.003 | -0.003 | 0 | 0.15 | 0.12 | 0.07 | 0.01 | 0.1 | 0.04 |
| 7 | 25229053 | rs1073578 | -0.382 | -1.706 | -0.427 | 0.852 | 0.014 | -0.02 | -0.02 | 0 | 0.26 | 0.14 | 0.24 | 0.44 | 0.13 | 0.3 |
| 7 | 25233396 | rs11768833 | -1.739 | -2.228 | 0.112 | 0.637 | 0.032 | 0.042 | 0.042 | 0 | 0.46 | 0.63 | 0.48 | 0.78 | 0.29 | 0.52 |
| 7 | 25235329 | rs11766477 | 0.382 | -2.26 | 0.416 | 0.866 | 0.003 | 0.02 | 0.02 | 0.325 | 0.08 | 0.14 | 0.14 | 0.13 | 0.08 | 0.1 |
| 7 | 25235462 | rs12535769 | 1.037 | -2.285 | 0.624 | 0.879 | -0.004 | 0.019 | 0.019 | 0 | 0.08 | 0.14 | 0.07 | 0.22 | 0.06 | 0.1 |
| 7 | 25239962 | rs10951056 | -1.206 | -2.301 | 0.686 | 0.87 | -0.008 | 0.044 | 0.044 | 0.447 | 0.19 | 0.36 | 0.21 | 0.35 | 0.14 | 0.21 |
| 7 | 25240087 | rs6959519 | -0.148 | -2.321 | 0.59 | 0.87 | 0.023 | -0.035 | -0.035 | 0 | 0.26 | 0.13 | 0.26 | 0.43 | 0.15 | 0.31 |
| 7 | 25245429 | rs11771762 | 0.372 | -1.957 | 0.53 | 0.45 | 0.003 | 0.02 | 0.02 | 0.353 | 0.08 | 0.14 | 0.14 | 0.14 | 0.08 | 0.1 |
| 7 | 25245779 | rs12673206 | -1.393 | -2.109 | 0.34 | 0.705 | -0.026 | 0.033 | 0.033 | 0 | 0.26 | 0.51 | 0.33 | 0.71 | 0.17 | 0.36 |
| 7 | 25246028 | rs10951060 | 0.034 | -1.901 | 0.187 | 0.737 | -0.001 | 0.035 | 0.035 | 0 | 0.15 | 0.16 | 0.18 | 0.57 | 0.08 | 0.23 |
| 7 | 25246674 | rs6967791 | -0.761 | -1.695 | 0.266 | 0.439 | -0.003 | -0.001 | -0.001 | 0.786 | 0.34 | 0.3 | 0.32 | 0.65 | 0.2 | 0.4 |
| 7 | 25258009 | rs2717888 | -0.377 | -1.594 | 0.461 | 0.026 | -0.015 | -0.088 | -0.088 | 0.153 | 0.13 | 0.29 | 0.08 | 0 | 0.12 | 0.08 |
| 7 | 25259224 | rs11764285 | -1.908 | -1.777 | -0.248 | -0.2 | -0.025 | 0.167 | 0.167 | 0 | 0.59 | 0.87 | 0.63 | 0.84 | 0.41 | 0.58 |
| 7 | 25262364 | rs739750 | -0.127 | -1.482 | 0.598 | 0.374 | 0.018 | 0.121 | 0.121 | 0.378 | 0.1 | 0.27 | 0.21 | 0.17 | 0.08 | 0.08 |
| 7 | 25262594 | rs739749 | -2.099 | -1.698 | 0.494 | 0.162 | -0.034 | 0.14 | 0.14 | 0 | 0.58 | 0.76 | 0.64 | 0.82 | 0.46 | 0.6 |
| 7 | 25264367 | rs6956128 | -1.124 | -1.799 | 0.701 | 0.105 | -0.007 | -0.023 | -0.023 | 0.734 | 0.43 | 0.54 | 0.47 | 0.83 | 0.29 | 0.49 |
| 7 | 25264556 | rs978095 | 0.421 | -1.769 | 0.65 | 0.104 | -0.001 | -0.056 | -0.056 | 0 | 0.1 | 0.19 | 0.13 | 0 | 0.09 | 0.04 |
| 7 | 25266400 | rs2717851 | -0.962 | -2.009 | 0.648 | 0.283 | -0.017 | -0.051 | -0.051 | 4.951 | 0.31 | 0.37 | 0.32 | 0.15 | 0.28 | 0.26 |
| 7 | 25267934 | rs877834 | -0.239 | -2.178 | 0.278 | 0.446 | 0.011 | -0.103 | -0.103 | 0.242 | 0.23 | 0.09 | 0.26 | 0.51 | 0.18 | 0.24 |
| 7 | 25271168 | rs1076221 | -0.571 | -1.881 | 0.578 | 0.583 | 0.015 | -0.072 | -0.072 | 0.016 | 0.24 | 0.13 | 0.26 | 0.51 | 0.18 | 0.23 |
| 7 | 25271524 | rs1076219 | -0.2 | -1.546 | 0.227 | 0.254 | 0 | 0 | 0 | 0 | 0.2 | 0.15 | 0.19 | 0.14 | 0.19 | 0.22 |
| 7 | 25272620 | rs929255 | -1.138 | -1.597 | 0.194 | 0.29 | 0.036 | 0.036 | 0.036 | 0 | 0.43 | 0.66 | 0.46 | 0.74 | 0.24 | 0.5 |
| 7 | 25275311 | rs2023857 | -1.352 | -1.067 | 0.389 | 0.303 | 0.017 | 0.029 | 0.029 | 0 | 0.48 | 0.62 | 0.5 | 0.74 | 0.38 | 0.54 |
| 7 | 25276424 | rs757202 | -2.173 | -1.248 | 0.095 | 0.108 | -0.033 | 0.054 | 0.054 | 0 | 0.62 | 0.75 | 0.66 | 0.77 | 0.51 | 0.62 |
| 7 | 25282158 | rs2158249 | -1.294 | -1.198 | 0.562 | 0.55 | 0.051 | 0.044 | 0.044 | 0 | 0.22 | 0.44 | 0.37 | 0.24 | 0.29 | 0.24 |
| 7 | 25285231 | rs11489352 | 0.069 | -1.508 | 0.277 | 0.691 | 0.021 | -0.052 | -0.052 | 0.006 | 0.2 | 0.14 | 0.15 | 0.41 | 0.12 | 0.2 |
| 7 | 25289582 | rs2391107 | 0.545 | -1.297 | 0.358 | 0.457 | -0.009 | 0.015 | 0.015 | 0 | 0.11 | 0.05 | 0.14 | 0.03 | 0.14 | 0.07 |
| 7 | 25290055 | rs1011967 | -0.647 | -1.128 | 0.712 | 0.621 | 0.062 | 0.164 | 0.164 | 0 | 0.09 | 0.26 | 0.21 | 0.21 | 0.14 | 0.16 |
| 7 | 25295346 | rs916893 | -1.279 | -1.493 | 0.486 | 0.473 | 0.036 | 0.015 | 0.015 | 0 | 0.49 | 0.34 | 0.44 | 0.56 | 0.36 | 0.45 |
| 7 | 25299989 | rs2107124 | 0.882 | -1.452 | 0.7 | 0.602 | -0.006 | -0.01 | -0.01 | 0 | 0.71 | 0.76 | 0.7 | 0.85 | 0.76 | 0.75 |
| 7 | 25303466 | rs2717907 | 0.224 | -1.473 | 0.717 | 0.64 | 0.007 | 0.005 | 0.005 | 0.737 | 0.78 | 0.85 | 0.81 | 0.87 | 0.84 | 0.79 |
| 7 | 25304791 | rs12672614 | -0.972 | -1.948 | 1.022 | 0.777 | 0.022 | 0.026 | 0.026 | 0 | 0.16 | 0.3 | 0.15 | 0.16 | 0.18 | 0.18 |
| 7 | 25306635 | rs11762781 | -0.828 | -1.844 | 0.842 | 0.733 | 0.002 | 0.004 | 0.004 | 0 | 0.26 | 0.18 | 0.29 | 0.16 | 0.21 | 0.24 |
| 7 | 25306852 | rs6961642 | -0.722 | -1.747 | 0.995 | 0.602 | 0.045 | -0.044 | -0.044 | 0 | 0.1 | 0.23 | 0.18 | 0.03 | 0.15 | 0.07 |
| 7 | 25307780 | rs2523090 | -0.774 | -1.839 | 0.621 | 0.673 | -0.019 | 0.017 | 0.017 | 0 | 0.26 | 0.27 | 0.26 | 0.32 | 0.18 | 0.36 |
| 7 | 25307974 | rs17151285 | -1.58 | -1.933 | 0.676 | 0.654 | 0.001 | -0.015 | -0.015 | 0 | 0.22 | 0.25 | 0.22 | 0.11 | 0.25 | 0.09 |
| 7 | 25308269 | rs757201 | -1.124 | -1.876 | 0.751 | 0.734 | 0.008 | -0.015 | -0.015 | 0.697 | 0.18 | 0.21 | 0.2 | 0.11 | 0.23 | 0.08 |
| 7 | 25312988 | rs2523084 | -1.751 | -1.921 | 0.684 | 0.768 | 0.002 | -0.001 | -0.001 | 0 | 0.46 | 0.53 | 0.45 | 0.43 | 0.4 | 0.44 |
| 7 | 25316325 | rs2717871 | -1.176 | -2.334 | 0.466 | 0.856 | -0.045 | 0.084 | 0.084 | 0.334 | 0.3 | 0.53 | 0.33 | 0.54 | 0.21 | 0.23 |
| 7 | 25316682 | rs765114 | -2.086 | -2.067 | -0.231 | 0.376 | -0.112 | 0.147 | 0.147 | 0 | 0.61 | 0.96 | 0.64 | 0.86 | 0.46 | 0.62 |
| 7 | 25325550 | rs7808695 | -0.206 | -1.718 | 0.744 | 1.018 | 0.018 | 0.037 | 0.037 | 0 | 0.07 | 0.22 | 0.12 | 0.11 | 0.07 | 0.02 |
| 7 | 25328060 | rs2188989 | -0.5 | -1.692 | 0.417 | 1.022 | -0.006 | 0.005 | 0.005 | 0 | 0.11 | 0.08 | 0.05 | 0 | 0.1 | 0.02 |
| 7 | 25330367 | rs17151313 | -1.787 | -1.989 | 0.551 | 1.363 | 0.049 | 0.022 | 0.022 | 0 | 0.36 | 0.79 | 0.39 | 0.69 | 0.23 | 0.54 |
| 7 | 25330508 | rs17151317 | -1.053 | -1.674 | 1.162 | 1.906 | 0.037 | 0.097 | 0.097 | 0 | 0.09 | 0.25 | 0.07 | 0.16 | 0.1 | 0.13 |
| 7 | 25331860 | rs6461855 | -1.103 | -1.775 | 0.573 | 1.621 | -0.05 | 0.055 | 0.055 | 1.364 | 0.25 | 0.55 | 0.27 | 0.59 | 0.14 | 0.3 |
| 7 | 25338512 | rs9691472 | -1.04 | -1.398 | 0.656 | 1.721 | -0.084 | 0.269 | 0.269 | 6.431 | 0.16 | 0.45 | 0.09 | 0.37 | 0.06 | 0.14 |
| 7 | 25342934 | rs7805812 | -0.519 | -0.959 | 0.229 | 0.553 | -0.015 | -0.099 | -0.099 | 0 | 0.3 | 0.48 | 0.17 | 0.1 | 0.28 | 0.29 |

| | | | | | | | | | | | | | | | | |
|---|---|---|---|---|---|---|---|---|---|---|---|---|---|---|---|---|
| 7 | 25343429 | rs916847 | -0.379 | -0.302 | -0.563 | 0.799 | -0.032 | -0.043 | -0.043 | 0 | 0.75 | 0.85 | 0.57 | 0.65 | 0.7 | 0.69 |
| 7 | 25344021 | rs11762772 | 0.543 | -0.325 | -0.43 | 0.326 | -0.001 | -0.017 | -0.017 | 0 | 0.15 | 0.18 | 0.12 | 0 | 0.19 | 0.15 |
| 7 | 25352809 | rs721316 | -0.495 | -0.38 | -0.395 | 0.671 | -0.03 | -0.037 | -0.037 | 0 | 0.76 | 0.82 | 0.59 | 0.65 | 0.7 | 0.7 |
| 7 | 25354077 | rs6966737 | 0.578 | -0.545 | 0.164 | 0.211 | -0.019 | -0.039 | -0.039 | 1.026 | 0.15 | 0.23 | 0.06 | 0.07 | 0.09 | 0.14 |
| 7 | 25354220 | rs11760705 | 0.032 | -0.392 | -0.151 | 0.069 | -0.001 | -0.008 | -0.008 | 0 | 0.18 | 0.12 | 0.12 | 0.01 | 0.21 | 0.15 |
| 7 | 25356802 | rs11772035 | 0.287 | -0.39 | -0.471 | 0.07 | 0.015 | 0.061 | 0.061 | 0.205 | 0.16 | 0.07 | 0.06 | 0.04 | 0.15 | 0.1 |
| 7 | 25357402 | rs1015510 | -0.963 | -0.165 | -0.707 | 0.469 | -0.027 | -0.042 | -0.042 | 0 | 0.8 | 0.85 | 0.63 | 0.59 | 0.72 | 0.67 |
| 7 | 25357772 | rs16873828 | -0.567 | -0.423 | -0.324 | 0.404 | -0.027 | -0.093 | -0.093 | 0 | 0.33 | 0.47 | 0.19 | 0.13 | 0.28 | 0.26 |
| 7 | 25358941 | rs10239513 | 1.252 | -0.913 | -0.845 | 0.185 | -0.022 | -0.013 | -0.013 | 0 | 0.24 | 0.3 | 0.21 | 0.22 | 0.17 | 0.24 |
| 7 | 25360909 | rs13228488 | 1.094 | -0.705 | -0.894 | 0.289 | -0.044 | -0.035 | -0.035 | 0.781 | 0.23 | 0.04 | 0.22 | 0.23 | 0.25 | 0.17 |
| 7 | 25363037 | rs10270076 | -0.327 | -0.775 | -0.73 | 0.263 | 0.002 | 0.05 | 0.05 | 0 | 0.57 | 0.71 | 0.4 | 0.35 | 0.46 | 0.5 |
| 7 | 25365227 | rs10249367 | 0.318 | -0.75 | -1.039 | 0.21 | 0.073 | 0.063 | 0.063 | 0 | 0.41 | 0.74 | 0.3 | 0.3 | 0.27 | 0.38 |
| 7 | 25365487 | rs17247294 | 0.373 | -0.105 | -0.577 | 0.537 | -0.004 | -0.009 | -0.009 | 0 | 0.4 | 0.5 | 0.3 | 0.31 | 0.35 | 0.33 |
| 7 | 25365550 | rs10252578 | -0.209 | 0.717 | -0.662 | 0.96 | -0.026 | -0.042 | -0.042 | 1.828 | 0.79 | 0.85 | 0.63 | 0.59 | 0.72 | 0.66 |
| 7 | 25368902 | rs11764873 | -0.368 | 0.484 | -0.193 | 0.119 | 0.023 | -0.05 | -0.05 | 0 | 0.29 | 0.43 | 0.45 | 0.17 | 0.36 | 0.31 |
| 7 | 25370035 | rs11765793 | -0.603 | 0.082 | -1.11 | 0.308 | 0.025 | -0.095 | -0.095 | 0 | 0.82 | 0.98 | 0.91 | 0.67 | 0.84 | 0.7 |
| 7 | 25377346 | rs9655257 | 0.443 | 0.027 | -0.308 | 0.127 | -0.002 | -0.014 | -0.014 | 0 | 0.47 | 0.18 | 0.41 | 0.48 | 0.47 | 0.39 |
| 7 | 25378585 | rs9655259 | 0.111 | 0.17 | 0.08 | 0.16 | 0.006 | -0.004 | -0.004 | 2.315 | 0.52 | 0.67 | 0.46 | 0.5 | 0.48 | 0.4 |
| 7 | 25380101 | rs12700609 | 0.627 | 1.077 | -0.313 | 0.023 | 0.001 | -0.003 | -0.003 | 0 | 0.09 | 0.07 | 0.07 | 0.02 | 0.15 | 0.07 |
| 7 | 25380703 | rs10241627 | -0.175 | 0.981 | -0.261 | -0.045 | -0.007 | -0.046 | -0.046 | 0.053 | 0.57 | 0.81 | 0.54 | 0.48 | 0.51 | 0.45 |
| 7 | 25382023 | rs17151393 | 0.35 | 0.551 | -0.155 | 0.025 | 0.002 | -0.001 | -0.001 | 0 | 0.42 | 0.36 | 0.43 | 0.45 | 0.34 | 0.38 |
| 7 | 25388946 | rs6964890 | 0.139 | 1.146 | -0.518 | -0.073 | -0.03 | 0.028 | 0.028 | 0 | 0.16 | 0.12 | 0.15 | 0.03 | 0.27 | 0.12 |
| 7 | 25393047 | rs11764103 | -0.009 | 1.124 | -0.316 | -0.101 | -0.005 | 0 | 0 | 0 | 0.17 | 0.17 | 0.17 | 0.03 | 0.27 | 0.12 |
| 7 | 25397118 | rs6461861 | -0.591 | 0.95 | -0.182 | -0.179 | 0.041 | -0.02 | -0.02 | 0 | 0.6 | 0.78 | 0.6 | 0.54 | 0.66 | 0.49 |
| 7 | 25402437 | rs11972231 | 0.518 | 1.018 | -0.183 | -0.173 | -0.012 | 0.007 | 0.007 | 0 | 0.16 | 0.1 | 0.16 | 0.02 | 0.27 | 0.12 |
| 7 | 25402947 | rs10275758 | 0.422 | 0.927 | 0.293 | -0.159 | 0 | -0.001 | -0.001 | 0 | 0.43 | 0.48 | 0.41 | 0.51 | 0.38 | 0.38 |
| 7 | 25404838 | rs17151430 | 0.391 | 0.927 | 0.36 | -0.172 | -0.001 | -0.003 | -0.003 | 0 | 0.43 | 0.48 | 0.41 | 0.52 | 0.38 | 0.38 |
| 7 | 25407621 | rs2893205 | 0.558 | 1.215 | 0.28 | -0.224 | 0 | 0.001 | 0.001 | 0 | 0.4 | 0.46 | 0.36 | 0.49 | 0.33 | 0.36 |
| 7 | 25414019 | rs2188994 | -0.329 | 1.063 | -0.055 | -0.159 | 0.025 | -0.003 | -0.003 | 0 | 0.57 | 0.75 | 0.54 | 0.54 | 0.59 | 0.48 |
| 7 | 25418864 | rs1015498 | -0.38 | 0.634 | -0.347 | -0.296 | 0 | -0.005 | -0.005 | 0 | 0.48 | 0.56 | 0.43 | 0.34 | 0.48 | 0.26 |
| 7 | 25420721 | rs9639507 | 1.166 | 0.485 | -0.545 | -0.379 | -0.005 | -0.023 | -0.023 | 0 | 0.22 | 0.07 | 0.24 | 0.3 | 0.19 | 0.16 |
| 7 | 25426044 | rs2074324 | 1.338 | 0.222 | -0.759 | 0.065 | -0.033 | -0.05 | -0.05 | 0 | 0.13 | 0.02 | 0.13 | 0.18 | 0.14 | 0.15 |
| 7 | 25428644 | rs4719809 | -0.407 | 0.174 | -0.383 | -0.279 | -0.005 | 0.076 | 0.076 | 0.814 | 0.48 | 0.78 | 0.38 | 0.34 | 0.47 | 0.27 |
| 7 | 25444631 | rs757970 | 0.325 | 0.202 | -0.964 | -0.357 | -0.053 | 0.035 | 0.035 | 0 | 0.36 | 0.17 | 0.3 | 0.3 | 0.44 | 0.25 |
| 7 | 25445253 | rs886800 | -0.327 | 0.166 | -0.738 | -0.376 | -0.006 | 0.024 | 0.024 | 0 | 0.44 | 0.39 | 0.33 | 0.31 | 0.47 | 0.27 |
| 7 | 25445972 | rs9648015 | 1.073 | 0.256 | -2.02 | -0.284 | -0.056 | 0.013 | 0.013 | 0 | 0.24 | 0.01 | 0.38 | 0.15 | 0.24 | 0.28 |
| 7 | 25446445 | rs882766 | -0.01 | 0.83 | -1.293 | -0.167 | -0.051 | 0.033 | 0.033 | 0 | 0.36 | 0.17 | 0.3 | 0.3 | 0.44 | 0.25 |
| 7 | 25447947 | rs1034836 | 0.067 | 1.67 | -1.139 | -0.118 | -0.039 | -0.028 | -0.028 | 0.044 | 0.81 | 0.98 | 0.78 | 0.81 | 0.71 | 0.77 |
| 7 | 25448939 | rs7807877 | -0.325 | 2.428 | -0.238 | -0.079 | 0.056 | 0.056 | 0.056 | 0.001 | 0.18 | 0.41 | 0.35 | 0.23 | 0.21 | 0.22 |
| 7 | 25449533 | rs7776769 | -0.325 | 2.428 | -0.238 | -0.074 | 0.056 | 0.059 | 0.059 | 0.001 | 0.18 | 0.41 | 0.35 | 0.23 | 0.21 | 0.22 |
| 7 | 25449584 | rs7812277 | -0.091 | 2.609 | -0.304 | -0.589 | -0.002 | -0.03 | -0.03 | 0 | 0.8 | 0.81 | 0.86 | 0.98 | 0.76 | 0.91 |
| 7 | 25450169 | rs1012151 | 1.608 | 2.725 | -0.506 | -0.864 | 0.075 | -0.021 | -0.021 | 0 | 0.43 | 0.13 | 0.33 | 0.59 | 0.35 | 0.53 |
| 7 | 25454923 | rs12531319 | 0.792 | 2.698 | -0.362 | -0.742 | -0.001 | 0.004 | 0.004 | 0 | 0.13 | 0.13 | 0.28 | 0.24 | 0.15 | 0.18 |
| 7 | 25455380 | rs729525 | -0.67 | 2.785 | -0.583 | -0.825 | -0.057 | -0.05 | -0.05 | 0.28 | 0.8 | 0.98 | 0.8 | 0.78 | 0.72 | 0.82 |
| 7 | 25462701 | rs4722467 | 0.931 | 2.814 | -0.241 | -0.944 | 0.014 | 0.006 | 0.006 | 0.247 | 0.55 | 0.66 | 0.41 | 0.6 | 0.43 | 0.6 |
| 7 | 25463604 | rs17151612 | 0.378 | 2.848 | 0.002 | -0.825 | 0.044 | -0.052 | -0.052 | 0 | 0.1 | 0.33 | 0.07 | 0.02 | 0.09 | 0.08 |
| 7 | 25466521 | rs10233491 | 0.271 | 2.736 | -0.395 | -0.462 | 0.033 | 0.032 | 0.032 | 0 | 0.14 | 0.31 | 0.3 | 0.18 | 0.15 | 0.18 |
| 7 | 25467288 | rs17310121 | 0.662 | 2.378 | -1.027 | -0.648 | -0.019 | 0.034 | 0.034 | 0 | 0.09 | 0.03 | 0.05 | 0 | 0.1 | 0.02 |
| 7 | 25470191 | rs6461867 | 0.5 | 2.288 | -1.014 | -0.589 | -0.044 | 0.023 | 0.023 | 0 | 0.7 | 0.89 | 0.75 | 0.79 | 0.61 | 0.78 |
| 7 | 25471651 | rs7798971 | -0.559 | 2.332 | -0.893 | -0.456 | -0.033 | -0.003 | -0.003 | 0 | 0.22 | 0.12 | 0.24 | 0.2 | 0.33 | 0.25 |
| 7 | 25472861 | rs7804262 | -0.149 | 2.575 | -0.517 | -0.699 | 0.069 | -0.026 | -0.026 | 1.013 | 0.1 | 0.17 | 0.25 | 0.06 | 0.25 | 0.19 |
| 7 | 25474286 | rs17151641 | 0.454 | 2.351 | -0.892 | -0.635 | -0.011 | -0.045 | -0.045 | 0 | 0.09 | 0 | 0.14 | 0.16 | 0.07 | 0.12 |
| 7 | 25474384 | rs720847 | 0.591 | 2.553 | -0.44 | -1.164 | -0.038 | 0.051 | 0.051 | 0 | 0.66 | 0.76 | 0.7 | 0.87 | 0.58 | 0.76 |
| 7 | 25474568 | rs17251040 | 0.533 | 2.625 | -0.531 | -1.146 | -0.05 | 0.016 | 0.016 | 0.961 | 0.08 | 0.02 | 0.23 | 0.06 | 0.25 | 0.19 |
| 7 | 25474649 | rs6461869 | -0.775 | 2.777 | -0.526 | -1.147 | -0.051 | -0.019 | -0.019 | 0 | 0.85 | 0.98 | 0.82 | 0.87 | 0.76 | 0.83 |
| 7 | 25476316 | rs1476697 | 1.128 | 2.09 | -0.004 | -1.039 | 0.118 | 0.161 | 0.161 | 0 | 0.52 | 0.16 | 0.4 | 0.81 | 0.33 | 0.51 |
| 7 | 25481005 | rs2108327 | 0.949 | 2.306 | -0.461 | -0.943 | -0.015 | -0.026 | -0.026 | 0 | 0.55 | 0.47 | 0.45 | 0.81 | 0.34 | 0.51 |
| 7 | 25482619 | rs7791700 | -0.62 | 2.825 | -0.343 | -0.352 | 0.115 | -0.03 | -0.03 | 0 | 0.1 | 0.25 | 0.23 | 0.06 | 0.24 | 0.24 |
| 7 | 25496143 | rs17151656 | -0.318 | 2.068 | -0.85 | -0.645 | -0.009 | 0.005 | 0.005 | 0 | 0.2 | 0.21 | 0.13 | 0.01 | 0.19 | 0.08 |
| 7 | 25501256 | rs4719814 | -0.932 | 1.549 | -0.902 | -1.218 | 0.055 | 0.085 | 0.085 | 0.191 | 0.52 | 0.72 | 0.59 | 0.19 | 0.64 | 0.49 |
| 7 | 25508637 | rs6948634 | -0.719 | 2.204 | -0.923 | -1.311 | -0.002 | 0.066 | 0.066 | 1.341 | 0.46 | 0.69 | 0.37 | 0.16 | 0.43 | 0.27 |
| 7 | 25509954 | rs17151696 | -0.566 | 1.944 | -0.457 | -1.508 | 0.007 | -0.043 | -0.043 | 0 | 0.34 | 0.43 | 0.24 | 0.1 | 0.38 | 0.19 |
| 7 | 25513733 | rs289988 | 1.819 | 1.282 | -0.966 | -1.308 | 0 | -0.001 | -0.001 | 0.073 | 0.33 | 0.27 | 0.23 | 0.61 | 0.19 | 0.39 |
| 7 | 25514053 | rs13243315 | 0.453 | 1.469 | -0.966 | -1.468 | -0.02 | 0.046 | 0.046 | 1.082 | 0.19 | 0.03 | 0.17 | 0.1 | 0.2 | 0.17 |
| 7 | 25514657 | rs10265861 | 0.999 | 1.619 | -0.906 | -1.441 | -0.013 | 0.013 | 0.013 | 0 | 0.1 | 0.11 | 0.08 | 0.06 | 0.16 | 0.11 |
| 7 | 25515926 | rs7782450 | 0.155 | 1.39 | -0.829 | -1.442 | 0.003 | -0.055 | -0.055 | 0 | 0.25 | 0.31 | 0.17 | 0.01 | 0.24 | 0.11 |
| 7 | 25519751 | rs10486458 | 0.904 | 1.218 | -1.044 | -1.071 | -0.023 | 0.059 | 0.059 | 0 | 0.2 | 0.08 | 0.13 | 0.05 | 0.19 | 0.08 |
| 7 | 25519829 | rs6979504 | -0.663 | 1.32 | -0.876 | -1.07 | 0.043 | -0.067 | -0.067 | 0 | 0.31 | 0.52 | 0.44 | 0.13 | 0.44 | 0.42 |
| 7 | 25525767 | rs10486459 | 1.406 | 0.082 | -1.654 | -0.18 | 0.014 | -0.035 | -0.035 | 0 | 0.13 | 0.03 | 0.1 | 0.25 | 0.11 | 0.13 |

| | | | | | | | | | | | | | | | | |
|---|---|---|---|---|---|---|---|---|---|---|---|---|---|---|---|---|
| 7 | 25527291 | rs371405 | -1.439 | -0.03 | -1.811 | -0.009 | -0.026 | -0.062 | -0.062 | 0 | 0.88 | 0.99 | 0.83 | 0.81 | 0.87 | 0.89 |
| 7 | 25529065 | rs17318255 | 1.655 | -0.098 | -1.38 | -0.161 | 0.014 | 0.013 | 0.013 | 0.211 | 0.12 | 0.07 | 0.11 | 0.06 | 0.08 | 0.09 |
| 7 | 25531396 | rs17252583 | 0.998 | -0.061 | -1.434 | -0.2 | -0.014 | 0.037 | 0.037 | 0 | 0.2 | 0.09 | 0.12 | 0.12 | 0.18 | 0.09 |
| 7 | 25533563 | rs11974986 | 0.212 | -0.017 | -1.39 | -0.124 | 0.01 | 0.036 | 0.036 | 0.198 | 0.32 | 0.21 | 0.21 | 0.17 | 0.25 | 0.15 |
| 7 | 25540744 | rs6959518 | 0.908 | 0.52 | -1.168 | 0.138 | -0.006 | 0.007 | 0.007 | 0 | 0.1 | 0.12 | 0.08 | 0.06 | 0.16 | 0.11 |
| 7 | 25545837 | rs12668343 | 0.457 | 0.541 | -1.147 | 0.245 | 0.008 | -0.032 | -0.032 | 0.816 | 0.16 | 0.08 | 0.14 | 0.23 | 0.15 | 0.15 |
| 7 | 25553702 | rs12667972 | 0.294 | 0.474 | -1.036 | 0.263 | 0.005 | -0.009 | -0.009 | 0 | 0.17 | 0.15 | 0.15 | 0.25 | 0.15 | 0.15 |
| 7 | 25554856 | rs9638826 | -0.721 | 0.51 | -0.849 | 0.226 | -0.004 | 0.01 | 0.01 | 0.888 | 0.28 | 0.38 | 0.24 | 0.3 | 0.3 | 0.28 |
| 7 | 25559106 | rs974340 | -0.289 | 0.379 | -1.036 | 0.048 | 0.001 | 0.012 | 0.012 | 0 | 0.26 | 0.2 | 0.17 | 0.11 | 0.22 | 0.14 |
| 7 | 25560512 | rs731008 | 0.685 | 0.249 | -0.697 | 0.654 | 0.004 | -0.014 | -0.014 | 0 | 0.1 | 0.04 | 0.09 | 0.25 | 0.09 | 0.09 |
| 7 | 25561256 | rs6972325 | -0.063 | 0.341 | -0.275 | 0.245 | 0.007 | -0.033 | -0.033 | 0.022 | 0.07 | 0.2 | 0.07 | 0 | 0.05 | 0.04 |
| 7 | 25563961 | rs11975146 | -0.373 | 0.231 | -0.337 | 0.432 | 0 | 0 | 0 | 0 | 0.43 | 0.55 | 0.38 | 0.48 | 0.44 | 0.45 |
| 7 | 25564438 | rs289999 | -0.479 | -0.33 | -0.375 | 0.364 | -0.007 | -0.004 | -0.004 | 0 | 0.61 | 0.65 | 0.75 | 0.6 | 0.67 | 0.61 |
| 7 | 25565151 | rs6956023 | -0.202 | -0.136 | -0.381 | -0.114 | -0.006 | 0.147 | 0.147 | 0 | 0.5 | 0.67 | 0.39 | 0.17 | 0.5 | 0.41 |
| 7 | 25565388 | rs11979740 | 1.643 | -0.203 | -0.208 | -0.1 | -0.005 | -0.021 | -0.021 | 0 | 0.06 | 0.15 | 0.11 | 0.02 | 0.07 | 0.11 |
| 7 | 25566545 | rs17151805 | 2.342 | -0.315 | 0.225 | -0.145 | -0.003 | -0.043 | -0.043 | 0 | 0.16 | 0.23 | 0.1 | 0.08 | 0.16 | 0.14 |
| 7 | 25569619 | rs11765117 | 2.04 | -0.337 | 0.526 | -0.129 | -0.008 | -0.039 | -0.039 | 0.389 | 0.18 | 0.23 | 0.13 | 0.11 | 0.16 | 0.2 |
| 7 | 25572091 | rs2999578 | -1.825 | 0.388 | 0.656 | 0.079 | 0.048 | -0.017 | -0.017 | 0 | 0.8 | 0.63 | 0.73 | 0.81 | 0.73 | 0.75 |
| 7 | 25572327 | rs1027670 | 1.012 | 0.139 | 0.316 | 0.181 | -0.022 | -0.101 | -0.101 | 0 | 0.14 | 0.01 | 0.17 | 0.34 | 0.1 | 0.25 |
| 7 | 25572668 | rs1027671 | 0.959 | 0.008 | 0.179 | -0.014 | -0.037 | 0.048 | 0.048 | 0 | 0.31 | 0.09 | 0.28 | 0.16 | 0.31 | 0.19 |
| 7 | 25574197 | rs981903 | 2.134 | -0.045 | 0.252 | -0.043 | -0.002 | -0.001 | -0.001 | 0.025 | 0.22 | 0.24 | 0.16 | 0.26 | 0.16 | 0.16 |
| 7 | 25575717 | rs17782178 | 2.237 | -0.008 | 0.225 | -0.084 | -0.075 | 0.112 | 0.112 | 0.015 | 0.17 | 0.01 | 0.11 | 0 | 0.18 | 0.04 |
| 7 | 25578353 | rs1502943 | 0.803 | -0.016 | 0.276 | -0.025 | 0.026 | -0.003 | -0.003 | 0 | 0.37 | 0.27 | 0.28 | 0.61 | 0.27 | 0.4 |
| 7 | 25578946 | rs1502945 | -1.264 | -0.073 | 0.376 | 0.041 | 0.037 | -0.023 | -0.023 | 0 | 0.75 | 0.53 | 0.62 | 0.75 | 0.65 | 0.65 |
| 7 | 25582362 | rs2999569 | 2.004 | -0.121 | 0.247 | 0.053 | -0.002 | -0.001 | -0.001 | 0 | 0.22 | 0.24 | 0.16 | 0.23 | 0.16 | 0.15 |
| 7 | 25582666 | rs11973020 | -0.196 | -0.062 | 0.222 | 0.062 | -0.004 | -0.016 | -0.016 | 3.245 | 0.39 | 0.34 | 0.4 | 0.45 | 0.35 | 0.42 |
| 7 | 25583417 | rs11766484 | 1.502 | -0.587 | 0.082 | 0.032 | 0.001 | -0.004 | -0.004 | 0.332 | 0.13 | 0.09 | 0.17 | 0.17 | 0.14 | 0.15 |
| 7 | 25585623 | rs11761708 | 1.502 | -0.587 | 0.082 | 0.006 | 0.001 | -0.004 | -0.004 | 1.733 | 0.13 | 0.09 | 0.17 | 0.15 | 0.13 | 0.16 |
| 7 | 25586747 | rs10951077 | 2.396 | -0.44 | 0.206 | -0.262 | 0 | -0.003 | -0.003 | 0.394 | 0.16 | 0.14 | 0.13 | 0.09 | 0.16 | 0.15 |
| 7 | 25588469 | rs2952519 | -0.636 | -0.211 | 0.037 | -0.405 | -0.001 | -0.006 | -0.006 | 0.015 | 0.73 | 0.76 | 0.71 | 0.91 | 0.65 | 0.84 |
| 7 | 25589591 | rs2938109 | 0.638 | -0.68 | -0.312 | 0.142 | 0.003 | -0.001 | -0.001 | 0 | 0.24 | 0.24 | 0.12 | 0.26 | 0.17 | 0.19 |
| 7 | 25590456 | rs3817077 | -0.35 | -0.411 | -0.26 | 0.332 | -0.007 | -0.006 | -0.006 | 0.36 | 0.39 | 0.5 | 0.44 | 0.33 | 0.37 | 0.5 |
| 7 | 25590559 | rs1393672 | 0.489 | -0.616 | -0.296 | -0.025 | 0.014 | 0.003 | 0.003 | 0.344 | 0.22 | 0.18 | 0.12 | 0.21 | 0.17 | 0.15 |
| 7 | 25593187 | rs2938111 | 0.382 | -0.365 | -0.117 | 0.013 | -0.022 | -0.016 | -0.016 | 0 | 0.66 | 0.45 | 0.77 | 0.76 | 0.72 | 0.69 |
| 7 | 25593469 | rs17726478 | -0.131 | 0.325 | -0.015 | 0.227 | -0.003 | -0.01 | -0.01 | 0 | 0.09 | 0.11 | 0.08 | 0.03 | 0.1 | 0.12 |
| 7 | 25595363 | rs2952520 | 0.142 | 1.148 | 0.126 | 0.462 | 0.043 | 0.015 | 0.015 | 0 | 0.83 | 0.72 | 0.65 | 0.82 | 0.74 | 0.79 |
| 7 | 25596056 | rs6943072 | 0.19 | 0.662 | -0.237 | 0.272 | 0.016 | 0.023 | 0.023 | 0.091 | 0.28 | 0.19 | 0.3 | 0.21 | 0.27 | 0.28 |
| 7 | 25598135 | rs2960818 | -0.911 | 0.731 | -0.221 | 0.273 | 0.001 | 0.024 | 0.024 | 0 | 0.55 | 0.77 | 0.51 | 0.69 | 0.5 | 0.6 |
| 7 | 25600612 | rs17726931 | 1.58 | 0.727 | -0.256 | 0.206 | -0.024 | 0.03 | 0.03 | 0.048 | 0.06 | 0 | 0.04 | 0 | 0.07 | 0.06 |
| 7 | 25601508 | rs2999575 | -0.423 | 0.517 | 0.133 | 0.29 | 0.059 | 0.002 | 0.002 | 0 | 0.47 | 0.75 | 0.45 | 0.64 | 0.39 | 0.48 |
| 7 | 25602231 | rs7780911 | -0.343 | 0.255 | 0.334 | 0.033 | -0.008 | -0.027 | -0.027 | 0 | 0.39 | 0.57 | 0.38 | 0.2 | 0.35 | 0.35 |
| 7 | 25604366 | rs7808374 | 0.006 | 0.419 | -0.106 | 0.054 | -0.007 | -0.058 | -0.058 | 0 | 0.33 | 0.43 | 0.34 | 0.19 | 0.34 | 0.27 |
| 7 | 25604598 | rs10235801 | 1.259 | 0.426 | -0.236 | -0.035 | -0.029 | 0.11 | 0.11 | 0 | 0.37 | 0.13 | 0.2 | 0.14 | 0.38 | 0.28 |
| 7 | 25605725 | rs10240053 | 1.274 | 0.421 | -0.239 | -0.042 | -0.03 | 0.106 | 0.106 | 0 | 0.37 | 0.13 | 0.2 | 0.14 | 0.38 | 0.27 |
| 7 | 25606273 | rs2999577 | -0.529 | 0.592 | -0.145 | 0.231 | -0.006 | 0.001 | 0.001 | 0 | 0.42 | 0.47 | 0.43 | 0.65 | 0.41 | 0.46 |
| 7 | 25607939 | rs11973587 | -1.356 | 0.319 | -0.122 | 0.472 | -0.018 | 0.072 | 0.072 | 0.983 | 0.26 | 0.39 | 0.21 | 0.52 | 0.23 | 0.35 |
| 7 | 25612661 | rs975369 | 0.271 | 0.241 | -0.376 | 0.482 | 0.007 | 0.001 | 0.001 | 0.061 | 0.1 | 0.08 | 0.09 | 0.47 | 0.09 | 0.21 |
| 7 | 25615221 | rs1354264 | 0.389 | 0.288 | -0.346 | 0.503 | 0.005 | 0.013 | 0.013 | 0.008 | 0.09 | 0.09 | 0.09 | 0.47 | 0.08 | 0.21 |
| 7 | 25617358 | rs755476 | 0.1 | 0.202 | -0.381 | 0.327 | 0.001 | 0.006 | 0.006 | 0 | 0.54 | 0.62 | 0.71 | 0.82 | 0.56 | 0.57 |
| 7 | 25620561 | rs208546 | -0.798 | 0.178 | -0.313 | 0.306 | -0.004 | 0.016 | 0.016 | 0 | 0.65 | 0.74 | 0.78 | 0.84 | 0.63 | 0.69 |
| 7 | 25621579 | rs208549 | -0.766 | 0.178 | -0.275 | 0.305 | -0.014 | 0.049 | 0.049 | 0 | 0.66 | 0.79 | 0.79 | 0.86 | 0.63 | 0.72 |
| 7 | 25622517 | rs208553 | 1.35 | 0.182 | -0.285 | 0.3 | 0.003 | -0.004 | -0.004 | 0.178 | 0.38 | 0.53 | 0.57 | 0.35 | 0.39 | 0.32 |
| 7 | 25622753 | rs1074503 | 0.125 | 0.21 | -0.353 | 0.14 | -0.008 | -0.031 | -0.031 | 0 | 0.11 | 0.12 | 0.07 | 0.02 | 0.07 | 0.12 |
| 7 | 25624089 | rs169642 | 0.067 | 0.222 | -0.343 | 0.146 | 0.002 | 0.048 | 0.048 | 0 | 0.55 | 0.67 | 0.71 | 0.84 | 0.55 | 0.6 |
| 7 | 25626542 | rs882101 | -0.123 | 0.195 | -0.316 | 0.149 | -0.008 | -0.031 | -0.031 | 0 | 0.12 | 0.16 | 0.08 | 0.02 | 0.07 | 0.13 |
| 7 | 25629089 | rs208563 | 1.233 | 0.222 | -0.295 | 0.202 | 0.002 | -0.009 | -0.009 | 0 | 0.38 | 0.46 | 0.57 | 0.32 | 0.39 | 0.31 |
| 7 | 25631509 | rs2521746 | 0.601 | 0.308 | -0.324 | 0.187 | -0.005 | 0.017 | 0.017 | 0.008 | 0.47 | 0.61 | 0.63 | 0.34 | 0.48 | 0.4 |
| 7 | 25634832 | rs1393677 | 1.237 | 0.446 | -0.37 | 0.25 | 0.001 | -0.004 | -0.004 | 0.21 | 0.4 | 0.44 | 0.57 | 0.32 | 0.4 | 0.32 |
| 7 | 25636379 | rs2813894 | -1.082 | 0.536 | -0.505 | 0.142 | -0.002 | 0.005 | 0.005 | 0 | 0.73 | 0.77 | 0.82 | 0.86 | 0.71 | 0.74 |
| 7 | 25637608 | rs17151997 | -1.566 | 0.513 | -0.508 | 0.18 | -0.001 | 0.006 | 0.006 | 0 | 0.32 | 0.31 | 0.25 | 0.54 | 0.31 | 0.42 |
| 7 | 25639031 | rs1882526 | -1.106 | 0.416 | -0.48 | 0.19 | -0.012 | 0.042 | 0.042 | 0 | 0.26 | 0.3 | 0.21 | 0.54 | 0.23 | 0.37 |
| 7 | 25653642 | rs1502948 | 0.324 | 0.383 | -0.585 | 0.272 | -0.012 | -0.073 | -0.073 | 0 | 0.14 | 0.04 | 0.18 | 0.5 | 0.17 | 0.21 |
| 7 | 25655407 | rs12700650 | -0.902 | 0.444 | -0.532 | 0.184 | -0.005 | 0.008 | 0.008 | 0 | 0.72 | 0.77 | 0.8 | 0.85 | 0.68 | 0.73 |
| 7 | 25661976 | rs10248149 | 1.071 | 0.153 | -0.765 | 0.385 | 0.002 | -0.003 | -0.003 | 0 | 0.39 | 0.4 | 0.51 | 0.32 | 0.41 | 0.34 |
| 7 | 25662742 | rs1476082 | -0.612 | -0.224 | -0.691 | 1.228 | -0.013 | -0.031 | -0.031 | 0.134 | 0.11 | 0.13 | 0.07 | 0.02 | 0.06 | 0.08 |
| 7 | 25668255 | rs10235998 | -1.004 | 0.369 | -0.756 | 1.935 | -0.005 | -0.058 | -0.058 | 0.076 | 0.14 | 0.05 | 0.18 | 0.51 | 0.16 | 0.23 |
| 7 | 25674531 | rs12540380 | -1.534 | 0.458 | -0.289 | 2.129 | 0.015 | 0.069 | 0.069 | 0 | 0.1 | 0.21 | 0.08 | 0.29 | 0.13 | 0.17 |
| 7 | 25676320 | rs12533343 | -1.684 | 0.056 | -0.783 | 2.441 | 0.004 | -0.029 | -0.029 | 1.249 | 0.25 | 0.22 | 0.15 | 0.56 | 0.24 | 0.37 |
| 7 | 25676557 | rs12540389 | -1.684 | 0.056 | -0.783 | 2.441 | 0.004 | -0.029 | -0.029 | 0.251 | 0.25 | 0.22 | 0.15 | 0.56 | 0.24 | 0.37 |

| | | | | | | | | | | | | | | | | |
|---|---|---|---|---|---|---|---|---|---|---|---|---|---|---|---|---|
| 7 | 25676893 | rs17152101 | -0.549 | 0.129 | -0.91 | 2.455 | -0.017 | -0.103 | -0.103 | 1.447 | 0.16 | 0.01 | 0.09 | 0.32 | 0.15 | 0.23 |
| 7 | 25679798 | rs10951080 | -1.323 | 0.279 | -0.41 | 1.855 | 0.006 | 0.039 | 0.039 | 0 | 0.09 | 0.17 | 0.05 | 0.25 | 0.11 | 0.14 |
| 7 | 25681464 | rs10951081 | 1.224 | -0.117 | -0.019 | 1.31 | -0.001 | 0 | 0 | 0 | 0.65 | 0.7 | 0.75 | 0.31 | 0.66 | 0.58 |
| 7 | 25681758 | rs10951082 | 1.118 | -0.265 | 0.115 | 1.288 | -0.001 | -0.001 | -0.001 | 0 | 0.65 | 0.68 | 0.72 | 0.32 | 0.64 | 0.59 |
| 7 | 25687574 | rs1859040 | 1.138 | -0.218 | 0.266 | 1.315 | 0 | 0 | 0 | 0 | 0.65 | 0.64 | 0.72 | 0.33 | 0.64 | 0.6 |
| 7 | 25702250 | rs442 | 0.971 | -0.803 | 1.378 | 1.572 | -0.032 | -0.014 | -0.014 | 0 | 0.74 | 0.54 | 0.75 | 0.2 | 0.78 | 0.6 |
| 7 | 25706712 | rs2108388 | -1.798 | -1.17 | 1.296 | 1.27 | -0.014 | -0.066 | -0.066 | 0 | 0.36 | 0.53 | 0.38 | 0.93 | 0.29 | 0.57 |
| 7 | 25718419 | rs740345 | -1.006 | -1.169 | 1.389 | 1.259 | -0.003 | 0.007 | 0.007 | 0 | 0.25 | 0.24 | 0.34 | 0.65 | 0.25 | 0.39 |
| 7 | 25722313 | rs2214976 | -1.172 | -1.171 | 1.421 | 1.222 | -0.017 | -0.012 | -0.012 | 0 | 0.26 | 0.39 | 0.4 | 0.79 | 0.25 | 0.41 |
| 7 | 25725601 | rs1544479 | -2.224 | -1.2 | 1.432 | 1.183 | 0 | -0.038 | -0.038 | 0.021 | 0.4 | 0.36 | 0.53 | 0.93 | 0.34 | 0.59 |
| 7 | 25753008 | rs6953294 | -1.017 | -1.3 | 1.525 | 1.261 | -0.003 | 0.006 | 0.006 | 0 | 0.25 | 0.25 | 0.33 | 0.65 | 0.25 | 0.38 |
| 7 | 25756356 | rs12536392 | -1.085 | -1.31 | 1.563 | 1.186 | 0.01 | -0.004 | -0.004 | 1.687 | 0.14 | 0.13 | 0.14 | 0.14 | 0.08 | 0.16 |
| 7 | 25762797 | rs17152404 | -1.11 | -1.326 | 1.779 | 1.282 | -0.001 | 0.005 | 0.005 | 0 | 0.24 | 0.21 | 0.31 | 0.63 | 0.24 | 0.4 |
| 7 | 25775690 | rs2058024 | -1.234 | -1.329 | 1.793 | 1.26 | -0.009 | 0.007 | 0.007 | 0 | 0.25 | 0.34 | 0.38 | 0.76 | 0.24 | 0.41 |
| 7 | 25780315 | rs12537117 | -1.112 | -1.306 | 1.747 | 1.28 | -0.041 | 0.064 | 0.064 | 3.774 | 0.15 | 0.28 | 0.2 | 0.28 | 0.08 | 0.18 |
| 7 | 25789508 | rs17152544 | -1.995 | -1.322 | 1.954 | 0.854 | -0.012 | -0.057 | -0.057 | 0 | 0.4 | 0.56 | 0.53 | 0.91 | 0.33 | 0.58 |
| 7 | 25792666 | rs4722520 | -0.693 | -1.203 | 1.574 | 1.058 | -0.001 | -0.023 | -0.023 | 0 | 0.21 | 0.14 | 0.3 | 0.68 | 0.23 | 0.38 |
| 7 | 25798864 | rs17152613 | -1.006 | -1.28 | 1.423 | 0.845 | -0.001 | 0.002 | 0.002 | 0 | 0.24 | 0.24 | 0.36 | 0.72 | 0.24 | 0.38 |
| 7 | 25802471 | rs4722522 | -1.862 | -1.395 | 0.943 | 0.651 | 0.053 | 0.095 | 0.095 | 0.209 | 0.42 | 0.71 | 0.52 | 0.83 | 0.33 | 0.57 |
| 7 | 25807078 | rs10480044 | -1.004 | -1.514 | 1.322 | 0.621 | -0.047 | -0.042 | -0.042 | 0 | 0.12 | 0.27 | 0.1 | 0.07 | 0.05 | 0.07 |
| 7 | 25807780 | rs11768829 | -1.666 | -1.585 | 0.823 | 0.706 | 0.034 | 0.019 | 0.019 | 0.315 | 0.34 | 0.71 | 0.34 | 0.77 | 0.26 | 0.46 |
| 7 | 25813918 | rs4719825 | -1.371 | -1.297 | 1.142 | 0.83 | -0.075 | 0.021 | 0.021 | 0.035 | 0.15 | 0.39 | 0.18 | 0.16 | 0.04 | 0.1 |
| 7 | 25814109 | rs4719826 | -1.32 | -1.32 | 1.114 | 0.84 | -0.063 | 0.017 | 0.017 | 0.103 | 0.14 | 0.29 | 0.16 | 0.16 | 0.04 | 0.1 |
| 7 | 25819209 | rs7350035 | 0.494 | -0.515 | 0.29 | 0.194 | -0.08 | 0.081 | 0.081 | 0 | 0.82 | 0.97 | 0.85 | 0.97 | 0.74 | 0.83 |
| 7 | 25823647 | rs960589 | 0.156 | -0.691 | 0.059 | 1.031 | -0.035 | -0.084 | -0.084 | 0 | 0.12 | 0.02 | 0.11 | 0.42 | 0.17 | 0.18 |
| 7 | 25823776 | rs1376526 | 1.781 | -0.668 | 0.871 | 0.587 | 0.004 | -0.012 | -0.012 | 0 | 0.72 | 0.36 | 0.71 | 0.84 | 0.72 | 0.78 |
| 7 | 25827799 | rs122505 | 0.656 | -0.649 | 0.835 | 0.52 | -0.029 | 0.036 | 0.036 | 0 | 0.81 | 0.87 | 0.83 | 0.98 | 0.75 | 0.83 |
| 7 | 25830647 | rs9639521 | 1.603 | -0.631 | 0.583 | 0.513 | 0.002 | -0.009 | -0.009 | 0 | 0.72 | 0.37 | 0.7 | 0.85 | 0.72 | 0.77 |
| 7 | 25845010 | rs1993094 | -1.809 | -0.924 | 0.753 | 0.753 | -0.037 | 0 | 0 | 0.273 | 0.21 | 0.57 | 0.27 | 0.21 | 0.13 | 0.28 |
| 7 | 25856295 | rs10226258 | -1.45 | -1.276 | 0.694 | 0.373 | -0.007 | -0.015 | -0.015 | 0 | 0.19 | 0.12 | 0.11 | 0.06 | 0.08 | 0.11 |
| 7 | 25861138 | rs12535837 | -1.414 | -1.642 | 0.659 | 0.001 | -0.028 | -0.056 | -0.056 | 0.748 | 0.32 | 0.44 | 0.34 | 0.83 | 0.27 | 0.53 |
| 7 | 25873110 | rs10282436 | -0.276 | -1.703 | 1.214 | 0.981 | 0.005 | -0.029 | -0.029 | 0.263 | 0.19 | 0.15 | 0.17 | 0.43 | 0.21 | 0.46 |
| 7 | 25875151 | rs12700664 | 2.066 | -1.47 | 1.249 | 0.324 | -0.045 | 0.153 | 0.153 | 0.14 | 0.43 | 0.17 | 0.42 | 0.13 | 0.48 | 0.37 |
| 7 | 25876612 | rs10232819 | -1.137 | -1.479 | 1.428 | 0.331 | -0.02 | -0.04 | -0.04 | 0.037 | 0.32 | 0.44 | 0.34 | 0.82 | 0.27 | 0.53 |
| 7 | 25887278 | rs2290263 | 0.576 | -1.487 | 1.228 | 0.365 | -0.009 | 0.028 | 0.028 | 1.091 | 0.75 | 0.8 | 0.78 | 0.95 | 0.74 | 0.9 |
| 7 | 25904794 | rs17152823 | 0.596 | -2.069 | 0.309 | -0.394 | 0.018 | 0.02 | 0.02 | 0.068 | 0.23 | 0.15 | 0.09 | 0.12 | 0.13 | 0.18 |
| 7 | 25904853 | rs3823944 | -0.6 | -1.326 | 0.562 | -0.493 | -0.002 | 0 | 0 | 0.051 | 0.61 | 0.41 | 0.45 | 0.62 | 0.55 | 0.74 |
| 7 | 25905599 | rs10235225 | -1.438 | -1.105 | 0.025 | -0.291 | 0.002 | 0.001 | 0.001 | 0.151 | 0.72 | 0.71 | 0.53 | 0.61 | 0.62 | 0.78 |
| 7 | 25908210 | rs10230872 | -0.693 | -1.009 | 0.129 | -0.412 | 0.014 | -0.096 | -0.096 | 0 | 0.3 | 0.38 | 0.28 | 0.08 | 0.32 | 0.14 |
| 7 | 25909208 | rs1451386 | 0.886 | -1.678 | 0.391 | 0.341 | -0.026 | 0.224 | 0.224 | 0.104 | 0.87 | 0.59 | 0.79 | 0.53 | 0.93 | 0.87 |
| 7 | 25911033 | rs10272701 | 0.206 | -1.67 | -0.099 | 0.28 | -0.012 | -0.027 | -0.027 | 0 | 0.18 | 0.12 | 0.16 | 0.25 | 0.2 | 0.33 |
| 7 | 25911327 | rs7785415 | -0.177 | -2.172 | 0.087 | -0.268 | -0.037 | -0.079 | -0.079 | 0.156 | 0.14 | 0.42 | 0.1 | 0.02 | 0.09 | 0.05 |
| 7 | 25912817 | rs1868889 | 1.986 | -2.269 | -0.699 | -0.943 | -0.142 | 0.153 | 0.153 | 0 | 0.27 | 0.04 | 0.35 | 0.04 | 0.38 | 0.18 |
| 7 | 25914286 | rs1451389 | -0.03 | -2.601 | 0.061 | -0.194 | 0.008 | -0.014 | -0.014 | 0 | 0.64 | 0.71 | 0.63 | 0.47 | 0.69 | 0.63 |
| 7 | 25918590 | rs6965388 | 1.005 | -2.481 | 0.269 | -0.169 | -0.014 | -0.033 | -0.033 | 1.3 | 0.11 | 0.17 | 0.07 | 0.02 | 0.09 | 0.05 |
| 7 | 25920438 | rs6461889 | -0.43 | -2.744 | 0.216 | -0.067 | -0.016 | -0.013 | -0.013 | 0 | 0.87 | 0.78 | 0.89 | 0.89 | 0.9 | 0.86 |
| 7 | 25921083 | rs4722536 | -0.476 | -2.703 | 0.27 | -0.016 | -0.025 | 0.012 | 0.012 | 0 | 0.37 | 0.47 | 0.36 | 0.44 | 0.31 | 0.29 |
| 7 | 25923381 | rs17152874 | 0.464 | -2.605 | 0.165 | 0.042 | -0.017 | 0.046 | 0.046 | 0 | 0.26 | 0.34 | 0.29 | 0.42 | 0.22 | 0.23 |
| 7 | 25930573 | rs6461892 | 0.135 | -2.008 | -0.257 | 0.202 | -0.02 | -0.017 | -0.017 | 0.339 | 0.85 | 0.69 | 0.86 | 0.87 | 0.9 | 0.86 |
| 7 | 25935030 | rs896312 | -0.389 | -1.646 | -0.295 | 0.514 | -0.023 | 0.013 | 0.013 | 0 | 0.38 | 0.51 | 0.37 | 0.45 | 0.31 | 0.31 |
| 7 | 25939723 | rs12672425 | -0.539 | -1.855 | -0.439 | 0.574 | -0.018 | 0.007 | 0.007 | 0 | 0.31 | 0.33 | 0.32 | 0.58 | 0.2 | 0.29 |
| 7 | 25945652 | rs7809764 | -0.257 | -1.826 | -0.956 | 0.126 | 0.002 | -0.005 | -0.005 | 0 | 0.39 | 0.36 | 0.27 | 0.53 | 0.35 | 0.56 |
| 7 | 25951345 | rs1966841 | -0.502 | -1.875 | -1.039 | -0.483 | -0.015 | 0.071 | 0.071 | 0.048 | 0.56 | 0.3 | 0.55 | 0.94 | 0.57 | 0.77 |
| 7 | 25955056 | rs878655 | -0.514 | -2.115 | -0.92 | -0.731 | -0.036 | -0.089 | -0.089 | 0.032 | 0.87 | 0.58 | 0.93 | 1 | 0.95 | 1 |
| 7 | 25955754 | rs12154905 | 0.703 | -2.336 | -1.473 | -0.479 | 0.012 | -0.179 | -0.179 | 0.016 | 0.27 | 0.03 | 0.31 | 0.52 | 0.25 | 0.31 |
| 7 | 25963672 | rs10280475 | 1.097 | -2.246 | -1.385 | -0.476 | -0.069 | -0.072 | -0.072 | 0 | 0.24 | 0.07 | 0.2 | 0.32 | 0.31 | 0.35 |
| 7 | 25982367 | rs2391175 | 1.609 | -1.768 | -1.839 | -0.585 | -0.144 | 0.157 | 0.157 | 0 | 0.26 | 0.04 | 0.34 | 0.02 | 0.37 | 0.19 |
| 7 | 25991826 | rs4722551 | 1.699 | -1.35 | -1.864 | -0.501 | -0.04 | 0.07 | 0.07 | 0 | 0.15 | 0.02 | 0.18 | 0.02 | 0.17 | 0.1 |
| 7 | 25991977 | rs9639523 | -0.164 | -1.646 | -1.544 | -0.057 | 0.044 | -0.032 | -0.032 | 0 | 0.36 | 0.2 | 0.34 | 0.59 | 0.33 | 0.46 |
| 7 | 25997536 | rs4719841 | -0.48 | -1.5 | -1.945 | -0.087 | 0.067 | 0.023 | 0.023 | 0 | 0.46 | 0.22 | 0.38 | 0.63 | 0.39 | 0.58 |
| 7 | 26007582 | rs7786527 | 0.577 | -2.21 | -2.007 | 0.096 | 0.028 | -0.029 | -0.029 | 0 | 0.19 | 0.09 | 0.07 | 0.31 | 0.11 | 0.26 |
| 7 | 26008052 | rs745580 | -0.463 | -2.242 | -2.373 | -0.448 | -0.001 | 0.065 | 0.065 | 0 | 0.55 | 0.75 | 0.47 | 0.32 | 0.43 | 0.39 |
| 7 | 26008165 | rs2012224 | 0.898 | -2.173 | -2.358 | -0.37 | 0.027 | -0.028 | -0.028 | 0 | 0.18 | 0.09 | 0.07 | 0.3 | 0.11 | 0.26 |
| 7 | 26008408 | rs4521656 | 1.069 | -2.161 | -2.616 | -0.365 | 0.034 | -0.074 | -0.074 | 0 | 0.18 | 0.03 | 0.07 | 0.3 | 0.11 | 0.26 |
| 7 | 26015963 | rs6958870 | -0.738 | -1.754 | -2.606 | -0.034 | -0.07 | -0.053 | -0.053 | 0 | 0.63 | 0.87 | 0.52 | 0.42 | 0.46 | 0.58 |
| 7 | 26019400 | rs10242720 | 0.694 | -1.945 | -2.661 | -0.215 | -0.04 | 0.069 | 0.069 | 0 | 0.28 | 0.16 | 0.35 | 0.12 | 0.3 | 0.31 |
| 7 | 26023090 | rs6947773 | -0.793 | -1.911 | -2.718 | -0.13 | -0.076 | -0.073 | -0.073 | 0.381 | 0.62 | 0.87 | 0.52 | 0.44 | 0.46 | 0.58 |
| 7 | 26029743 | rs6461901 | 0.149 | -2.023 | -2.518 | -0.136 | 0.035 | 0.005 | 0.005 | 0.019 | 0.34 | 0.67 | 0.17 | 0.32 | 0.16 | 0.28 |
| 7 | 26029862 | rs9639525 | 0.252 | -1.777 | -2.86 | -0.162 | -0.013 | 0.009 | 0.009 | 0 | 0.06 | 0.11 | 0.05 | 0.09 | 0.03 | 0.18 |

| | | | | | | | | | | | | | | | | |
|---|---|---|---|---|---|---|---|---|---|---|---|---|---|---|---|---|
| 7 | 26030325 | rs766839 | 0.844 | -1.586 | -2.794 | -0.111 | -0.03 | 0.06 | 0.06 | 0 | 0.27 | 0.13 | 0.35 | 0.12 | 0.29 | 0.3 |
| 7 | 26035630 | rs4719848 | -0.234 | -0.757 | -2.455 | 0.439 | -0.164 | -0.174 | -0.174 | 0.003 | 0.65 | 0.97 | 0.64 | 0.53 | 0.46 | 0.61 |
| 7 | 26038458 | rs2893236 | -0.631 | -1.032 | -2.156 | 0.788 | -0.046 | -0.072 | -0.072 | 0.006 | 0.79 | 0.99 | 0.85 | 0.77 | 0.8 | 0.8 |
| 7 | 26041879 | rs12112544 | -0.816 | -2.144 | -1.181 | 0.612 | -0.072 | 0.051 | 0.051 | 1.705 | 0.29 | 0.57 | 0.14 | 0.45 | 0.11 | 0.37 |
| 7 | 26059739 | rs11982959 | -0.293 | -2.197 | -0.551 | 0.534 | -0.081 | 0.116 | 0.116 | 2.191 | 0.21 | 0.42 | 0.09 | 0.37 | 0.08 | 0.2 |
| 7 | 26070505 | rs6948960 | -0.594 | -2.194 | -0.53 | 0.535 | -0.024 | 0.015 | 0.015 | 3.1 | 0.23 | 0.64 | 0.11 | 0.37 | 0.09 | 0.2 |
| 7 | 26075094 | rs10486491 | 1.796 | -2.205 | -0.763 | 0.427 | -0.026 | 0.071 | 0.071 | 0 | 0.12 | 0 | 0.04 | 0 | 0.09 | 0.06 |
| 7 | 26092684 | rs17153134 | -0.613 | -1.949 | -0.148 | 0.946 | -0.056 | 0.062 | 0.062 | 2.116 | 0.22 | 0.52 | 0.1 | 0.37 | 0.09 | 0.21 |
| 7 | 26102288 | rs17153154 | -1.129 | -1.893 | -0.257 | 0.966 | -0.026 | 0.029 | 0.029 | 0 | 0.08 | 0.19 | 0.16 | 0.19 | 0.05 | 0.15 |
| 7 | 26108984 | rs17322566 | 0.459 | -1.825 | -0.407 | 0.991 | 0.005 | -0.007 | -0.007 | 0 | 0.1 | 0.03 | 0.12 | 0.09 | 0.06 | 0.02 |
| 7 | 26116505 | rs10272549 | -1.021 | -1.962 | 0.029 | 1.085 | -0.081 | 0.16 | 0.16 | 2.923 | 0.2 | 0.42 | 0.17 | 0.44 | 0.07 | 0.22 |
| 7 | 26120798 | rs7805293 | -1.443 | -2.007 | -0.202 | 1.064 | -0.039 | 0.029 | 0.029 | 2.238 | 0.25 | 0.67 | 0.23 | 0.51 | 0.09 | 0.35 |
| 7 | 26122949 | rs12535874 | -0.447 | -1.745 | -0.504 | 1.022 | -0.008 | -0.001 | -0.001 | 0.4 | 0.14 | 0.19 | 0.15 | 0.16 | 0.06 | 0.15 |
| 7 | 26128382 | rs6945001 | -1.478 | -1.848 | -0.75 | 1.034 | 0.022 | -0.033 | -0.033 | 0 | 0.24 | 0.74 | 0.23 | 0.48 | 0.09 | 0.25 |
| 7 | 26131087 | rs7793753 | 0.754 | -0.641 | -1.559 | 0.105 | -0.026 | 0.145 | 0.145 | 0.559 | 0.49 | 0.11 | 0.44 | 0.33 | 0.49 | 0.48 |
| 7 | 26134253 | rs4722567 | -0.242 | -0.287 | -1.168 | 0.208 | -0.001 | 0 | 0 | 0.002 | 0.73 | 0.3 | 0.72 | 0.48 | 0.74 | 0.64 |
| 7 | 26147219 | rs3814095 | -0.483 | 0.035 | -1.264 | -0.001 | 0.001 | 0.022 | 0.022 | 0.11 | 0.25 | 0.21 | 0.23 | 0.14 | 0.21 | 0.14 |
| 7 | 26151190 | rs6973742 | 1.015 | -0.089 | -1.2 | -0.496 | 0.004 | 0.134 | 0.134 | 0.395 | 0.49 | 0.16 | 0.47 | 0.31 | 0.53 | 0.53 |
| 7 | 26153228 | rs4722576 | 1.083 | -0.087 | -1.378 | -0.495 | -0.01 | 0.155 | 0.155 | 0.488 | 0.48 | 0.13 | 0.47 | 0.31 | 0.53 | 0.52 |
| 7 | 26154110 | rs4722579 | 1.203 | 0.124 | -2.215 | -0.96 | -0.063 | 0.006 | 0.006 | 0.039 | 0.32 | 0.02 | 0.23 | 0.3 | 0.31 | 0.44 |
| 7 | 26155399 | rs2057763 | -0.09 | -0.142 | -1.652 | -0.669 | -0.013 | 0.065 | 0.065 | 0.014 | 0.22 | 0.13 | 0.13 | 0.05 | 0.21 | 0.12 |
| 7 | 26167109 | rs876421 | -0.26 | -0.681 | -0.831 | -0.32 | -0.008 | -0.001 | -0.001 | 0 | 0.74 | 0.37 | 0.63 | 0.35 | 0.74 | 0.74 |
| 7 | 26170112 | rs2057761 | -0.947 | -0.66 | -0.941 | -0.227 | -0.002 | -0.009 | -0.009 | 0 | 0.81 | 0.79 | 0.68 | 0.37 | 0.75 | 0.77 |
| 7 | 26171214 | rs11770984 | 0.874 | -0.563 | -1.678 | -1.006 | -0.154 | 0.133 | 0.133 | 0 | 0.25 | 0.02 | 0.17 | 0.04 | 0.33 | 0.24 |
| 7 | 26173027 | rs10248605 | 0.962 | -1.216 | -1.576 | -0.827 | 0.003 | 0.024 | 0.024 | 0.022 | 0.17 | 0 | 0.08 | 0.17 | 0.14 | 0.27 |
| 7 | 26175542 | rs2057765 | -1.08 | -1.077 | -1.224 | -0.568 | -0.001 | -0.008 | -0.008 | 0 | 0.8 | 0.79 | 0.68 | 0.37 | 0.75 | 0.77 |
| 7 | 26182214 | rs4259314 | -0.25 | -1.536 | -0.848 | -0.233 | -0.071 | 0.056 | 0.056 | 0.386 | 0.34 | 0.18 | 0.27 | 0.15 | 0.44 | 0.39 |
| 7 | 26183366 | rs4722585 | -0.916 | -1.897 | -0.801 | 0.365 | 0 | 0.034 | 0.034 | 2.72 | 0.53 | 0.27 | 0.33 | 0.6 | 0.52 | 0.62 |
| 7 | 26190193 | rs13228646 | -0.586 | -0.951 | 0.462 | 0.932 | -0.024 | 0.03 | 0.03 | 0.411 | 0.36 | 0.27 | 0.28 | 0.16 | 0.46 | 0.39 |
| 7 | 26191698 | rs2073536 | -0.85 | -0.905 | 1.01 | 0.993 | 0 | 0 | 0 | 0.317 | 0.37 | 0.31 | 0.35 | 0.5 | 0.44 | 0.46 |
| 7 | 26195594 | rs10270359 | -0.933 | -1.121 | 0.884 | 1.083 | -0.006 | -0.001 | -0.001 | 0 | 0.41 | 0.38 | 0.36 | 0.6 | 0.42 | 0.52 |
| 7 | 26198040 | rs3753098 | -0.403 | -1.063 | 0.873 | 1.026 | -0.029 | 0.032 | 0.032 | 0 | 0.37 | 0.26 | 0.26 | 0.22 | 0.43 | 0.38 |
| 7 | 26199207 | rs757371 | -1.512 | -0.902 | 0.777 | 1.2 | 0 | 0.041 | 0.041 | 0 | 0.55 | 0.7 | 0.51 | 0.69 | 0.51 | 0.57 |
| 7 | 26205980 | rs2237333 | -0.142 | -0.552 | 0.859 | 1.759 | -0.008 | 0.078 | 0.078 | 0 | 0.05 | 0.05 | 0.1 | 0.35 | 0.02 | 0.16 |
| 7 | 26210803 | rs10238831 | 0.142 | -0.537 | 1.108 | 1.571 | 0.006 | 0.007 | 0.007 | 11.539 | 0.09 | 0.26 | 0.04 | 0.09 | 0.05 | 0.05 |
| 7 | 26219687 | rs11971002 | 0.018 | -0.648 | 0.892 | 1.58 | -0.009 | -0.001 | -0.001 | 0 | 0.09 | 0.18 | 0.03 | 0.08 | 0.05 | 0.08 |
| 7 | 26223294 | rs12113404 | -2.417 | -0.834 | 1.15 | 1.695 | -0.063 | 0.097 | 0.097 | 4.519 | 0.24 | 0.5 | 0.25 | 0.55 | 0.12 | 0.24 |
| 7 | 26227647 | rs10236009 | -0.037 | -0.572 | -0.067 | 1.825 | -0.009 | -0.001 | -0.001 | 0 | 0.09 | 0.17 | 0.03 | 0.08 | 0.05 | 0.08 |
| 7 | 26248965 | rs7793815 | 0.254 | 0.176 | -0.535 | 1.53 | 0.004 | 0.002 | 0.002 | 0 | 0.06 | 0.03 | 0.02 | 0 | 0.04 | 0.01 |
| 7 | 26250940 | rs11972057 | 0.254 | 0.176 | -0.534 | 1.53 | 0.003 | -0.001 | -0.001 | 2.86 | 0.06 | 0.03 | 0.02 | 0 | 0.04 | 0.01 |
| 7 | 26282656 | rs10951129 | -0.295 | 0.897 | -0.582 | 1.784 | -0.017 | -0.024 | -0.024 | 0 | 0.17 | 0.09 | 0.28 | 0.49 | 0.24 | 0.22 |
| 7 | 26318396 | rs12700720 | -0.127 | 0.042 | -1.217 | 1.249 | -0.051 | -0.002 | -0.002 | 0 | 0.36 | 0.15 | 0.21 | 0.33 | 0.38 | 0.44 |
| 7 | 26335007 | rs972158 | 0.761 | -0.984 | -0.604 | 1.541 | 0.004 | 0.048 | 0.048 | 0.355 | 0.48 | 0.33 | 0.63 | 0.34 | 0.57 | 0.31 |
| 7 | 26335430 | rs12538280 | 0.58 | -0.95 | -0.515 | 1.744 | -0.003 | 0.029 | 0.029 | 0 | 0.06 | 0.04 | 0.08 | 0.27 | 0.01 | 0.1 |
| 7 | 26342980 | rs6975395 | 0.07 | -0.96 | 0.404 | 1.64 | 0.002 | -0.006 | -0.006 | 0 | 0.38 | 0.32 | 0.23 | 0.24 | 0.39 | 0.46 |
| 7 | 26343917 | rs2698723 | 0.873 | -0.899 | 0.208 | 1.818 | -0.028 | 0.133 | 0.133 | 0 | 0.9 | 0.77 | 0.84 | 0.72 | 0.94 | 0.83 |
| 7 | 26351206 | rs12700723 | 0.249 | -0.71 | -0.068 | 1.806 | 0.001 | 0.034 | 0.034 | 2.652 | 0.06 | 0.05 | 0.11 | 0.22 | 0.04 | 0.04 |
| 7 | 26352073 | rs2893245 | -0.533 | -0.647 | 0.113 | 1.07 | 0.004 | -0.048 | -0.048 | 0.015 | 0.58 | 0.43 | 0.49 | 0.79 | 0.53 | 0.74 |
| 7 | 26370190 | rs4722593 | -0.098 | -0.41 | -0.424 | 1.217 | -0.075 | -0.086 | -0.086 | 0 | 0.23 | 0.09 | 0.29 | 0.39 | 0.36 | 0.28 |
| 7 | 26371279 | rs6461929 | -0.11 | -1.44 | 0.099 | 1.173 | 0.001 | -0.001 | -0.001 | 0 | 0.4 | 0.49 | 0.28 | 0.42 | 0.21 | 0.51 |
| 7 | 26372025 | rs12668538 | 0.103 | -1.498 | 0.246 | 1.204 | -0.041 | -0.044 | -0.044 | 3.834 | 0.37 | 0.39 | 0.21 | 0.16 | 0.2 | 0.46 |
| 7 | 26374032 | rs7793945 | 0.278 | -1.998 | -0.167 | 0.747 | -0.006 | 0.008 | 0.008 | 0 | 0.16 | 0.19 | 0.19 | 0.34 | 0.1 | 0.13 |
| 7 | 26377088 | rs10270099 | -0.509 | -2.052 | -0.179 | 0.724 | 0.048 | -0.005 | -0.005 | 0 | 0.52 | 0.58 | 0.41 | 0.5 | 0.3 | 0.6 |
| 7 | 26385449 | rs17153451 | 1.011 | -2.051 | 0.245 | 0.627 | 0.025 | 0.02 | 0.02 | 0.051 | 0.31 | 0.21 | 0.15 | 0.05 | 0.16 | 0.44 |
| 7 | 26385615 | rs12537520 | -0.652 | -2.474 | 0.23 | 0.524 | -0.03 | 0.034 | 0.034 | 0 | 0.24 | 0.3 | 0.26 | 0.46 | 0.14 | 0.18 |
| 7 | 26385660 | rs12534835 | 0.421 | -2.417 | 0.05 | 0.527 | 0.025 | -0.027 | -0.027 | 0 | 0.15 | 0.06 | 0.16 | 0.19 | 0.11 | 0.11 |
| 7 | 26386812 | rs10230728 | -0.488 | -2.211 | 0.244 | 0.592 | -0.022 | 0.003 | 0.003 | 0 | 0.55 | 0.48 | 0.4 | 0.5 | 0.3 | 0.61 |
| 7 | 26388552 | rs10254065 | -0.399 | -2.249 | 0.297 | 0.626 | 0.031 | -0.007 | -0.007 | 0 | 0.56 | 0.61 | 0.42 | 0.51 | 0.3 | 0.61 |
| 7 | 26388832 | rs3801888 | 0.142 | -2.223 | 0.092 | 0.556 | 0.059 | 0.049 | 0.049 | 0.931 | 0.47 | 0.29 | 0.32 | 0.24 | 0.26 | 0.56 |
| 7 | 26388888 | rs3801887 | 0.824 | -2.414 | -0.417 | 0.322 | 0.019 | 0.021 | 0.021 | 0 | 0.12 | 0.03 | 0.15 | 0.07 | 0.11 | 0.15 |
| 7 | 26392337 | rs2698721 | -0.266 | -1.818 | -1.082 | -0.002 | -0.01 | 0.034 | 0.034 | 1.045 | 0.46 | 0.2 | 0.51 | 0.62 | 0.55 | 0.49 |
| 7 | 26392726 | rs6967109 | 0.932 | -1.528 | -1.08 | -0.299 | -0.013 | -0.035 | -0.035 | 0 | 0.29 | 0.26 | 0.11 | 0.04 | 0.11 | 0.35 |
| 7 | 26396198 | rs1406754 | 0.416 | -0.783 | -1.471 | 0.115 | -0.133 | -0.059 | -0.059 | 0 | 0.27 | 0.05 | 0.28 | 0.29 | 0.4 | 0.25 |
| 7 | 26401513 | rs7782538 | 1.135 | -1.007 | -1.26 | -0.155 | 0.03 | 0.036 | 0.036 | 0 | 0.31 | 0.17 | 0.13 | 0.04 | 0.18 | 0.38 |
| 7 | 26402948 | rs2698719 | 0.717 | 0.195 | -1.601 | -0.197 | -0.064 | -0.055 | -0.055 | 0.06 | 0.15 | 0.05 | 0.26 | 0.29 | 0.32 | 0.22 |
| 7 | 26403574 | rs2699810 | 0.283 | 0.038 | -1.531 | -0.064 | 0.027 | 0 | 0 | 0.213 | 0.5 | 0.28 | 0.45 | 0.46 | 0.53 | 0.62 |
| 7 | 26418709 | rs1468286 | 0.73 | 0.466 | -1.395 | 0.196 | 0.013 | 0.013 | 0.013 | 0 | 0.55 | 0.61 | 0.36 | 0.47 | 0.41 | 0.57 |
| 7 | 26427543 | rs6461936 | 0.442 | 0.687 | -1.493 | 0.276 | -0.008 | -0.025 | -0.025 | 0 | 0.12 | 0.04 | 0.17 | 0.14 | 0.12 | 0.06 |
| 7 | 26427889 | rs1406593 | -0.775 | 0.517 | -1.009 | 0.211 | 0.002 | 0 | 0 | 0 | 0.28 | 0.3 | 0.35 | 0.36 | 0.34 | 0.19 |

| | | | | | | | | | | | | | | | | |
|---|---|---|---|---|---|---|---|---|---|---|---|---|---|---|---|---|
| 7 | 26432924 | rs1229655 | -1.969 | -0.216 | -1.94 | -0.537 | -0.016 | 0.213 | 0.213 | 0 | 0.68 | 0.99 | 0.85 | 0.96 | 0.73 | 0.65 |
| 7 | 26439331 | rs1013688 | -0.105 | 0.895 | -1.23 | 0.342 | 0.044 | 0.027 | 0.027 | 0 | 0.22 | 0.34 | 0.47 | 0.38 | 0.4 | 0.31 |
| 7 | 26440503 | rs976346 | -0.966 | 0.224 | -0.963 | 0.071 | -0.003 | -0.002 | -0.002 | 0 | 0.43 | 0.53 | 0.67 | 0.58 | 0.58 | 0.52 |
| 7 | 26447021 | rs1229670 | 0.589 | 0.276 | -0.627 | 0.694 | -0.001 | 0.044 | 0.044 | 0 | 0.08 | 0.08 | 0.1 | 0.36 | 0.05 | 0.13 |
| 7 | 26450219 | rs2699818 | -0.243 | 0.524 | 0.108 | 0.308 | -0.009 | -0.041 | -0.041 | 0 | 0.17 | 0.18 | 0.07 | 0.03 | 0.1 | 0.09 |
| 7 | 26458246 | rs2698732 | -1.321 | 0.407 | 0.427 | 0.155 | 0.002 | -0.015 | -0.015 | 0 | 0.32 | 0.25 | 0.23 | 0.07 | 0.22 | 0.16 |
| 7 | 26460409 | rs1229678 | -0.282 | 0.815 | 0.864 | 0.44 | 0 | 0.053 | 0.053 | 0 | 0.15 | 0.24 | 0.14 | 0.44 | 0.17 | 0.19 |
| 7 | 26460646 | rs12669192 | 0.529 | 0.804 | 0.853 | 0.446 | -0.015 | -0.012 | -0.012 | 1.538 | 0.1 | 0.11 | 0.16 | 0.05 | 0.07 | 0.06 |
| 7 | 26463160 | rs10085659 | 0.021 | 1.236 | 0.962 | 0.385 | 0.069 | 0.124 | 0.124 | 0 | 0.14 | 0.23 | 0.39 | 0.44 | 0.26 | 0.35 |
| 7 | 26465811 | rs1534697 | -0.807 | 1.061 | 0.807 | 0.246 | -0.001 | -0.043 | -0.043 | 0 | 0.28 | 0.22 | 0.21 | 0.04 | 0.2 | 0.15 |
| 7 | 26470789 | rs2525762 | 0.102 | 1.302 | 0.882 | 0.429 | 0.02 | 0.065 | 0.065 | 0.417 | 0.21 | 0.09 | 0.07 | 0.03 | 0.15 | 0.11 |
| 7 | 26473995 | rs10248777 | 0.685 | 1.72 | 1.247 | 0.737 | 0.107 | 0.214 | 0.214 | 0 | 0.09 | 0.24 | 0.25 | 0.33 | 0.18 | 0.27 |
| 7 | 26474874 | rs10238880 | 0.527 | 1.717 | 1.294 | 0.731 | 0.132 | 0.251 | 0.251 | 0 | 0.1 | 0.28 | 0.26 | 0.33 | 0.19 | 0.26 |
| 7 | 26476051 | rs1997359 | -0.368 | 1.289 | 0.957 | 0.836 | 0.001 | 0.058 | 0.058 | 0 | 0.15 | 0.24 | 0.13 | 0.44 | 0.17 | 0.19 |
| 7 | 26478596 | rs2525764 | -0.47 | 1.342 | 0.812 | 0.718 | -0.004 | 0.059 | 0.059 | 0 | 0.26 | 0.12 | 0.23 | 0.13 | 0.24 | 0.2 |
| 7 | 26481804 | rs10951138 | -0.814 | 1.294 | 0.908 | 0.731 | -0.002 | 0.135 | 0.135 | 0 | 0.19 | 0.44 | 0.18 | 0.48 | 0.2 | 0.2 |
| 7 | 26482974 | rs2698727 | -0.643 | 1.317 | 0.632 | 0.271 | -0.004 | 0.058 | 0.058 | 0 | 0.26 | 0.12 | 0.23 | 0.13 | 0.24 | 0.19 |
| 7 | 26485860 | rs6461942 | -1.269 | 1.454 | 0.717 | 0.803 | 0.002 | 0 | 0 | 0 | 0.4 | 0.56 | 0.39 | 0.42 | 0.42 | 0.32 |
| 7 | 26497052 | rs4719866 | -0.204 | 1.009 | 0.659 | 0.483 | -0.01 | 0.028 | 0.028 | 1.748 | 0.25 | 0.2 | 0.15 | 0.07 | 0.27 | 0.21 |
| 7 | 26498747 | rs17153671 | 0.129 | 1.158 | 0.855 | 0.774 | -0.013 | 0.069 | 0.069 | 0 | 0.14 | 0.2 | 0.07 | 0.37 | 0.1 | 0.14 |
| 7 | 26501182 | rs4722598 | -2.003 | 1.382 | -0.078 | 0.528 | 0.048 | 0.087 | 0.087 | 0 | 0.62 | 0.87 | 0.76 | 0.81 | 0.67 | 0.72 |
| 7 | 26504359 | rs10235248 | -0.127 | 2.147 | 0.153 | 1.111 | -0.043 | -0.041 | -0.041 | 0 | 0.1 | 0.03 | 0.36 | 0.31 | 0.24 | 0.27 |
| 7 | 26508020 | rs6963302 | -0.21 | 1.939 | 0.48 | 0.806 | 0 | -0.005 | -0.005 | 0 | 0.22 | 0.29 | 0.13 | 0.15 | 0.23 | 0.19 |
| 7 | 26510916 | rs6979146 | -0.126 | 1.838 | 1.094 | 0.978 | 0.022 | 0.103 | 0.103 | 0.081 | 0.16 | 0.36 | 0.12 | 0.31 | 0.16 | 0.15 |
| 7 | 26513564 | rs9638838 | 0.085 | 1.98 | 1.06 | 0.983 | -0.023 | -0.022 | -0.022 | 0 | 0.11 | 0.08 | 0.35 | 0.31 | 0.24 | 0.3 |
| 7 | 26513728 | rs916969 | -1.367 | 2.033 | 1.128 | 0.818 | -0.003 | -0.013 | -0.013 | 0 | 0.44 | 0.69 | 0.37 | 0.48 | 0.41 | 0.42 |
| 7 | 26517555 | rs2214548 | -2.472 | 1.873 | 0.764 | 0.734 | 0.047 | 0.063 | 0.063 | 0.547 | 0.62 | 0.99 | 0.75 | 0.8 | 0.69 | 0.76 |
| 7 | 26520292 | rs17153720 | 0.035 | 1.648 | 0.629 | 0.809 | 0.022 | 0.106 | 0.106 | 0.081 | 0.16 | 0.36 | 0.12 | 0.31 | 0.16 | 0.15 |
| 7 | 26520694 | rs4722600 | 0.156 | 1.238 | -0.227 | 0.203 | 0.005 | 0.055 | 0.055 | 2.786 | 0.22 | 0.12 | 0.09 | 0.08 | 0.19 | 0.2 |
| 7 | 26526960 | rs12666574 | -1.549 | 0.867 | -0.574 | -0.051 | 0.018 | 0.071 | 0.071 | 0.577 | 0.58 | 0.99 | 0.74 | 0.76 | 0.63 | 0.71 |
| 7 | 26531720 | rs10951140 | -1.721 | 1.467 | 1.057 | 0.173 | 0.086 | 0 | 0 | 0 | 0.18 | 0.35 | 0.47 | 0.12 | 0.25 | 0.3 |
| 7 | 26532145 | rs757393 | -2.34 | 1.276 | 1.335 | 0.134 | -0.005 | 0.006 | 0.006 | 0 | 0.31 | 0.71 | 0.53 | 0.24 | 0.3 | 0.37 |
| 7 | 26532349 | rs17153742 | -0.298 | 1.517 | 0.663 | 0.244 | -0.014 | -0.042 | -0.042 | 0.51 | 0.12 | 0.16 | 0.04 | 0.02 | 0.05 | 0.06 |
| 7 | 26533803 | rs6969646 | -0.836 | 1.646 | 0.56 | 0.265 | -0.01 | -0.043 | -0.043 | 0 | 0.17 | 0.37 | 0.09 | 0.12 | 0.13 | 0.1 |
| 7 | 26536913 | rs3180361 | -0.91 | 1.843 | -0.016 | 0.541 | -0.015 | -0.011 | -0.011 | 0 | 0.1 | 0.1 | 0.29 | 0.12 | 0.16 | 0.25 |
| 7 | 26543316 | rs1859600 | -0.996 | 1.852 | 0.006 | 0.545 | -0.006 | -0.005 | -0.005 | 0 | 0.11 | 0.11 | 0.29 | 0.12 | 0.16 | 0.25 |
| 7 | 26545099 | rs4722604 | -0.872 | 1.755 | 0.005 | 0.545 | 0.004 | 0.001 | 0.001 | 0 | 0.1 | 0.15 | 0.27 | 0.12 | 0.15 | 0.24 |
| 7 | 26552429 | rs6461949 | -0.875 | 1.135 | 0.229 | 0.056 | -0.008 | -0.053 | -0.053 | 0 | 0.1 | 0.24 | 0.06 | 0 | 0.07 | 0.12 |
| 7 | 26553272 | rs6461951 | 0.206 | 1.038 | 0.365 | -0.22 | 0.008 | -0.044 | -0.044 | 0 | 0.08 | 0.16 | 0.05 | 0 | 0.06 | 0.02 |
| 7 | 26557421 | rs1541424 | 0.431 | 1.375 | 0.208 | 0.335 | -0.008 | -0.008 | -0.008 | 0 | 0.85 | 0.86 | 0.72 | 0.25 | 0.77 | 0.62 |
| 7 | 26557618 | rs886716 | -0.694 | 0.931 | 0.193 | -0.683 | -0.001 | -0.064 | -0.064 | 0 | 0.33 | 0.64 | 0.21 | 0.08 | 0.31 | 0.28 |
| 7 | 26559492 | rs10240247 | -0.27 | 0.961 | 0.638 | -0.597 | -0.01 | -0.019 | -0.019 | 0 | 0.08 | 0.25 | 0.05 | 0.01 | 0.03 | 0.08 |
| 7 | 26561804 | rs4719872 | -0.549 | 0.857 | 0.656 | -0.736 | 0.011 | -0.095 | -0.095 | 0.282 | 0.26 | 0.6 | 0.18 | 0.06 | 0.28 | 0.26 |
| 7 | 26566266 | rs6943220 | -1.101 | 0.86 | 0.662 | -0.728 | 0.001 | -0.062 | -0.062 | 0 | 0.32 | 0.61 | 0.21 | 0.07 | 0.31 | 0.34 |
| 7 | 26570142 | rs12537002 | -1.022 | 0.974 | 0.614 | -0.97 | 0.009 | 0.001 | 0.001 | 0 | 0.25 | 0.26 | 0.24 | 0.05 | 0.19 | 0.29 |
| 7 | 26570259 | rs2391283 | -0.319 | 1.283 | 0.684 | 0.33 | 0 | 0.028 | 0.028 | 0 | 0.87 | 0.86 | 0.73 | 0.52 | 0.77 | 0.67 |
| 7 | 26574973 | rs730956 | -0.395 | 1.264 | 0.929 | 0.16 | 0.114 | -0.056 | -0.056 | 0 | 0.11 | 0.38 | 0.08 | 0 | 0.14 | 0.06 |
| 7 | 26575513 | rs1076284 | -0.433 | 1.196 | 0.663 | 0.091 | 0.008 | 0.075 | 0.075 | 1.37 | 0.17 | 0.05 | 0.09 | 0.05 | 0.14 | 0.26 |
| 7 | 26576535 | rs740182 | 1.435 | 1.023 | 0.507 | 0.062 | -0.015 | 0.032 | 0.032 | 0.967 | 0.65 | 0.68 | 0.6 | 0.2 | 0.62 | 0.32 |
| 7 | 26577614 | rs17153818 | -1.117 | 1.224 | 0.81 | 0.131 | 0.104 | -0.061 | -0.061 | 0 | 0.14 | 0.45 | 0.21 | 0.02 | 0.17 | 0.07 |
| 7 | 26578105 | rs17153822 | -1.117 | 1.224 | 0.588 | 0.131 | 0.089 | -0.061 | -0.061 | 0 | 0.14 | 0.48 | 0.21 | 0.02 | 0.17 | 0.07 |
| 7 | 26580996 | rs4601210 | -1.12 | 1.152 | 0.581 | 0.172 | 0.1 | -0.033 | -0.033 | 0 | 0.11 | 0.48 | 0.21 | 0.02 | 0.13 | 0.03 |
| 7 | 26581796 | rs2391285 | -0.143 | 1.272 | 0.21 | 0.614 | -0.023 | -0.028 | -0.028 | 0.057 | 0.82 | 0.86 | 0.71 | 0.57 | 0.72 | 0.63 |
| 7 | 26583720 | rs17284017 | 0.188 | 0.979 | -0.086 | 0.289 | -0.007 | 0.023 | 0.023 | 0.092 | 0.12 | 0.05 | 0.09 | 0.07 | 0.12 | 0.2 |
| 7 | 26583781 | rs10215963 | 1.585 | 0.804 | 0.056 | 0.317 | 0.001 | 0.019 | 0.019 | 5.714 | 0.56 | 0.41 | 0.53 | 0.18 | 0.47 | 0.25 |
| 7 | 26586556 | rs17153906 | 0.185 | 0.335 | -0.694 | 0.452 | 0.007 | -0.101 | -0.101 | 3.148 | 0.21 | 0.02 | 0.11 | 0.3 | 0.17 | 0.25 |
| 7 | 26586734 | rs757808 | 1.715 | 0.306 | -0.015 | 0.216 | -0.002 | -0.007 | -0.007 | 0 | 0.41 | 0.43 | 0.37 | 0.2 | 0.38 | 0.22 |
| 7 | 26588410 | rs1029559 | 0.726 | 0.304 | 0.091 | 0.189 | 0.037 | 0.017 | 0.017 | 0.125 | 0.64 | 0.49 | 0.6 | 0.51 | 0.57 | 0.48 |
| 7 | 26588721 | rs735598 | -0.902 | 0.629 | 0.308 | 0.635 | 0 | -0.013 | -0.013 | 0.152 | 0.35 | 0.31 | 0.22 | 0.59 | 0.33 | 0.39 |
| 7 | 26590298 | rs882541 | 1.722 | 0.62 | 0.327 | 0.605 | -0.007 | -0.021 | -0.021 | 0 | 0.43 | 0.46 | 0.48 | 0.19 | 0.39 | 0.23 |
| 7 | 26590857 | rs1859601 | -0.854 | 0.73 | 0.279 | 0.602 | 0.009 | 0 | 0 | 0 | 0.85 | 0.81 | 0.75 | 0.85 | 0.81 | 0.76 |
| 7 | 26591929 | rs1557987 | 0.566 | 0.75 | 0.387 | 0.612 | 0.004 | 0.017 | 0.017 | 0 | 0.73 | 0.7 | 0.59 | 0.58 | 0.73 | 0.62 |
| 7 | 26592248 | rs6959388 | -0.87 | 0.793 | 0.271 | 0.614 | 0.013 | 0 | 0 | 0 | 0.85 | 0.81 | 0.75 | 0.85 | 0.81 | 0.76 |
| 7 | 26592617 | rs1029560 | 0.463 | 0.879 | 0.37 | 0.596 | 0.028 | -0.013 | -0.013 | 0.264 | 0.75 | 0.73 | 0.65 | 0.8 | 0.62 | 0.61 |
| 7 | 26593732 | rs12670740 | 0.509 | 0.701 | 0.463 | 0.866 | 0.031 | -0.024 | -0.024 | 0.012 | 0.21 | 0.08 | 0.12 | 0.26 | 0.18 | 0.25 |
| 7 | 26594315 | rs740183 | 0.57 | 0.815 | 0.465 | 0.887 | 0.022 | 0.004 | 0.004 | 0 | 0.74 | 0.73 | 0.65 | 0.71 | 0.62 | 0.61 |
| 7 | 26594823 | rs740184 | -0.093 | 0.792 | 0.359 | 0.895 | 0.002 | -0.001 | -0.001 | 0.324 | 0.12 | 0.08 | 0.1 | 0.14 | 0.19 | 0.15 |
| 7 | 26597211 | rs6957717 | -0.295 | 0.691 | 0.341 | 0.875 | 0.028 | 0 | 0 | 0 | 0.28 | 0.13 | 0.16 | 0.3 | 0.26 | 0.38 |
| 7 | 26600300 | rs12671326 | 0.441 | 0.687 | 0.275 | 0.88 | 0.032 | -0.022 | -0.022 | 0 | 0.21 | 0.08 | 0.12 | 0.26 | 0.18 | 0.25 |

| | | | | | | | | | | | | | | | | |
|---|---|---|---|---|---|---|---|---|---|---|---|---|---|---|---|---|
| 7 | 26600663 | rs3948698 | 1.664 | 0.619 | 0.409 | 0.815 | 0 | 0 | 0 | 0 | 0.45 | 0.47 | 0.4 | 0.23 | 0.44 | 0.35 |
| 7 | 26601316 | rs1557988 | 0.558 | 0.348 | 0.305 | 1.017 | 0.004 | -0.007 | -0.007 | 0.533 | 0.64 | 0.71 | 0.63 | 0.59 | 0.63 | 0.51 |
| 7 | 26605282 | rs17153945 | 0.88 | 0.337 | 0.258 | 0.842 | 0.001 | -0.022 | -0.022 | 2.565 | 0.07 | 0.2 | 0.06 | 0.05 | 0.08 | 0.13 |
| 7 | 26607065 | rs7789704 | -0.686 | 0.349 | 0.293 | 0.882 | 0.02 | 0.037 | 0.037 | 0 | 0.84 | 0.76 | 0.73 | 0.76 | 0.8 | 0.76 |
| 7 | 26607180 | rs7789471 | 0.658 | 0.376 | 0.378 | 0.718 | 0.006 | -0.013 | -0.013 | 0 | 0.09 | 0.05 | 0.03 | 0.27 | 0.05 | 0.13 |
| 7 | 26607413 | rs7776731 | 0.393 | 0.585 | 0.363 | 0.719 | 0.001 | -0.011 | -0.011 | 0 | 0.63 | 0.68 | 0.61 | 0.5 | 0.62 | 0.51 |
| 7 | 26607472 | rs7793934 | 0.393 | 0.563 | 0.363 | 0.719 | 0.001 | -0.011 | -0.011 | 0 | 0.63 | 0.68 | 0.61 | 0.5 | 0.62 | 0.51 |
| 7 | 26608284 | rs4624932 | 0.851 | 0.558 | 0.237 | 0.79 | -0.001 | 0 | 0 | 0 | 0.55 | 0.48 | 0.55 | 0.45 | 0.54 | 0.38 |
| 7 | 26611361 | rs6960339 | -0.547 | 0.33 | 0.469 | 0.925 | -0.02 | 0.122 | 0.122 | 0 | 0.14 | 0.4 | 0.09 | 0.27 | 0.06 | 0.14 |
| 7 | 26612333 | rs4722613 | -0.442 | 0.84 | 0.274 | 1.142 | -0.002 | -0.003 | -0.003 | 0 | 0.11 | 0.1 | 0.08 | 0.12 | 0.16 | 0.13 |
| 7 | 26614241 | rs4722614 | 1.103 | 0.741 | 0.229 | 1.143 | -0.008 | 0.008 | 0.008 | 0.507 | 0.13 | 0.18 | 0.13 | 0.16 | 0.12 | 0.05 |
| 7 | 26615293 | rs13311768 | 1.404 | 0.673 | 0.204 | 1.028 | 0.002 | 0.002 | 0.002 | 0.751 | 0.47 | 0.39 | 0.5 | 0.43 | 0.4 | 0.34 |
| 7 | 26617594 | rs2893255 | 0.265 | 0.676 | 0.132 | 0.958 | -0.001 | 0.009 | 0.009 | 0 | 0.63 | 0.53 | 0.6 | 0.54 | 0.62 | 0.53 |
| 7 | 26617908 | rs4557597 | -0.326 | 0.704 | -0.015 | 0.971 | -0.015 | 0.005 | 0.005 | 0 | 0.16 | 0.13 | 0.11 | 0.11 | 0.21 | 0.2 |
| 7 | 26619017 | rs731158 | 0.421 | 0.641 | -0.102 | 0.987 | 0.003 | -0.017 | -0.017 | 0.396 | 0.11 | 0.05 | 0.06 | 0.11 | 0.11 | 0.18 |
| 7 | 26619480 | rs11983370 | 1.473 | 0.695 | 0.046 | 0.849 | 0.009 | 0.007 | 0.007 | 1.362 | 0.44 | 0.29 | 0.38 | 0.43 | 0.38 | 0.33 |
| 7 | 26621872 | rs11975448 | 0.883 | 1.025 | 0.03 | 0.831 | -0.006 | 0.006 | 0.006 | 0.341 | 0.54 | 0.58 | 0.64 | 0.47 | 0.51 | 0.48 |
| 7 | 26622508 | rs740192 | 0.821 | 1.069 | 0.034 | 0.862 | 0.002 | -0.001 | -0.001 | 0 | 0.57 | 0.51 | 0.66 | 0.46 | 0.61 | 0.5 |
| 7 | 26624392 | rs2286182 | 1.089 | 0.832 | -0.093 | 0.895 | -0.001 | 0.002 | 0.002 | 0.888 | 0.51 | 0.41 | 0.61 | 0.47 | 0.49 | 0.49 |
| 7 | 26625527 | rs2190190 | 1.578 | 0.794 | -0.521 | 0.796 | 0.02 | 0.045 | 0.045 | 0.625 | 0.49 | 0.27 | 0.42 | 0.4 | 0.45 | 0.33 |
| 7 | 26626213 | rs17286397 | 2.548 | 0.696 | -0.787 | 0.636 | 0.001 | 0.021 | 0.021 | 2.536 | 0.31 | 0.21 | 0.32 | 0.13 | 0.32 | 0.21 |
| 7 | 26626743 | rs10242264 | -0.615 | 0.57 | -1.003 | 1.156 | 0.008 | -0.065 | -0.065 | 0.247 | 0.24 | 0.09 | 0.19 | 0.27 | 0.23 | 0.21 |
| 7 | 26627743 | rs12534493 | 0.897 | 1.271 | -0.703 | 1.056 | -0.043 | -0.026 | -0.026 | 0.063 | 0.07 | 0.01 | 0.22 | 0.07 | 0.13 | 0.17 |
| 7 | 26629336 | rs10486465 | -0.522 | 0.981 | -0.385 | 1.144 | -0.02 | 0.036 | 0.036 | 0 | 0.23 | 0.38 | 0.14 | 0.37 | 0.17 | 0.25 |
| 7 | 26629532 | rs10254487 | -0.869 | 1.446 | 0.163 | 1.23 | 0.089 | 0.132 | 0.132 | 0 | 0.15 | 0.43 | 0.35 | 0.25 | 0.18 | 0.22 |
| 7 | 26630540 | rs10464222 | -0.126 | 1.465 | -0.616 | 0.822 | -0.013 | 0.004 | 0.004 | 0 | 0.13 | 0.06 | 0.07 | 0.13 | 0.13 | 0.22 |
| 7 | 26633643 | rs17286978 | 0.274 | 1.467 | -0.887 | 0.742 | -0.041 | 0.025 | 0.025 | 0 | 0.24 | 0.06 | 0.15 | 0.15 | 0.27 | 0.24 |
| 7 | 26633863 | rs10226436 | 1.068 | 1.504 | -0.243 | 0.989 | 0 | 0.004 | 0.004 | 0.118 | 0.5 | 0.58 | 0.68 | 0.57 | 0.48 | 0.44 |
| 7 | 26637362 | rs10247053 | 0.138 | 1.508 | -0.041 | 0.877 | -0.013 | 0.047 | 0.047 | 0 | 0.31 | 0.21 | 0.19 | 0.14 | 0.32 | 0.41 |
| 7 | 26638405 | rs4722617 | 0.554 | 1.4 | 0.862 | 1.029 | -0.013 | 0.086 | 0.086 | 0 | 0.79 | 0.63 | 0.87 | 0.63 | 0.8 | 0.85 |
| 7 | 26639622 | rs4719875 | 1.625 | 1.682 | 0.957 | 0.958 | 0.002 | 0 | 0 | 0 | 0.41 | 0.32 | 0.43 | 0.43 | 0.34 | 0.27 |
| 7 | 26640101 | rs2214810 | 0.345 | 1.695 | 0.954 | 0.907 | -0.013 | 0.041 | 0.041 | 0 | 0.3 | 0.21 | 0.19 | 0.15 | 0.32 | 0.42 |
| 7 | 26640144 | rs13226306 | -0.762 | 1.731 | 0.995 | 1.108 | -0.005 | 0.012 | 0.012 | 0 | 0.21 | 0.34 | 0.31 | 0.25 | 0.19 | 0.25 |
| 7 | 26640272 | rs17154049 | 1.245 | 1.855 | 0.985 | 0.864 | 0.022 | -0.028 | -0.028 | 0 | 0.05 | 0.15 | 0.04 | 0 | 0.06 | 0.08 |
| 7 | 26642070 | rs6945115 | 0.122 | 1.836 | 0.952 | 0.875 | -0.001 | 0 | 0 | 0.035 | 0.32 | 0.34 | 0.21 | 0.15 | 0.32 | 0.42 |
| 7 | 26644642 | rs11763033 | 1.238 | 1.645 | 0.759 | 0.909 | 0.022 | 0.032 | 0.032 | 0.583 | 0.5 | 0.35 | 0.48 | 0.71 | 0.4 | 0.36 |
| 7 | 26645259 | rs10807833 | -0.507 | 1.738 | 0.496 | 0.924 | 0.026 | 0.009 | 0.009 | 0 | 0.92 | 0.9 | 0.95 | 0.92 | 0.86 | 0.94 |
| 7 | 26648786 | rs1005401 | -0.706 | 1.54 | 0.599 | 1.18 | 0 | -0.001 | -0.001 | 0.057 | 0.43 | 0.45 | 0.61 | 0.45 | 0.5 | 0.65 |
| 7 | 26648907 | rs2078118 | -0.814 | 1.23 | 0.202 | 0.664 | -0.008 | -0.019 | -0.019 | 0 | 0.93 | 0.98 | 0.97 | 0.92 | 0.92 | 0.95 |
| 7 | 26653227 | rs4640957 | 0.108 | 1.315 | 0.365 | 0.835 | -0.003 | 0.01 | 0.01 | 0 | 0.31 | 0.3 | 0.19 | 0.42 | 0.29 | 0.4 |
| 7 | 26654077 | rs7777206 | 0.657 | 1.318 | 0.129 | 0.847 | -0.005 | 0.014 | 0.014 | 0 | 0.26 | 0.27 | 0.16 | 0.34 | 0.25 | 0.27 |
| 7 | 26654392 | rs11763972 | 0.912 | 1.499 | 0.005 | 0.798 | 0.037 | -0.017 | -0.017 | 0.139 | 0.07 | 0.17 | 0.24 | 0.05 | 0.13 | 0.16 |
| 7 | 26654514 | rs10486467 | -0.256 | 1.436 | -0.619 | 0.736 | -0.011 | 0.022 | 0.022 | 0 | 0.07 | 0.17 | 0.04 | 0.09 | 0.04 | 0.13 |
| 7 | 26656114 | rs1557994 | 0.434 | 1.569 | -1.463 | 0.299 | -0.001 | 0.013 | 0.013 | 0 | 0.8 | 0.79 | 0.73 | 0.66 | 0.78 | 0.77 |
| 7 | 26656514 | rs17360198 | 1.152 | 1.838 | -1.524 | 0.293 | -0.017 | 0.116 | 0.116 | 0.78 | 0.39 | 0.14 | 0.28 | 0.17 | 0.35 | 0.2 |
| 7 | 26656707 | rs8180738 | 0.382 | 2.016 | -0.887 | 0.883 | 0.003 | 0.045 | 0.045 | 0 | 0.91 | 0.85 | 0.81 | 0.77 | 0.9 | 0.89 |
| 7 | 26657056 | rs1557995 | 0.405 | 2.189 | -1.296 | 0.383 | -0.037 | 0.031 | 0.031 | 0.994 | 0.06 | 0.01 | 0.15 | 0.01 | 0.14 | 0.12 |
| 7 | 26657771 | rs4722625 | -0.203 | 2.022 | -1.879 | 0.413 | -0.179 | -0.073 | -0.073 | 0 | 0.35 | 0.04 | 0.48 | 0.67 | 0.46 | 0.55 |
| 7 | 26657962 | rs4722626 | 0.584 | 1.746 | -0.847 | -0.277 | 0.011 | -0.043 | -0.043 | 0 | 0.62 | 0.44 | 0.73 | 0.8 | 0.59 | 0.75 |
| 7 | 26663492 | rs6967151 | -1.323 | 1.739 | 0.334 | 0.101 | 0.007 | 0.009 | 0.009 | 0 | 0.23 | 0.7 | 0.22 | 0.28 | 0.25 | 0.22 |
| 7 | 26685179 | rs17154189 | -1.563 | 1.91 | 0.339 | 0.224 | 0.004 | 0.005 | 0.005 | 0 | 0.26 | 0.69 | 0.25 | 0.33 | 0.28 | 0.22 |
| 7 | 26690417 | rs10242687 | -1.246 | 1.737 | 1.02 | 0.457 | 0.081 | 0.332 | 0.332 | 0.061 | 0.1 | 0.55 | 0.14 | 0.32 | 0.09 | 0.14 |
| 7 | 26695215 | rs212837 | 0.957 | 1.551 | 0.29 | -0.17 | -0.003 | -0.007 | -0.007 | 0.272 | 0.65 | 0.32 | 0.61 | 0.57 | 0.63 | 0.65 |
| 7 | 26699295 | rs7799519 | -0.247 | 1.546 | 0.994 | -0.428 | 0.04 | -0.023 | -0.023 | 0 | 0.08 | 0.27 | 0.12 | 0.01 | 0.08 | 0.04 |
| 7 | 26707657 | rs13186 | 1.048 | 0.803 | 0.773 | -1.061 | 0.031 | 0.095 | 0.095 | 0 | 0.51 | 0.32 | 0.61 | 0.86 | 0.55 | 0.66 |
| 7 | 26711817 | rs212842 | 0.769 | 0.722 | 0.622 | -1.091 | 0.014 | 0.068 | 0.068 | 0 | 0.55 | 0.7 | 0.64 | 0.85 | 0.56 | 0.67 |
| 7 | 26729740 | rs17291131 | 1.291 | 1.298 | 0.397 | -0.965 | -0.042 | 0.041 | 0.041 | 0 | 0.07 | 0.01 | 0.05 | 0 | 0.11 | 0.07 |
| 7 | 26734509 | rs10228105 | 0.351 | 0.836 | 0.675 | -0.898 | -0.037 | -0.116 | -0.116 | 0 | 0.24 | 0.43 | 0.2 | 0.03 | 0.2 | 0.13 |
| 7 | 26742154 | rs7795074 | 0.579 | 0.784 | 0.593 | -0.917 | -0.001 | 0.007 | 0.007 | 0 | 0.69 | 0.68 | 0.71 | 0.88 | 0.67 | 0.75 |
| 7 | 26747064 | rs2299102 | 1.003 | 0.741 | 0.586 | -0.943 | 0.007 | 0.004 | 0.004 | 0 | 0.51 | 0.5 | 0.62 | 0.85 | 0.55 | 0.66 |
| 7 | 26754682 | rs17310851 | 2.041 | 0.717 | 0.466 | -0.913 | 0.039 | 0.08 | 0.08 | 0 | 0.16 | 0.03 | 0.07 | 0.03 | 0.12 | 0.09 |
| 7 | 26763002 | rs1024453 | 0.088 | 0.901 | 0.715 | -0.794 | 0.041 | -0.023 | -0.023 | 0 | 0.08 | 0.37 | 0.12 | 0.01 | 0.08 | 0.04 |
| 7 | 26765596 | rs1129771 | 0.254 | 0.909 | 0.815 | -0.791 | 0.049 | -0.016 | -0.016 | 0 | 0.08 | 0.3 | 0.12 | 0.01 | 0.08 | 0.04 |
| 7 | 26774969 | rs11772673 | 0.156 | 0.879 | 0.846 | -0.847 | 0.028 | 0.087 | 0.087 | 0 | 0.6 | 0.76 | 0.74 | 0.86 | 0.64 | 0.69 |
| 7 | 26777367 | rs11973332 | 0.376 | 0.887 | 0.885 | -0.844 | 0.025 | -0.029 | -0.029 | 1.131 | 0.08 | 0.24 | 0.11 | 0.01 | 0.08 | 0.04 |
| 7 | 26778732 | rs11764558 | 1.07 | 1.096 | 0.959 | -0.503 | 0.001 | 0.012 | 0.012 | 0 | 0.47 | 0.26 | 0.53 | 0.56 | 0.5 | 0.58 |
| 7 | 26790969 | rs12700758 | 0.458 | 1.027 | 0.773 | -0.568 | 0.026 | 0.117 | 0.117 | 0 | 0.53 | 0.71 | 0.65 | 0.85 | 0.55 | 0.66 |
| 7 | 26793515 | rs12700761 | 0.644 | 1.061 | 0.576 | -0.553 | 0 | -0.02 | -0.02 | 0.006 | 0.52 | 0.52 | 0.63 | 0.85 | 0.55 | 0.66 |
| 7 | 26823963 | rs513387 | 2.159 | 1.36 | -0.348 | -0.352 | -0.069 | -0.15 | -0.15 | 0.085 | 0.33 | 0.03 | 0.4 | 0.44 | 0.35 | 0.41 |

| | | | | | | | | | | | | | | | | |
|---|---|---|---|---|---|---|---|---|---|---|---|---|---|---|---|---|
| 7 | 26833021 | rs3801839 | 1.317 | 1.394 | 0.221 | -0.475 | -0.036 | 0.003 | 0.003 | 0 | 0.46 | 0.19 | 0.52 | 0.55 | 0.47 | 0.6 |
| 7 | 26851805 | rs490366 | -0.635 | 1.119 | 0.55 | -0.881 | -0.005 | -0.055 | -0.055 | 0 | 0.77 | 0.69 | 0.81 | 0.95 | 0.74 | 0.89 |
| 7 | 26854266 | rs10262414 | -0.109 | 1.104 | 0.679 | -0.901 | 0.001 | 0.013 | 0.013 | 0 | 0.23 | 0.16 | 0.18 | 0.11 | 0.22 | 0.2 |
| 7 | 26862139 | rs570840 | 1.166 | 1.09 | 0.873 | -0.922 | 0.007 | 0.027 | 0.027 | 0 | 0.52 | 0.38 | 0.6 | 0.83 | 0.52 | 0.69 |
| 7 | 26872652 | rs10486480 | 1.941 | 1.003 | 0.25 | -0.955 | 0.02 | 0.069 | 0.069 | 0 | 0.15 | 0.01 | 0.06 | 0.04 | 0.12 | 0.07 |
| 7 | 26881302 | rs3801814 | -0.169 | 1.161 | 0.44 | -0.815 | 0.001 | 0.014 | 0.014 | 0 | 0.23 | 0.16 | 0.18 | 0.11 | 0.22 | 0.2 |
| 7 | 26938809 | rs213516 | -1.852 | 1.563 | 1.398 | -0.17 | 0.051 | 0.136 | 0.136 | 0 | 0.15 | 0.54 | 0.17 | 0.28 | 0.18 | 0.16 |
| 7 | 26954746 | rs13311400 | -1.323 | 1.67 | 1.314 | -0.416 | 0.015 | 0.014 | 0.014 | 0 | 0.48 | 0.7 | 0.39 | 0.44 | 0.48 | 0.42 |
| 7 | 27026872 | rs2140998 | -0.946 | 2.018 | 1.26 | -0.434 | 0.114 | -0.015 | -0.015 | 0 | 0.08 | 0.3 | 0.12 | 0.01 | 0.14 | 0.04 |
| 7 | 27051961 | rs2264803 | 0.168 | 1.821 | 0.98 | -0.45 | -0.071 | 0.094 | 0.094 | 0 | 0.32 | 0.03 | 0.22 | 0.12 | 0.36 | 0.23 |
| 7 | 27073731 | rs799234 | -0.633 | 1.962 | 1.225 | -0.49 | -0.002 | 0.003 | 0.003 | 0 | 0.42 | 0.37 | 0.45 | 0.22 | 0.48 | 0.33 |
| 7 | 27089764 | rs10081212 | -0.945 | 1.744 | 0.966 | -0.672 | 0.062 | -0.009 | -0.009 | 0 | 0.07 | 0.29 | 0.11 | 0.01 | 0.09 | 0.03 |
| 7 | 27108127 | rs2465274 | 0.125 | 1.721 | 0.017 | -0.711 | -0.033 | 0.089 | 0.089 | 0 | 0.3 | 0.01 | 0.2 | 0.11 | 0.29 | 0.22 |
| 7 | 27111195 | rs2428427 | -1.329 | 1.623 | 0.213 | -0.717 | 0.01 | 0.012 | 0.012 | 0 | 0.48 | 0.83 | 0.43 | 0.44 | 0.48 | 0.4 |
| 7 | 27118974 | rs17427421 | 0.96 | 1.636 | 0.585 | -0.87 | -0.013 | 0.039 | 0.039 | 0 | 0.07 | 0 | 0.03 | 0 | 0.07 | 0.02 |
| 7 | 27135314 | rs10951154 | 2.151 | 1.578 | 1.222 | -0.329 | 0.077 | -0.057 | -0.057 | 0 | 0.89 | 0.41 | 0.84 | 0.99 | 0.85 | 0.95 |
| 7 | 27143757 | rs2522828 | 1.059 | 1.454 | 0.507 | -0.173 | -0.001 | -0.004 | -0.004 | 0.1 | 0.94 | 0.93 | 0.94 | 1 | 0.91 | 0.96 |
| 7 | 27145024 | rs2428432 | 0.808 | 1.214 | 0.835 | -0.371 | -0.018 | -0.065 | -0.065 | 0 | 0.7 | 0.52 | 0.64 | 0.86 | 0.73 | 0.86 |
| 7 | 27145517 | rs2428433 | -1.385 | 1.354 | 1.146 | -0.263 | 0.054 | 0.046 | 0.046 | 0.122 | 0.1 | 0.28 | 0.26 | 0.14 | 0.12 | 0.07 |
| 7 | 27145717 | rs2465276 | -1.133 | 1.358 | 1.191 | -0.261 | 0.052 | 0.045 | 0.045 | 0 | 0.09 | 0.23 | 0.25 | 0.14 | 0.12 | 0.06 |
| 7 | 27167346 | rs2107063 | 1.892 | 1.43 | 1.254 | -0.174 | 0.075 | 0.056 | 0.056 | 0.444 | 0.89 | 0.65 | 0.75 | 0.86 | 0.87 | 0.88 |
| 7 | 27168590 | rs1801085 | -1.518 | 1.411 | 1.242 | -0.182 | 0.113 | 0.118 | 0.118 | 0.677 | 0.08 | 0.33 | 0.25 | 0.14 | 0.1 | 0.07 |
| 7 | 27171588 | rs1859166 | 2.301 | 1.616 | 1.05 | -0.37 | 0.011 | -0.017 | -0.017 | 0 | 0.81 | 0.39 | 0.65 | 0.86 | 0.8 | 0.83 |
| 7 | 27172571 | rs4719884 | 0.052 | 1.772 | 0.651 | -0.253 | 0 | 0.032 | 0.032 | 0 | 0.25 | 0.08 | 0.22 | 0.14 | 0.25 | 0.3 |
| 7 | 27182142 | rs3757639 | -0.569 | 2.699 | 1.484 | 0.609 | -0.033 | -0.016 | -0.016 | 0 | 0.05 | 0 | 0.12 | 0.26 | 0.14 | 0.06 |
| 7 | 27182245 | rs3757640 | 3.046 | 2.537 | 1.677 | 0.542 | 0.008 | 0.015 | 0.015 | 0.491 | 0.77 | 0.36 | 0.56 | 0.6 | 0.71 | 0.75 |
| 7 | 27189195 | rs3807597 | 1.164 | 2.17 | 1.765 | 0.472 | 0.054 | 0.053 | 0.053 | 0 | 0.92 | 0.77 | 0.76 | 0.86 | 0.87 | 0.89 |
| 7 | 27196069 | rs2301720 | 1.09 | 2.149 | 1.525 | 0.463 | 0.093 | 0.087 | 0.087 | 0.047 | 0.9 | 0.55 | 0.73 | 0.86 | 0.86 | 0.89 |
| 7 | 27196113 | rs2301721 | 2.655 | 2.094 | 1.403 | 0.43 | 0.004 | 0.028 | 0.028 | 4.241 | 0.79 | 0.1 | 0.65 | 0.86 | 0.83 | 0.83 |
| 7 | 27200905 | rs875898 | 1.952 | 2.103 | 1.487 | 0.398 | 0.009 | -0.018 | -0.018 | 0.001 | 0.81 | 0.32 | 0.68 | 0.86 | 0.82 | 0.85 |
| 7 | 27201532 | rs875896 | 2.179 | 2.057 | 0.927 | 0.384 | -0.027 | -0.052 | -0.052 | 0.016 | 0.73 | 0.1 | 0.6 | 0.62 | 0.75 | 0.79 |
| 7 | 27202289 | rs10259620 | 1.175 | 2.544 | 1.186 | 0.801 | 0.068 | 0.123 | 0.123 | 0 | 0.84 | 0.57 | 0.68 | 0.62 | 0.78 | 0.85 |
| 7 | 27203139 | rs7810502 | 1.512 | 2.647 | 1.085 | 0.811 | 0.138 | 0.128 | 0.128 | 0 | 0.82 | 0.49 | 0.59 | 0.61 | 0.7 | 0.8 |
| 7 | 27205282 | rs3801776 | 1.299 | 1.95 | 0.566 | 0.809 | 0.018 | 0.03 | 0.03 | 0 | 0.8 | 0.73 | 0.64 | 0.63 | 0.75 | 0.82 |
| 7 | 27206688 | rs11773804 | 0.51 | 1.57 | 0.084 | 0.604 | 0.005 | -0.015 | -0.015 | 0 | 0.42 | 0.2 | 0.47 | 0.51 | 0.42 | 0.5 |
| 7 | 27215041 | rs6949451 | 0.473 | 1.574 | 0.129 | 0.729 | -0.001 | -0.002 | -0.002 | 0.01 | 0.46 | 0.19 | 0.39 | 0.5 | 0.48 | 0.53 |
| 7 | 27223563 | rs6968828 | 0.347 | 1.093 | 0.422 | 0.46 | -0.003 | -0.006 | -0.006 | 0.498 | 0.53 | 0.46 | 0.55 | 0.53 | 0.57 | 0.59 |
| 7 | 27224963 | rs17437495 | -0.864 | 0.997 | 0.712 | 0.84 | -0.003 | -0.013 | -0.013 | 0 | 0.11 | 0.22 | 0.09 | 0.05 | 0.09 | 0.03 |
| 7 | 27229549 | rs4722670 | 0.557 | 1.067 | 0.773 | 0.737 | 0.014 | 0.154 | 0.154 | 0 | 0.93 | 0.86 | 0.84 | 0.58 | 0.91 | 0.9 |
| 7 | 27236958 | rs757181 | 1.072 | 1.155 | 0.797 | 0.702 | 0.008 | 0.137 | 0.137 | 0 | 0.92 | 0.81 | 0.83 | 0.58 | 0.91 | 0.88 |
| 7 | 27237453 | rs2189239 | -0.764 | 1.029 | 0.891 | 0.568 | 0.005 | -0.06 | -0.06 | 0 | 0.11 | 0.31 | 0.1 | 0 | 0.09 | 0.03 |
| 7 | 27243106 | rs10233387 | 1.014 | 0.956 | 0.126 | 0.46 | -0.076 | -0.073 | -0.073 | 0 | 0.38 | 0.07 | 0.29 | 0.48 | 0.45 | 0.43 |
| 7 | 27252246 | rs6958837 | 1.332 | 0.834 | 1.458 | 1.228 | -0.015 | 0.224 | 0.224 | 1.268 | 0.86 | 0.4 | 0.78 | 0.52 | 0.9 | 0.9 |
| 7 | 27260775 | rs10486494 | -0.485 | 0.826 | 0.721 | 0.768 | -0.008 | 0.032 | 0.032 | 0 | 0.15 | 0.07 | 0.17 | 0.07 | 0.15 | 0.17 |
| 7 | 27283498 | rs17437824 | -0.963 | 1.088 | 1.702 | 1.286 | 0.005 | -0.044 | -0.044 | 0 | 0.08 | 0.22 | 0.05 | 0 | 0.06 | 0.02 |
| 7 | 27286796 | rs6956598 | 1.671 | 0.998 | 2.162 | 1.435 | 0.02 | 0.247 | 0.247 | 0.422 | 0.88 | 0.39 | 0.78 | 0.57 | 0.89 | 0.9 |
| 7 | 27298264 | rs11764401 | 0.717 | 0.625 | 1.294 | 0.75 | -0.051 | 0.142 | 0.142 | 0 | 0.22 | 0.01 | 0.1 | 0.02 | 0.22 | 0.24 |
| 7 | 27300262 | rs6960464 | 1.903 | 0.561 | 2.409 | 1.343 | 0.015 | 0.202 | 0.202 | 0.665 | 0.86 | 0.42 | 0.79 | 0.57 | 0.87 | 0.9 |
| 7 | 27301441 | rs11564044 | -0.96 | 0.577 | 2.317 | 1.337 | -0.005 | -0.041 | -0.041 | 0 | 0.09 | 0.19 | 0.05 | 0 | 0.06 | 0.02 |
| 7 | 27303632 | rs6943291 | 2.078 | 0.531 | 2.112 | 1.316 | -0.009 | 0.04 | 0.04 | 0.768 | 0.82 | 0.3 | 0.78 | 0.57 | 0.85 | 0.89 |
| 7 | 27311559 | rs17437978 | -1.696 | 0.536 | 1.817 | 0.675 | -0.011 | -0.051 | -0.051 | 0 | 0.15 | 0.57 | 0.09 | 0.05 | 0.1 | 0.03 |
| 7 | 27322106 | rs10262979 | 0.398 | 0.473 | -0.329 | 0.762 | 0.01 | 0.154 | 0.154 | 0 | 0.48 | 0.09 | 0.37 | 0.27 | 0.39 | 0.52 |
| 7 | 27326739 | rs11564037 | -1.038 | 0.882 | 0.291 | 1.265 | 0.031 | 0.069 | 0.069 | 0 | 0.09 | 0.16 | 0.09 | 0.23 | 0.18 | 0.08 |
| 7 | 27328305 | rs1476658 | 0.369 | 1.076 | 0.802 | 1.103 | 0.047 | 0.006 | 0.006 | 0 | 0.9 | 0.57 | 0.78 | 0.96 | 0.89 | 0.96 |
| 7 | 27336004 | rs10254047 | -1.824 | 1.005 | 0.856 | 0.997 | 0.012 | 0.055 | 0.055 | 0 | 0.2 | 0.55 | 0.24 | 0.31 | 0.25 | 0.18 |
| 7 | 27336745 | rs6462003 | -0.358 | 1.023 | 0.82 | 1.043 | 0.01 | 0.025 | 0.025 | 0.269 | 0.69 | 0.64 | 0.61 | 0.58 | 0.64 | 0.69 |
| 7 | 27339901 | rs17438159 | -0.324 | 1.07 | 1.267 | 1.031 | 0.062 | 0.175 | 0.175 | 0 | 0.08 | 0.27 | 0.19 | 0.27 | 0.14 | 0.1 |
| 7 | 27341145 | rs6964636 | -0.976 | 0.979 | 0.983 | 1.007 | 0.018 | 0.048 | 0.048 | 0.009 | 0.74 | 0.66 | 0.68 | 0.62 | 0.71 | 0.72 |
| 7 | 27349479 | rs6462008 | -0.082 | 0.979 | 0.459 | 0.953 | -0.006 | -0.017 | -0.017 | 0.167 | 0.64 | 0.68 | 0.65 | 0.53 | 0.64 | 0.61 |
| 7 | 27351298 | rs1859669 | 1.12 | 0.871 | -0.027 | 0.845 | 0.017 | 0.145 | 0.145 | 0.091 | 0.46 | 0.09 | 0.36 | 0.26 | 0.37 | 0.5 |
| 7 | 27351650 | rs11563582 | 0.113 | 1.038 | 0.544 | 1.08 | 0.005 | -0.006 | -0.006 | 0.33 | 0.06 | 0.16 | 0.08 | 0.04 | 0.08 | 0.02 |
| 7 | 27352125 | rs6957104 | 0.355 | 1.041 | 0.108 | 1.232 | 0.002 | -0.01 | -0.01 | 0.755 | 0.65 | 0.57 | 0.7 | 0.75 | 0.65 | 0.65 |
| 7 | 27353904 | rs11564003 | -0.599 | 1.317 | -0.011 | 1.313 | 0.022 | 0.074 | 0.074 | 0 | 0.07 | 0.1 | 0.05 | 0.24 | 0.1 | 0.26 |
| 7 | 27356185 | rs11980136 | 0.444 | 1.367 | -0.038 | 0.38 | 0.023 | 0.008 | 0.008 | 0 | 0.63 | 0.46 | 0.53 | 0.6 | 0.53 | 0.67 |
| 7 | 27358452 | rs4722686 | 0.141 | 1.655 | 0.282 | 0.131 | -0.019 | -0.026 | -0.026 | 0 | 0.23 | 0.2 | 0.33 | 0.4 | 0.31 | 0.28 |
| 7 | 27359793 | rs2391415 | -1.774 | 0.521 | -0.154 | -0.946 | 0.02 | 0.047 | 0.047 | 0 | 0.64 | 0.8 | 0.81 | 0.99 | 0.68 | 0.72 |
| 7 | 27360015 | rs6953618 | 1.686 | 0.143 | -0.502 | -1.204 | 0.02 | 0.009 | 0.009 | 0 | 0.31 | 0.2 | 0.16 | 0.01 | 0.24 | 0.26 |
| 7 | 27362205 | rs1990120 | 1.352 | -0.184 | -0.576 | -1.323 | -0.027 | -0.022 | -0.022 | 0.168 | 0.07 | 0.12 | 0.03 | 0.04 | 0.02 | 0.05 |
| 7 | 27362214 | rs1990121 | -1.092 | -0.215 | -0.736 | -1.274 | -0.1 | -0.176 | -0.176 | 0.392 | 0.77 | 0.98 | 0.67 | 0.48 | 0.72 | 0.65 |

| | | | | | | | | | | | | | | | | |
|---|---|---|---|---|---|---|---|---|---|---|---|---|---|---|---|---|
| 7 | 27363392 | rs11772043 | -0.845 | -0.4 | -0.442 | -1.216 | -0.003 | 0.001 | 0.001 | 0 | 0.45 | 0.62 | 0.49 | 0.43 | 0.46 | 0.37 |
| 7 | 27363415 | rs11765116 | 0.961 | -0.329 | -0.374 | -1.064 | -0.017 | -0.024 | -0.024 | 0 | 0.23 | 0.2 | 0.33 | 0.39 | 0.3 | 0.28 |
| 7 | 27368442 | rs17473999 | 0.575 | -0.741 | -0.363 | -0.981 | -0.024 | -0.003 | -0.003 | 0 | 0.08 | 0.11 | 0.04 | 0.08 | 0.01 | 0.06 |
| 7 | 27372021 | rs10155834 | -0.192 | -0.624 | -0.824 | -1.093 | -0.102 | -0.175 | -0.175 | 0.487 | 0.66 | 0.98 | 0.56 | 0.46 | 0.6 | 0.68 |
| 7 | 27374565 | rs4722688 | 0.486 | -0.338 | -0.524 | -1.29 | -0.005 | -0.005 | -0.005 | 0 | 0.59 | 0.48 | 0.54 | 0.36 | 0.64 | 0.5 |
| 7 | 27378456 | rs4722689 | -1.075 | -0.342 | -0.445 | -1.244 | -0.002 | 0.023 | 0.023 | 0 | 0.75 | 0.7 | 0.63 | 0.5 | 0.75 | 0.73 |
| 7 | 27379499 | rs2158769 | -0.071 | -0.53 | -0.569 | -1.268 | -0.029 | -0.014 | -0.014 | 0 | 0.34 | 0.21 | 0.31 | 0.35 | 0.41 | 0.42 |
| 7 | 27386257 | rs6960719 | 1.223 | -0.278 | -0.294 | -1.086 | -0.004 | 0.007 | 0.007 | 0 | 0.47 | 0.39 | 0.44 | 0.41 | 0.49 | 0.45 |
| 7 | 27386618 | rs11979443 | 0.402 | -0.275 | -0.505 | -1.149 | -0.013 | 0.079 | 0.079 | 0.003 | 0.15 | 0.04 | 0.12 | 0.01 | 0.15 | 0.17 |
| 7 | 27389676 | rs6970737 | 1.222 | -0.306 | -0.184 | -1.075 | -0.003 | 0.004 | 0.004 | 0 | 0.46 | 0.42 | 0.43 | 0.36 | 0.49 | 0.45 |
| 7 | 27389942 | rs28357212 | -0.516 | -0.125 | -0.025 | -0.899 | -0.006 | -0.022 | -0.022 | 0 | 0.16 | 0.26 | 0.09 | 0.1 | 0.11 | 0.23 |
| 7 | 27391584 | rs4722692 | 1.014 | -0.312 | -0.384 | -1.027 | -0.004 | -0.008 | -0.008 | 0.038 | 0.26 | 0.27 | 0.33 | 0.38 | 0.3 | 0.32 |
| 7 | 27391639 | rs4719890 | -0.597 | -0.387 | -0.345 | -0.938 | 0 | -0.014 | -0.014 | 0 | 0.72 | 0.79 | 0.61 | 0.46 | 0.7 | 0.7 |
| 7 | 27392662 | rs12700801 | 0.72 | -0.308 | -0.538 | -0.954 | -0.084 | -0.002 | -0.002 | 0 | 0.34 | 0.08 | 0.35 | 0.26 | 0.44 | 0.35 |
| 7 | 27393144 | rs11981761 | 2.197 | -0.405 | -0.356 | -1.068 | 0.002 | -0.008 | -0.008 | 0.195 | 0.22 | 0.16 | 0.1 | 0 | 0.21 | 0.14 |
| 7 | 27393374 | rs10271302 | 0.799 | -0.39 | -0.431 | -1.042 | -0.079 | 0.001 | 0.001 | 0.315 | 0.34 | 0.08 | 0.35 | 0.26 | 0.44 | 0.35 |
| 7 | 27393518 | rs10274728 | -0.619 | -0.468 | -0.17 | -1.02 | 0.026 | 0.041 | 0.041 | 0 | 0.5 | 0.28 | 0.43 | 0.37 | 0.55 | 0.59 |
| 7 | 27397076 | rs10265434 | 0.304 | -0.434 | -0.045 | -0.83 | 0.002 | -0.006 | -0.006 | 0 | 0.14 | 0.14 | 0.08 | 0.11 | 0.11 | 0.12 |
| 7 | 27405091 | rs11563973 | 0.134 | -0.526 | -0.165 | -0.671 | -0.021 | 0.03 | 0.03 | 0 | 0.3 | 0.58 | 0.47 | 0.63 | 0.24 | 0.39 |
| 7 | 27408972 | rs11563971 | 1.166 | -0.299 | -0.695 | -0.578 | -0.019 | -0.093 | -0.093 | 0.02 | 0.21 | 0.08 | 0.32 | 0.52 | 0.24 | 0.26 |
| 7 | 27412812 | rs2189037 | -0.812 | -0.065 | -0.893 | -0.769 | -0.066 | -0.041 | -0.041 | 0.561 | 0.72 | 0.93 | 0.72 | 0.74 | 0.67 | 0.68 |
| 7 | 27414030 | rs6978648 | -0.34 | -0.203 | -0.332 | -1.008 | -0.009 | -0.048 | -0.048 | 0 | 0.24 | 0.33 | 0.16 | 0.11 | 0.21 | 0.16 |
| 7 | 27420533 | rs1055182 | 2.056 | -0.228 | -0.308 | -1.245 | -0.001 | -0.029 | -0.029 | 0 | 0.21 | 0.18 | 0.12 | 0 | 0.21 | 0.14 |
| 7 | 27422005 | rs9648340 | 1.537 | -0.23 | -0.618 | -1.01 | -0.052 | -0.156 | -0.156 | 0 | 0.21 | 0.02 | 0.32 | 0.5 | 0.24 | 0.28 |
| 7 | 27423066 | rs6961010 | 0.435 | -0.329 | -0.437 | -1.035 | 0.005 | 0.019 | 0.019 | 0 | 0.49 | 0.36 | 0.46 | 0.58 | 0.46 | 0.56 |
| 7 | 27425950 | rs16874272 | 0.421 | -0.286 | -0.217 | -1.204 | -0.001 | -0.049 | -0.049 | 0 | 0.19 | 0.19 | 0.14 | 0.03 | 0.2 | 0.15 |
| 7 | 27426466 | rs7805157 | 0.989 | -0.345 | -0.433 | -0.977 | -0.005 | -0.035 | -0.035 | 0 | 0.28 | 0.35 | 0.14 | 0.09 | 0.23 | 0.28 |
| 7 | 27431146 | rs7807747 | -0.489 | -0.055 | -0.123 | -0.942 | 0.002 | 0.005 | 0.005 | 0 | 0.5 | 0.48 | 0.42 | 0.51 | 0.45 | 0.59 |
| 7 | 27433363 | rs10240666 | -0.943 | -0.148 | -0.09 | -1.027 | 0.005 | 0.031 | 0.031 | 0.031 | 0.75 | 0.68 | 0.63 | 0.51 | 0.69 | 0.73 |
| 7 | 27436685 | rs2074337 | -0.536 | -0.13 | -0.072 | -0.939 | 0 | 0.001 | 0.001 | 0 | 0.51 | 0.53 | 0.5 | 0.51 | 0.47 | 0.59 |
| 7 | 27440061 | rs6957243 | -0.227 | -0.199 | -0.386 | -1.26 | -0.017 | 0.081 | 0.081 | 0 | 0.3 | 0.14 | 0.19 | 0.03 | 0.31 | 0.22 |
| 7 | 27446285 | rs11978918 | 1.395 | 0.067 | -0.069 | -0.873 | -0.003 | 0.062 | 0.062 | 0 | 0.17 | 0.26 | 0.29 | 0.47 | 0.15 | 0.26 |
| 7 | 27447332 | rs11764246 | 0.619 | 0.211 | -0.503 | -1.107 | -0.006 | 0 | 0 | 0 | 0.13 | 0.08 | 0.08 | 0 | 0.14 | 0.05 |
| 7 | 27447562 | rs11563953 | 1.795 | 0.029 | -0.586 | -1.398 | 0.012 | 0.027 | 0.027 | 0 | 0.27 | 0.14 | 0.15 | 0.03 | 0.23 | 0.22 |
| 7 | 27448316 | rs11761572 | 1.138 | 0.244 | -0.558 | -1.093 | -0.001 | -0.016 | -0.016 | 0.284 | 0.11 | 0.08 | 0.08 | 0 | 0.13 | 0.05 |
| 7 | 27453964 | rs17439381 | 1.708 | 0.309 | -0.949 | -0.955 | -0.038 | -0.082 | -0.082 | 0 | 0.15 | 0.02 | 0.16 | 0.3 | 0.19 | 0.16 |
| 7 | 27456500 | rs4722699 | 0.176 | -0.117 | -0.492 | -0.817 | -0.002 | -0.001 | -0.001 | 0.206 | 0.22 | 0.23 | 0.24 | 0.2 | 0.25 | 0.27 |
| 7 | 27461889 | rs10280925 | -1.052 | -0.223 | -0.382 | -0.854 | -0.004 | -0.014 | -0.014 | 0.666 | 0.73 | 0.81 | 0.73 | 0.7 | 0.69 | 0.77 |
| 7 | 27465607 | rs6963552 | -1.134 | -0.224 | -0.371 | -0.839 | -0.019 | -0.043 | -0.043 | 0 | 0.72 | 0.9 | 0.73 | 0.7 | 0.69 | 0.75 |
| 7 | 27470722 | rs757141 | -0.117 | -0.255 | -0.164 | -0.704 | 0 | -0.002 | -0.002 | 0 | 0.53 | 0.49 | 0.41 | 0.29 | 0.53 | 0.5 |
| 7 | 27471992 | rs1076210 | 1.189 | -0.541 | -0.05 | -0.555 | -0.002 | 0.078 | 0.078 | 0 | 0.18 | 0.29 | 0.31 | 0.42 | 0.16 | 0.26 |
| 7 | 27480649 | rs3915087 | -1.8 | -0.521 | -0.203 | -0.64 | 0.008 | 0.041 | 0.041 | 0 | 0.75 | 0.86 | 0.86 | 0.96 | 0.78 | 0.79 |
| 7 | 27482830 | rs10249757 | 0.207 | -0.661 | -0.017 | -0.312 | 0 | 0.016 | 0.016 | 0 | 0.17 | 0.28 | 0.2 | 0.25 | 0.18 | 0.25 |
| 7 | 27485395 | rs6957723 | 0.4 | -0.791 | 0.32 | 0.018 | -0.011 | 0.072 | 0.072 | 0 | 0.21 | 0.35 | 0.33 | 0.44 | 0.17 | 0.28 |
| 7 | 27496594 | rs12540319 | -0.773 | -0.572 | 0.047 | -0.022 | -0.015 | 0.173 | 0.173 | 0.921 | 0.07 | 0.29 | 0.12 | 0.26 | 0.02 | 0.17 |
| 7 | 27499198 | rs17743751 | 0.431 | -0.237 | -0.147 | -0.333 | -0.002 | 0.006 | 0.006 | 0 | 0.34 | 0.45 | 0.46 | 0.61 | 0.35 | 0.38 |
| 7 | 27504104 | rs9690475 | -0.896 | -0.03 | -0.881 | -0.501 | 0.003 | 0.019 | 0.019 | 0 | 0.63 | 0.71 | 0.73 | 0.99 | 0.63 | 0.67 |
| 7 | 27504332 | rs17155270 | -0.254 | -0.013 | -0.892 | 0.405 | -0.074 | -0.094 | -0.094 | 0 | 0.24 | 0.06 | 0.25 | 0.39 | 0.29 | 0.29 |
| 7 | 27508585 | rs7802070 | -0.812 | -0.718 | -0.156 | 0.387 | -0.038 | -0.019 | -0.019 | 0 | 0.2 | 0.58 | 0.24 | 0.16 | 0.13 | 0.16 |
| 7 | 27509208 | rs10271999 | -0.014 | -0.507 | -0.444 | 0.224 | 0.001 | -0.016 | -0.016 | 0 | 0.42 | 0.49 | 0.27 | 0.35 | 0.42 | 0.39 |
| 7 | 27509593 | rs9639549 | 0.147 | 0.367 | -0.72 | 0.828 | -0.021 | -0.015 | -0.015 | 0 | 0.26 | 0.19 | 0.21 | 0.36 | 0.35 | 0.34 |
| 7 | 27520326 | rs11763233 | 0.033 | -0.398 | -0.351 | 1.332 | 0 | -0.018 | -0.018 | 2.577 | 0.09 | 0.12 | 0.06 | 0 | 0.13 | 0.03 |
| 7 | 27520393 | rs10951168 | -1.52 | -0.662 | 0.277 | 1.638 | -0.024 | 0.256 | 0.256 | 0.541 | 0.17 | 0.54 | 0.38 | 0.43 | 0.16 | 0.16 |
| 7 | 27522629 | rs17155332 | -0.481 | -0.556 | -0.128 | 1.758 | -0.014 | 0.129 | 0.129 | 0 | 0.05 | 0.09 | 0.27 | 0.37 | 0.03 | 0.13 |
| 7 | 27526292 | rs11762112 | -0.459 | -0.768 | -0.795 | 0.91 | 0.038 | 0.073 | 0.073 | 0 | 0.53 | 0.73 | 0.55 | 0.66 | 0.43 | 0.42 |
| 7 | 27533292 | rs17744890 | 0.034 | -0.272 | -1.555 | 0.26 | -0.007 | 0.118 | 0.118 | 0 | 0.42 | 0.07 | 0.21 | 0.23 | 0.4 | 0.29 |
| 7 | 27559957 | rs11771265 | -0.288 | 0.202 | -0.767 | 0.225 | 0 | -0.019 | -0.019 | 0.005 | 0.11 | 0.08 | 0.08 | 0 | 0.15 | 0.02 |
| 7 | 27562660 | rs7779240 | 0.577 | 0.811 | -1.554 | -0.382 | -0.088 | 0.139 | 0.139 | 0 | 0.66 | 0.91 | 0.64 | 0.96 | 0.55 | 0.69 |
| 7 | 27564744 | rs13242238 | 1.071 | 0.27 | -2.34 | 0.109 | -0.069 | -0.171 | -0.171 | 0.036 | 0.24 | 0.01 | 0.24 | 0.51 | 0.25 | 0.35 |
| 7 | 27566505 | rs17155443 | -0.024 | 0.133 | -1.41 | -0.015 | 0.016 | 0.217 | 0.217 | 0.025 | 0.09 | 0.25 | 0.25 | 0.43 | 0.12 | 0.19 |
| 7 | 27578068 | rs17607303 | 0.866 | 0.115 | -1.51 | -0.238 | -0.008 | 0.014 | 0.014 | 0 | 0.1 | 0.05 | 0.08 | 0 | 0.15 | 0.02 |
| 7 | 27591093 | rs12700817 | -0.923 | 0.078 | -1.158 | -0.293 | -0.043 | 0.059 | 0.059 | 0 | 0.76 | 0.92 | 0.72 | 0.96 | 0.7 | 0.71 |
| 7 | 27591532 | rs12540276 | 0.081 | 0.072 | -1.156 | -0.302 | -0.05 | 0.09 | 0.09 | 0 | 0.66 | 0.86 | 0.64 | 0.96 | 0.55 | 0.69 |
| 7 | 27660013 | rs6966091 | -1.375 | 0.289 | -1.232 | -0.324 | -0.037 | 0.045 | 0.045 | 0 | 0.8 | 0.92 | 0.75 | 0.98 | 0.73 | 0.8 |
| 7 | 27662104 | rs1115346 | -0.379 | 0.131 | -0.217 | -0.395 | 0.028 | -0.072 | -0.072 | 0.31 | 0.73 | 0.68 | 0.69 | 0.96 | 0.7 | 0.71 |
| 7 | 27663276 | rs7778454 | -0.146 | 0.23 | 0.036 | -0.219 | 0.039 | 0.259 | 0.259 | 0.3 | 0.1 | 0.38 | 0.27 | 0.4 | 0.13 | 0.19 |
| 7 | 27672726 | rs12700826 | -0.582 | 0.231 | 0.048 | -0.451 | 0.031 | -0.062 | -0.062 | 0 | 0.75 | 0.71 | 0.7 | 0.96 | 0.7 | 0.71 |
| 7 | 27700389 | rs13227574 | 1.734 | 0.146 | -0.267 | -0.565 | 0.04 | 0.073 | 0.073 | 0 | 0.27 | 0.14 | 0.09 | 0 | 0.18 | 0.14 |
| 7 | 27701861 | rs42088 | 0.406 | 0.526 | -0.306 | -0.37 | -0.055 | 0.093 | 0.093 | 0 | 0.26 | 0.09 | 0.3 | 0.02 | 0.36 | 0.21 |

| | | | | | | | | | | | | | | | | |
|---|---|---|---|---|---|---|---|---|---|---|---|---|---|---|---|---|
| 7 | 27716345 | rs42101 | 0.022 | 0.542 | 0.071 | -0.014 | -0.002 | -0.001 | -0.001 | 0 | 0.48 | 0.49 | 0.51 | 0.51 | 0.47 | 0.5 |
| 7 | 27721129 | rs42105 | -0.024 | 0.522 | 0.073 | 0.118 | -0.001 | 0 | 0 | 0 | 0.48 | 0.49 | 0.51 | 0.51 | 0.47 | 0.5 |
| 7 | 27727849 | rs17150563 | 0.729 | 0.429 | -0.027 | 0.154 | 0.009 | -0.073 | -0.073 | 0 | 0.28 | 0.11 | 0.36 | 0.54 | 0.25 | 0.42 |
| 7 | 27730193 | rs12540821 | 1.31 | 0.393 | 0.345 | 0.238 | -0.002 | -0.003 | -0.003 | 0.017 | 0.3 | 0.37 | 0.24 | 0.31 | 0.28 | 0.26 |
| 7 | 27749846 | rs6957836 | -0.087 | 0.392 | 0.486 | 0.185 | 0.003 | -0.018 | -0.018 | 0.217 | 0.39 | 0.24 | 0.43 | 0.57 | 0.36 | 0.53 |
| 7 | 27754487 | rs9639563 | -0.147 | 0.365 | 0.77 | 0.172 | -0.007 | 0.009 | 0.009 | 0 | 0.4 | 0.42 | 0.44 | 0.57 | 0.36 | 0.53 |
| 7 | 27762576 | rs17155691 | 0.682 | 0.373 | 0.733 | 0.126 | 0.004 | -0.003 | -0.003 | 0 | 0.07 | 0.07 | 0.06 | 0.04 | 0.1 | 0.12 |
| 7 | 27762622 | rs10250333 | 1.835 | 0.319 | 0.689 | 0.256 | -0.065 | -0.042 | -0.042 | 0 | 0.17 | 0.03 | 0.3 | 0.19 | 0.22 | 0.17 |
| 7 | 27768492 | rs13247831 | 2.118 | 0.351 | 0.794 | 0.187 | 0.033 | 0.087 | 0.087 | 0 | 0.25 | 0.09 | 0.09 | 0 | 0.18 | 0.15 |
| 7 | 27770863 | rs9639564 | -0.177 | 0.352 | 0.898 | 0.201 | -0.006 | -0.015 | -0.015 | 0 | 0.4 | 0.46 | 0.45 | 0.66 | 0.37 | 0.54 |
| 7 | 27778885 | rs2237337 | 0.251 | 0.348 | 0.921 | 0.206 | -0.007 | 0.019 | 0.019 | 0 | 0.37 | 0.18 | 0.43 | 0.65 | 0.37 | 0.54 |
| 7 | 27784949 | rs10214930 | 1.595 | 0.381 | 0.56 | 0.117 | -0.012 | -0.004 | -0.004 | 0 | 0.19 | 0.14 | 0.31 | 0.19 | 0.22 | 0.17 |
| 7 | 27786969 | rs4722739 | 0.913 | 0.443 | 0.694 | 0.358 | 0.001 | 0.026 | 0.026 | 0 | 0.48 | 0.47 | 0.47 | 0.34 | 0.51 | 0.44 |
| 7 | 27804601 | rs11769157 | 0.422 | 0.652 | 0.501 | 0.52 | 0.004 | -0.003 | -0.003 | 0 | 0.07 | 0.09 | 0.06 | 0.04 | 0.1 | 0.13 |
| 7 | 27819650 | rs10155851 | -0.839 | 0.543 | 0.223 | 0.607 | 0.004 | -0.109 | -0.109 | 0.588 | 0.72 | 0.94 | 0.64 | 0.49 | 0.74 | 0.59 |
| 7 | 27823017 | rs3823946 | -0.626 | 0.573 | 0.118 | 0.639 | -0.018 | -0.033 | -0.033 | 0 | 0.71 | 0.76 | 0.6 | 0.34 | 0.65 | 0.56 |
| 7 | 27826523 | rs6944913 | -0.932 | 0.416 | -0.369 | 0.632 | -0.038 | -0.18 | -0.18 | 0.227 | 0.73 | 0.98 | 0.65 | 0.47 | 0.74 | 0.59 |
| 7 | 27857715 | rs7784038 | -1.214 | 0.443 | -0.361 | 0.525 | -0.055 | -0.182 | -0.182 | 3.347 | 0.75 | 0.99 | 0.67 | 0.51 | 0.75 | 0.66 |
| 7 | 27860762 | rs10225487 | -0.75 | 0.477 | -0.241 | 0.499 | 0.01 | -0.096 | -0.096 | 0 | 0.71 | 0.91 | 0.64 | 0.48 | 0.74 | 0.59 |
| 7 | 27869782 | rs6948467 | 0.948 | 0.504 | -0.171 | 0.183 | 0.001 | 0.093 | 0.093 | 0 | 0.43 | 0.25 | 0.38 | 0.19 | 0.41 | 0.31 |
| 7 | 27875454 | rs6979972 | -0.104 | 0.633 | 0.166 | 0.184 | 0.025 | -0.012 | -0.012 | 3.138 | 0.08 | 0.15 | 0.06 | 0.04 | 0.1 | 0.13 |
| 7 | 27879473 | rs6972821 | 0.684 | 0.325 | -0.398 | 0.398 | -0.023 | -0.162 | -0.162 | 0.368 | 0.29 | 0.02 | 0.34 | 0.52 | 0.25 | 0.41 |
| 7 | 27880888 | rs3944121 | -1.029 | 0.471 | -0.098 | 0.224 | -0.012 | -0.011 | -0.011 | 0 | 0.78 | 0.79 | 0.62 | 0.65 | 0.67 | 0.7 |
| 7 | 27888014 | rs11981911 | -0.13 | 0.304 | 0.297 | -0.133 | 0.028 | 0.081 | 0.081 | 0 | 0.49 | 0.58 | 0.25 | 0.07 | 0.42 | 0.36 |
| 7 | 27900596 | rs6462058 | -0.838 | 0.469 | 0.983 | 0.019 | 0.049 | 0.006 | 0.006 | 0 | 0.18 | 0.48 | 0.17 | 0.13 | 0.23 | 0.07 |
| 7 | 27903978 | rs12531244 | 1.547 | 0.081 | 0.718 | -0.438 | 0.046 | 0.073 | 0.073 | 0 | 0.3 | 0.13 | 0.12 | 0 | 0.2 | 0.15 |
| 7 | 27904674 | rs7790213 | -0.616 | 0.224 | 0.555 | -0.223 | -0.001 | 0.002 | 0.002 | 0 | 0.44 | 0.61 | 0.52 | 0.67 | 0.47 | 0.51 |
| 7 | 27907998 | rs12700848 | 1.208 | 0.238 | 0.61 | -0.195 | -0.008 | -0.018 | -0.018 | 0 | 0.34 | 0.36 | 0.15 | 0.09 | 0.21 | 0.16 |
| 7 | 27908326 | rs757396 | -1.07 | 0.335 | 0.206 | -0.146 | -0.046 | -0.042 | -0.042 | 0 | 0.77 | 0.89 | 0.65 | 0.67 | 0.67 | 0.66 |
| 7 | 27909037 | rs735664 | 0.486 | 0.148 | 0.329 | -0.113 | -0.001 | -0.001 | -0.001 | 0 | 0.58 | 0.51 | 0.48 | 0.6 | 0.44 | 0.51 |
| 7 | 27912757 | rs17155969 | 0.841 | 0.11 | -0.197 | 0.004 | -0.024 | -0.121 | -0.121 | 1.017 | 0.24 | 0.05 | 0.34 | 0.52 | 0.25 | 0.35 |
| 7 | 27916218 | rs720659 | 0.973 | 0.318 | 0.09 | -0.402 | -0.019 | -0.077 | -0.077 | 0 | 0.39 | 0.46 | 0.21 | 0.05 | 0.31 | 0.18 |
| 7 | 27917744 | rs17150572 | 0.932 | 0.436 | -0.107 | -0.155 | -0.051 | -0.146 | -0.146 | 0.042 | 0.2 | 0.01 | 0.21 | 0.5 | 0.2 | 0.33 |
| 7 | 27917884 | rs6943712 | -1.272 | 0.785 | 0.209 | 0.071 | -0.001 | -0.021 | -0.021 | 0 | 0.8 | 0.88 | 0.62 | 0.72 | 0.8 | 0.8 |
| 7 | 27918229 | rs10266923 | 0.799 | 0.593 | 0.368 | -0.104 | 0.007 | 0.007 | 0.007 | 0 | 0.58 | 0.43 | 0.42 | 0.55 | 0.51 | 0.51 |
| 7 | 27919667 | rs17155987 | -0.738 | 0.942 | 0.461 | -0.013 | 0.079 | 0.077 | 0.077 | 0 | 0.11 | 0.4 | 0.11 | 0.12 | 0.14 | 0.17 |
| 7 | 27922780 | rs763746 | -0.648 | 0.795 | -0.065 | -0.094 | -0.014 | -0.033 | -0.033 | 0 | 0.77 | 0.86 | 0.57 | 0.63 | 0.7 | 0.69 |
| 7 | 27929030 | rs10231578 | 1.04 | 0.327 | -0.076 | -0.84 | 0.008 | -0.005 | -0.005 | 0 | 0.34 | 0.19 | 0.19 | 0 | 0.32 | 0.16 |
| 7 | 27931860 | rs213 | 0.454 | 0.115 | 0.416 | -0.453 | -0.027 | -0.026 | -0.026 | 0 | 0.85 | 0.7 | 0.92 | 0.94 | 0.89 | 0.98 |
| 7 | 27932307 | rs10258132 | -1.609 | 0.483 | 0.784 | -0.353 | 0.039 | 0.052 | 0.052 | 0 | 0.11 | 0.42 | 0.12 | 0.1 | 0.09 | 0.11 |
| 7 | 27933232 | rs4719913 | -0.288 | 0.504 | -0.169 | -0.231 | -0.015 | -0.049 | -0.049 | 0 | 0.34 | 0.14 | 0.46 | 0.41 | 0.36 | 0.38 |
| 7 | 27936243 | rs916856 | -0.749 | 0.525 | 0.384 | -0.166 | 0.003 | 0.002 | 0.002 | 0 | 0.39 | 0.42 | 0.53 | 0.55 | 0.4 | 0.43 |
| 7 | 27937267 | rs12666447 | 0.677 | 0.557 | -0.22 | -0.384 | -0.048 | -0.144 | -0.144 | 0.156 | 0.2 | 0.02 | 0.23 | 0.48 | 0.22 | 0.3 |
| 7 | 27939039 | rs198 | -0.186 | 0.651 | 0.329 | -0.531 | 0.009 | -0.029 | -0.029 | 0.197 | 0.88 | 0.74 | 0.91 | 0.97 | 0.87 | 0.9 |
| 7 | 27943809 | rs193 | -0.28 | 0.522 | 0.585 | -0.59 | -0.003 | 0.027 | 0.027 | 0 | 0.35 | 0.41 | 0.49 | 0.48 | 0.33 | 0.34 |
| 7 | 27944385 | rs195 | 0.243 | 0.429 | 0.719 | -0.587 | -0.006 | -0.013 | -0.013 | 0 | 0.29 | 0.23 | 0.45 | 0.38 | 0.32 | 0.33 |
| 7 | 27945933 | rs12700849 | 0.359 | 0.725 | 0.615 | -0.545 | -0.025 | -0.004 | -0.004 | 0 | 0.07 | 0.02 | 0.07 | 0.09 | 0.12 | 0.11 |
| 7 | 27950576 | rs10278615 | 2.007 | 0.538 | 0.9 | -0.554 | 0.003 | 0.031 | 0.031 | 0 | 0.32 | 0.18 | 0.17 | 0.01 | 0.3 | 0.25 |
| 7 | 27951021 | rs217 | 0.849 | 0.679 | 1.12 | -0.255 | -0.002 | 0 | 0 | 0 | 0.62 | 0.4 | 0.62 | 0.38 | 0.63 | 0.57 |
| 7 | 27953981 | rs10275982 | 2.077 | 0.647 | 0.672 | -0.59 | 0.01 | 0.095 | 0.095 | 0.175 | 0.37 | 0.16 | 0.2 | 0.01 | 0.35 | 0.28 |
| 7 | 27957857 | rs1989909 | 0.062 | 0.461 | 0.691 | -0.337 | -0.004 | -0.009 | -0.009 | 0 | 0.25 | 0.22 | 0.41 | 0.36 | 0.27 | 0.3 |
| 7 | 27958750 | rs17621988 | -0.468 | 0.483 | 0.752 | -0.374 | -0.001 | 0.04 | 0.04 | 0.012 | 0.3 | 0.4 | 0.45 | 0.48 | 0.28 | 0.32 |
| 7 | 27962929 | rs7791093 | 1.346 | 0.609 | 0.422 | -0.597 | -0.007 | 0.107 | 0.107 | 0.385 | 0.42 | 0.24 | 0.26 | 0.04 | 0.42 | 0.39 |
| 7 | 27968193 | rs2040666 | -0.135 | 0.568 | 0.471 | -0.493 | -0.002 | 0.023 | 0.023 | 0.033 | 0.27 | 0.22 | 0.23 | 0.09 | 0.26 | 0.28 |
| 7 | 27972593 | rs4722745 | 2.02 | 0.632 | 0.52 | -0.609 | 0.008 | -0.005 | -0.005 | 0 | 0.37 | 0.29 | 0.21 | 0.01 | 0.35 | 0.27 |
| 7 | 27975745 | rs17156041 | 0.91 | 0.789 | 0.413 | -0.175 | -0.045 | -0.111 | -0.111 | 0 | 0.19 | 0.03 | 0.24 | 0.47 | 0.21 | 0.29 |
| 7 | 27985058 | rs11763181 | -0.107 | 0.311 | 0.482 | -0.188 | -0.019 | 0.014 | 0.014 | 0.064 | 0.89 | 0.73 | 0.92 | 0.86 | 0.92 | 0.98 |
| 7 | 27988689 | rs17156078 | -0.827 | 0.337 | -0.011 | -0.069 | 0.004 | 0.41 | 0.41 | 2.72 | 0.05 | 0.25 | 0.23 | 0.41 | 0.01 | 0.04 |
| 7 | 27988862 | rs10244948 | 1.829 | 0.408 | -0.898 | -0.928 | -0.007 | 0.106 | 0.106 | 0.024 | 0.38 | 0.16 | 0.21 | 0.02 | 0.4 | 0.29 |
| 7 | 27992659 | rs17622806 | -0.355 | 0.722 | -0.793 | -0.252 | -0.026 | 0.07 | 0.07 | 0 | 0.24 | 0.11 | 0.2 | 0.09 | 0.26 | 0.27 |
| 7 | 28004198 | rs4722750 | -0.991 | 0.922 | -0.226 | -0.022 | 0.039 | 0.09 | 0.09 | 0 | 0.07 | 0.2 | 0.09 | 0.14 | 0.1 | 0.15 |
| 7 | 28009233 | rs2106922 | 1.419 | 0.749 | -0.664 | -0.148 | 0.019 | 0.07 | 0.07 | 2.179 | 0.47 | 0.16 | 0.44 | 0.34 | 0.45 | 0.39 |
| 7 | 28009981 | rs253310 | -0.696 | 0.889 | -0.804 | 0.046 | -0.056 | -0.132 | -0.132 | 0 | 0.82 | 0.96 | 0.74 | 0.51 | 0.8 | 0.75 |
| 7 | 28016334 | rs38504 | -0.568 | 0.842 | -0.868 | 0.06 | -0.042 | -0.14 | -0.14 | 0 | 0.81 | 0.98 | 0.77 | 0.52 | 0.81 | 0.76 |
| 7 | 28018689 | rs6965213 | -0.191 | 0.702 | -0.803 | -0.084 | 0.006 | 0.049 | 0.049 | 0.228 | 0.17 | 0.2 | 0.34 | 0.48 | 0.23 | 0.31 |
| 7 | 28028227 | rs38513 | -0.609 | 0.744 | -0.921 | 0.016 | -0.054 | -0.133 | -0.133 | 0 | 0.81 | 0.98 | 0.74 | 0.53 | 0.8 | 0.76 |
| 7 | 28037886 | rs38519 | 0.144 | 0.377 | -1.245 | -1.749 | -0.038 | 0.064 | 0.064 | 0.87 | 0.68 | 0.79 | 0.46 | 0.03 | 0.6 | 0.51 |
| 7 | 28038375 | rs740127 | 0.722 | 0.535 | -1.28 | -1.551 | -0.009 | 0.115 | 0.115 | 0.584 | 0.42 | 0.21 | 0.27 | 0.01 | 0.43 | 0.32 |
| 7 | 28039590 | rs38522 | -0.269 | 0.349 | -1.285 | -1.512 | 0.004 | 0.024 | 0.024 | 0 | 0.36 | 0.25 | 0.26 | 0.08 | 0.36 | 0.23 |

| | | | | | | | | | | | | | | | | |
|---|---|---|---|---|---|---|---|---|---|---|---|---|---|---|---|---|
| 7 | 28039797 | rs38523 | -0.35 | 0.191 | -1.46 | -1.002 | -0.003 | -0.002 | -0.002 | 0.115 | 0.44 | 0.69 | 0.54 | 0.41 | 0.39 | 0.53 |
| 7 | 28039916 | rs6945134 | -0.46 | -0.563 | -0.316 | -1.287 | -0.048 | -0.147 | -0.147 | 0 | 0.25 | 0.55 | 0.15 | 0 | 0.15 | 0.07 |
| 7 | 28042363 | rs4400283 | 0.252 | 0.363 | -0.214 | -0.716 | 0 | -0.006 | -0.006 | 0 | 0.15 | 0.15 | 0.18 | 0.09 | 0.19 | 0.26 |
| 7 | 28051077 | rs38526 | -1.082 | -0.446 | -0.262 | 0.276 | 0.059 | -0.132 | -0.132 | 0 | 0.83 | 0.99 | 0.69 | 0.57 | 0.91 | 0.84 |
| 7 | 28060576 | rs10251664 | -0.112 | -0.591 | 0.64 | 0.308 | -0.053 | -0.034 | -0.034 | 0 | 0.77 | 0.63 | 0.66 | 0.17 | 0.85 | 0.72 |
| 7 | 28070311 | rs4722752 | 0.026 | -0.62 | 0.878 | 0.283 | -0.053 | -0.042 | -0.042 | 0 | 0.74 | 0.46 | 0.56 | 0.18 | 0.82 | 0.72 |
| 7 | 28080493 | rs10486573 | 1.363 | -0.111 | 0.411 | 0.037 | -0.008 | 0.035 | 0.035 | 1.078 | 0.18 | 0.07 | 0.12 | 0 | 0.17 | 0.04 |
| 7 | 28093980 | rs17156275 | -0.276 | -0.232 | 0.833 | 0.326 | 0.017 | 0.118 | 0.118 | 0.484 | 0.21 | 0.37 | 0.2 | 0.49 | 0.24 | 0.38 |
| 7 | 28096199 | rs7785730 | -0.151 | -0.249 | 0.825 | 0.324 | 0.007 | 0.131 | 0.131 | 0.331 | 0.2 | 0.39 | 0.15 | 0.49 | 0.21 | 0.38 |
| 7 | 28099570 | rs6462063 | -0.737 | -0.047 | 0.345 | 0.177 | 0 | -0.013 | -0.013 | 0.483 | 0.48 | 0.84 | 0.44 | 0.54 | 0.55 | 0.66 |
| 7 | 28103030 | rs6462064 | -1.397 | 0.04 | 1.101 | 1.2 | 0.054 | 0.436 | 0.436 | 4.297 | 0.08 | 0.43 | 0.06 | 0.35 | 0.06 | 0.12 |
| 7 | 28106847 | rs7798247 | -0.414 | 0.229 | 0.345 | 1.262 | 0.037 | 0.364 | 0.364 | 0.079 | 0.05 | 0.24 | 0.03 | 0.39 | 0.06 | 0.12 |
| 7 | 28111936 | rs7456467 | 0.517 | 0.025 | -0.559 | 1.066 | 0.037 | -0.062 | -0.062 | 0.311 | 0.18 | 0.05 | 0.36 | 0.42 | 0.11 | 0.15 |
| 7 | 28119070 | rs10268254 | 0.635 | 0.448 | -0.561 | 0.441 | -0.008 | 0.029 | 0.029 | 0.422 | 0.16 | 0.13 | 0.15 | 0.02 | 0.21 | 0.12 |
| 7 | 28119143 | rs10486576 | 0.659 | 0.235 | -0.765 | 0.839 | 0.024 | -0.036 | -0.036 | 0 | 0.17 | 0.09 | 0.36 | 0.42 | 0.11 | 0.18 |
| 7 | 28119713 | rs6968494 | -0.971 | 1.182 | 0.096 | 0.994 | 0.006 | 0.038 | 0.038 | 1.44 | 0.28 | 0.69 | 0.27 | 0.46 | 0.33 | 0.35 |
| 7 | 28122977 | rs10951191 | 1.179 | 1.099 | 0.522 | 0.834 | 0.079 | 0.155 | 0.155 | 0.17 | 0.83 | 0.49 | 0.84 | 0.66 | 0.76 | 0.68 |
| 7 | 28128410 | rs6950127 | -1.676 | -0.028 | 0.238 | 1.015 | -0.039 | 0.165 | 0.165 | 7.746 | 0.11 | 0.73 | 0.1 | 0.31 | 0.06 | 0.15 |
| 7 | 28131416 | rs11760530 | 0.151 | -0.127 | -0.294 | 0.326 | 0.008 | 0.017 | 0.017 | 0.141 | 0.17 | 0.1 | 0.11 | 0 | 0.14 | 0.04 |
| 7 | 28133980 | rs6977730 | -1.652 | 0.085 | 0.696 | 1.035 | 0.019 | 0.325 | 0.325 | 6.233 | 0.09 | 0.56 | 0.09 | 0.26 | 0.05 | 0.06 |
| 7 | 28138193 | rs6971086 | -0.348 | 0.111 | -0.104 | 0.577 | 0.006 | -0.025 | -0.025 | 0 | 0.34 | 0.28 | 0.28 | 0.6 | 0.28 | 0.38 |
| 7 | 28143490 | rs2158623 | 0.236 | 0.219 | -0.484 | 0.301 | -0.013 | 0.009 | 0.009 | 0 | 0.79 | 0.91 | 0.8 | 0.85 | 0.76 | 0.66 |
| 7 | 28145564 | rs2008296 | -0.55 | 0.052 | -0.428 | 0.287 | -0.021 | -0.043 | -0.043 | 0 | 0.89 | 0.98 | 0.92 | 0.76 | 0.85 | 0.85 |
| 7 | 28147452 | rs4722754 | 0.687 | -0.276 | -0.516 | 0.269 | 0.004 | 0.001 | 0.001 | 0 | 0.12 | 0.09 | 0.19 | 0.48 | 0.12 | 0.2 |
| 7 | 28150475 | rs10275834 | 1.214 | -0.588 | 0.46 | 0.017 | -0.02 | -0.026 | -0.026 | 2.741 | 0.68 | 0.49 | 0.65 | 0.21 | 0.7 | 0.55 |
| 7 | 28152095 | rs17156401 | 0.397 | -0.543 | 0.668 | 0.044 | -0.017 | 0.014 | 0.014 | 0.007 | 0.07 | 0.18 | 0.05 | 0.07 | 0.03 | 0.11 |
| 7 | 28161379 | rs6962289 | 1.513 | -0.721 | 0.772 | -0.118 | -0.012 | -0.029 | -0.029 | 0 | 0.61 | 0.52 | 0.47 | 0.23 | 0.62 | 0.35 |
| 7 | 28161983 | rs9648026 | 1.272 | -0.902 | 0.645 | -0.007 | 0.006 | -0.046 | -0.046 | 0 | 0.08 | 0 | 0.19 | 0.2 | 0.07 | 0.21 |
| 7 | 28176305 | rs917117 | -1.115 | -0.977 | 0.621 | -0.111 | -0.029 | -0.012 | -0.012 | 1.584 | 0.29 | 0.4 | 0.44 | 0.75 | 0.21 | 0.52 |
| 7 | 28177301 | rs11495981 | -1.115 | -0.977 | 0.621 | -0.111 | -0.03 | -0.013 | -0.013 | 0 | 0.29 | 0.4 | 0.44 | 0.75 | 0.21 | 0.52 |
| 7 | 28179396 | rs849139 | 0.431 | -0.84 | 0.418 | -0.298 | -0.02 | -0.037 | -0.037 | 0 | 0.61 | 0.82 | 0.65 | 0.5 | 0.61 | 0.55 |
| 7 | 28183702 | rs849140 | 1.027 | -0.686 | 0.248 | -0.363 | 0 | 0 | 0 | 0 | 0.54 | 0.53 | 0.61 | 0.71 | 0.64 | 0.45 |
| 7 | 28185091 | rs849141 | -0.24 | -0.757 | 0.195 | -0.391 | -0.017 | 0.104 | 0.104 | 0 | 0.71 | 0.89 | 0.84 | 0.98 | 0.74 | 0.76 |
| 7 | 28185891 | rs849142 | 1.981 | -0.689 | 0.23 | -0.343 | -0.071 | 0.145 | 0.145 | 0 | 0.38 | 0.15 | 0.35 | 0.01 | 0.51 | 0.21 |
| 7 | 28189411 | rs1635852 | 1.763 | -0.615 | 0.169 | -0.293 | -0.036 | 0.064 | 0.064 | 0 | 0.4 | 0.23 | 0.36 | 0.23 | 0.52 | 0.24 |
| 7 | 28190121 | rs10276381 | -0.639 | -0.942 | 0.575 | -0.218 | -0.047 | 0.071 | 0.071 | 0.04 | 0.18 | 0.36 | 0.23 | 0.28 | 0.09 | 0.31 |
| 7 | 28224053 | rs849336 | 1.488 | -0.912 | 0.016 | -0.369 | 0.052 | 0.176 | 0.176 | 0 | 0.47 | 0.2 | 0.43 | 0.03 | 0.66 | 0.35 |
| 7 | 28234599 | rs849329 | -0.499 | -0.228 | 0.425 | 0.005 | 0.001 | -0.031 | -0.031 | 0.583 | 0.79 | 0.82 | 0.89 | 1 | 0.87 | 0.89 |
| 7 | 28238860 | rs849315 | -0.47 | -0.278 | 0.426 | 0.005 | 0.002 | -0.029 | -0.029 | 0 | 0.78 | 0.83 | 0.89 | 1 | 0.87 | 0.89 |
| 7 | 28245295 | rs7801967 | -0.434 | 0.039 | 0.74 | 0.875 | 0.003 | 0.086 | 0.086 | 0 | 0.14 | 0.27 | 0.24 | 0.69 | 0.14 | 0.22 |
| 7 | 28250083 | rs10225945 | -1.34 | 0.034 | 0.434 | 0.683 | -0.028 | 0.001 | 0.001 | 0 | 0.25 | 0.57 | 0.3 | 0.77 | 0.19 | 0.34 |
| 7 | 28259131 | rs1513276 | -0.576 | 0.515 | 0.166 | 1.013 | 0.005 | 0.168 | 0.168 | 0 | 0.14 | 0.26 | 0.23 | 0.5 | 0.12 | 0.2 |
| 7 | 28260312 | rs849330 | -0.221 | 0.828 | -0.506 | 0.273 | -0.023 | -0.015 | -0.015 | 0 | 0.39 | 0.27 | 0.48 | 0.48 | 0.57 | 0.33 |
| 7 | 28267876 | rs488837 | 0.389 | 1.514 | 0.459 | -0.033 | 0.056 | 0.022 | 0.022 | 2.865 | 0.84 | 0.39 | 0.74 | 0.84 | 0.83 | 0.93 |
| 7 | 28269635 | rs2528486 | 0.989 | 1.391 | 0.159 | -0.057 | 0.024 | 0.021 | 0.021 | 0.089 | 0.73 | 0.4 | 0.55 | 0.7 | 0.68 | 0.79 |
| 7 | 28278258 | rs4722770 | 0.479 | 1.295 | 0.228 | -0.342 | 0.003 | 0.003 | 0.003 | 0 | 0.49 | 0.49 | 0.4 | 0.66 | 0.42 | 0.6 |
| 7 | 28279243 | rs4722771 | 0.383 | 1.403 | 0.282 | -0.385 | 0.017 | 0.168 | 0.168 | 0 | 0.51 | 0.75 | 0.57 | 0.83 | 0.47 | 0.6 |
| 7 | 28280505 | rs10486580 | -0.89 | 1.284 | 0.47 | -0.276 | 0.032 | 0.187 | 0.187 | 1.418 | 0.06 | 0.26 | 0.17 | 0.22 | 0.06 | 0.17 |
| 7 | 28285580 | rs849338 | 0.433 | 1.383 | 0.078 | -0.877 | -0.017 | 0.072 | 0.072 | 0 | 0.33 | 0.17 | 0.22 | 0.02 | 0.35 | 0.21 |
| 7 | 28285903 | rs4722772 | 0.895 | 1.125 | -0.028 | -1.306 | -0.007 | -0.058 | -0.058 | 1.762 | 0.34 | 0.65 | 0.27 | 0.04 | 0.25 | 0.42 |
| 7 | 28288174 | rs849341 | 0.424 | 1.016 | 0.309 | -1.076 | 0 | -0.042 | -0.042 | 0 | 0.37 | 0.39 | 0.25 | 0.05 | 0.34 | 0.39 |
| 7 | 28301970 | rs177464 | 0.622 | 1.176 | 0.012 | -1.045 | -0.008 | 0.045 | 0.045 | 0 | 0.34 | 0.22 | 0.22 | 0.02 | 0.34 | 0.21 |
| 7 | 28302072 | rs177465 | -0.082 | 1.404 | 0.003 | -0.45 | 0.004 | -0.012 | -0.012 | 0 | 0.58 | 0.51 | 0.44 | 0.7 | 0.53 | 0.63 |
| 7 | 28309468 | rs4719930 | -0.221 | 1.072 | -0.238 | -0.739 | 0.008 | 0.025 | 0.025 | 0 | 0.44 | 0.4 | 0.35 | 0.1 | 0.37 | 0.6 |
| 7 | 28316074 | rs13226529 | 0.531 | 0.569 | -0.174 | -0.958 | 0.012 | -0.016 | -0.016 | 0.282 | 0.29 | 0.24 | 0.18 | 0.01 | 0.18 | 0.23 |
| 7 | 28319962 | rs177485 | 0.139 | 1.237 | -1.034 | -0.481 | -0.054 | 0.086 | 0.086 | 0 | 0.56 | 0.85 | 0.63 | 0.71 | 0.55 | 0.67 |
| 7 | 28320034 | rs177486 | -0.12 | 1.297 | -1.141 | -0.639 | -0.066 | 0.103 | 0.103 | 0 | 0.61 | 0.89 | 0.66 | 0.83 | 0.57 | 0.7 |
| 7 | 28331586 | rs849322 | -0.266 | 0.986 | -0.646 | -0.085 | -0.001 | 0.02 | 0.02 | 0 | 0.52 | 0.73 | 0.51 | 0.62 | 0.51 | 0.56 |
| 7 | 28333077 | rs177505 | -0.243 | 0.483 | -0.15 | -0.718 | 0.011 | -0.034 | -0.034 | 1.064 | 0.33 | 0.52 | 0.32 | 0.16 | 0.36 | 0.27 |
| 7 | 28338080 | rs2175738 | 0.293 | -1.023 | -0.153 | -1.328 | 0.035 | 0.07 | 0.07 | 0.946 | 0.43 | 0.24 | 0.62 | 0.84 | 0.67 | 0.48 |
| 7 | 28340850 | rs4719932 | -0.553 | -1.68 | 0.099 | -1.903 | -0.031 | -0.077 | -0.077 | 0 | 0.76 | 0.67 | 0.91 | 0.99 | 0.88 | 0.81 |
| 7 | 28349720 | rs1860758 | -0.647 | -1.627 | 0.17 | -1.873 | 0.003 | -0.029 | -0.029 | 0 | 0.78 | 0.72 | 0.91 | 1 | 0.88 | 0.82 |
| 7 | 28355752 | rs17156573 | -0.504 | -1.123 | -0.109 | -1.745 | 0.041 | 0.051 | 0.051 | 0.114 | 0.52 | 0.62 | 0.32 | 0.05 | 0.31 | 0.44 |
| 7 | 28356889 | rs17156577 | -0.815 | -0.162 | 0.283 | -0.702 | -0.011 | -0.056 | -0.056 | 0 | 0.16 | 0.36 | 0.18 | 0.05 | 0.11 | 0.18 |
| 7 | 28360334 | rs6973453 | 0.22 | 0.511 | -0.209 | -0.527 | 0.016 | -0.048 | -0.048 | 0 | 0.62 | 0.57 | 0.41 | 0.24 | 0.46 | 0.48 |
| 7 | 28361117 | rs17156579 | 0.313 | 1.225 | -0.472 | 0.03 | -0.031 | -0.03 | -0.03 | 0 | 0.05 | 0 | 0.16 | 0.08 | 0.08 | 0.1 |
| 7 | 28366100 | rs17156585 | 0.219 | 0.879 | -0.54 | -0.76 | -0.001 | -0.004 | -0.004 | 0 | 0.2 | 0.21 | 0.15 | 0.09 | 0.23 | 0.23 |
| 7 | 28367949 | rs12700867 | -0.591 | 0.859 | 0.331 | -0.142 | -0.019 | -0.03 | -0.03 | 0 | 0.11 | 0.07 | 0.23 | 0.23 | 0.12 | 0.21 |
| 7 | 28370058 | rs6978238 | 0.288 | -0.151 | 0.529 | -1.055 | 0.005 | 0.045 | 0.045 | 0 | 0.43 | 0.35 | 0.33 | 0.09 | 0.41 | 0.4 |

| | | | | | | | | | | | | | | | | |
|---|---|---|---|---|---|---|---|---|---|---|---|---|---|---|---|---|
| 7 | 28370069 | rs13230543 | -0.787 | -0.126 | 0.921 | -0.522 | 0.002 | 0.011 | 0.011 | 0.146 | 0.12 | 0.16 | 0.24 | 0.25 | 0.12 | 0.21 |
| 7 | 28372747 | rs12673465 | 0.849 | 0.091 | 0.703 | 0.098 | -0.001 | -0.001 | -0.001 | 0 | 0.6 | 0.51 | 0.52 | 0.37 | 0.6 | 0.55 |
| 7 | 28374188 | rs10251129 | -0.298 | 0.149 | 0.466 | 0.474 | 0.008 | -0.009 | -0.009 | 0.394 | 0.73 | 0.84 | 0.79 | 0.68 | 0.73 | 0.75 |
| 7 | 28381283 | rs6971345 | -0.007 | 0.184 | 0.308 | 0.498 | -0.017 | -0.001 | -0.001 | 0 | 0.7 | 0.48 | 0.77 | 0.67 | 0.73 | 0.75 |
| 7 | 28382146 | rs17156603 | -0.549 | 0.834 | 0.06 | 0.914 | -0.013 | -0.011 | -0.011 | 0 | 0.11 | 0.08 | 0.26 | 0.34 | 0.14 | 0.21 |
| 7 | 28386703 | rs7806362 | -0.062 | 0.48 | 0.173 | 0.461 | 0.008 | -0.037 | -0.037 | 0 | 0.36 | 0.35 | 0.29 | 0.18 | 0.43 | 0.3 |
| 7 | 28389264 | rs17642145 | 1.091 | 0.468 | 0.146 | 0.302 | 0.001 | 0.053 | 0.053 | 0 | 0.19 | 0.04 | 0.1 | 0 | 0.15 | 0.16 |
| 7 | 28389465 | rs10229500 | -0.077 | 0.009 | 0.505 | 0.232 | -0.026 | -0.065 | -0.065 | 0 | 0.69 | 0.66 | 0.77 | 0.88 | 0.79 | 0.73 |
| 7 | 28389760 | rs10243659 | 0.605 | 0.07 | 0.454 | 0.421 | -0.021 | 0.077 | 0.077 | 0 | 0.8 | 0.68 | 0.72 | 0.6 | 0.85 | 0.69 |
| 7 | 28391047 | rs11772815 | 1.569 | 0.091 | 0.451 | 0.415 | 0 | -0.011 | -0.011 | 0.102 | 0.41 | 0.27 | 0.34 | 0.4 | 0.39 | 0.34 |
| 7 | 28391082 | rs6958133 | -1.084 | 0.114 | 0.545 | 0.164 | -0.045 | 0 | 0 | 0 | 0.9 | 0.8 | 0.95 | 0.89 | 0.97 | 0.92 |
| 7 | 28391142 | rs11765845 | 0.345 | 0.128 | 0.429 | 0.344 | 0 | 0.005 | 0.005 | 0 | 0.73 | 0.46 | 0.75 | 0.67 | 0.69 | 0.76 |
| 7 | 28392183 | rs7796544 | 0.632 | 0.157 | 0.425 | 0.151 | 0.031 | 0.004 | 0.004 | 0 | 0.17 | 0.26 | 0.2 | 0.19 | 0.28 | 0.14 |
| 7 | 28394531 | rs12700884 | 0.917 | 0.386 | -0.168 | 0.246 | -0.003 | 0.022 | 0.022 | 0 | 0.48 | 0.33 | 0.58 | 0.55 | 0.49 | 0.47 |
| 7 | 28395729 | rs17156635 | 0.223 | 0.414 | -0.29 | 0.265 | -0.011 | -0.016 | -0.016 | 0.627 | 0.05 | 0.02 | 0.06 | 0.13 | 0.05 | 0.14 |
| 7 | 28395910 | rs7799688 | 1.225 | 0.39 | -0.295 | 0.241 | -0.007 | 0.083 | 0.083 | 0 | 0.19 | 0.07 | 0.1 | 0 | 0.15 | 0.16 |
| 7 | 28397794 | rs7799687 | 0.242 | 0.317 | -0.237 | 0.29 | -0.008 | 0.039 | 0.039 | 0 | 0.27 | 0.13 | 0.26 | 0.14 | 0.24 | 0.32 |
| 7 | 28397908 | rs714218 | 0.041 | 0.408 | -0.252 | 0.013 | 0.003 | 0.003 | 0.003 | 0 | 0.73 | 0.5 | 0.75 | 0.69 | 0.69 | 0.76 |
| 7 | 28398460 | rs1860759 | 0.363 | 0.545 | -0.14 | 0.301 | 0 | 0.022 | 0.022 | 0 | 0.52 | 0.44 | 0.57 | 0.69 | 0.51 | 0.57 |
| 7 | 28408294 | rs10254657 | -0.646 | 0.556 | -0.556 | 0.5 | -0.01 | -0.014 | -0.014 | 0 | 0.56 | 0.73 | 0.6 | 0.58 | 0.54 | 0.57 |
| 7 | 28408943 | rs6953524 | -0.871 | 1.285 | -0.156 | 0.363 | 0.01 | 0.038 | 0.038 | 0 | 0.27 | 0.3 | 0.31 | 0.52 | 0.36 | 0.4 |
| 7 | 28415827 | rs6462079 | -0.705 | 1.138 | -0.441 | -0.126 | 0 | -0.052 | -0.052 | 0 | 0.76 | 0.69 | 0.76 | 1 | 0.79 | 0.85 |
| 7 | 28416417 | rs10239810 | 0.839 | 1.313 | -1.393 | 0.075 | -0.042 | 0.126 | 0.126 | 0 | 0.52 | 0.15 | 0.58 | 0.7 | 0.53 | 0.62 |
| 7 | 28421036 | rs1811248 | 0.04 | 1.019 | -1.309 | -0.092 | 0.01 | 0.083 | 0.083 | 0.422 | 0.63 | 0.21 | 0.64 | 0.87 | 0.69 | 0.67 |
| 7 | 28421983 | rs887620 | 0.099 | 1.348 | -1.486 | 0.022 | -0.067 | -0.042 | -0.042 | 0 | 0.17 | 0.01 | 0.11 | 0.18 | 0.23 | 0.14 |
| 7 | 28422083 | rs887622 | 0.134 | 1.589 | -0.942 | 0.264 | -0.036 | 0.151 | 0.151 | 0 | 0.29 | 0.07 | 0.27 | 0.03 | 0.3 | 0.25 |
| 7 | 28422134 | rs887623 | -0.726 | 1.859 | -0.881 | 0.477 | -0.015 | -0.014 | -0.014 | 0 | 0.13 | 0.09 | 0.15 | 0.23 | 0.24 | 0.23 |
| 7 | 28428391 | rs7794304 | 0.951 | 1.454 | -0.917 | 0.552 | -0.025 | -0.014 | -0.014 | 0.328 | 0.53 | 0.24 | 0.47 | 0.49 | 0.48 | 0.49 |
| 7 | 28429243 | rs42694 | 0.284 | 1.283 | -1.278 | 0.388 | -0.071 | 0.16 | 0.16 | 0 | 0.26 | 0.01 | 0.24 | 0.02 | 0.27 | 0.19 |
| 7 | 28430358 | rs6952227 | -0.122 | 1.125 | -0.74 | 1.082 | -0.008 | -0.022 | -0.022 | 0.938 | 0.09 | 0.03 | 0.11 | 0.24 | 0.15 | 0.13 |
| 7 | 28430860 | rs42695 | 1.063 | 1.116 | 0.181 | 1.061 | 0.116 | 0.378 | 0.378 | 0 | 0.92 | 0.57 | 0.81 | 0.74 | 0.91 | 0.83 |
| 7 | 28437116 | rs42699 | 0.011 | 1.171 | -0.753 | 0.2 | -0.017 | 0.082 | 0.082 | 0 | 0.24 | 0.1 | 0.24 | 0.02 | 0.25 | 0.19 |
| 7 | 28440320 | rs10233653 | 1.125 | 0.761 | -1.295 | 0.179 | -0.027 | 0.002 | 0.002 | 1.718 | 0.42 | 0.09 | 0.37 | 0.41 | 0.37 | 0.35 |
| 7 | 28446340 | rs1860757 | -0.725 | 0.819 | -0.302 | 0.674 | 0.001 | 0.017 | 0.017 | 0.374 | 0.25 | 0.28 | 0.3 | 0.51 | 0.31 | 0.41 |
| 7 | 28454580 | rs979915 | 0.239 | 0.314 | -1.092 | -0.239 | -0.038 | 0.202 | 0.202 | 0 | 0.32 | 0.02 | 0.27 | 0.02 | 0.28 | 0.29 |
| 7 | 28455233 | rs42714 | 0 | 0.531 | -0.961 | -0.214 | -0.008 | 0.008 | 0.008 | 0 | 0.92 | 0.96 | 0.93 | 0.98 | 0.91 | 0.9 |
| 7 | 28462344 | rs42733 | -0.371 | 0.504 | -0.944 | 0.241 | -0.006 | 0.009 | 0.009 | 0 | 0.93 | 0.97 | 0.95 | 0.98 | 0.92 | 0.89 |
| 7 | 28465434 | rs17156685 | 0.487 | 0.242 | -1.206 | 0.481 | -0.006 | -0.016 | -0.016 | 0 | 0.09 | 0 | 0.23 | 0.06 | 0.08 | 0.15 |
| 7 | 28465762 | rs174024 | 0.604 | 0.231 | -0.717 | 0.42 | 0.041 | 0.154 | 0.154 | 0 | 0.51 | 0.24 | 0.59 | 0.73 | 0.62 | 0.53 |
| 7 | 28468319 | rs2391665 | -0.696 | 0.23 | -0.753 | 0.496 | 0.029 | 0.113 | 0.113 | 0 | 0.51 | 0.76 | 0.41 | 0.34 | 0.39 | 0.51 |
| 7 | 28472224 | rs6949786 | -0.269 | 0.491 | -0.885 | 0.49 | 0.012 | 0.07 | 0.07 | 0 | 0.42 | 0.27 | 0.35 | 0.26 | 0.37 | 0.43 |
| 7 | 28484955 | rs17156694 | -1.572 | 0.565 | -0.145 | 1.628 | 0.034 | 0.317 | 0.317 | 0.025 | 0.07 | 0.25 | 0.06 | 0.58 | 0.07 | 0.22 |
| 7 | 28494295 | rs177572 | 0.348 | 0.999 | -0.735 | 0.904 | -0.01 | 0.022 | 0.022 | 0 | 0.58 | 0.31 | 0.52 | 0.63 | 0.52 | 0.52 |
| 7 | 28494445 | rs6977204 | 0.633 | 0.584 | -1.123 | 0.9 | 0.046 | -0.048 | -0.048 | 0 | 0.33 | 0.14 | 0.38 | 0.56 | 0.31 | 0.33 |
| 7 | 28497039 | rs177575 | -0.226 | 1.874 | -1.024 | 0.453 | -0.048 | 0.042 | 0.042 | 0 | 0.92 | 0.99 | 0.77 | 1 | 0.8 | 0.88 |
| 7 | 28498670 | rs177580 | -0.305 | 0.79 | -0.089 | 0.212 | -0.001 | -0.017 | -0.017 | 0.043 | 0.54 | 0.49 | 0.35 | 0.34 | 0.45 | 0.54 |
| 7 | 28499363 | rs177584 | -0.053 | 0.76 | 0.371 | 0.058 | -0.01 | -0.044 | -0.044 | 0 | 0.21 | 0.32 | 0.14 | 0.1 | 0.16 | 0.19 |
| 7 | 28500407 | rs177585 | -0.185 | 0.766 | 0.345 | 0.067 | -0.013 | -0.051 | -0.051 | 0 | 0.22 | 0.36 | 0.13 | 0.1 | 0.17 | 0.18 |
| 7 | 28504566 | rs6462088 | 0.931 | 1.016 | 0.256 | 0.377 | 0.02 | 0.054 | 0.054 | 1.032 | 0.46 | 0.2 | 0.46 | 0.62 | 0.44 | 0.4 |
| 7 | 28507628 | rs7796539 | 0.591 | 1.268 | 0.418 | 0.01 | -0.009 | -0.08 | -0.08 | 0.414 | 0.23 | 0.29 | 0.18 | 0.01 | 0.19 | 0.27 |
| 7 | 28508236 | rs1859020 | 0.826 | 1.572 | 0.311 | 0.488 | -0.015 | -0.004 | -0.004 | 0.178 | 0.72 | 0.69 | 0.64 | 0.63 | 0.64 | 0.68 |
| 7 | 28513002 | rs12671247 | -0.004 | 1.703 | -0.012 | 0.577 | -0.016 | -0.013 | -0.013 | 0 | 0.12 | 0.11 | 0.23 | 0.2 | 0.17 | 0.17 |
| 7 | 28513975 | rs4719935 | -0.027 | 1.697 | -0.015 | 0.564 | -0.016 | -0.012 | -0.012 | 0.331 | 0.12 | 0.1 | 0.23 | 0.2 | 0.17 | 0.17 |
| 7 | 28515179 | rs217505 | -0.43 | 1.905 | -0.141 | 0.542 | -0.03 | -0.009 | -0.009 | 0 | 0.87 | 0.95 | 0.88 | 0.84 | 0.81 | 0.87 |
| 7 | 28517022 | rs217508 | 0.313 | 1.917 | -0.023 | 0.875 | -0.009 | -0.009 | -0.009 | 0.388 | 0.8 | 0.85 | 0.73 | 0.43 | 0.73 | 0.78 |
| 7 | 28517072 | rs4719936 | 0.918 | 1.518 | -0.069 | 0.307 | 0.008 | -0.004 | -0.004 | 0 | 0.23 | 0.17 | 0.16 | 0.01 | 0.2 | 0.28 |
| 7 | 28517216 | rs217509 | 1.414 | 1.377 | -0.11 | 0.3 | 0.043 | 0.149 | 0.149 | 0.106 | 0.4 | 0.12 | 0.3 | 0.16 | 0.36 | 0.33 |
| 7 | 28518055 | rs217510 | -0.69 | 1.439 | -0.065 | 0.416 | -0.032 | -0.006 | -0.006 | 0.269 | 0.87 | 0.93 | 0.88 | 0.85 | 0.81 | 0.86 |
| 7 | 28523442 | rs217519 | 1.372 | 1.179 | -0.145 | 0.181 | 0.022 | 0.159 | 0.159 | 0.014 | 0.4 | 0.11 | 0.3 | 0.15 | 0.37 | 0.32 |
| 7 | 28523547 | rs4722801 | 0.666 | 1.265 | 0.396 | 0.142 | -0.017 | -0.111 | -0.111 | 0 | 0.25 | 0.27 | 0.18 | 0.01 | 0.2 | 0.29 |
| 7 | 28532464 | rs4722804 | -0.036 | 1.376 | 0.5 | 0.33 | 0.035 | 0.043 | 0.043 | 0 | 0.15 | 0.4 | 0.25 | 0.21 | 0.17 | 0.16 |
| 7 | 28538169 | rs217498 | 1.522 | 1.194 | -0.131 | 0.292 | -0.047 | 0.092 | 0.092 | 0.632 | 0.38 | 0.02 | 0.29 | 0.19 | 0.36 | 0.32 |
| 7 | 28538472 | rs11971533 | 0.797 | 1.174 | 0.053 | 0.174 | -0.03 | -0.136 | -0.136 | 0 | 0.26 | 0.33 | 0.18 | 0.01 | 0.19 | 0.28 |
| 7 | 28540618 | rs217503 | 0.529 | 1.221 | -0.01 | 0.227 | -0.003 | 0.01 | 0.01 | 0.014 | 0.53 | 0.43 | 0.53 | 0.41 | 0.53 | 0.47 |
| 7 | 28544875 | rs217513 | 1.259 | 1.25 | -0.556 | 0.345 | 0.011 | 0.101 | 0.101 | 0.332 | 0.39 | 0.1 | 0.31 | 0.2 | 0.36 | 0.33 |
| 7 | 28545611 | rs65264 | -0.019 | 1.336 | -0.892 | 0.327 | -0.036 | -0.046 | -0.046 | 1.215 | 0.79 | 0.82 | 0.72 | 0.43 | 0.72 | 0.76 |
| 7 | 28550394 | rs441355 | 1.211 | 1.062 | -0.944 | 0.228 | -0.047 | 0.139 | 0.139 | 0.564 | 0.38 | 0.02 | 0.29 | 0.15 | 0.36 | 0.32 |
| 7 | 28551960 | rs10215797 | -0.239 | 1.186 | -0.189 | 1.001 | 0.006 | 0.009 | 0.009 | 0.983 | 0.85 | 0.79 | 0.86 | 0.78 | 0.81 | 0.85 |
| 7 | 28552347 | rs2391670 | -0.001 | 0.918 | -0.174 | 0.896 | -0.001 | 0.001 | 0.001 | 0 | 0.31 | 0.25 | 0.32 | 0.37 | 0.28 | 0.38 |

| | | | | | | | | | | | | | | | | |
|---|---|---|---|---|---|---|---|---|---|---|---|---|---|---|---|---|
| 7 | 28556302 | rs216708 | 0.329 | 0.773 | -0.38 | 0.483 | -0.004 | 0.025 | 0.025 | 0 | 0.52 | 0.4 | 0.65 | 0.68 | 0.54 | 0.6 |
| 7 | 28558565 | rs148735 | 0.297 | 1.651 | 0.2 | 0.573 | 0.04 | -0.006 | -0.006 | 5.468 | 0.84 | 0.8 | 0.84 | 0.87 | 0.71 | 0.76 |
| 7 | 28558705 | rs11980665 | -0.427 | 0.698 | 0.486 | 0.063 | -0.071 | -0.01 | -0.01 | 0.007 | 0.19 | 0.45 | 0.12 | 0.15 | 0.08 | 0.13 |
| 7 | 28562743 | rs216712 | 1.417 | 0.569 | 0.42 | 0.033 | -0.003 | 0.004 | 0.004 | 0 | 0.64 | 0.32 | 0.72 | 0.71 | 0.63 | 0.63 |
| 7 | 28565641 | rs160346 | 1.423 | 0.518 | 0.423 | 0.035 | -0.001 | 0 | 0 | 0.022 | 0.66 | 0.32 | 0.73 | 0.71 | 0.66 | 0.63 |
| 7 | 28568804 | rs177594 | 0.307 | 1.523 | 0.511 | 0.313 | 0.002 | -0.017 | -0.017 | 0 | 0.1 | 0.14 | 0.06 | 0.06 | 0.09 | 0.13 |
| 7 | 28570687 | rs150610 | 0.262 | 1.6 | 0.333 | 0.326 | -0.006 | -0.007 | -0.007 | 0.017 | 0.94 | 0.94 | 0.9 | 0.92 | 0.8 | 0.9 |
| 7 | 28572882 | rs216715 | 0.845 | 1.385 | 0.774 | 1.126 | 0.062 | 0.091 | 0.091 | 1.069 | 0.83 | 0.58 | 0.82 | 0.77 | 0.78 | 0.83 |
| 7 | 28573565 | rs12539927 | 0.769 | 1.463 | -0.194 | 1.442 | -0.036 | -0.079 | -0.079 | 0 | 0.16 | 0.03 | 0.22 | 0.31 | 0.17 | 0.19 |
| 7 | 28573757 | rs170062 | 1.222 | 1.314 | 0.18 | 1.298 | 0.001 | 0 | 0 | 0.527 | 0.61 | 0.35 | 0.53 | 0.44 | 0.51 | 0.57 |
| 7 | 28575693 | rs216720 | 1.559 | 1.11 | 0.015 | 1.034 | 0 | -0.041 | -0.041 | 0 | 0.57 | 0.56 | 0.41 | 0.06 | 0.49 | 0.43 |
| 7 | 28577549 | rs2078980 | -0.516 | 0.554 | -0.293 | 1.525 | 0.001 | 0.023 | 0.023 | 0 | 0.13 | 0.1 | 0.19 | 0.41 | 0.09 | 0.26 |
| 7 | 28587817 | rs160335 | 0.516 | 0.854 | -0.482 | 1.237 | -0.031 | -0.028 | -0.028 | 0 | 0.9 | 0.97 | 0.86 | 0.6 | 0.83 | 0.85 |
| 7 | 28592608 | rs10951200 | 1.135 | 0.862 | -0.492 | -0.09 | 0.021 | 0.093 | 0.093 | 0.18 | 0.44 | 0.28 | 0.33 | 0.05 | 0.39 | 0.41 |
| 7 | 28594922 | rs10486588 | 0.557 | 1.371 | -1.108 | 0.371 | -0.008 | -0.03 | -0.03 | 0 | 0.18 | 0.09 | 0.17 | 0.47 | 0.21 | 0.25 |
| 7 | 28597194 | rs216735 | 1.077 | 1.138 | -1.007 | 0.52 | -0.006 | -0.073 | -0.073 | 0 | 0.15 | 0.06 | 0.27 | 0.42 | 0.14 | 0.21 |
| 7 | 28601707 | rs216743 | -0.197 | 1.598 | -0.419 | 0.736 | -0.006 | 0.004 | 0.004 | 0.309 | 0.92 | 0.94 | 0.89 | 0.95 | 0.83 | 0.97 |
| 7 | 28609796 | rs1976489 | 0.237 | 1.478 | -0.872 | 0.779 | -0.009 | -0.041 | -0.041 | 0 | 0.2 | 0.1 | 0.19 | 0.42 | 0.23 | 0.25 |
| 7 | 28616630 | rs150613 | 0.653 | 1.519 | -0.719 | 0.552 | -0.003 | 0.002 | 0.002 | 1.369 | 0.83 | 0.87 | 0.84 | 0.89 | 0.76 | 0.86 |
| 7 | 28619038 | rs17156863 | -0.013 | 1.087 | -0.756 | 0.966 | -0.007 | -0.034 | -0.034 | 1.378 | 0.2 | 0.1 | 0.2 | 0.47 | 0.23 | 0.25 |
| 7 | 28622352 | rs216750 | -0.305 | 2.028 | -0.52 | 0.344 | -0.011 | 0.01 | 0.01 | 0.13 | 0.83 | 0.84 | 0.8 | 0.92 | 0.6 | 0.93 |
| 7 | 28627789 | rs160337 | -0.508 | 2.035 | -0.342 | 0.346 | 0.024 | -0.01 | -0.01 | 0 | 0.84 | 0.82 | 0.79 | 0.94 | 0.6 | 0.93 |
| 7 | 28628202 | rs160338 | -0.618 | 2.048 | -0.357 | 0.359 | 0.016 | -0.002 | -0.002 | 0.168 | 0.85 | 0.85 | 0.79 | 0.92 | 0.6 | 0.93 |
| 7 | 28635741 | rs1008262 | 0.206 | 0.723 | -0.191 | 0.511 | 0.01 | -0.016 | -0.016 | 0.075 | 0.29 | 0.24 | 0.38 | 0.4 | 0.19 | 0.31 |
| 7 | 28641433 | rs13233942 | 0.347 | 0.666 | -0.3 | 0.738 | -0.006 | -0.101 | -0.101 | 0 | 0.28 | 0.05 | 0.36 | 0.43 | 0.21 | 0.33 |
| 7 | 28643045 | rs310338 | 0.498 | 0.626 | -0.012 | 0.239 | 0.003 | 0.008 | 0.008 | 0 | 0.54 | 0.33 | 0.46 | 0.38 | 0.59 | 0.46 |
| 7 | 28644857 | rs41273 | -0.726 | 0.491 | -0.053 | -0.06 | -0.009 | -0.014 | -0.014 | 0 | 0.13 | 0.21 | 0.09 | 0.09 | 0.11 | 0.16 |
| 7 | 28649438 | rs1637457 | -0.008 | 0.11 | -0.289 | -0.569 | -0.01 | -0.044 | -0.044 | 0.608 | 0.63 | 0.5 | 0.73 | 0.89 | 0.72 | 0.71 |
| 7 | 28650054 | rs41275 | 0.12 | -0.121 | -0.353 | -0.47 | -0.026 | -0.019 | -0.019 | 0 | 0.22 | 0.36 | 0.19 | 0.13 | 0.14 | 0.19 |
| 7 | 28650128 | rs17156919 | 1.277 | 0.132 | -0.775 | -0.012 | 0.006 | -0.087 | -0.087 | 0.337 | 0.24 | 0.05 | 0.34 | 0.37 | 0.17 | 0.4 |
| 7 | 28650263 | rs41276 | -0.024 | 0.351 | -0.549 | 0.302 | -0.033 | 0.083 | 0.083 | 0.234 | 0.24 | 0.38 | 0.2 | 0.4 | 0.15 | 0.24 |
| 7 | 28654701 | rs160375 | 0.853 | 1.384 | -1.041 | -0.064 | -0.072 | 0.039 | 0.039 | 0 | 0.18 | 0.05 | 0.18 | 0.12 | 0.34 | 0.07 |
| 7 | 28665767 | rs160343 | 0.388 | 1.004 | -0.817 | 0.01 | -0.015 | 0.008 | 0.008 | 0 | 0.19 | 0.15 | 0.2 | 0.12 | 0.25 | 0.2 |
| 7 | 28666636 | rs41295 | -0.634 | 0.761 | -0.949 | -0.394 | -0.034 | 0.117 | 0.117 | 0.006 | 0.61 | 0.8 | 0.71 | 0.88 | 0.52 | 0.78 |
| 7 | 28668502 | rs160357 | -0.191 | 1.282 | -0.46 | -0.024 | 0.032 | 0.002 | 0.002 | 0.064 | 0.32 | 0.5 | 0.32 | 0.34 | 0.44 | 0.23 |
| 7 | 28668780 | rs41298 | 0.671 | 1.576 | -0.504 | 0.188 | -0.003 | -0.009 | -0.009 | 0 | 0.28 | 0.27 | 0.33 | 0.37 | 0.26 | 0.34 |
| 7 | 28669836 | rs28234 | 0.751 | 1.12 | -0.476 | 0.244 | -0.006 | -0.014 | -0.014 | 0 | 0.27 | 0.22 | 0.32 | 0.37 | 0.27 | 0.34 |
| 7 | 28672354 | rs41304 | 0.739 | 1.135 | -0.496 | 0.354 | -0.003 | -0.005 | -0.005 | 0 | 0.27 | 0.26 | 0.33 | 0.36 | 0.27 | 0.33 |
| 7 | 28674854 | rs41307 | -0.524 | 1.214 | -0.775 | 0.388 | -0.015 | 0.041 | 0.041 | 0 | 0.55 | 0.69 | 0.62 | 0.65 | 0.49 | 0.66 |
| 7 | 28682081 | rs10228740 | -0.172 | 0.844 | -0.861 | 0.404 | -0.002 | 0 | 0 | 0 | 0.22 | 0.21 | 0.21 | 0.19 | 0.2 | 0.19 |
| 7 | 28683976 | rs3888613 | 1.05 | 1.848 | -1.615 | 0.601 | -0.107 | -0.095 | -0.095 | 0 | 0.19 | 0.02 | 0.28 | 0.34 | 0.35 | 0.17 |
| 7 | 28685819 | rs11543662 | 1.004 | 1.957 | -1.691 | 0.704 | -0.123 | -0.047 | -0.047 | 0.568 | 0.2 | 0.02 | 0.36 | 0.21 | 0.37 | 0.16 |
| 7 | 28686501 | rs41321 | -0.567 | -0.282 | -1.807 | 0.116 | 0.059 | 0.019 | 0.019 | 0 | 0.58 | 0.77 | 0.65 | 0.6 | 0.72 | 0.57 |
| 7 | 28688787 | rs7811703 | -0.106 | 0.095 | -0.177 | -0.605 | -0.024 | -0.049 | -0.049 | 0 | 0.21 | 0.31 | 0.1 | 0.08 | 0.12 | 0.26 |
| 7 | 28689587 | rs41327 | -1.312 | 0.022 | -0.194 | -0.634 | 0.04 | 0.005 | 0.005 | 0 | 0.36 | 0.84 | 0.3 | 0.34 | 0.24 | 0.36 |
| 7 | 28690496 | rs42322 | -1.678 | 0.29 | -0.154 | -0.555 | -0.021 | 0.06 | 0.06 | 0 | 0.47 | 0.92 | 0.44 | 0.33 | 0.39 | 0.55 |
| 7 | 28690735 | rs41328 | 0.164 | -0.329 | 0.3 | -0.341 | -0.013 | -0.082 | -0.082 | 0 | 0.14 | 0.32 | 0.16 | 0.01 | 0.11 | 0.17 |
| 7 | 28692444 | rs35506237 | -0.762 | -0.045 | 0.344 | 0.185 | 0.032 | 0.122 | 0.122 | 0 | 0.08 | 0.24 | 0.15 | 0.24 | 0.1 | 0.07 |
| 7 | 28693933 | rs9655280 | -0.628 | 0.153 | -0.175 | -0.79 | -0.017 | -0.046 | -0.046 | 0 | 0.24 | 0.29 | 0.12 | 0.08 | 0.16 | 0.31 |
| 7 | 28698698 | rs4719945 | -0.201 | 0.416 | -0.446 | -0.19 | 0.011 | -0.014 | -0.014 | 0 | 0.86 | 0.93 | 0.81 | 0.81 | 0.93 | 0.69 |
| 7 | 28703163 | rs10258405 | -0.047 | 0.988 | 0.183 | -0.42 | -0.009 | -0.05 | -0.05 | 0 | 0.4 | 0.5 | 0.24 | 0.14 | 0.38 | 0.23 |
| 7 | 28703795 | rs41334 | -0.236 | 1.617 | -0.127 | -0.073 | 0.086 | 0.245 | 0.245 | 0 | 0.06 | 0.13 | 0.26 | 0.53 | 0.21 | 0.14 |
| 7 | 28706813 | rs10245004 | 0.444 | 0.07 | -0.025 | -1.382 | -0.003 | -0.025 | -0.025 | 0 | 0.3 | 0.28 | 0.18 | 0.14 | 0.32 | 0.17 |
| 7 | 28707099 | rs978443 | 0.45 | 0.068 | -0.026 | -1.383 | -0.003 | -0.025 | -0.025 | 0 | 0.3 | 0.28 | 0.18 | 0.14 | 0.32 | 0.16 |
| 7 | 28708856 | rs982947 | -0.345 | 0.274 | -0.506 | -1.182 | 0.008 | 0.138 | 0.138 | 0 | 0.52 | 0.74 | 0.38 | 0.14 | 0.48 | 0.44 |
| 7 | 28709321 | rs982950 | -0.244 | 0.715 | -1.064 | -0.689 | 0.015 | 0.096 | 0.096 | 0 | 0.44 | 0.29 | 0.32 | 0.14 | 0.44 | 0.39 |
| 7 | 28715056 | rs16874653 | -0.424 | 0.841 | -0.89 | -0.351 | 0 | 0.005 | 0.005 | 0 | 0.37 | 0.31 | 0.3 | 0.26 | 0.33 | 0.33 |
| 7 | 28716154 | rs4719946 | -0.649 | 0.697 | -0.771 | -0.488 | -0.014 | 0.03 | 0.03 | 0.038 | 0.45 | 0.7 | 0.56 | 0.35 | 0.4 | 0.49 |
| 7 | 28719253 | rs9969149 | 0.488 | 0.94 | -1.224 | -1.239 | -0.002 | -0.002 | -0.002 | 1.099 | 0.57 | 0.45 | 0.45 | 0.63 | 0.6 | 0.43 |
| 7 | 28720609 | rs12113964 | -0.268 | 1.124 | -0.666 | -1.375 | 0.039 | -0.02 | -0.02 | 0 | 0.19 | 0.31 | 0.32 | 0.14 | 0.2 | 0.24 |
| 7 | 28722263 | rs6968464 | -0.26 | 0.914 | -0.714 | -1.45 | -0.001 | 0.001 | 0.001 | 0 | 0.88 | 0.87 | 0.86 | 0.96 | 0.84 | 0.83 |
| 7 | 28728440 | rs41351 | 1.054 | 1.189 | -0.157 | -1.567 | 0.008 | -0.04 | -0.04 | 0 | 0.9 | 0.77 | 0.85 | 0.95 | 0.91 | 0.89 |
| 7 | 28730009 | rs9691873 | 0.141 | 0.717 | -0.938 | -2.369 | -0.005 | 0.087 | 0.087 | 0 | 0.43 | 0.29 | 0.3 | 0.08 | 0.43 | 0.35 |
| 7 | 28730143 | rs10267387 | -0.319 | 0.992 | -0.813 | -2.383 | 0.023 | -0.031 | -0.031 | 0.375 | 0.08 | 0.09 | 0.11 | 0 | 0.12 | 0.1 |
| 7 | 28737298 | rs17157048 | -0.326 | 0.902 | -0.97 | -2.654 | -0.002 | -0.006 | -0.006 | 0 | 0.19 | 0.18 | 0.27 | 0.05 | 0.22 | 0.24 |
| 7 | 28745342 | rs10951201 | -0.185 | 0.653 | -0.629 | -2.934 | 0 | -0.025 | -0.025 | 0.009 | 0.47 | 0.56 | 0.45 | 0.11 | 0.42 | 0.46 |
| 7 | 28746913 | rs12540480 | 1.92 | 0.657 | -0.871 | -3.032 | -0.047 | 0.082 | 0.082 | 0 | 0.22 | 0.01 | 0.09 | 0.05 | 0.23 | 0.19 |
| 7 | 28747296 | rs10265166 | 1.438 | 0.657 | -0.725 | -2.927 | 0.008 | 0.023 | 0.023 | 0 | 0.11 | 0.05 | 0.05 | 0.04 | 0.09 | 0.1 |
| 7 | 28753387 | rs7806547 | -0.128 | 0.698 | -0.637 | -3.022 | 0.006 | 0.017 | 0.017 | 0 | 0.46 | 0.59 | 0.43 | 0.11 | 0.38 | 0.44 |

| | | | | | | | | | | | | | | | | |
|---|---|---|---|---|---|---|---|---|---|---|---|---|---|---|---|---|
| 7 | 28755781 | rs11771481 | 1.473 | 0.103 | -0.887 | -3.423 | 0.01 | 0.017 | 0.017 | 0 | 0.37 | 0.31 | 0.27 | 0.09 | 0.31 | 0.25 |
| 7 | 28757047 | rs9648352 | 0.918 | 0.416 | -0.84 | -3.251 | -0.018 | 0.035 | 0.035 | 0 | 0.13 | 0.05 | 0.09 | 0.03 | 0.16 | 0.12 |
| 7 | 28758638 | rs879593 | -0.303 | 0.863 | -0.999 | -2.665 | 0.005 | -0.017 | -0.017 | 0 | 0.23 | 0.3 | 0.43 | 0.15 | 0.29 | 0.31 |
| 7 | 28759014 | rs879591 | -0.939 | 0.563 | -1.025 | -2.631 | -0.005 | 0.007 | 0.007 | 0 | 0.37 | 0.36 | 0.5 | 0.16 | 0.43 | 0.43 |
| 7 | 28764109 | rs2066979 | 0.672 | -0.569 | -0.965 | -2.253 | 0.002 | -0.026 | -0.026 | 0 | 0.63 | 0.6 | 0.71 | 0.24 | 0.62 | 0.56 |
| 7 | 28766878 | rs10486591 | 0.339 | -0.167 | -0.994 | -1.979 | -0.001 | 0.034 | 0.034 | 0 | 0.6 | 0.63 | 0.68 | 0.24 | 0.62 | 0.63 |
| 7 | 28775381 | rs6462103 | -0.255 | 0.058 | -1.139 | -1.472 | 0.008 | 0.018 | 0.018 | 0.001 | 0.94 | 0.93 | 0.94 | 0.88 | 0.93 | 0.89 |
| 7 | 28794477 | rs740315 | -0.81 | -0.208 | -1.454 | -1.817 | 0.005 | 0.02 | 0.02 | 0.235 | 0.26 | 0.21 | 0.16 | 0.01 | 0.23 | 0.24 |
| 7 | 28796955 | rs1468447 | 0.24 | -0.081 | -1.285 | -1.807 | 0.038 | -0.028 | -0.028 | 0 | 0.79 | 0.52 | 0.7 | 0.82 | 0.74 | 0.74 |
| 7 | 28802524 | rs2073537 | 0.575 | -0.259 | -1.266 | -1.721 | 0.016 | -0.014 | -0.014 | 1.697 | 0.75 | 0.49 | 0.66 | 0.77 | 0.72 | 0.66 |
| 7 | 28809566 | rs4722844 | 0.547 | -0.242 | -1.29 | -1.777 | 0.014 | -0.062 | -0.062 | 0.731 | 0.76 | 0.53 | 0.67 | 0.84 | 0.74 | 0.58 |
| 7 | 28810310 | rs17730573 | 1.237 | -0.333 | -1.411 | -1.87 | -0.012 | 0.052 | 0.052 | 0 | 0.09 | 0 | 0.03 | 0 | 0.08 | 0.06 |
| 7 | 28811631 | rs17730621 | -0.577 | -0.548 | -1.692 | -2.05 | 0.014 | 0.039 | 0.039 | 0.11 | 0.45 | 0.41 | 0.4 | 0.73 | 0.37 | 0.32 |
| 7 | 28815239 | rs2299116 | 2.348 | -0.546 | -1.726 | -2.079 | -0.02 | 0.079 | 0.079 | 0 | 0.3 | 0.17 | 0.18 | 0.08 | 0.32 | 0.28 |
| 7 | 28818245 | rs10224680 | -0.623 | -0.581 | -1.753 | -2.101 | 0.021 | 0.063 | 0.063 | 0.05 | 0.47 | 0.4 | 0.49 | 0.78 | 0.39 | 0.44 |
| 7 | 28818730 | rs2299117 | 0.367 | -0.423 | -1.653 | -1.982 | 0.015 | -0.032 | -0.032 | 0 | 0.75 | 0.47 | 0.65 | 0.83 | 0.71 | 0.7 |
| 7 | 28828220 | rs1859683 | -2.164 | -0.522 | -1.811 | -2.009 | -0.01 | -0.074 | -0.074 | 0 | 0.82 | 0.95 | 0.76 | 0.3 | 0.83 | 0.78 |
| 7 | 28830609 | rs989438 | -1.298 | -0.623 | -1.825 | -2.054 | 0 | -0.018 | -0.018 | 0 | 0.77 | 0.95 | 0.66 | 0.22 | 0.81 | 0.65 |
| 7 | 28832297 | rs2237361 | -1.72 | -0.465 | -1.679 | -1.937 | -0.047 | 0.077 | 0.077 | 0.155 | 0.41 | 0.84 | 0.46 | 0.26 | 0.41 | 0.4 |
| 7 | 28834480 | rs28608003 | -1.204 | -0.297 | -1.38 | -1.801 | 0 | -0.033 | -0.033 | 0 | 0.22 | 0.38 | 0.18 | 0.14 | 0.2 | 0.22 |
| 7 | 28838253 | rs7791555 | 0.138 | -0.158 | -1.318 | -1.69 | 0.018 | 0.014 | 0.014 | 0.362 | 0.64 | 0.69 | 0.6 | 0.3 | 0.74 | 0.64 |
| 7 | 28842067 | rs2237364 | -1.577 | 0.086 | -2.313 | -1.598 | -0.049 | 0.035 | 0.035 | 0 | 0.4 | 0.8 | 0.47 | 0.31 | 0.43 | 0.44 |
| 7 | 28844815 | rs2282909 | -0.268 | 0.027 | -2.247 | -1.543 | 0.002 | -0.005 | -0.005 | 0 | 0.68 | 0.65 | 0.64 | 0.43 | 0.75 | 0.67 |
| 7 | 28848524 | rs12538892 | 0.223 | 0.053 | -2.22 | -1.587 | 0.046 | -0.018 | -0.018 | 0 | 0.2 | 0.46 | 0.3 | 0.16 | 0.25 | 0.21 |
| 7 | 28851986 | rs1964240 | 0.4 | 0.46 | -1.394 | -0.817 | 0.003 | 0 | 0 | 0.369 | 0.68 | 0.66 | 0.63 | 0.34 | 0.75 | 0.66 |
| 7 | 28852146 | rs10951204 | -0.912 | 0.445 | -1.43 | -0.861 | -0.009 | -0.066 | -0.066 | 0.654 | 0.8 | 0.96 | 0.69 | 0.31 | 0.84 | 0.84 |
| 7 | 28852462 | rs17669844 | -1.133 | 0.512 | -1.41 | -0.898 | 0 | -0.04 | -0.04 | 0.961 | 0.21 | 0.22 | 0.25 | 0.01 | 0.32 | 0.19 |
| 7 | 28857606 | rs12531253 | 1.14 | -0.55 | -0.971 | -0.369 | -0.009 | -0.012 | -0.012 | 0.264 | 0.63 | 0.54 | 0.55 | 0.3 | 0.72 | 0.63 |
| 7 | 28858456 | rs10951205 | -0.877 | -0.599 | -0.932 | -0.341 | 0.002 | -0.118 | -0.118 | 0.41 | 0.83 | 0.96 | 0.72 | 0.43 | 0.86 | 0.86 |
| 7 | 28859450 | rs2190305 | -0.811 | -0.563 | -0.927 | -0.345 | -0.02 | -0.119 | -0.119 | 0.085 | 0.82 | 0.96 | 0.71 | 0.43 | 0.84 | 0.86 |
| 7 | 28859859 | rs4722856 | -0.404 | 0.045 | -0.308 | -0.937 | 0.007 | -0.038 | -0.038 | 8.883 | 0.1 | 0.17 | 0.06 | 0 | 0.1 | 0.09 |
| 7 | 28867194 | rs11975539 | -1.752 | 0.044 | -0.314 | -0.909 | -0.016 | -0.103 | -0.103 | 0 | 0.26 | 0.41 | 0.19 | 0.02 | 0.2 | 0.26 |
| 7 | 28870741 | rs6462107 | -1.338 | 0.28 | 0 | -0.346 | -0.009 | -0.093 | -0.093 | 0 | 0.17 | 0.39 | 0.15 | 0.02 | 0.12 | 0.21 |
| 7 | 28884912 | rs4719955 | -1.149 | 0.579 | -0.714 | -0.209 | 0.001 | -0.074 | -0.074 | 0 | 0.15 | 0.31 | 0.15 | 0.02 | 0.12 | 0.2 |
| 7 | 28891257 | rs763809 | 0.39 | 0.81 | -1.567 | 0.336 | -0.017 | -0.033 | -0.033 | 0 | 0.69 | 0.85 | 0.62 | 0.39 | 0.72 | 0.67 |
| 7 | 28893354 | rs3735565 | -0.448 | 1.262 | -1.726 | -0.26 | -0.042 | 0.063 | 0.063 | 0 | 0.25 | 0.15 | 0.29 | 0.08 | 0.36 | 0.2 |
| 7 | 28896396 | rs12700916 | 0.135 | 0.557 | -1.363 | 0.047 | -0.008 | 0.005 | 0.005 | 0 | 0.38 | 0.46 | 0.31 | 0.42 | 0.33 | 0.39 |
| 7 | 28902397 | rs1859685 | -0.72 | 0.718 | -1.586 | 0.025 | -0.037 | -0.026 | -0.026 | 0.065 | 0.64 | 0.8 | 0.69 | 0.55 | 0.6 | 0.68 |
| 7 | 28902589 | rs4719956 | 0.311 | 0.426 | -0.785 | 0.238 | 0.027 | 0.068 | 0.068 | 0 | 0.12 | 0.23 | 0.26 | 0.2 | 0.18 | 0.13 |
| 7 | 28903158 | rs13235271 | 0.71 | 0.207 | -1.65 | 0.319 | 0 | -0.063 | -0.063 | 0.455 | 0.21 | 0.02 | 0.11 | 0.17 | 0.15 | 0.18 |
| 7 | 28903231 | rs10225559 | -0.9 | 0.646 | -0.772 | 0.108 | 0 | 0.003 | 0.003 | 0 | 0.21 | 0.56 | 0.44 | 0.21 | 0.25 | 0.22 |
| 7 | 28907630 | rs10261739 | -0.544 | 0.317 | -1.227 | -0.619 | 0.007 | -0.036 | -0.036 | 1.185 | 0.29 | 0.71 | 0.17 | 0.02 | 0.18 | 0.22 |
| 7 | 28907762 | rs2057954 | 0.776 | 2.118 | -1.532 | 0.524 | -0.034 | -0.045 | -0.045 | 0 | 0.77 | 0.82 | 0.73 | 0.55 | 0.7 | 0.62 |
| 7 | 28908683 | rs7777049 | 0.372 | 2.453 | -1.491 | 0.093 | -0.073 | -0.02 | -0.02 | 0.003 | 0.82 | 0.93 | 0.73 | 0.79 | 0.61 | 0.82 |
| 7 | 28915791 | rs6462110 | 0.488 | 0.779 | -1.062 | -0.605 | 0.025 | 0.054 | 0.054 | 0.004 | 0.56 | 0.36 | 0.62 | 0.8 | 0.65 | 0.6 |
| 7 | 28918565 | rs1024551 | 0.264 | 0.517 | -1.034 | -0.716 | 0.001 | -0.012 | -0.012 | 1.34 | 0.9 | 0.9 | 0.95 | 1 | 0.94 | 0.95 |
| 7 | 28926168 | rs1859681 | -0.359 | 0.386 | -0.942 | -0.803 | 0.005 | 0.032 | 0.032 | 0 | 0.4 | 0.18 | 0.45 | 0.38 | 0.38 | 0.47 |
| 7 | 28928171 | rs6951061 | -0.574 | 0.072 | -0.242 | -0.963 | -0.017 | -0.01 | -0.01 | 0.446 | 0.3 | 0.4 | 0.39 | 0.23 | 0.18 | 0.43 |
| 7 | 28933323 | rs1558031 | 0.443 | 0.302 | -0.512 | -0.86 | 0.041 | 0.036 | 0.036 | 0 | 0.2 | 0.1 | 0.2 | 0.12 | 0.12 | 0.35 |
| 7 | 28933628 | rs1558032 | 0.443 | 0.302 | -0.512 | -0.86 | 0.041 | 0.036 | 0.036 | 0 | 0.2 | 0.1 | 0.2 | 0.12 | 0.12 | 0.35 |
| 7 | 28934503 | rs4719959 | 0.856 | 0.759 | -0.607 | -0.525 | 0.009 | 0.013 | 0.013 | 0 | 0.14 | 0.14 | 0.22 | 0.29 | 0.24 | 0.12 |
| 7 | 28934828 | rs6950121 | 0.306 | 0.725 | -0.489 | -0.32 | -0.016 | 0.041 | 0.041 | 0 | 0.45 | 0.26 | 0.43 | 0.63 | 0.53 | 0.46 |
| 7 | 28939870 | rs929385 | 1.018 | 0.962 | -1.224 | -0.386 | -0.018 | 0.018 | 0.018 | 0 | 0.26 | 0.13 | 0.28 | 0.23 | 0.29 | 0.2 |
| 7 | 28941454 | rs10216225 | 0.524 | 0.421 | -1.043 | -0.672 | -0.047 | 0.041 | 0.041 | 0.812 | 0.43 | 0.19 | 0.42 | 0.62 | 0.5 | 0.46 |
| 7 | 28944145 | rs10242171 | 0.792 | -0.095 | 0.065 | -0.933 | -0.026 | -0.075 | -0.075 | 0 | 0.77 | 0.38 | 0.84 | 0.93 | 0.88 | 0.88 |
| 7 | 28961321 | rs1006514 | -1.183 | 0.395 | 0.21 | -0.793 | 0 | -0.078 | -0.078 | 4.206 | 0.22 | 0.46 | 0.27 | 0.09 | 0.22 | 0.24 |
| 7 | 28972863 | rs221127 | 1.129 | 0.629 | 0.482 | -0.826 | 0.026 | -0.036 | -0.036 | 0 | 0.81 | 0.65 | 0.72 | 0.9 | 0.77 | 0.76 |
| 7 | 28977011 | rs221134 | -0.194 | 0.293 | 0.227 | -0.889 | 0 | -0.002 | -0.002 | 0.003 | 0.23 | 0.28 | 0.27 | 0.61 | 0.23 | 0.32 |
| 7 | 28978148 | rs10479817 | 1.713 | -0.411 | 0.067 | -1.279 | 0.02 | 0.192 | 0.192 | 0.147 | 0.5 | 0.07 | 0.43 | 0.28 | 0.57 | 0.4 |
| 7 | 28980086 | rs221139 | 0.694 | -0.369 | 0.342 | -1.241 | -0.002 | -0.048 | -0.048 | 0 | 0.79 | 0.56 | 0.72 | 0.89 | 0.8 | 0.76 |
| 7 | 28991862 | rs221151 | 0.207 | -0.28 | -0.134 | -1.336 | 0.003 | -0.015 | -0.015 | 0.308 | 0.2 | 0.19 | 0.26 | 0.62 | 0.21 | 0.3 |
| 7 | 28996944 | rs740250 | 0.497 | -0.476 | -0.368 | -1.58 | -0.009 | 0.008 | 0.008 | 0 | 0.85 | 0.92 | 0.79 | 0.87 | 0.83 | 0.77 |
| 7 | 28998663 | rs2286219 | 1.093 | -0.595 | 0.474 | -1.62 | -0.008 | -0.028 | -0.028 | 0 | 0.63 | 0.55 | 0.51 | 0.26 | 0.62 | 0.44 |
| 7 | 29001210 | rs221159 | 0.156 | -0.507 | 0.44 | -1.614 | -0.001 | -0.022 | -0.022 | 0.325 | 0.22 | 0.17 | 0.29 | 0.61 | 0.25 | 0.34 |
| 7 | 29001891 | rs107542 | 0.359 | -0.737 | 0.514 | -1.542 | 0.002 | -0.016 | -0.016 | 0 | 0.2 | 0.17 | 0.26 | 0.61 | 0.21 | 0.34 |
| 7 | 29005441 | rs221170 | 1.91 | -1.094 | 0.75 | -1.799 | -0.012 | -0.063 | -0.063 | 1.59 | 0.71 | 0.44 | 0.7 | 0.89 | 0.77 | 0.76 |
| 7 | 29006197 | rs221173 | 0.682 | -1.104 | 0.79 | -1.776 | -0.001 | -0.016 | -0.016 | 0 | 0.84 | 0.74 | 0.78 | 0.89 | 0.83 | 0.79 |
| 7 | 29011102 | rs221181 | 0.574 | -1.168 | 0.724 | -1.689 | -0.001 | -0.007 | -0.007 | 0 | 0.84 | 0.8 | 0.79 | 0.89 | 0.83 | 0.79 |
| 7 | 29011448 | rs221182 | 0.493 | -1.171 | 0.684 | -1.69 | -0.001 | 0.009 | 0.009 | 0 | 0.85 | 0.87 | 0.79 | 0.89 | 0.83 | 0.79 |

| | | | | | | | | | | | | | | | | |
|---|---|---|---|---|---|---|---|---|---|---|---|---|---|---|---|---|
| 7 | 29014135 | rs122353 | 1.637 | -1.12 | 0.776 | -1.721 | -0.026 | -0.138 | -0.138 | 0 | 0.75 | 0.56 | 0.75 | 0.99 | 0.8 | 0.89 |
| 7 | 29017188 | rs13227954 | 1.035 | -1.068 | 0.826 | -1.483 | 0.01 | 0.025 | 0.025 | 0.729 | 0.64 | 0.31 | 0.67 | 0.29 | 0.69 | 0.53 |
| 7 | 29018593 | rs504453 | -1 | -1.058 | 0.806 | -1.335 | 0.023 | 0.041 | 0.041 | 0.877 | 0.11 | 0.39 | 0.1 | 0.11 | 0.06 | 0.13 |
| 7 | 29021336 | rs886834 | 0.727 | -0.661 | 0.832 | -1.277 | 0.002 | -0.005 | -0.005 | 1.474 | 0.7 | 0.45 | 0.69 | 0.29 | 0.7 | 0.53 |
| 7 | 29030246 | rs317766 | 0.686 | -1.168 | 0.3 | -1.285 | 0.006 | -0.008 | -0.008 | 0 | 0.73 | 0.79 | 0.81 | 0.47 | 0.81 | 0.69 |
| 7 | 29034918 | rs1034839 | -0.452 | -0.989 | 0.708 | -1.378 | 0.016 | 0.012 | 0.012 | 2.576 | 0.07 | 0.55 | 0.07 | 0.01 | 0.01 | 0.02 |
| 7 | 29037093 | rs317700 | 0.244 | -1.079 | 0.451 | -1.482 | 0.008 | 0.01 | 0.01 | 0 | 0.09 | 0.06 | 0.06 | 0.06 | 0.07 | 0.12 |
| 7 | 29039366 | rs317696 | -1.276 | -0.775 | 0.375 | -1.808 | -0.007 | -0.045 | -0.045 | 0.038 | 0.21 | 0.23 | 0.11 | 0.06 | 0.17 | 0.16 |
| 7 | 29041190 | rs2252521 | 1.109 | -1.167 | -0.378 | -1.873 | -0.004 | 0 | 0 | 0 | 0.63 | 0.37 | 0.71 | 0.48 | 0.7 | 0.65 |
| 7 | 29041980 | rs2529478 | 1.961 | -1.219 | -0.647 | -2.039 | -0.001 | 0.241 | 0.241 | 0.254 | 0.48 | 0.13 | 0.41 | 0.21 | 0.55 | 0.44 |
| 7 | 29049119 | rs642392 | -1.694 | -1.151 | -0.593 | -1.995 | -0.002 | -0.003 | -0.003 | 0.075 | 0.36 | 0.37 | 0.38 | 0.33 | 0.32 | 0.38 |
| 7 | 29051493 | rs673681 | -1.77 | -1.136 | -0.606 | -2.008 | -0.001 | -0.002 | -0.002 | 0 | 0.37 | 0.37 | 0.4 | 0.33 | 0.33 | 0.38 |
| 7 | 29052099 | rs686846 | -2.213 | -1.177 | -0.642 | -2.027 | 0.031 | 0.056 | 0.056 | 0 | 0.43 | 0.8 | 0.46 | 0.33 | 0.34 | 0.39 |
| 7 | 29059381 | rs2648661 | 0.861 | -1.179 | -0.685 | -2.024 | 0.016 | 0.02 | 0.02 | 0 | 0.1 | 0.05 | 0.05 | 0.06 | 0.07 | 0.12 |
| 7 | 29059435 | rs2529481 | -0.633 | -1.174 | -0.672 | -2.037 | 0.011 | 0.021 | 0.021 | 0 | 0.21 | 0.12 | 0.09 | 0.06 | 0.15 | 0.16 |
| 7 | 29059726 | rs2190531 | -1.067 | -1.187 | -0.64 | -2.037 | -0.006 | 0.021 | 0.021 | 1.784 | 0.21 | 0.64 | 0.34 | 0.27 | 0.18 | 0.22 |
| 7 | 29070209 | rs7313 | 2.475 | -1.026 | -0.923 | -1.99 | -0.036 | 0.091 | 0.091 | 0 | 0.39 | 0.23 | 0.36 | 0.15 | 0.46 | 0.32 |
| 7 | 29070320 | rs1052200 | -0.159 | -1.025 | -0.848 | -1.973 | -0.001 | 0.002 | 0.002 | 0.06 | 0.14 | 0.16 | 0.28 | 0.26 | 0.15 | 0.21 |
| 7 | 29071712 | rs17157410 | -0.953 | -1.114 | -0.811 | -1.97 | -0.025 | -0.077 | -0.077 | 0.068 | 0.28 | 0.57 | 0.17 | 0.07 | 0.19 | 0.18 |
| 7 | 29079657 | rs4141276 | 0.156 | -0.903 | -0.968 | -1.869 | -0.015 | 0.043 | 0.043 | 0.204 | 0.12 | 0.04 | 0.15 | 0.02 | 0.12 | 0.1 |
| 7 | 29079870 | rs4141275 | 1.526 | -0.732 | -0.72 | -1.629 | 0.079 | 0.094 | 0.094 | 0 | 0.51 | 0.18 | 0.62 | 0.42 | 0.61 | 0.52 |
| 7 | 29083262 | rs320088 | -0.383 | -0.854 | -0.753 | -1.439 | -0.025 | -0.13 | -0.13 | 0 | 0.8 | 0.98 | 0.81 | 0.49 | 0.79 | 0.7 |
| 7 | 29084392 | rs505532 | 0.337 | -0.389 | -0.328 | -1.29 | -0.026 | -0.009 | -0.009 | 0.339 | 0.62 | 0.52 | 0.68 | 0.42 | 0.69 | 0.56 |
| 7 | 29101160 | rs317710 | 0.837 | 0.061 | -1.57 | -1.585 | -0.123 | 0.288 | 0.288 | 0 | 0.42 | 0.06 | 0.35 | 0.11 | 0.53 | 0.38 |
| 7 | 29101891 | rs320101 | -1.709 | 0.178 | -0.478 | -1.166 | -0.001 | -0.062 | -0.062 | 0 | 0.11 | 0.39 | 0.06 | 0.01 | 0.05 | 0.05 |
| 7 | 29102596 | rs317712 | 0.587 | 0.487 | -1.407 | -1.456 | 0 | 0.123 | 0.123 | 0 | 0.58 | 0.27 | 0.53 | 0.21 | 0.6 | 0.49 |
| 7 | 29102661 | rs317713 | 1.016 | 0.788 | -1.511 | -0.903 | -0.003 | 0.001 | 0.001 | 0.244 | 0.8 | 0.82 | 0.86 | 0.84 | 0.76 | 0.91 |
| 7 | 29103126 | rs7777266 | -0.003 | 0.782 | -1.461 | -0.846 | 0.004 | 0.004 | 0.004 | 0 | 0.86 | 0.84 | 0.88 | 0.85 | 0.82 | 0.93 |
| 7 | 29111093 | rs2072237 | 0.875 | 0.758 | -1.637 | -0.128 | -0.011 | -0.135 | -0.135 | 0 | 0.2 | 0.02 | 0.15 | 0.43 | 0.17 | 0.3 |
| 7 | 29111578 | rs2074784 | -0.096 | 0.754 | -1.576 | -0.141 | -0.021 | -0.045 | -0.045 | 0.16 | 0.31 | 0.12 | 0.3 | 0.62 | 0.28 | 0.46 |
| 7 | 29113023 | rs160862 | 1.723 | 0.708 | -1.515 | -0.339 | -0.036 | 0.194 | 0.194 | 0.257 | 0.48 | 0.16 | 0.5 | 0.23 | 0.51 | 0.47 |
| 7 | 29117086 | rs17676174 | 0.041 | 0.718 | -1.437 | -0.299 | -0.014 | 0.031 | 0.031 | 0 | 0.14 | 0.08 | 0.11 | 0.02 | 0.17 | 0.06 |
| 7 | 29118083 | rs17748871 | 0.467 | 0.714 | -1.462 | -0.308 | -0.028 | 0.041 | 0.041 | 0 | 0.12 | 0.02 | 0.1 | 0.02 | 0.17 | 0.06 |
| 7 | 29118314 | rs17748895 | 0.467 | 0.714 | -1.462 | -0.308 | -0.028 | 0.041 | 0.041 | 0 | 0.12 | 0.02 | 0.1 | 0.02 | 0.17 | 0.06 |
| 7 | 29121211 | rs317726 | 1.498 | 0.994 | -1.352 | -0.312 | -0.001 | 0.07 | 0.07 | 0 | 0.52 | 0.6 | 0.55 | 0.23 | 0.51 | 0.47 |
| 7 | 29122176 | rs3735558 | -0.195 | 1.037 | -1.614 | 0.069 | -0.023 | -0.046 | -0.046 | 0 | 0.31 | 0.14 | 0.3 | 0.62 | 0.28 | 0.46 |
| 7 | 29132650 | rs17749450 | 0.276 | 1.129 | -1.458 | 0.076 | -0.008 | 0.029 | 0.029 | 0 | 0.11 | 0.03 | 0.09 | 0.01 | 0.12 | 0.05 |
| 7 | 29135569 | rs2286992 | -0.221 | 0.849 | -1.717 | 0.226 | -0.008 | -0.089 | -0.089 | 0 | 0.28 | 0.13 | 0.18 | 0.37 | 0.25 | 0.3 |
| 7 | 29136284 | rs11768067 | 0.54 | 0.825 | -1.754 | 0.202 | 0 | -0.09 | -0.09 | 0 | 0.22 | 0.09 | 0.16 | 0.38 | 0.18 | 0.3 |
| 7 | 29140213 | rs12670165 | 0.623 | 0.672 | -1.417 | 0.16 | -0.027 | -0.127 | -0.127 | 0 | 0.21 | 0.02 | 0.15 | 0.37 | 0.19 | 0.29 |
| 7 | 29141558 | rs1568658 | -0.705 | 0.432 | -1.34 | -0.097 | 0.004 | -0.002 | -0.002 | 0 | 0.36 | 0.7 | 0.27 | 0.56 | 0.29 | 0.4 |
| 7 | 29142663 | rs323199 | 0.548 | 1.04 | -1.874 | -0.483 | 0.054 | 0.119 | 0.119 | 0 | 0.5 | 0.7 | 0.55 | 0.27 | 0.44 | 0.44 |
| 7 | 29145731 | rs1522918 | -0.067 | 0.691 | -2.42 | -1.282 | 0.013 | 0.094 | 0.094 | 0 | 0.49 | 0.66 | 0.49 | 0.76 | 0.42 | 0.54 |
| 7 | 29148538 | rs17676979 | 0.716 | -0.043 | -2.366 | -0.824 | -0.029 | 0.142 | 0.142 | 0.14 | 0.26 | 0.06 | 0.24 | 0.08 | 0.25 | 0.19 |
| 7 | 29151397 | rs323183 | 0.313 | -0.51 | -1.551 | 0.19 | -0.004 | -0.013 | -0.013 | 0.363 | 0.19 | 0.11 | 0.15 | 0.33 | 0.18 | 0.21 |
| 7 | 29152371 | rs323182 | 0.128 | -1.178 | -0.443 | -0.583 | -0.024 | -0.039 | -0.039 | 0 | 0.11 | 0.15 | 0.08 | 0.03 | 0.1 | 0.01 |
| 7 | 29152416 | rs2011337 | -0.014 | -1.23 | -0.187 | -0.606 | -0.031 | -0.049 | -0.049 | 0 | 0.13 | 0.22 | 0.09 | 0.03 | 0.1 | 0.01 |
| 7 | 29152778 | rs11760588 | 0.384 | -1.125 | -0.261 | -0.685 | -0.019 | -0.003 | -0.003 | 0 | 0.1 | 0.06 | 0.1 | 0.09 | 0.16 | 0.08 |
| 7 | 29153237 | rs2021551 | -1.053 | -1.298 | 0.27 | -0.684 | -0.001 | -0.027 | -0.027 | 0 | 0.23 | 0.28 | 0.19 | 0.13 | 0.26 | 0.1 |
| 7 | 29154703 | rs1403408 | -1.462 | -1.676 | 0.346 | -0.848 | -0.021 | -0.059 | -0.059 | 0 | 0.35 | 0.42 | 0.22 | 0.17 | 0.31 | 0.27 |
| 7 | 29160852 | rs13246846 | -1.743 | -2.08 | 0.392 | -1.085 | 0.048 | 0.098 | 0.098 | 1.93 | 0.46 | 0.69 | 0.29 | 0.17 | 0.36 | 0.3 |
| 7 | 29171070 | rs6950695 | 0.453 | -1.759 | 0.583 | -0.913 | 0.01 | -0.003 | -0.003 | 0 | 0.12 | 0.16 | 0.12 | 0.09 | 0.16 | 0.08 |
| 7 | 29172256 | rs2687640 | -0.235 | -1.722 | 0.696 | -0.837 | 0.013 | -0.004 | -0.004 | 0 | 0.09 | 0.26 | 0.07 | 0.03 | 0.04 | 0.02 |
| 7 | 29173765 | rs6966319 | -0.829 | -1.706 | 0.703 | -0.826 | -0.026 | -0.037 | -0.037 | 1.398 | 0.14 | 0.31 | 0.09 | 0.08 | 0.06 | 0.18 |
| 7 | 29178492 | rs245866 | 0.834 | -1.639 | 0.564 | -0.731 | -0.009 | -0.035 | -0.035 | 0.169 | 0.66 | 0.59 | 0.79 | 0.91 | 0.7 | 0.87 |
| 7 | 29178843 | rs245867 | 0.415 | -1.579 | 0.468 | -0.487 | -0.037 | -0.063 | -0.063 | 0 | 0.7 | 0.62 | 0.84 | 0.92 | 0.73 | 0.87 |
| 7 | 29180335 | rs13238274 | 0.719 | -0.954 | 0.115 | -0.489 | -0.001 | -0.002 | -0.002 | 0.353 | 0.1 | 0.08 | 0.1 | 0.04 | 0.16 | 0.08 |
| 7 | 29181626 | rs13245225 | 0.719 | -0.954 | 0.115 | -0.489 | -0.001 | -0.002 | -0.002 | 0 | 0.1 | 0.08 | 0.1 | 0.04 | 0.16 | 0.08 |
| 7 | 29184489 | rs10486607 | 0.622 | -0.956 | 0.118 | -0.489 | -0.004 | 0.005 | 0.005 | 0 | 0.1 | 0.06 | 0.1 | 0.04 | 0.16 | 0.08 |
| 7 | 29184944 | rs245874 | 0.018 | -0.985 | -0.102 | -0.425 | 0.019 | 0.042 | 0.042 | 0.144 | 0.48 | 0.68 | 0.62 | 0.72 | 0.56 | 0.63 |
| 7 | 29185531 | rs17677737 | 1.174 | -1.009 | -0.271 | -0.522 | -0.005 | 0.018 | 0.018 | 0 | 0.07 | 0.03 | 0.06 | 0 | 0.08 | 0.03 |
| 7 | 29185604 | rs245878 | 1.26 | -1.15 | 0.081 | -0.15 | 0.011 | 0.007 | 0.007 | 0.003 | 0.31 | 0.38 | 0.37 | 0.38 | 0.38 | 0.37 |
| 7 | 29186256 | rs245880 | 0.275 | -1.291 | -0.137 | -0.213 | -0.006 | 0.008 | 0.008 | 0.102 | 0.43 | 0.64 | 0.56 | 0.69 | 0.41 | 0.56 |
| 7 | 29186576 | rs245881 | 0.116 | -1.451 | -0.094 | -0.467 | -0.006 | -0.067 | -0.067 | 0.409 | 0.68 | 0.71 | 0.8 | 0.99 | 0.79 | 0.9 |
| 7 | 29187251 | rs940041 | 0.19 | -1.124 | -0.301 | -0.007 | -0.033 | 0.143 | 0.143 | 0 | 0.09 | 0.29 | 0.18 | 0.26 | 0.01 | 0.07 |
| 7 | 29188554 | rs245885 | 0.01 | -1.076 | -0.699 | -0.158 | 0.004 | 0.038 | 0.038 | 0.665 | 0.51 | 0.72 | 0.62 | 0.67 | 0.49 | 0.6 |
| 7 | 29190881 | rs245891 | 1.243 | -1.127 | -0.912 | -0.169 | -0.011 | -0.006 | -0.006 | 0 | 0.31 | 0.28 | 0.37 | 0.38 | 0.38 | 0.37 |
| 7 | 29192357 | rs6948891 | 0.608 | -0.914 | -1.001 | -0.12 | -0.037 | 0.091 | 0.091 | 0.002 | 0.08 | 0.23 | 0.16 | 0.22 | 0 | 0.07 |
| 7 | 29192895 | rs245895 | 0.033 | -0.174 | -2.004 | -0.521 | 0.012 | 0.096 | 0.096 | 0 | 0.72 | 0.9 | 0.82 | 0.96 | 0.72 | 0.86 |

| | | | | | | | | | | | | | | | | |
|---|---|---|---|---|---|---|---|---|---|---|---|---|---|---|---|---|
| 7 | 29195455 | rs10256795 | 0.693 | -0.368 | -1.816 | -0.365 | -0.02 | -0.028 | -0.028 | 0 | 0.31 | 0.2 | 0.29 | 0.32 | 0.37 | 0.36 |
| 7 | 29196464 | rs245903 | 0.008 | 0.016 | -1.571 | -0.686 | 0.017 | 0.117 | 0.117 | 0 | 0.82 | 0.99 | 0.92 | 1 | 0.87 | 0.95 |
| 7 | 29197087 | rs245904 | -0.324 | 1.158 | -0.83 | -0.027 | 0.003 | -0.006 | -0.006 | 1.119 | 0.21 | 0.21 | 0.26 | 0.11 | 0.27 | 0.19 |
| 7 | 29199869 | rs245906 | 0.007 | 0.917 | -0.877 | -0.19 | 0.012 | 0.03 | 0.03 | 0.972 | 0.53 | 0.59 | 0.58 | 0.75 | 0.59 | 0.59 |
| 7 | 29209352 | rs245908 | 0.381 | 1.603 | -0.69 | 0.088 | 0.032 | -0.018 | -0.018 | 0 | 0.36 | 0.21 | 0.34 | 0.57 | 0.33 | 0.48 |
| 7 | 29209361 | rs6956877 | -0.363 | 1.957 | -0.007 | -0.08 | 0.021 | 0.079 | 0.079 | 1.051 | 0.15 | 0.51 | 0.25 | 0.2 | 0.12 | 0.04 |
| 7 | 29211267 | rs17157602 | 1.184 | 1.789 | -0.448 | 0.697 | -0.017 | -0.048 | -0.048 | 2.912 | 0.08 | 0.01 | 0.13 | 0.19 | 0.11 | 0.13 |
| 7 | 29213691 | rs6944596 | 0.133 | 1.236 | -0.472 | 1.135 | -0.004 | 0.014 | 0.014 | 0 | 0.15 | 0.19 | 0.2 | 0.44 | 0.07 | 0.2 |
| 7 | 29216460 | rs245912 | -1.358 | 1.085 | -0.743 | 0.898 | -0.003 | 0.133 | 0.133 | 2.479 | 0.47 | 0.81 | 0.62 | 0.79 | 0.49 | 0.51 |
| 7 | 29218159 | rs245914 | 1.124 | 1.166 | -0.817 | 0.961 | -0.002 | -0.002 | -0.002 | 0 | 0.17 | 0.17 | 0.17 | 0.13 | 0.16 | 0.16 |
| 7 | 29225521 | rs6962639 | 0.09 | 1.058 | -0.876 | 0.943 | -0.015 | -0.013 | -0.013 | 0 | 0.06 | 0.05 | 0.17 | 0.22 | 0.15 | 0.1 |
| 7 | 29232999 | rs245923 | 1.273 | 0.772 | -0.69 | 0.817 | -0.017 | -0.004 | -0.004 | 0 | 0.15 | 0.14 | 0.06 | 0.11 | 0.06 | 0.15 |
| 7 | 29233445 | rs39050 | -1.035 | 0.328 | -1.629 | 0.159 | 0.014 | 0.098 | 0.098 | 0 | 0.83 | 0.99 | 0.95 | 0.97 | 0.87 | 0.82 |
| 7 | 29237994 | rs3812398 | 0.016 | 0.478 | -1.227 | 0.738 | -0.005 | -0.002 | -0.002 | 0.444 | 0.15 | 0.17 | 0.18 | 0.28 | 0.07 | 0.18 |
| 7 | 29242220 | rs3812391 | -0.073 | 1.618 | -0.841 | 0.819 | 0.021 | 0.023 | 0.023 | 0 | 0.07 | 0.13 | 0.08 | 0.19 | 0.13 | 0.09 |
| 7 | 29244759 | rs3812389 | -0.124 | 1.863 | -0.872 | 0.786 | -0.027 | -0.024 | -0.024 | 0 | 0.14 | 0.07 | 0.22 | 0.19 | 0.24 | 0.19 |
| 7 | 29247409 | rs2269903 | -0.033 | 1.818 | -0.799 | 0.771 | -0.035 | -0.033 | -0.033 | 0 | 0.06 | 0.01 | 0.06 | 0.19 | 0.13 | 0.09 |
| 7 | 29249930 | rs245928 | -0.511 | 1.024 | -0.238 | 0.376 | 0.011 | -0.001 | -0.001 | 0 | 0.32 | 0.44 | 0.31 | 0.27 | 0.36 | 0.32 |
| 7 | 29250007 | rs17756941 | -0.222 | 1.226 | -0.077 | 0.313 | 0.035 | 0.009 | 0.009 | 0 | 0.15 | 0.19 | 0.22 | 0.15 | 0.28 | 0.15 |
| 7 | 29250335 | rs11981737 | 0.092 | 1.178 | -0.598 | -0.136 | 0.018 | -0.012 | -0.012 | 0.088 | 0.33 | 0.21 | 0.35 | 0.37 | 0.25 | 0.3 |
| 7 | 29251614 | rs39054 | 0.262 | 1.939 | -0.4 | -0.619 | -0.015 | 0.031 | 0.031 | 0.33 | 0.47 | 0.23 | 0.57 | 0.38 | 0.56 | 0.41 |
| 7 | 29252163 | rs13311144 | 0.426 | 2.172 | -0.386 | -0.594 | 0.001 | 0.01 | 0.01 | 0 | 0.06 | 0.04 | 0.15 | 0.02 | 0.19 | 0.03 |
| 7 | 29252965 | rs13235466 | 0.426 | 2.172 | -0.396 | -0.594 | -0.003 | 0.013 | 0.013 | 0 | 0.06 | 0.03 | 0.15 | 0.02 | 0.19 | 0.03 |
| 7 | 29253317 | rs13235885 | 0.426 | 2.172 | -0.396 | -0.594 | -0.003 | 0.013 | 0.013 | 0 | 0.06 | 0.03 | 0.15 | 0.02 | 0.19 | 0.03 |
| 7 | 29255135 | rs245930 | 0.484 | 2.092 | 0.157 | -0.707 | -0.001 | 0.078 | 0.078 | 6.475 | 0.23 | 0.32 | 0.3 | 0.55 | 0.19 | 0.22 |
| 7 | 29256264 | rs39061 | 0.508 | 1.977 | 0.038 | -0.844 | -0.004 | 0.018 | 0.018 | 0.998 | 0.46 | 0.2 | 0.57 | 0.38 | 0.58 | 0.4 |
| 7 | 29256568 | rs39062 | 0.999 | 1.888 | 0.03 | -0.827 | 0.023 | -0.011 | -0.011 | 0 | 0.39 | 0.16 | 0.42 | 0.35 | 0.39 | 0.37 |
| 7 | 29260018 | rs245934 | -0.243 | 1.934 | 0.149 | -0.769 | 0.057 | 0.058 | 0.058 | 7.407 | 0.29 | 0.41 | 0.46 | 0.56 | 0.38 | 0.25 |
| 7 | 29265915 | rs39067 | -0.743 | 1.663 | -0.005 | -0.832 | 0.019 | 0.083 | 0.083 | 0 | 0.72 | 0.85 | 0.91 | 0.93 | 0.77 | 0.63 |
| 7 | 29270958 | rs245945 | 0.662 | 1.717 | 0.058 | -0.54 | -0.003 | 0.077 | 0.077 | 5.611 | 0.22 | 0.3 | 0.3 | 0.54 | 0.18 | 0.21 |
| 7 | 29273986 | rs245952 | -0.35 | 1.771 | 0.004 | -0.572 | 0.048 | 0.05 | 0.05 | 5.688 | 0.29 | 0.4 | 0.46 | 0.56 | 0.38 | 0.25 |
| 7 | 29282337 | rs245960 | -0.354 | 1.657 | -0.162 | -0.608 | 0.095 | -0.057 | -0.057 | 1.152 | 0.11 | 0.22 | 0.2 | 0.02 | 0.22 | 0.05 |
| 7 | 29286334 | rs39092 | -0.627 | 0.93 | -0.845 | -0.67 | 0.006 | -0.006 | -0.006 | 0 | 0.72 | 0.75 | 0.9 | 0.92 | 0.8 | 0.64 |
| 7 | 29290153 | rs245962 | -0.105 | 1.312 | -1.499 | -0.243 | -0.019 | -0.031 | -0.031 | 0 | 0.32 | 0.77 | 0.48 | 0.56 | 0.34 | 0.25 |
| 7 | 29293095 | rs39099 | 0.677 | 1.215 | -1.581 | -0.291 | -0.025 | 0.023 | 0.023 | 0 | 0.33 | 0.07 | 0.34 | 0.25 | 0.34 | 0.25 |
| 7 | 29294462 | rs39101 | -0.126 | 0.623 | -0.688 | -0.816 | -0.012 | -0.022 | -0.022 | 0 | 0.6 | 0.41 | 0.84 | 0.86 | 0.76 | 0.56 |
| 7 | 29295025 | rs3812368 | 0.485 | 1.477 | 0.012 | -0.799 | 0.042 | -0.015 | -0.015 | 0 | 0.17 | 0.31 | 0.21 | 0.1 | 0.25 | 0.2 |
| 7 | 29295336 | rs7796223 | 0.284 | 1.415 | 0.394 | -0.826 | 0.063 | -0.023 | -0.023 | 0 | 0.18 | 0.39 | 0.23 | 0.1 | 0.25 | 0.2 |
| 7 | 29295856 | rs245974 | 0.513 | 1.125 | 0.108 | -1.736 | 0.015 | 0.106 | 0.106 | 0.315 | 0.55 | 0.25 | 0.52 | 0.89 | 0.41 | 0.57 |
| 7 | 29295894 | rs3812367 | 0.664 | 1.25 | -0.11 | -1.375 | 0.063 | -0.012 | -0.012 | 0 | 0.18 | 0.39 | 0.23 | 0.12 | 0.25 | 0.2 |
| 7 | 29300968 | rs245984 | 1.598 | 0.552 | -0.329 | -1.309 | -0.025 | -0.004 | -0.004 | 0 | 0.13 | 0.22 | 0.08 | 0.07 | 0.06 | 0.07 |
| 7 | 29304663 | rs245989 | -0.959 | 0.649 | -0.858 | -1.409 | -0.051 | 0.097 | 0.097 | 0.211 | 0.75 | 0.91 | 0.76 | 1 | 0.67 | 0.73 |
| 7 | 29314083 | rs3793306 | 0.667 | 0.696 | -0.621 | -0.058 | 0.011 | 0.092 | 0.092 | 0 | 0.1 | 0.11 | 0.35 | 0.44 | 0.14 | 0.16 |
| 7 | 29314959 | rs2158410 | 0.291 | -1.104 | -0.611 | -0.16 | 0.006 | 0 | 0 | 0 | 0.37 | 0.57 | 0.42 | 0.41 | 0.42 | 0.36 |
| 7 | 29319074 | rs3812354 | 0.874 | -1.746 | -0.932 | -0.403 | 0.055 | -0.021 | -0.021 | 0 | 0.18 | 0.05 | 0.04 | 0.18 | 0.06 | 0.17 |
| 7 | 29319249 | rs11980456 | -0.087 | -2.201 | -0.742 | -0.854 | 0.021 | -0.073 | -0.073 | 0.348 | 0.23 | 0.36 | 0.28 | 0.04 | 0.32 | 0.26 |
| 7 | 29319266 | rs39109 | 1.471 | -2.048 | -0.73 | 0.129 | -0.017 | -0.01 | -0.01 | 1.555 | 0.15 | 0.12 | 0.24 | 0.22 | 0.26 | 0.14 |
| 7 | 29322257 | rs3812351 | 1.55 | -1.958 | 0.07 | 0.076 | -0.094 | -0.004 | -0.004 | 0 | 0.22 | 0.03 | 0.2 | 0.16 | 0.28 | 0.2 |
| 7 | 29324232 | rs39112 | 0.093 | -2.105 | 0.746 | 0.235 | 0.004 | 0.013 | 0.013 | 0 | 0.33 | 0.62 | 0.44 | 0.52 | 0.4 | 0.25 |
| 7 | 29327526 | rs2023908 | 0.142 | -1.922 | 1.165 | 0.38 | -0.006 | -0.008 | -0.008 | 3.627 | 0.69 | 0.66 | 0.76 | 0.81 | 0.78 | 0.67 |
| 7 | 29331527 | rs2214569 | -0.135 | -1.893 | 1.199 | 0.415 | -0.002 | 0.003 | 0.003 | 0.97 | 0.56 | 0.57 | 0.61 | 0.65 | 0.51 | 0.6 |
| 7 | 29333089 | rs12533816 | -0.155 | -1.748 | 1.299 | 0.44 | -0.003 | 0.013 | 0.013 | 0.291 | 0.53 | 0.57 | 0.61 | 0.66 | 0.51 | 0.6 |
| 7 | 29336969 | rs4722898 | 1.086 | -1.766 | 1.08 | 0.578 | 0.003 | -0.005 | -0.005 | 0 | 0.17 | 0.18 | 0.22 | 0.32 | 0.14 | 0.3 |
| 7 | 29339617 | rs3793301 | 1.184 | -1.582 | 1.215 | 0.485 | 0.028 | 0.029 | 0.029 | 0 | 0.25 | 0.41 | 0.3 | 0.32 | 0.33 | 0.17 |
| 7 | 29339788 | rs3793300 | 1.494 | -1.274 | 0.405 | 0.183 | -0.065 | 0.01 | 0.01 | 0 | 0.21 | 0.05 | 0.23 | 0.18 | 0.31 | 0.2 |
| 7 | 29343177 | rs3812338 | 1.102 | -1.296 | 0.411 | 0.178 | -0.067 | 0.022 | 0.022 | 0 | 0.23 | 0.06 | 0.26 | 0.18 | 0.33 | 0.21 |
| 7 | 29345291 | rs11771116 | 1.203 | -1.662 | 0.847 | 0.36 | 0.005 | 0.011 | 0.011 | 0 | 0.24 | 0.29 | 0.3 | 0.36 | 0.32 | 0.16 |
| 7 | 29345604 | rs39754 | 1.215 | -1.848 | 1.016 | 0.335 | -0.003 | 0.004 | 0.004 | 0 | 0.18 | 0.23 | 0.21 | 0.28 | 0.15 | 0.32 |
| 7 | 29351988 | rs3793297 | 1.664 | -1.87 | 0.951 | 0.262 | -0.089 | 0.01 | 0.01 | 0 | 0.22 | 0.07 | 0.25 | 0.17 | 0.33 | 0.23 |
| 7 | 29353445 | rs3793295 | 1.458 | -1.894 | 1.254 | 0.252 | -0.017 | 0.008 | 0.008 | 0 | 0.23 | 0.17 | 0.26 | 0.17 | 0.33 | 0.22 |
| 7 | 29355811 | rs1004056 | -0.742 | -2.084 | 1.187 | 0.141 | 0.043 | 0.086 | 0.086 | 0 | 0.78 | 0.93 | 0.93 | 0.99 | 0.89 | 0.8 |
| 7 | 29356150 | rs1003536 | 1.613 | -2.087 | 1.216 | 0.204 | -0.021 | -0.032 | -0.032 | 0.587 | 0.22 | 0.15 | 0.29 | 0.36 | 0.32 | 0.16 |
| 7 | 29356403 | rs1016726 | 0.366 | -2.043 | 1.037 | 0.247 | -0.024 | 0.022 | 0.022 | 0 | 0.13 | 0.18 | 0.15 | 0.18 | 0.08 | 0.1 |
| 7 | 29358778 | rs40451 | 0.225 | -1.938 | 0.302 | 0.263 | 0.022 | -0.029 | -0.029 | 3.827 | 0.28 | 0.23 | 0.33 | 0.46 | 0.21 | 0.41 |
| 7 | 29359027 | rs39122 | 0.067 | -1.933 | 0.301 | 0.265 | 0 | -0.002 | -0.002 | 0.123 | 0.3 | 0.29 | 0.36 | 0.46 | 0.23 | 0.42 |
| 7 | 29365366 | rs3828985 | 0.347 | -1.832 | 0.113 | 0.268 | -0.017 | -0.003 | -0.003 | 0 | 0.44 | 0.3 | 0.48 | 0.45 | 0.51 | 0.55 |
| 7 | 29366116 | rs39128 | -1.777 | -1.21 | -0.033 | 0.291 | -0.01 | 0.049 | 0.049 | 0 | 0.91 | 1 | 0.99 | 1 | 0.94 | 0.89 |
| 7 | 29370447 | rs39129 | 0.017 | -0.933 | 0.404 | 0.552 | -0.003 | 0.013 | 0.013 | 0 | 0.48 | 0.58 | 0.57 | 0.67 | 0.53 | 0.49 |
| 7 | 29382710 | rs3793283 | 1.153 | -0.769 | -0.019 | 0.705 | -0.064 | -0.098 | -0.098 | 0.366 | 0.26 | 0.07 | 0.3 | 0.39 | 0.35 | 0.18 |

| | | | | | | | | | | | | | | | | |
|---|---|---|---|---|---|---|---|---|---|---|---|---|---|---|---|---|
| 7 | 29384157 | rs7801318 | 0.238 | -1.145 | 0.397 | 0.562 | -0.008 | 0.055 | 0.055 | 0 | 0.46 | 0.35 | 0.34 | 0.17 | 0.45 | 0.5 |
| 7 | 29384554 | rs6973226 | -1.031 | -1.175 | 0.313 | 0.419 | -0.003 | -0.048 | -0.048 | 0.293 | 0.77 | 0.79 | 0.89 | 1 | 0.86 | 0.78 |
| 7 | 29387383 | rs7789918 | 1.145 | -1.028 | 0.282 | 0.552 | -0.07 | 0.037 | 0.037 | 0 | 0.23 | 0.09 | 0.22 | 0.15 | 0.31 | 0.18 |
| 7 | 29388867 | rs6966495 | -0.081 | -1.028 | 0.38 | 0.599 | -0.032 | 0.041 | 0.041 | 0 | 0.28 | 0.58 | 0.33 | 0.44 | 0.19 | 0.4 |
| 7 | 29391611 | rs10253183 | 0.579 | -0.786 | 0.725 | 0.556 | 0.043 | -0.039 | -0.039 | 0.029 | 0.27 | 0.37 | 0.29 | 0.17 | 0.35 | 0.29 |
| 7 | 29397321 | rs6943806 | -0.423 | -0.781 | 0.544 | 0.622 | 0.014 | 0.106 | 0.106 | 0.004 | 0.51 | 0.75 | 0.63 | 0.72 | 0.54 | 0.61 |
| 7 | 29405979 | rs10244747 | -0.613 | -0.831 | 0.332 | 0.653 | 0.013 | 0.088 | 0.088 | 0 | 0.54 | 0.76 | 0.65 | 0.73 | 0.55 | 0.6 |
| 7 | 29407488 | rs2301776 | 1.409 | -0.848 | -0.13 | 0.893 | -0.018 | 0.03 | 0.03 | 0.019 | 0.07 | 0.08 | 0.1 | 0.16 | 0.02 | 0.07 |
| 7 | 29409596 | rs3793276 | 1.327 | -0.901 | -0.227 | 0.756 | -0.036 | -0.023 | -0.023 | 0 | 0.25 | 0.07 | 0.26 | 0.27 | 0.31 | 0.18 |
| 7 | 29410168 | rs7810666 | 0.234 | -0.084 | 0.459 | 1.62 | -0.007 | 0.01 | 0.01 | 0 | 0.74 | 0.7 | 0.68 | 0.58 | 0.81 | 0.69 |
| 7 | 29413388 | rs12700958 | -0.645 | 0.708 | 0.229 | 0.697 | -0.016 | -0.011 | -0.011 | 0 | 0.42 | 0.33 | 0.48 | 0.54 | 0.49 | 0.52 |
| 7 | 29413879 | rs7794652 | 0.758 | 0.821 | 0.037 | 0.763 | 0.001 | 0 | 0 | 0 | 0.56 | 0.43 | 0.49 | 0.46 | 0.51 | 0.49 |
| 7 | 29416396 | rs17157833 | -0.575 | 1.534 | -0.293 | 0.196 | 0.051 | -0.095 | -0.095 | 1.034 | 0.28 | 0.44 | 0.3 | 0.12 | 0.35 | 0.23 |
| 7 | 29421786 | rs12673506 | -0.799 | 1.501 | -0.728 | 1.403 | -0.014 | 0.007 | 0.007 | 0 | 0.1 | 0.06 | 0.28 | 0.47 | 0.22 | 0.25 |
| 7 | 29423773 | rs4719970 | -1.348 | 2.264 | -0.193 | 0.65 | 0.038 | 0.016 | 0.016 | 0 | 0.34 | 0.48 | 0.48 | 0.57 | 0.51 | 0.5 |
| 7 | 29427735 | rs3793272 | -0.058 | 1.618 | -0.364 | -0.256 | 0.004 | 0.007 | 0.007 | 0 | 0.27 | 0.25 | 0.14 | 0.1 | 0.24 | 0.25 |
| 7 | 29428832 | rs10253637 | 0.448 | 0.742 | -0.852 | -0.906 | 0.018 | 0.033 | 0.033 | 0.398 | 0.74 | 0.18 | 0.69 | 0.84 | 0.83 | 0.71 |
| 7 | 29432479 | rs4719971 | 0.696 | 0.312 | 0.118 | -1.142 | -0.038 | -0.073 | -0.073 | 1.032 | 0.75 | 0.47 | 0.76 | 0.88 | 0.85 | 0.79 |
| 7 | 29432885 | rs10256495 | 0.516 | 0.49 | 0.991 | -1.241 | -0.026 | -0.055 | -0.055 | 0.769 | 0.26 | 0.53 | 0.22 | 0.13 | 0.19 | 0.38 |
| 7 | 29434444 | rs1034716 | -0.501 | 0.075 | 0.631 | -1.59 | 0.014 | 0.016 | 0.016 | 0 | 0.43 | 0.67 | 0.34 | 0.3 | 0.34 | 0.54 |
| 7 | 29436519 | rs6979199 | -0.329 | 0.142 | 0.666 | -1.586 | 0.011 | -0.053 | -0.053 | 3.441 | 0.14 | 0.47 | 0.14 | 0.05 | 0.11 | 0.19 |
| 7 | 29436971 | rs1034718 | -1.14 | 0.194 | 0.49 | -1.576 | -0.018 | -0.014 | -0.014 | 0 | 0.88 | 0.8 | 0.84 | 0.89 | 0.93 | 0.9 |
| 7 | 29437136 | rs1034719 | 1.892 | 0.734 | -0.248 | -1.643 | -0.112 | 0.141 | 0.141 | 0 | 0.3 | 0.01 | 0.17 | 0.09 | 0.32 | 0.12 |
| 7 | 29437196 | rs1034720 | 0.983 | 0.074 | 0.031 | -1.386 | -0.058 | -0.02 | -0.02 | 0 | 0.43 | 0.08 | 0.47 | 0.57 | 0.57 | 0.37 |
| 7 | 29438109 | rs12112301 | 0.785 | 0.131 | 0.191 | -1.267 | 0 | -0.001 | -0.001 | 0.241 | 0.11 | 0.13 | 0.1 | 0.17 | 0.08 | 0.12 |
| 7 | 29447041 | rs7800059 | 1.205 | 0.166 | 0.329 | -1.291 | 0 | -0.015 | -0.015 | 0 | 0.07 | 0.06 | 0.04 | 0.01 | 0.08 | 0.03 |
| 7 | 29447142 | rs2189439 | -1.033 | 0.339 | 0.427 | -1.415 | -0.027 | 0.017 | 0.017 | 0 | 0.57 | 0.78 | 0.55 | 0.4 | 0.48 | 0.64 |
| 7 | 29447298 | rs12536100 | 0.479 | 0.303 | 0.351 | -1.422 | -0.009 | -0.032 | -0.032 | 0 | 0.14 | 0.34 | 0.19 | 0.1 | 0.08 | 0.1 |
| 7 | 29448420 | rs17157914 | 0.633 | 0.385 | -0.729 | -1.33 | 0.002 | -0.004 | -0.004 | 1.061 | 0.12 | 0.11 | 0.14 | 0.16 | 0.13 | 0.13 |
| 7 | 29448660 | rs6959646 | -1.168 | 0.362 | -0.813 | -1.324 | 0.019 | 0.057 | 0.057 | 0 | 0.39 | 0.81 | 0.47 | 0.31 | 0.32 | 0.42 |
| 7 | 29450895 | rs11981975 | 1.54 | 0.499 | -0.826 | -1.053 | 0.012 | 0.028 | 0.028 | 0 | 0.18 | 0.04 | 0.1 | 0.09 | 0.16 | 0.22 |
| 7 | 29454101 | rs2228688 | 0.27 | 0.609 | -0.694 | -0.979 | 0.007 | 0.029 | 0.029 | 0 | 0.33 | 0.16 | 0.26 | 0.26 | 0.31 | 0.35 |
| 7 | 29456005 | rs2057739 | 0.445 | 0.304 | -0.276 | -0.804 | 0.004 | -0.014 | -0.014 | 0.021 | 0.72 | 0.45 | 0.59 | 0.75 | 0.72 | 0.71 |
| 7 | 29456491 | rs12534213 | 0.953 | 0.123 | -0.912 | -0.575 | 0.047 | 0.073 | 0.073 | 0.565 | 0.26 | 0.12 | 0.1 | 0.1 | 0.21 | 0.11 |
| 7 | 29461916 | rs7777560 | -0.717 | 0.456 | -0.538 | -0.267 | 0 | -0.001 | -0.001 | 0 | 0.92 | 0.87 | 0.77 | 0.93 | 0.93 | 0.92 |
| 7 | 29462681 | rs12700961 | 1.133 | 1.126 | -1.491 | 0.241 | -0.084 | -0.085 | -0.085 | 0.002 | 0.15 | 0.01 | 0.25 | 0.29 | 0.26 | 0.26 |
| 7 | 29463193 | rs10486614 | 0.927 | 1.225 | -1.527 | 0.21 | -0.095 | -0.087 | -0.087 | 0 | 0.16 | 0.01 | 0.28 | 0.29 | 0.28 | 0.26 |
| 7 | 29464549 | rs9648362 | -0.393 | 0.502 | -1.763 | 0.278 | 0.017 | -0.032 | -0.032 | 0 | 0.71 | 0.8 | 0.65 | 0.57 | 0.79 | 0.63 |
| 7 | 29465310 | rs7793452 | -0.885 | 0.278 | -1.387 | 0.283 | -0.01 | -0.002 | -0.002 | 0 | 0.78 | 0.7 | 0.69 | 0.81 | 0.84 | 0.7 |
| 7 | 29467025 | rs6980280 | -0.389 | 0.287 | -0.957 | 0.209 | -0.011 | -0.098 | -0.098 | 0 | 0.24 | 0.52 | 0.2 | 0.07 | 0.23 | 0.12 |
| 7 | 29467634 | rs10254932 | -0.258 | 0.299 | -1.055 | 0.214 | -0.008 | 0.018 | 0.018 | 0 | 0.43 | 0.39 | 0.53 | 0.75 | 0.45 | 0.58 |
| 7 | 29468114 | rs7800248 | -0.07 | -0.186 | -0.319 | 0.113 | -0.006 | -0.117 | -0.117 | 0.373 | 0.2 | 0.33 | 0.14 | 0.01 | 0.2 | 0.09 |
| 7 | 29468698 | rs7785829 | -2.195 | -0.123 | -0.394 | 0.18 | -0.062 | 0.046 | 0.046 | 0 | 0.68 | 0.93 | 0.73 | 0.81 | 0.66 | 0.72 |
| 7 | 29469833 | rs11976491 | 0.531 | -0.758 | 0.187 | 0.232 | 0.003 | 0.025 | 0.025 | 0 | 0.06 | 0.2 | 0.07 | 0.05 | 0.02 | 0.04 |
| 7 | 29470739 | rs10281343 | 0.121 | -0.752 | 0.006 | 0.305 | -0.026 | 0.038 | 0.038 | 0 | 0.26 | 0.33 | 0.24 | 0.44 | 0.16 | 0.32 |
| 7 | 29470861 | rs10281477 | -0.691 | -0.772 | -0.026 | 0.29 | -0.002 | 0.002 | 0.002 | 0 | 0.38 | 0.41 | 0.35 | 0.46 | 0.36 | 0.39 |
| 7 | 29481940 | rs3828983 | 1.048 | -0.593 | -0.436 | 0.411 | 0.003 | -0.044 | -0.044 | 0 | 0.08 | 0.01 | 0.07 | 0.27 | 0.07 | 0.09 |
| 7 | 29482956 | rs4722905 | -1.439 | -0.765 | -0.358 | 0.284 | 0.06 | -0.033 | -0.033 | 0.256 | 0.51 | 0.87 | 0.48 | 0.5 | 0.39 | 0.46 |
| 7 | 29483759 | rs849910 | -0.195 | -0.714 | -0.09 | 0.154 | 0 | -0.011 | -0.011 | 0 | 0.16 | 0.18 | 0.12 | 0.01 | 0.19 | 0.08 |
| 7 | 29484027 | rs849911 | 0.899 | -0.733 | -0.096 | 0.161 | -0.029 | 0.028 | 0.028 | 0 | 0.11 | 0.07 | 0.1 | 0.01 | 0.2 | 0.07 |
| 7 | 29485257 | rs849914 | -0.604 | -0.846 | 0.008 | 0.184 | 0.01 | 0.02 | 0.02 | 0 | 0.35 | 0.71 | 0.37 | 0.23 | 0.31 | 0.34 |
| 7 | 29486638 | rs849916 | -1.978 | -0.699 | 0.166 | 0.235 | -0.006 | -0.009 | -0.009 | 1.349 | 0.33 | 0.6 | 0.3 | 0.32 | 0.31 | 0.24 |
| 7 | 29487391 | rs3909208 | -1.425 | -0.758 | 0.12 | 0.29 | -0.067 | 0.046 | 0.046 | 0 | 0.23 | 0.5 | 0.19 | 0.32 | 0.11 | 0.16 |
| 7 | 29491403 | rs702827 | -0.382 | -0.502 | -0.206 | 0.335 | -0.016 | -0.016 | -0.016 | 0.664 | 0.2 | 0.1 | 0.16 | 0.26 | 0.26 | 0.16 |
| 7 | 29493196 | rs3793263 | 1.39 | -0.835 | 0.109 | 0.804 | -0.007 | 0.003 | 0.003 | 0 | 0.15 | 0.18 | 0.16 | 0.18 | 0.09 | 0.22 |
| 7 | 29496163 | rs6462140 | -0.788 | -0.393 | 0.214 | 0.681 | -0.034 | 0.021 | 0.021 | 0 | 0.17 | 0.28 | 0.13 | 0.26 | 0.1 | 0.12 |
| 7 | 29498356 | rs3793260 | -0.322 | 0.235 | 0.075 | 0.47 | 0.021 | 0.016 | 0.016 | 0 | 0.06 | 0.17 | 0.08 | 0.05 | 0.04 | 0.06 |
| 7 | 29499866 | rs849935 | -0.796 | 0.164 | -0.843 | 0.086 | -0.005 | 0.074 | 0.074 | 0 | 0.71 | 0.9 | 0.74 | 0.85 | 0.73 | 0.78 |
| 7 | 29500126 | rs10951224 | 0.489 | 0.155 | -0.546 | 0.469 | 0.003 | 0.004 | 0.004 | 0 | 0.3 | 0.33 | 0.3 | 0.36 | 0.34 | 0.35 |
| 7 | 29500502 | rs3793258 | 1.66 | 0.011 | -0.774 | 0.663 | 0.005 | -0.025 | -0.025 | 0 | 0.14 | 0.08 | 0.14 | 0.18 | 0.12 | 0.21 |
| 7 | 29503078 | rs10499582 | 1.302 | 0.222 | -0.743 | 0.453 | -0.011 | -0.022 | -0.022 | 5.308 | 0.23 | 0.15 | 0.26 | 0.35 | 0.29 | 0.33 |
| 7 | 29503748 | rs6462141 | 1.246 | -0.164 | -0.889 | 0.422 | -0.031 | 0.04 | 0.04 | 0 | 0.36 | 0.09 | 0.38 | 0.25 | 0.35 | 0.28 |
| 7 | 29504650 | rs10951225 | 0.651 | -0.035 | -0.275 | 0.318 | 0.025 | 0.021 | 0.021 | 0 | 0.31 | 0.47 | 0.34 | 0.36 | 0.36 | 0.36 |
| 7 | 29505181 | rs11763818 | 0.651 | -0.035 | -0.273 | 0.318 | 0.026 | 0.023 | 0.023 | 0 | 0.31 | 0.47 | 0.34 | 0.36 | 0.36 | 0.36 |
| 7 | 29506756 | rs1362360 | 1.829 | -0.111 | -0.847 | 0.364 | -0.003 | -0.014 | -0.014 | 0 | 0.13 | 0.03 | 0.09 | 0.11 | 0.12 | 0.21 |
| 7 | 29508944 | rs10272160 | -0.652 | -0.166 | -0.868 | 0.416 | 0.026 | 0.021 | 0.021 | 0.303 | 0.4 | 0.72 | 0.26 | 0.34 | 0.34 | 0.38 |
| 7 | 29509169 | rs10258578 | -1.128 | 0.117 | -1.115 | 0.317 | -0.014 | -0.003 | -0.003 | 0 | 0.67 | 0.84 | 0.55 | 0.69 | 0.69 | 0.73 |
| 7 | 29509414 | rs10261846 | -0.26 | 0.195 | -0.842 | 0.342 | 0.01 | 0.016 | 0.016 | 0 | 0.52 | 0.72 | 0.44 | 0.59 | 0.57 | 0.51 |
| 7 | 29511652 | rs1362362 | -0.232 | 0.333 | -0.614 | 0.335 | -0.009 | 0.01 | 0.01 | 0 | 0.56 | 0.65 | 0.62 | 0.39 | 0.52 | 0.39 |

| | | | | | | | | | | | | | | | | |
|---|---|---|---|---|---|---|---|---|---|---|---|---|---|---|---|---|
| 7 | 29511791 | rs10255035 | 0.122 | 0.072 | -0.272 | -0.06 | -0.007 | -0.035 | -0.035 | 0.443 | 0.39 | 0.48 | 0.41 | 0.27 | 0.37 | 0.23 |
| 7 | 29512086 | rs1362364 | 0.23 | -0.01 | -0.453 | -0.054 | -0.001 | 0.011 | 0.011 | 0.373 | 0.63 | 0.61 | 0.63 | 0.77 | 0.65 | 0.68 |
| 7 | 29512612 | rs17158067 | 0.153 | -0.044 | -0.529 | 0.401 | 0 | 0.002 | 0.002 | 0 | 0.08 | 0.06 | 0.12 | 0.25 | 0.05 | 0.09 |
| 7 | 29513178 | rs12531664 | -0.153 | -0.185 | -0.186 | 0.167 | 0.001 | -0.002 | -0.002 | 0 | 0.72 | 0.67 | 0.6 | 0.76 | 0.67 | 0.69 |
| 7 | 29513395 | rs7792461 | 0.591 | -0.186 | -0.196 | 0.167 | -0.001 | -0.001 | -0.001 | 0 | 0.63 | 0.61 | 0.48 | 0.52 | 0.62 | 0.61 |
| 7 | 29514059 | rs722611 | -1.281 | -0.059 | -0.641 | 0.084 | -0.082 | -0.064 | -0.064 | 0 | 0.82 | 0.99 | 0.76 | 0.78 | 0.79 | 0.78 |
| 7 | 29517436 | rs3812328 | 1.078 | -0.197 | -0.949 | -0.033 | -0.002 | -0.039 | -0.039 | 0 | 0.39 | 0.12 | 0.31 | 0.59 | 0.37 | 0.47 |
| 7 | 29517909 | rs3812325 | 0.158 | -0.298 | -0.16 | -0.118 | -0.005 | -0.02 | -0.02 | 0 | 0.31 | 0.38 | 0.27 | 0.18 | 0.29 | 0.21 |
| 7 | 29518458 | rs3793252 | -0.49 | -0.167 | -0.476 | 0.074 | 0.006 | -0.003 | -0.003 | 0 | 0.17 | 0.14 | 0.26 | 0.25 | 0.17 | 0.16 |
| 7 | 29518841 | rs17752156 | -0.166 | -0.142 | -0.547 | 0.087 | 0.002 | -0.006 | -0.006 | 0.015 | 0.15 | 0.11 | 0.25 | 0.28 | 0.15 | 0.18 |
| 7 | 29519929 | rs3750103 | 0.312 | -0.602 | -0.492 | 0.021 | -0.011 | 0.004 | 0.004 | 0 | 0.1 | 0.14 | 0.12 | 0.22 | 0.05 | 0.09 |
| 7 | 29520042 | rs1476991 | 1.618 | -0.722 | -0.723 | -0.212 | -0.04 | -0.045 | -0.045 | 0 | 0.36 | 0.16 | 0.29 | 0.38 | 0.42 | 0.46 |
| 7 | 29521228 | rs1420132 | -0.52 | -0.385 | -0.598 | -0.345 | 0.024 | 0.02 | 0.02 | 0 | 0.48 | 0.65 | 0.46 | 0.4 | 0.44 | 0.43 |
| 7 | 29524353 | rs2024421 | 0.574 | -0.398 | -0.518 | -0.417 | 0.011 | -0.007 | -0.007 | 0 | 0.63 | 0.53 | 0.53 | 0.63 | 0.62 | 0.68 |
| 7 | 29525013 | rs2270311 | 2.081 | -0.068 | -1.384 | -0.234 | -0.024 | -0.052 | -0.052 | 0.206 | 0.29 | 0.12 | 0.2 | 0.36 | 0.32 | 0.4 |
| 7 | 29525267 | rs2270312 | 0.125 | 0.069 | -1.341 | -0.516 | -0.032 | 0.026 | 0.026 | 0.072 | 0.66 | 0.82 | 0.58 | 0.72 | 0.63 | 0.68 |
| 7 | 29527157 | rs1946133 | 0.49 | -0.445 | -1.708 | -0.406 | -0.015 | -0.026 | -0.026 | 0 | 0.27 | 0.07 | 0.28 | 0.27 | 0.27 | 0.27 |
| 7 | 29529073 | rs1420133 | 0.169 | -0.289 | -1.653 | -0.482 | 0.002 | 0.02 | 0.02 | 0 | 0.32 | 0.17 | 0.31 | 0.29 | 0.35 | 0.29 |
| 7 | 29530538 | rs1345305 | -1.956 | -0.291 | -1.989 | -0.295 | -0.064 | -0.154 | -0.154 | 0.909 | 0.71 | 0.99 | 0.85 | 0.61 | 0.71 | 0.63 |
| 7 | 29532418 | rs1420135 | 0.61 | -0.257 | -1.707 | -0.304 | 0.005 | 0.01 | 0.01 | 0 | 0.3 | 0.2 | 0.3 | 0.29 | 0.3 | 0.28 |
| 7 | 29535325 | rs1990593 | 1.174 | -0.167 | -1.566 | -0.324 | -0.003 | -0.002 | -0.002 | 0 | 0.11 | 0.12 | 0.22 | 0.01 | 0.19 | 0.11 |
| 7 | 29535998 | rs10951227 | -1.257 | -0.363 | -1.638 | -0.407 | -0.014 | -0.032 | -0.032 | 0 | 0.62 | 0.81 | 0.74 | 0.61 | 0.6 | 0.58 |
| 7 | 29536564 | rs10233803 | -0.458 | -0.535 | -1.474 | -0.461 | 0.029 | 0.056 | 0.056 | 0 | 0.51 | 0.41 | 0.58 | 0.38 | 0.55 | 0.47 |
| 7 | 29536707 | rs10085463 | 0.754 | -0.487 | -1.602 | -0.443 | -0.001 | -0.01 | -0.01 | 0 | 0.19 | 0.26 | 0.17 | 0.08 | 0.17 | 0.14 |
| 7 | 29538328 | rs10254629 | 1.152 | -0.481 | -1.444 | -0.183 | -0.044 | 0.041 | 0.041 | 0 | 0.11 | 0.1 | 0.22 | 0.01 | 0.2 | 0.11 |
| 7 | 29538924 | rs3793251 | 1.449 | -0.5 | -1.456 | -0.111 | 0.012 | -0.023 | -0.023 | 0 | 0.21 | 0.11 | 0.19 | 0.28 | 0.18 | 0.23 |
| 7 | 29540205 | rs10262722 | -0.785 | -0.329 | -1.284 | 0.028 | 0.014 | 0.008 | 0.008 | 0.129 | 0.23 | 0.53 | 0.39 | 0.25 | 0.25 | 0.22 |
| 7 | 29545501 | rs1420138 | -0.823 | -0.512 | -1.381 | 0.396 | -0.047 | -0.031 | -0.031 | 1.085 | 0.58 | 0.8 | 0.63 | 0.63 | 0.51 | 0.52 |
| 7 | 29547172 | rs11770859 | -0.649 | -0.395 | -0.932 | 0.194 | 0.01 | 0.009 | 0.009 | 0 | 0.37 | 0.61 | 0.43 | 0.37 | 0.33 | 0.3 |
| 7 | 29548324 | rs1420143 | 0.23 | -0.351 | -0.733 | -0.035 | -0.009 | 0.033 | 0.033 | 0 | 0.44 | 0.59 | 0.5 | 0.22 | 0.44 | 0.38 |
| 7 | 29550829 | rs7782730 | -0.716 | -0.316 | -0.564 | -0.041 | 0.004 | -0.004 | -0.004 | 0.586 | 0.93 | 0.96 | 0.96 | 1 | 0.97 | 0.97 |
| 7 | 29551124 | rs1055010 | -0.631 | -0.845 | -0.389 | 0.149 | -0.002 | -0.004 | -0.004 | 0 | 0.28 | 0.23 | 0.36 | 0.34 | 0.27 | 0.33 |
| 7 | 29552213 | rs3750099 | -0.38 | -1.221 | 0.458 | 0.472 | -0.023 | 0.338 | 0.338 | 0 | 0.09 | 0.48 | 0.15 | 0.31 | 0.02 | 0.15 |
| 7 | 29552671 | rs1059182 | 0.989 | -0.868 | 0.316 | 0.282 | -0.009 | 0.024 | 0.024 | 1.323 | 0.1 | 0.04 | 0.15 | 0 | 0.09 | 0.04 |
| 7 | 29552777 | rs12611 | 0.216 | -0.393 | 0.299 | 0.403 | 0.006 | 0.046 | 0.046 | 0 | 0.3 | 0.54 | 0.38 | 0.42 | 0.27 | 0.38 |
| 7 | 29552983 | rs1059185 | -1.042 | -0.084 | 1.051 | 0.314 | 0.021 | 0.022 | 0.022 | 0.249 | 0.25 | 0.51 | 0.34 | 0.24 | 0.27 | 0.22 |
| 7 | 29553339 | rs1558634 | -0.05 | -0.213 | 0.23 | 0.005 | 0.004 | -0.013 | -0.013 | 0 | 0.83 | 0.85 | 0.8 | 1 | 0.87 | 0.93 |
| 7 | 29557720 | rs13243100 | -0.267 | 0.197 | 0.237 | 0.107 | 0.082 | -0.005 | -0.005 | 0 | 0.17 | 0.18 | 0.29 | 0.14 | 0.29 | 0.31 |
| 7 | 29557754 | rs17809999 | 0.885 | 0.084 | -0.121 | 0.076 | -0.001 | -0.009 | -0.009 | 0 | 0.11 | 0.1 | 0.16 | 0 | 0.1 | 0.04 |
| 7 | 29577572 | rs4722917 | 0.995 | -0.207 | 0.024 | 0.24 | 0.003 | -0.001 | -0.001 | 0.153 | 0.45 | 0.48 | 0.38 | 0.54 | 0.45 | 0.42 |
| 7 | 29594554 | rs11765707 | 0.415 | -0.212 | 0.034 | 0.194 | -0.001 | -0.001 | -0.001 | 0.003 | 0.54 | 0.52 | 0.55 | 0.54 | 0.54 | 0.47 |
| 7 | 29594774 | rs1362352 | -0.254 | 0.366 | 0.243 | 0.181 | 0.088 | 0.006 | 0.006 | 0 | 0.14 | 0.18 | 0.27 | 0.15 | 0.27 | 0.26 |
| 7 | 29597497 | rs10237982 | 1.451 | -0.058 | 0.132 | 0.286 | 0.005 | 0.004 | 0.004 | 0 | 0.62 | 0.52 | 0.42 | 0.57 | 0.6 | 0.52 |
| 7 | 29601142 | rs7794697 | 0.446 | -0.224 | -0.055 | 0.156 | -0.011 | 0.001 | 0.001 | 0 | 0.32 | 0.36 | 0.22 | 0.65 | 0.27 | 0.34 |
| 7 | 29603850 | rs17754040 | 1.882 | -0.011 | -0.126 | 0.099 | 0 | 0.068 | 0.068 | 0 | 0.32 | 0.22 | 0.25 | 0.03 | 0.35 | 0.19 |
| 7 | 29605156 | rs1362355 | 1.14 | 0.584 | 0.1 | 0.688 | 0.034 | 0.06 | 0.06 | 0 | 0.74 | 0.7 | 0.61 | 0.56 | 0.69 | 0.58 |
| 7 | 29605481 | rs1420112 | -0.114 | 0.249 | 0.199 | 0.625 | -0.053 | 0.127 | 0.127 | 0 | 0.87 | 0.75 | 0.86 | 0.73 | 0.96 | 0.84 |
| 7 | 29606425 | rs2302619 | 0.517 | 1.178 | -0.335 | 0.47 | -0.009 | 0.013 | 0.013 | 0 | 0.05 | 0.01 | 0.15 | 0.02 | 0.12 | 0.12 |
| 7 | 29612042 | rs4722933 | 0.947 | 0.457 | -0.281 | 0.581 | -0.001 | 0.003 | 0.003 | 0.699 | 0.64 | 0.39 | 0.45 | 0.49 | 0.7 | 0.58 |
| 7 | 29612336 | rs1344478 | -0.519 | 0.268 | 0.079 | 0.696 | 0.005 | -0.001 | -0.001 | 0 | 0.22 | 0.23 | 0.35 | 0.24 | 0.26 | 0.29 |
| 7 | 29613298 | rs17810805 | 0.647 | 0.463 | -0.142 | 0.663 | -0.039 | -0.028 | -0.028 | 0.451 | 0.09 | 0.02 | 0.07 | 0.13 | 0.16 | 0.16 |
| 7 | 29614278 | rs6976604 | 0.573 | 0.386 | 0.07 | 0.705 | -0.012 | 0.054 | 0.054 | 0.032 | 0.1 | 0.2 | 0.05 | 0.22 | 0.07 | 0.09 |
| 7 | 29614832 | rs1362356 | 0.83 | 0.4 | 0.08 | 0.625 | 0.012 | 0.024 | 0.024 | 0.238 | 0.73 | 0.57 | 0.5 | 0.71 | 0.77 | 0.67 |
| 7 | 29618664 | rs2098346 | 1.842 | 0.052 | -0.279 | 0.217 | 0.009 | 0.008 | 0.008 | 0 | 0.39 | 0.31 | 0.26 | 0.03 | 0.37 | 0.21 |
| 7 | 29628434 | rs1420116 | -0.997 | 0.046 | -0.168 | 0.546 | -0.026 | 0.07 | 0.07 | 0 | 0.91 | 0.8 | 0.93 | 0.88 | 0.97 | 0.95 |
| 7 | 29631511 | rs12667343 | -0.172 | 0.128 | -0.186 | 0.373 | 0.004 | 0.03 | 0.03 | 0 | 0.82 | 0.78 | 0.85 | 0.76 | 0.79 | 0.79 |
| 7 | 29632309 | rs1420117 | -0.396 | -0.018 | -0.377 | 0.544 | -0.014 | 0.026 | 0.026 | 0 | 0.3 | 0.39 | 0.19 | 0.55 | 0.24 | 0.3 |
| 7 | 29633340 | rs10499586 | 0.365 | 0.233 | -0.956 | 0.181 | 0.014 | -0.032 | -0.032 | 0.044 | 0.2 | 0.16 | 0.14 | 0.33 | 0.17 | 0.22 |
| 7 | 29635526 | rs10499588 | 0.776 | 0.66 | -1.275 | -0.239 | 0.014 | 0.038 | 0.038 | 0 | 0.34 | 0.24 | 0.22 | 0.03 | 0.27 | 0.2 |
| 7 | 29636018 | rs11766467 | 0.23 | 1.272 | -1.118 | 0.551 | -0.02 | -0.08 | -0.08 | 0 | 0.18 | 0.08 | 0.25 | 0.5 | 0.25 | 0.33 |
| 7 | 29641278 | rs6942633 | -0.489 | 1.397 | -0.251 | 0.097 | 0.037 | 0.034 | 0.034 | 0.468 | 0.23 | 0.44 | 0.31 | 0.62 | 0.27 | 0.29 |
| 7 | 29641468 | rs10253443 | 0.603 | 1.04 | -0.87 | -0.777 | 0.002 | 0.058 | 0.058 | 0 | 0.24 | 0.11 | 0.13 | 0 | 0.21 | 0.1 |
| 7 | 29643665 | rs823364 | -0.969 | 0.749 | -1.393 | -1.048 | -0.039 | -0.019 | -0.019 | 4.084 | 0.72 | 0.98 | 0.7 | 0.7 | 0.68 | 0.58 |
| 7 | 29647662 | rs1420123 | -0.715 | 0.708 | -0.412 | -1.108 | 0.001 | 0.004 | 0.004 | 0 | 0.68 | 0.75 | 0.67 | 0.7 | 0.65 | 0.54 |
| 7 | 29648077 | rs17158276 | 0.808 | 0.587 | -0.007 | -1.236 | 0 | -0.02 | -0.02 | 0 | 0.24 | 0.26 | 0.22 | 0.03 | 0.16 | 0.14 |
| 7 | 29652263 | rs2080301 | 0.18 | 0.535 | 0.226 | -1.196 | -0.007 | -0.06 | -0.06 | 0.038 | 0.47 | 0.51 | 0.36 | 0.06 | 0.35 | 0.24 |
| 7 | 29659729 | rs1641192 | -1.052 | 0.412 | -0.061 | -1.128 | -0.04 | -0.024 | -0.024 | 0 | 0.71 | 0.94 | 0.71 | 0.68 | 0.68 | 0.55 |
| 7 | 29661486 | rs1641215 | -0.223 | 0.658 | 0.068 | -1.491 | 0.017 | 0.251 | 0.251 | 0 | 0.5 | 0.82 | 0.45 | 0.12 | 0.43 | 0.26 |
| 7 | 29663018 | rs17158286 | 0.071 | 0.661 | 0.319 | -1.511 | -0.003 | -0.044 | -0.044 | 1.657 | 0.46 | 0.55 | 0.38 | 0.07 | 0.35 | 0.19 |

| | | | | | | | | | | | | | | | | |
|---|---|---|---|---|---|---|---|---|---|---|---|---|---|---|---|---|
| 7 | 29663845 | rs1035149 | -0.286 | 0.593 | 0.071 | -1.506 | 0.022 | 0.177 | 0.177 | 0 | 0.49 | 0.75 | 0.44 | 0.12 | 0.43 | 0.26 |
| 7 | 29668519 | rs16875105 | 1.141 | 0.742 | -0.285 | -0.809 | -0.084 | -0.13 | -0.13 | 0 | 0.17 | 0.01 | 0.23 | 0.44 | 0.24 | 0.25 |
| 7 | 29668591 | rs850015 | 0.92 | 0.065 | 0.736 | -1.602 | -0.043 | -0.156 | -0.156 | 0 | 0.22 | 0.43 | 0.23 | 0 | 0.13 | 0.09 |
| 7 | 29685425 | rs1728635 | 1.1 | 0.09 | 0.701 | -1.616 | -0.024 | -0.082 | -0.082 | 0.243 | 0.2 | 0.29 | 0.21 | 0 | 0.11 | 0.09 |
| 7 | 29688860 | rs17150665 | 0.955 | 0.101 | 0.794 | -1.635 | 0.019 | -0.101 | -0.101 | 1.746 | 0.17 | 0.29 | 0.13 | 0 | 0.18 | 0.03 |
| 7 | 29690658 | rs1728537 | 0.136 | -0.064 | 0.822 | -1.682 | -0.012 | -0.081 | -0.081 | 2.379 | 0.29 | 0.41 | 0.3 | 0.07 | 0.24 | 0.2 |
| 7 | 29691083 | rs1728538 | -0.552 | -0.032 | 0.523 | -1.656 | 0.016 | 0.121 | 0.121 | 0 | 0.46 | 0.7 | 0.45 | 0.09 | 0.41 | 0.24 |
| 7 | 29691213 | rs1728539 | 0.975 | 0 | 0.427 | -1.688 | 0.001 | 0 | 0 | 0 | 0.23 | 0.2 | 0.26 | 0.05 | 0.21 | 0.17 |
| 7 | 29691438 | rs17604090 | 1.152 | 0.124 | 0.032 | -1.572 | -0.01 | 0.039 | 0.039 | 0 | 0.16 | 0.08 | 0.09 | 0 | 0.15 | 0.03 |
| 7 | 29698147 | rs1728605 | -0.667 | 0.075 | 0.017 | -1.563 | 0.022 | 0.161 | 0.161 | 0 | 0.51 | 0.71 | 0.43 | 0.13 | 0.43 | 0.27 |
| 7 | 29726287 | rs4000281 | 0.852 | 0.293 | -0.072 | -1.433 | -0.004 | 0.011 | 0.011 | 0 | 0.16 | 0.1 | 0.09 | 0 | 0.16 | 0.03 |
| 7 | 29745196 | rs10234834 | -0.095 | 0.307 | 0.226 | -1.497 | -0.004 | -0.112 | -0.112 | 0 | 0.39 | 0.47 | 0.36 | 0.06 | 0.36 | 0.2 |
| 7 | 29751459 | rs850074 | 0.395 | 0.124 | 0.282 | -1.303 | -0.012 | -0.117 | -0.117 | 0.998 | 0.27 | 0.46 | 0.19 | 0.07 | 0.24 | 0.15 |
| 7 | 29758161 | rs850079 | -0.085 | 0.289 | 0.027 | -0.849 | 0 | 0.002 | 0.002 | 0 | 0.24 | 0.25 | 0.42 | 0.3 | 0.26 | 0.35 |
| 7 | 29771597 | rs850084 | -0.156 | 0.11 | -0.076 | -0.955 | 0 | -0.001 | -0.001 | 3.476 | 0.27 | 0.26 | 0.42 | 0.31 | 0.26 | 0.36 |
| 7 | 29772138 | rs850086 | -0.043 | 0.122 | -0.071 | -0.95 | 0 | 0 | 0 | 0 | 0.26 | 0.26 | 0.48 | 0.31 | 0.26 | 0.36 |
| 7 | 29778251 | rs1100626 | -0.913 | 0.079 | -0.145 | -1.119 | -0.004 | -0.026 | -0.026 | 0.533 | 0.94 | 0.92 | 0.94 | 1 | 0.94 | 0.98 |
| 7 | 29791678 | rs741277 | 0.805 | -0.311 | -0.096 | -1.317 | 0.011 | 0.019 | 0.019 | 0 | 0.58 | 0.4 | 0.45 | 0.68 | 0.65 | 0.58 |
| 7 | 29791748 | rs741278 | 1.044 | 0.171 | -0.396 | -0.931 | -0.018 | 0.011 | 0.011 | 0 | 0.16 | 0.09 | 0.37 | 0.16 | 0.19 | 0.26 |
| 7 | 29794120 | rs13236218 | 1.13 | 0.623 | -0.883 | -0.624 | -0.068 | -0.116 | -0.116 | 0 | 0.16 | 0.01 | 0.15 | 0.55 | 0.21 | 0.26 |
| 7 | 29795025 | rs850139 | -0.136 | 0.39 | -0.206 | -0.99 | 0.006 | -0.019 | -0.019 | 0 | 0.33 | 0.45 | 0.48 | 0.3 | 0.35 | 0.31 |
| 7 | 29805403 | rs10253408 | 0.097 | -0.131 | -0.019 | -0.859 | -0.006 | 0.015 | 0.015 | 0 | 0.13 | 0.24 | 0.09 | 0.14 | 0.14 | 0.08 |
| 7 | 29809440 | rs10951246 | 0.785 | -0.574 | 0.249 | -1.223 | -0.026 | -0.057 | -0.057 | 0 | 0.81 | 0.41 | 0.82 | 0.92 | 0.85 | 0.9 |
| 7 | 29809569 | rs11971710 | 0.64 | -0.689 | -0.535 | -1.678 | 0.021 | 0.055 | 0.055 | 0 | 0.42 | 0.31 | 0.27 | 0.14 | 0.36 | 0.4 |
| 7 | 29810454 | rs4719982 | -0.294 | -0.689 | -0.04 | -1.661 | 0.003 | -0.036 | -0.036 | 0 | 0.92 | 0.74 | 0.93 | 1 | 0.95 | 0.98 |
| 7 | 29814047 | rs4722938 | 0.487 | -1.005 | -0.082 | -1.872 | 0.021 | 0.173 | 0.173 | 0 | 0.47 | 0.19 | 0.58 | 0.8 | 0.54 | 0.6 |
| 7 | 29817100 | rs12532595 | 1.281 | -0.88 | -0.113 | -1.921 | 0.024 | 0.044 | 0.044 | 1.924 | 0.34 | 0.25 | 0.17 | 0.07 | 0.25 | 0.2 |
| 7 | 29817834 | rs1861286 | 0.076 | -0.746 | -0.287 | -1.823 | 0.052 | 0.164 | 0.164 | 0.132 | 0.48 | 0.27 | 0.66 | 0.85 | 0.61 | 0.71 |
| 7 | 29817972 | rs2041755 | 0.827 | -0.246 | 0.096 | -1.323 | -0.007 | -0.017 | -0.017 | 0 | 0.11 | 0.23 | 0.1 | 0.07 | 0.1 | 0.08 |
| 7 | 29833841 | rs10224984 | -0.435 | -0.656 | -0.371 | -1.594 | -0.015 | -0.037 | -0.037 | 0 | 0.32 | 0.31 | 0.23 | 0.15 | 0.27 | 0.31 |
| 7 | 29836613 | rs10274061 | 1.377 | 0.202 | -0.248 | -1.2 | 0.01 | 0.025 | 0.025 | 0.068 | 0.21 | 0.25 | 0.2 | 0.57 | 0.26 | 0.27 |
| 7 | 29836717 | rs6953643 | 0.674 | -0.039 | -0.378 | -1.154 | -0.001 | -0.006 | -0.006 | 0 | 0.33 | 0.34 | 0.46 | 0.28 | 0.33 | 0.32 |
| 7 | 29842217 | rs767430 | 1.941 | 0.142 | -0.933 | -0.907 | -0.057 | -0.029 | -0.029 | 0 | 0.19 | 0.05 | 0.37 | 0.17 | 0.23 | 0.25 |
| 7 | 29844708 | rs713493 | 0.195 | -0.277 | -0.528 | -1.06 | 0.002 | 0.009 | 0.009 | 0 | 0.52 | 0.39 | 0.38 | 0.65 | 0.54 | 0.47 |
| 7 | 29849897 | rs174917 | 0.705 | -0.435 | 0.113 | -1.213 | 0.022 | 0.064 | 0.064 | 0 | 0.91 | 0.73 | 0.86 | 0.84 | 0.9 | 0.84 |
| 7 | 29850789 | rs10236126 | 1.499 | -0.519 | 0.196 | -1.307 | -0.009 | -0.044 | -0.044 | 0 | 0.12 | 0.18 | 0.11 | 0.04 | 0.12 | 0.08 |
| 7 | 29854684 | rs174924 | 2.022 | -0.203 | -0.207 | -0.629 | -0.083 | -0.137 | -0.137 | 1.261 | 0.26 | 0.08 | 0.41 | 0.36 | 0.29 | 0.29 |
| 7 | 29863135 | rs6979192 | -0.225 | -0.243 | -0.082 | -0.695 | 0 | 0.002 | 0.002 | 0.376 | 0.61 | 0.64 | 0.62 | 0.46 | 0.54 | 0.47 |
| 7 | 29864111 | rs6462178 | 0.064 | -0.247 | -0.019 | -0.707 | 0 | -0.006 | -0.006 | 0.974 | 0.58 | 0.42 | 0.59 | 0.46 | 0.53 | 0.46 |
| 7 | 29874756 | rs1548606 | 0.264 | -0.299 | -0.022 | -0.705 | 0.005 | -0.012 | -0.012 | 0 | 0.57 | 0.41 | 0.57 | 0.38 | 0.54 | 0.45 |
| 7 | 29876672 | rs10278201 | -1.779 | -0.293 | -0.049 | -0.706 | -0.011 | -0.029 | -0.029 | 1.04 | 0.73 | 0.83 | 0.73 | 0.5 | 0.66 | 0.56 |
| 7 | 29878542 | rs13224111 | 1.063 | -0.36 | -0.246 | -0.698 | -0.072 | -0.106 | -0.106 | 0 | 0.38 | 0.11 | 0.44 | 0.36 | 0.35 | 0.34 |
| 7 | 29880530 | rs174933 | -0.112 | -0.164 | 0.371 | -0.792 | 0.001 | -0.023 | -0.023 | 0.031 | 0.84 | 0.75 | 0.81 | 0.87 | 0.79 | 0.78 |
| 7 | 29882816 | rs174935 | 0.538 | -0.171 | 0.555 | -0.788 | -0.004 | 0.001 | 0.001 | 0 | 0.81 | 0.6 | 0.77 | 0.79 | 0.78 | 0.76 |
| 7 | 29884697 | rs10252599 | 1.498 | -0.152 | 0.4 | -0.785 | -0.01 | -0.04 | -0.04 | 0.266 | 0.12 | 0.17 | 0.11 | 0.04 | 0.12 | 0.08 |
| 7 | 29886947 | rs174945 | 0.843 | -0.198 | 0.878 | -0.804 | 0.021 | 0.015 | 0.015 | 0 | 0.93 | 0.7 | 0.88 | 0.91 | 0.9 | 0.87 |
| 7 | 29887612 | rs174946 | 0.429 | -0.301 | 1.014 | -0.849 | -0.007 | -0.029 | -0.029 | 0.528 | 0.81 | 0.58 | 0.79 | 0.86 | 0.79 | 0.78 |
| 7 | 29887961 | rs13438578 | 1.519 | -0.279 | 0.929 | -0.845 | -0.011 | -0.041 | -0.041 | 0 | 0.12 | 0.17 | 0.11 | 0.04 | 0.12 | 0.08 |
| 7 | 29889180 | rs174947 | 1.448 | -0.257 | 0.488 | -0.864 | -0.023 | -0.023 | -0.023 | 0.068 | 0.74 | 0.43 | 0.72 | 0.77 | 0.74 | 0.69 |
| 7 | 29889233 | rs174948 | 1.63 | -0.154 | 0.168 | -1.002 | 0.012 | 0.056 | 0.056 | 0 | 0.6 | 0.25 | 0.62 | 0.77 | 0.6 | 0.64 |
| 7 | 29893423 | rs9691880 | 2.132 | 0.272 | -0.207 | -0.604 | -0.034 | -0.055 | -0.055 | 0.407 | 0.24 | 0.13 | 0.42 | 0.49 | 0.27 | 0.3 |
| 7 | 29894775 | rs740142 | 0.824 | -0.046 | 0.212 | -2.168 | -0.001 | 0.005 | 0.005 | 0 | 0.14 | 0.08 | 0.07 | 0 | 0.14 | 0.05 |
| 7 | 29909284 | rs10236764 | -0.421 | -0.155 | 0.613 | -2.158 | -0.02 | -0.023 | -0.023 | 2.631 | 0.14 | 0.25 | 0.1 | 0.11 | 0.13 | 0.14 |
| 7 | 29910227 | rs174957 | 2.407 | 0.067 | 0.265 | -2.314 | -0.04 | 0.168 | 0.168 | 0.37 | 0.45 | 0.21 | 0.6 | 0.8 | 0.47 | 0.61 |
| 7 | 29916585 | rs10276961 | 0.671 | -0.248 | 0.429 | -2.307 | -0.007 | -0.034 | -0.034 | 0.01 | 0.11 | 0.14 | 0.07 | 0.04 | 0.11 | 0.09 |
| 7 | 29917385 | rs917145 | -0.654 | -0.336 | 0.245 | -2.383 | -0.003 | -0.062 | -0.062 | 0.058 | 0.36 | 0.31 | 0.2 | 0.02 | 0.29 | 0.18 |
| 7 | 29920967 | rs174962 | 1.424 | -0.071 | -0.345 | -2.273 | 0.024 | 0.181 | 0.181 | 0.977 | 0.48 | 0.29 | 0.66 | 0.87 | 0.51 | 0.64 |
| 7 | 29921360 | rs6979942 | -1.484 | -0.316 | -0.374 | -2.196 | 0.054 | 0.129 | 0.129 | 1.331 | 0.5 | 0.6 | 0.29 | 0.13 | 0.42 | 0.32 |
| 7 | 29921695 | rs740144 | 0.349 | 0.929 | -0.462 | -1.797 | -0.001 | 0.001 | 0.001 | 0 | 0.14 | 0.08 | 0.08 | 0 | 0.15 | 0.05 |
| 7 | 29924686 | rs174966 | -0.055 | 0.808 | -0.145 | -2.431 | -0.011 | -0.019 | -0.019 | 0 | 0.74 | 0.65 | 0.72 | 0.84 | 0.77 | 0.9 |
| 7 | 29928463 | rs2240382 | -0.477 | 0.815 | -0.01 | -2.891 | -0.007 | -0.045 | -0.045 | 0 | 0.21 | 0.25 | 0.13 | 0.08 | 0.19 | 0.2 |
| 7 | 29931848 | rs174986 | -0.252 | 0.782 | 0.052 | -2.862 | -0.017 | -0.043 | -0.043 | 0 | 0.76 | 0.62 | 0.75 | 0.89 | 0.76 | 0.91 |
| 7 | 29936632 | rs10227569 | -0.38 | 0.828 | -0.543 | -2.841 | -0.023 | -0.055 | -0.055 | 0 | 0.75 | 0.54 | 0.74 | 0.89 | 0.75 | 0.91 |
| 7 | 29936886 | rs10257272 | 0.412 | 0.859 | -0.733 | -2.941 | 0.005 | 0.044 | 0.044 | 1.551 | 0.09 | 0.03 | 0.04 | 0.05 | 0.04 | 0.09 |
| 7 | 29939989 | rs2269982 | 1.411 | 0.875 | -0.729 | -2.997 | 0.02 | 0.181 | 0.181 | 1.303 | 0.47 | 0.22 | 0.58 | 0.86 | 0.54 | 0.71 |
| 7 | 29940034 | rs2269981 | 1.46 | 0.875 | -0.727 | -2.996 | 0.016 | 0.176 | 0.176 | 1.292 | 0.47 | 0.21 | 0.58 | 0.81 | 0.53 | 0.67 |
| 7 | 29943208 | rs7797769 | -0.114 | 0.851 | -0.694 | -3.025 | -0.014 | -0.044 | -0.044 | 0 | 0.72 | 0.43 | 0.73 | 0.89 | 0.73 | 0.85 |
| 7 | 29944869 | rs10488082 | 0.171 | 0.862 | -0.695 | -3.026 | -0.011 | -0.045 | -0.045 | 0 | 0.16 | 0.19 | 0.12 | 0.03 | 0.15 | 0.1 |
| 7 | 29944926 | rs10231810 | -0.123 | 0.865 | -0.698 | -3.025 | -0.014 | -0.044 | -0.044 | 0 | 0.73 | 0.43 | 0.73 | 0.89 | 0.73 | 0.85 |

| | | | | | | | | | | | | | | | | |
|---|---|---|---|---|---|---|---|---|---|---|---|---|---|---|---|---|
| 7 | 29945282 | rs12532093 | 1.498 | 0.866 | -0.698 | -3.025 | 0.018 | 0.178 | 0.178 | 1.292 | 0.47 | 0.2 | 0.58 | 0.81 | 0.54 | 0.68 |
| 7 | 29945597 | rs10264876 | -0.084 | 1.145 | -0.579 | -2.9 | -0.013 | -0.045 | -0.045 | 0 | 0.73 | 0.43 | 0.73 | 0.89 | 0.73 | 0.85 |
| 7 | 29953766 | rs2072176 | -1.02 | 1.363 | -0.868 | -2.767 | -0.004 | -0.042 | -0.042 | 0 | 0.26 | 0.27 | 0.16 | 0.08 | 0.22 | 0.23 |
| 7 | 29953913 | rs2072175 | -0.91 | 1.357 | -0.874 | -2.764 | 0.003 | -0.011 | -0.011 | 2.183 | 0.24 | 0.19 | 0.15 | 0.08 | 0.2 | 0.17 |
| 7 | 29954611 | rs2057902 | 0.419 | 1.678 | -0.827 | -2.78 | -0.013 | -0.045 | -0.045 | 2.698 | 0.64 | 0.64 | 0.75 | 0.93 | 0.69 | 0.8 |
| 7 | 29960315 | rs1049395 | -1.574 | 2.585 | -0.587 | -2.201 | -0.002 | -0.03 | -0.03 | 0 | 0.35 | 0.42 | 0.26 | 0.15 | 0.34 | 0.26 |
| 7 | 29960462 | rs13834 | -1.588 | 2.583 | -0.588 | -2.202 | -0.002 | -0.03 | -0.03 | 0.854 | 0.36 | 0.42 | 0.27 | 0.15 | 0.34 | 0.26 |
| 7 | 29960804 | rs1049394 | -1.571 | 2.6 | -0.58 | -2.194 | -0.002 | -0.031 | -0.031 | 0 | 0.35 | 0.42 | 0.26 | 0.15 | 0.34 | 0.27 |
| 7 | 29962427 | rs11540586 | -1.588 | 2.601 | -0.579 | -2.188 | -0.002 | -0.027 | -0.027 | 0 | 0.36 | 0.42 | 0.27 | 0.16 | 0.34 | 0.26 |
| 7 | 29962934 | rs6976789 | -1.59 | 2.618 | -0.572 | -2.244 | -0.002 | -0.03 | -0.03 | 0 | 0.36 | 0.42 | 0.26 | 0.16 | 0.34 | 0.27 |
| 7 | 29980024 | rs2107898 | -0.601 | 2.001 | -0.406 | -2.215 | -0.006 | -0.057 | -0.057 | 1.115 | 0.81 | 0.74 | 0.82 | 0.97 | 0.82 | 0.94 |
| 7 | 29991995 | rs4722964 | 0.389 | 2.168 | -0.348 | -2.028 | 0.001 | 0.007 | 0.007 | 0 | 0.8 | 0.82 | 0.84 | 0.95 | 0.78 | 0.88 |
| 7 | 29993712 | rs11763228 | -0.776 | 2.089 | -0.165 | -1.846 | -0.001 | -0.018 | -0.018 | 0.101 | 0.2 | 0.25 | 0.17 | 0.07 | 0.19 | 0.17 |
| 7 | 30014068 | rs11978244 | 0.048 | 1.89 | -0.167 | -1.708 | -0.005 | -0.044 | -0.044 | 2.121 | 0.16 | 0.21 | 0.13 | 0.03 | 0.15 | 0.1 |
| 7 | 30044128 | rs17158577 | -0.552 | 1.94 | -0.155 | -1.686 | -0.002 | -0.017 | -0.017 | 0 | 0.18 | 0.23 | 0.17 | 0.07 | 0.16 | 0.12 |
| 7 | 30067626 | rs10275475 | -1.103 | 0.887 | -0.145 | -1.61 | 0.004 | -0.002 | -0.002 | 0.913 | 0.25 | 0.23 | 0.21 | 0.07 | 0.22 | 0.15 |
| 7 | 30083021 | rs10488084 | -0.112 | 0.838 | -0.179 | -1.594 | -0.006 | -0.025 | -0.025 | 4.623 | 0.09 | 0.15 | 0.06 | 0.05 | 0.08 | 0.04 |
| 7 | 30087381 | rs10488085 | -1.075 | 0.905 | -0.194 | -1.597 | -0.008 | -0.066 | -0.066 | 0.299 | 0.25 | 0.41 | 0.23 | 0.11 | 0.24 | 0.15 |
| 7 | 30101716 | rs17326411 | -0.454 | 1.09 | -0.846 | -1.612 | -0.003 | -0.019 | -0.019 | 0 | 0.18 | 0.21 | 0.17 | 0.07 | 0.16 | 0.12 |
| 7 | 30115881 | rs6976806 | -0.722 | 1.091 | -0.374 | -1.135 | -0.002 | -0.019 | -0.019 | 0.418 | 0.18 | 0.21 | 0.17 | 0.07 | 0.16 | 0.12 |
| 7 | 30117417 | rs10488086 | -0.722 | 1.091 | -0.372 | -1.135 | -0.005 | -0.032 | -0.032 | 1.267 | 0.18 | 0.25 | 0.17 | 0.07 | 0.16 | 0.12 |
| 7 | 30118508 | rs1109208 | -0.722 | 1.091 | -0.372 | -1.135 | -0.005 | -0.032 | -0.032 | 0 | 0.18 | 0.25 | 0.17 | 0.07 | 0.16 | 0.12 |
| 7 | 30119410 | rs876050 | -0.741 | 1.091 | -0.375 | -1.135 | -0.002 | -0.019 | -0.019 | 0 | 0.18 | 0.21 | 0.17 | 0.07 | 0.16 | 0.12 |
| 7 | 30125924 | rs17546537 | -0.726 | 1.104 | -0.369 | -1.129 | -0.002 | -0.019 | -0.019 | 0.953 | 0.18 | 0.21 | 0.17 | 0.07 | 0.16 | 0.12 |
| 7 | 30128903 | rs11982167 | -0.056 | 1.035 | -0.371 | -1.122 | -0.004 | -0.033 | -0.033 | 0.561 | 0.15 | 0.18 | 0.12 | 0.03 | 0.14 | 0.11 |
| 7 | 30129125 | rs11975522 | -0.056 | 1.035 | -0.368 | -1.122 | -0.004 | -0.035 | -0.035 | 0 | 0.15 | 0.19 | 0.13 | 0.03 | 0.14 | 0.11 |
| 7 | 30131289 | rs11773644 | -0.102 | 1.036 | -0.398 | -1.122 | -0.004 | -0.032 | -0.032 | 0.079 | 0.16 | 0.18 | 0.13 | 0.03 | 0.14 | 0.11 |
| 7 | 30133202 | rs7806238 | -0.693 | 1.052 | -0.389 | -1.115 | -0.006 | -0.032 | -0.032 | 0 | 0.18 | 0.24 | 0.17 | 0.07 | 0.16 | 0.12 |
| 7 | 30141435 | rs10275279 | 0.159 | 1.106 | -0.506 | -1.132 | -0.003 | -0.033 | -0.033 | 0 | 0.77 | 0.73 | 0.82 | 0.92 | 0.78 | 0.89 |
| 7 | 30154691 | rs11769513 | -0.861 | 0.645 | -0.567 | -0.157 | -0.004 | -0.032 | -0.032 | 0 | 0.16 | 0.18 | 0.13 | 0.03 | 0.14 | 0.1 |
| 7 | 30177035 | rs6973967 | -0.266 | 0.9 | -0.609 | 0.26 | 0.004 | -0.005 | -0.005 | 0 | 0.43 | 0.55 | 0.63 | 0.6 | 0.53 | 0.47 |
| 7 | 30177278 | rs6954379 | -0.78 | 0.928 | 0.055 | 0.437 | 0.014 | 0.082 | 0.082 | 0.554 | 0.25 | 0.56 | 0.37 | 0.38 | 0.31 | 0.33 |
| 7 | 30179394 | rs887883 | 0.194 | 0.016 | 0.329 | -0.652 | -0.039 | -0.094 | -0.094 | 0 | 0.71 | 0.51 | 0.75 | 0.94 | 0.81 | 0.8 |
| 7 | 30182444 | rs997349 | 0.391 | 0.005 | 0.477 | -0.863 | -0.011 | -0.031 | -0.031 | 0 | 0.32 | 0.56 | 0.28 | 0.17 | 0.33 | 0.34 |
| 7 | 30182813 | rs10488088 | 0.783 | -0.027 | 0.567 | -1.087 | 0.019 | -0.04 | -0.04 | 0.243 | 0.08 | 0.17 | 0.13 | 0 | 0.14 | 0.06 |
| 7 | 30187932 | rs10268447 | -0.941 | -0.83 | 0.496 | -1.499 | 0.01 | 0.009 | 0.009 | 0.251 | 0.75 | 0.82 | 0.8 | 0.92 | 0.83 | 0.86 |
| 7 | 30194317 | rs7811155 | 1.085 | -0.309 | 0.562 | -1.532 | 0.002 | -0.02 | -0.02 | 3.05 | 0.08 | 0.09 | 0.13 | 0 | 0.13 | 0.06 |
| 7 | 30195058 | rs7780166 | 0.25 | 0.205 | 0.438 | -1.062 | -0.003 | 0.011 | 0.011 | 0 | 0.57 | 0.58 | 0.47 | 0.68 | 0.48 | 0.73 |
| 7 | 30198259 | rs1558579 | 1.226 | 0.293 | 0.474 | -1.019 | 0.019 | -0.043 | -0.043 | 0.133 | 0.08 | 0.19 | 0.13 | 0 | 0.13 | 0.06 |
| 7 | 30198899 | rs10951262 | 1.103 | 0.299 | 0.318 | -1.12 | -0.004 | -0.085 | -0.085 | 0.394 | 0.27 | 0.11 | 0.29 | 0.51 | 0.25 | 0.4 |
| 7 | 30200599 | rs3750089 | -0.426 | 0.203 | 0.658 | -1.168 | -0.024 | 0.016 | 0.016 | 0 | 0.64 | 0.75 | 0.59 | 0.68 | 0.6 | 0.78 |
| 7 | 30201413 | rs2110394 | 1.007 | 0.205 | 0.664 | -1.166 | -0.015 | -0.108 | -0.108 | 0 | 0.27 | 0.08 | 0.3 | 0.51 | 0.27 | 0.4 |
| 7 | 30201663 | rs11156 | 1.719 | 0.133 | 0.879 | -1.245 | 0.015 | 0 | 0 | 0.803 | 0.21 | 0.12 | 0.13 | 0.17 | 0.2 | 0.29 |
| 7 | 30202836 | rs7793699 | 0.461 | 0.193 | 1.094 | -1.277 | 0.06 | -0.067 | -0.067 | 0.996 | 0.11 | 0.41 | 0.16 | 0 | 0.14 | 0.07 |
| 7 | 30210227 | rs12176611 | 1.707 | 0.08 | 0.524 | -1.245 | 0.013 | 0.054 | 0.054 | 0 | 0.21 | 0.03 | 0.11 | 0.07 | 0.2 | 0.24 |
| 7 | 30211222 | rs723654 | 0.587 | 0.041 | 1.019 | -0.99 | -0.002 | 0.022 | 0.022 | 0 | 0.3 | 0.35 | 0.34 | 0.62 | 0.28 | 0.45 |
| 7 | 30211788 | rs6953710 | 0.525 | 0.108 | 0.908 | -0.983 | -0.008 | 0.011 | 0.011 | 0 | 0.37 | 0.32 | 0.32 | 0.28 | 0.45 | 0.35 |
| 7 | 30213452 | rs3823765 | 0.882 | 0.278 | 0.724 | -0.805 | 0.022 | 0.079 | 0.079 | 0 | 0.05 | 0.13 | 0.07 | 0.21 | 0.1 | 0.04 |
| 7 | 30214178 | rs10224386 | -0.758 | 0.221 | 0.69 | -0.875 | -0.004 | 0.021 | 0.021 | 0 | 0.7 | 0.76 | 0.66 | 0.91 | 0.71 | 0.81 |
| 7 | 30220830 | rs12701008 | -0.695 | 0.158 | 0.758 | -0.844 | -0.001 | 0 | 0 | 0 | 0.7 | 0.76 | 0.67 | 0.91 | 0.73 | 0.81 |
| 7 | 30225119 | rs6462192 | 0.161 | 0.012 | 0.766 | -0.751 | 0.008 | -0.017 | -0.017 | 0 | 0.61 | 0.48 | 0.58 | 0.68 | 0.61 | 0.75 |
| 7 | 30233732 | rs2192399 | 0.147 | 0.045 | 0.745 | -0.766 | 0.006 | -0.014 | -0.014 | 1.21 | 0.61 | 0.51 | 0.58 | 0.69 | 0.61 | 0.73 |
| 7 | 30237983 | rs11768018 | 1.287 | 0.27 | 0.608 | -0.49 | -0.079 | -0.146 | -0.146 | 0.459 | 0.2 | 0.02 | 0.28 | 0.52 | 0.24 | 0.31 |
| 7 | 30239137 | rs7808333 | 0.151 | -0.279 | 1.113 | -1.219 | -0.038 | 0.037 | 0.037 | 0.185 | 0.39 | 0.23 | 0.37 | 0.73 | 0.42 | 0.46 |
| 7 | 30243357 | rs12535035 | 0.142 | -0.145 | 1.763 | -0.801 | 0.003 | -0.071 | -0.071 | 0 | 0.72 | 0.6 | 0.66 | 0.9 | 0.74 | 0.81 |
| 7 | 30244287 | rs17158705 | -0.217 | 0.008 | 1.264 | -0.659 | -0.002 | 0.004 | 0.004 | 0 | 0.25 | 0.23 | 0.32 | 0.1 | 0.24 | 0.18 |
| 7 | 30251254 | rs10246865 | -2.012 | -0.217 | 2.584 | -0.047 | -0.029 | -0.023 | -0.023 | 2.038 | 0.1 | 0.62 | 0.07 | 0.03 | 0.02 | 0.02 |
| 7 | 30255901 | rs10251654 | -1.991 | -0.236 | 2.773 | -0.056 | -0.035 | -0.03 | -0.03 | 0 | 0.1 | 0.67 | 0.07 | 0.03 | 0.02 | 0.02 |
| 7 | 30276161 | rs6462196 | -1.765 | -0.167 | 2.63 | -0.025 | -0.005 | 0.001 | 0.001 | 0.48 | 0.09 | 0.59 | 0.07 | 0.03 | 0.02 | 0.02 |
| 7 | 30293692 | rs17158763 | -0.578 | -0.278 | 2.391 | -0.037 | -0.001 | -0.016 | -0.016 | 0 | 0.05 | 0.2 | 0.03 | 0 | 0.02 | 0.01 |
| 7 | 30299645 | rs6462197 | -2.004 | -0.13 | 2.486 | 0.206 | 0.037 | 0.163 | 0.163 | 1.73 | 0.07 | 0.52 | 0.08 | 0.08 | 0.02 | 0.04 |
| 7 | 30347038 | rs9638864 | -1.867 | -0.26 | 2.015 | 0.035 | 0.006 | 0.092 | 0.092 | 2.301 | 0.07 | 0.57 | 0.06 | 0.06 | 0.01 | 0.04 |
| 7 | 30368541 | rs7805248 | -0.99 | -0.405 | 1.613 | -0.192 | -0.017 | -0.053 | -0.053 | 0 | 0.1 | 0.16 | 0.04 | 0 | 0.06 | 0.04 |
| 7 | 30370786 | rs11974360 | -1.762 | -0.267 | 1.538 | -0.016 | 0.013 | 0.102 | 0.102 | 1.185 | 0.06 | 0.43 | 0.05 | 0.06 | 0.01 | 0.04 |
| 7 | 30388409 | rs4722979 | 0.061 | 0.302 | -0.373 | -0.368 | -0.077 | 0.044 | 0.044 | 0 | 0.25 | 0.02 | 0.15 | 0.12 | 0.3 | 0.28 |
| 7 | 30396131 | rs42579 | -0.512 | -0.16 | 0.752 | -0.418 | -0.011 | -0.02 | -0.02 | 0.029 | 0.07 | 0.12 | 0.04 | 0.02 | 0.05 | 0.06 |
| 7 | 30405691 | rs42593 | -0.919 | -0.217 | 1.14 | -0.421 | -0.015 | -0.023 | -0.023 | 0 | 0.07 | 0.17 | 0.05 | 0.02 | 0.06 | 0.06 |
| 7 | 30406471 | rs42595 | -0.919 | -0.217 | 1.14 | -0.432 | -0.015 | -0.026 | -0.026 | 0 | 0.07 | 0.17 | 0.05 | 0.02 | 0.06 | 0.06 |
| 7 | 30406767 | rs42596 | -0.549 | -0.216 | 1.084 | -0.432 | -0.007 | -0.013 | -0.013 | 0 | 0.07 | 0.08 | 0.04 | 0.02 | 0.06 | 0.06 |

| | | | | | | | | | | | | | | | | |
|---|---|---|---|---|---|---|---|---|---|---|---|---|---|---|---|---|
| 7 | 30407445 | rs42597 | -1.672 | -0.353 | 1.948 | -0.493 | -0.005 | -0.026 | -0.026 | 0.003 | 0.1 | 0.45 | 0.09 | 0.02 | 0.06 | 0.06 |
| 7 | 30408353 | rs42599 | -1.501 | -0.321 | 1.511 | -0.587 | 0.004 | 0.009 | 0.009 | 0 | 0.14 | 0.1 | 0.13 | 0.12 | 0.13 | 0.18 |
| 7 | 30414662 | rs13221972 | 0.405 | 0.343 | 1.134 | -0.85 | -0.039 | 0.037 | 0.037 | 0 | 0.07 | 0.01 | 0.11 | 0 | 0.14 | 0.09 |
| 7 | 30415102 | rs17158939 | -0.726 | -0.354 | 1.918 | -0.834 | -0.026 | 0.003 | 0.003 | 1.757 | 0.09 | 0.28 | 0.05 | 0.05 | 0.02 | 0.04 |
| 7 | 30418224 | rs1029965 | -2.257 | -0.379 | 1.691 | -0.846 | -0.042 | -0.028 | -0.028 | 0 | 0.18 | 0.71 | 0.12 | 0.07 | 0.08 | 0.08 |
| 7 | 30429268 | rs6947237 | -2.235 | -0.409 | 1.486 | -0.734 | -0.029 | -0.01 | -0.01 | 0 | 0.23 | 0.67 | 0.2 | 0.22 | 0.13 | 0.18 |
| 7 | 30443534 | rs6942557 | -0.935 | 0.109 | 1.468 | -0.715 | 0.079 | 0.027 | 0.027 | 0 | 0.05 | 0.26 | 0.05 | 0.02 | 0.08 | 0.03 |
| 7 | 30444277 | rs6961625 | -1.825 | 0.044 | 1.419 | -0.806 | -0.006 | -0.053 | -0.053 | 0 | 0.16 | 0.48 | 0.11 | 0.06 | 0.14 | 0.17 |
| 7 | 30446814 | rs7782089 | -1.66 | -0.462 | 1.395 | -0.774 | -0.035 | -0.029 | -0.029 | 0.84 | 0.11 | 0.36 | 0.05 | 0.03 | 0.04 | 0.04 |
| 7 | 30460476 | rs6956830 | -1.693 | -0.466 | 1.346 | -0.794 | -0.04 | -0.043 | -0.043 | 0 | 0.11 | 0.34 | 0.04 | 0.02 | 0.04 | 0.04 |
| 7 | 30464961 | rs5743373 | -1.604 | -0.443 | 1.357 | -0.782 | -0.033 | -0.041 | -0.041 | 0.018 | 0.11 | 0.35 | 0.05 | 0.02 | 0.04 | 0.04 |
| 7 | 30465035 | rs5743372 | -1.604 | -0.451 | 1.343 | -0.782 | -0.038 | -0.041 | -0.041 | 0 | 0.11 | 0.34 | 0.04 | 0.02 | 0.04 | 0.04 |
| 7 | 30469270 | rs17159043 | -1.596 | -0.311 | 0.619 | -0.849 | 0.009 | 0.003 | 0.003 | 0.333 | 0.2 | 0.2 | 0.09 | 0.04 | 0.16 | 0.2 |
| 7 | 30473023 | rs2907748 | -0.235 | 0.159 | 0.064 | -0.066 | -0.057 | -0.072 | -0.072 | 0 | 0.18 | 0.06 | 0.28 | 0.25 | 0.24 | 0.22 |
| 7 | 30478056 | rs2970498 | -0.65 | 0.027 | 0.768 | -0.1 | 0.024 | 0.043 | 0.043 | 0 | 0.2 | 0.3 | 0.3 | 0.26 | 0.24 | 0.22 |
| 7 | 30491311 | rs2075821 | -0.993 | -1.152 | 0.576 | -0.162 | 0.002 | 0.001 | 0.001 | 0 | 0.35 | 0.33 | 0.16 | 0.38 | 0.24 | 0.42 |
| 7 | 30492237 | rs2075820 | -0.818 | -1.353 | 1.145 | -0.169 | -0.014 | -0.002 | -0.002 | 0.565 | 0.35 | 0.4 | 0.18 | 0.39 | 0.23 | 0.42 |
| 7 | 30494622 | rs2075819 | -1.296 | -1.467 | 1.314 | -0.251 | -0.038 | -0.015 | -0.015 | 0 | 0.41 | 0.52 | 0.21 | 0.39 | 0.27 | 0.44 |
| 7 | 30499575 | rs2906766 | -1.54 | -0.498 | 1.682 | 0.113 | -0.014 | -0.005 | -0.005 | 0 | 0.37 | 0.59 | 0.32 | 0.39 | 0.31 | 0.37 |
| 7 | 30499961 | rs3823773 | 0.246 | -0.666 | 1.468 | 0.245 | 0.006 | -0.006 | -0.006 | 0 | 0.14 | 0.02 | 0.06 | 0.13 | 0.09 | 0.14 |
| 7 | 30506644 | rs2906773 | -1.651 | -0.334 | 1.709 | 0.271 | 0.029 | 0.008 | 0.008 | 0 | 0.44 | 0.67 | 0.41 | 0.61 | 0.34 | 0.49 |
| 7 | 30510534 | rs2709803 | -0.51 | -0.337 | 1.516 | 0.279 | -0.022 | 0.003 | 0.003 | 0 | 0.28 | 0.34 | 0.17 | 0.38 | 0.17 | 0.3 |
| 7 | 30510853 | rs2893375 | 0.115 | -0.29 | 1.28 | 0.282 | -0.02 | -0.038 | -0.038 | 0.335 | 0.15 | 0.1 | 0.23 | 0.22 | 0.17 | 0.18 |
| 7 | 30512030 | rs4722988 | 0.009 | -0.574 | 1.286 | 0.287 | -0.005 | -0.037 | -0.037 | 0 | 0.15 | 0.1 | 0.23 | 0.22 | 0.15 | 0.18 |
| 7 | 30512771 | rs2709800 | -1.894 | -0.574 | 1.305 | 0.246 | 0.066 | 0.045 | 0.045 | 0 | 0.51 | 0.79 | 0.54 | 0.61 | 0.41 | 0.52 |
| 7 | 30513209 | rs2970503 | -0.373 | -0.502 | 1.237 | 0.265 | 0.024 | -0.03 | -0.03 | 0 | 0.07 | 0.18 | 0.14 | 0 | 0.1 | 0.03 |
| 7 | 30514991 | rs2256023 | -1.834 | -0.511 | 0.928 | 0.295 | 0.07 | 0.052 | 0.052 | 0 | 0.5 | 0.77 | 0.53 | 0.61 | 0.41 | 0.51 |
| 7 | 30519030 | rs2736726 | -1.455 | -0.36 | 1.346 | 0.223 | 0.035 | 0.007 | 0.007 | 0 | 0.35 | 0.73 | 0.32 | 0.39 | 0.27 | 0.33 |
| 7 | 30529411 | rs10257280 | -0.437 | -0.308 | 0.89 | 0.302 | 0.016 | -0.029 | -0.029 | 0 | 0.07 | 0.15 | 0.13 | 0 | 0.09 | 0.03 |
| 7 | 30529957 | rs2529445 | -0.713 | -0.386 | 1.022 | 0.395 | -0.033 | 0.007 | 0.007 | 0 | 0.27 | 0.36 | 0.17 | 0.39 | 0.17 | 0.3 |
| 7 | 30530872 | rs6955931 | -0.6 | 0.085 | 0.688 | 0.392 | 0.019 | -0.029 | -0.029 | 1.087 | 0.07 | 0.15 | 0.13 | 0 | 0.09 | 0.03 |
| 7 | 30536611 | rs4270 | -0.801 | -0.061 | 0.452 | 0.604 | -0.003 | 0 | 0 | 0 | 0.28 | 0.31 | 0.19 | 0.43 | 0.21 | 0.31 |
| 7 | 30536916 | rs10278906 | -0.703 | 0.389 | 0.522 | 0.289 | 0.004 | -0.029 | -0.029 | 0 | 0.13 | 0.16 | 0.25 | 0 | 0.18 | 0.09 |
| 7 | 30538651 | rs38409 | 0.985 | -0.34 | 1.113 | 0.919 | -0.031 | 0.087 | 0.087 | 1.838 | 0.74 | 0.52 | 0.76 | 0.7 | 0.79 | 0.65 |
| 7 | 30539595 | rs6462210 | 0.204 | -0.003 | 0.616 | 0.9 | -0.039 | -0.002 | -0.002 | 0 | 0.08 | 0.01 | 0.05 | 0.03 | 0.11 | 0.13 |
| 7 | 30541032 | rs2391870 | 1.865 | -0.627 | 0.251 | 0.914 | 0.063 | -0.025 | -0.025 | 0.53 | 0.52 | 0.06 | 0.51 | 0.43 | 0.63 | 0.4 |
| 7 | 30544360 | rs38414 | 1.159 | -0.617 | 1.031 | 0.834 | -0.019 | 0.044 | 0.044 | 1.637 | 0.7 | 0.48 | 0.62 | 0.69 | 0.7 | 0.62 |
| 7 | 30549884 | rs38430 | 0.877 | -0.758 | 1.419 | 0.758 | -0.018 | 0.048 | 0.048 | 0.001 | 0.76 | 0.67 | 0.76 | 0.69 | 0.81 | 0.65 |
| 7 | 30554997 | rs38434 | 0.806 | -0.779 | 1.393 | 0.756 | 0.017 | -0.047 | -0.047 | 0 | 0.77 | 0.86 | 0.79 | 0.69 | 0.81 | 0.65 |
| 7 | 30588091 | rs38468 | 1.455 | -0.722 | 1.656 | 0.852 | -0.017 | 0.053 | 0.053 | 1.267 | 0.72 | 0.46 | 0.69 | 0.67 | 0.73 | 0.63 |
| 7 | 30592856 | rs38471 | 1.77 | -0.754 | 1.869 | 0.832 | 0.03 | -0.069 | -0.069 | 9.69 | 0.69 | 0.21 | 0.65 | 0.67 | 0.72 | 0.62 |
| 7 | 30603540 | rs10234545 | -2.099 | -0.637 | 2.013 | 0.8 | 0.07 | 0.005 | 0.005 | 0.172 | 0.08 | 0.38 | 0.17 | 0 | 0.1 | 0.03 |
| 7 | 30625161 | rs38494 | 1.091 | -0.608 | 1.937 | 0.91 | -0.024 | 0.104 | 0.104 | 0.471 | 0.74 | 0.49 | 0.7 | 0.67 | 0.76 | 0.63 |
| 7 | 30634764 | rs2072236 | -0.066 | -0.7 | 1.676 | 0.886 | 0.015 | 0.008 | 0.008 | 0 | 0.16 | 0.12 | 0.1 | 0.18 | 0.07 | 0.23 |
| 7 | 30648051 | rs728539 | -1.14 | -0.655 | 1.685 | 0.879 | -0.054 | -0.059 | -0.059 | 0.036 | 0.22 | 0.48 | 0.15 | 0.18 | 0.12 | 0.24 |
| 7 | 30660938 | rs10951271 | 2.764 | -0.592 | 1.185 | 0.81 | -0.14 | -0.036 | -0.036 | 1.417 | 0.43 | 0.05 | 0.46 | 0.45 | 0.48 | 0.28 |
| 7 | 30667433 | rs2709778 | -0.305 | -0.585 | 1.131 | 1.058 | -0.014 | -0.06 | -0.06 | 0 | 0.87 | 0.99 | 0.95 | 0.85 | 0.9 | 0.85 |
| 7 | 30673345 | rs2240401 | 2.087 | -0.665 | 1.197 | 1.08 | 0.049 | -0.051 | -0.051 | 0 | 0.54 | 0.08 | 0.57 | 0.49 | 0.65 | 0.4 |
| 7 | 30674374 | rs7791663 | -0.935 | -0.739 | 1.222 | 1.031 | -0.034 | -0.037 | -0.037 | 0 | 0.2 | 0.36 | 0.13 | 0.18 | 0.12 | 0.24 |
| 7 | 30694961 | rs12701020 | 0.304 | -0.569 | 0.325 | 0.578 | -0.053 | 0.006 | 0.006 | 0 | 0.11 | 0 | 0.1 | 0.03 | 0.16 | 0.13 |
| 7 | 30696190 | rs2267710 | 1.839 | -0.87 | 0.979 | 0.956 | 0.029 | -0.009 | -0.009 | 0 | 0.61 | 0.71 | 0.64 | 0.47 | 0.68 | 0.4 |
| 7 | 30707302 | rs2240404 | -1.357 | -1.073 | 0.77 | 0.67 | -0.024 | -0.021 | -0.021 | 0 | 0.2 | 0.68 | 0.15 | 0.17 | 0.11 | 0.21 |
| 7 | 30709475 | rs2190242 | 1.744 | -1.209 | -0.008 | 0.972 | 0.055 | -0.031 | -0.031 | 0 | 0.66 | 0.82 | 0.77 | 0.47 | 0.79 | 0.49 |
| 7 | 30711961 | rs2284216 | -0.637 | -1.137 | 0.005 | 0.983 | -0.021 | -0.008 | -0.008 | 0 | 0.16 | 0.21 | 0.11 | 0.17 | 0.09 | 0.24 |
| 7 | 30714333 | rs2284218 | 2.145 | -0.998 | -0.252 | 0.94 | 0.059 | 0.037 | 0.037 | 0.129 | 0.53 | 0.12 | 0.53 | 0.4 | 0.61 | 0.44 |
| 7 | 30715573 | rs2014663 | 0.426 | -1.006 | -0.285 | 0.863 | -0.015 | -0.147 | -0.147 | 0 | 0.83 | 0.99 | 0.86 | 0.63 | 0.85 | 0.72 |
| 7 | 30716643 | rs2267716 | -1.79 | -0.761 | -0.016 | 1.093 | 0.037 | 0.076 | 0.076 | 0.277 | 0.29 | 0.86 | 0.33 | 0.17 | 0.24 | 0.28 |
| 7 | 30717043 | rs2267717 | -0.713 | -0.523 | 0.366 | 1.706 | -0.014 | 0.123 | 0.123 | 0 | 0.15 | 0.27 | 0.14 | 0.37 | 0.11 | 0.27 |
| 7 | 30725701 | rs4723002 | -0.912 | -0.119 | -0.088 | 0.966 | -0.018 | 0.002 | 0.002 | 0 | 0.13 | 0.2 | 0.09 | 0.18 | 0.08 | 0.23 |
| 7 | 30743011 | rs255125 | 0.747 | 0.403 | -0.06 | 1.209 | -0.002 | 0.066 | 0.066 | 0.077 | 0.7 | 0.53 | 0.51 | 0.62 | 0.68 | 0.51 |
| 7 | 30748323 | rs24003 | -0.287 | 0.133 | -0.185 | 0.589 | 0.004 | 0.006 | 0.006 | 0 | 0.5 | 0.5 | 0.51 | 0.26 | 0.61 | 0.56 |
| 7 | 30754949 | rs255142 | -0.737 | -0.475 | 0.411 | 1.556 | -0.005 | -0.046 | -0.046 | 0.202 | 0.58 | 0.96 | 0.57 | 0.58 | 0.63 | 0.73 |
| 7 | 30755583 | rs10267836 | 0.249 | -0.344 | 0.758 | 1.341 | -0.002 | 0 | 0 | 1.984 | 0.11 | 0.14 | 0.05 | 0.12 | 0.09 | 0.29 |
| 7 | 30759705 | rs255149 | -1.276 | -0.395 | 0.831 | 1.284 | 0.014 | -0.025 | -0.025 | 0 | 0.59 | 0.91 | 0.57 | 0.58 | 0.64 | 0.74 |
| 7 | 30761210 | rs11974150 | -0.91 | -0.399 | 1.372 | 1.236 | -0.01 | 0.083 | 0.083 | 0 | 0.17 | 0.56 | 0.1 | 0.3 | 0.12 | 0.42 |
| 7 | 30761852 | rs255152 | -0.356 | -0.449 | 1.453 | 1.294 | -0.001 | -0.001 | -0.001 | 0.035 | 0.43 | 0.6 | 0.49 | 0.29 | 0.53 | 0.34 |
| 7 | 30764503 | rs255156 | -0.339 | -0.29 | 1.586 | 1.12 | 0 | 0 | 0 | 0 | 0.39 | 0.42 | 0.4 | 0.29 | 0.45 | 0.28 |
| 7 | 30769508 | rs255165 | -1.25 | -0.33 | 1.274 | 1.024 | 0.008 | 0.01 | 0.01 | 0 | 0.53 | 0.88 | 0.5 | 0.58 | 0.58 | 0.68 |
| 7 | 30778815 | rs10239122 | -0.596 | -0.804 | 1.578 | 0.994 | -0.057 | 0.03 | 0.03 | 0.398 | 0.21 | 0.38 | 0.1 | 0.22 | 0.1 | 0.2 |

| | | | | | | | | | | | | | | | | |
|---|---|---|---|---|---|---|---|---|---|---|---|---|---|---|---|---|
| 7 | 30780217 | rs4723008 | 0.27 | 0.042 | 0.964 | 0.702 | -0.063 | 0.021 | 0.021 | 0 | 0.23 | 0.1 | 0.3 | 0.14 | 0.37 | 0.18 |
| 7 | 30780661 | rs255173 | -1.765 | -0.138 | 0.998 | 0.532 | 0.005 | 0.006 | 0.006 | 0.02 | 0.53 | 0.74 | 0.51 | 0.37 | 0.56 | 0.48 |
| 7 | 30782261 | rs10267520 | 0.41 | -0.194 | 1.052 | 0.324 | 0.041 | -0.02 | -0.02 | 0.002 | 0.77 | 0.92 | 0.91 | 0.79 | 0.87 | 0.78 |
| 7 | 30782394 | rs10270738 | 1.534 | -0.198 | 1.245 | 0.421 | -0.014 | 0.157 | 0.157 | 0.344 | 0.51 | 0.23 | 0.61 | 0.78 | 0.52 | 0.6 |
| 7 | 30782763 | rs11771217 | 1.824 | -0.409 | 1.197 | 0.582 | 0.054 | -0.012 | -0.012 | 0 | 0.58 | 0.22 | 0.73 | 0.58 | 0.7 | 0.4 |
| 7 | 30784694 | rs10279462 | -0.469 | -0.535 | 1.482 | 0.454 | -0.04 | -0.106 | -0.106 | 0 | 0.83 | 0.7 | 0.91 | 1 | 0.89 | 0.93 |
| 7 | 30784826 | rs4645485 | 1.805 | -0.306 | 1.195 | 0.526 | 0.004 | 0.112 | 0.112 | 0 | 0.45 | 0.18 | 0.52 | 0.77 | 0.42 | 0.51 |
| 7 | 30787798 | rs6966354 | -0.602 | -0.245 | 1.234 | 0.621 | -0.006 | 0.032 | 0.032 | 0 | 0.11 | 0.18 | 0.06 | 0.27 | 0.09 | 0.29 |
| 7 | 30788778 | rs4723009 | 0.702 | -0.197 | 1.207 | 0.474 | -0.013 | 0.004 | 0.004 | 0 | 0.2 | 0.18 | 0.23 | 0.03 | 0.31 | 0.14 |
| 7 | 30791848 | rs4723010 | -0.11 | -0.257 | 1.384 | 0.431 | 0.006 | -0.035 | -0.035 | 0 | 0.26 | 0.28 | 0.33 | 0.04 | 0.39 | 0.27 |
| 7 | 30795288 | rs2302339 | 0.112 | -0.314 | 1.322 | 0.722 | -0.055 | 0.023 | 0.023 | 0 | 0.24 | 0.13 | 0.25 | 0.17 | 0.35 | 0.38 |
| 7 | 30795331 | rs2302340 | 2.625 | -0.665 | 1.137 | 0.787 | -0.092 | -0.146 | -0.146 | 2.656 | 0.4 | 0.06 | 0.51 | 0.55 | 0.43 | 0.24 |
| 7 | 30795547 | rs6970396 | 0.466 | -0.744 | 1.363 | 0.787 | -0.079 | -0.008 | -0.008 | 0 | 0.77 | 0.54 | 0.89 | 0.84 | 0.9 | 0.79 |
| 7 | 30796530 | rs1061644 | 0.453 | -0.739 | 1.365 | 0.79 | -0.078 | -0.007 | -0.007 | 0 | 0.77 | 0.54 | 0.89 | 0.84 | 0.9 | 0.79 |
| 7 | 30800679 | rs10227423 | -0.726 | -0.531 | 1.263 | 0.864 | 0.015 | 0.015 | 0.015 | 0.044 | 0.11 | 0.29 | 0.14 | 0.13 | 0.11 | 0.16 |
| 7 | 30803158 | rs10265594 | -0.784 | -0.502 | 1.096 | 0.822 | -0.086 | 0.014 | 0.014 | 0 | 0.34 | 0.18 | 0.36 | 0.3 | 0.47 | 0.54 |
| 7 | 30804143 | rs12154580 | 1.235 | -0.447 | 1.063 | 0.811 | -0.016 | 0.019 | 0.019 | 0 | 0.05 | 0.03 | 0.09 | 0 | 0.08 | 0.08 |
| 7 | 30805595 | rs7810199 | -1.227 | -0.8 | 1.167 | 0.841 | -0.002 | -0.003 | -0.003 | 0 | 0.44 | 0.45 | 0.4 | 0.46 | 0.5 | 0.72 |
| 7 | 30806949 | rs7780659 | -0.686 | -1.241 | 1.297 | 1.296 | -0.012 | 0.163 | 0.163 | 0 | 0.08 | 0.18 | 0.04 | 0.2 | 0.02 | 0.15 |
| 7 | 30807671 | rs10807853 | -0.317 | -1.354 | 1.517 | 0.848 | -0.026 | -0.077 | -0.077 | 0.329 | 0.87 | 0.74 | 0.94 | 1 | 0.92 | 0.95 |
| 7 | 30809097 | rs6958083 | -1.423 | -1.205 | 1.464 | 1.061 | -0.022 | 0.054 | 0.054 | 0 | 0.1 | 0.21 | 0.06 | 0.26 | 0.06 | 0.27 |
| 7 | 30809567 | rs6962730 | -1.194 | -1.566 | 1.263 | 1.086 | -0.006 | 0.001 | 0.001 | 0 | 0.2 | 0.2 | 0.08 | 0.19 | 0.11 | 0.2 |
| 7 | 30810264 | rs12670137 | -1.731 | -1.577 | 1.334 | 1.156 | -0.066 | 0.05 | 0.05 | 0 | 0.25 | 0.45 | 0.12 | 0.3 | 0.16 | 0.39 |
| 7 | 30811746 | rs12666880 | -0.34 | -1.016 | 0.762 | 0.52 | -0.03 | 0.025 | 0.025 | 0 | 0.33 | 0.17 | 0.34 | 0.21 | 0.36 | 0.43 |
| 7 | 30819142 | rs12701028 | 1.053 | -1.068 | 0.675 | 0.48 | -0.022 | 0.019 | 0.019 | 0.136 | 0.2 | 0.13 | 0.28 | 0.02 | 0.29 | 0.22 |
| 7 | 30823758 | rs2107551 | -1.892 | -1.514 | 0.486 | 0.757 | -0.015 | 0.005 | 0.005 | 0 | 0.57 | 0.73 | 0.49 | 0.57 | 0.52 | 0.75 |
| 7 | 30825575 | rs12672184 | -0.117 | -1.562 | 0.572 | 0.656 | 0.018 | -0.022 | -0.022 | 0 | 0.08 | 0.03 | 0.02 | 0.17 | 0.02 | 0.16 |
| 7 | 30825762 | rs9986953 | -0.838 | -1.542 | 0.686 | 0.64 | -0.003 | -0.002 | -0.002 | 0 | 0.21 | 0.28 | 0.12 | 0.15 | 0.18 | 0.29 |
| 7 | 30828615 | rs10272887 | -1.947 | -1.539 | 0.692 | 0.637 | -0.015 | -0.003 | -0.003 | 0.11 | 0.57 | 0.72 | 0.47 | 0.54 | 0.52 | 0.75 |
| 7 | 30829039 | rs11768076 | 0.92 | -1.535 | 0.747 | 0.446 | 0.045 | -0.094 | -0.094 | 0 | 0.22 | 0.3 | 0.3 | 0.02 | 0.3 | 0.23 |
| 7 | 30831082 | rs12701034 | -1.148 | -1.563 | 0.716 | 0.435 | -0.027 | 0.165 | 0.165 | 0 | 0.45 | 0.8 | 0.44 | 0.16 | 0.48 | 0.51 |
| 7 | 30831144 | rs16875355 | -0.154 | -1.539 | 0.724 | 0.449 | 0.056 | 0.236 | 0.236 | 0 | 0.05 | 0.32 | 0.06 | 0.21 | 0.02 | 0.07 |
| 7 | 30831692 | rs12701035 | 1.028 | -1.535 | 0.734 | 0.44 | -0.006 | -0.012 | -0.012 | 3.993 | 0.22 | 0.17 | 0.29 | 0.02 | 0.3 | 0.23 |
| 7 | 30835567 | rs2391919 | -1.402 | -1.462 | 0.386 | 0.48 | -0.051 | 0.078 | 0.078 | 0 | 0.49 | 0.85 | 0.47 | 0.37 | 0.5 | 0.59 |
| 7 | 30835879 | rs739981 | -1.381 | -1.509 | 0.361 | 0.48 | -0.055 | 0.074 | 0.074 | 0 | 0.48 | 0.86 | 0.47 | 0.37 | 0.5 | 0.59 |
| 7 | 30836893 | rs6946714 | -0.201 | -1.677 | 0.436 | 0.51 | 0.006 | -0.004 | -0.004 | 0 | 0.15 | 0.1 | 0.07 | 0.21 | 0.11 | 0.12 |
| 7 | 30837003 | rs1017012 | -0.931 | -1.672 | 0.356 | 0.512 | -0.001 | -0.006 | -0.006 | 0 | 0.43 | 0.58 | 0.42 | 0.16 | 0.48 | 0.51 |
| 7 | 30837276 | rs7781249 | -0.923 | -1.667 | 0.358 | 0.514 | -0.001 | -0.006 | -0.006 | 0 | 0.43 | 0.58 | 0.42 | 0.16 | 0.48 | 0.51 |
| 7 | 30838631 | rs7790537 | -1.438 | -1.647 | 0.158 | 0.523 | -0.056 | 0.074 | 0.074 | 0 | 0.49 | 0.9 | 0.47 | 0.37 | 0.5 | 0.58 |
| 7 | 30841441 | rs17781398 | 2.248 | -2.033 | -0.349 | 0.115 | -0.014 | -0.016 | -0.016 | 0 | 0.13 | 0.02 | 0.07 | 0.1 | 0.09 | 0.04 |
| 7 | 30842662 | rs4723018 | -0.529 | -1.881 | 0.592 | 0.029 | 0.009 | -0.05 | -0.05 | 0.572 | 0.36 | 0.54 | 0.39 | 0.16 | 0.39 | 0.47 |
| 7 | 30842782 | rs10250368 | -0.1 | -1.884 | 0.658 | 0.026 | 0.008 | -0.032 | -0.032 | 0 | 0.3 | 0.38 | 0.33 | 0.12 | 0.35 | 0.42 |
| 7 | 30845619 | rs12701041 | -0.432 | -1.826 | 0.393 | 0.061 | 0.011 | -0.05 | -0.05 | 1.219 | 0.36 | 0.54 | 0.38 | 0.16 | 0.39 | 0.47 |
| 7 | 30852670 | rs11981431 | -1.778 | -2.076 | 0.282 | 0.374 | 0.051 | -0.031 | -0.031 | 0.481 | 0.36 | 0.76 | 0.21 | 0.52 | 0.22 | 0.51 |
| 7 | 30857341 | rs6960287 | -1.726 | -2.031 | 0.378 | 0.395 | 0.045 | -0.03 | -0.03 | 0 | 0.35 | 0.74 | 0.2 | 0.52 | 0.22 | 0.51 |
| 7 | 30860218 | rs10155856 | -1.769 | -2.038 | 0.396 | 0.391 | 0.039 | -0.025 | -0.025 | 0 | 0.35 | 0.74 | 0.2 | 0.52 | 0.22 | 0.51 |
| 7 | 30860705 | rs2107552 | -1.736 | -2.037 | 0.387 | 0.392 | 0.041 | -0.027 | -0.027 | 0 | 0.35 | 0.75 | 0.2 | 0.52 | 0.22 | 0.51 |
| 7 | 30865765 | rs10243139 | -0.989 | -2.03 | 0.594 | 0.387 | -0.028 | 0.126 | 0.126 | 0 | 0.2 | 0.53 | 0.14 | 0.4 | 0.1 | 0.29 |
| 7 | 30865976 | rs2040837 | -0.711 | -2.003 | 0.628 | 0.399 | -0.021 | 0.16 | 0.16 | 0 | 0.18 | 0.34 | 0.12 | 0.4 | 0.1 | 0.29 |
| 7 | 30868662 | rs17159487 | -1.256 | -2.13 | 0.504 | 0.202 | -0.055 | -0.039 | -0.039 | 0.021 | 0.19 | 0.36 | 0.1 | 0.1 | 0.16 | 0.23 |
| 7 | 30876432 | rs10258097 | -1.881 | -2.043 | -0.001 | 0.128 | -0.139 | -0.108 | -0.108 | 4.873 | 0.62 | 0.94 | 0.52 | 0.52 | 0.57 | 0.76 |
| 7 | 30876642 | rs10229281 | -1.136 | -2.501 | 1.471 | 1.153 | -0.025 | 0.131 | 0.131 | 0 | 0.18 | 0.29 | 0.11 | 0.39 | 0.09 | 0.23 |
| 7 | 30884273 | rs6972154 | -0.85 | -2.536 | 1.221 | 0.627 | -0.041 | -0.006 | -0.006 | 0.27 | 0.12 | 0.27 | 0.05 | 0.1 | 0.08 | 0.22 |
| 7 | 30887782 | rs1468313 | -1.408 | -2.493 | 1.199 | 0.709 | 0.021 | 0.005 | 0.005 | 0.055 | 0.2 | 0.14 | 0.11 | 0.13 | 0.14 | 0.16 |
| 7 | 30889617 | rs6957048 | -2.11 | -2.491 | 1.189 | 0.73 | -0.052 | 0.045 | 0.045 | 0 | 0.26 | 0.52 | 0.17 | 0.31 | 0.17 | 0.25 |
| 7 | 30894229 | rs10235270 | -2.11 | -2.491 | 1.189 | 0.73 | -0.052 | 0.045 | 0.045 | 0 | 0.26 | 0.52 | 0.17 | 0.31 | 0.17 | 0.25 |
| 7 | 30894983 | rs7780081 | -3.05 | -2.596 | 1.042 | 0.705 | 0.117 | -0.098 | -0.098 | 4.167 | 0.4 | 0.93 | 0.24 | 0.41 | 0.25 | 0.47 |
| 7 | 30895010 | rs7797330 | -2.439 | -2.549 | 1.17 | 0.711 | -0.032 | -0.01 | -0.01 | 0 | 0.33 | 0.41 | 0.16 | 0.24 | 0.23 | 0.38 |
| 7 | 30899543 | rs12674155 | -0.097 | -2.054 | 1.348 | 1.002 | 0.011 | -0.012 | -0.012 | 1.177 | 0.09 | 0.06 | 0.02 | 0.1 | 0.02 | 0.11 |
| 7 | 30905875 | rs10247175 | -2.052 | -2.037 | 1.51 | 0.921 | -0.039 | 0.046 | 0.046 | 0 | 0.17 | 0.48 | 0.14 | 0.22 | 0.13 | 0.14 |
| 7 | 30906193 | rs10277002 | -1.59 | -2.024 | 1.512 | 0.892 | -0.014 | 0.095 | 0.095 | 0 | 0.12 | 0.45 | 0.1 | 0.18 | 0.09 | 0.1 |
| 7 | 30910103 | rs763287 | -1.242 | -2.005 | 1.568 | 1.051 | 0.048 | 0.24 | 0.24 | 0 | 0.07 | 0.42 | 0.06 | 0.2 | 0.02 | 0.08 |
| 7 | 30911984 | rs3801323 | -2.796 | -2.142 | 1.24 | 0.988 | 0.038 | -0.009 | -0.009 | 0 | 0.25 | 0.79 | 0.16 | 0.32 | 0.13 | 0.34 |
| 7 | 30913359 | rs739979 | -1.836 | -2.125 | 1.376 | 0.998 | -0.061 | 0.062 | 0.062 | 0 | 0.16 | 0.62 | 0.09 | 0.29 | 0.06 | 0.28 |
| 7 | 30914408 | rs7782896 | -1.117 | -2.091 | 1.358 | 0.976 | -0.028 | -0.006 | -0.006 | 0.851 | 0.12 | 0.16 | 0.04 | 0.1 | 0.06 | 0.21 |
| 7 | 30914835 | rs7783861 | -2.557 | -1.985 | 0.739 | 0.983 | 0.106 | -0.069 | -0.069 | 0 | 0.34 | 0.87 | 0.18 | 0.42 | 0.16 | 0.44 |
| 7 | 30915478 | rs12669187 | -0.076 | -1.964 | 0.778 | 0.953 | -0.008 | 0.015 | 0.015 | 0 | 0.09 | 0.15 | 0.02 | 0.09 | 0.03 | 0.11 |
| 7 | 30916284 | rs12669292 | -0.043 | -1.963 | 0.633 | 0.954 | -0.006 | 0.007 | 0.007 | 0 | 0.09 | 0.11 | 0.02 | 0.08 | 0.03 | 0.11 |
| 7 | 30918363 | rs2107550 | -2.514 | -1.968 | 0.545 | 0.963 | 0.105 | -0.064 | -0.064 | 0 | 0.34 | 0.87 | 0.18 | 0.41 | 0.16 | 0.44 |

| | | | | | | | | | | | | | | | | |
|---|---|---|---|---|---|---|---|---|---|---|---|---|---|---|---|---|
| 7 | 30928268 | rs10281903 | -1.866 | -1.743 | 0.651 | 0.888 | -0.014 | 0 | 0 | 0 | 0.26 | 0.71 | 0.17 | 0.32 | 0.14 | 0.34 |
| 7 | 30931307 | rs1203181 | 1.756 | -1.719 | 0.402 | 0.809 | 0.053 | 0.016 | 0.016 | 0 | 0.66 | 0.22 | 0.79 | 0.67 | 0.79 | 0.58 |
| 7 | 30932526 | rs10229128 | -1.567 | -1.412 | 1.562 | 1.268 | -0.023 | 0.103 | 0.103 | 0 | 0.15 | 0.35 | 0.07 | 0.26 | 0.07 | 0.3 |
| 7 | 30934659 | rs4723021 | -0.357 | -1.308 | 1.04 | 1.3 | 0.002 | -0.003 | -0.003 | 0 | 0.1 | 0.08 | 0.03 | 0.11 | 0.06 | 0.17 |
| 7 | 30935748 | rs1465216 | -2.204 | -1.385 | 0.718 | 1.337 | 0.072 | -0.052 | -0.052 | 0.102 | 0.28 | 0.86 | 0.13 | 0.35 | 0.13 | 0.47 |
| 7 | 30935900 | rs12112380 | -0.984 | -0.991 | 0.824 | 0.924 | -0.024 | 0.01 | 0.01 | 0 | 0.1 | 0.31 | 0.04 | 0.09 | 0.04 | 0.22 |
| 7 | 30937178 | rs1000597 | -1.112 | -0.552 | 0.189 | 0.958 | -0.017 | -0.001 | -0.001 | 0 | 0.16 | 0.25 | 0.04 | 0.12 | 0.08 | 0.19 |
| 7 | 30938167 | rs1001733 | -1.01 | 0.477 | -0.035 | 1.231 | 0.005 | 0.025 | 0.025 | 0 | 0.16 | 0.21 | 0.19 | 0.31 | 0.17 | 0.12 |
| 7 | 30950744 | rs2075574 | -0.734 | 0.662 | -0.24 | 0.823 | 0 | 0 | 0 | 0 | 0.33 | 0.32 | 0.43 | 0.37 | 0.36 | 0.38 |
| 7 | 30955117 | rs1859838 | -0.809 | -0.607 | 0.564 | -0.27 | -0.052 | -0.144 | -0.144 | 0.002 | 0.23 | 0.43 | 0.17 | 0.01 | 0.16 | 0.13 |
| 7 | 30956451 | rs2267719 | -0.399 | -1.666 | 0.687 | 0.048 | -0.059 | 0.061 | 0.061 | 0 | 0.17 | 0.26 | 0.09 | 0.31 | 0.07 | 0.15 |
| 7 | 30956858 | rs1004317 | -1.695 | -1.692 | -0.164 | -0.675 | 0.03 | 0.03 | 0.03 | 0 | 0.57 | 0.91 | 0.38 | 0.35 | 0.42 | 0.56 |
| 7 | 30958408 | rs1004318 | -0.683 | -1.754 | 0.719 | -0.489 | -0.053 | 0.029 | 0.029 | 0.04 | 0.2 | 0.7 | 0.13 | 0.31 | 0.07 | 0.16 |
| 7 | 30958786 | rs10255904 | -0.587 | -1.713 | 0.879 | -0.341 | -0.076 | 0.089 | 0.089 | 0 | 0.19 | 0.63 | 0.18 | 0.37 | 0.07 | 0.15 |
| 7 | 30959002 | rs17159702 | -1.242 | -1.449 | 0.614 | -0.258 | 0.067 | -0.016 | -0.016 | 0 | 0.41 | 0.77 | 0.39 | 0.47 | 0.29 | 0.35 |
| 7 | 30963769 | rs11537660 | -0.799 | -1.263 | 0.748 | -0.096 | -0.032 | 0.013 | 0.013 | 0.123 | 0.21 | 0.73 | 0.2 | 0.31 | 0.08 | 0.17 |
| 7 | 30963889 | rs11537659 | -0.362 | -1.313 | 1.065 | -0.112 | -0.083 | 0.152 | 0.152 | 0 | 0.17 | 0.51 | 0.17 | 0.31 | 0.05 | 0.12 |
| 7 | 30964145 | rs11537661 | -0.006 | -1.263 | 1.125 | -0.158 | -0.055 | 0.083 | 0.083 | 0 | 0.14 | 0.31 | 0.09 | 0.22 | 0.05 | 0.11 |
| 7 | 30970989 | rs13222180 | -0.174 | -0.427 | 0.157 | -0.535 | -0.002 | 0.071 | 0.071 | 0 | 0.28 | 0.13 | 0.21 | 0.11 | 0.26 | 0.27 |
| 7 | 30971795 | rs10244884 | -1.247 | -0.267 | -0.165 | -0.764 | -0.001 | -0.032 | -0.032 | 0.046 | 0.49 | 0.55 | 0.51 | 0.11 | 0.5 | 0.53 |
| 7 | 30976064 | rs881499 | -0.036 | -0.178 | -0.94 | -0.872 | -0.014 | 0.114 | 0.114 | 0 | 0.28 | 0.11 | 0.22 | 0.04 | 0.29 | 0.24 |
| 7 | 30977442 | rs4723025 | -1.007 | -0.069 | -1.249 | -0.288 | -0.028 | -0.037 | -0.037 | 1.036 | 0.46 | 0.91 | 0.56 | 0.4 | 0.44 | 0.56 |
| 7 | 30989506 | rs4548071 | 0.038 | -0.15 | -1.878 | -0.239 | -0.002 | 0.001 | 0.001 | 0 | 0.11 | 0.1 | 0.25 | 0.09 | 0.14 | 0.2 |
| 7 | 30989577 | rs4507663 | -0.673 | -0.153 | -2.05 | -0.207 | -0.009 | -0.027 | -0.027 | 0 | 0.46 | 0.92 | 0.57 | 0.4 | 0.43 | 0.55 |
| 7 | 31001075 | rs10225302 | -0.492 | -0.858 | -2.339 | -0.763 | -0.121 | -0.021 | -0.021 | 0 | 0.66 | 0.96 | 0.39 | 0.29 | 0.43 | 0.64 |
| 7 | 31001860 | rs2075585 | 0.205 | -1.664 | -1.935 | -1.217 | -0.005 | -0.087 | -0.087 | 2.344 | 0.35 | 0.61 | 0.23 | 0.08 | 0.31 | 0.34 |
| 7 | 31003423 | rs2302019 | -1.08 | -1.714 | -2.093 | -1.086 | -0.131 | -0.049 | -0.049 | 0.363 | 0.66 | 0.96 | 0.39 | 0.31 | 0.43 | 0.64 |
| 7 | 31003819 | rs4988494 | -0.524 | -1.93 | -1.045 | -0.404 | -0.049 | -0.01 | -0.01 | 0 | 0.32 | 0.36 | 0.16 | 0.22 | 0.12 | 0.29 |
| 7 | 31006942 | rs2267723 | -0.95 | -1.003 | -1.652 | -0.738 | -0.031 | 0.126 | 0.126 | 3.859 | 0.54 | 0.89 | 0.43 | 0.25 | 0.47 | 0.54 |
| 7 | 31008004 | rs10227405 | -0.784 | -1.658 | -1.717 | -1.14 | 0.025 | 0.172 | 0.172 | 4.388 | 0.4 | 0.84 | 0.29 | 0.11 | 0.34 | 0.36 |
| 7 | 31008686 | rs4988496 | -0.426 | -1.754 | -0.304 | -1.208 | 0.033 | 0.01 | 0.01 | 0 | 0.1 | 0.34 | 0.05 | 0.06 | 0.06 | 0.03 |
| 7 | 31009576 | rs4988498 | -0.179 | -1.747 | 0.032 | -1.207 | 0.01 | -0.039 | -0.039 | 0 | 0.08 | 0.16 | 0.03 | 0 | 0.06 | 0.02 |
| 7 | 31009960 | rs10278927 | -0.385 | -1.797 | 0.009 | -1.227 | -0.002 | -0.013 | -0.013 | 0 | 0.1 | 0.17 | 0.03 | 0.06 | 0.06 | 0.03 |
| 7 | 31015565 | rs2741 | 0.086 | -1.78 | 0.279 | -1.389 | 0.022 | -0.006 | -0.006 | 0.047 | 0.15 | 0.08 | 0.14 | 0.17 | 0.12 | 0.17 |
| 7 | 31018138 | rs11761979 | 0.758 | -1.756 | 0.128 | -1.471 | -0.042 | 0.054 | 0.054 | 0.247 | 0.12 | 0.04 | 0.17 | 0.03 | 0.22 | 0.14 |
| 7 | 31019305 | rs6962369 | -0.885 | -1.918 | 0.822 | -1.475 | -0.036 | -0.015 | -0.015 | 0.071 | 0.26 | 0.3 | 0.18 | 0.2 | 0.15 | 0.27 |
| 7 | 31030833 | rs10233541 | -1.898 | -1.785 | 0.309 | -1.585 | 0.078 | 0.124 | 0.124 | 4.65 | 0.43 | 0.91 | 0.19 | 0.19 | 0.25 | 0.38 |
| 7 | 31035682 | rs10224932 | -0.81 | -1.585 | 0.902 | -1.878 | 0.026 | -0.074 | -0.074 | 0 | 0.13 | 0.39 | 0.06 | 0.01 | 0.08 | 0.1 |
| 7 | 31040481 | rs13438779 | 0.422 | -1.495 | 0.485 | -1.83 | 0.007 | -0.03 | -0.03 | 1.587 | 0.07 | 0.11 | 0.03 | 0 | 0.06 | 0.02 |
| 7 | 31040963 | rs10224073 | -1.603 | -1.555 | 0.406 | -1.907 | 0.028 | 0.059 | 0.059 | 0 | 0.47 | 0.68 | 0.32 | 0.15 | 0.42 | 0.48 |
| 7 | 31048898 | rs7794503 | -1.63 | -1.858 | -0.148 | -1.829 | 0.097 | 0.219 | 0.219 | 3.143 | 0.41 | 0.93 | 0.19 | 0.12 | 0.21 | 0.33 |
| 7 | 31049951 | rs6462246 | -1.418 | -1.948 | 0.17 | -1.872 | 0.024 | 0.025 | 0.025 | 5.427 | 0.31 | 0.75 | 0.15 | 0.06 | 0.18 | 0.22 |
| 7 | 31057567 | rs733317 | -1.719 | -1.945 | -0.057 | -1.907 | 0.111 | 0.204 | 0.204 | 1.924 | 0.41 | 0.82 | 0.19 | 0.1 | 0.22 | 0.37 |
| 7 | 31061611 | rs878014 | -1.035 | -1.946 | 0.634 | -1.735 | -0.021 | -0.046 | -0.046 | 0.029 | 0.22 | 0.28 | 0.1 | 0.06 | 0.13 | 0.28 |
| 7 | 31065393 | rs10224530 | 0.602 | -1.504 | 0.001 | -1.866 | -0.047 | 0.058 | 0.058 | 0.315 | 0.12 | 0.04 | 0.18 | 0.01 | 0.25 | 0.11 |
| 7 | 31066593 | rs1894847 | -1.444 | -1.503 | 0.365 | -1.825 | 0.005 | -0.013 | -0.013 | 0.018 | 0.39 | 0.7 | 0.33 | 0.07 | 0.38 | 0.41 |
| 7 | 31070366 | rs12701049 | -1.6 | -1.376 | 0.273 | -1.729 | 0.035 | 0.075 | 0.075 | 0 | 0.47 | 0.7 | 0.33 | 0.13 | 0.41 | 0.51 |
| 7 | 31074214 | rs10236961 | -0.69 | -1.211 | -0.085 | -1.644 | -0.009 | -0.063 | -0.063 | 0 | 0.31 | 0.39 | 0.26 | 0.11 | 0.35 | 0.41 |
| 7 | 31075721 | rs17159844 | -0.902 | -0.847 | -0.531 | -0.661 | 0.014 | 0.007 | 0.007 | 0 | 0.14 | 0.11 | 0.05 | 0.09 | 0.07 | 0.13 |
| 7 | 31084099 | rs7784067 | -0.931 | -0.079 | -0.945 | -0.846 | 0.01 | 0.133 | 0.133 | 0.267 | 0.51 | 0.83 | 0.42 | 0.29 | 0.44 | 0.5 |
| 7 | 31085162 | rs17159861 | -0.197 | -0.101 | -0.289 | -1.043 | -0.003 | -0.038 | -0.038 | 0.505 | 0.17 | 0.2 | 0.12 | 0.06 | 0.13 | 0.24 |
| 7 | 31089060 | rs1981701 | -1.417 | 0.536 | -0.925 | -0.907 | -0.001 | 0.098 | 0.098 | 0.186 | 0.51 | 0.84 | 0.49 | 0.32 | 0.52 | 0.52 |
| 7 | 31091235 | rs35413074 | -0.671 | 0.174 | -0.551 | -0.976 | -0.003 | -0.001 | -0.001 | 0 | 0.22 | 0.28 | 0.18 | 0.19 | 0.18 | 0.16 |
| 7 | 31093792 | rs741051 | -1.576 | -0.01 | -0.665 | -1.132 | -0.015 | 0.13 | 0.13 | 0.333 | 0.54 | 0.82 | 0.41 | 0.22 | 0.46 | 0.49 |
| 7 | 31096363 | rs10241138 | -0.759 | 0.112 | 0.311 | -0.752 | -0.009 | -0.068 | -0.068 | 0 | 0.18 | 0.31 | 0.14 | 0.07 | 0.14 | 0.22 |
| 7 | 31115225 | rs887703 | 1.127 | 0.6 | 0.729 | -0.18 | 0.006 | -0.004 | -0.004 | 0 | 0.71 | 0.39 | 0.7 | 0.73 | 0.63 | 0.63 |
| 7 | 31116168 | rs12668955 | 0.717 | -0.047 | 0.269 | 0.35 | -0.012 | 0.006 | 0.006 | 1.281 | 0.55 | 0.3 | 0.59 | 0.43 | 0.5 | 0.53 |
| 7 | 31116266 | rs2267727 | -1.225 | -0.328 | 0.667 | 0.525 | -0.005 | -0.008 | -0.008 | 0 | 0.13 | 0.2 | 0.11 | 0.08 | 0.09 | 0.12 |
| 7 | 31122542 | rs1541516 | -0.455 | -1.359 | 0.568 | 0.655 | -0.067 | -0.002 | -0.002 | 0 | 0.26 | 0.59 | 0.18 | 0.19 | 0.12 | 0.15 |
| 7 | 31122630 | rs2267730 | -0.265 | -0.761 | -0.115 | 1.065 | 0.041 | 0.037 | 0.037 | 0 | 0.34 | 0.47 | 0.48 | 0.49 | 0.45 | 0.34 |
| 7 | 31134317 | rs2267732 | -0.273 | -0.348 | -1.75 | 1.24 | -0.062 | -0.05 | -0.05 | 0 | 0.62 | 0.92 | 0.59 | 0.44 | 0.58 | 0.49 |
| 7 | 31134809 | rs2267733 | -0.727 | -0.332 | -1.804 | 0.902 | -0.093 | -0.094 | -0.094 | 0.034 | 0.69 | 0.97 | 0.69 | 0.64 | 0.62 | 0.59 |
| 7 | 31136610 | rs2267737 | -0.313 | -0.625 | -1.451 | 0.946 | -0.032 | -0.03 | -0.03 | 0 | 0.43 | 0.75 | 0.48 | 0.52 | 0.45 | 0.34 |
| 7 | 31136824 | rs2267738 | 1.149 | -1.79 | -0.836 | 0.93 | -0.004 | 0.006 | 0.006 | 0 | 0.71 | 0.69 | 0.83 | 0.54 | 0.79 | 0.56 |
| 7 | 31137003 | rs2267740 | -1.131 | -2.359 | -0.715 | 0.452 | -0.019 | -0.001 | -0.001 | 0 | 0.21 | 0.77 | 0.15 | 0.27 | 0.1 | 0.13 |
| 7 | 31141418 | rs2041568 | -0.868 | -2.94 | -0.467 | -0.362 | -0.045 | -0.033 | -0.033 | 0.604 | 0.27 | 0.67 | 0.1 | 0.12 | 0.07 | 0.17 |
| 7 | 31141534 | rs7457988 | -0.165 | -3.003 | 0.429 | -0.389 | -0.057 | -0.05 | -0.05 | 0 | 0.18 | 0.26 | 0.04 | 0.06 | 0.05 | 0.15 |
| 7 | 31147761 | rs4403321 | -0.005 | -2.416 | -0.382 | -0.435 | -0.023 | -0.031 | -0.031 | 0 | 0.17 | 0.1 | 0.37 | 0.61 | 0.22 | 0.4 |
| 7 | 31149140 | rs4503014 | -0.636 | -3.082 | -0.458 | -0.812 | -0.008 | -0.005 | -0.005 | 1.059 | 0.22 | 0.18 | 0.39 | 0.68 | 0.24 | 0.55 |

| | | | | | | | | | | | | | | | | |
|---|---|---|---|---|---|---|---|---|---|---|---|---|---|---|---|---|
| 7 | 31149283 | rs7793414 | -1.764 | -3.126 | -0.413 | -0.848 | 0.092 | 0.173 | 0.173 | 0 | 0.49 | 0.78 | 0.49 | 0.79 | 0.31 | 0.65 |
| 7 | 31151579 | rs1008385 | -1.216 | -2.947 | -0.057 | -0.767 | 0.011 | 0.019 | 0.019 | 0.356 | 0.41 | 0.41 | 0.44 | 0.75 | 0.3 | 0.62 |
| 7 | 31157200 | rs17159921 | 0.363 | -2.92 | -0.14 | -0.848 | -0.002 | 0.001 | 0.001 | 2.686 | 0.14 | 0.12 | 0.03 | 0.07 | 0.05 | 0.07 |
| 7 | 31158281 | rs4720028 | -0.663 | -2.728 | -0.532 | -1.22 | -0.027 | -0.011 | -0.011 | 0 | 0.25 | 0.19 | 0.45 | 0.66 | 0.32 | 0.51 |
| 7 | 31161159 | rs10262271 | -0.084 | -3.058 | -0.403 | -1.331 | -0.065 | -0.116 | -0.116 | 0 | 0.19 | 0.4 | 0.11 | 0.03 | 0.11 | 0.03 |
| 7 | 31161253 | rs7808175 | 0.203 | -2.7 | -1.006 | -1.204 | -0.018 | -0.033 | -0.033 | 0 | 0.17 | 0.14 | 0.38 | 0.65 | 0.21 | 0.48 |
| 7 | 31161945 | rs1012440 | 0.243 | -2.759 | -0.845 | -1.211 | -0.003 | 0.001 | 0.001 | 1.452 | 0.16 | 0.15 | 0.04 | 0.08 | 0.06 | 0.11 |
| 7 | 31164655 | rs9792050 | 0.226 | -2.233 | -0.461 | -0.976 | -0.021 | -0.034 | -0.034 | 0 | 0.82 | 0.8 | 0.9 | 0.95 | 0.89 | 0.97 |
| 7 | 31171531 | rs4723048 | -0.24 | -1.96 | -0.74 | -1.639 | 0.021 | 0.01 | 0.01 | 0 | 0.45 | 0.33 | 0.26 | 0.11 | 0.27 | 0.31 |
| 7 | 31175964 | rs7785534 | 0.848 | -1.422 | -1.198 | -0.77 | -0.008 | -0.018 | -0.018 | 0 | 0.12 | 0.05 | 0.28 | 0.59 | 0.14 | 0.37 |
| 7 | 31178749 | rs4291168 | -1.177 | -1.579 | -1.282 | -1.118 | -0.088 | 0.05 | 0.05 | 0.024 | 0.68 | 0.88 | 0.59 | 0.82 | 0.46 | 0.76 |
| 7 | 31191343 | rs1160169 | 0.971 | -1.765 | -0.958 | -0.979 | 0.003 | -0.002 | -0.002 | 0 | 0.09 | 0.04 | 0.09 | 0.08 | 0.05 | 0.11 |
| 7 | 31199267 | rs2041570 | -1.087 | -2.682 | -0.566 | -1.52 | -0.006 | 0.018 | 0.018 | 0 | 0.61 | 0.64 | 0.59 | 0.76 | 0.43 | 0.64 |
| 7 | 31200762 | rs17469030 | 0.869 | -3.129 | -0.787 | -1.752 | 0.007 | -0.002 | -0.002 | 0 | 0.17 | 0.14 | 0.1 | 0.01 | 0.13 | 0.02 |
| 7 | 31204503 | rs6945601 | -1.202 | -3.225 | -0.17 | -1.715 | 0.031 | 0.043 | 0.043 | 0.65 | 0.49 | 0.67 | 0.51 | 0.68 | 0.37 | 0.54 |
| 7 | 31205761 | rs7803859 | -0.612 | -3.133 | -0.386 | -1.85 | 0.025 | 0.035 | 0.035 | 4.731 | 0.35 | 0.29 | 0.25 | 0.09 | 0.23 | 0.16 |
| 7 | 31209019 | rs6462251 | -0.529 | -1.62 | -0.251 | -1.724 | 0.002 | 0.029 | 0.029 | 0 | 0.53 | 0.62 | 0.35 | 0.18 | 0.41 | 0.22 |
| 7 | 31213561 | rs7792372 | -0.577 | -1.435 | -0.181 | -1.617 | 0.004 | 0.048 | 0.048 | 0 | 0.54 | 0.66 | 0.34 | 0.18 | 0.39 | 0.24 |
| 7 | 31213647 | rs7809542 | -1.599 | -1.476 | -0.341 | -1.364 | -0.036 | 0.008 | 0.008 | 0 | 0.76 | 0.84 | 0.69 | 0.81 | 0.62 | 0.69 |
| 7 | 31228648 | rs878445 | 0.933 | -1.095 | -0.261 | -1.399 | 0 | -0.002 | -0.002 | 0 | 0.22 | 0.19 | 0.36 | 0.63 | 0.25 | 0.45 |
| 7 | 31230344 | rs6462256 | -1.572 | -1.096 | -0.36 | -1.389 | -0.024 | 0.025 | 0.025 | 0 | 0.76 | 0.82 | 0.71 | 0.93 | 0.63 | 0.7 |
| 7 | 31234875 | rs6462260 | 0.117 | -1.357 | -0.262 | -1.584 | -0.01 | -0.012 | -0.012 | 0.036 | 0.44 | 0.51 | 0.29 | 0.29 | 0.28 | 0.25 |
| 7 | 31236189 | rs4723062 | -1.854 | -1.319 | -0.304 | -1.578 | -0.084 | 0.072 | 0.072 | 0 | 0.77 | 0.92 | 0.72 | 0.92 | 0.64 | 0.72 |
| 7 | 31238061 | rs6965587 | 0.403 | -1.45 | -0.017 | -1.511 | 0.026 | 0.03 | 0.03 | 0 | 0.41 | 0.27 | 0.27 | 0.29 | 0.28 | 0.25 |
| 7 | 31240822 | rs177982 | 0.665 | -1.296 | -0.269 | -1.629 | -0.061 | 0.088 | 0.088 | 0 | 0.33 | 0.2 | 0.34 | 0.08 | 0.47 | 0.29 |
| 7 | 31247912 | rs177976 | -0.725 | -1.167 | 0.072 | -1.256 | -0.008 | 0.031 | 0.031 | 0 | 0.76 | 0.72 | 0.62 | 0.38 | 0.75 | 0.53 |
| 7 | 31259994 | rs177961 | -0.587 | -1.207 | -0.005 | -1.26 | -0.024 | 0.074 | 0.074 | 0 | 0.74 | 0.53 | 0.62 | 0.37 | 0.75 | 0.53 |
| 7 | 31271453 | rs2536534 | 1.81 | -1.253 | -0.282 | -1.432 | -0.084 | 0.067 | 0.067 | 0.156 | 0.22 | 0.07 | 0.28 | 0.07 | 0.36 | 0.28 |
| 7 | 31273711 | rs1011606 | 0.489 | -1.467 | -0.273 | -1.482 | 0.023 | 0.038 | 0.038 | 0.153 | 0.42 | 0.27 | 0.26 | 0.26 | 0.28 | 0.2 |
| 7 | 31273746 | rs1011605 | 0.627 | -1.313 | -0.417 | -1.669 | -0.06 | 0.067 | 0.067 | 0 | 0.31 | 0.23 | 0.35 | 0.11 | 0.46 | 0.33 |
| 7 | 31277761 | rs2690851 | 0.506 | -1.186 | -0.243 | -1.468 | 0.022 | 0.02 | 0.02 | 0.997 | 0.58 | 0.33 | 0.54 | 0.32 | 0.64 | 0.47 |
| 7 | 31282565 | rs881947 | -1.007 | -1.451 | -0.037 | -1.336 | 0.006 | -0.011 | -0.011 | 0.022 | 0.78 | 0.85 | 0.67 | 0.37 | 0.79 | 0.58 |
| 7 | 31290572 | rs2159766 | 0.745 | -1.394 | -0.2 | -1.712 | -0.007 | 0.009 | 0.009 | 0 | 0.29 | 0.24 | 0.24 | 0.23 | 0.27 | 0.19 |
| 7 | 31292900 | rs960434 | -1.191 | -1.415 | -0.179 | -1.713 | -0.01 | 0.013 | 0.013 | 0 | 0.77 | 0.79 | 0.67 | 0.36 | 0.8 | 0.57 |
| 7 | 31296238 | rs10281854 | -0.135 | -1.466 | -0.06 | -1.824 | -0.002 | 0.006 | 0.006 | 0 | 0.46 | 0.61 | 0.33 | 0.24 | 0.39 | 0.23 |
| 7 | 31299814 | rs2177553 | -0.118 | -1.409 | -0.117 | -1.698 | -0.01 | -0.004 | -0.004 | 0.027 | 0.62 | 0.61 | 0.58 | 0.33 | 0.68 | 0.51 |
| 7 | 31302734 | rs2690885 | 0.972 | -1.059 | -0.49 | -1.734 | -0.058 | 0.104 | 0.104 | 0 | 0.3 | 0.09 | 0.31 | 0.09 | 0.38 | 0.29 |
| 7 | 31303179 | rs2536523 | -0.047 | -1.306 | -0.057 | -1.759 | 0 | 0.055 | 0.055 | 0 | 0.48 | 0.66 | 0.44 | 0.12 | 0.53 | 0.35 |
| 7 | 31307585 | rs2690884 | 1.111 | -0.886 | -0.523 | -1.613 | -0.044 | 0.069 | 0.069 | 0 | 0.27 | 0.13 | 0.3 | 0.08 | 0.39 | 0.27 |
| 7 | 31313869 | rs1917011 | 0.584 | -0.736 | -0.438 | -1.387 | -0.015 | 0.014 | 0.014 | 0.005 | 0.29 | 0.18 | 0.22 | 0.24 | 0.27 | 0.19 |
| 7 | 31319422 | rs11762801 | 0.215 | -0.777 | 0.073 | -1.589 | 0.017 | -0.037 | -0.037 | 0 | 0.37 | 0.43 | 0.37 | 0.12 | 0.47 | 0.32 |
| 7 | 31323316 | rs1017118 | 0.196 | -0.781 | 0.072 | -1.59 | 0.016 | -0.036 | -0.036 | 0.071 | 0.37 | 0.43 | 0.37 | 0.12 | 0.47 | 0.32 |
| 7 | 31327300 | rs2690870 | -1.543 | -0.562 | -0.193 | -1.308 | -0.023 | -0.056 | -0.056 | 0 | 0.82 | 0.93 | 0.78 | 0.48 | 0.78 | 0.6 |
| 7 | 31331381 | rs7805619 | 0.431 | -1.275 | 0.24 | -1.55 | -0.033 | -0.022 | -0.022 | 0.341 | 0.37 | 0.47 | 0.37 | 0.36 | 0.27 | 0.24 |
| 7 | 31333947 | rs6950546 | 1.46 | -1.124 | -0.051 | -1.643 | -0.001 | -0.044 | -0.044 | 0.204 | 0.16 | 0.18 | 0.1 | 0 | 0.14 | 0.05 |
| 7 | 31335097 | rs10246694 | 0.52 | -0.932 | -0.02 | -1.497 | 0.011 | 0.002 | 0.002 | 0.742 | 0.36 | 0.28 | 0.37 | 0.4 | 0.27 | 0.27 |
| 7 | 31337504 | rs6942902 | -0.552 | -0.591 | -0.071 | -1.294 | -0.002 | 0 | 0 | 0 | 0.63 | 0.62 | 0.5 | 0.38 | 0.45 | 0.36 |
| 7 | 31344131 | rs17160070 | -0.214 | -0.673 | -0.283 | -1.48 | 0.001 | 0.002 | 0.002 | 0 | 0.46 | 0.47 | 0.4 | 0.53 | 0.43 | 0.52 |
| 7 | 31344595 | rs1860795 | -1.591 | -0.93 | -0.268 | -1.614 | -0.012 | 0.01 | 0.01 | 0 | 0.78 | 0.85 | 0.73 | 0.94 | 0.66 | 0.72 |
| 7 | 31346073 | rs10241207 | 0.869 | -1.48 | -0.197 | -1.731 | 0.036 | -0.008 | -0.008 | 0 | 0.31 | 0.14 | 0.31 | 0.41 | 0.23 | 0.2 |
| 7 | 31348947 | rs1880337 | 0.04 | -0.828 | -0.126 | -1.801 | 0.022 | 0.026 | 0.026 | 0 | 0.53 | 0.64 | 0.52 | 0.45 | 0.43 | 0.51 |
| 7 | 31349138 | rs1880336 | 0.514 | -1.017 | -0.132 | -1.736 | -0.033 | -0.004 | -0.004 | 0 | 0.34 | 0.39 | 0.33 | 0.41 | 0.23 | 0.21 |
| 7 | 31351245 | rs17160087 | 0.249 | -0.771 | -0.195 | -2.243 | -0.023 | -0.096 | -0.096 | 0 | 0.24 | 0.42 | 0.16 | 0.02 | 0.16 | 0.12 |
| 7 | 31354581 | rs9648374 | 0.908 | -0.772 | -0.856 | -2.113 | -0.005 | 0.028 | 0.028 | 0 | 0.32 | 0.25 | 0.31 | 0.29 | 0.3 | 0.21 |
| 7 | 31358060 | rs1558434 | 1.592 | -0.206 | -0.692 | -1.914 | -0.014 | 0.043 | 0.043 | 0 | 0.19 | 0.11 | 0.19 | 0.03 | 0.2 | 0.33 |
| 7 | 31361612 | rs10278661 | 0.409 | -0.086 | -1.036 | -1.901 | 0.015 | 0.123 | 0.123 | 0 | 0.44 | 0.23 | 0.36 | 0.29 | 0.36 | 0.23 |
| 7 | 31365758 | rs6462270 | -0.884 | -0.033 | -1.156 | -1.886 | -0.089 | -0.045 | -0.045 | 4.875 | 0.77 | 0.94 | 0.64 | 0.32 | 0.63 | 0.62 |
| 7 | 31371416 | rs10951290 | 1.026 | -0.175 | -1.246 | -1.957 | 0.012 | 0.072 | 0.072 | 1.43 | 0.36 | 0.11 | 0.28 | 0.29 | 0.27 | 0.2 |
| 7 | 31375757 | rs11767648 | -1.012 | 0.127 | -1.298 | -1.833 | -0.073 | -0.042 | -0.042 | 0 | 0.77 | 0.92 | 0.64 | 0.34 | 0.62 | 0.62 |
| 7 | 31377369 | rs10238918 | 0.531 | -0.541 | -1.052 | -1.734 | 0.033 | 0.063 | 0.063 | 1.367 | 0.39 | 0.21 | 0.3 | 0.29 | 0.27 | 0.24 |
| 7 | 31380817 | rs2233401 | 0.164 | -0.126 | -0.661 | -1.107 | -0.033 | -0.034 | -0.034 | 1.125 | 0.07 | 0.16 | 0.02 | 0 | 0.03 | 0.05 |
| 7 | 31382504 | rs2191888 | 0.98 | 0.636 | -1.029 | -0.787 | 0 | -0.014 | -0.014 | 0 | 0.12 | 0.1 | 0.11 | 0.01 | 0.14 | 0.03 |
| 7 | 31393552 | rs6949499 | -0.098 | 1.223 | -1.826 | -0.423 | -0.072 | 0.045 | 0.045 | 1.586 | 0.29 | 0.06 | 0.29 | 0.14 | 0.31 | 0.22 |
| 7 | 31395137 | rs962060 | 0.086 | 1.611 | -1.865 | 0.054 | -0.038 | -0.025 | -0.025 | 0.077 | 0.91 | 0.98 | 0.89 | 0.45 | 0.84 | 0.75 |
| 7 | 31399333 | rs7801993 | 0.124 | 1.519 | -1.946 | -0.161 | -0.096 | -0.014 | -0.014 | 0 | 0.67 | 0.91 | 0.61 | 0.35 | 0.55 | 0.58 |
| 7 | 31407901 | rs10265343 | -0.069 | 1.236 | -1.805 | -0.072 | -0.018 | -0.075 | -0.075 | 0 | 0.82 | 0.94 | 0.84 | 0.48 | 0.75 | 0.74 |
| 7 | 31411765 | rs12112223 | 0.942 | 0.644 | -1.339 | -1.138 | -0.001 | -0.056 | -0.056 | 0 | 0.28 | 0.38 | 0.21 | 0.07 | 0.29 | 0.27 |
| 7 | 31412771 | rs7801471 | -0.825 | 0.716 | -1.421 | -1.355 | -0.065 | -0.016 | -0.016 | 0 | 0.7 | 0.84 | 0.56 | 0.35 | 0.53 | 0.56 |
| 7 | 31413303 | rs1006229 | -0.365 | 0.55 | -1.051 | -1.301 | -0.02 | -0.03 | -0.03 | 1.934 | 0.42 | 0.45 | 0.34 | 0.28 | 0.25 | 0.3 |

| | | | | | | | | | | | | | | | | |
|---|---|---|---|---|---|---|---|---|---|---|---|---|---|---|---|---|
| 7 | 31415731 | rs12701079 | 0.308 | 0.299 | -1.372 | -1.303 | 0.071 | 0.032 | 0.032 | 0 | 0.3 | 0.12 | 0.27 | 0.26 | 0.15 | 0.21 |
| 7 | 31416548 | rs6462272 | 0.174 | 0.577 | -1.151 | -1.53 | 0 | -0.06 | -0.06 | 0.021 | 0.22 | 0.25 | 0.12 | 0 | 0.18 | 0.1 |
| 7 | 31419266 | rs17395568 | 0.141 | 0.242 | -1.491 | -1.217 | -0.018 | -0.021 | -0.021 | 0 | 0.23 | 0.07 | 0.15 | 0.27 | 0.2 | 0.2 |
| 7 | 31422667 | rs12113762 | -0.346 | 0.141 | -0.728 | -1.468 | -0.002 | -0.014 | -0.014 | 0 | 0.14 | 0.26 | 0.17 | 0.06 | 0.11 | 0.05 |
| 7 | 31434405 | rs6953761 | -0.888 | 0.151 | -1.506 | -1.853 | -0.037 | 0.035 | 0.035 | 0 | 0.7 | 0.97 | 0.75 | 0.16 | 0.67 | 0.62 |
| 7 | 31435744 | rs17160114 | 0.909 | 0.511 | -0.825 | -2.245 | 0.099 | -0.072 | -0.072 | 0 | 0.17 | 0.46 | 0.23 | 0.08 | 0.24 | 0.17 |
| 7 | 31438372 | rs11980453 | 0.772 | -0.176 | -0.664 | -2.225 | -0.046 | -0.106 | -0.106 | 0 | 0.14 | 0.53 | 0.21 | 0 | 0.06 | 0.13 |
| 7 | 31443359 | rs4492273 | -0.787 | 0.307 | -1.177 | -1.918 | -0.034 | 0.035 | 0.035 | 0 | 0.7 | 0.97 | 0.75 | 0.16 | 0.67 | 0.62 |
| 7 | 31444895 | rs1125984 | -0.796 | 0.306 | -1.175 | -1.916 | -0.038 | 0.034 | 0.034 | 0 | 0.7 | 0.97 | 0.75 | 0.16 | 0.67 | 0.62 |
| 7 | 31445815 | rs17160130 | -0.224 | 0.38 | -0.65 | -1.725 | 0 | -0.009 | -0.009 | 0 | 0.16 | 0.21 | 0.18 | 0.06 | 0.13 | 0.08 |
| 7 | 31452356 | rs12671007 | 0.932 | 0.311 | -0.948 | -1.605 | 0.007 | -0.006 | -0.006 | 0 | 0.06 | 0.05 | 0.1 | 0.06 | 0.04 | 0.05 |
| 7 | 31454564 | rs2108308 | 0.653 | 0.201 | -1.19 | -1.869 | 0.018 | 0.034 | 0.034 | 0 | 0.3 | 0.11 | 0.21 | 0.82 | 0.28 | 0.38 |
| 7 | 31454647 | rs10951292 | -0.604 | 0.721 | -0.6 | -1.763 | 0.021 | -0.046 | -0.046 | 0 | 0.2 | 0.43 | 0.25 | 0.08 | 0.2 | 0.13 |
| 7 | 31458772 | rs17474907 | 1.182 | 0.908 | -1.035 | -1.775 | -0.01 | 0.009 | 0.009 | 1.155 | 0.19 | 0.16 | 0.31 | 0.08 | 0.22 | 0.2 |
| 7 | 31459659 | rs12674467 | 0.394 | 0.836 | -1.09 | -1.814 | 0.01 | 0.005 | 0.005 | 0.013 | 0.11 | 0.05 | 0.15 | 0.06 | 0.09 | 0.07 |
| 7 | 31461564 | rs6957557 | -0.186 | 0.846 | -1.015 | -1.811 | -0.011 | 0.025 | 0.025 | 0.136 | 0.3 | 0.26 | 0.47 | 0.13 | 0.32 | 0.25 |
| 7 | 31462914 | rs2190442 | -0.941 | 0.578 | -1.112 | -2.079 | -0.007 | 0.066 | 0.066 | 0 | 0.58 | 0.72 | 0.69 | 0.96 | 0.57 | 0.67 |
| 7 | 31464032 | rs6462276 | 0.471 | 0.739 | -1.526 | -1.866 | 0.011 | 0.007 | 0.007 | 0.017 | 0.12 | 0.08 | 0.15 | 0.07 | 0.09 | 0.07 |
| 7 | 31465743 | rs11983505 | 0.399 | 0.763 | -1.397 | -1.576 | 0.004 | -0.007 | -0.007 | 0 | 0.26 | 0.31 | 0.42 | 0.24 | 0.27 | 0.24 |
| 7 | 31466377 | rs218151 | 0.099 | 0.77 | -2.053 | -1.546 | 0.014 | 0.039 | 0.039 | 0.476 | 0.36 | 0.05 | 0.27 | 0.71 | 0.35 | 0.45 |
| 7 | 31467955 | rs218149 | 0.228 | 0.811 | -1.665 | -0.452 | -0.031 | -0.042 | -0.042 | 0.011 | 0.36 | 0.21 | 0.47 | 0.54 | 0.41 | 0.39 |
| 7 | 31468306 | rs218148 | 0.315 | 0.535 | -1.449 | -0.687 | 0.046 | -0.002 | -0.002 | 0 | 0.47 | 0.25 | 0.55 | 0.62 | 0.59 | 0.47 |
| 7 | 31471205 | rs218141 | -0.829 | 1.11 | -0.858 | -0.583 | -0.014 | -0.01 | -0.01 | 0 | 0.61 | 0.67 | 0.65 | 0.5 | 0.53 | 0.52 |
| 7 | 31472061 | rs218137 | -1.025 | 0.862 | -0.957 | -0.602 | -0.009 | -0.026 | -0.026 | 0 | 0.63 | 0.76 | 0.71 | 0.5 | 0.58 | 0.55 |
| 7 | 31480332 | rs218122 | -1.06 | 0.704 | -0.991 | -0.538 | -0.007 | -0.037 | -0.037 | 0.743 | 0.63 | 0.77 | 0.7 | 0.54 | 0.57 | 0.55 |
| 7 | 31482533 | rs17160152 | 0.288 | 0.497 | -1.292 | -0.742 | -0.009 | -0.086 | -0.086 | 0 | 0.21 | 0.06 | 0.25 | 0.32 | 0.16 | 0.31 |
| 7 | 31486548 | rs218112 | 0.036 | 0.586 | -0.792 | -0.818 | 0.002 | -0.005 | -0.005 | 0.298 | 0.46 | 0.52 | 0.54 | 0.5 | 0.44 | 0.45 |
| 7 | 31492624 | rs218106 | -0.129 | 1.278 | -0.667 | -1.051 | -0.004 | 0.013 | 0.013 | 0.135 | 0.41 | 0.37 | 0.41 | 0.24 | 0.41 | 0.23 |
| 7 | 31492912 | rs6943112 | 0.057 | 1.153 | -0.465 | -0.43 | 0.001 | -0.021 | -0.021 | 0 | 0.09 | 0.06 | 0.05 | 0 | 0.11 | 0.05 |
| 7 | 31497220 | rs218098 | 0.895 | 1.072 | -0.078 | -0.45 | 0.017 | -0.048 | -0.048 | 0 | 0.74 | 0.53 | 0.65 | 0.89 | 0.75 | 0.77 |
| 7 | 31497237 | rs218097 | 1.272 | 0.454 | -0.167 | -0.795 | 0 | -0.015 | -0.015 | 0 | 0.67 | 0.64 | 0.65 | 0.9 | 0.71 | 0.71 |
| 7 | 31500579 | rs218094 | 0.723 | 0.046 | -0.6 | -0.955 | -0.006 | 0.056 | 0.056 | 1.499 | 0.34 | 0.18 | 0.35 | 0.3 | 0.34 | 0.3 |
| 7 | 31501174 | rs17160175 | 0.148 | 0.039 | -0.494 | -1.138 | -0.001 | -0.025 | -0.025 | 0.71 | 0.15 | 0.19 | 0.11 | 0 | 0.14 | 0.09 |
| 7 | 31504969 | rs218092 | -0.431 | 0.027 | -0.086 | -1.126 | 0 | -0.004 | -0.004 | 0 | 0.5 | 0.53 | 0.48 | 0.3 | 0.48 | 0.4 |
| 7 | 31512872 | rs11764304 | -0.421 | 0.157 | 0.282 | -0.668 | 0.004 | -0.078 | -0.078 | 1.611 | 0.15 | 0.28 | 0.11 | 0 | 0.14 | 0.1 |
| 7 | 31518967 | rs10228946 | -0.004 | -0.027 | -0.151 | -0.043 | -0.006 | -0.029 | -0.029 | 0 | 0.27 | 0.25 | 0.24 | 0.42 | 0.35 | 0.39 |
| 7 | 31520147 | rs740046 | 0.077 | -0.351 | 0.095 | -0.433 | 0.001 | -0.028 | -0.028 | 0 | 0.64 | 0.53 | 0.58 | 0.71 | 0.66 | 0.67 |
| 7 | 31522676 | rs4401747 | 1.025 | -0.233 | 0.259 | -0.478 | 0.019 | -0.025 | -0.025 | 0 | 0.65 | 0.52 | 0.49 | 0.73 | 0.63 | 0.56 |
| 7 | 31522993 | rs7788263 | 1.657 | -0.143 | 0.044 | -0.286 | -0.01 | -0.02 | -0.02 | 1.424 | 0.17 | 0.12 | 0.16 | 0.41 | 0.24 | 0.26 |
| 7 | 31526529 | rs16875548 | 1.639 | -0.147 | 0.042 | -0.273 | -0.01 | -0.021 | -0.021 | 0 | 0.17 | 0.12 | 0.16 | 0.41 | 0.24 | 0.27 |
| 7 | 31526824 | rs11763989 | 1.063 | -0.138 | -0.069 | -0.406 | -0.005 | 0.007 | 0.007 | 0 | 0.11 | 0.08 | 0.08 | 0 | 0.11 | 0.12 |
| 7 | 31528056 | rs6973071 | 2.107 | -0.358 | -0.392 | -0.445 | -0.077 | 0.007 | 0.007 | 0 | 0.24 | 0.01 | 0.15 | 0.26 | 0.24 | 0.21 |
| 7 | 31528906 | rs12673707 | -1.264 | -0.213 | -0.182 | -0.434 | 0.024 | -0.001 | -0.001 | 0.518 | 0.91 | 0.94 | 0.87 | 0.94 | 0.81 | 0.92 |
| 7 | 31533784 | rs882460 | 0.44 | -0.525 | 0.015 | -0.563 | 0.021 | 0.034 | 0.034 | 0 | 0.33 | 0.28 | 0.24 | 0.08 | 0.25 | 0.21 |
| 7 | 31534855 | rs6972352 | -1.529 | -0.449 | -0.008 | -0.557 | -0.052 | -0.01 | -0.01 | 0 | 0.92 | 0.99 | 0.88 | 0.94 | 0.81 | 0.92 |
| 7 | 31537510 | rs11773035 | 0.559 | -0.518 | 0.117 | -0.589 | -0.003 | 0.004 | 0.004 | 0 | 0.59 | 0.4 | 0.41 | 0.34 | 0.49 | 0.42 |
| 7 | 31551556 | rs719435 | 0.522 | -0.558 | 0.304 | -0.533 | 0.023 | -0.001 | -0.001 | 1.872 | 0.75 | 0.62 | 0.57 | 0.74 | 0.73 | 0.68 |
| 7 | 31552117 | rs2190443 | 1.814 | -0.667 | 0.104 | -0.579 | -0.003 | -0.038 | -0.038 | 0 | 0.07 | 0 | 0.25 | 0.15 | 0.08 | 0.18 |
| 7 | 31555490 | rs17398485 | 0.668 | -0.618 | 0.154 | -0.684 | -0.007 | 0.015 | 0.015 | 0 | 0.58 | 0.38 | 0.41 | 0.34 | 0.49 | 0.42 |
| 7 | 31562114 | rs217164 | -0.198 | -0.645 | 0.157 | -0.681 | -0.006 | -0.002 | -0.002 | 0.01 | 0.33 | 0.6 | 0.46 | 0.59 | 0.33 | 0.49 |
| 7 | 31563701 | rs217166 | -0.049 | -0.714 | 0.147 | -0.861 | -0.026 | 0.078 | 0.078 | 0 | 0.59 | 0.7 | 0.63 | 0.85 | 0.56 | 0.7 |
| 7 | 31567906 | rs2191336 | 1.556 | -0.744 | 0.584 | -0.769 | -0.017 | 0.052 | 0.052 | 0 | 0.09 | 0.18 | 0.27 | 0.21 | 0.08 | 0.2 |
| 7 | 31569547 | rs12701110 | 2.246 | -0.636 | -0.083 | -0.367 | -0.016 | -0.002 | -0.002 | 0.849 | 0.25 | 0.07 | 0.16 | 0.31 | 0.23 | 0.21 |
| 7 | 31573901 | rs10229918 | 0.222 | -0.757 | -0.08 | -0.257 | -0.006 | 0.004 | 0.004 | 0 | 0.66 | 0.4 | 0.53 | 0.45 | 0.67 | 0.51 |
| 7 | 31581104 | rs4720043 | -0.38 | -0.694 | -0.081 | -0.224 | -0.067 | 0.033 | 0.033 | 0 | 0.83 | 0.61 | 0.71 | 0.79 | 0.92 | 0.76 |
| 7 | 31584122 | rs719967 | 1.917 | -0.654 | -0.355 | -0.205 | 0.001 | 0.02 | 0.02 | 0 | 0.16 | 0.16 | 0.17 | 0.34 | 0.24 | 0.25 |
| 7 | 31587618 | rs10243603 | 1.318 | -0.512 | -0.374 | -0.264 | -0.108 | -0.004 | -0.004 | 1.05 | 0.33 | 0.09 | 0.29 | 0.37 | 0.42 | 0.3 |
| 7 | 31592733 | rs7811042 | 0.448 | -0.444 | 0.007 | 0.197 | -0.004 | 0.003 | 0.003 | 0 | 0.66 | 0.39 | 0.53 | 0.45 | 0.67 | 0.51 |
| 7 | 31594508 | rs2286711 | 1.139 | 0.456 | -0.594 | 0.47 | 0.002 | 0.021 | 0.021 | 0 | 0.16 | 0.16 | 0.17 | 0.33 | 0.24 | 0.25 |
| 7 | 31607134 | rs17160267 | 0.33 | -0.046 | -0.475 | -0.283 | 0.006 | 0.005 | 0.005 | 0 | 0.21 | 0.16 | 0.15 | 0.08 | 0.15 | 0.1 |
| 7 | 31608495 | rs4720044 | -0.227 | 0.251 | -0.688 | -0.176 | -0.001 | -0.001 | -0.001 | 0 | 0.42 | 0.57 | 0.59 | 0.41 | 0.43 | 0.39 |
| 7 | 31611522 | rs2041364 | 0.329 | -0.045 | -0.562 | -0.235 | -0.01 | -0.046 | -0.046 | 0.071 | 0.88 | 0.8 | 0.92 | 1 | 0.9 | 0.94 |
| 7 | 31615654 | rs6963296 | -0.606 | 0.007 | -1.084 | -0.384 | -0.011 | 0.017 | 0.017 | 0 | 0.49 | 0.7 | 0.62 | 0.44 | 0.51 | 0.45 |
| 7 | 31623746 | rs1014137 | -0.19 | -0.115 | -1.038 | -0.469 | -0.004 | -0.032 | -0.032 | 0.547 | 0.4 | 0.68 | 0.56 | 0.45 | 0.4 | 0.43 |
| 7 | 31629683 | rs1610058 | -0.244 | -0.119 | -1.075 | -0.462 | -0.003 | -0.033 | -0.033 | 0.28 | 0.41 | 0.67 | 0.56 | 0.46 | 0.4 | 0.44 |
| 7 | 31631594 | rs10274825 | 0.61 | -0.476 | -1.224 | -0.639 | 0.01 | 0.011 | 0.011 | 0 | 0.12 | 0.12 | 0.09 | 0 | 0.1 | 0.04 |
| 7 | 31633271 | rs12701126 | 1.036 | -0.572 | -1.289 | -0.689 | -0.065 | -0.07 | -0.07 | 0.999 | 0.43 | 0.12 | 0.33 | 0.53 | 0.48 | 0.49 |
| 7 | 31635011 | rs16875585 | 1.131 | -0.596 | -0.596 | -1.088 | -0.003 | -0.085 | -0.085 | 0 | 0.16 | 0.26 | 0.1 | 0 | 0.12 | 0.02 |
| 7 | 31638817 | rs7777851 | 0.338 | -0.623 | -0.922 | -0.926 | 0.008 | -0.093 | -0.093 | 0 | 0.17 | 0.03 | 0.36 | 0.35 | 0.14 | 0.35 |

| | | | | | | | | | | | | | | | | |
|---|---|---|---|---|---|---|---|---|---|---|---|---|---|---|---|---|
| 7 | 31640286 | rs6949643 | 1.372 | -0.736 | -1.064 | -1.049 | 0.051 | 0.025 | 0.025 | 0.194 | 0.45 | 0.19 | 0.4 | 0.64 | 0.6 | 0.53 |
| 7 | 31646038 | rs2109395 | -0.708 | -0.647 | -0.839 | -0.877 | 0 | -0.023 | -0.023 | 1.931 | 0.41 | 0.65 | 0.56 | 0.46 | 0.4 | 0.47 |
| 7 | 31652758 | rs38333 | -0.118 | -0.582 | -0.958 | -1.117 | 0.002 | 0.018 | 0.018 | 0 | 0.91 | 0.96 | 0.96 | 0.99 | 0.93 | 0.88 |
| 7 | 31653625 | rs763485 | -0.378 | -0.518 | -0.459 | -0.768 | 0.006 | 0.05 | 0.05 | 0.8 | 0.24 | 0.3 | 0.45 | 0.51 | 0.27 | 0.42 |
| 7 | 31654401 | rs10251898 | 0.384 | 0.224 | -0.637 | -1.259 | -0.041 | -0.027 | -0.027 | 0 | 0.07 | 0.02 | 0.09 | 0.11 | 0.16 | 0.1 |
| 7 | 31665800 | rs1003405 | 0.437 | 0.159 | -0.558 | -1.767 | 0.002 | 0.011 | 0.011 | 1.483 | 0.5 | 0.55 | 0.63 | 0.81 | 0.52 | 0.76 |
| 7 | 31667684 | rs8180867 | 0.706 | 0.124 | -0.588 | -1.783 | 0 | -0.015 | -0.015 | 0 | 0.48 | 0.44 | 0.62 | 0.81 | 0.52 | 0.75 |
| 7 | 31670531 | rs6462290 | 0.424 | 0.289 | -0.753 | -1.709 | -0.002 | 0.034 | 0.034 | 1.516 | 0.52 | 0.66 | 0.66 | 0.81 | 0.53 | 0.78 |
| 7 | 31672870 | rs38379 | 0.208 | 0.415 | -1.128 | -1.298 | -0.13 | 0.091 | 0.091 | 0 | 0.27 | 0.01 | 0.24 | 0.17 | 0.32 | 0.17 |
| 7 | 31682453 | rs4141001 | 0.12 | 0.918 | -0.226 | -0.883 | 0.026 | -0.055 | -0.055 | 0.172 | 0.76 | 0.57 | 0.78 | 0.81 | 0.75 | 0.81 |
| 7 | 31683410 | rs10247620 | -0.168 | 0.945 | -0.239 | -0.871 | 0.008 | -0.017 | -0.017 | 0 | 0.78 | 0.69 | 0.79 | 0.81 | 0.75 | 0.81 |
| 7 | 31685909 | rs27327 | 1.206 | 0.938 | -0.202 | -0.884 | -0.007 | 0.004 | 0.004 | 0.54 | 0.15 | 0.18 | 0.18 | 0.19 | 0.21 | 0.16 |
| 7 | 31691697 | rs3735424 | -0.217 | 0.964 | -0.138 | -0.882 | 0.014 | -0.029 | -0.029 | 0.036 | 0.78 | 0.67 | 0.79 | 0.8 | 0.75 | 0.81 |
| 7 | 31719569 | rs10258296 | 0.254 | 1.004 | -0.295 | -0.987 | -0.012 | 0.012 | 0.012 | 0 | 0.21 | 0.17 | 0.17 | 0.08 | 0.29 | 0.1 |
| 7 | 31720056 | rs1544573 | 0.346 | 1.144 | -0.303 | -0.596 | 0.027 | -0.058 | -0.058 | 0 | 0.75 | 0.6 | 0.76 | 0.8 | 0.71 | 0.79 |
| 7 | 31725977 | rs34156 | -0.939 | 0.241 | 0.216 | -0.026 | -0.022 | -0.041 | -0.041 | 0 | 0.09 | 0.36 | 0.07 | 0.01 | 0.03 | 0.03 |
| 7 | 31726465 | rs34158 | -0.413 | 0.714 | -0.888 | 0.508 | -0.035 | -0.049 | -0.049 | 3.949 | 0.32 | 0.21 | 0.41 | 0.47 | 0.43 | 0.33 |
| 7 | 31727749 | rs34161 | -1.276 | 0.345 | -0.732 | 0.935 | 0.026 | -0.03 | -0.03 | 0 | 0.34 | 0.8 | 0.29 | 0.35 | 0.27 | 0.33 |
| 7 | 31728980 | rs6953623 | -1.071 | 0.159 | 0.315 | 1.043 | -0.033 | -0.037 | -0.037 | 0.38 | 0.24 | 0.52 | 0.14 | 0.14 | 0.16 | 0.15 |
| 7 | 31730800 | rs2284227 | -1.891 | 0.467 | 1.539 | 1.917 | -0.009 | 0.204 | 0.204 | 0 | 0.17 | 0.34 | 0.19 | 0.47 | 0.15 | 0.14 |
| 7 | 31742050 | rs12534672 | -0.335 | 0.336 | 1.799 | 1.793 | -0.016 | -0.001 | -0.001 | 0 | 0.07 | 0.04 | 0.1 | 0.24 | 0.13 | 0.1 |
| 7 | 31744580 | rs2299909 | -0.38 | 0.333 | 1.793 | 1.812 | -0.024 | -0.014 | -0.014 | 1.199 | 0.07 | 0.03 | 0.1 | 0.25 | 0.13 | 0.1 |
| 7 | 31744814 | rs34182 | -2.317 | 0.283 | 2.083 | 1.576 | 0.003 | -0.008 | -0.008 | 0 | 0.3 | 0.45 | 0.35 | 0.66 | 0.34 | 0.37 |
| 7 | 31747249 | rs12669290 | -0.033 | 0.171 | 1.925 | 1.636 | -0.005 | -0.014 | -0.014 | 0.461 | 0.07 | 0.01 | 0.04 | 0.06 | 0.07 | 0.05 |
| 7 | 31755670 | rs34198 | -2.689 | 0.067 | 2.131 | 1.57 | 0.008 | 0.018 | 0.018 | 4.381 | 0.34 | 0.77 | 0.38 | 0.68 | 0.33 | 0.4 |
| 7 | 31756362 | rs2159688 | -2.046 | -0.036 | 1.877 | 1.36 | -0.004 | -0.006 | -0.006 | 0 | 0.37 | 0.56 | 0.39 | 0.62 | 0.38 | 0.42 |
| 7 | 31768673 | rs12532450 | -2.053 | 0.352 | 2.302 | 2.082 | 0.002 | 0.062 | 0.062 | 0.459 | 0.23 | 0.45 | 0.3 | 0.62 | 0.25 | 0.33 |
| 7 | 31772384 | rs10247756 | -1.369 | -0.172 | 2.092 | 1.744 | -0.015 | 0.227 | 0.227 | 0.08 | 0.16 | 0.45 | 0.22 | 0.36 | 0.13 | 0.2 |
| 7 | 31779145 | rs10254520 | 1.607 | 0.191 | 1.296 | 1.577 | 0.007 | -0.02 | -0.02 | 0 | 0.73 | 0.25 | 0.66 | 0.6 | 0.76 | 0.73 |
| 7 | 31786849 | rs7795363 | 1.026 | 1.046 | 1.208 | 1.903 | 0.051 | 0.202 | 0.202 | 0 | 0.87 | 0.64 | 0.79 | 0.67 | 0.84 | 0.87 |
| 7 | 31794740 | rs10239671 | -1.213 | 0.141 | 0.859 | 1.922 | -0.029 | 0.192 | 0.192 | 0 | 0.14 | 0.29 | 0.2 | 0.51 | 0.11 | 0.23 |
| 7 | 31794956 | rs10240123 | -0.001 | 0.461 | 0.473 | 1.67 | -0.001 | 0.003 | 0.003 | 0 | 0.64 | 0.66 | 0.77 | 0.58 | 0.62 | 0.68 |
| 7 | 31795155 | rs917749 | 0.693 | 1.299 | 1.115 | 2.009 | 0.017 | 0.085 | 0.085 | 0 | 0.78 | 0.68 | 0.61 | 0.47 | 0.73 | 0.73 |
| 7 | 31795168 | rs917750 | 0.897 | 1.466 | 1.079 | 2.087 | 0.015 | 0.039 | 0.039 | 0 | 0.77 | 0.67 | 0.59 | 0.36 | 0.72 | 0.68 |
| 7 | 31795856 | rs6952967 | -1.41 | 1.854 | 1.055 | 1.677 | 0.024 | 0.014 | 0.014 | 0 | 0.31 | 0.53 | 0.43 | 0.31 | 0.43 | 0.23 |
| 7 | 31804088 | rs10271159 | 0.618 | 1.889 | 0.96 | 1.634 | -0.022 | 0.033 | 0.033 | 0 | 0.1 | 0.05 | 0.11 | 0 | 0.18 | 0.07 |
| 7 | 31806797 | rs1978268 | 0.401 | 1.886 | 1.222 | 1.633 | 0.01 | -0.041 | -0.041 | 0 | 0.11 | 0.13 | 0.11 | 0 | 0.18 | 0.07 |
| 7 | 31808339 | rs4723102 | -0.53 | 2.763 | 1.29 | 1.664 | 0.05 | -0.009 | -0.009 | 0.575 | 0.18 | 0.28 | 0.36 | 0.12 | 0.27 | 0.12 |
| 7 | 31812263 | rs10275897 | -0.857 | 2.792 | 1.175 | 1.726 | 0.049 | 0.127 | 0.127 | 0.003 | 0.2 | 0.46 | 0.32 | 0.31 | 0.25 | 0.16 |
| 7 | 31812761 | rs28688395 | -0.921 | 2.703 | 1.177 | 1.728 | 0.096 | 0.121 | 0.121 | 0 | 0.21 | 0.46 | 0.41 | 0.31 | 0.28 | 0.17 |
| 7 | 31812836 | rs11980485 | -1.301 | 2.369 | 1.133 | 1.743 | 0.056 | 0.02 | 0.02 | 0 | 0.31 | 0.5 | 0.52 | 0.31 | 0.46 | 0.24 |
| 7 | 31815575 | rs10951303 | -0.596 | 1.572 | 0.752 | 2.023 | 0.012 | 0.001 | 0.001 | 0 | 0.14 | 0.21 | 0.13 | 0.11 | 0.18 | 0.07 |
| 7 | 31825276 | rs17160544 | -0.152 | 0.585 | -0.502 | 1.269 | 0.004 | -0.024 | -0.024 | 0 | 0.38 | 0.28 | 0.24 | 0.54 | 0.38 | 0.36 |
| 7 | 31825367 | rs11772988 | -1.213 | 0.129 | -0.859 | 0.459 | -0.005 | 0.046 | 0.046 | 0.886 | 0.13 | 0.04 | 0.11 | 0.02 | 0.14 | 0.07 |
| 7 | 31827180 | rs917754 | -0.448 | -0.873 | -1.294 | 0.32 | -0.001 | -0.035 | -0.035 | 0 | 0.6 | 0.76 | 0.35 | 0.51 | 0.4 | 0.63 |
| 7 | 31827417 | rs17160545 | 0.463 | -1.35 | -1.075 | 0.153 | 0.002 | 0.002 | 0.002 | 0 | 0.09 | 0.05 | 0.03 | 0.09 | 0.06 | 0.12 |
| 7 | 31829095 | rs2072423 | 0.75 | -2.337 | -0.462 | -0.036 | -0.055 | 0.008 | 0.008 | 0 | 0.74 | 0.53 | 0.82 | 0.71 | 0.86 | 0.72 |
| 7 | 31830095 | rs982857 | 1.384 | -2.789 | -0.514 | -0.344 | 0.05 | 0.033 | 0.033 | 0.139 | 0.48 | 0.26 | 0.66 | 0.59 | 0.64 | 0.5 |
| 7 | 31830457 | rs6961779 | -0.099 | -2.867 | -0.047 | -0.355 | -0.019 | -0.025 | -0.025 | 0.644 | 0.85 | 0.65 | 0.86 | 0.89 | 0.91 | 0.79 |
| 7 | 31834751 | rs3807626 | -1.647 | -2.87 | -0.437 | -0.338 | 0.03 | -0.013 | -0.013 | 0.081 | 0.55 | 0.8 | 0.37 | 0.47 | 0.38 | 0.61 |
| 7 | 31839327 | rs3807621 | -0.95 | -2.816 | -0.083 | -0.322 | -0.048 | 0.004 | 0.004 | 0 | 0.27 | 0.57 | 0.19 | 0.3 | 0.14 | 0.3 |
| 7 | 31844140 | rs1117825 | -1.542 | -2.899 | -0.299 | -0.381 | 0.05 | 0.029 | 0.029 | 0 | 0.5 | 0.66 | 0.33 | 0.42 | 0.36 | 0.51 |
| 7 | 31848706 | rs3213709 | 0.047 | -2.606 | -0.369 | -0.251 | -0.001 | -0.005 | -0.005 | 0 | 0.25 | 0.22 | 0.15 | 0.12 | 0.22 | 0.21 |
| 7 | 31849403 | rs6951050 | 0.072 | -2.6 | -0.354 | -0.248 | -0.002 | -0.007 | -0.007 | 1.663 | 0.24 | 0.21 | 0.15 | 0.12 | 0.22 | 0.21 |
| 7 | 31851037 | rs1035027 | 1.29 | -3.048 | -0.974 | -0.38 | 0.051 | -0.028 | -0.028 | 0 | 0.47 | 0.13 | 0.61 | 0.51 | 0.62 | 0.49 |
| 7 | 31854659 | rs7791212 | -0.354 | -3.26 | -0.989 | -0.641 | 0.006 | 0.007 | 0.007 | 0 | 0.22 | 0.09 | 0.15 | 0.17 | 0.21 | 0.19 |
| 7 | 31855569 | rs2302450 | -1.076 | -3.413 | -0.657 | -0.747 | 0.008 | 0.013 | 0.013 | 0 | 0.33 | 0.25 | 0.2 | 0.17 | 0.27 | 0.28 |
| 7 | 31855578 | rs1860790 | -0.947 | -3.452 | -0.646 | -0.733 | -0.004 | -0.001 | -0.001 | 0.667 | 0.93 | 0.93 | 0.98 | 0.92 | 0.98 | 0.96 |
| 7 | 31856070 | rs10951305 | -0.507 | -3.396 | -0.67 | -0.737 | 0.003 | 0.005 | 0.005 | 0.043 | 0.26 | 0.19 | 0.19 | 0.1 | 0.24 | 0.23 |
| 7 | 31861834 | rs10226190 | -0.177 | -3.386 | -0.592 | -0.607 | 0.002 | -0.011 | -0.011 | 0 | 0.9 | 0.92 | 0.86 | 0.77 | 0.94 | 0.91 |
| 7 | 31865695 | rs13238408 | -1.395 | -3.624 | -0.04 | -0.856 | -0.018 | -0.03 | -0.03 | 0 | 0.37 | 0.57 | 0.23 | 0.17 | 0.29 | 0.32 |
| 7 | 31867050 | rs1035028 | -0.665 | -3.606 | -0.54 | -0.805 | 0.009 | 0.023 | 0.023 | 0.302 | 0.28 | 0.17 | 0.16 | 0.1 | 0.24 | 0.23 |
| 7 | 31868064 | rs17160591 | 0.485 | -3.607 | -0.446 | -0.76 | -0.029 | -0.065 | -0.065 | 0 | 0.13 | 0.21 | 0.06 | 0 | 0.06 | 0.09 |
| 7 | 31868613 | rs4723103 | 0.551 | -3.495 | -0.486 | -0.717 | 0.004 | 0.004 | 0.004 | 0.256 | 0.68 | 0.74 | 0.81 | 0.84 | 0.76 | 0.74 |
| 7 | 31869225 | rs10951306 | -1.614 | -3.739 | -0.645 | -0.609 | 0.103 | 0 | 0 | 0 | 0.5 | 0.81 | 0.44 | 0.5 | 0.36 | 0.5 |
| 7 | 31869408 | rs3801352 | -0.261 | -3.66 | -0.219 | -0.785 | -0.016 | -0.003 | -0.003 | 0.51 | 0.15 | 0.21 | 0.1 | 0.15 | 0.11 | 0.15 |
| 7 | 31871153 | rs3729622 | -0.005 | -3.499 | -0.818 | -0.832 | -0.023 | -0.008 | -0.008 | 0 | 0.09 | 0.01 | 0.05 | 0.11 | 0.07 | 0.09 |
| 7 | 31874875 | rs12701142 | -1.065 | -3.236 | -0.834 | -1.002 | 0.021 | 0.073 | 0.073 | 2.148 | 0.4 | 0.22 | 0.33 | 0.23 | 0.32 | 0.33 |
| 7 | 31877261 | rs2270219 | 2.152 | -3.048 | -0.778 | -0.94 | -0.033 | -0.012 | -0.012 | 0 | 0.38 | 0.18 | 0.52 | 0.59 | 0.55 | 0.38 |

| | | | | | | | | | | | | | | | | |
|---|---|---|---|---|---|---|---|---|---|---|---|---|---|---|---|---|
| 7 | 31883140 | rs2191877 | -0.192 | -2.354 | -0.623 | -0.888 | -0.002 | -0.029 | -0.029 | 0 | 0.79 | 0.82 | 0.86 | 0.97 | 0.82 | 0.79 |
| 7 | 31888648 | rs7803484 | -0.717 | -2.405 | -0.672 | -0.777 | 0.004 | -0.017 | -0.017 | 0 | 0.14 | 0.07 | 0.15 | 0.2 | 0.08 | 0.09 |
| 7 | 31891598 | rs10487734 | -0.556 | -1.921 | -0.35 | -0.927 | 0.001 | -0.02 | -0.02 | 0 | 0.24 | 0.19 | 0.13 | 0.03 | 0.19 | 0.21 |
| 7 | 31892956 | rs3801344 | -0.501 | -1.986 | -0.043 | -0.856 | -0.013 | -0.004 | -0.004 | 0.125 | 0.84 | 0.69 | 0.88 | 0.82 | 0.9 | 0.82 |
| 7 | 31897241 | rs1016412 | -0.287 | -1.917 | 0.298 | -0.736 | -0.008 | 0.025 | 0.025 | 9.433 | 0.83 | 0.6 | 0.86 | 0.79 | 0.88 | 0.8 |
| 7 | 31899654 | rs4723104 | -0.396 | -1.918 | 0.333 | -0.756 | -0.005 | 0.023 | 0.023 | 9.433 | 0.83 | 0.63 | 0.87 | 0.79 | 0.88 | 0.81 |
| 7 | 31902703 | rs3922299 | -0.17 | -2.439 | 0.068 | -1.517 | -0.001 | 0.005 | 0.005 | 0.056 | 0.29 | 0.19 | 0.2 | 0.03 | 0.3 | 0.24 |
| 7 | 31903346 | rs10265932 | 0.66 | -2.57 | 0.157 | -1.586 | 0.018 | 0.046 | 0.046 | 0.71 | 0.59 | 0.36 | 0.73 | 0.76 | 0.68 | 0.59 |
| 7 | 31904027 | rs2270221 | -0.097 | -2.792 | -0.24 | -1.359 | 0.003 | 0.002 | 0.002 | 0 | 0.35 | 0.32 | 0.38 | 0.27 | 0.25 | 0.36 |
| 7 | 31912383 | rs2191869 | -0.887 | -2.955 | -0.45 | -1.479 | -0.085 | -0.027 | -0.027 | 0 | 0.33 | 0.55 | 0.34 | 0.24 | 0.19 | 0.33 |
| 7 | 31918879 | rs1860786 | 0.587 | -2.875 | -0.601 | -1.485 | 0.043 | -0.007 | -0.007 | 0 | 0.47 | 0.2 | 0.47 | 0.51 | 0.61 | 0.45 |
| 7 | 31919754 | rs3779243 | -0.247 | -2.734 | -0.387 | -1.435 | 0.001 | -0.016 | -0.016 | 0 | 0.59 | 0.6 | 0.69 | 0.76 | 0.6 | 0.61 |
| 7 | 31922902 | rs1860788 | 0.232 | -2.737 | -0.195 | -1.198 | -0.001 | 0 | 0 | 0 | 0.15 | 0.14 | 0.06 | 0.08 | 0.09 | 0.24 |
| 7 | 31933965 | rs7808892 | -1.052 | -2.855 | -0.064 | -1.309 | -0.098 | -0.045 | -0.045 | 0 | 0.32 | 0.49 | 0.33 | 0.24 | 0.19 | 0.33 |
| 7 | 31934481 | rs10252546 | 0.348 | -2.168 | -0.193 | -1.076 | -0.017 | 0.004 | 0.004 | 0 | 0.15 | 0.22 | 0.28 | 0.21 | 0.09 | 0.14 |
| 7 | 31943754 | rs12701156 | 0.711 | -1.882 | -0.371 | -1.146 | 0.03 | -0.015 | -0.015 | 0 | 0.13 | 0.07 | 0.27 | 0.21 | 0.08 | 0.15 |
| 7 | 31949763 | rs13223209 | -0.074 | -2.635 | 0.118 | -1.224 | -0.03 | -0.001 | -0.001 | 0 | 0.71 | 0.62 | 0.85 | 0.66 | 0.83 | 0.73 |
| 7 | 31953289 | rs10225593 | 1.743 | -2.63 | 0.153 | -1.527 | -0.014 | -0.029 | -0.029 | 0.271 | 0.35 | 0.27 | 0.44 | 0.4 | 0.52 | 0.3 |
| 7 | 31956217 | rs6962770 | 1.046 | -2.634 | 0.326 | -1.534 | -0.014 | 0.002 | 0.002 | 0 | 0.14 | 0.16 | 0.28 | 0.21 | 0.1 | 0.15 |
| 7 | 31965003 | rs4723113 | 1.947 | -2.289 | 0.521 | -1.63 | -0.079 | -0.077 | -0.077 | 0 | 0.3 | 0.13 | 0.38 | 0.42 | 0.42 | 0.28 |
| 7 | 31973974 | rs11770536 | 0.982 | -2.516 | 0.65 | -1.69 | -0.001 | 0.009 | 0.009 | 0 | 0.42 | 0.53 | 0.47 | 0.42 | 0.55 | 0.34 |
| 7 | 31977956 | rs6957512 | 0.918 | -2.237 | 0.361 | -1.566 | 0.015 | -0.068 | -0.068 | 0 | 0.21 | 0.03 | 0.1 | 0.34 | 0.15 | 0.22 |
| 7 | 31981713 | rs6462315 | -0.112 | -2.17 | 0.474 | -1.608 | 0.012 | 0.017 | 0.017 | 0.318 | 0.62 | 0.72 | 0.77 | 0.65 | 0.69 | 0.58 |
| 7 | 31986723 | rs7792024 | 0.392 | -1.697 | 0.426 | -1.572 | -0.025 | -0.034 | -0.034 | 0 | 0.07 | 0.15 | 0.02 | 0 | 0.03 | 0.05 |
| 7 | 32001658 | rs6952633 | -0.8 | -1.461 | 0.365 | -1.818 | 0.001 | -0.098 | -0.098 | 0.04 | 0.44 | 0.45 | 0.27 | 0.05 | 0.46 | 0.41 |
| 7 | 32009914 | rs6462318 | -0.631 | -0.572 | 0.304 | -1.733 | 0.013 | 0 | 0 | 0 | 0.44 | 0.62 | 0.53 | 0.55 | 0.36 | 0.42 |
| 7 | 32009927 | rs6961368 | -0.55 | -0.585 | 0.264 | -1.725 | 0.075 | -0.013 | -0.013 | 0 | 0.45 | 0.77 | 0.53 | 0.57 | 0.36 | 0.42 |
| 7 | 32012593 | rs7790003 | -1.372 | -0.498 | 0.327 | -1.74 | 0.003 | 0.023 | 0.023 | 0 | 0.63 | 0.56 | 0.65 | 0.6 | 0.61 | 0.61 |
| 7 | 32016539 | rs6949456 | -1.213 | -0.685 | 0.218 | -1.708 | -0.056 | -0.055 | -0.055 | 0.154 | 0.61 | 0.81 | 0.62 | 0.58 | 0.54 | 0.6 |
| 7 | 32018676 | rs12701165 | 1.727 | -0.581 | 0.222 | -1.692 | -0.023 | 0.043 | 0.043 | 0 | 0.15 | 0.08 | 0.09 | 0.03 | 0.17 | 0.17 |
| 7 | 32024231 | rs998413 | 1.234 | -0.789 | 0.536 | -1.79 | 0.01 | -0.002 | -0.002 | 0 | 0.16 | 0.08 | 0.09 | 0.02 | 0.15 | 0.15 |
| 7 | 32025543 | rs10951313 | -0.756 | -0.89 | 0.47 | -1.825 | 0.005 | 0.015 | 0.015 | 0 | 0.5 | 0.43 | 0.55 | 0.27 | 0.47 | 0.52 |
| 7 | 32026793 | rs17160741 | 1.96 | -0.926 | 0.344 | -1.736 | 0.008 | -0.011 | -0.011 | 0 | 0.09 | 0.09 | 0.05 | 0.32 | 0.05 | 0.08 |
| 7 | 32028025 | rs995267 | 0.808 | -0.938 | 0.451 | -1.824 | -0.033 | -0.097 | -0.097 | 0 | 0.19 | 0.27 | 0.11 | 0.02 | 0.15 | 0.15 |
| 7 | 32030184 | rs6954242 | 1.592 | -0.908 | -0.268 | -1.829 | 0.001 | -0.022 | -0.022 | 0.693 | 0.13 | 0.07 | 0.33 | 0.22 | 0.13 | 0.18 |
| 7 | 32030880 | rs2392015 | -0.601 | -0.99 | -0.238 | -1.927 | 0.037 | 0.015 | 0.015 | 0.423 | 0.36 | 0.68 | 0.21 | 0.35 | 0.24 | 0.3 |
| 7 | 32031510 | rs17160757 | 2.025 | -0.903 | -0.333 | -1.736 | 0.007 | -0.005 | -0.005 | 0 | 0.08 | 0.09 | 0.05 | 0.33 | 0.05 | 0.08 |
| 7 | 32032544 | rs12701169 | 1.098 | -0.399 | -0.103 | -2.407 | -0.031 | 0.063 | 0.063 | 0 | 0.17 | 0.05 | 0.11 | 0.03 | 0.2 | 0.19 |
| 7 | 32033738 | rs6462320 | -0.831 | -0.29 | 0.055 | -2.352 | 0.013 | 0.088 | 0.088 | 0 | 0.51 | 0.59 | 0.63 | 0.27 | 0.48 | 0.52 |
| 7 | 32034799 | rs4723120 | 0.662 | 0.069 | 0.143 | -1.791 | -0.001 | -0.014 | -0.014 | 0 | 0.16 | 0.07 | 0.43 | 0.22 | 0.16 | 0.2 |
| 7 | 32037234 | rs7796232 | 0.107 | 0.018 | 0.445 | -1.779 | -0.042 | 0.005 | 0.005 | 0 | 0.19 | 0.45 | 0.38 | 0.22 | 0.14 | 0.19 |
| 7 | 32037357 | rs13306873 | -0.065 | 0.596 | -0.52 | -2.062 | -0.049 | 0.094 | 0.094 | 0.382 | 0.26 | 0.09 | 0.17 | 0.03 | 0.3 | 0.26 |
| 7 | 32038104 | rs6968484 | -0.659 | 0.296 | -0.464 | -2.193 | 0.016 | 0.066 | 0.066 | 0 | 0.46 | 0.69 | 0.62 | 0.25 | 0.41 | 0.43 |
| 7 | 32038274 | rs12154278 | 0.504 | 0.379 | -0.393 | -1.644 | 0.002 | -0.017 | -0.017 | 0 | 0.16 | 0.08 | 0.35 | 0.2 | 0.15 | 0.18 |
| 7 | 32038718 | rs2192389 | -0.765 | 1.573 | 0.017 | -1.913 | -0.008 | -0.002 | -0.002 | 0 | 0.39 | 0.63 | 0.35 | 0.12 | 0.49 | 0.48 |
| 7 | 32043196 | rs2058545 | -0.658 | 1.84 | 0.167 | -1.269 | 0.057 | -0.071 | -0.071 | 1.496 | 0.2 | 0.39 | 0.19 | 0.09 | 0.28 | 0.19 |
| 7 | 32044030 | rs10251809 | -0.505 | 1.936 | -0.719 | -1.228 | 0.004 | -0.011 | -0.011 | 0 | 0.19 | 0.19 | 0.18 | 0.09 | 0.28 | 0.19 |
| 7 | 32048171 | rs17160788 | -0.098 | 1.447 | -0.64 | -1.043 | 0.016 | -0.029 | -0.029 | 0.41 | 0.16 | 0.23 | 0.15 | 0.09 | 0.22 | 0.18 |
| 7 | 32053711 | rs7783322 | 0.043 | 1.016 | -1.067 | -1.12 | -0.059 | 0.159 | 0.159 | 0 | 0.4 | 0.17 | 0.26 | 0.1 | 0.48 | 0.4 |
| 7 | 32057791 | rs30556 | -0.189 | 1.009 | -1.22 | -0.63 | -0.034 | -0.028 | -0.028 | 0.328 | 0.86 | 0.91 | 0.64 | 0.81 | 0.81 | 0.83 |
| 7 | 32061610 | rs2190127 | 0.235 | 0.309 | -0.428 | 0.269 | -0.037 | 0.233 | 0.233 | 0.288 | 0.14 | 0.47 | 0.11 | 0.33 | 0.08 | 0.12 |
| 7 | 32061966 | rs2214782 | 0.558 | 0.295 | -0.04 | 0.258 | -0.029 | 0.155 | 0.155 | 0.064 | 0.12 | 0.33 | 0.1 | 0.32 | 0.08 | 0.12 |
| 7 | 32067912 | rs12216602 | 0.185 | 0.15 | -0.304 | 0.102 | -0.003 | 0.008 | 0.008 | 0 | 0.12 | 0.07 | 0.12 | 0.05 | 0.13 | 0.07 |
| 7 | 32068771 | rs30565 | -0.858 | 0.135 | -0.356 | 0.14 | -0.01 | -0.021 | -0.021 | 5.44 | 0.87 | 0.92 | 0.65 | 0.81 | 0.85 | 0.84 |
| 7 | 32073838 | rs1557976 | -0.889 | 0.1 | -0.26 | 0.249 | -0.021 | 0.062 | 0.062 | 0 | 0.25 | 0.56 | 0.23 | 0.38 | 0.21 | 0.18 |
| 7 | 32075849 | rs30571 | 0.686 | 0.03 | -0.293 | 0.215 | 0.003 | -0.007 | -0.007 | 0 | 0.73 | 0.58 | 0.57 | 0.77 | 0.73 | 0.79 |
| 7 | 32077081 | rs10486499 | 1.48 | 0.021 | -1.15 | 0.318 | -0.078 | -0.127 | -0.127 | 0.108 | 0.18 | 0.02 | 0.2 | 0.34 | 0.24 | 0.26 |
| 7 | 32077460 | rs17160822 | 1.216 | 0.024 | -0.906 | 0.266 | 0.021 | -0.008 | -0.008 | 1.805 | 0.08 | 0.03 | 0.02 | 0.07 | 0.04 | 0.04 |
| 7 | 32081165 | rs10259038 | 0.52 | 0.397 | -0.771 | 0.193 | -0.026 | 0.074 | 0.074 | 0 | 0.14 | 0.32 | 0.11 | 0.3 | 0.11 | 0.14 |
| 7 | 32082198 | rs1024536 | 0.717 | 0.803 | -1.449 | -0.231 | -0.05 | 0.19 | 0.19 | 0.395 | 0.38 | 0.1 | 0.24 | 0.11 | 0.37 | 0.37 |
| 7 | 32093550 | rs30585 | -0.184 | 1.021 | -0.556 | 0.933 | 0 | 0.007 | 0.007 | 0 | 0.89 | 0.88 | 0.65 | 0.8 | 0.87 | 0.86 |
| 7 | 32105096 | rs30593 | 0.631 | 0.57 | -0.398 | 0.621 | -0.004 | 0.002 | 0.002 | 0.009 | 0.57 | 0.55 | 0.48 | 0.53 | 0.64 | 0.67 |
| 7 | 32111162 | rs30601 | 0.072 | 0.478 | -0.304 | 0.99 | -0.02 | 0.133 | 0.133 | 0 | 0.11 | 0.2 | 0.12 | 0.4 | 0.07 | 0.24 |
| 7 | 32119530 | rs2079373 | -0.128 | 0.561 | -0.633 | 0.979 | -0.01 | -0.032 | -0.032 | 0 | 0.88 | 0.94 | 0.66 | 0.81 | 0.86 | 0.86 |
| 7 | 32120897 | rs6462324 | 0.483 | 1.073 | -0.399 | 1.201 | -0.01 | -0.024 | -0.024 | 0 | 0.63 | 0.71 | 0.44 | 0.46 | 0.59 | 0.69 |
| 7 | 32133424 | rs11972318 | 0.523 | 0.943 | -0.265 | 0.982 | -0.004 | -0.012 | -0.012 | 0 | 0.63 | 0.64 | 0.43 | 0.46 | 0.61 | 0.69 |
| 7 | 32134056 | rs10246484 | 0.827 | 1.135 | -0.373 | 1.074 | 0.023 | 0.016 | 0.016 | 0.616 | 0.68 | 0.6 | 0.42 | 0.48 | 0.56 | 0.61 |
| 7 | 32134138 | rs11761910 | -0.802 | 0.809 | -0.495 | 0.788 | -0.001 | 0.002 | 0.002 | 5.948 | 0.29 | 0.31 | 0.26 | 0.46 | 0.27 | 0.33 |
| 7 | 32140526 | rs17423839 | 0.375 | 0.812 | -0.935 | -0.554 | -0.012 | 0.127 | 0.127 | 0 | 0.39 | 0.23 | 0.2 | 0.13 | 0.4 | 0.42 |

| | | | | | | | | | | | | | | | | |
|---|---|---|---|---|---|---|---|---|---|---|---|---|---|---|---|---|
| 7 | 32151370 | rs10486500 | 0.33 | 0.551 | -0.145 | -0.174 | 0.055 | 0.22 | 0.22 | 0 | 0.08 | 0.33 | 0.08 | 0.23 | 0.06 | 0.06 |
| 7 | 32155380 | rs11761940 | 0.733 | 0.591 | -0.36 | -0.282 | -0.013 | 0.131 | 0.131 | 0 | 0.37 | 0.18 | 0.19 | 0.13 | 0.37 | 0.41 |
| 7 | 32157711 | rs766856 | -0.112 | 0.541 | -0.544 | -0.738 | -0.005 | -0.019 | -0.019 | 3.272 | 0.39 | 0.51 | 0.3 | 0.27 | 0.36 | 0.22 |
| 7 | 32167619 | rs32304 | -0.243 | 0.964 | -0.229 | -0.275 | 0.003 | 0.011 | 0.011 | 0 | 0.76 | 0.75 | 0.5 | 0.4 | 0.72 | 0.63 |
| 7 | 32170646 | rs11763324 | 0.96 | 1.102 | -0.557 | -0.477 | -0.019 | 0.03 | 0.03 | 0 | 0.17 | 0.07 | 0.11 | 0.01 | 0.23 | 0.27 |
| 7 | 32171409 | rs32307 | -0.051 | 1.181 | 0.167 | 0.188 | 0.006 | 0.021 | 0.021 | 0.194 | 0.76 | 0.72 | 0.49 | 0.4 | 0.72 | 0.63 |
| 7 | 32179482 | rs6963935 | -0.127 | 0.724 | 0.162 | 0.701 | 0.039 | 0.179 | 0.179 | 0 | 0.08 | 0.26 | 0.07 | 0.23 | 0.06 | 0.05 |
| 7 | 32186706 | rs1029549 | 0.282 | 0.688 | -0.842 | 0.088 | -0.042 | 0.061 | 0.061 | 0 | 0.2 | 0.04 | 0.21 | 0.04 | 0.22 | 0.12 |
| 7 | 32187476 | rs4352745 | 0.83 | 0.1 | -0.539 | -0.334 | -0.005 | -0.012 | -0.012 | 0.003 | 0.33 | 0.33 | 0.17 | 0.19 | 0.21 | 0.38 |
| 7 | 32188096 | rs32322 | -1.511 | 0.552 | -0.667 | -0.714 | -0.032 | 0.058 | 0.058 | 0 | 0.79 | 0.93 | 0.65 | 0.99 | 0.74 | 0.85 |
| 7 | 32189295 | rs17160928 | 0.357 | 0.591 | -0.747 | -0.061 | -0.061 | -0.109 | -0.109 | 0 | 0.27 | 0.07 | 0.27 | 0.39 | 0.3 | 0.23 |
| 7 | 32191808 | rs1557975 | 0.935 | 0.249 | -0.41 | -0.211 | 0.002 | -0.004 | -0.004 | 0 | 0.33 | 0.29 | 0.22 | 0.52 | 0.24 | 0.29 |
| 7 | 32192596 | rs6415258 | -0.897 | 0.3 | 0.249 | 0.246 | 0.012 | 0.039 | 0.039 | 0 | 0.83 | 0.73 | 0.83 | 0.76 | 0.78 | 0.78 |
| 7 | 32199098 | rs996134 | -0.152 | 0.13 | 0.247 | 0.262 | 0.015 | 0.016 | 0.016 | 0.18 | 0.61 | 0.55 | 0.45 | 0.55 | 0.51 | 0.61 |
| 7 | 32201804 | rs11978519 | 1.487 | 0.011 | 0.038 | -0.108 | -0.006 | -0.01 | -0.01 | 0 | 0.26 | 0.25 | 0.16 | 0.17 | 0.16 | 0.18 |
| 7 | 32202786 | rs17425268 | 0.401 | -0.016 | -0.188 | -0.163 | 0 | 0.085 | 0.085 | 0 | 0.27 | 0.14 | 0.23 | 0.06 | 0.27 | 0.31 |
| 7 | 32206323 | rs10274609 | 0.194 | -0.059 | 0.291 | 0.131 | -0.01 | 0.083 | 0.083 | 0 | 0.14 | 0.21 | 0.09 | 0.35 | 0.1 | 0.29 |
| 7 | 32206419 | rs7792012 | 0.746 | -0.186 | 0.346 | 0.109 | -0.013 | 0.054 | 0.054 | 0 | 0.14 | 0.21 | 0.09 | 0.24 | 0.12 | 0.21 |
| 7 | 32209193 | rs12701178 | 1.475 | -0.055 | 0.187 | -0.21 | -0.032 | 0.049 | 0.049 | 0 | 0.22 | 0.08 | 0.35 | 0.01 | 0.27 | 0.17 |
| 7 | 32209283 | rs977624 | 0.608 | -0.238 | 0.28 | -0.14 | 0.031 | -0.013 | -0.013 | 0 | 0.38 | 0.34 | 0.23 | 0.41 | 0.27 | 0.39 |
| 7 | 32209459 | rs977626 | -0.229 | -0.155 | -0.036 | -0.415 | 0.008 | 0.01 | 0.01 | 0 | 0.52 | 0.55 | 0.31 | 0.76 | 0.38 | 0.69 |
| 7 | 32213806 | rs17160950 | 1.595 | -0.077 | -0.454 | -0.369 | 0.019 | 0.061 | 0.061 | 3.981 | 0.24 | 0.03 | 0.13 | 0.17 | 0.15 | 0.19 |
| 7 | 32217815 | rs7785525 | -0.581 | -0.082 | 0.015 | -0.333 | -0.03 | 0.019 | 0.019 | 0.06 | 0.3 | 0.52 | 0.21 | 0.59 | 0.24 | 0.49 |
| 7 | 32218044 | rs6972452 | -0.944 | 0.11 | -0.177 | -0.42 | -0.01 | -0.001 | -0.001 | 0 | 0.75 | 0.76 | 0.71 | 0.77 | 0.64 | 0.86 |
| 7 | 32220497 | rs987069 | -0.706 | 0.025 | -0.194 | -0.223 | -0.026 | 0.07 | 0.07 | 0 | 0.25 | 0.51 | 0.21 | 0.57 | 0.21 | 0.36 |
| 7 | 32220641 | rs17160951 | 1.085 | 0.098 | -0.195 | 0.02 | -0.001 | 0.06 | 0.06 | 0 | 0.07 | 0.11 | 0.07 | 0.35 | 0.06 | 0.1 |
| 7 | 32228302 | rs6978495 | -0.228 | 0.061 | -0.324 | -0.126 | 0.02 | 0.025 | 0.025 | 0.265 | 0.48 | 0.57 | 0.35 | 0.75 | 0.35 | 0.55 |
| 7 | 32228658 | rs6462339 | 0.467 | 0.155 | -0.11 | -0.125 | -0.026 | 0.016 | 0.016 | 4.21 | 0.34 | 0.55 | 0.25 | 0.52 | 0.21 | 0.29 |
| 7 | 32230177 | rs917454 | 0.389 | 0.684 | 0.304 | 0.41 | 0.003 | 0.001 | 0.001 | 0.785 | 0.54 | 0.53 | 0.62 | 0.53 | 0.54 | 0.46 |
| 7 | 32241867 | rs1860220 | -0.294 | 0.638 | 0.085 | 0.738 | -0.001 | 0.007 | 0.007 | 0 | 0.77 | 0.71 | 0.8 | 0.8 | 0.75 | 0.81 |
| 7 | 32242328 | rs12536286 | 0.364 | 0.625 | -0.303 | 0.672 | 0 | 0.085 | 0.085 | 0 | 0.07 | 0.1 | 0.08 | 0.37 | 0.06 | 0.1 |
| 7 | 32243360 | rs11765220 | 0.166 | 0.649 | -0.392 | 0.612 | 0 | 0.001 | 0.001 | 0 | 0.51 | 0.47 | 0.44 | 0.45 | 0.58 | 0.5 |
| 7 | 32244440 | rs4723128 | 0.089 | 0.679 | -1.361 | -0.42 | -0.019 | 0.174 | 0.174 | 0.014 | 0.41 | 0.2 | 0.48 | 0.18 | 0.37 | 0.4 |
| 7 | 32248311 | rs12534148 | -0.585 | 0.45 | -0.945 | 0.481 | -0.022 | -0.003 | -0.003 | 0 | 0.19 | 0.08 | 0.16 | 0.18 | 0.22 | 0.2 |
| 7 | 32248744 | rs2240674 | -0.783 | -0.176 | -1.047 | 0.386 | -0.017 | -0.005 | -0.005 | 4.374 | 0.61 | 0.75 | 0.47 | 0.6 | 0.53 | 0.7 |
| 7 | 32248809 | rs2240675 | 0.887 | -0.284 | -0.923 | 0.337 | -0.045 | -0.023 | -0.023 | 0 | 0.37 | 0.15 | 0.52 | 0.4 | 0.47 | 0.3 |
| 7 | 32248984 | rs10951324 | 0.068 | -0.497 | -0.127 | 0.231 | 0.002 | -0.001 | -0.001 | 0 | 0.47 | 0.61 | 0.36 | 0.53 | 0.39 | 0.56 |
| 7 | 32250148 | rs11977444 | 1.071 | -0.487 | -0.148 | 0.234 | -0.046 | -0.024 | -0.024 | 0.012 | 0.37 | 0.15 | 0.52 | 0.4 | 0.47 | 0.3 |
| 7 | 32250317 | rs7779445 | 0.95 | -0.97 | 0.507 | 0.396 | 0.013 | 0.03 | 0.03 | 0 | 0.34 | 0.16 | 0.26 | 0.28 | 0.29 | 0.48 |
| 7 | 32254014 | rs6966630 | 0.262 | -1.004 | 0.416 | 0.187 | -0.002 | 0.006 | 0.006 | 0 | 0.39 | 0.34 | 0.54 | 0.35 | 0.45 | 0.2 |
| 7 | 32255322 | rs4723134 | 1.601 | -1.143 | 0.143 | 0.078 | -0.092 | 0.115 | 0.115 | 0.48 | 0.23 | 0.04 | 0.43 | 0.07 | 0.34 | 0.1 |
| 7 | 32255642 | rs1860224 | -0.044 | -1.007 | 0.532 | 0.21 | 0.005 | 0.029 | 0.029 | 0 | 0.71 | 0.39 | 0.78 | 0.61 | 0.73 | 0.66 |
| 7 | 32256552 | rs6971522 | -0.686 | -0.987 | 0.472 | 0.181 | 0.008 | -0.034 | -0.034 | 0 | 0.85 | 0.67 | 0.89 | 0.94 | 0.87 | 0.87 |
| 7 | 32258435 | rs11763287 | 0.972 | -0.998 | 0.39 | 0.253 | -0.001 | 0.014 | 0.014 | 0 | 0.1 | 0.12 | 0.09 | 0.26 | 0.1 | 0.1 |
| 7 | 32258734 | rs12672267 | 1.093 | -1.099 | 0.422 | 0.129 | -0.009 | -0.02 | -0.02 | 0.214 | 0.35 | 0.45 | 0.26 | 0.26 | 0.28 | 0.46 |
| 7 | 32259273 | rs6945244 | -1.39 | -1.45 | 0.663 | 0.17 | 0.013 | 0.129 | 0.129 | 0 | 0.51 | 0.74 | 0.39 | 0.81 | 0.42 | 0.66 |
| 7 | 32261093 | rs13241693 | 0.797 | -1.878 | 0.806 | 0.149 | 0 | 0.012 | 0.012 | 0 | 0.11 | 0.12 | 0.09 | 0.24 | 0.11 | 0.1 |
| 7 | 32262377 | rs12669911 | 1.512 | -2 | 0.79 | 0.023 | 0.073 | 0.133 | 0.133 | 0 | 0.5 | 0.14 | 0.68 | 0.34 | 0.63 | 0.42 |
| 7 | 32266959 | rs4141108 | 0.334 | -1.858 | 1.111 | 0.036 | 0.004 | 0.097 | 0.097 | 0 | 0.25 | 0.42 | 0.21 | 0.4 | 0.24 | 0.3 |
| 7 | 32272342 | rs11767773 | 0.868 | -1.856 | 1.221 | 0.039 | -0.002 | -0.002 | -0.002 | 0 | 0.12 | 0.2 | 0.1 | 0.08 | 0.11 | 0.09 |
| 7 | 32273107 | rs7806224 | -0.91 | -1.933 | 1.379 | 0.078 | 0 | -0.004 | -0.004 | 0 | 0.37 | 0.62 | 0.3 | 0.52 | 0.34 | 0.37 |
| 7 | 32274979 | rs10216007 | 2.196 | -2.393 | 1.011 | -0.333 | -0.082 | -0.095 | -0.095 | 2.538 | 0.27 | 0.04 | 0.26 | 0.39 | 0.28 | 0.29 |
| 7 | 32278966 | rs6977493 | 1.473 | -1.813 | 1.405 | 0.027 | 0.006 | 0.013 | 0.013 | 0 | 0.07 | 0.11 | 0.08 | 0.13 | 0.09 | 0.06 |
| 7 | 32282457 | rs10263751 | -1.143 | -1.819 | 1.452 | 0.024 | 0.018 | 0.035 | 0.035 | 0 | 0.92 | 0.81 | 0.91 | 0.87 | 0.91 | 0.94 |
| 7 | 32282519 | rs10263673 | -0.587 | -1.821 | 1.424 | 0.023 | -0.011 | 0.054 | 0.054 | 0 | 0.3 | 0.51 | 0.22 | 0.39 | 0.25 | 0.31 |
| 7 | 32284238 | rs12538504 | 1.718 | -1.796 | 1.383 | 0.029 | 0.002 | 0.006 | 0.006 | 0 | 0.07 | 0.09 | 0.08 | 0.13 | 0.09 | 0.06 |
| 7 | 32290565 | rs17161076 | -0.463 | -1.798 | 1.358 | 0.019 | 0.006 | 0.016 | 0.016 | 0 | 0.19 | 0.34 | 0.18 | 0.21 | 0.2 | 0.15 |
| 7 | 32290730 | rs11975968 | 1.198 | -1.807 | 1.405 | 0.022 | -0.003 | 0.057 | 0.057 | 0 | 0.17 | 0.21 | 0.11 | 0.32 | 0.14 | 0.22 |
| 7 | 32293644 | rs929456 | 0.972 | -1.811 | 1.333 | 0.012 | 0.003 | -0.016 | -0.016 | 0 | 0.62 | 0.31 | 0.69 | 0.48 | 0.66 | 0.63 |
| 7 | 32295175 | rs6977468 | 0.656 | -1.784 | 1.342 | 0.034 | 0.019 | 0.052 | 0.052 | 0 | 0.09 | 0.25 | 0.1 | 0.14 | 0.1 | 0.06 |
| 7 | 32297310 | rs13438711 | 1.173 | -1.783 | 1.243 | 0.013 | -0.001 | -0.007 | -0.007 | 0.78 | 0.11 | 0.17 | 0.09 | 0.06 | 0.1 | 0.09 |
| 7 | 32299187 | rs17161105 | 0.886 | -1.784 | 1.262 | 0.012 | -0.002 | -0.015 | -0.015 | 0 | 0.12 | 0.26 | 0.1 | 0.06 | 0.1 | 0.09 |
| 7 | 32302338 | rs7357136 | 1.173 | -1.781 | 1.181 | 0.014 | -0.001 | -0.007 | -0.007 | 0 | 0.11 | 0.17 | 0.09 | 0.06 | 0.1 | 0.09 |
| 7 | 32305274 | rs7780515 | 0.915 | -1.779 | 0.962 | 0.013 | 0.004 | -0.021 | -0.021 | 0.004 | 0.62 | 0.26 | 0.69 | 0.47 | 0.66 | 0.62 |
| 7 | 32305391 | rs7780674 | 1.994 | -1.718 | 0.696 | -0.068 | -0.099 | 0.081 | 0.081 | 0.349 | 0.17 | 0.01 | 0.4 | 0.06 | 0.32 | 0.06 |
| 7 | 32308717 | rs17161127 | 1.616 | -2.158 | 1.592 | 0.714 | -0.04 | -0.078 | -0.078 | 8.008 | 0.12 | 0.03 | 0.08 | 0.32 | 0.13 | 0.22 |
| 7 | 32311672 | rs1450869 | 1.39 | -2.246 | 1.987 | 0.66 | 0.001 | -0.006 | -0.006 | 0.056 | 0.62 | 0.33 | 0.7 | 0.47 | 0.66 | 0.62 |
| 7 | 32315613 | rs10226228 | -1.285 | -2.243 | 2.004 | 0.662 | 0.001 | 0.011 | 0.011 | 0 | 0.36 | 0.43 | 0.28 | 0.53 | 0.34 | 0.38 |
| 7 | 32322496 | rs9771228 | -1.239 | -2.244 | 1.817 | 0.603 | 0.006 | 0.023 | 0.023 | 0 | 0.33 | 0.53 | 0.28 | 0.53 | 0.33 | 0.37 |

| | | | | | | | | | | | | | | | | |
|---|---|---|---|---|---|---|---|---|---|---|---|---|---|---|---|---|
| 7 | 32323579 | rs17161134 | 0.535 | -2.13 | 1.289 | 0.634 | -0.001 | 0.013 | 0.013 | 0 | 0.17 | 0.13 | 0.1 | 0.33 | 0.14 | 0.23 |
| 7 | 32326373 | rs215596 | 1.867 | -2.148 | 0.75 | 0.362 | 0.067 | 0.042 | 0.042 | 0 | 0.5 | 0.14 | 0.68 | 0.46 | 0.63 | 0.43 |
| 7 | 32333921 | rs215601 | 1.876 | -2.589 | 0.695 | 0.208 | 0.069 | 0.082 | 0.082 | 0 | 0.5 | 0.14 | 0.69 | 0.42 | 0.64 | 0.42 |
| 7 | 32336965 | rs215605 | 1.45 | -2.632 | 0.845 | 0.258 | 0.01 | 0.042 | 0.042 | 0 | 0.6 | 0.24 | 0.72 | 0.37 | 0.66 | 0.53 |
| 7 | 32344644 | rs12538119 | -0.106 | -2.566 | 0.892 | 0.221 | 0 | 0.001 | 0.001 | 0 | 0.1 | 0.12 | 0.09 | 0.19 | 0.1 | 0.14 |
| 7 | 32345283 | rs7779181 | -1.124 | -2.63 | 1.057 | 0.203 | -0.001 | -0.001 | -0.001 | 0 | 0.2 | 0.25 | 0.15 | 0.21 | 0.2 | 0.2 |
| 7 | 32347335 | rs215614 | 1.33 | -2.657 | 1.332 | 0.206 | -0.001 | 0.001 | 0.001 | 0 | 0.63 | 0.58 | 0.76 | 0.4 | 0.66 | 0.53 |
| 7 | 32354787 | rs11981809 | 0.98 | -2.566 | 1.039 | 0.272 | -0.006 | -0.009 | -0.009 | 1.3 | 0.07 | 0.03 | 0.07 | 0.12 | 0.08 | 0.06 |
| 7 | 32359489 | rs13235908 | 0.485 | -2.494 | 1.054 | 0.305 | -0.001 | -0.001 | -0.001 | 0 | 0.08 | 0.09 | 0.08 | 0.13 | 0.09 | 0.08 |
| 7 | 32360619 | rs1653876 | 0.382 | -2.53 | 1.211 | 0.206 | -0.002 | 0.06 | 0.06 | 0 | 0.74 | 0.5 | 0.81 | 0.48 | 0.77 | 0.66 |
| 7 | 32369148 | rs215634 | 1.87 | -2.596 | 0.784 | 0.11 | 0.059 | 0.091 | 0.091 | 0 | 0.49 | 0.19 | 0.71 | 0.39 | 0.63 | 0.32 |
| 7 | 32373104 | rs7804577 | -0.247 | -2.618 | 0.987 | 0.064 | -0.001 | -0.012 | -0.012 | 0 | 0.14 | 0.12 | 0.07 | 0.02 | 0.1 | 0.07 |
| 7 | 32376953 | rs10238006 | -0.187 | -2.523 | 0.563 | 0.149 | -0.001 | 0.067 | 0.067 | 0 | 0.27 | 0.5 | 0.19 | 0.51 | 0.23 | 0.34 |
| 7 | 32378979 | rs215669 | 1.194 | -2.152 | -0.469 | 0.276 | 0.051 | 0.073 | 0.073 | 0 | 0.48 | 0.11 | 0.7 | 0.4 | 0.64 | 0.35 |
| 7 | 32382872 | rs215674 | 0.225 | -2.008 | -0.199 | 0.579 | 0.031 | -0.044 | -0.044 | 0 | 0.7 | 0.8 | 0.88 | 0.61 | 0.82 | 0.53 |
| 7 | 32387493 | rs183347 | 0.22 | -2.016 | -0.205 | 0.582 | 0.031 | -0.047 | -0.047 | 0 | 0.7 | 0.8 | 0.88 | 0.6 | 0.81 | 0.53 |
| 7 | 32393467 | rs1562830 | 0.169 | -1.757 | 0.196 | 0.582 | 0.005 | 0.027 | 0.027 | 0 | 0.07 | 0.12 | 0.07 | 0.13 | 0.07 | 0.07 |
| 7 | 32397448 | rs4723146 | -0.721 | -1.8 | -0.019 | 0.61 | 0 | 0.003 | 0.003 | 0.422 | 0.12 | 0.15 | 0.11 | 0.25 | 0.16 | 0.17 |
| 7 | 32397633 | rs7778364 | -0.759 | -2.13 | 0.127 | 0.539 | -0.001 | 0 | 0 | 0 | 0.18 | 0.19 | 0.11 | 0.15 | 0.11 | 0.29 |
| 7 | 32398041 | rs215697 | 0.462 | -2.407 | 0.769 | 0.445 | -0.014 | 0.051 | 0.051 | 0 | 0.69 | 0.5 | 0.82 | 0.51 | 0.75 | 0.63 |
| 7 | 32399469 | rs1653892 | 1.205 | -2.489 | 1.081 | 0.469 | -0.006 | 0.001 | 0.001 | 0 | 0.57 | 0.41 | 0.79 | 0.51 | 0.71 | 0.43 |
| 7 | 32399833 | rs10486507 | -1.276 | -2.417 | 1.359 | 0.626 | -0.051 | 0.014 | 0.014 | 0 | 0.25 | 0.42 | 0.16 | 0.29 | 0.17 | 0.36 |
| 7 | 32403046 | rs451319 | 0.275 | -2.351 | 1.129 | 0.169 | -0.012 | -0.019 | -0.019 | 0.575 | 0.94 | 0.95 | 0.99 | 1 | 0.98 | 0.99 |
| 7 | 32415383 | rs399902 | 1.002 | -2.373 | 1.397 | 0.159 | -0.018 | -0.025 | -0.025 | 0.125 | 0.94 | 0.77 | 0.97 | 1 | 0.98 | 0.98 |
| 7 | 32423399 | rs215720 | 1.503 | -2.671 | 1.255 | 0.079 | -0.094 | -0.138 | -0.138 | 0 | 0.74 | 0.47 | 0.88 | 0.96 | 0.86 | 0.7 |
| 7 | 32428454 | rs2116669 | -1.048 | -2.594 | 1.167 | 0.113 | -0.059 | -0.081 | -0.081 | 0 | 0.18 | 0.39 | 0.09 | 0.04 | 0.1 | 0.24 |
| 7 | 32431246 | rs7456261 | 0.567 | -2.326 | 0.354 | -0.344 | -0.059 | 0.031 | 0.031 | 0 | 0.23 | 0.04 | 0.4 | 0.17 | 0.35 | 0.1 |
| 7 | 32440555 | rs369052 | 0.68 | -2.18 | 0.323 | -0.343 | -0.003 | -0.08 | -0.08 | 1.07 | 0.62 | 0.46 | 0.52 | 0.89 | 0.56 | 0.67 |
| 7 | 32442737 | rs452772 | 0.322 | -2.191 | 0.431 | -0.288 | -0.009 | -0.083 | -0.083 | 0 | 0.64 | 0.57 | 0.57 | 0.92 | 0.64 | 0.73 |
| 7 | 32444435 | rs215738 | 0.724 | -1.835 | 0.764 | 0.161 | -0.113 | -0.117 | -0.117 | 0 | 0.84 | 0.51 | 0.89 | 0.99 | 0.96 | 0.82 |
| 7 | 32446317 | rs170016 | 0.603 | -1.456 | 0.27 | 0.728 | 0.006 | -0.002 | -0.002 | 0 | 0.54 | 0.35 | 0.68 | 0.52 | 0.65 | 0.4 |
| 7 | 32446930 | rs1123893 | 1.068 | -0.946 | -0.043 | 0.725 | -0.05 | -0.023 | -0.023 | 0.112 | 0.16 | 0.06 | 0.2 | 0.21 | 0.2 | 0.22 |
| 7 | 32447262 | rs730725 | 1.192 | -1.837 | 0.846 | 1.03 | 0.039 | -0.02 | -0.02 | 0 | 0.57 | 0.26 | 0.66 | 0.53 | 0.67 | 0.4 |
| 7 | 32447350 | rs215743 | 0.894 | -1.79 | 1.032 | 1.138 | 0.013 | 0.08 | 0.08 | 0 | 0.92 | 0.64 | 0.9 | 0.9 | 0.96 | 0.95 |
| 7 | 32447684 | rs435584 | -0.763 | -1.784 | 0.632 | 1.138 | 0.001 | -0.001 | -0.001 | 0 | 0.35 | 0.39 | 0.25 | 0.37 | 0.29 | 0.55 |
| 7 | 32449568 | rs7792020 | -0.824 | -1.744 | -0.298 | 0.767 | -0.044 | -0.041 | -0.041 | 0 | 0.22 | 0.72 | 0.15 | 0.06 | 0.08 | 0.24 |
| 7 | 32451615 | rs2392055 | 0.08 | -1.061 | -0.323 | 0.992 | -0.009 | -0.07 | -0.07 | 0.073 | 0.74 | 0.88 | 0.67 | 0.56 | 0.72 | 0.62 |
| 7 | 32452397 | rs6975016 | -0.427 | -0.947 | 0.03 | 0.415 | -0.078 | -0.087 | -0.087 | 0 | 0.16 | 0.56 | 0.13 | 0.03 | 0.06 | 0.22 |
| 7 | 32456846 | rs215747 | 0.447 | 0.065 | -0.647 | 1.178 | -0.009 | -0.038 | -0.038 | 0 | 0.75 | 0.88 | 0.67 | 0.67 | 0.72 | 0.62 |
| 7 | 32457572 | rs215749 | -0.792 | 0.004 | 0.096 | 1.608 | 0 | 0.072 | 0.072 | 0 | 0.23 | 0.34 | 0.35 | 0.54 | 0.23 | 0.38 |
| 7 | 32457821 | rs215750 | 0.85 | 0.69 | 0.237 | 1.336 | 0.011 | 0.017 | 0.017 | 0 | 0.68 | 0.58 | 0.5 | 0.64 | 0.65 | 0.57 |
| 7 | 32460233 | rs10226019 | -0.405 | 1.138 | 0.203 | 0.893 | 0.019 | -0.007 | -0.007 | 0 | 0.19 | 0.3 | 0.11 | 0.19 | 0.23 | 0.12 |
| 7 | 32460840 | rs10273165 | -0.111 | 1.325 | 0.289 | 0.876 | -0.005 | -0.001 | -0.001 | 0 | 0.16 | 0.22 | 0.07 | 0.19 | 0.15 | 0.11 |
| 7 | 32461015 | rs7785659 | -0.129 | 1.318 | 0.286 | 0.872 | -0.005 | -0.001 | -0.001 | 0 | 0.16 | 0.22 | 0.07 | 0.19 | 0.15 | 0.11 |
| 7 | 32463933 | rs979355 | -0.705 | 1.128 | 0.305 | 1.21 | -0.009 | 0.008 | 0.008 | 2.774 | 0.19 | 0.27 | 0.18 | 0.3 | 0.16 | 0.14 |
| 7 | 32470370 | rs7804837 | -1.027 | 1.334 | 0.05 | 0.254 | -0.015 | 0.026 | 0.026 | 3.958 | 0.42 | 0.37 | 0.49 | 0.34 | 0.46 | 0.23 |
| 7 | 32472556 | rs6949694 | 0.142 | 0.801 | -0.227 | 0.204 | -0.008 | 0.097 | 0.097 | 1.488 | 0.44 | 0.16 | 0.41 | 0.21 | 0.4 | 0.32 |
| 7 | 32476929 | rs716500 | 0.228 | 0.823 | -0.296 | 0.257 | -0.019 | 0.066 | 0.066 | 3.374 | 0.34 | 0.16 | 0.33 | 0.24 | 0.36 | 0.19 |
| 7 | 32487376 | rs13223936 | 0.818 | 0.596 | 0.297 | 0.188 | -0.002 | -0.002 | -0.002 | 0 | 0.3 | 0.25 | 0.19 | 0.13 | 0.22 | 0.4 |
| 7 | 32493417 | rs958217 | 0.619 | 0.581 | 0.42 | 0.29 | -0.001 | -0.013 | -0.013 | 0 | 0.29 | 0.27 | 0.4 | 0.62 | 0.33 | 0.4 |
| 7 | 32496930 | rs3801328 | 0.461 | 0.565 | 0.465 | 0.282 | 0.001 | -0.004 | -0.004 | 0 | 0.32 | 0.34 | 0.42 | 0.62 | 0.33 | 0.4 |
| 7 | 32497039 | rs3801331 | 1.147 | 0.598 | 0.337 | 0.563 | -0.008 | -0.034 | -0.034 | 10.804 | 0.26 | 0.2 | 0.29 | 0.45 | 0.32 | 0.37 |
| 7 | 32512804 | rs4723159 | 1.1 | 0.599 | 0.386 | 0.446 | -0.017 | -0.072 | -0.072 | 0 | 0.26 | 0.17 | 0.29 | 0.51 | 0.32 | 0.37 |
| 7 | 32514580 | rs17362231 | 1.612 | 0.683 | 0.216 | 0.276 | -0.062 | 0.047 | 0.047 | 0 | 0.14 | 0.01 | 0.23 | 0.05 | 0.18 | 0.08 |
| 7 | 32514776 | rs16875962 | -0.15 | 0.511 | 0.473 | 0.299 | 0.013 | -0.03 | -0.03 | 0 | 0.37 | 0.47 | 0.37 | 0.25 | 0.42 | 0.21 |
| 7 | 32516370 | rs6462370 | 0.073 | 0.649 | 0.505 | 0.335 | 0.045 | -0.025 | -0.025 | 2.798 | 0.23 | 0.42 | 0.41 | 0.17 | 0.28 | 0.14 |
| 7 | 32524844 | rs2167274 | 0.998 | 0.583 | -0.146 | 0.387 | -0.015 | -0.07 | -0.07 | 0.794 | 0.31 | 0.17 | 0.33 | 0.5 | 0.38 | 0.36 |
| 7 | 32525631 | rs10272900 | 0.245 | 0.588 | -0.104 | 0.36 | -0.005 | 0.037 | 0.037 | 0.781 | 0.46 | 0.28 | 0.39 | 0.7 | 0.53 | 0.49 |
| 7 | 32534203 | rs10241303 | 1.136 | 0.711 | 0.057 | 0.538 | 0.003 | 0.002 | 0.002 | 0.905 | 0.16 | 0.1 | 0.06 | 0.2 | 0.15 | 0.13 |
| 7 | 32537679 | rs13244332 | 0.834 | 0.358 | -0.374 | 0.22 | 0.021 | -0.054 | -0.054 | 0 | 0.4 | 0.18 | 0.35 | 0.5 | 0.4 | 0.38 |
| 7 | 32546118 | rs1597555 | 0.597 | 0.302 | -0.265 | 0.197 | 0.042 | -0.03 | -0.03 | 0.01 | 0.42 | 0.25 | 0.36 | 0.5 | 0.4 | 0.38 |
| 7 | 32574243 | rs10260808 | -0.001 | 0.451 | 0.321 | 0.191 | -0.005 | 0.003 | 0.003 | 0 | 0.36 | 0.38 | 0.25 | 0.34 | 0.33 | 0.49 |
| 7 | 32575962 | rs17364223 | 0.963 | 0.523 | 0.221 | 0.182 | -0.093 | -0.01 | -0.01 | 0 | 0.18 | 0.05 | 0.36 | 0.16 | 0.27 | 0.13 |
| 7 | 32594431 | rs2290211 | 0.041 | 0.501 | 0.45 | 0.177 | -0.006 | 0.001 | 0.001 | 0 | 0.63 | 0.53 | 0.72 | 0.67 | 0.66 | 0.51 |
| 7 | 32598656 | rs2290214 | 0.343 | 0.52 | 0.647 | 0.164 | -0.004 | 0.017 | 0.017 | 0 | 0.19 | 0.33 | 0.17 | 0.13 | 0.17 | 0.36 |
| 7 | 32604222 | rs17161418 | -0.908 | 0.728 | 0.674 | 0.148 | 0.128 | 0.014 | 0.014 | 1 | 0.06 | 0.31 | 0.06 | 0 | 0.08 | 0.01 |
| 7 | 32613020 | rs1993050 | 0.158 | 0.747 | 0.659 | 0.149 | 0.006 | -0.001 | -0.001 | 0 | 0.63 | 0.39 | 0.71 | 0.67 | 0.66 | 0.51 |
| 7 | 32615553 | rs6462381 | -1.053 | 0.761 | 0.71 | 0.147 | 0.009 | -0.003 | -0.003 | 0.373 | 0.81 | 0.78 | 0.85 | 0.8 | 0.74 | 0.85 |
| 7 | 32625099 | rs3801332 | 0.033 | 0.763 | 0.753 | 0.156 | 0.04 | 0.007 | 0.007 | 0 | 0.22 | 0.37 | 0.21 | 0.13 | 0.24 | 0.37 |

| | | | | | | | | | | | | | | | | |
|---|---|---|---|---|---|---|---|---|---|---|---|---|---|---|---|---|
| 7 | 32626006 | rs3750083 | -0.372 | 0.976 | 0.759 | 0.558 | 0 | 0 | 0 | 0.685 | 0.28 | 0.26 | 0.37 | 0.31 | 0.35 | 0.49 |
| 7 | 32626781 | rs2290219 | 0.516 | 0.63 | 0.83 | 0.602 | -0.031 | 0.009 | 0.009 | 0.89 | 0.58 | 0.22 | 0.66 | 0.67 | 0.58 | 0.5 |
| 7 | 32629640 | rs12537908 | -1.411 | 0.626 | 0.81 | 0.61 | 0 | 0.001 | 0.001 | 2.223 | 0.81 | 0.8 | 0.89 | 0.8 | 0.82 | 0.87 |
| 7 | 32633021 | rs11983596 | 1.393 | 0.604 | 0.788 | 0.597 | 0.002 | 0.014 | 0.014 | 0 | 0.15 | 0.18 | 0.11 | 0.13 | 0.12 | 0.22 |
| 7 | 32645512 | rs17161469 | 0.376 | 0.635 | 0.463 | 0.638 | -0.003 | 0.03 | 0.03 | 0 | 0.2 | 0.41 | 0.18 | 0.13 | 0.16 | 0.35 |
| 7 | 32646096 | rs6945238 | 1.435 | 0.554 | 0.226 | 0.653 | -0.007 | -0.056 | -0.056 | 2.385 | 0.36 | 0.03 | 0.19 | 0.39 | 0.32 | 0.29 |
| 7 | 32665229 | rs6462391 | -1.894 | 0.583 | 0.263 | 0.61 | 0 | 0 | 0 | 1.015 | 0.83 | 0.85 | 0.89 | 0.8 | 0.82 | 0.87 |
| 7 | 32674289 | rs2122619 | -1.755 | 0.547 | 0.246 | 0.593 | 0.001 | 0.001 | 0.001 | 0.062 | 0.82 | 0.85 | 0.89 | 0.8 | 0.82 | 0.87 |
| 7 | 32679502 | rs4636106 | -1.099 | 0.612 | 0.299 | 0.618 | 0.029 | -0.017 | -0.017 | 0.194 | 0.79 | 0.62 | 0.84 | 0.8 | 0.74 | 0.86 |
| 7 | 32685774 | rs1839574 | 0.269 | 0.633 | 0.296 | 0.624 | -0.009 | 0.005 | 0.005 | 0.191 | 0.59 | 0.34 | 0.67 | 0.67 | 0.58 | 0.5 |
| 7 | 32694219 | rs7797763 | -1.032 | 0.579 | 0.314 | 0.599 | 0.028 | -0.018 | -0.018 | 0 | 0.78 | 0.62 | 0.84 | 0.8 | 0.74 | 0.86 |
| 7 | 32700515 | rs1839580 | 0.292 | 0.552 | 0.214 | 0.63 | -0.01 | 0.004 | 0.004 | 0.352 | 0.59 | 0.34 | 0.67 | 0.66 | 0.58 | 0.5 |
| 7 | 32711465 | rs13228793 | 1.358 | 0.483 | 0.05 | 0.721 | 0.021 | -0.007 | -0.007 | 0 | 0.36 | 0.04 | 0.19 | 0.35 | 0.32 | 0.29 |
| 7 | 32796927 | rs10085683 | 1.641 | 0.67 | 0.275 | 0.569 | 0 | 0 | 0 | 0 | 0.17 | 0.14 | 0.1 | 0.2 | 0.18 | 0.13 |
| 7 | 32800975 | rs13225132 | 1.006 | 0.649 | 0.224 | 0.597 | 0.029 | -0.031 | -0.031 | 0 | 0.41 | 0.19 | 0.25 | 0.46 | 0.38 | 0.39 |
| 7 | 32815508 | rs4723203 | 0.624 | 0.82 | 0.214 | 0.687 | -0.001 | 0.017 | 0.017 | 0 | 0.19 | 0.24 | 0.16 | 0.14 | 0.18 | 0.36 |
| 7 | 32818759 | rs17170516 | 1.511 | 0.867 | -0.212 | 0.674 | -0.048 | -0.028 | -0.028 | 0 | 0.16 | 0.04 | 0.41 | 0.2 | 0.18 | 0.12 |
| 7 | 32820437 | rs10243492 | 0.55 | 0.852 | -0.095 | 0.679 | -0.022 | 0.002 | 0.002 | 0 | 0.58 | 0.34 | 0.35 | 0.66 | 0.57 | 0.51 |
| 7 | 32824581 | rs13244487 | 0.945 | 0.809 | -0.305 | 0.697 | 0.03 | -0.019 | -0.019 | 0.203 | 0.39 | 0.2 | 0.22 | 0.41 | 0.35 | 0.36 |
| 7 | 32826172 | rs17136990 | -0.508 | 0.496 | -0.147 | 0.83 | -0.001 | -0.002 | -0.002 | 0.192 | 0.48 | 0.45 | 0.56 | 0.38 | 0.5 | 0.44 |
| 7 | 32829130 | rs4723212 | 0.222 | 0.634 | -0.327 | 0.729 | -0.02 | 0.039 | 0.039 | 0.235 | 0.3 | 0.21 | 0.41 | 0.19 | 0.35 | 0.2 |
| 7 | 32839266 | rs10264489 | 1.376 | 0.648 | -0.303 | 0.749 | -0.003 | 0.001 | 0.001 | 0 | 0.13 | 0.09 | 0.1 | 0.15 | 0.18 | 0.13 |
| 7 | 32843795 | rs4720078 | 0.888 | 0.542 | -0.584 | 0.653 | -0.011 | 0.038 | 0.038 | 0 | 0.2 | 0.15 | 0.35 | 0.04 | 0.22 | 0.19 |
| 7 | 32854040 | rs11975969 | 0.389 | 0.433 | -0.707 | 0.796 | 0.016 | 0.001 | 0.001 | 0.019 | 0.48 | 0.38 | 0.29 | 0.51 | 0.45 | 0.37 |
| 7 | 32857797 | rs11760370 | 0.406 | 0.442 | -0.711 | 0.8 | 0.014 | 0 | 0 | 0 | 0.48 | 0.39 | 0.3 | 0.51 | 0.45 | 0.37 |
| 7 | 32866723 | rs2392147 | -0.437 | 0.474 | -1.326 | 1.203 | 0.001 | -0.007 | -0.007 | 0 | 0.42 | 0.73 | 0.42 | 0.57 | 0.42 | 0.56 |
| 7 | 32890682 | rs2392151 | -0.572 | 0.458 | -1.61 | 1.25 | 0 | 0.013 | 0.013 | 0.102 | 0.38 | 0.84 | 0.28 | 0.35 | 0.34 | 0.39 |
| 7 | 32907344 | rs6462427 | 1.236 | 0.847 | -1.84 | 1.335 | -0.093 | -0.018 | -0.018 | 0.093 | 0.38 | 0.02 | 0.23 | 0.29 | 0.36 | 0.27 |
| 7 | 32908704 | rs1045477 | 1.236 | 0.847 | -1.84 | 1.335 | -0.093 | -0.018 | -0.018 | 0 | 0.38 | 0.02 | 0.23 | 0.29 | 0.37 | 0.27 |
| 7 | 32909182 | rs752730 | 1.22 | 0.933 | -1.882 | 1.586 | -0.032 | -0.028 | -0.028 | 0 | 0.18 | 0.08 | 0.42 | 0.22 | 0.2 | 0.15 |
| 7 | 32910615 | rs1545448 | 1.22 | 0.933 | -1.882 | 1.587 | -0.032 | -0.026 | -0.026 | 0.95 | 0.18 | 0.08 | 0.42 | 0.22 | 0.2 | 0.15 |
| 7 | 32912635 | rs9639668 | 0.075 | 0.765 | -0.841 | 1.667 | -0.016 | -0.045 | -0.045 | 0.407 | 0.29 | 0.39 | 0.21 | 0.19 | 0.25 | 0.42 |
| 7 | 32915204 | rs7785065 | -0.801 | 0.665 | -0.841 | 1.577 | 0.007 | -0.043 | -0.043 | 0.132 | 0.41 | 0.77 | 0.34 | 0.5 | 0.43 | 0.58 |
| 7 | 32953295 | rs3956705 | 1.529 | 0.813 | -1.057 | 1.475 | -0.07 | 0.034 | 0.034 | 0 | 0.4 | 0.02 | 0.22 | 0.26 | 0.36 | 0.26 |
| 7 | 32955265 | rs1376303 | 0.099 | 0.827 | -0.554 | 1.693 | 0.001 | 0.207 | 0.207 | 0 | 0.87 | 0.68 | 0.77 | 0.45 | 0.82 | 0.77 |
| 7 | 32965040 | rs12701264 | -1.236 | 0.873 | -0.664 | 1.755 | -0.001 | 0.078 | 0.078 | 0 | 0.94 | 0.86 | 0.94 | 0.81 | 0.9 | 0.92 |
| 7 | 32969419 | rs3750079 | -0.506 | 0.916 | -0.566 | 1.201 | -0.003 | -0.001 | -0.001 | 0 | 0.35 | 0.57 | 0.26 | 0.37 | 0.32 | 0.49 |
| 7 | 32972245 | rs17469596 | 1.283 | 1.044 | -1.084 | 0.874 | -0.04 | 0.054 | 0.054 | 0 | 0.16 | 0.1 | 0.31 | 0.03 | 0.19 | 0.08 |
| 7 | 32973818 | rs6945242 | 0.578 | 0.756 | -1.05 | 0.789 | -0.001 | 0.037 | 0.037 | 6.731 | 0.21 | 0.11 | 0.13 | 0.05 | 0.19 | 0.19 |
| 7 | 32976416 | rs7790246 | -0.718 | 0.694 | -1.324 | 1.086 | 0.037 | -0.05 | -0.05 | 1.14 | 0.38 | 0.79 | 0.39 | 0.54 | 0.31 | 0.55 |
| 7 | 32995331 | rs4720087 | 1.306 | 0.837 | -1.545 | 1.055 | -0.094 | 0.069 | 0.069 | 0 | 0.16 | 0.02 | 0.32 | 0.04 | 0.23 | 0.09 |
| 7 | 32999601 | rs13311608 | 0.906 | 0.479 | -1.394 | 1.133 | -0.018 | -0.029 | -0.029 | 0 | 0.46 | 0.12 | 0.29 | 0.42 | 0.47 | 0.35 |
| 7 | 33011535 | rs3109433 | -0.337 | 0.871 | -0.325 | 1.461 | 0.008 | 0.008 | 0.008 | 0 | 0.82 | 0.79 | 0.68 | 0.75 | 0.77 | 0.87 |
| 7 | 33040355 | rs6943296 | -0.372 | -0.053 | -0.885 | 0.461 | 0.011 | 0.023 | 0.023 | 0 | 0.44 | 0.61 | 0.54 | 0.18 | 0.59 | 0.3 |
| 7 | 33042945 | rs16881929 | 0.817 | -0.776 | -1.67 | 0.61 | 0.019 | 0.01 | 0.01 | 1.071 | 0.26 | 0.09 | 0.12 | 0.26 | 0.21 | 0.25 |
| 7 | 33053281 | rs6462446 | 0.071 | -0.969 | -1.585 | 0.663 | 0.054 | -0.037 | -0.037 | 0 | 0.62 | 0.81 | 0.72 | 0.46 | 0.71 | 0.6 |
| 7 | 33055164 | rs7790368 | -0.519 | -1.032 | -1.61 | 0.937 | 0.03 | -0.068 | -0.068 | 0 | 0.7 | 0.89 | 0.76 | 0.61 | 0.74 | 0.74 |
| 7 | 33060306 | rs16879126 | 1.638 | -1.031 | -1.675 | 1.279 | 0.002 | -0.008 | -0.008 | 0 | 0.05 | 0.02 | 0.07 | 0.18 | 0.05 | 0.1 |
| 7 | 33081018 | rs10230500 | 0.047 | -0.907 | -1.669 | 1.282 | 0.06 | -0.024 | -0.024 | 0 | 0.62 | 0.84 | 0.74 | 0.6 | 0.71 | 0.63 |
| 7 | 33086391 | rs7776847 | 1.419 | -0.257 | -1.585 | 1.019 | -0.012 | 0.009 | 0.009 | 0 | 0.15 | 0.12 | 0.3 | 0.05 | 0.23 | 0.06 |
| 7 | 33094982 | rs10251079 | 0.206 | -0.737 | -1.21 | 1.007 | 0.021 | -0.01 | -0.01 | 0.394 | 0.61 | 0.71 | 0.71 | 0.48 | 0.71 | 0.6 |
| 7 | 33127539 | rs7793131 | -0.961 | -0.031 | -1.583 | 1.074 | -0.042 | -0.056 | -0.056 | 0 | 0.81 | 0.99 | 0.85 | 0.81 | 0.79 | 0.84 |
| 7 | 33133742 | rs10245553 | 0.101 | 0.167 | -1.143 | 0.954 | -0.012 | 0.004 | 0.004 | 0 | 0.48 | 0.73 | 0.45 | 0.61 | 0.48 | 0.54 |
| 7 | 33152668 | rs9692508 | -0.144 | 0.084 | -0.562 | 0.652 | -0.007 | -0.005 | -0.005 | 0.677 | 0.12 | 0.18 | 0.11 | 0.1 | 0.09 | 0.16 |
| 7 | 33153065 | rs9690597 | 0.096 | -0.146 | -0.105 | 0.303 | -0.019 | -0.036 | -0.036 | 0.022 | 0.74 | 0.59 | 0.63 | 0.84 | 0.78 | 0.81 |
| 7 | 33153262 | rs10281493 | -0.548 | -0.053 | -0.188 | 0.464 | 0.004 | 0.005 | 0.005 | 0.471 | 0.3 | 0.62 | 0.34 | 0.4 | 0.31 | 0.24 |
| 7 | 33166070 | rs4720103 | 0.733 | -0.431 | -0.756 | 0.329 | 0.027 | 0.012 | 0.012 | 0 | 0.2 | 0.07 | 0.21 | 0.15 | 0.19 | 0.21 |
| 7 | 33167729 | rs10240165 | 0.718 | -0.439 | -0.759 | 0.346 | 0.028 | 0.01 | 0.01 | 0 | 0.2 | 0.07 | 0.21 | 0.15 | 0.19 | 0.21 |
| 7 | 33173159 | rs1420149 | 0.365 | -0.318 | -0.563 | 0.348 | -0.004 | 0.002 | 0.002 | 1.542 | 0.19 | 0.16 | 0.13 | 0.2 | 0.2 | 0.13 |
| 7 | 33193481 | rs12671333 | -0.556 | -0.267 | 0.257 | 0.492 | 0.026 | 0.078 | 0.078 | 0.358 | 0.3 | 0.45 | 0.26 | 0.51 | 0.3 | 0.3 |
| 7 | 33195732 | rs4720108 | 0.506 | -0.217 | 0.219 | 0.573 | -0.004 | -0.005 | -0.005 | 0 | 0.19 | 0.16 | 0.18 | 0.43 | 0.21 | 0.15 |
| 7 | 33197232 | rs1362368 | -0.136 | -0.267 | 0.23 | 0.534 | 0 | -0.005 | -0.005 | 0 | 0.39 | 0.38 | 0.4 | 0.58 | 0.4 | 0.37 |
| 7 | 33218763 | rs1009355 | -0.694 | -0.271 | 0.377 | 0.516 | 0.003 | 0.04 | 0.04 | 0.214 | 0.5 | 0.63 | 0.46 | 0.66 | 0.48 | 0.51 |
| 7 | 33237489 | rs10254116 | 0.796 | -0.262 | 0.315 | 0.189 | -0.011 | 0.001 | 0.001 | 0 | 0.29 | 0.3 | 0.47 | 0.31 | 0.35 | 0.21 |
| 7 | 33238200 | rs10486521 | 1.984 | -0.23 | 0.081 | 0.013 | -0.053 | 0.055 | 0.055 | 0.826 | 0.1 | 0.01 | 0.07 | 0 | 0.14 | 0.03 |
| 7 | 33241996 | rs10486523 | 0.942 | -0.307 | 0.313 | 0.019 | -0.003 | -0.004 | -0.004 | 0 | 0.1 | 0.06 | 0.06 | 0.08 | 0.09 | 0.15 |
| 7 | 33242474 | rs10486524 | 0.297 | -0.345 | 0.305 | 0.045 | -0.012 | 0.011 | 0.011 | 1.083 | 0.19 | 0.14 | 0.13 | 0.16 | 0.2 | 0.13 |
| 7 | 33243151 | rs986553 | -2.112 | -0.426 | 0.16 | 0.014 | -0.003 | 0.014 | 0.014 | 0.787 | 0.82 | 0.83 | 0.88 | 0.7 | 0.82 | 0.75 |
| 7 | 33256039 | rs10262453 | 1.057 | -0.302 | 0.042 | -0.075 | -0.037 | 0.094 | 0.094 | 0 | 0.28 | 0.13 | 0.4 | 0.04 | 0.32 | 0.19 |

| | | | | | | | | | | | | | | | | |
|---|---|---|---|---|---|---|---|---|---|---|---|---|---|---|---|---|
| 7 | 33277292 | rs2110767 | 0.917 | -0.347 | -0.006 | 0.037 | -0.039 | -0.059 | -0.059 | 0 | 0.14 | 0.06 | 0.16 | 0.43 | 0.19 | 0.14 |
| 7 | 33282577 | rs7793096 | 0.004 | -0.557 | 0.261 | -0.099 | 0.004 | 0.005 | 0.005 | 0 | 0.46 | 0.38 | 0.57 | 0.47 | 0.54 | 0.37 |
| 7 | 33284738 | rs6952877 | 2.152 | -0.666 | 0.165 | -0.224 | 0.005 | -0.008 | -0.008 | 0 | 0.17 | 0.02 | 0.2 | 0.15 | 0.17 | 0.18 |
| 7 | 33290931 | rs1420154 | 1.061 | -0.635 | 0.202 | -0.226 | -0.003 | -0.023 | -0.023 | 0.497 | 0.29 | 0.31 | 0.41 | 0.04 | 0.33 | 0.18 |
| 7 | 33304218 | rs7793862 | 0.658 | -0.759 | 0.233 | -0.097 | 0 | 0 | 0 | 0.785 | 0.27 | 0.24 | 0.27 | 0.23 | 0.26 | 0.32 |
| 7 | 33346932 | rs10253342 | 0.501 | -0.558 | 0.084 | -0.034 | -0.053 | -0.014 | -0.014 | 0 | 0.17 | 0.05 | 0.11 | 0.16 | 0.19 | 0.17 |
| 7 | 33354325 | rs7797739 | -0.262 | -0.776 | 0.42 | -0.265 | -0.007 | -0.081 | -0.081 | 0 | 0.13 | 0.33 | 0.09 | 0 | 0.1 | 0.04 |
| 7 | 33370588 | rs17170189 | 0.095 | -0.896 | 0.507 | -0.27 | -0.008 | -0.003 | -0.003 | 0 | 0.13 | 0.23 | 0.07 | 0.08 | 0.08 | 0.22 |
| 7 | 33373848 | rs6976311 | 2.392 | -0.847 | 0.096 | -0.224 | -0.04 | -0.097 | -0.097 | 0.143 | 0.17 | 0.01 | 0.11 | 0.29 | 0.17 | 0.19 |
| 7 | 33374005 | rs6977009 | 1.133 | -0.904 | 0.208 | -0.34 | -0.026 | 0.025 | 0.025 | 0.052 | 0.4 | 0.2 | 0.47 | 0.33 | 0.43 | 0.37 |
| 7 | 33377472 | rs11982012 | -0.299 | -0.903 | 0.387 | -0.473 | -0.004 | 0.002 | 0.002 | 0 | 0.28 | 0.22 | 0.2 | 0.08 | 0.29 | 0.21 |
| 7 | 33385262 | rs3884597 | 0.811 | -0.935 | 0.49 | -0.379 | -0.005 | -0.005 | -0.005 | 0 | 0.46 | 0.35 | 0.51 | 0.48 | 0.48 | 0.46 |
| 7 | 33400857 | rs17170204 | 0.43 | -0.93 | 0.38 | -0.415 | -0.003 | -0.007 | -0.007 | 0 | 0.14 | 0.18 | 0.07 | 0.08 | 0.1 | 0.16 |
| 7 | 33411267 | rs2109525 | 2.267 | -0.872 | 0.213 | -0.433 | -0.009 | -0.003 | -0.003 | 0 | 0.1 | 0.07 | 0.08 | 0 | 0.13 | 0.03 |
| 7 | 33423761 | rs17724206 | 1.96 | -0.942 | 0.129 | -0.454 | -0.044 | 0.081 | 0.081 | 0 | 0.21 | 0.07 | 0.34 | 0.04 | 0.26 | 0.1 |
| 7 | 33431364 | rs11972278 | -1.452 | -0.876 | 0.27 | -0.29 | -0.016 | 0.02 | 0.02 | 0.007 | 0.6 | 0.74 | 0.52 | 0.67 | 0.57 | 0.63 |
| 7 | 33453144 | rs17170220 | 1.862 | -0.89 | 0.158 | -0.31 | 0.007 | -0.005 | -0.005 | 0 | 0.17 | 0.1 | 0.2 | 0.19 | 0.19 | 0.21 |
| 7 | 33453281 | rs6977744 | 1.07 | -0.87 | 0.119 | -0.326 | -0.001 | -0.025 | -0.025 | 0 | 0.08 | 0.08 | 0.06 | 0 | 0.08 | 0.03 |
| 7 | 33456293 | rs6462477 | -1.511 | -0.85 | -0.167 | -0.27 | -0.025 | 0.024 | 0.024 | 0.033 | 0.6 | 0.8 | 0.53 | 0.67 | 0.57 | 0.63 |
| 7 | 33457501 | rs1419927 | -1.478 | -0.867 | -0.002 | -0.278 | -0.013 | 0.017 | 0.017 | 0.012 | 0.6 | 0.71 | 0.53 | 0.67 | 0.57 | 0.63 |
| 7 | 33459531 | rs6965729 | 1.825 | -0.911 | -0.088 | -0.287 | -0.019 | -0.037 | -0.037 | 0 | 0.17 | 0.01 | 0.2 | 0.19 | 0.19 | 0.2 |
| 7 | 33461577 | rs10951388 | -1.006 | -0.841 | 0.109 | -0.247 | -0.007 | 0.015 | 0.015 | 0.317 | 0.33 | 0.52 | 0.26 | 0.39 | 0.29 | 0.23 |
| 7 | 33469241 | rs4720118 | -1.471 | -0.841 | -0.117 | -0.26 | -0.045 | 0.023 | 0.023 | 0.154 | 0.6 | 0.85 | 0.53 | 0.66 | 0.57 | 0.64 |
| 7 | 33476376 | rs1419926 | -0.996 | -0.829 | -0.2 | -0.24 | -0.056 | -0.034 | -0.034 | 0 | 0.23 | 0.46 | 0.12 | 0.16 | 0.15 | 0.17 |
| 7 | 33484507 | rs1860601 | 0.3 | -0.13 | -0.963 | -0.061 | 0.006 | -0.007 | -0.007 | 0 | 0.2 | 0.22 | 0.15 | 0.08 | 0.23 | 0.18 |
| 7 | 33497548 | rs17170255 | 0.685 | -0.531 | -0.815 | 0.325 | -0.001 | 0.029 | 0.029 | 0 | 0.1 | 0.09 | 0.08 | 0.31 | 0.1 | 0.17 |
| 7 | 33499270 | rs13438256 | 0.016 | -0.648 | -0.769 | -0.335 | 0 | -0.008 | -0.008 | 0 | 0.31 | 0.3 | 0.31 | 0.5 | 0.33 | 0.45 |
| 7 | 33499822 | rs6973093 | 0.104 | -0.46 | -1.347 | -0.138 | -0.018 | 0.017 | 0.017 | 0 | 0.09 | 0.14 | 0.04 | 0.16 | 0.05 | 0.13 |
| 7 | 33505151 | rs1362229 | -1.694 | -0.299 | -1.609 | -0.348 | -0.108 | -0.08 | -0.08 | 3.296 | 0.73 | 0.98 | 0.82 | 0.71 | 0.7 | 0.78 |
| 7 | 33510163 | rs4637716 | -0.857 | -0.467 | -1.658 | -0.586 | 0.009 | 0.09 | 0.09 | 0.324 | 0.52 | 0.81 | 0.45 | 0.66 | 0.47 | 0.58 |
| 7 | 33511742 | rs2109526 | 2.078 | -0.545 | -1.805 | -0.534 | -0.025 | -0.084 | -0.084 | 3.001 | 0.13 | 0.05 | 0.2 | 0.46 | 0.17 | 0.23 |
| 7 | 33512125 | rs1419906 | -2.11 | -0.481 | -1.715 | -0.466 | -0.094 | -0.068 | -0.068 | 0 | 0.74 | 0.97 | 0.82 | 0.7 | 0.7 | 0.79 |
| 7 | 33512729 | rs10486535 | -0.033 | -0.595 | -1.505 | -0.519 | -0.004 | 0.004 | 0.004 | 0 | 0.2 | 0.23 | 0.12 | 0.19 | 0.17 | 0.25 |
| 7 | 33523792 | rs1419901 | -0.553 | -0.372 | -2.023 | -0.354 | 0.023 | 0.028 | 0.028 | 0 | 0.34 | 0.73 | 0.23 | 0.2 | 0.29 | 0.33 |
| 7 | 33526293 | rs7804723 | -1.506 | 0.201 | -2.214 | 0.072 | -0.11 | -0.077 | -0.077 | 0 | 0.74 | 0.98 | 0.82 | 0.7 | 0.7 | 0.79 |
| 7 | 33536947 | rs13227256 | 0.951 | 0.189 | -1.862 | 0.305 | -0.023 | -0.103 | -0.103 | 0 | 0.15 | 0.04 | 0.21 | 0.46 | 0.18 | 0.24 |
| 7 | 33537666 | rs11769528 | 1.365 | 0.254 | -1.839 | 0.334 | -0.033 | -0.091 | -0.091 | 0 | 0.13 | 0.01 | 0.2 | 0.46 | 0.17 | 0.23 |
| 7 | 33543478 | rs12536844 | 0.843 | 0.374 | -2.063 | -0.297 | -0.043 | 0.101 | 0.101 | 0 | 0.24 | 0.11 | 0.37 | 0.04 | 0.28 | 0.11 |
| 7 | 33543550 | rs2392241 | -1.535 | 0.502 | -2.052 | -0.113 | -0.038 | 0.031 | 0.031 | 0.906 | 0.76 | 0.88 | 0.85 | 0.81 | 0.73 | 0.82 |
| 7 | 33546619 | rs2392243 | -1.528 | 0.509 | -2.054 | -0.091 | -0.032 | 0 | 0 | 0.745 | 0.76 | 0.88 | 0.82 | 0.73 | 0.72 | 0.8 |
| 7 | 33549392 | rs11761619 | 0.587 | 0.934 | -1.036 | 0.193 | 0.091 | 0.041 | 0.041 | 0.565 | 0.07 | 0.31 | 0.1 | 0.07 | 0.11 | 0.04 |
| 7 | 33550041 | rs10262995 | 0.396 | 0.644 | -1.487 | 0.361 | -0.008 | -0.002 | -0.002 | 0 | 0.11 | 0.06 | 0.07 | 0.06 | 0.09 | 0.14 |
| 7 | 33552840 | rs10232000 | 0.466 | 0.435 | -1.303 | 0.53 | 0.006 | -0.071 | -0.071 | 0 | 0.19 | 0.11 | 0.23 | 0.47 | 0.2 | 0.3 |
| 7 | 33554162 | rs12674093 | -0.166 | 0.588 | -0.958 | 0.48 | -0.001 | 0.001 | 0.001 | 0 | 0.49 | 0.57 | 0.69 | 0.58 | 0.55 | 0.44 |
| 7 | 33554672 | rs10253978 | -0.353 | 0.824 | -0.831 | 0.754 | 0.006 | 0.011 | 0.011 | 0 | 0.07 | 0.08 | 0.04 | 0.12 | 0.09 | 0.13 |
| 7 | 33554950 | rs6966393 | 0.289 | 1.07 | -0.975 | 0.697 | 0.002 | 0 | 0 | 0 | 0.54 | 0.61 | 0.5 | 0.56 | 0.46 | 0.33 |
| 7 | 33555119 | rs2392244 | -1.22 | 0.897 | 0.028 | 1.072 | 0.014 | -0.012 | -0.012 | 0.095 | 0.27 | 0.39 | 0.2 | 0.28 | 0.31 | 0.29 |
| 7 | 33567912 | rs2392245 | -1.349 | 0.012 | 0.037 | 1.422 | 0.015 | -0.021 | -0.021 | 0.094 | 0.38 | 0.6 | 0.56 | 0.49 | 0.32 | 0.47 |
| 7 | 33568222 | rs7794579 | 0.066 | -0.066 | -0.14 | 1.242 | 0.001 | -0.013 | -0.013 | 0 | 0.12 | 0.15 | 0.08 | 0.05 | 0.11 | 0.06 |
| 7 | 33569954 | rs6971826 | -1.709 | -0.096 | -0.251 | 1.508 | 0.003 | 0.043 | 0.043 | 1.439 | 0.5 | 0.79 | 0.65 | 0.61 | 0.44 | 0.54 |
| 7 | 33571828 | rs2392246 | -2.083 | -0.278 | -0.321 | 1.365 | -0.105 | 0.014 | 0.014 | 3.18 | 0.56 | 0.96 | 0.67 | 0.69 | 0.46 | 0.75 |
| 7 | 33591485 | rs6974597 | -1.84 | -0.065 | 0.112 | 1.375 | -0.025 | -0.008 | -0.008 | 1.5 | 0.52 | 0.77 | 0.69 | 0.58 | 0.52 | 0.66 |
| 7 | 33595651 | rs17170286 | 0.361 | -0.027 | 0.262 | 1.244 | 0.001 | -0.039 | -0.039 | 0 | 0.11 | 0.14 | 0.07 | 0 | 0.09 | 0.05 |
| 7 | 33596515 | rs10486537 | -0.173 | -0.128 | 0.833 | 1.377 | -0.006 | -0.04 | -0.04 | 0 | 0.13 | 0.23 | 0.09 | 0.04 | 0.1 | 0.11 |
| 7 | 33597797 | rs1551098 | -1.555 | -0.131 | 0.535 | 1.366 | -0.001 | 0.003 | 0.003 | 2.758 | 0.39 | 0.55 | 0.61 | 0.52 | 0.4 | 0.57 |
| 7 | 33600759 | rs1451017 | -1.614 | -1.072 | 0.82 | 1.739 | -0.01 | 0.043 | 0.043 | 0.036 | 0.11 | 0.21 | 0.06 | 0.24 | 0.07 | 0.15 |
| 7 | 33603563 | rs1020519 | -2.097 | -1.193 | 0.799 | 1.721 | -0.003 | 0.02 | 0.02 | 0.167 | 0.23 | 0.33 | 0.13 | 0.41 | 0.19 | 0.22 |
| 7 | 33607320 | rs1451015 | -2.521 | -1.407 | 0.658 | 1.377 | 0.038 | -0.054 | -0.054 | 0 | 0.41 | 0.84 | 0.27 | 0.52 | 0.32 | 0.47 |
| 7 | 33607662 | rs2700695 | -2.149 | -1.573 | 1.063 | 1.296 | -0.039 | 0.038 | 0.038 | 1.091 | 0.28 | 0.6 | 0.18 | 0.42 | 0.18 | 0.39 |
| 7 | 33613560 | rs4590341 | -2.595 | -1.917 | 1.306 | 1.717 | -0.017 | 0.017 | 0.017 | 3.108 | 0.22 | 0.75 | 0.15 | 0.48 | 0.11 | 0.35 |
| 7 | 33618322 | rs2598395 | -2.306 | -1.806 | 0.797 | 1.761 | 0.047 | -0.051 | -0.051 | 0.58 | 0.32 | 0.83 | 0.2 | 0.48 | 0.21 | 0.4 |
| 7 | 33620899 | rs2598392 | -2.497 | -1.874 | 0.735 | 1.72 | 0.061 | -0.06 | -0.06 | 0 | 0.36 | 0.88 | 0.23 | 0.46 | 0.23 | 0.33 |
| 7 | 33622386 | rs1031347 | -1.954 | -1.88 | 0.815 | 1.717 | 0.014 | -0.02 | -0.02 | 0 | 0.28 | 0.76 | 0.19 | 0.47 | 0.19 | 0.27 |
| 7 | 33622588 | rs965571 | -2.827 | -1.868 | 0.754 | 1.721 | 0.075 | -0.058 | -0.058 | 0 | 0.39 | 0.89 | 0.24 | 0.49 | 0.25 | 0.46 |
| 7 | 33626739 | rs1985081 | -1.216 | -1.712 | 0.665 | 2.101 | 0.003 | -0.019 | -0.019 | 0.767 | 0.1 | 0.06 | 0.31 | 0.19 | 0.11 | 0.17 |
| 7 | 33631992 | rs13227571 | -0.959 | -2.17 | 1.182 | 2.437 | -0.018 | 0.118 | 0.118 | 2.719 | 0.2 | 0.52 | 0.13 | 0.42 | 0.15 | 0.23 |
| 7 | 33639870 | rs2700677 | -2.585 | -2.161 | 1.08 | 2.417 | 0.029 | -0.006 | -0.006 | 0.378 | 0.38 | 0.72 | 0.44 | 0.63 | 0.25 | 0.52 |
| 7 | 33644941 | rs1050719 | -2.499 | -1.961 | 1.077 | 2.35 | 0.035 | -0.021 | -0.021 | 0 | 0.37 | 0.79 | 0.49 | 0.6 | 0.27 | 0.52 |
| 7 | 33662618 | rs2034515 | -2.452 | -1.963 | 0.996 | 1.804 | 0.054 | -0.001 | -0.001 | 0 | 0.36 | 0.84 | 0.24 | 0.41 | 0.24 | 0.43 |

| | | | | | | | | | | | | | | | | |
|---|---|---|---|---|---|---|---|---|---|---|---|---|---|---|---|---|
| 7 | 33667314 | rs10486541 | -1.554 | -1.829 | 1.073 | 1.626 | -0.03 | 0.088 | 0.088 | 2.861 | 0.23 | 0.4 | 0.13 | 0.46 | 0.16 | 0.23 |
| 7 | 33671662 | rs10250216 | 2.212 | -1.831 | 0.975 | 1.718 | 0.047 | 0.074 | 0.074 | 1.701 | 0.6 | 0.27 | 0.54 | 0.18 | 0.72 | 0.45 |
| 7 | 33674311 | rs6954413 | -1.397 | -1.66 | 0.896 | 1.905 | -0.036 | 0.047 | 0.047 | 0.108 | 0.31 | 0.47 | 0.18 | 0.44 | 0.22 | 0.4 |
| 7 | 33675799 | rs10266903 | 1.554 | -1.174 | 0.381 | 0.646 | -0.016 | -0.029 | -0.029 | 0 | 0.66 | 0.39 | 0.55 | 0.19 | 0.76 | 0.58 |
| 7 | 33679634 | rs1451011 | -1.193 | -0.762 | 0.403 | 0.93 | -0.033 | -0.028 | -0.028 | 0 | 0.28 | 0.5 | 0.42 | 0.76 | 0.2 | 0.4 |
| 7 | 33683363 | rs6954592 | -0.995 | -0.293 | 0.487 | 1.166 | 0.031 | 0.007 | 0.007 | 0.235 | 0.22 | 0.42 | 0.29 | 0.74 | 0.23 | 0.32 |
| 7 | 33684775 | rs2392252 | 1.392 | -0.972 | 1.255 | 2.099 | -0.013 | 0.086 | 0.086 | 0 | 0.78 | 0.43 | 0.65 | 0.45 | 0.79 | 0.71 |
| 7 | 33684991 | rs10239102 | -1.429 | -1.279 | 1.458 | 2.52 | -0.004 | 0.097 | 0.097 | 0 | 0.08 | 0.14 | 0.19 | 0.31 | 0.08 | 0.23 |
| 7 | 33690186 | rs1451019 | 2.31 | -1.518 | 2.197 | 2.556 | 0.007 | 0.126 | 0.126 | 0.987 | 0.89 | 0.45 | 0.79 | 0.78 | 0.92 | 0.68 |
| 7 | 33690732 | rs1451020 | 1.452 | -2.241 | 1.673 | 2.05 | -0.035 | -0.014 | -0.014 | 3.495 | 0.74 | 0.34 | 0.74 | 0.76 | 0.87 | 0.64 |
| 7 | 33696721 | rs17170329 | -1.278 | -2.167 | 1.569 | 2.69 | 0.003 | -0.002 | -0.002 | 0.904 | 0.17 | 0.12 | 0.09 | 0.36 | 0.13 | 0.13 |
| 7 | 33701991 | rs17170333 | -1.109 | -2.136 | 1.479 | 2.636 | 0.027 | -0.011 | -0.011 | 0.48 | 0.11 | 0.02 | 0.03 | 0.12 | 0.05 | 0.11 |
| 7 | 33704883 | rs1376353 | 2.573 | -2.342 | 1.497 | 2.609 | 0.065 | 0.006 | 0.006 | 0.534 | 0.57 | 0.21 | 0.61 | 0.38 | 0.71 | 0.52 |
| 7 | 33708138 | rs7806997 | 1.261 | -2.314 | 2.405 | 2.733 | -0.014 | 0.132 | 0.132 | 0.114 | 0.79 | 0.54 | 0.77 | 0.57 | 0.84 | 0.73 |
| 7 | 33708637 | rs4723307 | -0.977 | -1.885 | 2.51 | 3.214 | -0.026 | -0.033 | -0.033 | 0 | 0.06 | 0 | 0.18 | 0.22 | 0.1 | 0.21 |
| 7 | 33712743 | rs1551099 | 1.094 | -1.955 | 3.063 | 3.172 | 0.012 | 0.008 | 0.008 | 0 | 0.89 | 0.52 | 0.89 | 0.94 | 0.92 | 0.98 |
| 7 | 33714770 | rs4463328 | -0.782 | -1.95 | 3.01 | 3.236 | -0.021 | -0.017 | -0.017 | 0 | 0.05 | 0 | 0.17 | 0.23 | 0.08 | 0.2 |
| 7 | 33718506 | rs6966497 | 2.033 | -1.995 | 3.17 | 3.111 | -0.043 | 0.011 | 0.011 | 0 | 0.84 | 0.42 | 0.86 | 0.85 | 0.89 | 0.94 |
| 7 | 33718644 | rs6966639 | -2.07 | -2 | 3.388 | 3.108 | -0.035 | 0.012 | 0.012 | 0 | 0.16 | 0.42 | 0.13 | 0.15 | 0.1 | 0.06 |
| 7 | 33719054 | rs6967252 | -0.907 | -1.962 | 2.956 | 3.07 | -0.019 | 0.014 | 0.014 | 0 | 0.06 | 0.1 | 0.03 | 0.08 | 0.02 | 0.04 |
| 7 | 33728199 | rs1545449 | 2.647 | -1.939 | 3.082 | 3.178 | -0.02 | 0.09 | 0.09 | 0.003 | 0.77 | 0.39 | 0.68 | 0.51 | 0.79 | 0.6 |
| 7 | 33733587 | rs4723316 | 2.393 | -1.983 | 2.837 | 2.762 | -0.033 | 0.032 | 0.032 | 0.052 | 0.77 | 0.29 | 0.83 | 0.62 | 0.86 | 0.76 |
| 7 | 33754092 | rs6956173 | -2.049 | -1.892 | 2.841 | 2.701 | -0.061 | -0.017 | -0.017 | 0.01 | 0.18 | 0.46 | 0.11 | 0.16 | 0.11 | 0.12 |
| 7 | 33754981 | rs7357093 | -1.053 | -1.837 | 2.326 | 3.306 | -0.022 | 0.025 | 0.025 | 0.156 | 0.06 | 0.01 | 0.27 | 0.35 | 0.11 | 0.22 |
| 7 | 33757751 | rs17170369 | -1.001 | -1.804 | 2.511 | 2.937 | -0.003 | -0.001 | -0.001 | 0.308 | 0.11 | 0.1 | 0.07 | 0.11 | 0.09 | 0.05 |
| 7 | 33759644 | rs10486546 | -0.92 | -1.726 | 2.421 | 2.903 | -0.015 | 0.073 | 0.073 | 0 | 0.06 | 0.28 | 0.04 | 0.11 | 0.03 | 0.07 |
| 7 | 33765010 | rs1421341 | -0.716 | -1.73 | 2.248 | 2.925 | -0.03 | 0.027 | 0.027 | 0 | 0.06 | 0.12 | 0.06 | 0.13 | 0.04 | 0.08 |
| 7 | 33766621 | rs3779237 | -2.493 | -1.861 | 2.4 | 3.184 | -0.05 | 0.137 | 0.137 | 0 | 0.24 | 0.48 | 0.32 | 0.48 | 0.21 | 0.17 |
| 7 | 33767012 | rs2232116 | -0.92 | -1.344 | 1.599 | 2.754 | -0.027 | 0.053 | 0.053 | 0 | 0.06 | 0.23 | 0.04 | 0.11 | 0.03 | 0.08 |
| 7 | 33770898 | rs10951391 | -1.432 | -1.199 | 1.243 | 2.895 | 0.003 | 0.106 | 0.106 | 0.255 | 0.06 | 0.1 | 0.28 | 0.31 | 0.1 | 0.19 |
| 7 | 33772241 | rs10278126 | -1.309 | -1.152 | 1.053 | 2.749 | -0.024 | 0.004 | 0.004 | 0.393 | 0.06 | 0.01 | 0.32 | 0.31 | 0.11 | 0.21 |
| 7 | 33774862 | rs17170389 | -1.823 | -1.529 | 1.861 | 2.661 | -0.047 | 0.087 | 0.087 | 0 | 0.17 | 0.38 | 0.15 | 0.3 | 0.12 | 0.15 |
| 7 | 33779883 | rs17811849 | -1.496 | -1.479 | 1.438 | 2.633 | -0.003 | -0.003 | -0.003 | 0.317 | 0.13 | 0.07 | 0.3 | 0.13 | 0.17 | 0.08 |
| 7 | 33787483 | rs10240704 | -1.539 | -1.472 | 1.522 | 2.752 | -0.016 | 0.089 | 0.089 | 0 | 0.15 | 0.23 | 0.17 | 0.46 | 0.11 | 0.28 |
| 7 | 33795876 | rs1421351 | 1.409 | -1.647 | 1.337 | 2.556 | -0.005 | 0.005 | 0.005 | 0 | 0.88 | 0.85 | 0.69 | 0.86 | 0.85 | 0.87 |
| 7 | 33798880 | rs4723328 | 1.298 | -1.92 | 0.263 | 1.756 | -0.097 | 0.161 | 0.161 | 0 | 0.36 | 0.09 | 0.38 | 0.07 | 0.49 | 0.23 |
| 7 | 33799093 | rs12055962 | -0.039 | -1.977 | 0.655 | 1.889 | 0.031 | 0.018 | 0.018 | 0.004 | 0.5 | 0.58 | 0.21 | 0.49 | 0.33 | 0.46 |
| 7 | 33799137 | rs724291 | 0.999 | -1.911 | 0.437 | 1.869 | -0.059 | -0.064 | -0.064 | 0 | 0.38 | 0.14 | 0.46 | 0.37 | 0.51 | 0.41 |
| 7 | 33803450 | rs4723329 | -1.753 | -1.585 | 1.343 | 2.533 | 0.039 | 0.389 | 0.389 | 0.021 | 0.05 | 0.28 | 0.12 | 0.3 | 0.05 | 0.2 |
| 7 | 33804485 | rs16879245 | -0.508 | -1.627 | 0.557 | 2.419 | -0.008 | 0.002 | 0.002 | 0 | 0.12 | 0.14 | 0.07 | 0.17 | 0.08 | 0.1 |
| 7 | 33811292 | rs17170408 | -1.581 | -1.113 | 0.333 | 2.554 | -0.008 | 0.037 | 0.037 | 0 | 0.12 | 0.19 | 0.31 | 0.26 | 0.15 | 0.14 |
| 7 | 33816709 | rs1362927 | 1.419 | -1.04 | -0.03 | 1.742 | -0.012 | 0.103 | 0.103 | 0.02 | 0.86 | 0.79 | 0.6 | 0.55 | 0.83 | 0.7 |
| 7 | 33819987 | rs1156238 | 0.049 | -0.734 | -0.802 | 0.264 | 0.015 | 0.063 | 0.063 | 0.48 | 0.69 | 0.83 | 0.81 | 0.84 | 0.75 | 0.8 |
| 7 | 33821645 | rs10265967 | 0.206 | -0.55 | -1.425 | -0.238 | 0.037 | 0.024 | 0.024 | 0.018 | 0.36 | 0.13 | 0.46 | 0.65 | 0.28 | 0.47 |
| 7 | 33824673 | rs11766089 | 0.699 | -0.608 | -0.592 | -0.449 | -0.031 | 0.022 | 0.022 | 0 | 0.3 | 0.25 | 0.31 | 0.19 | 0.48 | 0.3 |
| 7 | 33826589 | rs12671896 | 0.057 | -1.041 | -0.032 | -0.737 | 0.006 | 0.006 | 0.006 | 2.605 | 0.36 | 0.3 | 0.46 | 0.67 | 0.28 | 0.51 |
| 7 | 33827598 | rs10486619 | 0.552 | -1.08 | 0.407 | -0.74 | 0.019 | 0.038 | 0.038 | 2.885 | 0.33 | 0.23 | 0.21 | 0.13 | 0.26 | 0.19 |
| 7 | 33830585 | rs7795837 | 0.339 | -1.341 | -0.288 | -0.537 | 0.072 | 0.103 | 0.103 | 0.166 | 0.49 | 0.27 | 0.64 | 0.33 | 0.67 | 0.61 |
| 7 | 33845860 | rs759729 | -0.377 | -1.194 | -0.075 | -0.166 | -0.033 | 0.122 | 0.122 | 0.871 | 0.16 | 0.36 | 0.13 | 0.54 | 0.08 | 0.21 |
| 7 | 33846794 | rs7788027 | -0.346 | -1.187 | -0.121 | -0.162 | -0.031 | 0.113 | 0.113 | 1.004 | 0.16 | 0.33 | 0.13 | 0.54 | 0.08 | 0.21 |
| 7 | 33847892 | rs4723335 | 0.57 | -0.885 | -0.634 | -0.134 | -0.03 | 0.013 | 0.013 | 0.076 | 0.18 | 0.01 | 0.33 | 0.14 | 0.18 | 0.28 |
| 7 | 33853740 | rs1345517 | 0.058 | 0.499 | -0.333 | 0.554 | 0.001 | 0.049 | 0.049 | 0 | 0.12 | 0.13 | 0.15 | 0.44 | 0.17 | 0.22 |
| 7 | 33854696 | rs1421339 | -0.008 | -0.136 | -0.837 | -0.32 | 0.029 | 0.046 | 0.046 | 0 | 0.34 | 0.13 | 0.49 | 0.28 | 0.31 | 0.44 |
| 7 | 33857582 | rs17740845 | 0.981 | -0.36 | -0.645 | -0.507 | 0.025 | 0.004 | 0.004 | 0 | 0.19 | 0.02 | 0.23 | 0.15 | 0.13 | 0.21 |
| 7 | 33865741 | rs7785398 | 0.807 | -0.604 | -0.002 | -0.662 | -0.007 | 0.06 | 0.06 | 0.137 | 0.75 | 0.35 | 0.78 | 0.5 | 0.79 | 0.74 |
| 7 | 33869805 | rs10224342 | -1.559 | -0.421 | 0.057 | -0.809 | -0.018 | 0.038 | 0.038 | 0 | 0.59 | 0.75 | 0.71 | 0.79 | 0.53 | 0.71 |
| 7 | 33883098 | rs6948719 | 0.713 | 0.155 | -0.288 | -1.012 | 0.015 | -0.015 | -0.015 | 0.363 | 0.19 | 0.22 | 0.27 | 0.1 | 0.28 | 0.25 |
| 7 | 33883437 | rs4401748 | 1.8 | 0.044 | -0.335 | -1.07 | -0.051 | 0.128 | 0.128 | 0 | 0.29 | 0.05 | 0.2 | 0.11 | 0.31 | 0.11 |
| 7 | 33888495 | rs10243554 | 0.097 | -0.009 | -0.087 | -1.108 | -0.002 | 0.001 | 0.001 | 5.727 | 0.28 | 0.25 | 0.31 | 0.13 | 0.33 | 0.37 |
| 7 | 33897631 | rs888529 | 1.805 | -0.011 | -0.172 | -1.061 | -0.067 | 0.133 | 0.133 | 0 | 0.28 | 0.05 | 0.2 | 0.1 | 0.31 | 0.1 |
| 7 | 33899231 | rs7457120 | -0.248 | -0.112 | 0.317 | -0.893 | 0.011 | 0.033 | 0.033 | 0 | 0.94 | 0.71 | 0.92 | 0.92 | 0.98 | 0.95 |
| 7 | 33900465 | rs1421349 | 0.63 | 0.168 | -0.197 | -0.752 | -0.01 | 0.038 | 0.038 | 0 | 0.44 | 0.29 | 0.37 | 0.61 | 0.52 | 0.33 |
| 7 | 33902583 | rs16879278 | 0.637 | 0.443 | -0.035 | -0.303 | -0.005 | -0.019 | -0.019 | 0 | 0.12 | 0.14 | 0.05 | 0.03 | 0.07 | 0.12 |
| 7 | 33913048 | rs12540329 | 0.197 | 1.071 | 0.032 | 0.037 | 0.02 | 0.001 | 0.001 | 0 | 0.11 | 0.14 | 0.24 | 0.09 | 0.2 | 0.19 |
| 7 | 33913404 | rs3815652 | -0.318 | 1.145 | -0.102 | 1.011 | 0.017 | 0.1 | 0.1 | 0 | 0.1 | 0.16 | 0.15 | 0.42 | 0.18 | 0.22 |
| 7 | 33914560 | rs1362924 | 0.218 | 1.52 | -0.244 | 1.303 | -0.022 | -0.022 | -0.022 | 0 | 0.14 | 0.1 | 0.22 | 0.28 | 0.31 | 0.29 |
| 7 | 33924423 | rs10951401 | -0.268 | 0.396 | 0.041 | 0.887 | -0.01 | 0.009 | 0.009 | 0 | 0.37 | 0.41 | 0.53 | 0.47 | 0.36 | 0.45 |
| 7 | 33926351 | rs315860 | -0.82 | 0.022 | -0.103 | 0.728 | 0.004 | 0.008 | 0.008 | 0 | 0.49 | 0.41 | 0.54 | 0.57 | 0.52 | 0.63 |
| 7 | 33926669 | rs315862 | 0.325 | 0.16 | -0.197 | 0.785 | -0.022 | -0.02 | -0.02 | 0 | 0.2 | 0.18 | 0.25 | 0.31 | 0.35 | 0.36 |

| | | | | | | | | | | | | | | | | |
|---|---|---|---|---|---|---|---|---|---|---|---|---|---|---|---|---|
| 7 | 33930854 | rs7794587 | -0.621 | 0.157 | -0.339 | 0.784 | -0.059 | -0.051 | -0.051 | 0 | 0.81 | 0.89 | 0.76 | 0.69 | 0.65 | 0.64 |
| 7 | 33933854 | rs890602 | 2.11 | -0.1 | -0.62 | 0.63 | 0.002 | 0.023 | 0.023 | 0 | 0.32 | 0.04 | 0.16 | 0.22 | 0.24 | 0.17 |
| 7 | 33938388 | rs7787218 | 1.214 | -0.171 | -0.361 | 0.64 | -0.002 | -0.001 | -0.001 | 0 | 0.2 | 0.16 | 0.24 | 0.2 | 0.15 | 0.23 |
| 7 | 33943818 | rs4723342 | 0.259 | -0.046 | -0.442 | 0.697 | 0.003 | 0 | 0 | 0 | 0.56 | 0.53 | 0.46 | 0.42 | 0.5 | 0.38 |
| 7 | 33949959 | rs7801918 | -0.418 | 0.104 | -0.355 | 0.878 | 0.009 | 0.007 | 0.007 | 0 | 0.2 | 0.3 | 0.28 | 0.21 | 0.25 | 0.2 |
| 7 | 33967085 | rs13221600 | 1.213 | 0.088 | -1.163 | 0.95 | 0.023 | 0.027 | 0.027 | 0 | 0.35 | 0.1 | 0.17 | 0.26 | 0.26 | 0.18 |
| 7 | 33971904 | rs2016770 | -0.528 | 0.536 | -0.969 | 1.189 | 0.004 | 0.011 | 0.011 | 0 | 0.27 | 0.65 | 0.31 | 0.35 | 0.32 | 0.31 |
| 7 | 33973653 | rs10253162 | -0.04 | 0.522 | -0.656 | 1.423 | 0.05 | -0.004 | -0.004 | 0.031 | 0.24 | 0.49 | 0.26 | 0.23 | 0.31 | 0.3 |
| 7 | 33977757 | rs10247996 | -1.054 | 0.361 | -1.265 | 1.216 | -0.01 | -0.022 | -0.022 | 0 | 0.46 | 0.75 | 0.5 | 0.48 | 0.53 | 0.63 |
| 7 | 33984313 | rs1427481 | 0.234 | 0.734 | -0.671 | 1.208 | -0.002 | -0.004 | -0.004 | 0.041 | 0.54 | 0.53 | 0.48 | 0.63 | 0.54 | 0.47 |
| 7 | 33996948 | rs12155213 | 0.453 | 0.578 | -0.795 | 1.177 | 0.003 | -0.006 | -0.006 | 0 | 0.39 | 0.62 | 0.3 | 0.43 | 0.31 | 0.21 |
| 7 | 33998587 | rs4723343 | -0.411 | 0.614 | -0.727 | 0.85 | 0 | -0.002 | -0.002 | 0 | 0.41 | 0.62 | 0.61 | 0.53 | 0.38 | 0.54 |
| 7 | 33999889 | rs12672150 | 0.231 | 0.854 | -0.611 | 1.11 | -0.012 | 0.01 | 0.01 | 0 | 0.06 | 0.04 | 0.26 | 0.24 | 0.12 | 0.16 |
| 7 | 34012829 | rs1365888 | 0.079 | 0.839 | -0.479 | 1.104 | 0.001 | 0.035 | 0.035 | 0 | 0.07 | 0.06 | 0.26 | 0.24 | 0.12 | 0.18 |
| 7 | 34013178 | rs13243958 | 0.792 | 0.695 | -0.351 | 0.724 | 0.004 | 0.026 | 0.026 | 0 | 0.65 | 0.54 | 0.54 | 0.61 | 0.64 | 0.64 |
| 7 | 34017577 | rs10276432 | 0.742 | 0.67 | -0.106 | 0.659 | -0.006 | 0.004 | 0.004 | 0 | 0.63 | 0.54 | 0.54 | 0.62 | 0.63 | 0.61 |
| 7 | 34021755 | rs12701341 | 0.078 | 0.399 | -0.366 | 0.308 | 0.011 | 0.01 | 0.01 | 0 | 0.49 | 0.59 | 0.49 | 0.53 | 0.42 | 0.35 |
| 7 | 34035784 | rs6948650 | 1.335 | 0.247 | -0.557 | 0.255 | 0.049 | 0.022 | 0.022 | 0 | 0.18 | 0.12 | 0.07 | 0.09 | 0.09 | 0.08 |
| 7 | 34036618 | rs9986855 | -1.731 | 0.306 | -0.609 | 0.179 | -0.038 | 0.025 | 0.025 | 0 | 0.88 | 1 | 0.9 | 0.93 | 0.87 | 0.83 |
| 7 | 34038414 | rs17169617 | 0.152 | 0.05 | -0.347 | 0.328 | 0.014 | -0.015 | -0.015 | 0 | 0.25 | 0.24 | 0.32 | 0.31 | 0.2 | 0.24 |
| 7 | 34043348 | rs10951405 | 2.139 | 0.075 | -0.437 | 0.328 | 0.007 | -0.009 | -0.009 | 0.239 | 0.22 | 0.11 | 0.12 | 0.24 | 0.17 | 0.11 |
| 7 | 34046002 | rs974937 | 0.015 | 0.109 | -0.263 | 0.274 | -0.025 | 0.046 | 0.046 | 0 | 0.27 | 0.47 | 0.35 | 0.31 | 0.24 | 0.26 |
| 7 | 34047534 | rs10276214 | 0.274 | -0.221 | -0.402 | 0.31 | 0.033 | -0.04 | -0.04 | 0 | 0.24 | 0.17 | 0.31 | 0.31 | 0.19 | 0.25 |
| 7 | 34074661 | rs12670628 | 0.194 | -0.539 | -0.521 | 0.113 | 0.021 | 0.02 | 0.02 | 0.007 | 0.45 | 0.59 | 0.25 | 0.32 | 0.32 | 0.2 |
| 7 | 34087546 | rs17169630 | 0.793 | -0.491 | -0.326 | 0.067 | -0.077 | -0.048 | -0.048 | 0 | 0.2 | 0.41 | 0.09 | 0.09 | 0.1 | 0.08 |
| 7 | 34088690 | rs17823423 | 1.994 | -0.455 | -1.166 | -0.036 | -0.079 | -0.08 | -0.08 | 0.874 | 0.16 | 0.01 | 0.26 | 0.23 | 0.22 | 0.15 |
| 7 | 34089823 | rs7806522 | -0.075 | -1.349 | -0.598 | -0.56 | 0.017 | 0.017 | 0.017 | 0 | 0.44 | 0.6 | 0.24 | 0.32 | 0.32 | 0.2 |
| 7 | 34094687 | rs17169636 | 0.95 | -1.458 | -0.756 | -0.581 | -0.013 | -0.002 | -0.002 | 0.087 | 0.18 | 0.2 | 0.07 | 0.1 | 0.09 | 0.08 |
| 7 | 34103553 | rs1420328 | -0.667 | -0.763 | -1.184 | -0.997 | -0.016 | -0.064 | -0.064 | 0.214 | 0.88 | 0.77 | 0.92 | 1 | 0.91 | 0.93 |
| 7 | 34107393 | rs1362457 | 0.737 | 0.068 | -1.394 | -0.51 | -0.01 | 0.009 | 0.009 | 0 | 0.23 | 0.11 | 0.16 | 0.22 | 0.22 | 0.11 |
| 7 | 34111660 | rs961652 | 0 | 0.92 | -0.756 | 0.043 | -0.036 | 0.006 | 0.006 | 0 | 0.65 | 0.74 | 0.38 | 0.36 | 0.52 | 0.49 |
| 7 | 34121829 | rs2160308 | -0.487 | 1.117 | -0.464 | -0.229 | -0.015 | 0.085 | 0.085 | 0 | 0.3 | 0.2 | 0.21 | 0.08 | 0.32 | 0.18 |
| 7 | 34126782 | rs7790026 | -0.756 | 1.01 | 0.407 | -0.243 | -0.002 | -0.065 | -0.065 | 0 | 0.33 | 0.54 | 0.25 | 0.14 | 0.33 | 0.19 |
| 7 | 34132131 | rs1420331 | -0.154 | 1.109 | 1.072 | -0.225 | 0.02 | -0.06 | -0.06 | 0 | 0.22 | 0.32 | 0.17 | 0.07 | 0.26 | 0.11 |
| 7 | 34132727 | rs1420332 | 0.353 | 1.03 | 1.524 | -0.26 | 0.013 | -0.065 | -0.065 | 0 | 0.76 | 0.41 | 0.81 | 0.92 | 0.73 | 0.89 |
| 7 | 34133050 | rs973766 | 0.691 | 1.485 | 0.997 | -0.307 | -0.099 | 0.016 | 0.016 | 0 | 0.17 | 0.01 | 0.42 | 0.11 | 0.3 | 0.14 |
| 7 | 34137529 | rs6948894 | -0.805 | 0.967 | 1.901 | 0.234 | 0.023 | -0.084 | -0.084 | 0.353 | 0.24 | 0.37 | 0.17 | 0.08 | 0.27 | 0.11 |
| 7 | 34138717 | rs1946146 | -0.219 | 1.038 | 1.323 | 0.251 | -0.018 | 0.047 | 0.047 | 0.869 | 0.2 | 0.11 | 0.14 | 0.08 | 0.24 | 0.11 |
| 7 | 34140513 | rs1420335 | 1.139 | 0.906 | 1.161 | 0.886 | 0 | -0.025 | -0.025 | 0 | 0.68 | 0.31 | 0.74 | 0.54 | 0.69 | 0.68 |
| 7 | 34147964 | rs1362460 | 1.261 | 0.878 | 1.043 | 0.484 | -0.002 | 0.011 | 0.011 | 0.001 | 0.6 | 0.36 | 0.64 | 0.79 | 0.57 | 0.68 |
| 7 | 34151057 | rs757640 | 0.123 | 0.873 | 1.227 | 0.434 | 0.016 | -0.07 | -0.07 | 0 | 0.79 | 0.59 | 0.83 | 0.94 | 0.79 | 0.9 |
| 7 | 34153309 | rs3886609 | -0.178 | 0.91 | 1.265 | 0.456 | -0.015 | 0.011 | 0.011 | 0 | 0.16 | 0.07 | 0.17 | 0.12 | 0.22 | 0.12 |
| 7 | 34153768 | rs918038 | 1.19 | 0.903 | 1.272 | 0.462 | 0.011 | -0.043 | -0.043 | 0 | 0.63 | 0.5 | 0.66 | 0.83 | 0.58 | 0.78 |
| 7 | 34167361 | rs10486631 | -0.104 | 0.85 | 1.25 | 0.479 | -0.016 | 0.01 | 0.01 | 0 | 0.16 | 0.07 | 0.17 | 0.11 | 0.21 | 0.13 |
| 7 | 34168028 | rs6462533 | 1.494 | 0.849 | 1.304 | 0.466 | -0.002 | 0.005 | 0.005 | 0.02 | 0.6 | 0.38 | 0.64 | 0.79 | 0.57 | 0.68 |
| 7 | 34169964 | rs1861362 | -0.022 | 0.882 | 1.29 | 0.475 | 0.006 | -0.036 | -0.036 | 0.213 | 0.8 | 0.74 | 0.85 | 0.94 | 0.8 | 0.9 |
| 7 | 34173852 | rs10951406 | 0.288 | 0.887 | 1.316 | 0.49 | 0.005 | -0.062 | -0.062 | 0 | 0.77 | 0.45 | 0.81 | 0.9 | 0.78 | 0.86 |
| 7 | 34174173 | rs6961966 | -0.376 | 0.894 | 1.017 | 0.478 | 0 | 0 | 0 | 2.061 | 0.17 | 0.18 | 0.13 | 0.11 | 0.21 | 0.13 |
| 7 | 34178645 | rs10486632 | 0.181 | 1.033 | 0.527 | 0.363 | 0.007 | -0.051 | -0.051 | 0.531 | 0.77 | 0.63 | 0.83 | 0.9 | 0.79 | 0.86 |
| 7 | 34192011 | rs918037 | 0.406 | 0.864 | 0.051 | 0.652 | 0 | 0 | 0 | 0.001 | 0.6 | 0.56 | 0.7 | 0.66 | 0.55 | 0.75 |
| 7 | 34193589 | rs11972404 | -1.017 | 0.482 | 0.77 | 0.489 | 0.035 | 0.008 | 0.008 | 0.023 | 0.06 | 0.29 | 0.04 | 0 | 0.03 | 0.02 |
| 7 | 34193952 | rs17169667 | -1.716 | 0.082 | 1.157 | 0.097 | -0.032 | -0.091 | -0.091 | 0 | 0.17 | 0.58 | 0.23 | 0.02 | 0.1 | 0.05 |
| 7 | 34194102 | rs2058681 | -0.042 | 0.317 | -0.048 | 0.341 | -0.09 | -0.071 | -0.071 | 0.065 | 0.3 | 0.02 | 0.29 | 0.31 | 0.34 | 0.47 |
| 7 | 34197242 | rs2392263 | -0.069 | 0.241 | 0.837 | 0.374 | 0.051 | 0.017 | 0.017 | 0 | 0.06 | 0.28 | 0.17 | 0.02 | 0.03 | 0.05 |
| 7 | 34198270 | rs13223298 | -1.225 | 0.218 | 1.001 | 0.356 | -0.003 | -0.06 | -0.06 | 0 | 0.11 | 0.36 | 0.21 | 0.02 | 0.08 | 0.03 |
| 7 | 34200018 | rs11486838 | -0.578 | 0.663 | 0.542 | 0.794 | 0.004 | 0.009 | 0.009 | 0 | 0.1 | 0.14 | 0.19 | 0.17 | 0.11 | 0.12 |
| 7 | 34208711 | rs11768179 | -0.268 | 0.621 | 0.376 | 1.099 | -0.001 | 0.003 | 0.003 | 0 | 0.09 | 0.06 | 0.09 | 0.25 | 0.09 | 0.16 |
| 7 | 34221989 | rs7341527 | -0.042 | 0.597 | 0.464 | 0.322 | 0.01 | 0.129 | 0.129 | 0 | 0.57 | 0.73 | 0.39 | 0.14 | 0.55 | 0.42 |
| 7 | 34230370 | rs11768361 | 0.403 | 0.507 | 0.648 | 0.327 | 0.026 | 0.207 | 0.207 | 0.43 | 0.51 | 0.17 | 0.38 | 0.17 | 0.5 | 0.4 |
| 7 | 34239585 | rs13228877 | -1.453 | 0.583 | 1.163 | 0.335 | 0.012 | -0.027 | -0.027 | 0.386 | 0.08 | 0.57 | 0.09 | 0 | 0.06 | 0.01 |
| 7 | 34251359 | rs318628 | -1.82 | 0.573 | 0.902 | 0.331 | -0.018 | -0.053 | -0.053 | 0 | 0.11 | 0.69 | 0.11 | 0 | 0.1 | 0.01 |
| 7 | 34253747 | rs318623 | 0.41 | 0.708 | 0.725 | 0.674 | 0.026 | 0.125 | 0.125 | 0.29 | 0.51 | 0.21 | 0.41 | 0.17 | 0.52 | 0.4 |
| 7 | 34276627 | rs318571 | 0.073 | 0.868 | 0.41 | 1.193 | 0.001 | -0.001 | -0.001 | 0 | 0.62 | 0.55 | 0.56 | 0.64 | 0.63 | 0.5 |
| 7 | 34276765 | rs318572 | 0.121 | 1.015 | -0.972 | 1.213 | 0.016 | -0.028 | -0.028 | 0 | 0.6 | 0.21 | 0.52 | 0.6 | 0.62 | 0.49 |
| 7 | 34281928 | rs4720156 | -1.136 | 1.868 | -0.738 | 1.686 | -0.011 | -0.011 | -0.011 | 0 | 0.14 | 0.12 | 0.27 | 0.26 | 0.21 | 0.28 |
| 7 | 34283049 | rs17169692 | -1.416 | 1.863 | -0.783 | 1.979 | -0.019 | -0.037 | -0.037 | 0 | 0.17 | 0.11 | 0.24 | 0.29 | 0.21 | 0.19 |
| 7 | 34285221 | rs318576 | 1.779 | 1.185 | -0.386 | 2.145 | 0.003 | 0.007 | 0.007 | 1.612 | 0.63 | 0.29 | 0.5 | 0.33 | 0.68 | 0.49 |
| 7 | 34287258 | rs2108933 | -0.629 | 1.011 | 0.192 | 2.276 | -0.002 | 0.008 | 0.008 | 0.032 | 0.15 | 0.11 | 0.2 | 0.34 | 0.1 | 0.32 |
| 7 | 34293293 | rs2392265 | -1.06 | 1.471 | 0.15 | 1.947 | -0.052 | -0.048 | -0.048 | 0 | 0.21 | 0.1 | 0.3 | 0.27 | 0.28 | 0.2 |

| | | | | | | | | | | | | | | | | |
|---|---|---|---|---|---|---|---|---|---|---|---|---|---|---|---|---|
| 7 | 34293822 | rs17755229 | -1.686 | 1.368 | 0.65 | 2.028 | 0.014 | 0.023 | 0.023 | 0 | 0.33 | 0.39 | 0.44 | 0.44 | 0.38 | 0.27 |
| 7 | 34294321 | rs318583 | 1.286 | 0.659 | 0.947 | 2.028 | -0.029 | 0.053 | 0.053 | 0.654 | 0.86 | 0.6 | 0.89 | 0.71 | 0.91 | 0.71 |
| 7 | 34301101 | rs17169701 | -0.461 | 0.665 | 0.946 | 2.102 | -0.011 | 0.029 | 0.029 | 0 | 0.09 | 0.13 | 0.08 | 0.23 | 0.08 | 0.3 |
| 7 | 34326342 | rs995415 | -2.229 | 0.576 | 1.63 | 2.043 | -0.001 | 0.045 | 0.045 | 2.824 | 0.45 | 0.76 | 0.56 | 0.7 | 0.43 | 0.57 |
| 7 | 34330309 | rs12701349 | -1.29 | 0.544 | 1.714 | 2.031 | 0.025 | 0.037 | 0.037 | 0.73 | 0.3 | 0.33 | 0.46 | 0.41 | 0.35 | 0.26 |
| 7 | 34336057 | rs10282356 | -1.38 | 0.432 | 1.762 | 1.996 | -0.016 | 0.017 | 0.017 | 0.52 | 0.13 | 0.16 | 0.1 | 0.28 | 0.09 | 0.31 |
| 7 | 34338598 | rs13223800 | -2.151 | 0.699 | 1.695 | 1.907 | -0.006 | 0.038 | 0.038 | 3.042 | 0.46 | 0.68 | 0.56 | 0.7 | 0.47 | 0.58 |
| 7 | 34347206 | rs6946269 | -2.03 | 0.389 | 1.59 | 1.811 | 0 | -0.017 | -0.017 | 0.239 | 0.45 | 0.47 | 0.56 | 0.7 | 0.47 | 0.59 |
| 7 | 34350994 | rs6967834 | -0.065 | 0.25 | 1.029 | 1.516 | -0.013 | 0.043 | 0.043 | 0 | 0.07 | 0.11 | 0.09 | 0.22 | 0.04 | 0.26 |
| 7 | 34354465 | rs6462540 | -1.598 | -0.05 | 0.728 | 1.227 | 0.002 | 0.051 | 0.051 | 0 | 0.43 | 0.74 | 0.47 | 0.33 | 0.39 | 0.32 |
| 7 | 34371441 | rs17169743 | -0.804 | -0.076 | 0.99 | 1.306 | 0.003 | 0.022 | 0.022 | 0 | 0.15 | 0.29 | 0.17 | 0.18 | 0.16 | 0.12 |
| 7 | 34372314 | rs1419811 | -0.153 | -0.683 | 0.227 | 0.458 | -0.008 | 0.173 | 0.173 | 0 | 0.66 | 0.9 | 0.72 | 0.97 | 0.7 | 0.69 |
| 7 | 34374063 | rs10486639 | -0.345 | -0.876 | 1.265 | 1.461 | -0.018 | 0.225 | 0.225 | 0.096 | 0.09 | 0.31 | 0.09 | 0.31 | 0.04 | 0.26 |
| 7 | 34382289 | rs1419804 | -0.948 | -0.941 | 1.22 | 1.476 | -0.065 | 0.179 | 0.179 | 0 | 0.14 | 0.32 | 0.11 | 0.39 | 0.05 | 0.26 |
| 7 | 34388338 | rs6977125 | -2.386 | -0.863 | 0.649 | 1.328 | 0.027 | -0.044 | -0.044 | 0 | 0.33 | 0.81 | 0.33 | 0.57 | 0.26 | 0.4 |
| 7 | 34399494 | rs1419810 | -2.236 | -0.846 | 0.707 | 1.502 | 0.041 | -0.056 | -0.056 | 0 | 0.31 | 0.82 | 0.28 | 0.57 | 0.2 | 0.4 |
| 7 | 34401500 | rs17169769 | 0.412 | -0.629 | 0.758 | 1.362 | -0.02 | 0.006 | 0.006 | 0 | 0.06 | 0.07 | 0.03 | 0.08 | 0.02 | 0.02 |
| 7 | 34405042 | rs17169771 | -0.974 | -0.58 | 0.648 | 1.359 | -0.034 | 0.155 | 0.155 | 0 | 0.16 | 0.44 | 0.11 | 0.36 | 0.09 | 0.28 |
| 7 | 34412689 | rs4723357 | -1.116 | -0.755 | 0.29 | 1.37 | -0.043 | 0.035 | 0.035 | 0.043 | 0.26 | 0.48 | 0.21 | 0.44 | 0.14 | 0.42 |
| 7 | 34426320 | rs6975562 | 0.484 | -0.588 | 0.228 | 1.051 | -0.014 | 0.023 | 0.023 | 0 | 0.15 | 0.28 | 0.1 | 0.15 | 0.12 | 0.13 |
| 7 | 34428181 | rs7807919 | -1.299 | -0.966 | -0.433 | 0.815 | 0.059 | 0.016 | 0.016 | 0.06 | 0.46 | 0.87 | 0.37 | 0.62 | 0.32 | 0.58 |
| 7 | 34429228 | rs12537903 | -1.116 | -0.79 | 0.445 | 0.903 | -0.022 | 0.12 | 0.12 | 0 | 0.25 | 0.5 | 0.23 | 0.52 | 0.19 | 0.44 |
| 7 | 34435095 | rs7807040 | 1.784 | -0.467 | -0.716 | 0.393 | -0.09 | -0.117 | -0.117 | 0.33 | 0.35 | 0.06 | 0.36 | 0.44 | 0.42 | 0.26 |
| 7 | 34445471 | rs10225713 | -0.261 | -0.531 | -0.609 | 0.377 | -0.011 | 0.031 | 0.031 | 0.39 | 0.75 | 0.86 | 0.69 | 0.85 | 0.72 | 0.8 |
| 7 | 34448659 | rs1419840 | 0.502 | -0.815 | -0.027 | 0.612 | -0.019 | 0.028 | 0.028 | 0.886 | 0.67 | 0.43 | 0.63 | 0.57 | 0.68 | 0.65 |
| 7 | 34450880 | rs6947789 | 0.319 | -0.719 | 0.134 | 0.479 | -0.002 | -0.002 | -0.002 | 0 | 0.26 | 0.21 | 0.2 | 0.2 | 0.19 | 0.31 |
| 7 | 34451217 | rs736295 | -0.93 | -0.821 | -0.196 | 0.39 | -0.016 | 0.049 | 0.049 | 0.097 | 0.49 | 0.77 | 0.51 | 0.34 | 0.48 | 0.49 |
| 7 | 34451256 | rs4141226 | 0.343 | -0.775 | -0.083 | 0.435 | -0.002 | 0.003 | 0.003 | 0.04 | 0.2 | 0.22 | 0.27 | 0.15 | 0.24 | 0.17 |
| 7 | 34463295 | rs35114106 | -0.164 | -0.898 | -0.058 | 0.4 | 0.002 | -0.005 | -0.005 | 0 | 0.22 | 0.41 | 0.3 | 0.16 | 0.25 | 0.16 |
| 7 | 34463595 | rs2041176 | 0.077 | -0.966 | -0.044 | 0.528 | -0.007 | 0.019 | 0.019 | 0 | 0.87 | 0.85 | 0.9 | 0.67 | 0.91 | 0.82 |
| 7 | 34467317 | rs7790986 | 0.479 | -0.697 | -0.235 | 0.437 | -0.001 | 0.001 | 0.001 | 0 | 0.15 | 0.2 | 0.15 | 0.16 | 0.16 | 0.09 |
| 7 | 34477795 | rs11764303 | 1.684 | -0.649 | -0.282 | 0.412 | -0.027 | 0.034 | 0.034 | 0 | 0.06 | 0.01 | 0.13 | 0 | 0.08 | 0.08 |
| 7 | 34478120 | rs10227970 | 1.069 | -0.745 | -0.293 | 0.395 | 0.013 | -0.003 | -0.003 | 0 | 0.53 | 0.43 | 0.57 | 0.48 | 0.6 | 0.45 |
| 7 | 34481592 | rs9986800 | 0.044 | -0.209 | 0.154 | 0.474 | 0.016 | 0.039 | 0.039 | 1.289 | 0.16 | 0.3 | 0.15 | 0.2 | 0.12 | 0.21 |
| 7 | 34497585 | rs10250876 | -0.73 | 0.11 | -0.429 | 0.339 | -0.001 | -0.002 | -0.002 | 0 | 0.4 | 0.62 | 0.46 | 0.41 | 0.41 | 0.41 |
| 7 | 34507636 | rs1120768 | -0.753 | 0.098 | -0.465 | 0.35 | -0.002 | -0.005 | -0.005 | 0.17 | 0.41 | 0.66 | 0.47 | 0.42 | 0.41 | 0.41 |
| 7 | 34517895 | rs7781752 | 0.212 | 0.029 | -0.87 | 0.448 | -0.001 | -0.007 | -0.007 | 0 | 0.91 | 0.95 | 0.92 | 0.8 | 0.89 | 0.85 |
| 7 | 34519193 | rs17764093 | 0.8 | 0.029 | -1.023 | 0.342 | -0.001 | 0.005 | 0.005 | 0 | 0.07 | 0.04 | 0.08 | 0.07 | 0.1 | 0.05 |
| 7 | 34519793 | rs7792119 | 1.243 | 0.07 | -1.118 | 0.353 | 0.007 | 0.071 | 0.071 | 6.994 | 0.5 | 0.22 | 0.46 | 0.33 | 0.49 | 0.43 |
| 7 | 34520073 | rs7792581 | 1.168 | -0.325 | -1.031 | 0.143 | 0 | -0.021 | -0.021 | 3.533 | 0.17 | 0.11 | 0.06 | 0 | 0.07 | 0.14 |
| 7 | 34522166 | rs6462559 | -0.268 | -0.026 | -0.991 | 0.471 | 0.004 | -0.036 | -0.036 | 0 | 0.94 | 0.99 | 0.94 | 0.82 | 0.93 | 0.84 |
| 7 | 34523186 | rs1468559 | 1.144 | 0.002 | -0.167 | 0.728 | 0.019 | 0.222 | 0.222 | 0 | 0.88 | 0.68 | 0.89 | 0.68 | 0.88 | 0.83 |
| 7 | 34531750 | rs12333710 | 0.389 | 0.62 | -0.645 | 0.323 | 0.003 | -0.009 | -0.009 | 0.411 | 0.42 | 0.46 | 0.41 | 0.28 | 0.44 | 0.42 |
| 7 | 34532710 | rs4723373 | -0.232 | 0.538 | -0.848 | 0.21 | -0.018 | 0.044 | 0.044 | 0 | 0.92 | 0.84 | 0.93 | 0.82 | 0.93 | 0.87 |
| 7 | 34537408 | rs4723374 | 0.093 | 1.032 | -0.927 | 0.186 | -0.009 | -0.006 | -0.006 | 0.629 | 0.47 | 0.7 | 0.45 | 0.49 | 0.46 | 0.45 |
| 7 | 34539317 | rs10951418 | 0.723 | 0.505 | -1.445 | 0.21 | -0.064 | -0.033 | -0.033 | 0 | 0.4 | 0.06 | 0.44 | 0.32 | 0.42 | 0.41 |
| 7 | 34540082 | rs6971556 | 0.009 | 0.262 | -1.081 | 0.226 | -0.022 | 0.072 | 0.072 | 0 | 0.27 | 0.42 | 0.33 | 0.46 | 0.17 | 0.37 |
| 7 | 34540117 | rs6462561 | -1.605 | 0.797 | -1.724 | -0.215 | -0.101 | 0.029 | 0.029 | 0.043 | 0.78 | 0.98 | 0.83 | 0.81 | 0.69 | 0.76 |
| 7 | 34540560 | rs2041301 | 0.959 | 0.443 | -1.49 | -0.251 | 0.001 | -0.012 | -0.012 | 0 | 0.43 | 0.46 | 0.42 | 0.28 | 0.45 | 0.43 |
| 7 | 34541474 | rs1419865 | -0.311 | 0.114 | -1.098 | 0.01 | -0.023 | 0.037 | 0.037 | 0 | 0.23 | 0.31 | 0.33 | 0.51 | 0.11 | 0.31 |
| 7 | 34543087 | rs7802435 | -0.515 | 0.206 | -1.136 | -0.027 | -0.003 | 0.008 | 0.008 | 1.987 | 0.27 | 0.3 | 0.37 | 0.52 | 0.21 | 0.31 |
| 7 | 34543108 | rs10271519 | 0.259 | 0.11 | -0.91 | -0.203 | -0.003 | -0.055 | -0.055 | 0 | 0.1 | 0.17 | 0.07 | 0 | 0.11 | 0.03 |
| 7 | 34547662 | rs6947764 | -1.176 | 0.178 | -0.726 | -0.132 | 0.018 | -0.005 | -0.005 | 0 | 0.15 | 0.31 | 0.12 | 0.14 | 0.14 | 0.15 |
| 7 | 34561147 | rs10272610 | -1.558 | -0.358 | -0.861 | 0.115 | 0.01 | -0.011 | -0.011 | 0.006 | 0.35 | 0.7 | 0.39 | 0.45 | 0.24 | 0.33 |
| 7 | 34583827 | rs4723377 | -0.346 | -0.255 | -1.193 | -0.379 | -0.003 | 0.003 | 0.003 | 0 | 0.62 | 0.67 | 0.69 | 0.8 | 0.57 | 0.77 |
| 7 | 34607064 | rs10274492 | -0.17 | -0.507 | -1.029 | -0.468 | -0.055 | -0.116 | -0.116 | 0 | 0.21 | 0.31 | 0.09 | 0 | 0.11 | 0.03 |
| 7 | 34607206 | rs2108735 | -1.473 | -0.52 | -0.949 | -0.326 | -0.029 | -0.06 | -0.06 | 0 | 0.29 | 0.58 | 0.16 | 0.06 | 0.19 | 0.07 |
| 7 | 34607981 | rs1034876 | 0.18 | -0.19 | -0.92 | -0.049 | 0.025 | 0.022 | 0.022 | 2.342 | 0.08 | 0.27 | 0.07 | 0.06 | 0.08 | 0.04 |
| 7 | 34638272 | rs7792945 | 2.255 | -0.373 | -1.588 | -0.452 | -0.025 | 0.112 | 0.112 | 0 | 0.27 | 0.04 | 0.26 | 0.12 | 0.27 | 0.29 |
| 7 | 34638819 | rs1981606 | -1.1 | -0.477 | -1.325 | -0.195 | -0.015 | 0.101 | 0.101 | 0.551 | 0.21 | 0.59 | 0.33 | 0.45 | 0.16 | 0.31 |
| 7 | 34646059 | rs4452707 | -0.188 | -0.51 | -1.237 | -0.238 | -0.025 | -0.042 | -0.042 | 0 | 0.16 | 0.52 | 0.11 | 0.06 | 0.1 | 0.05 |
| 7 | 34652345 | rs1362166 | -2.143 | -0.511 | -1.534 | -0.44 | -0.136 | 0.029 | 0.029 | 0.027 | 0.76 | 0.97 | 0.76 | 0.81 | 0.64 | 0.8 |
| 7 | 34653161 | rs1894768 | -1.356 | -0.652 | -1.322 | -0.164 | -0.028 | 0.033 | 0.033 | 1.609 | 0.3 | 0.57 | 0.35 | 0.47 | 0.2 | 0.32 |
| 7 | 34660782 | rs10241719 | -1.252 | -0.592 | -1.165 | -0.2 | -0.02 | 0.111 | 0.111 | 0.415 | 0.24 | 0.46 | 0.32 | 0.46 | 0.18 | 0.31 |
| 7 | 34661743 | rs10951423 | 0.503 | -0.668 | -1.361 | -0.287 | 0.041 | -0.012 | -0.012 | 0.504 | 0.46 | 0.19 | 0.52 | 0.51 | 0.39 | 0.57 |
| 7 | 34661834 | rs12534368 | -0.067 | -0.671 | -1.519 | -0.283 | 0.011 | -0.13 | -0.13 | 0 | 0.2 | 0.05 | 0.27 | 0.4 | 0.15 | 0.29 |
| 7 | 34662712 | rs6960331 | -0.683 | -0.541 | -1.366 | -0.249 | 0.012 | -0.002 | -0.002 | 0.523 | 0.61 | 0.57 | 0.63 | 0.59 | 0.51 | 0.62 |
| 7 | 34662826 | rs12669756 | 0.759 | -0.491 | -1.367 | -0.157 | -0.015 | 0.061 | 0.061 | 0 | 0.06 | 0.01 | 0.21 | 0.4 | 0.06 | 0.27 |
| 7 | 34666534 | rs2067575 | 0.418 | -0.532 | -1.159 | -0.274 | 0.009 | 0.027 | 0.027 | 0.458 | 0.09 | 0.17 | 0.08 | 0.14 | 0.11 | 0.16 |

| | | | | | | | | | | | | | | | | |
|---|---|---|---|---|---|---|---|---|---|---|---|---|---|---|---|---|
| 7 | 34667275 | rs6974123 | -0.206 | -0.505 | -1.334 | -0.318 | 0 | -0.013 | -0.013 | 0 | 0.57 | 0.55 | 0.44 | 0.27 | 0.49 | 0.47 |
| 7 | 34667565 | rs17169924 | -0.962 | -0.635 | -1.264 | -0.323 | -0.051 | -0.062 | -0.062 | 0.149 | 0.31 | 0.51 | 0.17 | 0.14 | 0.22 | 0.19 |
| 7 | 34682881 | rs1819802 | -2.065 | -0.351 | -1.404 | -0.35 | -0.109 | 0.097 | 0.097 | 0.002 | 0.75 | 0.95 | 0.79 | 0.85 | 0.64 | 0.8 |
| 7 | 34684184 | rs2609240 | -0.999 | -0.774 | -1.028 | -0.22 | -0.008 | 0.004 | 0.004 | 1.269 | 0.63 | 0.68 | 0.67 | 0.64 | 0.53 | 0.63 |
| 7 | 34688425 | rs2609234 | -0.896 | -1.029 | -1.098 | -0.38 | -0.058 | -0.059 | -0.059 | 0 | 0.28 | 0.5 | 0.16 | 0.14 | 0.18 | 0.19 |
| 7 | 34693226 | rs714588 | -0.24 | -0.843 | -1.124 | -0.336 | 0 | -0.002 | -0.002 | 0 | 0.54 | 0.55 | 0.41 | 0.31 | 0.45 | 0.48 |
| 7 | 34695959 | rs2530543 | -1.342 | -0.89 | -0.855 | -0.179 | -0.035 | -0.047 | -0.047 | 0 | 0.26 | 0.48 | 0.18 | 0.17 | 0.21 | 0.19 |
| 7 | 34696545 | rs1023556 | -1.82 | -0.865 | -0.882 | -0.006 | -0.023 | 0.044 | 0.044 | 0 | 0.31 | 0.5 | 0.38 | 0.49 | 0.24 | 0.33 |
| 7 | 34697060 | rs2530544 | -1.046 | -0.595 | -0.776 | -0.608 | 0.003 | 0.013 | 0.013 | 0.102 | 0.2 | 0.43 | 0.16 | 0.2 | 0.22 | 0.16 |
| 7 | 34697224 | rs2530546 | -1.266 | -0.759 | -0.653 | -0.938 | -0.035 | -0.122 | -0.122 | 0 | 0.24 | 0.43 | 0.13 | 0.02 | 0.17 | 0.07 |
| 7 | 34697428 | rs2168891 | 0.268 | -0.688 | -0.411 | -0.872 | 0.011 | -0.018 | -0.018 | 0 | 0.07 | 0.16 | 0.05 | 0.01 | 0.08 | 0.03 |
| 7 | 34697460 | rs1963499 | -0.62 | -0.736 | -0.534 | -0.879 | -0.036 | -0.088 | -0.088 | 3.305 | 0.17 | 0.37 | 0.07 | 0.02 | 0.11 | 0.05 |
| 7 | 34697922 | rs2530547 | -0.111 | -0.692 | -1.124 | -0.965 | 0.044 | 0.121 | 0.121 | 0 | 0.57 | 0.77 | 0.42 | 0.19 | 0.41 | 0.5 |
| 7 | 34697982 | rs887020 | 0.324 | -0.977 | -1.519 | -1.243 | -0.025 | 0.133 | 0.133 | 2.033 | 0.4 | 0.11 | 0.57 | 0.8 | 0.53 | 0.58 |
| 7 | 34698210 | rs2530548 | -0.084 | -0.725 | -0.687 | -1.256 | -0.005 | -0.038 | -0.038 | 0 | 0.54 | 0.48 | 0.39 | 0.11 | 0.4 | 0.48 |
| 7 | 34698321 | rs2530551 | -0.774 | -0.702 | -0.691 | -1.251 | -0.081 | 0.1 | 0.1 | 2.658 | 0.63 | 0.88 | 0.47 | 0.21 | 0.5 | 0.58 |
| 7 | 34698376 | rs2530552 | 0.556 | -0.895 | -0.739 | -1.3 | -0.002 | -0.026 | -0.026 | 0 | 0.46 | 0.45 | 0.37 | 0.11 | 0.37 | 0.49 |
| 7 | 34699259 | rs13235228 | 0.999 | -1.166 | -0.933 | -1.44 | 0.05 | 0.087 | 0.087 | 0.123 | 0.37 | 0.23 | 0.26 | 0.06 | 0.28 | 0.37 |
| 7 | 34700389 | rs11761197 | 1.366 | -1.016 | -0.471 | -1.316 | 0.116 | 0.063 | 0.063 | 0 | 0.08 | 0.36 | 0.08 | 0.11 | 0.11 | 0.08 |
| 7 | 34701708 | rs411323 | -1.892 | -0.334 | -1.075 | -0.919 | -0.094 | -0.042 | -0.042 | 0 | 0.77 | 0.98 | 0.76 | 0.76 | 0.72 | 0.84 |
| 7 | 34702456 | rs2530561 | 0.351 | -0.657 | -0.642 | -0.851 | -0.001 | -0.003 | -0.003 | 0 | 0.13 | 0.16 | 0.09 | 0.05 | 0.1 | 0.12 |
| 7 | 34713727 | rs1458534 | -0.242 | -0.811 | -0.765 | -1.409 | 0.011 | 0.015 | 0.015 | 0.197 | 0.2 | 0.17 | 0.1 | 0.05 | 0.12 | 0.12 |
| 7 | 34717238 | rs1379923 | 1.48 | -1.1 | -0.894 | -1.698 | 0.022 | 0.022 | 0.022 | 0 | 0.3 | 0.22 | 0.27 | 0.06 | 0.24 | 0.37 |
| 7 | 34718523 | rs2531840 | 0.227 | -0.286 | -0.552 | -1.256 | 0.016 | 0.01 | 0.01 | 0.429 | 0.52 | 0.45 | 0.38 | 0.1 | 0.38 | 0.49 |
| 7 | 34719549 | rs2531841 | -0.221 | -0.238 | -0.726 | -1.226 | -0.024 | 0.14 | 0.14 | 1.865 | 0.59 | 0.8 | 0.45 | 0.21 | 0.47 | 0.57 |
| 7 | 34734251 | rs323928 | 0.259 | -0.339 | -0.685 | -1.261 | 0.032 | 0.047 | 0.047 | 0.733 | 0.51 | 0.46 | 0.37 | 0.1 | 0.37 | 0.49 |
| 7 | 34742051 | rs765023 | -0.654 | 0.079 | -0.727 | -0.738 | 0.018 | 0.022 | 0.022 | 0 | 0.3 | 0.53 | 0.29 | 0.44 | 0.37 | 0.21 |
| 7 | 34747017 | rs323920 | -1.218 | 0.009 | -0.839 | -0.909 | -0.014 | -0.009 | -0.009 | 0 | 0.38 | 0.68 | 0.38 | 0.5 | 0.45 | 0.3 |
| 7 | 34753046 | rs988367 | 0.251 | 0.22 | -0.774 | -0.849 | -0.004 | 0 | 0 | 0 | 0.17 | 0.17 | 0.22 | 0.37 | 0.27 | 0.19 |
| 7 | 34755872 | rs1419792 | 1.653 | 0.136 | -0.757 | -0.896 | -0.012 | 0.002 | 0.002 | 0 | 0.06 | 0.02 | 0.06 | 0.04 | 0.06 | 0.09 |
| 7 | 34758676 | rs17197848 | 0.422 | -0.317 | -1.165 | -1.265 | -0.024 | 0.056 | 0.056 | 0 | 0.21 | 0.03 | 0.15 | 0.09 | 0.16 | 0.16 |
| 7 | 34760208 | rs6961859 | 0.302 | -0.013 | -0.561 | -1.005 | -0.004 | -0.067 | -0.067 | 0 | 0.18 | 0.33 | 0.13 | 0.12 | 0.16 | 0.1 |
| 7 | 34770495 | rs324383 | -0.173 | -0.008 | -0.583 | -1.02 | -0.006 | -0.085 | -0.085 | 2.79 | 0.2 | 0.43 | 0.14 | 0.12 | 0.18 | 0.1 |
| 7 | 34777714 | rs324389 | -0.58 | 0.038 | -0.638 | -0.994 | -0.002 | -0.001 | -0.001 | 0 | 0.29 | 0.36 | 0.32 | 0.42 | 0.36 | 0.24 |
| 7 | 34783897 | rs324400 | -0.947 | -0.015 | -0.623 | -1.02 | 0.007 | 0.01 | 0.01 | 0 | 0.34 | 0.59 | 0.35 | 0.42 | 0.37 | 0.24 |
| 7 | 34786398 | rs10270663 | -0.42 | 0.046 | -0.756 | -0.98 | -0.039 | -0.045 | -0.045 | 0 | 0.28 | 0.2 | 0.31 | 0.41 | 0.36 | 0.24 |
| 7 | 34790123 | rs324396 | -1.995 | -0.3 | -1.223 | -1.263 | -0.115 | -0.189 | -0.189 | 0.11 | 0.74 | 0.97 | 0.65 | 0.56 | 0.68 | 0.69 |
| 7 | 34790349 | rs17777754 | -1.559 | 0.746 | -0.242 | -0.524 | -0.002 | -0.029 | -0.029 | 0 | 0.34 | 0.36 | 0.27 | 0.21 | 0.31 | 0.24 |
| 7 | 34791201 | rs33913770 | 0.892 | 0.754 | -0.4 | -0.473 | -0.025 | 0.019 | 0.019 | 0 | 0.12 | 0.04 | 0.09 | 0.13 | 0.15 | 0.1 |
| 7 | 34792221 | rs10243634 | -1.77 | 0.738 | -0.347 | -0.461 | 0 | 0.001 | 0.001 | 0 | 0.46 | 0.48 | 0.43 | 0.54 | 0.52 | 0.34 |
| 7 | 34802391 | rs324960 | -1.263 | 0.847 | -0.321 | -0.42 | -0.006 | -0.06 | -0.06 | 4.835 | 0.36 | 0.49 | 0.29 | 0.21 | 0.31 | 0.24 |
| 7 | 34803103 | rs10267134 | 1.368 | 0.98 | -0.46 | -0.291 | -0.054 | -0.061 | -0.061 | 0 | 0.11 | 0.01 | 0.15 | 0.32 | 0.2 | 0.1 |
| 7 | 34813308 | rs1419779 | -1.136 | 0.801 | -0.434 | -0.432 | -0.001 | -0.014 | -0.014 | 0 | 0.34 | 0.3 | 0.26 | 0.21 | 0.31 | 0.24 |
| 7 | 34820597 | rs324986 | -1.276 | 0.617 | -0.565 | -0.158 | -0.002 | -0.009 | -0.009 | 0 | 0.46 | 0.67 | 0.43 | 0.53 | 0.49 | 0.33 |
| 7 | 34852665 | rs17789420 | 0.417 | 0.714 | -0.411 | -0.331 | -0.018 | -0.001 | -0.001 | 0 | 0.08 | 0.02 | 0.06 | 0.13 | 0.09 | 0.08 |
| 7 | 34859868 | rs6977729 | -0.757 | 0.659 | -0.211 | -0.383 | 0.015 | -0.042 | -0.042 | 0 | 0.15 | 0.48 | 0.12 | 0.13 | 0.11 | 0.09 |
| 7 | 34867182 | rs17170011 | 0.24 | 0.966 | -1.14 | -0.184 | -0.011 | -0.004 | -0.004 | 0 | 0.08 | 0.03 | 0.06 | 0.14 | 0.1 | 0.08 |
| 7 | 34867557 | rs1419868 | -0.825 | 0.683 | -1.872 | -0.403 | -0.03 | 0.064 | 0.064 | 0 | 0.57 | 0.92 | 0.53 | 0.72 | 0.57 | 0.48 |
| 7 | 34867912 | rs17170012 | -0.961 | 1.225 | -0.591 | -0.367 | 0.007 | -0.003 | -0.003 | 0 | 0.12 | 0.17 | 0.11 | 0.09 | 0.1 | 0.19 |
| 7 | 34870001 | rs12111597 | 0.306 | 0.551 | -0.595 | -0.469 | -0.023 | 0.013 | 0.013 | 0 | 0.32 | 0.42 | 0.37 | 0.36 | 0.22 | 0.42 |
| 7 | 34871328 | rs17789834 | -0.483 | 1.41 | -1.039 | -0.516 | -0.055 | 0.012 | 0.012 | 0.161 | 0.12 | 0.03 | 0.08 | 0.07 | 0.18 | 0.19 |
| 7 | 34872332 | rs17170015 | 0.663 | 1.079 | -0.881 | -0.552 | 0.009 | -0.051 | -0.051 | 0 | 0.27 | 0.15 | 0.35 | 0.36 | 0.25 | 0.43 |
| 7 | 34873320 | rs1860369 | 0.663 | 0.912 | -0.425 | -0.531 | 0.001 | -0.002 | -0.002 | 1.931 | 0.31 | 0.27 | 0.35 | 0.35 | 0.21 | 0.41 |
| 7 | 34873745 | rs6947841 | 0.351 | 1.697 | 0.124 | -0.267 | 0.001 | -0.004 | -0.004 | 0.635 | 0.18 | 0.18 | 0.13 | 0.15 | 0.19 | 0.13 |
| 7 | 34873776 | rs33911118 | 0.683 | 1.606 | 0.005 | -0.285 | 0.011 | 0.05 | 0.05 | 0 | 0.26 | 0.39 | 0.34 | 0.43 | 0.29 | 0.22 |
| 7 | 34876014 | rs10249085 | -1.044 | 2.258 | 0.425 | -0.764 | 0.044 | -0.042 | -0.042 | 0 | 0.31 | 0.46 | 0.24 | 0.2 | 0.38 | 0.3 |
| 7 | 34877034 | rs10246825 | -1.44 | 2.018 | 0.598 | -1.086 | 0.003 | -0.038 | -0.038 | 0.132 | 0.32 | 0.42 | 0.22 | 0.16 | 0.31 | 0.19 |
| 7 | 34877996 | rs17200455 | 0.34 | 2.087 | 0.596 | -1.15 | -0.029 | 0.001 | 0.001 | 0 | 0.13 | 0.08 | 0.12 | 0.14 | 0.2 | 0.09 |
| 7 | 34879310 | rs740717 | 0.546 | 1.973 | 0.703 | -1.172 | -0.027 | -0.033 | -0.033 | 0.332 | 0.06 | 0.13 | 0.02 | 0 | 0.02 | 0 |
| 7 | 34880222 | rs11771697 | -0.078 | 1.993 | 0.642 | -1.165 | -0.004 | 0.013 | 0.013 | 0.938 | 0.09 | 0.03 | 0.06 | 0.01 | 0.1 | 0.04 |
| 7 | 34885132 | rs13231258 | -0.078 | 1.993 | 0.642 | -1.165 | -0.004 | 0.013 | 0.013 | 0.938 | 0.09 | 0.03 | 0.06 | 0.01 | 0.1 | 0.04 |
| 7 | 34885503 | rs10267935 | 0.232 | 2.216 | 0.664 | -1.161 | -0.034 | 0.004 | 0.004 | 0 | 0.13 | 0.08 | 0.13 | 0.14 | 0.23 | 0.09 |
| 7 | 34886141 | rs12154548 | 0.232 | 2.216 | 0.664 | -1.161 | -0.034 | 0.004 | 0.004 | 0 | 0.13 | 0.08 | 0.13 | 0.14 | 0.23 | 0.09 |
| 7 | 34886825 | rs10250709 | -2.075 | 2.142 | 0.749 | -1.213 | 0.008 | -0.042 | -0.042 | 0 | 0.33 | 0.41 | 0.23 | 0.15 | 0.35 | 0.13 |
| 7 | 34887170 | rs13245730 | -2.229 | 1.995 | 1.32 | -1.223 | -0.003 | 0.018 | 0.018 | 0.057 | 0.36 | 0.66 | 0.3 | 0.2 | 0.36 | 0.14 |
| 7 | 34887178 | rs13246028 | -2.043 | 2.005 | 1.297 | -1.218 | 0.007 | -0.042 | -0.042 | 0 | 0.33 | 0.4 | 0.23 | 0.15 | 0.35 | 0.13 |
| 7 | 34889182 | rs6972158 | -2.043 | 2.015 | 1.31 | -1.219 | 0.008 | -0.044 | -0.044 | 2.302 | 0.33 | 0.41 | 0.23 | 0.15 | 0.35 | 0.13 |
| 7 | 34889222 | rs10275028 | -2.043 | 2.015 | 1.31 | -1.219 | 0.008 | -0.044 | -0.044 | 0 | 0.33 | 0.4 | 0.23 | 0.15 | 0.35 | 0.13 |
| 7 | 34889552 | rs6958905 | -2.225 | 1.983 | 1.458 | -1.233 | -0.003 | 0.018 | 0.018 | 0.015 | 0.36 | 0.65 | 0.3 | 0.2 | 0.36 | 0.14 |

| | | | | | | | | | | | | | | | | |
|---|---|---|---|---|---|---|---|---|---|---|---|---|---|---|---|---|
| 7 | 34889675 | rs9655359 | -2.088 | 2.12 | 0.925 | -1.201 | 0.008 | -0.044 | -0.044 | 9.026 | 0.33 | 0.4 | 0.23 | 0.15 | 0.35 | 0.13 |
| 7 | 34899235 | rs10235648 | -1.238 | 2.278 | 0.488 | -1.105 | -0.009 | 0.008 | 0.008 | 0 | 0.23 | 0.24 | 0.2 | 0.15 | 0.33 | 0.13 |
| 7 | 34899573 | rs10238983 | -0.035 | 2.287 | 0.49 | -1.107 | 0.014 | -0.002 | -0.002 | 0 | 0.14 | 0.2 | 0.14 | 0.15 | 0.24 | 0.09 |
| 7 | 34901966 | rs11979570 | -1.167 | 2.262 | 0.486 | -1.096 | -0.009 | 0.008 | 0.008 | 0 | 0.23 | 0.24 | 0.2 | 0.15 | 0.33 | 0.13 |
| 7 | 34920075 | rs11771425 | -0.144 | 1.982 | 0.234 | -0.978 | 0.013 | 0.018 | 0.018 | 0.063 | 0.12 | 0.05 | 0.07 | 0.01 | 0.09 | 0.05 |
| 7 | 34924337 | rs7804942 | -1.465 | 2.423 | 0.439 | -1.019 | -0.014 | 0.015 | 0.015 | 0 | 0.36 | 0.31 | 0.28 | 0.22 | 0.46 | 0.31 |
| 7 | 34931893 | rs1807933 | 0.353 | 2.609 | 0.409 | -1.014 | -0.038 | 0.001 | 0.001 | 0 | 0.13 | 0.09 | 0.11 | 0.14 | 0.21 | 0.09 |
| 7 | 34932979 | rs995086 | -1.063 | 1.758 | 0.459 | -0.945 | -0.014 | 0.015 | 0.015 | 0 | 0.36 | 0.31 | 0.27 | 0.21 | 0.45 | 0.32 |
| 7 | 34938925 | rs2107578 | -1.214 | 1.82 | -0.556 | -0.87 | -0.052 | 0.069 | 0.069 | 0 | 0.38 | 0.77 | 0.36 | 0.27 | 0.48 | 0.31 |
| 7 | 34939884 | rs1859409 | 0.206 | 1.335 | -0.424 | -0.757 | 0.004 | -0.031 | -0.031 | 0 | 0.66 | 0.52 | 0.51 | 0.75 | 0.64 | 0.84 |
| 7 | 34940039 | rs10252228 | -0.022 | 1.13 | -0.813 | -0.907 | 0.009 | -0.023 | -0.023 | 0 | 0.35 | 0.39 | 0.27 | 0.19 | 0.41 | 0.16 |
| 7 | 34942820 | rs4723388 | -0.229 | 1.339 | -0.719 | -0.372 | 0.022 | 0.021 | 0.021 | 0.314 | 0.11 | 0.15 | 0.37 | 0.24 | 0.17 | 0.16 |
| 7 | 34944778 | rs17791600 | -0.944 | 1.242 | -0.652 | -0.502 | 0.017 | 0.009 | 0.009 | 0 | 0.15 | 0.2 | 0.38 | 0.24 | 0.19 | 0.16 |
| 7 | 34986082 | rs13234119 | 0.692 | 0.788 | -1.203 | -0.872 | 0.017 | 0.033 | 0.033 | 0.013 | 0.14 | 0 | 0.05 | 0.04 | 0.08 | 0.03 |
| 7 | 34986597 | rs1649220 | -0.569 | 0.957 | -1.343 | -1.292 | -0.009 | 0.01 | 0.01 | 0 | 0.41 | 0.37 | 0.64 | 0.37 | 0.47 | 0.38 |
| 7 | 34992635 | rs1649228 | -0.593 | 0.949 | -1.347 | -1.246 | -0.009 | 0.01 | 0.01 | 0 | 0.41 | 0.37 | 0.64 | 0.39 | 0.47 | 0.39 |
| 7 | 34995410 | rs2114733 | 0.796 | 0.901 | -1.429 | -1.267 | -0.069 | 0.037 | 0.037 | 0 | 0.24 | 0.03 | 0.23 | 0.14 | 0.26 | 0.22 |
| 7 | 35009754 | rs1637682 | -0.868 | 0.941 | -1.311 | -1.262 | 0.037 | 0.011 | 0.011 | 0 | 0.18 | 0.33 | 0.41 | 0.24 | 0.21 | 0.17 |
| 7 | 35020843 | rs328902 | 0.603 | 0.935 | -1.346 | -1.114 | -0.011 | 0.031 | 0.031 | 0 | 0.26 | 0.19 | 0.25 | 0.13 | 0.29 | 0.22 |
| 7 | 35023915 | rs328906 | -1.014 | 0.905 | -1.215 | -1.047 | 0.027 | 0.002 | 0.002 | 0 | 0.18 | 0.33 | 0.41 | 0.24 | 0.2 | 0.17 |
| 7 | 35031630 | rs2023328 | -0.459 | 0.896 | -1.285 | -1.015 | 0.008 | 0.002 | 0.002 | 0 | 0.16 | 0.18 | 0.38 | 0.24 | 0.19 | 0.16 |
| 7 | 35054045 | rs2893491 | 0.727 | 1.128 | -1.142 | -0.898 | 0.03 | 0.021 | 0.021 | 0 | 0.15 | 0.22 | 0.45 | 0.3 | 0.29 | 0.32 |
| 7 | 35080931 | rs329277 | -0.158 | 0.769 | -1.172 | -1.026 | -0.003 | 0.001 | 0.001 | 0.943 | 0.37 | 0.37 | 0.63 | 0.37 | 0.46 | 0.37 |
| 7 | 35095894 | rs329570 | 0.659 | 0.425 | -0.899 | -1.183 | 0.023 | 0.025 | 0.025 | 0.002 | 0.15 | 0.08 | 0.06 | 0.07 | 0.08 | 0.16 |
| 7 | 35141504 | rs34766665 | 0.309 | 0.398 | -0.876 | -1.187 | 0.016 | 0.017 | 0.017 | 0 | 0.17 | 0.15 | 0.08 | 0.07 | 0.1 | 0.15 |
| 7 | 35190409 | rs6972730 | -0.526 | 0.588 | -0.934 | -1.224 | 0.023 | 0.07 | 0.07 | 0 | 0.43 | 0.32 | 0.33 | 0.22 | 0.4 | 0.36 |
| 7 | 35225518 | rs10262841 | 0.634 | 0.604 | -0.963 | -1.205 | -0.02 | 0.041 | 0.041 | 0 | 0.26 | 0.11 | 0.24 | 0.19 | 0.29 | 0.21 |
| 7 | 35227297 | rs17136537 | 0.471 | 0.318 | -0.72 | -1.094 | 0 | -0.002 | -0.002 | 0 | 0.15 | 0.13 | 0.06 | 0.04 | 0.08 | 0.04 |
| 7 | 35269625 | rs2908049 | 0.19 | 0.532 | -0.787 | -0.754 | -0.006 | -0.033 | -0.033 | 0 | 0.34 | 0.44 | 0.46 | 0.28 | 0.38 | 0.49 |
| 7 | 35270988 | rs2109090 | 0.429 | 0.488 | -0.634 | -0.806 | 0.002 | 0.014 | 0.014 | 0 | 0.33 | 0.26 | 0.44 | 0.29 | 0.38 | 0.39 |
| 7 | 35273291 | rs2191883 | -0.498 | 0.42 | -0.84 | -1.041 | 0.002 | -0.015 | -0.015 | 1.479 | 0.56 | 0.48 | 0.58 | 0.35 | 0.54 | 0.44 |
| 7 | 35278424 | rs4723402 | 0.58 | 0.359 | -0.675 | -0.95 | 0.001 | 0.005 | 0.005 | 0 | 0.34 | 0.33 | 0.44 | 0.29 | 0.38 | 0.39 |
| 7 | 35280947 | rs2240994 | 0.623 | 0.382 | -0.735 | -0.94 | 0.004 | 0.018 | 0.018 | 0 | 0.34 | 0.26 | 0.44 | 0.29 | 0.38 | 0.39 |
| 7 | 35293972 | rs336284 | -0.191 | 0.458 | -0.658 | -0.428 | 0 | -0.001 | -0.001 | 0.206 | 0.48 | 0.52 | 0.53 | 0.53 | 0.52 | 0.54 |
| 7 | 35295491 | rs1003549 | 0.654 | 0.459 | -0.768 | -0.543 | 0.037 | 0.065 | 0.065 | 0 | 0.19 | 0.02 | 0.1 | 0.06 | 0.13 | 0.04 |
| 7 | 35305324 | rs1861095 | -0.317 | 0.549 | -0.558 | -0.517 | 0.001 | -0.019 | -0.019 | 0 | 0.56 | 0.51 | 0.58 | 0.36 | 0.53 | 0.38 |
| 7 | 35306983 | rs1419856 | 0.269 | 0.186 | -0.368 | -0.576 | 0.017 | 0.076 | 0.076 | 0 | 0.21 | 0.02 | 0.12 | 0.06 | 0.15 | 0.04 |
| 7 | 35312748 | rs4342481 | -0.189 | 0.234 | -0.309 | -0.477 | 0.04 | 0.066 | 0.066 | 0 | 0.25 | 0.09 | 0.13 | 0.09 | 0.16 | 0.15 |
| 7 | 35319577 | rs1425148 | 0.744 | 0.226 | -0.282 | -0.417 | -0.024 | -0.013 | -0.013 | 1.013 | 0.42 | 0.3 | 0.52 | 0.53 | 0.51 | 0.43 |
| 7 | 35326018 | rs6976419 | -0.87 | 0.271 | -0.267 | -0.391 | 0.018 | 0.046 | 0.046 | 0 | 0.68 | 0.56 | 0.67 | 0.62 | 0.67 | 0.56 |
| 7 | 35329471 | rs12538450 | -1.295 | 0.27 | -0.252 | -0.373 | -0.048 | -0.079 | -0.079 | 0.05 | 0.73 | 0.87 | 0.69 | 0.62 | 0.68 | 0.57 |
| 7 | 35332117 | rs10265222 | 0.221 | 0.357 | -0.157 | -0.074 | 0.002 | 0 | 0 | 0 | 0.43 | 0.55 | 0.53 | 0.53 | 0.51 | 0.42 |
| 7 | 35340998 | rs10237288 | 0.288 | 0.481 | -1.322 | -0.036 | -0.015 | -0.008 | -0.008 | 0.154 | 0.43 | 0.35 | 0.52 | 0.53 | 0.51 | 0.42 |
| 7 | 35353014 | rs11766153 | -0.172 | -0.172 | -1.504 | -0.445 | 0.034 | 0.061 | 0.061 | 0 | 0.18 | 0.02 | 0.09 | 0.06 | 0.11 | 0.03 |
| 7 | 35353294 | rs6942533 | -0.743 | -0.231 | -1.461 | -0.467 | 0.051 | 0.065 | 0.065 | 0 | 0.24 | 0.07 | 0.1 | 0.09 | 0.12 | 0.14 |
| 7 | 35354711 | rs235390 | -0.145 | 0.31 | -1.094 | -0.463 | 0.021 | 0.035 | 0.035 | 0 | 0.47 | 0.37 | 0.55 | 0.17 | 0.66 | 0.46 |
| 7 | 35356265 | rs6976218 | -0.684 | 1.138 | -0.718 | 0.133 | 0.002 | 0.009 | 0.009 | 0.004 | 0.34 | 0.41 | 0.33 | 0.57 | 0.3 | 0.38 |
| 7 | 35359729 | rs17675696 | -0.356 | 1.133 | -0.804 | -0.797 | -0.006 | 0.006 | 0.006 | 1.205 | 0.2 | 0.15 | 0.15 | 0.01 | 0.22 | 0.08 |
| 7 | 35361076 | rs11772478 | -0.429 | 1.294 | -0.571 | -1.445 | 0 | -0.034 | -0.034 | 0 | 0.49 | 0.52 | 0.41 | 0.1 | 0.54 | 0.39 |
| 7 | 35366205 | rs1125243 | 0.94 | 1.303 | -0.741 | -1.434 | 0.003 | 0.011 | 0.011 | 0 | 0.28 | 0.2 | 0.25 | 0.09 | 0.31 | 0.31 |
| 7 | 35367474 | rs235403 | 0.208 | 1.883 | -0.809 | -1.062 | -0.006 | -0.005 | -0.005 | 0 | 0.91 | 0.93 | 0.92 | 0.86 | 0.87 | 0.91 |
| 7 | 35369161 | rs6951055 | 0.037 | 1.313 | -0.822 | -1.132 | -0.006 | 0.012 | 0.012 | 1.215 | 0.2 | 0.14 | 0.15 | 0.01 | 0.22 | 0.08 |
| 7 | 35371417 | rs12532988 | 0.26 | 1.65 | -0.672 | -0.822 | 0 | 0.043 | 0.043 | 0 | 0.33 | 0.21 | 0.3 | 0.1 | 0.35 | 0.33 |
| 7 | 35372749 | rs235397 | -0.47 | 1.581 | -0.425 | -0.781 | 0.014 | 0.032 | 0.032 | 0.839 | 0.7 | 0.8 | 0.77 | 0.88 | 0.75 | 0.72 |
| 7 | 35374267 | rs9969168 | 0.374 | 1.468 | -0.275 | -0.658 | 0.001 | -0.003 | -0.003 | 0.272 | 0.34 | 0.41 | 0.34 | 0.67 | 0.29 | 0.36 |
| 7 | 35385397 | rs235413 | 0.018 | 1.342 | -0.527 | -0.729 | 0.013 | 0.033 | 0.033 | 0.167 | 0.3 | 0.25 | 0.36 | 0.09 | 0.3 | 0.29 |
| 7 | 35388122 | rs340370 | -0.106 | 1.483 | -0.161 | 0.046 | 0.001 | -0.018 | -0.018 | 0 | 0.68 | 0.61 | 0.68 | 0.77 | 0.68 | 0.83 |
| 7 | 35389499 | rs7800747 | -1.134 | 2.272 | -0.068 | 0.316 | 0.068 | 0.025 | 0.025 | 0 | 0.05 | 0.11 | 0.13 | 0.1 | 0.14 | 0.09 |
| 7 | 35392897 | rs10486663 | 0.201 | 1.897 | -0.388 | 0.325 | -0.019 | -0.022 | -0.022 | 0.28 | 0.28 | 0.17 | 0.2 | 0.38 | 0.31 | 0.19 |
| 7 | 35393412 | rs124557 | -0.329 | 1.263 | -0.731 | 0.02 | 0.029 | 0.016 | 0.016 | 0 | 0.86 | 0.92 | 0.9 | 0.92 | 0.92 | 0.89 |
| 7 | 35394051 | rs13239224 | -0.106 | 0.957 | -0.853 | -0.305 | -0.049 | 0.09 | 0.09 | 0 | 0.63 | 0.8 | 0.62 | 0.85 | 0.57 | 0.63 |
| 7 | 35394378 | rs340373 | 1.454 | 1.042 | -0.864 | -0.319 | -0.035 | 0.063 | 0.063 | 0 | 0.22 | 0.07 | 0.24 | 0.06 | 0.27 | 0.21 |
| 7 | 35394424 | rs7805453 | -0.414 | 1.096 | -0.787 | -0.26 | -0.003 | 0.095 | 0.095 | 0 | 0.64 | 0.85 | 0.67 | 0.86 | 0.65 | 0.68 |
| 7 | 35395048 | rs12701423 | 0.136 | 1.009 | -0.145 | -0.115 | -0.001 | 0.003 | 0.003 | 0 | 0.34 | 0.39 | 0.44 | 0.49 | 0.33 | 0.5 |
| 7 | 35397647 | rs6462603 | 0.392 | 0.871 | -0.123 | -0.111 | 0.024 | -0.029 | -0.029 | 7.14 | 0.32 | 0.27 | 0.39 | 0.47 | 0.25 | 0.45 |
| 7 | 35401570 | rs4445131 | 0.129 | 0.852 | -0.117 | -0.166 | -0.004 | 0.011 | 0.011 | 0 | 0.34 | 0.41 | 0.45 | 0.45 | 0.33 | 0.49 |
| 7 | 35404590 | rs340383 | 0.472 | 0.773 | 0.159 | -0.183 | 0.043 | 0.003 | 0.003 | 0 | 0.5 | 0.3 | 0.44 | 0.49 | 0.59 | 0.39 |
| 7 | 35408109 | rs13233781 | 0.453 | 0.493 | -0.241 | -0.264 | -0.018 | 0.013 | 0.013 | 0 | 0.31 | 0.36 | 0.4 | 0.42 | 0.26 | 0.44 |
| 7 | 35415420 | rs340402 | 1.288 | 0.776 | -0.317 | -0.352 | 0.017 | -0.053 | -0.053 | 0 | 0.23 | 0.26 | 0.24 | 0.07 | 0.28 | 0.22 |

| | | | | | | | | | | | | | | | | |
|---|---|---|---|---|---|---|---|---|---|---|---|---|---|---|---|---|
| 7 | 35416992 | rs11773862 | 0.478 | 1.114 | -0.642 | 0.062 | -0.022 | -0.03 | -0.03 | 19.42 | 0.29 | 0.19 | 0.21 | 0.42 | 0.33 | 0.19 |
| 7 | 35419743 | rs340407 | 0.361 | 0.982 | -0.156 | -0.116 | -0.001 | -0.014 | -0.014 | 0.28 | 0.82 | 0.73 | 0.86 | 0.91 | 0.85 | 0.84 |
| 7 | 35423720 | rs12113424 | -0.713 | 1.37 | -0.283 | -0.296 | 0.017 | -0.046 | -0.046 | 0 | 0.15 | 0.2 | 0.12 | 0.02 | 0.18 | 0.1 |
| 7 | 35423859 | rs2541899 | -0.679 | 0.486 | -0.562 | 0.021 | 0.017 | -0.041 | -0.041 | 0 | 0.76 | 0.82 | 0.84 | 0.59 | 0.79 | 0.83 |
| 7 | 35424928 | rs12672982 | -0.285 | 0.042 | -0.343 | -0.06 | -0.011 | -0.003 | -0.003 | 0.185 | 0.14 | 0.16 | 0.06 | 0.1 | 0.07 | 0.09 |
| 7 | 35428055 | rs4720172 | -0.076 | 0.537 | -0.415 | -0.263 | 0.032 | 0.033 | 0.033 | 0 | 0.39 | 0.18 | 0.37 | 0.34 | 0.35 | 0.47 |
| 7 | 35431359 | rs17170996 | 0.798 | 0.556 | -0.202 | -0.102 | -0.006 | -0.001 | -0.001 | 0 | 0.11 | 0.13 | 0.06 | 0.12 | 0.06 | 0.09 |
| 7 | 35437169 | rs2592857 | -0.136 | 0.717 | -0.423 | -0.467 | -0.007 | 0.029 | 0.029 | 0 | 0.37 | 0.69 | 0.44 | 0.24 | 0.36 | 0.3 |
| 7 | 35439366 | rs17676509 | 0.675 | 0.639 | -0.712 | -0.643 | -0.041 | 0.056 | 0.056 | 0.127 | 0.14 | 0.01 | 0.1 | 0.03 | 0.16 | 0.08 |
| 7 | 35443475 | rs2541911 | 0.857 | 0.821 | -0.374 | -0.429 | 0.013 | -0.023 | -0.023 | 0.517 | 0.26 | 0.61 | 0.38 | 0.11 | 0.29 | 0.22 |
| 7 | 35444548 | rs10236415 | 1.027 | 0.745 | -0.389 | -0.407 | -0.018 | 0.003 | 0.003 | 0 | 0.11 | 0.16 | 0.06 | 0.13 | 0.06 | 0.09 |
| 7 | 35449334 | rs998652 | -1.954 | 0.681 | -0.567 | -0.347 | -0.024 | -0.15 | -0.15 | 0 | 0.78 | 0.99 | 0.85 | 0.57 | 0.79 | 0.78 |
| 7 | 35449404 | rs4723424 | 0.422 | 0.711 | -0.569 | -0.371 | 0.015 | 0.021 | 0.021 | 0 | 0.38 | 0.12 | 0.37 | 0.34 | 0.35 | 0.47 |
| 7 | 35450900 | rs13235830 | -0.193 | 0.86 | -0.494 | -0.325 | 0.057 | 0.022 | 0.022 | 0 | 0.5 | 0.32 | 0.43 | 0.47 | 0.42 | 0.56 |
| 7 | 35451910 | rs2592851 | -1.69 | 1.014 | -0.762 | -0.186 | -0.003 | -0.128 | -0.128 | 0 | 0.77 | 0.93 | 0.84 | 0.57 | 0.79 | 0.78 |
| 7 | 35458934 | rs342976 | -1.606 | 1.046 | -0.79 | -0.185 | -0.066 | -0.118 | -0.118 | 0 | 0.77 | 0.94 | 0.81 | 0.57 | 0.71 | 0.79 |
| 7 | 35461773 | rs2551796 | 1.46 | 1.064 | -0.355 | -0.216 | 0.006 | -0.003 | -0.003 | 0.163 | 0.2 | 0.19 | 0.32 | 0.11 | 0.26 | 0.19 |
| 7 | 35463795 | rs10228514 | 0.682 | 0.948 | -0.57 | -0.077 | 0.014 | 0.005 | 0.005 | 0 | 0.12 | 0.21 | 0.09 | 0.12 | 0.14 | 0.09 |
| 7 | 35467026 | rs342988 | -1.379 | 0.642 | -0.7 | -0.057 | 0.009 | -0.047 | -0.047 | 0.166 | 0.76 | 0.88 | 0.82 | 0.57 | 0.76 | 0.77 |
| 7 | 35467896 | rs342989 | -1.692 | 0.628 | -0.978 | -0.042 | -0.026 | -0.149 | -0.149 | 0 | 0.78 | 0.98 | 0.84 | 0.57 | 0.79 | 0.78 |
| 7 | 35485135 | rs13223811 | 1.655 | 0.154 | -1.229 | -0.297 | -0.004 | 0.037 | 0.037 | 0 | 0.12 | 0.01 | 0.08 | 0.06 | 0.12 | 0.21 |
| 7 | 35486013 | rs4720174 | 0.109 | 0.343 | -1.002 | -0.231 | 0.046 | 0.037 | 0.037 | 0 | 0.38 | 0.14 | 0.35 | 0.36 | 0.33 | 0.42 |
| 7 | 35494821 | rs10951441 | 1.205 | 0.3 | -0.853 | -0.128 | -0.006 | 0.003 | 0.003 | 0 | 0.1 | 0.11 | 0.06 | 0.12 | 0.06 | 0.07 |
| 7 | 35501213 | rs343032 | 1.355 | 0.735 | -0.265 | -0.323 | 0.051 | -0.031 | -0.031 | 0 | 0.22 | 0.31 | 0.35 | 0.11 | 0.27 | 0.22 |
| 7 | 35501883 | rs343034 | -0.61 | 0.626 | -0.371 | -0.153 | -0.001 | -0.016 | -0.016 | 0.091 | 0.64 | 0.68 | 0.74 | 0.48 | 0.65 | 0.67 |
| 7 | 35506733 | rs343041 | 1.195 | 1.163 | 0.042 | -0.294 | 0.028 | -0.021 | -0.021 | 0 | 0.22 | 0.25 | 0.37 | 0.1 | 0.3 | 0.23 |
| 7 | 35506790 | rs343042 | -0.591 | 0.729 | -0.66 | -0.416 | 0.021 | -0.017 | -0.017 | 0 | 0.65 | 0.74 | 0.78 | 0.47 | 0.73 | 0.66 |
| 7 | 35508382 | rs343043 | 2.004 | 0.638 | -0.756 | -0.756 | -0.091 | 0.097 | 0.097 | 0 | 0.19 | 0.03 | 0.28 | 0.02 | 0.27 | 0.14 |
| 7 | 35509757 | rs13239463 | 0.254 | 0.579 | -0.502 | -0.363 | 0.04 | 0.031 | 0.031 | 0 | 0.42 | 0.26 | 0.38 | 0.73 | 0.38 | 0.48 |
| 7 | 35510477 | rs2592845 | 0.036 | 0.499 | -0.284 | -0.002 | 0.053 | 0.086 | 0.086 | 0 | 0.14 | 0.52 | 0.19 | 0.22 | 0.17 | 0.14 |
| 7 | 35518982 | rs12155020 | 1.227 | 0.526 | -0.59 | -0.042 | 0.01 | 0.012 | 0.012 | 0 | 0.1 | 0.14 | 0.09 | 0.14 | 0.14 | 0.06 |
| 7 | 35522553 | rs343053 | -1.617 | 0.412 | -0.802 | -0.198 | -0.007 | 0.13 | 0.13 | 0 | 0.79 | 0.99 | 0.89 | 0.98 | 0.83 | 0.82 |
| 7 | 35522899 | rs343054 | 0.909 | 0.214 | -0.926 | 0.482 | -0.003 | 0.074 | 0.074 | 0 | 0.46 | 0.3 | 0.5 | 0.29 | 0.54 | 0.55 |
| 7 | 35523030 | rs10262622 | -0.142 | 0.396 | -0.768 | 0.079 | 0.002 | -0.007 | -0.007 | 0 | 0.23 | 0.33 | 0.18 | 0.14 | 0.23 | 0.1 |
| 7 | 35525105 | rs10226053 | 1.088 | 0.438 | -0.773 | 0.299 | 0.001 | -0.002 | -0.002 | 0 | 0.12 | 0.12 | 0.1 | 0.14 | 0.1 | 0.03 |
| 7 | 35526390 | rs343056 | -1.651 | 0.476 | -0.839 | 0.264 | -0.014 | 0.112 | 0.112 | 0 | 0.79 | 0.98 | 0.88 | 0.98 | 0.82 | 0.79 |
| 7 | 35528049 | rs343058 | -0.919 | 0.859 | -0.166 | 0.88 | -0.002 | 0.001 | 0.001 | 0 | 0.88 | 0.89 | 0.91 | 0.86 | 0.9 | 0.97 |
| 7 | 35533096 | rs12701433 | 0.129 | 0.989 | -1.071 | 0.838 | -0.026 | -0.022 | -0.022 | 2.564 | 0.44 | 0.27 | 0.49 | 0.54 | 0.49 | 0.56 |
| 7 | 35533404 | rs13220973 | 0.025 | 0.93 | -1.028 | 0.811 | -0.001 | 0 | 0 | 0 | 0.29 | 0.29 | 0.24 | 0.4 | 0.25 | 0.39 |
| 7 | 35533597 | rs13236869 | 0.603 | 0.607 | -1.091 | 0.712 | -0.002 | 0 | 0 | 0 | 0.19 | 0.21 | 0.17 | 0.29 | 0.17 | 0.35 |
| 7 | 35533765 | rs6968864 | 0.902 | 0.458 | -1.026 | 0.682 | -0.018 | 0.009 | 0.009 | 0 | 0.31 | 0.25 | 0.45 | 0.23 | 0.36 | 0.28 |
| 7 | 35535511 | rs2052321 | -0.921 | 0.263 | -0.894 | 0.327 | 0.015 | 0.036 | 0.036 | 0 | 0.83 | 0.81 | 0.84 | 0.68 | 0.78 | 0.94 |
| 7 | 35535930 | rs1035218 | -0.903 | 0.259 | -0.717 | 0.325 | 0.025 | 0.055 | 0.055 | 0 | 0.83 | 0.77 | 0.84 | 0.68 | 0.78 | 0.94 |
| 7 | 35536505 | rs1035221 | -1.161 | 0.347 | -0.715 | 0.32 | -0.004 | 0.032 | 0.032 | 0 | 0.85 | 0.83 | 0.87 | 0.68 | 0.85 | 0.96 |
| 7 | 35536636 | rs6974145 | 2.316 | 0.309 | -0.947 | 0.11 | -0.05 | 0.051 | 0.051 | 0 | 0.19 | 0.1 | 0.27 | 0.02 | 0.28 | 0.16 |
| 7 | 35537950 | rs2392369 | 2.048 | -0.25 | -0.539 | -0.004 | 0.055 | 0.117 | 0.117 | 0 | 0.07 | 0.31 | 0.09 | 0.13 | 0.07 | 0.05 |
| 7 | 35538435 | rs2160621 | 0.722 | -0.816 | -0.873 | -0.293 | 0.046 | -0.06 | -0.06 | 0 | 0.3 | 0.18 | 0.3 | 0.5 | 0.22 | 0.44 |
| 7 | 35538760 | rs9886294 | 1.295 | -0.963 | -0.936 | -0.396 | 0.032 | -0.048 | -0.048 | 0 | 0.24 | 0.17 | 0.28 | 0.5 | 0.16 | 0.43 |
| 7 | 35543446 | rs2052322 | -1.457 | -0.604 | -0.698 | -0.125 | 0.003 | -0.02 | -0.02 | 0.13 | 0.85 | 0.92 | 0.88 | 0.69 | 0.85 | 0.96 |
| 7 | 35543469 | rs6969809 | 1.113 | -0.52 | -0.652 | -0.247 | 0.027 | 0.061 | 0.061 | 0 | 0.23 | 0.13 | 0.13 | 0.04 | 0.16 | 0.26 |
| 7 | 35545588 | rs16879846 | 2.136 | -0.174 | -0.333 | 0.081 | 0.059 | 0.135 | 0.135 | 0 | 0.07 | 0.31 | 0.09 | 0.13 | 0.07 | 0.05 |
| 7 | 35547956 | rs10486666 | 1.25 | -0.204 | -0.365 | 0.23 | 0.039 | 0.206 | 0.206 | 0 | 0.08 | 0.32 | 0.11 | 0.23 | 0.08 | 0.06 |
| 7 | 35548202 | rs4723430 | -1.525 | -0.185 | -0.79 | 0.22 | -0.006 | -0.041 | -0.041 | 0.167 | 0.86 | 0.97 | 0.9 | 0.77 | 0.87 | 0.96 |
| 7 | 35548431 | rs17698191 | 0.88 | -0.218 | -0.651 | 0.137 | 0.004 | 0.017 | 0.017 | 0.019 | 0.08 | 0.11 | 0.08 | 0.13 | 0.05 | 0.08 |
| 7 | 35551086 | rs10951443 | 1.076 | -0.601 | -1.007 | -0.171 | 0.009 | 0.033 | 0.033 | 0 | 0.18 | 0.13 | 0.13 | 0.04 | 0.16 | 0.2 |
| 7 | 35554788 | rs343064 | 0.216 | -0.759 | -1.212 | -0.327 | 0.059 | -0.047 | -0.047 | 0 | 0.44 | 0.1 | 0.4 | 0.48 | 0.38 | 0.49 |
| 7 | 35556427 | rs6972992 | 2.151 | -0.637 | -0.822 | -0.365 | 0.022 | -0.039 | -0.039 | 0 | 0.19 | 0.23 | 0.28 | 0.02 | 0.28 | 0.16 |
| 7 | 35557787 | rs343067 | 1.87 | -1.052 | -0.104 | -0.018 | -0.056 | 0.089 | 0.089 | 0 | 0.3 | 0.14 | 0.35 | 0.03 | 0.4 | 0.18 |
| 7 | 35562860 | rs343072 | 0.186 | -1.776 | 0.283 | 0.62 | 0.01 | -0.006 | -0.006 | 0 | 0.77 | 0.84 | 0.84 | 0.74 | 0.81 | 0.81 |
| 7 | 35564882 | rs343075 | 0.443 | -1.707 | 0.523 | 0.934 | -0.003 | 0.021 | 0.021 | 0.676 | 0.85 | 0.85 | 0.88 | 0.63 | 0.89 | 0.88 |
| 7 | 35566488 | rs13438729 | 0.918 | -1.699 | 0.583 | 0.914 | -0.008 | -0.027 | -0.027 | 0 | 0.07 | 0.12 | 0.04 | 0.01 | 0.04 | 0.03 |
| 7 | 35567145 | rs17765842 | -0.245 | -1.971 | 0.605 | 0.856 | 0.089 | -0.075 | -0.075 | 0.032 | 0.33 | 0.12 | 0.29 | 0.48 | 0.19 | 0.49 |
| 7 | 35572955 | rs12111746 | -0.501 | -1.589 | 1.189 | 0.64 | 0.001 | 0.057 | 0.057 | 0 | 0.18 | 0.38 | 0.2 | 0.26 | 0.17 | 0.25 |
| 7 | 35576505 | rs6462611 | -2.041 | -1.498 | 0.725 | 0.264 | -0.111 | 0.199 | 0.199 | 0 | 0.63 | 0.91 | 0.61 | 0.97 | 0.5 | 0.78 |
| 7 | 35576641 | rs7794457 | 0.282 | -1.528 | 1.547 | 0.586 | 0.013 | 0.056 | 0.056 | 0 | 0.11 | 0.39 | 0.13 | 0.12 | 0.11 | 0.17 |
| 7 | 35576991 | rs6962286 | -0.687 | -1.391 | 1.605 | 0.8 | 0 | 0 | 0 | 0.205 | 0.43 | 0.57 | 0.39 | 0.3 | 0.37 | 0.61 |
| 7 | 35581851 | rs17171472 | -1.141 | -0.734 | 1.993 | 0.618 | 0.051 | 0.033 | 0.033 | 0.034 | 0.16 | 0.39 | 0.17 | 0.21 | 0.17 | 0.1 |
| 7 | 35584937 | rs6967110 | -1.025 | -0.452 | 1.968 | 0.73 | 0.144 | 0.154 | 0.154 | 0.48 | 0.06 | 0.3 | 0.1 | 0.18 | 0.11 | 0.07 |
| 7 | 35585669 | rs17171480 | -0.697 | -0.443 | 1.921 | 0.732 | 0.105 | 0.118 | 0.118 | 0.064 | 0.06 | 0.21 | 0.09 | 0.18 | 0.11 | 0.07 |

| | | | | | | | | | | | | | | | | |
|---|---|---|---|---|---|---|---|---|---|---|---|---|---|---|---|---|
| 7 | 35586206 | rs6954895 | -0.355 | -0.947 | 2.004 | 0.449 | 0.001 | -0.015 | -0.015 | 0.124 | 0.37 | 0.37 | 0.28 | 0.06 | 0.25 | 0.36 |
| 7 | 35590308 | rs918880 | -0.712 | -0.922 | 1.395 | 0.489 | 0.001 | 0.007 | 0.007 | 0 | 0.43 | 0.57 | 0.36 | 0.24 | 0.36 | 0.43 |
| 7 | 35590780 | rs1610078 | -1.13 | -0.821 | 0.798 | 0.618 | -0.005 | 0.062 | 0.062 | 0 | 0.56 | 0.67 | 0.6 | 0.77 | 0.44 | 0.74 |
| 7 | 35592532 | rs10265961 | -0.72 | -0.865 | 0.534 | 0.576 | 0.021 | 0.109 | 0.109 | 0 | 0.44 | 0.79 | 0.36 | 0.21 | 0.34 | 0.4 |
| 7 | 35596241 | rs12540338 | -0.68 | -0.832 | 1.151 | 0.657 | -0.004 | 0.014 | 0.014 | 0 | 0.19 | 0.16 | 0.28 | 0.57 | 0.14 | 0.36 |
| 7 | 35596304 | rs10951449 | -0.993 | -0.822 | 1.06 | 0.657 | 0.001 | 0.152 | 0.152 | 0 | 0.21 | 0.49 | 0.33 | 0.57 | 0.21 | 0.37 |
| 7 | 35605025 | rs6979267 | 0.077 | -0.834 | 0.973 | 0.684 | 0.003 | 0.019 | 0.019 | 0 | 0.15 | 0.12 | 0.26 | 0.53 | 0.11 | 0.35 |
| 7 | 35608415 | rs7794286 | -0.247 | -0.862 | 0.998 | 0.589 | -0.013 | -0.023 | -0.023 | 0 | 0.38 | 0.44 | 0.29 | 0.16 | 0.25 | 0.39 |
| 7 | 35609340 | rs10238969 | -0.969 | -0.765 | 1.073 | 0.646 | 0.006 | 0.007 | 0.007 | 0 | 0.53 | 0.56 | 0.54 | 0.7 | 0.36 | 0.74 |
| 7 | 35610162 | rs1003247 | 0.029 | -0.714 | 0.417 | 0.537 | 0.029 | 0.046 | 0.046 | 0 | 0.34 | 0.23 | 0.23 | 0.17 | 0.24 | 0.42 |
| 7 | 35611107 | rs7456674 | -0.074 | -0.8 | -0.295 | 0.536 | -0.001 | -0.014 | -0.014 | 0 | 0.37 | 0.61 | 0.29 | 0.15 | 0.31 | 0.43 |
| 7 | 35613260 | rs10226256 | -1.024 | -0.329 | -1.122 | 0.557 | -0.097 | 0.05 | 0.05 | 0 | 0.59 | 0.92 | 0.61 | 0.72 | 0.45 | 0.77 |
| 7 | 35616053 | rs35977386 | -0.399 | -0.684 | -0.509 | 1.149 | 0.015 | -0.026 | -0.026 | 0 | 0.18 | 0.11 | 0.29 | 0.54 | 0.12 | 0.33 |
| 7 | 35616998 | rs34293086 | -0.17 | -0.663 | 0.26 | 0.618 | -0.014 | -0.038 | -0.038 | 0 | 0.3 | 0.42 | 0.24 | 0.15 | 0.23 | 0.37 |
| 7 | 35625326 | rs17171562 | -1.205 | -1.057 | 0.93 | 0.401 | -0.009 | -0.003 | -0.003 | 0 | 0.34 | 0.57 | 0.44 | 0.74 | 0.26 | 0.58 |
| 7 | 35626264 | rs12113684 | 0.255 | -1.033 | 0.922 | 0.401 | 0.01 | 0.069 | 0.069 | 0 | 0.07 | 0.19 | 0.1 | 0.14 | 0.06 | 0.23 |
| 7 | 35626559 | rs17678399 | 0.133 | -1.528 | 0.52 | 0.146 | 0.011 | 0.159 | 0.159 | 0 | 0.25 | 0.02 | 0.12 | 0.01 | 0.17 | 0.12 |
| 7 | 35640089 | rs12701439 | 0.33 | -1.509 | 0.917 | 0.343 | -0.033 | 0.051 | 0.051 | 0 | 0.71 | 0.61 | 0.66 | 0.42 | 0.8 | 0.64 |
| 7 | 35646526 | rs4640967 | 0.976 | -1.471 | 0.529 | 0.171 | 0.03 | 0.017 | 0.017 | 0.265 | 0.63 | 0.31 | 0.56 | 0.24 | 0.75 | 0.39 |
| 7 | 35650563 | rs11762428 | 1.939 | -1.538 | 0.076 | 0.112 | -0.057 | 0.033 | 0.033 | 0 | 0.38 | 0.09 | 0.43 | 0.24 | 0.57 | 0.26 |
| 7 | 35650705 | rs12701440 | 0.201 | -1.341 | 0.187 | 0.172 | 0.018 | 0.124 | 0.124 | 0 | 0.24 | 0.06 | 0.12 | 0 | 0.18 | 0.13 |
| 7 | 35654476 | rs10156082 | 1.575 | -1.097 | 0.299 | 0.307 | -0.029 | 0.06 | 0.06 | 0.355 | 0.4 | 0.13 | 0.43 | 0.23 | 0.57 | 0.26 |
| 7 | 35659575 | rs17616719 | 0.01 | -0.711 | 0.231 | 0.673 | -0.063 | -0.021 | -0.021 | 0.944 | 0.18 | 0.03 | 0.18 | 0.22 | 0.22 | 0.27 |
| 7 | 35660676 | rs4720180 | 0.885 | -0.308 | 0.286 | 0.605 | 0.067 | -0.004 | -0.004 | 0 | 0.54 | 0.2 | 0.55 | 0.45 | 0.65 | 0.53 |
| 7 | 35673799 | rs2305333 | -1.436 | -0.236 | 0.855 | 0.487 | -0.008 | 0.03 | 0.03 | 0 | 0.57 | 0.7 | 0.6 | 0.73 | 0.54 | 0.65 |
| 7 | 35678214 | rs3816422 | -1.653 | -0.253 | 0.861 | 0.479 | -0.049 | 0.043 | 0.043 | 0 | 0.59 | 0.87 | 0.61 | 0.73 | 0.54 | 0.67 |
| 7 | 35679425 | rs2060725 | 0.292 | -0.218 | 0.814 | 0.664 | -0.034 | -0.036 | -0.036 | 1.035 | 0.1 | 0.03 | 0.17 | 0.26 | 0.14 | 0.22 |
| 7 | 35682763 | rs1562836 | 0.292 | -0.236 | 0.914 | 0.656 | -0.037 | -0.04 | -0.04 | 0.076 | 0.1 | 0.03 | 0.17 | 0.26 | 0.14 | 0.22 |
| 7 | 35684511 | rs4142249 | -0.118 | -0.464 | 1.146 | 0.789 | -0.013 | 0.038 | 0.038 | 0 | 0.09 | 0.13 | 0.09 | 0.2 | 0.05 | 0.24 |
| 7 | 35686898 | rs13235590 | -0.307 | -0.591 | 1.08 | 0.679 | -0.066 | 0.056 | 0.056 | 0 | 0.34 | 0.09 | 0.31 | 0.26 | 0.37 | 0.33 |
| 7 | 35688370 | rs6462620 | -1.992 | -0.59 | 1.069 | 0.686 | -0.049 | 0.043 | 0.043 | 0 | 0.59 | 0.87 | 0.6 | 0.73 | 0.54 | 0.67 |
| 7 | 35690812 | rs2392376 | -1.261 | -0.494 | 1.027 | 0.733 | -0.001 | -0.001 | -0.001 | 0.112 | 0.48 | 0.57 | 0.5 | 0.54 | 0.49 | 0.42 |
| 7 | 35697108 | rs10275291 | -0.38 | -0.498 | 1.079 | 0.731 | -0.041 | 0.055 | 0.055 | 0 | 0.34 | 0.12 | 0.32 | 0.26 | 0.37 | 0.33 |
| 7 | 35706033 | rs10486674 | -0.338 | -0.498 | 1.08 | 0.731 | -0.052 | 0.056 | 0.056 | 0 | 0.34 | 0.09 | 0.31 | 0.26 | 0.37 | 0.33 |
| 7 | 35706058 | rs7779075 | 0.077 | -0.499 | 1.422 | 0.734 | -0.01 | 0.069 | 0.069 | 0 | 0.13 | 0.28 | 0.18 | 0.27 | 0.12 | 0.09 |
| 7 | 35709842 | rs3779234 | -0.366 | -0.489 | 1.235 | 0.73 | -0.041 | 0.055 | 0.055 | 0 | 0.34 | 0.12 | 0.32 | 0.26 | 0.37 | 0.33 |
| 7 | 35730645 | rs7793627 | -0.317 | -0.481 | 1.641 | 1.103 | -0.009 | 0.103 | 0.103 | 0 | 0.13 | 0.34 | 0.19 | 0.27 | 0.12 | 0.09 |
| 7 | 35732879 | rs2392382 | -1.128 | -0.472 | 1.599 | 1.069 | -0.007 | 0.004 | 0.004 | 0 | 0.43 | 0.27 | 0.4 | 0.46 | 0.42 | 0.57 |
| 7 | 35747325 | rs10266182 | -0.589 | -0.359 | 1.374 | 0.967 | -0.015 | 0.032 | 0.032 | 0 | 0.35 | 0.26 | 0.33 | 0.26 | 0.37 | 0.33 |
| 7 | 35751866 | rs10264379 | -1.201 | -0.311 | 0.994 | 0.912 | -0.001 | -0.001 | -0.001 | 0 | 0.48 | 0.53 | 0.51 | 0.53 | 0.49 | 0.42 |
| 7 | 35760229 | rs2060726 | -0.223 | -0.32 | 0.922 | 0.598 | -0.023 | 0.063 | 0.063 | 0 | 0.19 | 0.08 | 0.12 | 0 | 0.2 | 0.08 |
| 7 | 35764030 | rs6462623 | -0.556 | -0.451 | 1.687 | 0.833 | 0.028 | 0.217 | 0.217 | 0 | 0.11 | 0.46 | 0.13 | 0.26 | 0.1 | 0.08 |
| 7 | 35765932 | rs345397 | 1.932 | -0.643 | 1.666 | 0.892 | 0.013 | -0.028 | -0.028 | 0 | 0.6 | 0.27 | 0.65 | 0.51 | 0.61 | 0.63 |
| 7 | 35766263 | rs345395 | 1.114 | -0.626 | 1.535 | 0.855 | 0 | 0.034 | 0.034 | 0 | 0.71 | 0.76 | 0.77 | 0.87 | 0.72 | 0.71 |
| 7 | 35821504 | rs17767309 | -1.343 | -0.697 | 1.689 | 0.924 | 0.009 | -0.01 | -0.01 | 0 | 0.11 | 0.07 | 0.1 | 0.14 | 0.08 | 0.22 |
| 7 | 35830920 | rs7792246 | -1.319 | -0.691 | 1.8 | 0.919 | 0.005 | -0.008 | -0.008 | 0 | 0.32 | 0.75 | 0.26 | 0.35 | 0.31 | 0.16 |
| 7 | 35856339 | rs2893514 | -1.109 | -0.66 | 2.302 | 0.921 | 0.002 | 0.017 | 0.017 | 0.272 | 0.29 | 0.37 | 0.23 | 0.38 | 0.31 | 0.17 |
| 7 | 35860229 | rs10231322 | -1.867 | -0.728 | 2.317 | 0.884 | -0.029 | -0.053 | -0.053 | 0 | 0.13 | 0.3 | 0.12 | 0.01 | 0.08 | 0.21 |
| 7 | 35862309 | rs10214943 | -2.32 | -0.734 | 2.312 | 0.893 | 0.024 | -0.073 | -0.073 | 0 | 0.44 | 0.91 | 0.37 | 0.48 | 0.39 | 0.39 |
| 7 | 35863480 | rs12701460 | -0.2 | -0.64 | 2.416 | 0.935 | 0.004 | 0.107 | 0.107 | 0 | 0.09 | 0.23 | 0.11 | 0.25 | 0.1 | 0.08 |
| 7 | 35884997 | rs6462629 | -0.711 | -0.572 | 2.248 | 0.769 | 0.003 | 0.065 | 0.065 | 0.068 | 0.3 | 0.46 | 0.24 | 0.47 | 0.32 | 0.18 |
| 7 | 35920755 | rs9638914 | -0.738 | -0.56 | 2.237 | 0.758 | 0.002 | 0.016 | 0.016 | 0.074 | 0.29 | 0.37 | 0.23 | 0.38 | 0.31 | 0.17 |
| 7 | 35928645 | rs17172038 | -1.555 | -0.422 | 2.28 | 0.52 | -0.009 | 0.007 | 0.007 | 0 | 0.08 | 0.17 | 0.11 | 0.09 | 0.08 | 0.21 |
| 7 | 36004416 | rs6961681 | -2.07 | 0.229 | 2.072 | 0.445 | 0.085 | 0.022 | 0.022 | 0 | 0.06 | 0.33 | 0.04 | 0 | 0.03 | 0.03 |
| 7 | 36031972 | rs2717999 | -2.148 | 0.203 | 2.275 | 0.685 | 0.046 | 0.087 | 0.087 | 1.974 | 0.07 | 0.63 | 0.07 | 0.18 | 0.05 | 0.03 |
| 7 | 36053412 | rs6462639 | 0.528 | 0.991 | 1.091 | -0.39 | 0.014 | -0.108 | -0.108 | 0 | 0.71 | 0.37 | 0.69 | 1 | 0.65 | 0.79 |
| 7 | 36060956 | rs4723457 | 0.427 | 0.846 | 1.373 | -0.654 | 0.025 | -0.163 | -0.163 | 0 | 0.69 | 0.48 | 0.7 | 0.99 | 0.65 | 0.8 |
| 7 | 36061785 | rs7793728 | 0.238 | 0.872 | -0.185 | -0.274 | -0.014 | -0.034 | -0.034 | 0.322 | 0.24 | 0.16 | 0.2 | 0.37 | 0.28 | 0.37 |
| 7 | 36067466 | rs17172239 | 1.214 | 0.178 | -0.965 | -0.289 | -0.015 | -0.02 | -0.02 | 0 | 0.26 | 0.04 | 0.27 | 0.21 | 0.26 | 0.29 |
| 7 | 36070599 | rs13246428 | -0.275 | 0.63 | -0.032 | -0.339 | -0.007 | -0.006 | -0.006 | 3.014 | 0.75 | 0.82 | 0.74 | 1 | 0.71 | 0.85 |
| 7 | 36073321 | rs2718023 | 0.261 | 0.331 | 0.005 | 0.238 | -0.008 | 0.009 | 0.009 | 0 | 0.65 | 0.72 | 0.68 | 0.69 | 0.62 | 0.8 |
| 7 | 36073889 | rs2392412 | 1.099 | 0.398 | 0.18 | 0.194 | 0.005 | -0.008 | -0.008 | 0 | 0.33 | 0.25 | 0.41 | 0.46 | 0.25 | 0.55 |
| 7 | 36076406 | rs6970692 | 0.452 | 0.668 | 0.428 | 0.054 | -0.001 | 0.003 | 0.003 | 0 | 0.42 | 0.48 | 0.51 | 0.49 | 0.39 | 0.58 |
| 7 | 36079060 | rs6957831 | 1.145 | 0.637 | 0.234 | 0.323 | 0.003 | -0.004 | -0.004 | 0 | 0.34 | 0.28 | 0.41 | 0.43 | 0.25 | 0.55 |
| 7 | 36079709 | rs2718021 | -1.431 | 0.513 | 0.776 | 0.787 | 0.002 | -0.003 | -0.003 | 0 | 0.44 | 0.66 | 0.33 | 0.53 | 0.42 | 0.3 |
| 7 | 36083679 | rs2710803 | -0.301 | 0.358 | 0.493 | 0.643 | 0.009 | 0.026 | 0.026 | 0 | 0.25 | 0.16 | 0.16 | 0.22 | 0.23 | 0.27 |
| 7 | 36084529 | rs2710804 | -1.126 | 0.294 | 0.417 | 0.582 | -0.044 | 0.094 | 0.094 | 0 | 0.33 | 0.09 | 0.23 | 0.21 | 0.38 | 0.28 |
| 7 | 36084910 | rs2700936 | -1.608 | 0.467 | 0.773 | 0.716 | -0.004 | -0.003 | -0.003 | 0 | 0.43 | 0.39 | 0.34 | 0.54 | 0.46 | 0.35 |
| 7 | 36094146 | rs2710807 | -0.379 | 0.496 | 0.314 | 0.33 | 0.019 | 0.041 | 0.041 | 0 | 0.24 | 0.08 | 0.15 | 0.21 | 0.22 | 0.24 |

| | | | | | | | | | | | | | | | | |
|---|---|---|---|---|---|---|---|---|---|---|---|---|---|---|---|---|
| 7 | 36103289 | rs1806468 | 0.117 | 0.386 | 0.36 | 0.751 | 0.003 | 0.251 | 0.251 | 0 | 0.88 | 0.61 | 0.87 | 0.58 | 0.89 | 0.72 |
| 7 | 36110344 | rs12701466 | 1.737 | 0.377 | 0.604 | 0.668 | -0.023 | 0.049 | 0.049 | 0 | 0.52 | 0.11 | 0.61 | 0.34 | 0.5 | 0.41 |
| 7 | 36110383 | rs17263531 | -0.216 | 0.39 | 0.766 | 0.646 | 0.061 | -0.002 | -0.002 | 0 | 0.1 | 0.23 | 0.1 | 0.06 | 0.16 | 0.04 |
| 7 | 36112365 | rs3735405 | -0.393 | 0.498 | 0.669 | 0.715 | -0.01 | -0.023 | -0.023 | 0 | 0.27 | 0.27 | 0.15 | 0.17 | 0.23 | 0.28 |
| 7 | 36113173 | rs10243962 | 1.179 | 0.73 | -0.204 | 0.66 | 0.001 | 0.01 | 0.01 | 0 | 0.53 | 0.24 | 0.62 | 0.4 | 0.5 | 0.42 |
| 7 | 36114472 | rs10277914 | 1.194 | 0.735 | -0.202 | 0.546 | 0.002 | 0.044 | 0.044 | 0 | 0.53 | 0.24 | 0.62 | 0.34 | 0.5 | 0.41 |
| 7 | 36114743 | rs13246736 | 1.194 | 0.735 | -0.202 | 0.521 | 0.002 | 0.047 | 0.047 | 0 | 0.53 | 0.24 | 0.62 | 0.34 | 0.5 | 0.41 |
| 7 | 36116091 | rs2700945 | -0.158 | 0.161 | -0.099 | 0.559 | -0.06 | 0.107 | 0.107 | 0 | 0.26 | 0.47 | 0.2 | 0.57 | 0.18 | 0.38 |
| 7 | 36117089 | rs12701475 | 0.606 | 0.754 | -0.815 | 0.652 | 0.025 | 0.036 | 0.036 | 0.003 | 0.53 | 0.24 | 0.65 | 0.34 | 0.55 | 0.36 |
| 7 | 36117943 | rs11975227 | 0.581 | 0.9 | -1.072 | 0.675 | 0.018 | 0.053 | 0.053 | 0 | 0.5 | 0.21 | 0.63 | 0.34 | 0.53 | 0.35 |
| 7 | 36120302 | rs2718008 | 0.425 | 1.604 | -0.563 | 0.767 | 0.019 | -0.001 | -0.001 | 0.031 | 0.93 | 0.91 | 0.91 | 0.92 | 0.85 | 0.97 |
| 7 | 36122030 | rs2254363 | -0.799 | 1.323 | 0.155 | 1.551 | 0.034 | 0.329 | 0.329 | 0.017 | 0.15 | 0.43 | 0.26 | 0.51 | 0.15 | 0.17 |
| 7 | 36124292 | rs1986593 | 0.334 | 1.487 | -0.653 | 1.113 | -0.029 | 0.008 | 0.008 | 0.064 | 0.79 | 0.84 | 0.85 | 0.87 | 0.74 | 0.75 |
| 7 | 36124527 | rs12538868 | 0.701 | 0.854 | -1.194 | 0.236 | -0.005 | 0.006 | 0.006 | 0.073 | 0.38 | 0.34 | 0.35 | 0.31 | 0.39 | 0.37 |
| 7 | 36124622 | rs2718005 | -1.003 | 1.042 | -1.411 | 0.127 | -0.06 | -0.023 | -0.023 | 0.07 | 0.84 | 0.97 | 0.87 | 0.88 | 0.78 | 0.91 |
| 7 | 36124766 | rs2700947 | -0.994 | 1.001 | -1.415 | 0.149 | -0.079 | -0.029 | -0.029 | 0 | 0.84 | 0.99 | 0.87 | 0.88 | 0.77 | 0.91 |
| 7 | 36124844 | rs17263801 | 1.153 | 0.336 | -1.824 | -0.08 | -0.039 | 0.028 | 0.028 | 0 | 0.19 | 0.01 | 0.19 | 0.12 | 0.19 | 0.13 |
| 7 | 36124981 | rs1882059 | -1.081 | 0.497 | -1.574 | -0.015 | -0.079 | -0.032 | -0.032 | 0 | 0.84 | 0.99 | 0.87 | 0.88 | 0.77 | 0.9 |
| 7 | 36131138 | rs2718000 | -1.042 | 0.523 | -1.586 | -0.026 | -0.089 | -0.031 | -0.031 | 0 | 0.83 | 0.99 | 0.86 | 0.88 | 0.76 | 0.9 |
| 7 | 36132714 | rs17204544 | 1.21 | -0.079 | -1.695 | -0.187 | -0.048 | 0.036 | 0.036 | 0 | 0.39 | 0.18 | 0.42 | 0.3 | 0.42 | 0.43 |
| 7 | 36134316 | rs2700959 | -0.17 | -0.079 | -1.76 | -0.206 | -0.136 | -0.016 | -0.016 | 0 | 0.66 | 0.98 | 0.77 | 0.75 | 0.59 | 0.79 |
| 7 | 36134647 | rs12537099 | 0.712 | -0.218 | -1.535 | -0.066 | -0.001 | 0.051 | 0.051 | 0.098 | 0.51 | 0.29 | 0.55 | 0.64 | 0.5 | 0.68 |
| 7 | 36141552 | rs2718031 | 0.478 | -0.09 | -1.291 | -0.034 | 0.006 | -0.017 | -0.017 | 0.033 | 0.13 | 0.59 | 0.21 | 0.11 | 0.1 | 0.11 |
| 7 | 36143018 | rs7785609 | 0.223 | -0.063 | -1.354 | -0.058 | 0.016 | -0.013 | -0.013 | 0.116 | 0.71 | 0.4 | 0.65 | 0.77 | 0.65 | 0.8 |
| 7 | 36144754 | rs2893524 | 1.073 | -0.051 | -1.428 | -0.037 | -0.014 | -0.002 | -0.002 | 0 | 0.56 | 0.16 | 0.51 | 0.44 | 0.58 | 0.54 |
| 7 | 36147694 | rs1997412 | -0.807 | -0.052 | -1.361 | -0.036 | 0.015 | -0.01 | -0.01 | 0.251 | 0.88 | 0.45 | 0.79 | 0.89 | 0.9 | 0.89 |
| 7 | 36147815 | rs7783753 | 1.16 | -0.058 | -1.404 | -0.025 | -0.016 | 0.047 | 0.047 | 0 | 0.12 | 0.1 | 0.13 | 0.33 | 0.08 | 0.25 |
| 7 | 36148076 | rs4723477 | 1.05 | 0.004 | -1.49 | -0.021 | -0.008 | 0 | 0 | 0 | 0.56 | 0.17 | 0.52 | 0.44 | 0.58 | 0.54 |
| 7 | 36148652 | rs10246132 | 0.43 | 0.009 | -1.486 | -0.017 | 0.037 | -0.045 | -0.045 | 0 | 0.73 | 0.21 | 0.66 | 0.56 | 0.83 | 0.64 |
| 7 | 36149085 | rs9655368 | 0.952 | 0.114 | -1.45 | -0.025 | -0.076 | -0.015 | -0.015 | 0 | 0.17 | 0.04 | 0.14 | 0.12 | 0.25 | 0.1 |
| 7 | 36152783 | rs10951457 | 0.972 | 0.133 | -1.457 | -0.044 | -0.029 | -0.007 | -0.007 | 0.543 | 0.56 | 0.16 | 0.51 | 0.44 | 0.55 | 0.55 |
| 7 | 36155071 | rs2726051 | -0.337 | 0.267 | -1.318 | -0.09 | 0.04 | -0.032 | -0.032 | 0 | 0.86 | 0.44 | 0.77 | 0.89 | 0.85 | 0.89 |
| 7 | 36157988 | rs2392423 | 1.337 | 0.004 | -1.527 | -0.04 | -0.002 | -0.022 | -0.022 | 0 | 0.18 | 0 | 0.16 | 0.14 | 0.15 | 0.12 |
| 7 | 36167185 | rs2256964 | -0.863 | 0.012 | -0.9 | -0.04 | 0.058 | 0 | 0 | 0 | 0.88 | 0.61 | 0.8 | 0.89 | 0.88 | 0.89 |
| 7 | 36167974 | rs12540725 | 1.176 | 0.033 | -1.522 | 0.021 | -0.046 | 0.039 | 0.039 | 0.017 | 0.38 | 0.15 | 0.43 | 0.3 | 0.4 | 0.41 |
| 7 | 36169168 | rs7801365 | 0.59 | -0.021 | -1.327 | 0.21 | 0.012 | -0.015 | -0.015 | 0 | 0.14 | 0.13 | 0.13 | 0.33 | 0.09 | 0.25 |
| 7 | 36170199 | rs10951458 | 0.302 | 0.577 | -0.877 | -0.507 | 0.021 | -0.019 | -0.019 | 0.259 | 0.66 | 0.41 | 0.58 | 0.77 | 0.54 | 0.73 |
| 7 | 36170883 | rs2726070 | 0.315 | 0.618 | -1.287 | -0.831 | -0.028 | 0.032 | 0.032 | 0 | 0.41 | 0.24 | 0.45 | 0.31 | 0.43 | 0.33 |
| 7 | 36171685 | rs2700893 | 1.16 | 0.624 | -1.39 | -0.806 | -0.09 | -0.023 | -0.023 | 1.156 | 0.18 | 0.03 | 0.15 | 0.12 | 0.25 | 0.12 |
| 7 | 36171893 | rs17264903 | 0.249 | 0.54 | -1.308 | -0.776 | 0.044 | 0.026 | 0.026 | 0 | 0.44 | 0.23 | 0.37 | 0.59 | 0.36 | 0.51 |
| 7 | 36173508 | rs17264966 | 1.493 | 0.528 | -1.396 | -0.79 | -0.005 | -0.025 | -0.025 | 0 | 0.18 | 0 | 0.08 | 0.15 | 0.15 | 0.14 |
| 7 | 36177479 | rs11979422 | 0.996 | 0.527 | -1.149 | -0.691 | 0.008 | -0.067 | -0.067 | 0 | 0.24 | 0.08 | 0.29 | 0.45 | 0.21 | 0.37 |
| 7 | 36178054 | rs11975199 | 0.865 | 0.59 | -1.138 | -0.663 | -0.008 | 0.029 | 0.029 | 0.012 | 0.26 | 0.28 | 0.29 | 0.45 | 0.21 | 0.37 |
| 7 | 36178481 | rs2726115 | -0.757 | 0.638 | -1.118 | -0.707 | 0.042 | -0.055 | -0.055 | 1.231 | 0.85 | 0.46 | 0.82 | 0.9 | 0.8 | 0.84 |
| 7 | 36182488 | rs11979076 | 0.708 | 0.49 | -0.906 | -0.383 | 0.008 | -0.03 | -0.03 | 0 | 0.25 | 0.2 | 0.29 | 0.45 | 0.21 | 0.37 |
| 7 | 36183300 | rs4720196 | 0.8 | 0.506 | -0.953 | -0.422 | 0.001 | 0.004 | 0.004 | 0 | 0.23 | 0.18 | 0.29 | 0.18 | 0.18 | 0.22 |
| 7 | 36186593 | rs12669708 | 0.62 | 0.574 | -1.276 | -0.341 | 0.006 | -0.023 | -0.023 | 0.048 | 0.26 | 0.24 | 0.29 | 0.45 | 0.21 | 0.37 |
| 7 | 36188719 | rs1882062 | 0.529 | 0.842 | -1.3 | -0.367 | -0.001 | 0.034 | 0.034 | 0 | 0.22 | 0.1 | 0.28 | 0.18 | 0.18 | 0.22 |
| 7 | 36196971 | rs12154575 | 0.344 | 0.674 | -0.65 | -0.694 | -0.021 | -0.031 | -0.031 | 0 | 0.31 | 0.4 | 0.18 | 0.26 | 0.28 | 0.22 |
| 7 | 36198547 | rs2700908 | 0.162 | 0.871 | -0.804 | -1.141 | 0.003 | 0.07 | 0.07 | 0 | 0.42 | 0.33 | 0.37 | 0.11 | 0.4 | 0.3 |
| 7 | 36198976 | rs11760954 | 0.541 | 1.177 | -0.875 | -0.937 | 0.014 | 0.008 | 0.008 | 0 | 0.07 | 0.1 | 0.07 | 0.09 | 0.11 | 0.07 |
| 7 | 36199127 | rs2700909 | 0.311 | 1.122 | -0.819 | -0.966 | -0.022 | 0.165 | 0.165 | 0 | 0.4 | 0.16 | 0.36 | 0.11 | 0.39 | 0.3 |
| 7 | 36199170 | rs2726052 | 1.046 | 1.266 | -1.001 | -1.125 | -0.064 | 0.072 | 0.072 | 1.295 | 0.18 | 0.01 | 0.16 | 0.03 | 0.22 | 0.1 |
| 7 | 36201102 | rs4720197 | 0.339 | 1.321 | -0.886 | -1.031 | -0.008 | 0.018 | 0.018 | 0 | 0.3 | 0.27 | 0.27 | 0.17 | 0.28 | 0.27 |
| 7 | 36203128 | rs1420187 | -0.743 | 1.251 | -0.742 | -0.966 | 0.006 | 0.007 | 0.007 | 0 | 0.73 | 0.66 | 0.58 | 0.36 | 0.73 | 0.53 |
| 7 | 36210289 | rs10230457 | 1.033 | 1.174 | -0.887 | -1.008 | 0.013 | 0.076 | 0.076 | 0 | 0.23 | 0.09 | 0.19 | 0.08 | 0.18 | 0.2 |
| 7 | 36210485 | rs6973373 | 0.506 | 1.251 | -0.472 | -0.624 | -0.01 | -0.046 | -0.046 | 0 | 0.33 | 0.43 | 0.22 | 0.25 | 0.33 | 0.23 |
| 7 | 36211043 | rs11762552 | 0.589 | 1.676 | -0.422 | 0.096 | 0.022 | 0.039 | 0.039 | 0 | 0.05 | 0.12 | 0.03 | 0.09 | 0.05 | 0.07 |
| 7 | 36211110 | rs11769392 | 0.592 | 1.679 | -0.421 | 0.093 | 0.022 | 0.041 | 0.041 | 0 | 0.05 | 0.12 | 0.03 | 0.09 | 0.05 | 0.07 |
| 7 | 36211147 | rs1420195 | -0.474 | 1.216 | -0.703 | 0.116 | -0.001 | 0.011 | 0.011 | 0 | 0.78 | 0.8 | 0.62 | 0.45 | 0.78 | 0.6 |
| 7 | 36211749 | rs9648428 | -0.268 | 1.104 | -0.705 | -0.242 | 0.001 | -0.006 | -0.006 | 0 | 0.17 | 0.23 | 0.11 | 0.14 | 0.18 | 0.08 |
| 7 | 36216625 | rs2540659 | -0.464 | 1.262 | -0.898 | -0.437 | -0.016 | 0.016 | 0.016 | 2.289 | 0.15 | 0.08 | 0.17 | 0.01 | 0.21 | 0.06 |
| 7 | 36219439 | rs3021413 | -0.306 | 1.311 | -1.059 | -0.428 | -0.07 | 0.08 | 0.08 | 3.667 | 0.14 | 0.01 | 0.17 | 0.01 | 0.21 | 0.06 |
| 7 | 36219846 | rs16880939 | 0.39 | 0.935 | -0.838 | -0.693 | -0.001 | -0.023 | -0.023 | 0 | 0.11 | 0.12 | 0.16 | 0.01 | 0.09 | 0.11 |
| 7 | 36229407 | rs2726057 | 0.256 | 0.219 | -0.109 | -0.348 | -0.029 | 0.109 | 0.109 | 0.013 | 0.86 | 0.5 | 0.9 | 0.79 | 0.93 | 0.81 |
| 7 | 36230630 | rs17272724 | 0.943 | 0.092 | -1.25 | -0.514 | -0.11 | 0 | 0 | 0 | 0.33 | 0.08 | 0.3 | 0.27 | 0.44 | 0.29 |
| 7 | 36235099 | rs2726101 | -0.647 | -0.372 | -0.416 | -0.554 | -0.013 | -0.052 | -0.052 | 0 | 0.77 | 0.65 | 0.86 | 0.88 | 0.79 | 0.84 |
| 7 | 36241762 | rs888046 | 1.18 | -0.621 | -0.724 | -0.178 | -0.035 | 0.003 | 0.003 | 0 | 0.4 | 0.27 | 0.48 | 0.57 | 0.49 | 0.5 |
| 7 | 36243435 | rs10951463 | 1.438 | -0.595 | -0.695 | -0.183 | 0.005 | -0.027 | -0.027 | 0 | 0.09 | 0.04 | 0.2 | 0.2 | 0.08 | 0.07 |

| | | | | | | | | | | | | | | | | |
|---|---|---|---|---|---|---|---|---|---|---|---|---|---|---|---|---|
| 7 | 36244583 | rs6960355 | 0.223 | -0.632 | -0.207 | -0.297 | -0.006 | -0.074 | -0.074 | 1.141 | 0.58 | 0.49 | 0.72 | 0.87 | 0.6 | 0.71 |
| 7 | 36245577 | rs2726092 | 0.368 | -0.735 | -0.31 | -0.267 | -0.031 | 0.107 | 0.107 | 0.839 | 0.2 | 0.04 | 0.15 | 0.01 | 0.22 | 0.15 |
| 7 | 36252293 | rs17170531 | -0.098 | -0.811 | -0.099 | 0.037 | -0.029 | 0.017 | 0.017 | 0 | 0.68 | 0.47 | 0.79 | 0.6 | 0.77 | 0.65 |
| 7 | 36253012 | rs10277506 | 0.915 | -0.737 | -0.632 | 0.008 | 0.01 | -0.015 | -0.015 | 0 | 0.11 | 0.05 | 0.08 | 0.28 | 0.1 | 0.25 |
| 7 | 36254575 | rs4723496 | 1.252 | -0.725 | -0.487 | -0.256 | 0.018 | 0.004 | 0.004 | 0 | 0.2 | 0.12 | 0.09 | 0.22 | 0.13 | 0.12 |
| 7 | 36261771 | rs2726118 | 1.56 | -0.54 | -0.705 | -0.318 | -0.047 | 0.068 | 0.068 | 3.773 | 0.12 | 0.01 | 0.12 | 0.02 | 0.15 | 0.05 |
| 7 | 36265912 | rs13231204 | 1.463 | -0.452 | -0.663 | -0.162 | -0.016 | -0.047 | -0.047 | 1.117 | 0.09 | 0.01 | 0.19 | 0.18 | 0.09 | 0.08 |
| 7 | 36268815 | rs2392430 | 0.203 | -0.495 | -0.609 | -0.037 | 0.005 | -0.015 | -0.015 | 0 | 0.13 | 0.07 | 0.11 | 0.39 | 0.14 | 0.32 |
| 7 | 36269284 | rs2098368 | 0.691 | -0.595 | -0.63 | -0.23 | -0.01 | 0.046 | 0.046 | 0 | 0.45 | 0.19 | 0.6 | 0.35 | 0.54 | 0.42 |
| 7 | 36281897 | rs4723502 | 0.253 | -0.232 | -0.106 | -0.117 | 0.001 | 0.011 | 0.011 | 0.27 | 0.74 | 0.74 | 0.78 | 0.6 | 0.72 | 0.63 |
| 7 | 36285305 | rs196543 | -0.189 | -0.249 | -0.096 | -0.11 | -0.015 | 0.127 | 0.127 | 0.559 | 0.87 | 0.79 | 0.91 | 0.62 | 0.9 | 0.7 |
| 7 | 36287719 | rs4723504 | 0.267 | -0.496 | 0.065 | -0.252 | 0 | -0.001 | -0.001 | 0.06 | 0.29 | 0.26 | 0.3 | 0.4 | 0.22 | 0.2 |
| 7 | 36288575 | rs11762654 | 0.857 | -0.42 | 0.025 | -0.226 | -0.007 | 0.026 | 0.026 | 0 | 0.45 | 0.41 | 0.48 | 0.2 | 0.5 | 0.43 |
| 7 | 36301091 | rs986875 | 0.717 | -0.535 | -0.086 | 0.056 | -0.011 | 0.071 | 0.071 | 0.621 | 0.11 | 0.14 | 0.05 | 0.35 | 0.07 | 0.26 |
| 7 | 36304691 | rs17208592 | 1.375 | -0.538 | -0.103 | -0.159 | 0.014 | -0.003 | -0.003 | 1.497 | 0.17 | 0.12 | 0.1 | 0.22 | 0.12 | 0.13 |
| 7 | 36308589 | rs196575 | 0.393 | -0.582 | 0.03 | -0.233 | -0.03 | 0.004 | 0.004 | 0.064 | 0.63 | 0.54 | 0.66 | 0.58 | 0.74 | 0.68 |
| 7 | 36312929 | rs196579 | 1.393 | -0.492 | -0.049 | -0.337 | -0.027 | 0.037 | 0.037 | 4.571 | 0.14 | 0.08 | 0.13 | 0.02 | 0.18 | 0.07 |
| 7 | 36314442 | rs11765384 | 0.857 | -0.514 | 0.001 | -0.329 | 0.005 | 0.018 | 0.018 | 0 | 0.49 | 0.4 | 0.53 | 0.54 | 0.56 | 0.61 |
| 7 | 36316891 | rs196583 | -0.216 | -0.463 | -0.04 | -0.146 | -0.016 | 0.023 | 0.023 | 0 | 0.72 | 0.67 | 0.67 | 0.57 | 0.79 | 0.76 |
| 7 | 36317696 | rs196584 | -1.195 | -0.376 | -0.659 | -0.102 | 0 | 0.005 | 0.005 | 0 | 0.9 | 0.9 | 0.79 | 0.79 | 0.9 | 0.9 |
| 7 | 36319007 | rs11764972 | 1.787 | -0.324 | -0.935 | -0.482 | -0.102 | 0.005 | 0.005 | 0 | 0.23 | 0.03 | 0.23 | 0.16 | 0.31 | 0.11 |
| 7 | 36319026 | rs10244327 | 0.783 | -0.538 | -0.273 | -0.153 | -0.01 | -0.036 | -0.036 | 0.033 | 0.09 | 0.12 | 0.02 | 0 | 0.05 | 0.08 |
| 7 | 36319676 | rs17274649 | -0.104 | -0.562 | -0.27 | -0.151 | 0.018 | 0.013 | 0.013 | 0 | 0.26 | 0.23 | 0.11 | 0.22 | 0.16 | 0.21 |
| 7 | 36322615 | rs885638 | 0.873 | -0.256 | -0.69 | 0.124 | 0.026 | -0.008 | -0.008 | 0 | 0.17 | 0.1 | 0.09 | 0.23 | 0.12 | 0.14 |
| 7 | 36327317 | rs196594 | 1.576 | 0.233 | -0.699 | -0.038 | -0.04 | 0.028 | 0.028 | 7.893 | 0.06 | 0.01 | 0.07 | 0.02 | 0.1 | 0.04 |
| 7 | 36327410 | rs196595 | -0.725 | -0.172 | -0.331 | -0.096 | 0.001 | 0.001 | 0.001 | 0 | 0.44 | 0.46 | 0.42 | 0.79 | 0.33 | 0.53 |
| 7 | 36331782 | rs196604 | -0.439 | -0.067 | -0.528 | 0.022 | -0.001 | -0.007 | -0.007 | 0 | 0.34 | 0.27 | 0.32 | 0.42 | 0.27 | 0.25 |
| 7 | 36333687 | rs17275193 | 0.445 | 0.081 | -0.842 | -0.077 | 0.025 | 0.039 | 0.039 | 0 | 0.45 | 0.31 | 0.45 | 0.64 | 0.48 | 0.63 |
| 7 | 36335759 | rs10262791 | 0.542 | 0.503 | -0.61 | -0.274 | 0.01 | -0.059 | -0.059 | 0 | 0.31 | 0.41 | 0.27 | 0.12 | 0.31 | 0.16 |
| 7 | 36338589 | rs11773889 | 1.779 | 1.059 | -0.741 | -0.098 | -0.073 | -0.001 | -0.001 | 0 | 0.16 | 0.03 | 0.2 | 0.13 | 0.23 | 0.11 |
| 7 | 36338967 | rs2241142 | 1.036 | 1.08 | -0.289 | 0.233 | -0.025 | -0.04 | -0.04 | 0 | 0.07 | 0.01 | 0.16 | 0.19 | 0.06 | 0.07 |
| 7 | 36339126 | rs10848 | 1.054 | 0.864 | -0.419 | 0.339 | -0.019 | 0.037 | 0.037 | 0.586 | 0.51 | 0.2 | 0.43 | 0.43 | 0.48 | 0.3 |
| 7 | 36339630 | rs13224904 | 0.949 | 0.863 | -0.407 | 0.298 | -0.01 | -0.004 | -0.004 | 0.247 | 0.52 | 0.19 | 0.55 | 0.6 | 0.5 | 0.33 |
| 7 | 36339985 | rs7803174 | 0.896 | 0.992 | -0.195 | 0.38 | 0.024 | -0.047 | -0.047 | 0 | 0.26 | 0.14 | 0.27 | 0.33 | 0.27 | 0.51 |
| 7 | 36340134 | rs196618 | -0.559 | 1.047 | 0.143 | 0.291 | 0.044 | -0.057 | -0.057 | 0.007 | 0.88 | 0.63 | 0.91 | 0.97 | 0.87 | 0.89 |
| 7 | 36340308 | rs196619 | -0.559 | 1.047 | 0.143 | 0.291 | 0.044 | -0.057 | -0.057 | 0 | 0.88 | 0.63 | 0.91 | 0.97 | 0.87 | 0.89 |
| 7 | 36340675 | rs1046455 | 1.379 | 1.018 | -0.863 | 0.415 | -0.041 | 0.024 | 0.024 | 0.239 | 0.46 | 0.11 | 0.39 | 0.4 | 0.44 | 0.26 |
| 7 | 36340811 | rs3735402 | 0.212 | 0.68 | -0.53 | 0.643 | 0 | -0.001 | -0.001 | 0.394 | 0.28 | 0.29 | 0.33 | 0.35 | 0.33 | 0.52 |
| 7 | 36345194 | rs4723511 | 0.329 | 0.763 | -1.13 | 0.672 | 0.022 | -0.028 | -0.028 | 0 | 0.28 | 0.13 | 0.3 | 0.33 | 0.31 | 0.51 |
| 7 | 36345343 | rs196631 | -0.421 | 1.277 | -0.959 | 0.447 | 0.014 | -0.011 | -0.011 | 2.611 | 0.9 | 0.85 | 0.9 | 0.97 | 0.85 | 0.91 |
| 7 | 36352488 | rs741299 | -0.481 | 0.985 | -1.008 | 0.552 | 0.009 | 0.006 | 0.006 | 0.173 | 0.51 | 0.66 | 0.56 | 0.55 | 0.48 | 0.69 |
| 7 | 36353857 | rs10234940 | 0.221 | 0.785 | -2.311 | 0.304 | 0.019 | -0.001 | -0.001 | 0.034 | 0.49 | 0.24 | 0.41 | 0.5 | 0.48 | 0.39 |
| 7 | 36356324 | rs12531662 | 1.23 | 0.662 | -2.842 | 0.484 | -0.027 | -0.043 | -0.043 | 0 | 0.12 | 0.01 | 0.16 | 0.18 | 0.09 | 0.15 |
| 7 | 36356661 | rs196646 | 0.412 | 0.501 | -3.096 | 0.174 | 0.015 | 0.082 | 0.082 | 0 | 0.31 | 0.16 | 0.3 | 0.1 | 0.31 | 0.37 |
| 7 | 36374799 | rs3779230 | 1.625 | 0.475 | -2.966 | 0.18 | -0.028 | -0.037 | -0.037 | 0.168 | 0.11 | 0 | 0.16 | 0.17 | 0.09 | 0.14 |
| 7 | 36382183 | rs197318 | 0.141 | 0.366 | -2.947 | 0.434 | 0.017 | -0.007 | -0.007 | 0 | 0.33 | 0.17 | 0.31 | 0.33 | 0.33 | 0.54 |
| 7 | 36387944 | rs197324 | -0.531 | 0.401 | -3.075 | 0.447 | 0.022 | 0.007 | 0.007 | 0 | 0.46 | 0.3 | 0.5 | 0.52 | 0.46 | 0.65 |
| 7 | 36396358 | rs197328 | -0.586 | 0.443 | -3.038 | 0.493 | 0.031 | -0.001 | -0.001 | 1.474 | 0.47 | 0.23 | 0.5 | 0.49 | 0.45 | 0.64 |
| 7 | 36405894 | rs197334 | 0.333 | 0.444 | -3.03 | 0.464 | 0.015 | -0.016 | -0.016 | 0 | 0.33 | 0.12 | 0.3 | 0.31 | 0.32 | 0.53 |
| 7 | 36421494 | rs197345 | 0.495 | 0.553 | -3.125 | 0.573 | 0.008 | -0.041 | -0.041 | 1.516 | 0.31 | 0.1 | 0.29 | 0.31 | 0.3 | 0.52 |
| 7 | 36427461 | rs6976779 | 1.125 | 0.63 | -3.09 | 0.311 | 0.005 | -0.01 | -0.01 | 0 | 0.39 | 0.54 | 0.3 | 0.29 | 0.42 | 0.21 |
| 7 | 36430069 | rs197354 | -0.624 | 0.646 | -3.004 | 0.422 | 0.028 | -0.018 | -0.018 | 0 | 0.45 | 0.23 | 0.49 | 0.49 | 0.43 | 0.64 |
| 7 | 36436954 | rs1419985 | 0.379 | 0.963 | -2.579 | 0.825 | 0 | 0.013 | 0.013 | 0.641 | 0.15 | 0.16 | 0.21 | 0.4 | 0.15 | 0.27 |
| 7 | 36442113 | rs10235765 | 1.266 | 0.986 | -3.02 | 0.735 | -0.007 | -0.006 | -0.006 | 0 | 0.06 | 0.12 | 0.06 | 0.04 | 0.06 | 0.03 |
| 7 | 36445335 | rs1362269 | -0.573 | 1.306 | -3.259 | 0.647 | -0.003 | -0.032 | -0.032 | 0 | 0.25 | 0.2 | 0.28 | 0.4 | 0.26 | 0.31 |
| 7 | 36449637 | rs197368 | -1.539 | 1.283 | -3.591 | 0.386 | 0 | 0.005 | 0.005 | 0.01 | 0.62 | 0.41 | 0.63 | 0.72 | 0.63 | 0.79 |
| 7 | 36451360 | rs3801314 | -1.124 | 1.343 | -3.562 | 0.468 | -0.003 | 0.003 | 0.003 | 0 | 0.4 | 0.34 | 0.49 | 0.61 | 0.4 | 0.46 |
| 7 | 36454261 | rs3801310 | 1.165 | 1.439 | -3.589 | 0.605 | 0 | 0.072 | 0.072 | 0 | 0.11 | 0.12 | 0.19 | 0.4 | 0.12 | 0.28 |
| 7 | 36459076 | rs17213431 | 1.654 | 1.396 | -3.693 | 0.336 | -0.025 | 0.063 | 0.063 | 0 | 0.14 | 0 | 0.07 | 0.03 | 0.14 | 0.13 |
| 7 | 36466043 | rs2041632 | -1.209 | 1.478 | -3.746 | 0.351 | 0 | 0.04 | 0.04 | 0.134 | 0.53 | 0.39 | 0.55 | 0.66 | 0.53 | 0.6 |
| 7 | 36466655 | rs3815484 | 1.214 | 2.062 | -3.196 | 1.247 | -0.013 | -0.032 | -0.032 | 1.006 | 0.11 | 0.04 | 0.07 | 0.22 | 0.13 | 0.18 |
| 7 | 36467608 | rs17292369 | 1.668 | 2.198 | -3.347 | 1.147 | -0.022 | -0.042 | -0.042 | 1.688 | 0.17 | 0.01 | 0.23 | 0.19 | 0.19 | 0.06 |
| 7 | 36480876 | rs3801305 | -0.655 | 0.663 | -1.915 | 2.121 | 0.004 | 0 | 0 | 0 | 0.19 | 0.13 | 0.22 | 0.21 | 0.16 | 0.15 |
| 7 | 36488301 | rs9639730 | 0.084 | 1.202 | -1.514 | 2.426 | -0.026 | -0.033 | -0.033 | 1.17 | 0.13 | 0.03 | 0.16 | 0.17 | 0.11 | 0.11 |
| 7 | 36491811 | rs17170593 | 0.267 | 1.28 | -1.464 | 2.455 | 0 | 0.016 | 0.016 | 0 | 0.09 | 0.09 | 0.17 | 0.19 | 0.1 | 0.13 |
| 7 | 36492045 | rs3801302 | -1.177 | 1.282 | -1.465 | 2.463 | -0.003 | 0.001 | 0.001 | 0 | 0.2 | 0.19 | 0.25 | 0.41 | 0.23 | 0.31 |
| 7 | 36492212 | rs2052197 | 0.193 | 1.326 | -1.768 | 2.478 | -0.001 | 0.014 | 0.014 | 0 | 0.09 | 0.09 | 0.17 | 0.19 | 0.1 | 0.13 |
| 7 | 36498639 | rs1362273 | 0.177 | 1.401 | -2.189 | 2.344 | -0.013 | -0.032 | -0.032 | 0 | 0.11 | 0.04 | 0.07 | 0.22 | 0.13 | 0.18 |
| 7 | 36499921 | rs10227096 | -0.282 | 1.247 | -1.742 | 2.004 | -0.027 | -0.031 | -0.031 | 0 | 0.12 | 0.03 | 0.16 | 0.17 | 0.1 | 0.11 |

| | | | | | | | | | | | | | | | | |
|---|---|---|---|---|---|---|---|---|---|---|---|---|---|---|---|---|
| 7 | 36502623 | rs11763746 | -0.737 | 1.067 | -1.096 | 1.929 | -0.024 | -0.033 | -0.033 | 0 | 0.07 | 0 | 0.06 | 0.09 | 0.08 | 0.21 |
| 7 | 36503931 | rs1990506 | -0.869 | 0.905 | -0.698 | 1.722 | 0.005 | 0.042 | 0.042 | 0 | 0.1 | 0.14 | 0.2 | 0.23 | 0.13 | 0.17 |
| 7 | 36514874 | rs7808209 | -0.858 | 0.414 | -0.845 | 1.298 | 0.004 | -0.015 | -0.015 | 0 | 0.25 | 0.18 | 0.28 | 0.23 | 0.27 | 0.29 |
| 7 | 36514910 | rs11765557 | -0.07 | 0.467 | -0.602 | 0.494 | 0.002 | -0.022 | -0.022 | 0 | 0.14 | 0.17 | 0.1 | 0.01 | 0.17 | 0.04 |
| 7 | 36519651 | rs6954205 | 0.256 | 0.021 | -0.11 | 0.307 | -0.002 | -0.12 | -0.12 | 0 | 0.8 | 0.47 | 0.87 | 0.99 | 0.83 | 0.95 |
| 7 | 36522750 | rs10447562 | -0.876 | 0.122 | -0.412 | 1.095 | 0.007 | 0.01 | 0.01 | 0.045 | 0.28 | 0.23 | 0.32 | 0.75 | 0.28 | 0.45 |
| 7 | 36525157 | rs7777667 | -0.835 | 0.046 | -0.488 | 1.2 | 0.005 | -0.024 | -0.024 | 5.017 | 0.19 | 0.15 | 0.18 | 0.67 | 0.18 | 0.28 |
| 7 | 36529955 | rs12671968 | -1.059 | -0.167 | -0.54 | 1.136 | 0.014 | -0.024 | -0.024 | 0.022 | 0.23 | 0.18 | 0.19 | 0.66 | 0.17 | 0.34 |
| 7 | 36530132 | rs10235752 | -0.816 | -0.149 | -0.522 | 1.363 | 0.008 | 0.004 | 0.004 | 0.781 | 0.22 | 0.19 | 0.13 | 0.13 | 0.16 | 0.27 |
| 7 | 36530276 | rs6942586 | 0.929 | -0.087 | -0.556 | 0.817 | 0.022 | 0.053 | 0.053 | 0 | 0.53 | 0.3 | 0.64 | 0.29 | 0.58 | 0.53 |
| 7 | 36538581 | rs17170613 | -1.376 | -0.05 | -0.273 | 0.715 | -0.009 | -0.026 | -0.026 | 0 | 0.14 | 0.19 | 0.08 | 0.07 | 0.11 | 0.11 |
| 7 | 36546997 | rs4585649 | 0.245 | -0.108 | -1.149 | -0.24 | 0.024 | 0.091 | 0.091 | 0.293 | 0.41 | 0.19 | 0.35 | 0.78 | 0.33 | 0.4 |
| 7 | 36547245 | rs4489220 | 0.174 | -0.127 | -1.176 | -0.279 | 0.014 | 0.103 | 0.103 | 0.149 | 0.43 | 0.19 | 0.37 | 0.78 | 0.36 | 0.42 |
| 7 | 36547613 | rs7777022 | -0.319 | -0.269 | -0.79 | -0.347 | -0.003 | -0.028 | -0.028 | 0 | 0.09 | 0.18 | 0.07 | 0.03 | 0.08 | 0.07 |
| 7 | 36547667 | rs2392445 | 0.394 | -0.294 | -0.357 | -0.581 | -0.015 | -0.06 | -0.06 | 1.75 | 0.88 | 0.73 | 0.89 | 0.99 | 0.88 | 0.96 |
| 7 | 36547689 | rs3911827 | 0.735 | -0.477 | -1.216 | -0.573 | -0.01 | -0.036 | -0.036 | 1.298 | 0.21 | 0.12 | 0.26 | 0.61 | 0.2 | 0.22 |
| 7 | 36547889 | rs4723534 | 0.185 | -0.144 | -0.964 | -1.162 | -0.012 | 0.012 | 0.012 | 0 | 0.37 | 0.32 | 0.35 | 0.13 | 0.46 | 0.37 |
| 7 | 36547979 | rs13312060 | -0.139 | -0.037 | -1.116 | -1.033 | 0.006 | -0.029 | -0.029 | 0 | 0.6 | 0.54 | 0.63 | 0.86 | 0.52 | 0.58 |
| 7 | 36552656 | rs7986 | -0.299 | 0.475 | -1.601 | -0.631 | -0.114 | 0.094 | 0.094 | 0 | 0.3 | 0.03 | 0.32 | 0.15 | 0.37 | 0.3 |
| 7 | 36556478 | rs12531209 | -0.585 | 0.165 | -1.709 | -0.533 | -0.099 | 0.071 | 0.071 | 0 | 0.33 | 0.03 | 0.29 | 0.21 | 0.4 | 0.33 |
| 7 | 36556593 | rs17170619 | -0.91 | -0.225 | -0.938 | 0.089 | 0.009 | 0.021 | 0.021 | 0 | 0.45 | 0.29 | 0.49 | 0.39 | 0.45 | 0.49 |
| 7 | 36562244 | rs10273902 | -1.267 | -0.541 | 0.485 | 0.258 | 0.034 | 0.199 | 0.199 | 1.062 | 0.07 | 0.41 | 0.18 | 0.16 | 0.04 | 0.14 |
| 7 | 36567782 | rs3779222 | -1.338 | -0.551 | -0.034 | 0.392 | 0.01 | -0.02 | -0.02 | 1.714 | 0.08 | 0.03 | 0.12 | 0.14 | 0.07 | 0.07 |
| 7 | 36569019 | rs3801298 | 0.623 | -0.255 | 0.242 | 0.372 | -0.02 | 0.035 | 0.035 | 0 | 0.57 | 0.31 | 0.37 | 0.69 | 0.47 | 0.52 |
| 7 | 36569199 | rs3801299 | 1.058 | -0.39 | 0.016 | 0.352 | -0.016 | -0.009 | -0.009 | 0 | 0.48 | 0.1 | 0.37 | 0.56 | 0.52 | 0.49 |
| 7 | 36573313 | rs4236344 | 0.081 | -0.381 | 1.393 | 0.319 | -0.002 | 0.002 | 0.002 | 0 | 0.36 | 0.31 | 0.54 | 0.3 | 0.42 | 0.42 |
| 7 | 36576068 | rs4723537 | -0.99 | -0.307 | 1.422 | 0.33 | -0.022 | 0.001 | 0.001 | 0 | 0.44 | 0.72 | 0.65 | 0.43 | 0.52 | 0.52 |
| 7 | 36577047 | rs13307569 | 0.191 | -0.211 | 1.449 | 0.317 | -0.003 | 0.004 | 0.004 | 0 | 0.33 | 0.26 | 0.42 | 0.25 | 0.39 | 0.31 |
| 7 | 36588533 | rs10275819 | -0.587 | -0.532 | 1.185 | 0.284 | -0.025 | 0.104 | 0.104 | 0.372 | 0.42 | 0.26 | 0.49 | 0.26 | 0.45 | 0.33 |
| 7 | 36588659 | rs6973551 | -1.033 | -0.447 | 0.815 | 0.202 | -0.002 | 0.002 | 0.002 | 0 | 0.13 | 0.16 | 0.09 | 0.13 | 0.14 | 0.1 |
| 7 | 36590053 | rs12701509 | 0.265 | -0.396 | 0.476 | 0.188 | -0.151 | 0.017 | 0.017 | 0 | 0.33 | 0.02 | 0.44 | 0.25 | 0.43 | 0.33 |
| 7 | 36610493 | rs6948404 | 0.092 | -0.299 | 0.705 | 0.282 | 0.017 | 0.015 | 0.015 | 0.445 | 0.12 | 0.09 | 0.18 | 0.05 | 0.08 | 0.15 |
| 7 | 36612893 | rs11973927 | -0.248 | -0.311 | 0.78 | 0.24 | -0.007 | 0.023 | 0.023 | 0 | 0.11 | 0.16 | 0.05 | 0.12 | 0.08 | 0.09 |
| 7 | 36621071 | rs2392449 | -0.699 | -0.363 | 0.687 | 0.134 | -0.005 | 0.031 | 0.031 | 0.703 | 0.43 | 0.34 | 0.59 | 0.32 | 0.49 | 0.44 |
| 7 | 36622417 | rs12670273 | 0.05 | -0.247 | 0.635 | 0.274 | -0.024 | 0.016 | 0.016 | 0 | 0.32 | 0.23 | 0.41 | 0.26 | 0.42 | 0.3 |
| 7 | 36625156 | rs2392450 | -0.915 | -0.553 | 0.649 | 0.509 | -0.081 | 0.004 | 0.004 | 0.274 | 0.28 | 0.47 | 0.28 | 0.29 | 0.15 | 0.33 |
| 7 | 36625301 | rs2392452 | -0.898 | -0.543 | 0.653 | 0.519 | -0.08 | 0 | 0 | 0 | 0.28 | 0.47 | 0.28 | 0.29 | 0.15 | 0.33 |
| 7 | 36633202 | rs2727805 | -0.61 | 0.219 | 0.405 | 0.943 | -0.001 | -0.002 | -0.002 | 0.009 | 0.53 | 0.46 | 0.61 | 0.48 | 0.55 | 0.55 |
| 7 | 36635460 | rs2727810 | -1.002 | 0.822 | 0.275 | 1.174 | -0.002 | -0.018 | -0.018 | 0.061 | 0.31 | 0.76 | 0.44 | 0.58 | 0.33 | 0.32 |
| 7 | 36636449 | rs2727811 | -0.386 | 0.846 | -0.422 | 0.773 | -0.005 | 0.008 | 0.008 | 0 | 0.37 | 0.66 | 0.51 | 0.29 | 0.39 | 0.32 |
| 7 | 36637463 | rs2718177 | -1.105 | 0.976 | -0.584 | 0.867 | -0.001 | -0.008 | -0.008 | 0 | 0.53 | 0.79 | 0.61 | 0.44 | 0.53 | 0.43 |
| 7 | 36642252 | rs7808955 | 0.071 | 1.234 | 0.436 | 0.596 | 0.001 | -0.061 | -0.061 | 0 | 0.29 | 0.31 | 0.21 | 0.01 | 0.28 | 0.17 |
| 7 | 36643262 | rs2718179 | -0.385 | 1.216 | 0.521 | 0.725 | 0 | -0.019 | -0.019 | 0 | 0.34 | 0.56 | 0.44 | 0.23 | 0.33 | 0.28 |
| 7 | 36650296 | rs2718184 | -1.367 | 1.248 | 0.048 | 0.841 | 0.015 | 0 | 0 | 0 | 0.6 | 0.76 | 0.72 | 0.54 | 0.63 | 0.62 |
| 7 | 36651261 | rs2247298 | -0.289 | 1.361 | 0.268 | 0.92 | -0.003 | 0.081 | 0.081 | 0 | 0.19 | 0.31 | 0.32 | 0.4 | 0.2 | 0.2 |
| 7 | 36654238 | rs2001599 | -0.496 | 1.211 | 0.364 | 0.846 | 0.005 | -0.019 | -0.019 | 7.587 | 0.44 | 0.5 | 0.38 | 0.24 | 0.47 | 0.32 |
| 7 | 36656035 | rs3735386 | 0.422 | 1.338 | -0.521 | 0.841 | -0.038 | 0.024 | 0.024 | 0 | 0.11 | 0.01 | 0.13 | 0.03 | 0.13 | 0.09 |
| 7 | 36659892 | rs12537490 | -0.153 | 1.196 | -0.05 | 1.08 | -0.007 | 0.024 | 0.024 | 0.129 | 0.43 | 0.34 | 0.35 | 0.22 | 0.46 | 0.31 |
| 7 | 36669697 | rs12701517 | 0.168 | 1.147 | -0.153 | 1.278 | 0.005 | 0.027 | 0.027 | 0 | 0.83 | 0.75 | 0.82 | 0.78 | 0.85 | 0.71 |
| 7 | 36679794 | rs10243890 | -0.246 | 1.015 | -0.467 | 0.912 | -0.001 | -0.001 | -0.001 | 2.635 | 0.26 | 0.23 | 0.3 | 0.26 | 0.27 | 0.36 |
| 7 | 36680106 | rs12539267 | -0.524 | 0.952 | 0.024 | 0.689 | 0.006 | -0.006 | -0.006 | 0 | 0.15 | 0.31 | 0.16 | 0.07 | 0.13 | 0.13 |
| 7 | 36682895 | rs4504543 | -0.655 | 0.975 | -0.175 | 0.502 | 0.001 | -0.002 | -0.002 | 0 | 0.32 | 0.6 | 0.3 | 0.19 | 0.34 | 0.18 |
| 7 | 36684254 | rs11979472 | -0.516 | 0.778 | -0.184 | 0.187 | -0.012 | -0.046 | -0.046 | 0 | 0.22 | 0.52 | 0.22 | 0.07 | 0.18 | 0.1 |
| 7 | 36685212 | rs7794818 | -0.5 | 0.615 | -0.969 | -0.541 | 0 | -0.006 | -0.006 | 0.027 | 0.42 | 0.57 | 0.32 | 0.08 | 0.38 | 0.23 |
| 7 | 36686829 | rs11771672 | 0.235 | 0.528 | -0.921 | -0.334 | -0.022 | -0.001 | -0.001 | 0 | 0.19 | 0.13 | 0.17 | 0.12 | 0.24 | 0.12 |
| 7 | 36688836 | rs7804801 | 0.79 | 0.345 | -0.499 | -0.047 | 0 | -0.001 | -0.001 | 0 | 0.35 | 0.3 | 0.29 | 0.43 | 0.34 | 0.44 |
| 7 | 36690262 | rs11973236 | -0.672 | -0.044 | -0.648 | -0.01 | -0.025 | 0.065 | 0.065 | 0 | 0.64 | 0.78 | 0.59 | 0.85 | 0.59 | 0.65 |
| 7 | 36690449 | rs4570047 | -1.808 | -0.051 | -1.059 | -0.125 | -0.035 | 0.024 | 0.024 | 0 | 0.8 | 0.95 | 0.78 | 0.92 | 0.78 | 0.77 |
| 7 | 36691071 | rs11770435 | 1.006 | -0.055 | -0.946 | -0.081 | -0.03 | 0.047 | 0.047 | 0 | 0.16 | 0.03 | 0.18 | 0.07 | 0.2 | 0.11 |
| 7 | 36692185 | rs12690818 | -0.71 | -0.17 | -0.704 | 0.06 | -0.01 | 0.016 | 0.016 | 0 | 0.29 | 0.73 | 0.32 | 0.42 | 0.24 | 0.21 |
| 7 | 36694265 | rs12540585 | -0.118 | 0.089 | -0.49 | -0.202 | -0.002 | -0.048 | -0.048 | 0 | 0.22 | 0.35 | 0.23 | 0.09 | 0.22 | 0.08 |
| 7 | 36694799 | rs12530787 | -0.118 | 0.089 | -0.49 | -0.202 | -0.002 | -0.048 | -0.048 | 0 | 0.22 | 0.36 | 0.23 | 0.09 | 0.22 | 0.08 |
| 7 | 36695217 | rs4723551 | -0.455 | 0.207 | -0.816 | -0.243 | -0.042 | 0.08 | 0.08 | 0 | 0.64 | 0.87 | 0.6 | 0.85 | 0.59 | 0.65 |
| 7 | 36701694 | rs7807475 | 0.78 | 0.114 | -0.26 | 0.088 | -0.004 | 0.007 | 0.007 | 0 | 0.41 | 0.47 | 0.32 | 0.5 | 0.38 | 0.53 |
| 7 | 36706883 | rs11766973 | 0.369 | 0.183 | -0.422 | -0.044 | -0.045 | 0.046 | 0.046 | 0 | 0.17 | 0.06 | 0.21 | 0.1 | 0.24 | 0.11 |
| 7 | 36706968 | rs7797451 | 0.007 | 0.062 | -0.114 | -0.04 | 0 | -0.001 | -0.001 | 0.36 | 0.61 | 0.61 | 0.6 | 0.61 | 0.64 | 0.67 |
| 7 | 36709540 | rs12672188 | 0.515 | 0.101 | 0.068 | -0.016 | 0 | -0.002 | -0.002 | 0 | 0.56 | 0.38 | 0.48 | 0.61 | 0.55 | 0.65 |
| 7 | 36711493 | rs11978395 | 0.462 | -0.212 | 0.034 | -0.112 | 0 | 0.001 | 0.001 | 0.081 | 0.54 | 0.48 | 0.49 | 0.65 | 0.58 | 0.55 |
| 7 | 36715569 | rs10246586 | -0.53 | -0.276 | 0.01 | -0.068 | -0.019 | 0.067 | 0.067 | 0 | 0.22 | 0.38 | 0.26 | 0.31 | 0.17 | 0.13 |

| | | | | | | | | | | | | | | | | |
|---|---|---|---|---|---|---|---|---|---|---|---|---|---|---|---|---|
| 7 | 36721032 | rs4457232 | -1.039 | -0.388 | -0.473 | -0.36 | -0.046 | 0.101 | 0.101 | 0 | 0.75 | 0.97 | 0.76 | 0.95 | 0.75 | 0.71 |
| 7 | 36721093 | rs4304243 | -0.344 | -0.462 | 0.151 | -0.313 | -0.005 | 0.06 | 0.06 | 0 | 0.21 | 0.44 | 0.21 | 0.28 | 0.17 | 0.11 |
| 7 | 36721575 | rs4596555 | -0.923 | -0.509 | 0.293 | -0.333 | -0.021 | 0.039 | 0.039 | 0 | 0.23 | 0.52 | 0.29 | 0.29 | 0.18 | 0.13 |
| 7 | 36722365 | rs6961245 | 1.086 | -0.28 | 0.13 | -0.755 | 0 | -0.018 | -0.018 | 0 | 0.18 | 0.15 | 0.09 | 0 | 0.16 | 0.04 |
| 7 | 36723665 | rs12534128 | -0.107 | -0.312 | 0.262 | -0.585 | 0.001 | 0.004 | 0.004 | 0 | 0.08 | 0.07 | 0.11 | 0.18 | 0.06 | 0.07 |
| 7 | 36729102 | rs7802360 | -0.08 | -0.299 | 0.418 | -0.635 | -0.004 | 0.032 | 0.032 | 0 | 0.19 | 0.26 | 0.2 | 0.28 | 0.17 | 0.11 |
| 7 | 36729324 | rs4077337 | 0.316 | -0.325 | 0.262 | -0.692 | -0.002 | -0.006 | -0.006 | 0.243 | 0.54 | 0.53 | 0.55 | 0.65 | 0.58 | 0.61 |
| 7 | 36729951 | rs6462691 | 0.018 | -0.455 | 0.305 | -0.896 | 0.001 | -0.01 | -0.01 | 0 | 0.52 | 0.47 | 0.63 | 0.86 | 0.56 | 0.61 |
| 7 | 36729997 | rs4078285 | 0.466 | -0.398 | 0.418 | -0.807 | -0.014 | -0.071 | -0.071 | 0 | 0.22 | 0.41 | 0.13 | 0.09 | 0.19 | 0.11 |
| 7 | 36751049 | rs10275462 | 1.821 | -0.385 | -0.202 | -0.57 | -0.06 | -0.113 | -0.113 | 0 | 0.3 | 0.13 | 0.36 | 0.54 | 0.38 | 0.48 |
| 7 | 36757127 | rs4723562 | -0.899 | -0.15 | -0.002 | -0.653 | 0.002 | 0.045 | 0.045 | 9.191 | 0.75 | 0.88 | 0.88 | 1 | 0.77 | 0.95 |
| 7 | 36757463 | rs4723563 | -0.253 | -0.144 | 0.251 | -0.538 | -0.001 | -0.005 | -0.005 | 0 | 0.71 | 0.72 | 0.85 | 0.86 | 0.74 | 0.87 |
| 7 | 36762045 | rs13223177 | -0.65 | -0.061 | 0.411 | -0.224 | 0.009 | 0.051 | 0.051 | 0 | 0.23 | 0.37 | 0.24 | 0.32 | 0.22 | 0.13 |
| 7 | 36762883 | rs17170682 | 0.728 | -0.082 | 0.47 | -0.317 | 0 | -0.004 | -0.004 | 0 | 0.14 | 0.12 | 0.12 | 0.1 | 0.14 | 0.04 |
| 7 | 36763560 | rs6462703 | -0.443 | -0.116 | 0.628 | -0.398 | -0.003 | -0.019 | -0.019 | 0 | 0.3 | 0.53 | 0.22 | 0.25 | 0.26 | 0.16 |
| 7 | 36764123 | rs3807605 | -1.095 | -0.077 | 0.161 | -0.406 | 0 | 0.008 | 0.008 | 0.395 | 0.37 | 0.56 | 0.34 | 0.42 | 0.37 | 0.25 |
| 7 | 36765758 | rs3807609 | -0.928 | 0.506 | -0.245 | -0.415 | -0.008 | -0.02 | -0.02 | 0 | 0.33 | 0.7 | 0.33 | 0.39 | 0.32 | 0.23 |
| 7 | 36772392 | rs17405238 | 1.804 | 0.516 | -0.796 | -0.727 | -0.008 | 0.041 | 0.041 | 7.564 | 0.1 | 0.03 | 0.05 | 0.01 | 0.09 | 0.02 |
| 7 | 36775077 | rs17170686 | 0.251 | 0.491 | -0.714 | -0.708 | -0.021 | 0.021 | 0.021 | 0 | 0.16 | 0.1 | 0.19 | 0.11 | 0.18 | 0.07 |
| 7 | 36775215 | rs17379430 | -0.321 | 0.31 | -0.534 | -0.775 | 0.019 | 0.001 | 0.001 | 0 | 0.28 | 0.36 | 0.32 | 0.28 | 0.34 | 0.14 |
| 7 | 36775274 | rs17379465 | 0.12 | -0.127 | -1.401 | -1.411 | -0.034 | 0.16 | 0.16 | 0 | 0.31 | 0.06 | 0.27 | 0.05 | 0.32 | 0.34 |
| 7 | 36775650 | rs12532138 | -0.347 | -0.229 | -1.458 | -1.397 | -0.002 | 0.198 | 0.198 | 0 | 0.37 | 0.07 | 0.29 | 0.07 | 0.33 | 0.39 |
| 7 | 36777888 | rs1894910 | -0.275 | 0.296 | -0.792 | -1.079 | 0 | -0.005 | -0.005 | 0 | 0.72 | 0.69 | 0.75 | 0.93 | 0.71 | 0.68 |
| 7 | 36779910 | rs7801998 | 0.279 | 0.246 | -0.677 | -0.916 | -0.009 | -0.064 | -0.064 | 0 | 0.17 | 0.24 | 0.17 | 0.01 | 0.15 | 0.06 |
| 7 | 36782471 | rs17170695 | 0.827 | 0.454 | -1.391 | -0.124 | 0.004 | -0.056 | -0.056 | 0 | 0.24 | 0.12 | 0.19 | 0.4 | 0.24 | 0.12 |
| 7 | 36784023 | rs10951495 | -1.13 | 0.635 | -1.108 | -0.449 | 0.015 | -0.041 | -0.041 | 0 | 0.9 | 0.81 | 0.93 | 0.98 | 0.88 | 0.98 |
| 7 | 36784032 | rs13239851 | -0.382 | 0.883 | -1.088 | -0.706 | 0.083 | -0.027 | -0.027 | 0 | 0.09 | 0.5 | 0.16 | 0.01 | 0.11 | 0.06 |
| 7 | 36784390 | rs10229449 | 0.678 | 0.584 | -2.168 | -1.087 | -0.031 | 0.16 | 0.16 | 0.979 | 0.64 | 0.16 | 0.65 | 0.96 | 0.63 | 0.75 |
| 7 | 36786482 | rs4720217 | 0.788 | 1.021 | -1.522 | 0.074 | -0.017 | 0.032 | 0.032 | 0 | 0.46 | 0.31 | 0.51 | 0.63 | 0.48 | 0.31 |
| 7 | 36788357 | rs10239402 | -0.948 | 0.42 | -1.296 | 0.459 | -0.006 | 0.071 | 0.071 | 0.162 | 0.51 | 0.8 | 0.62 | 0.59 | 0.45 | 0.71 |
| 7 | 36789839 | rs2392461 | -0.358 | 1.058 | -1.163 | 0.972 | -0.064 | -0.105 | -0.105 | 0.659 | 0.33 | 0.02 | 0.32 | 0.4 | 0.32 | 0.48 |
| 7 | 36790954 | rs759416 | -1.046 | 0.665 | -0.821 | 1.231 | -0.009 | -0.035 | -0.035 | 0 | 0.39 | 0.74 | 0.56 | 0.52 | 0.38 | 0.65 |
| 7 | 36791806 | rs2160306 | -0.188 | 0.749 | -0.584 | 1.155 | -0.002 | 0.032 | 0.032 | 0.201 | 0.22 | 0.19 | 0.31 | 0.47 | 0.18 | 0.35 |
| 7 | 36793525 | rs6462706 | -0.651 | 0.543 | -0.984 | 0.516 | 0.009 | -0.005 | -0.005 | 0 | 0.37 | 0.68 | 0.41 | 0.37 | 0.34 | 0.62 |
| 7 | 36800555 | rs2392465 | 0.6 | 0.435 | -0.675 | 0.23 | 0.013 | 0.032 | 0.032 | 0 | 0.49 | 0.4 | 0.42 | 0.62 | 0.43 | 0.48 |
| 7 | 36800959 | rs888048 | -0.158 | 0.534 | -0.575 | -0.079 | -0.031 | 0.122 | 0.122 | 0 | 0.69 | 0.85 | 0.78 | 0.97 | 0.69 | 0.84 |
| 7 | 36802137 | rs2392466 | 1.195 | 0.566 | -0.614 | -0.322 | 0.011 | -0.009 | -0.009 | 0 | 0.63 | 0.52 | 0.55 | 0.65 | 0.59 | 0.42 |
| 7 | 36802963 | rs10215905 | 1.164 | -0.118 | -0.048 | -0.169 | 0.004 | 0.03 | 0.03 | 0.407 | 0.1 | 0.13 | 0.18 | 0.24 | 0.12 | 0.23 |
| 7 | 36803000 | rs741304 | 1.616 | -0.412 | 0.39 | -0.309 | -0.001 | 0.007 | 0.007 | 0 | 0.45 | 0.54 | 0.42 | 0.71 | 0.48 | 0.39 |
| 7 | 36804598 | rs11771320 | 2.324 | -0.464 | 0.494 | 0.036 | -0.01 | -0.027 | -0.027 | 0 | 0.37 | 0.21 | 0.34 | 0.54 | 0.39 | 0.3 |
| 7 | 36805549 | rs10276012 | 0.659 | -0.373 | 0.308 | 0.056 | 0.019 | 0.02 | 0.02 | 0.464 | 0.75 | 0.65 | 0.7 | 0.65 | 0.69 | 0.66 |
| 7 | 36807381 | rs10281171 | 1.049 | -0.338 | -0.11 | -0.077 | -0.011 | 0.212 | 0.212 | 0 | 0.56 | 0.8 | 0.61 | 0.94 | 0.6 | 0.62 |
| 7 | 36814860 | rs10271212 | -0.378 | -0.279 | -0.034 | 0.145 | 0.011 | -0.012 | -0.012 | 0.093 | 0.19 | 0.1 | 0.29 | 0.25 | 0.16 | 0.43 |
| 7 | 36819813 | rs4723572 | -1.22 | -0.058 | 0.332 | 0.202 | -0.002 | 0.006 | 0.006 | 0 | 0.54 | 0.45 | 0.58 | 0.39 | 0.51 | 0.66 |
| 7 | 36820963 | rs1420324 | -0.946 | -0.051 | 0.363 | 0.506 | 0.009 | -0.028 | -0.028 | 0 | 0.92 | 0.91 | 0.95 | 1 | 0.91 | 0.97 |
| 7 | 36823940 | rs10243286 | -0.608 | -0.46 | 0.665 | 1.103 | 0.002 | 0.095 | 0.095 | 0.122 | 0.18 | 0.3 | 0.26 | 0.54 | 0.21 | 0.34 |
| 7 | 36824655 | rs7806624 | -0.35 | -0.972 | 0.536 | 0.252 | 0.014 | 0.014 | 0.014 | 0.946 | 0.26 | 0.2 | 0.22 | 0.02 | 0.21 | 0.19 |
| 7 | 36825122 | rs7790608 | -1.447 | -1.283 | 1.268 | 0.484 | 0.005 | 0.002 | 0.002 | 0.135 | 0.44 | 0.56 | 0.51 | 0.37 | 0.35 | 0.55 |
| 7 | 36825290 | rs2392467 | -0.893 | -1.103 | 1.37 | 0.608 | -0.006 | -0.014 | -0.014 | 0 | 0.32 | 0.36 | 0.23 | 0.18 | 0.25 | 0.26 |
| 7 | 36825449 | rs2392469 | 0.438 | -0.935 | 1.67 | 0.847 | 0.003 | 0.084 | 0.084 | 0.059 | 0.11 | 0.24 | 0.3 | 0.26 | 0.14 | 0.28 |
| 7 | 36827835 | rs7795594 | -0.226 | -1.179 | 1.401 | 0.649 | -0.003 | -0.03 | -0.03 | 0 | 0.82 | 0.79 | 0.82 | 0.98 | 0.83 | 0.93 |
| 7 | 36829705 | rs10259514 | -1.022 | -1.344 | 1.06 | 0.429 | 0.008 | 0.011 | 0.011 | 0.03 | 0.29 | 0.15 | 0.32 | 0.24 | 0.25 | 0.47 |
| 7 | 36831723 | rs12668091 | -1.284 | -1.345 | 1.232 | 0.525 | -0.017 | 0.023 | 0.023 | 0 | 0.32 | 0.48 | 0.35 | 0.35 | 0.26 | 0.51 |
| 7 | 36838434 | rs6950956 | 0.543 | -1.299 | 0.999 | 0.469 | 0.004 | 0.002 | 0.002 | 0.048 | 0.73 | 0.81 | 0.61 | 0.76 | 0.7 | 0.67 |
| 7 | 36847304 | rs6960795 | 0.082 | -1.297 | 1.07 | 0.368 | 0.001 | -0.011 | -0.011 | 0.178 | 0.17 | 0.13 | 0.1 | 0.01 | 0.12 | 0.19 |
| 7 | 36849213 | rs10265806 | 0.906 | -1.312 | 1.187 | 0.466 | 0.001 | 0.106 | 0.106 | 0.082 | 0.11 | 0.17 | 0.24 | 0.34 | 0.14 | 0.3 |
| 7 | 36850396 | rs4723578 | -0.114 | -1.421 | 1.395 | 0.356 | 0.009 | 0.024 | 0.024 | 0 | 0.82 | 0.87 | 0.83 | 0.98 | 0.83 | 0.93 |
| 7 | 36851154 | rs7787226 | -1.149 | -1.403 | 1.504 | 0.5 | 0 | 0.037 | 0.037 | 0.012 | 0.31 | 0.37 | 0.36 | 0.49 | 0.3 | 0.53 |
| 7 | 36851426 | rs10951500 | 2.202 | -1.35 | 1.309 | 0.525 | -0.008 | -0.009 | -0.009 | 0.29 | 0.46 | 0.26 | 0.45 | 0.48 | 0.53 | 0.38 |
| 7 | 36851929 | rs17329669 | -0.282 | -1.295 | 1.205 | 0.455 | 0.014 | 0.07 | 0.07 | 0.009 | 0.2 | 0.06 | 0.1 | 0.01 | 0.12 | 0.21 |
| 7 | 36852287 | rs1592497 | -0.804 | -1.291 | 1.242 | 0.476 | 0 | 0 | 0 | 0 | 0.23 | 0.21 | 0.12 | 0.15 | 0.17 | 0.25 |
| 7 | 36867879 | rs13237538 | 1.864 | -1.314 | 1.212 | 0.617 | 0.008 | 0.002 | 0.002 | 0 | 0.47 | 0.35 | 0.47 | 0.48 | 0.53 | 0.39 |
| 7 | 36868010 | rs6969810 | 1.285 | -1.355 | 1.316 | 0.625 | 0.021 | 0.022 | 0.022 | 0 | 0.56 | 0.78 | 0.53 | 0.69 | 0.59 | 0.45 |
| 7 | 36869457 | rs6462713 | 0.956 | -1.315 | 1.258 | 0.665 | 0.017 | 0.176 | 0.176 | 0.146 | 0.1 | 0.26 | 0.25 | 0.29 | 0.13 | 0.28 |
| 7 | 36870228 | rs4723581 | 1.301 | -1.341 | 1.233 | 0.643 | -0.013 | -0.048 | -0.048 | 0.002 | 0.58 | 0.55 | 0.7 | 0.77 | 0.67 | 0.67 |
| 7 | 36871715 | rs759415 | 1.932 | -1.311 | 1.124 | 0.625 | 0.006 | -0.002 | -0.002 | 0 | 0.47 | 0.3 | 0.46 | 0.48 | 0.53 | 0.38 |
| 7 | 36877745 | rs756507 | 0.893 | -1.194 | 1.095 | 0.748 | 0.007 | 0.058 | 0.058 | 0.06 | 0.1 | 0.18 | 0.24 | 0.22 | 0.13 | 0.26 |
| 7 | 36890204 | rs6973243 | -1.525 | -1.331 | 0.981 | 0.685 | 0.023 | 0.008 | 0.008 | 0 | 0.5 | 0.58 | 0.51 | 0.51 | 0.42 | 0.58 |
| 7 | 36890612 | rs4140981 | 0.03 | -1.423 | 0.987 | 0.633 | 0 | 0.002 | 0.002 | 0 | 0.16 | 0.16 | 0.11 | 0.27 | 0.16 | 0.14 |

| | | | | | | | | | | | | | | | | |
|---|---|---|---|---|---|---|---|---|---|---|---|---|---|---|---|---|
| 7 | 36891841 | rs13234380 | 2.504 | -1.11 | 0.054 | 0.67 | -0.132 | -0.02 | -0.02 | 0 | 0.29 | 0.02 | 0.24 | 0.22 | 0.38 | 0.14 |
| 7 | 36893374 | rs6462719 | -1.755 | -1.708 | 0.922 | 0.895 | 0.022 | 0 | 0 | 0 | 0.5 | 0.57 | 0.5 | 0.47 | 0.42 | 0.58 |
| 7 | 36901579 | rs7785934 | 2.164 | -1.668 | 0.82 | 0.893 | -0.023 | 0.017 | 0.017 | 0 | 0.42 | 0.3 | 0.37 | 0.36 | 0.49 | 0.39 |
| 7 | 36905320 | rs6949576 | 1.87 | -1.865 | 1.264 | 1.317 | 0.003 | -0.002 | -0.002 | 0 | 0.56 | 0.44 | 0.54 | 0.61 | 0.57 | 0.58 |
| 7 | 36911493 | rs6965152 | -0.686 | -1.894 | 1.256 | 1.062 | -0.051 | -0.048 | -0.048 | 0.839 | 0.19 | 0.33 | 0.15 | 0.1 | 0.11 | 0.08 |
| 7 | 36913157 | rs7807884 | -0.437 | -1.958 | 1.267 | 1.034 | -0.006 | 0.021 | 0.021 | 0 | 0.09 | 0.29 | 0.05 | 0.08 | 0.05 | 0.03 |
| 7 | 36913619 | rs3778708 | 2.156 | -2.174 | 1.324 | 1.338 | 0.006 | 0.008 | 0.008 | 0 | 0.59 | 0.39 | 0.56 | 0.65 | 0.61 | 0.58 |
| 7 | 36915185 | rs1558688 | 1.488 | -2.151 | 1.473 | 1.366 | -0.001 | 0.005 | 0.005 | 0 | 0.7 | 0.43 | 0.61 | 0.66 | 0.69 | 0.65 |
| 7 | 36917995 | rs741301 | 1.917 | -2.147 | 1.476 | 1.361 | 0 | -0.001 | -0.001 | 0 | 0.67 | 0.37 | 0.6 | 0.65 | 0.68 | 0.61 |
| 7 | 36918688 | rs6960706 | -0.501 | -2.264 | 1.4 | 1.432 | 0.019 | 0.006 | 0.006 | 0 | 0.17 | 0.11 | 0.07 | 0.09 | 0.08 | 0.19 |
| 7 | 36920800 | rs3778709 | -1.643 | -2.24 | 1.522 | 1.554 | -0.003 | 0.007 | 0.007 | 0 | 0.3 | 0.48 | 0.38 | 0.33 | 0.3 | 0.35 |
| 7 | 36921670 | rs6956590 | -1.428 | -2.263 | 1.433 | 1.505 | 0.022 | 0.057 | 0.057 | 0 | 0.28 | 0.12 | 0.21 | 0.1 | 0.22 | 0.27 |
| 7 | 36922286 | rs17332226 | -0.166 | -2.261 | 1.41 | 1.512 | 0.048 | 0.005 | 0.005 | 0 | 0.15 | 0.04 | 0.05 | 0.07 | 0.07 | 0.17 |
| 7 | 36922825 | rs10281169 | -1.569 | -2.255 | 1.593 | 1.687 | -0.001 | -0.009 | -0.009 | 1.121 | 0.28 | 0.41 | 0.35 | 0.25 | 0.27 | 0.37 |
| 7 | 36923564 | rs6966983 | -1.497 | -2.252 | 1.517 | 1.718 | 0.013 | -0.011 | -0.011 | 0 | 0.3 | 0.19 | 0.28 | 0.33 | 0.25 | 0.3 |
| 7 | 36924415 | rs3778713 | -2.273 | -2.246 | 1.543 | 1.749 | 0.027 | -0.002 | -0.002 | 4.033 | 0.45 | 0.58 | 0.43 | 0.4 | 0.36 | 0.53 |
| 7 | 36926841 | rs2041801 | 2.599 | -2.246 | 1.518 | 1.745 | 0.026 | 0.09 | 0.09 | 0 | 0.49 | 0.2 | 0.46 | 0.34 | 0.56 | 0.4 |
| 7 | 36927501 | rs17170765 | -0.761 | -2.192 | 1.472 | 1.8 | -0.007 | 0.007 | 0.007 | 0 | 0.19 | 0.18 | 0.13 | 0.31 | 0.11 | 0.22 |
| 7 | 36928964 | rs7799004 | 1.205 | -1.938 | 1.345 | 1.726 | 0 | 0.066 | 0.066 | 0 | 0.81 | 0.6 | 0.7 | 0.75 | 0.81 | 0.72 |
| 7 | 36929344 | rs10225887 | 0.321 | -1.786 | 0.579 | 1.687 | -0.002 | -0.007 | -0.007 | 0 | 0.1 | 0.07 | 0.18 | 0.13 | 0.13 | 0.23 |
| 7 | 36936842 | rs4723589 | -0.169 | -1.751 | 0.393 | 1.701 | -0.013 | -0.047 | -0.047 | 0 | 0.12 | 0.04 | 0.2 | 0.23 | 0.16 | 0.25 |
| 7 | 36951044 | rs4723596 | 0.114 | -1.727 | 0.067 | 1.489 | -0.009 | -0.022 | -0.022 | 1.063 | 0.11 | 0.05 | 0.19 | 0.19 | 0.16 | 0.25 |
| 7 | 36951929 | rs4720228 | -1.774 | -1.75 | 0.307 | 1.283 | 0.032 | 0.013 | 0.013 | 0 | 0.49 | 0.65 | 0.51 | 0.56 | 0.4 | 0.57 |
| 7 | 36957908 | rs6942726 | -0.894 | -1.606 | 0.608 | 1.246 | -0.003 | -0.002 | -0.002 | 4.283 | 0.24 | 0.21 | 0.27 | 0.26 | 0.28 | 0.34 |
| 7 | 36959445 | rs17418220 | -1.01 | -1.641 | 0.593 | 1.283 | -0.005 | -0.002 | -0.002 | 0 | 0.26 | 0.21 | 0.34 | 0.27 | 0.3 | 0.37 |
| 7 | 36964059 | rs17418395 | -1.502 | -1.876 | 0.695 | 1.359 | -0.001 | -0.002 | -0.002 | 0 | 0.31 | 0.28 | 0.38 | 0.33 | 0.34 | 0.39 |
| 7 | 36966916 | rs1986619 | -0.708 | -2.289 | 0.564 | 1.304 | 0.045 | 0.015 | 0.015 | 0 | 0.17 | 0.05 | 0.06 | 0.08 | 0.08 | 0.2 |
| 7 | 36968412 | rs17170786 | -1.39 | -2.265 | 0.887 | 1.473 | 0.024 | -0.027 | -0.027 | 0.61 | 0.3 | 0.11 | 0.27 | 0.33 | 0.28 | 0.43 |
| 7 | 36970174 | rs1882083 | 1.064 | -2.442 | 1.493 | 1.509 | -0.014 | 0.046 | 0.046 | 0 | 0.77 | 0.47 | 0.74 | 0.69 | 0.84 | 0.86 |
| 7 | 36970474 | rs4720230 | -0.884 | -2.449 | 1.484 | 1.524 | 0.015 | 0.003 | 0.003 | 0 | 0.27 | 0.11 | 0.25 | 0.25 | 0.26 | 0.41 |
| 7 | 36970835 | rs10268319 | 1.764 | -2.262 | 1.297 | 1.333 | 0.035 | 0.024 | 0.024 | 0 | 0.5 | 0.36 | 0.47 | 0.44 | 0.57 | 0.46 |
| 7 | 36975080 | rs1882086 | 1.133 | -3.017 | 1.308 | 0.758 | -0.016 | 0.01 | 0.01 | 0.083 | 0.71 | 0.62 | 0.78 | 0.64 | 0.73 | 0.78 |
| 7 | 36979661 | rs6945781 | -1.25 | -3.29 | 1.769 | 0.226 | -0.053 | -0.1 | -0.1 | 0 | 0.21 | 0.29 | 0.09 | 0.01 | 0.08 | 0.08 |
| 7 | 36981343 | rs1986858 | -0.111 | -3.443 | 1.604 | 0.509 | 0.003 | -0.008 | -0.008 | 3.286 | 0.22 | 0.18 | 0.23 | 0.35 | 0.21 | 0.37 |
| 7 | 36985541 | rs2392477 | -2.14 | -3.543 | 1.637 | 0.439 | 0.088 | 0.066 | 0.066 | 0 | 0.49 | 0.76 | 0.49 | 0.64 | 0.36 | 0.61 |
| 7 | 36987889 | rs17170804 | -1.351 | -3.438 | 1.98 | 0.347 | -0.042 | 0.016 | 0.016 | 2.084 | 0.23 | 0.35 | 0.23 | 0.26 | 0.12 | 0.16 |
| 7 | 36989098 | rs4720231 | -0.901 | -3.232 | 2.071 | 0.47 | -0.017 | -0.008 | -0.008 | 0 | 0.35 | 0.38 | 0.29 | 0.27 | 0.29 | 0.43 |
| 7 | 36991362 | rs10233832 | -1.361 | -3.217 | 1.661 | 0.245 | 0.03 | 0.005 | 0.005 | 0 | 0.42 | 0.72 | 0.47 | 0.59 | 0.35 | 0.51 |
| 7 | 36996483 | rs7782153 | -1.857 | -3.139 | 2.304 | -0.341 | -0.071 | 0.012 | 0.012 | 0 | 0.25 | 0.5 | 0.25 | 0.26 | 0.14 | 0.23 |
| 7 | 37005260 | rs10226593 | -2.7 | -3.036 | 2.457 | -0.441 | 0.06 | 0.329 | 0.329 | 0.912 | 0.09 | 0.45 | 0.19 | 0.27 | 0.06 | 0.19 |
| 7 | 37007533 | rs17170812 | -1.593 | -3.098 | 2.36 | -0.792 | -0.027 | -0.006 | -0.006 | 0 | 0.24 | 0.29 | 0.27 | 0.22 | 0.16 | 0.21 |
| 7 | 37010772 | rs6962463 | -2.079 | -3.05 | 1.935 | -0.803 | 0.038 | 0.034 | 0.034 | 0.973 | 0.38 | 0.72 | 0.35 | 0.23 | 0.22 | 0.26 |
| 7 | 37043887 | rs10488027 | -1.856 | -2.681 | 1.742 | -1.119 | -0.025 | -0.055 | -0.055 | 0 | 0.36 | 0.41 | 0.22 | 0.05 | 0.2 | 0.17 |
| 7 | 37074776 | rs6965702 | -1.842 | -2.238 | 1.862 | -0.944 | -0.034 | -0.069 | -0.069 | 0 | 0.31 | 0.44 | 0.23 | 0.06 | 0.19 | 0.16 |
| 7 | 37076854 | rs10488031 | -0.826 | -2.331 | 1.823 | -1.015 | -0.006 | -0.037 | -0.037 | 0 | 0.13 | 0.18 | 0.05 | 0 | 0.07 | 0.05 |
| 7 | 37077758 | rs17170842 | -0.923 | -2.351 | 1.888 | -1.047 | -0.02 | -0.073 | -0.073 | 0 | 0.15 | 0.28 | 0.08 | 0 | 0.09 | 0.05 |
| 7 | 37088918 | rs17170854 | -1.363 | -2.348 | 1.802 | -1.118 | 0.001 | -0.001 | -0.001 | 0 | 0.19 | 0.16 | 0.18 | 0.05 | 0.13 | 0.12 |
| 7 | 37092810 | rs6967500 | -2.392 | -2.447 | 1.852 | -1.073 | -0.033 | -0.076 | -0.076 | 0 | 0.33 | 0.45 | 0.26 | 0.05 | 0.21 | 0.17 |
| 7 | 37097476 | rs7785575 | -0.939 | -2.435 | 1.863 | -1.067 | -0.025 | -0.085 | -0.085 | 0.779 | 0.16 | 0.3 | 0.08 | 0 | 0.09 | 0.05 |
| 7 | 37108104 | rs13233995 | -2.409 | -2.496 | 2.198 | -1.105 | -0.015 | -0.045 | -0.045 | 0.003 | 0.36 | 0.65 | 0.27 | 0.05 | 0.21 | 0.17 |
| 7 | 37110289 | rs880865 | -2.32 | -2.309 | 1.535 | -1.02 | -0.036 | -0.081 | -0.081 | 0.374 | 0.33 | 0.46 | 0.25 | 0.05 | 0.2 | 0.16 |
| 7 | 37114612 | rs17170863 | -1.469 | -1.744 | 1.366 | -0.588 | -0.004 | -0.008 | -0.008 | 0 | 0.17 | 0.16 | 0.17 | 0.04 | 0.11 | 0.1 |
| 7 | 37114721 | rs7455242 | -1.307 | -1.709 | 1.042 | -0.572 | 0.015 | 0.02 | 0.02 | 0 | 0.16 | 0.11 | 0.17 | 0.04 | 0.11 | 0.1 |
| 7 | 37135479 | rs6957997 | -1.779 | -1.303 | 1.129 | -0.745 | -0.053 | -0.145 | -0.145 | 0 | 0.24 | 0.59 | 0.14 | 0.01 | 0.13 | 0.09 |
| 7 | 37144066 | rs7809487 | -2.23 | -1.137 | 1.703 | -0.498 | -0.027 | -0.055 | -0.055 | 0 | 0.29 | 0.49 | 0.22 | 0.2 | 0.17 | 0.22 |
| 7 | 37144095 | rs10263072 | -1.575 | -1.107 | 1.769 | -0.474 | -0.023 | -0.039 | -0.039 | 0 | 0.22 | 0.37 | 0.18 | 0.12 | 0.13 | 0.17 |
| 7 | 37157641 | rs2111035 | -2.685 | -0.698 | 2.221 | -0.299 | 0.024 | 0.011 | 0.011 | 0.047 | 0.19 | 0.49 | 0.19 | 0.21 | 0.16 | 0.17 |
| 7 | 37158469 | rs1015909 | -2.685 | -0.698 | 2.221 | -0.299 | 0.024 | 0.011 | 0.011 | 0.047 | 0.19 | 0.49 | 0.19 | 0.21 | 0.16 | 0.17 |
| 7 | 37159141 | rs2541104 | -2.675 | -0.698 | 2.255 | -0.299 | 0.024 | 0.011 | 0.011 | 0.04 | 0.19 | 0.47 | 0.19 | 0.21 | 0.16 | 0.18 |
| 7 | 37166379 | rs2541072 | -2.64 | -0.714 | 2.657 | -0.306 | 0.026 | 0.013 | 0.013 | 0.092 | 0.19 | 0.48 | 0.2 | 0.22 | 0.16 | 0.17 |
| 7 | 37169563 | rs2541088 | -2.906 | -0.761 | 2.938 | -0.327 | -0.002 | -0.008 | -0.008 | 0.012 | 0.21 | 0.63 | 0.21 | 0.21 | 0.16 | 0.19 |
| 7 | 37170673 | rs2541094 | -1.138 | -0.742 | 3.133 | -0.321 | 0.012 | 0.029 | 0.029 | 0 | 0.08 | 0.18 | 0.12 | 0.1 | 0.07 | 0.08 |
| 7 | 37173048 | rs1420421 | -2.913 | -0.743 | 3.219 | -0.321 | 0.016 | 0.007 | 0.007 | 0 | 0.18 | 0.32 | 0.19 | 0.21 | 0.16 | 0.17 |
| 7 | 37187471 | rs10265823 | -2 | -0.685 | 3.228 | -0.363 | 0.05 | -0.02 | -0.02 | 0 | 0.05 | 0.17 | 0.05 | 0 | 0.06 | 0.04 |
| 7 | 37187498 | rs6942887 | 2.192 | -0.788 | 3.221 | -0.337 | 0.007 | 0.048 | 0.048 | 0.157 | 0.86 | 0.65 | 0.86 | 0.79 | 0.91 | 0.89 |
| 7 | 37190384 | rs6462733 | 2.718 | -0.993 | 3.02 | -0.187 | -0.014 | 0.018 | 0.018 | 0.081 | 0.78 | 0.36 | 0.82 | 0.72 | 0.85 | 0.82 |
| 7 | 37191340 | rs6956171 | -1.165 | -0.878 | 2.888 | -0.157 | 0.005 | 0.024 | 0.024 | 0.066 | 0.08 | 0.17 | 0.05 | 0.14 | 0.06 | 0.04 |
| 7 | 37199569 | rs11768811 | 2.833 | -0.868 | 2.887 | -0.38 | -0.006 | 0 | 0 | 0 | 0.75 | 0.31 | 0.78 | 0.75 | 0.82 | 0.81 |
| 7 | 37200613 | rs1345365 | 2.859 | -1.628 | 2.187 | -0.93 | 0.032 | 0.038 | 0.038 | 0.105 | 0.59 | 0.27 | 0.66 | 0.7 | 0.72 | 0.6 |

| | | | | | | | | | | | | | | | | |
|---|---|---|---|---|---|---|---|---|---|---|---|---|---|---|---|---|
| 7 | 37213108 | rs13225050 | -1.019 | -1.66 | 2.894 | -0.921 | -0.006 | -0.03 | -0.03 | 0 | 0.11 | 0.12 | 0.06 | 0 | 0.07 | 0.09 |
| 7 | 37213483 | rs10951509 | 2.933 | -1.689 | 2.845 | -0.936 | 0.036 | 0.033 | 0.033 | 0.282 | 0.61 | 0.26 | 0.67 | 0.7 | 0.75 | 0.6 |
| 7 | 37214704 | rs6980398 | -1.986 | -1.547 | 2.258 | -1.085 | -0.015 | -0.033 | -0.033 | 0 | 0.19 | 0.28 | 0.1 | 0.09 | 0.13 | 0.12 |
| 7 | 37220188 | rs12701548 | -1.728 | -1.541 | 2.086 | -0.724 | -0.04 | -0.033 | -0.033 | 0 | 0.28 | 0.52 | 0.29 | 0.19 | 0.18 | 0.29 |
| 7 | 37220932 | rs6946785 | 1.797 | -1.716 | 1.331 | -1.147 | 0.053 | 0.067 | 0.067 | 0.087 | 0.48 | 0.26 | 0.56 | 0.68 | 0.6 | 0.58 |
| 7 | 37223401 | rs2160430 | 0.371 | -1.719 | 1.615 | -1.116 | -0.018 | -0.008 | -0.008 | 0 | 0.75 | 0.63 | 0.78 | 0.77 | 0.78 | 0.86 |
| 7 | 37225235 | rs1420423 | -0.298 | -1.625 | 1.086 | -1.152 | -0.006 | 0.029 | 0.029 | 0 | 0.81 | 0.93 | 0.86 | 0.89 | 0.79 | 0.89 |
| 7 | 37234756 | rs2058730 | 1.521 | -1.961 | 0.851 | -1.389 | 0.038 | 0.058 | 0.058 | 0 | 0.46 | 0.27 | 0.48 | 0.66 | 0.63 | 0.49 |
| 7 | 37236103 | rs6462739 | 1.54 | -1.952 | 0.864 | -1.369 | 0.032 | 0.04 | 0.04 | 0 | 0.46 | 0.27 | 0.48 | 0.63 | 0.62 | 0.46 |
| 7 | 37241917 | rs6462740 | -0.064 | -1.948 | 1.196 | -1.534 | -0.015 | -0.038 | -0.038 | 1.131 | 0.28 | 0.42 | 0.16 | 0.13 | 0.22 | 0.16 |
| 7 | 37254465 | rs6963276 | 0.278 | -1.926 | 1.167 | -1.558 | 0.016 | 0.074 | 0.074 | 0 | 0.26 | 0.09 | 0.13 | 0.13 | 0.22 | 0.16 |
| 7 | 37260786 | rs10229046 | -0.018 | -0.735 | 1.141 | -1.57 | -0.001 | -0.044 | -0.044 | 0 | 0.29 | 0.44 | 0.18 | 0.13 | 0.27 | 0.17 |
| 7 | 37267938 | rs2111034 | 1.097 | -1.119 | 1.327 | -1.185 | 0.045 | 0.045 | 0.045 | 0 | 0.47 | 0.26 | 0.48 | 0.62 | 0.63 | 0.47 |
| 7 | 37286864 | rs2080410 | -1.11 | -1.252 | 1.209 | -1.313 | 0.052 | 0.056 | 0.056 | 0 | 0.53 | 0.76 | 0.52 | 0.38 | 0.36 | 0.53 |
| 7 | 37298800 | rs2717968 | -0.306 | -1.411 | 1.08 | -1.373 | 0.018 | 0.029 | 0.029 | 3.928 | 0.4 | 0.34 | 0.35 | 0.19 | 0.3 | 0.28 |
| 7 | 37303595 | rs2717972 | -0.521 | -1.386 | 1.034 | -1.373 | -0.023 | -0.027 | -0.027 | 0 | 0.43 | 0.53 | 0.41 | 0.3 | 0.31 | 0.29 |
| 7 | 37305527 | rs4723631 | 0.278 | -0.438 | 1.141 | -1.077 | 0.006 | 0.002 | 0.002 | 0 | 0.32 | 0.33 | 0.36 | 0.61 | 0.4 | 0.39 |
| 7 | 37306805 | rs2723974 | 0.499 | -0.432 | 1.152 | -1.08 | 0.001 | -0.008 | -0.008 | 0 | 0.58 | 0.51 | 0.53 | 0.29 | 0.55 | 0.38 |
| 7 | 37307515 | rs1076477 | -0.806 | -0.827 | 1.336 | -1.168 | -0.02 | 0.004 | 0.004 | 0 | 0.1 | 0.15 | 0.1 | 0.09 | 0.05 | 0.18 |
| 7 | 37313712 | rs10951513 | 0.068 | -0.77 | 1.205 | -1.234 | -0.011 | 0.005 | 0.005 | 0 | 0.07 | 0.17 | 0.08 | 0.09 | 0.07 | 0.05 |
| 7 | 37316506 | rs11981366 | 0.595 | -0.763 | 1.194 | -1.231 | -0.007 | 0.005 | 0.005 | 0 | 0.06 | 0.12 | 0.08 | 0.09 | 0.06 | 0.05 |
| 7 | 37326091 | rs2392487 | -0.95 | -0.75 | 1.183 | -1.205 | 0.004 | 0.022 | 0.022 | 1.271 | 0.49 | 0.5 | 0.69 | 0.78 | 0.54 | 0.75 |
| 7 | 37337853 | rs17198538 | 0.144 | -0.583 | 0.734 | -1.045 | 0.011 | 0.003 | 0.003 | 0.018 | 0.1 | 0.06 | 0.06 | 0.08 | 0.08 | 0.15 |
| 7 | 37341477 | rs2701004 | 0.205 | -0.598 | 0.78 | -1.043 | -0.022 | 0.008 | 0.008 | 0 | 0.88 | 0.83 | 0.94 | 0.83 | 0.92 | 0.84 |
| 7 | 37344369 | rs2723996 | 0.584 | -0.528 | 0.818 | -1.088 | -0.002 | -0.016 | -0.016 | 1.119 | 0.56 | 0.44 | 0.39 | 0.13 | 0.5 | 0.32 |
| 7 | 37353302 | rs12532146 | -0.648 | -0.25 | -0.026 | -0.722 | -0.02 | -0.003 | -0.003 | 0 | 0.2 | 0.26 | 0.16 | 0.18 | 0.13 | 0.23 |
| 7 | 37354091 | rs11773267 | -0.169 | 0.916 | -0.189 | -0.239 | 0.018 | 0 | 0 | 0 | 0.29 | 0.35 | 0.53 | 0.7 | 0.4 | 0.5 |
| 7 | 37356953 | rs11973001 | -0.793 | 0.054 | 0.108 | -0.828 | -0.019 | -0.125 | -0.125 | 0.09 | 0.29 | 0.51 | 0.19 | 0.05 | 0.26 | 0.18 |
| 7 | 37356965 | rs2724002 | 0.057 | -0.08 | 0.495 | -0.696 | 0 | -0.028 | -0.028 | 0 | 0.88 | 0.78 | 0.89 | 0.95 | 0.87 | 0.89 |
| 7 | 37357595 | rs2724003 | 0.765 | -0.499 | 0.664 | -0.201 | -0.034 | 0.012 | 0.012 | 0.258 | 0.75 | 0.49 | 0.76 | 0.78 | 0.83 | 0.81 |
| 7 | 37357655 | rs12666889 | -0.13 | -0.65 | 0.388 | -0.105 | -0.013 | 0.003 | 0.003 | 0 | 0.06 | 0.13 | 0.04 | 0.08 | 0.05 | 0.08 |
| 7 | 37359101 | rs10249209 | -1.144 | -0.983 | 0.847 | -0.082 | -0.05 | 0.025 | 0.025 | 0.598 | 0.25 | 0.44 | 0.18 | 0.28 | 0.16 | 0.2 |
| 7 | 37359548 | rs10282266 | -0.103 | -1.054 | 0.816 | -0.122 | -0.015 | 0.025 | 0.025 | 0 | 0.15 | 0.2 | 0.1 | 0.2 | 0.09 | 0.12 |
| 7 | 37360209 | rs6947058 | 0.854 | -0.629 | -0.4 | -0.71 | 0.026 | -0.007 | -0.007 | 0 | 0.43 | 0.21 | 0.44 | 0.58 | 0.35 | 0.38 |
| 7 | 37360358 | rs3847015 | -0.533 | -0.364 | -0.427 | -0.523 | -0.063 | 0.066 | 0.066 | 0.902 | 0.69 | 0.8 | 0.63 | 0.86 | 0.51 | 0.58 |
| 7 | 37361023 | rs17170988 | 0.741 | -0.912 | -0.611 | -0.175 | 0.031 | -0.078 | -0.078 | 0 | 0.26 | 0.12 | 0.34 | 0.51 | 0.2 | 0.28 |
| 7 | 37363581 | rs2723978 | 0.336 | -0.47 | -0.097 | -0.33 | 0.017 | -0.02 | -0.02 | 0 | 0.93 | 0.76 | 0.89 | 0.99 | 0.94 | 0.89 |
| 7 | 37365234 | rs10245068 | -0.392 | -1.085 | -0.61 | -0.257 | 0.009 | 0.002 | 0.002 | 0 | 0.18 | 0.14 | 0.1 | 0.2 | 0.12 | 0.2 |
| 7 | 37365422 | rs2723980 | 1.128 | -0.919 | -1.459 | -0.02 | -0.026 | -0.033 | -0.033 | 8.273 | 0.16 | 0.03 | 0.18 | 0.14 | 0.18 | 0.15 |
| 7 | 37368161 | rs12667592 | -0.274 | -1.13 | -1.461 | 0.693 | 0.004 | -0.046 | -0.046 | 0.022 | 0.21 | 0.08 | 0.23 | 0.37 | 0.16 | 0.23 |
| 7 | 37369189 | rs2700996 | 1.13 | -1.092 | -1.715 | 0.321 | 0.001 | 0 | 0 | 0 | 0.44 | 0.23 | 0.48 | 0.41 | 0.57 | 0.48 |
| 7 | 37369557 | rs10242920 | -0.039 | -0.603 | -1.519 | 0.193 | 0.006 | 0.001 | 0.001 | 0 | 0.25 | 0.2 | 0.2 | 0.23 | 0.23 | 0.26 |
| 7 | 37371598 | rs11532670 | 0.824 | -0.377 | -1.432 | 0.049 | -0.047 | -0.007 | -0.007 | 0 | 0.09 | 0 | 0.06 | 0.07 | 0.1 | 0.06 |
| 7 | 37374510 | rs6974491 | 0.682 | -0.623 | -1.5 | -0.357 | -0.001 | 0.014 | 0.014 | 0 | 0.15 | 0.07 | 0.14 | 0 | 0.15 | 0.09 |
| 7 | 37375317 | rs2717947 | 1.113 | -0.789 | -2.237 | -0.548 | -0.083 | 0.073 | 0.073 | 0 | 0.37 | 0.09 | 0.39 | 0.23 | 0.53 | 0.42 |
| 7 | 37381220 | rs2700991 | -0.005 | -0.917 | -1.572 | -0.316 | 0.012 | -0.001 | -0.001 | 0 | 0.72 | 0.73 | 0.72 | 0.69 | 0.83 | 0.82 |
| 7 | 37382520 | rs17259252 | 0.801 | 0.08 | -1 | -0.618 | 0.034 | -0.058 | -0.058 | 0 | 0.24 | 0.29 | 0.27 | 0.09 | 0.37 | 0.34 |
| 7 | 37386098 | rs2717952 | -0.019 | -0.38 | -1.631 | -0.432 | -0.006 | 0.102 | 0.102 | 0 | 0.43 | 0.13 | 0.4 | 0.22 | 0.41 | 0.26 |
| 7 | 37388634 | rs2724012 | -0.043 | -0.889 | -1.596 | -0.583 | 0.023 | 0.046 | 0.046 | 0 | 0.39 | 0.32 | 0.47 | 0.79 | 0.36 | 0.47 |
| 7 | 37389804 | rs2700986 | 1.164 | -0.943 | -1.568 | -0.434 | 0.007 | -0.002 | -0.002 | 0 | 0.21 | 0.17 | 0.23 | 0.2 | 0.2 | 0.15 |
| 7 | 37390464 | rs2724017 | -1.342 | -1.158 | -1.05 | -0.549 | -0.013 | 0.005 | 0.005 | 0 | 0.9 | 0.85 | 0.93 | 0.93 | 0.95 | 0.92 |
| 7 | 37391266 | rs1882079 | 0.424 | -1.263 | -0.773 | -0.695 | 0 | 0.008 | 0.008 | 0 | 0.47 | 0.57 | 0.52 | 0.7 | 0.54 | 0.63 |
| 7 | 37391820 | rs2700984 | -0.003 | -1.205 | -0.871 | -0.722 | 0.01 | 0.067 | 0.067 | 0 | 0.43 | 0.27 | 0.4 | 0.22 | 0.41 | 0.29 |
| 7 | 37395373 | rs2717954 | 0.809 | -0.867 | -1.068 | -0.861 | -0.022 | 0.011 | 0.011 | 0 | 0.26 | 0.14 | 0.23 | 0.21 | 0.29 | 0.2 |
| 7 | 37397262 | rs13247953 | 0.794 | -1.027 | -0.839 | -0.628 | 0.012 | -0.04 | -0.04 | 0 | 0.19 | 0.16 | 0.3 | 0.6 | 0.16 | 0.32 |
| 7 | 37400106 | rs2717955 | 0.239 | -0.9 | -0.81 | -0.91 | 0.002 | 0.009 | 0.009 | 0 | 0.7 | 0.74 | 0.8 | 0.92 | 0.76 | 0.8 |
| 7 | 37401277 | rs2700975 | -0.707 | -0.867 | -0.798 | -0.869 | -0.003 | 0.008 | 0.008 | 0 | 0.82 | 0.86 | 0.87 | 0.93 | 0.86 | 0.88 |
| 7 | 37406352 | rs1882070 | -0.026 | -1.065 | -0.407 | -0.849 | -0.029 | 0.04 | 0.04 | 0 | 0.25 | 0.38 | 0.31 | 0.61 | 0.16 | 0.32 |
| 7 | 37410782 | rs10246492 | 0.03 | -0.793 | -0.746 | -1.039 | -0.003 | 0.015 | 0.015 | 0 | 0.44 | 0.43 | 0.52 | 0.73 | 0.53 | 0.63 |
| 7 | 37412056 | rs4723645 | -0.238 | -0.731 | -0.7 | -0.997 | 0.004 | 0.01 | 0.01 | 0 | 0.56 | 0.45 | 0.59 | 0.74 | 0.64 | 0.7 |
| 7 | 37413914 | rs17260207 | 1.849 | -0.239 | -0.842 | -0.763 | 0.007 | -0.011 | -0.011 | 0 | 0.18 | 0.17 | 0.17 | 0.12 | 0.28 | 0.26 |
| 7 | 37419801 | rs1962234 | 1.862 | -0.231 | -0.85 | -0.76 | 0.006 | -0.01 | -0.01 | 0 | 0.17 | 0.17 | 0.17 | 0.12 | 0.28 | 0.26 |
| 7 | 37424821 | rs13235125 | 0.421 | -0.389 | -0.621 | 0.159 | -0.018 | 0.139 | 0.139 | 0 | 0.1 | 0.24 | 0.21 | 0.44 | 0.05 | 0.12 |
| 7 | 37425376 | rs2701011 | 0.354 | -0.572 | -1.032 | -0.289 | -0.043 | -0.009 | -0.009 | 0 | 0.29 | 0.08 | 0.33 | 0.25 | 0.31 | 0.24 |
| 7 | 37431858 | rs9648447 | -0.808 | -1.322 | -0.401 | -0.143 | 0.014 | -0.038 | -0.038 | 0 | 0.89 | 0.82 | 0.92 | 0.98 | 0.89 | 0.91 |
| 7 | 37435223 | rs1882074 | -0.715 | -1.35 | -0.418 | -0.141 | 0.004 | -0.036 | -0.036 | 0.867 | 0.85 | 0.82 | 0.91 | 0.98 | 0.87 | 0.89 |
| 7 | 37436388 | rs4720246 | 1.101 | -1.189 | -1.147 | -0.08 | 0.019 | -0.006 | -0.006 | 0 | 0.18 | 0.06 | 0.18 | 0.18 | 0.14 | 0.16 |
| 7 | 37436929 | rs1882077 | 0.509 | -0.935 | -0.893 | 0.443 | 0.056 | -0.012 | -0.012 | 0 | 0.49 | 0.19 | 0.48 | 0.48 | 0.61 | 0.58 |
| 7 | 37437877 | rs11979905 | 0.506 | -0.718 | -0.186 | 0.555 | -0.005 | -0.034 | -0.034 | 0 | 0.11 | 0.15 | 0.07 | 0.01 | 0.11 | 0.07 |

| | | | | | | | | | | | | | | | | |
|---|---|---|---|---|---|---|---|---|---|---|---|---|---|---|---|---|
| 7 | 37437956 | rs12671707 | -0.268 | -0.752 | -0.316 | 0.993 | -0.018 | 0.043 | 0.043 | 0 | 0.11 | 0.2 | 0.2 | 0.29 | 0.04 | 0.17 |
| 7 | 37438596 | rs6964474 | -0.627 | -0.789 | -0.411 | 0.902 | -0.026 | 0.016 | 0.016 | 0 | 0.33 | 0.62 | 0.28 | 0.41 | 0.21 | 0.34 |
| 7 | 37440489 | rs7803682 | 0.245 | -0.172 | -0.951 | 0.658 | -0.012 | -0.026 | -0.026 | 0 | 0.22 | 0.09 | 0.23 | 0.34 | 0.22 | 0.39 |
| 7 | 37440628 | rs13241143 | 0.525 | -0.91 | -0.989 | 0.548 | 0 | 0.003 | 0.003 | 0 | 0.23 | 0.25 | 0.34 | 0.43 | 0.18 | 0.18 |
| 7 | 37440684 | rs7788477 | -0.468 | -0.856 | -1.026 | -0.005 | -0.003 | 0.008 | 0.008 | 0 | 0.56 | 0.43 | 0.64 | 0.78 | 0.49 | 0.62 |
| 7 | 37444728 | rs17171026 | 0.642 | -0.957 | -0.8 | 0.06 | 0.005 | -0.008 | -0.008 | 0 | 0.24 | 0.24 | 0.34 | 0.4 | 0.18 | 0.2 |
| 7 | 37445253 | rs6977706 | 0.771 | 0.054 | -0.567 | -0.541 | -0.005 | 0.008 | 0.008 | 0 | 0.28 | 0.22 | 0.25 | 0.21 | 0.33 | 0.35 |
| 7 | 37445669 | rs1425125 | 0.345 | -0.297 | -0.309 | -0.614 | -0.017 | 0.02 | 0.02 | 0 | 0.37 | 0.32 | 0.32 | 0.21 | 0.47 | 0.39 |
| 7 | 37445820 | rs4565373 | -0.087 | -0.301 | -0.193 | -0.644 | 0.011 | 0.04 | 0.04 | 0 | 0.54 | 0.63 | 0.6 | 0.77 | 0.41 | 0.55 |
| 7 | 37451713 | rs1647801 | 0.734 | 0.125 | -0.253 | -0.704 | -0.035 | -0.001 | -0.001 | 0 | 0.18 | 0.1 | 0.19 | 0.19 | 0.32 | 0.27 |
| 7 | 37452489 | rs987702 | -0.301 | 0.008 | -0.081 | -0.589 | 0.011 | -0.016 | -0.016 | 0.141 | 0.63 | 0.59 | 0.68 | 0.78 | 0.53 | 0.61 |
| 7 | 37455031 | rs10253164 | 1.927 | -0.129 | -0.276 | -0.438 | -0.015 | 0.01 | 0.01 | 0 | 0.21 | 0.07 | 0.14 | 0.21 | 0.2 | 0.27 |
| 7 | 37460441 | rs12540876 | -0.008 | -0.168 | -0.134 | -0.423 | -0.006 | 0 | 0 | 0 | 0.39 | 0.43 | 0.53 | 0.58 | 0.33 | 0.35 |
| 7 | 37461214 | rs12531123 | 0.369 | -0.203 | -0.501 | -0.35 | 0.02 | -0.041 | -0.041 | 0.834 | 0.33 | 0.26 | 0.42 | 0.43 | 0.3 | 0.25 |
| 7 | 37466564 | rs17202203 | 0.543 | -0.624 | -0.731 | -0.625 | -0.004 | 0.029 | 0.029 | 0 | 0.36 | 0.25 | 0.23 | 0.25 | 0.33 | 0.33 |
| 7 | 37466643 | rs10280643 | 0.349 | -0.805 | -1.112 | -0.584 | -0.005 | 0.051 | 0.051 | 0 | 0.52 | 0.22 | 0.36 | 0.38 | 0.53 | 0.5 |
| 7 | 37466970 | rs10267165 | -0.236 | -0.388 | -0.618 | -0.082 | 0.025 | -0.014 | -0.014 | 0 | 0.43 | 0.71 | 0.54 | 0.5 | 0.38 | 0.46 |
| 7 | 37469073 | rs11770274 | 1.184 | -0.287 | -0.974 | -0.266 | -0.006 | -0.018 | -0.018 | 0 | 0.07 | 0.03 | 0.08 | 0.11 | 0.09 | 0.09 |
| 7 | 37470284 | rs918979 | 0.334 | 0.298 | -0.903 | -0.539 | -0.083 | 0.046 | 0.046 | 0.256 | 0.36 | 0.11 | 0.26 | 0.26 | 0.43 | 0.4 |
| 7 | 37472124 | rs7777258 | 0.605 | -0.608 | -0.697 | -0.382 | 0 | -0.001 | -0.001 | 0 | 0.28 | 0.27 | 0.17 | 0.26 | 0.24 | 0.38 |
| 7 | 37473071 | rs1647791 | 0.435 | -0.498 | -0.846 | -0.355 | 0.044 | 0.014 | 0.014 | 0 | 0.28 | 0.41 | 0.49 | 0.3 | 0.38 | 0.19 |
| 7 | 37475868 | rs7797162 | 0.463 | -0.478 | -0.898 | -0.329 | 0.041 | -0.027 | -0.027 | 0 | 0.34 | 0.21 | 0.27 | 0.42 | 0.23 | 0.37 |
| 7 | 37477863 | rs17171046 | -0.196 | 0.313 | -0.444 | -0.651 | 0.001 | -0.083 | -0.083 | 6.966 | 0.83 | 0.62 | 0.89 | 0.98 | 0.86 | 0.95 |
| 7 | 37480974 | rs6962537 | 0.67 | 0.754 | -1.389 | -0.43 | -0.003 | 0.03 | 0.03 | 0 | 0.53 | 0.6 | 0.74 | 0.74 | 0.55 | 0.52 |
| 7 | 37483175 | rs9655382 | -0.521 | 0.788 | -2.019 | 0.477 | -0.065 | -0.057 | -0.057 | 0 | 0.23 | 0.01 | 0.1 | 0.24 | 0.22 | 0.4 |
| 7 | 37489588 | rs1882071 | -0.535 | 0.488 | -1.413 | 0.604 | 0 | -0.001 | -0.001 | 0.174 | 0.3 | 0.67 | 0.42 | 0.33 | 0.29 | 0.3 |
| 7 | 37490003 | rs1882072 | -0.556 | 0.594 | -1.834 | 0.099 | 0.009 | 0.053 | 0.053 | 0.791 | 0.22 | 0.08 | 0.16 | 0.12 | 0.18 | 0.11 |
| 7 | 37493679 | rs1625531 | 0.453 | 1.695 | -1.514 | 0.261 | 0.036 | 0.002 | 0.002 | 0 | 0.91 | 0.77 | 0.89 | 0.87 | 0.87 | 0.75 |
| 7 | 37493692 | rs7811413 | 0.018 | 0.896 | -1.266 | -0.053 | 0.001 | -0.036 | -0.036 | 0 | 0.09 | 0.12 | 0.07 | 0 | 0.1 | 0.04 |
| 7 | 37493995 | rs1731989 | -0.065 | 0.852 | -1.391 | 0.186 | 0.007 | 0.003 | 0.003 | 0 | 0.94 | 0.78 | 0.92 | 0.92 | 0.94 | 0.93 |
| 7 | 37494415 | rs4548075 | 0.214 | 0.584 | -1.694 | 0.398 | 0 | 0.004 | 0.004 | 0 | 0.39 | 0.42 | 0.42 | 0.38 | 0.38 | 0.31 |
| 7 | 37494968 | rs1425129 | -0.722 | 0.185 | -1.965 | 0.258 | 0.006 | 0.055 | 0.055 | 1.414 | 0.16 | 0 | 0.15 | 0.06 | 0.11 | 0.05 |
| 7 | 37501026 | rs12539581 | -0.544 | 0.452 | -1.688 | 0.831 | 0.014 | -0.034 | -0.034 | 0 | 0.32 | 0.19 | 0.3 | 0.55 | 0.28 | 0.39 |
| 7 | 37501378 | rs10215851 | -0.67 | 0.504 | -1.77 | 0.718 | 0.02 | -0.03 | -0.03 | 0 | 0.37 | 0.23 | 0.33 | 0.51 | 0.34 | 0.4 |
| 7 | 37505367 | rs6977033 | 0.54 | 2.391 | -1.533 | 1.139 | -0.003 | -0.011 | -0.011 | 0 | 0.89 | 0.9 | 0.71 | 0.76 | 0.71 | 0.88 |
| 7 | 37509747 | rs11980720 | 0.828 | 1.189 | -1.296 | 1.379 | -0.029 | -0.029 | -0.029 | 0 | 0.64 | 0.76 | 0.5 | 0.54 | 0.52 | 0.57 |
| 7 | 37524894 | rs11763394 | 0.974 | 0.878 | -0.228 | 1.33 | 0.015 | 0.008 | 0.008 | 0.6 | 0.62 | 0.48 | 0.47 | 0.45 | 0.52 | 0.5 |
| 7 | 37529157 | rs8180866 | 0.611 | -0.173 | -1.13 | 0.912 | 0.024 | -0.026 | -0.026 | 0.213 | 0.22 | 0.04 | 0.08 | 0.16 | 0.15 | 0.19 |
| 7 | 37534044 | rs10237799 | -0.968 | 0.312 | -0.945 | 1.54 | 0.071 | 0.476 | 0.476 | 0 | 0.07 | 0.37 | 0.25 | 0.46 | 0.05 | 0.17 |
| 7 | 37534885 | rs2052680 | 0.588 | -0.2 | -0.376 | 1.457 | 0.025 | 0.019 | 0.019 | 0 | 0.49 | 0.73 | 0.63 | 0.64 | 0.42 | 0.49 |
| 7 | 37536367 | rs12671530 | 1.001 | -0.113 | -0.451 | 1.45 | -0.007 | -0.074 | -0.074 | 0 | 0.2 | 0.02 | 0.08 | 0.22 | 0.16 | 0.19 |
| 7 | 37541438 | rs1425119 | -0.74 | -0.109 | -0.165 | 1.455 | 0.068 | 0.455 | 0.455 | 0 | 0.07 | 0.35 | 0.24 | 0.46 | 0.05 | 0.18 |
| 7 | 37542051 | rs7791991 | -0.313 | -0.091 | -0.242 | 1.378 | -0.002 | -0.001 | -0.001 | 0 | 0.68 | 0.72 | 0.7 | 0.87 | 0.58 | 0.68 |
| 7 | 37547290 | rs7782150 | 1.382 | -0.177 | -0.237 | 1.305 | 0.006 | 0.119 | 0.119 | 0 | 0.4 | 0.16 | 0.37 | 0.18 | 0.37 | 0.32 |
| 7 | 37552669 | rs11770769 | 0.35 | -0.13 | -0.11 | 1.421 | 0.002 | 0 | 0 | 0 | 0.62 | 0.47 | 0.47 | 0.4 | 0.53 | 0.51 |
| 7 | 37552703 | rs10951521 | 0.35 | -0.13 | -0.11 | 1.418 | 0.002 | 0 | 0 | 0.312 | 0.62 | 0.47 | 0.47 | 0.4 | 0.53 | 0.51 |
| 7 | 37553149 | rs10951525 | -0.305 | -0.159 | 0.128 | 1.192 | 0.007 | -0.012 | -0.012 | 0 | 0.69 | 0.71 | 0.71 | 0.83 | 0.58 | 0.68 |
| 7 | 37553647 | rs2893542 | 0.129 | -0.17 | 0.1 | 1.328 | -0.006 | 0.012 | 0.012 | 3.266 | 0.25 | 0.29 | 0.31 | 0.64 | 0.21 | 0.37 |
| 7 | 37556237 | rs1364731 | 0.592 | -0.385 | 0.116 | 1.083 | -0.012 | -0.026 | -0.026 | 0 | 0.41 | 0.51 | 0.51 | 0.31 | 0.36 | 0.3 |
| 7 | 37562368 | rs1425132 | 0.39 | 0.045 | -0.088 | 1.12 | -0.02 | 0.069 | 0.069 | 0 | 0.55 | 0.33 | 0.42 | 0.29 | 0.49 | 0.4 |
| 7 | 37562888 | rs724489 | -0.207 | -0.036 | 0.15 | 1.46 | -0.002 | 0.007 | 0.007 | 0 | 0.62 | 0.73 | 0.71 | 0.79 | 0.59 | 0.69 |
| 7 | 37574668 | rs1469231 | -0.696 | 0.167 | 0.05 | 1.782 | -0.001 | 0.001 | 0.001 | 0 | 0.27 | 0.3 | 0.38 | 0.59 | 0.28 | 0.41 |
| 7 | 37581187 | rs1972684 | -0.354 | 0.159 | 0.075 | 1.554 | -0.001 | 0 | 0 | 0 | 0.24 | 0.24 | 0.18 | 0.28 | 0.23 | 0.18 |
| 7 | 37584705 | rs17171090 | -0.462 | 0.235 | -0.028 | 1.561 | 0.002 | -0.03 | -0.03 | 0 | 0.25 | 0.22 | 0.32 | 0.38 | 0.22 | 0.38 |
| 7 | 37591968 | rs6462772 | 0.296 | 0.221 | 0.231 | 1.506 | 0.014 | 0.02 | 0.02 | 0 | 0.45 | 0.64 | 0.61 | 0.7 | 0.41 | 0.58 |
| 7 | 37592231 | rs6971768 | 0.625 | 0.234 | 0.117 | 1.301 | -0.008 | -0.017 | -0.017 | 0.197 | 0.4 | 0.52 | 0.39 | 0.37 | 0.36 | 0.31 |
| 7 | 37594860 | rs11770352 | 1.25 | 0.239 | 0.049 | 1.296 | 0.006 | 0.014 | 0.014 | 0 | 0.35 | 0.29 | 0.32 | 0.28 | 0.31 | 0.29 |
| 7 | 37601546 | rs2033043 | 1.178 | 0.795 | -0.393 | 0.908 | -0.001 | 0.102 | 0.102 | 0 | 0.39 | 0.12 | 0.35 | 0.28 | 0.37 | 0.34 |
| 7 | 37603073 | rs6462776 | 0.487 | 0.795 | -0.105 | 0.973 | -0.001 | -0.004 | -0.004 | 0 | 0.47 | 0.52 | 0.45 | 0.36 | 0.45 | 0.38 |
| 7 | 37604528 | rs10257469 | 0.524 | 0.789 | -0.046 | 0.955 | 0.006 | 0.032 | 0.032 | 0 | 0.47 | 0.4 | 0.43 | 0.36 | 0.45 | 0.38 |
| 7 | 37608503 | rs4723673 | 2.622 | 0.764 | -0.109 | 0.947 | -0.026 | -0.043 | -0.043 | 0.266 | 0.07 | 0.01 | 0.11 | 0.2 | 0.1 | 0.08 |
| 7 | 37612307 | rs6957814 | 0.302 | 0.703 | 0.423 | 1.756 | 0.002 | 0.028 | 0.028 | 0 | 0.51 | 0.61 | 0.65 | 0.7 | 0.5 | 0.65 |
| 7 | 37618173 | rs1364729 | 0.596 | 0.718 | 0.245 | 1.68 | -0.015 | 0.088 | 0.088 | 0 | 0.2 | 0.09 | 0.11 | 0.04 | 0.18 | 0.12 |
| 7 | 37619922 | rs17265593 | -0.851 | 0.733 | 0.308 | 1.748 | 0.007 | 0.004 | 0.004 | 0 | 0.71 | 0.71 | 0.76 | 0.74 | 0.67 | 0.76 |
| 7 | 37620921 | rs13229754 | 1.812 | 0.775 | 0.023 | 1.733 | -0.002 | -0.081 | -0.081 | 0.23 | 0.2 | 0.01 | 0.13 | 0.2 | 0.15 | 0.2 |
| 7 | 37626839 | rs13222756 | 1.812 | 0.721 | 0.021 | 1.918 | -0.002 | -0.096 | -0.096 | 0 | 0.2 | 0.01 | 0.13 | 0.22 | 0.15 | 0.2 |
| 7 | 37663495 | rs10232731 | 1.977 | 0.946 | 0.132 | 1.626 | 0.014 | 0.103 | 0.103 | 0 | 0.35 | 0.1 | 0.29 | 0.22 | 0.32 | 0.33 |
| 7 | 37675572 | rs6462780 | 1.002 | 0.993 | 0.242 | 1.658 | -0.009 | -0.038 | -0.038 | 0.159 | 0.43 | 0.52 | 0.53 | 0.29 | 0.39 | 0.4 |
| 7 | 37684187 | rs11772261 | 1.083 | 0.986 | 0.264 | 1.637 | 0.017 | 0.069 | 0.069 | 0 | 0.42 | 0.3 | 0.35 | 0.24 | 0.38 | 0.35 |

| | | | | | | | | | | | | | | | | |
|---|---|---|---|---|---|---|---|---|---|---|---|---|---|---|---|---|
| 7 | 37689170 | rs2718073 | -1.162 | 1.073 | 0.347 | 1.72 | 0.007 | 0.008 | 0.008 | 0 | 0.71 | 0.71 | 0.76 | 0.7 | 0.67 | 0.76 |
| 7 | 37689851 | rs3847016 | 1.229 | 1.207 | 0.354 | 1.706 | 0.023 | 0.063 | 0.063 | 0 | 0.41 | 0.29 | 0.34 | 0.23 | 0.36 | 0.33 |
| 7 | 37690507 | rs2597283 | -1.158 | 1.213 | 0.255 | 1.723 | 0.006 | 0.012 | 0.012 | 0 | 0.71 | 0.72 | 0.59 | 0.63 | 0.66 | 0.7 |
| 7 | 37693767 | rs2597282 | -0.166 | 1.207 | 0.22 | 1.729 | -0.002 | 0.003 | 0.003 | 0 | 0.07 | 0.14 | 0.07 | 0.08 | 0.05 | 0.02 |
| 7 | 37699089 | rs2717937 | -0.908 | 1.191 | 0.213 | 1.884 | -0.007 | -0.041 | -0.041 | 2.132 | 0.3 | 0.21 | 0.23 | 0.5 | 0.3 | 0.44 |
| 7 | 37702069 | rs2718075 | -0.65 | 1.191 | 0.213 | 1.883 | -0.009 | -0.034 | -0.034 | 0 | 0.29 | 0.21 | 0.23 | 0.4 | 0.3 | 0.37 |
| 7 | 37705787 | rs3916058 | 2.664 | 1.048 | 0.088 | 1.64 | -0.045 | 0.064 | 0.064 | 0 | 0.25 | 0.03 | 0.2 | 0.16 | 0.25 | 0.23 |
| 7 | 37711525 | rs2718079 | -1.101 | 1.61 | 0.559 | 2.036 | 0.009 | -0.002 | -0.002 | 0 | 0.71 | 0.71 | 0.76 | 0.8 | 0.67 | 0.83 |
| 7 | 37712109 | rs4723693 | 1.184 | 1.599 | 0.573 | 2.021 | -0.009 | -0.037 | -0.037 | 0 | 0.42 | 0.48 | 0.36 | 0.23 | 0.36 | 0.33 |
| 7 | 37714079 | rs2717925 | 0.281 | 1.6 | 0.54 | 2.021 | -0.001 | 0.002 | 0.002 | 0 | 0.06 | 0.11 | 0.06 | 0.07 | 0.05 | 0.02 |
| 7 | 37723413 | rs2597277 | -1.08 | 1.625 | 0.556 | 2.012 | 0.008 | -0.002 | -0.002 | 0 | 0.71 | 0.72 | 0.76 | 0.79 | 0.67 | 0.83 |
| 7 | 37729431 | rs9942693 | 1.011 | 1.23 | 0.395 | 1.861 | -0.008 | -0.028 | -0.028 | 0 | 0.42 | 0.51 | 0.53 | 0.3 | 0.38 | 0.39 |
| 7 | 37736164 | rs10272013 | 1.039 | 1.231 | 0.393 | 1.861 | -0.007 | -0.025 | -0.025 | 0 | 0.42 | 0.48 | 0.53 | 0.3 | 0.38 | 0.39 |
| 7 | 37741303 | rs11764256 | 0.724 | 1.372 | 0.375 | 1.911 | -0.001 | -0.001 | -0.001 | 0 | 0.47 | 0.56 | 0.59 | 0.63 | 0.47 | 0.63 |
| 7 | 37741319 | rs11771135 | 0.724 | 1.372 | 0.374 | 2.115 | -0.001 | -0.001 | -0.001 | 0.022 | 0.47 | 0.56 | 0.59 | 0.53 | 0.47 | 0.63 |
| 7 | 37742031 | rs17171134 | -0.401 | 1.389 | 0.345 | 2.142 | 0.053 | 0.156 | 0.156 | 0 | 0.06 | 0.22 | 0.24 | 0.26 | 0.09 | 0.3 |
| 7 | 37746569 | rs2392510 | 0.678 | 1.453 | 0.164 | 2.073 | -0.001 | 0 | 0 | 0 | 0.47 | 0.47 | 0.58 | 0.53 | 0.46 | 0.63 |
| 7 | 37747276 | rs11535191 | 0.484 | 1.408 | -0.161 | 2.174 | -0.002 | -0.03 | -0.03 | 0 | 0.2 | 0.07 | 0.11 | 0.25 | 0.18 | 0.19 |
| 7 | 37755081 | rs1450178 | -1.061 | 1.545 | 0.268 | 2.037 | 0.005 | 0.016 | 0.016 | 0 | 0.31 | 0.35 | 0.49 | 0.49 | 0.35 | 0.37 |
| 7 | 37757216 | rs2709105 | -1.522 | 1.617 | 0.205 | 1.54 | -0.03 | -0.002 | -0.002 | 0 | 0.76 | 0.91 | 0.79 | 0.85 | 0.71 | 0.88 |
| 7 | 37758187 | rs1562767 | -1.241 | 1.554 | 0.65 | 1.513 | 0.017 | 0.003 | 0.003 | 0 | 0.73 | 0.63 | 0.76 | 0.78 | 0.7 | 0.84 |
| 7 | 37760902 | rs2257053 | 0.426 | 1.458 | 0.527 | 1.571 | 0.001 | -0.002 | -0.002 | 0.681 | 0.09 | 0.08 | 0.07 | 0.1 | 0.07 | 0.03 |
| 7 | 37763133 | rs2717931 | -1.031 | 1.468 | 0.604 | 1.626 | -0.004 | -0.016 | -0.016 | 0 | 0.28 | 0.52 | 0.22 | 0.23 | 0.26 | 0.21 |
| 7 | 37764184 | rs966053 | -0.889 | 1.705 | 0.726 | 1.796 | 0.003 | 0.041 | 0.041 | 0 | 0.26 | 0.48 | 0.21 | 0.31 | 0.24 | 0.21 |
| 7 | 37765688 | rs7779566 | 0.05 | 1.721 | 0.584 | 1.876 | 0.045 | 0.136 | 0.136 | 0 | 0.06 | 0.18 | 0.14 | 0.23 | 0.09 | 0.27 |
| 7 | 37774026 | rs2247244 | 0.268 | 1.636 | 0.335 | 1.8 | -0.003 | -0.009 | -0.009 | 0.48 | 0.77 | 0.82 | 0.82 | 0.69 | 0.76 | 0.78 |
| 7 | 37783669 | rs10269442 | -0.002 | 1.471 | 0.578 | 1.563 | 0.017 | 0.019 | 0.019 | 0 | 0.07 | 0.25 | 0.06 | 0.05 | 0.04 | 0.01 |
| 7 | 37786514 | rs2718042 | -0.866 | 1.565 | 0.201 | 1.557 | 0 | -0.002 | -0.002 | 0 | 0.84 | 0.85 | 0.89 | 0.9 | 0.84 | 0.8 |
| 7 | 37789556 | rs10243140 | 1.511 | 1.272 | 0.011 | 1.399 | 0.034 | 0.143 | 0.143 | 0 | 0.4 | 0.1 | 0.27 | 0.2 | 0.35 | 0.34 |
| 7 | 37792902 | rs10274478 | -1.409 | 1.383 | 0.063 | 1.496 | -0.031 | 0.008 | 0.008 | 0 | 0.75 | 0.87 | 0.8 | 0.86 | 0.73 | 0.89 |
| 7 | 37794937 | rs13233670 | 0.199 | 1.413 | 0.18 | 1.641 | 0.004 | 0.021 | 0.021 | 0.619 | 0.19 | 0.2 | 0.22 | 0.42 | 0.24 | 0.43 |
| 7 | 37798537 | rs6462787 | -0.364 | 1.398 | 0.305 | 1.533 | 0.032 | -0.009 | -0.009 | 0 | 0.09 | 0.31 | 0.08 | 0.06 | 0.1 | 0.02 |
| 7 | 37800529 | rs12155159 | -0.118 | 1.343 | 0.141 | 1.536 | 0 | -0.001 | -0.001 | 0.185 | 0.62 | 0.38 | 0.52 | 0.7 | 0.62 | 0.78 |
| 7 | 37808298 | rs2722273 | 1.637 | 1.171 | -0.057 | 1.538 | -0.069 | -0.024 | -0.024 | 0 | 0.23 | 0.04 | 0.19 | 0.13 | 0.25 | 0.1 |
| 7 | 37823995 | rs17171163 | -0.448 | 1.179 | -0.006 | 1.571 | 0.006 | 0.025 | 0.025 | 0.03 | 0.25 | 0.24 | 0.27 | 0.5 | 0.28 | 0.47 |
| 7 | 37833185 | rs10257984 | 1.76 | 1.146 | -0.154 | 1.496 | 0.019 | 0.069 | 0.069 | 0 | 0.38 | 0.12 | 0.43 | 0.31 | 0.37 | 0.41 |
| 7 | 37835603 | rs2722363 | 1.593 | 1.192 | -0.193 | 1.465 | -0.069 | -0.023 | -0.023 | 0 | 0.23 | 0.04 | 0.19 | 0.13 | 0.25 | 0.1 |
| 7 | 37843740 | rs13243936 | 0.218 | 1.285 | -0.125 | 1.692 | -0.025 | 0.003 | 0.003 | 0 | 0.05 | 0.01 | 0.27 | 0.29 | 0.08 | 0.28 |
| 7 | 37845286 | rs2722248 | 1.625 | 1.29 | -0.139 | 1.438 | -0.073 | -0.027 | -0.027 | 0 | 0.23 | 0.03 | 0.19 | 0.13 | 0.25 | 0.1 |
| 7 | 37858969 | rs4346913 | -0.175 | 1.228 | 0.032 | 1.341 | 0.021 | -0.011 | -0.011 | 0 | 0.09 | 0.3 | 0.09 | 0.06 | 0.1 | 0.04 |
| 7 | 37859423 | rs10241576 | 1.782 | 1.197 | -0.445 | 1.278 | 0.018 | 0.143 | 0.143 | 0 | 0.37 | 0.08 | 0.25 | 0.2 | 0.35 | 0.32 |
| 7 | 37863043 | rs1597550 | -1.266 | 1.194 | -0.413 | 1.277 | 0.039 | 0.133 | 0.133 | 0.089 | 0.58 | 0.8 | 0.72 | 0.8 | 0.63 | 0.67 |
| 7 | 37867036 | rs2722334 | 0.93 | 1.281 | -0.447 | 1.472 | -0.008 | -0.017 | -0.017 | 0.288 | 0.28 | 0.24 | 0.32 | 0.48 | 0.32 | 0.34 |
| 7 | 37867052 | rs2122624 | 0.674 | 1.29 | -0.292 | 1.443 | 0.009 | -0.008 | -0.008 | 1.137 | 0.43 | 0.35 | 0.39 | 0.54 | 0.42 | 0.57 |
| 7 | 37872545 | rs2722246 | 0.331 | 1.395 | -0.314 | 1.67 | 0 | 0.033 | 0.033 | 0 | 0.9 | 0.86 | 0.85 | 0.65 | 0.86 | 0.74 |
| 7 | 37872650 | rs17171172 | 0.522 | 1.572 | -0.074 | 1.663 | 0.063 | 0.042 | 0.042 | 0.119 | 0.14 | 0.3 | 0.14 | 0.15 | 0.19 | 0.19 |
| 7 | 37877695 | rs2598068 | 0.588 | 1.105 | -0.223 | 1.67 | 0.001 | -0.021 | -0.021 | 0 | 0.29 | 0.38 | 0.24 | 0.16 | 0.27 | 0.08 |
| 7 | 37879092 | rs2598067 | -0.676 | 1.047 | -0.249 | 1.746 | -0.003 | 0.026 | 0.026 | 0 | 0.92 | 0.89 | 0.88 | 0.73 | 0.95 | 0.78 |
| 7 | 37885121 | rs10261983 | 0.341 | 0.943 | -0.214 | 1.898 | 0.017 | -0.029 | -0.029 | 0 | 0.6 | 0.22 | 0.45 | 0.46 | 0.65 | 0.61 |
| 7 | 37888118 | rs16879722 | 1.222 | 0.922 | 0.064 | 1.432 | 0.001 | -0.006 | -0.006 | 0 | 0.23 | 0.25 | 0.21 | 0.1 | 0.24 | 0.06 |
| 7 | 37888174 | rs2598048 | -0.692 | 0.924 | 0.057 | 1.498 | -0.007 | 0.086 | 0.086 | 0 | 0.92 | 0.86 | 0.88 | 0.65 | 0.94 | 0.78 |
| 7 | 37888354 | rs4723713 | 1.175 | 0.936 | -0.187 | 1.245 | -0.024 | 0.084 | 0.084 | 0 | 0.44 | 0.1 | 0.31 | 0.32 | 0.44 | 0.41 |
| 7 | 37888627 | rs4720262 | 2.189 | 0.687 | -0.576 | 0.968 | -0.065 | 0.095 | 0.095 | 0 | 0.28 | 0.02 | 0.16 | 0.13 | 0.3 | 0.25 |
| 7 | 37890081 | rs2722371 | 0.667 | 0.981 | 0.165 | 1.523 | 0.003 | -0.019 | -0.019 | 0.015 | 0.27 | 0.41 | 0.22 | 0.18 | 0.25 | 0.11 |
| 7 | 37896896 | rs3213976 | 2.055 | 1.114 | 0.029 | 1.471 | -0.012 | -0.003 | -0.003 | 0 | 0.17 | 0.01 | 0.13 | 0.08 | 0.12 | 0.04 |
| 7 | 37897198 | rs6954013 | -0.289 | 0.846 | 0.192 | 1.353 | 0 | 0.007 | 0.007 | 0 | 0.68 | 0.65 | 0.68 | 0.47 | 0.7 | 0.66 |
| 7 | 37898279 | rs12536302 | 0.661 | 0.833 | -0.106 | 1.385 | -0.003 | 0 | 0 | 0 | 0.08 | 0.1 | 0.21 | 0.09 | 0.06 | 0.05 |
| 7 | 37901002 | rs17171184 | 0.902 | 1.048 | -0.538 | 1.448 | 0.016 | -0.006 | -0.006 | 0 | 0.12 | 0.13 | 0.13 | 0.07 | 0.19 | 0.19 |
| 7 | 37901490 | rs17235118 | 0.457 | 0.407 | -0.899 | 1.443 | -0.003 | 0 | 0 | 0 | 0.08 | 0.1 | 0.21 | 0.09 | 0.06 | 0.05 |
| 7 | 37906477 | rs2198050 | 0.056 | 0.61 | -1.021 | 1.41 | 0.001 | 0.03 | 0.03 | 0 | 0.92 | 0.92 | 0.96 | 0.66 | 0.95 | 0.77 |
| 7 | 37908779 | rs3807171 | 0.087 | 0.776 | -1.093 | 0.339 | 0.008 | 0.07 | 0.07 | 0 | 0.49 | 0.36 | 0.35 | 0.25 | 0.49 | 0.41 |
| 7 | 37912535 | rs2722309 | -0.005 | 0.517 | -1.256 | 0.592 | 0.003 | 0.007 | 0.007 | 0 | 0.62 | 0.32 | 0.68 | 0.69 | 0.64 | 0.66 |
| 7 | 37917888 | rs2598018 | -0.649 | 0.404 | -1.583 | 0.684 | -0.007 | 0.023 | 0.023 | 0.839 | 0.43 | 0.41 | 0.55 | 0.74 | 0.48 | 0.64 |
| 7 | 37920114 | rs6965805 | -0.657 | 0.347 | -1.806 | 0.772 | -0.014 | 0.019 | 0.019 | 0 | 0.12 | 0.26 | 0.13 | 0.22 | 0.1 | 0.24 |
| 7 | 37924737 | rs1530822 | -1.016 | 0.339 | -1.814 | 0.795 | -0.01 | 0.032 | 0.032 | 0.912 | 0.44 | 0.38 | 0.55 | 0.74 | 0.48 | 0.67 |
| 7 | 37926758 | rs2722366 | 0.074 | 0.33 | -1.847 | 1.12 | -0.064 | -0.084 | -0.084 | 0.278 | 0.31 | 0.11 | 0.26 | 0.42 | 0.37 | 0.36 |
| 7 | 37927702 | rs2722365 | -0.746 | 0.421 | -1.808 | 1.077 | -0.013 | 0.005 | 0.005 | 0.576 | 0.43 | 0.36 | 0.39 | 0.65 | 0.47 | 0.58 |
| 7 | 37932255 | rs10268173 | 0.378 | 0.269 | -1.579 | 1.178 | 0.011 | -0.04 | -0.04 | 0 | 0.12 | 0.06 | 0.06 | 0.32 | 0.08 | 0.18 |
| 7 | 37932888 | rs2598124 | -0.773 | 0.602 | -1.551 | 0.902 | -0.01 | 0.012 | 0.012 | 0.216 | 0.38 | 0.35 | 0.6 | 0.75 | 0.45 | 0.63 |

| | | | | | | | | | | | | | | | | |
|---|---|---|---|---|---|---|---|---|---|---|---|---|---|---|---|---|
| 7 | 37933059 | rs10488616 | 0.157 | 0.556 | -1.189 | 1.038 | 0.012 | -0.023 | -0.023 | 0.045 | 0.4 | 0.18 | 0.21 | 0.37 | 0.35 | 0.28 |
| 7 | 37934673 | rs7807193 | -0.049 | 0.56 | -0.492 | 0.999 | 0.01 | 0.013 | 0.013 | 0.472 | 0.27 | 0.33 | 0.45 | 0.32 | 0.28 | 0.38 |
| 7 | 37936530 | rs3213975 | 0.531 | 0.495 | 0.173 | 1.059 | -0.002 | -0.002 | -0.002 | 0 | 0.2 | 0.17 | 0.2 | 0.23 | 0.24 | 0.21 |
| 7 | 37937033 | rs10488617 | 0.514 | 0.494 | 0.173 | 1.058 | -0.002 | -0.002 | -0.002 | 0 | 0.2 | 0.18 | 0.2 | 0.23 | 0.24 | 0.2 |
| 7 | 37939840 | rs10276139 | 1.393 | 0.146 | -0.099 | 0.798 | 0.008 | 0.072 | 0.072 | 6.469 | 0.25 | 0.08 | 0.13 | 0.07 | 0.2 | 0.07 |
| 7 | 37940324 | rs12671838 | 0.516 | 0.144 | -0.08 | 0.8 | -0.01 | 0.002 | 0.002 | 0 | 0.07 | 0.15 | 0.24 | 0.09 | 0.04 | 0.17 |
| 7 | 37940824 | rs7811420 | 0.248 | -0.004 | -0.031 | 0.991 | 0.007 | 0 | 0 | 1.269 | 0.38 | 0.59 | 0.44 | 0.43 | 0.29 | 0.4 |
| 7 | 37941252 | rs1530820 | 1.474 | 0.06 | -0.123 | 1.05 | -0.05 | -0.018 | -0.018 | 0 | 0.17 | 0.05 | 0.18 | 0.13 | 0.23 | 0.2 |
| 7 | 37944375 | rs17236800 | 1.542 | 0.125 | -0.167 | 1.07 | 0.008 | 0.071 | 0.071 | 6.319 | 0.25 | 0.08 | 0.13 | 0.07 | 0.2 | 0.07 |
| 7 | 37944746 | rs4720265 | 0.974 | 0.144 | -0.129 | 1.08 | 0 | 0 | 0 | 0.144 | 0.21 | 0.18 | 0.2 | 0.23 | 0.24 | 0.21 |
| 7 | 37946348 | rs2598116 | 1.115 | -0.099 | -0.3 | 1.036 | 0.009 | -0.014 | -0.014 | 0 | 0.16 | 0.13 | 0.13 | 0.21 | 0.13 | 0.25 |
| 7 | 37947103 | rs1802074 | 1.08 | -0.085 | -0.145 | 1.01 | -0.023 | -0.015 | -0.015 | 0 | 0.3 | 0.4 | 0.18 | 0.26 | 0.22 | 0.13 |
| 7 | 37948289 | rs2722325 | 0.023 | -0.099 | -0.174 | 1.005 | 0.023 | 0.002 | 0.002 | 0.487 | 0.51 | 0.45 | 0.54 | 0.56 | 0.39 | 0.55 |
| 7 | 37951097 | rs1450857 | 0.177 | -0.075 | -0.162 | 1.026 | -0.002 | 0.001 | 0.001 | 0 | 0.22 | 0.25 | 0.38 | 0.3 | 0.17 | 0.42 |
| 7 | 37951726 | rs1132553 | -0.073 | 0.042 | -0.182 | 0.966 | 0.001 | 0.001 | 0.001 | 0.226 | 0.49 | 0.45 | 0.45 | 0.37 | 0.58 | 0.34 |
| 7 | 37952346 | rs3807193 | -0.092 | 0.072 | -0.167 | 0.98 | 0.001 | 0.001 | 0.001 | 0.275 | 0.49 | 0.45 | 0.45 | 0.37 | 0.58 | 0.34 |
| 7 | 37953840 | rs1132552 | 0.038 | 0.095 | -0.247 | 0.99 | 0.009 | 0.009 | 0.009 | 0.25 | 0.5 | 0.49 | 0.55 | 0.63 | 0.42 | 0.65 |
| 7 | 37955001 | rs2598111 | -0.056 | 0.142 | -0.24 | 0.992 | 0.002 | 0.009 | 0.009 | 0.13 | 0.51 | 0.5 | 0.56 | 0.73 | 0.44 | 0.66 |
| 7 | 37958289 | rs1376264 | 1.023 | 0.318 | -1.073 | 0.947 | -0.029 | 0.055 | 0.055 | 2.056 | 0.32 | 0.12 | 0.2 | 0.18 | 0.32 | 0.19 |
| 7 | 37961669 | rs1450852 | 0.434 | 0.167 | -0.759 | 1.179 | 0 | 0.001 | 0.001 | 0 | 0.18 | 0.18 | 0.37 | 0.3 | 0.16 | 0.42 |
| 7 | 37962818 | rs6964434 | 0.227 | 0.206 | -1.525 | 0.958 | 0.001 | 0.002 | 0.002 | 0.351 | 0.49 | 0.47 | 0.36 | 0.37 | 0.47 | 0.39 |
| 7 | 37962897 | rs6968251 | 0.227 | 0.206 | -1.525 | 0.958 | 0.001 | 0.002 | 0.002 | 0.351 | 0.49 | 0.47 | 0.36 | 0.37 | 0.47 | 0.39 |
| 7 | 37964903 | rs1349394 | 0.745 | 0.672 | -1.73 | 0.732 | -0.07 | 0.059 | 0.059 | 0 | 0.27 | 0.05 | 0.22 | 0.14 | 0.34 | 0.15 |
| 7 | 37965165 | rs3734952 | -0.451 | 0.796 | -0.601 | 1.174 | 0.012 | 0.009 | 0.009 | 0.496 | 0.24 | 0.35 | 0.22 | 0.23 | 0.24 | 0.21 |
| 7 | 37973014 | rs2598107 | -0.331 | 0.782 | -1.053 | 0.728 | -0.007 | -0.011 | -0.011 | 0.598 | 0.36 | 0.24 | 0.32 | 0.34 | 0.35 | 0.45 |
| 7 | 37984865 | rs6949704 | -0.252 | 0.697 | -0.44 | 0.374 | 0.001 | -0.024 | -0.024 | 0 | 0.22 | 0.37 | 0.35 | 0.17 | 0.26 | 0.1 |
| 7 | 37987519 | rs2290222 | -0.107 | 0.727 | -0.505 | 0.496 | -0.001 | -0.002 | -0.002 | 0.409 | 0.12 | 0.11 | 0.07 | 0.09 | 0.12 | 0.32 |
| 7 | 37988291 | rs1545442 | -0.858 | 0.418 | -0.873 | 0.118 | 0.001 | 0.008 | 0.008 | 0 | 0.18 | 0.11 | 0.1 | 0.09 | 0.16 | 0.35 |
| 7 | 37988311 | rs3816415 | 0.861 | 0.456 | -1.067 | -0.211 | -0.002 | 0.006 | 0.006 | 0 | 0.13 | 0.12 | 0.15 | 0.11 | 0.14 | 0.11 |
| 7 | 37988589 | rs2044831 | 0.007 | -0.426 | -0.8 | -0.038 | -0.045 | 0.094 | 0.094 | 0 | 0.75 | 0.51 | 0.75 | 0.59 | 0.78 | 0.74 |
| 7 | 37989095 | rs16879765 | 1.729 | -0.707 | -0.135 | 0.308 | -0.018 | 0.01 | 0.01 | 0 | 0.09 | 0.17 | 0.15 | 0.1 | 0.1 | 0.15 |
| 7 | 37989178 | rs2722276 | 1.424 | -1.271 | -0.243 | 0.047 | -0.015 | 0.034 | 0.034 | 0 | 0.44 | 0.12 | 0.51 | 0.39 | 0.53 | 0.31 |
| 7 | 37990222 | rs3734953 | 1.101 | -1.226 | -0.254 | 0.075 | -0.001 | -0.001 | -0.001 | 0 | 0.1 | 0.1 | 0.05 | 0.05 | 0.06 | 0.17 |
| 7 | 37990917 | rs2722278 | -1.29 | -1.218 | 0.394 | 0.096 | -0.003 | 0.148 | 0.148 | 0.771 | 0.91 | 0.77 | 0.93 | 0.67 | 0.93 | 0.93 |
| 7 | 37991129 | rs2722279 | -1.316 | -1.218 | 0.49 | 0.097 | 0 | 0.165 | 0.165 | 0.567 | 0.91 | 0.75 | 0.93 | 0.67 | 0.93 | 0.93 |
| 7 | 37991186 | rs1047785 | 0.996 | -1.229 | 0.274 | -0.086 | 0.038 | 0.014 | 0.014 | 0 | 0.55 | 0.2 | 0.54 | 0.33 | 0.59 | 0.48 |
| 7 | 37991389 | rs1376261 | -1.42 | -1.106 | 0.279 | 0.034 | 0 | 0.162 | 0.162 | 0.567 | 0.91 | 0.75 | 0.93 | 0.67 | 0.93 | 0.93 |
| 7 | 37991450 | rs1823522 | -1.42 | -1.106 | 0.279 | 0.034 | 0 | 0.162 | 0.162 | 0.567 | 0.91 | 0.75 | 0.93 | 0.67 | 0.93 | 0.93 |
| 7 | 37991523 | rs1376260 | -1.42 | -1.106 | 0.279 | 0.034 | 0 | 0.162 | 0.162 | 0.567 | 0.91 | 0.75 | 0.93 | 0.67 | 0.93 | 0.93 |
| 7 | 37992041 | rs2722281 | -0.057 | -1.144 | 0.098 | -0.045 | -0.002 | -0.003 | -0.003 | 0 | 0.13 | 0.15 | 0.09 | 0.13 | 0.1 | 0.21 |
| 7 | 37998848 | rs12701600 | -1.511 | -1 | 0.157 | 0.106 | -0.003 | 0.072 | 0.072 | 0.362 | 0.91 | 0.83 | 0.94 | 0.66 | 0.93 | 0.92 |
| 7 | 38003850 | rs1717740 | 1.042 | -1.001 | 0.018 | -0.115 | 0.007 | 0.01 | 0.01 | 0 | 0.16 | 0.12 | 0.08 | 0.05 | 0.11 | 0.21 |
| 7 | 38009269 | rs1668349 | 0.741 | -0.702 | 0.41 | -0.107 | 0.001 | 0.001 | 0.001 | 0 | 0.57 | 0.39 | 0.54 | 0.3 | 0.57 | 0.46 |
| 7 | 38027816 | rs2598097 | 0.268 | -0.656 | -0.314 | -0.617 | 0.005 | 0.039 | 0.039 | 0 | 0.23 | 0.05 | 0.3 | 0.15 | 0.16 | 0.28 |
| 7 | 38029651 | rs9986874 | 0.62 | -0.808 | -0.26 | -0.838 | 0.007 | -0.017 | -0.017 | 0 | 0.42 | 0.52 | 0.48 | 0.36 | 0.55 | 0.39 |
| 7 | 38038678 | rs7799502 | -1.9 | -0.985 | -0.225 | -0.918 | 0.005 | 0.007 | 0.007 | 0 | 0.81 | 0.86 | 0.88 | 0.95 | 0.87 | 0.79 |
| 7 | 38040271 | rs1574005 | 1.076 | -0.962 | -0.216 | -0.932 | 0.005 | -0.026 | -0.026 | 0.141 | 0.08 | 0.09 | 0.06 | 0 | 0.1 | 0.04 |
| 7 | 38043677 | rs1524053 | 1.018 | -1.104 | -0.149 | -1.017 | 0.024 | -0.043 | -0.043 | 0 | 0.35 | 0.46 | 0.43 | 0.22 | 0.49 | 0.27 |
| 7 | 38046212 | rs952368 | 2.909 | -1.143 | -0.222 | -0.994 | 0.007 | -0.029 | -0.029 | 0 | 0.07 | 0 | 0.04 | 0.12 | 0.07 | 0.1 |
| 7 | 38051822 | rs1721405 | 0.88 | -1.25 | -0.181 | -1.048 | 0.013 | 0.009 | 0.009 | 0 | 0.28 | 0.23 | 0.17 | 0.18 | 0.2 | 0.32 |
| 7 | 38052205 | rs1464717 | 0.446 | -1.116 | -0.244 | -0.987 | 0.001 | 0 | 0 | 0 | 0.62 | 0.65 | 0.6 | 0.4 | 0.68 | 0.59 |
| 7 | 38053602 | rs1717758 | -0.246 | -1.067 | -0.49 | -0.969 | 0.018 | -0.036 | -0.036 | 0 | 0.71 | 0.84 | 0.67 | 0.4 | 0.79 | 0.63 |
| 7 | 38056445 | rs1524078 | 0.743 | -1.05 | -0.487 | -1.003 | 0.015 | 0.068 | 0.068 | 0 | 0.55 | 0.74 | 0.57 | 0.27 | 0.62 | 0.47 |
| 7 | 38060785 | rs1721400 | -0.797 | -0.971 | -0.523 | -0.962 | 0.005 | 0.044 | 0.044 | 0 | 0.78 | 0.95 | 0.71 | 0.85 | 0.84 | 0.72 |
| 7 | 38063117 | rs7800054 | -0.523 | -0.986 | -0.088 | -0.746 | 0.01 | 0.014 | 0.014 | 0 | 0.59 | 0.65 | 0.59 | 0.79 | 0.73 | 0.52 |
| 7 | 38064005 | rs17171260 | 1.291 | -0.6 | -0.029 | -0.344 | -0.006 | 0.155 | 0.155 | 0 | 0.07 | 0.16 | 0.04 | 0.46 | 0.05 | 0.08 |
| 7 | 38064082 | rs6462801 | 0.293 | -0.463 | -0.127 | -0.637 | 0.004 | 0.003 | 0.003 | 0 | 0.33 | 0.6 | 0.34 | 0.58 | 0.42 | 0.3 |
| 7 | 38064636 | rs6462802 | 0.291 | -0.354 | -0.306 | -0.253 | -0.022 | -0.043 | -0.043 | 0 | 0.13 | 0.09 | 0.07 | 0.44 | 0.15 | 0.11 |
| 7 | 38065562 | rs7810404 | 0.443 | -0.347 | 0.252 | -0.34 | 0.059 | 0.009 | 0.009 | 0.008 | 0.08 | 0.34 | 0.07 | 0.04 | 0.11 | 0.13 |
| 7 | 38067723 | rs1524072 | 0.156 | -0.122 | 0.204 | -0.23 | -0.009 | 0.003 | 0.003 | 0 | 0.15 | 0.12 | 0.09 | 0.45 | 0.18 | 0.12 |
| 7 | 38077008 | rs17171271 | 0.137 | -0.12 | 0.27 | -0.229 | 0.004 | 0.04 | 0.04 | 0 | 0.15 | 0.15 | 0.1 | 0.45 | 0.18 | 0.12 |
| 7 | 38082249 | rs1357653 | 0.464 | 0.056 | 0.309 | -0.499 | 0.011 | -0.05 | -0.05 | 0 | 0.19 | 0.34 | 0.21 | 0.09 | 0.24 | 0.19 |
| 7 | 38093233 | rs1524071 | 0.228 | -0.419 | -0.025 | -0.769 | 0 | 0 | 0 | 0 | 0.47 | 0.5 | 0.38 | 0.29 | 0.47 | 0.6 |
| 7 | 38097548 | rs1980313 | 0.735 | -0.343 | -0.11 | -0.65 | -0.004 | -0.004 | -0.004 | 0 | 0.31 | 0.33 | 0.34 | 0.5 | 0.41 | 0.21 |
| 7 | 38098280 | rs7781364 | 0.878 | -0.217 | -0.186 | -0.62 | -0.013 | -0.012 | -0.012 | 0 | 0.15 | 0.1 | 0.1 | 0.4 | 0.17 | 0.14 |
| 7 | 38103949 | rs1474346 | -1.511 | -0.371 | -0.388 | -0.645 | 0.008 | -0.026 | -0.026 | 0 | 0.83 | 0.97 | 0.73 | 0.78 | 0.88 | 0.78 |
| 7 | 38109048 | rs1014842 | 0.021 | -0.431 | -0.452 | -0.762 | 0.004 | -0.002 | -0.002 | 0 | 0.43 | 0.53 | 0.4 | 0.61 | 0.46 | 0.38 |
| 7 | 38110394 | rs939666 | 2.228 | -0.49 | -0.462 | -0.836 | -0.001 | 0.021 | 0.021 | 0.179 | 0.22 | 0.1 | 0.13 | 0.09 | 0.18 | 0.25 |
| 7 | 38110425 | rs940347 | -0.128 | -0.405 | -0.476 | -0.801 | 0 | 0 | 0 | 0.738 | 0.64 | 0.66 | 0.53 | 0.69 | 0.64 | 0.63 |

| | | | | | | | | | | | | | | | | |
|---|---|---|---|---|---|---|---|---|---|---|---|---|---|---|---|---|
| 7 | 38123209 | rs17408199 | 2.576 | 0.144 | -0.744 | -0.617 | -0.058 | 0.045 | 0.045 | 0 | 0.15 | 0.03 | 0.23 | 0.1 | 0.23 | 0.07 |
| 7 | 38123440 | rs1524066 | 0.999 | -0.09 | -0.5 | -0.616 | 0 | -0.006 | -0.006 | 0.11 | 0.18 | 0.17 | 0.19 | 0.09 | 0.24 | 0.15 |
| 7 | 38140307 | rs1524063 | 0.289 | 0.391 | -0.598 | -0.248 | -0.02 | -0.027 | -0.027 | 0 | 0.3 | 0.28 | 0.33 | 0.5 | 0.39 | 0.21 |
| 7 | 38149932 | rs10280461 | 0.218 | 0.368 | -0.503 | -0.15 | 0.004 | 0.013 | 0.013 | 0 | 0.47 | 0.45 | 0.44 | 0.59 | 0.51 | 0.44 |
| 7 | 38153390 | rs4576333 | 1.04 | 0.234 | -0.986 | -0.381 | 0 | 0.003 | 0.003 | 0.431 | 0.22 | 0.2 | 0.14 | 0.08 | 0.18 | 0.26 |
| 7 | 38153444 | rs4346915 | 0.349 | 0.102 | -1.029 | -0.444 | -0.011 | 0.037 | 0.037 | 0 | 0.41 | 0.3 | 0.4 | 0.25 | 0.42 | 0.36 |
| 7 | 38155563 | rs2893552 | 1.087 | -0.087 | -1.411 | 0.093 | -0.062 | -0.046 | -0.046 | 0 | 0.24 | 0.09 | 0.39 | 0.31 | 0.38 | 0.17 |
| 7 | 38155667 | rs1524056 | -1.212 | -0.109 | -1.451 | -0.305 | 0.003 | 0.004 | 0.004 | 0 | 0.89 | 0.88 | 0.95 | 0.92 | 0.92 | 0.91 |
| 7 | 38156102 | rs10246805 | 1.259 | 0.063 | -1.621 | -0.171 | 0.008 | 0.019 | 0.019 | 0.081 | 0.21 | 0.11 | 0.16 | 0.1 | 0.16 | 0.24 |
| 7 | 38156645 | rs9791663 | 0.475 | -0.062 | -0.845 | -0.035 | 0.014 | 0.033 | 0.033 | 0 | 0.12 | 0.11 | 0.09 | 0.27 | 0.14 | 0.16 |
| 7 | 38157881 | rs13246622 | 0.881 | -0.173 | -1.371 | -0.044 | 0.026 | 0.02 | 0.02 | 0 | 0.12 | 0.01 | 0.05 | 0.1 | 0.07 | 0.16 |
| 7 | 38159445 | rs6944227 | 0.645 | 0.042 | -1.146 | 0.084 | 0.004 | 0.005 | 0.005 | 3.841 | 0.47 | 0.26 | 0.65 | 0.51 | 0.57 | 0.51 |
| 7 | 38161518 | rs6975425 | 0.451 | 0.186 | -0.536 | 0.435 | -0.008 | 0.011 | 0.011 | 1.005 | 0.67 | 0.46 | 0.79 | 0.63 | 0.75 | 0.73 |
| 7 | 38163538 | rs4720268 | -0.269 | 0.175 | -0.534 | 0.341 | -0.018 | -0.013 | -0.013 | 0 | 0.36 | 0.43 | 0.33 | 0.31 | 0.24 | 0.45 |
| 7 | 38176373 | rs6969314 | -0.723 | 1.079 | -0.147 | 0.245 | 0.018 | -0.022 | -0.022 | 0 | 0.32 | 0.51 | 0.3 | 0.23 | 0.41 | 0.31 |
| 7 | 38178196 | rs6462811 | -0.535 | 1.334 | 0.398 | 0.174 | 0.06 | 0.033 | 0.033 | 0 | 0.11 | 0.35 | 0.14 | 0.08 | 0.16 | 0.16 |
| 7 | 38178319 | rs6462813 | -1.136 | 1.324 | 0.355 | 0.138 | 0.042 | -0.027 | -0.027 | 0 | 0.26 | 0.44 | 0.26 | 0.17 | 0.38 | 0.26 |
| 7 | 38183787 | rs1357647 | 0.003 | 1.404 | -0.007 | 0.472 | -0.004 | -0.015 | -0.015 | 0 | 0.13 | 0.1 | 0.18 | 0.32 | 0.12 | 0.25 |
| 7 | 38184046 | rs1357648 | 0.13 | 1.218 | 0.333 | 0.473 | 0 | 0.012 | 0.012 | 0 | 0.3 | 0.35 | 0.39 | 0.27 | 0.33 | 0.26 |
| 7 | 38209837 | rs7801794 | -0.255 | 1.601 | 0.102 | 0.435 | -0.002 | -0.002 | -0.002 | 0 | 0.15 | 0.18 | 0.16 | 0.19 | 0.23 | 0.2 |
| 7 | 38217295 | rs10259650 | -0.985 | 1.583 | 0.411 | 0.314 | -0.016 | -0.025 | -0.025 | 1.3 | 0.62 | 0.56 | 0.74 | 0.7 | 0.67 | 0.66 |
| 7 | 38217555 | rs10230858 | -0.976 | 1.611 | 0.459 | 0.312 | -0.02 | -0.032 | -0.032 | 0.396 | 0.62 | 0.53 | 0.74 | 0.7 | 0.67 | 0.66 |
| 7 | 38218976 | rs1019179 | 1.199 | 1.767 | 0.371 | 0.207 | 0.018 | -0.037 | -0.037 | 1.595 | 0.26 | 0.12 | 0.33 | 0.27 | 0.22 | 0.3 |
| 7 | 38228480 | rs17171306 | 0.843 | 1.862 | 0.347 | 0.249 | -0.009 | -0.043 | -0.043 | 0.039 | 0.3 | 0.12 | 0.38 | 0.31 | 0.34 | 0.35 |
| 7 | 38228902 | rs2159499 | 0.036 | 1.86 | 0.376 | 0.224 | -0.009 | -0.003 | -0.003 | 0 | 0.41 | 0.32 | 0.49 | 0.48 | 0.45 | 0.5 |
| 7 | 38229107 | rs1014136 | -0.151 | 1.868 | 0.322 | 0.228 | 0.001 | -0.003 | -0.003 | 0 | 0.21 | 0.21 | 0.25 | 0.22 | 0.22 | 0.16 |
| 7 | 38259354 | rs2392540 | 0.789 | 1.818 | 0.386 | 0.21 | -0.016 | -0.011 | -0.011 | 0 | 0.13 | 0.08 | 0.19 | 0.22 | 0.15 | 0.15 |
| 7 | 38267754 | rs4720271 | -2.027 | 1.733 | 0.501 | 0.125 | 0.01 | 0.048 | 0.048 | 0.023 | 0.73 | 0.91 | 0.82 | 0.84 | 0.8 | 0.73 |
| 7 | 38268395 | rs7795499 | 0.102 | 1.687 | 0.743 | 0.17 | -0.01 | -0.005 | -0.005 | 0 | 0.41 | 0.32 | 0.49 | 0.48 | 0.45 | 0.5 |
| 7 | 38268811 | rs10252428 | 0.982 | 1.706 | 0.784 | 0.159 | 0.008 | -0.03 | -0.03 | 0 | 0.07 | 0.1 | 0.06 | 0 | 0.07 | 0.01 |
| 7 | 38269292 | rs10226230 | -0.289 | 1.727 | 0.6 | 0.163 | 0.001 | 0.022 | 0.022 | 0 | 0.51 | 0.43 | 0.55 | 0.62 | 0.58 | 0.56 |
| 7 | 38271544 | rs6978770 | 0.582 | 1.414 | 0.362 | -0.054 | -0.024 | -0.028 | -0.028 | 0 | 0.06 | 0.01 | 0.09 | 0.2 | 0.08 | 0.1 |
| 7 | 38273305 | rs6462820 | -0.69 | 1.357 | 0.343 | -0.17 | -0.017 | -0.006 | -0.006 | 0.145 | 0.61 | 0.86 | 0.71 | 0.64 | 0.68 | 0.52 |
| 7 | 38273611 | rs4720273 | 0.326 | 1.367 | 0.333 | -0.135 | 0.003 | 0.021 | 0.021 | 0.01 | 0.46 | 0.28 | 0.57 | 0.61 | 0.49 | 0.45 |
| 7 | 38277792 | rs2534596 | 1.136 | 1.364 | -0.027 | -0.077 | -0.036 | -0.1 | -0.1 | 0 | 0.32 | 0.11 | 0.49 | 0.48 | 0.34 | 0.38 |
| 7 | 38278789 | rs2159502 | 1.15 | 1.374 | 0.03 | -0.077 | -0.021 | -0.075 | -0.075 | 0.591 | 0.31 | 0.15 | 0.48 | 0.47 | 0.34 | 0.39 |
| 7 | 38288270 | rs2240825 | -2.265 | 1.004 | 0.741 | 0.267 | -0.019 | -0.009 | -0.009 | 0 | 0.45 | 0.87 | 0.36 | 0.48 | 0.46 | 0.45 |
| 7 | 38291851 | rs6954071 | -1.997 | 0.982 | 1.255 | 0.287 | -0.002 | 0.003 | 0.003 | 2.972 | 0.37 | 0.6 | 0.29 | 0.48 | 0.38 | 0.39 |
| 7 | 38292663 | rs13244147 | 0.949 | 1.013 | 1.244 | 0.308 | -0.012 | -0.023 | -0.023 | 0.112 | 0.09 | 0 | 0.06 | 0.14 | 0.13 | 0.07 |
| 7 | 38297145 | rs11971259 | -2.118 | 1.014 | 1.247 | 0.308 | -0.002 | 0.004 | 0.004 | 3.037 | 0.37 | 0.6 | 0.29 | 0.48 | 0.38 | 0.39 |
| 7 | 38298285 | rs28677126 | -1.726 | 1.018 | 1.236 | 0.3 | 0 | 0.001 | 0.001 | 2.385 | 0.25 | 0.27 | 0.2 | 0.34 | 0.25 | 0.33 |
| 7 | 38319339 | rs2240826 | -0.607 | 1.087 | 0.769 | 0.334 | -0.013 | -0.016 | -0.016 | 0.003 | 0.15 | 0.03 | 0.12 | 0.19 | 0.13 | 0.19 |
| 7 | 38319436 | rs2240827 | 0.992 | 1.125 | 0.28 | 0.35 | -0.016 | -0.031 | -0.031 | 0 | 0.09 | 0 | 0.06 | 0.15 | 0.13 | 0.06 |
| 7 | 38323485 | rs1546834 | -1.887 | 1.078 | 0.515 | 0.372 | 0.003 | 0.013 | 0.013 | 3.243 | 0.27 | 0.64 | 0.23 | 0.34 | 0.25 | 0.31 |
| 7 | 38330810 | rs1860517 | -1.97 | 0.999 | -0.199 | 0.395 | -0.041 | -0.034 | -0.034 | 0 | 0.45 | 0.91 | 0.35 | 0.49 | 0.44 | 0.41 |
| 7 | 38331271 | rs1860520 | -1.408 | 0.63 | 0.688 | 0.35 | -0.042 | 0.065 | 0.065 | 3.59 | 0.09 | 0.68 | 0.06 | 0.14 | 0.01 | 0.08 |
| 7 | 38331504 | rs1860521 | -1.387 | 0.64 | 0.558 | 0.355 | -0.043 | 0.059 | 0.059 | 3.643 | 0.09 | 0.7 | 0.06 | 0.14 | 0.01 | 0.09 |
| 7 | 38334818 | rs10441090 | -0.58 | 1.118 | -0.04 | -0.017 | -0.001 | 0.001 | 0.001 | 0 | 0.16 | 0.12 | 0.11 | 0.14 | 0.13 | 0.19 |
| 7 | 38335520 | rs2191311 | -1.062 | 1.148 | -0.321 | 0.018 | 0.004 | -0.034 | -0.034 | 0 | 0.26 | 0.62 | 0.19 | 0.16 | 0.24 | 0.23 |
| 7 | 38341226 | rs2248839 | -0.104 | 1.287 | -1.26 | -0.104 | -0.051 | 0.034 | 0.034 | 0 | 0.84 | 0.99 | 0.85 | 0.88 | 0.79 | 0.86 |
| 7 | 38341615 | rs2736973 | -0.128 | 1.341 | -1.261 | -0.105 | -0.033 | 0.033 | 0.033 | 0 | 0.84 | 0.99 | 0.87 | 0.88 | 0.81 | 0.87 |
| 7 | 38341925 | rs2240839 | 0.245 | 1.253 | -1.283 | -0.025 | 0.015 | 0.001 | 0.001 | 1.418 | 0.48 | 0.64 | 0.52 | 0.59 | 0.37 | 0.34 |
| 7 | 38343085 | rs2392546 | 1.11 | 0.774 | -1.402 | -0.748 | 0.001 | 0.026 | 0.026 | 0.171 | 0.17 | 0.12 | 0.15 | 0.06 | 0.16 | 0.1 |
| 7 | 38343670 | rs2736970 | -0.52 | 0.885 | -1.515 | -0.099 | 0.002 | -0.001 | -0.001 | 0 | 0.44 | 0.58 | 0.53 | 0.44 | 0.37 | 0.36 |
| 7 | 38343956 | rs2736969 | 0.727 | 0.312 | -1.228 | -0.146 | 0.012 | 0.006 | 0.006 | 0 | 0.16 | 0.08 | 0.14 | 0.09 | 0.11 | 0.08 |
| 7 | 38344769 | rs2736964 | -0.663 | 0.957 | -1.368 | -0.26 | -0.003 | 0.008 | 0.008 | 0.774 | 0.48 | 0.63 | 0.58 | 0.44 | 0.46 | 0.43 |
| 7 | 38348812 | rs12533564 | 0.694 | 0.976 | -1.232 | -0.235 | 0.007 | 0.029 | 0.029 | 0 | 0.3 | 0.55 | 0.41 | 0.39 | 0.3 | 0.36 |
| 7 | 38356415 | rs6953248 | 0.417 | 1.213 | -1.277 | 0.008 | -0.009 | -0.079 | -0.079 | 0 | 0.17 | 0.06 | 0.12 | 0.34 | 0.17 | 0.24 |
| 7 | 38357075 | rs11769443 | -0.266 | 1.226 | -0.741 | -0.204 | 0.007 | -0.002 | -0.002 | 0.678 | 0.33 | 0.53 | 0.44 | 0.34 | 0.37 | 0.37 |
| 7 | 38357323 | rs11765884 | 1.065 | 1.414 | -1.538 | -0.232 | -0.018 | 0.019 | 0.019 | 0 | 0.2 | 0.15 | 0.19 | 0.05 | 0.26 | 0.18 |
| 7 | 38358412 | rs2240845 | -0.255 | 1.298 | -1.486 | -0.119 | -0.025 | -0.028 | -0.028 | 0 | 0.32 | 0.25 | 0.46 | 0.47 | 0.4 | 0.42 |
| 7 | 38358956 | rs2534567 | -0.142 | 1.147 | -1.436 | 0.395 | -0.007 | 0.013 | 0.013 | 0.031 | 0.12 | 0.09 | 0.29 | 0.42 | 0.17 | 0.25 |
| 7 | 38359720 | rs2534569 | -1.245 | 0.712 | -1.576 | -0.022 | -0.026 | 0.034 | 0.034 | 0 | 0.84 | 0.99 | 0.87 | 0.88 | 0.82 | 0.88 |
| 7 | 38361999 | rs718880 | 0.294 | 0.8 | -1.307 | 0.543 | -0.008 | -0.013 | -0.013 | 0 | 0.1 | 0.05 | 0.25 | 0.31 | 0.11 | 0.21 |
| 7 | 38362957 | rs17171331 | -0.412 | 0.466 | -0.232 | 0.329 | 0.055 | 0.026 | 0.026 | 0 | 0.07 | 0.38 | 0.04 | 0 | 0.02 | 0.03 |
| 7 | 38363748 | rs2240850 | 0.462 | 0.252 | -0.746 | -0.168 | 0.009 | -0.014 | -0.014 | 0 | 0.4 | 0.18 | 0.48 | 0.38 | 0.35 | 0.43 |
| 7 | 38363929 | rs2240851 | -0.675 | 0.747 | -1.085 | -0.228 | -0.012 | -0.032 | -0.032 | 0 | 0.81 | 0.86 | 0.78 | 0.55 | 0.73 | 0.71 |
| 7 | 38367259 | rs2191312 | 0.405 | 0.474 | -0.819 | -0.894 | -0.062 | 0.119 | 0.119 | 0 | 0.31 | 0.08 | 0.23 | 0.16 | 0.36 | 0.25 |
| 7 | 38371474 | rs2534575 | -0.251 | 0.561 | -0.607 | -0.84 | 0 | -0.039 | -0.039 | 0 | 0.37 | 0.6 | 0.28 | 0.16 | 0.36 | 0.27 |

| | | | | | | | | | | | | | | | |
|---|---|---|---|---|---|---|---|---|---|---|---|---|---|---|---|
| 7 | 38374697 | rs2240853 | -0.081 | 0.75 | -0.114 | -0.901 | 0.042 | -0.121 | -0.121 | 0 | 0.25 | 0.55 | 0.24 | 0.06 | 0.3 | 0.15 |
| 7 | 38380446 | rs733905 | -0.181 | -0.286 | -0.055 | -0.917 | -0.066 | -0.077 | -0.077 | 0 | 0.24 | 0.47 | 0.12 | 0.13 | 0.13 | 0.07 |
| 7 | 38383243 | rs2735143 | -1.55 | -0.243 | -0.797 | -1.189 | 0.003 | 0.035 | 0.035 | 0 | 0.82 | 0.93 | 0.89 | 0.96 | 0.83 | 0.87 |
| 7 | 38402496 | rs2735118 | -0.771 | -0.096 | -0.952 | -0.314 | -0.003 | -0.039 | -0.039 | 0 | 0.73 | 0.82 | 0.8 | 0.56 | 0.73 | 0.62 |
| 7 | 38412022 | rs10235885 | 1.12 | -0.798 | -0.577 | -0.617 | -0.021 | -0.01 | -0.01 | 0.665 | 0.18 | 0.3 | 0.07 | 0.14 | 0.08 | 0.07 |
| 7 | 38412988 | rs4373430 | 1.168 | -0.593 | -1.409 | -0.698 | -0.04 | -0.048 | -0.048 | 1.049 | 0.17 | 0.04 | 0.33 | 0.26 | 0.24 | 0.26 |
| 7 | 38413859 | rs11761049 | 2.287 | 0.132 | -1.464 | -0.837 | -0.015 | 0.01 | 0.01 | 0 | 0.09 | 0.06 | 0.12 | 0.07 | 0.13 | 0.05 |
| 7 | 38417556 | rs2080284 | -0.358 | -0.028 | -1.096 | -1.168 | -0.009 | 0.02 | 0.02 | 0.584 | 0.54 | 0.43 | 0.54 | 0.38 | 0.5 | 0.51 |
| 7 | 38420184 | rs17158 | 2.145 | 0.107 | -1.408 | -1.061 | -0.004 | -0.017 | -0.017 | 6.902 | 0.15 | 0 | 0.07 | 0.14 | 0.11 | 0.05 |
| 7 | 38420403 | rs17497509 | 2.217 | 0.061 | -1.463 | -1.199 | -0.035 | 0.06 | 0.06 | 0 | 0.13 | 0.03 | 0.08 | 0 | 0.14 | 0.14 |
| 7 | 38424039 | rs2072505 | 1.87 | 0.046 | -0.999 | -1.105 | 0.004 | 0.004 | 0.004 | 0.324 | 0.16 | 0.23 | 0.21 | 0.16 | 0.21 | 0.12 |
| 7 | 38424065 | rs6960550 | 0.533 | -0.155 | -0.532 | -1.172 | 0.038 | -0.006 | -0.006 | 0 | 0.08 | 0.41 | 0.07 | 0 | 0.04 | 0.01 |
| 7 | 38428814 | rs6972549 | -1.098 | 0.032 | -0.994 | -1.206 | 0.002 | -0.022 | -0.022 | 0.303 | 0.68 | 0.76 | 0.79 | 0.55 | 0.69 | 0.57 |
| 7 | 38429370 | rs6977263 | 0.888 | 0.13 | -1.084 | -0.981 | -0.05 | 0.081 | 0.081 | 0.773 | 0.28 | 0.04 | 0.16 | 0.14 | 0.26 | 0.19 |
| 7 | 38431436 | rs1058656 | 1.011 | -0.24 | -0.565 | -0.691 | -0.002 | -0.007 | -0.007 | 0 | 0.17 | 0.23 | 0.22 | 0.13 | 0.18 | 0.13 |
| 7 | 38432937 | rs2024408 | 0.046 | -0.72 | -0.2 | -0.341 | 0 | 0 | 0 | 0 | 0.5 | 0.52 | 0.54 | 0.45 | 0.5 | 0.44 |
| 7 | 38432978 | rs3823598 | 1.929 | -0.72 | -0.417 | -0.262 | 0.022 | -0.065 | -0.065 | 0 | 0.14 | 0 | 0.05 | 0.18 | 0.07 | 0.06 |
| 7 | 38433361 | rs17413296 | 1.112 | -0.544 | -0.449 | -0.288 | -0.033 | -0.044 | -0.044 | 0 | 0.08 | 0.01 | 0.28 | 0.24 | 0.14 | 0.25 |
| 7 | 38433986 | rs2072507 | 0.622 | -0.652 | -0.28 | -0.205 | -0.034 | 0.036 | 0.036 | 0 | 0.36 | 0.21 | 0.46 | 0.27 | 0.44 | 0.39 |
| 7 | 38435447 | rs6953928 | 1.787 | -0.721 | 0.001 | -0.352 | -0.023 | 0.011 | 0.011 | 1.151 | 0.15 | 0.16 | 0.06 | 0.18 | 0.07 | 0.06 |
| 7 | 38439572 | rs1030016 | -0.963 | -0.722 | 0.155 | -0.546 | 0 | -0.039 | -0.039 | 0 | 0.67 | 0.67 | 0.79 | 0.97 | 0.68 | 0.85 |
| 7 | 38441643 | rs2299945 | 1.157 | -0.683 | 0.091 | -0.524 | -0.004 | 0.003 | 0.003 | 0 | 0.31 | 0.32 | 0.28 | 0.14 | 0.36 | 0.28 |
| 7 | 38450598 | rs2267806 | 0.984 | -0.791 | 0.2 | -0.237 | -0.003 | 0.064 | 0.064 | 0 | 0.07 | 0.07 | 0.05 | 0.35 | 0.02 | 0.16 |
| 7 | 38452648 | rs2267807 | 1.259 | -0.734 | 0.29 | -0.36 | 0 | -0.058 | -0.058 | 0 | 0.28 | 0.39 | 0.19 | 0.03 | 0.29 | 0.14 |
| 7 | 38454797 | rs2299946 | 0.471 | -0.678 | 0.083 | -0.349 | -0.005 | -0.005 | -0.005 | 0.473 | 0.42 | 0.39 | 0.41 | 0.47 | 0.45 | 0.45 |
| 7 | 38460351 | rs1019286 | -0.537 | -0.814 | -0.169 | -0.577 | -0.001 | 0.092 | 0.092 | 0 | 0.64 | 0.77 | 0.78 | 0.95 | 0.64 | 0.75 |
| 7 | 38468463 | rs2267808 | -0.313 | -0.807 | 0.071 | -0.398 | -0.056 | -0.002 | -0.002 | 0 | 0.75 | 0.61 | 0.86 | 0.79 | 0.86 | 0.82 |
| 7 | 38469039 | rs2215955 | 0.414 | -0.678 | -0.121 | -0.144 | 0.006 | -0.011 | -0.011 | 0 | 0.35 | 0.29 | 0.22 | 0.38 | 0.32 | 0.3 |
| 7 | 38474057 | rs2267809 | -0.39 | -0.68 | -0.004 | -0.138 | -0.005 | 0.023 | 0.023 | 0.289 | 0.43 | 0.65 | 0.59 | 0.7 | 0.44 | 0.63 |
| 7 | 38474750 | rs973527 | 1.348 | -0.772 | -0.102 | -0.245 | -0.023 | 0.044 | 0.044 | 0 | 0.28 | 0.23 | 0.26 | 0.09 | 0.35 | 0.18 |
| 7 | 38476474 | rs4720279 | 0.176 | -0.829 | -0.096 | -0.058 | 0.004 | 0.008 | 0.008 | 0 | 0.5 | 0.53 | 0.63 | 0.56 | 0.54 | 0.4 |
| 7 | 38479220 | rs3778884 | 0.714 | -0.664 | -0.228 | -0.104 | 0.004 | 0.002 | 0.002 | 0 | 0.29 | 0.29 | 0.43 | 0.36 | 0.36 | 0.32 |
| 7 | 38479434 | rs3778883 | -0.64 | -0.58 | -0.579 | -0.384 | -0.025 | 0.082 | 0.082 | 0 | 0.63 | 0.76 | 0.7 | 0.88 | 0.58 | 0.71 |
| 7 | 38488638 | rs2284248 | 1.225 | -0.338 | -0.584 | -0.23 | -0.005 | 0.003 | 0.003 | 0.375 | 0.22 | 0.21 | 0.13 | 0.03 | 0.25 | 0.11 |
| 7 | 38491495 | rs4723754 | -0.739 | -0.534 | -0.396 | 0.009 | -0.005 | 0.033 | 0.033 | 0 | 0.18 | 0.24 | 0.36 | 0.66 | 0.16 | 0.51 |
| 7 | 38493602 | rs4723756 | -0.929 | -0.528 | -0.518 | -0.339 | -0.009 | 0.032 | 0.032 | 0 | 0.73 | 0.79 | 0.85 | 0.97 | 0.72 | 0.88 |
| 7 | 38497295 | rs6943730 | 0.125 | -0.775 | -0.593 | -0.024 | 0.015 | -0.058 | -0.058 | 0.12 | 0.31 | 0.16 | 0.42 | 0.52 | 0.26 | 0.38 |
| 7 | 38501065 | rs2302611 | 1.336 | -0.624 | -0.692 | 0.073 | -0.031 | 0.002 | 0.002 | 0 | 0.06 | 0.01 | 0.3 | 0.34 | 0.12 | 0.2 |
| 7 | 38503023 | rs1019288 | -0.421 | -0.791 | -0.372 | 0.06 | -0.023 | 0.009 | 0.009 | 1.307 | 0.36 | 0.46 | 0.53 | 0.58 | 0.31 | 0.51 |
| 7 | 38510787 | rs1014570 | -1.119 | -0.813 | -0.302 | 0.139 | 0.004 | 0.224 | 0.224 | 0 | 0.17 | 0.4 | 0.4 | 0.42 | 0.16 | 0.43 |
| 7 | 38510842 | rs6969360 | -0.686 | -0.861 | -0.279 | 0.134 | -0.028 | 0.024 | 0.024 | 0 | 0.11 | 0.4 | 0.11 | 0.06 | 0.06 | 0.24 |
| 7 | 38512542 | rs7811329 | -0.094 | -0.901 | -0.325 | 0.093 | -0.058 | -0.016 | -0.016 | 0 | 0.3 | 0.4 | 0.22 | 0.22 | 0.19 | 0.31 |
| 7 | 38515584 | rs1861239 | 1.67 | -0.861 | -0.421 | 0.108 | 0.008 | -0.03 | -0.03 | 0 | 0.19 | 0 | 0.11 | 0.16 | 0.14 | 0.07 |
| 7 | 38530568 | rs2043786 | -0.917 | -0.664 | -0.322 | -0.051 | -0.017 | 0.004 | 0.004 | 0 | 0.64 | 0.74 | 0.69 | 0.63 | 0.6 | 0.72 |
| 7 | 38531418 | rs2196679 | -0.668 | -0.674 | -0.168 | 0.009 | -0.027 | 0.021 | 0.021 | 0 | 0.11 | 0.4 | 0.11 | 0.06 | 0.06 | 0.24 |
| 7 | 38531899 | rs3778880 | -0.593 | -0.565 | -0.638 | 0.095 | -0.002 | 0.052 | 0.052 | 0 | 0.14 | 0.16 | 0.32 | 0.42 | 0.12 | 0.43 |
| 7 | 38532657 | rs3778878 | 0.026 | -0.261 | -0.746 | -0.027 | 0.005 | 0.042 | 0.042 | 0 | 0.51 | 0.59 | 0.37 | 0.21 | 0.48 | 0.29 |
| 7 | 38533101 | rs10248732 | 0.083 | 0.088 | -0.383 | 0.016 | 0.003 | -0.072 | -0.072 | 0 | 0.32 | 0.58 | 0.26 | 0.05 | 0.34 | 0.22 |
| 7 | 38534409 | rs2028210 | -0.348 | 0.062 | -0.359 | 0.08 | 0.002 | 0 | 0 | 0 | 0.38 | 0.59 | 0.55 | 0.42 | 0.44 | 0.42 |
| 7 | 38534816 | rs10269341 | 0.257 | 0.098 | -0.259 | 0.182 | 0.031 | 0.033 | 0.033 | 0 | 0.3 | 0.52 | 0.45 | 0.39 | 0.35 | 0.3 |
| 7 | 38540061 | rs1971809 | 0.809 | -0.265 | -0.215 | 0.535 | 0.006 | 0.074 | 0.074 | 0 | 0.85 | 0.8 | 0.61 | 0.66 | 0.83 | 0.76 |
| 7 | 38541139 | rs12673437 | 1.219 | -0.79 | -0.778 | 0.347 | -0.001 | -0.041 | -0.041 | 0 | 0.2 | 0.01 | 0.11 | 0.17 | 0.13 | 0.08 |
| 7 | 38543957 | rs2196674 | -0.408 | -0.783 | -0.707 | 0.3 | -0.006 | -0.043 | -0.043 | 0 | 0.92 | 0.81 | 0.89 | 1 | 0.93 | 0.98 |
| 7 | 38545379 | rs10233502 | 0.372 | -0.711 | -0.511 | -0.174 | -0.004 | 0.027 | 0.027 | 0.065 | 0.66 | 0.53 | 0.62 | 0.4 | 0.63 | 0.53 |
| 7 | 38545499 | rs2043787 | 0.564 | -0.723 | -0.368 | -0.178 | 0.002 | 0.004 | 0.004 | 0 | 0.47 | 0.44 | 0.53 | 0.38 | 0.51 | 0.46 |
| 7 | 38546848 | rs3807411 | 1.686 | -0.762 | -0.368 | -0.015 | -0.005 | -0.003 | -0.003 | 0 | 0.15 | 0.13 | 0.18 | 0.19 | 0.24 | 0.23 |
| 7 | 38548727 | rs17171390 | 0.822 | -0.789 | -0.279 | 0.024 | -0.01 | 0.074 | 0.074 | 0 | 0.05 | 0.1 | 0.05 | 0.25 | 0.03 | 0.14 |
| 7 | 38550586 | rs10246472 | 0.801 | -0.976 | -0.079 | -0.179 | -0.007 | 0.012 | 0.012 | 0 | 0.42 | 0.35 | 0.24 | 0.05 | 0.42 | 0.31 |
| 7 | 38552036 | rs2028209 | 0.331 | -0.716 | -0.274 | -0.059 | -0.009 | 0.087 | 0.087 | 0 | 0.48 | 0.69 | 0.38 | 0.29 | 0.48 | 0.39 |
| 7 | 38557432 | rs2083146 | -0.119 | -0.551 | -0.202 | 0.081 | 0.002 | 0.037 | 0.037 | 0 | 0.88 | 0.88 | 0.69 | 0.67 | 0.86 | 0.77 |
| 7 | 38563611 | rs730582 | 0.849 | -0.46 | -0.397 | 0.152 | -0.023 | 0.004 | 0.004 | 0 | 0.07 | 0.01 | 0.29 | 0.33 | 0.11 | 0.22 |
| 7 | 38572120 | rs3800795 | 0.002 | -0.649 | -0.387 | -0.235 | 0.022 | 0.026 | 0.026 | 0 | 0.55 | 0.77 | 0.67 | 0.63 | 0.58 | 0.61 |
| 7 | 38572694 | rs3807393 | 0.527 | -0.72 | -0.452 | -0.31 | 0.038 | 0.075 | 0.075 | 0 | 0.53 | 0.71 | 0.43 | 0.22 | 0.64 | 0.43 |
| 7 | 38573234 | rs2392581 | 0.495 | -0.484 | -0.497 | -0.569 | 0.002 | 0 | 0 | 8.265 | 0.49 | 0.44 | 0.36 | 0.19 | 0.4 | 0.24 |
| 7 | 38581467 | rs10951562 | 1.265 | -0.526 | -0.913 | -0.293 | 0 | -0.005 | -0.005 | 0 | 0.22 | 0.01 | 0.17 | 0.15 | 0.13 | 0.07 |
| 7 | 38581530 | rs2118458 | -0.459 | -0.73 | -0.65 | -0.501 | -0.016 | -0.052 | -0.052 | 0 | 0.89 | 0.83 | 0.92 | 1 | 0.92 | 0.97 |
| 7 | 38582081 | rs1561308 | 0.111 | -0.746 | -0.445 | -0.142 | 0.018 | -0.008 | -0.008 | 0 | 0.61 | 0.66 | 0.68 | 0.55 | 0.73 | 0.72 |
| 7 | 38584734 | rs17171395 | 0.35 | -0.474 | -0.386 | 0.161 | -0.005 | 0.098 | 0.098 | 0 | 0.06 | 0.12 | 0.07 | 0.3 | 0.05 | 0.17 |
| 7 | 38586223 | rs893412 | 0.434 | -0.371 | -0.785 | 0.06 | -0.03 | -0.038 | -0.038 | 0.224 | 0.09 | 0.01 | 0.26 | 0.37 | 0.12 | 0.24 |

| | | | | | | | | | | | | | | | | |
|---|---|---|---|---|---|---|---|---|---|---|---|---|---|---|---|---|
| 7 | 38586401 | rs893413 | -0.083 | -0.356 | -0.672 | -0.184 | -0.002 | -0.001 | -0.001 | 0 | 0.44 | 0.47 | 0.57 | 0.51 | 0.49 | 0.43 |
| 7 | 38590127 | rs10265223 | 0.917 | -0.405 | -0.808 | -0.27 | -0.013 | 0.004 | 0.004 | 0 | 0.33 | 0.29 | 0.21 | 0.01 | 0.36 | 0.2 |
| 7 | 38590558 | rs6942910 | 0.411 | -0.5 | -0.719 | -0.033 | -0.011 | 0.002 | 0.002 | 0 | 0.42 | 0.39 | 0.47 | 0.38 | 0.48 | 0.43 |
| 7 | 38591623 | rs2196678 | -0.064 | -0.288 | -1.733 | -0.613 | -0.006 | 0.174 | 0.174 | 0 | 0.51 | 0.86 | 0.56 | 0.73 | 0.52 | 0.66 |
| 7 | 38593862 | rs10215533 | 0.057 | -0.321 | -0.733 | -0.198 | -0.006 | -0.035 | -0.035 | 0 | 0.91 | 0.88 | 0.93 | 1 | 0.91 | 0.97 |
| 7 | 38595629 | rs7785499 | 0.134 | -0.002 | -1.002 | 0.111 | -0.001 | -0.001 | -0.001 | 0 | 0.48 | 0.57 | 0.4 | 0.41 | 0.47 | 0.54 |
| 7 | 38595908 | rs6974116 | 0.58 | -0.11 | -1.738 | -0.39 | -0.014 | 0.078 | 0.078 | 0 | 0.39 | 0.23 | 0.32 | 0.22 | 0.41 | 0.37 |
| 7 | 38596612 | rs4427071 | 1.194 | -0.3 | -1.661 | -0.207 | -0.008 | 0.139 | 0.139 | 0.162 | 0.5 | 0.18 | 0.38 | 0.3 | 0.49 | 0.35 |
| 7 | 38600176 | rs6967370 | 0.739 | -0.503 | -1.28 | -0.472 | 0.01 | -0.009 | -0.009 | 0 | 0.78 | 0.8 | 0.84 | 0.97 | 0.84 | 0.8 |
| 7 | 38600741 | rs10226831 | -0.13 | -0.178 | -1.323 | -0.257 | 0.011 | 0.019 | 0.019 | 0 | 0.85 | 0.92 | 0.92 | 0.97 | 0.89 | 0.86 |
| 7 | 38603499 | rs12701634 | -0.399 | 0.368 | -1.432 | 0.655 | -0.063 | -0.122 | -0.122 | 0 | 0.2 | 0.04 | 0.23 | 0.45 | 0.29 | 0.36 |
| 7 | 38613717 | rs11770392 | 0.598 | 0.25 | -0.814 | -0.523 | 0.009 | -0.051 | -0.051 | 0 | 0.3 | 0.33 | 0.23 | 0.06 | 0.35 | 0.18 |
| 7 | 38617022 | rs6965722 | -0.356 | -0.403 | -0.873 | -0.315 | 0.041 | -0.074 | -0.074 | 0.002 | 0.61 | 0.78 | 0.67 | 0.53 | 0.68 | 0.47 |
| 7 | 38617573 | rs4146042 | -1.166 | -0.691 | -0.944 | -0.361 | 0.065 | -0.13 | -0.13 | 0 | 0.76 | 0.93 | 0.78 | 0.58 | 0.86 | 0.62 |
| 7 | 38619348 | rs6948732 | -0.268 | -0.75 | -1.042 | -0.446 | 0.057 | -0.007 | -0.007 | 0 | 0.6 | 0.82 | 0.5 | 0.46 | 0.68 | 0.5 |
| 7 | 38623588 | rs1030654 | 1.906 | -0.549 | -1.005 | -0.581 | -0.006 | -0.035 | -0.035 | 0 | 0.28 | 0.23 | 0.23 | 0.01 | 0.35 | 0.14 |
| 7 | 38624970 | rs6942412 | -0.695 | -0.275 | -0.436 | -0.229 | -0.002 | -0.021 | -0.021 | 0 | 0.27 | 0.48 | 0.2 | 0.23 | 0.24 | 0.27 |
| 7 | 38634185 | rs6962805 | -0.71 | -0.128 | -0.48 | -0.152 | 0.008 | 0.048 | 0.048 | 0 | 0.31 | 0.47 | 0.43 | 0.46 | 0.33 | 0.28 |
| 7 | 38634578 | rs6945114 | -0.79 | 0.127 | -0.592 | -0.852 | -0.006 | 0.009 | 0.009 | 0 | 0.37 | 0.31 | 0.46 | 0.23 | 0.44 | 0.37 |
| 7 | 38636891 | rs10263645 | -0.667 | -0.067 | -0.706 | -0.821 | -0.003 | 0 | 0 | 0 | 0.67 | 0.61 | 0.67 | 0.3 | 0.78 | 0.55 |
| 7 | 38637841 | rs7811508 | -1.228 | 0.53 | -0.61 | -0.102 | -0.004 | 0.007 | 0.007 | 0.077 | 0.33 | 0.32 | 0.39 | 0.18 | 0.39 | 0.31 |
| 7 | 38640752 | rs2116667 | 0.092 | 0.97 | -1.515 | -0.122 | -0.054 | 0.015 | 0.015 | 0 | 0.16 | 0.01 | 0.27 | 0.06 | 0.2 | 0.12 |
| 7 | 38641095 | rs13225949 | -0.051 | 0.79 | -1.009 | 0.102 | -0.044 | 0.049 | 0.049 | 0.47 | 0.13 | 0.04 | 0.12 | 0.05 | 0.19 | 0.15 |
| 7 | 38641366 | rs17171412 | 0.009 | 0.256 | -0.353 | 0.507 | 0.004 | 0.053 | 0.053 | 0.116 | 0.11 | 0.24 | 0.12 | 0.16 | 0.08 | 0.1 |
| 7 | 38641903 | rs10278233 | -0.465 | 0.712 | -1.084 | 0.335 | -0.065 | -0.023 | -0.023 | 0 | 0.13 | 0.01 | 0.28 | 0.12 | 0.21 | 0.2 |
| 7 | 38652088 | rs10464364 | 0.275 | -0.998 | -1.154 | -0.158 | 0.048 | 0.016 | 0.016 | 0 | 0.71 | 0.87 | 0.76 | 0.26 | 0.82 | 0.53 |
| 7 | 38652687 | rs11760351 | 0.622 | -1.121 | -1.081 | -0.702 | -0.002 | -0.006 | -0.006 | 0.063 | 0.16 | 0.19 | 0.1 | 0.13 | 0.14 | 0.15 |
| 7 | 38655758 | rs17500182 | 0.61 | -1.116 | -1.321 | -0.823 | -0.015 | 0.078 | 0.078 | 0 | 0.14 | 0 | 0.08 | 0.01 | 0.13 | 0.18 |
| 7 | 38657224 | rs2043260 | 0.831 | -1.121 | -0.874 | -0.975 | 0.005 | 0.008 | 0.008 | 0.116 | 0.49 | 0.56 | 0.53 | 0.23 | 0.57 | 0.29 |
| 7 | 38664199 | rs10267034 | -0.089 | -1.079 | -0.581 | -0.8 | -0.007 | 0.032 | 0.032 | 0.015 | 0.18 | 0.1 | 0.12 | 0.01 | 0.21 | 0.22 |
| 7 | 38673833 | rs9918553 | -0.804 | -1.155 | -0.288 | 0.118 | -0.006 | 0.035 | 0.035 | 0.622 | 0.25 | 0.52 | 0.27 | 0.38 | 0.24 | 0.43 |
| 7 | 38677137 | rs7781415 | 0.157 | -1.087 | -1.024 | -0.728 | 0.047 | 0.085 | 0.085 | 0 | 0.52 | 0.64 | 0.65 | 0.13 | 0.69 | 0.38 |
| 7 | 38678273 | rs10265891 | 0.773 | -0.45 | -0.754 | -0.351 | 0.042 | -0.09 | -0.09 | 0.089 | 0.33 | 0.49 | 0.46 | 0.1 | 0.45 | 0.13 |
| 7 | 38682411 | rs4723777 | 0.25 | -0.593 | -1.05 | 0.854 | 0.01 | -0.038 | -0.038 | 0 | 0.83 | 0.88 | 0.84 | 0.35 | 0.88 | 0.58 |
| 7 | 38692014 | rs6951440 | 0.501 | -0.707 | -0.564 | 0.446 | -0.032 | 0.077 | 0.077 | 0 | 0.77 | 0.48 | 0.64 | 0.64 | 0.84 | 0.7 |
| 7 | 38695210 | rs2392591 | 0.381 | -0.86 | -1.254 | 0.019 | 0.025 | 0.12 | 0.12 | 0.435 | 0.48 | 0.25 | 0.33 | 0.32 | 0.57 | 0.4 |
| 7 | 38695584 | rs10226586 | 0.981 | -0.767 | -1.133 | 0.049 | -0.008 | -0.047 | -0.047 | 0 | 0.27 | 0.1 | 0.31 | 0.32 | 0.27 | 0.3 |
| 7 | 38706384 | rs10279772 | 0.9 | -0.864 | -1.082 | 0.045 | 0.008 | -0.012 | -0.012 | 0 | 0.28 | 0.18 | 0.31 | 0.31 | 0.26 | 0.29 |
| 7 | 38713672 | rs17171431 | 0.29 | -0.558 | -1.001 | -1.036 | -0.002 | 0 | 0 | 0.008 | 0.14 | 0.16 | 0.08 | 0.19 | 0.11 | 0.18 |
| 7 | 38714438 | rs16879916 | 0.688 | -0.657 | -0.559 | -1.357 | 0.012 | -0.088 | -0.088 | 0 | 0.13 | 0.34 | 0.24 | 0.01 | 0.11 | 0.1 |
| 7 | 38717897 | rs11979904 | 0.399 | -0.519 | -0.712 | -1.347 | 0.015 | -0.151 | -0.151 | 0 | 0.4 | 0.46 | 0.46 | 0.01 | 0.48 | 0.21 |
| 7 | 38721690 | rs2116666 | -0.181 | -0.303 | -1 | -1.031 | 0.015 | 0.028 | 0.028 | 0 | 0.55 | 0.64 | 0.4 | 0.02 | 0.58 | 0.38 |
| 7 | 38721867 | rs13242754 | -0.541 | 0.179 | -0.53 | 0.199 | -0.006 | 0.02 | 0.02 | 0.009 | 0.05 | 0 | 0.02 | 0.01 | 0.05 | 0.06 |
| 7 | 38722499 | rs17171441 | -0.723 | 0.319 | -0.295 | 0.499 | 0.014 | 0.062 | 0.062 | 0 | 0.1 | 0.15 | 0.05 | 0.31 | 0.12 | 0.15 |
| 7 | 38724491 | rs10259635 | -1.015 | 0.004 | 0.148 | -0.611 | -0.013 | -0.061 | -0.061 | 0 | 0.12 | 0.34 | 0.05 | 0.02 | 0.08 | 0.04 |
| 7 | 38725804 | rs11980087 | -0.772 | -0.056 | -0.992 | -0.777 | 0.029 | -0.016 | -0.016 | 0 | 0.56 | 0.75 | 0.36 | 0.56 | 0.62 | 0.65 |
| 7 | 38725883 | rs12701651 | -0.139 | 0.017 | -0.141 | -1.101 | -0.03 | 0.045 | 0.045 | 0 | 0.29 | 0.42 | 0.12 | 0.54 | 0.19 | 0.44 |
| 7 | 38727427 | rs17171446 | 1.014 | 0.123 | -0.053 | -1.389 | 0.012 | -0.026 | -0.026 | 0 | 0.06 | 0.09 | 0.04 | 0 | 0.09 | 0.03 |
| 7 | 38729405 | rs12701653 | 0.833 | 0.21 | 0.071 | -1.427 | -0.003 | -0.035 | -0.035 | 0.052 | 0.11 | 0.11 | 0.06 | 0.01 | 0.09 | 0.17 |
| 7 | 38738905 | rs9692361 | -1.562 | 0.034 | 0.054 | -1.374 | 0.026 | -0.044 | -0.044 | 0 | 0.6 | 0.81 | 0.39 | 0.48 | 0.65 | 0.73 |
| 7 | 38740878 | rs6964269 | -1.493 | -0.096 | 0.093 | -1.443 | 0.029 | -0.005 | -0.005 | 0 | 0.59 | 0.74 | 0.4 | 0.42 | 0.68 | 0.73 |
| 7 | 38741626 | rs10227687 | -0.978 | -0.102 | 0.062 | -1.474 | 0.005 | 0.004 | 0.004 | 0 | 0.49 | 0.57 | 0.34 | 0.41 | 0.61 | 0.58 |
| 7 | 38749008 | rs7778570 | 0.029 | 0.147 | 0.118 | -1.53 | -0.006 | 0.02 | 0.02 | 0 | 0.24 | 0.34 | 0.14 | 0.43 | 0.19 | 0.58 |
| 7 | 38753575 | rs1859493 | 1.027 | 0.149 | 0.121 | -1.522 | -0.017 | 0.112 | 0.112 | 0 | 0.15 | 0.34 | 0.09 | 0.43 | 0.11 | 0.43 |
| 7 | 38758326 | rs7797871 | -1.124 | 0.068 | 0.197 | -1.589 | 0.02 | -0.005 | -0.005 | 3.142 | 0.4 | 0.7 | 0.21 | 0.43 | 0.29 | 0.62 |
| 7 | 38759848 | rs11978898 | 0.097 | -0.044 | 0.235 | -1.629 | -0.025 | -0.083 | -0.083 | 0 | 0.16 | 0.25 | 0.07 | 0 | 0.1 | 0.05 |
| 7 | 38761510 | rs12701660 | -1.986 | -0.227 | -0.021 | -1.63 | 0.02 | -0.029 | -0.029 | 0.169 | 0.65 | 0.81 | 0.44 | 0.56 | 0.72 | 0.77 |
| 7 | 38764168 | rs13231320 | 0.266 | -0.128 | 0.062 | -1.785 | -0.02 | -0.071 | -0.071 | 0 | 0.16 | 0.22 | 0.07 | 0 | 0.1 | 0.05 |
| 7 | 38764676 | rs11773094 | 0.226 | 0.283 | -0.092 | -1.553 | -0.101 | 0.039 | 0.039 | 0 | 0.22 | 0.07 | 0.2 | 0.13 | 0.4 | 0.12 |
| 7 | 38764855 | rs17767818 | 1.026 | 0.237 | 0.116 | -1.339 | -0.012 | 0.143 | 0.143 | 0 | 0.15 | 0.37 | 0.1 | 0.43 | 0.12 | 0.43 |
| 7 | 38765476 | rs12701661 | 1.198 | 0.234 | 0.116 | -1.341 | -0.003 | -0.032 | -0.032 | 0.23 | 0.1 | 0.11 | 0.06 | 0 | 0.08 | 0.16 |
| 7 | 38765527 | rs3735347 | -0.118 | 0.242 | 0.12 | -1.338 | -0.021 | 0.082 | 0.082 | 0 | 0.25 | 0.48 | 0.16 | 0.43 | 0.2 | 0.58 |
| 7 | 38765785 | rs13235997 | 0.906 | 0.705 | 0.234 | -1.135 | -0.003 | -0.032 | -0.032 | 0.055 | 0.1 | 0.11 | 0.06 | 0 | 0.08 | 0.16 |
| 7 | 38782897 | rs10224856 | -0.467 | 0.69 | 0.132 | -1.15 | -0.004 | -0.047 | -0.047 | 0 | 0.87 | 0.84 | 0.93 | 1 | 0.9 | 0.82 |
| 7 | 38784481 | rs2240353 | 0.982 | 0.558 | 0.24 | -1.2 | -0.015 | 0.045 | 0.045 | 0 | 0.24 | 0.38 | 0.12 | 0.44 | 0.19 | 0.46 |
| 7 | 38791112 | rs7795874 | -0.065 | 0.571 | 0.239 | -1.184 | -0.024 | 0.022 | 0.022 | 0 | 0.63 | 0.45 | 0.81 | 0.56 | 0.72 | 0.36 |
| 7 | 38793243 | rs3779127 | -0.007 | 0.648 | 0.208 | -1.335 | -0.024 | 0.017 | 0.017 | 0.105 | 0.22 | 0.14 | 0.2 | 0.13 | 0.37 | 0.08 |
| 7 | 38793549 | rs3779126 | 1.093 | 0.642 | 0.18 | -1.33 | 0.022 | -0.008 | -0.008 | 0 | 0.4 | 0.23 | 0.59 | 0.42 | 0.34 | 0.26 |
| 7 | 38796570 | rs2240350 | 1.006 | 0.641 | 0.138 | -1.327 | 0.019 | -0.005 | -0.005 | 0 | 0.41 | 0.25 | 0.6 | 0.42 | 0.35 | 0.28 |

| | | | | | | | | | | | | | | | | |
|---|---|---|---|---|---|---|---|---|---|---|---|---|---|---|---|---|
| 7 | 38803741 | rs2240352 | -0.263 | 0.626 | 0.184 | -1.287 | -0.002 | 0.001 | 0.001 | 0 | 0.64 | 0.56 | 0.67 | 0.55 | 0.71 | 0.35 |
| 7 | 38823574 | rs10486680 | 0.532 | 0.82 | 0.091 | -1.19 | 0.005 | -0.055 | -0.055 | 0.779 | 0.1 | 0.26 | 0.05 | 0 | 0.07 | 0.02 |
| 7 | 38825187 | rs4720303 | 0.07 | 0.88 | -0.03 | -1.228 | -0.147 | 0.047 | 0.047 | 0.157 | 0.75 | 0.97 | 0.78 | 0.86 | 0.6 | 0.75 |
| 7 | 38829347 | rs1557941 | 0.233 | 0.841 | 0.09 | -1.222 | 0.001 | -0.066 | -0.066 | 0 | 0.11 | 0.35 | 0.06 | 0 | 0.07 | 0.02 |
| 7 | 38829465 | rs1001476 | 0.206 | 0.84 | 0.088 | -1.223 | 0.001 | -0.067 | -0.067 | 0.38 | 0.11 | 0.35 | 0.06 | 0 | 0.07 | 0.02 |
| 7 | 38849694 | rs2392601 | 1.043 | 0.887 | -0.067 | -1.241 | 0.019 | -0.057 | -0.057 | 0.082 | 0.63 | 0.56 | 0.7 | 0.87 | 0.52 | 0.73 |
| 7 | 38853772 | rs17680408 | 0.913 | 0.984 | -0.112 | -1.22 | -0.077 | -0.016 | -0.016 | 0 | 0.13 | 0.03 | 0.16 | 0.13 | 0.3 | 0.07 |
| 7 | 38858256 | rs3801135 | 1.072 | 0.755 | -0.048 | -1.229 | 0.012 | -0.006 | -0.006 | 0 | 0.4 | 0.26 | 0.6 | 0.44 | 0.34 | 0.26 |
| 7 | 38861809 | rs4723800 | 0.936 | 0.77 | -0.118 | -1.217 | 0.008 | -0.033 | -0.033 | 0 | 0.63 | 0.6 | 0.7 | 0.87 | 0.52 | 0.73 |
| 7 | 38862911 | rs7785161 | 1.541 | 0.617 | -0.433 | -1.407 | 0.011 | -0.096 | -0.096 | 0.024 | 0.29 | 0.05 | 0.32 | 0.43 | 0.24 | 0.16 |
| 7 | 38870444 | rs10085898 | 0.951 | 1.262 | -0.121 | -0.959 | -0.055 | 0.079 | 0.079 | 0.036 | 0.65 | 0.82 | 0.74 | 0.87 | 0.53 | 0.74 |
| 7 | 38873937 | rs12533017 | 1.043 | 1.274 | -0.285 | -0.953 | -0.017 | 0.02 | 0.02 | 0 | 0.64 | 0.7 | 0.72 | 0.87 | 0.53 | 0.74 |
| 7 | 38893322 | rs3801127 | 0.715 | 0.874 | -0.422 | -0.557 | -0.09 | 0.104 | 0.104 | 1.144 | 0.68 | 0.87 | 0.7 | 0.88 | 0.5 | 0.73 |
| 7 | 38919395 | rs2190666 | 1.355 | 0.7 | -0.577 | -0.458 | 0.032 | 0.001 | 0.001 | 0.599 | 0.41 | 0.25 | 0.47 | 0.41 | 0.31 | 0.25 |
| 7 | 38931930 | rs11980882 | 0.517 | 0.69 | -0.433 | -0.487 | -0.03 | 0.092 | 0.092 | 0 | 0.26 | 0.52 | 0.19 | 0.44 | 0.19 | 0.47 |
| 7 | 38933942 | rs6955677 | 0.452 | 0.716 | -0.478 | -0.588 | -0.064 | 0.062 | 0.062 | 1.358 | 0.7 | 0.82 | 0.72 | 0.88 | 0.51 | 0.76 |
| 7 | 38946736 | rs917321 | 0.181 | 0.822 | -0.533 | -0.6 | -0.139 | 0.01 | 0.01 | 0.042 | 0.19 | 0.02 | 0.21 | 0.12 | 0.38 | 0.08 |
| 7 | 38949201 | rs2074853 | 0.826 | 0.634 | -0.514 | -0.617 | -0.076 | 0.089 | 0.089 | 1.124 | 0.67 | 0.87 | 0.71 | 0.86 | 0.5 | 0.73 |
| 7 | 38953696 | rs2108507 | 0.78 | 0.595 | -0.443 | -0.592 | 0.004 | -0.013 | -0.013 | 0 | 0.23 | 0.25 | 0.15 | 0.44 | 0.19 | 0.47 |
| 7 | 38954251 | rs4723807 | 0.356 | 0.836 | -1.353 | -0.513 | 0.016 | -0.015 | -0.015 | 0.381 | 0.61 | 0.28 | 0.74 | 0.56 | 0.69 | 0.34 |
| 7 | 38955367 | rs6973977 | 0.661 | 2.103 | -1.438 | -0.591 | 0 | -0.014 | -0.014 | 0 | 0.87 | 0.9 | 0.93 | 1 | 0.88 | 0.82 |
| 7 | 38955773 | rs6974798 | -0.735 | 2.13 | -1.496 | -0.592 | -0.144 | 0.003 | 0.003 | 0.06 | 0.2 | 0.02 | 0.22 | 0.13 | 0.38 | 0.08 |
| 7 | 38956526 | rs4720309 | -0.15 | 2.176 | -1.441 | -0.541 | -0.047 | 0.102 | 0.102 | 0 | 0.84 | 0.99 | 0.9 | 1 | 0.83 | 0.84 |
| 7 | 38958276 | rs6961729 | 1.37 | 2.18 | -1.385 | -0.494 | -0.041 | 0.057 | 0.057 | 0.745 | 0.67 | 0.75 | 0.71 | 0.86 | 0.5 | 0.73 |
| 7 | 38959842 | rs10255854 | -0.149 | 2.319 | -2.154 | -0.408 | -0.095 | -0.03 | -0.03 | 1.737 | 0.13 | 0.02 | 0.16 | 0.13 | 0.31 | 0.07 |
| 7 | 38961388 | rs6978093 | 0.575 | 0.496 | -1.391 | -0.288 | -0.037 | 0.114 | 0.114 | 0 | 0.81 | 0.98 | 0.88 | 1 | 0.81 | 0.8 |
| 7 | 38969045 | rs17171483 | -0.609 | 0.107 | -0.614 | 0.364 | -0.017 | 0.013 | 0.013 | 0 | 0.08 | 0.13 | 0.1 | 0.07 | 0.05 | 0.07 |
| 7 | 38970305 | rs17171484 | -1.299 | -0.287 | -0.496 | 1.794 | -0.033 | 0.16 | 0.16 | 0 | 0.11 | 0.22 | 0.11 | 0.39 | 0.05 | 0.17 |
| 7 | 38972696 | rs859520 | 1.934 | -1.639 | 0.394 | 2.048 | -0.079 | 0.054 | 0.054 | 0 | 0.77 | 0.4 | 0.82 | 0.34 | 0.91 | 0.72 |
| 7 | 38982655 | rs12234473 | -1.165 | -1.924 | 0.3 | 1.567 | 0.047 | -0.029 | -0.029 | 0.274 | 0.12 | 0.04 | 0.04 | 0.28 | 0.05 | 0.16 |
| 7 | 38987911 | rs859526 | 2.049 | -1.895 | 0.832 | 1.562 | -0.068 | -0.001 | -0.001 | 0.423 | 0.77 | 0.65 | 0.85 | 0.69 | 0.9 | 0.78 |
| 7 | 38990205 | rs859524 | 0.365 | -1.797 | 0.88 | 1.573 | -0.019 | -0.022 | -0.022 | 0 | 0.9 | 0.73 | 0.89 | 0.96 | 0.95 | 0.93 |
| 7 | 38995788 | rs2116640 | 1.134 | -1.604 | -0.031 | 0.89 | 0.015 | 0 | 0 | 0.044 | 0.46 | 0.17 | 0.59 | 0.52 | 0.59 | 0.65 |
| 7 | 39000551 | rs1229052 | -0.156 | -1.679 | 0.369 | 0.491 | 0.011 | 0.017 | 0.017 | 0.017 | 0.32 | 0.25 | 0.21 | 0.27 | 0.24 | 0.17 |
| 7 | 39004902 | rs1918727 | 1.022 | -1.646 | 0.307 | 0.497 | 0.102 | 0.142 | 0.142 | 0.897 | 0.52 | 0.24 | 0.65 | 0.7 | 0.69 | 0.72 |
| 7 | 39007380 | rs7792493 | 0.82 | -1.609 | 0.443 | 0.514 | 0.038 | 0.051 | 0.051 | 0.444 | 0.54 | 0.37 | 0.68 | 0.7 | 0.69 | 0.72 |
| 7 | 39027459 | rs17171509 | 0.116 | -1.208 | 0.211 | 0.401 | -0.033 | -0.034 | -0.034 | 0.275 | 0.06 | 0.02 | 0.05 | 0.13 | 0.11 | 0.09 |
| 7 | 39030401 | rs17171511 | -0.83 | -1.27 | 0.356 | 0.385 | -0.005 | -0.005 | -0.005 | 0.218 | 0.08 | 0.08 | 0.09 | 0.13 | 0.13 | 0.11 |
| 7 | 39036864 | rs17176480 | 0.643 | -1.222 | -0.255 | -0.402 | 0.027 | 0.084 | 0.084 | 0 | 0.37 | 0.19 | 0.43 | 0.15 | 0.29 | 0.48 |
| 7 | 39039285 | rs1229014 | -0.191 | -0.801 | -0.166 | -0.719 | -0.015 | 0.037 | 0.037 | 0 | 0.34 | 0.29 | 0.28 | 0.14 | 0.37 | 0.22 |
| 7 | 39046768 | rs940876 | 0.392 | -1.214 | -0.427 | -0.835 | -0.003 | 0.017 | 0.017 | 0.04 | 0.32 | 0.23 | 0.33 | 0.27 | 0.33 | 0.47 |
| 7 | 39054538 | rs1229057 | 1.411 | -1.251 | -0.634 | -1.283 | 0.002 | 0.018 | 0.018 | 0.183 | 0.14 | 0.1 | 0.09 | 0 | 0.12 | 0.07 |
| 7 | 39057062 | rs35787066 | 1.323 | -1.502 | -0.724 | -1.388 | 0.017 | 0.067 | 0.067 | 0 | 0.26 | 0.11 | 0.29 | 0.13 | 0.24 | 0.37 |
| 7 | 39058757 | rs859548 | -0.508 | -1.808 | -0.249 | -1.625 | -0.046 | -0.086 | -0.086 | 0.125 | 0.85 | 0.61 | 0.86 | 0.96 | 0.92 | 0.91 |
| 7 | 39060267 | rs17255994 | 0.609 | -1.556 | -0.502 | -1.643 | -0.076 | 0.013 | 0.013 | 0.021 | 0.16 | 0.03 | 0.14 | 0.11 | 0.23 | 0.12 |
| 7 | 39061882 | rs17171521 | 1.641 | -1.625 | -0.404 | -1.569 | 0.014 | 0.066 | 0.066 | 0 | 0.25 | 0.09 | 0.28 | 0.13 | 0.24 | 0.38 |
| 7 | 39072473 | rs7801660 | 1.057 | -1.542 | -0.349 | -1.495 | -0.002 | 0 | 0 | 0.392 | 0.25 | 0.24 | 0.33 | 0.72 | 0.32 | 0.29 |
| 7 | 39086869 | rs10085472 | -0.086 | -1.531 | -0.267 | -1.751 | -0.007 | -0.101 | -0.101 | 0.096 | 0.65 | 0.54 | 0.78 | 0.95 | 0.7 | 0.78 |
| 7 | 39090698 | rs12531458 | -0.409 | -1.368 | -0.253 | -1.807 | 0.001 | -0.031 | -0.031 | 0 | 0.42 | 0.47 | 0.51 | 0.89 | 0.46 | 0.44 |
| 7 | 39096388 | rs1949880 | -0.661 | -1.047 | -0.013 | -1.687 | -0.009 | -0.067 | -0.067 | 0 | 0.32 | 0.5 | 0.23 | 0.18 | 0.26 | 0.15 |
| 7 | 39116798 | rs10258862 | 0.591 | -1.25 | -0.03 | -2.21 | -0.018 | -0.057 | -0.057 | 0 | 0.43 | 0.37 | 0.37 | 0.11 | 0.31 | 0.46 |
| 7 | 39118242 | rs1534415 | -1.511 | -1.181 | -0.105 | -2.314 | -0.017 | 0.026 | 0.026 | 0 | 0.81 | 0.88 | 0.87 | 1 | 0.77 | 0.85 |
| 7 | 39118314 | rs11761423 | 1.757 | -1.209 | -0.069 | -2.286 | -0.007 | -0.002 | -0.002 | 0 | 0.11 | 0.09 | 0.1 | 0.13 | 0.12 | 0.04 |
| 7 | 39119320 | rs1527952 | 0.447 | -1.137 | -0.034 | -2.217 | 0.004 | -0.002 | -0.002 | 3.557 | 0.25 | 0.32 | 0.36 | 0.74 | 0.28 | 0.33 |
| 7 | 39121465 | rs1404999 | 0.655 | -1.198 | 0 | -2.191 | 0.003 | -0.016 | -0.016 | 0 | 0.42 | 0.33 | 0.35 | 0.11 | 0.31 | 0.46 |
| 7 | 39123165 | rs6959715 | -0.085 | -1.019 | -0.203 | -2.089 | -0.029 | 0.039 | 0.039 | 0 | 0.67 | 0.71 | 0.71 | 0.86 | 0.58 | 0.79 |
| 7 | 39124426 | rs17171532 | 1.617 | -1.022 | 0.012 | -1.972 | -0.026 | -0.028 | -0.028 | 0 | 0.1 | 0.22 | 0.09 | 0.04 | 0.03 | 0.06 |
| 7 | 39132816 | rs12672089 | 0.267 | 0.016 | -0.518 | -1.379 | -0.031 | -0.079 | -0.079 | 0 | 0.25 | 0.13 | 0.31 | 0.54 | 0.29 | 0.31 |
| 7 | 39136566 | rs1525800 | -0.21 | -0.036 | -0.49 | -1.487 | -0.019 | 0.013 | 0.013 | 0 | 0.36 | 0.34 | 0.43 | 0.7 | 0.42 | 0.36 |
| 7 | 39141221 | rs12216590 | 1.918 | -0.251 | -0.404 | -1.541 | 0.021 | 0.045 | 0.045 | 0 | 0.28 | 0.15 | 0.3 | 0.11 | 0.25 | 0.37 |
| 7 | 39145862 | rs11763728 | 0.245 | 0 | -0.402 | -1.382 | -0.003 | -0.011 | -0.011 | 0 | 0.26 | 0.29 | 0.33 | 0.54 | 0.29 | 0.32 |
| 7 | 39147916 | rs953874 | 0.653 | -0.008 | -0.456 | -1.433 | 0 | 0.008 | 0.008 | 0 | 0.2 | 0.24 | 0.3 | 0.54 | 0.27 | 0.32 |
| 7 | 39151039 | rs10951595 | -0.402 | 0.033 | -0.489 | -1.497 | -0.028 | 0.017 | 0.017 | 0 | 0.35 | 0.31 | 0.44 | 0.71 | 0.44 | 0.37 |
| 7 | 39172228 | rs2280667 | 0.82 | 0.046 | -0.45 | -1.476 | -0.02 | -0.038 | -0.038 | 0 | 0.19 | 0.12 | 0.29 | 0.54 | 0.27 | 0.31 |
| 7 | 39186030 | rs12701698 | -0.378 | 0.008 | -0.483 | -1.501 | -0.051 | 0.021 | 0.021 | 0 | 0.34 | 0.19 | 0.43 | 0.71 | 0.44 | 0.36 |
| 7 | 39194416 | rs10273694 | 1.887 | 0.098 | -0.18 | -1.655 | 0.023 | 0.043 | 0.043 | 0 | 0.31 | 0.2 | 0.32 | 0.1 | 0.27 | 0.36 |
| 7 | 39197354 | rs10242528 | -0.26 | 1 | 0.096 | -1.21 | 0.023 | -0.021 | -0.021 | 0.274 | 0.29 | 0.41 | 0.41 | 0.83 | 0.39 | 0.37 |
| 7 | 39198960 | rs6462895 | 0.614 | 0.717 | -0.101 | -1.408 | 0.026 | 0.039 | 0.039 | 0 | 0.55 | 0.56 | 0.48 | 0.23 | 0.37 | 0.47 |
| 7 | 39211106 | rs17171546 | 2.245 | 0.877 | -0.283 | -1.38 | 0.007 | 0.003 | 0.003 | 0 | 0.09 | 0.12 | 0.11 | 0.16 | 0.14 | 0.04 |

| | | | | | | | | | | | | | | | | |
|---|---|---|---|---|---|---|---|---|---|---|---|---|---|---|---|---|
| 7 | 39240060 | rs4323393 | 1.004 | 0.78 | 0.097 | -1.34 | 0.023 | 0.039 | 0.039 | 0 | 0.21 | 0.27 | 0.31 | 0.69 | 0.27 | 0.3 |
| 7 | 39242508 | rs4472423 | 0.603 | 0.561 | -0.068 | -1.39 | 0.033 | 0.052 | 0.052 | 0 | 0.53 | 0.39 | 0.46 | 0.15 | 0.36 | 0.48 |
| 7 | 39243058 | rs12534084 | 0.603 | 0.561 | -0.068 | -1.39 | 0.033 | 0.052 | 0.052 | 0 | 0.53 | 0.39 | 0.45 | 0.15 | 0.36 | 0.48 |
| 7 | 39243189 | rs12531437 | 0.603 | 0.561 | -0.068 | -1.39 | 0.033 | 0.052 | 0.052 | 0 | 0.53 | 0.39 | 0.45 | 0.15 | 0.36 | 0.48 |
| 7 | 39245404 | rs7777684 | 0.287 | 0.735 | -0.496 | -1.164 | -0.028 | 0.007 | 0.007 | 0 | 0.62 | 0.77 | 0.59 | 0.33 | 0.49 | 0.59 |
| 7 | 39268195 | rs4236356 | 0.173 | 1.051 | -0.184 | -0.722 | 0.03 | 0.018 | 0.018 | 0 | 0.25 | 0.34 | 0.36 | 0.7 | 0.35 | 0.34 |
| 7 | 39269010 | rs4291171 | 0.173 | 1.051 | -0.232 | -0.725 | 0.025 | 0.014 | 0.014 | 0 | 0.25 | 0.33 | 0.36 | 0.7 | 0.35 | 0.34 |
| 7 | 39276953 | rs11771925 | 0.133 | 1.128 | -0.547 | -0.709 | -0.052 | -0.019 | -0.019 | 2.43 | 0.29 | 0.19 | 0.38 | 0.68 | 0.45 | 0.38 |
| 7 | 39279545 | rs6974806 | 0.581 | 0.376 | -0.404 | -1.233 | -0.021 | 0.021 | 0.021 | 1.08 | 0.26 | 0.45 | 0.22 | 0.29 | 0.17 | 0.15 |
| 7 | 39282408 | rs4574755 | 0.47 | 0.516 | -0.429 | -0.498 | -0.016 | 0.01 | 0.01 | 0.128 | 0.28 | 0.41 | 0.26 | 0.29 | 0.2 | 0.17 |
| 7 | 39282503 | rs4404830 | 0.389 | 0.513 | -0.435 | -0.499 | -0.018 | 0.011 | 0.011 | 0.202 | 0.28 | 0.42 | 0.27 | 0.29 | 0.2 | 0.17 |
| 7 | 39285620 | rs4629763 | -1.384 | 0.541 | -0.602 | -0.528 | 0.046 | 0.095 | 0.095 | 0 | 0.57 | 0.68 | 0.65 | 0.97 | 0.64 | 0.57 |
| 7 | 39287246 | rs11762823 | 2.186 | 0.461 | -0.508 | -0.492 | 0.041 | 0.031 | 0.031 | 0 | 0.1 | 0.16 | 0.11 | 0.16 | 0.12 | 0.05 |
| 7 | 39287422 | rs10081326 | -1.44 | 0.541 | -1.177 | -0.518 | 0.052 | 0.148 | 0.148 | 0 | 0.58 | 0.77 | 0.67 | 0.98 | 0.64 | 0.6 |
| 7 | 39292111 | rs9655034 | 0.409 | 0.867 | -1.122 | -0.231 | -0.051 | -0.02 | -0.02 | 0 | 0.29 | 0.17 | 0.38 | 0.68 | 0.45 | 0.38 |
| 7 | 39297668 | rs6967316 | -0.782 | 0.194 | -1.334 | -0.606 | 0.032 | 0.092 | 0.092 | 0 | 0.59 | 0.74 | 0.63 | 0.97 | 0.62 | 0.58 |
| 7 | 39298546 | rs4294089 | -0.189 | -0.15 | -1.276 | 0.344 | 0.005 | -0.01 | -0.01 | 0 | 0.25 | 0.23 | 0.24 | 0.43 | 0.19 | 0.32 |
| 7 | 39302577 | rs12701705 | -0.015 | -0.028 | -1.324 | 0.119 | -0.044 | -0.06 | -0.06 | 0 | 0.37 | 0.23 | 0.6 | 0.51 | 0.48 | 0.34 |
| 7 | 39303142 | rs10257516 | -1.129 | 0.293 | -0.807 | 0.07 | 0.083 | 0.034 | 0.034 | 0 | 0.86 | 0.72 | 0.85 | 0.8 | 0.76 | 0.89 |
| 7 | 39304125 | rs4302750 | -0.122 | 0.163 | -0.802 | -0.098 | 0.013 | -0.024 | -0.024 | 0 | 0.82 | 0.65 | 0.79 | 0.89 | 0.81 | 0.85 |
| 7 | 39308521 | rs3923295 | 0.996 | -0.418 | -1.066 | -0.324 | -0.046 | -0.056 | -0.056 | 0 | 0.37 | 0.44 | 0.18 | 0.2 | 0.23 | 0.37 |
| 7 | 39308930 | rs4524676 | 1 | -0.029 | -1.269 | -0.145 | -0.029 | -0.044 | -0.044 | 0.305 | 0.19 | 0.08 | 0.14 | 0.25 | 0.2 | 0.22 |
| 7 | 39309284 | rs4723834 | -0.237 | -0.976 | -0.9 | -0.809 | 0 | -0.023 | -0.023 | 0 | 0.55 | 0.52 | 0.73 | 0.88 | 0.71 | 0.59 |
| 7 | 39318130 | rs4723836 | 0.202 | -0.633 | -1.05 | -0.618 | 0.003 | 0.096 | 0.096 | 0.551 | 0.45 | 0.24 | 0.6 | 0.7 | 0.57 | 0.47 |
| 7 | 39323778 | rs4318977 | -0.692 | -0.552 | -0.494 | -0.576 | 0.027 | 0.006 | 0.006 | 0.202 | 0.8 | 0.67 | 0.83 | 0.78 | 0.78 | 0.83 |
| 7 | 39324470 | rs13246263 | 0.937 | -0.891 | -0.561 | -0.628 | 0.013 | 0.016 | 0.016 | 0 | 0.19 | 0.13 | 0.17 | 0.08 | 0.15 | 0.11 |
| 7 | 39324983 | rs6462901 | -0.263 | -0.835 | -0.547 | -0.511 | -0.003 | 0 | 0 | 0 | 0.68 | 0.63 | 0.76 | 0.69 | 0.72 | 0.64 |
| 7 | 39327012 | rs4551241 | 2.779 | -0.37 | -0.703 | -0.343 | -0.047 | -0.041 | -0.041 | 0.296 | 0.07 | 0.01 | 0.09 | 0.16 | 0.16 | 0.05 |
| 7 | 39331323 | rs12701716 | 0.776 | -0.819 | -0.593 | -0.498 | -0.007 | -0.017 | -0.017 | 3.553 | 0.47 | 0.25 | 0.58 | 0.48 | 0.52 | 0.46 |
| 7 | 39332266 | rs4330591 | -1.427 | -0.729 | -0.486 | -0.48 | -0.002 | -0.003 | -0.003 | 0.402 | 0.8 | 0.78 | 0.87 | 0.86 | 0.85 | 0.75 |
| 7 | 39341026 | rs4377848 | 1.921 | -0.91 | -0.536 | -0.405 | 0.001 | 0.01 | 0.01 | 0 | 0.12 | 0.14 | 0.13 | 0.17 | 0.13 | 0.1 |
| 7 | 39345684 | rs6962816 | -0.643 | -0.772 | -0.549 | -0.317 | 0.013 | 0.009 | 0.009 | 0 | 0.72 | 0.66 | 0.76 | 0.66 | 0.69 | 0.64 |
| 7 | 39348637 | rs6968121 | 0.651 | -0.812 | -0.628 | -0.583 | -0.016 | -0.019 | -0.019 | 0 | 0.24 | 0.3 | 0.17 | 0.17 | 0.18 | 0.19 |
| 7 | 39355178 | rs13222559 | 0.491 | -0.438 | -1.27 | -0.339 | -0.066 | -0.093 | -0.093 | 1.394 | 0.33 | 0.16 | 0.54 | 0.52 | 0.47 | 0.26 |
| 7 | 39356036 | rs10447664 | 0.1 | 0.188 | -0.689 | -0.199 | 0.01 | 0.002 | 0.002 | 0 | 0.71 | 0.64 | 0.76 | 0.7 | 0.7 | 0.65 |
| 7 | 39363081 | rs12701732 | 0.447 | 0.549 | -0.897 | -0.132 | -0.001 | 0.001 | 0.001 | 0 | 0.11 | 0.12 | 0.2 | 0.08 | 0.14 | 0.04 |
| 7 | 39364993 | rs4723858 | -0.855 | 0.987 | -0.836 | 0.011 | -0.003 | 0.01 | 0.01 | 1.745 | 0.32 | 0.31 | 0.32 | 0.26 | 0.33 | 0.26 |
| 7 | 39366946 | rs4723859 | -0.134 | 0.892 | -0.785 | -0.149 | 0.006 | 0.005 | 0.005 | 1.979 | 0.2 | 0.18 | 0.2 | 0.08 | 0.15 | 0.12 |
| 7 | 39370539 | rs7786697 | -0.615 | 1.304 | -0.533 | 0.047 | -0.003 | 0.022 | 0.022 | 0 | 0.22 | 0.29 | 0.29 | 0.69 | 0.21 | 0.3 |
| 7 | 39374823 | rs12701735 | -0.627 | 2.381 | -0.652 | -0.16 | -0.043 | 0.027 | 0.027 | 0 | 0.24 | 0.12 | 0.32 | 0.15 | 0.37 | 0.16 |
| 7 | 39376593 | rs12533451 | 1.835 | 2.579 | -0.683 | -0.143 | -0.039 | -0.032 | -0.032 | 0 | 0.07 | 0.01 | 0.09 | 0.12 | 0.16 | 0.03 |
| 7 | 39378149 | rs2237413 | -0.553 | 2.449 | -0.399 | 0.116 | 0.003 | 0.092 | 0.092 | 0 | 0.21 | 0.44 | 0.23 | 0.6 | 0.19 | 0.27 |
| 7 | 39378982 | rs2074937 | 2.002 | 2.457 | -0.464 | 0.098 | -0.039 | -0.031 | -0.031 | 0 | 0.07 | 0.01 | 0.09 | 0.12 | 0.16 | 0.03 |
| 7 | 39383090 | rs2329388 | -1.827 | 2.314 | -0.407 | -0.002 | -0.002 | -0.021 | -0.021 | 0.06 | 0.44 | 0.56 | 0.55 | 0.74 | 0.56 | 0.41 |
| 7 | 39384830 | rs6962623 | 2.023 | 2.245 | -0.537 | -0.225 | 0.007 | 0.007 | 0.007 | 0 | 0.42 | 0.32 | 0.37 | 0.26 | 0.28 | 0.43 |
| 7 | 39402851 | rs6946515 | 0.051 | 2.431 | -0.406 | 0.138 | -0.012 | 0.058 | 0.058 | 0 | 0.17 | 0.07 | 0.22 | 0.03 | 0.2 | 0.12 |
| 7 | 39409900 | rs12672997 | 0.149 | 2.354 | -0.221 | 0.252 | 0.021 | 0.118 | 0.118 | 0.152 | 0.13 | 0.21 | 0.19 | 0.6 | 0.16 | 0.23 |
| 7 | 39414410 | rs17686832 | 0.114 | 2.372 | -0.374 | 0.162 | 0.005 | 0.055 | 0.055 | 0 | 0.16 | 0.07 | 0.2 | 0.03 | 0.17 | 0.11 |
| 7 | 39419783 | rs6462914 | 0.314 | 2.471 | -0.08 | 0.804 | 0.018 | 0.168 | 0.168 | 0.099 | 0.11 | 0.18 | 0.19 | 0.48 | 0.14 | 0.19 |
| 7 | 39424553 | rs6942833 | -0.756 | 2.718 | -0.048 | 0.722 | 0.007 | 0.006 | 0.006 | 0 | 0.21 | 0.3 | 0.3 | 0.27 | 0.32 | 0.23 |
| 7 | 39427296 | rs1990243 | -1.535 | 2.66 | -0.218 | 0.631 | 0.047 | -0.045 | -0.045 | 4.167 | 0.33 | 0.5 | 0.49 | 0.76 | 0.47 | 0.42 |
| 7 | 39431315 | rs10256992 | -1.378 | 2.591 | -0.092 | 0.74 | -0.012 | 0.009 | 0.009 | 0 | 0.3 | 0.57 | 0.4 | 0.63 | 0.3 | 0.39 |
| 7 | 39431588 | rs10227915 | -0.248 | 2.592 | -0.059 | 0.808 | 0.01 | 0.204 | 0.204 | 1.405 | 0.16 | 0.3 | 0.31 | 0.48 | 0.18 | 0.25 |
| 7 | 39432975 | rs10951603 | -1.911 | 2.759 | 0.177 | 0.379 | -0.008 | 0.027 | 0.027 | 0 | 0.41 | 0.68 | 0.48 | 0.76 | 0.53 | 0.46 |
| 7 | 39434312 | rs2190893 | -1.064 | 2.348 | 0.405 | 0.56 | 0.053 | 0.01 | 0.01 | 0.113 | 0.25 | 0.39 | 0.34 | 0.7 | 0.35 | 0.36 |
| 7 | 39436094 | rs10248165 | -1.884 | 2.338 | 0.389 | 0.531 | 0.001 | -0.005 | -0.005 | 1.012 | 0.37 | 0.63 | 0.44 | 0.75 | 0.47 | 0.43 |
| 7 | 39437717 | rs917400 | 0.054 | 2.195 | 0.394 | 0.659 | -0.002 | 0.054 | 0.054 | 0.169 | 0.13 | 0.36 | 0.1 | 0.17 | 0.12 | 0.16 |
| 7 | 39440893 | rs7789243 | -0.391 | 2.302 | 0.054 | 0.787 | 0.016 | 0.09 | 0.09 | 0.114 | 0.07 | 0.26 | 0.06 | 0.17 | 0.09 | 0.14 |
| 7 | 39444551 | rs10259786 | -0.928 | 2.454 | 0.033 | 0.597 | 0.014 | 0.025 | 0.025 | 0 | 0.23 | 0.25 | 0.33 | 0.58 | 0.34 | 0.27 |
| 7 | 39449983 | rs2237400 | -1.465 | 2.407 | -0.038 | 0.599 | 0.021 | 0.003 | 0.003 | 0 | 0.28 | 0.35 | 0.38 | 0.69 | 0.42 | 0.37 |
| 7 | 39455473 | rs2024101 | -1.366 | 2.302 | 0.05 | 0.67 | 0.035 | 0.048 | 0.048 | 0 | 0.24 | 0.34 | 0.34 | 0.62 | 0.35 | 0.29 |
| 7 | 39457714 | rs17619647 | 1.803 | 1.997 | -0.165 | 0.285 | 0.013 | 0.027 | 0.027 | 0 | 0.52 | 0.33 | 0.54 | 0.31 | 0.42 | 0.48 |
| 7 | 39462495 | rs3819426 | -0.798 | 2.301 | 0.109 | 0.806 | 0.053 | 0.172 | 0.172 | 0 | 0.17 | 0.33 | 0.29 | 0.52 | 0.25 | 0.23 |
| 7 | 39464732 | rs17712867 | 1.53 | 2.415 | -0.087 | 0.974 | -0.036 | -0.035 | -0.035 | 0 | 0.06 | 0 | 0.07 | 0.17 | 0.14 | 0.02 |
| 7 | 39472708 | rs2302123 | -0.869 | 1.918 | -0.034 | 0.571 | 0.026 | 0.134 | 0.134 | 0.073 | 0.21 | 0.54 | 0.28 | 0.58 | 0.24 | 0.3 |
| 7 | 39488248 | rs1155269 | -0.731 | 1.924 | -0.044 | 0.563 | 0.033 | 0.135 | 0.135 | 0 | 0.2 | 0.41 | 0.27 | 0.58 | 0.24 | 0.29 |
| 7 | 39498090 | rs17619856 | 0.444 | 1.911 | -0.217 | 0.25 | -0.002 | 0.012 | 0.012 | 0 | 0.1 | 0.08 | 0.18 | 0 | 0.13 | 0.07 |
| 7 | 39503427 | rs12533673 | -0.291 | 1.798 | -0.265 | 0.274 | -0.005 | 0 | 0 | 0 | 0.9 | 0.9 | 0.94 | 0.93 | 0.91 | 0.83 |
| 7 | 39505059 | rs6977512 | 0.766 | 1.895 | -0.212 | 0.346 | 0.027 | 0.029 | 0.029 | 0 | 0.7 | 0.43 | 0.65 | 0.38 | 0.65 | 0.61 |

| | | | | | | | | | | | | | | | | |
|---|---|---|---|---|---|---|---|---|---|---|---|---|---|---|---|---|
| 7 | 39505737 | rs2052017 | -0.428 | 1.789 | -0.263 | 0.301 | -0.002 | 0 | 0 | 0 | 0.9 | 0.87 | 0.94 | 0.93 | 0.91 | 0.83 |
| 7 | 39508462 | rs2329395 | 0.059 | 1.837 | -0.163 | 0.336 | 0.048 | 0.16 | 0.16 | 0 | 0.2 | 0.42 | 0.29 | 0.56 | 0.26 | 0.24 |
| 7 | 39510623 | rs1558162 | 1.073 | 1.773 | -0.481 | 0.107 | -0.037 | 0.044 | 0.044 | 0 | 0.72 | 0.81 | 0.83 | 0.84 | 0.69 | 0.69 |
| 7 | 39513782 | rs17713251 | -0.136 | 1.811 | -0.613 | -0.193 | -0.073 | 0.039 | 0.039 | 0.064 | 0.18 | 0.03 | 0.11 | 0.09 | 0.22 | 0.14 |
| 7 | 39513998 | rs3937443 | 0.76 | 1.794 | -0.606 | -0.192 | -0.085 | 0.108 | 0.108 | 0 | 0.74 | 0.92 | 0.84 | 0.91 | 0.7 | 0.74 |
| 7 | 39515179 | rs2015739 | 0.139 | 1.824 | -0.211 | 0.081 | 0.03 | 0.13 | 0.13 | 0 | 0.21 | 0.39 | 0.3 | 0.52 | 0.27 | 0.24 |
| 7 | 39516722 | rs4723866 | 0.283 | 1.657 | -0.404 | -0.087 | 0.01 | 0.018 | 0.018 | 0.044 | 0.63 | 0.5 | 0.6 | 0.45 | 0.61 | 0.61 |
| 7 | 39521010 | rs6962246 | -0.293 | 1.321 | -0.204 | 0.107 | 0.001 | 0.002 | 0.002 | 0 | 0.51 | 0.44 | 0.44 | 0.54 | 0.51 | 0.45 |
| 7 | 39525669 | rs1024595 | 0.257 | 1.313 | -0.318 | -0.192 | 0.002 | -0.045 | -0.045 | 0 | 0.08 | 0.19 | 0.05 | 0 | 0.07 | 0.1 |
| 7 | 39526246 | rs17770101 | -0.486 | 1.317 | -0.685 | 0.168 | 0 | -0.001 | -0.001 | 0.168 | 0.09 | 0.13 | 0.07 | 0.09 | 0.1 | 0.12 |
| 7 | 39533774 | rs4618604 | -0.371 | 0.893 | -0.738 | 0.342 | 0.001 | -0.019 | -0.019 | 0 | 0.61 | 0.79 | 0.49 | 0.58 | 0.57 | 0.55 |
| 7 | 39539545 | rs11980909 | -0.374 | 0.675 | -0.724 | 0.169 | -0.003 | -0.012 | -0.012 | 0 | 0.05 | 0.07 | 0.02 | 0 | 0.02 | 0.05 |
| 7 | 39540140 | rs6949167 | -0.401 | 0.61 | -0.89 | 0.181 | -0.003 | -0.045 | -0.045 | 0 | 0.63 | 0.81 | 0.5 | 0.57 | 0.6 | 0.56 |
| 7 | 39544787 | rs6972153 | 0.348 | 0.525 | -0.405 | -0.112 | -0.003 | 0.001 | 0.001 | 0 | 0.46 | 0.52 | 0.39 | 0.57 | 0.44 | 0.46 |
| 7 | 39544838 | rs6971762 | -0.131 | 0.735 | -0.475 | -0.022 | -0.019 | -0.021 | -0.021 | 0 | 0.57 | 0.75 | 0.45 | 0.6 | 0.49 | 0.51 |
| 7 | 39553810 | rs2329401 | 0.062 | 0.427 | -0.221 | -0.094 | 0.017 | -0.006 | -0.006 | 0.626 | 0.53 | 0.27 | 0.39 | 0.54 | 0.46 | 0.52 |
| 7 | 39554212 | rs11772773 | 0.403 | -0.179 | -0.613 | -0.255 | -0.015 | -0.02 | -0.02 | 0 | 0.4 | 0.12 | 0.36 | 0.44 | 0.41 | 0.45 |
| 7 | 39561066 | rs4723875 | -0.796 | 0.06 | -0.111 | -0.26 | -0.022 | -0.034 | -0.034 | 0.01 | 0.67 | 0.72 | 0.54 | 0.59 | 0.59 | 0.6 |
| 7 | 39573216 | rs2876855 | 0.624 | -0.085 | -0.614 | -0.246 | -0.111 | -0.146 | -0.146 | 0 | 0.3 | 0.07 | 0.42 | 0.41 | 0.39 | 0.4 |
| 7 | 39576753 | rs17770427 | 0.48 | -0.719 | -0.52 | -0.363 | 0.011 | 0.004 | 0.004 | 0 | 0.17 | 0.13 | 0.08 | 0.22 | 0.06 | 0.1 |
| 7 | 39587912 | rs2329449 | -0.735 | 0.014 | -0.474 | -0.377 | -0.034 | -0.018 | -0.018 | 0.022 | 0.61 | 0.81 | 0.61 | 0.58 | 0.54 | 0.55 |
| 7 | 39588232 | rs715283 | -0.034 | -0.031 | -0.302 | -0.387 | 0.03 | 0.02 | 0.02 | 0.978 | 0.49 | 0.31 | 0.47 | 0.55 | 0.42 | 0.5 |
| 7 | 39595948 | rs4723876 | -0.643 | -0.012 | -0.641 | -0.275 | -0.062 | -0.044 | -0.044 | 0 | 0.61 | 0.89 | 0.59 | 0.55 | 0.54 | 0.52 |
| 7 | 39605204 | rs10232859 | -0.496 | 0.133 | 0.14 | -0.506 | -0.024 | -0.049 | -0.049 | 0.811 | 0.2 | 0.3 | 0.07 | 0.11 | 0.1 | 0.12 |
| 7 | 39606107 | rs1051162 | -0.484 | 0.217 | -0.066 | -0.094 | -0.024 | -0.048 | -0.048 | 0 | 0.2 | 0.3 | 0.07 | 0.11 | 0.1 | 0.12 |
| 7 | 39607399 | rs6462932 | -0.987 | -0.303 | -0.365 | -0.396 | 0.003 | -0.037 | -0.037 | 0 | 0.6 | 0.85 | 0.68 | 0.55 | 0.59 | 0.6 |
| 7 | 39608877 | rs17171592 | -0.205 | -0.302 | -0.384 | -0.393 | -0.023 | -0.048 | -0.048 | 0.017 | 0.19 | 0.3 | 0.07 | 0.11 | 0.1 | 0.12 |
| 7 | 39609805 | rs2288743 | 1.156 | -0.219 | -0.187 | -0.417 | -0.028 | -0.049 | -0.049 | 0 | 0.3 | 0.15 | 0.38 | 0.34 | 0.36 | 0.38 |
| 7 | 39610177 | rs6947660 | -0.175 | -0.182 | -0.187 | -0.401 | -0.023 | -0.048 | -0.048 | 0 | 0.19 | 0.3 | 0.07 | 0.11 | 0.1 | 0.12 |
| 7 | 39611153 | rs17171597 | 0.635 | -0.285 | -0.14 | -0.464 | -0.036 | -0.057 | -0.057 | 3.982 | 0.11 | 0.19 | 0.04 | 0 | 0.03 | 0 |
| 7 | 39615376 | rs17171603 | 0.632 | -0.213 | 0.083 | -0.434 | 0.065 | 0.007 | 0.007 | 0 | 0.08 | 0.24 | 0.13 | 0.1 | 0.12 | 0.07 |
| 7 | 39622353 | rs17687978 | 1.651 | -0.064 | -0.384 | -0.542 | -0.031 | -0.047 | -0.047 | 0 | 0.1 | 0.02 | 0.23 | 0.17 | 0.13 | 0.05 |
| 7 | 39622502 | rs4720328 | -1.872 | -0.445 | -0.537 | -0.891 | -0.019 | -0.06 | -0.06 | 0 | 0.64 | 0.89 | 0.68 | 0.55 | 0.6 | 0.62 |
| 7 | 39623495 | rs10272782 | -1.266 | -0.068 | -0.056 | -0.751 | -0.007 | -0.07 | -0.07 | 0.557 | 0.3 | 0.5 | 0.2 | 0.22 | 0.22 | 0.21 |
| 7 | 39646505 | rs17714061 | 1.55 | 0.192 | -0.495 | -0.582 | -0.031 | -0.015 | -0.015 | 0.011 | 0.07 | 0 | 0.04 | 0.11 | 0.08 | 0.11 |
| 7 | 39648977 | rs4723882 | -0.67 | 0.7 | -0.214 | -0.539 | 0.044 | -0.062 | -0.062 | 3.624 | 0.15 | 0.56 | 0.25 | 0.1 | 0.17 | 0.12 |
| 7 | 39652633 | rs1113117 | -1.248 | 0.318 | -0.316 | -0.475 | 0.025 | -0.021 | -0.021 | 0 | 0.57 | 0.8 | 0.68 | 0.54 | 0.6 | 0.62 |
| 7 | 39656686 | rs11767632 | -0.921 | 0.259 | -0.122 | -0.424 | 0.033 | -0.003 | -0.003 | 0 | 0.52 | 0.42 | 0.56 | 0.44 | 0.59 | 0.55 |
| 7 | 39661524 | rs10254562 | -0.343 | 0.332 | -0.106 | -0.445 | -0.009 | -0.006 | -0.006 | 0 | 0.41 | 0.3 | 0.49 | 0.35 | 0.5 | 0.46 |
| 7 | 39662268 | rs2008819 | -0.612 | 0.33 | -0.107 | -0.445 | -0.004 | 0 | 0 | 0 | 0.44 | 0.3 | 0.49 | 0.35 | 0.5 | 0.46 |
| 7 | 39666267 | rs11982634 | -1.035 | 0.236 | 0.009 | -0.39 | 0.009 | -0.002 | -0.002 | 0 | 0.53 | 0.48 | 0.56 | 0.41 | 0.59 | 0.55 |
| 7 | 39695873 | rs2107710 | -1.142 | 0.376 | -0.589 | -0.347 | 0.072 | -0.029 | -0.029 | 0 | 0.54 | 0.83 | 0.67 | 0.41 | 0.62 | 0.58 |
| 7 | 39723768 | rs10486802 | -0.7 | 0.912 | -0.429 | -0.415 | -0.003 | -0.014 | -0.014 | 0 | 0.4 | 0.39 | 0.3 | 0.09 | 0.45 | 0.45 |
| 7 | 39724464 | rs17171612 | 0.554 | 0.871 | -0.357 | -0.298 | 0.039 | 0.212 | 0.212 | 0.097 | 0.15 | 0.43 | 0.38 | 0.3 | 0.16 | 0.14 |
| 7 | 39733688 | rs6956681 | -1.736 | 0.824 | -1.047 | -0.252 | 0.02 | -0.086 | -0.086 | 0.544 | 0.6 | 0.94 | 0.73 | 0.53 | 0.63 | 0.64 |
| 7 | 39736796 | rs3779203 | 1.373 | 1.528 | -0.696 | -0.652 | -0.036 | -0.003 | -0.003 | 0 | 0.08 | 0 | 0.06 | 0.06 | 0.09 | 0.09 |
| 7 | 39737269 | rs3779200 | 0.367 | 1.554 | -0.636 | -0.675 | 0 | -0.007 | -0.007 | 0 | 0.23 | 0.18 | 0.13 | 0.04 | 0.21 | 0.32 |
| 7 | 39740487 | rs7778955 | -0.138 | 1.312 | -0.575 | -0.589 | -0.014 | 0.013 | 0.013 | 0 | 0.31 | 0.22 | 0.22 | 0.04 | 0.36 | 0.36 |
| 7 | 39748116 | rs17508585 | -0.71 | 1.257 | -0.531 | -0.715 | -0.009 | 0.02 | 0.02 | 0 | 0.42 | 0.36 | 0.29 | 0.09 | 0.45 | 0.45 |
| 7 | 39787109 | rs7458148 | -0.828 | 1.036 | -0.419 | 0.025 | 0.003 | 0.005 | 0.005 | 0 | 0.42 | 0.55 | 0.49 | 0.46 | 0.43 | 0.54 |
| 7 | 39792651 | rs6950742 | -0.589 | 1.097 | -0.395 | 0.169 | 0.001 | 0.013 | 0.013 | 0 | 0.33 | 0.35 | 0.42 | 0.44 | 0.34 | 0.44 |
| 7 | 39796229 | rs6951760 | -0.271 | 0.8 | -0.424 | 0.506 | -0.001 | 0.007 | 0.007 | 0.056 | 0.17 | 0.15 | 0.25 | 0.39 | 0.16 | 0.12 |
| 7 | 39849482 | rs10252318 | 0.051 | 1.357 | -0.69 | -0.263 | -0.056 | 0.062 | 0.062 | 0 | 0.11 | 0.01 | 0.12 | 0 | 0.17 | 0.03 |
| 7 | 39849931 | rs4720341 | -1.03 | 0.647 | -0.107 | -0.907 | 0.015 | -0.134 | -0.134 | 0.649 | 0.33 | 0.51 | 0.42 | 0.02 | 0.36 | 0.14 |
| 7 | 39854141 | rs4720342 | -1.063 | 0.173 | -0.149 | -0.713 | 0.004 | 0.01 | 0.01 | 0 | 0.45 | 0.35 | 0.45 | 0.29 | 0.39 | 0.42 |
| 7 | 39858378 | rs917199 | 0.464 | 0.287 | -0.269 | -0.709 | 0 | -0.001 | -0.001 | 0 | 0.2 | 0.16 | 0.27 | 0.02 | 0.19 | 0.11 |
| 7 | 39863558 | rs11972197 | 0.464 | 0.287 | -0.269 | -0.709 | 0 | -0.001 | -0.001 | 0 | 0.2 | 0.16 | 0.27 | 0.02 | 0.19 | 0.11 |
| 7 | 39874986 | rs10272635 | -1.323 | 0.23 | 0.306 | 0.043 | -0.001 | -0.003 | -0.003 | 0 | 0.35 | 0.45 | 0.32 | 0.3 | 0.33 | 0.38 |
| 7 | 39875108 | rs6969532 | 0.101 | 0.249 | 0.332 | 0.029 | -0.025 | 0.022 | 0.022 | 0 | 0.08 | 0.04 | 0.1 | 0 | 0.14 | 0.04 |
| 7 | 39880053 | rs1005311 | -1.387 | 0.242 | 0.333 | 0.03 | -0.001 | -0.003 | -0.003 | 0 | 0.35 | 0.45 | 0.32 | 0.3 | 0.33 | 0.38 |
| 7 | 39885541 | rs11561862 | -0.127 | 0.096 | 0.328 | 0.13 | 0.002 | -0.006 | -0.006 | 0 | 0.22 | 0.13 | 0.19 | 0.29 | 0.18 | 0.34 |
| 7 | 39894883 | rs12532428 | 0.265 | 0.205 | 0.05 | 0.163 | -0.032 | 0.033 | 0.033 | 0 | 0.74 | 0.63 | 0.78 | 0.71 | 0.81 | 0.66 |
| 7 | 39913870 | rs1880129 | 0.846 | 0.296 | -0.094 | 0.145 | 0 | -0.004 | -0.004 | 0.127 | 0.66 | 0.59 | 0.68 | 0.7 | 0.67 | 0.63 |
| 7 | 39928599 | rs7777774 | 0.641 | 0.28 | -0.441 | 0.78 | -0.005 | 0.018 | 0.018 | 0 | 0.86 | 0.84 | 0.76 | 0.56 | 0.8 | 0.85 |
| 7 | 39930518 | rs12701773 | 1.186 | -0.558 | -0.968 | 0.098 | 0.015 | 0.263 | 0.263 | 0 | 0.54 | 0.13 | 0.49 | 0.15 | 0.62 | 0.5 |
| 7 | 39939960 | rs4723915 | 0.577 | -0.545 | -0.301 | 0.186 | 0.028 | 0.184 | 0.184 | 0.012 | 0.84 | 0.56 | 0.71 | 0.46 | 0.79 | 0.85 |
| 7 | 39942647 | rs4313070 | 0.904 | -0.527 | -0.841 | -0.295 | 0.023 | 0.173 | 0.173 | 0 | 0.55 | 0.23 | 0.49 | 0.15 | 0.62 | 0.5 |
| 7 | 39962864 | rs10240222 | -0.271 | -0.311 | -1.558 | 0.158 | -0.097 | -0.071 | -0.071 | 2.147 | 0.39 | 0.09 | 0.48 | 0.6 | 0.51 | 0.51 |
| 7 | 39968050 | rs2329614 | 0.382 | -0.196 | -1.528 | 0.166 | -0.006 | -0.007 | -0.007 | 0 | 0.82 | 0.78 | 0.76 | 0.58 | 0.75 | 0.83 |

| | | | | | | | | | | | | | | | | |
|---|---|---|---|---|---|---|---|---|---|---|---|---|---|---|---|---|
| 7 | 39972125 | rs10243593 | 0.486 | -0.526 | -1.206 | -0.008 | -0.013 | 0.028 | 0.028 | 0 | 0.56 | 0.31 | 0.44 | 0.31 | 0.49 | 0.51 |
| 7 | 39991673 | rs9655407 | 0.83 | -0.93 | -2.019 | -0.742 | -0.063 | 0.268 | 0.268 | 0 | 0.37 | 0.07 | 0.27 | 0.01 | 0.35 | 0.23 |
| 7 | 40006195 | rs1082023 | -0.594 | -1.066 | -1.854 | -0.799 | 0.01 | 0.128 | 0.128 | 0 | 0.6 | 0.75 | 0.71 | 0.99 | 0.64 | 0.76 |
| 7 | 40023364 | rs17171639 | -0.906 | -1.176 | -1.682 | -0.859 | -0.006 | -0.008 | -0.008 | 0 | 0.06 | 0.58 | 0.05 | 0 | 0.01 | 0 |
| 7 | 40027484 | rs3735135 | 1.39 | -1.362 | -1.834 | -0.584 | 0.046 | -0.074 | -0.074 | 0 | 0.16 | 0.05 | 0.12 | 0.34 | 0.09 | 0.3 |
| 7 | 40030488 | rs4723924 | 0.736 | -1.154 | -1.937 | -0.555 | -0.1 | -0.082 | -0.082 | 0.044 | 0.32 | 0.11 | 0.43 | 0.5 | 0.42 | 0.39 |
| 7 | 40054784 | rs1403381 | -0.043 | -1.202 | -1.734 | -0.623 | -0.037 | -0.017 | -0.017 | 0.54 | 0.6 | 0.84 | 0.45 | 0.32 | 0.45 | 0.56 |
| 7 | 40056118 | rs773411 | -0.611 | -1.228 | -1.731 | -0.632 | -0.023 | 0.012 | 0.012 | 0 | 0.78 | 0.35 | 0.83 | 0.69 | 0.91 | 0.68 |
| 7 | 40058992 | rs10225492 | 1.205 | -1.231 | -1.776 | -0.647 | -0.059 | 0.268 | 0.268 | 0 | 0.37 | 0.07 | 0.27 | 0.01 | 0.35 | 0.23 |
| 7 | 40060180 | rs17526699 | 1.205 | -1.231 | -1.776 | -0.647 | -0.059 | 0.268 | 0.268 | 0 | 0.37 | 0.07 | 0.27 | 0.01 | 0.35 | 0.23 |
| 7 | 40062866 | rs7797041 | -0.033 | -1.012 | -1.693 | -0.517 | -0.048 | -0.023 | -0.023 | 0 | 0.61 | 0.85 | 0.45 | 0.32 | 0.45 | 0.56 |
| 7 | 40067021 | rs6971502 | 1.753 | -0.917 | -1.765 | -0.54 | -0.06 | 0.069 | 0.069 | 0 | 0.22 | 0.07 | 0.27 | 0.13 | 0.27 | 0.28 |
| 7 | 40075461 | rs6966917 | -0.114 | -1.231 | -1.648 | -0.488 | -0.039 | -0.019 | -0.019 | 0.128 | 0.61 | 0.84 | 0.45 | 0.32 | 0.45 | 0.56 |
| 7 | 40096299 | rs12533272 | 0.017 | -1.217 | -1.682 | -0.47 | -0.008 | 0.008 | 0.008 | 0 | 0.59 | 0.72 | 0.44 | 0.32 | 0.44 | 0.56 |
| 7 | 40124137 | rs1146042 | 0.582 | -1.074 | -1.618 | -0.267 | -0.034 | -0.038 | -0.038 | 0 | 0.07 | 0.01 | 0.12 | 0.15 | 0.14 | 0.06 |
| 7 | 40127767 | rs2302341 | 1.359 | -1.407 | -1.618 | -0.37 | 0.044 | -0.068 | -0.068 | 0.062 | 0.16 | 0.05 | 0.12 | 0.31 | 0.09 | 0.3 |
| 7 | 40133044 | rs2077124 | -0.022 | -1.747 | -2.009 | -0.873 | 0.012 | -0.004 | -0.004 | 0.485 | 0.34 | 0.35 | 0.24 | 0.32 | 0.22 | 0.41 |
| 7 | 40134678 | rs17538328 | 1.493 | -1.745 | -2.039 | -0.878 | -0.017 | 0.083 | 0.083 | 0 | 0.16 | 0.04 | 0.11 | 0.01 | 0.13 | 0.09 |
| 7 | 40135669 | rs3801238 | 0.778 | -1.8 | -1.335 | -0.805 | -0.086 | 0.118 | 0.118 | 0 | 0.2 | 0.51 | 0.14 | 0.31 | 0.09 | 0.32 |
| 7 | 40155624 | rs7786147 | -1.315 | -1.728 | -1.3 | -0.745 | -0.061 | -0.087 | -0.087 | 0 | 0.18 | 0.26 | 0.09 | 0 | 0.05 | 0.1 |
| 7 | 40157040 | rs7795333 | 0.824 | -0.923 | -1.085 | -0.445 | -0.005 | 0.073 | 0.073 | 0 | 0.16 | 0.07 | 0.11 | 0.01 | 0.13 | 0.09 |
| 7 | 40158527 | rs17171658 | -0.051 | -0.731 | -1.133 | -0.432 | -0.046 | -0.026 | -0.026 | 0 | 0.6 | 0.88 | 0.45 | 0.32 | 0.44 | 0.57 |
| 7 | 40162586 | rs13233333 | 1.502 | -0.764 | -1.132 | -1.01 | 0.004 | -0.148 | -0.148 | 0.03 | 0.36 | 0.04 | 0.33 | 0.46 | 0.31 | 0.47 |
| 7 | 40169196 | rs12534074 | 0.976 | 0.131 | -0.099 | -0.204 | 0 | -0.001 | -0.001 | 0 | 0.56 | 0.43 | 0.48 | 0.47 | 0.44 | 0.55 |
| 7 | 40193535 | rs11760739 | 1.079 | 0.419 | -0.124 | -0.098 | -0.045 | -0.061 | -0.061 | 0.159 | 0.54 | 0.14 | 0.44 | 0.47 | 0.46 | 0.57 |
| 7 | 40194260 | rs2024200 | 0.69 | 0.412 | -0.107 | -0.101 | -0.064 | -0.076 | -0.076 | 0.558 | 0.61 | 0.86 | 0.49 | 0.47 | 0.46 | 0.61 |
| 7 | 40197168 | rs419576 | -0.641 | 0.535 | -0.245 | -0.038 | -0.031 | -0.065 | -0.065 | 0 | 0.9 | 0.99 | 0.84 | 0.61 | 0.88 | 0.9 |
| 7 | 40203356 | rs6954096 | 0.452 | 0.304 | 0.064 | -0.111 | -0.006 | -0.009 | -0.009 | 0.031 | 0.59 | 0.66 | 0.48 | 0.47 | 0.46 | 0.6 |
| 7 | 40225129 | rs6462962 | 0.49 | 0.257 | -0.046 | -0.167 | -0.003 | -0.004 | -0.004 | 1.95 | 0.59 | 0.6 | 0.48 | 0.47 | 0.46 | 0.61 |
| 7 | 40227307 | rs2056775 | 0.306 | 0.259 | -0.273 | -0.166 | -0.079 | -0.088 | -0.088 | 0 | 0.61 | 0.88 | 0.49 | 0.47 | 0.46 | 0.61 |
| 7 | 40242537 | rs10269349 | 0.597 | 0.369 | -0.271 | -0.127 | -0.001 | -0.003 | -0.003 | 0 | 0.57 | 0.39 | 0.47 | 0.47 | 0.46 | 0.58 |
| 7 | 40247794 | rs11768838 | 0.623 | 0.463 | -1.495 | -0.082 | -0.05 | -0.061 | -0.061 | 0 | 0.54 | 0.14 | 0.44 | 0.47 | 0.46 | 0.57 |
| 7 | 40248683 | rs6951375 | 0.612 | 0.481 | -1.487 | -0.094 | -0.051 | -0.066 | -0.066 | 0.525 | 0.54 | 0.14 | 0.44 | 0.47 | 0.46 | 0.57 |
| 7 | 40255122 | rs6950489 | 0.346 | 1.096 | -1.588 | 0.138 | -0.034 | 0.027 | 0.027 | 0 | 0.93 | 0.99 | 0.92 | 0.99 | 0.86 | 0.95 |
| 7 | 40265510 | rs6462965 | 0.39 | 0.869 | -2.052 | 0.201 | -0.083 | -0.095 | -0.095 | 0.422 | 0.61 | 0.88 | 0.49 | 0.46 | 0.47 | 0.61 |
| 7 | 40267017 | rs6948247 | 0.856 | 1.238 | -1.335 | -0.958 | 0.055 | -0.136 | -0.136 | 0 | 0.74 | 0.52 | 0.62 | 0.98 | 0.65 | 0.88 |
| 7 | 40271224 | rs4723936 | 0.673 | 0.626 | -1.088 | -1.251 | -0.004 | 0 | 0 | 0.041 | 0.45 | 0.47 | 0.4 | 0.46 | 0.38 | 0.42 |
| 7 | 40284166 | rs6964864 | -0.173 | 0.685 | -1.24 | -1.497 | -0.059 | -0.055 | -0.055 | 0 | 0.36 | 0.18 | 0.44 | 0.52 | 0.46 | 0.44 |
| 7 | 40290983 | rs4723938 | 1.248 | 0.357 | -0.566 | -1.688 | 0.007 | -0.01 | -0.01 | 0.014 | 0.4 | 0.36 | 0.34 | 0.45 | 0.37 | 0.4 |
| 7 | 40298878 | rs4296945 | -0.32 | 0.329 | -0.418 | -1.702 | -0.013 | -0.087 | -0.087 | 0 | 0.77 | 0.71 | 0.79 | 0.98 | 0.82 | 0.85 |
| 7 | 40305112 | rs6462968 | -0.594 | 0.024 | -0.618 | -1.846 | 0.016 | -0.002 | -0.002 | 0.212 | 0.32 | 0.72 | 0.26 | 0.03 | 0.2 | 0.16 |
| 7 | 40307178 | rs4129802 | 0.581 | 0.514 | -0.621 | -1.654 | -0.006 | -0.025 | -0.025 | 0 | 0.26 | 0.21 | 0.35 | 0.5 | 0.27 | 0.4 |
| 7 | 40314165 | rs12669315 | 1.802 | 0.492 | -0.537 | -1.679 | -0.014 | -0.103 | -0.103 | 0.665 | 0.37 | 0.11 | 0.33 | 0.45 | 0.36 | 0.4 |
| 7 | 40314243 | rs4637704 | 0.421 | 0.579 | -0.585 | -1.687 | -0.007 | -0.029 | -0.029 | 0 | 0.27 | 0.21 | 0.36 | 0.5 | 0.29 | 0.4 |
| 7 | 40318967 | rs952750 | -0.951 | 0.486 | -0.55 | -1.613 | 0.054 | -0.012 | -0.012 | 0 | 0.56 | 0.23 | 0.61 | 0.54 | 0.65 | 0.56 |
| 7 | 40320493 | rs6960480 | -1.697 | 0.497 | -0.453 | -1.603 | -0.013 | 0.013 | 0.013 | 0 | 0.63 | 0.5 | 0.68 | 0.55 | 0.67 | 0.57 |
| 7 | 40338553 | rs6962341 | 1.128 | 0.551 | -0.427 | -1.578 | -0.014 | 0.003 | 0.003 | 0 | 0.42 | 0.46 | 0.36 | 0.46 | 0.34 | 0.41 |
| 7 | 40344782 | rs10247300 | 0.217 | 0.656 | -0.635 | -1.61 | -0.01 | -0.033 | -0.033 | 0 | 0.29 | 0.22 | 0.37 | 0.5 | 0.28 | 0.4 |
| 7 | 40353713 | rs10278370 | -0.907 | 0.454 | -0.686 | -1.455 | 0.039 | -0.008 | -0.008 | 0.016 | 0.56 | 0.31 | 0.62 | 0.54 | 0.65 | 0.59 |
| 7 | 40356836 | rs10951633 | 1.586 | 0.636 | -0.787 | -1.354 | 0.018 | -0.036 | -0.036 | 0 | 0.36 | 0.21 | 0.3 | 0.44 | 0.33 | 0.39 |
| 7 | 40360502 | rs12701816 | -0.479 | 1.179 | -1.122 | -1.585 | -0.022 | 0.027 | 0.027 | 0 | 0.3 | 0.24 | 0.33 | 0.06 | 0.36 | 0.28 |
| 7 | 40364588 | rs10256660 | 0.018 | 0.877 | -0.918 | -1.244 | 0.001 | -0.058 | -0.058 | 0 | 0.73 | 0.68 | 0.74 | 0.99 | 0.64 | 0.81 |
| 7 | 40364726 | rs10256791 | 1.091 | 0.741 | -0.852 | -0.585 | 0.007 | 0.008 | 0.008 | 0 | 0.6 | 0.5 | 0.6 | 0.58 | 0.54 | 0.64 |
| 7 | 40368926 | rs17171686 | 0.47 | 0.744 | -0.868 | -0.581 | -0.005 | 0.04 | 0.04 | 3.913 | 0.09 | 0.06 | 0.13 | 0.43 | 0.1 | 0.16 |
| 7 | 40381031 | rs4613890 | -0.248 | 0.729 | -0.828 | -0.61 | -0.009 | -0.028 | -0.028 | 0.021 | 0.73 | 0.79 | 0.75 | 1 | 0.64 | 0.9 |
| 7 | 40386835 | rs10951634 | 0.767 | 0.606 | -0.843 | -0.58 | -0.007 | -0.008 | -0.008 | 0.396 | 0.63 | 0.66 | 0.62 | 0.56 | 0.54 | 0.65 |
| 7 | 40391181 | rs11761602 | 1.978 | 0.437 | -1.449 | -0.385 | -0.062 | -0.101 | -0.101 | 0 | 0.34 | 0.04 | 0.32 | 0.39 | 0.37 | 0.37 |
| 7 | 40394497 | rs17171693 | 0.747 | 0.926 | -0.33 | 0.375 | -0.029 | -0.011 | -0.011 | 0.302 | 0.08 | 0 | 0.12 | 0.43 | 0.08 | 0.16 |
| 7 | 40394987 | rs17171694 | 0.552 | 0.931 | -0.319 | 0.377 | -0.011 | 0.012 | 0.012 | 3.937 | 0.08 | 0.04 | 0.12 | 0.43 | 0.08 | 0.16 |
| 7 | 40409406 | rs6965764 | 1.445 | 0.913 | -0.266 | 0.354 | 0.002 | 0.003 | 0.003 | 0 | 0.61 | 0.51 | 0.6 | 0.56 | 0.54 | 0.64 |
| 7 | 40418721 | rs12533531 | 0.118 | 1.037 | -0.301 | 0.326 | 0 | 0.095 | 0.095 | 4.369 | 0.09 | 0.18 | 0.14 | 0.48 | 0.09 | 0.17 |
| 7 | 40420032 | rs9648486 | -0.404 | 1.026 | -0.334 | 0.255 | -0.093 | 0.148 | 0.148 | 0 | 0.26 | 0.07 | 0.25 | 0.01 | 0.38 | 0.1 |
| 7 | 40421458 | rs7779860 | -0.404 | 1.026 | -0.331 | 0.255 | -0.089 | 0.142 | 0.142 | 0 | 0.26 | 0.07 | 0.25 | 0 | 0.38 | 0.1 |
| 7 | 40427000 | rs17171705 | 0.481 | 1.089 | -0.262 | 0.326 | -0.004 | 0.037 | 0.037 | 3.898 | 0.08 | 0.06 | 0.13 | 0.48 | 0.09 | 0.17 |
| 7 | 40427617 | rs10234636 | 0.434 | 1.079 | -0.204 | 0.342 | 0.014 | 0.292 | 0.292 | 2.348 | 0.89 | 0.64 | 0.86 | 0.53 | 0.9 | 0.75 |
| 7 | 40427634 | rs12532479 | 0.159 | 1.082 | -0.226 | 0.343 | 0.001 | 0.106 | 0.106 | 4.693 | 0.09 | 0.17 | 0.14 | 0.45 | 0.09 | 0.17 |
| 7 | 40432219 | rs12673516 | -0.184 | 1.096 | -0.223 | 0.315 | 0 | 0.083 | 0.083 | 3.891 | 0.1 | 0.18 | 0.14 | 0.48 | 0.09 | 0.25 |
| 7 | 40436624 | rs10263744 | 0.388 | 1.275 | -0.469 | 0.351 | 0.001 | -0.06 | -0.06 | 1.522 | 0.73 | 0.73 | 0.75 | 1 | 0.62 | 0.9 |
| 7 | 40440233 | rs17171710 | 0.248 | 1.456 | -0.38 | 0.494 | -0.004 | 0.037 | 0.037 | 3.898 | 0.08 | 0.06 | 0.13 | 0.48 | 0.09 | 0.16 |

| | | | | | | | | | | | | | | | | |
|---|---|---|---|---|---|---|---|---|---|---|---|---|---|---|---|---|
| 7 | 40449904 | rs12234460 | 0.248 | 1.484 | -0.51 | 0.516 | -0.003 | 0.044 | 0.044 | 3.898 | 0.08 | 0.07 | 0.13 | 0.47 | 0.09 | 0.16 |
| 7 | 40458304 | rs4364550 | 0.296 | 1.49 | -0.519 | 0.426 | 0 | -0.053 | -0.053 | 0.65 | 0.75 | 0.75 | 0.75 | 1 | 0.63 | 0.9 |
| 7 | 40474936 | rs2190261 | -0.274 | 0.398 | -0.172 | 1.315 | -0.003 | 0.043 | 0.043 | 3.898 | 0.08 | 0.07 | 0.13 | 0.48 | 0.09 | 0.18 |
| 7 | 40485949 | rs10229414 | -0.605 | 0.414 | 0.209 | 0.339 | 0.045 | 0.078 | 0.078 | 0.07 | 0.07 | 0.23 | 0.06 | 0.12 | 0.12 | 0.17 |
| 7 | 40489083 | rs7809620 | -0.931 | -0.055 | -0.064 | -0.258 | 0.005 | 0.035 | 0.035 | 0 | 0.18 | 0.12 | 0.19 | 0.03 | 0.17 | 0.1 |
| 7 | 40510686 | rs17433827 | -0.6 | 0.093 | -0.231 | -0.274 | -0.032 | 0.095 | 0.095 | 0 | 0.15 | 0 | 0.1 | 0 | 0.16 | 0.07 |
| 7 | 40515627 | rs10248808 | -1.473 | 0.066 | 0.763 | 0.073 | 0.049 | 0.012 | 0.012 | 0.223 | 0.07 | 0.38 | 0.09 | 0 | 0.04 | 0.04 |
| 7 | 40525965 | rs4320458 | 0.23 | 0.322 | -0.086 | 0.298 | -0.026 | -0.019 | -0.019 | 0 | 0.08 | 0.01 | 0.18 | 0.06 | 0.12 | 0.17 |
| 7 | 40586674 | rs11982075 | -1.904 | -0.214 | 1.321 | 0.084 | -0.019 | -0.052 | -0.052 | 0 | 0.14 | 0.59 | 0.14 | 0.05 | 0.12 | 0.11 |
| 7 | 40607390 | rs17171734 | -1.257 | -0.224 | 1.399 | 0.06 | 0.002 | 0.005 | 0.005 | 0 | 0.1 | 0.1 | 0.11 | 0.03 | 0.11 | 0.1 |
| 7 | 40612023 | rs740214 | -0.109 | -0.8 | 0.949 | -0.034 | 0.017 | 0 | 0 | 6.702 | 0.17 | 0.05 | 0.09 | 0.15 | 0.12 | 0.1 |
| 7 | 40612250 | rs10263677 | -1.962 | -1.026 | 1.351 | 0.049 | -0.027 | -0.057 | -0.057 | 0 | 0.15 | 0.65 | 0.15 | 0.05 | 0.12 | 0.11 |
| 7 | 40631826 | rs1019018 | -1.485 | -1.03 | 1.515 | 0.047 | -0.01 | -0.039 | -0.039 | 0 | 0.12 | 0.36 | 0.13 | 0.05 | 0.11 | 0.11 |
| 7 | 40638365 | rs11973006 | -2.296 | -1.212 | 0.918 | -0.169 | -0.035 | -0.097 | -0.097 | 0.094 | 0.23 | 0.59 | 0.2 | 0.05 | 0.19 | 0.15 |
| 7 | 40685811 | rs10282661 | -2.095 | -0.809 | 0.536 | -0.255 | 0.031 | 0.178 | 0.178 | 1.29 | 0.49 | 0.97 | 0.34 | 0.22 | 0.36 | 0.26 |
| 7 | 40687150 | rs6954401 | -2.205 | 0.18 | 1.674 | 0.349 | -0.025 | -0.057 | -0.057 | 0.087 | 0.14 | 0.64 | 0.1 | 0.05 | 0.09 | 0.1 |
| 7 | 40697882 | rs10251707 | -1.622 | 0.474 | 1.621 | 0.268 | 0.021 | -0.131 | -0.131 | 0 | 0.2 | 0.5 | 0.18 | 0.04 | 0.19 | 0.17 |
| 7 | 40699189 | rs10280713 | -2.038 | -0.026 | 1.599 | 0.51 | -0.012 | 0.003 | 0.003 | 0.117 | 0.29 | 0.59 | 0.23 | 0.28 | 0.24 | 0.2 |
| 7 | 40716701 | rs6952969 | -2.052 | -0.003 | 1.369 | 0.52 | -0.006 | 0.001 | 0.001 | 0.054 | 0.29 | 0.63 | 0.24 | 0.29 | 0.25 | 0.19 |
| 7 | 40726304 | rs6977963 | -0.988 | 0.096 | 1.093 | 0.282 | -0.038 | 0.078 | 0.078 | 0 | 0.19 | 0.06 | 0.18 | 0.02 | 0.23 | 0.18 |
| 7 | 40727223 | rs6965055 | -1.827 | 0.063 | 1.234 | 0.401 | -0.004 | -0.003 | -0.003 | 0 | 0.31 | 0.44 | 0.3 | 0.31 | 0.28 | 0.22 |
| 7 | 40728563 | rs17171762 | 0.199 | 0.061 | 0.479 | 0.431 | -0.009 | 0.033 | 0.033 | 0 | 0.1 | 0.13 | 0.1 | 0.28 | 0.05 | 0.04 |
| 7 | 40728868 | rs10250246 | -1.176 | 0.198 | 0.094 | 0.504 | -0.002 | -0.001 | -0.001 | 0 | 0.32 | 0.67 | 0.31 | 0.31 | 0.28 | 0.22 |
| 7 | 40748650 | rs1203739 | -1.237 | 0.198 | -1.121 | 0.352 | -0.044 | -0.136 | -0.136 | 0 | 0.38 | 0.94 | 0.35 | 0.53 | 0.32 | 0.32 |
| 7 | 40756923 | rs1203731 | -1.228 | 0.161 | -1.106 | 0.294 | -0.029 | -0.114 | -0.114 | 0 | 0.38 | 0.92 | 0.35 | 0.52 | 0.32 | 0.32 |
| 7 | 40760874 | rs1921744 | -1.017 | 0.44 | -0.642 | -0.569 | -0.027 | 0.043 | 0.043 | 0 | 0.2 | 0.12 | 0.2 | 0 | 0.25 | 0.09 |
| 7 | 40787104 | rs17438725 | -0.493 | -0.038 | -1.251 | -0.479 | -0.003 | 0.083 | 0.083 | 0 | 0.18 | 0.03 | 0.09 | 0 | 0.12 | 0.09 |
| 7 | 40790257 | rs6970061 | 0.905 | 0.061 | -2.485 | -0.235 | -0.014 | -0.06 | -0.06 | 0 | 0.46 | 0.1 | 0.47 | 0.47 | 0.49 | 0.6 |
| 7 | 40790382 | rs6970230 | -0.358 | 0.321 | -2.33 | -0.374 | -0.004 | 0.038 | 0.038 | 0 | 0.23 | 0.12 | 0.13 | 0.01 | 0.16 | 0.11 |
| 7 | 40795112 | rs17529882 | 0.854 | 0.301 | -2.305 | -0.35 | -0.009 | 0.072 | 0.072 | 0 | 0.17 | 0.03 | 0.09 | 0 | 0.11 | 0.08 |
| 7 | 40796618 | rs1921752 | -0.234 | 0.332 | -2.006 | 0.006 | -0.012 | -0.036 | -0.036 | 0.94 | 0.68 | 0.25 | 0.72 | 0.54 | 0.65 | 0.7 |
| 7 | 40809143 | rs2876978 | -0.214 | 0.294 | -2.018 | 0.039 | -0.011 | -0.039 | -0.039 | 0 | 0.68 | 0.26 | 0.73 | 0.55 | 0.66 | 0.7 |
| 7 | 40812079 | rs12701832 | -0.498 | -1.265 | -2.876 | -1.154 | 0.057 | 0.162 | 0.162 | 0 | 0.45 | 0.24 | 0.43 | 0.13 | 0.34 | 0.44 |
| 7 | 40815025 | rs17530165 | -0.358 | -1.736 | -3.373 | -1.3 | 0.004 | -0.027 | -0.027 | 0 | 0.42 | 0.15 | 0.3 | 0.45 | 0.38 | 0.52 |
| 7 | 40816465 | rs2329877 | -1.584 | -1.781 | -3.212 | -1.337 | -0.045 | -0.041 | -0.041 | 0 | 0.61 | 0.25 | 0.52 | 0.56 | 0.49 | 0.65 |
| 7 | 40816580 | rs17530234 | 1.142 | -1.909 | -3.131 | -1.296 | 0.012 | 0.013 | 0.013 | 0.254 | 0.18 | 0.11 | 0.09 | 0.02 | 0.11 | 0.09 |
| 7 | 40820783 | rs12155167 | 1.373 | -1.887 | -3.226 | -1.286 | 0.014 | 0.052 | 0.052 | 0.555 | 0.17 | 0.06 | 0.08 | 0.02 | 0.1 | 0.09 |
| 7 | 40824133 | rs1921743 | 0.783 | -1.991 | -3.29 | -1.323 | -0.052 | -0.1 | -0.1 | 0 | 0.22 | 0.06 | 0.14 | 0.43 | 0.22 | 0.23 |
| 7 | 40826041 | rs7807596 | -0.56 | -2.032 | -2.842 | -1.341 | 0.022 | 0.004 | 0.004 | 0 | 0.43 | 0.31 | 0.3 | 0.45 | 0.39 | 0.52 |
| 7 | 40836244 | rs1921748 | -0.526 | -1.858 | -3.034 | -1.271 | -0.007 | -0.038 | -0.038 | 0 | 0.39 | 0.15 | 0.29 | 0.45 | 0.38 | 0.48 |
| 7 | 40847909 | rs2302025 | -1.4 | -1.794 | -2.95 | -1.226 | -0.046 | -0.006 | -0.006 | 0.053 | 0.57 | 0.22 | 0.37 | 0.47 | 0.48 | 0.57 |
| 7 | 40866663 | rs17688601 | -0.294 | -1.736 | -2.921 | -1.323 | 0.038 | 0.215 | 0.215 | 0.718 | 0.34 | 0.03 | 0.23 | 0.05 | 0.25 | 0.35 |
| 7 | 40868141 | rs4723979 | 1.064 | -1.547 | -2.922 | -1.165 | -0.013 | -0.056 | -0.056 | 0 | 0.21 | 0.15 | 0.26 | 0.51 | 0.21 | 0.3 |
| 7 | 40870771 | rs10247275 | 1.046 | -1.559 | -2.888 | -1.161 | -0.01 | -0.036 | -0.036 | 0 | 0.21 | 0.16 | 0.13 | 0.42 | 0.21 | 0.22 |
| 7 | 40871858 | rs13246232 | -0.912 | -1.572 | -2.861 | -1.266 | 0.016 | -0.057 | -0.057 | 0.082 | 0.38 | 0.21 | 0.42 | 0.53 | 0.37 | 0.56 |
| 7 | 40873221 | rs1733483 | -1.915 | -1.719 | -2.6 | -1.276 | -0.046 | -0.04 | -0.04 | 3.645 | 0.65 | 0.76 | 0.56 | 0.61 | 0.52 | 0.78 |
| 7 | 40875192 | rs17621345 | -0.39 | -2.103 | -2.69 | -2.098 | 0.04 | 0.181 | 0.181 | 1.25 | 0.33 | 0.1 | 0.21 | 0.03 | 0.24 | 0.36 |
| 7 | 40886576 | rs7808121 | -0.152 | -1.748 | -2.209 | -1.416 | -0.036 | -0.001 | -0.001 | 0 | 0.23 | 0.11 | 0.28 | 0.8 | 0.24 | 0.3 |
| 7 | 40892728 | rs4723983 | -0.788 | -1.586 | -2.109 | -1.4 | -0.017 | 0.11 | 0.11 | 0.172 | 0.42 | 0.26 | 0.4 | 0.82 | 0.37 | 0.38 |
| 7 | 40900175 | rs699484 | -0.696 | -1.501 | -2.151 | -1.362 | -0.031 | 0.12 | 0.12 | 0.013 | 0.41 | 0.22 | 0.4 | 0.83 | 0.37 | 0.37 |
| 7 | 40906697 | rs728585 | -0.284 | -1.451 | -1.273 | -0.871 | 0.002 | 0.015 | 0.015 | 0 | 0.29 | 0.18 | 0.17 | 0.75 | 0.23 | 0.32 |
| 7 | 40918070 | rs7801303 | -1.163 | -1.294 | -0.63 | -0.596 | -0.017 | 0.062 | 0.062 | 0.086 | 0.59 | 0.7 | 0.55 | 0.9 | 0.43 | 0.69 |
| 7 | 40919389 | rs6974783 | 0.117 | -0.936 | -0.739 | -0.421 | -0.014 | 0.098 | 0.098 | 0 | 0.27 | 0.09 | 0.23 | 0.09 | 0.23 | 0.18 |
| 7 | 40922984 | rs758008 | -1.034 | -1.042 | 0.218 | 0.798 | -0.038 | -0.038 | -0.038 | 0.12 | 0.28 | 0.45 | 0.28 | 0.8 | 0.18 | 0.5 |
| 7 | 40925373 | rs17714729 | -0.779 | -0.134 | -0.501 | 1.135 | 0.003 | -0.011 | -0.011 | 0 | 0.06 | 0.04 | 0.04 | 0.17 | 0.04 | 0.03 |
| 7 | 40926906 | rs992223 | -0.505 | -0.282 | -0.268 | 0.737 | -0.031 | -0.012 | -0.012 | 0.29 | 0.36 | 0.48 | 0.19 | 0.67 | 0.24 | 0.36 |
| 7 | 40928113 | rs10276497 | -0.021 | 0.566 | -0.423 | 0.933 | 0.009 | 0.03 | 0.03 | 0 | 0.51 | 0.58 | 0.39 | 0.65 | 0.45 | 0.48 |
| 7 | 40928658 | rs6957465 | -1.148 | 0.562 | -0.002 | 1.121 | -0.009 | 0.046 | 0.046 | 0.605 | 0.19 | 0.45 | 0.16 | 0.22 | 0.15 | 0.33 |
| 7 | 40939687 | rs11767956 | 0.257 | 0.842 | -0.37 | 1.009 | 0.001 | 0.006 | 0.006 | 0.16 | 0.6 | 0.48 | 0.43 | 0.4 | 0.44 | 0.55 |
| 7 | 40939791 | rs10486695 | -1.241 | 1.278 | 0.256 | 1.626 | 0.008 | 0.047 | 0.047 | 0.218 | 0.12 | 0.23 | 0.09 | 0.19 | 0.15 | 0.14 |
| 7 | 40939856 | rs17171789 | -0.757 | 1.112 | 0.858 | 1.53 | 0.029 | 0.052 | 0.052 | 0 | 0.09 | 0.34 | 0.07 | 0.07 | 0.06 | 0.06 |
| 7 | 40944385 | rs258076 | 0.562 | 1.615 | -0.406 | 1.612 | 0.02 | 0.076 | 0.076 | 0 | 0.42 | 0.21 | 0.28 | 0.35 | 0.36 | 0.46 |
| 7 | 40944971 | rs12672085 | 0.462 | 1.639 | -0.184 | 1.575 | -0.038 | 0.017 | 0.017 | 0.161 | 0.31 | 0.15 | 0.48 | 0.22 | 0.41 | 0.15 |
| 7 | 40945186 | rs29871 | 0.405 | 0.916 | 0.471 | 1.126 | 0.004 | 0.004 | 0.004 | 0.071 | 0.51 | 0.59 | 0.35 | 0.54 | 0.43 | 0.69 |
| 7 | 40946513 | rs29873 | -0.094 | -0.187 | 0.441 | 0.677 | 0 | 0 | 0 | 0 | 0.5 | 0.55 | 0.35 | 0.58 | 0.44 | 0.68 |
| 7 | 40947301 | rs29874 | -0.334 | -0.129 | 0.485 | 0.667 | -0.011 | -0.023 | -0.023 | 0 | 0.52 | 0.8 | 0.36 | 0.58 | 0.44 | 0.67 |
| 7 | 40948817 | rs40463 | -0.568 | -0.243 | 0.382 | 0.62 | -0.007 | -0.09 | -0.09 | 0 | 0.83 | 0.99 | 0.85 | 0.76 | 0.86 | 0.85 |
| 7 | 40950856 | rs6942942 | 1.246 | -0.52 | 0.625 | 0.405 | -0.001 | -0.009 | -0.009 | 0 | 0.26 | 0.35 | 0.22 | 0.29 | 0.23 | 0.33 |
| 7 | 40953754 | rs12701839 | 1.381 | -0.505 | 0.555 | 0.369 | -0.043 | 0.062 | 0.062 | 1.185 | 0.35 | 0.15 | 0.49 | 0.18 | 0.43 | 0.16 |

| | | | | | | | | | | | | | | | | |
|---|---|---|---|---|---|---|---|---|---|---|---|---|---|---|---|---|
| 7 | 40954481 | rs29880 | -0.302 | -0.451 | 0.726 | 0.44 | -0.008 | -0.018 | -0.018 | 0 | 0.17 | 0.27 | 0.1 | 0.11 | 0.14 | 0.15 |
| 7 | 40955755 | rs29881 | -0.652 | -0.463 | 0.711 | 0.444 | 0.02 | -0.037 | -0.037 | 0 | 0.82 | 0.94 | 0.84 | 0.76 | 0.86 | 0.84 |
| 7 | 40957669 | rs17171796 | -0.445 | -0.416 | 0.884 | 0.453 | -0.012 | -0.025 | -0.025 | 0.003 | 0.18 | 0.35 | 0.11 | 0.11 | 0.14 | 0.15 |
| 7 | 40958343 | rs17689310 | 2.12 | 0.09 | -0.001 | -0.023 | -0.071 | -0.028 | -0.028 | 0 | 0.17 | 0.01 | 0.25 | 0.13 | 0.21 | 0.06 |
| 7 | 40959796 | rs17771654 | 0.955 | -0.173 | 0.679 | 0.084 | 0.001 | 0.008 | 0.008 | 0 | 0.33 | 0.39 | 0.26 | 0.47 | 0.3 | 0.53 |
| 7 | 40961856 | rs11765424 | 1.038 | -0.208 | 0.458 | 0.073 | -0.009 | -0.033 | -0.033 | 0.619 | 0.33 | 0.2 | 0.24 | 0.47 | 0.29 | 0.53 |
| 7 | 40962660 | rs10234656 | 0.74 | -0.221 | 0.417 | 0.02 | -0.012 | 0.002 | 0.002 | 0 | 0.07 | 0.01 | 0.05 | 0.03 | 0.06 | 0.08 |
| 7 | 40963206 | rs9648489 | 0.706 | -0.25 | 0.492 | 0.027 | -0.002 | -0.003 | -0.003 | 0.685 | 0.07 | 0.07 | 0.05 | 0.03 | 0.06 | 0.08 |
| 7 | 40963295 | rs29888 | 0.994 | -0.257 | 0.417 | 0.028 | 0 | -0.004 | -0.004 | 1.366 | 0.08 | 0.13 | 0.04 | 0.08 | 0.09 | 0.07 |
| 7 | 40965127 | rs7789197 | -1.082 | -0.261 | 0.429 | 0.112 | 0.001 | 0.042 | 0.042 | 0.617 | 0.22 | 0.55 | 0.14 | 0.28 | 0.2 | 0.34 |
| 7 | 40985798 | rs917231 | 0.859 | -0.144 | -0.023 | -0.05 | -0.026 | 0.047 | 0.047 | 0.033 | 0.34 | 0.19 | 0.49 | 0.2 | 0.42 | 0.15 |
| 7 | 41007730 | rs6974676 | -0.771 | -0.123 | 0.067 | 0.318 | 0.008 | -0.034 | -0.034 | 0 | 0.86 | 0.96 | 0.86 | 0.76 | 0.87 | 0.84 |
| 7 | 41007872 | rs6954727 | 0.205 | -0.119 | 0.156 | 0.312 | 0.003 | -0.016 | -0.016 | 0 | 0.78 | 0.84 | 0.81 | 0.69 | 0.77 | 0.75 |
| 7 | 41011305 | rs4720392 | -0.638 | -0.098 | 0.336 | 0.914 | -0.002 | 0.017 | 0.017 | 0 | 0.88 | 0.88 | 0.86 | 0.79 | 0.87 | 0.87 |
| 7 | 41013284 | rs17622266 | 2.043 | -0.363 | -0.134 | 0.718 | -0.029 | 0.02 | 0.02 | 0.103 | 0.31 | 0.05 | 0.2 | 0.27 | 0.29 | 0.33 |
| 7 | 41016411 | rs17771935 | 1.787 | -0.017 | 0 | 0.887 | -0.029 | -0.034 | -0.034 | 0 | 0.06 | 0 | 0.03 | 0.16 | 0.07 | 0.23 |
| 7 | 41016442 | rs2214880 | -0.072 | -0.261 | 0.354 | 0.784 | -0.022 | -0.002 | -0.002 | 0 | 0.45 | 0.15 | 0.56 | 0.42 | 0.51 | 0.43 |
| 7 | 41018143 | rs1990142 | -2.326 | -0.25 | 0.32 | 0.797 | -0.016 | -0.063 | -0.063 | 0 | 0.9 | 0.99 | 0.86 | 0.78 | 0.87 | 0.87 |
| 7 | 41019212 | rs12701851 | 1.679 | -0.264 | 0.977 | 0.781 | 0.005 | 0.014 | 0.014 | 0 | 0.39 | 0.31 | 0.25 | 0.36 | 0.36 | 0.44 |
| 7 | 41020157 | rs6960159 | 1.546 | -0.259 | 0.867 | 0.784 | 0 | 0 | 0 | 0.065 | 0.4 | 0.37 | 0.26 | 0.36 | 0.36 | 0.44 |
| 7 | 41024666 | rs2024032 | -0.219 | -0.248 | 1.004 | 0.639 | 0.005 | 0.181 | 0.181 | 0.501 | 0.79 | 0.51 | 0.79 | 0.62 | 0.79 | 0.64 |
| 7 | 41025298 | rs2024033 | -1.095 | -0.431 | 1.07 | 0.804 | -0.013 | 0.089 | 0.089 | 0 | 0.86 | 0.72 | 0.84 | 0.78 | 0.87 | 0.87 |
| 7 | 41025898 | rs10229451 | 1.445 | -0.311 | 0.405 | 0.607 | -0.011 | 0.005 | 0.005 | 0 | 0.06 | 0 | 0.05 | 0.02 | 0.06 | 0.07 |
| 7 | 41042665 | rs6964018 | 0.883 | -0.32 | 0.434 | 0.62 | -0.048 | 0.082 | 0.082 | 0 | 0.37 | 0.15 | 0.53 | 0.24 | 0.42 | 0.2 |
| 7 | 41051036 | rs10486701 | -0.967 | -0.21 | 0.566 | 0.67 | 0.033 | 0.172 | 0.172 | 2.419 | 0.05 | 0.23 | 0.02 | 0.18 | 0.02 | 0.21 |
| 7 | 41065647 | rs1859699 | 1.464 | -0.297 | 0.428 | 0.593 | -0.011 | 0.027 | 0.027 | 0 | 0.39 | 0.23 | 0.28 | 0.36 | 0.42 | 0.45 |
| 7 | 41065688 | rs10275188 | -0.918 | -0.363 | 0.506 | 0.584 | -0.019 | 0.069 | 0.069 | 0.184 | 0.15 | 0.38 | 0.08 | 0.21 | 0.08 | 0.3 |
| 7 | 41065849 | rs1859700 | -0.538 | -0.304 | 0.263 | 0.606 | -0.029 | -0.023 | -0.023 | 0.851 | 0.55 | 0.74 | 0.36 | 0.57 | 0.5 | 0.76 |
| 7 | 41067781 | rs1548630 | 0.18 | -0.286 | -0.365 | 0.565 | -0.001 | -0.003 | -0.003 | 0 | 0.51 | 0.49 | 0.34 | 0.39 | 0.48 | 0.55 |
| 7 | 41067917 | rs2214876 | 1.214 | -0.103 | -1.855 | 0.234 | -0.12 | 0.109 | 0.109 | 0.023 | 0.33 | 0.04 | 0.44 | 0.19 | 0.37 | 0.12 |
| 7 | 41071575 | rs1990136 | -1.463 | 0.24 | -1.273 | 0.646 | -0.006 | -0.071 | -0.071 | 0 | 0.89 | 0.99 | 0.82 | 0.76 | 0.87 | 0.87 |
| 7 | 41071890 | rs1990138 | 0.488 | 0.427 | -0.811 | 0.331 | -0.001 | -0.001 | -0.001 | 0 | 0.41 | 0.37 | 0.29 | 0.49 | 0.43 | 0.46 |
| 7 | 41075387 | rs10280625 | -0.192 | 0.536 | -1.044 | -0.181 | -0.004 | 0.075 | 0.075 | 0 | 0.26 | 0.13 | 0.2 | 0.06 | 0.28 | 0.12 |
| 7 | 41075449 | rs2108185 | 0.515 | 0.333 | -0.429 | 0.453 | -0.002 | 0.003 | 0.003 | 0 | 0.71 | 0.71 | 0.61 | 0.68 | 0.73 | 0.69 |
| 7 | 41075484 | rs1990139 | 1.092 | 0.324 | -0.04 | 0.723 | -0.001 | -0.004 | -0.004 | 0 | 0.74 | 0.69 | 0.57 | 0.55 | 0.68 | 0.72 |
| 7 | 41075786 | rs4724018 | -1.43 | 0.044 | 0.394 | 0.425 | -0.008 | 0.004 | 0.004 | 0 | 0.22 | 0.38 | 0.23 | 0.21 | 0.2 | 0.13 |
| 7 | 41085469 | rs10225807 | -1.398 | -0.262 | 0.463 | -0.088 | -0.014 | -0.082 | -0.082 | 1.478 | 0.21 | 0.32 | 0.26 | 0.01 | 0.15 | 0.17 |
| 7 | 41086119 | rs17622724 | 0.054 | -0.221 | -0.144 | -0.083 | -0.01 | 0.059 | 0.059 | 0 | 0.1 | 0.01 | 0.07 | 0.01 | 0.09 | 0.14 |
| 7 | 41094298 | rs11974873 | 1.82 | -0.246 | -0.005 | 0.095 | 0.013 | 0.035 | 0.035 | 0 | 0.66 | 0.23 | 0.51 | 0.8 | 0.69 | 0.72 |
| 7 | 41104111 | rs10085715 | -0.054 | -0.012 | 0.036 | 0.29 | 0 | 0.001 | 0.001 | 0 | 0.07 | 0.07 | 0.17 | 0.01 | 0.06 | 0.03 |
| 7 | 41110311 | rs2190345 | 1.126 | -0.023 | 0.413 | 0.343 | -0.001 | 0.005 | 0.005 | 0.004 | 0.78 | 0.46 | 0.59 | 0.8 | 0.78 | 0.85 |
| 7 | 41113263 | rs2190346 | 1.639 | -0.058 | -0.069 | 0.321 | 0 | 0.005 | 0.005 | 0 | 0.62 | 0.36 | 0.46 | 0.82 | 0.62 | 0.66 |
| 7 | 41118061 | rs17690319 | 1.213 | -0.237 | -0.114 | 0.248 | -0.002 | -0.03 | -0.03 | 0 | 0.05 | 0 | 0.03 | 0.08 | 0.05 | 0.06 |
| 7 | 41124590 | rs2108187 | -1.622 | -0.29 | 0.013 | 0.012 | -0.01 | -0.129 | -0.129 | 0 | 0.27 | 0.57 | 0.31 | 0.01 | 0.22 | 0.24 |
| 7 | 41125447 | rs2214877 | -0.896 | -0.149 | 0.454 | 0.253 | 0.003 | -0.11 | -0.11 | 0 | 0.16 | 0.54 | 0.23 | 0.01 | 0.13 | 0.08 |
| 7 | 41136770 | rs17690456 | 0.987 | -0.059 | 0.054 | 0.608 | -0.009 | -0.028 | -0.028 | 0 | 0.05 | 0 | 0.11 | 0.11 | 0.05 | 0.05 |
| 7 | 41141268 | rs12701863 | -1.004 | 0.161 | -0.147 | 0.355 | -0.002 | 0 | 0 | 0 | 0.42 | 0.57 | 0.49 | 0.42 | 0.41 | 0.27 |
| 7 | 41142944 | rs12701864 | -1.028 | 0.112 | 0.355 | 0.351 | -0.001 | 0 | 0 | 0 | 0.4 | 0.32 | 0.47 | 0.42 | 0.41 | 0.27 |
| 7 | 41149228 | rs17623158 | 0.941 | 0.314 | 0.149 | 0.301 | -0.014 | 0.052 | 0.052 | 0 | 0.09 | 0.01 | 0.08 | 0.01 | 0.09 | 0.11 |
| 7 | 41161681 | rs10256962 | 0.203 | 0.055 | 0.1 | 0.078 | -0.047 | -0.068 | -0.068 | 1.55 | 0.76 | 0.92 | 0.76 | 0.58 | 0.68 | 0.84 |
| 7 | 41162499 | rs6975061 | 1.429 | -0.182 | 0.37 | -0.022 | 0 | -0.001 | -0.001 | 5.933 | 0.55 | 0.47 | 0.63 | 0.56 | 0.57 | 0.7 |
| 7 | 41166649 | rs6958423 | 0.751 | -0.062 | 0.252 | 0.069 | -0.029 | -0.054 | -0.054 | 0 | 0.7 | 0.87 | 0.74 | 0.57 | 0.68 | 0.81 |
| 7 | 41173925 | rs6463008 | 0.763 | -0.097 | 0.218 | 0.105 | -0.043 | -0.061 | -0.061 | 0.247 | 0.7 | 0.89 | 0.74 | 0.59 | 0.67 | 0.81 |
| 7 | 41176030 | rs2108157 | 1.133 | -0.16 | 0.624 | 0.049 | -0.001 | -0.002 | -0.002 | 0 | 0.67 | 0.63 | 0.72 | 0.59 | 0.67 | 0.8 |
| 7 | 41207483 | rs29510 | 2.332 | -0.389 | 0.381 | -0.15 | -0.004 | -0.015 | -0.015 | 0.43 | 0.3 | 0.17 | 0.27 | 0.27 | 0.29 | 0.46 |
| 7 | 41228193 | rs6951284 | 1.399 | -0.317 | 0.354 | -0.258 | -0.002 | -0.032 | -0.032 | 0 | 0.1 | 0.14 | 0.09 | 0.01 | 0.09 | 0.11 |
| 7 | 41229231 | rs255013 | 1.078 | -0.361 | 0.388 | -0.023 | -0.002 | 0.026 | 0.026 | 0 | 0.58 | 0.62 | 0.69 | 0.81 | 0.59 | 0.75 |
| 7 | 41237230 | rs17772990 | 1.271 | -0.461 | 0.173 | 0.08 | -0.051 | -0.015 | -0.015 | 1.134 | 0.16 | 0.02 | 0.2 | 0.14 | 0.21 | 0.13 |
| 7 | 41241212 | rs10245608 | -1.208 | -0.52 | 0.656 | -0.132 | 0.006 | -0.001 | -0.001 | 0 | 0.42 | 0.49 | 0.36 | 0.43 | 0.47 | 0.29 |
| 7 | 41242205 | rs17716257 | 2.085 | -0.411 | 0.587 | -0.055 | -0.008 | -0.031 | -0.031 | 0.017 | 0.05 | 0 | 0.11 | 0.15 | 0.05 | 0.11 |
| 7 | 41242283 | rs13311231 | -0.7 | -0.193 | 0.909 | -0.156 | -0.023 | 0.04 | 0.04 | 0 | 0.28 | 0.13 | 0.22 | 0.18 | 0.32 | 0.14 |
| 7 | 41245268 | rs6976413 | -1.349 | -0.262 | 0.896 | -0.085 | -0.041 | -0.004 | -0.004 | 0 | 0.27 | 0.48 | 0.17 | 0.25 | 0.18 | 0.22 |
| 7 | 41249901 | rs10260526 | 1.018 | -0.191 | 0.65 | -0.183 | -0.006 | 0.066 | 0.066 | 0 | 0.11 | 0.01 | 0.08 | 0.01 | 0.1 | 0.11 |
| 7 | 41252255 | rs7781291 | -1.634 | 0.137 | 0.901 | -0.182 | 0.012 | 0 | 0 | 0.061 | 0.43 | 0.6 | 0.28 | 0.43 | 0.3 | 0.31 |
| 7 | 41258750 | rs6971930 | -0.439 | 0.003 | 1.219 | 0.11 | -0.038 | -0.065 | -0.065 | 0 | 0.11 | 0.3 | 0.06 | 0 | 0.04 | 0.03 |
| 7 | 41265555 | rs7784685 | -0.231 | 0.394 | 0.519 | 0.255 | -0.029 | -0.034 | -0.034 | 0 | 0.27 | 0.19 | 0.23 | 0.42 | 0.35 | 0.18 |
| 7 | 41286287 | rs10242681 | -1.065 | 0.264 | 0.546 | -0.087 | -0.005 | 0.134 | 0.134 | 0.136 | 0.46 | 0.75 | 0.36 | 0.2 | 0.43 | 0.26 |
| 7 | 41287859 | rs10951648 | -0.928 | 0.237 | 0.902 | -0.111 | 0.001 | -0.036 | -0.036 | 0 | 0.43 | 0.48 | 0.32 | 0.17 | 0.43 | 0.26 |
| 7 | 41292492 | rs6955172 | -0.872 | 0.238 | 1.031 | -0.099 | -0.002 | 0 | 0 | 0 | 0.43 | 0.41 | 0.33 | 0.18 | 0.43 | 0.26 |

| | | | | | | | | | | | | | | | | |
|---|---|---|---|---|---|---|---|---|---|---|---|---|---|---|---|---|
| 7 | 41294684 | rs10238779 | -1.062 | 0.231 | 0.672 | -0.102 | -0.002 | 0.027 | 0.027 | 0 | 0.44 | 0.56 | 0.33 | 0.18 | 0.44 | 0.26 |
| 7 | 41294895 | rs12668579 | 0.45 | 0.185 | 0.654 | -0.181 | -0.025 | 0.012 | 0.012 | 0 | 0.22 | 0.14 | 0.19 | 0.17 | 0.3 | 0.11 |
| 7 | 41295663 | rs6951537 | -2.681 | 0.143 | 0.268 | -0.08 | -0.057 | -0.094 | -0.094 | 0 | 0.69 | 0.94 | 0.62 | 0.63 | 0.7 | 0.56 |
| 7 | 41295903 | rs11765463 | 0.507 | 0.061 | 0.481 | -0.127 | -0.029 | 0.013 | 0.013 | 0 | 0.22 | 0.14 | 0.19 | 0.17 | 0.3 | 0.11 |
| 7 | 41295987 | rs11765437 | -0.996 | 0.142 | 0.49 | -0.055 | -0.008 | 0.104 | 0.104 | 0.044 | 0.45 | 0.69 | 0.36 | 0.21 | 0.44 | 0.27 |
| 7 | 41296236 | rs12701879 | -1.001 | 0.136 | 0.326 | -0.087 | -0.001 | -0.003 | -0.003 | 0.154 | 0.44 | 0.53 | 0.33 | 0.18 | 0.43 | 0.26 |
| 7 | 41301734 | rs12155345 | -0.43 | 0.043 | -0.646 | -0.085 | 0.004 | -0.031 | -0.031 | 0 | 0.32 | 0.42 | 0.25 | 0.16 | 0.32 | 0.15 |
| 7 | 41301819 | rs10246308 | -1.029 | -0.002 | -1.034 | -0.127 | 0 | 0.006 | 0.006 | 0 | 0.44 | 0.58 | 0.35 | 0.18 | 0.43 | 0.26 |
| 7 | 41304710 | rs11768187 | -0.345 | 0.053 | -2.464 | -0.096 | -0.014 | 0.037 | 0.037 | 0 | 0.3 | 0.2 | 0.24 | 0.16 | 0.32 | 0.15 |
| 7 | 41306172 | rs6943312 | -0.866 | 0.017 | -2.515 | 0.052 | 0.03 | -0.039 | -0.039 | 0 | 0.37 | 0.77 | 0.37 | 0.45 | 0.32 | 0.39 |
| 7 | 41308698 | rs2158559 | 0.865 | 0.303 | -2.029 | -0.649 | 0.004 | 0.03 | 0.03 | 0 | 0.11 | 0.03 | 0.08 | 0.01 | 0.1 | 0.11 |
| 7 | 41311029 | rs13226725 | -0.523 | 0.493 | -2.032 | -0.457 | -0.057 | 0.049 | 0.049 | 0 | 0.37 | 0.17 | 0.3 | 0.24 | 0.47 | 0.24 |
| 7 | 41312992 | rs718224 | -0.656 | 0.459 | -1.391 | -0.139 | -0.004 | 0.027 | 0.027 | 0 | 0.28 | 0.19 | 0.23 | 0.16 | 0.28 | 0.15 |
| 7 | 41313851 | rs17171832 | -0.732 | -0.037 | -1.33 | -0.077 | 0.008 | 0.003 | 0.003 | 0 | 0.18 | 0.16 | 0.12 | 0.03 | 0.12 | 0.16 |
| 7 | 41314465 | rs417916 | 0.369 | 0.537 | -1.76 | 0.567 | 0.005 | 0.022 | 0.022 | 0 | 0.52 | 0.17 | 0.62 | 0.58 | 0.46 | 0.71 |
| 7 | 41319241 | rs10255776 | -0.007 | 0.713 | -1.155 | 1.268 | -0.002 | 0.001 | 0.001 | 0.868 | 0.27 | 0.24 | 0.21 | 0.19 | 0.35 | 0.27 |
| 7 | 41320138 | rs377204 | -0.927 | 0.503 | -0.471 | 1.646 | 0.012 | 0.11 | 0.11 | 0 | 0.2 | 0.34 | 0.22 | 0.5 | 0.23 | 0.2 |
| 7 | 41332894 | rs37262 | 0.235 | -0.261 | -0.782 | 0.765 | 0.019 | 0.033 | 0.033 | 0.163 | 0.45 | 0.32 | 0.59 | 0.37 | 0.44 | 0.46 |
| 7 | 41337044 | rs37267 | -0.817 | -1.536 | -0.457 | 1.249 | 0.021 | -0.008 | -0.008 | 0.377 | 0.2 | 0.15 | 0.11 | 0.26 | 0.13 | 0.09 |
| 7 | 41338980 | rs37268 | -1.755 | -1.83 | -0.674 | 0.889 | 0.081 | -0.105 | -0.105 | 0 | 0.33 | 0.92 | 0.24 | 0.46 | 0.21 | 0.17 |
| 7 | 41345488 | rs37277 | -1.419 | -1.729 | -0.088 | 0.453 | -0.019 | 0 | 0 | 0 | 0.24 | 0.33 | 0.16 | 0.26 | 0.17 | 0.18 |
| 7 | 41350221 | rs11768528 | 1.282 | -1.999 | -0.969 | -0.065 | -0.167 | 0.095 | 0.095 | 0.072 | 0.34 | 0.04 | 0.35 | 0.17 | 0.52 | 0.25 |
| 7 | 41352338 | rs6946001 | 0.205 | -1.794 | 0.087 | -0.116 | 0.028 | -0.067 | -0.067 | 0 | 0.11 | 0.37 | 0.08 | 0 | 0.12 | 0.07 |
| 7 | 41360478 | rs37312 | -1.947 | -1.906 | 0.12 | -0.078 | 0.035 | 0.041 | 0.041 | 2.025 | 0.38 | 0.8 | 0.26 | 0.29 | 0.31 | 0.25 |
| 7 | 41370672 | rs17171847 | 0.292 | -1.875 | 0.114 | -0.081 | 0.03 | -0.061 | -0.061 | 0 | 0.11 | 0.42 | 0.07 | 0 | 0.11 | 0.07 |
| 7 | 41374447 | rs1020147 | -1.313 | -1.928 | -0.345 | -0.113 | -0.008 | -0.111 | -0.111 | 0 | 0.2 | 0.55 | 0.13 | 0.02 | 0.19 | 0.14 |
| 7 | 41375616 | rs28811 | -2.188 | -1.918 | -0.318 | -0.062 | 0.043 | 0.053 | 0.053 | 3.652 | 0.38 | 0.87 | 0.26 | 0.29 | 0.31 | 0.25 |
| 7 | 41375956 | rs37326 | -0.764 | -1.702 | -0.231 | 0.195 | 0.009 | -0.007 | -0.007 | 0 | 0.19 | 0.17 | 0.11 | 0.27 | 0.12 | 0.13 |
| 7 | 41382433 | rs7779436 | -0.763 | -1.567 | -0.297 | 0.266 | -0.023 | 0.016 | 0.016 | 1.66 | 0.19 | 0.31 | 0.13 | 0.27 | 0.11 | 0.11 |
| 7 | 41389199 | rs920414 | -1.017 | -1.535 | -0.266 | 0.441 | -0.024 | 0.056 | 0.056 | 0 | 0.25 | 0.52 | 0.2 | 0.33 | 0.2 | 0.16 |
| 7 | 41407459 | rs7780802 | -1.791 | -1.669 | -1.017 | 0.126 | 0.025 | -0.013 | -0.013 | 0.081 | 0.33 | 0.9 | 0.26 | 0.31 | 0.27 | 0.24 |
| 7 | 41412956 | rs10251186 | -0.311 | -1.729 | -1.44 | 0.165 | -0.004 | -0.116 | -0.116 | 0.067 | 0.61 | 0.98 | 0.56 | 0.45 | 0.69 | 0.36 |
| 7 | 41412994 | rs10280469 | -0.621 | -1.184 | -0.023 | 0.796 | 0.001 | 0.061 | 0.061 | 0 | 0.14 | 0.25 | 0.12 | 0.29 | 0.14 | 0.11 |
| 7 | 41416071 | rs7809878 | -1.031 | -1.08 | -0.39 | 0.815 | -0.007 | 0.089 | 0.089 | 1.147 | 0.19 | 0.49 | 0.17 | 0.29 | 0.15 | 0.11 |
| 7 | 41417008 | rs12701888 | 0.635 | -1.469 | -1.857 | 0.914 | 0.101 | -0.051 | -0.051 | 0 | 0.56 | 0.07 | 0.47 | 0.37 | 0.74 | 0.53 |
| 7 | 41417267 | rs10267966 | 0.741 | -0.763 | -0.943 | 1.173 | -0.01 | 0.003 | 0.003 | 2.753 | 0.57 | 0.47 | 0.72 | 0.55 | 0.69 | 0.75 |
| 7 | 41431684 | rs273188 | 1.221 | 0.311 | 0.084 | 0.966 | 0 | -0.031 | -0.031 | 0 | 0.58 | 0.4 | 0.53 | 0.09 | 0.59 | 0.46 |
| 7 | 41441549 | rs378367 | 1.061 | 0.205 | 0.63 | 1.745 | -0.001 | 0 | 0 | 0 | 0.61 | 0.62 | 0.66 | 0.33 | 0.59 | 0.48 |
| 7 | 41461200 | rs273172 | 1.087 | 0.231 | 0.741 | 2.18 | 0.019 | 0.022 | 0.022 | 0 | 0.62 | 0.48 | 0.65 | 0.53 | 0.58 | 0.57 |
| 7 | 41464183 | rs273200 | 1.408 | 0.772 | 0.868 | 1.561 | -0.006 | 0.019 | 0.019 | 0 | 0.58 | 0.61 | 0.52 | 0.09 | 0.51 | 0.49 |
| 7 | 41468649 | rs273207 | 1.311 | 0.761 | 1.491 | 2.152 | -0.003 | 0.043 | 0.043 | 0 | 0.74 | 0.46 | 0.64 | 0.28 | 0.77 | 0.66 |
| 7 | 41469325 | rs273209 | 0.767 | 0.743 | 1.624 | 2.449 | -0.002 | -0.001 | -0.001 | 0 | 0.79 | 0.77 | 0.79 | 0.54 | 0.78 | 0.67 |
| 7 | 41471164 | rs273211 | 1.169 | 0.741 | 1.714 | 2.411 | 0.011 | 0.055 | 0.055 | 0 | 0.77 | 0.55 | 0.65 | 0.28 | 0.77 | 0.66 |
| 7 | 41474901 | rs273105 | 0.982 | 0.862 | 1.475 | 2.165 | 0.002 | 0.032 | 0.032 | 0 | 0.79 | 0.74 | 0.8 | 0.5 | 0.78 | 0.68 |
| 7 | 41477327 | rs273114 | 1.65 | 0.904 | 1.393 | 2.077 | -0.001 | 0 | 0 | 0 | 0.7 | 0.68 | 0.76 | 0.5 | 0.68 | 0.65 |
| 7 | 41478897 | rs10253670 | -2.232 | 0.904 | 1.739 | 1.974 | 0.039 | 0.076 | 0.076 | 0.122 | 0.14 | 0.41 | 0.1 | 0.18 | 0.12 | 0.04 |
| 7 | 41485619 | rs273123 | 1.059 | 1 | 1.55 | 2.206 | 0.037 | 0.059 | 0.059 | 0 | 0.75 | 0.53 | 0.75 | 0.63 | 0.68 | 0.69 |
| 7 | 41491762 | rs1482548 | -1.582 | 0.541 | 1.431 | 2.161 | -0.015 | 0.187 | 0.187 | 0 | 0.19 | 0.41 | 0.24 | 0.48 | 0.14 | 0.16 |
| 7 | 41492780 | rs10243538 | -1.484 | 0.843 | 1.671 | 2.2 | 0.047 | 0.016 | 0.016 | 0 | 0.1 | 0.58 | 0.08 | 0.09 | 0.12 | 0.11 |
| 7 | 41494434 | rs6962297 | -0.701 | 0.908 | 1.297 | 2.229 | 0.074 | 0.041 | 0.041 | 0 | 0.08 | 0.33 | 0.07 | 0.09 | 0.12 | 0.11 |
| 7 | 41494840 | rs273129 | 0.179 | 0.614 | 0.361 | 2.377 | -0.01 | -0.019 | -0.019 | 0 | 0.83 | 0.88 | 0.82 | 0.71 | 0.8 | 0.8 |
| 7 | 41503620 | rs273136 | 0.398 | 0.628 | 0.389 | 2.478 | 0.021 | 0.101 | 0.101 | 0 | 0.81 | 0.7 | 0.79 | 0.53 | 0.78 | 0.79 |
| 7 | 41505256 | rs17774910 | -1.203 | 0.698 | 0.331 | 2.438 | 0.007 | -0.032 | -0.032 | 0 | 0.15 | 0.07 | 0.21 | 0.3 | 0.12 | 0.15 |
| 7 | 41509518 | rs12701914 | -0.003 | 1.512 | -0.853 | 1.013 | -0.108 | -0.034 | -0.034 | 0 | 0.18 | 0.02 | 0.16 | 0.16 | 0.33 | 0.14 |
| 7 | 41511739 | rs273148 | 0.286 | 0.66 | -0.799 | 0.965 | -0.007 | -0.023 | -0.023 | 0 | 0.67 | 0.78 | 0.68 | 0.58 | 0.65 | 0.76 |
| 7 | 41516677 | rs273156 | 0.509 | -0.689 | -0.179 | 0.615 | 0.001 | 0.002 | 0.002 | 0 | 0.46 | 0.56 | 0.35 | 0.41 | 0.27 | 0.57 |
| 7 | 41516733 | rs1482556 | -0.197 | -1.246 | -0.052 | 0.608 | -0.011 | 0.002 | 0.002 | 0 | 0.08 | 0.11 | 0.07 | 0.11 | 0.02 | 0.06 |
| 7 | 41517598 | rs273157 | 1.174 | -1.307 | 0.272 | 0.46 | 0 | 0 | 0 | 0 | 0.34 | 0.27 | 0.28 | 0.28 | 0.26 | 0.51 |
| 7 | 41519437 | rs568520 | 1.183 | -1.311 | 0.27 | 0.458 | 0 | 0 | 0 | 0 | 0.33 | 0.27 | 0.29 | 0.28 | 0.26 | 0.51 |
| 7 | 41519812 | rs1482553 | -0.124 | -1.395 | 0.605 | 0.412 | 0.077 | 0.03 | 0.03 | 0 | 0.52 | 0.32 | 0.62 | 0.59 | 0.71 | 0.44 |
| 7 | 41524795 | rs4724050 | 0.089 | -0.621 | 0.562 | 0.392 | -0.051 | -0.046 | -0.046 | 3.09 | 0.33 | 0.19 | 0.48 | 0.42 | 0.52 | 0.33 |
| 7 | 41531573 | rs569972 | 0.328 | -0.49 | 1.063 | 0.516 | -0.013 | 0.004 | 0.004 | 0 | 0.93 | 0.91 | 0.94 | 0.88 | 0.98 | 0.94 |
| 7 | 41537374 | rs487838 | 0.394 | -0.534 | 1.162 | 0.496 | -0.019 | 0.007 | 0.007 | 0 | 0.92 | 0.89 | 0.94 | 0.88 | 0.98 | 0.94 |
| 7 | 41542150 | rs580514 | 1.693 | -0.521 | 1.052 | 0.516 | -0.081 | -0.002 | -0.002 | 0 | 0.85 | 0.53 | 0.91 | 0.88 | 0.97 | 0.93 |
| 7 | 41572437 | rs510073 | 0.451 | -0.262 | 1.098 | 0.52 | -0.009 | 0.005 | 0.005 | 0 | 0.93 | 0.91 | 0.94 | 0.91 | 0.98 | 0.94 |
| 7 | 41577967 | rs2035425 | -0.07 | -0.729 | 0.926 | 0.168 | 0.083 | 0.023 | 0.023 | 0 | 0.49 | 0.29 | 0.53 | 0.57 | 0.65 | 0.44 |
| 7 | 41581421 | rs17704722 | 0.733 | -0.496 | 0.58 | -0.071 | 0.013 | 0.066 | 0.066 | 0 | 0.3 | 0.12 | 0.23 | 0.13 | 0.3 | 0.19 |
| 7 | 41582896 | rs519654 | 0.313 | -0.482 | 0.644 | -0.001 | -0.012 | 0.007 | 0.007 | 0 | 0.93 | 0.9 | 0.93 | 0.89 | 0.98 | 0.93 |
| 7 | 41606407 | rs473147 | 1.572 | -0.791 | 0.792 | -0.156 | -0.061 | -0.013 | -0.013 | 0 | 0.86 | 0.65 | 0.9 | 0.89 | 0.96 | 0.92 |

| | | | | | | | | | | | | | | | | |
|---|---|---|---|---|---|---|---|---|---|---|---|---|---|---|---|---|
| 7 | 41612820 | rs4724053 | 0.305 | -0.648 | 0.586 | -0.061 | 0.047 | 0.057 | 0.057 | 0 | 0.21 | 0.31 | 0.33 | 0.46 | 0.37 | 0.29 |
| 7 | 41615988 | rs516810 | 1.505 | -0.678 | 0.003 | -0.117 | -0.024 | -0.039 | -0.039 | 0.492 | 0.34 | 0.04 | 0.33 | 0.3 | 0.29 | 0.47 |
| 7 | 41617294 | rs549847 | 1.141 | -0.631 | 0.098 | -0.077 | -0.053 | -0.044 | -0.044 | 0.002 | 0.63 | 0.16 | 0.55 | 0.43 | 0.59 | 0.63 |
| 7 | 41645139 | rs10256711 | -0.023 | -0.864 | 0.011 | 0.142 | -0.012 | 0.022 | 0.022 | 0 | 0.93 | 0.9 | 0.94 | 0.82 | 0.98 | 0.93 |
| 7 | 41649281 | rs1115129 | 1.11 | -0.899 | -0.219 | 0.113 | -0.023 | -0.003 | -0.003 | 0 | 0.34 | 0.04 | 0.33 | 0.23 | 0.3 | 0.46 |
| 7 | 41659417 | rs10486718 | 1.013 | -0.807 | 0.581 | 0.445 | -0.058 | 0.052 | 0.052 | 0 | 0.88 | 0.69 | 0.91 | 0.81 | 0.98 | 0.92 |
| 7 | 41662627 | rs11971117 | 0.463 | 0.041 | 0.103 | 0.55 | 0.001 | 0.028 | 0.028 | 0.002 | 0.15 | 0.2 | 0.28 | 0.45 | 0.21 | 0.24 |
| 7 | 41663805 | rs4724061 | 1.017 | -0.584 | 0.16 | 0.21 | -0.071 | 0.013 | 0.013 | 0.281 | 0.83 | 0.42 | 0.84 | 0.83 | 0.93 | 0.9 |
| 7 | 41665687 | rs2218068 | -1.7 | -0.553 | 0.389 | 0.158 | -0.025 | -0.023 | -0.023 | 0.382 | 0.1 | 0.42 | 0.05 | 0.02 | 0.01 | 0.01 |
| 7 | 41671911 | rs4724063 | 1.492 | -0.293 | 0.17 | 0.32 | -0.049 | 0.028 | 0.028 | 0 | 0.8 | 0.4 | 0.71 | 0.78 | 0.88 | 0.88 |
| 7 | 41679114 | rs17719161 | -0.349 | -0.306 | 0.158 | 0.33 | -0.038 | 0.051 | 0.051 | 0 | 0.1 | 0.19 | 0.18 | 0.21 | 0.04 | 0.09 |
| 7 | 41681228 | rs6463035 | -1.104 | 0.049 | -1.235 | 0.073 | -0.179 | 0.013 | 0.013 | 0.132 | 0.64 | 0.98 | 0.69 | 0.78 | 0.62 | 0.53 |
| 7 | 41683647 | rs7455525 | -0.966 | 0.094 | -0.753 | 0.252 | -0.048 | 0.031 | 0.031 | 0.131 | 0.19 | 0.64 | 0.3 | 0.22 | 0.13 | 0.12 |
| 7 | 41691876 | rs6961184 | -0.909 | 0.163 | -1.064 | 0.224 | -0.032 | 0.01 | 0.01 | 0.204 | 0.2 | 0.67 | 0.3 | 0.22 | 0.13 | 0.12 |
| 7 | 41696061 | rs2877084 | 0.965 | 1.375 | -1.493 | 0.195 | -0.027 | 0.028 | 0.028 | 0 | 0.05 | 0.01 | 0.04 | 0 | 0.15 | 0.04 |
| 7 | 41700245 | rs4724065 | 0.199 | 0.664 | -1.659 | 0.496 | -0.036 | -0.13 | -0.13 | 0 | 0.17 | 0.02 | 0.19 | 0.44 | 0.21 | 0.2 |
| 7 | 41706132 | rs1476665 | -0.26 | 0.299 | -1.294 | 0.379 | -0.005 | 0.006 | 0.006 | 0.128 | 0.64 | 0.42 | 0.5 | 0.59 | 0.7 | 0.63 |
| 7 | 41718754 | rs17705333 | 0.565 | 0.326 | -1.95 | 0.296 | -0.019 | 0.036 | 0.036 | 0 | 0.25 | 0.05 | 0.18 | 0.15 | 0.25 | 0.2 |
| 7 | 41743294 | rs2877098 | -0.202 | 0.247 | -0.632 | 0.968 | -0.001 | -0.02 | -0.02 | 0.066 | 0.69 | 0.73 | 0.57 | 0.59 | 0.69 | 0.69 |
| 7 | 41746557 | rs2007475 | 0.663 | -0.028 | -0.007 | 0.966 | 0.003 | 0 | 0 | 0.194 | 0.5 | 0.48 | 0.37 | 0.46 | 0.56 | 0.51 |
| 7 | 41747403 | rs6977571 | 0.745 | -0.086 | 0.389 | 0.94 | 0.028 | -0.003 | -0.003 | 0 | 0.49 | 0.3 | 0.35 | 0.46 | 0.56 | 0.51 |
| 7 | 41750087 | rs1003291 | -1.296 | -0.13 | 0.133 | 1.102 | -0.062 | -0.062 | -0.062 | 0 | 0.82 | 0.96 | 0.62 | 0.74 | 0.8 | 0.72 |
| 7 | 41757878 | rs17776182 | 0.418 | 0.106 | 0.04 | 1.035 | -0.049 | -0.1 | -0.1 | 0 | 0.14 | 0.02 | 0.13 | 0.44 | 0.16 | 0.27 |
| 7 | 41762747 | rs2108174 | 0.72 | -0.188 | 0.41 | 0.984 | 0.009 | -0.004 | -0.004 | 0 | 0.55 | 0.63 | 0.38 | 0.5 | 0.57 | 0.53 |
| 7 | 41764992 | rs995124 | -0.247 | -0.246 | 0.057 | 0.945 | 0.016 | -0.025 | -0.025 | 2.016 | 0.24 | 0.05 | 0.21 | 0.24 | 0.23 | 0.19 |
| 7 | 41775475 | rs3886608 | 1.059 | -0.203 | 0.224 | 0.882 | 0.033 | 0.001 | 0.001 | 0.367 | 0.5 | 0.31 | 0.36 | 0.47 | 0.56 | 0.52 |
| 7 | 41781911 | rs12537065 | -0.374 | -0.101 | 0.845 | 0.887 | -0.002 | 0.005 | 0.005 | 0 | 0.77 | 0.59 | 0.58 | 0.73 | 0.8 | 0.71 |
| 7 | 41783359 | rs4724070 | -0.773 | -0.109 | 0.88 | 0.884 | -0.007 | -0.012 | -0.012 | 0.166 | 0.79 | 0.86 | 0.61 | 0.73 | 0.8 | 0.71 |
| 7 | 41793043 | rs2214875 | 0.696 | -0.175 | 0.908 | 0.865 | -0.001 | 0.002 | 0.002 | 0 | 0.15 | 0.16 | 0.17 | 0.19 | 0.17 | 0.14 |
| 7 | 41800176 | rs1990133 | 0.007 | -0.21 | 0.939 | 0.866 | -0.015 | 0.002 | 0.002 | 0 | 0.73 | 0.57 | 0.57 | 0.71 | 0.79 | 0.7 |
| 7 | 41806349 | rs4720405 | -0.397 | -0.206 | 0.892 | 0.867 | 0.001 | -0.005 | -0.005 | 0 | 0.76 | 0.76 | 0.6 | 0.71 | 0.79 | 0.71 |
| 7 | 41806538 | rs4720406 | 1.014 | -0.171 | 0.689 | 0.88 | 0.009 | -0.005 | -0.005 | 1.33 | 0.58 | 0.33 | 0.4 | 0.52 | 0.62 | 0.57 |
| 7 | 41810755 | rs10951661 | -0.131 | -0.154 | 0.703 | 0.933 | -0.011 | 0.006 | 0.006 | 0 | 0.74 | 0.55 | 0.58 | 0.72 | 0.79 | 0.71 |
| 7 | 41811908 | rs10486722 | 0.395 | -0.084 | 0.566 | 0.951 | -0.015 | 0.021 | 0.021 | 0 | 0.64 | 0.48 | 0.44 | 0.48 | 0.67 | 0.57 |
| 7 | 41812483 | rs10216044 | -0.994 | 0.026 | -0.109 | 0.916 | 0.028 | 0.04 | 0.04 | 0 | 0.44 | 0.75 | 0.35 | 0.32 | 0.39 | 0.49 |
| 7 | 41821917 | rs12701937 | 1.665 | 0.011 | -0.339 | 1.052 | -0.132 | -0.055 | -0.055 | 0.777 | 0.36 | 0.03 | 0.25 | 0.31 | 0.42 | 0.28 |
| 7 | 41822335 | rs2190339 | 0.64 | -0.236 | 0.359 | 1.046 | -0.002 | 0 | 0 | 0 | 0.56 | 0.44 | 0.39 | 0.52 | 0.54 | 0.41 |
| 7 | 41823010 | rs17776768 | 0.441 | -0.271 | 0.173 | 1.009 | 0.03 | -0.047 | -0.047 | 0 | 0.16 | 0.04 | 0.11 | 0.21 | 0.12 | 0.13 |
| 7 | 41825744 | rs7805201 | 0.651 | -0.171 | 0.356 | 0.881 | -0.001 | 0 | 0 | 0 | 0.09 | 0.07 | 0.08 | 0.06 | 0.11 | 0.06 |
| 7 | 41826934 | rs11773475 | 0.81 | -0.13 | 0.278 | 1.01 | -0.004 | 0.006 | 0.006 | 0 | 0.52 | 0.38 | 0.38 | 0.49 | 0.53 | 0.4 |
| 7 | 41828527 | rs10234444 | 0.844 | -0.054 | -0.168 | 0.994 | -0.002 | -0.006 | -0.006 | 0 | 0.17 | 0.15 | 0.32 | 0.37 | 0.17 | 0.24 |
| 7 | 41830184 | rs257114 | 0.545 | -0.425 | 0.248 | 0.421 | -0.001 | -0.025 | -0.025 | 0 | 0.17 | 0.26 | 0.21 | 0.05 | 0.19 | 0.29 |
| 7 | 41839311 | rs29694 | 0.159 | -0.2 | -0.762 | 0.707 | -0.003 | -0.053 | -0.053 | 0 | 0.32 | 0.61 | 0.32 | 0.14 | 0.31 | 0.36 |
| 7 | 41846628 | rs42513 | -0.155 | 0.408 | -1.284 | 0.923 | 0.036 | 0.051 | 0.051 | 0.137 | 0.46 | 0.79 | 0.44 | 0.33 | 0.42 | 0.46 |
| 7 | 41850294 | rs29688 | -0.112 | 0.394 | -1.308 | 0.999 | 0.034 | 0.004 | 0.004 | 0.035 | 0.46 | 0.77 | 0.47 | 0.43 | 0.43 | 0.48 |
| 7 | 41851037 | rs10226747 | 0.179 | 0.586 | -0.866 | 1.004 | -0.023 | -0.046 | -0.046 | 0 | 0.11 | 0 | 0.13 | 0.14 | 0.09 | 0.08 |
| 7 | 41854681 | rs6947337 | -0.598 | 0.808 | -0.514 | 0.763 | -0.021 | -0.001 | -0.001 | 0 | 0.36 | 0.23 | 0.51 | 0.64 | 0.43 | 0.4 |
| 7 | 41862682 | rs1544471 | 0.589 | 0.058 | -0.496 | 0.187 | -0.026 | -0.03 | -0.03 | 0 | 0.67 | 0.57 | 0.72 | 0.85 | 0.76 | 0.82 |
| 7 | 41866396 | rs11763434 | 1.65 | 0.061 | -0.453 | -0.056 | -0.039 | 0.121 | 0.121 | 0 | 0.35 | 0.18 | 0.23 | 0.23 | 0.42 | 0.5 |
| 7 | 41866769 | rs4720408 | -0.302 | -0.259 | -0.387 | -0.136 | 0.036 | 0.033 | 0.033 | 0.003 | 0.22 | 0.09 | 0.17 | 0.07 | 0.13 | 0.14 |
| 7 | 41871611 | rs6943781 | 0.843 | -0.262 | 0.413 | -0.201 | 0.001 | 0.013 | 0.013 | 0 | 0.49 | 0.62 | 0.34 | 0.31 | 0.52 | 0.55 |
| 7 | 41876783 | rs7803239 | -0.409 | 0.204 | 0.225 | 0.082 | -0.027 | -0.03 | -0.03 | 0 | 0.72 | 0.87 | 0.53 | 0.39 | 0.66 | 0.69 |
| 7 | 41888455 | rs6463053 | -0.309 | 0.056 | 0.41 | -0.082 | 0.031 | 0.012 | 0.012 | 0.068 | 0.12 | 0.44 | 0.11 | 0.09 | 0.09 | 0.05 |
| 7 | 41891246 | rs2024031 | 0.848 | -0.169 | 0.39 | -0.217 | 0.001 | -0.019 | -0.019 | 0 | 0.56 | 0.58 | 0.34 | 0.32 | 0.53 | 0.58 |
| 7 | 41891383 | rs4724078 | 0.627 | -0.243 | 0.262 | -0.302 | -0.006 | 0.027 | 0.027 | 0 | 0.07 | 0.01 | 0.1 | 0.01 | 0.08 | 0.05 |
| 7 | 41899191 | rs718939 | 1 | -0.204 | 0.431 | -0.26 | 0.002 | -0.027 | -0.027 | 0.11 | 0.56 | 0.56 | 0.34 | 0.32 | 0.53 | 0.58 |
| 7 | 41907182 | rs988329 | -0.428 | -0.223 | 0.437 | -0.234 | -0.021 | -0.028 | -0.028 | 0 | 0.18 | 0.27 | 0.19 | 0.06 | 0.13 | 0.12 |
| 7 | 41911549 | rs13232007 | 1.514 | -0.228 | -0.121 | -0.247 | -0.034 | 0.213 | 0.213 | 1.442 | 0.44 | 0.14 | 0.24 | 0.24 | 0.45 | 0.53 |
| 7 | 41916923 | rs17706596 | 1.718 | -0.287 | -0.279 | -0.275 | -0.094 | 0.201 | 0.201 | 1.815 | 0.42 | 0.05 | 0.22 | 0.23 | 0.44 | 0.53 |
| 7 | 41922310 | rs6954585 | -0.405 | -0.022 | 0.019 | -0.063 | -0.008 | -0.005 | -0.005 | 0 | 0.73 | 0.8 | 0.52 | 0.39 | 0.66 | 0.7 |
| 7 | 41926152 | rs17171966 | -1.221 | -0.098 | 0.338 | 0.095 | -0.019 | -0.021 | -0.021 | 0 | 0.23 | 0.45 | 0.25 | 0.15 | 0.18 | 0.17 |
| 7 | 41926703 | rs17706679 | -0.35 | 0.052 | -0.305 | 0.25 | 0.012 | 0.014 | 0.014 | 0 | 0.16 | 0.09 | 0.18 | 0.07 | 0.13 | 0.13 |
| 7 | 41929342 | rs17639438 | 0.001 | 0.795 | -0.355 | 0.603 | -0.011 | -0.012 | -0.012 | 0 | 0.21 | 0.15 | 0.43 | 0.62 | 0.29 | 0.31 |
| 7 | 41946930 | rs10246037 | -0.96 | -0.256 | -0.831 | 0.266 | 0.014 | 0.022 | 0.022 | 6.056 | 0.33 | 0.74 | 0.37 | 0.2 | 0.24 | 0.12 |
| 7 | 41951210 | rs4724081 | 1.504 | 0.135 | -1.989 | 0.457 | -0.05 | -0.005 | -0.005 | 0 | 0.36 | 0.05 | 0.22 | 0.35 | 0.38 | 0.49 |
| 7 | 41969521 | rs6951077 | 0.566 | 0.529 | -1.813 | 0.506 | -0.031 | 0.01 | 0.01 | 1.577 | 0.14 | 0.02 | 0.19 | 0.06 | 0.19 | 0.1 |
| 7 | 41971266 | rs7811368 | 0.38 | 0.38 | -1.007 | 0.894 | 0.023 | 0.064 | 0.064 | 0.597 | 0.12 | 0.43 | 0.12 | 0.15 | 0.07 | 0.03 |
| 7 | 41973509 | rs6952496 | 0.564 | 0.68 | -1.392 | 1.018 | -0.047 | -0.094 | -0.094 | 0.571 | 0.69 | 0.88 | 0.57 | 0.55 | 0.64 | 0.63 |
| 7 | 41973787 | rs7805291 | 0.935 | 0.688 | -1.478 | 1.035 | -0.001 | 0 | 0 | 0.105 | 0.63 | 0.65 | 0.54 | 0.55 | 0.62 | 0.62 |

| | | | | | | | | | | | | | | | | |
|---|---|---|---|---|---|---|---|---|---|---|---|---|---|---|---|---|
| 7 | 41975256 | rs10251717 | -0.211 | 0.946 | -1.545 | 1.2 | -0.051 | -0.043 | -0.043 | 0 | 0.81 | 0.96 | 0.78 | 0.84 | 0.77 | 0.8 |
| 7 | 41975571 | rs2286234 | 0.353 | 0.774 | -1.343 | 0.958 | 0 | 0 | 0 | 0.003 | 0.64 | 0.65 | 0.54 | 0.63 | 0.63 | 0.64 |
| 7 | 41983613 | rs10234024 | 0.177 | 0.952 | -2 | 0.535 | -0.139 | -0.119 | -0.119 | 0.332 | 0.3 | 0.02 | 0.36 | 0.37 | 0.36 | 0.35 |
| 7 | 41988488 | rs6463079 | -0.601 | 0.595 | -0.627 | 0.477 | -0.011 | -0.013 | -0.013 | 0 | 0.06 | 0.64 | 0.03 | 0 | 0 | 0.01 |
| 7 | 41989362 | rs11763154 | -0.092 | 0.459 | -1.149 | 0.439 | -0.088 | -0.138 | -0.138 | 0.177 | 0.25 | 0.04 | 0.21 | 0.44 | 0.31 | 0.3 |
| 7 | 41993594 | rs6965030 | -0.892 | 0.201 | -0.996 | 0.16 | -0.034 | -0.061 | -0.061 | 0 | 0.15 | 0.65 | 0.14 | 0.01 | 0.07 | 0.03 |
| 7 | 42000101 | rs7805604 | -0.773 | 0.78 | -1.548 | 0.4 | -0.036 | -0.064 | -0.064 | 0.203 | 0.34 | 0.15 | 0.33 | 0.46 | 0.38 | 0.33 |
| 7 | 42000924 | rs3735361 | -0.433 | 0.946 | -1.628 | 0.782 | -0.108 | -0.124 | -0.124 | 0.9 | 0.26 | 0.05 | 0.35 | 0.56 | 0.33 | 0.35 |
| 7 | 42001171 | rs3823720 | -0.561 | 0.9 | -1.576 | 0.496 | -0.089 | -0.137 | -0.137 | 1.863 | 0.25 | 0.05 | 0.28 | 0.41 | 0.33 | 0.34 |
| 7 | 42003500 | rs2051935 | -0.379 | 0.533 | -0.971 | -0.69 | 0.022 | 0.111 | 0.111 | 0.387 | 0.41 | 0.22 | 0.46 | 0.76 | 0.35 | 0.54 |
| 7 | 42015020 | rs7794242 | -0.96 | 0.247 | -0.679 | -0.588 | 0 | -0.029 | -0.029 | 1.53 | 0.08 | 0.13 | 0.05 | 0.01 | 0.07 | 0.07 |
| 7 | 42017972 | rs4724086 | -0.83 | 0.243 | -0.6 | -0.545 | 0.028 | 0.087 | 0.087 | 1.861 | 0.48 | 0.67 | 0.51 | 0.78 | 0.38 | 0.59 |
| 7 | 42020130 | rs4364531 | -0.415 | 0.398 | -1.298 | -0.411 | 0.017 | 0.138 | 0.138 | 0.365 | 0.41 | 0.18 | 0.46 | 0.77 | 0.36 | 0.55 |
| 7 | 42030686 | rs9648515 | -0.556 | 0.517 | -0.965 | -0.338 | 0.005 | 0.012 | 0.012 | 1.462 | 0.46 | 0.58 | 0.5 | 0.78 | 0.39 | 0.57 |
| 7 | 42031107 | rs9648517 | -0.406 | 0.556 | -1.606 | 0.862 | -0.049 | -0.1 | -0.1 | 0 | 0.14 | 0.01 | 0.11 | 0.39 | 0.18 | 0.22 |
| 7 | 42031224 | rs2237429 | -0.429 | 0.39 | -1.473 | 0.637 | 0.018 | -0.044 | -0.044 | 0 | 0.25 | 0.18 | 0.32 | 0.38 | 0.19 | 0.36 |
| 7 | 42032565 | rs2237427 | 0.698 | -0.165 | -1.89 | 0.058 | 0.043 | -0.04 | -0.04 | 0 | 0.62 | 0.8 | 0.69 | 0.58 | 0.71 | 0.52 |
| 7 | 42033195 | rs2299146 | 0.723 | -0.139 | -1.95 | 0.019 | 0.041 | -0.06 | -0.06 | 0 | 0.62 | 0.87 | 0.7 | 0.58 | 0.69 | 0.52 |
| 7 | 42034728 | rs10486731 | 1.181 | -0.318 | -1.988 | -0.156 | 0.014 | 0.058 | 0.058 | 3.933 | 0.11 | 0.02 | 0.05 | 0 | 0.07 | 0.02 |
| 7 | 42034831 | rs6969239 | -0.362 | 0.362 | -2.863 | -0.731 | -0.077 | -0.087 | -0.087 | 0 | 0.46 | 0.91 | 0.38 | 0.52 | 0.54 | 0.43 |
| 7 | 42035623 | rs11772482 | 0.177 | 0.382 | -2.697 | -0.69 | -0.04 | -0.068 | -0.068 | 0.085 | 0.38 | 0.82 | 0.32 | 0.52 | 0.42 | 0.33 |
| 7 | 42036345 | rs846272 | 0.28 | 0.305 | -2.67 | -0.869 | -0.017 | -0.046 | -0.046 | 0.302 | 0.36 | 0.8 | 0.28 | 0.51 | 0.34 | 0.32 |
| 7 | 42055725 | rs1029589 | 0.287 | 1.505 | -2.077 | -0.434 | 0.014 | 0.051 | 0.051 | 0 | 0.12 | 0.11 | 0.12 | 0.4 | 0.2 | 0.17 |
| 7 | 42058291 | rs7806689 | -1.622 | 1.308 | -2.155 | -0.618 | 0.005 | 0.034 | 0.034 | 0.137 | 0.94 | 0.98 | 0.98 | 0.99 | 0.97 | 0.93 |
| 7 | 42061502 | rs846277 | -0.346 | 1.308 | -2.076 | -0.642 | -0.001 | -0.015 | -0.015 | 0 | 0.9 | 0.92 | 0.94 | 0.99 | 0.87 | 0.92 |
| 7 | 42061523 | rs2299142 | 0.948 | 1.217 | -2.325 | -0.744 | -0.024 | -0.075 | -0.075 | 0.05 | 0.48 | 0.05 | 0.61 | 0.45 | 0.42 | 0.51 |
| 7 | 42066881 | rs846288 | -0.543 | 1.608 | -1.705 | -0.402 | -0.018 | -0.017 | -0.017 | 1.866 | 0.38 | 0.74 | 0.31 | 0.55 | 0.43 | 0.39 |
| 7 | 42071698 | rs846335 | -0.902 | 1.591 | -1.911 | -0.406 | -0.057 | -0.053 | -0.053 | 0 | 0.43 | 0.9 | 0.37 | 0.56 | 0.55 | 0.44 |
| 7 | 42073962 | rs846329 | 0.922 | 1.448 | -1.079 | -0.221 | 0.001 | -0.019 | -0.019 | 0 | 0.21 | 0.37 | 0.17 | 0.16 | 0.23 | 0.21 |
| 7 | 42076893 | rs846327 | -0.523 | 1.405 | -1.535 | -0.341 | -0.073 | -0.033 | -0.033 | 0.249 | 0.41 | 0.86 | 0.34 | 0.57 | 0.46 | 0.42 |
| 7 | 42088222 | rs846266 | -0.849 | 1.155 | -1.574 | -0.297 | -0.017 | -0.033 | -0.033 | 0 | 0.44 | 0.89 | 0.38 | 0.57 | 0.58 | 0.43 |
| 7 | 42090528 | rs846264 | -0.85 | 1.145 | -1.605 | -0.298 | -0.027 | -0.035 | -0.035 | 0 | 0.44 | 0.9 | 0.38 | 0.57 | 0.57 | 0.43 |
| 7 | 42095836 | rs2282920 | -0.86 | 1.318 | -1.197 | 0.025 | -0.016 | -0.036 | -0.036 | 0 | 0.43 | 0.78 | 0.35 | 0.49 | 0.58 | 0.44 |
| 7 | 42098880 | rs917230 | -0.533 | 1.195 | -0.636 | -0.342 | 0.005 | 0.003 | 0.003 | 0.001 | 0.4 | 0.54 | 0.29 | 0.53 | 0.47 | 0.52 |
| 7 | 42098976 | rs917229 | -1.385 | 1.081 | -0.613 | -0.302 | 0.04 | -0.006 | -0.006 | 0 | 0.5 | 0.81 | 0.37 | 0.53 | 0.6 | 0.54 |
| 7 | 42101820 | rs1527499 | -1.385 | 1.06 | -0.594 | -0.346 | 0.04 | 0 | 0 | 0 | 0.5 | 0.81 | 0.37 | 0.53 | 0.6 | 0.54 |
| 7 | 42101972 | rs2237419 | 0.527 | 1.227 | 0.188 | -0.244 | 0.004 | -0.023 | -0.023 | 0 | 0.25 | 0.26 | 0.17 | 0.14 | 0.25 | 0.31 |
| 7 | 42103319 | rs2282919 | -0.305 | 1.027 | 0.314 | -0.397 | -0.004 | -0.01 | -0.01 | 0 | 0.34 | 0.26 | 0.23 | 0.51 | 0.38 | 0.43 |
| 7 | 42106599 | rs6966658 | -0.642 | 1.351 | 0.492 | -0.377 | 0.019 | -0.028 | -0.028 | 0.009 | 0.06 | 0.13 | 0.06 | 0.01 | 0.08 | 0.08 |
| 7 | 42108145 | rs6959829 | -1.911 | 1.366 | 0.539 | -0.411 | 0.046 | -0.061 | -0.061 | 1.211 | 0.11 | 0.34 | 0.08 | 0.01 | 0.11 | 0.1 |
| 7 | 42112533 | rs10282353 | -1.317 | 1.106 | 0.275 | -0.416 | -0.011 | 0.006 | 0.006 | 0.435 | 0.48 | 0.82 | 0.37 | 0.54 | 0.52 | 0.55 |
| 7 | 42113146 | rs17172002 | -0.796 | 1.146 | 0.326 | -0.378 | 0.064 | -0.017 | -0.017 | 0 | 0.07 | 0.32 | 0.09 | 0 | 0.08 | 0.08 |
| 7 | 42115888 | rs10951666 | 0.202 | 1.117 | 0.201 | -0.441 | -0.133 | -0.168 | -0.168 | 2.914 | 0.28 | 0.02 | 0.21 | 0.54 | 0.39 | 0.37 |
| 7 | 42118825 | rs4724094 | 0.764 | 1.125 | 0.277 | -0.488 | 0.053 | -0.047 | -0.047 | 0 | 0.9 | 0.67 | 0.89 | 0.99 | 0.88 | 0.91 |
| 7 | 42121553 | rs865860 | 1.083 | 0.904 | 0.045 | -0.393 | -0.018 | 0.017 | 0.017 | 1.301 | 0.52 | 0.18 | 0.63 | 0.45 | 0.5 | 0.44 |
| 7 | 42122193 | rs10256288 | -0.401 | 1.085 | 0.101 | -0.57 | 0.011 | -0.094 | -0.094 | 0.094 | 0.2 | 0.39 | 0.35 | 0.09 | 0.18 | 0.17 |
| 7 | 42123955 | rs846293 | -0.295 | 1.042 | -0.624 | -0.434 | -0.054 | -0.028 | -0.028 | 0 | 0.66 | 0.78 | 0.74 | 0.41 | 0.59 | 0.59 |
| 7 | 42125900 | rs12539142 | 1.434 | 0.663 | -0.539 | -0.45 | 0.053 | 0.008 | 0.008 | 0.977 | 0.39 | 0.21 | 0.32 | 0.33 | 0.33 | 0.35 |
| 7 | 42126604 | rs3779144 | -0.227 | 1.022 | -0.555 | -0.668 | -0.016 | -0.008 | -0.008 | 0 | 0.67 | 0.76 | 0.78 | 0.38 | 0.62 | 0.59 |
| 7 | 42128896 | rs846312 | -1.013 | 0.965 | 0.023 | -0.893 | 0.014 | -0.08 | -0.08 | 0 | 0.23 | 0.36 | 0.43 | 0.08 | 0.21 | 0.17 |
| 7 | 42131390 | rs17640804 | -0.909 | 0.998 | -0.084 | -0.88 | -0.009 | 0.013 | 0.013 | 0 | 0.23 | 0.21 | 0.43 | 0.08 | 0.22 | 0.18 |
| 7 | 42151599 | rs6463088 | 0.875 | 0.492 | 0.256 | -0.656 | -0.003 | -0.005 | -0.005 | 0 | 0.47 | 0.5 | 0.37 | 0.27 | 0.41 | 0.4 |
| 7 | 42151721 | rs10951667 | -0.314 | 0.471 | 0.274 | -0.693 | 0.041 | 0.097 | 0.097 | 3.496 | 0.45 | 0.16 | 0.59 | 0.73 | 0.56 | 0.58 |
| 7 | 42152794 | rs10951669 | -0.484 | 0.57 | 0.298 | -0.648 | 0.054 | 0.104 | 0.104 | 2.827 | 0.47 | 0.21 | 0.6 | 0.72 | 0.56 | 0.58 |
| 7 | 42152856 | rs6463089 | -0.469 | 0.804 | 1.399 | -0.606 | 0.015 | -0.04 | -0.04 | 0 | 0.07 | 0.18 | 0.08 | 0.01 | 0.09 | 0.07 |
| 7 | 42156363 | rs3801187 | -2.139 | 0.825 | 1.073 | -0.6 | 0.027 | -0.1 | -0.1 | 0 | 0.14 | 0.52 | 0.13 | 0.01 | 0.13 | 0.09 |
| 7 | 42157147 | rs17172014 | -0.751 | 0.772 | 0.888 | -0.459 | 0.043 | -0.001 | -0.001 | 0 | 0.07 | 0.31 | 0.05 | 0 | 0.04 | 0.02 |
| 7 | 42160492 | rs846399 | -1.43 | 1.01 | 0.531 | -0.659 | 0.022 | -0.073 | -0.073 | 1.425 | 0.29 | 0.44 | 0.5 | 0.1 | 0.33 | 0.19 |
| 7 | 42160703 | rs846398 | -0.322 | 0.864 | 0.286 | -0.49 | 0.019 | 0.013 | 0.013 | 0 | 0.67 | 0.59 | 0.71 | 0.36 | 0.65 | 0.65 |
| 7 | 42160959 | rs10242174 | 1.54 | 0.68 | -0.189 | -0.5 | 0.084 | 0.026 | 0.026 | 0.59 | 0.38 | 0.12 | 0.2 | 0.28 | 0.31 | 0.43 |
| 7 | 42161527 | rs3801192 | -0.354 | 1.356 | 0.3 | -0.827 | 0.013 | -0.035 | -0.035 | 0 | 0.07 | 0.11 | 0.08 | 0.01 | 0.09 | 0.06 |
| 7 | 42164049 | rs3801198 | -0.598 | 1.045 | -0.115 | -0.632 | 0.007 | 0 | 0 | 0.386 | 0.05 | 0.03 | 0.02 | 0.05 | 0.03 | 0.07 |
| 7 | 42171041 | rs846381 | -1.107 | 1.253 | -0.004 | -0.745 | -0.004 | 0.005 | 0.005 | 0 | 0.22 | 0.22 | 0.44 | 0.07 | 0.26 | 0.18 |
| 7 | 42174596 | rs6974655 | 0.101 | 1.449 | -0.23 | -0.256 | 0.002 | 0.003 | 0.003 | 0 | 0.77 | 0.72 | 0.83 | 0.76 | 0.74 | 0.72 |
| 7 | 42177469 | rs12668912 | 1.31 | 1.595 | -0.734 | -0.282 | -0.048 | -0.025 | -0.025 | 0 | 0.54 | 0.19 | 0.49 | 0.66 | 0.55 | 0.63 |
| 7 | 42179377 | rs6463092 | 0.168 | 1.426 | -0.35 | -0.13 | -0.002 | 0.032 | 0.032 | 0 | 0.73 | 0.48 | 0.84 | 0.76 | 0.78 | 0.71 |
| 7 | 42179545 | rs17753324 | -0.658 | 1.362 | -0.953 | -0.313 | -0.054 | -0.019 | -0.019 | 0 | 0.12 | 0 | 0.28 | 0.08 | 0.17 | 0.07 |
| 7 | 42188538 | rs3801204 | -0.454 | 1.023 | -0.899 | -0.339 | 0.002 | 0.013 | 0.013 | 0 | 0.2 | 0.18 | 0.12 | 0.15 | 0.2 | 0.26 |
| 7 | 42188856 | rs10230715 | -1.603 | 1.32 | -0.596 | -0.758 | 0.002 | 0.012 | 0.012 | 1.16 | 0.46 | 0.47 | 0.42 | 0.29 | 0.52 | 0.47 |

| | | | | | | | | | | | | | | | | |
|---|---|---|---|---|---|---|---|---|---|---|---|---|---|---|---|---|
| 7 | 42192247 | rs3779159 | 0.38 | 1.041 | -0.642 | -0.744 | -0.017 | -0.008 | -0.008 | 0 | 0.16 | 0.06 | 0.12 | 0.14 | 0.19 | 0.21 |
| 7 | 42198560 | rs3801210 | -0.942 | 1.048 | -0.61 | -0.846 | 0 | -0.011 | -0.011 | 0 | 0.28 | 0.32 | 0.27 | 0.16 | 0.32 | 0.27 |
| 7 | 42201874 | rs3801211 | -0.666 | 1.032 | -0.912 | -0.858 | -0.068 | 0.037 | 0.037 | 0 | 0.26 | 0.05 | 0.24 | 0.14 | 0.32 | 0.24 |
| 7 | 42203353 | rs846401 | -1.139 | 0.75 | -0.814 | -1.036 | -0.024 | -0.005 | -0.005 | 1.96 | 0.41 | 0.72 | 0.5 | 0.38 | 0.41 | 0.48 |
| 7 | 42207032 | rs3801212 | 0.347 | 0.583 | -0.102 | -0.974 | 0.01 | 0.294 | 0.294 | 0.881 | 0.07 | 0.46 | 0.13 | 0.2 | 0.03 | 0.05 |
| 7 | 42209050 | rs6963356 | 0.039 | 0.577 | -0.105 | -0.984 | -0.004 | 0.301 | 0.301 | 0.882 | 0.08 | 0.46 | 0.14 | 0.22 | 0.03 | 0.08 |
| 7 | 42209334 | rs2190513 | -2.228 | 0.455 | -0.583 | -1.123 | -0.038 | -0.059 | -0.059 | 0 | 0.59 | 0.93 | 0.67 | 0.38 | 0.6 | 0.56 |
| 7 | 42210825 | rs3801213 | 1.503 | 0.713 | -0.333 | -1.425 | -0.013 | -0.011 | -0.011 | 0.722 | 0.12 | 0.03 | 0.09 | 0.14 | 0.13 | 0.19 |
| 7 | 42211849 | rs2286294 | -1.092 | 0.655 | -0.065 | -1.596 | -0.007 | 0.004 | 0.004 | 1.28 | 0.4 | 0.57 | 0.42 | 0.17 | 0.45 | 0.44 |
| 7 | 42213071 | rs3801214 | 0.585 | 0.868 | -0.598 | -1.648 | -0.001 | 0.012 | 0.012 | 0 | 0.11 | 0.06 | 0.13 | 0.01 | 0.11 | 0.06 |
| 7 | 42217027 | rs9886211 | -0.335 | 0.998 | -0.671 | -1.469 | -0.039 | 0.058 | 0.058 | 0.07 | 0.27 | 0.13 | 0.24 | 0.15 | 0.31 | 0.35 |
| 7 | 42221305 | rs3779173 | 1.129 | 1.078 | -0.907 | -1.46 | -0.012 | 0.01 | 0.01 | 0 | 0.13 | 0.03 | 0.08 | 0.14 | 0.13 | 0.23 |
| 7 | 42221813 | rs12540671 | 0.565 | 1.152 | -0.776 | -1.241 | -0.075 | 0.082 | 0.082 | 0 | 0.13 | 0.01 | 0.14 | 0.01 | 0.18 | 0.12 |
| 7 | 42228900 | rs2108368 | 0.48 | 1.192 | -0.869 | -1.575 | -0.091 | 0.102 | 0.102 | 0 | 0.16 | 0.01 | 0.15 | 0.01 | 0.2 | 0.13 |
| 7 | 42229946 | rs10951671 | 0.48 | 1.199 | -0.869 | -1.575 | -0.093 | 0.102 | 0.102 | 0 | 0.16 | 0.01 | 0.15 | 0.01 | 0.2 | 0.13 |
| 7 | 42232422 | rs10261063 | -0.012 | 1.013 | -0.547 | -1.665 | -0.006 | -0.042 | -0.042 | 0 | 0.36 | 0.47 | 0.34 | 0.23 | 0.35 | 0.37 |
| 7 | 42233283 | rs3801216 | -1.315 | 1.135 | -1.154 | -1.299 | -0.026 | 0.068 | 0.068 | 1.724 | 0.65 | 0.89 | 0.57 | 0.26 | 0.61 | 0.61 |
| 7 | 42236334 | rs6976209 | -0.429 | 1.34 | -0.05 | -0.807 | 0.047 | -0.127 | -0.127 | 0.042 | 0.19 | 0.36 | 0.18 | 0.02 | 0.19 | 0.13 |
| 7 | 42240694 | rs1019046 | -0.057 | 1.072 | -0.103 | -0.492 | 0.012 | -0.017 | -0.017 | 0 | 0.85 | 0.75 | 0.81 | 0.86 | 0.85 | 0.74 |
| 7 | 42245499 | rs17172023 | 0.145 | 1.779 | -1.046 | -0.315 | -0.06 | -0.05 | -0.05 | 0.141 | 0.12 | 0.02 | 0.12 | 0.18 | 0.18 | 0.14 |
| 7 | 42247876 | rs3801220 | -0.292 | 1.279 | -1.82 | -0.794 | -0.12 | 0.047 | 0.047 | 0 | 0.36 | 0.12 | 0.48 | 0.72 | 0.49 | 0.52 |
| 7 | 42253313 | rs3801232 | -0.118 | 1.35 | -1.897 | -0.771 | -0.11 | -0.012 | -0.012 | 0.134 | 0.32 | 0.08 | 0.42 | 0.68 | 0.42 | 0.47 |
| 7 | 42278116 | rs7778774 | -0.174 | 0.803 | -1.199 | -0.741 | -0.016 | 0.014 | 0.014 | 0 | 0.36 | 0.29 | 0.46 | 0.68 | 0.44 | 0.48 |
| 7 | 42281481 | rs6463094 | -0.664 | 0.237 | -0.931 | -1.026 | -0.025 | 0.037 | 0.037 | 1.208 | 0.43 | 0.75 | 0.51 | 0.68 | 0.5 | 0.51 |
| 7 | 42286475 | rs6463096 | -1.319 | 0.536 | -0.851 | -1.163 | 0.043 | -0.002 | -0.002 | 0.573 | 0.87 | 0.71 | 0.81 | 0.88 | 0.87 | 0.77 |
| 7 | 42292179 | rs17755180 | 0.296 | 0.531 | -0.973 | -1.131 | -0.022 | -0.005 | -0.005 | 0.036 | 0.29 | 0.25 | 0.45 | 0.67 | 0.41 | 0.43 |
| 7 | 42304214 | rs10229660 | -0.063 | 0.183 | -1.022 | -1.048 | 0 | 0.003 | 0.003 | 2.202 | 0.58 | 0.57 | 0.42 | 0.31 | 0.47 | 0.41 |
| 7 | 42305237 | rs10277168 | 0.245 | 0.549 | -0.814 | -0.978 | -0.044 | 0.068 | 0.068 | 0.02 | 0.56 | 0.2 | 0.42 | 0.3 | 0.53 | 0.44 |
| 7 | 42307527 | rs6950755 | 0.76 | -0.426 | -1.183 | -0.962 | -0.034 | -0.078 | -0.078 | 0.133 | 0.23 | 0.01 | 0.19 | 0.25 | 0.22 | 0.28 |
| 7 | 42309380 | rs4720420 | 0.824 | -0.682 | -1.217 | -0.895 | -0.039 | -0.075 | -0.075 | 0 | 0.4 | 0.19 | 0.54 | 0.47 | 0.56 | 0.42 |
| 7 | 42310981 | rs17172032 | 0.207 | -0.571 | -1.29 | -0.723 | 0.071 | -0.004 | -0.004 | 0 | 0.35 | 0.05 | 0.23 | 0.3 | 0.25 | 0.31 |
| 7 | 42311183 | rs10275082 | -1.221 | -0.519 | -1.068 | -0.7 | 0.01 | 0.016 | 0.016 | 0 | 0.84 | 0.75 | 0.83 | 0.78 | 0.87 | 0.79 |
| 7 | 42314044 | rs12701960 | -0.192 | -0.557 | -1.416 | -0.509 | 0.071 | 0.018 | 0.018 | 2.762 | 0.35 | 0.12 | 0.23 | 0.3 | 0.25 | 0.31 |
| 7 | 42318852 | rs886829 | 0.128 | 0.48 | -0.674 | -0.662 | 0.014 | 0.128 | 0.128 | 0.52 | 0.51 | 0.29 | 0.57 | 0.76 | 0.63 | 0.62 |
| 7 | 42319056 | rs886827 | -0.035 | 0.284 | -0.276 | -0.395 | 0.008 | 0.014 | 0.014 | 0.256 | 0.57 | 0.47 | 0.6 | 0.7 | 0.71 | 0.63 |
| 7 | 42319972 | rs1006369 | -1.405 | 0.426 | 0.403 | -0.392 | 0.04 | 0.004 | 0.004 | 0.344 | 0.05 | 0.28 | 0.05 | 0 | 0.04 | 0.01 |
| 7 | 42323532 | rs10488183 | -0.682 | 0.738 | -0.062 | -0.492 | -0.036 | 0.005 | 0.005 | 0 | 0.07 | 0.01 | 0.13 | 0.03 | 0.11 | 0.05 |
| 7 | 42324551 | rs2190521 | 0.807 | 0.499 | -0.643 | -0.758 | -0.044 | 0.052 | 0.052 | 1.68 | 0.28 | 0.08 | 0.3 | 0.21 | 0.28 | 0.36 |
| 7 | 42333628 | rs7792073 | -2.221 | 0.007 | -0.039 | -0.953 | -0.007 | -0.037 | -0.037 | 0.381 | 0.19 | 0.76 | 0.22 | 0.03 | 0.16 | 0.07 |
| 7 | 42344694 | rs7785123 | -0.123 | 0.08 | 0.407 | -0.886 | 0.01 | 0.024 | 0.024 | 0.119 | 0.39 | 0.69 | 0.38 | 0.24 | 0.33 | 0.39 |
| 7 | 42348227 | rs13233071 | -0.654 | 0.089 | 0.454 | -0.889 | 0.016 | 0.141 | 0.141 | 0 | 0.47 | 0.81 | 0.51 | 0.26 | 0.43 | 0.43 |
| 7 | 42356814 | rs10271006 | 0.472 | 0.085 | 0.732 | -0.89 | 0 | -0.006 | -0.006 | 0 | 0.33 | 0.38 | 0.34 | 0.21 | 0.31 | 0.37 |
| 7 | 42364924 | rs17172040 | -1.205 | 0.16 | 0.841 | -0.869 | 0.036 | -0.002 | -0.002 | 0.013 | 0.11 | 0.3 | 0.15 | 0.06 | 0.12 | 0.04 |
| 7 | 42377219 | rs2330441 | 0.366 | 0.132 | 0.506 | -0.84 | -0.007 | -0.045 | -0.045 | 0.022 | 0.36 | 0.52 | 0.36 | 0.21 | 0.32 | 0.38 |
| 7 | 42377974 | rs2190517 | -0.899 | 0.147 | 0.232 | -0.761 | 0.037 | 0.016 | 0.016 | 0 | 0.1 | 0.32 | 0.15 | 0.06 | 0.1 | 0.04 |
| 7 | 42379168 | rs917285 | -0.272 | 0.199 | -0.604 | -0.685 | -0.048 | 0.114 | 0.114 | 0.102 | 0.46 | 0.97 | 0.49 | 0.27 | 0.4 | 0.42 |
| 7 | 42384620 | rs7791745 | -0.019 | 0.19 | -0.174 | -0.163 | -0.002 | 0.016 | 0.016 | 0.094 | 0.76 | 0.7 | 0.82 | 0.66 | 0.74 | 0.66 |
| 7 | 42412407 | rs12540120 | -1.062 | 0.557 | 0.241 | -0.574 | 0.086 | -0.051 | -0.051 | 0 | 0.15 | 0.52 | 0.24 | 0.04 | 0.2 | 0.05 |
| 7 | 42420811 | rs6954249 | 0.094 | 0.313 | -0.118 | -0.998 | -0.015 | -0.09 | -0.09 | 0.063 | 0.17 | 0.29 | 0.09 | 0.01 | 0.12 | 0.08 |
| 7 | 42428579 | rs12530988 | -0.593 | -0.028 | -0.363 | -0.585 | 0.003 | 0.105 | 0.105 | 0.102 | 0.22 | 0.41 | 0.35 | 0.37 | 0.21 | 0.2 |
| 7 | 42434846 | rs1073164 | 1.716 | 0.324 | -0.572 | -1.379 | -0.051 | 0.005 | 0.005 | 0.75 | 0.29 | 0.06 | 0.31 | 0.29 | 0.31 | 0.33 |
| 7 | 42440761 | rs6966512 | -0.378 | 0.399 | -0.317 | -1.272 | 0.054 | -0.009 | -0.009 | 0 | 0.1 | 0.25 | 0.2 | 0.04 | 0.15 | 0.05 |
| 7 | 42445655 | rs9942705 | -0.74 | 0.322 | -0.871 | -1.138 | -0.023 | -0.095 | -0.095 | 0 | 0.23 | 0.31 | 0.11 | 0 | 0.16 | 0.09 |
| 7 | 42446415 | rs6959987 | -0.714 | 0.35 | -0.865 | -1.123 | -0.024 | -0.098 | -0.098 | 0 | 0.23 | 0.33 | 0.11 | 0 | 0.16 | 0.09 |
| 7 | 42456634 | rs10240250 | -1.401 | 0.342 | -0.642 | -0.644 | -0.026 | -0.058 | -0.058 | 0.039 | 0.15 | 0.34 | 0.13 | 0.02 | 0.09 | 0.04 |
| 7 | 42461966 | rs10257638 | -0.669 | 0.263 | -0.564 | -0.594 | -0.014 | -0.033 | -0.033 | 0.029 | 0.11 | 0.28 | 0.12 | 0.02 | 0.08 | 0.01 |
| 7 | 42464159 | rs4601201 | -0.03 | 0.245 | -0.903 | -0.752 | -0.021 | 0.05 | 0.05 | 0 | 0.4 | 0.15 | 0.43 | 0.71 | 0.43 | 0.58 |
| 7 | 42466264 | rs10274882 | -2.033 | 0.646 | -0.218 | -0.369 | -0.025 | -0.064 | -0.064 | 0 | 0.16 | 0.36 | 0.14 | 0.02 | 0.11 | 0.04 |
| 7 | 42489711 | rs4050289 | -1.302 | 0.765 | -0.525 | -0.359 | 0.007 | -0.033 | -0.033 | 0 | 0.12 | 0.21 | 0.16 | 0.02 | 0.15 | 0.02 |
| 7 | 42494497 | rs730485 | -1.791 | 0.598 | -0.491 | -0.357 | -0.007 | -0.063 | -0.063 | 0.865 | 0.17 | 0.34 | 0.15 | 0.02 | 0.13 | 0.04 |
| 7 | 42494790 | rs17172062 | -0.824 | 0.488 | -0.572 | -0.177 | -0.01 | -0.056 | -0.056 | 1.226 | 0.1 | 0.24 | 0.05 | 0 | 0.05 | 0.01 |
| 7 | 42496091 | rs17172063 | -0.684 | 0.979 | -1.383 | 0.279 | -0.067 | -0.054 | -0.054 | 5.567 | 0.15 | 0.01 | 0.11 | 0.2 | 0.21 | 0.3 |
| 7 | 42496317 | rs1465682 | 0.177 | 0.267 | -1.081 | -0.393 | -0.007 | -0.035 | -0.035 | 0.022 | 0.44 | 0.39 | 0.38 | 0.16 | 0.35 | 0.34 |
| 7 | 42496495 | rs1123227 | 0.532 | 0.316 | -1.27 | 0.077 | -0.004 | -0.011 | -0.011 | 0 | 0.25 | 0.14 | 0.29 | 0.3 | 0.25 | 0.36 |
| 7 | 42497247 | rs6977301 | 0.586 | -0.576 | -0.676 | 0.531 | -0.03 | 0.059 | 0.059 | 0.173 | 0.79 | 0.73 | 0.83 | 0.55 | 0.91 | 0.79 |
| 7 | 42499307 | rs4724107 | -0.475 | -0.376 | -0.678 | 0.516 | 0.004 | -0.026 | -0.026 | 0 | 0.94 | 0.91 | 0.97 | 1 | 0.95 | 1 |
| 7 | 42501585 | rs2877149 | 1.163 | -0.375 | -0.595 | 0.501 | -0.003 | 0.039 | 0.039 | 0 | 0.85 | 0.84 | 0.86 | 0.54 | 0.92 | 0.79 |
| 7 | 42503057 | rs6943150 | 1.088 | -0.748 | -0.591 | 0.042 | -0.003 | -0.006 | -0.006 | 0.352 | 0.57 | 0.55 | 0.68 | 0.25 | 0.73 | 0.48 |
| 7 | 42504691 | rs10085826 | 1.046 | -0.689 | -0.722 | 0.054 | 0.002 | -0.015 | -0.015 | 0 | 0.05 | 0.06 | 0.03 | 0 | 0.05 | 0.01 |

| | | | | | | | | | | | | | | | | |
|---|---|---|---|---|---|---|---|---|---|---|---|---|---|---|---|---|
| 7 | 42505060 | rs7806030 | 0.881 | -0.439 | -1.015 | -0.49 | -0.01 | -0.01 | -0.01 | 0.84 | 0.26 | 0.27 | 0.3 | 0.04 | 0.43 | 0.13 |
| 7 | 42505123 | rs10244840 | -0.294 | -0.79 | -0.987 | 0.02 | 0 | -0.001 | -0.001 | 0 | 0.29 | 0.28 | 0.21 | 0.28 | 0.22 | 0.3 |
| 7 | 42513491 | rs4143614 | -1.017 | -0.229 | -0.848 | 0.416 | 0.011 | -0.027 | -0.027 | 0 | 0.18 | 0.12 | 0.14 | 0.47 | 0.09 | 0.22 |
| 7 | 42522398 | rs934405 | 0.669 | 0.737 | -0.621 | 0.019 | 0.013 | -0.024 | -0.024 | 0 | 0.76 | 0.65 | 0.79 | 0.85 | 0.68 | 0.9 |
| 7 | 42523690 | rs17172071 | 0.58 | 0.042 | -1.11 | 0.089 | -0.005 | -0.08 | -0.08 | 2.558 | 0.22 | 0.05 | 0.15 | 0.34 | 0.19 | 0.36 |
| 7 | 42524501 | rs17172073 | 0.229 | 0.146 | -0.453 | -0.269 | -0.004 | -0.068 | -0.068 | 0 | 0.12 | 0.24 | 0.09 | 0.01 | 0.11 | 0.04 |
| 7 | 42525576 | rs12154826 | 0.886 | 0.641 | -0.676 | -0.564 | -0.044 | 0.041 | 0.041 | 0 | 0.15 | 0.07 | 0.19 | 0.05 | 0.28 | 0.04 |
| 7 | 42530615 | rs934404 | 2.144 | 0.101 | -0.733 | -0.432 | 0.003 | 0.002 | 0.002 | 0 | 0.29 | 0.25 | 0.47 | 0.26 | 0.26 | 0.31 |
| 7 | 42534499 | rs6463117 | 0.063 | 0.25 | -0.362 | -0.627 | -0.018 | 0.005 | 0.005 | 1.625 | 0.64 | 0.55 | 0.76 | 0.66 | 0.71 | 0.61 |
| 7 | 42541434 | rs4724111 | 1.24 | 0.137 | -0.48 | -0.67 | 0.032 | 0.003 | 0.003 | 0.468 | 0.43 | 0.22 | 0.52 | 0.61 | 0.37 | 0.53 |
| 7 | 42542632 | rs10261329 | 0.343 | -0.056 | -0.115 | -0.54 | -0.001 | -0.018 | -0.018 | 0 | 0.28 | 0.19 | 0.2 | 0.33 | 0.28 | 0.38 |
| 7 | 42547000 | rs10480181 | 0.735 | -0.173 | 0.128 | -0.466 | -0.01 | -0.069 | -0.069 | 0 | 0.12 | 0.28 | 0.07 | 0.01 | 0.08 | 0.04 |
| 7 | 42552131 | rs10237075 | 0.561 | -0.176 | 0.148 | -0.468 | -0.006 | -0.074 | -0.074 | 0 | 0.12 | 0.35 | 0.08 | 0.01 | 0.08 | 0.04 |
| 7 | 42564537 | rs10279276 | 0.039 | -0.214 | 0.138 | -0.485 | -0.027 | -0.099 | -0.099 | 0 | 0.14 | 0.47 | 0.08 | 0.01 | 0.08 | 0.04 |
| 7 | 42565284 | rs11767154 | -0.954 | -0.215 | 0.013 | -0.537 | 0.029 | 0.01 | 0.01 | 0.834 | 0.41 | 0.7 | 0.29 | 0.34 | 0.34 | 0.42 |
| 7 | 42566664 | rs12701985 | 0.94 | -0.07 | -0.279 | -0.532 | 0.012 | -0.034 | -0.034 | 0 | 0.07 | 0.12 | 0.06 | 0 | 0.09 | 0.03 |
| 7 | 42575126 | rs10229847 | 0.743 | -0.156 | -0.096 | -0.527 | 0.07 | 0.003 | 0.003 | 0.059 | 0.08 | 0.35 | 0.07 | 0.01 | 0.06 | 0.04 |
| 7 | 42584265 | rs6972060 | 1.333 | -0.18 | -0.342 | -0.463 | 0.002 | -0.072 | -0.072 | 0 | 0.19 | 0.05 | 0.14 | 0.32 | 0.17 | 0.34 |
| 7 | 42586866 | rs1880406 | 1.003 | -0.166 | -0.294 | -0.512 | 0.012 | -0.034 | -0.034 | 0 | 0.07 | 0.12 | 0.06 | 0 | 0.09 | 0.03 |
| 7 | 42594033 | rs4551244 | 0.449 | -0.223 | -0.326 | -0.553 | -0.012 | -0.092 | -0.092 | 0 | 0.13 | 0.43 | 0.08 | 0.01 | 0.08 | 0.04 |
| 7 | 42597593 | rs6976333 | -0.908 | -0.229 | -0.578 | -0.542 | 0.034 | 0.014 | 0.014 | 1.385 | 0.4 | 0.72 | 0.29 | 0.33 | 0.34 | 0.41 |
| 7 | 42604257 | rs10239387 | 1.358 | -0.127 | -0.701 | -0.397 | -0.025 | -0.095 | -0.095 | 0.309 | 0.18 | 0.04 | 0.14 | 0.33 | 0.17 | 0.31 |
| 7 | 42605354 | rs7785714 | 0.51 | -0.228 | -0.34 | -0.507 | -0.008 | -0.082 | -0.082 | 0 | 0.12 | 0.43 | 0.08 | 0.01 | 0.07 | 0.04 |
| 7 | 42605660 | rs4367431 | -1.252 | 0.065 | -0.639 | -0.688 | -0.055 | 0.053 | 0.053 | 0 | 0.85 | 0.93 | 0.79 | 0.95 | 0.73 | 0.95 |
| 7 | 42612690 | rs4634534 | 1.461 | -0.007 | -0.578 | -0.591 | 0.019 | 0.02 | 0.02 | 0.357 | 0.45 | 0.14 | 0.5 | 0.63 | 0.39 | 0.57 |
| 7 | 42613720 | rs7811252 | 1.059 | 0.083 | -0.332 | -0.457 | 0.101 | 0.031 | 0.031 | 0 | 0.06 | 0.35 | 0.06 | 0.01 | 0.06 | 0.04 |
| 7 | 42614219 | rs1524594 | 0.049 | 0.033 | -0.802 | -0.482 | -0.091 | -0.031 | -0.031 | 0.297 | 0.65 | 0.83 | 0.64 | 0.63 | 0.55 | 0.63 |
| 7 | 42615429 | rs4400272 | -0.631 | 0.286 | -0.522 | -0.823 | -0.023 | 0.02 | 0.02 | 0 | 0.84 | 0.88 | 0.79 | 0.95 | 0.73 | 0.95 |
| 7 | 42615554 | rs1917215 | 2.054 | -0.124 | -0.976 | -0.61 | -0.035 | -0.098 | -0.098 | 0 | 0.31 | 0.03 | 0.45 | 0.35 | 0.28 | 0.4 |
| 7 | 42615974 | rs4724115 | 1.786 | -0.023 | -0.777 | -0.571 | 0.013 | -0.038 | -0.038 | 0 | 0.33 | 0.15 | 0.45 | 0.36 | 0.28 | 0.4 |
| 7 | 42617751 | rs13312158 | 0.499 | -0.049 | -0.657 | -0.566 | 0.013 | -0.03 | -0.03 | 0 | 0.35 | 0.25 | 0.24 | 0.55 | 0.32 | 0.49 |
| 7 | 42636452 | rs1524583 | 1.751 | -0.029 | -0.697 | -0.561 | -0.002 | -0.024 | -0.024 | 0 | 0.36 | 0.18 | 0.47 | 0.37 | 0.34 | 0.41 |
| 7 | 42642443 | rs952889 | 0.979 | 0.523 | -0.089 | -0.713 | 0.02 | -0.08 | -0.08 | 0.898 | 0.71 | 0.45 | 0.71 | 0.92 | 0.66 | 0.9 |
| 7 | 42643421 | rs9639861 | -0.935 | 0.739 | -0.318 | -0.454 | -0.014 | 0.001 | 0.001 | 0 | 0.88 | 0.91 | 0.81 | 0.99 | 0.75 | 0.96 |
| 7 | 42657424 | rs1880403 | 0.454 | 0.294 | -0.113 | -0.41 | 0.048 | -0.098 | -0.098 | 0 | 0.72 | 0.51 | 0.71 | 0.92 | 0.62 | 0.93 |
| 7 | 42662187 | rs17172094 | 0.591 | -0.761 | -0.278 | 0.017 | -0.009 | 0.003 | 0.003 | 0.055 | 0.41 | 0.54 | 0.44 | 0.44 | 0.29 | 0.44 |
| 7 | 42665355 | rs17172095 | 0.673 | -0.976 | 0.337 | -0.053 | -0.007 | 0.017 | 0.017 | 0 | 0.38 | 0.49 | 0.49 | 0.47 | 0.33 | 0.37 |
| 7 | 42666722 | rs2204350 | -0.335 | -0.765 | -0.127 | -0.129 | -0.116 | -0.035 | -0.035 | 0 | 0.67 | 0.91 | 0.7 | 0.68 | 0.53 | 0.73 |
| 7 | 42668807 | rs1880404 | 0.435 | -0.752 | 0.015 | -0.16 | 0.01 | -0.005 | -0.005 | 0 | 0.48 | 0.59 | 0.54 | 0.5 | 0.4 | 0.43 |
| 7 | 42671386 | rs13222816 | 2.14 | -0.939 | 0.162 | -0.274 | -0.003 | -0.038 | -0.038 | 0 | 0.3 | 0.15 | 0.41 | 0.42 | 0.28 | 0.35 |
| 7 | 42684548 | rs2656505 | 1.555 | -0.804 | -0.295 | -0.304 | -0.108 | 0 | 0 | 0 | 0.16 | 0.01 | 0.18 | 0.12 | 0.31 | 0.12 |
| 7 | 42690554 | rs6974906 | 0.987 | -0.98 | 0.302 | -0.246 | -0.005 | -0.002 | -0.002 | 0 | 0.19 | 0.22 | 0.16 | 0.18 | 0.13 | 0.29 |
| 7 | 42704102 | rs2583879 | -0.064 | -0.744 | 0.384 | -0.218 | 0.021 | -0.005 | -0.005 | 0 | 0.52 | 0.34 | 0.49 | 0.5 | 0.6 | 0.58 |
| 7 | 42704271 | rs3913703 | 1.079 | -0.711 | 0.492 | -0.281 | -0.006 | 0.028 | 0.028 | 0 | 0.33 | 0.38 | 0.43 | 0.46 | 0.3 | 0.33 |
| 7 | 42732842 | rs13234515 | 1.364 | -0.702 | 0.16 | -0.984 | 0 | -0.026 | -0.026 | 0 | 0.31 | 0.19 | 0.39 | 0.41 | 0.29 | 0.33 |
| 7 | 42745436 | rs17543692 | 0.255 | -1.115 | 0.486 | -2.052 | 0.019 | 0.022 | 0.022 | 0 | 0.21 | 0.16 | 0.18 | 0.17 | 0.16 | 0.29 |
| 7 | 42745578 | rs2583893 | 1.134 | -1.171 | 0.569 | -2.149 | 0.026 | 0.048 | 0.048 | 2.835 | 0.62 | 0.24 | 0.68 | 0.75 | 0.7 | 0.63 |
| 7 | 42748204 | rs1039590 | 0.187 | -1.177 | 0.768 | -2.184 | -0.035 | -0.023 | -0.023 | 0 | 0.21 | 0.4 | 0.2 | 0.21 | 0.15 | 0.31 |
| 7 | 42756339 | rs965739 | 0.704 | -1.292 | 0.674 | -2.178 | -0.009 | -0.112 | -0.112 | 0 | 0.81 | 0.48 | 0.85 | 0.94 | 0.84 | 0.91 |
| 7 | 42759430 | rs6463139 | -0.118 | -1.451 | 0.559 | -1.86 | -0.012 | -0.012 | -0.012 | 0.332 | 0.22 | 0.29 | 0.18 | 0.15 | 0.15 | 0.32 |
| 7 | 42762246 | rs719910 | -0.022 | -0.991 | 0.658 | -1.613 | -0.012 | -0.042 | -0.042 | 0 | 0.57 | 0.5 | 0.66 | 0.78 | 0.65 | 0.45 |
| 7 | 42789194 | rs6966201 | -0.048 | -0.805 | 0.248 | -1.353 | -0.072 | 0.046 | 0.046 | 11.103 | 0.2 | 0.04 | 0.23 | 0.14 | 0.31 | 0.26 |
| 7 | 42800841 | rs2583890 | 0.432 | -1.049 | 0.516 | -1.279 | 0.015 | -0.035 | -0.035 | 2.011 | 0.72 | 0.45 | 0.7 | 0.81 | 0.65 | 0.62 |
| 7 | 42810378 | rs2656510 | 1.115 | -1.337 | -0.087 | -1.391 | 0.019 | 0.106 | 0.106 | 3.881 | 0.54 | 0.16 | 0.6 | 0.75 | 0.58 | 0.44 |
| 7 | 42811567 | rs11984213 | -1 | -1.965 | 1.06 | -1.054 | -0.045 | -0.042 | -0.042 | 0 | 0.18 | 0.31 | 0.1 | 0.06 | 0.08 | 0.16 |
| 7 | 42811744 | rs11977440 | -0.557 | -1.826 | 0.504 | -0.991 | 0.004 | 0.005 | 0.005 | 0.077 | 0.15 | 0.12 | 0.07 | 0.04 | 0.07 | 0.16 |
| 7 | 42823075 | rs1504837 | 0.738 | -0.878 | 1.031 | -0.906 | 0.047 | 0.082 | 0.082 | 0.267 | 0.62 | 0.23 | 0.74 | 0.9 | 0.77 | 0.74 |
| 7 | 42831789 | rs11769164 | -0.322 | -0.823 | 1.01 | -0.889 | 0.031 | 0.121 | 0.121 | 0 | 0.3 | 0.11 | 0.18 | 0.04 | 0.21 | 0.25 |
| 7 | 42835506 | rs9691053 | -0.546 | -0.584 | 0.468 | -0.775 | -0.013 | -0.052 | -0.052 | 0 | 0.36 | 0.59 | 0.23 | 0.09 | 0.22 | 0.26 |
| 7 | 42869090 | rs621013 | 1.605 | -0.461 | 2.554 | -0.001 | 0.023 | 0.112 | 0.112 | 0.476 | 0.93 | 0.47 | 0.92 | 0.91 | 0.97 | 0.99 |
| 7 | 42884961 | rs17172123 | -0.604 | -0.213 | 2.151 | -0.047 | -0.002 | 0 | 0 | 0 | 0.05 | 0.06 | 0.07 | 0.07 | 0.08 | 0.15 |
| 7 | 42906793 | rs607148 | 1.985 | -0.26 | 2.839 | -0.038 | -0.034 | 0.021 | 0.021 | 3.137 | 0.92 | 0.3 | 0.91 | 0.91 | 0.97 | 0.99 |
| 7 | 42927961 | rs595541 | 1.309 | 0.049 | 3.009 | -0.377 | -0.007 | -0.031 | -0.031 | 0 | 0.83 | 0.45 | 0.85 | 0.9 | 0.89 | 0.91 |
| 7 | 42941854 | rs602524 | 1.489 | -0.303 | 2.99 | -0.477 | -0.01 | -0.038 | -0.038 | 0.003 | 0.81 | 0.72 | 0.75 | 0.91 | 0.87 | 0.92 |
| 7 | 42948900 | rs1130010 | -0.64 | -0.316 | 2.843 | -0.478 | 0.005 | -0.028 | -0.028 | 0.348 | 0.05 | 0 | 0.14 | 0.09 | 0.04 | 0.07 |
| 7 | 42949038 | rs1048434 | -0.864 | -0.332 | 3.086 | -0.489 | 0.001 | -0.03 | -0.03 | 0 | 0.11 | 0.09 | 0.09 | 0 | 0.08 | 0.01 |
| 7 | 42949101 | rs1048423 | -0.864 | -0.332 | 3.086 | -0.489 | 0.001 | -0.03 | -0.03 | 0 | 0.11 | 0.09 | 0.09 | 0 | 0.08 | 0.01 |
| 7 | 42951936 | rs639459 | 2.549 | -0.372 | 3.538 | -0.478 | -0.023 | -0.047 | -0.047 | 0.322 | 0.79 | 0.48 | 0.74 | 0.91 | 0.87 | 0.92 |
| 7 | 42952486 | rs661592 | 1.891 | -0.344 | 3.369 | -0.466 | -0.005 | -0.02 | -0.02 | 0 | 0.83 | 0.78 | 0.77 | 0.91 | 0.87 | 0.92 |

| | | | | | | | | | | | | | | | | |
|---|---|---|---|---|---|---|---|---|---|---|---|---|---|---|---|---|
| 7 | 42956696 | rs17722119 | -0.645 | -0.342 | 3.336 | -0.46 | 0.005 | -0.029 | -0.029 | 0 | 0.05 | 0 | 0.14 | 0.09 | 0.04 | 0.07 |
| 7 | 42972411 | rs647618 | 2.106 | -0.318 | 3.09 | -0.34 | -0.004 | -0.003 | -0.003 | 0.073 | 0.78 | 0.78 | 0.71 | 0.81 | 0.82 | 0.8 |
| 7 | 42979942 | rs10486743 | 0.219 | -1.014 | 2.429 | -0.846 | 0.022 | 0.03 | 0.03 | 0.451 | 0.11 | 0.06 | 0.04 | 0.01 | 0.04 | 0.03 |
| 7 | 42988394 | rs6463157 | -0.175 | -1.321 | 1.89 | -1.581 | -0.017 | -0.037 | -0.037 | 2.732 | 0.25 | 0.41 | 0.21 | 0.15 | 0.23 | 0.28 |
| 7 | 42991650 | rs584184 | 0.012 | -1.322 | 1.981 | -1.565 | -0.021 | -0.023 | -0.023 | 0 | 0.75 | 0.53 | 0.78 | 0.82 | 0.77 | 0.72 |
| 7 | 43009602 | rs1896487 | -0.112 | -1.017 | 1.257 | -1.402 | -0.006 | -0.028 | -0.028 | 0 | 0.3 | 0.45 | 0.19 | 0.21 | 0.3 | 0.38 |
| 7 | 43010631 | rs10245441 | 1.355 | -1.195 | 1.021 | -1.514 | -0.034 | -0.037 | -0.037 | 0.015 | 0.08 | 0.03 | 0.08 | 0.18 | 0.19 | 0.29 |
| 7 | 43012510 | rs677442 | 2.144 | -1.223 | 0.945 | -1.46 | -0.102 | -0.155 | -0.155 | 0 | 0.29 | 0.04 | 0.32 | 0.55 | 0.35 | 0.26 |
| 7 | 43013834 | rs596167 | -0.84 | -1.254 | 1.119 | -1.391 | 0.011 | 0.007 | 0.007 | 0.431 | 0.37 | 0.67 | 0.39 | 0.3 | 0.34 | 0.47 |
| 7 | 43022660 | rs636860 | 1.257 | -1.255 | 1.111 | -1.394 | -0.077 | -0.067 | -0.067 | 1.125 | 0.41 | 0.1 | 0.41 | 0.55 | 0.43 | 0.27 |
| 7 | 43025181 | rs10256012 | -0.805 | -1.234 | 1.127 | -1.386 | -0.009 | -0.006 | -0.006 | 0.109 | 0.36 | 0.56 | 0.39 | 0.29 | 0.34 | 0.45 |
| 7 | 43026461 | rs799084 | 0.064 | -1.311 | 1.02 | -1.287 | -0.019 | -0.003 | -0.003 | 0.711 | 0.71 | 0.55 | 0.62 | 0.76 | 0.73 | 0.63 |
| 7 | 43030978 | rs986413 | -0.368 | -1.315 | 1.21 | -1.435 | -0.009 | -0.057 | -0.057 | 0.616 | 0.76 | 0.64 | 0.8 | 0.9 | 0.77 | 0.71 |
| 7 | 43032827 | rs12537581 | -0.837 | -1.973 | 0.796 | -1.809 | -0.046 | -0.067 | -0.067 | 0.447 | 0.25 | 0.51 | 0.3 | 0.11 | 0.13 | 0.1 |
| 7 | 43045905 | rs11763965 | -0.134 | -1.689 | 0.136 | -2.047 | -0.008 | -0.031 | -0.031 | 0 | 0.28 | 0.36 | 0.19 | 0.22 | 0.27 | 0.29 |
| 7 | 43052388 | rs7785548 | -0.124 | -1.695 | 0.133 | -2.05 | -0.007 | -0.031 | -0.031 | 0 | 0.28 | 0.36 | 0.19 | 0.22 | 0.27 | 0.29 |
| 7 | 43053715 | rs880976 | -0.723 | -1.534 | 0.177 | -2.046 | -0.01 | -0.026 | -0.026 | 0.122 | 0.73 | 0.52 | 0.63 | 0.79 | 0.74 | 0.63 |
| 7 | 43057398 | rs887060 | 0.299 | -1.627 | 0.358 | -2.127 | -0.005 | -0.032 | -0.032 | 0 | 0.58 | 0.46 | 0.59 | 0.79 | 0.64 | 0.59 |
| 7 | 43077116 | rs7786719 | 1.853 | -1.865 | 0.201 | -2.191 | -0.119 | -0.115 | -0.115 | 0 | 0.28 | 0.04 | 0.35 | 0.57 | 0.38 | 0.28 |
| 7 | 43077850 | rs2215170 | -0.071 | -1.859 | 0.293 | -2.251 | -0.008 | -0.017 | -0.017 | 0 | 0.28 | 0.32 | 0.24 | 0.22 | 0.26 | 0.31 |
| 7 | 43083245 | rs1972543 | -1.635 | -1.771 | 0.407 | -2.142 | -0.044 | 0.059 | 0.059 | 0 | 0.8 | 0.93 | 0.82 | 0.9 | 0.77 | 0.71 |
| 7 | 43125022 | rs10224910 | 0.407 | -1.68 | -0.564 | -2.197 | 0.001 | 0.007 | 0.007 | 1.384 | 0.51 | 0.54 | 0.57 | 0.76 | 0.47 | 0.4 |
| 7 | 43128524 | rs1012033 | 1.461 | -1.688 | -1.056 | -1.759 | -0.041 | -0.031 | -0.031 | 0 | 0.3 | 0.17 | 0.38 | 0.59 | 0.39 | 0.26 |
| 7 | 43140213 | rs10273260 | -1.101 | -1.579 | -1.072 | -1.714 | 0.011 | 0.01 | 0.01 | 0 | 0.76 | 0.77 | 0.77 | 0.8 | 0.8 | 0.62 |
| 7 | 43147965 | rs7798775 | -0.316 | -1.247 | -0.888 | -1.665 | 0.023 | -0.058 | -0.058 | 13.503 | 0.3 | 0.44 | 0.29 | 0.18 | 0.35 | 0.32 |
| 7 | 43149715 | rs4724175 | 1.292 | -1.117 | -1.276 | -1.463 | -0.097 | -0.075 | -0.075 | 0 | 0.29 | 0.08 | 0.39 | 0.6 | 0.39 | 0.25 |
| 7 | 43155250 | rs10951716 | 0.226 | -0.689 | -0.322 | -1.433 | 0.039 | -0.061 | -0.061 | 0 | 0.11 | 0.21 | 0.1 | 0.03 | 0.12 | 0.05 |
| 7 | 43157968 | rs2190947 | -0.43 | 0.293 | -1.187 | -0.959 | -0.073 | 0.058 | 0.058 | 1.177 | 0.7 | 0.87 | 0.63 | 0.82 | 0.63 | 0.68 |
| 7 | 43158219 | rs7787143 | -0.301 | 0.21 | -0.24 | -0.906 | 0.081 | -0.042 | -0.042 | 0 | 0.09 | 0.37 | 0.1 | 0 | 0.11 | 0.01 |
| 7 | 43158477 | rs2024125 | 0.058 | -0.291 | -0.159 | -0.666 | -0.001 | -0.001 | -0.001 | 0 | 0.59 | 0.66 | 0.51 | 0.74 | 0.49 | 0.56 |
| 7 | 43162969 | rs1006107 | 0.298 | -0.405 | -0.025 | -1.039 | -0.023 | -0.076 | -0.076 | 0 | 0.22 | 0.27 | 0.09 | 0.02 | 0.09 | 0.07 |
| 7 | 43163707 | rs4724177 | 0.533 | -0.023 | -0.026 | -0.157 | 0.032 | 0.054 | 0.054 | 0.005 | 0.19 | 0.32 | 0.34 | 0.32 | 0.26 | 0.24 |
| 7 | 43166061 | rs6978743 | -0.559 | -0.115 | -0.195 | -0.037 | -0.001 | 0.002 | 0.002 | 1.243 | 0.43 | 0.42 | 0.51 | 0.59 | 0.54 | 0.53 |
| 7 | 43166443 | rs6959165 | -0.865 | -0.146 | -0.245 | -0.039 | 0.009 | 0.038 | 0.038 | 0.938 | 0.47 | 0.75 | 0.53 | 0.59 | 0.54 | 0.53 |
| 7 | 43171955 | rs17708960 | 0.882 | -0.497 | -0.798 | -0.117 | -0.005 | 0.151 | 0.151 | 4.355 | 0.25 | 0.02 | 0.24 | 0.02 | 0.18 | 0.08 |
| 7 | 43173610 | rs12690832 | -0.701 | 0.106 | -0.407 | 0.298 | 0.001 | -0.001 | -0.001 | 0.417 | 0.42 | 0.42 | 0.43 | 0.58 | 0.52 | 0.52 |
| 7 | 43174414 | rs17795651 | 0.887 | 0.382 | -0.435 | 0.493 | -0.023 | -0.039 | -0.039 | 0 | 0.07 | 0 | 0.03 | 0.18 | 0.06 | 0.05 |
| 7 | 43186431 | rs6463166 | -0.567 | 0.38 | -0.086 | 0.294 | 0.025 | 0.038 | 0.038 | 0.878 | 0.35 | 0.46 | 0.39 | 0.45 | 0.46 | 0.46 |
| 7 | 43186655 | rs7341456 | 0.535 | 0.588 | -0.29 | 0.328 | 0.036 | 0.122 | 0.122 | 0 | 0.18 | 0.33 | 0.26 | 0.37 | 0.25 | 0.23 |
| 7 | 43192377 | rs7807551 | -0.398 | 0.101 | -0.718 | -0.004 | -0.004 | -0.016 | -0.016 | 0 | 0.62 | 0.58 | 0.69 | 0.81 | 0.75 | 0.87 |
| 7 | 43193772 | rs2107270 | -0.496 | 0.118 | -0.689 | 0.003 | 0.011 | 0.022 | 0.022 | 2.085 | 0.63 | 0.67 | 0.7 | 0.81 | 0.75 | 0.87 |
| 7 | 43194078 | rs10269585 | -0.75 | 0.583 | -0.496 | 0.42 | 0.01 | 0.013 | 0.013 | 0.211 | 0.36 | 0.53 | 0.39 | 0.42 | 0.46 | 0.47 |
| 7 | 43194619 | rs1029479 | -0.144 | 0.712 | -0.291 | 0.988 | 0.021 | 0.111 | 0.111 | 0 | 0.18 | 0.29 | 0.25 | 0.34 | 0.25 | 0.21 |
| 7 | 43195455 | rs2330745 | 0.417 | 0.6 | -0.914 | 1.053 | -0.006 | -0.013 | -0.013 | 0 | 0.06 | 0.02 | 0.03 | 0.15 | 0.05 | 0.05 |
| 7 | 43195668 | rs4636109 | -0.019 | 0.031 | -1.098 | 0.591 | 0.043 | -0.021 | -0.021 | 0 | 0.39 | 0.81 | 0.37 | 0.46 | 0.33 | 0.44 |
| 7 | 43196387 | rs2107272 | 0.879 | 0.382 | -1.343 | 0.627 | -0.043 | -0.079 | -0.079 | 4.995 | 0.26 | 0.07 | 0.3 | 0.38 | 0.3 | 0.35 |
| 7 | 43200551 | rs6972287 | 0.059 | -0.266 | -1.103 | 0.215 | 0.021 | 0.005 | 0.005 | 0.013 | 0.35 | 0.19 | 0.31 | 0.35 | 0.27 | 0.26 |
| 7 | 43201282 | rs6957566 | 0.277 | 0.028 | -0.43 | -0.086 | -0.002 | -0.025 | -0.025 | 0.147 | 0.47 | 0.52 | 0.46 | 0.19 | 0.59 | 0.48 |
| 7 | 43201742 | rs721965 | 1.361 | 0.409 | -0.303 | -0.046 | -0.052 | 0.071 | 0.071 | 5.108 | 0.31 | 0.12 | 0.35 | 0.18 | 0.44 | 0.34 |
| 7 | 43202662 | rs3757564 | 0.242 | 0.111 | -0.102 | 0.436 | 0.034 | 0.059 | 0.059 | 0.008 | 0.09 | 0.32 | 0.05 | 0.14 | 0.05 | 0.05 |
| 7 | 43207653 | rs6463169 | -0.364 | -0.265 | -0.21 | 0.386 | 0.016 | -0.012 | -0.012 | 0 | 0.7 | 0.76 | 0.77 | 0.68 | 0.79 | 0.63 |
| 7 | 43207862 | rs6463170 | -0.503 | 0.138 | -0.137 | 0.659 | -0.007 | 0.047 | 0.047 | 1.928 | 0.29 | 0.39 | 0.33 | 0.51 | 0.23 | 0.21 |
| 7 | 43212606 | rs12674179 | 0.908 | 0.114 | -0.444 | 0.916 | -0.009 | 0.068 | 0.068 | 0 | 0.07 | 0.09 | 0.07 | 0.27 | 0.04 | 0.08 |
| 7 | 43214842 | rs9655043 | -1.264 | 0.136 | -0.091 | 0.413 | -0.001 | 0.002 | 0.002 | 2.524 | 0.43 | 0.48 | 0.45 | 0.48 | 0.39 | 0.3 |
| 7 | 43218933 | rs7802446 | -1.007 | 0.157 | -0.093 | 0.338 | -0.004 | -0.006 | -0.006 | 0 | 0.36 | 0.41 | 0.42 | 0.32 | 0.34 | 0.26 |
| 7 | 43222805 | rs7783147 | -0.058 | 0.122 | -0.002 | 0.281 | 0.004 | -0.037 | -0.037 | 0 | 0.22 | 0.26 | 0.3 | 0.07 | 0.26 | 0.18 |
| 7 | 43222995 | rs7787227 | -1.043 | 0.081 | 0.265 | 0.27 | 0.005 | -0.03 | -0.03 | 0 | 0.35 | 0.46 | 0.39 | 0.23 | 0.34 | 0.22 |
| 7 | 43228198 | rs10243801 | -0.083 | 0.156 | 0.147 | 0.244 | 0.004 | -0.037 | -0.037 | 0.243 | 0.22 | 0.26 | 0.3 | 0.07 | 0.26 | 0.18 |
| 7 | 43229414 | rs4437556 | 1.804 | 0.032 | -0.242 | 0.038 | -0.051 | 0.06 | 0.06 | 5.234 | 0.32 | 0.15 | 0.34 | 0.21 | 0.41 | 0.34 |
| 7 | 43230281 | rs13246891 | 0.886 | 0.219 | -0.001 | 0.146 | -0.04 | 0.058 | 0.058 | 0.436 | 0.14 | 0.01 | 0.23 | 0.06 | 0.16 | 0.09 |
| 7 | 43230504 | rs17172171 | 0.485 | 0.232 | 0.261 | 0.264 | 0.03 | -0.031 | -0.031 | 0.79 | 0.07 | 0.16 | 0.06 | 0.01 | 0.1 | 0.08 |
| 7 | 43235298 | rs4724184 | 0.75 | 0.066 | 0.058 | 0.523 | -0.015 | 0.091 | 0.091 | 0 | 0.08 | 0.14 | 0.08 | 0.26 | 0.05 | 0.08 |
| 7 | 43243719 | rs6949342 | 0.173 | 0.081 | -0.12 | 0.305 | -0.002 | 0 | 0 | 1.343 | 0.6 | 0.62 | 0.56 | 0.64 | 0.64 | 0.55 |
| 7 | 43248626 | rs10951717 | 0.018 | 0.594 | -0.189 | 0.029 | -0.004 | 0.006 | 0.006 | 0.67 | 0.71 | 0.68 | 0.65 | 0.41 | 0.75 | 0.58 |
| 7 | 43254610 | rs1029482 | 0.298 | 0.572 | -0.712 | -0.944 | 0.017 | 0.102 | 0.102 | 0.944 | 0.62 | 0.76 | 0.75 | 0.87 | 0.66 | 0.64 |
| 7 | 43254706 | rs12540272 | -0.359 | -0.029 | -0.58 | -0.942 | 0.029 | 0.042 | 0.042 | 1.879 | 0.3 | 0.24 | 0.16 | 0.12 | 0.19 | 0.28 |
| 7 | 43256590 | rs13225071 | 0.058 | 0.502 | -0.393 | -0.956 | 0.014 | 0.001 | 0.001 | 0 | 0.16 | 0.15 | 0.15 | 0.19 | 0.3 | 0.11 |
| 7 | 43262476 | rs12702025 | -1.349 | 0.184 | -0.147 | -0.979 | -0.01 | 0.027 | 0.027 | 0.624 | 0.57 | 0.75 | 0.48 | 0.4 | 0.54 | 0.41 |
| 7 | 43266557 | rs13243213 | -0.751 | 0.025 | 0.129 | -0.981 | 0.001 | 0.007 | 0.007 | 0.086 | 0.46 | 0.47 | 0.31 | 0.28 | 0.45 | 0.39 |

| | | | | | | | | | | | | | | | | |
|---|---|---|---|---|---|---|---|---|---|---|---|---|---|---|---|---|
| 7 | 43268470 | rs4460277 | -0.801 | -0.025 | -0.121 | -1.038 | -0.061 | 0.027 | 0.027 | 0.054 | 0.27 | 0.14 | 0.23 | 0.19 | 0.4 | 0.14 |
| 7 | 43273124 | rs10486733 | -1.244 | 0.052 | 0.108 | -0.923 | -0.001 | -0.013 | -0.013 | 0 | 0.54 | 0.52 | 0.45 | 0.32 | 0.54 | 0.43 |
| 7 | 43274439 | rs2040840 | -0.961 | 0.158 | -0.153 | -0.873 | 0.018 | 0.059 | 0.059 | 0 | 0.49 | 0.73 | 0.4 | 0.39 | 0.46 | 0.42 |
| 7 | 43276428 | rs2330785 | -0.501 | 0.263 | -0.029 | -0.832 | 0.006 | -0.001 | -0.001 | 0 | 0.23 | 0.23 | 0.21 | 0.18 | 0.35 | 0.12 |
| 7 | 43278848 | rs12530692 | -0.481 | -0.758 | -0.015 | -0.841 | -0.041 | -0.073 | -0.073 | 3.182 | 0.29 | 0.43 | 0.17 | 0.1 | 0.14 | 0.27 |
| 7 | 43282331 | rs4304246 | -0.677 | -0.592 | -0.392 | -0.496 | 0.025 | 0.032 | 0.032 | 1.82 | 0.37 | 0.78 | 0.31 | 0.31 | 0.31 | 0.27 |
| 7 | 43283115 | rs17172181 | -0.307 | -0.835 | 0.075 | -0.646 | -0.028 | -0.115 | -0.115 | 0.009 | 0.32 | 0.48 | 0.28 | 0.09 | 0.28 | 0.29 |
| 7 | 43286129 | rs17172184 | -0.73 | -0.915 | -0.053 | -0.637 | -0.001 | 0 | 0 | 0.439 | 0.41 | 0.57 | 0.34 | 0.3 | 0.3 | 0.39 |
| 7 | 43286839 | rs17172185 | -0.893 | -0.692 | 0.154 | -0.311 | 0.022 | 0.187 | 0.187 | 0.239 | 0.09 | 0.36 | 0.06 | 0.25 | 0.07 | 0.05 |
| 7 | 43286893 | rs17723330 | -0.826 | -0.686 | -0.002 | -0.309 | 0.009 | 0.148 | 0.148 | 0.121 | 0.09 | 0.31 | 0.05 | 0.25 | 0.07 | 0.05 |
| 7 | 43288484 | rs11771692 | -0.904 | -0.725 | -1.398 | -0.234 | -0.106 | -0.16 | -0.16 | 0.056 | 0.59 | 0.97 | 0.58 | 0.52 | 0.49 | 0.53 |
| 7 | 43291972 | rs6972797 | -0.396 | -1.503 | -0.037 | -0.392 | -0.049 | -0.039 | -0.039 | 0 | 0.13 | 0.43 | 0.08 | 0.08 | 0.05 | 0.14 |
| 7 | 43292359 | rs2240270 | -1.172 | -1.61 | -0.767 | -0.653 | 0.065 | 0.103 | 0.103 | 6.268 | 0.52 | 0.8 | 0.4 | 0.35 | 0.38 | 0.44 |
| 7 | 43298065 | rs2301871 | -0.851 | -1.656 | -0.603 | -1.013 | 0.036 | 0.051 | 0.051 | 6.134 | 0.46 | 0.65 | 0.34 | 0.17 | 0.29 | 0.26 |
| 7 | 43299455 | rs11763488 | -0.029 | -0.853 | -1.333 | -0.759 | 0.017 | 0.132 | 0.132 | 5.481 | 0.31 | 0.11 | 0.25 | 0.09 | 0.24 | 0.12 |
| 7 | 43307817 | rs7784053 | -0.153 | -1.098 | -0.503 | 0.213 | -0.034 | 0.039 | 0.039 | 0 | 0.16 | 0.27 | 0.21 | 0.28 | 0.09 | 0.29 |
| 7 | 43319101 | rs17172190 | 0.317 | -1.251 | -0.329 | -0.002 | -0.004 | -0.025 | -0.025 | 0 | 0.06 | 0.1 | 0.05 | 0 | 0.04 | 0.01 |
| 7 | 43320413 | rs9648078 | 1.46 | -1.495 | -0.945 | -0.404 | -0.009 | 0.056 | 0.056 | 0 | 0.11 | 0 | 0.03 | 0.01 | 0.09 | 0.09 |
| 7 | 43320431 | rs9648079 | -1.47 | -2.069 | -0.718 | -0.599 | 0.094 | 0.043 | 0.043 | 1.11 | 0.38 | 0.85 | 0.32 | 0.3 | 0.22 | 0.43 |
| 7 | 43321192 | rs10250916 | -0.043 | -1.887 | -0.079 | -0.395 | 0.036 | 0.032 | 0.032 | 2.71 | 0.05 | 0.38 | 0.03 | 0 | 0.01 | 0.07 |
| 7 | 43325451 | rs2214610 | -0.972 | -1.198 | -1.427 | -0.558 | -0.136 | -0.127 | -0.127 | 2 | 0.66 | 0.98 | 0.59 | 0.4 | 0.59 | 0.59 |
| 7 | 43336243 | rs886550 | -0.756 | -1.731 | -0.233 | 0.323 | -0.067 | 0.163 | 0.163 | 0 | 0.22 | 0.45 | 0.29 | 0.48 | 0.13 | 0.28 |
| 7 | 43346016 | rs2079263 | -0.327 | -1.116 | -0.432 | 0.049 | -0.005 | 0.043 | 0.043 | 2.244 | 0.26 | 0.32 | 0.34 | 0.53 | 0.23 | 0.42 |
| 7 | 43346678 | rs12532870 | -0.116 | -1.534 | -1.247 | -0.41 | 0.022 | 0.071 | 0.071 | 1.364 | 0.23 | 0.11 | 0.1 | 0.04 | 0.17 | 0.14 |
| 7 | 43350189 | rs6463180 | -0.885 | -1.642 | -1.236 | -0.416 | 0.042 | -0.044 | -0.044 | 0 | 0.39 | 0.84 | 0.43 | 0.53 | 0.3 | 0.44 |
| 7 | 43356791 | rs6968910 | -1.394 | -1.618 | -1.338 | -0.405 | -0.049 | -0.075 | -0.075 | 0 | 0.55 | 0.94 | 0.48 | 0.55 | 0.4 | 0.55 |
| 7 | 43357543 | rs6463182 | -0.816 | -1.486 | -0.486 | -0.337 | -0.004 | -0.007 | -0.007 | 0.648 | 0.07 | 0.6 | 0.05 | 0 | 0.01 | 0.01 |
| 7 | 43361008 | rs4509217 | 0.96 | -1.255 | -1.11 | -0.259 | -0.013 | 0.082 | 0.082 | 0 | 0.14 | 0.01 | 0.04 | 0.01 | 0.1 | 0.11 |
| 7 | 43365014 | rs917017 | -0.919 | -1.076 | -1.008 | -0.086 | -0.077 | -0.071 | -0.071 | 0.438 | 0.61 | 0.92 | 0.55 | 0.54 | 0.56 | 0.54 |
| 7 | 43365819 | rs2057809 | -0.44 | -0.629 | 0.275 | 0.516 | -0.027 | 0.218 | 0.218 | 0.011 | 0.15 | 0.43 | 0.25 | 0.36 | 0.09 | 0.13 |
| 7 | 43371852 | rs17723745 | -0.343 | -0.409 | 0.486 | 0.579 | 0.001 | 0.001 | 0.001 | 1.459 | 0.41 | 0.58 | 0.33 | 0.44 | 0.47 | 0.44 |
| 7 | 43373648 | rs17172198 | 0.22 | -0.285 | 0.703 | 0.591 | 0.087 | 0.149 | 0.149 | 0.409 | 0.15 | 0.43 | 0.17 | 0.24 | 0.19 | 0.17 |
| 7 | 43376210 | rs13232367 | -0.173 | -1.165 | 0.953 | 0.569 | -0.001 | -0.004 | -0.004 | 0 | 0.3 | 0.28 | 0.17 | 0.19 | 0.28 | 0.28 |
| 7 | 43377308 | rs17723805 | 1.001 | -1.438 | 0.605 | 0.605 | 0.007 | 0.005 | 0.005 | 0 | 0.08 | 0.05 | 0.17 | 0.27 | 0.05 | 0.19 |
| 7 | 43393057 | rs7808837 | -0.723 | -1.108 | 0.724 | 0.097 | -0.001 | 0 | 0 | 0.047 | 0.43 | 0.42 | 0.38 | 0.42 | 0.45 | 0.47 |
| 7 | 43393858 | rs12702035 | 1.214 | -1.131 | 0.371 | -0.049 | -0.078 | 0.108 | 0.108 | 0 | 0.17 | 0.02 | 0.12 | 0.01 | 0.24 | 0.11 |
| 7 | 43397700 | rs17172202 | -0.561 | -1.081 | 0.616 | 0.094 | 0.01 | 0.143 | 0.143 | 0 | 0.06 | 0.21 | 0.1 | 0.21 | 0.06 | 0.04 |
| 7 | 43398963 | rs2052148 | -0.013 | -1.233 | 0.551 | -0.163 | -0.001 | 0.013 | 0.013 | 3.212 | 0.55 | 0.61 | 0.46 | 0.32 | 0.47 | 0.32 |
| 7 | 43417909 | rs7794076 | -0.816 | -1.075 | -0.126 | -0.063 | 0.003 | 0.002 | 0.002 | 0 | 0.69 | 0.69 | 0.51 | 0.54 | 0.55 | 0.61 |
| 7 | 43424112 | rs10246620 | 0.81 | -1.011 | -0.288 | -0.021 | -0.003 | 0.001 | 0.001 | 8.598 | 0.16 | 0.14 | 0.11 | 0.21 | 0.12 | 0.28 |
| 7 | 43427551 | rs12702040 | 0.048 | -1.073 | -0.251 | -0.094 | 0.01 | -0.002 | -0.002 | 0.38 | 0.27 | 0.33 | 0.26 | 0.26 | 0.34 | 0.19 |
| 7 | 43428499 | rs10257003 | -0.203 | -1.042 | -0.919 | -0.218 | -0.001 | 0.007 | 0.007 | 0.661 | 0.34 | 0.29 | 0.23 | 0.22 | 0.35 | 0.4 |
| 7 | 43431056 | rs1860756 | 0.872 | -2.06 | -0.356 | 0.434 | 0.075 | 0.102 | 0.102 | 0.862 | 0.26 | 0.06 | 0.17 | 0.08 | 0.13 | 0.15 |
| 7 | 43431159 | rs10224012 | -0.225 | -1.746 | -0.198 | 0.561 | 0.087 | 0.057 | 0.057 | 0.612 | 0.43 | 0.23 | 0.28 | 0.28 | 0.26 | 0.43 |
| 7 | 43433870 | rs10951730 | -0.952 | -0.923 | -0.234 | 0.674 | 0.014 | 0.088 | 0.088 | 0 | 0.08 | 0.19 | 0.14 | 0.21 | 0.1 | 0.07 |
| 7 | 43444666 | rs13238475 | -0.806 | -0.769 | -0.898 | 0.73 | 0.007 | 0.055 | 0.055 | 0 | 0.1 | 0.2 | 0.18 | 0.21 | 0.12 | 0.08 |
| 7 | 43446761 | rs6980106 | -0.013 | -1.367 | -1.773 | -0.003 | 0.079 | 0.065 | 0.065 | 0.372 | 0.26 | 0.01 | 0.16 | 0.09 | 0.13 | 0.16 |
| 7 | 43447790 | rs10255565 | -0.673 | -1.386 | -1.084 | 0.491 | -0.019 | 0.007 | 0.007 | 1.079 | 0.28 | 0.32 | 0.15 | 0.33 | 0.16 | 0.26 |
| 7 | 43448803 | rs7350018 | -0.152 | -0.727 | -1.335 | 0.189 | -0.009 | -0.005 | -0.005 | 0 | 0.12 | 0.04 | 0.16 | 0.13 | 0.18 | 0.17 |
| 7 | 43451327 | rs11766157 | 0.634 | -0.945 | -2.242 | -0.607 | -0.005 | 0.127 | 0.127 | 0.007 | 0.24 | 0.03 | 0.14 | 0.05 | 0.22 | 0.21 |
| 7 | 43451710 | rs4724203 | 0.097 | 0.104 | -1.661 | -0.576 | 0.017 | 0.031 | 0.031 | 0 | 0.17 | 0.22 | 0.29 | 0.71 | 0.2 | 0.22 |
| 7 | 43451778 | rs10951731 | 0.125 | -0.172 | -1.661 | -0.711 | 0.018 | 0.124 | 0.124 | 0 | 0.42 | 0.18 | 0.32 | 0.19 | 0.43 | 0.39 |
| 7 | 43458861 | rs6463190 | 0.005 | -0.29 | -1.009 | -0.763 | 0.015 | 0.038 | 0.038 | 0 | 0.43 | 0.39 | 0.34 | 0.22 | 0.43 | 0.4 |
| 7 | 43463317 | rs4724205 | -0.734 | -0.453 | -0.85 | -0.897 | -0.006 | 0.026 | 0.026 | 0.206 | 0.62 | 0.73 | 0.63 | 0.93 | 0.63 | 0.62 |
| 7 | 43474285 | rs1860754 | -0.396 | -0.028 | -1.355 | -0.616 | -0.007 | 0.01 | 0.01 | 0 | 0.55 | 0.39 | 0.48 | 0.35 | 0.52 | 0.55 |
| 7 | 43475901 | rs9655442 | 0.967 | 0.073 | -1.436 | -0.595 | -0.006 | 0.041 | 0.041 | 0 | 0.26 | 0.14 | 0.16 | 0.08 | 0.25 | 0.29 |
| 7 | 43480199 | rs2240984 | -0.031 | 0.046 | -1.977 | -0.32 | 0 | -0.041 | -0.041 | 0.072 | 0.22 | 0.08 | 0.25 | 0.22 | 0.23 | 0.25 |
| 7 | 43485248 | rs6960889 | 0.009 | -0.971 | -1.838 | -0.059 | 0.012 | 0.004 | 0.004 | 0 | 0.77 | 0.85 | 0.82 | 0.87 | 0.8 | 0.75 |
| 7 | 43486413 | rs13222216 | 0.312 | -1.412 | -1.864 | -0.208 | 0 | 0 | 0 | 0.156 | 0.48 | 0.49 | 0.55 | 0.6 | 0.57 | 0.45 |
| 7 | 43488660 | rs7783180 | 0.971 | -1.215 | -2.158 | -0.683 | -0.036 | 0.159 | 0.159 | 3.521 | 0.41 | 0.11 | 0.26 | 0.18 | 0.39 | 0.42 |
| 7 | 43490137 | rs4724206 | 0.189 | -1.307 | -2.051 | -0.516 | 0.011 | 0.006 | 0.006 | 0 | 0.22 | 0.16 | 0.26 | 0.15 | 0.2 | 0.22 |
| 7 | 43491105 | rs10253450 | -0.166 | -1.355 | -2.273 | -0.549 | 0.049 | 0.067 | 0.067 | 0.136 | 0.51 | 0.35 | 0.39 | 0.35 | 0.43 | 0.51 |
| 7 | 43492179 | rs6463191 | -0.391 | -1.233 | -2.327 | -0.564 | 0.019 | 0.075 | 0.075 | 0 | 0.48 | 0.31 | 0.38 | 0.24 | 0.42 | 0.46 |
| 7 | 43493657 | rs10232587 | -0.587 | -1.221 | -2.305 | -0.549 | 0.005 | 0.006 | 0.006 | 2.779 | 0.7 | 0.39 | 0.62 | 0.43 | 0.63 | 0.71 |
| 7 | 43495098 | rs10270105 | -0.154 | -1.223 | -2.291 | -0.773 | 0.03 | 0.012 | 0.012 | 0 | 0.27 | 0.16 | 0.21 | 0.21 | 0.21 | 0.27 |
| 7 | 43496138 | rs2304331 | 1.64 | -1.199 | -2.297 | -0.802 | 0.013 | 0.033 | 0.033 | 1.488 | 0.23 | 0.17 | 0.18 | 0.15 | 0.21 | 0.25 |
| 7 | 43502217 | rs6957396 | -0.869 | -1.169 | -2.255 | -0.782 | 0.057 | 0.053 | 0.053 | 2.204 | 0.72 | 0.56 | 0.55 | 0.39 | 0.63 | 0.71 |
| 7 | 43506158 | rs2304328 | -0.475 | -0.989 | -2.069 | -0.59 | 0.005 | 0.007 | 0.007 | 0.155 | 0.53 | 0.52 | 0.4 | 0.35 | 0.43 | 0.51 |
| 7 | 43512070 | rs13241883 | -0.184 | -1.133 | -2.067 | -0.59 | -0.031 | -0.026 | -0.026 | 0 | 0.24 | 0.39 | 0.22 | 0.16 | 0.18 | 0.13 |

| | | | | | | | | | | | | | | | | |
|---|---|---|---|---|---|---|---|---|---|---|---|---|---|---|---|---|
| 7 | 43517171 | rs6965038 | 1.346 | -0.917 | -2.174 | -0.587 | 0.025 | 0.041 | 0.041 | 0.855 | 0.25 | 0.16 | 0.18 | 0.15 | 0.2 | 0.25 |
| 7 | 43519337 | rs2304327 | 1.097 | -0.737 | -2.536 | -0.406 | 0.009 | 0.017 | 0.017 | 0 | 0.14 | 0.06 | 0.13 | 0.08 | 0.14 | 0.08 |
| 7 | 43520811 | rs10229477 | 0.556 | -0.727 | -2.589 | -0.473 | -0.036 | 0.22 | 0.22 | 0.352 | 0.5 | 0.09 | 0.35 | 0.2 | 0.44 | 0.51 |
| 7 | 43522684 | rs4433042 | 0.612 | -0.731 | -2.576 | -0.473 | -0.008 | 0.198 | 0.198 | 0.252 | 0.51 | 0.15 | 0.35 | 0.2 | 0.45 | 0.57 |
| 7 | 43536753 | rs7786662 | 0.108 | -0.668 | -2.456 | -0.331 | 0.019 | 0.02 | 0.02 | 0.074 | 0.34 | 0.24 | 0.23 | 0.22 | 0.26 | 0.38 |
| 7 | 43537812 | rs17802356 | 1.502 | -0.686 | -2.711 | -0.378 | 0.003 | 0.055 | 0.055 | 0.635 | 0.25 | 0.05 | 0.18 | 0.16 | 0.21 | 0.3 |
| 7 | 43540108 | rs17172216 | 1.009 | -0.141 | -2.417 | -0.268 | 0.008 | 0.029 | 0.029 | 0 | 0.14 | 0.07 | 0.13 | 0.07 | 0.14 | 0.08 |
| 7 | 43541932 | rs10246139 | 0.199 | -0.133 | -2.392 | -0.305 | -0.018 | 0.003 | 0.003 | 0.077 | 0.6 | 0.32 | 0.42 | 0.38 | 0.48 | 0.68 |
| 7 | 43544507 | rs17151028 | 0.723 | -0.07 | -2.429 | -0.318 | 0.018 | -0.002 | -0.002 | 0 | 0.37 | 0.33 | 0.28 | 0.38 | 0.28 | 0.5 |
| 7 | 43549041 | rs11767004 | -0.616 | 0.228 | -2.116 | -0.013 | 0.043 | 0.055 | 0.055 | 0.733 | 0.74 | 0.43 | 0.56 | 0.47 | 0.63 | 0.75 |
| 7 | 43549242 | rs17172220 | 0.077 | 0.047 | -2.218 | 0.07 | 0.01 | 0.029 | 0.029 | 1.315 | 0.26 | 0.16 | 0.17 | 0.09 | 0.21 | 0.29 |
| 7 | 43559539 | rs1025540 | 0.117 | -0.225 | -2.096 | 0.465 | 0.034 | -0.004 | -0.004 | 0.169 | 0.36 | 0.17 | 0.28 | 0.36 | 0.29 | 0.49 |
| 7 | 43560154 | rs6463197 | 0.204 | -0.433 | -1.937 | 0.961 | 0.031 | -0.028 | -0.028 | 7.587 | 0.68 | 0.27 | 0.57 | 0.58 | 0.77 | 0.78 |
| 7 | 43561814 | rs10487674 | -0.112 | -0.304 | -1.505 | 1.457 | 0.006 | -0.001 | -0.001 | 0 | 0.13 | 0.12 | 0.13 | 0.36 | 0.1 | 0.11 |
| 7 | 43562910 | rs10260214 | 0.556 | -0.279 | -1.812 | 0.943 | 0.021 | 0.022 | 0.022 | 0.087 | 0.51 | 0.64 | 0.55 | 0.43 | 0.4 | 0.48 |
| 7 | 43564142 | rs17209297 | 0.441 | 0.883 | -2.09 | 0.663 | -0.133 | 0.048 | 0.048 | 1.957 | 0.25 | 0.02 | 0.25 | 0.06 | 0.34 | 0.26 |
| 7 | 43566266 | rs17172223 | -0.62 | 0.94 | -1.66 | 1.02 | 0 | 0.038 | 0.038 | 0.873 | 0.35 | 0.23 | 0.3 | 0.26 | 0.35 | 0.29 |
| 7 | 43574713 | rs10226241 | 0.164 | 0.775 | -1.07 | 0.952 | -0.02 | 0.005 | 0.005 | 0 | 0.17 | 0.22 | 0.14 | 0.25 | 0.1 | 0.13 |
| 7 | 43575577 | rs4724213 | 0.419 | 0.805 | -1.026 | 1.03 | -0.009 | 0.01 | 0.01 | 0.965 | 0.2 | 0.17 | 0.25 | 0.06 | 0.24 | 0.16 |
| 7 | 43577715 | rs13234419 | 0.122 | 1.171 | -0.953 | 1.038 | -0.005 | -0.002 | -0.002 | 0 | 0.12 | 0.09 | 0.14 | 0.17 | 0.27 | 0.16 |
| 7 | 43579724 | rs4720449 | 0.294 | 1.115 | -0.96 | 1.039 | -0.026 | 0.033 | 0.033 | 0 | 0.17 | 0.24 | 0.22 | 0.3 | 0.1 | 0.14 |
| 7 | 43580166 | rs6952467 | -1.487 | 1.138 | -0.88 | 1.043 | -0.003 | -0.001 | -0.001 | 0 | 0.49 | 0.49 | 0.53 | 0.48 | 0.62 | 0.45 |
| 7 | 43583428 | rs11764232 | -0.496 | 1.065 | -0.891 | 1.038 | 0.001 | 0.007 | 0.007 | 0.616 | 0.35 | 0.25 | 0.38 | 0.31 | 0.34 | 0.29 |
| 7 | 43584521 | rs10245574 | -0.64 | 1.077 | -0.933 | 1.043 | -0.001 | -0.003 | -0.003 | 0.998 | 0.36 | 0.39 | 0.39 | 0.31 | 0.34 | 0.29 |
| 7 | 43592502 | rs9648531 | -0.942 | 1.132 | -0.953 | 1.03 | -0.005 | -0.003 | -0.003 | 0 | 0.29 | 0.29 | 0.29 | 0.39 | 0.35 | 0.29 |
| 7 | 43594471 | rs9638972 | 0.279 | 1.157 | -0.937 | 1.07 | -0.065 | -0.028 | -0.028 | 0 | 0.11 | 0.02 | 0.13 | 0.15 | 0.25 | 0.15 |
| 7 | 43594560 | rs2304319 | -1.205 | 0.996 | -0.95 | 1.013 | 0.028 | 0.009 | 0.009 | 0.842 | 0.47 | 0.32 | 0.53 | 0.45 | 0.6 | 0.44 |
| 7 | 43595400 | rs4720450 | -1.266 | 1.005 | -0.966 | 0.967 | 0.008 | 0.004 | 0.004 | 0.882 | 0.47 | 0.44 | 0.53 | 0.45 | 0.6 | 0.44 |
| 7 | 43598222 | rs11768401 | 1.744 | 1.044 | -1.043 | 1.324 | 0.006 | -0.023 | -0.023 | 0.002 | 0.07 | 0.03 | 0.08 | 0.16 | 0.08 | 0.07 |
| 7 | 43600227 | rs2304316 | -0.702 | 1.071 | -0.758 | 1.114 | 0.025 | 0.007 | 0.007 | 1.63 | 0.33 | 0.4 | 0.39 | 0.39 | 0.5 | 0.32 |
| 7 | 43604107 | rs12533509 | 0.254 | 0.944 | -0.421 | 0.751 | 0.026 | -0.046 | -0.046 | 0 | 0.21 | 0.36 | 0.26 | 0.05 | 0.24 | 0.15 |
| 7 | 43607828 | rs12538259 | 0.284 | 0.951 | -0.437 | 0.757 | 0.024 | -0.04 | -0.04 | 1.453 | 0.21 | 0.35 | 0.26 | 0.06 | 0.24 | 0.15 |
| 7 | 43622806 | rs10251937 | -0.823 | 1.167 | -0.063 | 0.858 | 0.012 | 0.001 | 0.001 | 0.466 | 0.34 | 0.44 | 0.48 | 0.29 | 0.38 | 0.27 |
| 7 | 43625858 | rs10248844 | 1.2 | 1.256 | -0.484 | 0.752 | -0.071 | 0.069 | 0.069 | 0.971 | 0.15 | 0.01 | 0.14 | 0.01 | 0.22 | 0.12 |
| 7 | 43643835 | rs10279856 | -1.307 | 1.144 | -0.473 | 0.866 | 0.004 | 0.004 | 0.004 | 2.641 | 0.51 | 0.47 | 0.64 | 0.62 | 0.64 | 0.44 |
| 7 | 43655211 | rs10215478 | 0.002 | 1.146 | -0.549 | 0.837 | -0.034 | 0.013 | 0.013 | 0.52 | 0.25 | 0.11 | 0.31 | 0.11 | 0.33 | 0.21 |
| 7 | 43664198 | rs10233697 | 1.606 | 1.151 | -0.56 | 0.814 | -0.071 | 0.068 | 0.068 | 1.045 | 0.15 | 0.02 | 0.14 | 0.01 | 0.22 | 0.12 |
| 7 | 43665500 | rs1044220 | 1.606 | 1.151 | -0.56 | 0.814 | -0.071 | 0.068 | 0.068 | 1.045 | 0.15 | 0.02 | 0.14 | 0.01 | 0.22 | 0.12 |
| 7 | 43665808 | rs2108411 | 0.409 | 1.187 | -0.596 | 0.969 | -0.043 | -0.079 | -0.079 | 0 | 0.74 | 0.87 | 0.68 | 0.58 | 0.63 | 0.77 |
| 7 | 43670323 | rs1057523 | -0.512 | 1.17 | -0.474 | 0.988 | 0.003 | 0.002 | 0.002 | 0.342 | 0.31 | 0.35 | 0.47 | 0.29 | 0.35 | 0.27 |
| 7 | 43670562 | rs11984341 | 1.553 | 0.983 | 0.116 | 0.9 | 0.017 | 0.016 | 0.016 | 3.303 | 0.47 | 0.39 | 0.35 | 0.38 | 0.36 | 0.56 |
| 7 | 43675674 | rs11768098 | 0.532 | 1.145 | -0.123 | 0.947 | -0.002 | -0.021 | -0.021 | 1.863 | 0.14 | 0.1 | 0.32 | 0.28 | 0.14 | 0.15 |
| 7 | 43687296 | rs2240516 | 1.283 | 1.149 | -0.099 | 0.944 | 0.006 | 0.003 | 0.003 | 0 | 0.65 | 0.62 | 0.5 | 0.39 | 0.57 | 0.69 |
| 7 | 43690266 | rs12669280 | 1.655 | 1.179 | -0.354 | 0.936 | 0.002 | 0.011 | 0.011 | 0 | 0.06 | 0.09 | 0.15 | 0.18 | 0.06 | 0.08 |
| 7 | 43700346 | rs10281917 | 1.644 | 1.168 | -0.384 | 0.891 | -0.071 | 0.068 | 0.068 | 1.045 | 0.15 | 0.01 | 0.14 | 0.01 | 0.22 | 0.12 |
| 7 | 43709735 | rs10231164 | 1.644 | 1.168 | -0.384 | 0.891 | -0.071 | 0.068 | 0.068 | 1.045 | 0.15 | 0.02 | 0.14 | 0.01 | 0.21 | 0.12 |
| 7 | 43713856 | rs9639884 | -0.682 | 1.194 | -0.403 | 0.938 | -0.061 | -0.07 | -0.07 | 1.298 | 0.83 | 0.96 | 0.85 | 0.67 | 0.75 | 0.84 |
| 7 | 43738921 | rs3807517 | 0.846 | 0.869 | -1.047 | 0.627 | -0.01 | -0.005 | -0.005 | 0.052 | 0.12 | 0.09 | 0.19 | 0.18 | 0.11 | 0.16 |
| 7 | 43755241 | rs10245456 | 1.56 | 0.791 | -0.841 | 0.824 | 0.001 | -0.005 | -0.005 | 0.01 | 0.09 | 0.09 | 0.17 | 0.1 | 0.12 | 0.06 |
| 7 | 43765667 | rs849172 | 0.765 | 0.783 | -0.848 | 0.814 | -0.033 | -0.081 | -0.081 | 0 | 0.58 | 0.85 | 0.54 | 0.57 | 0.41 | 0.64 |
| 7 | 43781905 | rs17724202 | 1.638 | 0.785 | -0.809 | 0.829 | 0.001 | -0.014 | -0.014 | 0 | 0.09 | 0.06 | 0.17 | 0.1 | 0.12 | 0.06 |
| 7 | 43789148 | rs1181597 | 0.128 | 0.656 | -0.916 | 0.747 | -0.136 | -0.041 | -0.041 | 0 | 0.68 | 0.91 | 0.71 | 0.66 | 0.53 | 0.71 |
| 7 | 43797453 | rs1637530 | -0.731 | 0.768 | -0.902 | 0.794 | -0.063 | -0.05 | -0.05 | 2.038 | 0.83 | 0.96 | 0.85 | 0.71 | 0.75 | 0.84 |
| 7 | 43801983 | rs10486752 | 0.41 | 0.267 | -1.196 | 0.359 | 0.014 | -0.007 | -0.007 | 0.228 | 0.16 | 0.11 | 0.2 | 0.23 | 0.12 | 0.18 |
| 7 | 43804695 | rs1181574 | 0.254 | 0.431 | -1.101 | 0.458 | -0.052 | -0.07 | -0.07 | 2.157 | 0.76 | 0.87 | 0.69 | 0.51 | 0.69 | 0.75 |
| 7 | 43805932 | rs10229363 | 1.708 | 0.443 | -1.176 | 0.421 | -0.069 | 0.068 | 0.068 | 1.045 | 0.15 | 0.01 | 0.13 | 0.01 | 0.21 | 0.13 |
| 7 | 43810764 | rs699512 | -0.866 | 0.451 | -1.112 | 0.477 | -0.063 | -0.067 | -0.067 | 0 | 0.83 | 0.96 | 0.85 | 0.68 | 0.75 | 0.84 |
| 7 | 43821453 | rs849161 | 1.033 | 0.202 | -0.726 | 0.463 | 0.048 | 0.034 | 0.034 | 1.944 | 0.5 | 0.63 | 0.37 | 0.41 | 0.36 | 0.56 |
| 7 | 43827414 | rs849165 | 0.372 | 0.203 | -0.703 | 0.468 | -0.006 | -0.016 | -0.016 | 0 | 0.59 | 0.68 | 0.54 | 0.49 | 0.48 | 0.62 |
| 7 | 43838444 | rs17246016 | 1.371 | 0.238 | -0.725 | 0.504 | 0.001 | -0.005 | -0.005 | 0 | 0.1 | 0.08 | 0.17 | 0.08 | 0.12 | 0.06 |
| 7 | 43843423 | rs1131372 | 1.612 | 0.274 | -0.759 | 0.521 | 0 | -0.023 | -0.023 | 1.052 | 0.09 | 0.05 | 0.17 | 0.08 | 0.12 | 0.06 |
| 7 | 43847709 | rs589570 | 0.757 | 0.2 | -0.649 | 0.485 | 0.043 | 0.031 | 0.031 | 2.841 | 0.5 | 0.62 | 0.37 | 0.4 | 0.35 | 0.57 |
| 7 | 43863684 | rs2528366 | 1.944 | -0.031 | -0.9 | 0.358 | -0.013 | -0.031 | -0.031 | 0 | 0.36 | 0.13 | 0.3 | 0.37 | 0.31 | 0.45 |
| 7 | 43864699 | rs623108 | 0.962 | 0.507 | -0.433 | 0.51 | 0.057 | 0.031 | 0.031 | 2.297 | 0.49 | 0.65 | 0.36 | 0.43 | 0.33 | 0.56 |
| 7 | 43865459 | rs10250233 | 0.621 | 0.816 | -0.341 | 0.447 | 0 | 0.007 | 0.007 | 0 | 0.07 | 0.14 | 0.15 | 0.18 | 0.06 | 0.09 |
| 7 | 43877983 | rs586475 | -0.523 | 1.094 | -0.549 | 0.48 | -0.035 | -0.05 | -0.05 | 10.021 | 0.83 | 0.93 | 0.85 | 0.7 | 0.78 | 0.84 |
| 7 | 43879274 | rs1181531 | 1.098 | 0.58 | -0.388 | 0.352 | 0 | 0 | 0 | 0.009 | 0.51 | 0.49 | 0.51 | 0.51 | 0.49 | 0.59 |
| 7 | 43890379 | rs604060 | 0.015 | 1.054 | -0.729 | 0.677 | -0.036 | -0.039 | -0.039 | 2.652 | 0.82 | 0.93 | 0.83 | 0.71 | 0.74 | 0.84 |
| 7 | 43905193 | rs10224322 | 0.067 | 1.016 | -0.864 | 0.15 | -0.028 | 0.052 | 0.052 | 0.02 | 0.12 | 0.01 | 0.09 | 0.01 | 0.13 | 0.11 |

| | | | | | | | | | | | | | | | | |
|---|---|---|---|---|---|---|---|---|---|---|---|---|---|---|---|---|
| 7 | 43910682 | rs2730610 | 0.97 | 0.829 | -0.778 | 0.146 | -0.027 | 0.026 | 0.026 | 0 | 0.62 | 0.79 | 0.62 | 0.71 | 0.55 | 0.66 |
| 7 | 43916491 | rs2232109 | 0.239 | 0.652 | 0.207 | 0.06 | -0.02 | 0.061 | 0.061 | 0.037 | 0.13 | 0.03 | 0.09 | 0.01 | 0.13 | 0.12 |
| 7 | 43916727 | rs2232108 | 1.505 | 0.793 | 0.631 | 0.093 | 0.003 | 0.012 | 0.012 | 0 | 0.52 | 0.39 | 0.52 | 0.58 | 0.46 | 0.65 |
| 7 | 43917013 | rs2232105 | 1.526 | 0.757 | 0.892 | 0.085 | 0.006 | 0.024 | 0.024 | 0 | 0.52 | 0.33 | 0.52 | 0.58 | 0.46 | 0.65 |
| 7 | 43917274 | rs2232103 | -0.162 | 0.83 | 0.818 | 0.114 | -0.007 | -0.003 | -0.003 | 0 | 0.75 | 0.8 | 0.71 | 0.73 | 0.68 | 0.78 |
| 7 | 43930775 | rs4724254 | -0.217 | 0.811 | 0.786 | 0.254 | -0.007 | -0.007 | -0.007 | 0 | 0.05 | 0.06 | 0.16 | 0.06 | 0.14 | 0.06 |
| 7 | 43932337 | rs2528392 | 1.492 | 0.757 | 0.892 | 0.232 | 0.004 | -0.001 | -0.001 | 0 | 0.5 | 0.48 | 0.37 | 0.51 | 0.32 | 0.6 |
| 7 | 43932910 | rs2956425 | 1.615 | 0.74 | 0.931 | 0.224 | 0.059 | -0.01 | -0.01 | 0.793 | 0.49 | 0.27 | 0.36 | 0.51 | 0.32 | 0.6 |
| 7 | 43935301 | rs10248914 | 0.96 | 0.838 | 0.783 | 0.306 | 0 | 0.004 | 0.004 | 0.944 | 0.07 | 0.14 | 0.07 | 0.12 | 0.07 | 0.01 |
| 7 | 43943717 | rs11768707 | -0.729 | 0.848 | 0.757 | 0.309 | 0.042 | 0.039 | 0.039 | 0 | 0.06 | 0.18 | 0.18 | 0.06 | 0.14 | 0.06 |
| 7 | 43965747 | rs2528382 | 0.569 | 0.807 | 0.775 | 0.302 | 0.018 | 0.027 | 0.027 | 0.722 | 0.08 | 0.25 | 0.09 | 0.12 | 0.07 | 0.01 |
| 7 | 43969408 | rs2079390 | -0.088 | 0.594 | 0.418 | 0.141 | -0.003 | -0.001 | -0.001 | 0 | 0.7 | 0.74 | 0.7 | 0.69 | 0.65 | 0.78 |
| 7 | 43969657 | rs17724605 | 1.707 | 0.491 | 0.47 | 0.093 | -0.003 | 0 | 0 | 0.294 | 0.47 | 0.48 | 0.37 | 0.51 | 0.32 | 0.59 |
| 7 | 43981034 | rs2527811 | 1.35 | 0.588 | 0.456 | 0.095 | 0.002 | 0.001 | 0.001 | 0.267 | 0.51 | 0.56 | 0.44 | 0.6 | 0.37 | 0.6 |
| 7 | 43984254 | rs2527813 | 0.951 | 0.649 | 0.443 | 0.09 | -0.009 | 0.02 | 0.02 | 0.111 | 0.57 | 0.68 | 0.61 | 0.66 | 0.52 | 0.66 |
| 7 | 43985345 | rs10229021 | 0.598 | 0.893 | 0.416 | 0.152 | -0.03 | 0.059 | 0.059 | 0.036 | 0.13 | 0.01 | 0.11 | 0.01 | 0.13 | 0.06 |
| 7 | 43989377 | rs11770228 | -0.125 | 0.909 | 0.493 | 0.184 | 0.017 | 0.016 | 0.016 | 0 | 0.05 | 0.09 | 0.16 | 0.06 | 0.14 | 0.06 |
| 7 | 43989395 | rs6463211 | 1.557 | 0.11 | 0.182 | -0.022 | 0.004 | 0.005 | 0.005 | 0.896 | 0.41 | 0.52 | 0.51 | 0.52 | 0.45 | 0.55 |
| 7 | 43997075 | rs1894752 | 0.635 | -0.045 | 0.364 | 0.117 | 0.003 | 0.002 | 0.002 | 4.147 | 0.58 | 0.62 | 0.7 | 0.65 | 0.64 | 0.61 |
| 7 | 44082242 | rs10228823 | -1.615 | 0.54 | 0.938 | -0.387 | -0.01 | -0.021 | -0.021 | 3.276 | 0.24 | 0.35 | 0.28 | 0.11 | 0.24 | 0.2 |
| 7 | 44083774 | rs17551738 | -1.053 | 0.535 | 0.946 | -0.391 | -0.041 | -0.048 | -0.048 | 0 | 0.19 | 0.3 | 0.12 | 0.05 | 0.1 | 0.14 |
| 7 | 44092723 | rs10951755 | -0.153 | 0.74 | 1.027 | -0.212 | -0.007 | 0.001 | 0.001 | 14.194 | 0.15 | 0.34 | 0.19 | 0.1 | 0.15 | 0.17 |
| 7 | 44095999 | rs11766481 | -0.122 | 0.739 | 0.933 | -0.213 | -0.001 | -0.001 | -0.001 | 0 | 0.14 | 0.17 | 0.18 | 0.1 | 0.15 | 0.18 |
| 7 | 44102037 | rs11767165 | -1.169 | 0.735 | 0.993 | -0.252 | -0.013 | -0.028 | -0.028 | 0 | 0.22 | 0.43 | 0.28 | 0.11 | 0.23 | 0.2 |
| 7 | 44111189 | rs28745596 | 0.824 | 0.725 | 0.953 | -0.252 | -0.015 | 0.019 | 0.019 | 0 | 0.07 | 0.01 | 0.08 | 0.01 | 0.08 | 0.03 |
| 7 | 44112529 | rs7778745 | -1.155 | 0.728 | 0.965 | -0.251 | -0.011 | -0.028 | -0.028 | 0 | 0.22 | 0.43 | 0.28 | 0.11 | 0.23 | 0.21 |
| 7 | 44112873 | rs3218660 | -1.056 | 0.729 | 0.711 | -0.251 | -0.007 | -0.017 | -0.017 | 0 | 0.21 | 0.31 | 0.27 | 0.11 | 0.23 | 0.21 |
| 7 | 44114251 | rs6955679 | -0.114 | 0.746 | 0.055 | -0.244 | -0.001 | -0.001 | -0.001 | 0.331 | 0.14 | 0.17 | 0.18 | 0.1 | 0.15 | 0.18 |
| 7 | 44115429 | rs1139270 | -0.034 | 0.903 | -0.695 | -0.203 | -0.003 | -0.003 | -0.003 | 1.545 | 0.25 | 0.21 | 0.2 | 0.1 | 0.32 | 0.24 |
| 7 | 44121115 | rs4640970 | 1.431 | 0.79 | -0.823 | -0.026 | -0.005 | 0.082 | 0.082 | 1.97 | 0.45 | 0.14 | 0.44 | 0.68 | 0.37 | 0.54 |
| 7 | 44127371 | rs4724284 | 0.53 | 1.396 | -0.527 | 0.122 | 0.001 | -0.01 | -0.01 | 0 | 0.7 | 0.68 | 0.74 | 0.8 | 0.63 | 0.8 |
| 7 | 44134848 | rs4236374 | -0.682 | 1.301 | -0.919 | 0.862 | -0.08 | -0.081 | -0.081 | 0.87 | 0.16 | 0.02 | 0.26 | 0.22 | 0.26 | 0.08 |
| 7 | 44134894 | rs9784990 | -0.772 | 1.155 | -0.448 | 0.794 | -0.014 | 0.04 | 0.04 | 3.846 | 0.35 | 0.53 | 0.42 | 0.5 | 0.33 | 0.42 |
| 7 | 44143341 | rs10271244 | -0.53 | 1.118 | -0.491 | 1.019 | -0.081 | -0.076 | -0.076 | 0 | 0.17 | 0.02 | 0.29 | 0.21 | 0.28 | 0.07 |
| 7 | 44155252 | rs2075070 | -1.41 | 0.825 | 0.367 | 0.711 | -0.003 | 0.012 | 0.012 | 0.239 | 0.4 | 0.37 | 0.39 | 0.26 | 0.46 | 0.31 |
| 7 | 44159224 | rs2908280 | -1.938 | 0.698 | 0.886 | 0.986 | -0.003 | 0.053 | 0.053 | 0.656 | 0.48 | 0.76 | 0.65 | 0.62 | 0.52 | 0.5 |
| 7 | 44161809 | rs3087370 | -0.469 | 0.742 | 1.161 | 1.385 | -0.008 | -0.006 | -0.006 | 0 | 0.19 | 0.19 | 0.3 | 0.22 | 0.28 | 0.08 |
| 7 | 44162355 | rs2979422 | -0.703 | 0.591 | 1.216 | 1.281 | -0.021 | -0.004 | -0.004 | 0 | 0.22 | 0.31 | 0.21 | 0.19 | 0.12 | 0.2 |
| 7 | 44171471 | rs2971684 | -1.928 | 0.67 | 1.131 | 1.219 | 0.001 | 0.005 | 0.005 | 0.693 | 0.44 | 0.66 | 0.62 | 0.6 | 0.46 | 0.43 |
| 7 | 44177191 | rs17136076 | -0.677 | 0.927 | 1.309 | 1.345 | 0.016 | 0.012 | 0.012 | 0 | 0.19 | 0.29 | 0.32 | 0.22 | 0.28 | 0.08 |
| 7 | 44177412 | rs17769733 | -0.293 | 0.938 | 1.026 | 1.35 | -0.085 | -0.076 | -0.076 | 0 | 0.17 | 0.02 | 0.3 | 0.22 | 0.28 | 0.08 |
| 7 | 44178829 | rs882019 | -1.724 | 0.72 | 1.194 | 1.232 | -0.001 | 0 | 0 | 1.776 | 0.4 | 0.66 | 0.62 | 0.61 | 0.46 | 0.41 |
| 7 | 44183433 | rs2908277 | -0.348 | 0.659 | 1.194 | 1.405 | -0.008 | 0.007 | 0.007 | 0 | 0.15 | 0.29 | 0.19 | 0.17 | 0.12 | 0.18 |
| 7 | 44189023 | rs2268576 | -1.694 | 1.088 | 0.98 | 1.407 | -0.03 | 0.037 | 0.037 | 1.037 | 0.44 | 0.78 | 0.62 | 0.62 | 0.51 | 0.43 |
| 7 | 44189274 | rs2268575 | -0.344 | 1.027 | 0.839 | 1.275 | 0.009 | 0.007 | 0.007 | 3.906 | 0.23 | 0.16 | 0.25 | 0.18 | 0.19 | 0.22 |
| 7 | 44193404 | rs2908297 | -1.578 | 1.062 | 0.959 | 1.356 | 0.019 | 0.088 | 0.088 | 1.987 | 0.15 | 0.61 | 0.21 | 0.25 | 0.17 | 0.2 |
| 7 | 44204322 | rs2080033 | -1.472 | 0.858 | 0.259 | 1.28 | -0.004 | 0.094 | 0.094 | 1.744 | 0.48 | 0.82 | 0.62 | 0.65 | 0.54 | 0.41 |
| 7 | 44211337 | rs2971671 | -0.269 | 1.127 | 0.49 | 1.281 | -0.004 | 0.006 | 0.006 | 7.484 | 0.24 | 0.2 | 0.26 | 0.16 | 0.26 | 0.22 |
| 7 | 44216137 | rs2908290 | -1.269 | 1.146 | 0.603 | 1.341 | 0.013 | 0.033 | 0.033 | 5.143 | 0.34 | 0.57 | 0.43 | 0.45 | 0.41 | 0.35 |
| 7 | 44218856 | rs2300586 | -1.189 | 1.175 | 0.329 | 1.504 | 0.05 | 0.159 | 0.159 | 0 | 0.09 | 0.28 | 0.16 | 0.29 | 0.15 | 0.12 |
| 7 | 44221581 | rs758988 | 0.283 | 0.454 | -0.676 | 0.633 | 0.014 | 0.029 | 0.029 | 0 | 0.82 | 0.86 | 0.88 | 0.93 | 0.86 | 0.87 |
| 7 | 44222471 | rs758985 | 0.328 | 0.507 | -0.902 | 0.783 | 0.003 | 0.008 | 0.008 | 0.018 | 0.81 | 0.92 | 0.85 | 0.84 | 0.82 | 0.85 |
| 7 | 44222802 | rs1990458 | 1.185 | 0.649 | -1.278 | 0.753 | -0.028 | 0.002 | 0.002 | 0.589 | 0.33 | 0.18 | 0.41 | 0.34 | 0.39 | 0.46 |
| 7 | 44227020 | rs2268569 | 0.091 | 0.797 | -0.584 | 0.95 | 0.055 | 0.13 | 0.13 | 0 | 0.07 | 0.22 | 0.07 | 0.18 | 0.08 | 0.12 |
| 7 | 44229232 | rs12702070 | 1.674 | 0.492 | -0.943 | 0.922 | 0.004 | -0.026 | -0.026 | 0 | 0.09 | 0.01 | 0.04 | 0.07 | 0.06 | 0.05 |
| 7 | 44231216 | rs3757840 | 0.865 | 0.447 | -0.68 | 0.915 | -0.02 | -0.011 | -0.011 | 0 | 0.45 | 0.26 | 0.46 | 0.48 | 0.47 | 0.53 |
| 7 | 44235668 | rs4607517 | 0.919 | 0.481 | -0.89 | 0.747 | -0.008 | -0.007 | -0.007 | 0 | 0.21 | 0.07 | 0.18 | 0.19 | 0.18 | 0.13 |
| 7 | 44240407 | rs1004558 | 0.656 | 0.508 | -0.789 | 0.758 | -0.003 | -0.003 | -0.003 | 0 | 0.23 | 0.3 | 0.29 | 0.19 | 0.18 | 0.13 |
| 7 | 44243953 | rs1476892 | 0.468 | 0.445 | -1.093 | 0.536 | 0.004 | 0.025 | 0.025 | 0.078 | 0.76 | 0.81 | 0.73 | 0.84 | 0.73 | 0.78 |
| 7 | 44250826 | rs2072439 | 0.404 | 0.582 | -1.198 | 0.557 | -0.01 | 0.012 | 0.012 | 0.152 | 0.76 | 0.91 | 0.75 | 0.83 | 0.76 | 0.77 |
| 7 | 44263028 | rs2075067 | -0.149 | 0.536 | -0.785 | 0.568 | 0.063 | 0.078 | 0.078 | 0.14 | 0.09 | 0.24 | 0.09 | 0.17 | 0.16 | 0.15 |
| 7 | 44263456 | rs1003573 | 0.659 | 0.341 | -1.042 | 0.428 | -0.004 | 0.001 | 0.001 | 0.05 | 0.6 | 0.57 | 0.63 | 0.6 | 0.59 | 0.65 |
| 7 | 44276691 | rs10253419 | -0.096 | 0.399 | -2.087 | -0.322 | -0.068 | 0.127 | 0.127 | 0 | 0.27 | 0.02 | 0.19 | 0.04 | 0.29 | 0.15 |
| 7 | 44277705 | rs3757836 | -0.473 | 0.21 | -1.915 | -0.283 | -0.007 | -0.002 | -0.002 | 0 | 0.1 | 0.13 | 0.08 | 0.1 | 0.05 | 0.11 |
| 7 | 44280948 | rs2075074 | -0.899 | -0.226 | -2.177 | -0.409 | 0.047 | 0.148 | 0.148 | 0 | 0.48 | 0.15 | 0.29 | 0.22 | 0.41 | 0.39 |
| 7 | 44288665 | rs10441113 | -0.421 | 0.257 | -2.169 | -0.571 | -0.001 | 0.162 | 0.162 | 0 | 0.39 | 0.14 | 0.25 | 0.09 | 0.4 | 0.26 |
| 7 | 44296307 | rs4526269 | -0.332 | 0.202 | -2.24 | -0.853 | -0.034 | 0.244 | 0.244 | 0.073 | 0.38 | 0.06 | 0.24 | 0.04 | 0.4 | 0.27 |
| 7 | 44301371 | rs35397936 | -0.426 | 0.06 | -1.976 | -0.768 | 0.006 | 0.126 | 0.126 | 0.105 | 0.39 | 0.2 | 0.25 | 0.06 | 0.39 | 0.27 |
| 7 | 44306081 | rs7796267 | 0.022 | -0.047 | -2.019 | -0.805 | 0.007 | 0.088 | 0.088 | 0 | 0.29 | 0.16 | 0.18 | 0.06 | 0.28 | 0.17 |

| | | | | | | | | | | | | | | | | |
|---|---|---|---|---|---|---|---|---|---|---|---|---|---|---|---|---|
| 7 | 44310945 | rs10230538 | 0.365 | -0.036 | -2.018 | -0.807 | -0.006 | 0.102 | 0.102 | 0.049 | 0.27 | 0.12 | 0.17 | 0.03 | 0.27 | 0.17 |
| 7 | 44313930 | rs12702074 | -0.414 | 0.083 | -1.823 | -0.82 | 0.023 | 0.154 | 0.154 | 0 | 0.43 | 0.25 | 0.26 | 0.07 | 0.4 | 0.28 |
| 7 | 44315541 | rs4642534 | 0.832 | -0.015 | -1.452 | -0.167 | -0.004 | 0.008 | 0.008 | 0 | 0.45 | 0.61 | 0.49 | 0.61 | 0.52 | 0.57 |
| 7 | 44316168 | rs10267121 | 0.389 | 0.083 | -1.384 | -0.039 | -0.016 | 0.135 | 0.135 | 0.181 | 0.9 | 0.79 | 0.78 | 0.67 | 0.95 | 0.85 |
| 7 | 44320434 | rs4724298 | 1.154 | 1.187 | -2.217 | -0.758 | -0.061 | -0.052 | -0.052 | 0 | 0.09 | 0.01 | 0.15 | 0.21 | 0.2 | 0.18 |
| 7 | 44327055 | rs10231886 | -0.184 | 0.336 | -1.917 | -1.358 | 0.015 | 0.219 | 0.219 | 0 | 0.43 | 0.19 | 0.26 | 0.06 | 0.4 | 0.29 |
| 7 | 44339406 | rs13246801 | -0.672 | 0.125 | -1.587 | -1.342 | 0.007 | 0.028 | 0.028 | 0 | 0.52 | 0.46 | 0.33 | 0.1 | 0.45 | 0.3 |
| 7 | 44342440 | rs10215216 | 0.134 | 0.016 | -1.671 | -1.435 | -0.022 | 0.149 | 0.149 | 0 | 0.34 | 0.16 | 0.23 | 0.06 | 0.36 | 0.26 |
| 7 | 44342969 | rs13234819 | -0.442 | 0.081 | -1.654 | -1.378 | 0.001 | -0.028 | -0.028 | 0 | 0.42 | 0.46 | 0.29 | 0.06 | 0.4 | 0.27 |
| 7 | 44353609 | rs7804804 | -0.619 | 0.34 | -1.028 | -0.653 | -0.003 | 0.022 | 0.022 | 0 | 0.07 | 0.2 | 0.07 | 0.05 | 0.04 | 0.01 |
| 7 | 44361910 | rs9918696 | 0.505 | 0.459 | -1.362 | -1.086 | 0.003 | 0.013 | 0.013 | 0 | 0.49 | 0.55 | 0.65 | 0.9 | 0.54 | 0.59 |
| 7 | 44367292 | rs35705461 | 0.494 | 0.33 | -1.908 | -1.21 | 0.016 | 0.126 | 0.126 | 0.124 | 0.31 | 0.12 | 0.2 | 0.05 | 0.26 | 0.3 |
| 7 | 44369065 | rs6962696 | -0.263 | 0.19 | -1.611 | -1.085 | -0.007 | -0.031 | -0.031 | 0 | 0.16 | 0.21 | 0.17 | 0.03 | 0.14 | 0.11 |
| 7 | 44371637 | rs6974322 | 0.538 | 0.424 | -1.942 | -0.683 | -0.006 | 0.005 | 0.005 | 0 | 0.4 | 0.64 | 0.36 | 0.62 | 0.4 | 0.39 |
| 7 | 44374331 | rs11764828 | 0.43 | 0.285 | -1.745 | -0.781 | 0 | 0.057 | 0.057 | 0 | 0.11 | 0.01 | 0.11 | 0.01 | 0.1 | 0.1 |
| 7 | 44376833 | rs12535503 | 0.227 | 0.581 | -1.889 | -1.451 | -0.018 | 0.042 | 0.042 | 0 | 0.4 | 0.27 | 0.32 | 0.07 | 0.44 | 0.3 |
| 7 | 44377943 | rs7797047 | -0.708 | 0.193 | -1.033 | -1.264 | -0.008 | -0.055 | -0.055 | 0 | 0.2 | 0.37 | 0.15 | 0.09 | 0.19 | 0.1 |
| 7 | 44412735 | rs11762603 | 0.466 | 0.161 | -1.374 | -1.325 | -0.008 | 0.028 | 0.028 | 0 | 0.13 | 0.07 | 0.08 | 0.01 | 0.16 | 0.08 |
| 7 | 44420901 | rs11772858 | 0.73 | 0.121 | -1.437 | -1.358 | 0.017 | 0.028 | 0.028 | 0 | 0.07 | 0.01 | 0.02 | 0 | 0.06 | 0.05 |
| 7 | 44424983 | rs17711306 | -0.136 | -0.041 | -1.551 | -1.474 | -0.024 | 0.084 | 0.084 | 0 | 0.15 | 0.02 | 0.08 | 0.01 | 0.17 | 0.11 |
| 7 | 44430251 | rs2331071 | 1.441 | -0.212 | -2.398 | -1.465 | -0.12 | -0.027 | -0.027 | 0 | 0.22 | 0.01 | 0.23 | 0.26 | 0.35 | 0.28 |
| 7 | 44442296 | rs1473114 | -2.053 | -0.464 | -2.458 | -1.759 | -0.108 | 0.117 | 0.117 | 0.762 | 0.75 | 0.95 | 0.79 | 0.93 | 0.71 | 0.77 |
| 7 | 44443455 | rs11770266 | 0.514 | -0.059 | -2.237 | -1.503 | 0.024 | 0.046 | 0.046 | 0 | 0.1 | 0.01 | 0.03 | 0.01 | 0.06 | 0.08 |
| 7 | 44444122 | rs11550029 | 0.661 | -0.016 | -2.19 | -1.45 | -0.024 | 0.084 | 0.084 | 0 | 0.15 | 0.02 | 0.08 | 0.01 | 0.17 | 0.11 |
| 7 | 44466729 | rs2240100 | 0.755 | -0.368 | -1.791 | -0.8 | -0.007 | -0.021 | -0.021 | 0.101 | 0.37 | 0.29 | 0.53 | 0.52 | 0.43 | 0.46 |
| 7 | 44552209 | rs3187907 | -0.538 | -0.013 | -1.427 | -1.199 | -0.026 | 0.105 | 0.105 | 0 | 0.18 | 0.02 | 0.1 | 0.01 | 0.18 | 0.13 |
| 7 | 44553238 | rs217434 | -0.913 | -0.017 | -1.25 | -1.201 | 0 | -0.027 | -0.027 | 0 | 0.19 | 0.15 | 0.12 | 0.01 | 0.18 | 0.12 |
| 7 | 44553826 | rs217432 | -1.74 | -0.037 | -1.231 | -1.219 | -0.007 | -0.082 | -0.082 | 0 | 0.24 | 0.28 | 0.15 | 0.01 | 0.2 | 0.14 |
| 7 | 44555081 | rs217430 | -0.99 | 0.304 | -1.381 | -1.166 | -0.003 | -0.026 | -0.026 | 0 | 0.19 | 0.15 | 0.14 | 0.01 | 0.22 | 0.12 |
| 7 | 44555406 | rs10264715 | -0.059 | 0.446 | -1.488 | -1.043 | -0.06 | 0.082 | 0.082 | 0 | 0.15 | 0.03 | 0.1 | 0.01 | 0.21 | 0.09 |
| 7 | 44561600 | rs217421 | -0.543 | 0.221 | -1.362 | -0.968 | 0.006 | -0.068 | -0.068 | 0.269 | 0.21 | 0.25 | 0.14 | 0.01 | 0.23 | 0.12 |
| 7 | 44570067 | rs11763759 | 0.772 | 0.468 | -1.742 | -0.494 | -0.07 | 0.125 | 0.125 | 1.874 | 0.25 | 0.02 | 0.18 | 0.04 | 0.29 | 0.22 |
| 7 | 44580991 | rs17655652 | 0.96 | 0.572 | -1.667 | -0.485 | -0.061 | 0.165 | 0.165 | 2.706 | 0.26 | 0.03 | 0.24 | 0.01 | 0.29 | 0.23 |
| 7 | 44581986 | rs17725246 | -0.869 | 1.092 | -0.811 | 0.603 | 0.025 | 0.096 | 0.096 | 0 | 0.15 | 0.24 | 0.28 | 0.38 | 0.2 | 0.19 |
| 7 | 44599468 | rs217388 | -1.011 | 0.072 | -1.326 | 0.225 | -0.098 | 0.098 | 0.098 | 0.544 | 0.68 | 0.94 | 0.64 | 0.84 | 0.59 | 0.71 |
| 7 | 44600376 | rs17798255 | -0.108 | -0.306 | -0.517 | 0.444 | -0.027 | 0.3 | 0.3 | 0 | 0.09 | 0.31 | 0.05 | 0.45 | 0.03 | 0.28 |
| 7 | 44602187 | rs217385 | 0.854 | -0.354 | -0.406 | 0.335 | -0.002 | -0.084 | -0.084 | 0 | 0.45 | 0.43 | 0.31 | 0.02 | 0.41 | 0.28 |
| 7 | 44602589 | rs7791240 | 0.527 | -0.344 | -0.466 | 0.513 | -0.026 | -0.025 | -0.025 | 0 | 0.1 | 0 | 0.08 | 0.12 | 0.08 | 0.08 |
| 7 | 44604017 | rs217384 | -1.146 | -0.405 | -0.445 | 0.533 | -0.116 | 0.106 | 0.106 | 0.377 | 0.67 | 0.93 | 0.63 | 0.84 | 0.57 | 0.7 |
| 7 | 44606184 | rs2304372 | 0.443 | -0.474 | -0.289 | 0.559 | -0.023 | 0.199 | 0.199 | 0 | 0.07 | 0.18 | 0.05 | 0.44 | 0.03 | 0.28 |
| 7 | 44617748 | rs1055967 | 0.431 | -0.433 | -0.295 | 0.564 | -0.022 | 0.187 | 0.187 | 0 | 0.07 | 0.17 | 0.05 | 0.45 | 0.03 | 0.28 |
| 7 | 44619302 | rs217361 | -1.274 | -0.394 | -0.358 | 0.464 | -0.098 | 0.098 | 0.098 | 0.544 | 0.68 | 0.94 | 0.64 | 0.84 | 0.59 | 0.71 |
| 7 | 44620836 | rs8580 | -1.295 | -0.398 | -0.405 | 0.462 | -0.126 | 0.092 | 0.092 | 0.465 | 0.69 | 0.96 | 0.64 | 0.84 | 0.59 | 0.71 |
| 7 | 44626377 | rs217355 | 0.501 | -0.181 | -0.222 | 0.434 | 0 | -0.098 | -0.098 | 0 | 0.44 | 0.43 | 0.32 | 0.02 | 0.44 | 0.27 |
| 7 | 44643802 | rs10236105 | -0.522 | -0.214 | -0.233 | 0.52 | 0 | -0.003 | -0.003 | 0.184 | 0.54 | 0.49 | 0.51 | 0.73 | 0.52 | 0.66 |
| 7 | 44646185 | rs17551563 | 0.481 | -0.179 | -0.302 | 0.556 | -0.011 | -0.026 | -0.026 | 0 | 0.1 | 0 | 0.07 | 0.12 | 0.07 | 0.08 |
| 7 | 44669698 | rs740094 | 0.178 | -0.164 | -0.209 | 0.457 | -0.005 | 0.003 | 0.003 | 0.47 | 0.43 | 0.37 | 0.42 | 0.61 | 0.45 | 0.57 |
| 7 | 44680228 | rs17133676 | 0.839 | -0.55 | -0.285 | 0.672 | -0.003 | 0.35 | 0.35 | 0 | 0.05 | 0.23 | 0.06 | 0.4 | 0.02 | 0.2 |
| 7 | 44686497 | rs757702 | -1.272 | -0.672 | 0.094 | 0.617 | -0.041 | 0.316 | 0.316 | 1.914 | 0.1 | 0.6 | 0.09 | 0.4 | 0.02 | 0.2 |
| 7 | 44693897 | rs10951768 | -1.679 | -0.665 | 0.035 | 0.708 | -0.033 | 0.23 | 0.23 | 1.823 | 0.13 | 0.61 | 0.14 | 0.45 | 0.07 | 0.22 |
| 7 | 44705721 | rs12155014 | 0.362 | -0.634 | -0.21 | 0.734 | -0.005 | -0.002 | -0.002 | 2.392 | 0.1 | 0 | 0.07 | 0.08 | 0.07 | 0.08 |
| 7 | 44705980 | rs10247064 | 1.029 | -0.748 | -0.102 | 0.733 | 0.005 | 0.072 | 0.072 | 0 | 0.06 | 0.12 | 0.19 | 0.24 | 0.1 | 0.09 |
| 7 | 44709372 | rs710887 | -1.659 | -0.93 | -0.463 | 0.282 | 0.005 | 0.001 | 0.001 | 2.107 | 0.7 | 0.76 | 0.78 | 1 | 0.7 | 0.78 |
| 7 | 44713691 | rs740040 | 0.142 | -0.636 | -0.508 | 1.082 | 0.025 | 0.474 | 0.474 | 0.045 | 0.07 | 0.42 | 0.07 | 0.38 | 0.02 | 0.2 |
| 7 | 44714512 | rs3735477 | 0.433 | -0.637 | -1.029 | 1.043 | -0.177 | 0.02 | 0.02 | 0.078 | 0.34 | 0.02 | 0.36 | 0.23 | 0.42 | 0.31 |
| 7 | 44730284 | rs7805156 | -0.821 | -0.806 | 0.421 | 1.592 | 0 | 0.451 | 0.451 | 0.829 | 0.08 | 0.51 | 0.08 | 0.38 | 0.02 | 0.2 |
| 7 | 44747578 | rs2070607 | -0.64 | -0.788 | -0.388 | 1.433 | 0.027 | 0.008 | 0.008 | 0 | 0.1 | 0.01 | 0.05 | 0.08 | 0.04 | 0.07 |
| 7 | 44750079 | rs917098 | -1.326 | -0.698 | -0.32 | 1.371 | -0.071 | 0.087 | 0.087 | 0.362 | 0.2 | 0.62 | 0.13 | 0.45 | 0.06 | 0.27 |
| 7 | 44769869 | rs4720475 | 0.412 | 0.378 | -0.969 | 0.636 | -0.027 | 0.021 | 0.021 | 0 | 0.62 | 0.73 | 0.59 | 0.66 | 0.53 | 0.76 |
| 7 | 44784697 | rs799449 | 0.372 | 0.464 | -1.58 | -0.157 | -0.055 | 0.081 | 0.081 | 1.213 | 0.63 | 0.84 | 0.68 | 0.83 | 0.52 | 0.71 |
| 7 | 44790988 | rs799451 | 0.41 | 0.451 | -1.586 | -0.083 | -0.054 | 0.075 | 0.075 | 1.153 | 0.63 | 0.84 | 0.68 | 0.8 | 0.52 | 0.71 |
| 7 | 44800176 | rs3735478 | 0.606 | 0.501 | -1.667 | -0.117 | -0.111 | 0.017 | 0.017 | 0.066 | 0.21 | 0.01 | 0.19 | 0.14 | 0.3 | 0.16 |
| 7 | 44803286 | rs4724319 | 1.177 | 0.359 | -0.979 | -0.108 | 0.012 | 0.035 | 0.035 | 0.392 | 0.48 | 0.61 | 0.51 | 0.77 | 0.41 | 0.65 |
| 7 | 44804225 | rs2289379 | -0.402 | 0.266 | -1.023 | -0.102 | -0.052 | 0.069 | 0.069 | 1.35 | 0.35 | 0.13 | 0.28 | 0.2 | 0.43 | 0.27 |
| 7 | 44808091 | rs1050331 | 1.074 | 0.306 | -0.473 | -0.137 | 0.001 | 0.004 | 0.004 | 0.379 | 0.49 | 0.54 | 0.5 | 0.75 | 0.47 | 0.64 |
| 7 | 44808223 | rs1050338 | -0.564 | 0.352 | -0.523 | -0.153 | -0.031 | 0.051 | 0.051 | 0.207 | 0.39 | 0.2 | 0.41 | 0.22 | 0.47 | 0.3 |
| 7 | 44808416 | rs3812257 | -0.242 | 0.353 | -0.525 | -0.156 | -0.054 | 0.063 | 0.063 | 1.024 | 0.36 | 0.13 | 0.37 | 0.23 | 0.43 | 0.29 |
| 7 | 44809273 | rs1065646 | 1.107 | 0.232 | -0.38 | -0.159 | 0.003 | 0.019 | 0.019 | 0.362 | 0.49 | 0.57 | 0.52 | 0.75 | 0.47 | 0.64 |
| 7 | 44809276 | rs1065647 | -1.058 | 0.233 | -0.542 | -0.158 | 0.009 | 0.061 | 0.061 | 0.551 | 0.5 | 0.65 | 0.53 | 0.75 | 0.47 | 0.64 |

| | | | | | | | | | | | | | | | | |
|---|---|---|---|---|---|---|---|---|---|---|---|---|---|---|---|---|
| 7 | 44822385 | rs4724321 | -1.142 | -0.683 | -1.085 | -0.571 | 0.005 | 0.004 | 0.004 | 1.225 | 0.33 | 0.68 | 0.38 | 0.76 | 0.19 | 0.42 |
| 7 | 44837298 | rs6970925 | -1.756 | -0.337 | -1.034 | -0.527 | -0.028 | 0.149 | 0.149 | 2.021 | 0.64 | 0.87 | 0.77 | 0.98 | 0.61 | 0.61 |
| 7 | 44841646 | rs11984372 | 1.822 | -0.383 | -1.043 | -0.1 | -0.04 | 0.174 | 0.174 | 2.581 | 0.35 | 0.09 | 0.22 | 0.02 | 0.39 | 0.38 |
| 7 | 44845032 | rs17134462 | 0.805 | -0.115 | -0.925 | 0.053 | -0.001 | 0.014 | 0.014 | 0 | 0.09 | 0 | 0.06 | 0.03 | 0.07 | 0.06 |
| 7 | 44866832 | rs7016 | 1.659 | 0.019 | -0.543 | 0.006 | -0.006 | 0.079 | 0.079 | 1.673 | 0.37 | 0.19 | 0.23 | 0.02 | 0.39 | 0.37 |
| 7 | 44917040 | rs8840 | -0.214 | 0.13 | -0.855 | 0.027 | -0.051 | 0.054 | 0.054 | 0 | 0.35 | 0.15 | 0.38 | 0.23 | 0.44 | 0.24 |
| 7 | 44925896 | rs1990053 | -0.192 | 0.164 | -0.84 | 0.043 | -0.051 | 0.052 | 0.052 | 0 | 0.35 | 0.15 | 0.38 | 0.23 | 0.44 | 0.24 |
| 7 | 44932538 | rs757694 | -0.063 | 0.168 | -0.852 | 0.013 | -0.061 | 0.036 | 0.036 | 0.003 | 0.34 | 0.14 | 0.37 | 0.24 | 0.44 | 0.22 |
| 7 | 44934521 | rs35735865 | 1.438 | 0.113 | -0.944 | -0.287 | -0.025 | 0.203 | 0.203 | 1.305 | 0.37 | 0.1 | 0.22 | 0 | 0.37 | 0.36 |
| 7 | 44944893 | rs11763472 | 0.945 | -0.373 | 0.145 | 0.637 | -0.049 | 0.066 | 0.066 | 0.025 | 0.72 | 0.55 | 0.71 | 0.58 | 0.81 | 0.73 |
| 7 | 44959923 | rs2331175 | -0.356 | 0.037 | -0.157 | 0.125 | -0.006 | -0.058 | -0.058 | 0 | 0.84 | 0.74 | 0.76 | 0.96 | 0.86 | 0.87 |
| 7 | 44959936 | rs4724334 | 1.18 | 0.142 | -0.465 | 0.056 | -0.001 | 0.042 | 0.042 | 1.237 | 0.5 | 0.43 | 0.47 | 0.71 | 0.54 | 0.66 |
| 7 | 44964910 | rs10951777 | 1.209 | -0.026 | -0.256 | 0.293 | -0.029 | -0.063 | -0.063 | 0 | 0.74 | 0.64 | 0.69 | 0.9 | 0.8 | 0.81 |
| 7 | 44968196 | rs4236376 | 0.541 | 0.478 | 0.194 | 0.493 | -0.012 | -0.074 | -0.074 | 0.02 | 0.83 | 0.65 | 0.75 | 0.96 | 0.86 | 0.87 |
| 7 | 44976349 | rs1990042 | 0.848 | 0.931 | -0.028 | 1.044 | 0.012 | 0.057 | 0.057 | 0.319 | 0.75 | 0.63 | 0.57 | 0.6 | 0.75 | 0.7 |
| 7 | 44978151 | rs7803214 | -0.054 | 0.403 | -0.787 | 0.915 | -0.011 | -0.015 | -0.015 | 0.671 | 0.09 | 0 | 0.09 | 0.11 | 0.06 | 0.04 |
| 7 | 44978350 | rs4724335 | 0.575 | 0.32 | -1.065 | 0.594 | -0.038 | -0.009 | -0.009 | 0.312 | 0.32 | 0.27 | 0.31 | 0.35 | 0.46 | 0.44 |
| 7 | 44989199 | rs6463255 | -0.06 | 0.751 | -0.549 | 0.793 | 0.029 | 0.048 | 0.048 | 0 | 0.1 | 0.2 | 0.25 | 0.24 | 0.15 | 0.21 |
| 7 | 44995327 | rs2877301 | 0.455 | 0.478 | -1.149 | -0.437 | 0.018 | 0.069 | 0.069 | 2.841 | 0.51 | 0.33 | 0.63 | 0.78 | 0.59 | 0.56 |
| 7 | 44998714 | rs6946581 | -0.05 | -0.242 | -0.909 | -0.799 | -0.016 | 0.063 | 0.063 | 4.913 | 0.66 | 0.75 | 0.71 | 0.81 | 0.64 | 0.81 |
| 7 | 44999863 | rs3812259 | 0.309 | -0.3 | -0.292 | -0.79 | -0.049 | -0.048 | -0.048 | 0 | 0.13 | 0.29 | 0.06 | 0.03 | 0.05 | 0.25 |
| 7 | 45004063 | rs7792760 | -0.718 | -0.305 | -0.92 | -1.186 | -0.013 | 0.062 | 0.062 | 0 | 0.81 | 0.98 | 0.86 | 0.89 | 0.85 | 0.88 |
| 7 | 45014304 | rs6976664 | 0.899 | -0.344 | -0.953 | -1.159 | -0.058 | -0.003 | -0.003 | 0 | 0.13 | 0.04 | 0.17 | 0.1 | 0.23 | 0.07 |
| 7 | 45018263 | rs10951781 | 1.601 | -0.565 | -1.058 | -0.894 | -0.049 | -0.042 | -0.042 | 0 | 0.08 | 0.01 | 0.22 | 0.15 | 0.15 | 0.16 |
| 7 | 45025696 | rs10228004 | 0.836 | -0.756 | -0.786 | -1.02 | 0.04 | 0.022 | 0.022 | 0 | 0.05 | 0.29 | 0.03 | 0 | 0.02 | 0.09 |
| 7 | 45028992 | rs7795102 | 1.586 | -1.257 | -0.916 | -1.24 | 0.026 | 0.094 | 0.094 | 0 | 0.48 | 0.27 | 0.51 | 0.73 | 0.58 | 0.42 |
| 7 | 45038881 | rs2158564 | 1.601 | -1.267 | -0.919 | -1.255 | 0.026 | 0.095 | 0.095 | 0 | 0.48 | 0.27 | 0.51 | 0.73 | 0.58 | 0.42 |
| 7 | 45055410 | rs10227754 | 1.005 | -1.221 | -0.502 | -1.203 | 0.023 | 0.006 | 0.006 | 0 | 0.05 | 0.28 | 0.03 | 0 | 0.02 | 0.09 |
| 7 | 45067017 | rs1294962 | -1.255 | -1.424 | -0.794 | -1.371 | -0.023 | 0.065 | 0.065 | 0.068 | 0.79 | 0.98 | 0.83 | 0.89 | 0.83 | 0.88 |
| 7 | 45069584 | rs10271311 | 0.055 | -1.545 | -0.559 | -1.428 | -0.063 | -0.089 | -0.089 | 0.814 | 0.15 | 0.45 | 0.05 | 0.01 | 0.05 | 0.25 |
| 7 | 45073799 | rs1294966 | -1.262 | -1.428 | -0.825 | -1.369 | -0.023 | 0.064 | 0.064 | 0.093 | 0.79 | 0.98 | 0.83 | 0.89 | 0.83 | 0.88 |
| 7 | 45084056 | rs4724353 | -2.226 | -1.592 | -0.774 | -1.416 | 0.005 | 0.071 | 0.071 | 0 | 0.55 | 0.76 | 0.5 | 0.31 | 0.44 | 0.58 |
| 7 | 45084525 | rs17451462 | 1.1 | -1.637 | -0.843 | -1.445 | -0.02 | 0.069 | 0.069 | 0.2 | 0.21 | 0.02 | 0.17 | 0.11 | 0.17 | 0.12 |
| 7 | 45085936 | rs1866581 | -2.237 | -1.605 | -0.493 | -1.421 | 0.001 | 0.004 | 0.004 | 0 | 0.54 | 0.64 | 0.48 | 0.31 | 0.44 | 0.58 |
| 7 | 45086238 | rs11765574 | 1.218 | -1.578 | -0.441 | -1.363 | -0.007 | -0.011 | -0.011 | 0 | 0.07 | 0.1 | 0.04 | 0.01 | 0.05 | 0.05 |
| 7 | 45100586 | rs7788562 | -3.101 | -1.391 | -0.675 | -1.396 | -0.017 | -0.009 | -0.009 | 0.25 | 0.67 | 0.81 | 0.66 | 0.37 | 0.64 | 0.63 |
| 7 | 45104131 | rs11552377 | 1.158 | -1.42 | -0.622 | -1.37 | -0.019 | 0.064 | 0.064 | 0.093 | 0.21 | 0.02 | 0.17 | 0.11 | 0.17 | 0.12 |
| 7 | 45105117 | rs3735486 | -0.798 | -0.889 | -0.535 | -1.396 | 0.006 | 0.028 | 0.028 | 0.611 | 0.41 | 0.25 | 0.44 | 0.3 | 0.38 | 0.34 |
| 7 | 45112471 | rs11977608 | 1.091 | -0.865 | -0.369 | -1.376 | 0.032 | 0.015 | 0.015 | 0 | 0.05 | 0.3 | 0.03 | 0 | 0.02 | 0.09 |
| 7 | 45113170 | rs2289367 | -0.137 | -0.725 | -0.589 | -1.328 | 0 | -0.002 | -0.002 | 0 | 0.2 | 0.23 | 0.28 | 0.19 | 0.21 | 0.21 |
| 7 | 45117538 | rs6953826 | -0.137 | -0.704 | -0.58 | -1.322 | 0.002 | 0 | 0 | 0 | 0.2 | 0.23 | 0.27 | 0.19 | 0.21 | 0.21 |
| 7 | 45122324 | rs3735493 | 1.066 | -0.596 | -0.927 | -1.248 | -0.019 | 0.038 | 0.038 | 0.021 | 0.21 | 0.02 | 0.17 | 0.12 | 0.17 | 0.12 |
| 7 | 45124286 | rs3735494 | -0.518 | -0.676 | -0.75 | -1.251 | -0.003 | -0.01 | -0.01 | 0 | 0.36 | 0.35 | 0.42 | 0.29 | 0.34 | 0.29 |
| 7 | 45131853 | rs11765226 | 1.003 | -0.745 | -1.257 | -1.233 | -0.006 | 0.037 | 0.037 | 0.876 | 0.23 | 0.02 | 0.16 | 0.12 | 0.17 | 0.12 |
| 7 | 45139163 | rs6967748 | -1.164 | -0.883 | -1.223 | -1.132 | 0.07 | 0.086 | 0.086 | 0 | 0.45 | 0.67 | 0.27 | 0.19 | 0.27 | 0.44 |
| 7 | 45139892 | rs4553 | -1.241 | -0.877 | -1.243 | -1.129 | 0.082 | 0.1 | 0.1 | 0 | 0.46 | 0.7 | 0.27 | 0.19 | 0.27 | 0.44 |
| 7 | 45143892 | rs6973982 | 1.453 | -0.281 | -1.212 | -0.831 | -0.054 | -0.055 | -0.055 | 0 | 0.12 | 0.01 | 0.22 | 0.18 | 0.16 | 0.14 |
| 7 | 45146656 | rs3757572 | -2.706 | -0.721 | -1.548 | -1.208 | -0.001 | -0.004 | -0.004 | 0 | 0.69 | 0.68 | 0.67 | 0.42 | 0.71 | 0.67 |
| 7 | 45148019 | rs3757574 | 1.106 | -1.149 | -1.569 | -1.248 | -0.008 | 0.002 | 0.002 | 0 | 0.22 | 0.02 | 0.16 | 0.15 | 0.16 | 0.14 |
| 7 | 45148667 | rs2304693 | 2.524 | -1.094 | -1.563 | -1.358 | -0.054 | -0.053 | -0.053 | 0 | 0.12 | 0.01 | 0.22 | 0.18 | 0.16 | 0.13 |
| 7 | 45148773 | rs2304694 | 2.527 | -1.093 | -1.562 | -1.358 | -0.054 | -0.053 | -0.053 | 0 | 0.12 | 0.01 | 0.22 | 0.18 | 0.16 | 0.13 |
| 7 | 45150331 | rs7810512 | -1.372 | -1.214 | -1.392 | -1.971 | 0.009 | 0.009 | 0.009 | 3.543 | 0.35 | 0.64 | 0.32 | 0.26 | 0.27 | 0.44 |
| 7 | 45155888 | rs10258996 | 1.021 | -1.056 | -1.353 | -1.856 | -0.01 | 0.028 | 0.028 | 0 | 0.22 | 0.02 | 0.17 | 0.11 | 0.16 | 0.12 |
| 7 | 45161049 | rs4720497 | -0.106 | -0.739 | -1.091 | -1.857 | -0.01 | -0.006 | -0.006 | 0 | 0.24 | 0.53 | 0.29 | 0.24 | 0.21 | 0.25 |
| 7 | 45165116 | rs1866569 | -0.97 | -0.65 | -0.518 | -1.697 | -0.017 | -0.022 | -0.022 | 9.126 | 0.32 | 0.57 | 0.29 | 0.25 | 0.25 | 0.5 |
| 7 | 45167831 | rs7799075 | -0.365 | -0.497 | -0.599 | -1.714 | -0.01 | -0.026 | -0.026 | 0 | 0.36 | 0.4 | 0.17 | 0.12 | 0.19 | 0.46 |
| 7 | 45170510 | rs10244707 | 1.237 | -0.287 | -1.215 | -1.65 | 0.007 | 0.044 | 0.044 | 0.838 | 0.21 | 0.12 | 0.14 | 0.1 | 0.14 | 0.08 |
| 7 | 45171410 | rs10278144 | -1.228 | -0.318 | -1.273 | -1.659 | -0.014 | 0.021 | 0.021 | 0 | 0.57 | 0.73 | 0.45 | 0.36 | 0.44 | 0.6 |
| 7 | 45173384 | rs1866571 | -0.985 | 0.377 | -0.816 | -1.419 | -0.005 | 0.071 | 0.071 | 0.655 | 0.47 | 0.76 | 0.45 | 0.34 | 0.41 | 0.38 |
| 7 | 45178748 | rs12536178 | 0.122 | 0.437 | -0.649 | -1.264 | 0.03 | 0.017 | 0.017 | 6.142 | 0.2 | 0.47 | 0.25 | 0.22 | 0.2 | 0.25 |
| 7 | 45182591 | rs10229398 | 1.132 | 0.267 | -0.879 | -0.941 | -0.007 | 0.025 | 0.025 | 0.849 | 0.17 | 0.04 | 0.12 | 0.1 | 0.13 | 0.08 |
| 7 | 45184555 | rs1866572 | -0.284 | 0.407 | -0.037 | -0.197 | -0.019 | 0.013 | 0.013 | 1.965 | 0.42 | 0.25 | 0.51 | 0.61 | 0.5 | 0.66 |
| 7 | 45186727 | rs10230614 | 0.1 | 0.854 | 0.231 | -0.41 | -0.008 | -0.01 | -0.01 | 0 | 0.95 | 0.98 | 0.93 | 0.89 | 0.93 | 0.93 |
| 7 | 45190414 | rs12666020 | 1.658 | 0.79 | 0.153 | -0.238 | -0.109 | -0.041 | -0.041 | 0.514 | 0.41 | 0.04 | 0.48 | 0.38 | 0.41 | 0.19 |
| 7 | 45190502 | rs2119053 | 1.384 | 0.248 | 0.435 | -0.402 | -0.009 | -0.032 | -0.032 | 0 | 0.78 | 0.41 | 0.84 | 0.89 | 0.83 | 0.72 |
| 7 | 45192586 | rs4720500 | 0.801 | 0.354 | 0.584 | -0.231 | -0.001 | 0.005 | 0.005 | 1.828 | 0.69 | 0.36 | 0.77 | 0.84 | 0.72 | 0.67 |
| 7 | 45196053 | rs6463265 | -0.695 | 0.356 | 0.48 | -0.332 | 0.001 | -0.04 | -0.04 | 0.319 | 0.29 | 0.64 | 0.17 | 0.02 | 0.23 | 0.24 |
| 7 | 45201045 | rs3801408 | 1.602 | 0.432 | 0.463 | -0.258 | -0.146 | 0.003 | 0.003 | 4.098 | 0.41 | 0.04 | 0.51 | 0.35 | 0.48 | 0.21 |
| 7 | 45209186 | rs11763766 | -0.385 | 0.577 | 0.799 | -0.173 | -0.006 | -0.13 | -0.13 | 0 | 0.26 | 0.49 | 0.14 | 0.02 | 0.22 | 0.23 |

| | | | | | | | | | | | | | | | | |
|---|---|---|---|---|---|---|---|---|---|---|---|---|---|---|---|---|
| 7 | 45211941 | rs1294947 | -0.481 | 0.601 | 0.679 | -0.09 | 0.007 | -0.044 | -0.044 | 0.197 | 0.76 | 0.83 | 0.76 | 0.51 | 0.77 | 0.52 |
| 7 | 45212736 | rs1294946 | 0.872 | 0.649 | 0.541 | -0.062 | 0.019 | 0.018 | 0.018 | 1.509 | 0.49 | 0.3 | 0.61 | 0.49 | 0.55 | 0.29 |
| 7 | 45213469 | rs3757577 | -0.271 | 0.745 | 0.433 | 0.08 | -0.004 | -0.002 | -0.002 | 0.091 | 0.05 | 0.04 | 0.08 | 0.12 | 0.08 | 0.08 |
| 7 | 45215088 | rs1294945 | -0.381 | 0.672 | 0.641 | 0.019 | -0.004 | 0.027 | 0.027 | 0 | 0.76 | 0.69 | 0.76 | 0.51 | 0.76 | 0.53 |
| 7 | 45216785 | rs17172483 | -0.177 | 0.698 | 0.779 | 0.057 | -0.007 | -0.034 | -0.034 | 1.053 | 0.07 | 0.16 | 0.05 | 0 | 0.05 | 0.03 |
| 7 | 45218366 | rs1294935 | 1.239 | 0.743 | 0.704 | 0.04 | -0.03 | 0.058 | 0.058 | 4.012 | 0.45 | 0.24 | 0.54 | 0.39 | 0.49 | 0.22 |
| 7 | 45223259 | rs12702121 | -1.198 | 1.004 | 1.615 | -0.158 | 0.008 | -0.061 | -0.061 | 0 | 0.26 | 0.53 | 0.19 | 0.12 | 0.24 | 0.28 |
| 7 | 45223331 | rs1046292 | -1.165 | 1.019 | 1.613 | -0.151 | 0.011 | -0.065 | -0.065 | 0 | 0.26 | 0.51 | 0.19 | 0.12 | 0.24 | 0.28 |
| 7 | 45223486 | rs3173757 | -1.787 | 1.013 | 1.573 | -0.156 | -0.001 | -0.015 | -0.015 | 0.154 | 0.33 | 0.64 | 0.24 | 0.12 | 0.29 | 0.31 |
| 7 | 45225708 | rs13222185 | -1.798 | 1.025 | 1.536 | -0.151 | -0.001 | -0.019 | -0.019 | 0 | 0.33 | 0.62 | 0.24 | 0.12 | 0.29 | 0.31 |
| 7 | 45225822 | rs11760688 | -0.881 | 1.059 | 1.486 | -0.135 | -0.001 | -0.031 | -0.031 | 0 | 0.07 | 0.19 | 0.05 | 0 | 0.06 | 0.03 |
| 7 | 45225854 | rs11767365 | -2.017 | 1.074 | 1.487 | -0.131 | 0.011 | 0.032 | 0.032 | 0 | 0.12 | 0.29 | 0.14 | 0.1 | 0.11 | 0.1 |
| 7 | 45225883 | rs12702124 | -0.466 | 1.077 | 1.201 | -0.061 | 0.001 | -0.088 | -0.088 | 0 | 0.2 | 0.29 | 0.11 | 0.01 | 0.18 | 0.21 |
| 7 | 45228095 | rs1866576 | -1.478 | 1.042 | 0.786 | 0.979 | 0.018 | 0.059 | 0.059 | 0 | 0.05 | 0.13 | 0.08 | 0.1 | 0.06 | 0.07 |
| 7 | 45231308 | rs11761143 | -1.184 | 1.01 | 0.28 | 0.94 | 0 | -0.026 | -0.026 | 0 | 0.06 | 0.18 | 0.05 | 0 | 0.05 | 0.03 |
| 7 | 45232842 | rs4724374 | -1.537 | 1.035 | 0.201 | 1.005 | 0.005 | 0.031 | 0.031 | 0 | 0.11 | 0.27 | 0.13 | 0.1 | 0.1 | 0.08 |
| 7 | 45234630 | rs1294911 | 1.403 | 1.624 | -0.417 | 1.213 | -0.009 | -0.009 | -0.009 | 0.003 | 0.93 | 0.95 | 0.77 | 0.89 | 0.86 | 0.92 |
| 7 | 45234851 | rs1294910 | -0.414 | 1.252 | -0.295 | 1.326 | -0.007 | 0.005 | 0.005 | 0 | 0.26 | 0.35 | 0.2 | 0.38 | 0.23 | 0.24 |
| 7 | 45235120 | rs1294908 | 1.407 | 1.295 | -0.336 | 1.585 | 0.002 | 0.018 | 0.018 | 0 | 0.82 | 0.77 | 0.72 | 0.72 | 0.79 | 0.82 |
| 7 | 45235862 | rs1294906 | 1.504 | 1.597 | -0.287 | 1.81 | 0.033 | 0.087 | 0.087 | 0.268 | 0.92 | 0.84 | 0.75 | 0.76 | 0.84 | 0.93 |
| 7 | 45235891 | rs1294905 | 1.228 | 1.682 | -0.287 | 1.81 | 0.036 | 0.116 | 0.116 | 0.041 | 0.93 | 0.85 | 0.77 | 0.76 | 0.87 | 0.93 |
| 7 | 45237407 | rs1294897 | 1.29 | 1.649 | -0.54 | 1.825 | 0.011 | 0.042 | 0.042 | 0.187 | 0.93 | 0.89 | 0.75 | 0.76 | 0.85 | 0.93 |
| 7 | 45243877 | rs1294889 | 0.223 | 1.534 | -1.034 | 1.323 | -0.023 | 0.026 | 0.026 | 0.664 | 0.57 | 0.76 | 0.5 | 0.67 | 0.53 | 0.74 |
| 7 | 45245316 | rs6463268 | 0.598 | 1.941 | -1.66 | 1.789 | -0.036 | 0.006 | 0.006 | 0 | 0.74 | 0.84 | 0.78 | 0.78 | 0.68 | 0.72 |
| 7 | 45246984 | rs1294884 | -0.758 | 1.696 | -2.139 | 1.551 | -0.008 | -0.023 | -0.023 | 2.126 | 0.21 | 0.13 | 0.39 | 0.71 | 0.19 | 0.29 |
| 7 | 45247913 | rs4724376 | 1.202 | 2.341 | -2.611 | 0.811 | -0.041 | 0.001 | 0.001 | 1.553 | 0.11 | 0.03 | 0.15 | 0.1 | 0.33 | 0.3 |
| 7 | 45248202 | rs1294881 | -0.646 | 2.471 | -2.754 | 0.435 | -0.009 | 0.002 | 0.002 | 0 | 0.84 | 0.84 | 0.86 | 1 | 0.79 | 0.95 |
| 7 | 45248653 | rs11767371 | 1.116 | 2.303 | -2.621 | 0.402 | -0.046 | 0.006 | 0.006 | 1.026 | 0.11 | 0.03 | 0.14 | 0.09 | 0.32 | 0.26 |
| 7 | 45251293 | rs17172499 | 1.729 | 0.612 | -2.411 | 0.195 | 0.144 | 0.064 | 0.064 | 1.241 | 0.45 | 0.18 | 0.42 | 0.36 | 0.25 | 0.33 |
| 7 | 45251443 | rs1294875 | 0.346 | 0.543 | -1.776 | 0.089 | -0.014 | 0.028 | 0.028 | 0 | 0.67 | 0.73 | 0.69 | 0.89 | 0.45 | 0.58 |
| 7 | 45255940 | rs975349 | -0.351 | 0.71 | -1.484 | 0.202 | 0.03 | 0.16 | 0.16 | 0.027 | 0.21 | 0.49 | 0.26 | 0.53 | 0.22 | 0.25 |
| 7 | 45256975 | rs1294865 | -0.57 | -0.601 | -2.113 | -0.917 | 0.006 | 0.008 | 0.008 | 0 | 0.39 | 0.62 | 0.5 | 0.71 | 0.29 | 0.44 |
| 7 | 45262913 | rs757680 | -1.464 | -0.372 | -2.229 | -1.103 | -0.044 | 0.051 | 0.051 | 0 | 0.91 | 1 | 0.92 | 0.99 | 0.84 | 0.87 |
| 7 | 45263026 | rs1294857 | -0.519 | 0.077 | -2.163 | -1.038 | 0.005 | 0.007 | 0.007 | 0 | 0.39 | 0.63 | 0.51 | 0.71 | 0.3 | 0.43 |
| 7 | 45263634 | rs17230937 | 1.693 | 0.084 | -2.229 | -1.185 | 0.024 | 0.053 | 0.053 | 0 | 0.17 | 0.08 | 0.08 | 0.01 | 0.08 | 0.15 |
| 7 | 45266404 | rs10248863 | 2.36 | 0.213 | -2.328 | -1.258 | -0.038 | 0 | 0 | 2.329 | 0.11 | 0.03 | 0.14 | 0.09 | 0.29 | 0.26 |
| 7 | 45267645 | rs917088 | 0.491 | -0.166 | -1.781 | -1.852 | -0.001 | -0.019 | -0.019 | 0 | 0.62 | 0.59 | 0.68 | 0.89 | 0.45 | 0.51 |
| 7 | 45268637 | rs757678 | -0.462 | -0.279 | -1.816 | -1.613 | -0.005 | 0.016 | 0.016 | 0 | 0.78 | 0.81 | 0.84 | 0.99 | 0.77 | 0.84 |
| 7 | 45269903 | rs1294847 | 0.049 | -0.469 | -1.359 | -1.148 | 0.05 | 0.089 | 0.089 | 0 | 0.49 | 0.61 | 0.62 | 0.89 | 0.4 | 0.44 |
| 7 | 45273264 | rs17172506 | 1.806 | -1.092 | -1.766 | -1.065 | 0.046 | -0.038 | -0.038 | 0 | 0.19 | 0.01 | 0.15 | 0.19 | 0.14 | 0.13 |
| 7 | 45277431 | rs10272379 | 0.452 | -0.892 | -1.218 | -1.473 | 0.05 | 0.058 | 0.058 | 0.602 | 0.35 | 0.19 | 0.24 | 0.16 | 0.18 | 0.22 |
| 7 | 45280440 | rs6976907 | 0.785 | -0.655 | -0.903 | -2.013 | -0.016 | 0.027 | 0.027 | 0.045 | 0.37 | 0.3 | 0.56 | 0.77 | 0.49 | 0.6 |
| 7 | 45281716 | rs10224119 | 0.378 | -0.153 | -1.113 | -2.281 | 0.05 | 0.083 | 0.083 | 0.063 | 0.39 | 0.22 | 0.28 | 0.16 | 0.26 | 0.28 |
| 7 | 45283115 | rs2040864 | 0.44 | -0.45 | -0.771 | -2.276 | 0.029 | 0.035 | 0.035 | 0.105 | 0.38 | 0.27 | 0.28 | 0.16 | 0.24 | 0.27 |
| 7 | 45287160 | rs2331224 | 0.341 | -0.304 | -0.967 | -2.398 | 0.059 | 0.118 | 0.118 | 0 | 0.39 | 0.17 | 0.27 | 0.16 | 0.26 | 0.28 |
| 7 | 45289092 | rs2331226 | 1.791 | -0.409 | -1.113 | -2.404 | 0.038 | 0.069 | 0.069 | 0 | 0.26 | 0.08 | 0.21 | 0.12 | 0.17 | 0.22 |
| 7 | 45293723 | rs10236372 | -0.184 | -0.375 | -0.154 | -2.124 | 0 | -0.053 | -0.053 | 0 | 0.76 | 0.76 | 0.85 | 0.99 | 0.73 | 0.87 |
| 7 | 45297878 | rs2040863 | 0.417 | -0.381 | -0.205 | -2.137 | 0.009 | -0.182 | -0.182 | 0 | 0.69 | 0.5 | 0.82 | 0.99 | 0.69 | 0.86 |
| 7 | 45299755 | rs6946436 | 1.53 | -0.323 | -1.004 | -2.247 | 0.067 | 0.151 | 0.151 | 0 | 0.36 | 0.08 | 0.26 | 0.14 | 0.24 | 0.33 |
| 7 | 45310492 | rs1527969 | 1.389 | -0.387 | -0.953 | -2.21 | 0.057 | 0.131 | 0.131 | 0 | 0.37 | 0.1 | 0.28 | 0.17 | 0.27 | 0.35 |
| 7 | 45310510 | rs799714 | -1.54 | -0.08 | -0.924 | -2.23 | -0.037 | 0.036 | 0.036 | 0 | 0.9 | 0.96 | 0.9 | 0.99 | 0.82 | 0.9 |
| 7 | 45318389 | rs799703 | 0.241 | -0.394 | -0.214 | -2.251 | 0.056 | -0.064 | -0.064 | 0 | 0.85 | 0.68 | 0.87 | 0.95 | 0.79 | 0.88 |
| 7 | 45324117 | rs6978685 | -0.466 | -0.322 | -0.183 | -2.223 | 0.022 | -0.043 | -0.043 | 0 | 0.88 | 0.83 | 0.89 | 0.99 | 0.81 | 0.92 |
| 7 | 45325658 | rs7797477 | 1.892 | -0.109 | -0.132 | -1.678 | -0.024 | -0.031 | -0.031 | 0 | 0.2 | 0.15 | 0.41 | 0.55 | 0.31 | 0.42 |
| 7 | 45326476 | rs9784917 | 0.777 | -0.364 | 0.015 | -1.848 | 0.055 | 0.077 | 0.077 | 0.764 | 0.48 | 0.3 | 0.34 | 0.17 | 0.32 | 0.37 |
| 7 | 45334883 | rs799707 | -0.349 | -1.737 | 0.114 | -1.626 | 0.018 | -0.062 | -0.062 | 0 | 0.4 | 0.53 | 0.55 | 0.82 | 0.49 | 0.55 |
| 7 | 45336250 | rs4724385 | -0.962 | -1.486 | -0.541 | -1.508 | -0.038 | 0.226 | 0.226 | 0 | 0.56 | 0.91 | 0.63 | 0.87 | 0.58 | 0.64 |
| 7 | 45349609 | rs12539253 | 1.072 | -1.63 | 0.497 | -1.049 | 0.013 | 0.083 | 0.083 | 0 | 0.06 | 0.11 | 0.1 | 0.27 | 0.1 | 0.1 |
| 7 | 45351309 | rs1852621 | 1.204 | -1.755 | 0.586 | -1.125 | -0.004 | 0.001 | 0.001 | 0 | 0.73 | 0.3 | 0.83 | 0.69 | 0.77 | 0.82 |
| 7 | 45367036 | rs1830264 | 0.657 | -1.535 | 0.597 | -1.04 | 0.017 | -0.026 | -0.026 | 0.705 | 0.06 | 0.17 | 0.02 | 0 | 0.04 | 0.01 |
| 7 | 45370510 | rs12702136 | 1.018 | -1.504 | 0.358 | -1.026 | 0.007 | -0.027 | -0.027 | 0 | 0.05 | 0.12 | 0.02 | 0 | 0.04 | 0.01 |
| 7 | 45380052 | rs12667007 | 1.002 | -1.439 | 0.159 | -0.929 | 0.021 | 0.108 | 0.108 | 0 | 0.06 | 0.12 | 0.1 | 0.27 | 0.1 | 0.1 |
| 7 | 45380502 | rs10435170 | 0.912 | -1.654 | -0.07 | -1.061 | -0.004 | 0.003 | 0.003 | 0 | 0.73 | 0.3 | 0.82 | 0.65 | 0.76 | 0.78 |
| 7 | 45386297 | rs4361689 | 1.415 | -2.564 | -0.584 | -1.598 | 0.071 | 0.021 | 0.021 | 0 | 0.13 | 0.02 | 0.12 | 0.06 | 0.05 | 0.09 |
| 7 | 45386995 | rs11976218 | -1.167 | -3.549 | -0.477 | -2.133 | 0.087 | 0.09 | 0.09 | 1.934 | 0.49 | 0.64 | 0.26 | 0.12 | 0.23 | 0.27 |
| 7 | 45390815 | rs1969510 | -0.753 | -3.308 | 0.004 | -2.02 | -0.046 | -0.082 | -0.082 | 0 | 0.29 | 0.41 | 0.12 | 0.08 | 0.18 | 0.14 |
| 7 | 45404603 | rs1852613 | 1.206 | -3.088 | 0.236 | -1.863 | 0.01 | 0.002 | 0.002 | 0.523 | 0.1 | 0.09 | 0.02 | 0 | 0.07 | 0.02 |
| 7 | 45405129 | rs1819034 | -0.702 | -2.964 | -0.033 | -2.162 | 0.01 | 0.008 | 0.008 | 3.204 | 0.51 | 0.53 | 0.35 | 0.16 | 0.41 | 0.26 |
| 7 | 45407140 | rs6949064 | -1.129 | -2.817 | -0.149 | -2.097 | 0.012 | 0.055 | 0.055 | 3.546 | 0.57 | 0.65 | 0.36 | 0.17 | 0.42 | 0.32 |

| | | | | | | | | | | | | | | | | |
|---|---|---|---|---|---|---|---|---|---|---|---|---|---|---|---|---|
| 7 | 45428369 | rs1969508 | -0.669 | -3.143 | 0.361 | -1.735 | -0.015 | -0.088 | -0.088 | 2.368 | 0.28 | 0.4 | 0.22 | 0.08 | 0.26 | 0.14 |
| 7 | 45434192 | rs2091304 | -0.587 | -3.164 | 0.261 | -1.72 | -0.017 | -0.075 | -0.075 | 1.004 | 0.27 | 0.38 | 0.21 | 0.08 | 0.23 | 0.14 |
| 7 | 45437234 | rs6960713 | -1.23 | -3.686 | 0.117 | -1.779 | 0.054 | 0.06 | 0.06 | 1.491 | 0.47 | 0.63 | 0.33 | 0.16 | 0.27 | 0.31 |
| 7 | 45450413 | rs7805779 | -0.687 | -3.486 | 0.21 | -1.686 | -0.033 | -0.098 | -0.098 | 0 | 0.29 | 0.48 | 0.22 | 0.11 | 0.24 | 0.18 |
| 7 | 45464222 | rs1917716 | 1.598 | -3.26 | -0.465 | -2.316 | -0.01 | 0.032 | 0.032 | 7.957 | 0.17 | 0.08 | 0.16 | 0.06 | 0.17 | 0.13 |
| 7 | 45476338 | rs10499634 | 0.676 | -2.94 | 0.059 | -1.778 | -0.002 | -0.019 | -0.019 | 0.288 | 0.11 | 0.12 | 0.03 | 0 | 0.06 | 0.02 |
| 7 | 45481896 | rs2103184 | 1.614 | -3.038 | -0.03 | -1.833 | 0.009 | 0.005 | 0.005 | 0.364 | 0.56 | 0.41 | 0.66 | 0.84 | 0.74 | 0.68 |
| 7 | 45489306 | rs1589881 | -1.184 | -2.373 | -0.153 | -1.565 | -0.028 | -0.082 | -0.082 | 0.115 | 0.29 | 0.45 | 0.22 | 0.11 | 0.24 | 0.17 |
| 7 | 45489331 | rs1589880 | 0.312 | -2.333 | -0.222 | -1.559 | -0.003 | 0.024 | 0.024 | 0 | 0.16 | 0.11 | 0.17 | 0.07 | 0.17 | 0.12 |
| 7 | 45494555 | rs17566076 | -0.276 | -2.031 | -0.194 | -1.364 | -0.013 | -0.089 | -0.089 | 0.45 | 0.28 | 0.4 | 0.22 | 0.07 | 0.24 | 0.15 |
| 7 | 45495784 | rs17488020 | -0.072 | -1.973 | -0.418 | -1.294 | 0 | -0.018 | -0.018 | 2.053 | 0.26 | 0.28 | 0.22 | 0.08 | 0.25 | 0.14 |
| 7 | 45498400 | rs885244 | 0.58 | -1.229 | -0.714 | -0.961 | -0.004 | -0.018 | -0.018 | 0 | 0.48 | 0.47 | 0.64 | 0.82 | 0.65 | 0.67 |
| 7 | 45518854 | rs1949873 | -0.123 | -0.496 | -1.357 | -0.094 | 0.036 | -0.003 | -0.003 | 1.247 | 0.88 | 0.97 | 0.92 | 0.88 | 0.94 | 0.93 |
| 7 | 45523448 | rs1534131 | -0.479 | 0.209 | -1.217 | -0.543 | 0 | 0.025 | 0.025 | 0.6 | 0.39 | 0.36 | 0.28 | 0.16 | 0.39 | 0.25 |
| 7 | 45526173 | rs4720510 | 0.389 | -0.127 | -1.541 | -0.712 | 0.006 | 0.036 | 0.036 | 7.033 | 0.79 | 0.87 | 0.89 | 0.91 | 0.83 | 0.87 |
| 7 | 45527477 | rs10229543 | 0.515 | 0.007 | -1.21 | -0.353 | 0.001 | 0.039 | 0.039 | 1.548 | 0.5 | 0.43 | 0.54 | 0.31 | 0.51 | 0.43 |
| 7 | 45527612 | rs10229769 | -0.045 | -0.169 | -1.193 | -0.295 | -0.012 | 0.101 | 0.101 | 0.22 | 0.67 | 0.89 | 0.81 | 0.83 | 0.71 | 0.79 |
| 7 | 45539170 | rs4724403 | -0.162 | -0.88 | -0.968 | -0.537 | 0.023 | 0.155 | 0.155 | 0 | 0.59 | 0.89 | 0.79 | 0.81 | 0.65 | 0.7 |
| 7 | 45551710 | rs1525661 | -0.796 | -0.858 | -0.714 | 0.285 | 0.02 | 0.203 | 0.203 | 0.014 | 0.17 | 0.47 | 0.25 | 0.39 | 0.18 | 0.32 |
| 7 | 45553681 | rs17488993 | 0.048 | -0.857 | -0.679 | 0.239 | 0.064 | 0.033 | 0.033 | 0 | 0.06 | 0.19 | 0.06 | 0.07 | 0.09 | 0.12 |
| 7 | 45554986 | rs7793106 | -0.256 | -0.972 | -0.752 | 0.214 | 0.017 | 0.073 | 0.073 | 0 | 0.59 | 0.9 | 0.75 | 0.69 | 0.64 | 0.67 |
| 7 | 45565811 | rs1525655 | 0.949 | -0.964 | -0.643 | 0.2 | -0.003 | 0.003 | 0.003 | 0 | 0.41 | 0.44 | 0.51 | 0.29 | 0.46 | 0.34 |
| 7 | 45581268 | rs10951822 | 0.235 | -0.998 | -0.662 | 0.232 | 0.006 | 0.037 | 0.037 | 0 | 0.52 | 0.7 | 0.69 | 0.62 | 0.55 | 0.54 |
| 7 | 45598469 | rs1525645 | -1.231 | -0.954 | -0.753 | 0.302 | 0.003 | 0.186 | 0.186 | 0.044 | 0.18 | 0.5 | 0.28 | 0.52 | 0.19 | 0.36 |
| 7 | 45605998 | rs7357339 | 1.031 | -0.273 | -1.931 | 0.228 | -0.11 | -0.091 | -0.091 | 0.026 | 0.24 | 0.02 | 0.37 | 0.28 | 0.28 | 0.31 |
| 7 | 45615783 | rs12702149 | -0.086 | -0.428 | -1.626 | 0.424 | 0.017 | 0.035 | 0.035 | 0.098 | 0.47 | 0.2 | 0.36 | 0.66 | 0.4 | 0.51 |
| 7 | 45624449 | rs7456393 | -0.335 | -0.495 | -1.427 | 0.361 | 0.007 | 0.019 | 0.019 | 0.178 | 0.51 | 0.37 | 0.41 | 0.66 | 0.47 | 0.61 |
| 7 | 45629845 | rs11771815 | -0.169 | -0.452 | -1.426 | 0.46 | 0.041 | 0.011 | 0.011 | 0.369 | 0.48 | 0.26 | 0.36 | 0.48 | 0.41 | 0.46 |
| 7 | 45632622 | rs17567824 | -0.784 | -0.425 | -1.441 | 0.397 | 0 | -0.015 | -0.015 | 0 | 0.08 | 0.1 | 0.06 | 0 | 0.08 | 0.02 |
| 7 | 45644419 | rs11770994 | -0.345 | -0.416 | -1.474 | 0.473 | 0.034 | 0.019 | 0.019 | 0.382 | 0.48 | 0.31 | 0.4 | 0.62 | 0.43 | 0.54 |
| 7 | 45648803 | rs11766192 | 0.627 | -0.466 | -1.545 | 0.475 | -0.001 | -0.082 | -0.082 | 0 | 0.37 | 0.06 | 0.28 | 0.46 | 0.3 | 0.32 |
| 7 | 45656924 | rs12112953 | -1.827 | -0.117 | -1.24 | 0.421 | 0.001 | 0.023 | 0.023 | 0 | 0.1 | 0.2 | 0.11 | 0.16 | 0.13 | 0.22 |
| 7 | 45664592 | rs6961503 | -0.686 | -0.352 | -1.359 | 0.718 | 0.019 | 0.058 | 0.058 | 0.102 | 0.5 | 0.35 | 0.28 | 0.69 | 0.42 | 0.61 |
| 7 | 45679021 | rs34154333 | -2.173 | -0.079 | -1.481 | 0.621 | -0.016 | 0.032 | 0.032 | 0 | 0.13 | 0.09 | 0.09 | 0.04 | 0.15 | 0.25 |
| 7 | 45680258 | rs9639924 | 0.718 | -0.238 | -1.44 | 0.797 | 0.004 | -0.11 | -0.11 | 0.724 | 0.33 | 0.07 | 0.16 | 0.49 | 0.26 | 0.31 |
| 7 | 45684544 | rs1357111 | -1.373 | -0.032 | -1.265 | -0.143 | 0 | 0 | 0 | 0.058 | 0.11 | 0.14 | 0.09 | 0.13 | 0.13 | 0.17 |
| 7 | 45685196 | rs9638987 | 0.541 | -0.104 | -1.419 | -0.154 | 0.039 | 0.148 | 0.148 | 0.905 | 0.35 | 0.08 | 0.15 | 0.13 | 0.24 | 0.3 |
| 7 | 45685250 | rs870010 | -0.598 | -0.188 | -0.981 | -0.103 | 0.008 | 0.028 | 0.028 | 0.171 | 0.51 | 0.43 | 0.27 | 0.33 | 0.41 | 0.54 |
| 7 | 45688233 | rs3735672 | -1.683 | 0.359 | -0.328 | 0.587 | -0.007 | 0.006 | 0.006 | 0.004 | 0.1 | 0.16 | 0.06 | 0.16 | 0.05 | 0.09 |
| 7 | 45689929 | rs868362 | -1.77 | 0.453 | -0.352 | 0.562 | 0.011 | 0.022 | 0.022 | 0 | 0.15 | 0.32 | 0.11 | 0.19 | 0.13 | 0.13 |
| 7 | 45695437 | rs2471226 | -0.439 | 0.865 | -0.72 | -0.262 | 0 | -0.023 | -0.023 | 0.095 | 0.32 | 0.35 | 0.2 | 0.11 | 0.26 | 0.24 |
| 7 | 45695468 | rs10268339 | -0.084 | 1.493 | -1.188 | 0.105 | -0.049 | -0.043 | -0.043 | 0 | 0.14 | 0.01 | 0.12 | 0.16 | 0.18 | 0.07 |
| 7 | 45698253 | rs2960288 | 0.128 | 1.17 | -0.501 | 0.382 | 0.006 | 0.04 | 0.04 | 6.88 | 0.78 | 0.45 | 0.84 | 0.72 | 0.79 | 0.84 |
| 7 | 45703971 | rs1042009 | 1.304 | 0.903 | -0.853 | 0.65 | -0.002 | 0.018 | 0.018 | 4.429 | 0.63 | 0.16 | 0.63 | 0.33 | 0.68 | 0.71 |
| 7 | 45705017 | rs1521469 | -1.183 | 0.95 | -0.189 | 0.759 | 0.059 | -0.018 | -0.018 | 1.266 | 0.86 | 0.6 | 0.88 | 0.92 | 0.84 | 0.96 |
| 7 | 45721919 | rs4724425 | 1.154 | 0.918 | -0.499 | 0.865 | 0.001 | -0.032 | -0.032 | 0 | 0.67 | 0.22 | 0.64 | 0.42 | 0.7 | 0.73 |
| 7 | 45722930 | rs7778387 | 0.385 | 0.903 | -0.225 | 1.042 | -0.016 | 0.166 | 0.166 | 0 | 0.11 | 0.25 | 0.18 | 0.35 | 0.06 | 0.1 |
| 7 | 45723834 | rs2461108 | 0.508 | 1.657 | -0.343 | 0.835 | 0.038 | -0.001 | -0.001 | 0.151 | 0.76 | 0.42 | 0.68 | 0.78 | 0.59 | 0.81 |
| 7 | 45723950 | rs10241950 | -0.241 | 1.667 | -0.38 | 0.893 | -0.013 | 0.088 | 0.088 | 0 | 0.17 | 0.25 | 0.2 | 0.44 | 0.12 | 0.13 |
| 7 | 45724696 | rs2461106 | 1.176 | 1.184 | -1.283 | 0.527 | 0.017 | -0.024 | -0.024 | 0 | 0.23 | 0.02 | 0.28 | 0.19 | 0.17 | 0.32 |
| 7 | 45732690 | rs12670400 | -1.179 | 2.187 | -0.096 | 1.219 | -0.01 | 0.096 | 0.096 | 0 | 0.2 | 0.32 | 0.22 | 0.44 | 0.16 | 0.17 |
| 7 | 45741525 | rs17172602 | 0.67 | 2.137 | 0.374 | 1.166 | 0.023 | -0.022 | -0.022 | 0 | 0.14 | 0.26 | 0.12 | 0.09 | 0.18 | 0.05 |
| 7 | 45748520 | rs2248284 | 2.129 | 2.187 | 0.022 | 1.187 | -0.025 | 0.053 | 0.053 | 1.076 | 0.56 | 0.12 | 0.46 | 0.35 | 0.44 | 0.66 |
| 7 | 45754582 | rs3735669 | 0.162 | 2.296 | 0.172 | 1.307 | -0.006 | 0.154 | 0.154 | 0 | 0.11 | 0.23 | 0.2 | 0.33 | 0.08 | 0.09 |
| 7 | 45757343 | rs2280495 | -0.175 | 2.353 | -0.166 | 1.282 | -0.048 | -0.034 | -0.034 | 0.002 | 0.09 | 0.02 | 0.19 | 0.12 | 0.23 | 0.09 |
| 7 | 45757806 | rs2461127 | 2.101 | 2.302 | -0.241 | 1.228 | 0.037 | 0.092 | 0.092 | 0.472 | 0.51 | 0.11 | 0.45 | 0.35 | 0.41 | 0.65 |
| 7 | 45757847 | rs10225980 | 1.703 | 2.344 | -0.244 | 1.215 | -0.003 | -0.002 | -0.002 | 0.608 | 0.07 | 0.08 | 0.03 | 0.08 | 0.08 | 0.03 |
| 7 | 45759262 | rs2471267 | 2.1 | 2.336 | -0.261 | 1.21 | 0.033 | 0.093 | 0.093 | 0.28 | 0.51 | 0.09 | 0.44 | 0.34 | 0.41 | 0.6 |
| 7 | 45761351 | rs10249706 | 1.855 | 2.299 | -0.231 | 1.229 | -0.003 | -0.002 | -0.002 | 0.003 | 0.07 | 0.08 | 0.03 | 0.08 | 0.08 | 0.03 |
| 7 | 45761952 | rs3735666 | -0.055 | 2.034 | 0.387 | 1.246 | 0.014 | -0.043 | -0.043 | 0 | 0.19 | 0.38 | 0.12 | 0.12 | 0.18 | 0.05 |
| 7 | 45764099 | rs1554465 | -0.589 | 1.958 | -0.536 | 1.099 | -0.049 | -0.034 | -0.034 | 0 | 0.09 | 0.02 | 0.18 | 0.12 | 0.23 | 0.1 |
| 7 | 45765234 | rs2960287 | 1.044 | 1.949 | -0.488 | 1.102 | -0.038 | -0.003 | -0.003 | 0.018 | 0.59 | 0.23 | 0.48 | 0.43 | 0.49 | 0.64 |
| 7 | 45868309 | rs4720522 | -1.212 | 2.296 | 0.027 | 1.105 | -0.047 | -0.034 | -0.034 | 0.051 | 0.09 | 0.02 | 0.18 | 0.12 | 0.23 | 0.1 |
| 7 | 45894105 | rs10275903 | -0.474 | 1.208 | -0.355 | 1.208 | -0.01 | -0.023 | -0.023 | 0 | 0.06 | 0.01 | 0.09 | 0.19 | 0.09 | 0.04 |
| 7 | 45898114 | rs13222310 | -0.877 | 2.082 | -0.389 | 1.288 | -0.013 | -0.007 | -0.007 | 0 | 0.09 | 0.04 | 0.16 | 0.1 | 0.15 | 0.09 |
| 7 | 45899405 | rs4724442 | -1.054 | 2.086 | -0.04 | 1.48 | 0.013 | 0.018 | 0.018 | 0.817 | 0.26 | 0.27 | 0.36 | 0.36 | 0.31 | 0.19 |
| 7 | 45901549 | rs2173035 | -0.552 | 1.767 | 0.41 | 1.367 | -0.01 | -0.002 | -0.002 | 0.014 | 0.06 | 0.2 | 0.04 | 0.05 | 0.04 | 0.12 |
| 7 | 45904498 | rs7794738 | 0.085 | 1.881 | 0.205 | 1.424 | 0.003 | -0.004 | -0.004 | 0.06 | 0.07 | 0.05 | 0.16 | 0.09 | 0.14 | 0.08 |
| 7 | 45906594 | rs12666800 | 0.085 | 1.881 | 0.205 | 1.424 | 0.003 | -0.004 | -0.004 | 1.239 | 0.07 | 0.05 | 0.16 | 0.09 | 0.14 | 0.08 |

| | | | | | | | | | | | | | | | | |
|---|---|---|---|---|---|---|---|---|---|---|---|---|---|---|---|---|
| 7 | 45906903 | rs12667301 | 0.467 | 1.858 | 0.283 | 1.467 | 0 | 0.004 | 0.004 | 1.886 | 0.15 | 0.12 | 0.18 | 0.26 | 0.15 | 0.09 |
| 7 | 45908689 | rs1995050 | -1.694 | 1.655 | 0.72 | 1.338 | 0.027 | 0.046 | 0.046 | 0 | 0.34 | 0.53 | 0.41 | 0.42 | 0.37 | 0.32 |
| 7 | 45908915 | rs10228265 | -1.798 | 1.855 | 0.643 | 1.2 | 0.058 | 0.057 | 0.057 | 0 | 0.23 | 0.43 | 0.32 | 0.35 | 0.32 | 0.27 |
| 7 | 45908994 | rs1553009 | 0.567 | 2.034 | 0.513 | 1.234 | -0.02 | -0.046 | -0.046 | 0 | 0.17 | 0.06 | 0.2 | 0.25 | 0.18 | 0.09 |
| 7 | 45909998 | rs1027438 | -1.726 | 1.883 | 0.546 | 1.009 | 0.003 | -0.003 | -0.003 | 0.432 | 0.41 | 0.56 | 0.52 | 0.61 | 0.51 | 0.36 |
| 7 | 45912764 | rs10241749 | -1.176 | 1.291 | -0.568 | 0.385 | -0.013 | 0.08 | 0.08 | 0 | 0.76 | 0.98 | 0.8 | 0.9 | 0.78 | 0.59 |
| 7 | 45915907 | rs10243669 | 0.505 | 1.904 | 0.207 | 0.941 | -0.032 | -0.037 | -0.037 | 0 | 0.07 | 0.01 | 0.11 | 0.2 | 0.16 | 0.05 |
| 7 | 45920582 | rs6964374 | -0.349 | 1.541 | 0.377 | 0.823 | -0.005 | 0.004 | 0.004 | 0 | 0.06 | 0.18 | 0.04 | 0.06 | 0.04 | 0.13 |
| 7 | 45927677 | rs1065780 | -1.195 | 1.989 | 0.42 | 0.808 | 0.038 | 0.064 | 0.064 | 0.721 | 0.29 | 0.41 | 0.38 | 0.52 | 0.39 | 0.18 |
| 7 | 45931135 | rs2854843 | 0.536 | 1.748 | 0.627 | 0.942 | -0.013 | -0.037 | -0.037 | 0.019 | 0.14 | 0.08 | 0.19 | 0.32 | 0.16 | 0.11 |
| 7 | 45932669 | rs4619 | -1.143 | 1.858 | 0.736 | 0.944 | 0.043 | 0.123 | 0.123 | 0 | 0.25 | 0.44 | 0.36 | 0.53 | 0.33 | 0.18 |
| 7 | 45934139 | rs9658238 | -0.266 | 1.853 | 0.638 | 0.933 | 0.003 | 0.002 | 0.002 | 0 | 0.09 | 0.1 | 0.15 | 0.2 | 0.18 | 0.08 |
| 7 | 45934159 | rs9658239 | 0.66 | 1.832 | 0.422 | 0.937 | -0.025 | -0.035 | -0.035 | 0 | 0.07 | 0.01 | 0.09 | 0.2 | 0.12 | 0.04 |
| 7 | 45940903 | rs1496496 | -1.266 | 1.825 | 0.471 | 0.905 | 0.04 | 0.118 | 0.118 | 0 | 0.26 | 0.44 | 0.36 | 0.53 | 0.34 | 0.18 |
| 7 | 45945469 | rs12702181 | 0.579 | 1.436 | 0.368 | 0.534 | -0.001 | 0.007 | 0.007 | 0.796 | 0.4 | 0.38 | 0.35 | 0.33 | 0.39 | 0.38 |
| 7 | 45949570 | rs2270628 | -1.418 | 1.963 | 0.62 | 0.903 | 0.084 | 0.069 | 0.069 | 0.191 | 0.12 | 0.38 | 0.18 | 0.19 | 0.2 | 0.08 |
| 7 | 45950446 | rs10282088 | -0.114 | 1.533 | 0.68 | 0.452 | 0.001 | 0.02 | 0.02 | 0 | 0.13 | 0.14 | 0.19 | 0.32 | 0.16 | 0.11 |
| 7 | 45951230 | rs13223993 | -0.836 | 1.442 | 0.526 | 0.465 | 0.057 | 0.119 | 0.119 | 0 | 0.19 | 0.47 | 0.27 | 0.33 | 0.22 | 0.14 |
| 7 | 45953573 | rs2453839 | -0.804 | 0.943 | 0.295 | -0.131 | -0.004 | -0.003 | -0.003 | 0 | 0.21 | 0.41 | 0.15 | 0.21 | 0.19 | 0.18 |
| 7 | 45955029 | rs3110697 | -0.154 | 0.884 | -0.611 | -0.86 | -0.002 | 0.055 | 0.055 | 0 | 0.53 | 0.63 | 0.38 | 0.75 | 0.57 | 0.5 |
| 7 | 45963114 | rs924140 | 0.145 | 0.918 | -0.578 | -1.095 | 0.014 | 0.038 | 0.038 | 0.007 | 0.47 | 0.58 | 0.33 | 0.76 | 0.46 | 0.46 |
| 7 | 45965795 | rs2453836 | 0.912 | 1.267 | -0.948 | -1.38 | 0.026 | 0.057 | 0.057 | 0 | 0.32 | 0.21 | 0.18 | 0.02 | 0.21 | 0.27 |
| 7 | 45977173 | rs2471552 | 0.008 | 1.395 | -0.936 | -1.282 | -0.027 | -0.015 | -0.015 | 0.016 | 0.86 | 0.93 | 0.58 | 0.79 | 0.79 | 0.76 |
| 7 | 45979023 | rs1496499 | 0.357 | 1.407 | -0.899 | -1.468 | 0.03 | 0.102 | 0.102 | 0 | 0.49 | 0.64 | 0.49 | 0.91 | 0.46 | 0.57 |
| 7 | 45984645 | rs1534151 | 0.355 | 1.466 | -0.947 | -1.378 | -0.001 | 0.046 | 0.046 | 0 | 0.38 | 0.24 | 0.25 | 0.02 | 0.33 | 0.33 |
| 7 | 45984820 | rs2949833 | 1.322 | 1.237 | -1.525 | -1.47 | -0.067 | -0.074 | -0.074 | 0 | 0.25 | 0.01 | 0.18 | 0.71 | 0.3 | 0.26 |
| 7 | 45997370 | rs11764490 | 0.619 | 1.688 | -0.816 | -1.224 | 0.014 | -0.015 | -0.015 | 0 | 0.05 | 0.04 | 0.05 | 0 | 0.11 | 0.04 |
| 7 | 46006544 | rs788718 | 0.679 | 1.239 | -0.987 | -1.408 | 0.03 | 0.101 | 0.101 | 0 | 0.31 | 0.13 | 0.19 | 0.03 | 0.21 | 0.26 |
| 7 | 46007868 | rs788720 | 0.358 | 2.174 | -0.835 | -1.239 | -0.02 | 0.004 | 0.004 | 0.043 | 0.86 | 0.91 | 0.68 | 1 | 0.66 | 0.84 |
| 7 | 46008110 | rs11977526 | 0.674 | 1.314 | -0.154 | -1.553 | 0.014 | 0.034 | 0.034 | 0.578 | 0.44 | 0.33 | 0.28 | 0.79 | 0.4 | 0.39 |
| 7 | 46016304 | rs1723957 | -0.336 | 1.95 | 0.171 | -1.289 | -0.009 | -0.002 | -0.002 | 0 | 0.89 | 0.93 | 0.73 | 1 | 0.76 | 0.87 |
| 7 | 46016662 | rs1722141 | -0.275 | 2.04 | 0.152 | -1.296 | -0.01 | 0.006 | 0.006 | 0.036 | 0.9 | 0.94 | 0.73 | 1 | 0.78 | 0.87 |
| 7 | 46018146 | rs2965069 | 0.82 | 1.087 | 0.104 | -1.447 | -0.018 | -0.039 | -0.039 | 0 | 0.4 | 0.43 | 0.25 | 0.78 | 0.33 | 0.42 |
| 7 | 46020785 | rs7795962 | 0.683 | 1.273 | -0.397 | -1.889 | 0.015 | 0.154 | 0.154 | 0 | 0.33 | 0.12 | 0.21 | 0.02 | 0.29 | 0.25 |
| 7 | 46021684 | rs2949307 | 1.157 | 1.494 | -0.133 | -1.484 | 0.07 | -0.1 | -0.1 | 0 | 0.75 | 0.59 | 0.61 | 0.96 | 0.62 | 0.76 |
| 7 | 46026181 | rs788748 | 0.102 | 1.238 | -0.459 | -1.286 | -0.041 | 0.147 | 0.147 | 0 | 0.57 | 0.76 | 0.5 | 0.92 | 0.49 | 0.63 |
| 7 | 46027518 | rs6962288 | -0.128 | 1.119 | 0.18 | -1.319 | -0.011 | -0.012 | -0.012 | 0.141 | 0.06 | 0.13 | 0.03 | 0.04 | 0.02 | 0.09 |
| 7 | 46028577 | rs1723939 | 0.454 | 1.059 | 0.48 | -1.307 | 0 | -0.017 | -0.017 | 0.278 | 0.48 | 0.51 | 0.46 | 0.88 | 0.47 | 0.54 |
| 7 | 46033433 | rs4278081 | 0.211 | 1.247 | 0.417 | -0.958 | 0.02 | 0.039 | 0.039 | 0 | 0.38 | 0.26 | 0.24 | 0.17 | 0.31 | 0.35 |
| 7 | 46034024 | rs2091148 | 1.038 | 1.288 | 0.4 | -0.988 | 0.016 | 0.045 | 0.045 | 1.005 | 0.41 | 0.27 | 0.38 | 0.82 | 0.35 | 0.5 |
| 7 | 46041100 | rs788740 | 0.355 | 1.232 | 0.388 | -1.056 | 0.025 | -0.042 | -0.042 | 0 | 0.85 | 0.76 | 0.69 | 0.98 | 0.77 | 0.89 |
| 7 | 46044387 | rs2965083 | 0.939 | 1.015 | 0.123 | -0.387 | 0.014 | -0.004 | -0.004 | 0 | 0.4 | 0.25 | 0.22 | 0.62 | 0.34 | 0.42 |
| 7 | 46049342 | rs879965 | 1.077 | 0.98 | 0.133 | -0.002 | 0.013 | 0.023 | 0.023 | 0 | 0.52 | 0.26 | 0.31 | 0.52 | 0.45 | 0.47 |
| 7 | 46049467 | rs879966 | 0.102 | 1.085 | 0.496 | 0.009 | 0.001 | -0.003 | -0.003 | 0 | 0.68 | 0.64 | 0.41 | 0.81 | 0.59 | 0.74 |
| 7 | 46059924 | rs1722135 | 0.712 | 1.165 | 0.442 | -0.049 | -0.015 | -0.093 | -0.093 | 2.051 | 0.21 | 0.31 | 0.14 | 0 | 0.18 | 0.06 |
| 7 | 46066152 | rs788745 | -0.23 | 1.22 | 0.392 | 0.044 | -0.027 | -0.022 | -0.022 | 0 | 0.88 | 0.94 | 0.54 | 0.81 | 0.76 | 0.8 |
| 7 | 46066786 | rs17172622 | -0.319 | 1.142 | 0.423 | 0.056 | -0.007 | 0 | 0 | 0.113 | 0.06 | 0.1 | 0.02 | 0.07 | 0.02 | 0.12 |
| 7 | 46068448 | rs788744 | -0.212 | 1.337 | 0.37 | 0.076 | -0.011 | -0.01 | -0.01 | 0 | 0.88 | 0.89 | 0.55 | 0.81 | 0.76 | 0.8 |
| 7 | 46072124 | rs7780838 | -0.003 | 1.016 | 0.453 | 0.089 | 0.013 | 0.018 | 0.018 | 0 | 0.09 | 0.15 | 0.08 | 0.22 | 0.12 | 0.16 |
| 7 | 46074194 | rs10277696 | 0.228 | 1.021 | 0.441 | 0.107 | 0.01 | 0.012 | 0.012 | 0 | 0.08 | 0.12 | 0.07 | 0.22 | 0.12 | 0.16 |
| 7 | 46076308 | rs812570 | -0.807 | 0.945 | 0.46 | 0.017 | -0.004 | -0.016 | -0.016 | 3.453 | 0.36 | 0.67 | 0.25 | 0.29 | 0.31 | 0.34 |
| 7 | 46089002 | rs4439027 | -1.864 | 1.215 | 0.301 | 0.738 | -0.006 | 0.029 | 0.029 | 0 | 0.33 | 0.39 | 0.36 | 0.56 | 0.3 | 0.48 |
| 7 | 46093628 | rs12111606 | -1.342 | 1.169 | 0.288 | 0.773 | -0.011 | 0.014 | 0.014 | 0 | 0.24 | 0.26 | 0.28 | 0.36 | 0.17 | 0.32 |
| 7 | 46113834 | rs13226164 | -0.061 | 1.208 | 0.223 | 0.729 | 0.004 | 0.006 | 0.006 | 0 | 0.16 | 0.08 | 0.1 | 0.09 | 0.14 | 0.12 |
| 7 | 46116722 | rs6968126 | -0.248 | 1.211 | 0.286 | 0.887 | 0.008 | 0.014 | 0.014 | 0 | 0.09 | 0.13 | 0.09 | 0.21 | 0.14 | 0.17 |
| 7 | 46117485 | rs12666823 | -0.813 | 1.168 | 0.333 | 0.88 | -0.007 | -0.002 | -0.002 | 3.564 | 0.06 | 0.1 | 0.02 | 0.07 | 0.02 | 0.12 |
| 7 | 46117498 | rs1404161 | 2.428 | 1.516 | 0.291 | 0.855 | -0.001 | 0 | 0 | 0 | 0.56 | 0.56 | 0.32 | 0.42 | 0.45 | 0.38 |
| 7 | 46118303 | rs1525826 | -1.552 | 1.559 | 0.232 | 0.771 | 0.003 | 0.014 | 0.014 | 0 | 0.15 | 0.24 | 0.11 | 0.27 | 0.15 | 0.3 |
| 7 | 46122350 | rs2464923 | 0.949 | 1.599 | 0.064 | 0.723 | -0.049 | 0.038 | 0.038 | 0 | 0.73 | 0.87 | 0.58 | 0.89 | 0.61 | 0.76 |
| 7 | 46149011 | rs12667006 | -0.738 | 1.598 | 0.185 | 0.991 | -0.004 | -0.002 | -0.002 | 0.373 | 0.06 | 0.09 | 0.03 | 0.06 | 0.02 | 0.14 |
| 7 | 46155974 | rs1085924 | -0.128 | 1.982 | 0.056 | 0.96 | -0.062 | 0.023 | 0.023 | 0 | 0.91 | 0.99 | 0.69 | 0.98 | 0.76 | 0.89 |
| 7 | 46157057 | rs1581571 | 2.094 | 1.594 | 0.179 | 1.04 | 0.003 | 0.004 | 0.004 | 0 | 0.58 | 0.49 | 0.34 | 0.43 | 0.48 | 0.42 |
| 7 | 46170512 | rs10499642 | 0.033 | 1.068 | 0.059 | 1.041 | -0.043 | 0.009 | 0.009 | 0 | 0.07 | 0.01 | 0.28 | 0.02 | 0.16 | 0.1 |
| 7 | 46170767 | rs17580669 | -2.159 | 0.851 | 0.222 | 1.265 | 0.005 | 0.022 | 0.022 | 0 | 0.31 | 0.42 | 0.56 | 0.41 | 0.32 | 0.43 |
| 7 | 46171884 | rs6944635 | -1.939 | 0.811 | 0.327 | 1.221 | 0.002 | 0.011 | 0.011 | 0 | 0.22 | 0.35 | 0.49 | 0.28 | 0.19 | 0.34 |
| 7 | 46172466 | rs1486160 | -2.208 | 0.41 | 0.502 | 1.504 | 0 | 0.034 | 0.034 | 0 | 0.24 | 0.33 | 0.16 | 0.4 | 0.24 | 0.37 |
| 7 | 46175077 | rs7794166 | -2.642 | -0.143 | 0.464 | 1.148 | 0.023 | 0.024 | 0.024 | 0 | 0.5 | 0.66 | 0.49 | 0.62 | 0.42 | 0.61 |
| 7 | 46176155 | rs1586360 | -0.843 | 0.141 | 0.773 | 1.348 | -0.016 | -0.019 | -0.019 | 0 | 0.12 | 0.16 | 0.07 | 0.04 | 0.06 | 0.13 |
| 7 | 46186950 | rs11772890 | -2.899 | 0.104 | 0.592 | 1.34 | 0.001 | 0.001 | 0.001 | 0 | 0.45 | 0.58 | 0.45 | 0.66 | 0.41 | 0.62 |

| | | | | | | | | | | | | | | | | |
|---|---|---|---|---|---|---|---|---|---|---|---|---|---|---|---|---|
| 7 | 46190860 | rs17580738 | 0.053 | 0.184 | 0.455 | 1.299 | 0.002 | -0.003 | -0.003 | 0 | 0.06 | 0.05 | 0.07 | 0.02 | 0.09 | 0.09 |
| 7 | 46199753 | rs6968071 | -1.05 | 0.194 | 0.306 | 1.363 | -0.011 | -0.002 | -0.002 | 0.435 | 0.19 | 0.3 | 0.15 | 0.15 | 0.14 | 0.1 |
| 7 | 46206780 | rs1486170 | -2.564 | 0.329 | 0.294 | 1.352 | -0.001 | 0 | 0 | 0 | 0.43 | 0.5 | 0.44 | 0.61 | 0.41 | 0.56 |
| 7 | 46210835 | rs11768394 | -0.485 | 0.586 | -0.009 | 1.473 | -0.022 | -0.028 | -0.028 | 0.503 | 0.07 | 0.03 | 0.03 | 0.17 | 0.11 | 0.15 |
| 7 | 46212371 | rs10266929 | -1.935 | 0.503 | 0.322 | 1.39 | 0.001 | 0.018 | 0.018 | 0 | 0.33 | 0.43 | 0.4 | 0.55 | 0.35 | 0.45 |
| 7 | 46212931 | rs12672756 | -1.149 | 0.783 | 0.315 | 1.677 | -0.012 | -0.028 | -0.028 | 0 | 0.15 | 0.08 | 0.15 | 0.25 | 0.17 | 0.2 |
| 7 | 46213818 | rs4559153 | -2.212 | 0.373 | 0.685 | 1.4 | -0.008 | 0.002 | 0.002 | 0 | 0.35 | 0.56 | 0.21 | 0.35 | 0.28 | 0.42 |
| 7 | 46216799 | rs17151092 | -1.179 | 0.612 | 0.67 | 1.421 | -0.008 | -0.001 | -0.001 | 0 | 0.07 | 0.03 | 0.1 | 0.07 | 0.08 | 0.03 |
| 7 | 46218147 | rs11763946 | -2.938 | 0.606 | 1.302 | 1.7 | -0.001 | 0.013 | 0.013 | 0 | 0.29 | 0.6 | 0.23 | 0.43 | 0.25 | 0.28 |
| 7 | 46221810 | rs7797417 | -2.274 | 0.571 | 1.839 | 1.74 | 0.016 | 0.034 | 0.034 | 0 | 0.11 | 0.3 | 0.11 | 0.14 | 0.1 | 0.11 |
| 7 | 46223986 | rs11762916 | -1.603 | 0.555 | 1.959 | 1.745 | 0.002 | 0.013 | 0.013 | 0 | 0.15 | 0.19 | 0.11 | 0.29 | 0.15 | 0.17 |
| 7 | 46226952 | rs10951847 | -3.306 | 0.56 | 1.873 | 1.744 | -0.001 | 0.014 | 0.014 | 0 | 0.28 | 0.6 | 0.23 | 0.43 | 0.25 | 0.28 |
| 7 | 46233664 | rs7795327 | -3.368 | 0.533 | 1.884 | 1.707 | -0.001 | 0.005 | 0.005 | 0 | 0.29 | 0.67 | 0.24 | 0.42 | 0.25 | 0.28 |
| 7 | 46234083 | rs11978068 | -1.966 | 0.733 | 1.414 | 1.597 | 0.006 | 0.051 | 0.051 | 0 | 0.17 | 0.29 | 0.12 | 0.29 | 0.15 | 0.17 |
| 7 | 46237040 | rs7778867 | -0.948 | 0.719 | 0.979 | 1.349 | 0.012 | 0.001 | 0.001 | 0 | 0.06 | 0.07 | 0.1 | 0.07 | 0.08 | 0.03 |
| 7 | 46241570 | rs6976880 | -1.277 | 0.661 | 0.942 | 1.452 | 0 | 0.022 | 0.022 | 0 | 0.12 | 0.19 | 0.09 | 0.14 | 0.11 | 0.02 |
| 7 | 46246580 | rs9691259 | -2.094 | 0.649 | 1.089 | 1.429 | 0.003 | 0.028 | 0.028 | 0 | 0.2 | 0.37 | 0.21 | 0.22 | 0.2 | 0.04 |
| 7 | 46248354 | rs936856 | 1.155 | 0.901 | 0.317 | 0.944 | -0.024 | 0.048 | 0.048 | 0 | 0.64 | 0.2 | 0.62 | 0.8 | 0.63 | 0.65 |
| 7 | 46251052 | rs10281778 | 1.338 | 0.794 | 0.24 | 0.967 | -0.03 | 0.049 | 0.049 | 0 | 0.62 | 0.2 | 0.61 | 0.77 | 0.61 | 0.65 |
| 7 | 46251743 | rs7457604 | 1.559 | 0.535 | 0.045 | 1.267 | -0.053 | -0.012 | -0.012 | 0 | 0.59 | 0.14 | 0.56 | 0.67 | 0.57 | 0.51 |
| 7 | 46259005 | rs17583337 | 0.204 | 0.733 | 1.046 | 1.072 | -0.02 | 0.04 | 0.04 | 0 | 0.14 | 0.06 | 0.09 | 0.01 | 0.16 | 0.07 |
| 7 | 46259925 | rs7778803 | 1.058 | 0.722 | 1.117 | 1.069 | 0.008 | 0.004 | 0.004 | 0 | 0.76 | 0.69 | 0.84 | 0.78 | 0.72 | 0.64 |
| 7 | 46261337 | rs10258322 | 2.435 | 0.71 | 1.03 | 1.063 | -0.045 | 0.011 | 0.011 | 0 | 0.57 | 0.14 | 0.56 | 0.69 | 0.56 | 0.51 |
| 7 | 46264195 | rs17583463 | 0.245 | 0.805 | 0.851 | 1.045 | -0.02 | 0.037 | 0.037 | 0 | 0.14 | 0.06 | 0.09 | 0 | 0.16 | 0.07 |
| 7 | 46264235 | rs4720535 | 0.361 | 0.771 | 0.889 | 1.178 | 0.026 | -0.03 | -0.03 | 0 | 0.8 | 0.69 | 0.88 | 0.88 | 0.77 | 0.78 |
| 7 | 46290701 | rs13234885 | 2.134 | 0.689 | 0.984 | 1.084 | -0.027 | 0.014 | 0.014 | 0 | 0.58 | 0.18 | 0.55 | 0.7 | 0.56 | 0.51 |
| 7 | 46294240 | rs10270664 | 1.458 | 0.684 | 0.992 | 1.082 | 0.01 | -0.036 | -0.036 | 1.359 | 0.66 | 0.61 | 0.79 | 0.87 | 0.61 | 0.72 |
| 7 | 46295363 | rs10229794 | 2.139 | 0.68 | 1.075 | 1.123 | -0.007 | 0.003 | 0.003 | 0 | 0.58 | 0.31 | 0.56 | 0.7 | 0.56 | 0.52 |
| 7 | 46297747 | rs11981109 | -0.47 | 0.627 | 0.838 | 1.159 | -0.025 | 0.11 | 0.11 | 0 | 0.07 | 0.21 | 0.22 | 0.17 | 0.05 | 0.21 |
| 7 | 46310166 | rs17584269 | -0.108 | 1.069 | 0.431 | 0.962 | -0.025 | 0.038 | 0.038 | 0 | 0.15 | 0.08 | 0.11 | 0 | 0.18 | 0.08 |
| 7 | 46310371 | rs11767887 | 1.582 | 0.235 | -0.908 | 0.632 | -0.075 | 0.045 | 0.045 | 0.876 | 0.28 | 0.01 | 0.22 | 0.14 | 0.33 | 0.21 |
| 7 | 46311210 | rs13228308 | 1.154 | 0.536 | -0.322 | 0.784 | -0.022 | 0.012 | 0.012 | 0 | 0.57 | 0.22 | 0.55 | 0.7 | 0.54 | 0.56 |
| 7 | 46320115 | rs1903674 | -0.144 | 1.043 | -0.237 | 1.197 | 0.005 | 0 | 0 | 0.01 | 0.22 | 0.22 | 0.19 | 0.29 | 0.35 | 0.22 |
| 7 | 46322635 | rs997490 | 1.028 | 0.487 | -0.135 | 1.173 | 0.013 | 0.001 | 0.001 | 4.972 | 0.54 | 0.37 | 0.53 | 0.4 | 0.48 | 0.43 |
| 7 | 46323681 | rs7805659 | 0.028 | 0.459 | 0.039 | 1.096 | -0.011 | 0.006 | 0.006 | 0 | 0.76 | 0.67 | 0.72 | 0.69 | 0.83 | 0.65 |
| 7 | 46331207 | rs13237404 | 1.5 | 0.112 | -0.475 | 0.648 | 0.023 | -0.016 | -0.016 | 2.897 | 0.1 | 0.01 | 0.07 | 0.07 | 0.08 | 0.1 |
| 7 | 46331712 | rs1966956 | -0.749 | 0.156 | -0.324 | 0.704 | 0.016 | 0.023 | 0.023 | 1.031 | 0.34 | 0.42 | 0.36 | 0.45 | 0.43 | 0.35 |
| 7 | 46338819 | rs6959818 | 0.247 | 0.815 | 0.307 | 1.106 | 0.012 | 0.013 | 0.013 | 0.028 | 0.14 | 0.17 | 0.1 | 0.3 | 0.24 | 0.19 |
| 7 | 46344069 | rs6952576 | -0.785 | 0.729 | 0.319 | 1.059 | 0.017 | 0.029 | 0.029 | 0.041 | 0.24 | 0.32 | 0.18 | 0.37 | 0.33 | 0.29 |
| 7 | 46350602 | rs7790440 | 0.278 | 0.747 | 0.338 | 1.063 | 0.031 | 0.032 | 0.032 | 0.025 | 0.14 | 0.19 | 0.11 | 0.3 | 0.24 | 0.19 |
| 7 | 46360746 | rs12538141 | 1.334 | 0.601 | 0.17 | 0.974 | 0.031 | -0.01 | -0.01 | 0 | 0.1 | 0.01 | 0.07 | 0.07 | 0.08 | 0.1 |
| 7 | 46368995 | rs10252568 | 1.002 | 0.203 | 0.249 | 0.802 | -0.016 | 0.001 | 0.001 | 0 | 0.62 | 0.42 | 0.63 | 0.59 | 0.77 | 0.54 |
| 7 | 46369680 | rs7457578 | 1.142 | 0.183 | 0.262 | 0.796 | 0.04 | -0.002 | -0.002 | 0 | 0.61 | 0.29 | 0.62 | 0.59 | 0.76 | 0.51 |
| 7 | 46384262 | rs6977956 | 2.302 | 0.276 | -0.164 | 0.572 | -0.148 | 0.08 | 0.08 | 0.886 | 0.32 | 0.02 | 0.36 | 0.13 | 0.41 | 0.18 |
| 7 | 46384534 | rs6960609 | 1.945 | 0.423 | 0.096 | 0.834 | -0.051 | 0.107 | 0.107 | 0.047 | 0.48 | 0.11 | 0.51 | 0.29 | 0.52 | 0.33 |
| 7 | 46386508 | rs4570054 | 1.128 | 0.223 | 0.256 | 0.845 | 0.017 | -0.002 | -0.002 | 0 | 0.61 | 0.35 | 0.62 | 0.59 | 0.76 | 0.51 |
| 7 | 46386613 | rs7458399 | 0.207 | 0.203 | 0.307 | 0.846 | -0.041 | 0.021 | 0.021 | 0 | 0.74 | 0.57 | 0.7 | 0.66 | 0.85 | 0.64 |
| 7 | 46387679 | rs1350927 | 0.92 | 0.228 | 0.307 | 0.85 | -0.034 | 0.004 | 0.004 | 0 | 0.63 | 0.55 | 0.63 | 0.59 | 0.77 | 0.53 |
| 7 | 46390040 | rs753667 | 0.909 | 0.487 | 0.314 | 0.973 | 0.014 | -0.013 | -0.013 | 1.132 | 0.65 | 0.32 | 0.55 | 0.51 | 0.77 | 0.49 |
| 7 | 46390788 | rs898930 | -0.102 | 0.672 | 0.326 | 0.926 | -0.019 | 0.056 | 0.056 | 0 | 0.79 | 0.61 | 0.73 | 0.66 | 0.86 | 0.65 |
| 7 | 46396881 | rs13244035 | 0.562 | 0.659 | 0.247 | 0.97 | -0.041 | 0.028 | 0.028 | 0 | 0.13 | 0.23 | 0.17 | 0.14 | 0.09 | 0.14 |
| 7 | 46402100 | rs7794677 | 0.865 | 1.446 | 0.137 | 1.125 | 0.058 | 0.077 | 0.077 | 0.01 | 0.1 | 0.18 | 0.08 | 0.32 | 0.22 | 0.16 |
| 7 | 46402263 | rs7794838 | 0.865 | 1.446 | 0.137 | 1.125 | 0.058 | 0.077 | 0.077 | 0.01 | 0.1 | 0.18 | 0.09 | 0.32 | 0.22 | 0.16 |
| 7 | 46402269 | rs7811653 | 1.308 | 1.151 | -0.112 | 0.964 | 0.024 | 0.026 | 0.026 | 0 | 0.1 | 0.01 | 0.07 | 0.03 | 0.08 | 0.11 |
| 7 | 46403343 | rs7800108 | 0.008 | 1.102 | 0.049 | 0.897 | -0.004 | 0.029 | 0.029 | 0.001 | 0.77 | 0.65 | 0.71 | 0.63 | 0.82 | 0.59 |
| 7 | 46418650 | rs7778674 | 1.778 | 0.989 | -0.799 | 0.581 | -0.121 | 0.043 | 0.043 | 1.446 | 0.36 | 0.02 | 0.4 | 0.18 | 0.42 | 0.18 |
| 7 | 46432488 | rs17586544 | 1.889 | 1.978 | -0.412 | 0.996 | -0.064 | 0.065 | 0.065 | 0.415 | 0.11 | 0.01 | 0.14 | 0 | 0.22 | 0.05 |
| 7 | 46432603 | rs1462278 | 1.386 | 1.101 | 0.699 | 1.708 | -0.009 | 0.006 | 0.006 | 0.026 | 0.67 | 0.36 | 0.55 | 0.57 | 0.75 | 0.48 |
| 7 | 46450230 | rs10275461 | -0.376 | 0.667 | 0.787 | 1.353 | 0.039 | 0.073 | 0.073 | 0 | 0.15 | 0.24 | 0.09 | 0.35 | 0.22 | 0.18 |
| 7 | 46451098 | rs6959948 | -1.05 | 0.684 | 0.796 | 1.342 | 0.009 | 0.038 | 0.038 | 0 | 0.25 | 0.34 | 0.16 | 0.38 | 0.29 | 0.28 |
| 7 | 46462742 | rs260370 | -1.191 | 0.489 | 1.015 | 1.296 | 0.033 | 0.233 | 0.233 | 0 | 0.14 | 0.52 | 0.33 | 0.31 | 0.11 | 0.34 |
| 7 | 46463342 | rs260368 | -1.864 | 0.262 | 1.081 | 1.13 | 0.016 | -0.02 | -0.02 | 0 | 0.28 | 0.8 | 0.42 | 0.37 | 0.21 | 0.44 |
| 7 | 46465996 | rs260362 | -1.378 | 0.273 | 1.359 | 1.006 | 0.043 | -0.024 | -0.024 | 0.113 | 0.1 | 0.38 | 0.07 | 0.03 | 0.08 | 0.13 |
| 7 | 46472363 | rs12537559 | -0.577 | 0.248 | 1.812 | 1.006 | -0.05 | -0.068 | -0.068 | 0 | 0.16 | 0.35 | 0.11 | 0.03 | 0.11 | 0.14 |
| 7 | 46489309 | rs4470909 | -0.14 | 0.292 | 1.685 | 1.017 | 0.049 | 0.004 | 0.004 | 0 | 0.06 | 0.22 | 0.03 | 0.03 | 0.06 | 0.1 |
| 7 | 46491184 | rs2345016 | -2.044 | 0.242 | 1.549 | 1.104 | 0 | 0 | 0 | 0 | 0.27 | 0.74 | 0.42 | 0.37 | 0.21 | 0.43 |
| 7 | 46501859 | rs11972406 | 0.281 | 0.317 | 1.475 | 0.997 | 0.014 | -0.01 | -0.01 | 0 | 0.05 | 0.12 | 0.03 | 0.03 | 0.06 | 0.1 |
| 7 | 46511393 | rs3890267 | -1.388 | 0.232 | 1.425 | 0.974 | -0.053 | -0.1 | -0.1 | 0 | 0.22 | 0.55 | 0.14 | 0.06 | 0.16 | 0.24 |
| 7 | 46525612 | rs10236600 | 0.173 | 0.709 | 0.298 | 1.319 | -0.016 | -0.014 | -0.014 | 0.235 | 0.11 | 0.06 | 0.07 | 0.3 | 0.21 | 0.16 |

| | | | | | | | | | | | | | | | | |
|---|---|---|---|---|---|---|---|---|---|---|---|---|---|---|---|---|
| 7 | 46530146 | rs970605 | 0.589 | 0.297 | 0.094 | 1.27 | 0.024 | 0.031 | 0.031 | 0 | 0.11 | 0.01 | 0.07 | 0.02 | 0.1 | 0.13 |
| 7 | 46532584 | rs10243723 | -0.408 | 0.76 | 0.474 | 1.731 | 0.018 | 0.236 | 0.236 | 0 | 0.06 | 0.21 | 0.28 | 0.31 | 0.05 | 0.19 |
| 7 | 46535632 | rs9648087 | -1.202 | 0.785 | -0.442 | 0.651 | -0.046 | 0.076 | 0.076 | 1.632 | 0.63 | 0.93 | 0.6 | 0.82 | 0.61 | 0.83 |
| 7 | 46570611 | rs12535902 | -1.075 | 1.05 | 0.554 | 1.227 | 0.02 | 0.016 | 0.016 | 1.089 | 0.17 | 0.34 | 0.11 | 0.22 | 0.19 | 0.29 |
| 7 | 46577056 | rs723149 | -0.959 | 0.587 | 0.175 | 1.701 | -0.078 | 0.077 | 0.077 | 1.088 | 0.54 | 0.95 | 0.43 | 0.72 | 0.54 | 0.69 |
| 7 | 46591617 | rs6946252 | -1.474 | 0.687 | 0.831 | 1.057 | 0.023 | 0.058 | 0.058 | 0.016 | 0.1 | 0.3 | 0.07 | 0.15 | 0.08 | 0.14 |
| 7 | 46611204 | rs10807935 | -0.116 | 0.522 | 0.212 | 0.599 | -0.046 | 0.027 | 0.027 | 0 | 0.32 | 0.24 | 0.42 | 0.21 | 0.51 | 0.42 |
| 7 | 46611926 | rs6974838 | -0.664 | 0.538 | 1.393 | 0.717 | 0.112 | 0.086 | 0.086 | 0 | 0.07 | 0.3 | 0.06 | 0.1 | 0.08 | 0.14 |
| 7 | 46617526 | rs6964695 | 0.79 | -0.171 | 1.395 | 0.364 | 0.002 | 0.093 | 0.093 | 0 | 0.53 | 0.23 | 0.5 | 0.76 | 0.42 | 0.42 |
| 7 | 46619758 | rs7781043 | 0.441 | -0.231 | 1.473 | 0.337 | -0.018 | -0.011 | -0.011 | 0 | 0.94 | 0.86 | 0.99 | 0.98 | 0.98 | 0.98 |
| 7 | 46620145 | rs7784776 | 0.702 | -0.215 | 1.393 | 0.342 | -0.001 | -0.007 | -0.007 | 1.917 | 0.54 | 0.41 | 0.51 | 0.76 | 0.42 | 0.42 |
| 7 | 46622308 | rs17172724 | 1.174 | -0.048 | 1.037 | 0.483 | 0.098 | 0.085 | 0.085 | 0.05 | 0.5 | 0.12 | 0.46 | 0.66 | 0.33 | 0.3 |
| 7 | 46623843 | rs10951858 | 0.331 | -0.152 | 1.096 | 0.501 | -0.049 | 0.02 | 0.02 | 0 | 0.3 | 0.22 | 0.43 | 0.22 | 0.52 | 0.39 |
| 7 | 46640168 | rs17172727 | -0.873 | -0.131 | 1.491 | 0.437 | -0.001 | -0.027 | -0.027 | 0.003 | 0.08 | 0.29 | 0.03 | 0 | 0.03 | 0.04 |
| 7 | 46640784 | rs6947204 | 0.194 | 0.028 | 1.387 | 0.418 | -0.042 | 0.024 | 0.024 | 0 | 0.3 | 0.2 | 0.42 | 0.19 | 0.49 | 0.39 |
| 7 | 46640900 | rs6968945 | -0.824 | -0.003 | 1.544 | 0.409 | -0.033 | 0.075 | 0.075 | 0 | 0.42 | 0.76 | 0.49 | 0.23 | 0.55 | 0.47 |
| 7 | 46641358 | rs17257433 | 0.121 | 0.252 | 1.222 | 0.506 | -0.041 | 0.022 | 0.022 | 0 | 0.29 | 0.2 | 0.42 | 0.19 | 0.49 | 0.39 |
| 7 | 46642397 | rs983398 | 0.176 | 0.269 | 1.177 | 0.611 | -0.057 | -0.042 | -0.042 | 0.708 | 0.29 | 0.18 | 0.42 | 0.44 | 0.5 | 0.43 |
| 7 | 46642993 | rs11773468 | -0.215 | 0.255 | 1.207 | 0.581 | -0.024 | -0.018 | -0.018 | 0 | 0.69 | 0.74 | 0.57 | 0.54 | 0.5 | 0.55 |
| 7 | 46649625 | rs17198174 | -0.154 | 0.147 | 0.797 | 0.142 | -0.04 | 0.032 | 0.032 | 0 | 0.16 | 0.09 | 0.16 | 0.08 | 0.26 | 0.24 |
| 7 | 46655415 | rs7810635 | 0.351 | 0.135 | 0.629 | 0.168 | -0.056 | -0.049 | -0.049 | 0 | 0.29 | 0.17 | 0.42 | 0.46 | 0.5 | 0.44 |
| 7 | 46656869 | rs9648555 | -0.165 | 0.375 | 0.089 | 0.29 | -0.005 | -0.004 | -0.004 | 0 | 0.62 | 0.31 | 0.53 | 0.51 | 0.47 | 0.53 |
| 7 | 46665658 | rs12535934 | 0.358 | -0.533 | -1.274 | 0.647 | 0.124 | -0.011 | -0.011 | 0 | 0.45 | 0.13 | 0.36 | 0.42 | 0.21 | 0.26 |
| 7 | 46670743 | rs993183 | -0.243 | 0.229 | -1.441 | 0.472 | -0.026 | -0.02 | -0.02 | 0.241 | 0.45 | 0.29 | 0.59 | 0.47 | 0.47 | 0.26 |
| 7 | 46678255 | rs767395 | -0.154 | 0.15 | -1.438 | 0.368 | -0.049 | -0.052 | -0.052 | 0 | 0.43 | 0.14 | 0.54 | 0.46 | 0.45 | 0.23 |
| 7 | 46689571 | rs17172739 | -0.031 | 0.78 | -1.717 | 0.568 | -0.099 | -0.09 | -0.09 | 0.508 | 0.2 | 0.04 | 0.28 | 0.33 | 0.26 | 0.08 |
| 7 | 46691762 | rs12112847 | -0.043 | 0.965 | -1.71 | 0.612 | -0.056 | -0.052 | -0.052 | 0.696 | 0.2 | 0.07 | 0.29 | 0.35 | 0.3 | 0.11 |
| 7 | 46692667 | rs13237848 | -0.336 | 0.135 | -2.473 | 0.011 | -0.035 | -0.062 | -0.062 | 0.295 | 0.61 | 0.11 | 0.69 | 0.42 | 0.62 | 0.39 |
| 7 | 46695819 | rs2082364 | 0.427 | -0.268 | -1.481 | -0.217 | 0.039 | 0.058 | 0.058 | 1.095 | 0.28 | 0.4 | 0.27 | 0.58 | 0.36 | 0.56 |
| 7 | 46696175 | rs10257870 | 0.377 | 0.186 | -1.839 | 0.085 | 0 | 0.024 | 0.024 | 0 | 0.08 | 0.07 | 0.07 | 0.26 | 0.1 | 0.21 |
| 7 | 46699079 | rs4724485 | 0.597 | -0.441 | -2.033 | -0.756 | 0.034 | 0.125 | 0.125 | 0 | 0.33 | 0.06 | 0.34 | 0.07 | 0.26 | 0.17 |
| 7 | 46700168 | rs12540816 | 0.441 | -0.445 | -1.996 | -0.729 | 0.033 | 0.146 | 0.146 | 0 | 0.36 | 0.07 | 0.39 | 0.08 | 0.3 | 0.22 |
| 7 | 46708387 | rs856540 | -0.271 | 0.176 | -1.159 | -0.876 | -0.005 | -0.031 | -0.031 | 0 | 0.46 | 0.49 | 0.52 | 0.89 | 0.64 | 0.71 |
| 7 | 46713060 | rs11973070 | 0.124 | 0.986 | -1.182 | -0.141 | -0.005 | 0 | 0 | 0 | 0.18 | 0.16 | 0.26 | 0.49 | 0.29 | 0.47 |
| 7 | 46717899 | rs6975578 | 0.124 | 0.986 | -1.182 | -0.141 | -0.005 | 0 | 0 | 0 | 0.18 | 0.17 | 0.26 | 0.49 | 0.29 | 0.47 |
| 7 | 46717956 | rs6975330 | 0.342 | 0.407 | -1.798 | -0.874 | -0.03 | 0 | 0 | 0.098 | 0.58 | 0.24 | 0.72 | 0.58 | 0.6 | 0.74 |
| 7 | 46721237 | rs17259858 | 1.699 | 0.347 | -1.867 | -0.904 | 0.014 | 0.115 | 0.115 | 0.016 | 0.33 | 0.02 | 0.38 | 0.08 | 0.24 | 0.16 |
| 7 | 46721854 | rs856565 | 0.637 | 0.603 | -1.557 | -1.075 | 0.008 | 0.015 | 0.015 | 0 | 0.22 | 0.24 | 0.2 | 0.41 | 0.27 | 0.16 |
| 7 | 46723510 | rs856563 | -0.703 | 0.557 | -1.528 | -1.065 | 0.01 | 0.008 | 0.008 | 0.037 | 0.46 | 0.41 | 0.54 | 0.91 | 0.63 | 0.74 |
| 7 | 46728937 | rs1404727 | 0.779 | 0.698 | -1.581 | -0.99 | -0.005 | 0 | 0 | 0 | 0.18 | 0.16 | 0.26 | 0.49 | 0.29 | 0.47 |
| 7 | 46733850 | rs878024 | 2.076 | 0.604 | -1.706 | -1.005 | 0.014 | 0.105 | 0.105 | 0.016 | 0.33 | 0.02 | 0.38 | 0.1 | 0.24 | 0.16 |
| 7 | 46734307 | rs856589 | 0.078 | 0.621 | -1.484 | -0.975 | -0.023 | -0.025 | -0.025 | 0 | 0.27 | 0.21 | 0.28 | 0.4 | 0.34 | 0.27 |
| 7 | 46745813 | rs10242889 | 0.07 | 0.342 | -0.65 | -0.98 | -0.032 | -0.151 | -0.151 | 0.527 | 0.2 | 0.49 | 0.08 | 0.01 | 0.13 | 0.1 |
| 7 | 46751036 | rs969951 | 0.824 | 0.603 | -0.568 | -0.861 | 0.001 | 0.02 | 0.02 | 0 | 0.16 | 0.17 | 0.25 | 0.5 | 0.32 | 0.48 |
| 7 | 46752880 | rs1030640 | 0.161 | 0.445 | 0.027 | -0.979 | -0.041 | -0.11 | -0.11 | 0.078 | 0.19 | 0.32 | 0.07 | 0 | 0.09 | 0.09 |
| 7 | 46754686 | rs1525242 | 0.477 | 0.472 | -0.032 | -0.967 | -0.029 | -0.091 | -0.091 | 0 | 0.18 | 0.27 | 0.07 | 0 | 0.09 | 0.09 |
| 7 | 46754744 | rs970316 | 1.007 | 0.894 | -0.22 | -0.2 | -0.008 | -0.009 | -0.009 | 0 | 0.12 | 0.1 | 0.15 | 0.26 | 0.28 | 0.33 |
| 7 | 46760129 | rs856554 | 0.442 | 0.699 | -0.175 | -0.012 | 0.007 | 0.012 | 0.012 | 0 | 0.22 | 0.24 | 0.18 | 0.32 | 0.25 | 0.15 |
| 7 | 46778653 | rs957755 | 2.061 | -0.424 | -0.828 | -0.664 | 0.016 | 0.042 | 0.042 | 2.378 | 0.25 | 0.04 | 0.25 | 0.16 | 0.19 | 0.22 |
| 7 | 46780614 | rs6463347 | -0.942 | -0.252 | -0.652 | -0.522 | 0.016 | 0.014 | 0.014 | 0 | 0.75 | 0.69 | 0.7 | 0.68 | 0.71 | 0.84 |
| 7 | 46786796 | rs12672158 | -0.268 | -0.27 | -0.603 | -0.515 | -0.001 | -0.005 | -0.005 | 0 | 0.46 | 0.42 | 0.43 | 0.52 | 0.51 | 0.6 |
| 7 | 46790462 | rs6956126 | -1.72 | 0.282 | -0.621 | -0.289 | -0.054 | 0.053 | 0.053 | 0 | 0.89 | 0.99 | 0.93 | 1 | 0.84 | 0.97 |
| 7 | 46793982 | rs7789758 | -0.002 | -0.053 | -0.417 | -0.531 | -0.013 | -0.005 | -0.005 | 0 | 0.6 | 0.75 | 0.56 | 0.44 | 0.57 | 0.67 |
| 7 | 46795322 | rs11974105 | 0.797 | -0.252 | -0.047 | -0.617 | -0.043 | -0.049 | -0.049 | 0 | 0.1 | 0.15 | 0.03 | 0 | 0.01 | 0 |
| 7 | 46796371 | rs6967755 | -0.39 | -0.185 | 0.172 | -0.766 | -0.038 | -0.104 | -0.104 | 0 | 0.16 | 0.38 | 0.07 | 0.01 | 0.08 | 0.04 |
| 7 | 46796519 | rs17587111 | 0.463 | 1.061 | -0.499 | -0.913 | -0.029 | 0.021 | 0.021 | 0 | 0.13 | 0.05 | 0.08 | 0 | 0.18 | 0.04 |
| 7 | 46796544 | rs11767621 | 1.015 | 0.296 | -1.055 | -0.805 | 0.002 | 0.026 | 0.026 | 1.744 | 0.23 | 0.07 | 0.13 | 0.17 | 0.21 | 0.17 |
| 7 | 46801428 | rs1464841 | -0.122 | 0.762 | -0.393 | -1.123 | 0.009 | -0.036 | -0.036 | 0 | 0.19 | 0.25 | 0.17 | 0.03 | 0.26 | 0.15 |
| 7 | 46802560 | rs10229581 | 1.203 | 0.567 | -0.271 | -0.918 | 0.005 | 0.006 | 0.006 | 1.074 | 0.25 | 0.19 | 0.27 | 0.23 | 0.23 | 0.23 |
| 7 | 46803846 | rs13230502 | -0.058 | 0.756 | -0.374 | -1.022 | 0.023 | -0.073 | -0.073 | 0 | 0.2 | 0.35 | 0.18 | 0.03 | 0.26 | 0.15 |
| 7 | 46804913 | rs1525240 | -0.747 | 0.307 | -1.447 | -1.115 | -0.008 | 0.088 | 0.088 | 0.631 | 0.75 | 0.97 | 0.87 | 0.88 | 0.74 | 0.79 |
| 7 | 46807301 | rs13245231 | 0.895 | 0.439 | -1.177 | -0.947 | -0.017 | 0.039 | 0.039 | 0 | 0.15 | 0.06 | 0.15 | 0.03 | 0.22 | 0.21 |
| 7 | 46811178 | rs12668204 | 0.896 | 0.267 | -0.775 | 0.004 | 0 | 0.003 | 0.003 | 0 | 0.23 | 0.21 | 0.17 | 0.37 | 0.22 | 0.31 |
| 7 | 46813079 | rs10232103 | -1.37 | -0.178 | -1.236 | 0.15 | 0.004 | -0.001 | -0.001 | 0.212 | 0.79 | 0.97 | 0.74 | 0.8 | 0.78 | 0.85 |
| 7 | 46815537 | rs1404719 | 0.018 | -0.23 | -0.657 | -0.049 | -0.001 | -0.001 | -0.001 | 0 | 0.43 | 0.55 | 0.53 | 0.48 | 0.44 | 0.54 |
| 7 | 46822686 | rs1525229 | -0.689 | -0.226 | -0.528 | 0.073 | -0.001 | -0.003 | -0.003 | 0.839 | 0.72 | 0.71 | 0.73 | 0.68 | 0.7 | 0.8 |
| 7 | 46822880 | rs2346399 | -0.736 | -0.229 | -0.516 | 0.071 | -0.001 | -0.004 | -0.004 | 0 | 0.72 | 0.74 | 0.74 | 0.68 | 0.7 | 0.8 |
| 7 | 46831835 | rs12155095 | -0.671 | -0.131 | -0.32 | -0.063 | -0.003 | -0.002 | -0.002 | 0 | 0.72 | 0.63 | 0.72 | 0.73 | 0.71 | 0.77 |
| 7 | 46844783 | rs1917612 | -0.592 | -0.102 | -0.205 | -0.135 | -0.006 | -0.003 | -0.003 | 0 | 0.71 | 0.6 | 0.72 | 0.73 | 0.71 | 0.77 |

| | | | | | | | | | | | | | | | | |
|---|---|---|---|---|---|---|---|---|---|---|---|---|---|---|---|---|
| 7 | 46850069 | rs10951872 | 1.005 | -0.271 | -0.399 | -0.282 | 0 | 0.006 | 0.006 | 0 | 0.29 | 0.26 | 0.32 | 0.53 | 0.27 | 0.29 |
| 7 | 46851363 | rs1534103 | 0.877 | -0.384 | -0.554 | -0.418 | -0.028 | 0.045 | 0.045 | 1.173 | 0.25 | 0.08 | 0.2 | 0.16 | 0.26 | 0.26 |
| 7 | 46853914 | rs11760799 | -0.982 | -0.567 | -0.321 | -0.22 | -0.001 | -0.001 | -0.001 | 0.576 | 0.72 | 0.71 | 0.75 | 0.7 | 0.71 | 0.77 |
| 7 | 46856454 | rs1525233 | 0.982 | -0.564 | -0.445 | -0.369 | 0.001 | -0.045 | -0.045 | 0 | 0.19 | 0.23 | 0.24 | 0.03 | 0.22 | 0.16 |
| 7 | 46865930 | rs9791386 | 1.107 | -0.337 | -0.417 | -0.257 | 0.007 | 0.034 | 0.034 | 0 | 0.3 | 0.37 | 0.35 | 0.48 | 0.3 | 0.35 |
| 7 | 46875405 | rs11770936 | 1.766 | -0.912 | -0.525 | -0.306 | -0.028 | -0.027 | -0.027 | 0.364 | 0.13 | 0.03 | 0.13 | 0.17 | 0.15 | 0.18 |
| 7 | 46875785 | rs1525235 | -0.449 | -0.983 | -0.482 | -0.363 | 0.004 | 0.043 | 0.043 | 0 | 0.43 | 0.34 | 0.41 | 0.21 | 0.41 | 0.41 |
| 7 | 46878747 | rs12702255 | 0.962 | -0.997 | -0.454 | -0.368 | -0.001 | -0.006 | -0.006 | 0.102 | 0.21 | 0.19 | 0.24 | 0.03 | 0.23 | 0.16 |
| 7 | 46879078 | rs7794050 | -1.355 | -0.959 | -0.454 | -0.335 | -0.008 | 0.01 | 0.01 | 0 | 0.75 | 0.69 | 0.77 | 0.68 | 0.76 | 0.79 |
| 7 | 46880743 | rs4354226 | 1.18 | -0.794 | -0.349 | -0.281 | 0.014 | 0.032 | 0.032 | 0 | 0.31 | 0.35 | 0.36 | 0.48 | 0.33 | 0.35 |
| 7 | 46883650 | rs4415228 | -0.032 | -0.793 | -0.418 | -0.271 | 0.012 | -0.006 | -0.006 | 0 | 0.54 | 0.61 | 0.6 | 0.51 | 0.57 | 0.54 |
| 7 | 46885232 | rs4724511 | 1.154 | -0.612 | -0.369 | -0.24 | -0.006 | -0.011 | -0.011 | 0 | 0.3 | 0.27 | 0.35 | 0.49 | 0.33 | 0.36 |
| 7 | 46889411 | rs4236382 | -0.071 | -0.74 | -0.378 | -0.312 | 0.012 | 0.023 | 0.023 | 0 | 0.52 | 0.4 | 0.52 | 0.65 | 0.52 | 0.59 |
| 7 | 46892201 | rs4580949 | -0.45 | -0.834 | -0.422 | -0.371 | 0.002 | 0.084 | 0.084 | 0 | 0.45 | 0.3 | 0.42 | 0.21 | 0.44 | 0.42 |
| 7 | 46893288 | rs12668566 | -1.428 | -0.834 | -0.35 | -0.338 | -0.012 | 0.034 | 0.034 | 0 | 0.78 | 0.69 | 0.78 | 0.68 | 0.77 | 0.78 |
| 7 | 46894870 | rs6976404 | -0.647 | -0.758 | -0.4 | -0.589 | 0.006 | 0.033 | 0.033 | 0 | 0.48 | 0.43 | 0.43 | 0.21 | 0.44 | 0.43 |
| 7 | 46899150 | rs4349889 | 0.42 | -0.764 | -0.385 | -0.631 | -0.003 | -0.058 | -0.058 | 0 | 0.25 | 0.32 | 0.27 | 0.04 | 0.25 | 0.19 |
| 7 | 46901914 | rs12702263 | -0.538 | -0.748 | -0.353 | -0.607 | 0.004 | 0.042 | 0.042 | 0 | 0.46 | 0.4 | 0.43 | 0.21 | 0.44 | 0.43 |
| 7 | 46902535 | rs7785858 | -0.636 | -0.738 | -0.345 | -0.623 | 0.01 | 0.028 | 0.028 | 0 | 0.44 | 0.41 | 0.4 | 0.22 | 0.38 | 0.42 |
| 7 | 46902668 | rs7803146 | 0.351 | -0.7 | -0.422 | -0.575 | 0.007 | 0.005 | 0.005 | 0.596 | 0.21 | 0.19 | 0.14 | 0.18 | 0.14 | 0.24 |
| 7 | 46915752 | rs12702267 | 0.204 | -0.737 | -0.288 | -0.334 | 0.001 | 0 | 0 | 0 | 0.43 | 0.4 | 0.63 | 0.42 | 0.47 | 0.4 |
| 7 | 46919134 | rs6977455 | -0.029 | -0.773 | -0.384 | -0.335 | -0.002 | 0 | 0 | 10.479 | 0.42 | 0.55 | 0.51 | 0.57 | 0.36 | 0.44 |
| 7 | 46919715 | rs7810453 | -0.85 | -0.422 | -0.449 | -0.077 | -0.029 | -0.019 | -0.019 | 0.779 | 0.67 | 0.76 | 0.78 | 0.61 | 0.61 | 0.63 |
| 7 | 46939896 | rs740140 | 2.069 | -0.379 | -0.566 | -0.02 | 0.004 | -0.036 | -0.036 | 0.29 | 0.1 | 0.03 | 0.1 | 0.19 | 0.09 | 0.12 |
| 7 | 46940630 | rs2016094 | 0.875 | -0.49 | -0.368 | 0.002 | -0.006 | 0.012 | 0.012 | 0 | 0.24 | 0.26 | 0.39 | 0.39 | 0.24 | 0.23 |
| 7 | 46942076 | rs10951880 | 0.298 | 0.268 | -0.314 | 0.147 | -0.003 | 0.039 | 0.039 | 0 | 0.09 | 0.07 | 0.07 | 0.39 | 0.1 | 0.29 |
| 7 | 46942387 | rs10256313 | -0.935 | 0.143 | -0.302 | -0.299 | 0.004 | -0.053 | -0.053 | 0.22 | 0.76 | 0.77 | 0.85 | 1 | 0.7 | 0.91 |
| 7 | 46947466 | rs12702272 | -0.884 | 0.013 | -0.274 | -0.253 | 0.01 | -0.078 | -0.078 | 0 | 0.76 | 0.73 | 0.85 | 1 | 0.73 | 0.93 |
| 7 | 46950860 | rs11767243 | -1.308 | 0.134 | 0.154 | 0.679 | -0.001 | 0.313 | 0.313 | 0.148 | 0.15 | 0.46 | 0.1 | 0.54 | 0.09 | 0.35 |
| 7 | 46951569 | rs10951881 | -0.869 | -0.048 | -0.27 | 0.295 | 0.001 | 0 | 0 | 0 | 0.63 | 0.68 | 0.74 | 0.84 | 0.67 | 0.76 |
| 7 | 46951703 | rs11975289 | 0.229 | 0.072 | -0.393 | 0.436 | 0.008 | -0.018 | -0.018 | 0 | 0.25 | 0.44 | 0.24 | 0.23 | 0.24 | 0.23 |
| 7 | 46953695 | rs12672191 | 0.216 | 0.072 | -0.404 | 0.436 | 0.007 | -0.019 | -0.019 | 0 | 0.26 | 0.44 | 0.24 | 0.23 | 0.24 | 0.23 |
| 7 | 46959491 | rs917132 | -1.039 | 0.076 | -0.605 | 0.371 | 0.002 | 0.026 | 0.026 | 0 | 0.63 | 0.72 | 0.75 | 0.84 | 0.67 | 0.76 |
| 7 | 46968452 | rs10269378 | 0.806 | 0.426 | -0.757 | 0.368 | -0.027 | -0.016 | -0.016 | 0 | 0.24 | 0.07 | 0.35 | 0.22 | 0.31 | 0.23 |
| 7 | 46975314 | rs4724520 | 0.797 | 0.431 | -0.757 | 0.372 | -0.03 | -0.029 | -0.029 | 0 | 0.25 | 0.07 | 0.44 | 0.25 | 0.32 | 0.23 |
| 7 | 46978309 | rs2346406 | -0.939 | 0.667 | 0.033 | 0.545 | 0.01 | -0.018 | -0.018 | 0 | 0.33 | 0.58 | 0.37 | 0.25 | 0.34 | 0.27 |
| 7 | 46980422 | rs11767105 | -1.426 | 0.526 | 0.311 | 0.628 | 0.011 | -0.037 | -0.037 | 1.481 | 0.14 | 0.4 | 0.08 | 0.1 | 0.12 | 0.13 |
| 7 | 46983183 | rs17172792 | -1.375 | 0.351 | 0.264 | 1.139 | -0.028 | 0.063 | 0.063 | 0 | 0.27 | 0.52 | 0.15 | 0.38 | 0.19 | 0.38 |
| 7 | 46983232 | rs17172793 | -0.625 | 0.556 | 0.359 | 0.685 | 0.003 | -0.051 | -0.051 | 0 | 0.16 | 0.29 | 0.1 | 0.09 | 0.14 | 0.14 |
| 7 | 46986218 | rs9639966 | -0.932 | 0.752 | 0.003 | 1.013 | 0 | -0.016 | -0.016 | 0 | 0.36 | 0.7 | 0.43 | 0.47 | 0.35 | 0.34 |
| 7 | 46988142 | rs4436029 | -0.182 | 0.715 | 0.231 | 1.397 | 0.001 | 0.095 | 0.095 | 0.492 | 0.13 | 0.18 | 0.26 | 0.46 | 0.17 | 0.23 |
| 7 | 46990237 | rs4724521 | -1.548 | 0.604 | 0.317 | 0.699 | -0.002 | 0.032 | 0.032 | 1.081 | 0.36 | 0.5 | 0.44 | 0.52 | 0.39 | 0.37 |
| 7 | 46994345 | rs6953890 | -1.113 | 0.735 | 0.841 | 1.001 | 0.001 | 0.098 | 0.098 | 1.304 | 0.25 | 0.43 | 0.34 | 0.52 | 0.3 | 0.31 |
| 7 | 47005740 | rs10239638 | 0.659 | 1.448 | 0.533 | 1.014 | 0.03 | 0.011 | 0.011 | 0 | 0.15 | 0.22 | 0.25 | 0.17 | 0.22 | 0.33 |
| 7 | 47007215 | rs6956956 | 0.41 | 1.417 | 0.622 | 0.992 | 0.05 | 0.017 | 0.017 | 0 | 0.16 | 0.28 | 0.26 | 0.17 | 0.22 | 0.33 |
| 7 | 47008390 | rs929341 | -0.446 | 1.109 | 0.402 | 1.122 | 0 | 0.039 | 0.039 | 1.216 | 0.21 | 0.25 | 0.32 | 0.52 | 0.26 | 0.26 |
| 7 | 47022589 | rs6954915 | 0.166 | 1.601 | 0.7 | 0.884 | 0.032 | 0.017 | 0.017 | 0 | 0.16 | 0.24 | 0.25 | 0.18 | 0.22 | 0.33 |
| 7 | 47022767 | rs6955374 | -1.503 | 1.506 | 1.082 | 0.764 | -0.005 | 0.001 | 0.001 | 0 | 0.29 | 0.71 | 0.35 | 0.24 | 0.34 | 0.42 |
| 7 | 47023177 | rs6463372 | -1.065 | 1.475 | 1.178 | 0.703 | 0.028 | -0.003 | -0.003 | 0 | 0.25 | 0.33 | 0.31 | 0.21 | 0.28 | 0.34 |
| 7 | 47023504 | rs10487768 | -1.473 | 1.121 | 1.539 | 0.957 | -0.003 | -0.08 | -0.08 | 0.032 | 0.13 | 0.32 | 0.09 | 0 | 0.1 | 0.07 |
| 7 | 47023761 | rs6942503 | -0.014 | 0.796 | 0.531 | 1.37 | 0 | -0.004 | -0.004 | 0.433 | 0.27 | 0.23 | 0.27 | 0.36 | 0.26 | 0.21 |
| 7 | 47032225 | rs2346855 | 0.004 | 1.131 | 0.976 | 1.333 | 0.001 | 0.01 | 0.01 | 0.587 | 0.2 | 0.28 | 0.2 | 0.24 | 0.22 | 0.11 |
| 7 | 47033404 | rs12672541 | 2.284 | 1.167 | 1.098 | 1.4 | -0.007 | 0.084 | 0.084 | 0 | 0.46 | 0.26 | 0.42 | 0.26 | 0.44 | 0.51 |
| 7 | 47034331 | rs7804773 | -1.282 | 1.208 | 1.024 | 1.622 | -0.035 | -0.048 | -0.048 | 1.944 | 0.92 | 0.99 | 0.82 | 0.74 | 0.88 | 0.85 |
| 7 | 47034472 | rs10951886 | 0.371 | 1.199 | 1.182 | 1.413 | 0.001 | 0.008 | 0.008 | 0.659 | 0.2 | 0.24 | 0.2 | 0.25 | 0.23 | 0.11 |
| 7 | 47036388 | rs12540758 | -0.013 | 1.178 | 1.315 | 1.407 | 0.007 | 0.012 | 0.012 | 1.464 | 0.12 | 0.18 | 0.11 | 0.17 | 0.13 | 0.1 |
| 7 | 47037783 | rs17443746 | 2.5 | 1.304 | 1.307 | 1.364 | 0.008 | 0.068 | 0.068 | 0 | 0.43 | 0.23 | 0.38 | 0.26 | 0.39 | 0.45 |
| 7 | 47039263 | rs17172801 | 0.012 | 1.308 | 1.292 | 1.366 | 0.007 | 0.012 | 0.012 | 1.464 | 0.12 | 0.18 | 0.11 | 0.17 | 0.13 | 0.1 |
| 7 | 47039741 | rs13238599 | 0.029 | 1.3 | 1.374 | 1.363 | 0.002 | -0.016 | -0.016 | 0 | 0.8 | 0.81 | 0.76 | 0.67 | 0.8 | 0.72 |
| 7 | 47040175 | rs1859524 | 2.569 | 1.297 | 1.31 | 1.361 | 0 | 0.082 | 0.082 | 0 | 0.42 | 0.16 | 0.37 | 0.26 | 0.39 | 0.45 |
| 7 | 47042254 | rs6966942 | 1.387 | 1.318 | 1.18 | 1.391 | 0 | 0.012 | 0.012 | 0.384 | 0.68 | 0.63 | 0.65 | 0.5 | 0.67 | 0.62 |
| 7 | 47054582 | rs12670333 | -0.093 | 1.314 | 0.838 | 1.386 | 0.001 | -0.035 | -0.035 | 0 | 0.81 | 0.88 | 0.76 | 0.67 | 0.8 | 0.72 |
| 7 | 47059736 | rs10282172 | 1.632 | 1.27 | 0.599 | 1.264 | 0 | 0.058 | 0.058 | 0 | 0.47 | 0.32 | 0.43 | 0.26 | 0.44 | 0.51 |
| 7 | 47064694 | rs4724530 | -1.124 | 1.19 | 0.717 | 1.765 | 0.003 | 0.011 | 0.011 | 0.578 | 0.24 | 0.26 | 0.26 | 0.3 | 0.28 | 0.17 |
| 7 | 47070010 | rs1008364 | -0.25 | 1.251 | 0.132 | 1.509 | -0.011 | -0.024 | -0.024 | 0 | 0.18 | 0.11 | 0.21 | 0.25 | 0.22 | 0.11 |
| 7 | 47075092 | rs17172808 | -1.29 | 0.938 | 0.801 | 1.068 | 0.025 | 0.079 | 0.079 | 0 | 0.09 | 0.32 | 0.11 | 0.16 | 0.07 | 0.13 |
| 7 | 47077430 | rs2286117 | 1.278 | 1.135 | 0.08 | 0.561 | -0.005 | 0.03 | 0.03 | 0.321 | 0.56 | 0.34 | 0.46 | 0.28 | 0.53 | 0.45 |
| 7 | 47079999 | rs12537881 | 0.883 | 1.23 | 0.568 | 0.35 | 0.006 | -0.02 | -0.02 | 0 | 0.67 | 0.45 | 0.61 | 0.23 | 0.65 | 0.63 |
| 7 | 47081588 | rs6948884 | 0.98 | 0.834 | 0.727 | 0.417 | 0.003 | 0.012 | 0.012 | 0 | 0.49 | 0.39 | 0.44 | 0.23 | 0.41 | 0.47 |

| | | | | | | | | | | | | | | | | |
|---|---|---|---|---|---|---|---|---|---|---|---|---|---|---|---|---|
| 7 | 47088439 | rs6463379 | -0.397 | 0.904 | 0.483 | 0.665 | 0.013 | 0.108 | 0.108 | 0.057 | 0.09 | 0.33 | 0.07 | 0.16 | 0.06 | 0.1 |
| 7 | 47091670 | rs10276554 | -0.378 | 0.918 | 0.351 | 0.516 | 0.033 | 0.108 | 0.108 | 0.092 | 0.09 | 0.39 | 0.08 | 0.14 | 0.06 | 0.1 |
| 7 | 47093009 | rs7799055 | 1.238 | 0.887 | 0.372 | 0.477 | 0.002 | 0.127 | 0.127 | 0.021 | 0.46 | 0.18 | 0.43 | 0.23 | 0.4 | 0.47 |
| 7 | 47094523 | rs11771745 | -0.974 | 0.918 | 0.819 | 0.829 | 0.031 | 0.106 | 0.106 | 0.147 | 0.1 | 0.41 | 0.08 | 0.14 | 0.06 | 0.1 |
| 7 | 47103211 | rs11761505 | 0.363 | 1.452 | 0.039 | 0.987 | 0.006 | 0.012 | 0.012 | 0.005 | 0.73 | 0.68 | 0.61 | 0.34 | 0.68 | 0.63 |
| 7 | 47112988 | rs10951888 | -1.078 | 1.569 | 0.78 | 0.46 | 0.038 | -0.013 | -0.013 | 0.352 | 0.07 | 0.29 | 0.06 | 0 | 0.06 | 0.02 |
| 7 | 47117617 | rs6954800 | 0.702 | 2.004 | 0.915 | 0.581 | 0.004 | -0.005 | -0.005 | 3.303 | 0.68 | 0.38 | 0.59 | 0.69 | 0.62 | 0.77 |
| 7 | 47118424 | rs3747528 | -0.66 | 1.524 | 0.836 | 1.204 | -0.006 | 0.058 | 0.058 | 0 | 0.08 | 0.08 | 0.03 | 0.32 | 0.05 | 0.16 |
| 7 | 47118677 | rs3747527 | -0.66 | 1.524 | 0.836 | 1.204 | -0.006 | 0.058 | 0.058 | 0 | 0.08 | 0.08 | 0.03 | 0.32 | 0.05 | 0.16 |
| 7 | 47119135 | rs2013523 | 1.301 | 0.679 | 0.168 | 0.489 | 0.024 | -0.005 | -0.005 | 0 | 0.39 | 0.24 | 0.42 | 0.39 | 0.32 | 0.54 |
| 7 | 47119768 | rs886658 | -0.667 | 0.752 | 0.492 | 0.446 | 0.018 | -0.003 | -0.003 | 0.472 | 0.75 | 0.68 | 0.65 | 0.72 | 0.68 | 0.79 |
| 7 | 47119985 | rs874321 | 0.624 | 0.771 | 0.211 | 0.474 | 0.006 | 0.104 | 0.104 | 0.359 | 0.5 | 0.32 | 0.51 | 0.71 | 0.47 | 0.72 |
| 7 | 47128635 | rs17446835 | 0.619 | 0.434 | 0.435 | 0.54 | 0.016 | 0.04 | 0.04 | 0 | 0.18 | 0.02 | 0.09 | 0.06 | 0.17 | 0.07 |
| 7 | 47128879 | rs6957383 | -1.449 | -0.004 | 0.545 | 0.272 | 0.005 | 0.003 | 0.003 | 0 | 0.46 | 0.54 | 0.31 | 0.35 | 0.43 | 0.36 |
| 7 | 47129358 | rs17172814 | 0.053 | 0.175 | 0.004 | 0.406 | 0.022 | -0.034 | -0.034 | 0 | 0.16 | 0.08 | 0.09 | 0.29 | 0.11 | 0.23 |
| 7 | 47133202 | rs1866361 | -0.949 | 0.193 | 0.03 | 0.365 | 0.012 | -0.034 | -0.034 | 0.042 | 0.38 | 0.14 | 0.23 | 0.35 | 0.38 | 0.34 |
| 7 | 47137770 | rs10499651 | 0.729 | 0.29 | -0.032 | 0.193 | 0.014 | 0.068 | 0.068 | 0 | 0.18 | 0.01 | 0.09 | 0.04 | 0.17 | 0.07 |
| 7 | 47141339 | rs7792919 | -0.655 | 0.421 | 0.678 | 0.321 | 0.018 | 0.05 | 0.05 | 0.005 | 0.2 | 0.29 | 0.17 | 0.28 | 0.24 | 0.28 |
| 7 | 47145414 | rs1866356 | -0.957 | 0.304 | 0.797 | 0.201 | -0.002 | -0.003 | -0.003 | 0 | 0.4 | 0.39 | 0.27 | 0.31 | 0.44 | 0.36 |
| 7 | 47145824 | rs10499653 | 1.214 | 0.294 | 0.508 | 0.17 | -0.003 | 0.101 | 0.101 | 0 | 0.16 | 0 | 0.09 | 0 | 0.16 | 0.03 |
| 7 | 47145978 | rs17172817 | -0.154 | 0.41 | 0.664 | 0.217 | -0.004 | 0.013 | 0.013 | 0 | 0.18 | 0.19 | 0.11 | 0.28 | 0.17 | 0.27 |
| 7 | 47146035 | rs6463390 | -1.376 | 0.362 | 0.798 | 0.157 | 0.017 | 0.039 | 0.039 | 0 | 0.47 | 0.68 | 0.32 | 0.31 | 0.48 | 0.38 |
| 7 | 47154615 | rs2881495 | -1.03 | 0.345 | 0.868 | 0.269 | 0.004 | 0.004 | 0.004 | 0 | 0.4 | 0.37 | 0.24 | 0.21 | 0.37 | 0.3 |
| 7 | 47154807 | rs4557608 | 0.105 | 0.256 | 0.22 | 0.246 | 0.027 | 0.069 | 0.069 | 0 | 0.23 | 0.12 | 0.13 | 0.03 | 0.21 | 0.08 |
| 7 | 47154985 | rs6968013 | -0.839 | 0.452 | 0.149 | 0.485 | 0.016 | 0.034 | 0.034 | 0 | 0.46 | 0.67 | 0.32 | 0.31 | 0.48 | 0.38 |
| 7 | 47157710 | rs11773471 | -0.677 | 0.371 | 0.878 | 0.747 | 0.045 | 0.163 | 0.163 | 0 | 0.08 | 0.36 | 0.06 | 0.12 | 0.05 | 0.07 |
| 7 | 47160224 | rs10255142 | 0.848 | 0.029 | 0.979 | 0.838 | -0.04 | -0.002 | -0.002 | 5.501 | 0.36 | 0.52 | 0.37 | 0.35 | 0.19 | 0.42 |
| 7 | 47161018 | rs10258830 | 0.793 | 0.022 | 1.184 | 0.834 | 0 | 0 | 0 | 0 | 0.35 | 0.38 | 0.36 | 0.35 | 0.19 | 0.42 |
| 7 | 47163189 | rs11770341 | 0.393 | 0.078 | 0.812 | 0.777 | 0 | -0.014 | -0.014 | 0.194 | 0.16 | 0.11 | 0.11 | 0.2 | 0.2 | 0.19 |
| 7 | 47163756 | rs1898663 | 0.396 | 0.583 | 1.017 | 0.725 | 0.004 | -0.004 | -0.004 | 0.016 | 0.44 | 0.62 | 0.48 | 0.45 | 0.35 | 0.53 |
| 7 | 47166274 | rs876922 | 1.171 | 0.566 | 1.057 | 0.63 | -0.002 | 0.004 | 0.004 | 2.006 | 0.35 | 0.28 | 0.4 | 0.33 | 0.29 | 0.44 |
| 7 | 47185886 | rs6973264 | 1.016 | 0.536 | 1.226 | 0.593 | -0.006 | -0.009 | -0.009 | 0 | 0.38 | 0.49 | 0.42 | 0.34 | 0.29 | 0.44 |
| 7 | 47185952 | rs4144076 | -0.614 | 0.572 | 1.21 | 0.639 | -0.005 | 0.037 | 0.037 | 0 | 0.24 | 0.38 | 0.19 | 0.29 | 0.28 | 0.29 |
| 7 | 47186412 | rs2918194 | -0.591 | 0.579 | 1.055 | 0.558 | -0.052 | -0.002 | -0.002 | 0.002 | 0.61 | 0.87 | 0.61 | 0.64 | 0.56 | 0.73 |
| 7 | 47188064 | rs1438603 | 1.181 | 0.575 | 1.121 | 0.794 | -0.002 | -0.007 | -0.007 | 0.001 | 0.37 | 0.43 | 0.41 | 0.31 | 0.29 | 0.44 |
| 7 | 47188122 | rs768904 | 0.151 | 0.798 | 1.36 | 1.051 | 0.032 | 0.068 | 0.068 | 0 | 0.07 | 0.23 | 0.06 | 0.1 | 0.07 | 0.08 |
| 7 | 47195053 | rs2728484 | -0.904 | 1.035 | 0.56 | 0.87 | -0.029 | -0.04 | -0.04 | 0 | 0.78 | 0.88 | 0.69 | 0.64 | 0.73 | 0.77 |
| 7 | 47198606 | rs2578435 | -0.734 | 1.037 | 0.612 | 0.862 | -0.017 | 0.037 | 0.037 | 0.077 | 0.25 | 0.55 | 0.2 | 0.33 | 0.26 | 0.27 |
| 7 | 47199047 | rs2578436 | -1.037 | 1.15 | 0.155 | 0.939 | -0.04 | -0.051 | -0.051 | 0 | 0.8 | 0.91 | 0.69 | 0.64 | 0.73 | 0.77 |
| 7 | 47206621 | rs10241085 | 0.159 | 0.866 | 0.308 | 0.787 | -0.004 | 0.015 | 0.015 | 0 | 0.16 | 0.2 | 0.12 | 0.23 | 0.18 | 0.18 |
| 7 | 47208813 | rs1370917 | -0.227 | 0.698 | -0.239 | 0.33 | -0.064 | 0.014 | 0.014 | 1.935 | 0.7 | 0.85 | 0.58 | 0.21 | 0.58 | 0.53 |
| 7 | 47209007 | rs2728487 | 0.294 | 0.575 | -0.197 | 0.424 | -0.001 | -0.001 | -0.001 | 0.44 | 0.57 | 0.61 | 0.5 | 0.46 | 0.5 | 0.55 |
| 7 | 47209261 | rs2578445 | 0.508 | 0.354 | -0.221 | 0.169 | 0.016 | 0.034 | 0.034 | 0.142 | 0.49 | 0.43 | 0.44 | 0.36 | 0.42 | 0.4 |
| 7 | 47211732 | rs2728491 | -0.561 | 0.281 | -0.745 | 0.088 | -0.015 | 0.014 | 0.014 | 0 | 0.72 | 0.84 | 0.79 | 0.72 | 0.69 | 0.76 |
| 7 | 47214236 | rs12531862 | 2.37 | -0.463 | -1.14 | -0.08 | 0.038 | 0.019 | 0.019 | 4.706 | 0.16 | 0.02 | 0.2 | 0.07 | 0.08 | 0.11 |
| 7 | 47214444 | rs1347801 | -0.797 | 0.237 | -0.604 | -0.311 | 0.069 | -0.021 | -0.021 | 0 | 0.25 | 0.47 | 0.34 | 0.26 | 0.33 | 0.34 |
| 7 | 47217923 | rs1020112 | -0.006 | -0.358 | -0.92 | -0.303 | 0.02 | 0.049 | 0.049 | 0.349 | 0.33 | 0.17 | 0.17 | 0.14 | 0.29 | 0.28 |
| 7 | 47224680 | rs12702310 | 1.399 | -0.458 | -1.121 | -0.38 | 0.003 | 0.101 | 0.101 | 0 | 0.17 | 0.01 | 0.08 | 0 | 0.16 | 0.04 |
| 7 | 47225542 | rs1530071 | 0.422 | -0.612 | -0.759 | -0.191 | -0.001 | -0.001 | -0.001 | 0.316 | 0.15 | 0.15 | 0.09 | 0.14 | 0.13 | 0.24 |
| 7 | 47226179 | rs1530074 | 0.422 | -0.612 | -0.759 | -0.191 | -0.001 | -0.001 | -0.001 | 0 | 0.15 | 0.15 | 0.09 | 0.14 | 0.13 | 0.24 |
| 7 | 47233161 | rs2578390 | -0.975 | -1.277 | -0.525 | -0.009 | 0.012 | 0.002 | 0.002 | 0 | 0.7 | 0.65 | 0.58 | 0.65 | 0.58 | 0.71 |
| 7 | 47234555 | rs7782928 | 0.483 | -1.416 | -0.407 | -0.1 | -0.012 | -0.019 | -0.019 | 1 | 0.3 | 0.23 | 0.26 | 0.43 | 0.35 | 0.34 |
| 7 | 47238707 | rs1370916 | -0.948 | -1.464 | -0.603 | -0.102 | -0.004 | 0.008 | 0.008 | 0.244 | 0.53 | 0.63 | 0.39 | 0.56 | 0.5 | 0.62 |
| 7 | 47242308 | rs12702316 | 0.223 | -1.501 | -1.091 | -0.106 | -0.008 | -0.014 | -0.014 | 0.801 | 0.31 | 0.25 | 0.26 | 0.42 | 0.34 | 0.34 |
| 7 | 47246076 | rs2118664 | 0.294 | -0.858 | -0.459 | 0.013 | -0.003 | 0 | 0 | 0 | 0.17 | 0.18 | 0.1 | 0.14 | 0.14 | 0.24 |
| 7 | 47248297 | rs12702317 | 0.771 | -0.854 | -0.557 | 0.068 | 0.012 | -0.017 | -0.017 | 0.009 | 0.46 | 0.26 | 0.52 | 0.49 | 0.39 | 0.34 |
| 7 | 47250136 | rs2578398 | 0.321 | -0.823 | -1.038 | 0.041 | -0.005 | 0 | 0 | 0 | 0.17 | 0.19 | 0.09 | 0.14 | 0.14 | 0.23 |
| 7 | 47250231 | rs2728464 | -0.398 | -0.71 | -1.209 | 0.093 | 0.011 | -0.002 | -0.002 | 1.606 | 0.64 | 0.42 | 0.61 | 0.63 | 0.54 | 0.59 |
| 7 | 47251761 | rs12668145 | 0.366 | -0.104 | -1.811 | 0.285 | -0.056 | -0.094 | -0.094 | 0 | 0.42 | 0.09 | 0.51 | 0.48 | 0.39 | 0.33 |
| 7 | 47260060 | rs2728468 | -1.387 | -0.122 | -1.233 | 0.245 | -0.089 | -0.019 | -0.019 | 0.272 | 0.61 | 0.9 | 0.6 | 0.6 | 0.58 | 0.76 |
| 7 | 47261581 | rs893525 | -0.891 | -0.486 | -0.794 | 0.046 | 0.057 | 0.064 | 0.064 | 0 | 0.4 | 0.68 | 0.33 | 0.19 | 0.24 | 0.44 |
| 7 | 47263856 | rs2728470 | 0.321 | -0.84 | -0.293 | -0.014 | -0.042 | -0.014 | -0.014 | 0 | 0.2 | 0.28 | 0.1 | 0.13 | 0.12 | 0.22 |
| 7 | 47266834 | rs753063 | -1.031 | -0.528 | -1.102 | 0.124 | 0.094 | 0.102 | 0.102 | 0.001 | 0.43 | 0.83 | 0.4 | 0.26 | 0.3 | 0.45 |
| 7 | 47271381 | rs2578414 | 0.117 | -0.348 | -0.409 | -0.01 | -0.022 | -0.009 | -0.009 | 2.289 | 0.28 | 0.36 | 0.2 | 0.23 | 0.22 | 0.26 |
| 7 | 47272813 | rs17520335 | 0.924 | -0.22 | -1.073 | 0.038 | -0.021 | 0.041 | 0.041 | 0 | 0.09 | 0.02 | 0.17 | 0.03 | 0.08 | 0.07 |
| 7 | 47274398 | rs987792 | -1.779 | 0.013 | -1.057 | 0.123 | -0.046 | 0.013 | 0.013 | 0 | 0.62 | 0.83 | 0.65 | 0.64 | 0.6 | 0.77 |
| 7 | 47275737 | rs10265057 | 0.283 | 0.008 | -1.101 | 0.124 | -0.012 | -0.048 | -0.048 | 0 | 0.14 | 0.26 | 0.2 | 0.05 | 0.09 | 0.18 |
| 7 | 47276175 | rs11491841 | -0.042 | 0.618 | -1.788 | 0.467 | -0.024 | -0.045 | -0.045 | 0 | 0.18 | 0.13 | 0.24 | 0.36 | 0.29 | 0.33 |
| 7 | 47278232 | rs2578419 | -0.553 | 0.796 | -1.459 | 0.188 | -0.07 | 0.005 | 0.005 | 0 | 0.63 | 0.88 | 0.65 | 0.64 | 0.6 | 0.77 |

| | | | | | | | | | | | | | | | | |
|---|---|---|---|---|---|---|---|---|---|---|---|---|---|---|---|---|
| 7 | 47282889 | rs6463401 | -0.522 | 0.754 | -0.658 | -0.223 | 0.001 | -0.035 | -0.035 | 0 | 0.31 | 0.37 | 0.36 | 0.18 | 0.34 | 0.34 |
| 7 | 47283472 | rs7808403 | -0.417 | 0.47 | -0.473 | -0.996 | -0.003 | -0.012 | -0.012 | 0.003 | 0.38 | 0.3 | 0.29 | 0.08 | 0.43 | 0.32 |
| 7 | 47283974 | rs2728474 | 1.179 | -0.028 | -0.987 | -0.67 | 0.067 | -0.048 | -0.048 | 0 | 0.3 | 0.12 | 0.36 | 0.56 | 0.22 | 0.35 |
| 7 | 47285131 | rs4512292 | -0.496 | -0.038 | -0.417 | -0.765 | -0.009 | -0.004 | -0.004 | 0 | 0.79 | 0.81 | 0.8 | 0.65 | 0.68 | 0.71 |
| 7 | 47285454 | rs3935078 | 1.094 | -0.125 | -0.363 | -0.836 | 0.073 | -0.017 | -0.017 | 0 | 0.34 | 0.16 | 0.42 | 0.55 | 0.25 | 0.35 |
| 7 | 47288211 | rs4272275 | -0.185 | 0.13 | -0.036 | -0.66 | 0.015 | 0.002 | 0.002 | 1.392 | 0.77 | 0.71 | 0.79 | 0.65 | 0.68 | 0.71 |
| 7 | 47291954 | rs6463403 | -1.119 | 0.499 | -0.257 | -0.625 | -0.046 | -0.018 | -0.018 | 0 | 0.84 | 0.93 | 0.87 | 0.74 | 0.74 | 0.83 |
| 7 | 47305969 | rs4724556 | -0.411 | 0.166 | -0.105 | -0.49 | -0.018 | -0.012 | -0.012 | 0.593 | 0.78 | 0.8 | 0.78 | 0.64 | 0.68 | 0.71 |
| 7 | 47308750 | rs4724557 | 0.737 | -0.41 | 0.386 | -0.75 | 0.014 | -0.002 | -0.002 | 0 | 0.39 | 0.38 | 0.41 | 0.52 | 0.25 | 0.39 |
| 7 | 47315291 | rs9876 | 1.511 | -0.449 | 0.211 | -0.458 | 0.073 | -0.096 | -0.096 | 0 | 0.32 | 0.11 | 0.39 | 0.48 | 0.24 | 0.35 |
| 7 | 47316586 | rs3184329 | -0.17 | -0.498 | 0.34 | -0.504 | -0.001 | -0.018 | -0.018 | 0.12 | 0.43 | 0.5 | 0.38 | 0.12 | 0.43 | 0.34 |
| 7 | 47317046 | rs3750161 | -0.684 | -0.498 | 0.342 | -0.498 | 0.009 | 0.003 | 0.003 | 1.628 | 0.78 | 0.73 | 0.78 | 0.64 | 0.68 | 0.7 |
| 7 | 47317324 | rs3210894 | 1.561 | -0.448 | 0.308 | -0.468 | 0.072 | -0.096 | -0.096 | 3.492 | 0.32 | 0.11 | 0.39 | 0.48 | 0.24 | 0.35 |
| 7 | 47317510 | rs3750163 | 0.167 | -0.01 | 0.724 | -0.29 | -0.014 | 0.02 | 0.02 | 0.958 | 0.35 | 0.23 | 0.36 | 0.11 | 0.43 | 0.31 |
| 7 | 47319852 | rs6971754 | 0.545 | 0.251 | 0.757 | -0.4 | -0.025 | 0.073 | 0.073 | 0.61 | 0.31 | 0.14 | 0.33 | 0.05 | 0.35 | 0.21 |
| 7 | 47321050 | rs12702333 | 0.098 | 0.377 | 0.363 | -0.319 | 0.05 | 0.017 | 0.017 | 1.429 | 0.73 | 0.45 | 0.76 | 0.63 | 0.68 | 0.69 |
| 7 | 47323456 | rs3750164 | -0.358 | 0.575 | -0.327 | -0.231 | -0.002 | -0.002 | -0.002 | 0 | 0.77 | 0.74 | 0.78 | 0.63 | 0.68 | 0.69 |
| 7 | 47328384 | rs2347785 | -0.325 | 0.268 | -0.28 | -0.146 | 0.008 | 0.006 | 0.006 | 2.329 | 0.77 | 0.72 | 0.78 | 0.59 | 0.67 | 0.68 |
| 7 | 47332238 | rs884824 | 0.202 | 0.242 | -0.313 | -0.147 | 0.012 | -0.044 | -0.044 | 0 | 0.36 | 0.38 | 0.37 | 0.14 | 0.43 | 0.32 |
| 7 | 47333101 | rs4512293 | 0.253 | -0.635 | -0.78 | 0.3 | 0.039 | -0.019 | -0.019 | 0 | 0.4 | 0.3 | 0.41 | 0.45 | 0.24 | 0.37 |
| 7 | 47339892 | rs11764487 | 0.312 | 0.646 | -1.194 | 0.443 | 0 | 0.046 | 0.046 | 0 | 0.48 | 0.34 | 0.47 | 0.32 | 0.43 | 0.51 |
| 7 | 47345002 | rs6959554 | 0.816 | 1.056 | -1.2 | 0.277 | 0.005 | 0.036 | 0.036 | 0 | 0.39 | 0.19 | 0.37 | 0.24 | 0.34 | 0.35 |
| 7 | 47356861 | rs12702335 | 0.104 | 1.2 | -1.611 | 0.927 | -0.058 | -0.07 | -0.07 | 0 | 0.16 | 0.01 | 0.25 | 0.22 | 0.16 | 0.17 |
| 7 | 47364673 | rs7781509 | 0.44 | 1.416 | -1.306 | 0.555 | 0.002 | 0.025 | 0.025 | 0 | 0.44 | 0.36 | 0.41 | 0.31 | 0.39 | 0.47 |
| 7 | 47367818 | rs11766862 | 0.395 | 1.423 | -1.354 | 0.558 | 0.004 | 0.033 | 0.033 | 0 | 0.45 | 0.35 | 0.41 | 0.31 | 0.39 | 0.48 |
| 7 | 47370119 | rs6950527 | 1.053 | 1.427 | -1.393 | 0.55 | 0.018 | 0.033 | 0.033 | 0 | 0.38 | 0.14 | 0.35 | 0.24 | 0.32 | 0.34 |
| 7 | 47370958 | rs7788911 | -0.163 | 1.437 | -1.39 | 0.554 | 0.053 | -0.012 | -0.012 | 0 | 0.06 | 0.15 | 0.06 | 0.07 | 0.07 | 0.14 |
| 7 | 47374770 | rs1464921 | 0.905 | 1.23 | -1.61 | 0.612 | 0.021 | 0.035 | 0.035 | 0 | 0.38 | 0.16 | 0.35 | 0.24 | 0.32 | 0.34 |
| 7 | 47382518 | rs12375125 | 0.956 | 1.206 | -1.357 | 1.102 | -0.017 | -0.014 | -0.014 | 0 | 0.61 | 0.31 | 0.52 | 0.38 | 0.55 | 0.44 |
| 7 | 47386422 | rs2462645 | 1.153 | 0.749 | -1.246 | 1.16 | 0.018 | -0.003 | -0.003 | 0 | 0.58 | 0.23 | 0.54 | 0.35 | 0.6 | 0.43 |
| 7 | 47399398 | rs3323 | -0.23 | 1.116 | -1.171 | 1.421 | -0.034 | -0.019 | -0.019 | 0 | 0.09 | 0.03 | 0.12 | 0.12 | 0.19 | 0.17 |
| 7 | 47408443 | rs2293362 | 0.611 | 0.114 | -1.291 | 1.57 | 0.033 | -0.06 | -0.06 | 0.099 | 0.1 | 0 | 0.05 | 0.2 | 0.04 | 0.01 |
| 7 | 47416733 | rs6964472 | 0.777 | 0.407 | -0.075 | 1.683 | -0.03 | 0.131 | 0.131 | 0 | 0.77 | 0.59 | 0.69 | 0.48 | 0.83 | 0.65 |
| 7 | 47422049 | rs11496038 | -0.756 | 0.866 | -0.412 | 1.283 | 0.035 | -0.008 | -0.008 | 3.126 | 0.12 | 0.24 | 0.18 | 0.11 | 0.21 | 0.18 |
| 7 | 47424086 | rs4724568 | -1.195 | 0.371 | -0.437 | 1.67 | 0.002 | -0.003 | -0.003 | 3.33 | 0.24 | 0.22 | 0.34 | 0.56 | 0.21 | 0.36 |
| 7 | 47427643 | rs12537130 | -0.889 | 0.388 | -0.779 | 1.803 | -0.015 | -0.063 | -0.063 | 0.072 | 0.16 | 0.05 | 0.25 | 0.32 | 0.15 | 0.12 |
| 7 | 47429054 | rs6976292 | 0.19 | 0.297 | -0.591 | 1.116 | 0.001 | -0.011 | -0.011 | 0 | 0.68 | 0.63 | 0.83 | 0.82 | 0.72 | 0.85 |
| 7 | 47429406 | rs6956779 | 0.085 | -0.074 | -0.595 | 1.067 | -0.077 | 0.016 | 0.016 | 1.124 | 0.86 | 0.64 | 0.97 | 0.85 | 0.99 | 0.94 |
| 7 | 47431511 | rs4724573 | -0.43 | 0.416 | -0.755 | 1.242 | -0.027 | -0.036 | -0.036 | 0 | 0.16 | 0.1 | 0.33 | 0.29 | 0.21 | 0.11 |
| 7 | 47432669 | rs2470996 | -1.406 | 0.127 | -0.345 | 1.068 | 0.04 | 0.086 | 0.086 | 0.028 | 0.52 | 0.68 | 0.65 | 0.82 | 0.66 | 0.5 |
| 7 | 47433750 | rs10279872 | -0.485 | -0.069 | 0.03 | 0.77 | 0 | 0.01 | 0.01 | 0.592 | 0.14 | 0.19 | 0.2 | 0.31 | 0.21 | 0.29 |
| 7 | 47435954 | rs2462637 | 0.962 | -0.269 | 0.423 | 0.315 | -0.081 | -0.034 | -0.034 | 0.108 | 0.78 | 0.57 | 0.92 | 0.86 | 0.94 | 0.84 |
| 7 | 47436497 | rs17544028 | 0.226 | 0.083 | -0.028 | 0.183 | -0.016 | -0.003 | -0.003 | 0 | 0.07 | 0.03 | 0.15 | 0.08 | 0.16 | 0.17 |
| 7 | 47436558 | rs2177795 | 0.36 | -0.191 | 0.255 | 0.195 | -0.024 | -0.064 | -0.064 | 0.143 | 0.86 | 0.77 | 0.96 | 1 | 0.97 | 0.89 |
| 7 | 47436838 | rs17626611 | -0.263 | 0.676 | -0.125 | 0.33 | 0.035 | -0.017 | -0.017 | 1.683 | 0.09 | 0.16 | 0.1 | 0.03 | 0.18 | 0.05 |
| 7 | 47437150 | rs12540482 | 0.117 | 0.34 | -0.665 | 0.242 | -0.068 | 0.028 | 0.028 | 0.417 | 0.22 | 0.01 | 0.22 | 0.11 | 0.26 | 0.2 |
| 7 | 47446215 | rs834602 | -0.951 | 0.239 | -0.382 | 0.518 | 0.012 | 0.007 | 0.007 | 0 | 0.38 | 0.26 | 0.3 | 0.27 | 0.36 | 0.29 |
| 7 | 47447541 | rs11764365 | 1.859 | 0.179 | -0.702 | 0.669 | -0.141 | -0.056 | -0.056 | 1.217 | 0.41 | 0.05 | 0.55 | 0.42 | 0.48 | 0.45 |
| 7 | 47450271 | rs834604 | -0.924 | 0.074 | -0.598 | 0.677 | 0.046 | 0.066 | 0.066 | 0 | 0.43 | 0.25 | 0.3 | 0.27 | 0.35 | 0.41 |
| 7 | 47455916 | rs12216631 | -0.882 | 0.058 | -0.177 | 0.62 | -0.028 | -0.03 | -0.03 | 0 | 0.21 | 0.38 | 0.12 | 0.16 | 0.14 | 0.27 |
| 7 | 47457466 | rs865603 | -0.124 | 0.397 | 0.597 | 0.712 | 0.023 | 0.096 | 0.096 | 0.552 | 0.92 | 0.75 | 0.96 | 0.84 | 0.97 | 0.92 |
| 7 | 47479377 | rs17172860 | -0.032 | 0.407 | -0.23 | 0.621 | -0.012 | -0.055 | -0.055 | 0 | 0.11 | 0.27 | 0.07 | 0.01 | 0.07 | 0.1 |
| 7 | 47479583 | rs1799376 | -0.576 | 0.296 | -0.393 | 0.6 | 0.028 | 0.079 | 0.079 | 0 | 0.45 | 0.33 | 0.35 | 0.26 | 0.41 | 0.4 |
| 7 | 47481602 | rs11763044 | 1.871 | 0.395 | -0.454 | 0.628 | -0.151 | -0.075 | -0.075 | 0 | 0.38 | 0.03 | 0.53 | 0.43 | 0.47 | 0.44 |
| 7 | 47481628 | rs16881490 | -0.456 | 0.641 | 0.449 | 0.655 | -0.013 | -0.059 | -0.059 | 0 | 0.12 | 0.29 | 0.07 | 0.01 | 0.07 | 0.1 |
| 7 | 47483997 | rs1704974 | 1.238 | 1.475 | 0.701 | 1.166 | 0.002 | 0.106 | 0.106 | 0.486 | 0.83 | 0.47 | 0.89 | 0.7 | 0.87 | 0.84 |
| 7 | 47485965 | rs1799384 | 1.473 | 1.692 | 0.333 | 1.045 | 0.071 | 0.043 | 0.043 | 0 | 0.74 | 0.49 | 0.76 | 0.67 | 0.63 | 0.78 |
| 7 | 47488901 | rs13241485 | -0.549 | 1.964 | -0.413 | 1.157 | -0.041 | -0.038 | -0.038 | 0 | 0.07 | 0 | 0.1 | 0.16 | 0.18 | 0.06 |
| 7 | 47494033 | rs7778450 | -1.149 | 1.873 | -0.018 | 0.598 | 0.023 | -0.037 | -0.037 | 0 | 0.11 | 0.14 | 0.11 | 0 | 0.18 | 0.05 |
| 7 | 47494959 | rs2941528 | -0.324 | 1.477 | -0.028 | 0.383 | 0.008 | 0.008 | 0.008 | 2.776 | 0.32 | 0.29 | 0.21 | 0.21 | 0.25 | 0.27 |
| 7 | 47496258 | rs4724581 | 1.842 | 1.301 | -0.134 | 0.316 | 0.016 | -0.032 | -0.032 | 0 | 0.37 | 0.18 | 0.52 | 0.58 | 0.32 | 0.46 |
| 7 | 47498364 | rs2941549 | -0.01 | 1.244 | -0.05 | 0.315 | 0.006 | 0.021 | 0.021 | 3.921 | 0.31 | 0.21 | 0.22 | 0.17 | 0.29 | 0.24 |
| 7 | 47499395 | rs6943731 | 0.154 | 1.357 | -0.293 | 0.326 | -0.031 | -0.017 | -0.017 | 0 | 0.09 | 0.03 | 0.08 | 0.09 | 0.14 | 0.08 |
| 7 | 47500360 | rs12702340 | 0.302 | 1.386 | 0.621 | 0.323 | -0.008 | -0.003 | -0.003 | 0.223 | 0.66 | 0.57 | 0.77 | 0.69 | 0.69 | 0.62 |
| 7 | 47502291 | rs7793087 | 1.168 | 1.254 | 0.473 | 0.303 | 0.011 | 0.023 | 0.023 | 1.402 | 0.51 | 0.2 | 0.65 | 0.6 | 0.53 | 0.51 |
| 7 | 47511161 | rs3801081 | 0.672 | 1.106 | 0.736 | -0.022 | 0.014 | -0.024 | -0.024 | 0 | 0.71 | 0.62 | 0.76 | 0.89 | 0.7 | 0.83 |
| 7 | 47515257 | rs868107 | -0.771 | 0.743 | 0.838 | 0.026 | 0.007 | -0.024 | -0.024 | 0 | 0.05 | 0.08 | 0.05 | 0.01 | 0.07 | 0.03 |
| 7 | 47518446 | rs2941539 | 2.052 | 0.587 | 0.666 | 0.1 | -0.002 | 0.013 | 0.013 | 0.074 | 0.63 | 0.24 | 0.7 | 0.67 | 0.65 | 0.59 |
| 7 | 47520850 | rs1543035 | -0.383 | 0.563 | 1.083 | 0.141 | -0.032 | 0.05 | 0.05 | 0.58 | 0.14 | 0.31 | 0.06 | 0.24 | 0.08 | 0.22 |

| | | | | | | | | | | | | | | | | |
|---|---|---|---|---|---|---|---|---|---|---|---|---|---|---|---|---|
| 7 | 47521863 | rs2964955 | 1.821 | 0.609 | 0.834 | 0.134 | 0.007 | 0 | 0 | 0 | 0.68 | 0.25 | 0.75 | 0.67 | 0.73 | 0.65 |
| 7 | 47524798 | rs2941542 | 0.666 | 0.525 | 0.809 | 0.105 | -0.04 | 0.023 | 0.023 | 0 | 0.83 | 0.88 | 0.81 | 0.91 | 0.74 | 0.82 |
| 7 | 47525903 | rs2964959 | 1.854 | 0.583 | 0.66 | 0.221 | -0.009 | 0.004 | 0.004 | 1.366 | 0.65 | 0.19 | 0.7 | 0.65 | 0.66 | 0.6 |
| 7 | 47526059 | rs2941544 | 1.854 | 0.583 | 0.66 | 0.221 | -0.009 | 0.004 | 0.004 | 1.366 | 0.65 | 0.19 | 0.7 | 0.65 | 0.66 | 0.6 |
| 7 | 47532716 | rs7803908 | -1.481 | 0.913 | 0.933 | 0.211 | 0.043 | -0.028 | -0.028 | 0 | 0.21 | 0.43 | 0.23 | 0.12 | 0.26 | 0.2 |
| 7 | 47537679 | rs884972 | 0.096 | 0.11 | 0.249 | 0.27 | 0 | 0.001 | 0.001 | 2.293 | 0.54 | 0.52 | 0.76 | 0.59 | 0.69 | 0.48 |
| 7 | 47538546 | rs2271311 | 0.832 | 0.129 | 0.299 | 0.603 | -0.01 | 0.025 | 0.025 | 2.011 | 0.22 | 0.25 | 0.27 | 0.29 | 0.17 | 0.19 |
| 7 | 47539322 | rs10265492 | 0.75 | -0.118 | 0.541 | 0.585 | -0.037 | 0.048 | 0.048 | 1.691 | 0.22 | 0.37 | 0.27 | 0.28 | 0.14 | 0.19 |
| 7 | 47543793 | rs1009701 | 0.6 | -0.49 | 0.759 | 0.462 | -0.027 | -0.024 | -0.024 | 0 | 0.38 | 0.52 | 0.26 | 0.33 | 0.28 | 0.46 |
| 7 | 47545479 | rs6974901 | 0.562 | -0.5 | 0.434 | 0.473 | -0.003 | 0.002 | 0.002 | 0 | 0.08 | 0.12 | 0.07 | 0.1 | 0.07 | 0.06 |
| 7 | 47546375 | rs334496 | -0.606 | -0.463 | 0.742 | 0.322 | -0.014 | -0.016 | -0.016 | 0.557 | 0.61 | 0.57 | 0.73 | 0.67 | 0.71 | 0.54 |
| 7 | 47548893 | rs12668436 | 0.438 | -0.032 | 0.164 | 0.338 | 0.01 | -0.033 | -0.033 | 0 | 0.31 | 0.16 | 0.27 | 0.37 | 0.28 | 0.27 |
| 7 | 47549170 | rs182838 | 1.022 | 0.08 | -0.022 | 0.352 | -0.11 | -0.058 | -0.058 | 0 | 0.28 | 0.08 | 0.43 | 0.3 | 0.43 | 0.27 |
| 7 | 47552208 | rs586005 | 0.984 | 0.089 | 0.071 | 0.355 | -0.059 | -0.026 | -0.026 | 0 | 0.28 | 0.12 | 0.43 | 0.3 | 0.42 | 0.27 |
| 7 | 47554879 | rs17658891 | 0.274 | 0.107 | -0.138 | 0.458 | -0.002 | 0.002 | 0.002 | 3.84 | 0.31 | 0.34 | 0.29 | 0.38 | 0.28 | 0.27 |
| 7 | 47560950 | rs11977711 | 0.658 | 0.174 | -0.784 | 0.343 | 0.026 | 0.019 | 0.019 | 3.629 | 0.34 | 0.14 | 0.23 | 0.32 | 0.27 | 0.46 |
| 7 | 47564745 | rs334521 | -0.114 | -0.14 | -0.751 | 0.415 | 0.072 | -0.014 | -0.014 | 0.573 | 0.54 | 0.35 | 0.67 | 0.52 | 0.74 | 0.56 |
| 7 | 47564942 | rs334522 | 0.53 | 0.987 | -1.163 | 0.55 | -0.061 | -0.056 | -0.056 | 0 | 0.19 | 0.07 | 0.32 | 0.37 | 0.29 | 0.25 |
| 7 | 47565504 | rs334523 | 0.309 | 0.505 | -0.813 | 0.053 | 0.001 | 0.007 | 0.007 | 0.472 | 0.55 | 0.45 | 0.62 | 0.74 | 0.6 | 0.51 |
| 7 | 47566295 | rs334526 | 0.261 | 0.785 | -0.926 | 0.446 | -0.018 | -0.002 | -0.002 | 0 | 0.87 | 0.69 | 0.9 | 0.89 | 0.91 | 0.92 |
| 7 | 47567227 | rs334527 | -0.049 | 1.49 | -1.581 | 0.806 | -0.124 | -0.139 | -0.139 | 2.25 | 0.19 | 0.01 | 0.38 | 0.44 | 0.35 | 0.25 |
| 7 | 47574318 | rs10242770 | 0.051 | 1.061 | -1.427 | 0.114 | -0.022 | 0.137 | 0.137 | 0 | 0.48 | 0.23 | 0.34 | 0.2 | 0.47 | 0.46 |
| 7 | 47577952 | rs592048 | 0.139 | 1.404 | -1.6 | 0.584 | -0.071 | -0.082 | -0.082 | 3.008 | 0.2 | 0.07 | 0.35 | 0.44 | 0.35 | 0.34 |
| 7 | 47578201 | rs334508 | -0.82 | 1.061 | -0.688 | 0.504 | -0.002 | 0 | 0 | 0 | 0.69 | 0.67 | 0.77 | 0.66 | 0.8 | 0.75 |
| 7 | 47580445 | rs10245886 | -0.088 | 1.282 | -0.505 | -0.025 | 0.051 | -0.077 | -0.077 | 0 | 0.22 | 0.5 | 0.23 | 0.09 | 0.27 | 0.28 |
| 7 | 47587877 | rs613485 | -0.128 | 1.334 | -0.609 | 0.241 | -0.014 | -0.031 | -0.031 | 0 | 0.77 | 0.79 | 0.66 | 0.56 | 0.68 | 0.65 |
| 7 | 47589807 | rs675242 | -0.022 | 0.995 | -0.521 | -0.016 | -0.008 | 0.08 | 0.08 | 0 | 0.45 | 0.24 | 0.43 | 0.22 | 0.48 | 0.4 |
| 7 | 47596775 | rs615300 | -0.038 | 0.784 | -0.78 | 0.116 | 0 | 0.039 | 0.039 | 0 | 0.47 | 0.38 | 0.42 | 0.27 | 0.45 | 0.41 |
| 7 | 47598606 | rs7807054 | -0.268 | 0.917 | -0.771 | 0.081 | -0.002 | 0 | 0 | 0.301 | 0.55 | 0.49 | 0.61 | 0.48 | 0.63 | 0.48 |
| 7 | 47601082 | rs12155302 | 0.959 | 0.756 | -1.102 | -0.008 | 0.015 | 0.032 | 0.032 | 0 | 0.27 | 0.17 | 0.21 | 0.1 | 0.22 | 0.12 |
| 7 | 47607478 | rs11763932 | -0.235 | 0.723 | -0.961 | 0.16 | 0.007 | -0.003 | -0.003 | 0 | 0.56 | 0.37 | 0.6 | 0.45 | 0.63 | 0.47 |
| 7 | 47619815 | rs6463424 | 0.171 | 1.532 | -0.6 | 0.436 | 0.005 | 0 | 0 | 0 | 0.69 | 0.33 | 0.59 | 0.65 | 0.59 | 0.65 |
| 7 | 47628498 | rs17710080 | -0.087 | 1.396 | -1.111 | 0.816 | -0.042 | -0.051 | -0.051 | 0.039 | 0.2 | 0.06 | 0.28 | 0.29 | 0.25 | 0.37 |
| 7 | 47629367 | rs17710092 | -0.84 | 1.556 | -1.127 | 0.958 | -0.015 | -0.032 | -0.032 | 0.017 | 0.28 | 0.22 | 0.36 | 0.42 | 0.34 | 0.48 |
| 7 | 47629909 | rs6463425 | -0.166 | 1.119 | -0.82 | 0.718 | 0.004 | 0.002 | 0.002 | 0 | 0.07 | 0.15 | 0.08 | 0.07 | 0.09 | 0.07 |
| 7 | 47633187 | rs6974363 | -0.545 | 0.824 | -1.249 | 0.305 | 0.01 | 0.032 | 0.032 | 0.045 | 0.58 | 0.65 | 0.66 | 0.82 | 0.64 | 0.81 |
| 7 | 47636122 | rs12702357 | 0.463 | 0.994 | -1.592 | 0.437 | -0.068 | 0.008 | 0.008 | 0.796 | 0.19 | 0.02 | 0.2 | 0.18 | 0.27 | 0.3 |
| 7 | 47636610 | rs10807947 | -0.493 | 0.847 | -1.172 | 0.848 | -0.001 | 0.036 | 0.036 | 0 | 0.34 | 0.48 | 0.34 | 0.53 | 0.37 | 0.48 |
| 7 | 47639999 | rs10224591 | -0.981 | 1.081 | -1.052 | 0.482 | 0.012 | 0.012 | 0.012 | 1.251 | 0.5 | 0.54 | 0.58 | 0.47 | 0.59 | 0.72 |
| 7 | 47641594 | rs10272973 | 0.331 | 0.926 | -1.548 | 0.425 | -0.102 | 0.007 | 0.007 | 1.166 | 0.24 | 0.03 | 0.33 | 0.17 | 0.31 | 0.35 |
| 7 | 47650640 | rs11765122 | -0.525 | 0.858 | -0.438 | 0.528 | 0.003 | -0.007 | -0.007 | 0.04 | 0.27 | 0.53 | 0.26 | 0.29 | 0.3 | 0.37 |
| 7 | 47665372 | rs13235604 | -0.687 | 1.148 | -0.297 | 0.674 | 0.046 | 0.04 | 0.04 | 0.084 | 0.24 | 0.52 | 0.26 | 0.29 | 0.34 | 0.31 |
| 7 | 47676143 | rs4724603 | -0.029 | 0.268 | -0.77 | 0.235 | 0.005 | 0.013 | 0.013 | 0.208 | 0.68 | 0.76 | 0.61 | 0.79 | 0.69 | 0.63 |
| 7 | 47686368 | rs7797887 | -0.08 | 0.359 | -0.364 | 0.064 | -0.004 | -0.056 | -0.056 | 0.064 | 0.24 | 0.31 | 0.31 | 0.1 | 0.22 | 0.34 |
| 7 | 47687938 | rs965924 | 0.267 | 0.561 | -0.527 | 0.216 | -0.001 | -0.002 | -0.002 | 0 | 0.23 | 0.24 | 0.19 | 0.26 | 0.26 | 0.19 |
| 7 | 47690398 | rs2347990 | -0.678 | 0.503 | -0.635 | 0.191 | 0.006 | 0.018 | 0.018 | 0.001 | 0.48 | 0.59 | 0.51 | 0.36 | 0.47 | 0.53 |
| 7 | 47698685 | rs12531737 | 0.653 | 0.606 | -0.678 | 0.618 | -0.056 | -0.051 | -0.051 | 0 | 0.16 | 0.03 | 0.26 | 0.19 | 0.23 | 0.17 |
| 7 | 47700779 | rs11770543 | 0.549 | 0.467 | -0.388 | 0.594 | -0.001 | 0.006 | 0.006 | 0 | 0.26 | 0.32 | 0.31 | 0.27 | 0.28 | 0.19 |
| 7 | 47702761 | rs7790476 | 0.669 | 0.504 | -0.727 | 0.373 | -0.006 | 0.016 | 0.016 | 0 | 0.25 | 0.12 | 0.3 | 0.19 | 0.27 | 0.17 |
| 7 | 47704843 | rs6463429 | -0.397 | 0.554 | -0.947 | 0.486 | -0.002 | -0.001 | -0.001 | 0 | 0.65 | 0.62 | 0.7 | 0.65 | 0.67 | 0.68 |
| 7 | 47706121 | rs10233927 | 0.4 | 0.929 | -1.263 | 0.554 | -0.026 | 0.033 | 0.033 | 0.787 | 0.3 | 0.11 | 0.31 | 0.28 | 0.32 | 0.42 |
| 7 | 47706667 | rs12532470 | 1.022 | 1.047 | -1.51 | 0.674 | -0.068 | -0.001 | -0.001 | 0 | 0.25 | 0.02 | 0.32 | 0.17 | 0.24 | 0.27 |
| 7 | 47708525 | rs3934868 | -0.256 | 0.988 | -1.114 | 0.801 | 0.032 | 0.176 | 0.176 | 0.26 | 0.85 | 0.65 | 0.8 | 0.58 | 0.79 | 0.88 |
| 7 | 47710987 | rs4720605 | 0.239 | 0.99 | -1.234 | 0.79 | -0.008 | 0.01 | 0.01 | 2.173 | 0.58 | 0.58 | 0.54 | 0.28 | 0.49 | 0.44 |
| 7 | 47713313 | rs7792269 | 0.545 | 1.411 | -0.645 | 1.184 | 0.011 | -0.006 | -0.006 | 0 | 0.62 | 0.48 | 0.48 | 0.41 | 0.57 | 0.62 |
| 7 | 47716264 | rs6463430 | 0.39 | 1.453 | -0.726 | 1.561 | 0 | 0.012 | 0.012 | 0.116 | 0.88 | 0.88 | 0.85 | 0.74 | 0.83 | 0.88 |
| 7 | 47727607 | rs10227002 | -0.385 | 1.401 | -0.838 | 1.435 | -0.015 | -0.026 | -0.026 | 0 | 0.11 | 0.05 | 0.24 | 0.37 | 0.18 | 0.13 |
| 7 | 47729511 | rs4720608 | -1.176 | 1.342 | 0.212 | 1.224 | 0.035 | 0.118 | 0.118 | 0 | 0.16 | 0.37 | 0.29 | 0.37 | 0.21 | 0.14 |
| 7 | 47730673 | rs11772364 | -0.249 | 1.37 | -0.146 | 1.264 | -0.019 | -0.044 | -0.044 | 2.734 | 0.11 | 0.05 | 0.24 | 0.37 | 0.17 | 0.13 |
| 7 | 47732411 | rs4724613 | 0.014 | 1.682 | -0.901 | 0.812 | -0.065 | -0.09 | -0.09 | 0.11 | 0.71 | 0.88 | 0.63 | 0.53 | 0.62 | 0.5 |
| 7 | 47741157 | rs6947796 | -0.071 | 0.853 | -0.605 | 0.229 | -0.001 | -0.002 | -0.002 | 0 | 0.55 | 0.53 | 0.49 | 0.52 | 0.55 | 0.42 |
| 7 | 47743362 | rs7792409 | 0.144 | 0.713 | -1.314 | 0.098 | -0.047 | -0.089 | -0.089 | 0 | 0.21 | 0.03 | 0.28 | 0.39 | 0.23 | 0.23 |
| 7 | 47747268 | rs13232026 | 0.654 | 0.453 | -0.544 | -0.166 | 0.007 | 0.032 | 0.032 | 3.297 | 0.32 | 0.21 | 0.18 | 0.13 | 0.32 | 0.19 |
| 7 | 47749358 | rs7795529 | 0.333 | 0.386 | 0.107 | -0.271 | -0.011 | -0.055 | -0.055 | 0.045 | 0.34 | 0.46 | 0.21 | 0.11 | 0.32 | 0.18 |
| 7 | 47750051 | rs10255531 | 0.632 | 0.231 | -0.14 | -0.422 | 0 | -0.006 | -0.006 | 0 | 0.29 | 0.27 | 0.19 | 0.11 | 0.3 | 0.16 |
| 7 | 47750095 | rs6967974 | -0.724 | 0.469 | -0.859 | -0.267 | 0 | 0.004 | 0.004 | 0 | 0.78 | 0.82 | 0.74 | 0.79 | 0.81 | 0.68 |
| 7 | 47750184 | rs6968120 | -0.731 | 0.468 | -0.859 | -0.255 | -0.001 | 0.014 | 0.014 | 0 | 0.78 | 0.82 | 0.74 | 0.82 | 0.81 | 0.69 |
| 7 | 47751799 | rs6956508 | -0.576 | 0.551 | -0.923 | -0.275 | -0.01 | 0.013 | 0.013 | 0 | 0.76 | 0.82 | 0.73 | 0.81 | 0.79 | 0.69 |
| 7 | 47751893 | rs4278082 | -0.117 | 0.743 | -0.935 | -0.301 | -0.008 | -0.011 | -0.011 | 0 | 0.34 | 0.15 | 0.42 | 0.69 | 0.36 | 0.4 |

| | | | | | | | | | | | | | | | | |
|---|---|---|---|---|---|---|---|---|---|---|---|---|---|---|---|---|
| 7 | 47751960 | rs4364533 | 0.385 | 0.775 | -1.158 | 0.106 | -0.031 | -0.089 | -0.089 | 0 | 0.25 | 0.04 | 0.25 | 0.41 | 0.29 | 0.33 |
| 7 | 47756822 | rs10281623 | -0.572 | 0.309 | -1.221 | -0.326 | 0.011 | 0.075 | 0.075 | 0 | 0.8 | 0.89 | 0.86 | 0.98 | 0.87 | 0.8 |
| 7 | 47758976 | rs4724614 | 0.231 | 0.281 | -0.594 | -0.069 | -0.012 | -0.003 | -0.003 | 0 | 0.72 | 0.55 | 0.66 | 0.72 | 0.79 | 0.74 |
| 7 | 47759777 | rs10499664 | 0.596 | 0.809 | -0.917 | -0.097 | -0.05 | -0.074 | -0.074 | 0 | 0.25 | 0.09 | 0.3 | 0.42 | 0.33 | 0.33 |
| 7 | 47764757 | rs6977529 | 1.078 | 0.748 | -0.769 | -0.378 | 0.002 | 0.001 | 0.001 | 4.24 | 0.38 | 0.32 | 0.26 | 0.3 | 0.35 | 0.28 |
| 7 | 47770509 | rs6942837 | -0.151 | 0.79 | -0.519 | -0.108 | -0.005 | 0.016 | 0.016 | 0 | 0.74 | 0.62 | 0.67 | 0.68 | 0.79 | 0.73 |
| 7 | 47775298 | rs12702377 | 0.671 | 0.552 | -0.863 | -0.164 | 0.002 | -0.004 | -0.004 | 0 | 0.45 | 0.34 | 0.39 | 0.49 | 0.49 | 0.45 |
| 7 | 47775355 | rs11761091 | 0.238 | 0.553 | -0.686 | -0.168 | -0.004 | -0.012 | -0.012 | 0 | 0.64 | 0.54 | 0.57 | 0.68 | 0.68 | 0.63 |
| 7 | 47777082 | rs1078236 | 0.332 | 0.599 | -0.672 | -0.113 | -0.012 | -0.01 | -0.01 | 0 | 0.67 | 0.42 | 0.6 | 0.69 | 0.71 | 0.66 |
| 7 | 47777685 | rs17544941 | 0.753 | 0.655 | -0.985 | -0.154 | 0 | -0.006 | -0.006 | 0.742 | 0.11 | 0.07 | 0.09 | 0.01 | 0.12 | 0.11 |
| 7 | 47778326 | rs17172896 | 0.668 | 0.644 | -0.985 | -0.16 | 0 | -0.019 | -0.019 | 0 | 0.11 | 0.09 | 0.1 | 0.01 | 0.12 | 0.11 |
| 7 | 47779633 | rs882160 | 0.902 | 0.687 | -1.202 | -0.098 | -0.032 | -0.047 | -0.047 | 0 | 0.26 | 0.13 | 0.3 | 0.4 | 0.33 | 0.37 |
| 7 | 47779969 | rs7800244 | 1.833 | 0.673 | -1.193 | -0.17 | -0.026 | -0.022 | -0.022 | 0 | 0.17 | 0.06 | 0.17 | 0.18 | 0.18 | 0.18 |
| 7 | 47781538 | rs7805780 | 0.597 | 0.744 | -0.841 | -0.248 | 0.001 | -0.057 | -0.057 | 0 | 0.14 | 0.17 | 0.11 | 0.01 | 0.14 | 0.12 |
| 7 | 47781978 | rs6463435 | -0.192 | 0.788 | -0.978 | 0.049 | -0.012 | 0.023 | 0.023 | 0.513 | 0.65 | 0.71 | 0.62 | 0.68 | 0.64 | 0.55 |
| 7 | 47782640 | rs4724619 | 0.936 | 0.972 | -0.891 | 0.3 | -0.002 | -0.004 | -0.004 | 3.961 | 0.21 | 0.2 | 0.25 | 0.27 | 0.23 | 0.28 |
| 7 | 47784580 | rs4720613 | 1.334 | 0.892 | -0.816 | 0.638 | -0.019 | 0.002 | 0.002 | 1.381 | 0.05 | 0.01 | 0.18 | 0.31 | 0.07 | 0.04 |
| 7 | 47789918 | rs7787946 | -0.382 | 0.761 | -0.642 | -0.138 | 0.006 | 0.025 | 0.025 | 2.839 | 0.5 | 0.43 | 0.61 | 0.66 | 0.58 | 0.64 |
| 7 | 47790912 | rs4724620 | -1.665 | 0.795 | -0.872 | -0.19 | 0.006 | 0.018 | 0.018 | 0 | 0.84 | 0.89 | 0.88 | 0.97 | 0.9 | 0.84 |
| 7 | 47791977 | rs6965341 | -0.439 | 0.804 | -0.784 | -0.132 | -0.001 | 0.001 | 0.001 | 0.315 | 0.67 | 0.67 | 0.7 | 0.74 | 0.72 | 0.68 |
| 7 | 47795058 | rs921630 | 0.143 | 0.918 | -0.866 | 0.092 | -0.003 | -0.006 | -0.006 | 0.027 | 0.4 | 0.38 | 0.56 | 0.59 | 0.46 | 0.45 |
| 7 | 47795664 | rs4724626 | -0.253 | 0.752 | -0.809 | 0.001 | -0.004 | -0.014 | -0.014 | 0 | 0.57 | 0.49 | 0.72 | 0.82 | 0.64 | 0.62 |
| 7 | 47798316 | rs718077 | -0.255 | 0.744 | -0.78 | 0.039 | 0 | 0 | 0 | 0 | 0.57 | 0.49 | 0.73 | 0.79 | 0.65 | 0.62 |
| 7 | 47798901 | rs17627589 | 0.421 | 0.757 | -0.932 | -0.205 | -0.002 | -0.008 | -0.008 | 0 | 0.29 | 0.4 | 0.25 | 0.21 | 0.29 | 0.26 |
| 7 | 47802084 | rs10807948 | -0.077 | 1.022 | -1.107 | 0.046 | -0.011 | 0.022 | 0.022 | 0 | 0.47 | 0.35 | 0.48 | 0.42 | 0.52 | 0.48 |
| 7 | 47802472 | rs6953403 | -0.077 | 0.986 | -1.107 | 0.046 | -0.012 | 0.022 | 0.022 | 0 | 0.47 | 0.35 | 0.48 | 0.41 | 0.52 | 0.48 |
| 7 | 47804264 | rs4724639 | -1.289 | 0.992 | -1.355 | -0.84 | -0.016 | 0.04 | 0.04 | 0 | 0.78 | 0.89 | 0.82 | 0.94 | 0.76 | 0.79 |
| 7 | 47805513 | rs13438587 | 1.887 | 0.9 | -1.081 | -0.686 | -0.012 | -0.038 | -0.038 | 0 | 0.17 | 0.24 | 0.09 | 0.07 | 0.14 | 0.09 |
| 7 | 47806077 | rs10276437 | 1.075 | 0.905 | -1.206 | -0.684 | -0.035 | -0.019 | -0.019 | 0.112 | 0.19 | 0.04 | 0.25 | 0.22 | 0.22 | 0.26 |
| 7 | 47808059 | rs11971419 | 1.649 | 0.876 | -0.816 | -0.725 | -0.013 | -0.039 | -0.039 | 0.118 | 0.17 | 0.24 | 0.09 | 0.07 | 0.14 | 0.09 |
| 7 | 47811443 | rs6463443 | -0.419 | 1.181 | -1.208 | -0.395 | -0.019 | 0.023 | 0.023 | 0 | 0.73 | 0.89 | 0.8 | 0.86 | 0.72 | 0.65 |
| 7 | 47817391 | rs17627715 | 0.633 | 1.025 | -0.964 | -0.342 | -0.001 | -0.004 | -0.004 | 0 | 0.13 | 0.09 | 0.09 | 0.01 | 0.16 | 0.13 |
| 7 | 47818966 | rs1453288 | -0.28 | 1.186 | -0.93 | -0.285 | -0.001 | 0 | 0 | 0.563 | 0.21 | 0.19 | 0.25 | 0.21 | 0.24 | 0.26 |
| 7 | 47821926 | rs4724641 | 0.766 | 1.638 | -0.582 | -0.26 | -0.005 | -0.009 | -0.009 | 0 | 0.23 | 0.26 | 0.37 | 0.57 | 0.26 | 0.27 |
| 7 | 47823298 | rs7797545 | -0.388 | 1.506 | -0.476 | -0.254 | 0.013 | -0.015 | -0.015 | 0 | 0.39 | 0.5 | 0.56 | 0.78 | 0.48 | 0.45 |
| 7 | 47837117 | rs965143 | -0.534 | 1.022 | -0.054 | -0.235 | -0.007 | 0.022 | 0.022 | 0 | 0.71 | 0.64 | 0.66 | 0.65 | 0.74 | 0.63 |
| 7 | 47839736 | rs4720616 | -1.357 | 0.935 | 0.079 | -0.231 | -0.011 | -0.019 | -0.019 | 0 | 0.78 | 0.8 | 0.81 | 0.92 | 0.84 | 0.83 |
| 7 | 47839846 | rs6463448 | 0.706 | 0.759 | -0.023 | -0.599 | -0.021 | 0.008 | 0.008 | 0 | 0.46 | 0.29 | 0.48 | 0.61 | 0.47 | 0.43 |
| 7 | 47840387 | rs13231277 | 0.373 | 0.66 | 0.295 | -0.548 | -0.005 | -0.032 | -0.032 | 0 | 0.13 | 0.22 | 0.09 | 0.04 | 0.12 | 0.07 |
| 7 | 47847183 | rs3887211 | -0.112 | 0.98 | -0.08 | -0.353 | -0.001 | 0 | 0 | 0 | 0.63 | 0.69 | 0.59 | 0.63 | 0.64 | 0.7 |
| 7 | 47847679 | rs1074780 | -0.823 | 0.551 | 0.32 | -0.305 | -0.002 | -0.001 | -0.001 | 2.567 | 0.7 | 0.71 | 0.66 | 0.71 | 0.74 | 0.76 |
| 7 | 47847751 | rs3734931 | 1.442 | 0.656 | 0.156 | -0.228 | 0 | 0 | 0 | 0 | 0.05 | 0.06 | 0.03 | 0.03 | 0.05 | 0.14 |
| 7 | 47852837 | rs2290386 | 1.325 | 0.553 | -0.149 | -0.427 | -0.013 | -0.029 | -0.029 | 0.875 | 0.16 | 0.02 | 0.21 | 0.22 | 0.18 | 0.19 |
| 7 | 47853413 | rs6944477 | 1.219 | 0.466 | -0.008 | -0.535 | 0.01 | 0.018 | 0.018 | 0 | 0.27 | 0.19 | 0.23 | 0.08 | 0.23 | 0.19 |
| 7 | 47854956 | rs17545564 | 1.398 | 0.485 | -0.072 | -0.529 | -0.009 | -0.029 | -0.029 | 0 | 0.16 | 0.02 | 0.2 | 0.22 | 0.17 | 0.19 |
| 7 | 47857982 | rs3800568 | 0.791 | 0.523 | 0.487 | -0.604 | -0.019 | -0.006 | -0.006 | 0 | 0.37 | 0.32 | 0.41 | 0.63 | 0.41 | 0.42 |
| 7 | 47858097 | rs13246591 | 0.947 | 0.524 | 0.653 | -0.432 | 0.001 | -0.025 | -0.025 | 0 | 0.13 | 0.21 | 0.1 | 0.04 | 0.12 | 0.06 |
| 7 | 47860510 | rs17659820 | 0.955 | 0.517 | 0.599 | -0.418 | -0.002 | -0.002 | -0.002 | 0 | 0.07 | 0.09 | 0.03 | 0.03 | 0.05 | 0.14 |
| 7 | 47867350 | rs7786516 | -1.219 | 0.521 | 0.71 | -0.423 | -0.017 | -0.041 | -0.041 | 0 | 0.72 | 0.68 | 0.76 | 0.92 | 0.77 | 0.81 |
| 7 | 47869038 | rs7788583 | -0.059 | 0.516 | 0.476 | -0.446 | -0.003 | -0.03 | -0.03 | 0.305 | 0.19 | 0.34 | 0.13 | 0.07 | 0.17 | 0.21 |
| 7 | 47872845 | rs921634 | -0.66 | 0.501 | 0.375 | -0.45 | -0.007 | -0.022 | -0.022 | 0 | 0.66 | 0.65 | 0.73 | 0.9 | 0.73 | 0.66 |
| 7 | 47877939 | rs10245949 | 0.725 | 0.617 | 0.212 | -0.353 | -0.052 | -0.03 | -0.03 | 0.571 | 0.34 | 0.23 | 0.38 | 0.62 | 0.38 | 0.38 |
| 7 | 47878993 | rs6960394 | 1.078 | 0.533 | 0.232 | -0.355 | 0.026 | 0.052 | 0.052 | 0 | 0.28 | 0.13 | 0.23 | 0.07 | 0.23 | 0.21 |
| 7 | 47880403 | rs4724650 | 1.334 | 0.624 | -0.2 | -0.311 | -0.025 | -0.041 | -0.041 | 0 | 0.17 | 0.02 | 0.23 | 0.23 | 0.21 | 0.2 |
| 7 | 47882306 | rs10429127 | 1.093 | 0.582 | -0.035 | -0.298 | 0.002 | -0.011 | -0.011 | 0 | 0.13 | 0.18 | 0.1 | 0.05 | 0.12 | 0.06 |
| 7 | 47886707 | rs10280228 | 0.112 | 0.632 | -0.18 | -0.257 | 0.004 | 0.001 | 0.001 | 0.325 | 0.42 | 0.55 | 0.34 | 0.12 | 0.35 | 0.27 |
| 7 | 47894326 | rs13228166 | -0.329 | 0.971 | -0.223 | -0.069 | -0.006 | 0.003 | 0.003 | 0 | 0.61 | 0.63 | 0.6 | 0.35 | 0.61 | 0.49 |
| 7 | 47896158 | rs6967514 | -0.336 | 0.92 | -0.18 | 0.027 | -0.005 | 0.001 | 0.001 | 0 | 0.61 | 0.63 | 0.6 | 0.36 | 0.61 | 0.5 |
| 7 | 47898597 | rs2053985 | 0.47 | 1.252 | -0.012 | 0.095 | 0.008 | -0.006 | -0.006 | 0.042 | 0.13 | 0.17 | 0.11 | 0.07 | 0.15 | 0.09 |
| 7 | 47898734 | rs2053986 | 0.21 | 0.962 | -0.065 | 0.618 | -0.006 | 0.003 | 0.003 | 0 | 0.56 | 0.51 | 0.59 | 0.56 | 0.58 | 0.63 |
| 7 | 47898751 | rs2053987 | 0.21 | 0.938 | -0.065 | 0.618 | -0.005 | 0.003 | 0.003 | 0 | 0.56 | 0.51 | 0.58 | 0.56 | 0.58 | 0.63 |
| 7 | 47901119 | rs11763656 | -0.263 | 1.462 | -0.015 | 1.494 | -0.005 | 0.014 | 0.014 | 0 | 0.11 | 0.14 | 0.15 | 0.35 | 0.16 | 0.18 |
| 7 | 47902551 | rs1546740 | -0.214 | 0.716 | 0.272 | 1.378 | -0.005 | 0.104 | 0.104 | 0 | 0.06 | 0.1 | 0.1 | 0.29 | 0.05 | 0.14 |
| 7 | 47903141 | rs17710983 | -0.263 | 0.723 | 0.681 | 0.423 | 0.027 | 0.025 | 0.025 | 0 | 0.15 | 0.4 | 0.22 | 0.14 | 0.16 | 0.22 |
| 7 | 47904737 | rs1554495 | 0.588 | 0.61 | 0.429 | 0.101 | -0.009 | 0.015 | 0.015 | 0 | 0.43 | 0.31 | 0.41 | 0.63 | 0.45 | 0.43 |
| 7 | 47905044 | rs7782964 | 0.499 | 0.956 | 0.5 | -0.531 | -0.017 | -0.009 | -0.009 | 0 | 0.12 | 0.22 | 0.19 | 0.08 | 0.09 | 0.18 |
| 7 | 47906367 | rs6972918 | -0.309 | 0.894 | 0.384 | -0.831 | -0.012 | -0.063 | -0.063 | 1.368 | 0.34 | 0.43 | 0.34 | 0.1 | 0.29 | 0.31 |
| 7 | 47906665 | rs2167877 | -0.2 | 0.908 | -0.01 | -0.825 | -0.011 | -0.056 | -0.056 | 0 | 0.35 | 0.6 | 0.35 | 0.1 | 0.29 | 0.31 |
| 7 | 47913579 | rs1470859 | -1.191 | 0.831 | -0.085 | -0.872 | -0.045 | 0.056 | 0.056 | 0 | 0.86 | 0.98 | 0.91 | 0.97 | 0.83 | 0.87 |

| | | | | | | | | | | | | | | | | |
|---|---|---|---|---|---|---|---|---|---|---|---|---|---|---|---|---|
| 7 | 47914774 | rs6943832 | -1.062 | 0.456 | 0.643 | -0.321 | 0.005 | 0.033 | 0.033 | 0 | 0.39 | 0.71 | 0.38 | 0.16 | 0.37 | 0.37 |
| 7 | 47915917 | rs4724657 | -0.923 | -0.093 | 0.392 | -0.539 | -0.011 | -0.057 | -0.057 | 0 | 0.35 | 0.59 | 0.35 | 0.1 | 0.29 | 0.32 |
| 7 | 47916770 | rs2348659 | -0.74 | -0.097 | 0.39 | -0.54 | -0.011 | -0.056 | -0.056 | 0.109 | 0.35 | 0.59 | 0.35 | 0.1 | 0.29 | 0.32 |
| 7 | 47917495 | rs10951934 | -0.137 | 0.475 | -0.336 | -0.307 | 0 | -0.032 | -0.032 | 0 | 0.22 | 0.23 | 0.16 | 0.01 | 0.2 | 0.14 |
| 7 | 47936850 | rs7780754 | -0.533 | 0.386 | -0.369 | 0.487 | -0.012 | -0.039 | -0.039 | 0.227 | 0.42 | 0.77 | 0.54 | 0.55 | 0.43 | 0.31 |
| 7 | 47939661 | rs6463453 | -0.602 | 0.568 | -0.537 | 0.528 | -0.019 | -0.021 | -0.021 | 0 | 0.42 | 0.78 | 0.53 | 0.6 | 0.42 | 0.32 |
| 7 | 47941714 | rs6945365 | -0.308 | 0.254 | -0.667 | 1.008 | 0.044 | -0.039 | -0.039 | 2.94 | 0.19 | 0.08 | 0.15 | 0.25 | 0.11 | 0.21 |
| 7 | 47943953 | rs11766212 | 0.192 | -0.184 | -1.309 | 0.892 | -0.15 | -0.147 | -0.147 | 0.048 | 0.24 | 0.04 | 0.41 | 0.44 | 0.37 | 0.22 |
| 7 | 47951430 | rs7802908 | -1.356 | -0.91 | -1.566 | 0.508 | -0.024 | 0.03 | 0.03 | 0 | 0.62 | 0.92 | 0.69 | 0.73 | 0.65 | 0.62 |
| 7 | 47968927 | rs2686817 | -0.181 | -0.977 | -0.526 | 0.594 | 0.018 | 0.02 | 0.02 | 0.118 | 0.38 | 0.51 | 0.51 | 0.51 | 0.51 | 0.38 |
| 7 | 47969644 | rs17711336 | 0.602 | -1.334 | -0.609 | 0.789 | 0.003 | -0.012 | -0.012 | 0 | 0.07 | 0.07 | 0.1 | 0.13 | 0.07 | 0.15 |
| 7 | 47973034 | rs6955251 | -0.726 | -1.802 | -0.729 | 0.455 | -0.005 | 0.023 | 0.023 | 4.589 | 0.21 | 0.23 | 0.28 | 0.4 | 0.17 | 0.37 |
| 7 | 47975082 | rs2686821 | 0.075 | -1.794 | -0.559 | 0.406 | 0.006 | 0.013 | 0.013 | 0.053 | 0.4 | 0.52 | 0.53 | 0.51 | 0.52 | 0.39 |
| 7 | 47981235 | rs2686828 | -1.707 | -1.917 | -0.111 | 0.205 | 0.007 | 0.003 | 0.003 | 0 | 0.67 | 0.69 | 0.8 | 0.96 | 0.72 | 0.77 |
| 7 | 47984753 | rs2686829 | -1.227 | -1.86 | -0.266 | 0.198 | 0.048 | 0.155 | 0.155 | 1.171 | 0.62 | 0.86 | 0.81 | 0.91 | 0.69 | 0.76 |
| 7 | 47991245 | rs11973038 | 1.789 | -1.762 | -0.054 | 0.392 | -0.032 | 0.016 | 0.016 | 1.207 | 0.07 | 0.12 | 0.01 | 0.1 | 0.03 | 0.02 |
| 7 | 48003665 | rs2686835 | 0.454 | -2.326 | 0.049 | 0.385 | -0.013 | -0.017 | -0.017 | 0 | 0.4 | 0.28 | 0.49 | 0.51 | 0.52 | 0.38 |
| 7 | 48006649 | rs2686838 | 0.454 | -2.345 | 0.174 | 0.376 | -0.013 | -0.017 | -0.017 | 0 | 0.4 | 0.28 | 0.49 | 0.51 | 0.52 | 0.38 |
| 7 | 48008738 | rs3176581 | 0.815 | -2.393 | 0.228 | 0.548 | -0.006 | 0.069 | 0.069 | 0.93 | 0.1 | 0.14 | 0.14 | 0.23 | 0.07 | 0.2 |
| 7 | 48010184 | rs3176565 | 0.059 | -2.408 | 0.117 | 0.522 | 0.01 | -0.011 | -0.011 | 0 | 0.12 | 0.07 | 0.15 | 0.12 | 0.1 | 0.22 |
| 7 | 48014111 | rs2708862 | 0.351 | -2.236 | 0.262 | 0.653 | -0.013 | -0.017 | -0.017 | 1.302 | 0.4 | 0.28 | 0.49 | 0.51 | 0.52 | 0.38 |
| 7 | 48016353 | rs2307254 | 2.94 | -2.292 | 0.267 | 0.638 | -0.011 | 0.008 | 0.008 | 0.907 | 0.05 | 0.03 | 0 | 0.1 | 0.02 | 0.02 |
| 7 | 48016442 | rs2307252 | 0.102 | -2.297 | 0.289 | 0.661 | -0.007 | 0.136 | 0.136 | 1.581 | 0.12 | 0.33 | 0.16 | 0.23 | 0.07 | 0.2 |
| 7 | 48016717 | rs2708861 | 0.361 | -2.259 | 0.21 | 0.663 | -0.013 | -0.018 | -0.018 | 0.992 | 0.4 | 0.28 | 0.49 | 0.51 | 0.52 | 0.38 |
| 7 | 48018677 | rs3176499 | 1.653 | -2.142 | 0.097 | 0.522 | -0.032 | 0.016 | 0.016 | 1.207 | 0.07 | 0.12 | 0.01 | 0.1 | 0.03 | 0.02 |
| 7 | 48023754 | rs2708906 | -0.611 | -1.979 | 0.114 | 0.529 | 0.042 | 0.065 | 0.065 | 0.016 | 0.51 | 0.65 | 0.64 | 0.75 | 0.57 | 0.55 |
| 7 | 48035893 | rs2708899 | -0.126 | -1.499 | 0.263 | 1.23 | -0.021 | 0.087 | 0.087 | 0.026 | 0.89 | 0.84 | 0.86 | 0.7 | 0.95 | 0.81 |
| 7 | 48037815 | rs2708885 | 0.449 | -1.708 | 0.298 | 1.336 | -0.005 | -0.001 | -0.001 | 0 | 0.33 | 0.27 | 0.49 | 0.41 | 0.51 | 0.36 |
| 7 | 48045367 | rs12702405 | 0.232 | -1.595 | 0.219 | 1.144 | -0.007 | -0.013 | -0.013 | 0.842 | 0.59 | 0.6 | 0.38 | 0.29 | 0.45 | 0.45 |
| 7 | 48050570 | rs7779252 | 0.191 | -1.618 | 0.205 | 1.233 | -0.003 | -0.001 | -0.001 | 0 | 0.59 | 0.66 | 0.38 | 0.33 | 0.45 | 0.45 |
| 7 | 48085802 | rs2708851 | 0.495 | -1.499 | 0.331 | 1.276 | -0.011 | -0.005 | -0.005 | 0 | 0.31 | 0.29 | 0.49 | 0.44 | 0.5 | 0.36 |
| 7 | 48086077 | rs2708912 | -0.064 | -1.787 | 0.766 | 1.359 | -0.035 | 0.115 | 0.115 | 0 | 0.89 | 0.77 | 0.86 | 0.75 | 0.95 | 0.81 |
| 7 | 48089571 | rs2708890 | -0.39 | -1.757 | 0.77 | 1.372 | -0.025 | 0.132 | 0.132 | 0.132 | 0.9 | 0.78 | 0.86 | 0.75 | 0.95 | 0.8 |
| 7 | 48090640 | rs2686789 | 1.089 | -1.905 | 0.748 | 1.334 | -0.041 | 0.069 | 0.069 | 0.827 | 0.81 | 0.42 | 0.73 | 0.68 | 0.88 | 0.67 |
| 7 | 48092486 | rs2686791 | -0.49 | -1.887 | 0.737 | 1.342 | -0.022 | 0.144 | 0.144 | 0.223 | 0.9 | 0.78 | 0.86 | 0.75 | 0.95 | 0.81 |
| 7 | 48098949 | rs6463459 | -0.537 | -1.697 | 0.594 | 1.029 | 0.013 | 0.019 | 0.019 | 0 | 0.09 | 0.37 | 0.13 | 0.06 | 0.07 | 0.13 |
| 7 | 48110103 | rs17132076 | 0.962 | -2.014 | 0.385 | 0.87 | -0.035 | -0.041 | -0.041 | 0 | 0.09 | 0.17 | 0.02 | 0 | 0.01 | 0 |
| 7 | 48117684 | rs9639018 | 0.994 | -1.345 | -0.017 | 0.868 | -0.052 | 0.055 | 0.055 | 0 | 0.48 | 0.15 | 0.36 | 0.29 | 0.5 | 0.49 |
| 7 | 48127194 | rs7793905 | -0.415 | -1.215 | 0.657 | 0.729 | 0.016 | 0.003 | 0.003 | 0 | 0.39 | 0.48 | 0.48 | 0.48 | 0.45 | 0.35 |
| 7 | 48134719 | rs11768083 | -0.66 | -0.843 | 0.034 | 0.858 | -0.052 | -0.027 | -0.027 | 0.112 | 0.72 | 0.81 | 0.55 | 0.56 | 0.57 | 0.67 |
| 7 | 48137413 | rs7341420 | -0.584 | -0.76 | 0.006 | 0.824 | -0.036 | -0.024 | -0.024 | 0 | 0.71 | 0.8 | 0.57 | 0.54 | 0.62 | 0.69 |
| 7 | 48142194 | rs6463462 | 0.021 | -0.808 | 0.199 | 0.786 | -0.003 | 0.009 | 0.009 | 0.272 | 0.59 | 0.61 | 0.56 | 0.52 | 0.6 | 0.66 |
| 7 | 48153838 | rs10279431 | 0.502 | -0.948 | 0.14 | 0.825 | -0.018 | 0.056 | 0.056 | 0 | 0.7 | 0.64 | 0.63 | 0.56 | 0.75 | 0.54 |
| 7 | 48155507 | rs7784208 | 0.64 | -0.975 | 0.155 | 0.813 | -0.022 | 0.063 | 0.063 | 0 | 0.69 | 0.62 | 0.63 | 0.56 | 0.76 | 0.52 |
| 7 | 48168692 | rs10238679 | 1.759 | -0.804 | -0.543 | 0.197 | -0.092 | 0.119 | 0.119 | 0.011 | 0.23 | 0.03 | 0.18 | 0.03 | 0.32 | 0.13 |
| 7 | 48182649 | rs12531591 | 1.292 | -0.958 | -0.234 | 0.579 | -0.031 | 0.103 | 0.103 | 0 | 0.08 | 0.14 | 0.12 | 0.22 | 0.03 | 0.15 |
| 7 | 48183283 | rs11769253 | 1.652 | -0.9 | -0.022 | 0.404 | 0.006 | -0.044 | -0.044 | 0 | 0.24 | 0.27 | 0.19 | 0.03 | 0.33 | 0.13 |
| 7 | 48185689 | rs7384280 | -1.123 | -1.223 | -0.482 | 0.656 | -0.102 | -0.05 | -0.05 | 0.754 | 0.66 | 0.92 | 0.66 | 0.56 | 0.58 | 0.61 |
| 7 | 48194935 | rs10215134 | -0.356 | -1.19 | -0.335 | 0.611 | -0.007 | -0.008 | -0.008 | 0 | 0.54 | 0.63 | 0.64 | 0.47 | 0.55 | 0.55 |
| 7 | 48196475 | rs4130215 | -0.677 | -1.505 | -0.063 | 0.523 | -0.069 | 0.082 | 0.082 | 0.325 | 0.2 | 0.42 | 0.21 | 0.29 | 0.11 | 0.25 |
| 7 | 48196627 | rs4130214 | 0.04 | -1.221 | -0.248 | 0.45 | 0.026 | 0.031 | 0.031 | 0 | 0.43 | 0.62 | 0.38 | 0.3 | 0.39 | 0.39 |
| 7 | 48237721 | rs28637820 | -0.232 | -1.19 | -0.253 | 0.57 | -0.043 | 0.07 | 0.07 | 0.048 | 0.17 | 0.27 | 0.22 | 0.29 | 0.11 | 0.19 |
| 7 | 48237802 | rs2188 | 1.094 | -0.703 | -1.11 | 0.442 | -0.094 | -0.05 | -0.05 | 0.039 | 0.34 | 0.13 | 0.34 | 0.48 | 0.46 | 0.42 |
| 7 | 48244414 | rs6583476 | 1.564 | -0.375 | -1.394 | 0.576 | -0.138 | -0.011 | -0.011 | 0 | 0.28 | 0.02 | 0.19 | 0.21 | 0.34 | 0.33 |
| 7 | 48257014 | rs1526106 | 0.826 | -0.71 | -1.057 | 0.367 | -0.092 | -0.04 | -0.04 | 0 | 0.37 | 0.13 | 0.35 | 0.47 | 0.47 | 0.45 |
| 7 | 48260869 | rs10252745 | 1.123 | 0.081 | -0.606 | 0.03 | -0.008 | 0.01 | 0.01 | 0 | 0.24 | 0.19 | 0.17 | 0.02 | 0.3 | 0.13 |
| 7 | 48263829 | rs10255965 | 1.119 | 0.091 | -0.607 | 0.034 | -0.008 | 0.01 | 0.01 | 0 | 0.24 | 0.18 | 0.17 | 0.02 | 0.3 | 0.13 |
| 7 | 48263899 | rs17712031 | 0.028 | 0.031 | -0.333 | 0.898 | -0.018 | 0.007 | 0.007 | 0 | 0.05 | 0.1 | 0.05 | 0.09 | 0.03 | 0.04 |
| 7 | 48266195 | rs17712049 | -0.107 | 0.213 | -0.433 | 0.992 | 0 | 0.013 | 0.013 | 0 | 0.88 | 0.9 | 0.75 | 0.81 | 0.89 | 0.79 |
| 7 | 48266601 | rs6957787 | 0.774 | 0.053 | -0.481 | 0.197 | 0 | 0.01 | 0.01 | 2.216 | 0.51 | 0.45 | 0.32 | 0.21 | 0.52 | 0.28 |
| 7 | 48266876 | rs17132152 | 1.733 | -0.074 | -0.734 | -0.103 | 0.009 | 0.006 | 0.006 | 2.103 | 0.08 | 0.03 | 0 | 0 | 0.01 | 0.01 |
| 7 | 48269820 | rs7779575 | 1.094 | -0.166 | -0.45 | -0.518 | -0.001 | -0.042 | -0.042 | 0 | 0.43 | 0.34 | 0.24 | 0.02 | 0.45 | 0.2 |
| 7 | 48272716 | rs1526105 | 0.169 | 0.003 | -0.22 | 0.047 | -0.008 | -0.001 | -0.001 | 0 | 0.67 | 0.63 | 0.51 | 0.24 | 0.57 | 0.43 |
| 7 | 48275852 | rs7777880 | 0.708 | -0.114 | 0.063 | 0.032 | 0.005 | 0.078 | 0.078 | 0 | 0.12 | 0.21 | 0.18 | 0.15 | 0.11 | 0.19 |
| 7 | 48277751 | rs11185610 | 0.817 | 0.211 | -0.365 | 0.493 | -0.077 | -0.083 | -0.083 | 0 | 0.18 | 0.05 | 0.24 | 0.42 | 0.28 | 0.39 |
| 7 | 48281178 | rs10224277 | 1.503 | -0.177 | -0.064 | 0.083 | 0 | 0.1 | 0.1 | 4.717 | 0.46 | 0.31 | 0.32 | 0.09 | 0.45 | 0.23 |
| 7 | 48285664 | rs1405127 | -0.373 | -0.185 | 0.28 | 0.244 | -0.052 | 0.028 | 0.028 | 1.262 | 0.22 | 0.44 | 0.2 | 0.15 | 0.13 | 0.21 |
| 7 | 48292302 | rs6583461 | 0.246 | 0.012 | 0.15 | 0.264 | -0.078 | -0.086 | -0.086 | 0 | 0.28 | 0.16 | 0.44 | 0.62 | 0.41 | 0.5 |
| 7 | 48310167 | rs1405126 | 1.258 | -0.022 | 0.241 | 0.297 | 0.002 | 0.043 | 0.043 | 0 | 0.11 | 0.13 | 0.17 | 0.15 | 0.11 | 0.19 |

| | | | | | | | | | | | | | | | | |
|---|---|---|---|---|---|---|---|---|---|---|---|---|---|---|---|---|
| 7 | 48311563 | rs17712293 | 0.445 | -0.038 | 0.228 | 0.248 | 0.002 | 0.035 | 0.035 | 0 | 0.14 | 0.15 | 0.27 | 0.22 | 0.13 | 0.24 |
| 7 | 48313881 | rs17712299 | 0.898 | -0.169 | 0.32 | 0.17 | -0.011 | 0.019 | 0.019 | 0.01 | 0.07 | 0.14 | 0.05 | 0.15 | 0.05 | 0.06 |
| 7 | 48314330 | rs17132196 | 2.557 | -0.183 | 0.259 | -0.157 | 0.008 | 0.005 | 0.005 | 2.438 | 0.08 | 0.03 | 0 | 0 | 0.01 | 0.01 |
| 7 | 48314798 | rs13233225 | 1.787 | -0.195 | 0.314 | -0.202 | 0.011 | 0.095 | 0.095 | 4.314 | 0.41 | 0.26 | 0.21 | 0.01 | 0.39 | 0.19 |
| 7 | 48316055 | rs17132201 | 2.547 | -0.162 | 0.291 | -0.151 | 0.008 | 0.005 | 0.005 | 2.438 | 0.08 | 0.03 | 0 | 0 | 0.01 | 0.01 |
| 7 | 48318098 | rs17132206 | 2.547 | -0.162 | 0.291 | -0.151 | 0.008 | 0.005 | 0.005 | 2.438 | 0.08 | 0.03 | 0 | 0 | 0.01 | 0.01 |
| 7 | 48318400 | rs17132208 | 2.547 | -0.162 | 0.291 | -0.151 | 0.008 | 0.005 | 0.005 | 2.438 | 0.08 | 0.03 | 0 | 0 | 0.01 | 0.01 |
| 7 | 48318811 | rs2222648 | -2.055 | -0.415 | -0.145 | -0.748 | 0 | -0.042 | -0.042 | 2.373 | 0.76 | 0.83 | 0.84 | 0.99 | 0.72 | 0.88 |
| 7 | 48325832 | rs9692496 | -1.081 | 0.159 | 0.224 | -0.173 | 0 | -0.002 | -0.002 | 0 | 0.46 | 0.6 | 0.51 | 0.77 | 0.48 | 0.56 |
| 7 | 48330287 | rs6970910 | 0.23 | 0.38 | -0.24 | -0.068 | -0.012 | 0.003 | 0.003 | 6.319 | 0.64 | 0.63 | 0.5 | 0.23 | 0.54 | 0.44 |
| 7 | 48336844 | rs6583546 | -1.731 | 0.439 | -0.337 | -0.281 | 0.011 | 0.076 | 0.076 | 3.031 | 0.58 | 0.7 | 0.78 | 0.99 | 0.59 | 0.81 |
| 7 | 48339730 | rs6943725 | -0.757 | 1.752 | -0.534 | 0.429 | -0.018 | 0.01 | 0.01 | 8.543 | 0.36 | 0.33 | 0.59 | 0.79 | 0.5 | 0.6 |
| 7 | 48349647 | rs3931814 | 0.744 | 2.026 | -0.496 | 0.719 | -0.004 | 0.001 | 0.001 | 0.803 | 0.13 | 0.04 | 0.15 | 0.1 | 0.12 | 0.17 |
| 7 | 48350983 | rs4260852 | 0.498 | 1.903 | -0.309 | 0.52 | -0.006 | -0.003 | -0.003 | 0 | 0.11 | 0.11 | 0.19 | 0.17 | 0.16 | 0.18 |
| 7 | 48354820 | rs10248040 | -0.95 | 1.851 | -0.05 | 1.002 | 0.011 | -0.005 | -0.005 | 0.033 | 0.35 | 0.47 | 0.48 | 0.78 | 0.42 | 0.56 |
| 7 | 48360337 | rs6949736 | -0.56 | 1.906 | -0.286 | 1.197 | -0.023 | -0.03 | -0.03 | 0 | 0.29 | 0.25 | 0.37 | 0.64 | 0.37 | 0.5 |
| 7 | 48361784 | rs17712647 | -0.311 | 1.347 | -0.108 | 1.272 | -0.007 | 0.033 | 0.033 | 0.505 | 0.06 | 0.14 | 0.11 | 0.15 | 0.04 | 0.05 |
| 7 | 48364245 | rs4024044 | 0.011 | 1.657 | -0.269 | 1.115 | 0.002 | -0.002 | -0.002 | 0 | 0.84 | 0.88 | 0.88 | 0.89 | 0.8 | 0.85 |
| 7 | 48367798 | rs1568946 | 1.088 | 1.545 | 0.048 | 1.684 | 0.004 | 0.064 | 0.064 | 0 | 0.71 | 0.55 | 0.76 | 0.58 | 0.69 | 0.7 |
| 7 | 48373487 | rs713061 | 0.179 | 1.917 | 0.28 | 2.048 | 0.027 | 0.073 | 0.073 | 1.9 | 0.13 | 0.23 | 0.23 | 0.37 | 0.14 | 0.33 |
| 7 | 48378802 | rs12056015 | 0.915 | 1.95 | 0.185 | 1.763 | 0.006 | 0.06 | 0.06 | 0 | 0.71 | 0.57 | 0.65 | 0.59 | 0.66 | 0.69 |
| 7 | 48385877 | rs4917078 | -0.85 | 1.978 | 0.08 | 1.859 | -0.01 | -0.004 | -0.004 | 0 | 0.88 | 0.95 | 0.82 | 0.9 | 0.83 | 0.86 |
| 7 | 48392514 | rs11772384 | 1.701 | 1.517 | -0.76 | 0.873 | -0.002 | 0.077 | 0.077 | 1.282 | 0.29 | 0.14 | 0.14 | 0 | 0.27 | 0.16 |
| 7 | 48392771 | rs11983883 | -0.439 | 1.408 | -0.58 | 1.068 | 0.012 | 0.088 | 0.088 | 0 | 0.11 | 0.24 | 0.14 | 0.28 | 0.14 | 0.12 |
| 7 | 48403751 | rs17661843 | 1.576 | 1.164 | -0.927 | 0.97 | 0.026 | 0.025 | 0.025 | 0.959 | 0.16 | 0.05 | 0.11 | 0.13 | 0.13 | 0.11 |
| 7 | 48407217 | rs1526093 | -1.393 | 1.462 | -0.497 | 1.18 | 0.013 | 0.124 | 0.124 | 0 | 0.17 | 0.4 | 0.17 | 0.31 | 0.18 | 0.17 |
| 7 | 48420261 | rs17661929 | -0.459 | 1.487 | -0.559 | 1.181 | -0.004 | 0.013 | 0.013 | 0 | 0.09 | 0.14 | 0.13 | 0.25 | 0.14 | 0.12 |
| 7 | 48428181 | rs17132286 | -1.434 | 1.453 | -0.449 | 1.157 | 0.016 | 0.103 | 0.103 | 0 | 0.17 | 0.4 | 0.17 | 0.27 | 0.18 | 0.17 |
| 7 | 48428981 | rs4917118 | 0.759 | 1.766 | -0.53 | 1.174 | 0.016 | 0.064 | 0.064 | 0 | 0.73 | 0.57 | 0.67 | 0.63 | 0.66 | 0.69 |
| 7 | 48429429 | rs717707 | -0.999 | 1.75 | -0.343 | 1.203 | -0.005 | 0.082 | 0.082 | 0 | 0.12 | 0.28 | 0.09 | 0.15 | 0.1 | 0.14 |
| 7 | 48434008 | rs6957855 | 1.395 | 1.789 | -0.683 | 0.702 | 0.003 | 0.072 | 0.072 | 1.379 | 0.27 | 0.13 | 0.14 | 0 | 0.25 | 0.15 |
| 7 | 48436309 | rs1560792 | 1.078 | 2.118 | -0.209 | 1.847 | 0.007 | 0.035 | 0.035 | 0 | 0.72 | 0.6 | 0.67 | 0.62 | 0.66 | 0.71 |
| 7 | 48439043 | rs10278745 | -1.822 | 3.205 | 0.317 | 2.571 | 0.035 | 0.037 | 0.037 | 0 | 0.26 | 0.37 | 0.48 | 0.43 | 0.42 | 0.21 |
| 7 | 48443116 | rs10236551 | -1.097 | 3.334 | 0.34 | 2.676 | -0.006 | 0.001 | 0.001 | 0 | 0.19 | 0.16 | 0.28 | 0.22 | 0.22 | 0.12 |
| 7 | 48444995 | rs11489714 | -2.335 | 3.339 | 0.347 | 2.688 | 0.08 | 0.095 | 0.095 | 0 | 0.14 | 0.32 | 0.27 | 0.26 | 0.26 | 0.13 |
| 7 | 48445174 | rs7780793 | -2.118 | 3.347 | 0.237 | 2.691 | 0.046 | 0.056 | 0.056 | 0 | 0.13 | 0.23 | 0.26 | 0.26 | 0.26 | 0.13 |
| 7 | 48445268 | rs10248739 | 0.459 | 3.364 | 0.099 | 2.625 | -0.029 | -0.022 | -0.022 | 0 | 0.11 | 0.04 | 0.21 | 0.16 | 0.16 | 0.08 |
| 7 | 48448916 | rs10242905 | 0.351 | 3.364 | 0.099 | 2.644 | -0.028 | -0.024 | -0.024 | 0 | 0.12 | 0.04 | 0.21 | 0.16 | 0.16 | 0.08 |
| 7 | 48450157 | rs17548783 | -1.63 | 3.195 | 0.271 | 2.518 | 0.07 | 0.005 | 0.005 | 0 | 0.3 | 0.46 | 0.52 | 0.58 | 0.47 | 0.3 |
| 7 | 48451939 | rs1865195 | -1.565 | 2.711 | 0.039 | 2.433 | 0.07 | 0.108 | 0.108 | 0 | 0.06 | 0.15 | 0.2 | 0.21 | 0.2 | 0.09 |
| 7 | 48452200 | rs7789493 | -1.565 | 2.711 | 0.039 | 2.433 | 0.07 | 0.108 | 0.108 | 0 | 0.06 | 0.15 | 0.2 | 0.21 | 0.2 | 0.09 |
| 7 | 48455084 | rs17662118 | -1.521 | 2.207 | 0.045 | 2.42 | 0.045 | 0.114 | 0.114 | 0 | 0.06 | 0.14 | 0.18 | 0.22 | 0.17 | 0.09 |
| 7 | 48457088 | rs2060330 | 0.521 | 2.055 | -0.195 | 2.125 | 0.005 | 0.014 | 0.014 | 0 | 0.78 | 0.72 | 0.7 | 0.56 | 0.64 | 0.77 |
| 7 | 48457344 | rs6945363 | -1.303 | 1.97 | 0.551 | 2.537 | 0.042 | 0.203 | 0.203 | 0.44 | 0.18 | 0.48 | 0.17 | 0.41 | 0.19 | 0.2 |
| 7 | 48457425 | rs6973815 | -1.588 | 1.858 | 1.07 | 2.875 | 0.018 | 0.154 | 0.154 | 0 | 0.2 | 0.4 | 0.2 | 0.43 | 0.22 | 0.24 |
| 7 | 48457486 | rs6943378 | -1.19 | 1.934 | 1.084 | 2.71 | 0.047 | 0.071 | 0.071 | 0.145 | 0.16 | 0.35 | 0.19 | 0.25 | 0.22 | 0.12 |
| 7 | 48457628 | rs4917133 | 0.923 | 1.365 | 0.688 | 2.159 | -0.007 | -0.007 | -0.007 | 6.874 | 0.54 | 0.37 | 0.57 | 0.56 | 0.5 | 0.65 |
| 7 | 48457808 | rs4917136 | 0.603 | 1.318 | 0.649 | 2.236 | 0 | 0.001 | 0.001 | 0 | 0.66 | 0.7 | 0.73 | 0.7 | 0.64 | 0.64 |
| 7 | 48459516 | rs1433515 | -0.604 | 0.515 | 0.285 | 1.827 | -0.005 | -0.001 | -0.001 | 0 | 0.38 | 0.58 | 0.48 | 0.34 | 0.32 | 0.31 |
| 7 | 48462044 | rs1109443 | 0.025 | 0.611 | 0.47 | 1.924 | 0.05 | 0.103 | 0.103 | 1.013 | 0.06 | 0.24 | 0.16 | 0.12 | 0.08 | 0.05 |
| 7 | 48462563 | rs875812 | -0.08 | 0.532 | 0.278 | 1.896 | 0 | 0.004 | 0.004 | 0 | 0.24 | 0.23 | 0.26 | 0.28 | 0.26 | 0.27 |
| 7 | 48467355 | rs13233421 | -0.149 | 0.517 | 0.255 | 1.896 | 0.001 | 0.007 | 0.007 | 0 | 0.25 | 0.24 | 0.27 | 0.35 | 0.29 | 0.24 |
| 7 | 48479620 | rs12154722 | 0.033 | 0.224 | 0.094 | 1.77 | 0 | -0.019 | -0.019 | 0.658 | 0.12 | 0.08 | 0.19 | 0.16 | 0.11 | 0.15 |
| 7 | 48479836 | rs6959861 | -0.688 | -0.044 | 0.192 | 1.681 | 0.013 | -0.016 | -0.016 | 0 | 0.22 | 0.16 | 0.26 | 0.25 | 0.17 | 0.2 |
| 7 | 48481370 | rs10499681 | -0.072 | 0.156 | 0.316 | 1.715 | 0.001 | 0.028 | 0.028 | 0 | 0.26 | 0.33 | 0.27 | 0.36 | 0.29 | 0.24 |
| 7 | 48499310 | rs4083233 | -0.892 | 0.154 | 0.122 | 1.702 | 0.008 | -0.015 | -0.015 | 0 | 0.38 | 0.34 | 0.36 | 0.44 | 0.37 | 0.33 |
| 7 | 48506566 | rs4917152 | -0.824 | 0.091 | 0.102 | 1.72 | -0.027 | 0.029 | 0.029 | 0 | 0.2 | 0.35 | 0.26 | 0.22 | 0.15 | 0.14 |
| 7 | 48506707 | rs4917154 | -0.824 | 0.091 | 0.102 | 1.72 | -0.027 | 0.029 | 0.029 | 0 | 0.2 | 0.35 | 0.27 | 0.22 | 0.15 | 0.14 |
| 7 | 48507428 | rs7781930 | -1.634 | 0.023 | -0.193 | 1.369 | -0.046 | 0.071 | 0.071 | 0 | 0.56 | 0.84 | 0.61 | 0.65 | 0.53 | 0.58 |
| 7 | 48526625 | rs6958584 | -1.392 | -0.074 | 0.078 | 1.447 | 0.02 | -0.027 | -0.027 | 0 | 0.36 | 0.74 | 0.49 | 0.5 | 0.31 | 0.31 |
| 7 | 48527244 | rs11762388 | -0.378 | -0.117 | 0.412 | 1.604 | -0.006 | 0.009 | 0.009 | 0 | 0.19 | 0.23 | 0.25 | 0.28 | 0.15 | 0.14 |
| 7 | 48528081 | rs17662629 | -0.951 | -0.136 | 0.737 | 1.608 | -0.018 | 0.033 | 0.033 | 0 | 0.23 | 0.32 | 0.27 | 0.36 | 0.18 | 0.2 |
| 7 | 48537543 | rs1918613 | 0.904 | -0.145 | 0.912 | 1.576 | 0.059 | 0.14 | 0.14 | 0 | 0.06 | 0.25 | 0.17 | 0.14 | 0.08 | 0.06 |
| 7 | 48537588 | rs1918612 | -1.273 | 0.126 | 0.263 | 1.433 | -0.036 | 0.009 | 0.009 | 0.13 | 0.87 | 0.96 | 0.87 | 0.92 | 0.82 | 0.93 |
| 7 | 48538770 | rs1997238 | 2.392 | 0.105 | 0.257 | 1.452 | -0.123 | 0.022 | 0.022 | 0 | 0.46 | 0.04 | 0.34 | 0.41 | 0.48 | 0.48 |
| 7 | 48540424 | rs4144066 | 1.859 | 0.099 | 0.221 | 1.435 | -0.046 | -0.022 | -0.022 | 0 | 0.16 | 0.01 | 0.09 | 0.15 | 0.19 | 0.13 |
| 7 | 48545789 | rs12669930 | 1.031 | 0.04 | 0.394 | 1.443 | 0.092 | 0.188 | 0.188 | 0 | 0.06 | 0.29 | 0.18 | 0.14 | 0.08 | 0.06 |
| 7 | 48549633 | rs7776966 | -0.073 | 0.037 | 0.362 | 1.44 | 0.066 | 0.137 | 0.137 | 0 | 0.09 | 0.34 | 0.21 | 0.14 | 0.1 | 0.2 |
| 7 | 48553615 | rs4917021 | -1.145 | 0.02 | 0.322 | 1.442 | -0.036 | 0.032 | 0.032 | 0 | 0.31 | 0.52 | 0.31 | 0.37 | 0.24 | 0.25 |

| | | | | | | | | | | | | | | | | |
|---|---|---|---|---|---|---|---|---|---|---|---|---|---|---|---|---|
| 7 | 48554584 | rs5025818 | 2.233 | 0.09 | 0.123 | 1.367 | -0.101 | 0.037 | 0.037 | 0 | 0.46 | 0.07 | 0.35 | 0.41 | 0.48 | 0.48 |
| 7 | 48557141 | rs17132432 | 1.65 | 0.011 | 0.485 | 1.295 | -0.001 | -0.033 | -0.033 | 0 | 0.07 | 0.12 | 0.04 | 0 | 0.05 | 0.05 |
| 7 | 48561095 | rs4917161 | 0.316 | -0.065 | 0.266 | 1.399 | -0.01 | -0.045 | -0.045 | 0.032 | 0.09 | 0.01 | 0.17 | 0.18 | 0.08 | 0.12 |
| 7 | 48568990 | rs17132444 | 1.578 | -0.02 | 0.423 | 1.038 | 0.001 | -0.04 | -0.04 | 0 | 0.07 | 0.13 | 0.04 | 0 | 0.05 | 0.05 |
| 7 | 48579066 | rs12112361 | 0.623 | -0.032 | 0.624 | 1.136 | 0.066 | 0.174 | 0.174 | 0.011 | 0.07 | 0.28 | 0.18 | 0.15 | 0.08 | 0.07 |
| 7 | 48587910 | rs1534393 | 0.053 | -0.109 | 0.588 | 1.149 | -0.012 | 0.031 | 0.031 | 0 | 0.15 | 0.23 | 0.23 | 0.28 | 0.13 | 0.13 |
| 7 | 48591660 | rs10252100 | -1.787 | -0.092 | 0.501 | 1.087 | 0.037 | -0.081 | -0.081 | 0 | 0.36 | 0.86 | 0.5 | 0.44 | 0.3 | 0.36 |
| 7 | 48596337 | rs10231745 | -1.177 | -0.097 | 0.587 | 1.103 | -0.021 | 0.114 | 0.114 | 0 | 0.25 | 0.5 | 0.43 | 0.43 | 0.22 | 0.2 |
| 7 | 48599764 | rs12113960 | 0.293 | -0.031 | 0.05 | 1.101 | 0.035 | 0.069 | 0.069 | 0 | 0.14 | 0.41 | 0.22 | 0.15 | 0.13 | 0.12 |
| 7 | 48614305 | rs7783364 | -0.163 | 0.067 | -0.479 | 1.052 | 0.012 | -0.012 | -0.012 | 0 | 0.06 | 0.24 | 0.04 | 0 | 0.04 | 0.11 |
| 7 | 48623327 | rs6974375 | 2.201 | 0.22 | -1.196 | 1.091 | -0.066 | 0.046 | 0.046 | 0 | 0.3 | 0.05 | 0.25 | 0.24 | 0.28 | 0.32 |
| 7 | 48626500 | rs7789709 | -0.344 | 0.308 | -1.268 | 1.193 | -0.053 | -0.044 | -0.044 | 0 | 0.68 | 0.93 | 0.77 | 0.68 | 0.63 | 0.68 |
| 7 | 48627470 | rs1527837 | -0.602 | 0.313 | -1.321 | 1.164 | -0.071 | -0.029 | -0.029 | 1.019 | 0.71 | 0.96 | 0.79 | 0.74 | 0.66 | 0.73 |
| 7 | 48628104 | rs2178089 | -1.401 | 0.279 | -1.134 | 1.13 | 0.009 | -0.094 | -0.094 | 0 | 0.38 | 0.91 | 0.53 | 0.37 | 0.35 | 0.36 |
| 7 | 48633056 | rs10238373 | -0.169 | 0.249 | -0.855 | 1.026 | 0.002 | -0.011 | -0.011 | 0 | 0.14 | 0.1 | 0.23 | 0.21 | 0.14 | 0.13 |
| 7 | 48638428 | rs12536486 | 1.654 | 0.376 | -1.208 | 0.66 | -0.028 | 0.041 | 0.041 | 0 | 0.34 | 0.09 | 0.27 | 0.26 | 0.29 | 0.29 |
| 7 | 48652085 | rs4917024 | -0.471 | 0.425 | -1.02 | 0.754 | -0.038 | -0.036 | -0.036 | 0.042 | 0.73 | 0.84 | 0.79 | 0.62 | 0.66 | 0.7 |
| 7 | 48652671 | rs1918604 | -0.971 | 0.28 | -0.447 | 0.717 | -0.001 | 0 | 0 | 0 | 0.37 | 0.59 | 0.51 | 0.34 | 0.36 | 0.4 |
| 7 | 48657541 | rs4917165 | 2.068 | 0.207 | -0.534 | 0.679 | -0.027 | 0.038 | 0.038 | 0 | 0.33 | 0.09 | 0.27 | 0.28 | 0.29 | 0.3 |
| 7 | 48660363 | rs1318024 | -1.064 | 0.203 | -0.303 | 0.683 | -0.001 | 0 | 0 | 0.057 | 0.38 | 0.61 | 0.49 | 0.33 | 0.35 | 0.4 |
| 7 | 48665024 | rs10216298 | 0.137 | 0.118 | -0.167 | 0.768 | 0 | 0.003 | 0.003 | 0 | 0.14 | 0.15 | 0.12 | 0.21 | 0.13 | 0.12 |
| 7 | 48673941 | rs1527842 | -0.435 | 0.15 | -0.255 | 0.748 | -0.018 | -0.015 | -0.015 | 0 | 0.71 | 0.79 | 0.77 | 0.61 | 0.64 | 0.78 |
| 7 | 48678081 | rs1918583 | -0.703 | 0.188 | -0.306 | 0.765 | -0.095 | -0.085 | -0.085 | 0 | 0.73 | 0.95 | 0.78 | 0.61 | 0.64 | 0.78 |
| 7 | 48686032 | rs3923511 | 0.66 | 0.149 | -0.024 | 0.752 | -0.016 | 0.008 | 0.008 | 0 | 0.69 | 0.43 | 0.56 | 0.71 | 0.75 | 0.66 |
| 7 | 48686600 | rs6955686 | -0.051 | 0.321 | -0.139 | 0.654 | -0.006 | -0.007 | -0.007 | 0.164 | 0.33 | 0.32 | 0.25 | 0.43 | 0.4 | 0.26 |
| 7 | 48698562 | rs7807488 | 0.754 | 0.471 | 0.38 | 0.668 | -0.001 | 0.013 | 0.013 | 0 | 0.06 | 0.13 | 0.24 | 0.07 | 0.06 | 0.07 |
| 7 | 48700374 | rs3812250 | 1.094 | 0.559 | 0.046 | 0.51 | 0.025 | 0.004 | 0.004 | 0 | 0.48 | 0.3 | 0.46 | 0.64 | 0.54 | 0.46 |
| 7 | 48709903 | rs12718295 | 0.294 | 0.526 | 0.279 | 0.085 | 0.003 | 0.013 | 0.013 | 0.037 | 0.31 | 0.39 | 0.29 | 0.4 | 0.33 | 0.24 |
| 7 | 48718598 | rs4540359 | 0.655 | 0.463 | -0.197 | -0.05 | -0.001 | 0.002 | 0.002 | 0 | 0.12 | 0.13 | 0.06 | 0.19 | 0.14 | 0.1 |
| 7 | 48722168 | rs7809837 | 1.237 | 0.513 | -0.119 | -0.016 | -0.004 | 0.026 | 0.026 | 0 | 0.43 | 0.23 | 0.38 | 0.45 | 0.44 | 0.41 |
| 7 | 48722484 | rs11771415 | 0.519 | 0.677 | -0.393 | 0.155 | -0.015 | -0.06 | -0.06 | 0.378 | 0.26 | 0.18 | 0.23 | 0.4 | 0.26 | 0.15 |
| 7 | 48724541 | rs9886262 | 0.053 | 0.425 | -0.105 | -0.069 | -0.003 | -0.062 | -0.062 | 0 | 0.85 | 0.81 | 0.93 | 0.99 | 0.87 | 0.78 |
| 7 | 48733874 | rs4360243 | 0.07 | 0.654 | -0.202 | 0.233 | -0.005 | -0.003 | -0.003 | 0 | 0.58 | 0.6 | 0.58 | 0.52 | 0.58 | 0.65 |
| 7 | 48743676 | rs6958356 | 0.059 | 0.65 | -0.185 | 0.231 | -0.006 | -0.003 | -0.003 | 0 | 0.58 | 0.62 | 0.58 | 0.52 | 0.58 | 0.65 |
| 7 | 48744571 | rs4363147 | 0.068 | 0.652 | -0.213 | 0.233 | -0.005 | -0.003 | -0.003 | 0.052 | 0.58 | 0.6 | 0.58 | 0.52 | 0.58 | 0.65 |
| 7 | 48747909 | rs11971128 | -0.211 | 0.652 | -0.528 | 0.16 | 0.003 | -0.001 | -0.001 | 0 | 0.14 | 0.12 | 0.09 | 0.14 | 0.15 | 0.13 |
| 7 | 48760256 | rs10262957 | 1.106 | 0.626 | -0.346 | -0.14 | 0.002 | -0.023 | -0.023 | 0 | 0.07 | 0.13 | 0.06 | 0.01 | 0.07 | 0.09 |
| 7 | 48784835 | rs7799291 | -1.3 | 0.216 | -0.629 | -0.268 | -0.006 | -0.007 | -0.007 | 1.315 | 0.68 | 0.9 | 0.63 | 0.64 | 0.68 | 0.64 |
| 7 | 48787644 | rs2082825 | -1.132 | 0.16 | -0.165 | -0.293 | 0.012 | 0.011 | 0.011 | 0 | 0.66 | 0.77 | 0.63 | 0.64 | 0.68 | 0.64 |
| 7 | 48789559 | rs4917171 | -0.584 | 0.13 | -0.133 | -0.3 | 0.007 | 0.017 | 0.017 | 0 | 0.6 | 0.63 | 0.57 | 0.63 | 0.61 | 0.55 |
| 7 | 48801250 | rs1865569 | 0.298 | 0.389 | 0.028 | -0.519 | 0 | -0.014 | -0.014 | 0 | 0.16 | 0.14 | 0.13 | 0.07 | 0.16 | 0.11 |
| 7 | 48809570 | rs17132613 | -1.774 | 0.343 | -0.174 | -0.562 | 0.003 | -0.008 | -0.008 | 6.809 | 0.72 | 0.9 | 0.76 | 0.68 | 0.74 | 0.75 |
| 7 | 48809644 | rs17330072 | 0.153 | 0.272 | -0.089 | -0.645 | 0.001 | -0.057 | -0.057 | 0.007 | 0.2 | 0.19 | 0.13 | 0.01 | 0.18 | 0.15 |
| 7 | 48810202 | rs12674192 | 0.319 | 0.284 | -0.021 | -0.571 | 0.007 | -0.004 | -0.004 | 0 | 0.06 | 0.19 | 0.15 | 0.03 | 0.06 | 0.11 |
| 7 | 48815027 | rs1529867 | 0.482 | 0.22 | 0.344 | -0.578 | -0.006 | 0.008 | 0.008 | 0 | 0.42 | 0.35 | 0.43 | 0.57 | 0.46 | 0.44 |
| 7 | 48817965 | rs11767191 | -0.07 | 0.237 | 0.229 | -0.58 | 0 | 0.022 | 0.022 | 0 | 0.48 | 0.63 | 0.59 | 0.61 | 0.52 | 0.55 |
| 7 | 48840395 | rs6979016 | 0.78 | 0.283 | -0.164 | -0.337 | -0.007 | -0.017 | -0.017 | 0 | 0.35 | 0.29 | 0.38 | 0.44 | 0.38 | 0.37 |
| 7 | 48850787 | rs4557646 | 0.579 | 0.532 | -0.697 | -0.275 | 0 | -0.006 | -0.006 | 0 | 0.05 | 0.1 | 0.14 | 0.03 | 0.07 | 0.11 |
| 7 | 48850987 | rs2363131 | 1.733 | 0.589 | -1.012 | -0.29 | -0.065 | -0.084 | -0.084 | 0.096 | 0.26 | 0.01 | 0.23 | 0.4 | 0.27 | 0.25 |
| 7 | 48852331 | rs6583466 | -0.874 | 0.573 | -0.494 | -0.416 | 0.003 | -0.055 | -0.055 | 2.422 | 0.32 | 0.42 | 0.32 | 0.1 | 0.33 | 0.34 |
| 7 | 48853719 | rs2190013 | 1.36 | 0.51 | -1.249 | -0.192 | -0.058 | -0.085 | -0.085 | 0 | 0.23 | 0.01 | 0.15 | 0.35 | 0.21 | 0.24 |
| 7 | 48855974 | rs17331155 | -0.416 | 0.094 | -0.757 | -0.774 | -0.002 | -0.022 | -0.022 | 0.857 | 0.2 | 0.2 | 0.15 | 0.08 | 0.18 | 0.19 |
| 7 | 48857113 | rs7786596 | -0.168 | 0.695 | -0.502 | -0.918 | 0.023 | -0.014 | -0.014 | 0 | 0.92 | 0.87 | 0.8 | 0.97 | 0.88 | 0.85 |
| 7 | 48857278 | rs17390477 | -0.047 | 0.712 | -0.638 | -0.915 | 0.002 | 0.014 | 0.014 | 0 | 0.19 | 0.14 | 0.15 | 0.08 | 0.18 | 0.19 |
| 7 | 48859514 | rs2158642 | -0.871 | 0.793 | -0.599 | -0.866 | 0.001 | 0 | 0 | 0 | 0.95 | 0.94 | 0.81 | 0.99 | 0.89 | 0.89 |
| 7 | 48866316 | rs12531263 | 1.315 | 0.559 | -0.94 | -0.759 | -0.038 | -0.014 | -0.014 | 0 | 0.38 | 0.29 | 0.44 | 0.5 | 0.49 | 0.37 |
| 7 | 48872127 | rs17132679 | 0.624 | 0.441 | -0.932 | -0.682 | -0.003 | -0.063 | -0.063 | 0 | 0.15 | 0.2 | 0.08 | 0 | 0.11 | 0.08 |
| 7 | 48872760 | rs10272096 | 0.149 | 0.956 | -0.861 | -0.574 | 0.011 | -0.008 | -0.008 | 0 | 0.42 | 0.31 | 0.2 | 0.37 | 0.33 | 0.22 |
| 7 | 48878120 | rs4917028 | 0.201 | 0.768 | -1.011 | -0.575 | 0.042 | -0.016 | -0.016 | 0 | 0.42 | 0.26 | 0.17 | 0.37 | 0.29 | 0.21 |
| 7 | 48895330 | rs2190012 | 0.281 | 0.797 | -1.484 | -0.643 | -0.003 | 0.091 | 0.091 | 0 | 0.39 | 0.1 | 0.29 | 0.25 | 0.37 | 0.3 |
| 7 | 48906680 | rs4917030 | 0.654 | 1.085 | -0.619 | -0.886 | -0.003 | -0.01 | -0.01 | 0 | 0.33 | 0.39 | 0.26 | 0.4 | 0.35 | 0.26 |
| 7 | 48907718 | rs10232860 | 1.803 | 1.084 | -0.711 | -0.92 | 0.008 | -0.026 | -0.026 | 0 | 0.2 | 0.17 | 0.12 | 0.4 | 0.16 | 0.12 |
| 7 | 48908477 | rs17333102 | 1.004 | 0.974 | -0.928 | -0.922 | 0.008 | 0.018 | 0.018 | 0 | 0.23 | 0.07 | 0.18 | 0.2 | 0.2 | 0.26 |
| 7 | 48914927 | rs12668449 | 2.301 | 0.803 | -1.125 | -0.998 | -0.039 | -0.082 | -0.082 | 0 | 0.17 | 0.01 | 0.19 | 0.3 | 0.19 | 0.3 |
| 7 | 48915248 | rs17132757 | 1.286 | 0.782 | -0.71 | -1.031 | -0.006 | 0.005 | 0.005 | 0 | 0.07 | 0.13 | 0.15 | 0.08 | 0.04 | 0.09 |
| 7 | 48917108 | rs10261356 | 2.268 | 0.786 | -0.52 | -1.028 | -0.01 | -0.046 | -0.046 | 0 | 0.18 | 0.09 | 0.19 | 0.29 | 0.19 | 0.3 |
| 7 | 48918669 | rs6946623 | 0.345 | 0.792 | -0.187 | -1.076 | -0.01 | -0.063 | -0.063 | 0 | 0.23 | 0.32 | 0.25 | 0.09 | 0.2 | 0.11 |
| 7 | 48920484 | rs2158641 | 0.289 | 0.816 | -0.194 | -1.048 | -0.003 | -0.005 | -0.005 | 0 | 0.41 | 0.41 | 0.44 | 0.38 | 0.39 | 0.41 |
| 7 | 48925603 | rs17092976 | 1.38 | 0.756 | -0.188 | -0.901 | -0.005 | 0.005 | 0.005 | 0 | 0.07 | 0.11 | 0.15 | 0.08 | 0.04 | 0.09 |

| | | | | | | | | | | | | | | | | |
|---|---|---|---|---|---|---|---|---|---|---|---|---|---|---|---|---|
| 7 | 48926725 | rs7809618 | -0.946 | 0.814 | -0.165 | -0.91 | 0 | -0.005 | -0.005 | 0 | 0.74 | 0.77 | 0.71 | 0.77 | 0.74 | 0.67 |
| 7 | 48927065 | rs17132771 | -0.055 | 0.873 | 0.058 | -0.913 | -0.001 | 0 | 0 | 0 | 0.48 | 0.52 | 0.35 | 0.39 | 0.51 | 0.28 |
| 7 | 48928297 | rs885685 | -0.159 | 0.973 | 0.107 | -0.658 | -0.001 | -0.003 | -0.003 | 0 | 0.43 | 0.49 | 0.44 | 0.47 | 0.41 | 0.36 |
| 7 | 48930260 | rs2190009 | 0.058 | 0.893 | 0.105 | -0.984 | -0.003 | -0.008 | -0.008 | 0.535 | 0.46 | 0.48 | 0.45 | 0.37 | 0.4 | 0.46 |
| 7 | 48930555 | rs12539546 | 1.22 | 0.744 | 0.138 | -1.027 | 0.009 | 0.013 | 0.013 | 0 | 0.22 | 0.19 | 0.1 | 0.13 | 0.16 | 0.18 |
| 7 | 48941456 | rs10266289 | 0.042 | 0.806 | 0.305 | -1.029 | 0.001 | 0.002 | 0.002 | 0 | 0.52 | 0.51 | 0.55 | 0.66 | 0.59 | 0.54 |
| 7 | 48941748 | rs10228895 | 0.153 | 0.823 | 0.579 | -1.046 | 0.011 | 0.035 | 0.035 | 0 | 0.51 | 0.39 | 0.53 | 0.65 | 0.57 | 0.49 |
| 7 | 48944233 | rs12531347 | 1.302 | 0.853 | 0.488 | -1.039 | 0.013 | 0.019 | 0.019 | 0 | 0.24 | 0.19 | 0.1 | 0.13 | 0.17 | 0.19 |
| 7 | 48964255 | rs6968293 | 0.323 | 0.844 | 0.348 | -1.415 | 0.012 | -0.017 | -0.017 | 3.872 | 0.4 | 0.48 | 0.5 | 0.69 | 0.48 | 0.53 |
| 7 | 48973578 | rs2215361 | 1.449 | 0.955 | 0.366 | -1.553 | 0.001 | -0.011 | -0.011 | 0 | 0.25 | 0.19 | 0.23 | 0.68 | 0.24 | 0.34 |
| 7 | 48976637 | rs7803161 | 1.555 | 0.893 | 0.373 | -1.483 | 0.013 | 0.024 | 0.024 | 0 | 0.23 | 0.18 | 0.09 | 0.1 | 0.15 | 0.19 |
| 7 | 48978276 | rs6953545 | -0.391 | 0.917 | 0.242 | -1.468 | 0.001 | 0.014 | 0.014 | 0 | 0.64 | 0.67 | 0.59 | 0.78 | 0.63 | 0.72 |
| 7 | 48981329 | rs7802229 | -0.382 | 0.895 | 0.24 | -1.47 | 0 | 0.014 | 0.014 | 0 | 0.64 | 0.67 | 0.6 | 0.78 | 0.62 | 0.71 |
| 7 | 48983994 | rs6942473 | -0.26 | 1.051 | 0.226 | -1.484 | -0.004 | 0.019 | 0.019 | 0 | 0.63 | 0.66 | 0.57 | 0.78 | 0.57 | 0.71 |
| 7 | 48984046 | rs6958208 | -1.258 | 0.676 | 0.32 | -1.422 | 0.002 | -0.006 | -0.006 | 0.061 | 0.76 | 0.73 | 0.66 | 0.79 | 0.75 | 0.74 |
| 7 | 48985673 | rs7779027 | 0.413 | 0.667 | 0.316 | -1.433 | -0.015 | 0.015 | 0.015 | 0 | 0.38 | 0.31 | 0.48 | 0.68 | 0.45 | 0.48 |
| 7 | 48987444 | rs6583481 | -1.355 | 0.737 | 0.21 | -1.33 | -0.004 | 0.006 | 0.006 | 0 | 0.77 | 0.83 | 0.68 | 0.79 | 0.76 | 0.74 |
| 7 | 48993937 | rs993871 | -1.419 | 0.934 | 0.834 | -0.961 | 0.003 | 0.043 | 0.043 | 0 | 0.6 | 0.74 | 0.59 | 0.74 | 0.64 | 0.64 |
| 7 | 48999108 | rs17132823 | 0.213 | 0.74 | 0.988 | -0.946 | -0.011 | -0.003 | -0.003 | 0.042 | 0.1 | 0.26 | 0.04 | 0.07 | 0.06 | 0.15 |
| 7 | 49001350 | rs6977535 | -0.78 | 0.791 | 1.043 | -0.945 | 0 | -0.01 | -0.01 | 0.052 | 0.47 | 0.54 | 0.51 | 0.74 | 0.51 | 0.62 |
| 7 | 49002811 | rs10282410 | 0.169 | 0.846 | 0.81 | -0.991 | -0.04 | 0 | 0 | 0 | 0.35 | 0.25 | 0.5 | 0.66 | 0.43 | 0.43 |
| 7 | 49009677 | rs957814 | -2.242 | 0.8 | 0.742 | -1.056 | -0.004 | 0.096 | 0.096 | 0.423 | 0.71 | 0.9 | 0.83 | 0.97 | 0.75 | 0.81 |
| 7 | 49010701 | rs2191297 | -1.371 | 0.869 | 0.694 | -0.997 | 0.003 | 0.03 | 0.03 | 0 | 0.62 | 0.76 | 0.63 | 0.73 | 0.64 | 0.64 |
| 7 | 49014630 | rs7780870 | -1.202 | 0.91 | 0.793 | -1.045 | 0 | -0.016 | -0.016 | 0 | 0.6 | 0.64 | 0.61 | 0.74 | 0.64 | 0.58 |
| 7 | 49016312 | rs12534189 | 1.939 | 0.88 | 0.448 | -0.971 | -0.033 | -0.095 | -0.095 | 0 | 0.16 | 0.01 | 0.18 | 0.39 | 0.16 | 0.26 |
| 7 | 49017001 | rs2364008 | -1.01 | 0.69 | 0.864 | -2.079 | 0.001 | 0.039 | 0.039 | 0 | 0.6 | 0.75 | 0.63 | 0.73 | 0.64 | 0.64 |
| 7 | 49026535 | rs10229530 | 1.58 | 0.863 | 0.629 | -1.951 | -0.007 | -0.019 | -0.019 | 0 | 0.08 | 0.05 | 0.18 | 0.23 | 0.11 | 0.16 |
| 7 | 49040143 | rs6583489 | 0.422 | 0.875 | 1.371 | -1.6 | 0.004 | -0.003 | -0.003 | 0 | 0.61 | 0.54 | 0.49 | 0.33 | 0.53 | 0.51 |
| 7 | 49041510 | rs10227381 | 0.129 | 0.991 | 1.36 | -1.55 | -0.036 | 0.008 | 0.008 | 0 | 0.64 | 0.76 | 0.52 | 0.33 | 0.54 | 0.52 |
| 7 | 49050763 | rs4256549 | 0.141 | 1.019 | 1.349 | -1.541 | -0.037 | 0.008 | 0.008 | 0 | 0.64 | 0.77 | 0.52 | 0.33 | 0.54 | 0.5 |
| 7 | 49060792 | rs10258925 | -0.906 | 1.031 | 0.97 | -1.507 | -0.002 | -0.022 | -0.022 | 0 | 0.77 | 0.82 | 0.75 | 0.34 | 0.76 | 0.71 |
| 7 | 49063613 | rs7791409 | 0.773 | 1.032 | 0.95 | -1.557 | -0.001 | -0.003 | -0.003 | 0 | 0.52 | 0.48 | 0.44 | 0.33 | 0.43 | 0.53 |
| 7 | 49069994 | rs7781172 | 0.314 | 0.826 | 0.042 | -1.505 | -0.03 | -0.026 | -0.026 | 1.142 | 0.11 | 0.27 | 0.06 | 0.05 | 0.05 | 0.17 |
| 7 | 49071509 | rs7804817 | 1.424 | 1.235 | -1.006 | -1.294 | 0.008 | 0.04 | 0.04 | 0 | 0.37 | 0.22 | 0.36 | 0.29 | 0.34 | 0.34 |
| 7 | 49103406 | rs4244231 | 1.038 | 1.749 | -0.938 | -0.534 | -0.039 | 0.01 | 0.01 | 0 | 0.64 | 0.74 | 0.52 | 0.34 | 0.54 | 0.54 |
| 7 | 49116140 | rs12718322 | -0.143 | 1.552 | -1.358 | -0.482 | -0.062 | 0.069 | 0.069 | 0.666 | 0.12 | 0.01 | 0.22 | 0 | 0.2 | 0.16 |
| 7 | 49127859 | rs7796330 | 0.455 | 0.93 | -0.661 | 0.109 | -0.011 | -0.037 | -0.037 | 0 | 0.77 | 0.85 | 0.76 | 0.38 | 0.75 | 0.75 |
| 7 | 49145008 | rs7795076 | -0.947 | 0.592 | 0.121 | -0.527 | 0.009 | 0.006 | 0.006 | 0 | 0.17 | 0.36 | 0.32 | 0.15 | 0.18 | 0.23 |
| 7 | 49145538 | rs6977474 | -0.523 | 0.704 | 0.189 | -1.362 | 0 | -0.025 | -0.025 | 0 | 0.24 | 0.22 | 0.17 | 0.03 | 0.25 | 0.26 |
| 7 | 49158700 | rs11185630 | 1.833 | -0.238 | 1.141 | -1.228 | -0.015 | -0.045 | -0.045 | 0.124 | 0.7 | 0.4 | 0.8 | 0.91 | 0.75 | 0.85 |
| 7 | 49161069 | rs4278110 | -0.318 | -0.217 | 1.306 | -1.219 | -0.031 | -0.047 | -0.047 | 1.536 | 0.09 | 0.31 | 0.03 | 0 | 0.01 | 0 |
| 7 | 49170764 | rs4917199 | 1.883 | -0.275 | 1.36 | -1.278 | 0.003 | -0.004 | -0.004 | 1.028 | 0.69 | 0.33 | 0.79 | 0.91 | 0.74 | 0.85 |
| 7 | 49177628 | rs4397348 | -0.029 | -0.207 | 1.266 | -1.226 | -0.026 | -0.041 | -0.041 | 0.042 | 0.08 | 0.24 | 0.03 | 0 | 0.01 | 0 |
| 7 | 49200835 | rs4323425 | -0.19 | -0.171 | 1.126 | -1.191 | -0.017 | -0.048 | -0.048 | 0 | 0.91 | 0.83 | 0.96 | 1 | 0.92 | 0.98 |
| 7 | 49203447 | rs12155275 | -0.461 | -0.002 | 0.651 | -1.113 | 0.004 | -0.005 | -0.005 | 0.212 | 0.13 | 0.16 | 0.13 | 0.09 | 0.17 | 0.13 |
| 7 | 49208805 | rs6980397 | 0.76 | -0.346 | 0.814 | -1.049 | -0.066 | -0.14 | -0.14 | 0 | 0.81 | 0.6 | 0.8 | 1 | 0.89 | 0.95 |
| 7 | 49227996 | rs6962014 | -0.501 | -0.023 | 0.273 | -1.364 | -0.016 | 0.041 | 0.041 | 0 | 0.19 | 0.04 | 0.2 | 0.14 | 0.22 | 0.2 |
| 7 | 49231841 | rs4610694 | -0.267 | 0.822 | 0.694 | -1.621 | 0.016 | -0.029 | -0.029 | 0 | 0.11 | 0.16 | 0.25 | 0.02 | 0.2 | 0.11 |
| 7 | 49234401 | rs4917209 | 0.918 | 0.78 | 0.624 | -1.638 | 0.051 | -0.087 | -0.087 | 0 | 0.83 | 0.66 | 0.69 | 0.97 | 0.73 | 0.77 |
| 7 | 49236711 | rs7792833 | 1.161 | 0.704 | 0.992 | -1.732 | 0.043 | -0.115 | -0.115 | 0 | 0.8 | 0.54 | 0.68 | 0.97 | 0.71 | 0.72 |
| 7 | 49245689 | rs7800355 | 1.131 | 0.677 | 0.915 | -1.745 | 0.03 | -0.105 | -0.105 | 1.098 | 0.8 | 0.51 | 0.67 | 0.97 | 0.72 | 0.71 |
| 7 | 49282513 | rs4566981 | 0.687 | 0.676 | 0.32 | -1.683 | 0.042 | -0.119 | -0.119 | 0 | 0.82 | 0.55 | 0.72 | 0.98 | 0.77 | 0.78 |
| 7 | 49284123 | rs6958192 | 0.671 | 0.682 | 0.323 | -1.68 | 0.043 | -0.118 | -0.118 | 0 | 0.82 | 0.55 | 0.72 | 0.98 | 0.77 | 0.78 |
| 7 | 49284679 | rs12718160 | -0.616 | 0.33 | -0.712 | -1.24 | -0.008 | 0.076 | 0.076 | 0 | 0.31 | 0.11 | 0.3 | 0.18 | 0.37 | 0.36 |
| 7 | 49286618 | rs992880 | 0.688 | 0.793 | -0.198 | -0.098 | 0.013 | 0.14 | 0.14 | 0 | 0.77 | 0.5 | 0.72 | 0.53 | 0.78 | 0.73 |
| 7 | 49286949 | rs17309873 | 0.472 | 0.462 | -1.152 | -0.256 | 0.001 | 0.019 | 0.019 | 0 | 0.18 | 0.04 | 0.15 | 0.17 | 0.17 | 0.16 |
| 7 | 49287008 | rs2195666 | 0.746 | 0.327 | -1.495 | -0.945 | -0.001 | -0.003 | -0.003 | 0.438 | 0.43 | 0.6 | 0.48 | 0.16 | 0.41 | 0.37 |
| 7 | 49295013 | rs12718161 | 0.417 | 0.678 | -1.59 | -0.742 | -0.027 | 0.028 | 0.028 | 0 | 0.24 | 0.07 | 0.19 | 0.23 | 0.24 | 0.24 |
| 7 | 49295246 | rs714741 | -0.162 | 0.734 | -1.296 | 0.079 | -0.007 | -0.003 | -0.003 | 0 | 0.31 | 0.21 | 0.24 | 0.37 | 0.32 | 0.34 |
| 7 | 49300291 | rs6949245 | 1.505 | 1.087 | -1.535 | -0.048 | -0.002 | -0.019 | -0.019 | 0 | 0.39 | 0.63 | 0.37 | 0.15 | 0.3 | 0.33 |
| 7 | 49308651 | rs6955167 | 0.613 | 1.077 | -1.615 | -0.013 | -0.027 | 0.038 | 0.038 | 0 | 0.24 | 0.07 | 0.19 | 0.21 | 0.24 | 0.24 |
| 7 | 49312703 | rs12718168 | -0.398 | 1.02 | -1.358 | 0.2 | -0.049 | -0.128 | -0.128 | 1.697 | 0.29 | 0.08 | 0.38 | 0.48 | 0.37 | 0.33 |
| 7 | 49313567 | rs10251159 | -0.356 | 0.993 | -1.292 | 0.784 | -0.044 | -0.088 | -0.088 | 0 | 0.28 | 0.09 | 0.37 | 0.4 | 0.36 | 0.33 |
| 7 | 49315026 | rs1155875 | -1.17 | 0.766 | -0.877 | 0.465 | -0.002 | -0.007 | -0.007 | 0.447 | 0.42 | 0.38 | 0.39 | 0.66 | 0.41 | 0.52 |
| 7 | 49317234 | rs6961065 | 0.675 | 0.733 | -0.858 | 0.791 | 0 | -0.003 | -0.003 | 3.246 | 0.07 | 0.07 | 0.03 | 0.04 | 0.06 | 0.11 |
| 7 | 49320983 | rs9649707 | 0.211 | 0.75 | -0.909 | 0.82 | -0.026 | -0.133 | -0.133 | 0 | 0.24 | 0.06 | 0.3 | 0.45 | 0.27 | 0.29 |
| 7 | 49334542 | rs10235620 | -0.83 | 0.837 | -0.872 | 0.772 | -0.006 | 0.016 | 0.016 | 0 | 0.35 | 0.36 | 0.35 | 0.49 | 0.33 | 0.4 |
| 7 | 49338202 | rs718555 | -0.499 | 1.048 | -1.011 | 0.844 | -0.014 | -0.01 | -0.01 | 0 | 0.31 | 0.14 | 0.31 | 0.59 | 0.36 | 0.4 |
| 7 | 49339156 | rs10223898 | -0.078 | 1.091 | -0.597 | 0.817 | 0.01 | -0.014 | -0.014 | 0 | 0.08 | 0.12 | 0.09 | 0.06 | 0.1 | 0.09 |

| | | | | | | | | | | | | | | | | |
|---|---|---|---|---|---|---|---|---|---|---|---|---|---|---|---|---|
| 7 | 49348892 | rs12718172 | 1.471 | 0.944 | -0.076 | 0.831 | 0.027 | 0.031 | 0.031 | 0.133 | 0.81 | 0.52 | 0.85 | 0.76 | 0.81 | 0.8 |
| 7 | 49359804 | rs10253384 | -1.487 | 0.779 | -0.333 | 1.092 | -0.002 | -0.011 | -0.011 | 0 | 0.27 | 0.2 | 0.2 | 0.34 | 0.31 | 0.31 |
| 7 | 49361783 | rs11562081 | -0.29 | 0.803 | -0.464 | 1.066 | -0.027 | -0.017 | -0.017 | 0 | 0.17 | 0.06 | 0.15 | 0.17 | 0.21 | 0.2 |
| 7 | 49365151 | rs10277837 | -0.754 | 0.797 | -0.451 | 1.063 | -0.011 | -0.001 | -0.001 | 0.438 | 0.2 | 0.07 | 0.15 | 0.17 | 0.22 | 0.2 |
| 7 | 49366759 | rs2116020 | -0.559 | 0.826 | -0.863 | 0.908 | -0.08 | -0.049 | -0.049 | 0.203 | 0.3 | 0.07 | 0.4 | 0.38 | 0.39 | 0.34 |
| 7 | 49371689 | rs11769954 | 1.443 | 0.949 | -0.885 | 0.931 | -0.056 | -0.019 | -0.019 | 0.102 | 0.15 | 0.02 | 0.27 | 0.2 | 0.19 | 0.14 |
| 7 | 49376156 | rs13235335 | -0.329 | 0.839 | -0.792 | 0.886 | -0.094 | -0.049 | -0.049 | 0 | 0.3 | 0.07 | 0.42 | 0.38 | 0.4 | 0.33 |
| 7 | 49403143 | rs12666784 | 1.093 | 0.869 | -0.36 | 0.988 | 0.001 | 0.018 | 0.018 | 0 | 0.06 | 0.06 | 0.04 | 0.17 | 0.08 | 0.11 |
| 7 | 49404098 | rs6968712 | 0.985 | 0.806 | -0.28 | 0.916 | -0.001 | -0.002 | -0.002 | 0.423 | 0.59 | 0.45 | 0.5 | 0.45 | 0.5 | 0.56 |
| 7 | 49407335 | rs10257060 | -0.576 | 1.107 | -1.08 | 0.686 | -0.087 | -0.051 | -0.051 | 0.246 | 0.33 | 0.05 | 0.36 | 0.38 | 0.4 | 0.38 |
| 7 | 49408840 | rs10242803 | -0.584 | 1.079 | -0.923 | 0.695 | -0.042 | -0.002 | -0.002 | 0 | 0.36 | 0.13 | 0.39 | 0.39 | 0.43 | 0.39 |
| 7 | 49412633 | rs7803133 | 1.597 | 0.599 | -1.895 | 0.094 | -0.029 | 0.043 | 0.043 | 3.202 | 0.28 | 0.07 | 0.12 | 0.13 | 0.23 | 0.19 |
| 7 | 49414950 | rs10485944 | -0.061 | 0.691 | -1.828 | 0.044 | -0.106 | -0.046 | -0.046 | 0 | 0.35 | 0.05 | 0.39 | 0.39 | 0.44 | 0.39 |
| 7 | 49416593 | rs6954263 | 1.252 | 0.432 | -1.901 | -0.557 | 0.006 | 0.059 | 0.059 | 3.499 | 0.26 | 0.14 | 0.31 | 0.02 | 0.24 | 0.09 |
| 7 | 49422750 | rs10250376 | -0.864 | 0.336 | -1.56 | -0.47 | 0.005 | -0.028 | -0.028 | 0 | 0.94 | 0.85 | 0.76 | 1 | 0.92 | 0.95 |
| 7 | 49425806 | rs10278222 | 1.812 | 0.402 | -1.643 | -0.645 | 0.001 | -0.013 | -0.013 | 0.711 | 0.07 | 0.1 | 0.06 | 0 | 0.08 | 0.04 |
| 7 | 49426506 | rs1988494 | 0.342 | 0.046 | -2.011 | -0.512 | -0.038 | -0.047 | -0.047 | 0 | 0.4 | 0.11 | 0.38 | 0.43 | 0.37 | 0.59 |
| 7 | 49426729 | rs1988495 | 1.519 | 0.105 | -2.323 | -0.554 | -0.056 | -0.008 | -0.008 | 2.598 | 0.19 | 0.02 | 0.2 | 0.19 | 0.2 | 0.17 |
| 7 | 49438926 | rs13246472 | 1.223 | 0.029 | -1.087 | -0.568 | -0.008 | 0.003 | 0.003 | 0 | 0.43 | 0.45 | 0.3 | 0.56 | 0.47 | 0.32 |
| 7 | 49449367 | rs7782678 | 0.407 | 0.045 | -1.041 | -0.398 | 0.003 | 0.003 | 0.003 | 0 | 0.54 | 0.64 | 0.36 | 0.58 | 0.57 | 0.51 |
| 7 | 49454996 | rs13221062 | 1.685 | 0.739 | -1.031 | 0.351 | -0.006 | -0.006 | -0.006 | 0 | 0.17 | 0.15 | 0.16 | 0.26 | 0.24 | 0.18 |
| 7 | 49460153 | rs2023965 | 0.312 | 0.277 | -1.032 | 0.374 | 0.002 | -0.022 | -0.022 | 0 | 0.24 | 0.13 | 0.26 | 0.27 | 0.22 | 0.22 |
| 7 | 49461592 | rs10263065 | 0.841 | 0.088 | -0.885 | 0.138 | -0.009 | -0.037 | -0.037 | 0 | 0.31 | 0.47 | 0.17 | 0.23 | 0.29 | 0.2 |
| 7 | 49462209 | rs2365053 | 0.974 | 0.073 | -0.726 | 0.14 | -0.002 | -0.007 | -0.007 | 0.085 | 0.3 | 0.33 | 0.15 | 0.24 | 0.29 | 0.2 |
| 7 | 49465995 | rs1158257 | -0.021 | 0.272 | -0.809 | 0.275 | -0.005 | -0.002 | -0.002 | 0 | 0.55 | 0.58 | 0.44 | 0.52 | 0.51 | 0.42 |
| 7 | 49469003 | rs740083 | 1.2 | 0.379 | -1.851 | 0.234 | 0.011 | -0.002 | -0.002 | 0 | 0.21 | 0.26 | 0.12 | 0.22 | 0.25 | 0.16 |
| 7 | 49480294 | rs10236064 | 0.084 | 0.117 | -2.244 | 0.134 | 0.012 | -0.009 | -0.009 | 0.541 | 0.33 | 0.2 | 0.31 | 0.3 | 0.31 | 0.31 |
| 7 | 49481903 | rs7778595 | 1.436 | 0.223 | -2.632 | -0.436 | 0.001 | 0.014 | 0.014 | 1.416 | 0.11 | 0.07 | 0.06 | 0 | 0.08 | 0.05 |
| 7 | 49486765 | rs2253433 | -0.174 | 0.446 | -2.893 | -0.038 | -0.076 | 0.102 | 0.102 | 0.549 | 0.45 | 0.86 | 0.49 | 0.27 | 0.47 | 0.47 |
| 7 | 49499488 | rs6965339 | -0.15 | 0.568 | -2.536 | -0.007 | 0.006 | 0.007 | 0.007 | 0 | 0.08 | 0.09 | 0.04 | 0.01 | 0.07 | 0.04 |
| 7 | 49506248 | rs4917060 | -0.267 | 0.61 | -2.748 | 0.095 | -0.068 | 0.1 | 0.1 | 0.336 | 0.49 | 0.86 | 0.51 | 0.3 | 0.51 | 0.48 |
| 7 | 49508231 | rs6961632 | -1.433 | 0.336 | -2.779 | 0.354 | 0.032 | -0.089 | -0.089 | 0.768 | 0.85 | 1 | 0.87 | 0.77 | 0.92 | 0.75 |
| 7 | 49509028 | rs10228342 | -0.13 | 0.598 | -2.793 | 0.281 | 0.005 | -0.036 | -0.036 | 0 | 0.65 | 0.8 | 0.53 | 0.41 | 0.67 | 0.54 |
| 7 | 49518688 | rs2886259 | 1.573 | 0.754 | -2.828 | 0.337 | -0.038 | -0.078 | -0.078 | 0 | 0.15 | 0.01 | 0.27 | 0.26 | 0.14 | 0.15 |
| 7 | 49525399 | rs12536558 | 1.094 | 0.83 | -2.883 | 0.173 | -0.062 | 0.04 | 0.04 | 0.585 | 0.24 | 0.02 | 0.11 | 0.12 | 0.25 | 0.11 |
| 7 | 49536308 | rs3904031 | 0.806 | 0.818 | -2.441 | 0.167 | 0.007 | -0.015 | -0.015 | 0 | 0.38 | 0.49 | 0.4 | 0.29 | 0.42 | 0.41 |
| 7 | 49542874 | rs17394967 | 1.664 | 0.813 | -2.892 | 0.198 | -0.035 | 0.035 | 0.035 | 0.714 | 0.21 | 0.02 | 0.08 | 0.09 | 0.19 | 0.1 |
| 7 | 49557766 | rs6970228 | -1.17 | 0.813 | -2.846 | 0.196 | -0.005 | 0.051 | 0.051 | 0.323 | 0.75 | 0.88 | 0.88 | 0.87 | 0.75 | 0.89 |
| 7 | 49562655 | rs17133366 | 1.797 | 0.735 | -3.02 | 0.065 | -0.028 | -0.081 | -0.081 | 0 | 0.16 | 0.01 | 0.12 | 0.28 | 0.16 | 0.16 |
| 7 | 49564504 | rs4475431 | -0.274 | 0.816 | -2.199 | 0.776 | -0.037 | -0.066 | -0.066 | 0.193 | 0.34 | 0.13 | 0.43 | 0.59 | 0.35 | 0.46 |
| 7 | 49573825 | rs12112354 | 1.2 | 0.852 | -2.191 | 0.792 | -0.003 | 0.062 | 0.062 | 0.131 | 0.41 | 0.75 | 0.45 | 0.3 | 0.38 | 0.43 |
| 7 | 49581956 | rs6979977 | 1.207 | 0.858 | -2.188 | 0.795 | -0.003 | 0.061 | 0.061 | 0.126 | 0.41 | 0.75 | 0.45 | 0.3 | 0.38 | 0.43 |
| 7 | 49602718 | rs7792658 | -0.3 | 0.87 | -2.321 | 0.799 | -0.036 | -0.066 | -0.066 | 0 | 0.34 | 0.13 | 0.43 | 0.59 | 0.35 | 0.46 |
| 7 | 49603238 | rs4917063 | -1.373 | 0.903 | -2.309 | 0.812 | -0.045 | 0.041 | 0.041 | 0.648 | 0.79 | 0.99 | 0.92 | 0.93 | 0.79 | 0.9 |
| 7 | 49604838 | rs7802832 | -0.501 | 0.945 | -2.204 | 0.847 | -0.002 | -0.025 | -0.025 | 0 | 0.72 | 0.67 | 0.86 | 0.89 | 0.73 | 0.89 |
| 7 | 49613268 | rs1859478 | 0.538 | 1.04 | -2.791 | 1.062 | -0.008 | -0.061 | -0.061 | 0 | 0.18 | 0.12 | 0.16 | 0.31 | 0.2 | 0.29 |
| 7 | 49614427 | rs10251209 | 0.964 | 1.067 | -2.778 | 1.053 | -0.035 | -0.072 | -0.072 | 0 | 0.16 | 0.02 | 0.27 | 0.28 | 0.15 | 0.18 |
| 7 | 49615332 | rs10265687 | -0.133 | 1.177 | -2.6 | 0.54 | -0.037 | -0.067 | -0.067 | 0 | 0.34 | 0.13 | 0.43 | 0.59 | 0.35 | 0.46 |
| 7 | 49617533 | rs10245006 | -1.073 | 1.371 | -2.383 | 0.658 | -0.019 | 0.025 | 0.025 | 0.927 | 0.81 | 0.97 | 0.93 | 0.93 | 0.83 | 0.92 |
| 7 | 49620107 | rs10264162 | -0.132 | 1.153 | -2.047 | 0.736 | -0.006 | -0.015 | -0.015 | 0 | 0.39 | 0.32 | 0.48 | 0.55 | 0.42 | 0.48 |
| 7 | 49621833 | rs2886379 | 0.425 | 1.069 | -1.522 | 0.44 | -0.001 | -0.001 | -0.001 | 0 | 0.37 | 0.6 | 0.42 | 0.31 | 0.34 | 0.42 |
| 7 | 49629291 | rs9642306 | -0.485 | 1.158 | -1.058 | 0.583 | 0 | 0.001 | 0.001 | 0 | 0.27 | 0.3 | 0.24 | 0.32 | 0.34 | 0.33 |
| 7 | 49633802 | rs4492324 | 0.061 | 0.989 | -1.636 | -0.786 | -0.132 | 0.153 | 0.153 | 0 | 0.36 | 0.02 | 0.26 | 0.15 | 0.4 | 0.19 |
| 7 | 49637607 | rs12112531 | -0.304 | 0.448 | -1.454 | -1.022 | 0.034 | 0.054 | 0.054 | 0 | 0.42 | 0.79 | 0.45 | 0.69 | 0.35 | 0.56 |
| 7 | 49638772 | rs4546608 | 0.665 | 0.873 | -1.825 | -1.403 | -0.002 | 0.008 | 0.008 | 0 | 0.2 | 0.14 | 0.18 | 0.07 | 0.24 | 0.28 |
| 7 | 49639378 | rs13244520 | 1.385 | 0.79 | -2.001 | -1.532 | -0.021 | 0.099 | 0.099 | 1.086 | 0.27 | 0.04 | 0.15 | 0.15 | 0.23 | 0.06 |
| 7 | 49640912 | rs9649728 | 0.847 | 0.161 | -0.984 | -1.919 | 0.006 | 0.011 | 0.011 | 0 | 0.24 | 0.44 | 0.35 | 0.69 | 0.25 | 0.44 |
| 7 | 49644622 | rs4421315 | -0.046 | -0.161 | -0.016 | -2.03 | -0.061 | -0.109 | -0.109 | 0.014 | 0.2 | 0.36 | 0.11 | 0 | 0.11 | 0.12 |
| 7 | 49651943 | rs10241433 | -0.559 | -0.137 | -0.184 | -2.046 | -0.055 | 0.052 | 0.052 | 1.301 | 0.69 | 0.86 | 0.61 | 0.84 | 0.58 | 0.62 |
| 7 | 49653422 | rs6975043 | -0.188 | -0.197 | -0.093 | -2.012 | -0.071 | -0.123 | -0.123 | 0 | 0.21 | 0.52 | 0.12 | 0 | 0.11 | 0.12 |
| 7 | 49653598 | rs6950454 | 0.456 | -0.15 | -0.176 | -1.991 | -0.004 | -0.028 | -0.028 | 0.088 | 0.17 | 0.15 | 0.09 | 0 | 0.1 | 0.12 |
| 7 | 49657446 | rs7796855 | 0.847 | 0.061 | -0.507 | -1.943 | -0.005 | 0.135 | 0.135 | 0.145 | 0.48 | 0.32 | 0.49 | 0.84 | 0.47 | 0.5 |
| 7 | 49657549 | rs12533512 | 1.848 | 0.067 | -0.575 | -1.979 | -0.02 | 0.101 | 0.101 | 1.022 | 0.28 | 0.04 | 0.15 | 0.16 | 0.23 | 0.06 |
| 7 | 49659033 | rs4916988 | -1.767 | -0.087 | -0.379 | -1.84 | -0.064 | -0.009 | -0.009 | 0 | 0.89 | 0.99 | 0.79 | 0.91 | 0.82 | 0.9 |
| 7 | 49659214 | rs12718209 | 0.982 | -0.062 | -0.289 | -1.792 | -0.004 | 0.129 | 0.129 | 0.145 | 0.48 | 0.34 | 0.49 | 0.84 | 0.47 | 0.5 |
| 7 | 49665976 | rs4540358 | 0.805 | -0.103 | -0.393 | -1.718 | -0.008 | 0.029 | 0.029 | 0 | 0.19 | 0.12 | 0.18 | 0.07 | 0.23 | 0.27 |
| 7 | 49670813 | rs6971984 | -0.377 | -0.321 | 0.321 | -1.527 | -0.074 | -0.134 | -0.134 | 0.215 | 0.2 | 0.42 | 0.11 | 0 | 0.11 | 0.12 |
| 7 | 49672367 | rs6953677 | 0.91 | -0.071 | -0.116 | -1.419 | 0.01 | 0.09 | 0.09 | 0 | 0.51 | 0.26 | 0.4 | 0.25 | 0.55 | 0.5 |
| 7 | 49676497 | rs10282536 | -0.735 | -0.129 | 0.472 | -0.42 | -0.023 | 0.122 | 0.122 | 0.229 | 0.25 | 0.57 | 0.33 | 0.49 | 0.2 | 0.3 |

| | | | | | | | | | | | | | | | | |
|---|---|---|---|---|---|---|---|---|---|---|---|---|---|---|---|---|
| 7 | 49681873 | rs3923403 | 0.366 | -0.103 | 0.933 | -0.331 | 0.002 | 0.173 | 0.173 | 0 | 0.15 | 0.31 | 0.26 | 0.48 | 0.13 | 0.23 |
| 7 | 49686888 | rs4294139 | -0.578 | -0.39 | 0.671 | -0.867 | 0.006 | 0.008 | 0.008 | 0 | 0.44 | 0.42 | 0.48 | 0.56 | 0.41 | 0.59 |
| 7 | 49696467 | rs4464911 | 1.699 | -0.093 | 0.334 | -0.746 | -0.014 | 0.037 | 0.037 | 0.074 | 0.35 | 0.16 | 0.24 | 0.3 | 0.35 | 0.28 |
| 7 | 49696734 | rs10263034 | 1.56 | -0.131 | 0.594 | -0.732 | -0.001 | 0.008 | 0.008 | 0.01 | 0.37 | 0.31 | 0.25 | 0.3 | 0.35 | 0.28 |
| 7 | 49697140 | rs10278891 | 0.826 | -0.5 | 0.46 | -0.789 | 0.001 | 0.046 | 0.046 | 0 | 0.49 | 0.41 | 0.49 | 0.78 | 0.48 | 0.51 |
| 7 | 49706591 | rs6961746 | -1.736 | -0.445 | 0.355 | -0.755 | -0.065 | -0.035 | -0.035 | 0 | 0.88 | 0.99 | 0.79 | 0.86 | 0.81 | 0.89 |
| 7 | 49710785 | rs10236813 | 0.702 | -0.398 | 0.463 | -0.547 | -0.019 | 0.105 | 0.105 | 0 | 0.15 | 0.27 | 0.24 | 0.47 | 0.09 | 0.23 |
| 7 | 49711297 | rs12718221 | -0.141 | -0.439 | 0.557 | -0.607 | -0.007 | -0.031 | -0.031 | 0 | 0.37 | 0.43 | 0.29 | 0.08 | 0.33 | 0.38 |
| 7 | 49711428 | rs2059544 | 1.599 | -0.483 | 0.655 | -0.51 | -0.003 | 0.002 | 0.002 | 0.017 | 0.36 | 0.27 | 0.26 | 0.31 | 0.39 | 0.28 |
| 7 | 49717680 | rs12668897 | 0.757 | -0.411 | 0.201 | -0.449 | -0.009 | 0.069 | 0.069 | 0 | 0.14 | 0.23 | 0.24 | 0.47 | 0.09 | 0.23 |
| 7 | 49728176 | rs4917079 | 0.639 | -0.437 | 0.115 | -0.608 | 0.001 | 0.026 | 0.026 | 0 | 0.5 | 0.44 | 0.49 | 0.78 | 0.48 | 0.51 |
| 7 | 49730525 | rs10257300 | -0.318 | -0.355 | 0.338 | -0.293 | -0.027 | 0.116 | 0.116 | 0 | 0.2 | 0.39 | 0.28 | 0.51 | 0.12 | 0.23 |
| 7 | 49733584 | rs4557649 | 0.066 | -0.526 | -0.297 | -0.35 | 0.001 | -0.04 | -0.04 | 0.169 | 0.36 | 0.2 | 0.53 | 0.51 | 0.34 | 0.45 |
| 7 | 49738741 | rs10266625 | -0.423 | -0.176 | 0.013 | -0.328 | 0.001 | 0.001 | 0.001 | 0 | 0.52 | 0.53 | 0.58 | 0.63 | 0.41 | 0.54 |
| 7 | 49739058 | rs10263787 | -0.677 | -0.066 | -0.508 | -0.224 | 0.001 | 0.031 | 0.031 | 0 | 0.57 | 0.71 | 0.64 | 0.63 | 0.45 | 0.57 |
| 7 | 49739731 | rs17663429 | 0.814 | 0.043 | -0.318 | 0.085 | -0.006 | 0.04 | 0.04 | 0 | 0.08 | 0.04 | 0.23 | 0.42 | 0.08 | 0.19 |
| 7 | 49742459 | rs1346111 | 0.883 | 0.297 | -0.774 | -0.52 | -0.048 | 0.038 | 0.038 | 0 | 0.31 | 0.1 | 0.22 | 0.27 | 0.37 | 0.31 |
| 7 | 49746554 | rs727051 | -0.216 | 0.433 | -0.113 | -0.615 | -0.001 | -0.033 | -0.033 | 0 | 0.27 | 0.35 | 0.19 | 0.15 | 0.25 | 0.19 |
| 7 | 49754752 | rs1543902 | -0.642 | 0.619 | -0.111 | 0.009 | 0.009 | 0.06 | 0.06 | 1.438 | 0.5 | 0.83 | 0.42 | 0.4 | 0.47 | 0.48 |
| 7 | 49759036 | rs1365086 | 0.12 | 1.066 | 0.268 | -0.07 | -0.007 | 0.008 | 0.008 | 0 | 0.58 | 0.37 | 0.68 | 0.65 | 0.51 | 0.78 |
| 7 | 49760185 | rs10261805 | -0.355 | 1.021 | 0.125 | -0.069 | -0.005 | 0 | 0 | 1.568 | 0.94 | 0.96 | 0.94 | 0.93 | 0.89 | 0.96 |
| 7 | 49761362 | rs11185584 | 0.96 | 0.939 | 0.372 | -0.072 | 0.01 | -0.013 | -0.013 | 0 | 0.44 | 0.18 | 0.49 | 0.52 | 0.41 | 0.48 |
| 7 | 49762093 | rs1425881 | 0.419 | 0.901 | 0.942 | -0.095 | -0.001 | 0 | 0 | 0.165 | 0.35 | 0.35 | 0.25 | 0.28 | 0.38 | 0.18 |
| 7 | 49762598 | rs13240380 | 0.932 | 0.92 | 0.88 | -0.052 | 0.011 | -0.004 | -0.004 | 1.131 | 0.45 | 0.22 | 0.5 | 0.53 | 0.41 | 0.48 |
| 7 | 49775431 | rs12532230 | 0.143 | 0.873 | 0.962 | -0.072 | -0.017 | 0.021 | 0.021 | 0 | 0.58 | 0.26 | 0.68 | 0.65 | 0.51 | 0.78 |
| 7 | 49787859 | rs7799604 | 0.865 | 0.928 | 1.004 | -0.046 | -0.002 | 0.009 | 0.009 | 0.268 | 0.31 | 0.3 | 0.22 | 0.16 | 0.36 | 0.12 |
| 7 | 49803095 | rs1477578 | 1.115 | 0.746 | 0.227 | -0.025 | 0.003 | 0.019 | 0.019 | 0.364 | 0.84 | 0.36 | 0.87 | 0.79 | 0.86 | 0.89 |
| 7 | 49808395 | rs1477579 | 0.865 | 0.784 | -0.223 | 0.125 | -0.008 | -0.038 | -0.038 | 0 | 0.43 | 0.12 | 0.49 | 0.52 | 0.41 | 0.48 |
| 7 | 49810966 | rs12673518 | 0.617 | 1.027 | -0.128 | 0.007 | 0.03 | 0.002 | 0.002 | 0 | 0.46 | 0.33 | 0.54 | 0.51 | 0.37 | 0.46 |
| 7 | 49818127 | rs482968 | -0.8 | 0.141 | -0.67 | -0.055 | 0.021 | 0.055 | 0.055 | 1.209 | 0.51 | 0.73 | 0.43 | 0.4 | 0.48 | 0.47 |
| 7 | 49819656 | rs2162179 | 0.122 | 0.616 | -1.49 | 0.12 | 0.021 | -0.012 | -0.012 | 0 | 0.05 | 0.01 | 0.02 | 0.1 | 0.02 | 0.06 |
| 7 | 49819713 | rs557626 | -0.326 | 1.117 | -1.299 | -0.237 | -0.018 | 0.004 | 0.004 | 0 | 0.13 | 0.06 | 0.16 | 0.13 | 0.14 | 0.33 |
| 7 | 49820719 | rs17632348 | -0.056 | 0.758 | -0.731 | -0.155 | 0 | 0.001 | 0.001 | 1.08 | 0.29 | 0.26 | 0.21 | 0.17 | 0.32 | 0.09 |
| 7 | 49825108 | rs900647 | -0.164 | 1.396 | -0.956 | 0.15 | -0.067 | -0.074 | -0.074 | 0 | 0.11 | 0.02 | 0.25 | 0.63 | 0.24 | 0.09 |
| 7 | 49826804 | rs10280152 | -0.816 | 0.896 | -0.357 | -0.748 | -0.006 | -0.019 | -0.019 | 0.17 | 0.21 | 0.57 | 0.14 | 0.16 | 0.18 | 0.13 |
| 7 | 49833688 | rs10499687 | -0.443 | 0.863 | -0.465 | -1.046 | -0.001 | -0.023 | -0.023 | 0 | 0.1 | 0.16 | 0.11 | 0 | 0.1 | 0.09 |
| 7 | 49836460 | rs10237193 | 1.173 | -0.206 | -1.401 | -1.857 | -0.024 | 0.143 | 0.143 | 0 | 0.46 | 0.08 | 0.28 | 0.27 | 0.53 | 0.46 |
| 7 | 49841119 | rs691785 | -0.059 | -0.135 | -0.891 | -1.835 | 0.003 | 0.012 | 0.012 | 0 | 0.64 | 0.71 | 0.42 | 0.27 | 0.64 | 0.53 |
| 7 | 49845694 | rs13228679 | 0.035 | 0.15 | -1.177 | -1.646 | -0.001 | -0.025 | -0.025 | 0 | 0.1 | 0.16 | 0.11 | 0 | 0.1 | 0.09 |
| 7 | 49848808 | rs12375076 | 0.865 | 0.276 | -1.295 | -1.4 | -0.063 | -0.063 | -0.063 | 5.76 | 0.1 | 0.01 | 0.24 | 0.64 | 0.22 | 0.08 |
| 7 | 49852730 | rs10239501 | 1.102 | 0.065 | -0.717 | -1.893 | -0.014 | 0.077 | 0.077 | 0 | 0.38 | 0.2 | 0.2 | 0.09 | 0.38 | 0.35 |
| 7 | 49859810 | rs10260065 | -0.62 | -1.043 | -0.486 | -1.596 | -0.056 | -0.035 | -0.035 | 6.969 | 0.81 | 0.74 | 0.93 | 0.94 | 0.92 | 0.92 |
| 7 | 49860530 | rs6957449 | -0.062 | -0.779 | -0.234 | -1.605 | 0.021 | -0.018 | -0.018 | 0 | 0.22 | 0.36 | 0.17 | 0.16 | 0.24 | 0.19 |
| 7 | 49862947 | rs1483797 | -0.34 | -0.91 | -0.481 | -1.542 | -0.07 | -0.045 | -0.045 | 0.786 | 0.77 | 0.66 | 0.91 | 0.93 | 0.91 | 0.77 |
| 7 | 49869567 | rs12718224 | 0.384 | 0.924 | -0.91 | -0.937 | -0.069 | -0.084 | -0.084 | 0.293 | 0.12 | 0.01 | 0.24 | 0.65 | 0.25 | 0.08 |
| 7 | 49876993 | rs1384888 | -0.839 | 0.502 | -0.628 | -1.38 | 0.02 | -0.033 | -0.033 | 0 | 0.35 | 0.39 | 0.66 | 0.84 | 0.52 | 0.34 |
| 7 | 49883749 | rs12536329 | -0.012 | -0.31 | -1.536 | -1.773 | 0.037 | 0.158 | 0.158 | 0.969 | 0.4 | 0.12 | 0.51 | 0.85 | 0.66 | 0.27 |
| 7 | 49907824 | rs692039 | -0.283 | -1.663 | -1.576 | -2.457 | 0.05 | 0.087 | 0.087 | 0 | 0.68 | 0.84 | 0.63 | 0.88 | 0.76 | 0.69 |
| 7 | 49909451 | rs6963083 | 0.224 | -2.078 | -0.381 | -2.641 | -0.026 | -0.051 | -0.051 | 0 | 0.6 | 0.46 | 0.59 | 0.88 | 0.76 | 0.52 |
| 7 | 49928919 | rs17664027 | 1.376 | -1.212 | -0.909 | -2.253 | -0.058 | -0.066 | -0.066 | 0.916 | 0.11 | 0.03 | 0.24 | 0.64 | 0.24 | 0.08 |
| 7 | 49931275 | rs6968993 | 0.799 | -1.158 | -0.232 | -2.425 | 0.002 | -0.021 | -0.021 | 1.721 | 0.45 | 0.44 | 0.33 | 0.24 | 0.47 | 0.44 |
| 7 | 49949984 | rs13236056 | 1.52 | -1.263 | -0.264 | -2.458 | -0.022 | -0.053 | -0.053 | 0 | 0.19 | 0.27 | 0.09 | 0.05 | 0.12 | 0.24 |
| 7 | 49953313 | rs1483071 | -0.005 | -1.337 | -0.171 | -2.473 | -0.003 | -0.025 | -0.025 | 0 | 0.6 | 0.56 | 0.58 | 0.89 | 0.71 | 0.67 |
| 7 | 49973902 | rs6583402 | -0.534 | -1.399 | -0.305 | -2.468 | 0.008 | 0.003 | 0.003 | 0 | 0.66 | 0.67 | 0.62 | 0.88 | 0.79 | 0.69 |
| 7 | 49998624 | rs7785151 | -0.79 | -1.429 | -0.287 | -2.468 | 0.045 | 0.073 | 0.073 | 0.044 | 0.68 | 0.81 | 0.64 | 0.88 | 0.78 | 0.69 |
| 7 | 50010988 | rs1396277 | -0.86 | -1.48 | -0.311 | -2.487 | 0.043 | 0.076 | 0.076 | 0 | 0.69 | 0.82 | 0.84 | 0.92 | 0.78 | 0.78 |
| 7 | 50032637 | rs1476240 | -0.691 | -1.658 | -0.499 | -2.62 | 0.046 | 0.058 | 0.058 | 0.167 | 0.71 | 0.81 | 0.65 | 0.88 | 0.81 | 0.72 |
| 7 | 50043518 | rs11980286 | 1.131 | -1.654 | -0.435 | -2.589 | -0.02 | -0.069 | -0.069 | 0.082 | 0.23 | 0.3 | 0.11 | 0.05 | 0.14 | 0.27 |
| 7 | 50052934 | rs2128663 | 1.06 | -1.497 | -0.485 | -2.52 | -0.043 | 0.022 | 0.022 | 0 | 0.28 | 0.13 | 0.25 | 0.2 | 0.37 | 0.21 |
| 7 | 50087159 | rs3807318 | 1.225 | -1.508 | -0.524 | -2.592 | -0.055 | 0.012 | 0.012 | 0.008 | 0.25 | 0.11 | 0.22 | 0.2 | 0.34 | 0.2 |
| 7 | 50087970 | rs11983726 | 1.151 | -1.796 | -0.279 | -2.66 | -0.018 | -0.058 | -0.058 | 0 | 0.2 | 0.29 | 0.1 | 0.05 | 0.13 | 0.26 |
| 7 | 50099404 | rs2221656 | -0.857 | -2 | -0.911 | -2.498 | 0.066 | 0.064 | 0.064 | 0.53 | 0.79 | 0.92 | 0.91 | 0.93 | 0.89 | 0.85 |
| 7 | 50119502 | rs1911754 | 0.28 | -1.902 | -0.697 | -2.49 | 0.025 | 0.03 | 0.03 | 0 | 0.65 | 0.72 | 0.83 | 0.88 | 0.77 | 0.72 |
| 7 | 50122310 | rs1567278 | -0.265 | -1.897 | -0.692 | -2.492 | 0.045 | 0.049 | 0.049 | 0.319 | 0.7 | 0.8 | 0.85 | 0.88 | 0.83 | 0.73 |
| 7 | 50143938 | rs17133685 | -0.332 | -1.756 | -0.631 | -2.387 | -0.001 | -0.02 | -0.02 | 0 | 0.07 | 0.13 | 0.03 | 0.05 | 0.05 | 0.09 |
| 7 | 50160189 | rs1379175 | -0.37 | -2.021 | -0.674 | -2.395 | 0.05 | 0.059 | 0.059 | 0.129 | 0.72 | 0.82 | 0.86 | 0.93 | 0.82 | 0.8 |
| 7 | 50171491 | rs11760445 | 0.746 | -1.768 | -1.151 | -2.483 | -0.173 | 0.055 | 0.055 | 0.857 | 0.3 | 0.01 | 0.25 | 0.15 | 0.4 | 0.23 |
| 7 | 50172468 | rs17633571 | 0.575 | -1.765 | -0.96 | -2.476 | -0.054 | 0.058 | 0.058 | 0.007 | 0.32 | 0.15 | 0.26 | 0.2 | 0.4 | 0.33 |
| 7 | 50180112 | rs11185600 | 0.472 | -1.878 | -0.353 | -2.5 | 0.018 | -0.017 | -0.017 | 0.983 | 0.33 | 0.45 | 0.59 | 0.73 | 0.42 | 0.32 |

| | | | | | | | | | | | | | | | | |
|---|---|---|---|---|---|---|---|---|---|---|---|---|---|---|---|---|
| 7 | 50182079 | rs10234768 | -0.61 | -1.97 | -0.343 | -2.602 | 0.021 | 0.022 | 0.022 | 0.257 | 0.71 | 0.76 | 0.85 | 0.93 | 0.82 | 0.79 |
| 7 | 50191216 | rs10260654 | 0.612 | -1.828 | -0.206 | -2.513 | 0.011 | -0.009 | -0.009 | 0 | 0.33 | 0.39 | 0.58 | 0.73 | 0.42 | 0.32 |
| 7 | 50193102 | rs4917010 | -0.129 | -1.78 | -0.704 | -2.414 | 0.025 | 0.025 | 0.025 | 0 | 0.67 | 0.71 | 0.84 | 0.93 | 0.81 | 0.78 |
| 7 | 50205435 | rs6965975 | -0.062 | -1.693 | -0.627 | -2.464 | -0.003 | -0.035 | -0.035 | 0 | 0.34 | 0.38 | 0.35 | 0.05 | 0.29 | 0.27 |
| 7 | 50207040 | rs10246184 | 0.448 | -1.797 | -1.42 | -2.494 | 0.092 | 0.181 | 0.181 | 0.653 | 0.56 | 0.17 | 0.62 | 0.88 | 0.7 | 0.37 |
| 7 | 50207532 | rs12531206 | 0.817 | -1.538 | -1.919 | -2.457 | -0.015 | -0.011 | -0.011 | 1.744 | 0.27 | 0.13 | 0.36 | 0.7 | 0.3 | 0.19 |
| 7 | 50211199 | rs882034 | 0.807 | -1.055 | -2.012 | -2.245 | -0.158 | 0.016 | 0.016 | 0.002 | 0.28 | 0.01 | 0.25 | 0.17 | 0.41 | 0.17 |
| 7 | 50222500 | rs897694 | 0.311 | -1.477 | -1.659 | -2.326 | 0.019 | 0.112 | 0.112 | 0.165 | 0.31 | 0.17 | 0.3 | 0.03 | 0.26 | 0.26 |
| 7 | 50225391 | rs7805803 | 0.702 | -0.64 | -1.291 | -2.292 | -0.001 | 0.023 | 0.023 | 2.39 | 0.44 | 0.34 | 0.39 | 0.18 | 0.44 | 0.18 |
| 7 | 50234164 | rs6955221 | 0.709 | -0.52 | -1.278 | -0.806 | 0.048 | 0.023 | 0.023 | 0.708 | 0.28 | 0.16 | 0.14 | 0.2 | 0.19 | 0.24 |
| 7 | 50234363 | rs4917122 | 1.19 | -0.268 | -1.343 | -0.794 | -0.046 | -0.064 | -0.064 | 0 | 0.13 | 0.04 | 0.23 | 0.53 | 0.24 | 0.12 |
| 7 | 50234801 | rs874355 | 1.21 | -0.267 | -1.343 | -0.794 | -0.046 | -0.063 | -0.063 | 0 | 0.13 | 0.04 | 0.23 | 0.53 | 0.24 | 0.12 |
| 7 | 50236464 | rs1379183 | 0.311 | -0.272 | -1.08 | -1.715 | 0.002 | 0.002 | 0.002 | 1.276 | 0.53 | 0.38 | 0.6 | 0.2 | 0.56 | 0.32 |
| 7 | 50237083 | rs6957965 | 1.119 | -0.455 | -1.734 | -1.962 | -0.134 | 0.041 | 0.041 | 0 | 0.31 | 0.03 | 0.33 | 0.17 | 0.44 | 0.17 |
| 7 | 50237496 | rs13231384 | 1.15 | -0.483 | -1.728 | -1.956 | -0.129 | 0.039 | 0.039 | 0 | 0.31 | 0.03 | 0.33 | 0.17 | 0.44 | 0.17 |
| 7 | 50241536 | rs921908 | -1.009 | -0.2 | -1.432 | -1.577 | 0.074 | 0.012 | 0.012 | 0 | 0.62 | 0.79 | 0.81 | 0.32 | 0.77 | 0.52 |
| 7 | 50241812 | rs921911 | 0.626 | 0.69 | -1.316 | -1.512 | 0.01 | 0.027 | 0.027 | 0.204 | 0.23 | 0.16 | 0.28 | 0.02 | 0.16 | 0.12 |
| 7 | 50243488 | rs1037676 | 0.266 | 1.284 | -0.765 | -0.618 | 0.064 | -0.012 | -0.012 | 0 | 0.09 | 0.16 | 0.17 | 0.03 | 0.17 | 0.06 |
| 7 | 50249013 | rs11761913 | 0.24 | 0.89 | -1.446 | -0.9 | 0.02 | 0.225 | 0.225 | 0 | 0.45 | 0.07 | 0.63 | 0.13 | 0.58 | 0.4 |
| 7 | 50249571 | rs4917125 | 0.515 | 1.235 | -1.3 | -0.455 | -0.018 | -0.023 | -0.023 | 0 | 0.06 | 0.01 | 0.08 | 0.13 | 0.09 | 0.04 |
| 7 | 50250354 | rs17133733 | 0.249 | 0.51 | -0.834 | -0.443 | 0.028 | -0.005 | -0.005 | 0 | 0.29 | 0.27 | 0.18 | 0.65 | 0.22 | 0.13 |
| 7 | 50251216 | rs10257997 | -0.394 | 1.702 | 0.064 | -0.052 | 0.091 | 0.008 | 0.008 | 0 | 0.1 | 0.21 | 0.25 | 0.05 | 0.18 | 0.08 |
| 7 | 50256069 | rs2168587 | -0.543 | 0.683 | 0.09 | -0.12 | -0.02 | -0.066 | -0.066 | 0.738 | 0.12 | 0.28 | 0.07 | 0.06 | 0.08 | 0.41 |
| 7 | 50269672 | rs1456893 | 0.412 | 2.229 | 0.032 | 0.37 | 0.021 | 0.017 | 0.017 | 0.389 | 0.77 | 0.63 | 0.78 | 0.41 | 0.69 | 0.85 |
| 7 | 50281539 | rs981132 | 0.412 | 2.229 | 0.029 | 0.397 | 0.019 | 0.016 | 0.016 | 0.389 | 0.77 | 0.63 | 0.78 | 0.41 | 0.69 | 0.85 |
| 7 | 50305863 | rs4917014 | 0.495 | 1.622 | -1.051 | -0.111 | -0.054 | -0.059 | -0.059 | 0.105 | 0.31 | 0.04 | 0.51 | 0.35 | 0.35 | 0.28 |
| 7 | 50313352 | rs10276619 | 0.018 | 1.616 | -0.502 | -0.103 | 0.091 | 0.069 | 0.069 | 0.138 | 0.73 | 0.42 | 0.71 | 0.45 | 0.62 | 0.76 |
| 7 | 50318308 | rs17552904 | 0.397 | 1.426 | -0.879 | -0.123 | -0.009 | -0.025 | -0.025 | 0 | 0.29 | 0.05 | 0.45 | 0.3 | 0.29 | 0.26 |
| 7 | 50324652 | rs4917131 | 0.116 | 2.153 | -0.177 | 0.987 | -0.005 | 0.103 | 0.103 | 1.607 | 0.77 | 0.34 | 0.81 | 0.52 | 0.81 | 0.74 |
| 7 | 50325717 | rs716719 | -0.205 | 2.39 | -0.775 | 1.366 | -0.072 | -0.12 | -0.12 | 0.757 | 0.22 | 0.06 | 0.4 | 0.48 | 0.29 | 0.25 |
| 7 | 50332196 | rs10239000 | -0.137 | 2.602 | -0.216 | 1.405 | 0.01 | 0.007 | 0.007 | 0.223 | 0.81 | 0.74 | 0.84 | 0.79 | 0.75 | 0.9 |
| 7 | 50333463 | rs4917016 | 0.244 | 2.945 | -0.276 | 1.134 | 0.015 | -0.006 | -0.006 | 0 | 0.84 | 0.8 | 0.82 | 0.89 | 0.71 | 0.91 |
| 7 | 50335232 | rs4917017 | -1.125 | 1.366 | -0.361 | 1.481 | 0.002 | 0.017 | 0.017 | 0.17 | 0.27 | 0.26 | 0.43 | 0.55 | 0.29 | 0.38 |
| 7 | 50336551 | rs9886239 | -1.338 | 1.4 | -0.294 | 1.512 | 0.011 | 0.046 | 0.046 | 0.253 | 0.29 | 0.36 | 0.45 | 0.59 | 0.31 | 0.4 |
| 7 | 50344544 | rs11506039 | -0.272 | 1.184 | -0.643 | 1.449 | -0.044 | -0.032 | -0.032 | 0 | 0.1 | 0.16 | 0.02 | 0.07 | 0.01 | 0.23 |
| 7 | 50346134 | rs7781977 | -0.919 | 1.365 | -0.606 | 1.569 | 0.001 | 0.014 | 0.014 | 0.242 | 0.28 | 0.27 | 0.44 | 0.5 | 0.29 | 0.35 |
| 7 | 50350267 | rs6583437 | -0.823 | 1.232 | -0.671 | 1.434 | -0.001 | -0.011 | -0.011 | 0 | 0.35 | 0.27 | 0.47 | 0.57 | 0.34 | 0.38 |
| 7 | 50361683 | rs6583440 | -0.915 | 1.13 | -0.806 | 1.184 | -0.006 | -0.027 | -0.027 | 3.889 | 0.34 | 0.18 | 0.48 | 0.5 | 0.37 | 0.39 |
| 7 | 50364034 | rs6583442 | -0.741 | 0.906 | -0.543 | 1.539 | 0 | 0.027 | 0.027 | 0 | 0.06 | 0.03 | 0.09 | 0.19 | 0.04 | 0.04 |
| 7 | 50364626 | rs10230193 | -0.397 | 0.707 | -0.393 | 1.161 | -0.014 | 0.035 | 0.035 | 3.339 | 0.36 | 0.57 | 0.43 | 0.5 | 0.33 | 0.44 |
| 7 | 50366637 | rs10251980 | -1.221 | 1.311 | 0.379 | 0.774 | 0.116 | 0.062 | 0.062 | 0.053 | 0.09 | 0.34 | 0.17 | 0.06 | 0.13 | 0.05 |
| 7 | 50417522 | rs11765988 | -0.131 | 1.233 | -0.153 | 0.14 | -0.008 | 0.021 | 0.021 | 4.564 | 0.3 | 0.17 | 0.28 | 0.18 | 0.33 | 0.22 |
| 7 | 50418242 | rs7785321 | -0.182 | 1.625 | -0.009 | -0.19 | 0.056 | -0.035 | -0.035 | 1.079 | 0.08 | 0.35 | 0.08 | 0 | 0.09 | 0.05 |
| 7 | 50418625 | rs7800411 | -0.039 | 1.433 | -0.046 | -0.074 | 0.003 | -0.017 | -0.017 | 0.12 | 0.3 | 0.24 | 0.28 | 0.18 | 0.33 | 0.22 |
| 7 | 50424061 | rs7790846 | 0.002 | 0.538 | -0.857 | 0.23 | 0.028 | -0.034 | -0.034 | 1.812 | 0.66 | 0.86 | 0.74 | 0.53 | 0.73 | 0.51 |
| 7 | 50428445 | rs12718598 | 0.346 | 0.937 | -0.452 | 0.051 | -0.001 | 0 | 0 | 0 | 0.48 | 0.51 | 0.6 | 0.44 | 0.5 | 0.49 |
| 7 | 50435777 | rs12669559 | 0.791 | 0.835 | -0.363 | -0.133 | -0.006 | 0.077 | 0.077 | 0 | 0.37 | 0.23 | 0.38 | 0.19 | 0.37 | 0.34 |
| 7 | 50438720 | rs7780012 | -0.689 | 0.455 | -0.113 | -0.036 | 0.05 | 0.001 | 0.001 | 0.058 | 0.15 | 0.44 | 0.11 | 0.11 | 0.2 | 0.07 |
| 7 | 50442726 | rs7809377 | 0.634 | -0.776 | -0.729 | -0.135 | 0.005 | 0.061 | 0.061 | 0.037 | 0.16 | 0.05 | 0.14 | 0.05 | 0.13 | 0.07 |
| 7 | 50452552 | rs10230385 | -0.555 | -0.823 | -0.024 | 0.061 | -0.035 | 0.006 | 0.006 | 0 | 0.7 | 0.59 | 0.61 | 0.31 | 0.79 | 0.64 |
| 7 | 50454736 | rs6962370 | 0.227 | -1.187 | -0.265 | -0.304 | 0.015 | 0.023 | 0.023 | 0 | 0.16 | 0.07 | 0.12 | 0.01 | 0.15 | 0.22 |
| 7 | 50460096 | rs6964823 | 1.122 | -0.797 | 0.113 | -0.119 | -0.006 | 0.01 | 0.01 | 0 | 0.4 | 0.43 | 0.38 | 0.19 | 0.47 | 0.35 |
| 7 | 50460208 | rs7797772 | 0.5 | -0.795 | 0.119 | -0.137 | 0.003 | -0.007 | -0.007 | 0.864 | 0.08 | 0.05 | 0.03 | 0.06 | 0.09 | 0.09 |
| 7 | 50462138 | rs11763774 | 1.016 | -0.82 | 0.216 | 0.188 | -0.003 | 0.001 | 0.001 | 0 | 0.42 | 0.47 | 0.57 | 0.39 | 0.48 | 0.37 |
| 7 | 50462935 | rs6952409 | 0.223 | -0.768 | 0.019 | 0.177 | -0.002 | 0.029 | 0.029 | 0 | 0.21 | 0.09 | 0.2 | 0.05 | 0.23 | 0.21 |
| 7 | 50469981 | rs11980379 | -0.717 | -0.766 | 0.196 | 0.177 | 0.001 | 0.021 | 0.021 | 0 | 0.29 | 0.17 | 0.23 | 0.11 | 0.32 | 0.29 |
| 7 | 50470604 | rs4132601 | -0.729 | -0.793 | 0.192 | 0.172 | 0.002 | 0.022 | 0.022 | 0 | 0.29 | 0.17 | 0.23 | 0.11 | 0.32 | 0.29 |
| 7 | 50474223 | rs12538151 | -1.489 | -0.869 | 0.31 | 0.379 | 0.006 | -0.008 | -0.008 | 0.796 | 0.76 | 0.76 | 0.83 | 0.61 | 0.83 | 0.72 |
| 7 | 50476139 | rs12719019 | 1.205 | -0.82 | 0.401 | 0.13 | -0.052 | 0.099 | 0.099 | 0 | 0.38 | 0.24 | 0.35 | 0.19 | 0.47 | 0.34 |
| 7 | 50479414 | rs6959427 | -0.293 | -0.584 | 0.951 | 0.365 | 0.024 | 0.015 | 0.015 | 0 | 0.1 | 0.22 | 0.11 | 0.07 | 0.13 | 0.12 |
| 7 | 50486309 | rs7792486 | 0.658 | -0.665 | 0.818 | 0.413 | -0.006 | 0.013 | 0.013 | 0.171 | 0.06 | 0.08 | 0.03 | 0.12 | 0.04 | 0.06 |
| 7 | 50493729 | rs10499691 | 0.666 | -0.662 | 0.735 | 0.395 | -0.019 | -0.029 | -0.029 | 0 | 0.07 | 0 | 0.14 | 0.07 | 0.11 | 0.1 |
| 7 | 50494207 | rs11976368 | 0.435 | -0.669 | 0.945 | 0.391 | -0.001 | 0.005 | 0.005 | 0.015 | 0.08 | 0.09 | 0.15 | 0.07 | 0.11 | 0.1 |
| 7 | 50497131 | rs17634488 | 0.684 | -0.694 | 0.942 | 0.401 | -0.002 | 0.005 | 0.005 | 0.161 | 0.06 | 0.06 | 0.03 | 0.12 | 0.04 | 0.06 |
| 7 | 50507893 | rs4947437 | -0.994 | -0.771 | 1.187 | 0.406 | -0.022 | 0.047 | 0.047 | 0 | 0.74 | 0.57 | 0.87 | 0.59 | 0.82 | 0.71 |
| 7 | 50510269 | rs10243511 | -0.029 | -0.781 | 1.308 | 0.401 | -0.011 | 0.022 | 0.022 | 0 | 0.66 | 0.43 | 0.72 | 0.52 | 0.71 | 0.61 |
| 7 | 50512284 | rs2305358 | -0.029 | -0.781 | 1.308 | 0.424 | -0.011 | 0.021 | 0.021 | 0 | 0.66 | 0.43 | 0.72 | 0.52 | 0.71 | 0.61 |
| 7 | 50512929 | rs10230343 | -0.029 | -0.781 | 1.308 | 0.42 | -0.011 | 0.022 | 0.022 | 0 | 0.66 | 0.43 | 0.72 | 0.52 | 0.71 | 0.61 |
| 7 | 50517271 | rs3735275 | -1.194 | -0.782 | 1.372 | 0.439 | 0.014 | -0.021 | -0.021 | 3.432 | 0.76 | 0.76 | 0.89 | 0.59 | 0.84 | 0.72 |

| | | | | | | | | | | | | | | | | |
|---|---|---|---|---|---|---|---|---|---|---|---|---|---|---|---|---|
| 7 | 50525420 | rs4947510 | -0.147 | -0.778 | 1.295 | 0.438 | -0.011 | 0.025 | 0.025 | 0 | 0.67 | 0.57 | 0.73 | 0.52 | 0.71 | 0.61 |
| 7 | 50526219 | rs11575553 | 0.59 | -0.62 | 0.972 | 0.466 | -0.011 | -0.022 | -0.022 | 0 | 0.07 | 0.01 | 0.14 | 0.07 | 0.11 | 0.1 |
| 7 | 50529384 | rs2060762 | -1.145 | -0.757 | 0.784 | 0.522 | 0.017 | -0.025 | -0.025 | 0.008 | 0.76 | 0.78 | 0.89 | 0.59 | 0.84 | 0.72 |
| 7 | 50530309 | rs11575548 | 0.435 | -0.086 | 1.155 | 0.584 | -0.035 | -0.028 | -0.028 | 0 | 0.08 | 0.01 | 0.14 | 0.07 | 0.13 | 0.11 |
| 7 | 50530762 | rs11575544 | 1.256 | -0.058 | 1.055 | 0.549 | -0.023 | 0.024 | 0.024 | 0 | 0.1 | 0.02 | 0.07 | 0.04 | 0.09 | 0.04 |
| 7 | 50538589 | rs11238131 | -0.198 | -0.339 | 0.934 | 0.559 | 0.017 | 0 | 0 | 0 | 0.54 | 0.55 | 0.77 | 0.52 | 0.7 | 0.62 |
| 7 | 50542843 | rs11772714 | -1.325 | -0.403 | 0.848 | 0.556 | 0.035 | -0.013 | -0.013 | 0 | 0.68 | 0.74 | 0.86 | 0.62 | 0.79 | 0.7 |
| 7 | 50542914 | rs11238133 | -0.427 | -0.354 | 0.802 | 0.556 | 0.028 | -0.034 | -0.034 | 0 | 0.61 | 0.73 | 0.72 | 0.55 | 0.67 | 0.59 |
| 7 | 50542927 | rs11238134 | -0.427 | -0.354 | 0.802 | 0.556 | 0.028 | -0.034 | -0.034 | 0 | 0.61 | 0.73 | 0.72 | 0.55 | 0.67 | 0.59 |
| 7 | 50544314 | rs4490786 | -0.463 | -0.333 | 1.099 | 0.432 | 0.004 | -0.007 | -0.007 | 1.492 | 0.21 | 0.22 | 0.24 | 0.13 | 0.22 | 0.18 |
| 7 | 50547865 | rs11575453 | 0.989 | -0.311 | 1.205 | 0.44 | 0 | -0.001 | -0.001 | 0 | 0.13 | 0.14 | 0.09 | 0.07 | 0.09 | 0.07 |
| 7 | 50547922 | rs2329367 | 0.896 | -0.298 | 1.189 | 0.367 | -0.035 | -0.028 | -0.028 | 0.379 | 0.08 | 0.01 | 0.15 | 0.07 | 0.13 | 0.11 |
| 7 | 50550144 | rs12718541 | 0.324 | -0.296 | 1.288 | 0.386 | 0 | 0 | 0 | 0 | 0.47 | 0.52 | 0.62 | 0.46 | 0.56 | 0.51 |
| 7 | 50564204 | rs3807566 | 1.145 | 0.05 | 1.114 | 0.241 | -0.02 | -0.016 | -0.016 | 0 | 0.34 | 0.29 | 0.41 | 0.36 | 0.4 | 0.35 |
| 7 | 50565404 | rs1037351 | -1.052 | 0.004 | 1.257 | 0.183 | -0.003 | 0.005 | 0.005 | 0 | 0.3 | 0.23 | 0.32 | 0.1 | 0.37 | 0.34 |
| 7 | 50567435 | rs11575387 | 1.198 | -0.006 | 1.267 | 0.173 | -0.003 | -0.046 | -0.046 | 0 | 0.13 | 0.2 | 0.08 | 0.03 | 0.08 | 0.07 |
| 7 | 50567493 | rs6951648 | -0.152 | 0.003 | 1.258 | 0.164 | 0.001 | -0.045 | -0.045 | 0 | 0.21 | 0.21 | 0.16 | 0.04 | 0.22 | 0.22 |
| 7 | 50570888 | rs7791875 | 1.067 | -0.079 | 1.102 | 0.241 | 0.001 | -0.001 | -0.001 | 0 | 0.13 | 0.12 | 0.09 | 0.03 | 0.09 | 0.07 |
| 7 | 50571022 | rs3807558 | -1.657 | -0.127 | 1.147 | 0.251 | 0.033 | -0.027 | -0.027 | 0 | 0.68 | 0.73 | 0.86 | 0.56 | 0.79 | 0.68 |
| 7 | 50572890 | rs6592961 | -0.56 | -0.006 | 1.082 | 0.199 | 0.024 | -0.02 | -0.02 | 0.143 | 0.23 | 0.36 | 0.35 | 0.19 | 0.23 | 0.2 |
| 7 | 50573325 | rs12718572 | 1.592 | -0.055 | 0.938 | 0.198 | -0.135 | -0.045 | -0.045 | 7.594 | 0.29 | 0.02 | 0.36 | 0.25 | 0.36 | 0.23 |
| 7 | 50573333 | rs7809758 | -1.341 | 0.154 | 1.335 | 0.044 | 0.009 | -0.023 | -0.023 | 0 | 0.33 | 0.34 | 0.45 | 0.2 | 0.4 | 0.35 |
| 7 | 50581845 | rs6592963 | 0.941 | 0.142 | 1.568 | 0.359 | 0.006 | -0.007 | -0.007 | 0 | 0.78 | 0.74 | 0.59 | 0.81 | 0.72 | 0.69 |
| 7 | 50598703 | rs10244632 | 1.319 | 0.177 | 1.879 | 0.276 | 0.027 | -0.015 | -0.015 | 0.692 | 0.77 | 0.55 | 0.57 | 0.81 | 0.7 | 0.67 |
| 7 | 50610071 | rs10268819 | -0.605 | 0.191 | 1.539 | 0.254 | 0.004 | -0.033 | -0.033 | 0 | 0.91 | 0.89 | 0.92 | 0.99 | 0.86 | 0.87 |
| 7 | 50610613 | rs3807553 | 0.836 | 0.237 | 1.322 | 0.307 | -0.033 | -0.032 | -0.032 | 1.211 | 0.08 | 0.01 | 0.15 | 0.07 | 0.12 | 0.15 |
| 7 | 50614173 | rs10499694 | 0.601 | 0.014 | 1.432 | 0.313 | -0.004 | 0.013 | 0.013 | 0 | 0.46 | 0.35 | 0.43 | 0.41 | 0.48 | 0.34 |
| 7 | 50615440 | rs2329371 | 1.391 | 0.025 | 1.383 | 0.175 | 0.03 | -0.023 | -0.023 | 0.055 | 0.31 | 0.22 | 0.14 | 0.39 | 0.22 | 0.32 |
| 7 | 50618876 | rs4947644 | 0.647 | 0.038 | 1.252 | 0.198 | -0.02 | -0.009 | -0.009 | 0 | 0.45 | 0.27 | 0.57 | 0.5 | 0.46 | 0.36 |
| 7 | 50627009 | rs1470747 | 1.2 | -0.085 | 0.494 | 0.192 | 0.035 | -0.037 | -0.037 | 0.536 | 0.32 | 0.19 | 0.15 | 0.4 | 0.24 | 0.32 |
| 7 | 50627101 | rs1966839 | 0.293 | 0.052 | 0.515 | 0.124 | 0.022 | -0.047 | -0.047 | 0 | 0.41 | 0.2 | 0.25 | 0.47 | 0.36 | 0.47 |
| 7 | 50632796 | rs6593011 | -0.245 | 0.188 | 0.679 | -0.398 | 0.03 | -0.076 | -0.076 | 0.81 | 0.15 | 0.43 | 0.17 | 0.03 | 0.17 | 0.16 |
| 7 | 50637429 | rs978784 | 1.01 | 0.079 | 0.142 | -0.048 | 0.021 | -0.037 | -0.037 | 0 | 0.33 | 0.25 | 0.24 | 0.51 | 0.24 | 0.32 |
| 7 | 50646280 | rs1839740 | 0.058 | 0.105 | 0.079 | -0.08 | 0.012 | 0.01 | 0.01 | 0 | 0.42 | 0.35 | 0.34 | 0.57 | 0.35 | 0.47 |
| 7 | 50656334 | rs12669770 | 0.201 | 0.118 | 0.035 | -0.053 | 0.01 | 0.007 | 0.007 | 0 | 0.41 | 0.31 | 0.34 | 0.57 | 0.35 | 0.48 |
| 7 | 50660787 | rs3807550 | 0.471 | 0.498 | -0.167 | 0.157 | -0.032 | -0.033 | -0.033 | 0 | 0.08 | 0.01 | 0.1 | 0.06 | 0.11 | 0.15 |
| 7 | 50661409 | rs4245555 | 0.806 | 0.394 | -0.632 | -0.016 | -0.082 | -0.106 | -0.106 | 0 | 0.32 | 0.04 | 0.4 | 0.46 | 0.39 | 0.36 |
| 7 | 50661661 | rs4245556 | 0.787 | 0.467 | -0.568 | 0.001 | -0.089 | -0.111 | -0.111 | 0 | 0.29 | 0.04 | 0.4 | 0.46 | 0.37 | 0.36 |
| 7 | 50661735 | rs4947406 | 1.689 | 0.893 | 0.249 | -0.158 | 0.005 | -0.025 | -0.025 | 0 | 0.72 | 0.37 | 0.68 | 0.92 | 0.65 | 0.7 |
| 7 | 50671350 | rs12718729 | -1.083 | 0.523 | 0.381 | -0.273 | 0.021 | -0.003 | -0.003 | 0 | 0.47 | 0.41 | 0.67 | 0.52 | 0.68 | 0.62 |
| 7 | 50672907 | rs4947709 | -0.975 | 0.335 | 0.264 | -0.319 | 0.008 | 0.004 | 0.004 | 0 | 0.45 | 0.7 | 0.63 | 0.49 | 0.62 | 0.5 |
| 7 | 50673029 | rs4947710 | 0.622 | 0.707 | 0.719 | -0.343 | -0.003 | 0.001 | 0.001 | 0.03 | 0.88 | 0.83 | 0.89 | 0.94 | 0.89 | 0.85 |
| 7 | 50681256 | rs2715129 | -1.769 | 0.858 | 0.666 | -0.546 | 0.029 | -0.113 | -0.113 | 4.978 | 0.18 | 0.36 | 0.14 | 0.02 | 0.23 | 0.14 |
| 7 | 50685602 | rs11770199 | -0.464 | 1.144 | -0.25 | -0.396 | -0.035 | 0.034 | 0.034 | 0.625 | 0.08 | 0.03 | 0.09 | 0.01 | 0.19 | 0.09 |
| 7 | 50688054 | rs17544225 | 0.322 | 0.95 | -0.27 | -0.308 | -0.032 | -0.02 | -0.02 | 0 | 0.08 | 0.01 | 0.1 | 0.05 | 0.11 | 0.14 |
| 7 | 50688132 | rs2250152 | -1.262 | 1.401 | -0.113 | -0.336 | 0.011 | -0.004 | -0.004 | 0 | 0.37 | 0.58 | 0.42 | 0.38 | 0.45 | 0.46 |
| 7 | 50689284 | rs2237440 | -1.854 | 1.341 | -0.932 | -0.256 | -0.058 | -0.056 | -0.056 | 0 | 0.48 | 0.97 | 0.57 | 0.43 | 0.58 | 0.61 |
| 7 | 50689952 | rs2299150 | 0.009 | 1.447 | -0.73 | 0.277 | -0.029 | -0.053 | -0.053 | 0 | 0.18 | 0.12 | 0.3 | 0.36 | 0.25 | 0.31 |
| 7 | 50696094 | rs980716 | 0.849 | 1.042 | -0.381 | -0.21 | 0 | 0 | 0 | 2.227 | 0.8 | 0.8 | 0.9 | 0.92 | 0.8 | 0.69 |
| 7 | 50697051 | rs6948959 | 2.071 | 1.156 | 0.224 | -0.042 | 0.001 | -0.018 | -0.018 | 0 | 0.76 | 0.32 | 0.8 | 0.89 | 0.75 | 0.71 |
| 7 | 50697279 | rs2529393 | 0.728 | 1.366 | 0.064 | 0.04 | -0.025 | 0.037 | 0.037 | 0 | 0.9 | 0.99 | 0.95 | 0.94 | 0.87 | 0.82 |
| 7 | 50699818 | rs17544971 | -0.593 | 1.332 | 0.123 | 0.037 | -0.005 | -0.004 | -0.004 | 0 | 0.06 | 0.05 | 0.1 | 0.05 | 0.12 | 0.11 |
| 7 | 50700153 | rs2237444 | -0.346 | 1.412 | 0.332 | 0.331 | 0.038 | 0.027 | 0.027 | 1.561 | 0.35 | 0.48 | 0.51 | 0.45 | 0.51 | 0.57 |
| 7 | 50705770 | rs6593077 | 1.1 | 1.374 | 0.702 | 0.325 | 0.022 | -0.006 | -0.006 | 1.927 | 0.53 | 0.31 | 0.43 | 0.49 | 0.36 | 0.25 |
| 7 | 50707844 | rs2237447 | 0.014 | 1.443 | 0.675 | 0.286 | 0.015 | 0.012 | 0.012 | 0.087 | 0.3 | 0.4 | 0.4 | 0.4 | 0.4 | 0.46 |
| 7 | 50709650 | rs17133917 | 0.626 | 1.48 | 0.702 | 0.289 | 0.011 | -0.004 | -0.004 | 0.312 | 0.6 | 0.5 | 0.54 | 0.54 | 0.48 | 0.36 |
| 7 | 50724555 | rs2237456 | 1.197 | 1.318 | 0.537 | 0.461 | 0.056 | 0.032 | 0.032 | 5.458 | 0.49 | 0.31 | 0.31 | 0.34 | 0.31 | 0.22 |
| 7 | 50728691 | rs2237458 | 0.102 | 1.421 | 0.477 | 0.396 | 0.02 | 0.015 | 0.015 | 0.074 | 0.3 | 0.42 | 0.4 | 0.4 | 0.39 | 0.45 |
| 7 | 50729681 | rs2237461 | 0.708 | 1.171 | 0.455 | 0.78 | -0.02 | 0.035 | 0.035 | 0 | 0.91 | 0.98 | 0.95 | 0.93 | 0.88 | 0.82 |
| 7 | 50730452 | rs2715094 | 1.324 | 1.143 | 0.474 | 0.792 | -0.032 | 0.012 | 0.012 | 0.437 | 0.84 | 0.92 | 0.84 | 0.89 | 0.76 | 0.71 |
| 7 | 50736662 | rs2529415 | -0.188 | 0.42 | 0.51 | 0.675 | 0.033 | 0.003 | 0.003 | 0.629 | 0.53 | 0.74 | 0.63 | 0.43 | 0.64 | 0.63 |
| 7 | 50739833 | rs2529410 | 0.288 | 0.456 | 0.618 | 0.666 | 0.002 | 0.016 | 0.016 | 0 | 0.47 | 0.66 | 0.53 | 0.4 | 0.53 | 0.51 |
| 7 | 50742180 | rs1800504 | 0.315 | 0.455 | 0.613 | 0.662 | 0 | 0.017 | 0.017 | 0 | 0.47 | 0.66 | 0.53 | 0.4 | 0.53 | 0.49 |
| 7 | 50742617 | rs2190503 | 0.757 | 0.372 | 0.633 | 0.605 | -0.003 | -0.027 | -0.027 | 0 | 0.89 | 0.86 | 0.92 | 0.96 | 0.9 | 0.84 |
| 7 | 50745307 | rs2237477 | 0.392 | 0.381 | 0.636 | 0.623 | 0.01 | 0.002 | 0.002 | 0.892 | 0.42 | 0.27 | 0.4 | 0.55 | 0.35 | 0.33 |
| 7 | 50745794 | rs2237478 | 1.051 | 0.384 | 0.5 | 0.632 | 0.029 | -0.046 | -0.046 | 0.006 | 0.35 | 0.18 | 0.31 | 0.51 | 0.25 | 0.21 |
| 7 | 50748639 | rs2237482 | -0.217 | 0.496 | 0.606 | 0.659 | 0.039 | 0.03 | 0.03 | 0.156 | 0.06 | 0.13 | 0.1 | 0.04 | 0.1 | 0.12 |
| 7 | 50751090 | rs10248619 | 1.732 | 0.436 | 1.007 | 0.621 | 0.02 | -0.043 | -0.043 | 0 | 0.82 | 0.52 | 0.82 | 0.92 | 0.79 | 0.72 |
| 7 | 50754679 | rs2282930 | 0.884 | 0.387 | 0.898 | 0.619 | 0.001 | -0.002 | -0.002 | 1.839 | 0.36 | 0.3 | 0.3 | 0.51 | 0.25 | 0.2 |

| | | | | | | | | | | | | | | | | |
|---|---|---|---|---|---|---|---|---|---|---|---|---|---|---|---|---|
| 7 | 50755208 | rs1004402 | 0.838 | 0.532 | 1.064 | 0.549 | 0 | 0.03 | 0.03 | 0 | 0.89 | 0.92 | 0.93 | 0.96 | 0.9 | 0.84 |
| 7 | 50758245 | rs933360 | 2.582 | 0.473 | 1.232 | 0.617 | 0.002 | 0.003 | 0.003 | 1.03 | 0.76 | 0.25 | 0.68 | 0.75 | 0.75 | 0.68 |
| 7 | 50760209 | rs2299155 | -0.544 | 0.576 | 1.039 | 0.655 | 0.001 | -0.005 | -0.005 | 0 | 0.08 | 0.07 | 0.13 | 0.16 | 0.06 | 0.05 |
| 7 | 50768087 | rs17152102 | -1.675 | 0.52 | 1.06 | 0.658 | 0 | -0.005 | -0.005 | 0.083 | 0.14 | 0.15 | 0.22 | 0.2 | 0.15 | 0.16 |
| 7 | 50786663 | rs2108349 | 2.304 | 0.468 | 0.969 | 0.619 | -0.001 | 0.015 | 0.015 | 5.82 | 0.7 | 0.23 | 0.55 | 0.71 | 0.71 | 0.65 |
| 7 | 50789189 | rs10216232 | 0.022 | 0.5 | 0.897 | 0.596 | -0.031 | -0.005 | -0.005 | 0.409 | 0.94 | 0.98 | 0.94 | 0.92 | 0.9 | 0.87 |
| 7 | 50791579 | rs6943153 | 2.198 | 0.497 | 0.957 | 0.429 | 0.002 | 0.03 | 0.03 | 5.314 | 0.7 | 0.25 | 0.56 | 0.76 | 0.7 | 0.67 |
| 7 | 50802434 | rs6968827 | 1.596 | 0.05 | 0.794 | 0.473 | -0.003 | -0.012 | -0.012 | 0 | 0.78 | 0.7 | 0.79 | 0.8 | 0.81 | 0.79 |
| 7 | 50809085 | rs13236710 | -0.947 | -0.009 | 0.332 | 0.482 | 0.022 | 0.007 | 0.007 | 7.976 | 0.14 | 0.06 | 0.13 | 0.17 | 0.09 | 0.09 |
| 7 | 50831871 | rs6979369 | 1.788 | 0.294 | 0.583 | 0.773 | 0.003 | 0.009 | 0.009 | 2.428 | 0.8 | 0.31 | 0.58 | 0.74 | 0.76 | 0.7 |
| 7 | 50835893 | rs4947861 | 1.036 | 0.299 | 0.732 | 0.778 | 0.025 | 0.031 | 0.031 | 1.153 | 0.88 | 0.78 | 0.81 | 0.76 | 0.83 | 0.81 |
| 7 | 50837004 | rs7805310 | 0.166 | 0.339 | -0.526 | 0.791 | 0.011 | 0.06 | 0.06 | 6.589 | 0.05 | 0.09 | 0.13 | 0.2 | 0.06 | 0.04 |
| 7 | 50839798 | rs6976572 | 0.196 | 0.232 | -1.577 | 0.851 | 0.008 | 0.071 | 0.071 | 0 | 0.49 | 0.26 | 0.38 | 0.34 | 0.54 | 0.51 |
| 7 | 50856792 | rs7791286 | 1.055 | 0.851 | -1.15 | 0.384 | -0.011 | -0.025 | -0.025 | 0.13 | 0.8 | 0.32 | 0.7 | 0.92 | 0.83 | 0.71 |
| 7 | 50858645 | rs6593185 | 0.715 | 0.834 | -1.272 | 0.398 | -0.196 | 0.11 | 0.11 | 0 | 0.33 | 0.02 | 0.28 | 0.13 | 0.46 | 0.22 |
| 7 | 50871909 | rs6943901 | 1.264 | 0.788 | -0.307 | 0.767 | 0.004 | 0.012 | 0.012 | 2.084 | 0.8 | 0.34 | 0.58 | 0.72 | 0.76 | 0.67 |
| 7 | 50881795 | rs1015573 | 1.15 | 0.999 | -1.089 | 0.835 | -0.006 | -0.021 | -0.021 | 0 | 0.73 | 0.12 | 0.65 | 0.73 | 0.73 | 0.66 |
| 7 | 50889126 | rs10270964 | 0.676 | 0.102 | -0.395 | 0.733 | -0.036 | -0.003 | -0.003 | 0 | 0.83 | 0.3 | 0.8 | 0.86 | 0.93 | 0.82 |
| 7 | 50894054 | rs4947923 | 0.499 | 0.896 | -0.801 | 0.925 | -0.005 | 0.03 | 0.03 | 1.812 | 0.83 | 0.34 | 0.77 | 0.79 | 0.84 | 0.79 |
| 7 | 50897276 | rs4947475 | -0.098 | 1.164 | -1.986 | 0.324 | -0.124 | 0.124 | 0.124 | 0 | 0.37 | 0.05 | 0.34 | 0.23 | 0.47 | 0.4 |
| 7 | 50901533 | rs1548608 | -1.056 | 0.915 | -1.553 | 1.013 | -0.003 | -0.018 | -0.018 | 0 | 0.13 | 0.08 | 0.28 | 0.26 | 0.18 | 0.33 |
| 7 | 50902340 | rs9649850 | -1.25 | 0.355 | -1.538 | 1.325 | 0.006 | -0.015 | -0.015 | 0 | 0.33 | 0.28 | 0.41 | 0.42 | 0.33 | 0.48 |
| 7 | 50903714 | rs13226406 | -1.223 | -0.171 | -1.757 | 1.118 | 0.027 | -0.062 | -0.062 | 1.411 | 0.36 | 0.15 | 0.4 | 0.42 | 0.35 | 0.51 |
| 7 | 50904256 | rs11238357 | 1.204 | -0.414 | -1.345 | 0.619 | -0.015 | -0.001 | -0.001 | 0 | 0.33 | 0.35 | 0.21 | 0.31 | 0.18 | 0.16 |
| 7 | 50906752 | rs4947936 | -1.245 | -0.288 | -1.584 | 0.405 | -0.174 | 0.026 | 0.026 | 0 | 0.79 | 0.98 | 0.74 | 0.82 | 0.57 | 0.79 |
| 7 | 50914179 | rs12718959 | -0.88 | -0.322 | -0.902 | 0.549 | 0 | 0 | 0 | 0 | 0.77 | 0.8 | 0.64 | 0.74 | 0.56 | 0.74 |
| 7 | 50915776 | rs17454517 | -0.403 | -1.534 | -1.448 | -0.057 | 0.024 | -0.029 | -0.029 | 0 | 0.65 | 0.47 | 0.59 | 0.74 | 0.53 | 0.72 |
| 7 | 50921069 | rs12532788 | 0.73 | -2.468 | -1.797 | -0.152 | 0.019 | -0.01 | -0.01 | 0 | 0.25 | 0.19 | 0.23 | 0.32 | 0.14 | 0.19 |
| 7 | 50924695 | rs2876900 | 0.962 | -2.552 | -1.844 | -0.145 | 0.032 | -0.056 | -0.056 | 0 | 0.22 | 0.03 | 0.13 | 0.28 | 0.13 | 0.17 |
| 7 | 50929594 | rs6593252 | -1.818 | -2.297 | -1.599 | -0.405 | -0.101 | 0.037 | 0.037 | 0 | 0.79 | 0.91 | 0.73 | 0.82 | 0.57 | 0.8 |
| 7 | 50932580 | rs2161825 | -1.14 | -2.528 | -0.757 | -0.452 | 0.055 | -0.014 | -0.014 | 0 | 0.73 | 0.72 | 0.58 | 0.74 | 0.54 | 0.72 |
| 7 | 50933120 | rs17152188 | 0.531 | -3.032 | -0.63 | -0.2 | 0.004 | -0.021 | -0.021 | 0 | 0.35 | 0.1 | 0.2 | 0.31 | 0.29 | 0.41 |
| 7 | 50933370 | rs10264737 | -1.247 | -2.898 | -0.455 | -0.239 | -0.014 | 0.009 | 0.009 | 0.296 | 0.69 | 0.69 | 0.68 | 0.82 | 0.55 | 0.79 |
| 7 | 50933678 | rs10274003 | -0.011 | -2.877 | -0.457 | -0.205 | -0.012 | 0.055 | 0.055 | 0 | 0.1 | 0.33 | 0.16 | 0.08 | 0.03 | 0.09 |
| 7 | 50938948 | rs11531636 | -0.689 | -2.821 | -0.31 | -0.17 | 0.018 | -0.014 | -0.014 | 0 | 0.65 | 0.5 | 0.49 | 0.68 | 0.45 | 0.67 |
| 7 | 50948855 | rs10249897 | -1.405 | -2.985 | -0.453 | -0.243 | -0.049 | -0.014 | -0.014 | 0.148 | 0.76 | 0.79 | 0.54 | 0.68 | 0.48 | 0.69 |
| 7 | 50954338 | rs10258680 | 0.381 | -3.092 | -0.388 | -0.274 | -0.022 | 0.014 | 0.014 | 0 | 0.09 | 0.19 | 0.15 | 0.08 | 0.04 | 0.1 |
| 7 | 50981344 | rs6977927 | 1.64 | -3.054 | -0.348 | -0.359 | -0.039 | -0.056 | -0.056 | 0.478 | 0.1 | 0.21 | 0.05 | 0 | 0.03 | 0.01 |
| 7 | 50984237 | rs7800732 | -1.601 | -2.765 | -0.583 | -0.218 | -0.11 | -0.044 | -0.044 | 0 | 0.76 | 0.88 | 0.55 | 0.67 | 0.48 | 0.69 |
| 7 | 50989383 | rs4948034 | 1.261 | -2.935 | -0.717 | -0.196 | 0.048 | -0.067 | -0.067 | 2.676 | 0.22 | 0.04 | 0.12 | 0.3 | 0.13 | 0.17 |
| 7 | 50995578 | rs9642409 | -0.384 | -2.5 | -0.494 | -0.069 | 0.002 | 0.001 | 0.001 | 0.225 | 0.67 | 0.66 | 0.39 | 0.59 | 0.44 | 0.59 |
| 7 | 51012673 | rs2189820 | -0.464 | -2.841 | -0.315 | 0.01 | 0.02 | -0.001 | -0.001 | 0 | 0.37 | 0.62 | 0.43 | 0.37 | 0.24 | 0.33 |
| 7 | 51012828 | rs7458079 | -1.375 | -2.563 | -0.737 | -0.141 | -0.063 | 0.025 | 0.025 | 0 | 0.66 | 0.89 | 0.59 | 0.71 | 0.43 | 0.71 |
| 7 | 51024118 | rs6593312 | -0.126 | -1.816 | -0.327 | -0.025 | 0 | 0.009 | 0.009 | 0 | 0.69 | 0.65 | 0.57 | 0.63 | 0.72 | 0.61 |
| 7 | 51025937 | rs10486748 | -0.261 | -1.668 | -0.14 | -0.001 | 0.021 | 0.011 | 0.011 | 0 | 0.9 | 0.77 | 0.76 | 0.92 | 0.93 | 0.85 |
| 7 | 51029101 | rs6942987 | 0.658 | -1.789 | -0.074 | -0.009 | 0.006 | 0.016 | 0.016 | 0 | 0.08 | 0.19 | 0.16 | 0.08 | 0.04 | 0.09 |
| 7 | 51029371 | rs4948092 | 0.823 | -1.902 | -0.152 | -0.001 | -0.011 | -0.036 | -0.036 | 0 | 0.21 | 0.09 | 0.18 | 0.29 | 0.22 | 0.24 |
| 7 | 51041748 | rs7788787 | 0.96 | -1.719 | 0.145 | 0.168 | -0.007 | 0 | 0 | 0.155 | 0.08 | 0.18 | 0.15 | 0.06 | 0.04 | 0.09 |
| 7 | 51043153 | rs17656156 | 0.673 | -1.213 | -0.092 | -0.01 | -0.011 | -0.028 | -0.028 | 1.915 | 0.26 | 0.06 | 0.21 | 0.3 | 0.25 | 0.29 |
| 7 | 51046552 | rs11971137 | 0.769 | -1.854 | 0.371 | -0.369 | -0.082 | -0.1 | -0.1 | 0 | 0.15 | 0.37 | 0.04 | 0 | 0.03 | 0.05 |
| 7 | 51049803 | rs17656198 | 2.336 | -1.642 | -0.019 | -0.292 | -0.155 | 0.003 | 0.003 | 0 | 0.28 | 0.03 | 0.29 | 0.23 | 0.42 | 0.21 |
| 7 | 51051491 | rs12216580 | 0.924 | -2.233 | 0.522 | -0.346 | 0.058 | 0.051 | 0.051 | 0.899 | 0.49 | 0.3 | 0.51 | 0.63 | 0.66 | 0.52 |
| 7 | 51053326 | rs916936 | -0.888 | -1.797 | 0.562 | -0.291 | 0.005 | 0.002 | 0.002 | 0 | 0.9 | 0.75 | 0.76 | 0.93 | 0.95 | 0.87 |
| 7 | 51057161 | rs12537931 | 0.566 | -1.846 | 0.668 | -0.322 | -0.056 | -0.065 | -0.065 | 1.553 | 0.78 | 0.63 | 0.73 | 0.93 | 0.91 | 0.81 |
| 7 | 51058134 | rs2329572 | 1.407 | -1.493 | 0.666 | -0.151 | -0.001 | 0.011 | 0.011 | 0 | 0.08 | 0.16 | 0.16 | 0.07 | 0.05 | 0.1 |
| 7 | 51061987 | rs11765623 | 0.171 | -1.577 | 0.495 | -0.176 | -0.025 | -0.022 | -0.022 | 0 | 0.14 | 0.21 | 0.17 | 0.07 | 0.07 | 0.15 |
| 7 | 51062256 | rs17798871 | 0.972 | -1.717 | -0.176 | -0.131 | -0.047 | -0.055 | -0.055 | 0 | 0.25 | 0.01 | 0.2 | 0.29 | 0.24 | 0.28 |
| 7 | 51063910 | rs2051820 | -0.104 | -1.748 | 0.177 | -0.154 | -0.033 | -0.047 | -0.047 | 0.265 | 0.83 | 0.55 | 0.71 | 0.93 | 0.91 | 0.81 |
| 7 | 51065188 | rs7802321 | 0.817 | -1.782 | 0.391 | -0.169 | -0.001 | -0.001 | -0.001 | 0 | 0.26 | 0.26 | 0.22 | 0.28 | 0.24 | 0.29 |
| 7 | 51068073 | rs970907 | 0.767 | -1.719 | -0.445 | -0.141 | 0.037 | 0.036 | 0.036 | 0 | 0.57 | 0.28 | 0.49 | 0.64 | 0.67 | 0.53 |
| 7 | 51074440 | rs17725940 | 0.721 | -1.222 | -0.767 | 0.062 | -0.003 | -0.014 | -0.014 | 0 | 0.25 | 0.05 | 0.21 | 0.28 | 0.24 | 0.29 |
| 7 | 51077711 | rs10486749 | -0.328 | -1.288 | -0.523 | 0.298 | -0.037 | 0.027 | 0.027 | 1.014 | 0.34 | 0.49 | 0.32 | 0.39 | 0.21 | 0.4 |
| 7 | 51077759 | rs757323 | 1.068 | -0.754 | -0.562 | 0.229 | -0.021 | 0.002 | 0.002 | 0 | 0.33 | 0.3 | 0.37 | 0.33 | 0.52 | 0.27 |
| 7 | 51081097 | rs17134124 | 0.676 | -0.805 | 0.123 | 0.442 | -0.007 | 0.001 | 0.001 | 0.02 | 0.11 | 0.28 | 0.21 | 0.08 | 0.07 | 0.11 |
| 7 | 51082475 | rs6956095 | 1.139 | -0.58 | -0.481 | 0.112 | -0.002 | 0.108 | 0.108 | 0 | 0.48 | 0.16 | 0.43 | 0.34 | 0.53 | 0.41 |
| 7 | 51084321 | rs1046964 | -0.546 | -0.076 | -0.173 | 0.783 | 0.001 | 0.028 | 0.028 | 0 | 0.9 | 0.92 | 0.85 | 0.72 | 0.85 | 0.8 |
| 7 | 51090065 | rs999605 | -0.685 | -0.542 | 0.235 | 0.443 | -0.041 | -0.061 | -0.061 | 0 | 0.23 | 0.46 | 0.27 | 0.11 | 0.16 | 0.23 |
| 7 | 51090935 | rs2240086 | -0.269 | -0.574 | 0.251 | 0.682 | -0.001 | 0.001 | 0.001 | 0.195 | 0.22 | 0.19 | 0.26 | 0.31 | 0.22 | 0.3 |
| 7 | 51096974 | rs2240090 | 0.354 | -0.578 | 0.135 | 0.834 | 0.002 | -0.003 | -0.003 | 0.099 | 0.63 | 0.66 | 0.59 | 0.46 | 0.69 | 0.6 |

| | | | | | | | | | | | | | | | | |
|---|---|---|---|---|---|---|---|---|---|---|---|---|---|---|---|---|
| 7 | 51102820 | rs2240093 | -0.902 | -1.11 | 0.155 | 0.867 | -0.012 | 0.008 | 0.008 | 0 | 0.39 | 0.53 | 0.38 | 0.46 | 0.29 | 0.49 |
| 7 | 51104236 | rs2285843 | -0.859 | -0.736 | 0.499 | 1.179 | -0.012 | 0.077 | 0.077 | 0 | 0.28 | 0.47 | 0.36 | 0.46 | 0.23 | 0.41 |
| 7 | 51121341 | rs1898449 | -0.867 | -0.777 | 0.32 | 1.16 | -0.013 | 0.066 | 0.066 | 0 | 0.29 | 0.51 | 0.36 | 0.46 | 0.24 | 0.38 |
| 7 | 51130351 | rs6970066 | -0.938 | -0.893 | -0.398 | 0.817 | 0.015 | 0.009 | 0.009 | 0 | 0.3 | 0.76 | 0.34 | 0.22 | 0.2 | 0.38 |
| 7 | 51143605 | rs1437490 | -0.206 | -0.662 | -0.472 | 1.114 | -0.002 | 0.003 | 0.003 | 0 | 0.57 | 0.54 | 0.45 | 0.25 | 0.48 | 0.46 |
| 7 | 51145573 | rs17554286 | 0.612 | -1.31 | -0.709 | 0.494 | 0.002 | 0.038 | 0.038 | 3.184 | 0.26 | 0.15 | 0.22 | 0.12 | 0.27 | 0.24 |
| 7 | 51146550 | rs6950779 | 1.686 | -0.98 | -1.002 | 0.144 | -0.045 | 0.042 | 0.042 | 0 | 0.23 | 0.2 | 0.23 | 0.15 | 0.36 | 0.3 |
| 7 | 51148028 | rs7803708 | -1.679 | -0.976 | -0.881 | 0.292 | 0.015 | -0.024 | -0.024 | 0 | 0.84 | 0.9 | 0.76 | 0.7 | 0.88 | 0.79 |
| 7 | 51150108 | rs17152220 | 1.12 | -1.046 | -0.565 | -0.057 | -0.011 | 0.023 | 0.023 | 0.545 | 0.22 | 0.12 | 0.13 | 0.02 | 0.2 | 0.09 |
| 7 | 51151231 | rs1545125 | 0.059 | -1.403 | -0.583 | 0.266 | 0.001 | 0.011 | 0.011 | 0 | 0.44 | 0.56 | 0.37 | 0.26 | 0.39 | 0.48 |
| 7 | 51151661 | rs2289175 | -1.225 | -1.689 | -0.762 | 0.537 | -0.027 | -0.014 | -0.014 | 0 | 0.73 | 0.77 | 0.7 | 0.61 | 0.62 | 0.7 |
| 7 | 51152063 | rs2289177 | -0.662 | -2.089 | -0.655 | 0.552 | -0.008 | -0.005 | -0.005 | 0.461 | 0.67 | 0.65 | 0.56 | 0.42 | 0.54 | 0.65 |
| 7 | 51156296 | rs6593325 | 0.524 | -2.101 | -0.655 | 0.334 | 0 | 0.001 | 0.001 | 0 | 0.41 | 0.56 | 0.36 | 0.27 | 0.35 | 0.35 |
| 7 | 51158612 | rs10156067 | -0.588 | -2.112 | -0.634 | 0.33 | -0.031 | 0.003 | 0.003 | 0.766 | 0.66 | 0.71 | 0.51 | 0.3 | 0.54 | 0.5 |
| 7 | 51165130 | rs10899772 | -1.003 | -2.101 | -0.637 | 0.381 | -0.048 | -0.02 | -0.02 | 0 | 0.7 | 0.8 | 0.61 | 0.59 | 0.59 | 0.53 |
| 7 | 51167232 | rs4948190 | 0.432 | -2.005 | -0.573 | 0.255 | 0 | 0 | 0 | 0 | 0.4 | 0.56 | 0.36 | 0.27 | 0.37 | 0.36 |
| 7 | 51170630 | rs6957736 | 1.359 | -1.827 | -0.652 | 0.031 | 0.004 | -0.017 | -0.017 | 0.003 | 0.15 | 0.13 | 0.08 | 0 | 0.09 | 0.03 |
| 7 | 51182498 | rs4947572 | 0.17 | -1.983 | -0.315 | 0.126 | 0 | -0.002 | -0.002 | 0 | 0.44 | 0.42 | 0.43 | 0.57 | 0.4 | 0.29 |
| 7 | 51187785 | rs4416783 | 0.957 | -1.651 | -0.277 | 0.171 | -0.004 | 0.002 | 0.002 | 0.094 | 0.27 | 0.26 | 0.23 | 0.16 | 0.38 | 0.38 |
| 7 | 51196639 | rs4948194 | -0.637 | -1.68 | -0.228 | 0.196 | 0.015 | 0.008 | 0.008 | 0.486 | 0.7 | 0.62 | 0.58 | 0.59 | 0.58 | 0.55 |
| 7 | 51201878 | rs6593335 | -0.775 | -1.784 | 0.037 | 0.458 | -0.023 | 0.004 | 0.004 | 0 | 0.61 | 0.7 | 0.54 | 0.59 | 0.51 | 0.48 |
| 7 | 51207720 | rs4948195 | 1.796 | -1.594 | 0.507 | 0.818 | 0.002 | 0.001 | 0.001 | 0 | 0.08 | 0.06 | 0.17 | 0.07 | 0.09 | 0.04 |
| 7 | 51208903 | rs10256402 | 1.622 | -1.6 | 0.571 | 0.82 | -0.006 | -0.003 | -0.003 | 0 | 0.09 | 0.09 | 0.17 | 0.07 | 0.09 | 0.07 |
| 7 | 51213607 | rs10231416 | 0.53 | -1.658 | 0.791 | 0.903 | -0.037 | 0.036 | 0.036 | 0.054 | 0.12 | 0.36 | 0.19 | 0.14 | 0.09 | 0.18 |
| 7 | 51230648 | rs2118306 | 0.136 | -1.793 | 0.611 | 0.801 | -0.046 | -0.021 | -0.021 | 0.138 | 0.15 | 0.29 | 0.09 | 0.06 | 0.08 | 0.25 |
| 7 | 51240364 | rs10236935 | 0.886 | -1.78 | 0.528 | 0.725 | -0.024 | -0.049 | -0.049 | 0 | 0.12 | 0.19 | 0.08 | 0 | 0.08 | 0.06 |
| 7 | 51251544 | rs10280348 | 0.837 | -1.784 | 0.462 | 0.723 | -0.012 | -0.032 | -0.032 | 0.084 | 0.12 | 0.15 | 0.07 | 0 | 0.08 | 0.06 |
| 7 | 51252719 | rs2043733 | 1.077 | -1.92 | 0.138 | 0.771 | 0.038 | 0.024 | 0.024 | 0 | 0.52 | 0.28 | 0.43 | 0.4 | 0.59 | 0.4 |
| 7 | 51277376 | rs801139 | -0.141 | -1.466 | 0.371 | 0.73 | 0 | 0.002 | 0.002 | 0 | 0.72 | 0.76 | 0.8 | 0.79 | 0.74 | 0.72 |
| 7 | 51287470 | rs801142 | -0.146 | -1.459 | 0.336 | 0.765 | 0 | 0.002 | 0.002 | 0 | 0.72 | 0.76 | 0.8 | 0.78 | 0.74 | 0.71 |
| 7 | 51289518 | rs1089285 | -1.635 | -1.402 | 0.839 | 1.193 | -0.027 | 0.172 | 0.172 | 0.134 | 0.13 | 0.42 | 0.22 | 0.34 | 0.07 | 0.29 |
| 7 | 51291926 | rs801144 | -0.479 | -1.228 | 0.814 | 1.143 | 0.014 | 0.207 | 0.207 | 0.057 | 0.14 | 0.31 | 0.3 | 0.54 | 0.12 | 0.11 |
| 7 | 51292344 | rs9649865 | 0.902 | -1.108 | 0.318 | 0.705 | -0.042 | 0.15 | 0.15 | 0 | 0.43 | 0.23 | 0.41 | 0.17 | 0.5 | 0.37 |
| 7 | 51295543 | rs887058 | -0.06 | -0.985 | 0.352 | 0.715 | 0.001 | 0.018 | 0.018 | 0 | 0.6 | 0.67 | 0.72 | 0.77 | 0.61 | 0.65 |
| 7 | 51300075 | rs3815336 | -0.286 | -1 | 0.342 | 0.666 | -0.024 | 0.08 | 0.08 | 0 | 0.63 | 0.84 | 0.74 | 0.78 | 0.63 | 0.68 |
| 7 | 51300561 | rs4948204 | 0.259 | -1.092 | 0.647 | 0.888 | 0.003 | 0.014 | 0.014 | 0 | 0.57 | 0.66 | 0.68 | 0.65 | 0.61 | 0.4 |
| 7 | 51303534 | rs2215162 | -0.382 | -1.176 | 0.41 | 0.806 | -0.096 | 0.102 | 0.102 | 0 | 0.65 | 0.98 | 0.76 | 0.82 | 0.63 | 0.67 |
| 7 | 51309769 | rs4948209 | -0.28 | -1.373 | 0.814 | 0.73 | -0.029 | 0.112 | 0.112 | 1.902 | 0.22 | 0.47 | 0.45 | 0.62 | 0.18 | 0.13 |
| 7 | 51315142 | rs12673454 | 0.032 | -1.333 | 0.694 | 0.742 | -0.011 | 0.088 | 0.088 | 0.897 | 0.19 | 0.28 | 0.41 | 0.55 | 0.18 | 0.1 |
| 7 | 51315315 | rs1860160 | 0.969 | -1.3 | 0.22 | 0.486 | -0.069 | 0.068 | 0.068 | 0.17 | 0.26 | 0.01 | 0.15 | 0.12 | 0.26 | 0.26 |
| 7 | 51319328 | rs2215164 | -0.598 | -1.218 | 0.269 | 0.837 | -0.031 | 0.121 | 0.121 | 0 | 0.62 | 0.86 | 0.74 | 0.82 | 0.62 | 0.62 |
| 7 | 51319875 | rs1001148 | -0.608 | -1.216 | 0.27 | 0.835 | -0.031 | 0.118 | 0.118 | 0 | 0.62 | 0.87 | 0.74 | 0.82 | 0.62 | 0.62 |
| 7 | 51327575 | rs11767820 | 0.898 | -0.692 | 0.319 | 1.017 | -0.136 | 0.095 | 0.095 | 0.676 | 0.34 | 0.04 | 0.25 | 0.15 | 0.42 | 0.3 |
| 7 | 51341309 | rs917430 | -0.219 | -0.737 | 0.549 | 1.33 | -0.007 | 0.002 | 0.002 | 0.112 | 0.47 | 0.67 | 0.51 | 0.55 | 0.49 | 0.58 |
| 7 | 51342600 | rs13222665 | 0.795 | -0.491 | -0.241 | 1.124 | -0.147 | 0.089 | 0.089 | 0.676 | 0.34 | 0.03 | 0.25 | 0.15 | 0.42 | 0.3 |
| 7 | 51343606 | rs740545 | 0.761 | -0.491 | -0.208 | 1.124 | -0.102 | 0.107 | 0.107 | 0.654 | 0.34 | 0.06 | 0.25 | 0.15 | 0.42 | 0.3 |
| 7 | 51375461 | rs17756186 | 0.611 | -0.378 | -0.258 | 1.13 | -0.044 | 0.049 | 0.049 | 0.607 | 0.34 | 0.15 | 0.26 | 0.21 | 0.42 | 0.47 |
| 7 | 51375641 | rs12673911 | 0.942 | -0.288 | -0.445 | 1.088 | -0.128 | 0.078 | 0.078 | 0.943 | 0.32 | 0.05 | 0.25 | 0.16 | 0.42 | 0.28 |
| 7 | 51376184 | rs12719052 | 0.942 | -0.276 | -0.445 | 1.088 | -0.131 | 0.078 | 0.078 | 0.943 | 0.32 | 0.05 | 0.25 | 0.16 | 0.42 | 0.28 |
| 7 | 51377189 | rs12532786 | 0.927 | -0.301 | -0.434 | 1.089 | -0.086 | 0.086 | 0.086 | 0.955 | 0.32 | 0.08 | 0.25 | 0.16 | 0.42 | 0.28 |
| 7 | 51393774 | rs11981993 | 0.574 | -0.145 | -0.138 | 1.506 | -0.001 | 0.056 | 0.056 | 0 | 0.92 | 0.91 | 0.81 | 0.7 | 0.93 | 0.92 |
| 7 | 51400718 | rs2329727 | 0.948 | -0.161 | -0.144 | 1.501 | -0.002 | 0.063 | 0.063 | 0.007 | 0.91 | 0.9 | 0.81 | 0.65 | 0.93 | 0.75 |
| 7 | 51413456 | rs873209 | -0.139 | -0.297 | -0.287 | 1.216 | -0.044 | -0.042 | -0.042 | 0.256 | 0.59 | 0.8 | 0.55 | 0.51 | 0.49 | 0.43 |
| 7 | 51416721 | rs10253920 | -0.815 | -0.312 | 0.601 | 1.246 | 0.077 | 0.148 | 0.148 | 0.098 | 0.06 | 0.44 | 0.07 | 0.08 | 0.02 | 0.07 |
| 7 | 51423526 | rs6969097 | 0.179 | -0.359 | 0.455 | 1.212 | -0.002 | 0.001 | 0.001 | 1.294 | 0.54 | 0.41 | 0.49 | 0.42 | 0.47 | 0.38 |
| 7 | 51425884 | rs10254410 | -0.917 | -0.353 | 0.663 | 1.551 | 0.018 | 0.256 | 0.256 | 0 | 0.1 | 0.37 | 0.16 | 0.35 | 0.06 | 0.11 |
| 7 | 51428391 | rs1880435 | 0.534 | -0.465 | 0.24 | 1.137 | 0.012 | 0.043 | 0.043 | 0.232 | 0.47 | 0.34 | 0.3 | 0.15 | 0.38 | 0.38 |
| 7 | 51428616 | rs930363 | -1.186 | -0.402 | -0.148 | 1.323 | -0.052 | 0.047 | 0.047 | 0.592 | 0.71 | 0.85 | 0.76 | 0.8 | 0.57 | 0.74 |
| 7 | 51435507 | rs7349963 | 0.746 | -0.243 | 0.054 | 1.63 | -0.006 | 0.059 | 0.059 | 0 | 0.1 | 0.13 | 0.3 | 0.3 | 0.09 | 0.08 |
| 7 | 51436113 | rs1288312 | -1.251 | -0.16 | -0.106 | 1.462 | 0 | 0.041 | 0.041 | 0.235 | 0.27 | 0.38 | 0.46 | 0.52 | 0.25 | 0.2 |
| 7 | 51442799 | rs581221 | -0.155 | -0.491 | -0.346 | 1.155 | 0.002 | -0.001 | -0.001 | 0 | 0.53 | 0.51 | 0.46 | 0.53 | 0.62 | 0.6 |
| 7 | 51498968 | rs4507720 | 0.935 | 0.156 | -1.085 | 0.785 | -0.058 | 0.068 | 0.068 | 2.206 | 0.28 | 0.15 | 0.34 | 0.11 | 0.38 | 0.19 |
| 7 | 51500101 | rs12056030 | 0.616 | -0.315 | -1.34 | 1.02 | -0.009 | 0.07 | 0.07 | 0.041 | 0.47 | 0.11 | 0.36 | 0.37 | 0.43 | 0.47 |
| 7 | 51516977 | rs12533007 | -0.403 | -0.252 | -1 | 1.252 | -0.035 | 0.02 | 0.02 | 0 | 0.6 | 0.75 | 0.56 | 0.68 | 0.5 | 0.7 |
| 7 | 51519404 | rs12718496 | 0.205 | -0.288 | -0.897 | 1.377 | -0.031 | 0.001 | 0.001 | 2.798 | 0.4 | 0.73 | 0.54 | 0.38 | 0.46 | 0.3 |
| 7 | 51519839 | rs10250139 | -0.731 | -0.359 | -0.498 | 1.369 | -0.018 | 0.229 | 0.229 | 0.036 | 0.12 | 0.5 | 0.2 | 0.32 | 0.08 | 0.22 |
| 7 | 51520250 | rs12540310 | 0.607 | -0.32 | -0.855 | 1.35 | 0.019 | 0.088 | 0.088 | 0 | 0.48 | 0.21 | 0.36 | 0.36 | 0.43 | 0.48 |
| 7 | 51522441 | rs6962123 | -0.396 | -0.082 | -0.523 | 1.302 | -0.046 | 0.022 | 0.022 | 0 | 0.61 | 0.84 | 0.57 | 0.68 | 0.5 | 0.7 |
| 7 | 51528902 | rs12537855 | 1.196 | 0.152 | -0.565 | 1.205 | -0.046 | 0.06 | 0.06 | 1.909 | 0.28 | 0.1 | 0.34 | 0.11 | 0.35 | 0.2 |

| | | | | | | | | | | | | | | | | |
|---|---|---|---|---|---|---|---|---|---|---|---|---|---|---|---|---|
| 7 | 51532411 | rs1880428 | -0.584 | 0.17 | -0.561 | 1.463 | -0.01 | -0.015 | -0.015 | 0.394 | 0.9 | 0.91 | 0.92 | 0.8 | 0.89 | 0.9 |
| 7 | 51537887 | rs1949829 | 0.326 | -0.292 | -0.408 | 1.278 | -0.007 | 0.169 | 0.169 | 0.01 | 0.09 | 0.26 | 0.16 | 0.27 | 0.05 | 0.1 |
| 7 | 51542081 | rs1316471 | 0.914 | -0.295 | -1.372 | 0.615 | -0.013 | -0.011 | -0.011 | 0 | 0.09 | 0.01 | 0.13 | 0.06 | 0.11 | 0.12 |
| 7 | 51542925 | rs10236091 | 1.344 | -0.238 | -1.375 | 0.471 | -0.002 | -0.001 | -0.001 | 2.779 | 0.28 | 0.24 | 0.33 | 0.11 | 0.34 | 0.21 |
| 7 | 51544056 | rs17734557 | -0.002 | -0.614 | -1.847 | 0.745 | 0.003 | -0.04 | -0.04 | 0 | 0.14 | 0.02 | 0.15 | 0.11 | 0.13 | 0.24 |
| 7 | 51546823 | rs7807708 | 0.852 | -0.614 | -2.035 | 0.674 | 0.006 | -0.1 | -0.1 | 0 | 0.24 | 0.06 | 0.26 | 0.41 | 0.2 | 0.28 |
| 7 | 51547147 | rs6592926 | -0.106 | -0.105 | -1.396 | 0.876 | 0.001 | 0.019 | 0.019 | 0.235 | 0.89 | 0.85 | 0.9 | 0.73 | 0.88 | 0.78 |
| 7 | 51550923 | rs17555162 | 0.563 | -0.112 | -1.412 | 0.6 | -0.009 | -0.007 | -0.007 | 0 | 0.08 | 0.01 | 0.05 | 0.12 | 0.07 | 0.07 |
| 7 | 51553380 | rs4947604 | -0.665 | -0.259 | -1.435 | 0.514 | 0.001 | -0.004 | -0.004 | 0 | 0.82 | 0.86 | 0.87 | 0.77 | 0.83 | 0.89 |
| 7 | 51553397 | rs4947605 | -0.079 | -0.301 | -1.54 | 0.488 | 0.001 | -0.015 | -0.015 | 0 | 0.78 | 0.85 | 0.81 | 0.71 | 0.79 | 0.82 |
| 7 | 51553479 | rs4947606 | -0.057 | -0.315 | -1.585 | 0.633 | 0.007 | -0.039 | -0.039 | 0 | 0.78 | 0.85 | 0.82 | 0.66 | 0.82 | 0.71 |
| 7 | 51555658 | rs12718499 | 0.318 | -0.676 | -1.229 | 0.246 | 0.008 | 0.075 | 0.075 | 0.006 | 0.48 | 0.39 | 0.41 | 0.24 | 0.52 | 0.4 |
| 7 | 51557339 | rs17134279 | 0.849 | -0.641 | -0.848 | 0.367 | -0.008 | 0.022 | 0.022 | 0 | 0.28 | 0.3 | 0.39 | 0.42 | 0.27 | 0.29 |
| 7 | 51567327 | rs1917283 | 0.45 | -0.503 | -1.345 | -0.167 | -0.03 | 0.101 | 0.101 | 0 | 0.25 | 0.04 | 0.17 | 0.13 | 0.27 | 0.22 |
| 7 | 51567805 | rs7812201 | -0.71 | -0.535 | -1.174 | -0.062 | 0.001 | -0.007 | -0.007 | 0 | 0.76 | 0.73 | 0.79 | 0.71 | 0.77 | 0.82 |
| 7 | 51569418 | rs1917286 | 1.184 | -0.576 | -1.013 | -0.051 | -0.013 | 0.04 | 0.04 | 0 | 0.25 | 0.29 | 0.36 | 0.41 | 0.21 | 0.26 |
| 7 | 51577465 | rs1574717 | 0.491 | -0.551 | -1.12 | -0.065 | -0.029 | 0.098 | 0.098 | 0 | 0.25 | 0.04 | 0.17 | 0.13 | 0.27 | 0.23 |
| 7 | 51578632 | rs4947614 | -0.548 | -0.503 | -0.895 | 0.087 | 0.002 | -0.011 | -0.011 | 0 | 0.75 | 0.72 | 0.78 | 0.66 | 0.77 | 0.69 |
| 7 | 51582644 | rs7798551 | -0.796 | -0.447 | -1.064 | 0.077 | -0.003 | -0.013 | -0.013 | 0 | 0.77 | 0.82 | 0.8 | 0.71 | 0.77 | 0.83 |
| 7 | 51582989 | rs7776713 | 0.695 | -0.423 | -0.995 | 0.079 | -0.003 | 0.026 | 0.026 | 0 | 0.3 | 0.33 | 0.41 | 0.41 | 0.31 | 0.3 |
| 7 | 51612031 | rs11238125 | 1.523 | -0.086 | -1.055 | 0.149 | -0.049 | 0.035 | 0.035 | 0 | 0.2 | 0.07 | 0.22 | 0.11 | 0.26 | 0.2 |
| 7 | 51613687 | rs7778403 | 0.589 | -0.316 | -1.202 | 0.121 | -0.025 | 0.089 | 0.089 | 0 | 0.24 | 0.04 | 0.16 | 0.12 | 0.26 | 0.22 |
| 7 | 51615505 | rs4557653 | 1.121 | -0.269 | -0.696 | 0.14 | -0.002 | 0.001 | 0.001 | 0 | 0.25 | 0.21 | 0.26 | 0.15 | 0.29 | 0.31 |
| 7 | 51618352 | rs12533589 | 1.578 | -0.26 | -0.772 | 0.142 | -0.109 | 0.02 | 0.02 | 0 | 0.22 | 0.01 | 0.24 | 0.12 | 0.28 | 0.18 |
| 7 | 51619613 | rs10081251 | 0.703 | -0.292 | -0.556 | 0.391 | -0.007 | 0.004 | 0.004 | 0 | 0.25 | 0.16 | 0.19 | 0.26 | 0.26 | 0.23 |
| 7 | 51629362 | rs4947617 | 1.115 | -0.352 | -0.331 | 0.453 | -0.018 | 0.077 | 0.077 | 0.673 | 0.23 | 0.29 | 0.35 | 0.44 | 0.19 | 0.27 |
| 7 | 51633217 | rs1860464 | -1.231 | -0.172 | -0.656 | 0.328 | -0.014 | 0.025 | 0.025 | 0 | 0.82 | 0.91 | 0.84 | 0.85 | 0.81 | 0.88 |
| 7 | 51637202 | rs4947375 | -0.272 | -0.17 | -0.249 | 0.299 | 0.011 | 0.02 | 0.02 | 0.106 | 0.56 | 0.41 | 0.47 | 0.37 | 0.59 | 0.48 |
| 7 | 51637583 | rs1159835 | 1.464 | -0.134 | -0.327 | 0.252 | -0.042 | 0.035 | 0.035 | 0.314 | 0.22 | 0.06 | 0.23 | 0.12 | 0.28 | 0.18 |
| 7 | 51651873 | rs2191253 | -0.403 | -0.175 | -0.3 | 0.284 | 0.019 | 0.031 | 0.031 | 0.281 | 0.54 | 0.32 | 0.46 | 0.39 | 0.59 | 0.48 |
| 7 | 51654040 | rs2215331 | -0.375 | -0.164 | -0.222 | 0.289 | -0.003 | -0.004 | -0.004 | 0 | 0.56 | 0.51 | 0.48 | 0.39 | 0.59 | 0.48 |
| 7 | 51657242 | rs2329828 | 0.424 | 0.101 | -0.24 | 0.356 | -0.003 | -0.02 | -0.02 | 2.603 | 0.08 | 0.1 | 0.03 | 0 | 0.05 | 0.07 |
| 7 | 51658069 | rs6953923 | 0.151 | 0.096 | -0.384 | 0.382 | -0.008 | 0.004 | 0.004 | 0 | 0.25 | 0.19 | 0.19 | 0.26 | 0.27 | 0.23 |
| 7 | 51660226 | rs758671 | -0.948 | 0.129 | -0.437 | 0.397 | -0.01 | -0.015 | -0.015 | 0 | 0.39 | 0.5 | 0.26 | 0.3 | 0.35 | 0.43 |
| 7 | 51668274 | rs4628217 | 0.009 | 0.773 | -0.825 | 0.945 | -0.025 | -0.009 | -0.009 | 0 | 0.23 | 0.1 | 0.18 | 0.26 | 0.27 | 0.22 |
| 7 | 51673344 | rs4947623 | -0.215 | 0.763 | -0.657 | 1.272 | -0.003 | 0.013 | 0.013 | 0 | 0.69 | 0.52 | 0.62 | 0.67 | 0.69 | 0.67 |
| 7 | 51673984 | rs12718516 | 0.811 | 0.869 | -0.871 | 1.19 | 0.001 | 0.004 | 0.004 | 0 | 0.46 | 0.46 | 0.44 | 0.42 | 0.41 | 0.46 |
| 7 | 51674137 | rs6979000 | 0.305 | 1.107 | -0.784 | 1.222 | 0.003 | 0.002 | 0.002 | 0 | 0.71 | 0.63 | 0.64 | 0.71 | 0.69 | 0.8 |
| 7 | 51674838 | rs1860466 | -0.014 | 1.143 | -1.262 | 1.361 | 0.003 | 0.001 | 0.001 | 0 | 0.49 | 0.46 | 0.31 | 0.56 | 0.41 | 0.52 |
| 7 | 51675218 | rs13241700 | -0.994 | 1.393 | -1.528 | 1.713 | -0.044 | -0.059 | -0.059 | 0 | 0.09 | 0.01 | 0.09 | 0.25 | 0.14 | 0.11 |
| 7 | 51679299 | rs17578222 | -0.409 | 1.304 | -1.723 | 1.577 | -0.031 | -0.064 | -0.064 | 0 | 0.15 | 0.03 | 0.22 | 0.22 | 0.18 | 0.24 |
| 7 | 51680653 | rs11765942 | 0.114 | 0.114 | -1.297 | 0.642 | -0.035 | -0.068 | -0.068 | 2.673 | 0.21 | 0.33 | 0.16 | 0.05 | 0.14 | 0.06 |
| 7 | 51680811 | rs13226225 | -0.235 | -0.034 | -1.601 | 0.943 | -0.004 | -0.02 | -0.02 | 0 | 0.14 | 0.04 | 0.08 | 0.12 | 0.12 | 0.13 |
| 7 | 51682201 | rs964804 | 0.835 | 0.285 | -1.535 | 0.967 | -0.007 | 0.003 | 0.003 | 0.256 | 0.27 | 0.38 | 0.15 | 0.3 | 0.23 | 0.25 |
| 7 | 51682401 | rs4947627 | -0.236 | 0.544 | -1.288 | 0.961 | 0.006 | 0.011 | 0.011 | 0 | 0.17 | 0.3 | 0.16 | 0.18 | 0.22 | 0.28 |
| 7 | 51683067 | rs10239205 | 0.483 | -0.137 | -0.566 | 0.539 | -0.001 | -0.008 | -0.008 | 0.179 | 0.05 | 0.23 | 0.02 | 0 | 0 | 0.03 |
| 7 | 51685174 | rs1990344 | 0.233 | -0.203 | -0.552 | 0.592 | 0 | -0.001 | -0.001 | 0 | 0.26 | 0.26 | 0.37 | 0.33 | 0.22 | 0.27 |
| 7 | 51690383 | rs17134352 | 0.999 | 0.097 | -0.629 | 0.684 | -0.027 | -0.033 | -0.033 | 2.546 | 0.11 | 0.03 | 0.06 | 0.14 | 0.14 | 0.15 |
| 7 | 51691587 | rs12669969 | 1.79 | 0.037 | -0.609 | 0.703 | -0.008 | 0.017 | 0.017 | 2.804 | 0.19 | 0.09 | 0.1 | 0.15 | 0.18 | 0.17 |
| 7 | 51692766 | rs987252 | -0.797 | 0.074 | -0.65 | 0.735 | -0.003 | -0.002 | -0.002 | 0.874 | 0.63 | 0.64 | 0.56 | 0.62 | 0.55 | 0.63 |
| 7 | 51693151 | rs6952304 | 1.003 | 0.197 | -0.759 | 0.408 | -0.007 | -0.041 | -0.041 | 0.107 | 0.24 | 0.33 | 0.12 | 0.15 | 0.18 | 0.21 |
| 7 | 51694472 | rs17134353 | 1.277 | 0.348 | -1.152 | 0.524 | -0.017 | -0.064 | -0.064 | 0 | 0.16 | 0.02 | 0.22 | 0.22 | 0.14 | 0.19 |
| 7 | 51697118 | rs4947376 | -1.208 | 0.644 | -1.088 | 0.148 | -0.036 | -0.009 | -0.009 | 0 | 0.84 | 0.9 | 0.8 | 0.86 | 0.75 | 0.75 |
| 7 | 51700020 | rs12718523 | 0.436 | 0.421 | -0.725 | 0.076 | 0.003 | 0.013 | 0.013 | 0 | 0.3 | 0.2 | 0.24 | 0.24 | 0.25 | 0.22 |
| 7 | 51703164 | rs6952787 | -0.996 | 1.001 | -0.597 | 0.251 | -0.027 | -0.014 | -0.014 | 0 | 0.82 | 0.85 | 0.76 | 0.81 | 0.69 | 0.69 |
| 7 | 51703325 | rs6973521 | 0.234 | 0.368 | -0.454 | -0.017 | -0.004 | -0.048 | -0.048 | 0 | 0.38 | 0.42 | 0.21 | 0.14 | 0.33 | 0.21 |
| 7 | 51706830 | rs7789538 | -0.115 | 0.385 | -0.267 | 0.884 | 0.002 | 0.041 | 0.041 | 0 | 0.14 | 0.14 | 0.14 | 0.35 | 0.14 | 0.2 |
| 7 | 51709304 | rs10263016 | -0.055 | 0.02 | 0.15 | 0.002 | -0.023 | -0.105 | -0.105 | 0 | 0.16 | 0.29 | 0.08 | 0 | 0.08 | 0.07 |
| 7 | 51715681 | rs17500908 | 0.815 | -0.217 | -0.538 | -0.459 | -0.027 | 0.084 | 0.084 | 1.601 | 0.19 | 0.01 | 0.18 | 0.08 | 0.18 | 0.1 |
| 7 | 51716684 | rs10234700 | 0.997 | -0.289 | -0.314 | -0.506 | -0.026 | -0.067 | -0.067 | 0 | 0.12 | 0.2 | 0.06 | 0 | 0.05 | 0.04 |
| 7 | 51718636 | rs10261309 | -0.05 | -0.326 | -0.234 | -0.532 | -0.025 | -0.103 | -0.103 | 0 | 0.17 | 0.29 | 0.08 | 0 | 0.08 | 0.08 |
| 7 | 51719970 | rs11238132 | -0.737 | -0.359 | 0.155 | -0.523 | -0.004 | -0.05 | -0.05 | 2.649 | 0.37 | 0.59 | 0.28 | 0.09 | 0.28 | 0.19 |
| 7 | 51721558 | rs7786795 | -1.538 | -0.195 | -0.286 | -0.142 | -0.025 | -0.01 | -0.01 | 0 | 0.65 | 0.72 | 0.67 | 0.63 | 0.54 | 0.64 |
| 7 | 51722990 | rs6963263 | -0.81 | -0.263 | -0.293 | -0.204 | 0.024 | 0.054 | 0.054 | 0.429 | 0.53 | 0.62 | 0.5 | 0.33 | 0.4 | 0.41 |
| 7 | 51728439 | rs6948705 | -1.087 | 0.18 | -0.487 | -0.067 | -0.033 | 0.091 | 0.091 | 1.054 | 0.81 | 0.99 | 0.82 | 0.93 | 0.81 | 0.92 |
| 7 | 51730710 | rs10267195 | 0.578 | 0.148 | -0.418 | -0.043 | 0 | 0.001 | 0.001 | 0 | 0.24 | 0.23 | 0.22 | 0.27 | 0.22 | 0.27 |
| 7 | 51730792 | rs10249138 | 1.2 | 0.027 | -0.592 | 0.052 | 0.008 | -0.021 | -0.021 | 0.361 | 0.15 | 0.13 | 0.21 | 0.24 | 0.12 | 0.23 |
| 7 | 51733827 | rs12537160 | 1.407 | 0.037 | -0.577 | 0.075 | -0.019 | -0.015 | -0.015 | 0.022 | 0.36 | 0.27 | 0.35 | 0.41 | 0.44 | 0.36 |
| 7 | 51749659 | rs7803183 | -1.288 | 0.372 | -1.087 | 0.067 | -0.038 | 0.086 | 0.086 | 0.731 | 0.82 | 1 | 0.82 | 0.93 | 0.82 | 0.91 |

| | | | | | | | | | | | | | | | | |
|---|---|---|---|---|---|---|---|---|---|---|---|---|---|---|---|---|
| 7 | 51751994 | rs7786998 | 0.497 | 0.385 | -0.595 | 0.344 | -0.005 | 0.014 | 0.014 | 0 | 0.18 | 0.21 | 0.23 | 0.25 | 0.14 | 0.29 |
| 7 | 51756063 | rs6979413 | 0.418 | 0.454 | -0.96 | 0.162 | -0.001 | 0.022 | 0.022 | 0 | 0.57 | 0.69 | 0.6 | 0.65 | 0.58 | 0.65 |
| 7 | 51765420 | rs11773632 | 0.606 | 0.522 | -1.181 | -0.095 | -0.046 | 0.077 | 0.077 | 0 | 0.17 | 0 | 0.11 | 0.02 | 0.18 | 0.06 |
| 7 | 51766831 | rs12673228 | 0.453 | 0.252 | -0.345 | 0.111 | -0.002 | 0.073 | 0.073 | 0 | 0.15 | 0.35 | 0.21 | 0.27 | 0.11 | 0.13 |
| 7 | 51767142 | rs9784925 | 0.398 | 0.596 | -0.544 | 0.169 | -0.006 | -0.006 | -0.006 | 0 | 0.27 | 0.28 | 0.37 | 0.38 | 0.36 | 0.4 |
| 7 | 51774342 | rs12718533 | 0.511 | 0.798 | -0.323 | 0.164 | 0.001 | 0.001 | 0.001 | 0.112 | 0.26 | 0.28 | 0.37 | 0.38 | 0.36 | 0.39 |
| 7 | 51774799 | rs10244676 | 0.505 | 0.793 | -0.326 | 0.161 | 0.001 | 0.001 | 0.001 | 0.123 | 0.26 | 0.28 | 0.37 | 0.38 | 0.36 | 0.4 |
| 7 | 51779695 | rs11765595 | 1.083 | 0.455 | -0.575 | 0.111 | 0.01 | 0.04 | 0.04 | 0 | 0.28 | 0.18 | 0.16 | 0.09 | 0.2 | 0.2 |
| 7 | 51785689 | rs4947636 | -0.443 | 0.523 | -0.171 | 0.135 | 0.004 | -0.051 | -0.051 | 0 | 0.72 | 0.83 | 0.66 | 0.5 | 0.72 | 0.69 |
| 7 | 51798888 | rs965277 | -0.764 | 0.658 | -0.024 | 0.157 | 0.023 | 0.047 | 0.047 | 0 | 0.58 | 0.76 | 0.67 | 0.73 | 0.69 | 0.69 |
| 7 | 51803145 | rs2329810 | 0.752 | 0.624 | -0.048 | 0.041 | -0.009 | -0.04 | -0.04 | 1.013 | 0.25 | 0.37 | 0.18 | 0.11 | 0.26 | 0.16 |
| 7 | 51806112 | rs17152331 | 1.147 | 0.592 | -0.334 | 0.039 | 0.025 | 0.063 | 0.063 | 0.322 | 0.08 | 0.3 | 0.07 | 0.08 | 0.08 | 0.08 |
| 7 | 51807027 | rs13244843 | -0.667 | 0.974 | -0.691 | 0.133 | 0.046 | 0.063 | 0.063 | 0 | 0.5 | 0.31 | 0.58 | 0.66 | 0.61 | 0.59 |
| 7 | 51809260 | rs17134469 | 0.17 | 0.975 | -0.838 | 0.211 | -0.128 | 0.011 | 0.011 | 0 | 0.35 | 0.06 | 0.4 | 0.33 | 0.44 | 0.43 |
| 7 | 51809901 | rs12718545 | -0.719 | 0.99 | -0.864 | 0.228 | 0.018 | 0.024 | 0.024 | 0 | 0.51 | 0.46 | 0.6 | 0.66 | 0.61 | 0.6 |
| 7 | 51810301 | rs12718546 | 0.82 | 1.311 | -1.744 | 0.455 | -0.099 | -0.071 | -0.071 | 0 | 0.18 | 0.02 | 0.29 | 0.29 | 0.27 | 0.35 |
| 7 | 51813707 | rs721114 | 0.358 | 0.637 | -1.108 | 0.593 | -0.008 | -0.052 | -0.052 | 6.19 | 0.15 | 0.1 | 0.17 | 0.33 | 0.17 | 0.17 |
| 7 | 51816358 | rs2329812 | 0.219 | 0.593 | -1.01 | 0.549 | 0.005 | -0.017 | -0.017 | 0 | 0.35 | 0.28 | 0.25 | 0.36 | 0.35 | 0.28 |
| 7 | 51824061 | rs628653 | 0.119 | 0.594 | -0.988 | 0.408 | 0 | -0.001 | -0.001 | 0 | 0.64 | 0.65 | 0.54 | 0.67 | 0.67 | 0.56 |
| 7 | 51824545 | rs641117 | 0.993 | 1.354 | -1.512 | 0.606 | -0.054 | -0.065 | -0.065 | 0 | 0.11 | 0.01 | 0.13 | 0.28 | 0.17 | 0.21 |
| 7 | 51824832 | rs662083 | -0.368 | 1.247 | -2.272 | 0.729 | -0.086 | -0.076 | -0.076 | 0.285 | 0.73 | 0.97 | 0.64 | 0.71 | 0.65 | 0.7 |
| 7 | 51825899 | rs1008386 | 1.027 | 1.331 | -2.275 | 0.163 | -0.028 | 0.026 | 0.026 | 0 | 0.13 | 0.06 | 0.12 | 0 | 0.19 | 0.05 |
| 7 | 51827021 | rs631671 | -0.24 | 1.028 | -2.091 | 0.411 | -0.046 | -0.075 | -0.075 | 0.748 | 0.64 | 0.88 | 0.6 | 0.59 | 0.58 | 0.57 |
| 7 | 51827355 | rs4947638 | 1.936 | 0.56 | -1.733 | 0.292 | 0.035 | 0.002 | 0.002 | 0 | 0.3 | 0.21 | 0.29 | 0.27 | 0.21 | 0.26 |
| 7 | 51829861 | rs2537505 | 1.769 | 0.636 | -1.715 | 0.309 | -0.041 | -0.054 | -0.054 | 0 | 0.12 | 0.03 | 0.12 | 0.25 | 0.15 | 0.21 |
| 7 | 51833487 | rs682792 | 0.023 | 0.463 | -1.171 | -0.003 | -0.017 | -0.012 | -0.012 | 0 | 0.57 | 0.71 | 0.42 | 0.57 | 0.51 | 0.47 |
| 7 | 51833835 | rs6954126 | 1.045 | 0.806 | -1.169 | -0.22 | -0.018 | 0.009 | 0.009 | 0 | 0.13 | 0.07 | 0.11 | 0 | 0.18 | 0.05 |
| 7 | 51835276 | rs6960015 | 0.95 | -0.064 | -0.983 | 0.133 | 0.008 | -0.037 | -0.037 | 0 | 0.07 | 0.14 | 0.18 | 0.01 | 0.07 | 0.1 |
| 7 | 51845731 | rs736248 | 1.353 | -0.487 | -0.949 | 0.405 | 0 | -0.002 | -0.002 | 0 | 0.24 | 0.22 | 0.22 | 0.25 | 0.17 | 0.17 |
| 7 | 51846147 | rs673982 | -0.402 | 0.281 | -1.301 | 0.41 | -0.049 | -0.018 | -0.018 | 0.005 | 0.75 | 0.92 | 0.71 | 0.71 | 0.64 | 0.73 |
| 7 | 51848170 | rs595846 | 0.602 | 0.113 | -0.673 | 0.426 | -0.001 | 0 | 0 | 0 | 0.53 | 0.51 | 0.48 | 0.55 | 0.52 | 0.46 |
| 7 | 51849478 | rs13225022 | -0.044 | 0.206 | -1.124 | 0.726 | -0.02 | -0.027 | -0.027 | 0 | 0.31 | 0.19 | 0.32 | 0.4 | 0.37 | 0.38 |
| 7 | 51865608 | rs4947642 | -1.145 | 0.487 | -1.197 | 0.916 | -0.063 | -0.051 | -0.051 | 0 | 0.84 | 0.92 | 0.81 | 0.72 | 0.73 | 0.73 |
| 7 | 51869701 | rs1473654 | 0.002 | -0.682 | -0.66 | 0.486 | 0.002 | 0.011 | 0.011 | 0.005 | 0.56 | 0.64 | 0.52 | 0.32 | 0.43 | 0.37 |
| 7 | 51873129 | rs627860 | -0.545 | -1.22 | -0.535 | 0.913 | -0.001 | -0.031 | -0.031 | 0 | 0.28 | 0.15 | 0.27 | 0.35 | 0.27 | 0.29 |
| 7 | 51876412 | rs1005032 | 1.085 | -1.577 | -0.235 | 0.97 | -0.045 | 0.076 | 0.076 | 0 | 0.18 | 0.29 | 0.19 | 0.36 | 0.1 | 0.18 |
| 7 | 51886164 | rs10264309 | 1.062 | -1.518 | -0.257 | 0.997 | -0.053 | 0.106 | 0.106 | 0 | 0.18 | 0.37 | 0.2 | 0.38 | 0.1 | 0.23 |
| 7 | 51886597 | rs603745 | -0.375 | -1.495 | -0.337 | 0.96 | -0.015 | -0.074 | -0.074 | 0 | 0.28 | 0.07 | 0.25 | 0.34 | 0.26 | 0.23 |
| 7 | 51887620 | rs11771485 | 0.709 | -1.551 | -0.326 | 0.947 | 0.002 | -0.001 | -0.001 | 0.068 | 0.11 | 0.08 | 0.07 | 0 | 0.1 | 0.05 |
| 7 | 51896753 | rs759002 | 0.023 | -1.671 | -0.255 | 0.948 | -0.06 | 0.054 | 0.054 | 0 | 0.26 | 0.54 | 0.25 | 0.5 | 0.13 | 0.31 |
| 7 | 51907832 | rs10215689 | 1.091 | -1.642 | -0.302 | 0.757 | -0.054 | 0.111 | 0.111 | 0 | 0.18 | 0.4 | 0.21 | 0.38 | 0.1 | 0.23 |
| 7 | 51916445 | rs654525 | -0.98 | -1.336 | -0.382 | 0.217 | 0.027 | 0.088 | 0.088 | 0 | 0.55 | 0.65 | 0.52 | 0.95 | 0.39 | 0.58 |
| 7 | 51916740 | rs4401799 | 0.712 | -1.344 | -0.372 | 0.25 | 0.017 | 0.019 | 0.019 | 0 | 0.28 | 0.22 | 0.17 | 0.28 | 0.2 | 0.28 |
| 7 | 51919827 | rs619342 | -0.42 | -1.195 | -0.654 | 0.245 | 0.041 | 0.062 | 0.062 | 0 | 0.44 | 0.23 | 0.45 | 0.71 | 0.36 | 0.47 |
| 7 | 51930023 | rs670754 | 0.359 | -0.497 | -1.166 | 0.023 | -0.025 | -0.13 | -0.13 | 0.013 | 0.17 | 0.02 | 0.28 | 0.43 | 0.16 | 0.19 |
| 7 | 51932604 | rs605705 | -0.674 | -0.438 | -1.132 | 0.005 | 0.004 | 0.125 | 0.125 | 0 | 0.58 | 0.73 | 0.66 | 0.95 | 0.42 | 0.62 |
| 7 | 51936695 | rs665087 | -0.643 | -0.295 | -2.597 | 0.074 | 0.002 | 0.136 | 0.136 | 0 | 0.58 | 0.75 | 0.66 | 0.95 | 0.42 | 0.62 |
| 7 | 51943403 | rs581988 | -0.859 | 0.729 | -2.063 | 1.511 | -0.029 | -0.128 | -0.128 | 0.484 | 0.19 | 0.06 | 0.4 | 0.45 | 0.19 | 0.24 |
| 7 | 51949727 | rs674648 | 0.361 | 0.718 | -2.269 | 0.994 | -0.078 | -0.094 | -0.094 | 0.693 | 0.69 | 0.95 | 0.54 | 0.38 | 0.64 | 0.57 |
| 7 | 51951935 | rs597467 | -0.715 | 0.146 | -2.527 | 1.208 | -0.063 | -0.135 | -0.135 | 0.009 | 0.33 | 0.06 | 0.5 | 0.49 | 0.36 | 0.39 |
| 7 | 51952293 | rs801447 | -0.519 | -0.455 | -1.806 | 0.227 | -0.029 | -0.093 | -0.093 | 0 | 0.64 | 0.55 | 0.8 | 0.88 | 0.74 | 0.68 |
| 7 | 51953172 | rs10250712 | 0.082 | -0.676 | -1.001 | 0.415 | -0.006 | 0.2 | 0.2 | 0 | 0.06 | 0.26 | 0.02 | 0.2 | 0 | 0.04 |
| 7 | 51953844 | rs10254486 | 0.031 | -0.694 | -0.986 | 0.407 | -0.019 | 0.149 | 0.149 | 0 | 0.05 | 0.21 | 0.02 | 0.2 | 0 | 0.05 |
| 7 | 51954436 | rs10240034 | 0.049 | -0.875 | -0.196 | -0.162 | 0.01 | -0.034 | -0.034 | 0 | 0.33 | 0.46 | 0.27 | 0.2 | 0.37 | 0.25 |
| 7 | 51954824 | rs11238145 | 0.283 | -1.151 | -0.102 | -0.272 | 0.012 | -0.018 | -0.018 | 0 | 0.31 | 0.38 | 0.52 | 0.7 | 0.35 | 0.36 |
| 7 | 51970823 | rs1993956 | 0.381 | -1.249 | 0.258 | -0.209 | 0.003 | 0.017 | 0.017 | 0 | 0.32 | 0.38 | 0.53 | 0.51 | 0.35 | 0.34 |
| 7 | 51972865 | rs17202853 | 0.614 | -1.257 | 0.054 | -0.304 | -0.077 | 0.039 | 0.039 | 0.353 | 0.26 | 0.1 | 0.25 | 0.2 | 0.36 | 0.25 |
| 7 | 51976812 | rs7801225 | -0.572 | -1.291 | 0.589 | -0.3 | 0.014 | 0.011 | 0.011 | 0 | 0.6 | 0.61 | 0.78 | 0.71 | 0.71 | 0.59 |
| 7 | 51978391 | rs10255174 | -0.73 | -1.347 | 0.484 | -0.34 | 0.034 | 0.027 | 0.027 | 0 | 0.62 | 0.74 | 0.79 | 0.72 | 0.72 | 0.62 |
| 7 | 51984019 | rs432995 | 0.046 | -1.446 | 0.493 | -0.403 | -0.002 | 0.01 | 0.01 | 0.317 | 0.64 | 0.33 | 0.68 | 0.72 | 0.62 | 0.55 |
| 7 | 51987979 | rs631999 | 0.482 | -1.378 | 0.519 | -0.375 | -0.007 | -0.01 | -0.01 | 0 | 0.36 | 0.39 | 0.19 | 0.2 | 0.27 | 0.18 |
| 7 | 51991339 | rs7783471 | 1.06 | -1.421 | 0.059 | -0.603 | 0.001 | -0.014 | -0.014 | 0 | 0.1 | 0.09 | 0.04 | 0 | 0.09 | 0.06 |
| 7 | 51999888 | rs372475 | 0.558 | -1.57 | -0.186 | -1.102 | -0.041 | 0.064 | 0.064 | 0 | 0.36 | 0.16 | 0.28 | 0.24 | 0.42 | 0.35 |
| 7 | 52006246 | rs1600385 | 0.546 | -2.017 | -0.308 | -1.18 | -0.012 | 0.014 | 0.014 | 0 | 0.47 | 0.22 | 0.61 | 0.57 | 0.55 | 0.58 |
| 7 | 52006415 | rs406001 | -0.484 | -1.997 | 0.442 | -1.194 | -0.029 | -0.035 | -0.035 | 0 | 0.93 | 0.78 | 0.96 | 1 | 0.99 | 0.98 |
| 7 | 52017424 | rs11521104 | 0.299 | -1.729 | 0.3 | -1.25 | 0 | -0.023 | -0.023 | 0.325 | 0.17 | 0.19 | 0.1 | 0.03 | 0.13 | 0.27 |
| 7 | 52018577 | rs17204439 | 1.617 | -1.654 | 0.101 | -1.308 | -0.064 | -0.038 | -0.038 | 0 | 0.15 | 0.03 | 0.17 | 0.23 | 0.24 | 0.15 |
| 7 | 52018700 | rs4947661 | 1.166 | -1.484 | 0.201 | -0.895 | -0.002 | -0.123 | -0.123 | 0 | 0.29 | 0.12 | 0.41 | 0.47 | 0.22 | 0.24 |
| 7 | 52020592 | rs4947392 | 0.145 | -1.493 | 0.222 | -0.864 | 0.036 | -0.013 | -0.013 | 0 | 0.46 | 0.31 | 0.51 | 0.5 | 0.34 | 0.5 |

| | | | | | | | | | | | | | | | | |
|---|---|---|---|---|---|---|---|---|---|---|---|---|---|---|---|---|
| 7 | 52029895 | rs7804217 | 0.873 | -1.295 | -0.322 | -0.916 | -0.041 | 0.009 | 0.009 | 0 | 0.1 | 0.04 | 0.16 | 0.09 | 0.25 | 0.12 |
| 7 | 52030913 | rs454066 | 0.342 | -1.524 | 0.234 | -0.859 | 0.024 | 0.053 | 0.053 | 0.01 | 0.45 | 0.27 | 0.43 | 0.35 | 0.64 | 0.46 |
| 7 | 52035335 | rs4292640 | -0.301 | -1.216 | 0.398 | -0.852 | -0.039 | -0.01 | -0.01 | 0 | 0.22 | 0.43 | 0.12 | 0.18 | 0.13 | 0.28 |
| 7 | 52050366 | rs17134720 | 1.029 | -1.114 | 0.435 | -0.537 | -0.009 | 0.034 | 0.034 | 0 | 0.26 | 0.29 | 0.42 | 0.52 | 0.21 | 0.23 |
| 7 | 52058887 | rs13225954 | -0.067 | -1.177 | 0.53 | -0.801 | -0.006 | 0.006 | 0.006 | 0 | 0.16 | 0.16 | 0.16 | 0.11 | 0.26 | 0.24 |
| 7 | 52062583 | rs6955285 | 1.285 | -1.443 | 0.597 | -0.96 | -0.011 | 0.045 | 0.045 | 0 | 0.37 | 0.27 | 0.33 | 0.22 | 0.4 | 0.32 |
| 7 | 52064705 | rs7801309 | -0.056 | -1.33 | 0.504 | -0.905 | -0.01 | -0.006 | -0.006 | 2.551 | 0.79 | 0.72 | 0.9 | 0.86 | 0.87 | 0.78 |
| 7 | 52070744 | rs13231221 | 0.233 | -1.186 | 0.381 | -0.773 | -0.007 | 0.006 | 0.006 | 0 | 0.14 | 0.12 | 0.15 | 0.1 | 0.25 | 0.18 |
| 7 | 52072257 | rs2113636 | 0.001 | -1.237 | 0.461 | -0.789 | 0 | -0.001 | -0.001 | 0 | 0.4 | 0.4 | 0.57 | 0.63 | 0.45 | 0.41 |
| 7 | 52073758 | rs17273555 | 0.158 | -1.105 | 0.127 | -0.702 | 0 | 0 | 0 | 0 | 0.14 | 0.15 | 0.16 | 0.11 | 0.25 | 0.18 |
| 7 | 52074136 | rs13243210 | -0.526 | -1.245 | 0.082 | -0.78 | 0.011 | -0.013 | -0.013 | 0 | 0.21 | 0.3 | 0.18 | 0.16 | 0.27 | 0.19 |
| 7 | 52076060 | rs13237752 | -0.765 | -1.461 | -0.096 | -0.822 | -0.003 | 0 | 0 | 0 | 0.71 | 0.73 | 0.57 | 0.39 | 0.79 | 0.74 |
| 7 | 52084694 | rs10046510 | 0.132 | -1.358 | -0.419 | -0.875 | -0.008 | -0.014 | -0.014 | 0.077 | 0.54 | 0.56 | 0.52 | 0.38 | 0.66 | 0.62 |
| 7 | 52088946 | rs2194579 | 1.302 | -1.134 | -1.444 | -0.782 | -0.017 | -0.13 | -0.13 | 0 | 0.24 | 0.05 | 0.38 | 0.51 | 0.19 | 0.22 |
| 7 | 52089353 | rs10277728 | 1.355 | -0.692 | -1.057 | -0.785 | -0.003 | -0.009 | -0.009 | 0 | 0.1 | 0.08 | 0.05 | 0 | 0.11 | 0.06 |
| 7 | 52090123 | rs6953592 | 0.679 | -0.41 | -1.138 | -0.572 | -0.031 | 0.049 | 0.049 | 0 | 0.36 | 0.17 | 0.32 | 0.3 | 0.39 | 0.43 |
| 7 | 52094719 | rs2329939 | -0.363 | 0.806 | -0.818 | 0.161 | -0.012 | -0.008 | -0.008 | 1.895 | 0.15 | 0.11 | 0.14 | 0.19 | 0.23 | 0.13 |
| 7 | 52098740 | rs1978512 | 1.114 | 0.514 | -0.943 | 0.185 | 0.001 | -0.032 | -0.032 | 0 | 0.24 | 0.21 | 0.41 | 0.53 | 0.26 | 0.29 |
| 7 | 52099373 | rs10278980 | 1.411 | 0.482 | -1.085 | 0.309 | -0.01 | -0.063 | -0.063 | 0 | 0.22 | 0.11 | 0.38 | 0.44 | 0.25 | 0.26 |
| 7 | 52102546 | rs4947685 | 0.024 | 0.588 | -1.023 | 0.291 | -0.012 | -0.009 | -0.009 | 0 | 0.15 | 0.11 | 0.14 | 0.2 | 0.23 | 0.13 |
| 7 | 52107816 | rs10249902 | -0.49 | 0.14 | -0.44 | -0.13 | -0.001 | -0.003 | -0.003 | 0.15 | 0.44 | 0.44 | 0.56 | 0.76 | 0.5 | 0.49 |
| 7 | 52110788 | rs2216883 | -0.693 | -0.352 | -0.213 | -0.48 | -0.001 | 0.028 | 0.028 | 0 | 0.55 | 0.6 | 0.61 | 0.76 | 0.55 | 0.53 |
| 7 | 52122844 | rs4947688 | -0.123 | -0.369 | 0.173 | -0.345 | -0.002 | -0.01 | -0.01 | 0.479 | 0.41 | 0.49 | 0.52 | 0.64 | 0.4 | 0.42 |
| 7 | 52124519 | rs1863081 | -0.466 | -0.373 | -0.06 | -0.401 | 0.023 | 0.124 | 0.124 | 0 | 0.47 | 0.76 | 0.56 | 0.76 | 0.42 | 0.51 |
| 7 | 52129322 | rs6946992 | -0.466 | -0.373 | -0.069 | -0.401 | 0.022 | 0.128 | 0.128 | 0 | 0.47 | 0.77 | 0.56 | 0.76 | 0.42 | 0.51 |
| 7 | 52130600 | rs10232268 | -0.457 | -0.374 | -0.088 | -0.401 | 0.02 | 0.131 | 0.131 | 0 | 0.46 | 0.8 | 0.56 | 0.76 | 0.42 | 0.51 |
| 7 | 52135471 | rs10279635 | -1.351 | -0.007 | -0.545 | -0.597 | 0.001 | 0.002 | 0.002 | 0 | 0.92 | 0.93 | 0.95 | 0.98 | 0.94 | 0.94 |
| 7 | 52142647 | rs12533618 | 1.026 | -0.19 | -0.591 | -0.35 | 0 | 0.014 | 0.014 | 0 | 0.11 | 0.11 | 0.22 | 0.26 | 0.1 | 0.12 |
| 7 | 52148307 | rs11979834 | 1.793 | -0.258 | -0.48 | -0.396 | -0.013 | -0.05 | -0.05 | 0 | 0.19 | 0.11 | 0.25 | 0.4 | 0.25 | 0.3 |
| 7 | 52158949 | rs10899740 | 0.616 | -0.363 | -0.219 | -0.399 | -0.003 | 0.007 | 0.007 | 0 | 0.31 | 0.3 | 0.29 | 0.39 | 0.3 | 0.34 |
| 7 | 52160643 | rs2464949 | -0.134 | -0.287 | -0.061 | -0.463 | -0.016 | 0.014 | 0.014 | 0 | 0.93 | 0.78 | 0.93 | 0.89 | 0.95 | 0.92 |
| 7 | 52160893 | rs7779581 | 0.726 | -0.182 | -0.546 | -0.646 | -0.014 | 0.01 | 0.01 | 0.01 | 0.27 | 0.12 | 0.36 | 0.23 | 0.29 | 0.21 |
| 7 | 52161151 | rs12718586 | 0.219 | -0.277 | -0.29 | -0.764 | -0.001 | 0.007 | 0.007 | 0 | 0.41 | 0.32 | 0.3 | 0.27 | 0.4 | 0.4 |
| 7 | 52162898 | rs2952763 | 0.215 | -0.278 | -0.614 | -0.692 | 0.008 | 0.043 | 0.043 | 0.633 | 0.53 | 0.36 | 0.53 | 0.72 | 0.58 | 0.65 |
| 7 | 52167385 | rs6965611 | 1.556 | -0.015 | -1.272 | -0.423 | -0.061 | -0.16 | -0.16 | 0 | 0.24 | 0.03 | 0.24 | 0.41 | 0.26 | 0.31 |
| 7 | 52167885 | rs2465786 | 0.702 | -0.025 | -1.004 | -0.532 | -0.015 | 0.019 | 0.019 | 0.107 | 0.28 | 0.15 | 0.3 | 0.26 | 0.31 | 0.34 |
| 7 | 52168088 | rs2464946 | -0.625 | -0.218 | -0.347 | -0.396 | -0.002 | 0.01 | 0.01 | 0 | 0.67 | 0.4 | 0.69 | 0.51 | 0.7 | 0.63 |
| 7 | 52176891 | rs2952754 | -0.238 | -0.245 | -0.036 | -0.547 | -0.016 | -0.011 | -0.011 | 0 | 0.93 | 0.8 | 0.97 | 0.97 | 0.98 | 0.97 |
| 7 | 52178844 | rs2464957 | 0.14 | -0.169 | -0.724 | -0.87 | 0.006 | 0.014 | 0.014 | 0 | 0.32 | 0.17 | 0.41 | 0.32 | 0.27 | 0.42 |
| 7 | 52181844 | rs4947694 | 1.177 | -0.193 | -0.917 | -0.94 | -0.021 | -0.029 | -0.029 | 0 | 0.11 | 0.01 | 0.12 | 0.11 | 0.09 | 0.19 |
| 7 | 52183846 | rs10236557 | -1.298 | -0.132 | -0.329 | -0.97 | -0.002 | 0.002 | 0.002 | 0 | 0.54 | 0.54 | 0.54 | 0.44 | 0.45 | 0.54 |
| 7 | 52185600 | rs2465791 | 0.189 | 0.055 | -0.576 | -0.834 | -0.002 | 0.01 | 0.01 | 0 | 0.32 | 0.17 | 0.42 | 0.32 | 0.29 | 0.42 |
| 7 | 52188555 | rs2953432 | 0.252 | 0.044 | -0.464 | -0.86 | -0.002 | 0.005 | 0.005 | 0 | 0.31 | 0.22 | 0.4 | 0.32 | 0.28 | 0.41 |
| 7 | 52190733 | rs2465796 | 0.168 | -0.024 | -0.572 | -1.005 | 0 | -0.004 | -0.004 | 0.044 | 0.52 | 0.49 | 0.64 | 0.77 | 0.55 | 0.72 |
| 7 | 52201059 | rs2465798 | -0.459 | -0.036 | -0.926 | -0.958 | -0.007 | 0.044 | 0.044 | 0 | 0.45 | 0.26 | 0.46 | 0.36 | 0.41 | 0.46 |
| 7 | 52202071 | rs7808716 | 1.822 | -0.159 | -0.834 | -0.855 | -0.032 | -0.098 | -0.098 | 0 | 0.22 | 0.07 | 0.23 | 0.48 | 0.28 | 0.29 |
| 7 | 52203796 | rs13240590 | -0.746 | -0.316 | -0.444 | -1.022 | -0.023 | -0.021 | -0.021 | 0 | 0.92 | 0.79 | 0.95 | 0.97 | 0.98 | 0.98 |
| 7 | 52208909 | rs4526312 | -0.094 | -0.429 | -0.775 | -1.173 | -0.038 | -0.046 | -0.046 | 0.022 | 0.8 | 0.55 | 0.83 | 0.85 | 0.88 | 0.84 |
| 7 | 52209695 | rs4273801 | 0.497 | -0.31 | -1.09 | -1.085 | -0.009 | -0.009 | -0.009 | 0 | 0.3 | 0.18 | 0.39 | 0.36 | 0.28 | 0.42 |
| 7 | 52211550 | rs12718604 | -0.297 | -0.353 | -1.058 | -1.056 | 0.021 | 0.052 | 0.052 | 0 | 0.56 | 0.2 | 0.59 | 0.36 | 0.59 | 0.52 |
| 7 | 52217141 | rs6951196 | 1.789 | -0.118 | -1.357 | -0.776 | -0.092 | -0.184 | -0.184 | 0 | 0.25 | 0.01 | 0.24 | 0.45 | 0.3 | 0.31 |
| 7 | 52218507 | rs7790298 | 0.305 | -0.345 | -0.876 | -0.953 | -0.028 | -0.034 | -0.034 | 0.798 | 0.81 | 0.61 | 0.85 | 0.89 | 0.88 | 0.86 |
| 7 | 52222322 | rs10256183 | 0.586 | -0.039 | -1.657 | -0.883 | -0.035 | 0.123 | 0.123 | 0 | 0.35 | 0.13 | 0.33 | 0.13 | 0.38 | 0.24 |
| 7 | 52222952 | rs12671238 | 1.104 | 0.305 | -1.258 | -0.187 | -0.013 | -0.057 | -0.057 | 0 | 0.26 | 0.15 | 0.27 | 0.43 | 0.29 | 0.42 |
| 7 | 52236264 | rs3923846 | 1.071 | 0.483 | -1.428 | -0.382 | -0.044 | 0.032 | 0.032 | 0.548 | 0.19 | 0.03 | 0.18 | 0.1 | 0.24 | 0.15 |
| 7 | 52237443 | rs4319038 | 1.009 | 0.41 | -0.832 | -0.276 | 0.017 | 0.07 | 0.07 | 0 | 0.16 | 0.26 | 0.25 | 0.36 | 0.17 | 0.14 |
| 7 | 52240990 | rs4413723 | 1.05 | 0.524 | -0.769 | -0.234 | 0.007 | 0.024 | 0.024 | 0 | 0.09 | 0.13 | 0.17 | 0.2 | 0.08 | 0.12 |
| 7 | 52244143 | rs10231296 | -0.303 | -0.022 | -0.342 | -0.426 | -0.032 | -0.022 | -0.022 | 0 | 0.9 | 0.79 | 0.95 | 0.94 | 0.94 | 0.9 |
| 7 | 52245441 | rs10238949 | 0.398 | -0.175 | -0.429 | -0.705 | -0.015 | 0.064 | 0.064 | 0 | 0.41 | 0.25 | 0.39 | 0.18 | 0.45 | 0.33 |
| 7 | 52246309 | rs7793296 | 0.686 | -0.272 | 0.104 | -0.445 | -0.037 | 0.004 | 0.004 | 0 | 0.79 | 0.48 | 0.76 | 0.74 | 0.86 | 0.78 |
| 7 | 52252958 | rs4349938 | 1.032 | -0.193 | -0.163 | -0.454 | -0.007 | -0.007 | -0.007 | 0 | 0.48 | 0.33 | 0.47 | 0.49 | 0.48 | 0.59 |
| 7 | 52256544 | rs4947699 | 0.817 | 0.243 | 0.19 | -0.32 | -0.002 | 0.013 | 0.013 | 0 | 0.12 | 0.19 | 0.12 | 0.17 | 0.11 | 0.08 |
| 7 | 52265380 | rs12673349 | 0.745 | 0.066 | 0.048 | -0.504 | -0.01 | 0.009 | 0.009 | 0 | 0.52 | 0.31 | 0.48 | 0.53 | 0.52 | 0.63 |
| 7 | 52269884 | rs11238186 | -0.515 | 0.337 | -0.463 | -0.412 | -0.004 | 0.029 | 0.029 | 0 | 0.72 | 0.79 | 0.72 | 0.89 | 0.7 | 0.75 |
| 7 | 52273284 | rs10230289 | 0.312 | 0.208 | -0.74 | -0.298 | -0.001 | -0.001 | -0.001 | 0 | 0.56 | 0.45 | 0.52 | 0.54 | 0.56 | 0.68 |
| 7 | 52278157 | rs4557656 | 0.296 | 0.502 | -0.96 | -0.307 | -0.084 | 0.046 | 0.046 | 0 | 0.16 | 0.02 | 0.3 | 0.07 | 0.23 | 0.18 |
| 7 | 52282461 | rs6964146 | -0.025 | 0.375 | -0.927 | 0.114 | -0.017 | -0.017 | -0.017 | 0 | 0.91 | 0.94 | 0.95 | 0.89 | 0.9 | 0.9 |
| 7 | 52282513 | rs12533876 | 1.539 | -0.377 | -0.365 | -0.198 | -0.022 | -0.018 | -0.018 | 0 | 0.78 | 0.33 | 0.88 | 0.87 | 0.86 | 0.85 |
| 7 | 52284537 | rs4596601 | 0.227 | -0.348 | 0.011 | -0.186 | 0.01 | 0.011 | 0.011 | 0 | 0.9 | 0.85 | 0.94 | 0.89 | 0.9 | 0.89 |

| | | | | | | | | | | | | | | | | |
|---|---|---|---|---|---|---|---|---|---|---|---|---|---|---|---|---|
| 7 | 52287973 | rs4523204 | 0.309 | -0.698 | -0.468 | -0.367 | 0.059 | 0.085 | 0.085 | 0.039 | 0.51 | 0.28 | 0.63 | 0.73 | 0.62 | 0.69 |
| 7 | 52294105 | rs4245554 | -0.38 | -0.507 | -0.519 | -0.269 | -0.01 | -0.01 | -0.01 | 0 | 0.91 | 0.91 | 0.95 | 0.89 | 0.9 | 0.9 |
| 7 | 52295068 | rs6948352 | 1.441 | -0.481 | -0.487 | -0.275 | 0.018 | 0.041 | 0.041 | 1.418 | 0.28 | 0.14 | 0.26 | 0.14 | 0.24 | 0.15 |
| 7 | 52305206 | rs4286900 | 0.868 | -0.461 | -0.699 | 0.087 | -0.129 | -0.088 | -0.088 | 0 | 0.37 | 0.07 | 0.44 | 0.44 | 0.48 | 0.55 |
| 7 | 52306798 | rs12718620 | 1.461 | -0.457 | -0.566 | -0.077 | 0 | 0.067 | 0.067 | 1.308 | 0.28 | 0.06 | 0.27 | 0.14 | 0.25 | 0.16 |
| 7 | 52316977 | rs6963554 | 0.207 | -0.621 | -0.321 | -0.039 | 0.043 | 0.06 | 0.06 | 0.841 | 0.48 | 0.3 | 0.63 | 0.73 | 0.62 | 0.71 |
| 7 | 52326551 | rs2191714 | -1.501 | -0.525 | -0.38 | -0.048 | 0.013 | 0.058 | 0.058 | 1.475 | 0.74 | 0.94 | 0.75 | 0.86 | 0.78 | 0.85 |
| 7 | 52330420 | rs2041481 | -1.288 | -0.544 | 0.015 | -0.048 | -0.005 | -0.012 | -0.012 | 0.182 | 0.72 | 0.77 | 0.74 | 0.86 | 0.78 | 0.85 |
| 7 | 52336399 | rs1860689 | 0.132 | -0.417 | 0.249 | 0.353 | 0.034 | 0.177 | 0.177 | 2.224 | 0.16 | 0.48 | 0.17 | 0.34 | 0.16 | 0.16 |
| 7 | 52347667 | rs11768415 | 0.551 | -0.673 | -0.336 | 0.107 | -0.129 | -0.076 | -0.076 | 0 | 0.4 | 0.07 | 0.46 | 0.43 | 0.48 | 0.54 |
| 7 | 52350879 | rs17134954 | 0.704 | -0.751 | -0.229 | 0.009 | -0.009 | -0.023 | -0.023 | 0.846 | 0.1 | 0.2 | 0.08 | 0.04 | 0.08 | 0.09 |
| 7 | 52356515 | rs11764734 | 0.602 | -0.667 | -0.462 | 0.065 | -0.133 | -0.08 | -0.08 | 0.512 | 0.4 | 0.06 | 0.46 | 0.43 | 0.48 | 0.54 |
| 7 | 52386376 | rs9886124 | -0.142 | -0.736 | -0.025 | 0.196 | -0.004 | 0.046 | 0.046 | 2.071 | 0.19 | 0.29 | 0.14 | 0.29 | 0.19 | 0.21 |
| 7 | 52424318 | rs7802586 | -0.661 | -0.762 | 0.004 | 0.184 | -0.018 | 0.034 | 0.034 | 2.471 | 0.23 | 0.57 | 0.18 | 0.29 | 0.19 | 0.22 |
| 7 | 52426894 | rs2041562 | 1.732 | -0.591 | -0.412 | 0.125 | -0.045 | -0.125 | -0.125 | 0 | 0.21 | 0 | 0.14 | 0.36 | 0.23 | 0.28 |
| 7 | 52427003 | rs6971252 | 0.241 | -0.7 | -0.127 | 0.172 | -0.055 | -0.044 | -0.044 | 0 | 0.1 | 0.32 | 0.05 | 0.02 | 0.04 | 0.07 |
| 7 | 52428432 | rs12673852 | 1.209 | -0.688 | -0.11 | 0.179 | 0.034 | 0.101 | 0.101 | 1.644 | 0.13 | 0.25 | 0.13 | 0.27 | 0.15 | 0.15 |
| 7 | 52428669 | rs7791231 | -0.306 | -0.551 | -0.163 | 0.223 | -0.019 | 0.032 | 0.032 | 2.378 | 0.23 | 0.57 | 0.18 | 0.29 | 0.19 | 0.22 |
| 7 | 52432938 | rs296307 | 0.326 | -0.407 | -0.41 | 0.035 | 0.021 | -0.097 | -0.097 | 0 | 0.71 | 0.53 | 0.59 | 0.9 | 0.68 | 0.67 |
| 7 | 52433121 | rs296308 | 1.54 | -0.249 | -1.126 | 0.18 | -0.018 | -0.02 | -0.02 | 0 | 0.27 | 0.06 | 0.27 | 0.27 | 0.24 | 0.15 |
| 7 | 52439967 | rs296316 | -0.854 | 0.047 | -1.234 | -0.024 | -0.054 | 0.056 | 0.056 | 0 | 0.84 | 0.95 | 0.69 | 0.95 | 0.77 | 0.82 |
| 7 | 52443808 | rs296328 | 1.165 | -0.089 | -1.287 | 0.001 | -0.03 | 0.05 | 0.05 | 0 | 0.52 | 0.14 | 0.43 | 0.63 | 0.49 | 0.47 |
| 7 | 52452724 | rs1035042 | -0.966 | -0.209 | -1.279 | 0.194 | 0.041 | -0.019 | -0.019 | 3.866 | 0.34 | 0.88 | 0.27 | 0.32 | 0.28 | 0.35 |
| 7 | 52468999 | rs4947712 | -0.56 | 0.381 | -1.08 | 0.067 | -0.062 | 0.058 | 0.058 | 0 | 0.84 | 0.95 | 0.67 | 0.97 | 0.72 | 0.77 |
| 7 | 52469741 | rs9642474 | 1.018 | -0.058 | -0.131 | 0.271 | 0.017 | -0.022 | -0.022 | 0.027 | 0.6 | 0.46 | 0.5 | 0.75 | 0.51 | 0.53 |
| 7 | 52470990 | rs6965881 | 0.772 | -0.356 | 0.596 | 0.636 | -0.023 | -0.021 | -0.021 | 0 | 0.64 | 0.68 | 0.51 | 0.56 | 0.51 | 0.51 |
| 7 | 52471137 | rs12718633 | 0.78 | -0.44 | 1.099 | 0.595 | 0.015 | -0.066 | -0.066 | 0 | 0.7 | 0.49 | 0.59 | 0.89 | 0.64 | 0.57 |
| 7 | 52472450 | rs1012036 | 1.36 | -0.826 | 0.457 | 0.511 | -0.022 | -0.075 | -0.075 | 0 | 0.26 | 0.04 | 0.2 | 0.32 | 0.23 | 0.34 |
| 7 | 52473094 | rs2215662 | 1.246 | -0.799 | 0.722 | 0.546 | -0.014 | -0.013 | -0.013 | 0 | 0.57 | 0.3 | 0.47 | 0.56 | 0.5 | 0.49 |
| 7 | 52473969 | rs9642477 | 0.508 | -0.798 | 0.916 | 0.579 | 0.013 | 0.012 | 0.012 | 0 | 0.69 | 0.5 | 0.57 | 0.69 | 0.62 | 0.55 |
| 7 | 52475793 | rs1121768 | 1.136 | -0.666 | 0.475 | 0.597 | -0.02 | -0.019 | -0.019 | 2.562 | 0.57 | 0.26 | 0.47 | 0.56 | 0.5 | 0.5 |
| 7 | 52481425 | rs11238202 | -0.592 | -0.665 | 0.506 | 0.483 | 0.015 | 0.088 | 0.088 | 0 | 0.52 | 0.71 | 0.39 | 0.74 | 0.46 | 0.62 |
| 7 | 52482227 | rs996561 | 0.921 | -0.722 | 0.587 | 0.402 | -0.007 | 0.063 | 0.063 | 0.373 | 0.54 | 0.28 | 0.49 | 0.77 | 0.51 | 0.52 |
| 7 | 52495428 | rs4947713 | 0.891 | -0.543 | 0.409 | 0.654 | -0.002 | 0.006 | 0.006 | 0.022 | 0.07 | 0.26 | 0.07 | 0.06 | 0.07 | 0.08 |
| 7 | 52497012 | rs17135029 | 1.162 | -0.523 | 0.332 | 0.657 | 0.001 | 0.003 | 0.003 | 0 | 0.06 | 0.2 | 0.06 | 0.06 | 0.07 | 0.07 |
| 7 | 52507626 | rs12718642 | 1.353 | -0.688 | -0.168 | 0.52 | -0.053 | -0.022 | -0.022 | 0 | 0.5 | 0.1 | 0.44 | 0.46 | 0.49 | 0.49 |
| 7 | 52510591 | rs10226649 | 1.05 | -0.721 | -0.138 | 0.504 | 0.016 | 0.006 | 0.006 | 3.173 | 0.17 | 0.07 | 0.09 | 0.14 | 0.11 | 0.07 |
| 7 | 52516374 | rs10277927 | 0.279 | -0.606 | 0.056 | 0.512 | 0.038 | 0.044 | 0.044 | 0.388 | 0.75 | 0.56 | 0.57 | 0.61 | 0.64 | 0.67 |
| 7 | 52517562 | rs1547473 | -0.958 | -0.691 | -0.036 | 0.011 | -0.063 | 0.048 | 0.048 | 0.984 | 0.83 | 0.95 | 0.67 | 0.96 | 0.72 | 0.76 |
| 7 | 52517718 | rs6962552 | 1.787 | -0.902 | -0.281 | -0.126 | -0.007 | 0.075 | 0.075 | 0.387 | 0.27 | 0.06 | 0.25 | 0.14 | 0.25 | 0.17 |
| 7 | 52520976 | rs6950637 | -0.034 | -0.702 | 0.099 | -0.076 | 0.034 | -0.035 | -0.035 | 3.962 | 0.77 | 0.58 | 0.6 | 0.89 | 0.67 | 0.7 |
| 7 | 52521444 | rs12538005 | 1.151 | -0.729 | -0.354 | -0.03 | -0.024 | 0.125 | 0.125 | 1.185 | 0.52 | 0.16 | 0.47 | 0.74 | 0.52 | 0.53 |
| 7 | 52525336 | rs6959593 | 0.034 | -0.852 | 0.048 | 0.057 | -0.036 | -0.023 | -0.023 | 0.047 | 0.22 | 0.34 | 0.11 | 0.16 | 0.12 | 0.07 |
| 7 | 52526486 | rs17815318 | 0.502 | -0.174 | -0.478 | 0.065 | 0 | -0.003 | -0.003 | 0.437 | 0.08 | 0.06 | 0.04 | 0 | 0.08 | 0.09 |
| 7 | 52530601 | rs7794935 | 0.425 | -0.536 | -0.718 | 0.086 | 0.031 | 0.079 | 0.079 | 0.479 | 0.46 | 0.16 | 0.36 | 0.36 | 0.42 | 0.31 |
| 7 | 52538830 | rs7776642 | 0.475 | -0.598 | -0.765 | 0.114 | 0.022 | 0.101 | 0.101 | 1.081 | 0.44 | 0.13 | 0.35 | 0.31 | 0.39 | 0.24 |
| 7 | 52543480 | rs10251994 | -0.366 | -0.691 | -0.349 | 0.097 | -0.016 | 0.004 | 0.004 | 0 | 0.06 | 0.14 | 0.03 | 0.05 | 0.05 | 0.09 |
| 7 | 52544356 | rs4269485 | -0.143 | -0.472 | 0.033 | 0.336 | -0.001 | 0.001 | 0.001 | 0 | 0.61 | 0.33 | 0.49 | 0.51 | 0.65 | 0.41 |
| 7 | 52548691 | rs984197 | -1.014 | -0.609 | 0.496 | 0.288 | 0.003 | -0.001 | -0.001 | 0.217 | 0.27 | 0.66 | 0.2 | 0.26 | 0.31 | 0.27 |
| 7 | 52548888 | rs12718665 | 1.595 | -0.838 | 0.511 | 0.102 | 0.005 | 0.011 | 0.011 | 0.316 | 0.29 | 0.2 | 0.29 | 0.29 | 0.3 | 0.27 |
| 7 | 52554862 | rs6972820 | -1.369 | -0.834 | 0.458 | 0.171 | 0.004 | 0.044 | 0.044 | 0 | 0.44 | 0.72 | 0.28 | 0.31 | 0.43 | 0.33 |
| 7 | 52557596 | rs12056006 | 2.108 | -0.761 | 0.384 | 0.137 | -0.006 | 0.028 | 0.028 | 0.612 | 0.25 | 0.07 | 0.27 | 0.23 | 0.26 | 0.18 |
| 7 | 52557887 | rs2159813 | 1.203 | -0.264 | 0.56 | 0.18 | 0.025 | 0.019 | 0.019 | 0 | 0.1 | 0.14 | 0.09 | 0.15 | 0.15 | 0.12 |
| 7 | 52557903 | rs2159814 | -0.893 | -0.293 | 0.519 | 0.179 | 0 | -0.011 | -0.011 | 1.251 | 0.66 | 0.65 | 0.53 | 0.54 | 0.68 | 0.5 |
| 7 | 52569318 | rs17152615 | -0.152 | -0.284 | 0.532 | 0.165 | -0.001 | -0.005 | -0.005 | 0.056 | 0.3 | 0.34 | 0.16 | 0.16 | 0.26 | 0.2 |
| 7 | 52575764 | rs10276896 | -0.913 | -0.296 | 0.584 | 0.171 | -0.001 | -0.004 | -0.004 | 0 | 0.67 | 0.65 | 0.53 | 0.49 | 0.67 | 0.41 |
| 7 | 52575811 | rs10248355 | 0.278 | -0.282 | 0.487 | 0.181 | 0.001 | 0.004 | 0.004 | 1.622 | 0.43 | 0.56 | 0.4 | 0.43 | 0.47 | 0.39 |
| 7 | 52577868 | rs12539002 | 1.438 | -0.348 | 0.246 | 0.36 | 0.036 | -0.023 | -0.023 | 2.596 | 0.56 | 0.13 | 0.52 | 0.58 | 0.62 | 0.56 |
| 7 | 52579738 | rs12537723 | 1.343 | -0.062 | 0.387 | 0.474 | -0.011 | -0.019 | -0.019 | 0.177 | 0.67 | 0.26 | 0.56 | 0.6 | 0.63 | 0.56 |
| 7 | 52580826 | rs989023 | -0.025 | -0.086 | 0.589 | 0.545 | 0.035 | 0.086 | 0.086 | 0.017 | 0.82 | 0.62 | 0.65 | 0.7 | 0.79 | 0.72 |
| 7 | 52582500 | rs10280700 | 0.901 | -0.079 | 0.332 | 0.53 | 0.002 | 0.056 | 0.056 | 0.568 | 0.76 | 0.44 | 0.62 | 0.65 | 0.73 | 0.62 |
| 7 | 52592662 | rs1549868 | 2.181 | -0.033 | -0.181 | 0.347 | 0.001 | 0.015 | 0.015 | 0.253 | 0.08 | 0.05 | 0.06 | 0.02 | 0.09 | 0.02 |
| 7 | 52594753 | rs6945518 | 1.539 | -0.065 | -0.308 | 0.382 | -0.036 | -0.069 | -0.069 | 0 | 0.23 | 0.05 | 0.31 | 0.31 | 0.24 | 0.33 |
| 7 | 52595134 | rs11975368 | 0.901 | -0.052 | 0.028 | 0.398 | 0.005 | 0.07 | 0.07 | 0.006 | 0.08 | 0.24 | 0.15 | 0.14 | 0.06 | 0.16 |
| 7 | 52596669 | rs6969276 | 1.256 | -0.011 | -0.14 | 0.416 | 0.004 | 0.061 | 0.061 | 0 | 0.07 | 0.21 | 0.15 | 0.14 | 0.06 | 0.17 |
| 7 | 52607698 | rs10226087 | 0.141 | 0.044 | -0.245 | 0.252 | 0 | -0.003 | -0.003 | 0 | 0.2 | 0.22 | 0.14 | 0.13 | 0.22 | 0.1 |
| 7 | 52608767 | rs10260554 | 0.455 | -0.013 | -0.385 | 0.377 | -0.043 | -0.062 | -0.062 | 0 | 0.35 | 0.11 | 0.38 | 0.4 | 0.36 | 0.35 |
| 7 | 52612123 | rs17135127 | 1.458 | 0.168 | -0.261 | 0.484 | 0.012 | -0.016 | -0.016 | 0 | 0.09 | 0.13 | 0.05 | 0.05 | 0.11 | 0.05 |
| 7 | 52612575 | rs804755 | -0.124 | 0.176 | -0.258 | 0.486 | 0.009 | -0.001 | -0.001 | 0 | 0.19 | 0.23 | 0.14 | 0.19 | 0.25 | 0.17 |

| | | | | | | | | | | | | | | | | |
|---|---|---|---|---|---|---|---|---|---|---|---|---|---|---|---|---|
| 7 | 52622294 | rs10250763 | 1.051 | -0.03 | -0.546 | 0.343 | -0.004 | 0.006 | 0.006 | 6.178 | 0.12 | 0.06 | 0.08 | 0.09 | 0.11 | 0.07 |
| 7 | 52623937 | rs804751 | 0.252 | -0.272 | -0.587 | 0.188 | 0.007 | -0.006 | -0.006 | 0 | 0.71 | 0.72 | 0.84 | 0.58 | 0.79 | 0.64 |
| 7 | 52626048 | rs2076888 | 1.299 | 0.141 | -0.661 | 0.124 | -0.016 | 0.019 | 0.019 | 0 | 0.09 | 0.07 | 0.06 | 0.04 | 0.1 | 0.04 |
| 7 | 52630218 | rs810834 | 0.169 | 0.312 | -0.947 | 0 | 0.002 | 0 | 0 | 1.621 | 0.66 | 0.69 | 0.75 | 0.63 | 0.69 | 0.65 |
| 7 | 52643487 | rs10253778 | 1.166 | 0.654 | -0.956 | -0.043 | -0.02 | 0.02 | 0.02 | 0 | 0.09 | 0.06 | 0.07 | 0.04 | 0.12 | 0.06 |
| 7 | 52647593 | rs1442243 | 0.504 | 0.613 | -1.046 | 0.05 | -0.002 | -0.004 | -0.004 | 0 | 0.07 | 0.03 | 0.04 | 0.09 | 0.06 | 0.05 |
| 7 | 52650637 | rs9642489 | 1.402 | 0.445 | -1.168 | 0.246 | 0.008 | -0.005 | -0.005 | 0 | 0.13 | 0.12 | 0.18 | 0.37 | 0.09 | 0.15 |
| 7 | 52652332 | rs9642365 | 0.653 | 0.348 | -1.43 | -0.005 | 0.013 | -0.035 | -0.035 | 0.09 | 0.18 | 0.13 | 0.32 | 0.37 | 0.15 | 0.21 |
| 7 | 52653399 | rs17831541 | 1.067 | 0.237 | -1.251 | -0.128 | -0.001 | 0 | 0 | 0 | 0.06 | 0.03 | 0.04 | 0.07 | 0.06 | 0.04 |
| 7 | 52655552 | rs17135206 | 0.916 | 0.282 | -0.602 | -0.116 | 0.002 | 0.048 | 0.048 | 0 | 0.22 | 0.24 | 0.25 | 0.42 | 0.21 | 0.22 |
| 7 | 52655888 | rs10488323 | 2.337 | 0.45 | -0.987 | -0.328 | -0.047 | 0.047 | 0.047 | 0 | 0.08 | 0 | 0.11 | 0.01 | 0.13 | 0.14 |
| 7 | 52656778 | rs213185 | 0.223 | 0.146 | -0.956 | -0.478 | 0 | 0.001 | 0.001 | 0 | 0.23 | 0.25 | 0.22 | 0.34 | 0.26 | 0.35 |
| 7 | 52657920 | rs1816790 | 1.745 | 0.249 | -1.232 | -0.87 | -0.013 | 0.073 | 0.073 | 0.102 | 0.21 | 0.08 | 0.27 | 0.02 | 0.23 | 0.18 |
| 7 | 52659243 | rs2960632 | -0.236 | 0.314 | -0.795 | -0.618 | -0.001 | -0.009 | -0.009 | 0 | 0.44 | 0.44 | 0.42 | 0.76 | 0.43 | 0.56 |
| 7 | 52659340 | rs9791698 | 0.559 | -0.052 | -0.233 | -0.315 | -0.007 | 0.053 | 0.053 | 0.416 | 0.26 | 0.42 | 0.3 | 0.36 | 0.25 | 0.27 |
| 7 | 52660581 | rs1822228 | 2.097 | -0.113 | -0.613 | -1.019 | -0.023 | 0.074 | 0.074 | 0 | 0.21 | 0.07 | 0.18 | 0.02 | 0.23 | 0.15 |
| 7 | 52664036 | rs1348246 | 1.13 | -0.07 | -0.018 | -0.472 | 0.006 | -0.027 | -0.027 | 0.463 | 0.22 | 0.15 | 0.23 | 0.42 | 0.18 | 0.23 |
| 7 | 52671158 | rs10241795 | 0.481 | -0.093 | 0.217 | -0.488 | -0.015 | 0.081 | 0.081 | 0.026 | 0.28 | 0.4 | 0.28 | 0.46 | 0.24 | 0.26 |
| 7 | 52671586 | rs4483100 | 0.883 | -0.123 | 0.464 | -0.558 | 0.01 | 0.007 | 0.007 | 0.001 | 0.05 | 0.08 | 0.03 | 0.08 | 0.06 | 0.04 |
| 7 | 52673183 | rs9649808 | 1.329 | -0.131 | 0.427 | -0.53 | 0.038 | 0.031 | 0.031 | 0 | 0.05 | 0.23 | 0.04 | 0.05 | 0.05 | 0.03 |
| 7 | 52679045 | rs1442256 | 1.432 | -0.115 | -0.027 | -0.482 | 0.004 | -0.078 | -0.078 | 0 | 0.2 | 0.06 | 0.22 | 0.39 | 0.17 | 0.2 |
| 7 | 52682027 | rs1442252 | -0.209 | -0.295 | 0.106 | -0.73 | 0 | -0.004 | -0.004 | 0 | 0.71 | 0.7 | 0.73 | 0.82 | 0.67 | 0.76 |
| 7 | 52684353 | rs10274203 | 1.528 | -0.268 | -0.001 | -0.694 | 0.01 | -0.054 | -0.054 | 3.031 | 0.21 | 0.12 | 0.23 | 0.39 | 0.17 | 0.2 |
| 7 | 52684685 | rs2920017 | -1.785 | -0.352 | -0.213 | -0.834 | -0.011 | -0.008 | -0.008 | 0.075 | 0.89 | 0.93 | 0.81 | 0.88 | 0.78 | 0.88 |
| 7 | 52685141 | rs1442251 | 1.095 | -0.385 | -0.077 | -0.782 | -0.076 | 0.059 | 0.059 | 0 | 0.38 | 0.17 | 0.43 | 0.31 | 0.43 | 0.39 |
| 7 | 52685663 | rs2961030 | 1.085 | -0.384 | -0.066 | -0.781 | -0.071 | 0.059 | 0.059 | 0 | 0.38 | 0.18 | 0.43 | 0.31 | 0.43 | 0.39 |
| 7 | 52693587 | rs2920014 | -0.125 | -0.354 | 0.656 | -0.375 | 0.015 | -0.018 | -0.018 | 0 | 0.6 | 0.54 | 0.62 | 0.74 | 0.49 | 0.63 |
| 7 | 52696431 | rs10228124 | 0.856 | -0.196 | 0.787 | -0.139 | 0.032 | 0.025 | 0.025 | 0 | 0.05 | 0.23 | 0.05 | 0.05 | 0.06 | 0.02 |
| 7 | 52698527 | rs12671939 | 1.394 | -0.321 | 0.549 | -0.137 | -0.011 | -0.109 | -0.109 | 0 | 0.17 | 0.03 | 0.21 | 0.38 | 0.14 | 0.15 |
| 7 | 52706218 | rs1661632 | 0.997 | -0.194 | 0.624 | -0.167 | -0.015 | 0.066 | 0.066 | 0 | 0.35 | 0.19 | 0.36 | 0.28 | 0.33 | 0.34 |
| 7 | 52708424 | rs12670614 | 1.017 | -0.191 | 0.708 | -0.145 | 0.003 | -0.001 | -0.001 | 0 | 0.2 | 0.19 | 0.24 | 0.42 | 0.15 | 0.18 |
| 7 | 52710740 | rs1442260 | 0.972 | -0.329 | 0.431 | -0.222 | -0.041 | 0.078 | 0.078 | 0 | 0.32 | 0.12 | 0.28 | 0.11 | 0.38 | 0.27 |
| 7 | 52718129 | rs1620792 | 1.034 | -0.281 | 0.378 | -0.197 | -0.03 | 0.063 | 0.063 | 0 | 0.33 | 0.13 | 0.35 | 0.28 | 0.32 | 0.33 |
| 7 | 52722167 | rs10499703 | 0.8 | -0.333 | 0.797 | -0.13 | 0.016 | 0.088 | 0.088 | 0 | 0.84 | 0.6 | 0.88 | 0.79 | 0.84 | 0.76 |
| 7 | 52722258 | rs10249489 | -0.239 | -0.308 | 0.858 | -0.153 | 0 | -0.002 | -0.002 | 0.066 | 0.56 | 0.6 | 0.57 | 0.55 | 0.57 | 0.46 |
| 7 | 52722432 | rs7810220 | 0.89 | -0.305 | 0.806 | -0.175 | -0.012 | 0.036 | 0.036 | 0 | 0.32 | 0.22 | 0.29 | 0.11 | 0.38 | 0.27 |
| 7 | 52727932 | rs1661606 | 0.29 | -0.237 | 0.57 | -0.138 | 0.06 | 0.105 | 0.105 | 0 | 0.9 | 0.66 | 0.92 | 0.83 | 0.89 | 0.88 |
| 7 | 52733291 | rs11762211 | 0.85 | -0.226 | 0.099 | -0.169 | -0.055 | 0.08 | 0.08 | 0 | 0.31 | 0.09 | 0.28 | 0.11 | 0.38 | 0.27 |
| 7 | 52734328 | rs13243026 | 1.076 | -0.054 | -0.158 | -0.04 | 0.007 | -0.021 | -0.021 | 0.036 | 0.18 | 0.15 | 0.24 | 0.4 | 0.14 | 0.15 |
| 7 | 52736727 | rs11238221 | -0.159 | 0.025 | -0.082 | -0.28 | -0.003 | 0.008 | 0.008 | 0 | 0.55 | 0.4 | 0.55 | 0.55 | 0.56 | 0.55 |
| 7 | 52743776 | rs1442239 | -0.338 | 0.008 | -0.346 | -0.09 | -0.007 | 0.02 | 0.02 | 0 | 0.61 | 0.62 | 0.65 | 0.82 | 0.54 | 0.61 |
| 7 | 52743803 | rs12672075 | 0.288 | 0.427 | -0.128 | 0.63 | -0.007 | 0.026 | 0.026 | 0 | 0.2 | 0.22 | 0.25 | 0.42 | 0.13 | 0.18 |
| 7 | 52748225 | rs10272941 | 0.691 | 0.862 | -0.292 | 0.33 | 0 | -0.002 | -0.002 | 0 | 0.73 | 0.74 | 0.81 | 0.71 | 0.72 | 0.71 |
| 7 | 52750565 | rs2330173 | 0.352 | 1.153 | -0.426 | 0.314 | -0.036 | -0.03 | -0.03 | 0 | 0.06 | 0.01 | 0.19 | 0.1 | 0.13 | 0.18 |
| 7 | 52751348 | rs12534294 | 0.402 | 1.104 | -0.243 | 0.013 | 0.001 | -0.005 | -0.005 | 0.509 | 0.4 | 0.39 | 0.39 | 0.58 | 0.37 | 0.31 |
| 7 | 52751510 | rs11238224 | 0.055 | 1.124 | -0.231 | 0.009 | -0.017 | -0.018 | -0.018 | 0 | 0.85 | 0.87 | 0.73 | 0.71 | 0.75 | 0.72 |
| 7 | 52752010 | rs6973042 | 0.949 | 1.106 | -0.178 | 0.004 | -0.002 | 0 | 0 | 0.375 | 0.72 | 0.71 | 0.67 | 0.7 | 0.68 | 0.58 |
| 7 | 52752234 | rs6952975 | 1.4 | 1.149 | -0.214 | 0.017 | -0.003 | -0.023 | -0.023 | 0 | 0.32 | 0.31 | 0.27 | 0.12 | 0.31 | 0.26 |
| 7 | 52754700 | rs1372573 | 0.203 | 1.246 | -0.392 | 0.164 | -0.018 | -0.019 | -0.019 | 0.756 | 0.85 | 0.87 | 0.73 | 0.71 | 0.76 | 0.72 |
| 7 | 52757076 | rs1661612 | -0.799 | 0.451 | -0.251 | 0.085 | 0 | 0.003 | 0.003 | 0 | 0.91 | 0.88 | 0.92 | 0.82 | 0.88 | 0.9 |
| 7 | 52757480 | rs1358603 | 0.135 | 0.496 | -0.38 | 0.079 | 0 | 0.021 | 0.021 | 0 | 0.45 | 0.42 | 0.53 | 0.69 | 0.5 | 0.49 |
| 7 | 52759429 | rs1881658 | 0.185 | 0.372 | -0.242 | 0.131 | -0.022 | -0.023 | -0.023 | 1.163 | 0.77 | 0.63 | 0.85 | 0.81 | 0.81 | 0.76 |
| 7 | 52760671 | rs6958725 | 0.293 | 0.717 | -0.393 | -0.011 | -0.012 | -0.009 | -0.009 | 0 | 0.56 | 0.43 | 0.66 | 0.39 | 0.64 | 0.56 |
| 7 | 52762420 | rs1867018 | 0.19 | 1.037 | -0.364 | 0.265 | 0.005 | -0.003 | -0.003 | 0 | 0.25 | 0.25 | 0.33 | 0.26 | 0.32 | 0.29 |
| 7 | 52762734 | rs6965436 | 0.327 | -0.207 | -0.109 | 0.387 | -0.029 | 0.002 | 0.002 | 0.076 | 0.64 | 0.47 | 0.71 | 0.39 | 0.74 | 0.58 |
| 7 | 52763452 | rs1852796 | 0.641 | -0.075 | 0.393 | 0.878 | -0.002 | -0.004 | -0.004 | 0 | 0.7 | 0.6 | 0.65 | 0.69 | 0.71 | 0.73 |
| 7 | 52763685 | rs1661631 | 1.082 | -0.29 | 0.501 | 0.946 | -0.001 | 0.022 | 0.022 | 0 | 0.66 | 0.56 | 0.67 | 0.52 | 0.65 | 0.66 |
| 7 | 52764676 | rs1729830 | -1.319 | -0.606 | 0.866 | 0.932 | -0.008 | 0.018 | 0.018 | 0 | 0.35 | 0.53 | 0.32 | 0.48 | 0.33 | 0.34 |
| 7 | 52764733 | rs1729831 | -1.953 | -0.943 | 1.097 | 0.817 | 0.067 | 0.135 | 0.135 | 0 | 0.49 | 0.82 | 0.47 | 0.77 | 0.38 | 0.48 |
| 7 | 52774490 | rs1529005 | -0.312 | -1.017 | 1.165 | 1.024 | -0.01 | -0.044 | -0.044 | 0 | 0.25 | 0.34 | 0.22 | 0.09 | 0.24 | 0.19 |
| 7 | 52774983 | rs10259939 | 0.198 | -1.006 | 1.013 | 1.098 | -0.006 | 0.027 | 0.027 | 0.086 | 0.08 | 0.05 | 0.09 | 0.38 | 0.09 | 0.15 |
| 7 | 52777420 | rs890635 | -1.503 | -0.997 | 1.254 | 0.995 | 0.019 | 0.015 | 0.015 | 0 | 0.41 | 0.65 | 0.45 | 0.64 | 0.36 | 0.41 |
| 7 | 52780923 | rs1729851 | -1.52 | -0.986 | 1.193 | 1 | 0.029 | 0.023 | 0.023 | 0 | 0.41 | 0.71 | 0.45 | 0.64 | 0.36 | 0.41 |
| 7 | 52781324 | rs1529001 | -1.417 | -0.971 | 0.985 | 1.007 | 0.002 | 0.002 | 0.002 | 0 | 0.4 | 0.61 | 0.44 | 0.64 | 0.35 | 0.41 |
| 7 | 52781527 | rs1649704 | -1.417 | -0.941 | 1.02 | 0.997 | 0.002 | 0.002 | 0.002 | 0 | 0.4 | 0.61 | 0.44 | 0.64 | 0.35 | 0.41 |
| 7 | 52782466 | rs2330177 | 0.037 | -0.941 | 0.374 | 1.3 | -0.024 | 0.048 | 0.048 | 0 | 0.06 | 0.12 | 0.12 | 0.16 | 0.02 | 0.07 |
| 7 | 52784785 | rs13438347 | -0.378 | -0.869 | -0.216 | 1.246 | 0.013 | -0.066 | -0.066 | 0 | 0.15 | 0.05 | 0.12 | 0.42 | 0.12 | 0.2 |
| 7 | 52788234 | rs9690428 | -0.847 | -0.628 | -0.124 | 1.228 | 0.012 | -0.004 | -0.004 | 0 | 0.42 | 0.6 | 0.36 | 0.61 | 0.29 | 0.37 |
| 7 | 52790428 | rs1528997 | -0.91 | -0.669 | -0.2 | 0.838 | 0.029 | 0.022 | 0.022 | 0.119 | 0.44 | 0.64 | 0.39 | 0.75 | 0.29 | 0.38 |

| | | | | | | | | | | | | | | | | |
|---|---|---|---|---|---|---|---|---|---|---|---|---|---|---|---|---|
| 7 | 52793270 | rs1230545 | -0.345 | -0.346 | -0.005 | 0.791 | -0.001 | -0.003 | -0.003 | 0 | 0.2 | 0.26 | 0.15 | 0.17 | 0.2 | 0.2 |
| 7 | 52793371 | rs1230544 | -1.076 | -0.447 | -0.888 | 0.288 | -0.081 | 0.075 | 0.075 | 0.628 | 0.66 | 0.85 | 0.64 | 0.81 | 0.52 | 0.5 |
| 7 | 52794073 | rs12532211 | 1.439 | -0.964 | -0.95 | 0.464 | -0.001 | -0.001 | -0.001 | 0.033 | 0.16 | 0.12 | 0.14 | 0.15 | 0.11 | 0.03 |
| 7 | 52794572 | rs1311637 | -1.535 | -0.933 | -1.168 | 0.455 | -0.133 | 0.059 | 0.059 | 0.164 | 0.67 | 0.9 | 0.64 | 0.8 | 0.53 | 0.51 |
| 7 | 52794612 | rs1528995 | 0.522 | -1.153 | -0.33 | 0.704 | -0.015 | 0.031 | 0.031 | 0 | 0.22 | 0.28 | 0.13 | 0.36 | 0.15 | 0.14 |
| 7 | 52796887 | rs1230531 | -0.121 | -1.093 | -0.271 | 0.673 | -0.004 | 0.002 | 0.002 | 0.02 | 0.36 | 0.36 | 0.44 | 0.37 | 0.33 | 0.3 |
| 7 | 52804569 | rs6977717 | -0.897 | -0.518 | -0.065 | 0.178 | -0.002 | -0.016 | -0.016 | 0.493 | 0.3 | 0.6 | 0.25 | 0.21 | 0.3 | 0.34 |
| 7 | 52807643 | rs2330178 | -0.908 | -0.541 | 0.186 | 0.142 | -0.002 | -0.016 | -0.016 | 0 | 0.3 | 0.6 | 0.25 | 0.21 | 0.3 | 0.34 |
| 7 | 52810926 | rs17682796 | 1.51 | -0.423 | -0.199 | 0.195 | -0.015 | -0.003 | -0.003 | 0 | 0.07 | 0.01 | 0.05 | 0.06 | 0.06 | 0.08 |
| 7 | 52811025 | rs17135293 | 1.244 | -0.428 | -0.072 | 0.193 | 0.001 | 0.002 | 0.002 | 1.079 | 0.08 | 0.08 | 0.05 | 0.06 | 0.06 | 0.08 |
| 7 | 52826003 | rs11515006 | 1.732 | 0.447 | -0.81 | -0.12 | -0.099 | -0.017 | -0.017 | 3.57 | 0.17 | 0.02 | 0.24 | 0.12 | 0.29 | 0.23 |
| 7 | 52839549 | rs6976719 | -0.99 | -0.085 | 0.084 | 0.274 | 0.021 | 0.003 | 0.003 | 0.076 | 0.18 | 0.42 | 0.17 | 0.17 | 0.16 | 0.18 |
| 7 | 52840889 | rs11975075 | 0.574 | -0.274 | -0.586 | 0.356 | 0.008 | -0.017 | -0.017 | 0.475 | 0.23 | 0.06 | 0.2 | 0.19 | 0.18 | 0.11 |
| 7 | 52846014 | rs2965507 | 0.224 | -0.068 | -0.438 | 0.11 | 0.048 | 0.034 | 0.034 | 0 | 0.08 | 0.21 | 0.11 | 0.11 | 0.1 | 0.1 |
| 7 | 52855306 | rs2965511 | 0.224 | -0.068 | -0.438 | 0.11 | 0.048 | 0.034 | 0.034 | 0 | 0.08 | 0.21 | 0.11 | 0.11 | 0.1 | 0.1 |
| 7 | 52857194 | rs971648 | -1.492 | -0.277 | -0.574 | 0.017 | -0.061 | 0.131 | 0.131 | 0 | 0.58 | 0.89 | 0.64 | 0.81 | 0.49 | 0.54 |
| 7 | 52859106 | rs1528989 | -1.258 | -0.091 | -0.115 | 0.029 | 0.002 | -0.03 | -0.03 | 0.258 | 0.28 | 0.4 | 0.24 | 0.2 | 0.29 | 0.33 |
| 7 | 52865546 | rs2912902 | -0.539 | 0.032 | -0.13 | 0.122 | 0.003 | -0.02 | -0.02 | 0 | 0.2 | 0.29 | 0.18 | 0.14 | 0.23 | 0.25 |
| 7 | 52866542 | rs1919543 | -0.701 | 0.034 | -0.129 | 0.123 | 0.003 | -0.02 | -0.02 | 0.396 | 0.2 | 0.29 | 0.18 | 0.14 | 0.23 | 0.25 |
| 7 | 52868222 | rs17135330 | 0.733 | 0.016 | -0.364 | 0.113 | 0.008 | 0.031 | 0.031 | 0 | 0.13 | 0.09 | 0.07 | 0.03 | 0.13 | 0.15 |
| 7 | 52875730 | rs2912913 | -1.185 | -0.115 | -0.094 | 0.166 | -0.004 | -0.021 | -0.021 | 0.017 | 0.31 | 0.45 | 0.28 | 0.24 | 0.33 | 0.36 |
| 7 | 52878284 | rs11773689 | 0.986 | -0.136 | -0.103 | 0.147 | 0.006 | 0.025 | 0.025 | 0 | 0.12 | 0.09 | 0.07 | 0.04 | 0.12 | 0.15 |
| 7 | 52882629 | rs7785023 | -1.188 | -0.144 | 0.031 | 0.172 | -0.015 | -0.011 | -0.011 | 0 | 0.3 | 0.37 | 0.24 | 0.23 | 0.28 | 0.33 |
| 7 | 52893743 | rs2965587 | -1.141 | -0.133 | 0.051 | 0.153 | -0.004 | -0.003 | -0.003 | 0 | 0.29 | 0.32 | 0.23 | 0.23 | 0.28 | 0.33 |
| 7 | 52897504 | rs11976451 | 1.454 | -0.119 | 0.095 | 0.168 | -0.009 | -0.001 | -0.001 | 0.019 | 0.07 | 0.12 | 0.06 | 0.04 | 0.05 | 0.05 |
| 7 | 52899187 | rs1863786 | -0.453 | -0.155 | -0.106 | 0.177 | 0.003 | 0.004 | 0.004 | 1.404 | 0.21 | 0.19 | 0.17 | 0.19 | 0.23 | 0.28 |
| 7 | 52899925 | rs2912967 | -0.479 | -0.163 | -0.097 | 0.173 | -0.002 | -0.002 | -0.002 | 0 | 0.21 | 0.23 | 0.17 | 0.19 | 0.23 | 0.28 |
| 7 | 52903807 | rs11238232 | 0.568 | -0.182 | -0.08 | 0.13 | 0.016 | 0 | 0 | 0.868 | 0.1 | 0.22 | 0.11 | 0.06 | 0.11 | 0.21 |
| 7 | 52905808 | rs2965634 | -0.491 | -0.185 | -0.172 | 0.128 | -0.001 | -0.002 | -0.002 | 0 | 0.22 | 0.23 | 0.17 | 0.2 | 0.23 | 0.29 |
| 7 | 52905841 | rs2912974 | -0.49 | -0.188 | -0.173 | 0.127 | -0.001 | -0.002 | -0.002 | 0.815 | 0.22 | 0.23 | 0.17 | 0.2 | 0.23 | 0.29 |
| 7 | 52907463 | rs990236 | 0.969 | -0.154 | -0.53 | 0.154 | -0.001 | 0.014 | 0.014 | 0 | 0.06 | 0.08 | 0.07 | 0.1 | 0.07 | 0.07 |
| 7 | 52910684 | rs13246506 | 1.292 | -0.027 | -1.249 | -0.03 | -0.05 | 0.017 | 0.017 | 0.03 | 0.18 | 0.01 | 0.16 | 0.09 | 0.2 | 0.08 |
| 7 | 52917246 | rs2913002 | -0.576 | 0.207 | -1.015 | -0.078 | 0.01 | 0.003 | 0.003 | 0.253 | 0.21 | 0.32 | 0.22 | 0.21 | 0.23 | 0.32 |
| 7 | 52922213 | rs11765529 | 2.425 | 0.244 | -1.208 | -0.081 | -0.023 | -0.007 | -0.007 | 0.574 | 0.05 | 0.01 | 0.04 | 0.05 | 0.06 | 0.07 |
| 7 | 52929758 | rs2965656 | -1.402 | -0.07 | -1.343 | 0.028 | -0.064 | 0.082 | 0.082 | 0 | 0.61 | 0.88 | 0.64 | 0.77 | 0.56 | 0.66 |
| 7 | 52930883 | rs2195139 | 0.395 | 0.134 | -0.782 | 0.017 | -0.011 | 0.022 | 0.022 | 2.063 | 0.16 | 0.06 | 0.11 | 0.1 | 0.16 | 0.22 |
| 7 | 52936160 | rs11764533 | 0.944 | 0.099 | -0.756 | 0.02 | 0.018 | 0.029 | 0.029 | 0 | 0.11 | 0.05 | 0.05 | 0.05 | 0.1 | 0.14 |
| 7 | 52941674 | rs11767372 | 1.33 | 0.133 | -0.74 | 0.036 | 0.011 | 0.022 | 0.022 | 0 | 0.1 | 0.04 | 0.05 | 0.05 | 0.09 | 0.14 |
| 7 | 52944424 | rs7805401 | 0.841 | 0.184 | -0.555 | 0.103 | -0.001 | -0.003 | -0.003 | 0 | 0.67 | 0.64 | 0.72 | 0.73 | 0.67 | 0.53 |
| 7 | 52945935 | rs7798662 | 1.779 | 0.39 | -0.819 | 0.355 | -0.023 | -0.007 | -0.007 | 0.46 | 0.05 | 0.01 | 0.04 | 0.05 | 0.06 | 0.08 |
| 7 | 52947719 | rs7789851 | 0.302 | 0.307 | -0.689 | 0.173 | 0 | 0.006 | 0.006 | 0 | 0.43 | 0.34 | 0.41 | 0.63 | 0.47 | 0.42 |
| 7 | 52948897 | rs12718708 | 0.965 | 0.069 | -0.851 | 0.129 | -0.019 | 0.029 | 0.029 | 0 | 0.19 | 0.05 | 0.15 | 0.1 | 0.2 | 0.1 |
| 7 | 52950472 | rs6967460 | -0.774 | 0.152 | -0.314 | 0.419 | 0.003 | 0.049 | 0.049 | 0 | 0.93 | 0.88 | 0.74 | 0.86 | 0.92 | 0.92 |
| 7 | 52952144 | rs6974056 | -0.016 | 0.038 | -0.476 | 0.08 | 0.007 | -0.022 | -0.022 | 0 | 0.23 | 0.18 | 0.13 | 0 | 0.18 | 0.08 |
| 7 | 52954110 | rs4476958 | -0.764 | 0.475 | -0.093 | 0.59 | -0.001 | 0.011 | 0.011 | 0 | 0.94 | 0.94 | 0.75 | 0.86 | 0.92 | 0.92 |
| 7 | 52957973 | rs7800684 | 0.514 | 0.38 | -0.138 | 0.407 | 0.004 | -0.003 | -0.003 | 0 | 0.14 | 0.11 | 0.07 | 0 | 0.11 | 0.04 |
| 7 | 52965371 | rs10248576 | 0.117 | 0.562 | -0.167 | 0.678 | 0 | -0.001 | -0.001 | 0 | 0.81 | 0.85 | 0.67 | 0.84 | 0.81 | 0.88 |
| 7 | 52967922 | rs6955298 | 0.231 | 0.516 | 0.33 | 1.464 | -0.006 | 0.069 | 0.069 | 0 | 0.16 | 0.24 | 0.21 | 0.38 | 0.13 | 0.23 |
| 7 | 52968984 | rs7806124 | -0.135 | 0.496 | 0.296 | 1.485 | 0.001 | 0.002 | 0.002 | 0 | 0.93 | 0.9 | 0.94 | 0.91 | 0.92 | 0.89 |
| 7 | 52971229 | rs12535352 | 0.231 | 0.544 | 0.014 | 1.432 | -0.027 | -0.031 | -0.031 | 0 | 0.14 | 0.05 | 0.16 | 0.16 | 0.17 | 0.1 |
| 7 | 52971254 | rs7791066 | 0.71 | 0.245 | 0.793 | 1.355 | 0.014 | 0.067 | 0.067 | 0.042 | 0.89 | 0.69 | 0.88 | 0.84 | 0.9 | 0.85 |
| 7 | 52975364 | rs10257778 | 0.286 | 0.483 | 0.075 | 1.263 | -0.001 | 0.027 | 0.027 | 7.053 | 0.07 | 0.15 | 0.03 | 0.13 | 0.05 | 0.14 |
| 7 | 52989552 | rs10276114 | -1.012 | 0.686 | 0.406 | 1.571 | 0.035 | 0.177 | 0.177 | 0 | 0.16 | 0.42 | 0.19 | 0.36 | 0.15 | 0.17 |
| 7 | 52990209 | rs10249461 | -0.332 | 0.112 | -0.047 | 1.3 | 0.003 | -0.003 | -0.003 | 0 | 0.4 | 0.39 | 0.37 | 0.54 | 0.37 | 0.35 |
| 7 | 52990398 | rs4947756 | 0.742 | -0.25 | 0.128 | 1.18 | 0 | 0.029 | 0.029 | 0.595 | 0.47 | 0.39 | 0.49 | 0.3 | 0.52 | 0.58 |
| 7 | 52990517 | rs4947417 | 1.247 | -0.975 | 0.207 | 0.862 | -0.01 | -0.015 | -0.015 | 0 | 0.08 | 0.12 | 0.14 | 0.02 | 0.07 | 0.04 |
| 7 | 52990869 | rs6951369 | 0.031 | -1.494 | -0.124 | 0.995 | -0.037 | 0.04 | 0.04 | 0.232 | 0.66 | 0.5 | 0.64 | 0.53 | 0.74 | 0.77 |
| 7 | 52992974 | rs10239546 | 0.375 | -1.395 | -0.197 | 0.977 | -0.006 | 0.004 | 0.004 | 0.028 | 0.6 | 0.43 | 0.57 | 0.51 | 0.67 | 0.72 |
| 7 | 52994281 | rs7804465 | 0.446 | -1.311 | -0.756 | 0.569 | 0.002 | 0.001 | 0.001 | 0.008 | 0.35 | 0.38 | 0.38 | 0.38 | 0.37 | 0.22 |
| 7 | 52995247 | rs10248341 | 0.644 | -1.587 | -0.483 | 0.621 | -0.017 | 0.194 | 0.194 | 0.101 | 0.13 | 0.36 | 0.17 | 0.33 | 0.08 | 0.07 |
| 7 | 52995409 | rs6593033 | -0.866 | -1.624 | -0.5 | 0.595 | 0.005 | 0.004 | 0.004 | 0 | 0.78 | 0.8 | 0.79 | 0.86 | 0.81 | 0.83 |
| 7 | 52995901 | rs12718724 | 0.127 | -1.46 | -0.537 | 0.564 | -0.009 | 0.11 | 0.11 | 0 | 0.18 | 0.37 | 0.22 | 0.35 | 0.14 | 0.11 |
| 7 | 52996013 | rs4326334 | 1.376 | -1.363 | -1.254 | 0.22 | -0.155 | 0.06 | 0.06 | 1.07 | 0.32 | 0.03 | 0.29 | 0.24 | 0.42 | 0.45 |
| 7 | 53000794 | rs4245557 | 0.033 | -1.095 | -0.229 | 0.495 | -0.01 | 0.02 | 0.02 | 0 | 0.66 | 0.47 | 0.63 | 0.49 | 0.7 | 0.71 |
| 7 | 53001439 | rs12718725 | 0.474 | -0.962 | -0.388 | 0.563 | 0.035 | 0.043 | 0.043 | 1.282 | 0.5 | 0.29 | 0.45 | 0.47 | 0.54 | 0.64 |
| 7 | 53006704 | rs7806786 | 0.408 | -0.7 | -0.27 | 0.514 | -0.012 | 0.011 | 0.011 | 0 | 0.4 | 0.31 | 0.41 | 0.37 | 0.5 | 0.4 |
| 7 | 53008596 | rs11771058 | 0.975 | -0.66 | -0.285 | 0.456 | -0.023 | 0.072 | 0.072 | 0.612 | 0.3 | 0.2 | 0.27 | 0.08 | 0.35 | 0.29 |
| 7 | 53008761 | rs1689295 | 0.035 | -0.976 | -0.017 | 0.344 | -0.006 | -0.008 | -0.008 | 0 | 0.44 | 0.46 | 0.39 | 0.61 | 0.32 | 0.53 |
| 7 | 53009493 | rs1689297 | -1.174 | -0.878 | -0.001 | 0.356 | -0.012 | -0.047 | -0.047 | 1.158 | 0.83 | 0.75 | 0.8 | 0.98 | 0.84 | 0.93 |

| | | | | | | | | | | | | | | | | |
|---|---|---|---|---|---|---|---|---|---|---|---|---|---|---|---|---|
| 7 | 53011628 | rs1689304 | -0.461 | -0.894 | -0.153 | 0.461 | 0.001 | 0 | 0 | 0 | 0.77 | 0.76 | 0.73 | 0.8 | 0.79 | 0.88 |
| 7 | 53011910 | rs11760600 | 1.07 | -0.898 | -0.207 | 0.38 | -0.029 | 0.043 | 0.043 | 0.285 | 0.31 | 0.24 | 0.33 | 0.18 | 0.45 | 0.32 |
| 7 | 53012513 | rs7795100 | 1.102 | -0.983 | -0.312 | 0.67 | -0.026 | 0.047 | 0.047 | 0.515 | 0.29 | 0.18 | 0.27 | 0.15 | 0.36 | 0.29 |
| 7 | 53015990 | rs1320888 | 1.529 | -0.975 | -0.316 | 0.373 | -0.012 | 0.043 | 0.043 | 0.02 | 0.24 | 0.17 | 0.23 | 0.05 | 0.31 | 0.24 |
| 7 | 53016462 | rs1689310 | 0.014 | -1.001 | -0.249 | 0.447 | 0.004 | 0.006 | 0.006 | 0 | 0.46 | 0.51 | 0.4 | 0.61 | 0.34 | 0.56 |
| 7 | 53025804 | rs6968538 | 1.105 | -0.868 | 0.058 | 0.645 | 0.011 | 0.091 | 0.091 | 0 | 0.16 | 0.26 | 0.25 | 0.34 | 0.25 | 0.13 |
| 7 | 53031792 | rs6955255 | -0.06 | -1.149 | 0.097 | 0.584 | -0.006 | 0.089 | 0.089 | 0 | 0.44 | 0.28 | 0.37 | 0.26 | 0.34 | 0.54 |
| 7 | 53031812 | rs6955269 | -0.578 | -1.117 | -0.23 | 0.653 | -0.002 | 0.003 | 0.003 | 0.257 | 0.6 | 0.55 | 0.62 | 0.61 | 0.59 | 0.67 |
| 7 | 53034197 | rs10499707 | 0.843 | -1.157 | -0.446 | 0.519 | -0.002 | 0.027 | 0.027 | 0 | 0.28 | 0.16 | 0.26 | 0.24 | 0.22 | 0.4 |
| 7 | 53035166 | rs11763192 | -0.549 | -0.836 | -0.153 | 0.652 | -0.019 | 0.058 | 0.058 | 0 | 0.84 | 0.75 | 0.85 | 0.73 | 0.9 | 0.91 |
| 7 | 53036443 | rs4947763 | -0.489 | -0.51 | -0.277 | 0.51 | 0.001 | -0.001 | -0.001 | 0 | 0.29 | 0.27 | 0.23 | 0.27 | 0.18 | 0.3 |
| 7 | 53037887 | rs2172510 | 0.936 | -0.279 | -0.299 | 0.441 | -0.012 | 0.029 | 0.029 | 0 | 0.38 | 0.3 | 0.49 | 0.22 | 0.55 | 0.31 |
| 7 | 53039588 | rs4947764 | -0.083 | -0.244 | -0.257 | 0.526 | -0.012 | 0.004 | 0.004 | 0 | 0.24 | 0.24 | 0.19 | 0.25 | 0.12 | 0.22 |
| 7 | 53049631 | rs11238241 | 1.103 | 0.679 | 0.232 | 0.031 | 0.016 | 0.003 | 0.003 | 0 | 0.13 | 0.23 | 0.26 | 0.16 | 0.24 | 0.09 |
| 7 | 53062116 | rs4947770 | 0.762 | 0.576 | 0.227 | 0.002 | -0.001 | 0.088 | 0.088 | 0 | 0.5 | 0.34 | 0.61 | 0.29 | 0.6 | 0.42 |
| 7 | 53063823 | rs17135475 | 1.616 | 0.897 | 0.326 | 0.133 | 0.029 | -0.021 | -0.021 | 0.203 | 0.05 | 0.15 | 0.05 | 0.02 | 0.06 | 0.08 |
| 7 | 53064379 | rs17735647 | 1.351 | 1.01 | 0.068 | 0.23 | -0.003 | -0.011 | -0.011 | 0 | 0.08 | 0.04 | 0.08 | 0.13 | 0.07 | 0.06 |
| 7 | 53066952 | rs923823 | -0.063 | 0.897 | 0.05 | 0.229 | -0.002 | 0.003 | 0.003 | 0 | 0.62 | 0.45 | 0.73 | 0.45 | 0.71 | 0.56 |
| 7 | 53068819 | rs17668139 | 0.443 | 1.23 | -0.217 | 0.268 | 0.002 | 0 | 0 | 0 | 0.13 | 0.17 | 0.13 | 0.15 | 0.13 | 0.14 |
| 7 | 53068973 | rs11238242 | 0.093 | 0.912 | -0.259 | 0.598 | -0.001 | 0.03 | 0.03 | 0 | 0.67 | 0.59 | 0.79 | 0.55 | 0.74 | 0.56 |
| 7 | 53075827 | rs2087912 | -0.388 | 0.771 | -0.944 | 1.13 | 0.001 | -0.015 | -0.015 | 2.577 | 0.76 | 0.76 | 0.87 | 0.55 | 0.82 | 0.62 |
| 7 | 53080551 | rs12673285 | -0.393 | 0.904 | -0.825 | 0.936 | -0.003 | 0 | 0 | 0 | 0.13 | 0.07 | 0.25 | 0.14 | 0.23 | 0.11 |
| 7 | 53081669 | rs12673084 | -0.393 | 0.904 | -0.825 | 0.936 | -0.003 | 0 | 0 | 0.782 | 0.13 | 0.07 | 0.25 | 0.14 | 0.23 | 0.11 |
| 7 | 53100319 | rs2330339 | -0.328 | 0.718 | -0.84 | 1.017 | 0 | -0.01 | -0.01 | 0 | 0.08 | 0.06 | 0.08 | 0.13 | 0.07 | 0.07 |
| 7 | 53102259 | rs4947776 | -0.408 | 0.476 | -0.791 | 1.132 | -0.001 | -0.001 | -0.001 | 0 | 0.13 | 0.08 | 0.12 | 0.11 | 0.12 | 0.06 |
| 7 | 53102360 | rs4947777 | -1.684 | -0.012 | -0.252 | 1.096 | -0.015 | 0.006 | 0.006 | 0 | 0.25 | 0.26 | 0.22 | 0.27 | 0.19 | 0.14 |
| 7 | 53103257 | rs10229649 | 0.17 | -0.297 | 0.353 | 0.992 | 0.008 | 0.008 | 0.008 | 0.939 | 0.09 | 0.22 | 0.21 | 0.1 | 0.15 | 0.05 |
| 7 | 53103267 | rs10248048 | -0.105 | -0.221 | 0.061 | 0.626 | 0.004 | 0.018 | 0.018 | 0 | 0.51 | 0.41 | 0.65 | 0.4 | 0.63 | 0.39 |
| 7 | 53104700 | rs9770361 | 0.444 | -0.277 | -0.313 | 0.538 | -0.019 | 0.046 | 0.046 | 0 | 0.4 | 0.19 | 0.42 | 0.3 | 0.46 | 0.28 |
| 7 | 53105335 | rs977663 | 0.308 | -0.37 | -0.096 | 0.507 | 0.007 | -0.025 | -0.025 | 0 | 0.1 | 0.03 | 0.09 | 0.15 | 0.07 | 0.08 |
| 7 | 53105925 | rs13226989 | 0.335 | -0.346 | -0.07 | 0.543 | -0.013 | 0.07 | 0.07 | 0 | 0.42 | 0.21 | 0.43 | 0.27 | 0.49 | 0.34 |
| 7 | 53111699 | rs17643382 | 0.881 | -0.219 | -0.154 | 0.646 | -0.022 | -0.058 | -0.058 | 0.151 | 0.36 | 0.22 | 0.25 | 0.47 | 0.3 | 0.47 |
| 7 | 53119005 | rs1174760 | -0.084 | -0.179 | 0.117 | 0.853 | -0.005 | 0.029 | 0.029 | 0 | 0.92 | 0.92 | 0.92 | 0.86 | 0.95 | 0.93 |
| 7 | 53120437 | rs880048 | -0.954 | 0.778 | -0.588 | 1.346 | -0.071 | -0.122 | -0.122 | 0 | 0.21 | 0.06 | 0.4 | 0.5 | 0.32 | 0.24 |
| 7 | 53127559 | rs1174864 | -0.447 | 0.239 | -0.407 | 0.479 | 0 | 0.001 | 0.001 | 0 | 0.48 | 0.48 | 0.47 | 0.19 | 0.58 | 0.29 |
| 7 | 53131432 | rs6967541 | 0.604 | 0.335 | -0.565 | 0.402 | -0.045 | -0.036 | -0.036 | 0 | 0.12 | 0.01 | 0.23 | 0.14 | 0.18 | 0.11 |
| 7 | 53134593 | rs1174749 | 0.537 | 0.145 | -0.243 | 0.287 | -0.001 | -0.05 | -0.05 | 0 | 0.31 | 0.28 | 0.22 | 0.05 | 0.4 | 0.17 |
| 7 | 53142268 | rs12535625 | -1.079 | 0.201 | 0.034 | 0.908 | 0.006 | 0.059 | 0.059 | 0.135 | 0.22 | 0.26 | 0.43 | 0.45 | 0.27 | 0.21 |
| 7 | 53150728 | rs1021465 | 0.046 | 0.212 | 0.058 | 0.907 | -0.001 | -0.003 | -0.003 | 0 | 0.15 | 0.15 | 0.25 | 0.14 | 0.18 | 0.11 |
| 7 | 53154944 | rs7782134 | -1.054 | 0.062 | -0.254 | 0.977 | -0.015 | -0.032 | -0.032 | 0 | 0.93 | 1 | 0.9 | 0.91 | 0.94 | 0.93 |
| 7 | 53155331 | rs1464888 | -0.948 | 0.039 | 0.026 | 0.847 | 0.006 | 0.098 | 0.098 | 0.446 | 0.23 | 0.52 | 0.43 | 0.45 | 0.27 | 0.19 |
| 7 | 53156188 | rs6967345 | -0.466 | -0.233 | -1.389 | 1.142 | 0.042 | 0.129 | 0.129 | 0 | 0.53 | 0.73 | 0.52 | 0.28 | 0.65 | 0.33 |
| 7 | 53167501 | rs6593048 | -0.609 | -0.303 | -1.719 | 0.833 | 0.007 | -0.024 | -0.024 | 0.65 | 0.55 | 0.81 | 0.64 | 0.5 | 0.64 | 0.38 |
| 7 | 53168727 | rs17557291 | 0.391 | -0.351 | -2.116 | 0.733 | 0.034 | -0.041 | -0.041 | 1.113 | 0.43 | 0.12 | 0.23 | 0.44 | 0.29 | 0.54 |
| 7 | 53168912 | rs17557298 | 1 | 0.015 | -1.833 | 0.036 | 0.013 | 0.041 | 0.041 | 0 | 0.32 | 0.1 | 0.24 | 0.25 | 0.23 | 0.45 |
| 7 | 53171595 | rs10249125 | 0.153 | 0.43 | -1.332 | 0.046 | -0.003 | 0.047 | 0.047 | 0 | 0.21 | 0.14 | 0.15 | 0.02 | 0.22 | 0.09 |
| 7 | 53174688 | rs6973717 | -0.545 | 0.325 | -0.656 | 0.341 | -0.002 | -0.071 | -0.071 | 1.584 | 0.59 | 0.51 | 0.68 | 0.89 | 0.64 | 0.61 |
| 7 | 53175158 | rs10236661 | -0.51 | -0.006 | -0.451 | 1.702 | -0.004 | 0.067 | 0.067 | 0 | 0.07 | 0.13 | 0.15 | 0.28 | 0.04 | 0.16 |
| 7 | 53177183 | rs877940 | -0.678 | -0.265 | 0.08 | 1.797 | -0.017 | 0.209 | 0.209 | 0 | 0.11 | 0.38 | 0.19 | 0.39 | 0.06 | 0.12 |
| 7 | 53177456 | rs2222614 | -0.87 | 0.018 | -0.477 | 0.9 | -0.007 | 0.028 | 0.028 | 0 | 0.16 | 0.08 | 0.11 | 0.05 | 0.2 | 0.12 |
| 7 | 53182000 | rs6970154 | 1.14 | -0.242 | -0.798 | 1.159 | -0.01 | -0.013 | -0.013 | 0 | 0.77 | 0.85 | 0.74 | 0.77 | 0.7 | 0.78 |
| 7 | 53186040 | rs6954000 | -0.288 | -0.026 | -0.509 | 1.074 | 0.008 | 0.099 | 0.099 | 0 | 0.21 | 0.39 | 0.31 | 0.53 | 0.22 | 0.24 |
| 7 | 53187490 | rs7793024 | 0.81 | 0.267 | -0.692 | 1.196 | -0.007 | -0.009 | -0.009 | 0 | 0.8 | 0.88 | 0.76 | 0.78 | 0.73 | 0.81 |
| 7 | 53198269 | rs6952331 | 0.273 | -0.007 | -0.831 | 0.778 | -0.022 | 0.027 | 0.027 | 0 | 0.85 | 0.93 | 0.85 | 0.94 | 0.77 | 0.84 |
| 7 | 53204653 | rs13231130 | 1.402 | -0.63 | -1.327 | 0.113 | -0.039 | 0.079 | 0.079 | 0 | 0.34 | 0.19 | 0.28 | 0.18 | 0.38 | 0.34 |
| 7 | 53206225 | rs2330341 | 0.049 | -0.82 | -0.855 | 0.55 | 0.043 | 0.014 | 0.014 | 0.021 | 0.38 | 0.2 | 0.35 | 0.3 | 0.26 | 0.33 |
| 7 | 53214203 | rs1525574 | -0.371 | -0.678 | -0.962 | 0.455 | -0.053 | 0.053 | 0.053 | 0 | 0.87 | 0.99 | 0.86 | 0.96 | 0.78 | 0.85 |
| 7 | 53214419 | rs1357961 | 1.051 | -0.844 | -0.184 | 0.683 | 0 | -0.003 | -0.003 | 0 | 0.45 | 0.45 | 0.44 | 0.5 | 0.51 | 0.41 |
| 7 | 53216299 | rs2877103 | -0.209 | -0.382 | -0.366 | 0.291 | -0.053 | 0.053 | 0.053 | 0 | 0.87 | 0.99 | 0.86 | 0.96 | 0.78 | 0.85 |
| 7 | 53243789 | rs1608852 | 0.732 | -0.151 | 0.261 | 0.192 | 0.066 | 0 | 0 | 0 | 0.75 | 0.57 | 0.61 | 0.72 | 0.59 | 0.7 |
| 7 | 53244852 | rs4947421 | 0.296 | -0.673 | 0.365 | 0.06 | 0.018 | -0.016 | -0.016 | 0 | 0.31 | 0.24 | 0.33 | 0.51 | 0.16 | 0.27 |
| 7 | 53249376 | rs7809635 | -0.211 | -0.546 | 0.218 | 0.063 | 0.026 | 0.04 | 0.04 | 0 | 0.19 | 0.41 | 0.26 | 0.24 | 0.21 | 0.21 |
| 7 | 53251092 | rs7808774 | -1.049 | -0.535 | 0.179 | 0.063 | -0.036 | 0.025 | 0.025 | 0 | 0.93 | 0.98 | 0.87 | 0.96 | 0.8 | 0.91 |
| 7 | 53252768 | rs7805945 | -0.345 | -0.682 | 0.334 | 0.051 | 0.077 | 0.184 | 0.184 | 0 | 0.08 | 0.33 | 0.13 | 0.23 | 0.12 | 0.11 |
| 7 | 53253372 | rs1404880 | 0.592 | -1.29 | -0.348 | -0.164 | -0.021 | 0.034 | 0.034 | 0 | 0.1 | 0 | 0.05 | 0.02 | 0.08 | 0.07 |
| 7 | 53254366 | rs1357959 | 0.412 | -1.332 | -0.219 | -0.1 | 0.076 | -0.117 | -0.117 | 0 | 0.3 | 0.05 | 0.32 | 0.51 | 0.16 | 0.27 |
| 7 | 53256063 | rs11978966 | 0.584 | -0.706 | 0.318 | 0.1 | -0.004 | -0.043 | -0.043 | 0 | 0.11 | 0.15 | 0.14 | 0 | 0.1 | 0.1 |
| 7 | 53260331 | rs10276168 | 1.739 | -0.87 | 0.069 | -0.16 | -0.047 | 0.027 | 0.027 | 0 | 0.35 | 0.2 | 0.27 | 0.29 | 0.44 | 0.41 |
| 7 | 53262051 | rs17558136 | 0.443 | -0.981 | 0.241 | -0.263 | 0.001 | 0.008 | 0.008 | 0 | 0.11 | 0.04 | 0.05 | 0.02 | 0.08 | 0.07 |

| | | | | | | | | | | | | | | | | |
|---|---|---|---|---|---|---|---|---|---|---|---|---|---|---|---|---|
| 7 | 53262113 | rs10231635 | 0.655 | -1.045 | 0.832 | -0.283 | -0.001 | -0.008 | -0.008 | 0 | 0.48 | 0.49 | 0.33 | 0.3 | 0.51 | 0.48 |
| 7 | 53263554 | rs10282010 | 0.856 | -1.048 | 0.909 | -0.284 | 0.004 | 0.044 | 0.044 | 0 | 0.47 | 0.3 | 0.32 | 0.3 | 0.51 | 0.48 |
| 7 | 53265599 | rs1525577 | -0.227 | -1.073 | 0.968 | -0.345 | -0.011 | 0.005 | 0.005 | 0 | 0.81 | 0.86 | 0.73 | 0.96 | 0.7 | 0.81 |
| 7 | 53270914 | rs6952158 | 0.277 | -1.157 | 0.987 | -0.345 | -0.016 | 0.001 | 0.001 | 0 | 0.31 | 0.35 | 0.4 | 0.65 | 0.18 | 0.3 |
| 7 | 53272719 | rs7795436 | 1.731 | -1.123 | 0.921 | -0.358 | -0.007 | 0.009 | 0.009 | 0 | 0.39 | 0.41 | 0.28 | 0.28 | 0.44 | 0.43 |
| 7 | 53276952 | rs1554713 | 0.604 | -1.125 | 0.747 | -0.197 | 0.037 | -0.075 | -0.075 | 0 | 0.27 | 0.1 | 0.35 | 0.51 | 0.17 | 0.26 |
| 7 | 53277006 | rs17558317 | 1.362 | -0.945 | 0.714 | -0.365 | 0.002 | -0.001 | -0.001 | 0 | 0.11 | 0.08 | 0.14 | 0 | 0.1 | 0.09 |
| 7 | 53283134 | rs10246618 | 1.178 | -0.643 | 0.965 | -0.057 | -0.011 | 0.018 | 0.018 | 0 | 0.39 | 0.36 | 0.28 | 0.29 | 0.44 | 0.44 |
| 7 | 53290896 | rs6651136 | 1.017 | -0.566 | 1.112 | -0.255 | 0.021 | -0.075 | -0.075 | 0 | 0.8 | 0.66 | 0.83 | 0.94 | 0.76 | 0.85 |
| 7 | 53292132 | rs6956230 | 1.341 | -0.593 | 0.828 | 0.185 | -0.048 | 0.001 | 0.001 | 0 | 0.4 | 0.24 | 0.4 | 0.41 | 0.47 | 0.56 |
| 7 | 53292859 | rs4147413 | -1.196 | -0.489 | 0.661 | 0.211 | -0.037 | 0.012 | 0.012 | 0 | 0.94 | 0.99 | 0.89 | 0.96 | 0.84 | 0.93 |
| 7 | 53294765 | rs12718755 | 0.093 | -0.517 | 0.863 | 0.215 | 0.037 | -0.022 | -0.022 | 0.152 | 0.4 | 0.3 | 0.42 | 0.51 | 0.28 | 0.29 |
| 7 | 53294958 | rs10233599 | 0.683 | -0.437 | 0.859 | 0.128 | 0.005 | -0.021 | -0.021 | 0 | 0.82 | 0.78 | 0.83 | 0.94 | 0.76 | 0.86 |
| 7 | 53299136 | rs10232192 | 0.387 | -0.486 | 0.971 | 0.221 | 0.066 | -0.061 | -0.061 | 0 | 0.87 | 0.58 | 0.86 | 0.96 | 0.82 | 0.92 |
| 7 | 53303291 | rs12666375 | -0.041 | -0.691 | 0.733 | 0.375 | 0.045 | -0.028 | -0.028 | 0 | 0.39 | 0.24 | 0.42 | 0.53 | 0.29 | 0.3 |
| 7 | 53306008 | rs7793602 | -0.444 | -0.291 | 0.944 | 0.209 | 0.013 | -0.036 | -0.036 | 0 | 0.1 | 0.25 | 0.04 | 0.01 | 0.07 | 0.03 |
| 7 | 53308306 | rs10247908 | -0.031 | -0.277 | 0.843 | 0.215 | -0.001 | -0.025 | -0.025 | 0 | 0.08 | 0.13 | 0.04 | 0.01 | 0.06 | 0.03 |
| 7 | 53311527 | rs6971101 | -0.627 | -0.615 | 0.828 | 0.344 | -0.001 | -0.001 | -0.001 | 0.004 | 0.44 | 0.52 | 0.41 | 0.3 | 0.41 | 0.32 |
| 7 | 53314226 | rs4299959 | -0.5 | -0.561 | 0.676 | 0.295 | -0.005 | 0.007 | 0.007 | 0 | 0.91 | 0.94 | 0.89 | 0.93 | 0.89 | 0.94 |
| 7 | 53316102 | rs7808809 | 0.328 | -0.592 | 0.788 | 0.298 | -0.003 | -0.004 | -0.004 | 0 | 0.35 | 0.36 | 0.37 | 0.28 | 0.35 | 0.28 |
| 7 | 53327665 | rs6956897 | -0.582 | -0.001 | 0.597 | 0.521 | -0.041 | -0.001 | -0.001 | 0 | 0.91 | 0.97 | 0.88 | 0.93 | 0.87 | 0.92 |
| 7 | 53330504 | rs12718775 | 0.973 | -0.252 | 0.743 | 0.581 | 0.009 | -0.043 | -0.043 | 0 | 0.43 | 0.18 | 0.39 | 0.53 | 0.35 | 0.58 |
| 7 | 53331061 | rs6945046 | 0.167 | -0.13 | 0.782 | 0.621 | -0.003 | 0.012 | 0.012 | 0.043 | 0.57 | 0.64 | 0.5 | 0.65 | 0.52 | 0.65 |
| 7 | 53345258 | rs6971358 | -1.276 | -0.113 | 0.822 | 0.502 | 0.038 | 0.029 | 0.029 | 0 | 0.14 | 0.4 | 0.09 | 0.13 | 0.12 | 0.09 |
| 7 | 53349479 | rs4131439 | 0.261 | 0.261 | 0.155 | 0.602 | 0.009 | 0.021 | 0.021 | 0.009 | 0.49 | 0.3 | 0.47 | 0.44 | 0.41 | 0.52 |
| 7 | 53349912 | rs13222080 | 0.639 | -0.093 | 0.478 | 0.487 | -0.001 | -0.002 | -0.002 | 0 | 0.2 | 0.15 | 0.16 | 0.1 | 0.21 | 0.11 |
| 7 | 53356343 | rs11768111 | -0.235 | 0.072 | 0.479 | 0.577 | 0.007 | 0.013 | 0.013 | 0 | 0.06 | 0.11 | 0.04 | 0.07 | 0.07 | 0.03 |
| 7 | 53357384 | rs4498495 | -1.489 | -0.111 | 0.412 | 0.563 | -0.032 | -0.024 | -0.024 | 0.008 | 0.58 | 0.81 | 0.51 | 0.5 | 0.58 | 0.4 |
| 7 | 53360479 | rs9690458 | -0.081 | 0.012 | 0.636 | 0.625 | 0.024 | 0.035 | 0.035 | 0 | 0.06 | 0.18 | 0.05 | 0.07 | 0.07 | 0.03 |
| 7 | 53362473 | rs12673526 | 0.39 | 0.081 | 0.491 | 0.731 | -0.002 | -0.023 | -0.023 | 0 | 0.09 | 0.05 | 0.1 | 0.23 | 0.1 | 0.07 |
| 7 | 53365317 | rs6593078 | -1.465 | 0.003 | 0.58 | 0.673 | -0.031 | -0.02 | -0.02 | 0 | 0.58 | 0.84 | 0.51 | 0.51 | 0.57 | 0.4 |
| 7 | 53373482 | rs12691388 | -0.759 | -0.073 | 0.833 | 0.614 | -0.001 | 0.013 | 0.013 | 0.417 | 0.41 | 0.46 | 0.36 | 0.44 | 0.43 | 0.22 |
| 7 | 53376491 | rs9691859 | -0.479 | 0.033 | 0.659 | 0.771 | -0.007 | -0.018 | -0.018 | 0 | 0.14 | 0.13 | 0.14 | 0.26 | 0.15 | 0.08 |
| 7 | 53381584 | rs6958437 | -1.252 | 0.188 | 0.637 | 0.77 | -0.043 | -0.033 | -0.033 | 0.223 | 0.57 | 0.87 | 0.51 | 0.52 | 0.57 | 0.4 |
| 7 | 53385379 | rs11981254 | -1.303 | 0.181 | 0.644 | 0.709 | -0.039 | -0.057 | -0.057 | 0 | 0.58 | 0.86 | 0.51 | 0.51 | 0.57 | 0.39 |
| 7 | 53405036 | rs7786939 | -1.087 | 0.056 | 0.813 | 0.365 | -0.017 | 0.023 | 0.023 | 0.157 | 0.55 | 0.69 | 0.41 | 0.41 | 0.5 | 0.38 |
| 7 | 53412123 | rs10266948 | 0.513 | 0.025 | 0.941 | 0.373 | 0.008 | 0.041 | 0.041 | 0 | 0.08 | 0.13 | 0.05 | 0.19 | 0.09 | 0.06 |
| 7 | 53420481 | rs10252217 | 0.439 | 0.022 | 1 | 0.372 | 0.016 | 0.059 | 0.059 | 0 | 0.08 | 0.17 | 0.05 | 0.19 | 0.09 | 0.06 |
| 7 | 53424191 | rs13229298 | -0.94 | -0.035 | 1.266 | 0.323 | -0.002 | 0.014 | 0.014 | 0.209 | 0.53 | 0.56 | 0.39 | 0.4 | 0.49 | 0.38 |
| 7 | 53436568 | rs6973726 | -0.368 | -0.019 | 1.085 | 0.232 | 0.002 | 0.005 | 0.005 | 0.305 | 0.45 | 0.42 | 0.34 | 0.2 | 0.4 | 0.31 |
| 7 | 53438252 | rs1012690 | -0.147 | -0.046 | 0.964 | 0.243 | 0.017 | 0.052 | 0.052 | 0.67 | 0.39 | 0.27 | 0.3 | 0.19 | 0.36 | 0.29 |
| 7 | 53448643 | rs3750141 | -1.356 | 0.052 | 0.621 | 0.252 | -0.016 | -0.024 | -0.024 | 0 | 0.59 | 0.78 | 0.57 | 0.48 | 0.59 | 0.45 |
| 7 | 53451179 | rs17135696 | -0.379 | 0.077 | 0.778 | 0.252 | 0.002 | -0.008 | -0.008 | 0.019 | 0.12 | 0.46 | 0.08 | 0.08 | 0.09 | 0.02 |
| 7 | 53454043 | rs1158661 | -0.063 | 0.04 | 0.688 | 0.18 | -0.016 | -0.04 | -0.04 | 0 | 0.34 | 0.43 | 0.25 | 0.19 | 0.33 | 0.17 |
| 7 | 53454892 | rs11983950 | 1.295 | -0.012 | 0.722 | 0.135 | 0.002 | -0.018 | -0.018 | 0.041 | 0.15 | 0.21 | 0.25 | 0.09 | 0.14 | 0.17 |
| 7 | 53455603 | rs2877132 | -0.962 | -0.059 | 0.251 | 0.159 | -0.003 | -0.014 | -0.014 | 0 | 0.33 | 0.61 | 0.35 | 0.16 | 0.26 | 0.22 |
| 7 | 53456941 | rs9642511 | -1.431 | -0.012 | -0.164 | 0.165 | -0.041 | -0.053 | -0.053 | 0 | 0.6 | 0.85 | 0.57 | 0.48 | 0.59 | 0.43 |
| 7 | 53457866 | rs10233940 | -0.134 | 0.013 | -0.082 | 0.136 | -0.005 | -0.062 | -0.062 | 0.099 | 0.24 | 0.39 | 0.28 | 0.1 | 0.18 | 0.21 |
| 7 | 53465744 | rs6968186 | -1.096 | 0.136 | -1.136 | 0.334 | -0.035 | -0.053 | -0.053 | 1.587 | 0.59 | 0.85 | 0.57 | 0.49 | 0.59 | 0.42 |
| 7 | 53467605 | rs7810981 | 0.502 | 0.211 | -1.044 | 0.2 | 0.006 | 0.047 | 0.047 | 0 | 0.25 | 0.1 | 0.18 | 0.13 | 0.25 | 0.16 |
| 7 | 53481902 | rs4947802 | -0.674 | 0.741 | -1.1 | 0.299 | -0.019 | 0.028 | 0.028 | 0 | 0.87 | 0.94 | 0.91 | 0.92 | 0.85 | 0.91 |
| 7 | 53490161 | rs7779192 | 1.046 | 0.849 | -0.923 | 0.111 | -0.007 | 0.022 | 0.022 | 0.011 | 0.72 | 0.72 | 0.82 | 0.83 | 0.7 | 0.86 |
| 7 | 53500844 | rs2111443 | -0.813 | 0.995 | -0.931 | 0.194 | -0.005 | 0.029 | 0.029 | 0 | 0.87 | 0.94 | 0.93 | 0.92 | 0.87 | 0.9 |
| 7 | 53503144 | rs7777919 | 0.998 | 1.186 | -0.907 | 0.998 | -0.014 | -0.007 | -0.007 | 0 | 0.77 | 0.76 | 0.8 | 0.67 | 0.66 | 0.84 |
| 7 | 53513975 | rs6943713 | -0.835 | 1.114 | -0.974 | 0.927 | -0.005 | 0.029 | 0.029 | 0 | 0.87 | 0.94 | 0.93 | 0.92 | 0.87 | 0.9 |
| 7 | 53536783 | rs6971950 | -0.349 | 1.165 | -0.969 | 0.934 | -0.018 | 0.016 | 0.016 | 0 | 0.85 | 0.9 | 0.9 | 0.86 | 0.8 | 0.89 |
| 7 | 53537843 | rs17692753 | 1.777 | 1.003 | -1.552 | 0.583 | -0.05 | -0.005 | -0.005 | 0 | 0.2 | 0.04 | 0.19 | 0.18 | 0.21 | 0.27 |
| 7 | 53540887 | rs10271450 | 0.835 | 0.971 | -0.448 | 0.602 | 0.034 | 0.012 | 0.012 | 0 | 0.76 | 0.61 | 0.78 | 0.66 | 0.66 | 0.84 |
| 7 | 53541479 | rs2330522 | -0.322 | 1.021 | -0.589 | 0.653 | -0.02 | -0.018 | -0.018 | 0 | 0.83 | 0.88 | 0.84 | 0.68 | 0.72 | 0.86 |
| 7 | 53549482 | rs7458373 | -0.393 | 1.037 | -0.613 | 0.654 | -0.044 | -0.038 | -0.038 | 0.084 | 0.84 | 0.9 | 0.85 | 0.68 | 0.72 | 0.86 |
| 7 | 53551618 | rs7803407 | 0.458 | 0.811 | 0.116 | 0.44 | 0.03 | -0.016 | -0.016 | 0 | 0.08 | 0.28 | 0.06 | 0.02 | 0.06 | 0.03 |
| 7 | 53553734 | rs10231055 | 0.568 | 0.828 | 0.085 | 0.448 | 0.017 | -0.05 | -0.05 | 0 | 0.77 | 0.6 | 0.83 | 0.85 | 0.74 | 0.86 |
| 7 | 53571744 | rs2194649 | -0.443 | 0.837 | 0.17 | 0.501 | 0.02 | -0.027 | -0.027 | 1.107 | 0.84 | 0.69 | 0.89 | 0.86 | 0.8 | 0.88 |
| 7 | 53580706 | rs2042536 | -1.185 | 0.796 | -0.314 | 0.486 | -0.003 | 0.031 | 0.031 | 0 | 0.87 | 0.94 | 0.93 | 0.93 | 0.87 | 0.9 |
| 7 | 53590250 | rs11764319 | 1.511 | 0.382 | -0.965 | 0.201 | -0.039 | 0.044 | 0.044 | 0 | 0.25 | 0.01 | 0.18 | 0.11 | 0.26 | 0.2 |
| 7 | 53603254 | rs10279984 | 0.974 | 0.196 | -0.277 | 0.154 | -0.001 | 0.003 | 0.003 | 1.276 | 0.65 | 0.37 | 0.76 | 0.64 | 0.64 | 0.82 |
| 7 | 53613811 | rs12718797 | 2.134 | 0.281 | -0.354 | 0.116 | -0.037 | 0.013 | 0.013 | 0 | 0.2 | 0.08 | 0.19 | 0.16 | 0.22 | 0.25 |
| 7 | 53619262 | rs4947815 | 1.036 | 0.054 | -0.015 | -0.117 | 0.002 | 0.023 | 0.023 | 2.31 | 0.63 | 0.32 | 0.75 | 0.68 | 0.64 | 0.8 |
| 7 | 53623518 | rs2194654 | 0.014 | 0.462 | -0.128 | 0.11 | 0.004 | 0 | 0 | 0 | 0.78 | 0.78 | 0.81 | 0.7 | 0.67 | 0.84 |

| | | | | | | | | | | | | | | | | |
|---|---|---|---|---|---|---|---|---|---|---|---|---|---|---|---|---|
| 7 | 53634725 | rs4947818 | 0.404 | 0.094 | -0.053 | -0.503 | -0.016 | -0.073 | -0.073 | 0 | 0.73 | 0.72 | 0.86 | 0.95 | 0.82 | 0.87 |
| 7 | 53635159 | rs9656687 | 0.262 | -0.131 | 0.149 | -0.258 | 0.042 | 0.069 | 0.069 | 0 | 0.07 | 0.13 | 0.08 | 0.26 | 0.17 | 0.08 |
| 7 | 53635424 | rs17622844 | 1.766 | -0.13 | 0.069 | -0.083 | -0.028 | -0.015 | -0.015 | 0.02 | 0.06 | 0 | 0.04 | 0.05 | 0.1 | 0.02 |
| 7 | 53637372 | rs2194637 | 1.943 | -0.365 | -0.254 | -0.213 | -0.022 | 0.042 | 0.042 | 0 | 0.23 | 0.03 | 0.18 | 0.11 | 0.23 | 0.21 |
| 7 | 53638569 | rs2330424 | 0.453 | -0.326 | -0.128 | -0.127 | 0.043 | 0.07 | 0.07 | 0 | 0.07 | 0.13 | 0.08 | 0.26 | 0.17 | 0.08 |
| 7 | 53639058 | rs1030305 | -0.305 | -0.481 | -0.099 | -0.272 | -0.042 | -0.089 | -0.089 | 0 | 0.78 | 0.7 | 0.9 | 0.95 | 0.88 | 0.89 |
| 7 | 53639105 | rs1030306 | 2.399 | -0.603 | -0.449 | -0.403 | -0.005 | 0.027 | 0.027 | 0.168 | 0.08 | 0.04 | 0.19 | 0 | 0.08 | 0.12 |
| 7 | 53640749 | rs6943826 | 1.056 | -0.608 | -0.317 | -0.338 | 0.003 | 0.05 | 0.05 | 0 | 0.32 | 0.21 | 0.25 | 0.16 | 0.33 | 0.22 |
| 7 | 53652231 | rs10235430 | 0.511 | -0.741 | -0.346 | -0.386 | 0.025 | 0.086 | 0.086 | 0 | 0.38 | 0.23 | 0.26 | 0.16 | 0.35 | 0.23 |
| 7 | 53652608 | rs2059407 | -0.227 | -0.704 | -0.085 | -0.314 | 0.016 | 0.014 | 0.014 | 0 | 0.78 | 0.75 | 0.84 | 0.68 | 0.73 | 0.81 |
| 7 | 53654002 | rs7803594 | 0.92 | -0.679 | -0.067 | -0.223 | -0.006 | -0.001 | -0.001 | 0 | 0.39 | 0.37 | 0.57 | 0.56 | 0.38 | 0.58 |
| 7 | 53656227 | rs6958765 | -0.139 | -0.697 | -0.271 | -0.389 | 0.009 | 0.054 | 0.054 | 0 | 0.56 | 0.64 | 0.65 | 0.79 | 0.57 | 0.68 |
| 7 | 53657870 | rs918944 | -1.754 | -0.531 | -0.299 | -0.262 | -0.002 | -0.011 | -0.011 | 0 | 0.89 | 0.87 | 0.92 | 0.93 | 0.9 | 0.91 |
| 7 | 53657896 | rs918945 | -1.407 | -0.549 | -0.316 | -0.27 | -0.006 | -0.024 | -0.024 | 0 | 0.87 | 0.79 | 0.92 | 0.93 | 0.9 | 0.91 |
| 7 | 53659747 | rs12535918 | 1.698 | -0.668 | -0.668 | 0.262 | -0.067 | -0.037 | -0.037 | 0.064 | 0.24 | 0.05 | 0.27 | 0.29 | 0.27 | 0.32 |
| 7 | 53661665 | rs2161966 | 1.383 | -1.086 | -0.57 | -0.056 | 0.018 | 0.057 | 0.057 | 0.462 | 0.25 | 0.05 | 0.17 | 0.11 | 0.23 | 0.21 |
| 7 | 53661764 | rs11975558 | 1.617 | -0.859 | 0.009 | 0.057 | -0.015 | -0.029 | -0.029 | 0 | 0.06 | 0.16 | 0.02 | 0 | 0.02 | 0.01 |
| 7 | 53664058 | rs11238275 | 0.327 | -1.159 | -0.603 | -0.157 | -0.001 | -0.005 | -0.005 | 0.024 | 0.63 | 0.43 | 0.72 | 0.69 | 0.66 | 0.75 |
| 7 | 53667614 | rs11238276 | 1.016 | -1.229 | -1.21 | -0.364 | -0.006 | 0.06 | 0.06 | 0.064 | 0.29 | 0.02 | 0.21 | 0.1 | 0.25 | 0.21 |
| 7 | 53668078 | rs6955484 | 0.29 | -0.969 | -0.27 | 0.205 | 0.016 | 0.025 | 0.025 | 0 | 0.35 | 0.52 | 0.5 | 0.55 | 0.48 | 0.53 |
| 7 | 53668289 | rs6593090 | -0.813 | -0.97 | 0.044 | 0.464 | -0.034 | 0.059 | 0.059 | 0 | 0.89 | 0.8 | 0.87 | 0.83 | 0.94 | 0.89 |
| 7 | 53684368 | rs17624890 | 1.274 | -1.345 | -0.619 | 0.079 | 0.021 | 0.116 | 0.116 | 0 | 0.29 | 0.01 | 0.15 | 0.09 | 0.2 | 0.15 |
| 7 | 53688656 | rs10235116 | -0.032 | -1.264 | -0.243 | 0.175 | -0.003 | 0.006 | 0.006 | 0 | 0.56 | 0.32 | 0.42 | 0.39 | 0.51 | 0.44 |
| 7 | 53706347 | rs6593097 | -0.877 | -0.995 | -0.726 | 0.358 | -0.036 | -0.014 | -0.014 | 0 | 0.66 | 0.74 | 0.55 | 0.57 | 0.56 | 0.5 |
| 7 | 53706549 | rs13228939 | -0.683 | -1 | -0.797 | 0.397 | -0.028 | -0.022 | -0.022 | 1.275 | 0.65 | 0.71 | 0.53 | 0.5 | 0.51 | 0.49 |
| 7 | 53707699 | rs1012889 | -2.223 | -0.633 | -0.984 | 0.374 | -0.092 | -0.076 | -0.076 | 0.034 | 0.79 | 0.96 | 0.62 | 0.64 | 0.68 | 0.65 |
| 7 | 53711214 | rs13226109 | 0.221 | -1.12 | -0.524 | 0.047 | -0.008 | -0.059 | -0.059 | 0.052 | 0.3 | 0.44 | 0.27 | 0.14 | 0.28 | 0.28 |
| 7 | 53712973 | rs7785632 | 0.992 | -1.243 | -0.578 | 0.036 | 0.031 | -0.012 | -0.012 | 0 | 0.31 | 0.25 | 0.25 | 0.32 | 0.23 | 0.18 |
| 7 | 53715983 | rs10261178 | -0.202 | -1.111 | -0.61 | 0.113 | 0 | 0.003 | 0.003 | 0 | 0.57 | 0.54 | 0.39 | 0.24 | 0.44 | 0.39 |
| 7 | 53718191 | rs2330583 | -0.788 | -1.039 | -0.794 | 0.094 | -0.012 | -0.02 | -0.02 | 0.184 | 0.61 | 0.69 | 0.52 | 0.46 | 0.51 | 0.46 |
| 7 | 53724916 | rs766069 | -0.595 | -1.03 | -0.824 | 0.024 | -0.005 | -0.01 | -0.01 | 0.003 | 0.6 | 0.62 | 0.49 | 0.38 | 0.46 | 0.45 |
| 7 | 53725263 | rs10277106 | 1.662 | -1.144 | -0.898 | -0.052 | 0.059 | 0.084 | 0.084 | 0 | 0.27 | 0.06 | 0.12 | 0.11 | 0.16 | 0.12 |
| 7 | 53726826 | rs10225959 | -0.308 | -0.954 | -0.795 | 0.082 | 0.001 | -0.002 | -0.002 | 0.429 | 0.34 | 0.61 | 0.38 | 0.3 | 0.3 | 0.33 |
| 7 | 53729051 | rs17135936 | 0.832 | -1.15 | -0.756 | 0.082 | 0.043 | 0.005 | 0.005 | 0 | 0.33 | 0.24 | 0.22 | 0.26 | 0.18 | 0.18 |
| 7 | 53761339 | rs13232315 | -0.676 | -0.485 | 0.118 | 0.456 | -0.035 | 0.114 | 0.114 | 1.629 | 0.15 | 0.58 | 0.23 | 0.28 | 0.12 | 0.13 |
| 7 | 53772035 | rs13229596 | -1.346 | -0.797 | 0.143 | 0.553 | -0.01 | 0.008 | 0.008 | 0 | 0.32 | 0.6 | 0.4 | 0.34 | 0.3 | 0.27 |
| 7 | 53776356 | rs17632216 | 0.519 | -0.63 | -0.193 | 0.513 | -0.034 | 0.012 | 0.012 | 0 | 0.17 | 0.01 | 0.16 | 0.06 | 0.18 | 0.15 |
| 7 | 53786903 | rs2193600 | 0.185 | -0.655 | 0.415 | 0.576 | -0.025 | 0.1 | 0.1 | 0.94 | 0.1 | 0.28 | 0.15 | 0.16 | 0.05 | 0.05 |
| 7 | 53787288 | rs7795256 | -0.111 | -0.443 | 0.195 | 0.403 | -0.004 | -0.021 | -0.021 | 0 | 0.21 | 0.31 | 0.23 | 0.11 | 0.22 | 0.2 |
| 7 | 53789410 | rs1861894 | -1.116 | -0.525 | 0.03 | 0.686 | -0.014 | -0.005 | -0.005 | 0 | 0.64 | 0.74 | 0.51 | 0.57 | 0.55 | 0.48 |
| 7 | 53795117 | rs17135974 | -0.54 | -0.275 | 0.293 | 0.941 | 0.005 | 0.053 | 0.053 | 0 | 0.09 | 0.15 | 0.21 | 0.18 | 0.12 | 0.16 |
| 7 | 53807652 | rs2330589 | -0.406 | -0.274 | 0.142 | 1.017 | -0.001 | 0.016 | 0.016 | 0 | 0.09 | 0.08 | 0.22 | 0.23 | 0.14 | 0.16 |
| 7 | 53827111 | rs1264108 | -0.048 | -0.296 | 0.113 | 1 | -0.011 | -0.034 | -0.034 | 0.231 | 0.08 | 0.03 | 0.21 | 0.18 | 0.11 | 0.16 |
| 7 | 53836224 | rs6942701 | 0.726 | -0.923 | 0.303 | 0.743 | -0.016 | -0.017 | -0.017 | 0.064 | 0.32 | 0.43 | 0.15 | 0.28 | 0.25 | 0.2 |
| 7 | 53836381 | rs6966615 | -0.763 | -1.062 | 0.315 | 0.637 | 0.013 | 0.023 | 0.023 | 1.556 | 0.55 | 0.65 | 0.29 | 0.34 | 0.41 | 0.31 |
| 7 | 53837397 | rs1569251 | -0.118 | -1.077 | 0.183 | 0.61 | -0.017 | -0.051 | -0.051 | 0 | 0.89 | 0.97 | 0.79 | 0.82 | 0.86 | 0.78 |
| 7 | 53841647 | rs6965214 | -0.358 | -1.177 | 0.296 | 0.425 | 0.016 | 0.03 | 0.03 | 0 | 0.21 | 0.11 | 0.13 | 0.06 | 0.16 | 0.1 |
| 7 | 53844931 | rs2091615 | -1.639 | -1.099 | 0.31 | 0.529 | -0.015 | -0.002 | -0.002 | 0.055 | 0.63 | 0.73 | 0.48 | 0.58 | 0.53 | 0.46 |
| 7 | 53865681 | rs1534734 | -1.688 | -1.171 | 0.419 | 0.518 | -0.027 | -0.009 | -0.009 | 0 | 0.64 | 0.79 | 0.48 | 0.58 | 0.53 | 0.46 |
| 7 | 53866525 | rs4144070 | 0.3 | -1.216 | 0.176 | 0.573 | -0.006 | -0.024 | -0.024 | 0.577 | 0.07 | 0.03 | 0.18 | 0.26 | 0.1 | 0.13 |
| 7 | 53870862 | rs1406637 | 0.831 | -1.225 | 0.315 | 0.461 | -0.039 | -0.055 | -0.055 | 0 | 0.32 | 0.46 | 0.14 | 0.21 | 0.22 | 0.21 |
| 7 | 53872885 | rs12216690 | 0.899 | -1.288 | 0.283 | 0.464 | -0.022 | -0.03 | -0.03 | 0 | 0.32 | 0.42 | 0.14 | 0.21 | 0.22 | 0.21 |
| 7 | 53909342 | rs7778779 | -0.703 | -1.265 | 0.255 | 0.488 | -0.008 | -0.016 | -0.016 | 0 | 0.22 | 0.23 | 0.14 | 0.06 | 0.16 | 0.1 |
| 7 | 53913543 | rs1919798 | -0.78 | -1.104 | 0.119 | 0.549 | -0.012 | -0.08 | -0.08 | 0 | 0.22 | 0.08 | 0.27 | 0.32 | 0.23 | 0.24 |
| 7 | 53915632 | rs7797182 | -2.219 | -0.96 | -0.169 | 0.797 | -0.165 | -0.042 | -0.042 | 0 | 0.7 | 0.98 | 0.54 | 0.73 | 0.6 | 0.61 |
| 7 | 53915850 | rs12536631 | -0.806 | -1.157 | 0.019 | 0.753 | -0.004 | -0.073 | -0.073 | 0.102 | 0.22 | 0.1 | 0.28 | 0.37 | 0.26 | 0.25 |
| 7 | 53920963 | rs6963388 | 1.087 | -1.088 | 0.03 | 0.796 | -0.009 | -0.013 | -0.013 | 0 | 0.08 | 0.04 | 0.18 | 0.26 | 0.11 | 0.15 |
| 7 | 53920981 | rs1919799 | 0.093 | -1.131 | 0.205 | 0.745 | -0.012 | 0.006 | 0.006 | 0 | 0.41 | 0.51 | 0.34 | 0.53 | 0.37 | 0.37 |
| 7 | 53924589 | rs17559045 | -0.769 | -1.088 | 0.059 | 0.728 | -0.015 | -0.01 | -0.01 | 0 | 0.23 | 0.25 | 0.15 | 0.12 | 0.18 | 0.1 |
| 7 | 53926458 | rs7787516 | 0.46 | -0.869 | -0.24 | 0.832 | 0.001 | 0 | 0 | 0 | 0.1 | 0.13 | 0.08 | 0.06 | 0.11 | 0.09 |
| 7 | 53941475 | rs2222806 | -1.675 | -0.728 | -0.527 | 1.017 | -0.092 | -0.001 | -0.001 | 0.078 | 0.67 | 0.89 | 0.51 | 0.64 | 0.55 | 0.53 |
| 7 | 53950851 | rs1569247 | 0.242 | -0.545 | -0.382 | 1.11 | 0.002 | 0.076 | 0.076 | 0 | 0.09 | 0.12 | 0.2 | 0.32 | 0.14 | 0.15 |
| 7 | 53967836 | rs11532826 | -1.288 | -0.676 | -0.732 | 0.776 | -0.031 | -0.027 | -0.027 | 0 | 0.62 | 0.79 | 0.47 | 0.53 | 0.48 | 0.47 |
| 7 | 53970911 | rs10464789 | 1.648 | -1.009 | -0.984 | 0.468 | 0.028 | -0.004 | -0.004 | 0 | 0.2 | 0.01 | 0.09 | 0.16 | 0.13 | 0.13 |
| 7 | 53980226 | rs1529250 | -0.52 | -0.565 | -0.633 | 0.714 | 0.037 | 0.082 | 0.082 | 4.088 | 0.54 | 0.75 | 0.29 | 0.28 | 0.36 | 0.28 |
| 7 | 53982626 | rs17736906 | 0.248 | 0.022 | -0.493 | 0.937 | -0.011 | -0.002 | -0.002 | 0.731 | 0.08 | 0.05 | 0.2 | 0.25 | 0.12 | 0.15 |
| 7 | 53993722 | rs4615525 | 0.838 | -0.137 | -0.17 | 0.581 | 0.002 | -0.007 | -0.007 | 0.295 | 0.12 | 0.13 | 0.1 | 0.06 | 0.12 | 0.04 |
| 7 | 53996432 | rs1526780 | -1.01 | -0.27 | -0.328 | 0.667 | -0.052 | -0.017 | -0.017 | 0.107 | 0.61 | 0.79 | 0.45 | 0.37 | 0.51 | 0.42 |
| 7 | 54009270 | rs12668378 | 0.861 | -0.325 | -0.443 | 0.717 | -0.071 | -0.063 | -0.063 | 0.256 | 0.42 | 0.12 | 0.5 | 0.49 | 0.5 | 0.45 |

| | | | | | | | | | | | | | | | | |
|---|---|---|---|---|---|---|---|---|---|---|---|---|---|---|---|---|
| 7 | 54015461 | rs7791859 | -0.025 | -0.435 | 0.305 | 0.675 | 0.039 | 0.138 | 0.138 | 0 | 0.92 | 0.73 | 0.93 | 0.86 | 0.92 | 0.92 |
| 7 | 54018828 | rs1526768 | 0.62 | -0.446 | 0.305 | 0.715 | 0.007 | -0.028 | -0.028 | 0 | 0.48 | 0.18 | 0.52 | 0.53 | 0.53 | 0.53 |
| 7 | 54019032 | rs6942618 | -0.42 | -0.59 | 0.632 | 0.72 | -0.009 | 0.035 | 0.035 | 0 | 0.14 | 0.29 | 0.15 | 0.21 | 0.14 | 0.24 |
| 7 | 54034670 | rs6956800 | -0.466 | -0.594 | 0.629 | 0.721 | -0.01 | 0.047 | 0.047 | 0 | 0.14 | 0.3 | 0.15 | 0.26 | 0.14 | 0.24 |
| 7 | 54044068 | rs7785867 | 1.85 | -0.509 | 0.347 | 0.498 | -0.023 | 0.025 | 0.025 | 0 | 0.05 | 0 | 0.1 | 0.01 | 0.08 | 0.07 |
| 7 | 54048025 | rs7789754 | 0.851 | -0.535 | 0.445 | 0.545 | -0.002 | 0 | 0 | 0.144 | 0.07 | 0.06 | 0.12 | 0.21 | 0.09 | 0.15 |
| 7 | 54051462 | rs12666229 | 1.784 | -0.614 | 0.28 | 0.477 | -0.12 | -0.1 | -0.1 | 0.706 | 0.32 | 0.04 | 0.45 | 0.59 | 0.43 | 0.45 |
| 7 | 54051865 | rs11979191 | -2 | -0.676 | 0.341 | 0.49 | -0.027 | -0.023 | -0.023 | 0.048 | 0.69 | 0.82 | 0.58 | 0.55 | 0.61 | 0.56 |
| 7 | 54054334 | rs1526755 | 0.521 | -1.338 | 0.731 | 0.389 | 0.007 | 0.037 | 0.037 | 2.676 | 0.29 | 0.18 | 0.24 | 0.13 | 0.25 | 0.22 |
| 7 | 54058897 | rs17153040 | -0.496 | -0.77 | 1.061 | 0.97 | -0.002 | 0.022 | 0.022 | 0 | 0.12 | 0.14 | 0.13 | 0.22 | 0.12 | 0.21 |
| 7 | 54059050 | rs17136044 | -0.088 | -0.738 | 0.983 | 0.402 | -0.013 | -0.033 | -0.033 | 0 | 0.06 | 0.19 | 0.03 | 0 | 0.05 | 0.08 |
| 7 | 54075091 | rs1983477 | 0.902 | -0.992 | 0.698 | 0.138 | -0.042 | -0.126 | -0.126 | 0 | 0.76 | 0.42 | 0.87 | 0.99 | 0.83 | 0.8 |
| 7 | 54076129 | rs4947860 | -0.885 | -1.021 | 0.637 | 0.322 | -0.002 | 0.027 | 0.027 | 0 | 0.09 | 0.13 | 0.13 | 0.18 | 0.09 | 0.05 |
| 7 | 54076794 | rs1526770 | 1.368 | -0.939 | 0.474 | 0.378 | 0.011 | 0.008 | 0.008 | 0.001 | 0.67 | 0.3 | 0.74 | 0.8 | 0.74 | 0.75 |
| 7 | 54081192 | rs6962838 | -0.872 | -0.604 | 0.359 | -0.048 | -0.028 | -0.082 | -0.082 | 0 | 0.25 | 0.38 | 0.22 | 0.05 | 0.18 | 0.15 |
| 7 | 54095256 | rs1405428 | 1.601 | 0.538 | 0.407 | 0.678 | 0.025 | 0.034 | 0.034 | 0 | 0.7 | 0.5 | 0.72 | 0.49 | 0.66 | 0.56 |
| 7 | 54095935 | rs11238290 | 0.852 | 0.357 | -0.144 | 0.134 | -0.088 | 0.016 | 0.016 | 0.884 | 0.35 | 0.08 | 0.43 | 0.38 | 0.42 | 0.48 |
| 7 | 54097204 | rs1526760 | 0.675 | 0.363 | -0.029 | 0.385 | -0.045 | 0.268 | 0.268 | 0.042 | 0.83 | 0.55 | 0.89 | 0.53 | 0.9 | 0.78 |
| 7 | 54097645 | rs2222689 | 1.199 | 0.497 | -0.245 | 0.394 | -0.073 | -0.007 | -0.007 | 0 | 0.31 | 0.07 | 0.42 | 0.38 | 0.38 | 0.42 |
| 7 | 54098497 | rs10899757 | 0.579 | 0.439 | 0.009 | 0.333 | -0.039 | 0.08 | 0.08 | 0.268 | 0.38 | 0.16 | 0.47 | 0.84 | 0.44 | 0.56 |
| 7 | 54108381 | rs11760550 | 0.558 | 0.498 | 0.033 | 0.357 | -0.039 | 0.078 | 0.078 | 0.905 | 0.37 | 0.16 | 0.47 | 0.84 | 0.44 | 0.56 |
| 7 | 54126697 | rs1916967 | 1.202 | 0.583 | -0.197 | 0.264 | -0.113 | -0.041 | -0.041 | 0 | 0.31 | 0.03 | 0.42 | 0.38 | 0.38 | 0.42 |
| 7 | 54127555 | rs13235686 | 0 | 0.837 | 0.127 | -0.226 | -0.013 | 0 | 0 | 0 | 0.1 | 0.07 | 0.17 | 0.05 | 0.21 | 0.16 |
| 7 | 54127780 | rs17560050 | -0.746 | 0.775 | 0.104 | -0.225 | -0.038 | 0.013 | 0.013 | 0 | 0.14 | 0.07 | 0.18 | 0.05 | 0.24 | 0.22 |
| 7 | 54131575 | rs13240898 | -0.746 | 0.775 | 0.104 | -0.225 | -0.038 | 0.013 | 0.013 | 0 | 0.14 | 0.07 | 0.18 | 0.05 | 0.24 | 0.22 |
| 7 | 54134080 | rs17560147 | -0.746 | 0.775 | 0.104 | -0.225 | -0.038 | 0.013 | 0.013 | 0 | 0.14 | 0.07 | 0.18 | 0.05 | 0.24 | 0.22 |
| 7 | 54135851 | rs11766546 | -0.02 | 0.6 | 0.382 | -0.215 | -0.024 | -0.056 | -0.056 | 2.665 | 0.9 | 0.64 | 0.94 | 0.99 | 0.96 | 0.94 |
| 7 | 54145081 | rs6976840 | -1.045 | 0.975 | 0.222 | -0.065 | 0.06 | -0.028 | -0.028 | 0.017 | 0.17 | 0.33 | 0.21 | 0.05 | 0.24 | 0.22 |
| 7 | 54146242 | rs2222535 | 0.38 | 0.927 | -0.465 | 0.358 | -0.042 | -0.001 | -0.001 | 0 | 0.37 | 0.15 | 0.46 | 0.74 | 0.45 | 0.6 |
| 7 | 54148237 | rs7788615 | 1.674 | 1.238 | 0.258 | 0.125 | 0.007 | -0.018 | -0.018 | 1.705 | 0.7 | 0.33 | 0.66 | 0.9 | 0.64 | 0.69 |
| 7 | 54150785 | rs7793672 | -0.167 | 1.419 | 0.141 | 0.097 | -0.003 | -0.004 | -0.004 | 0 | 0.08 | 0.08 | 0.1 | 0.01 | 0.11 | 0.08 |
| 7 | 54165227 | rs1405443 | 1.484 | 1.033 | 1.177 | 0.596 | -0.038 | 0.047 | 0.047 | 0.046 | 0.79 | 0.49 | 0.84 | 0.76 | 0.86 | 0.72 |
| 7 | 54176406 | rs3851234 | -1.16 | 1.088 | 1.283 | 0.615 | -0.019 | 0.051 | 0.051 | 0.289 | 0.13 | 0.25 | 0.13 | 0.21 | 0.07 | 0.18 |
| 7 | 54177773 | rs17646326 | 0.921 | 1.259 | 0.462 | 0.749 | -0.105 | -0.091 | -0.091 | 0 | 0.26 | 0.01 | 0.38 | 0.4 | 0.34 | 0.36 |
| 7 | 54182781 | rs10232172 | -1.087 | 1.221 | 1.37 | 0.651 | -0.016 | 0.054 | 0.054 | 0.195 | 0.12 | 0.23 | 0.12 | 0.21 | 0.07 | 0.16 |
| 7 | 54185285 | rs10230947 | -1.834 | 1.471 | 1.496 | 0.59 | 0.045 | 0.027 | 0.027 | 0 | 0.23 | 0.37 | 0.32 | 0.25 | 0.27 | 0.31 |
| 7 | 54188503 | rs10225470 | -1.294 | 1.779 | 1.632 | 0.724 | 0.116 | 0.101 | 0.101 | 0 | 0.16 | 0.38 | 0.3 | 0.24 | 0.25 | 0.23 |
| 7 | 54189089 | rs2330668 | 1.655 | 1.561 | 1.459 | 0.649 | 0.021 | 0.007 | 0.007 | 0 | 0.74 | 0.41 | 0.67 | 0.76 | 0.69 | 0.7 |
| 7 | 54198501 | rs1524395 | -1.815 | 1.578 | 1.336 | 0.656 | 0.038 | 0.015 | 0.015 | 0 | 0.25 | 0.54 | 0.32 | 0.24 | 0.31 | 0.3 |
| 7 | 54208630 | rs10224185 | -1.112 | 1.247 | 1.037 | 0.321 | 0.02 | -0.056 | -0.056 | 3.581 | 0.14 | 0.28 | 0.17 | 0.02 | 0.16 | 0.16 |
| 7 | 54210427 | rs9642543 | 0.683 | 1.269 | 0.271 | 0.492 | -0.09 | -0.099 | -0.099 | 0 | 0.23 | 0.01 | 0.31 | 0.42 | 0.3 | 0.37 |
| 7 | 54213764 | rs2330676 | 0.296 | 1.442 | 0.411 | 0.202 | 0.009 | -0.018 | -0.018 | 0.089 | 0.11 | 0.12 | 0.19 | 0.02 | 0.18 | 0.14 |
| 7 | 54219207 | rs11238298 | 1.944 | 1.15 | 0.415 | 0.149 | -0.021 | 0.004 | 0.004 | 0.017 | 0.63 | 0.17 | 0.57 | 0.7 | 0.61 | 0.61 |
| 7 | 54219462 | rs6959483 | 1.305 | 1.116 | 0.938 | 0.13 | 0.018 | 0.02 | 0.02 | 0 | 0.75 | 0.61 | 0.74 | 0.73 | 0.7 | 0.74 |
| 7 | 54221842 | rs1524393 | 0.158 | 1.106 | 1.099 | 0.126 | -0.003 | -0.01 | -0.01 | 0 | 0.09 | 0.08 | 0.12 | 0.03 | 0.09 | 0.11 |
| 7 | 54225078 | rs12718846 | 1.872 | 0.956 | 0.691 | 0.252 | -0.019 | 0.053 | 0.053 | 0 | 0.37 | 0.08 | 0.25 | 0.2 | 0.31 | 0.21 |
| 7 | 54227617 | rs1524405 | 1.783 | 0.833 | 1.53 | 0.304 | 0.006 | 0.006 | 0.006 | 0.054 | 0.73 | 0.41 | 0.72 | 0.73 | 0.7 | 0.72 |
| 7 | 54228492 | rs12530576 | -0.249 | 0.884 | 1.53 | 0.328 | 0.014 | -0.028 | -0.028 | 0 | 0.14 | 0.15 | 0.2 | 0.02 | 0.21 | 0.11 |
| 7 | 54238625 | rs1581515 | 0.686 | 0.741 | 1.069 | 0.489 | 0.037 | 0.31 | 0.31 | 0.064 | 0.93 | 0.66 | 0.94 | 0.75 | 0.96 | 0.9 |
| 7 | 54247600 | rs7799840 | 1.546 | 0.739 | 0.807 | 0.477 | 0 | 0.179 | 0.179 | 0 | 0.87 | 0.58 | 0.91 | 0.75 | 0.91 | 0.84 |
| 7 | 54249748 | rs2049409 | 0.95 | 0.696 | 0.897 | 0.569 | 0.042 | 0.344 | 0.344 | 0.384 | 0.92 | 0.59 | 0.93 | 0.75 | 0.96 | 0.9 |
| 7 | 54254685 | rs4498497 | 1.074 | 0.859 | -0.646 | 0.507 | -0.017 | 0.004 | 0.004 | 0 | 0.64 | 0.19 | 0.57 | 0.71 | 0.61 | 0.6 |
| 7 | 54265587 | rs4425715 | -0.683 | 0.729 | -0.32 | 0.401 | 0.003 | -0.001 | -0.001 | 0 | 0.57 | 0.59 | 0.72 | 0.52 | 0.67 | 0.78 |
| 7 | 54266710 | rs4418304 | -0.869 | 0.763 | -0.227 | 0.351 | 0.003 | -0.019 | -0.019 | 0 | 0.59 | 0.69 | 0.7 | 0.55 | 0.64 | 0.76 |
| 7 | 54313171 | rs13437953 | -0.033 | 0.568 | 0.318 | 0.143 | -0.014 | -0.039 | -0.039 | 0 | 0.09 | 0.15 | 0.07 | 0.01 | 0.08 | 0.14 |
| 7 | 54316072 | rs7791494 | 0.107 | 1.06 | 0.714 | -0.157 | 0.006 | -0.035 | -0.035 | 0 | 0.15 | 0.16 | 0.23 | 0.02 | 0.18 | 0.1 |
| 7 | 54317563 | rs10273428 | -0.574 | 1.2 | 0.626 | -0.198 | -0.002 | -0.039 | -0.039 | 0 | 0.13 | 0.16 | 0.09 | 0.01 | 0.13 | 0.21 |
| 7 | 54318757 | rs4338053 | -0.499 | 1.123 | 0.834 | -0.2 | 0.001 | 0 | 0 | 0 | 0.43 | 0.44 | 0.58 | 0.5 | 0.51 | 0.5 |
| 7 | 54394095 | rs10253714 | -0.211 | 1.017 | 0.865 | 0.006 | -0.014 | -0.008 | -0.008 | 0 | 0.11 | 0.15 | 0.09 | 0.11 | 0.09 | 0.2 |
| 7 | 54396110 | rs12534887 | 0.912 | 0.949 | 0.983 | -0.045 | -0.022 | -0.063 | -0.063 | 0 | 0.26 | 0.13 | 0.34 | 0.48 | 0.32 | 0.4 |
| 7 | 54412203 | rs17670957 | -1.21 | 0.933 | 1.048 | -0.028 | 0.002 | 0 | 0 | 0 | 0.54 | 0.6 | 0.67 | 0.63 | 0.6 | 0.72 |
| 7 | 54412608 | rs4628218 | -0.485 | 0.94 | 1.092 | -0.026 | 0.002 | 0.001 | 0.001 | 0 | 0.43 | 0.4 | 0.57 | 0.52 | 0.51 | 0.5 |
| 7 | 54419111 | rs1917576 | -1.616 | 0.898 | 1.046 | -0.045 | 0.005 | -0.009 | -0.009 | 0 | 0.58 | 0.68 | 0.7 | 0.64 | 0.66 | 0.75 |
| 7 | 54423086 | rs11764910 | -1.026 | 1.016 | 0.771 | -0.014 | 0 | -0.001 | -0.001 | 0 | 0.52 | 0.48 | 0.66 | 0.54 | 0.6 | 0.71 |
| 7 | 54430725 | rs1525164 | -0.669 | 1.073 | 0.73 | 0.015 | -0.002 | 0.013 | 0.013 | 0 | 0.44 | 0.34 | 0.59 | 0.43 | 0.55 | 0.53 |
| 7 | 54430805 | rs13234631 | 0.404 | 1.176 | 0.635 | 0.026 | -0.03 | -0.02 | -0.02 | 0.159 | 0.29 | 0.13 | 0.34 | 0.41 | 0.37 | 0.42 |
| 7 | 54439565 | rs1080136 | 0.829 | 1.276 | 0.656 | 0.046 | -0.01 | -0.024 | -0.024 | 0 | 0.26 | 0.13 | 0.32 | 0.41 | 0.31 | 0.39 |
| 7 | 54444505 | rs17647441 | 0.234 | 1.299 | 0.49 | 0.058 | 0.001 | -0.017 | -0.017 | 0.113 | 0.14 | 0.12 | 0.25 | 0.02 | 0.19 | 0.1 |
| 7 | 54467507 | rs10234994 | -1.075 | 0.522 | -0.091 | -0.387 | 0 | 0.017 | 0.017 | 0 | 0.66 | 0.46 | 0.7 | 0.53 | 0.69 | 0.74 |

| | | | | | | | | | | | | | | | | |
|---|---|---|---|---|---|---|---|---|---|---|---|---|---|---|---|---|
| 7 | 54488242 | rs7806139 | -0.934 | 0.675 | 0.052 | -0.127 | 0.002 | 0.001 | 0.001 | 0 | 0.55 | 0.46 | 0.68 | 0.52 | 0.64 | 0.7 |
| 7 | 54497328 | rs10499738 | 0.228 | 0.97 | -0.457 | -0.207 | -0.001 | -0.021 | -0.021 | 2.358 | 0.14 | 0.13 | 0.26 | 0 | 0.25 | 0.15 |
| 7 | 54523070 | rs4480061 | -0.854 | 0.635 | -0.448 | -0.172 | 0.021 | 0.015 | 0.015 | 0.006 | 0.52 | 0.32 | 0.66 | 0.52 | 0.63 | 0.68 |
| 7 | 54535622 | rs6948197 | -1.365 | 0.757 | -0.003 | -0.338 | -0.002 | -0.003 | -0.003 | 0.511 | 0.23 | 0.24 | 0.34 | 0.11 | 0.29 | 0.29 |
| 7 | 54552899 | rs526461 | -1.83 | 0.591 | 0.407 | 0.049 | 0.008 | 0.018 | 0.018 | 0.489 | 0.29 | 0.48 | 0.42 | 0.3 | 0.34 | 0.34 |
| 7 | 54558397 | rs480777 | -1.091 | 0.53 | 0.569 | 0.049 | -0.03 | 0.077 | 0.077 | 0 | 0.12 | 0.23 | 0.15 | 0.3 | 0.09 | 0.2 |
| 7 | 54561418 | rs11767914 | 0.197 | 0.615 | 0.541 | -0.009 | -0.014 | 0 | 0 | 0 | 0.13 | 0.08 | 0.26 | 0 | 0.23 | 0.12 |
| 7 | 54563190 | rs11765590 | -0.296 | 0.606 | 0.635 | -0.013 | 0.024 | -0.064 | -0.064 | 1.107 | 0.15 | 0.2 | 0.26 | 0 | 0.23 | 0.12 |
| 7 | 54564662 | rs2462578 | -1.72 | 0.599 | 0.694 | 0.066 | 0.007 | 0.028 | 0.028 | 0.242 | 0.26 | 0.4 | 0.41 | 0.3 | 0.32 | 0.3 |
| 7 | 54574605 | rs479006 | -1.445 | 0.547 | 0.693 | 0.091 | -0.025 | 0.099 | 0.099 | 0 | 0.12 | 0.24 | 0.25 | 0.3 | 0.14 | 0.25 |
| 7 | 54576747 | rs507047 | -1.825 | 0.535 | 0.771 | 0.071 | -0.022 | 0.106 | 0.106 | 0 | 0.14 | 0.3 | 0.27 | 0.3 | 0.15 | 0.26 |
| 7 | 54582342 | rs681510 | -0.42 | 0.515 | 0.805 | 0.108 | -0.003 | 0.096 | 0.096 | 0 | 0.05 | 0.13 | 0.18 | 0.19 | 0.09 | 0.1 |
| 7 | 54595330 | rs671767 | -1.699 | 0.589 | 0.779 | 0.084 | -0.018 | 0.069 | 0.069 | 0 | 0.13 | 0.24 | 0.26 | 0.31 | 0.14 | 0.25 |
| 7 | 54600112 | rs577570 | -1.47 | 0.683 | 0.669 | 0.073 | 0.013 | 0.018 | 0.018 | 0.1 | 0.26 | 0.38 | 0.43 | 0.31 | 0.34 | 0.31 |
| 7 | 54610293 | rs513529 | -1.665 | 0.699 | 0.523 | 0.097 | 0.012 | 0.022 | 0.022 | 1.339 | 0.28 | 0.52 | 0.43 | 0.3 | 0.34 | 0.31 |
| 7 | 54618619 | rs3862849 | 0.333 | 0.675 | -0.458 | -0.2 | -0.056 | -0.109 | -0.109 | 1.312 | 0.29 | 0.07 | 0.31 | 0.57 | 0.34 | 0.49 |
| 7 | 54619136 | rs2204133 | 1.386 | 1.398 | 0.246 | -0.016 | 0.01 | 0.009 | 0.009 | 0.224 | 0.68 | 0.35 | 0.5 | 0.6 | 0.59 | 0.67 |
| 7 | 54619264 | rs2204134 | 0.067 | 1.455 | -0.107 | 0.143 | 0.013 | 0.028 | 0.028 | 0.116 | 0.82 | 0.74 | 0.73 | 0.72 | 0.77 | 0.87 |
| 7 | 54621134 | rs964823 | 0.595 | 1.393 | -0.189 | 0.078 | 0.008 | 0.007 | 0.007 | 0.322 | 0.78 | 0.71 | 0.69 | 0.72 | 0.74 | 0.85 |
| 7 | 54634567 | rs1403237 | -1.497 | 1.216 | 0.536 | -0.043 | 0.053 | -0.008 | -0.008 | 2.905 | 0.12 | 0.44 | 0.24 | 0.08 | 0.15 | 0.21 |
| 7 | 54640917 | rs1851438 | 1.921 | 1.105 | 0.693 | 0.667 | 0.042 | 0.225 | 0.225 | 0 | 0.85 | 0.47 | 0.67 | 0.63 | 0.84 | 0.72 |
| 7 | 54646788 | rs12533009 | 0.653 | 1.414 | -0.2 | 0.533 | -0.103 | 0.027 | 0.027 | 0 | 0.23 | 0.02 | 0.3 | 0.14 | 0.35 | 0.25 |
| 7 | 54651526 | rs1031490 | -1.027 | 0.789 | 0.365 | 0.879 | 0.023 | 0.163 | 0.163 | 0 | 0.07 | 0.44 | 0.17 | 0.12 | 0.05 | 0.18 |
| 7 | 54652511 | rs10250847 | 1.533 | 0.135 | 0.103 | 0.021 | 0.039 | 0.058 | 0.058 | 0.373 | 0.29 | 0.15 | 0.13 | 0.05 | 0.18 | 0.15 |
| 7 | 54655459 | rs1153922 | -1.554 | 0.771 | 0.867 | 0.88 | 0.09 | -0.033 | -0.033 | 0 | 0.09 | 0.39 | 0.18 | 0.01 | 0.11 | 0.09 |
| 7 | 54658511 | rs331824 | -2.29 | 0.793 | 0.783 | 0.844 | 0.041 | -0.046 | -0.046 | 0.442 | 0.16 | 0.51 | 0.25 | 0.08 | 0.18 | 0.24 |
| 7 | 54667620 | rs1454524 | -0.893 | 0.527 | 0.554 | 0.899 | 0.013 | -0.035 | -0.035 | 0 | 0.07 | 0.24 | 0.14 | 0 | 0.05 | 0.08 |
| 7 | 54674873 | rs679830 | 1.451 | 0.55 | 0.515 | 1.066 | 0.006 | 0.111 | 0.111 | 1.393 | 0.92 | 0.72 | 0.87 | 0.83 | 0.95 | 0.92 |
| 7 | 54675049 | rs6970726 | -0.405 | 0.606 | 0.307 | 0.422 | -0.013 | 0.032 | 0.032 | 0.986 | 0.33 | 0.43 | 0.39 | 0.47 | 0.32 | 0.47 |
| 7 | 54676738 | rs4947924 | 0.329 | 0.892 | 0.287 | -0.991 | 0.008 | -0.006 | -0.006 | 0 | 0.9 | 0.85 | 0.88 | 0.95 | 0.87 | 0.92 |
| 7 | 54678150 | rs10227226 | 0.422 | 1.169 | -0.21 | -1.357 | -0.05 | 0.047 | 0.047 | 3.387 | 0.33 | 0.14 | 0.5 | 0.77 | 0.43 | 0.62 |
| 7 | 54687869 | rs7782796 | 0.22 | 1.002 | -0.679 | -2.116 | 0.012 | -0.145 | -0.145 | 0.525 | 0.28 | 0.46 | 0.2 | 0.02 | 0.29 | 0.12 |
| 7 | 54690500 | rs1454511 | -0.033 | 0.8 | -0.752 | -2.089 | 0.001 | -0.019 | -0.019 | 0.352 | 0.54 | 0.54 | 0.34 | 0.06 | 0.44 | 0.27 |
| 7 | 54690641 | rs13232607 | 1.48 | 0.977 | -1.711 | -2.003 | 0.005 | -0.027 | -0.027 | 0 | 0.27 | 0.24 | 0.15 | 0.03 | 0.15 | 0.14 |
| 7 | 54703359 | rs7806988 | 1.405 | 1.36 | -1.545 | -1.758 | -0.029 | 0.035 | 0.035 | 0 | 0.13 | 0.04 | 0.12 | 0.02 | 0.19 | 0.07 |
| 7 | 54705006 | rs10793421 | 0.561 | 0.401 | -1.375 | -1.356 | 0.029 | 0.013 | 0.013 | 0 | 0.38 | 0.28 | 0.19 | 0.04 | 0.24 | 0.2 |
| 7 | 54706018 | rs648890 | -0.13 | 1.285 | -1.113 | -0.25 | -0.006 | 0.002 | 0.002 | 0 | 0.07 | 0.04 | 0.09 | 0.03 | 0.08 | 0.06 |
| 7 | 54708056 | rs17563093 | 0.054 | 1.05 | -1.527 | -0.61 | -0.02 | 0.071 | 0.071 | 0 | 0.28 | 0.09 | 0.25 | 0.18 | 0.34 | 0.31 |
| 7 | 54708311 | rs582252 | 0.049 | 0.361 | -1.508 | -1.377 | -0.006 | 0.199 | 0.199 | 1.123 | 0.5 | 0.26 | 0.57 | 0.88 | 0.51 | 0.54 |
| 7 | 54709978 | rs616709 | 0.205 | 0.362 | -1.457 | -1.219 | -0.006 | 0.176 | 0.176 | 1.392 | 0.5 | 0.26 | 0.57 | 0.85 | 0.52 | 0.54 |
| 7 | 54711414 | rs535535 | 0.053 | 0.284 | -1.267 | -1.472 | -0.009 | -0.028 | -0.028 | 0 | 0.69 | 0.44 | 0.72 | 0.85 | 0.75 | 0.6 |
| 7 | 54711923 | rs9692211 | 0.232 | 0.223 | -1.034 | -1.509 | -0.022 | -0.039 | -0.039 | 0 | 0.24 | 0.47 | 0.25 | 0.15 | 0.2 | 0.32 |
| 7 | 54715735 | rs678243 | -0.066 | 0.28 | -0.931 | -1.699 | -0.002 | -0.085 | -0.085 | 0.05 | 0.69 | 0.47 | 0.74 | 0.9 | 0.69 | 0.67 |
| 7 | 54716920 | rs662865 | -0.032 | 0.267 | -0.917 | -1.68 | -0.019 | -0.03 | -0.03 | 0 | 0.88 | 0.65 | 0.89 | 0.93 | 0.92 | 0.76 |
| 7 | 54717472 | rs10499747 | 1.447 | 0.2 | -0.954 | -1.794 | -0.002 | 0.018 | 0.018 | 0 | 0.14 | 0.05 | 0.08 | 0 | 0.14 | 0.08 |
| 7 | 54717529 | rs568201 | 0.121 | 0.202 | -0.952 | -1.793 | -0.021 | -0.102 | -0.102 | 0 | 0.74 | 0.6 | 0.81 | 0.92 | 0.78 | 0.67 |
| 7 | 54718300 | rs1569261 | -0.386 | 0.804 | -1.928 | -1.395 | 0.01 | 0.007 | 0.007 | 0 | 0.6 | 0.39 | 0.53 | 0.65 | 0.71 | 0.49 |
| 7 | 54724015 | rs1569267 | -0.404 | 1.222 | -1.765 | -0.842 | 0.001 | 0.032 | 0.032 | 0 | 0.46 | 0.41 | 0.45 | 0.65 | 0.57 | 0.4 |
| 7 | 54733732 | rs6971934 | 0.831 | 1.362 | -1.947 | -0.634 | -0.005 | 0.01 | 0.01 | 0 | 0.21 | 0.18 | 0.15 | 0.03 | 0.23 | 0.15 |
| 7 | 54741478 | rs518709 | 0.762 | 1.311 | -1.829 | -0.113 | -0.042 | -0.059 | -0.059 | 0 | 0.67 | 0.85 | 0.68 | 0.39 | 0.59 | 0.6 |
| 7 | 54741870 | rs17672458 | -0.444 | 1.196 | -1.616 | 0.243 | -0.04 | -0.084 | -0.084 | 0 | 0.27 | 0.11 | 0.29 | 0.61 | 0.34 | 0.32 |
| 7 | 54751771 | rs10238349 | 0.383 | 1.174 | -0.54 | -0.288 | -0.022 | -0.028 | -0.028 | 0 | 0.74 | 0.89 | 0.57 | 0.41 | 0.7 | 0.75 |
| 7 | 54752386 | rs7788658 | 0.421 | 0.468 | -0.408 | -1.052 | -0.012 | -0.016 | -0.016 | 0 | 0.22 | 0.37 | 0.17 | 0.13 | 0.18 | 0.17 |
| 7 | 54757128 | rs17563764 | 1.18 | 0.211 | -1.042 | -1.505 | -0.024 | 0.072 | 0.072 | 0 | 0.13 | 0.03 | 0.08 | 0 | 0.14 | 0.07 |
| 7 | 54773399 | rs10282532 | 0.19 | -0.013 | -0.944 | -1.766 | 0.002 | 0.001 | 0.001 | 0 | 0.48 | 0.5 | 0.34 | 0.27 | 0.47 | 0.55 |
| 7 | 54779296 | rs4518612 | 1.164 | -0.279 | -0.894 | -1.948 | 0.002 | 0.014 | 0.014 | 0 | 0.28 | 0.17 | 0.2 | 0.22 | 0.3 | 0.38 |
| 7 | 54780365 | rs7805129 | 0.368 | -0.252 | -0.964 | -1.921 | -0.017 | -0.085 | -0.085 | 0 | 0.2 | 0.3 | 0.13 | 0.05 | 0.17 | 0.17 |
| 7 | 54781060 | rs6948049 | -0.583 | -0.247 | -1.257 | -1.975 | -0.104 | -0.098 | -0.098 | 0 | 0.7 | 0.96 | 0.51 | 0.41 | 0.66 | 0.72 |
| 7 | 54783915 | rs17172374 | 1.555 | -0.247 | -1.142 | -1.907 | 0.015 | 0.044 | 0.044 | 0 | 0.14 | 0.07 | 0.07 | 0 | 0.11 | 0.06 |
| 7 | 54787317 | rs11768499 | 1.466 | -0.411 | -1.483 | -2.12 | -0.074 | -0.059 | -0.059 | 0 | 0.18 | 0.02 | 0.15 | 0.22 | 0.26 | 0.32 |
| 7 | 54795837 | rs17564002 | -0.059 | -0.569 | -1.356 | -2.224 | -0.002 | -0.01 | -0.01 | 0 | 0.36 | 0.38 | 0.29 | 0.27 | 0.38 | 0.48 |
| 7 | 54796443 | rs10215012 | 0.321 | -0.567 | -1.254 | -2.222 | -0.001 | 0.002 | 0.002 | 0 | 0.32 | 0.29 | 0.28 | 0.27 | 0.36 | 0.45 |
| 7 | 54801714 | rs10499749 | 1.45 | -0.961 | -1.075 | -2.21 | -0.001 | -0.018 | -0.018 | 0.005 | 0.21 | 0.38 | 0.17 | 0.14 | 0.19 | 0.17 |
| 7 | 54819127 | rs2141539 | -1.141 | -1.121 | -1.8 | -2.309 | -0.143 | -0.137 | -0.137 | 0 | 0.77 | 0.98 | 0.49 | 0.42 | 0.66 | 0.67 |
| 7 | 54825019 | rs2256906 | 0.519 | -1.299 | -1.525 | -2.034 | -0.003 | -0.006 | -0.006 | 0 | 0.29 | 0.31 | 0.22 | 0.27 | 0.31 | 0.4 |
| 7 | 54826323 | rs2699215 | 1.862 | -1.305 | -1.528 | -2.037 | 0.025 | 0.05 | 0.05 | 0 | 0.1 | 0.03 | 0.07 | 0 | 0.06 | 0.03 |
| 7 | 54836484 | rs2699236 | -0.7 | -1.343 | -1.419 | -2.056 | -0.087 | -0.022 | -0.022 | 0.002 | 0.61 | 0.9 | 0.42 | 0.41 | 0.54 | 0.61 |
| 7 | 54841341 | rs2699237 | 0.313 | -1.308 | -1.395 | -2.048 | -0.002 | -0.004 | -0.004 | 0 | 0.31 | 0.31 | 0.23 | 0.27 | 0.33 | 0.43 |
| 7 | 54857816 | rs6946833 | 0.099 | -1.334 | -1.403 | -2.043 | 0.008 | 0.013 | 0.013 | 0 | 0.47 | 0.58 | 0.31 | 0.36 | 0.45 | 0.52 |

| | | | | | | | | | | | | | | | | |
|---|---|---|---|---|---|---|---|---|---|---|---|---|---|---|---|---|
| 7 | 54859417 | rs1113443 | -0.675 | -1.284 | -1.507 | -2.025 | -0.088 | -0.02 | -0.02 | 0 | 0.61 | 0.9 | 0.42 | 0.41 | 0.54 | 0.61 |
| 7 | 54862831 | rs6968982 | 1.361 | -1.183 | -1.453 | -1.934 | -0.063 | -0.056 | -0.056 | 0 | 0.19 | 0.02 | 0.14 | 0.22 | 0.27 | 0.35 |
| 7 | 54864072 | rs1112565 | 0.5 | -1.072 | -1.228 | -1.824 | -0.04 | -0.064 | -0.064 | 0.883 | 0.28 | 0.49 | 0.17 | 0.14 | 0.18 | 0.18 |
| 7 | 54871940 | rs17172391 | 1.614 | -1.06 | -1.284 | -1.84 | 0 | -0.019 | -0.019 | 0 | 0.11 | 0.08 | 0.1 | 0 | 0.08 | 0.03 |
| 7 | 54873006 | rs2293589 | 0.138 | -1.095 | -1.385 | -1.876 | 0.014 | 0.021 | 0.021 | 0 | 0.46 | 0.68 | 0.31 | 0.42 | 0.44 | 0.49 |
| 7 | 54874029 | rs2330769 | -0.031 | -1.336 | -1.162 | -1.9 | 0.006 | 0.002 | 0.002 | 0.11 | 0.34 | 0.74 | 0.22 | 0.35 | 0.33 | 0.4 |
| 7 | 54880969 | rs2699212 | 0.823 | -1.697 | -0.626 | -1.721 | -0.005 | -0.028 | -0.028 | 0 | 0.12 | 0.15 | 0.11 | 0.01 | 0.08 | 0.03 |
| 7 | 54893417 | rs10499751 | -0.419 | -1.427 | -0.515 | -1.178 | -0.022 | -0.022 | -0.022 | 0 | 0.15 | 0.41 | 0.06 | 0.08 | 0.07 | 0.08 |
| 7 | 54896159 | rs13236867 | -0.385 | -0.908 | -1.386 | -0.988 | 0.022 | 0.087 | 0.087 | 0 | 0.17 | 0.02 | 0.16 | 0.01 | 0.11 | 0.1 |
| 7 | 54905350 | rs17746029 | -0.341 | -0.589 | -1.157 | -0.061 | 0.024 | -0.035 | -0.035 | 1.4 | 0.32 | 0.25 | 0.16 | 0.44 | 0.21 | 0.3 |
| 7 | 54905511 | rs12672832 | -0.25 | -0.425 | -1.417 | -0.849 | 0.007 | 0.059 | 0.059 | 0 | 0.18 | 0.06 | 0.17 | 0.01 | 0.16 | 0.12 |
| 7 | 54926166 | rs11238330 | -0.008 | -0.215 | -0.98 | -0.651 | -0.003 | 0.031 | 0.031 | 0 | 0.53 | 0.57 | 0.51 | 0.75 | 0.48 | 0.66 |
| 7 | 54930362 | rs2877212 | -0.079 | -0.34 | -0.835 | -0.738 | -0.037 | 0.088 | 0.088 | 0 | 0.45 | 0.2 | 0.46 | 0.25 | 0.52 | 0.33 |
| 7 | 54930470 | rs11238333 | 0.46 | -0.577 | -0.451 | -0.622 | 0.011 | 0.023 | 0.023 | 0 | 0.3 | 0.33 | 0.42 | 0.61 | 0.32 | 0.4 |
| 7 | 54941222 | rs7778465 | 0.67 | -0.378 | -0.239 | -0.79 | 0.004 | 0.004 | 0.004 | 0 | 0.13 | 0.21 | 0.1 | 0.13 | 0.12 | 0.17 |
| 7 | 54946332 | rs11982432 | 0.4 | -0.576 | -0.293 | -0.871 | -0.004 | -0.008 | -0.008 | 0.62 | 0.2 | 0.15 | 0.32 | 0.73 | 0.19 | 0.52 |
| 7 | 54950551 | rs10899766 | -0.398 | -0.724 | -0.208 | -0.989 | 0.008 | 0.02 | 0.02 | 0 | 0.4 | 0.29 | 0.41 | 0.75 | 0.35 | 0.63 |
| 7 | 54958458 | rs6945082 | 0.412 | -0.663 | -0.179 | -1.08 | 0.012 | 0.056 | 0.056 | 0 | 0.33 | 0.18 | 0.19 | 0.02 | 0.3 | 0.14 |
| 7 | 54961755 | rs17565126 | -0.293 | -0.582 | -0.076 | -0.83 | -0.007 | 0.03 | 0.03 | 0 | 0.51 | 0.28 | 0.5 | 0.62 | 0.49 | 0.51 |
| 7 | 54962084 | rs17650312 | 1.459 | -0.589 | -0.518 | -1.154 | 0.006 | 0.047 | 0.047 | 0 | 0.12 | 0.03 | 0.1 | 0.04 | 0.13 | 0.15 |
| 7 | 54968003 | rs10488134 | -0.105 | -0.595 | -0.046 | -1.047 | -0.003 | 0.004 | 0.004 | 0 | 0.54 | 0.42 | 0.51 | 0.62 | 0.5 | 0.53 |
| 7 | 54970589 | rs12055922 | -0.076 | -0.599 | -0.194 | -1.023 | -0.006 | 0.01 | 0.01 | 0 | 0.54 | 0.36 | 0.5 | 0.61 | 0.5 | 0.53 |
| 7 | 54973910 | rs2538066 | 0.212 | -0.901 | -0.175 | -0.999 | 0.014 | 0.016 | 0.016 | 0 | 0.43 | 0.63 | 0.23 | 0.14 | 0.31 | 0.32 |
| 7 | 54976715 | rs10277377 | 0.488 | 0.158 | -0.199 | -0.609 | -0.001 | 0.011 | 0.011 | 0 | 0.08 | 0.05 | 0.06 | 0.19 | 0.11 | 0.07 |
| 7 | 55002632 | rs2330915 | -0.01 | 0.486 | -0.148 | -0.717 | -0.001 | 0 | 0 | 0 | 0.23 | 0.23 | 0.35 | 0.65 | 0.27 | 0.37 |
| 7 | 55004546 | rs2005868 | 0.203 | 0.521 | -0.7 | -0.695 | -0.077 | -0.102 | -0.102 | 0 | 0.21 | 0.03 | 0.33 | 0.65 | 0.26 | 0.37 |
| 7 | 55059669 | rs17584903 | 0.202 | 0.338 | -1.062 | -1.37 | -0.013 | 0.083 | 0.083 | 0 | 0.16 | 0.05 | 0.13 | 0 | 0.19 | 0.12 |
| 7 | 55082418 | rs6969537 | 0.273 | 1.186 | -0.602 | -1.146 | 0.024 | -0.02 | -0.02 | 0 | 0.92 | 0.84 | 0.76 | 1 | 0.86 | 0.96 |
| 7 | 55090379 | rs11770506 | -0.16 | 1.242 | -0.98 | -0.834 | -0.053 | -0.041 | -0.041 | 0 | 0.24 | 0.13 | 0.41 | 0.68 | 0.35 | 0.4 |
| 7 | 55114316 | rs1015793 | -0.87 | 0.621 | -0.288 | -1.508 | 0.05 | -0.041 | -0.041 | 0 | 0.14 | 0.35 | 0.15 | 0.03 | 0.14 | 0.08 |
| 7 | 55123968 | rs11773818 | 0.326 | 1.385 | -0.91 | -1.711 | -0.067 | 0.009 | 0.009 | 0.056 | 0.19 | 0.06 | 0.36 | 0.86 | 0.28 | 0.41 |
| 7 | 55128207 | rs729969 | 0.307 | 1.562 | -0.604 | -1.81 | 0.051 | -0.032 | -0.032 | 0.031 | 0.87 | 0.69 | 0.88 | 0.97 | 0.86 | 0.92 |
| 7 | 55128731 | rs1024750 | -0.925 | 0.958 | -1.362 | -2.631 | 0.011 | 0.078 | 0.078 | 0 | 0.78 | 0.91 | 0.9 | 0.98 | 0.82 | 0.84 |
| 7 | 55134272 | rs1558542 | 0.852 | 0.857 | -1.487 | -2.833 | -0.016 | 0.054 | 0.054 | 0.507 | 0.14 | 0.04 | 0.15 | 0 | 0.22 | 0.05 |
| 7 | 55142435 | rs17172433 | 0.209 | 0.939 | -1.085 | -2.601 | 0.133 | -0.024 | -0.024 | 0 | 0.08 | 0.27 | 0.12 | 0 | 0.14 | 0.04 |
| 7 | 55144156 | rs984654 | 1.055 | 0.905 | -1.485 | -2.716 | -0.06 | 0.027 | 0.027 | 0.089 | 0.2 | 0.06 | 0.36 | 0.89 | 0.27 | 0.42 |
| 7 | 55147478 | rs10245472 | -0.347 | 1.099 | -0.792 | -2.567 | 0.169 | -0.041 | -0.041 | 0 | 0.1 | 0.42 | 0.17 | 0 | 0.17 | 0.05 |
| 7 | 55151537 | rs17172438 | -0.599 | 1.024 | -0.971 | -2.603 | 0.077 | -0.04 | -0.04 | 0.64 | 0.14 | 0.36 | 0.16 | 0.03 | 0.18 | 0.08 |
| 7 | 55152337 | rs10244108 | 0.307 | 0.981 | -1.023 | -3.059 | -0.005 | 0.015 | 0.015 | 0.17 | 0.29 | 0.17 | 0.2 | 0.01 | 0.35 | 0.2 |
| 7 | 55154636 | rs759169 | -0.603 | 1.217 | -0.841 | -2.51 | 0.017 | 0.08 | 0.08 | 0 | 0.81 | 0.96 | 0.93 | 0.98 | 0.86 | 0.9 |
| 7 | 55160538 | rs4947490 | 0.23 | 1.131 | -0.801 | -2.422 | 0.023 | -0.113 | -0.113 | 0 | 0.73 | 0.63 | 0.77 | 0.98 | 0.69 | 0.85 |
| 7 | 55162011 | rs917880 | -0.511 | 1.215 | -1.597 | -2.122 | -0.011 | 0.021 | 0.021 | 1.505 | 0.41 | 0.63 | 0.45 | 0.17 | 0.49 | 0.4 |
| 7 | 55162305 | rs917881 | 0.211 | 1.378 | -1.844 | -2.211 | -0.015 | 0.063 | 0.063 | 0 | 0.83 | 0.95 | 0.73 | 1 | 0.83 | 0.85 |
| 7 | 55162336 | rs11977660 | 0.789 | 1.116 | -2.166 | -1.786 | -0.017 | 0.06 | 0.06 | 0 | 0.48 | 0.22 | 0.49 | 0.68 | 0.51 | 0.43 |
| 7 | 55169291 | rs4947491 | 0.015 | 1.081 | -1.48 | -1.94 | -0.002 | -0.011 | -0.011 | 0 | 0.7 | 0.75 | 0.62 | 0.92 | 0.65 | 0.7 |
| 7 | 55171190 | rs6954351 | 0.729 | 1.15 | -2.155 | -2.102 | -0.015 | 0.07 | 0.07 | 0 | 0.15 | 0.03 | 0.12 | 0 | 0.17 | 0.07 |
| 7 | 55172768 | rs2110290 | 1.412 | 1.219 | -2.07 | -2.002 | -0.018 | -0.002 | -0.002 | 0 | 0.31 | 0.23 | 0.37 | 0.72 | 0.33 | 0.37 |
| 7 | 55174342 | rs2330951 | -0.439 | 1.444 | -2.249 | -2.392 | 0.021 | -0.03 | -0.03 | 0 | 0.9 | 0.89 | 0.72 | 1 | 0.77 | 0.89 |
| 7 | 55175876 | rs7796139 | -0.505 | 0.509 | -2.287 | -2.238 | 0.03 | 0.155 | 0.155 | 0 | 0.33 | 0.08 | 0.17 | 0 | 0.29 | 0.14 |
| 7 | 55177869 | rs2058502 | 0.159 | 0.813 | -1.833 | -1.945 | 0.003 | 0.151 | 0.151 | 0 | 0.53 | 0.79 | 0.53 | 0.91 | 0.52 | 0.62 |
| 7 | 55179844 | rs7796872 | 0.137 | 1.475 | -1.727 | -1.71 | -0.039 | 0.033 | 0.033 | 0 | 0.06 | 0.01 | 0.22 | 0 | 0.13 | 0.08 |
| 7 | 55180110 | rs6593206 | 0.254 | 0.399 | -1.302 | -2.727 | 0 | -0.08 | -0.08 | 0 | 0.59 | 0.65 | 0.73 | 1 | 0.65 | 0.82 |
| 7 | 55181442 | rs759160 | -0.032 | 0.003 | -0.837 | -2.961 | 0.002 | -0.132 | -0.132 | 0.003 | 0.78 | 0.63 | 0.79 | 1 | 0.75 | 0.9 |
| 7 | 55186456 | rs4947978 | 0.398 | -0.616 | -1.213 | -2.419 | -0.009 | -0.002 | -0.002 | 0 | 0.58 | 0.53 | 0.69 | 0.67 | 0.64 | 0.59 |
| 7 | 55187053 | rs12667668 | 0.524 | -0.663 | -1.189 | -2.439 | 0.016 | 0.182 | 0.182 | 0 | 0.12 | 0.23 | 0.16 | 0.32 | 0.16 | 0.26 |
| 7 | 55187138 | rs9649847 | 1.346 | -0.619 | -1.09 | -2.49 | 0.002 | -0.04 | -0.04 | 0.761 | 0.09 | 0.16 | 0.1 | 0 | 0.12 | 0.06 |
| 7 | 55190489 | rs7786831 | 1.05 | -0.734 | -1.226 | -2.549 | 0.05 | 0.066 | 0.066 | 0.003 | 0.19 | 0.07 | 0.05 | 0 | 0.1 | 0.08 |
| 7 | 55192256 | rs13244925 | 1.111 | -0.632 | -0.7 | -2.477 | -0.009 | -0.004 | -0.004 | 0 | 0.37 | 0.33 | 0.63 | 0.69 | 0.53 | 0.51 |
| 7 | 55195625 | rs4947979 | -0.057 | -0.889 | -0.582 | -2.488 | -0.018 | -0.09 | -0.09 | 0 | 0.69 | 0.64 | 0.85 | 0.99 | 0.79 | 0.88 |
| 7 | 55196855 | rs6593207 | 1.017 | -0.866 | -0.646 | -2.286 | 0.01 | -0.016 | -0.016 | 0.151 | 0.19 | 0.13 | 0.1 | 0.31 | 0.1 | 0.28 |
| 7 | 55202598 | rs17172446 | 0.786 | -0.489 | -0.628 | -2.455 | 0.005 | -0.061 | -0.061 | 0 | 0.21 | 0.28 | 0.3 | 0.02 | 0.23 | 0.19 |
| 7 | 55206861 | rs887824 | 0.744 | -0.431 | -0.889 | -2.149 | -0.017 | -0.048 | -0.048 | 0.413 | 0.19 | 0.14 | 0.05 | 0 | 0.08 | 0.06 |
| 7 | 55209637 | rs6964705 | 0.319 | 0.566 | -1.079 | -1.651 | -0.048 | -0.03 | -0.03 | 0 | 0.31 | 0.22 | 0.45 | 0.67 | 0.48 | 0.41 |
| 7 | 55211879 | rs13234622 | 0.865 | 0.807 | -1.311 | -1.48 | -0.08 | -0.086 | -0.086 | 0 | 0.22 | 0.11 | 0.37 | 0.66 | 0.36 | 0.35 |
| 7 | 55214348 | rs2072454 | 0.28 | 0.284 | -0.968 | -1.742 | 0 | -0.001 | -0.001 | 0.485 | 0.52 | 0.49 | 0.49 | 0.35 | 0.48 | 0.56 |
| 7 | 55220177 | rs11506105 | -0.287 | 0.473 | -0.974 | -1.609 | -0.007 | -0.007 | -0.007 | 0 | 0.66 | 0.68 | 0.55 | 0.35 | 0.55 | 0.59 |
| 7 | 55221167 | rs11760524 | 1.318 | 0.121 | -1.647 | -1.896 | 0.024 | 0.089 | 0.089 | 0.209 | 0.19 | 0.05 | 0.11 | 0 | 0.12 | 0.09 |
| 7 | 55229543 | rs3752651 | 1.72 | 0.765 | -1.451 | -1.14 | -0.051 | 0.006 | 0.006 | 0 | 0.15 | 0.07 | 0.16 | 0.14 | 0.2 | 0.11 |
| 7 | 55230840 | rs7781264 | 0.227 | 0.318 | -0.953 | -1.787 | -0.008 | -0.114 | -0.114 | 0 | 0.66 | 0.41 | 0.79 | 1 | 0.68 | 0.88 |

| | | | | | | | | | | | | | | | | |
|---|---|---|---|---|---|---|---|---|---|---|---|---|---|---|---|---|
| 7 | 55232333 | rs11976696 | 0.111 | 0.609 | -0.661 | -0.994 | -0.001 | 0.02 | 0.02 | 0 | 0.22 | 0.24 | 0.31 | 0.35 | 0.2 | 0.41 |
| 7 | 55237283 | rs845551 | 0.004 | -0.111 | -0.135 | -0.864 | -0.018 | 0.12 | 0.12 | 0 | 0.75 | 0.53 | 0.81 | 0.63 | 0.77 | 0.64 |
| 7 | 55238268 | rs10228436 | 0.573 | -0.232 | -0.639 | -1.112 | 0.061 | -0.008 | -0.008 | 0 | 0.47 | 0.22 | 0.39 | 0.54 | 0.35 | 0.44 |
| 7 | 55238464 | rs10277413 | 0.573 | -0.292 | -0.639 | -1.112 | 0.061 | -0.008 | -0.008 | 0.88 | 0.47 | 0.22 | 0.39 | 0.53 | 0.36 | 0.44 |
| 7 | 55242609 | rs2017000 | 1.057 | -0.299 | -0.917 | -1.053 | 0.011 | -0.109 | -0.109 | 0 | 0.35 | 0.06 | 0.37 | 0.62 | 0.3 | 0.39 |
| 7 | 55243754 | rs9692301 | 0.312 | 0.348 | -0.151 | -1.293 | -0.012 | -0.001 | -0.001 | 0.166 | 0.25 | 0.17 | 0.33 | 0.01 | 0.29 | 0.22 |
| 7 | 55247588 | rs845558 | -0.153 | 0.337 | 0.539 | -1.258 | 0.029 | -0.119 | -0.119 | 0 | 0.32 | 0.55 | 0.42 | 0.02 | 0.4 | 0.23 |
| 7 | 55247960 | rs845559 | -0.533 | 0.261 | 0.944 | -0.995 | 0.031 | -0.044 | -0.044 | 0 | 0.11 | 0.3 | 0.13 | 0.02 | 0.14 | 0.05 |
| 7 | 55250130 | rs7795743 | 0.729 | -0.113 | 0.465 | -0.993 | 0.002 | -0.039 | -0.039 | 0 | 0.64 | 0.57 | 0.57 | 0.18 | 0.61 | 0.45 |
| 7 | 55251593 | rs13222385 | 0.928 | -0.218 | 0.222 | -1.195 | -0.112 | 0.153 | 0.153 | 1.075 | 0.31 | 0.08 | 0.36 | 0.11 | 0.37 | 0.36 |
| 7 | 55253425 | rs2075106 | 0.632 | -0.203 | 0.328 | -1.165 | 0.001 | -0.028 | -0.028 | 0.012 | 0.63 | 0.59 | 0.57 | 0.18 | 0.61 | 0.45 |
| 7 | 55254186 | rs6593210 | 1.535 | -0.23 | 0.072 | -1.169 | 0.011 | 0.02 | 0.02 | 0.223 | 0.29 | 0.2 | 0.18 | 0.08 | 0.23 | 0.09 |
| 7 | 55254805 | rs845562 | -1.129 | -0.266 | -0.022 | -0.684 | 0.002 | -0.071 | -0.071 | 4.905 | 0.84 | 0.92 | 0.86 | 0.59 | 0.84 | 0.74 |
| 7 | 55255963 | rs1554718 | 0.807 | -0.193 | 0.007 | -1.073 | 0.003 | 0.014 | 0.014 | 0 | 0.59 | 0.35 | 0.55 | 0.2 | 0.61 | 0.45 |
| 7 | 55255986 | rs2740761 | 1.029 | -0.117 | 0.114 | -0.71 | 0.006 | 0.006 | 0.006 | 0.185 | 0.3 | 0.21 | 0.18 | 0.08 | 0.23 | 0.09 |
| 7 | 55259763 | rs6970262 | -0.496 | 0.018 | 0.186 | -0.776 | 0.002 | -0.008 | -0.008 | 0 | 0.61 | 0.66 | 0.71 | 0.87 | 0.64 | 0.87 |
| 7 | 55266417 | rs1140475 | -0.009 | 0.092 | 0.011 | -0.341 | -0.007 | 0.004 | 0.004 | 0 | 0.92 | 0.93 | 0.9 | 0.92 | 0.89 | 0.94 |
| 7 | 55266757 | rs2293348 | 0.382 | 0.012 | -0.229 | -0.85 | -0.03 | 0.08 | 0.08 | 0.041 | 0.28 | 0.14 | 0.38 | 0.13 | 0.32 | 0.28 |
| 7 | 55268916 | rs2293347 | 1.041 | -0.03 | -0.396 | -0.355 | -0.018 | -0.052 | -0.052 | 0 | 0.09 | 0.01 | 0.15 | 0.25 | 0.11 | 0.23 |
| 7 | 55277387 | rs940810 | 1.333 | -0.138 | 0.208 | -0.525 | -0.083 | 0.091 | 0.091 | 0.161 | 0.21 | 0.04 | 0.34 | 0.08 | 0.27 | 0.26 |
| 7 | 55277964 | rs6593211 | -0.914 | -0.301 | 0.852 | -0.494 | -0.01 | -0.019 | -0.019 | 0 | 0.34 | 0.59 | 0.31 | 0.72 | 0.28 | 0.53 |
| 7 | 55279494 | rs940807 | 1.105 | -0.274 | 0.617 | -0.374 | -0.027 | -0.058 | -0.058 | 0.334 | 0.09 | 0.01 | 0.16 | 0.25 | 0.12 | 0.28 |
| 7 | 55279745 | rs940806 | -1.323 | -0.317 | 0.713 | -0.431 | -0.025 | 0.097 | 0.097 | 0.005 | 0.64 | 0.82 | 0.7 | 0.86 | 0.6 | 0.81 |
| 7 | 55280152 | rs1107616 | -0.3 | -0.559 | 0.986 | -0.383 | 0.016 | 0.021 | 0.021 | 0.328 | 0.25 | 0.16 | 0.27 | 0.72 | 0.23 | 0.45 |
| 7 | 55280268 | rs1107617 | 0.555 | -0.552 | 1.118 | -0.371 | 0 | 0.014 | 0.014 | 0 | 0.17 | 0.19 | 0.12 | 0.48 | 0.11 | 0.22 |
| 7 | 55280283 | rs1107618 | -0.894 | -0.564 | 1.261 | -0.398 | -0.015 | -0.02 | -0.02 | 0 | 0.29 | 0.62 | 0.3 | 0.72 | 0.23 | 0.45 |
| 7 | 55281096 | rs11761161 | 1.529 | -0.556 | 1.129 | -0.396 | -0.006 | -0.092 | -0.092 | 0 | 0.32 | 0.53 | 0.22 | 0.06 | 0.3 | 0.14 |
| 7 | 55282677 | rs1111650 | 0.247 | -0.545 | 1.062 | -0.372 | 0.054 | 0.045 | 0.045 | 0.371 | 0.5 | 0.78 | 0.35 | 0.54 | 0.41 | 0.35 |
| 7 | 55284254 | rs7776830 | 0.3 | -0.403 | 1.062 | -0.325 | 0.026 | 0.169 | 0.169 | 0 | 0.12 | 0.39 | 0.18 | 0.25 | 0.12 | 0.23 |
| 7 | 55286337 | rs2331014 | 1.308 | -0.31 | 0.222 | -0.555 | 0 | 0.059 | 0.059 | 0 | 0.3 | 0.17 | 0.19 | 0.06 | 0.29 | 0.14 |
| 7 | 55292028 | rs10233099 | 0.152 | -0.399 | 0.656 | 0.281 | -0.036 | 0.074 | 0.074 | 0.191 | 0.17 | 0.28 | 0.09 | 0.29 | 0.08 | 0.22 |
| 7 | 55292321 | rs12718947 | 1.675 | -0.503 | 0.616 | 0.322 | 0.003 | 0.03 | 0.03 | 0 | 0.3 | 0.15 | 0.28 | 0.21 | 0.32 | 0.26 |
| 7 | 55292883 | rs12536061 | 0.481 | -0.33 | 0.926 | 0.151 | -0.008 | 0.021 | 0.021 | 0.389 | 0.35 | 0.27 | 0.41 | 0.14 | 0.36 | 0.35 |
| 7 | 55293361 | rs7809332 | 0.519 | -0.231 | 1.078 | 0.259 | -0.02 | 0.089 | 0.089 | 0.073 | 0.74 | 0.48 | 0.78 | 0.63 | 0.78 | 0.73 |
| 7 | 55301151 | rs17746319 | 0.672 | -0.106 | 0.749 | 0.35 | -0.026 | -0.066 | -0.066 | 2.987 | 0.11 | 0.03 | 0.09 | 0.29 | 0.12 | 0.13 |
| 7 | 55301435 | rs11769590 | 1.878 | -0.16 | 0.773 | 0.208 | 0.004 | 0.032 | 0.032 | 0 | 0.31 | 0.15 | 0.28 | 0.21 | 0.32 | 0.26 |
| 7 | 55302777 | rs6945081 | -0.381 | -0.181 | 1.028 | 0.275 | -0.022 | 0.08 | 0.08 | 1.025 | 0.19 | 0.34 | 0.13 | 0.29 | 0.15 | 0.3 |
| 7 | 55305966 | rs4947993 | 1.126 | -0.097 | 1.002 | 0.329 | 0.02 | 0.035 | 0.035 | 0 | 0.77 | 0.68 | 0.78 | 0.64 | 0.74 | 0.72 |
| 7 | 55306579 | rs2037701 | -0.212 | -0.528 | 0.516 | 0.297 | 0.01 | -0.051 | -0.051 | 0 | 0.59 | 0.53 | 0.46 | 0.77 | 0.56 | 0.71 |
| 7 | 55308630 | rs4947995 | 0.515 | -0.364 | -0.012 | 0.28 | -0.012 | -0.045 | -0.045 | 0 | 0.42 | 0.19 | 0.36 | 0.48 | 0.45 | 0.45 |
| 7 | 55313228 | rs11982525 | -0.83 | -0.586 | 0.183 | 0.237 | 0.001 | -0.002 | -0.002 | 0.006 | 0.23 | 0.25 | 0.14 | 0.3 | 0.19 | 0.17 |
| 7 | 55313350 | rs17673576 | 0.708 | -0.483 | -0.043 | 0.332 | -0.043 | -0.07 | -0.07 | 0.242 | 0.11 | 0.01 | 0.09 | 0.3 | 0.14 | 0.12 |
| 7 | 55313378 | rs11767730 | 1.523 | -0.613 | 0.183 | 0.024 | 0 | 0.031 | 0.031 | 0 | 0.31 | 0.15 | 0.27 | 0.2 | 0.31 | 0.28 |
| 7 | 55323098 | rs6948867 | -1.575 | -1.019 | 1.217 | 0.285 | -0.047 | -0.034 | -0.034 | 0.152 | 0.28 | 0.5 | 0.17 | 0.23 | 0.19 | 0.23 |
| 7 | 55326587 | rs10215769 | -1.694 | -1.033 | 0.827 | 0.279 | -0.018 | -0.013 | -0.013 | 0 | 0.29 | 0.65 | 0.18 | 0.23 | 0.19 | 0.23 |
| 7 | 55327452 | rs7803459 | -0.978 | -1.029 | 1.042 | 0.01 | -0.063 | -0.109 | -0.109 | 0.449 | 0.17 | 0.58 | 0.09 | 0 | 0.07 | 0.12 |
| 7 | 55327748 | rs1525642 | -1.263 | -0.803 | 0.874 | 0.651 | -0.039 | 0.058 | 0.058 | 0 | 0.28 | 0.49 | 0.18 | 0.57 | 0.21 | 0.43 |
| 7 | 55334240 | rs10255892 | 0.266 | -0.378 | 0.306 | 0.018 | 0.001 | -0.006 | -0.006 | 0 | 0.37 | 0.59 | 0.33 | 0.21 | 0.39 | 0.28 |
| 7 | 55339948 | rs6974790 | 0.586 | -0.61 | -0.92 | 0.292 | -0.01 | -0.021 | -0.021 | 0 | 0.35 | 0.06 | 0.45 | 0.36 | 0.31 | 0.42 |
| 7 | 55340022 | rs6593217 | -0.686 | -0.651 | -0.398 | 0.19 | 0.013 | 0.067 | 0.067 | 0 | 0.75 | 0.6 | 0.77 | 0.57 | 0.71 | 0.7 |
| 7 | 55342649 | rs12538798 | 1.248 | -0.778 | -0.746 | -0.099 | -0.014 | -0.014 | -0.014 | 0.42 | 0.21 | 0.02 | 0.36 | 0.16 | 0.18 | 0.21 |
| 7 | 55343650 | rs13438690 | 0.507 | -0.561 | -0.314 | -0.243 | 0.002 | -0.037 | -0.037 | 0 | 0.4 | 0.55 | 0.32 | 0.16 | 0.4 | 0.28 |
| 7 | 55345282 | rs10215194 | 1.182 | -0.199 | -0.233 | -0.276 | -0.002 | 0.005 | 0.005 | 0 | 0.32 | 0.28 | 0.27 | 0.16 | 0.35 | 0.24 |
| 7 | 55347512 | rs10232489 | 0.734 | -0.354 | -0.484 | -0.198 | 0.007 | -0.05 | -0.05 | 0.479 | 0.38 | 0.52 | 0.31 | 0.16 | 0.39 | 0.27 |
| 7 | 55370212 | rs4948007 | 0.931 | -0.241 | -1.202 | 0.045 | -0.071 | -0.114 | -0.114 | 0 | 0.25 | 0.02 | 0.44 | 0.32 | 0.27 | 0.2 |
| 7 | 55371250 | rs1880625 | 0.391 | -0.487 | -0.876 | -0.114 | 0.004 | -0.004 | -0.004 | 0.75 | 0.51 | 0.69 | 0.37 | 0.48 | 0.52 | 0.51 |
| 7 | 55374827 | rs7805062 | 1.538 | -0.902 | -0.914 | -0.251 | 0.008 | -0.016 | -0.016 | 1.71 | 0.27 | 0.34 | 0.19 | 0.13 | 0.36 | 0.26 |
| 7 | 55388586 | rs12672167 | 1.486 | -0.576 | -1.372 | -0.073 | -0.067 | -0.114 | -0.114 | 0 | 0.25 | 0.02 | 0.44 | 0.33 | 0.27 | 0.2 |
| 7 | 55390990 | rs2177808 | 1.231 | -0.548 | -1.244 | -0.058 | 0.001 | -0.01 | -0.01 | 0 | 0.28 | 0.22 | 0.45 | 0.33 | 0.28 | 0.2 |
| 7 | 55393032 | rs6956733 | -1.033 | -0.455 | -1.515 | 0.035 | -0.048 | -0.034 | -0.034 | 0 | 0.86 | 0.97 | 0.85 | 0.8 | 0.82 | 0.74 |
| 7 | 55398692 | rs9649851 | -1.182 | -0.278 | -1.374 | 0.155 | -0.035 | 0 | 0 | 0 | 0.78 | 0.97 | 0.84 | 0.8 | 0.79 | 0.71 |
| 7 | 55416964 | rs6943047 | 0.109 | -0.46 | -0.669 | 0.451 | -0.015 | 0.015 | 0.015 | 0 | 0.24 | 0.28 | 0.18 | 0.35 | 0.15 | 0.24 |
| 7 | 55423050 | rs10243871 | -0.103 | 0.204 | -0.654 | -0.311 | 0.017 | 0.054 | 0.054 | 0 | 0.53 | 0.68 | 0.65 | 0.67 | 0.43 | 0.45 |
| 7 | 55427182 | rs7787709 | -0.093 | 0.432 | -0.682 | -0.219 | -0.01 | 0.041 | 0.041 | 0 | 0.54 | 0.66 | 0.65 | 0.67 | 0.45 | 0.47 |
| 7 | 55433884 | rs2272263 | 0.973 | 0.666 | -0.444 | -0.091 | -0.014 | -0.001 | -0.001 | 0 | 0.17 | 0.14 | 0.18 | 0.16 | 0.33 | 0.24 |
| 7 | 55442775 | rs11974368 | 0.674 | 0.419 | -0.76 | -0.308 | -0.01 | 0.039 | 0.039 | 0 | 0.21 | 0.14 | 0.13 | 0.08 | 0.2 | 0.16 |
| 7 | 55442803 | rs10282700 | -0.312 | 0.125 | -0.211 | -0.33 | -0.001 | 0.004 | 0.004 | 0.198 | 0.43 | 0.58 | 0.33 | 0.23 | 0.53 | 0.41 |
| 7 | 55456524 | rs6593246 | -1.401 | -0.15 | -0.475 | -0.324 | 0.032 | -0.031 | -0.031 | 0 | 0.81 | 0.99 | 0.9 | 0.8 | 0.87 | 0.74 |
| 7 | 55459563 | rs2049497 | 0.635 | -0.077 | -0.267 | -0.278 | 0.002 | -0.005 | -0.005 | 0.275 | 0.38 | 0.31 | 0.56 | 0.57 | 0.34 | 0.33 |

| | | | | | | | | | | | | | | | | |
|---|---|---|---|---|---|---|---|---|---|---|---|---|---|---|---|---|
| 7 | 55481389 | rs1568824 | 0.177 | -0.118 | -0.246 | -0.339 | -0.01 | -0.014 | -0.014 | 0 | 0.55 | 0.53 | 0.73 | 0.7 | 0.65 | 0.6 |
| 7 | 55489494 | rs815970 | -1.295 | -0.21 | -0.314 | -0.369 | 0.039 | -0.033 | -0.033 | 0 | 0.8 | 0.97 | 0.89 | 0.77 | 0.86 | 0.76 |
| 7 | 55490914 | rs3807357 | 1.484 | 0.235 | -0.268 | 0.047 | -0.024 | -0.033 | -0.033 | 1.19 | 0.06 | 0 | 0.11 | 0.26 | 0.07 | 0.13 |
| 7 | 55495109 | rs3778864 | -0.086 | 0.261 | -0.057 | -0.179 | -0.008 | -0.011 | -0.011 | 4.469 | 0.55 | 0.57 | 0.73 | 0.69 | 0.65 | 0.6 |
| 7 | 55498417 | rs12718964 | 1.53 | 0.39 | 0.118 | -0.203 | 0.003 | 0 | 0 | 0 | 0.17 | 0.17 | 0.17 | 0.16 | 0.31 | 0.23 |
| 7 | 55499591 | rs815989 | -0.359 | 0.353 | 0.097 | -0.189 | 0.032 | 0.051 | 0.051 | 1.321 | 0.58 | 0.85 | 0.76 | 0.69 | 0.65 | 0.6 |
| 7 | 55499723 | rs3823584 | 0.511 | 0.346 | 0.283 | -0.17 | -0.001 | -0.031 | -0.031 | 0 | 0.35 | 0.24 | 0.55 | 0.54 | 0.34 | 0.35 |
| 7 | 55507580 | rs1722981 | -0.19 | -0.069 | -0.574 | -0.279 | 0.032 | 0.067 | 0.067 | 1.195 | 0.59 | 0.85 | 0.76 | 0.71 | 0.64 | 0.6 |
| 7 | 55514819 | rs815946 | 0.275 | 0.187 | -0.534 | -0.096 | 0.003 | -0.01 | -0.01 | 0 | 0.34 | 0.31 | 0.52 | 0.55 | 0.28 | 0.36 |
| 7 | 55515871 | rs6972560 | 1.368 | 0.068 | -0.417 | -0.141 | -0.004 | -0.001 | -0.001 | 0 | 0.19 | 0.18 | 0.21 | 0.16 | 0.36 | 0.25 |
| 7 | 55533847 | rs6961353 | 0.058 | -0.17 | -0.478 | -0.272 | 0.012 | 0.036 | 0.036 | 0.264 | 0.47 | 0.61 | 0.36 | 0.22 | 0.56 | 0.42 |
| 7 | 55537497 | rs940633 | 0.211 | -0.279 | -0.508 | -0.268 | 0.034 | 0.033 | 0.033 | 0.78 | 0.55 | 0.35 | 0.74 | 0.67 | 0.69 | 0.6 |
| 7 | 55547694 | rs815964 | 0.587 | -0.284 | -0.589 | -0.234 | -0.003 | -0.087 | -0.087 | 0 | 0.33 | 0.12 | 0.53 | 0.51 | 0.31 | 0.37 |
| 7 | 55550693 | rs6957545 | 1.408 | -1.049 | -0.307 | -0.209 | 0.03 | -0.005 | -0.005 | 0.26 | 0.22 | 0.28 | 0.21 | 0.16 | 0.37 | 0.23 |
| 7 | 55551832 | rs815965 | 0.567 | -1.131 | -0.376 | -0.196 | 0.001 | -0.065 | -0.065 | 0.158 | 0.33 | 0.14 | 0.53 | 0.51 | 0.31 | 0.37 |
| 7 | 55554182 | rs17565689 | 0.332 | -1.127 | -0.623 | -0.131 | 0.012 | 0.004 | 0.004 | 1.717 | 0.24 | 0.21 | 0.19 | 0.24 | 0.2 | 0.29 |
| 7 | 55556628 | rs815966 | 0.689 | -0.952 | -1.206 | -0.079 | 0.003 | -0.01 | -0.01 | 0.002 | 0.35 | 0.24 | 0.54 | 0.51 | 0.32 | 0.37 |
| 7 | 55560395 | rs12718968 | 1.328 | -0.859 | -1.281 | -0.039 | -0.012 | 0.002 | 0.002 | 0 | 0.22 | 0.2 | 0.2 | 0.16 | 0.37 | 0.22 |
| 7 | 55561884 | rs815952 | -0.852 | -1.379 | -0.447 | 0.175 | 0.041 | -0.034 | -0.034 | 0 | 0.83 | 0.92 | 0.91 | 0.72 | 0.89 | 0.8 |
| 7 | 55592518 | rs7798349 | 0.383 | -1.375 | -0.394 | 0.077 | 0.005 | -0.081 | -0.081 | 0.018 | 0.21 | 0.38 | 0.16 | 0.06 | 0.2 | 0.19 |
| 7 | 55597235 | rs28642833 | 0.554 | -1.143 | -1.681 | 0.308 | 0.04 | -0.036 | -0.036 | 0 | 0.17 | 0.08 | 0.1 | 0.28 | 0.11 | 0.2 |
| 7 | 55603590 | rs10251825 | -0.133 | -0.086 | -1.313 | 0.716 | 0.058 | 0.067 | 0.067 | 0.534 | 0.55 | 0.26 | 0.72 | 0.65 | 0.66 | 0.58 |
| 7 | 55610012 | rs4948031 | -0.139 | -0.054 | -1.298 | 0.729 | 0.057 | 0.071 | 0.071 | 0.643 | 0.55 | 0.26 | 0.72 | 0.67 | 0.66 | 0.59 |
| 7 | 55612169 | rs17673975 | -0.09 | -0.035 | -1.363 | 0.889 | -0.006 | -0.061 | -0.061 | 0.125 | 0.1 | 0 | 0.14 | 0.23 | 0.1 | 0.14 |
| 7 | 55624401 | rs2030734 | 0.243 | 0.088 | -0.857 | 0.312 | 0.021 | 0.026 | 0.026 | 0.281 | 0.57 | 0.38 | 0.73 | 0.67 | 0.66 | 0.59 |
| 7 | 55640771 | rs6943773 | 0.291 | -0.054 | -1.609 | 0.339 | 0.07 | 0.054 | 0.054 | 0.437 | 0.56 | 0.23 | 0.74 | 0.65 | 0.68 | 0.59 |
| 7 | 55651352 | rs4947514 | 0.321 | 0.02 | -1.597 | 0.376 | 0.06 | 0.039 | 0.039 | 0.526 | 0.55 | 0.25 | 0.74 | 0.63 | 0.66 | 0.58 |
| 7 | 55654391 | rs6968800 | 0.713 | -0.79 | -1.516 | -0.318 | 0.003 | 0.003 | 0.003 | 0 | 0.73 | 0.32 | 0.78 | 0.86 | 0.77 | 0.84 |
| 7 | 55654430 | rs6948102 | 0.996 | -1.161 | -1.048 | -0.492 | -0.008 | -0.011 | -0.011 | 0 | 0.69 | 0.43 | 0.76 | 0.83 | 0.73 | 0.81 |
| 7 | 55654767 | rs11769548 | 0.663 | -1.342 | -0.93 | -1.06 | 0.01 | -0.024 | -0.024 | 0.687 | 0.74 | 0.6 | 0.64 | 0.86 | 0.69 | 0.86 |
| 7 | 55655081 | rs11772826 | 0.978 | -0.38 | -1.407 | -1.153 | 0.004 | -0.008 | -0.008 | 2.896 | 0.3 | 0.31 | 0.23 | 0.25 | 0.32 | 0.24 |
| 7 | 55656981 | rs4245568 | 0.161 | -0.421 | -1.653 | -1.232 | 0.039 | -0.014 | -0.014 | 0 | 0.39 | 0.19 | 0.26 | 0.6 | 0.34 | 0.57 |
| 7 | 55657127 | rs4245569 | 0.073 | -0.187 | -1.667 | -1.218 | 0.025 | 0 | 0 | 0.361 | 0.4 | 0.28 | 0.29 | 0.61 | 0.36 | 0.61 |
| 7 | 55657718 | rs7802222 | 0.329 | 0.306 | -1.282 | -0.976 | 0.045 | -0.023 | -0.023 | 0 | 0.78 | 0.58 | 0.61 | 0.87 | 0.71 | 0.91 |
| 7 | 55661653 | rs12718974 | -0.663 | 0.873 | -1.86 | -1.02 | -0.003 | -0.015 | -0.015 | 0.047 | 0.12 | 0.11 | 0.07 | 0 | 0.14 | 0.05 |
| 7 | 55662586 | rs7795911 | 0.28 | 1.136 | -2.452 | -0.615 | -0.07 | -0.007 | -0.007 | 0.048 | 0.37 | 0.15 | 0.32 | 0.34 | 0.49 | 0.41 |
| 7 | 55668163 | rs4298454 | 0.623 | 0.689 | -2.437 | -0.64 | -0.04 | 0.023 | 0.023 | 0 | 0.3 | 0.09 | 0.23 | 0.22 | 0.33 | 0.37 |
| 7 | 55680064 | rs35520500 | -0.423 | 0.08 | -1.852 | -0.083 | 0.006 | 0.026 | 0.026 | 0 | 0.09 | 0.04 | 0.04 | 0 | 0.08 | 0.04 |
| 7 | 55688100 | rs4948051 | -0.071 | -0.554 | -1.991 | 0.093 | 0.092 | -0.099 | -0.099 | 0.755 | 0.23 | 0.03 | 0.37 | 0.69 | 0.16 | 0.44 |
| 7 | 55700256 | rs9986837 | -0.268 | 0.307 | -1.078 | -0.296 | 0.013 | -0.058 | -0.058 | 4.056 | 0.21 | 0.33 | 0.15 | 0.09 | 0.24 | 0.1 |
| 7 | 55703387 | rs10250208 | -0.257 | 0.607 | -1.309 | -0.474 | -0.018 | 0.065 | 0.065 | 0.214 | 0.54 | 0.79 | 0.34 | 0.31 | 0.55 | 0.35 |
| 7 | 55705566 | rs10245396 | 0.491 | 0.758 | -1.148 | -0.498 | 0.002 | -0.087 | -0.087 | 0 | 0.19 | 0.26 | 0.13 | 0 | 0.2 | 0.05 |
| 7 | 55714282 | rs10239854 | -0.14 | 0.662 | -1.055 | -0.486 | -0.003 | 0.077 | 0.077 | 0 | 0.52 | 0.76 | 0.34 | 0.31 | 0.55 | 0.38 |
| 7 | 55715405 | rs3935165 | -0.132 | 0.855 | -0.517 | -0.597 | 0.042 | -0.078 | -0.078 | 0.016 | 0.23 | 0.51 | 0.19 | 0.09 | 0.29 | 0.12 |
| 7 | 55715496 | rs3935167 | 1.135 | 0.719 | -0.628 | -0.568 | 0.013 | 0.007 | 0.007 | 0 | 0.29 | 0.25 | 0.16 | 0.22 | 0.27 | 0.25 |
| 7 | 55716138 | rs10261389 | 0.099 | 1.031 | -0.333 | -0.569 | 0.019 | -0.056 | -0.056 | 3.298 | 0.2 | 0.33 | 0.15 | 0.09 | 0.24 | 0.1 |
| 7 | 55716877 | rs4621761 | -0.462 | 1.204 | -1.022 | -0.146 | -0.024 | -0.113 | -0.113 | 0 | 0.69 | 0.89 | 0.51 | 0.47 | 0.71 | 0.57 |
| 7 | 55718702 | rs10244823 | -0.835 | 1.372 | -0.285 | 0.378 | 0.019 | -0.02 | -0.02 | 0.384 | 0.33 | 0.49 | 0.26 | 0.23 | 0.39 | 0.25 |
| 7 | 55728418 | rs4073869 | -0.391 | 1.06 | -0.776 | 0.069 | -0.004 | 0.002 | 0.002 | 0.009 | 0.37 | 0.42 | 0.5 | 0.72 | 0.44 | 0.62 |
| 7 | 55736508 | rs4948060 | -0.745 | 0.609 | -1.345 | 0.096 | 0.041 | 0.14 | 0.14 | 0.106 | 0.55 | 0.89 | 0.67 | 0.77 | 0.65 | 0.66 |
| 7 | 55762611 | rs4947521 | -0.724 | 0.561 | -0.623 | 0.859 | -0.002 | 0.01 | 0.01 | 0 | 0.22 | 0.28 | 0.29 | 0.42 | 0.2 | 0.37 |
| 7 | 55778416 | rs4397354 | -0.495 | 0.195 | -0.318 | 1.109 | -0.014 | 0.05 | 0.05 | 0.051 | 0.13 | 0.23 | 0.18 | 0.41 | 0.1 | 0.33 |
| 7 | 55798357 | rs10280445 | -0.192 | 0.334 | -0.686 | 0.95 | 0.006 | 0 | 0 | 0.09 | 0.28 | 0.54 | 0.35 | 0.23 | 0.34 | 0.19 |
| 7 | 55838545 | rs11766051 | -0.31 | -0.032 | -0.479 | 0.932 | -0.034 | 0.097 | 0.097 | 0.195 | 0.15 | 0.34 | 0.2 | 0.42 | 0.1 | 0.33 |
| 7 | 55841188 | rs6700 | -0.597 | -0.139 | 0.554 | 0.884 | -0.055 | 0.149 | 0.149 | 0.37 | 0.17 | 0.51 | 0.21 | 0.42 | 0.1 | 0.33 |
| 7 | 55848790 | rs6966061 | -0.377 | -0.121 | 0.633 | 0.915 | -0.043 | 0.115 | 0.115 | 0.296 | 0.16 | 0.41 | 0.2 | 0.42 | 0.1 | 0.33 |
| 7 | 55856799 | rs9642404 | -0.664 | -0.034 | 0.889 | 0.886 | -0.031 | 0.091 | 0.091 | 1.003 | 0.18 | 0.61 | 0.24 | 0.42 | 0.14 | 0.33 |
| 7 | 55866443 | rs34616126 | 1.037 | -1.252 | -0.6 | -0.025 | 0.068 | 0.142 | 0.142 | 0 | 0.28 | 0.01 | 0.15 | 0.04 | 0.15 | 0.15 |
| 7 | 55880211 | rs28803557 | 0.497 | -1.59 | -0.32 | -0.044 | -0.055 | 0.05 | 0.05 | 0 | 0.21 | 0.39 | 0.22 | 0.43 | 0.12 | 0.33 |
| 7 | 55880449 | rs12538049 | 0.023 | -1.632 | -0.824 | -0.217 | 0.008 | 0.004 | 0.004 | 0 | 0.46 | 0.45 | 0.49 | 0.65 | 0.37 | 0.49 |
| 7 | 55881666 | rs10233558 | 0.844 | -1.533 | -0.772 | -0.346 | -0.055 | 0.05 | 0.05 | 0 | 0.21 | 0.4 | 0.22 | 0.43 | 0.12 | 0.33 |
| 7 | 55885674 | rs7787574 | -0.876 | -1.471 | -0.755 | -0.222 | -0.029 | 0.132 | 0.132 | 0 | 0.59 | 0.89 | 0.63 | 0.87 | 0.51 | 0.63 |
| 7 | 55889109 | rs4588807 | -0.391 | -1.32 | -0.709 | -0.366 | 0.018 | -0.013 | -0.013 | 0.059 | 0.33 | 0.75 | 0.34 | 0.48 | 0.26 | 0.45 |
| 7 | 55889334 | rs1113765 | -0.142 | -1.202 | -0.588 | -0.263 | 0.009 | -0.008 | -0.008 | 0 | 0.29 | 0.75 | 0.33 | 0.48 | 0.22 | 0.4 |
| 7 | 55899496 | rs6593284 | -1.209 | -0.061 | -1.037 | -0.537 | -0.046 | 0.194 | 0.194 | 0 | 0.64 | 0.97 | 0.76 | 0.96 | 0.68 | 0.76 |
| 7 | 55905896 | rs10230845 | 0.886 | -0.319 | -0.408 | 0.066 | -0.048 | 0.049 | 0.049 | 0 | 0.2 | 0.38 | 0.21 | 0.43 | 0.11 | 0.32 |
| 7 | 55908211 | rs10239528 | 0.183 | -0.254 | -0.604 | 0.069 | -0.057 | 0.049 | 0.049 | 0 | 0.26 | 0.5 | 0.29 | 0.46 | 0.17 | 0.38 |
| 7 | 55944185 | rs6593287 | -0.224 | 0.288 | -0.636 | 0.425 | 0.012 | 0.012 | 0.012 | 0 | 0.38 | 0.47 | 0.47 | 0.5 | 0.53 | 0.36 |
| 7 | 55944254 | rs6593288 | 0.982 | 0.983 | -0.752 | 0.494 | -0.022 | -0.014 | -0.014 | 0 | 0.07 | 0.02 | 0.15 | 0.25 | 0.21 | 0.13 |

| | | | | | | | | | | | | | | | | |
|---|---|---|---|---|---|---|---|---|---|---|---|---|---|---|---|---|
| 7 | 55950947 | rs11761352 | 0.561 | -0.249 | -0.973 | -0.129 | -0.049 | 0.01 | 0.01 | 0.087 | 0.29 | 0.41 | 0.25 | 0.42 | 0.16 | 0.35 |
| 7 | 55961981 | rs10252442 | 0.816 | 0.244 | -1.034 | -0.123 | -0.007 | -0.003 | -0.003 | 0 | 0.37 | 0.4 | 0.32 | 0.31 | 0.24 | 0.4 |
| 7 | 55968447 | rs6421738 | -1.773 | 0.596 | -1.247 | -0.292 | -0.06 | 0.073 | 0.073 | 0 | 0.77 | 0.99 | 0.79 | 0.95 | 0.75 | 0.78 |
| 7 | 55988679 | rs28828764 | 1.009 | 0.502 | -1.137 | -0.282 | -0.005 | -0.002 | -0.002 | 0 | 0.38 | 0.4 | 0.33 | 0.31 | 0.25 | 0.4 |
| 7 | 55991144 | rs2135116 | 1.009 | 0.502 | -1.137 | -0.282 | -0.005 | -0.002 | -0.002 | 0.163 | 0.38 | 0.4 | 0.33 | 0.31 | 0.25 | 0.4 |
| 7 | 55999702 | rs13222366 | -0.161 | 0.568 | -0.92 | -0.257 | 0.047 | 0.025 | 0.025 | 0.018 | 0.31 | 0.4 | 0.39 | 0.58 | 0.44 | 0.3 |
| 7 | 56018666 | rs12671091 | -0.155 | 0.596 | -1.015 | -0.241 | 0.041 | 0.023 | 0.023 | 0.025 | 0.32 | 0.38 | 0.39 | 0.58 | 0.44 | 0.3 |
| 7 | 56022981 | rs15892 | -1.578 | 0.514 | -1.203 | -0.176 | -0.059 | 0.065 | 0.065 | 0 | 0.77 | 0.98 | 0.79 | 0.93 | 0.75 | 0.78 |
| 7 | 56024287 | rs4948093 | -1.619 | 0.527 | -1.197 | -0.17 | -0.06 | 0.063 | 0.063 | 0 | 0.77 | 0.99 | 0.79 | 0.93 | 0.75 | 0.78 |
| 7 | 56033141 | rs11238386 | -0.009 | 0.708 | -0.879 | 0.045 | -0.004 | -0.001 | -0.001 | 0 | 0.37 | 0.4 | 0.35 | 0.33 | 0.36 | 0.36 |
| 7 | 56036983 | rs12668532 | 0.82 | 1.154 | -1.052 | 0.123 | -0.071 | -0.077 | -0.077 | 0 | 0.13 | 0.05 | 0.17 | 0.35 | 0.2 | 0.14 |
| 7 | 56047215 | rs4543497 | -1.326 | 1.019 | -0.976 | -0.029 | -0.006 | -0.004 | -0.004 | 0.21 | 0.64 | 0.61 | 0.67 | 0.7 | 0.74 | 0.6 |
| 7 | 56049019 | rs2304376 | 1.667 | 0.98 | -1.042 | -0.07 | -0.022 | 0.072 | 0.072 | 0 | 0.23 | 0.05 | 0.2 | 0.07 | 0.24 | 0.22 |
| 7 | 56066079 | rs11974040 | 0.259 | 1.046 | -0.904 | -0.021 | -0.024 | -0.015 | -0.015 | 0.238 | 0.27 | 0.21 | 0.32 | 0.36 | 0.38 | 0.24 |
| 7 | 56067016 | rs4689 | -1.199 | 0.937 | -1.005 | -0.126 | -0.006 | -0.004 | -0.004 | 0.935 | 0.64 | 0.61 | 0.67 | 0.7 | 0.74 | 0.6 |
| 7 | 56074118 | rs12155441 | 0.269 | 0.969 | -0.89 | 0.023 | 0.03 | 0.07 | 0.07 | 0 | 0.15 | 0.21 | 0.18 | 0.39 | 0.21 | 0.16 |
| 7 | 56077235 | rs6970842 | 0.414 | 1.152 | -0.862 | 0.024 | 0.017 | 0.047 | 0.047 | 0 | 0.14 | 0.18 | 0.18 | 0.39 | 0.2 | 0.15 |
| 7 | 56088001 | rs4496923 | -0.7 | 1.114 | -0.881 | -0.018 | 0.008 | 0.018 | 0.018 | 0 | 0.23 | 0.25 | 0.27 | 0.43 | 0.3 | 0.21 |
| 7 | 56104388 | rs4275190 | -0.788 | 0.821 | -0.865 | -0.182 | 0.01 | 0.022 | 0.022 | 2.088 | 0.59 | 0.63 | 0.67 | 0.71 | 0.67 | 0.58 |
| 7 | 56106397 | rs6965614 | 1.444 | 0.826 | -0.989 | -0.226 | 0.021 | 0.07 | 0.07 | 0 | 0.23 | 0.07 | 0.21 | 0.07 | 0.22 | 0.23 |
| 7 | 56118293 | rs6593296 | 0.902 | 0.828 | -1.025 | -0.222 | 0.013 | -0.028 | -0.028 | 0.907 | 0.27 | 0.1 | 0.27 | 0.27 | 0.29 | 0.29 |
| 7 | 56120881 | rs7793921 | -0.472 | 0.837 | -0.868 | -0.212 | -0.009 | -0.016 | -0.016 | 0 | 0.57 | 0.52 | 0.6 | 0.68 | 0.65 | 0.54 |
| 7 | 56146310 | rs11765364 | 1.291 | 0.805 | -1.096 | -0.208 | 0.011 | 0.079 | 0.079 | 0 | 0.24 | 0.06 | 0.21 | 0.07 | 0.23 | 0.22 |
| 7 | 56147589 | rs10800 | 0.42 | 1.084 | -0.907 | -0.132 | 0.022 | 0.057 | 0.057 | 0 | 0.15 | 0.19 | 0.18 | 0.4 | 0.22 | 0.17 |
| 7 | 56151489 | rs2242508 | 0.685 | 1.011 | -1.004 | -0.352 | 0.033 | 0.101 | 0.101 | 0 | 0.51 | 0.69 | 0.46 | 0.3 | 0.41 | 0.47 |
| 7 | 56158581 | rs12673255 | 0.369 | 1.029 | -0.843 | -0.124 | 0.022 | 0.056 | 0.056 | 0 | 0.15 | 0.18 | 0.18 | 0.4 | 0.21 | 0.15 |
| 7 | 56167595 | rs4948106 | 0.34 | 0.755 | -0.614 | -0.219 | -0.008 | 0.018 | 0.018 | 0 | 0.57 | 0.61 | 0.54 | 0.34 | 0.51 | 0.56 |
| 7 | 56171489 | rs816411 | 0.349 | 0.757 | -0.613 | -0.218 | -0.008 | 0.018 | 0.018 | 0 | 0.57 | 0.61 | 0.54 | 0.33 | 0.51 | 0.56 |
| 7 | 56184492 | rs2270244 | -0.161 | 1.279 | -0.01 | 0.403 | 0.023 | 0.092 | 0.092 | 0 | 0.14 | 0.21 | 0.16 | 0.46 | 0.18 | 0.15 |
| 7 | 56223751 | rs698648 | -1.079 | 0.346 | 0.034 | -0.266 | 0.004 | -0.07 | -0.07 | 0 | 0.37 | 0.55 | 0.41 | 0.9 | 0.35 | 0.57 |
| 7 | 56291170 | rs1357741 | -0.308 | 0.024 | 0.086 | 0.519 | -0.058 | 0.082 | 0.082 | 0.833 | 0.15 | 0.41 | 0.09 | 0.22 | 0.06 | 0.22 |
| 7 | 56294281 | rs1722282 | -0.986 | 0.151 | -0.232 | 0.245 | -0.011 | 0.067 | 0.067 | 0 | 0.25 | 0.57 | 0.24 | 0.61 | 0.21 | 0.34 |
| 7 | 63695082 | rs11763256 | -0.171 | -0.651 | -0.42 | -0.375 | -0.01 | 0.022 | 0.022 | 0 | 0.69 | 0.77 | 0.74 | 0.9 | 0.67 | 0.65 |
| 7 | 63699812 | rs10085751 | 0.965 | -0.258 | -0.734 | -0.168 | -0.077 | -0.088 | -0.088 | 0 | 0.32 | 0.1 | 0.34 | 0.47 | 0.39 | 0.34 |
| 7 | 63704031 | rs11980921 | 0.966 | -0.248 | -0.718 | -0.163 | -0.074 | -0.085 | -0.085 | 0 | 0.32 | 0.11 | 0.34 | 0.47 | 0.39 | 0.34 |
| 7 | 63725595 | rs1852105 | -0.019 | -0.189 | 0.208 | -0.212 | 0.086 | 0.104 | 0.104 | 0.183 | 0.94 | 0.68 | 0.95 | 0.9 | 0.95 | 0.87 |
| 7 | 63744350 | rs651934 | 0.41 | -0.45 | -0.019 | -0.323 | 0.004 | -0.009 | -0.009 | 0 | 0.74 | 0.34 | 0.75 | 0.83 | 0.69 | 0.66 |
| 7 | 63750368 | rs679883 | 0.725 | -0.509 | -0.148 | -0.144 | -0.002 | 0.001 | 0.001 | 0 | 0.71 | 0.32 | 0.68 | 0.73 | 0.66 | 0.62 |
| 7 | 63751201 | rs1467328 | -0.666 | -0.745 | -0.177 | -0.252 | 0.003 | 0.05 | 0.05 | 0.209 | 0.5 | 0.4 | 0.56 | 0.28 | 0.54 | 0.44 |
| 7 | 63752903 | rs604013 | 1.407 | -0.959 | -0.467 | -0.25 | -0.076 | -0.035 | -0.035 | 2.659 | 0.41 | 0.05 | 0.37 | 0.62 | 0.41 | 0.44 |
| 7 | 63756245 | rs660228 | 0.747 | -0.716 | -0.304 | -0.067 | -0.001 | 0.018 | 0.018 | 0 | 0.7 | 0.39 | 0.69 | 0.48 | 0.71 | 0.62 |
| 7 | 63758482 | rs599522 | 0.634 | -0.716 | -0.31 | -0.067 | -0.001 | 0.016 | 0.016 | 0.005 | 0.7 | 0.38 | 0.69 | 0.48 | 0.71 | 0.62 |
| 7 | 63772707 | rs7790689 | -1.156 | -0.867 | -0.063 | -0.16 | -0.052 | 0.092 | 0.092 | 0 | 0.24 | 0.43 | 0.3 | 0.36 | 0.15 | 0.16 |
| 7 | 63778717 | rs6460137 | -0.947 | -0.794 | -0.567 | -0.165 | -0.017 | 0.067 | 0.067 | 1.015 | 0.53 | 0.88 | 0.56 | 0.62 | 0.58 | 0.42 |
| 7 | 63796171 | rs619102 | 0.525 | -0.584 | -0.433 | -0.164 | -0.001 | -0.001 | -0.001 | 0 | 0.6 | 0.39 | 0.55 | 0.54 | 0.65 | 0.61 |
| 7 | 63812551 | rs649475 | 1.874 | -0.487 | -1.123 | -0.106 | -0.104 | 0.02 | 0.02 | 0.019 | 0.35 | 0.04 | 0.27 | 0.28 | 0.36 | 0.37 |
| 7 | 63824928 | rs2127047 | -0.698 | -0.341 | -1.007 | -0.199 | -0.021 | 0.133 | 0.133 | 0.293 | 0.48 | 0.88 | 0.47 | 0.27 | 0.54 | 0.35 |
| 7 | 63847894 | rs649761 | 0.804 | -0.473 | -0.764 | -0.218 | 0.01 | 0.079 | 0.079 | 0 | 0.51 | 0.14 | 0.51 | 0.64 | 0.44 | 0.51 |
| 7 | 63852980 | rs4718070 | -0.258 | -0.576 | -0.767 | -0.313 | -0.001 | 0.007 | 0.007 | 0 | 0.26 | 0.2 | 0.32 | 0.08 | 0.26 | 0.28 |
| 7 | 63860590 | rs608294 | 0.814 | -0.488 | -0.679 | -0.242 | 0.01 | 0.079 | 0.079 | 0 | 0.51 | 0.14 | 0.51 | 0.64 | 0.44 | 0.51 |
| 7 | 63870365 | rs4454181 | -1.049 | -0.515 | -0.764 | -0.243 | 0.012 | 0.116 | 0.116 | 0.22 | 0.53 | 0.81 | 0.58 | 0.7 | 0.6 | 0.54 |
| 7 | 63881238 | rs6974833 | -0.877 | -0.448 | -0.735 | -0.245 | -0.028 | 0.137 | 0.137 | 0.392 | 0.51 | 0.84 | 0.55 | 0.7 | 0.55 | 0.55 |
| 7 | 63886568 | rs17771603 | 1.092 | -0.418 | -0.647 | -0.254 | -0.006 | 0.006 | 0.006 | 0 | 0.16 | 0.13 | 0.17 | 0.08 | 0.21 | 0.18 |
| 7 | 63899114 | rs10247383 | -2.042 | -0.412 | -0.808 | -0.229 | -0.091 | 0.001 | 0.001 | 0 | 0.65 | 0.96 | 0.73 | 0.7 | 0.65 | 0.63 |
| 7 | 63910447 | rs10271255 | -0.341 | -0.562 | -0.56 | -0.079 | 0.001 | -0.004 | -0.004 | 0 | 0.3 | 0.71 | 0.31 | 0.44 | 0.3 | 0.3 |
| 7 | 63939340 | rs13233386 | 0.471 | -0.422 | -0.42 | -0.19 | 0.036 | -0.154 | -0.154 | 0.081 | 0.23 | 0.47 | 0.22 | 0.01 | 0.28 | 0.13 |
| 7 | 63952472 | rs7796427 | -0.875 | -0.506 | -0.696 | -0.157 | -0.025 | -0.029 | -0.029 | 0.026 | 0.51 | 0.86 | 0.59 | 0.46 | 0.55 | 0.54 |
| 7 | 63962963 | rs1885912 | -1.253 | -0.435 | -0.7 | -0.175 | -0.033 | 0.067 | 0.067 | 0 | 0.57 | 0.86 | 0.67 | 0.64 | 0.6 | 0.58 |
| 7 | 63975912 | rs13239925 | 0.868 | -0.272 | -0.221 | -0.305 | 0.014 | -0.064 | -0.064 | 2.83 | 0.2 | 0.25 | 0.21 | 0.01 | 0.28 | 0.13 |
| 7 | 63980233 | rs4717195 | 1.6 | -0.511 | -0.485 | -0.113 | -0.03 | 0.119 | 0.119 | 2.666 | 0.08 | 0.13 | 0.14 | 0.37 | 0.03 | 0.08 |
| 7 | 64019592 | rs6962076 | -0.049 | -0.054 | -0.404 | -0.61 | -0.007 | 0.01 | 0.01 | 0 | 0.27 | 0.2 | 0.32 | 0.08 | 0.32 | 0.37 |
| 7 | 64030116 | rs1446886 | 0.75 | -0.042 | -0.357 | -0.568 | 0.008 | -0.026 | -0.026 | 0.011 | 0.19 | 0.22 | 0.22 | 0.08 | 0.25 | 0.33 |
| 7 | 64054804 | rs10279234 | 0.603 | -0.053 | -0.341 | -0.623 | 0.057 | -0.127 | -0.127 | 1.414 | 0.21 | 0.51 | 0.23 | 0.02 | 0.28 | 0.13 |
| 7 | 64059230 | rs4626471 | -1.505 | -0.148 | -0.349 | -0.563 | 0.028 | 0.091 | 0.091 | 0 | 0.57 | 0.83 | 0.65 | 0.25 | 0.65 | 0.55 |
| 7 | 64060424 | rs6415206 | -1.505 | -0.148 | -0.349 | -0.563 | 0.028 | 0.091 | 0.091 | 0 | 0.57 | 0.83 | 0.64 | 0.25 | 0.65 | 0.55 |
| 7 | 64060585 | rs10949928 | 0.711 | -0.02 | -0.337 | -0.557 | -0.001 | -0.016 | -0.016 | 0 | 0.1 | 0.07 | 0.11 | 0 | 0.09 | 0.05 |
| 7 | 64072452 | rs13226579 | -0.632 | 0.024 | -0.632 | -0.532 | 0.004 | -0.019 | -0.019 | 0.276 | 0.36 | 0.59 | 0.43 | 0.17 | 0.42 | 0.22 |
| 7 | 64075291 | rs11763777 | 0.525 | 0.216 | -0.454 | -0.092 | 0.006 | -0.001 | -0.001 | 0 | 0.18 | 0.2 | 0.21 | 0.15 | 0.23 | 0.33 |
| 7 | 64101034 | rs2862876 | -0.608 | 0.256 | -1.684 | 0.347 | -0.05 | -0.001 | -0.001 | 0 | 0.55 | 0.95 | 0.67 | 0.63 | 0.45 | 0.55 |

| | | | | | | | | | | | | | | | | |
|---|---|---|---|---|---|---|---|---|---|---|---|---|---|---|---|---|
| 7 | 64102495 | rs13223688 | -0.012 | -0.16 | -1.156 | -0.022 | 0 | -0.018 | -0.018 | 2.2 | 0.15 | 0.16 | 0.22 | 0.02 | 0.17 | 0.14 |
| 7 | 64105180 | rs7811223 | 0.936 | -0.154 | -1.12 | -0.007 | -0.003 | -0.004 | -0.004 | 0 | 0.26 | 0.24 | 0.19 | 0.35 | 0.26 | 0.38 |
| 7 | 64105197 | rs7776508 | 0.468 | -0.981 | -0.866 | -0.193 | -0.015 | -0.005 | -0.005 | 0.7 | 0.56 | 0.51 | 0.5 | 0.47 | 0.67 | 0.71 |
| 7 | 64105259 | rs7798993 | -0.236 | -1.061 | -0.751 | -0.012 | -0.017 | 0.053 | 0.053 | 0 | 0.3 | 0.42 | 0.41 | 0.45 | 0.27 | 0.21 |
| 7 | 64110593 | rs10807682 | -1.043 | -1.089 | -0.765 | -0.024 | 0.004 | 0.028 | 0.028 | 0 | 0.62 | 0.8 | 0.76 | 0.65 | 0.68 | 0.56 |
| 7 | 64118662 | rs10234472 | -1.855 | -1.456 | -0.497 | 0.159 | 0.031 | 0.03 | 0.03 | 0 | 0.85 | 0.93 | 0.9 | 0.92 | 0.93 | 0.92 |
| 7 | 64128448 | rs10085911 | -1.85 | -1.428 | -0.519 | 0.173 | 0.02 | 0.019 | 0.019 | 0 | 0.85 | 0.89 | 0.9 | 0.92 | 0.93 | 0.92 |
| 7 | 64132105 | rs10085484 | 1.632 | -1.644 | -0.311 | 0.673 | -0.017 | 0.052 | 0.052 | 1.181 | 0.1 | 0.14 | 0.08 | 0.3 | 0.05 | 0.04 |
| 7 | 64135796 | rs6460176 | 1.045 | -1.458 | -0.426 | 0.174 | -0.002 | -0.006 | -0.006 | 0 | 0.29 | 0.27 | 0.31 | 0.12 | 0.4 | 0.34 |
| 7 | 64137087 | rs2862793 | -0.233 | -1.264 | -1.044 | 0.141 | 0.001 | 0.01 | 0.01 | 0 | 0.53 | 0.42 | 0.57 | 0.68 | 0.53 | 0.58 |
| 7 | 64141291 | rs6965811 | -1.818 | -1.247 | -1.062 | 0.15 | 0.024 | 0.023 | 0.023 | 0 | 0.85 | 0.9 | 0.91 | 0.92 | 0.93 | 0.92 |
| 7 | 64150010 | rs2862798 | 0.669 | -1.024 | -0.95 | 0.069 | -0.033 | 0.054 | 0.054 | 0.642 | 0.39 | 0.23 | 0.39 | 0.25 | 0.44 | 0.51 |
| 7 | 64150849 | rs6460178 | 0.259 | -0.097 | -0.597 | 0.252 | 0.019 | -0.029 | -0.029 | 0.29 | 0.32 | 0.45 | 0.33 | 0.24 | 0.4 | 0.34 |
| 7 | 64211341 | rs4585620 | -0.442 | -0.368 | -1.581 | 0.409 | 0.029 | 0.002 | 0.002 | 0 | 0.67 | 0.91 | 0.79 | 0.69 | 0.77 | 0.59 |
| 7 | 64225274 | rs1581570 | 0.558 | -0.547 | -1.445 | -0.083 | -0.028 | 0.036 | 0.036 | 0 | 0.4 | 0.18 | 0.36 | 0.3 | 0.52 | 0.44 |
| 7 | 64226341 | rs1113255 | 0.008 | -1.019 | -0.551 | 0.389 | -0.026 | 0.157 | 0.157 | 0.011 | 0.09 | 0.26 | 0.17 | 0.26 | 0.04 | 0.12 |
| 7 | 64226499 | rs1589895 | -0.683 | -0.837 | -0.33 | 0.231 | 0.016 | 0.011 | 0.011 | 0 | 0.93 | 0.82 | 0.91 | 0.9 | 0.95 | 0.97 |
| 7 | 64231596 | rs10807684 | 0.229 | -1.091 | -0.347 | 0.257 | 0.001 | -0.004 | -0.004 | 0 | 0.39 | 0.37 | 0.34 | 0.22 | 0.36 | 0.38 |
| 7 | 64247483 | rs6972180 | 0.365 | -1.053 | -0.391 | 0.551 | -0.005 | 0.012 | 0.012 | 0 | 0.87 | 0.87 | 0.82 | 0.73 | 0.93 | 0.78 |
| 7 | 64254060 | rs11760270 | 0.463 | -0.992 | -0.69 | 0.399 | -0.062 | 0.149 | 0.149 | 0 | 0.38 | 0.05 | 0.32 | 0.16 | 0.36 | 0.29 |
| 7 | 64254791 | rs10949938 | -0.428 | -1.015 | -0.527 | 0.388 | -0.017 | 0.165 | 0.165 | 0 | 0.61 | 0.89 | 0.67 | 0.84 | 0.63 | 0.71 |
| 7 | 64256482 | rs4718112 | -0.091 | -0.947 | -0.447 | 0.681 | 0.002 | 0.074 | 0.074 | 0 | 0.07 | 0.05 | 0.19 | 0.38 | 0.08 | 0.11 |
| 7 | 64267756 | rs10155953 | -0.049 | -0.999 | -0.023 | 0.381 | 0.011 | 0.056 | 0.056 | 0 | 0.57 | 0.75 | 0.62 | 0.72 | 0.59 | 0.68 |
| 7 | 64284959 | rs7792444 | -0.17 | -0.966 | -0.054 | 0.403 | 0.002 | 0.053 | 0.053 | 0 | 0.59 | 0.8 | 0.63 | 0.72 | 0.59 | 0.69 |
| 7 | 64292695 | rs6460195 | 0.453 | -0.954 | -0.138 | 0.253 | -0.003 | 0.129 | 0.129 | 0 | 0.38 | 0.12 | 0.33 | 0.16 | 0.37 | 0.29 |
| 7 | 64312707 | rs7804955 | 0.232 | -0.753 | 0.727 | 0.447 | -0.008 | 0.02 | 0.02 | 0 | 0.44 | 0.29 | 0.43 | 0.38 | 0.5 | 0.49 |
| 7 | 64314296 | rs2863984 | 0.645 | -0.696 | 0.801 | 0.417 | -0.04 | 0.031 | 0.031 | 0 | 0.38 | 0.19 | 0.36 | 0.28 | 0.44 | 0.45 |
| 7 | 64323479 | rs6960979 | 0.055 | -0.675 | 0.891 | 0.408 | 0 | -0.002 | -0.002 | 0 | 0.45 | 0.48 | 0.45 | 0.38 | 0.5 | 0.49 |
| 7 | 64325935 | rs1568910 | 0.508 | -0.658 | 0.985 | 0.375 | 0.006 | -0.009 | -0.009 | 0 | 0.39 | 0.38 | 0.36 | 0.28 | 0.44 | 0.44 |
| 7 | 64340294 | rs4717222 | 0.136 | -0.67 | 1.311 | 0.507 | 0.021 | -0.006 | -0.006 | 0 | 0.51 | 0.33 | 0.51 | 0.49 | 0.47 | 0.51 |
| 7 | 64342398 | rs4718154 | 0.117 | -0.705 | 1.23 | 0.51 | 0.003 | -0.01 | -0.01 | 0 | 0.44 | 0.44 | 0.44 | 0.38 | 0.49 | 0.48 |
| 7 | 64343985 | rs7788090 | 1.26 | -0.98 | 0.647 | 0.258 | 0.008 | 0.001 | 0.001 | 0 | 0.17 | 0.04 | 0.12 | 0.15 | 0.1 | 0.19 |
| 7 | 64347479 | rs6460206 | 0.877 | -0.931 | 0.828 | 0.379 | 0.015 | -0.02 | -0.02 | 2.335 | 0.39 | 0.43 | 0.37 | 0.29 | 0.44 | 0.44 |
| 7 | 64364381 | rs4718163 | 0.637 | -0.988 | 0.795 | 0.357 | 0.006 | -0.003 | -0.003 | 0 | 0.42 | 0.45 | 0.47 | 0.55 | 0.45 | 0.49 |
| 7 | 64393948 | rs13236444 | 0.058 | -0.899 | 0.695 | 0.347 | -0.004 | -0.017 | -0.017 | 0 | 0.49 | 0.13 | 0.5 | 0.43 | 0.47 | 0.49 |
| 7 | 64397931 | rs12534485 | -0.817 | -0.837 | 0.792 | 0.335 | 0.006 | -0.012 | -0.012 | 0 | 0.6 | 0.52 | 0.63 | 0.71 | 0.56 | 0.56 |
| 7 | 64399736 | rs12535862 | 0.38 | -1.182 | 0.675 | 0.435 | 0.001 | 0.002 | 0.002 | 0 | 0.05 | 0.1 | 0.07 | 0.1 | 0.05 | 0.03 |
| 7 | 64402765 | rs10236729 | -0.521 | -1.183 | 0.672 | 0.444 | 0.004 | 0.047 | 0.047 | 0 | 0.09 | 0.17 | 0.12 | 0.22 | 0.09 | 0.05 |
| 7 | 64409809 | rs2091228 | 0.408 | -1.082 | 0.457 | 0.064 | -0.023 | 0.021 | 0.021 | 0 | 0.38 | 0.17 | 0.55 | 0.28 | 0.47 | 0.34 |
| 7 | 64416284 | rs7800787 | 0.587 | -1.208 | 0.87 | 0.092 | -0.002 | 0 | 0 | 0 | 0.38 | 0.42 | 0.27 | 0.4 | 0.4 | 0.43 |
| 7 | 64426173 | rs2049443 | -0.247 | -1.242 | 1.018 | 0.138 | 0.022 | 0.129 | 0.129 | 0.117 | 0.94 | 0.75 | 0.94 | 0.87 | 0.96 | 0.97 |
| 7 | 64436980 | rs1852015 | -0.129 | -1.244 | 1.087 | 0.137 | 0.028 | 0.141 | 0.141 | 0.143 | 0.94 | 0.73 | 0.94 | 0.87 | 0.97 | 0.97 |
| 7 | 64439701 | rs3807069 | 0.616 | -1.261 | 1.092 | 0.125 | -0.004 | 0 | 0 | 0 | 0.38 | 0.4 | 0.26 | 0.41 | 0.4 | 0.43 |
| 7 | 64442904 | rs7780100 | -0.055 | -1.259 | 1.073 | 0.124 | -0.007 | -0.009 | -0.009 | 0 | 0.56 | 0.33 | 0.67 | 0.47 | 0.56 | 0.54 |
| 7 | 64450556 | rs10807685 | -0.024 | -1.253 | 1.044 | 0.133 | 0.025 | 0.135 | 0.135 | 0.221 | 0.94 | 0.73 | 0.94 | 0.87 | 0.97 | 0.97 |
| 7 | 64451699 | rs4717229 | -0.251 | -1.232 | 0.992 | 0.142 | 0.018 | 0.121 | 0.121 | 0.065 | 0.94 | 0.77 | 0.95 | 0.87 | 0.97 | 0.97 |
| 7 | 64456549 | rs2140724 | 0.598 | -1.284 | 0.904 | 0.013 | -0.019 | 0.01 | 0.01 | 0 | 0.38 | 0.21 | 0.55 | 0.3 | 0.47 | 0.36 |
| 7 | 64462326 | rs4718180 | -0.102 | -1.295 | 0.937 | 0.032 | -0.008 | -0.013 | -0.013 | 0.096 | 0.55 | 0.31 | 0.67 | 0.46 | 0.56 | 0.54 |
| 7 | 64566009 | rs1709517 | 1.156 | -1.302 | 0.427 | 0.071 | 0.026 | -0.011 | -0.011 | 0 | 0.17 | 0.04 | 0.13 | 0.16 | 0.09 | 0.2 |
| 7 | 64679561 | rs2900856 | 0.576 | -1.549 | 0.493 | 0.055 | -0.026 | 0.01 | 0.01 | 0 | 0.38 | 0.23 | 0.56 | 0.3 | 0.52 | 0.37 |
| 7 | 64696330 | rs34739196 | 0.894 | -1.65 | 0.524 | 0.233 | -0.032 | -0.03 | -0.03 | 4.516 | 0.29 | 0.18 | 0.23 | 0.44 | 0.37 | 0.39 |
| 7 | 64698500 | rs495336 | 0.188 | -1.588 | 0.354 | 0.193 | -0.008 | -0.015 | -0.015 | 1.42 | 0.56 | 0.3 | 0.68 | 0.46 | 0.58 | 0.57 |
| 7 | 64700434 | rs657560 | -0.573 | -1.502 | 0.512 | 0.208 | 0.021 | 0.023 | 0.023 | 0 | 0.66 | 0.43 | 0.74 | 0.55 | 0.63 | 0.6 |
| 7 | 64703012 | rs575433 | -0.603 | -1.827 | 0.475 | 0.057 | 0.001 | -0.002 | -0.002 | 0 | 0.6 | 0.62 | 0.43 | 0.7 | 0.49 | 0.64 |
| 7 | 64727862 | rs17710097 | 0.937 | -1.609 | 0.214 | 0.253 | -0.066 | -0.041 | -0.041 | 0 | 0.29 | 0.12 | 0.23 | 0.39 | 0.37 | 0.38 |
| 7 | 64739969 | rs663732 | 1.224 | -1.551 | 0.201 | 0.364 | 0.05 | -0.019 | -0.019 | 0 | 0.16 | 0.04 | 0.12 | 0.16 | 0.06 | 0.19 |
| 7 | 64745413 | rs6460232 | 0.784 | -1.537 | 0.33 | 0.327 | -0.027 | -0.003 | -0.003 | 0.86 | 0.35 | 0.23 | 0.25 | 0.38 | 0.38 | 0.39 |
| 7 | 64751148 | rs6956221 | 0.263 | -1.229 | 0.533 | 0.33 | -0.015 | 0.013 | 0.013 | 0 | 0.41 | 0.33 | 0.57 | 0.3 | 0.52 | 0.39 |
| 7 | 64767457 | rs7796960 | -0.099 | -1.286 | 0.435 | 0.332 | 0 | 0 | 0 | 0 | 0.46 | 0.52 | 0.62 | 0.42 | 0.56 | 0.42 |
| 7 | 64768219 | rs11765030 | -0.14 | -1.351 | 0.6 | 0.287 | -0.019 | -0.016 | -0.016 | 0 | 0.45 | 0.3 | 0.59 | 0.45 | 0.56 | 0.42 |
| 7 | 64770261 | rs4718223 | -1.116 | -1.67 | 0.726 | 0.261 | -0.015 | -0.002 | -0.002 | 0 | 0.82 | 0.79 | 0.87 | 0.84 | 0.95 | 0.81 |
| 7 | 64782426 | rs6963801 | 0.064 | -1.728 | 0.749 | 0.3 | -0.035 | -0.028 | -0.028 | 0 | 0.44 | 0.25 | 0.59 | 0.46 | 0.56 | 0.42 |
| 7 | 64786318 | rs12536316 | 0.714 | -1.755 | 0.939 | 0.294 | -0.044 | -0.007 | -0.007 | 0 | 0.35 | 0.19 | 0.24 | 0.37 | 0.39 | 0.39 |
| 7 | 64794391 | rs13244870 | 0.62 | -1.811 | 0.78 | 0.18 | -0.066 | -0.006 | -0.006 | 2.448 | 0.19 | 0.35 | 0.15 | 0.17 | 0.06 | 0.19 |
| 7 | 64799139 | rs11765265 | 0.229 | -1.643 | 0.643 | 0.179 | -0.034 | 0.03 | 0.03 | 0 | 0.41 | 0.25 | 0.56 | 0.31 | 0.52 | 0.4 |
| 7 | 64827987 | rs10259348 | -0.099 | -1.625 | 0.047 | -0.158 | -0.038 | 0.078 | 0.078 | 0 | 0.22 | 0.51 | 0.42 | 0.31 | 0.17 | 0.22 |
| 7 | 64830594 | rs2418470 | -1.095 | -1.727 | -0.714 | -0.343 | -0.043 | -0.004 | -0.004 | 0 | 0.64 | 0.83 | 0.73 | 0.62 | 0.61 | 0.59 |
| 7 | 64840402 | rs7799794 | 0.12 | -1.314 | -0.998 | -0.28 | -0.019 | -0.038 | -0.038 | 0.561 | 0.07 | 0 | 0.07 | 0.13 | 0.09 | 0.04 |
| 7 | 64844720 | rs10230566 | 0.353 | -1.632 | -0.857 | -0.4 | -0.013 | -0.095 | -0.095 | 0 | 0.15 | 0.52 | 0.16 | 0.02 | 0.12 | 0.03 |

| | | | | | | | | | | | | | | | | |
|---|---|---|---|---|---|---|---|---|---|---|---|---|---|---|---|---|
| 7 | 64864800 | rs4122883 | -0.063 | -1.592 | -0.818 | -0.387 | -0.025 | 0.074 | 0.074 | 0 | 0.21 | 0.54 | 0.42 | 0.32 | 0.16 | 0.21 |
| 7 | 64864912 | rs3846972 | -0.101 | -1.594 | -0.88 | -0.388 | -0.023 | 0.056 | 0.056 | 0.004 | 0.21 | 0.58 | 0.43 | 0.31 | 0.16 | 0.21 |
| 7 | 64875899 | rs10232533 | -0.511 | -1.439 | -1.196 | -0.557 | -0.064 | 0.094 | 0.094 | 0.951 | 0.77 | 0.93 | 0.82 | 0.99 | 0.63 | 0.82 |
| 7 | 64891546 | rs6460248 | -0.911 | -1.624 | -1.128 | -0.462 | -0.023 | -0.033 | -0.033 | 0 | 0.61 | 0.73 | 0.72 | 0.48 | 0.59 | 0.59 |
| 7 | 64896694 | rs11760982 | 0.547 | -1.622 | -1.246 | -0.574 | -0.096 | 0.136 | 0.136 | 0.934 | 0.22 | 0.03 | 0.17 | 0.01 | 0.37 | 0.18 |
| 7 | 64900821 | rs11771033 | -0.794 | -1.816 | -1.075 | -0.485 | -0.007 | -0.005 | -0.005 | 0 | 0.47 | 0.62 | 0.64 | 0.46 | 0.57 | 0.42 |
| 7 | 64902831 | rs17139634 | 1.472 | -1.896 | -1.131 | -0.472 | 0.019 | 0.004 | 0.004 | 0 | 0.17 | 0.16 | 0.12 | 0.15 | 0.06 | 0.18 |
| 7 | 64904905 | rs17146452 | 1.624 | -1.571 | -1.842 | -0.749 | -0.026 | -0.084 | -0.084 | 0.033 | 0.15 | 0.02 | 0.09 | 0.3 | 0.15 | 0.17 |
| 7 | 64907357 | rs9638226 | 0.402 | -1.582 | -1.599 | -0.791 | -0.014 | 0.022 | 0.022 | 3.273 | 0.28 | 0.21 | 0.23 | 0.15 | 0.41 | 0.21 |
| 7 | 64916237 | rs4718244 | -0.076 | -1.587 | -1.466 | -0.536 | -0.01 | -0.007 | -0.007 | 0.162 | 0.46 | 0.59 | 0.63 | 0.46 | 0.57 | 0.42 |
| 7 | 64924420 | rs4718245 | 1.184 | -1.454 | -1.719 | -0.462 | 0.005 | 0.008 | 0.008 | 0 | 0.06 | 0.01 | 0.26 | 0.3 | 0.05 | 0.17 |
| 7 | 64925829 | rs1812771 | -0.553 | -1.503 | -1.736 | -0.528 | -0.046 | -0.018 | -0.018 | 0 | 0.66 | 0.83 | 0.77 | 0.61 | 0.62 | 0.61 |
| 7 | 64926823 | rs12333375 | -0.564 | -1.502 | -1.747 | -0.534 | -0.049 | -0.018 | -0.018 | 0.329 | 0.66 | 0.84 | 0.77 | 0.61 | 0.62 | 0.61 |
| 7 | 64941887 | rs11762103 | 0.173 | -1.174 | -1.876 | -0.794 | 0.004 | 0.031 | 0.031 | 0 | 0.67 | 0.75 | 0.55 | 0.17 | 0.71 | 0.5 |
| 7 | 64956181 | rs10274919 | 0.707 | -1.194 | -1.609 | -0.2 | 0.017 | -0.028 | -0.028 | 0 | 0.37 | 0.48 | 0.47 | 0.21 | 0.5 | 0.35 |
| 7 | 64957497 | rs1964363 | -0.118 | -0.493 | -1.899 | 0.06 | -0.045 | -0.012 | -0.012 | 0 | 0.64 | 0.83 | 0.76 | 0.61 | 0.61 | 0.61 |
| 7 | 65063274 | rs7786206 | -0.238 | -0.162 | -2.216 | 0.057 | -0.115 | -0.093 | -0.093 | 0.834 | 0.7 | 0.93 | 0.74 | 0.56 | 0.58 | 0.62 |
| 7 | 65070538 | rs4236206 | -0.198 | -0.273 | -2.144 | 0.054 | -0.046 | -0.051 | -0.051 | 0 | 0.7 | 0.85 | 0.75 | 0.56 | 0.63 | 0.62 |
| 7 | 65087974 | rs4718262 | -0.1 | -0.166 | -2.316 | 0.053 | -0.086 | -0.101 | -0.101 | 0 | 0.7 | 0.92 | 0.76 | 0.55 | 0.59 | 0.58 |
| 7 | 65201335 | rs10950010 | -0.234 | -0.25 | -2.252 | 0.08 | -0.052 | -0.071 | -0.071 | 0.879 | 0.7 | 0.88 | 0.75 | 0.53 | 0.6 | 0.6 |
| 7 | 65202326 | rs6460259 | -0.113 | -0.297 | -2.218 | 0.283 | -0.042 | -0.051 | -0.051 | 0 | 0.69 | 0.85 | 0.71 | 0.44 | 0.6 | 0.58 |
| 7 | 65203587 | rs1568780 | -0.244 | -0.292 | -2.221 | 0.273 | -0.052 | -0.071 | -0.071 | 0.694 | 0.7 | 0.88 | 0.75 | 0.53 | 0.6 | 0.6 |
| 7 | 65215455 | rs6460260 | -1.332 | -0.674 | -2.437 | 0.015 | 0.044 | 0.012 | 0.012 | 0 | 0.8 | 0.92 | 0.78 | 0.86 | 0.89 | 0.8 |
| 7 | 65216099 | rs4718273 | -1.332 | -0.672 | -2.436 | 0.016 | 0.044 | 0.012 | 0.012 | 0 | 0.8 | 0.91 | 0.78 | 0.86 | 0.89 | 0.8 |
| 7 | 65300423 | rs35735127 | 1.003 | -0.649 | -2.374 | 0.012 | 0.024 | 0.011 | 0.011 | 0 | 0.21 | 0.16 | 0.22 | 0.14 | 0.11 | 0.21 |
| 7 | 65326821 | rs11763147 | -0.855 | -0.857 | -2.566 | -0.108 | -0.017 | 0.041 | 0.041 | 0 | 0.61 | 0.31 | 0.55 | 0.79 | 0.56 | 0.65 |
| 7 | 65350058 | rs13236109 | 1.094 | -0.842 | -2.553 | -0.103 | 0.027 | 0.012 | 0.012 | 0 | 0.21 | 0.15 | 0.22 | 0.15 | 0.11 | 0.2 |
| 7 | 65360829 | rs35391607 | 1.095 | -0.837 | -2.55 | -0.103 | 0.024 | 0.011 | 0.011 | 0 | 0.21 | 0.16 | 0.22 | 0.15 | 0.11 | 0.2 |
| 7 | 65376456 | rs10950022 | 0.182 | -0.854 | -2.57 | -0.044 | -0.091 | -0.116 | -0.116 | 0.908 | 0.3 | 0.06 | 0.24 | 0.47 | 0.39 | 0.4 |
| 7 | 65387101 | rs6949812 | 1.1 | -0.84 | -2.563 | -0.093 | 0.027 | 0.012 | 0.012 | 0 | 0.21 | 0.15 | 0.22 | 0.15 | 0.11 | 0.2 |
| 7 | 65424658 | rs2961102 | 1.198 | -0.759 | -2.498 | -0.107 | 0.044 | 0.012 | 0.012 | 0 | 0.2 | 0.08 | 0.21 | 0.15 | 0.11 | 0.2 |
| 7 | 65425894 | rs9530 | -1.07 | -0.852 | -2.439 | -0.012 | -0.011 | 0.021 | 0.021 | 0 | 0.62 | 0.35 | 0.56 | 0.83 | 0.56 | 0.67 |
| 7 | 65437774 | rs12540589 | 0.187 | -0.796 | -2.522 | 0.081 | -0.054 | -0.078 | -0.078 | 0 | 0.3 | 0.12 | 0.25 | 0.48 | 0.39 | 0.41 |
| 7 | 65449716 | rs1039664 | 1.136 | -0.764 | -2.581 | -0.07 | 0.039 | 0.013 | 0.013 | 0 | 0.2 | 0.12 | 0.22 | 0.15 | 0.11 | 0.2 |
| 7 | 65453292 | rs34466769 | 0.282 | -0.537 | -2.418 | -0.039 | 0.08 | 0.021 | 0.021 | 0 | 0.29 | 0.12 | 0.26 | 0.22 | 0.16 | 0.23 |
| 7 | 65509258 | rs420880 | 0.609 | 0.186 | -2.276 | -0.054 | 0.02 | 0.009 | 0.009 | 0 | 0.59 | 0.79 | 0.68 | 0.36 | 0.61 | 0.56 |
| 7 | 65515276 | rs7811214 | -0.446 | 0.152 | -2.203 | 0.174 | 0.002 | -0.034 | -0.034 | 0 | 0.33 | 0.22 | 0.24 | 0.52 | 0.34 | 0.37 |
| 7 | 65546062 | rs2220626 | 0.623 | -0.297 | -2.106 | 0.124 | 0.021 | 0.06 | 0.06 | 0 | 0.46 | 0.69 | 0.55 | 0.34 | 0.58 | 0.43 |
| 7 | 65556280 | rs160646 | 1.11 | -0.378 | -2.179 | 0.183 | 0.049 | 0.011 | 0.011 | 0 | 0.21 | 0.06 | 0.21 | 0.15 | 0.11 | 0.21 |
| 7 | 65560561 | rs709609 | 0.242 | -0.297 | -2.096 | 0.153 | 0.027 | 0.033 | 0.033 | 0.65 | 0.56 | 0.75 | 0.6 | 0.4 | 0.63 | 0.48 |
| 7 | 65579722 | rs6946143 | 0.526 | -0.276 | -2.069 | 0.133 | 0.02 | 0.055 | 0.055 | 0 | 0.46 | 0.69 | 0.55 | 0.34 | 0.58 | 0.43 |
| 7 | 65584318 | rs160637 | 1.129 | -0.513 | -2.22 | 0.163 | 0.049 | 0.011 | 0.011 | 0 | 0.21 | 0.06 | 0.21 | 0.15 | 0.11 | 0.21 |
| 7 | 65597341 | rs419603 | 1.184 | -0.521 | -2.223 | 0.159 | 0.047 | 0.009 | 0.009 | 0 | 0.21 | 0.06 | 0.21 | 0.15 | 0.11 | 0.21 |
| 7 | 65599196 | rs2243480 | 1.12 | -0.522 | -2.224 | 0.159 | 0.049 | 0.011 | 0.011 | 0 | 0.21 | 0.06 | 0.21 | 0.15 | 0.11 | 0.21 |
| 7 | 65617971 | rs316305 | -0.451 | -0.58 | -2.29 | 0.482 | -0.006 | -0.015 | -0.015 | 0.624 | 0.78 | 0.82 | 0.85 | 0.65 | 0.74 | 0.69 |
| 7 | 65618674 | rs316306 | -0.451 | -0.58 | -2.29 | 0.482 | -0.006 | -0.015 | -0.015 | 0.624 | 0.78 | 0.82 | 0.85 | 0.65 | 0.74 | 0.69 |
| 7 | 65665056 | rs6968534 | 1.963 | -0.426 | -2.748 | 0.095 | -0.13 | -0.013 | -0.013 | 0 | 0.25 | 0.03 | 0.34 | 0.2 | 0.37 | 0.32 |
| 7 | 65683317 | rs6952182 | 0.194 | -0.867 | -2.451 | 0.361 | 0.021 | -0.017 | -0.017 | 1.397 | 0.57 | 0.76 | 0.64 | 0.5 | 0.64 | 0.49 |
| 7 | 65706814 | rs12532675 | 1.504 | -1.437 | -1.682 | 0.44 | 0.026 | 0.007 | 0.007 | 2.19 | 0.09 | 0 | 0.05 | 0.07 | 0.05 | 0.04 |
| 7 | 65709988 | rs17138154 | 0.249 | -1.348 | -1.695 | 0.426 | -0.035 | -0.024 | -0.024 | 0 | 0.08 | 0.01 | 0.1 | 0.11 | 0.12 | 0.09 |
| 7 | 65720516 | rs11769702 | 0.437 | -1.346 | -1.65 | 0.612 | -0.042 | -0.059 | -0.059 | 0 | 0.21 | 0.07 | 0.14 | 0.31 | 0.25 | 0.31 |
| 7 | 65723846 | rs1547529 | 0.342 | -1.426 | -1.661 | 0.616 | -0.006 | -0.01 | -0.01 | 1.725 | 0.22 | 0.18 | 0.15 | 0.31 | 0.26 | 0.31 |
| 7 | 65731855 | rs1875057 | 0.199 | -1.451 | -1.669 | 0.599 | 0.02 | -0.029 | -0.029 | 0 | 0.57 | 0.76 | 0.64 | 0.54 | 0.64 | 0.49 |
| 7 | 65734783 | rs6945843 | 0.674 | -1.497 | -1.553 | 0.589 | 0.017 | 0.029 | 0.029 | 0 | 0.46 | 0.64 | 0.54 | 0.38 | 0.57 | 0.43 |
| 7 | 65750967 | rs4149463 | 0.679 | -1.387 | -1.453 | 0.608 | -0.014 | -0.027 | -0.027 | 0.069 | 0.06 | 0 | 0.07 | 0.11 | 0.08 | 0.08 |
| 7 | 65773859 | rs11984115 | 0.277 | -1.424 | -1.44 | 0.61 | 0.018 | -0.029 | -0.029 | 0.416 | 0.57 | 0.76 | 0.64 | 0.54 | 0.64 | 0.49 |
| 7 | 65796626 | rs7783779 | 0.283 | -1.497 | -1.433 | 0.617 | 0.019 | -0.03 | -0.03 | 0 | 0.57 | 0.76 | 0.64 | 0.54 | 0.64 | 0.49 |
| 7 | 65822360 | rs778732 | 0.692 | -1.547 | -1.465 | 0.593 | 0.066 | 0.014 | 0.014 | 0 | 0.23 | 0.06 | 0.21 | 0.15 | 0.11 | 0.2 |
| 7 | 65825257 | rs3757417 | 0.814 | -1.359 | -1.393 | 0.548 | -0.035 | -0.028 | -0.028 | 0.06 | 0.08 | 0.01 | 0.1 | 0.12 | 0.12 | 0.09 |
| 7 | 65846275 | rs778720 | -0.514 | -1.795 | -1.376 | 0.535 | 0.025 | 0.005 | 0.005 | 1.186 | 0.44 | 0.25 | 0.4 | 0.56 | 0.37 | 0.52 |
| 7 | 65891461 | rs6976714 | -0.439 | -1.815 | -1.385 | 0.527 | 0.022 | 0.002 | 0.002 | 0.851 | 0.43 | 0.24 | 0.4 | 0.56 | 0.37 | 0.52 |
| 7 | 65938980 | rs7789768 | -0.897 | -1.844 | -1.38 | 0.515 | 0.022 | 0.038 | 0.038 | 0 | 0.54 | 0.3 | 0.45 | 0.62 | 0.42 | 0.57 |
| 7 | 65988610 | rs10950043 | -0.436 | -1.784 | -1.345 | 0.572 | 0.02 | -0.029 | -0.029 | 0.448 | 0.43 | 0.24 | 0.36 | 0.45 | 0.36 | 0.51 |
| 7 | 66026115 | rs801195 | 0.388 | -1.784 | -1.343 | 0.572 | 0.02 | -0.03 | -0.03 | 0 | 0.57 | 0.77 | 0.64 | 0.55 | 0.64 | 0.49 |
| 7 | 66029429 | rs3757419 | 0.434 | -1.753 | -1.338 | 0.578 | 0.002 | -0.021 | -0.021 | 0 | 0.1 | 0.06 | 0.14 | 0.21 | 0.13 | 0.12 |
| 7 | 66067032 | rs6971059 | 0.427 | -1.799 | -1.323 | 0.577 | 0.02 | -0.029 | -0.029 | 0 | 0.57 | 0.76 | 0.64 | 0.55 | 0.64 | 0.49 |
| 7 | 66068129 | rs13311962 | -0.298 | -1.859 | -1.362 | 0.599 | -0.006 | -0.012 | -0.012 | 0 | 0.78 | 0.82 | 0.79 | 0.68 | 0.74 | 0.7 |
| 7 | 66089165 | rs2707851 | -0.348 | -1.895 | -1.351 | 0.61 | -0.006 | -0.012 | -0.012 | 0.117 | 0.78 | 0.82 | 0.85 | 0.69 | 0.74 | 0.7 |
| 7 | 66092308 | rs1638736 | 1.463 | -1.998 | -1.416 | 0.55 | 0.049 | -0.018 | -0.018 | 0.039 | 0.17 | 0.01 | 0.17 | 0.14 | 0.06 | 0.19 |

| | | | | | | | | | | | | | | | | |
|---|---|---|---|---|---|---|---|---|---|---|---|---|---|---|---|---|
| 7 | 66105198 | rs9791713 | 0.403 | -1.806 | -1.283 | 0.68 | 0.02 | -0.029 | -0.029 | 0 | 0.57 | 0.76 | 0.64 | 0.54 | 0.64 | 0.5 |
| 7 | 66107024 | rs1267818 | 1.251 | -1.782 | -1.275 | 0.688 | 0.044 | 0.015 | 0.015 | 0.049 | 0.2 | 0.06 | 0.21 | 0.14 | 0.11 | 0.2 |
| 7 | 66110040 | rs1267817 | 1.236 | -1.787 | -1.277 | 0.685 | 0.045 | 0.016 | 0.016 | 0.06 | 0.2 | 0.06 | 0.21 | 0.14 | 0.11 | 0.2 |
| 7 | 66120035 | rs3846973 | 0.434 | -1.766 | -1.338 | 0.611 | 0.024 | -0.027 | -0.027 | 0 | 0.57 | 0.76 | 0.64 | 0.55 | 0.63 | 0.49 |
| 7 | 66141303 | rs9654843 | 0.976 | -1.691 | -1.335 | 0.601 | -0.003 | -0.004 | -0.004 | 0 | 0.08 | 0.05 | 0.1 | 0.11 | 0.11 | 0.1 |
| 7 | 66147110 | rs3800823 | 1.946 | -1.842 | -1.149 | 0.54 | -0.04 | 0.011 | 0.011 | 0 | 0.32 | 0.24 | 0.42 | 0.27 | 0.46 | 0.34 |
| 7 | 66158420 | rs2659911 | 1.271 | -1.814 | -1.224 | 0.526 | 0.044 | 0.015 | 0.015 | 0.049 | 0.2 | 0.06 | 0.21 | 0.14 | 0.11 | 0.2 |
| 7 | 66197799 | rs2178742 | 1.271 | -1.838 | -1.219 | 0.536 | 0.044 | 0.015 | 0.015 | 0.049 | 0.2 | 0.06 | 0.21 | 0.14 | 0.11 | 0.2 |
| 7 | 66242593 | rs4717319 | -1.55 | -1.673 | -2.029 | 0.57 | -0.021 | 0.018 | 0.018 | 0 | 0.63 | 0.33 | 0.55 | 0.73 | 0.54 | 0.66 |
| 7 | 66251369 | rs10234747 | 0.259 | -2.725 | -1.808 | 0.863 | 0.002 | -0.021 | -0.021 | 1.103 | 0.1 | 0.06 | 0.14 | 0.22 | 0.13 | 0.12 |
| 7 | 66263994 | rs10479729 | -0.055 | -2.809 | -1.854 | 0.845 | 0.028 | -0.038 | -0.038 | 0.059 | 0.19 | 0.08 | 0.18 | 0.28 | 0.18 | 0.15 |
| 7 | 66265445 | rs2077180 | -0.055 | -2.809 | -1.854 | 0.853 | 0.028 | -0.04 | -0.04 | 0.544 | 0.19 | 0.08 | 0.18 | 0.28 | 0.18 | 0.15 |
| 7 | 66274955 | rs1064795 | 0.253 | -2.801 | -1.849 | 0.848 | 0.002 | -0.021 | -0.021 | 0 | 0.1 | 0.06 | 0.14 | 0.22 | 0.13 | 0.11 |
| 7 | 66276217 | rs11028 | -0.506 | -2.968 | -1.882 | 0.818 | 0.065 | 0.001 | 0.001 | 0 | 0.41 | 0.26 | 0.4 | 0.42 | 0.29 | 0.36 |
| 7 | 66278207 | rs4717322 | 0.409 | -2.933 | -1.981 | 0.829 | 0.015 | -0.061 | -0.061 | 0 | 0.16 | 0.06 | 0.1 | 0.31 | 0.16 | 0.24 |
| 7 | 66281040 | rs1984651 | -0.007 | -2.804 | -1.912 | 0.859 | 0.028 | -0.038 | -0.038 | 0.02 | 0.19 | 0.08 | 0.18 | 0.28 | 0.18 | 0.15 |
| 7 | 66315976 | rs10231411 | 0.311 | -2.761 | -1.887 | 0.885 | 0.002 | -0.024 | -0.024 | 0.188 | 0.1 | 0.06 | 0.14 | 0.21 | 0.13 | 0.12 |
| 7 | 66323500 | rs2016325 | 1.962 | -2.909 | -1.891 | 0.823 | -0.038 | 0.076 | 0.076 | 0 | 0.34 | 0.1 | 0.44 | 0.21 | 0.53 | 0.39 |
| 7 | 66328454 | rs17747927 | 0.67 | -2.531 | -1.405 | 1.307 | -0.02 | -0.02 | -0.02 | 0.161 | 0.08 | 0.01 | 0.1 | 0.11 | 0.11 | 0.1 |
| 7 | 66329634 | rs17747939 | 0.526 | -2.613 | -1.451 | 1.359 | 0.002 | -0.021 | -0.021 | 0.686 | 0.09 | 0.04 | 0.07 | 0.29 | 0.08 | 0.1 |
| 7 | 66343304 | rs6974046 | 0.399 | -2.613 | -1.446 | 1.359 | 0.004 | -0.004 | -0.004 | 1.081 | 0.09 | 0.06 | 0.07 | 0.29 | 0.08 | 0.11 |
| 7 | 66346373 | rs10234640 | 0.598 | -2.588 | -1.485 | 1.065 | -0.013 | -0.013 | -0.013 | 0 | 0.08 | 0.04 | 0.1 | 0.11 | 0.11 | 0.1 |
| 7 | 66350300 | rs4718421 | -1.699 | -2.637 | -1.433 | 1.039 | 0.065 | 0.106 | 0.106 | 0 | 0.56 | 0.77 | 0.52 | 0.77 | 0.38 | 0.46 |
| 7 | 66377234 | rs12532805 | -0.502 | -1.837 | -2 | 1.03 | 0.028 | -0.038 | -0.038 | 1.22 | 0.19 | 0.08 | 0.18 | 0.28 | 0.18 | 0.15 |
| 7 | 66384315 | rs10235423 | -0.47 | -1.831 | -1.996 | 1.033 | 0.027 | -0.036 | -0.036 | 0 | 0.19 | 0.08 | 0.18 | 0.28 | 0.18 | 0.15 |
| 7 | 66399017 | rs17748115 | 0.352 | -1.821 | -1.992 | 1.032 | -0.002 | -0.003 | -0.003 | 0.203 | 0.09 | 0.06 | 0.1 | 0.11 | 0.11 | 0.09 |
| 7 | 66404770 | rs10264873 | 0.001 | -1.706 | -1.924 | 1.03 | 0.027 | -0.038 | -0.038 | 0 | 0.19 | 0.07 | 0.18 | 0.28 | 0.18 | 0.15 |
| 7 | 66415069 | rs2901311 | -0.12 | -1.12 | -1.573 | 1.216 | 0.097 | 0.018 | 0.018 | 0 | 0.4 | 0.14 | 0.39 | 0.35 | 0.28 | 0.35 |
| 7 | 66421446 | rs4718428 | -0.85 | -1.025 | -1.541 | 1.098 | 0.084 | 0.085 | 0.085 | 0 | 0.5 | 0.32 | 0.48 | 0.71 | 0.37 | 0.47 |
| 7 | 66448791 | rs7784393 | -0.253 | -0.338 | -1.853 | 1.255 | -0.002 | 0.019 | 0.019 | 0 | 0.09 | 0.06 | 0.08 | 0.36 | 0.09 | 0.12 |
| 7 | 66452054 | rs10273658 | -0.411 | -0.403 | -1.81 | 0.76 | 0.081 | 0.03 | 0.03 | 0 | 0.48 | 0.2 | 0.43 | 0.61 | 0.37 | 0.44 |
| 7 | 66464764 | rs7790407 | -0.301 | 0.077 | -1.004 | 0.995 | 0.003 | 0.056 | 0.056 | 0 | 0.1 | 0.15 | 0.09 | 0.31 | 0.12 | 0.14 |
| 7 | 66466119 | rs13340529 | -0.365 | 0.272 | -0.49 | 0.218 | 0.026 | -0.11 | -0.11 | 0.276 | 0.18 | 0.33 | 0.2 | 0.01 | 0.2 | 0.21 |
| 7 | 66472198 | rs11766656 | 0.184 | 0.165 | -1.045 | -0.341 | 0.021 | 0.052 | 0.052 | 1.488 | 0.51 | 0.23 | 0.36 | 0.43 | 0.47 | 0.41 |
| 7 | 66472231 | rs11766621 | -0.903 | 0.114 | -0.979 | 0.231 | 0.014 | 0.035 | 0.035 | 0.088 | 0.71 | 0.59 | 0.74 | 0.61 | 0.68 | 0.63 |
| 7 | 66479228 | rs3748107 | 0.181 | -0.505 | -0.984 | -0.021 | 0.008 | -0.008 | -0.008 | 0.035 | 0.38 | 0.33 | 0.38 | 0.54 | 0.32 | 0.28 |
| 7 | 66487445 | rs6944726 | 1.084 | -0.296 | -1.219 | 0.122 | -0.009 | -0.034 | -0.034 | 0.874 | 0.08 | 0.02 | 0.07 | 0.3 | 0.11 | 0.1 |
| 7 | 66506086 | rs6974864 | -0.319 | -0.37 | -1.026 | -0.568 | 0.01 | -0.085 | -0.085 | 0 | 0.22 | 0.38 | 0.18 | 0.07 | 0.22 | 0.3 |
| 7 | 66515614 | rs10228308 | -0.145 | -0.311 | -0.947 | -0.536 | 0.015 | -0.069 | -0.069 | 0 | 0.21 | 0.32 | 0.18 | 0.07 | 0.22 | 0.3 |
| 7 | 66530404 | rs7780458 | -0.692 | -0.341 | -1.006 | -0.601 | 0.005 | -0.069 | -0.069 | 1.461 | 0.27 | 0.33 | 0.26 | 0.07 | 0.26 | 0.34 |
| 7 | 66533356 | rs10240542 | 0.66 | -0.243 | -0.632 | -0.289 | 0.039 | -0.08 | -0.08 | 0 | 0.12 | 0.29 | 0.09 | 0.01 | 0.14 | 0.17 |
| 7 | 66535742 | rs4718450 | 1.259 | -0.551 | -1.257 | -0.146 | 0.029 | 0.022 | 0.022 | 1.391 | 0.1 | 0 | 0.04 | 0.05 | 0.05 | 0.06 |
| 7 | 66546046 | rs13225388 | -0.555 | -0.597 | -1.132 | -0.261 | -0.003 | 0.038 | 0.038 | 0.048 | 0.57 | 0.69 | 0.54 | 0.3 | 0.44 | 0.49 |
| 7 | 66575738 | rs6959635 | -0.33 | -0.977 | -0.95 | -0.574 | -0.01 | -0.107 | -0.107 | 0 | 0.28 | 0.48 | 0.23 | 0.07 | 0.24 | 0.32 |
| 7 | 66637001 | rs4433018 | -1.469 | -0.378 | -0.896 | -0.246 | 0.004 | 0.008 | 0.008 | 0 | 0.67 | 0.62 | 0.66 | 0.6 | 0.63 | 0.61 |
| 7 | 66667774 | rs6967217 | -0.025 | -0.636 | -0.628 | -0.466 | 0 | -0.048 | -0.048 | 0 | 0.25 | 0.3 | 0.21 | 0.07 | 0.24 | 0.31 |
| 7 | 66668500 | rs4718475 | -0.025 | -0.636 | -0.628 | -0.466 | 0 | -0.048 | -0.048 | 0.051 | 0.25 | 0.3 | 0.21 | 0.07 | 0.24 | 0.31 |
| 7 | 66671807 | rs4718480 | 2.062 | -0.802 | -0.763 | -0.419 | 0.023 | -0.003 | -0.003 | 0 | 0.09 | 0 | 0.05 | 0.07 | 0.04 | 0.06 |
| 7 | 66672143 | rs6951443 | -1.512 | -0.663 | -0.694 | -0.081 | 0.003 | 0.007 | 0.007 | 0 | 0.66 | 0.65 | 0.65 | 0.59 | 0.61 | 0.6 |
| 7 | 66688206 | rs12155193 | -0.575 | -0.705 | -0.743 | -0.068 | -0.007 | -0.004 | -0.004 | 0 | 0.38 | 0.57 | 0.48 | 0.33 | 0.3 | 0.39 |
| 7 | 66689729 | rs11761805 | 1.574 | -0.621 | -0.936 | -0.183 | -0.005 | 0.068 | 0.068 | 0 | 0.16 | 0.05 | 0.06 | 0 | 0.14 | 0.1 |
| 7 | 66726274 | rs12531759 | -0.139 | -0.909 | -1.103 | 0.171 | 0.047 | -0.099 | -0.099 | 0 | 0.28 | 0.13 | 0.24 | 0.47 | 0.24 | 0.33 |
| 7 | 66728987 | rs17144562 | 0.903 | -0.51 | -1.001 | 0.399 | -0.042 | -0.078 | -0.078 | 0 | 0.13 | 0.02 | 0.29 | 0.43 | 0.18 | 0.17 |
| 7 | 66765880 | rs7804288 | 1.21 | -0.707 | -0.518 | 0.197 | 0.031 | -0.071 | -0.071 | 0 | 0.12 | 0.28 | 0.08 | 0.01 | 0.13 | 0.14 |
| 7 | 66768879 | rs9647886 | -0.86 | -0.647 | -0.496 | 0.362 | -0.01 | 0.01 | 0.01 | 0 | 0.6 | 0.66 | 0.57 | 0.69 | 0.52 | 0.57 |
| 7 | 66773745 | rs7791853 | 0.286 | -0.72 | -0.542 | 0.427 | 0.018 | 0.027 | 0.027 | 1.596 | 0.19 | 0.31 | 0.34 | 0.26 | 0.21 | 0.26 |
| 7 | 66774157 | rs2041575 | -0.629 | -0.509 | -0.667 | 0.373 | -0.01 | 0.009 | 0.009 | 0 | 0.6 | 0.66 | 0.57 | 0.69 | 0.52 | 0.57 |
| 7 | 66785714 | rs1808293 | -0.244 | -0.419 | -0.28 | 0.242 | -0.001 | -0.001 | -0.001 | 0.811 | 0.51 | 0.51 | 0.59 | 0.34 | 0.54 | 0.44 |
| 7 | 66786374 | rs2548080 | -0.182 | -0.462 | -0.138 | 0.277 | 0.005 | 0.059 | 0.059 | 0.641 | 0.5 | 0.39 | 0.41 | 0.21 | 0.53 | 0.41 |
| 7 | 66793316 | rs11772331 | 0.341 | -0.253 | 0.138 | 0.13 | 0.022 | -0.017 | -0.017 | 0 | 0.16 | 0.25 | 0.16 | 0.11 | 0.23 | 0.23 |
| 7 | 66796109 | rs11760843 | -0.197 | -0.38 | 0.198 | 0.062 | 0.016 | -0.024 | -0.024 | 0 | 0.19 | 0.3 | 0.21 | 0.1 | 0.26 | 0.24 |
| 7 | 66798043 | rs4717369 | -1.044 | -1.127 | 0.064 | 0.064 | 0.006 | 0.038 | 0.038 | 0 | 0.45 | 0.66 | 0.57 | 0.65 | 0.48 | 0.48 |
| 7 | 66805000 | rs4718532 | -1.044 | -1.127 | 0.064 | 0.064 | 0.006 | 0.038 | 0.038 | 0 | 0.45 | 0.67 | 0.58 | 0.66 | 0.48 | 0.48 |
| 7 | 66814900 | rs7357162 | 0.158 | -1.184 | 0.18 | 0.179 | -0.013 | 0.027 | 0.027 | 0 | 0.25 | 0.28 | 0.23 | 0.41 | 0.21 | 0.21 |
| 7 | 66821067 | rs13230948 | -0.131 | -1.2 | 0.192 | -0.008 | 0.009 | -0.022 | -0.022 | 3.81 | 0.35 | 0.43 | 0.41 | 0.25 | 0.41 | 0.36 |
| 7 | 66823022 | rs12698576 | -2.084 | -0.877 | -0.234 | -0.179 | -0.025 | 0.052 | 0.052 | 0 | 0.74 | 0.87 | 0.73 | 0.9 | 0.74 | 0.72 |
| 7 | 66824704 | rs7792207 | 1.7 | -0.77 | -0.074 | -0.054 | 0.017 | -0.034 | -0.034 | 2.734 | 0.09 | 0.1 | 0.07 | 0 | 0.16 | 0.15 |
| 7 | 66832228 | rs11765724 | 1.848 | -0.828 | 0.026 | -0.103 | -0.001 | -0.036 | -0.036 | 0 | 0.16 | 0.15 | 0.07 | 0 | 0.13 | 0.1 |
| 7 | 66832426 | rs956523 | -0.113 | -0.991 | 0.192 | -0.064 | 0.008 | -0.055 | -0.055 | 0.534 | 0.34 | 0.56 | 0.24 | 0.11 | 0.37 | 0.32 |

| | | | | | | | | | | | | | | | | |
|---|---|---|---|---|---|---|---|---|---|---|---|---|---|---|---|---|
| 7 | 66844011 | rs10085684 | 0.151 | -1.066 | 0.156 | 0.23 | -0.023 | 0.14 | 0.14 | 0.36 | 0.11 | 0.24 | 0.1 | 0.31 | 0.08 | 0.13 |
| 7 | 66845239 | rs4718544 | -1.295 | -1.098 | 0.063 | 0.16 | -0.002 | -0.008 | -0.008 | 0 | 0.6 | 0.64 | 0.5 | 0.51 | 0.6 | 0.55 |
| 7 | 66853624 | rs7804093 | 0.497 | -1.278 | -0.049 | -0.04 | 0.011 | -0.029 | -0.029 | 0 | 0.27 | 0.3 | 0.22 | 0.11 | 0.35 | 0.25 |
| 7 | 66853648 | rs4717380 | 1.515 | -1.3 | -0.079 | 0.285 | 0.007 | -0.001 | -0.001 | 0 | 0.14 | 0.05 | 0.13 | 0.09 | 0.13 | 0.08 |
| 7 | 66854619 | rs17145211 | 1.515 | -1.3 | -0.079 | 0.285 | 0.007 | -0.001 | -0.001 | 0 | 0.14 | 0.05 | 0.13 | 0.09 | 0.13 | 0.08 |
| 7 | 66855248 | rs6967566 | -0.068 | -1.311 | -0.043 | 0.351 | -0.022 | 0.032 | 0.032 | 0 | 0.26 | 0.32 | 0.36 | 0.55 | 0.22 | 0.24 |
| 7 | 66855925 | rs17145225 | -0.213 | -1.151 | 0.131 | 0.513 | -0.026 | 0.242 | 0.242 | 0.319 | 0.11 | 0.27 | 0.23 | 0.46 | 0.1 | 0.16 |
| 7 | 66858450 | rs2110004 | -0.155 | -1.185 | 0.11 | 0.812 | -0.031 | 0.046 | 0.046 | 0.013 | 0.14 | 0.25 | 0.11 | 0.23 | 0.1 | 0.21 |
| 7 | 66859973 | rs4281012 | -0.648 | -1.216 | 0.132 | 0.849 | -0.039 | 0.154 | 0.154 | 0 | 0.17 | 0.29 | 0.28 | 0.48 | 0.13 | 0.25 |
| 7 | 66861725 | rs11976226 | 0.626 | -1.286 | 0.076 | 0.118 | 0.001 | 0.001 | 0.001 | 3.083 | 0.28 | 0.26 | 0.19 | 0.09 | 0.25 | 0.11 |
| 7 | 66863024 | rs4559132 | -0.374 | -1.27 | -0.299 | 0.15 | -0.016 | -0.011 | -0.011 | 0 | 0.29 | 0.52 | 0.23 | 0.24 | 0.24 | 0.26 |
| 7 | 66864120 | rs10252540 | -0.601 | -1.195 | -0.292 | 0.194 | 0.018 | 0.016 | 0.016 | 0.148 | 0.46 | 0.55 | 0.35 | 0.33 | 0.37 | 0.36 |
| 7 | 66865643 | rs12698588 | 0.309 | -0.924 | -0.567 | -0.166 | -0.005 | -0.034 | -0.034 | 0 | 0.27 | 0.18 | 0.23 | 0.45 | 0.32 | 0.47 |
| 7 | 66877880 | rs4266531 | 1.568 | -0.889 | 0.006 | -0.648 | 0 | -0.027 | -0.027 | 0.185 | 0.12 | 0.12 | 0.06 | 0 | 0.13 | 0.04 |
| 7 | 66878782 | rs4718560 | -0.778 | -0.855 | -0.209 | -0.581 | -0.01 | -0.025 | -0.025 | 0 | 0.92 | 0.98 | 0.92 | 0.89 | 0.92 | 0.92 |
| 7 | 66883410 | rs4507650 | 0.567 | -0.891 | -0.042 | -0.626 | 0.041 | 0.078 | 0.078 | 4.257 | 0.54 | 0.3 | 0.63 | 0.78 | 0.65 | 0.74 |
| 7 | 66887996 | rs4993699 | 0.723 | -0.277 | 0.021 | -0.753 | 0.005 | 0.016 | 0.016 | 1.758 | 0.52 | 0.44 | 0.66 | 0.8 | 0.58 | 0.75 |
| 7 | 66888014 | rs11977025 | 0.991 | -0.165 | -0.058 | -0.83 | 0.002 | -0.034 | -0.034 | 0 | 0.15 | 0.14 | 0.07 | 0.01 | 0.14 | 0.06 |
| 7 | 66888441 | rs11982022 | -0.456 | -0.264 | -0.065 | -0.789 | -0.004 | -0.003 | -0.003 | 0 | 0.9 | 0.86 | 0.9 | 0.91 | 0.91 | 0.91 |
| 7 | 66901285 | rs4644126 | 1.069 | -0.279 | -0.035 | -0.774 | -0.004 | -0.014 | -0.014 | 0.427 | 0.14 | 0.11 | 0.07 | 0.01 | 0.13 | 0.06 |
| 7 | 66901317 | rs12666354 | 0.96 | -0.442 | 0.063 | -0.816 | -0.001 | 0.02 | 0.02 | 2.632 | 0.47 | 0.58 | 0.65 | 0.79 | 0.48 | 0.73 |
| 7 | 66901528 | rs12669143 | -0.074 | -0.169 | 0.128 | -0.708 | -0.012 | 0.022 | 0.022 | 0 | 0.7 | 0.77 | 0.81 | 0.9 | 0.67 | 0.83 |
| 7 | 66904395 | rs6948216 | 0.538 | -0.098 | 0.172 | -0.494 | 0.004 | 0.051 | 0.051 | 0.323 | 0.5 | 0.59 | 0.64 | 0.79 | 0.48 | 0.73 |
| 7 | 66908992 | rs4618582 | 0.623 | -0.052 | -0.238 | -0.489 | 0.001 | 0.022 | 0.022 | 0.257 | 0.49 | 0.43 | 0.63 | 0.8 | 0.48 | 0.73 |
| 7 | 66924597 | rs10279017 | -0.227 | -0.937 | -0.336 | 0.13 | 0.014 | -0.003 | -0.003 | 0 | 0.46 | 0.55 | 0.48 | 0.48 | 0.28 | 0.45 |
| 7 | 66931015 | rs7794580 | -0.094 | -1.183 | 0.031 | 0.087 | -0.034 | -0.013 | -0.013 | 0 | 0.68 | 0.59 | 0.77 | 0.82 | 0.83 | 0.79 |
| 7 | 66934791 | rs4717388 | -0.211 | -1.134 | 0.185 | 0.162 | 0.002 | 0.012 | 0.012 | 0.082 | 0.95 | 0.8 | 0.97 | 0.95 | 0.97 | 0.99 |
| 7 | 66945510 | rs7810927 | 0.701 | -1.191 | -0.11 | 0.268 | 0.045 | 0.014 | 0.014 | 0 | 0.51 | 0.38 | 0.51 | 0.39 | 0.65 | 0.5 |
| 7 | 66945831 | rs4579412 | 1.282 | -1.353 | -0.356 | 0.376 | 0.032 | -0.089 | -0.089 | 0 | 0.21 | 0.02 | 0.18 | 0.24 | 0.15 | 0.21 |
| 7 | 66950818 | rs6953207 | 0.405 | -1.287 | -0.086 | 0.349 | -0.023 | 0.01 | 0.01 | 0.807 | 0.72 | 0.43 | 0.66 | 0.58 | 0.77 | 0.69 |
| 7 | 66954848 | rs4718584 | 0.446 | -1.303 | -0.012 | 0.272 | -0.018 | 0.009 | 0.009 | 0.747 | 0.72 | 0.38 | 0.66 | 0.58 | 0.77 | 0.69 |
| 7 | 66956421 | rs12666509 | 1.383 | -1.301 | 0.106 | 0.266 | -0.015 | 0.093 | 0.093 | 0.341 | 0.15 | 0.33 | 0.24 | 0.32 | 0.13 | 0.25 |
| 7 | 66962579 | rs10249596 | -0.27 | -1.37 | 0.071 | 0.232 | -0.04 | 0.051 | 0.051 | 0.331 | 0.78 | 0.51 | 0.72 | 0.62 | 0.83 | 0.72 |
| 7 | 66962936 | rs12698593 | 1.338 | -0.336 | -0.225 | 0.033 | -0.023 | 0.003 | 0.003 | 0 | 0.35 | 0.26 | 0.26 | 0.32 | 0.41 | 0.29 |
| 7 | 66965868 | rs11533888 | 1.48 | -0.343 | 0.014 | 0.117 | -0.015 | 0.093 | 0.093 | 0.341 | 0.15 | 0.33 | 0.24 | 0.33 | 0.14 | 0.25 |
| 7 | 66971934 | rs4335032 | 0.199 | -0.224 | -0.497 | 0.057 | -0.021 | 0.008 | 0.008 | 0.989 | 0.71 | 0.41 | 0.66 | 0.57 | 0.76 | 0.68 |
| 7 | 66987667 | rs6460366 | 1.518 | 0.383 | -0.672 | -0.056 | 0.004 | -0.016 | -0.016 | 0.566 | 0.05 | 0.06 | 0.12 | 0.01 | 0.08 | 0.12 |
| 7 | 66988316 | rs11763265 | 1.04 | 0.439 | -0.805 | -0.074 | -0.027 | 0 | 0 | 0.191 | 0.12 | 0.07 | 0.16 | 0.09 | 0.21 | 0.12 |
| 7 | 66990895 | rs4718591 | 0.637 | 0.151 | -0.764 | -0.038 | -0.007 | 0.005 | 0.005 | 0 | 0.4 | 0.39 | 0.38 | 0.37 | 0.49 | 0.41 |
| 7 | 66998246 | rs10950078 | 1.648 | 0.049 | -0.893 | -0.159 | 0.002 | 0.004 | 0.004 | 0 | 0.12 | 0.07 | 0.08 | 0 | 0.1 | 0.1 |
| 7 | 66998857 | rs4495312 | 1.243 | -0.136 | -0.445 | 0.344 | -0.014 | 0.071 | 0.071 | 0.435 | 0.15 | 0.22 | 0.23 | 0.36 | 0.13 | 0.26 |
| 7 | 67002797 | rs4502996 | 0.419 | -0.053 | -1.197 | 0.043 | -0.06 | -0.085 | -0.085 | 3.405 | 0.3 | 0.03 | 0.29 | 0.27 | 0.32 | 0.28 |
| 7 | 67003386 | rs6953181 | 1.126 | 0.115 | -1.047 | 0.169 | -0.023 | 0.023 | 0.023 | 0 | 0.36 | 0.29 | 0.25 | 0.3 | 0.41 | 0.3 |
| 7 | 67004528 | rs6460372 | -0.349 | -0.358 | -1.02 | 0.64 | 0.054 | -0.005 | -0.005 | 0.019 | 0.76 | 0.89 | 0.83 | 0.82 | 0.88 | 0.82 |
| 7 | 67004828 | rs4718593 | 0.182 | 0.738 | -0.188 | 0.958 | 0.002 | 0.054 | 0.054 | 0.661 | 0.74 | 0.64 | 0.59 | 0.56 | 0.75 | 0.61 |
| 7 | 67017462 | rs10236205 | -0.514 | 1.024 | -0.044 | 1.117 | 0.038 | 0.138 | 0.138 | 0.017 | 0.19 | 0.5 | 0.22 | 0.33 | 0.23 | 0.26 |
| 7 | 67026320 | rs4576304 | -0.141 | 1.013 | -1.203 | 0.148 | -0.022 | 0.107 | 0.107 | 0.684 | 0.75 | 0.94 | 0.88 | 0.95 | 0.73 | 0.89 |
| 7 | 67033390 | rs1019080 | 1.057 | 1.029 | -1.089 | 0.261 | 0.032 | 0.136 | 0.136 | 0 | 0.13 | 0.4 | 0.14 | 0.25 | 0.17 | 0.2 |
| 7 | 67035262 | rs758245 | 1.217 | 0.885 | -1.684 | -0.355 | -0.006 | -0.038 | -0.038 | 0.477 | 0.29 | 0.16 | 0.35 | 0.49 | 0.29 | 0.34 |
| 7 | 67037628 | rs4236215 | 0.912 | 0.916 | -1.762 | -0.516 | -0.017 | -0.035 | -0.035 | 0 | 0.34 | 0.21 | 0.49 | 0.5 | 0.38 | 0.41 |
| 7 | 67042696 | rs17633215 | 2.76 | 0.456 | -2.024 | -0.777 | -0.031 | -0.011 | -0.011 | 0.138 | 0.12 | 0 | 0.11 | 0.11 | 0.1 | 0.22 |
| 7 | 67052528 | rs716578 | 0.48 | 0.329 | -1.969 | -0.811 | 0.001 | 0.019 | 0.019 | 0.188 | 0.48 | 0.26 | 0.62 | 0.61 | 0.49 | 0.63 |
| 7 | 67054054 | rs10261724 | 1.303 | 0.367 | -1.924 | -0.746 | 0.006 | 0.012 | 0.012 | 0 | 0.06 | 0.1 | 0.08 | 0.09 | 0.06 | 0.05 |
| 7 | 67057015 | rs4718600 | 2.172 | 0.405 | -2.39 | -0.422 | -0.053 | -0.115 | -0.115 | 0 | 0.18 | 0.01 | 0.25 | 0.35 | 0.19 | 0.31 |
| 7 | 67058023 | rs11763487 | 1.297 | 0.183 | -2.18 | -0.685 | -0.005 | -0.038 | -0.038 | 0 | 0.29 | 0.17 | 0.35 | 0.5 | 0.3 | 0.34 |
| 7 | 67059296 | rs1109786 | 2.226 | 0.172 | -2.071 | -0.696 | 0.034 | 0.128 | 0.128 | 0 | 0.13 | 0.4 | 0.14 | 0.25 | 0.17 | 0.2 |
| 7 | 67060423 | rs10267997 | -0.746 | 0.037 | -1.305 | -0.796 | 0.001 | 0.045 | 0.045 | 0.075 | 0.63 | 0.74 | 0.74 | 0.84 | 0.63 | 0.67 |
| 7 | 67062333 | rs7782163 | 2.516 | 0.044 | -1.203 | -0.63 | -0.027 | -0.036 | -0.036 | 0 | 0.08 | 0.13 | 0.03 | 0 | 0.01 | 0 |
| 7 | 67066278 | rs10950082 | 0.962 | 0.1 | -1.092 | -0.613 | 0 | 0.005 | 0.005 | 0.017 | 0.24 | 0.26 | 0.24 | 0.35 | 0.27 | 0.42 |
| 7 | 67068878 | rs7779951 | -0.756 | 0.158 | -1.165 | -0.675 | -0.023 | 0.042 | 0.042 | 0 | 0.77 | 0.86 | 0.73 | 0.91 | 0.75 | 0.85 |
| 7 | 67073968 | rs12667041 | 0.561 | 0.148 | -1.121 | -0.612 | -0.009 | -0.014 | -0.014 | 0 | 0.44 | 0.23 | 0.42 | 0.53 | 0.47 | 0.45 |
| 7 | 67074643 | rs17633687 | 2.139 | 0.457 | -1.095 | -0.515 | 0.012 | 0.06 | 0.06 | 0 | 0.13 | 0.25 | 0.12 | 0.25 | 0.17 | 0.2 |
| 7 | 67076373 | rs13238909 | 1.946 | 0.252 | -1.313 | -0.649 | -0.001 | 0.033 | 0.033 | 0 | 0.15 | 0.06 | 0.08 | 0.05 | 0.17 | 0.08 |
| 7 | 67077181 | rs4717400 | -1.832 | 0.342 | -1.225 | -0.592 | -0.007 | 0.015 | 0.015 | 0 | 0.84 | 0.9 | 0.88 | 0.91 | 0.85 | 0.9 |
| 7 | 67079810 | rs13245296 | 0.305 | 0.312 | -1.144 | -0.537 | 0 | 0 | 0 | 0 | 0.4 | 0.42 | 0.34 | 0.4 | 0.44 | 0.5 |
| 7 | 67084772 | rs11971450 | 1.502 | 0.431 | -1.245 | -0.543 | -0.017 | 0.018 | 0.018 | 0 | 0.07 | 0.04 | 0.16 | 0 | 0.11 | 0.04 |
| 7 | 67088079 | rs11772426 | 1.009 | 0.007 | -1.292 | -0.562 | -0.016 | -0.058 | -0.058 | 0 | 0.28 | 0.15 | 0.33 | 0.48 | 0.28 | 0.38 |
| 7 | 67093906 | rs7798818 | 0.17 | -0.409 | -0.814 | -0.793 | 0.002 | 0.007 | 0.007 | 1.356 | 0.25 | 0.32 | 0.3 | 0.26 | 0.29 | 0.24 |
| 7 | 67094001 | rs6460386 | -0.143 | -0.409 | -0.796 | -0.716 | 0 | 0 | 0 | 0 | 0.4 | 0.36 | 0.37 | 0.26 | 0.44 | 0.3 |

| | | | | | | | | | | | | | | | | |
|---|---|---|---|---|---|---|---|---|---|---|---|---|---|---|---|---|
| 7 | 67098945 | rs11974840 | -0.324 | -0.356 | -0.792 | -0.72 | -0.001 | -0.004 | -0.004 | 1.502 | 0.47 | 0.52 | 0.42 | 0.26 | 0.49 | 0.34 |
| 7 | 67102353 | rs12539606 | -0.381 | -0.521 | -1.013 | -0.674 | 0.024 | 0.134 | 0.134 | 0 | 0.51 | 0.71 | 0.43 | 0.21 | 0.48 | 0.32 |
| 7 | 67107174 | rs6460388 | -2.232 | -0.168 | -1.035 | -0.268 | -0.026 | -0.036 | -0.036 | 0 | 0.92 | 0.97 | 0.82 | 0.81 | 0.88 | 0.97 |
| 7 | 67109115 | rs6952180 | 1.096 | -0.361 | -0.801 | -0.598 | -0.005 | -0.042 | -0.042 | 0.063 | 0.14 | 0.17 | 0.19 | 0 | 0.11 | 0.04 |
| 7 | 67114978 | rs4601196 | -0.215 | -0.775 | -0.697 | -0.601 | -0.008 | -0.002 | -0.002 | 0 | 0.36 | 0.61 | 0.34 | 0.35 | 0.3 | 0.33 |
| 7 | 67120558 | rs6972789 | -0.467 | -0.629 | -0.899 | -0.852 | 0.007 | 0.025 | 0.025 | 0 | 0.48 | 0.56 | 0.4 | 0.22 | 0.44 | 0.35 |
| 7 | 67122981 | rs17142110 | 2.458 | -0.564 | -1.085 | -0.652 | 0.008 | -0.015 | -0.015 | 0 | 0.14 | 0.04 | 0.06 | 0.17 | 0.11 | 0.18 |
| 7 | 67125244 | rs887010 | -2.886 | -0.477 | -1.191 | -0.663 | -0.026 | -0.053 | -0.053 | 0 | 0.87 | 0.96 | 0.81 | 0.83 | 0.85 | 0.96 |
| 7 | 67125882 | rs1029705 | -1.145 | -0.544 | -1.124 | -0.679 | 0.003 | -0.04 | -0.04 | 0 | 0.71 | 0.9 | 0.75 | 0.68 | 0.74 | 0.77 |
| 7 | 67126466 | rs11770692 | 0.539 | -0.21 | -0.251 | -0.355 | 0.015 | 0.194 | 0.194 | 0 | 0.08 | 0.31 | 0.08 | 0.21 | 0.06 | 0.13 |
| 7 | 67129310 | rs4718606 | 1.772 | -0.023 | -0.625 | -0.405 | 0.029 | -0.027 | -0.027 | 0 | 0.08 | 0.15 | 0.2 | 0.01 | 0.13 | 0.04 |
| 7 | 67129638 | rs996034 | 1.421 | -0.157 | -0.794 | -0.412 | -0.026 | -0.048 | -0.048 | 0 | 0.24 | 0.17 | 0.33 | 0.45 | 0.3 | 0.41 |
| 7 | 67140419 | rs2714614 | 0.68 | -0.067 | -0.516 | -0.383 | -0.023 | 0.068 | 0.068 | 0 | 0.15 | 0.41 | 0.12 | 0.2 | 0.11 | 0.2 |
| 7 | 67142683 | rs17712825 | 1.228 | 0.05 | -1.14 | -0.711 | -0.013 | 0.091 | 0.091 | 0 | 0.25 | 0.06 | 0.12 | 0.02 | 0.24 | 0.19 |
| 7 | 67143694 | rs2530416 | -0.139 | -0.043 | -1.176 | -0.602 | 0.003 | 0.06 | 0.06 | 0 | 0.48 | 0.67 | 0.47 | 0.8 | 0.47 | 0.72 |
| 7 | 67143897 | rs12669927 | 1.626 | 0.445 | -1.483 | 0.009 | -0.035 | -0.056 | -0.056 | 0.003 | 0.17 | 0.13 | 0.29 | 0.45 | 0.25 | 0.34 |
| 7 | 67144217 | rs7797358 | -1.511 | 0.126 | -1.077 | -0.152 | 0.013 | -0.017 | -0.017 | 0 | 0.81 | 0.91 | 0.8 | 0.83 | 0.84 | 0.96 |
| 7 | 67146381 | rs1024590 | 0.312 | 0.598 | -0.659 | 0.021 | 0.005 | 0.006 | 0.006 | 0 | 0.35 | 0.37 | 0.27 | 0.4 | 0.43 | 0.33 |
| 7 | 67146459 | rs1024591 | 0.423 | 0.512 | -0.501 | 0 | -0.004 | -0.005 | -0.005 | 0 | 0.33 | 0.27 | 0.25 | 0.4 | 0.41 | 0.33 |
| 7 | 67159881 | rs2530427 | 0.199 | 0.452 | -0.612 | 0.013 | 0.004 | 0.017 | 0.017 | 0.038 | 0.33 | 0.41 | 0.25 | 0.4 | 0.4 | 0.33 |
| 7 | 67165048 | rs12698617 | -0.858 | 0.487 | -0.589 | -0.042 | -0.014 | -0.012 | -0.012 | 0.328 | 0.44 | 0.63 | 0.46 | 0.48 | 0.53 | 0.39 |
| 7 | 67172209 | rs12674230 | 1.66 | 0.48 | -0.706 | 0.015 | -0.032 | -0.004 | -0.004 | 0 | 0.2 | 0.11 | 0.28 | 0.19 | 0.21 | 0.4 |
| 7 | 67172420 | rs2714606 | -1.788 | 0.243 | -1.154 | -0.372 | -0.001 | 0 | 0 | 4.729 | 0.77 | 0.78 | 0.75 | 0.83 | 0.72 | 0.93 |
| 7 | 67174442 | rs2530445 | -0.85 | 0.238 | -1.057 | -0.385 | -0.013 | 0.001 | 0.001 | 0 | 0.57 | 0.68 | 0.48 | 0.63 | 0.51 | 0.53 |
| 7 | 67176832 | rs2530451 | 1.691 | 0.222 | -1.027 | -0.307 | -0.011 | -0.001 | -0.001 | 0 | 0.22 | 0.17 | 0.24 | 0.17 | 0.28 | 0.07 |
| 7 | 67176975 | rs2530452 | 0.682 | -0.537 | -0.935 | -0.278 | 0.015 | -0.011 | -0.011 | 0 | 0.57 | 0.39 | 0.58 | 0.52 | 0.6 | 0.62 |
| 7 | 67177415 | rs10269693 | 1.538 | -0.371 | -1.017 | -0.255 | -0.002 | -0.012 | -0.012 | 0 | 0.15 | 0.17 | 0.21 | 0.08 | 0.14 | 0.05 |
| 7 | 67187678 | rs11762504 | -1.138 | -0.507 | -1.107 | -0.086 | 0.022 | -0.061 | -0.061 | 0.238 | 0.76 | 0.93 | 0.83 | 0.63 | 0.77 | 0.68 |
| 7 | 67188788 | rs7807441 | 0.457 | -0.47 | -0.834 | -0.217 | -0.006 | 0.009 | 0.009 | 0 | 0.59 | 0.52 | 0.59 | 0.51 | 0.58 | 0.61 |
| 7 | 67189026 | rs7806722 | 0.904 | -0.342 | -0.916 | -0.253 | -0.006 | 0.006 | 0.006 | 0 | 0.37 | 0.34 | 0.37 | 0.34 | 0.36 | 0.56 |
| 7 | 67195038 | rs4566929 | -0.903 | -0.055 | -0.852 | -0.036 | -0.008 | 0.026 | 0.026 | 0 | 0.75 | 0.74 | 0.8 | 0.59 | 0.74 | 0.67 |
| 7 | 67203277 | rs12673339 | 1.422 | 0.069 | -0.923 | -0.106 | -0.021 | 0.023 | 0.023 | 0 | 0.3 | 0.2 | 0.35 | 0.19 | 0.33 | 0.45 |
| 7 | 67204151 | rs9647899 | 0.7 | -0.115 | -0.896 | -0.195 | 0.008 | 0.016 | 0.016 | 0 | 0.29 | 0.21 | 0.26 | 0.23 | 0.24 | 0.2 |
| 7 | 67206709 | rs6948905 | 0.155 | 0.199 | -0.484 | 0.117 | -0.002 | -0.001 | -0.001 | 0.164 | 0.59 | 0.41 | 0.61 | 0.42 | 0.57 | 0.65 |
| 7 | 67212001 | rs6952046 | -1.038 | 0.107 | -0.463 | 0.23 | 0.006 | 0.104 | 0.104 | 0 | 0.82 | 0.73 | 0.84 | 0.59 | 0.8 | 0.7 |
| 7 | 67214927 | rs4415205 | -1.156 | 0.09 | -0.478 | 0.241 | -0.003 | 0.108 | 0.108 | 0 | 0.84 | 0.75 | 0.85 | 0.58 | 0.81 | 0.77 |
| 7 | 67216428 | rs6973730 | -2.193 | 0.015 | -0.639 | 0.256 | -0.038 | -0.072 | -0.072 | 0 | 0.89 | 0.98 | 0.89 | 0.65 | 0.84 | 0.79 |
| 7 | 67223669 | rs4613871 | 1.15 | 0.013 | -0.511 | 0.105 | -0.002 | 0 | 0 | 0.067 | 0.22 | 0.32 | 0.23 | 0.18 | 0.22 | 0.05 |
| 7 | 67227790 | rs10263656 | 0.186 | 0.048 | -0.432 | 0.089 | -0.004 | -0.004 | -0.004 | 0.051 | 0.61 | 0.42 | 0.62 | 0.4 | 0.58 | 0.7 |
| 7 | 67235252 | rs6946808 | 0.748 | 0.369 | -0.006 | 0.155 | 0.004 | 0.028 | 0.028 | 0 | 0.67 | 0.49 | 0.65 | 0.52 | 0.64 | 0.71 |
| 7 | 67236758 | rs10232026 | 0.249 | 0.4 | -0.09 | 0.058 | -0.001 | -0.026 | -0.026 | 0 | 0.18 | 0.21 | 0.21 | 0.06 | 0.16 | 0.05 |
| 7 | 67237875 | rs6944468 | 0.569 | 0.387 | -0.37 | 0.134 | 0.002 | 0.031 | 0.031 | 0 | 0.66 | 0.5 | 0.64 | 0.51 | 0.65 | 0.69 |
| 7 | 67249701 | rs7798608 | 1.17 | 0.123 | -0.439 | -1.108 | -0.003 | -0.017 | -0.017 | 0 | 0.12 | 0.11 | 0.09 | 0 | 0.2 | 0.04 |
| 7 | 67255748 | rs7805713 | 0.135 | -0.036 | 0.036 | -1.15 | 0.007 | 0.012 | 0.012 | 0 | 0.07 | 0.21 | 0.14 | 0.06 | 0.06 | 0.07 |
| 7 | 67258286 | rs9691392 | 0.531 | -0.251 | -0.399 | -1.307 | -0.037 | 0.045 | 0.045 | 0 | 0.26 | 0.16 | 0.22 | 0.13 | 0.37 | 0.16 |
| 7 | 67261526 | rs10279927 | 0.398 | -0.554 | -0.274 | -1.231 | -0.001 | 0.006 | 0.006 | 0.551 | 0.52 | 0.48 | 0.54 | 0.41 | 0.46 | 0.66 |
| 7 | 67262760 | rs1564596 | 1.542 | -0.777 | -0.557 | -1.466 | -0.022 | 0.029 | 0.029 | 0.279 | 0.29 | 0.12 | 0.34 | 0.24 | 0.32 | 0.46 |
| 7 | 67265685 | rs6943261 | -0.869 | -0.731 | -0.278 | -1.261 | -0.029 | 0.119 | 0.119 | 1.058 | 0.78 | 0.64 | 0.77 | 0.54 | 0.83 | 0.81 |
| 7 | 67279358 | rs6972474 | 0.677 | 0.03 | 0.149 | -1.29 | -0.005 | 0.007 | 0.007 | 0 | 0.63 | 0.43 | 0.68 | 0.44 | 0.69 | 0.63 |
| 7 | 67282132 | rs2865107 | -1.242 | 0.165 | 0.265 | -0.948 | 0 | 0.049 | 0.049 | 0 | 0.87 | 0.87 | 0.8 | 0.58 | 0.84 | 0.82 |
| 7 | 67283756 | rs4718628 | 1.054 | -0.216 | 0.247 | -0.983 | -0.015 | -0.001 | -0.001 | 0 | 0.24 | 0.31 | 0.28 | 0.25 | 0.21 | 0.18 |
| 7 | 67291013 | rs6946643 | -0.278 | -0.313 | -0.046 | -0.921 | 0.006 | 0.018 | 0.018 | 0 | 0.71 | 0.67 | 0.73 | 0.58 | 0.64 | 0.77 |
| 7 | 67291581 | rs13244158 | 1.5 | -0.273 | -0.277 | -0.814 | -0.019 | -0.001 | -0.001 | 0 | 0.12 | 0 | 0.04 | 0.11 | 0.07 | 0.13 |
| 7 | 67305177 | rs12673877 | -0.209 | -0.094 | -0.064 | -0.7 | -0.008 | -0.014 | -0.014 | 0 | 0.71 | 0.74 | 0.73 | 0.59 | 0.64 | 0.78 |
| 7 | 67307712 | rs7776685 | -0.206 | 0.015 | 0.061 | -1.525 | 0.016 | 0.048 | 0.048 | 0 | 0.44 | 0.28 | 0.39 | 0.32 | 0.36 | 0.57 |
| 7 | 67319114 | rs6973904 | 0.23 | -0.179 | 0.85 | -0.788 | 0.006 | 0.004 | 0.004 | 0 | 0.9 | 0.77 | 0.95 | 0.88 | 0.94 | 0.88 |
| 7 | 67320928 | rs4718639 | 0.336 | -0.241 | 0.655 | -1.007 | 0.004 | 0.045 | 0.045 | 0 | 0.47 | 0.4 | 0.52 | 0.67 | 0.49 | 0.42 |
| 7 | 67323044 | rs3846975 | 0.038 | -0.204 | 0.741 | -0.997 | 0.014 | 0.008 | 0.008 | 0 | 0.89 | 0.73 | 0.94 | 0.88 | 0.94 | 0.88 |
| 7 | 67329328 | rs10241584 | 0.342 | -0.203 | 0.716 | -0.997 | 0.004 | 0.045 | 0.045 | 0 | 0.47 | 0.4 | 0.49 | 0.67 | 0.49 | 0.38 |
| 7 | 67335673 | rs10266956 | 0.347 | 0.389 | 0.76 | -0.269 | 0.015 | 0.008 | 0.008 | 0 | 0.89 | 0.73 | 0.94 | 0.88 | 0.94 | 0.87 |
| 7 | 67349749 | rs900278 | 0.08 | 0.063 | 0.474 | -0.256 | -0.001 | 0.032 | 0.032 | 0.011 | 0.9 | 0.66 | 0.95 | 0.88 | 0.95 | 0.89 |
| 7 | 67352771 | rs1600879 | 0.628 | -0.037 | 0.189 | -0.327 | -0.02 | 0.003 | 0.003 | 0 | 0.87 | 0.49 | 0.92 | 0.88 | 0.93 | 0.89 |
| 7 | 67361027 | rs6971526 | 0.499 | -0.089 | -0.434 | -0.739 | -0.021 | 0.201 | 0.201 | 0 | 0.47 | 0.18 | 0.43 | 0.17 | 0.5 | 0.51 |
| 7 | 67365305 | rs6460410 | -0.045 | 0.248 | -0.664 | -0.44 | -0.002 | 0.04 | 0.04 | 0 | 0.41 | 0.29 | 0.49 | 0.7 | 0.43 | 0.37 |
| 7 | 67373364 | rs10272439 | -0.221 | -0.387 | -0.803 | -0.316 | 0.022 | 0.015 | 0.015 | 0.789 | 0.6 | 0.71 | 0.69 | 0.67 | 0.65 | 0.64 |
| 7 | 67386514 | rs13240999 | 0.301 | -0.305 | -0.609 | -0.371 | -0.003 | -0.005 | -0.005 | 0 | 0.08 | 0.09 | 0.03 | 0.06 | 0.04 | 0.1 |
| 7 | 67396184 | rs7791037 | -0.132 | -0.342 | -0.128 | -0.773 | -0.012 | -0.002 | -0.002 | 0.278 | 0.42 | 0.38 | 0.4 | 0.53 | 0.5 | 0.36 |
| 7 | 67398107 | rs17144954 | -0.745 | -0.585 | -0.166 | -0.856 | 0 | -0.005 | -0.005 | 0 | 0.15 | 0.23 | 0.19 | 0.17 | 0.13 | 0.38 |
| 7 | 67399606 | rs10950102 | 0.685 | -0.395 | -0.194 | -0.834 | 0.008 | 0.003 | 0.003 | 0 | 0.08 | 0.11 | 0.16 | 0.12 | 0.09 | 0.27 |

| | | | | | | | | | | | | | | | | |
|---|---|---|---|---|---|---|---|---|---|---|---|---|---|---|---|---|
| 7 | 67401021 | rs10263643 | 0.128 | -0.423 | -0.119 | -0.826 | -0.037 | -0.032 | -0.032 | 0 | 0.34 | 0.27 | 0.37 | 0.48 | 0.46 | 0.26 |
| 7 | 67405642 | rs6958698 | -0.716 | -0.511 | -0.314 | -0.927 | -0.018 | -0.076 | -0.076 | 0 | 0.12 | 0.28 | 0.08 | 0 | 0.08 | 0.05 |
| 7 | 67408820 | rs6946431 | -0.202 | -0.487 | -0.676 | -0.688 | 0.001 | -0.005 | -0.005 | 0 | 0.45 | 0.65 | 0.46 | 0.48 | 0.54 | 0.29 |
| 7 | 67416726 | rs6460415 | 0.549 | -0.95 | -0.795 | -0.885 | -0.009 | -0.011 | -0.011 | 0 | 0.57 | 0.41 | 0.66 | 0.64 | 0.72 | 0.47 |
| 7 | 67424221 | rs11763596 | 0.493 | -1.029 | -0.809 | -0.945 | -0.024 | -0.028 | -0.028 | 0 | 0.58 | 0.47 | 0.67 | 0.65 | 0.72 | 0.46 |
| 7 | 67426093 | rs6971706 | 0.377 | -1.196 | -0.634 | -0.888 | -0.016 | 0.002 | 0.002 | 0 | 0.09 | 0.12 | 0.14 | 0.17 | 0.02 | 0.13 |
| 7 | 67426323 | rs6961611 | 0.377 | -1.196 | -0.634 | -0.888 | -0.016 | 0.002 | 0.002 | 0 | 0.09 | 0.12 | 0.14 | 0.17 | 0.02 | 0.13 |
| 7 | 67428498 | rs11973776 | 1.459 | -1.2 | -0.629 | -0.905 | -0.018 | -0.019 | -0.019 | 1.071 | 0.06 | 0.08 | 0.01 | 0 | 0 | 0 |
| 7 | 67430718 | rs7802158 | 0.721 | -1.778 | -0.688 | -1.402 | -0.014 | -0.059 | -0.059 | 9.062 | 0.62 | 0.46 | 0.8 | 0.81 | 0.71 | 0.65 |
| 7 | 67434847 | rs12673611 | -0.732 | -1.886 | -0.658 | -1.457 | 0.01 | -0.045 | -0.045 | 0 | 0.8 | 0.67 | 0.88 | 0.87 | 0.81 | 0.76 |
| 7 | 67435457 | rs11973463 | 0.345 | -1.85 | -0.68 | -1.442 | -0.016 | -0.065 | -0.065 | 0.946 | 0.7 | 0.51 | 0.85 | 0.82 | 0.78 | 0.66 |
| 7 | 67438274 | rs6969045 | 0.797 | -1.821 | -0.686 | -1.437 | -0.014 | -0.062 | -0.062 | 0 | 0.62 | 0.46 | 0.8 | 0.82 | 0.71 | 0.65 |
| 7 | 67448407 | rs4717427 | 1.121 | -0.816 | -1.597 | -1.354 | -0.131 | -0.147 | -0.147 | 0 | 0.22 | 0.01 | 0.32 | 0.51 | 0.33 | 0.22 |
| 7 | 67462375 | rs13308082 | 1.166 | -1.2 | -0.833 | -1.134 | -0.002 | -0.019 | -0.019 | 0.049 | 0.08 | 0.05 | 0.05 | 0 | 0.07 | 0.01 |
| 7 | 67464503 | rs4717428 | -0.766 | -1.123 | -0.617 | -1.122 | 0.003 | -0.036 | -0.036 | 0.464 | 0.81 | 0.75 | 0.89 | 0.87 | 0.82 | 0.76 |
| 7 | 67470070 | rs4718691 | 0.358 | -0.749 | -0.73 | -0.913 | -0.007 | -0.01 | -0.01 | 0.441 | 0.58 | 0.48 | 0.68 | 0.64 | 0.69 | 0.52 |
| 7 | 67482110 | rs12665945 | 0 | -0.318 | -0.801 | -0.777 | 0.001 | -0.028 | -0.028 | 0 | 0.83 | 0.82 | 0.91 | 1 | 0.88 | 0.93 |
| 7 | 67490321 | rs12535497 | 0.079 | 0.286 | -0.242 | -0.51 | 0.021 | 0.02 | 0.02 | 0.055 | 0.24 | 0.28 | 0.33 | 0.29 | 0.3 | 0.38 |
| 7 | 67494709 | rs4292583 | -0.277 | 0.257 | -0.53 | -0.635 | 0.006 | -0.004 | -0.004 | 0 | 0.8 | 0.83 | 0.9 | 1 | 0.84 | 0.88 |
| 7 | 67498336 | rs6942436 | 0.605 | 0.243 | -0.957 | -0.529 | -0.042 | -0.04 | -0.04 | 0 | 0.37 | 0.11 | 0.39 | 0.54 | 0.41 | 0.3 |
| 7 | 67514001 | rs4349878 | -0.082 | 0.168 | -0.18 | -0.401 | -0.021 | 0.037 | 0.037 | 0 | 0.91 | 0.62 | 0.89 | 0.84 | 0.95 | 0.87 |
| 7 | 67515801 | rs4718701 | -0.2 | 0.2 | -0.177 | -0.48 | -0.011 | 0.015 | 0.015 | 0 | 0.92 | 0.63 | 0.89 | 0.93 | 0.95 | 0.87 |
| 7 | 67523020 | rs7809564 | 0.235 | 0.432 | -0.905 | -0.415 | 0.001 | 0.077 | 0.077 | 0 | 0.41 | 0.2 | 0.32 | 0.17 | 0.39 | 0.31 |
| 7 | 67537424 | rs2642148 | 0.107 | 0.349 | -1.275 | -0.42 | 0.019 | -0.025 | -0.025 | 0 | 0.57 | 0.18 | 0.59 | 0.57 | 0.58 | 0.62 |
| 7 | 67541376 | rs13437667 | -0.228 | 0.404 | -0.865 | -1.248 | -0.009 | -0.104 | -0.104 | 0 | 0.32 | 0.52 | 0.26 | 0.03 | 0.28 | 0.3 |
| 7 | 67544285 | rs11773133 | 0.552 | 0.653 | -1.662 | -1.059 | -0.067 | 0.153 | 0.153 | 0 | 0.23 | 0.01 | 0.14 | 0 | 0.21 | 0.18 |
| 7 | 67550831 | rs2642143 | 0.03 | 0.728 | -1.174 | -0.613 | -0.001 | -0.019 | -0.019 | 0 | 0.48 | 0.5 | 0.38 | 0.22 | 0.44 | 0.49 |
| 7 | 67556056 | rs6968714 | -0.178 | 0.539 | -2.133 | 0.292 | 0.012 | -0.016 | -0.016 | 0 | 0.32 | 0.13 | 0.43 | 0.6 | 0.32 | 0.44 |
| 7 | 67559054 | rs2019539 | 0.264 | 0.73 | -2.606 | 0.265 | -0.095 | -0.081 | -0.081 | 0 | 0.38 | 0.06 | 0.33 | 0.52 | 0.47 | 0.43 |
| 7 | 67565499 | rs17636163 | 0.441 | 0.838 | -2.735 | -0.103 | -0.032 | 0.082 | 0.082 | 0.065 | 0.13 | 0.01 | 0.09 | 0 | 0.1 | 0.11 |
| 7 | 67567429 | rs4718709 | 0.181 | 0.651 | -2.134 | 0.404 | 0.009 | 0.013 | 0.013 | 0 | 0.55 | 0.19 | 0.43 | 0.53 | 0.58 | 0.58 |
| 7 | 67572507 | rs9654877 | -0.046 | 0.584 | -1.54 | 0.292 | -0.001 | 0 | 0 | 0 | 0.14 | 0.14 | 0.16 | 0.09 | 0.12 | 0.09 |
| 7 | 67577782 | rs2159481 | 0.287 | 0.175 | -2.312 | 0.399 | 0.043 | -0.081 | -0.081 | 0 | 0.4 | 0.11 | 0.43 | 0.51 | 0.34 | 0.58 |
| 7 | 67583762 | rs2642124 | 0.031 | 0.539 | -1.292 | 0.114 | 0.044 | -0.035 | -0.035 | 0 | 0.14 | 0.35 | 0.13 | 0.1 | 0.13 | 0.2 |
| 7 | 67584032 | rs11772145 | 1.289 | 0.216 | -1.683 | 0.134 | -0.015 | -0.02 | -0.02 | 0 | 0.13 | 0.01 | 0.14 | 0.16 | 0.13 | 0.25 |
| 7 | 67587520 | rs10238581 | 1.237 | 0.228 | -1.608 | 0.104 | -0.003 | 0.006 | 0.006 | 0.952 | 0.14 | 0.12 | 0.16 | 0.08 | 0.12 | 0.07 |
| 7 | 67587945 | rs2642142 | 0.209 | 0.133 | -1.399 | 0.19 | 0.044 | -0.035 | -0.035 | 0 | 0.14 | 0.35 | 0.13 | 0.1 | 0.13 | 0.2 |
| 7 | 67590433 | rs2642123 | 0.296 | 0.182 | -1.357 | 0.139 | 0.003 | -0.046 | -0.046 | 0 | 0.21 | 0.26 | 0.25 | 0.08 | 0.17 | 0.14 |
| 7 | 67590626 | rs6977786 | 1.051 | 0.082 | -1.849 | 0.515 | 0.01 | -0.038 | -0.038 | 0 | 0.25 | 0.07 | 0.24 | 0.29 | 0.2 | 0.25 |
| 7 | 67590850 | rs6979453 | 0.152 | 0.17 | -1.718 | 0.558 | 0.038 | -0.028 | -0.028 | 0 | 0.4 | 0.14 | 0.35 | 0.45 | 0.34 | 0.51 |
| 7 | 67594586 | rs7777871 | -0.092 | 0.524 | -1.222 | 1.042 | 0.012 | 0.035 | 0.035 | 0.067 | 0.08 | 0.27 | 0.1 | 0.06 | 0.06 | 0.09 |
| 7 | 67598227 | rs1990352 | -0.728 | 0.719 | -2.042 | 1.095 | -0.002 | -0.004 | -0.004 | 0 | 0.87 | 0.96 | 0.82 | 0.82 | 0.87 | 0.75 |
| 7 | 67604314 | rs17146301 | -0.506 | 0.9 | -2.007 | 1.17 | 0 | 0 | 0 | 0.006 | 0.06 | 0.07 | 0.05 | 0.15 | 0.04 | 0.14 |
| 7 | 67606033 | rs10233260 | 0.365 | 0.889 | -2.147 | 1.064 | -0.055 | -0.072 | -0.072 | 0.373 | 0.23 | 0.07 | 0.23 | 0.25 | 0.34 | 0.12 |
| 7 | 67610420 | rs9654878 | -0.469 | 0.738 | -2.109 | 0.676 | -0.026 | 0.011 | 0.011 | 0.075 | 0.81 | 0.98 | 0.79 | 0.82 | 0.82 | 0.74 |
| 7 | 67610939 | rs10950111 | 0.397 | 0.664 | -2.128 | 0.795 | 0 | -0.011 | -0.011 | 0.051 | 0.76 | 0.91 | 0.75 | 0.68 | 0.79 | 0.6 |
| 7 | 67611708 | rs12672105 | -0.919 | 0.765 | -1.784 | 0.852 | -0.001 | 0.011 | 0.011 | 0 | 0.07 | 0.09 | 0.06 | 0.15 | 0.05 | 0.14 |
| 7 | 67612719 | rs17715059 | 0.187 | 0.694 | -2.246 | 0.851 | -0.021 | 0.003 | 0.003 | 0 | 0.2 | 0.13 | 0.24 | 0.14 | 0.24 | 0.17 |
| 7 | 67613518 | rs6955185 | 0.894 | 0.693 | -1.863 | 0.96 | 0.046 | 0.019 | 0.019 | 0 | 0.21 | 0.38 | 0.28 | 0.2 | 0.28 | 0.11 |
| 7 | 67618911 | rs4140683 | 0.847 | 0.638 | -1.828 | 0.999 | 0.044 | 0.019 | 0.019 | 0 | 0.21 | 0.41 | 0.28 | 0.2 | 0.28 | 0.11 |
| 7 | 67619952 | rs1990351 | -0.594 | 0.466 | -1.976 | 1.016 | -0.032 | -0.055 | -0.055 | 3.145 | 0.46 | 0.89 | 0.59 | 0.44 | 0.55 | 0.34 |
| 7 | 67620455 | rs2052095 | -0.421 | 0.485 | -1.978 | 0.985 | -0.06 | -0.04 | -0.04 | 0 | 0.42 | 0.82 | 0.54 | 0.39 | 0.52 | 0.3 |
| 7 | 67623433 | rs2079833 | -0.89 | 0.462 | -2.089 | 0.942 | -0.013 | -0.007 | -0.007 | 0 | 0.63 | 0.9 | 0.69 | 0.6 | 0.68 | 0.57 |
| 7 | 67625408 | rs10243130 | 0.479 | 0.247 | -1.525 | 0.771 | 0.048 | 0.022 | 0.022 | 0 | 0.21 | 0.42 | 0.28 | 0.2 | 0.28 | 0.11 |
| 7 | 67628115 | rs12234472 | -0.341 | -0.166 | -2.372 | 0.621 | 0.028 | 0 | 0 | 0 | 0.28 | 0.07 | 0.19 | 0.2 | 0.22 | 0.32 |
| 7 | 67631003 | rs2903502 | -0.012 | 0.544 | -2.461 | 0.785 | -0.043 | -0.055 | -0.055 | 1.394 | 0.6 | 0.87 | 0.54 | 0.48 | 0.6 | 0.63 |
| 7 | 67632557 | rs1014383 | -0.122 | 0.068 | -2.525 | 0.647 | 0.025 | -0.04 | -0.04 | 0 | 0.35 | 0.15 | 0.32 | 0.44 | 0.28 | 0.48 |
| 7 | 67633440 | rs4717437 | 0.323 | 0.235 | -1.825 | -0.146 | -0.003 | -0.011 | -0.011 | 0.852 | 0.32 | 0.57 | 0.33 | 0.22 | 0.29 | 0.31 |
| 7 | 67637342 | rs2867161 | 0.97 | 0.203 | -1.133 | -0.219 | 0.003 | -0.003 | -0.003 | 0 | 0.1 | 0.17 | 0.05 | 0.07 | 0.09 | 0.13 |
| 7 | 67637981 | rs12535104 | 0.37 | 0.112 | -0.778 | -0.488 | -0.001 | -0.016 | -0.016 | 0 | 0.41 | 0.44 | 0.28 | 0.09 | 0.4 | 0.29 |
| 7 | 67638578 | rs17145165 | -0.424 | 0.001 | -0.618 | -0.319 | -0.001 | -0.001 | -0.001 | 0 | 0.63 | 0.67 | 0.6 | 0.39 | 0.6 | 0.53 |
| 7 | 67641115 | rs17145184 | 0.902 | 0.012 | -0.555 | -0.414 | 0.012 | 0.107 | 0.107 | 0 | 0.11 | 0.31 | 0.11 | 0.23 | 0.1 | 0.16 |
| 7 | 67642170 | rs10950112 | -1.147 | 0.037 | -0.997 | -0.346 | -0.015 | -0.075 | -0.075 | 0.972 | 0.73 | 0.88 | 0.66 | 0.46 | 0.7 | 0.64 |
| 7 | 67642280 | rs17643158 | 0.786 | -0.335 | -0.82 | -1.356 | 0.014 | 0.049 | 0.049 | 0 | 0.31 | 0.19 | 0.2 | 0.06 | 0.24 | 0.24 |
| 7 | 67644074 | rs17643180 | 0.705 | 0.036 | -0.67 | -1.016 | -0.004 | -0.002 | -0.002 | 0 | 0.19 | 0.11 | 0.24 | 0.15 | 0.19 | 0.18 |
| 7 | 67645786 | rs10242225 | -0.858 | 0.22 | -0.713 | -1.695 | -0.042 | 0.015 | 0.015 | 0 | 0.74 | 0.87 | 0.52 | 0.19 | 0.66 | 0.7 |
| 7 | 67647531 | rs11761240 | -0.157 | -0.07 | 0.047 | -1.942 | -0.007 | -0.08 | -0.08 | 0 | 0.33 | 0.53 | 0.31 | 0.09 | 0.32 | 0.25 |
| 7 | 67647907 | rs6962996 | 0.111 | -0.021 | 0.231 | -1.983 | 0 | -0.004 | -0.004 | 0 | 0.3 | 0.32 | 0.29 | 0.09 | 0.3 | 0.25 |
| 7 | 67649847 | rs10236956 | -0.667 | -0.143 | 0.215 | -2.133 | 0.017 | 0.102 | 0.102 | 0 | 0.45 | 0.74 | 0.37 | 0.17 | 0.42 | 0.44 |

| | | | | | | | | | | | | | | | | |
|---|---|---|---|---|---|---|---|---|---|---|---|---|---|---|---|---|
| 7 | 67656907 | rs4718716 | -1.005 | -0.112 | 0.222 | -2.229 | 0 | 0 | 0 | 0 | 0.28 | 0.71 | 0.21 | 0.15 | 0.26 | 0.27 |
| 7 | 67661167 | rs10155935 | 0.53 | 0.161 | 0.117 | -2.26 | 0.002 | -0.024 | -0.024 | 0 | 0.08 | 0.14 | 0.06 | 0.01 | 0.09 | 0.06 |
| 7 | 67665731 | rs4718718 | 0.45 | 0.253 | -0.014 | -2.314 | 0.018 | 0.09 | 0.09 | 0.934 | 0.35 | 0.15 | 0.3 | 0.22 | 0.31 | 0.4 |
| 7 | 67666173 | rs7807405 | 1.153 | 0.09 | -0.039 | -2.318 | 0.017 | 0.039 | 0.039 | 0.073 | 0.15 | 0.06 | 0.1 | 0.07 | 0.11 | 0.32 |
| 7 | 67668651 | rs17145208 | 1.212 | 0.05 | 0.019 | -2.31 | 0.004 | -0.001 | -0.001 | 0.949 | 0.22 | 0.17 | 0.23 | 0.04 | 0.21 | 0.19 |
| 7 | 67676780 | rs4718720 | -0.808 | 0.144 | -0.199 | -2.29 | 0.023 | 0.219 | 0.219 | 0.752 | 0.5 | 0.93 | 0.39 | 0.19 | 0.43 | 0.59 |
| 7 | 67686930 | rs12698678 | -0.288 | 0.238 | 0.225 | -2.159 | 0.072 | 0.054 | 0.054 | 0 | 0.11 | 0.6 | 0.11 | 0.09 | 0.11 | 0.07 |
| 7 | 67690924 | rs3108019 | -0.387 | 0.039 | 0.128 | -2.299 | 0.031 | 0.058 | 0.058 | 1.326 | 0.66 | 0.8 | 0.76 | 0.77 | 0.74 | 0.61 |
| 7 | 67690962 | rs3113158 | -0.398 | 0.04 | 0.083 | -2.298 | 0.037 | 0.076 | 0.076 | 1.326 | 0.66 | 0.82 | 0.76 | 0.78 | 0.74 | 0.61 |
| 7 | 67692747 | rs6950238 | 0.914 | 0.134 | 0.083 | -2.239 | -0.007 | -0.043 | -0.043 | 0.485 | 0.23 | 0.26 | 0.23 | 0.04 | 0.21 | 0.19 |
| 7 | 67699174 | rs10248573 | 0.927 | 0.133 | 0.083 | -2.239 | -0.006 | -0.041 | -0.041 | 0 | 0.23 | 0.25 | 0.23 | 0.04 | 0.21 | 0.19 |
| 7 | 67700504 | rs3113099 | -1.616 | 0.122 | -0.103 | -2.286 | -0.045 | 0.008 | 0.008 | 0.05 | 0.81 | 0.98 | 0.8 | 0.84 | 0.81 | 0.93 |
| 7 | 67702235 | rs6960380 | 0.508 | 0.23 | 0.337 | -2.27 | 0.14 | 0.134 | 0.134 | 0 | 0.07 | 0.4 | 0.08 | 0.08 | 0.13 | 0.06 |
| 7 | 67703646 | rs6979688 | 1.288 | 0.151 | 0.244 | -2.342 | 0.006 | 0.005 | 0.005 | 0.025 | 0.22 | 0.15 | 0.22 | 0.04 | 0.21 | 0.19 |
| 7 | 67705446 | rs3108021 | -1.642 | 0.188 | 0.121 | -2.302 | -0.03 | 0.02 | 0.02 | 0.044 | 0.81 | 0.96 | 0.8 | 0.84 | 0.81 | 0.93 |
| 7 | 67708317 | rs3113123 | 0.118 | 0.363 | 0.322 | -2.254 | 0 | -0.031 | -0.031 | 0.611 | 0.31 | 0.59 | 0.32 | 0.12 | 0.35 | 0.26 |
| 7 | 67715094 | rs4718727 | 1.3 | 0.333 | 0.011 | -2.249 | -0.141 | -0.08 | -0.08 | 0.297 | 0.31 | 0.01 | 0.41 | 0.66 | 0.38 | 0.34 |
| 7 | 67716291 | rs3113126 | 0.119 | 0.141 | 0.329 | -2.268 | 0 | 0 | 0 | 0 | 0.43 | 0.6 | 0.52 | 0.74 | 0.53 | 0.41 |
| 7 | 67725587 | rs1524860 | -0.104 | -0.098 | 0.349 | -2.282 | -0.025 | -0.055 | -0.055 | 0 | 0.62 | 0.55 | 0.72 | 0.78 | 0.74 | 0.61 |
| 7 | 67728772 | rs3108040 | 0.203 | -0.007 | -0.413 | -2.313 | 0.026 | 0.162 | 0.162 | 0 | 0.52 | 0.27 | 0.51 | 0.8 | 0.57 | 0.71 |
| 7 | 67730559 | rs12154600 | 0.226 | -0.053 | -0.364 | -2.307 | 0.009 | 0.009 | 0.009 | 1.625 | 0.29 | 0.72 | 0.29 | 0.04 | 0.24 | 0.2 |
| 7 | 67742611 | rs10434956 | -0.13 | 0.128 | -0.544 | -2.142 | 0.016 | 0.054 | 0.054 | 0 | 0.54 | 0.31 | 0.52 | 0.73 | 0.55 | 0.7 |
| 7 | 67749723 | rs6954186 | 0.43 | 0.118 | -0.695 | -2.108 | -0.043 | 0.095 | 0.095 | 0 | 0.45 | 0.1 | 0.44 | 0.72 | 0.44 | 0.64 |
| 7 | 67763464 | rs13438777 | 1.216 | -0.041 | -0.447 | -2.045 | 0.018 | 0.018 | 0.018 | 0.026 | 0.14 | 0.07 | 0.08 | 0.06 | 0.06 | 0.31 |
| 7 | 67765455 | rs1917406 | 1.632 | 0.001 | -0.915 | -2.089 | 0.018 | 0.045 | 0.045 | 0.234 | 0.15 | 0.07 | 0.11 | 0.02 | 0.13 | 0.16 |
| 7 | 67769546 | rs9638255 | 0.118 | 0.487 | -0.559 | -1.811 | -0.002 | 0.001 | 0.001 | 0 | 0.66 | 0.31 | 0.67 | 0.74 | 0.65 | 0.85 |
| 7 | 67775112 | rs940597 | 0.809 | 1.129 | -0.366 | -1.911 | -0.003 | 0.005 | 0.005 | 0 | 0.64 | 0.38 | 0.62 | 0.77 | 0.65 | 0.6 |
| 7 | 67777535 | rs3113173 | 0.221 | 1.285 | -0.34 | -1.963 | -0.021 | 0.086 | 0.086 | 0.062 | 0.72 | 0.86 | 0.8 | 0.91 | 0.74 | 0.64 |
| 7 | 67778319 | rs3113176 | -0.936 | 1.66 | -0.038 | -1.816 | 0.107 | 0.136 | 0.136 | 0 | 0.08 | 0.49 | 0.18 | 0.14 | 0.09 | 0.04 |
| 7 | 67778529 | rs3108087 | -0.103 | 1.513 | -0.681 | -1.758 | -0.074 | 0.072 | 0.072 | 0 | 0.86 | 0.96 | 0.89 | 0.97 | 0.8 | 0.89 |
| 7 | 67788551 | rs12154587 | 0.427 | 1.373 | -0.836 | -1.975 | 0 | 0.19 | 0.19 | 0 | 0.45 | 0.11 | 0.41 | 0.17 | 0.42 | 0.29 |
| 7 | 67796007 | rs1524865 | 0.156 | 0.904 | -0.182 | -1.705 | 0 | 0 | 0 | 0 | 0.34 | 0.29 | 0.42 | 0.8 | 0.36 | 0.55 |
| 7 | 67797507 | rs3108004 | 0.164 | 0.94 | -0.084 | -1.672 | 0 | -0.001 | -0.001 | 0 | 0.34 | 0.29 | 0.43 | 0.8 | 0.36 | 0.55 |
| 7 | 67810679 | rs6460447 | 0.041 | 0.937 | -0.429 | -1.453 | 0.006 | 0.011 | 0.011 | 0 | 0.59 | 0.37 | 0.53 | 0.27 | 0.62 | 0.43 |
| 7 | 67812679 | rs994086 | 0.022 | 0.841 | -0.559 | -1.672 | 0.002 | 0.005 | 0.005 | 0.356 | 0.72 | 0.24 | 0.73 | 0.89 | 0.76 | 0.76 |
| 7 | 67817108 | rs7792400 | 0.182 | 1.41 | -0.935 | -1.752 | -0.05 | 0.119 | 0.119 | 0.309 | 0.23 | 0.06 | 0.16 | 0.02 | 0.24 | 0.13 |
| 7 | 67828081 | rs1874243 | 0.752 | 1.484 | -0.599 | -1.746 | -0.035 | 0.054 | 0.054 | 0 | 0.62 | 0.3 | 0.63 | 0.81 | 0.58 | 0.67 |
| 7 | 67836835 | rs4717452 | -0.33 | 1.486 | 0.922 | -1.7 | 0.007 | -0.026 | -0.026 | 0 | 0.74 | 0.66 | 0.79 | 0.91 | 0.72 | 0.74 |
| 7 | 67844301 | rs1874247 | 0.283 | 1.469 | 1.221 | -1.708 | 0.004 | -0.013 | -0.013 | 0 | 0.69 | 0.4 | 0.7 | 0.81 | 0.67 | 0.71 |
| 7 | 67845582 | rs2056611 | -0.376 | 1.493 | 1.259 | -1.701 | 0.004 | -0.017 | -0.017 | 4.845 | 0.74 | 0.7 | 0.8 | 0.91 | 0.72 | 0.74 |
| 7 | 67848496 | rs33978605 | 0.081 | 1.475 | 1.287 | -1.687 | 0.023 | -0.036 | -0.036 | 0.347 | 0.71 | 0.55 | 0.71 | 0.82 | 0.68 | 0.73 |
| 7 | 67856395 | rs1554651 | 0.097 | 1.423 | 1.281 | -1.592 | 0.021 | -0.028 | -0.028 | 0 | 0.7 | 0.51 | 0.71 | 0.8 | 0.68 | 0.72 |
| 7 | 67882598 | rs13223163 | 1.258 | 1.321 | 0.737 | -1.507 | 0.029 | 0.137 | 0.137 | 0.011 | 0.51 | 0.22 | 0.49 | 0.75 | 0.47 | 0.63 |
| 7 | 67885035 | rs17137439 | -1.291 | 1.262 | 1.374 | -1.378 | 0.099 | 0.139 | 0.139 | 0 | 0.09 | 0.42 | 0.18 | 0.11 | 0.11 | 0.06 |
| 7 | 67887396 | rs12532549 | 0.175 | 1.261 | 1.297 | -1.369 | 0.026 | 0.002 | 0.002 | 0 | 0.05 | 0.12 | 0.13 | 0.04 | 0.08 | 0.03 |
| 7 | 67889919 | rs6955144 | -1.234 | 1.109 | 0.943 | -1.569 | 0.056 | 0.119 | 0.119 | 0 | 0.53 | 0.76 | 0.52 | 0.26 | 0.57 | 0.4 |
| 7 | 67909312 | rs10269153 | -0.29 | 1.305 | 1.316 | -1.609 | 0.057 | -0.036 | -0.036 | 0 | 0.17 | 0.44 | 0.13 | 0.07 | 0.21 | 0.14 |
| 7 | 67912083 | rs12540706 | 0.222 | 1.297 | 0.917 | -1.736 | -0.035 | 0.102 | 0.102 | 0 | 0.3 | 0.12 | 0.29 | 0.09 | 0.32 | 0.26 |
| 7 | 67920764 | rs17137443 | -1.07 | 1.149 | 1.08 | -1.752 | 0.018 | 0.029 | 0.029 | 0 | 0.5 | 0.52 | 0.43 | 0.09 | 0.51 | 0.37 |
| 7 | 67921441 | rs2868540 | -1.441 | 1.269 | 0.86 | -1.734 | 0.073 | 0.104 | 0.104 | 0 | 0.55 | 0.75 | 0.54 | 0.24 | 0.59 | 0.43 |
| 7 | 67929825 | rs10256053 | -0.22 | 1.572 | 0.973 | -1.546 | 0.026 | -0.102 | -0.102 | 0.094 | 0.17 | 0.32 | 0.13 | 0.01 | 0.17 | 0.11 |
| 7 | 67934228 | rs7796950 | -1.346 | 1.589 | 0.628 | -1.267 | 0.036 | 0.084 | 0.084 | 0 | 0.08 | 0.24 | 0.13 | 0.15 | 0.09 | 0.07 |
| 7 | 67938007 | rs11770329 | 0.403 | 0.991 | -0.291 | -1.296 | -0.063 | 0.147 | 0.147 | 0 | 0.33 | 0.07 | 0.29 | 0.12 | 0.32 | 0.25 |
| 7 | 67946161 | rs1474285 | -0.527 | 0.671 | 0.164 | -0.648 | 0.027 | 0.087 | 0.087 | 0 | 0.21 | 0.54 | 0.2 | 0.28 | 0.21 | 0.18 |
| 7 | 67946524 | rs1554496 | 0.674 | 0.344 | -0.611 | 0.158 | 0.031 | 0.034 | 0.034 | 0 | 0.36 | 0.18 | 0.33 | 0.28 | 0.32 | 0.41 |
| 7 | 67947961 | rs17139782 | -0.235 | 0.243 | -0.298 | 0.113 | -0.002 | 0 | 0 | 0 | 0.53 | 0.55 | 0.66 | 0.58 | 0.65 | 0.61 |
| 7 | 67952586 | rs903527 | -0.517 | 1.123 | 0.116 | 0.446 | 0.037 | 0.123 | 0.123 | 0 | 0.26 | 0.48 | 0.32 | 0.48 | 0.28 | 0.39 |
| 7 | 67953848 | rs4717457 | 0.717 | 1.185 | 0.386 | 0.508 | 0.015 | 0.097 | 0.097 | 0 | 0.12 | 0.21 | 0.22 | 0.3 | 0.13 | 0.2 |
| 7 | 67957981 | rs2868538 | -0.812 | 1.248 | 0.151 | 0.471 | 0.025 | 0.16 | 0.16 | 0 | 0.23 | 0.5 | 0.3 | 0.48 | 0.25 | 0.39 |
| 7 | 67968501 | rs17139792 | -0.492 | 0.938 | 0.432 | 0.436 | 0.055 | 0.056 | 0.056 | 0 | 0.16 | 0.52 | 0.12 | 0.18 | 0.16 | 0.2 |
| 7 | 67973884 | rs1403157 | -0.545 | 0.455 | 0.168 | 0.447 | -0.026 | 0.068 | 0.068 | 0.188 | 0.8 | 0.71 | 0.73 | 0.6 | 0.86 | 0.84 |
| 7 | 67996983 | rs6979790 | -0.697 | 0.728 | -0.216 | 0.343 | -0.001 | -0.003 | -0.003 | 0 | 0.38 | 0.39 | 0.55 | 0.58 | 0.44 | 0.47 |
| 7 | 68002819 | rs4718769 | 1.332 | 0.751 | -0.306 | 0.387 | -0.014 | -0.035 | -0.035 | 0 | 0.12 | 0.04 | 0.25 | 0.38 | 0.13 | 0.21 |
| 7 | 68003277 | rs4718771 | 0.621 | 0.72 | -0.162 | 0.372 | 0.005 | 0.075 | 0.075 | 0 | 0.15 | 0.3 | 0.28 | 0.38 | 0.14 | 0.21 |
| 7 | 68004011 | rs17670419 | 0.227 | 0.681 | -0.248 | 0.247 | -0.007 | 0.042 | 0.042 | 0 | 0.22 | 0.13 | 0.28 | 0.12 | 0.26 | 0.21 |
| 7 | 68007956 | rs10950134 | -0.956 | 0.693 | -0.144 | 0.194 | 0 | -0.001 | -0.001 | 0 | 0.47 | 0.59 | 0.61 | 0.63 | 0.51 | 0.62 |
| 7 | 68015028 | rs7790867 | -0.535 | 0.722 | -0.268 | 0.258 | -0.001 | -0.002 | -0.002 | 0 | 0.38 | 0.43 | 0.55 | 0.58 | 0.44 | 0.47 |
| 7 | 68022241 | rs9986974 | 1.006 | 0.706 | -0.273 | 0.137 | 0.004 | -0.002 | -0.002 | 0 | 0.08 | 0.13 | 0.05 | 0.04 | 0.07 | 0.15 |
| 7 | 68022415 | rs9986976 | -0.094 | 0.402 | -0.183 | 0.062 | -0.004 | 0.015 | 0.015 | 0.154 | 0.75 | 0.68 | 0.58 | 0.27 | 0.79 | 0.66 |

| | | | | | | | | | | | | | | | | |
|---|---|---|---|---|---|---|---|---|---|---|---|---|---|---|---|---|
| 7 | 68023074 | rs1357305 | 0.541 | 0.366 | -0.304 | -0.448 | -0.001 | 0.016 | 0.016 | 0 | 0.62 | 0.69 | 0.53 | 0.18 | 0.61 | 0.54 |
| 7 | 68025258 | rs10226192 | 0.371 | 0.243 | -0.199 | -0.725 | 0.002 | 0.003 | 0.003 | 0 | 0.47 | 0.38 | 0.28 | 0.1 | 0.54 | 0.29 |
| 7 | 68025407 | rs10282048 | 0.375 | -0.532 | -0.472 | -1.049 | -0.028 | -0.094 | -0.094 | 0.549 | 0.22 | 0.28 | 0.1 | 0.01 | 0.14 | 0.1 |
| 7 | 68025695 | rs10226722 | -0.396 | -0.603 | -0.615 | -0.271 | 0.004 | -0.003 | -0.003 | 0 | 0.6 | 0.6 | 0.43 | 0.44 | 0.63 | 0.43 |
| 7 | 68030164 | rs11760460 | 0.914 | -0.714 | -0.53 | -0.046 | -0.027 | -0.023 | -0.023 | 0 | 0.14 | 0.14 | 0.05 | 0.06 | 0.05 | 0.04 |
| 7 | 68031011 | rs11763698 | -0.614 | -0.887 | -1.054 | -0.07 | 0.04 | -0.045 | -0.045 | 2.118 | 0.36 | 0.19 | 0.38 | 0.52 | 0.26 | 0.34 |
| 7 | 68031324 | rs17750149 | 0.203 | -0.922 | -1.124 | 0.049 | -0.032 | -0.099 | -0.099 | 0 | 0.2 | 0.04 | 0.22 | 0.37 | 0.18 | 0.21 |
| 7 | 68037531 | rs10242635 | -1.103 | -0.765 | -0.683 | -0.31 | -0.005 | 0.003 | 0.003 | 0 | 0.68 | 0.67 | 0.62 | 0.65 | 0.71 | 0.67 |
| 7 | 68037870 | rs10276315 | 0.108 | -0.771 | -0.721 | -0.316 | 0.035 | -0.036 | -0.036 | 0 | 0.36 | 0.22 | 0.38 | 0.52 | 0.27 | 0.34 |
| 7 | 68042545 | rs10250062 | -0.978 | -0.704 | -0.673 | -0.269 | 0.001 | -0.002 | -0.002 | 0 | 0.69 | 0.75 | 0.63 | 0.65 | 0.71 | 0.67 |
| 7 | 68045874 | rs1357308 | -0.914 | -0.33 | -0.306 | -0.744 | 0.029 | -0.05 | -0.05 | 0 | 0.72 | 0.63 | 0.78 | 0.95 | 0.62 | 0.75 |
| 7 | 68047178 | rs10248387 | 1.46 | -0.372 | -0.311 | -0.794 | 0.006 | 0.023 | 0.023 | 0 | 0.09 | 0.32 | 0.06 | 0.05 | 0.05 | 0.14 |
| 7 | 68048592 | rs17670645 | 0.592 | -0.939 | -0.765 | -0.713 | 0.013 | 0.07 | 0.07 | 0 | 0.2 | 0.06 | 0.12 | 0.07 | 0.16 | 0.16 |
| 7 | 68050119 | rs1522499 | 0.089 | -1.003 | -0.809 | -0.564 | 0 | -0.006 | -0.006 | 0 | 0.58 | 0.4 | 0.6 | 0.7 | 0.49 | 0.55 |
| 7 | 68051008 | rs2090934 | -2.112 | -0.674 | -0.905 | -0.6 | -0.08 | 0.081 | 0.081 | 0 | 0.82 | 0.93 | 0.84 | 1 | 0.69 | 0.9 |
| 7 | 68051712 | rs10265554 | 1.41 | -0.976 | -0.555 | -0.229 | 0.012 | 0.006 | 0.006 | 0 | 0.1 | 0.28 | 0.08 | 0.05 | 0.08 | 0.15 |
| 7 | 68058102 | rs6946893 | 0.832 | -1.149 | -0.942 | 0.231 | 0.001 | -0.017 | -0.017 | 4.774 | 0.32 | 0.22 | 0.38 | 0.4 | 0.3 | 0.33 |
| 7 | 68058887 | rs17139865 | 1.657 | -1.583 | -0.913 | 0.415 | -0.025 | -0.01 | -0.01 | 0 | 0.06 | 0.08 | 0.02 | 0.06 | 0.02 | 0.04 |
| 7 | 68059881 | rs9638547 | -0.429 | -1.473 | -0.873 | 0.385 | 0.006 | -0.001 | -0.001 | 0 | 0.26 | 0.19 | 0.2 | 0.3 | 0.18 | 0.23 |
| 7 | 68068890 | rs13243699 | -0.099 | -1.285 | -1.163 | 0.28 | 0.019 | 0.024 | 0.024 | 0 | 0.33 | 0.24 | 0.16 | 0.16 | 0.24 | 0.25 |
| 7 | 68070039 | rs4718793 | 0.739 | -1.27 | -1.212 | 0.316 | -0.002 | -0.022 | -0.022 | 0 | 0.26 | 0.22 | 0.51 | 0.58 | 0.31 | 0.35 |
| 7 | 68071508 | rs11974253 | 1.046 | -1.296 | -1.21 | 0.5 | -0.001 | -0.029 | -0.029 | 0 | 0.22 | 0.16 | 0.39 | 0.41 | 0.25 | 0.27 |
| 7 | 68073390 | rs10428951 | -0.675 | -1.676 | -0.99 | 0.285 | -0.046 | 0.041 | 0.041 | 0 | 0.81 | 0.54 | 0.86 | 0.74 | 0.89 | 0.83 |
| 7 | 68083568 | rs4535613 | 0.275 | -1.754 | -1.269 | 0.433 | -0.001 | 0.001 | 0.001 | 0 | 0.33 | 0.36 | 0.39 | 0.58 | 0.32 | 0.39 |
| 7 | 68084026 | rs4499981 | 0.802 | -1.804 | -1.309 | 0.373 | 0.019 | 0.065 | 0.065 | 0 | 0.45 | 0.23 | 0.46 | 0.31 | 0.57 | 0.48 |
| 7 | 68087967 | rs4262230 | -1.554 | -1.893 | -1.052 | 0.283 | -0.021 | -0.035 | -0.035 | 0 | 0.87 | 0.76 | 0.93 | 0.95 | 0.93 | 0.91 |
| 7 | 68088133 | rs4507644 | 1.652 | -1.77 | -1.752 | 0.181 | -0.001 | 0.002 | 0.002 | 0 | 0.19 | 0.18 | 0.21 | 0.3 | 0.19 | 0.25 |
| 7 | 68088138 | rs4419706 | 0.674 | -1.76 | -1.712 | 0.223 | 0.019 | 0.072 | 0.072 | 0 | 0.45 | 0.23 | 0.39 | 0.3 | 0.57 | 0.48 |
| 7 | 68090345 | rs4717464 | 1.178 | -1.406 | -2.193 | 0.136 | -0.005 | -0.008 | -0.008 | 0 | 0.18 | 0.05 | 0.25 | 0.17 | 0.15 | 0.16 |
| 7 | 68092473 | rs7794902 | 0.623 | -0.901 | -1.991 | 0.053 | -0.059 | 0.124 | 0.124 | 0 | 0.34 | 0.18 | 0.27 | 0.06 | 0.42 | 0.26 |
| 7 | 68099251 | rs17750362 | 1.578 | -0.895 | -1.827 | 0.722 | -0.026 | -0.042 | -0.042 | 0 | 0.08 | 0.01 | 0.12 | 0.13 | 0.09 | 0.16 |
| 7 | 68099467 | rs17682856 | 1.877 | -0.781 | -1.716 | 0.995 | -0.003 | -0.04 | -0.04 | 2.91 | 0.07 | 0.01 | 0.04 | 0.16 | 0.06 | 0.06 |
| 7 | 68100045 | rs17139879 | 1.426 | -0.643 | -1.506 | 0.751 | -0.002 | 0 | 0 | 0 | 0.08 | 0.07 | 0.08 | 0.06 | 0.04 | 0.04 |
| 7 | 68101280 | rs4718795 | 0.174 | -0.078 | -1.499 | 0.848 | -0.001 | -0.001 | -0.001 | 0.841 | 0.07 | 0.1 | 0.21 | 0.1 | 0.08 | 0.04 |
| 7 | 68104188 | rs10280848 | 0.393 | 0.153 | -1.342 | 0.867 | 0.011 | 0.012 | 0.012 | 0 | 0.47 | 0.61 | 0.55 | 0.69 | 0.42 | 0.5 |
| 7 | 68111061 | rs7782570 | 0.811 | 0.292 | -1.481 | 0.309 | -0.105 | 0.032 | 0.032 | 0 | 0.21 | 0.03 | 0.16 | 0.1 | 0.34 | 0.14 |
| 7 | 68113660 | rs12672160 | -0.034 | -0.402 | -0.467 | 0.479 | -0.013 | -0.002 | -0.002 | 0 | 0.42 | 0.45 | 0.31 | 0.43 | 0.35 | 0.41 |
| 7 | 68127655 | rs2868923 | 0.411 | -0.481 | 0.046 | 0.4 | -0.032 | 0.009 | 0.009 | 0.01 | 0.86 | 0.71 | 0.79 | 0.9 | 0.93 | 0.87 |
| 7 | 68128716 | rs6969932 | 0.238 | -0.306 | -0.055 | 0.221 | 0.001 | 0.001 | 0.001 | 0 | 0.4 | 0.62 | 0.26 | 0.38 | 0.34 | 0.42 |
| 7 | 68132028 | rs4718801 | -0.171 | -0.529 | -0.287 | 0.021 | -0.005 | -0.003 | -0.003 | 0.02 | 0.79 | 0.78 | 0.69 | 0.79 | 0.84 | 0.88 |
| 7 | 68136274 | rs11768279 | 0.87 | -0.373 | 0.462 | 0.083 | 0.007 | -0.05 | -0.05 | 3.476 | 0.31 | 0.13 | 0.19 | 0.39 | 0.27 | 0.34 |
| 7 | 68138053 | rs12531895 | 0.462 | -0.378 | 0.829 | 0.058 | -0.001 | 0 | 0 | 0 | 0.36 | 0.38 | 0.22 | 0.4 | 0.31 | 0.41 |
| 7 | 68146505 | rs11981119 | 0.835 | -0.306 | 0.793 | 0.088 | 0.014 | -0.021 | -0.021 | 0 | 0.32 | 0.21 | 0.2 | 0.38 | 0.27 | 0.36 |
| 7 | 68149915 | rs10807719 | 0.21 | -0.332 | 0.94 | 0.049 | -0.002 | 0 | 0 | 0 | 0.39 | 0.61 | 0.25 | 0.4 | 0.31 | 0.42 |
| 7 | 68154951 | rs10234474 | 0.629 | -0.235 | 0.786 | 0.108 | -0.013 | 0.01 | 0.01 | 0 | 0.34 | 0.4 | 0.21 | 0.39 | 0.27 | 0.35 |
| 7 | 68166590 | rs4269418 | 0.75 | 0.023 | 0.639 | 0.156 | 0.009 | -0.001 | -0.001 | 0 | 0.38 | 0.28 | 0.21 | 0.39 | 0.31 | 0.35 |
| 7 | 68166849 | rs4632950 | 0.901 | 0.024 | 0.562 | 0.157 | 0.026 | -0.01 | -0.01 | 0 | 0.36 | 0.17 | 0.2 | 0.39 | 0.31 | 0.35 |
| 7 | 68185001 | rs11982155 | 0.864 | 0.608 | 0.193 | -0.025 | -0.001 | 0.062 | 0.062 | 0 | 0.22 | 0.1 | 0.12 | 0.11 | 0.21 | 0.19 |
| 7 | 68191264 | rs1002606 | 0.333 | 0.603 | 0.497 | 0.176 | -0.003 | -0.001 | -0.001 | 0 | 0.39 | 0.43 | 0.21 | 0.39 | 0.31 | 0.4 |
| 7 | 68196278 | rs17139956 | -0.38 | 1.371 | 0.522 | 0.265 | 0.052 | -0.024 | -0.024 | 0.026 | 0.21 | 0.37 | 0.39 | 0.18 | 0.29 | 0.18 |
| 7 | 68198661 | rs990446 | 0.352 | 1.443 | -0.331 | -0.046 | -0.061 | -0.072 | -0.072 | 0 | 0.41 | 0.12 | 0.35 | 0.51 | 0.43 | 0.46 |
| 7 | 68214762 | rs2527672 | -0.622 | 1.455 | -0.343 | -0.049 | 0 | 0.015 | 0.015 | 0 | 0.61 | 0.52 | 0.59 | 0.57 | 0.61 | 0.57 |
| 7 | 68218637 | rs2527670 | -0.78 | 1.524 | -0.774 | -0.071 | 0.001 | 0.004 | 0.004 | 0 | 0.62 | 0.65 | 0.61 | 0.58 | 0.61 | 0.57 |
| 7 | 68229868 | rs2057917 | -0.327 | 1.851 | -1.563 | 0.529 | -0.038 | -0.046 | -0.046 | 0.194 | 0.42 | 0.15 | 0.35 | 0.51 | 0.43 | 0.51 |
| 7 | 68230246 | rs2057916 | -0.151 | 0.526 | -1.383 | 0.691 | 0.036 | -0.057 | -0.057 | 0 | 0.38 | 0.15 | 0.29 | 0.51 | 0.31 | 0.41 |
| 7 | 68234449 | rs17341408 | -0.973 | 1.213 | -1.041 | 0.791 | -0.004 | -0.003 | -0.003 | 0 | 0.25 | 0.25 | 0.36 | 0.28 | 0.25 | 0.3 |
| 7 | 68235087 | rs757784 | 0.308 | 0.867 | -1.479 | 0.568 | 0.03 | -0.009 | -0.009 | 0 | 0.45 | 0.29 | 0.46 | 0.45 | 0.44 | 0.41 |
| 7 | 68235512 | rs2428834 | 0.171 | 1.076 | -1.311 | -0.045 | -0.012 | 0.031 | 0.031 | 0 | 0.73 | 0.8 | 0.65 | 0.93 | 0.69 | 0.69 |
| 7 | 68243122 | rs2711516 | 0.395 | 0.984 | -1.296 | -0.289 | -0.02 | 0.074 | 0.074 | 0 | 0.67 | 0.79 | 0.64 | 0.92 | 0.62 | 0.67 |
| 7 | 68244215 | rs2711483 | 0.553 | 0.969 | -0.714 | -0.156 | -0.008 | 0.02 | 0.02 | 0 | 0.64 | 0.67 | 0.62 | 0.88 | 0.6 | 0.64 |
| 7 | 68247814 | rs2711481 | 0.348 | 1.008 | -0.743 | -0.126 | -0.02 | 0.068 | 0.068 | 0 | 0.67 | 0.78 | 0.63 | 0.92 | 0.62 | 0.67 |
| 7 | 68253974 | rs2869615 | 0.364 | 0.835 | 0.19 | 0.103 | -0.021 | 0.102 | 0.102 | 0 | 0.27 | 0.4 | 0.18 | 0.5 | 0.21 | 0.31 |
| 7 | 68254550 | rs1859578 | 0.364 | 0.835 | 0.19 | 0.103 | -0.021 | 0.102 | 0.102 | 0 | 0.27 | 0.4 | 0.18 | 0.5 | 0.21 | 0.31 |
| 7 | 68262350 | rs2527636 | 0.388 | 1.171 | 0.395 | 0.266 | 0.01 | -0.044 | -0.044 | 0 | 0.66 | 0.55 | 0.52 | 0.85 | 0.6 | 0.57 |
| 7 | 68270995 | rs9654886 | 1.21 | 1.055 | -0.353 | 0.12 | -0.002 | -0.038 | -0.038 | 0 | 0.37 | 0.12 | 0.32 | 0.39 | 0.35 | 0.32 |
| 7 | 68285442 | rs2107945 | 0.184 | 1.294 | -0.372 | -0.02 | -0.015 | 0.116 | 0.116 | 0 | 0.28 | 0.08 | 0.31 | 0.05 | 0.28 | 0.31 |
| 7 | 68286270 | rs10265677 | -0.455 | 1.234 | 0.404 | -0.067 | -0.002 | -0.032 | -0.032 | 0 | 0.35 | 0.6 | 0.36 | 0.08 | 0.3 | 0.39 |
| 7 | 68287168 | rs6460468 | -0.22 | 1.443 | 0.26 | 0.74 | -0.019 | -0.058 | -0.058 | 0 | 0.77 | 0.92 | 0.79 | 0.57 | 0.72 | 0.82 |
| 7 | 68295086 | rs757783 | -1.752 | 1.286 | 1.14 | 0.716 | 0.105 | 0.184 | 0.184 | 1.751 | 0.07 | 0.56 | 0.14 | 0.1 | 0.05 | 0.06 |

| | | | | | | | | | | | | | | | | |
|---|---|---|---|---|---|---|---|---|---|---|---|---|---|---|---|---|
| 7 | 68319863 | rs4327750 | 0.012 | 1.528 | 0.279 | 0.43 | -0.01 | 0.038 | 0.038 | 0 | 0.27 | 0.15 | 0.34 | 0.14 | 0.27 | 0.23 |
| 7 | 68320522 | rs10950153 | -0.046 | 1.185 | 0.334 | 0.73 | -0.012 | 0.05 | 0.05 | 0 | 0.2 | 0.31 | 0.32 | 0.42 | 0.14 | 0.25 |
| 7 | 68326660 | rs10237067 | -1.125 | 1.891 | 0.81 | 0.721 | 0.014 | 0.036 | 0.036 | 0 | 0.55 | 0.74 | 0.8 | 0.68 | 0.6 | 0.58 |
| 7 | 68337786 | rs2215558 | -1.176 | 1.828 | 0.74 | 1.041 | 0.019 | 0.034 | 0.034 | 0 | 0.57 | 0.74 | 0.81 | 0.7 | 0.62 | 0.6 |
| 7 | 68345599 | rs10271980 | -0.873 | 1.855 | 0.734 | 1.058 | -0.002 | -0.003 | -0.003 | 0 | 0.53 | 0.55 | 0.73 | 0.58 | 0.57 | 0.54 |
| 7 | 68349402 | rs11977562 | 0.483 | 1.721 | 0.805 | 1.071 | -0.023 | 0.03 | 0.03 | 0 | 0.15 | 0.23 | 0.14 | 0.28 | 0.07 | 0.14 |
| 7 | 68352718 | rs13245506 | 0.004 | 1.789 | 0.652 | 1.073 | -0.016 | -0.012 | -0.012 | 0.112 | 0.08 | 0.07 | 0.23 | 0.14 | 0.2 | 0.15 |
| 7 | 68354674 | rs12698742 | -0.588 | 1.748 | 0.68 | 1.048 | 0.005 | 0.037 | 0.037 | 0 | 0.23 | 0.35 | 0.37 | 0.42 | 0.27 | 0.29 |
| 7 | 68356415 | rs2191749 | -0.403 | 1.661 | 0.355 | 0.944 | 0.025 | 0.035 | 0.035 | 0 | 0.05 | 0.13 | 0.09 | 0.15 | 0.08 | 0.09 |
| 7 | 68358970 | rs10226583 | -0.566 | 1.431 | 0.165 | 0.711 | 0.001 | 0.001 | 0.001 | 0 | 0.51 | 0.54 | 0.73 | 0.56 | 0.55 | 0.51 |
| 7 | 68365245 | rs6978595 | 0.598 | 1.522 | -0.299 | 0.873 | 0.013 | 0.019 | 0.019 | 0 | 0.06 | 0.12 | 0.2 | 0.15 | 0.1 | 0.12 |
| 7 | 68372748 | rs2079975 | -1.016 | 1.357 | -0.557 | 0.539 | 0.001 | -0.017 | -0.017 | 0 | 0.62 | 0.86 | 0.8 | 0.59 | 0.63 | 0.57 |
| 7 | 68379401 | rs7786650 | -0.031 | 1.753 | -0.356 | 0.144 | -0.032 | 0.028 | 0.028 | 0 | 0.05 | 0 | 0.05 | 0 | 0.13 | 0.06 |
| 7 | 68384522 | rs2869902 | -0.592 | 1.586 | -0.422 | -0.095 | -0.017 | 0.068 | 0.068 | 1.44 | 0.4 | 0.7 | 0.52 | 0.13 | 0.43 | 0.31 |
| 7 | 68387791 | rs10950156 | -0.305 | 1.619 | -0.164 | -0.03 | 0.014 | -0.023 | -0.023 | 0.551 | 0.07 | 0.1 | 0.07 | 0 | 0.13 | 0.06 |
| 7 | 68405136 | rs2869905 | -0.519 | 1.183 | -0.123 | 0.033 | -0.016 | 0.057 | 0.057 | 0.656 | 0.4 | 0.7 | 0.52 | 0.13 | 0.43 | 0.3 |
| 7 | 68409806 | rs11773125 | -0.164 | 1.23 | 0.163 | 0.103 | -0.007 | 0.009 | 0.009 | 1.67 | 0.34 | 0.27 | 0.43 | 0.13 | 0.41 | 0.29 |
| 7 | 68412653 | rs7806002 | -0.663 | 1.196 | 0.098 | 0.303 | -0.001 | -0.001 | -0.001 | 0 | 0.43 | 0.43 | 0.63 | 0.28 | 0.53 | 0.42 |
| 7 | 68413318 | rs13228923 | 0.685 | 0.75 | -0.102 | 0.496 | -0.005 | -0.012 | -0.012 | 0 | 0.35 | 0.29 | 0.4 | 0.53 | 0.37 | 0.47 |
| 7 | 68415653 | rs17671261 | -0.238 | 0.6 | 0.362 | 0.192 | -0.018 | 0.045 | 0.045 | 0.996 | 0.32 | 0.23 | 0.28 | 0.09 | 0.39 | 0.28 |
| 7 | 68417790 | rs6951384 | -0.307 | 0.625 | 0.317 | 0.203 | 0.015 | -0.074 | -0.074 | 0 | 0.33 | 0.43 | 0.29 | 0.09 | 0.39 | 0.28 |
| 7 | 68421575 | rs7784905 | 0.931 | 0.453 | -0.417 | 0.615 | -0.024 | 0.004 | 0.004 | 0.41 | 0.42 | 0.28 | 0.39 | 0.41 | 0.48 | 0.28 |
| 7 | 68423676 | rs1476865 | -0.746 | 0.467 | -0.24 | 0.858 | 0 | 0 | 0 | 0.634 | 0.52 | 0.53 | 0.56 | 0.51 | 0.47 | 0.68 |
| 7 | 68424896 | rs10464154 | -0.308 | 0.332 | 0.006 | 1.4 | 0.007 | -0.002 | -0.002 | 0 | 0.16 | 0.18 | 0.28 | 0.48 | 0.1 | 0.31 |
| 7 | 68425037 | rs10464189 | -0.395 | 0.306 | 0.255 | 1.354 | -0.011 | 0.038 | 0.038 | 0 | 0.16 | 0.24 | 0.28 | 0.44 | 0.1 | 0.29 |
| 7 | 68425573 | rs7805210 | 0.888 | 0.448 | 0.755 | 0.52 | 0.005 | -0.084 | -0.084 | 0 | 0.79 | 0.54 | 0.72 | 0.95 | 0.81 | 0.87 |
| 7 | 68429563 | rs6969978 | 1.527 | 0.875 | 1.171 | 0.929 | 0.09 | 0.211 | 0.211 | 0 | 0.89 | 0.64 | 0.88 | 0.72 | 0.84 | 0.88 |
| 7 | 68433598 | rs1017911 | 0.083 | 0.827 | 1.402 | 0.852 | 0.014 | -0.01 | -0.01 | 0 | 0.1 | 0.22 | 0.17 | 0.06 | 0.11 | 0.11 |
| 7 | 68434641 | rs11505255 | 0.108 | 0.827 | 1.395 | 0.852 | 0.014 | -0.01 | -0.01 | 0 | 0.1 | 0.21 | 0.16 | 0.06 | 0.11 | 0.11 |
| 7 | 68438954 | rs1010779 | 0.861 | 0.838 | 1.253 | 0.851 | 0.001 | -0.059 | -0.059 | 0.207 | 0.83 | 0.52 | 0.79 | 0.94 | 0.85 | 0.83 |
| 7 | 68446151 | rs4718837 | 2.085 | 0.897 | 1.035 | 0.716 | -0.018 | -0.019 | -0.019 | 0.473 | 0.72 | 0.32 | 0.68 | 0.65 | 0.65 | 0.72 |
| 7 | 68448569 | rs7805438 | 2.541 | 0.669 | 0.617 | 0.495 | -0.075 | -0.043 | -0.043 | 0 | 0.58 | 0.17 | 0.59 | 0.6 | 0.53 | 0.58 |
| 7 | 68455500 | rs1485693 | 1.36 | 0.976 | 0.992 | 0.681 | 0.016 | -0.088 | -0.088 | 0.329 | 0.79 | 0.52 | 0.76 | 0.94 | 0.77 | 0.77 |
| 7 | 68457986 | rs17140203 | -0.415 | 0.359 | 0.925 | 0.604 | 0.002 | -0.028 | -0.028 | 0 | 0.12 | 0.24 | 0.17 | 0.05 | 0.11 | 0.11 |
| 7 | 68458474 | rs7810725 | 1.38 | 0.161 | 0.442 | 0.624 | 0.017 | 0.005 | 0.005 | 0 | 0.5 | 0.34 | 0.39 | 0.43 | 0.57 | 0.37 |
| 7 | 68458785 | rs11771792 | -0.985 | 0.383 | 0.697 | 0.618 | 0.016 | -0.025 | -0.025 | 0.492 | 0.15 | 0.24 | 0.22 | 0.1 | 0.17 | 0.12 |
| 7 | 68459585 | rs4143504 | 0.978 | 0.003 | 0.321 | 0.567 | -0.023 | -0.071 | -0.071 | 0 | 0.84 | 0.47 | 0.79 | 0.95 | 0.89 | 0.85 |
| 7 | 68461525 | rs6460477 | 0.811 | 0 | 0.405 | 0.578 | -0.014 | -0.065 | -0.065 | 0.121 | 0.85 | 0.53 | 0.79 | 0.96 | 0.89 | 0.85 |
| 7 | 68462060 | rs16868315 | -0.336 | -0.168 | 0.22 | 0.853 | 0.013 | -0.024 | -0.024 | 0 | 0.16 | 0.13 | 0.25 | 0.29 | 0.1 | 0.27 |
| 7 | 68464812 | rs10950162 | 0.941 | -0.263 | 0.249 | 0.654 | -0.029 | -0.081 | -0.081 | 0 | 0.82 | 0.48 | 0.82 | 0.95 | 0.88 | 0.85 |
| 7 | 68473340 | rs2024237 | 0.351 | -0.278 | 0.381 | 0.642 | 0.006 | -0.028 | -0.028 | 0 | 0.94 | 0.87 | 0.96 | 0.98 | 0.95 | 0.92 |
| 7 | 68479862 | rs7793273 | 1.467 | -0.175 | 0.286 | 0.75 | -0.003 | -0.021 | -0.021 | 0.012 | 0.74 | 0.44 | 0.73 | 0.78 | 0.77 | 0.73 |
| 7 | 68486634 | rs2869738 | 2.343 | -0.084 | -0.209 | 0.687 | -0.141 | 0.119 | 0.119 | 0 | 0.4 | 0.02 | 0.3 | 0.18 | 0.48 | 0.36 |
| 7 | 68504050 | rs2904188 | 0.22 | -0.393 | 0.676 | 0.609 | 0.02 | 0.014 | 0.014 | 0 | 0.9 | 0.79 | 0.89 | 0.83 | 0.88 | 0.89 |
| 7 | 68505785 | rs9632553 | 2.346 | -0.583 | 0.622 | 0.629 | 0.037 | -0.06 | -0.06 | 0 | 0.61 | 0.16 | 0.49 | 0.47 | 0.71 | 0.51 |
| 7 | 68508054 | rs11561744 | 1.509 | -0.515 | 0.8 | 0.676 | -0.013 | 0.064 | 0.064 | 0.997 | 0.78 | 0.59 | 0.66 | 0.53 | 0.83 | 0.67 |
| 7 | 68508645 | rs2062403 | 2.133 | -0.517 | 0.839 | 0.674 | 0.011 | -0.013 | -0.013 | 0 | 0.67 | 0.3 | 0.53 | 0.51 | 0.76 | 0.59 |
| 7 | 68514117 | rs767638 | -0.28 | -0.485 | 0.69 | 0.708 | 0.004 | 0.001 | 0.001 | 0 | 0.11 | 0.14 | 0.24 | 0.3 | 0.06 | 0.22 |
| 7 | 68523481 | rs12540557 | -0.519 | -0.31 | 0.701 | 0.709 | 0.011 | -0.013 | -0.013 | 0 | 0.22 | 0.18 | 0.32 | 0.3 | 0.18 | 0.24 |
| 7 | 68526895 | rs923135 | -0.162 | -0.314 | 0.654 | 0.683 | -0.009 | -0.034 | -0.034 | 0.351 | 0.11 | 0.3 | 0.13 | 0.02 | 0.07 | 0.08 |
| 7 | 68531960 | rs764049 | -0.934 | -0.312 | 0.443 | 0.692 | -0.055 | -0.042 | -0.042 | 0 | 0.16 | 0.49 | 0.15 | 0.06 | 0.07 | 0.11 |
| 7 | 68532146 | rs3113288 | -2.364 | -0.262 | -0.141 | 0.727 | 0.021 | -0.045 | -0.045 | 2.107 | 0.49 | 0.86 | 0.59 | 0.54 | 0.41 | 0.49 |
| 7 | 68535404 | rs17683343 | -0.658 | 0.016 | -0.18 | 0.791 | 0.015 | -0.039 | -0.039 | 0.142 | 0.21 | 0.11 | 0.32 | 0.3 | 0.18 | 0.24 |
| 7 | 68539977 | rs2869745 | -0.912 | 0.536 | 0.321 | 0.666 | 0.013 | 0.007 | 0.007 | 0 | 0.09 | 0.11 | 0.1 | 0.17 | 0.12 | 0.09 |
| 7 | 68543159 | rs17140325 | -0.908 | 0.123 | 0.372 | 0.447 | -0.021 | -0.016 | -0.016 | 0 | 0.11 | 0.34 | 0.22 | 0.04 | 0.06 | 0.07 |
| 7 | 68544052 | rs3102817 | -1.071 | 0.336 | -1.035 | 0.431 | -0.011 | 0.019 | 0.019 | 0 | 0.44 | 0.95 | 0.53 | 0.33 | 0.39 | 0.41 |
| 7 | 68548693 | rs17683378 | 0.196 | 0.25 | -0.989 | 0.528 | 0.006 | 0.012 | 0.012 | 0 | 0.06 | 0.03 | 0.08 | 0.3 | 0.04 | 0.12 |
| 7 | 68549573 | rs3102809 | -0.376 | 0.243 | -0.201 | 0.3 | 0.002 | -0.082 | -0.082 | 0 | 0.3 | 0.39 | 0.4 | 0.05 | 0.32 | 0.22 |
| 7 | 68549627 | rs11767085 | 0.004 | 0.465 | -0.523 | 0.249 | -0.041 | -0.002 | -0.002 | 0.054 | 0.08 | 0 | 0.09 | 0.11 | 0.11 | 0.11 |
| 7 | 68549929 | rs3113304 | -1.433 | 0.22 | -0.114 | 0.335 | 0.017 | 0.028 | 0.028 | 0 | 0.44 | 0.87 | 0.53 | 0.37 | 0.41 | 0.4 |
| 7 | 68553958 | rs3102796 | -0.533 | 0.123 | -0.257 | 0.243 | -0.001 | -0.014 | -0.014 | 0.365 | 0.3 | 0.29 | 0.39 | 0.05 | 0.32 | 0.22 |
| 7 | 68561321 | rs3113316 | -1.024 | 0.36 | -0.858 | 0.587 | 0.01 | -0.008 | -0.008 | 0 | 0.44 | 0.87 | 0.62 | 0.41 | 0.43 | 0.42 |
| 7 | 68564589 | rs10215141 | -1.089 | 0.401 | 0.052 | 0.89 | -0.006 | 0.09 | 0.09 | 0 | 0.23 | 0.41 | 0.37 | 0.51 | 0.22 | 0.24 |
| 7 | 68567914 | rs10277599 | -1.543 | 0.293 | 0.446 | 1.764 | -0.002 | 0.233 | 0.233 | 0.18 | 0.05 | 0.2 | 0.2 | 0.51 | 0.05 | 0.06 |
| 7 | 68569292 | rs2103036 | -0.462 | 0.171 | -0.265 | 0.39 | -0.007 | 0.047 | 0.047 | 0 | 0.56 | 0.38 | 0.67 | 0.96 | 0.55 | 0.74 |
| 7 | 68569986 | rs12539135 | -0.019 | 0.535 | -0.297 | 0.595 | 0.019 | 0.141 | 0.141 | 0 | 0.49 | 0.16 | 0.44 | 0.4 | 0.44 | 0.66 |
| 7 | 68573396 | rs10229812 | -0.602 | -0.256 | 0.391 | -0.109 | 0.004 | -0.055 | -0.055 | 0.128 | 0.4 | 0.55 | 0.61 | 0.88 | 0.44 | 0.42 |
| 7 | 68574359 | rs12698769 | -0.037 | 0.359 | 0.652 | 0.256 | 0.017 | 0.141 | 0.141 | 0 | 0.49 | 0.16 | 0.43 | 0.4 | 0.44 | 0.66 |
| 7 | 68574861 | rs4717488 | -0.201 | 0.357 | 0.645 | 0.238 | -0.008 | 0.035 | 0.035 | 0 | 0.52 | 0.37 | 0.48 | 0.48 | 0.49 | 0.68 |

| | | | | | | | | | | | | | | | | |
|---|---|---|---|---|---|---|---|---|---|---|---|---|---|---|---|---|
| 7 | 68593216 | rs6976718 | -0.496 | 0.078 | 0.37 | -0.115 | -0.026 | 0.124 | 0.124 | 0 | 0.58 | 0.75 | 0.66 | 0.94 | 0.49 | 0.73 |
| 7 | 68594912 | rs10464191 | -0.242 | 0.272 | -0.288 | -0.031 | -0.014 | 0.201 | 0.201 | 0 | 0.53 | 0.2 | 0.63 | 0.92 | 0.49 | 0.72 |
| 7 | 68610860 | rs12234865 | -1.323 | 0.848 | 0.479 | 1.105 | 0.052 | 0.341 | 0.341 | 0 | 0.06 | 0.21 | 0.19 | 0.5 | 0.09 | 0.08 |
| 7 | 68624342 | rs1195218 | 0.611 | 0.996 | -0.221 | 0.498 | -0.021 | 0.034 | 0.034 | 0 | 0.06 | 0 | 0.04 | 0 | 0.08 | 0.02 |
| 7 | 68625525 | rs2870105 | 0.195 | 0.676 | -0.356 | 0.82 | 0.005 | -0.067 | -0.067 | 1.701 | 0.14 | 0.04 | 0.2 | 0.47 | 0.14 | 0.07 |
| 7 | 68636019 | rs10238378 | -0.951 | 1.263 | -0.19 | 0.367 | -0.063 | -0.011 | -0.011 | 0 | 0.3 | 0.11 | 0.53 | 0.7 | 0.38 | 0.39 |
| 7 | 68649968 | rs1195261 | 1.245 | 0.331 | 0.189 | 0.123 | -0.007 | 0.024 | 0.024 | 0 | 0.57 | 0.34 | 0.57 | 0.38 | 0.54 | 0.6 |
| 7 | 68653387 | rs3846983 | 0.319 | 0.343 | 0.436 | 0.084 | -0.001 | -0.013 | -0.013 | 0 | 0.18 | 0.21 | 0.29 | 0.56 | 0.2 | 0.11 |
| 7 | 68656780 | rs1195265 | 1.02 | 0.131 | 0.016 | -0.348 | -0.061 | 0.081 | 0.081 | 0.004 | 0.29 | 0.13 | 0.4 | 0.2 | 0.35 | 0.31 |
| 7 | 68659880 | rs1915973 | 1.282 | 0.286 | 0.011 | -0.347 | -0.012 | 0.016 | 0.016 | 0 | 0.08 | 0.03 | 0.05 | 0.04 | 0.11 | 0.15 |
| 7 | 68664061 | rs1718582 | 1.062 | 0.259 | 0.255 | -0.361 | -0.047 | 0.069 | 0.069 | 0.002 | 0.29 | 0.16 | 0.4 | 0.19 | 0.35 | 0.31 |
| 7 | 68664836 | rs4717494 | 0.932 | -0.033 | 0.261 | -0.36 | 0 | 0.018 | 0.018 | 0 | 0.17 | 0.2 | 0.25 | 0.48 | 0.15 | 0.1 |
| 7 | 68665692 | rs2049250 | -0.57 | 0.092 | 0.054 | -0.596 | 0.004 | -0.026 | -0.026 | 0 | 0.23 | 0.27 | 0.13 | 0.06 | 0.26 | 0.3 |
| 7 | 68670227 | rs35207662 | 0.835 | 0.052 | -0.027 | -0.59 | 0.002 | -0.009 | -0.009 | 0.284 | 0.09 | 0.12 | 0.04 | 0.02 | 0.09 | 0.1 |
| 7 | 68673096 | rs1543239 | 1.337 | 0.016 | -0.127 | -0.545 | 0.022 | 0.018 | 0.018 | 0 | 0.28 | 0.19 | 0.18 | 0.19 | 0.19 | 0.29 |
| 7 | 68673806 | rs10228728 | -0.519 | 0.511 | -0.262 | -0.194 | 0 | -0.007 | -0.007 | 0 | 0.14 | 0.12 | 0.08 | 0.04 | 0.17 | 0.19 |
| 7 | 68675461 | rs12333528 | 0.346 | 0.593 | -0.214 | -0.115 | 0.001 | -0.009 | -0.009 | 0.23 | 0.09 | 0.07 | 0.06 | 0.03 | 0.11 | 0.15 |
| 7 | 68681198 | rs17603178 | 0.013 | 0.133 | -0.32 | 0.199 | -0.001 | -0.003 | -0.003 | 0.057 | 0.09 | 0.11 | 0.04 | 0.02 | 0.08 | 0.1 |
| 7 | 68696421 | rs17751099 | -0.493 | -0.787 | -1.059 | 0.824 | 0.014 | -0.105 | -0.105 | 0.005 | 0.24 | 0.07 | 0.26 | 0.42 | 0.23 | 0.16 |
| 7 | 68704394 | rs6953746 | 0.544 | -0.782 | -0.933 | 0.437 | 0.03 | -0.012 | -0.012 | 0.328 | 0.24 | 0.13 | 0.1 | 0.23 | 0.12 | 0.3 |
| 7 | 68712121 | rs12056012 | 0.769 | -0.794 | -1.042 | 0.427 | 0.072 | 0.002 | 0.002 | 0 | 0.23 | 0.08 | 0.07 | 0.18 | 0.08 | 0.29 |
| 7 | 68718752 | rs10250140 | 0.338 | -0.936 | -0.958 | 0.329 | 0.022 | -0.001 | -0.001 | 1.896 | 0.58 | 0.36 | 0.61 | 0.57 | 0.66 | 0.71 |
| 7 | 68722152 | rs7802012 | -0.223 | -0.78 | -1.233 | 0.405 | 0.007 | -0.009 | -0.009 | 0 | 0.68 | 0.73 | 0.71 | 0.57 | 0.73 | 0.76 |
| 7 | 68724055 | rs7811944 | 0.247 | 0.125 | -1.265 | 0.534 | -0.079 | -0.01 | -0.01 | 0 | 0.34 | 0.14 | 0.36 | 0.32 | 0.53 | 0.43 |
| 7 | 68724740 | rs10807722 | 0.159 | -0.05 | -1.152 | 0.641 | -0.013 | 0.012 | 0.012 | 0 | 0.38 | 0.26 | 0.5 | 0.34 | 0.56 | 0.43 |
| 7 | 68729490 | rs4717496 | 0.276 | -0.167 | -0.829 | 0.96 | -0.001 | 0.005 | 0.005 | 0 | 0.73 | 0.71 | 0.68 | 0.54 | 0.73 | 0.75 |
| 7 | 68732517 | rs1718768 | -0.117 | -0.202 | -1.587 | 0.702 | 0.006 | 0.003 | 0.003 | 0.233 | 0.26 | 0.08 | 0.19 | 0.22 | 0.22 | 0.21 |
| 7 | 68734969 | rs1504274 | -0.689 | -0.097 | -1.292 | 1.121 | -0.004 | -0.004 | -0.004 | 0.091 | 0.19 | 0.17 | 0.32 | 0.22 | 0.23 | 0.12 |
| 7 | 68736093 | rs1195215 | 1.031 | -0.838 | -1.57 | 0.511 | 0.049 | 0.187 | 0.187 | 1.527 | 0.5 | 0.16 | 0.41 | 0.24 | 0.59 | 0.37 |
| 7 | 68738072 | rs1879873 | 0.326 | -0.891 | -0.593 | 0.955 | -0.018 | 0.034 | 0.034 | 0 | 0.14 | 0.31 | 0.09 | 0.16 | 0.08 | 0.19 |
| 7 | 68744504 | rs2001927 | 0.584 | -0.908 | -0.339 | 0.979 | -0.005 | 0.06 | 0.06 | 0.205 | 0.72 | 0.54 | 0.6 | 0.42 | 0.75 | 0.68 |
| 7 | 68748090 | rs1718607 | 1.666 | -1.007 | -0.213 | 0.902 | 0.01 | 0.062 | 0.062 | 0.41 | 0.57 | 0.26 | 0.47 | 0.25 | 0.62 | 0.47 |
| 7 | 68751088 | rs10258649 | -0.258 | -0.919 | -0.165 | 1.006 | -0.017 | 0.002 | 0.002 | 0 | 0.16 | 0.31 | 0.11 | 0.14 | 0.12 | 0.2 |
| 7 | 68772902 | rs12540889 | 0.418 | -0.882 | -0.463 | 0.923 | -0.018 | 0.042 | 0.042 | 0 | 0.08 | 0 | 0.17 | 0 | 0.09 | 0.06 |
| 7 | 68777454 | rs2062856 | 0.939 | -0.98 | -0.157 | 1.183 | -0.015 | 0.027 | 0.027 | 0 | 0.08 | 0.12 | 0.02 | 0.22 | 0.01 | 0.13 |
| 7 | 68784336 | rs2965445 | 1.568 | -1.045 | -0.175 | 1.042 | 0.02 | 0.087 | 0.087 | 1.192 | 0.55 | 0.21 | 0.47 | 0.24 | 0.62 | 0.47 |
| 7 | 68785411 | rs2938485 | -0.613 | -1.165 | 0.562 | 1.135 | -0.044 | 0.105 | 0.105 | 0 | 0.91 | 0.75 | 0.97 | 0.78 | 0.98 | 0.87 |
| 7 | 68790622 | rs12667664 | 1.598 | -1.11 | 0.499 | 0.927 | 0.006 | 0.017 | 0.017 | 1.356 | 0.59 | 0.34 | 0.51 | 0.25 | 0.66 | 0.47 |
| 7 | 68814378 | rs7786926 | -1.073 | -1.092 | 0.614 | 1.379 | 0.004 | -0.014 | -0.014 | 0.019 | 0.26 | 0.24 | 0.36 | 0.41 | 0.23 | 0.31 |
| 7 | 68824095 | rs2271032 | -0.058 | -1.11 | 0.659 | 1.508 | -0.022 | -0.059 | -0.059 | 0 | 0.15 | 0.02 | 0.24 | 0.21 | 0.15 | 0.13 |
| 7 | 68830773 | rs1317000 | 0.014 | -0.945 | 0.841 | 1.255 | 0 | 0.025 | 0.025 | 0 | 0.83 | 0.73 | 0.85 | 0.71 | 0.84 | 0.8 |
| 7 | 68833315 | rs2965441 | 0.58 | -0.992 | 1.034 | 1.233 | -0.004 | 0.058 | 0.058 | 0.215 | 0.81 | 0.5 | 0.84 | 0.71 | 0.84 | 0.8 |
| 7 | 68836921 | rs4457215 | -0.739 | -0.982 | 0.929 | 1.247 | -0.009 | 0.08 | 0.08 | 0 | 0.89 | 0.75 | 0.9 | 0.71 | 0.92 | 0.81 |
| 7 | 68838972 | rs2965447 | 2.112 | -1.018 | 0.731 | 1.166 | 0.024 | 0.063 | 0.063 | 0 | 0.53 | 0.23 | 0.47 | 0.28 | 0.61 | 0.47 |
| 7 | 68865799 | rs6957450 | 2.1 | -1.099 | 0.676 | 1.355 | 0.024 | 0.017 | 0.017 | 0 | 0.57 | 0.22 | 0.52 | 0.39 | 0.66 | 0.53 |
| 7 | 68884077 | rs10447529 | 1.952 | -1.114 | 0.768 | 1.234 | 0.015 | 0.041 | 0.041 | 0 | 0.54 | 0.29 | 0.47 | 0.28 | 0.62 | 0.44 |
| 7 | 68887938 | rs10243719 | 1.529 | -0.421 | 0.335 | 1.169 | -0.003 | -0.011 | -0.011 | 0 | 0.36 | 0.44 | 0.35 | 0.23 | 0.44 | 0.39 |
| 7 | 68889217 | rs4718885 | 0.322 | -0.325 | 0.383 | 1.098 | -0.011 | 0.023 | 0.023 | 0 | 0.81 | 0.7 | 0.84 | 0.7 | 0.84 | 0.75 |
| 7 | 68891086 | rs11978315 | 0.516 | -0.335 | 0.306 | 0.988 | -0.015 | 0.035 | 0.035 | 0 | 0.8 | 0.66 | 0.81 | 0.63 | 0.84 | 0.75 |
| 7 | 68895339 | rs11771719 | -1.044 | 0.447 | 0.193 | 1.011 | 0.011 | -0.002 | -0.002 | 0 | 0.43 | 0.35 | 0.39 | 0.36 | 0.38 | 0.34 |
| 7 | 68895393 | rs11760432 | 1.705 | -0.088 | 0.161 | 0.974 | 0.024 | 0.078 | 0.078 | 0 | 0.48 | 0.25 | 0.49 | 0.26 | 0.62 | 0.45 |
| 7 | 68898029 | rs12698800 | -0.345 | -0.225 | -0.153 | 1.206 | 0.012 | -0.017 | -0.017 | 0 | 0.3 | 0.25 | 0.32 | 0.33 | 0.23 | 0.29 |
| 7 | 68898379 | rs6460512 | -0.199 | -0.157 | 0.026 | 1.294 | 0.01 | 0.111 | 0.111 | 0 | 0.14 | 0.31 | 0.17 | 0.4 | 0.11 | 0.22 |
| 7 | 68905642 | rs6969183 | -1.276 | 0.027 | -0.541 | 0.41 | 0.025 | 0.042 | 0.042 | 0 | 0.47 | 0.58 | 0.48 | 0.74 | 0.35 | 0.51 |
| 7 | 68905738 | rs6959964 | -1.153 | 0.069 | -0.521 | 0.394 | 0.019 | 0.029 | 0.029 | 0 | 0.46 | 0.58 | 0.48 | 0.72 | 0.35 | 0.51 |
| 7 | 68908066 | rs6957234 | -1.06 | -0.088 | -0.606 | -0.193 | 0.003 | 0.001 | 0.001 | 0.958 | 0.41 | 0.64 | 0.51 | 0.85 | 0.34 | 0.6 |
| 7 | 68910326 | rs17140564 | -0.997 | 0.166 | -0.485 | 0.117 | -0.003 | -0.003 | -0.003 | 0 | 0.22 | 0.25 | 0.29 | 0.22 | 0.28 | 0.21 |
| 7 | 68913086 | rs16869103 | -0.763 | -0.286 | -0.738 | -0.205 | -0.025 | 0.044 | 0.044 | 0 | 0.25 | 0.05 | 0.25 | 0.13 | 0.19 | 0.2 |
| 7 | 68913353 | rs6460519 | 0.296 | -0.319 | -0.174 | 0.061 | -0.022 | 0.097 | 0.097 | 0 | 0.16 | 0.23 | 0.17 | 0.63 | 0.1 | 0.29 |
| 7 | 68917040 | rs763391 | -1.778 | -0.387 | 0.152 | -0.175 | 0.007 | -0.001 | -0.001 | 0 | 0.28 | 0.47 | 0.34 | 0.2 | 0.26 | 0.26 |
| 7 | 68927579 | rs6950764 | -0.022 | -0.495 | -0.012 | -0.256 | -0.01 | -0.026 | -0.026 | 0 | 0.05 | 0.19 | 0.03 | 0.01 | 0.02 | 0.08 |
| 7 | 68963429 | rs2865170 | -1.112 | -0.441 | -0.058 | -0.214 | 0.004 | 0.006 | 0.006 | 0.042 | 0.19 | 0.26 | 0.28 | 0.18 | 0.2 | 0.19 |
| 7 | 68968759 | rs17349245 | 0.864 | -0.532 | -0.28 | -0.307 | 0.019 | 0.01 | 0.01 | 0 | 0.14 | 0.04 | 0.05 | 0.14 | 0.06 | 0.04 |
| 7 | 68995564 | rs17140651 | -0.205 | -0.425 | -0.213 | -0.266 | -0.022 | 0.01 | 0.01 | 0 | 0.11 | 0.06 | 0.21 | 0.07 | 0.14 | 0.14 |
| 7 | 69004448 | rs12698804 | -1.14 | -0.522 | 0.044 | -0.285 | 0.003 | 0.002 | 0.002 | 0.042 | 0.19 | 0.25 | 0.29 | 0.15 | 0.2 | 0.21 |
| 7 | 69006274 | rs10265054 | -0.868 | -0.644 | 0.066 | -0.595 | 0.006 | -0.039 | -0.039 | 0 | 0.91 | 0.84 | 0.86 | 0.99 | 0.86 | 0.89 |
| 7 | 69006876 | rs41859 | -2.096 | -0.824 | -0.27 | -0.54 | -0.004 | -0.006 | -0.006 | 0 | 0.43 | 0.53 | 0.37 | 0.31 | 0.31 | 0.35 |
| 7 | 69007901 | rs41858 | -1.299 | -0.804 | -0.546 | -0.571 | -0.009 | 0.024 | 0.024 | 0 | 0.26 | 0.2 | 0.28 | 0.22 | 0.25 | 0.28 |
| 7 | 69031759 | rs10950191 | -1.172 | -0.709 | -0.463 | -0.504 | 0.008 | 0.05 | 0.05 | 0 | 0.31 | 0.2 | 0.27 | 0.2 | 0.25 | 0.28 |

| | | | | | | | | | | | | | | | | |
|---|---|---|---|---|---|---|---|---|---|---|---|---|---|---|---|---|
| 7 | 69041229 | rs10487945 | 1.267 | -0.722 | -0.459 | -0.435 | 0.019 | 0.011 | 0.011 | 0 | 0.13 | 0.04 | 0.03 | 0.12 | 0.07 | 0.04 |
| 7 | 69048337 | rs11773581 | 0.227 | -0.52 | -0.261 | -0.342 | -0.012 | 0.014 | 0.014 | 0.241 | 0.09 | 0.21 | 0.08 | 0.08 | 0.05 | 0.06 |
| 7 | 69064637 | rs3735260 | -0.177 | -0.54 | -0.237 | -0.371 | -0.026 | 0.023 | 0.023 | 0.048 | 0.11 | 0.29 | 0.09 | 0.08 | 0.06 | 0.08 |
| 7 | 69163557 | rs628885 | 0.008 | -0.237 | -0.288 | -0.385 | -0.032 | 0.019 | 0.019 | 0 | 0.87 | 0.95 | 0.77 | 0.93 | 0.82 | 0.84 |
| 7 | 69171104 | rs6460525 | 0.99 | -0.25 | -0.243 | -0.351 | -0.012 | 0.043 | 0.043 | 0.111 | 0.08 | 0.23 | 0.08 | 0.08 | 0.04 | 0.06 |
| 7 | 69210523 | rs7793984 | 1.37 | -0.029 | -0.639 | -0.69 | -0.019 | 0.08 | 0.08 | 0.481 | 0.62 | 0.75 | 0.66 | 0.78 | 0.65 | 0.69 |
| 7 | 69235712 | rs210605 | -0.119 | -0.072 | -0.605 | -0.625 | -0.029 | 0.019 | 0.019 | 0 | 0.13 | 0.05 | 0.23 | 0.07 | 0.18 | 0.15 |
| 7 | 69238158 | rs11765886 | 1.246 | -0.153 | -0.659 | -0.721 | 0 | 0.065 | 0.065 | 0 | 0.65 | 0.74 | 0.66 | 0.78 | 0.64 | 0.71 |
| 7 | 69247003 | rs210591 | -1.89 | -0.367 | -0.4 | -0.635 | 0.002 | 0.003 | 0.003 | 0 | 0.43 | 0.49 | 0.38 | 0.3 | 0.32 | 0.36 |
| 7 | 69266221 | rs17353815 | -0.993 | -0.217 | -0.323 | -0.548 | 0.006 | 0.034 | 0.034 | 0 | 0.3 | 0.25 | 0.19 | 0.21 | 0.25 | 0.27 |
| 7 | 69266730 | rs10278978 | 2.155 | -0.339 | -0.315 | -0.52 | -0.007 | 0.021 | 0.021 | 0.754 | 0.06 | 0.15 | 0.07 | 0.08 | 0.04 | 0.06 |
| 7 | 69282553 | rs3800979 | 1.312 | -0.339 | -0.317 | -0.519 | -0.015 | 0.036 | 0.036 | 0.351 | 0.08 | 0.23 | 0.08 | 0.08 | 0.04 | 0.06 |
| 7 | 69298131 | rs10237984 | 1.508 | -0.761 | -0.161 | -0.478 | -0.026 | 0.098 | 0.098 | 0 | 0.16 | 0.2 | 0.22 | 0.54 | 0.08 | 0.32 |
| 7 | 69300968 | rs12698810 | 0.605 | -0.731 | -0.479 | -0.459 | -0.001 | 0.004 | 0.004 | 0 | 0.09 | 0.05 | 0.13 | 0.06 | 0.11 | 0.11 |
| 7 | 69329347 | rs1014416 | 2.519 | -0.599 | -0.563 | -0.879 | -0.111 | 0.129 | 0.129 | 0.043 | 0.33 | 0.04 | 0.32 | 0.14 | 0.44 | 0.28 |
| 7 | 69352617 | rs7790458 | 1.463 | -0.567 | -0.355 | -0.716 | -0.024 | 0.004 | 0.004 | 0 | 0.13 | 0.04 | 0.03 | 0.13 | 0.1 | 0.06 |
| 7 | 69375390 | rs10266126 | -0.059 | -0.76 | -0.383 | -0.8 | -0.001 | 0 | 0 | 0 | 0.23 | 0.27 | 0.12 | 0.21 | 0.14 | 0.14 |
| 7 | 69403890 | rs17141032 | 1.611 | -0.779 | -0.368 | -0.817 | -0.011 | 0.009 | 0.009 | 0.71 | 0.08 | 0.16 | 0.07 | 0.08 | 0.04 | 0.08 |
| 7 | 69418905 | rs12698823 | -0.465 | -0.802 | -0.43 | -0.876 | 0.005 | 0.012 | 0.012 | 0 | 0.15 | 0.13 | 0.17 | 0.09 | 0.13 | 0.18 |
| 7 | 69425893 | rs12698826 | -1.057 | -0.835 | -0.42 | -0.744 | 0.011 | 0.02 | 0.02 | 0 | 0.3 | 0.17 | 0.2 | 0.26 | 0.23 | 0.26 |
| 7 | 69425994 | rs6958981 | -1.691 | -0.807 | -0.315 | -0.724 | -0.008 | -0.003 | -0.003 | 0.698 | 0.42 | 0.52 | 0.29 | 0.34 | 0.28 | 0.35 |
| 7 | 69445526 | rs13238398 | -0.939 | -0.769 | -0.323 | -0.739 | -0.001 | -0.003 | -0.003 | 0 | 0.17 | 0.2 | 0.17 | 0.09 | 0.14 | 0.2 |
| 7 | 69455154 | rs12698829 | -0.244 | -0.715 | -0.329 | -0.697 | 0.003 | 0 | 0 | 0 | 0.13 | 0.12 | 0.15 | 0.11 | 0.13 | 0.18 |
| 7 | 69470248 | rs11764911 | 1.894 | -0.668 | -0.249 | -0.646 | 0 | -0.001 | -0.001 | 0 | 0.08 | 0.12 | 0.07 | 0.08 | 0.04 | 0.08 |
| 7 | 69475046 | rs1035013 | -1.574 | -0.678 | -0.265 | -0.688 | -0.009 | -0.009 | -0.009 | 0 | 0.41 | 0.52 | 0.29 | 0.3 | 0.29 | 0.34 |
| 7 | 69482645 | rs7800759 | -1.01 | -0.68 | -0.268 | -0.691 | 0.013 | 0.041 | 0.041 | 0 | 0.32 | 0.25 | 0.21 | 0.22 | 0.24 | 0.26 |
| 7 | 69505703 | rs6415223 | -1.568 | -0.682 | -0.259 | -0.69 | -0.009 | -0.009 | -0.009 | 0 | 0.41 | 0.51 | 0.29 | 0.3 | 0.29 | 0.34 |
| 7 | 69509686 | rs6951909 | -1.007 | -0.687 | -0.257 | -0.684 | 0.013 | 0.021 | 0.021 | 0 | 0.32 | 0.25 | 0.21 | 0.26 | 0.25 | 0.26 |
| 7 | 69516075 | rs6950485 | -1.568 | -0.683 | -0.259 | -0.685 | -0.009 | -0.009 | -0.009 | 0 | 0.41 | 0.52 | 0.29 | 0.3 | 0.29 | 0.34 |
| 7 | 69523434 | rs917717 | -0.944 | -0.633 | -0.281 | -0.684 | 0.021 | 0.02 | 0.02 | 0 | 0.31 | 0.25 | 0.21 | 0.26 | 0.24 | 0.26 |
| 7 | 69530888 | rs13242485 | -1.062 | -0.76 | -0.298 | -0.704 | 0.015 | 0.043 | 0.043 | 0 | 0.32 | 0.25 | 0.21 | 0.22 | 0.25 | 0.26 |
| 7 | 69534070 | rs17141181 | 1.85 | -0.704 | -0.252 | -0.671 | -0.007 | 0.006 | 0.006 | 0.457 | 0.08 | 0.13 | 0.07 | 0.08 | 0.04 | 0.08 |
| 7 | 69541182 | rs7794623 | -1.432 | -0.715 | -0.245 | -0.68 | 0.015 | 0.016 | 0.016 | 0 | 0.4 | 0.41 | 0.28 | 0.3 | 0.29 | 0.34 |
| 7 | 69546617 | rs735060 | -1.022 | -0.716 | -0.252 | -0.681 | 0.011 | 0.031 | 0.031 | 0 | 0.32 | 0.27 | 0.21 | 0.22 | 0.25 | 0.26 |
| 7 | 69562563 | rs1476105 | -0.988 | -0.717 | -0.253 | -0.678 | 0.012 | 0.02 | 0.02 | 0 | 0.32 | 0.25 | 0.21 | 0.26 | 0.25 | 0.26 |
| 7 | 69567938 | rs10279392 | -1.421 | -0.709 | -0.254 | -0.675 | 0.014 | 0.005 | 0.005 | 0 | 0.4 | 0.41 | 0.28 | 0.34 | 0.29 | 0.34 |
| 7 | 69570127 | rs6953170 | -1.534 | -0.709 | -0.242 | -0.678 | -0.008 | -0.008 | -0.008 | 1.125 | 0.41 | 0.51 | 0.29 | 0.3 | 0.29 | 0.34 |
| 7 | 69579024 | rs2058382 | 2.601 | -0.74 | -0.447 | -0.8 | -0.117 | 0.107 | 0.107 | 0 | 0.31 | 0.04 | 0.34 | 0.15 | 0.44 | 0.25 |
| 7 | 69593167 | rs12698854 | 0.319 | -1.089 | -0.314 | -0.711 | 0.002 | 0.008 | 0.008 | 0 | 0.1 | 0.06 | 0.14 | 0.07 | 0.11 | 0.11 |
| 7 | 69630394 | rs7799192 | -0.834 | -1.022 | -0.273 | -0.684 | 0.048 | 0.02 | 0.02 | 0 | 0.31 | 0.23 | 0.19 | 0.26 | 0.19 | 0.25 |
| 7 | 69641784 | rs10256186 | -1.51 | -0.999 | -0.289 | -0.673 | 0.024 | 0.004 | 0.004 | 0 | 0.39 | 0.4 | 0.27 | 0.34 | 0.23 | 0.33 |
| 7 | 69647225 | rs10268924 | -1.405 | -0.997 | -0.292 | -0.689 | -0.029 | -0.011 | -0.011 | 0 | 0.41 | 0.51 | 0.28 | 0.3 | 0.23 | 0.33 |
| 7 | 69649683 | rs6975279 | -1.396 | -0.997 | -0.289 | -0.689 | -0.021 | -0.007 | -0.007 | 0 | 0.4 | 0.51 | 0.28 | 0.3 | 0.23 | 0.33 |
| 7 | 69682576 | rs10256093 | 1.804 | -0.851 | -0.626 | -0.607 | -0.007 | 0.006 | 0.006 | 0.457 | 0.08 | 0.13 | 0.07 | 0.08 | 0.04 | 0.08 |
| 7 | 69688862 | rs4718937 | 0.346 | -0.862 | -0.519 | -0.655 | -0.024 | 0.048 | 0.048 | 1.01 | 0.23 | 0.34 | 0.26 | 0.58 | 0.12 | 0.4 |
| 7 | 69690408 | rs4475377 | -1.109 | -0.547 | -1.114 | -0.707 | 0.037 | 0.035 | 0.035 | 0 | 0.31 | 0.25 | 0.2 | 0.22 | 0.2 | 0.25 |
| 7 | 69732592 | rs6975814 | -1.121 | -1.892 | -1.07 | -0.663 | 0.032 | 0.015 | 0.015 | 0 | 0.31 | 0.27 | 0.2 | 0.26 | 0.19 | 0.25 |
| 7 | 69739690 | rs12698887 | -1.098 | -1.904 | -1.043 | -0.669 | 0.018 | 0.017 | 0.017 | 0 | 0.31 | 0.29 | 0.24 | 0.22 | 0.19 | 0.25 |
| 7 | 69741327 | rs13246213 | 0.359 | -1.825 | -1.178 | -0.65 | -0.002 | -0.01 | -0.01 | 0.558 | 0.09 | 0.06 | 0.18 | 0.11 | 0.12 | 0.11 |
| 7 | 69759833 | rs2866277 | -1.418 | -2.003 | -1.059 | -1.082 | -0.011 | -0.003 | -0.003 | 0 | 0.4 | 0.47 | 0.28 | 0.3 | 0.23 | 0.33 |
| 7 | 69777353 | rs6955333 | 0.22 | -1.884 | -1.577 | -1.094 | 0.006 | 0.007 | 0.007 | 0 | 0.1 | 0.07 | 0.16 | 0.07 | 0.11 | 0.11 |
| 7 | 69783020 | rs10085696 | -0.775 | -1.75 | -1.509 | -1.031 | 0.042 | 0.034 | 0.034 | 0 | 0.31 | 0.26 | 0.22 | 0.22 | 0.18 | 0.25 |
| 7 | 69783198 | rs12672930 | 1.329 | -1.131 | -1.724 | -0.916 | -0.13 | 0.114 | 0.114 | 0 | 0.32 | 0.04 | 0.38 | 0.15 | 0.5 | 0.26 |
| 7 | 69786742 | rs10255669 | -0.102 | -1.484 | -0.622 | -0.313 | 0.016 | 0.008 | 0.008 | 0 | 0.14 | 0.11 | 0.02 | 0.13 | 0.04 | 0.05 |
| 7 | 69811212 | rs10234844 | -1.126 | -1.686 | -0.389 | -0.4 | -0.035 | -0.013 | -0.013 | 4.397 | 0.41 | 0.55 | 0.28 | 0.31 | 0.22 | 0.34 |
| 7 | 69828729 | rs1557920 | 0.139 | -1.481 | 0.221 | -0.342 | -0.005 | 0.044 | 0.044 | 1.249 | 0.09 | 0.25 | 0.05 | 0.08 | 0.04 | 0.07 |
| 7 | 69836148 | rs12698901 | -0.571 | -1.054 | -0.199 | -0.53 | -0.037 | 0.007 | 0.007 | 0 | 0.1 | 0.02 | 0.16 | 0.06 | 0.2 | 0.09 |
| 7 | 69837319 | rs12698902 | -2.101 | -1.564 | 0.074 | -0.655 | -0.026 | -0.03 | -0.03 | 0.108 | 0.31 | 0.51 | 0.28 | 0.19 | 0.28 | 0.28 |
| 7 | 69840168 | rs10486876 | -0.993 | -1.647 | 0.316 | -0.647 | -0.039 | 0.019 | 0.019 | 0.524 | 0.13 | 0.39 | 0.06 | 0.08 | 0.05 | 0.1 |
| 7 | 69872379 | rs7804672 | -1.24 | -2.237 | 0.089 | -1.18 | 0.028 | 0.024 | 0.024 | 0 | 0.37 | 0.33 | 0.15 | 0.22 | 0.19 | 0.24 |
| 7 | 69877261 | rs11972782 | -2.173 | -2.091 | 0.427 | -1.278 | 0.056 | 0.067 | 0.067 | 0 | 0.5 | 0.71 | 0.33 | 0.33 | 0.38 | 0.31 |
| 7 | 69879914 | rs10486878 | 0.559 | -2.244 | 0.315 | -1.428 | -0.008 | 0.001 | 0.001 | 0 | 0.09 | 0.13 | 0.03 | 0.14 | 0.04 | 0.05 |
| 7 | 69887085 | rs11766624 | -1.061 | -2.236 | 0.44 | -1.43 | 0.001 | -0.003 | -0.003 | 0 | 0.28 | 0.27 | 0.13 | 0.09 | 0.15 | 0.2 |
| 7 | 69893262 | rs17488553 | -0.686 | -2.26 | 0.401 | -1.473 | 0.037 | 0.066 | 0.066 | 0 | 0.21 | 0.09 | 0.09 | 0.01 | 0.11 | 0.12 |
| 7 | 69897826 | rs10275122 | -1.423 | -2.088 | 0.495 | -1.366 | -0.005 | -0.003 | -0.003 | 0 | 0.38 | 0.4 | 0.16 | 0.22 | 0.19 | 0.24 |
| 7 | 69908696 | rs6977656 | -0.189 | -1.567 | 0.476 | -1.166 | -0.026 | 0.003 | 0.003 | 0 | 0.08 | 0.03 | 0.12 | 0.07 | 0.19 | 0.05 |
| 7 | 69925030 | rs11760753 | -0.023 | -1.635 | 0.424 | -1.196 | 0.005 | 0.007 | 0.007 | 0.267 | 0.2 | 0.16 | 0.11 | 0.08 | 0.15 | 0.14 |
| 7 | 69928976 | rs17141669 | -0.64 | -1.753 | 0.456 | -1.247 | -0.013 | -0.026 | -0.026 | 0 | 0.07 | 0.12 | 0.02 | 0 | 0.01 | 0.02 |
| 7 | 69930151 | rs10950203 | -0.029 | -1.687 | 0.401 | -1.212 | 0.004 | 0.006 | 0.006 | 0.28 | 0.2 | 0.16 | 0.11 | 0.08 | 0.15 | 0.14 |

| | | | | | | | | | | | | | | | | |
|---|---|---|---|---|---|---|---|---|---|---|---|---|---|---|---|---|
| 7 | 69935641 | rs4717530 | 1.496 | -1.848 | 0.165 | -1.707 | -0.038 | 0.017 | 0.017 | 1.016 | 0.39 | 0.2 | 0.54 | 0.67 | 0.49 | 0.6 |
| 7 | 69935986 | rs4717531 | -0.158 | -1.944 | 0.352 | -1.769 | -0.012 | -0.021 | -0.021 | 0 | 0.74 | 0.69 | 0.88 | 0.91 | 0.83 | 0.84 |
| 7 | 69945340 | rs13232100 | -1.051 | -1.862 | 0.107 | -1.837 | -0.001 | -0.015 | -0.015 | 0 | 0.49 | 0.5 | 0.38 | 0.17 | 0.42 | 0.27 |
| 7 | 69946964 | rs10486873 | 0.251 | -1.517 | 0.036 | -1.552 | -0.001 | 0.006 | 0.006 | 3.052 | 0.09 | 0.13 | 0.07 | 0.16 | 0.09 | 0.12 |
| 7 | 69954583 | rs11760622 | -0.026 | -1.638 | -0.03 | -1.925 | 0.023 | 0.053 | 0.053 | 0 | 0.33 | 0.22 | 0.2 | 0.09 | 0.26 | 0.22 |
| 7 | 69957025 | rs3113281 | -1.463 | -1.795 | 0.063 | -2.1 | -0.01 | 0.056 | 0.056 | 0.142 | 0.57 | 0.75 | 0.39 | 0.22 | 0.51 | 0.38 |
| 7 | 69966192 | rs10486872 | -0.183 | -1.543 | 0.272 | -1.938 | 0.001 | -0.017 | -0.017 | 0 | 0.16 | 0.24 | 0.15 | 0.08 | 0.19 | 0.08 |
| 7 | 69971020 | rs3094893 | -0.942 | -1.572 | 0.392 | -1.952 | 0.003 | -0.057 | -0.057 | 0 | 0.22 | 0.45 | 0.18 | 0.13 | 0.26 | 0.16 |
| 7 | 69985843 | rs3113269 | -1.457 | -1.439 | 0.078 | -1.737 | -0.026 | 0.027 | 0.027 | 0.599 | 0.63 | 0.79 | 0.5 | 0.22 | 0.54 | 0.42 |
| 7 | 69991859 | rs3094903 | -1.41 | -1.435 | 0.079 | -1.736 | -0.014 | 0.014 | 0.014 | 0.69 | 0.63 | 0.73 | 0.49 | 0.22 | 0.54 | 0.42 |
| 7 | 69997575 | rs10224955 | -0.327 | -1.248 | 0.195 | -1.318 | -0.001 | -0.011 | -0.011 | 0 | 0.07 | 0.11 | 0.03 | 0 | 0.02 | 0.02 |
| 7 | 70000870 | rs17141722 | -0.171 | -1.343 | 0.328 | -1.373 | -0.003 | 0.002 | 0.002 | 0.569 | 0.06 | 0.11 | 0.04 | 0.05 | 0.07 | 0.09 |
| 7 | 70003835 | rs10268257 | -1.073 | -1.388 | 0.356 | -1.398 | -0.019 | -0.039 | -0.039 | 0 | 0.09 | 0.22 | 0.03 | 0 | 0.02 | 0.02 |
| 7 | 70004665 | rs6980457 | -1.386 | -1.328 | 0.299 | -1.359 | -0.046 | -0.06 | -0.06 | 0 | 0.15 | 0.37 | 0.07 | 0.05 | 0.09 | 0.11 |
| 7 | 70013591 | rs4718958 | -0.104 | -0.483 | -0.044 | -0.568 | -0.003 | 0.126 | 0.126 | 0.6 | 0.81 | 0.7 | 0.8 | 0.45 | 0.85 | 0.62 |
| 7 | 70014509 | rs6967298 | -0.108 | 0.275 | -0.074 | -0.586 | 0.03 | 0.106 | 0.106 | 0.397 | 0.81 | 0.7 | 0.78 | 0.43 | 0.8 | 0.62 |
| 7 | 70017937 | rs6946285 | 0.084 | 0.299 | -0.549 | -0.986 | 0.01 | 0.024 | 0.024 | 0 | 0.3 | 0.26 | 0.18 | 0.15 | 0.25 | 0.23 |
| 7 | 70018135 | rs11523675 | 0.084 | 0.299 | -0.549 | -0.99 | 0.01 | 0.024 | 0.024 | 0 | 0.3 | 0.26 | 0.18 | 0.15 | 0.25 | 0.23 |
| 7 | 70020160 | rs734930 | 0.328 | 0.443 | -0.637 | -0.818 | -0.002 | 0.017 | 0.017 | 0 | 0.53 | 0.44 | 0.6 | 0.28 | 0.56 | 0.39 |
| 7 | 70022011 | rs17491493 | 1.492 | 0.351 | -0.861 | -1.012 | 0.008 | 0.137 | 0.137 | 0 | 0.36 | 0.19 | 0.46 | 0.11 | 0.31 | 0.27 |
| 7 | 70034730 | rs757726 | 0.537 | 0.482 | -0.664 | -0.821 | 0.015 | 0.029 | 0.029 | 0.009 | 0.45 | 0.41 | 0.5 | 0.17 | 0.37 | 0.34 |
| 7 | 70035509 | rs740114 | 0.694 | 0.499 | -0.754 | -0.822 | 0.017 | 0.038 | 0.038 | 0.133 | 0.43 | 0.35 | 0.5 | 0.18 | 0.37 | 0.34 |
| 7 | 70037996 | rs1013870 | 0.946 | 0.681 | -0.594 | -0.799 | -0.014 | 0.001 | 0.001 | 0 | 0.09 | 0.06 | 0.09 | 0.1 | 0.18 | 0.05 |
| 7 | 70040448 | rs4718965 | -0.297 | 0.899 | -0.356 | -0.898 | 0.004 | -0.04 | -0.04 | 0 | 0.9 | 0.86 | 0.88 | 1 | 0.89 | 0.98 |
| 7 | 70040558 | rs4718966 | 1.461 | 0.266 | -0.637 | -1.168 | -0.012 | 0.03 | 0.03 | 0 | 0.42 | 0.25 | 0.45 | 0.68 | 0.41 | 0.6 |
| 7 | 70043551 | rs12533226 | 0.062 | -0.125 | -0.294 | -0.952 | -0.009 | -0.049 | -0.049 | 0.351 | 0.19 | 0.16 | 0.18 | 0.01 | 0.17 | 0.14 |
| 7 | 70043670 | rs12698910 | -0.116 | 0.406 | -0.205 | -0.329 | -0.014 | 0.012 | 0.012 | 0 | 0.18 | 0.31 | 0.15 | 0.21 | 0.13 | 0.18 |
| 7 | 70046675 | rs17141782 | -0.482 | 0.879 | -0.431 | -0.329 | 0 | -0.003 | -0.003 | 0 | 0.22 | 0.26 | 0.2 | 0.15 | 0.26 | 0.22 |
| 7 | 70047583 | rs719114 | 0.383 | 1.325 | -0.13 | 0.583 | 0.025 | 0.2 | 0.2 | 0 | 0.08 | 0.14 | 0.26 | 0.55 | 0.09 | 0.22 |
| 7 | 70050442 | rs2079337 | -0.152 | 1.305 | -0.211 | -0.575 | -0.002 | -0.016 | -0.016 | 0 | 0.25 | 0.18 | 0.24 | 0 | 0.27 | 0.12 |
| 7 | 70051715 | rs38321 | 0.84 | 1.8 | -0.532 | -0.209 | -0.06 | -0.06 | -0.06 | 1.062 | 0.12 | 0.01 | 0.1 | 0.19 | 0.18 | 0.1 |
| 7 | 70052522 | rs13236715 | -0.058 | 1.589 | -0.327 | -0.608 | -0.033 | -0.038 | -0.038 | 0 | 0.08 | 0.2 | 0.04 | 0.02 | 0.04 | 0.04 |
| 7 | 70053106 | rs38319 | 0.883 | 1.711 | -0.553 | -0.546 | -0.06 | -0.018 | -0.018 | 0.02 | 0.12 | 0.01 | 0.1 | 0.13 | 0.18 | 0.1 |
| 7 | 70067679 | rs38313 | -0.607 | 1.089 | -0.227 | -0.434 | 0.002 | -0.03 | -0.03 | 0 | 0.43 | 0.42 | 0.32 | 0.16 | 0.49 | 0.29 |
| 7 | 70068614 | rs38307 | -0.73 | 1.424 | -0.407 | -0.205 | 0.002 | 0.031 | 0.031 | 0 | 0.51 | 0.56 | 0.4 | 0.17 | 0.53 | 0.33 |
| 7 | 70071416 | rs723980 | 0.801 | 1.861 | -0.138 | 0.547 | 0.005 | 0.006 | 0.006 | 0 | 0.78 | 0.75 | 0.78 | 0.74 | 0.71 | 0.79 |
| 7 | 70071920 | rs10486867 | -1.201 | 1.749 | -0.229 | 0.589 | 0.004 | -0.004 | -0.004 | 0 | 0.44 | 0.64 | 0.46 | 0.55 | 0.45 | 0.27 |
| 7 | 70071979 | rs38300 | 0.612 | 2.257 | -0.287 | 0.94 | -0.044 | -0.046 | -0.046 | 0 | 0.1 | 0.03 | 0.14 | 0.24 | 0.17 | 0.1 |
| 7 | 70081436 | rs11768511 | 0.271 | 1.957 | -0.221 | 0.629 | 0.019 | 0.003 | 0.003 | 0 | 0.14 | 0.05 | 0.15 | 0.14 | 0.13 | 0.09 |
| 7 | 70081456 | rs17493336 | 0.276 | 1.678 | -0.093 | 0.115 | -0.007 | -0.102 | -0.102 | 0 | 0.39 | 0.48 | 0.23 | 0.07 | 0.38 | 0.23 |
| 7 | 70082106 | rs10233063 | -0.519 | 1.893 | 0.017 | 0.438 | 0.006 | 0.049 | 0.049 | 0 | 0.54 | 0.66 | 0.45 | 0.26 | 0.49 | 0.32 |
| 7 | 70082913 | rs10267303 | -0.332 | 1.86 | -0.012 | -0.057 | 0.02 | 0.082 | 0.082 | 0 | 0.52 | 0.66 | 0.37 | 0.21 | 0.46 | 0.31 |
| 7 | 70083527 | rs7780941 | -0.984 | 2.174 | -0.036 | 0.277 | -0.021 | -0.022 | -0.022 | 6.623 | 0.89 | 0.93 | 0.85 | 0.74 | 0.84 | 0.91 |
| 7 | 70085453 | rs11762238 | -0.104 | 1.879 | 0.5 | 0.111 | 0.006 | 0.019 | 0.019 | 0 | 0.51 | 0.56 | 0.37 | 0.15 | 0.5 | 0.28 |
| 7 | 70085934 | rs7791651 | 0.023 | 1.86 | 0.529 | 0.118 | 0.009 | 0.035 | 0.035 | 0 | 0.49 | 0.36 | 0.35 | 0.21 | 0.5 | 0.28 |
| 7 | 70088113 | rs3922333 | -0.689 | 1.952 | 0.544 | 0.125 | 0.003 | 0.014 | 0.014 | 0 | 0.88 | 0.84 | 0.84 | 0.69 | 0.82 | 0.91 |
| 7 | 70088696 | rs7459368 | 0.086 | 1.995 | 0.187 | 0.114 | 0 | -0.022 | -0.022 | 0 | 0.5 | 0.48 | 0.37 | 0.15 | 0.5 | 0.28 |
| 7 | 70094620 | rs4718971 | 0.977 | 1.519 | -0.047 | 0.31 | 0 | -0.015 | -0.015 | 0 | 0.34 | 0.28 | 0.46 | 0.53 | 0.3 | 0.58 |
| 7 | 70096901 | rs4718972 | 0.04 | 1.362 | 0.061 | 0.044 | 0.009 | 0.02 | 0.02 | 0 | 0.51 | 0.45 | 0.55 | 0.86 | 0.43 | 0.67 |
| 7 | 70097265 | rs6965211 | 0.126 | 1.387 | 0.28 | 0.525 | -0.011 | -0.003 | -0.003 | 0 | 0.42 | 0.53 | 0.44 | 0.61 | 0.35 | 0.6 |
| 7 | 70099488 | rs17141917 | 0.245 | 1.664 | -0.461 | 0.469 | -0.006 | -0.003 | -0.003 | 0 | 0.41 | 0.46 | 0.43 | 0.61 | 0.33 | 0.6 |
| 7 | 70100347 | rs7809475 | -0.408 | 1.861 | -0.949 | 0.312 | -0.021 | 0.007 | 0.007 | 0 | 0.85 | 0.94 | 0.81 | 0.87 | 0.77 | 0.89 |
| 7 | 70106061 | rs2866719 | 0.128 | 1.202 | -0.769 | 0.685 | 0.006 | -0.021 | -0.021 | 2.019 | 0.36 | 0.3 | 0.39 | 0.51 | 0.33 | 0.57 |
| 7 | 70111420 | rs11979094 | 0.124 | 1.037 | -0.866 | 0.2 | -0.015 | 0.016 | 0.016 | 0 | 0.62 | 0.65 | 0.59 | 0.28 | 0.52 | 0.51 |
| 7 | 70113274 | rs4718975 | 0.356 | 0.851 | -1.492 | -0.456 | -0.042 | 0.082 | 0.082 | 1.177 | 0.22 | 0.07 | 0.18 | 0.05 | 0.3 | 0.16 |
| 7 | 70113409 | rs4718976 | 0.369 | 0.517 | -0.96 | -0.073 | 0.053 | 0.037 | 0.037 | 0.148 | 0.77 | 0.59 | 0.66 | 0.67 | 0.66 | 0.77 |
| 7 | 70115244 | rs10276134 | 0.582 | 0.524 | -1.213 | -0.197 | -0.039 | 0.09 | 0.09 | 0 | 0.26 | 0.09 | 0.2 | 0.08 | 0.31 | 0.17 |
| 7 | 70116973 | rs11773370 | -0.558 | 0.723 | -0.698 | 0.17 | 0.097 | 0.093 | 0.093 | 0.47 | 0.87 | 0.73 | 0.78 | 0.69 | 0.69 | 0.85 |
| 7 | 70117018 | rs17141930 | 0.955 | 0.02 | -1.484 | -0.067 | -0.005 | -0.15 | -0.15 | 0 | 0.26 | 0.03 | 0.31 | 0.38 | 0.2 | 0.34 |
| 7 | 70117140 | rs6973425 | 0.099 | 0.209 | -0.916 | -0.029 | 0.083 | 0.102 | 0.102 | 0 | 0.8 | 0.45 | 0.7 | 0.61 | 0.64 | 0.76 |
| 7 | 70117254 | rs6973599 | 0.954 | -0.364 | -1.463 | 0.474 | -0.028 | -0.091 | -0.091 | 0 | 0.2 | 0.03 | 0.27 | 0.29 | 0.18 | 0.32 |
| 7 | 70124648 | rs1011780 | 0.33 | -0.802 | -0.945 | -0.257 | 0 | -0.066 | -0.066 | 0.044 | 0.27 | 0.36 | 0.19 | 0.02 | 0.31 | 0.14 |
| 7 | 70125364 | rs7791363 | -0.635 | -0.841 | -1.716 | -0.021 | -0.107 | -0.018 | -0.018 | 0.113 | 0.8 | 0.93 | 0.79 | 0.79 | 0.56 | 0.75 |
| 7 | 70126146 | rs4718978 | 0.902 | -1.708 | -1.576 | 0.228 | 0.063 | -0.083 | -0.083 | 0 | 0.19 | 0.09 | 0.27 | 0.46 | 0.06 | 0.28 |
| 7 | 70127527 | rs10282280 | 0.42 | -0.505 | -0.939 | 0.088 | 0.01 | 0.01 | 0.01 | 0.574 | 0.27 | 0.2 | 0.22 | 0.13 | 0.13 | 0.19 |
| 7 | 70131300 | rs6961823 | 0.252 | -0.317 | -0.911 | 0.177 | -0.001 | -0.002 | -0.002 | 0 | 0.28 | 0.26 | 0.23 | 0.14 | 0.14 | 0.21 |
| 7 | 70133502 | rs17141982 | -0.328 | 1.049 | -0.91 | 0.968 | 0.001 | 0.004 | 0.004 | 0.031 | 0.24 | 0.25 | 0.3 | 0.35 | 0.32 | 0.35 |
| 7 | 70135261 | rs4718980 | 0.57 | 2.619 | -1.385 | 1.03 | -0.09 | -0.05 | -0.05 | 0 | 0.87 | 0.98 | 0.84 | 0.81 | 0.68 | 0.91 |
| 7 | 70135447 | rs11772110 | 0.665 | 1.78 | -1.228 | 0.646 | 0.001 | 0.004 | 0.004 | 0 | 0.52 | 0.43 | 0.52 | 0.62 | 0.45 | 0.63 |

| | | | | | | | | | | | | | | | | |
|---|---|---|---|---|---|---|---|---|---|---|---|---|---|---|---|---|
| 7 | 70136082 | rs6460550 | -0.017 | 1.933 | -0.857 | 0.291 | 0.017 | -0.037 | -0.037 | 2.051 | 0.85 | 0.8 | 0.83 | 1 | 0.67 | 0.88 |
| 7 | 70136765 | rs17141991 | 0.362 | 1.354 | -1.061 | 0.191 | 0.091 | -0.088 | -0.088 | 0.087 | 0.26 | 0.04 | 0.25 | 0.38 | 0.11 | 0.25 |
| 7 | 70137117 | rs10257489 | 1.069 | 1.334 | -0.924 | 0.451 | -0.018 | 0.081 | 0.081 | 0 | 0.52 | 0.78 | 0.47 | 0.31 | 0.48 | 0.4 |
| 7 | 70137165 | rs11771201 | 1.459 | 1.23 | -0.183 | 0.566 | 0.008 | 0.03 | 0.03 | 0 | 0.44 | 0.36 | 0.36 | 0.23 | 0.39 | 0.34 |
| 7 | 70137481 | rs740166 | 1.393 | 1.242 | 0.184 | 0.558 | 0.013 | 0.047 | 0.047 | 0 | 0.46 | 0.61 | 0.38 | 0.23 | 0.39 | 0.34 |
| 7 | 70141539 | rs6949410 | 1.583 | 1.467 | -0.071 | 0.43 | 0.008 | 0.016 | 0.016 | 0 | 0.41 | 0.35 | 0.33 | 0.22 | 0.31 | 0.33 |
| 7 | 70143677 | rs12533012 | -0.742 | 0.728 | -0.372 | 0.803 | 0.009 | -0.048 | -0.048 | 0 | 0.18 | 0.09 | 0.32 | 0.57 | 0.15 | 0.45 |
| 7 | 70144848 | rs10234816 | 1.178 | -0.216 | -0.182 | 0.201 | -0.044 | 0.019 | 0.019 | 0 | 0.26 | 0.39 | 0.3 | 0.3 | 0.15 | 0.26 |
| 7 | 70144987 | rs10234983 | 0.308 | -0.269 | -0.285 | -0.358 | -0.01 | -0.025 | -0.025 | 0 | 0.24 | 0.4 | 0.18 | 0.12 | 0.19 | 0.11 |
| 7 | 70145442 | rs17502134 | 1.342 | 0.097 | -0.638 | -0.528 | -0.014 | 0.057 | 0.057 | 0 | 0.28 | 0.15 | 0.15 | 0.01 | 0.31 | 0.19 |
| 7 | 70148486 | rs4718982 | 0.35 | -0.378 | -0.552 | 0.053 | -0.009 | 0.007 | 0.007 | 0 | 0.36 | 0.62 | 0.54 | 0.52 | 0.25 | 0.5 |
| 7 | 70153068 | rs6946693 | 0.29 | 0.208 | -0.593 | -0.102 | -0.092 | 0.013 | 0.013 | 0 | 0.72 | 0.94 | 0.83 | 0.73 | 0.67 | 0.75 |
| 7 | 70155165 | rs6978392 | 0.522 | 0.266 | 0.105 | 0.029 | 0.099 | 0.029 | 0.029 | 0.123 | 0.81 | 0.64 | 0.79 | 0.73 | 0.66 | 0.7 |
| 7 | 70161221 | rs11768039 | 0.053 | 0.032 | -0.478 | 0.086 | 0.035 | -0.022 | -0.022 | 0 | 0.21 | 0.1 | 0.08 | 0.3 | 0.08 | 0.23 |
| 7 | 70169449 | rs4075496 | 0.981 | 0.453 | -0.406 | 0.009 | -0.027 | -0.001 | -0.001 | 1.344 | 0.05 | 0.01 | 0.06 | 0.02 | 0.08 | 0.02 |
| 7 | 70171159 | rs3923606 | 1.058 | 0.095 | -0.015 | -0.629 | 0.008 | -0.011 | -0.011 | 1.422 | 0.42 | 0.37 | 0.23 | 0.08 | 0.32 | 0.25 |
| 7 | 70173197 | rs4610627 | -0.375 | 0.311 | 0.384 | -0.588 | 0.027 | -0.035 | -0.035 | 0 | 0.82 | 0.75 | 0.83 | 0.91 | 0.71 | 0.86 |
| 7 | 70174823 | rs11970891 | 0.073 | -0.084 | 0.855 | -0.095 | 0.019 | 0.297 | 0.297 | 0.02 | 0.09 | 0.25 | 0.33 | 0.36 | 0.08 | 0.16 |
| 7 | 70176751 | rs7796173 | 1.66 | -0.163 | 0.783 | -0.192 | 0.003 | -0.002 | -0.002 | 0 | 0.18 | 0.19 | 0.13 | 0.23 | 0.14 | 0.18 |
| 7 | 70178152 | rs8180759 | -0.923 | -0.111 | 0.894 | -0.208 | 0.111 | -0.098 | -0.098 | 0.098 | 0.85 | 0.66 | 0.86 | 1 | 0.73 | 0.93 |
| 7 | 70179714 | rs12698917 | 0.485 | -0.118 | 0.459 | -0.166 | -0.029 | -0.039 | -0.039 | 0 | 0.09 | 0.02 | 0.09 | 0.19 | 0.11 | 0.22 |
| 7 | 70183274 | rs6959578 | -0.518 | -0.131 | 0.553 | -0.229 | 0.027 | -0.03 | -0.03 | 0.922 | 0.82 | 0.77 | 0.83 | 0.91 | 0.71 | 0.86 |
| 7 | 70184785 | rs12532316 | 0.614 | -0.15 | 0.04 | -0.224 | 0.028 | 0.039 | 0.039 | 0 | 0.73 | 0.67 | 0.49 | 0.57 | 0.58 | 0.72 |
| 7 | 70186701 | rs17604300 | 2.448 | -0.169 | -0.747 | -0.841 | 0.025 | 0.118 | 0.118 | 0 | 0.29 | 0.15 | 0.18 | 0.01 | 0.24 | 0.22 |
| 7 | 70186936 | rs10279132 | -1.095 | 0.105 | -0.818 | -0.468 | -0.027 | 0.007 | 0.007 | 0 | 0.31 | 0.12 | 0.47 | 0.69 | 0.29 | 0.46 |
| 7 | 70192653 | rs10282512 | -0.413 | 0.335 | -0.382 | -0.041 | -0.014 | -0.032 | -0.032 | 0 | 0.17 | 0.12 | 0.41 | 0.57 | 0.21 | 0.4 |
| 7 | 70193768 | rs7807160 | 0.812 | 0.25 | -0.179 | -0.101 | -0.004 | -0.001 | -0.001 | 0 | 0.19 | 0.23 | 0.16 | 0.22 | 0.19 | 0.18 |
| 7 | 70199714 | rs7783704 | 0.374 | 0.153 | -0.333 | -0.042 | -0.004 | 0.001 | 0.001 | 0 | 0.72 | 0.78 | 0.77 | 0.79 | 0.65 | 0.68 |
| 7 | 70201815 | rs10270627 | 0.566 | 0.341 | -0.45 | -0.028 | -0.053 | 0.005 | 0.005 | 0 | 0.71 | 0.82 | 0.75 | 0.78 | 0.57 | 0.66 |
| 7 | 70203536 | rs1534344 | -0.897 | 0.391 | -0.475 | -0.056 | -0.03 | 0.017 | 0.017 | 0 | 0.83 | 0.89 | 0.88 | 0.99 | 0.71 | 0.93 |
| 7 | 70209637 | rs723340 | -0.731 | -1.381 | -0.205 | 0.061 | 0.097 | 0.094 | 0.094 | 0 | 0.46 | 0.84 | 0.59 | 0.77 | 0.31 | 0.46 |
| 7 | 70209816 | rs6942467 | 0.206 | -1.208 | 0.527 | 0.122 | 0.042 | -0.019 | -0.019 | 0 | 0.09 | 0.2 | 0.09 | 0.04 | 0.12 | 0.06 |
| 7 | 70213618 | rs17142037 | -0.278 | -1.557 | 0.656 | 0.232 | -0.038 | 0.138 | 0.138 | 0 | 0.19 | 0.41 | 0.44 | 0.59 | 0.14 | 0.29 |
| 7 | 70215224 | rs6460553 | -1.083 | -1.504 | 0.051 | 0.366 | -0.002 | 0.007 | 0.007 | 0 | 0.26 | 0.73 | 0.43 | 0.45 | 0.19 | 0.23 |
| 7 | 70216522 | rs6957531 | -1.186 | -1.465 | 0.06 | 0.077 | -0.001 | 0 | 0 | 0 | 0.28 | 0.72 | 0.46 | 0.49 | 0.27 | 0.24 |
| 7 | 70217058 | rs4717551 | -1.903 | -1.537 | -0.337 | -0.326 | -0.061 | 0.167 | 0.167 | 0 | 0.59 | 0.84 | 0.71 | 0.98 | 0.49 | 0.69 |
| 7 | 70217631 | rs1881010 | 1.238 | -1.476 | -0.31 | -0.037 | -0.006 | 0 | 0 | 0 | 0.12 | 0.1 | 0.04 | 0.1 | 0.03 | 0.1 |
| 7 | 70221259 | rs1544009 | 0.608 | -1.409 | -0.403 | 0.184 | -0.032 | -0.023 | -0.023 | 0 | 0.09 | 0.05 | 0.3 | 0.39 | 0.14 | 0.17 |
| 7 | 70223193 | rs2178044 | 1.67 | -1.659 | -0.366 | 0.018 | 0.027 | 0.184 | 0.184 | 0.011 | 0.48 | 0.12 | 0.39 | 0.19 | 0.6 | 0.52 |
| 7 | 70228020 | rs2293507 | 0.888 | -2.06 | -0.128 | 0.266 | -0.015 | -0.003 | -0.003 | 0 | 0.13 | 0.15 | 0.04 | 0.1 | 0.03 | 0.07 |
| 7 | 70230863 | rs6959436 | 1.06 | -1.703 | 0.285 | 0.503 | -0.052 | 0.055 | 0.055 | 0.45 | 0.69 | 0.56 | 0.73 | 0.53 | 0.8 | 0.62 |
| 7 | 70231649 | rs2178045 | -0.435 | -1.072 | -0.317 | 0.857 | 0.009 | -0.011 | -0.011 | 0 | 0.87 | 0.89 | 0.87 | 0.78 | 0.9 | 0.87 |
| 7 | 70233139 | rs3750170 | -0.142 | -0.552 | -0.203 | 1.042 | -0.005 | 0.026 | 0.026 | 0 | 0.11 | 0.13 | 0.31 | 0.37 | 0.14 | 0.19 |
| 7 | 70236787 | rs2293505 | -0.05 | -0.559 | -0.253 | 0.893 | -0.002 | 0.001 | 0.001 | 0 | 0.07 | 0.05 | 0.1 | 0.18 | 0.08 | 0.2 |
| 7 | 70237585 | rs6947003 | -1.319 | -1.147 | 0.088 | 0.346 | 0.001 | 0 | 0 | 0 | 0.33 | 0.36 | 0.46 | 0.65 | 0.26 | 0.49 |
| 7 | 70241056 | rs6971633 | -0.944 | -1.326 | 0.659 | 0.598 | -0.009 | 0.147 | 0.147 | 0 | 0.17 | 0.33 | 0.27 | 0.37 | 0.14 | 0.17 |
| 7 | 70241298 | rs6460557 | -0.678 | -1.369 | 0.655 | 0.589 | -0.002 | 0.178 | 0.178 | 0 | 0.16 | 0.33 | 0.27 | 0.37 | 0.13 | 0.17 |
| 7 | 70242821 | rs6460558 | 0.251 | -0.993 | 0.663 | 0.279 | 0.001 | -0.005 | -0.005 | 0 | 0.87 | 0.85 | 0.92 | 0.95 | 0.84 | 0.89 |
| 7 | 70244289 | rs2049635 | 1.069 | -1.509 | 0.436 | 0.025 | 0.042 | 0.025 | 0.025 | 0 | 0.15 | 0.03 | 0.05 | 0.09 | 0.03 | 0.07 |
| 7 | 70246898 | rs951392 | 2.136 | -1.716 | 0.155 | -0.436 | -0.069 | 0.097 | 0.097 | 0 | 0.34 | 0.19 | 0.29 | 0.01 | 0.48 | 0.33 |
| 7 | 70248469 | rs9886351 | -1.045 | -1.694 | 0.355 | -0.306 | 0.027 | 0.095 | 0.095 | 0 | 0.26 | 0.47 | 0.33 | 0.42 | 0.28 | 0.28 |
| 7 | 70249234 | rs6948548 | 0.076 | -1.613 | 0.477 | -0.227 | 0.004 | 0.201 | 0.201 | 1.641 | 0.13 | 0.29 | 0.25 | 0.37 | 0.12 | 0.21 |
| 7 | 70249868 | rs3817575 | 0.443 | -1.589 | 0.36 | -0.232 | 0.004 | 0.152 | 0.152 | 0.044 | 0.12 | 0.23 | 0.25 | 0.37 | 0.12 | 0.16 |
| 7 | 70251651 | rs10499822 | 1.4 | -1.555 | 0.021 | -0.193 | 0.009 | -0.004 | -0.004 | 0 | 0.46 | 0.41 | 0.55 | 0.38 | 0.6 | 0.49 |
| 7 | 70251923 | rs2293501 | 0.778 | -1.768 | 0.146 | -0.103 | 0.007 | -0.002 | -0.002 | 0 | 0.58 | 0.58 | 0.63 | 0.42 | 0.76 | 0.61 |
| 7 | 70252185 | rs2293500 | 0.27 | -0.718 | -0.233 | 0.471 | -0.017 | 0.019 | 0.019 | 0 | 0.78 | 0.73 | 0.7 | 0.53 | 0.81 | 0.67 |
| 7 | 70260049 | rs13246944 | 0.755 | -0.276 | -0.436 | -0.043 | 0 | -0.02 | -0.02 | 0.023 | 0.57 | 0.51 | 0.44 | 0.3 | 0.56 | 0.53 |
| 7 | 70266515 | rs1105953 | 0.597 | -0.188 | -1.012 | -0.53 | 0.042 | 0.148 | 0.148 | 0 | 0.5 | 0.28 | 0.43 | 0.16 | 0.62 | 0.48 |
| 7 | 70270337 | rs4717554 | -1.011 | 0.561 | -1.019 | 0.469 | -0.014 | -0.017 | -0.017 | 0.006 | 0.2 | 0.19 | 0.29 | 0.4 | 0.33 | 0.26 |
| 7 | 70270672 | rs2903400 | -0.241 | -0.025 | -1.438 | -0.533 | 0.006 | -0.005 | -0.005 | 0.321 | 0.26 | 0.22 | 0.21 | 0.41 | 0.21 | 0.3 |
| 7 | 70270987 | rs12667834 | -0.149 | 0.343 | -0.666 | -0.358 | 0.023 | 0.046 | 0.046 | 0 | 0.85 | 0.79 | 0.61 | 0.49 | 0.74 | 0.75 |
| 7 | 70271908 | rs1918427 | -0.186 | 0.352 | -0.685 | -0.46 | 0.019 | 0.038 | 0.038 | 0 | 0.85 | 0.81 | 0.61 | 0.49 | 0.74 | 0.74 |
| 7 | 70274757 | rs1881013 | 0.167 | -0.194 | -0.875 | -1.198 | -0.049 | -0.109 | -0.109 | 0 | 0.25 | 0.45 | 0.22 | 0.06 | 0.17 | 0.26 |
| 7 | 70286089 | rs1949979 | 1.18 | -0.594 | -1.409 | -1.412 | -0.126 | 0.154 | 0.154 | 0 | 0.41 | 0.08 | 0.31 | 0.27 | 0.49 | 0.27 |
| 7 | 70295629 | rs10263077 | 0.204 | -0.914 | -0.68 | -1.122 | -0.036 | -0.085 | -0.085 | 0 | 0.23 | 0.36 | 0.21 | 0.06 | 0.17 | 0.26 |
| 7 | 70316365 | rs6460575 | 1.064 | -0.533 | -1.498 | -1.199 | -0.064 | 0.111 | 0.111 | 0.121 | 0.37 | 0.09 | 0.26 | 0.26 | 0.41 | 0.25 |
| 7 | 70317727 | rs7784844 | 0.543 | -0.702 | -1.236 | -1.197 | 0.071 | 0.095 | 0.095 | 0 | 0.57 | 0.31 | 0.68 | 0.76 | 0.72 | 0.52 |
| 7 | 70318843 | rs1590016 | 0.117 | -1.231 | -0.444 | -1.123 | -0.051 | -0.026 | -0.026 | 0 | 0.79 | 0.45 | 0.89 | 0.81 | 0.92 | 0.82 |
| 7 | 70321643 | rs1527588 | -0.099 | -0.93 | 0.165 | -0.985 | -0.016 | -0.052 | -0.052 | 0.019 | 0.9 | 0.77 | 0.95 | 0.99 | 0.95 | 0.94 |

| | | | | | | | | | | | | | | | | |
|---|---|---|---|---|---|---|---|---|---|---|---|---|---|---|---|---|
| 7 | 70325633 | rs1527592 | -0.099 | -0.93 | 0.165 | -0.985 | -0.016 | -0.052 | -0.052 | 0 | 0.9 | 0.77 | 0.95 | 0.99 | 0.95 | 0.94 |
| 7 | 70326475 | rs1997226 | -0.148 | -0.901 | -0.125 | -0.98 | -0.012 | 0.013 | 0.013 | 0 | 0.41 | 0.37 | 0.55 | 0.55 | 0.51 | 0.51 |
| 7 | 70337942 | rs12698956 | 0.654 | -0.988 | 0.205 | -0.6 | -0.031 | -0.025 | -0.025 | 1.065 | 0.81 | 0.77 | 0.93 | 0.93 | 0.9 | 0.82 |
| 7 | 70343081 | rs6942810 | 0.131 | -0.611 | 0.023 | -0.194 | 0.003 | 0.007 | 0.007 | 0 | 0.08 | 0.06 | 0.03 | 0.19 | 0.04 | 0.12 |
| 7 | 70350082 | rs4717561 | 1.305 | -0.9 | 0.04 | -0.507 | -0.003 | 0 | 0 | 0.162 | 0.77 | 0.77 | 0.89 | 0.82 | 0.85 | 0.74 |
| 7 | 70354372 | rs1534346 | -0.585 | -1.035 | -0.048 | -0.612 | 0.009 | 0.086 | 0.086 | 0 | 0.52 | 0.35 | 0.59 | 0.66 | 0.57 | 0.65 |
| 7 | 70364102 | rs6460597 | 0.8 | -1.132 | 0.471 | -0.67 | -0.001 | 0.013 | 0.013 | 0 | 0.44 | 0.35 | 0.39 | 0.34 | 0.42 | 0.33 |
| 7 | 70368805 | rs2866951 | 0.238 | -1.153 | 0.303 | -0.685 | -0.037 | 0.04 | 0.04 | 0 | 0.39 | 0.27 | 0.53 | 0.65 | 0.51 | 0.54 |
| 7 | 70370469 | rs7808758 | 0.461 | -1.033 | 0.419 | -0.614 | 0.004 | 0.029 | 0.029 | 0 | 0.48 | 0.56 | 0.41 | 0.34 | 0.43 | 0.35 |
| 7 | 70371194 | rs1918425 | 0.461 | -1.033 | 0.419 | -0.614 | 0.004 | 0.029 | 0.029 | 0 | 0.48 | 0.56 | 0.41 | 0.34 | 0.43 | 0.35 |
| 7 | 70378407 | rs7786437 | 0.842 | -0.972 | 0.275 | -0.52 | -0.046 | -0.012 | -0.012 | 0 | 0.74 | 0.57 | 0.87 | 0.82 | 0.85 | 0.71 |
| 7 | 70383718 | rs940731 | 0.438 | -0.335 | 0.691 | -0.7 | 0.002 | 0.008 | 0.008 | 0 | 0.42 | 0.39 | 0.36 | 0.26 | 0.37 | 0.26 |
| 7 | 70387953 | rs940728 | 0.148 | -0.391 | 0.927 | -0.512 | -0.046 | -0.042 | -0.042 | 0.3 | 0.32 | 0.47 | 0.3 | 0.27 | 0.19 | 0.25 |
| 7 | 70397051 | rs4717565 | -0.184 | 0.076 | 1.089 | -0.426 | -0.003 | -0.019 | -0.019 | 0 | 0.42 | 0.49 | 0.41 | 0.34 | 0.35 | 0.34 |
| 7 | 70404146 | rs13312126 | 0.446 | -0.122 | 1.039 | -0.507 | 0.027 | 0.09 | 0.09 | 0 | 0.53 | 0.22 | 0.56 | 0.66 | 0.62 | 0.61 |
| 7 | 70405492 | rs10950229 | 0.479 | -0.178 | 1.096 | -0.454 | 0.038 | 0.08 | 0.08 | 0 | 0.53 | 0.24 | 0.56 | 0.65 | 0.63 | 0.6 |
| 7 | 70407537 | rs12113420 | -0.239 | -0.179 | 1.456 | -0.441 | -0.002 | -0.012 | -0.012 | 0.065 | 0.42 | 0.45 | 0.42 | 0.29 | 0.36 | 0.4 |
| 7 | 70413659 | rs1525292 | 1.836 | -0.179 | 1.844 | -0.337 | 0.038 | 0.094 | 0.094 | 0.245 | 0.92 | 0.58 | 0.93 | 0.92 | 0.95 | 0.9 |
| 7 | 70419818 | rs1464853 | 0.588 | -0.237 | 1.631 | -0.373 | 0.019 | 0.037 | 0.037 | 0 | 0.55 | 0.35 | 0.57 | 0.63 | 0.64 | 0.6 |
| 7 | 70426380 | rs4719035 | 1.963 | -0.111 | 1.944 | -0.118 | 0.041 | 0.095 | 0.095 | 0 | 0.93 | 0.59 | 0.93 | 0.93 | 0.95 | 0.9 |
| 7 | 70449956 | rs1568868 | 1.495 | 0.267 | 1.747 | -0.208 | 0.018 | -0.04 | -0.04 | 0 | 0.89 | 0.63 | 0.93 | 0.98 | 0.9 | 0.92 |
| 7 | 70453467 | rs10253135 | 0.301 | 0.164 | 1.545 | -0.42 | 0 | -0.003 | -0.003 | 0 | 0.35 | 0.34 | 0.32 | 0.26 | 0.27 | 0.36 |
| 7 | 70455873 | rs4719045 | 0.863 | 0.165 | 1.65 | -0.395 | 0.025 | -0.026 | -0.026 | 0.022 | 0.9 | 0.69 | 0.94 | 0.99 | 0.9 | 0.96 |
| 7 | 70462013 | rs2177725 | 0.419 | 0.155 | 1.475 | -0.402 | -0.004 | -0.014 | -0.014 | 0 | 0.35 | 0.39 | 0.34 | 0.25 | 0.27 | 0.36 |
| 7 | 70465729 | rs12698968 | 0.615 | 0.266 | 0.809 | 0.157 | 0.039 | 0.09 | 0.09 | 0 | 0.52 | 0.14 | 0.54 | 0.64 | 0.62 | 0.55 |
| 7 | 70470196 | rs7806679 | 0.778 | 0.374 | 0.882 | -0.058 | -0.01 | 0.008 | 0.008 | 0.29 | 0.05 | 0.04 | 0.02 | 0.01 | 0.08 | 0.03 |
| 7 | 70470837 | rs2049475 | 0.545 | 0.273 | 0.902 | 0.201 | 0.036 | 0.149 | 0.149 | 0 | 0.53 | 0.16 | 0.58 | 0.75 | 0.63 | 0.59 |
| 7 | 70480631 | rs10950235 | 0.501 | 0.152 | 0.625 | 0.785 | 0.024 | -0.037 | -0.037 | 0 | 0.42 | 0.14 | 0.44 | 0.51 | 0.36 | 0.31 |
| 7 | 70491251 | rs10950237 | 0.468 | 0.08 | 0.645 | 0.982 | 0.032 | -0.065 | -0.065 | 0.7 | 0.42 | 0.14 | 0.4 | 0.46 | 0.36 | 0.31 |
| 7 | 70493364 | rs4717577 | 0.077 | -0.012 | 1.056 | 0.741 | 0.014 | 0.002 | 0.002 | 0 | 0.51 | 0.39 | 0.55 | 0.51 | 0.57 | 0.62 |
| 7 | 70500643 | rs13227328 | 0.614 | -0.267 | 0.74 | -0.162 | 0.028 | 0.062 | 0.062 | 0 | 0.36 | 0.24 | 0.29 | 0.07 | 0.29 | 0.29 |
| 7 | 70501199 | rs12698976 | 1.114 | -0.264 | 0.552 | -0.17 | -0.017 | 0.205 | 0.205 | 0 | 0.31 | 0.02 | 0.27 | 0.06 | 0.29 | 0.26 |
| 7 | 70511536 | rs7777882 | -0.066 | -0.192 | 0.974 | -0.088 | -0.012 | -0.111 | -0.111 | 0 | 0.58 | 0.51 | 0.67 | 0.92 | 0.65 | 0.69 |
| 7 | 70524233 | rs4479997 | -0.086 | -0.53 | 0.752 | 0.662 | -0.012 | 0.048 | 0.048 | 0.37 | 0.15 | 0.19 | 0.38 | 0.66 | 0.14 | 0.24 |
| 7 | 70525507 | rs10260200 | 0.565 | -0.323 | 0.594 | 0.183 | 0.004 | -0.015 | -0.015 | 0 | 0.37 | 0.4 | 0.28 | 0.2 | 0.39 | 0.43 |
| 7 | 70533310 | rs12698994 | 0.002 | -0.184 | 0.402 | -0.01 | 0.009 | -0.012 | -0.012 | 0 | 0.82 | 0.78 | 0.85 | 0.88 | 0.82 | 0.91 |
| 7 | 70537478 | rs10429063 | 0.438 | -0.402 | 0.042 | 0.516 | 0.021 | -0.035 | -0.035 | 0 | 0.15 | 0.12 | 0.35 | 0.4 | 0.12 | 0.12 |
| 7 | 70546596 | rs6460638 | 1.12 | -0.16 | -0.002 | 0.189 | 0.001 | -0.011 | -0.011 | 0 | 0.1 | 0.02 | 0.06 | 0.11 | 0.07 | 0.07 |
| 7 | 70558485 | rs10950240 | -0.286 | -0.167 | 0.082 | -0.078 | -0.001 | -0.009 | -0.009 | 0 | 0.55 | 0.38 | 0.63 | 0.53 | 0.5 | 0.42 |
| 7 | 70562841 | rs12538806 | 1.185 | 0.189 | 0.499 | 0.048 | -0.019 | 0.124 | 0.124 | 0 | 0.28 | 0.1 | 0.2 | 0.01 | 0.31 | 0.24 |
| 7 | 70572236 | rs7779440 | -0.161 | 0.117 | 0.612 | -0.633 | -0.014 | 0.113 | 0.113 | 0 | 0.14 | 0.22 | 0.27 | 0.53 | 0.09 | 0.24 |
| 7 | 70572266 | rs10265996 | 0.163 | 0.191 | 0.356 | -0.785 | 0.067 | -0.029 | -0.029 | 0 | 0.07 | 0.19 | 0.09 | 0.02 | 0.11 | 0.04 |
| 7 | 70573387 | rs7798323 | 0.548 | 0.175 | -0.252 | -0.806 | -0.02 | 0.025 | 0.025 | 0 | 0.48 | 0.23 | 0.42 | 0.38 | 0.49 | 0.47 |
| 7 | 70577861 | rs847724 | -1.006 | 0.626 | -0.377 | -1.436 | -0.011 | 0.099 | 0.099 | 0 | 0.71 | 0.88 | 0.8 | 0.99 | 0.69 | 0.75 |
| 7 | 70585797 | rs11762189 | 1.437 | 0.977 | -0.698 | -1.458 | -0.031 | -0.03 | -0.03 | 0.008 | 0.11 | 0.06 | 0.15 | 0.18 | 0.22 | 0.17 |
| 7 | 70597522 | rs10275799 | -0.758 | 0.174 | -0.149 | -2.169 | -0.014 | 0.124 | 0.124 | 0 | 0.72 | 0.9 | 0.81 | 0.99 | 0.72 | 0.75 |
| 7 | 70601412 | rs11762382 | -1.602 | 0.641 | 0.896 | -1.112 | 0.038 | -0.031 | -0.031 | 0 | 0.18 | 0.32 | 0.16 | 0.13 | 0.2 | 0.13 |
| 7 | 70604381 | rs11769535 | -1.29 | 1.156 | 1.171 | -1.151 | 0.11 | -0.023 | -0.023 | 0 | 0.08 | 0.24 | 0.11 | 0.02 | 0.14 | 0.05 |
| 7 | 70617004 | rs2040970 | -0.055 | 0.425 | 0.725 | -1.392 | -0.006 | 0.028 | 0.028 | 0 | 0.3 | 0.22 | 0.22 | 0.06 | 0.33 | 0.15 |
| 7 | 70619098 | rs2867041 | -0.475 | 0.428 | 1.156 | -1.442 | 0.005 | -0.061 | -0.061 | 0 | 0.34 | 0.36 | 0.24 | 0.07 | 0.37 | 0.17 |
| 7 | 70619951 | rs17142542 | -0.405 | 0.439 | 1.167 | -1.444 | 0.005 | -0.063 | -0.063 | 0 | 0.33 | 0.36 | 0.24 | 0.07 | 0.37 | 0.17 |
| 7 | 70620003 | rs12665914 | -0.484 | 0.411 | 1.162 | -1.444 | 0.006 | -0.064 | -0.064 | 0.01 | 0.34 | 0.36 | 0.24 | 0.07 | 0.37 | 0.17 |
| 7 | 70624620 | rs728136 | -1.177 | -0.034 | 0.833 | -1.703 | 0.026 | 0.131 | 0.131 | 0 | 0.49 | 0.68 | 0.36 | 0.08 | 0.41 | 0.3 |
| 7 | 70628972 | rs4717582 | -0.362 | -0.365 | 0.497 | -1.527 | -0.065 | -0.087 | -0.087 | 0 | 0.17 | 0.29 | 0.12 | 0.01 | 0.05 | 0.13 |
| 7 | 70630136 | rs7804724 | -0.242 | 0.611 | -0.025 | -1.14 | -0.008 | 0.035 | 0.035 | 0 | 0.31 | 0.22 | 0.23 | 0.07 | 0.34 | 0.12 |
| 7 | 70631020 | rs10261209 | -1.518 | 0.929 | 0.603 | -0.283 | 0.027 | -0.034 | -0.034 | 0 | 0.18 | 0.65 | 0.23 | 0.11 | 0.18 | 0.09 |
| 7 | 70638282 | rs10215771 | -1.478 | 0.576 | 0.656 | -0.595 | 0.011 | -0.135 | -0.135 | 0 | 0.29 | 0.53 | 0.25 | 0.08 | 0.23 | 0.25 |
| 7 | 70641761 | rs10271414 | -1.449 | -0.388 | 0.523 | -0.026 | -0.019 | -0.004 | -0.004 | 1.628 | 0.23 | 0.78 | 0.27 | 0.2 | 0.1 | 0.2 |
| 7 | 70644029 | rs1796057 | -1.3 | -0.354 | 0.538 | -0.194 | -0.022 | -0.024 | -0.024 | 0 | 0.22 | 0.76 | 0.17 | 0.09 | 0.09 | 0.2 |
| 7 | 70655941 | rs4339520 | -0.635 | 0.125 | 0.625 | 0.636 | -0.001 | 0.069 | 0.069 | 0 | 0.06 | 0.12 | 0.05 | 0.14 | 0.05 | 0.04 |
| 7 | 70659219 | rs1922520 | -0.506 | -0.837 | 0.838 | 0.316 | -0.037 | -0.017 | -0.017 | 0 | 0.21 | 0.24 | 0.19 | 0.15 | 0.1 | 0.15 |
| 7 | 70659343 | rs1202657 | -1.42 | -0.582 | 0.495 | 0.967 | -0.009 | 0.084 | 0.084 | 0.88 | 0.23 | 0.66 | 0.25 | 0.45 | 0.24 | 0.21 |
| 7 | 70669190 | rs11766740 | -0.176 | -0.445 | 0.883 | 1.311 | 0.006 | 0.043 | 0.043 | 0 | 0.1 | 0.16 | 0.14 | 0.2 | 0.15 | 0.14 |
| 7 | 70673908 | rs17592881 | 1.379 | 0.092 | 0.265 | 0.798 | -0.071 | 0.046 | 0.046 | 0.83 | 0.45 | 0.05 | 0.3 | 0.42 | 0.36 | 0.56 |
| 7 | 70675645 | rs7807224 | -0.854 | 0.122 | 1.015 | 0.928 | 0.065 | 0.307 | 0.307 | 0.02 | 0.07 | 0.43 | 0.06 | 0.21 | 0.06 | 0.07 |
| 7 | 70676108 | rs6979579 | 0.888 | 0.079 | 0.943 | 0.885 | 0.049 | 0.024 | 0.024 | 0 | 0.67 | 0.46 | 0.44 | 0.47 | 0.45 | 0.71 |
| 7 | 70679318 | rs9638288 | 0.533 | 0.136 | 1.344 | 0.978 | -0.016 | 0.026 | 0.026 | 0 | 0.55 | 0.67 | 0.44 | 0.74 | 0.43 | 0.63 |
| 7 | 70684291 | rs7788558 | 0.54 | 0.135 | 1.242 | 0.95 | -0.023 | 0.002 | 0.002 | 0 | 0.55 | 0.73 | 0.38 | 0.64 | 0.43 | 0.63 |
| 7 | 70686526 | rs10950250 | 1.295 | 0.182 | 0.731 | 0.867 | -0.019 | 0.074 | 0.074 | 0.595 | 0.46 | 0.12 | 0.3 | 0.42 | 0.36 | 0.56 |

| | | | | | | | | | | | | | | | | |
|---|---|---|---|---|---|---|---|---|---|---|---|---|---|---|---|---|
| 7 | 70688504 | rs12539391 | 1.226 | 0.184 | 0.742 | 0.868 | -0.001 | 0.067 | 0.067 | 0.273 | 0.46 | 0.19 | 0.28 | 0.42 | 0.36 | 0.56 |
| 7 | 70693915 | rs17142940 | -1.925 | 0.099 | 1.088 | 1.216 | -0.013 | 0.188 | 0.188 | 2.611 | 0.1 | 0.67 | 0.17 | 0.32 | 0.07 | 0.07 |
| 7 | 70695280 | rs11975543 | -1.506 | 0.194 | 0.949 | 1.114 | 0.057 | 0.322 | 0.322 | 0.636 | 0.08 | 0.51 | 0.07 | 0.23 | 0.07 | 0.07 |
| 7 | 70698028 | rs7806416 | -0.619 | 0.064 | 0.001 | 0.243 | 0.014 | 0.019 | 0.019 | 2.11 | 0.28 | 0.22 | 0.23 | 0.06 | 0.2 | 0.18 |
| 7 | 70701596 | rs10260768 | -0.624 | 0.392 | 0.245 | 0.42 | 0.001 | -0.055 | -0.055 | 0.227 | 0.1 | 0.24 | 0.09 | 0 | 0.09 | 0.01 |
| 7 | 70703896 | rs2867046 | -0.664 | 1.6 | -0.111 | 0.74 | -0.075 | -0.018 | -0.018 | 0 | 0.83 | 0.92 | 0.64 | 0.85 | 0.58 | 0.8 |
| 7 | 70705454 | rs6974582 | 0.546 | 1.561 | -0.055 | 1.02 | 0.011 | 0 | 0 | 0.018 | 0.73 | 0.67 | 0.52 | 0.82 | 0.44 | 0.77 |
| 7 | 70706501 | rs7811880 | 0.913 | 1.437 | -0.283 | 0.788 | -0.009 | 0.012 | 0.012 | 0 | 0.64 | 0.67 | 0.5 | 0.82 | 0.38 | 0.7 |
| 7 | 70713870 | rs16869397 | 0.325 | 1.27 | 0.172 | 1.285 | 0.007 | -0.011 | -0.011 | 0 | 0.11 | 0.22 | 0.06 | 0.04 | 0.11 | 0.09 |
| 7 | 70715414 | rs4717586 | 0.366 | 2.215 | -0.154 | 1.254 | -0.056 | -0.015 | -0.015 | 0.021 | 0.17 | 0.08 | 0.32 | 0.14 | 0.36 | 0.21 |
| 7 | 70719222 | rs11772787 | -0.961 | 2.209 | -0.326 | 1.063 | 0.019 | 0.036 | 0.036 | 0 | 0.24 | 0.33 | 0.14 | 0.32 | 0.26 | 0.4 |
| 7 | 70720859 | rs4719095 | -0.757 | 1.865 | -0.633 | 0.643 | 0.01 | 0.02 | 0.02 | 0 | 0.75 | 0.79 | 0.77 | 0.96 | 0.79 | 0.83 |
| 7 | 70721939 | rs12666964 | 1.192 | 1.948 | -0.661 | 0.692 | -0.023 | -0.007 | -0.007 | 0 | 0.1 | 0.03 | 0.07 | 0.15 | 0.13 | 0.2 |
| 7 | 70722342 | rs17594016 | 1.143 | 1.951 | -0.646 | 0.719 | -0.01 | -0.003 | -0.003 | 0.789 | 0.1 | 0.04 | 0.07 | 0.15 | 0.13 | 0.2 |
| 7 | 70726179 | rs737214 | -0.364 | 1.96 | -0.674 | 0.739 | -0.048 | -0.066 | -0.066 | 0.148 | 0.22 | 0.06 | 0.12 | 0.32 | 0.26 | 0.4 |
| 7 | 70726379 | rs2014978 | 1.82 | 1.505 | -0.627 | 0.742 | -0.011 | 0.007 | 0.007 | 0.039 | 0.36 | 0.38 | 0.31 | 0.49 | 0.23 | 0.33 |
| 7 | 70729787 | rs11764495 | -0.898 | 1.54 | -0.813 | 0.515 | -0.005 | -0.007 | -0.007 | 0 | 0.37 | 0.31 | 0.43 | 0.4 | 0.57 | 0.51 |
| 7 | 70736168 | rs7793970 | -1.009 | 1.521 | -0.812 | 0.41 | -0.001 | -0.001 | -0.001 | 0.022 | 0.38 | 0.37 | 0.52 | 0.46 | 0.57 | 0.51 |
| 7 | 70738028 | rs10238468 | -0.228 | 1.543 | -0.868 | 0.41 | -0.083 | -0.085 | -0.085 | 0.006 | 0.28 | 0.14 | 0.48 | 0.46 | 0.52 | 0.44 |
| 7 | 70738518 | rs7795083 | -0.307 | 1.765 | -1.122 | 0.334 | 0.001 | -0.037 | -0.037 | 0 | 0.22 | 0.29 | 0.11 | 0.14 | 0.19 | 0.28 |
| 7 | 70738575 | rs7794178 | 0.415 | 1.726 | -1.826 | 0.652 | 0.017 | 0.101 | 0.101 | 0 | 0.51 | 0.38 | 0.66 | 0.82 | 0.59 | 0.52 |
| 7 | 70738729 | rs7794356 | 0.731 | 0.365 | -1.231 | 0.758 | 0.007 | -0.005 | -0.005 | 0 | 0.35 | 0.35 | 0.25 | 0.49 | 0.21 | 0.29 |
| 7 | 70740155 | rs17594632 | 0.404 | 0.913 | -1.236 | 0.454 | -0.007 | 0.039 | 0.039 | 0 | 0.09 | 0 | 0.02 | 0.01 | 0.05 | 0.07 |
| 7 | 70743033 | rs1202640 | -0.074 | 0.912 | -1.207 | 0.522 | -0.009 | 0.001 | 0.001 | 0.969 | 0.59 | 0.67 | 0.38 | 0.61 | 0.46 | 0.68 |
| 7 | 70743371 | rs12536849 | -0.23 | 0.5 | -1.495 | -0.193 | 0.002 | 0.035 | 0.035 | 0.558 | 0.17 | 0.12 | 0.12 | 0.08 | 0.14 | 0.13 |
| 7 | 70747828 | rs9638614 | -0.425 | 0.295 | -1.859 | 0.427 | -0.039 | -0.111 | -0.111 | 0 | 0.28 | 0.11 | 0.28 | 0.46 | 0.3 | 0.34 |
| 7 | 70750493 | rs9638616 | -1.405 | -0.462 | -0.745 | 0.091 | -0.006 | 0.011 | 0.011 | 0 | 0.4 | 0.5 | 0.32 | 0.48 | 0.32 | 0.39 |
| 7 | 70755998 | rs4446620 | 1.27 | -0.278 | -0.414 | -0.04 | -0.012 | 0.059 | 0.059 | 0.236 | 0.53 | 0.21 | 0.59 | 0.36 | 0.56 | 0.5 |
| 7 | 70757913 | rs4719103 | 0.282 | -0.193 | 0.1 | 0.033 | 0.021 | 0.023 | 0.023 | 0 | 0.85 | 0.54 | 0.88 | 0.84 | 0.86 | 0.87 |
| 7 | 70761953 | rs6976905 | -1.266 | 0.143 | -0.158 | -0.204 | 0.003 | -0.03 | -0.03 | 0 | 0.6 | 0.53 | 0.56 | 0.83 | 0.51 | 0.72 |
| 7 | 70764328 | rs1317290 | -0.267 | 0.083 | -0.105 | -0.183 | 0.002 | -0.007 | -0.007 | 0 | 0.38 | 0.38 | 0.31 | 0.49 | 0.32 | 0.39 |
| 7 | 70764516 | rs10272997 | 0.282 | 0.185 | -0.072 | -0.043 | 0.001 | -0.005 | -0.005 | 0 | 0.23 | 0.16 | 0.25 | 0.33 | 0.18 | 0.33 |
| 7 | 70769692 | rs9638618 | 0.332 | 0.255 | 0.312 | -0.131 | 0.01 | 0.061 | 0.061 | 0 | 0.7 | 0.6 | 0.67 | 0.53 | 0.71 | 0.65 |
| 7 | 70778156 | rs1950148 | 0.32 | 0.168 | 0.318 | -0.209 | -0.027 | -0.101 | -0.101 | 0.095 | 0.31 | 0.13 | 0.28 | 0.47 | 0.3 | 0.39 |
| 7 | 70778579 | rs17628870 | 1.713 | 0.179 | 0.417 | -0.313 | 0.006 | 0.065 | 0.065 | 0 | 0.34 | 0.23 | 0.35 | 0.04 | 0.38 | 0.26 |
| 7 | 70788156 | rs1574440 | 0.494 | 0.201 | 0.522 | -0.146 | 0 | -0.032 | -0.032 | 0 | 0.23 | 0.09 | 0.19 | 0.31 | 0.19 | 0.3 |
| 7 | 70798935 | rs6460653 | 0 | 0.158 | 0.615 | -0.123 | -0.041 | -0.073 | -0.073 | 0 | 0.64 | 0.78 | 0.66 | 0.54 | 0.61 | 0.6 |
| 7 | 70799863 | rs6944999 | 0.511 | 0.146 | 0.574 | -0.117 | 0.007 | -0.01 | -0.01 | 0 | 0.6 | 0.46 | 0.56 | 0.36 | 0.59 | 0.56 |
| 7 | 70800407 | rs13240200 | 1.804 | 0.174 | 0.26 | -0.185 | -0.002 | -0.067 | -0.067 | 0 | 0.35 | 0.4 | 0.38 | 0.04 | 0.39 | 0.26 |
| 7 | 70801406 | rs9638622 | 0.556 | 0.457 | -0.317 | 0.438 | -0.018 | 0.031 | 0.031 | 0.068 | 0.23 | 0 | 0.11 | 0.13 | 0.16 | 0.2 |
| 7 | 70811708 | rs16869429 | 0.403 | 1.033 | 0.395 | 1.091 | -0.043 | -0.013 | -0.013 | 0 | 0.07 | 0.01 | 0.16 | 0.33 | 0.14 | 0.19 |
| 7 | 70815845 | rs10240080 | -0.142 | 0.002 | 0.736 | 0.744 | 0.07 | 0.041 | 0.041 | 0.076 | 0.07 | 0.34 | 0.14 | 0.06 | 0.09 | 0.03 |
| 7 | 70828718 | rs2527298 | 0.041 | 0.086 | -0.467 | 0.824 | -0.003 | -0.029 | -0.029 | 0 | 0.21 | 0.1 | 0.17 | 0.31 | 0.18 | 0.29 |
| 7 | 70829578 | rs17143419 | 0.752 | -0.012 | -0.234 | 0.744 | 0.01 | 0.002 | 0.002 | 0 | 0.06 | 0.13 | 0.12 | 0.05 | 0.09 | 0.02 |
| 7 | 70833344 | rs4719118 | -0.741 | -0.578 | -0.07 | 1.567 | 0 | 0.066 | 0.066 | 0 | 0.08 | 0.13 | 0.21 | 0.4 | 0.12 | 0.09 |
| 7 | 70837798 | rs7792469 | 1.339 | -0.795 | -0.518 | 0.961 | -0.071 | 0.118 | 0.118 | 0 | 0.33 | 0.18 | 0.42 | 0.05 | 0.46 | 0.19 |
| 7 | 70838143 | rs929342 | 0.045 | -1.378 | -0.717 | 0.903 | 0.006 | -0.075 | -0.075 | 0.647 | 0.71 | 0.85 | 0.69 | 0.51 | 0.72 | 0.52 |
| 7 | 70847554 | rs4719119 | 1.257 | -1.464 | -0.107 | 0.63 | -0.007 | 0.007 | 0.007 | 0 | 0.39 | 0.62 | 0.47 | 0.19 | 0.48 | 0.27 |
| 7 | 70848821 | rs12699037 | 1.033 | -1.589 | -0.016 | 0.715 | 0.018 | 0.1 | 0.1 | 0.332 | 0.57 | 0.27 | 0.54 | 0.15 | 0.59 | 0.39 |
| 7 | 70851287 | rs4719120 | 0.497 | -1.552 | 0.153 | 0.788 | 0.002 | 0.011 | 0.011 | 0 | 0.63 | 0.65 | 0.58 | 0.29 | 0.61 | 0.44 |
| 7 | 70859491 | rs2189930 | 1.269 | -1.888 | -0.085 | 1.033 | 0.018 | -0.028 | -0.028 | 2.781 | 0.42 | 0.26 | 0.65 | 0.42 | 0.63 | 0.32 |
| 7 | 70860252 | rs6964977 | -0.559 | -1.953 | 0.625 | 1.05 | 0.063 | 0.18 | 0.18 | 0 | 0.06 | 0.38 | 0.05 | 0.13 | 0.02 | 0.03 |
| 7 | 70864261 | rs2960881 | -0.101 | -2.07 | 0.394 | 1.131 | 0.02 | -0.014 | -0.014 | 0 | 0.73 | 0.83 | 0.79 | 0.65 | 0.78 | 0.52 |
| 7 | 70870965 | rs1012127 | -0.107 | -1.945 | 0.264 | 1.195 | 0.02 | -0.032 | -0.032 | 0.026 | 0.76 | 0.86 | 0.73 | 0.62 | 0.78 | 0.53 |
| 7 | 70880650 | rs2066984 | 1.441 | -2.144 | 0.265 | 1.153 | 0.019 | -0.004 | -0.004 | 0 | 0.42 | 0.25 | 0.6 | 0.38 | 0.63 | 0.3 |
| 7 | 70883515 | rs734053 | 0.022 | -3.075 | 0.517 | 1.316 | 0.025 | -0.038 | -0.038 | 0 | 0.76 | 0.87 | 0.74 | 0.62 | 0.79 | 0.54 |
| 7 | 70885536 | rs2960888 | 0.754 | -3.141 | 0.433 | 1.198 | 0.069 | 0.038 | 0.038 | 1.373 | 0.17 | 0.04 | 0.03 | 0.09 | 0.04 | 0.12 |
| 7 | 70890039 | rs1859526 | 0.672 | -3.06 | 0.352 | 1.212 | 0.094 | -0.032 | -0.032 | 0.022 | 0.59 | 0.81 | 0.71 | 0.52 | 0.75 | 0.42 |
| 7 | 70899397 | rs740099 | 0.264 | -3.157 | 0.327 | 0.294 | -0.01 | -0.018 | -0.018 | 0 | 0.14 | 0.17 | 0.14 | 0.08 | 0.14 | 0.21 |
| 7 | 70908346 | rs4388345 | -1.18 | -3.26 | 0.108 | 0.375 | 0.058 | 0.011 | 0.011 | 0.415 | 0.84 | 0.96 | 0.97 | 0.9 | 0.96 | 0.89 |
| 7 | 70911123 | rs4236247 | 1.678 | -3.358 | 0.131 | 0.403 | 0.104 | 0.144 | 0.144 | 0 | 0.47 | 0.26 | 0.59 | 0.25 | 0.66 | 0.3 |
| 7 | 70929635 | rs4521648 | 0.491 | -3.129 | 0.158 | 0.608 | -0.027 | -0.072 | -0.072 | 0 | 0.11 | 0.01 | 0.18 | 0.56 | 0.12 | 0.3 |
| 7 | 70932550 | rs4535614 | 1.271 | -3.219 | 0.484 | 0.544 | 0.03 | 0.016 | 0.016 | 0 | 0.6 | 0.39 | 0.72 | 0.34 | 0.79 | 0.52 |
| 7 | 70935693 | rs10261379 | -0.937 | -2.63 | 0.912 | 0.716 | -0.019 | 0.079 | 0.079 | 0 | 0.21 | 0.29 | 0.22 | 0.57 | 0.17 | 0.36 |
| 7 | 70946807 | rs11561753 | 1.043 | -2.605 | 0.535 | 0.728 | -0.03 | 0.068 | 0.068 | 0 | 0.77 | 0.44 | 0.76 | 0.44 | 0.83 | 0.64 |
| 7 | 70954231 | rs7808818 | 1.232 | -2.148 | 0.078 | 0.202 | 0.005 | 0.137 | 0.137 | 0 | 0.4 | 0.25 | 0.49 | 0.11 | 0.58 | 0.27 |
| 7 | 70970026 | rs4717599 | 0.737 | -2.081 | 0.457 | 0.98 | 0.029 | -0.008 | -0.008 | 0 | 0.57 | 0.37 | 0.69 | 0.51 | 0.73 | 0.58 |
| 7 | 70972962 | rs10279551 | -0.019 | -2.236 | 0.645 | 1.09 | 0.056 | 0.05 | 0.05 | 0 | 0.31 | 0.17 | 0.19 | 0.19 | 0.18 | 0.18 |
| 7 | 70973447 | rs4605942 | 1.658 | -2.385 | 0.495 | 1.038 | 0.038 | 0.089 | 0.089 | 0.03 | 0.46 | 0.31 | 0.53 | 0.11 | 0.64 | 0.31 |

| | | | | | | | | | | | | | | | | |
|---|---|---|---|---|---|---|---|---|---|---|---|---|---|---|---|---|
| 7 | 70973959 | rs6943641 | -0.386 | -2.445 | 0.655 | 1.03 | 0.019 | 0.019 | 0.019 | 0 | 0.36 | 0.29 | 0.2 | 0.2 | 0.21 | 0.22 |
| 7 | 70976393 | rs6955186 | -0.423 | -1.863 | 0.337 | 0.725 | 0.021 | 0.219 | 0.219 | 0 | 0.08 | 0.26 | 0.1 | 0.28 | 0.07 | 0.21 |
| 7 | 70982773 | rs7796112 | 1.424 | -1.882 | 0.064 | 0.672 | -0.002 | 0.216 | 0.216 | 0.045 | 0.85 | 0.52 | 0.87 | 0.71 | 0.89 | 0.74 |
| 7 | 70984103 | rs7783170 | 0.659 | -1.58 | -0.585 | 0.74 | 0.083 | -0.018 | -0.018 | 0 | 0.55 | 0.76 | 0.65 | 0.41 | 0.72 | 0.55 |
| 7 | 70994375 | rs7807431 | -0.832 | -0.831 | -0.272 | 0.156 | -0.012 | -0.036 | -0.036 | 0 | 0.33 | 0.38 | 0.4 | 0.83 | 0.24 | 0.5 |
| 7 | 70996732 | rs6947549 | 0.529 | -0.136 | -0.352 | 0.861 | -0.006 | 0.003 | 0.003 | 0 | 0.87 | 0.88 | 0.84 | 0.53 | 0.86 | 0.68 |
| 7 | 71000614 | rs6460664 | 0.015 | 0.055 | -0.532 | -0.17 | -0.001 | 0 | 0 | 0 | 0.36 | 0.63 | 0.42 | 0.23 | 0.36 | 0.23 |
| 7 | 71006669 | rs6460666 | 0.627 | -0.285 | -0.997 | -0.611 | -0.002 | 0.003 | 0.003 | 0 | 0.68 | 0.66 | 0.71 | 0.48 | 0.73 | 0.48 |
| 7 | 71007374 | rs6973852 | 1.127 | -0.394 | -0.813 | -0.93 | 0.023 | -0.002 | -0.002 | 0 | 0.56 | 0.28 | 0.64 | 0.48 | 0.65 | 0.41 |
| 7 | 71019833 | rs12699052 | 1.643 | -0.059 | -0.816 | -0.88 | -0.069 | 0.065 | 0.065 | 0 | 0.34 | 0.1 | 0.32 | 0.25 | 0.4 | 0.27 |
| 7 | 71020307 | rs6460671 | 0.059 | 0.108 | -0.184 | -0.726 | 0 | -0.003 | -0.003 | 1.7 | 0.32 | 0.32 | 0.36 | 0.24 | 0.32 | 0.21 |
| 7 | 71020953 | rs6977973 | -0.03 | 0.116 | 0.076 | -0.722 | -0.002 | -0.021 | -0.021 | 0.079 | 0.33 | 0.42 | 0.37 | 0.24 | 0.32 | 0.21 |
| 7 | 71021037 | rs6460672 | -0.443 | 0.171 | -0.083 | -0.524 | -0.006 | 0.006 | 0.006 | 0 | 0.87 | 0.87 | 0.79 | 0.52 | 0.84 | 0.66 |
| 7 | 71023877 | rs7806531 | -0.637 | -0.182 | 0.431 | -0.891 | 0 | 0.001 | 0.001 | 0 | 0.37 | 0.6 | 0.45 | 0.24 | 0.41 | 0.25 |
| 7 | 71026463 | rs7780636 | 1.32 | -0.238 | 0.371 | -0.943 | 0.008 | 0.025 | 0.025 | 0 | 0.11 | 0.07 | 0.05 | 0.03 | 0.07 | 0.11 |
| 7 | 71031469 | rs4719152 | -1.195 | -0.186 | 0.41 | -0.951 | 0.012 | 0.052 | 0.052 | 0 | 0.48 | 0.67 | 0.49 | 0.27 | 0.48 | 0.37 |
| 7 | 71035420 | rs10232345 | -1.252 | -0.144 | 0.987 | -0.968 | 0.009 | -0.058 | -0.058 | 0.15 | 0.11 | 0.38 | 0.07 | 0 | 0.08 | 0.03 |
| 7 | 71037932 | rs11773571 | 0.213 | 0.116 | 0.83 | -0.908 | 0.026 | 0.003 | 0.003 | 0 | 0.26 | 0.39 | 0.37 | 0.24 | 0.34 | 0.17 |
| 7 | 71042418 | rs11976847 | -1.309 | -0.023 | 0.987 | -0.688 | 0.009 | -0.036 | -0.036 | 0 | 0.09 | 0.33 | 0.06 | 0 | 0.06 | 0.07 |
| 7 | 71043615 | rs10950267 | 1.365 | -0.021 | 0.541 | -0.827 | -0.053 | 0.092 | 0.092 | 0.306 | 0.52 | 0.06 | 0.52 | 0.72 | 0.58 | 0.64 |
| 7 | 71049656 | rs2903582 | 0.209 | 0.033 | 0.944 | -0.761 | 0.003 | -0.002 | -0.002 | 0.502 | 0.74 | 0.35 | 0.62 | 0.78 | 0.7 | 0.76 |
| 7 | 71050482 | rs10807732 | 0.587 | 0.072 | 1.141 | -0.782 | 0.016 | 0.012 | 0.012 | 0 | 0.2 | 0.3 | 0.33 | 0.21 | 0.25 | 0.18 |
| 7 | 71054650 | rs6961201 | 0.549 | -0.167 | 1.126 | -0.79 | 0.013 | 0.042 | 0.042 | 0.018 | 0.13 | 0.05 | 0.07 | 0.03 | 0.09 | 0.14 |
| 7 | 71057795 | rs1464904 | 1.226 | 0.035 | 1.344 | -0.59 | 0.066 | 0.018 | 0.018 | 0 | 0.93 | 0.64 | 0.95 | 1 | 0.95 | 0.94 |
| 7 | 71065519 | rs10273024 | 0.124 | 0.139 | 1.356 | -0.569 | 0.033 | -0.038 | -0.038 | 0 | 0.1 | 0.32 | 0.06 | 0.01 | 0.07 | 0.03 |
| 7 | 71066926 | rs10237977 | 0.63 | 0.167 | 1.2 | -0.563 | 0.006 | -0.043 | -0.043 | 0 | 0.08 | 0.16 | 0.05 | 0 | 0.08 | 0.03 |
| 7 | 71071646 | rs6944213 | 0.535 | 0.125 | 1.131 | -0.574 | 0 | -0.034 | -0.034 | 0 | 0.09 | 0.13 | 0.04 | 0 | 0.07 | 0.04 |
| 7 | 71081245 | rs6460677 | 0.616 | 0.094 | 0.744 | -0.91 | -0.009 | 0.083 | 0.083 | 0 | 0.15 | 0.02 | 0.1 | 0.02 | 0.15 | 0.14 |
| 7 | 71087923 | rs6977420 | 0.464 | -0.128 | 1.56 | -0.974 | -0.004 | -0.041 | -0.041 | 0.478 | 0.1 | 0.16 | 0.04 | 0 | 0.07 | 0.04 |
| 7 | 71091521 | rs11762927 | 1.319 | 0.628 | 1.122 | -1.043 | -0.039 | 0.149 | 0.149 | 0.011 | 0.54 | 0.12 | 0.63 | 0.77 | 0.49 | 0.64 |
| 7 | 71107224 | rs4719155 | -0.81 | 0.856 | 1.38 | -0.818 | 0.053 | 0.028 | 0.028 | 0 | 0.84 | 0.71 | 0.8 | 0.76 | 0.7 | 0.84 |
| 7 | 71110648 | rs11972325 | 0.738 | 0.745 | 1.479 | -0.911 | 0 | -0.05 | -0.05 | 0.022 | 0.1 | 0.18 | 0.04 | 0 | 0.06 | 0.04 |
| 7 | 71110740 | rs17763663 | 0.78 | 0.348 | 0.937 | -0.55 | -0.003 | -0.061 | -0.061 | 0.393 | 0.1 | 0.01 | 0.16 | 0.25 | 0.06 | 0.12 |
| 7 | 71110900 | rs10243782 | 0.109 | 0.316 | 1.138 | -0.978 | -0.009 | 0.005 | 0.005 | 0 | 0.68 | 0.68 | 0.7 | 0.74 | 0.56 | 0.69 |
| 7 | 71114444 | rs2056647 | 1.179 | 0.262 | 0.999 | -0.967 | -0.042 | 0.012 | 0.012 | 0 | 0.69 | 0.13 | 0.72 | 0.73 | 0.64 | 0.72 |
| 7 | 71115769 | rs1525720 | -0.072 | 0.164 | 1.235 | -0.994 | -0.002 | -0.048 | -0.048 | 0.559 | 0.23 | 0.21 | 0.12 | 0.01 | 0.17 | 0.18 |
| 7 | 71122193 | rs2140890 | 0.863 | 0.446 | 1.486 | -0.67 | 0.014 | -0.02 | -0.02 | 0 | 0.66 | 0.52 | 0.7 | 0.73 | 0.61 | 0.72 |
| 7 | 71125248 | rs605586 | -0.71 | 0.472 | 1.458 | -0.659 | 0.007 | -0.006 | -0.006 | 0 | 0.32 | 0.37 | 0.28 | 0.27 | 0.39 | 0.25 |
| 7 | 71125622 | rs1548763 | 0.969 | 0.497 | 1.211 | -0.619 | 0.002 | -0.009 | -0.009 | 0 | 0.64 | 0.42 | 0.67 | 0.72 | 0.61 | 0.72 |
| 7 | 71126019 | rs619096 | 0.337 | 0.826 | 0.68 | -0.238 | -0.006 | -0.006 | -0.006 | 0 | 0.16 | 0.16 | 0.19 | 0.26 | 0.27 | 0.18 |
| 7 | 71128401 | rs11975195 | 0.45 | 0.247 | 0.769 | -0.047 | -0.012 | 0.17 | 0.17 | 0 | 0.1 | 0.21 | 0.15 | 0.58 | 0.05 | 0.29 |
| 7 | 71129699 | rs567591 | -1.507 | 0.079 | 0.127 | -0.945 | -0.013 | 0.106 | 0.106 | 0 | 0.58 | 0.77 | 0.61 | 0.99 | 0.54 | 0.8 |
| 7 | 71133902 | rs995497 | 1.711 | 0.098 | -0.209 | -0.953 | -0.098 | 0.28 | 0.28 | 0 | 0.38 | 0.03 | 0.37 | 0.01 | 0.44 | 0.19 |
| 7 | 71135192 | rs2302440 | 0.424 | 0.132 | 0.21 | -0.938 | 0 | -0.001 | -0.001 | 0.554 | 0.57 | 0.5 | 0.57 | 0.58 | 0.55 | 0.54 |
| 7 | 71138339 | rs10231680 | 0.207 | 0.173 | 0.801 | -0.096 | 0.002 | 0.243 | 0.243 | 0 | 0.1 | 0.25 | 0.14 | 0.58 | 0.05 | 0.33 |
| 7 | 71148103 | rs664112 | -0.196 | 0.427 | -0.076 | -0.253 | -0.037 | 0.007 | 0.007 | 0 | 0.17 | 0.07 | 0.18 | 0.15 | 0.27 | 0.13 |
| 7 | 71152622 | rs6944414 | 1.221 | 0.198 | 0.55 | -0.129 | 0.009 | -0.009 | -0.009 | 0 | 0.71 | 0.56 | 0.65 | 0.31 | 0.68 | 0.49 |
| 7 | 71154079 | rs1608875 | 0.289 | 0.347 | 0.361 | -1.118 | -0.003 | 0.002 | 0.002 | 0 | 0.81 | 0.8 | 0.8 | 0.86 | 0.73 | 0.84 |
| 7 | 71165833 | rs10275110 | 0.4 | -0.288 | 0.801 | -1.564 | -0.008 | 0.115 | 0.115 | 0 | 0.1 | 0.25 | 0.15 | 0.29 | 0.06 | 0.19 |
| 7 | 71168010 | rs10256029 | 0.731 | -0.44 | 0.198 | -2.005 | 0.009 | -0.064 | -0.064 | 0 | 0.13 | 0.22 | 0.08 | 0 | 0.15 | 0.18 |
| 7 | 71168582 | rs1525727 | 0.078 | -0.473 | -0.12 | -2.138 | 0.005 | -0.071 | -0.071 | 0.257 | 0.67 | 0.51 | 0.78 | 0.92 | 0.63 | 0.69 |
| 7 | 71178166 | rs1043987 | 1.49 | -0.442 | -0.36 | -2.033 | -0.005 | -0.041 | -0.041 | 0 | 0.08 | 0.03 | 0.14 | 0.28 | 0.06 | 0.18 |
| 7 | 71179316 | rs6943205 | 0.254 | -0.419 | -0.435 | -2.092 | 0.002 | -0.005 | -0.005 | 0 | 0.27 | 0.24 | 0.27 | 0.63 | 0.22 | 0.45 |
| 7 | 71184181 | rs605065 | -0.6 | 0.343 | -0.293 | -2.029 | 0 | -0.057 | -0.057 | 0.557 | 0.61 | 0.58 | 0.62 | 0.98 | 0.63 | 0.8 |
| 7 | 71185237 | rs620269 | -0.843 | -0.057 | -0.266 | -1.869 | 0.001 | -0.01 | -0.01 | 0 | 0.39 | 0.52 | 0.41 | 0.72 | 0.39 | 0.44 |
| 7 | 71189112 | rs678798 | -0.2 | 0.14 | -0.418 | -1.841 | 0.003 | 0.013 | 0.013 | 0 | 0.55 | 0.63 | 0.57 | 0.97 | 0.57 | 0.76 |
| 7 | 71189138 | rs17143909 | 0.947 | 0.173 | -1.254 | -1.692 | 0 | 0 | 0 | 0 | 0.29 | 0.32 | 0.23 | 0.27 | 0.24 | 0.4 |
| 7 | 71194861 | rs12154827 | -0.098 | 0.642 | -0.542 | 0.219 | -0.03 | -0.099 | -0.099 | 0.436 | 0.18 | 0.04 | 0.24 | 0.56 | 0.2 | 0.14 |
| 7 | 71196604 | rs12699065 | 0.98 | 0.309 | 0.536 | -0.259 | 0.017 | -0.058 | -0.058 | 0.266 | 0.82 | 0.5 | 0.87 | 0.93 | 0.8 | 0.91 |
| 7 | 71200151 | rs677014 | -1.066 | 0.398 | 0.235 | -0.092 | -0.004 | -0.04 | -0.04 | 0 | 0.27 | 0.33 | 0.18 | 0.1 | 0.25 | 0.13 |
| 7 | 71211570 | rs10273681 | 0.91 | 0.695 | -1.025 | 0.568 | -0.011 | 0.01 | 0.01 | 0 | 0.88 | 0.99 | 0.9 | 0.93 | 0.84 | 0.86 |
| 7 | 71214274 | rs10245217 | 1.423 | 0.659 | -1.052 | 0.937 | -0.002 | -0.021 | -0.021 | 0.29 | 0.83 | 0.96 | 0.86 | 0.8 | 0.79 | 0.8 |
| 7 | 71228862 | rs2191740 | -1.219 | 0.809 | -1.293 | 1.516 | -0.005 | -0.056 | -0.056 | 0 | 0.19 | 0.11 | 0.24 | 0.57 | 0.16 | 0.38 |
| 7 | 71232992 | rs992125 | -1.339 | 0.597 | -0.197 | 1.42 | -0.005 | 0.082 | 0.082 | 0.197 | 0.21 | 0.39 | 0.25 | 0.57 | 0.16 | 0.4 |
| 7 | 71254570 | rs7786834 | -0.395 | 0.928 | -1.511 | 0.013 | -0.151 | 0.162 | 0.162 | 0 | 0.66 | 0.99 | 0.76 | 0.92 | 0.56 | 0.73 |
| 7 | 71255458 | rs6977083 | -0.503 | 0.309 | -0.683 | 0.307 | -0.006 | 0.026 | 0.026 | 0 | 0.32 | 0.43 | 0.32 | 0.5 | 0.33 | 0.31 |
| 7 | 71266533 | rs6955664 | -1.179 | 0.168 | 0.179 | 0.741 | 0.021 | 0.23 | 0.23 | 0.107 | 0.1 | 0.27 | 0.18 | 0.61 | 0.09 | 0.21 |
| 7 | 71272257 | rs12699091 | -1.042 | -0.121 | 0.836 | 0.429 | -0.006 | 0.043 | 0.043 | 0 | 0.28 | 0.51 | 0.29 | 0.61 | 0.28 | 0.25 |
| 7 | 71273602 | rs17423790 | 0.029 | -0.142 | 0.613 | 0.128 | 0 | -0.034 | -0.034 | 0 | 0.19 | 0.17 | 0.11 | 0 | 0.19 | 0.05 |

| | | | | | | | | | | | | | | | | |
|---|---|---|---|---|---|---|---|---|---|---|---|---|---|---|---|---|
| 7 | 71297516 | rs7777622 | 0.584 | 0.237 | 0.422 | 0.077 | 0.048 | 0.026 | 0.026 | 0 | 0.5 | 0.68 | 0.46 | 0.48 | 0.42 | 0.55 |
| 7 | 71302421 | rs10226380 | 0.642 | 0.002 | 0.936 | -0.131 | 0.046 | 0.029 | 0.029 | 0 | 0.51 | 0.31 | 0.44 | 0.48 | 0.44 | 0.55 |
| 7 | 71304661 | rs7790027 | 0.574 | 0.449 | 0.891 | 0.147 | -0.008 | 0.006 | 0.006 | 0.095 | 0.14 | 0.1 | 0.12 | 0.12 | 0.16 | 0.1 |
| 7 | 71304749 | rs7794787 | 1.271 | 0.039 | 0.489 | 0.104 | -0.02 | -0.013 | -0.013 | 0 | 0.36 | 0.59 | 0.35 | 0.31 | 0.29 | 0.3 |
| 7 | 71307372 | rs987034 | 0.833 | 0.703 | 0.522 | 0.246 | 0.062 | 0.064 | 0.064 | 0 | 0.45 | 0.2 | 0.38 | 0.35 | 0.36 | 0.45 |
| 7 | 71314281 | rs2023988 | 0.401 | 1.185 | 0.761 | 0.104 | -0.005 | 0.16 | 0.16 | 0 | 0.55 | 0.81 | 0.61 | 0.87 | 0.46 | 0.65 |
| 7 | 71336068 | rs7349979 | -0.143 | 1.656 | 1.383 | 0.46 | 0.038 | -0.038 | -0.038 | 0.54 | 0.18 | 0.34 | 0.17 | 0.12 | 0.22 | 0.11 |
| 7 | 71338530 | rs17340639 | 0.344 | 1.66 | 1.191 | 0.462 | -0.016 | 0.012 | 0.012 | 0.16 | 0.16 | 0.07 | 0.14 | 0.12 | 0.22 | 0.11 |
| 7 | 71341808 | rs7785844 | -1.075 | 1.618 | 1.04 | 0.316 | -0.027 | -0.007 | -0.007 | 0.108 | 0.42 | 0.76 | 0.47 | 0.53 | 0.53 | 0.26 |
| 7 | 71360451 | rs6973743 | 0.873 | 1.514 | 0.821 | 0.233 | -0.023 | 0.115 | 0.115 | 0 | 0.19 | 0.03 | 0.1 | 0 | 0.23 | 0.06 |
| 7 | 71361261 | rs726469 | -0.124 | 1.736 | 1.377 | 0.418 | 0.082 | -0.058 | -0.058 | 0 | 0.18 | 0.48 | 0.18 | 0.1 | 0.21 | 0.1 |
| 7 | 71366745 | rs2214747 | -0.573 | 1.592 | 0.912 | 0.447 | 0.065 | 0.137 | 0.137 | 0 | 0.22 | 0.49 | 0.2 | 0.5 | 0.25 | 0.18 |
| 7 | 71386614 | rs13229300 | -0.422 | 1.717 | 0.705 | 0.541 | 0.061 | 0.14 | 0.14 | 0 | 0.21 | 0.53 | 0.2 | 0.48 | 0.25 | 0.17 |
| 7 | 71388036 | rs6460695 | -0.681 | 1.195 | 0.06 | 0.368 | 0.006 | 0.003 | 0.003 | 0.115 | 0.4 | 0.47 | 0.33 | 0.51 | 0.52 | 0.23 |
| 7 | 71389353 | rs17341778 | -0.177 | 0.953 | -0.564 | -0.447 | -0.027 | 0.062 | 0.062 | 0 | 0.39 | 0.24 | 0.44 | 0.18 | 0.44 | 0.2 |
| 7 | 71395150 | rs1075042 | -0.35 | 0.598 | -0.592 | -0.803 | -0.004 | -0.081 | -0.081 | 0 | 0.66 | 0.41 | 0.77 | 0.86 | 0.61 | 0.49 |
| 7 | 71395521 | rs12537492 | 0.716 | 1.438 | -0.583 | -0.595 | -0.002 | 0 | 0 | 0.387 | 0.16 | 0.12 | 0.22 | 0.09 | 0.21 | 0.1 |
| 7 | 71400441 | rs1859552 | 0.707 | 1.08 | -0.329 | -0.57 | 0.001 | -0.023 | -0.023 | 0 | 0.24 | 0.3 | 0.24 | 0.09 | 0.21 | 0.1 |
| 7 | 71406392 | rs10271976 | -0.461 | 0.734 | -0.81 | -0.271 | 0.01 | -0.028 | -0.028 | 0.025 | 0.72 | 0.8 | 0.78 | 0.53 | 0.79 | 0.82 |
| 7 | 71408360 | rs10227914 | 1.005 | 0.944 | -0.809 | -0.966 | -0.011 | 0.008 | 0.008 | 0.009 | 0.16 | 0.15 | 0.18 | 0.1 | 0.18 | 0.14 |
| 7 | 71411789 | rs974134 | -0.489 | 0.991 | -0.675 | -0.929 | -0.005 | -0.001 | -0.001 | 0 | 0.71 | 0.7 | 0.54 | 0.3 | 0.75 | 0.69 |
| 7 | 71413328 | rs17426915 | 1.494 | 0.951 | -0.864 | -1.084 | -0.075 | 0.134 | 0.134 | 0 | 0.23 | 0.02 | 0.13 | 0.03 | 0.23 | 0.1 |
| 7 | 71414285 | rs10252758 | -0.047 | 0.957 | -0.706 | -1.013 | 0 | 0.027 | 0.027 | 2.55 | 0.47 | 0.63 | 0.4 | 0.26 | 0.5 | 0.59 |
| 7 | 71418458 | rs1076303 | 0.14 | 0.706 | -0.709 | -1.017 | 0.001 | 0.029 | 0.029 | 3.523 | 0.48 | 0.38 | 0.58 | 0.73 | 0.44 | 0.4 |
| 7 | 71424080 | rs6460700 | 1.04 | 0.721 | -0.678 | -1.069 | -0.014 | -0.047 | -0.047 | 0.29 | 0.3 | 0.36 | 0.21 | 0.16 | 0.32 | 0.45 |
| 7 | 71424171 | rs6460701 | -0.924 | 0.852 | -1.278 | -1.05 | 0.006 | -0.003 | -0.003 | 0 | 0.74 | 0.8 | 0.56 | 0.29 | 0.79 | 0.72 |
| 7 | 71424467 | rs13238682 | 1.107 | 0.642 | -1.039 | -1.023 | -0.04 | 0.112 | 0.112 | 0 | 0.23 | 0.05 | 0.13 | 0.03 | 0.23 | 0.1 |
| 7 | 71425392 | rs10276706 | -0.017 | 0.688 | -0.912 | -0.731 | -0.008 | 0.055 | 0.055 | 0.031 | 0.54 | 0.64 | 0.36 | 0.19 | 0.54 | 0.56 |
| 7 | 71437062 | rs7795273 | 0.493 | 0.486 | -1.066 | -0.545 | -0.087 | 0.159 | 0.159 | 0 | 0.25 | 0.02 | 0.11 | 0.03 | 0.26 | 0.1 |
| 7 | 71439807 | rs12530737 | -0.886 | 0.481 | -0.15 | 0.245 | 0.023 | 0.082 | 0.082 | 0.665 | 0.26 | 0.52 | 0.36 | 0.56 | 0.28 | 0.33 |
| 7 | 71443057 | rs17137604 | -1.029 | 0.483 | 0.249 | 0.969 | 0.028 | 0.242 | 0.242 | 0 | 0.06 | 0.19 | 0.17 | 0.38 | 0.09 | 0.16 |
| 7 | 71470001 | rs1099451 | -1.913 | 0.459 | 0.927 | 0.989 | 0.108 | 0.501 | 0.501 | 1.28 | 0.07 | 0.47 | 0.1 | 0.38 | 0.07 | 0.13 |
| 7 | 71482224 | rs10273495 | 0.189 | 0.076 | 0.938 | 0.85 | -0.011 | 0.257 | 0.257 | 0 | 0.1 | 0.32 | 0.07 | 0.35 | 0.06 | 0.21 |
| 7 | 71482823 | rs844796 | -1.236 | 0.033 | 0.881 | 0.808 | -0.02 | 0.123 | 0.123 | 0.076 | 0.16 | 0.66 | 0.15 | 0.4 | 0.13 | 0.27 |
| 7 | 71489191 | rs10280568 | 0.043 | 0.063 | 0.727 | 0.914 | -0.014 | 0.24 | 0.24 | 0 | 0.1 | 0.34 | 0.07 | 0.35 | 0.06 | 0.23 |
| 7 | 71504260 | rs844669 | -0.833 | -0.132 | -0.09 | 0.568 | 0.006 | 0.029 | 0.029 | 0 | 0.61 | 0.72 | 0.57 | 0.75 | 0.61 | 0.7 |
| 7 | 71508645 | rs4717625 | 0.574 | 0.161 | -0.426 | 0.46 | -0.069 | -0.059 | -0.059 | 0.072 | 0.14 | 0.01 | 0.15 | 0.16 | 0.17 | 0.15 |
| 7 | 71512298 | rs4997672 | 0.282 | 0.142 | -0.396 | 0.462 | -0.068 | -0.057 | -0.057 | 0 | 0.16 | 0.02 | 0.19 | 0.2 | 0.22 | 0.18 |
| 7 | 71513381 | rs10277807 | 0.319 | 0.03 | 0.26 | 0.585 | 0.001 | 0.208 | 0.208 | 0 | 0.11 | 0.34 | 0.08 | 0.33 | 0.08 | 0.25 |
| 7 | 71521799 | rs844762 | 0.804 | -0.139 | -0.011 | -0.637 | -0.002 | 0.014 | 0.014 | 0 | 0.32 | 0.24 | 0.22 | 0.04 | 0.31 | 0.17 |
| 7 | 71524672 | rs844759 | -1.495 | -0.2 | -0.323 | -0.905 | -0.013 | 0.128 | 0.128 | 2.349 | 0.68 | 0.91 | 0.83 | 1 | 0.7 | 0.88 |
| 7 | 71533900 | rs4717626 | -0.001 | -0.197 | 0.101 | -0.586 | 0 | 0.001 | 0.001 | 0 | 0.47 | 0.55 | 0.5 | 0.67 | 0.46 | 0.63 |
| 7 | 71546947 | rs12699121 | -1.333 | -0.641 | -0.274 | -0.978 | -0.065 | 0.195 | 0.195 | 1.387 | 0.7 | 0.97 | 0.84 | 1 | 0.7 | 0.88 |
| 7 | 71550079 | rs4719207 | 0.142 | -0.884 | -0.162 | -1.166 | 0.005 | -0.017 | -0.017 | 4.355 | 0.24 | 0.18 | 0.3 | 0.26 | 0.22 | 0.21 |
| 7 | 71557399 | rs6953054 | 1.057 | -0.798 | 0.159 | -1.319 | 0.008 | -0.05 | -0.05 | 0 | 0.24 | 0.27 | 0.19 | 0.03 | 0.31 | 0.17 |
| 7 | 71578708 | rs7798356 | 1.089 | -0.834 | -0.187 | -1.171 | 0.001 | 0.066 | 0.066 | 1.368 | 0.35 | 0.18 | 0.18 | 0 | 0.31 | 0.13 |
| 7 | 71603773 | rs10950298 | -0.372 | -0.714 | -0.362 | -0.857 | 0.018 | 0.076 | 0.076 | 0.135 | 0.5 | 0.69 | 0.51 | 0.71 | 0.48 | 0.61 |
| 7 | 71625867 | rs7789624 | 1.022 | -0.756 | -0.136 | -0.954 | 0 | -0.065 | -0.065 | 1.439 | 0.32 | 0.26 | 0.2 | 0 | 0.32 | 0.12 |
| 7 | 71626255 | rs10267684 | -1.52 | -0.756 | -0.426 | -1.063 | -0.044 | 0.139 | 0.139 | 0.029 | 0.68 | 0.87 | 0.8 | 0.98 | 0.66 | 0.79 |
| 7 | 71633053 | rs7798609 | 1.267 | -0.672 | -0.369 | -1.021 | -0.005 | 0.025 | 0.025 | 2.957 | 0.31 | 0.16 | 0.18 | 0 | 0.32 | 0.12 |
| 7 | 71637229 | rs7808523 | -1.064 | -0.356 | -0.57 | -0.661 | -0.024 | 0.097 | 0.097 | 0 | 0.66 | 0.82 | 0.79 | 0.98 | 0.66 | 0.79 |
| 7 | 71683507 | rs12112510 | -0.404 | -0.078 | 0.191 | 0.535 | -0.01 | 0.144 | 0.144 | 0 | 0.09 | 0.13 | 0.26 | 0.42 | 0.09 | 0.23 |
| 7 | 71688190 | rs11972942 | 0.149 | -0.381 | -0.062 | -0.218 | 0.001 | -0.002 | -0.002 | 0 | 0.6 | 0.63 | 0.49 | 0.58 | 0.58 | 0.65 |
| 7 | 71711227 | rs496808 | -0.767 | -0.396 | 0.084 | -0.307 | 0.002 | 0.008 | 0.008 | 0.044 | 0.68 | 0.75 | 0.76 | 1 | 0.67 | 0.88 |
| 7 | 71712651 | rs2677277 | -0.744 | -0.407 | 0.078 | -0.312 | 0.003 | 0.009 | 0.009 | 0.036 | 0.68 | 0.75 | 0.76 | 1 | 0.67 | 0.88 |
| 7 | 71714102 | rs491413 | -0.926 | -0.415 | 0.06 | -0.315 | -0.012 | 0.109 | 0.109 | 0.26 | 0.69 | 0.89 | 0.77 | 1 | 0.67 | 0.88 |
| 7 | 71727921 | rs548405 | -0.261 | -0.592 | -0.079 | -0.448 | 0.004 | -0.012 | -0.012 | 0.425 | 0.63 | 0.65 | 0.51 | 0.58 | 0.58 | 0.65 |
| 7 | 71732674 | rs477387 | -0.572 | -0.532 | -0.036 | -0.582 | 0.002 | 0.257 | 0.257 | 0.081 | 0.15 | 0.45 | 0.15 | 0.43 | 0.1 | 0.24 |
| 7 | 71751316 | rs12699131 | 0.397 | -0.493 | -0.173 | -0.567 | 0.013 | 0.012 | 0.012 | 0.22 | 0.54 | 0.29 | 0.6 | 0.57 | 0.57 | 0.69 |
| 7 | 71769515 | rs10256020 | 0.162 | -0.114 | 0.03 | -0.273 | 0.038 | 0.296 | 0.296 | 0 | 0.11 | 0.34 | 0.14 | 0.42 | 0.09 | 0.19 |
| 7 | 71797644 | rs10229537 | -0.275 | -0.486 | -0.288 | -0.466 | 0.01 | 0.188 | 0.188 | 0 | 0.13 | 0.37 | 0.16 | 0.4 | 0.09 | 0.22 |
| 7 | 71808745 | rs10274751 | -0.203 | -0.35 | -0.281 | -0.449 | 0.025 | 0.195 | 0.195 | 0 | 0.13 | 0.37 | 0.16 | 0.4 | 0.1 | 0.23 |
| 7 | 71815170 | rs11764286 | -0.097 | -0.343 | -0.18 | -0.44 | 0.06 | 0.319 | 0.319 | 0 | 0.06 | 0.26 | 0.13 | 0.38 | 0.07 | 0.12 |
| 7 | 71847345 | rs1574549 | 0.666 | -0.574 | -0.798 | -0.221 | -0.061 | 0.133 | 0.133 | 0.117 | 0.22 | 0.03 | 0.22 | 0 | 0.29 | 0.1 |
| 7 | 71854104 | rs6460721 | -0.994 | -0.566 | -0.162 | 0.123 | -0.004 | -0.017 | -0.017 | 0 | 0.07 | 0.31 | 0.03 | 0 | 0.01 | 0 |
| 7 | 73069245 | rs4377835 | -0.025 | 0.868 | -2.062 | -0.894 | 0.044 | 0.299 | 0.299 | 0 | 0.48 | 0.84 | 0.32 | 0.07 | 0.36 | 0.23 |
| 7 | 73108924 | rs10479665 | 0.868 | 0.12 | -0.46 | 0.052 | 0.038 | 0.43 | 0.43 | 0.359 | 0.93 | 0.63 | 0.81 | 0.56 | 0.96 | 0.8 |
| 7 | 73113483 | rs6956879 | 0.965 | -0.124 | -0.426 | -0.263 | -0.005 | 0.016 | 0.016 | 0 | 0.53 | 0.37 | 0.28 | 0.32 | 0.45 | 0.3 |
| 7 | 73139762 | rs6959538 | -0.949 | -0.055 | -0.858 | -0.384 | 0.006 | 0.024 | 0.024 | 0 | 0.13 | 0.09 | 0.06 | 0 | 0.09 | 0.08 |

| | | | | | | | | | | | | | | | | |
|---|---|---|---|---|---|---|---|---|---|---|---|---|---|---|---|---|
| 7 | 73140386 | rs9654749 | -0.594 | -0.169 | -0.172 | 0.61 | -0.003 | 0.228 | 0.228 | 0 | 0.14 | 0.39 | 0.22 | 0.49 | 0.16 | 0.4 |
| 7 | 73141464 | rs2030922 | 0.363 | -0.825 | -1.146 | -0.101 | 0.1 | 0.157 | 0.157 | 0 | 0.5 | 0.26 | 0.56 | 0.23 | 0.61 | 0.34 |
| 7 | 73146039 | rs4717104 | -0.577 | 0.387 | -1.261 | -0.243 | -0.032 | 0.091 | 0.091 | 0 | 0.57 | 0.66 | 0.56 | 0.76 | 0.54 | 0.66 |
| 7 | 73150511 | rs1059779 | -0.666 | 0.704 | -1.334 | 0.111 | -0.039 | -0.014 | -0.014 | 0.31 | 0.11 | 0.04 | 0.13 | 0.09 | 0.22 | 0.09 |
| 7 | 73151644 | rs6460052 | -0.758 | -0.455 | -1.914 | -0.059 | -0.036 | -0.022 | -0.022 | 0 | 0.61 | 0.83 | 0.54 | 0.52 | 0.49 | 0.55 |
| 7 | 73155017 | rs10232827 | -0.057 | -0.498 | -1.9 | -0.059 | 0.159 | 0.02 | 0.02 | 0 | 0.5 | 0.79 | 0.41 | 0.43 | 0.27 | 0.46 |
| 7 | 73156211 | rs10266412 | -0.754 | -0.472 | -1.905 | -0.059 | -0.033 | -0.022 | -0.022 | 0.426 | 0.61 | 0.83 | 0.54 | 0.52 | 0.49 | 0.55 |
| 7 | 73157315 | rs34062071 | 0.082 | -0.404 | -1.883 | -0.015 | -0.038 | -0.014 | -0.014 | 0.218 | 0.11 | 0.04 | 0.12 | 0.09 | 0.21 | 0.09 |
| 7 | 73178435 | rs11764572 | -0.581 | -0.71 | -1.113 | -0.147 | 0 | 0 | 0 | 0 | 0.58 | 0.58 | 0.51 | 0.51 | 0.48 | 0.55 |
| 7 | 73182017 | rs12536661 | 0.538 | -1.591 | -0.677 | -0.311 | -0.03 | 0 | 0 | 0 | 0.32 | 0.37 | 0.32 | 0.27 | 0.2 | 0.25 |
| 7 | 73182941 | rs11979807 | -0.998 | -1.455 | -0.713 | -0.293 | 0 | 0 | 0 | 0 | 0.58 | 0.57 | 0.51 | 0.51 | 0.48 | 0.55 |
| 7 | 73187517 | rs6460055 | -0.205 | -1.48 | -0.725 | -0.303 | -0.006 | 0 | 0 | 0 | 0.46 | 0.53 | 0.39 | 0.42 | 0.27 | 0.45 |
| 7 | 73187651 | rs4484568 | -0.902 | -1.368 | -0.677 | -0.261 | 0 | 0 | 0 | 0 | 0.58 | 0.57 | 0.51 | 0.51 | 0.48 | 0.55 |
| 7 | 73188318 | rs4131376 | -0.903 | -1.369 | -0.641 | -0.262 | 0 | 0 | 0 | 4.429 | 0.58 | 0.57 | 0.51 | 0.51 | 0.48 | 0.55 |
| 7 | 73198616 | rs6979031 | -0.914 | -1.203 | -0.628 | -0.085 | -0.001 | -0.002 | -0.002 | 0 | 0.58 | 0.58 | 0.51 | 0.48 | 0.48 | 0.54 |
| 7 | 73219508 | rs13232536 | 0.482 | -1.814 | -0.444 | -0.077 | 0.094 | 0.049 | 0.049 | 0 | 0.51 | 0.29 | 0.61 | 0.65 | 0.73 | 0.55 |
| 7 | 73226807 | rs9647713 | -0.32 | -1.902 | -0.276 | -0.178 | -0.024 | -0.015 | -0.015 | 0 | 0.67 | 0.66 | 0.72 | 0.8 | 0.81 | 0.77 |
| 7 | 73239618 | rs4255023 | 0.204 | -1.619 | 0.042 | 0.199 | 0.052 | 0.005 | 0.005 | 0 | 0.53 | 0.34 | 0.61 | 0.58 | 0.72 | 0.57 |
| 7 | 73241386 | rs7455225 | 0.689 | -1.024 | -0.114 | 0.61 | 0.069 | 0.029 | 0.029 | 0 | 0.46 | 0.15 | 0.5 | 0.39 | 0.65 | 0.35 |
| 7 | 73245519 | rs8629 | 0.051 | -1.056 | -0.014 | 0.564 | -0.009 | 0 | 0 | 0 | 0.54 | 0.46 | 0.61 | 0.58 | 0.73 | 0.57 |
| 7 | 73246461 | rs1127155 | -0.228 | -1.172 | -0.058 | 0.549 | -0.021 | 0.003 | 0.003 | 0 | 0.59 | 0.55 | 0.62 | 0.58 | 0.73 | 0.57 |
| 7 | 73247204 | rs9691413 | 0.12 | -1.174 | -0.058 | 0.548 | -0.006 | 0 | 0 | 3.793 | 0.54 | 0.45 | 0.61 | 0.58 | 0.73 | 0.57 |
| 7 | 73247732 | rs6968360 | -0.228 | -1.172 | -0.06 | 0.55 | -0.021 | 0.003 | 0.003 | 0 | 0.59 | 0.55 | 0.62 | 0.58 | 0.73 | 0.57 |
| 7 | 73253772 | rs13223593 | 0.086 | -1.163 | -0.007 | 0.554 | 0.001 | 0 | 0 | 0 | 0.54 | 0.46 | 0.61 | 0.58 | 0.73 | 0.57 |
| 7 | 73269937 | rs6949053 | -0.189 | -1.15 | -0.108 | 0.564 | -0.034 | -0.002 | -0.002 | 0 | 0.58 | 0.51 | 0.61 | 0.62 | 0.73 | 0.57 |
| 7 | 73275501 | rs11770024 | 0.103 | -1.04 | -0.25 | 0.607 | 0.028 | 0.009 | 0.009 | 0 | 0.53 | 0.4 | 0.61 | 0.63 | 0.72 | 0.58 |
| 7 | 73275769 | rs11767129 | 0.062 | -1.021 | -0.631 | 0.616 | 0.085 | 0.021 | 0.021 | 0 | 0.53 | 0.29 | 0.6 | 0.63 | 0.73 | 0.58 |
| 7 | 73278177 | rs7796838 | 0.061 | -0.951 | -0.883 | 0.49 | 0.08 | 0.021 | 0.021 | 0.077 | 0.46 | 0.25 | 0.57 | 0.63 | 0.66 | 0.58 |
| 7 | 73280691 | rs13244770 | 0.257 | -0.652 | -0.844 | 0.225 | 0 | -0.006 | -0.006 | 0.03 | 0.41 | 0.26 | 0.49 | 0.43 | 0.57 | 0.42 |
| 7 | 73282030 | rs11767926 | 0.249 | -0.644 | -0.843 | 0.226 | 0.003 | -0.006 | -0.006 | 0 | 0.42 | 0.26 | 0.49 | 0.43 | 0.58 | 0.42 |
| 7 | 73284148 | rs11763011 | 0.034 | -0.741 | -1.026 | 0.118 | 0.015 | 0.002 | 0.002 | 0.007 | 0.45 | 0.24 | 0.49 | 0.44 | 0.58 | 0.46 |
| 7 | 73301489 | rs4717827 | -0.883 | -0.75 | -0.442 | -0.082 | -0.053 | 0 | 0 | 0 | 0.27 | 0.48 | 0.28 | 0.3 | 0.16 | 0.3 |
| 7 | 73301666 | rs4717829 | -0.406 | -0.797 | -0.454 | -0.077 | 0.028 | -0.023 | -0.023 | 0 | 0.19 | 0.09 | 0.23 | 0.29 | 0.13 | 0.29 |
| 7 | 73304636 | rs12669267 | -0.426 | -0.81 | -0.338 | -0.06 | -0.015 | 0.012 | 0.012 | 0 | 0.19 | 0.26 | 0.25 | 0.3 | 0.14 | 0.3 |
| 7 | 73313856 | rs11769827 | -1.136 | -0.051 | -0.854 | -0.416 | 0 | -0.011 | -0.011 | 0 | 0.37 | 0.42 | 0.4 | 0.33 | 0.36 | 0.43 |
| 7 | 73320111 | rs12538362 | 1.133 | -0.034 | -2.159 | -1.159 | 0 | 0.024 | 0.024 | 0 | 0.21 | 0.09 | 0.15 | 0 | 0.19 | 0.16 |
| 7 | 73326629 | rs4717839 | -0.807 | -0.03 | -1.97 | -0.605 | -0.013 | 0.018 | 0.018 | 0.252 | 0.36 | 0.16 | 0.38 | 0.32 | 0.38 | 0.47 |
| 7 | 73359124 | rs12530707 | -0.209 | -0.851 | -2.177 | -0.377 | 0.048 | -0.02 | -0.02 | 0 | 0.39 | 0.76 | 0.37 | 0.51 | 0.17 | 0.49 |
| 7 | 73376167 | rs6460065 | -0.406 | 0.325 | -2.506 | -0.21 | 0.007 | -0.015 | -0.015 | 0.424 | 0.6 | 0.89 | 0.56 | 0.62 | 0.37 | 0.65 |
| 7 | 73387168 | rs6460066 | -0.064 | 0.573 | -1.922 | -0.721 | -0.004 | 0.027 | 0.027 | 0 | 0.81 | 0.87 | 0.81 | 0.92 | 0.78 | 0.81 |
| 7 | 73403193 | rs6964149 | 0.663 | 1.408 | -2.033 | -0.659 | -0.101 | 0.102 | 0.102 | 5.795 | 0.27 | 0.04 | 0.2 | 0.09 | 0.35 | 0.17 |
| 7 | 73405560 | rs6975518 | 0.262 | 0.458 | -1.871 | -0.52 | -0.017 | 0.155 | 0.155 | 2.061 | 0.4 | 0.1 | 0.3 | 0.17 | 0.54 | 0.29 |
| 7 | 73410408 | rs6973541 | -0.216 | -0.108 | -0.449 | -0.23 | -0.015 | 0.01 | 0.01 | 0 | 0.07 | 0.28 | 0.02 | 0.04 | 0.01 | 0.07 |
| 7 | 73416168 | rs4717860 | 1.212 | -0.467 | -0.661 | -0.295 | 0.046 | 0.043 | 0.043 | 0.022 | 0.61 | 0.26 | 0.54 | 0.31 | 0.76 | 0.51 |
| 7 | 73423846 | rs10224499 | 0.154 | -0.34 | -0.649 | -0.476 | 0.004 | -0.001 | -0.001 | 0 | 0.34 | 0.35 | 0.52 | 0.6 | 0.39 | 0.48 |
| 7 | 73425238 | rs6948963 | 0.606 | -0.158 | -0.715 | -0.286 | 0.001 | 0.017 | 0.017 | 0.762 | 0.45 | 0.43 | 0.37 | 0.31 | 0.49 | 0.39 |
| 7 | 73428671 | rs6460068 | 1.379 | -0.17 | -0.646 | -0.298 | -0.043 | 0.086 | 0.086 | 0.922 | 0.37 | 0.17 | 0.34 | 0.27 | 0.47 | 0.29 |
| 7 | 73428879 | rs4717863 | -0.537 | -0.122 | -0.522 | -0.229 | 0 | 0.001 | 0.001 | 0.663 | 0.48 | 0.49 | 0.6 | 0.61 | 0.49 | 0.54 |
| 7 | 73436066 | rs6979609 | -0.541 | 0.056 | -0.128 | 0.296 | 0.021 | 0.097 | 0.097 | 0 | 0.08 | 0.32 | 0.11 | 0.23 | 0.05 | 0.12 |
| 7 | 73443132 | rs7784595 | -0.51 | 0.056 | -0.227 | 0.296 | 0.012 | 0.083 | 0.083 | 0 | 0.08 | 0.28 | 0.11 | 0.23 | 0.05 | 0.12 |
| 7 | 73447506 | rs873647 | -0.472 | 0.069 | -0.451 | 0.288 | -0.004 | 0.051 | 0.051 | 0 | 0.08 | 0.22 | 0.1 | 0.23 | 0.05 | 0.12 |
| 7 | 73447658 | rs870424 | -0.472 | 0.069 | -0.451 | 0.288 | -0.004 | 0.051 | 0.051 | 0 | 0.08 | 0.22 | 0.1 | 0.24 | 0.05 | 0.12 |
| 7 | 73454199 | rs4717865 | 0.39 | 0.31 | -1.12 | 1.005 | -0.032 | -0.044 | -0.044 | 0.029 | 0.09 | 0.01 | 0.28 | 0.38 | 0.09 | 0.28 |
| 7 | 73470714 | rs2071307 | 1.107 | 0.011 | -1.322 | -0.292 | -0.028 | 0.132 | 0.132 | 1.316 | 0.36 | 0.13 | 0.25 | 0.11 | 0.42 | 0.23 |
| 7 | 73470782 | rs2856728 | 0.963 | -0.542 | -0.903 | -0.362 | -0.004 | 0 | 0 | 2.251 | 0.78 | 0.75 | 0.81 | 0.68 | 0.83 | 0.73 |
| 7 | 73473494 | rs2528795 | -1.301 | -0.728 | -0.445 | -0.975 | -0.037 | -0.123 | -0.123 | 0 | 0.26 | 0.47 | 0.18 | 0.08 | 0.2 | 0.16 |
| 7 | 73484224 | rs10233395 | -2.027 | 0.093 | 0.062 | -0.689 | 0.022 | -0.001 | -0.001 | 0 | 0.08 | 0.31 | 0.05 | 0.06 | 0.06 | 0.08 |
| 7 | 73489736 | rs1091813 | 2 | 0.029 | 0.023 | -0.612 | 0.019 | 0.039 | 0.039 | 0 | 0.91 | 0.6 | 0.9 | 0.89 | 0.92 | 0.9 |
| 7 | 73491212 | rs1091811 | 1.619 | 0.043 | -0.054 | -0.596 | 0.115 | 0.113 | 0.113 | 0 | 0.89 | 0.48 | 0.85 | 0.87 | 0.86 | 0.87 |
| 7 | 73493533 | rs810549 | 1.532 | -0.419 | -0.698 | -0.808 | 0.018 | 0.017 | 0.017 | 3.53 | 0.27 | 0.04 | 0.28 | 0.25 | 0.22 | 0.13 |
| 7 | 73497196 | rs6460071 | -1.656 | 0.913 | 0.402 | -0.576 | 0.173 | 0.124 | 0.124 | 0 | 0.08 | 0.51 | 0.1 | 0.08 | 0.12 | 0.11 |
| 7 | 73504233 | rs810536 | 2.487 | 0.759 | 0.277 | -0.66 | -0.003 | -0.026 | -0.026 | 0 | 0.71 | 0.43 | 0.73 | 0.84 | 0.74 | 0.69 |
| 7 | 73524207 | rs150862 | 2.336 | 1.17 | 0.619 | -0.343 | 0.022 | 0.041 | 0.041 | 0 | 0.8 | 0.46 | 0.8 | 0.79 | 0.8 | 0.65 |
| 7 | 73529976 | rs2285025 | -0.83 | 1.198 | 0.614 | -0.339 | -0.001 | -0.001 | -0.001 | 0 | 0.11 | 0.09 | 0.09 | 0.11 | 0.13 | 0.24 |
| 7 | 73538326 | rs178413 | 0.122 | 0.724 | 0.53 | -0.444 | -0.016 | 0.018 | 0.018 | 0.288 | 0.09 | 0.04 | 0.08 | 0 | 0.12 | 0.05 |
| 7 | 73554035 | rs444297 | 2.703 | 0.706 | 0.68 | -0.109 | 0.001 | -0.021 | -0.021 | 3.196 | 0.65 | 0.45 | 0.68 | 0.78 | 0.67 | 0.6 |
| 7 | 73568191 | rs170323 | 1.718 | 0.936 | 0.897 | 0.03 | 0.021 | 0.083 | 0.083 | 0 | 0.92 | 0.59 | 0.91 | 0.87 | 0.95 | 0.91 |
| 7 | 73568425 | rs12539781 | -1.117 | 0.908 | 0.718 | -0.025 | -0.007 | 0.006 | 0.006 | 0.193 | 0.2 | 0.12 | 0.15 | 0.11 | 0.22 | 0.28 |
| 7 | 73579794 | rs223014 | 1.832 | 0.945 | 0.87 | 0.124 | 0.003 | -0.014 | -0.014 | 0 | 0.66 | 0.54 | 0.59 | 0.74 | 0.66 | 0.58 |

| | | | | | | | | | | | | | | | | |
|---|---|---|---|---|---|---|---|---|---|---|---|---|---|---|---|---|
| 7 | 73584129 | rs6953474 | -1.163 | 0.93 | 0.604 | 0.035 | -0.025 | 0.02 | 0.02 | 0.139 | 0.27 | 0.13 | 0.31 | 0.14 | 0.3 | 0.34 |
| 7 | 73584226 | rs222999 | 1.514 | 0.796 | 1.01 | 0.183 | 0.008 | 0.03 | 0.03 | 0 | 0.93 | 0.76 | 0.92 | 0.89 | 0.95 | 0.92 |
| 7 | 73599917 | rs150875 | 2.179 | 0.82 | 1.148 | 0.267 | 0.006 | 0.065 | 0.065 | 0.12 | 0.93 | 0.66 | 0.92 | 0.93 | 0.97 | 0.92 |
| 7 | 73624615 | rs7810996 | -1.014 | 1.066 | 1.432 | 0.176 | 0.005 | -0.009 | -0.009 | 0 | 0.27 | 0.32 | 0.32 | 0.12 | 0.3 | 0.28 |
| 7 | 73630155 | rs2293757 | -1.482 | 1.17 | 1.355 | 0.226 | 0.011 | 0.01 | 0.01 | 0 | 0.06 | 0.08 | 0.05 | 0.08 | 0.06 | 0.1 |
| 7 | 73649825 | rs3135698 | -1.485 | 1.275 | 1.515 | 0.273 | 0.077 | 0.072 | 0.072 | 0 | 0.07 | 0.38 | 0.07 | 0.08 | 0.07 | 0.1 |
| 7 | 73650292 | rs3135696 | -1.485 | 1.275 | 1.515 | 0.273 | 0.077 | 0.072 | 0.072 | 0 | 0.07 | 0.38 | 0.07 | 0.08 | 0.07 | 0.1 |
| 7 | 73673643 | rs12538007 | -1.335 | 1.464 | 0.54 | 0.402 | 0 | -0.001 | -0.001 | 0 | 0.36 | 0.66 | 0.41 | 0.21 | 0.38 | 0.38 |
| 7 | 73695848 | rs17145359 | -1.535 | 0.82 | 1.164 | 0.562 | 0.058 | -0.027 | -0.027 | 0 | 0.1 | 0.39 | 0.09 | 0.03 | 0.08 | 0.04 |
| 7 | 73697327 | rs17145361 | -1.747 | 0.612 | 0.97 | 0.546 | -0.014 | -0.057 | -0.057 | 0 | 0.21 | 0.46 | 0.14 | 0.07 | 0.15 | 0.26 |
| 7 | 73703236 | rs10236516 | -1.765 | 0.556 | 0.965 | 0.541 | -0.02 | -0.06 | -0.06 | 0 | 0.21 | 0.48 | 0.14 | 0.07 | 0.15 | 0.26 |
| 7 | 73724910 | rs229878 | 1.029 | 0.694 | 0.531 | 1.411 | 0.01 | -0.039 | -0.039 | 0 | 0.69 | 0.25 | 0.58 | 0.62 | 0.76 | 0.6 |
| 7 | 73728109 | rs229874 | 0.969 | 0.686 | 0.559 | 1.408 | 0.007 | -0.034 | -0.034 | 0.015 | 0.7 | 0.27 | 0.58 | 0.62 | 0.76 | 0.6 |
| 7 | 73732583 | rs229872 | 0.951 | 0.677 | 0.703 | 1.404 | 0 | -0.001 | -0.001 | 0 | 0.71 | 0.34 | 0.6 | 0.62 | 0.76 | 0.6 |
| 7 | 73738067 | rs519462 | 0.625 | 0.658 | 1.401 | 1.383 | -0.001 | -0.004 | -0.004 | 0 | 0.75 | 0.71 | 0.78 | 0.85 | 0.78 | 0.69 |
| 7 | 73742424 | rs573296 | 0.816 | 0.717 | 0.889 | 1.608 | -0.004 | 0.014 | 0.014 | 0.325 | 0.71 | 0.4 | 0.59 | 0.51 | 0.76 | 0.6 |
| 7 | 73752699 | rs574637 | 0.955 | 0.723 | 0.692 | 1.54 | 0 | -0.003 | -0.003 | 0 | 0.7 | 0.33 | 0.57 | 0.5 | 0.76 | 0.61 |
| 7 | 73753250 | rs539518 | 0.823 | 0.818 | 0.696 | 1.552 | -0.001 | 0.001 | 0.001 | 0.389 | 0.71 | 0.35 | 0.58 | 0.49 | 0.76 | 0.61 |
| 7 | 73789730 | rs35705083 | 0.637 | 1.047 | 0.118 | 1.287 | 0.011 | 0.019 | 0.019 | 0.08 | 0.56 | 0.26 | 0.5 | 0.44 | 0.63 | 0.59 |
| 7 | 73803303 | rs7790414 | -0.168 | 1.328 | -0.015 | 0.641 | -0.002 | -0.022 | -0.022 | 0.525 | 0.17 | 0.18 | 0.1 | 0.02 | 0.18 | 0.14 |
| 7 | 73810096 | rs10949838 | -0.399 | 1.113 | -0.202 | 0.797 | 0.015 | 0.015 | 0.015 | 0 | 0.61 | 0.53 | 0.5 | 0.52 | 0.51 | 0.46 |
| 7 | 73811146 | rs1468417 | 1.031 | 1.037 | 0.785 | 1.122 | 0.088 | 0.182 | 0.182 | 0.642 | 0.91 | 0.58 | 0.72 | 0.76 | 0.9 | 0.89 |
| 7 | 73819191 | rs2240435 | 0.269 | 1.173 | 0.394 | 0.75 | 0.001 | -0.003 | -0.003 | 0 | 0.71 | 0.7 | 0.6 | 0.77 | 0.66 | 0.66 |
| 7 | 73820636 | rs4717890 | 0.252 | 1.176 | 0.395 | 0.752 | 0.001 | -0.003 | -0.003 | 0 | 0.71 | 0.7 | 0.6 | 0.77 | 0.66 | 0.66 |
| 7 | 73822588 | rs4717138 | 0.144 | 1.18 | 0.299 | 0.773 | -0.002 | 0.001 | 0.001 | 0 | 0.71 | 0.75 | 0.61 | 0.77 | 0.66 | 0.66 |
| 7 | 73824052 | rs6974767 | -0.217 | 0.91 | 0.238 | 1.11 | -0.011 | 0.039 | 0.039 | 0 | 0.29 | 0.4 | 0.35 | 0.44 | 0.23 | 0.21 |
| 7 | 73862412 | rs2158797 | 0.11 | 0.63 | -0.239 | 0.62 | -0.003 | 0.051 | 0.051 | 0 | 0.48 | 0.61 | 0.48 | 0.24 | 0.46 | 0.5 |
| 7 | 75235549 | rs17149227 | -1.807 | -0.804 | 1.097 | 0.247 | 0.02 | 0.14 | 0.14 | 0 | 0.09 | 0.32 | 0.26 | 0.27 | 0.05 | 0.22 |
| 7 | 75237158 | rs6963510 | -1.146 | -1.242 | 0.829 | -0.402 | -0.072 | -0.121 | -0.121 | 0 | 0.19 | 0.43 | 0.05 | 0 | 0.07 | 0.1 |
| 7 | 75240915 | rs7794510 | -1.246 | -1.656 | 0.597 | 0.205 | -0.058 | -0.008 | -0.008 | 0 | 0.25 | 0.67 | 0.08 | 0.19 | 0.07 | 0.2 |
| 7 | 75243991 | rs1179605 | -1.358 | -1.503 | 1.244 | 0.037 | -0.028 | 0.003 | 0.003 | 0 | 0.17 | 0.54 | 0.09 | 0.18 | 0.08 | 0.13 |
| 7 | 75246126 | rs1179610 | -1.55 | -1.45 | 1.374 | -0.148 | 0.02 | 0.023 | 0.023 | 0 | 0.19 | 0.54 | 0.14 | 0.3 | 0.16 | 0.18 |
| 7 | 75255491 | rs1179625 | -0.824 | -1.074 | -0.045 | -0.96 | -0.012 | 0.043 | 0.043 | 0.218 | 0.44 | 0.65 | 0.36 | 0.35 | 0.43 | 0.4 |
| 7 | 75255633 | rs7810980 | -0.516 | -1.002 | 0.522 | -1.096 | 0.039 | -0.091 | -0.091 | 0 | 0.16 | 0.42 | 0.09 | 0.03 | 0.14 | 0.1 |
| 7 | 75263792 | rs12534422 | -0.309 | -0.961 | 0.822 | -1.165 | 0.006 | -0.043 | -0.043 | 0 | 0.29 | 0.37 | 0.25 | 0.1 | 0.29 | 0.28 |
| 7 | 75264463 | rs4731231 | 0.729 | -0.952 | 0.676 | -1.153 | -0.034 | 0.071 | 0.071 | 0 | 0.21 | 0.06 | 0.2 | 0.07 | 0.23 | 0.2 |
| 7 | 75274068 | rs1179594 | -1.446 | -1.12 | 0.704 | -1.198 | 0.001 | 0.076 | 0.076 | 0 | 0.49 | 0.67 | 0.36 | 0.33 | 0.41 | 0.35 |
| 7 | 75274161 | rs1179595 | -0.348 | -1.245 | 0.843 | -1.126 | 0.022 | 0.017 | 0.017 | 0 | 0.14 | 0.28 | 0.11 | 0.23 | 0.14 | 0.08 |
| 7 | 75279583 | rs12154679 | -0.022 | -1.683 | 0.245 | -1.4 | 0.019 | 0.061 | 0.061 | 0 | 0.22 | 0.07 | 0.17 | 0.07 | 0.15 | 0.16 |
| 7 | 75280922 | rs6960110 | -0.017 | -1.925 | 0.09 | -1.531 | -0.009 | -0.056 | -0.056 | 0 | 0.33 | 0.43 | 0.23 | 0.11 | 0.25 | 0.31 |
| 7 | 75283383 | rs17353131 | 0.084 | -2.118 | -0.412 | -1.721 | -0.054 | 0.116 | 0.116 | 1.003 | 0.23 | 0.04 | 0.11 | 0.06 | 0.22 | 0.15 |
| 7 | 75284941 | rs2867803 | -0.465 | -1.881 | -0.086 | -1.595 | -0.003 | 0.087 | 0.087 | 0 | 0.52 | 0.36 | 0.32 | 0.17 | 0.44 | 0.44 |
| 7 | 75287410 | rs757364 | -0.97 | -1.781 | -0.231 | -1.501 | -0.012 | 0.035 | 0.035 | 0.206 | 0.59 | 0.76 | 0.41 | 0.33 | 0.49 | 0.49 |
| 7 | 75294275 | rs2525467 | -1.788 | -1.95 | -0.714 | -0.588 | 0.006 | 0.008 | 0.008 | 0 | 0.76 | 0.77 | 0.7 | 0.54 | 0.55 | 0.59 |
| 7 | 75304219 | rs4728147 | -0.761 | -1.754 | -0.938 | -0.218 | -0.003 | -0.003 | -0.003 | 0 | 0.66 | 0.67 | 0.62 | 0.48 | 0.53 | 0.55 |
| 7 | 75309746 | rs10247961 | 1.001 | -1.756 | -0.887 | -0.12 | 0.006 | -0.024 | -0.024 | 0 | 0.12 | 0.09 | 0.13 | 0.21 | 0.13 | 0.15 |
| 7 | 75317218 | rs2240133 | -2.247 | -1.873 | -0.872 | -0.051 | -0.02 | -0.024 | -0.024 | 2.501 | 0.79 | 0.89 | 0.75 | 0.68 | 0.65 | 0.69 |
| 7 | 75317281 | rs2240134 | 0.817 | -2.094 | -0.882 | -0.264 | 0 | -0.011 | -0.011 | 0.091 | 0.35 | 0.37 | 0.29 | 0.22 | 0.35 | 0.41 |
| 7 | 75317583 | rs10244441 | 0.578 | -1.561 | -0.909 | 0.083 | 0.005 | -0.01 | -0.01 | 0.115 | 0.13 | 0.13 | 0.14 | 0.2 | 0.13 | 0.15 |
| 7 | 75329671 | rs2240135 | 0.695 | -0.943 | -0.675 | 0.451 | -0.002 | 0.002 | 0.002 | 0.019 | 0.31 | 0.27 | 0.3 | 0.24 | 0.35 | 0.34 |
| 7 | 75342453 | rs2240139 | -0.248 | -0.778 | -1.126 | 0.562 | -0.009 | -0.012 | -0.012 | 0 | 0.56 | 0.67 | 0.62 | 0.56 | 0.49 | 0.51 |
| 7 | 75351138 | rs4392824 | -0.078 | -0.581 | -2.254 | 0.406 | 0.002 | 0.049 | 0.049 | 0 | 0.2 | 0.1 | 0.1 | 0.01 | 0.18 | 0.08 |
| 7 | 75367376 | rs757359 | -1.028 | -0.161 | -1.903 | 1.043 | -0.028 | -0.012 | -0.012 | 0 | 0.7 | 0.82 | 0.75 | 0.74 | 0.6 | 0.59 |
| 7 | 75369103 | rs4385415 | 0.17 | 0.483 | -1.636 | 0.647 | 0.002 | -0.01 | -0.01 | 0 | 0.38 | 0.57 | 0.34 | 0.25 | 0.4 | 0.28 |
| 7 | 75370368 | rs6978786 | -0.14 | 1.103 | -1.814 | 1.111 | -0.023 | -0.046 | -0.046 | 0 | 0.62 | 0.83 | 0.67 | 0.53 | 0.58 | 0.55 |
| 7 | 75377439 | rs6945301 | 0.195 | 0.883 | -1.739 | 0.74 | -0.028 | 0.039 | 0.039 | 0 | 0.55 | 0.82 | 0.59 | 0.43 | 0.44 | 0.44 |
| 7 | 75386669 | rs4620231 | -0.357 | 0.934 | -1.522 | 1.062 | -0.003 | -0.018 | -0.018 | 0 | 0.21 | 0.15 | 0.43 | 0.25 | 0.2 | 0.21 |
| 7 | 75388031 | rs12538253 | 0.236 | 1.154 | -2.141 | 0.887 | -0.017 | -0.015 | -0.015 | 0 | 0.25 | 0.14 | 0.2 | 0.3 | 0.3 | 0.25 |
| 7 | 75388355 | rs7800985 | 0.974 | 0.508 | -1.666 | 0.239 | 0.024 | -0.06 | -0.06 | 3.008 | 0.25 | 0.41 | 0.26 | 0.14 | 0.3 | 0.23 |
| 7 | 75389831 | rs10275942 | 0.923 | 0.466 | -1.685 | 0.225 | 0.022 | -0.06 | -0.06 | 0 | 0.26 | 0.41 | 0.26 | 0.14 | 0.3 | 0.23 |
| 7 | 75398722 | rs11465352 | 0.918 | 0.198 | -2.193 | -0.101 | -0.033 | -0.076 | -0.076 | 0 | 0.2 | 0.01 | 0.09 | 0.3 | 0.19 | 0.28 |
| 7 | 75398998 | rs2302009 | 0.923 | 0.39 | -1.624 | -0.077 | 0.015 | -0.081 | -0.081 | 1.382 | 0.24 | 0.46 | 0.25 | 0.14 | 0.27 | 0.22 |
| 7 | 75400673 | rs10275464 | 0.923 | 0.374 | -1.447 | -0.077 | 0.015 | -0.081 | -0.081 | 0.801 | 0.24 | 0.46 | 0.25 | 0.14 | 0.27 | 0.22 |
| 7 | 75401418 | rs2240478 | 0.975 | 0.431 | -1.478 | -0.025 | 0 | 0.008 | 0.008 | 0 | 0.21 | 0.19 | 0.43 | 0.25 | 0.2 | 0.2 |
| 7 | 75404761 | rs7787623 | 0.563 | 0.581 | -1.097 | -0.211 | 0.017 | -0.085 | -0.085 | 0 | 0.24 | 0.48 | 0.25 | 0.14 | 0.27 | 0.22 |
| 7 | 75415801 | rs7811372 | -0.369 | 0.649 | -1.471 | 0.222 | -0.004 | -0.003 | -0.003 | 0.116 | 0.81 | 0.8 | 0.8 | 0.73 | 0.71 | 0.81 |
| 7 | 75421326 | rs13237353 | -0.59 | 0.26 | -1.918 | -0.066 | -0.013 | 0.05 | 0.05 | 0 | 0.44 | 0.3 | 0.38 | 0.68 | 0.42 | 0.57 |
| 7 | 75427565 | rs2108329 | -0.115 | 0.239 | -1.624 | 0.732 | 0.008 | -0.017 | -0.017 | 0.811 | 0.18 | 0.14 | 0.21 | 0.5 | 0.16 | 0.31 |
| 7 | 75432999 | rs7784660 | -0.719 | 0.029 | -1.371 | 0.526 | -0.007 | 0.028 | 0.028 | 0 | 0.29 | 0.36 | 0.38 | 0.47 | 0.25 | 0.38 |

| | | | | | | | | | | | | | | | | |
|---|---|---|---|---|---|---|---|---|---|---|---|---|---|---|---|---|
| 7 | 75437107 | rs7807085 | 0.195 | 0.044 | -1.728 | 0.42 | 0.013 | -0.023 | -0.023 | 0.011 | 0.18 | 0.15 | 0.21 | 0.54 | 0.15 | 0.31 |
| 7 | 75437685 | rs13340508 | -0.325 | 0.102 | -1.727 | 0.425 | 0 | -0.001 | -0.001 | 0 | 0.21 | 0.21 | 0.35 | 0.35 | 0.21 | 0.25 |
| 7 | 75439671 | rs13340490 | -1.052 | -0.017 | -1.582 | 0.459 | -0.004 | -0.017 | -0.017 | 0 | 0.32 | 0.33 | 0.53 | 0.89 | 0.3 | 0.54 |
| 7 | 75439696 | rs13340504 | -0.038 | 0.375 | -1.495 | 1.012 | 0 | 0.007 | 0.007 | 0 | 0.14 | 0.15 | 0.32 | 0.35 | 0.15 | 0.23 |
| 7 | 75441762 | rs4732416 | -1.017 | 0.764 | -1.795 | 0.548 | -0.014 | 0.008 | 0.008 | 0.014 | 0.31 | 0.24 | 0.37 | 0.33 | 0.36 | 0.39 |
| 7 | 75441794 | rs11465307 | -0.319 | 0.998 | -1.801 | 0.493 | -0.026 | 0.027 | 0.027 | 0 | 0.18 | 0.12 | 0.14 | 0.07 | 0.28 | 0.16 |
| 7 | 75442730 | rs2302006 | -1.112 | 0.763 | -1.798 | 1.089 | 0.002 | -0.004 | -0.004 | 0.03 | 0.25 | 0.27 | 0.41 | 0.56 | 0.18 | 0.35 |
| 7 | 75442759 | rs2302005 | -0.622 | 0.76 | -2.117 | 0.782 | 0.017 | -0.012 | -0.012 | 0.035 | 0.32 | 0.23 | 0.42 | 0.63 | 0.23 | 0.32 |
| 7 | 75442855 | rs2302004 | -1.61 | 0.857 | -2.123 | 0.653 | -0.004 | 0.035 | 0.035 | 0.641 | 0.47 | 0.39 | 0.54 | 0.7 | 0.45 | 0.44 |
| 7 | 75445377 | rs3757598 | -0.202 | 1 | -2.621 | 0.761 | -0.031 | -0.051 | -0.051 | 0 | 0.21 | 0.1 | 0.24 | 0.29 | 0.23 | 0.15 |
| 7 | 75451657 | rs7802169 | -0.976 | 1.048 | -2.426 | -0.109 | 0.008 | 0.183 | 0.183 | 0.172 | 0.51 | 0.22 | 0.57 | 0.8 | 0.53 | 0.6 |
| 7 | 75452584 | rs4279521 | -1.081 | 1.055 | -2.423 | -0.105 | 0.009 | 0.129 | 0.129 | 0.017 | 0.52 | 0.31 | 0.58 | 0.8 | 0.53 | 0.59 |
| 7 | 75466308 | rs6966161 | -0.736 | 0.828 | -2.787 | -0.425 | -0.021 | 0.171 | 0.171 | 0.126 | 0.48 | 0.26 | 0.53 | 0.81 | 0.46 | 0.57 |
| 7 | 75475297 | rs4732439 | 0.036 | 0.664 | -2.556 | -0.641 | 0.007 | -0.028 | -0.028 | 0 | 0.3 | 0.1 | 0.41 | 0.68 | 0.23 | 0.3 |
| 7 | 75479448 | rs6946822 | 0.035 | 0.678 | -1.8 | -0.482 | 0.003 | 0.105 | 0.105 | 0 | 0.44 | 0.7 | 0.42 | 0.23 | 0.47 | 0.4 |
| 7 | 75496097 | rs2024049 | 0.069 | 0.699 | -1.594 | -0.408 | -0.007 | 0.021 | 0.021 | 0.144 | 0.42 | 0.6 | 0.43 | 0.23 | 0.49 | 0.39 |
| 7 | 75509452 | rs42184 | -0.155 | 0.722 | -1.826 | 0.291 | -0.042 | -0.125 | -0.125 | 0.002 | 0.85 | 0.99 | 0.79 | 0.55 | 0.83 | 0.83 |
| 7 | 75510561 | rs7807392 | -0.459 | 0.48 | -1.28 | -0.38 | -0.019 | -0.088 | -0.088 | 0 | 0.19 | 0.39 | 0.16 | 0.03 | 0.18 | 0.31 |
| 7 | 75511134 | rs10255642 | -0.439 | 0.481 | -1.279 | -0.38 | -0.018 | -0.087 | -0.087 | 0 | 0.18 | 0.39 | 0.16 | 0.02 | 0.18 | 0.25 |
| 7 | 75529137 | rs10280802 | 0.647 | 0.313 | -1.612 | -1.048 | -0.002 | -0.007 | -0.007 | 0 | 0.25 | 0.19 | 0.14 | 0.01 | 0.23 | 0.19 |
| 7 | 75536736 | rs28737229 | -0.389 | 0.152 | -1.55 | -0.967 | -0.005 | 0.009 | 0.009 | 0.158 | 0.46 | 0.41 | 0.33 | 0.03 | 0.48 | 0.43 |
| 7 | 75546892 | rs3898649 | -1.511 | -0.239 | -0.476 | 0.353 | 0.006 | -0.031 | -0.031 | 0 | 0.28 | 0.93 | 0.42 | 0.28 | 0.27 | 0.34 |
| 7 | 75553319 | rs6967208 | -0.581 | -0.457 | 0.107 | 0.924 | -0.008 | -0.01 | -0.01 | 0 | 0.06 | 0.31 | 0.03 | 0.01 | 0.01 | 0.03 |
| 7 | 75565740 | rs239960 | -0.158 | -0.436 | -0.199 | 0.955 | 0.059 | 0.257 | 0.257 | 0 | 0.11 | 0.49 | 0.24 | 0.25 | 0.09 | 0.1 |
| 7 | 75576170 | rs1860702 | -0.761 | -0.359 | -0.489 | 0.916 | -0.005 | -0.008 | -0.008 | 0 | 0.06 | 0.3 | 0.03 | 0 | 0.01 | 0.02 |
| 7 | 75579490 | rs239953 | 0.163 | -0.248 | -0.866 | 1.066 | -0.012 | -0.131 | -0.131 | 0.351 | 0.84 | 0.99 | 0.8 | 0.54 | 0.86 | 0.85 |
| 7 | 75583325 | rs10262966 | -1.164 | -0.03 | 0.098 | 0.585 | -0.008 | -0.011 | -0.011 | 0 | 0.06 | 0.3 | 0.03 | 0.01 | 0.01 | 0.1 |
| 7 | 75586536 | rs4728533 | 0.802 | 0.136 | 0.2 | 0.991 | 0.021 | 0.191 | 0.191 | 0 | 0.88 | 0.48 | 0.76 | 0.74 | 0.91 | 0.84 |
| 7 | 75588704 | rs12537277 | 1.054 | -0.04 | 0.38 | 0.787 | -0.012 | -0.007 | -0.007 | 0 | 0.71 | 0.6 | 0.76 | 0.77 | 0.73 | 0.59 |
| 7 | 75589903 | rs2868177 | 0.928 | 0.281 | -0.257 | 0.671 | 0.002 | -0.002 | -0.002 | 0 | 0.4 | 0.45 | 0.5 | 0.46 | 0.34 | 0.27 |
| 7 | 75590100 | rs2868178 | 1.176 | 0.104 | -0.11 | 0.676 | 0.009 | -0.029 | -0.029 | 0 | 0.37 | 0.26 | 0.48 | 0.46 | 0.34 | 0.25 |
| 7 | 75590808 | rs2868180 | 0.703 | 0.029 | 0.948 | 0.647 | 0.001 | -0.001 | -0.001 | 0 | 0.42 | 0.63 | 0.51 | 0.46 | 0.34 | 0.3 |
| 7 | 75592925 | rs6965343 | -0.302 | 0.103 | 1.079 | 0.6 | -0.002 | -0.003 | -0.003 | 1.114 | 0.66 | 0.72 | 0.67 | 0.5 | 0.59 | 0.61 |
| 7 | 75595682 | rs7804806 | -1.148 | 0.045 | 1.642 | 0.532 | 0.007 | -0.008 | -0.008 | 3.084 | 0.62 | 0.66 | 0.58 | 0.57 | 0.69 | 0.75 |
| 7 | 75595970 | rs7796654 | 0.455 | -0.073 | 1.691 | 0.478 | -0.017 | 0.018 | 0.018 | 0 | 0.76 | 0.5 | 0.76 | 0.75 | 0.76 | 0.62 |
| 7 | 75597545 | rs10954724 | -0.208 | -0.103 | 1.646 | 0.644 | -0.003 | -0.004 | -0.004 | 0.091 | 0.63 | 0.41 | 0.6 | 0.47 | 0.58 | 0.54 |
| 7 | 75601169 | rs6953065 | 0.535 | -0.084 | 1.466 | 0.578 | -0.056 | 0.019 | 0.019 | 0 | 0.32 | 0.15 | 0.27 | 0.3 | 0.38 | 0.34 |
| 7 | 75601867 | rs17148944 | 0.776 | 0.347 | 1.068 | 0.29 | -0.016 | 0.044 | 0.044 | 0 | 0.12 | 0.03 | 0.04 | 0 | 0.1 | 0.17 |
| 7 | 75607608 | rs4732513 | -1.136 | 0.489 | 1.149 | 0.311 | 0.019 | 0.023 | 0.023 | 0 | 0.55 | 0.35 | 0.43 | 0.39 | 0.64 | 0.69 |
| 7 | 75608986 | rs10239977 | -0.117 | 0.826 | 0.871 | 0.076 | -0.038 | 0.106 | 0.106 | 0 | 0.26 | 0.08 | 0.21 | 0.04 | 0.33 | 0.27 |
| 7 | 75611149 | rs10954732 | 0.909 | 0.773 | 0.998 | 0.183 | 0 | 0 | 0 | 0 | 0.65 | 0.38 | 0.61 | 0.46 | 0.65 | 0.53 |
| 7 | 75615287 | rs1057870 | -0.214 | 0.698 | 0.945 | 0.07 | -0.019 | 0.056 | 0.056 | 0 | 0.27 | 0.13 | 0.21 | 0.04 | 0.34 | 0.29 |
| 7 | 75616105 | rs17685 | -0.148 | 0.568 | 0.928 | 0.09 | -0.019 | -0.03 | -0.03 | 0 | 0.26 | 0.17 | 0.21 | 0.35 | 0.29 | 0.34 |
| 7 | 75616577 | rs8509 | 0.874 | 0.604 | 0.877 | 0.288 | -0.007 | -0.008 | -0.008 | 0 | 0.11 | 0.1 | 0.04 | 0.12 | 0.1 | 0.16 |
| 7 | 75619650 | rs7788926 | -0.27 | 0.513 | 1.111 | 0.48 | 0 | 0.01 | 0.01 | 0 | 0.86 | 0.77 | 0.89 | 0.82 | 0.89 | 0.77 |
| 7 | 75620641 | rs7794040 | -0.27 | 0.513 | 1.111 | 0.479 | 0 | 0.009 | 0.009 | 0 | 0.86 | 0.77 | 0.89 | 0.82 | 0.89 | 0.77 |
| 7 | 75622811 | rs6952753 | 2.105 | 0.459 | 1.039 | 0.464 | 0.009 | -0.03 | -0.03 | 0 | 0.38 | 0.22 | 0.36 | 0.36 | 0.32 | 0.24 |
| 7 | 75628101 | rs7785025 | 1.251 | 0.544 | 1.164 | 0.565 | 0.005 | -0.044 | -0.044 | 0 | 0.64 | 0.23 | 0.58 | 0.39 | 0.67 | 0.55 |
| 7 | 75629778 | rs3801472 | -0.804 | 0.556 | 1.116 | 0.547 | 0.005 | 0.014 | 0.014 | 0 | 0.88 | 0.83 | 0.9 | 0.82 | 0.89 | 0.82 |
| 7 | 75634687 | rs1044484 | 2.008 | 0.317 | 0.37 | -0.15 | -0.061 | 0.073 | 0.073 | 0 | 0.12 | 0.01 | 0.11 | 0 | 0.16 | 0.02 |
| 7 | 75636240 | rs6978677 | 0.353 | 0.092 | 0.785 | 0.207 | -0.004 | 0.016 | 0.016 | 0 | 0.69 | 0.58 | 0.67 | 0.56 | 0.71 | 0.61 |
| 7 | 75651085 | rs6467975 | 0.016 | 0.069 | 0.795 | 0.229 | 0.085 | 0.255 | 0.255 | 0.779 | 0.06 | 0.34 | 0.09 | 0.17 | 0.04 | 0.06 |
| 7 | 75657850 | rs11983987 | 0.652 | 0.06 | 0.676 | 0.209 | -0.001 | 0 | 0 | 0 | 0.17 | 0.16 | 0.22 | 0.26 | 0.18 | 0.16 |
| 7 | 75685593 | rs1104878 | -1.273 | -0.015 | 0.559 | 0.16 | 0.017 | -0.044 | -0.044 | 0.398 | 0.38 | 0.81 | 0.41 | 0.61 | 0.33 | 0.46 |
| 7 | 75695081 | rs3779419 | 0.196 | -0.04 | 0.706 | 0.17 | -0.015 | -0.002 | -0.002 | 0 | 0.2 | 0.3 | 0.23 | 0.26 | 0.17 | 0.17 |
| 7 | 75695908 | rs3815457 | 0.302 | -0.021 | 0.519 | 0.181 | 0.108 | 0.29 | 0.29 | 0.088 | 0.05 | 0.34 | 0.08 | 0.16 | 0.04 | 0.06 |
| 7 | 75696420 | rs10476 | -1.314 | -0.039 | 0.459 | 0.185 | 0.019 | -0.043 | -0.043 | 0 | 0.39 | 0.81 | 0.42 | 0.61 | 0.33 | 0.46 |
| 7 | 75702719 | rs7804781 | 0.218 | -0.006 | 0.458 | 0.201 | 0.036 | 0.158 | 0.158 | 0 | 0.16 | 0.41 | 0.18 | 0.34 | 0.15 | 0.23 |
| 7 | 75762619 | rs9649679 | 0.293 | -0.017 | 0.439 | 0.124 | 0.034 | 0.039 | 0.039 | 0 | 0.16 | 0.39 | 0.13 | 0.17 | 0.15 | 0.22 |
| 7 | 75769049 | rs6953665 | -0.958 | 0.071 | -0.315 | 0.036 | 0.016 | -0.04 | -0.04 | 0.893 | 0.38 | 0.79 | 0.41 | 0.61 | 0.33 | 0.46 |
| 7 | 75772878 | rs10952840 | -0.982 | 0.079 | -0.311 | 0.04 | 0.017 | -0.04 | -0.04 | 0 | 0.38 | 0.79 | 0.41 | 0.61 | 0.33 | 0.46 |
| 7 | 75790405 | rs9691174 | -0.982 | 0.079 | -0.311 | 0.04 | 0.017 | -0.04 | -0.04 | 0 | 0.38 | 0.79 | 0.41 | 0.61 | 0.33 | 0.46 |
| 7 | 75802674 | rs10277259 | -0.989 | 0.069 | -0.315 | 0.035 | 0.018 | -0.039 | -0.039 | 0.741 | 0.38 | 0.79 | 0.41 | 0.61 | 0.33 | 0.46 |
| 7 | 75803132 | rs4728587 | 0.343 | 0.051 | -0.109 | 0.205 | 0.005 | -0.004 | -0.004 | 0.003 | 0.18 | 0.14 | 0.22 | 0.26 | 0.17 | 0.17 |
| 7 | 75812003 | rs7788837 | 0.182 | 0.277 | -0.453 | -0.069 | -0.105 | 0.145 | 0.145 | 0 | 0.25 | 0.02 | 0.23 | 0.03 | 0.36 | 0.29 |
| 7 | 75841589 | rs4728614 | 0.122 | -0.04 | -0.073 | 0.513 | 0.006 | -0.012 | -0.012 | 0.013 | 0.16 | 0.12 | 0.21 | 0.28 | 0.16 | 0.15 |
| 7 | 75846453 | rs10952869 | -1.129 | -0.041 | 0.225 | 0.319 | -0.014 | 0.042 | 0.042 | 0 | 0.25 | 0.66 | 0.36 | 0.62 | 0.21 | 0.29 |
| 7 | 75876754 | rs1894761 | -0.988 | -0.311 | 0.021 | 0.378 | -0.02 | 0.054 | 0.054 | 0 | 0.29 | 0.6 | 0.35 | 0.62 | 0.27 | 0.27 |
| 7 | 75879334 | rs11984438 | -0.319 | -0.062 | -0.339 | 0.575 | 0.005 | -0.024 | -0.024 | 0 | 0.22 | 0.21 | 0.27 | 0.44 | 0.25 | 0.18 |

| | | | | | | | | | | | | | | | | |
|---|---|---|---|---|---|---|---|---|---|---|---|---|---|---|---|---|
| 7 | 75896535 | rs6974346 | 0.658 | 0.08 | -0.184 | 0.814 | -0.002 | 0.021 | 0.021 | 0 | 0.12 | 0.16 | 0.06 | 0.26 | 0.11 | 0.06 |
| 7 | 75906486 | rs13239419 | -0.331 | 0.131 | -0.366 | 0.695 | -0.01 | -0.009 | -0.009 | 0 | 0.63 | 0.72 | 0.6 | 0.41 | 0.66 | 0.67 |
| 7 | 75908947 | rs7808886 | -0.317 | 0.139 | -0.27 | 0.679 | -0.004 | -0.004 | -0.004 | 0 | 0.63 | 0.68 | 0.61 | 0.41 | 0.66 | 0.67 |
| 7 | 75927125 | rs2908201 | -0.464 | -0.212 | 0.079 | 0.278 | 0.015 | 0.033 | 0.033 | 0 | 0.38 | 0.73 | 0.43 | 0.76 | 0.29 | 0.41 |
| 7 | 75930789 | rs2903813 | 1.267 | 0.049 | -0.063 | 0.138 | -0.003 | -0.005 | -0.005 | 0.149 | 0.1 | 0.09 | 0.22 | 0.16 | 0.18 | 0.11 |
| 7 | 75931160 | rs2009836 | 0.337 | -0.106 | -0.366 | -0.141 | 0.007 | 0.041 | 0.041 | 0 | 0.6 | 0.27 | 0.56 | 0.24 | 0.66 | 0.62 |
| 7 | 75933712 | rs2070804 | 0.992 | -0.096 | 0.127 | 0.514 | 0.008 | 0.085 | 0.085 | 0 | 0.12 | 0.26 | 0.05 | 0.25 | 0.11 | 0.06 |
| 7 | 75937049 | rs2908197 | -0.747 | -0.903 | -0.333 | -0.653 | -0.012 | 0.081 | 0.081 | 0 | 0.57 | 0.73 | 0.68 | 0.93 | 0.51 | 0.57 |
| 7 | 75946763 | rs2961040 | -1.099 | -0.879 | -0.537 | -0.631 | -0.064 | 0.192 | 0.192 | 0 | 0.59 | 0.92 | 0.69 | 0.93 | 0.51 | 0.59 |
| 7 | 75953297 | rs758944 | 0.533 | -1.226 | -0.269 | -0.322 | -0.005 | 0.031 | 0.031 | 0 | 0.24 | 0.33 | 0.29 | 0.45 | 0.3 | 0.18 |
| 7 | 75954657 | rs12334209 | 0.659 | -1.583 | 0.119 | -0.249 | -0.049 | 0.031 | 0.031 | 0.033 | 0.13 | 0.28 | 0.09 | 0.18 | 0.05 | 0.15 |
| 7 | 75957644 | rs1046304 | 2.23 | -1.549 | -0.092 | -0.366 | -0.068 | -0.084 | -0.084 | 0 | 0.19 | 0.02 | 0.31 | 0.31 | 0.2 | 0.25 |
| 7 | 75963169 | rs2109857 | -1.671 | -1.555 | -0.075 | -0.402 | -0.09 | 0.182 | 0.182 | 0 | 0.61 | 0.92 | 0.72 | 0.93 | 0.55 | 0.61 |
| 7 | 75975427 | rs10241401 | 0.991 | -1.458 | 0.255 | -0.112 | -0.019 | 0.009 | 0.009 | 0 | 0.12 | 0.15 | 0.09 | 0.18 | 0.05 | 0.15 |
| 7 | 75984532 | rs917424 | 1.154 | -0.929 | 0.11 | -0.089 | -0.008 | -0.016 | -0.016 | 0.491 | 0.23 | 0.16 | 0.32 | 0.31 | 0.21 | 0.28 |
| 7 | 75985552 | rs17149177 | 1.586 | -0.868 | -0.232 | -0.062 | -0.056 | -0.075 | -0.075 | 0 | 0.21 | 0.03 | 0.31 | 0.31 | 0.21 | 0.25 |
| 7 | 76004709 | rs10224066 | -0.287 | -0.991 | -0.447 | -0.488 | -0.009 | 0.017 | 0.017 | 0 | 0.55 | 0.68 | 0.53 | 0.57 | 0.45 | 0.6 |
| 7 | 76005696 | rs10231308 | -0.197 | -0.989 | -0.291 | -0.368 | -0.02 | -0.004 | -0.004 | 0 | 0.35 | 0.45 | 0.21 | 0.27 | 0.25 | 0.33 |
| 7 | 76006522 | rs10272095 | 2.763 | -0.862 | -0.414 | -0.268 | 0.005 | -0.004 | -0.004 | 1.275 | 0.07 | 0.04 | 0.07 | 0.16 | 0.02 | 0.07 |
| 7 | 76014581 | rs11767818 | 1.758 | -1.061 | -0.452 | -0.43 | -0.026 | 0.038 | 0.038 | 0 | 0.2 | 0.05 | 0.1 | 0.05 | 0.2 | 0.19 |
| 7 | 76016222 | rs11771651 | 0.044 | -1.15 | -0.294 | -0.361 | -0.007 | 0 | 0 | 0 | 0.47 | 0.32 | 0.5 | 0.51 | 0.42 | 0.53 |
| 7 | 76018346 | rs7790894 | 3.177 | -1.183 | -0.506 | -0.161 | 0 | 0 | 0 | 0 | 0.06 | 0.04 | 0.07 | 0.15 | 0.01 | 0.05 |
| 7 | 76019015 | rs999096 | 2.189 | -1.204 | -0.548 | -0.139 | -0.083 | -0.092 | -0.092 | 4.21 | 0.18 | 0.01 | 0.3 | 0.3 | 0.21 | 0.27 |
| 7 | 76019590 | rs10227141 | 1.228 | -1.216 | -0.056 | -0.261 | -0.03 | 0.013 | 0.013 | 1.099 | 0.1 | 0.15 | 0.07 | 0.15 | 0.02 | 0.07 |
| 7 | 76037583 | rs17149199 | 1.905 | -1.332 | -0.285 | -0.609 | -0.011 | -0.034 | -0.034 | 0.135 | 0.11 | 0.15 | 0.05 | 0 | 0.07 | 0.02 |
| 7 | 76038848 | rs12540545 | 1.84 | -1.337 | -0.214 | -0.611 | -0.023 | -0.053 | -0.053 | 0 | 0.11 | 0.18 | 0.05 | 0 | 0.07 | 0.02 |
| 7 | 76040527 | rs7799638 | -0.633 | -1.416 | 0.076 | -0.533 | -0.085 | 0.077 | 0.077 | 0.011 | 0.16 | 0.52 | 0.18 | 0.25 | 0.05 | 0.13 |
| 7 | 76048535 | rs750103 | 0.287 | -1.212 | 0.029 | -0.527 | -0.007 | 0.006 | 0.006 | 0 | 0.07 | 0.08 | 0.07 | 0.13 | 0.03 | 0.09 |
| 7 | 76048676 | rs10241243 | 2.139 | -1.065 | 0.385 | -0.395 | -0.015 | 0.086 | 0.086 | 0.653 | 0.07 | 0.24 | 0.1 | 0.12 | 0.02 | 0.06 |
| 7 | 76050132 | rs2868366 | -0.25 | -0.886 | -0.879 | -0.565 | -0.007 | 0.002 | 0.002 | 0 | 0.58 | 0.62 | 0.56 | 0.54 | 0.47 | 0.58 |
| 7 | 76054195 | rs13232187 | 2.17 | -0.606 | -1.034 | -0.677 | -0.002 | -0.027 | -0.027 | 0 | 0.11 | 0.15 | 0.03 | 0 | 0.09 | 0.04 |
| 7 | 76054372 | rs2286428 | 1.714 | -0.522 | -1.363 | -0.692 | -0.018 | 0.053 | 0.053 | 0 | 0.23 | 0.05 | 0.14 | 0.05 | 0.21 | 0.18 |
| 7 | 76059561 | rs3789833 | 1.632 | -0.068 | -1.12 | -0.243 | -0.043 | -0.022 | -0.022 | 0.045 | 0.23 | 0.06 | 0.28 | 0.23 | 0.24 | 0.29 |
| 7 | 76134203 | rs1465314 | 0.924 | -0.092 | -1.107 | -0.409 | -0.004 | -0.013 | -0.013 | 0.063 | 0.28 | 0.2 | 0.33 | 0.37 | 0.25 | 0.4 |
| 7 | 76137375 | rs2110464 | 0.182 | 0.175 | -1.326 | -0.396 | -0.015 | -0.026 | -0.026 | 0 | 0.3 | 0.21 | 0.38 | 0.45 | 0.41 | 0.31 |
| 7 | 76143705 | rs887911 | 0.068 | 0.253 | -1.357 | -0.38 | -0.011 | -0.023 | -0.023 | 0 | 0.32 | 0.21 | 0.38 | 0.44 | 0.4 | 0.32 |
| 7 | 76185628 | rs10952955 | 1.169 | 0.584 | -1.152 | -0.953 | -0.005 | 0.012 | 0.012 | 0 | 0.13 | 0.1 | 0.12 | 0.02 | 0.14 | 0.14 |
| 7 | 76209459 | rs17718122 | 0.1 | 0.483 | -1.153 | -0.685 | -0.007 | 0.11 | 0.11 | 0 | 0.34 | 0.15 | 0.16 | 0.05 | 0.3 | 0.21 |
| 7 | 76288272 | rs10247269 | -0.794 | 0.577 | -1.123 | -0.114 | 0 | -0.04 | -0.04 | 0 | 0.63 | 0.86 | 0.65 | 0.4 | 0.59 | 0.71 |
| 7 | 76289033 | rs17645907 | -0.06 | 0.713 | -0.169 | -0.624 | -0.005 | -0.078 | -0.078 | 0 | 0.26 | 0.33 | 0.23 | 0.01 | 0.24 | 0.24 |
| 7 | 76307066 | rs10268548 | -0.017 | 0.603 | 0.024 | -0.677 | -0.011 | -0.11 | -0.11 | 0 | 0.22 | 0.62 | 0.15 | 0.01 | 0.17 | 0.2 |
| 7 | 76309668 | rs7456697 | 1.054 | 0.819 | -0.986 | -0.598 | -0.102 | -0.067 | -0.067 | 0 | 0.21 | 0.02 | 0.3 | 0.25 | 0.26 | 0.32 |
| 7 | 76309742 | rs10280733 | 0.419 | 0.793 | -0.65 | -0.781 | -0.021 | -0.123 | -0.123 | 0 | 0.19 | 0.41 | 0.12 | 0 | 0.14 | 0.11 |
| 7 | 76336888 | rs34856074 | -0.109 | 1.067 | -1.46 | -0.719 | -0.017 | 0.043 | 0.043 | 0 | 0.34 | 0.26 | 0.35 | 0.19 | 0.35 | 0.39 |
| 7 | 76341486 | rs3909148 | 1.257 | 1.192 | -1.879 | -0.539 | -0.08 | -0.037 | -0.037 | 1.475 | 0.17 | 0.02 | 0.26 | 0.19 | 0.22 | 0.28 |
| 7 | 76342312 | rs13227110 | 0.129 | 0.916 | -1.536 | -0.209 | -0.051 | 0.169 | 0.169 | 0.179 | 0.45 | 0.13 | 0.48 | 0.2 | 0.48 | 0.53 |
| 7 | 76347197 | rs10273087 | -0.591 | 1.125 | -1.535 | 0.384 | -0.032 | -0.074 | -0.074 | 0.614 | 0.62 | 0.9 | 0.68 | 0.47 | 0.56 | 0.74 |
| 7 | 76414540 | rs10241304 | -0.461 | 2.003 | -0.464 | 0.703 | 0.116 | 0.054 | 0.054 | 0.118 | 0.06 | 0.18 | 0.11 | 0.11 | 0.15 | 0.2 |
| 7 | 76418140 | rs11765301 | -0.368 | 1.845 | -1.13 | 0.736 | 0 | 0.001 | 0.001 | 0 | 0.28 | 0.26 | 0.44 | 0.27 | 0.28 | 0.38 |
| 7 | 76429959 | rs10215877 | 0.123 | 1.644 | -2.185 | 1.05 | 0.031 | -0.076 | -0.076 | 0 | 0.31 | 0.07 | 0.33 | 0.44 | 0.27 | 0.27 |
| 7 | 76432653 | rs12540927 | 0.344 | 1.917 | -1.927 | 0.791 | -0.069 | 0.028 | 0.028 | 0.144 | 0.37 | 0.06 | 0.19 | 0.25 | 0.36 | 0.25 |
| 7 | 76476970 | rs9691954 | 0.069 | 1.151 | -0.412 | 1.044 | 0.001 | -0.005 | -0.005 | 0.042 | 0.35 | 0.31 | 0.44 | 0.56 | 0.37 | 0.42 |
| 7 | 76499312 | rs7798157 | -0.472 | 0.938 | -0.398 | 0.568 | -0.005 | -0.02 | -0.02 | 0 | 0.64 | 0.62 | 0.84 | 0.81 | 0.71 | 0.76 |
| 7 | 76505270 | rs3864610 | 0.51 | 1.099 | -0.42 | 0.646 | 0 | -0.001 | -0.001 | 0 | 0.31 | 0.32 | 0.36 | 0.3 | 0.35 | 0.37 |
| 7 | 76510219 | rs2868248 | -0.158 | 1.22 | -0.722 | 0.592 | 0 | 0.055 | 0.055 | 0 | 0.49 | 0.36 | 0.6 | 0.41 | 0.48 | 0.45 |
| 7 | 76524381 | rs7777346 | 0.429 | 1.125 | -0.144 | 0.433 | 0 | -0.022 | -0.022 | 0 | 0.32 | 0.43 | 0.36 | 0.3 | 0.35 | 0.33 |
| 7 | 76533023 | rs17803197 | -0.478 | 1.392 | -0.062 | 0.698 | 0.066 | 0.04 | 0.04 | 0.116 | 0.11 | 0.26 | 0.14 | 0.09 | 0.21 | 0.13 |
| 7 | 76538443 | rs7793752 | -1.08 | 1.041 | -0.086 | 0.736 | -0.001 | -0.001 | -0.001 | 0.125 | 0.74 | 0.78 | 0.89 | 0.81 | 0.79 | 0.81 |
| 7 | 76548003 | rs7781966 | 0.195 | 0.954 | -0.579 | 0.84 | 0.001 | -0.001 | -0.001 | 0.122 | 0.39 | 0.41 | 0.4 | 0.44 | 0.4 | 0.42 |
| 7 | 76550045 | rs17148511 | 0.695 | 0.903 | -1.244 | 0.496 | -0.037 | 0.003 | 0.003 | 0 | 0.24 | 0.02 | 0.1 | 0.19 | 0.21 | 0.18 |
| 7 | 76557047 | rs2539499 | 0.06 | 1.296 | -0.858 | 0.266 | -0.041 | 0.053 | 0.053 | 0.221 | 0.36 | 0.26 | 0.4 | 0.24 | 0.55 | 0.28 |
| 7 | 76637441 | rs10278476 | -0.194 | 1.557 | -0.433 | 0.353 | 0.002 | 0.011 | 0.011 | 0 | 0.36 | 0.7 | 0.3 | 0.24 | 0.32 | 0.31 |
| 7 | 76720449 | rs3930016 | 0.098 | 1.188 | -0.532 | -0.042 | -0.002 | 0.006 | 0.006 | 0 | 0.65 | 0.7 | 0.8 | 0.87 | 0.58 | 0.79 |
| 7 | 76738591 | rs3864639 | -0.082 | 1.19 | -1.104 | 0.583 | -0.024 | -0.054 | -0.054 | 0.078 | 0.81 | 0.96 | 0.88 | 0.78 | 0.79 | 0.74 |
| 7 | 76759712 | rs6962391 | -1.141 | 1.519 | -0.004 | 0.865 | 0.135 | 0.138 | 0.138 | 0.012 | 0.07 | 0.27 | 0.12 | 0.18 | 0.22 | 0.07 |
| 7 | 76788477 | rs3972417 | 0.059 | 0.997 | -0.176 | 0.559 | -0.009 | -0.003 | -0.003 | 0.451 | 0.57 | 0.66 | 0.51 | 0.49 | 0.52 | 0.35 |
| 7 | 76812454 | rs3095464 | -0.498 | 0.046 | 0.069 | 0.199 | 0.001 | 0.019 | 0.019 | 0 | 0.34 | 0.42 | 0.46 | 0.5 | 0.35 | 0.3 |
| 7 | 76820800 | rs10252204 | -0.778 | -0.601 | -0.291 | 0.176 | -0.016 | -0.014 | -0.014 | 0 | 0.59 | 0.83 | 0.48 | 0.57 | 0.4 | 0.36 |
| 7 | 76821842 | rs41534948 | 0.389 | -1.376 | -0.289 | -0.034 | 0.028 | 0.013 | 0.013 | 0 | 0.18 | 0.13 | 0.03 | 0.06 | 0.06 | 0.12 |

| | | | | | | | | | | | | | | | | |
|---|---|---|---|---|---|---|---|---|---|---|---|---|---|---|---|---|
| 7 | 76826438 | rs3093277 | -0.017 | -1.186 | 1.027 | 0.263 | -0.026 | 0.053 | 0.053 | 2.974 | 0.07 | 0.24 | 0.01 | 0.06 | 0.01 | 0.05 |
| 7 | 76828953 | rs2075761 | -0.001 | -1.397 | 0.738 | 0.286 | 0.018 | -0.046 | -0.046 | 0 | 0.13 | 0.06 | 0.26 | 0.26 | 0.09 | 0.15 |
| 7 | 76834066 | rs10488499 | 1.09 | -1.611 | 0.728 | 0.033 | -0.004 | 0.105 | 0.105 | 0.061 | 0.21 | 0.01 | 0.07 | 0.05 | 0.16 | 0.08 |
| 7 | 76841972 | rs3095471 | -1.188 | -1.653 | 1.023 | 0.08 | 0.063 | -0.007 | -0.007 | 0 | 0.45 | 0.75 | 0.42 | 0.54 | 0.31 | 0.29 |
| 7 | 76845398 | rs7788171 | -1.338 | -1.668 | 1.16 | 0.087 | -0.04 | 0.063 | 0.063 | 0.627 | 0.2 | 0.39 | 0.3 | 0.31 | 0.09 | 0.17 |
| 7 | 76851323 | rs7810066 | -0.513 | -1.778 | 1.192 | 0.148 | 0 | 0 | 0 | 0 | 0.74 | 0.73 | 0.86 | 0.77 | 0.74 | 0.87 |
| 7 | 76853725 | rs3108406 | 1.524 | -1.767 | 1.319 | 0.174 | -0.008 | 0.172 | 0.172 | 0.35 | 0.88 | 0.56 | 0.91 | 0.77 | 0.9 | 0.9 |
| 7 | 76856975 | rs6943246 | -0.034 | -1.754 | 0.878 | 0.144 | -0.035 | 0.038 | 0.038 | 1.327 | 0.08 | 0.22 | 0.02 | 0.06 | 0.01 | 0.06 |
| 7 | 76863337 | rs3114306 | 1.258 | -1.741 | 0.734 | 0.154 | 0.073 | -0.007 | -0.007 | 0 | 0.55 | 0.21 | 0.57 | 0.46 | 0.69 | 0.71 |
| 7 | 76871113 | rs17230060 | 1.046 | -1.694 | 0.6 | 0.112 | -0.007 | 0.111 | 0.111 | 0.036 | 0.21 | 0.01 | 0.07 | 0.05 | 0.16 | 0.08 |
| 7 | 76880223 | rs4729246 | 0.593 | -1.571 | 0.746 | 0.232 | 0.019 | -0.03 | -0.03 | 0 | 0.1 | 0.06 | 0.25 | 0.26 | 0.04 | 0.12 |
| 7 | 76881171 | rs12704841 | 0.849 | -1.57 | 1.009 | 0.248 | 0.138 | 0.271 | 0.271 | 0 | 0.94 | 0.52 | 0.91 | 0.82 | 0.91 | 0.96 |
| 7 | 76881983 | rs3114315 | 0.797 | -1.576 | 0.987 | -0.125 | -0.019 | 0.184 | 0.184 | 0.009 | 0.83 | 0.41 | 0.66 | 0.56 | 0.86 | 0.84 |
| 7 | 76883110 | rs3114317 | 0.389 | -1.573 | 0.988 | -0.124 | 0.001 | 0.241 | 0.241 | 0 | 0.85 | 0.46 | 0.66 | 0.56 | 0.86 | 0.84 |
| 7 | 76884226 | rs2056708 | 1.687 | -1.682 | 0.937 | -0.175 | -0.016 | 0.03 | 0.03 | 0.054 | 0.77 | 0.26 | 0.65 | 0.51 | 0.86 | 0.78 |
| 7 | 76885769 | rs3108428 | 1.086 | -1.89 | 0.757 | -0.302 | 0.098 | 0.002 | 0.002 | 0 | 0.53 | 0.21 | 0.57 | 0.46 | 0.69 | 0.68 |
| 7 | 76887189 | rs3108430 | -0.238 | -1.851 | 0.883 | -0.352 | -0.031 | -0.135 | -0.135 | 1.527 | 0.7 | 0.51 | 0.83 | 0.93 | 0.74 | 0.86 |
| 7 | 76891485 | rs1109968 | 0.365 | -1.8 | 0.682 | -0.229 | 0.005 | 0.016 | 0.016 | 0 | 0.11 | 0.11 | 0.25 | 0.42 | 0.05 | 0.12 |
| 7 | 76891835 | rs1111665 | 1.263 | -1.715 | 0.586 | -0.32 | 0.088 | 0.013 | 0.013 | 0 | 0.51 | 0.22 | 0.55 | 0.45 | 0.66 | 0.64 |
| 7 | 76900052 | rs3114325 | 0.542 | -1.07 | 0.426 | -0.913 | -0.028 | -0.055 | -0.055 | 0 | 0.63 | 0.46 | 0.81 | 0.87 | 0.71 | 0.78 |
| 7 | 76903217 | rs7810561 | -1.261 | -0.377 | 0.462 | -0.191 | -0.024 | 0.012 | 0.012 | 0.846 | 0.09 | 0.19 | 0.04 | 0.06 | 0.03 | 0.08 |
| 7 | 76908199 | rs3828947 | -1.236 | -0.235 | 0.029 | -0.745 | 0.047 | 0.087 | 0.087 | 0 | 0.44 | 0.7 | 0.27 | 0.13 | 0.32 | 0.46 |
| 7 | 76909035 | rs6958464 | -0.837 | -0.507 | 0.52 | -0.884 | -0.012 | -0.046 | -0.046 | 3.858 | 0.33 | 0.54 | 0.24 | 0.13 | 0.28 | 0.39 |
| 7 | 76911470 | rs10247327 | -0.157 | -0.631 | 0.39 | -0.998 | -0.019 | -0.065 | -0.065 | 0.066 | 0.15 | 0.19 | 0.07 | 0 | 0.1 | 0.06 |
| 7 | 76914470 | rs6971515 | -2.252 | -0.657 | 0.616 | -1.148 | 0.022 | 0.033 | 0.033 | 0.148 | 0.38 | 0.7 | 0.21 | 0.1 | 0.29 | 0.32 |
| 7 | 76916010 | rs6953266 | -1.334 | -0.578 | 0.721 | -1.106 | -0.018 | -0.047 | -0.047 | 0 | 0.22 | 0.51 | 0.14 | 0.09 | 0.17 | 0.25 |
| 7 | 76916819 | rs3763402 | -1.145 | -0.593 | 0.837 | -1.112 | -0.009 | -0.032 | -0.032 | 0 | 0.21 | 0.41 | 0.12 | 0.09 | 0.17 | 0.25 |
| 7 | 76917194 | rs6958942 | 0.849 | -0.555 | 0.741 | -0.982 | -0.003 | -0.001 | -0.001 | 0.282 | 0.09 | 0.1 | 0.19 | 0.11 | 0.14 | 0.19 |
| 7 | 76917430 | rs6465532 | -2.384 | -0.564 | 0.762 | -0.976 | 0.031 | 0.11 | 0.11 | 0 | 0.47 | 0.8 | 0.39 | 0.2 | 0.42 | 0.5 |
| 7 | 76919068 | rs7778518 | -0.936 | -0.599 | 0.833 | -0.804 | -0.046 | -0.02 | -0.02 | 0.053 | 0.18 | 0.39 | 0.07 | 0.08 | 0.07 | 0.18 |
| 7 | 76920048 | rs17151162 | 0.312 | -0.552 | 0.05 | -0.806 | 0.006 | -0.001 | -0.001 | 0 | 0.11 | 0.19 | 0.2 | 0.11 | 0.14 | 0.19 |
| 7 | 76921114 | rs17151202 | -0.577 | -0.585 | -0.17 | -0.791 | 0.02 | -0.008 | -0.008 | 0 | 0.16 | 0.01 | 0.05 | 0.07 | 0.07 | 0.18 |
| 7 | 76923550 | rs17151246 | -1.193 | -0.247 | -0.133 | -0.809 | -0.01 | 0.008 | 0.008 | 0 | 0.27 | 0.12 | 0.28 | 0.19 | 0.24 | 0.41 |
| 7 | 76928011 | rs10273411 | -2.219 | -0.316 | -0.099 | -0.938 | 0.011 | 0.162 | 0.162 | 0 | 0.48 | 0.82 | 0.36 | 0.16 | 0.44 | 0.52 |
| 7 | 76940613 | rs1139802 | -0.129 | -0.006 | 0.456 | -0.768 | -0.048 | -0.112 | -0.112 | 1.731 | 0.16 | 0.51 | 0.08 | 0 | 0.09 | 0.05 |
| 7 | 76941830 | rs6973950 | -2.115 | -0.032 | 0.299 | -0.621 | -0.005 | 0.085 | 0.085 | 0 | 0.44 | 0.81 | 0.5 | 0.22 | 0.43 | 0.52 |
| 7 | 76942455 | rs17805954 | -0.327 | 0.096 | 0.354 | -0.499 | -0.006 | 0.004 | 0.004 | 0 | 0.16 | 0.16 | 0.21 | 0.1 | 0.27 | 0.35 |
| 7 | 76943517 | rs6942520 | 0.191 | -0.071 | 0.293 | -0.488 | -0.013 | 0.013 | 0.013 | 0 | 0.13 | 0.1 | 0.16 | 0.07 | 0.16 | 0.31 |
| 7 | 76950035 | rs6950870 | 0.313 | -0.04 | 0.333 | -0.497 | -0.01 | 0.012 | 0.012 | 0 | 0.12 | 0.11 | 0.16 | 0.05 | 0.16 | 0.27 |
| 7 | 76950590 | rs17151683 | 0.346 | -0.228 | 0.592 | -0.548 | -0.009 | -0.023 | -0.023 | 0.007 | 0.13 | 0.11 | 0.04 | 0.01 | 0.09 | 0.07 |
| 7 | 76950686 | rs17151689 | -0.192 | -0.336 | 0.656 | -0.541 | -0.003 | -0.001 | -0.001 | 0 | 0.1 | 0.12 | 0.04 | 0.07 | 0.04 | 0.1 |
| 7 | 76950699 | rs17151692 | 0.151 | -0.316 | 0.759 | -0.545 | -0.019 | -0.039 | -0.039 | 0 | 0.14 | 0.16 | 0.04 | 0.01 | 0.08 | 0.07 |
| 7 | 76952866 | rs12155404 | 0.634 | -0.331 | 0.737 | -0.554 | 0.02 | 0.027 | 0.027 | 0 | 0.12 | 0.05 | 0.04 | 0 | 0.09 | 0.06 |
| 7 | 76956000 | rs11980285 | -0.309 | -0.169 | 0.924 | -0.552 | -0.001 | -0.037 | -0.037 | 0 | 0.17 | 0.2 | 0.1 | 0.03 | 0.2 | 0.11 |
| 7 | 76956597 | rs11766346 | 0.262 | -0.17 | 0.925 | -0.563 | -0.007 | -0.027 | -0.027 | 0 | 0.14 | 0.14 | 0.04 | 0.01 | 0.08 | 0.07 |
| 7 | 76956639 | rs2293452 | -0.909 | -0.41 | 0.703 | 0.09 | 0.001 | 0.089 | 0.089 | 0 | 0.26 | 0.46 | 0.43 | 0.48 | 0.25 | 0.35 |
| 7 | 76956774 | rs10271991 | -2.433 | -0.48 | 0.697 | 0.097 | -0.008 | 0.014 | 0.014 | 0 | 0.54 | 0.81 | 0.57 | 0.58 | 0.49 | 0.57 |
| 7 | 76958980 | rs10262175 | 0.401 | -0.534 | 0.706 | 0.127 | -0.03 | 0.189 | 0.189 | 0 | 0.15 | 0.35 | 0.27 | 0.42 | 0.09 | 0.09 |
| 7 | 76959815 | rs3750046 | -1.341 | -0.531 | 0.678 | 0.118 | -0.014 | 0.015 | 0.015 | 0 | 0.32 | 0.55 | 0.37 | 0.45 | 0.29 | 0.2 |
| 7 | 76960884 | rs17806432 | 0.844 | -0.68 | 0.646 | 0.076 | -0.002 | 0.013 | 0.013 | 0 | 0.11 | 0.1 | 0.12 | 0.05 | 0.09 | 0.26 |
| 7 | 76967750 | rs17151944 | -0.076 | -0.751 | 0.635 | 0.106 | -0.001 | -0.06 | -0.06 | 0 | 0.17 | 0.22 | 0.1 | 0.02 | 0.2 | 0.11 |
| 7 | 76978907 | rs4729343 | -2.127 | -0.838 | 0.342 | 0.31 | 0.022 | 0.016 | 0.016 | 0 | 0.52 | 0.84 | 0.55 | 0.59 | 0.44 | 0.57 |
| 7 | 76983314 | rs1181904 | 1.604 | -0.857 | 0.741 | 0.368 | -0.004 | -0.03 | -0.03 | 0.016 | 0.62 | 0.46 | 0.7 | 0.83 | 0.63 | 0.51 |
| 7 | 76983515 | rs6979188 | 0.932 | -0.851 | 0.578 | 0.343 | 0.01 | 0.016 | 0.016 | 0 | 0.11 | 0.09 | 0.12 | 0.06 | 0.06 | 0.26 |
| 7 | 76984572 | rs2037753 | -1.613 | -0.81 | 0.493 | 0.375 | 0.073 | -0.036 | -0.036 | 0 | 0.41 | 0.83 | 0.48 | 0.5 | 0.3 | 0.44 |
| 7 | 76990126 | rs4727365 | -0.84 | -0.706 | 0.58 | 0.229 | -0.015 | -0.09 | -0.09 | 0 | 0.27 | 0.54 | 0.21 | 0.09 | 0.23 | 0.32 |
| 7 | 76991149 | rs1181900 | -0.648 | -0.63 | 0.511 | 0.244 | -0.041 | -0.037 | -0.037 | 0.063 | 0.9 | 0.98 | 0.93 | 0.92 | 0.86 | 0.88 |
| 7 | 76992278 | rs1859573 | -1.019 | -0.629 | 0.539 | 0.208 | -0.017 | -0.086 | -0.086 | 0 | 0.28 | 0.55 | 0.22 | 0.09 | 0.23 | 0.35 |
| 7 | 77001201 | rs1726576 | 2.21 | -0.648 | 0.164 | 0.332 | 0.02 | 0.049 | 0.049 | 0 | 0.48 | 0.19 | 0.37 | 0.37 | 0.54 | 0.43 |
| 7 | 77012042 | rs6972651 | 0.615 | -0.596 | -0.091 | 0.111 | -0.001 | -0.001 | -0.001 | 0 | 0.11 | 0.14 | 0.19 | 0.09 | 0.06 | 0.24 |
| 7 | 77025766 | rs11772568 | 1.168 | -0.609 | -0.248 | 0.02 | 0.02 | 0.068 | 0.068 | 0 | 0.12 | 0 | 0.03 | 0 | 0.07 | 0.06 |
| 7 | 77026040 | rs2286158 | 1.989 | -0.543 | -0.186 | 0.33 | 0.043 | -0.006 | -0.006 | 0.108 | 0.5 | 0.19 | 0.46 | 0.43 | 0.57 | 0.48 |
| 7 | 77026460 | rs2286156 | -0.323 | -0.864 | 0.278 | 0.286 | -0.001 | -0.012 | -0.012 | 0 | 0.23 | 0.24 | 0.11 | 0.07 | 0.13 | 0.27 |
| 7 | 77027076 | rs10267936 | 0.417 | -0.793 | -0.393 | 0.357 | -0.03 | 0.075 | 0.075 | 0 | 0.13 | 0.21 | 0.24 | 0.41 | 0.07 | 0.09 |
| 7 | 77032997 | rs1859572 | -0.655 | -0.407 | -0.958 | 0.463 | 0.07 | -0.003 | -0.003 | 0.912 | 0.37 | 0.77 | 0.38 | 0.48 | 0.2 | 0.37 |
| 7 | 77036025 | rs6465628 | -0.943 | -0.37 | -0.86 | 0.536 | 0.064 | 0.005 | 0.005 | 0.04 | 0.42 | 0.76 | 0.44 | 0.5 | 0.31 | 0.44 |
| 7 | 77043863 | rs6465633 | -0.07 | 0.194 | -0.901 | 0.336 | -0.006 | 0.013 | 0.013 | 0 | 0.46 | 0.65 | 0.46 | 0.37 | 0.46 | 0.71 |
| 7 | 77044448 | rs9641218 | 0.714 | -0.394 | -0.623 | 0.055 | -0.008 | -0.008 | -0.008 | 0 | 0.74 | 0.52 | 0.58 | 0.26 | 0.74 | 0.81 |
| 7 | 77046222 | rs10953236 | -0.436 | -0.632 | -0.625 | -0.516 | 0.01 | 0.027 | 0.027 | 0 | 0.58 | 0.67 | 0.64 | 0.65 | 0.65 | 0.37 |

| | | | | | | | | | | | | | | | | |
|---|---|---|---|---|---|---|---|---|---|---|---|---|---|---|---|---|
| 7 | 77047056 | rs6970927 | -1.274 | -0.608 | -0.973 | -0.63 | -0.025 | -0.067 | -0.067 | 0 | 0.85 | 0.96 | 0.79 | 0.71 | 0.83 | 0.67 |
| 7 | 77051490 | rs10278905 | -0.401 | -0.61 | -0.877 | -0.628 | 0.003 | 0.016 | 0.016 | 0 | 0.61 | 0.72 | 0.65 | 0.63 | 0.66 | 0.37 |
| 7 | 77055836 | rs740158 | 0.577 | -0.553 | -1.675 | -0.253 | -0.175 | -0.049 | -0.049 | 1.591 | 0.37 | 0.04 | 0.36 | 0.22 | 0.49 | 0.24 |
| 7 | 77055885 | rs740157 | -0.186 | -0.447 | -1.242 | -0.502 | 0.026 | 0.085 | 0.085 | 0 | 0.5 | 0.26 | 0.57 | 0.63 | 0.57 | 0.35 |
| 7 | 77059517 | rs5004556 | -0.472 | -0.12 | -1.107 | -1.025 | -0.008 | -0.078 | -0.078 | 0 | 0.75 | 0.9 | 0.6 | 0.32 | 0.75 | 0.57 |
| 7 | 77059616 | rs12704962 | 0.865 | 0.267 | -1.539 | -1.338 | -0.091 | -0.022 | -0.022 | 0 | 0.23 | 0.01 | 0.21 | 0.15 | 0.28 | 0.14 |
| 7 | 77061342 | rs1978181 | 0.89 | 0.223 | -0.813 | -1.519 | 0.004 | -0.009 | -0.009 | 0 | 0.38 | 0.4 | 0.21 | 0.11 | 0.28 | 0.31 |
| 7 | 77062679 | rs998994 | 0.044 | 0.229 | -0.493 | -1.497 | -0.003 | 0.102 | 0.102 | 0.54 | 0.51 | 0.7 | 0.37 | 0.17 | 0.47 | 0.43 |
| 7 | 77068965 | rs17153477 | 1.312 | 0.047 | -0.671 | -1.516 | -0.009 | -0.032 | -0.032 | 0 | 0.29 | 0.34 | 0.16 | 0.1 | 0.19 | 0.3 |
| 7 | 77069360 | rs10953244 | 0.502 | 0.399 | -0.487 | -1.288 | 0.005 | -0.011 | -0.011 | 0 | 0.61 | 0.44 | 0.41 | 0.26 | 0.55 | 0.45 |
| 7 | 77070195 | rs11773184 | 0.56 | 0.635 | -0.468 | -1.069 | 0.013 | 0.003 | 0.003 | 0 | 0.09 | 0.09 | 0.12 | 0.06 | 0.14 | 0.09 |
| 7 | 77075756 | rs6963259 | -0.303 | 0.029 | -0.296 | -1.206 | 0.01 | -0.013 | -0.013 | 0.161 | 0.6 | 0.37 | 0.77 | 0.93 | 0.72 | 0.71 |
| 7 | 77082896 | rs12704984 | 0.404 | 0.258 | -0.76 | -1.213 | -0.062 | 0.041 | 0.041 | 0.463 | 0.37 | 0.15 | 0.32 | 0.21 | 0.52 | 0.22 |
| 7 | 77086107 | rs7783632 | 1.015 | 0.256 | -0.529 | -1.087 | 0.013 | -0.062 | -0.062 | 0 | 0.2 | 0.11 | 0.38 | 0.67 | 0.17 | 0.4 |
| 7 | 77090958 | rs4729435 | -0.025 | 0.29 | -0.407 | -1.215 | 0.01 | 0.062 | 0.062 | 0 | 0.54 | 0.73 | 0.63 | 0.79 | 0.41 | 0.8 |
| 7 | 77093557 | rs10257454 | -1.019 | 0.429 | -0.569 | -1.205 | -0.009 | -0.038 | -0.038 | 3.361 | 0.9 | 0.86 | 0.94 | 1 | 0.92 | 0.99 |
| 7 | 77095463 | rs17153801 | 1.109 | 0.403 | -1.145 | -1.2 | -0.004 | -0.036 | -0.036 | 0 | 0.28 | 0.3 | 0.16 | 0.08 | 0.2 | 0.26 |
| 7 | 77098235 | rs2107913 | -0.913 | 0.842 | -1.826 | -0.732 | 0.004 | -0.002 | -0.002 | 0 | 0.61 | 0.4 | 0.7 | 0.88 | 0.68 | 0.69 |
| 7 | 77098772 | rs2868784 | -0.033 | 0.827 | -1.855 | -0.73 | -0.019 | 0.027 | 0.027 | 0 | 0.39 | 0.29 | 0.31 | 0.19 | 0.51 | 0.18 |
| 7 | 77101532 | rs10225158 | -0.465 | 0.964 | -1.944 | 0.305 | -0.054 | -0.04 | -0.04 | 0.102 | 0.33 | 0.13 | 0.49 | 0.69 | 0.4 | 0.56 |
| 7 | 77107465 | rs6465689 | -0.382 | 0.346 | -1.72 | -0.041 | 0.081 | 0.089 | 0.089 | 0.635 | 0.49 | 0.79 | 0.59 | 0.74 | 0.38 | 0.45 |
| 7 | 77114154 | rs6967577 | 0.143 | -0.086 | -0.914 | 0.461 | -0.005 | 0.09 | 0.09 | 0 | 0.08 | 0.23 | 0.11 | 0.17 | 0.06 | 0.2 |
| 7 | 77117524 | rs6465694 | -1.297 | 0.763 | -1.632 | -0.04 | -0.072 | 0.058 | 0.058 | 0.792 | 0.72 | 0.97 | 0.82 | 0.91 | 0.69 | 0.75 |
| 7 | 77122188 | rs2097892 | 0.611 | -0.115 | -0.705 | -0.336 | -0.014 | 0.036 | 0.036 | 0 | 0.32 | 0.37 | 0.43 | 0.58 | 0.29 | 0.24 |
| 7 | 77131251 | rs6465722 | -0.337 | -0.303 | -0.479 | -0.248 | -0.031 | 0.025 | 0.025 | 0 | 0.16 | 0.3 | 0.14 | 0.17 | 0.09 | 0.21 |
| 7 | 77165919 | rs3763404 | 0.101 | -0.782 | -0.665 | -0.685 | -0.015 | 0.009 | 0.009 | 0.405 | 0.23 | 0.27 | 0.42 | 0.69 | 0.14 | 0.35 |
| 7 | 77171617 | rs2052162 | -1.787 | -0.511 | -0.972 | -1.084 | -0.068 | 0.053 | 0.053 | 1.878 | 0.73 | 0.96 | 0.8 | 0.9 | 0.65 | 0.77 |
| 7 | 77171954 | rs12533502 | -1.701 | -0.586 | -0.836 | -1.141 | -0.043 | 0.118 | 0.118 | 1.174 | 0.72 | 0.91 | 0.79 | 0.99 | 0.65 | 0.75 |
| 7 | 77174497 | rs9886084 | 0.449 | -0.315 | -0.44 | -1.332 | 0.054 | -0.072 | -0.072 | 0 | 0.13 | 0.33 | 0.09 | 0.04 | 0.15 | 0.06 |
| 7 | 77176390 | rs12113982 | 0.642 | -0.55 | -0.399 | -1.383 | -0.007 | 0.005 | 0.005 | 0 | 0.23 | 0.26 | 0.24 | 0.18 | 0.32 | 0.28 |
| 7 | 77222886 | rs10270482 | -0.83 | -0.648 | -0.134 | -1.409 | 0.044 | 0.042 | 0.042 | 0.753 | 0.48 | 0.6 | 0.55 | 0.73 | 0.34 | 0.47 |
| 7 | 77233827 | rs1972568 | -1.939 | -0.593 | -0.185 | -1.425 | -0.026 | 0.051 | 0.051 | 0.512 | 0.71 | 0.86 | 0.79 | 0.91 | 0.65 | 0.74 |
| 7 | 77237390 | rs7776973 | -0.997 | -0.773 | -0.213 | -1.475 | 0.001 | 0.082 | 0.082 | 0 | 0.45 | 0.72 | 0.35 | 0.22 | 0.48 | 0.35 |
| 7 | 77239456 | rs10237942 | 0.875 | -0.683 | 0.141 | -1.507 | 0.028 | -0.051 | -0.051 | 0.496 | 0.12 | 0.21 | 0.08 | 0.04 | 0.14 | 0.04 |
| 7 | 77239735 | rs17381884 | 1.199 | -0.266 | -0.153 | -1.664 | -0.079 | 0.082 | 0.082 | 0 | 0.16 | 0.01 | 0.1 | 0.01 | 0.22 | 0.09 |
| 7 | 77255941 | rs17467232 | 0.427 | -0.391 | 0.091 | -1.617 | -0.028 | 0.015 | 0.015 | 0 | 0.23 | 0.17 | 0.23 | 0.18 | 0.31 | 0.28 |
| 7 | 77256713 | rs3750050 | 0.6 | -0.466 | 0.109 | -1.608 | 0.026 | -0.013 | -0.013 | 0 | 0.27 | 0.22 | 0.44 | 0.69 | 0.16 | 0.39 |
| 7 | 77297415 | rs9641271 | -1.452 | -0.401 | 0.2 | -1.87 | -0.019 | 0.029 | 0.029 | 0.279 | 0.62 | 0.73 | 0.71 | 0.83 | 0.51 | 0.69 |
| 7 | 77297541 | rs17156247 | 0.246 | -0.424 | 0.125 | -1.88 | -0.01 | 0.01 | 0.01 | 0 | 0.24 | 0.26 | 0.24 | 0.13 | 0.31 | 0.31 |
| 7 | 77298752 | rs4729594 | -2.326 | -0.319 | 0.04 | -1.846 | -0.054 | 0.055 | 0.055 | 1.407 | 0.73 | 0.93 | 0.8 | 0.91 | 0.65 | 0.74 |
| 7 | 77302059 | rs11514912 | 0.508 | -0.505 | 0.18 | -1.756 | 0 | -0.003 | -0.003 | 0.055 | 0.27 | 0.25 | 0.44 | 0.7 | 0.16 | 0.35 |
| 7 | 77308295 | rs17807185 | -1.608 | 0.663 | 1.315 | -1.292 | -0.001 | 0.038 | 0.038 | 0 | 0.43 | 0.67 | 0.37 | 0.23 | 0.4 | 0.53 |
| 7 | 77314818 | rs12113119 | -1.616 | 0.663 | 1.393 | -1.306 | -0.003 | 0.051 | 0.051 | 0 | 0.43 | 0.69 | 0.38 | 0.24 | 0.4 | 0.53 |
| 7 | 77325349 | rs2301938 | 0.22 | 0.581 | 1.384 | -1.269 | -0.026 | 0.006 | 0.006 | 0 | 0.09 | 0.2 | 0.09 | 0.1 | 0.05 | 0.2 |
| 7 | 77327333 | rs35216166 | 0.663 | 0.737 | 1.267 | -1.296 | -0.033 | 0.001 | 0.001 | 0 | 0.07 | 0 | 0.05 | 0.06 | 0.09 | 0.04 |
| 7 | 77357274 | rs11765986 | -0.271 | 0.772 | 1.418 | -1.307 | -0.032 | 0.032 | 0.032 | 0 | 0.24 | 0.18 | 0.23 | 0.14 | 0.3 | 0.29 |
| 7 | 77393739 | rs4729710 | 0.289 | 0.736 | 1.259 | -1.292 | -0.018 | 0.003 | 0.003 | 0 | 0.09 | 0.17 | 0.09 | 0.1 | 0.05 | 0.2 |
| 7 | 77405741 | rs2868813 | -2.364 | 0.734 | 1.127 | -1.397 | -0.009 | 0.128 | 0.128 | 0 | 0.53 | 0.87 | 0.44 | 0.32 | 0.51 | 0.57 |
| 7 | 77407556 | rs11970824 | -0.751 | 0.785 | 1.363 | -1.368 | 0.036 | -0.049 | -0.049 | 0 | 0.14 | 0.4 | 0.09 | 0.08 | 0.15 | 0.07 |
| 7 | 77416439 | rs6465825 | -1.67 | 0.786 | 1.359 | -1.372 | 0.025 | -0.047 | -0.047 | 0.756 | 0.37 | 0.51 | 0.31 | 0.22 | 0.43 | 0.36 |
| 7 | 77423152 | rs1045463 | -0.262 | 0.807 | 1.388 | -1.369 | -0.057 | 0.037 | 0.037 | 0 | 0.22 | 0.08 | 0.21 | 0.14 | 0.29 | 0.29 |
| 7 | 77423574 | rs7953 | -2.397 | 0.777 | 1.393 | -1.381 | -0.009 | 0.131 | 0.131 | 0 | 0.52 | 0.86 | 0.44 | 0.32 | 0.51 | 0.57 |
| 7 | 77426482 | rs6970595 | -1.693 | 0.791 | 1.44 | -1.37 | 0.018 | -0.035 | -0.035 | 0 | 0.37 | 0.53 | 0.32 | 0.22 | 0.43 | 0.36 |
| 7 | 77432914 | rs11761324 | -0.422 | 1.013 | 1.244 | -1.465 | -0.033 | 0.03 | 0.03 | 0 | 0.23 | 0.13 | 0.21 | 0.14 | 0.29 | 0.3 |
| 7 | 77433484 | rs11972096 | -1.549 | 1.009 | 1.096 | -1.467 | 0.026 | -0.049 | -0.049 | 0 | 0.37 | 0.5 | 0.31 | 0.22 | 0.44 | 0.36 |
| 7 | 77434215 | rs6975196 | -0.072 | 1.059 | 1.241 | -1.421 | -0.007 | 0.043 | 0.043 | 0 | 0.1 | 0.39 | 0.11 | 0.1 | 0.05 | 0.2 |
| 7 | 77449598 | rs10953356 | -0.365 | 1.106 | 1.225 | -1.424 | -0.039 | 0.029 | 0.029 | 0 | 0.22 | 0.13 | 0.21 | 0.14 | 0.3 | 0.3 |
| 7 | 77450358 | rs6966446 | 0.285 | 1.095 | 1.235 | -1.421 | -0.015 | 0.026 | 0.026 | 0 | 0.09 | 0.27 | 0.1 | 0.1 | 0.05 | 0.2 |
| 7 | 77459325 | rs17158527 | -0.562 | 1.089 | 1.234 | -1.425 | -0.047 | -0.03 | -0.03 | 0.041 | 0.16 | 0.36 | 0.14 | 0.1 | 0.08 | 0.21 |
| 7 | 77467459 | rs17807736 | -0.415 | 1.115 | 1.225 | -1.415 | -0.035 | 0.031 | 0.031 | 0 | 0.22 | 0.13 | 0.21 | 0.14 | 0.29 | 0.3 |
| 7 | 77468437 | rs17158641 | 0.273 | 1.088 | 1.227 | -1.404 | -0.015 | 0.026 | 0.026 | 0 | 0.09 | 0.27 | 0.1 | 0.1 | 0.05 | 0.2 |
| 7 | 77489783 | rs10215731 | -0.501 | 1.211 | 1.208 | -1.413 | 0.003 | -0.004 | -0.004 | 0.139 | 0.23 | 0.24 | 0.21 | 0.14 | 0.28 | 0.29 |
| 7 | 77496555 | rs4291201 | -0.548 | 1.114 | 1.223 | -1.403 | -0.003 | 0.002 | 0.002 | 0 | 0.23 | 0.23 | 0.23 | 0.14 | 0.3 | 0.3 |
| 7 | 77501962 | rs2471585 | -0.751 | 1.112 | 1.268 | -1.41 | 0.036 | -0.047 | -0.047 | 0 | 0.14 | 0.38 | 0.09 | 0.08 | 0.15 | 0.06 |
| 7 | 77525199 | rs2471603 | -2.471 | 1.056 | 1.153 | -1.434 | -0.002 | 0.122 | 0.122 | 0 | 0.54 | 0.89 | 0.45 | 0.32 | 0.53 | 0.57 |
| 7 | 77533397 | rs12531239 | -0.484 | 0.99 | 1.234 | -1.43 | 0.024 | -0.043 | -0.043 | 0 | 0.13 | 0.3 | 0.08 | 0.08 | 0.15 | 0.07 |
| 7 | 77537964 | rs848494 | 0.436 | 0.996 | 1.243 | -1.424 | -0.004 | 0.003 | 0.003 | 0 | 0.77 | 0.77 | 0.77 | 0.85 | 0.71 | 0.71 |
| 7 | 77545076 | rs12536643 | -0.523 | 0.956 | 1.095 | -1.413 | -0.041 | -0.027 | -0.027 | 0.671 | 0.16 | 0.3 | 0.13 | 0.1 | 0.08 | 0.21 |
| 7 | 77549906 | rs12234571 | 0.248 | 1.039 | 1.139 | -1.331 | -0.02 | 0.015 | 0.015 | 0 | 0.09 | 0.21 | 0.1 | 0.1 | 0.05 | 0.2 |

| | | | | | | | | | | | | | | | | |
|---|---|---|---|---|---|---|---|---|---|---|---|---|---|---|---|---|
| 7 | 77557014 | rs17159512 | 0.197 | 1.044 | 1.035 | -1.329 | -0.017 | 0.023 | 0.023 | 0 | 0.09 | 0.25 | 0.1 | 0.1 | 0.05 | 0.2 |
| 7 | 77558095 | rs13245154 | -0.557 | 1.194 | 0.785 | -1.263 | 0.026 | -0.042 | -0.042 | 0 | 0.13 | 0.27 | 0.08 | 0.07 | 0.15 | 0.07 |
| 7 | 77562294 | rs2428939 | 0.865 | 1.247 | 0.216 | -1.253 | 0.002 | -0.017 | -0.017 | 0.053 | 0.62 | 0.59 | 0.65 | 0.76 | 0.63 | 0.5 |
| 7 | 77582265 | rs848461 | 1.633 | 1.149 | -0.941 | -1.051 | -0.041 | 0.109 | 0.109 | 0.137 | 0.46 | 0.08 | 0.56 | 0.68 | 0.49 | 0.42 |
| 7 | 77592652 | rs848467 | 0.874 | 1.497 | -0.679 | -0.486 | -0.032 | -0.064 | -0.064 | 0 | 0.71 | 0.85 | 0.68 | 0.59 | 0.69 | 0.48 |
| 7 | 77593973 | rs1211911 | 1.42 | 0.978 | -0.31 | -0.951 | 0.019 | 0.02 | 0.02 | 0.212 | 0.61 | 0.28 | 0.66 | 0.65 | 0.68 | 0.47 |
| 7 | 77594628 | rs848472 | 0.878 | 1.012 | 0.369 | -1.232 | 0.008 | -0.05 | -0.05 | 0 | 0.74 | 0.57 | 0.73 | 0.82 | 0.76 | 0.61 |
| 7 | 77595063 | rs1205283 | 1.404 | 0.983 | 0.419 | -1.262 | -0.006 | -0.012 | -0.012 | 1.571 | 0.72 | 0.41 | 0.73 | 0.75 | 0.76 | 0.61 |
| 7 | 77595308 | rs1205284 | 2.206 | 1.04 | 0.233 | -1.177 | -0.005 | 0.091 | 0.091 | 0.009 | 0.51 | 0.27 | 0.57 | 0.65 | 0.54 | 0.42 |
| 7 | 77596047 | rs848451 | 2.206 | 1.04 | 0.24 | -1.297 | -0.006 | 0.114 | 0.114 | 0.009 | 0.51 | 0.25 | 0.57 | 0.67 | 0.54 | 0.42 |
| 7 | 77596796 | rs6971524 | -1.05 | 1.18 | 0.168 | -1.204 | -0.002 | -0.036 | -0.036 | 0 | 0.23 | 0.35 | 0.16 | 0.16 | 0.22 | 0.19 |
| 7 | 77596812 | rs848452 | 0.379 | 1.298 | 0.421 | -1.17 | -0.006 | 0.029 | 0.029 | 0 | 0.88 | 0.73 | 0.87 | 0.83 | 0.9 | 0.7 |
| 7 | 77602832 | rs848458 | 1.006 | 1.272 | -0.122 | -1.191 | 0.001 | 0.038 | 0.038 | 0.13 | 0.56 | 0.39 | 0.61 | 0.81 | 0.6 | 0.56 |
| 7 | 77610479 | rs17160037 | -0.81 | 1.42 | -0.138 | -0.88 | 0.001 | -0.017 | -0.017 | 0 | 0.08 | 0.13 | 0.12 | 0.06 | 0.07 | 0.06 |
| 7 | 77610528 | rs17160040 | -0.889 | 1.697 | -0.21 | -1.16 | 0.003 | -0.035 | -0.035 | 0.035 | 0.44 | 0.46 | 0.35 | 0.16 | 0.51 | 0.36 |
| 7 | 77614334 | rs6465932 | -0.212 | 1.859 | -0.557 | -1.255 | -0.024 | 0.074 | 0.074 | 0 | 0.37 | 0.25 | 0.24 | 0.06 | 0.44 | 0.29 |
| 7 | 77616400 | rs13244959 | -0.56 | 1.587 | -0.652 | -1.033 | -0.005 | -0.005 | -0.005 | 0 | 0.43 | 0.4 | 0.36 | 0.12 | 0.52 | 0.36 |
| 7 | 77622712 | rs719313 | -0.036 | 1.562 | -1.469 | -0.953 | -0.064 | 0.153 | 0.153 | 0.051 | 0.39 | 0.11 | 0.25 | 0.11 | 0.45 | 0.39 |
| 7 | 77623551 | rs7789223 | -0.496 | 1.294 | -0.407 | -0.383 | 0.004 | 0.003 | 0.003 | 0 | 0.58 | 0.57 | 0.44 | 0.36 | 0.68 | 0.54 |
| 7 | 77625851 | rs10953392 | -0.823 | 1.754 | 0.038 | -0.509 | 0.018 | -0.025 | -0.025 | 0 | 0.19 | 0.32 | 0.2 | 0.12 | 0.31 | 0.15 |
| 7 | 77628137 | rs9986939 | -0.901 | 1.594 | 0.016 | -0.504 | 0.008 | -0.027 | -0.027 | 0 | 0.29 | 0.55 | 0.24 | 0.22 | 0.35 | 0.38 |
| 7 | 77634980 | rs2906510 | -0.525 | 1.558 | -0.11 | -0.604 | -0.006 | 0.058 | 0.058 | 0.271 | 0.44 | 0.65 | 0.32 | 0.18 | 0.48 | 0.41 |
| 7 | 77635119 | rs7805185 | -0.433 | 1.548 | -0.221 | -0.173 | -0.003 | 0.012 | 0.012 | 0.204 | 0.45 | 0.58 | 0.46 | 0.32 | 0.53 | 0.48 |
| 7 | 77639656 | rs12673147 | -0.892 | 1.598 | -0.178 | 0.479 | -0.029 | -0.044 | -0.044 | 0 | 0.12 | 0.03 | 0.17 | 0.17 | 0.15 | 0.11 |
| 7 | 77645588 | rs2108082 | 0.817 | 1.361 | 0.46 | 0.409 | -0.004 | -0.03 | -0.03 | 0 | 0.92 | 0.87 | 0.92 | 1 | 0.92 | 0.96 |
| 7 | 77645731 | rs2428918 | -1.092 | 1.395 | 0.772 | 0.462 | 0.021 | 0.114 | 0.114 | 0 | 0.22 | 0.44 | 0.41 | 0.58 | 0.24 | 0.39 |
| 7 | 77654514 | rs2471597 | 0.842 | 1.043 | 0.387 | 0.8 | -0.016 | -0.047 | -0.047 | 0 | 0.91 | 0.82 | 0.92 | 1 | 0.93 | 0.97 |
| 7 | 77661138 | rs2471577 | -0.636 | 1.349 | 0.04 | 1.169 | 0.001 | 0.127 | 0.127 | 0 | 0.11 | 0.18 | 0.33 | 0.55 | 0.1 | 0.3 |
| 7 | 77663087 | rs3779348 | -0.975 | 1.689 | -0.22 | 0.749 | -0.012 | -0.011 | -0.011 | 0 | 0.09 | 0.05 | 0.16 | 0.17 | 0.17 | 0.06 |
| 7 | 77663180 | rs3779347 | -1.945 | 1.343 | 0.436 | 0.612 | 0.023 | 0.013 | 0.013 | 0 | 0.19 | 0.4 | 0.23 | 0.18 | 0.23 | 0.07 |
| 7 | 77663486 | rs2471614 | 1.098 | 1.679 | 0.303 | 0.315 | 0.009 | -0.042 | -0.042 | 0 | 0.9 | 0.85 | 0.91 | 0.99 | 0.85 | 0.96 |
| 7 | 77663791 | rs17160602 | -0.867 | 1.846 | 0.473 | 0.109 | 0.05 | -0.041 | -0.041 | 0 | 0.08 | 0.19 | 0.1 | 0.01 | 0.17 | 0.06 |
| 7 | 77664569 | rs2428927 | -0.414 | 1.074 | -0.536 | 0.399 | -0.025 | -0.074 | -0.074 | 0.598 | 0.67 | 0.9 | 0.55 | 0.5 | 0.69 | 0.69 |
| 7 | 77664661 | rs3779344 | -0.756 | 0.495 | 0.177 | 0.377 | -0.041 | 0.039 | 0.039 | 0.306 | 0.2 | 0.44 | 0.18 | 0.21 | 0.13 | 0.27 |
| 7 | 77665201 | rs2428929 | 0.109 | 0.668 | 0.252 | 0.231 | -0.003 | 0.011 | 0.011 | 0 | 0.45 | 0.44 | 0.42 | 0.31 | 0.54 | 0.49 |
| 7 | 77665925 | rs2471619 | 0.452 | 0.825 | -0.063 | -0.116 | 0.01 | -0.041 | -0.041 | 0 | 0.33 | 0.41 | 0.32 | 0.15 | 0.41 | 0.29 |
| 7 | 77666501 | rs757583 | 0.931 | 0.612 | -0.142 | -0.127 | -0.011 | -0.021 | -0.021 | 1.925 | 0.12 | 0.03 | 0.11 | 0.16 | 0.14 | 0.21 |
| 7 | 77668115 | rs3779341 | -0.528 | 0.59 | 0.144 | 0.023 | -0.036 | 0.03 | 0.03 | 1.948 | 0.18 | 0.39 | 0.17 | 0.21 | 0.12 | 0.17 |
| 7 | 77668741 | rs3807786 | 1.541 | 0.909 | 0.285 | -0.259 | -0.005 | 0 | 0 | 0 | 0.21 | 0.18 | 0.23 | 0.1 | 0.28 | 0.11 |
| 7 | 77668942 | rs2428936 | 0.934 | 0.844 | 0.332 | -0.163 | 0.002 | -0.01 | -0.01 | 0.274 | 0.06 | 0.13 | 0.06 | 0.01 | 0.09 | 0.04 |
| 7 | 77671200 | rs6465973 | -0.93 | 0.39 | 0.356 | 0.498 | -0.045 | 0.279 | 0.279 | 0.045 | 0.15 | 0.42 | 0.35 | 0.55 | 0.11 | 0.2 |
| 7 | 77671381 | rs757863 | -0.212 | 0.48 | 0.221 | 0.023 | 0.004 | -0.02 | -0.02 | 0 | 0.41 | 0.48 | 0.35 | 0.21 | 0.45 | 0.32 |
| 7 | 77671455 | rs3807779 | 0.423 | 0.388 | 0.278 | 0.038 | 0.004 | -0.036 | -0.036 | 0 | 0.32 | 0.48 | 0.37 | 0.25 | 0.36 | 0.25 |
| 7 | 77671786 | rs735406 | -0.312 | 0.439 | 0.201 | -0.022 | -0.001 | -0.001 | -0.001 | 0 | 0.41 | 0.52 | 0.43 | 0.31 | 0.45 | 0.35 |
| 7 | 77671832 | rs757865 | -1.912 | 0.449 | -0.119 | -0.023 | -0.018 | -0.082 | -0.082 | 0 | 0.65 | 0.86 | 0.62 | 0.52 | 0.68 | 0.66 |
| 7 | 77672995 | rs11763565 | 0.332 | 0.534 | -0.049 | -0.007 | -0.019 | -0.029 | -0.029 | 0 | 0.08 | 0 | 0.04 | 0.06 | 0.08 | 0.09 |
| 7 | 77675129 | rs10277160 | -1.109 | 0.498 | -0.036 | -0.026 | -0.032 | 0.066 | 0.066 | 0 | 0.53 | 0.88 | 0.5 | 0.34 | 0.54 | 0.48 |
| 7 | 77676395 | rs11768469 | -0.006 | 0.568 | 0.135 | 0.164 | -0.003 | -0.008 | -0.008 | 3.698 | 0.08 | 0.04 | 0.05 | 0.07 | 0.08 | 0.09 |
| 7 | 77677220 | rs3779330 | 1.724 | 0.604 | 0.355 | 0.115 | -0.002 | 0.026 | 0.026 | 0 | 0.77 | 0.52 | 0.78 | 0.77 | 0.78 | 0.65 |
| 7 | 77677237 | rs3779329 | -0.008 | 0.583 | 0.355 | 0.177 | 0.011 | 0.073 | 0.073 | 0 | 0.09 | 0.25 | 0.15 | 0.15 | 0.09 | 0.14 |
| 7 | 77679834 | rs1019021 | 0.272 | 0.634 | 0.141 | 0.207 | -0.009 | -0.03 | -0.03 | 0.008 | 0.07 | 0 | 0.02 | 0.06 | 0.05 | 0.09 |
| 7 | 77684816 | rs3779325 | -0.378 | 0.665 | 0.276 | 0.04 | 0.013 | -0.014 | -0.014 | 0 | 0.68 | 0.6 | 0.67 | 0.73 | 0.66 | 0.68 |
| 7 | 77685345 | rs7792331 | 0.273 | 0.634 | 0.18 | 0.075 | 0.014 | -0.01 | -0.01 | 0 | 0.6 | 0.5 | 0.59 | 0.69 | 0.56 | 0.56 |
| 7 | 77691587 | rs3779320 | -0.625 | 0.289 | -0.321 | -0.128 | -0.034 | -0.033 | -0.033 | 0.018 | 0.24 | 0.06 | 0.15 | 0.24 | 0.25 | 0.29 |
| 7 | 77692114 | rs3807763 | 1.65 | 0.155 | -0.103 | -0.241 | 0.008 | -0.064 | -0.064 | 0 | 0.35 | 0.15 | 0.39 | 0.49 | 0.29 | 0.31 |
| 7 | 77693091 | rs6977382 | -1.111 | 0.108 | 0.029 | -0.367 | -0.013 | 0.016 | 0.016 | 0.046 | 0.74 | 0.77 | 0.73 | 0.88 | 0.68 | 0.77 |
| 7 | 77693385 | rs6977897 | 0.093 | 0.305 | -0.256 | -0.279 | -0.008 | 0.027 | 0.027 | 0 | 0.59 | 0.35 | 0.55 | 0.73 | 0.54 | 0.6 |
| 7 | 77694502 | rs3779317 | -0.834 | 0.404 | -0.366 | -0.233 | -0.002 | 0.005 | 0.005 | 0 | 0.23 | 0.21 | 0.19 | 0.2 | 0.29 | 0.24 |
| 7 | 77696285 | rs3807758 | 0.502 | 0.421 | -0.643 | -0.076 | -0.032 | 0.007 | 0.007 | 0.139 | 0.13 | 0.01 | 0.09 | 0.1 | 0.16 | 0.13 |
| 7 | 77706847 | rs2868838 | 0.474 | 0.348 | -0.939 | -0.088 | 0.005 | -0.072 | -0.072 | 0.007 | 0.23 | 0.08 | 0.33 | 0.48 | 0.21 | 0.21 |
| 7 | 77707557 | rs4590377 | 0.013 | 0.351 | -0.585 | -0.418 | 0.012 | -0.053 | -0.053 | 0 | 0.29 | 0.49 | 0.21 | 0.19 | 0.27 | 0.38 |
| 7 | 77711290 | rs7788132 | -0.457 | 0.245 | -0.906 | -0.454 | -0.02 | 0.067 | 0.067 | 0 | 0.45 | 0.29 | 0.31 | 0.29 | 0.49 | 0.48 |
| 7 | 77711756 | rs3779312 | 1.254 | 0.264 | -0.96 | -0.359 | -0.018 | 0.017 | 0.017 | 0 | 0.19 | 0.1 | 0.15 | 0.15 | 0.2 | 0.24 |
| 7 | 77714115 | rs3779309 | 1.149 | 0.261 | -0.789 | -0.399 | 0.001 | -0.004 | -0.004 | 0.003 | 0.19 | 0.21 | 0.16 | 0.15 | 0.2 | 0.24 |
| 7 | 77717443 | rs3807745 | -1.652 | 0.162 | -1.15 | -0.42 | -0.001 | 0 | 0 | 0 | 0.72 | 0.74 | 0.67 | 0.79 | 0.73 | 0.77 |
| 7 | 77721473 | rs3807743 | -0.537 | 0.124 | -1.404 | -0.502 | -0.001 | 0.047 | 0.047 | 0 | 0.55 | 0.71 | 0.59 | 0.7 | 0.56 | 0.61 |
| 7 | 77728411 | rs6976139 | -0.674 | 0.397 | -1.13 | -0.284 | 0.002 | 0.031 | 0.031 | 0 | 0.45 | 0.63 | 0.29 | 0.23 | 0.45 | 0.49 |
| 7 | 77729487 | rs3807737 | -0.03 | 0.378 | -0.919 | -0.172 | -0.001 | -0.027 | -0.027 | 0 | 0.29 | 0.61 | 0.21 | 0.14 | 0.28 | 0.33 |
| 7 | 77740168 | rs3823793 | 1.797 | 0.501 | -1.488 | -0.136 | -0.038 | 0.021 | 0.021 | 0 | 0.14 | 0.02 | 0.1 | 0.07 | 0.15 | 0.19 |

| | | | | | | | | | | | | | | | | |
|---|---|---|---|---|---|---|---|---|---|---|---|---|---|---|---|---|
| 7 | 77740606 | rs3807735 | 1.178 | 0.424 | -1.693 | -0.208 | -0.022 | 0.009 | 0.009 | 0.719 | 0.16 | 0 | 0.08 | 0.1 | 0.16 | 0.16 |
| 7 | 77740909 | rs3807733 | 0.591 | 0.63 | -1.16 | 0.022 | 0.002 | 0.001 | 0.001 | 0 | 0.07 | 0.13 | 0.06 | 0.06 | 0.09 | 0.12 |
| 7 | 77741395 | rs3807730 | 0.023 | 0.931 | -1.365 | 0.052 | -0.107 | 0.041 | 0.041 | 0 | 0.28 | 0.1 | 0.23 | 0.19 | 0.36 | 0.33 |
| 7 | 77743622 | rs3807728 | 0.232 | 0.865 | -1.084 | 0.073 | -0.006 | 0.009 | 0.009 | 0 | 0.45 | 0.57 | 0.46 | 0.36 | 0.49 | 0.46 |
| 7 | 77744459 | rs3807723 | 0.181 | 0.988 | -1.189 | 0.147 | -0.024 | 0.042 | 0.042 | 0 | 0.3 | 0.18 | 0.22 | 0.13 | 0.34 | 0.29 |
| 7 | 77747902 | rs10246111 | -1.811 | 0.878 | -1.165 | 0.2 | 0.043 | -0.075 | -0.075 | 0 | 0.79 | 0.93 | 0.72 | 0.56 | 0.87 | 0.81 |
| 7 | 77751791 | rs3807721 | 0.137 | 0.939 | -1.149 | 0.244 | 0.004 | 0.023 | 0.023 | 0 | 0.49 | 0.74 | 0.5 | 0.45 | 0.53 | 0.53 |
| 7 | 77752282 | rs3779300 | -0.601 | 1.014 | -1.181 | 0.22 | 0.024 | -0.02 | -0.02 | 0 | 0.6 | 0.75 | 0.59 | 0.49 | 0.69 | 0.6 |
| 7 | 77756275 | rs2074640 | -0.378 | 0.923 | -1.424 | 0.554 | 0.008 | -0.096 | -0.096 | 0.001 | 0.67 | 0.91 | 0.63 | 0.53 | 0.7 | 0.71 |
| 7 | 77756580 | rs2074641 | -1.456 | 0.788 | -1.364 | 0.575 | 0.042 | -0.06 | -0.06 | 0 | 0.78 | 0.92 | 0.72 | 0.57 | 0.87 | 0.79 |
| 7 | 77756724 | rs7812015 | -1.456 | 0.788 | -1.364 | 0.575 | 0.042 | -0.06 | -0.06 | 0.433 | 0.78 | 0.92 | 0.72 | 0.57 | 0.87 | 0.79 |
| 7 | 77762013 | rs2074646 | -0.075 | 1.038 | -1.172 | 0.437 | -0.002 | -0.009 | -0.009 | 0 | 0.61 | 0.78 | 0.56 | 0.46 | 0.6 | 0.59 |
| 7 | 77764079 | rs11975124 | 0.074 | 1.04 | -0.753 | 0.502 | 0.008 | -0.005 | -0.005 | 0.903 | 0.25 | 0.35 | 0.18 | 0.2 | 0.27 | 0.33 |
| 7 | 77764253 | rs1465221 | 0.105 | 1.417 | -0.714 | 0.265 | 0.048 | -0.012 | -0.012 | 0 | 0.23 | 0.3 | 0.31 | 0.21 | 0.33 | 0.22 |
| 7 | 77766520 | rs987493 | -1.308 | 0.311 | -1.374 | 0.761 | 0.055 | -0.022 | -0.022 | 0 | 0.77 | 0.92 | 0.79 | 0.73 | 0.88 | 0.8 |
| 7 | 77767434 | rs7778303 | 1.342 | 0.824 | -1.58 | 0.642 | -0.059 | 0.011 | 0.011 | 0 | 0.11 | 0.01 | 0.09 | 0.04 | 0.16 | 0.08 |
| 7 | 77768217 | rs2868895 | -0.4 | 0.746 | -1.383 | 0.637 | 0.016 | 0.005 | 0.005 | 0 | 0.65 | 0.84 | 0.7 | 0.69 | 0.72 | 0.73 |
| 7 | 77769423 | rs3807707 | -0.087 | 0.552 | -0.949 | 0.632 | 0 | 0 | 0 | 0 | 0.44 | 0.49 | 0.38 | 0.47 | 0.48 | 0.5 |
| 7 | 77770521 | rs3807703 | 1.621 | 1.114 | -0.965 | 0.764 | -0.037 | -0.012 | -0.012 | 0 | 0.16 | 0.03 | 0.1 | 0.12 | 0.17 | 0.18 |
| 7 | 77770699 | rs3807702 | 0.292 | 1.384 | -0.378 | 1.001 | 0.028 | 0.097 | 0.097 | 0 | 0.08 | 0.24 | 0.15 | 0.17 | 0.11 | 0.1 |
| 7 | 77771239 | rs10228748 | 0.292 | 1.384 | -0.405 | 1.001 | 0.027 | 0.094 | 0.094 | 0 | 0.08 | 0.24 | 0.15 | 0.17 | 0.11 | 0.1 |
| 7 | 77771752 | rs3823789 | 0.899 | 1.534 | -0.754 | 1.076 | 0.028 | 0.007 | 0.007 | 0 | 0.21 | 0.33 | 0.32 | 0.22 | 0.24 | 0.23 |
| 7 | 77771770 | rs3807701 | 1.1 | 1.268 | -0.682 | 1.126 | -0.029 | -0.027 | -0.027 | 6.101 | 0.06 | 0.01 | 0.12 | 0.17 | 0.1 | 0.1 |
| 7 | 77773132 | rs3823787 | 0.11 | 1.197 | -0.615 | 0.712 | 0.02 | -0.009 | -0.009 | 0 | 0.33 | 0.35 | 0.4 | 0.25 | 0.4 | 0.3 |
| 7 | 77774487 | rs6963746 | -0.301 | 1.108 | -0.713 | 0.994 | 0.007 | 0.108 | 0.108 | 0 | 0.09 | 0.16 | 0.14 | 0.32 | 0.11 | 0.09 |
| 7 | 77776110 | rs17524120 | 0.478 | 1.459 | -1.146 | 0.582 | -0.075 | 0.01 | 0.01 | 0 | 0.13 | 0.01 | 0.11 | 0.08 | 0.22 | 0.17 |
| 7 | 77778668 | rs3807694 | -1.376 | 1.205 | -1.289 | 0.296 | 0.023 | 0.009 | 0.009 | 0 | 0.47 | 0.39 | 0.39 | 0.51 | 0.6 | 0.46 |
| 7 | 77779144 | rs3807692 | -0.268 | 1.364 | -1.282 | 0.476 | -0.007 | -0.031 | -0.031 | 0 | 0.2 | 0.14 | 0.17 | 0.33 | 0.22 | 0.16 |
| 7 | 77782780 | rs3807689 | -0.376 | 1.041 | -1.658 | 0.225 | 0.01 | 0.014 | 0.014 | 1.382 | 0.67 | 0.87 | 0.7 | 0.71 | 0.7 | 0.69 |
| 7 | 77783782 | rs3807688 | 0.284 | 0.99 | -1.676 | 0.259 | -0.027 | -0.004 | -0.004 | 0.018 | 0.19 | 0.12 | 0.14 | 0.15 | 0.27 | 0.19 |
| 7 | 77785591 | rs3779289 | 1.909 | 0.893 | -1.3 | 0.165 | -0.008 | -0.009 | -0.009 | 1.301 | 0.33 | 0.54 | 0.42 | 0.31 | 0.29 | 0.37 |
| 7 | 77786274 | rs2868865 | -1.102 | 0.539 | -1.07 | 0.294 | 0.025 | 0.049 | 0.049 | 0.303 | 0.77 | 0.91 | 0.84 | 0.9 | 0.82 | 0.82 |
| 7 | 77791253 | rs7796631 | 0.709 | 0.71 | -0.938 | 0.449 | -0.005 | 0.061 | 0.061 | 0 | 0.09 | 0.13 | 0.09 | 0.27 | 0.05 | 0.08 |
| 7 | 77793321 | rs1524742 | 1.441 | 0.842 | -0.682 | 0.406 | -0.001 | -0.009 | -0.009 | 0 | 0.08 | 0.06 | 0.13 | 0.19 | 0.09 | 0.1 |
| 7 | 77793413 | rs1524743 | 1.002 | 0.821 | -0.505 | 0.396 | -0.002 | 0.017 | 0.017 | 0 | 0.09 | 0.15 | 0.14 | 0.19 | 0.09 | 0.1 |
| 7 | 77795921 | rs740027 | 1.415 | 0.843 | -0.457 | 0.366 | 0 | 0.025 | 0.025 | 0 | 0.08 | 0.12 | 0.14 | 0.2 | 0.09 | 0.1 |
| 7 | 77796919 | rs2072156 | 1.447 | 0.843 | -0.457 | 0.365 | 0 | 0.026 | 0.026 | 0 | 0.08 | 0.12 | 0.13 | 0.2 | 0.09 | 0.1 |
| 7 | 77798673 | rs2049434 | -0.343 | 0.901 | -0.498 | 0.334 | 0 | 0 | 0 | 0.25 | 0.23 | 0.25 | 0.24 | 0.29 | 0.3 | 0.26 |
| 7 | 77801671 | rs1524745 | 1.065 | 0.624 | -0.471 | 0.385 | -0.001 | 0 | 0 | 0.293 | 0.47 | 0.5 | 0.53 | 0.34 | 0.51 | 0.48 |
| 7 | 77802070 | rs3807676 | 1.055 | 0.655 | -0.921 | 0.378 | -0.036 | 0.012 | 0.012 | 0.572 | 0.11 | 0.04 | 0.08 | 0.08 | 0.15 | 0.16 |
| 7 | 77803275 | rs2691529 | -0.075 | 0.452 | -0.807 | 0.465 | 0.016 | -0.01 | -0.01 | 0.062 | 0.67 | 0.72 | 0.75 | 0.63 | 0.76 | 0.73 |
| 7 | 77809425 | rs3807671 | 0.087 | 0.505 | -0.506 | 0.58 | -0.004 | 0.193 | 0.193 | 0 | 0.82 | 0.45 | 0.74 | 0.52 | 0.85 | 0.83 |
| 7 | 77810868 | rs3807669 | 0.376 | 0.569 | -1.154 | 0.383 | -0.035 | 0.09 | 0.09 | 0.048 | 0.35 | 0.05 | 0.23 | 0.18 | 0.33 | 0.35 |
| 7 | 77817115 | rs3807663 | 0.496 | 0.844 | -0.844 | 0.341 | -0.024 | 0.057 | 0.057 | 0 | 0.45 | 0.23 | 0.5 | 0.35 | 0.52 | 0.48 |
| 7 | 77818152 | rs3807660 | 0.412 | 0.892 | -0.864 | 0.354 | -0.013 | 0.057 | 0.057 | 0 | 0.46 | 0.3 | 0.5 | 0.35 | 0.52 | 0.48 |
| 7 | 77821780 | rs3823778 | 0.391 | 0.679 | -0.845 | 0.362 | 0.004 | -0.022 | -0.022 | 0 | 0.4 | 0.31 | 0.52 | 0.5 | 0.4 | 0.47 |
| 7 | 77825705 | rs1965530 | -0.45 | 0.922 | -0.444 | 0.042 | 0.035 | -0.045 | -0.045 | 0 | 0.8 | 0.75 | 0.76 | 0.95 | 0.67 | 0.9 |
| 7 | 77825707 | rs1965529 | 1.118 | 0.049 | -0.402 | 0.137 | 0.017 | -0.015 | -0.015 | 0 | 0.28 | 0.21 | 0.33 | 0.39 | 0.21 | 0.37 |
| 7 | 77826131 | rs1965528 | 0.358 | -0.211 | -0.601 | 0.143 | 0.001 | 0 | 0 | 0 | 0.45 | 0.52 | 0.41 | 0.63 | 0.32 | 0.51 |
| 7 | 77828434 | rs3807648 | -0.648 | 0.034 | -0.602 | -0.171 | -0.006 | -0.004 | -0.004 | 0.151 | 0.54 | 0.48 | 0.61 | 0.54 | 0.6 | 0.38 |
| 7 | 77828940 | rs3779273 | 0.662 | 0.018 | -0.731 | -0.178 | 0.023 | 0.015 | 0.015 | 0 | 0.45 | 0.38 | 0.38 | 0.46 | 0.4 | 0.62 |
| 7 | 77829071 | rs1997017 | -0.395 | -0.049 | -0.707 | -0.097 | -0.037 | 0.1 | 0.1 | 0 | 0.81 | 0.54 | 0.84 | 0.7 | 0.89 | 0.87 |
| 7 | 77830091 | rs3807645 | 1.846 | -0.124 | -1.431 | -0.062 | 0.018 | 0.028 | 0.028 | 0 | 0.27 | 0.15 | 0.26 | 0.18 | 0.22 | 0.27 |
| 7 | 77830751 | rs1829989 | -0.479 | 0.124 | -1.071 | 0.177 | 0.007 | 0.013 | 0.013 | 0 | 0.47 | 0.39 | 0.58 | 0.53 | 0.53 | 0.34 |
| 7 | 77831075 | rs17809684 | 1.393 | 0.653 | -1.507 | 0.196 | -0.09 | 0.019 | 0.019 | 0 | 0.14 | 0.02 | 0.26 | 0.07 | 0.28 | 0.08 |
| 7 | 77832716 | rs10499837 | 1.433 | 0.554 | -1.382 | 0.227 | 0.02 | 0.067 | 0.067 | 0 | 0.34 | 0.15 | 0.29 | 0.19 | 0.29 | 0.31 |
| 7 | 77836239 | rs6963699 | 0.892 | 0.783 | -1.721 | 0.493 | -0.021 | -0.014 | -0.014 | 0 | 0.19 | 0.07 | 0.2 | 0.17 | 0.19 | 0.16 |
| 7 | 77836723 | rs10256412 | -0.098 | 0.916 | -1.527 | 0.123 | 0.008 | 0.013 | 0.013 | 0.012 | 0.48 | 0.43 | 0.58 | 0.53 | 0.55 | 0.34 |
| 7 | 77837131 | rs17149920 | -0.129 | 1.302 | -1.705 | 0.521 | -0.004 | -0.034 | -0.034 | 1.868 | 0.13 | 0.06 | 0.09 | 0.28 | 0.14 | 0.33 |
| 7 | 77837900 | rs17809941 | -0.049 | 1.55 | -1.481 | 0.133 | -0.079 | 0.016 | 0.016 | 0 | 0.13 | 0.01 | 0.24 | 0.07 | 0.25 | 0.08 |
| 7 | 77841333 | rs17149959 | -0.671 | 0.374 | -1.148 | -0.155 | -0.008 | -0.035 | -0.035 | 0 | 0.11 | 0.2 | 0.02 | 0.02 | 0.03 | 0.04 |
| 7 | 77841529 | rs6976643 | 0.003 | 0.977 | -1.763 | -0.367 | -0.058 | 0.012 | 0.012 | 0 | 0.23 | 0.12 | 0.38 | 0.21 | 0.38 | 0.2 |
| 7 | 77845098 | rs6959540 | -0.685 | 0.22 | -1.516 | -0.101 | 0 | -0.001 | -0.001 | 0 | 0.7 | 0.72 | 0.76 | 0.62 | 0.72 | 0.54 |
| 7 | 77846676 | rs12672053 | 0.748 | 0.282 | -1.554 | -0.064 | -0.079 | 0.008 | 0.008 | 0.083 | 0.21 | 0.08 | 0.37 | 0.19 | 0.38 | 0.21 |
| 7 | 77850352 | rs1107560 | 0.824 | -0.387 | -0.815 | -0.038 | -0.024 | 0.083 | 0.083 | 0 | 0.14 | 0.24 | 0.17 | 0.32 | 0.08 | 0.2 |
| 7 | 77850398 | rs1107561 | 0.578 | -0.403 | -0.749 | -0.042 | -0.034 | 0.093 | 0.093 | 0 | 0.16 | 0.31 | 0.18 | 0.33 | 0.08 | 0.2 |
| 7 | 77857381 | rs1965526 | 1.208 | -0.63 | -1.071 | 0.078 | -0.01 | -0.049 | -0.049 | 0 | 0.14 | 0.04 | 0.18 | 0.33 | 0.16 | 0.31 |
| 7 | 77859083 | rs6954324 | 1.031 | -0.787 | -1.228 | -0.085 | -0.096 | 0.01 | 0.01 | 0.954 | 0.25 | 0.07 | 0.39 | 0.19 | 0.42 | 0.2 |
| 7 | 77860274 | rs1829993 | 0.09 | -0.831 | -0.973 | -0.124 | -0.004 | -0.005 | -0.005 | 0 | 0.41 | 0.37 | 0.56 | 0.53 | 0.5 | 0.41 |

| | | | | | | | | | | | | | | | | |
|---|---|---|---|---|---|---|---|---|---|---|---|---|---|---|---|---|
| 7 | 77866711 | rs1852008 | -0.171 | -0.954 | -0.944 | -0.304 | -0.012 | -0.027 | -0.027 | 0.02 | 0.75 | 0.67 | 0.92 | 0.9 | 0.88 | 0.81 |
| 7 | 77868102 | rs10953499 | 1.391 | -0.96 | -0.989 | -0.216 | -0.013 | -0.057 | -0.057 | 0.514 | 0.14 | 0.04 | 0.18 | 0.33 | 0.16 | 0.31 |
| 7 | 77870658 | rs4730150 | 0.38 | -0.947 | -0.964 | -0.233 | 0.006 | -0.005 | -0.005 | 0 | 0.61 | 0.62 | 0.74 | 0.57 | 0.73 | 0.5 |
| 7 | 77870975 | rs4730151 | 0.914 | -0.977 | -0.803 | -0.249 | -0.005 | 0 | 0 | 0 | 0.36 | 0.37 | 0.35 | 0.38 | 0.33 | 0.31 |
| 7 | 77871607 | rs10256416 | 1.118 | -1.057 | -0.764 | -0.222 | -0.032 | 0.095 | 0.095 | 0 | 0.14 | 0.32 | 0.17 | 0.31 | 0.08 | 0.2 |
| 7 | 77872067 | rs12536887 | 2.055 | -0.999 | -1.138 | -0.313 | -0.056 | 0.044 | 0.044 | 0.345 | 0.21 | 0.01 | 0.18 | 0.06 | 0.25 | 0.09 |
| 7 | 77874485 | rs798355 | -0.538 | -1.07 | -0.704 | -0.23 | -0.014 | -0.014 | -0.014 | 0 | 0.4 | 0.5 | 0.37 | 0.7 | 0.28 | 0.59 |
| 7 | 77874688 | rs798356 | -1.1 | -1.055 | -0.861 | -0.26 | -0.004 | 0.082 | 0.082 | 0 | 0.55 | 0.92 | 0.45 | 0.74 | 0.35 | 0.72 |
| 7 | 77875299 | rs13239642 | 1.983 | -1.02 | -0.818 | -0.269 | -0.012 | 0.059 | 0.059 | 0.078 | 0.22 | 0.06 | 0.17 | 0.06 | 0.24 | 0.09 |
| 7 | 77875410 | rs699332 | -0.056 | -0.841 | -0.652 | -0.14 | -0.026 | 0.038 | 0.038 | 0 | 0.26 | 0.45 | 0.2 | 0.37 | 0.14 | 0.27 |
| 7 | 77878100 | rs798361 | -1.166 | -0.511 | -1.327 | -0.149 | -0.094 | 0.006 | 0.006 | 0 | 0.76 | 0.93 | 0.63 | 0.8 | 0.6 | 0.81 |
| 7 | 77886264 | rs798319 | -0.363 | -0.821 | -0.749 | 0.223 | -0.03 | 0.016 | 0.016 | 0 | 0.36 | 0.43 | 0.24 | 0.45 | 0.19 | 0.36 |
| 7 | 77888356 | rs798320 | -0.524 | -0.784 | -1.09 | 0.231 | -0.003 | 0 | 0 | 0 | 0.37 | 0.6 | 0.25 | 0.45 | 0.2 | 0.36 |
| 7 | 77896360 | rs3915115 | 0.402 | -0.789 | -1.177 | 0.242 | 0.002 | -0.014 | -0.014 | 0 | 0.18 | 0.15 | 0.25 | 0.32 | 0.18 | 0.33 |
| 7 | 77896555 | rs798329 | -0.384 | -0.754 | -1.074 | 0.256 | 0.025 | -0.018 | -0.018 | 0 | 0.36 | 0.66 | 0.25 | 0.45 | 0.19 | 0.34 |
| 7 | 77911947 | rs798334 | -0.219 | -0.769 | -1.2 | 0.24 | -0.026 | 0.006 | 0.006 | 0.006 | 0.29 | 0.61 | 0.23 | 0.35 | 0.13 | 0.25 |
| 7 | 77912876 | rs11770919 | 1.012 | -0.245 | -1.397 | 0.527 | -0.031 | -0.02 | -0.02 | 2.315 | 0.08 | 0.01 | 0.03 | 0.1 | 0.09 | 0.11 |
| 7 | 77913918 | rs11761364 | 0.209 | -0.321 | -1.372 | 0.509 | -0.007 | -0.052 | -0.052 | 0 | 0.17 | 0.06 | 0.25 | 0.33 | 0.18 | 0.33 |
| 7 | 77916282 | rs798337 | -0.671 | -0.157 | -1.428 | 0.333 | 0.034 | 0.024 | 0.024 | 0 | 0.39 | 0.78 | 0.5 | 0.73 | 0.29 | 0.5 |
| 7 | 77931730 | rs798276 | -0.748 | -0.099 | -1.604 | 0.343 | 0.027 | -0.005 | -0.005 | 0.597 | 0.42 | 0.81 | 0.41 | 0.67 | 0.34 | 0.62 |
| 7 | 77939996 | rs798285 | 0.015 | 0.096 | -1.703 | 0.299 | 0 | 0 | 0 | 0 | 0.59 | 0.36 | 0.52 | 0.39 | 0.57 | 0.59 |
| 7 | 77944878 | rs798292 | 0.419 | 0.233 | -1.566 | 0.374 | 0.001 | 0.049 | 0.049 | 0 | 0.23 | 0.36 | 0.31 | 0.32 | 0.21 | 0.25 |
| 7 | 77952630 | rs11971642 | -0.299 | 0.525 | -0.811 | 0.827 | -0.032 | 0.021 | 0.021 | 0 | 0.25 | 0.5 | 0.12 | 0.37 | 0.15 | 0.4 |
| 7 | 77953166 | rs798299 | 0.212 | 1.603 | -1.216 | 1.146 | -0.034 | -0.023 | -0.023 | 0.363 | 0.76 | 0.87 | 0.62 | 0.73 | 0.65 | 0.77 |
| 7 | 77954342 | rs798302 | 0.503 | 1.434 | -0.677 | 1.503 | 0.027 | 0.088 | 0.088 | 0 | 0.88 | 0.74 | 0.81 | 0.64 | 0.83 | 0.78 |
| 7 | 77954675 | rs17404237 | -0.5 | 2.071 | -1.041 | 1.022 | -0.026 | -0.012 | -0.012 | 0 | 0.22 | 0.1 | 0.34 | 0.25 | 0.3 | 0.27 |
| 7 | 77955134 | rs798305 | 0.889 | 1.246 | -1.437 | 0.347 | 0.002 | 0.065 | 0.065 | 0.647 | 0.62 | 0.84 | 0.76 | 0.81 | 0.65 | 0.49 |
| 7 | 77955456 | rs798306 | 0.252 | 1.18 | -1.567 | 0.237 | -0.032 | 0.013 | 0.013 | 0.118 | 0.78 | 0.96 | 0.88 | 0.86 | 0.74 | 0.69 |
| 7 | 77958122 | rs798309 | 1.617 | 0.725 | -1.495 | 0.27 | 0.008 | 0.038 | 0.038 | 0 | 0.5 | 0.6 | 0.43 | 0.24 | 0.56 | 0.3 |
| 7 | 77958559 | rs798311 | -1.218 | 0.815 | -1.518 | 0.289 | -0.007 | -0.008 | -0.008 | 1.178 | 0.87 | 0.97 | 0.92 | 0.88 | 0.83 | 0.81 |
| 7 | 77958823 | rs798313 | 0.68 | 0.617 | -1.049 | 0.203 | 0 | 0 | 0 | 0 | 0.7 | 0.7 | 0.78 | 0.83 | 0.73 | 0.61 |
| 7 | 77959140 | rs798315 | -0.957 | 0.786 | -1.037 | 0.342 | -0.001 | -0.001 | -0.001 | 0 | 0.87 | 0.87 | 0.91 | 0.89 | 0.83 | 0.81 |
| 7 | 77964528 | rs2691542 | -0.633 | 0.584 | -1.359 | 0.07 | -0.007 | -0.001 | -0.001 | 0 | 0.83 | 0.96 | 0.9 | 0.88 | 0.8 | 0.82 |
| 7 | 77966572 | rs2691546 | 0.509 | 0.58 | -1.238 | -0.238 | 0.001 | -0.006 | -0.006 | 0.016 | 0.7 | 0.85 | 0.56 | 0.29 | 0.69 | 0.59 |
| 7 | 77972921 | rs13244251 | 2.374 | 0.519 | -1.187 | -0.944 | -0.008 | 0.02 | 0.02 | 0 | 0.21 | 0.15 | 0.13 | 0.06 | 0.26 | 0.1 |
| 7 | 77973787 | rs2075014 | 0.222 | 0.632 | -0.99 | -0.778 | -0.001 | 0.012 | 0.012 | 0 | 0.62 | 0.75 | 0.54 | 0.29 | 0.63 | 0.48 |
| 7 | 77976040 | rs17150562 | 1.171 | 0.529 | -0.826 | -0.824 | -0.02 | -0.031 | -0.031 | 0 | 0.15 | 0.29 | 0.12 | 0.05 | 0.08 | 0.19 |
| 7 | 77977855 | rs2159511 | 0.163 | 0.497 | -0.992 | -0.844 | -0.002 | -0.01 | -0.01 | 0 | 0.6 | 0.55 | 0.51 | 0.28 | 0.62 | 0.45 |
| 7 | 77979258 | rs13233399 | 1.491 | 0.583 | -0.745 | -0.971 | 0.014 | 0.039 | 0.039 | 2.529 | 0.37 | 0.35 | 0.21 | 0.11 | 0.35 | 0.24 |
| 7 | 77979388 | rs17347135 | 0.629 | 1.153 | -0.19 | -0.492 | 0.072 | 0.037 | 0.037 | 0 | 0.11 | 0.26 | 0.19 | 0.12 | 0.17 | 0.06 |
| 7 | 77980534 | rs2361378 | 0.842 | 0.482 | -0.198 | -0.539 | -0.001 | -0.007 | -0.007 | 0.076 | 0.61 | 0.54 | 0.44 | 0.19 | 0.64 | 0.49 |
| 7 | 77982085 | rs2191323 | 0.076 | 0.361 | -0.212 | -0.364 | 0.003 | 0.002 | 0.002 | 0.039 | 0.74 | 0.74 | 0.8 | 0.75 | 0.74 | 0.68 |
| 7 | 77982497 | rs714437 | 1.551 | 0.783 | -0.165 | -0.559 | -0.001 | -0.023 | -0.023 | 0 | 0.57 | 0.53 | 0.34 | 0.1 | 0.54 | 0.4 |
| 7 | 77985864 | rs4730234 | 0.318 | 0.764 | 0.283 | -0.452 | 0.034 | 0.092 | 0.092 | 0 | 0.14 | 0.19 | 0.46 | 0.63 | 0.2 | 0.26 |
| 7 | 77987710 | rs6977674 | -0.482 | 0.786 | 0.213 | -0.502 | -0.003 | 0.013 | 0.013 | 0 | 0.79 | 0.81 | 0.89 | 0.84 | 0.8 | 0.79 |
| 7 | 77990305 | rs7802036 | -0.713 | 0.747 | 0.219 | -0.009 | 0.021 | 0.103 | 0.103 | 0 | 0.21 | 0.29 | 0.44 | 0.59 | 0.27 | 0.38 |
| 7 | 77992996 | rs4476926 | 0.542 | 0.602 | 0.094 | 0.164 | -0.002 | 0.009 | 0.009 | 0 | 0.12 | 0.08 | 0.33 | 0.45 | 0.13 | 0.24 |
| 7 | 77993077 | rs12539285 | 1.744 | 0.727 | -0.22 | 0.018 | -0.036 | 0.053 | 0.053 | 0 | 0.09 | 0 | 0.03 | 0 | 0.08 | 0.09 |
| 7 | 77993705 | rs9641383 | -0.196 | 0.695 | -0.091 | 0.118 | -0.002 | 0.01 | 0.01 | 0 | 0.79 | 0.82 | 0.89 | 0.83 | 0.8 | 0.79 |
| 7 | 77994606 | rs2191324 | 0.803 | 0.796 | 0.012 | 0.106 | 0.002 | 0.018 | 0.018 | 0 | 0.63 | 0.52 | 0.51 | 0.37 | 0.64 | 0.53 |
| 7 | 77996712 | rs2215377 | -0.369 | 0.806 | -0.089 | 0.084 | -0.011 | 0.025 | 0.025 | 0 | 0.8 | 0.88 | 0.9 | 0.85 | 0.81 | 0.79 |
| 7 | 77998838 | rs7783086 | 1.322 | 0.742 | -0.028 | 0.05 | 0.002 | 0.012 | 0.012 | 0 | 0.56 | 0.39 | 0.42 | 0.26 | 0.57 | 0.41 |
| 7 | 77999020 | rs12539788 | 1.659 | 0.762 | -0.735 | -0.65 | -0.042 | 0.045 | 0.045 | 0 | 0.19 | 0.06 | 0.16 | 0.07 | 0.28 | 0.14 |
| 7 | 78014174 | rs2012593 | 1.474 | 0.542 | -0.569 | -0.637 | -0.02 | 0.098 | 0.098 | 0 | 0.32 | 0.13 | 0.2 | 0.07 | 0.35 | 0.17 |
| 7 | 78019346 | rs13223430 | 0.851 | 0.461 | -0.204 | -0.421 | 0 | -0.019 | -0.019 | 1.806 | 0.61 | 0.45 | 0.39 | 0.2 | 0.63 | 0.51 |
| 7 | 78019476 | rs1468596 | 0.494 | 0.455 | -0.064 | -0.466 | 0.024 | 0.114 | 0.114 | 0.033 | 0.17 | 0.3 | 0.49 | 0.62 | 0.17 | 0.27 |
| 7 | 78021500 | rs16886004 | 1.777 | 0.006 | -0.749 | -0.699 | -0.001 | -0.009 | -0.009 | 0 | 0.18 | 0.21 | 0.09 | 0.06 | 0.18 | 0.22 |
| 7 | 78022368 | rs881447 | 1.023 | 0.23 | -0.838 | -0.617 | -0.012 | 0.104 | 0.104 | 0 | 0.39 | 0.22 | 0.22 | 0.01 | 0.38 | 0.18 |
| 7 | 78023516 | rs4730257 | -0.565 | 0.777 | -0.079 | 0.309 | 0.001 | -0.028 | -0.028 | 0 | 0.89 | 0.97 | 0.94 | 0.77 | 0.88 | 0.88 |
| 7 | 78024012 | rs9641392 | -0.646 | 0.774 | -0.076 | 0.303 | 0.003 | -0.027 | -0.027 | 0 | 0.89 | 0.97 | 0.94 | 0.77 | 0.89 | 0.88 |
| 7 | 78026633 | rs2191327 | 1.073 | 0.781 | -0.545 | -0.643 | -0.001 | -0.024 | -0.024 | 0 | 0.61 | 0.44 | 0.39 | 0.14 | 0.64 | 0.52 |
| 7 | 78035433 | rs6466202 | 0.61 | 1.786 | -1.198 | -0.107 | -0.049 | -0.071 | -0.071 | 1.344 | 0.73 | 0.87 | 0.63 | 0.36 | 0.69 | 0.61 |
| 7 | 78038652 | rs7804000 | 0.629 | 1.919 | -1.15 | -0.136 | -0.066 | -0.059 | -0.059 | 0 | 0.73 | 0.87 | 0.63 | 0.34 | 0.66 | 0.6 |
| 7 | 78038797 | rs7807814 | 0.462 | 1.431 | -1.202 | -0.329 | -0.071 | -0.023 | -0.023 | 0 | 0.64 | 0.85 | 0.5 | 0.31 | 0.48 | 0.58 |
| 7 | 78040841 | rs6967983 | 1.368 | 0.449 | -0.431 | -1.019 | 0.003 | -0.006 | -0.006 | 0 | 0.63 | 0.6 | 0.78 | 0.91 | 0.69 | 0.67 |
| 7 | 78045248 | rs10953556 | -1.045 | 0.589 | -0.709 | -0.898 | -0.017 | -0.009 | -0.009 | 0 | 0.18 | 0.06 | 0.17 | 0.17 | 0.22 | 0.13 |
| 7 | 78045567 | rs2191330 | 1.963 | 0.578 | -0.584 | -0.939 | 0.003 | 0.059 | 0.059 | 0.08 | 0.54 | 0.65 | 0.65 | 0.74 | 0.54 | 0.61 |
| 7 | 78047073 | rs17417090 | -1.45 | 0.659 | -0.645 | -0.77 | 0.005 | 0.012 | 0.012 | 0 | 0.3 | 0.24 | 0.2 | 0.18 | 0.28 | 0.28 |
| 7 | 78048441 | rs1860537 | 1.153 | 0.678 | -0.513 | -0.937 | -0.001 | -0.009 | -0.009 | 0 | 0.73 | 0.75 | 0.85 | 0.91 | 0.8 | 0.76 |

| | | | | | | | | | | | | | | | | |
|---|---|---|---|---|---|---|---|---|---|---|---|---|---|---|---|---|
| 7 | 78048626 | rs17150696 | -0.599 | 0.75 | -0.512 | -0.945 | -0.014 | -0.062 | -0.062 | 0.021 | 0.2 | 0.23 | 0.08 | 0.01 | 0.14 | 0.14 |
| 7 | 78050158 | rs2191331 | 0.79 | 1.456 | -0.148 | -0.143 | -0.008 | 0.003 | 0.003 | 0.773 | 0.84 | 0.79 | 0.81 | 0.84 | 0.85 | 0.81 |
| 7 | 78050595 | rs2215379 | 1.211 | 1.81 | -0.077 | -0.041 | -0.002 | 0 | 0 | 0 | 0.76 | 0.79 | 0.75 | 0.79 | 0.7 | 0.76 |
| 7 | 78055031 | rs6466213 | -1.063 | 1.967 | 0.449 | -0.015 | 0.066 | -0.075 | -0.075 | 0 | 0.12 | 0.4 | 0.1 | 0 | 0.13 | 0.1 |
| 7 | 78057840 | rs10485886 | -1.509 | 1.893 | 0.468 | 0.041 | 0.077 | -0.059 | -0.059 | 0.207 | 0.19 | 0.41 | 0.22 | 0.08 | 0.26 | 0.17 |
| 7 | 78060486 | rs12154832 | -1.556 | 0.983 | 0.087 | 0.707 | 0.012 | 0.049 | 0.049 | 0.013 | 0.08 | 0.24 | 0.16 | 0.15 | 0.09 | 0.11 |
| 7 | 78066797 | rs6961988 | -0.126 | 0.852 | -0.863 | 0.546 | -0.005 | 0.034 | 0.034 | 0.076 | 0.49 | 0.29 | 0.51 | 0.57 | 0.42 | 0.58 |
| 7 | 78066878 | rs10266582 | -0.83 | 0.69 | -0.108 | 0.651 | 0.003 | 0.01 | 0.01 | 0 | 0.13 | 0.13 | 0.2 | 0.16 | 0.14 | 0.21 |
| 7 | 78068055 | rs10485887 | 0.971 | 0.787 | -1.184 | 0.719 | -0.107 | -0.034 | -0.034 | 0.263 | 0.47 | 0.03 | 0.35 | 0.42 | 0.49 | 0.41 |
| 7 | 78068384 | rs11981822 | -0.484 | 0.618 | -0.051 | 0.683 | -0.019 | 0.049 | 0.049 | 0 | 0.29 | 0.63 | 0.47 | 0.48 | 0.2 | 0.36 |
| 7 | 78070628 | rs2216076 | 0.233 | 0.379 | 0.344 | 0.592 | 0.012 | -0.02 | -0.02 | 0 | 0.18 | 0.15 | 0.25 | 0.39 | 0.14 | 0.32 |
| 7 | 78071641 | rs2885552 | 0.676 | 0.354 | 0.885 | 0.558 | 0.097 | 0.093 | 0.093 | 0.105 | 0.95 | 0.68 | 0.96 | 0.98 | 0.98 | 0.96 |
| 7 | 78072702 | rs2362598 | 0.978 | 0.552 | 1.04 | 0.517 | 0.054 | 0.037 | 0.037 | 0.171 | 0.91 | 0.74 | 0.93 | 0.95 | 0.88 | 0.89 |
| 7 | 78072970 | rs7802215 | 1.666 | 0.661 | 1.214 | 0.485 | 0.078 | 0.02 | 0.02 | 0 | 0.84 | 0.57 | 0.86 | 0.85 | 0.79 | 0.8 |
| 7 | 78072997 | rs13243127 | 2.048 | 0.646 | 1.074 | 0.474 | -0.089 | -0.1 | -0.1 | 0 | 0.55 | 0.04 | 0.5 | 0.45 | 0.56 | 0.45 |
| 7 | 78073633 | rs17430936 | -0.098 | 0.677 | 1.366 | 0.46 | 0.022 | 0.011 | 0.011 | 0 | 0.06 | 0.15 | 0.05 | 0.04 | 0.1 | 0.06 |
| 7 | 78073861 | rs2110868 | 1.64 | 0.68 | 1.405 | 0.458 | 0.067 | -0.005 | -0.005 | 0.451 | 0.84 | 0.63 | 0.89 | 0.89 | 0.79 | 0.8 |
| 7 | 78074957 | rs1157211 | 1.096 | 0.638 | 1.5 | 0.453 | 0.048 | -0.01 | -0.01 | 0 | 0.9 | 0.74 | 0.94 | 0.92 | 0.88 | 0.87 |
| 7 | 78077635 | rs7784064 | -1.59 | 0.636 | 1.201 | 0.502 | 0.002 | 0.014 | 0.014 | 0 | 0.39 | 0.58 | 0.46 | 0.52 | 0.42 | 0.49 |
| 7 | 78079216 | rs17431866 | 0.26 | 0.681 | 0.557 | 0.529 | -0.025 | -0.016 | -0.016 | 0 | 0.05 | 0 | 0.04 | 0.04 | 0.1 | 0.06 |
| 7 | 78079675 | rs6957462 | -1.328 | 0.62 | 0.794 | 0.51 | 0.002 | -0.014 | -0.014 | 0 | 0.33 | 0.37 | 0.41 | 0.47 | 0.33 | 0.4 |
| 7 | 78080548 | rs2110871 | 1.929 | 0.329 | 1.012 | 0.38 | 0.033 | -0.091 | -0.091 | 0.161 | 0.56 | 0.08 | 0.53 | 0.5 | 0.67 | 0.5 |
| 7 | 78080669 | rs2885553 | 0.928 | 0.325 | 1.161 | 0.375 | 0.019 | -0.1 | -0.1 | 1.496 | 0.83 | 0.48 | 0.89 | 0.94 | 0.84 | 0.87 |
| 7 | 78080820 | rs10485891 | -1.034 | 0.438 | 0.82 | 0.571 | 0.002 | -0.02 | -0.02 | 0 | 0.28 | 0.22 | 0.36 | 0.41 | 0.25 | 0.38 |
| 7 | 78081149 | rs10485892 | 0.558 | 0.068 | 0.742 | 0.7 | 0.02 | -0.023 | -0.023 | 0 | 0.13 | 0.09 | 0.13 | 0.33 | 0.08 | 0.25 |
| 7 | 78081804 | rs4623346 | 1.18 | 0.335 | 1.323 | 0.836 | 0.008 | 0.149 | 0.149 | 0.191 | 0.86 | 0.4 | 0.88 | 0.67 | 0.9 | 0.74 |
| 7 | 78082607 | rs2160321 | 1.133 | 0.219 | 0.675 | 0.361 | -0.004 | -0.001 | -0.001 | 0 | 0.7 | 0.36 | 0.78 | 0.63 | 0.75 | 0.67 |
| 7 | 78083514 | rs7810530 | -1.855 | 0.348 | 0.176 | 0.381 | -0.005 | 0.003 | 0.003 | 1.053 | 0.39 | 0.76 | 0.28 | 0.39 | 0.34 | 0.41 |
| 7 | 78085219 | rs2302640 | -1.312 | 0.168 | 0.067 | 0.569 | -0.005 | 0.004 | 0.004 | 1.992 | 0.28 | 0.74 | 0.21 | 0.39 | 0.21 | 0.34 |
| 7 | 78086846 | rs10261909 | -1.04 | 0.298 | -0.48 | 0.795 | 0.001 | -0.002 | -0.002 | 0.026 | 0.36 | 0.42 | 0.44 | 0.51 | 0.32 | 0.51 |
| 7 | 78089202 | rs17150825 | -1.191 | 0.2 | -0.755 | 0.703 | 0.014 | 0.017 | 0.017 | 0 | 0.34 | 0.32 | 0.26 | 0.18 | 0.26 | 0.23 |
| 7 | 78090357 | rs2110872 | -0.657 | 0.258 | -0.743 | 1.047 | 0.048 | 0.089 | 0.089 | 0 | 0.12 | 0.26 | 0.32 | 0.29 | 0.17 | 0.29 |
| 7 | 78092244 | rs17150832 | -0.512 | -0.126 | -1.86 | 0.933 | 0.05 | 0.006 | 0.006 | 0 | 0.26 | 0.1 | 0.15 | 0.15 | 0.17 | 0.17 |
| 7 | 78092652 | rs1558699 | 1.788 | 0.442 | -1.855 | 1.256 | -0.096 | -0.018 | -0.018 | 0.615 | 0.55 | 0.07 | 0.42 | 0.45 | 0.54 | 0.43 |
| 7 | 78092725 | rs9649305 | -0.552 | -0.35 | -1.027 | 1.165 | 0.024 | -0.013 | -0.013 | 0.016 | 0.22 | 0.19 | 0.17 | 0.2 | 0.14 | 0.18 |
| 7 | 78093859 | rs6466236 | -0.82 | 0.051 | -0.871 | 1.08 | -0.001 | 0 | 0 | 0 | 0.08 | 0.11 | 0.03 | 0.12 | 0.07 | 0.06 |
| 7 | 78094640 | rs7803705 | 0.012 | 0.279 | -1.802 | 0.624 | -0.041 | 0.042 | 0.042 | 0 | 0.28 | 0.14 | 0.33 | 0.19 | 0.3 | 0.3 |
| 7 | 78095593 | rs7789215 | 0.918 | -0.25 | -1.78 | 1.042 | 0.008 | -0.027 | -0.027 | 0 | 0.52 | 0.16 | 0.37 | 0.51 | 0.58 | 0.53 |
| 7 | 78095872 | rs7803276 | 1.097 | -0.019 | -1.493 | 1.077 | -0.029 | 0.191 | 0.191 | 0 | 0.46 | 0.14 | 0.29 | 0.16 | 0.47 | 0.43 |
| 7 | 78096656 | rs2192883 | 0.686 | -0.123 | -1.586 | 0.67 | -0.011 | 0.059 | 0.059 | 0 | 0.32 | 0.16 | 0.23 | 0.18 | 0.34 | 0.24 |
| 7 | 78098152 | rs11972334 | 1.309 | -0.079 | -1.214 | 0.776 | 0.002 | -0.001 | -0.001 | 0.614 | 0.17 | 0.11 | 0.32 | 0.19 | 0.16 | 0.23 |
| 7 | 78103473 | rs10485896 | 1.08 | -0.067 | -0.143 | 0.878 | -0.022 | -0.088 | -0.088 | 0 | 0.21 | 0.42 | 0.12 | 0.07 | 0.15 | 0.24 |
| 7 | 78104278 | rs7808641 | 0.211 | -0.087 | -0.052 | 0.996 | -0.013 | -0.009 | -0.009 | 0.2 | 0.26 | 0.54 | 0.17 | 0.24 | 0.23 | 0.33 |
| 7 | 78104517 | rs7807844 | 1.116 | -0.046 | -0.109 | 0.92 | -0.021 | -0.031 | -0.031 | 0 | 0.21 | 0.42 | 0.13 | 0.14 | 0.15 | 0.26 |
| 7 | 78105184 | rs10485898 | 0.296 | -0.022 | -0.593 | 0.744 | -0.008 | 0.024 | 0.024 | 0 | 0.27 | 0.63 | 0.18 | 0.57 | 0.23 | 0.39 |
| 7 | 78108303 | rs6951193 | 0.141 | 0.065 | -1 | 0.487 | -0.006 | -0.048 | -0.048 | 0.018 | 0.32 | 0.2 | 0.43 | 0.52 | 0.31 | 0.4 |
| 7 | 78108724 | rs7789155 | -0.855 | 0.217 | -1.684 | -0.061 | -0.127 | 0.041 | 0.041 | 0.042 | 0.81 | 0.98 | 0.84 | 0.9 | 0.59 | 0.85 |
| 7 | 78109304 | rs12667712 | 0.495 | 0.008 | -0.542 | -0.011 | -0.007 | 0.004 | 0.004 | 0 | 0.35 | 0.43 | 0.46 | 0.6 | 0.31 | 0.41 |
| 7 | 78111984 | rs7784896 | 0.108 | -0.201 | -0.5 | 0.008 | -0.009 | -0.014 | -0.014 | 0.018 | 0.39 | 0.47 | 0.61 | 0.66 | 0.32 | 0.47 |
| 7 | 78112731 | rs2215934 | -0.221 | -0.558 | -0.62 | -0.128 | 0.047 | 0.048 | 0.048 | 0.792 | 0.41 | 0.68 | 0.45 | 0.66 | 0.26 | 0.34 |
| 7 | 78115241 | rs13227908 | 0.905 | -0.415 | -0.551 | -0.245 | 0.017 | 0.07 | 0.07 | 0 | 0.41 | 0.36 | 0.38 | 0.23 | 0.34 | 0.5 |
| 7 | 78116572 | rs7780361 | 1.983 | -0.568 | -0.708 | -0.336 | -0.002 | 0.023 | 0.023 | 0 | 0.15 | 0.03 | 0.23 | 0.13 | 0.14 | 0.35 |
| 7 | 78117090 | rs6974799 | 0.88 | -0.63 | -0.649 | -0.294 | 0 | -0.009 | -0.009 | 2.167 | 0.19 | 0.19 | 0.33 | 0.46 | 0.16 | 0.2 |
| 7 | 78117548 | rs17435216 | 1.805 | -0.696 | -0.751 | -0.373 | -0.035 | 0.056 | 0.056 | 0.571 | 0.23 | 0.01 | 0.12 | 0.09 | 0.19 | 0.14 |
| 7 | 78118216 | rs759333 | -0.798 | -0.688 | -0.423 | -0.394 | 0.053 | -0.02 | -0.02 | 0.258 | 0.78 | 0.75 | 0.8 | 0.82 | 0.59 | 0.83 |
| 7 | 78118506 | rs759334 | -0.271 | -0.741 | -0.409 | -0.371 | 0.003 | -0.005 | -0.005 | 0 | 0.63 | 0.71 | 0.56 | 0.68 | 0.46 | 0.48 |
| 7 | 78118846 | rs1548867 | 1.421 | -0.8 | -0.307 | -0.402 | -0.02 | -0.014 | -0.014 | 0 | 0.18 | 0.2 | 0.09 | 0.14 | 0.09 | 0.14 |
| 7 | 78118982 | rs1548868 | 0.428 | -0.755 | -0.509 | -0.385 | -0.002 | -0.007 | -0.007 | 0.133 | 0.44 | 0.49 | 0.23 | 0.23 | 0.3 | 0.29 |
| 7 | 78119109 | rs3735442 | 0.433 | -0.761 | -0.512 | -0.387 | -0.002 | -0.008 | -0.008 | 0.036 | 0.44 | 0.49 | 0.23 | 0.23 | 0.3 | 0.29 |
| 7 | 78119931 | rs7782195 | -1.118 | -0.065 | 0.004 | -0.354 | 0 | 0.013 | 0.013 | 0.427 | 0.64 | 0.72 | 0.77 | 0.82 | 0.63 | 0.78 |
| 7 | 78121458 | rs7805441 | -0.837 | 0.203 | -0.459 | -0.565 | -0.076 | -0.052 | -0.052 | 0 | 0.61 | 0.89 | 0.45 | 0.5 | 0.53 | 0.72 |
| 7 | 78122206 | rs7809117 | 0.681 | -0.178 | -0.373 | -0.978 | -0.051 | -0.048 | -0.048 | 0 | 0.21 | 0.55 | 0.13 | 0.13 | 0.1 | 0.16 |
| 7 | 78122616 | rs10268479 | 0.155 | -0.076 | -0.614 | -0.995 | -0.022 | 0.056 | 0.056 | 0.554 | 0.37 | 0.24 | 0.31 | 0.28 | 0.4 | 0.49 |
| 7 | 78125648 | rs7789112 | -0.429 | 0.142 | -0.942 | -0.803 | -0.027 | 0.046 | 0.046 | 1.073 | 0.55 | 0.89 | 0.4 | 0.41 | 0.4 | 0.64 |
| 7 | 78136192 | rs2110630 | -1.385 | 0.068 | -0.881 | -0.871 | -0.057 | -0.063 | -0.063 | 0 | 0.68 | 0.91 | 0.51 | 0.47 | 0.64 | 0.75 |
| 7 | 78138542 | rs4730345 | -0.788 | 0.096 | -0.892 | -0.876 | -0.074 | -0.043 | -0.043 | 0.041 | 0.62 | 0.9 | 0.49 | 0.48 | 0.58 | 0.71 |
| 7 | 78138977 | rs1030012 | 0.863 | 0.051 | -0.863 | -1.037 | 0.006 | 0.017 | 0.017 | 0.106 | 0.22 | 0.14 | 0.22 | 0.25 | 0.21 | 0.42 |
| 7 | 78139581 | rs1030015 | -0.693 | -0.146 | -1.244 | -1.347 | -0.023 | 0.172 | 0.172 | 0.404 | 0.48 | 0.85 | 0.43 | 0.34 | 0.53 | 0.59 |
| 7 | 78141660 | rs6947977 | 0.145 | 0.039 | -0.991 | -1.335 | -0.005 | 0.021 | 0.021 | 0.012 | 0.34 | 0.29 | 0.3 | 0.26 | 0.36 | 0.48 |

| | | | | | | | | | | | | | | | | |
|---|---|---|---|---|---|---|---|---|---|---|---|---|---|---|---|---|
| 7 | 78145656 | rs2362630 | 0.303 | -0.078 | -1.207 | -1.172 | 0.01 | -0.023 | -0.023 | 0 | 0.26 | 0.3 | 0.27 | 0.26 | 0.33 | 0.48 |
| 7 | 78147225 | rs3801362 | 1.38 | -0.05 | -1.558 | -0.981 | 0.046 | -0.003 | -0.003 | 0 | 0.13 | 0.04 | 0.06 | 0.12 | 0.06 | 0.12 |
| 7 | 78148980 | rs9691998 | -0.094 | 0.004 | -1.436 | -0.993 | -0.007 | -0.003 | -0.003 | 0 | 0.22 | 0.2 | 0.14 | 0.23 | 0.19 | 0.24 |
| 7 | 78154019 | rs7806320 | -1.064 | 0.215 | -2.331 | -0.96 | -0.098 | -0.048 | -0.048 | 0 | 0.64 | 0.87 | 0.45 | 0.44 | 0.52 | 0.67 |
| 7 | 78158472 | rs12666870 | -0.066 | 0.616 | -2.164 | -0.755 | 0.049 | 0.01 | 0.01 | 0 | 0.35 | 0.15 | 0.27 | 0.36 | 0.28 | 0.56 |
| 7 | 78158609 | rs10271370 | 0.023 | 1.319 | -1.548 | -0.991 | 0.041 | -0.095 | -0.095 | 0 | 0.2 | 0.44 | 0.15 | 0.06 | 0.21 | 0.13 |
| 7 | 78163301 | rs2885563 | -0.108 | 1.152 | -2.264 | -0.283 | -0.009 | 0.023 | 0.023 | 0.015 | 0.22 | 0.09 | 0.21 | 0.22 | 0.23 | 0.43 |
| 7 | 78170588 | rs727200 | 0.012 | 1.906 | -2.286 | 0.05 | -0.073 | 0.007 | 0.007 | 0 | 0.82 | 0.95 | 0.77 | 0.87 | 0.73 | 0.89 |
| 7 | 78173595 | rs17437602 | -0.411 | 1.469 | -2.141 | 0.087 | -0.081 | 0.024 | 0.024 | 0 | 0.23 | 0.04 | 0.28 | 0.19 | 0.32 | 0.26 |
| 7 | 78175596 | rs1978326 | 0.239 | 0.965 | -1.291 | 0.486 | 0.003 | 0.037 | 0.037 | 0.378 | 0.94 | 0.84 | 0.94 | 0.94 | 0.95 | 0.98 |
| 7 | 78175862 | rs2192651 | 0.446 | 0.557 | -1.996 | 0.509 | -0.007 | 0.006 | 0.006 | 2.747 | 0.51 | 0.81 | 0.49 | 0.56 | 0.46 | 0.56 |
| 7 | 78176756 | rs12705567 | -0.444 | -0.209 | -1.559 | 0.469 | -0.022 | -0.018 | -0.018 | 0 | 0.59 | 0.69 | 0.61 | 0.54 | 0.53 | 0.65 |
| 7 | 78176966 | rs763452 | -0.84 | -0.409 | -0.672 | 0.337 | -0.001 | -0.001 | -0.001 | 0.187 | 0.69 | 0.63 | 0.63 | 0.59 | 0.58 | 0.76 |
| 7 | 78176997 | rs763451 | 0.033 | -1.006 | -1.19 | -0.671 | -0.05 | 0.087 | 0.087 | 1.075 | 0.21 | 0.06 | 0.12 | 0.03 | 0.3 | 0.17 |
| 7 | 78182068 | rs10262180 | -0.116 | -1.674 | -0.016 | -0.377 | -0.004 | -0.034 | -0.034 | 0 | 0.81 | 0.69 | 0.81 | 0.85 | 0.78 | 0.65 |
| 7 | 78182572 | rs4730378 | -1.223 | -1.911 | 0.122 | -0.463 | 0.005 | 0.004 | 0.004 | 0 | 0.44 | 0.57 | 0.49 | 0.62 | 0.29 | 0.25 |
| 7 | 78184222 | rs12705579 | 0.503 | -1.853 | 0.363 | -0.363 | -0.005 | -0.025 | -0.025 | 0 | 0.12 | 0.1 | 0.04 | 0.01 | 0.06 | 0.07 |
| 7 | 78185888 | rs2192656 | 0.482 | -1.949 | 0.802 | -0.327 | 0.008 | 0.01 | 0.01 | 0 | 0.72 | 0.6 | 0.56 | 0.35 | 0.67 | 0.67 |
| 7 | 78185955 | rs2192658 | 0.206 | -1.581 | 0.237 | -0.036 | 0.031 | 0.259 | 0.259 | 0.489 | 0.08 | 0.22 | 0.29 | 0.52 | 0.08 | 0.04 |
| 7 | 78190213 | rs13242509 | 0.25 | -1.791 | -0.337 | -0.261 | -0.006 | -0.036 | -0.036 | 0 | 0.12 | 0.13 | 0.06 | 0.01 | 0.11 | 0.08 |
| 7 | 78190606 | rs12705589 | 0.894 | -1.839 | -0.375 | -0.258 | -0.003 | 0.106 | 0.106 | 3.287 | 0.46 | 0.23 | 0.33 | 0.23 | 0.52 | 0.46 |
| 7 | 78191370 | rs12705591 | 0.294 | -1.794 | -0.43 | -0.235 | -0.007 | -0.036 | -0.036 | 0 | 0.12 | 0.14 | 0.05 | 0.01 | 0.09 | 0.07 |
| 7 | 78192343 | rs17438461 | 1.482 | -1.414 | -1.021 | 0.062 | -0.105 | 0.047 | 0.047 | 0 | 0.34 | 0.09 | 0.29 | 0.23 | 0.44 | 0.35 |
| 7 | 78213223 | rs10269564 | -0.455 | -1.138 | -0.203 | 0.692 | -0.028 | 0.263 | 0.263 | 0 | 0.08 | 0.19 | 0.31 | 0.53 | 0.03 | 0.27 |
| 7 | 78214830 | rs2362640 | 0.935 | -1.006 | -0.224 | 0.149 | -0.007 | 0.046 | 0.046 | 0 | 0.55 | 0.35 | 0.52 | 0.77 | 0.44 | 0.69 |
| 7 | 78215346 | rs11979133 | -0.44 | -1.517 | -0.192 | -0.085 | -0.001 | 0.001 | 0.001 | 0.059 | 0.85 | 0.82 | 0.74 | 0.9 | 0.77 | 0.84 |
| 7 | 78216837 | rs319864 | -0.048 | -1.35 | -0.46 | -0.079 | 0.005 | -0.005 | -0.005 | 0 | 0.1 | 0.12 | 0.27 | 0.09 | 0.2 | 0.17 |
| 7 | 78217826 | rs17439102 | 2.172 | -1.62 | -0.738 | -0.136 | -0.108 | 0.048 | 0.048 | 0.272 | 0.35 | 0.08 | 0.33 | 0.29 | 0.43 | 0.35 |
| 7 | 78218753 | rs6955962 | 0.41 | -0.709 | 0.13 | 0.266 | -0.007 | -0.048 | -0.048 | 0 | 0.09 | 0.37 | 0.05 | 0 | 0.06 | 0.05 |
| 7 | 78219309 | rs17439220 | 0.836 | -0.765 | -0.942 | 0.478 | -0.065 | -0.043 | -0.043 | 0 | 0.1 | 0.01 | 0.11 | 0.13 | 0.2 | 0.04 |
| 7 | 78219577 | rs619747 | 1.287 | -0.919 | -0.864 | 0.406 | 0.003 | 0.006 | 0.006 | 0 | 0.6 | 0.35 | 0.62 | 0.38 | 0.66 | 0.64 |
| 7 | 78223217 | rs7777792 | 0.731 | -0.74 | -1.27 | 0.742 | 0.01 | -0.026 | -0.026 | 0.015 | 0.74 | 0.75 | 0.54 | 0.5 | 0.74 | 0.76 |
| 7 | 78225633 | rs319866 | -0.517 | -0.384 | -1.829 | 0.571 | 0.043 | -0.072 | -0.072 | 3.592 | 0.84 | 0.94 | 0.8 | 0.57 | 0.93 | 0.93 |
| 7 | 78228963 | rs38093 | -0.361 | 0.281 | -1.574 | 0.246 | -0.036 | -0.016 | -0.016 | 0 | 0.93 | 0.99 | 0.77 | 0.9 | 0.82 | 0.84 |
| 7 | 78229176 | rs319870 | 0.061 | 0.335 | -1.406 | 0.114 | -0.083 | -0.031 | -0.031 | 0.491 | 0.13 | 0.01 | 0.13 | 0.13 | 0.22 | 0.07 |
| 7 | 78230243 | rs319872 | -0.37 | 0.358 | -1.115 | 0.229 | 0.012 | -0.038 | -0.038 | 0 | 0.28 | 0.62 | 0.23 | 0.13 | 0.31 | 0.15 |
| 7 | 78232484 | rs22536 | 0.574 | -0.079 | -1.281 | 0.968 | 0.025 | -0.045 | -0.045 | 0 | 0.73 | 0.85 | 0.57 | 0.52 | 0.76 | 0.76 |
| 7 | 78233434 | rs6466310 | -1.04 | 0.704 | -1.306 | 0.391 | -0.009 | 0.017 | 0.017 | 1.411 | 0.23 | 0.18 | 0.22 | 0.08 | 0.22 | 0.12 |
| 7 | 78233755 | rs9640728 | -1.39 | 0.221 | -0.703 | 1.197 | -0.014 | 0.039 | 0.039 | 0 | 0.24 | 0.29 | 0.32 | 0.49 | 0.19 | 0.24 |
| 7 | 78234264 | rs38095 | 0.801 | 0.025 | -0.157 | 0.89 | -0.018 | 0.016 | 0.016 | 0 | 0.73 | 0.5 | 0.63 | 0.28 | 0.74 | 0.74 |
| 7 | 78236116 | rs38096 | 1.203 | 1.089 | 0.403 | 0.741 | 0.019 | 0.01 | 0.01 | 0.323 | 0.7 | 0.53 | 0.63 | 0.31 | 0.68 | 0.7 |
| 7 | 78238766 | rs38098 | 0.872 | 1.22 | 0.869 | 1.036 | -0.005 | 0.054 | 0.054 | 0.313 | 0.77 | 0.71 | 0.72 | 0.4 | 0.76 | 0.72 |
| 7 | 78241011 | rs38100 | 1.64 | 1.221 | 0.945 | 1.008 | 0.021 | -0.001 | -0.001 | 0 | 0.71 | 0.61 | 0.62 | 0.3 | 0.66 | 0.7 |
| 7 | 78242398 | rs38101 | 0.884 | 1.125 | 1.057 | 1.54 | -0.021 | -0.019 | -0.019 | 0.797 | 0.87 | 0.94 | 0.86 | 0.55 | 0.81 | 0.81 |
| 7 | 78247172 | rs38109 | 1.747 | 1.037 | 1.178 | 1.511 | 0.003 | 0.04 | 0.04 | 0.596 | 0.76 | 0.7 | 0.72 | 0.38 | 0.72 | 0.72 |
| 7 | 78247295 | rs38110 | 2.455 | 0.673 | 1.095 | 0.892 | 0 | 0.052 | 0.052 | 0.91 | 0.54 | 0.34 | 0.52 | 0.25 | 0.55 | 0.63 |
| 7 | 78249193 | rs38111 | 1.693 | 0.672 | 1.09 | 0.871 | 0.014 | -0.002 | -0.002 | 0 | 0.72 | 0.62 | 0.62 | 0.3 | 0.67 | 0.71 |
| 7 | 78256342 | rs38116 | 2.435 | 0.67 | 0.985 | 0.888 | 0 | 0.06 | 0.06 | 1.368 | 0.54 | 0.33 | 0.51 | 0.23 | 0.55 | 0.63 |
| 7 | 78256643 | rs38117 | 1.945 | 0.807 | 0.991 | 1.135 | 0.018 | -0.001 | -0.001 | 0 | 0.71 | 0.53 | 0.62 | 0.3 | 0.67 | 0.71 |
| 7 | 78257163 | rs38118 | 0.411 | 0.761 | 0.806 | 1.233 | -0.014 | -0.007 | -0.007 | 0 | 0.88 | 0.93 | 0.85 | 0.55 | 0.82 | 0.81 |
| 7 | 78259702 | rs38121 | -0.589 | 0.014 | 0.068 | 0.074 | -0.018 | -0.07 | -0.07 | 0 | 0.59 | 0.51 | 0.64 | 0.87 | 0.66 | 0.62 |
| 7 | 78264474 | rs34692025 | -0.045 | 0.103 | 0.027 | 0.408 | 0.027 | -0.036 | -0.036 | 0 | 0.13 | 0.07 | 0.15 | 0.18 | 0.11 | 0.1 |
| 7 | 78269375 | rs38127 | -0.486 | 0.113 | 0.05 | -0.086 | 0.02 | 0.019 | 0.019 | 2.586 | 0.54 | 0.61 | 0.65 | 0.93 | 0.65 | 0.63 |
| 7 | 78277679 | rs12705674 | 2.048 | 0.052 | -0.081 | 0.293 | -0.061 | 0.007 | 0.007 | 0 | 0.24 | 0.07 | 0.27 | 0.17 | 0.3 | 0.26 |
| 7 | 78279749 | rs322006 | -0.183 | -0.057 | 0.202 | 0.212 | 0.004 | 0.028 | 0.028 | 1.298 | 0.47 | 0.58 | 0.53 | 0.76 | 0.55 | 0.59 |
| 7 | 78285960 | rs4730427 | 0.11 | 0.34 | 0.018 | 0.576 | -0.038 | -0.029 | -0.029 | 0 | 0.07 | 0.01 | 0.19 | 0.24 | 0.13 | 0.06 |
| 7 | 78287193 | rs12673114 | 1.713 | 0.418 | 0.32 | 0.593 | -0.021 | -0.005 | -0.005 | 0 | 0.26 | 0.17 | 0.3 | 0.25 | 0.33 | 0.31 |
| 7 | 78289970 | rs6466333 | 0.949 | 0.398 | 0.636 | 0.587 | 0.042 | 0.017 | 0.017 | 0 | 0.3 | 0.35 | 0.38 | 0.32 | 0.41 | 0.32 |
| 7 | 78290844 | rs1496783 | 1.668 | 0.329 | 0.096 | 0.566 | -0.018 | -0.004 | -0.004 | 1.717 | 0.26 | 0.18 | 0.3 | 0.25 | 0.33 | 0.3 |
| 7 | 78294834 | rs321991 | -0.169 | 0.179 | 0.332 | 1.036 | 0.001 | 0.021 | 0.021 | 0.04 | 0.1 | 0.07 | 0.12 | 0.39 | 0.11 | 0.24 |
| 7 | 78295714 | rs321989 | -0.008 | 0.202 | 0.343 | 1.046 | 0 | 0.032 | 0.032 | 0.01 | 0.09 | 0.07 | 0.12 | 0.39 | 0.11 | 0.24 |
| 7 | 78302547 | rs321983 | -1.921 | -0.063 | 0.735 | 0.63 | 0.002 | 0.001 | 0.001 | 0.317 | 0.39 | 0.58 | 0.43 | 0.66 | 0.37 | 0.42 |
| 7 | 78303261 | rs321982 | -1.488 | -0.579 | 1.193 | 0.474 | -0.02 | 0.094 | 0.094 | 0.781 | 0.21 | 0.3 | 0.27 | 0.54 | 0.19 | 0.37 |
| 7 | 78303699 | rs9886142 | 0.505 | -0.566 | 1.37 | 0.461 | -0.01 | -0.06 | -0.06 | 0 | 0.16 | 0.19 | 0.08 | 0.01 | 0.11 | 0.03 |
| 7 | 78303886 | rs321980 | -1.083 | -0.322 | 1.3 | 0.904 | -0.01 | 0.061 | 0.061 | 0.145 | 0.15 | 0.25 | 0.14 | 0.4 | 0.14 | 0.31 |
| 7 | 78303931 | rs321979 | -1.473 | -0.33 | 1.351 | 0.91 | -0.006 | 0.095 | 0.095 | 0.352 | 0.15 | 0.15 | 0.25 | 0.53 | 0.15 | 0.32 |
| 7 | 78304299 | rs321978 | -0.811 | -0.282 | 0.965 | 0.942 | -0.012 | 0.078 | 0.078 | 0.174 | 0.15 | 0.27 | 0.16 | 0.41 | 0.14 | 0.31 |
| 7 | 78304878 | rs17161332 | 0.358 | -0.087 | 0.464 | 0.598 | 0 | -0.033 | -0.033 | 0 | 0.06 | 0 | 0.13 | 0.14 | 0.05 | 0.06 |
| 7 | 78313673 | rs321969 | -1.04 | -0.299 | 0.487 | -0.193 | -0.001 | -0.064 | -0.064 | 0 | 0.73 | 0.68 | 0.74 | 0.96 | 0.75 | 0.72 |

| | | | | | | | | | | | | | | | | |
|---|---|---|---|---|---|---|---|---|---|---|---|---|---|---|---|---|
| 7 | 78315591 | rs321964 | -0.963 | -0.273 | 0.535 | -0.234 | 0 | -0.079 | -0.079 | 0.21 | 0.73 | 0.67 | 0.75 | 0.97 | 0.74 | 0.72 |
| 7 | 78315924 | rs2159017 | 0.311 | -0.534 | 0.19 | -0.144 | 0.025 | 0.04 | 0.04 | 1.025 | 0.25 | 0.16 | 0.11 | 0.11 | 0.17 | 0.09 |
| 7 | 78319728 | rs7791589 | 0.527 | -0.613 | 0.907 | 0.668 | -0.003 | -0.003 | -0.003 | 0 | 0.53 | 0.52 | 0.43 | 0.38 | 0.56 | 0.37 |
| 7 | 78320848 | rs9640732 | 1.195 | -0.607 | 0.752 | 0.581 | -0.06 | 0.064 | 0.064 | 0.028 | 0.41 | 0.15 | 0.36 | 0.21 | 0.47 | 0.27 |
| 7 | 78323029 | rs321999 | -0.239 | -0.69 | 0.913 | 0.68 | 0.017 | 0.096 | 0.096 | 0 | 0.63 | 0.78 | 0.72 | 0.87 | 0.67 | 0.64 |
| 7 | 78324890 | rs322000 | -0.432 | -0.465 | 1.054 | 1.138 | 0.001 | 0.121 | 0.121 | 0 | 0.09 | 0.08 | 0.24 | 0.45 | 0.1 | 0.27 |
| 7 | 78327299 | rs10485911 | 0.667 | -0.466 | 1.173 | 0.939 | -0.001 | -0.019 | -0.019 | 0 | 0.05 | 0.08 | 0.14 | 0 | 0.05 | 0.04 |
| 7 | 78330515 | rs10953668 | -0.686 | -0.294 | 1.698 | 1.005 | 0.024 | 0.036 | 0.036 | 0 | 0.17 | 0.44 | 0.24 | 0.21 | 0.2 | 0.15 |
| 7 | 78331066 | rs13246026 | 1.422 | -0.317 | 1.73 | 0.859 | -0.008 | 0.035 | 0.035 | 0.258 | 0.41 | 0.27 | 0.33 | 0.21 | 0.41 | 0.25 |
| 7 | 78332900 | rs322005 | -0.211 | -0.304 | 1.674 | 1.131 | 0.011 | 0.088 | 0.088 | 0.142 | 0.62 | 0.79 | 0.71 | 0.8 | 0.66 | 0.61 |
| 7 | 78335301 | rs16886265 | 0.477 | -0.243 | 1.783 | 1.219 | 0.028 | 0.013 | 0.013 | 0 | 0.05 | 0.18 | 0.14 | 0.04 | 0.05 | 0.09 |
| 7 | 78336093 | rs323166 | -0.802 | -0.239 | 1.769 | 1.236 | -0.01 | 0.187 | 0.187 | 0 | 0.18 | 0.42 | 0.36 | 0.59 | 0.19 | 0.35 |
| 7 | 78341360 | rs323135 | -0.563 | -0.203 | 1.761 | 1.514 | -0.001 | 0.128 | 0.128 | 0 | 0.17 | 0.43 | 0.31 | 0.36 | 0.18 | 0.23 |
| 7 | 78344269 | rs10485912 | 1.418 | -0.36 | 1.339 | 1.349 | -0.03 | -0.029 | -0.029 | 0.874 | 0.22 | 0.08 | 0.25 | 0.27 | 0.24 | 0.39 |
| 7 | 78344389 | rs7793494 | 1.134 | -0.297 | 1.55 | 1.298 | -0.037 | 0.055 | 0.055 | 0 | 0.76 | 0.55 | 0.67 | 0.63 | 0.78 | 0.72 |
| 7 | 78344461 | rs12665877 | 0.398 | -0.354 | 1.491 | 1.291 | -0.005 | -0.031 | -0.031 | 0 | 0.33 | 0.19 | 0.3 | 0.41 | 0.3 | 0.45 |
| 7 | 78346008 | rs10255125 | 1.683 | -0.404 | 1.436 | 1.289 | -0.019 | 0.025 | 0.025 | 0 | 0.39 | 0.25 | 0.34 | 0.22 | 0.42 | 0.26 |
| 7 | 78346314 | rs17151352 | -0.037 | -0.552 | 1.408 | 1.208 | -0.018 | 0.019 | 0.019 | 0 | 0.14 | 0.25 | 0.18 | 0.21 | 0.13 | 0.15 |
| 7 | 78346879 | rs7779874 | 0.262 | -0.522 | 1.412 | 1.222 | -0.005 | 0.012 | 0.012 | 0 | 0.91 | 0.9 | 0.86 | 0.84 | 0.91 | 0.87 |
| 7 | 78348879 | rs17151376 | -0.118 | -0.631 | 1.575 | 1.059 | -0.017 | 0.012 | 0.012 | 0 | 0.14 | 0.22 | 0.19 | 0.2 | 0.13 | 0.15 |
| 7 | 78352199 | rs17151383 | 1.443 | -0.949 | 1.092 | 1.123 | 0.004 | -0.011 | -0.011 | 0 | 0.08 | 0.05 | 0.05 | 0.18 | 0.03 | 0.06 |
| 7 | 78358514 | rs4370470 | 0.723 | -0.82 | 1.085 | 1.017 | 0.001 | -0.012 | -0.012 | 0 | 0.1 | 0.08 | 0.19 | 0.24 | 0.09 | 0.11 |
| 7 | 78359355 | rs10242163 | 1.301 | -0.709 | 1.269 | 0.931 | -0.023 | 0.026 | 0.026 | 0 | 0.39 | 0.26 | 0.36 | 0.29 | 0.45 | 0.24 |
| 7 | 78366575 | rs10808167 | 0.863 | -0.939 | 1.627 | 1.083 | -0.002 | -0.006 | -0.006 | 0.173 | 0.45 | 0.46 | 0.4 | 0.29 | 0.54 | 0.27 |
| 7 | 78377750 | rs1207881 | 0.679 | -1.05 | 1.814 | 0.685 | 0.005 | 0.016 | 0.016 | 0.094 | 0.31 | 0.32 | 0.25 | 0.41 | 0.3 | 0.52 |
| 7 | 78380016 | rs4730487 | 0.943 | -1.084 | 1.732 | 0.688 | -0.04 | -0.002 | -0.002 | 0.518 | 0.75 | 0.64 | 0.64 | 0.75 | 0.86 | 0.8 |
| 7 | 78380442 | rs1207870 | 0.749 | -1.037 | 1.396 | 0.699 | -0.016 | -0.037 | -0.037 | 0.014 | 0.3 | 0.18 | 0.24 | 0.4 | 0.3 | 0.52 |
| 7 | 78390300 | rs1888238 | 1.075 | -0.934 | 1.201 | 0.743 | -0.012 | 0.024 | 0.024 | 0 | 0.44 | 0.34 | 0.38 | 0.34 | 0.5 | 0.26 |
| 7 | 78396047 | rs16886321 | 0.902 | -0.972 | 1.264 | 0.181 | -0.02 | 0.029 | 0.029 | 0.07 | 0.1 | 0.15 | 0.19 | 0.24 | 0.06 | 0.1 |
| 7 | 78401699 | rs1207868 | 0.453 | -1.029 | 1.215 | 0.179 | 0.003 | -0.003 | -0.003 | 0 | 0.33 | 0.35 | 0.26 | 0.33 | 0.32 | 0.52 |
| 7 | 78403188 | rs12705771 | 0.405 | -1.208 | 0.838 | -0.183 | -0.027 | -0.056 | -0.056 | 0.462 | 0.87 | 0.77 | 0.83 | 1 | 0.91 | 0.91 |
| 7 | 78407722 | rs10242583 | 0.559 | -0.841 | 0.672 | 0.087 | -0.039 | 0.078 | 0.078 | 0.372 | 0.1 | 0.28 | 0.2 | 0.24 | 0.07 | 0.09 |
| 7 | 78416056 | rs37852 | 0.117 | -0.063 | 0.213 | -0.177 | -0.011 | 0.032 | 0.032 | 0 | 0.65 | 0.76 | 0.74 | 0.73 | 0.67 | 0.48 |
| 7 | 78416820 | rs37854 | 0.111 | -0.538 | -0.095 | -0.88 | 0.02 | 0.034 | 0.034 | 0 | 0.79 | 0.91 | 0.91 | 0.98 | 0.86 | 0.87 |
| 7 | 78420458 | rs37859 | 0.744 | -0.453 | 0.267 | -0.379 | 0.01 | -0.05 | -0.05 | 0 | 0.36 | 0.2 | 0.27 | 0.55 | 0.39 | 0.29 |
| 7 | 78424993 | rs37867 | 0.817 | -0.442 | 0.201 | -0.389 | -0.01 | -0.081 | -0.081 | 0 | 0.36 | 0.15 | 0.28 | 0.55 | 0.4 | 0.3 |
| 7 | 78434965 | rs1207887 | 0.472 | -0.488 | 0.653 | -0.475 | 0.027 | 0.016 | 0.016 | 0 | 0.47 | 0.3 | 0.45 | 0.42 | 0.51 | 0.62 |
| 7 | 78438407 | rs37887 | 1.122 | -0.478 | 0.609 | -0.452 | -0.025 | -0.062 | -0.062 | 0 | 0.86 | 0.78 | 0.83 | 1 | 0.9 | 0.92 |
| 7 | 78438634 | rs1207890 | 0.455 | -0.398 | 0.004 | -0.418 | 0.027 | 0.013 | 0.013 | 0.002 | 0.47 | 0.31 | 0.45 | 0.42 | 0.51 | 0.63 |
| 7 | 78441787 | rs17151525 | 0.629 | -0.813 | 0.319 | -0.874 | -0.001 | -0.014 | -0.014 | 0 | 0.05 | 0.07 | 0.13 | 0 | 0.06 | 0.04 |
| 7 | 78445408 | rs246464 | 0.493 | -0.842 | 0.012 | -0.857 | -0.056 | -0.073 | -0.073 | 8.06 | 0.84 | 0.6 | 0.74 | 0.98 | 0.91 | 0.91 |
| 7 | 78456148 | rs519514 | -0.671 | -0.644 | -1.019 | -0.721 | -0.09 | 0.02 | 0.02 | 0 | 0.61 | 0.87 | 0.68 | 0.34 | 0.51 | 0.64 |
| 7 | 78456168 | rs508733 | -0.187 | 0.478 | -0.478 | -0.105 | -0.01 | 0.014 | 0.014 | 0 | 0.38 | 0.32 | 0.51 | 0.32 | 0.42 | 0.51 |
| 7 | 78463648 | rs510604 | 0.647 | -0.003 | 0.007 | -0.034 | -0.023 | -0.009 | -0.009 | 0 | 0.95 | 0.77 | 0.97 | 0.97 | 1 | 0.95 |
| 7 | 78467217 | rs2065198 | 0.33 | 0.017 | -0.051 | -0.839 | -0.008 | 0.043 | 0.043 | 0 | 0.42 | 0.35 | 0.52 | 0.74 | 0.49 | 0.36 |
| 7 | 78468373 | rs472023 | 0.79 | -0.221 | 0.128 | -1.778 | 0.001 | 0.018 | 0.018 | 0.526 | 0.56 | 0.38 | 0.53 | 0.72 | 0.61 | 0.43 |
| 7 | 78469638 | rs17455327 | 1.487 | -0.19 | -0.137 | -2.018 | -0.027 | 0.056 | 0.056 | 0 | 0.1 | 0.01 | 0.04 | 0.01 | 0.09 | 0.03 |
| 7 | 78470511 | rs554380 | 1.116 | -0.288 | 0.043 | -2.04 | 0.001 | 0.064 | 0.064 | 1.724 | 0.23 | 0.09 | 0.08 | 0 | 0.2 | 0.08 |
| 7 | 78471483 | rs563887 | -0.19 | -0.309 | 0.412 | -1.397 | -0.001 | -0.011 | -0.011 | 0 | 0.76 | 0.65 | 0.64 | 0.74 | 0.78 | 0.47 |
| 7 | 78472436 | rs10242235 | 1.187 | -0.323 | 0.458 | -1.421 | 0.016 | -0.03 | -0.03 | 0 | 0.08 | 0.25 | 0.05 | 0 | 0.04 | 0.01 |
| 7 | 78473591 | rs961928 | 1.814 | -0.33 | 0.186 | -1.428 | -0.023 | 0.044 | 0.044 | 0 | 0.33 | 0.18 | 0.44 | 0.71 | 0.41 | 0.35 |
| 7 | 78474408 | rs1330495 | 2.106 | -0.358 | 0.24 | -1.447 | -0.03 | -0.074 | -0.074 | 0 | 0.28 | 0.11 | 0.31 | 0.52 | 0.35 | 0.23 |
| 7 | 78479542 | rs6950191 | 1.071 | -0.349 | 0.302 | -1.537 | -0.01 | 0.003 | 0.003 | 0 | 0.18 | 0.11 | 0.33 | 0.23 | 0.22 | 0.48 |
| 7 | 78479740 | rs576744 | 0.264 | -0.269 | 0.085 | -1.507 | -0.005 | -0.028 | -0.028 | 0.553 | 0.69 | 0.4 | 0.6 | 0.74 | 0.74 | 0.46 |
| 7 | 78483211 | rs571376 | -0.348 | -0.247 | 0.087 | -1.484 | 0 | -0.022 | -0.022 | 0 | 0.76 | 0.57 | 0.65 | 0.74 | 0.78 | 0.47 |
| 7 | 78497786 | rs4730552 | 1.698 | -0.185 | -0.314 | -1.356 | -0.004 | 0.014 | 0.014 | 0 | 0.31 | 0.26 | 0.42 | 0.73 | 0.36 | 0.29 |
| 7 | 78504119 | rs508990 | 0.899 | -0.184 | -0.4 | -1.272 | 0.001 | 0.006 | 0.006 | 2.034 | 0.54 | 0.24 | 0.41 | 0.53 | 0.59 | 0.31 |
| 7 | 78504355 | rs5000370 | 0.43 | -0.178 | -0.38 | -1.248 | 0.003 | -0.015 | -0.015 | 0.02 | 0.64 | 0.26 | 0.45 | 0.53 | 0.68 | 0.34 |
| 7 | 78507181 | rs1799025 | -0.353 | -0.021 | -0.25 | -0.987 | 0.004 | 0.081 | 0.081 | 0.163 | 0.36 | 0.18 | 0.14 | 0.01 | 0.33 | 0.11 |
| 7 | 78514140 | rs7789885 | -0.2 | 0.051 | -0.243 | -0.433 | -0.001 | -0.004 | -0.004 | 0 | 0.14 | 0.14 | 0.17 | 0.21 | 0.14 | 0.1 |
| 7 | 78524614 | rs1799017 | -0.847 | -0.196 | -0.436 | -0.668 | 0.005 | -0.01 | -0.01 | 0 | 0.58 | 0.23 | 0.6 | 0.46 | 0.58 | 0.7 |
| 7 | 78529329 | rs1799011 | -0.007 | -0.207 | -0.648 | -0.895 | -0.061 | 0.176 | 0.176 | 0.133 | 0.43 | 0.09 | 0.44 | 0.24 | 0.45 | 0.59 |
| 7 | 78532094 | rs17151643 | -0.369 | -0.029 | 0.01 | -0.906 | 0.006 | -0.055 | -0.055 | 0 | 0.15 | 0.3 | 0.08 | 0.04 | 0.14 | 0.03 |
| 7 | 78536207 | rs12705833 | 0.353 | -0.265 | 0.142 | -0.768 | 0.001 | 0.017 | 0.017 | 0.106 | 0.52 | 0.61 | 0.52 | 0.78 | 0.54 | 0.44 |
| 7 | 78537682 | rs7791106 | -0.943 | -0.454 | 0.312 | -0.155 | -0.037 | 0.068 | 0.068 | 0 | 0.18 | 0.53 | 0.19 | 0.23 | 0.1 | 0.2 |
| 7 | 78537751 | rs918285 | -0.484 | 0.2 | -0.618 | -0.422 | -0.03 | 0.039 | 0.039 | 0.971 | 0.1 | 0 | 0.1 | 0.07 | 0.1 | 0.18 |
| 7 | 78544172 | rs1799022 | -0.058 | -0.031 | -0.572 | -0.398 | 0.009 | 0.03 | 0.03 | 0 | 0.38 | 0.23 | 0.28 | 0.25 | 0.37 | 0.44 |
| 7 | 78545424 | rs17151659 | 0.149 | -0.021 | -0.752 | -0.482 | 0.026 | 0.014 | 0.014 | 0.014 | 0.33 | 0.12 | 0.24 | 0.25 | 0.31 | 0.39 |
| 7 | 78547202 | rs12705836 | 1.225 | -0.132 | -1.307 | -0.419 | -0.128 | -0.156 | -0.156 | 1.674 | 0.34 | 0.04 | 0.34 | 0.49 | 0.44 | 0.3 |

| | | | | | | | | | | | | | | | | |
|---|---|---|---|---|---|---|---|---|---|---|---|---|---|---|---|---|
| 7 | 78547842 | rs4730569 | 0.723 | -0.321 | -1.161 | -0.481 | -0.08 | -0.082 | -0.082 | 2.004 | 0.43 | 0.07 | 0.42 | 0.57 | 0.51 | 0.41 |
| 7 | 78550410 | rs9656198 | -0.059 | -0.723 | 0.462 | -0.189 | -0.026 | 0.107 | 0.107 | 7.228 | 0.08 | 0.28 | 0.22 | 0.18 | 0.01 | 0.05 |
| 7 | 78554551 | rs10230752 | -0.533 | -0.74 | 0.971 | -0.2 | -0.019 | 0.144 | 0.144 | 5.754 | 0.09 | 0.39 | 0.22 | 0.18 | 0.01 | 0.07 |
| 7 | 78573030 | rs6943858 | 0.312 | -0.779 | 1.042 | -0.322 | 0.002 | 0.002 | 0.002 | 0 | 0.35 | 0.27 | 0.24 | 0.25 | 0.28 | 0.37 |
| 7 | 78575481 | rs1642888 | -0.181 | -0.757 | 1.021 | -0.274 | 0.003 | -0.004 | -0.004 | 0.541 | 0.39 | 0.44 | 0.48 | 0.43 | 0.36 | 0.47 |
| 7 | 78578763 | rs7797351 | 0.52 | -0.773 | 1.37 | -0.282 | -0.009 | 0.001 | 0.001 | 0 | 0.56 | 0.46 | 0.51 | 0.58 | 0.64 | 0.52 |
| 7 | 78586678 | rs12705859 | 0.373 | -0.488 | 1.335 | -0.283 | -0.002 | 0 | 0 | 0 | 0.54 | 0.47 | 0.45 | 0.57 | 0.6 | 0.53 |
| 7 | 78595857 | rs1073416 | -0.575 | -0.526 | 1.332 | -0.357 | 0.024 | 0.015 | 0.015 | 0 | 0.66 | 0.85 | 0.7 | 0.74 | 0.66 | 0.64 |
| 7 | 78601619 | rs2191724 | -0.672 | -0.757 | 1.379 | -0.591 | 0.034 | 0.062 | 0.062 | 0 | 0.67 | 0.89 | 0.79 | 0.82 | 0.69 | 0.7 |
| 7 | 78604974 | rs10249654 | 0.195 | -0.745 | 1.453 | -0.582 | -0.031 | -0.033 | -0.033 | 0.016 | 0.06 | 0.18 | 0.02 | 0 | 0 | 0.01 |
| 7 | 78608574 | rs10225728 | -1.556 | -0.738 | 1.518 | -0.591 | -0.003 | -0.067 | -0.067 | 0 | 0.11 | 0.33 | 0.06 | 0 | 0.07 | 0.07 |
| 7 | 78615062 | rs7793390 | -0.668 | -0.729 | 1.484 | -0.59 | 0.033 | 0.057 | 0.057 | 0 | 0.67 | 0.87 | 0.79 | 0.82 | 0.69 | 0.7 |
| 7 | 78616461 | rs12537793 | 0.272 | -0.72 | 1.527 | -0.583 | 0.004 | 0.003 | 0.003 | 0 | 0.57 | 0.6 | 0.73 | 0.82 | 0.63 | 0.62 |
| 7 | 78628456 | rs4730596 | -0.276 | -0.696 | 1.359 | -0.552 | 0.044 | 0.062 | 0.062 | 0.065 | 0.62 | 0.81 | 0.78 | 0.83 | 0.69 | 0.7 |
| 7 | 78632489 | rs4730599 | -0.079 | -0.697 | 1.402 | -0.443 | 0.009 | 0 | 0 | 0.03 | 0.6 | 0.65 | 0.58 | 0.64 | 0.68 | 0.65 |
| 7 | 78641909 | rs12536821 | 0.269 | -0.698 | 1.341 | -0.442 | -0.005 | -0.003 | -0.003 | 0 | 0.55 | 0.49 | 0.54 | 0.64 | 0.62 | 0.59 |
| 7 | 78644077 | rs1476106 | -0.275 | -0.666 | 1.329 | -0.428 | 0.044 | 0.061 | 0.061 | 0.07 | 0.62 | 0.8 | 0.78 | 0.83 | 0.69 | 0.7 |
| 7 | 78651208 | rs6945664 | 0.365 | -0.622 | 1.074 | -0.442 | -0.004 | -0.002 | -0.002 | 0 | 0.56 | 0.5 | 0.54 | 0.64 | 0.62 | 0.6 |
| 7 | 78659092 | rs10224582 | -0.225 | 0.067 | 1.123 | -0.128 | 0.015 | 0 | 0 | 0 | 0.41 | 0.69 | 0.44 | 0.37 | 0.37 | 0.4 |
| 7 | 78664320 | rs17151887 | -1.42 | 0.368 | 0.83 | -0.603 | 0.139 | 0.037 | 0.037 | 0 | 0.05 | 0.34 | 0.05 | 0 | 0.06 | 0.06 |
| 7 | 78664486 | rs10485927 | -1.42 | 0.368 | 0.83 | -0.603 | 0.139 | 0.037 | 0.037 | 0 | 0.05 | 0.34 | 0.05 | 0 | 0.06 | 0.06 |
| 7 | 78664922 | rs17406935 | -0.018 | 0.28 | -0.365 | -0.488 | -0.017 | 0.001 | 0.001 | 0 | 0.08 | 0 | 0.08 | 0.08 | 0.06 | 0.1 |
| 7 | 78678354 | rs4437570 | -0.57 | 0.365 | 0.033 | -0.367 | 0.013 | 0.064 | 0.064 | 0 | 0.42 | 0.72 | 0.36 | 0.19 | 0.41 | 0.38 |
| 7 | 78680726 | rs17407075 | -1.732 | 1.117 | 0.351 | 0.148 | 0.001 | -0.016 | -0.016 | 0.39 | 0.05 | 0.08 | 0.03 | 0 | 0.05 | 0.05 |
| 7 | 78687119 | rs7809986 | -1.571 | 0.983 | 0.683 | 0.053 | 0.001 | -0.059 | -0.059 | 0.277 | 0.21 | 0.63 | 0.18 | 0.06 | 0.19 | 0.22 |
| 7 | 78688292 | rs17407267 | -1.458 | 1.347 | 0.832 | -0.043 | 0.023 | -0.041 | -0.041 | 0.259 | 0.09 | 0.19 | 0.06 | 0 | 0.11 | 0.02 |
| 7 | 78691005 | rs12530795 | 1.928 | 0.77 | 0.051 | 0.659 | -0.007 | -0.111 | -0.111 | 0 | 0.64 | 0.06 | 0.53 | 0.46 | 0.76 | 0.55 |
| 7 | 78693250 | rs2191719 | 1.282 | 0.218 | 1.512 | 0.883 | -0.037 | 0.1 | 0.1 | 0.006 | 0.76 | 0.59 | 0.71 | 0.48 | 0.87 | 0.58 |
| 7 | 78703278 | rs6942475 | 0.757 | 0.24 | 2.143 | 0.935 | 0.014 | -0.027 | -0.027 | 0.477 | 0.81 | 0.85 | 0.73 | 0.49 | 0.89 | 0.59 |
| 7 | 78726778 | rs1013289 | 1.492 | 0.255 | 2.514 | 0.919 | -0.032 | 0.07 | 0.07 | 0.106 | 0.75 | 0.6 | 0.61 | 0.44 | 0.85 | 0.58 |
| 7 | 78727984 | rs17151948 | -0.138 | 0.28 | 2.359 | 0.886 | -0.005 | -0.027 | -0.027 | 0.046 | 0.05 | 0.15 | 0.02 | 0 | 0.02 | 0 |
| 7 | 78729245 | rs6977105 | 2.382 | 0.285 | 2.227 | 0.899 | 0.068 | -0.12 | -0.12 | 0.514 | 0.66 | 0.06 | 0.56 | 0.44 | 0.83 | 0.57 |
| 7 | 78732358 | rs12674038 | 1.919 | 0.258 | 2.614 | 0.959 | -0.02 | 0.026 | 0.026 | 0.442 | 0.7 | 0.35 | 0.58 | 0.45 | 0.83 | 0.56 |
| 7 | 78740625 | rs1016189 | 2.193 | 0.273 | 2.215 | 1.431 | 0.073 | -0.025 | -0.025 | 0 | 0.68 | 0.13 | 0.67 | 0.63 | 0.84 | 0.56 |
| 7 | 78742376 | rs2886053 | 1.871 | 0.296 | 2.355 | 1.435 | -0.022 | 0.005 | 0.005 | 0.144 | 0.71 | 0.4 | 0.7 | 0.63 | 0.85 | 0.57 |
| 7 | 78748641 | rs2215585 | 1.38 | 0.377 | 2.276 | 1.674 | -0.031 | 0.028 | 0.028 | 0.083 | 0.74 | 0.43 | 0.6 | 0.58 | 0.85 | 0.53 |
| 7 | 78750133 | rs10280055 | 1.555 | 0.351 | 2.279 | 1.68 | -0.022 | 0.021 | 0.021 | 0.04 | 0.73 | 0.43 | 0.59 | 0.58 | 0.83 | 0.52 |
| 7 | 78751706 | rs17151993 | -0.305 | 1.46 | 1.661 | 1.735 | -0.073 | -0.072 | -0.072 | 0.008 | 0.11 | 0.01 | 0.18 | 0.24 | 0.27 | 0.14 |
| 7 | 78767182 | rs10255742 | 1.916 | 0.073 | 2.316 | 1.38 | -0.003 | -0.002 | -0.002 | 1.556 | 0.64 | 0.3 | 0.62 | 0.63 | 0.75 | 0.49 |
| 7 | 78775199 | rs7787597 | 2.099 | 0.132 | 2.6 | 1.696 | 0.057 | -0.048 | -0.048 | 0.396 | 0.68 | 0.16 | 0.57 | 0.5 | 0.83 | 0.51 |
| 7 | 78781075 | rs6957950 | 0.918 | 0.107 | 2.803 | 1.708 | -0.02 | 0.022 | 0.022 | 0 | 0.78 | 0.64 | 0.7 | 0.68 | 0.86 | 0.53 |
| 7 | 78783115 | rs1990433 | 1.457 | 0.132 | 2.667 | 1.67 | -0.045 | 0.023 | 0.023 | 0.452 | 0.73 | 0.47 | 0.69 | 0.64 | 0.84 | 0.51 |
| 7 | 78793083 | rs10953783 | 1.985 | 0.183 | 2.222 | 1.808 | 0.054 | -0.062 | -0.062 | 0 | 0.69 | 0.18 | 0.58 | 0.49 | 0.83 | 0.5 |
| 7 | 78801465 | rs4573181 | 2.022 | 0.188 | 2.387 | 1.851 | 0.062 | -0.076 | -0.076 | 0 | 0.68 | 0.19 | 0.59 | 0.48 | 0.83 | 0.51 |
| 7 | 78808948 | rs13222189 | 1.666 | 0.184 | 2.432 | 1.832 | -0.007 | 0.002 | 0.002 | 1.115 | 0.72 | 0.34 | 0.68 | 0.63 | 0.84 | 0.52 |
| 7 | 78814410 | rs2024254 | 0.6 | 0.157 | 2.672 | 1.528 | -0.01 | 0.009 | 0.009 | 0 | 0.8 | 0.77 | 0.71 | 0.7 | 0.86 | 0.52 |
| 7 | 78814643 | rs7783481 | -2.199 | 0.299 | 2.377 | 1.622 | -0.033 | 0.089 | 0.089 | 0.689 | 0.1 | 0.38 | 0.11 | 0.19 | 0.04 | 0.03 |
| 7 | 78818348 | rs2191813 | -2.788 | 0.292 | 2.212 | 1.619 | -0.054 | 0.062 | 0.062 | 2.674 | 0.12 | 0.58 | 0.13 | 0.19 | 0.04 | 0.03 |
| 7 | 78821030 | rs17152038 | -2.009 | 0.341 | 1.498 | 1.616 | 0.006 | 0.123 | 0.123 | 0.07 | 0.06 | 0.31 | 0.09 | 0.15 | 0.01 | 0.01 |
| 7 | 78830766 | rs7798423 | -1.273 | 0.401 | 1.054 | 1.543 | -0.012 | 0.07 | 0.07 | 0.18 | 0.09 | 0.36 | 0.04 | 0.13 | 0.03 | 0.02 |
| 7 | 78833734 | rs7809989 | -0.916 | 0.479 | 0.15 | 1.593 | -0.021 | 0.064 | 0.064 | 0 | 0.08 | 0.26 | 0.03 | 0.15 | 0.03 | 0.03 |
| 7 | 78839317 | rs12535987 | 1.227 | 0.831 | -0.954 | 1.338 | 0.03 | -0.128 | -0.128 | 0 | 0.65 | 0.07 | 0.57 | 0.48 | 0.77 | 0.55 |
| 7 | 78843289 | rs12706063 | 1.244 | 0.874 | -1.134 | 1.284 | -0.026 | -0.128 | -0.128 | 0 | 0.58 | 0.03 | 0.52 | 0.48 | 0.69 | 0.54 |
| 7 | 78858392 | rs2056753 | -1.479 | 2.36 | 0.166 | 1.458 | 0.136 | 0.049 | 0.049 | 0 | 0.08 | 0.22 | 0.23 | 0.2 | 0.2 | 0.15 |
| 7 | 78865618 | rs10953812 | 1.078 | 2.282 | 0.638 | 1.851 | -0.011 | 0.037 | 0.037 | 1.211 | 0.67 | 0.56 | 0.6 | 0.5 | 0.69 | 0.58 |
| 7 | 78869490 | rs717067 | 1.202 | 2.159 | 0.631 | 1.822 | -0.003 | 0.003 | 0.003 | 0.981 | 0.64 | 0.36 | 0.58 | 0.49 | 0.69 | 0.57 |
| 7 | 78871941 | rs1528272 | 1.06 | 2.146 | 0.673 | 1.815 | -0.011 | 0.033 | 0.033 | 1.793 | 0.67 | 0.58 | 0.6 | 0.5 | 0.69 | 0.58 |
| 7 | 78872516 | rs1406160 | 2.048 | 2.246 | 0.559 | 1.796 | 0.014 | 0.026 | 0.026 | 1.302 | 0.56 | 0.27 | 0.4 | 0.26 | 0.61 | 0.42 |
| 7 | 78890691 | rs6953483 | 1.315 | 2.175 | 0.785 | 1.785 | -0.007 | 0.016 | 0.016 | 0.795 | 0.66 | 0.4 | 0.58 | 0.5 | 0.69 | 0.58 |
| 7 | 78894075 | rs10253536 | -1.059 | 2.221 | 1.21 | 2.038 | -0.017 | 0.111 | 0.111 | 0 | 0.19 | 0.46 | 0.2 | 0.34 | 0.14 | 0.13 |
| 7 | 78903813 | rs6946286 | 1.937 | 2.147 | 1.557 | 2.065 | 0.005 | -0.011 | -0.011 | 0 | 0.66 | 0.31 | 0.7 | 0.48 | 0.7 | 0.55 |
| 7 | 78910974 | rs1406157 | 2.162 | 2.009 | 1.5 | 2.015 | 0.029 | -0.024 | -0.024 | 0 | 0.61 | 0.2 | 0.68 | 0.45 | 0.68 | 0.53 |
| 7 | 78914631 | rs10234403 | 1.854 | 2.01 | 1.582 | 1.993 | -0.009 | 0.019 | 0.019 | 0.731 | 0.66 | 0.37 | 0.7 | 0.48 | 0.7 | 0.55 |
| 7 | 78933657 | rs1528265 | -2.045 | 2.018 | 1.599 | 1.989 | -0.01 | 0.02 | 0.02 | 0.436 | 0.34 | 0.6 | 0.3 | 0.51 | 0.3 | 0.43 |
| 7 | 78947104 | rs13230579 | 1.794 | 1.886 | 1.833 | 1.955 | -0.015 | 0.021 | 0.021 | 0.995 | 0.64 | 0.47 | 0.7 | 0.45 | 0.68 | 0.55 |
| 7 | 78947912 | rs11763567 | -1.186 | 2.165 | 1.672 | 2.012 | 0.068 | 0.018 | 0.018 | 0 | 0.09 | 0.16 | 0.24 | 0.22 | 0.2 | 0.14 |
| 7 | 78954604 | rs12216581 | 2.059 | 2.139 | 1.702 | 2.035 | 0.021 | -0.016 | -0.016 | 0.017 | 0.61 | 0.25 | 0.69 | 0.45 | 0.68 | 0.54 |
| 7 | 78963773 | rs7811833 | 0.619 | 2.195 | 1.708 | 2.029 | 0.002 | -0.004 | -0.004 | 0 | 0.84 | 0.86 | 0.8 | 0.83 | 0.85 | 0.71 |
| 7 | 78971036 | rs10499842 | -0.866 | 2.448 | 1.995 | 2.08 | 0.078 | 0.029 | 0.029 | 0 | 0.08 | 0.15 | 0.23 | 0.2 | 0.2 | 0.15 |

| | | | | | | | | | | | | | | | | |
|---|---|---|---|---|---|---|---|---|---|---|---|---|---|---|---|---|
| 7 | 78973208 | rs4730769 | 2.476 | 2.086 | 1.929 | 2.049 | -0.006 | 0.016 | 0.016 | 0 | 0.59 | 0.29 | 0.49 | 0.57 | 0.6 | 0.52 |
| 7 | 78974072 | rs1918930 | 0.321 | 2.021 | 1.992 | 1.958 | -0.002 | 0 | 0 | 0 | 0.85 | 0.85 | 0.92 | 0.96 | 0.86 | 0.92 |
| 7 | 78987060 | rs1918935 | -0.743 | 1.906 | 1.815 | 1.952 | 0.001 | -0.003 | -0.003 | 0 | 0.14 | 0.12 | 0.2 | 0.16 | 0.13 | 0.26 |
| 7 | 79023504 | rs2364913 | -1.724 | 1.797 | 2.037 | 2.068 | -0.033 | 0.12 | 0.12 | 0.059 | 0.19 | 0.45 | 0.23 | 0.39 | 0.14 | 0.34 |
| 7 | 79029377 | rs10261439 | -1.416 | 1.719 | 2.037 | 1.945 | 0 | -0.049 | -0.049 | 0.2 | 0.22 | 0.23 | 0.12 | 0.04 | 0.22 | 0.1 |
| 7 | 79030922 | rs4730806 | -2.248 | 1.76 | 2.068 | 2.052 | 0.021 | -0.014 | -0.014 | 0 | 0.36 | 0.71 | 0.32 | 0.43 | 0.27 | 0.41 |
| 7 | 79036238 | rs10254950 | -1.417 | 1.713 | 2.085 | 2.069 | -0.044 | 0.133 | 0.133 | 0.386 | 0.19 | 0.46 | 0.23 | 0.39 | 0.15 | 0.33 |
| 7 | 79043802 | rs9641589 | -0.856 | 1.73 | 1.773 | 2.088 | 0.021 | 0.048 | 0.048 | 0 | 0.09 | 0.16 | 0.22 | 0.29 | 0.13 | 0.13 |
| 7 | 79048593 | rs10214949 | -1.41 | 1.691 | 1.966 | 2.139 | -0.035 | 0.003 | 0.003 | 0 | 0.22 | 0.52 | 0.1 | 0.22 | 0.14 | 0.16 |
| 7 | 79048739 | rs10224698 | -2.085 | 1.771 | 1.64 | 2.11 | 0.01 | -0.03 | -0.03 | 0 | 0.32 | 0.72 | 0.32 | 0.52 | 0.27 | 0.29 |
| 7 | 79057761 | rs10485959 | 2.251 | 2.323 | 0.926 | 1.609 | 0.047 | 0.03 | 0.03 | 1.43 | 0.43 | 0.11 | 0.36 | 0.25 | 0.35 | 0.33 |
| 7 | 79061837 | rs10259665 | 1.315 | 2.352 | 1.686 | 1.645 | 0.059 | 0.059 | 0.059 | 1.389 | 0.76 | 0.54 | 0.67 | 0.68 | 0.68 | 0.66 |
| 7 | 79063644 | rs10953862 | 0.315 | 2.362 | 1.229 | 1.609 | -0.008 | -0.01 | -0.01 | 0 | 0.83 | 0.87 | 0.75 | 0.78 | 0.77 | 0.71 |
| 7 | 79067580 | rs17152506 | -0.948 | 2.42 | 1.352 | 1.629 | 0.014 | 0.033 | 0.033 | 0 | 0.1 | 0.16 | 0.21 | 0.29 | 0.14 | 0.15 |
| 7 | 79070452 | rs4730838 | 1.69 | 2.25 | 1.468 | 1.538 | -0.012 | -0.009 | -0.009 | 0 | 0.6 | 0.67 | 0.48 | 0.48 | 0.54 | 0.51 |
| 7 | 79078161 | rs10265155 | -2.206 | 2.506 | 1.82 | 1.491 | 0.131 | 0.094 | 0.094 | 0 | 0.19 | 0.48 | 0.32 | 0.39 | 0.3 | 0.19 |
| 7 | 79084800 | rs10231884 | 1.583 | 2.288 | 1.796 | 1.567 | 0.001 | 0.002 | 0.002 | 0.889 | 0.59 | 0.61 | 0.47 | 0.4 | 0.57 | 0.46 |
| 7 | 79085543 | rs4730857 | 2.118 | 2.292 | 1.89 | 1.585 | 0.027 | 0.054 | 0.054 | 1.828 | 0.51 | 0.27 | 0.39 | 0.29 | 0.44 | 0.34 |
| 7 | 79086119 | rs17152533 | -1.03 | 2.387 | 1.054 | 1.605 | 0.073 | -0.024 | -0.024 | 0 | 0.05 | 0.2 | 0.07 | 0 | 0.13 | 0.04 |
| 7 | 79086317 | rs6964700 | -0.157 | 2.375 | 1.035 | 1.628 | 0.005 | -0.015 | -0.015 | 0 | 0.84 | 0.91 | 0.76 | 0.75 | 0.84 | 0.69 |
| 7 | 79097483 | rs10252126 | 0.99 | 2.237 | 1.197 | 1.42 | -0.012 | -0.007 | -0.007 | 0 | 0.59 | 0.64 | 0.58 | 0.54 | 0.5 | 0.5 |
| 7 | 79110792 | rs1017039 | 0.563 | 2.056 | 1.195 | 1.435 | -0.002 | -0.021 | -0.021 | 0 | 0.63 | 0.76 | 0.64 | 0.54 | 0.63 | 0.52 |
| 7 | 79115715 | rs4388396 | 0.663 | 2.095 | 1.012 | 1.454 | 0 | -0.006 | -0.006 | 0.071 | 0.62 | 0.67 | 0.64 | 0.54 | 0.63 | 0.48 |
| 7 | 79115832 | rs4286869 | 1.353 | 2.036 | 0.7 | 1.338 | 0 | 0.001 | 0.001 | 0.582 | 0.52 | 0.47 | 0.41 | 0.28 | 0.44 | 0.33 |
| 7 | 79122018 | rs2079385 | -1.276 | 1.977 | 0.448 | 1.282 | 0 | -0.001 | -0.001 | 1.432 | 0.1 | 0.13 | 0.32 | 0.25 | 0.2 | 0.15 |
| 7 | 79136577 | rs10254114 | -1.228 | 1.087 | 0.718 | 0.917 | -0.019 | 0.025 | 0.025 | 0 | 0.2 | 0.58 | 0.44 | 0.25 | 0.12 | 0.21 |
| 7 | 79139403 | rs10485961 | -0.854 | 0.818 | 0.612 | 0.302 | 0.008 | -0.055 | -0.055 | 0 | 0.14 | 0.2 | 0.11 | 0 | 0.17 | 0.07 |
| 7 | 79168005 | rs10480737 | -1.511 | -0.228 | 0.713 | 0.611 | -0.033 | -0.01 | -0.01 | 0.237 | 0.19 | 0.26 | 0.08 | 0.18 | 0.07 | 0.19 |
| 7 | 79173769 | rs4730917 | -2.011 | -0.389 | 1.65 | 1.12 | -0.051 | 0.024 | 0.024 | 0 | 0.24 | 0.58 | 0.3 | 0.34 | 0.1 | 0.26 |
| 7 | 79174185 | rs7788571 | -1.581 | -0.019 | 0.024 | 0.878 | -0.137 | -0.146 | -0.146 | 2.588 | 0.56 | 0.98 | 0.56 | 0.51 | 0.48 | 0.6 |
| 7 | 79243030 | rs6961362 | -0.739 | -0.378 | 1.217 | 1.001 | -0.024 | 0.002 | 0.002 | 0 | 0.15 | 0.19 | 0.07 | 0.17 | 0.06 | 0.19 |
| 7 | 79248120 | rs6955922 | -1.675 | -0.42 | 1.369 | 0.995 | -0.07 | 0.091 | 0.091 | 0.17 | 0.2 | 0.51 | 0.3 | 0.33 | 0.09 | 0.26 |
| 7 | 79268520 | rs4398836 | -0.781 | -0.377 | 0.69 | 0.915 | 0.008 | -0.003 | -0.003 | 0 | 0.14 | 0.12 | 0.06 | 0.17 | 0.06 | 0.18 |
| 7 | 79284609 | rs6979533 | -0.594 | -0.41 | 0.652 | 0.929 | 0.02 | 0.006 | 0.006 | 0 | 0.13 | 0.09 | 0.05 | 0.08 | 0.06 | 0.08 |
| 7 | 79288947 | rs2465511 | -1.692 | -0.301 | 0.161 | 1.091 | -0.08 | -0.083 | -0.083 | 0.69 | 0.53 | 0.95 | 0.61 | 0.55 | 0.52 | 0.56 |
| 7 | 79293845 | rs2714676 | -0.314 | -0.183 | 0.754 | 0.995 | 0.046 | -0.022 | -0.022 | 0 | 0.29 | 0.53 | 0.29 | 0.2 | 0.42 | 0.33 |
| 7 | 79294183 | rs2714679 | -2.027 | -0.354 | 0.234 | 0.932 | -0.093 | -0.1 | -0.1 | 0.076 | 0.55 | 0.98 | 0.62 | 0.58 | 0.53 | 0.59 |
| 7 | 79296074 | rs2365693 | -1.115 | -0.324 | 0.38 | 1.009 | 0.02 | -0.037 | -0.037 | 0 | 0.18 | 0.1 | 0.26 | 0.27 | 0.1 | 0.16 |
| 7 | 79297997 | rs6959961 | -1.161 | -0.127 | 0.606 | 0.83 | -0.009 | 0.008 | 0.008 | 0.898 | 0.42 | 0.62 | 0.34 | 0.36 | 0.44 | 0.44 |
| 7 | 79303403 | rs1403875 | -1.628 | -0.091 | 0.032 | 1.539 | -0.09 | -0.004 | -0.004 | 0 | 0.5 | 0.96 | 0.58 | 0.56 | 0.48 | 0.63 |
| 7 | 79305142 | rs1403874 | -0.352 | -0.029 | 0.732 | 1.607 | -0.016 | 0.008 | 0.008 | 0 | 0.27 | 0.29 | 0.29 | 0.21 | 0.37 | 0.39 |
| 7 | 79314722 | rs16886601 | -1.293 | -0.047 | 0.937 | 1.936 | -0.003 | 0.052 | 0.052 | 0.039 | 0.08 | 0.11 | 0.09 | 0.22 | 0.08 | 0.17 |
| 7 | 79315761 | rs2140493 | 0.722 | 0.364 | 1.164 | 1.757 | -0.007 | 0.034 | 0.034 | 0 | 0.84 | 0.52 | 0.77 | 0.87 | 0.87 | 0.93 |
| 7 | 79316164 | rs10270681 | -0.315 | 0.414 | 1.081 | 1.642 | 0.01 | -0.006 | -0.006 | 0 | 0.28 | 0.36 | 0.3 | 0.2 | 0.36 | 0.36 |
| 7 | 79319930 | rs1403881 | -1.181 | 0.387 | 1.017 | 1.54 | 0.023 | 0.015 | 0.015 | 1.095 | 0.37 | 0.5 | 0.39 | 0.42 | 0.44 | 0.53 |
| 7 | 79323090 | rs11975512 | -0.372 | 0.32 | 0.811 | 1.03 | -0.032 | 0.057 | 0.057 | 0 | 0.29 | 0.2 | 0.29 | 0.09 | 0.35 | 0.35 |
| 7 | 79324680 | rs2462129 | 1.509 | 0.189 | 1.462 | 1.197 | -0.003 | 0.035 | 0.035 | 0.559 | 0.52 | 0.27 | 0.45 | 0.69 | 0.52 | 0.51 |
| 7 | 79334324 | rs2689787 | 1.582 | 0.136 | 1.432 | 1.139 | -0.004 | 0.058 | 0.058 | 0.85 | 0.51 | 0.2 | 0.44 | 0.69 | 0.52 | 0.51 |
| 7 | 79337748 | rs4419754 | -0.183 | 0.236 | 1.41 | 1.117 | -0.039 | 0.067 | 0.067 | 0 | 0.28 | 0.18 | 0.26 | 0.09 | 0.34 | 0.35 |
| 7 | 79338211 | rs2689790 | 0.337 | 0.188 | 1.652 | 1.095 | 0.001 | -0.012 | -0.012 | 0 | 0.86 | 0.83 | 0.94 | 0.99 | 0.9 | 0.96 |
| 7 | 79339063 | rs2465513 | 1.423 | 0.208 | 1.804 | 1.207 | -0.007 | 0.017 | 0.017 | 0.119 | 0.55 | 0.37 | 0.47 | 0.74 | 0.53 | 0.57 |
| 7 | 79341676 | rs2714703 | 1.249 | 0.203 | 1.672 | 1.197 | -0.013 | 0.026 | 0.026 | 0.211 | 0.54 | 0.29 | 0.46 | 0.69 | 0.53 | 0.51 |
| 7 | 79347341 | rs2714707 | 1.13 | 0.126 | 1.451 | 1.419 | -0.016 | 0.049 | 0.049 | 0 | 0.82 | 0.6 | 0.83 | 0.77 | 0.84 | 0.84 |
| 7 | 79347456 | rs1403880 | 1.335 | 0.141 | 1.52 | 1.422 | -0.026 | 0.048 | 0.048 | 0.169 | 0.81 | 0.42 | 0.81 | 0.82 | 0.85 | 0.91 |
| 7 | 79351368 | rs4288313 | -0.945 | 0.108 | 1.58 | 1.39 | -0.008 | 0.014 | 0.014 | 0.02 | 0.14 | 0.39 | 0.08 | 0.15 | 0.09 | 0.13 |
| 7 | 79362257 | rs2689813 | 0.548 | 0.134 | 1.666 | 1.39 | -0.002 | 0.012 | 0.012 | 0 | 0.86 | 0.68 | 0.93 | 0.85 | 0.9 | 0.87 |
| 7 | 79366631 | rs17832522 | 0.414 | 0.061 | 1.344 | 1.011 | 0.017 | 0.023 | 0.023 | 0 | 0.18 | 0.12 | 0.11 | 0.07 | 0.12 | 0.38 |
| 7 | 79367667 | rs2714700 | 1.914 | 0.78 | 1.281 | 1.683 | -0.028 | -0.007 | -0.007 | 0 | 0.57 | 0.28 | 0.44 | 0.51 | 0.53 | 0.44 |
| 7 | 79371206 | rs2714702 | 0.051 | 0.782 | 1.413 | 1.655 | -0.007 | -0.045 | -0.045 | 0.351 | 0.87 | 0.78 | 0.94 | 0.99 | 0.9 | 0.95 |
| 7 | 79389666 | rs1950158 | -0.126 | 0.781 | 1.397 | 1.655 | -0.003 | -0.026 | -0.026 | 0.233 | 0.88 | 0.82 | 0.94 | 0.99 | 0.9 | 0.95 |
| 7 | 79396718 | rs2689805 | 0.376 | 0.771 | 1.402 | 1.67 | 0.002 | 0.019 | 0.019 | 0.018 | 0.86 | 0.61 | 0.92 | 0.85 | 0.9 | 0.87 |
| 7 | 79402551 | rs10499847 | 0.386 | 0.935 | 1.279 | 1.654 | -0.041 | 0.043 | 0.043 | 0.449 | 0.08 | 0.01 | 0.13 | 0 | 0.22 | 0.05 |
| 7 | 79430869 | rs2714682 | 1.906 | 1.072 | 1.29 | 1.716 | -0.038 | -0.018 | -0.018 | 1.131 | 0.57 | 0.24 | 0.45 | 0.49 | 0.56 | 0.45 |
| 7 | 79437535 | rs12535819 | -0.081 | 1.138 | 1.19 | 1.609 | 0.228 | -0.037 | -0.037 | 0 | 0.09 | 0.21 | 0.14 | 0 | 0.22 | 0.05 |
| 7 | 79438813 | rs2714664 | 1.197 | 1.101 | 1.147 | 1.61 | 0.088 | -0.038 | -0.038 | 0 | 0.79 | 0.67 | 0.8 | 0.85 | 0.68 | 0.83 |
| 7 | 79443551 | rs2714657 | 1.197 | 1.101 | 1.147 | 1.61 | 0.088 | -0.038 | -0.038 | 0 | 0.79 | 0.67 | 0.8 | 0.85 | 0.68 | 0.83 |
| 7 | 79452761 | rs2365696 | 2.387 | 0.733 | 0.116 | 1.319 | -0.026 | -0.003 | -0.003 | 0.793 | 0.39 | 0.02 | 0.29 | 0.29 | 0.38 | 0.35 |
| 7 | 79455233 | rs1922725 | -1.528 | 1.499 | 0.821 | 1.495 | 0.091 | -0.055 | -0.055 | 0.044 | 0.22 | 0.45 | 0.2 | 0.15 | 0.29 | 0.17 |
| 7 | 79495978 | rs11975054 | -0.27 | 1.605 | 0.822 | 1.562 | 0.201 | 0.085 | 0.085 | 0.123 | 0.07 | 0.22 | 0.11 | 0.14 | 0.17 | 0.11 |

| | | | | | | | | | | | | | | | | |
|---|---|---|---|---|---|---|---|---|---|---|---|---|---|---|---|---|
| 7 | 79530882 | rs2707916 | -0.679 | 1.477 | 0.767 | 0.792 | 0.005 | 0.001 | 0.001 | 0 | 0.14 | 0.16 | 0.07 | 0.01 | 0.12 | 0.05 |
| 7 | 79532641 | rs2707908 | -1.857 | 1.471 | 0.935 | 0.785 | 0.057 | -0.05 | -0.05 | 0.744 | 0.23 | 0.53 | 0.2 | 0.14 | 0.29 | 0.17 |
| 7 | 79550607 | rs7776691 | -0.618 | 1.508 | 0.722 | 0.83 | 0.227 | 0.097 | 0.097 | 0 | 0.09 | 0.31 | 0.1 | 0.14 | 0.17 | 0.11 |
| 7 | 79553908 | rs333211 | 0.436 | 1.427 | 0.608 | 0.81 | -0.004 | -0.002 | -0.002 | 2.609 | 0.86 | 0.76 | 0.92 | 0.86 | 0.9 | 0.88 |
| 7 | 79556915 | rs10276215 | 1.596 | 1.081 | 0.115 | 0.477 | -0.022 | 0.049 | 0.049 | 1.999 | 0.37 | 0.11 | 0.32 | 0.27 | 0.42 | 0.31 |
| 7 | 79585451 | rs10229206 | 1.694 | 1.491 | 0.707 | 1.011 | 0.063 | 0.061 | 0.061 | 0.215 | 0.76 | 0.53 | 0.6 | 0.6 | 0.68 | 0.8 |
| 7 | 79589187 | rs9640814 | -0.421 | 1.555 | 0.561 | 0.996 | -0.007 | -0.006 | -0.006 | 0 | 0.08 | 0.03 | 0.31 | 0.28 | 0.19 | 0.07 |
| 7 | 79594834 | rs10229737 | 1.465 | 1.53 | 0.62 | 0.789 | 0.057 | -0.032 | -0.032 | 0 | 0.76 | 0.43 | 0.79 | 0.83 | 0.69 | 0.83 |
| 7 | 79604156 | rs4731111 | -1.187 | 1.786 | 0.544 | 1.298 | 0.039 | 0.04 | 0.04 | 0 | 0.11 | 0.23 | 0.25 | 0.28 | 0.17 | 0.07 |
| 7 | 79619012 | rs6466884 | -1.256 | 1.671 | 0.423 | 0.887 | 0.087 | 0.089 | 0.089 | 0.076 | 0.12 | 0.59 | 0.11 | 0.16 | 0.16 | 0.07 |
| 7 | 79630265 | rs10244217 | 0.088 | 1.035 | -0.858 | 0.559 | 0.041 | -0.036 | -0.036 | 0.044 | 0.51 | 0.85 | 0.5 | 0.46 | 0.59 | 0.38 |
| 7 | 79632588 | rs17802112 | -0.153 | 1.409 | -0.189 | 0.675 | 0.051 | 0.053 | 0.053 | 0 | 0.07 | 0.12 | 0.09 | 0.14 | 0.15 | 0.07 |
| 7 | 79636682 | rs17802148 | 0.462 | 1.431 | -0.184 | 0.661 | 0.037 | -0.028 | -0.028 | 0 | 0.05 | 0.06 | 0.06 | 0 | 0.13 | 0.03 |
| 7 | 79644412 | rs728459 | 0.137 | 1.499 | -0.009 | 0.481 | -0.017 | -0.016 | -0.016 | 0 | 0.42 | 0.68 | 0.39 | 0.44 | 0.45 | 0.35 |
| 7 | 79645196 | rs2040759 | 0.998 | 1.438 | 0.391 | 0.469 | -0.002 | -0.003 | -0.003 | 0 | 0.32 | 0.28 | 0.28 | 0.29 | 0.3 | 0.28 |
| 7 | 79657365 | rs6466888 | -0.013 | 1.441 | 0.188 | 0.472 | -0.062 | -0.048 | -0.048 | 0 | 0.43 | 0.86 | 0.43 | 0.43 | 0.45 | 0.36 |
| 7 | 79659914 | rs17153268 | 0.627 | 1.257 | 0.128 | 0.457 | 0.014 | 0.021 | 0.021 | 0 | 0.36 | 0.76 | 0.33 | 0.29 | 0.3 | 0.29 |
| 7 | 79663989 | rs17153296 | -0.217 | 1.233 | -0.133 | 0.668 | -0.031 | -0.045 | -0.045 | 0 | 0.08 | 0.02 | 0.19 | 0.31 | 0.14 | 0.07 |
| 7 | 79676787 | rs6466904 | -0.053 | 0.99 | 0.521 | -0.174 | -0.001 | 0.006 | 0.006 | 0 | 0.48 | 0.54 | 0.41 | 0.34 | 0.49 | 0.4 |
| 7 | 79679154 | rs7778809 | -0.338 | 0.97 | 0.077 | -0.174 | -0.081 | -0.079 | -0.079 | 0 | 0.52 | 0.93 | 0.47 | 0.46 | 0.51 | 0.44 |
| 7 | 79694753 | rs10486920 | -0.538 | 1.141 | 0.131 | -0.226 | -0.002 | -0.001 | -0.001 | 0 | 0.15 | 0.15 | 0.14 | 0.14 | 0.22 | 0.14 |
| 7 | 79701003 | rs4731190 | -0.257 | 0.722 | -0.034 | -0.124 | -0.001 | -0.003 | -0.003 | 0.31 | 0.51 | 0.78 | 0.46 | 0.47 | 0.51 | 0.46 |
| 7 | 79701190 | rs10250164 | 0.063 | 0.837 | -0.32 | -0.253 | -0.035 | 0.055 | 0.055 | 1.271 | 0.45 | 0.2 | 0.37 | 0.34 | 0.49 | 0.41 |
| 7 | 79723943 | rs6945349 | 0.663 | 1.229 | 0.201 | -0.246 | 0.066 | -0.071 | -0.071 | 0.106 | 0.88 | 0.41 | 0.87 | 1 | 0.84 | 0.97 |
| 7 | 79725465 | rs12531497 | 0.876 | 1.175 | 0.158 | -0.083 | 0.059 | 0.072 | 0.072 | 0.677 | 0.87 | 0.42 | 0.85 | 0.87 | 0.83 | 0.93 |
| 7 | 79726578 | rs886364 | 0.519 | 1.076 | -1.099 | -0.306 | -0.091 | -0.07 | -0.07 | 0 | 0.5 | 0.08 | 0.53 | 0.55 | 0.49 | 0.61 |
| 7 | 79729075 | rs10238400 | 0.515 | 1.163 | -0.974 | -0.269 | -0.014 | 0.006 | 0.006 | 0.127 | 0.35 | 0.12 | 0.3 | 0.31 | 0.33 | 0.32 |
| 7 | 79736039 | rs10248649 | 0.514 | 1.389 | -1.008 | -0.044 | 0.025 | 0.037 | 0.037 | 1.889 | 0.88 | 0.71 | 0.88 | 0.86 | 0.83 | 0.93 |
| 7 | 79740668 | rs12673522 | 0.899 | 1.272 | -2.005 | -0.142 | -0.091 | -0.101 | -0.101 | 0 | 0.25 | 0.01 | 0.27 | 0.31 | 0.28 | 0.27 |
| 7 | 79750012 | rs10236183 | 0.482 | 0.574 | -1.511 | 0.181 | 0.06 | -0.096 | -0.096 | 1.93 | 0.37 | 0.11 | 0.54 | 0.56 | 0.27 | 0.56 |
| 7 | 79750613 | rs1011971 | 0.271 | 0.656 | -1.407 | -0.334 | 0.057 | 0.031 | 0.031 | 0 | 0.39 | 0.21 | 0.58 | 0.69 | 0.3 | 0.6 |
| 7 | 79755875 | rs6979946 | 0.813 | 1.256 | -0.571 | -0.168 | 0.008 | -0.034 | -0.034 | 0 | 0.9 | 0.83 | 0.91 | 1 | 0.84 | 0.97 |
| 7 | 79759832 | rs6948782 | -0.863 | 0.999 | -0.628 | -0.466 | 0 | -0.005 | -0.005 | 0.413 | 0.4 | 0.7 | 0.36 | 0.25 | 0.39 | 0.44 |
| 7 | 79760082 | rs2079162 | 0.785 | 1.002 | -0.424 | -0.364 | 0 | 0.016 | 0.016 | 1.533 | 0.7 | 0.52 | 0.66 | 0.44 | 0.68 | 0.58 |
| 7 | 79763302 | rs2714465 | 1.536 | 1.046 | -0.704 | -0.45 | -0.033 | 0.197 | 0.197 | 3.118 | 0.51 | 0.11 | 0.48 | 0.24 | 0.48 | 0.37 |
| 7 | 79765622 | rs2428469 | 1.411 | 0.519 | -0.082 | 0.035 | 0.01 | -0.001 | -0.001 | 0.011 | 0.65 | 0.32 | 0.72 | 0.36 | 0.7 | 0.73 |
| 7 | 79766351 | rs2714458 | 1.182 | 0.142 | -0.279 | 0.062 | -0.055 | 0.053 | 0.053 | 0 | 0.72 | 0.48 | 0.8 | 0.43 | 0.82 | 0.76 |
| 7 | 79770436 | rs1034702 | 0.508 | 0.224 | -0.53 | 0.401 | -0.011 | 0.307 | 0.307 | 0 | 0.95 | 0.81 | 0.89 | 0.48 | 0.96 | 0.9 |
| 7 | 79772027 | rs12706644 | 0.042 | 0.258 | -0.89 | 0.007 | 0.012 | 0.01 | 0.01 | 0 | 0.09 | 0.05 | 0.03 | 0.07 | 0.06 | 0.04 |
| 7 | 79774364 | rs2523186 | 1.425 | 0.325 | -1.638 | -0.254 | -0.022 | 0.057 | 0.057 | 0 | 0.39 | 0.06 | 0.42 | 0.24 | 0.35 | 0.32 |
| 7 | 79776428 | rs2523189 | 1.698 | -0.043 | -2.034 | -0.524 | -0.063 | -0.047 | -0.047 | 0.348 | 0.31 | 0.02 | 0.41 | 0.28 | 0.31 | 0.28 |
| 7 | 79779284 | rs17153496 | 0.775 | -0.207 | -1.638 | -0.693 | -0.02 | -0.002 | -0.002 | 0 | 0.06 | 0.11 | 0.02 | 0.06 | 0.01 | 0.03 |
| 7 | 79780292 | rs10274707 | -0.377 | -0.045 | -2.053 | -0.881 | -0.128 | 0.018 | 0.018 | 0 | 0.27 | 0.03 | 0.21 | 0.18 | 0.36 | 0.39 |
| 7 | 79786628 | rs2714457 | -1.479 | 0.108 | -1.54 | -1.016 | -0.005 | 0.015 | 0.015 | 0 | 0.37 | 0.65 | 0.33 | 0.25 | 0.42 | 0.35 |
| 7 | 79789180 | rs2714454 | -0.688 | 0.083 | -1.364 | -1.038 | -0.003 | -0.062 | -0.062 | 0 | 0.25 | 0.56 | 0.16 | 0.11 | 0.24 | 0.08 |
| 7 | 79792606 | rs17806203 | 0.055 | 0.149 | -1.362 | -0.974 | -0.047 | -0.023 | -0.023 | 0 | 0.11 | 0.03 | 0.17 | 0.13 | 0.18 | 0.26 |
| 7 | 79799613 | rs11980616 | 0.502 | 0.133 | -1.024 | -1.032 | 0.039 | -0.01 | -0.01 | 0 | 0.12 | 0.38 | 0.1 | 0.08 | 0.12 | 0.05 |
| 7 | 79800347 | rs916905 | -0.141 | -0.154 | -1.22 | -0.707 | -0.002 | 0.008 | 0.008 | 0.159 | 0.72 | 0.38 | 0.74 | 0.55 | 0.74 | 0.66 |
| 7 | 79809984 | rs10230368 | -2.01 | -0.158 | -0.225 | -1.17 | 0.01 | -0.024 | -0.024 | 0.03 | 0.29 | 0.53 | 0.28 | 0.24 | 0.33 | 0.31 |
| 7 | 79814927 | rs6956455 | 2.11 | -0.21 | -0.257 | -1.201 | -0.004 | 0.008 | 0.008 | 0 | 0.63 | 0.37 | 0.68 | 0.76 | 0.59 | 0.65 |
| 7 | 79817949 | rs4512317 | 2.33 | -0.446 | -0.117 | -1.239 | -0.001 | 0.054 | 0.054 | 0.08 | 0.6 | 0.34 | 0.68 | 0.8 | 0.61 | 0.64 |
| 7 | 79832633 | rs17245731 | -1.141 | -0.01 | 0.099 | -1.017 | -0.018 | 0.025 | 0.025 | 0 | 0.2 | 0.11 | 0.2 | 0.09 | 0.26 | 0.26 |
| 7 | 79834048 | rs12155252 | 0.823 | 0.027 | -0.071 | -1.045 | -0.007 | -0.03 | -0.03 | 0.083 | 0.92 | 0.86 | 0.97 | 1 | 0.93 | 0.96 |
| 7 | 79838358 | rs13222403 | -1.307 | 0.048 | -0.168 | -1.069 | 0.001 | -0.005 | -0.005 | 1.46 | 0.26 | 0.23 | 0.15 | 0.08 | 0.25 | 0.12 |
| 7 | 79840111 | rs3801357 | 0.156 | 0.106 | -0.777 | -1.078 | -0.031 | -0.017 | -0.017 | 0.127 | 0.06 | 0.01 | 0.12 | 0.08 | 0.12 | 0.23 |
| 7 | 79841313 | rs10237963 | 0.392 | 0.091 | -0.771 | -1.088 | -0.004 | -0.022 | -0.022 | 0 | 0.93 | 0.88 | 0.98 | 1 | 0.93 | 0.97 |
| 7 | 79841954 | rs10238438 | -1.27 | 0.108 | -0.784 | -1.087 | -0.046 | 0.033 | 0.033 | 0 | 0.22 | 0.08 | 0.22 | 0.17 | 0.29 | 0.34 |
| 7 | 79843305 | rs2886611 | -1.518 | 0.11 | -0.78 | -1.058 | -0.009 | -0.043 | -0.043 | 1.106 | 0.28 | 0.52 | 0.18 | 0.11 | 0.25 | 0.13 |
| 7 | 79847565 | rs17153599 | -0.865 | 0.123 | -0.773 | -1.045 | -0.006 | -0.044 | -0.044 | 0.308 | 0.23 | 0.53 | 0.16 | 0.11 | 0.2 | 0.12 |
| 7 | 79850207 | rs6973427 | 1.963 | 0.201 | -1.225 | -0.981 | -0.007 | 0.074 | 0.074 | 0 | 0.59 | 0.32 | 0.68 | 0.83 | 0.57 | 0.62 |
| 7 | 79854751 | rs2886612 | 0.413 | 0.375 | -1.114 | -0.977 | -0.001 | -0.051 | -0.051 | 0.592 | 0.82 | 0.4 | 0.9 | 0.96 | 0.83 | 0.94 |
| 7 | 79856411 | rs2886609 | 1.349 | 0.653 | -1.94 | -0.756 | -0.008 | 0.047 | 0.047 | 0.019 | 0.6 | 0.34 | 0.68 | 0.81 | 0.55 | 0.63 |
| 7 | 79857872 | rs7458619 | 0.618 | -0.493 | -1.83 | -0.803 | -0.001 | -0.011 | -0.011 | 0.058 | 0.7 | 0.37 | 0.82 | 0.89 | 0.72 | 0.86 |
| 7 | 79872721 | rs13243329 | 0.951 | -0.084 | -2.7 | -0.885 | 0.056 | 0.287 | 0.287 | 0.802 | 0.56 | 0.1 | 0.69 | 0.93 | 0.64 | 0.63 |
| 7 | 79872964 | rs4728040 | -0.982 | 0.705 | -1.685 | 0.418 | -0.006 | 0.01 | 0.01 | 0.473 | 0.09 | 0.04 | 0.1 | 0.06 | 0.11 | 0.25 |
| 7 | 79887271 | rs306699 | 0.541 | 1.177 | -1.626 | 1.153 | -0.042 | -0.047 | -0.047 | 0 | 0.91 | 1 | 0.72 | 0.57 | 0.84 | 0.86 |
| 7 | 79888510 | rs2077759 | -0.364 | 1.743 | -2.093 | 0.001 | -0.075 | 0.044 | 0.044 | 0 | 0.25 | 0.1 | 0.18 | 0.16 | 0.33 | 0.27 |
| 7 | 79892337 | rs306705 | -0.666 | 0.388 | -1.776 | -0.65 | 0.003 | 0.075 | 0.075 | 0 | 0.49 | 0.44 | 0.29 | 0.21 | 0.45 | 0.36 |
| 7 | 79893373 | rs515693 | 0.144 | 0.435 | -1.883 | -0.895 | 0.035 | 0.054 | 0.054 | 0 | 0.29 | 0.18 | 0.13 | 0.05 | 0.19 | 0.11 |

| | | | | | | | | | | | | | | | | |
|---|---|---|---|---|---|---|---|---|---|---|---|---|---|---|---|---|
| 7 | 79894529 | rs306710 | -1.198 | 0.589 | -1.024 | -1.221 | -0.004 | 0.085 | 0.085 | 0 | 0.55 | 0.74 | 0.34 | 0.21 | 0.5 | 0.37 |
| 7 | 79895883 | rs306713 | 0.107 | 0.598 | -1.163 | -1.223 | -0.026 | 0.016 | 0.016 | 0 | 0.22 | 0.24 | 0.19 | 0.16 | 0.31 | 0.26 |
| 7 | 79898931 | rs1828522 | 0.309 | 0.536 | -0.57 | -1.165 | 0.022 | 0.028 | 0.028 | 0 | 0.29 | 0.23 | 0.13 | 0.05 | 0.19 | 0.1 |
| 7 | 79900518 | rs182524 | 0.327 | 0.84 | -0.618 | -1.177 | -0.028 | 0.011 | 0.011 | 1.704 | 0.19 | 0.15 | 0.16 | 0.16 | 0.3 | 0.26 |
| 7 | 79902287 | rs306721 | 0.225 | 0.694 | -0.547 | -1.177 | 0.008 | -0.011 | -0.011 | 0 | 0.21 | 0.28 | 0.14 | 0.16 | 0.25 | 0.24 |
| 7 | 79903870 | rs4728044 | 0.442 | 0.206 | -1.091 | -1.595 | 0.053 | 0.159 | 0.159 | 0 | 0.53 | 0.32 | 0.72 | 0.93 | 0.61 | 0.61 |
| 7 | 79904808 | rs306694 | -0.342 | 0.174 | -0.82 | -1.131 | -0.008 | -0.03 | -0.03 | 1.016 | 0.85 | 0.74 | 0.87 | 0.93 | 0.84 | 0.72 |
| 7 | 79911122 | rs10246790 | -0.052 | 0.498 | -1.516 | -1.208 | -0.067 | 0.032 | 0.032 | 2.291 | 0.23 | 0.04 | 0.1 | 0.18 | 0.24 | 0.23 |
| 7 | 79912682 | rs4731318 | -0.356 | -0.355 | -1.205 | -1.339 | 0.048 | 0.055 | 0.055 | 0 | 0.48 | 0.61 | 0.21 | 0.27 | 0.29 | 0.33 |
| 7 | 79924081 | rs17153690 | 1.442 | -0.136 | -1.734 | -1.376 | -0.024 | 0.036 | 0.036 | 0.169 | 0.22 | 0.02 | 0.19 | 0.14 | 0.19 | 0.36 |
| 7 | 79926885 | rs673193 | 0.294 | -0.217 | -0.9 | -1.548 | 0.033 | 0.062 | 0.062 | 0 | 0.43 | 0.19 | 0.73 | 0.73 | 0.63 | 0.52 |
| 7 | 79930984 | rs11767908 | 0.905 | -0.876 | -0.906 | -1.76 | -0.004 | -0.008 | -0.008 | 0 | 0.1 | 0.12 | 0.01 | 0 | 0.02 | 0.06 |
| 7 | 79934744 | rs504540 | 0.627 | -0.378 | -0.809 | -1.579 | -0.003 | 0.045 | 0.045 | 0 | 0.4 | 0.16 | 0.71 | 0.73 | 0.58 | 0.49 |
| 7 | 79937894 | rs4731336 | -0.04 | -0.352 | -0.72 | -1.59 | 0.004 | 0.005 | 0.005 | 0 | 0.52 | 0.58 | 0.23 | 0.27 | 0.37 | 0.37 |
| 7 | 79945112 | rs37920 | 0.254 | -0.277 | -0.614 | -1.506 | -0.005 | -0.046 | -0.046 | 0.222 | 0.09 | 0.25 | 0.06 | 0 | 0.06 | 0.15 |
| 7 | 79952755 | rs304732 | -0.632 | -0.296 | -0.708 | -1.527 | -0.007 | 0.036 | 0.036 | 0 | 0.61 | 0.77 | 0.29 | 0.26 | 0.44 | 0.52 |
| 7 | 79953523 | rs304733 | -0.281 | -0.027 | -0.107 | -1.098 | 0.004 | 0.053 | 0.053 | 0 | 0.09 | 0.32 | 0.09 | 0.09 | 0.07 | 0.15 |
| 7 | 79955724 | rs304735 | -1.119 | 0.479 | -0.567 | -0.758 | 0.046 | 0.042 | 0.042 | 0 | 0.82 | 0.87 | 0.49 | 0.43 | 0.62 | 0.88 |
| 7 | 79961091 | rs304737 | -1.005 | 0.641 | -0.728 | -0.046 | -0.038 | -0.06 | -0.06 | 0.001 | 0.81 | 0.9 | 0.53 | 0.57 | 0.7 | 0.85 |
| 7 | 79968083 | rs304739 | 0.254 | -0.22 | -0.573 | -0.374 | 0.001 | -0.022 | -0.022 | 0 | 0.18 | 0.14 | 0.09 | 0 | 0.15 | 0.13 |
| 7 | 79973826 | rs10227851 | 1.054 | -0.198 | -0.673 | -0.305 | 0.009 | 0.005 | 0.005 | 0.282 | 0.17 | 0.06 | 0.06 | 0.18 | 0.12 | 0.22 |
| 7 | 79982423 | rs1511682 | -0.566 | -0.381 | -0.463 | -0.366 | -0.018 | -0.038 | -0.038 | 0.2 | 0.33 | 0.4 | 0.15 | 0.2 | 0.26 | 0.35 |
| 7 | 79991029 | rs304761 | -1.823 | -0.082 | -0.628 | -0.217 | -0.053 | -0.055 | -0.055 | 0 | 0.81 | 0.9 | 0.51 | 0.61 | 0.65 | 0.87 |
| 7 | 79991379 | rs6467126 | -0.869 | -1.014 | -0.034 | -0.537 | 0.05 | 0.033 | 0.033 | 0 | 0.45 | 0.67 | 0.21 | 0.34 | 0.32 | 0.39 |
| 7 | 79993965 | rs6467133 | -0.276 | -1.039 | 0.087 | -0.653 | -0.037 | -0.041 | -0.041 | 0 | 0.28 | 0.55 | 0.14 | 0.14 | 0.19 | 0.16 |
| 7 | 79997202 | rs304768 | 0.317 | -0.847 | 0.223 | -0.432 | -0.003 | -0.031 | -0.031 | 0.715 | 0.11 | 0.18 | 0.06 | 0 | 0.11 | 0.08 |
| 7 | 79997946 | rs2906199 | -0.582 | -0.86 | 0.335 | -0.419 | -0.003 | 0.007 | 0.007 | 0.439 | 0.13 | 0.23 | 0.08 | 0.14 | 0.14 | 0.13 |
| 7 | 79998853 | rs10265717 | 0.902 | -0.896 | 0.331 | -0.447 | -0.005 | 0.002 | 0.002 | 0.079 | 0.17 | 0.12 | 0.06 | 0.2 | 0.13 | 0.22 |
| 7 | 79999975 | rs10270183 | 0.369 | -0.997 | 0.116 | -0.748 | -0.033 | -0.105 | -0.105 | 1.845 | 0.22 | 0.34 | 0.08 | 0 | 0.12 | 0.11 |
| 7 | 80002614 | rs304774 | -0.43 | -0.916 | 0.155 | -0.663 | 0.015 | 0.044 | 0.044 | 0.489 | 0.28 | 0.23 | 0.11 | 0.17 | 0.24 | 0.3 |
| 7 | 80003498 | rs7793055 | 0.203 | -0.895 | 0.097 | -0.693 | -0.012 | -0.078 | -0.078 | 0.609 | 0.24 | 0.29 | 0.08 | 0 | 0.15 | 0.16 |
| 7 | 80004495 | rs304776 | -1.405 | -0.807 | -0.231 | -0.612 | 0.04 | 0.083 | 0.083 | 0 | 0.53 | 0.8 | 0.23 | 0.31 | 0.39 | 0.47 |
| 7 | 80016757 | rs304782 | 0.245 | -0.783 | -0.253 | -0.589 | -0.007 | -0.047 | -0.047 | 0 | 0.11 | 0.2 | 0.06 | 0.01 | 0.11 | 0.08 |
| 7 | 80018114 | rs304783 | -1.008 | -0.433 | -0.377 | -0.361 | 0.037 | 0.087 | 0.087 | 0 | 0.53 | 0.84 | 0.24 | 0.31 | 0.39 | 0.46 |
| 7 | 80025778 | rs304797 | -0.474 | -0.483 | -0.74 | -0.207 | 0.034 | 0.049 | 0.049 | 0 | 0.36 | 0.75 | 0.17 | 0.16 | 0.26 | 0.24 |
| 7 | 80030601 | rs573316 | -0.186 | -0.069 | -0.127 | 0.324 | -0.017 | -0.013 | -0.013 | 0 | 0.16 | 0.43 | 0.1 | 0.14 | 0.08 | 0.14 |
| 7 | 80040648 | rs471683 | -0.284 | -0.094 | -0.117 | 0.519 | -0.021 | 0.057 | 0.057 | 0.202 | 0.16 | 0.46 | 0.11 | 0.23 | 0.08 | 0.13 |
| 7 | 80042917 | rs534997 | -0.08 | -0.094 | 0.028 | 0.501 | -0.005 | -0.03 | -0.03 | 0 | 0.15 | 0.35 | 0.09 | 0.13 | 0.08 | 0.14 |
| 7 | 80048871 | rs10246082 | 0.05 | -0.085 | 0.063 | 0.499 | -0.004 | -0.015 | -0.015 | 1.152 | 0.14 | 0.26 | 0.08 | 0.13 | 0.08 | 0.13 |
| 7 | 80051323 | rs17153898 | 0.022 | -0.082 | -0.081 | 0.544 | -0.003 | 0.087 | 0.087 | 0.421 | 0.14 | 0.28 | 0.1 | 0.23 | 0.08 | 0.14 |
| 7 | 80053909 | rs7785386 | -0.032 | -0.012 | -0.324 | 0.555 | -0.008 | 0.085 | 0.085 | 0.062 | 0.16 | 0.35 | 0.1 | 0.23 | 0.08 | 0.14 |
| 7 | 80054945 | rs12535593 | 0.466 | 0.433 | -1.335 | 0.905 | -0.026 | -0.017 | -0.017 | 0 | 0.25 | 0.06 | 0.25 | 0.24 | 0.23 | 0.37 |
| 7 | 80055910 | rs6467158 | -0.067 | 0.789 | -1.06 | 0.687 | -0.014 | -0.006 | -0.006 | 0 | 0.59 | 0.62 | 0.45 | 0.6 | 0.5 | 0.7 |
| 7 | 80060495 | rs10441382 | -0.301 | 0.947 | -1.115 | 0.699 | -0.028 | -0.007 | -0.007 | 0 | 0.4 | 0.31 | 0.4 | 0.42 | 0.45 | 0.55 |
| 7 | 80065677 | rs704871 | 0.103 | 0.557 | -0.366 | 0.456 | -0.002 | -0.001 | -0.001 | 0 | 0.71 | 0.65 | 0.73 | 0.72 | 0.78 | 0.75 |
| 7 | 80073310 | rs813348 | 0.218 | 0.592 | -0.244 | 0.405 | -0.001 | -0.001 | -0.001 | 0.003 | 0.66 | 0.6 | 0.71 | 0.72 | 0.72 | 0.75 |
| 7 | 80074216 | rs808956 | 0.003 | 0.596 | -0.366 | 0.451 | 0.009 | 0.021 | 0.021 | 0 | 0.69 | 0.88 | 0.75 | 0.75 | 0.74 | 0.77 |
| 7 | 80075157 | rs799964 | 0.303 | 0.572 | 0.199 | 0.427 | -0.016 | -0.016 | -0.016 | 0 | 0.66 | 0.57 | 0.71 | 0.72 | 0.72 | 0.75 |
| 7 | 80082966 | rs799974 | -0.011 | 0.781 | -0.469 | 0.349 | 0 | 0.019 | 0.019 | 0.781 | 0.72 | 0.82 | 0.78 | 0.79 | 0.74 | 0.79 |
| 7 | 80088592 | rs17147968 | -0.601 | 1.635 | -0.484 | 1.085 | -0.066 | -0.073 | -0.073 | 0 | 0.18 | 0.12 | 0.4 | 0.59 | 0.3 | 0.33 |
| 7 | 80098487 | rs799930 | 0.5 | 1.601 | -0.278 | 1.068 | -0.024 | -0.027 | -0.027 | 0 | 0.81 | 0.8 | 0.59 | 0.42 | 0.7 | 0.66 |
| 7 | 80098568 | rs12671431 | -0.725 | 0.579 | 0.215 | 0.831 | -0.02 | 0.017 | 0.017 | 0.906 | 0.06 | 0.11 | 0.03 | 0.07 | 0.02 | 0.02 |
| 7 | 80104891 | rs6944302 | -0.932 | 0.765 | 1.091 | 0.644 | 0.02 | -0.042 | -0.042 | 0.605 | 0.1 | 0.31 | 0.06 | 0 | 0.07 | 0.04 |
| 7 | 80107798 | rs6467192 | -2.103 | 0.488 | 0.905 | 0.416 | 0 | 0.001 | 0.001 | 0.19 | 0.19 | 0.73 | 0.11 | 0.13 | 0.13 | 0.13 |
| 7 | 80119690 | rs11978272 | -1.264 | 0.482 | 1.5 | 0.401 | 0.054 | 0.003 | 0.003 | 0.419 | 0.07 | 0.38 | 0.05 | 0 | 0.03 | 0 |
| 7 | 80121054 | rs10234980 | -0.373 | 0.443 | 1.523 | 0.38 | -0.02 | -0.048 | -0.048 | 0 | 0.09 | 0.15 | 0.04 | 0 | 0.05 | 0.03 |
| 7 | 80123999 | rs6975345 | -2.378 | 0.291 | 1.383 | 0.355 | 0.006 | 0.006 | 0.006 | 0.788 | 0.2 | 0.77 | 0.12 | 0.13 | 0.13 | 0.14 |
| 7 | 80134355 | rs1404314 | -0.789 | 0.352 | 1.434 | 0.429 | 0.028 | 0.057 | 0.057 | 0 | 0.1 | 0.19 | 0.05 | 0.13 | 0.1 | 0.06 |
| 7 | 80135050 | rs7800673 | -0.778 | 0.356 | 1.484 | 0.45 | 0.022 | 0.064 | 0.064 | 0 | 0.1 | 0.2 | 0.06 | 0.13 | 0.09 | 0.11 |
| 7 | 80137548 | rs10276559 | -0.716 | 0.326 | 1.469 | 0.448 | 0.012 | 0.019 | 0.019 | 0 | 0.1 | 0.16 | 0.06 | 0.09 | 0.1 | 0.06 |
| 7 | 80138074 | rs6970109 | -1.615 | 0.3 | 1.695 | 0.439 | 0.007 | 0.048 | 0.048 | 0 | 0.13 | 0.48 | 0.08 | 0.13 | 0.1 | 0.11 |
| 7 | 80141069 | rs10230573 | -0.787 | 0.17 | 0.764 | 0.185 | 0.01 | 0.028 | 0.028 | 0 | 0.53 | 0.54 | 0.66 | 0.7 | 0.6 | 0.5 |
| 7 | 80142245 | rs7805711 | -0.177 | 0.493 | 0.623 | 0.207 | 0.003 | -0.007 | -0.007 | 0 | 0.42 | 0.53 | 0.6 | 0.65 | 0.49 | 0.51 |
| 7 | 80143428 | rs17260769 | -0.068 | 0.418 | 0.502 | 0.202 | -0.008 | -0.001 | -0.001 | 0 | 0.41 | 0.36 | 0.59 | 0.61 | 0.49 | 0.46 |
| 7 | 80146139 | rs10232443 | -1 | 0.356 | 0.077 | 0.121 | -0.01 | 0.006 | 0.006 | 0 | 0.31 | 0.43 | 0.33 | 0.68 | 0.27 | 0.3 |
| 7 | 80150131 | rs1107660 | -1.159 | 0.313 | -0.408 | 0.228 | -0.044 | 0.043 | 0.043 | 0 | 0.6 | 0.9 | 0.69 | 0.73 | 0.63 | 0.56 |
| 7 | 80150594 | rs940541 | -1.223 | 0.259 | -0.451 | 0.229 | -0.053 | 0.057 | 0.057 | 0 | 0.61 | 0.89 | 0.7 | 0.75 | 0.63 | 0.58 |
| 7 | 80156314 | rs2366735 | -0.821 | 0.532 | 0.041 | 0.955 | 0.012 | 0.073 | 0.073 | 0 | 0.51 | 0.65 | 0.64 | 0.7 | 0.57 | 0.44 |
| 7 | 80157564 | rs6959775 | -0.239 | 0.392 | 0.131 | 1.043 | -0.018 | -0.014 | -0.014 | 0.219 | 0.39 | 0.34 | 0.6 | 0.6 | 0.55 | 0.36 |

| | | | | | | | | | | | | | | | | |
|---|---|---|---|---|---|---|---|---|---|---|---|---|---|---|---|---|
| 7 | 80158861 | rs7455906 | -1.442 | 0.537 | -0.445 | 1.608 | 0.009 | -0.036 | -0.036 | 4.71 | 0.26 | 0.72 | 0.15 | 0.44 | 0.2 | 0.21 |
| 7 | 80160041 | rs6974934 | -0.848 | 0.18 | -0.378 | 1.637 | -0.059 | 0.01 | 0.01 | 0 | 0.56 | 0.93 | 0.67 | 0.7 | 0.6 | 0.43 |
| 7 | 80164380 | rs10282644 | 0.195 | 0.415 | 0.268 | 1.765 | 0 | 0.022 | 0.022 | 0 | 0.54 | 0.58 | 0.49 | 0.67 | 0.46 | 0.73 |
| 7 | 80167254 | rs4112274 | -1.02 | 0.398 | 0.496 | 2.085 | 0.041 | 0.261 | 0.261 | 0.185 | 0.06 | 0.26 | 0.12 | 0.38 | 0.06 | 0.26 |
| 7 | 80169404 | rs10215825 | 0.722 | 0.347 | 0.413 | 1.945 | 0.034 | 0.105 | 0.105 | 0 | 0.47 | 0.15 | 0.36 | 0.25 | 0.4 | 0.4 |
| 7 | 80177084 | rs819454 | 0.414 | 0.398 | 0.637 | 2.102 | 0.001 | 0.001 | 0.001 | 0 | 0.5 | 0.41 | 0.46 | 0.37 | 0.44 | 0.44 |
| 7 | 80181115 | rs1464794 | 0.753 | 0.421 | 0.338 | 2.167 | 0.033 | 0.077 | 0.077 | 0 | 0.47 | 0.15 | 0.36 | 0.27 | 0.41 | 0.41 |
| 7 | 80197286 | rs819457 | -0.021 | 0.397 | 0.288 | 1.928 | -0.099 | 0.028 | 0.028 | 0 | 0.58 | 0.84 | 0.52 | 0.69 | 0.5 | 0.73 |
| 7 | 80211423 | rs7779873 | 0.405 | 0.47 | 0.595 | 2.039 | 0.015 | 0.04 | 0.04 | 0 | 0.51 | 0.58 | 0.42 | 0.58 | 0.45 | 0.69 |
| 7 | 80220608 | rs2065668 | 0.748 | 0.114 | 0.559 | 2.046 | 0.049 | 0.041 | 0.041 | 0 | 0.45 | 0.3 | 0.33 | 0.34 | 0.36 | 0.43 |
| 7 | 80223996 | rs819437 | 0.004 | 0.163 | 0.615 | 2.294 | -0.01 | 0.066 | 0.066 | 0 | 0.54 | 0.73 | 0.47 | 0.69 | 0.46 | 0.71 |
| 7 | 80227837 | rs10499857 | -0.493 | 0.198 | 0.876 | 2.337 | 0.047 | 0.259 | 0.259 | 0 | 0.05 | 0.2 | 0.11 | 0.34 | 0.06 | 0.28 |
| 7 | 80236014 | rs13236689 | 0.291 | 0.19 | 0.937 | 2.337 | 0.015 | 0.014 | 0.014 | 0 | 0.5 | 0.45 | 0.36 | 0.6 | 0.42 | 0.68 |
| 7 | 80239418 | rs1761663 | 0.356 | 0.159 | 0.932 | 2.198 | 0.042 | 0.048 | 0.048 | 0 | 0.49 | 0.52 | 0.36 | 0.35 | 0.4 | 0.43 |
| 7 | 80252518 | rs4545029 | -0.199 | 0.188 | 0.849 | 2.134 | 0 | 0.003 | 0.003 | 0.001 | 0.45 | 0.29 | 0.45 | 0.31 | 0.53 | 0.29 |
| 7 | 80262830 | rs10216184 | 0.741 | -0.044 | 0.81 | 2.162 | 0.002 | 0.002 | 0.002 | 0 | 0.43 | 0.27 | 0.32 | 0.25 | 0.37 | 0.4 |
| 7 | 80266068 | rs1334511 | -0.626 | -0.013 | 0.72 | 2.345 | 0.072 | 0.298 | 0.298 | 0 | 0.05 | 0.27 | 0.12 | 0.35 | 0.07 | 0.27 |
| 7 | 80271108 | rs1534314 | -0.731 | -0.016 | 0.669 | 2.363 | 0.056 | 0.307 | 0.307 | 0.066 | 0.06 | 0.26 | 0.19 | 0.44 | 0.07 | 0.3 |
| 7 | 80273308 | rs3212160 | -0.042 | -0.041 | 0.62 | 2.296 | 0 | 0.07 | 0.07 | 0 | 0.47 | 0.22 | 0.46 | 0.31 | 0.54 | 0.3 |
| 7 | 80275455 | rs1049654 | -0.229 | -0.221 | 0.599 | 2.201 | 0.02 | 0.06 | 0.06 | 0 | 0.49 | 0.37 | 0.47 | 0.31 | 0.54 | 0.3 |
| 7 | 80278465 | rs3211820 | -0.656 | -0.142 | 0.313 | 2.225 | -0.012 | -0.012 | -0.012 | 0 | 0.57 | 0.6 | 0.59 | 0.64 | 0.61 | 0.57 |
| 7 | 80279460 | rs3211827 | 0.585 | -0.117 | -0.379 | 2.294 | 0.042 | 0.082 | 0.082 | 0 | 0.43 | 0.1 | 0.3 | 0.25 | 0.36 | 0.4 |
| 7 | 80285850 | rs3173798 | -0.776 | 0.159 | -0.217 | 2.436 | 0.05 | 0.282 | 0.282 | 0 | 0.06 | 0.23 | 0.19 | 0.42 | 0.07 | 0.3 |
| 7 | 80288854 | rs3211881 | 0.327 | -0.017 | -0.543 | 1.802 | 0.025 | 0.007 | 0.007 | 0 | 0.11 | 0.06 | 0.05 | 0.07 | 0.06 | 0.07 |
| 7 | 80289402 | rs3211885 | 0.167 | 0.26 | -0.398 | 1.864 | 0.024 | 0.009 | 0.009 | 0 | 0.48 | 0.38 | 0.42 | 0.52 | 0.42 | 0.67 |
| 7 | 80298173 | rs3211931 | 0.308 | 0.544 | -1.304 | 1.783 | -0.011 | -0.051 | -0.051 | 0 | 0.45 | 0.2 | 0.38 | 0.51 | 0.44 | 0.48 |
| 7 | 80304072 | rs3211958 | 0.25 | 0.539 | -1.286 | 1.651 | -0.057 | -0.065 | -0.065 | 0.104 | 0.44 | 0.12 | 0.37 | 0.44 | 0.44 | 0.46 |
| 7 | 80307224 | rs12706949 | 0.325 | 0.561 | -1.282 | 1.647 | -0.042 | -0.046 | -0.046 | 0 | 0.43 | 0.13 | 0.37 | 0.43 | 0.44 | 0.45 |
| 7 | 80341163 | rs11971792 | -0.679 | 0.278 | -1.012 | 2.121 | -0.012 | -0.027 | -0.027 | 0.07 | 0.1 | 0.11 | 0.19 | 0.42 | 0.11 | 0.16 |
| 7 | 80351673 | rs954410 | -1.611 | -0.132 | -0.934 | 1.908 | 0.004 | -0.013 | -0.013 | 0 | 0.16 | 0.19 | 0.2 | 0.42 | 0.12 | 0.21 |
| 7 | 80357963 | rs16886849 | -0.588 | -0.119 | -0.932 | 2.05 | -0.006 | -0.012 | -0.012 | 0 | 0.1 | 0.13 | 0.12 | 0.2 | 0.11 | 0.15 |
| 7 | 80370829 | rs1527470 | 1.105 | -0.066 | -0.834 | 1.115 | -0.068 | 0.211 | 0.211 | 0 | 0.41 | 0.11 | 0.32 | 0.08 | 0.44 | 0.34 |
| 7 | 80373728 | rs12113860 | -0.131 | 0.074 | -0.72 | 1.25 | 0.013 | -0.001 | -0.001 | 0 | 0.93 | 0.73 | 0.99 | 1 | 0.99 | 0.95 |
| 7 | 80374438 | rs1949972 | 0.161 | 0.07 | -0.718 | 1.322 | 0.006 | 0.215 | 0.215 | 1.278 | 0.92 | 0.69 | 0.9 | 0.77 | 0.99 | 0.94 |
| 7 | 80375186 | rs1358340 | 0.191 | -0.075 | -0.332 | 1.112 | 0 | 0.01 | 0.01 | 0 | 0.58 | 0.36 | 0.44 | 0.3 | 0.56 | 0.65 |
| 7 | 80382592 | rs3778674 | -0.264 | 0.19 | -0.892 | 1.202 | -0.002 | -0.005 | -0.005 | 0 | 0.13 | 0.08 | 0.1 | 0.15 | 0.11 | 0.2 |
| 7 | 80384050 | rs12706974 | -0.162 | 0.217 | -0.838 | 1.385 | -0.023 | -0.041 | -0.041 | 0 | 0.37 | 0.27 | 0.45 | 0.52 | 0.43 | 0.33 |
| 7 | 80384926 | rs3807093 | 0.006 | 0.716 | -0.893 | 1.733 | -0.017 | -0.02 | -0.02 | 0 | 0.13 | 0.1 | 0.31 | 0.16 | 0.21 | 0.1 |
| 7 | 80385604 | rs10247306 | -1.593 | 0.214 | -1.219 | 1.41 | 0.022 | 0.138 | 0.138 | 0 | 0.53 | 0.77 | 0.78 | 0.74 | 0.58 | 0.53 |
| 7 | 80385987 | rs6978637 | -0.963 | -0.523 | -0.609 | 2.244 | 0.01 | -0.016 | -0.016 | 0 | 0.24 | 0.76 | 0.21 | 0.51 | 0.11 | 0.23 |
| 7 | 80387360 | rs10233114 | -0.05 | -0.486 | 0.008 | 1.948 | -0.008 | 0.012 | 0.012 | 0 | 0.07 | 0.17 | 0.04 | 0.05 | 0.02 | 0.07 |
| 7 | 80387766 | rs1058425 | -2.209 | -0.597 | -0.324 | 2.269 | 0.016 | -0.01 | -0.01 | 0 | 0.34 | 0.65 | 0.33 | 0.43 | 0.19 | 0.36 |
| 7 | 80387949 | rs2272353 | -0.119 | -0.307 | -0.194 | 2.477 | -0.009 | 0.003 | 0.003 | 0 | 0.11 | 0.06 | 0.15 | 0.11 | 0.12 | 0.11 |
| 7 | 80388136 | rs3807094 | -0.993 | -0.25 | -0.354 | 2.516 | -0.02 | -0.042 | -0.042 | 0 | 0.2 | 0.43 | 0.21 | 0.16 | 0.15 | 0.19 |
| 7 | 80389412 | rs3807097 | 0.015 | -0.021 | -0.459 | 2.544 | -0.026 | 0.055 | 0.055 | 1.4 | 0.11 | 0.17 | 0.12 | 0.27 | 0.04 | 0.17 |
| 7 | 80401470 | rs10266458 | -1.436 | 0.536 | -0.497 | 1.852 | -0.023 | 0.057 | 0.057 | 0 | 0.24 | 0.52 | 0.25 | 0.43 | 0.16 | 0.3 |
| 7 | 80403185 | rs12671126 | -0.795 | 1.165 | -1.169 | 2.045 | -0.009 | -0.065 | -0.065 | 0 | 0.16 | 0.15 | 0.21 | 0.46 | 0.17 | 0.32 |
| 7 | 80407801 | rs13224811 | 1.277 | 1.555 | -1.781 | 1.314 | -0.007 | 0.137 | 0.137 | 0.376 | 0.44 | 0.12 | 0.28 | 0.25 | 0.39 | 0.38 |
| 7 | 80411104 | rs3778677 | -1.586 | 2.508 | -1.344 | 1.859 | -0.003 | -0.002 | -0.002 | 0 | 0.26 | 0.32 | 0.4 | 0.53 | 0.32 | 0.39 |
| 7 | 80418689 | rs17275986 | 0.34 | 2.577 | -1.358 | 1.735 | -0.02 | -0.007 | -0.007 | 2.969 | 0.08 | 0.06 | 0.11 | 0.11 | 0.18 | 0.11 |
| 7 | 80425858 | rs13242982 | 0.035 | 2.71 | -1.339 | 1.303 | -0.009 | -0.004 | -0.004 | 0 | 0.07 | 0.02 | 0.1 | 0.06 | 0.11 | 0.09 |
| 7 | 80427495 | rs1880959 | 0.115 | 2.601 | -1.44 | 1.24 | -0.017 | 0.019 | 0.019 | 0 | 0.07 | 0.07 | 0.05 | 0 | 0.09 | 0.02 |
| 7 | 80444576 | rs12668734 | -1.336 | 2.713 | -1.325 | 1.373 | 0.005 | 0.012 | 0.012 | 0.17 | 0.25 | 0.34 | 0.4 | 0.55 | 0.32 | 0.4 |
| 7 | 80451440 | rs17154508 | -0.384 | 2.708 | -1.149 | 1.345 | 0.025 | -0.026 | -0.026 | 0 | 0.09 | 0.25 | 0.19 | 0.06 | 0.1 | 0.07 |
| 7 | 80451815 | rs12535655 | 0.373 | 2.831 | -1.615 | 1.195 | -0.025 | 0.038 | 0.038 | 0 | 0.14 | 0 | 0.09 | 0.03 | 0.13 | 0.09 |
| 7 | 80453333 | rs2030925 | -0.149 | 2.679 | -1.357 | 1.339 | -0.02 | -0.03 | -0.03 | 0 | 0.15 | 0.1 | 0.23 | 0.35 | 0.23 | 0.27 |
| 7 | 80460016 | rs2040908 | 0.489 | 2.087 | -1.324 | 0.821 | 0.018 | 0.041 | 0.041 | 0.224 | 0.74 | 0.6 | 0.59 | 0.44 | 0.68 | 0.6 |
| 7 | 80468127 | rs11767508 | -0.845 | 1.484 | -1.514 | 0.378 | -0.039 | 0.033 | 0.033 | 0 | 0.22 | 0.05 | 0.19 | 0.13 | 0.25 | 0.16 |
| 7 | 80468801 | rs17228184 | -0.58 | 1.454 | -1.42 | 0.257 | -0.02 | 0.054 | 0.054 | 0 | 0.19 | 0.06 | 0.18 | 0.07 | 0.21 | 0.13 |
| 7 | 80471535 | rs4731861 | 0.786 | 2.065 | -0.657 | 0.487 | 0.056 | 0.006 | 0.006 | 0 | 0.87 | 0.76 | 0.84 | 0.89 | 0.78 | 0.89 |
| 7 | 80474046 | rs1016840 | -0.77 | -0.084 | -0.819 | -0.217 | -0.021 | -0.029 | -0.029 | 0 | 0.42 | 0.48 | 0.2 | 0.21 | 0.21 | 0.31 |
| 7 | 80476677 | rs11980158 | -0.681 | -0.19 | -0.699 | -0.239 | 0.01 | -0.01 | -0.01 | 0 | 0.4 | 0.38 | 0.18 | 0.07 | 0.21 | 0.27 |
| 7 | 80476920 | rs12707036 | -0.967 | -0.29 | -0.692 | -0.085 | -0.028 | -0.032 | -0.032 | 0 | 0.43 | 0.51 | 0.2 | 0.22 | 0.22 | 0.31 |
| 7 | 80478992 | rs12707038 | -0.147 | -0.825 | -0.831 | -0.414 | 0.021 | 0.038 | 0.038 | 0 | 0.22 | 0.16 | 0.12 | 0.07 | 0.16 | 0.16 |
| 7 | 80479894 | rs12707039 | -1.293 | -0.811 | -0.836 | -0.411 | 0.011 | -0.009 | -0.009 | 0 | 0.4 | 0.38 | 0.19 | 0.08 | 0.21 | 0.27 |
| 7 | 80488068 | rs10954378 | -0.736 | -0.559 | -0.355 | -0.049 | -0.021 | -0.024 | -0.024 | 0 | 0.19 | 0.19 | 0.06 | 0.03 | 0.06 | 0.1 |
| 7 | 80494049 | rs11764532 | 0.257 | 0.401 | -0.498 | 0.24 | 0.218 | 0.147 | 0.147 | 0.608 | 0.86 | 0.6 | 0.68 | 0.75 | 0.74 | 0.81 |
| 7 | 80508422 | rs765764 | 1.524 | 0.438 | -0.618 | 0.327 | -0.029 | -0.018 | -0.018 | 1.169 | 0.66 | 0.23 | 0.59 | 0.61 | 0.69 | 0.69 |
| 7 | 80522906 | rs17154583 | -1.106 | 0.171 | -0.324 | 0.393 | -0.04 | -0.002 | -0.002 | 0 | 0.15 | 0.36 | 0.08 | 0.15 | 0.05 | 0.1 |

| | | | | | | | | | | | | | | | | |
|---|---|---|---|---|---|---|---|---|---|---|---|---|---|---|---|---|
| 7 | 80541773 | rs9641941 | -0.502 | 0.243 | -0.35 | 0.443 | -0.004 | 0.03 | 0.03 | 0 | 0.09 | 0.23 | 0.05 | 0.14 | 0.04 | 0.1 |
| 7 | 80544531 | rs12707079 | 1.656 | 0.301 | -0.767 | 0.391 | -0.033 | -0.023 | -0.023 | 0.289 | 0.63 | 0.18 | 0.61 | 0.62 | 0.7 | 0.67 |
| 7 | 80546745 | rs7793555 | 0.55 | 0.67 | -0.316 | 0.501 | 0.1 | 0.043 | 0.043 | 0 | 0.81 | 0.62 | 0.7 | 0.77 | 0.75 | 0.78 |
| 7 | 80553950 | rs6964151 | 0.741 | 0.335 | -0.461 | 0.669 | -0.003 | 0.014 | 0.014 | 0 | 0.44 | 0.22 | 0.61 | 0.36 | 0.57 | 0.53 |
| 7 | 80563385 | rs723581 | 0.145 | 0.046 | -0.221 | 1.096 | 0.001 | 0.008 | 0.008 | 0 | 0.86 | 0.84 | 0.89 | 0.81 | 0.85 | 0.89 |
| 7 | 80566789 | rs17154598 | 0.174 | -0.47 | 0.015 | 1.081 | -0.035 | -0.025 | -0.025 | 0 | 0.17 | 0.33 | 0.05 | 0.13 | 0.06 | 0.11 |
| 7 | 80569893 | rs10243928 | -0.6 | 0.064 | 0.176 | 1.207 | 0.001 | 0 | 0 | 0 | 0.25 | 0.24 | 0.19 | 0.3 | 0.29 | 0.24 |
| 7 | 80570562 | rs1019016 | 1.591 | -0.373 | 0.183 | 1.048 | -0.008 | -0.01 | -0.01 | 0 | 0.58 | 0.36 | 0.69 | 0.48 | 0.59 | 0.65 |
| 7 | 80583280 | rs10232696 | 0.228 | -0.346 | 0.121 | 1.078 | 0.013 | 0.03 | 0.03 | 0 | 0.12 | 0.19 | 0.11 | 0.19 | 0.13 | 0.1 |
| 7 | 80583482 | rs7783868 | 1.729 | -0.514 | -0.004 | 0.97 | -0.003 | -0.003 | -0.003 | 0 | 0.57 | 0.44 | 0.68 | 0.48 | 0.58 | 0.65 |
| 7 | 80587737 | rs740210 | 1.255 | -0.048 | 0.091 | 1.296 | 0.006 | 0.04 | 0.04 | 0 | 0.72 | 0.63 | 0.77 | 0.62 | 0.71 | 0.79 |
| 7 | 80588716 | rs853061 | 0.528 | 0.237 | -0.104 | 1.022 | -0.008 | 0.01 | 0.01 | 0 | 0.9 | 0.93 | 0.92 | 1 | 0.88 | 0.97 |
| 7 | 80590518 | rs853057 | 1.329 | 0.024 | -0.452 | 1.377 | -0.013 | 0.016 | 0.016 | 0.039 | 0.4 | 0.22 | 0.56 | 0.25 | 0.52 | 0.53 |
| 7 | 80598261 | rs7806898 | 1.256 | 0.315 | -1.096 | 1.866 | 0 | -0.001 | -0.001 | 0 | 0.45 | 0.37 | 0.55 | 0.41 | 0.51 | 0.52 |
| 7 | 80599142 | rs1358503 | 1.118 | 0.318 | -1.068 | 1.978 | -0.002 | 0.002 | 0.002 | 0 | 0.45 | 0.33 | 0.55 | 0.22 | 0.51 | 0.51 |
| 7 | 80599357 | rs4236644 | 0.443 | 0.251 | -1.226 | 1.964 | -0.013 | 0.006 | 0.006 | 0 | 0.63 | 0.44 | 0.8 | 0.58 | 0.75 | 0.73 |
| 7 | 80600158 | rs1528388 | 1.347 | 0.282 | -1.22 | 1.958 | 0 | -0.001 | -0.001 | 0 | 0.46 | 0.37 | 0.55 | 0.41 | 0.51 | 0.52 |
| 7 | 80606705 | rs6950903 | -0.029 | -0.293 | -1.346 | 1.552 | -0.009 | 0.004 | 0.004 | 0 | 0.18 | 0.11 | 0.25 | 0.16 | 0.26 | 0.21 |
| 7 | 80610569 | rs7788299 | 0.9 | -0.324 | -1.342 | 1.26 | 0.016 | 0.006 | 0.006 | 0 | 0.53 | 0.34 | 0.61 | 0.43 | 0.59 | 0.59 |
| 7 | 80611355 | rs1208092 | -0.687 | -0.589 | -1.012 | 0.945 | -0.059 | -0.028 | -0.028 | 1.421 | 0.28 | 0.49 | 0.13 | 0.24 | 0.13 | 0.21 |
| 7 | 80618038 | rs701325 | 0.296 | -0.241 | -1.926 | 1.527 | 0.022 | -0.098 | -0.098 | 0 | 0.78 | 0.92 | 0.82 | 0.57 | 0.81 | 0.71 |
| 7 | 80619575 | rs853051 | 0.099 | 0.456 | -2.589 | 0.821 | -0.034 | 0.051 | 0.051 | 1.027 | 0.23 | 0.13 | 0.16 | 0.11 | 0.26 | 0.12 |
| 7 | 80619607 | rs853050 | -0.02 | 0.09 | -2.35 | 1.62 | -0.002 | -0.033 | -0.033 | 0 | 0.79 | 0.97 | 0.84 | 0.82 | 0.81 | 0.84 |
| 7 | 80620397 | rs853048 | 0.352 | 0.24 | -2.352 | 1.42 | 0.027 | -0.122 | -0.122 | 0 | 0.74 | 0.92 | 0.69 | 0.57 | 0.76 | 0.69 |
| 7 | 80621583 | rs853045 | 0.063 | 0.677 | -2.411 | 1.514 | -0.091 | -0.123 | -0.123 | 1.533 | 0.27 | 0.04 | 0.24 | 0.35 | 0.35 | 0.25 |
| 7 | 80636021 | rs701326 | 0.151 | -0.218 | -1.35 | 1.322 | -0.022 | -0.004 | -0.004 | 0.112 | 0.69 | 0.43 | 0.69 | 0.7 | 0.76 | 0.68 |
| 7 | 80642423 | rs7801859 | -0.93 | -0.403 | 0.065 | 1.323 | 0.149 | 0.319 | 0.319 | 0.326 | 0.06 | 0.49 | 0.21 | 0.19 | 0.06 | 0.09 |
| 7 | 80648073 | rs4019352 | -0.136 | -0.565 | 0.046 | 1.215 | -0.005 | -0.011 | -0.011 | 0 | 0.75 | 0.79 | 0.9 | 0.89 | 0.82 | 0.78 |
| 7 | 80652059 | rs853107 | 1.207 | -1.052 | -0.435 | 1.55 | 0.099 | 0.185 | 0.185 | 0 | 0.5 | 0.1 | 0.4 | 0.28 | 0.66 | 0.35 |
| 7 | 80654259 | rs2108056 | -0.263 | -1.056 | 0.041 | 2.047 | -0.01 | 0.021 | 0.021 | 0 | 0.19 | 0.22 | 0.25 | 0.29 | 0.1 | 0.33 |
| 7 | 80660357 | rs10262336 | -0.237 | -1.101 | 0.371 | 2.225 | -0.024 | 0.098 | 0.098 | 0 | 0.11 | 0.2 | 0.24 | 0.3 | 0.07 | 0.29 |
| 7 | 80668458 | rs853072 | 0.494 | -1.083 | 0.48 | 2.141 | -0.033 | 0.022 | 0.022 | 0 | 0.64 | 0.61 | 0.64 | 0.59 | 0.74 | 0.49 |
| 7 | 80679694 | rs17298913 | 1.356 | -0.705 | -0.322 | 1.757 | -0.056 | 0.013 | 0.013 | 0 | 0.21 | 0.02 | 0.15 | 0.19 | 0.23 | 0.07 |
| 7 | 80681879 | rs1636593 | -0.976 | -1.023 | 0.034 | 1.818 | 0.001 | 0.001 | 0.001 | 0 | 0.77 | 0.83 | 0.88 | 0.89 | 0.81 | 0.78 |
| 7 | 80682961 | rs728106 | 0.49 | -0.834 | 0.088 | 1.579 | 0.1 | 0.032 | 0.032 | 0 | 0.59 | 0.2 | 0.6 | 0.45 | 0.73 | 0.5 |
| 7 | 80683308 | rs1636579 | 0.381 | -0.79 | 0.106 | 1.192 | 0.042 | -0.025 | -0.025 | 0 | 0.61 | 0.3 | 0.61 | 0.59 | 0.75 | 0.51 |
| 7 | 80699883 | rs6944056 | 0.699 | -0.249 | 0.358 | 0.861 | 0.033 | 0.028 | 0.028 | 0 | 0.42 | 0.21 | 0.39 | 0.36 | 0.33 | 0.36 |
| 7 | 80702178 | rs7788645 | -0.236 | -0.143 | 0.643 | 0.831 | -0.016 | 0.091 | 0.091 | 0 | 0.57 | 0.74 | 0.74 | 0.93 | 0.51 | 0.65 |
| 7 | 80703957 | rs13240198 | -0.132 | -0.132 | 0.824 | 0.997 | 0.001 | 0.087 | 0.087 | 0 | 0.13 | 0.22 | 0.31 | 0.48 | 0.18 | 0.29 |
| 7 | 80709513 | rs10277023 | -1.437 | -0.306 | 0.599 | 0.755 | -0.018 | 0.032 | 0.032 | 0 | 0.26 | 0.65 | 0.36 | 0.57 | 0.21 | 0.38 |
| 7 | 80713204 | rs17277762 | 1.006 | -0.309 | 0.57 | 0.756 | 0.027 | 0.004 | 0.004 | 0 | 0.41 | 0.21 | 0.39 | 0.42 | 0.33 | 0.36 |
| 7 | 80718358 | rs2697949 | -0.632 | -0.641 | 0.662 | 1.041 | -0.004 | 0.109 | 0.109 | 0 | 0.16 | 0.35 | 0.32 | 0.57 | 0.18 | 0.3 |
| 7 | 80739951 | rs12113822 | 0.646 | -0.713 | -0.348 | 0.739 | -0.014 | -0.041 | -0.041 | 0 | 0.08 | 0.14 | 0.02 | 0 | 0.03 | 0.01 |
| 7 | 80743902 | rs4096939 | 0.047 | -0.37 | -0.574 | 0.695 | -0.011 | 0.058 | 0.058 | 0.037 | 0.62 | 0.75 | 0.83 | 0.84 | 0.63 | 0.71 |
| 7 | 80747605 | rs6942673 | -0.081 | -0.38 | -0.582 | 0.68 | -0.015 | 0.076 | 0.076 | 0.013 | 0.63 | 0.79 | 0.84 | 0.93 | 0.63 | 0.79 |
| 7 | 80748070 | rs3914079 | 0.274 | -0.299 | -0.813 | 0.729 | 0 | 0.004 | 0.004 | 0.345 | 0.58 | 0.6 | 0.81 | 0.84 | 0.63 | 0.68 |
| 7 | 80749360 | rs10277984 | -1.354 | -0.403 | -0.847 | 2.001 | 0.037 | -0.055 | -0.055 | 0 | 0.28 | 0.79 | 0.29 | 0.55 | 0.21 | 0.32 |
| 7 | 80763989 | rs1528736 | -1.268 | -0.34 | -0.816 | 1.711 | 0.034 | -0.051 | -0.051 | 0 | 0.28 | 0.79 | 0.29 | 0.46 | 0.2 | 0.32 |
| 7 | 80769699 | rs7791061 | -1.324 | 0.141 | -0.027 | 2.621 | 0 | 0.104 | 0.104 | 0 | 0.15 | 0.31 | 0.1 | 0.49 | 0.13 | 0.12 |
| 7 | 80780608 | rs327673 | 1.003 | 0.03 | -0.014 | 2.394 | -0.003 | 0.017 | 0.017 | 0 | 0.83 | 0.75 | 0.89 | 0.73 | 0.83 | 0.89 |
| 7 | 80781016 | rs327674 | 0.823 | -0.198 | -0.702 | 1.41 | -0.069 | 0.044 | 0.044 | 0 | 0.39 | 0.15 | 0.6 | 0.29 | 0.47 | 0.59 |
| 7 | 80793932 | rs478703 | 1.271 | -0.166 | 0.379 | 1.425 | 0.003 | 0.037 | 0.037 | 0 | 0.78 | 0.57 | 0.83 | 0.36 | 0.78 | 0.85 |
| 7 | 80797311 | rs565429 | 0.295 | -0.188 | 0.278 | 1.38 | 0.002 | -0.003 | -0.003 | 0.094 | 0.83 | 0.78 | 0.9 | 0.71 | 0.83 | 0.89 |
| 7 | 80802406 | rs327678 | 0.29 | -0.188 | 0.278 | 1.38 | 0.002 | -0.003 | -0.003 | 0.034 | 0.83 | 0.78 | 0.9 | 0.71 | 0.83 | 0.89 |
| 7 | 80805518 | rs11977048 | 0.586 | -0.467 | 0.72 | 1.173 | -0.014 | -0.051 | -0.051 | 0.011 | 0.39 | 0.48 | 0.23 | 0.18 | 0.31 | 0.26 |
| 7 | 80805828 | rs327683 | 0.238 | -0.472 | 0.711 | 1.203 | 0.002 | -0.003 | -0.003 | 2.683 | 0.83 | 0.78 | 0.9 | 0.71 | 0.83 | 0.89 |
| 7 | 80805984 | rs7798421 | 0.319 | -0.323 | 1.152 | 1.309 | -0.019 | -0.125 | -0.125 | 4.747 | 0.39 | 0.44 | 0.23 | 0.09 | 0.31 | 0.26 |
| 7 | 80812355 | rs7797263 | -0.043 | -0.462 | 1.06 | 1.494 | 0.013 | 0.013 | 0.013 | 0 | 0.43 | 0.56 | 0.42 | 0.34 | 0.33 | 0.48 |
| 7 | 80812398 | rs7797400 | -0.246 | -0.339 | 1.102 | 1.523 | 0.017 | -0.004 | -0.004 | 3.413 | 0.47 | 0.58 | 0.48 | 0.59 | 0.37 | 0.53 |
| 7 | 80827037 | rs1859678 | 0.493 | -0.348 | 0.897 | 1.261 | 0.002 | -0.04 | -0.04 | 0 | 0.35 | 0.24 | 0.18 | 0.05 | 0.3 | 0.16 |
| 7 | 80827086 | rs327701 | 0.386 | -0.072 | 0.295 | 1.425 | 0.027 | -0.021 | -0.021 | 2.845 | 0.49 | 0.19 | 0.48 | 0.41 | 0.58 | 0.4 |
| 7 | 80831501 | rs327708 | 0.225 | -0.431 | 0.336 | 0.701 | 0.08 | 0.136 | 0.136 | 0 | 0.53 | 0.24 | 0.55 | 0.74 | 0.62 | 0.47 |
| 7 | 80831733 | rs327710 | 0.166 | -0.318 | 0.009 | 0.545 | -0.008 | -0.021 | -0.021 | 0 | 0.56 | 0.4 | 0.57 | 0.77 | 0.67 | 0.5 |
| 7 | 80832457 | rs1912364 | 0.334 | 0.409 | -0.744 | 0.57 | 0.009 | 0.035 | 0.035 | 0 | 0.56 | 0.37 | 0.66 | 0.25 | 0.61 | 0.67 |
| 7 | 80832856 | rs12707261 | -0.754 | 1.482 | -1.065 | 0.89 | -0.039 | -0.047 | -0.047 | 0 | 0.2 | 0.1 | 0.23 | 0.3 | 0.3 | 0.21 |
| 7 | 80834923 | rs17154908 | -0.499 | 0.21 | -0.806 | 0.297 | 0.001 | -0.002 | -0.002 | 0 | 0.14 | 0.12 | 0.23 | 0.14 | 0.09 | 0.2 |
| 7 | 80837411 | rs2190296 | 0.417 | -0.13 | -0.881 | -0.164 | 0 | 0.004 | 0.004 | 0.695 | 0.08 | 0.07 | 0.13 | 0.21 | 0.03 | 0.09 |
| 7 | 80839653 | rs327718 | -0.808 | -0.025 | -1.274 | -1.366 | -0.04 | 0.169 | 0.169 | 0 | 0.59 | 0.81 | 0.69 | 0.92 | 0.43 | 0.64 |
| 7 | 80856984 | rs1912367 | 0.468 | -0.314 | 0.205 | -1.447 | 0 | 0.031 | 0.031 | 3.754 | 0.34 | 0.19 | 0.45 | 0.67 | 0.31 | 0.45 |

| | | | | | | | | | | | | | | | | |
|---|---|---|---|---|---|---|---|---|---|---|---|---|---|---|---|---|
| 7 | 80857196 | rs327649 | -0.914 | -0.251 | 0.222 | -1.48 | -0.037 | 0.17 | 0.17 | 0 | 0.59 | 0.81 | 0.69 | 0.92 | 0.43 | 0.65 |
| 7 | 80857451 | rs186609 | -0.906 | -0.248 | 0.288 | -1.479 | -0.022 | 0.116 | 0.116 | 0 | 0.59 | 0.75 | 0.68 | 0.92 | 0.43 | 0.65 |
| 7 | 80860413 | rs1397015 | 0.483 | 0.112 | -0.021 | -1.477 | -0.016 | 0.027 | 0.027 | 5.981 | 0.33 | 0.19 | 0.5 | 0.67 | 0.33 | 0.45 |
| 7 | 80863837 | rs17154948 | -0.008 | -0.28 | 0.059 | -1.323 | 0.023 | 0.026 | 0.026 | 4.884 | 0.22 | 0.1 | 0.09 | 0.18 | 0.12 | 0.16 |
| 7 | 80866153 | rs2158376 | 0.27 | -0.159 | 0.354 | -1.676 | -0.018 | 0.01 | 0.01 | 0 | 0.39 | 0.65 | 0.32 | 0.08 | 0.54 | 0.38 |
| 7 | 80867976 | rs17154957 | -0.113 | -0.567 | 0.91 | -1.46 | -0.001 | -0.021 | -0.021 | 0 | 0.46 | 0.46 | 0.38 | 0.25 | 0.6 | 0.42 |
| 7 | 80869802 | rs17154968 | -0.288 | -0.76 | 0.998 | -1.512 | 0.02 | 0.017 | 0.017 | 0 | 0.06 | 0.31 | 0.03 | 0.01 | 0.02 | 0.02 |
| 7 | 80880969 | rs325972 | -0.493 | -0.794 | 0.605 | -1.54 | 0.013 | 0.188 | 0.188 | 0.11 | 0.49 | 0.84 | 0.46 | 0.26 | 0.61 | 0.44 |
| 7 | 80886497 | rs161828 | -0.547 | -0.779 | 0.598 | -1.533 | 0.015 | 0.199 | 0.199 | 0.056 | 0.5 | 0.86 | 0.46 | 0.26 | 0.61 | 0.44 |
| 7 | 80904549 | rs1509914 | -1.069 | -0.764 | 1.094 | -1.469 | -0.029 | -0.032 | -0.032 | 0.144 | 0.09 | 0.54 | 0.05 | 0.01 | 0.02 | 0.02 |
| 7 | 80910329 | rs245433 | -0.57 | -0.671 | 0.739 | -1.553 | 0.028 | 0.055 | 0.055 | 1.439 | 0.57 | 0.68 | 0.5 | 0.28 | 0.7 | 0.46 |
| 7 | 80921505 | rs11771361 | 0.886 | -0.761 | 0.857 | -1.527 | 0.001 | -0.016 | -0.016 | 0 | 0.13 | 0.12 | 0.07 | 0.01 | 0.11 | 0.04 |
| 7 | 80933229 | rs17626401 | 0.315 | -0.709 | 0.852 | -1.576 | -0.018 | 0.08 | 0.08 | 1.43 | 0.4 | 0.21 | 0.41 | 0.26 | 0.61 | 0.42 |
| 7 | 80933327 | rs7790316 | 0.803 | -0.736 | 0.89 | -1.594 | -0.005 | -0.036 | -0.036 | 1.577 | 0.13 | 0.17 | 0.08 | 0.01 | 0.11 | 0.05 |
| 7 | 80934172 | rs6943816 | 0.202 | -0.699 | 0.9 | -1.556 | -0.002 | 0.018 | 0.018 | 0.657 | 0.41 | 0.33 | 0.42 | 0.26 | 0.61 | 0.42 |
| 7 | 80938045 | rs765619 | -0.449 | -1.019 | 0.716 | -1.27 | 0.002 | -0.026 | -0.026 | 0 | 0.92 | 0.9 | 0.94 | 1 | 0.9 | 0.92 |
| 7 | 80945405 | rs10215927 | 0.144 | -0.86 | 0.37 | -1.215 | 0.018 | 0.028 | 0.028 | 1.285 | 0.52 | 0.38 | 0.46 | 0.34 | 0.7 | 0.45 |
| 7 | 80947517 | rs10486820 | 0.78 | -1.064 | 0.303 | -1.202 | 0.023 | 0.009 | 0.009 | 0 | 0.35 | 0.3 | 0.46 | 0.63 | 0.18 | 0.44 |
| 7 | 80952495 | rs6946105 | -0.237 | -0.961 | -0.462 | -1.148 | 0.003 | -0.048 | -0.048 | 3.21 | 0.87 | 0.69 | 0.92 | 0.96 | 0.89 | 0.89 |
| 7 | 80953472 | rs6969877 | 0.517 | -0.274 | -1.567 | -0.907 | -0.055 | 0.136 | 0.136 | 0 | 0.36 | 0.16 | 0.34 | 0.05 | 0.61 | 0.38 |
| 7 | 80957136 | rs12539379 | 0.801 | -0.271 | -1.565 | -0.875 | 0.012 | -0.033 | -0.033 | 0 | 0.15 | 0.08 | 0.12 | 0.29 | 0.09 | 0.08 |
| 7 | 80962598 | rs1014079 | -0.094 | 0.502 | -1.298 | -1.046 | -0.045 | 0.124 | 0.124 | 0 | 0.37 | 0.18 | 0.34 | 0.05 | 0.62 | 0.38 |
| 7 | 80968055 | rs6952065 | 0.196 | -0.095 | -1.036 | -0.17 | 0.036 | 0.036 | 0.036 | 0 | 0.51 | 0.58 | 0.26 | 0.31 | 0.29 | 0.3 |
| 7 | 80970635 | rs17155092 | -0.23 | -0.898 | -0.741 | 0.74 | 0.006 | -0.008 | -0.008 | 0 | 0.17 | 0.09 | 0.17 | 0.45 | 0.09 | 0.43 |
| 7 | 80971145 | rs10230861 | -0.837 | -0.823 | -0.358 | 0.238 | -0.001 | 0.002 | 0.002 | 0 | 0.48 | 0.56 | 0.63 | 0.84 | 0.53 | 0.74 |
| 7 | 80975390 | rs12534756 | 0.959 | -0.634 | -0.486 | 0.44 | -0.094 | 0.023 | 0.023 | 0 | 0.2 | 0.07 | 0.2 | 0.12 | 0.37 | 0.2 |
| 7 | 80984831 | rs10263457 | -0.643 | -0.719 | 0.082 | 0.533 | -0.042 | 0.022 | 0.022 | 0 | 0.28 | 0.44 | 0.38 | 0.66 | 0.16 | 0.54 |
| 7 | 80992634 | rs10226237 | -0.879 | -0.675 | -0.093 | 0.347 | 0.001 | -0.002 | -0.002 | 0 | 0.51 | 0.53 | 0.66 | 0.83 | 0.58 | 0.73 |
| 7 | 80998755 | rs10229228 | -0.485 | -0.623 | -0.412 | 0.446 | 0.008 | 0.001 | 0.001 | 0 | 0.29 | 0.22 | 0.45 | 0.69 | 0.21 | 0.53 |
| 7 | 81001744 | rs2189915 | 0.918 | -0.298 | -0.51 | 0.303 | -0.096 | 0.045 | 0.045 | 0 | 0.2 | 0.06 | 0.2 | 0.11 | 0.37 | 0.2 |
| 7 | 81010851 | rs6955019 | -0.439 | -1.076 | 0.265 | 0.507 | -0.01 | 0.042 | 0.042 | 0 | 0.27 | 0.3 | 0.29 | 0.55 | 0.2 | 0.46 |
| 7 | 81026186 | rs6955708 | -0.897 | -0.791 | 0.129 | 0.126 | -0.111 | 0.124 | 0.124 | 0 | 0.77 | 0.93 | 0.8 | 0.99 | 0.62 | 0.77 |
| 7 | 81027425 | rs17629690 | 0.978 | -1.473 | 0.44 | -0.08 | 0.004 | -0.008 | -0.008 | 0.086 | 0.32 | 0.2 | 0.38 | 0.31 | 0.25 | 0.21 |
| 7 | 81029279 | rs10278362 | -0.477 | -1.648 | 0.791 | -0.144 | -0.003 | 0 | 0 | 0 | 0.4 | 0.41 | 0.33 | 0.55 | 0.32 | 0.48 |
| 7 | 81044104 | rs17155159 | 0.857 | -1.508 | 0.91 | 0.141 | 0.008 | -0.044 | -0.044 | 0.093 | 0.31 | 0.17 | 0.37 | 0.37 | 0.24 | 0.27 |
| 7 | 81045962 | rs2967926 | -0.049 | -1.75 | 0.998 | 0.205 | -0.015 | -0.02 | -0.02 | 0 | 0.21 | 0.29 | 0.11 | 0.11 | 0.19 | 0.12 |
| 7 | 81046460 | rs2967925 | -0.356 | -1.582 | 0.579 | 0.167 | 0.012 | -0.001 | -0.001 | 0 | 0.54 | 0.72 | 0.51 | 0.48 | 0.43 | 0.39 |
| 7 | 81046710 | rs2967924 | -0.368 | -1.585 | 0.578 | 0.019 | 0.011 | 0.018 | 0.018 | 0 | 0.54 | 0.74 | 0.55 | 0.55 | 0.44 | 0.39 |
| 7 | 81047852 | rs2906234 | -0.726 | -1.351 | 0.482 | -0.288 | -0.029 | -0.126 | -0.126 | 0.209 | 0.25 | 0.45 | 0.15 | 0.01 | 0.23 | 0.11 |
| 7 | 81056006 | rs2967917 | -0.031 | -1.585 | 0.442 | 0.539 | -0.001 | -0.003 | -0.003 | 0 | 0.52 | 0.5 | 0.7 | 0.64 | 0.43 | 0.53 |
| 7 | 81060070 | rs2906207 | -0.127 | -1.641 | 0.293 | 0.624 | 0.01 | -0.032 | -0.032 | 0 | 0.56 | 0.78 | 0.46 | 0.49 | 0.4 | 0.54 |
| 7 | 81060188 | rs2906208 | -0.106 | -1.654 | 0.387 | 0.633 | 0.011 | -0.025 | -0.025 | 0 | 0.56 | 0.69 | 0.45 | 0.49 | 0.4 | 0.54 |
| 7 | 81070971 | rs2906226 | 0.262 | -1.796 | 0.074 | 0.55 | -0.02 | 0.074 | 0.074 | 2.796 | 0.33 | 0.09 | 0.28 | 0.21 | 0.33 | 0.35 |
| 7 | 81075055 | rs2967907 | -0.306 | -1.435 | -0.559 | 0.583 | 0.017 | -0.013 | -0.013 | 0 | 0.54 | 0.62 | 0.41 | 0.5 | 0.38 | 0.55 |
| 7 | 81075188 | rs2906229 | -0.253 | -1.449 | -0.648 | 0.576 | 0.011 | -0.006 | -0.006 | 0 | 0.53 | 0.58 | 0.41 | 0.5 | 0.38 | 0.55 |
| 7 | 81076550 | rs2189911 | -1.037 | -1.312 | -1.807 | 0.244 | -0.125 | 0.105 | 0.105 | 0 | 0.71 | 0.92 | 0.76 | 0.91 | 0.5 | 0.78 |
| 7 | 81084663 | rs2967903 | -0.598 | -0.967 | -1.256 | 0.459 | 0.037 | 0.014 | 0.014 | 0.002 | 0.52 | 0.78 | 0.36 | 0.44 | 0.37 | 0.53 |
| 7 | 81087389 | rs7791326 | -0.247 | -0.266 | -1.164 | 0.175 | -0.006 | 0.041 | 0.041 | 0.041 | 0.25 | 0.09 | 0.2 | 0.17 | 0.24 | 0.28 |
| 7 | 81092376 | rs10273273 | -0.051 | -0.901 | -0.86 | -0.107 | -0.011 | -0.018 | -0.018 | 0 | 0.45 | 0.51 | 0.46 | 0.21 | 0.32 | 0.44 |
| 7 | 81100527 | rs12666154 | -0.541 | -1.072 | -0.473 | 1.22 | -0.017 | 0.136 | 0.136 | 0 | 0.06 | 0.11 | 0.02 | 0.5 | 0.02 | 0.2 |
| 7 | 81122819 | rs10808295 | 0.258 | -0.172 | -0.63 | 0.925 | -0.015 | -0.003 | -0.003 | 0.229 | 0.89 | 0.93 | 0.95 | 0.9 | 0.86 | 0.88 |
| 7 | 81146818 | rs6956927 | -1.336 | -0.84 | 0.844 | 1.083 | -0.04 | 0.231 | 0.231 | 0 | 0.09 | 0.26 | 0.09 | 0.47 | 0.03 | 0.25 |
| 7 | 81166295 | rs17155218 | -1.3 | -0.925 | 0.944 | 1.044 | -0.043 | 0.208 | 0.208 | 0 | 0.1 | 0.24 | 0.09 | 0.46 | 0.03 | 0.25 |
| 7 | 81171725 | rs11972009 | -1.403 | -1.135 | 0.82 | 0.85 | -0.052 | 0.128 | 0.128 | 0 | 0.16 | 0.31 | 0.11 | 0.46 | 0.07 | 0.25 |
| 7 | 81176804 | rs2909580 | 0.594 | -0.891 | -0.492 | -0.047 | -0.076 | 0.087 | 0.087 | 0 | 0.33 | 0.01 | 0.32 | 0.15 | 0.36 | 0.3 |
| 7 | 81190100 | rs4141453 | 0.712 | -1.575 | 0.368 | 0.402 | -0.033 | 0.349 | 0.349 | 0 | 0.89 | 0.58 | 0.88 | 0.59 | 0.97 | 0.71 |
| 7 | 81195596 | rs12532699 | 0.928 | -1.015 | -0.211 | 0.061 | -0.137 | 0.028 | 0.028 | 0 | 0.27 | 0.01 | 0.32 | 0.15 | 0.38 | 0.23 |
| 7 | 81207444 | rs10486831 | 0.438 | -1.445 | 0.481 | 0.872 | -0.023 | -0.034 | -0.034 | 0 | 0.1 | 0.03 | 0.06 | 0.18 | 0.1 | 0.15 |
| 7 | 81231258 | rs11560402 | -0.352 | -1.275 | 0.847 | 1.018 | -0.005 | 0.343 | 0.343 | 0 | 0.06 | 0.25 | 0.07 | 0.42 | 0.01 | 0.28 |
| 7 | 81234408 | rs7790727 | -0.03 | -1.53 | 0.715 | 0.849 | 0.021 | -0.015 | -0.015 | 0.198 | 0.21 | 0.13 | 0.08 | 0.22 | 0.12 | 0.15 |
| 7 | 81242237 | rs7786648 | -1.142 | -1.627 | 0.737 | 0.946 | -0.032 | 0.208 | 0.208 | 0 | 0.13 | 0.29 | 0.13 | 0.46 | 0.07 | 0.28 |
| 7 | 81266952 | rs2114736 | -1.593 | -1.685 | 0.836 | 0.839 | 0.012 | 0.004 | 0.004 | 2.11 | 0.37 | 0.64 | 0.24 | 0.69 | 0.2 | 0.47 |
| 7 | 81273423 | rs10276241 | -1.075 | -0.995 | 1.029 | 1.354 | 0.008 | -0.022 | -0.022 | 0 | 0.07 | 0.17 | 0.06 | 0.03 | 0.05 | 0.06 |
| 7 | 81273756 | rs4464906 | 0.125 | -0.736 | 0.772 | 1.291 | -0.049 | -0.039 | -0.039 | 0 | 0.18 | 0.01 | 0.32 | 0.17 | 0.2 | 0.17 |
| 7 | 81276415 | rs2367533 | -1.035 | -0.815 | 1.74 | 1.578 | -0.032 | 0.236 | 0.236 | 0.442 | 0.08 | 0.21 | 0.09 | 0.43 | 0.04 | 0.26 |
| 7 | 81285767 | rs982904 | 0.728 | -0.769 | 1.626 | 1.497 | -0.03 | 0.229 | 0.229 | 0.2 | 0.92 | 0.81 | 0.91 | 0.57 | 0.96 | 0.77 |
| 7 | 81290522 | rs7799610 | 1.473 | -0.788 | 1.69 | 1.421 | -0.032 | 0.303 | 0.303 | 0 | 0.86 | 0.58 | 0.86 | 0.53 | 0.93 | 0.72 |
| 7 | 81297342 | rs10266050 | -1.349 | -0.349 | 1.192 | 0.662 | -0.019 | 0.039 | 0.039 | 0.881 | 0.09 | 0.51 | 0.05 | 0.05 | 0.02 | 0.07 |
| 7 | 81306540 | rs17155352 | -1.401 | -0.586 | 0.96 | 0.337 | 0.002 | -0.007 | -0.007 | 0 | 0.12 | 0.13 | 0.08 | 0.02 | 0.09 | 0.19 |

| | | | | | | | | | | | | | | | | |
|---|---|---|---|---|---|---|---|---|---|---|---|---|---|---|---|---|
| 7 | 81307464 | rs2190218 | -0.247 | 0.315 | 0.195 | 0.722 | -0.073 | -0.069 | -0.069 | 0 | 0.12 | 0.01 | 0.1 | 0.25 | 0.18 | 0.18 |
| 7 | 81308396 | rs17155363 | 0.432 | -0.47 | 0.263 | 0.907 | 0.003 | -0.007 | -0.007 | 0 | 0.17 | 0.15 | 0.23 | 0.32 | 0.14 | 0.19 |
| 7 | 81309224 | rs1029567 | -0.61 | -0.392 | -0.307 | 0.591 | -0.001 | 0.006 | 0.006 | 0 | 0.88 | 0.92 | 0.87 | 0.89 | 0.86 | 0.86 |
| 7 | 81309505 | rs1029568 | -0.954 | -0.531 | 0.203 | 0.858 | -0.026 | 0.018 | 0.018 | 1.066 | 0.32 | 0.6 | 0.34 | 0.38 | 0.23 | 0.38 |
| 7 | 81317508 | rs757828 | -0.623 | 0.158 | 0.376 | 0.989 | -0.012 | 0.048 | 0.048 | 1.286 | 0.27 | 0.6 | 0.32 | 0.37 | 0.22 | 0.36 |
| 7 | 81322168 | rs4732401 | 0.819 | 0.338 | 1.062 | 0.288 | 0.036 | -0.051 | -0.051 | 0 | 0.83 | 0.5 | 0.84 | 0.88 | 0.85 | 0.87 |
| 7 | 81330627 | rs5745774 | 0.132 | 0.12 | 1.133 | 0.356 | -0.01 | 0.002 | 0.002 | 0 | 0.09 | 0.16 | 0.11 | 0.13 | 0.07 | 0.11 |
| 7 | 81335460 | rs5745752 | -0.389 | 0.085 | 0.911 | 0.311 | -0.003 | -0.028 | -0.028 | 0 | 0.3 | 0.13 | 0.24 | 0.33 | 0.28 | 0.4 |
| 7 | 81339304 | rs2074725 | 0.61 | -0.019 | 1.352 | 0.035 | 0.089 | -0.021 | -0.021 | 0 | 0.85 | 0.58 | 0.84 | 0.88 | 0.8 | 0.86 |
| 7 | 81343093 | rs6467865 | 0.54 | -0.022 | 1.419 | 0.018 | 0.085 | -0.03 | -0.03 | 0.279 | 0.85 | 0.57 | 0.83 | 0.88 | 0.8 | 0.85 |
| 7 | 81346925 | rs5745720 | 0.048 | 0.09 | 0.642 | 0.064 | -0.006 | 0.004 | 0.004 | 0 | 0.88 | 0.94 | 0.86 | 0.88 | 0.8 | 0.85 |
| 7 | 81358060 | rs5745696 | 0.123 | -0.207 | 0.6 | 0.036 | 0.002 | 0.001 | 0.001 | 0 | 0.11 | 0.1 | 0.11 | 0.11 | 0.08 | 0.1 |
| 7 | 81359051 | rs5745687 | 1.503 | 0.106 | 0.203 | -0.008 | -0.009 | 0.041 | 0.041 | 0 | 0.08 | 0 | 0.04 | 0 | 0.06 | 0.01 |
| 7 | 81363897 | rs4732404 | 0.024 | 0.322 | 0.2 | 0.043 | -0.037 | 0.001 | 0.001 | 0 | 0.88 | 0.96 | 0.85 | 0.86 | 0.76 | 0.84 |
| 7 | 81364187 | rs12540393 | 0.495 | 0.293 | 0.426 | 0.031 | 0.066 | -0.009 | -0.009 | 0 | 0.86 | 0.76 | 0.84 | 0.86 | 0.76 | 0.84 |
| 7 | 81369419 | rs12707453 | -0.081 | 0.207 | -0.183 | 0.203 | -0.02 | -0.101 | -0.101 | 0 | 0.19 | 0.01 | 0.17 | 0.3 | 0.16 | 0.26 |
| 7 | 81375051 | rs5745661 | 0.345 | 0.446 | 0.038 | 0.144 | -0.016 | 0.003 | 0.003 | 0 | 0.87 | 0.93 | 0.85 | 0.86 | 0.76 | 0.84 |
| 7 | 81381326 | rs5745646 | 0.228 | 0.463 | -0.126 | 0.152 | -0.072 | -0.015 | -0.015 | 0 | 0.88 | 0.98 | 0.85 | 0.86 | 0.76 | 0.84 |
| 7 | 81388042 | rs5745635 | -1.493 | -0.401 | 0.638 | 0.12 | 0.011 | 0.023 | 0.023 | 0.891 | 0.06 | 0.31 | 0.03 | 0.01 | 0.01 | 0.07 |
| 7 | 81398356 | rs5745616 | -0.662 | -0.541 | -0.197 | 0.073 | 0.015 | -0.017 | -0.017 | 0 | 0.29 | 0.2 | 0.22 | 0.32 | 0.21 | 0.39 |
| 7 | 81400939 | rs3735520 | -0.018 | -0.621 | -0.416 | -0.359 | 0 | 0.002 | 0.002 | 1.372 | 0.59 | 0.63 | 0.42 | 0.39 | 0.43 | 0.57 |
| 7 | 81405985 | rs17155457 | 0.656 | -0.526 | -0.782 | -0.263 | -0.002 | -0.027 | -0.027 | 3.122 | 0.06 | 0.01 | 0.02 | 0.13 | 0.03 | 0.1 |
| 7 | 81407080 | rs1558001 | -0.825 | 0.603 | -0.685 | -0.356 | -0.023 | -0.045 | -0.045 | 0 | 0.7 | 0.79 | 0.5 | 0.54 | 0.59 | 0.75 |
| 7 | 81409040 | rs10277195 | -0.368 | 0.659 | -0.681 | -0.328 | -0.017 | -0.023 | -0.023 | 0 | 0.65 | 0.68 | 0.44 | 0.52 | 0.45 | 0.66 |
| 7 | 81411028 | rs975360 | -0.026 | 0.461 | -0.814 | -0.574 | -0.011 | 0.061 | 0.061 | 0 | 0.43 | 0.32 | 0.23 | 0.19 | 0.43 | 0.39 |
| 7 | 81415783 | rs17501560 | 1.534 | 0.674 | -1.206 | -0.485 | -0.077 | 0.006 | 0.006 | 0 | 0.12 | 0.01 | 0.09 | 0.05 | 0.21 | 0.08 |
| 7 | 81428307 | rs1019617 | 0.365 | -0.246 | -0.828 | -0.676 | 0.046 | 0.015 | 0.015 | 0.107 | 0.51 | 0.43 | 0.7 | 0.53 | 0.65 | 0.41 |
| 7 | 81429590 | rs6951168 | 1.052 | -0.4 | -0.791 | -0.742 | 0.006 | 0.006 | 0.006 | 0 | 0.24 | 0.18 | 0.09 | 0.13 | 0.16 | 0.23 |
| 7 | 81430939 | rs7456691 | 0.316 | 0.158 | -0.868 | -0.518 | 0.013 | 0.006 | 0.006 | 0.09 | 0.46 | 0.36 | 0.66 | 0.53 | 0.6 | 0.34 |
| 7 | 81448657 | rs17155547 | -0.395 | 0.367 | -1.138 | -0.268 | -0.019 | -0.027 | -0.027 | 0 | 0.14 | 0.08 | 0.14 | 0.24 | 0.15 | 0.31 |
| 7 | 81460489 | rs12707462 | -0.92 | -0.252 | -1.018 | -0.601 | 0.002 | 0.001 | 0.001 | 0 | 0.69 | 0.72 | 0.84 | 0.81 | 0.78 | 0.71 |
| 7 | 81466742 | rs17155560 | 1.384 | 0.161 | -1.223 | -0.726 | -0.014 | -0.045 | -0.045 | 0 | 0.3 | 0.21 | 0.54 | 0.48 | 0.38 | 0.24 |
| 7 | 81476599 | rs12707465 | -0.312 | -0.673 | -0.898 | -0.76 | 0.015 | 0.014 | 0.014 | 0.207 | 0.58 | 0.43 | 0.79 | 0.77 | 0.73 | 0.67 |
| 7 | 81486442 | rs5002653 | 0.537 | -0.412 | -1.129 | -0.694 | -0.003 | 0.006 | 0.006 | 0 | 0.09 | 0.1 | 0.05 | 0.06 | 0.08 | 0.19 |
| 7 | 81486758 | rs4307284 | 0.451 | -0.301 | -1.195 | -0.773 | -0.001 | 0.038 | 0.038 | 0.059 | 0.21 | 0.13 | 0.11 | 0.11 | 0.15 | 0.32 |
| 7 | 81492276 | rs11769657 | 0.786 | -0.118 | -1.181 | -0.644 | -0.041 | -0.047 | -0.047 | 0 | 0.34 | 0.23 | 0.64 | 0.48 | 0.54 | 0.28 |
| 7 | 81498151 | rs17155595 | 0.392 | 0.37 | -1.434 | -0.758 | -0.032 | -0.031 | -0.031 | 0 | 0.38 | 0.3 | 0.53 | 0.52 | 0.47 | 0.41 |
| 7 | 81501546 | rs12155465 | 0.993 | 0.349 | -1.058 | -0.778 | -0.017 | -0.027 | -0.027 | 0.024 | 0.3 | 0.24 | 0.49 | 0.48 | 0.39 | 0.23 |
| 7 | 81507279 | rs1922073 | 2.018 | -0.054 | -1.078 | -0.773 | -0.001 | -0.006 | -0.006 | 0.698 | 0.14 | 0.13 | 0.06 | 0.06 | 0.07 | 0.12 |
| 7 | 81516978 | rs9986815 | 0.681 | -0.367 | -1.101 | -0.939 | -0.038 | -0.076 | -0.076 | 0 | 0.24 | 0.31 | 0.11 | 0.06 | 0.13 | 0.16 |
| 7 | 81517801 | rs4732415 | 0.004 | 0.356 | -1.387 | -0.354 | -0.02 | -0.017 | -0.017 | 0 | 0.1 | 0.06 | 0.22 | 0.17 | 0.2 | 0.13 |
| 7 | 81521295 | rs1733623 | -0.587 | 0.477 | -1.3 | -0.293 | 0.041 | 0.015 | 0.015 | 0.131 | 0.49 | 0.68 | 0.41 | 0.46 | 0.46 | 0.46 |
| 7 | 81534837 | rs1229453 | 0.571 | 0.544 | -1.437 | 0.022 | -0.006 | -0.032 | -0.032 | 0.414 | 0.17 | 0.05 | 0.09 | 0.22 | 0.17 | 0.27 |
| 7 | 81535350 | rs4728489 | -0.18 | 0.809 | -1.422 | -0.205 | -0.039 | 0.006 | 0.006 | 0 | 0.26 | 0.18 | 0.33 | 0.26 | 0.36 | 0.34 |
| 7 | 81538507 | rs1229460 | -0.59 | 1.239 | -0.99 | 0.686 | 0.001 | 0.002 | 0.002 | 0.176 | 0.62 | 0.76 | 0.63 | 0.63 | 0.62 | 0.67 |
| 7 | 81543626 | rs1229469 | 0.513 | 1.261 | -0.935 | 1.049 | -0.025 | -0.049 | -0.049 | 0 | 0.19 | 0.07 | 0.2 | 0.27 | 0.21 | 0.28 |
| 7 | 81543886 | rs1229470 | -0.843 | 1.457 | -1.052 | 1.066 | -0.034 | 0.003 | 0.003 | 0 | 0.89 | 0.98 | 0.92 | 0.92 | 0.85 | 0.88 |
| 7 | 81550157 | rs1229476 | 0.015 | 1.55 | -0.372 | 1.443 | -0.012 | -0.03 | -0.03 | 0 | 0.2 | 0.11 | 0.21 | 0.27 | 0.21 | 0.27 |
| 7 | 81555317 | rs1229484 | 0.12 | 1.421 | 0.051 | 1.321 | -0.002 | -0.003 | -0.003 | 0 | 0.23 | 0.18 | 0.24 | 0.24 | 0.23 | 0.25 |
| 7 | 81558785 | rs11982413 | 0.376 | 1.68 | 0.491 | 1.233 | 0.005 | -0.026 | -0.026 | 0.1 | 0.28 | 0.35 | 0.23 | 0.18 | 0.28 | 0.25 |
| 7 | 81559428 | rs1229486 | -0.559 | 1.678 | 0.612 | 1.223 | 0.003 | 0.008 | 0.008 | 0.757 | 0.49 | 0.56 | 0.46 | 0.43 | 0.49 | 0.46 |
| 7 | 81560832 | rs11983646 | 0.388 | 1.601 | 0.64 | 1.134 | -0.002 | 0.002 | 0.002 | 1.28 | 0.28 | 0.28 | 0.23 | 0.19 | 0.28 | 0.25 |
| 7 | 81562516 | rs11971148 | 0.312 | 1.601 | 0.645 | 1.134 | 0.005 | -0.025 | -0.025 | 0 | 0.28 | 0.35 | 0.23 | 0.19 | 0.28 | 0.25 |
| 7 | 81564122 | rs1229492 | -1.209 | 1.567 | 0.141 | 1.294 | 0.054 | 0.047 | 0.047 | 0 | 0.61 | 0.83 | 0.69 | 0.74 | 0.75 | 0.73 |
| 7 | 81565355 | rs1229494 | 0.566 | 1.636 | -0.11 | 1.054 | -0.01 | -0.014 | -0.014 | 0 | 0.22 | 0.12 | 0.22 | 0.27 | 0.23 | 0.23 |
| 7 | 81566723 | rs9769000 | -0.443 | 1.315 | 0.218 | 0.73 | 0 | -0.034 | -0.034 | 0 | 0.3 | 0.45 | 0.24 | 0.19 | 0.28 | 0.26 |
| 7 | 81569820 | rs1922077 | -0.894 | 1.846 | 0.436 | 1.427 | 0.073 | 0.141 | 0.141 | 0 | 0.1 | 0.29 | 0.24 | 0.31 | 0.25 | 0.23 |
| 7 | 81571561 | rs4732418 | -0.865 | 1.844 | 0.511 | 1.433 | 0.027 | 0.058 | 0.058 | 0 | 0.1 | 0.17 | 0.22 | 0.3 | 0.24 | 0.23 |
| 7 | 81573465 | rs17155669 | -0.16 | 1.345 | 0.95 | 1.203 | 0 | -0.007 | -0.007 | 0.061 | 0.25 | 0.24 | 0.21 | 0.2 | 0.24 | 0.24 |
| 7 | 81575366 | rs966584 | -0.689 | 1.475 | 1.008 | 1.244 | 0.06 | 0.162 | 0.162 | 0 | 0.09 | 0.29 | 0.21 | 0.31 | 0.19 | 0.22 |
| 7 | 81584569 | rs1558371 | -0.903 | 1.418 | 1.229 | 1.198 | 0.017 | 0.02 | 0.02 | 0.054 | 0.33 | 0.4 | 0.42 | 0.5 | 0.42 | 0.46 |
| 7 | 81585921 | rs12668403 | -1.003 | 1.403 | 1.082 | 1.219 | 0.016 | 0.023 | 0.023 | 0.116 | 0.34 | 0.52 | 0.43 | 0.51 | 0.42 | 0.46 |
| 7 | 81588636 | rs1229502 | 0.71 | 1.193 | 0.522 | 1.117 | -0.039 | -0.017 | -0.017 | 0 | 0.26 | 0.11 | 0.26 | 0.24 | 0.3 | 0.25 |
| 7 | 81591716 | rs1229503 | 0.698 | 1.178 | 0.525 | 1.158 | -0.032 | -0.015 | -0.015 | 0 | 0.26 | 0.13 | 0.27 | 0.24 | 0.3 | 0.25 |
| 7 | 81593454 | rs3735517 | 0.179 | 0.847 | 0.686 | 1.035 | 0 | -0.029 | -0.029 | 0 | 0.26 | 0.35 | 0.23 | 0.2 | 0.23 | 0.24 |
| 7 | 81600197 | rs37067 | -0.299 | 0.48 | 0.433 | 0.865 | 0.004 | 0.002 | 0.002 | 0.667 | 0.45 | 0.53 | 0.58 | 0.6 | 0.64 | 0.6 |
| 7 | 81600267 | rs1229504 | 0.824 | 0.806 | 0.335 | 0.978 | -0.067 | -0.003 | -0.003 | 0 | 0.27 | 0.1 | 0.32 | 0.23 | 0.35 | 0.27 |
| 7 | 81600547 | rs37070 | -1.385 | 0.547 | 0.351 | 0.919 | 0.036 | 0.034 | 0.034 | 0 | 0.64 | 0.78 | 0.71 | 0.73 | 0.73 | 0.66 |
| 7 | 81601759 | rs10486940 | 0.889 | 0.546 | 0.358 | 0.922 | 0.035 | 0.022 | 0.022 | 2.984 | 0.19 | 0.1 | 0.12 | 0.12 | 0.09 | 0.06 |

| | | | | | | | | | | | | | | | | |
|---|---|---|---|---|---|---|---|---|---|---|---|---|---|---|---|---|
| 7 | 81609774 | rs37087 | -0.524 | 0.837 | 0.443 | 1.017 | 0.043 | 0.028 | 0.028 | 0.94 | 0.46 | 0.73 | 0.6 | 0.62 | 0.64 | 0.6 |
| 7 | 81610724 | rs37089 | -0.611 | 1.039 | 0.569 | 1.208 | 0.079 | 0.14 | 0.14 | 0.023 | 0.19 | 0.49 | 0.27 | 0.35 | 0.29 | 0.34 |
| 7 | 81617296 | rs37093 | -0.323 | 0.578 | -0.256 | 0.881 | -0.04 | -0.031 | -0.031 | 0.791 | 0.6 | 0.88 | 0.5 | 0.42 | 0.43 | 0.5 |
| 7 | 81646013 | rs11984173 | 1.412 | 0.053 | -0.11 | 0.755 | 0.061 | -0.015 | -0.015 | 0 | 0.27 | 0.12 | 0.23 | 0.26 | 0.17 | 0.2 |
| 7 | 81656066 | rs7797314 | 0.303 | 0.176 | 0.326 | 0.804 | 0.042 | -0.005 | -0.005 | 0.029 | 0.13 | 0.31 | 0.19 | 0.09 | 0.17 | 0.16 |
| 7 | 81657357 | rs2019073 | 0.337 | 0.11 | 0.445 | 0.759 | 0.024 | 0.018 | 0.018 | 0.1 | 0.45 | 0.39 | 0.29 | 0.32 | 0.26 | 0.3 |
| 7 | 81663275 | rs258728 | 0.306 | 0.066 | 0.307 | 0.74 | -0.004 | -0.001 | -0.001 | 0.102 | 0.45 | 0.56 | 0.3 | 0.32 | 0.26 | 0.3 |
| 7 | 81663428 | rs10231240 | 0.004 | 0.302 | 0.452 | 0.733 | 0.036 | -0.003 | -0.003 | 0.225 | 0.14 | 0.31 | 0.19 | 0.11 | 0.17 | 0.16 |
| 7 | 81668717 | rs3779452 | -0.313 | 0.341 | 0.178 | 1.05 | -0.006 | -0.007 | -0.007 | 3.813 | 0.85 | 0.86 | 0.82 | 0.69 | 0.79 | 0.76 |
| 7 | 81671035 | rs17155782 | 0.733 | -0.204 | 0.396 | 0.841 | -0.028 | -0.028 | -0.028 | 0 | 0.18 | 0.3 | 0.06 | 0.07 | 0.09 | 0.1 |
| 7 | 81671265 | rs6970802 | 0.663 | -0.221 | 0.263 | 0.843 | -0.014 | -0.023 | -0.023 | 0 | 0.36 | 0.4 | 0.32 | 0.25 | 0.27 | 0.31 |
| 7 | 81675161 | rs6467882 | -0.816 | 0.068 | 0.263 | 1.052 | -0.015 | -0.025 | -0.025 | 0 | 0.87 | 0.91 | 0.78 | 0.62 | 0.81 | 0.79 |
| 7 | 81684061 | rs2282936 | 0.325 | -0.221 | 0.41 | 0.821 | -0.063 | -0.06 | -0.06 | 0.091 | 0.21 | 0.46 | 0.08 | 0.1 | 0.09 | 0.1 |
| 7 | 81690700 | rs2237504 | -0.235 | -0.283 | 0.008 | 0.879 | 0.027 | 0.029 | 0.029 | 0 | 0.53 | 0.65 | 0.44 | 0.35 | 0.37 | 0.42 |
| 7 | 81691091 | rs10242428 | 0.791 | -0.667 | -0.093 | 0.666 | -0.003 | -0.02 | -0.02 | 0.183 | 0.16 | 0.21 | 0.12 | 0.07 | 0.14 | 0.19 |
| 7 | 81697666 | rs7802058 | -0.232 | -0.162 | -0.583 | 0.954 | 0.024 | -0.004 | -0.004 | 0 | 0.55 | 0.75 | 0.45 | 0.42 | 0.37 | 0.42 |
| 7 | 81702259 | rs6467883 | 1.049 | -0.167 | -0.719 | 0.884 | 0.004 | 0.004 | 0.004 | 0 | 0.19 | 0.18 | 0.26 | 0.16 | 0.15 | 0.13 |
| 7 | 81702854 | rs3823923 | 0.76 | 0.54 | -0.781 | 0.981 | -0.01 | 0.003 | 0.003 | 0 | 0.31 | 0.31 | 0.38 | 0.29 | 0.41 | 0.33 |
| 7 | 81708318 | rs1011696 | 0.099 | 0.59 | -0.921 | 0.952 | 0.033 | 0.024 | 0.024 | 0 | 0.5 | 0.56 | 0.41 | 0.35 | 0.35 | 0.44 |
| 7 | 81710361 | rs10247422 | 0.06 | 1.019 | -1.14 | 1.193 | -0.039 | -0.057 | -0.057 | 0 | 0.83 | 0.93 | 0.79 | 0.64 | 0.77 | 0.77 |
| 7 | 81713692 | rs2059037 | 0.105 | 0.448 | -0.863 | 0.695 | 0.003 | 0.002 | 0.002 | 0 | 0.19 | 0.18 | 0.14 | 0.18 | 0.15 | 0.13 |
| 7 | 81721513 | rs258651 | 0.454 | 0.635 | 0.012 | 0.504 | 0.04 | 0.051 | 0.051 | 0 | 0.73 | 0.55 | 0.65 | 0.59 | 0.67 | 0.64 |
| 7 | 81722247 | rs258653 | -0.107 | 0.911 | 0.07 | 0.551 | 0.037 | 0.153 | 0.153 | 0 | 0.83 | 0.61 | 0.74 | 0.59 | 0.81 | 0.8 |
| 7 | 81723042 | rs10250522 | 0.547 | 0.344 | 0.266 | 0.264 | -0.022 | 0.014 | 0.014 | 0 | 0.14 | 0.29 | 0.12 | 0.14 | 0.06 | 0.11 |
| 7 | 81723050 | rs7787882 | -0.203 | 0.614 | 0.776 | 0.302 | 0.019 | -0.035 | -0.035 | 0.358 | 0.11 | 0.25 | 0.12 | 0.03 | 0.15 | 0.19 |
| 7 | 81723112 | rs10265730 | 0.201 | 0.228 | 0.078 | -0.383 | -0.001 | -0.008 | -0.008 | 0 | 0.54 | 0.5 | 0.31 | 0.24 | 0.39 | 0.48 |
| 7 | 81724717 | rs10225295 | 0.209 | 0.282 | -0.032 | -0.36 | -0.002 | 0.015 | 0.015 | 0 | 0.57 | 0.63 | 0.33 | 0.23 | 0.39 | 0.45 |
| 7 | 81725241 | rs258659 | 0.913 | 0.314 | -0.156 | -0.313 | 0.007 | 0.008 | 0.008 | 0 | 0.27 | 0.27 | 0.44 | 0.38 | 0.41 | 0.31 |
| 7 | 81727778 | rs10808296 | 0.758 | 0.238 | -0.366 | -0.318 | 0.021 | 0.025 | 0.025 | 0 | 0.48 | 0.49 | 0.23 | 0.21 | 0.26 | 0.33 |
| 7 | 81732714 | rs1963788 | 0.852 | 0.307 | -0.179 | -0.297 | -0.025 | -0.034 | -0.034 | 0 | 0.24 | 0.16 | 0.43 | 0.38 | 0.4 | 0.3 |
| 7 | 81735015 | rs258684 | -0.301 | 0.006 | -0.257 | -0.232 | 0.014 | 0.002 | 0.002 | 0 | 0.38 | 0.48 | 0.55 | 0.41 | 0.57 | 0.51 |
| 7 | 81737327 | rs6960102 | 0.969 | -1.045 | -0.467 | -0.378 | 0.017 | 0.01 | 0.01 | 0 | 0.21 | 0.15 | 0.08 | 0.13 | 0.14 | 0.2 |
| 7 | 81737700 | rs258685 | -0.606 | -1.142 | -0.393 | -0.558 | -0.012 | -0.015 | -0.015 | 0 | 0.58 | 0.65 | 0.4 | 0.52 | 0.43 | 0.59 |
| 7 | 81738903 | rs2097914 | 0.56 | -1.079 | -0.424 | -0.569 | -0.009 | 0.011 | 0.011 | 0 | 0.09 | 0.04 | 0.06 | 0.03 | 0.09 | 0.16 |
| 7 | 81739031 | rs4732424 | 0.306 | -1.207 | -0.323 | -0.581 | -0.024 | 0.012 | 0.012 | 0 | 0.28 | 0.43 | 0.26 | 0.36 | 0.2 | 0.22 |
| 7 | 81741904 | rs12533683 | 1.408 | -1.052 | -0.405 | -0.53 | -0.004 | -0.026 | -0.026 | 0.192 | 0.19 | 0.13 | 0.21 | 0.37 | 0.18 | 0.16 |
| 7 | 81751207 | rs258688 | -1.543 | -1.267 | -1.056 | -0.731 | -0.158 | -0.136 | -0.136 | 0 | 0.74 | 0.99 | 0.53 | 0.61 | 0.52 | 0.72 |
| 7 | 81754449 | rs7341478 | -0.02 | -1.306 | -0.797 | -0.779 | -0.002 | -0.006 | -0.006 | 0 | 0.39 | 0.58 | 0.19 | 0.16 | 0.24 | 0.43 |
| 7 | 81755870 | rs258690 | -1.05 | -1.383 | -0.557 | -0.794 | 0.008 | 0.003 | 0.003 | 0 | 0.69 | 0.7 | 0.48 | 0.61 | 0.49 | 0.65 |
| 7 | 81758397 | rs258694 | 0.334 | -1.317 | -0.543 | -0.709 | 0.009 | -0.013 | -0.013 | 0 | 0.34 | 0.3 | 0.31 | 0.45 | 0.25 | 0.29 |
| 7 | 81759508 | rs3801748 | 0.527 | -0.953 | -0.3 | -0.547 | 0.004 | -0.001 | -0.001 | 0 | 0.47 | 0.46 | 0.32 | 0.5 | 0.35 | 0.41 |
| 7 | 81764719 | rs10234372 | 0.402 | -1.037 | -0.149 | -0.536 | -0.029 | -0.024 | -0.024 | 0 | 0.23 | 0.35 | 0.1 | 0.13 | 0.16 | 0.24 |
| 7 | 81768894 | rs258710 | 0.763 | -0.874 | -0.559 | -0.43 | -0.019 | -0.017 | -0.017 | 0 | 0.16 | 0.23 | 0.1 | 0.08 | 0.07 | 0.13 |
| 7 | 81771456 | rs258716 | 0.863 | -0.853 | -0.561 | -0.421 | -0.016 | -0.015 | -0.015 | 0 | 0.15 | 0.22 | 0.1 | 0.08 | 0.07 | 0.1 |
| 7 | 81771742 | rs258719 | 0.863 | -0.853 | -0.561 | -0.421 | -0.016 | -0.015 | -0.015 | 0 | 0.15 | 0.22 | 0.1 | 0.08 | 0.07 | 0.1 |
| 7 | 81772396 | rs258723 | -0.08 | -0.287 | -0.761 | -0.387 | 0.014 | 0.007 | 0.007 | 0.155 | 0.49 | 0.41 | 0.4 | 0.48 | 0.36 | 0.45 |
| 7 | 81773149 | rs10486945 | 0.586 | 0.107 | -1.594 | -0.134 | -0.038 | -0.059 | -0.059 | 0.051 | 0.33 | 0.09 | 0.29 | 0.41 | 0.28 | 0.35 |
| 7 | 81775555 | rs10954664 | -0.496 | 0.747 | -1.1 | 0.355 | -0.039 | -0.029 | -0.029 | 0 | 0.76 | 0.84 | 0.52 | 0.67 | 0.55 | 0.67 |
| 7 | 81781051 | rs3801740 | -0.742 | -0.021 | -0.466 | -0.123 | 0.001 | 0.005 | 0.005 | 0 | 0.39 | 0.62 | 0.3 | 0.24 | 0.34 | 0.44 |
| 7 | 81782514 | rs6942458 | -0.3 | -0.058 | -0.085 | -0.215 | -0.017 | -0.071 | -0.071 | 0 | 0.26 | 0.48 | 0.2 | 0.16 | 0.23 | 0.29 |
| 7 | 81785166 | rs11984254 | 0.212 | -0.412 | 0.218 | -0.008 | -0.024 | -0.006 | -0.006 | 0.027 | 0.13 | 0.29 | 0.07 | 0.11 | 0.08 | 0.17 |
| 7 | 81786628 | rs3801737 | 1.421 | -0.273 | -0.787 | 0.022 | -0.17 | -0.166 | -0.166 | 0 | 0.34 | 0.04 | 0.35 | 0.52 | 0.45 | 0.29 |
| 7 | 81788703 | rs3801730 | -0.235 | -0.815 | -0.366 | -0.106 | -0.014 | -0.012 | -0.012 | 0 | 0.42 | 0.5 | 0.46 | 0.3 | 0.34 | 0.36 |
| 7 | 81791527 | rs38536 | -0.908 | -1 | -0.248 | -0.107 | 0.041 | 0 | 0 | 0 | 0.53 | 0.78 | 0.53 | 0.39 | 0.41 | 0.48 |
| 7 | 81792001 | rs3801727 | 0.743 | -1.2 | 0.078 | -0.063 | -0.031 | 0.026 | 0.026 | 1.089 | 0.09 | 0.22 | 0.13 | 0.14 | 0.03 | 0.08 |
| 7 | 81792390 | rs38537 | 0.931 | -1.09 | 0.123 | -0.23 | -0.019 | 0.001 | 0.001 | 0.108 | 0.13 | 0.26 | 0.07 | 0.1 | 0.08 | 0.17 |
| 7 | 81793397 | rs17246425 | 0.191 | -0.889 | -0.117 | -0.251 | 0.008 | 0.022 | 0.022 | 0 | 0.31 | 0.19 | 0.32 | 0.15 | 0.29 | 0.21 |
| 7 | 81796853 | rs3801725 | 0.13 | -0.889 | -0.055 | -0.235 | 0.004 | 0.01 | 0.01 | 0 | 0.31 | 0.22 | 0.31 | 0.16 | 0.29 | 0.22 |
| 7 | 81802385 | rs17247184 | 1.606 | -0.698 | -0.488 | -0.009 | -0.146 | -0.109 | -0.109 | 0.774 | 0.35 | 0.04 | 0.36 | 0.46 | 0.48 | 0.3 |
| 7 | 81802908 | rs3801721 | 0.368 | -0.821 | -0.018 | -0.03 | -0.043 | 0.031 | 0.031 | 0 | 0.13 | 0.3 | 0.15 | 0.2 | 0.06 | 0.14 |
| 7 | 81804220 | rs38545 | 0.018 | -0.816 | -0.204 | -0.163 | -0.001 | 0 | 0 | 0.36 | 0.83 | 0.8 | 0.86 | 0.94 | 0.83 | 0.75 |
| 7 | 81805657 | rs1156450 | -0.576 | -1.148 | -0.205 | -0.141 | 0.006 | -0.002 | -0.002 | 0 | 0.73 | 0.78 | 0.69 | 0.76 | 0.73 | 0.7 |
| 7 | 81809550 | rs2237514 | 0.243 | -0.855 | -0.222 | -0.22 | -0.018 | 0.005 | 0.005 | 0 | 0.6 | 0.48 | 0.53 | 0.58 | 0.68 | 0.57 |
| 7 | 81810590 | rs6948895 | 0.552 | -0.113 | -0.645 | -0.286 | 0.027 | -0.004 | -0.004 | 0 | 0.48 | 0.3 | 0.41 | 0.53 | 0.52 | 0.48 |
| 7 | 81812996 | rs7796947 | 0.859 | 0.192 | -0.785 | -0.148 | -0.019 | -0.024 | -0.024 | 0 | 0.42 | 0.26 | 0.29 | 0.5 | 0.39 | 0.31 |
| 7 | 81821061 | rs2367912 | -0.779 | -0.346 | -0.493 | -0.552 | -0.013 | -0.029 | -0.029 | 0 | 0.35 | 0.46 | 0.24 | 0.19 | 0.25 | 0.36 |
| 7 | 81821589 | rs13235513 | 1.086 | -0.739 | -1.021 | -0.642 | 0.061 | 0.064 | 0.064 | 0.495 | 0.55 | 0.2 | 0.51 | 0.71 | 0.69 | 0.49 |
| 7 | 81823590 | rs1990071 | 0.211 | -0.12 | -1.268 | -0.62 | -0.047 | 0.01 | 0.01 | 0 | 0.71 | 0.92 | 0.72 | 0.78 | 0.63 | 0.67 |
| 7 | 81823930 | rs12707478 | 0.646 | -0.063 | -1.31 | -0.617 | -0.08 | -0.061 | -0.061 | 0.277 | 0.18 | 0.01 | 0.22 | 0.21 | 0.31 | 0.23 |

| | | | | | | | | | | | | | | | | |
|---|---|---|---|---|---|---|---|---|---|---|---|---|---|---|---|---|
| 7 | 81824126 | rs4732432 | 0.197 | -0.209 | -0.523 | -0.726 | -0.015 | -0.027 | -0.027 | 0 | 0.83 | 0.68 | 0.72 | 0.88 | 0.89 | 0.86 |
| 7 | 81824403 | rs4728500 | -0.658 | -0.274 | -0.736 | -0.867 | -0.003 | -0.03 | -0.03 | 0 | 0.87 | 0.85 | 0.92 | 1 | 0.91 | 0.84 |
| 7 | 81824570 | rs4732433 | 1.075 | -0.391 | -0.702 | -0.923 | 0 | 0.007 | 0.007 | 0 | 0.4 | 0.34 | 0.49 | 0.7 | 0.37 | 0.35 |
| 7 | 81825346 | rs1859541 | 0.347 | -0.71 | -0.11 | -0.784 | -0.011 | 0.147 | 0.147 | 0 | 0.1 | 0.43 | 0.26 | 0.17 | 0.06 | 0.11 |
| 7 | 81827878 | rs6950804 | -0.273 | -0.597 | -0.185 | -0.863 | 0.039 | 0.096 | 0.096 | 0 | 0.16 | 0.48 | 0.35 | 0.21 | 0.18 | 0.22 |
| 7 | 81830424 | rs3801714 | 0.362 | -0.901 | -0.063 | -0.861 | 0.015 | 0.115 | 0.115 | 0 | 0.1 | 0.39 | 0.26 | 0.17 | 0.08 | 0.06 |
| 7 | 81843391 | rs9692165 | -0.588 | -1.026 | -0.636 | -1.178 | -0.005 | -0.048 | -0.048 | 1.057 | 0.8 | 0.8 | 0.91 | 0.99 | 0.91 | 0.86 |
| 7 | 81848326 | rs2367914 | 0.343 | -0.646 | -0.876 | -1.245 | -0.004 | 0.086 | 0.086 | 0 | 0.55 | 0.76 | 0.66 | 0.8 | 0.55 | 0.57 |
| 7 | 81851811 | rs10229296 | 0.188 | -1.564 | -0.972 | -1.348 | -0.092 | -0.01 | -0.01 | 0 | 0.27 | 0.47 | 0.29 | 0.25 | 0.12 | 0.15 |
| 7 | 81853237 | rs11514983 | 0.817 | -0.766 | -1.248 | -1.173 | -0.075 | -0.006 | -0.006 | 0 | 0.25 | 0.07 | 0.26 | 0.19 | 0.36 | 0.32 |
| 7 | 81855594 | rs10954669 | -0.536 | -1.12 | -0.926 | -1.084 | 0.007 | -0.009 | -0.009 | 0 | 0.8 | 0.81 | 0.9 | 0.99 | 0.91 | 0.84 |
| 7 | 81868051 | rs2214726 | -0.881 | -0.678 | -1.005 | -1.075 | 0 | -0.003 | -0.003 | 0.883 | 0.83 | 0.87 | 0.92 | 1 | 0.89 | 0.88 |
| 7 | 81868200 | rs10275620 | 1.177 | -1.065 | -0.712 | -1.109 | -0.045 | 0.001 | 0.001 | 0 | 0.11 | 0.26 | 0.04 | 0.09 | 0.04 | 0.08 |
| 7 | 81868265 | rs10275727 | 0.964 | -1.09 | -0.763 | -1.12 | -0.051 | -0.004 | -0.004 | 1.065 | 0.12 | 0.3 | 0.04 | 0.09 | 0.04 | 0.08 |
| 7 | 81877715 | rs38553 | -0.242 | -0.662 | -1.289 | -1.271 | -0.045 | 0.094 | 0.094 | 0 | 0.27 | 0.13 | 0.24 | 0.04 | 0.35 | 0.25 |
| 7 | 81877993 | rs3801706 | 0.875 | -0.85 | -1.417 | -1.203 | -0.021 | 0.013 | 0.013 | 0 | 0.27 | 0.09 | 0.21 | 0.2 | 0.29 | 0.31 |
| 7 | 81880361 | rs38554 | -1.418 | -0.677 | -1.35 | -1.014 | -0.071 | -0.099 | -0.099 | 0 | 0.62 | 0.9 | 0.46 | 0.44 | 0.43 | 0.56 |
| 7 | 81880696 | rs38555 | -2.285 | -0.651 | -1.213 | -0.849 | -0.094 | -0.125 | -0.125 | 0.407 | 0.73 | 0.96 | 0.64 | 0.48 | 0.67 | 0.72 |
| 7 | 81883891 | rs17156069 | 1.009 | -0.661 | -0.612 | -0.866 | 0.018 | 0.083 | 0.083 | 0.239 | 0.06 | 0.31 | 0.03 | 0.05 | 0.02 | 0.02 |
| 7 | 81892962 | rs40634 | -0.772 | -0.581 | -1.57 | -0.986 | 0.001 | 0.053 | 0.053 | 0 | 0.34 | 0.17 | 0.26 | 0.19 | 0.35 | 0.35 |
| 7 | 81897925 | rs3801703 | 0.664 | -0.202 | -1.538 | -0.679 | -0.029 | 0.004 | 0.004 | 0 | 0.26 | 0.07 | 0.2 | 0.23 | 0.3 | 0.31 |
| 7 | 81908393 | rs1010344 | -0.174 | -0.458 | -1.549 | -0.742 | -0.03 | 0.018 | 0.018 | 0 | 0.19 | 0.07 | 0.26 | 0.11 | 0.25 | 0.21 |
| 7 | 81908725 | rs2299166 | -0.348 | -0.743 | -1.05 | -0.789 | -0.037 | 0.002 | 0.002 | 0 | 0.17 | 0.38 | 0.12 | 0.14 | 0.1 | 0.09 |
| 7 | 81912148 | rs3757631 | 0.066 | -0.386 | -1.12 | -0.691 | 0.002 | 0 | 0 | 0 | 0.31 | 0.36 | 0.18 | 0.27 | 0.25 | 0.3 |
| 7 | 81914169 | rs17177211 | 0.599 | -0.419 | -1.165 | -0.602 | -0.032 | 0 | 0 | 0 | 0.17 | 0.05 | 0.24 | 0.12 | 0.25 | 0.21 |
| 7 | 81915186 | rs10280428 | -0.232 | -0.882 | -0.495 | -0.823 | -0.03 | -0.082 | -0.082 | 0.013 | 0.15 | 0.48 | 0.07 | 0.01 | 0.06 | 0.09 |
| 7 | 81915620 | rs10225267 | -0.233 | -0.756 | -0.315 | -0.747 | 0.058 | 0.059 | 0.059 | 2.753 | 0.06 | 0.44 | 0.03 | 0.01 | 0 | 0.02 |
| 7 | 81923441 | rs2299169 | 1.087 | -0.335 | -1.096 | -0.662 | -0.007 | 0.033 | 0.033 | 0 | 0.21 | 0.07 | 0.14 | 0.16 | 0.2 | 0.3 |
| 7 | 81923913 | rs740340 | -0.993 | -0.636 | -1.201 | -0.666 | 0.045 | 0.04 | 0.04 | 0 | 0.51 | 0.72 | 0.51 | 0.34 | 0.44 | 0.44 |
| 7 | 81926808 | rs6942428 | 0.478 | -0.639 | -1.618 | -0.454 | -0.055 | -0.015 | -0.015 | 0 | 0.25 | 0.05 | 0.29 | 0.19 | 0.34 | 0.29 |
| 7 | 81942310 | rs10954673 | 1.123 | -0.87 | -1.08 | -0.593 | 0 | 0.001 | 0.001 | 0 | 0.2 | 0.14 | 0.14 | 0.11 | 0.19 | 0.32 |
| 7 | 81943935 | rs929416 | -0.297 | -1.223 | -0.824 | -0.449 | 0.025 | 0.015 | 0.015 | 0.073 | 0.35 | 0.68 | 0.32 | 0.25 | 0.25 | 0.53 |
| 7 | 81944479 | rs12673548 | 1.244 | -1.084 | -0.917 | -0.526 | 0 | 0.001 | 0.001 | 0 | 0.2 | 0.14 | 0.14 | 0.11 | 0.19 | 0.32 |
| 7 | 81951013 | rs2041047 | 0.397 | -1.07 | -1.388 | -0.461 | -0.002 | 0.007 | 0.007 | 0 | 0.27 | 0.22 | 0.3 | 0.2 | 0.34 | 0.25 |
| 7 | 81952939 | rs42048 | -0.805 | -1.111 | -1.432 | -0.461 | -0.054 | -0.016 | -0.016 | 0 | 0.74 | 0.95 | 0.71 | 0.8 | 0.66 | 0.75 |
| 7 | 81957411 | rs17156126 | 0.863 | -1.124 | -0.79 | -0.457 | -0.037 | -0.062 | -0.062 | 0.492 | 0.12 | 0.22 | 0.05 | 0.01 | 0.05 | 0.19 |
| 7 | 81958387 | rs42054 | -0.069 | -1.149 | -0.914 | -0.486 | -0.009 | 0.054 | 0.054 | 0 | 0.53 | 0.72 | 0.56 | 0.7 | 0.47 | 0.43 |
| 7 | 81960231 | rs42236 | 1.613 | -0.807 | -1.148 | -0.247 | -0.145 | -0.168 | -0.168 | 0 | 0.26 | 0.01 | 0.32 | 0.55 | 0.35 | 0.2 |
| 7 | 81968032 | rs2299173 | 0.146 | -0.014 | -0.407 | -0.015 | -0.018 | -0.004 | -0.004 | 0 | 0.14 | 0.3 | 0.06 | 0.07 | 0.07 | 0.19 |
| 7 | 81981195 | rs2368020 | -0.385 | 0.078 | -0.718 | 0.058 | -0.029 | -0.005 | -0.005 | 0 | 0.77 | 0.88 | 0.71 | 0.8 | 0.66 | 0.75 |
| 7 | 81984104 | rs724118 | -0.477 | 0.133 | -0.843 | 0.133 | -0.031 | -0.032 | -0.032 | 0.737 | 0.76 | 0.89 | 0.72 | 0.75 | 0.67 | 0.76 |
| 7 | 81987870 | rs2190235 | -0.918 | 0.423 | -0.824 | 0.085 | -0.046 | -0.047 | -0.047 | 0.789 | 0.92 | 1 | 0.87 | 0.83 | 0.84 | 0.87 |
| 7 | 81993260 | rs2018982 | -0.29 | -0.041 | -0.296 | -0.21 | -0.002 | 0.006 | 0.006 | 0 | 0.68 | 0.35 | 0.66 | 0.67 | 0.66 | 0.62 |
| 7 | 82004280 | rs11768220 | 0.881 | -0.087 | -0.767 | -0.514 | 0.009 | 0.027 | 0.027 | 2.648 | 0.11 | 0.03 | 0.03 | 0.01 | 0.05 | 0.08 |
| 7 | 82005470 | rs1544461 | 0.36 | -0.112 | -0.664 | -0.653 | 0 | -0.003 | -0.003 | 0.008 | 0.45 | 0.3 | 0.5 | 0.58 | 0.42 | 0.3 |
| 7 | 82011353 | rs2299184 | 0.39 | -1.199 | -0.774 | -1.855 | -0.022 | -0.056 | -0.056 | 0 | 0.38 | 0.41 | 0.23 | 0.11 | 0.23 | 0.17 |
| 7 | 82011505 | rs2299185 | 0.134 | -1.174 | -0.814 | -1.843 | 0.062 | 0.088 | 0.088 | 12.087 | 0.41 | 0.71 | 0.27 | 0.15 | 0.24 | 0.23 |
| 7 | 82012494 | rs2237526 | -0.107 | -0.884 | -0.994 | -1.519 | 0 | -0.014 | -0.014 | 0 | 0.43 | 0.63 | 0.25 | 0.13 | 0.35 | 0.43 |
| 7 | 82013724 | rs10274659 | 0.17 | -0.199 | -1.45 | -0.58 | -0.018 | -0.016 | -0.016 | 0.252 | 0.38 | 0.45 | 0.24 | 0.23 | 0.25 | 0.19 |
| 7 | 82014592 | rs3801682 | -0.689 | 0.366 | -1.88 | -0.497 | -0.072 | 0.155 | 0.155 | 0 | 0.3 | 0.04 | 0.22 | 0.1 | 0.31 | 0.39 |
| 7 | 82024922 | rs3801671 | -0.622 | 0.019 | -1.268 | 0.118 | 0.017 | 0.029 | 0.029 | 0 | 0.11 | 0.04 | 0.04 | 0.01 | 0.06 | 0.01 |
| 7 | 82035021 | rs17156246 | -1.142 | -0.336 | -1.051 | -0.351 | -0.001 | -0.025 | -0.025 | 0 | 0.22 | 0.14 | 0.08 | 0.02 | 0.15 | 0.1 |
| 7 | 82052351 | rs763448 | -0.067 | -1.04 | -2.236 | 0.611 | -0.036 | -0.147 | -0.147 | 2.247 | 0.2 | 0.02 | 0.32 | 0.44 | 0.18 | 0.29 |
| 7 | 82057406 | rs4732461 | -0.214 | -1.323 | -2.28 | 0.572 | -0.018 | -0.095 | -0.095 | 0 | 0.19 | 0.07 | 0.33 | 0.43 | 0.19 | 0.26 |
| 7 | 82058986 | rs757847 | 0.501 | -0.243 | -2.615 | -0.022 | -0.065 | -0.069 | -0.069 | 0 | 0.7 | 0.86 | 0.51 | 0.56 | 0.59 | 0.65 |
| 7 | 82059917 | rs731160 | 1.144 | -0.786 | -3.07 | -0.687 | -0.035 | -0.027 | -0.027 | 0 | 0.46 | 0.07 | 0.31 | 0.43 | 0.44 | 0.56 |
| 7 | 82066387 | rs10234381 | 0.783 | -0.889 | -1.746 | -0.736 | -0.011 | -0.101 | -0.101 | 0 | 0.17 | 0.37 | 0.1 | 0.01 | 0.14 | 0.07 |
| 7 | 82066634 | rs17148126 | 0.878 | -0.873 | -1.928 | -0.872 | 0.022 | 0.04 | 0.04 | 0.536 | 0.14 | 0.05 | 0.04 | 0.01 | 0.08 | 0.02 |
| 7 | 82067309 | rs3801664 | -0.473 | -0.805 | -1.898 | -0.815 | -0.024 | -0.037 | -0.037 | 1.981 | 0.34 | 0.41 | 0.22 | 0.13 | 0.23 | 0.12 |
| 7 | 82075560 | rs12707492 | 1.909 | -0.735 | -2.165 | -0.773 | -0.072 | -0.099 | -0.099 | 1.996 | 0.33 | 0.04 | 0.27 | 0.44 | 0.36 | 0.55 |
| 7 | 82079601 | rs6977797 | -0.178 | -0.771 | -1.937 | -0.824 | 0.036 | 0.054 | 0.054 | 0 | 0.32 | 0.78 | 0.16 | 0.02 | 0.2 | 0.08 |
| 7 | 82084287 | rs6965042 | -0.191 | -0.768 | -1.943 | -0.823 | 0.038 | 0.059 | 0.059 | 0 | 0.32 | 0.79 | 0.16 | 0.02 | 0.2 | 0.08 |
| 7 | 82088388 | rs10486960 | 1.231 | -0.436 | -1.981 | -0.054 | -0.032 | -0.058 | -0.058 | 0 | 0.14 | 0.03 | 0.31 | 0.22 | 0.15 | 0.17 |
| 7 | 82089478 | rs7780221 | -0.033 | -0.386 | -1.934 | -0.13 | -0.084 | -0.092 | -0.092 | 3.624 | 0.72 | 0.89 | 0.52 | 0.57 | 0.59 | 0.66 |
| 7 | 82095415 | rs17148130 | 1.688 | -1.108 | -2.436 | -0.139 | 0.037 | -0.062 | -0.062 | 1.812 | 0.16 | 0.07 | 0.07 | 0.33 | 0.09 | 0.17 |
| 7 | 82101994 | rs6967410 | 0.208 | -0.85 | -2.549 | -0.407 | -0.09 | -0.06 | -0.06 | 0.651 | 0.58 | 0.85 | 0.38 | 0.56 | 0.51 | 0.6 |
| 7 | 82103053 | rs10262899 | 1.516 | -0.783 | -2.14 | -0.303 | -0.015 | 0.014 | 0.014 | 0 | 0.17 | 0.23 | 0.09 | 0.32 | 0.09 | 0.17 |
| 7 | 82123522 | rs12113700 | 1.366 | -0.527 | -1.767 | -0.437 | 0.039 | -0.055 | -0.055 | 0 | 0.22 | 0.42 | 0.11 | 0.12 | 0.25 | 0.31 |
| 7 | 82125194 | rs6467900 | -0.495 | -0.61 | -1.965 | -0.454 | -0.093 | -0.087 | -0.087 | 0.457 | 0.69 | 0.88 | 0.48 | 0.59 | 0.58 | 0.57 |

| | | | | | | | | | | | | | | | | |
|---|---|---|---|---|---|---|---|---|---|---|---|---|---|---|---|---|
| 7 | 82132710 | rs6467901 | 1.077 | -0.678 | -1.921 | -0.469 | -0.003 | 0.009 | 0.009 | 0.018 | 0.34 | 0.43 | 0.17 | 0.44 | 0.32 | 0.47 |
| 7 | 82136552 | rs2158636 | 0.446 | -0.54 | -2.034 | -0.534 | -0.012 | -0.02 | -0.02 | 2.921 | 0.32 | 0.33 | 0.28 | 0.14 | 0.24 | 0.1 |
| 7 | 82137236 | rs6947580 | -1.553 | 0.117 | -1.94 | -0.214 | -0.054 | 0.03 | 0.03 | 0 | 0.92 | 0.99 | 0.81 | 0.99 | 0.79 | 0.92 |
| 7 | 82138216 | rs10233240 | 0.349 | 0.148 | -1.768 | 0.012 | -0.023 | 0.061 | 0.061 | 0 | 0.28 | 0.58 | 0.2 | 0.41 | 0.22 | 0.38 |
| 7 | 82138942 | rs2107850 | 0.06 | 0.29 | -2.118 | -0.015 | -0.089 | -0.005 | -0.005 | 0 | 0.76 | 0.91 | 0.49 | 0.76 | 0.61 | 0.64 |
| 7 | 82139481 | rs7792064 | 0.071 | 0.236 | -2.122 | -0.019 | -0.085 | -0.005 | -0.005 | 0 | 0.76 | 0.91 | 0.5 | 0.76 | 0.62 | 0.68 |
| 7 | 82143020 | rs757740 | -0.291 | 0.433 | -2.116 | 0.098 | -0.033 | -0.042 | -0.042 | 0 | 0.25 | 0.13 | 0.48 | 0.32 | 0.3 | 0.26 |
| 7 | 82143888 | rs740133 | 0.415 | -1.281 | -2.133 | -0.256 | 0.009 | 0.038 | 0.038 | 0 | 0.38 | 0.28 | 0.31 | 0.31 | 0.34 | 0.57 |
| 7 | 82144312 | rs9642136 | -0.205 | -1.428 | -1.535 | 0.039 | -0.023 | 0.033 | 0.033 | 0 | 0.92 | 0.98 | 0.87 | 0.99 | 0.9 | 0.91 |
| 7 | 82147302 | rs10486963 | 1.622 | -2.454 | -1.02 | 0.583 | -0.007 | -0.005 | -0.005 | 0.023 | 0.58 | 0.4 | 0.44 | 0.43 | 0.61 | 0.53 |
| 7 | 82148144 | rs10252807 | 0.966 | -2.561 | -0.67 | 0.235 | 0.005 | -0.036 | -0.036 | 0.911 | 0.11 | 0.29 | 0.2 | 0.02 | 0.11 | 0.03 |
| 7 | 82149580 | rs6950390 | -2.142 | -2.529 | -0.707 | 0.227 | -0.029 | 0.026 | 0.026 | 0 | 0.95 | 1 | 0.97 | 1 | 0.92 | 0.92 |
| 7 | 82150331 | rs10486965 | 0.323 | -2.639 | -0.675 | 0.389 | 0.001 | 0.005 | 0.005 | 0 | 0.1 | 0.09 | 0.06 | 0.2 | 0.15 | 0.25 |
| 7 | 82153240 | rs6971119 | 1.875 | -2.622 | -0.745 | 0.504 | -0.005 | 0.002 | 0.002 | 0.117 | 0.65 | 0.4 | 0.61 | 0.65 | 0.66 | 0.61 |
| 7 | 82158582 | rs7802743 | 0.242 | -2.467 | -1.123 | 0.429 | 0 | 0.008 | 0.008 | 0 | 0.79 | 0.87 | 0.7 | 0.86 | 0.81 | 0.86 |
| 7 | 82168867 | rs11972597 | 1.023 | -2.459 | -1.133 | 0.425 | 0.009 | 0.074 | 0.074 | 0 | 0.27 | 0.12 | 0.22 | 0.12 | 0.23 | 0.34 |
| 7 | 82169556 | rs1859545 | 0.156 | -2.516 | -1.164 | 0.55 | 0.001 | 0.001 | 0.001 | 0 | 0.15 | 0.14 | 0.1 | 0.2 | 0.2 | 0.17 |
| 7 | 82170046 | rs7788561 | 1.741 | -2.514 | -1.198 | 0.573 | 0.014 | 0.075 | 0.075 | 0.341 | 0.21 | 0.05 | 0.19 | 0.12 | 0.15 | 0.33 |
| 7 | 82170371 | rs12668197 | 0.635 | -2.788 | -0.962 | 0.764 | 0.06 | -0.047 | -0.047 | 0.737 | 0.33 | 0.16 | 0.23 | 0.46 | 0.18 | 0.42 |
| 7 | 82172769 | rs17156397 | 0.991 | -2.583 | -0.877 | 0.942 | -0.013 | -0.044 | -0.044 | 0 | 0.07 | 0 | 0.17 | 0.23 | 0.07 | 0.06 |
| 7 | 82175885 | rs10486969 | 1.427 | -2.611 | -0.871 | 0.942 | 0.006 | -0.041 | -0.041 | 0 | 0.07 | 0 | 0.17 | 0.18 | 0.05 | 0.05 |
| 7 | 82177900 | rs13438136 | 0.648 | -2.826 | -0.91 | 0.847 | 0.046 | -0.042 | -0.042 | 0 | 0.31 | 0.17 | 0.22 | 0.46 | 0.18 | 0.43 |
| 7 | 82178561 | rs17156412 | -0.625 | -2.715 | -0.859 | 0.88 | -0.01 | -0.037 | -0.037 | 0 | 0.21 | 0.13 | 0.27 | 0.4 | 0.24 | 0.22 |
| 7 | 82178889 | rs7793652 | 0.954 | -2.967 | -0.294 | 0.688 | 0.004 | 0.057 | 0.057 | 0 | 0.45 | 0.26 | 0.42 | 0.2 | 0.55 | 0.21 |
| 7 | 82178978 | rs10954681 | 1.007 | -2.965 | -0.293 | 0.689 | 0.003 | 0.053 | 0.053 | 0 | 0.44 | 0.26 | 0.42 | 0.2 | 0.54 | 0.21 |
| 7 | 82183483 | rs7810918 | -0.601 | -2.928 | -0.086 | 0.824 | -0.006 | 0.022 | 0.022 | 0 | 0.2 | 0.38 | 0.28 | 0.38 | 0.17 | 0.21 |
| 7 | 82183868 | rs11760881 | 1.455 | -3.032 | -0.081 | 0.831 | -0.01 | 0.008 | 0.008 | 0 | 0.43 | 0.26 | 0.25 | 0.35 | 0.46 | 0.24 |
| 7 | 82184977 | rs2189990 | -0.357 | -2.979 | -0.051 | 0.825 | 0.003 | 0.034 | 0.034 | 0 | 0.64 | 0.78 | 0.54 | 0.73 | 0.64 | 0.45 |
| 7 | 82187231 | rs10265943 | 0.503 | -2.982 | -0.037 | 0.764 | 0.01 | 0.01 | 0.01 | 0 | 0.33 | 0.43 | 0.37 | 0.32 | 0.29 | 0.51 |
| 7 | 82188970 | rs6467906 | 0.466 | -3.003 | -0.104 | 0.771 | 0 | -0.011 | -0.011 | 0.152 | 0.52 | 0.77 | 0.46 | 0.52 | 0.51 | 0.28 |
| 7 | 82190666 | rs4335074 | -0.024 | -2.933 | -0.072 | 0.782 | -0.014 | 0.004 | 0.004 | 0 | 0.4 | 0.57 | 0.55 | 0.49 | 0.33 | 0.56 |
| 7 | 82193853 | rs2189993 | 1.215 | -2.776 | -0.177 | 0.835 | -0.001 | -0.001 | -0.001 | 0.049 | 0.47 | 0.46 | 0.44 | 0.53 | 0.51 | 0.29 |
| 7 | 82193993 | rs17156423 | 1.41 | -2.196 | -0.091 | 1.054 | -0.038 | -0.056 | -0.056 | 13.771 | 0.09 | 0.01 | 0.08 | 0.21 | 0.12 | 0.15 |
| 7 | 82194982 | rs17156429 | -0.065 | -2.256 | 0.007 | 1.056 | 0.009 | -0.066 | -0.066 | 0 | 0.21 | 0.08 | 0.11 | 0.39 | 0.16 | 0.21 |
| 7 | 82197469 | rs10486972 | 0.014 | -2.232 | 0.021 | 1.062 | 0.008 | -0.052 | -0.052 | 0 | 0.21 | 0.12 | 0.12 | 0.39 | 0.16 | 0.21 |
| 7 | 82199173 | rs2214733 | -1.215 | -1.941 | -0.05 | 1.092 | -0.012 | -0.036 | -0.036 | 0 | 0.93 | 1 | 0.89 | 0.88 | 0.93 | 0.9 |
| 7 | 82204358 | rs2368084 | 0.793 | -2.008 | -0.343 | 1.162 | 0.014 | 0.015 | 0.015 | 0 | 0.5 | 0.37 | 0.35 | 0.38 | 0.6 | 0.35 |
| 7 | 82204878 | rs4728516 | -1.244 | -1.669 | -1.084 | 1.52 | -0.017 | -0.033 | -0.033 | 0 | 0.94 | 1 | 0.89 | 0.88 | 0.93 | 0.91 |
| 7 | 82206178 | rs7457005 | -0.458 | -1.738 | -1.103 | 1.512 | 0.004 | 0.017 | 0.017 | 0.081 | 0.73 | 0.91 | 0.62 | 0.78 | 0.76 | 0.57 |
| 7 | 82207785 | rs2189984 | 0.233 | -1.333 | -0.748 | 1.652 | 0.008 | 0.069 | 0.069 | 0 | 0.83 | 0.74 | 0.8 | 0.64 | 0.8 | 0.75 |
| 7 | 82214954 | rs12531738 | -0.087 | -1.347 | -1.426 | 1.173 | 0.017 | -0.033 | -0.033 | 0 | 0.07 | 0.02 | 0.05 | 0.21 | 0.05 | 0.08 |
| 7 | 82218457 | rs10486973 | 0.491 | -1.369 | -1.549 | 0.911 | -0.001 | 0.048 | 0.048 | 1.421 | 0.23 | 0.06 | 0.27 | 0.14 | 0.17 | 0.33 |
| 7 | 82219425 | rs1981576 | -0.687 | -1.136 | -1.506 | 0.889 | -0.013 | -0.039 | -0.039 | 0 | 0.93 | 1 | 0.88 | 0.86 | 0.93 | 0.89 |
| 7 | 82222805 | rs10231433 | 0.144 | -1.163 | -0.909 | 0.801 | 0.006 | -0.001 | -0.001 | 0 | 0.27 | 0.28 | 0.41 | 0.27 | 0.19 | 0.4 |
| 7 | 82223696 | rs4732466 | -0.102 | -0.42 | -1.085 | 0.158 | 0.004 | 0.121 | 0.121 | 0.437 | 0.47 | 0.78 | 0.44 | 0.27 | 0.53 | 0.29 |
| 7 | 82224180 | rs1034784 | 0.982 | -0.197 | -0.298 | 0.664 | -0.001 | 0.002 | 0.002 | 0 | 0.92 | 0.94 | 0.91 | 0.88 | 0.91 | 0.9 |
| 7 | 82227551 | rs10267460 | -1.53 | -0.901 | 0.07 | 0.628 | -0.029 | 0.009 | 0.009 | 0 | 0.24 | 0.67 | 0.19 | 0.37 | 0.12 | 0.41 |
| 7 | 82229179 | rs17278657 | -0.313 | -0.587 | 0.136 | 0.117 | 0.009 | -0.027 | -0.027 | 0 | 0.06 | 0.09 | 0.05 | 0 | 0.1 | 0.01 |
| 7 | 82230985 | rs17156492 | -1.103 | -0.828 | 1.373 | 0.02 | -0.009 | -0.008 | -0.008 | 0 | 0.06 | 0.14 | 0.03 | 0.04 | 0.04 | 0.07 |
| 7 | 82245831 | rs11768819 | -1.672 | -0.762 | 1.435 | -0.054 | -0.005 | -0.032 | -0.032 | 0.031 | 0.09 | 0.19 | 0.06 | 0.03 | 0.09 | 0.15 |
| 7 | 82247882 | rs7788705 | -2.774 | -1.01 | 1.929 | 0.106 | -0.023 | 0.015 | 0.015 | 0 | 0.21 | 0.48 | 0.23 | 0.21 | 0.2 | 0.26 |
| 7 | 82251633 | rs2368088 | -1.878 | -0.796 | 1.858 | -0.08 | -0.009 | -0.022 | -0.022 | 0 | 0.1 | 0.26 | 0.06 | 0.05 | 0.08 | 0.15 |
| 7 | 82251883 | rs10267783 | -0.526 | -0.683 | 1.447 | -0.019 | 0.026 | -0.036 | -0.036 | 0 | 0.07 | 0.23 | 0.07 | 0 | 0.11 | 0.01 |
| 7 | 82256206 | rs10281234 | -0.961 | -0.693 | 1.414 | -0.005 | 0.025 | 0.031 | 0.031 | 0 | 0.08 | 0.23 | 0.17 | 0.12 | 0.12 | 0.06 |
| 7 | 82285637 | rs757133 | -1.085 | -0.781 | 1.269 | -0.104 | 0.026 | -0.074 | -0.074 | 0 | 0.13 | 0.45 | 0.11 | 0.02 | 0.16 | 0.09 |
| 7 | 82286106 | rs757134 | 1.423 | -0.368 | 0.263 | -0.772 | -0.041 | 0.155 | 0.155 | 0 | 0.55 | 0.16 | 0.42 | 0.81 | 0.5 | 0.56 |
| 7 | 82299789 | rs17156564 | -1.639 | -0.598 | 0.661 | -0.421 | -0.003 | -0.006 | -0.006 | 0 | 0.26 | 0.67 | 0.29 | 0.14 | 0.22 | 0.24 |
| 7 | 82303636 | rs12707512 | -1.357 | -0.25 | 0.534 | 0.005 | 0.01 | 0.061 | 0.061 | 0 | 0.06 | 0.19 | 0.12 | 0.1 | 0.05 | 0.14 |
| 7 | 82306895 | rs10954686 | 0.501 | -0.242 | -0.559 | 0.126 | -0.023 | 0.03 | 0.03 | 0.083 | 0.45 | 0.12 | 0.36 | 0.37 | 0.43 | 0.45 |
| 7 | 82313975 | rs4732474 | -1.072 | -0.332 | 0.304 | 0.375 | -0.027 | 0.064 | 0.064 | 0 | 0.28 | 0.56 | 0.22 | 0.43 | 0.23 | 0.33 |
| 7 | 82317559 | rs2106920 | 0.682 | -0.613 | 0.257 | -0.051 | -0.006 | -0.037 | -0.037 | 0 | 0.13 | 0.16 | 0.08 | 0 | 0.09 | 0.02 |
| 7 | 82320160 | rs7785156 | 0.574 | -0.335 | 0.246 | -0.055 | 0.003 | -0.024 | -0.024 | 0.825 | 0.13 | 0.19 | 0.3 | 0.04 | 0.13 | 0.1 |
| 7 | 82327115 | rs1989907 | 0.387 | -0.309 | 0.5 | -0.043 | 0.001 | -0.037 | -0.037 | 2.33 | 0.14 | 0.22 | 0.3 | 0.04 | 0.13 | 0.1 |
| 7 | 82328200 | rs10229392 | -1.335 | -0.355 | 0.602 | -0.051 | 0.028 | -0.03 | -0.03 | 0 | 0.42 | 0.8 | 0.52 | 0.47 | 0.35 | 0.43 |
| 7 | 82331576 | rs997166 | 0.233 | -0.473 | 0.88 | 0.132 | -0.022 | 0.108 | 0.108 | 0 | 0.06 | 0.22 | 0.11 | 0.3 | 0.03 | 0.07 |
| 7 | 82337806 | rs2189029 | -1.445 | -0.456 | 0.795 | 0.107 | 0.02 | -0.018 | -0.018 | 0.678 | 0.42 | 0.73 | 0.52 | 0.47 | 0.35 | 0.43 |
| 7 | 82337859 | rs2371208 | -0.985 | -0.453 | 0.789 | 0.113 | -0.032 | 0.082 | 0.082 | 0 | 0.28 | 0.5 | 0.22 | 0.43 | 0.23 | 0.34 |
| 7 | 82342570 | rs985020 | -0.898 | -0.232 | 0.912 | 0.324 | -0.002 | 0 | 0 | 0 | 0.4 | 0.63 | 0.3 | 0.43 | 0.32 | 0.36 |
| 7 | 82342937 | rs2214330 | 0.358 | -0.23 | 0.9 | 0.308 | 0.009 | 0.053 | 0.053 | 0 | 0.24 | 0.15 | 0.12 | 0.04 | 0.22 | 0.11 |

| | | | | | | | | | | | | | | | | |
|---|---|---|---|---|---|---|---|---|---|---|---|---|---|---|---|---|
| 7 | 82347290 | rs10246264 | -0.864 | -0.224 | 1.051 | 0.372 | -0.035 | 0.206 | 0.206 | 0 | 0.15 | 0.45 | 0.17 | 0.39 | 0.1 | 0.25 |
| 7 | 82348028 | rs7806760 | -1.153 | -0.233 | 0.684 | 0.342 | 0.009 | -0.009 | -0.009 | 0 | 0.44 | 0.66 | 0.53 | 0.49 | 0.39 | 0.44 |
| 7 | 82348139 | rs970867 | -1.254 | -0.445 | -0.336 | 0.167 | -0.029 | 0.02 | 0.02 | 0.083 | 0.66 | 0.98 | 0.81 | 0.74 | 0.67 | 0.67 |
| 7 | 82348481 | rs6975360 | -0.323 | -0.388 | 0.838 | 0.278 | -0.001 | -0.027 | -0.027 | 0 | 0.27 | 0.33 | 0.34 | 0.06 | 0.25 | 0.16 |
| 7 | 82350584 | rs6467910 | -1.321 | 0.233 | 1.183 | 1.277 | 0.048 | -0.001 | -0.001 | 14.77 | 0.12 | 0.59 | 0.28 | 0.09 | 0.12 | 0.26 |
| 7 | 82357284 | rs969015 | -1.321 | 0.463 | 1.287 | 0.287 | 0.047 | -0.001 | -0.001 | 0.364 | 0.12 | 0.59 | 0.28 | 0.09 | 0.12 | 0.26 |
| 7 | 82358006 | rs17156615 | -0.858 | 0.469 | 1.367 | 0.287 | 0.028 | 0.01 | 0.01 | 0 | 0.11 | 0.4 | 0.27 | 0.08 | 0.1 | 0.2 |
| 7 | 82361634 | rs9642144 | 0.03 | 0.403 | 0.787 | 0.148 | -0.001 | -0.004 | -0.004 | 0.57 | 0.05 | 0.07 | 0.04 | 0.02 | 0.05 | 0.05 |
| 7 | 82371376 | rs6956744 | -1.207 | 0.361 | 0.078 | 0.182 | 0.006 | -0.033 | -0.033 | 0 | 0.18 | 0.67 | 0.32 | 0.1 | 0.17 | 0.32 |
| 7 | 82375808 | rs2106919 | 0.373 | 0.07 | -0.242 | 0.179 | -0.013 | -0.015 | -0.015 | 0 | 0.67 | 0.6 | 0.66 | 0.76 | 0.73 | 0.78 |
| 7 | 82376124 | rs10236324 | -0.422 | -1.045 | 0.132 | 0.476 | -0.054 | 0.071 | 0.071 | 0.985 | 0.27 | 0.48 | 0.32 | 0.42 | 0.14 | 0.09 |
| 7 | 82376148 | rs4732480 | -0.336 | -0.302 | -0.833 | -0.713 | -0.096 | 0.158 | 0.158 | 0.008 | 0.64 | 0.88 | 0.66 | 0.9 | 0.47 | 0.62 |
| 7 | 82376952 | rs7780196 | -0.22 | -0.603 | -0.067 | -0.948 | -0.002 | -0.028 | -0.028 | 0 | 0.55 | 0.58 | 0.6 | 0.81 | 0.41 | 0.53 |
| 7 | 82377645 | rs10244323 | -0.243 | -0.668 | -0.186 | -0.86 | 0.014 | 0.157 | 0.157 | 0.297 | 0.56 | 0.84 | 0.6 | 0.74 | 0.41 | 0.5 |
| 7 | 82378880 | rs6953162 | -0.104 | -1.142 | 0.512 | -0.967 | -0.024 | -0.054 | -0.054 | 0 | 0.1 | 0.33 | 0.03 | 0.01 | 0.03 | 0.03 |
| 7 | 82385732 | rs1859174 | -0.388 | -0.794 | 0.395 | -0.837 | -0.002 | 0.095 | 0.095 | 0.217 | 0.57 | 0.8 | 0.6 | 0.72 | 0.46 | 0.54 |
| 7 | 82386082 | rs10486985 | -0.188 | -1.106 | 0.461 | -0.475 | 0.019 | 0.035 | 0.035 | 0 | 0.5 | 0.39 | 0.52 | 0.57 | 0.46 | 0.53 |
| 7 | 82389249 | rs6959361 | 0.008 | -1.095 | 0.341 | -0.516 | 0.022 | 0.026 | 0.026 | 0 | 0.45 | 0.35 | 0.5 | 0.57 | 0.4 | 0.51 |
| 7 | 82395322 | rs17210438 | 0.001 | -0.716 | -0.296 | -0.87 | -0.021 | 0.064 | 0.064 | 0 | 0.56 | 0.3 | 0.52 | 0.72 | 0.5 | 0.64 |
| 7 | 82399191 | rs2158219 | -0.604 | -0.624 | -0.211 | -0.917 | -0.033 | 0.043 | 0.043 | 0 | 0.65 | 0.77 | 0.6 | 0.85 | 0.5 | 0.64 |
| 7 | 82413325 | rs12707521 | -1.015 | 0.487 | -0.123 | -1.147 | 0.006 | -0.029 | -0.029 | 0 | 0.84 | 0.79 | 0.82 | 0.91 | 0.86 | 0.78 |
| 7 | 82413699 | rs12672552 | 1.294 | 0.226 | -0.571 | -1.186 | -0.033 | -0.034 | -0.034 | 0 | 0.13 | 0 | 0.03 | 0.14 | 0.11 | 0.09 |
| 7 | 82414933 | rs9656532 | 0.796 | -0.372 | -0.031 | -0.788 | 0.009 | -0.052 | -0.052 | 0.005 | 0.28 | 0.37 | 0.25 | 0.13 | 0.37 | 0.15 |
| 7 | 82422405 | rs17156675 | 1.915 | -0.641 | -0.757 | -1.004 | -0.04 | -0.114 | -0.114 | 0 | 0.31 | 0.04 | 0.46 | 0.53 | 0.3 | 0.47 |
| 7 | 82422937 | rs6979066 | 1.221 | -0.587 | -0.684 | -0.852 | -0.043 | -0.112 | -0.112 | 0.145 | 0.35 | 0.04 | 0.39 | 0.51 | 0.35 | 0.52 |
| 7 | 82428871 | rs6965452 | -2.11 | -0.567 | -0.607 | -0.862 | -0.001 | -0.005 | -0.005 | 0 | 0.85 | 0.85 | 0.82 | 0.91 | 0.86 | 0.78 |
| 7 | 82429778 | rs7806943 | 0.174 | -0.539 | -0.927 | -0.699 | -0.071 | 0.112 | 0.112 | 0 | 0.49 | 0.04 | 0.42 | 0.65 | 0.46 | 0.61 |
| 7 | 82430654 | rs12667348 | 0.19 | -0.543 | -0.929 | -0.701 | -0.072 | 0.11 | 0.11 | 0 | 0.48 | 0.04 | 0.42 | 0.65 | 0.46 | 0.61 |
| 7 | 82435033 | rs12668093 | 0.596 | -0.468 | -0.396 | -0.549 | -0.008 | 0.021 | 0.021 | 0 | 0.21 | 0.34 | 0.1 | 0.28 | 0.14 | 0.11 |
| 7 | 82438734 | rs17156710 | 0.64 | -0.451 | -0.302 | -0.493 | -0.003 | 0.046 | 0.046 | 0 | 0.06 | 0.19 | 0.06 | 0.11 | 0.04 | 0.02 |
| 7 | 82448100 | rs2522831 | 0.021 | -0.534 | -0.601 | -0.776 | 0.014 | 0.2 | 0.2 | 0 | 0.51 | 0.22 | 0.55 | 0.81 | 0.47 | 0.6 |
| 7 | 82450364 | rs2522832 | 0.021 | -0.534 | -0.601 | -0.743 | 0.014 | 0.198 | 0.198 | 0 | 0.51 | 0.22 | 0.55 | 0.81 | 0.47 | 0.6 |
| 7 | 82451558 | rs1044639 | 0.406 | -0.542 | -0.681 | -0.724 | -0.047 | 0.102 | 0.102 | 0 | 0.45 | 0.04 | 0.42 | 0.68 | 0.43 | 0.58 |
| 7 | 82453708 | rs2522833 | 0.403 | -0.56 | -0.688 | -0.732 | -0.047 | 0.102 | 0.102 | 0 | 0.45 | 0.04 | 0.42 | 0.69 | 0.43 | 0.56 |
| 7 | 82461293 | rs2023847 | 1.874 | -0.586 | -0.828 | -0.874 | -0.098 | 0.064 | 0.064 | 0 | 0.22 | 0.02 | 0.19 | 0.1 | 0.36 | 0.14 |
| 7 | 82485050 | rs7795945 | 0.44 | -0.776 | -0.717 | -0.145 | 0.043 | 0.031 | 0.031 | 0 | 0.38 | 0.17 | 0.16 | 0.3 | 0.29 | 0.22 |
| 7 | 82486615 | rs12707524 | 0.853 | -0.872 | -0.795 | -0.188 | 0.036 | 0.052 | 0.052 | 0 | 0.28 | 0.1 | 0.13 | 0.14 | 0.21 | 0.14 |
| 7 | 82495412 | rs13229774 | 0.379 | -0.809 | -0.739 | -0.14 | 0.045 | 0.029 | 0.029 | 0 | 0.39 | 0.17 | 0.15 | 0.31 | 0.29 | 0.22 |
| 7 | 82500328 | rs13241689 | 0.01 | -0.777 | -0.163 | -0.121 | -0.006 | 0 | 0 | 0 | 0.43 | 0.42 | 0.3 | 0.45 | 0.33 | 0.23 |
| 7 | 82506874 | rs2371364 | -1.888 | -0.607 | -0.12 | 0.095 | -0.002 | 0.009 | 0.009 | 0 | 0.9 | 0.94 | 0.75 | 0.67 | 0.88 | 0.66 |
| 7 | 82508211 | rs10243348 | -0.156 | -0.767 | 0.33 | -0.053 | 0.055 | 0.008 | 0.008 | 0.131 | 0.48 | 0.71 | 0.35 | 0.46 | 0.36 | 0.27 |
| 7 | 82509435 | rs10250881 | 0.915 | -0.731 | 0.323 | -0.191 | -0.024 | 0.119 | 0.119 | 0.234 | 0.42 | 0.21 | 0.4 | 0.21 | 0.52 | 0.39 |
| 7 | 82514138 | rs13237603 | 0.501 | -0.938 | 0.563 | -0.034 | 0.024 | 0.032 | 0.032 | 0 | 0.33 | 0.17 | 0.28 | 0.24 | 0.27 | 0.19 |
| 7 | 82516932 | rs17282616 | 1.006 | -1.226 | 0.108 | -0.762 | -0.042 | 0.134 | 0.134 | 0 | 0.23 | 0.02 | 0.08 | 0.01 | 0.23 | 0.14 |
| 7 | 82524066 | rs2189971 | -1.591 | -1.184 | 0.104 | -0.927 | -0.003 | -0.003 | -0.003 | 0 | 0.82 | 0.84 | 0.79 | 0.92 | 0.84 | 0.74 |
| 7 | 82524663 | rs2189972 | 0.575 | -0.954 | 0.279 | -0.643 | 0.006 | -0.024 | -0.024 | 0 | 0.2 | 0.14 | 0.48 | 0.56 | 0.2 | 0.38 |
| 7 | 82534715 | rs10808302 | -1.249 | -0.76 | 0.209 | -0.607 | -0.032 | 0.098 | 0.098 | 0.504 | 0.58 | 0.79 | 0.64 | 0.77 | 0.52 | 0.59 |
| 7 | 82544987 | rs17156844 | -0.166 | -0.774 | 0.41 | -0.313 | -0.041 | 0.113 | 0.113 | 0.217 | 0.24 | 0.5 | 0.5 | 0.55 | 0.2 | 0.38 |
| 7 | 82557161 | rs10954694 | 0.332 | -0.597 | 0.049 | -0.37 | -0.004 | -0.034 | -0.034 | 0 | 0.32 | 0.18 | 0.42 | 0.44 | 0.36 | 0.41 |
| 7 | 82557228 | rs10954695 | 0.134 | -0.701 | 0.012 | -0.415 | 0 | -0.02 | -0.02 | 0.776 | 0.34 | 0.24 | 0.42 | 0.45 | 0.36 | 0.42 |
| 7 | 82559291 | rs10487645 | 0.412 | -0.818 | 0.334 | -0.526 | -0.05 | -0.009 | -0.009 | 0 | 0.15 | 0.43 | 0.17 | 0.11 | 0.06 | 0.11 |
| 7 | 82561324 | rs9690648 | 0.208 | -0.843 | 0.455 | -0.541 | -0.062 | -0.023 | -0.023 | 0 | 0.16 | 0.6 | 0.19 | 0.11 | 0.06 | 0.12 |
| 7 | 82562340 | rs6959723 | -0.929 | -0.611 | 0.095 | -1.263 | -0.016 | 0.075 | 0.075 | 0.161 | 0.76 | 0.91 | 0.78 | 0.91 | 0.81 | 0.76 |
| 7 | 82563111 | rs7781142 | -0.676 | -0.805 | 0.265 | -1.061 | -0.001 | -0.002 | -0.002 | 0 | 0.56 | 0.59 | 0.62 | 0.76 | 0.51 | 0.61 |
| 7 | 82569707 | rs6467919 | -0.647 | -0.797 | 0.265 | -1.057 | 0.005 | -0.035 | -0.035 | 0 | 0.56 | 0.52 | 0.61 | 0.76 | 0.51 | 0.61 |
| 7 | 82581859 | rs976714 | -0.088 | -0.35 | -0.188 | -0.409 | -0.027 | -0.09 | -0.09 | 1.249 | 0.31 | 0.1 | 0.39 | 0.44 | 0.36 | 0.4 |
| 7 | 82582846 | rs10954696 | -0.088 | -0.35 | -0.188 | -0.409 | -0.027 | -0.09 | -0.09 | 0 | 0.31 | 0.1 | 0.39 | 0.44 | 0.36 | 0.4 |
| 7 | 82583609 | rs10487648 | 1 | -0.247 | 0.007 | -1.014 | -0.021 | 0.071 | 0.071 | 0 | 0.14 | 0.01 | 0.05 | 0.01 | 0.15 | 0.09 |
| 7 | 82630706 | rs1001594 | -0.593 | -0.105 | 0.137 | -1.116 | -0.005 | 0.051 | 0.051 | 0 | 0.56 | 0.66 | 0.66 | 0.82 | 0.51 | 0.7 |
| 7 | 82638965 | rs10435449 | 0.006 | -0.287 | -0.255 | -1.033 | -0.049 | -0.128 | -0.128 | 0 | 0.25 | 0.04 | 0.37 | 0.55 | 0.32 | 0.34 |
| 7 | 82645155 | rs17817131 | 0.679 | -0.328 | 0.212 | -1.058 | 0.001 | 0.001 | 0.001 | 0 | 0.28 | 0.27 | 0.26 | 0.27 | 0.18 | 0.3 |
| 7 | 82646797 | rs41601 | 0.751 | -0.236 | -0.244 | -1.278 | -0.047 | 0.134 | 0.134 | 0 | 0.4 | 0.06 | 0.2 | 0.19 | 0.46 | 0.3 |
| 7 | 82651558 | rs41608 | 1.83 | -0.233 | -0.25 | -1.284 | -0.04 | 0.045 | 0.045 | 0 | 0.27 | 0.05 | 0.16 | 0.18 | 0.33 | 0.21 |
| 7 | 82653249 | rs6977045 | 1.936 | -0.01 | -0.175 | -1.317 | -0.001 | 0.034 | 0.034 | 0 | 0.08 | 0.01 | 0.04 | 0 | 0.09 | 0.03 |
| 7 | 82659856 | rs12707538 | 1.002 | 0.052 | -0.135 | -1.276 | -0.007 | 0.063 | 0.063 | 0 | 0.13 | 0.01 | 0.05 | 0.01 | 0.13 | 0.09 |
| 7 | 82662097 | rs7810390 | -0.014 | 0.029 | 0.28 | -1.274 | 0 | -0.012 | -0.012 | 0 | 0.43 | 0.58 | 0.62 | 0.8 | 0.4 | 0.61 |
| 7 | 82665830 | rs10274504 | 0.485 | 0.177 | -0.2 | -1.026 | 0.001 | -0.036 | -0.036 | 0 | 0.14 | 0.15 | 0.16 | 0 | 0.14 | 0.09 |
| 7 | 82673172 | rs6948626 | 0.577 | 0.183 | -0.285 | -1.024 | -0.005 | 0 | 0 | 0 | 0.13 | 0.07 | 0.15 | 0 | 0.14 | 0.08 |
| 7 | 82688580 | rs1978177 | -1.671 | -0.308 | -0.404 | -0.883 | -0.05 | 0.01 | 0.01 | 0 | 0.73 | 0.97 | 0.85 | 0.81 | 0.68 | 0.79 |

| | | | | | | | | | | | | | | | | |
|---|---|---|---|---|---|---|---|---|---|---|---|---|---|---|---|---|
| 7 | 82689901 | rs4732493 | 0.077 | -0.114 | 0.192 | -0.503 | 0.023 | 0.221 | 0.221 | 0 | 0.14 | 0.34 | 0.34 | 0.51 | 0.19 | 0.3 |
| 7 | 82698998 | rs7797507 | 0.892 | 0.205 | 0.07 | -0.481 | 0.013 | 0.047 | 0.047 | 1.122 | 0.05 | 0.14 | 0.21 | 0.17 | 0.06 | 0.13 |
| 7 | 82713148 | rs10487657 | -0.661 | 0.14 | 0.099 | -0.471 | -0.001 | -0.013 | -0.013 | 0.095 | 0.54 | 0.52 | 0.49 | 0.29 | 0.47 | 0.48 |
| 7 | 82721742 | rs11562069 | -1.27 | -0.187 | 0.621 | 0.211 | 0.002 | -0.002 | -0.002 | 0 | 0.35 | 0.44 | 0.54 | 0.67 | 0.42 | 0.42 |
| 7 | 82732583 | rs17819684 | -0.889 | -0.129 | 0.659 | 0.231 | -0.013 | -0.027 | -0.027 | 0 | 0.32 | 0.19 | 0.48 | 0.6 | 0.4 | 0.32 |
| 7 | 82734845 | rs7791742 | -1.004 | -0.07 | 0.403 | 0.225 | 0.004 | -0.006 | -0.006 | 0 | 0.35 | 0.47 | 0.6 | 0.68 | 0.44 | 0.45 |
| 7 | 82746437 | rs10954710 | -0.567 | 0.781 | -0.216 | 0.708 | -0.029 | -0.06 | -0.06 | 1.349 | 0.22 | 0.1 | 0.37 | 0.57 | 0.38 | 0.31 |
| 7 | 82760225 | rs6946469 | -0.567 | 0.557 | -1.446 | 0.401 | -0.016 | 0.024 | 0.024 | 0.744 | 0.47 | 0.85 | 0.48 | 0.65 | 0.5 | 0.54 |
| 7 | 82761117 | rs10954712 | -0.248 | 0.282 | -1.382 | 0.215 | 0.01 | 0.018 | 0.018 | 0 | 0.48 | 0.6 | 0.55 | 0.39 | 0.45 | 0.51 |
| 7 | 82761250 | rs6950738 | -0.152 | -0.04 | -1.458 | -0.056 | -0.027 | -0.041 | -0.041 | 1.719 | 0.69 | 0.8 | 0.69 | 0.44 | 0.64 | 0.59 |
| 7 | 82767699 | rs11978731 | -0.125 | -0.252 | -1.248 | -0.709 | -0.001 | 0.002 | 0.002 | 0 | 0.17 | 0.19 | 0.12 | 0.06 | 0.21 | 0.16 |
| 7 | 82771395 | rs7783211 | 0.132 | -0.58 | -0.742 | -0.826 | -0.01 | 0.003 | 0.003 | 0.225 | 0.37 | 0.39 | 0.44 | 0.67 | 0.45 | 0.61 |
| 7 | 82775328 | rs10081314 | -1.742 | -1.33 | -0.352 | -1.265 | 0.001 | -0.024 | -0.024 | 0.597 | 0.69 | 0.81 | 0.78 | 0.99 | 0.69 | 0.92 |
| 7 | 82781096 | rs10954714 | 0.73 | -1.557 | -0.489 | -1.145 | -0.023 | -0.024 | -0.024 | 0 | 0.29 | 0.21 | 0.29 | 0.61 | 0.34 | 0.42 |
| 7 | 82783846 | rs7807790 | 1.638 | -1.395 | -0.245 | -1.296 | 0.014 | 0.002 | 0.002 | 1.421 | 0.31 | 0.23 | 0.18 | 0.01 | 0.26 | 0.07 |
| 7 | 82789419 | rs4361713 | -0.799 | -1.244 | -0.131 | -1.078 | 0.021 | 0.025 | 0.025 | 0 | 0.73 | 0.58 | 0.71 | 0.38 | 0.68 | 0.62 |
| 7 | 82791402 | rs4341113 | 0.103 | -1.336 | -0.159 | -1.154 | -0.001 | -0.028 | -0.028 | 0 | 0.61 | 0.55 | 0.5 | 0.13 | 0.65 | 0.49 |
| 7 | 82796409 | rs4732502 | -0.447 | -0.699 | 0.069 | -0.571 | 0.007 | 0.008 | 0.008 | 0 | 0.68 | 0.57 | 0.71 | 0.38 | 0.67 | 0.57 |
| 7 | 82811218 | rs11773347 | 0.24 | -0.644 | -0.096 | -0.691 | -0.005 | -0.004 | -0.004 | 0 | 0.73 | 0.87 | 0.64 | 0.72 | 0.71 | 0.7 |
| 7 | 82817384 | rs4728528 | -0.537 | -0.708 | 0.021 | -0.794 | 0.01 | 0.005 | 0.005 | 0 | 0.67 | 0.53 | 0.7 | 0.39 | 0.64 | 0.56 |
| 7 | 82818143 | rs11761272 | 1.586 | -0.823 | -0.207 | -0.824 | 0.041 | 0.152 | 0.152 | 0.21 | 0.33 | 0.11 | 0.17 | 0.07 | 0.26 | 0.09 |
| 7 | 82823924 | rs7795829 | -0.092 | -0.69 | -0.066 | -0.716 | 0.001 | -0.002 | -0.002 | 0.18 | 0.62 | 0.39 | 0.54 | 0.36 | 0.55 | 0.43 |
| 7 | 82830700 | rs11768582 | -0.337 | -0.579 | -0.441 | -0.569 | -0.005 | -0.011 | -0.011 | 0 | 0.26 | 0.09 | 0.34 | 0.29 | 0.27 | 0.31 |
| 7 | 82834944 | rs17157258 | 0.677 | -0.507 | -0.529 | -0.536 | 0.01 | -0.056 | -0.056 | 0 | 0.09 | 0 | 0.2 | 0.26 | 0.05 | 0.13 |
| 7 | 82835999 | rs1159635 | -0.636 | -0.818 | -0.251 | -0.15 | -0.041 | 0.056 | 0.056 | 0 | 0.81 | 0.67 | 0.89 | 0.64 | 0.91 | 0.79 |
| 7 | 82837056 | rs11769322 | 1.296 | -0.755 | -0.33 | -0.221 | -0.06 | 0.079 | 0.079 | 0 | 0.13 | 0.01 | 0.13 | 0 | 0.21 | 0.15 |
| 7 | 82837298 | rs12707570 | 1.94 | -0.848 | -0.169 | -0.139 | 0.001 | 0.014 | 0.014 | 0.577 | 0.47 | 0.35 | 0.4 | 0.31 | 0.53 | 0.39 |
| 7 | 82841213 | rs11773318 | 1.138 | -0.725 | -0.307 | -0.16 | -0.003 | 0.092 | 0.092 | 0 | 0.48 | 0.23 | 0.43 | 0.08 | 0.53 | 0.32 |
| 7 | 82844374 | rs1557785 | -0.734 | -0.591 | -0.221 | 0.004 | -0.029 | 0.064 | 0.064 | 0 | 0.17 | 0.35 | 0.25 | 0.31 | 0.1 | 0.17 |
| 7 | 82848330 | rs7779384 | -0.095 | -0.561 | -0.31 | -0.023 | 0.013 | 0.015 | 0.015 | 0 | 0.32 | 0.56 | 0.46 | 0.56 | 0.38 | 0.48 |
| 7 | 82861350 | rs6947770 | 0.926 | -0.811 | -0.251 | -0.01 | -0.027 | -0.007 | -0.007 | 0 | 0.64 | 0.52 | 0.56 | 0.33 | 0.78 | 0.61 |
| 7 | 82865056 | rs6968758 | 1.219 | -0.852 | -0.154 | -0.005 | 0.067 | 0.02 | 0.02 | 0 | 0.62 | 0.22 | 0.55 | 0.33 | 0.78 | 0.61 |
| 7 | 82869498 | rs13232363 | 2.291 | -0.855 | -0.308 | -0.159 | 0.032 | 0.126 | 0.126 | 0.103 | 0.32 | 0.07 | 0.19 | 0.08 | 0.26 | 0.09 |
| 7 | 82870122 | rs1123203 | 0.578 | -0.869 | -0.054 | 0.039 | -0.076 | 0.026 | 0.026 | 0 | 0.7 | 0.49 | 0.68 | 0.38 | 0.86 | 0.67 |
| 7 | 82872150 | rs2709913 | -0.708 | -0.793 | -0.073 | 0.004 | -0.031 | 0.037 | 0.037 | 0.013 | 0.29 | 0.68 | 0.24 | 0.41 | 0.18 | 0.3 |
| 7 | 82880350 | rs2709887 | 0.094 | -0.564 | -0.11 | -0.034 | 0.026 | 0.017 | 0.017 | 0 | 0.66 | 0.53 | 0.45 | 0.44 | 0.57 | 0.5 |
| 7 | 82881692 | rs10264381 | 0.368 | -0.758 | 0.019 | 0.132 | -0.045 | 0.06 | 0.06 | 0 | 0.2 | 0.33 | 0.11 | 0.37 | 0.09 | 0.25 |
| 7 | 82893679 | rs2244639 | 0.62 | -0.612 | -0.143 | 0.043 | 0.018 | -0.004 | -0.004 | 0 | 0.56 | 0.67 | 0.34 | 0.5 | 0.37 | 0.34 |
| 7 | 82911915 | rs17289948 | -0.775 | -0.549 | -0.266 | 0.193 | -0.04 | -0.023 | -0.023 | 0 | 0.29 | 0.14 | 0.46 | 0.31 | 0.37 | 0.3 |
| 7 | 82915430 | rs2189103 | 2.06 | -0.017 | -0.385 | 0.253 | -0.086 | -0.07 | -0.07 | 0 | 0.14 | 0.05 | 0.21 | 0.25 | 0.29 | 0.38 |
| 7 | 82918301 | rs2522346 | 0.202 | -0.336 | -0.121 | 0.156 | -0.072 | 0.082 | 0.082 | 0 | 0.21 | 0.43 | 0.12 | 0.38 | 0.09 | 0.23 |
| 7 | 82918349 | rs11761032 | 0.154 | -0.326 | -0.228 | 0.086 | -0.029 | -0.054 | -0.054 | 0 | 0.22 | 0.08 | 0.34 | 0.29 | 0.26 | 0.23 |
| 7 | 82918369 | rs13224075 | 1.613 | -0.276 | -0.196 | 0.085 | 0.023 | 0.089 | 0.089 | 0.693 | 0.5 | 0.29 | 0.43 | 0.32 | 0.55 | 0.47 |
| 7 | 82919330 | rs2214355 | 1.422 | -0.386 | -0.273 | 0.115 | 0.087 | 0.038 | 0.038 | 1.524 | 0.42 | 0.2 | 0.41 | 0.34 | 0.27 | 0.15 |
| 7 | 82920114 | rs2214353 | -1.452 | -0.856 | -0.068 | 0.387 | 0.002 | -0.001 | -0.001 | 0 | 0.42 | 0.72 | 0.45 | 0.44 | 0.41 | 0.45 |
| 7 | 82923367 | rs17157388 | 0.176 | -0.862 | 0.028 | 0.386 | 0.03 | 0.01 | 0.01 | 0 | 0.07 | 0.23 | 0.19 | 0.05 | 0.08 | 0.06 |
| 7 | 82925773 | rs2522343 | -0.675 | -0.867 | -0.122 | 0.372 | -0.039 | 0.05 | 0.05 | 0.016 | 0.28 | 0.61 | 0.3 | 0.44 | 0.17 | 0.29 |
| 7 | 82926352 | rs10486907 | 1.423 | -0.909 | -0.139 | 0.381 | 0.032 | -0.026 | -0.026 | 0.226 | 0.06 | 0.01 | 0.13 | 0.24 | 0.01 | 0.05 |
| 7 | 82926427 | rs2522342 | -0.769 | -0.556 | 0.036 | 0.155 | -0.035 | 0.05 | 0.05 | 0.016 | 0.28 | 0.61 | 0.3 | 0.44 | 0.18 | 0.29 |
| 7 | 82929501 | rs2522339 | -1.352 | -0.583 | -0.014 | 0.236 | 0.002 | -0.001 | -0.001 | 0.006 | 0.42 | 0.71 | 0.45 | 0.44 | 0.41 | 0.45 |
| 7 | 82932149 | rs2522333 | -0.203 | -1.451 | 0.294 | 0.677 | -0.052 | 0.07 | 0.07 | 0 | 0.2 | 0.38 | 0.11 | 0.39 | 0.09 | 0.24 |
| 7 | 82933022 | rs2709915 | -1.135 | -1.632 | 0.104 | -0.167 | 0.023 | 0.017 | 0.017 | 0.771 | 0.66 | 0.79 | 0.85 | 0.92 | 0.8 | 0.88 |
| 7 | 82937146 | rs2709911 | -0.093 | -1.437 | 0.588 | 0.224 | -0.002 | 0.017 | 0.017 | 0 | 0.09 | 0.15 | 0.08 | 0.12 | 0.09 | 0.08 |
| 7 | 82939626 | rs2522368 | -0.631 | -1.582 | 0.771 | 0.547 | 0.001 | 0.007 | 0.007 | 0 | 0.63 | 0.56 | 0.53 | 0.57 | 0.4 | 0.39 |
| 7 | 82942970 | rs17157433 | 0.867 | -1.47 | -0.095 | 0.668 | -0.107 | -0.116 | -0.116 | 0.788 | 0.33 | 0.08 | 0.42 | 0.43 | 0.58 | 0.58 |
| 7 | 82950337 | rs9656547 | -0.365 | -1.668 | 0.033 | 0.558 | 0 | -0.003 | -0.003 | 0 | 0.61 | 0.62 | 0.54 | 0.55 | 0.38 | 0.38 |
| 7 | 82957777 | rs2371662 | -0.613 | -1.595 | -0.281 | 0.484 | -0.013 | 0.032 | 0.032 | 0 | 0.62 | 0.79 | 0.47 | 0.32 | 0.41 | 0.4 |
| 7 | 82958464 | rs12707578 | -0.505 | -1.593 | -0.378 | 0.485 | -0.011 | 0.031 | 0.031 | 0 | 0.62 | 0.79 | 0.47 | 0.32 | 0.4 | 0.41 |
| 7 | 82960004 | rs10239446 | -1.537 | -0.993 | 0.266 | 0.755 | 0.009 | -0.01 | -0.01 | 0 | 0.32 | 0.79 | 0.44 | 0.36 | 0.25 | 0.3 |
| 7 | 82965322 | rs2371667 | -0.981 | -0.227 | 0.286 | 0.452 | -0.004 | -0.032 | -0.032 | 0 | 0.28 | 0.56 | 0.34 | 0.2 | 0.25 | 0.3 |
| 7 | 82968338 | rs9690908 | 1.357 | 0.293 | 0.913 | 0.979 | 0.153 | 0.282 | 0.282 | 0.359 | 0.93 | 0.58 | 0.81 | 0.8 | 0.91 | 0.87 |
| 7 | 82968487 | rs11768410 | 1.033 | -0.8 | 0.215 | 0.524 | 0.014 | -0.011 | -0.011 | 0 | 0.58 | 0.29 | 0.52 | 0.45 | 0.58 | 0.61 |
| 7 | 82977626 | rs2371660 | -0.936 | -1.497 | 0.639 | 0.417 | -0.026 | 0.001 | 0.001 | 0 | 0.32 | 0.48 | 0.35 | 0.39 | 0.24 | 0.19 |
| 7 | 82982193 | rs10236360 | -0.329 | -2.044 | 0.737 | -0.435 | -0.035 | -0.034 | -0.034 | 0 | 0.28 | 0.69 | 0.17 | 0.13 | 0.13 | 0.13 |
| 7 | 82984827 | rs10234926 | 0.816 | -2.074 | 0.72 | -0.448 | 0.047 | 0.028 | 0.028 | 0.2 | 0.2 | 0.12 | 0.12 | 0.13 | 0.1 | 0.08 |
| 7 | 82992227 | rs2709977 | 0.381 | -2.134 | 0.825 | -0.496 | 0.033 | 0.032 | 0.032 | 0.16 | 0.7 | 0.22 | 0.83 | 0.87 | 0.87 | 0.88 |
| 7 | 82999141 | rs2723028 | 0.901 | -2.185 | 0.767 | -0.665 | 0.092 | 0.136 | 0.136 | 0 | 0.59 | 0.19 | 0.76 | 0.87 | 0.76 | 0.8 |
| 7 | 83009942 | rs10273103 | 0.168 | -1.995 | 1.048 | -0.662 | -0.037 | -0.016 | -0.016 | 0 | 0.21 | 0.5 | 0.14 | 0.15 | 0.12 | 0.04 |
| 7 | 83021583 | rs10252196 | 0.471 | -2.016 | 1.114 | -0.659 | -0.013 | -0.006 | -0.006 | 0 | 0.2 | 0.26 | 0.13 | 0.15 | 0.12 | 0.05 |

| | | | | | | | | | | | | | | | | |
|---|---|---|---|---|---|---|---|---|---|---|---|---|---|---|---|---|
| 7 | 83025318 | rs2709922 | 1.253 | -1.983 | 0.93 | -0.809 | 0.036 | 0.136 | 0.136 | 0 | 0.59 | 0.14 | 0.72 | 0.84 | 0.69 | 0.82 |
| 7 | 83027345 | rs7777731 | 0.565 | -1.942 | 1.053 | -0.8 | -0.001 | -0.01 | -0.01 | 0 | 0.13 | 0.09 | 0.1 | 0 | 0.15 | 0.09 |
| 7 | 83033641 | rs17285647 | 0.522 | -1.75 | 0.992 | -0.849 | 0.012 | 0.06 | 0.06 | 0.261 | 0.24 | 0.09 | 0.14 | 0.01 | 0.25 | 0.21 |
| 7 | 83037751 | rs2722985 | 1.276 | -1.72 | 0.947 | -0.792 | 0.039 | 0.203 | 0.203 | 0 | 0.48 | 0.13 | 0.68 | 0.84 | 0.6 | 0.7 |
| 7 | 83041344 | rs2709963 | 1.329 | -1.609 | 1.51 | -0.741 | 0.067 | 0.063 | 0.063 | 1.019 | 0.95 | 0.51 | 0.97 | 1 | 1 | 1 |
| 7 | 83057126 | rs12112469 | 0.392 | -1.507 | 1.053 | -0.866 | 0.007 | 0.012 | 0.012 | 0.356 | 0.25 | 0.13 | 0.15 | 0.01 | 0.25 | 0.21 |
| 7 | 83058827 | rs2722964 | -0.079 | -1.482 | 1.052 | -0.855 | -0.03 | -0.164 | -0.164 | 1.584 | 0.71 | 0.57 | 0.83 | 1 | 0.75 | 0.79 |
| 7 | 83061378 | rs2247921 | -0.079 | -1.482 | 1.052 | -0.855 | -0.03 | -0.164 | -0.164 | 0 | 0.71 | 0.57 | 0.83 | 1 | 0.75 | 0.79 |
| 7 | 83085727 | rs2723023 | 1.558 | -1.104 | -0.146 | -0.151 | -0.113 | -0.12 | -0.12 | 1.871 | 0.3 | 0.06 | 0.37 | 0.47 | 0.4 | 0.39 |
| 7 | 83087972 | rs2466440 | 0.392 | -1.266 | -0.063 | -0.368 | 0.049 | 0.231 | 0.231 | 0 | 0.49 | 0.17 | 0.69 | 0.86 | 0.6 | 0.7 |
| 7 | 83101090 | rs2722989 | 0.575 | -1.433 | 0.405 | 0.024 | -0.01 | 0 | 0 | 0.099 | 0.61 | 0.42 | 0.73 | 0.66 | 0.79 | 0.65 |
| 7 | 83101484 | rs2722990 | -1.249 | -1.146 | 0.714 | 0.343 | -0.028 | 0.059 | 0.059 | 0 | 0.86 | 0.7 | 0.86 | 0.8 | 0.94 | 0.78 |
| 7 | 83109037 | rs10270844 | -0.655 | -0.946 | 0.453 | 0.163 | -0.036 | 0.03 | 0.03 | 0 | 0.57 | 0.76 | 0.55 | 0.39 | 0.54 | 0.38 |
| 7 | 83113683 | rs2255872 | 0.578 | -0.946 | 0.531 | 0.164 | -0.034 | 0.03 | 0.03 | 0 | 0.43 | 0.27 | 0.46 | 0.62 | 0.46 | 0.62 |
| 7 | 83116386 | rs2255614 | 0.331 | -0.852 | 0.989 | 0.224 | 0.055 | 0.152 | 0.152 | 0 | 0.92 | 0.66 | 0.75 | 0.77 | 0.9 | 0.94 |
| 7 | 83121863 | rs2723006 | 0.179 | -0.99 | 0.911 | 0.107 | 0.007 | 0.038 | 0.038 | 0 | 0.49 | 0.41 | 0.49 | 0.61 | 0.51 | 0.7 |
| 7 | 83126859 | rs3801516 | 0.332 | -1.045 | 0.809 | -0.037 | 0.013 | 0.07 | 0.07 | 0 | 0.42 | 0.22 | 0.27 | 0.16 | 0.39 | 0.25 |
| 7 | 83128484 | rs2535376 | 0.087 | -1.212 | 1.037 | 0.27 | -0.053 | 0.029 | 0.029 | 0 | 0.19 | 0.45 | 0.16 | 0.18 | 0.09 | 0.26 |
| 7 | 83134047 | rs13221248 | 1.639 | -0.926 | 0.108 | 0.457 | -0.006 | 0.01 | 0.01 | 0 | 0.12 | 0.15 | 0.12 | 0.18 | 0.06 | 0.22 |
| 7 | 83134459 | rs11982066 | 0.745 | -0.856 | -0.215 | 0.478 | -0.041 | -0.035 | -0.035 | 0 | 0.34 | 0.2 | 0.35 | 0.42 | 0.43 | 0.44 |
| 7 | 83140212 | rs10485857 | 1.292 | -0.945 | -0.149 | 0.537 | 0.014 | -0.013 | -0.013 | 0 | 0.13 | 0.09 | 0.12 | 0.16 | 0.05 | 0.21 |
| 7 | 83141662 | rs3801519 | 1.274 | -0.94 | -0.15 | 0.536 | 0.012 | -0.013 | -0.013 | 0 | 0.13 | 0.09 | 0.12 | 0.16 | 0.06 | 0.21 |
| 7 | 83144112 | rs2535372 | 0.175 | -0.419 | -0.247 | 0.379 | -0.001 | -0.001 | -0.001 | 0 | 0.23 | 0.22 | 0.15 | 0.22 | 0.16 | 0.1 |
| 7 | 83147251 | rs2535371 | 0.14 | -0.797 | 0.038 | 0.376 | -0.015 | -0.003 | -0.003 | 2.397 | 0.23 | 0.26 | 0.14 | 0.22 | 0.13 | 0.1 |
| 7 | 83147705 | rs2535370 | 0.056 | -0.567 | -0.053 | 0.434 | -0.007 | 0.095 | 0.095 | 0.001 | 0.78 | 0.39 | 0.6 | 0.6 | 0.81 | 0.74 |
| 7 | 83149564 | rs1860610 | -0.603 | -0.219 | -0.206 | 0.366 | 0.058 | 0.137 | 0.137 | 0 | 0.93 | 0.72 | 0.74 | 0.76 | 0.88 | 0.96 |
| 7 | 83152236 | rs17157704 | 1.236 | -0.21 | -1.326 | -0.365 | -0.092 | -0.071 | -0.071 | 0 | 0.31 | 0.07 | 0.35 | 0.38 | 0.45 | 0.46 |
| 7 | 83158344 | rs17505926 | 0.579 | -0.862 | -0.948 | 0.047 | 0.033 | 0.022 | 0.022 | 2.37 | 0.56 | 0.19 | 0.47 | 0.38 | 0.68 | 0.64 |
| 7 | 83167328 | rs7810691 | -0.023 | -1.342 | -0.335 | -0.691 | -0.028 | -0.128 | -0.128 | 0 | 0.38 | 0.49 | 0.17 | 0.01 | 0.26 | 0.22 |
| 7 | 83171681 | rs17447180 | 0.383 | -1.566 | -0.91 | 0.04 | 0.02 | -0.069 | -0.069 | 0 | 0.16 | 0 | 0.12 | 0.21 | 0.08 | 0.09 |
| 7 | 83173289 | rs10248786 | -0.164 | -1.215 | -0.692 | 0.327 | 0.03 | -0.044 | -0.044 | 0 | 0.31 | 0.15 | 0.27 | 0.41 | 0.22 | 0.32 |
| 7 | 83173734 | rs3801537 | -0.411 | -1.276 | -0.628 | 0.277 | 0.038 | -0.024 | -0.024 | 0.578 | 0.37 | 0.2 | 0.3 | 0.41 | 0.24 | 0.3 |
| 7 | 83173932 | rs1693381 | -0.441 | -1.425 | -0.015 | 0.307 | 0.001 | -0.002 | -0.002 | 0 | 0.37 | 0.37 | 0.31 | 0.42 | 0.23 | 0.29 |
| 7 | 83174363 | rs1639309 | -1.112 | -1.362 | -0.164 | 0.5 | -0.062 | -0.045 | -0.045 | 0 | 0.7 | 0.87 | 0.6 | 0.59 | 0.49 | 0.54 |
| 7 | 83178817 | rs1693387 | -0.578 | -2.023 | 0.229 | 0.364 | 0.007 | 0 | 0 | 0.052 | 0.42 | 0.64 | 0.33 | 0.41 | 0.21 | 0.31 |
| 7 | 83178951 | rs2041427 | 1.306 | -2.072 | 0.208 | 0.341 | 0.003 | 0.009 | 0.009 | 0.022 | 0.27 | 0.22 | 0.26 | 0.17 | 0.25 | 0.23 |
| 7 | 83179622 | rs1639304 | -0.641 | -2.022 | 0.279 | 0.398 | 0.007 | 0 | 0 | 0.063 | 0.42 | 0.64 | 0.33 | 0.41 | 0.21 | 0.31 |
| 7 | 83185739 | rs6952103 | 0.257 | -2.181 | 0.193 | 0.446 | 0.011 | -0.009 | -0.009 | 0.627 | 0.15 | 0.13 | 0.15 | 0.19 | 0.08 | 0.1 |
| 7 | 83191301 | rs17157828 | 1.616 | -2.209 | 0.036 | 0.413 | -0.017 | -0.054 | -0.054 | 0.423 | 0.16 | 0.02 | 0.14 | 0.22 | 0.12 | 0.17 |
| 7 | 83204964 | rs215289 | -0.295 | -2.339 | 0.575 | 0.305 | 0.054 | 0.032 | 0.032 | 0 | 0.5 | 0.68 | 0.44 | 0.37 | 0.34 | 0.36 |
| 7 | 83211117 | rs6966242 | -0.79 | -2.382 | 0.752 | 0.297 | -0.057 | -0.004 | -0.004 | 0 | 0.23 | 0.36 | 0.17 | 0.19 | 0.08 | 0.14 |
| 7 | 83219267 | rs215311 | -1.294 | -2.133 | 0.743 | 0.437 | -0.043 | -0.038 | -0.038 | 0 | 0.69 | 0.86 | 0.59 | 0.58 | 0.46 | 0.54 |
| 7 | 83226044 | rs215318 | 1.307 | -2.182 | 0.867 | 0.364 | -0.002 | -0.006 | -0.006 | 0 | 0.27 | 0.3 | 0.28 | 0.17 | 0.25 | 0.22 |
| 7 | 83226255 | rs3801563 | 0.354 | -2.203 | 0.785 | 0.382 | 0.009 | -0.008 | -0.008 | 0 | 0.15 | 0.13 | 0.15 | 0.19 | 0.08 | 0.1 |
| 7 | 83229565 | rs3823906 | -0.447 | -2.13 | 0.787 | 0.383 | -0.007 | -0.001 | -0.001 | 0 | 0.39 | 0.44 | 0.31 | 0.38 | 0.21 | 0.31 |
| 7 | 83233110 | rs3801568 | -0.451 | -2.132 | 0.786 | 0.383 | -0.007 | -0.001 | -0.001 | 0 | 0.39 | 0.44 | 0.31 | 0.38 | 0.21 | 0.31 |
| 7 | 83242503 | rs42027 | -0.939 | -1.417 | -0.082 | 0.601 | -0.002 | 0.048 | 0.048 | 0 | 0.56 | 0.82 | 0.57 | 0.63 | 0.54 | 0.68 |
| 7 | 83248171 | rs42019 | 1.704 | -1.365 | -0.285 | 0.61 | 0.042 | 0.018 | 0.018 | 0.121 | 0.14 | 0.14 | 0.24 | 0.26 | 0.33 | 0.37 |
| 7 | 83248206 | rs42018 | -1.03 | -1.435 | -0.407 | 0.553 | -0.054 | 0.026 | 0.026 | 0.038 | 0.57 | 0.93 | 0.58 | 0.63 | 0.54 | 0.69 |
| 7 | 83261703 | rs42001 | -1.871 | -1.451 | -0.377 | 0.581 | -0.034 | -0.025 | -0.025 | 0 | 0.83 | 0.99 | 0.84 | 0.8 | 0.8 | 0.91 |
| 7 | 83266000 | rs10246477 | -0.912 | -1.145 | 0.647 | 0.954 | 0.026 | 0.001 | 0.001 | 0.026 | 0.41 | 0.68 | 0.33 | 0.37 | 0.2 | 0.3 |
| 7 | 83267983 | rs536299 | -1.29 | -0.955 | 0.247 | 1.012 | 0.023 | 0.007 | 0.007 | 0.07 | 0.86 | 0.88 | 0.76 | 0.74 | 0.67 | 0.62 |
| 7 | 83269003 | rs17157933 | 0.126 | -1.035 | 0.234 | 0.898 | 0.002 | -0.004 | -0.004 | 0 | 0.15 | 0.15 | 0.16 | 0.19 | 0.08 | 0.1 |
| 7 | 83269806 | rs302124 | 0.733 | -0.535 | 0.489 | 0.812 | -0.02 | -0.012 | -0.012 | 0.057 | 0.59 | 0.33 | 0.59 | 0.56 | 0.55 | 0.42 |
| 7 | 83280198 | rs302113 | -0.406 | 0.425 | 0.758 | 0.961 | 0.017 | 0.005 | 0.005 | 0.025 | 0.86 | 0.88 | 0.77 | 0.72 | 0.68 | 0.67 |
| 7 | 83287607 | rs1972459 | 0.105 | 0.341 | 0.532 | 0.983 | 0.074 | 0.026 | 0.026 | 0 | 0.8 | 0.73 | 0.72 | 0.72 | 0.65 | 0.61 |
| 7 | 83290740 | rs10274738 | -1.069 | -0.86 | 0.797 | 0.81 | -0.036 | -0.044 | -0.044 | 0 | 0.09 | 0.38 | 0.04 | 0 | 0.01 | 0.04 |
| 7 | 83295674 | rs7807686 | -1.288 | -0.926 | 0.288 | 1.091 | -0.09 | 0.06 | 0.06 | 0 | 0.17 | 0.5 | 0.24 | 0.19 | 0.06 | 0.15 |
| 7 | 83297241 | rs2158725 | -0.969 | -0.207 | -0.686 | 1.273 | -0.034 | -0.081 | -0.081 | 0 | 0.13 | 0.01 | 0.21 | 0.28 | 0.14 | 0.13 |
| 7 | 83297975 | rs595787 | 1.142 | -0.775 | -0.408 | 1.965 | -0.024 | 0.128 | 0.128 | 1.17 | 0.05 | 0.08 | 0.09 | 0.31 | 0.01 | 0.09 |
| 7 | 83298203 | rs533887 | 0.462 | -0.847 | -0.279 | 1.407 | 0.005 | -0.001 | -0.001 | 0 | 0.68 | 0.75 | 0.57 | 0.66 | 0.73 | 0.74 |
| 7 | 83306889 | rs567663 | 0.536 | -0.918 | 0.132 | 1.234 | 0.001 | -0.015 | -0.015 | 0 | 0.64 | 0.72 | 0.49 | 0.44 | 0.66 | 0.47 |
| 7 | 83307600 | rs540157 | 0.986 | -0.219 | -0.492 | -0.025 | -0.008 | 0.116 | 0.116 | 0.35 | 0.39 | 0.21 | 0.2 | 0.02 | 0.36 | 0.23 |
| 7 | 83318052 | rs2527837 | 1.267 | -0.537 | -1.201 | -0.412 | -0.025 | 0.103 | 0.103 | 0 | 0.19 | 0 | 0.11 | 0.02 | 0.17 | 0.14 |
| 7 | 83318378 | rs17458455 | 1.231 | -0.453 | -1.278 | -0.591 | -0.095 | 0.138 | 0.138 | 0.12 | 0.21 | 0.01 | 0.11 | 0 | 0.24 | 0.13 |
| 7 | 83323789 | rs17458902 | 0.234 | -0.656 | -0.899 | -0.795 | -0.02 | 0.021 | 0.021 | 0 | 0.36 | 0.26 | 0.23 | 0.01 | 0.5 | 0.21 |
| 7 | 83326168 | rs1181370 | -0.347 | -0.822 | -0.572 | -0.749 | 0.007 | 0.001 | 0.001 | 0 | 0.59 | 0.63 | 0.39 | 0.11 | 0.67 | 0.47 |
| 7 | 83326808 | rs1181369 | 1.5 | -0.765 | -0.521 | -0.707 | 0.014 | 0.037 | 0.037 | 0 | 0.21 | 0.09 | 0.13 | 0.1 | 0.17 | 0.26 |
| 7 | 83332518 | rs12113156 | 1.68 | -0.575 | -0.563 | -0.738 | -0.078 | 0.103 | 0.103 | 0.158 | 0.17 | 0.01 | 0.09 | 0 | 0.19 | 0.11 |

| | | | | | | | | | | | | | | | | |
|---|---|---|---|---|---|---|---|---|---|---|---|---|---|---|---|---|
| 7 | 83332892 | rs17459332 | 1.533 | -0.538 | -0.535 | -0.704 | -0.05 | 0.049 | 0.049 | 0 | 0.15 | 0.05 | 0.12 | 0.03 | 0.3 | 0.09 |
| 7 | 83334604 | rs532597 | -1.721 | -0.834 | -0.536 | -0.439 | 0.003 | -0.012 | -0.012 | 0 | 0.77 | 0.82 | 0.64 | 0.65 | 0.78 | 0.82 |
| 7 | 83338753 | rs679423 | -0.357 | -0.886 | -0.514 | -0.517 | 0 | -0.007 | -0.007 | 0 | 0.35 | 0.69 | 0.35 | 0.49 | 0.27 | 0.44 |
| 7 | 83347858 | rs9640943 | 0.876 | -0.9 | -0.523 | -0.363 | 0 | -0.029 | -0.029 | 0 | 0.1 | 0.03 | 0.15 | 0.21 | 0.1 | 0.26 |
| 7 | 83349603 | rs1181361 | -1.036 | -0.937 | -0.567 | -0.392 | 0.016 | -0.024 | -0.024 | 0.391 | 0.7 | 0.82 | 0.6 | 0.62 | 0.73 | 0.75 |
| 7 | 83352427 | rs1921595 | -0.972 | -1.046 | 0.013 | 0.118 | 0.006 | -0.012 | -0.012 | 0 | 0.41 | 0.69 | 0.39 | 0.49 | 0.32 | 0.46 |
| 7 | 83359224 | rs1921598 | 0.268 | -1.014 | 0.349 | 0.011 | 0.02 | 0.107 | 0.107 | 0.358 | 0.06 | 0.22 | 0.13 | 0.21 | 0.06 | 0.22 |
| 7 | 83369234 | rs17158034 | -0.063 | -1.022 | 0.489 | -0.287 | 0.039 | 0.009 | 0.009 | 0.342 | 0.07 | 0.39 | 0.03 | 0 | 0.01 | 0.02 |
| 7 | 83370845 | rs2141780 | 0.588 | -1.205 | 0.217 | -0.25 | 0.058 | -0.035 | -0.035 | 0 | 0.58 | 0.23 | 0.57 | 0.45 | 0.73 | 0.42 |
| 7 | 83381426 | rs17158062 | 1.144 | -0.903 | -0.047 | -0.221 | -0.003 | 0.004 | 0.004 | 0 | 0.06 | 0.03 | 0.04 | 0.03 | 0.09 | 0.05 |
| 7 | 83385765 | rs10954734 | -0.643 | -0.906 | -0.035 | -0.163 | 0.017 | -0.053 | -0.053 | 0 | 0.73 | 0.85 | 0.67 | 0.46 | 0.79 | 0.47 |
| 7 | 83400104 | rs1882344 | -0.704 | -0.91 | -0.738 | -0.164 | 0.014 | -0.062 | -0.062 | 0 | 0.74 | 0.87 | 0.66 | 0.45 | 0.8 | 0.52 |
| 7 | 83403157 | rs6949364 | 0.13 | -0.794 | -0.82 | -0.254 | -0.063 | 0.125 | 0.125 | 0 | 0.53 | 0.89 | 0.63 | 0.72 | 0.54 | 0.37 |
| 7 | 83405454 | rs10224170 | 0.2 | -0.351 | -0.37 | -0.372 | 0.002 | -0.002 | -0.002 | 0 | 0.58 | 0.43 | 0.56 | 0.45 | 0.67 | 0.43 |
| 7 | 83406694 | rs1526484 | 0.223 | -0.324 | -0.509 | -0.356 | 0.016 | -0.009 | -0.009 | 0 | 0.58 | 0.39 | 0.55 | 0.45 | 0.67 | 0.43 |
| 7 | 83412551 | rs6467963 | -0.75 | 0.108 | -0.39 | 0.241 | 0.015 | -0.043 | -0.043 | 0 | 0.73 | 0.87 | 0.76 | 0.55 | 0.8 | 0.56 |
| 7 | 83413999 | rs1465005 | 0.945 | 0.128 | -0.432 | 0.214 | -0.014 | -0.023 | -0.023 | 0 | 0.41 | 0.22 | 0.48 | 0.42 | 0.48 | 0.3 |
| 7 | 83415942 | rs6947649 | -0.454 | 0.507 | -0.597 | 0.266 | 0.014 | -0.048 | -0.048 | 0 | 0.73 | 0.88 | 0.76 | 0.55 | 0.8 | 0.61 |
| 7 | 83421885 | rs6980017 | -0.655 | 1.008 | 0.398 | -0.206 | 0.03 | -0.117 | -0.117 | 0.592 | 0.26 | 0.45 | 0.15 | 0.03 | 0.28 | 0.18 |
| 7 | 83422832 | rs17158185 | -0.45 | 1.999 | 0.557 | 0.403 | 0.148 | 0.029 | 0.029 | 0 | 0.12 | 0.33 | 0.21 | 0.11 | 0.27 | 0.22 |
| 7 | 83422849 | rs2141030 | -1.05 | 0.368 | 0.227 | 0.718 | -0.017 | 0.039 | 0.039 | 0 | 0.12 | 0.18 | 0.04 | 0.21 | 0.07 | 0.07 |
| 7 | 83423307 | rs11764330 | -0.619 | 0.64 | -0.178 | -0.018 | 0.002 | 0.012 | 0.012 | 0 | 0.1 | 0.06 | 0.06 | 0.01 | 0.07 | 0.06 |
| 7 | 83424108 | rs6953094 | -0.78 | 0.396 | 0.012 | -0.471 | -0.002 | -0.052 | -0.052 | 0.132 | 0.39 | 0.46 | 0.26 | 0.13 | 0.34 | 0.42 |
| 7 | 83429355 | rs12666664 | 0.141 | -0.078 | -0.332 | -0.508 | 0.01 | -0.059 | -0.059 | 0 | 0.17 | 0.07 | 0.32 | 0.69 | 0.18 | 0.24 |
| 7 | 83431203 | rs6467967 | -1.193 | -0.329 | 0.141 | -0.776 | -0.015 | 0.048 | 0.048 | 0 | 0.58 | 0.67 | 0.6 | 0.85 | 0.52 | 0.71 |
| 7 | 83431290 | rs2030861 | -1.189 | -0.334 | 0.158 | -0.78 | -0.014 | 0.046 | 0.046 | 0 | 0.58 | 0.67 | 0.6 | 0.84 | 0.52 | 0.71 |
| 7 | 83433436 | rs1526490 | 0.207 | -0.641 | -0.305 | -1.225 | -0.027 | 0.137 | 0.137 | 0 | 0.3 | 0.04 | 0.17 | 0.07 | 0.24 | 0.38 |
| 7 | 83433484 | rs1526491 | -0.469 | -0.608 | 0.687 | -1.048 | 0.005 | 0.005 | 0.005 | 0 | 0.5 | 0.49 | 0.53 | 0.78 | 0.42 | 0.64 |
| 7 | 83437224 | rs6942510 | -1.21 | -0.637 | 0.904 | -1.081 | -0.028 | 0.074 | 0.074 | 0 | 0.6 | 0.78 | 0.64 | 0.85 | 0.56 | 0.69 |
| 7 | 83437463 | rs10255815 | 0.621 | -0.406 | 0.632 | -1.153 | 0.035 | 0.004 | 0.004 | 0 | 0.09 | 0.24 | 0.1 | 0.06 | 0.14 | 0.04 |
| 7 | 83444771 | rs11761508 | -0.25 | -0.557 | -0.039 | -0.914 | -0.006 | 0.195 | 0.195 | 0.546 | 0.43 | 0.09 | 0.32 | 0.11 | 0.41 | 0.55 |
| 7 | 83452312 | rs701271 | 0.218 | -0.547 | 0.065 | -0.91 | 0.01 | 0.161 | 0.161 | 0.563 | 0.57 | 0.86 | 0.67 | 0.89 | 0.59 | 0.52 |
| 7 | 83474010 | rs10248603 | -0.051 | -0.446 | 1.208 | -0.779 | 0.044 | 0.002 | 0.002 | 0.557 | 0.13 | 0.45 | 0.12 | 0.07 | 0.14 | 0.04 |
| 7 | 83479375 | rs797648 | 0.126 | -0.819 | 0.686 | -0.985 | -0.042 | 0.169 | 0.169 | 1.363 | 0.58 | 0.94 | 0.66 | 0.83 | 0.57 | 0.42 |
| 7 | 83491815 | rs9690506 | 0.115 | -0.796 | 1.362 | -0.842 | 0.025 | -0.008 | -0.008 | 0 | 0.12 | 0.34 | 0.12 | 0.07 | 0.14 | 0.04 |
| 7 | 83502159 | rs797635 | -0.128 | -1 | 0.994 | -0.86 | 0.008 | 0.063 | 0.063 | 0.385 | 0.63 | 0.81 | 0.76 | 0.8 | 0.67 | 0.5 |
| 7 | 83504609 | rs11769587 | 1.336 | -0.592 | 0.769 | -0.72 | -0.051 | 0.165 | 0.165 | 0 | 0.39 | 0.08 | 0.46 | 0.11 | 0.4 | 0.26 |
| 7 | 83505392 | rs11981075 | 0.433 | -0.477 | 0.645 | -0.835 | -0.01 | 0.032 | 0.032 | 2.963 | 0.27 | 0.12 | 0.2 | 0.2 | 0.29 | 0.44 |
| 7 | 83506643 | rs1526483 | -0.117 | -0.518 | 0.888 | -0.443 | -0.001 | 0.056 | 0.056 | 0 | 0.86 | 0.85 | 0.83 | 0.44 | 0.85 | 0.82 |
| 7 | 83507541 | rs1524332 | 0.293 | -0.731 | 1.126 | -0.64 | 0.037 | -0.025 | -0.025 | 0 | 0.1 | 0.41 | 0.1 | 0.05 | 0.13 | 0.07 |
| 7 | 83515095 | rs12666591 | 0.482 | -0.771 | 0.764 | -0.46 | 0.039 | 0.006 | 0.006 | 1.319 | 0.1 | 0.43 | 0.1 | 0.07 | 0.12 | 0.06 |
| 7 | 83518131 | rs10487867 | 1.091 | -0.61 | -0.004 | -0.622 | 0.001 | 0.143 | 0.143 | 0 | 0.44 | 0.16 | 0.48 | 0.13 | 0.42 | 0.31 |
| 7 | 83519060 | rs7795229 | 0.411 | -0.336 | -0.405 | -0.278 | -0.023 | 0.036 | 0.036 | 0.018 | 0.6 | 0.74 | 0.62 | 0.26 | 0.56 | 0.4 |
| 7 | 83530065 | rs701276 | -0.037 | 0.005 | -0.556 | -0.065 | -0.018 | -0.008 | -0.008 | 0 | 0.25 | 0.2 | 0.26 | 0.29 | 0.34 | 0.42 |
| 7 | 83531098 | rs701277 | -0.242 | -0.432 | -1.093 | 0.156 | -0.009 | -0.126 | -0.126 | 0 | 0.85 | 0.99 | 0.87 | 0.49 | 0.88 | 0.76 |
| 7 | 83533954 | rs1404151 | 0.949 | -0.598 | -1.713 | -1.128 | -0.008 | 0.136 | 0.136 | 0 | 0.34 | 0.12 | 0.16 | 0.01 | 0.29 | 0.2 |
| 7 | 83549032 | rs2888206 | 0.312 | -0.637 | -0.99 | -1.451 | 0.001 | -0.048 | -0.048 | 0 | 0.49 | 0.45 | 0.3 | 0.02 | 0.5 | 0.25 |
| 7 | 83555527 | rs701303 | 0.744 | -0.798 | -1 | -0.924 | -0.028 | -0.033 | -0.033 | 0 | 0.3 | 0.18 | 0.53 | 0.56 | 0.38 | 0.49 |
| 7 | 83555894 | rs1068156 | -0.629 | -0.951 | -1.135 | -1.497 | 0.004 | -0.163 | -0.163 | 1.476 | 0.67 | 0.55 | 0.76 | 1 | 0.59 | 0.79 |
| 7 | 83558010 | rs6944013 | 2.245 | -0.709 | -1.311 | -1.344 | -0.027 | 0.047 | 0.047 | 0 | 0.11 | 0.03 | 0.13 | 0 | 0.21 | 0.04 |
| 7 | 83562164 | rs701308 | 0.135 | -0.844 | -1 | -1.175 | 0.005 | 0.073 | 0.073 | 0 | 0.49 | 0.25 | 0.61 | 0.63 | 0.47 | 0.6 |
| 7 | 83566026 | rs696874 | -0.898 | -0.7 | -0.782 | -1.73 | 0.003 | -0.094 | -0.094 | 0.532 | 0.73 | 0.69 | 0.81 | 1 | 0.66 | 0.86 |
| 7 | 83566585 | rs10232913 | 0.833 | -0.498 | -1.148 | -1.28 | 0.019 | 0.036 | 0.036 | 0.772 | 0.2 | 0.08 | 0.08 | 0.09 | 0.1 | 0.06 |
| 7 | 83566638 | rs701309 | -0.115 | -0.287 | -0.817 | -1.224 | -0.003 | 0.039 | 0.039 | 0 | 0.55 | 0.34 | 0.65 | 0.87 | 0.48 | 0.69 |
| 7 | 83568746 | rs16887591 | 0.187 | -0.799 | -0.568 | -0.903 | -0.028 | -0.011 | -0.011 | 0.007 | 0.22 | 0.3 | 0.12 | 0.23 | 0.07 | 0.23 |
| 7 | 83570506 | rs17158353 | 0.207 | -0.809 | -0.592 | -0.937 | -0.024 | -0.012 | -0.012 | 1.199 | 0.22 | 0.29 | 0.11 | 0.22 | 0.07 | 0.24 |
| 7 | 83571278 | rs10215636 | 0.495 | -0.547 | -1.304 | -1.731 | -0.045 | 0.103 | 0.103 | 1.851 | 0.29 | 0.11 | 0.22 | 0.02 | 0.4 | 0.1 |
| 7 | 83580714 | rs797831 | -1.311 | -0.454 | -1.828 | -1.864 | -0.039 | 0.041 | 0.041 | 0 | 0.74 | 0.97 | 0.83 | 0.89 | 0.76 | 0.75 |
| 7 | 83581709 | rs17158388 | 0.718 | -0.369 | -2.008 | -2.129 | 0 | -0.031 | -0.031 | 3.155 | 0.36 | 0.33 | 0.2 | 0.04 | 0.29 | 0.16 |
| 7 | 83581975 | rs701319 | 1.004 | -0.459 | -1.687 | -2.185 | -0.025 | -0.03 | -0.03 | 0 | 0.22 | 0.14 | 0.47 | 0.62 | 0.32 | 0.42 |
| 7 | 83583757 | rs797827 | -0.408 | -0.68 | -1.55 | -2.164 | -0.013 | -0.027 | -0.027 | 1.16 | 0.62 | 0.48 | 0.69 | 0.69 | 0.64 | 0.62 |
| 7 | 83586686 | rs797824 | 0.417 | -0.681 | -1.13 | -2.172 | 0 | -0.001 | -0.001 | 0 | 0.27 | 0.27 | 0.5 | 0.63 | 0.35 | 0.46 |
| 7 | 83589142 | rs1052343 | 2.216 | -0.719 | -1.59 | -2.177 | -0.015 | 0.038 | 0.038 | 0.275 | 0.17 | 0.08 | 0.15 | 0.02 | 0.24 | 0.05 |
| 7 | 83590231 | rs10239654 | 2.216 | -0.719 | -1.59 | -2.161 | -0.015 | 0.038 | 0.038 | 0.275 | 0.17 | 0.08 | 0.15 | 0.02 | 0.24 | 0.05 |
| 7 | 83590852 | rs797821 | 0.127 | 0.048 | -1.277 | -1.665 | -0.011 | -0.028 | -0.028 | 0 | 0.62 | 0.48 | 0.69 | 0.69 | 0.64 | 0.63 |
| 7 | 83599335 | rs740948 | 0.872 | -0.043 | -0.887 | -1.148 | 0.003 | -0.001 | -0.001 | 0 | 0.46 | 0.4 | 0.26 | 0.06 | 0.43 | 0.2 |
| 7 | 83606518 | rs701320 | -0.746 | 0.435 | -1.737 | -0.774 | -0.021 | 0.018 | 0.018 | 0 | 0.77 | 0.99 | 0.85 | 0.89 | 0.79 | 0.79 |
| 7 | 83629442 | rs3801593 | 0.04 | 0.811 | -1.006 | -0.151 | -0.046 | -0.043 | -0.043 | 1.339 | 0.77 | 0.91 | 0.54 | 0.38 | 0.69 | 0.56 |
| 7 | 83631923 | rs6978182 | -0.443 | -0.529 | -0.611 | -0.179 | 0.043 | 0.048 | 0.048 | 0.784 | 0.35 | 0.27 | 0.18 | 0.15 | 0.22 | 0.3 |

| | | | | | | | | | | | | | | | | |
|---|---|---|---|---|---|---|---|---|---|---|---|---|---|---|---|---|
| 7 | 83632549 | rs3801599 | -0.412 | -0.507 | -0.601 | -0.17 | 0.04 | 0.046 | 0.046 | 0.603 | 0.34 | 0.27 | 0.18 | 0.15 | 0.22 | 0.3 |
| 7 | 83634713 | rs7804122 | -0.293 | -0.535 | -0.608 | -0.176 | 0.036 | 0.037 | 0.037 | 0 | 0.33 | 0.27 | 0.17 | 0.15 | 0.21 | 0.3 |
| 7 | 83642555 | rs17158505 | -0.479 | -0.301 | -0.103 | -0.063 | -0.009 | 0.006 | 0.006 | 0.337 | 0.15 | 0.22 | 0.08 | 0.24 | 0.09 | 0.21 |
| 7 | 83644954 | rs4732531 | -0.85 | -0.292 | 0.37 | 0.503 | -0.04 | 0.102 | 0.102 | 0 | 0.27 | 0.48 | 0.3 | 0.44 | 0.16 | 0.26 |
| 7 | 83647970 | rs7788378 | -0.765 | -0.291 | 0.48 | 0.489 | 0 | 0.002 | 0.002 | 0 | 0.24 | 0.26 | 0.26 | 0.43 | 0.16 | 0.28 |
| 7 | 83651854 | rs6976211 | 0.898 | -0.172 | 0.777 | 0.439 | 0.015 | 0.1 | 0.1 | 0.043 | 0.51 | 0.3 | 0.4 | 0.18 | 0.53 | 0.36 |
| 7 | 83652875 | rs11974894 | -0.898 | -0.231 | 0.807 | 0.43 | 0.003 | -0.005 | -0.005 | 0.005 | 0.76 | 0.82 | 0.69 | 0.79 | 0.74 | 0.67 |
| 7 | 83653553 | rs6944702 | 1.433 | -0.112 | 0.605 | 0.453 | -0.038 | 0.111 | 0.111 | 0 | 0.41 | 0.19 | 0.37 | 0.14 | 0.5 | 0.26 |
| 7 | 83656800 | rs10235789 | 1.425 | -0.095 | 0.614 | 0.461 | -0.038 | 0.112 | 0.112 | 0 | 0.41 | 0.19 | 0.37 | 0.14 | 0.5 | 0.26 |
| 7 | 83659577 | rs17158529 | 0.634 | -0.082 | 0.636 | 0.474 | 0.045 | 0.09 | 0.09 | 0 | 0.06 | 0.21 | 0.03 | 0.19 | 0.06 | 0.12 |
| 7 | 83661339 | rs17158533 | -0.4 | -0.09 | 0.666 | 0.474 | 0.007 | -0.019 | -0.019 | 0.05 | 0.24 | 0.22 | 0.26 | 0.43 | 0.16 | 0.28 |
| 7 | 83664129 | rs17158535 | -0.708 | -0.039 | 0.354 | 0.402 | -0.013 | 0.039 | 0.039 | 0 | 0.3 | 0.43 | 0.3 | 0.62 | 0.22 | 0.4 |
| 7 | 83667173 | rs1404473 | -0.195 | -0.056 | 0.233 | 0.36 | -0.007 | 0.001 | 0.001 | 0 | 0.72 | 0.65 | 0.69 | 0.77 | 0.73 | 0.66 |
| 7 | 83678952 | rs4355720 | 0.531 | -0.014 | 0.034 | 0.257 | -0.016 | -0.017 | -0.017 | 0 | 0.57 | 0.49 | 0.69 | 0.65 | 0.68 | 0.52 |
| 7 | 83686148 | rs2527040 | -0.422 | -0.152 | 0.064 | 0.151 | 0.051 | 0.075 | 0.075 | 0.11 | 0.39 | 0.73 | 0.3 | 0.15 | 0.26 | 0.36 |
| 7 | 83689614 | rs2527039 | -0.117 | -0.331 | 0.264 | 0.057 | -0.034 | -0.051 | -0.051 | 0.18 | 0.36 | 0.43 | 0.28 | 0.13 | 0.22 | 0.35 |
| 7 | 83691571 | rs2189849 | -0.694 | -0.286 | 0.211 | 0.146 | 0.053 | 0.048 | 0.048 | 0.007 | 0.43 | 0.69 | 0.33 | 0.33 | 0.29 | 0.47 |
| 7 | 83698177 | rs2527031 | 0.556 | -0.345 | 0.014 | 0.058 | 0.015 | 0.026 | 0.026 | 0 | 0.23 | 0.17 | 0.22 | 0.09 | 0.18 | 0.24 |
| 7 | 83699733 | rs17296151 | 0.847 | 0.675 | -0.682 | 0.16 | -0.082 | -0.055 | -0.055 | 0 | 0.14 | 0.01 | 0.14 | 0.14 | 0.23 | 0.22 |
| 7 | 83701677 | rs10488266 | 0.361 | 0.237 | -0.304 | 0.202 | -0.006 | -0.007 | -0.007 | 0.257 | 0.13 | 0.08 | 0.1 | 0.13 | 0.13 | 0.15 |
| 7 | 83702476 | rs2691694 | -0.698 | -0.068 | -0.307 | 0.244 | -0.005 | 0.004 | 0.004 | 0 | 0.91 | 0.93 | 0.92 | 0.93 | 0.89 | 0.91 |
| 7 | 83717020 | rs2691699 | 0.031 | -0.385 | -0.1 | 0.204 | -0.025 | -0.004 | -0.004 | 0 | 0.3 | 0.48 | 0.23 | 0.31 | 0.22 | 0.37 |
| 7 | 83722691 | rs7804363 | 1.268 | -0.48 | -0.698 | 0.155 | -0.103 | -0.082 | -0.082 | 0.267 | 0.41 | 0.1 | 0.53 | 0.47 | 0.52 | 0.31 |
| 7 | 83723672 | rs2057845 | 1.268 | -0.48 | -0.677 | 0.159 | -0.092 | -0.08 | -0.08 | 3.453 | 0.41 | 0.12 | 0.54 | 0.48 | 0.52 | 0.31 |
| 7 | 83726605 | rs7801600 | -1.15 | -0.563 | -0.661 | 0.115 | -0.005 | 0.026 | 0.026 | 0 | 0.79 | 0.97 | 0.81 | 0.9 | 0.82 | 0.76 |
| 7 | 83734989 | rs3801631 | 0.804 | -0.521 | -0.373 | 0.119 | 0 | 0.021 | 0.021 | 0 | 0.07 | 0.09 | 0.03 | 0.22 | 0.07 | 0.13 |
| 7 | 83737144 | rs3801637 | 1.345 | -0.605 | -0.309 | 0.04 | -0.054 | -0.039 | -0.039 | 0 | 0.42 | 0.16 | 0.54 | 0.47 | 0.51 | 0.31 |
| 7 | 83745375 | rs13222995 | 0.735 | -0.582 | 0.082 | -0.07 | -0.023 | 0 | 0 | 0 | 0.23 | 0.32 | 0.21 | 0.19 | 0.18 | 0.25 |
| 7 | 83745474 | rs12707617 | 0.416 | -0.559 | 0.103 | -0.071 | -0.016 | -0.05 | -0.05 | 3.249 | 0.72 | 0.59 | 0.79 | 0.9 | 0.75 | 0.69 |
| 7 | 83747200 | rs7781923 | -0.127 | -0.694 | 0.233 | -0.145 | 0 | -0.005 | -0.005 | 0 | 0.8 | 0.8 | 0.84 | 0.92 | 0.78 | 0.76 |
| 7 | 83747642 | rs1990046 | 0.91 | -0.697 | 0.239 | -0.145 | -0.021 | -0.033 | -0.033 | 0.746 | 0.09 | 0.21 | 0.05 | 0.02 | 0.02 | 0.08 |
| 7 | 83759108 | rs7793598 | -0.102 | -0.631 | -0.229 | -0.112 | 0 | -0.005 | -0.005 | 7.443 | 0.8 | 0.8 | 0.84 | 0.92 | 0.78 | 0.76 |
| 7 | 83770348 | rs10488270 | 1.033 | -0.537 | -0.579 | -0.024 | -0.003 | -0.041 | -0.041 | 0.234 | 0.21 | 0.03 | 0.19 | 0.2 | 0.18 | 0.25 |
| 7 | 83774499 | rs7781126 | 0.062 | -0.406 | -0.362 | 0.01 | -0.014 | -0.007 | -0.007 | 0.553 | 0.32 | 0.53 | 0.27 | 0.23 | 0.2 | 0.33 |
| 7 | 83775796 | rs723997 | 1.078 | -0.565 | -0.506 | -0.049 | 0.023 | 0.036 | 0.036 | 0.501 | 0.46 | 0.25 | 0.56 | 0.68 | 0.56 | 0.43 |
| 7 | 83782435 | rs7781152 | -1.148 | -0.704 | -1.021 | -0.134 | 0.008 | 0.007 | 0.007 | 0 | 0.9 | 0.96 | 0.93 | 0.94 | 0.89 | 0.92 |
| 7 | 83782839 | rs17158675 | 1.031 | -0.563 | -0.521 | -0.027 | -0.007 | -0.02 | -0.02 | 0.004 | 0.08 | 0.19 | 0.05 | 0.02 | 0.03 | 0.08 |
| 7 | 83786460 | rs464 | -0.263 | -0.022 | -0.474 | 0.04 | -0.013 | 0.044 | 0.044 | 0 | 0.74 | 0.84 | 0.81 | 0.9 | 0.71 | 0.76 |
| 7 | 83794756 | rs17158686 | 0.472 | 0.488 | -0.558 | 0.482 | -0.001 | 0.002 | 0.002 | 0 | 0.07 | 0.08 | 0.03 | 0.2 | 0.07 | 0.14 |
| 7 | 83795479 | rs6946868 | 0.272 | 0.705 | -0.615 | 0.381 | -0.017 | -0.016 | -0.016 | 0.003 | 0.43 | 0.28 | 0.55 | 0.5 | 0.49 | 0.36 |
| 7 | 83801357 | rs12666837 | -0.637 | 1.138 | -0.34 | 0.01 | 0.01 | -0.018 | -0.018 | 0 | 0.23 | 0.32 | 0.19 | 0.16 | 0.35 | 0.3 |
| 7 | 83801652 | rs10261069 | 0.565 | 1.277 | -0.376 | -0.158 | -0.004 | -0.009 | -0.009 | 0 | 0.87 | 0.85 | 0.89 | 0.99 | 0.84 | 0.95 |
| 7 | 83802951 | rs9642175 | 0.048 | 1.222 | 0.005 | 0.016 | 0.006 | -0.026 | -0.026 | 0.017 | 0.71 | 0.62 | 0.61 | 0.79 | 0.68 | 0.74 |
| 7 | 83805716 | rs7808707 | 0.398 | 0.954 | -0.046 | 0.988 | 0.005 | -0.004 | -0.004 | 0 | 0.07 | 0.02 | 0.05 | 0.27 | 0.07 | 0.17 |
| 7 | 83808712 | rs17302032 | 1.695 | 0.928 | -0.583 | 0.708 | -0.072 | 0.053 | 0.053 | 0 | 0.24 | 0.02 | 0.2 | 0.08 | 0.28 | 0.08 |
| 7 | 83809111 | rs13224698 | 0.6 | 0.643 | 0.377 | 1.001 | 0.014 | -0.035 | -0.035 | 0 | 0.07 | 0.11 | 0.05 | 0 | 0.05 | 0.01 |
| 7 | 83814352 | rs9642178 | 0.618 | 0.54 | 0.488 | 1.424 | -0.004 | -0.019 | -0.019 | 0 | 0.06 | 0.01 | 0.04 | 0.29 | 0.07 | 0.17 |
| 7 | 83819515 | rs11976488 | 0.871 | 0.587 | 0.484 | 1.614 | -0.01 | -0.015 | -0.015 | 0.331 | 0.06 | 0.02 | 0.09 | 0.14 | 0.08 | 0.08 |
| 7 | 83821946 | rs11979170 | -0.465 | 0.455 | 0.543 | 1.489 | 0.006 | -0.022 | -0.022 | 0 | 0.69 | 0.61 | 0.51 | 0.79 | 0.65 | 0.71 |
| 7 | 83837045 | rs7804323 | 0.053 | 0.418 | 0.209 | 1.718 | -0.004 | 0.126 | 0.126 | 0 | 0.07 | 0.12 | 0.08 | 0.44 | 0.08 | 0.18 |
| 7 | 83846895 | rs1015629 | -0.446 | 0.422 | 0.374 | 1.371 | -0.001 | 0.024 | 0.024 | 0.06 | 0.7 | 0.75 | 0.55 | 0.8 | 0.69 | 0.68 |
| 7 | 83847064 | rs1015628 | 0.507 | 0.132 | 0.584 | 1.263 | -0.01 | -0.018 | -0.018 | 0 | 0.07 | 0.14 | 0.04 | 0.02 | 0.05 | 0.08 |
| 7 | 83857204 | rs7802925 | 0.192 | 0.133 | 0.586 | 1.541 | -0.006 | 0.147 | 0.147 | 0 | 0.07 | 0.11 | 0.07 | 0.41 | 0.07 | 0.16 |
| 7 | 83864172 | rs1989963 | -0.055 | 0.241 | 0.296 | 1.47 | -0.013 | 0.055 | 0.055 | 0.265 | 0.63 | 0.74 | 0.52 | 0.76 | 0.58 | 0.63 |
| 7 | 83880644 | rs10954751 | 1.602 | 0.154 | 0.341 | 1.419 | -0.077 | 0.127 | 0.127 | 0.631 | 0.42 | 0.12 | 0.35 | 0.17 | 0.42 | 0.32 |
| 7 | 83884705 | rs10233152 | -0.667 | 0.019 | 0.351 | 1.544 | 0 | 0.002 | 0.002 | 0.173 | 0.69 | 0.68 | 0.59 | 0.69 | 0.69 | 0.67 |
| 7 | 83885441 | rs1356804 | 0.929 | 0.026 | 0.118 | 1.542 | -0.003 | 0.01 | 0.01 | 0 | 0.06 | 0.02 | 0.05 | 0.28 | 0.07 | 0.15 |
| 7 | 83905380 | rs2372167 | -0.76 | -0.032 | 0.174 | 1.449 | -0.016 | 0.063 | 0.063 | 0.505 | 0.71 | 0.9 | 0.61 | 0.81 | 0.69 | 0.68 |
| 7 | 83934959 | rs6949774 | 0.413 | 0.119 | 0.088 | 1.275 | -0.001 | 0.06 | 0.06 | 0 | 0.07 | 0.07 | 0.06 | 0.28 | 0.07 | 0.15 |
| 7 | 83936564 | rs594533 | 1.632 | 0.147 | -0.033 | 0.903 | -0.066 | 0.115 | 0.115 | 0.278 | 0.41 | 0.16 | 0.38 | 0.18 | 0.42 | 0.32 |
| 7 | 83939664 | rs9986714 | -0.724 | 0.441 | 0.364 | 0.763 | 0 | -0.059 | -0.059 | 0 | 0.14 | 0.32 | 0.11 | 0.07 | 0.14 | 0.11 |
| 7 | 83941201 | rs2040696 | 0.015 | 0.616 | -0.504 | 1.1 | 0 | 0.002 | 0.002 | 0.213 | 0.7 | 0.73 | 0.6 | 0.71 | 0.69 | 0.67 |
| 7 | 83954737 | rs6467991 | -1.015 | 0.468 | -0.502 | 1.352 | -0.009 | 0.049 | 0.049 | 0 | 0.24 | 0.62 | 0.16 | 0.4 | 0.19 | 0.27 |
| 7 | 83958436 | rs515139 | -0.594 | 0.483 | -1.04 | 1.218 | 0.025 | -0.021 | -0.021 | 0.123 | 0.39 | 0.75 | 0.47 | 0.63 | 0.33 | 0.53 |
| 7 | 83971334 | rs1468213 | -0.016 | 0.953 | -1.063 | 1.264 | -0.027 | -0.03 | -0.03 | 12.801 | 0.05 | 0.01 | 0.08 | 0.15 | 0.07 | 0.06 |
| 7 | 83982343 | rs611954 | -0.808 | 1.109 | -0.745 | 1.564 | -0.052 | 0.04 | 0.04 | 1.446 | 0.58 | 0.87 | 0.62 | 0.68 | 0.53 | 0.67 |
| 7 | 83984606 | rs11972641 | 0.003 | 0.764 | -0.159 | 1.495 | 0 | 0.068 | 0.068 | 0.404 | 0.27 | 0.52 | 0.38 | 0.34 | 0.24 | 0.23 |
| 7 | 83986207 | rs7806920 | 0.05 | 0.737 | -0.077 | 1.413 | -0.017 | -0.029 | -0.029 | 0 | 0.26 | 0.34 | 0.16 | 0.19 | 0.22 | 0.38 |
| 7 | 83996546 | rs1557788 | -0.493 | 0.844 | 0.093 | 1.148 | -0.001 | -0.006 | -0.006 | 0.013 | 0.35 | 0.41 | 0.53 | 0.78 | 0.28 | 0.41 |

| | | | | | | | | | | | | | | | | |
|---|---|---|---|---|---|---|---|---|---|---|---|---|---|---|---|---|
| 7 | 84010005 | rs656300 | -0.116 | 0.432 | 0.001 | 1.092 | -0.02 | 0.25 | 0.25 | 0 | 0.09 | 0.3 | 0.25 | 0.34 | 0.03 | 0.12 |
| 7 | 84014620 | rs2522370 | -0.587 | 0.377 | 0.055 | 1.102 | -0.01 | 0.033 | 0.033 | 0 | 0.23 | 0.35 | 0.44 | 0.63 | 0.14 | 0.32 |
| 7 | 84016382 | rs12673557 | 0.885 | 0.403 | -0.018 | 0.88 | -0.028 | -0.061 | -0.061 | 0 | 0.14 | 0.04 | 0.19 | 0.29 | 0.11 | 0.2 |
| 7 | 84016715 | rs520178 | 0.328 | 0.335 | 0.185 | 0.79 | -0.011 | 0.137 | 0.137 | 0 | 0.07 | 0.24 | 0.16 | 0.18 | 0.02 | 0.11 |
| 7 | 84020366 | rs1228875 | 0.454 | 0.505 | 0.05 | 0.553 | -0.001 | -0.035 | -0.035 | 0 | 0.21 | 0.2 | 0.13 | 0 | 0.21 | 0.1 |
| 7 | 84039765 | rs1634605 | -0.541 | 0.412 | 0.17 | 0.685 | -0.011 | -0.02 | -0.02 | 0 | 0.3 | 0.62 | 0.3 | 0.18 | 0.24 | 0.21 |
| 7 | 84051368 | rs1228963 | 0.214 | 0.178 | 0.184 | 0.572 | 0 | -0.021 | -0.021 | 0 | 0.23 | 0.2 | 0.13 | 0 | 0.21 | 0.11 |
| 7 | 84081186 | rs2706903 | -0.649 | 0.333 | 0.329 | 0.593 | -0.019 | 0.164 | 0.164 | 0 | 0.1 | 0.42 | 0.17 | 0.18 | 0.02 | 0.11 |
| 7 | 84089388 | rs1228935 | -1.418 | 0.62 | 0.524 | 0.414 | -0.005 | 0.201 | 0.201 | 0.13 | 0.51 | 0.81 | 0.7 | 0.92 | 0.43 | 0.48 |
| 7 | 84098378 | rs2715036 | -0.467 | 0.627 | 0.492 | 0.684 | -0.003 | 0.145 | 0.145 | 0 | 0.21 | 0.44 | 0.36 | 0.46 | 0.13 | 0.3 |
| 7 | 84101774 | rs1236923 | -1.576 | 0.689 | 0.457 | 0.594 | 0.009 | 0.135 | 0.135 | 0 | 0.5 | 0.72 | 0.68 | 0.92 | 0.43 | 0.48 |
| 7 | 84104060 | rs1228970 | -2.498 | 0.528 | 0.325 | 0.32 | -0.023 | 0.112 | 0.112 | 0.072 | 0.74 | 0.93 | 0.82 | 0.97 | 0.65 | 0.85 |
| 7 | 84104530 | rs1228971 | -0.958 | 0.541 | 1.026 | 0.946 | 0.002 | 0.001 | 0.001 | 0 | 0.38 | 0.44 | 0.35 | 0.45 | 0.34 | 0.33 |
| 7 | 84131879 | rs10954754 | 0.002 | 0.545 | 0.94 | 0.984 | -0.023 | -0.063 | -0.063 | 5.367 | 0.15 | 0.04 | 0.19 | 0.3 | 0.11 | 0.21 |
| 7 | 84136556 | rs1228925 | -1.372 | 0.549 | 1.007 | 0.85 | 0.001 | 0.004 | 0.004 | 0 | 0.43 | 0.67 | 0.51 | 0.62 | 0.35 | 0.41 |
| 7 | 84139004 | rs1029541 | -0.051 | 0.6 | 1.118 | 0.85 | 0.004 | -0.049 | -0.049 | 0 | 0.24 | 0.4 | 0.16 | 0.16 | 0.22 | 0.11 |
| 7 | 84146330 | rs1922342 | -0.62 | 0.679 | 0.833 | 1.077 | 0 | 0.05 | 0.05 | 0 | 0.18 | 0.25 | 0.35 | 0.46 | 0.12 | 0.29 |
| 7 | 84168615 | rs4732558 | -1.473 | 0.46 | 0.848 | 0.798 | 0.001 | 0.019 | 0.019 | 0 | 0.52 | 0.62 | 0.65 | 0.81 | 0.42 | 0.59 |
| 7 | 84178141 | rs3815067 | -0.553 | 0.253 | 1.078 | 1.065 | -0.009 | -0.06 | -0.06 | 0 | 0.24 | 0.04 | 0.25 | 0.32 | 0.18 | 0.34 |
| 7 | 84243643 | rs9642186 | -1.625 | 0.378 | 1.564 | 0.913 | 0.002 | 0.018 | 0.018 | 0 | 0.51 | 0.46 | 0.63 | 0.81 | 0.42 | 0.59 |
| 7 | 84250535 | rs12669783 | -1.301 | 0.373 | 1.485 | 0.927 | -0.032 | -0.06 | -0.06 | 0.439 | 0.46 | 0.15 | 0.52 | 0.5 | 0.41 | 0.57 |
| 7 | 84251379 | rs4132667 | -1.8 | 0.377 | 1.604 | 0.705 | 0.003 | 0.065 | 0.065 | 0 | 0.53 | 0.68 | 0.65 | 0.81 | 0.42 | 0.59 |
| 7 | 84267724 | rs4391370 | 0.204 | 0.461 | 1.413 | 0.757 | -0.035 | 0.047 | 0.047 | 0 | 0.14 | 0 | 0.19 | 0.05 | 0.13 | 0.25 |
| 7 | 84273418 | rs4422706 | -1.688 | 0.604 | 1.616 | 0.726 | 0.002 | 0.019 | 0.019 | 0 | 0.52 | 0.62 | 0.65 | 0.81 | 0.42 | 0.59 |
| 7 | 84277134 | rs12707665 | -1.187 | 0.841 | 1.691 | 0.917 | -0.018 | 0.007 | 0.007 | 0 | 0.29 | 0.47 | 0.51 | 0.63 | 0.19 | 0.39 |
| 7 | 84284168 | rs7795440 | -1.759 | 0.828 | 1.753 | 0.87 | 0.004 | 0.026 | 0.026 | 0 | 0.49 | 0.62 | 0.65 | 0.81 | 0.41 | 0.55 |
| 7 | 84289915 | rs4128405 | -0.905 | 0.818 | 1.814 | 1.039 | -0.001 | 0.003 | 0.003 | 0 | 0.28 | 0.33 | 0.32 | 0.41 | 0.19 | 0.38 |
| 7 | 84295995 | rs11765801 | -1.736 | 0.84 | 1.635 | 1.02 | 0.003 | 0.022 | 0.022 | 0 | 0.49 | 0.62 | 0.66 | 0.81 | 0.42 | 0.55 |
| 7 | 84305487 | rs10230448 | 0.612 | 0.865 | 1.641 | 1.03 | -0.03 | 0.062 | 0.062 | 0 | 0.18 | 0.05 | 0.13 | 0.03 | 0.22 | 0.16 |
| 7 | 84309621 | rs11971107 | -0.002 | 0.843 | 1.617 | 1.141 | -0.022 | -0.097 | -0.097 | 0.031 | 0.2 | 0.04 | 0.25 | 0.35 | 0.17 | 0.3 |
| 7 | 84311765 | rs6952175 | -1.68 | 0.762 | 1.686 | 1.07 | 0.003 | 0.015 | 0.015 | 0 | 0.49 | 0.62 | 0.66 | 0.81 | 0.41 | 0.55 |
| 7 | 84344258 | rs12154270 | 0.637 | 0.98 | 1.616 | 0.776 | -0.056 | 0.056 | 0.056 | 0 | 0.17 | 0.02 | 0.12 | 0.05 | 0.2 | 0.14 |
| 7 | 84346026 | rs11761247 | -1.012 | 0.928 | 1.818 | 0.818 | -0.009 | 0.088 | 0.088 | 0.121 | 0.28 | 0.58 | 0.4 | 0.49 | 0.23 | 0.23 |
| 7 | 84395443 | rs12707676 | 0.139 | 0.956 | 1.885 | 0.795 | -0.037 | 0.064 | 0.064 | 0.787 | 0.19 | 0.05 | 0.13 | 0.06 | 0.22 | 0.16 |
| 7 | 84405422 | rs4529405 | -1.893 | 1.021 | 1.785 | 0.731 | 0.004 | 0.042 | 0.042 | 0.134 | 0.54 | 0.68 | 0.73 | 0.96 | 0.49 | 0.57 |
| 7 | 84429366 | rs7458400 | -0.415 | 1.198 | 1.428 | 0.859 | -0.037 | -0.075 | -0.075 | 0 | 0.26 | 0.09 | 0.34 | 0.44 | 0.27 | 0.33 |
| 7 | 84460354 | rs1835401 | -0.333 | 1.232 | 1.395 | 0.805 | -0.01 | 0.032 | 0.032 | 0.459 | 0.24 | 0.12 | 0.17 | 0.14 | 0.22 | 0.23 |
| 7 | 84470000 | rs1594392 | -0.346 | 1.179 | 1.454 | 0.806 | 0.001 | 0.023 | 0.023 | 0.427 | 0.26 | 0.2 | 0.18 | 0.14 | 0.22 | 0.23 |
| 7 | 84480679 | rs4518607 | -0.852 | 1.135 | 1.743 | 0.855 | -0.017 | -0.011 | -0.011 | 0 | 0.31 | 0.59 | 0.23 | 0.29 | 0.23 | 0.25 |
| 7 | 84499593 | rs1819081 | -0.766 | 1.154 | 1.67 | 0.879 | -0.012 | 0.029 | 0.029 | 0.018 | 0.3 | 0.46 | 0.39 | 0.36 | 0.23 | 0.25 |
| 7 | 84504645 | rs10499882 | -0.384 | 1.234 | 1.414 | 0.892 | -0.066 | -0.115 | -0.115 | 0.206 | 0.26 | 0.05 | 0.33 | 0.43 | 0.27 | 0.33 |
| 7 | 84542156 | rs12707690 | -0.685 | 1.148 | 1.451 | 0.805 | -0.004 | 0.022 | 0.022 | 0.009 | 0.28 | 0.2 | 0.21 | 0.14 | 0.25 | 0.25 |
| 7 | 84544961 | rs11767793 | 0.128 | 1.18 | 1.315 | 0.756 | -0.036 | 0.065 | 0.065 | 0.115 | 0.2 | 0.05 | 0.15 | 0.05 | 0.22 | 0.17 |
| 7 | 84559637 | rs4560742 | -1.036 | 0.952 | 1.482 | 0.95 | 0.005 | -0.007 | -0.007 | 0 | 0.36 | 0.45 | 0.3 | 0.33 | 0.34 | 0.28 |
| 7 | 84561292 | rs2372504 | -2.096 | 0.962 | 1.38 | 0.997 | 0.006 | 0.024 | 0.024 | 1.398 | 0.58 | 0.7 | 0.61 | 0.76 | 0.54 | 0.61 |
| 7 | 84577411 | rs13234914 | -0.554 | 1.046 | 1.494 | 0.864 | -0.048 | 0.089 | 0.089 | 1.305 | 0.28 | 0.1 | 0.21 | 0.05 | 0.32 | 0.2 |
| 7 | 84578002 | rs10256889 | -1.413 | 1.042 | 1.4 | 0.902 | 0.001 | -0.004 | -0.004 | 0 | 0.38 | 0.61 | 0.32 | 0.33 | 0.34 | 0.29 |
| 7 | 84591966 | rs1881480 | -1.253 | 1.098 | 1.259 | 0.91 | 0.003 | -0.004 | -0.004 | 0 | 0.37 | 0.45 | 0.3 | 0.33 | 0.34 | 0.29 |
| 7 | 84597933 | rs7794976 | -0.718 | 1.424 | 1.003 | 0.717 | -0.054 | 0.097 | 0.097 | 1.35 | 0.29 | 0.09 | 0.21 | 0.06 | 0.32 | 0.2 |
| 7 | 84612468 | rs7780675 | -1.051 | 1.491 | 0.83 | 0.731 | 0.019 | -0.029 | -0.029 | 0 | 0.33 | 0.53 | 0.26 | 0.27 | 0.33 | 0.21 |
| 7 | 84626388 | rs2040957 | -1.102 | 1.493 | 0.758 | 0.732 | 0.007 | -0.015 | -0.015 | 0 | 0.34 | 0.59 | 0.27 | 0.27 | 0.33 | 0.21 |
| 7 | 84627231 | rs10261725 | -0.745 | 1.409 | 1.248 | 0.721 | -0.014 | 0.016 | 0.016 | 1.4 | 0.29 | 0.21 | 0.22 | 0.06 | 0.32 | 0.2 |
| 7 | 84628989 | rs7800072 | -0.95 | 1.508 | 1.48 | 0.714 | 0.005 | -0.009 | -0.009 | 0 | 0.31 | 0.35 | 0.24 | 0.27 | 0.32 | 0.21 |
| 7 | 84630516 | rs2286184 | -0.17 | 1.464 | 1.236 | 0.931 | -0.016 | -0.079 | -0.079 | 0 | 0.2 | 0.08 | 0.29 | 0.47 | 0.2 | 0.32 |
| 7 | 84643764 | rs1468396 | -1.041 | 1.448 | 1.487 | 0.744 | 0.014 | -0.022 | -0.022 | 0 | 0.33 | 0.55 | 0.27 | 0.27 | 0.33 | 0.21 |
| 7 | 84644500 | rs17559084 | -0.623 | 1.372 | 1.454 | 0.722 | -0.113 | 0.151 | 0.151 | 0.799 | 0.29 | 0.03 | 0.2 | 0.06 | 0.32 | 0.2 |
| 7 | 84652988 | rs1016027 | -1.071 | 1.402 | 1.436 | 0.726 | 0.008 | -0.017 | -0.017 | 0 | 0.33 | 0.58 | 0.27 | 0.27 | 0.33 | 0.21 |
| 7 | 84677860 | rs17559978 | -0.968 | 1.496 | 1.057 | 0.767 | 0.019 | -0.026 | -0.026 | 0 | 0.31 | 0.54 | 0.25 | 0.27 | 0.3 | 0.19 |
| 7 | 84689251 | rs2214820 | -0.846 | 1.626 | 0.181 | 0.831 | -0.017 | 0.012 | 0.012 | 0.036 | 0.29 | 0.19 | 0.22 | 0.27 | 0.3 | 0.19 |
| 7 | 84692941 | rs2240394 | -0.886 | 2.16 | 0.533 | 1.066 | 0.138 | 0.092 | 0.092 | 0.07 | 0.08 | 0.29 | 0.12 | 0.2 | 0.14 | 0.02 |
| 7 | 84699387 | rs6966472 | -0.971 | 2.118 | 0.467 | 1.064 | 0.158 | 0.115 | 0.115 | 0.198 | 0.09 | 0.37 | 0.12 | 0.2 | 0.14 | 0.02 |
| 7 | 84700633 | rs12673176 | -0.806 | 1.654 | -0.805 | 1.122 | -0.041 | -0.147 | -0.147 | 1.036 | 0.23 | 0.05 | 0.31 | 0.49 | 0.23 | 0.37 |
| 7 | 84707925 | rs6962019 | -0.485 | 1.322 | -0.694 | 1.248 | 0.005 | -0.112 | -0.112 | 0 | 0.18 | 0.06 | 0.28 | 0.45 | 0.16 | 0.27 |
| 7 | 84719338 | rs10954763 | -0.694 | 1.608 | -0.526 | 1.142 | -0.003 | -0.031 | -0.031 | 0 | 0.31 | 0.17 | 0.29 | 0.39 | 0.28 | 0.35 |
| 7 | 84721127 | rs12536409 | -0.542 | 1.616 | -0.668 | 1.148 | -0.022 | -0.062 | -0.062 | 0 | 0.3 | 0.14 | 0.29 | 0.39 | 0.28 | 0.35 |
| 7 | 84737683 | rs10486848 | -0.065 | 2.214 | -0.706 | 0.803 | -0.045 | 0.046 | 0.046 | 0 | 0.1 | 0.01 | 0.09 | 0 | 0.2 | 0.07 |
| 7 | 84738495 | rs1029563 | -1.178 | 2.182 | 0.082 | 1.256 | 0.032 | 0.03 | 0.03 | 0 | 0.16 | 0.27 | 0.12 | 0.24 | 0.24 | 0.16 |
| 7 | 84751951 | rs2190208 | -1.23 | 1.948 | 0.309 | 1.049 | 0.001 | 0.019 | 0.019 | 0.02 | 0.47 | 0.62 | 0.31 | 0.64 | 0.5 | 0.5 |
| 7 | 84754151 | rs17647464 | -1.348 | 1.847 | 0.15 | 1.011 | -0.001 | 0.049 | 0.049 | 0.245 | 0.48 | 0.8 | 0.32 | 0.64 | 0.51 | 0.5 |

| | | | | | | | | | | | | | | | | |
|---|---|---|---|---|---|---|---|---|---|---|---|---|---|---|---|---|
| 7 | 84764394 | rs764077 | -1.333 | 1.717 | -0.056 | 1.027 | -0.017 | 0.072 | 0.072 | 0.45 | 0.51 | 0.86 | 0.36 | 0.66 | 0.51 | 0.6 |
| 7 | 84785117 | rs2191483 | -1.156 | 1.243 | -0.152 | 0.647 | 0.022 | 0.19 | 0.19 | 0.604 | 0.55 | 0.8 | 0.57 | 0.89 | 0.59 | 0.56 |
| 7 | 84802050 | rs7780423 | -1.465 | 1.254 | -0.207 | 0.611 | 0.001 | 0.225 | 0.225 | 0.804 | 0.58 | 0.91 | 0.61 | 0.91 | 0.61 | 0.65 |
| 7 | 84810210 | rs2732757 | 0.157 | 1.531 | -0.111 | 0.92 | -0.026 | -0.025 | -0.025 | 0 | 0.13 | 0.09 | 0.12 | 0.21 | 0.24 | 0.09 |
| 7 | 84820483 | rs1013264 | -0.715 | 1.4 | 0.674 | 0.503 | 0.014 | 0.034 | 0.034 | 0 | 0.44 | 0.67 | 0.47 | 0.7 | 0.36 | 0.56 |
| 7 | 84845546 | rs2159579 | -1.487 | 1.377 | 0.863 | 0.782 | 0.049 | 0.067 | 0.067 | 0.286 | 0.21 | 0.53 | 0.2 | 0.23 | 0.26 | 0.18 |
| 7 | 84861335 | rs11973517 | -1.322 | 0.764 | 1.095 | 0.967 | 0.053 | 0.305 | 0.305 | 1.285 | 0.06 | 0.37 | 0.05 | 0.16 | 0.01 | 0 |
| 7 | 84866308 | rs2732736 | 1.046 | 0.839 | -0.412 | -0.204 | -0.036 | 0.155 | 0.155 | 0.218 | 0.41 | 0.16 | 0.27 | 0.01 | 0.51 | 0.25 |
| 7 | 84873886 | rs2732734 | 1.016 | 0.711 | -0.049 | 0.166 | -0.004 | 0.102 | 0.102 | 0.897 | 0.46 | 0.28 | 0.34 | 0.17 | 0.49 | 0.26 |
| 7 | 84877814 | rs12668415 | -0.272 | 0.995 | 0.347 | 0.854 | 0.003 | 0.034 | 0.034 | 0.3 | 0.14 | 0.19 | 0.14 | 0.38 | 0.15 | 0.35 |
| 7 | 84885923 | rs2534848 | 1.593 | 0.801 | 0.547 | 1.151 | -0.013 | 0.071 | 0.071 | 2.533 | 0.45 | 0.17 | 0.4 | 0.28 | 0.48 | 0.29 |
| 7 | 84890498 | rs4370460 | -0.099 | 0.562 | 0.951 | 1.15 | 0.004 | 0.01 | 0.01 | 0 | 0.24 | 0.24 | 0.17 | 0.13 | 0.2 | 0.2 |
| 7 | 84892114 | rs1527930 | 0.972 | 0.715 | 1.341 | 1.15 | 0.017 | 0.037 | 0.037 | 0.753 | 0.75 | 0.65 | 0.6 | 0.41 | 0.71 | 0.49 |
| 7 | 84896786 | rs2696169 | 2.191 | 0.556 | 1.056 | 1.161 | -0.024 | 0.075 | 0.075 | 2.352 | 0.45 | 0.18 | 0.41 | 0.28 | 0.53 | 0.3 |
| 7 | 84905855 | rs2178115 | 0.952 | 0.596 | 0.981 | 1.186 | -0.028 | -0.005 | -0.005 | 0 | 0.06 | 0.01 | 0.2 | 0.05 | 0.07 | 0.13 |
| 7 | 84914691 | rs6944323 | -0.342 | 0.644 | 1.139 | 1.145 | 0.005 | 0.008 | 0.008 | 0 | 0.1 | 0.11 | 0.3 | 0.18 | 0.11 | 0.21 |
| 7 | 84916291 | rs10252813 | -1.347 | 0.608 | 1.206 | 1.146 | 0.008 | 0.05 | 0.05 | 0 | 0.08 | 0.21 | 0.11 | 0.13 | 0.04 | 0.08 |
| 7 | 84935694 | rs2696166 | 2.215 | 0.489 | 1.045 | 1.07 | -0.006 | 0.058 | 0.058 | 0.669 | 0.46 | 0.26 | 0.42 | 0.28 | 0.53 | 0.29 |
| 7 | 84946665 | rs1429743 | -0.851 | 0.469 | 0.772 | 1.22 | -0.01 | 0.021 | 0.021 | 0 | 0.3 | 0.17 | 0.31 | 0.22 | 0.27 | 0.3 |
| 7 | 84953373 | rs12704123 | -1.558 | 0.631 | 0.944 | 1.137 | -0.007 | -0.008 | -0.008 | 0.378 | 0.4 | 0.36 | 0.42 | 0.58 | 0.39 | 0.61 |
| 7 | 84955066 | rs13438056 | -1.489 | 0.649 | 0.87 | 1.145 | -0.001 | -0.001 | -0.001 | 0 | 0.42 | 0.45 | 0.43 | 0.58 | 0.4 | 0.61 |
| 7 | 84968982 | rs1540847 | -2.015 | 0.624 | 0.932 | 1.132 | 0 | 0.001 | 0.001 | 1.527 | 0.48 | 0.56 | 0.53 | 0.71 | 0.43 | 0.68 |
| 7 | 84978077 | rs7781046 | -1.456 | 0.351 | 0.771 | 1.03 | -0.002 | -0.006 | -0.006 | 0.941 | 0.41 | 0.47 | 0.38 | 0.3 | 0.35 | 0.59 |
| 7 | 84987757 | rs6968184 | 0.44 | 0.33 | 0.625 | 0.966 | 0.018 | 0.03 | 0.03 | 0 | 0.18 | 0.13 | 0.06 | 0 | 0.14 | 0.15 |
| 7 | 84988303 | rs7805482 | -2.129 | 0.267 | 0.849 | 1.014 | 0.019 | 0.015 | 0.015 | 2.225 | 0.47 | 0.76 | 0.51 | 0.43 | 0.39 | 0.66 |
| 7 | 84991442 | rs1851704 | -0.613 | 0.347 | 0.279 | 0.827 | 0.003 | 0.076 | 0.076 | 0 | 0.25 | 0.17 | 0.27 | 0.04 | 0.22 | 0.27 |
| 7 | 84993669 | rs2082074 | -1.096 | 0.397 | 0.216 | 0.962 | -0.001 | -0.002 | -0.002 | 0 | 0.38 | 0.44 | 0.38 | 0.3 | 0.35 | 0.59 |
| 7 | 85004375 | rs1916667 | 0.887 | 0.126 | 0.026 | 0.897 | -0.015 | 0.013 | 0.013 | 0.174 | 0.63 | 0.36 | 0.71 | 0.44 | 0.68 | 0.57 |
| 7 | 85005259 | rs1429746 | -0.847 | 0.39 | -0.109 | 0.874 | -0.014 | -0.031 | -0.031 | 0 | 0.25 | 0.22 | 0.25 | 0.58 | 0.3 | 0.31 |
| 7 | 85005940 | rs993900 | 0.42 | 0.88 | 0.29 | 0.228 | 0.006 | -0.062 | -0.062 | 2.607 | 0.76 | 0.75 | 0.71 | 0.95 | 0.76 | 0.85 |
| 7 | 85009349 | rs2463468 | 1.836 | 0.722 | 0.399 | 0.653 | 0.007 | 0.012 | 0.012 | 0 | 0.5 | 0.42 | 0.45 | 0.32 | 0.46 | 0.52 |
| 7 | 85014355 | rs1403786 | 0.198 | 0.742 | 0.277 | 0.715 | -0.025 | -0.055 | -0.055 | 0.761 | 0.19 | 0.14 | 0.17 | 0.45 | 0.23 | 0.29 |
| 7 | 85026991 | rs12666995 | 0.268 | 0.631 | 0.285 | 0.719 | -0.038 | -0.077 | -0.077 | 0 | 0.18 | 0.1 | 0.17 | 0.45 | 0.24 | 0.3 |
| 7 | 85030592 | rs2463478 | 1.844 | 0.566 | 0.303 | 0.716 | 0.021 | 0.044 | 0.044 | 0.02 | 0.5 | 0.34 | 0.34 | 0.29 | 0.46 | 0.52 |
| 7 | 85031803 | rs9649073 | -1.05 | 0.689 | -0.181 | 0.719 | 0.001 | -0.003 | -0.003 | 0 | 0.29 | 0.31 | 0.36 | 0.23 | 0.3 | 0.19 |
| 7 | 85033062 | rs13236638 | -0.05 | 0.757 | -0.236 | 0.753 | 0.002 | 0.02 | 0.02 | 0.047 | 0.21 | 0.31 | 0.19 | 0.45 | 0.24 | 0.3 |
| 7 | 85034506 | rs10245800 | -1.085 | 0.828 | -0.254 | 0.622 | -0.021 | 0.008 | 0.008 | 0 | 0.26 | 0.14 | 0.34 | 0.23 | 0.3 | 0.19 |
| 7 | 85047858 | rs2498572 | 0.82 | 0.903 | 0.008 | 0.865 | -0.007 | -0.005 | -0.005 | 0.105 | 0.81 | 0.86 | 0.7 | 0.76 | 0.79 | 0.85 |
| 7 | 85055535 | rs11768400 | 0.866 | 0.563 | 0.417 | 0.055 | -0.008 | -0.111 | -0.111 | 0 | 0.43 | 0.46 | 0.26 | 0.04 | 0.41 | 0.4 |
| 7 | 85055617 | rs2435279 | 0.725 | 0.44 | 0.357 | 0.487 | -0.014 | 0.106 | 0.106 | 0 | 0.92 | 0.84 | 0.85 | 0.67 | 0.94 | 0.97 |
| 7 | 85056906 | rs6465004 | 1.307 | 0.324 | 0.633 | 0.389 | -0.009 | 0.174 | 0.174 | 0.837 | 0.39 | 0.12 | 0.22 | 0.04 | 0.4 | 0.4 |
| 7 | 85079830 | rs2498551 | 1.074 | 0.673 | 1.002 | 0.809 | -0.016 | 0.134 | 0.134 | 0 | 0.91 | 0.81 | 0.85 | 0.67 | 0.94 | 0.97 |
| 7 | 85079960 | rs2463664 | 1.363 | 0.666 | 1.014 | 0.806 | -0.012 | 0.192 | 0.192 | 0 | 0.91 | 0.74 | 0.85 | 0.67 | 0.94 | 0.97 |
| 7 | 85082956 | rs1357549 | 2.194 | 0.687 | 1.024 | 0.871 | 0.027 | 0.238 | 0.238 | 0 | 0.87 | 0.65 | 0.72 | 0.48 | 0.88 | 0.8 |
| 7 | 85097007 | rs1852697 | 1.722 | 0.609 | 1.047 | 0.794 | 0.007 | 0.192 | 0.192 | 0 | 0.86 | 0.65 | 0.69 | 0.61 | 0.9 | 0.88 |
| 7 | 85099770 | rs2462051 | -0.444 | 0.619 | 1.05 | 0.551 | -0.012 | 0.008 | 0.008 | 1.072 | 0.46 | 0.41 | 0.45 | 0.57 | 0.5 | 0.49 |
| 7 | 85103671 | rs2091086 | 0.332 | 0.442 | 1.118 | -0.302 | 0.008 | 0.002 | 0.002 | 0 | 0.93 | 0.87 | 0.82 | 0.94 | 0.94 | 0.92 |
| 7 | 85111529 | rs2140461 | -0.219 | 0.552 | 1.206 | -0.349 | -0.007 | -0.014 | -0.014 | 0 | 0.42 | 0.49 | 0.39 | 0.71 | 0.37 | 0.34 |
| 7 | 85117732 | rs4728584 | -0.134 | 0.555 | 1.039 | -0.348 | 0.005 | 0.009 | 0.009 | 0.294 | 0.41 | 0.42 | 0.39 | 0.71 | 0.37 | 0.34 |
| 7 | 85124635 | rs2463678 | -0.624 | 0.681 | 0.675 | -0.209 | 0.002 | 0 | 0 | 0 | 0.13 | 0.15 | 0.17 | 0.19 | 0.15 | 0.19 |
| 7 | 85137782 | rs17160014 | 0.206 | 0.983 | 0.525 | 0.275 | -0.029 | -0.179 | -0.179 | 0 | 0.29 | 0.01 | 0.24 | 0.53 | 0.22 | 0.28 |
| 7 | 85143014 | rs2463686 | -0.631 | 1.214 | 0.879 | -0.103 | 0.069 | 0.05 | 0.05 | 0.677 | 0.09 | 0.21 | 0.19 | 0.22 | 0.17 | 0.16 |
| 7 | 85143724 | rs16887926 | -0.07 | 1.259 | 0.648 | -0.124 | 0.004 | 0.001 | 0.001 | 0 | 0.06 | 0.1 | 0.18 | 0.06 | 0.05 | 0.12 |
| 7 | 85147535 | rs10254290 | 2.169 | 1.016 | 0.696 | -0.24 | 0.021 | 0.006 | 0.006 | 0 | 0.77 | 0.31 | 0.59 | 0.72 | 0.77 | 0.71 |
| 7 | 85147760 | rs17328908 | 0.107 | 0.993 | 0.702 | -0.301 | -0.007 | 0.016 | 0.016 | 1.045 | 0.36 | 0.07 | 0.35 | 0.68 | 0.35 | 0.37 |
| 7 | 85148963 | rs7807771 | -1.479 | 1.098 | 0.777 | -0.311 | 0.01 | 0.045 | 0.045 | 0 | 0.11 | 0.65 | 0.22 | 0.13 | 0.05 | 0.13 |
| 7 | 85155842 | rs1980389 | 0.203 | 1.297 | 0.022 | -0.429 | -0.03 | -0.005 | -0.005 | 0 | 0.89 | 0.96 | 0.81 | 0.86 | 0.83 | 0.84 |
| 7 | 85160755 | rs6951823 | 1.002 | 1.023 | 0.139 | -0.614 | -0.021 | 0.233 | 0.233 | 0.584 | 0.42 | 0.11 | 0.23 | 0.04 | 0.43 | 0.35 |
| 7 | 85194414 | rs10499891 | -0.212 | 0.79 | 0.738 | 0.3 | -0.025 | -0.004 | -0.004 | 0.129 | 0.06 | 0.01 | 0.1 | 0.06 | 0.09 | 0.12 |
| 7 | 85203213 | rs17160047 | -1.108 | 0.73 | 1.509 | 0.273 | 0.029 | -0.003 | -0.003 | 0 | 0.09 | 0.22 | 0.1 | 0.06 | 0.09 | 0.12 |
| 7 | 85205095 | rs2372800 | 0.41 | 0.455 | 1.53 | 0.109 | -0.009 | -0.013 | -0.013 | 2.176 | 0.65 | 0.6 | 0.6 | 0.22 | 0.77 | 0.55 |
| 7 | 85213231 | rs6970690 | 0.463 | 0.427 | 1.383 | 0.123 | -0.03 | -0.037 | -0.037 | 0 | 0.64 | 0.52 | 0.6 | 0.24 | 0.77 | 0.55 |
| 7 | 85216272 | rs10264125 | 0.448 | 0.439 | 1.388 | 0.128 | -0.03 | -0.038 | -0.038 | 0.575 | 0.64 | 0.52 | 0.6 | 0.24 | 0.77 | 0.55 |
| 7 | 85217250 | rs4728594 | 0.448 | 0.439 | 1.388 | 0.128 | -0.03 | -0.038 | -0.038 | 0.274 | 0.64 | 0.52 | 0.6 | 0.24 | 0.77 | 0.55 |
| 7 | 85221926 | rs4147416 | 0.195 | 0.42 | 1.264 | 0.114 | 0.041 | 0.051 | 0.051 | 1.094 | 0.68 | 0.82 | 0.63 | 0.24 | 0.78 | 0.55 |
| 7 | 85222453 | rs6953137 | -0.849 | 0.581 | 0.939 | 0.122 | 0.065 | 0.004 | 0.004 | 0 | 0.11 | 0.21 | 0.18 | 0.14 | 0.16 | 0.21 |
| 7 | 85233643 | rs7780046 | 1.335 | 0.561 | 0.92 | 0.11 | 0.02 | 0.143 | 0.143 | 0.038 | 0.52 | 0.27 | 0.42 | 0.09 | 0.62 | 0.34 |
| 7 | 85240205 | rs4279515 | 0.196 | 0.814 | 0.461 | 0.608 | 0.005 | -0.049 | -0.049 | 0 | 0.08 | 0 | 0.08 | 0.19 | 0.06 | 0.08 |
| 7 | 85250155 | rs7794764 | 0.854 | 1.06 | 0.669 | 0.112 | 0.071 | 0.008 | 0.008 | 0 | 0.89 | 0.78 | 0.82 | 0.85 | 0.84 | 0.79 |

| | | | | | | | | | | | | | | | | |
|---|---|---|---|---|---|---|---|---|---|---|---|---|---|---|---|---|
| 7 | 85269460 | rs12704156 | 1.107 | 0.71 | 0.029 | -0.138 | 0.028 | 0.215 | 0.215 | 0.071 | 0.52 | 0.19 | 0.42 | 0.09 | 0.62 | 0.34 |
| 7 | 85275522 | rs4466324 | -0.229 | 0.431 | 0.057 | 0.092 | 0.046 | 0.023 | 0.023 | 1.129 | 0.31 | 0.08 | 0.26 | 0.71 | 0.22 | 0.46 |
| 7 | 85280071 | rs12534172 | 0.128 | 0.591 | 0.036 | 0.011 | -0.022 | -0.031 | -0.031 | 0 | 0.06 | 0.01 | 0.18 | 0.15 | 0.07 | 0.11 |
| 7 | 85291871 | rs4376434 | -0.87 | 0.473 | 0.687 | -0.117 | -0.003 | -0.025 | -0.025 | 0.349 | 0.39 | 0.31 | 0.45 | 0.85 | 0.28 | 0.56 |
| 7 | 85297352 | rs4543471 | 0.267 | 1.125 | 0.864 | -0.11 | 0.122 | 0.012 | 0.012 | 0 | 0.94 | 0.81 | 0.89 | 0.93 | 0.89 | 0.89 |
| 7 | 85303815 | rs4291195 | 0.081 | 1.131 | 0.766 | -0.107 | 0.087 | 0.003 | 0.003 | 0 | 0.94 | 0.86 | 0.9 | 0.93 | 0.89 | 0.89 |
| 7 | 85335684 | rs10261522 | -0.22 | 0.889 | 0.605 | -0.133 | 0.051 | 0.046 | 0.046 | 1.071 | 0.32 | 0.14 | 0.34 | 0.74 | 0.21 | 0.45 |
| 7 | 85338042 | rs10235777 | 1.344 | 0.865 | 0.877 | -0.238 | 0.011 | 0.213 | 0.213 | 0.064 | 0.51 | 0.22 | 0.41 | 0.07 | 0.61 | 0.34 |
| 7 | 85338112 | rs4323400 | -0.21 | 0.909 | 0.664 | -0.213 | 0.052 | 0.057 | 0.057 | 1.024 | 0.32 | 0.14 | 0.37 | 0.76 | 0.21 | 0.45 |
| 7 | 85340656 | rs10244696 | 0.925 | 0.946 | 0.732 | -0.334 | 0 | -0.06 | -0.06 | 0 | 0.55 | 0.61 | 0.45 | 0.07 | 0.61 | 0.34 |
| 7 | 85345855 | rs12534577 | 0.31 | 0.534 | -0.353 | 0.207 | 0.026 | -0.055 | -0.055 | 0.053 | 0.22 | 0.1 | 0.18 | 0.51 | 0.15 | 0.37 |
| 7 | 85347424 | rs13246206 | 0.556 | 0.758 | -0.496 | -0.631 | 0.012 | 0.179 | 0.179 | 0.043 | 0.51 | 0.25 | 0.41 | 0.07 | 0.6 | 0.34 |
| 7 | 85356773 | rs6948502 | 0.272 | 0.478 | -0.59 | -0.736 | 0.006 | -0.007 | -0.007 | 0 | 0.38 | 0.34 | 0.3 | 0.07 | 0.38 | 0.31 |
| 7 | 85365197 | rs884128 | -0.141 | 0.185 | -1.057 | 0.241 | 0.015 | -0.081 | -0.081 | 0.003 | 0.26 | 0.06 | 0.27 | 0.45 | 0.2 | 0.39 |
| 7 | 85381614 | rs10952857 | -0.695 | -0.166 | -0.516 | 0.079 | -0.017 | 0.121 | 0.121 | 0 | 0.22 | 0.54 | 0.23 | 0.43 | 0.2 | 0.29 |
| 7 | 85383292 | rs9649088 | -0.197 | -0.223 | -0.039 | 0.612 | -0.024 | 0.035 | 0.035 | 0.384 | 0.16 | 0.3 | 0.11 | 0.22 | 0.15 | 0.11 |
| 7 | 85405337 | rs10273682 | -0.6 | -0.192 | 0.16 | 0.616 | 0.108 | -0.021 | -0.021 | 0.189 | 0.39 | 0.17 | 0.3 | 0.48 | 0.19 | 0.4 |
| 7 | 85407839 | rs6465020 | -1.106 | -0.153 | 0.288 | 0.519 | -0.003 | -0.003 | -0.003 | 0 | 0.47 | 0.51 | 0.45 | 0.68 | 0.3 | 0.58 |
| 7 | 85415440 | rs10952858 | -1.343 | -0.758 | 0.318 | 0.527 | 0.027 | 0.025 | 0.025 | 0.018 | 0.49 | 0.55 | 0.46 | 0.89 | 0.33 | 0.68 |
| 7 | 85419011 | rs9640563 | -1.004 | -0.759 | 0.053 | 0.528 | -0.002 | 0.001 | 0.001 | 0.057 | 0.46 | 0.44 | 0.41 | 0.69 | 0.25 | 0.58 |
| 7 | 85431893 | rs17160166 | -1.152 | -0.623 | -0.005 | 0.543 | -0.015 | -0.011 | -0.011 | 0.174 | 0.48 | 0.47 | 0.42 | 0.71 | 0.27 | 0.61 |
| 7 | 85440774 | rs10246682 | 0.389 | -0.698 | -0.138 | 0.699 | 0.027 | 0.027 | 0.027 | 0.881 | 0.11 | 0.04 | 0.05 | 0.02 | 0.07 | 0.02 |
| 7 | 85443602 | rs10226311 | -0.664 | -0.685 | -0.076 | 0.71 | 0.065 | -0.007 | -0.007 | 0 | 0.41 | 0.24 | 0.31 | 0.51 | 0.2 | 0.43 |
| 7 | 85458197 | rs10273712 | -0.317 | -0.567 | -0.316 | 0.553 | 0.016 | 0.009 | 0.009 | 0.119 | 0.18 | 0.14 | 0.1 | 0.02 | 0.14 | 0.02 |
| 7 | 85461399 | rs1396587 | -0.134 | -0.596 | -0.255 | 0.324 | 0.006 | 0.104 | 0.104 | 0 | 0.07 | 0.08 | 0.14 | 0.38 | 0.14 | 0.28 |
| 7 | 85470182 | rs9640974 | 0.341 | -0.748 | -0.292 | 0.257 | 0.055 | -0.053 | -0.053 | 0 | 0.28 | 0.09 | 0.24 | 0.48 | 0.13 | 0.41 |
| 7 | 85485663 | rs7350007 | 0.046 | -0.727 | -0.308 | 0.212 | 0.027 | 0.068 | 0.068 | 1.496 | 0.16 | 0.05 | 0.09 | 0.02 | 0.14 | 0.02 |
| 7 | 85495837 | rs12537524 | 0.343 | -0.725 | -0.281 | 0.259 | 0.044 | -0.043 | -0.043 | 0 | 0.27 | 0.11 | 0.24 | 0.48 | 0.13 | 0.41 |
| 7 | 85514562 | rs6971791 | 0.847 | -0.706 | -0.256 | 0.254 | 0.002 | -0.014 | -0.014 | 0.597 | 0.74 | 0.76 | 0.75 | 0.61 | 0.73 | 0.71 |
| 7 | 85516921 | rs4336535 | 0.087 | -0.587 | -0.292 | 0.319 | -0.004 | 0 | 0 | 0 | 0.29 | 0.25 | 0.26 | 0.48 | 0.15 | 0.4 |
| 7 | 85525999 | rs4461822 | 0.112 | -0.575 | -0.585 | 0.217 | 0.013 | 0.093 | 0.093 | 0 | 0.67 | 0.91 | 0.62 | 0.17 | 0.73 | 0.34 |
| 7 | 85533208 | rs4464877 | -1.166 | -0.66 | -0.559 | 0.127 | 0.048 | 0.136 | 0.136 | 0 | 0.52 | 0.26 | 0.49 | 0.88 | 0.4 | 0.7 |
| 7 | 85555944 | rs17160342 | 0.506 | -0.61 | -0.895 | 0.283 | 0.048 | -0.141 | -0.141 | 0.252 | 0.26 | 0 | 0.25 | 0.46 | 0.13 | 0.39 |
| 7 | 85556475 | rs7457191 | 0.484 | -0.714 | -0.889 | 0.302 | 0.082 | -0.115 | -0.115 | 0 | 0.26 | 0.03 | 0.23 | 0.44 | 0.12 | 0.39 |
| 7 | 85558378 | rs717708 | -0.452 | -0.217 | -0.503 | 0.461 | -0.001 | 0.001 | 0.001 | 0.35 | 0.47 | 0.43 | 0.34 | 0.48 | 0.26 | 0.42 |
| 7 | 85558996 | rs2726001 | -0.773 | -0.076 | -0.472 | -0.263 | -0.009 | -0.046 | -0.046 | 0 | 0.54 | 0.49 | 0.48 | 0.85 | 0.39 | 0.69 |
| 7 | 85571719 | rs13222653 | 0.057 | -0.059 | -0.769 | -0.436 | 0.018 | 0.04 | 0.04 | 0.063 | 0.44 | 0.3 | 0.34 | 0.2 | 0.34 | 0.45 |
| 7 | 85573660 | rs2107256 | 0.216 | 0.48 | -0.684 | -0.35 | -0.003 | -0.018 | -0.018 | 0.493 | 0.58 | 0.58 | 0.68 | 0.78 | 0.58 | 0.67 |
| 7 | 85574619 | rs10237969 | -0.361 | 0.704 | -0.79 | -0.528 | -0.006 | 0.034 | 0.034 | 4.851 | 0.47 | 0.66 | 0.35 | 0.29 | 0.51 | 0.29 |
| 7 | 85582101 | rs763637 | -0.226 | 1.113 | -0.428 | -0.586 | -0.003 | -0.003 | -0.003 | 0 | 0.44 | 0.43 | 0.3 | 0.3 | 0.42 | 0.27 |
| 7 | 85585601 | rs13228601 | 0.287 | 1.019 | -0.191 | -0.703 | -0.003 | -0.003 | -0.003 | 0.219 | 0.16 | 0.18 | 0.13 | 0.12 | 0.17 | 0.09 |
| 7 | 85593562 | rs7785694 | 1.076 | 0.518 | -0.467 | -0.841 | -0.025 | -0.019 | -0.019 | 0 | 0.17 | 0.25 | 0.1 | 0.11 | 0.11 | 0.1 |
| 7 | 85593608 | rs10952871 | -0.194 | 0.477 | -0.522 | -0.868 | 0.023 | 0.023 | 0.023 | 5.141 | 0.46 | 0.67 | 0.33 | 0.29 | 0.42 | 0.27 |
| 7 | 85594984 | rs17222177 | 0.864 | 0.568 | -0.815 | -0.82 | -0.003 | -0.003 | -0.003 | 0 | 0.16 | 0.18 | 0.13 | 0.12 | 0.17 | 0.09 |
| 7 | 85608916 | rs10235951 | -1.158 | 1.007 | -1.152 | -0.688 | -0.075 | -0.101 | -0.101 | 0 | 0.64 | 0.92 | 0.47 | 0.41 | 0.62 | 0.4 |
| 7 | 85611483 | rs10245061 | -0.693 | 0.861 | -1.173 | -0.668 | -0.061 | -0.04 | -0.04 | 0 | 0.61 | 0.92 | 0.42 | 0.34 | 0.51 | 0.34 |
| 7 | 85612834 | rs10252288 | -0.505 | 0.849 | -1.1 | -0.673 | -0.003 | 0.005 | 0.005 | 1.326 | 0.58 | 0.69 | 0.39 | 0.34 | 0.5 | 0.34 |
| 7 | 85617211 | rs6465034 | -0.895 | 0.879 | -1.281 | -0.681 | -0.084 | -0.095 | -0.095 | 0 | 0.62 | 0.92 | 0.43 | 0.41 | 0.53 | 0.38 |
| 7 | 85618492 | rs6944937 | 0.809 | 1.137 | -0.803 | -0.807 | 0.008 | 0.003 | 0.003 | 0 | 0.25 | 0.11 | 0.18 | 0.17 | 0.25 | 0.19 |
| 7 | 85624839 | rs2373019 | -0.727 | 1.624 | -0.951 | -0.721 | -0.056 | 0.034 | 0.034 | 0 | 0.84 | 0.96 | 0.76 | 0.94 | 0.78 | 0.83 |
| 7 | 85640225 | rs17160554 | 1.003 | 0.869 | -0.738 | -0.609 | 0.023 | -0.01 | -0.01 | 0 | 0.19 | 0.09 | 0.13 | 0.15 | 0.16 | 0.15 |
| 7 | 85643340 | rs2158318 | -0.36 | 1.108 | -0.405 | -0.592 | 0.095 | 0.033 | 0.033 | 0 | 0.07 | 0.28 | 0.06 | 0.07 | 0.11 | 0.06 |
| 7 | 85672109 | rs17160578 | 1.338 | 0.881 | -0.643 | -0.514 | -0.005 | -0.054 | -0.054 | 0.271 | 0.17 | 0.02 | 0.12 | 0.15 | 0.15 | 0.12 |
| 7 | 85699781 | rs12540083 | 0.997 | 0.472 | -0.524 | -0.894 | 0.032 | -0.056 | -0.056 | 0 | 0.33 | 0.15 | 0.37 | 0.6 | 0.27 | 0.47 |
| 7 | 85711326 | rs10243282 | -0.184 | 0.648 | -0.526 | -1.014 | -0.1 | 0.062 | 0.062 | 0.913 | 0.64 | 0.9 | 0.6 | 0.78 | 0.54 | 0.66 |
| 7 | 85737242 | rs1117120 | -0.416 | 1.107 | -0.565 | -1.24 | -0.077 | 0.094 | 0.094 | 0.794 | 0.67 | 0.9 | 0.62 | 0.85 | 0.59 | 0.7 |
| 7 | 85774632 | rs41441 | 0.741 | 0.982 | -0.07 | -1.22 | -0.006 | -0.019 | -0.019 | 0.045 | 0.17 | 0.3 | 0.15 | 0.11 | 0.17 | 0.07 |
| 7 | 85813923 | rs6465042 | 0.478 | 1.086 | -0.086 | -1.346 | 0.012 | -0.034 | -0.034 | 0 | 0.18 | 0.26 | 0.17 | 0.05 | 0.22 | 0.06 |
| 7 | 85814138 | rs10223926 | 1.38 | 0.646 | 0.024 | -1.442 | 0.008 | -0.001 | -0.001 | 0 | 0.22 | 0.16 | 0.13 | 0.18 | 0.16 | 0.13 |
| 7 | 85815656 | rs12531407 | 1.252 | 0.837 | -0.16 | -1.721 | -0.002 | -0.007 | -0.007 | 0.081 | 0.14 | 0.17 | 0.12 | 0.05 | 0.12 | 0.04 |
| 7 | 85841693 | rs7782603 | 1.19 | 0.803 | -0.192 | -1.781 | -0.015 | -0.04 | -0.04 | 0 | 0.14 | 0.3 | 0.13 | 0.05 | 0.13 | 0.04 |
| 7 | 85849534 | rs6465047 | 0.424 | 0.698 | -0.253 | -1.826 | -0.004 | -0.014 | -0.014 | 0 | 0.36 | 0.41 | 0.27 | 0.13 | 0.32 | 0.17 |
| 7 | 85857978 | rs13233795 | 0.401 | 0.809 | -0.481 | -1.892 | -0.091 | 0.096 | 0.096 | 1.798 | 0.36 | 0.07 | 0.43 | 0.24 | 0.45 | 0.34 |
| 7 | 85866217 | rs7458310 | 0.533 | 0.845 | -0.344 | -1.825 | -0.006 | -0.015 | -0.015 | 0 | 0.35 | 0.41 | 0.25 | 0.13 | 0.28 | 0.16 |
| 7 | 85867258 | rs10262028 | -0.184 | 0.892 | -0.307 | -1.834 | -0.001 | -0.024 | -0.024 | 0 | 0.61 | 0.54 | 0.7 | 0.87 | 0.62 | 0.82 |
| 7 | 85871095 | rs1860781 | 1.42 | 0.677 | -0.367 | -1.784 | -0.011 | -0.03 | -0.03 | 0 | 0.14 | 0.22 | 0.13 | 0.05 | 0.13 | 0.04 |
| 7 | 85890669 | rs2708580 | 1.632 | 0.666 | -0.578 | -1.72 | 0.019 | 0.042 | 0.042 | 0 | 0.2 | 0.04 | 0.11 | 0.08 | 0.16 | 0.13 |
| 7 | 85893683 | rs2708582 | 1.549 | 0.664 | -0.543 | -1.721 | 0.023 | 0.036 | 0.036 | 0 | 0.21 | 0.11 | 0.12 | 0.08 | 0.16 | 0.13 |
| 7 | 85901134 | rs17226867 | 1.221 | 0.843 | -0.727 | -1.834 | -0.029 | 0.013 | 0.013 | 0.095 | 0.22 | 0.04 | 0.28 | 0.18 | 0.25 | 0.18 |

| | | | | | | | | | | | | | | | | |
|---|---|---|---|---|---|---|---|---|---|---|---|---|---|---|---|---|
| 7 | 85927182 | rs2041511 | -0.62 | 1.06 | -0.267 | -1.811 | 0.045 | -0.079 | -0.079 | 0.009 | 0.81 | 0.56 | 0.82 | 0.95 | 0.78 | 0.94 |
| 7 | 85944686 | rs6963410 | 0.52 | 1.116 | -0.185 | -1.81 | 0.032 | -0.049 | -0.049 | 0.012 | 0.19 | 0.32 | 0.2 | 0.05 | 0.26 | 0.06 |
| 7 | 85947103 | rs1978266 | -0.454 | 1.113 | -0.238 | -1.812 | 0.062 | -0.084 | -0.084 | 0.6 | 0.8 | 0.53 | 0.79 | 0.95 | 0.74 | 0.94 |
| 7 | 85952083 | rs6977312 | 1.315 | 1.049 | -0.377 | -1.979 | -0.026 | 0.042 | 0.042 | 0 | 0.27 | 0.46 | 0.36 | 0.64 | 0.21 | 0.5 |
| 7 | 85962908 | rs11767843 | 0.077 | 0.998 | -0.545 | -2.251 | -0.079 | 0.094 | 0.094 | 2.5 | 0.36 | 0.06 | 0.35 | 0.23 | 0.42 | 0.35 |
| 7 | 85982270 | rs2373210 | 1.378 | 0.964 | -0.456 | -2.236 | -0.03 | 0.047 | 0.047 | 0.002 | 0.26 | 0.46 | 0.35 | 0.63 | 0.19 | 0.48 |
| 7 | 85990367 | rs9969120 | -0.857 | 0.84 | -0.685 | -2.315 | 0.013 | 0.017 | 0.017 | 0 | 0.53 | 0.46 | 0.55 | 0.3 | 0.65 | 0.42 |
| 7 | 85994103 | rs4236502 | 0.234 | 0.849 | -0.743 | -2.318 | -0.088 | 0.084 | 0.084 | 2.779 | 0.37 | 0.07 | 0.37 | 0.25 | 0.45 | 0.37 |
| 7 | 86019034 | rs10235241 | -0.531 | 0.795 | -0.611 | -2.278 | 0.003 | 0.045 | 0.045 | 0 | 0.49 | 0.41 | 0.51 | 0.31 | 0.56 | 0.4 |
| 7 | 86036570 | rs1320780 | 1.943 | 0.813 | -0.709 | -2.186 | -0.044 | -0.095 | -0.095 | 0 | 0.22 | 0.05 | 0.32 | 0.62 | 0.22 | 0.48 |
| 7 | 86059778 | rs10239986 | 1.475 | 0.501 | -0.341 | -2.171 | -0.01 | 0.018 | 0.018 | 0 | 0.25 | 0.32 | 0.33 | 0.63 | 0.19 | 0.48 |
| 7 | 86066709 | rs2037877 | -1.729 | 0.348 | -0.473 | -2.351 | 0.027 | 0.042 | 0.042 | 0.913 | 0.78 | 0.9 | 0.9 | 0.93 | 0.84 | 0.9 |
| 7 | 86072337 | rs1881817 | 1.244 | 0.559 | -0.366 | -2.319 | 0.014 | -0.021 | -0.021 | 0 | 0.11 | 0.22 | 0.15 | 0.05 | 0.15 | 0.03 |
| 7 | 86074295 | rs2110391 | 0.1 | 0.032 | -0.409 | -2.301 | 0.013 | -0.03 | -0.03 | 0 | 0.27 | 0.31 | 0.28 | 0.06 | 0.39 | 0.16 |
| 7 | 86079467 | rs1534755 | 0.111 | 0.134 | -0.394 | -2.26 | 0.001 | 0.005 | 0.005 | 0.728 | 0.46 | 0.4 | 0.35 | 0.13 | 0.47 | 0.24 |
| 7 | 86081848 | rs956030 | 1.575 | -0.239 | -0.635 | -2.31 | -0.058 | -0.087 | -0.087 | 0 | 0.27 | 0.04 | 0.4 | 0.62 | 0.31 | 0.47 |
| 7 | 86085024 | rs6946717 | -0.195 | -0.076 | -0.524 | -2.248 | 0.004 | -0.003 | -0.003 | 0 | 0.49 | 0.46 | 0.39 | 0.13 | 0.55 | 0.25 |
| 7 | 86097013 | rs2056800 | 0.779 | -0.067 | -0.503 | -2.132 | -0.011 | 0.002 | 0.002 | 0 | 0.34 | 0.43 | 0.46 | 0.63 | 0.3 | 0.5 |
| 7 | 86103402 | rs1358684 | -0.362 | -0.199 | -0.675 | -2.267 | 0.048 | 0.047 | 0.047 | 0 | 0.6 | 0.73 | 0.81 | 0.72 | 0.72 | 0.67 |
| 7 | 86104673 | rs12113079 | 0.47 | 0.003 | -0.558 | -2.375 | 0.008 | -0.03 | -0.03 | 0 | 0.26 | 0.29 | 0.29 | 0.07 | 0.36 | 0.16 |
| 7 | 86106885 | rs1881812 | -0.019 | 0.116 | -0.557 | -2.394 | 0.019 | -0.038 | -0.038 | 0 | 0.29 | 0.34 | 0.34 | 0.07 | 0.45 | 0.18 |
| 7 | 86107652 | rs1395892 | -1.52 | -0.114 | -0.691 | -2.457 | -0.007 | 0.002 | 0.002 | 0 | 0.83 | 0.81 | 0.91 | 0.78 | 0.88 | 0.78 |
| 7 | 86115537 | rs6965526 | -0.208 | -0.146 | -0.799 | -2.542 | 0.016 | 0.045 | 0.045 | 0 | 0.6 | 0.73 | 0.79 | 0.72 | 0.68 | 0.68 |
| 7 | 86115922 | rs1881819 | 1.797 | -0.26 | -0.903 | -2.628 | -0.075 | -0.082 | -0.082 | 0.407 | 0.27 | 0.04 | 0.39 | 0.64 | 0.31 | 0.48 |
| 7 | 86118757 | rs10952885 | -0.152 | -0.071 | -0.825 | -2.543 | 0.014 | 0.036 | 0.036 | 0 | 0.6 | 0.68 | 0.78 | 0.72 | 0.68 | 0.68 |
| 7 | 86121316 | rs1507591 | 0.602 | -0.023 | -0.912 | -2.474 | 0.008 | -0.03 | -0.03 | 0.413 | 0.26 | 0.29 | 0.3 | 0.07 | 0.36 | 0.16 |
| 7 | 86126902 | rs1320794 | -0.782 | -0.383 | -0.744 | -2.344 | 0.01 | -0.004 | -0.004 | 0 | 0.8 | 0.74 | 0.86 | 0.77 | 0.8 | 0.77 |
| 7 | 86129956 | rs1395891 | -1.299 | -0.499 | -0.714 | -2.317 | -0.018 | 0.005 | 0.005 | 0 | 0.82 | 0.75 | 0.9 | 0.77 | 0.88 | 0.78 |
| 7 | 86134197 | rs3897749 | 1.62 | -0.629 | -1.093 | -2.35 | 0.052 | 0.057 | 0.057 | 0.282 | 0.19 | 0.06 | 0.08 | 0.06 | 0.11 | 0.06 |
| 7 | 86144698 | rs3913932 | 0.849 | -0.351 | -0.987 | -2.321 | 0.012 | 0 | 0 | 0.104 | 0.37 | 0.49 | 0.51 | 0.6 | 0.38 | 0.53 |
| 7 | 86146600 | rs3913931 | 0.688 | -0.498 | -0.895 | -2.423 | -0.001 | -0.009 | -0.009 | 0 | 0.25 | 0.22 | 0.3 | 0.07 | 0.39 | 0.18 |
| 7 | 86150075 | rs3847033 | 1.25 | -0.169 | -0.375 | -2.067 | -0.005 | 0.02 | 0.02 | 0 | 0.29 | 0.4 | 0.4 | 0.6 | 0.28 | 0.48 |
| 7 | 86166392 | rs781011 | 0.892 | -0.226 | -0.394 | -2.076 | -0.018 | 0.01 | 0.01 | 0 | 0.19 | 0.28 | 0.11 | 0.27 | 0.12 | 0.22 |
| 7 | 86171821 | rs10278499 | 0.145 | 0.05 | -0.634 | -2.541 | 0.001 | -0.014 | -0.014 | 0 | 0.08 | 0.09 | 0.11 | 0.01 | 0.1 | 0.05 |
| 7 | 86173228 | rs1405879 | -0.149 | -0.134 | -0.735 | -2.388 | -0.012 | 0.11 | 0.11 | 0.399 | 0.44 | 0.23 | 0.39 | 0.13 | 0.5 | 0.25 |
| 7 | 86222259 | rs1507599 | -0.569 | -0.543 | -0.346 | -2.618 | 0.003 | -0.042 | -0.042 | 0 | 0.3 | 0.57 | 0.22 | 0.09 | 0.32 | 0.15 |
| 7 | 86227662 | rs12704277 | -0.16 | -1.072 | -0.876 | -2.671 | 0.002 | 0.115 | 0.115 | 0.176 | 0.38 | 0.11 | 0.22 | 0.19 | 0.32 | 0.29 |
| 7 | 86238294 | rs12704279 | 0.545 | -1.1 | -1.013 | -2.709 | -0.066 | 0.062 | 0.062 | 1.482 | 0.23 | 0.02 | 0.11 | 0.1 | 0.21 | 0.1 |
| 7 | 86244405 | rs13243345 | -0.208 | -1.129 | -0.982 | -2.704 | -0.005 | 0.104 | 0.104 | 0.082 | 0.37 | 0.13 | 0.24 | 0.2 | 0.31 | 0.29 |
| 7 | 86252828 | rs1830249 | 0.931 | -1.149 | -0.559 | -2.87 | -0.032 | -0.028 | -0.028 | 0 | 0.16 | 0.31 | 0.13 | 0.1 | 0.1 | 0.2 |
| 7 | 86254166 | rs274638 | 1.974 | -1.16 | -1.083 | -2.977 | -0.077 | -0.04 | -0.04 | 0 | 0.28 | 0.06 | 0.52 | 0.72 | 0.32 | 0.56 |
| 7 | 86257517 | rs701330 | 0.516 | -0.919 | -0.609 | -2.848 | -0.046 | 0.09 | 0.09 | 2.59 | 0.23 | 0.03 | 0.1 | 0.09 | 0.21 | 0.09 |
| 7 | 86264311 | rs274627 | 1.718 | -0.934 | -0.505 | -2.786 | -0.044 | 0.008 | 0.008 | 0.039 | 0.32 | 0.09 | 0.53 | 0.72 | 0.34 | 0.62 |
| 7 | 86284433 | rs802432 | 0.038 | -1.011 | 0.021 | -2.823 | -0.005 | -0.044 | -0.044 | 0.19 | 0.28 | 0.27 | 0.14 | 0.08 | 0.2 | 0.08 |
| 7 | 86293693 | rs802422 | -0.778 | -1.068 | 0.117 | -2.859 | -0.016 | -0.053 | -0.053 | 0 | 0.44 | 0.49 | 0.27 | 0.18 | 0.3 | 0.27 |
| 7 | 86307788 | rs12669901 | 2 | -1.004 | -0.526 | -2.9 | -0.06 | 0.001 | 0.001 | 0.509 | 0.31 | 0.06 | 0.53 | 0.72 | 0.34 | 0.63 |
| 7 | 86308600 | rs1581638 | 1.001 | -0.961 | -0.493 | -2.82 | 0.02 | 0.018 | 0.018 | 0.013 | 0.15 | 0.1 | 0.08 | 0.1 | 0.1 | 0.17 |
| 7 | 86325382 | rs724225 | -0.605 | -0.886 | -0.374 | -2.839 | 0.019 | -0.018 | -0.018 | 0.909 | 0.78 | 0.8 | 0.81 | 0.9 | 0.64 | 0.9 |
| 7 | 86325620 | rs724224 | 0.751 | -0.879 | -1.342 | -2.895 | 0.003 | 0.224 | 0.224 | 3.376 | 0.5 | 0.21 | 0.64 | 0.82 | 0.47 | 0.83 |
| 7 | 86335589 | rs17607448 | 0.84 | -1.643 | -1.266 | -2.62 | 0.029 | 0.022 | 0.022 | 0 | 0.15 | 0.08 | 0.07 | 0.11 | 0.1 | 0.17 |
| 7 | 86335812 | rs2299218 | -0.411 | -1.14 | -0.874 | -2.449 | 0.007 | 0.022 | 0.022 | 2.139 | 0.28 | 0.21 | 0.18 | 0.08 | 0.2 | 0.1 |
| 7 | 86345495 | rs2214653 | 1.128 | -0.743 | -1.285 | -2.132 | -0.142 | 0.002 | 0.002 | 0.11 | 0.37 | 0.05 | 0.57 | 0.71 | 0.44 | 0.68 |
| 7 | 86348037 | rs2237547 | -0.86 | -1.048 | -0.515 | -2.229 | 0.083 | 0.115 | 0.115 | 0 | 0.43 | 0.77 | 0.26 | 0.23 | 0.29 | 0.25 |
| 7 | 86351217 | rs10239714 | -0.725 | -1.121 | 0.399 | -2.256 | 0.004 | 0.002 | 0.002 | 0 | 0.42 | 0.6 | 0.25 | 0.23 | 0.29 | 0.25 |
| 7 | 86361467 | rs12671181 | 0.9 | -1.373 | 0.7 | -2.356 | -0.007 | -0.004 | -0.004 | 0.091 | 0.14 | 0.11 | 0.08 | 0.05 | 0.09 | 0.12 |
| 7 | 86374533 | rs2237553 | -1.21 | -1.43 | 0.676 | -2.41 | 0.091 | 0.129 | 0.129 | 0.002 | 0.44 | 0.79 | 0.26 | 0.22 | 0.29 | 0.26 |
| 7 | 86386989 | rs2158785 | -0.941 | -1.434 | 1.078 | -2.412 | -0.02 | -0.039 | -0.039 | 0 | 0.39 | 0.49 | 0.22 | 0.15 | 0.26 | 0.21 |
| 7 | 86395926 | rs6943659 | -1.374 | -1.279 | 0.822 | -2.412 | 0.075 | 0.125 | 0.125 | 0.031 | 0.45 | 0.77 | 0.27 | 0.22 | 0.31 | 0.25 |
| 7 | 86401275 | rs723630 | -0.648 | -1.431 | 1.3 | -2.437 | 0 | -0.004 | -0.004 | 0 | 0.32 | 0.31 | 0.17 | 0.14 | 0.26 | 0.21 |
| 7 | 86401804 | rs2282965 | -1.372 | -1.485 | 1.32 | -2.418 | 0.053 | 0.091 | 0.091 | 0 | 0.39 | 0.8 | 0.25 | 0.21 | 0.3 | 0.29 |
| 7 | 86403263 | rs6943762 | 0.773 | -1.376 | 1.144 | -2.408 | -0.031 | 0.035 | 0.035 | 0.162 | 0.13 | 0.02 | 0.06 | 0.08 | 0.13 | 0.09 |
| 7 | 86403475 | rs6465084 | -0.804 | -1.338 | 1.173 | -2.377 | 0 | -0.004 | -0.004 | 1.865 | 0.32 | 0.31 | 0.17 | 0.14 | 0.26 | 0.21 |
| 7 | 86415987 | rs2228595 | 1.4 | -1.165 | 0.356 | -2.079 | 0.025 | -0.005 | -0.005 | 0 | 0.12 | 0 | 0.07 | 0.05 | 0.07 | 0.12 |
| 7 | 86417845 | rs2299219 | -0.756 | -1.378 | 0.347 | -2.287 | 0.064 | 0.088 | 0.088 | 0.006 | 0.35 | 0.1 | 0.18 | 0.2 | 0.23 | 0.25 |
| 7 | 86420217 | rs11772755 | -0.59 | -1.575 | 1.056 | -2.448 | -0.012 | -0.048 | -0.048 | 0 | 0.09 | 0.31 | 0.06 | 0 | 0.06 | 0.04 |
| 7 | 86420377 | rs10234440 | -0.095 | -1.602 | 0.81 | -2.464 | 0.037 | 0.082 | 0.082 | 0.141 | 0.31 | 0.05 | 0.13 | 0.14 | 0.2 | 0.19 |
| 7 | 86422232 | rs2237562 | -1.215 | -1.532 | 1.179 | -2.418 | 0.07 | 0.09 | 0.09 | 0 | 0.46 | 0.75 | 0.26 | 0.21 | 0.3 | 0.3 |
| 7 | 86443168 | rs2237563 | 1.037 | -0.881 | 0.445 | -2.485 | -0.02 | 0.017 | 0.017 | 0 | 0.09 | 0.05 | 0.13 | 0.02 | 0.21 | 0.03 |
| 7 | 86443385 | rs17608250 | 1.297 | -1.532 | 0.624 | -2.236 | 0.039 | 0.003 | 0.003 | 0 | 0.14 | 0.01 | 0.07 | 0.06 | 0.07 | 0.12 |

| | | | | | | | | | | | | | | | |
|---|---|---|---|---|---|---|---|---|---|---|---|---|---|---|---|
| 7 | 86455536 | rs2237565 | -1.355 | -1.403 | 0.792 | -2.494 | 0.098 | 0.193 | 0.193 | 0 | 0.49 | 0.79 | 0.25 | 0.19 | 0.33 | 0.32 |
| 7 | 86467251 | rs10952891 | -0.861 | -0.347 | 1.537 | -1.856 | 0.004 | -0.038 | -0.038 | 0 | 0.1 | 0.17 | 0.06 | 0 | 0.1 | 0.06 |
| 7 | 86475667 | rs2282966 | -1.489 | -0.515 | 1.835 | -1.953 | -0.022 | -0.072 | -0.072 | 0 | 0.33 | 0.51 | 0.18 | 0.17 | 0.26 | 0.2 |
| 7 | 86476311 | rs6967992 | -1.082 | -0.221 | 1.772 | -1.777 | 0.023 | -0.036 | -0.036 | 0 | 0.08 | 0.32 | 0.05 | 0 | 0.05 | 0.05 |
| 7 | 86502733 | rs7788695 | -1.685 | -0.347 | 1.305 | -1.963 | 0.014 | 0.041 | 0.041 | 0.515 | 0.35 | 0.76 | 0.21 | 0.16 | 0.27 | 0.2 |
| 7 | 86512637 | rs10236047 | -0.875 | -0.351 | 1.419 | -1.907 | -0.001 | -0.002 | -0.002 | 0 | 0.09 | 0.08 | 0.09 | 0.13 | 0.09 | 0.11 |
| 7 | 86535446 | rs10258008 | -0.945 | -0.355 | 1.516 | -1.909 | 0 | 0.006 | 0.006 | 0 | 0.1 | 0.14 | 0.09 | 0.12 | 0.09 | 0.11 |
| 7 | 86536344 | rs7781178 | -1.852 | -0.377 | 1.512 | -1.934 | 0.003 | 0.003 | 0.003 | 0.269 | 0.35 | 0.67 | 0.2 | 0.19 | 0.27 | 0.2 |
| 7 | 86542996 | rs11981220 | -1.728 | -0.345 | 1.649 | -1.91 | 0.002 | 0.019 | 0.019 | 0 | 0.13 | 0.53 | 0.12 | 0.12 | 0.09 | 0.12 |
| 7 | 86544667 | rs2519713 | -2.061 | 0.013 | 0.436 | -2.251 | -0.018 | 0.183 | 0.183 | 1.739 | 0.53 | 0.9 | 0.38 | 0.23 | 0.54 | 0.26 |
| 7 | 86549011 | rs2519709 | -0.391 | -0.216 | 0.947 | -1.87 | 0.016 | -0.025 | -0.025 | 0 | 0.05 | 0.12 | 0.03 | 0 | 0.04 | 0 |
| 7 | 86567364 | rs7797095 | -0.234 | -0.168 | 0.365 | -1.837 | -0.014 | 0.072 | 0.072 | 0.372 | 0.18 | 0.05 | 0.08 | 0.06 | 0.17 | 0.08 |
| 7 | 86593170 | rs1999947 | -0.874 | -0.096 | 0.493 | -1.639 | -0.005 | -0.03 | -0.03 | 1.766 | 0.08 | 0.12 | 0.05 | 0 | 0.06 | 0.08 |
| 7 | 86607231 | rs1767721 | -1.468 | -0.238 | 0.411 | -1.846 | 0.014 | 0.065 | 0.065 | 4.368 | 0.37 | 0.67 | 0.23 | 0.11 | 0.29 | 0.17 |
| 7 | 86608786 | rs1006248 | -1.75 | -0.092 | 0.754 | -1.682 | 0.005 | -0.042 | -0.042 | 0 | 0.34 | 0.57 | 0.24 | 0.19 | 0.29 | 0.13 |
| 7 | 86633357 | rs1767698 | -1.857 | -0.032 | 0.797 | -1.624 | 0.004 | -0.029 | -0.029 | 0.05 | 0.34 | 0.57 | 0.31 | 0.27 | 0.3 | 0.15 |
| 7 | 86654355 | rs9886155 | -0.961 | -0.025 | 0.816 | -1.574 | -0.003 | -0.038 | -0.038 | 0 | 0.25 | 0.28 | 0.14 | 0.15 | 0.19 | 0.1 |
| 7 | 86661802 | rs6971856 | -1.019 | -0.034 | 0.812 | -1.578 | -0.004 | -0.038 | -0.038 | 0.59 | 0.26 | 0.28 | 0.14 | 0.14 | 0.19 | 0.14 |
| 7 | 86662659 | rs6953148 | -1.846 | 0.102 | 1.05 | -1.479 | 0.002 | -0.041 | -0.041 | 0 | 0.33 | 0.47 | 0.27 | 0.28 | 0.26 | 0.16 |
| 7 | 86663809 | rs4728664 | -0.159 | 0.093 | 0.585 | -1.47 | 0.031 | 0.031 | 0.031 | 0 | 0.06 | 0.18 | 0.13 | 0.14 | 0.07 | 0.03 |
| 7 | 86665092 | rs10271876 | -1.715 | 0.044 | 0.184 | -1.373 | -0.002 | 0.007 | 0.007 | 3.223 | 0.35 | 0.67 | 0.3 | 0.3 | 0.29 | 0.17 |
| 7 | 86678176 | rs4727139 | 0.55 | 0.43 | 0.008 | -0.981 | 0.009 | 0.025 | 0.025 | 0.596 | 0.58 | 0.38 | 0.68 | 0.9 | 0.69 | 0.78 |
| 7 | 86678206 | rs4728666 | 0.55 | 0.485 | 0.01 | -0.981 | 0.009 | 0.028 | 0.028 | 0.596 | 0.58 | 0.38 | 0.68 | 0.9 | 0.68 | 0.78 |
| 7 | 86678967 | rs4728669 | 0.755 | 0.496 | 0.11 | -0.915 | 0.002 | 0.021 | 0.021 | 0.311 | 0.53 | 0.41 | 0.64 | 0.75 | 0.61 | 0.65 |
| 7 | 86681714 | rs11768321 | 0.887 | 0.732 | 0.374 | -0.446 | 0.001 | 0.001 | 0.001 | 0 | 0.51 | 0.47 | 0.52 | 0.53 | 0.49 | 0.54 |
| 7 | 86691254 | rs13243613 | -0.672 | 0.57 | 1.427 | 0.296 | 0.002 | 0.028 | 0.028 | 0 | 0.08 | 0.12 | 0.13 | 0.35 | 0.1 | 0.17 |
| 7 | 86702671 | rs17161290 | -1.554 | 0.548 | 1.768 | 0.235 | 0.02 | -0.022 | -0.022 | 4.893 | 0.1 | 0.24 | 0.06 | 0.02 | 0.08 | 0.14 |
| 7 | 86706918 | rs6948837 | -2.495 | 0.525 | 1.863 | 0.241 | 0.046 | 0.103 | 0.103 | 0.442 | 0.18 | 0.49 | 0.2 | 0.37 | 0.18 | 0.31 |
| 7 | 86714309 | rs12704346 | -2.93 | 0.455 | 1.773 | 0.184 | 0.009 | 0.037 | 0.037 | 1.124 | 0.23 | 0.56 | 0.23 | 0.37 | 0.19 | 0.31 |
| 7 | 86715547 | rs10231949 | -2.833 | 0.406 | 1.924 | 0.325 | 0.014 | 0.067 | 0.067 | 0.985 | 0.23 | 0.5 | 0.32 | 0.44 | 0.2 | 0.18 |
| 7 | 86716102 | rs10257010 | -1.832 | 0.633 | 2.107 | 0.428 | 0.038 | 0.094 | 0.094 | 0 | 0.16 | 0.29 | 0.28 | 0.43 | 0.18 | 0.17 |
| 7 | 86719524 | rs4728679 | -1.919 | 0.635 | 2.076 | 0.429 | 0.037 | 0.093 | 0.093 | 0 | 0.16 | 0.31 | 0.28 | 0.43 | 0.18 | 0.17 |
| 7 | 86719708 | rs4728681 | -2.449 | 0.61 | 1.922 | 0.43 | 0.032 | 0.092 | 0.092 | 0.144 | 0.21 | 0.49 | 0.32 | 0.44 | 0.2 | 0.18 |
| 7 | 86719916 | rs11975476 | -3.124 | 0.606 | 1.925 | 0.424 | 0.016 | 0.054 | 0.054 | 1.595 | 0.25 | 0.57 | 0.34 | 0.46 | 0.22 | 0.21 |
| 7 | 86729268 | rs13226884 | -0.92 | 0.79 | 2.04 | 0.487 | 0.044 | -0.05 | -0.05 | 1.658 | 0.09 | 0.28 | 0.08 | 0 | 0.09 | 0.01 |
| 7 | 86734235 | rs13245768 | -1.265 | 0.779 | 1.863 | 0.512 | 0.023 | 0.154 | 0.154 | 0 | 0.1 | 0.22 | 0.23 | 0.43 | 0.1 | 0.17 |
| 7 | 86735343 | rs2158156 | -2.642 | 0.752 | 1.765 | 0.49 | 0.021 | 0.05 | 0.05 | 0.52 | 0.22 | 0.63 | 0.34 | 0.45 | 0.22 | 0.21 |
| 7 | 86741039 | rs2214351 | -1.865 | 0.501 | 1.541 | 1.055 | 0.029 | 0.187 | 0.187 | 0.313 | 0.12 | 0.29 | 0.25 | 0.51 | 0.12 | 0.22 |
| 7 | 86751525 | rs6958626 | 2.328 | 0.69 | 1.604 | 1.148 | 0.065 | 0.237 | 0.237 | 0 | 0.9 | 0.74 | 0.73 | 0.48 | 0.85 | 0.78 |
| 7 | 86755158 | rs734607 | -1.462 | 0.454 | 1.861 | 0.977 | 0.004 | -0.023 | -0.023 | 0 | 0.11 | 0.23 | 0.12 | 0.03 | 0.09 | 0.04 |
| 7 | 86759238 | rs17161323 | -1.408 | 0.439 | 1.877 | 1.05 | 0.001 | -0.013 | -0.013 | 0 | 0.11 | 0.18 | 0.12 | 0.04 | 0.09 | 0.04 |
| 7 | 86764877 | rs6465093 | 1.357 | 0.403 | 2.057 | 1.163 | 0.065 | 0.261 | 0.261 | 0 | 0.93 | 0.82 | 0.73 | 0.5 | 0.89 | 0.8 |
| 7 | 86765760 | rs11978231 | -1.336 | 0.396 | 2.094 | 1.435 | -0.001 | 0.006 | 0.006 | 0 | 0.1 | 0.17 | 0.11 | 0.08 | 0.09 | 0.05 |
| 7 | 86767689 | rs7812191 | 2.487 | 0.375 | 2.308 | 1.425 | 0.039 | 0.243 | 0.243 | 0.39 | 0.82 | 0.53 | 0.62 | 0.42 | 0.82 | 0.75 |
| 7 | 86775235 | rs6964225 | 1.936 | 0.376 | 2.358 | 1.568 | 0.136 | 0.183 | 0.183 | 0 | 0.92 | 0.74 | 0.87 | 0.77 | 0.86 | 0.86 |
| 7 | 86779262 | rs6957785 | 2.086 | 0.367 | 2.358 | 1.442 | 0.041 | 0.142 | 0.142 | 0 | 0.84 | 0.72 | 0.68 | 0.54 | 0.8 | 0.76 |
| 7 | 86781423 | rs2286267 | 2.65 | 0.349 | 2.373 | 1.434 | 0.058 | 0.219 | 0.219 | 0.358 | 0.81 | 0.52 | 0.66 | 0.54 | 0.79 | 0.76 |
| 7 | 86789426 | rs1058972 | 2.261 | 0.362 | 2.511 | 1.53 | 0.063 | 0.22 | 0.22 | 0 | 0.83 | 0.61 | 0.67 | 0.53 | 0.8 | 0.75 |
| 7 | 86793303 | rs3789251 | 1.147 | 0.366 | 2.486 | 1.532 | 0.068 | 0.213 | 0.213 | 0 | 0.93 | 0.82 | 0.79 | 0.61 | 0.89 | 0.81 |
| 7 | 86794121 | rs2074757 | 1.09 | 0.366 | 2.501 | 1.535 | 0.069 | 0.217 | 0.217 | 0 | 0.94 | 0.83 | 0.79 | 0.61 | 0.89 | 0.81 |
| 7 | 86800351 | rs3747807 | 1.179 | 0.366 | 2.496 | 1.535 | 0.068 | 0.216 | 0.216 | 0 | 0.93 | 0.82 | 0.79 | 0.61 | 0.89 | 0.81 |
| 7 | 86802008 | rs7800541 | 2.276 | 0.362 | 2.524 | 1.534 | 0.062 | 0.219 | 0.219 | 0 | 0.83 | 0.61 | 0.67 | 0.53 | 0.8 | 0.76 |
| 7 | 86823619 | rs11971131 | -0.454 | 0.364 | 2.444 | 1.5 | 0.001 | 0.002 | 0.002 | 0 | 0.08 | 0.06 | 0.09 | 0.02 | 0.06 | 0.04 |
| 7 | 86830190 | rs17766292 | 2.794 | -0.123 | 2.308 | 1.149 | 0.002 | 0.009 | 0.009 | 0.901 | 0.57 | 0.17 | 0.51 | 0.39 | 0.66 | 0.57 |
| 7 | 86847321 | rs10245036 | 2.481 | 0.215 | 2.409 | 1.066 | 0.044 | 0.203 | 0.203 | 0.085 | 0.8 | 0.52 | 0.66 | 0.53 | 0.79 | 0.75 |
| 7 | 86851097 | rs4728689 | 1.143 | 0.508 | 1.825 | 1.136 | 0.065 | 0.205 | 0.205 | 0 | 0.93 | 0.82 | 0.78 | 0.61 | 0.88 | 0.8 |
| 7 | 86861313 | rs6942407 | 2.183 | 0.142 | 1.508 | 0.851 | 0.004 | -0.036 | -0.036 | 0 | 0.74 | 0.22 | 0.68 | 0.57 | 0.8 | 0.7 |
| 7 | 86863060 | rs10273853 | 1.185 | 0.507 | 1.522 | 1.05 | 0.036 | 0.148 | 0.148 | 0 | 0.89 | 0.76 | 0.76 | 0.65 | 0.87 | 0.78 |
| 7 | 86875826 | rs2888556 | -1.853 | 0.226 | 1.349 | 0.728 | -0.004 | -0.07 | -0.07 | 0.347 | 0.27 | 0.48 | 0.17 | 0.1 | 0.21 | 0.13 |
| 7 | 86878081 | rs2214931 | 0.447 | 0.335 | 1.007 | 0.636 | -0.006 | -0.006 | -0.006 | 0 | 0.12 | 0.01 | 0.1 | 0.1 | 0.12 | 0.06 |
| 7 | 86885119 | rs1990171 | -1.838 | 0.52 | 1.468 | 0.744 | -0.001 | -0.069 | -0.069 | 1.419 | 0.26 | 0.48 | 0.17 | 0.1 | 0.21 | 0.13 |
| 7 | 86896568 | rs802101 | -0.849 | 0.561 | 1.439 | 0.716 | -0.004 | -0.004 | -0.004 | 0.822 | 0.13 | 0.1 | 0.05 | 0 | 0.12 | 0.06 |
| 7 | 86923179 | rs1089519 | 0.884 | 0.586 | 1.419 | 0.968 | 0.007 | 0.094 | 0.094 | 0 | 0.9 | 0.82 | 0.78 | 0.62 | 0.89 | 0.72 |
| 7 | 86941669 | rs1358064 | 1.343 | 0.579 | 1.492 | 0.74 | -0.001 | -0.058 | -0.058 | 0.489 | 0.78 | 0.73 | 0.85 | 0.98 | 0.82 | 0.88 |
| 7 | 86944642 | rs7778976 | 1.94 | 0.599 | 1.042 | 0.871 | -0.004 | -0.013 | -0.013 | 0 | 0.65 | 0.35 | 0.62 | 0.52 | 0.69 | 0.6 |
| 7 | 86951809 | rs802046 | -1.079 | 0.317 | 1.471 | 0.388 | -0.011 | -0.059 | -0.059 | 0 | 0.11 | 0.28 | 0.03 | 0 | 0.04 | 0 |
| 7 | 86965599 | rs996745 | 2.209 | 0.401 | 1.569 | 0.623 | -0.001 | 0.004 | 0.004 | 0 | 0.61 | 0.42 | 0.64 | 0.67 | 0.64 | 0.52 |
| 7 | 86969438 | rs17149389 | -0.256 | 0.549 | 1.797 | 0.627 | -0.004 | -0.007 | -0.007 | 0 | 0.07 | 0.1 | 0.08 | 0.04 | 0.05 | 0.06 |
| 7 | 86969559 | rs802035 | 1.434 | 0.51 | 2.076 | 0.605 | 0.04 | 0.027 | 0.027 | 0 | 0.78 | 0.64 | 0.81 | 0.71 | 0.74 | 0.64 |

| | | | | | | | | | | | | | | | | |
|---|---|---|---|---|---|---|---|---|---|---|---|---|---|---|---|---|
| 7 | 86977894 | rs802036 | -1.596 | 0.578 | 2.486 | 0.58 | -0.009 | -0.083 | -0.083 | 0 | 0.13 | 0.3 | 0.06 | 0 | 0.07 | 0.03 |
| 7 | 86978347 | rs802037 | 0.821 | 0.589 | 2.481 | 0.57 | -0.02 | -0.02 | -0.02 | 0.152 | 0.84 | 0.9 | 0.83 | 0.72 | 0.76 | 0.67 |
| 7 | 86979519 | rs6954473 | 2.668 | 0.588 | 2.367 | 0.57 | 0.004 | -0.009 | -0.009 | 0 | 0.64 | 0.49 | 0.7 | 0.68 | 0.64 | 0.59 |
| 7 | 86983715 | rs7808249 | 2.36 | 0.629 | 2.261 | 0.552 | 0.025 | -0.014 | -0.014 | 0.01 | 0.7 | 0.6 | 0.77 | 0.72 | 0.69 | 0.65 |
| 7 | 86985568 | rs802023 | 0.904 | 0.751 | 1.753 | 0.64 | -0.019 | -0.019 | -0.019 | 0 | 0.84 | 0.9 | 0.83 | 0.72 | 0.76 | 0.67 |
| 7 | 86989039 | rs802026 | -1.626 | 0.238 | 2.109 | 0.467 | -0.016 | -0.078 | -0.078 | 3.015 | 0.16 | 0.25 | 0.12 | 0.01 | 0.1 | 0.16 |
| 7 | 86995922 | rs2051950 | -1.756 | 0.701 | 2.365 | 0.503 | 0.031 | -0.05 | -0.05 | 0.348 | 0.1 | 0.3 | 0.1 | 0.01 | 0.08 | 0.17 |
| 7 | 86995999 | rs7802658 | 2.605 | 0.828 | 2.537 | 0.865 | 0.001 | 0.008 | 0.008 | 0.05 | 0.81 | 0.37 | 0.76 | 0.76 | 0.83 | 0.71 |
| 7 | 86996489 | rs802054 | 0.579 | 0.826 | 2.574 | 0.897 | -0.016 | -0.019 | -0.019 | 0 | 0.93 | 0.99 | 0.89 | 0.76 | 0.91 | 0.88 |
| 7 | 86997163 | rs701339 | 0.579 | 0.826 | 2.587 | 0.897 | -0.012 | -0.012 | -0.012 | 0.493 | 0.93 | 0.98 | 0.89 | 0.76 | 0.91 | 0.88 |
| 7 | 86997694 | rs10247098 | 2.57 | 0.82 | 2.843 | 0.888 | 0.006 | 0.019 | 0.019 | 0.029 | 0.81 | 0.41 | 0.77 | 0.75 | 0.83 | 0.71 |
| 7 | 86998335 | rs3789250 | 2.57 | 0.82 | 2.832 | 0.888 | 0.007 | 0.02 | 0.02 | 0.029 | 0.81 | 0.42 | 0.77 | 0.75 | 0.83 | 0.71 |
| 7 | 86998842 | rs6966140 | 2.791 | 0.866 | 2.758 | 0.821 | -0.001 | -0.002 | -0.002 | 0 | 0.74 | 0.36 | 0.73 | 0.75 | 0.78 | 0.71 |
| 7 | 86999865 | rs7786781 | -0.899 | 0.912 | 2.821 | 0.937 | 0.003 | -0.029 | -0.029 | 0 | 0.07 | 0.13 | 0.09 | 0.01 | 0.08 | 0.17 |
| 7 | 87000825 | rs802057 | 1.271 | 0.864 | 2.921 | 0.963 | 0.036 | 0.121 | 0.121 | 0.375 | 0.92 | 0.8 | 0.88 | 0.75 | 0.91 | 0.88 |
| 7 | 87004281 | rs802062 | 1.109 | 0.844 | 2.935 | 1.004 | 0.041 | 0.143 | 0.143 | 0.011 | 0.92 | 0.8 | 0.87 | 0.74 | 0.91 | 0.89 |
| 7 | 87007179 | rs258962 | 1.305 | 0.843 | 2.935 | 1.004 | 0.043 | 0.135 | 0.135 | 0.015 | 0.92 | 0.8 | 0.87 | 0.74 | 0.9 | 0.89 |
| 7 | 87007955 | rs258965 | 0.475 | 0.842 | 2.939 | 1.004 | -0.018 | -0.01 | -0.01 | 0 | 0.93 | 0.99 | 0.88 | 0.74 | 0.9 | 0.89 |
| 7 | 87010204 | rs31627 | 1.174 | 0.816 | 2.925 | 1.153 | 0.043 | 0.167 | 0.167 | 0.112 | 0.92 | 0.8 | 0.86 | 0.72 | 0.9 | 0.88 |
| 7 | 87010888 | rs31629 | 1.174 | 0.816 | 2.938 | 1.156 | 0.045 | 0.167 | 0.167 | 0.112 | 0.92 | 0.8 | 0.86 | 0.72 | 0.9 | 0.88 |
| 7 | 87012415 | rs31632 | 1.174 | 0.819 | 2.922 | 1.153 | 0.043 | 0.167 | 0.167 | 0.112 | 0.92 | 0.8 | 0.86 | 0.72 | 0.9 | 0.88 |
| 7 | 87022596 | rs2072207 | 2.665 | 0.835 | 2.935 | 1.164 | 0.009 | 0.048 | 0.048 | 0.336 | 0.82 | 0.41 | 0.76 | 0.71 | 0.82 | 0.72 |
| 7 | 87022981 | rs31645 | 0.596 | 0.86 | 2.781 | 1.206 | -0.016 | -0.001 | -0.001 | 0 | 0.93 | 0.99 | 0.87 | 0.72 | 0.9 | 0.88 |
| 7 | 87029287 | rs31651 | 1.116 | 0.867 | 2.82 | 1.212 | 0.038 | 0.149 | 0.149 | 0.135 | 0.92 | 0.81 | 0.87 | 0.73 | 0.9 | 0.88 |
| 7 | 87029687 | rs31652 | 1.249 | 0.984 | 3.015 | 1.262 | 0.06 | 0.192 | 0.192 | 0.141 | 0.91 | 0.77 | 0.85 | 0.72 | 0.87 | 0.85 |
| 7 | 87030903 | rs2097937 | 2.645 | 1.002 | 3.104 | 1.206 | 0.027 | 0.07 | 0.07 | 0.433 | 0.84 | 0.42 | 0.75 | 0.71 | 0.81 | 0.72 |
| 7 | 87031809 | rs1526090 | 2.607 | 1.002 | 3.104 | 1.206 | 0.029 | 0.07 | 0.07 | 0.368 | 0.84 | 0.42 | 0.75 | 0.71 | 0.81 | 0.72 |
| 7 | 87032613 | rs31653 | 0.568 | 1.001 | 3.104 | 1.206 | -0.036 | -0.029 | -0.029 | 0 | 0.93 | 0.99 | 0.87 | 0.73 | 0.89 | 0.88 |
| 7 | 87033948 | rs12154399 | 2.719 | 0.992 | 3.102 | 1.2 | 0.026 | 0.069 | 0.069 | 0.712 | 0.83 | 0.42 | 0.75 | 0.71 | 0.81 | 0.71 |
| 7 | 87036942 | rs2373593 | 1.651 | 0.949 | 3.163 | 1.18 | 0.054 | 0.203 | 0.203 | 0.12 | 0.91 | 0.72 | 0.86 | 0.72 | 0.89 | 0.88 |
| 7 | 87038074 | rs17149539 | -1.584 | 0.949 | 3.16 | 1.18 | 0.084 | -0.005 | -0.005 | 0 | 0.07 | 0.3 | 0.11 | 0.01 | 0.08 | 0.15 |
| 7 | 87040629 | rs31659 | 0.674 | 0.941 | 3.199 | 1.168 | 0.018 | 0.094 | 0.094 | 0.199 | 0.93 | 0.88 | 0.87 | 0.72 | 0.89 | 0.88 |
| 7 | 87041856 | rs11761050 | -1.603 | 0.94 | 3.175 | 1.158 | 0.082 | -0.007 | -0.007 | 0 | 0.07 | 0.3 | 0.11 | 0.01 | 0.08 | 0.16 |
| 7 | 87043583 | rs7807638 | -1.639 | 0.942 | 3.168 | 1.159 | 0.082 | -0.007 | -0.007 | 0.359 | 0.07 | 0.31 | 0.11 | 0.01 | 0.08 | 0.16 |
| 7 | 87043708 | rs4148830 | 2.682 | 0.942 | 3.064 | 1.169 | 0.028 | 0.069 | 0.069 | 0.41 | 0.84 | 0.42 | 0.75 | 0.71 | 0.81 | 0.72 |
| 7 | 87044443 | rs31660 | 0.554 | 0.946 | 3.023 | 1.173 | -0.036 | -0.028 | -0.028 | 0.108 | 0.93 | 0.99 | 0.87 | 0.73 | 0.89 | 0.88 |
| 7 | 87045584 | rs31662 | 1.171 | 0.948 | 3.039 | 1.174 | 0.077 | 0.241 | 0.241 | 0.017 | 0.92 | 0.71 | 0.86 | 0.73 | 0.89 | 0.88 |
| 7 | 87046470 | rs4148829 | -1.591 | 0.678 | 2.697 | 1.014 | 0.046 | -0.026 | -0.026 | 0 | 0.07 | 0.21 | 0.1 | 0.01 | 0.08 | 0.16 |
| 7 | 87047813 | rs7788404 | -1.588 | 0.679 | 2.683 | 1.014 | 0.042 | -0.028 | -0.028 | 0 | 0.07 | 0.2 | 0.1 | 0.01 | 0.08 | 0.16 |
| 7 | 87048530 | rs4148828 | -0.014 | 0.643 | 1.848 | 1.222 | -0.025 | -0.04 | -0.04 | 0 | 0.12 | 0.01 | 0.1 | 0.19 | 0.13 | 0.05 |
| 7 | 87052268 | rs31666 | 1.851 | 0.403 | 2.451 | 1.446 | 0.013 | 0.126 | 0.126 | 0.239 | 0.87 | 0.66 | 0.81 | 0.69 | 0.85 | 0.73 |
| 7 | 87053150 | rs31667 | 2.21 | 0.282 | 2.726 | 1.422 | 0.01 | 0.158 | 0.158 | 0.389 | 0.85 | 0.5 | 0.79 | 0.69 | 0.84 | 0.73 |
| 7 | 87053206 | rs31668 | 0.572 | 0.285 | 2.609 | 1.425 | 0.004 | 0.019 | 0.019 | 0.464 | 0.93 | 0.94 | 0.9 | 0.71 | 0.93 | 0.88 |
| 7 | 87054247 | rs31669 | 0.797 | 0.28 | 2.623 | 1.418 | 0.004 | 0.015 | 0.015 | 0.099 | 0.92 | 0.93 | 0.9 | 0.72 | 0.92 | 0.88 |
| 7 | 87056011 | rs1017054 | -2.429 | 0.29 | 2.738 | 1.316 | 0.087 | 0.014 | 0.014 | 1.806 | 0.08 | 0.39 | 0.11 | 0.03 | 0.08 | 0.14 |
| 7 | 87059164 | rs31670 | 2.833 | 0.202 | 2.603 | 1.249 | -0.004 | 0.005 | 0.005 | 0 | 0.76 | 0.36 | 0.76 | 0.69 | 0.79 | 0.72 |
| 7 | 87068464 | rs31674 | 2.848 | 0.488 | 2.464 | 1.159 | -0.004 | 0.006 | 0.006 | 0 | 0.76 | 0.37 | 0.76 | 0.69 | 0.79 | 0.72 |
| 7 | 87069758 | rs31675 | 0.38 | 0.484 | 2.48 | 1.156 | 0.006 | 0.007 | 0.007 | 0.463 | 0.93 | 0.95 | 0.91 | 0.72 | 0.93 | 0.87 |
| 7 | 87069880 | rs31676 | 2.808 | 0.493 | 2.46 | 1.162 | -0.004 | 0.006 | 0.006 | 0 | 0.76 | 0.37 | 0.76 | 0.69 | 0.79 | 0.72 |
| 7 | 87072505 | rs17149606 | -1.653 | 0.547 | 2.374 | 1.126 | 0.08 | 0.017 | 0.017 | 0.335 | 0.07 | 0.24 | 0.1 | 0.03 | 0.08 | 0.14 |
| 7 | 87073594 | rs17149608 | -1.569 | 0.443 | 2.129 | 1.299 | 0 | 0.001 | 0.001 | 0 | 0.15 | 0.14 | 0.13 | 0.23 | 0.13 | 0.12 |
| 7 | 87074419 | rs4148826 | -2.723 | 0.302 | 2.37 | 1.337 | -0.026 | 0.005 | 0.005 | 0 | 0.23 | 0.53 | 0.23 | 0.27 | 0.19 | 0.25 |
| 7 | 87074964 | rs1468615 | -2.237 | 0.306 | 2.357 | 1.339 | 0.003 | -0.004 | -0.004 | 3.26 | 0.2 | 0.2 | 0.2 | 0.27 | 0.19 | 0.25 |
| 7 | 87075362 | rs4148824 | -2.787 | 0.293 | 2.498 | 1.333 | -0.02 | 0.001 | 0.001 | 0 | 0.24 | 0.59 | 0.23 | 0.28 | 0.19 | 0.25 |
| 7 | 87076587 | rs1014283 | -2.747 | 0.296 | 2.524 | 1.335 | -0.026 | 0.005 | 0.005 | 0 | 0.23 | 0.53 | 0.23 | 0.27 | 0.19 | 0.25 |
| 7 | 87077996 | rs4148822 | -0.321 | 0.296 | 2.451 | 1.339 | -0.001 | -0.036 | -0.036 | 0.459 | 0.07 | 0.01 | 0.09 | 0.2 | 0.07 | 0.14 |
| 7 | 87079069 | rs1558375 | -2.509 | 0.308 | 2.521 | 1.353 | -0.03 | 0.007 | 0.007 | 0 | 0.23 | 0.51 | 0.22 | 0.27 | 0.19 | 0.25 |
| 7 | 87079103 | rs1558376 | -2.509 | 0.308 | 2.521 | 1.353 | -0.03 | 0.007 | 0.007 | 0 | 0.23 | 0.51 | 0.22 | 0.27 | 0.19 | 0.25 |
| 7 | 87080350 | rs1473152 | -2.515 | 0.364 | 2.51 | 1.341 | -0.028 | 0.006 | 0.006 | 0 | 0.24 | 0.51 | 0.24 | 0.27 | 0.19 | 0.25 |
| 7 | 87081349 | rs2072383 | -1.89 | 0.361 | 2.635 | 1.337 | -0.027 | -0.006 | -0.006 | 0 | 0.21 | 0.32 | 0.17 | 0.2 | 0.15 | 0.14 |
| 7 | 87086931 | rs17149647 | -2.233 | 0.351 | 2.656 | 1.364 | -0.046 | -0.007 | -0.007 | 0 | 0.22 | 0.41 | 0.17 | 0.2 | 0.15 | 0.14 |
| 7 | 87087731 | rs4148816 | -2.536 | 0.355 | 2.669 | 1.367 | -0.031 | 0.009 | 0.009 | 0 | 0.24 | 0.51 | 0.24 | 0.27 | 0.19 | 0.25 |
| 7 | 87087834 | rs4148815 | -2.3 | 0.355 | 2.669 | 1.367 | -0.049 | -0.009 | -0.009 | 0 | 0.22 | 0.49 | 0.17 | 0.2 | 0.15 | 0.14 |
| 7 | 87091543 | rs4148814 | -1.889 | 0.353 | 2.672 | 1.38 | -0.029 | -0.002 | -0.002 | 0 | 0.21 | 0.32 | 0.17 | 0.2 | 0.15 | 0.14 |
| 7 | 87091826 | rs2302385 | -1.889 | 0.353 | 2.672 | 1.375 | -0.029 | -0.004 | -0.004 | 0 | 0.21 | 0.32 | 0.17 | 0.2 | 0.15 | 0.14 |
| 7 | 87092185 | rs2302387 | -2.268 | 0.354 | 2.665 | 1.375 | -0.049 | -0.009 | -0.009 | 0 | 0.22 | 0.49 | 0.17 | 0.2 | 0.15 | 0.14 |
| 7 | 87101486 | rs4148811 | -2.489 | 0.354 | 2.647 | 1.375 | -0.031 | 0.009 | 0.009 | 0 | 0.24 | 0.51 | 0.24 | 0.27 | 0.19 | 0.25 |
| 7 | 87103263 | rs4148809 | -1.858 | -0.729 | 1.213 | 1.24 | -0.11 | -0.026 | -0.026 | 0 | 0.61 | 0.98 | 0.57 | 0.62 | 0.45 | 0.56 |
| 7 | 87103670 | rs2888611 | -1.77 | -0.606 | 1.365 | 1.262 | -0.03 | 0.01 | 0.01 | 0 | 0.23 | 0.5 | 0.24 | 0.28 | 0.19 | 0.25 |

| | | | | | | | | | | | | | | | | |
|---|---|---|---|---|---|---|---|---|---|---|---|---|---|---|---|---|
| 7 | 87105795 | rs4148808 | -1.023 | -0.602 | 1.303 | 1.269 | -0.025 | 0.04 | 0.04 | 0 | 0.2 | 0.34 | 0.16 | 0.28 | 0.14 | 0.13 |
| 7 | 87106365 | rs4148805 | -1.032 | -0.6 | 1.3 | 1.331 | -0.025 | 0.038 | 0.038 | 0 | 0.2 | 0.33 | 0.16 | 0.28 | 0.14 | 0.13 |
| 7 | 87110017 | rs12539936 | -1.744 | -0.595 | 1.073 | 1.338 | -0.023 | 0.026 | 0.026 | 0 | 0.23 | 0.57 | 0.25 | 0.29 | 0.19 | 0.24 |
| 7 | 87122022 | rs2178658 | -0.105 | -0.702 | 0.706 | 1.403 | 0.029 | -0.033 | -0.033 | 0 | 0.33 | 0.15 | 0.3 | 0.33 | 0.26 | 0.21 |
| 7 | 87124822 | rs17209837 | -1.081 | -0.375 | 0.496 | 1.07 | -0.017 | 0.023 | 0.023 | 0 | 0.19 | 0.28 | 0.16 | 0.28 | 0.14 | 0.14 |
| 7 | 87128865 | rs6946119 | -0.049 | -0.78 | -0.086 | 0.949 | -0.041 | 0.064 | 0.064 | 0 | 0.25 | 0.02 | 0.25 | 0.12 | 0.24 | 0.17 |
| 7 | 87132916 | rs1055302 | -1.279 | -0.899 | 0.554 | 1.186 | -0.024 | 0.022 | 0.022 | 0 | 0.21 | 0.31 | 0.16 | 0.29 | 0.14 | 0.15 |
| 7 | 87137018 | rs1882478 | -1.105 | -0.924 | 0.159 | 0.711 | -0.014 | 0.008 | 0.008 | 0 | 0.34 | 0.62 | 0.41 | 0.56 | 0.26 | 0.24 |
| 7 | 87138532 | rs2235047 | -0.958 | -0.693 | 0.382 | 0.98 | -0.007 | 0.378 | 0.378 | 0 | 0.05 | 0.19 | 0.1 | 0.42 | 0.02 | 0.11 |
| 7 | 87138645 | rs1045642 | -2.186 | -0.552 | -0.396 | 0.292 | -0.032 | 0.004 | 0.004 | 1.276 | 0.58 | 0.85 | 0.57 | 0.6 | 0.48 | 0.43 |
| 7 | 87140808 | rs10808071 | -0.603 | -0.5 | 0.087 | -0.102 | 0.005 | -0.052 | -0.052 | 0 | 0.22 | 0.25 | 0.17 | 0.09 | 0.21 | 0.19 |
| 7 | 87141497 | rs1002204 | -2.013 | -0.482 | -0.144 | -0.144 | -0.038 | 0.005 | 0.005 | 0.993 | 0.57 | 0.84 | 0.58 | 0.6 | 0.49 | 0.41 |
| 7 | 87141751 | rs17149699 | -1.363 | -0.751 | 0.474 | 0.011 | -0.014 | 0.324 | 0.324 | 0.046 | 0.08 | 0.32 | 0.12 | 0.37 | 0.05 | 0.11 |
| 7 | 87141814 | rs6949448 | -2.259 | 0.195 | 0.199 | -0.176 | -0.03 | -0.005 | -0.005 | 0.203 | 0.59 | 0.84 | 0.6 | 0.61 | 0.58 | 0.44 |
| 7 | 87143153 | rs4148751 | -0.751 | 0.519 | 0.512 | -0.163 | -0.012 | 0.291 | 0.291 | 0 | 0.07 | 0.22 | 0.11 | 0.37 | 0.05 | 0.12 |
| 7 | 87143504 | rs1922243 | -1.883 | 0.482 | 0.855 | -0.18 | -0.005 | 0.457 | 0.457 | 0.918 | 0.1 | 0.47 | 0.14 | 0.37 | 0.05 | 0.07 |
| 7 | 87147147 | rs4148747 | 0.019 | 0.621 | 0.805 | -0.302 | 0.006 | -0.043 | -0.043 | 0 | 0.13 | 0.22 | 0.11 | 0.07 | 0.14 | 0.16 |
| 7 | 87149922 | rs2235067 | 0.316 | 0.554 | 0.887 | -0.066 | 0.008 | -0.034 | -0.034 | 0.103 | 0.12 | 0.18 | 0.11 | 0.07 | 0.13 | 0.15 |
| 7 | 87153160 | rs2373588 | -0.504 | 0.535 | 0.922 | 0.026 | -0.015 | 0.292 | 0.292 | 0 | 0.07 | 0.21 | 0.11 | 0.37 | 0.04 | 0.06 |
| 7 | 87153585 | rs10280101 | 0.071 | 0.545 | 0.943 | -0.023 | 0.006 | -0.039 | -0.039 | 0 | 0.13 | 0.2 | 0.11 | 0.07 | 0.13 | 0.18 |
| 7 | 87154646 | rs10225473 | 0.151 | 0.54 | 0.935 | -0.025 | 0.005 | -0.034 | -0.034 | 0 | 0.13 | 0.18 | 0.11 | 0.07 | 0.13 | 0.18 |
| 7 | 87160561 | rs2032583 | 0.021 | 0.532 | 0.928 | -0.046 | 0.006 | -0.04 | -0.04 | 0 | 0.13 | 0.2 | 0.11 | 0.07 | 0.13 | 0.18 |
| 7 | 87161049 | rs4148739 | 0.021 | 0.539 | 0.917 | -0.046 | 0.006 | -0.04 | -0.04 | 3.075 | 0.13 | 0.2 | 0.11 | 0.07 | 0.13 | 0.18 |
| 7 | 87161520 | rs11983225 | 0.021 | 0.539 | 0.917 | -0.046 | 0.006 | -0.04 | -0.04 | 0 | 0.13 | 0.2 | 0.11 | 0.07 | 0.13 | 0.18 |
| 7 | 87163016 | rs11760837 | 0.021 | 0.539 | 0.917 | -0.046 | 0.006 | -0.04 | -0.04 | 0 | 0.13 | 0.2 | 0.11 | 0.07 | 0.13 | 0.18 |
| 7 | 87164483 | rs10274587 | -0.07 | 0.525 | 0.909 | -0.052 | 0.003 | -0.035 | -0.035 | 0.713 | 0.13 | 0.18 | 0.11 | 0.07 | 0.13 | 0.18 |
| 7 | 87164986 | rs10248420 | -1.666 | 0.517 | 1.043 | -0.015 | -0.021 | 0.117 | 0.117 | 0.607 | 0.22 | 0.54 | 0.24 | 0.43 | 0.18 | 0.24 |
| 7 | 87165750 | rs2235040 | -0.002 | 0.563 | 1.021 | -0.424 | 0.003 | -0.035 | -0.035 | 0 | 0.13 | 0.18 | 0.11 | 0.07 | 0.13 | 0.18 |
| 7 | 87169669 | rs10268314 | -0.002 | 1.039 | 0.727 | -0.537 | 0.005 | -0.039 | -0.039 | 0 | 0.13 | 0.2 | 0.11 | 0.07 | 0.13 | 0.18 |
| 7 | 87171152 | rs4148737 | -1.095 | 1.356 | 0.975 | -0.525 | 0 | -0.005 | -0.005 | 0 | 0.41 | 0.47 | 0.46 | 0.3 | 0.44 | 0.22 |
| 7 | 87171383 | rs4148736 | -1.095 | 1.356 | 0.975 | -0.563 | 0 | -0.005 | -0.005 | 0 | 0.41 | 0.47 | 0.46 | 0.3 | 0.44 | 0.22 |
| 7 | 87172136 | rs6961419 | -1.095 | 1.065 | 1.228 | -0.378 | 0 | -0.005 | -0.005 | 0 | 0.41 | 0.47 | 0.46 | 0.3 | 0.44 | 0.22 |
| 7 | 87172881 | rs4148735 | -1.095 | 1.065 | 1.228 | -0.378 | 0 | -0.005 | -0.005 | 0 | 0.41 | 0.47 | 0.46 | 0.3 | 0.44 | 0.22 |
| 7 | 87174066 | rs2235046 | -1.973 | 1.056 | 1.256 | -0.382 | -0.008 | 0.066 | 0.066 | 0.016 | 0.55 | 0.8 | 0.59 | 0.37 | 0.57 | 0.39 |
| 7 | 87174504 | rs2091766 | -1.116 | 1.117 | 1.244 | -0.415 | 0.001 | -0.011 | -0.011 | 0 | 0.38 | 0.47 | 0.45 | 0.3 | 0.41 | 0.21 |
| 7 | 87178626 | rs2235013 | -1.593 | 1.195 | 1.186 | -0.462 | -0.001 | -0.012 | -0.012 | 0 | 0.45 | 0.52 | 0.55 | 0.38 | 0.49 | 0.4 |
| 7 | 87179086 | rs2235035 | -0.567 | 1.206 | 1.192 | -0.467 | 0.004 | 0.009 | 0.009 | 0 | 0.29 | 0.21 | 0.41 | 0.3 | 0.32 | 0.2 |
| 7 | 87179143 | rs2235033 | -1.619 | 1.209 | 1.192 | -0.465 | -0.001 | -0.012 | -0.012 | 0 | 0.45 | 0.52 | 0.55 | 0.38 | 0.49 | 0.4 |
| 7 | 87179601 | rs1128503 | -2.395 | 1.278 | 1.142 | -0.562 | -0.013 | 0.072 | 0.072 | 0.247 | 0.56 | 0.86 | 0.6 | 0.37 | 0.58 | 0.41 |
| 7 | 87180198 | rs10276036 | -2.369 | 1.293 | 1.065 | -0.562 | 0 | 0.074 | 0.074 | 0.025 | 0.55 | 0.81 | 0.59 | 0.37 | 0.58 | 0.41 |
| 7 | 87183354 | rs1922240 | -0.452 | 1.245 | 0.926 | -0.508 | 0.001 | 0.008 | 0.008 | 0 | 0.28 | 0.2 | 0.41 | 0.29 | 0.32 | 0.2 |
| 7 | 87185894 | rs1922241 | -0.575 | 1.314 | 0.941 | -0.496 | 0.003 | 0.012 | 0.012 | 0 | 0.29 | 0.2 | 0.41 | 0.29 | 0.32 | 0.2 |
| 7 | 87186018 | rs11772987 | 0.414 | 1.223 | 0.88 | -0.44 | 0.008 | -0.032 | -0.032 | 0 | 0.12 | 0.18 | 0.11 | 0.07 | 0.13 | 0.18 |
| 7 | 87189930 | rs868755 | -2.606 | 1.165 | 0.493 | -0.357 | -0.108 | -0.138 | -0.138 | 0 | 0.6 | 0.99 | 0.64 | 0.58 | 0.61 | 0.42 |
| 7 | 87190452 | rs2235023 | -0.301 | 1.292 | 0.926 | -0.643 | 0.052 | -0.053 | -0.053 | 0 | 0.1 | 0.3 | 0.04 | 0 | 0.09 | 0.01 |
| 7 | 87191669 | rs13237132 | -0.579 | 1.226 | 0.996 | -0.653 | 0.004 | 0.01 | 0.01 | 0 | 0.29 | 0.27 | 0.42 | 0.29 | 0.32 | 0.19 |
| 7 | 87192731 | rs11975994 | -2.422 | 1.234 | 0.893 | -0.663 | -0.014 | 0.065 | 0.065 | 0.152 | 0.56 | 0.86 | 0.6 | 0.37 | 0.58 | 0.4 |
| 7 | 87193597 | rs4148734 | -0.088 | 1.232 | 0.84 | -0.634 | -0.056 | 0.079 | 0.079 | 0 | 0.26 | 0.03 | 0.33 | 0.16 | 0.3 | 0.15 |
| 7 | 87195106 | rs1202170 | 1.45 | 1.235 | 0.995 | -0.592 | -0.001 | -0.006 | -0.006 | 0 | 0.54 | 0.5 | 0.45 | 0.63 | 0.51 | 0.61 |
| 7 | 87195850 | rs1202169 | 2.011 | 1.156 | 0.544 | -0.541 | -0.002 | 0.068 | 0.068 | 0 | 0.45 | 0.19 | 0.41 | 0.63 | 0.41 | 0.6 |
| 7 | 87195962 | rs1202168 | 2.011 | 1.156 | 0.544 | -0.541 | -0.002 | 0.068 | 0.068 | 0 | 0.45 | 0.19 | 0.41 | 0.63 | 0.41 | 0.6 |
| 7 | 87197059 | rs1202167 | 2.023 | 1.16 | 0.545 | -0.526 | -0.002 | 0.059 | 0.059 | 0 | 0.45 | 0.19 | 0.4 | 0.63 | 0.41 | 0.6 |
| 7 | 87199564 | rs2235015 | -0.662 | 1.245 | 0.879 | -0.686 | 0.034 | -0.108 | -0.108 | 1.81 | 0.2 | 0.39 | 0.15 | 0.07 | 0.21 | 0.18 |
| 7 | 87200842 | rs10259849 | 0.069 | 1.205 | 0.737 | -0.775 | -0.014 | 0.081 | 0.081 | 0 | 0.73 | 0.91 | 0.65 | 0.84 | 0.7 | 0.85 |
| 7 | 87201086 | rs2520464 | 2.145 | 1.226 | 0.797 | -0.759 | -0.001 | 0.07 | 0.07 | 0.124 | 0.44 | 0.18 | 0.4 | 0.63 | 0.41 | 0.59 |
| 7 | 87201482 | rs10260862 | -0.758 | 1.141 | 0.926 | -0.855 | 0.031 | -0.109 | -0.109 | 0 | 0.21 | 0.39 | 0.15 | 0.07 | 0.21 | 0.18 |
| 7 | 87202615 | rs10264990 | 0.717 | 1.309 | 0.926 | -0.788 | 0.001 | 0.059 | 0.059 | 0 | 0.68 | 0.83 | 0.63 | 0.87 | 0.66 | 0.82 |
| 7 | 87205663 | rs1989830 | 1.135 | 1.348 | 1.102 | -0.835 | 0.044 | -0.098 | -0.098 | 1.62 | 0.76 | 0.68 | 0.8 | 0.97 | 0.64 | 0.82 |
| 7 | 87209150 | rs1202175 | 1.169 | 1.322 | 1.101 | -0.836 | 0.044 | -0.097 | -0.097 | 0.107 | 0.75 | 0.68 | 0.8 | 0.97 | 0.63 | 0.82 |
| 7 | 87213232 | rs4148733 | 0.331 | 1.193 | 1.111 | -0.854 | -0.01 | -0.018 | -0.018 | 0 | 0.17 | 0.2 | 0.28 | 0.13 | 0.14 | 0.12 |
| 7 | 87213258 | rs1202186 | 1.1 | 1.227 | 0.967 | -0.854 | 0.046 | -0.098 | -0.098 | 0.885 | 0.76 | 0.67 | 0.8 | 0.97 | 0.64 | 0.82 |
| 7 | 87213384 | rs1202185 | 1.1 | 1.227 | 0.967 | -0.854 | 0.046 | -0.098 | -0.098 | 0 | 0.76 | 0.68 | 0.8 | 0.97 | 0.63 | 0.82 |
| 7 | 87213901 | rs1202184 | 2.274 | 1.16 | 0.929 | -0.848 | -0.065 | 0.169 | 0.169 | 0.504 | 0.51 | 0.06 | 0.43 | 0.67 | 0.48 | 0.66 |
| 7 | 87215304 | rs1202182 | 1.178 | 1.369 | 1.041 | -0.999 | 0.045 | -0.098 | -0.098 | 1.319 | 0.76 | 0.68 | 0.8 | 0.97 | 0.63 | 0.82 |
| 7 | 87216150 | rs1202181 | 1.191 | 1.368 | 1.04 | -0.999 | 0.044 | -0.097 | -0.097 | 0 | 0.75 | 0.68 | 0.8 | 0.97 | 0.63 | 0.82 |
| 7 | 87216817 | rs17327624 | -0.171 | 1.257 | 0.67 | -1.087 | 0.007 | 0.01 | 0.01 | 0 | 0.21 | 0.16 | 0.3 | 0.11 | 0.19 | 0.14 |
| 7 | 87220334 | rs12535512 | 2.454 | 0.87 | 0.347 | -0.755 | -0.096 | -0.115 | -0.115 | 0 | 0.43 | 0.01 | 0.35 | 0.48 | 0.41 | 0.62 |
| 7 | 87220562 | rs2188526 | 2.18 | 0.86 | 0.397 | -0.762 | -0.046 | -0.035 | -0.035 | 0.259 | 0.48 | 0.08 | 0.38 | 0.48 | 0.45 | 0.65 |
| 7 | 87220886 | rs3789243 | 1.865 | 0.823 | 0.763 | -0.753 | -0.003 | 0.011 | 0.011 | 0 | 0.55 | 0.46 | 0.48 | 0.61 | 0.47 | 0.68 |

| | | | | | | | | | | | | | | | | |
|---|---|---|---|---|---|---|---|---|---|---|---|---|---|---|---|---|
| 7 | 87221216 | rs1858923 | 2.178 | 0.835 | 0.72 | -0.748 | -0.046 | -0.035 | -0.035 | 0 | 0.48 | 0.08 | 0.38 | 0.48 | 0.45 | 0.65 |
| 7 | 87224429 | rs2214104 | -0.297 | 0.864 | 1.111 | -0.501 | -0.011 | 0.003 | 0.003 | 0.042 | 0.07 | 0.13 | 0.05 | 0.05 | 0.04 | 0.04 |
| 7 | 87224772 | rs2188525 | 0.806 | 0.865 | 1.104 | -0.502 | 0.004 | 0.018 | 0.018 | 0.013 | 0.05 | 0.02 | 0.03 | 0 | 0.04 | 0.03 |
| 7 | 87225046 | rs2235074 | -0.339 | 0.891 | 1.19 | -0.49 | -0.012 | 0.001 | 0.001 | 0.038 | 0.07 | 0.12 | 0.05 | 0.05 | 0.04 | 0.03 |
| 7 | 87230193 | rs3213619 | 0.068 | 1.153 | 1.245 | -0.486 | -0.001 | 0 | 0 | 0 | 0.06 | 0.09 | 0.04 | 0.05 | 0.04 | 0.04 |
| 7 | 87233602 | rs4728709 | -1.925 | 1.097 | 1.552 | -0.522 | -0.028 | 0.035 | 0.035 | 0 | 0.12 | 0.41 | 0.14 | 0.13 | 0.06 | 0.06 |
| 7 | 87233989 | rs17149810 | -1.925 | 1.097 | 1.552 | -0.522 | -0.028 | 0.035 | 0.035 | 0 | 0.12 | 0.41 | 0.14 | 0.13 | 0.06 | 0.06 |
| 7 | 87234049 | rs4148732 | 0.542 | 1.037 | 1.365 | -0.595 | 0.011 | 0.025 | 0.025 | 0 | 0.14 | 0.03 | 0.26 | 0.14 | 0.12 | 0.12 |
| 7 | 87237759 | rs28381787 | 0.448 | -0.152 | 1.453 | -0.556 | 0.006 | 0.021 | 0.021 | 0.23 | 0.06 | 0.02 | 0.03 | 0 | 0.04 | 0.03 |
| 7 | 87244960 | rs13233308 | 2.677 | -0.069 | 1.474 | -0.6 | -0.121 | -0.098 | -0.098 | 0 | 0.48 | 0.02 | 0.37 | 0.48 | 0.47 | 0.66 |
| 7 | 87246983 | rs28381779 | 0.859 | -0.214 | 1.51 | -0.571 | 0.004 | 0.025 | 0.025 | 0 | 0.13 | 0.03 | 0.26 | 0.14 | 0.11 | 0.11 |
| 7 | 87261736 | rs10276499 | -1.769 | -0.365 | 1.754 | -0.618 | -0.027 | 0.041 | 0.041 | 0.018 | 0.12 | 0.39 | 0.14 | 0.13 | 0.06 | 0.06 |
| 7 | 87262581 | rs10264856 | -1.769 | -0.365 | 1.754 | -0.615 | -0.027 | 0.029 | 0.029 | 0.018 | 0.12 | 0.39 | 0.14 | 0.12 | 0.06 | 0.06 |
| 7 | 87269497 | rs2214101 | -0.733 | -0.361 | 1.747 | -0.624 | -0.011 | -0.003 | -0.003 | 4.611 | 0.08 | 0.19 | 0.11 | 0.08 | 0.06 | 0.06 |
| 7 | 87275641 | rs10246878 | 0.29 | -0.326 | 1.553 | -0.815 | -0.081 | 0.092 | 0.092 | 0.012 | 0.85 | 0.99 | 0.86 | 0.98 | 0.75 | 0.87 |
| 7 | 87278760 | rs10267099 | 0.243 | -0.309 | 1.642 | -0.807 | 0.014 | -0.053 | -0.053 | 0 | 0.84 | 0.84 | 0.85 | 0.98 | 0.75 | 0.87 |
| 7 | 87279856 | rs12539395 | -1.873 | -0.2 | 1.218 | -0.711 | -0.027 | 0.041 | 0.041 | 0.018 | 0.12 | 0.39 | 0.14 | 0.13 | 0.06 | 0.06 |
| 7 | 87288324 | rs6951067 | -1.873 | -0.301 | 1.127 | -0.633 | -0.027 | 0.041 | 0.041 | 0.018 | 0.12 | 0.39 | 0.14 | 0.14 | 0.06 | 0.06 |
| 7 | 87330423 | rs6465118 | -0.772 | 0.33 | 1.089 | -0.248 | -0.011 | -0.003 | -0.003 | 0 | 0.08 | 0.17 | 0.11 | 0.08 | 0.06 | 0.06 |
| 7 | 87370183 | rs17251003 | 0.632 | 0.271 | 0.933 | -0.271 | 0.017 | 0.031 | 0.031 | 0 | 0.14 | 0.03 | 0.25 | 0.14 | 0.1 | 0.12 |
| 7 | 87397345 | rs992422 | -1.561 | 0.3 | 0.977 | -0.251 | -0.023 | 0.038 | 0.038 | 0 | 0.11 | 0.36 | 0.13 | 0.14 | 0.06 | 0.06 |
| 7 | 87418861 | rs1029421 | -2.608 | 0.206 | 0.866 | -0.255 | -0.124 | -0.142 | -0.142 | 0 | 0.55 | 0.99 | 0.62 | 0.53 | 0.54 | 0.35 |
| 7 | 87430669 | rs12704370 | -2.605 | 0.35 | 0.952 | -0.356 | -0.088 | -0.081 | -0.081 | 0 | 0.54 | 0.94 | 0.62 | 0.52 | 0.53 | 0.35 |
| 7 | 87466971 | rs2285647 | -1.074 | 0.013 | 0.841 | 0.059 | 0.023 | 0.001 | 0.001 | 0 | 0.24 | 0.28 | 0.19 | 0.25 | 0.32 | 0.16 |
| 7 | 87493807 | rs10278291 | -1.986 | -0.116 | 0.401 | -0.218 | -0.026 | 0.038 | 0.038 | 0 | 0.1 | 0.35 | 0.13 | 0.14 | 0.06 | 0.06 |
| 7 | 87511396 | rs10246786 | -2.44 | -0.991 | 0.865 | 0.264 | -0.03 | 0.043 | 0.043 | 0 | 0.11 | 0.35 | 0.13 | 0.14 | 0.06 | 0.06 |
| 7 | 87514832 | rs12669619 | -1.835 | -0.987 | 0.846 | 0.266 | -0.019 | 0.034 | 0.034 | 0 | 0.09 | 0.22 | 0.12 | 0.14 | 0.06 | 0.06 |
| 7 | 87538027 | rs6977687 | -1.586 | -0.873 | 0.726 | 0.267 | -0.068 | -0.011 | -0.011 | 0.08 | 0.25 | 0.54 | 0.4 | 0.3 | 0.16 | 0.17 |
| 7 | 87550234 | rs2158929 | -0.768 | -0.718 | 0.73 | 0.087 | -0.004 | -0.007 | -0.007 | 0.058 | 0.18 | 0.2 | 0.3 | 0.16 | 0.14 | 0.15 |
| 7 | 87564277 | rs10239100 | -0.976 | -0.639 | 0.318 | 0.086 | -0.068 | -0.039 | -0.039 | 0 | 0.22 | 0.54 | 0.33 | 0.21 | 0.14 | 0.16 |
| 7 | 87573647 | rs12019361 | -0.904 | 0.256 | 0.077 | 0.171 | 0.015 | -0.024 | -0.024 | 0 | 0.07 | 0.12 | 0.03 | 0.01 | 0.07 | 0.01 |
| 7 | 87594188 | rs1637500 | -0.2 | -0.355 | -0.141 | -0.906 | 0.011 | 0.075 | 0.075 | 0 | 0.51 | 0.41 | 0.37 | 0.08 | 0.53 | 0.37 |
| 7 | 87605618 | rs1688888 | -0.486 | -0.488 | 0.403 | 0.15 | 0 | 0.001 | 0.001 | 0 | 0.55 | 0.49 | 0.47 | 0.36 | 0.56 | 0.44 |
| 7 | 87606246 | rs1688890 | -0.54 | -0.629 | 0.999 | 0.23 | -0.001 | 0.007 | 0.007 | 0 | 0.57 | 0.71 | 0.49 | 0.41 | 0.56 | 0.44 |
| 7 | 87623250 | rs1688899 | -0.641 | -0.706 | 1.092 | 0.224 | -0.003 | 0.006 | 0.006 | 0 | 0.58 | 0.79 | 0.5 | 0.41 | 0.56 | 0.44 |
| 7 | 87624805 | rs1637509 | 0.222 | -0.607 | 1.011 | 0.058 | -0.013 | 0.082 | 0.082 | 0.063 | 0.46 | 0.34 | 0.32 | 0.06 | 0.47 | 0.32 |
| 7 | 87633527 | rs2282950 | -1.136 | -0.605 | 1.001 | 0.094 | -0.001 | 0.08 | 0.08 | 0 | 0.09 | 0.16 | 0.15 | 0.3 | 0.08 | 0.12 |
| 7 | 87644091 | rs2022163 | 0.491 | -0.586 | 0.992 | 0.17 | 0.001 | 0.003 | 0.003 | 0 | 0.43 | 0.29 | 0.51 | 0.56 | 0.44 | 0.56 |
| 7 | 87651567 | rs10235694 | 0.443 | -0.561 | 1.007 | 0.21 | 0.002 | 0.01 | 0.01 | 0 | 0.45 | 0.54 | 0.53 | 0.63 | 0.45 | 0.56 |
| 7 | 87681827 | rs4728725 | -0.861 | -1.172 | 0.293 | -0.148 | -0.133 | 0.06 | 0.06 | 0.03 | 0.77 | 0.95 | 0.8 | 0.9 | 0.6 | 0.86 |
| 7 | 87695787 | rs6958024 | 0.717 | -1.016 | 0.592 | 0.307 | 0.007 | -0.005 | -0.005 | 0.792 | 0.42 | 0.16 | 0.48 | 0.58 | 0.41 | 0.58 |
| 7 | 87699736 | rs2282954 | 1.29 | -1.744 | 0.1 | 0.803 | -0.091 | -0.098 | -0.098 | 3.177 | 0.28 | 0.02 | 0.21 | 0.31 | 0.3 | 0.39 |
| 7 | 87717611 | rs4728731 | 0.218 | -1.627 | 0.242 | 0.615 | 0.012 | 0.014 | 0.014 | 0.002 | 0.43 | 0.18 | 0.49 | 0.62 | 0.41 | 0.58 |
| 7 | 87717762 | rs917205 | -0.96 | -1.713 | 0.589 | 0.622 | 0.005 | 0.167 | 0.167 | 0 | 0.08 | 0.28 | 0.2 | 0.25 | 0.06 | 0.14 |
| 7 | 87721274 | rs10234048 | -0.34 | -1.747 | 0.601 | 0.285 | 0.001 | -0.013 | -0.013 | 0 | 0.52 | 0.51 | 0.69 | 0.87 | 0.46 | 0.72 |
| 7 | 87721928 | rs10238083 | -0.824 | -1.738 | 0.599 | 0.306 | 0.031 | 0.005 | 0.005 | 2.224 | 0.05 | 0.27 | 0.05 | 0 | 0.03 | 0.03 |
| 7 | 87731692 | rs2237541 | 0.184 | -1.742 | 0.461 | 0.288 | 0.013 | 0.116 | 0.116 | 0.067 | 0.44 | 0.16 | 0.51 | 0.83 | 0.42 | 0.68 |
| 7 | 87739123 | rs726328 | -0.996 | -2.15 | 0.325 | 0.073 | -0.016 | 0.03 | 0.03 | 0.767 | 0.63 | 0.68 | 0.73 | 0.91 | 0.49 | 0.78 |
| 7 | 87746839 | rs6955696 | 0.169 | -2.063 | -0.134 | 0.12 | 0.019 | 0.128 | 0.128 | 0.095 | 0.45 | 0.17 | 0.53 | 0.87 | 0.42 | 0.68 |
| 7 | 87754915 | rs17255978 | 0.227 | -2.075 | 0.508 | 0.742 | 0.001 | -0.002 | -0.002 | 0 | 0.12 | 0.12 | 0.15 | 0.04 | 0.04 | 0.06 |
| 7 | 87755471 | rs12538010 | 0.227 | -2.075 | 0.508 | 0.742 | 0.001 | -0.002 | -0.002 | 0 | 0.12 | 0.12 | 0.15 | 0.04 | 0.04 | 0.06 |
| 7 | 87763101 | rs4728732 | -0.936 | -2.16 | 0.59 | 0.726 | -0.082 | -0.086 | -0.086 | 0.506 | 0.19 | 0.44 | 0.22 | 0.07 | 0.08 | 0.1 |
| 7 | 87763113 | rs2041009 | -0.421 | -2.362 | 0.371 | 0.969 | 0.036 | -0.086 | -0.086 | 0.532 | 0.15 | 0.03 | 0.25 | 0.49 | 0.11 | 0.29 |
| 7 | 87768174 | rs10268574 | -2.028 | -2.375 | 0.452 | 0.964 | -0.026 | 0.006 | 0.006 | 0 | 0.35 | 0.56 | 0.48 | 0.56 | 0.19 | 0.38 |
| 7 | 87774727 | rs3761806 | 0.335 | -2.29 | 0.494 | 1.168 | 0.035 | -0.047 | -0.047 | 0.855 | 0.11 | 0.03 | 0.11 | 0.24 | 0.09 | 0.18 |
| 7 | 87775157 | rs10238607 | -0.423 | -2.33 | 0.585 | 1.142 | -0.09 | -0.065 | -0.065 | 0.548 | 0.17 | 0.44 | 0.2 | 0.07 | 0.05 | 0.09 |
| 7 | 87781111 | rs10487008 | -0.58 | -2.317 | 0.532 | 1.239 | 0.016 | -0.015 | -0.015 | 0 | 0.15 | 0.1 | 0.26 | 0.49 | 0.12 | 0.28 |
| 7 | 87794707 | rs2299205 | 0.373 | -2.275 | 0.087 | 1.217 | 0.035 | -0.047 | -0.047 | 0 | 0.11 | 0.03 | 0.11 | 0.24 | 0.09 | 0.17 |
| 7 | 87812968 | rs7792913 | -1.435 | -2.142 | 0.129 | 0.785 | -0.042 | 0.013 | 0.013 | 0 | 0.33 | 0.54 | 0.46 | 0.56 | 0.16 | 0.38 |
| 7 | 87820419 | rs3789236 | -0.735 | -1.152 | -0.197 | 1.208 | -0.011 | 0.07 | 0.07 | 0.027 | 0.13 | 0.14 | 0.22 | 0.5 | 0.08 | 0.31 |
| 7 | 87826957 | rs7810426 | 0.232 | -0.647 | -0.459 | 0.486 | 0.014 | 0.01 | 0.01 | 0 | 0.44 | 0.39 | 0.39 | 0.38 | 0.34 | 0.34 |
| 7 | 87835469 | rs1063964 | -0.266 | 0.621 | -0.692 | -0.096 | 0.061 | -0.083 | -0.083 | 0 | 0.22 | 0.49 | 0.2 | 0.07 | 0.31 | 0.07 |
| 7 | 87839690 | rs17150369 | 1.012 | 1.078 | -1.023 | -0.447 | -0.003 | 0.048 | 0.048 | 0 | 0.17 | 0.08 | 0.12 | 0 | 0.17 | 0.06 |
| 7 | 87840263 | rs6964163 | 1.136 | 1.035 | -1.077 | -0.554 | 0.005 | 0.004 | 0.004 | 0 | 0.47 | 0.4 | 0.38 | 0.18 | 0.45 | 0.29 |
| 7 | 87856515 | rs43111 | -1.067 | 1.057 | -1.408 | -0.446 | 0.013 | 0.023 | 0.023 | 0 | 0.49 | 0.45 | 0.63 | 0.81 | 0.57 | 0.66 |
| 7 | 87876090 | rs43123 | -2.285 | 0.988 | -1.578 | -0.5 | 0.004 | 0.151 | 0.151 | 0.253 | 0.67 | 0.99 | 0.78 | 0.89 | 0.73 | 0.78 |
| 7 | 87906171 | rs8122 | -1.015 | 1.021 | -1.421 | -0.507 | 0.02 | 0.043 | 0.043 | 0 | 0.48 | 0.41 | 0.62 | 0.84 | 0.57 | 0.66 |
| 7 | 87908388 | rs9655959 | 1.016 | 0.961 | -2.016 | -0.644 | -0.001 | 0.018 | 0.018 | 0 | 0.18 | 0.1 | 0.14 | 0 | 0.19 | 0.06 |
| 7 | 87913361 | rs1981529 | -2.252 | 0.918 | -2.231 | -0.604 | -0.005 | 0.153 | 0.153 | 0.229 | 0.67 | 0.99 | 0.78 | 0.91 | 0.73 | 0.78 |

| | | | | | | | | | | | | | | | | |
|---|---|---|---|---|---|---|---|---|---|---|---|---|---|---|---|---|
| 7 | 87939356 | rs12534694 | 0.906 | 1.484 | -1.548 | 0.218 | -0.014 | 0 | 0 | 1.277 | 0.07 | 0.01 | 0.07 | 0.07 | 0.06 | 0.09 |
| 7 | 87941709 | rs12704392 | 0.539 | 1.59 | -1.651 | 0.43 | -0.064 | -0.136 | -0.136 | 0 | 0.17 | 0.02 | 0.29 | 0.56 | 0.2 | 0.44 |
| 7 | 87959102 | rs240011 | -0.483 | 1.568 | -1.212 | -0.072 | -0.028 | 0.088 | 0.088 | 0 | 0.24 | 0.08 | 0.22 | 0.08 | 0.25 | 0.16 |
| 7 | 87971692 | rs6959795 | -0.516 | 1.542 | -1.077 | 0.01 | -0.027 | 0.07 | 0.07 | 0 | 0.83 | 0.94 | 0.85 | 1 | 0.8 | 0.94 |
| 7 | 87977805 | rs7800039 | 0.705 | 1.394 | -0.694 | 0.411 | -0.001 | -0.011 | -0.011 | 0.232 | 0.65 | 0.6 | 0.55 | 0.23 | 0.66 | 0.51 |
| 7 | 87981432 | rs17260153 | -0.183 | 1.727 | -0.174 | 0.278 | -0.006 | 0.006 | 0.006 | 0 | 0.19 | 0.2 | 0.25 | 0.1 | 0.24 | 0.24 |
| 7 | 87983941 | rs7777957 | -1.499 | 1.285 | 0.344 | 0.767 | 0.012 | -0.008 | -0.008 | 1.183 | 0.32 | 0.61 | 0.32 | 0.27 | 0.38 | 0.31 |
| 7 | 87992766 | rs11762499 | 0.525 | 1.526 | 0.313 | 0.754 | -0.022 | -0.036 | -0.036 | 0 | 0.07 | 0 | 0.03 | 0.13 | 0.1 | 0.07 |
| 7 | 87997384 | rs733004 | -1.483 | 1.456 | 0.378 | 0.761 | 0.014 | -0.019 | -0.019 | 0 | 0.32 | 0.61 | 0.32 | 0.22 | 0.39 | 0.31 |
| 7 | 87999503 | rs6948731 | -1.238 | 1.465 | 0.387 | 0.741 | 0.024 | -0.06 | -0.06 | 0 | 0.28 | 0.61 | 0.31 | 0.1 | 0.35 | 0.24 |
| 7 | 88016450 | rs10487012 | -0.422 | 1.677 | 0.366 | 0.76 | -0.003 | -0.053 | -0.053 | 0.014 | 0.77 | 0.87 | 0.61 | 0.4 | 0.79 | 0.54 |
| 7 | 88019109 | rs11773152 | 1.673 | 1.766 | 0.444 | 0.704 | 0.03 | 0.076 | 0.076 | 0 | 0.45 | 0.27 | 0.3 | 0.17 | 0.41 | 0.24 |
| 7 | 88023006 | rs2191472 | 0.619 | 1.585 | 0.472 | 1.317 | 0.003 | -0.012 | -0.012 | 0 | 0.66 | 0.68 | 0.63 | 0.44 | 0.69 | 0.47 |
| 7 | 88039884 | rs2024195 | -0.181 | 1.891 | 0.693 | 1.327 | 0.033 | 0.073 | 0.073 | 0 | 0.1 | 0.21 | 0.16 | 0.19 | 0.11 | 0.16 |
| 7 | 88054557 | rs41654 | 0.324 | 1.761 | 0.156 | 1.31 | -0.054 | -0.15 | -0.15 | 0 | 0.2 | 0.02 | 0.41 | 0.55 | 0.2 | 0.43 |
| 7 | 88054931 | rs11972558 | 0.68 | 1.928 | 0.657 | 1.1 | 0.011 | 0.001 | 0.001 | 0.152 | 0.62 | 0.4 | 0.42 | 0.38 | 0.6 | 0.47 |
| 7 | 88059907 | rs41657 | -0.291 | 2.177 | 0.704 | 0.962 | 0.018 | -0.011 | -0.011 | 1.654 | 0.05 | 0.06 | 0.09 | 0 | 0.14 | 0.02 |
| 7 | 88062343 | rs17353068 | -0.094 | 1.938 | 0.74 | 1.076 | 0.01 | 0.027 | 0.027 | 0 | 0.09 | 0.1 | 0.16 | 0.19 | 0.11 | 0.21 |
| 7 | 88067616 | rs181123 | 0.784 | 1.476 | 1.011 | 0.479 | -0.001 | 0.096 | 0.096 | 0 | 0.93 | 0.72 | 0.97 | 0.89 | 0.98 | 0.98 |
| 7 | 88075888 | rs10487017 | 0.244 | 1.2 | 0.81 | 0.269 | 0.024 | 0.013 | 0.013 | 0 | 0.35 | 0.19 | 0.45 | 0.32 | 0.27 | 0.37 |
| 7 | 88076132 | rs10487018 | 1.353 | 1.297 | 0.725 | 0.125 | 0 | 0.001 | 0.001 | 0 | 0.16 | 0.14 | 0.12 | 0.07 | 0.17 | 0.12 |
| 7 | 88077195 | rs10487019 | 1.38 | 1.432 | 0.666 | 0.128 | -0.011 | 0.002 | 0.002 | 0 | 0.05 | 0.03 | 0.05 | 0.05 | 0.06 | 0.1 |
| 7 | 88077426 | rs6961325 | 1.646 | 1.605 | 0.795 | 0.029 | -0.004 | -0.016 | -0.016 | 0.218 | 0.32 | 0.23 | 0.28 | 0.44 | 0.35 | 0.37 |
| 7 | 88080623 | rs181106 | -1.005 | 1.577 | 0.804 | 0.013 | -0.005 | 0.011 | 0.011 | 0 | 0.81 | 0.77 | 0.87 | 0.82 | 0.81 | 0.87 |
| 7 | 88082373 | rs1476452 | 0.312 | 1.481 | 0.443 | -0.039 | 0.026 | 0.005 | 0.005 | 0 | 0.35 | 0.18 | 0.45 | 0.34 | 0.27 | 0.38 |
| 7 | 88090950 | rs7801811 | 0.726 | 1.314 | 0.578 | -0.546 | 0.014 | -0.018 | -0.018 | 1.546 | 0.67 | 0.4 | 0.73 | 0.78 | 0.62 | 0.75 |
| 7 | 88091162 | rs181086 | 0.43 | 1.466 | 0.675 | -0.619 | 0.019 | -0.022 | -0.022 | 0 | 0.22 | 0.32 | 0.21 | 0.07 | 0.32 | 0.13 |
| 7 | 88092563 | rs10232253 | 0.736 | 1.457 | 0.371 | -0.599 | 0.016 | -0.022 | -0.022 | 0.003 | 0.67 | 0.41 | 0.72 | 0.79 | 0.62 | 0.75 |
| 7 | 88095957 | rs1468106 | 1.113 | 1.414 | 0.087 | -0.643 | 0.026 | -0.045 | -0.045 | 0 | 0.23 | 0.03 | 0.32 | 0.29 | 0.16 | 0.36 |
| 7 | 88098737 | rs2373452 | 0.453 | 1.389 | 0.214 | -0.648 | 0.032 | -0.045 | -0.045 | 0.033 | 0.23 | 0.43 | 0.22 | 0.08 | 0.31 | 0.14 |
| 7 | 88099701 | rs181073 | 0.577 | 1.368 | 0.217 | -0.64 | 0.01 | -0.018 | -0.018 | 0 | 0.66 | 0.37 | 0.71 | 0.8 | 0.61 | 0.75 |
| 7 | 88100188 | rs10952920 | 0.389 | 1.333 | 0.281 | -0.698 | 0.039 | 0.034 | 0.034 | 0 | 0.34 | 0.12 | 0.38 | 0.3 | 0.26 | 0.39 |
| 7 | 88102745 | rs10238377 | 0.289 | 1.331 | 0.306 | -0.699 | 0.022 | 0.019 | 0.019 | 0 | 0.35 | 0.22 | 0.39 | 0.3 | 0.27 | 0.39 |
| 7 | 88105166 | rs4728750 | 1.613 | 1.307 | 0.171 | -0.729 | -0.017 | -0.058 | -0.058 | 0 | 0.29 | 0.16 | 0.3 | 0.51 | 0.33 | 0.36 |
| 7 | 88109823 | rs2213987 | -0.148 | 1.135 | 0.222 | -0.732 | 0 | 0 | 0 | 0 | 0.6 | 0.66 | 0.57 | 0.63 | 0.68 | 0.59 |
| 7 | 88115537 | rs4727171 | 1.637 | 1.147 | 0.066 | -0.699 | -0.01 | -0.062 | -0.062 | 0 | 0.28 | 0.15 | 0.29 | 0.5 | 0.31 | 0.35 |
| 7 | 88117619 | rs181055 | 0.484 | 1.245 | 0.351 | -0.622 | 0.097 | -0.031 | -0.031 | 0.147 | 0.07 | 0.25 | 0.09 | 0 | 0.15 | 0.02 |
| 7 | 88120640 | rs1011444 | 1.44 | 1.184 | -0.06 | -0.637 | 0.001 | 0.002 | 0.002 | 0 | 0.17 | 0.17 | 0.14 | 0.07 | 0.17 | 0.11 |
| 7 | 88122950 | rs181049 | 0.189 | 1.187 | -0.034 | -0.649 | 0.002 | 0.001 | 0.001 | 0 | 0.47 | 0.57 | 0.47 | 0.55 | 0.55 | 0.48 |
| 7 | 88123226 | rs181048 | -0.407 | 1.234 | -0.032 | -0.693 | 0.071 | -0.05 | -0.05 | 0 | 0.14 | 0.34 | 0.14 | 0.04 | 0.21 | 0.12 |
| 7 | 88130171 | rs13225805 | 1.427 | 1.341 | -0.025 | -0.736 | -0.02 | -0.066 | -0.066 | 0.007 | 0.31 | 0.15 | 0.32 | 0.51 | 0.34 | 0.36 |
| 7 | 88144473 | rs4728755 | 0.572 | 1.411 | -0.072 | -0.742 | -0.005 | 0.031 | 0.031 | 0 | 0.48 | 0.37 | 0.46 | 0.58 | 0.52 | 0.46 |
| 7 | 88148465 | rs181027 | -0.395 | 1.576 | -0.331 | -0.518 | 0.079 | -0.063 | -0.063 | 0 | 0.11 | 0.37 | 0.1 | 0 | 0.15 | 0.02 |
| 7 | 88161734 | rs6977770 | 0.477 | 1.234 | -1.185 | -0.102 | 0.001 | -0.035 | -0.035 | 0.126 | 0.39 | 0.57 | 0.3 | 0.08 | 0.43 | 0.16 |
| 7 | 88166231 | rs2078799 | 0.647 | 1.225 | -1.234 | -0.106 | 0.003 | -0.04 | -0.04 | 0.04 | 0.38 | 0.57 | 0.3 | 0.08 | 0.43 | 0.16 |
| 7 | 88173578 | rs1119075 | -0.411 | 1.513 | -2.008 | 0.076 | 0.006 | -0.064 | -0.064 | 0 | 0.7 | 0.93 | 0.63 | 0.58 | 0.76 | 0.5 |
| 7 | 88178616 | rs2519526 | -0.15 | 1.545 | -1.376 | 0.695 | -0.013 | -0.054 | -0.054 | 0 | 0.29 | 0.18 | 0.31 | 0.5 | 0.33 | 0.34 |
| 7 | 88190053 | rs6962353 | -0.569 | 1.866 | -1.24 | 0.305 | -0.066 | 0.003 | 0.003 | 0 | 0.22 | 0.1 | 0.28 | 0.22 | 0.32 | 0.44 |
| 7 | 88191618 | rs2519523 | 1.34 | 1.534 | -1.001 | -0.153 | -0.024 | 0.032 | 0.032 | 0 | 0.65 | 0.76 | 0.65 | 0.77 | 0.58 | 0.52 |
| 7 | 88197858 | rs4728764 | 1.337 | 1.48 | -0.836 | -0.168 | -0.026 | 0.032 | 0.032 | 0 | 0.65 | 0.76 | 0.63 | 0.77 | 0.58 | 0.48 |
| 7 | 88199328 | rs2718315 | 1.302 | 1.408 | -0.639 | -0.209 | -0.005 | 0.005 | 0.005 | 0 | 0.63 | 0.65 | 0.62 | 0.76 | 0.58 | 0.48 |
| 7 | 88201442 | rs717655 | 0.452 | 1.535 | -0.699 | -0.121 | 0.039 | -0.022 | -0.022 | 0 | 0.94 | 0.84 | 0.94 | 1 | 0.91 | 0.99 |
| 7 | 88203866 | rs2188253 | -0.931 | 1.394 | -0.579 | -0.097 | -0.019 | 0.017 | 0.017 | 0 | 0.28 | 0.2 | 0.32 | 0.23 | 0.33 | 0.48 |
| 7 | 88204888 | rs2519515 | -1.338 | 0.844 | -1.302 | 0.131 | -0.061 | -0.03 | -0.03 | 0 | 0.39 | 0.93 | 0.33 | 0.58 | 0.37 | 0.28 |
| 7 | 88206152 | rs2718320 | -1.069 | 0.655 | -0.058 | 0.225 | -0.012 | 0.083 | 0.083 | 0 | 0.22 | 0.54 | 0.2 | 0.57 | 0.15 | 0.18 |
| 7 | 88208465 | rs2263064 | -0.218 | 0.271 | 0.521 | 0.539 | -0.014 | 0.1 | 0.1 | 0 | 0.11 | 0.16 | 0.14 | 0.56 | 0.07 | 0.14 |
| 7 | 88209768 | rs12531431 | 1.802 | 0.345 | 0.548 | 0.684 | -0.009 | -0.067 | -0.067 | 0.139 | 0.64 | 0.16 | 0.68 | 0.51 | 0.63 | 0.74 |
| 7 | 88210663 | rs2519533 | 1.289 | 0.508 | 1.551 | 0.598 | 0.172 | -0.003 | -0.003 | 0 | 0.88 | 0.58 | 0.84 | 0.95 | 0.74 | 0.87 |
| 7 | 88213062 | rs2188256 | 1.266 | 0.513 | 1.657 | 0.571 | 0.139 | -0.019 | -0.019 | 0 | 0.89 | 0.67 | 0.85 | 0.95 | 0.74 | 0.87 |
| 7 | 88217901 | rs12112417 | 0.105 | 0.509 | 1.382 | 0.554 | 0.002 | -0.013 | -0.013 | 0 | 0.1 | 0.12 | 0.04 | 0 | 0.05 | 0.05 |
| 7 | 88222707 | rs2538913 | 1.123 | 0.569 | 1.415 | 0.652 | 0.035 | -0.009 | -0.009 | 0 | 0.89 | 0.83 | 0.84 | 0.95 | 0.72 | 0.85 |
| 7 | 88233444 | rs2718359 | 1.205 | -0.611 | 1.878 | 0.745 | -0.006 | -0.067 | -0.067 | 0 | 0.73 | 0.47 | 0.73 | 0.88 | 0.76 | 0.78 |
| 7 | 88239231 | rs12533988 | 0.449 | -0.947 | 1.649 | 0.54 | -0.001 | 0.11 | 0.11 | 0 | 0.73 | 0.93 | 0.85 | 0.92 | 0.77 | 0.74 |
| 7 | 88247365 | rs2538898 | 1.223 | -1.06 | 2.116 | 0.435 | 0.004 | -0.007 | -0.007 | 1.102 | 0.65 | 0.66 | 0.81 | 0.89 | 0.74 | 0.72 |
| 7 | 88248135 | rs2718293 | -1.559 | -1.384 | 1.659 | 0.858 | -0.053 | 0.213 | 0.213 | 0 | 0.19 | 0.42 | 0.3 | 0.48 | 0.18 | 0.15 |
| 7 | 88259445 | rs2718296 | 0.185 | -1.436 | 0.893 | 0.805 | 0.002 | 0.11 | 0.11 | 0 | 0.73 | 0.93 | 0.85 | 0.92 | 0.76 | 0.74 |
| 7 | 88263202 | rs10487028 | 2.005 | -1.345 | 0.877 | 0.895 | 0.083 | 0.01 | 0.01 | 0 | 0.49 | 0.08 | 0.53 | 0.41 | 0.58 | 0.58 |
| 7 | 88265803 | rs10487029 | 2.013 | -1.345 | 0.877 | 0.892 | 0.083 | 0.013 | 0.013 | 0 | 0.49 | 0.08 | 0.52 | 0.41 | 0.58 | 0.58 |
| 7 | 88268399 | rs2718387 | -0.794 | -1.253 | 1.012 | 1.086 | -0.01 | 0.093 | 0.093 | 0 | 0.12 | 0.15 | 0.26 | 0.47 | 0.15 | 0.14 |

| | | | | | | | | | | | | | | | | |
|---|---|---|---|---|---|---|---|---|---|---|---|---|---|---|---|---|
| 7 | 88271503 | rs2538909 | 0.365 | -1.195 | 0.823 | 0.691 | 0.001 | 0.116 | 0.116 | 0.102 | 0.71 | 0.92 | 0.82 | 0.92 | 0.73 | 0.74 |
| 7 | 88272436 | rs2538910 | -0.059 | -1.616 | 1.068 | 1.376 | -0.002 | 0.009 | 0.009 | 0 | 0.11 | 0.1 | 0.13 | 0.22 | 0.13 | 0.09 |
| 7 | 88280047 | rs10156041 | 1.288 | -1.745 | 1.57 | 1.281 | 0.044 | 0.014 | 0.014 | 0 | 0.59 | 0.75 | 0.59 | 0.66 | 0.61 | 0.61 |
| 7 | 88283826 | rs2538912 | 0.155 | -1.724 | 1.366 | 1.276 | 0.03 | 0.103 | 0.103 | 0 | 0.73 | 0.92 | 0.82 | 0.92 | 0.75 | 0.75 |
| 7 | 88287698 | rs7779756 | 1.58 | -1.729 | 1.441 | 1.286 | 0.065 | 0.03 | 0.03 | 0 | 0.55 | 0.29 | 0.56 | 0.65 | 0.61 | 0.6 |
| 7 | 88295001 | rs12531232 | 1.391 | -1.728 | 1.384 | 1.298 | 0.031 | 0.014 | 0.014 | 0 | 0.59 | 0.68 | 0.59 | 0.67 | 0.61 | 0.61 |
| 7 | 88301914 | rs6465149 | -0.488 | -1.696 | 1.351 | 1.277 | -0.007 | 0.112 | 0.112 | 0 | 0.1 | 0.25 | 0.22 | 0.25 | 0.12 | 0.13 |
| 7 | 88302772 | rs6959780 | 1.5 | -1.611 | 1.411 | 1.555 | 0.008 | -0.004 | -0.004 | 0.384 | 0.55 | 0.56 | 0.53 | 0.48 | 0.58 | 0.58 |
| 7 | 88302926 | rs10252643 | -0.86 | -1.545 | 0.979 | 1.521 | -0.022 | 0.057 | 0.057 | 0 | 0.14 | 0.25 | 0.23 | 0.25 | 0.14 | 0.14 |
| 7 | 88303672 | rs2098278 | 1.453 | -1.289 | 0.297 | 1.383 | 0.112 | -0.006 | -0.006 | 0 | 0.51 | 0.12 | 0.5 | 0.44 | 0.58 | 0.57 |
| 7 | 88311428 | rs2538918 | -2.082 | -0.291 | 0.151 | 1.738 | 0 | 0.151 | 0.151 | 0.062 | 0.44 | 0.91 | 0.48 | 0.75 | 0.37 | 0.51 |
| 7 | 88311477 | rs10235198 | -1.268 | -0.093 | 1.257 | 2.315 | 0.005 | 0.158 | 0.158 | 0 | 0.19 | 0.58 | 0.21 | 0.53 | 0.12 | 0.32 |
| 7 | 88325106 | rs10241505 | -0.347 | -0.096 | 1.643 | 2.871 | 0.001 | 0.051 | 0.051 | 0 | 0.15 | 0.26 | 0.15 | 0.35 | 0.11 | 0.25 |
| 7 | 88327568 | rs17360725 | 0.434 | -0.1 | 1.412 | 2.952 | -0.039 | -0.015 | -0.015 | 0 | 0.11 | 0 | 0.05 | 0.08 | 0.09 | 0.21 |
| 7 | 88368532 | rs9641030 | -1.452 | -0.156 | 1.581 | 2.903 | 0.005 | 0.211 | 0.211 | 0 | 0.19 | 0.48 | 0.27 | 0.51 | 0.13 | 0.34 |
| 7 | 88370585 | rs2519908 | -0.392 | -0.172 | 1.746 | 2.746 | 0.02 | 0.015 | 0.015 | 0.052 | 0.13 | 0.24 | 0.17 | 0.13 | 0.18 | 0.07 |
| 7 | 88371952 | rs17164033 | -0.923 | -0.194 | 1.774 | 2.767 | 0.006 | 0.062 | 0.062 | 0 | 0.17 | 0.36 | 0.11 | 0.24 | 0.1 | 0.29 |
| 7 | 88387997 | rs2888567 | -1.949 | -0.241 | 1.659 | 2.745 | -0.003 | -0.015 | -0.015 | 0 | 0.31 | 0.74 | 0.29 | 0.35 | 0.28 | 0.35 |
| 7 | 88396505 | rs2519945 | -2.187 | -0.382 | 1.66 | 2.736 | 0 | -0.001 | -0.001 | 0.141 | 0.36 | 0.63 | 0.33 | 0.47 | 0.31 | 0.44 |
| 7 | 88402485 | rs2519954 | -2.264 | -0.413 | 1.54 | 2.724 | 0.013 | -0.013 | -0.013 | 0 | 0.37 | 0.81 | 0.4 | 0.54 | 0.31 | 0.44 |
| 7 | 88408427 | rs3847035 | -2.501 | -0.474 | 1.512 | 2.699 | 0.021 | -0.007 | -0.007 | 0.143 | 0.39 | 0.81 | 0.4 | 0.54 | 0.31 | 0.44 |
| 7 | 88423881 | rs2293583 | -0.214 | -0.53 | 1.689 | 2.852 | 0.002 | 0.006 | 0.006 | 0 | 0.14 | 0.11 | 0.06 | 0.12 | 0.09 | 0.22 |
| 7 | 88424291 | rs4363132 | -0.294 | -0.566 | 1.93 | 2.886 | 0.016 | 0 | 0 | 0.022 | 0.13 | 0.22 | 0.16 | 0.11 | 0.18 | 0.08 |
| 7 | 88425772 | rs17164406 | 0.345 | -0.608 | 2.052 | 2.931 | -0.022 | 0.13 | 0.13 | 0 | 0.08 | 0.19 | 0.16 | 0.32 | 0.03 | 0.15 |
| 7 | 88433768 | rs7779337 | -1.902 | -0.78 | 1.66 | 2.734 | 0.008 | -0.001 | -0.001 | 0 | 0.28 | 0.27 | 0.31 | 0.68 | 0.19 | 0.43 |
| 7 | 88434794 | rs17164472 | -0.312 | -0.678 | 1.716 | 2.699 | 0.026 | 0.004 | 0.004 | 0.086 | 0.13 | 0.26 | 0.17 | 0.11 | 0.18 | 0.07 |
| 7 | 88435045 | rs10487038 | -2.336 | -0.684 | 1.679 | 2.734 | -0.006 | -0.002 | -0.002 | 0 | 0.39 | 0.5 | 0.4 | 0.52 | 0.34 | 0.46 |
| 7 | 88447161 | rs17164532 | -1.417 | -0.687 | 1.663 | 2.658 | 0.009 | -0.028 | -0.028 | 0 | 0.25 | 0.22 | 0.23 | 0.43 | 0.17 | 0.39 |
| 7 | 88447187 | rs3915191 | -2.465 | -0.722 | 1.684 | 2.655 | -0.001 | -0.015 | -0.015 | 0 | 0.42 | 0.55 | 0.5 | 0.81 | 0.38 | 0.53 |
| 7 | 88448121 | rs6968172 | -0.34 | -0.759 | 1.628 | 2.789 | 0.025 | 0.042 | 0.042 | 0.049 | 0.14 | 0.27 | 0.19 | 0.15 | 0.18 | 0.07 |
| 7 | 88448157 | rs17164545 | -1.295 | -0.799 | 1.478 | 2.767 | 0.003 | -0.009 | -0.009 | 0 | 0.08 | 0.06 | 0.13 | 0.25 | 0.08 | 0.11 |
| 7 | 88450383 | rs3915192 | -2.246 | -0.785 | 1.729 | 2.682 | -0.004 | -0.002 | -0.002 | 0 | 0.32 | 0.37 | 0.4 | 0.71 | 0.26 | 0.46 |
| 7 | 88450572 | rs17164562 | -1.413 | -0.765 | 1.649 | 2.712 | -0.006 | 0.01 | 0.01 | 0 | 0.24 | 0.32 | 0.27 | 0.65 | 0.17 | 0.42 |
| 7 | 88451505 | rs13221086 | -0.886 | -0.764 | 1.658 | 2.634 | 0.006 | -0.013 | -0.013 | 0 | 0.2 | 0.21 | 0.2 | 0.44 | 0.12 | 0.36 |
| 7 | 88453043 | rs6974679 | -2.655 | -0.74 | 1.79 | 2.588 | 0.026 | 0.122 | 0.122 | 0 | 0.44 | 0.76 | 0.52 | 0.81 | 0.4 | 0.53 |
| 7 | 88454633 | rs3843539 | -2.492 | -0.739 | 1.838 | 2.588 | 0.004 | 0.01 | 0.01 | 0 | 0.43 | 0.6 | 0.51 | 0.81 | 0.4 | 0.53 |
| 7 | 88461461 | rs7804445 | -1.383 | -0.735 | 1.815 | 2.492 | 0.002 | -0.004 | -0.004 | 0 | 0.24 | 0.27 | 0.3 | 0.68 | 0.17 | 0.42 |
| 7 | 88470006 | rs1969551 | -1.396 | -0.656 | 1.906 | 2.454 | 0.006 | -0.011 | -0.011 | 1.132 | 0.19 | 0.29 | 0.2 | 0.13 | 0.23 | 0.12 |
| 7 | 88490822 | rs6962263 | -1.561 | -0.65 | 1.323 | 2.436 | -0.002 | 0.004 | 0.004 | 1.022 | 0.2 | 0.57 | 0.23 | 0.18 | 0.23 | 0.12 |
| 7 | 88491230 | rs17164834 | -1.228 | -0.639 | 1.198 | 2.376 | 0.015 | -0.004 | -0.004 | 0 | 0.06 | 0.2 | 0.05 | 0.03 | 0.05 | 0.04 |
| 7 | 88508355 | rs1526251 | -2.234 | -0.677 | 1.076 | 2.482 | 0.017 | 0.021 | 0.021 | 0 | 0.41 | 0.69 | 0.43 | 0.6 | 0.35 | 0.46 |
| 7 | 88513041 | rs2214632 | -1.459 | -0.683 | 0.87 | 2.544 | -0.013 | 0.042 | 0.042 | 0 | 0.26 | 0.39 | 0.25 | 0.43 | 0.17 | 0.36 |
| 7 | 88543687 | rs1528934 | -2.088 | -0.672 | 0.612 | 2.214 | 0.015 | 0.013 | 0.013 | 0 | 0.4 | 0.71 | 0.43 | 0.57 | 0.35 | 0.44 |
| 7 | 88547866 | rs17301014 | -2.252 | -0.74 | 0.175 | 2.243 | 0.042 | -0.045 | -0.045 | 0 | 0.33 | 0.83 | 0.4 | 0.56 | 0.28 | 0.28 |
| 7 | 88550219 | rs7790821 | -1.702 | -0.6 | 0.198 | 1.902 | 0.009 | 0.014 | 0.014 | 0 | 0.27 | 0.34 | 0.23 | 0.26 | 0.27 | 0.3 |
| 7 | 88550492 | rs10248254 | 0.045 | -0.51 | -0.499 | 1.611 | -0.028 | -0.001 | -0.001 | 0 | 0.11 | 0.01 | 0.05 | 0.07 | 0.09 | 0.19 |
| 7 | 88556361 | rs17165493 | -0.698 | -0.758 | -0.343 | 1.833 | 0.002 | -0.004 | -0.004 | 0 | 0.11 | 0.14 | 0.12 | 0.18 | 0.04 | 0.19 |
| 7 | 88556835 | rs12672637 | -0.415 | -0.68 | -0.433 | 1.756 | 0.016 | -0.022 | -0.022 | 0.153 | 0.11 | 0.1 | 0.11 | 0.18 | 0.05 | 0.18 |
| 7 | 88558183 | rs1528932 | -1.005 | -0.239 | -0.309 | 1.647 | 0.001 | 0.006 | 0.006 | 0.505 | 0.21 | 0.23 | 0.09 | 0.13 | 0.16 | 0.27 |
| 7 | 88558912 | rs2189749 | -0.813 | -0.295 | -0.254 | 1.777 | -0.001 | 0.01 | 0.01 | 0 | 0.07 | 0.12 | 0.15 | 0.2 | 0.05 | 0.15 |
| 7 | 88560427 | rs10267374 | -1.023 | -0.239 | -0.229 | 1.521 | 0.001 | 0.004 | 0.004 | 0.46 | 0.21 | 0.24 | 0.09 | 0.13 | 0.15 | 0.26 |
| 7 | 88568549 | rs10258416 | -0.731 | -0.165 | 0.03 | 2.116 | 0 | 0.022 | 0.022 | 1.37 | 0.16 | 0.25 | 0.16 | 0.32 | 0.16 | 0.23 |
| 7 | 88571862 | rs1024508 | -1.319 | 0.507 | -0.789 | 1.909 | 0.067 | 0.027 | 0.027 | 0 | 0.49 | 0.32 | 0.49 | 0.46 | 0.56 | 0.58 |
| 7 | 88573567 | rs13231899 | 1.422 | 0.341 | -0.675 | 2.27 | 0.01 | 0.006 | 0.006 | 0.107 | 0.06 | 0.04 | 0.01 | 0.04 | 0.03 | 0.06 |
| 7 | 88581859 | rs4727195 | -1.307 | -0.641 | -0.972 | 1.413 | -0.024 | -0.015 | -0.015 | 0 | 0.71 | 0.68 | 0.78 | 0.83 | 0.78 | 0.84 |
| 7 | 88582676 | rs1990020 | -1.296 | -0.648 | -0.975 | 1.387 | -0.024 | -0.015 | -0.015 | 0 | 0.71 | 0.68 | 0.78 | 0.83 | 0.78 | 0.84 |
| 7 | 88588037 | rs10486887 | -0.363 | 0.427 | -1.414 | 1.417 | -0.053 | -0.005 | -0.005 | 2.478 | 0.37 | 0.25 | 0.39 | 0.37 | 0.46 | 0.39 |
| 7 | 88591502 | rs6943877 | -0.364 | 0.426 | -1.478 | 1.313 | -0.057 | 0.029 | 0.029 | 0 | 0.37 | 0.23 | 0.35 | 0.26 | 0.46 | 0.39 |
| 7 | 88595238 | rs10486888 | -0.321 | 0.461 | -1.48 | 1.299 | -0.061 | 0.034 | 0.034 | 0 | 0.37 | 0.23 | 0.34 | 0.26 | 0.45 | 0.38 |
| 7 | 88595651 | rs17165847 | 1.074 | 0.336 | -1.45 | 1.268 | -0.027 | -0.04 | -0.04 | 0 | 0.07 | 0 | 0.08 | 0.16 | 0.06 | 0.17 |
| 7 | 88596054 | rs10249495 | -0.647 | 0.278 | -1.348 | 0.967 | 0.01 | 0.006 | 0.006 | 0 | 0.51 | 0.5 | 0.58 | 0.65 | 0.6 | 0.57 |
| 7 | 88596385 | rs6959888 | -0.022 | 0.266 | -1.135 | 0.961 | -0.016 | -0.025 | -0.025 | 0 | 0.12 | 0.06 | 0.13 | 0.17 | 0.11 | 0.22 |
| 7 | 88596821 | rs6964420 | -0.339 | 0.464 | -0.952 | 0.569 | -0.022 | -0.052 | -0.052 | 0 | 0.75 | 0.86 | 0.59 | 0.5 | 0.65 | 0.67 |
| 7 | 88601192 | rs17393952 | 1.472 | 0.125 | -0.877 | 0.143 | -0.001 | -0.045 | -0.045 | 0.019 | 0.24 | 0.18 | 0.11 | 0 | 0.21 | 0.21 |
| 7 | 88604651 | rs10224451 | 0.497 | -0.057 | -0.719 | 0.27 | 0.01 | 0.004 | 0.004 | 0 | 0.34 | 0.69 | 0.24 | 0.27 | 0.26 | 0.36 |
| 7 | 88607314 | rs10243825 | 0.977 | 0.085 | -0.824 | 0.286 | -0.001 | -0.001 | -0.001 | 0 | 0.06 | 0.06 | 0.1 | 0.09 | 0.12 | 0.03 |
| 7 | 88613111 | rs2214628 | -0.932 | 0.041 | -0.885 | 0.215 | -0.047 | -0.071 | -0.071 | 5.481 | 0.76 | 0.91 | 0.67 | 0.62 | 0.69 | 0.72 |
| 7 | 88615862 | rs6971362 | 1.608 | 0.06 | -0.855 | 0.138 | -0.001 | -0.045 | -0.045 | 1.07 | 0.24 | 0.18 | 0.11 | 0 | 0.22 | 0.22 |
| 7 | 88637780 | rs756749 | 1.193 | -0.32 | -0.376 | 1.379 | 0 | 0 | 0 | 0.071 | 0.61 | 0.34 | 0.53 | 0.32 | 0.63 | 0.55 |

| | | | | | | | | | | | | | | | | |
|---|---|---|---|---|---|---|---|---|---|---|---|---|---|---|---|---|
| 7 | 88653885 | rs16868583 | -1.117 | -0.289 | -0.136 | 1.385 | -0.051 | 0.186 | 0.186 | 0 | 0.14 | 0.45 | 0.14 | 0.3 | 0.06 | 0.17 |
| 7 | 88654133 | rs10262711 | -1.143 | -0.297 | -0.121 | 1.381 | -0.051 | 0.173 | 0.173 | 0 | 0.15 | 0.43 | 0.13 | 0.3 | 0.06 | 0.17 |
| 7 | 88657022 | rs2040524 | -0.254 | -0.303 | -0.131 | 1.369 | -0.001 | 0 | 0 | 0 | 0.22 | 0.22 | 0.26 | 0.3 | 0.2 | 0.15 |
| 7 | 88657093 | rs10253454 | 0.787 | -0.288 | -0.161 | 1.373 | -0.006 | -0.032 | -0.032 | 0.133 | 0.16 | 0.06 | 0.24 | 0.28 | 0.19 | 0.12 |
| 7 | 88662774 | rs13438042 | -0.971 | -0.26 | -0.118 | 1.395 | -0.023 | 0.025 | 0.025 | 0 | 0.3 | 0.43 | 0.36 | 0.58 | 0.25 | 0.29 |
| 7 | 88676411 | rs12538305 | 1.406 | -0.264 | -0.141 | 1.248 | 0.02 | 0.053 | 0.053 | 0 | 0.5 | 0.38 | 0.38 | 0.15 | 0.56 | 0.46 |
| 7 | 88677955 | rs6975830 | -0.124 | -0.252 | -0.143 | 1.198 | 0.002 | -0.005 | -0.005 | 0 | 0.76 | 0.79 | 0.74 | 0.68 | 0.79 | 0.72 |
| 7 | 88679791 | rs7782027 | -1.102 | -0.26 | -0.267 | 1.196 | -0.068 | -0.099 | -0.099 | 0.82 | 0.82 | 0.99 | 0.76 | 0.7 | 0.78 | 0.75 |
| 7 | 88685040 | rs7788652 | -0.086 | -0.253 | -0.26 | 1.2 | 0.004 | -0.008 | -0.008 | 0.154 | 0.75 | 0.79 | 0.73 | 0.68 | 0.79 | 0.72 |
| 7 | 88686705 | rs10486896 | 0.104 | -0.254 | -0.254 | 1.229 | -0.018 | 0.169 | 0.169 | 0 | 0.08 | 0.26 | 0.1 | 0.28 | 0.05 | 0.14 |
| 7 | 88688729 | rs2189053 | 1 | -0.257 | -0.275 | 1.273 | -0.006 | 0.103 | 0.103 | 0 | 0.06 | 0.22 | 0.02 | 0.16 | 0.02 | 0.08 |
| 7 | 88689707 | rs2158143 | -0.391 | -0.258 | -0.281 | 1.198 | -0.017 | 0.069 | 0.069 | 0 | 0.25 | 0.4 | 0.35 | 0.52 | 0.23 | 0.25 |
| 7 | 88690800 | rs10249100 | -0.391 | -0.258 | -0.293 | 1.198 | -0.02 | 0.081 | 0.081 | 0 | 0.25 | 0.42 | 0.35 | 0.52 | 0.23 | 0.26 |
| 7 | 88692519 | rs10254094 | 0.991 | -0.254 | -0.339 | 1.196 | -0.009 | 0.081 | 0.081 | 0 | 0.06 | 0.22 | 0.02 | 0.15 | 0.02 | 0.07 |
| 7 | 88695596 | rs13233091 | 1.418 | -0.233 | -0.877 | 1.188 | 0.026 | 0.085 | 0.085 | 0 | 0.5 | 0.33 | 0.38 | 0.16 | 0.56 | 0.47 |
| 7 | 88700594 | rs2189055 | 1.182 | -0.032 | -1.176 | 1.166 | -0.042 | -0.037 | -0.037 | 0 | 0.08 | 0.01 | 0.05 | 0.12 | 0.1 | 0.17 |
| 7 | 88700818 | rs2158142 | 0.479 | -0.777 | -0.555 | 1.711 | -0.005 | -0.044 | -0.044 | 0 | 0.16 | 0.06 | 0.23 | 0.37 | 0.19 | 0.12 |
| 7 | 88704998 | rs2373836 | 1.101 | -0.486 | -0.77 | 1.191 | -0.005 | -0.034 | -0.034 | 0 | 0.55 | 0.49 | 0.41 | 0.18 | 0.56 | 0.51 |
| 7 | 88709142 | rs2189057 | -0.019 | -0.51 | -0.755 | 1.237 | -0.028 | 0.071 | 0.071 | 0 | 0.25 | 0.39 | 0.34 | 0.52 | 0.21 | 0.24 |
| 7 | 88711537 | rs2189058 | -0.611 | -0.389 | -1.261 | 1.291 | -0.039 | 0.037 | 0.037 | 0 | 0.32 | 0.57 | 0.37 | 0.54 | 0.23 | 0.27 |
| 7 | 88713746 | rs17166630 | 0.161 | -0.472 | -1.584 | 1.299 | -0.014 | -0.012 | -0.012 | 1.685 | 0.05 | 0.09 | 0.02 | 0.02 | 0.01 | 0.03 |
| 7 | 88722433 | rs6465182 | -0.373 | 0.017 | -1.768 | 0.654 | -0.017 | -0.14 | -0.14 | 0.292 | 0.8 | 0.97 | 0.69 | 0.59 | 0.79 | 0.71 |
| 7 | 88723903 | rs12704428 | -0.024 | 0.551 | -1.196 | -0.378 | 0.007 | 0.007 | 0.007 | 0.045 | 0.49 | 0.44 | 0.39 | 0.16 | 0.55 | 0.48 |
| 7 | 88737192 | rs10242213 | 0.145 | 0.731 | -1.718 | -0.113 | -0.02 | 0.084 | 0.084 | 0 | 0.56 | 0.84 | 0.61 | 0.65 | 0.54 | 0.51 |
| 7 | 88737228 | rs1029439 | -0.058 | 1.018 | -1.216 | -0.237 | 0.013 | -0.013 | -0.013 | 0 | 0.13 | 0.21 | 0.1 | 0.08 | 0.18 | 0.13 |
| 7 | 88737610 | rs10272493 | -0.226 | 0.679 | -2.74 | -1.587 | -0.021 | 0.197 | 0.197 | 0 | 0.61 | 0.96 | 0.63 | 0.83 | 0.7 | 0.57 |
| 7 | 88738077 | rs4141350 | 0.426 | 1.722 | -2.699 | -1.759 | 0.026 | 0.073 | 0.073 | 0 | 0.46 | 0.71 | 0.43 | 0.69 | 0.39 | 0.4 |
| 7 | 88738358 | rs10276350 | 0.219 | 1.718 | -2.72 | -1.787 | 0.06 | -0.001 | -0.001 | 0 | 0.16 | 0.24 | 0.21 | 0.16 | 0.32 | 0.2 |
| 7 | 88739100 | rs2189059 | -0.839 | 1.513 | -2.96 | -2.044 | 0.002 | 0.14 | 0.14 | 0 | 0.69 | 0.96 | 0.76 | 0.84 | 0.78 | 0.63 |
| 7 | 88750473 | rs7787455 | -0.353 | 1.493 | -2.974 | -2.057 | -0.001 | 0.19 | 0.19 | 0 | 0.63 | 0.95 | 0.65 | 0.84 | 0.72 | 0.6 |
| 7 | 88751636 | rs2373842 | -0.873 | 1.492 | -2.929 | -2.05 | 0.003 | 0.137 | 0.137 | 0 | 0.69 | 0.96 | 0.76 | 0.84 | 0.78 | 0.63 |
| 7 | 88751897 | rs10268458 | -0.45 | 0.818 | -3.311 | -2.425 | 0.012 | 0.219 | 0.219 | 0 | 0.55 | 0.92 | 0.63 | 0.81 | 0.66 | 0.53 |
| 7 | 88753614 | rs7803285 | 0.362 | 0.632 | -2.294 | -1.951 | 0.053 | 0.045 | 0.045 | 1.843 | 0.18 | 0.31 | 0.27 | 0.6 | 0.32 | 0.31 |
| 7 | 88756032 | rs17166916 | 1.153 | 0.321 | -1.967 | -2.121 | -0.001 | -0.007 | -0.007 | 0 | 0.24 | 0.22 | 0.2 | 0.18 | 0.25 | 0.17 |
| 7 | 88759421 | rs10260551 | 0.703 | 0.485 | -1.946 | -2.052 | 0.007 | 0.007 | 0.007 | 2.215 | 0.17 | 0.19 | 0.26 | 0.59 | 0.31 | 0.3 |
| 7 | 88760129 | rs17166931 | 1.252 | 0.437 | -1.717 | -2.064 | 0.002 | -0.01 | -0.01 | 0.131 | 0.07 | 0.29 | 0.04 | 0.03 | 0.04 | 0.02 |
| 7 | 88760360 | rs12704439 | 0.538 | 0.449 | -1.773 | -2.062 | 0.001 | 0.009 | 0.009 | 0 | 0.29 | 0.23 | 0.31 | 0.18 | 0.31 | 0.21 |
| 7 | 88761835 | rs6951573 | 0.148 | 0.414 | -1.778 | -2.072 | -0.031 | -0.084 | -0.084 | 0.493 | 0.22 | 0.54 | 0.19 | 0.06 | 0.15 | 0.12 |
| 7 | 88764400 | rs17307450 | 0.722 | 0.237 | -1.853 | -2.217 | 0.01 | 0.048 | 0.048 | 0 | 0.17 | 0.08 | 0.17 | 0.01 | 0.14 | 0.12 |
| 7 | 88768424 | rs2189069 | 0.68 | 0.256 | -1.853 | -2.159 | 0.001 | 0.008 | 0.008 | 0 | 0.29 | 0.23 | 0.32 | 0.19 | 0.31 | 0.2 |
| 7 | 88769118 | rs7803663 | 2.585 | 0.752 | -1.902 | -2.193 | -0.033 | -0.011 | -0.011 | 0 | 0.06 | 0 | 0.07 | 0.05 | 0.09 | 0.05 |
| 7 | 88770627 | rs10233510 | -1.173 | 0.666 | -1.893 | -2.229 | 0.021 | 0.106 | 0.106 | 0 | 0.5 | 0.74 | 0.49 | 0.65 | 0.54 | 0.52 |
| 7 | 88775048 | rs1014759 | -0.325 | 1.07 | -1.042 | -2.2 | 0.063 | 0.02 | 0.02 | 0.19 | 0.9 | 0.72 | 0.87 | 0.92 | 0.82 | 0.89 |
| 7 | 88775551 | rs12111622 | 0.778 | 0.915 | -1.088 | -2.434 | 0 | 0 | 0 | 0.468 | 0.27 | 0.24 | 0.31 | 0.19 | 0.29 | 0.2 |
| 7 | 88778601 | rs2015523 | 0.659 | 0.708 | -0.915 | -2.433 | -0.008 | -0.083 | -0.083 | 2.007 | 0.17 | 0.27 | 0.2 | 0.01 | 0.15 | 0.14 |
| 7 | 88778734 | rs2015535 | 0.924 | 0.734 | -1.04 | -2.421 | 0 | -0.031 | -0.031 | 0 | 0.16 | 0.16 | 0.19 | 0.01 | 0.15 | 0.14 |
| 7 | 88780559 | rs722002 | 1.311 | 0.629 | -1.182 | -2.144 | 0.002 | 0.01 | 0.01 | 0 | 0.14 | 0.05 | 0.13 | 0.06 | 0.12 | 0.09 |
| 7 | 88782822 | rs17308009 | 1.783 | 0.446 | -1.195 | -2.334 | 0.014 | 0.076 | 0.076 | 0 | 0.22 | 0.04 | 0.12 | 0.12 | 0.16 | 0.27 |
| 7 | 88784356 | rs7781087 | 0.019 | 0.626 | -1.176 | -2.543 | 0.036 | -0.031 | -0.031 | 0 | 0.3 | 0.41 | 0.45 | 0.22 | 0.42 | 0.29 |
| 7 | 88794199 | rs2189052 | 1.031 | 0.174 | -1.166 | -1.86 | -0.034 | -0.086 | -0.086 | 0 | 0.12 | 0.01 | 0.3 | 0.59 | 0.12 | 0.2 |
| 7 | 88804940 | rs2888659 | -0.272 | 0.322 | -1.133 | -1.972 | 0.012 | 0.007 | 0.007 | 0 | 0.78 | 0.72 | 0.6 | 0.34 | 0.67 | 0.75 |
| 7 | 88809649 | rs9640583 | 3.068 | -0.29 | -1.731 | -2.195 | -0.002 | 0.086 | 0.086 | 0 | 0.21 | 0.01 | 0.11 | 0.07 | 0.15 | 0.14 |
| 7 | 88809850 | rs7779011 | 2.078 | 0.553 | -1.258 | -2.023 | 0.009 | -0.004 | -0.004 | 0 | 0.14 | 0.18 | 0.24 | 0.13 | 0.18 | 0.16 |
| 7 | 88811466 | rs7784997 | -0.861 | 0.592 | -1.308 | -2.101 | -0.095 | -0.092 | -0.092 | 0 | 0.81 | 0.96 | 0.64 | 0.37 | 0.69 | 0.79 |
| 7 | 88813289 | rs2214339 | 2.327 | 0.147 | -1.202 | -2.364 | 0.063 | 0.141 | 0.141 | 0.214 | 0.32 | 0.13 | 0.14 | 0.08 | 0.2 | 0.24 |
| 7 | 88815895 | rs6465186 | 0.594 | 1.297 | -0.693 | -1.829 | 0.006 | -0.051 | -0.051 | 0.164 | 0.62 | 0.52 | 0.45 | 0.22 | 0.42 | 0.53 |
| 7 | 88817483 | rs10275638 | 0.979 | 1.744 | -0.811 | -1.379 | 0.004 | -0.009 | -0.009 | 0 | 0.16 | 0.17 | 0.25 | 0.11 | 0.2 | 0.19 |
| 7 | 88818755 | rs10085897 | 0.1 | 1.748 | -0.427 | -0.893 | 0.007 | 0.003 | 0.003 | 0 | 0.8 | 0.75 | 0.61 | 0.31 | 0.69 | 0.78 |
| 7 | 88821660 | rs10277089 | 0.329 | 1.382 | -0.378 | -0.99 | 0.041 | -0.021 | -0.021 | 0 | 0.07 | 0.16 | 0.06 | 0.03 | 0.11 | 0.08 |
| 7 | 88830659 | rs757153 | 0.398 | 0.32 | -0.824 | -1.605 | 0.001 | -0.027 | -0.027 | 0 | 0.19 | 0.15 | 0.1 | 0.03 | 0.12 | 0.14 |
| 7 | 88848044 | rs10279101 | 1.475 | 0.475 | -0.957 | -1.958 | 0.01 | 0.047 | 0.047 | 0 | 0.55 | 0.6 | 0.51 | 0.25 | 0.57 | 0.55 |
| 7 | 88850732 | rs17167557 | 2.258 | 0.006 | -1.487 | -2.432 | 0.018 | 0.081 | 0.081 | 0 | 0.13 | 0 | 0.04 | 0 | 0.06 | 0.1 |
| 7 | 88852488 | rs1008594 | -0.401 | 0.72 | -0.966 | -1.933 | -0.006 | -0.004 | -0.004 | 0.191 | 0.78 | 0.78 | 0.64 | 0.29 | 0.74 | 0.72 |
| 7 | 88868129 | rs11984375 | 1.982 | 0.158 | -0.557 | -2.01 | 0.003 | -0.001 | -0.001 | 0.157 | 0.37 | 0.35 | 0.21 | 0.13 | 0.3 | 0.35 |
| 7 | 88868981 | rs13223868 | 0.497 | 0.026 | -0.516 | -1.85 | -0.039 | -0.096 | -0.096 | 0.04 | 0.22 | 0.34 | 0.11 | 0.03 | 0.12 | 0.14 |
| 7 | 88890500 | rs1208835 | -0.3 | 0.301 | -0.674 | -1.768 | -0.031 | -0.005 | -0.005 | 0.919 | 0.77 | 0.86 | 0.61 | 0.28 | 0.69 | 0.69 |
| 7 | 88890591 | rs7783629 | 1.145 | 0.237 | -0.569 | -1.964 | -0.002 | 0.031 | 0.031 | 0.182 | 0.59 | 0.69 | 0.32 | 0.15 | 0.42 | 0.49 |
| 7 | 88895753 | rs2708619 | 1.454 | 0.398 | -0.743 | -1.709 | -0.023 | -0.061 | -0.061 | 3.081 | 0.11 | 0.03 | 0.28 | 0.62 | 0.11 | 0.22 |
| 7 | 88904674 | rs1208853 | -0.982 | 0.283 | -0.537 | -2.029 | -0.003 | -0.08 | -0.08 | 0.229 | 0.39 | 0.46 | 0.4 | 0.16 | 0.4 | 0.35 |

| | | | | | | | | | | | | | | | | |
|---|---|---|---|---|---|---|---|---|---|---|---|---|---|---|---|---|
| 7 | 88904777 | rs1208854 | 0.645 | 0.687 | -0.594 | -1.756 | 0.016 | -0.012 | -0.012 | 0.241 | 0.18 | 0.19 | 0.29 | 0.13 | 0.27 | 0.21 |
| 7 | 88919393 | rs7778127 | 1.581 | 0.291 | -0.809 | -1.904 | 0.001 | -0.002 | -0.002 | 0 | 0.35 | 0.32 | 0.21 | 0.12 | 0.29 | 0.31 |
| 7 | 88919955 | rs1208888 | 0.829 | 0.867 | -0.894 | -1.168 | -0.037 | 0.063 | 0.063 | 0 | 0.69 | 0.82 | 0.39 | 0.17 | 0.54 | 0.55 |
| 7 | 88924014 | rs2030662 | 0.305 | 1.181 | -0.709 | -0.835 | 0.008 | -0.007 | -0.007 | 1.264 | 0.16 | 0.14 | 0.33 | 0.1 | 0.27 | 0.2 |
| 7 | 88931661 | rs17168262 | 0.827 | 1.034 | -0.761 | -0.925 | 0.006 | 0.023 | 0.023 | 0 | 0.12 | 0.03 | 0.06 | 0 | 0.08 | 0.07 |
| 7 | 88933320 | rs13239763 | -0.079 | 1.475 | -0.444 | -0.054 | -0.009 | 0.021 | 0.021 | 0 | 0.05 | 0.04 | 0.21 | 0.67 | 0.06 | 0.21 |
| 7 | 88935494 | rs38930 | -0.894 | 1.484 | -0.418 | -0.162 | 0.009 | -0.014 | -0.014 | 0 | 0.22 | 0.25 | 0.38 | 0.11 | 0.38 | 0.24 |
| 7 | 88945484 | rs38948 | -0.155 | 0.491 | -0.41 | 0.356 | 0.003 | -0.014 | -0.014 | 0 | 0.05 | 0.11 | 0.05 | 0 | 0.11 | 0.04 |
| 7 | 88969883 | rs801844 | -0.695 | -0.97 | -0.793 | 0.889 | 0.056 | 0.027 | 0.027 | 0 | 0.39 | 0.78 | 0.36 | 0.72 | 0.23 | 0.43 |
| 7 | 88974907 | rs1013540 | -0.564 | -1.062 | 0.277 | 0.525 | -0.011 | -0.015 | -0.015 | 0 | 0.34 | 0.41 | 0.16 | 0.23 | 0.22 | 0.25 |
| 7 | 88981054 | rs6965360 | 0.691 | -0.945 | -0.146 | 0.059 | -0.008 | 0.033 | 0.033 | 0 | 0.08 | 0.01 | 0.06 | 0 | 0.08 | 0.03 |
| 7 | 88995297 | rs801873 | -0.204 | -1.367 | 0.902 | 0.115 | -0.043 | -0.019 | -0.019 | 3.793 | 0.12 | 0.28 | 0.06 | 0.05 | 0.06 | 0.05 |
| 7 | 89011745 | rs6966330 | 1.404 | -1.718 | 0.692 | 0.142 | 0.067 | 0.028 | 0.028 | 0.244 | 0.56 | 0.19 | 0.57 | 0.31 | 0.69 | 0.57 |
| 7 | 89016595 | rs801900 | 0.518 | -1.154 | 0.911 | 0.441 | -0.008 | 0.035 | 0.035 | 0 | 0.71 | 0.65 | 0.65 | 0.37 | 0.75 | 0.63 |
| 7 | 89020306 | rs801904 | 0.479 | -1.098 | 0.438 | 0.433 | 0 | -0.004 | -0.004 | 1.017 | 0.72 | 0.75 | 0.66 | 0.37 | 0.74 | 0.62 |
| 7 | 89031107 | rs6960193 | 0.468 | -1.089 | 0.411 | 0.444 | 0.002 | -0.027 | -0.027 | 0 | 0.72 | 0.81 | 0.66 | 0.39 | 0.74 | 0.61 |
| 7 | 89034943 | rs929192 | 0.276 | -0.996 | 0.364 | 0.511 | 0 | 0.013 | 0.013 | 0.028 | 0.77 | 0.75 | 0.69 | 0.39 | 0.77 | 0.61 |
| 7 | 89060931 | rs10233998 | -0.914 | -2.255 | 0.442 | -0.237 | -0.102 | -0.04 | -0.04 | 0 | 0.24 | 0.39 | 0.09 | 0.13 | 0.08 | 0.13 |
| 7 | 89061690 | rs39000 | -0.254 | -2.188 | 0.879 | -0.408 | 0.057 | 0.001 | 0.001 | 0.124 | 0.06 | 0.25 | 0.07 | 0 | 0.06 | 0.03 |
| 7 | 89077204 | rs39011 | -0.341 | -2.358 | 0.193 | -0.957 | 0.004 | 0.001 | 0.001 | 0 | 0.69 | 0.75 | 0.72 | 0.73 | 0.74 | 0.8 |
| 7 | 89086638 | rs2373919 | 1.76 | -1.825 | -1.135 | -1.505 | -0.024 | -0.079 | -0.079 | 0 | 0.26 | 0.08 | 0.37 | 0.59 | 0.27 | 0.49 |
| 7 | 89098791 | rs39031 | -2.211 | -1.798 | -1.187 | -1.533 | -0.047 | -0.017 | -0.017 | 0 | 0.82 | 0.96 | 0.76 | 0.82 | 0.77 | 0.89 |
| 7 | 89099843 | rs39033 | -0.654 | -1.703 | -0.847 | -1.606 | 0.003 | -0.039 | -0.039 | 8.952 | 0.4 | 0.41 | 0.33 | 0.14 | 0.48 | 0.3 |
| 7 | 89103561 | rs17161528 | -0.212 | -1.551 | -0.85 | -1.477 | 0.036 | -0.071 | -0.071 | 0 | 0.31 | 0.37 | 0.29 | 0.11 | 0.43 | 0.27 |
| 7 | 89104802 | rs17617686 | 0.383 | -1.217 | -1.242 | -1.401 | -0.069 | 0.07 | 0.07 | 0.799 | 0.24 | 0.09 | 0.22 | 0.11 | 0.35 | 0.23 |
| 7 | 89107084 | rs12535393 | 1.373 | -1.031 | -1.102 | -1.234 | -0.032 | -0.07 | -0.07 | 0.645 | 0.26 | 0.09 | 0.37 | 0.6 | 0.28 | 0.51 |
| 7 | 89111561 | rs2028008 | 0.37 | -0.432 | -0.893 | -1.229 | -0.035 | 0.04 | 0.04 | 0.257 | 0.19 | 0.09 | 0.19 | 0.11 | 0.28 | 0.19 |
| 7 | 89115474 | rs39043 | -0.678 | -0.696 | -0.388 | -1.129 | -0.002 | 0.103 | 0.103 | 0 | 0.48 | 0.69 | 0.37 | 0.23 | 0.5 | 0.35 |
| 7 | 89116491 | rs6465197 | 0.015 | -0.364 | -0.266 | -0.373 | 0.003 | 0 | 0 | 0 | 0.41 | 0.62 | 0.5 | 0.58 | 0.35 | 0.55 |
| 7 | 89116966 | rs7799830 | 0.294 | -0.642 | 0.222 | -0.5 | -0.003 | -0.014 | -0.014 | 0 | 0.56 | 0.53 | 0.58 | 0.67 | 0.55 | 0.72 |
| 7 | 89118409 | rs4727203 | -2.183 | -0.142 | 1.149 | 0.02 | -0.03 | 0.028 | 0.028 | 4.469 | 0.13 | 0.74 | 0.2 | 0.18 | 0.05 | 0.14 |
| 7 | 89129100 | rs6465198 | -0.558 | 0.131 | 1.585 | -0.162 | 0.026 | 0.009 | 0.009 | 0 | 0.14 | 0.39 | 0.17 | 0.13 | 0.13 | 0.14 |
| 7 | 89137422 | rs983218 | -1.803 | 0.093 | 1.141 | 0.047 | 0.073 | 0.052 | 0.052 | 0.087 | 0.57 | 0.85 | 0.65 | 0.74 | 0.63 | 0.69 |
| 7 | 89150909 | rs11975137 | -0.288 | 1.127 | 1.213 | 0.568 | 0.006 | -0.016 | -0.016 | 0.016 | 0.12 | 0.22 | 0.15 | 0.09 | 0.12 | 0.09 |
| 7 | 89161129 | rs4489249 | -0.288 | 1.142 | 0.828 | 0.58 | 0.008 | -0.016 | -0.016 | 0 | 0.12 | 0.22 | 0.15 | 0.09 | 0.12 | 0.09 |
| 7 | 89184131 | rs12056158 | -0.271 | 1.131 | 0.652 | 0.757 | -0.116 | -0.103 | -0.103 | 0.241 | 0.3 | 0.07 | 0.43 | 0.6 | 0.48 | 0.47 |
| 7 | 89190648 | rs17161956 | -0.632 | 1.11 | 0.992 | 0.713 | 0.019 | -0.041 | -0.041 | 0 | 0.15 | 0.45 | 0.17 | 0.09 | 0.12 | 0.09 |
| 7 | 89195515 | rs1072107 | 0.739 | 1.37 | 0.929 | 0.971 | -0.013 | -0.017 | -0.017 | 0.462 | 0.07 | 0.03 | 0.14 | 0.21 | 0.09 | 0.09 |
| 7 | 89198032 | rs3922360 | 0.042 | 0.412 | 1.102 | 0.423 | -0.116 | -0.101 | -0.101 | 0 | 0.3 | 0.07 | 0.43 | 0.6 | 0.48 | 0.48 |
| 7 | 89202570 | rs10236388 | -1.752 | 0.434 | 2.115 | 0.542 | 0.006 | -0.008 | -0.008 | 0 | 0.16 | 0.64 | 0.19 | 0.14 | 0.13 | 0.15 |
| 7 | 89254213 | rs2106740 | -1.121 | 0.76 | 0.591 | 0.373 | 0.022 | -0.036 | -0.036 | 0 | 0.13 | 0.29 | 0.15 | 0.09 | 0.12 | 0.07 |
| 7 | 89271899 | rs10263640 | -1.269 | 0.66 | 0.362 | 0.404 | 0.019 | -0.032 | -0.032 | 0 | 0.14 | 0.33 | 0.15 | 0.11 | 0.13 | 0.07 |
| 7 | 89295268 | rs11764742 | -0.369 | -0.222 | -0.337 | 0.709 | 0.021 | -0.048 | -0.048 | 0 | 0.29 | 0.14 | 0.25 | 0.45 | 0.2 | 0.29 |
| 7 | 89311159 | rs10250195 | -0.557 | -0.301 | 0.141 | 0.673 | -0.017 | 0.017 | 0.017 | 0.173 | 0.32 | 0.61 | 0.28 | 0.45 | 0.2 | 0.29 |
| 7 | 89321267 | rs2214219 | -0.881 | -0.408 | -0.901 | 0.713 | 0.017 | -0.113 | -0.113 | 0 | 0.75 | 0.99 | 0.75 | 0.7 | 0.84 | 0.73 |
| 7 | 89332591 | rs916761 | 0.888 | 0.908 | 0.433 | 0.877 | 0.034 | 0.033 | 0.033 | 0.482 | 0.77 | 0.42 | 0.67 | 0.66 | 0.63 | 0.69 |
| 7 | 89333043 | rs2521319 | 0.03 | 1.024 | 1.022 | 0.945 | 0.028 | 0.018 | 0.018 | 0 | 0.85 | 0.75 | 0.76 | 0.81 | 0.72 | 0.75 |
| 7 | 89334668 | rs2724463 | 0.855 | 1.018 | 1.367 | 0.94 | 0.069 | 0.067 | 0.067 | 0.037 | 0.79 | 0.6 | 0.68 | 0.66 | 0.63 | 0.71 |
| 7 | 89335357 | rs10247416 | 0.005 | 0.942 | 1.103 | 0.914 | 0.031 | 0.053 | 0.053 | 0.031 | 0.07 | 0.17 | 0.07 | 0.14 | 0.08 | 0.05 |
| 7 | 89338128 | rs2724452 | 1.65 | 0.874 | 0.585 | 0.736 | -0.009 | -0.01 | -0.01 | 0 | 0.46 | 0.16 | 0.52 | 0.37 | 0.37 | 0.39 |
| 7 | 89338551 | rs6944957 | -0.57 | 1.012 | 0.417 | 0.653 | -0.025 | -0.015 | -0.015 | 0 | 0.87 | 0.92 | 0.77 | 0.81 | 0.71 | 0.75 |
| 7 | 89340308 | rs1014736 | -0.454 | 0.534 | 0.607 | 0.384 | -0.022 | -0.134 | -0.134 | 0 | 0.19 | 0.55 | 0.1 | 0 | 0.16 | 0.09 |
| 7 | 89341627 | rs968907 | -0.857 | 0.658 | -0.197 | 0.606 | 0.01 | 0.001 | 0.001 | 0 | 0.32 | 0.27 | 0.16 | 0.33 | 0.26 | 0.32 |
| 7 | 89341720 | rs968908 | 0.74 | 1.253 | -0.242 | 0.437 | -0.046 | 0.021 | 0.021 | 1.334 | 0.64 | 0.19 | 0.59 | 0.67 | 0.47 | 0.62 |
| 7 | 89345351 | rs2188679 | 0.821 | 1.267 | 0.172 | 0.106 | 0.011 | -0.049 | -0.049 | 0 | 0.65 | 0.48 | 0.65 | 0.82 | 0.52 | 0.66 |
| 7 | 89345537 | rs6944565 | -1.431 | 1.196 | 0.475 | 0.189 | 0.008 | -0.019 | -0.019 | 0 | 0.13 | 0.22 | 0.15 | 0.04 | 0.16 | 0.14 |
| 7 | 89352860 | rs2521310 | -1.516 | 1.57 | 1.082 | 0.144 | 0.029 | -0.016 | -0.016 | 0.107 | 0.17 | 0.37 | 0.12 | 0.16 | 0.15 | 0.2 |
| 7 | 89353960 | rs10281928 | 1.195 | 1.803 | 0.933 | 0.277 | 0.029 | 0.027 | 0.027 | 0.145 | 0.63 | 0.5 | 0.47 | 0.55 | 0.53 | 0.46 |
| 7 | 89365778 | rs1024424 | -2.121 | 1.466 | 1.454 | 0.56 | -0.006 | 0.072 | 0.072 | 0.358 | 0.24 | 0.48 | 0.25 | 0.4 | 0.17 | 0.47 |
| 7 | 89372040 | rs1548440 | -0.144 | 1.785 | 1.561 | 0.531 | 0.007 | 0.014 | 0.014 | 0 | 0.17 | 0.26 | 0.3 | 0.19 | 0.21 | 0.15 |
| 7 | 89373249 | rs12667937 | 2.454 | 1.808 | 1.664 | 0.492 | 0.021 | 0.025 | 0.025 | 0.809 | 0.69 | 0.48 | 0.71 | 0.59 | 0.69 | 0.44 |
| 7 | 89375225 | rs4520094 | -1.189 | 1.777 | 1.802 | 0.525 | -0.004 | 0.325 | 0.325 | 0.318 | 0.08 | 0.36 | 0.15 | 0.4 | 0.02 | 0.27 |
| 7 | 89377947 | rs6980440 | 0.044 | 1.807 | 1.704 | 0.512 | -0.021 | 0.022 | 0.022 | 0 | 0.92 | 0.97 | 0.95 | 1 | 0.86 | 0.9 |
| 7 | 89382342 | rs4728833 | 1.607 | 1.69 | 1.787 | 0.473 | 0.027 | 0.198 | 0.198 | 0 | 0.84 | 0.52 | 0.78 | 0.6 | 0.84 | 0.63 |
| 7 | 89384704 | rs6958119 | -0.251 | 1.758 | 1.686 | 0.444 | 0.027 | -0.026 | -0.026 | 4.415 | 0.05 | 0.12 | 0.05 | 0 | 0.09 | 0.04 |
| 7 | 89386626 | rs17618989 | -0.352 | 1.749 | 1.667 | 0.439 | 0.015 | -0.021 | -0.021 | 0.024 | 0.05 | 0.09 | 0.05 | 0 | 0.09 | 0.04 |
| 7 | 89387363 | rs9942677 | 2.144 | 1.707 | 1.705 | 0.49 | 0 | 0 | 0 | 0 | 0.65 | 0.33 | 0.46 | 0.56 | 0.56 | 0.47 |
| 7 | 89387626 | rs7350025 | 0.059 | 1.673 | 1.759 | 0.409 | 0.008 | 0.014 | 0.014 | 0 | 0.16 | 0.2 | 0.29 | 0.19 | 0.2 | 0.14 |
| 7 | 89392021 | rs13437729 | -0.502 | 1.702 | 1.797 | 0.379 | -0.009 | -0.07 | -0.07 | 0 | 0.11 | 0.26 | 0.07 | 0 | 0.06 | 0.15 |

| | | | | | | | | | | | | | | | | |
|---|---|---|---|---|---|---|---|---|---|---|---|---|---|---|---|---|
| 7 | 89397701 | rs10272254 | 1.572 | 1.725 | 1.713 | 0.409 | 0.006 | -0.014 | -0.014 | 0 | 0.65 | 0.59 | 0.6 | 0.8 | 0.59 | 0.6 |
| 7 | 89409167 | rs6947709 | 1.977 | 1.76 | 1.9 | 0.394 | 0.002 | -0.003 | -0.003 | 0 | 0.6 | 0.45 | 0.55 | 0.65 | 0.55 | 0.58 |
| 7 | 89409488 | rs6948429 | -0.412 | 1.617 | 1.734 | 0.349 | 0 | -0.002 | -0.002 | 0 | 0.2 | 0.19 | 0.29 | 0.19 | 0.21 | 0.15 |
| 7 | 89410098 | rs12704484 | 2.2 | 1.821 | 1.837 | 0.549 | 0.027 | 0.009 | 0.009 | 1.834 | 0.76 | 0.35 | 0.69 | 0.73 | 0.71 | 0.49 |
| 7 | 89432925 | rs6465211 | 1.759 | 1.867 | 1.819 | 0.563 | 0.008 | -0.021 | -0.021 | 0 | 0.6 | 0.51 | 0.56 | 0.81 | 0.5 | 0.57 |
| 7 | 89441028 | rs10248450 | 1.69 | 1.818 | 1.822 | 0.546 | 0.011 | -0.029 | -0.029 | 0 | 0.6 | 0.47 | 0.55 | 0.8 | 0.49 | 0.57 |
| 7 | 89444167 | rs1858966 | -0.783 | 1.93 | 1.826 | 0.497 | -0.004 | -0.002 | -0.002 | 0 | 0.25 | 0.27 | 0.27 | 0.18 | 0.21 | 0.24 |
| 7 | 89449367 | rs10487085 | -0.402 | 1.791 | 1.875 | 0.537 | 0.007 | -0.02 | -0.02 | 0.052 | 0.1 | 0.15 | 0.13 | 0.01 | 0.17 | 0.11 |
| 7 | 89449405 | rs7805617 | 1.641 | 1.785 | 1.877 | 0.543 | 0.01 | -0.028 | -0.028 | 0 | 0.6 | 0.49 | 0.56 | 0.81 | 0.49 | 0.57 |
| 7 | 89453630 | rs916759 | 1.642 | 1.79 | 1.762 | 0.546 | 0.009 | -0.024 | -0.024 | 0 | 0.6 | 0.5 | 0.56 | 0.8 | 0.49 | 0.57 |
| 7 | 89459030 | rs6960013 | 2.373 | 1.54 | 1.318 | 0.29 | -0.004 | -0.007 | -0.007 | 0.125 | 0.44 | 0.08 | 0.31 | 0.4 | 0.4 | 0.28 |
| 7 | 89460928 | rs2023739 | 2.228 | 1.554 | 1.37 | 0.271 | 0.039 | 0.046 | 0.046 | 0 | 0.47 | 0.27 | 0.46 | 0.65 | 0.4 | 0.52 |
| 7 | 89474743 | rs10215711 | 0.704 | 1.39 | 1.278 | 0.3 | 0.004 | -0.012 | -0.012 | 0 | 0.69 | 0.62 | 0.69 | 0.8 | 0.66 | 0.68 |
| 7 | 89482839 | rs2188677 | 0.83 | 1.444 | 1.215 | 0.163 | 0 | 0.01 | 0.01 | 0.284 | 0.68 | 0.66 | 0.57 | 0.56 | 0.66 | 0.44 |
| 7 | 89489616 | rs12671169 | 2.289 | 1.424 | 1.018 | 0.155 | 0.014 | 0.019 | 0.019 | 0 | 0.44 | 0.13 | 0.31 | 0.39 | 0.4 | 0.28 |
| 7 | 89509761 | rs12668274 | 0.922 | 1.273 | 1.119 | 0.3 | -0.002 | -0.022 | -0.022 | 0 | 0.68 | 0.71 | 0.57 | 0.55 | 0.66 | 0.44 |
| 7 | 89512049 | rs12535395 | 0.051 | 1.283 | 0.938 | 0.317 | 0.001 | -0.002 | -0.002 | 0 | 0.12 | 0.1 | 0.09 | 0.15 | 0.1 | 0.05 |
| 7 | 89516662 | rs2079890 | 1.744 | 1.298 | 0.964 | 0.244 | 0 | 0.002 | 0.002 | 0 | 0.55 | 0.44 | 0.47 | 0.4 | 0.56 | 0.39 |
| 7 | 89532912 | rs11487032 | -0.071 | 1.174 | 0.543 | 0.318 | 0.001 | 0.003 | 0.003 | 0.71 | 0.12 | 0.1 | 0.08 | 0.09 | 0.1 | 0.04 |
| 7 | 89534606 | rs7781928 | 0.814 | 1.175 | 0.951 | 0.27 | 0 | 0.024 | 0.024 | 0 | 0.72 | 0.65 | 0.57 | 0.49 | 0.7 | 0.51 |
| 7 | 89536358 | rs10255948 | 1.342 | 1.246 | 0.875 | 0.244 | -0.002 | -0.004 | -0.004 | 0.674 | 0.6 | 0.54 | 0.49 | 0.4 | 0.6 | 0.47 |
| 7 | 89538107 | rs1981681 | 2.097 | 1.474 | 0.89 | 0.105 | -0.002 | -0.002 | -0.002 | 1.124 | 0.47 | 0.44 | 0.35 | 0.39 | 0.4 | 0.28 |
| 7 | 89538623 | rs17686777 | -0.282 | 1.616 | 0.759 | 0.189 | -0.019 | 0.009 | 0.009 | 0.529 | 0.13 | 0.1 | 0.14 | 0.01 | 0.2 | 0.19 |
| 7 | 89544278 | rs6972809 | -1.358 | 1.388 | 1.169 | 0.426 | 0.013 | 0.04 | 0.04 | 0 | 0.26 | 0.56 | 0.38 | 0.41 | 0.28 | 0.47 |
| 7 | 89544313 | rs6972943 | -1.361 | 1.423 | 1.205 | 0.472 | 0.005 | 0.004 | 0.004 | 0 | 0.25 | 0.53 | 0.34 | 0.26 | 0.25 | 0.45 |
| 7 | 89553014 | rs9640589 | -0.227 | 1.322 | 1.478 | 0.57 | -0.001 | 0.002 | 0.002 | 0 | 0.17 | 0.2 | 0.24 | 0.18 | 0.17 | 0.11 |
| 7 | 89556164 | rs4728843 | 0.394 | 1.139 | 1.522 | 0.555 | 0.006 | -0.007 | -0.007 | 0 | 0.1 | 0.07 | 0.17 | 0.25 | 0.06 | 0.24 |
| 7 | 89557548 | rs7801279 | -1.667 | 1.047 | 1.723 | 0.412 | -0.009 | 0.01 | 0.01 | 0 | 0.31 | 0.62 | 0.47 | 0.59 | 0.26 | 0.37 |
| 7 | 89560253 | rs6946219 | -0.198 | 1.062 | 1.566 | 0.425 | -0.001 | 0.005 | 0.005 | 0 | 0.16 | 0.2 | 0.24 | 0.19 | 0.16 | 0.11 |
| 7 | 89564164 | rs1990394 | -0.091 | 0.529 | 1.345 | 0.324 | -0.017 | -0.034 | -0.034 | 0 | 0.06 | 0.16 | 0.03 | 0 | 0.02 | 0.07 |
| 7 | 89570624 | rs17769458 | 0.411 | 0.542 | 0.863 | 0.464 | 0.001 | 0.014 | 0.014 | 1.36 | 0.09 | 0.08 | 0.17 | 0.29 | 0.03 | 0.22 |
| 7 | 89572966 | rs1990395 | 0.81 | 0.684 | 0.387 | -0.224 | 0.008 | -0.006 | -0.006 | 0 | 0.61 | 0.43 | 0.61 | 0.68 | 0.53 | 0.7 |
| 7 | 89578317 | rs6465218 | -1.202 | 0.883 | 0.25 | -0.327 | -0.075 | 0.076 | 0.076 | 0 | 0.86 | 0.99 | 0.91 | 0.99 | 0.76 | 0.88 |
| 7 | 89580280 | rs2374079 | -0.364 | 0.684 | 0.739 | -0.228 | -0.001 | 0.033 | 0.033 | 0.216 | 0.08 | 0.23 | 0.15 | 0.13 | 0.05 | 0.09 |
| 7 | 89582420 | rs6465220 | -1.09 | 0.67 | 0.54 | -0.383 | -0.092 | 0.108 | 0.108 | 0 | 0.82 | 0.99 | 0.9 | 0.99 | 0.71 | 0.87 |
| 7 | 89584264 | rs6971802 | 0.712 | 0.19 | 0.708 | -0.225 | -0.023 | 0.055 | 0.055 | 1.575 | 0.11 | 0.15 | 0.18 | 0.31 | 0.05 | 0.25 |
| 7 | 89595954 | rs10487089 | 0.529 | -0.094 | 0.489 | -0.595 | 0 | -0.03 | -0.03 | 0 | 0.17 | 0.17 | 0.17 | 0.01 | 0.22 | 0.2 |
| 7 | 89601247 | rs6968049 | 1.263 | -0.108 | 0.583 | -0.578 | 0.018 | 0.003 | 0.003 | 0 | 0.44 | 0.23 | 0.39 | 0.55 | 0.39 | 0.34 |
| 7 | 89611841 | rs6959993 | -0.17 | 0.209 | 0.394 | -0.687 | 0.022 | -0.055 | -0.055 | 0 | 0.73 | 0.59 | 0.74 | 0.87 | 0.66 | 0.78 |
| 7 | 89621311 | rs12704506 | 0.03 | 0.12 | 0.296 | -0.435 | 0.008 | -0.052 | -0.052 | 0 | 0.17 | 0.2 | 0.17 | 0.01 | 0.22 | 0.2 |
| 7 | 89622961 | rs17162736 | 0.016 | 0.033 | -0.119 | -0.107 | -0.003 | -0.038 | -0.038 | 0.014 | 0.15 | 0.06 | 0.09 | 0.19 | 0.15 | 0.11 |
| 7 | 89623271 | rs11975637 | 0.452 | -0.087 | 0.314 | -0.215 | 0.006 | -0.014 | -0.014 | 0 | 0.05 | 0.2 | 0.03 | 0 | 0.01 | 0.07 |
| 7 | 89623326 | rs10487092 | 0.693 | -0.335 | 0.278 | -0.071 | -0.008 | 0.023 | 0.023 | 1.909 | 0.11 | 0.11 | 0.18 | 0.3 | 0.05 | 0.24 |
| 7 | 89624173 | rs13230930 | -0.012 | -0.334 | 0.344 | -0.288 | 0.034 | -0.114 | -0.114 | 0.515 | 0.18 | 0.39 | 0.19 | 0.02 | 0.23 | 0.21 |
| 7 | 89626246 | rs9987049 | 0.437 | -0.549 | -0.501 | -0.042 | -0.024 | 0.036 | 0.036 | 2.645 | 0.13 | 0.17 | 0.19 | 0.3 | 0.05 | 0.24 |
| 7 | 89626611 | rs259152 | -0.774 | -0.028 | -0.145 | 0.263 | -0.001 | 0.019 | 0.019 | 1.427 | 0.27 | 0.68 | 0.4 | 0.47 | 0.25 | 0.36 |
| 7 | 89629174 | rs17769931 | -0.368 | 0.163 | -0.454 | 0.345 | 0.007 | -0.033 | -0.033 | 0 | 0.16 | 0.06 | 0.11 | 0.19 | 0.13 | 0.1 |
| 7 | 89635264 | rs10487094 | -0.746 | 0.452 | -0.146 | 0.323 | -0.001 | 0.004 | 0.004 | 0 | 0.18 | 0.22 | 0.12 | 0.17 | 0.17 | 0.13 |
| 7 | 89635741 | rs10259230 | -1.436 | 0.846 | 0.229 | -0.041 | 0.027 | -0.022 | -0.022 | 0.078 | 0.32 | 0.47 | 0.32 | 0.2 | 0.37 | 0.26 |
| 7 | 89635799 | rs12670363 | 1.281 | -0.114 | 0.069 | -0.099 | 0.003 | -0.005 | -0.005 | 0 | 0.25 | 0.21 | 0.33 | 0.43 | 0.19 | 0.13 |
| 7 | 89636224 | rs7798970 | -0.699 | -0.072 | 0.098 | -0.094 | -0.009 | 0.028 | 0.028 | 0 | 0.55 | 0.68 | 0.63 | 0.63 | 0.53 | 0.38 |
| 7 | 89636404 | rs7799411 | 0.298 | -0.146 | 0.162 | -0.139 | -0.011 | -0.023 | -0.023 | 0 | 0.38 | 0.23 | 0.51 | 0.45 | 0.39 | 0.25 |
| 7 | 89641818 | rs17162755 | 0.246 | -0.145 | 0.174 | -0.131 | -0.012 | 0.02 | 0.02 | 0.143 | 0.17 | 0.29 | 0.11 | 0.18 | 0.14 | 0.12 |
| 7 | 89645073 | rs4466326 | 0.277 | -0.177 | 0.169 | -0.17 | -0.009 | -0.02 | -0.02 | 0 | 0.37 | 0.26 | 0.51 | 0.45 | 0.38 | 0.26 |
| 7 | 89645264 | rs10452943 | 1.667 | -0.231 | 0.112 | -0.221 | 0.018 | 0.099 | 0.099 | 0 | 0.22 | 0.06 | 0.15 | 0 | 0.18 | 0.09 |
| 7 | 89647264 | rs17162758 | 0.1 | -0.213 | 0.326 | -0.156 | -0.003 | -0.006 | -0.006 | 3.099 | 0.39 | 0.43 | 0.26 | 0.19 | 0.32 | 0.21 |
| 7 | 89647400 | rs740291 | 0.214 | -0.039 | 0.249 | 0.029 | 0.011 | 0.049 | 0.049 | 0 | 0.15 | 0.19 | 0.36 | 0.44 | 0.21 | 0.17 |
| 7 | 89653769 | rs10245062 | 0.116 | -0.049 | 0.074 | 0.014 | 0.045 | 0.141 | 0.141 | 0 | 0.16 | 0.29 | 0.37 | 0.45 | 0.21 | 0.17 |
| 7 | 89673676 | rs7793198 | -1.168 | -0.471 | -0.015 | -0.24 | -0.007 | 0.03 | 0.03 | 1.142 | 0.4 | 0.78 | 0.51 | 0.64 | 0.36 | 0.37 |
| 7 | 89680075 | rs1024557 | 0.087 | -1.036 | -0.003 | -0.124 | -0.004 | 0.001 | 0.001 | 0 | 0.36 | 0.37 | 0.42 | 0.41 | 0.28 | 0.21 |
| 7 | 89681721 | rs12671408 | 1.522 | -1.003 | 0.017 | -0.102 | 0.018 | -0.066 | -0.066 | 0 | 0.21 | 0.11 | 0.29 | 0.4 | 0.18 | 0.1 |
| 7 | 89682186 | rs10264568 | 0.659 | -0.803 | 0.138 | -0.1 | -0.005 | -0.071 | -0.071 | 0 | 0.12 | 0.26 | 0.13 | 0.01 | 0.1 | 0.07 |
| 7 | 89683536 | rs4380852 | -0.07 | -0.67 | 0.096 | 0.095 | -0.003 | 0.002 | 0.002 | 0.061 | 0.21 | 0.24 | 0.17 | 0.21 | 0.17 | 0.17 |
| 7 | 89684293 | rs6952332 | -0.262 | -0.681 | 0.094 | -0.046 | -0.004 | 0.008 | 0.008 | 0 | 0.22 | 0.27 | 0.31 | 0.6 | 0.18 | 0.22 |
| 7 | 89684555 | rs6952664 | -0.549 | -0.348 | 0.069 | 0.176 | 0 | 0.011 | 0.011 | 0 | 0.57 | 0.64 | 0.59 | 0.62 | 0.45 | 0.39 |
| 7 | 89690275 | rs757951 | -0.765 | -0.861 | 0.287 | 0.671 | -0.002 | -0.001 | -0.001 | 0 | 0.34 | 0.64 | 0.49 | 0.76 | 0.23 | 0.54 |
| 7 | 89717332 | rs2190399 | -1.045 | -0.865 | 0.233 | 0.697 | 0.063 | 0.236 | 0.236 | 0 | 0.51 | 0.79 | 0.67 | 0.9 | 0.39 | 0.64 |
| 7 | 89724651 | rs717922 | -0.107 | -0.477 | 0.321 | 0.632 | -0.016 | -0.062 | -0.062 | 0 | 0.11 | 0.21 | 0.07 | 0 | 0.07 | 0.2 |
| 7 | 89724772 | rs17770685 | -0.303 | -0.239 | -0.229 | 0.945 | 0.001 | -0.029 | -0.029 | 0 | 0.16 | 0.12 | 0.28 | 0.37 | 0.16 | 0.3 |

| | | | | | | | | | | | | | | | | |
|---|---|---|---|---|---|---|---|---|---|---|---|---|---|---|---|---|
| 7 | 89734282 | rs39203 | 0.83 | -1.167 | 0.413 | 0.517 | -0.063 | 0.001 | 0.001 | 0 | 0.72 | 0.53 | 0.61 | 0.28 | 0.85 | 0.51 |
| 7 | 89735315 | rs39206 | 0.619 | -1.187 | 0.57 | 0.804 | -0.041 | 0.037 | 0.037 | 0 | 0.74 | 0.71 | 0.68 | 0.35 | 0.85 | 0.65 |
| 7 | 89793393 | rs2051959 | -0.913 | -0.819 | 0.618 | 0.472 | -0.005 | 0.013 | 0.013 | 0 | 0.26 | 0.27 | 0.37 | 0.65 | 0.22 | 0.35 |
| 7 | 89793641 | rs2286254 | 0.029 | -0.831 | 0.79 | 0.464 | -0.026 | 0.047 | 0.047 | 0 | 0.18 | 0.21 | 0.28 | 0.65 | 0.09 | 0.32 |
| 7 | 89807586 | rs6465236 | -0.922 | -0.222 | 0.184 | -0.366 | 0.005 | 0.018 | 0.018 | 0.023 | 0.67 | 0.73 | 0.78 | 0.84 | 0.72 | 0.7 |
| 7 | 89813943 | rs6465237 | -1.329 | -0.263 | 0.184 | -0.385 | 0.003 | 0.057 | 0.057 | 0.048 | 0.69 | 0.83 | 0.84 | 0.91 | 0.72 | 0.84 |
| 7 | 89816334 | rs11563944 | -0.297 | 0.123 | 0.175 | -0.392 | -0.001 | 0.011 | 0.011 | 0 | 0.27 | 0.27 | 0.38 | 0.65 | 0.24 | 0.36 |
| 7 | 89816701 | rs11563564 | 0.774 | 0.114 | 0.125 | -0.357 | 0.001 | -0.014 | -0.014 | 0 | 0.4 | 0.46 | 0.41 | 0.19 | 0.51 | 0.35 |
| 7 | 89838707 | rs194499 | 1.067 | 0.105 | -0.008 | -0.371 | 0.006 | 0.045 | 0.045 | 0.027 | 0.32 | 0.2 | 0.16 | 0.09 | 0.28 | 0.17 |
| 7 | 89841084 | rs194503 | 0.123 | 0.106 | 0.099 | -0.347 | -0.005 | 0.02 | 0.02 | 0 | 0.74 | 0.7 | 0.61 | 0.35 | 0.78 | 0.65 |
| 7 | 89843560 | rs194507 | 0.557 | 0.208 | -0.048 | -0.361 | 0.001 | -0.006 | -0.006 | 0.208 | 0.42 | 0.49 | 0.45 | 0.26 | 0.51 | 0.49 |
| 7 | 89845903 | rs11563440 | 0.737 | 0.155 | -0.014 | -0.31 | -0.019 | 0.041 | 0.041 | 0 | 0.18 | 0.21 | 0.28 | 0.65 | 0.09 | 0.31 |
| 7 | 89848991 | rs7780514 | 1.792 | 0.137 | -0.065 | -0.357 | -0.004 | -0.028 | -0.028 | 0.089 | 0.27 | 0.25 | 0.14 | 0.08 | 0.22 | 0.15 |
| 7 | 89856644 | rs2016903 | 1.831 | 0.405 | -0.821 | -0.771 | -0.07 | 0.065 | 0.065 | 0.493 | 0.13 | 0.01 | 0.08 | 0.01 | 0.21 | 0.09 |
| 7 | 89857750 | rs10487105 | 0.977 | 0.37 | -0.741 | -0.766 | -0.022 | 0.048 | 0.048 | 0 | 0.18 | 0.23 | 0.28 | 0.65 | 0.09 | 0.31 |
| 7 | 89861832 | rs194524 | 0.255 | 0.608 | -0.984 | -1.011 | -0.036 | 0.089 | 0.089 | 0.02 | 0.4 | 0.27 | 0.39 | 0.18 | 0.51 | 0.34 |
| 7 | 89861890 | rs194525 | -1.43 | 0.967 | -1.066 | -0.773 | -0.036 | 0.037 | 0.037 | 0 | 0.92 | 0.97 | 0.9 | 1 | 0.87 | 0.96 |
| 7 | 89861957 | rs17621350 | 1.265 | 0.947 | -1.065 | -0.783 | -0.065 | 0.071 | 0.071 | 0.354 | 0.13 | 0.02 | 0.08 | 0.01 | 0.21 | 0.09 |
| 7 | 89862684 | rs194527 | -1.366 | 0.91 | -0.949 | -0.66 | -0.036 | 0.037 | 0.037 | 0 | 0.92 | 0.97 | 0.9 | 1 | 0.87 | 0.96 |
| 7 | 89862707 | rs11563439 | 1.219 | 0.747 | -0.874 | -0.581 | 0 | 0.029 | 0.029 | 0 | 0.06 | 0.01 | 0.02 | 0 | 0.04 | 0.01 |
| 7 | 89864417 | rs11563931 | 0.954 | 0.736 | -0.886 | -0.594 | 0 | 0.033 | 0.033 | 0 | 0.06 | 0.01 | 0.03 | 0 | 0.04 | 0.02 |
| 7 | 89864573 | rs1060194 | 1.215 | 0.995 | -0.941 | -0.57 | -0.072 | 0.038 | 0.038 | 0.313 | 0.13 | 0.01 | 0.08 | 0.02 | 0.21 | 0.11 |
| 7 | 89865661 | rs194531 | -1.02 | 0.777 | -0.558 | -0.346 | -0.034 | 0.033 | 0.033 | 0 | 0.92 | 0.95 | 0.9 | 1 | 0.87 | 0.96 |
| 7 | 89871132 | rs194534 | -0.413 | 0.498 | -0.613 | -0.523 | -0.02 | 0.054 | 0.054 | 0 | 0.83 | 0.93 | 0.86 | 1 | 0.83 | 0.82 |
| 7 | 89880949 | rs10240790 | 1.398 | -0.055 | -0.11 | -0.367 | -0.012 | -0.012 | -0.012 | 0.078 | 0.64 | 0.45 | 0.49 | 0.27 | 0.73 | 0.52 |
| 7 | 89884167 | rs11563921 | 0.455 | -0.1 | -0.147 | -0.383 | 0 | 0.04 | 0.04 | 0.093 | 0.11 | 0.05 | 0.06 | 0 | 0.09 | 0.06 |
| 7 | 89893945 | rs6951894 | -0.178 | -0.204 | 0.072 | -0.8 | -0.011 | 0.041 | 0.041 | 0 | 0.55 | 0.68 | 0.7 | 0.92 | 0.55 | 0.74 |
| 7 | 89895262 | rs10248104 | -1.009 | -0.205 | 0.062 | -0.805 | -0.004 | -0.005 | -0.005 | 0 | 0.65 | 0.73 | 0.76 | 0.92 | 0.65 | 0.8 |
| 7 | 89897703 | rs17863057 | 1.902 | -0.134 | -0.166 | -0.903 | -0.004 | -0.021 | -0.021 | 0 | 0.1 | 0.1 | 0.04 | 0 | 0.04 | 0.14 |
| 7 | 89899975 | rs17862139 | 0.812 | -0.11 | -0.194 | -0.889 | -0.013 | 0.058 | 0.058 | 0.088 | 0.1 | 0.01 | 0.06 | 0 | 0.09 | 0.06 |
| 7 | 89902533 | rs10226014 | 0.75 | -0.14 | -0.12 | -0.832 | -0.025 | 0.073 | 0.073 | 0.027 | 0.42 | 0.29 | 0.37 | 0.2 | 0.52 | 0.36 |
| 7 | 89912301 | rs1029365 | 0.44 | -0.1 | -0.15 | -0.834 | -0.007 | 0.003 | 0.003 | 0 | 0.8 | 0.88 | 0.85 | 1 | 0.79 | 0.9 |
| 7 | 89913269 | rs1015402 | 0.251 | -0.118 | -0.1 | -0.843 | 0 | 0.092 | 0.092 | 0.082 | 0.51 | 0.35 | 0.4 | 0.2 | 0.56 | 0.5 |
| 7 | 89925561 | rs2040514 | 0.737 | -0.138 | -0.022 | -0.471 | -0.012 | -0.007 | -0.007 | 0 | 0.36 | 0.48 | 0.44 | 0.69 | 0.31 | 0.34 |
| 7 | 89935888 | rs10487111 | 0.357 | -0.132 | -0.111 | -0.468 | 0.005 | 0.004 | 0.004 | 0 | 0.4 | 0.57 | 0.44 | 0.69 | 0.31 | 0.34 |
| 7 | 89946003 | rs10225744 | 0.58 | -0.114 | -0.101 | -0.462 | -0.009 | 0.014 | 0.014 | 0 | 0.43 | 0.4 | 0.43 | 0.3 | 0.52 | 0.48 |
| 7 | 89949597 | rs42608 | 0.681 | 0.027 | 0.136 | -0.337 | -0.003 | 0.008 | 0.008 | 0 | 0.79 | 0.83 | 0.79 | 0.89 | 0.79 | 0.78 |
| 7 | 89961876 | rs42627 | 0.348 | 0.068 | 0.016 | -0.392 | -0.003 | 0.144 | 0.144 | 0.266 | 0.5 | 0.27 | 0.4 | 0.2 | 0.55 | 0.48 |
| 7 | 89973355 | rs2106272 | 1.371 | 0.044 | 0.213 | -0.17 | -0.004 | 0.028 | 0.028 | 0 | 0.28 | 0.34 | 0.38 | 0.67 | 0.24 | 0.3 |
| 7 | 89979227 | rs12704526 | 1.393 | 0.072 | 0.05 | -0.291 | -0.001 | 0.018 | 0.018 | 0.036 | 0.05 | 0.07 | 0.09 | 0.11 | 0.05 | 0.17 |
| 7 | 89981958 | rs42659 | 0.073 | 0.06 | 0.047 | 0.041 | -0.009 | 0.027 | 0.027 | 0.006 | 0.53 | 0.34 | 0.53 | 0.64 | 0.55 | 0.6 |
| 7 | 89982132 | rs6972561 | 1.593 | 0.061 | 0.12 | 0.094 | -0.002 | -0.008 | -0.008 | 0 | 0.27 | 0.4 | 0.25 | 0.24 | 0.24 | 0.19 |
| 7 | 89983808 | rs42663 | -0.128 | 0.067 | 0.154 | 0.052 | 0.011 | -0.004 | -0.004 | 0.018 | 0.83 | 0.73 | 0.8 | 0.89 | 0.82 | 0.79 |
| 7 | 89984409 | rs42664 | 0.141 | 0.055 | 0.151 | 0.058 | -0.007 | 0.038 | 0.038 | 0.026 | 0.52 | 0.31 | 0.53 | 0.65 | 0.55 | 0.59 |
| 7 | 89992918 | rs174073 | 0.293 | 0.149 | -0.098 | -0.237 | 0.004 | 0.03 | 0.03 | 0 | 0.47 | 0.28 | 0.49 | 0.64 | 0.48 | 0.59 |
| 7 | 90014408 | rs17863999 | 0.135 | 0.324 | -0.062 | -0.081 | 0.02 | 0.005 | 0.005 | 0.578 | 0.14 | 0.22 | 0.2 | 0.11 | 0.19 | 0.21 |
| 7 | 90015825 | rs6965126 | -0.565 | 0.335 | -0.21 | -0.064 | 0.003 | -0.002 | -0.002 | 0 | 0.18 | 0.19 | 0.21 | 0.11 | 0.21 | 0.22 |
| 7 | 90023079 | rs6945128 | 1.023 | 0.21 | -0.58 | -0.43 | -0.013 | -0.062 | -0.062 | 0 | 0.27 | 0.19 | 0.3 | 0.54 | 0.31 | 0.39 |
| 7 | 90037937 | rs11563429 | 0.84 | 0.176 | -0.447 | -0.557 | -0.001 | 0.006 | 0.006 | 0.148 | 0.06 | 0.07 | 0.1 | 0.1 | 0.06 | 0.18 |
| 7 | 90042857 | rs3810874 | 0.216 | -0.004 | -0.433 | -0.576 | -0.001 | -0.01 | -0.01 | 0.496 | 0.18 | 0.13 | 0.34 | 0.56 | 0.2 | 0.16 |
| 7 | 90043182 | rs1017104 | -0.024 | -0.022 | -0.359 | -0.584 | 0 | 0.016 | 0.016 | 0 | 0.18 | 0.16 | 0.34 | 0.56 | 0.2 | 0.16 |
| 7 | 90043314 | rs1017106 | -1.85 | -0.311 | -0.428 | -0.919 | 0.01 | 0.001 | 0.001 | 0 | 0.9 | 0.93 | 0.96 | 0.99 | 0.96 | 0.87 |
| 7 | 90044945 | rs9731 | 1.225 | -0.321 | -0.065 | -1.009 | -0.014 | -0.007 | -0.007 | 0 | 0.59 | 0.57 | 0.46 | 0.32 | 0.66 | 0.47 |
| 7 | 90052434 | rs17862178 | 2.127 | 0.006 | -0.805 | -1.168 | -0.04 | 0.055 | 0.055 | 0 | 0.23 | 0.04 | 0.12 | 0.07 | 0.2 | 0.16 |
| 7 | 90068856 | rs10268943 | -0.862 | -0.024 | -0.337 | -1.088 | -0.001 | 0.001 | 0.001 | 0 | 0.28 | 0.29 | 0.48 | 0.68 | 0.3 | 0.35 |
| 7 | 90075626 | rs17867660 | 1.141 | 0.102 | -0.21 | -1.195 | -0.01 | 0.011 | 0.011 | 0 | 0.28 | 0.25 | 0.21 | 0.09 | 0.39 | 0.26 |
| 7 | 90076713 | rs10277213 | 0.996 | -0.221 | 0.006 | -0.89 | -0.006 | 0.074 | 0.074 | 0 | 0.07 | 0.21 | 0.11 | 0.11 | 0.05 | 0.18 |
| 7 | 90078799 | rs12111838 | 1.166 | -0.272 | -0.175 | -1.078 | -0.01 | -0.01 | -0.01 | 1.945 | 0.55 | 0.54 | 0.46 | 0.31 | 0.62 | 0.42 |
| 7 | 90082176 | rs11764579 | 1.173 | -0.045 | -0.521 | -0.936 | 0.005 | 0.002 | 0.002 | 0 | 0.06 | 0.03 | 0.04 | 0 | 0.04 | 0.03 |
| 7 | 90084456 | rs177690 | 1.418 | 0.095 | -0.564 | -0.875 | 0.023 | 0.074 | 0.074 | 0 | 0.13 | 0.01 | 0.05 | 0.01 | 0.06 | 0.13 |
| 7 | 90095070 | rs6954511 | 1.963 | -0.019 | -0.377 | -1.01 | 0.001 | 0.011 | 0.011 | 0 | 0.23 | 0.21 | 0.2 | 0.16 | 0.19 | 0.13 |
| 7 | 90098903 | rs10268928 | 0.666 | 0.033 | -0.189 | -0.92 | 0.003 | 0.048 | 0.048 | 0 | 0.17 | 0.2 | 0.33 | 0.62 | 0.21 | 0.21 |
| 7 | 90100606 | rs177667 | -0.552 | -0.124 | -0.101 | -1.062 | -0.024 | 0.06 | 0.06 | 0 | 0.6 | 0.71 | 0.7 | 0.91 | 0.49 | 0.63 |
| 7 | 90101081 | rs11266823 | 0.769 | 0.013 | -0.364 | -0.802 | -0.014 | -0.061 | -0.061 | 0 | 0.16 | 0.08 | 0.33 | 0.56 | 0.2 | 0.2 |
| 7 | 90109586 | rs177659 | 1.679 | -0.068 | -0.335 | -0.916 | 0.016 | 0.07 | 0.07 | 0 | 0.13 | 0.01 | 0.04 | 0.01 | 0.06 | 0.13 |
| 7 | 90117365 | rs11563535 | 1.713 | 0.061 | -0.188 | -0.666 | 0.001 | -0.002 | -0.002 | 0 | 0.24 | 0.22 | 0.2 | 0.22 | 0.19 | 0.13 |
| 7 | 90123400 | rs11563423 | 1.875 | 0.071 | -0.197 | -0.651 | -0.001 | -0.004 | -0.004 | 0 | 0.22 | 0.2 | 0.2 | 0.22 | 0.19 | 0.13 |
| 7 | 90136655 | rs12113813 | 1.298 | 0.019 | -0.08 | -0.67 | 0.001 | 0.001 | 0.001 | 0 | 0.28 | 0.26 | 0.24 | 0.22 | 0.23 | 0.16 |
| 7 | 90141855 | rs11563856 | 1.153 | -0.017 | -0.164 | -0.478 | -0.002 | 0.151 | 0.151 | 0 | 0.05 | 0.08 | 0.25 | 0.51 | 0.05 | 0.18 |

| | | | | | | | | | | | | | | | | |
|---|---|---|---|---|---|---|---|---|---|---|---|---|---|---|---|---|
| 7 | 90141903 | rs12704547 | 0.499 | -0.01 | -0.33 | -0.768 | -0.016 | 0.031 | 0.031 | 0 | 0.34 | 0.27 | 0.25 | 0.12 | 0.46 | 0.32 |
| 7 | 90148372 | rs11563848 | 1.529 | -0.021 | -0.414 | -0.751 | 0.001 | 0.003 | 0.003 | 0 | 0.28 | 0.22 | 0.23 | 0.22 | 0.23 | 0.17 |
| 7 | 90151034 | rs11563419 | 1.976 | 0.132 | -0.744 | -0.922 | -0.013 | 0.053 | 0.053 | 0 | 0.14 | 0.04 | 0.07 | 0.02 | 0.17 | 0.18 |
| 7 | 90153821 | rs7800844 | 0.721 | 0.151 | -0.571 | -0.805 | 0.005 | -0.032 | -0.032 | 0 | 0.28 | 0.18 | 0.37 | 0.52 | 0.27 | 0.34 |
| 7 | 90157105 | rs6956471 | 1.577 | 0.179 | -1.134 | -1.079 | -0.067 | -0.006 | -0.006 | 0 | 0.13 | 0.03 | 0.13 | 0.1 | 0.25 | 0.09 |
| 7 | 90157692 | rs17864056 | 1.75 | -0.582 | -0.736 | -0.721 | -0.021 | 0.073 | 0.073 | 0 | 0.26 | 0.08 | 0.15 | 0.08 | 0.23 | 0.15 |
| 7 | 90161308 | rs12154771 | 0.745 | -0.439 | -0.915 | -0.726 | -0.052 | 0.082 | 0.082 | 0 | 0.34 | 0.18 | 0.24 | 0.12 | 0.48 | 0.32 |
| 7 | 90163753 | rs11563368 | 1.368 | -0.392 | -0.848 | -0.587 | 0.002 | 0.004 | 0.004 | 0 | 0.28 | 0.25 | 0.24 | 0.22 | 0.22 | 0.17 |
| 7 | 90166905 | rs13226727 | 0.812 | -0.331 | -1.246 | -0.638 | 0.049 | 0.085 | 0.085 | 0 | 0.56 | 0.25 | 0.36 | 0.2 | 0.67 | 0.44 |
| 7 | 90182290 | rs6949622 | -1.363 | -0.054 | -1.455 | -0.51 | -0.016 | 0.083 | 0.083 | 0 | 0.72 | 0.93 | 0.86 | 0.92 | 0.78 | 0.83 |
| 7 | 90186171 | rs11563840 | 0.467 | -0.262 | -1.16 | 0.182 | 0.001 | 0.114 | 0.114 | 0 | 0.06 | 0.07 | 0.25 | 0.53 | 0.05 | 0.19 |
| 7 | 90187804 | rs10273323 | 0.463 | -0.441 | -0.798 | -0.049 | -0.015 | -0.038 | -0.038 | 0 | 0.24 | 0.36 | 0.24 | 0.15 | 0.21 | 0.17 |
| 7 | 90190462 | rs993357 | 0.755 | -0.423 | -0.753 | -0.003 | -0.017 | -0.032 | -0.032 | 0 | 0.22 | 0.35 | 0.22 | 0.15 | 0.18 | 0.17 |
| 7 | 90196305 | rs17865923 | 0.236 | -0.16 | -1.238 | 0.491 | 0.001 | 0.115 | 0.115 | 0 | 0.06 | 0.05 | 0.25 | 0.53 | 0.05 | 0.19 |
| 7 | 90199232 | rs17867705 | 1.413 | -0.066 | -1.5 | -0.052 | -0.015 | 0.001 | 0.001 | 0 | 0.22 | 0.04 | 0.16 | 0.14 | 0.18 | 0.13 |
| 7 | 90205434 | rs17862237 | 0.564 | 0.105 | -1.213 | -0.238 | 0.001 | -0.003 | -0.003 | 2.644 | 0.06 | 0.06 | 0.1 | 0.14 | 0.04 | 0.11 |
| 7 | 90214383 | rs9690523 | 1.04 | -0.703 | -0.715 | -0.378 | -0.012 | 0.008 | 0.008 | 0 | 0.29 | 0.37 | 0.46 | 0.66 | 0.24 | 0.38 |
| 7 | 90216277 | rs2888807 | 0.255 | -0.53 | -1.051 | -1.063 | -0.038 | 0.037 | 0.037 | 0 | 0.3 | 0.21 | 0.24 | 0.14 | 0.45 | 0.33 |
| 7 | 90218346 | rs9690273 | 1.054 | -0.774 | -0.913 | -1.224 | -0.004 | -0.008 | -0.008 | 0 | 0.52 | 0.47 | 0.58 | 0.63 | 0.43 | 0.51 |
| 7 | 90229038 | rs13234217 | -0.211 | -0.671 | -0.735 | -1.306 | -0.016 | 0.007 | 0.007 | 0 | 0.37 | 0.33 | 0.33 | 0.34 | 0.49 | 0.45 |
| 7 | 90229239 | rs6945415 | -1.296 | -0.727 | -0.722 | -1.321 | -0.016 | -0.004 | -0.004 | 0 | 0.67 | 0.54 | 0.47 | 0.34 | 0.72 | 0.59 |
| 7 | 90233618 | rs6465273 | 0.032 | -0.089 | -1.054 | -1.661 | 0.018 | 0.168 | 0.168 | 4.113 | 0.41 | 0.19 | 0.21 | 0.05 | 0.37 | 0.29 |
| 7 | 90247308 | rs2106730 | -1.421 | 0.064 | -1.069 | -1.656 | 0.014 | -0.008 | -0.008 | 0 | 0.85 | 0.82 | 0.85 | 0.91 | 0.72 | 0.86 |
| 7 | 90247442 | rs17865021 | 0.535 | 0.01 | -1.086 | -1.604 | -0.003 | 0.028 | 0.028 | 0.605 | 0.17 | 0.13 | 0.09 | 0.05 | 0.17 | 0.15 |
| 7 | 90255237 | rs17867002 | 0.877 | 0.005 | -1.047 | -1.49 | 0.002 | 0.007 | 0.007 | 0 | 0.07 | 0.07 | 0.1 | 0.2 | 0.05 | 0.12 |
| 7 | 90261374 | rs10247140 | -0.29 | -0.18 | -1.24 | -1.71 | 0.01 | -0.015 | -0.015 | 0 | 0.84 | 0.9 | 0.86 | 0.8 | 0.87 | 0.78 |
| 7 | 90269865 | rs7782861 | 2.603 | -0.259 | -1.304 | -1.797 | 0.014 | 0.025 | 0.025 | 0.32 | 0.16 | 0.08 | 0.08 | 0 | 0.12 | 0.03 |
| 7 | 90269990 | rs7801922 | 0.555 | -0.188 | -1.239 | -1.789 | -0.013 | 0.005 | 0.005 | 0.691 | 0.67 | 0.72 | 0.71 | 0.71 | 0.57 | 0.59 |
| 7 | 90270664 | rs12672866 | 1.739 | -0.114 | -1.121 | -1.74 | -0.02 | 0.007 | 0.007 | 0 | 0.3 | 0.39 | 0.49 | 0.66 | 0.24 | 0.37 |
| 7 | 90272877 | rs7797824 | -0.067 | 0.06 | -1.088 | -1.554 | 0.004 | 0.001 | 0.001 | 0.035 | 0.25 | 0.28 | 0.26 | 0.29 | 0.34 | 0.3 |
| 7 | 90280817 | rs17866339 | 0.619 | -1.396 | -0.582 | -1.425 | -0.012 | 0.061 | 0.061 | 0 | 0.07 | 0.14 | 0.1 | 0.2 | 0.04 | 0.11 |
| 7 | 90282798 | rs11563414 | 1.154 | -1.394 | -0.591 | -1.424 | 0.003 | 0.006 | 0.006 | 0 | 0.06 | 0.05 | 0.1 | 0.2 | 0.04 | 0.11 |
| 7 | 90283594 | rs1011429 | -2.11 | -1.407 | -0.64 | -1.466 | -0.007 | 0.019 | 0.019 | 4.348 | 0.94 | 0.98 | 0.97 | 1 | 0.93 | 0.99 |
| 7 | 90285329 | rs6964141 | -0.578 | -1.353 | -0.556 | -1.437 | -0.014 | 0.01 | 0.01 | 0 | 0.16 | 0.25 | 0.15 | 0.2 | 0.11 | 0.16 |
| 7 | 90301209 | rs7807078 | 0.767 | -1.453 | -0.536 | -1.369 | 0.004 | 0.023 | 0.023 | 0.769 | 0.48 | 0.59 | 0.66 | 0.75 | 0.55 | 0.56 |
| 7 | 90305331 | rs7808418 | -1.215 | -1.476 | -0.684 | -1.454 | 0 | 0.159 | 0.159 | 4.214 | 0.76 | 0.96 | 0.89 | 1 | 0.8 | 0.84 |
| 7 | 90305548 | rs6976601 | 1.702 | -1.064 | -0.571 | -1.448 | 0.011 | 0.002 | 0.002 | 0 | 0.32 | 0.38 | 0.54 | 0.66 | 0.38 | 0.4 |
| 7 | 90315383 | rs6963621 | -0.469 | -1.046 | -0.657 | -1.313 | -0.001 | 0.001 | 0.001 | 0 | 0.25 | 0.3 | 0.22 | 0.25 | 0.23 | 0.27 |
| 7 | 90329724 | rs17716105 | 0.83 | -0.882 | -0.703 | -1.414 | -0.001 | -0.011 | -0.011 | 0.027 | 0.15 | 0.17 | 0.09 | 0.09 | 0.16 | 0.16 |
| 7 | 90334930 | rs4728931 | -0.69 | -0.845 | -0.823 | -1.486 | -0.014 | -0.02 | -0.02 | 0 | 0.84 | 0.9 | 0.83 | 0.8 | 0.83 | 0.84 |
| 7 | 90335099 | rs17716111 | 1.041 | -0.896 | -0.878 | -2.038 | 0 | -0.003 | -0.003 | 0 | 0.14 | 0.14 | 0.09 | 0.09 | 0.16 | 0.17 |
| 7 | 90338850 | rs11767691 | 0.862 | -0.832 | -0.858 | -2.183 | 0 | -0.003 | -0.003 | 0.111 | 0.14 | 0.16 | 0.09 | 0.09 | 0.16 | 0.16 |
| 7 | 90346227 | rs994511 | 0.966 | -0.766 | -0.92 | -2.197 | 0.004 | -0.052 | -0.052 | 0.037 | 0.39 | 0.46 | 0.65 | 0.85 | 0.42 | 0.5 |
| 7 | 90365828 | rs7796430 | -1.658 | -0.405 | -0.994 | -1.859 | -0.011 | 0.008 | 0.008 | 0 | 0.91 | 0.95 | 0.93 | 1 | 0.87 | 0.94 |
| 7 | 90399859 | rs11763871 | 0.099 | -0.652 | -0.929 | -1.932 | 0 | -0.001 | -0.001 | 0 | 0.19 | 0.18 | 0.17 | 0.12 | 0.21 | 0.24 |
| 7 | 90401125 | rs6972909 | 0.966 | -0.644 | -0.786 | -1.951 | 0.007 | -0.006 | -0.006 | 0.292 | 0.05 | 0.06 | 0.06 | 0.05 | 0.08 | 0.13 |
| 7 | 90404676 | rs3808227 | 0.153 | -0.657 | -0.824 | -1.96 | 0 | 0 | 0 | 0 | 0.19 | 0.18 | 0.17 | 0.12 | 0.21 | 0.24 |
| 7 | 90405834 | rs3779554 | 1.318 | -0.904 | -0.856 | -2.324 | 0.004 | -0.034 | -0.034 | 0 | 0.37 | 0.45 | 0.58 | 0.82 | 0.39 | 0.41 |
| 7 | 90407237 | rs11983021 | 2.145 | -0.952 | -0.866 | -2.554 | -0.05 | 0.103 | 0.103 | 0 | 0.26 | 0.12 | 0.17 | 0.02 | 0.3 | 0.16 |
| 7 | 90417457 | rs17163201 | 0.878 | -0.793 | -0.552 | -2.29 | 0.018 | 0.076 | 0.076 | 3.814 | 0.24 | 0.12 | 0.11 | 0.01 | 0.19 | 0.16 |
| 7 | 90425217 | rs17394921 | 0.957 | -0.799 | -0.58 | -2.295 | -0.003 | 0.154 | 0.154 | 4.479 | 0.24 | 0.05 | 0.11 | 0.01 | 0.19 | 0.16 |
| 7 | 90449362 | rs10245696 | 0.631 | -0.768 | -0.546 | -2.28 | 0.002 | -0.025 | -0.025 | 0 | 0.57 | 0.63 | 0.75 | 0.94 | 0.6 | 0.65 |
| 7 | 90452976 | rs17163266 | 0.755 | -0.658 | -0.692 | -2.088 | 0.009 | 0.111 | 0.111 | 4.361 | 0.24 | 0.08 | 0.11 | 0.01 | 0.19 | 0.16 |
| 7 | 90461628 | rs7786646 | 0.483 | -0.546 | -0.554 | -1.977 | 0.024 | -0.011 | -0.011 | 0 | 0.06 | 0.12 | 0.06 | 0.05 | 0.08 | 0.13 |
| 7 | 90476833 | rs1978068 | 0.573 | -0.719 | -0.544 | -2.106 | 0 | -0.034 | -0.034 | 0.588 | 0.48 | 0.52 | 0.66 | 0.9 | 0.5 | 0.54 |
| 7 | 90481699 | rs7809531 | -0.533 | -0.715 | -0.386 | -1.903 | 0.011 | -0.053 | -0.053 | 0 | 0.32 | 0.37 | 0.29 | 0.15 | 0.38 | 0.43 |
| 7 | 90483905 | rs12534295 | -0.414 | -0.799 | -0.366 | -1.974 | 0.004 | 0.015 | 0.015 | 0.083 | 0.27 | 0.22 | 0.2 | 0.11 | 0.25 | 0.31 |
| 7 | 90494919 | rs758706 | 1.385 | -0.712 | 0.021 | -1.472 | -0.004 | 0.001 | 0.001 | 0 | 0.37 | 0.34 | 0.49 | 0.67 | 0.43 | 0.41 |
| 7 | 90500326 | rs7780162 | -0.688 | -0.574 | 0.041 | -1.386 | 0.009 | -0.005 | -0.005 | 0 | 0.94 | 0.92 | 0.94 | 0.95 | 0.9 | 0.89 |
| 7 | 90518027 | rs17476172 | 0.772 | -0.864 | -0.132 | -1.534 | 0.043 | 0.133 | 0.133 | 2.335 | 0.27 | 0.07 | 0.16 | 0.04 | 0.2 | 0.3 |
| 7 | 90520097 | rs3802038 | -0.135 | -0.814 | -0.057 | -1.531 | 0.023 | -0.012 | -0.012 | 0 | 0.93 | 0.79 | 0.93 | 0.95 | 0.9 | 0.9 |
| 7 | 90522559 | rs3779568 | 0.871 | -0.84 | -0.099 | -1.567 | 0.003 | 0.003 | 0.003 | 0 | 0.5 | 0.5 | 0.59 | 0.74 | 0.57 | 0.49 |
| 7 | 90523546 | rs6957439 | 0.911 | -0.285 | -0.322 | -1.356 | -0.01 | 0.03 | 0.03 | 0 | 0.12 | 0.05 | 0.08 | 0 | 0.13 | 0.1 |
| 7 | 90543678 | rs7783741 | 1.127 | -0.305 | -0.277 | -1.332 | -0.001 | 0.001 | 0.001 | 0 | 0.14 | 0.13 | 0.09 | 0.06 | 0.15 | 0.1 |
| 7 | 90545729 | rs3808242 | -0.972 | -0.464 | -0.352 | -1.448 | 0.038 | 0.086 | 0.086 | 2.288 | 0.74 | 0.87 | 0.83 | 0.96 | 0.81 | 0.71 |
| 7 | 90545844 | rs3808243 | -0.972 | -0.464 | -0.352 | -1.448 | 0.038 | 0.086 | 0.086 | 2.288 | 0.74 | 0.87 | 0.83 | 0.96 | 0.81 | 0.71 |
| 7 | 90546764 | rs2286696 | 0.653 | -0.47 | -0.235 | -1.406 | 0.013 | 0.039 | 0.039 | 0.063 | 0.5 | 0.61 | 0.57 | 0.73 | 0.55 | 0.53 |
| 7 | 90549300 | rs17398589 | 1.444 | -0.464 | -0.202 | -1.402 | -0.005 | -0.012 | -0.012 | 0 | 0.09 | 0.05 | 0.16 | 0.17 | 0.11 | 0.07 |
| 7 | 90550191 | rs10953026 | 0.148 | -0.49 | -0.147 | -1.343 | 0.002 | 0.003 | 0.003 | 0.729 | 0.59 | 0.61 | 0.64 | 0.73 | 0.65 | 0.59 |

| | | | | | | | | | | | | | | | | |
|---|---|---|---|---|---|---|---|---|---|---|---|---|---|---|---|---|
| 7 | 90561832 | rs11976125 | 0.485 | -0.264 | -0.132 | -1.238 | 0.003 | -0.031 | -0.031 | 0 | 0.18 | 0.22 | 0.11 | 0.06 | 0.19 | 0.11 |
| 7 | 90581572 | rs7807704 | 1.111 | -0.554 | -0.075 | -0.765 | -0.001 | 0.001 | 0.001 | 0 | 0.14 | 0.13 | 0.09 | 0.06 | 0.17 | 0.1 |
| 7 | 90588732 | rs3808261 | 1.317 | -0.602 | 0.064 | -0.777 | -0.001 | 0.027 | 0.027 | 0 | 0.06 | 0.07 | 0.09 | 0.19 | 0.05 | 0.09 |
| 7 | 90589751 | rs732956 | 0.916 | -0.713 | 0.244 | -0.946 | -0.001 | 0.009 | 0.009 | 0.025 | 0.44 | 0.39 | 0.59 | 0.73 | 0.54 | 0.43 |
| 7 | 90600629 | rs975004 | 1.31 | -0.618 | 0.256 | -0.837 | -0.002 | 0.041 | 0.041 | 0 | 0.06 | 0.09 | 0.09 | 0.19 | 0.05 | 0.09 |
| 7 | 90601518 | rs3824047 | -0.164 | -0.73 | 0.168 | -0.927 | 0.024 | 0.018 | 0.018 | 0.788 | 0.64 | 0.7 | 0.74 | 0.77 | 0.75 | 0.63 |
| 7 | 90608720 | rs802394 | 1.171 | -0.692 | 0.052 | -0.99 | 0.011 | -0.011 | -0.011 | 0 | 0.4 | 0.43 | 0.57 | 0.71 | 0.48 | 0.47 |
| 7 | 90610105 | rs802396 | 1.932 | -0.644 | -0.071 | -0.946 | -0.004 | -0.007 | -0.007 | 0.263 | 0.32 | 0.29 | 0.4 | 0.48 | 0.36 | 0.29 |
| 7 | 90611107 | rs17479582 | 0.92 | -0.479 | 0.008 | -0.896 | 0.03 | 0.122 | 0.122 | 2.808 | 0.25 | 0.06 | 0.15 | 0.04 | 0.18 | 0.27 |
| 7 | 90629849 | rs1208898 | 1.04 | -0.425 | -0.041 | -0.812 | 0.006 | -0.009 | -0.009 | 0 | 0.42 | 0.48 | 0.58 | 0.7 | 0.47 | 0.47 |
| 7 | 90630558 | rs802382 | -0.393 | -0.443 | -0.03 | -0.86 | -0.002 | 0.075 | 0.075 | 0 | 0.89 | 0.72 | 0.89 | 0.81 | 0.93 | 0.91 |
| 7 | 90630720 | rs802383 | 1.62 | -0.38 | 0.003 | -0.804 | -0.005 | -0.007 | -0.007 | 0.031 | 0.37 | 0.33 | 0.44 | 0.53 | 0.44 | 0.39 |
| 7 | 90632954 | rs802387 | 0.506 | -0.231 | -0.584 | -1.003 | 0.007 | 0.011 | 0.011 | 0.291 | 0.53 | 0.53 | 0.66 | 0.71 | 0.61 | 0.53 |
| 7 | 90639136 | rs11773740 | 0.564 | -0.149 | -1.149 | -1.173 | -0.004 | 0 | 0 | 0 | 0.23 | 0.2 | 0.16 | 0.06 | 0.28 | 0.17 |
| 7 | 90650256 | rs7789771 | -0.649 | -0.72 | -1.126 | -0.386 | 0.02 | 0.066 | 0.066 | 1.292 | 0.29 | 0.24 | 0.2 | 0.11 | 0.25 | 0.37 |
| 7 | 90656040 | rs10488001 | 0.447 | -0.702 | -1.427 | -0.203 | 0.021 | 0.101 | 0.101 | 0.249 | 0.16 | 0.02 | 0.05 | 0 | 0.08 | 0.06 |
| 7 | 90660299 | rs705346 | -0.25 | -0.4 | -1.118 | 0.489 | 0.009 | -0.015 | -0.015 | 0.12 | 0.15 | 0.1 | 0.26 | 0.41 | 0.08 | 0.17 |
| 7 | 90660684 | rs1358101 | -0.274 | -0.433 | -0.696 | -0.788 | -0.018 | -0.059 | -0.059 | 0 | 0.35 | 0.47 | 0.24 | 0.11 | 0.29 | 0.42 |
| 7 | 90671675 | rs7782262 | -0.589 | -0.089 | -0.572 | -1.073 | -0.004 | -0.09 | -0.09 | 0.532 | 0.25 | 0.59 | 0.18 | 0.06 | 0.21 | 0.19 |
| 7 | 90674192 | rs962281 | -0.076 | -0.242 | -0.058 | -1.213 | 0.002 | -0.037 | -0.037 | 2.09 | 0.14 | 0.17 | 0.11 | 0.01 | 0.16 | 0.13 |
| 7 | 90674492 | rs697393 | 0.77 | -0.15 | -1.334 | -2.234 | -0.003 | 0.071 | 0.071 | 0 | 0.43 | 0.32 | 0.63 | 0.86 | 0.48 | 0.53 |
| 7 | 90679137 | rs17163534 | 1.377 | -0.522 | -1.834 | -2.332 | 0.007 | 0.001 | 0.001 | 0 | 0.1 | 0.02 | 0.24 | 0.06 | 0.06 | 0.13 |
| 7 | 90679837 | rs17163541 | 1.173 | -0.517 | -1.077 | -2.328 | -0.016 | -0.025 | -0.025 | 0 | 0.09 | 0.18 | 0.06 | 0.04 | 0.05 | 0.06 |
| 7 | 90692002 | rs696271 | -0.11 | -0.508 | -0.898 | -2.352 | 0.001 | -0.124 | -0.124 | 0 | 0.24 | 0.47 | 0.17 | 0.06 | 0.2 | 0.19 |
| 7 | 90698364 | rs705352 | 1.571 | 0.337 | -1.462 | -1.316 | -0.059 | -0.06 | -0.06 | 0 | 0.1 | 0.01 | 0.21 | 0.55 | 0.21 | 0.23 |
| 7 | 90699073 | rs705353 | -1.601 | -0.189 | -1.699 | -1.893 | -0.028 | 0.044 | 0.044 | 1.206 | 0.81 | 0.9 | 0.86 | 0.91 | 0.78 | 0.73 |
| 7 | 90713562 | rs696267 | -0.575 | 0.22 | -1.695 | -1.899 | -0.005 | 0.17 | 0.17 | 0.663 | 0.63 | 0.86 | 0.74 | 0.91 | 0.68 | 0.69 |
| 7 | 90718286 | rs705350 | 0.337 | 0.406 | -1.23 | -1.647 | -0.006 | 0.005 | 0.005 | 0 | 0.3 | 0.26 | 0.4 | 0.8 | 0.44 | 0.42 |
| 7 | 90726315 | rs705357 | 0.042 | 0.074 | -1.127 | -0.983 | 0.003 | -0.001 | -0.001 | 0 | 0.65 | 0.71 | 0.77 | 0.88 | 0.73 | 0.67 |
| 7 | 90739777 | rs803157 | 0.163 | 0.251 | -0.984 | -0.03 | -0.001 | 0 | 0 | 0.301 | 0.37 | 0.33 | 0.48 | 0.37 | 0.4 | 0.44 |
| 7 | 90742248 | rs17163593 | 0.411 | -1.308 | -1.154 | 1.078 | -0.039 | -0.101 | -0.101 | 5.136 | 0.13 | 0.01 | 0.2 | 0.6 | 0.18 | 0.19 |
| 7 | 90742807 | rs803160 | 1.142 | -1.564 | -1.386 | 0.513 | 0.01 | 0.159 | 0.159 | 0 | 0.45 | 0.21 | 0.33 | 0.1 | 0.55 | 0.46 |
| 7 | 90744442 | rs810640 | 0.619 | -1.675 | -0.902 | 0.978 | 0.048 | 0.018 | 0.018 | 0 | 0.54 | 0.22 | 0.49 | 0.44 | 0.67 | 0.44 |
| 7 | 90746617 | rs809132 | 1.45 | -1.586 | -1.29 | 0.804 | -0.095 | 0.273 | 0.273 | 0.006 | 0.4 | 0.04 | 0.29 | 0.01 | 0.52 | 0.26 |
| 7 | 90747682 | rs10488003 | -0.606 | -1.189 | -0.438 | 0.917 | 0.012 | 0.15 | 0.15 | 0 | 0.12 | 0.26 | 0.34 | 0.53 | 0.16 | 0.3 |
| 7 | 90752135 | rs42999 | -0.134 | -1.354 | -0.723 | 0.096 | -0.015 | -0.025 | -0.025 | 0 | 0.76 | 0.76 | 0.85 | 0.84 | 0.84 | 0.73 |
| 7 | 90755496 | rs6944536 | 0.443 | -1.216 | -1.048 | 0.319 | -0.043 | -0.119 | -0.119 | 1.107 | 0.15 | 0.02 | 0.19 | 0.45 | 0.17 | 0.17 |
| 7 | 90756548 | rs43002 | 0.106 | -1.447 | -0.619 | 0.434 | -0.033 | -0.058 | -0.058 | 0.9 | 0.72 | 0.66 | 0.84 | 0.86 | 0.82 | 0.61 |
| 7 | 90758236 | rs10488004 | 0.48 | -1.231 | -1.584 | 0.31 | 0.097 | -0.042 | -0.042 | 0.281 | 0.58 | 0.15 | 0.52 | 0.47 | 0.74 | 0.47 |
| 7 | 90764892 | rs803163 | -0.664 | -0.376 | -1.397 | 0.436 | -0.005 | -0.135 | -0.135 | 0 | 0.22 | 0.04 | 0.25 | 0.53 | 0.21 | 0.21 |
| 7 | 90768739 | rs803165 | -0.756 | -0.542 | -1.003 | -0.083 | -0.005 | 0.169 | 0.169 | 3.346 | 0.52 | 0.79 | 0.7 | 0.97 | 0.46 | 0.62 |
| 7 | 90770662 | rs43007 | 1.751 | -0.838 | -0.78 | -0.14 | -0.014 | 0.024 | 0.024 | 0 | 0.77 | 0.7 | 0.72 | 0.43 | 0.87 | 0.64 |
| 7 | 90771000 | rs11763415 | 0.761 | -0.866 | -0.165 | -0.225 | 0 | 0 | 0 | 0.026 | 0.3 | 0.27 | 0.25 | 0.37 | 0.32 | 0.34 |
| 7 | 90771789 | rs43010 | -0.228 | -0.85 | -0.141 | -0.283 | 0.011 | 0.022 | 0.022 | 0 | 0.91 | 0.96 | 0.96 | 1 | 0.94 | 0.88 |
| 7 | 90773196 | rs803172 | 0.183 | -0.853 | -0.01 | -0.221 | -0.001 | -0.005 | -0.005 | 0.311 | 0.37 | 0.35 | 0.4 | 0.87 | 0.36 | 0.5 |
| 7 | 90773360 | rs803173 | 0.082 | -0.871 | 0.011 | -0.231 | -0.002 | -0.058 | -0.058 | 0.039 | 0.38 | 0.46 | 0.4 | 0.87 | 0.36 | 0.5 |
| 7 | 90774840 | rs43013 | 1.298 | -0.621 | -0.046 | -0.143 | -0.094 | 0.257 | 0.257 | 4.284 | 0.4 | 0.08 | 0.25 | 0.02 | 0.47 | 0.27 |
| 7 | 90781987 | rs813379 | -0.141 | -0.445 | 0.03 | -0.045 | 0.006 | 0.022 | 0.022 | 0 | 0.91 | 0.96 | 0.95 | 0.99 | 0.93 | 0.88 |
| 7 | 90787226 | rs10953033 | 0.533 | -0.454 | 0.058 | -0.052 | -0.013 | 0.035 | 0.035 | 2.501 | 0.87 | 0.96 | 0.77 | 0.97 | 0.86 | 0.85 |
| 7 | 90788633 | rs803169 | 0.773 | -0.5 | 0.134 | -0.012 | 0 | 0 | 0 | 0 | 0.32 | 0.29 | 0.25 | 0.38 | 0.32 | 0.36 |
| 7 | 90789353 | rs813003 | 0.916 | -0.34 | -0.659 | -0.034 | -0.016 | 0.07 | 0.07 | 3.235 | 0.43 | 0.36 | 0.35 | 0.1 | 0.49 | 0.35 |
| 7 | 90792408 | rs803171 | 1.359 | -0.281 | -0.747 | -0.026 | -0.067 | 0.198 | 0.198 | 5.893 | 0.39 | 0.1 | 0.26 | 0.02 | 0.48 | 0.26 |
| 7 | 90794795 | rs803174 | -0.009 | -0.268 | -0.723 | 0.008 | 0.008 | 0.025 | 0.025 | 0 | 0.91 | 0.97 | 0.95 | 1 | 0.93 | 0.88 |
| 7 | 90800787 | rs2518782 | 1.025 | -0.285 | -0.716 | 0.049 | 0.015 | -0.051 | -0.051 | 0 | 0.85 | 0.93 | 0.81 | 0.5 | 0.9 | 0.74 |
| 7 | 90801714 | rs7779721 | 1.252 | -0.229 | -0.779 | 0.02 | -0.067 | 0.206 | 0.206 | 2.11 | 0.4 | 0.1 | 0.25 | 0.02 | 0.48 | 0.26 |
| 7 | 90802001 | rs766544 | 0.63 | -0.107 | -0.671 | 0.081 | -0.009 | 0.016 | 0.016 | 0.037 | 0.35 | 0.44 | 0.27 | 0.38 | 0.32 | 0.35 |
| 7 | 90802565 | rs4276614 | 0.823 | 0.239 | -0.748 | 0.411 | -0.068 | 0.193 | 0.193 | 6.036 | 0.39 | 0.1 | 0.25 | 0.02 | 0.47 | 0.26 |
| 7 | 90812330 | rs2724355 | 0.511 | 0.367 | -0.658 | 0.524 | -0.024 | -0.024 | -0.024 | 0 | 0.43 | 0.24 | 0.37 | 0.51 | 0.48 | 0.39 |
| 7 | 90818888 | rs6465305 | -0.299 | 0.749 | -1.109 | -0.504 | 0.015 | 0.063 | 0.063 | 1.25 | 0.54 | 0.33 | 0.56 | 0.12 | 0.61 | 0.42 |
| 7 | 90820630 | rs2063390 | -0.098 | 0.691 | -1.789 | -0.445 | -0.02 | 0.236 | 0.236 | 10.703 | 0.48 | 0.15 | 0.35 | 0.1 | 0.53 | 0.39 |
| 7 | 90825228 | rs4727250 | -0.143 | 0.747 | -1.903 | -0.42 | -0.021 | 0.25 | 0.25 | 8.259 | 0.49 | 0.15 | 0.36 | 0.1 | 0.53 | 0.39 |
| 7 | 90828977 | rs4236518 | -0.915 | 1.102 | -1.11 | 0.682 | -0.009 | -0.019 | -0.019 | 0.043 | 0.06 | 0.02 | 0.1 | 0.07 | 0.06 | 0.1 |
| 7 | 90833184 | rs12704597 | -1.121 | 1.738 | -1.293 | 1.166 | -0.039 | -0.05 | -0.05 | 0 | 0.12 | 0.05 | 0.16 | 0.29 | 0.19 | 0.14 |
| 7 | 90833383 | rs6975786 | -0.931 | 1.802 | -0.385 | 1.268 | 0.058 | 0.069 | 0.069 | 0 | 0.06 | 0.27 | 0.03 | 0.08 | 0.06 | 0.05 |
| 7 | 90835846 | rs4554397 | -1.485 | 2.25 | -0.706 | 1.192 | 0.006 | 0.003 | 0.003 | 0 | 0.41 | 0.49 | 0.58 | 0.63 | 0.49 | 0.56 |
| 7 | 90837216 | rs11771034 | -0.722 | 2.202 | -0.766 | 1.304 | -0.027 | -0.033 | -0.033 | 0.95 | 0.11 | 0.08 | 0.15 | 0.28 | 0.18 | 0.14 |
| 7 | 90840353 | rs11970859 | -1.699 | 1.468 | -0.697 | 1.014 | -0.002 | 0 | 0 | 0.018 | 0.08 | 0.08 | 0.05 | 0.07 | 0.08 | 0.09 |
| 7 | 90851322 | rs11976442 | -1.416 | 1.83 | -0.8 | 1.335 | -0.011 | -0.037 | -0.037 | 0 | 0.13 | 0.1 | 0.23 | 0.38 | 0.16 | 0.18 |
| 7 | 90852632 | rs6963218 | -1.416 | 1.912 | -0.8 | 1.341 | -0.014 | -0.037 | -0.037 | 0.15 | 0.13 | 0.1 | 0.23 | 0.39 | 0.16 | 0.18 |

| | | | | | | | | | | | | | | | | |
|---|---|---|---|---|---|---|---|---|---|---|---|---|---|---|---|---|
| 7 | 90854065 | rs10259757 | 2.227 | 2.243 | 0.152 | 1.24 | 0.017 | 0.083 | 0.083 | 0 | 0.73 | 0.53 | 0.66 | 0.49 | 0.68 | 0.71 |
| 7 | 90860351 | rs7809443 | -2.098 | 2.13 | 0.744 | 1.235 | 0.003 | 0.09 | 0.09 | 0 | 0.19 | 0.34 | 0.26 | 0.39 | 0.19 | 0.23 |
| 7 | 90862407 | rs12538957 | -2.014 | 1.987 | 0.804 | 1.234 | 0.051 | 0.031 | 0.031 | 0.658 | 0.06 | 0.32 | 0.05 | 0.02 | 0.03 | 0.06 |
| 7 | 90866589 | rs6955473 | -0.615 | 1.638 | -0.047 | 0.234 | 0 | 0 | 0 | 0 | 0.47 | 0.48 | 0.53 | 0.41 | 0.58 | 0.49 |
| 7 | 90866693 | rs4455763 | 0.973 | 1.487 | -0.057 | 0.232 | -0.007 | -0.001 | -0.001 | 0 | 0.47 | 0.46 | 0.39 | 0.49 | 0.3 | 0.46 |
| 7 | 90870373 | rs10277895 | -0.448 | 1.437 | 0.792 | 1.022 | 0.062 | 0.371 | 0.371 | 0 | 0.07 | 0.31 | 0.19 | 0.33 | 0.06 | 0.16 |
| 7 | 90871330 | rs4374920 | -0.909 | 1.216 | 1.013 | 0.818 | 0.036 | 0.06 | 0.06 | 0 | 0.52 | 0.67 | 0.7 | 0.72 | 0.64 | 0.63 |
| 7 | 90871515 | rs17351398 | 0.415 | 1.277 | 0.972 | 0.722 | -0.067 | 0.083 | 0.083 | 0 | 0.32 | 0.21 | 0.41 | 0.08 | 0.46 | 0.31 |
| 7 | 90872656 | rs6969413 | -1.06 | 1.278 | 0.965 | 0.726 | 0.036 | 0.06 | 0.06 | 0 | 0.52 | 0.67 | 0.7 | 0.72 | 0.64 | 0.63 |
| 7 | 90872756 | rs11771236 | -0.11 | 1.345 | 1.048 | 0.885 | 0.012 | -0.034 | -0.034 | 0.032 | 0.12 | 0.07 | 0.09 | 0.31 | 0.12 | 0.16 |
| 7 | 90872842 | rs10244545 | -1.372 | 1.349 | 1.056 | 0.799 | -0.019 | 0.111 | 0.111 | 0 | 0.2 | 0.43 | 0.28 | 0.64 | 0.18 | 0.32 |
| 7 | 90878249 | rs6465306 | -1.516 | 1.344 | 1.054 | 0.787 | -0.016 | 0.064 | 0.064 | 0.029 | 0.22 | 0.55 | 0.29 | 0.64 | 0.19 | 0.32 |
| 7 | 90880142 | rs12540728 | -0.287 | 1.523 | 0.621 | 0.994 | 0.003 | -0.029 | -0.029 | 0 | 0.13 | 0.07 | 0.1 | 0.21 | 0.16 | 0.14 |
| 7 | 90888070 | rs12537425 | 0.451 | 1.585 | 0.969 | 1.21 | 0.012 | -0.002 | -0.002 | 1.104 | 0.29 | 0.32 | 0.45 | 0.22 | 0.36 | 0.25 |
| 7 | 90888413 | rs6977978 | 0.118 | 1.714 | 0.505 | 1.307 | -0.042 | -0.037 | -0.037 | 0 | 0.91 | 0.97 | 0.85 | 0.78 | 0.83 | 0.82 |
| 7 | 90897731 | rs1052015 | -0.183 | 1.687 | 0.516 | 1.323 | -0.003 | -0.036 | -0.036 | 0 | 0.13 | 0.06 | 0.1 | 0.22 | 0.16 | 0.14 |
| 7 | 90904938 | rs10252817 | -0.655 | 1.494 | 0.873 | 1.211 | -0.01 | -0.008 | -0.008 | 0.082 | 0.44 | 0.67 | 0.56 | 0.45 | 0.52 | 0.4 |
| 7 | 90905041 | rs10252923 | 0.375 | 1.568 | 0.897 | 1.167 | 0.04 | -0.009 | -0.009 | 0 | 0.3 | 0.46 | 0.46 | 0.22 | 0.36 | 0.26 |
| 7 | 90910024 | rs10281519 | -0.937 | 1.463 | 0.32 | 1.177 | -0.034 | -0.02 | -0.02 | 1.096 | 0.94 | 0.99 | 0.94 | 0.91 | 0.88 | 0.95 |
| 7 | 90915354 | rs2888830 | 1.185 | 1.196 | 0.444 | 1.118 | 0.005 | 0.045 | 0.045 | 0.295 | 0.73 | 0.54 | 0.73 | 0.61 | 0.68 | 0.62 |
| 7 | 90919567 | rs1346665 | 0.276 | 1.14 | 0.429 | 0.993 | 0.002 | 0 | 0 | 0.069 | 0.59 | 0.54 | 0.4 | 0.62 | 0.49 | 0.53 |
| 7 | 90922544 | rs10250915 | -0.399 | -0.235 | -0.04 | 0.748 | -0.003 | -0.001 | -0.001 | 0 | 0.35 | 0.37 | 0.38 | 0.61 | 0.28 | 0.32 |
| 7 | 90934346 | rs2163086 | -0.703 | -0.207 | 0.264 | 0.668 | -0.013 | -0.007 | -0.007 | 0 | 0.4 | 0.52 | 0.38 | 0.63 | 0.29 | 0.35 |
| 7 | 90936457 | rs10265137 | -0.648 | -0.26 | 0.381 | 0.645 | -0.017 | -0.008 | -0.008 | 0 | 0.4 | 0.44 | 0.38 | 0.63 | 0.29 | 0.35 |
| 7 | 90949831 | rs930110 | -0.615 | 0.475 | 0.068 | 0.792 | 0 | -0.001 | -0.001 | 0 | 0.42 | 0.58 | 0.41 | 0.54 | 0.47 | 0.3 |
| 7 | 90952930 | rs2163087 | -0.82 | 0.368 | 0.08 | 0.76 | -0.008 | 0.051 | 0.051 | 0 | 0.46 | 0.68 | 0.49 | 0.68 | 0.49 | 0.35 |
| 7 | 90955960 | rs1429535 | -0.485 | 0.629 | -0.489 | 0.435 | -0.024 | 0.004 | 0.004 | 0.162 | 0.85 | 0.95 | 0.8 | 0.83 | 0.81 | 0.78 |
| 7 | 90956438 | rs11486955 | 0.919 | -0.114 | -0.536 | 1.097 | 0.017 | -0.026 | -0.026 | 0 | 0.15 | 0.15 | 0.2 | 0.31 | 0.09 | 0.1 |
| 7 | 90956958 | rs1126204 | -0.195 | -0.128 | -0.271 | 1.056 | 0.041 | 0.024 | 0.024 | 0 | 0.3 | 0.44 | 0.28 | 0.34 | 0.4 | 0.25 |
| 7 | 90958247 | rs1344708 | -0.871 | -0.004 | -0.526 | 0.911 | -0.007 | 0.051 | 0.051 | 0 | 0.46 | 0.68 | 0.48 | 0.69 | 0.49 | 0.35 |
| 7 | 90963727 | rs6954427 | 0.802 | -0.822 | -1.285 | 0.664 | -0.049 | -0.002 | -0.002 | 0 | 0.42 | 0.11 | 0.3 | 0.35 | 0.54 | 0.38 |
| 7 | 90969050 | rs997852 | 1.483 | -0.408 | -0.973 | 1.062 | -0.001 | -0.002 | -0.002 | 0 | 0.12 | 0.13 | 0.16 | 0.22 | 0.11 | 0.09 |
| 7 | 90974367 | rs7791011 | 0.28 | -0.851 | -0.183 | 1.151 | -0.006 | -0.005 | -0.005 | 0 | 0.75 | 0.74 | 0.69 | 0.71 | 0.8 | 0.76 |
| 7 | 90986046 | rs1429529 | -0.52 | -0.747 | 0.27 | 1.303 | 0.041 | 0.23 | 0.23 | 0 | 0.09 | 0.32 | 0.15 | 0.3 | 0.1 | 0.12 |
| 7 | 90986376 | rs4728968 | -0.485 | -0.741 | 0.314 | 1.306 | 0.035 | 0.212 | 0.212 | 0 | 0.09 | 0.3 | 0.14 | 0.3 | 0.1 | 0.12 |
| 7 | 90988012 | rs6975598 | 0.238 | -0.853 | 0.298 | 1.184 | -0.009 | -0.002 | -0.002 | 3.748 | 0.76 | 0.71 | 0.69 | 0.68 | 0.79 | 0.77 |
| 7 | 90989868 | rs1429520 | 0.411 | -0.855 | 0.331 | 1.206 | -0.013 | 0.01 | 0.01 | 0.112 | 0.74 | 0.66 | 0.69 | 0.6 | 0.77 | 0.74 |
| 7 | 90990111 | rs1429519 | 1.192 | -0.89 | 0.28 | 1.169 | 0.001 | -0.005 | -0.005 | 0 | 0.66 | 0.34 | 0.53 | 0.38 | 0.69 | 0.64 |
| 7 | 90992321 | rs10249530 | 0.794 | -1.003 | 0.066 | 1.281 | 0.003 | -0.007 | -0.007 | 0.021 | 0.71 | 0.69 | 0.66 | 0.52 | 0.73 | 0.65 |
| 7 | 90992609 | rs7793152 | 0.813 | -1.001 | 0.07 | 1.282 | 0.003 | -0.006 | -0.006 | 0 | 0.7 | 0.68 | 0.66 | 0.52 | 0.73 | 0.65 |
| 7 | 90993299 | rs10253828 | 0.285 | -1.003 | 0.068 | 1.356 | 0.003 | -0.002 | -0.002 | 1.087 | 0.76 | 0.76 | 0.7 | 0.65 | 0.83 | 0.75 |
| 7 | 90999109 | rs10953045 | 0.725 | -0.98 | 0.098 | 1.335 | -0.006 | 0.008 | 0.008 | 0 | 0.7 | 0.63 | 0.66 | 0.52 | 0.73 | 0.66 |
| 7 | 91010280 | rs10236943 | 0.433 | -1.174 | 0.164 | 1.451 | -0.022 | 0.137 | 0.137 | 0 | 0.19 | 0.35 | 0.29 | 0.37 | 0.16 | 0.28 |
| 7 | 91029468 | rs2564445 | 0.189 | -0.967 | 0.074 | 1.39 | 0.032 | 0.058 | 0.058 | 0 | 0.92 | 0.67 | 0.93 | 0.86 | 0.89 | 0.87 |
| 7 | 91033751 | rs2564443 | 0.515 | -0.982 | 0.058 | 1.497 | 0.033 | 0.091 | 0.091 | 0.002 | 0.91 | 0.63 | 0.92 | 0.84 | 0.89 | 0.87 |
| 7 | 91038222 | rs2709204 | -0.639 | -1.067 | -0.067 | 1.68 | -0.002 | 0.001 | 0.001 | 0 | 0.41 | 0.52 | 0.58 | 0.58 | 0.36 | 0.54 |
| 7 | 91043217 | rs720487 | 0.95 | -1.157 | -0.495 | 1.253 | 0.019 | -0.033 | -0.033 | 0.434 | 0.6 | 0.1 | 0.54 | 0.4 | 0.65 | 0.47 |
| 7 | 91086794 | rs4577890 | -0.326 | -0.793 | 0.039 | 0.903 | 0.004 | 0 | 0 | 0 | 0.39 | 0.32 | 0.57 | 0.61 | 0.33 | 0.49 |
| 7 | 91090780 | rs4076028 | 0.389 | -0.755 | -0.402 | 0.728 | 0.026 | 0.007 | 0.007 | 0 | 0.59 | 0.41 | 0.43 | 0.37 | 0.69 | 0.51 |
| 7 | 91098469 | rs4427099 | -0.179 | -0.496 | -1.19 | 0.722 | 0.053 | 0.029 | 0.029 | 0 | 0.43 | 0.24 | 0.56 | 0.64 | 0.33 | 0.5 |
| 7 | 91102306 | rs11981000 | 0.774 | -0.367 | -1.191 | 0.72 | -0.042 | 0.176 | 0.176 | 0 | 0.47 | 0.1 | 0.31 | 0.2 | 0.52 | 0.36 |
| 7 | 91102443 | rs10224650 | 0.856 | -0.365 | -1.021 | 0.925 | -0.015 | 0.136 | 0.136 | 0 | 0.07 | 0.17 | 0.13 | 0.16 | 0.04 | 0.05 |
| 7 | 91104171 | rs10276095 | -0.116 | 0.377 | -1.069 | 0.845 | 0.001 | 0.005 | 0.005 | 0 | 0.63 | 0.33 | 0.78 | 0.77 | 0.69 | 0.67 |
| 7 | 91113318 | rs7791302 | 1.134 | 0.376 | -1.538 | 0.486 | 0.016 | 0.1 | 0.1 | 0 | 0.26 | 0.07 | 0.11 | 0.08 | 0.18 | 0.17 |
| 7 | 91113955 | rs4626524 | -0.402 | 0.407 | -1.433 | 0.526 | 0.002 | -0.014 | -0.014 | 0.787 | 0.63 | 0.32 | 0.69 | 0.62 | 0.68 | 0.66 |
| 7 | 91142400 | rs11761294 | 1.223 | 0.696 | -1.49 | 0.505 | -0.093 | 0.017 | 0.017 | 0 | 0.2 | 0.04 | 0.21 | 0.13 | 0.34 | 0.17 |
| 7 | 91145137 | rs2709220 | -0.335 | 0.12 | -0.699 | 0.783 | 0.004 | -0.015 | -0.015 | 0 | 0.38 | 0.68 | 0.31 | 0.39 | 0.32 | 0.34 |
| 7 | 91149325 | rs6465323 | -0.604 | -0.216 | -0.549 | 0.282 | 0.026 | 0.03 | 0.03 | 0 | 0.39 | 0.69 | 0.24 | 0.14 | 0.26 | 0.4 |
| 7 | 91155612 | rs6956521 | -0.697 | -0.42 | -0.1 | 0.366 | -0.026 | 0.044 | 0.044 | 0.758 | 0.1 | 0.5 | 0.09 | 0.1 | 0.03 | 0.21 |
| 7 | 91155850 | rs17163952 | 0.085 | -0.386 | 0.058 | 0.341 | 0.032 | 0.045 | 0.045 | 0.187 | 0.07 | 0.37 | 0.04 | 0.04 | 0.03 | 0.1 |
| 7 | 91167397 | rs13241074 | 0.377 | 0.039 | -1.009 | 0.805 | -0.02 | -0.029 | -0.029 | 0 | 0.12 | 0.06 | 0.35 | 0.38 | 0.18 | 0.15 |
| 7 | 91168595 | rs6942835 | -0.937 | -0.745 | -0.609 | 0.193 | 0.024 | 0.028 | 0.028 | 0 | 0.38 | 0.69 | 0.31 | 0.16 | 0.25 | 0.4 |
| 7 | 91170941 | rs4727257 | -0.933 | -0.738 | -0.606 | 0.197 | 0.023 | 0.028 | 0.028 | 0 | 0.38 | 0.69 | 0.31 | 0.16 | 0.25 | 0.38 |
| 7 | 91172008 | rs17691195 | -0.005 | -0.471 | -0.866 | 0.206 | 0.026 | 0.122 | 0.122 | 0 | 0.27 | 0.07 | 0.14 | 0.05 | 0.2 | 0.17 |
| 7 | 91173580 | rs12538853 | -0.891 | 0.063 | -0.045 | 1.23 | -0.002 | -0.002 | -0.002 | 0 | 0.07 | 0.06 | 0.07 | 0.13 | 0.1 | 0.17 |
| 7 | 91174263 | rs7787692 | -1.334 | -0.375 | 0.385 | 1.064 | -0.021 | 0.108 | 0.108 | 0.117 | 0.1 | 0.54 | 0.16 | 0.13 | 0.04 | 0.24 |
| 7 | 91176242 | rs2540590 | -0.76 | -0.249 | -0.322 | 0.999 | 0.002 | -0.002 | -0.002 | 0 | 0.18 | 0.18 | 0.26 | 0.26 | 0.17 | 0.19 |
| 7 | 91178624 | rs17691277 | 0.525 | 0.008 | -1.265 | 0.572 | -0.059 | 0.051 | 0.051 | 2.633 | 0.28 | 0.08 | 0.22 | 0.18 | 0.32 | 0.22 |
| 7 | 91180083 | rs17716299 | 0.997 | -0.64 | -1.684 | 0.697 | 0.029 | 0.003 | 0.003 | 0 | 0.24 | 0.13 | 0.15 | 0.2 | 0.17 | 0.29 |

| | | | | | | | | | | | | | | | | |
|---|---|---|---|---|---|---|---|---|---|---|---|---|---|---|---|---|
| 7 | 91181056 | rs2710941 | 0.696 | -0.576 | -1.649 | 0.608 | -0.008 | -0.034 | -0.034 | 0 | 0.16 | 0.07 | 0.25 | 0.28 | 0.17 | 0.19 |
| 7 | 91181705 | rs2710942 | 0.122 | -0.666 | -0.647 | 0.58 | -0.001 | 0.004 | 0.004 | 0.235 | 0.47 | 0.45 | 0.5 | 0.45 | 0.49 | 0.41 |
| 7 | 91183628 | rs2465153 | -0.59 | -0.439 | -0.59 | 0.885 | 0.002 | -0.037 | -0.037 | 0 | 0.78 | 0.58 | 0.87 | 0.89 | 0.8 | 0.78 |
| 7 | 91184357 | rs4140877 | 0.886 | -0.684 | -1.174 | 0.601 | -0.067 | 0.078 | 0.078 | 0.222 | 0.34 | 0.06 | 0.23 | 0.21 | 0.34 | 0.34 |
| 7 | 91186352 | rs1076465 | -0.126 | -0.698 | -0.523 | 0.743 | 0.001 | -0.001 | -0.001 | 0 | 0.69 | 0.67 | 0.63 | 0.73 | 0.61 | 0.78 |
| 7 | 91186474 | rs1076464 | -0.031 | -1.302 | -0.212 | 0.968 | 0.011 | -0.014 | -0.014 | 0.147 | 0.33 | 0.31 | 0.38 | 0.47 | 0.28 | 0.44 |
| 7 | 91187387 | rs12530936 | 0.893 | -1.195 | -0.523 | 0.762 | 0.025 | -0.006 | -0.006 | 0 | 0.22 | 0.14 | 0.15 | 0.22 | 0.15 | 0.22 |
| 7 | 91189624 | rs2710952 | 0.211 | -0.965 | -0.176 | 0.81 | 0.012 | 0.015 | 0.015 | 0 | 0.49 | 0.38 | 0.65 | 0.6 | 0.57 | 0.63 |
| 7 | 91191512 | rs1017144 | -0.509 | -1.069 | -0.242 | 0.73 | 0.005 | 0.004 | 0.004 | 0 | 0.79 | 0.59 | 0.87 | 0.77 | 0.79 | 0.85 |
| 7 | 91191664 | rs12539100 | 0.821 | -0.949 | -1.185 | 0.759 | 0.042 | 0.055 | 0.055 | 0 | 0.29 | 0.12 | 0.21 | 0.17 | 0.22 | 0.22 |
| 7 | 91191928 | rs2430449 | 0.28 | -0.495 | -1.276 | 0.924 | -0.024 | 0.004 | 0.004 | 0.555 | 0.44 | 0.13 | 0.64 | 0.58 | 0.55 | 0.55 |
| 7 | 91195485 | rs2160256 | -0.423 | -0.218 | -0.984 | 1.357 | -0.001 | 0.027 | 0.027 | 0.012 | 0.1 | 0.07 | 0.11 | 0.34 | 0.12 | 0.17 |
| 7 | 91200083 | rs2540574 | -0.489 | -0.465 | -1.061 | 1.36 | 0 | 0.05 | 0.05 | 0 | 0.1 | 0.12 | 0.1 | 0.36 | 0.11 | 0.17 |
| 7 | 91206783 | rs2540571 | -0.909 | -0.739 | -1.793 | 0.96 | -0.13 | 0.182 | 0.182 | 0.007 | 0.58 | 0.97 | 0.53 | 0.85 | 0.5 | 0.6 |
| 7 | 91207699 | rs2540569 | -0.842 | -0.736 | -1.559 | 0.96 | -0.057 | 0.169 | 0.169 | 0 | 0.57 | 0.9 | 0.52 | 0.85 | 0.5 | 0.6 |
| 7 | 91208600 | rs2188148 | 1.046 | -1.441 | -1.91 | 1.004 | -0.006 | 0.013 | 0.013 | 0 | 0.24 | 0.02 | 0.19 | 0.14 | 0.18 | 0.16 |
| 7 | 91208821 | rs2430445 | 0.112 | -0.841 | -1.533 | 1.045 | -0.007 | -0.007 | -0.007 | 0 | 0.08 | 0.04 | 0.1 | 0.37 | 0.11 | 0.17 |
| 7 | 91210770 | rs992769 | -0.588 | -0.815 | -1.738 | 0.797 | -0.018 | 0.192 | 0.192 | 0 | 0.56 | 0.93 | 0.48 | 0.84 | 0.41 | 0.63 |
| 7 | 91226820 | rs2710976 | -0.222 | -0.917 | -1.665 | 0.902 | -0.005 | -0.008 | -0.008 | 0 | 0.09 | 0.08 | 0.14 | 0.35 | 0.12 | 0.15 |
| 7 | 91233469 | rs2540592 | -0.441 | -0.763 | -1.655 | 0.63 | -0.003 | 0.189 | 0.189 | 0 | 0.56 | 0.94 | 0.48 | 0.84 | 0.4 | 0.62 |
| 7 | 91246651 | rs6962300 | -1.376 | -0.545 | -0.244 | 1.197 | -0.031 | 0.032 | 0.032 | 0.418 | 0.17 | 0.66 | 0.17 | 0.18 | 0.09 | 0.23 |
| 7 | 91257948 | rs10249071 | -0.586 | 0.338 | -0.338 | 1.058 | 0.012 | 0.005 | 0.005 | 1.039 | 0.23 | 0.43 | 0.2 | 0.22 | 0.18 | 0.21 |
| 7 | 91263039 | rs4728985 | 0.395 | 0.434 | -0.88 | 1.439 | -0.021 | 0.002 | 0.002 | 0 | 0.06 | 0.01 | 0.11 | 0.37 | 0.1 | 0.15 |
| 7 | 91272344 | rs1476442 | -0.731 | 0.17 | -0.311 | 0.629 | -0.007 | -0.017 | -0.017 | 0.158 | 0.2 | 0.48 | 0.18 | 0.15 | 0.14 | 0.14 |
| 7 | 91280470 | rs12673563 | -0.924 | 0.153 | -0.368 | 0.74 | 0.015 | 0.046 | 0.046 | 0 | 0.08 | 0.27 | 0.12 | 0.12 | 0.08 | 0.06 |
| 7 | 91285428 | rs12668101 | -0.612 | 0.189 | -0.582 | 0.753 | 0.007 | 0.024 | 0.024 | 0 | 0.07 | 0.19 | 0.12 | 0.12 | 0.08 | 0.06 |
| 7 | 91289228 | rs6976523 | 0.046 | 0.001 | -1.448 | 0.254 | 0.04 | 0.113 | 0.113 | 0 | 0.49 | 0.2 | 0.45 | 0.31 | 0.56 | 0.38 |
| 7 | 91299161 | rs2974129 | 0.631 | -1.023 | -0.782 | 0.794 | 0.011 | -0.002 | -0.002 | 0 | 0.44 | 0.35 | 0.58 | 0.45 | 0.61 | 0.39 |
| 7 | 91303198 | rs12668203 | -0.048 | -1.328 | -1.008 | 0.802 | 0.003 | -0.005 | -0.005 | 0 | 0.17 | 0.17 | 0.12 | 0.25 | 0.1 | 0.16 |
| 7 | 91303535 | rs12672616 | -0.048 | -1.328 | -1.008 | 0.802 | 0.003 | -0.005 | -0.005 | 0 | 0.17 | 0.17 | 0.12 | 0.25 | 0.1 | 0.16 |
| 7 | 91320854 | rs2157998 | -0.616 | -1.248 | -0.878 | 0.689 | -0.006 | -0.004 | -0.004 | 0 | 0.67 | 0.66 | 0.72 | 0.73 | 0.73 | 0.62 |
| 7 | 91325649 | rs10953051 | -0.877 | -1.094 | -0.972 | 0.919 | 0.033 | -0.01 | -0.01 | 0 | 0.22 | 0.16 | 0.13 | 0.25 | 0.1 | 0.24 |
| 7 | 91345077 | rs12667992 | -0.929 | -1.107 | -0.967 | 0.994 | 0.033 | -0.017 | -0.017 | 0 | 0.22 | 0.17 | 0.14 | 0.29 | 0.1 | 0.24 |
| 7 | 91375638 | rs803269 | -0.321 | -1.184 | -0.657 | 0.774 | -0.014 | -0.015 | -0.015 | 0 | 0.65 | 0.61 | 0.73 | 0.77 | 0.75 | 0.62 |
| 7 | 91379996 | rs2521455 | -0.438 | -1.238 | -0.396 | 0.75 | 0.012 | 0.012 | 0.012 | 0 | 0.66 | 0.73 | 0.73 | 0.77 | 0.75 | 0.62 |
| 7 | 91380845 | rs803282 | 0.597 | -1.466 | -0.286 | 0.698 | 0.04 | 0.011 | 0.011 | 0 | 0.49 | 0.34 | 0.59 | 0.44 | 0.65 | 0.43 |
| 7 | 91398179 | rs1209000 | 1.246 | -0.673 | -0.386 | 0.904 | -0.005 | 0.003 | 0.003 | 0 | 0.35 | 0.29 | 0.27 | 0.23 | 0.44 | 0.25 |
| 7 | 91398541 | rs42985 | -0.359 | -0.804 | -0.097 | 0.738 | 0.075 | 0.013 | 0.013 | 0 | 0.48 | 0.73 | 0.42 | 0.56 | 0.36 | 0.54 |
| 7 | 91408398 | rs11763307 | -0.508 | -1.082 | 0.468 | 1.368 | -0.014 | 0.093 | 0.093 | 0 | 0.11 | 0.22 | 0.14 | 0.38 | 0.09 | 0.13 |
| 7 | 91429600 | rs4729006 | -0.478 | -1.082 | 0.468 | 1.368 | -0.013 | 0.087 | 0.087 | 0 | 0.11 | 0.21 | 0.13 | 0.38 | 0.09 | 0.14 |
| 7 | 91433827 | rs6949881 | 0.861 | -0.759 | 0.313 | 0.962 | 0.004 | 0.067 | 0.067 | 0.375 | 0.45 | 0.3 | 0.35 | 0.16 | 0.37 | 0.41 |
| 7 | 91440359 | rs1209001 | 1.417 | -0.768 | 0.309 | 1.07 | -0.031 | 0.001 | 0.001 | 0.406 | 0.23 | 0.15 | 0.19 | 0.18 | 0.34 | 0.23 |
| 7 | 91444679 | rs13229505 | 0.747 | -0.789 | 0.443 | 0.995 | 0.004 | 0.095 | 0.095 | 0.406 | 0.45 | 0.26 | 0.34 | 0.16 | 0.37 | 0.41 |
| 7 | 91467435 | rs6949700 | 0.646 | -0.798 | 0.458 | 0.991 | 0.003 | 0.017 | 0.017 | 0.845 | 0.46 | 0.37 | 0.35 | 0.16 | 0.38 | 0.41 |
| 7 | 91475007 | rs9656002 | -0.672 | -0.784 | 0.504 | 0.996 | 0.002 | 0.101 | 0.101 | 0.364 | 0.55 | 0.77 | 0.66 | 0.84 | 0.63 | 0.59 |
| 7 | 91492103 | rs10242792 | -0.82 | -0.764 | 0.56 | 1.08 | 0.007 | 0.093 | 0.093 | 0.214 | 0.55 | 0.75 | 0.64 | 0.84 | 0.61 | 0.59 |
| 7 | 91503228 | rs10266424 | 0.262 | -0.747 | 0.61 | 1.203 | -0.064 | -0.039 | -0.039 | 0 | 0.4 | 0.19 | 0.5 | 0.46 | 0.52 | 0.43 |
| 7 | 91509551 | rs2269811 | -0.511 | -0.764 | 0.705 | 1.203 | -0.008 | 0.064 | 0.064 | 0 | 0.12 | 0.22 | 0.13 | 0.36 | 0.09 | 0.14 |
| 7 | 91509867 | rs1011372 | 0.262 | -0.748 | 0.607 | 1.201 | -0.064 | -0.039 | -0.039 | 0.771 | 0.4 | 0.19 | 0.5 | 0.46 | 0.52 | 0.43 |
| 7 | 91539717 | rs7781767 | -1.043 | -0.64 | 0.819 | 1.3 | -0.006 | 0.278 | 0.278 | 0 | 0.14 | 0.46 | 0.14 | 0.37 | 0.09 | 0.19 |
| 7 | 91561592 | rs4729015 | -0.534 | -0.77 | 0.688 | 1.173 | 0.002 | 0.003 | 0.003 | 0.543 | 0.55 | 0.6 | 0.64 | 0.84 | 0.61 | 0.59 |
| 7 | 91568272 | rs1859037 | -0.534 | -0.772 | 0.761 | 1.179 | 0.002 | 0.003 | 0.003 | 0.543 | 0.55 | 0.6 | 0.64 | 0.84 | 0.61 | 0.59 |
| 7 | 91570615 | rs7797369 | -0.497 | -0.805 | 0.747 | 1.163 | -0.003 | -0.041 | -0.041 | 0.109 | 0.55 | 0.5 | 0.64 | 0.84 | 0.61 | 0.59 |
| 7 | 91571074 | rs10227225 | 0.172 | -0.835 | 0.733 | 1.187 | -0.068 | -0.031 | -0.031 | 0.009 | 0.44 | 0.17 | 0.51 | 0.47 | 0.52 | 0.45 |
| 7 | 91572749 | rs2961019 | -1.379 | -1.07 | 0.768 | 1.138 | -0.025 | 0.094 | 0.094 | 0 | 0.64 | 0.83 | 0.73 | 0.92 | 0.63 | 0.63 |
| 7 | 91574110 | rs12666179 | -0.561 | -0.934 | 0.773 | 1.187 | 0.012 | 0.096 | 0.096 | 0.165 | 0.55 | 0.34 | 0.63 | 0.82 | 0.61 | 0.57 |
| 7 | 91575417 | rs6953256 | 0.226 | -0.934 | 0.772 | 1.201 | -0.068 | -0.028 | -0.028 | 0 | 0.44 | 0.17 | 0.51 | 0.47 | 0.52 | 0.45 |
| 7 | 91580490 | rs415 | -0.546 | -0.934 | 0.774 | 1.2 | -0.001 | -0.018 | -0.018 | 0 | 0.56 | 0.56 | 0.64 | 0.84 | 0.61 | 0.59 |
| 7 | 91583334 | rs418 | 0.224 | -0.93 | 0.821 | 1.202 | -0.072 | -0.035 | -0.035 | 0 | 0.44 | 0.18 | 0.51 | 0.47 | 0.52 | 0.45 |
| 7 | 91584356 | rs2282971 | 0.526 | -0.931 | 0.832 | 1.195 | 0.01 | 0.082 | 0.082 | 0.115 | 0.45 | 0.63 | 0.37 | 0.18 | 0.39 | 0.43 |
| 7 | 91588036 | rs12530646 | 0.458 | -0.923 | 0.766 | 1.198 | -0.001 | -0.023 | -0.023 | 1.305 | 0.44 | 0.44 | 0.36 | 0.16 | 0.39 | 0.41 |
| 7 | 91588194 | rs7781597 | 0.454 | -0.84 | 0.769 | 1.259 | 0 | -0.007 | -0.007 | 0 | 0.44 | 0.44 | 0.35 | 0.18 | 0.38 | 0.44 |
| 7 | 91590408 | rs17164275 | 0.464 | -0.954 | 0.784 | 1.175 | -0.001 | -0.021 | -0.021 | 2.328 | 0.44 | 0.44 | 0.35 | 0.16 | 0.39 | 0.41 |
| 7 | 91594510 | rs6942649 | 0.397 | -0.953 | 0.762 | 1.175 | -0.003 | -0.039 | -0.039 | 0.113 | 0.45 | 0.5 | 0.36 | 0.16 | 0.39 | 0.41 |
| 7 | 91610111 | rs10228800 | 0.399 | -0.961 | 0.718 | 1.158 | -0.003 | -0.041 | -0.041 | 0.057 | 0.45 | 0.5 | 0.36 | 0.16 | 0.39 | 0.41 |
| 7 | 91616891 | rs10279210 | 0.498 | -0.954 | 0.72 | 1.161 | 0.013 | 0.101 | 0.101 | 0.165 | 0.45 | 0.66 | 0.37 | 0.18 | 0.39 | 0.43 |
| 7 | 91626699 | rs2299232 | 0.624 | -0.961 | 0.721 | 1.158 | 0.002 | 0.069 | 0.069 | 0.011 | 0.43 | 0.26 | 0.35 | 0.16 | 0.38 | 0.41 |
| 7 | 91626953 | rs2299233 | -0.273 | -0.931 | 0.51 | 1.178 | -0.072 | -0.031 | -0.031 | 0 | 0.56 | 0.83 | 0.49 | 0.54 | 0.48 | 0.55 |
| 7 | 91629984 | rs3753108 | 0.501 | -0.927 | 0.514 | 1.167 | 0.013 | 0.102 | 0.102 | 0.128 | 0.46 | 0.66 | 0.37 | 0.18 | 0.39 | 0.43 |

| | | | | | | | | | | | | | | | | |
|---|---|---|---|---|---|---|---|---|---|---|---|---|---|---|---|---|
| 7 | 91630620 | rs6964587 | 0.41 | -0.927 | 0.515 | 1.166 | -0.003 | -0.041 | -0.041 | 0 | 0.45 | 0.5 | 0.36 | 0.16 | 0.39 | 0.41 |
| 7 | 91633213 | rs12540565 | 0.452 | -0.918 | 0.546 | 1.17 | -0.002 | -0.035 | -0.035 | 0 | 0.44 | 0.46 | 0.36 | 0.16 | 0.39 | 0.41 |
| 7 | 91640273 | rs6465344 | 0.505 | -0.919 | 0.537 | 1.171 | 0.013 | 0.102 | 0.102 | 0.128 | 0.46 | 0.66 | 0.37 | 0.18 | 0.39 | 0.43 |
| 7 | 91641928 | rs13245393 | 0.561 | -0.916 | 0.422 | 1.157 | -0.003 | -0.035 | -0.035 | 3.569 | 0.45 | 0.51 | 0.36 | 0.16 | 0.39 | 0.41 |
| 7 | 91644553 | rs7805077 | 0.403 | -0.914 | 0.423 | 1.158 | -0.003 | -0.041 | -0.041 | 0 | 0.45 | 0.5 | 0.36 | 0.16 | 0.39 | 0.41 |
| 7 | 91644613 | rs4729018 | -0.269 | -0.918 | 0.42 | 1.168 | -0.072 | -0.035 | -0.035 | 0.027 | 0.56 | 0.84 | 0.49 | 0.53 | 0.48 | 0.55 |
| 7 | 91648341 | rs2888851 | 0.358 | -0.835 | 0.391 | 0.983 | -0.002 | -0.03 | -0.03 | 1.759 | 0.44 | 0.46 | 0.36 | 0.16 | 0.39 | 0.41 |
| 7 | 91654517 | rs2299238 | 0.377 | -0.835 | 0.39 | 0.983 | -0.001 | -0.025 | -0.025 | 0.105 | 0.44 | 0.45 | 0.36 | 0.16 | 0.39 | 0.41 |
| 7 | 91657994 | rs6952389 | 0.295 | -0.822 | 0.321 | 0.989 | -0.003 | -0.035 | -0.035 | 0.001 | 0.45 | 0.51 | 0.36 | 0.16 | 0.39 | 0.41 |
| 7 | 91663364 | rs7811564 | 0.356 | -0.824 | 0.325 | 0.669 | -0.002 | -0.03 | -0.03 | 0 | 0.44 | 0.46 | 0.36 | 0.16 | 0.39 | 0.41 |
| 7 | 91664645 | rs7781221 | 0.356 | -0.824 | 0.325 | 0.669 | -0.002 | -0.03 | -0.03 | 0 | 0.44 | 0.46 | 0.36 | 0.16 | 0.39 | 0.41 |
| 7 | 91666369 | rs2282972 | 0.356 | -0.824 | 0.325 | 0.669 | -0.002 | -0.03 | -0.03 | 0 | 0.44 | 0.46 | 0.36 | 0.16 | 0.39 | 0.41 |
| 7 | 91666661 | rs2282973 | -0.19 | -0.898 | 0.398 | 0.799 | -0.005 | 0.064 | 0.064 | 0 | 0.11 | 0.17 | 0.13 | 0.35 | 0.09 | 0.12 |
| 7 | 91668854 | rs7789492 | 0.483 | -0.907 | 0.489 | 0.811 | 0.014 | 0.109 | 0.109 | 0.149 | 0.46 | 0.67 | 0.37 | 0.18 | 0.39 | 0.43 |
| 7 | 91671853 | rs926192 | 0.483 | -0.907 | 0.51 | 0.811 | 0.014 | 0.109 | 0.109 | 0.149 | 0.46 | 0.67 | 0.37 | 0.18 | 0.39 | 0.43 |
| 7 | 91672403 | rs9784993 | 0.562 | -0.946 | 0.493 | 0.793 | -0.003 | -0.035 | -0.035 | 0 | 0.45 | 0.51 | 0.36 | 0.16 | 0.39 | 0.41 |
| 7 | 91674302 | rs9785013 | 0.489 | -0.908 | 0.523 | 0.81 | 0.015 | 0.118 | 0.118 | 0.15 | 0.46 | 0.68 | 0.37 | 0.18 | 0.39 | 0.43 |
| 7 | 91679887 | rs1859113 | 0.399 | -0.907 | 0.521 | 0.809 | -0.002 | -0.032 | -0.032 | 0 | 0.45 | 0.51 | 0.36 | 0.16 | 0.39 | 0.41 |
| 7 | 91680974 | rs10275125 | -1.928 | -0.98 | 0.35 | 0.782 | -0.086 | -0.033 | -0.033 | 0.113 | 0.8 | 0.98 | 0.82 | 0.82 | 0.69 | 0.81 |
| 7 | 91691070 | rs2961024 | -1.94 | -0.984 | 0.318 | 0.78 | -0.086 | -0.03 | -0.03 | 0.905 | 0.8 | 0.98 | 0.82 | 0.82 | 0.69 | 0.81 |
| 7 | 91726927 | rs1063243 | 0.596 | -0.977 | 0.346 | 0.779 | -0.002 | -0.027 | -0.027 | 0 | 0.45 | 0.52 | 0.36 | 0.16 | 0.39 | 0.41 |
| 7 | 91733829 | rs4512319 | 1.501 | -1.083 | 0.444 | 0.74 | -0.027 | 0.104 | 0.104 | 0 | 0.36 | 0.16 | 0.28 | 0.08 | 0.37 | 0.37 |
| 7 | 91736464 | rs12538678 | -1.687 | -1.073 | 0.122 | 0.757 | -0.038 | -0.001 | -0.001 | 0 | 0.78 | 0.88 | 0.81 | 0.81 | 0.7 | 0.78 |
| 7 | 91736874 | rs7802788 | 0.611 | -1.073 | 0.166 | 0.757 | 0.015 | 0.113 | 0.113 | 0.137 | 0.46 | 0.67 | 0.37 | 0.18 | 0.39 | 0.43 |
| 7 | 91738437 | rs2075882 | -0.11 | -1.078 | 0.155 | 0.761 | -0.005 | 0.067 | 0.067 | 0 | 0.11 | 0.17 | 0.12 | 0.35 | 0.09 | 0.13 |
| 7 | 91740796 | rs10274756 | 0.706 | -1.067 | 0.171 | 0.734 | -0.002 | -0.032 | -0.032 | 0 | 0.45 | 0.51 | 0.36 | 0.16 | 0.39 | 0.41 |
| 7 | 91746727 | rs2301559 | 0.584 | -1.067 | 0.178 | 0.734 | -0.002 | -0.026 | -0.026 | 0 | 0.45 | 0.45 | 0.36 | 0.16 | 0.39 | 0.41 |
| 7 | 91750016 | rs6956431 | 0.706 | -1.067 | 0.178 | 0.734 | -0.002 | -0.032 | -0.032 | 0 | 0.45 | 0.51 | 0.36 | 0.16 | 0.39 | 0.41 |
| 7 | 91751120 | rs7811873 | 0.712 | -1.081 | 0.172 | 0.728 | -0.001 | -0.024 | -0.024 | 0 | 0.45 | 0.45 | 0.36 | 0.16 | 0.39 | 0.41 |
| 7 | 91752690 | rs2269727 | 0.57 | -1.05 | 0.188 | 0.769 | -0.002 | -0.026 | -0.026 | 0 | 0.45 | 0.45 | 0.36 | 0.16 | 0.39 | 0.41 |
| 7 | 91753579 | rs10488512 | 0.692 | -1.05 | 0.187 | 0.769 | -0.002 | -0.032 | -0.032 | 0 | 0.45 | 0.51 | 0.36 | 0.16 | 0.39 | 0.41 |
| 7 | 91758111 | rs7784248 | 0.692 | -1.05 | 0.187 | 0.769 | -0.002 | -0.032 | -0.032 | 0.659 | 0.45 | 0.51 | 0.36 | 0.16 | 0.39 | 0.41 |
| 7 | 91759597 | rs4644173 | 0.603 | -1.05 | 0.17 | 0.77 | 0.015 | 0.116 | 0.116 | 0.137 | 0.46 | 0.67 | 0.37 | 0.18 | 0.39 | 0.43 |
| 7 | 91766322 | rs9656003 | 0.64 | -1.091 | 0.194 | 0.773 | 0.011 | 0.116 | 0.116 | 1.389 | 0.46 | 0.67 | 0.37 | 0.19 | 0.39 | 0.37 |
| 7 | 91768615 | rs6465350 | 0.607 | -1.091 | 0.199 | 0.773 | -0.002 | -0.033 | -0.033 | 0 | 0.44 | 0.45 | 0.36 | 0.17 | 0.39 | 0.35 |
| 7 | 91773213 | rs6465351 | -0.256 | -1.093 | -0.228 | 0.785 | -0.071 | -0.039 | -0.039 | 0 | 0.57 | 0.83 | 0.49 | 0.53 | 0.48 | 0.55 |
| 7 | 91773359 | rs7807490 | -0.256 | -1.093 | -0.253 | 0.785 | -0.073 | -0.04 | -0.04 | 0 | 0.57 | 0.83 | 0.49 | 0.53 | 0.48 | 0.55 |
| 7 | 91779971 | rs6465353 | 0.648 | -1.092 | -0.253 | 0.784 | 0.011 | 0.116 | 0.116 | 1.389 | 0.46 | 0.67 | 0.37 | 0.19 | 0.39 | 0.36 |
| 7 | 91785970 | rs10271174 | 0.706 | -1.094 | -0.255 | 0.784 | 0.008 | 0.077 | 0.077 | 0.822 | 0.45 | 0.6 | 0.37 | 0.19 | 0.39 | 0.36 |
| 7 | 91797296 | rs10236483 | -0.239 | -1.072 | -0.484 | 0.786 | -0.073 | -0.04 | -0.04 | 0 | 0.57 | 0.83 | 0.49 | 0.53 | 0.48 | 0.55 |
| 7 | 91842554 | rs11542682 | -0.811 | -1.073 | -0.405 | 0.851 | -0.002 | 0.031 | 0.031 | 0 | 0.12 | 0.16 | 0.12 | 0.34 | 0.08 | 0.19 |
| 7 | 91856816 | rs11769780 | 0.248 | -1.077 | -0.401 | 0.849 | -0.012 | 0.046 | 0.046 | 0 | 0.08 | 0.01 | 0.06 | 0 | 0.08 | 0.03 |
| 7 | 91861656 | rs1034575 | -0.637 | -1.06 | -0.394 | 0.858 | -0.003 | 0.034 | 0.034 | 0 | 0.12 | 0.16 | 0.12 | 0.34 | 0.08 | 0.19 |
| 7 | 91874686 | rs12704637 | 1.518 | -1.023 | -0.504 | 0.758 | -0.109 | 0.199 | 0.199 | 1.844 | 0.35 | 0.03 | 0.25 | 0.08 | 0.38 | 0.32 |
| 7 | 91901716 | rs7796370 | -0.977 | -0.907 | -0.5 | 0.729 | -0.051 | -0.061 | -0.061 | 2.853 | 0.8 | 0.97 | 0.68 | 0.71 | 0.79 | 0.77 |
| 7 | 91925256 | rs721015 | 0.072 | -0.421 | -0.748 | 0.938 | 0.005 | 0.02 | 0.02 | 0.39 | 0.5 | 0.72 | 0.47 | 0.53 | 0.44 | 0.51 |
| 7 | 91998495 | rs17164519 | -0.308 | -0.342 | -0.67 | 1.015 | 0.009 | 0 | 0 | 0.004 | 0.1 | 0.06 | 0.11 | 0.34 | 0.07 | 0.16 |
| 7 | 92008220 | rs12536421 | 1.111 | -0.333 | -0.81 | 0.847 | -0.005 | 0.024 | 0.024 | 1.386 | 0.37 | 0.28 | 0.34 | 0.17 | 0.35 | 0.31 |
| 7 | 92028039 | rs38794 | -0.392 | 0.146 | -1.832 | 0.284 | -0.005 | 0.011 | 0.011 | 0.366 | 0.49 | 0.24 | 0.52 | 0.47 | 0.56 | 0.49 |
| 7 | 92031948 | rs38795 | 1.107 | 0.268 | -1.69 | 0.435 | -0.072 | -0.022 | -0.022 | 0 | 0.2 | 0.02 | 0.19 | 0.16 | 0.32 | 0.19 |
| 7 | 92043016 | rs39744 | -0.096 | 0.034 | -1.094 | 0.552 | 0.001 | 0.004 | 0.004 | 1.088 | 0.55 | 0.31 | 0.56 | 0.45 | 0.59 | 0.48 |
| 7 | 92055272 | rs6977036 | -0.404 | -0.321 | -0.597 | 0.955 | 0.03 | -0.014 | -0.014 | 0 | 0.41 | 0.68 | 0.34 | 0.55 | 0.29 | 0.54 |
| 7 | 92057299 | rs4729040 | -1.158 | 0.302 | 0.708 | 1.383 | 0.032 | 0.138 | 0.138 | 0 | 0.15 | 0.38 | 0.2 | 0.35 | 0.15 | 0.21 |
| 7 | 92057320 | rs10278819 | -1.152 | 0.271 | 0.729 | 0.747 | 0.011 | -0.07 | -0.07 | 2.165 | 0.13 | 0.47 | 0.1 | 0.03 | 0.12 | 0.15 |
| 7 | 92063500 | rs4729041 | -1.417 | 0.594 | 0.013 | 1.677 | -0.001 | 0.024 | 0.024 | 0 | 0.11 | 0.1 | 0.18 | 0.31 | 0.15 | 0.1 |
| 7 | 92068289 | rs10281972 | -1.993 | 0.416 | 0.774 | 1.732 | -0.013 | 0.204 | 0.204 | 0.178 | 0.09 | 0.31 | 0.12 | 0.29 | 0.05 | 0.06 |
| 7 | 92068523 | rs7777732 | -1.517 | 0.298 | 0.654 | 1.439 | 0.003 | -0.008 | -0.008 | 0 | 0.12 | 0.19 | 0.09 | 0.04 | 0.15 | 0.07 |
| 7 | 92069057 | rs7781900 | -0.891 | 0.327 | 0.563 | 1.475 | -0.028 | -0.019 | -0.019 | 0 | 0.05 | 0.01 | 0.06 | 0.04 | 0.1 | 0.05 |
| 7 | 92069164 | rs4729042 | -1.686 | 0.271 | 0.763 | 1.502 | -0.015 | 0.141 | 0.141 | 0 | 0.08 | 0.22 | 0.12 | 0.29 | 0.05 | 0.06 |
| 7 | 92080395 | rs2287179 | -1.278 | 0.254 | 0.768 | 1.505 | -0.023 | 0.123 | 0.123 | 0 | 0.05 | 0.1 | 0.1 | 0.27 | 0.02 | 0.03 |
| 7 | 92096133 | rs10257545 | -1.042 | 0.346 | 0.741 | 1.445 | -0.003 | -0.001 | -0.001 | 0.547 | 0.05 | 0.07 | 0.06 | 0.03 | 0.1 | 0.05 |
| 7 | 92097166 | rs10232215 | -2.759 | 0.294 | 0.901 | 1.491 | -0.001 | 0.078 | 0.078 | 0 | 0.2 | 0.4 | 0.2 | 0.34 | 0.2 | 0.13 |
| 7 | 92110467 | rs10235602 | 1.22 | 0.307 | 0.939 | 1.462 | 0.005 | 0 | 0 | 0 | 0.94 | 0.89 | 0.94 | 0.97 | 0.89 | 0.95 |
| 7 | 92136754 | rs7791858 | -0.462 | 1.375 | 1.302 | 1.374 | 0.071 | -0.001 | -0.001 | 0 | 0.14 | 0.27 | 0.35 | 0.11 | 0.23 | 0.22 |
| 7 | 92140397 | rs2111200 | -2.546 | 1.238 | 0.968 | 1.383 | 0.001 | 0.083 | 0.083 | 0 | 0.2 | 0.41 | 0.21 | 0.33 | 0.2 | 0.13 |
| 7 | 92144751 | rs10252324 | -1.792 | 1.447 | 0.76 | 1.284 | 0.023 | -0.001 | -0.001 | 0 | 0.07 | 0.16 | 0.1 | 0.05 | 0.16 | 0.08 |
| 7 | 92155775 | rs6976142 | -1.528 | 1.3 | 0.754 | 1.29 | 0.008 | 0 | 0 | 0 | 0.07 | 0.13 | 0.08 | 0.05 | 0.12 | 0.08 |
| 7 | 92157886 | rs12386601 | -1.552 | 1.301 | 0.767 | 1.29 | 0.021 | 0.001 | 0.001 | 0 | 0.07 | 0.15 | 0.08 | 0.05 | 0.12 | 0.08 |
| 7 | 92167829 | rs10488516 | -1.371 | 1.723 | 0.594 | 0.705 | 0.012 | 0.001 | 0.001 | 0 | 0.06 | 0.11 | 0.08 | 0.05 | 0.12 | 0.08 |

| | | | | | | | | | | | | | | | | |
|---|---|---|---|---|---|---|---|---|---|---|---|---|---|---|---|---|
| 7 | 92182674 | rs6950934 | -1.749 | 1.715 | 0.609 | 0.621 | 0.025 | -0.007 | -0.007 | 0 | 0.07 | 0.17 | 0.08 | 0.04 | 0.12 | 0.09 |
| 7 | 92216059 | rs10488519 | 0.345 | 1.517 | 0.494 | 0.596 | -0.008 | 0.026 | 0.026 | 0 | 0.06 | 0 | 0.02 | 0 | 0.05 | 0.02 |
| 7 | 92223192 | rs17164630 | -1.199 | 1.091 | 0.499 | 0.404 | 0.004 | -0.025 | -0.025 | 0 | 0.07 | 0.27 | 0.03 | 0.01 | 0.04 | 0.04 |
| 7 | 92230162 | rs740047 | 2.197 | 1.468 | 0.464 | 0.541 | 0.066 | -0.01 | -0.01 | 0.09 | 0.88 | 0.72 | 0.8 | 0.89 | 0.79 | 0.91 |
| 7 | 92236164 | rs8179 | 1.148 | 1.262 | 0.34 | 0.436 | -0.025 | 0.028 | 0.028 | 0 | 0.85 | 0.89 | 0.86 | 0.98 | 0.78 | 0.92 |
| 7 | 92236829 | rs4272 | -1.422 | 1.154 | 0.582 | 0.423 | -0.003 | 0.002 | 0.002 | 0 | 0.16 | 0.19 | 0.21 | 0.1 | 0.22 | 0.09 |
| 7 | 92243672 | rs42377 | -2.167 | 1.21 | 0.563 | 0.274 | 0.018 | -0.063 | -0.063 | 0.128 | 0.27 | 0.41 | 0.27 | 0.1 | 0.32 | 0.16 |
| 7 | 92244422 | rs42039 | -1.048 | 1.123 | 0.541 | 0.287 | -0.042 | 0.052 | 0.052 | 0 | 0.18 | 0.09 | 0.17 | 0.02 | 0.27 | 0.11 |
| 7 | 92250354 | rs42045 | -1.876 | 0.627 | 0.149 | 0.236 | 0.008 | 0.133 | 0.133 | 3.302 | 0.51 | 0.8 | 0.76 | 0.8 | 0.55 | 0.44 |
| 7 | 92253233 | rs3731348 | -0.146 | 0.599 | 0.118 | 0.211 | 0 | 0.011 | 0.011 | 0 | 0.06 | 0.02 | 0.03 | 0.01 | 0.05 | 0.04 |
| 7 | 92264410 | rs2282978 | -1.445 | 0.489 | -0.065 | 0.179 | 0.003 | -0.057 | -0.057 | 3.735 | 0.36 | 0.49 | 0.4 | 0.11 | 0.36 | 0.18 |
| 7 | 92264993 | rs2282979 | -0.52 | 0.486 | -0.092 | 0.179 | -0.042 | 0.074 | 0.074 | 0.479 | 0.26 | 0.12 | 0.35 | 0.1 | 0.31 | 0.14 |
| 7 | 92273536 | rs3731343 | -2.27 | 0.47 | -0.094 | 0.163 | -0.003 | 0.153 | 0.153 | 4.101 | 0.52 | 0.88 | 0.77 | 0.83 | 0.55 | 0.44 |
| 7 | 92279363 | rs2282983 | -1.741 | -0.373 | -0.1 | -0.001 | 0 | 0.003 | 0.003 | 0.208 | 0.42 | 0.59 | 0.43 | 0.26 | 0.39 | 0.43 |
| 7 | 92279693 | rs2282984 | 0.42 | -0.261 | -0.221 | -0.201 | -0.015 | 0.075 | 0.075 | 0 | 0.08 | 0.17 | 0.03 | 0.15 | 0.03 | 0.25 |
| 7 | 92280695 | rs11765954 | -0.424 | -0.195 | -0.31 | -0.243 | -0.048 | 0.101 | 0.101 | 0.633 | 0.26 | 0.11 | 0.21 | 0.02 | 0.31 | 0.13 |
| 7 | 92303554 | rs3757823 | -1.721 | -1.203 | 0.338 | 0.54 | -0.042 | 0.183 | 0.183 | 0 | 0.2 | 0.53 | 0.14 | 0.5 | 0.11 | 0.34 |
| 7 | 92315330 | rs6964803 | -1.326 | -1.071 | -0.08 | 0.482 | -0.012 | 0.182 | 0.182 | 0.553 | 0.08 | 0.35 | 0.08 | 0.34 | 0.03 | 0.06 |
| 7 | 92317752 | rs7781436 | -1.591 | -1.424 | -0.62 | -0.032 | -0.054 | 0.101 | 0.101 | 0.77 | 0.75 | 0.99 | 0.89 | 0.96 | 0.78 | 0.79 |
| 7 | 92326468 | rs2282990 | -0.883 | -1.356 | 0.635 | 0.854 | -0.009 | 0.006 | 0.006 | 0 | 0.12 | 0.11 | 0.06 | 0.12 | 0.08 | 0.28 |
| 7 | 92328956 | rs1004051 | -1.449 | -1.271 | 0.664 | 0.906 | -0.009 | 0.186 | 0.186 | 0.401 | 0.08 | 0.36 | 0.08 | 0.34 | 0.03 | 0.05 |
| 7 | 92353332 | rs10234981 | -1.513 | -1.138 | 0.596 | 0.434 | -0.03 | 0.014 | 0.014 | 0 | 0.15 | 0.41 | 0.07 | 0.13 | 0.08 | 0.27 |
| 7 | 92377183 | rs4729049 | -1.393 | -1.453 | 0.555 | 0.502 | -0.013 | 0.013 | 0.013 | 0 | 0.12 | 0.17 | 0.06 | 0.15 | 0.1 | 0.33 |
| 7 | 92380438 | rs17164772 | -1.992 | -1.541 | 0.898 | 0.455 | -0.032 | 0.027 | 0.027 | 0 | 0.15 | 0.44 | 0.09 | 0.15 | 0.1 | 0.33 |
| 7 | 92402480 | rs3802073 | -2.03 | -1.535 | 0.902 | 0.451 | -0.026 | 0.015 | 0.015 | 0 | 0.14 | 0.34 | 0.07 | 0.14 | 0.1 | 0.33 |
| 7 | 92408370 | rs445 | -0.498 | -2.032 | 0.727 | 0.489 | 0.008 | -0.007 | -0.007 | 2.426 | 0.23 | 0.18 | 0.15 | 0.36 | 0.11 | 0.1 |
| 7 | 92411623 | rs2282995 | -1.442 | -1.084 | 0.865 | 0.289 | -0.004 | 0.002 | 0.002 | 0 | 0.12 | 0.09 | 0.05 | 0.15 | 0.1 | 0.33 |
| 7 | 92435186 | rs6975474 | -2.769 | -0.722 | 1.062 | 0.256 | 0.003 | 0.022 | 0.022 | 0.003 | 0.11 | 0.4 | 0.07 | 0.08 | 0.06 | 0.3 |
| 7 | 92449293 | rs10246604 | -2.587 | -0.627 | 0.891 | 0.3 | 0.016 | 0.036 | 0.036 | 0 | 0.09 | 0.35 | 0.07 | 0.08 | 0.06 | 0.3 |
| 7 | 92470345 | rs10269570 | -0.686 | -0.76 | 0.374 | 0.161 | 0.011 | 0.032 | 0.032 | 0 | 0.16 | 0.1 | 0.09 | 0.04 | 0.14 | 0.19 |
| 7 | 92477082 | rs4605988 | -1.259 | -0.291 | 0.557 | 0.424 | -0.017 | -0.022 | -0.022 | 0 | 0.19 | 0.24 | 0.11 | 0.11 | 0.14 | 0.4 |
| 7 | 92527551 | rs10272533 | -2.416 | -0.232 | 1.726 | 0.867 | -0.04 | 0.012 | 0.012 | 1.728 | 0.12 | 0.69 | 0.08 | 0.07 | 0.04 | 0.21 |
| 7 | 92567471 | rs10255859 | -2.319 | 1.015 | 1.617 | 2.247 | 0.096 | 0.351 | 0.351 | 1.192 | 0.08 | 0.48 | 0.25 | 0.25 | 0.09 | 0.14 |
| 7 | 92577760 | rs6955627 | -2.372 | 0.955 | 1.764 | 2.246 | 0.085 | 0.332 | 0.332 | 3.762 | 0.08 | 0.5 | 0.25 | 0.25 | 0.08 | 0.12 |
| 7 | 92611456 | rs4727283 | -1.241 | 0.484 | 1.095 | 1.92 | 0.001 | 0 | 0 | 0 | 0.16 | 0.16 | 0.35 | 0.6 | 0.17 | 0.37 |
| 7 | 92615875 | rs10230949 | -0.647 | 0.763 | 0.842 | 1.693 | 0 | 0.023 | 0.023 | 0 | 0.05 | 0.05 | 0.03 | 0.15 | 0.05 | 0.17 |
| 7 | 92616315 | rs17164910 | 0.561 | 0.028 | 0.389 | 1.125 | 0.02 | 0.019 | 0.019 | 0 | 0.11 | 0.04 | 0.05 | 0.08 | 0.05 | 0.08 |
| 7 | 92623141 | rs1013803 | -2.268 | 0.115 | 0.917 | 0.767 | 0.003 | -0.004 | -0.004 | 0 | 0.46 | 0.57 | 0.53 | 0.9 | 0.41 | 0.69 |
| 7 | 92635659 | rs4729057 | -2.279 | 0.114 | 0.629 | 0.798 | 0.003 | -0.012 | -0.012 | 0 | 0.45 | 0.57 | 0.53 | 0.9 | 0.4 | 0.68 |
| 7 | 92636769 | rs6949790 | -1.887 | -0.034 | 0.673 | 0.886 | -0.01 | -0.027 | -0.027 | 0 | 0.38 | 0.49 | 0.48 | 0.76 | 0.35 | 0.49 |
| 7 | 92637108 | rs17778801 | -1.01 | 0.298 | 0.218 | 1.2 | -0.002 | 0 | 0 | 0 | 0.08 | 0.09 | 0.1 | 0.14 | 0.14 | 0.19 |
| 7 | 92646033 | rs7806426 | -0.947 | -0.359 | 0.464 | 1.597 | -0.049 | 0.168 | 0.168 | 0 | 0.21 | 0.46 | 0.23 | 0.48 | 0.16 | 0.13 |
| 7 | 92648340 | rs723668 | -0.05 | 0.074 | 0.372 | 1.757 | -0.005 | 0.035 | 0.035 | 0.074 | 0.06 | 0.05 | 0.26 | 0.27 | 0.12 | 0.27 |
| 7 | 92650743 | rs10488528 | 0.22 | 0.112 | 0.172 | 1.356 | -0.027 | -0.006 | -0.006 | 0 | 0.06 | 0 | 0.03 | 0.07 | 0.07 | 0.07 |
| 7 | 92653334 | rs10488529 | 0.099 | -0.417 | -0.037 | 1.04 | 0.001 | -0.008 | -0.008 | 0 | 0.14 | 0.08 | 0.16 | 0.16 | 0.11 | 0.21 |
| 7 | 92653680 | rs11979483 | -0.764 | -0.439 | 0.087 | 0.497 | -0.034 | -0.099 | -0.099 | 1.315 | 0.23 | 0.45 | 0.16 | 0.07 | 0.17 | 0.11 |
| 7 | 92654839 | rs4294124 | -0.465 | 0.079 | 0.191 | 0.657 | 0.001 | -0.003 | -0.003 | 3.431 | 0.07 | 0.03 | 0.04 | 0 | 0.06 | 0.01 |
| 7 | 92657985 | rs10215082 | -1.16 | 0.015 | 0.074 | 0.416 | -0.06 | 0.142 | 0.142 | 2.269 | 0.61 | 0.86 | 0.74 | 0.92 | 0.56 | 0.69 |
| 7 | 92659160 | rs2188177 | -1.143 | -0.196 | 0.108 | 0.514 | -0.12 | 0.272 | 0.272 | 2.705 | 0.6 | 0.94 | 0.72 | 0.92 | 0.52 | 0.7 |
| 7 | 92666717 | rs4729062 | -0.54 | -0.061 | 0.42 | 0.788 | 0.01 | 0.006 | 0.006 | 0 | 0.49 | 0.38 | 0.57 | 0.58 | 0.45 | 0.65 |
| 7 | 92667211 | rs11981412 | -0.981 | -0.058 | 0.357 | 0.791 | -0.002 | -0.008 | -0.008 | 0 | 0.54 | 0.82 | 0.6 | 0.58 | 0.47 | 0.66 |
| 7 | 92674917 | rs12704649 | -0.704 | 0.294 | -0.082 | 1.017 | 0.004 | 0.001 | 0.001 | 0 | 0.49 | 0.44 | 0.57 | 0.58 | 0.45 | 0.65 |
| 7 | 92682609 | rs739385 | -1.015 | 0.074 | 0.509 | 1.334 | -0.028 | 0.012 | 0.012 | 2.868 | 0.19 | 0.6 | 0.19 | 0.19 | 0.16 | 0.03 |
| 7 | 92685201 | rs10248180 | -1.381 | -0.131 | -0.568 | 1.764 | 0.021 | -0.081 | -0.081 | 0.299 | 0.41 | 0.93 | 0.58 | 0.56 | 0.38 | 0.34 |
| 7 | 92695002 | rs10267557 | -0.959 | 0.211 | 0.525 | 2.021 | -0.006 | 0.015 | 0.015 | 0.355 | 0.19 | 0.26 | 0.15 | 0.25 | 0.2 | 0.04 |
| 7 | 92704801 | rs10257148 | -0.954 | 0.216 | 0.382 | 1.99 | 0.003 | -0.007 | -0.007 | 0 | 0.19 | 0.18 | 0.14 | 0.26 | 0.19 | 0.04 |
| 7 | 92727142 | rs7798225 | -2.299 | 0.046 | 1.347 | 2.366 | 0.002 | 0.291 | 0.291 | 4.926 | 0.09 | 0.39 | 0.3 | 0.33 | 0.08 | 0.1 |
| 7 | 92728427 | rs10953089 | -1.59 | -0.03 | 1.925 | 2.541 | -0.014 | 0.13 | 0.13 | 1.372 | 0.2 | 0.59 | 0.43 | 0.52 | 0.2 | 0.13 |
| 7 | 92733766 | rs10279499 | -0.428 | -0.072 | 2.115 | 2.56 | 0.008 | -0.005 | -0.005 | 0.133 | 0.13 | 0.12 | 0.1 | 0.14 | 0.1 | 0.02 |
| 7 | 92734983 | rs6969691 | -0.899 | -0.14 | 2.225 | 2.495 | -0.021 | 0.005 | 0.005 | 0.822 | 0.15 | 0.29 | 0.13 | 0.14 | 0.13 | 0.03 |
| 7 | 92736179 | rs12668221 | -0.785 | -0.141 | 2.188 | 2.494 | 0.009 | -0.003 | -0.003 | 0 | 0.14 | 0.12 | 0.11 | 0.13 | 0.12 | 0.03 |
| 7 | 92759462 | rs4267 | -2.278 | -0.17 | 2.466 | 2.574 | 0.04 | 0.389 | 0.389 | 2.81 | 0.08 | 0.4 | 0.31 | 0.34 | 0.07 | 0.12 |
| 7 | 92764489 | rs10488532 | -0.865 | -0.57 | 1.653 | 1.877 | 0.046 | 0.097 | 0.097 | 0 | 0.19 | 0.04 | 0.11 | 0.04 | 0.13 | 0.2 |
| 7 | 92775019 | rs28662 | 0.714 | 0.334 | 1.581 | 1.793 | -0.007 | 0.09 | 0.09 | 0 | 0.8 | 0.95 | 0.89 | 0.97 | 0.81 | 0.81 |
| 7 | 92812699 | rs988995 | -1.785 | 0.467 | 2.623 | 2.426 | 0.039 | 0.329 | 0.329 | 0.244 | 0.06 | 0.34 | 0.26 | 0.3 | 0.05 | 0.13 |
| 7 | 92895930 | rs12539635 | -2.023 | 1.335 | 2.804 | 2.435 | 0.059 | 0.469 | 0.469 | 0 | 0.05 | 0.31 | 0.28 | 0.38 | 0.06 | 0.16 |
| 7 | 92900159 | rs10488537 | -0.018 | 1.329 | 2.196 | 2.417 | -0.028 | -0.023 | -0.023 | 0 | 0.09 | 0 | 0.05 | 0.06 | 0.08 | 0.03 |
| 7 | 92941747 | rs13224723 | -0.851 | 1.143 | 2.279 | 1.892 | -0.001 | -0.04 | -0.04 | 0 | 0.2 | 0.19 | 0.13 | 0.03 | 0.19 | 0.2 |
| 7 | 92943760 | rs2188404 | -0.873 | 1.557 | 1.554 | 1.377 | 0.019 | 0.092 | 0.092 | 0.196 | 0.2 | 0.35 | 0.42 | 0.39 | 0.27 | 0.27 |

| | | | | | | | | | | | | | | | | |
|---|---|---|---|---|---|---|---|---|---|---|---|---|---|---|---|---|
| 7 | 92969716 | rs4729080 | -0.642 | 1.415 | 0.583 | 1.38 | -0.029 | -0.09 | -0.09 | 0.692 | 0.2 | 0.06 | 0.39 | 0.34 | 0.24 | 0.31 |
| 7 | 92985252 | rs2374639 | -1.339 | 1.508 | 0.924 | 1.048 | 0 | 0.007 | 0.007 | 3.494 | 0.27 | 0.39 | 0.32 | 0.28 | 0.29 | 0.35 |
| 7 | 92987445 | rs2285505 | -1.613 | 1.456 | 0.961 | 0.304 | -0.009 | -0.025 | -0.025 | 0 | 0.34 | 0.53 | 0.31 | 0.27 | 0.36 | 0.36 |
| 7 | 92989228 | rs2285507 | -1.238 | 1.4 | 0.99 | 0.3 | -0.01 | -0.013 | -0.013 | 17.925 | 0.29 | 0.35 | 0.26 | 0.23 | 0.26 | 0.31 |
| 7 | 92991487 | rs6956125 | 1.599 | 1.906 | 0.618 | 0.206 | 0.015 | -0.024 | -0.024 | 0 | 0.62 | 0.47 | 0.6 | 0.71 | 0.5 | 0.61 |
| 7 | 92991879 | rs6960300 | 1.982 | 1.871 | 0.758 | -0.003 | 0.055 | 0.01 | 0.01 | 0.293 | 0.74 | 0.44 | 0.66 | 0.74 | 0.6 | 0.76 |
| 7 | 92995752 | rs10233804 | -1.297 | 1.087 | -0.34 | -1.323 | 0.005 | 0.11 | 0.11 | 0.881 | 0.58 | 0.79 | 0.69 | 0.99 | 0.44 | 0.74 |
| 7 | 93002194 | rs9649173 | 2.148 | 1.872 | 0.305 | -0.261 | 0.042 | -0.015 | -0.015 | 0.587 | 0.74 | 0.44 | 0.66 | 0.79 | 0.6 | 0.74 |
| 7 | 93009475 | rs11765782 | 1.273 | 1.913 | -0.185 | -0.236 | -0.082 | -0.024 | -0.024 | 0 | 0.82 | 0.98 | 0.78 | 0.85 | 0.7 | 0.82 |
| 7 | 93009888 | rs10251417 | -2.221 | 1.062 | 0.111 | -0.004 | 0.026 | 0.026 | 0.026 | 0.087 | 0.2 | 0.51 | 0.24 | 0.21 | 0.25 | 0.24 |
| 7 | 93013606 | rs17785143 | -1.004 | 0.808 | -1.145 | 0.15 | -0.02 | -0.042 | -0.042 | 0 | 0.12 | 0.01 | 0.14 | 0.16 | 0.15 | 0.16 |
| 7 | 93019948 | rs2106431 | -0.102 | 0.632 | -1.366 | -0.366 | 0.016 | 0.06 | 0.06 | 5.373 | 0.38 | 0.16 | 0.45 | 0.78 | 0.33 | 0.54 |
| 7 | 93022840 | rs11767067 | -0.151 | 1.174 | -1.532 | -0.985 | -0.034 | 0.054 | 0.054 | 0.267 | 0.3 | 0.18 | 0.27 | 0.06 | 0.4 | 0.26 |
| 7 | 93023568 | rs11764153 | -0.495 | 1.04 | -1.275 | -0.139 | -0.006 | -0.013 | -0.013 | 0.179 | 0.26 | 0.22 | 0.43 | 0.51 | 0.29 | 0.38 |
| 7 | 93039164 | rs4276616 | -0.41 | 0.248 | -0.943 | -0.464 | -0.02 | -0.038 | -0.038 | 0.555 | 0.37 | 0.43 | 0.26 | 0.81 | 0.24 | 0.46 |
| 7 | 93040470 | rs2023778 | -0.738 | 0.188 | -0.767 | -0.656 | 0.003 | -0.01 | -0.01 | 0 | 0.42 | 0.61 | 0.43 | 0.86 | 0.3 | 0.54 |
| 7 | 93046845 | rs17165377 | -0.393 | 0.039 | -0.758 | -0.248 | -0.005 | 0.17 | 0.17 | 0 | 0.17 | 0.36 | 0.14 | 0.38 | 0.13 | 0.25 |
| 7 | 93055650 | rs1042138 | 1.077 | -0.048 | -1.055 | -0.303 | 0.041 | -0.093 | -0.093 | 0 | 0.21 | 0.05 | 0.13 | 0.43 | 0.12 | 0.23 |
| 7 | 93055753 | rs1801197 | -0.962 | 0.019 | -0.934 | -0.407 | -0.014 | -0.053 | -0.053 | 0 | 0.43 | 0.54 | 0.43 | 0.86 | 0.31 | 0.55 |
| 7 | 93061227 | rs2283004 | -0.831 | 0.247 | -1.355 | -0.319 | -0.028 | -0.075 | -0.075 | 3.71 | 0.21 | 0.12 | 0.38 | 0.4 | 0.25 | 0.25 |
| 7 | 93063207 | rs3815058 | 0.489 | 0.341 | -1.454 | 0.06 | -0.028 | -0.049 | -0.049 | 0 | 0.09 | 0 | 0.05 | 0.14 | 0.08 | 0.13 |
| 7 | 93066173 | rs10243420 | -0.723 | 0.302 | -1.075 | -0.693 | -0.003 | -0.002 | -0.002 | 0 | 0.69 | 0.7 | 0.77 | 0.89 | 0.62 | 0.7 |
| 7 | 93078060 | rs2214213 | -0.671 | 0.855 | -1.318 | 0.017 | -0.001 | -0.02 | -0.02 | 1.835 | 0.56 | 0.86 | 0.69 | 0.53 | 0.59 | 0.58 |
| 7 | 93078245 | rs7783961 | 0.147 | 0.981 | -0.697 | -0.436 | 0.002 | -0.056 | -0.056 | 0 | 0.23 | 0.26 | 0.24 | 0.03 | 0.25 | 0.21 |
| 7 | 93079627 | rs6465381 | -0.827 | 1.046 | -0.686 | 0.026 | -0.001 | 0 | 0 | 0 | 0.13 | 0.12 | 0.06 | 0.11 | 0.18 | 0.07 |
| 7 | 93079738 | rs6976450 | -0.164 | 1.157 | -1.565 | 0.697 | -0.065 | -0.136 | -0.136 | 0.096 | 0.22 | 0.01 | 0.17 | 0.3 | 0.22 | 0.37 |
| 7 | 93095867 | rs2237579 | 1.102 | 0.645 | -0.058 | 0.608 | 0.003 | 0.006 | 0.006 | 2.357 | 0.56 | 0.29 | 0.3 | 0.4 | 0.46 | 0.48 |
| 7 | 93101080 | rs10250228 | 0.818 | 0.642 | 0.004 | 0.607 | 0 | -0.001 | -0.001 | 0 | 0.59 | 0.54 | 0.33 | 0.4 | 0.46 | 0.57 |
| 7 | 93107093 | rs2074120 | 0.659 | 0.453 | 0.453 | 0.921 | -0.03 | 0.032 | 0.032 | 1.978 | 0.63 | 0.84 | 0.41 | 0.71 | 0.48 | 0.57 |
| 7 | 93116299 | rs2301680 | 0.937 | 0.463 | 0.403 | 0.926 | 0.002 | -0.006 | -0.006 | 0.077 | 0.61 | 0.64 | 0.39 | 0.71 | 0.48 | 0.57 |
| 7 | 93125921 | rs17788636 | 0.254 | 1.185 | -0.035 | 0.749 | -0.093 | 0.065 | 0.065 | 0.193 | 0.16 | 0.01 | 0.13 | 0.05 | 0.25 | 0.12 |
| 7 | 93129686 | rs17165500 | 0.675 | 0.696 | 0.183 | 0.911 | 0.036 | 0.388 | 0.388 | 0.372 | 0.05 | 0.35 | 0.09 | 0.31 | 0.02 | 0.09 |
| 7 | 93131162 | rs2023772 | 0.936 | 0.701 | 0.135 | 0.919 | -0.057 | 0.042 | 0.042 | 1.93 | 0.64 | 0.89 | 0.42 | 0.72 | 0.48 | 0.66 |
| 7 | 93142429 | rs17727433 | -1.059 | 0.888 | 0.613 | 0.972 | 0.04 | 0.054 | 0.054 | 0.32 | 0.28 | 0.56 | 0.3 | 0.42 | 0.36 | 0.23 |
| 7 | 93143761 | rs2188799 | -0.586 | 1.05 | 0.126 | 0.406 | 0.007 | -0.013 | -0.013 | 0 | 0.23 | 0.23 | 0.21 | 0.11 | 0.34 | 0.14 |
| 7 | 93147883 | rs989506 | -0.006 | 0.787 | -0.209 | 0.556 | -0.009 | -0.003 | -0.003 | 1.406 | 0.84 | 0.91 | 0.61 | 0.83 | 0.83 | 0.8 |
| 7 | 93159402 | rs17165541 | -0.415 | -0.001 | 0.317 | 0.732 | -0.052 | 0.233 | 0.233 | 0.418 | 0.12 | 0.4 | 0.21 | 0.37 | 0.05 | 0.18 |
| 7 | 93159655 | rs13223739 | 0.313 | -0.081 | 0.234 | 0.574 | 0.001 | -0.003 | -0.003 | 0 | 0.1 | 0.1 | 0.07 | 0.04 | 0.09 | 0.09 |
| 7 | 93163719 | rs10488551 | 1.494 | -0.443 | -0.641 | 0.144 | -0.037 | 0.041 | 0.041 | 3.721 | 0.4 | 0.04 | 0.23 | 0.29 | 0.38 | 0.37 |
| 7 | 93175689 | rs6961603 | 0.46 | -0.35 | -0.217 | 0.326 | -0.044 | 0.192 | 0.192 | 0.044 | 0.11 | 0.29 | 0.19 | 0.38 | 0.05 | 0.13 |
| 7 | 93186244 | rs1544360 | -0.31 | -0.306 | -0.673 | 0.131 | -0.137 | -0.054 | -0.054 | 0.511 | 0.69 | 0.94 | 0.59 | 0.7 | 0.59 | 0.67 |
| 7 | 93194590 | rs7801223 | -0.968 | -0.275 | 0.105 | 0.277 | -0.034 | 0.082 | 0.082 | 0.947 | 0.19 | 0.66 | 0.26 | 0.38 | 0.12 | 0.21 |
| 7 | 93194944 | rs17080 | 0.614 | 0.082 | -0.155 | 0.113 | 0.016 | -0.029 | -0.029 | 0 | 0.06 | 0.07 | 0.04 | 0 | 0.07 | 0.08 |
| 7 | 93201902 | rs10237272 | 0.224 | -0.889 | -0.831 | -0.097 | -0.045 | 0.099 | 0.099 | 0.544 | 0.5 | 0.93 | 0.53 | 0.7 | 0.51 | 0.65 |
| 7 | 93207873 | rs13236112 | 1.231 | -0.793 | -0.578 | 0.019 | -0.001 | 0.001 | 0.001 | 0 | 0.08 | 0.07 | 0.06 | 0.08 | 0.08 | 0.08 |
| 7 | 93222426 | rs17165596 | 0.792 | -0.946 | -0.052 | 0.075 | -0.022 | 0.291 | 0.291 | 0.573 | 0.07 | 0.3 | 0.19 | 0.39 | 0.03 | 0.14 |
| 7 | 93222805 | rs2528522 | 0.202 | -0.944 | -0.294 | -0.102 | -0.045 | 0.164 | 0.164 | 0.578 | 0.5 | 0.93 | 0.52 | 0.75 | 0.51 | 0.66 |
| 7 | 93276888 | rs2528550 | -1.508 | -0.87 | -0.508 | -0.047 | 0.012 | -0.045 | -0.045 | 0 | 0.84 | 0.97 | 0.75 | 0.78 | 0.89 | 0.83 |
| 7 | 93297506 | rs2677039 | 0.285 | -0.834 | -0.101 | 0.118 | -0.005 | 0.012 | 0.012 | 0 | 0.44 | 0.55 | 0.33 | 0.36 | 0.48 | 0.52 |
| 7 | 93309656 | rs1557659 | 0.72 | -0.696 | 0.004 | 0.352 | -0.018 | 0.318 | 0.318 | 0.847 | 0.06 | 0.27 | 0.18 | 0.4 | 0.02 | 0.14 |
| 7 | 93311711 | rs2677031 | -0.948 | -1.038 | -0.411 | -0.053 | -0.007 | 0.003 | 0.003 | 0.239 | 0.27 | 0.72 | 0.27 | 0.52 | 0.15 | 0.28 |
| 7 | 93312679 | rs6968642 | 0.995 | -0.541 | -0.346 | -0.296 | -0.015 | 0.02 | 0.02 | 0 | 0.31 | 0.21 | 0.25 | 0.22 | 0.39 | 0.35 |
| 7 | 93321008 | rs17165665 | 0.514 | -0.898 | -0.049 | -0.117 | -0.033 | 0.373 | 0.373 | 0.86 | 0.09 | 0.41 | 0.14 | 0.48 | 0.03 | 0.19 |
| 7 | 93322256 | rs2214001 | 0.389 | -0.078 | -0.36 | -1.003 | -0.006 | 0.031 | 0.031 | 0.459 | 0.45 | 0.74 | 0.42 | 0.7 | 0.49 | 0.59 |
| 7 | 93334446 | rs2078032 | 0.162 | 0.384 | -0.087 | -0.7 | 0.023 | -0.077 | -0.077 | 0.534 | 0.18 | 0.32 | 0.18 | 0.02 | 0.23 | 0.17 |
| 7 | 93336473 | rs9655749 | 0.992 | 0.17 | -0.275 | -0.808 | -0.017 | 0.009 | 0.009 | 5.774 | 0.14 | 0.19 | 0.06 | 0.13 | 0.09 | 0.09 |
| 7 | 93338280 | rs10254794 | 0.129 | 0.548 | -0.485 | -0.645 | 0.013 | -0.055 | -0.055 | 0 | 0.18 | 0.25 | 0.17 | 0.02 | 0.23 | 0.17 |
| 7 | 93339775 | rs8180847 | -0.147 | 0.29 | -0.918 | -0.12 | 0.004 | 0.004 | 0.004 | 0 | 0.4 | 0.32 | 0.32 | 0.69 | 0.36 | 0.48 |
| 7 | 93340580 | rs2677061 | -0.296 | 0.623 | -0.675 | -0.492 | 0 | -0.039 | -0.039 | 0 | 0.45 | 0.48 | 0.5 | 0.14 | 0.47 | 0.29 |
| 7 | 93341409 | rs10281294 | -0.389 | 0.391 | -0.302 | 0.056 | -0.007 | -0.01 | -0.01 | 0 | 0.3 | 0.39 | 0.33 | 0.23 | 0.26 | 0.24 |
| 7 | 93347267 | rs2677059 | 0.379 | 1.064 | 0.192 | 0.86 | 0.07 | 0.153 | 0.153 | 0 | 0.93 | 0.78 | 0.89 | 0.78 | 0.9 | 0.93 |
| 7 | 93349015 | rs1858826 | 0.105 | 1.098 | 0.192 | 0.571 | 0.038 | -0.031 | -0.031 | 0.23 | 0.94 | 0.88 | 0.94 | 1 | 0.9 | 0.99 |
| 7 | 93350257 | rs1557660 | 0.052 | 1.13 | 0.446 | 0.564 | 0.061 | -0.028 | -0.028 | 0 | 0.94 | 0.83 | 0.94 | 1 | 0.91 | 0.99 |
| 7 | 93350440 | rs2188274 | 0.46 | -0.314 | 0.597 | 0.213 | -0.03 | -0.058 | -0.058 | 0 | 0.71 | 0.5 | 0.83 | 0.85 | 0.81 | 0.78 |
| 7 | 93352716 | rs17165691 | 1.239 | 0.321 | -0.293 | 0.254 | -0.113 | -0.045 | -0.045 | 0 | 0.25 | 0.02 | 0.25 | 0.26 | 0.36 | 0.26 |
| 7 | 93353042 | rs2157814 | 1.055 | 0.137 | -0.34 | 0.221 | -0.003 | -0.009 | -0.009 | 0 | 0.27 | 0.34 | 0.27 | 0.14 | 0.25 | 0.2 |
| 7 | 93358149 | rs2157813 | -1.254 | 0.197 | -0.34 | 0.254 | 0.061 | -0.028 | -0.028 | 0 | 0.94 | 0.83 | 0.94 | 1 | 0.9 | 0.99 |
| 7 | 93361351 | rs2677055 | -1.254 | 0.197 | -0.322 | 0.254 | 0.051 | -0.03 | -0.03 | 0 | 0.94 | 0.84 | 0.94 | 1 | 0.9 | 0.99 |
| 7 | 93364926 | rs2677049 | -1.254 | 0.197 | -0.326 | 0.254 | 0.061 | -0.028 | -0.028 | 2.237 | 0.94 | 0.83 | 0.94 | 1 | 0.9 | 0.99 |

| | | | | | | | | | | | | | | | | |
|---|---|---|---|---|---|---|---|---|---|---|---|---|---|---|---|---|
| 7 | 93369643 | rs2188273 | -0.544 | 0.085 | -0.425 | 0.286 | 0.021 | 0.005 | 0.005 | 0 | 0.83 | 0.77 | 0.83 | 0.84 | 0.76 | 0.81 |
| 7 | 93387256 | rs2677071 | 0.496 | -0.075 | -0.153 | 0.287 | -0.014 | -0.012 | -0.012 | 0.009 | 0.29 | 0.25 | 0.21 | 0.15 | 0.15 | 0.3 |
| 7 | 93396080 | rs4729093 | 1.206 | -0.058 | -0.322 | 0.214 | -0.022 | -0.015 | -0.015 | 0.1 | 0.15 | 0.09 | 0.25 | 0.14 | 0.2 | 0.16 |
| 7 | 93401210 | rs2724063 | -1.21 | -0.061 | -0.164 | 0.251 | -0.018 | 0.004 | 0.004 | 0.232 | 0.69 | 0.74 | 0.69 | 0.7 | 0.55 | 0.74 |
| 7 | 93401719 | rs2724062 | -0.079 | -0.82 | 0.009 | 0.006 | -0.004 | 0.04 | 0.04 | 0 | 0.84 | 0.64 | 0.79 | 0.75 | 0.88 | 0.89 |
| 7 | 93403501 | rs17165733 | 1.347 | -0.902 | -0.084 | -0.215 | -0.011 | -0.048 | -0.048 | 0 | 0.15 | 0.22 | 0.06 | 0 | 0.09 | 0.05 |
| 7 | 93403525 | rs12670534 | 1.581 | -0.542 | -0.322 | -0.064 | -0.104 | -0.093 | -0.093 | 0 | 0.32 | 0.07 | 0.48 | 0.62 | 0.46 | 0.41 |
| 7 | 93404220 | rs2724121 | -0.333 | -0.607 | -0.127 | -0.113 | 0.003 | -0.091 | -0.091 | 4.549 | 0.86 | 0.68 | 0.9 | 1 | 0.89 | 0.96 |
| 7 | 93404574 | rs11982215 | 0.962 | -0.654 | -0.215 | -0.101 | -0.004 | -0.012 | -0.012 | 0 | 0.23 | 0.3 | 0.15 | 0.07 | 0.17 | 0.11 |
| 7 | 93405173 | rs13242459 | 1.289 | -0.251 | -0.568 | 0.114 | -0.167 | -0.108 | -0.108 | 0 | 0.31 | 0.02 | 0.38 | 0.37 | 0.45 | 0.35 |
| 7 | 93406587 | rs10259426 | 0.441 | -0.287 | -0.386 | 0.127 | -0.031 | -0.058 | -0.058 | 0.015 | 0.31 | 0.45 | 0.2 | 0.07 | 0.19 | 0.14 |
| 7 | 93407398 | rs2519609 | -0.633 | -0.191 | -0.496 | 0.199 | -0.016 | -0.015 | -0.015 | 0 | 0.62 | 0.73 | 0.46 | 0.38 | 0.45 | 0.57 |
| 7 | 93407706 | rs16868514 | 1.131 | 0.115 | -0.585 | 0.277 | -0.008 | -0.04 | -0.04 | 0.083 | 0.07 | 0 | 0.12 | 0.14 | 0.06 | 0.16 |
| 7 | 93414482 | rs2724111 | -0.225 | 0.197 | -0.542 | 0.183 | -0.009 | 0.017 | 0.017 | 0 | 0.32 | 0.33 | 0.36 | 0.56 | 0.27 | 0.5 |
| 7 | 93415545 | rs7778691 | 0.355 | 0.125 | -0.62 | 0.247 | 0 | 0.018 | 0.018 | 0 | 0.62 | 0.59 | 0.59 | 0.44 | 0.63 | 0.49 |
| 7 | 93416051 | rs11505801 | 0.248 | 0.225 | -0.92 | -0.281 | 0 | -0.001 | -0.001 | 0 | 0.63 | 0.67 | 0.69 | 0.69 | 0.64 | 0.55 |
| 7 | 93429143 | rs2519637 | 0.386 | 0.461 | -0.407 | -0.483 | 0.04 | -0.027 | -0.027 | 0 | 0.09 | 0.37 | 0.05 | 0 | 0.05 | 0.02 |
| 7 | 93435604 | rs13243204 | 1.052 | 0.19 | -0.998 | -0.843 | -0.017 | 0.053 | 0.053 | 2.358 | 0.12 | 0 | 0.07 | 0.02 | 0.1 | 0.09 |
| 7 | 93435670 | rs2724099 | -0.303 | 0.025 | -0.86 | -0.576 | -0.018 | 0.017 | 0.017 | 0.939 | 0.14 | 0.56 | 0.19 | 0.15 | 0.08 | 0.11 |
| 7 | 93435967 | rs10464585 | 0.319 | 0.032 | -1.487 | -0.634 | -0.012 | -0.031 | -0.031 | 0.203 | 0.29 | 0.16 | 0.38 | 0.63 | 0.28 | 0.4 |
| 7 | 93448448 | rs2519631 | -0.918 | -0.076 | -2.007 | -0.797 | -0.096 | 0.154 | 0.154 | 3.337 | 0.56 | 0.96 | 0.64 | 0.82 | 0.44 | 0.6 |
| 7 | 93451083 | rs2519630 | 0.422 | -0.219 | -2.037 | -0.878 | -0.013 | -0.041 | -0.041 | 0.169 | 0.67 | 0.52 | 0.67 | 0.79 | 0.73 | 0.69 |
| 7 | 93452850 | rs2724117 | -0.909 | -0.156 | -2.051 | -0.61 | -0.095 | 0.039 | 0.039 | 3.232 | 0.56 | 0.96 | 0.64 | 0.72 | 0.43 | 0.6 |
| 7 | 93455006 | rs10243038 | 1.388 | -0.163 | -1.994 | -0.763 | 0.012 | 0.011 | 0.011 | 0.431 | 0.13 | 0.16 | 0.05 | 0.04 | 0.07 | 0.13 |
| 7 | 93460205 | rs17165797 | 1.537 | -0.059 | -2.133 | -0.635 | -0.025 | -0.078 | -0.078 | 1.335 | 0.15 | 0.06 | 0.18 | 0.44 | 0.15 | 0.12 |
| 7 | 93468542 | rs2519662 | 0.419 | -0.012 | -1.497 | -0.993 | 0.011 | 0.104 | 0.104 | 0.146 | 0.42 | 0.72 | 0.58 | 0.78 | 0.36 | 0.48 |
| 7 | 93471420 | rs2724085 | 0.519 | 0.033 | -1.62 | -0.973 | 0.005 | 0.032 | 0.032 | 0 | 0.42 | 0.63 | 0.58 | 0.78 | 0.36 | 0.47 |
| 7 | 93475293 | rs2519658 | 0.515 | 0.026 | -1.615 | -0.968 | 0.006 | 0.033 | 0.033 | 0 | 0.42 | 0.63 | 0.58 | 0.78 | 0.36 | 0.47 |
| 7 | 93510823 | rs2724112 | -1.021 | 0.435 | -2.059 | -0.825 | -0.149 | 0.046 | 0.046 | 1.327 | 0.71 | 0.97 | 0.72 | 0.84 | 0.54 | 0.7 |
| 7 | 93513413 | rs6970118 | -0.221 | 0.174 | -0.662 | -0.463 | -0.028 | -0.046 | -0.046 | 0 | 0.14 | 0.27 | 0.06 | 0.04 | 0.07 | 0.13 |
| 7 | 93514647 | rs34914495 | -0.356 | 1.208 | -0.877 | 0.195 | -0.023 | 0.054 | 0.054 | 2.546 | 0.12 | 0 | 0.07 | 0.02 | 0.1 | 0.09 |
| 7 | 93528312 | rs2519649 | -0.772 | 0.746 | 0.477 | 0.014 | -0.007 | -0.037 | -0.037 | 1.667 | 0.09 | 0.36 | 0.04 | 0 | 0.03 | 0.02 |
| 7 | 93530510 | rs180283 | 0.598 | 0.951 | 0.147 | -0.35 | 0.005 | 0.028 | 0.028 | 0 | 0.51 | 0.35 | 0.65 | 0.76 | 0.57 | 0.62 |
| 7 | 93531908 | rs2724113 | 0.744 | 0.999 | 0.907 | -0.408 | -0.001 | -0.001 | -0.001 | 0.019 | 0.47 | 0.37 | 0.54 | 0.67 | 0.5 | 0.52 |
| 7 | 93533463 | rs180280 | -0.084 | 0.953 | 0.873 | -0.398 | 0.007 | -0.003 | -0.003 | 0 | 0.05 | 0.09 | 0.14 | 0.09 | 0.07 | 0.11 |
| 7 | 93535185 | rs7788636 | 0.816 | 1.017 | 0.86 | -0.392 | -0.008 | 0.041 | 0.041 | 0 | 0.46 | 0.27 | 0.52 | 0.67 | 0.5 | 0.51 |
| 7 | 93535432 | rs10246567 | 0.751 | 1.002 | 0.985 | -0.376 | 0.001 | -0.011 | -0.011 | 0 | 0.47 | 0.4 | 0.54 | 0.67 | 0.5 | 0.51 |
| 7 | 93537158 | rs180274 | 0.406 | 1.27 | 0.957 | -0.473 | 0.01 | 0.018 | 0.018 | 0 | 0.59 | 0.34 | 0.6 | 0.68 | 0.66 | 0.56 |
| 7 | 93538075 | rs17165876 | 0.16 | 0.948 | 1.117 | -0.796 | -0.005 | -0.032 | -0.032 | 0 | 0.12 | 0.15 | 0.03 | 0.01 | 0.06 | 0.06 |
| 7 | 93538295 | rs180273 | -0.878 | 0.832 | 1.205 | -0.908 | -0.002 | -0.005 | -0.005 | 0 | 0.84 | 0.89 | 0.86 | 0.84 | 0.81 | 0.77 |
| 7 | 93542743 | rs180262 | 1.32 | 0.78 | 1.641 | -1.019 | -0.01 | -0.034 | -0.034 | 0 | 0.64 | 0.46 | 0.63 | 0.74 | 0.68 | 0.59 |
| 7 | 93549656 | rs180241 | 0.069 | 0.695 | 1.653 | -1.052 | 0.003 | 0.01 | 0.01 | 0 | 0.8 | 0.86 | 0.84 | 0.88 | 0.78 | 0.72 |
| 7 | 93550447 | rs180238 | 1.653 | 0.684 | 1.642 | -1.027 | -0.018 | -0.04 | -0.04 | 0.676 | 0.62 | 0.49 | 0.64 | 0.74 | 0.68 | 0.58 |
| 7 | 93551428 | rs4262 | 1.956 | 1.495 | 1.476 | -1.19 | 0.004 | 0.029 | 0.029 | 0 | 0.58 | 0.34 | 0.59 | 0.74 | 0.6 | 0.55 |
| 7 | 93551606 | rs17165909 | -0.114 | 1.395 | 1.515 | -1.23 | -0.02 | -0.012 | -0.012 | 0.346 | 0.09 | 0.19 | 0.14 | 0.12 | 0.03 | 0.09 |
| 7 | 93552275 | rs17303650 | 0.498 | 1.452 | 1.343 | -1.245 | -0.001 | -0.004 | -0.004 | 0 | 0.05 | 0.05 | 0.05 | 0 | 0.05 | 0.04 |
| 7 | 93553341 | rs180236 | 1.394 | 1.461 | 1.395 | -1.266 | -0.005 | 0.025 | 0.025 | 0 | 0.68 | 0.67 | 0.75 | 0.89 | 0.63 | 0.64 |
| 7 | 93554976 | rs180235 | 1.15 | 1.451 | 0.975 | -1.057 | -0.088 | 0.095 | 0.095 | 0 | 0.64 | 0.89 | 0.76 | 0.85 | 0.55 | 0.58 |
| 7 | 93555311 | rs180233 | 0.636 | 1.465 | 0.961 | -1.058 | -0.075 | 0.06 | 0.06 | 0.004 | 0.68 | 0.91 | 0.8 | 0.85 | 0.61 | 0.62 |
| 7 | 93557588 | rs180231 | 0.931 | 1.451 | 0.456 | -1.025 | -0.067 | 0.083 | 0.083 | 0 | 0.64 | 0.85 | 0.76 | 0.85 | 0.56 | 0.58 |
| 7 | 93564883 | rs2191879 | -0.172 | 1.565 | 0.312 | -0.93 | -0.032 | 0.059 | 0.059 | 0 | 0.24 | 0.1 | 0.17 | 0.08 | 0.29 | 0.26 |
| 7 | 93565361 | rs6955457 | -1.149 | 1.198 | 0.385 | -0.661 | -0.026 | 0.105 | 0.105 | 0.004 | 0.54 | 0.81 | 0.44 | 0.27 | 0.51 | 0.51 |
| 7 | 93568355 | rs2374692 | -0.998 | 1.312 | 0.081 | -0.465 | 0.002 | 0.013 | 0.013 | 0.024 | 0.56 | 0.57 | 0.48 | 0.35 | 0.55 | 0.52 |
| 7 | 93577154 | rs13241960 | -1.243 | 1.75 | 0.412 | -0.171 | 0.038 | -0.05 | -0.05 | 0 | 0.2 | 0.37 | 0.17 | 0.05 | 0.24 | 0.2 |
| 7 | 93592846 | rs1014424 | -0.493 | 1 | 0.149 | 0.956 | 0.003 | 0.032 | 0.032 | 0.621 | 0.06 | 0.04 | 0.05 | 0.31 | 0.04 | 0.11 |
| 7 | 93601549 | rs17245644 | -0.747 | 0.982 | -0.158 | 0.751 | -0.044 | 0.014 | 0.014 | 0 | 0.08 | 0.02 | 0.1 | 0.05 | 0.16 | 0.08 |
| 7 | 93606881 | rs4729105 | -1.251 | 0.283 | 0.084 | 0.971 | 0.005 | -0.018 | -0.018 | 2.02 | 0.36 | 0.65 | 0.31 | 0.56 | 0.32 | 0.42 |
| 7 | 93613738 | rs2041518 | -1.419 | 0.231 | 0.47 | 0.908 | -0.009 | 0.018 | 0.018 | 0.001 | 0.38 | 0.57 | 0.32 | 0.56 | 0.32 | 0.41 |
| 7 | 93618626 | rs17245818 | -0.26 | -0.764 | 0.712 | 0.684 | 0.047 | 0.079 | 0.079 | 1.232 | 0.25 | 0.58 | 0.31 | 0.5 | 0.35 | 0.34 |
| 7 | 93621393 | rs6945362 | 0.35 | -0.81 | 0.798 | 0.702 | -0.057 | -0.072 | -0.072 | 0.206 | 0.2 | 0.09 | 0.26 | 0.46 | 0.33 | 0.34 |
| 7 | 93622247 | rs10953105 | 0.678 | -1.208 | 0.706 | 0.639 | 0.035 | -0.034 | -0.034 | 0 | 0.11 | 0.06 | 0.03 | 0.14 | 0.03 | 0.08 |
| 7 | 93629242 | rs11973524 | -0.64 | -1.185 | 0.87 | 0.629 | 0.079 | 0.057 | 0.057 | 1.189 | 0.06 | 0.51 | 0.05 | 0 | 0.02 | 0 |
| 7 | 93637625 | rs10260824 | -0.628 | -1.213 | 0.957 | 0.636 | -0.068 | 0.034 | 0.034 | 10.009 | 0.16 | 0.61 | 0.08 | 0.15 | 0.05 | 0.09 |
| 7 | 93642391 | rs9649181 | -1.123 | -1.097 | -0.851 | 0.375 | -0.033 | 0.017 | 0.017 | 2.411 | 0.37 | 0.77 | 0.36 | 0.61 | 0.38 | 0.43 |
| 7 | 93654752 | rs7794610 | -1.566 | -1.166 | -1.133 | 0.257 | -0.076 | 0.263 | 0.263 | 0.775 | 0.51 | 0.91 | 0.44 | 0.76 | 0.47 | 0.5 |
| 7 | 93662801 | rs4729107 | -1.929 | -1.115 | -1.326 | 0.022 | -0.12 | 0.313 | 0.313 | 0 | 0.61 | 0.98 | 0.55 | 0.99 | 0.58 | 0.78 |
| 7 | 93668153 | rs9641129 | 0.105 | -0.054 | -0.441 | 0.926 | -0.05 | -0.064 | -0.064 | 0 | 0.2 | 0.12 | 0.27 | 0.52 | 0.33 | 0.34 |
| 7 | 93670529 | rs10487243 | -0.629 | 0.075 | -0.491 | 0.447 | 0.052 | 0.025 | 0.025 | 0 | 0.06 | 0.52 | 0.05 | 0 | 0.02 | 0 |
| 7 | 93673611 | rs11973636 | -0.61 | 0.459 | -1.399 | 0.689 | -0.067 | -0.02 | -0.02 | 0 | 0.3 | 0.14 | 0.28 | 0.63 | 0.36 | 0.41 |

| | | | | | | | | | | | | | | | | |
|---|---|---|---|---|---|---|---|---|---|---|---|---|---|---|---|---|
| 7 | 93682230 | rs10215958 | -0.701 | -0.443 | -0.877 | 0.567 | -0.034 | 0.013 | 0.013 | 0.726 | 0.29 | 0.61 | 0.25 | 0.31 | 0.15 | 0.24 |
| 7 | 93682407 | rs41522 | -0.556 | 0.274 | -1.356 | -0.607 | -0.013 | 0.09 | 0.09 | 0 | 0.6 | 0.73 | 0.54 | 0.94 | 0.56 | 0.7 |
| 7 | 93683737 | rs2188876 | 0.255 | 0.368 | -1.238 | 0.056 | 0 | 0.006 | 0.006 | 0 | 0.47 | 0.66 | 0.57 | 0.39 | 0.43 | 0.34 |
| 7 | 93686991 | rs41528 | -0.243 | 0.553 | -1.877 | 0.181 | -0.034 | 0.006 | 0.006 | 0 | 0.73 | 0.93 | 0.79 | 0.74 | 0.71 | 0.67 |
| 7 | 93688964 | rs41532 | -0.239 | 0.464 | -1.794 | 0.093 | -0.005 | -0.023 | -0.023 | 0 | 0.27 | 0.18 | 0.23 | 0.36 | 0.31 | 0.34 |
| 7 | 93689452 | rs1159548 | 0.656 | 0.04 | -0.574 | -0.066 | -0.001 | -0.003 | -0.003 | 0 | 0.43 | 0.51 | 0.54 | 0.38 | 0.4 | 0.34 |
| 7 | 93691744 | rs9918668 | -0.496 | -0.693 | -0.477 | -0.381 | 0.002 | 0.022 | 0.022 | 1.475 | 0.38 | 0.73 | 0.26 | 0.68 | 0.34 | 0.4 |
| 7 | 93693901 | rs3286 | -1.576 | -1.33 | -0.051 | -0.237 | -0.006 | -0.033 | -0.033 | 0 | 0.39 | 0.61 | 0.23 | 0.82 | 0.3 | 0.24 |
| 7 | 93694611 | rs41539 | -0.187 | -0.719 | -0.679 | 0.066 | 0.016 | -0.01 | -0.01 | 0.215 | 0.2 | 0.1 | 0.13 | 0.19 | 0.2 | 0.12 |
| 7 | 93698376 | rs10273701 | -0.899 | -0.66 | 0.337 | 0.456 | -0.001 | 0.05 | 0.05 | 0.057 | 0.25 | 0.38 | 0.18 | 0.64 | 0.23 | 0.16 |
| 7 | 93698571 | rs12375149 | 0.143 | -1.49 | 0.373 | -0.251 | 0.001 | 0.075 | 0.075 | 0.064 | 0.27 | 0.15 | 0.17 | 0.04 | 0.28 | 0.34 |
| 7 | 93700162 | rs7795786 | -0.82 | -1.827 | 0.827 | -0.182 | 0.033 | 0.086 | 0.086 | 2.862 | 0.4 | 0.72 | 0.33 | 0.2 | 0.33 | 0.45 |
| 7 | 93705033 | rs11975386 | 0.181 | -1.739 | 1.199 | -0.035 | -0.013 | 0.015 | 0.015 | 0 | 0.08 | 0.11 | 0.13 | 0.15 | 0.04 | 0.11 |
| 7 | 93714157 | rs10259011 | -1.131 | -1.77 | 1.677 | -0.165 | 0.037 | 0.034 | 0.034 | 0.196 | 0.05 | 0.36 | 0.04 | 0 | 0 | 0.01 |
| 7 | 93714505 | rs10259273 | -0.134 | -1.789 | 1.648 | -0.247 | -0.019 | -0.132 | -0.132 | 0 | 0.33 | 0.46 | 0.19 | 0.04 | 0.29 | 0.35 |
| 7 | 93722036 | rs41542 | 0.423 | -1.555 | 1.433 | 0.682 | 0.004 | 0.107 | 0.107 | 0 | 0.92 | 0.91 | 0.92 | 0.42 | 0.89 | 0.91 |
| 7 | 93740752 | rs4257948 | 1.176 | -1.855 | 1.634 | 0.338 | -0.001 | 0.011 | 0.011 | 2.068 | 0.47 | 0.45 | 0.58 | 0.25 | 0.51 | 0.41 |
| 7 | 93746499 | rs10245760 | -1.475 | -1.958 | 2.004 | 0.491 | -0.042 | 0.006 | 0.006 | 1.851 | 0.11 | 0.64 | 0.15 | 0.1 | 0.03 | 0.11 |
| 7 | 93766448 | rs11973087 | -1.628 | -1.937 | 1.766 | 0.433 | -0.046 | -0.022 | -0.022 | 2.135 | 0.14 | 0.63 | 0.17 | 0.12 | 0.06 | 0.12 |
| 7 | 93787413 | rs7357193 | -0.783 | -1.836 | 1.209 | 0.53 | 0.017 | 0.032 | 0.032 | 2.489 | 0.38 | 0.69 | 0.31 | 0.16 | 0.31 | 0.42 |
| 7 | 93812586 | rs1990543 | -2.391 | -1.544 | 1.977 | 1.805 | -0.042 | 0.173 | 0.173 | 0 | 0.16 | 0.65 | 0.22 | 0.48 | 0.08 | 0.11 |
| 7 | 93865697 | rs2052213 | -1.068 | -1.489 | 2.085 | 1.221 | -0.001 | 0.109 | 0.109 | 0 | 0.09 | 0.17 | 0.09 | 0.66 | 0.05 | 0.16 |
| 7 | 93878287 | rs949524 | -1.812 | -1.454 | 1.601 | 0.779 | -0.038 | 0.005 | 0.005 | 0.436 | 0.11 | 0.58 | 0.11 | 0.1 | 0.02 | 0.07 |
| 7 | 93883599 | rs1861114 | -1.176 | -1.634 | 1.197 | 0.692 | -0.044 | -0.02 | -0.02 | 0 | 0.17 | 0.39 | 0.13 | 0.16 | 0.08 | 0.09 |
| 7 | 93890183 | rs10228755 | -1.858 | -1.818 | 1.776 | 0.657 | -0.069 | -0.036 | -0.036 | 0 | 0.21 | 0.57 | 0.15 | 0.17 | 0.09 | 0.09 |
| 7 | 93892286 | rs7777958 | -1.988 | -1.917 | 1.029 | 0.509 | 0.024 | 0.184 | 0.184 | 0 | 0.46 | 0.92 | 0.37 | 0.82 | 0.35 | 0.56 |
| 7 | 93903748 | rs6465406 | -1.762 | -1.885 | 0.977 | 0.621 | -0.01 | -0.022 | -0.022 | 0 | 0.36 | 0.6 | 0.28 | 0.26 | 0.29 | 0.4 |
| 7 | 93904262 | rs1029999 | -1.78 | -1.745 | 0.94 | 0.621 | -0.003 | -0.005 | -0.005 | 0.01 | 0.37 | 0.64 | 0.28 | 0.29 | 0.3 | 0.4 |
| 7 | 93904860 | rs2024362 | -1.026 | -1.396 | 1.025 | 0.621 | 0.012 | -0.021 | -0.021 | 1.339 | 0.21 | 0.57 | 0.18 | 0.21 | 0.22 | 0.32 |
| 7 | 93905952 | rs11983945 | -1.544 | -1.458 | 1.049 | 0.589 | 0.045 | 0.109 | 0.109 | 1.462 | 0.06 | 0.52 | 0.05 | 0.09 | 0.01 | 0.01 |
| 7 | 93910179 | rs983894 | -0.214 | -1.351 | 0.394 | 0.565 | -0.008 | 0.019 | 0.019 | 0 | 0.17 | 0.14 | 0.14 | 0.1 | 0.2 | 0.31 |
| 7 | 93923144 | rs7782469 | -2.059 | -2.01 | -0.362 | 0.981 | -0.014 | 0.016 | 0.016 | 2.296 | 0.54 | 0.96 | 0.38 | 0.66 | 0.39 | 0.61 |
| 7 | 93926316 | rs12669399 | -1.063 | -2.08 | 0.185 | 1.013 | -0.018 | -0.013 | -0.013 | 0 | 0.29 | 0.43 | 0.18 | 0.27 | 0.26 | 0.37 |
| 7 | 93928050 | rs735392 | -0.738 | -2.028 | -0.588 | 0.844 | 0.006 | 0.015 | 0.015 | 0 | 0.27 | 0.2 | 0.16 | 0.23 | 0.25 | 0.37 |
| 7 | 93930713 | rs12704714 | -1.457 | -1.409 | -1.421 | 0.762 | -0.013 | 0.032 | 0.032 | 0 | 0.47 | 0.83 | 0.39 | 0.63 | 0.41 | 0.61 |
| 7 | 93935077 | rs7780548 | -1.378 | -1.472 | -1.109 | 1.068 | 0.018 | 0.009 | 0.009 | 0 | 0.42 | 0.7 | 0.37 | 0.63 | 0.34 | 0.57 |
| 7 | 93945455 | rs10953111 | -1.487 | -0.877 | -1.213 | 1.197 | 0.025 | 0.032 | 0.032 | 0 | 0.44 | 0.7 | 0.37 | 0.63 | 0.37 | 0.63 |
| 7 | 93951884 | rs6465409 | -0.199 | -0.803 | -1.139 | 1.445 | -0.056 | -0.044 | -0.044 | 0.066 | 0.16 | 0.06 | 0.15 | 0.27 | 0.24 | 0.38 |
| 7 | 93959918 | rs17166182 | -1.301 | -0.891 | -1.095 | 1.283 | 0.026 | 0.022 | 0.022 | 0.01 | 0.42 | 0.69 | 0.34 | 0.63 | 0.33 | 0.63 |
| 7 | 93963296 | rs2157811 | -1.06 | -1.638 | -0.574 | 1.487 | -0.039 | 0.023 | 0.023 | 0.007 | 0.27 | 0.63 | 0.19 | 0.41 | 0.09 | 0.28 |
| 7 | 93966036 | rs1557658 | -1.145 | -1.359 | -0.362 | 1.335 | -0.076 | 0.06 | 0.06 | 6.044 | 0.3 | 0.48 | 0.17 | 0.6 | 0.12 | 0.33 |
| 7 | 93967861 | rs13234589 | -1.289 | -0.029 | -0.912 | 0.7 | -0.017 | 0.041 | 0.041 | 0 | 0.18 | 0.06 | 0.28 | 0.08 | 0.15 | 0.11 |
| 7 | 93972820 | rs732584 | -1.425 | -0.088 | -0.712 | 0.696 | 0 | 0.01 | 0.01 | 0 | 0.14 | 0.1 | 0.26 | 0.09 | 0.1 | 0.09 |
| 7 | 93982039 | rs10464587 | -1.717 | 0.452 | -0.355 | 1.638 | -0.002 | 0 | 0 | 0 | 0.2 | 0.23 | 0.38 | 0.43 | 0.2 | 0.29 |
| 7 | 93999375 | rs7784854 | 0.177 | 0.533 | -0.21 | 1.536 | -0.003 | 0.001 | 0.001 | 0 | 0.05 | 0.03 | 0.03 | 0 | 0.05 | 0.02 |
| 7 | 94004197 | rs17166206 | -0.593 | 0.641 | -0.085 | 1.791 | 0.008 | 0.104 | 0.104 | 0 | 0.08 | 0.19 | 0.13 | 0.29 | 0.11 | 0.19 |
| 7 | 94008535 | rs4729127 | -1.269 | 0.638 | -0.094 | 1.79 | 0.007 | 0.126 | 0.126 | 0 | 0.09 | 0.27 | 0.14 | 0.29 | 0.11 | 0.19 |
| 7 | 94021475 | rs3763469 | -0.034 | 1.539 | -0.124 | 1.119 | 0.006 | -0.004 | -0.004 | 0 | 0.15 | 0.14 | 0.17 | 0.11 | 0.21 | 0.21 |
| 7 | 94021546 | rs3763468 | -0.594 | 1.584 | -0.289 | 1.097 | -0.01 | 0.012 | 0.012 | 0 | 0.13 | 0.05 | 0.28 | 0.11 | 0.11 | 0.09 |
| 7 | 94023646 | rs390201 | 0.177 | 1.319 | -0.267 | 1.046 | -0.006 | -0.022 | -0.022 | 0 | 0.13 | 0.06 | 0.13 | 0.25 | 0.18 | 0.26 |
| 7 | 94028427 | rs4729131 | 0.12 | 1.037 | 0.76 | 1.597 | -0.007 | -0.02 | -0.02 | 0.183 | 0.13 | 0.1 | 0.13 | 0.26 | 0.18 | 0.26 |
| 7 | 94030899 | rs1800222 | 0.002 | 0.687 | 0.437 | 0.947 | -0.044 | 0.146 | 0.146 | 0.097 | 0.19 | 0.57 | 0.17 | 0.39 | 0.15 | 0.1 |
| 7 | 94031457 | rs11770203 | -0.448 | 1.014 | 0.433 | 0.981 | -0.013 | -0.017 | -0.017 | 0 | 0.23 | 0.16 | 0.28 | 0.38 | 0.36 | 0.46 |
| 7 | 94032753 | rs1858822 | -0.448 | 1.014 | 0.433 | 0.981 | -0.013 | -0.017 | -0.017 | 0 | 0.23 | 0.16 | 0.28 | 0.38 | 0.36 | 0.46 |
| 7 | 94033031 | rs411717 | -1.095 | 1.265 | 0.127 | 0.631 | -0.01 | -0.01 | -0.01 | 0.189 | 0.36 | 0.29 | 0.61 | 0.51 | 0.54 | 0.56 |
| 7 | 94033800 | rs406226 | 0.216 | 1.015 | 0.071 | 0.659 | 0 | 0 | 0 | 0 | 0.13 | 0.13 | 0.34 | 0.13 | 0.18 | 0.1 |
| 7 | 94039187 | rs2521206 | 0.182 | 1.012 | 0.537 | 0.725 | 0.061 | 0.223 | 0.223 | 0.567 | 0.14 | 0.36 | 0.2 | 0.48 | 0.21 | 0.26 |
| 7 | 94043274 | rs421587 | -0.333 | 0.982 | 0.245 | 0.506 | -0.005 | 0.005 | 0.005 | 0 | 0.24 | 0.25 | 0.27 | 0.17 | 0.32 | 0.24 |
| 7 | 94046199 | rs10487254 | 0.293 | 1.196 | 0.318 | 0.512 | -0.004 | 0.001 | 0.001 | 1.058 | 0.13 | 0.12 | 0.33 | 0.12 | 0.17 | 0.1 |
| 7 | 94046541 | rs2521205 | -1.027 | 1.175 | 0.331 | 0.51 | -0.01 | 0.007 | 0.007 | 0.183 | 0.37 | 0.37 | 0.6 | 0.29 | 0.49 | 0.34 |
| 7 | 94048482 | rs369982 | -0.557 | 1.822 | 0.147 | 0.457 | -0.031 | -0.025 | -0.025 | 0 | 0.27 | 0.19 | 0.5 | 0.37 | 0.43 | 0.36 |
| 7 | 94049356 | rs42528 | -1.159 | 1.3 | 0.12 | 0.444 | 0.029 | 0.006 | 0.006 | 1.66 | 0.51 | 0.44 | 0.77 | 0.54 | 0.74 | 0.61 |
| 7 | 94051795 | rs10235102 | 0.283 | 1.306 | -0.463 | 0.466 | 0.029 | 0.064 | 0.064 | 1.315 | 0.19 | 0.08 | 0.12 | 0.05 | 0.14 | 0.04 |
| 7 | 94052495 | rs42530 | 0.544 | 1.888 | -0.85 | 0.461 | -0.111 | -0.092 | -0.092 | 0 | 0.2 | 0.02 | 0.28 | 0.25 | 0.32 | 0.29 |
| 7 | 94056006 | rs400218 | -0.303 | 1.046 | -0.378 | 0.463 | 0.007 | -0.037 | -0.037 | 0 | 0.38 | 0.24 | 0.66 | 0.49 | 0.62 | 0.58 |
| 7 | 94056044 | rs2621215 | -1.168 | 0.87 | -0.399 | 0.001 | 0.028 | 0.032 | 0.032 | 1.323 | 0.62 | 0.71 | 0.84 | 0.89 | 0.77 | 0.68 |
| 7 | 94060509 | rs1062394 | 0.151 | 0.805 | -0.284 | 0.674 | 0.001 | 0 | 0 | 0 | 0.13 | 0.1 | 0.1 | 0.04 | 0.1 | 0.02 |
| 7 | 94075973 | rs17166289 | 0.126 | 0.797 | -0.427 | 1.411 | 0.007 | -0.061 | -0.061 | 0 | 0.14 | 0.02 | 0.13 | 0.21 | 0.11 | 0.04 |
| 7 | 94101318 | rs2188646 | 0.053 | 0.669 | -0.093 | 1.779 | -0.007 | 0.039 | 0.039 | 0.286 | 0.09 | 0.19 | 0.11 | 0.15 | 0.1 | 0.19 |

| | | | | | | | | | | | | | | | | |
|---|---|---|---|---|---|---|---|---|---|---|---|---|---|---|---|---|
| 7 | 94104205 | rs10278176 | -0.289 | 0.106 | -0.776 | 1.611 | -0.028 | 0.012 | 0.012 | 0 | 0.29 | 0.11 | 0.3 | 0.28 | 0.29 | 0.39 |
| 7 | 94106403 | rs12704720 | -0.876 | 0.006 | 0.059 | 1.567 | -0.003 | -0.005 | -0.005 | 0.328 | 0.37 | 0.28 | 0.41 | 0.43 | 0.38 | 0.58 |
| 7 | 94110725 | rs12704721 | -0.728 | 0.068 | 0.154 | 1.544 | -0.001 | 0 | 0 | 0 | 0.27 | 0.26 | 0.28 | 0.41 | 0.3 | 0.34 |
| 7 | 94111956 | rs2188648 | -0.119 | 0.059 | 0.141 | 1.528 | -0.066 | -0.014 | -0.014 | 0.45 | 0.28 | 0.04 | 0.3 | 0.28 | 0.28 | 0.39 |
| 7 | 94114619 | rs7797075 | -0.184 | 0.083 | 0.178 | 1.55 | 0.002 | 0.03 | 0.03 | 0 | 0.21 | 0.33 | 0.18 | 0.26 | 0.21 | 0.15 |
| 7 | 94133800 | rs10277520 | -1.224 | 0.082 | -0.211 | 1.535 | -0.011 | -0.009 | -0.009 | 0 | 0.47 | 0.24 | 0.48 | 0.54 | 0.5 | 0.54 |
| 7 | 94136402 | rs10248931 | -1.708 | 0.011 | -0.211 | 1.212 | 0.004 | 0.025 | 0.025 | 0 | 0.56 | 0.34 | 0.58 | 0.7 | 0.59 | 0.74 |
| 7 | 94137554 | rs10268766 | -1.842 | -0.001 | -0.256 | 1.206 | 0 | 0 | 0 | 0 | 0.57 | 0.56 | 0.6 | 0.7 | 0.6 | 0.74 |
| 7 | 94141226 | rs2374735 | -1.691 | 0.007 | -1.372 | 1.219 | 0.007 | 0.043 | 0.043 | 0 | 0.56 | 0.29 | 0.58 | 0.69 | 0.59 | 0.74 |
| 7 | 94146975 | rs2301646 | -0.446 | -0.025 | -1.389 | 1.237 | -0.089 | -0.008 | -0.008 | 0 | 0.31 | 0.07 | 0.34 | 0.34 | 0.39 | 0.49 |
| 7 | 94152988 | rs10267474 | -1.261 | 0.009 | -1.365 | 1.347 | -0.007 | 0.011 | 0.011 | 0 | 0.47 | 0.29 | 0.45 | 0.49 | 0.5 | 0.54 |
| 7 | 94174485 | rs10261761 | -1.586 | 0.046 | -1.293 | 1.289 | 0.007 | 0.043 | 0.043 | 0 | 0.56 | 0.29 | 0.58 | 0.69 | 0.59 | 0.74 |
| 7 | 94174854 | rs3793326 | -1.586 | 0.046 | -1.293 | 1.289 | 0.007 | 0.043 | 0.043 | 0 | 0.56 | 0.29 | 0.58 | 0.69 | 0.59 | 0.74 |
| 7 | 94196161 | rs10243372 | 0.02 | 0.267 | -1.398 | 1.393 | -0.001 | -0.005 | -0.005 | 0 | 0.19 | 0.15 | 0.15 | 0.29 | 0.19 | 0.15 |
| 7 | 94200797 | rs12539995 | 1.117 | 0.291 | -1.576 | 1.419 | -0.023 | -0.031 | -0.031 | 0.489 | 0.08 | 0 | 0.13 | 0.12 | 0.05 | 0.08 |
| 7 | 94201152 | rs1987104 | -0.22 | -0.007 | -1.556 | 1.344 | -0.021 | 0.038 | 0.038 | 4.596 | 0.16 | 0.66 | 0.19 | 0.21 | 0.15 | 0.25 |
| 7 | 94209932 | rs17400379 | 1.017 | -0.081 | -1.749 | 1.333 | -0.041 | -0.032 | -0.032 | 0 | 0.12 | 0.01 | 0.09 | 0.14 | 0.15 | 0.19 |
| 7 | 94213193 | rs10247056 | 1.232 | -0.177 | -1.516 | 1.237 | 0.016 | -0.02 | -0.02 | 2.797 | 0.13 | 0.03 | 0.09 | 0.16 | 0.09 | 0.04 |
| 7 | 94216110 | rs1851422 | -0.247 | -0.187 | -1.439 | 1.233 | -0.024 | 0.052 | 0.052 | 4.872 | 0.17 | 0.62 | 0.19 | 0.21 | 0.15 | 0.26 |
| 7 | 94227509 | rs10232398 | -0.108 | 0.272 | -1.702 | 1.393 | -0.066 | -0.04 | -0.04 | 0 | 0.74 | 0.96 | 0.7 | 0.71 | 0.72 | 0.65 |
| 7 | 94253617 | rs7778237 | 2.181 | 0.267 | -1.727 | 1.371 | -0.053 | 0.09 | 0.09 | 0 | 0.39 | 0.12 | 0.38 | 0.21 | 0.4 | 0.24 |
| 7 | 94262067 | rs10953114 | -0.095 | 0.361 | -1.694 | 1.415 | -0.067 | -0.04 | -0.04 | 0.339 | 0.74 | 0.96 | 0.7 | 0.71 | 0.72 | 0.65 |
| 7 | 94274361 | rs13241489 | 0.815 | 0.361 | -1.664 | 1.438 | -0.001 | -0.041 | -0.041 | 0 | 0.12 | 0.03 | 0.16 | 0.21 | 0.15 | 0.25 |
| 7 | 94281335 | rs1568214 | 2.131 | 0.334 | -1.703 | 1.438 | -0.041 | 0.095 | 0.095 | 0 | 0.39 | 0.15 | 0.38 | 0.21 | 0.4 | 0.24 |
| 7 | 94294682 | rs3750105 | 0.353 | 1.118 | -1.458 | 1.676 | -0.016 | -0.075 | -0.075 | 0 | 0.12 | 0.01 | 0.16 | 0.21 | 0.15 | 0.25 |
| 7 | 94296769 | rs13073 | 0.296 | 1.162 | -1.448 | 1.7 | -0.067 | -0.04 | -0.04 | 0 | 0.74 | 0.96 | 0.7 | 0.72 | 0.72 | 0.64 |
| 7 | 94296793 | rs13226637 | -0.326 | 0.981 | -0.465 | 1.706 | 0.003 | -0.013 | -0.013 | 0.527 | 0.12 | 0.1 | 0.16 | 0.21 | 0.15 | 0.25 |
| 7 | 94299429 | rs10239188 | -0.156 | 1.267 | -0.34 | 1.135 | 0.033 | 0.036 | 0.036 | 0 | 0.05 | 0.1 | 0.04 | 0.13 | 0.08 | 0.11 |
| 7 | 94312206 | rs1357317 | 0.578 | 1.516 | -0.802 | 0.925 | -0.026 | 0.016 | 0.016 | 0 | 0.93 | 0.97 | 0.93 | 0.97 | 0.89 | 0.9 |
| 7 | 94314873 | rs1533831 | 1.676 | 0.633 | -0.018 | 1.356 | 0.002 | 0.052 | 0.052 | 1.692 | 0.71 | 0.44 | 0.79 | 0.6 | 0.69 | 0.51 |
| 7 | 94329411 | rs17166434 | -0.036 | 0.529 | -0.108 | 1.458 | 0.014 | -0.077 | -0.077 | 0 | 0.13 | 0.01 | 0.09 | 0.2 | 0.12 | 0.28 |
| 7 | 94338286 | rs1879857 | 1.33 | 0.598 | 0.568 | 1.481 | 0.044 | 0.016 | 0.016 | 0.178 | 0.85 | 0.69 | 0.88 | 0.79 | 0.81 | 0.79 |
| 7 | 94338863 | rs10808087 | 0.341 | 0.607 | 0.189 | 1.116 | 0 | -0.015 | -0.015 | 0 | 0.32 | 0.22 | 0.34 | 0.41 | 0.33 | 0.31 |
| 7 | 94344023 | rs10279084 | 0.402 | 0.527 | 0.155 | 1.082 | -0.001 | -0.015 | -0.015 | 1.201 | 0.31 | 0.22 | 0.34 | 0.41 | 0.33 | 0.31 |
| 7 | 94349839 | rs1522545 | 0.642 | 0.563 | 0.066 | 1.118 | 0.044 | 0.018 | 0.018 | 0.178 | 0.85 | 0.69 | 0.88 | 0.79 | 0.81 | 0.79 |
| 7 | 94356849 | rs12704740 | 0.406 | 0.532 | 0.253 | 1.119 | -0.012 | 0.016 | 0.016 | 0.79 | 0.14 | 0.16 | 0.09 | 0.2 | 0.12 | 0.28 |
| 7 | 94359748 | rs7799573 | 1.691 | 0.605 | 0.428 | 0.95 | -0.028 | 0.115 | 0.115 | 0.906 | 0.39 | 0.12 | 0.44 | 0.17 | 0.36 | 0.2 |
| 7 | 94364780 | rs6973800 | 0.333 | 0.686 | 0.589 | 1.052 | 0.07 | 0.102 | 0.102 | 0.01 | 0.07 | 0.3 | 0.05 | 0.19 | 0.08 | 0.11 |
| 7 | 94368896 | rs9641147 | -0.556 | 0.638 | 0.685 | 1.026 | 0.003 | 0.004 | 0.004 | 0.062 | 0.46 | 0.41 | 0.44 | 0.61 | 0.45 | 0.59 |
| 7 | 94375808 | rs6953258 | -1.18 | 0.643 | 0.56 | 0.928 | -0.009 | 0.121 | 0.121 | 1.115 | 0.54 | 0.79 | 0.48 | 0.8 | 0.51 | 0.7 |
| 7 | 94406425 | rs4376436 | 0.577 | 0.703 | 0.808 | 1.4 | 0.028 | 0.051 | 0.051 | 0 | 0.71 | 0.58 | 0.58 | 0.58 | 0.64 | 0.75 |
| 7 | 94408558 | rs6965194 | -0.415 | 0.788 | 0.877 | 1.413 | 0 | 0.03 | 0.03 | 0 | 0.26 | 0.3 | 0.4 | 0.42 | 0.28 | 0.24 |
| 7 | 94420238 | rs10281073 | -0.006 | 0.71 | 0.831 | 1.872 | -0.019 | -0.037 | -0.037 | 0 | 0.22 | 0.07 | 0.25 | 0.26 | 0.23 | 0.21 |
| 7 | 94423720 | rs13231561 | -0.875 | 0.583 | 1.008 | 1.572 | -0.01 | 0.044 | 0.044 | 1.494 | 0.2 | 0.32 | 0.13 | 0.38 | 0.18 | 0.46 |
| 7 | 94444750 | rs1926095 | -1.343 | 0.799 | 1.079 | 1.047 | 0.007 | 0.041 | 0.041 | 0.009 | 0.46 | 0.63 | 0.52 | 0.79 | 0.46 | 0.7 |
| 7 | 94463888 | rs10755864 | -0.922 | 0.815 | 1.059 | 1.134 | -0.012 | -0.009 | -0.009 | 0 | 0.4 | 0.45 | 0.49 | 0.61 | 0.38 | 0.6 |
| 7 | 94481267 | rs10280013 | -0.782 | 0.804 | 0.962 | 1.161 | 0.001 | 0 | 0 | 0 | 0.37 | 0.36 | 0.49 | 0.61 | 0.35 | 0.58 |
| 7 | 94489098 | rs10499920 | -1.343 | 1.266 | 1.465 | 0.992 | 0.147 | 0.104 | 0.104 | 0 | 0.11 | 0.43 | 0.11 | 0.2 | 0.19 | 0.13 |
| 7 | 94518128 | rs1569091 | -0.989 | 0.247 | 0.888 | 0.26 | -0.005 | -0.063 | -0.063 | 0.067 | 0.46 | 0.5 | 0.61 | 0.93 | 0.45 | 0.73 |
| 7 | 94534886 | rs1527680 | -1.371 | -0.188 | 0.907 | 0.974 | -0.041 | 0.05 | 0.05 | 1.435 | 0.21 | 0.44 | 0.31 | 0.37 | 0.16 | 0.28 |
| 7 | 94535888 | rs35429726 | -0.639 | -0.194 | 0.727 | 1.052 | 0.012 | -0.014 | -0.014 | 0 | 0.15 | 0.09 | 0.16 | 0.27 | 0.1 | 0.19 |
| 7 | 94539536 | rs12536183 | -0.492 | -0.245 | 0.622 | 1.022 | 0.016 | -0.007 | -0.007 | 0 | 0.14 | 0.02 | 0.22 | 0.14 | 0.1 | 0.13 |
| 7 | 94540416 | rs10230714 | -1.609 | -0.228 | 1.129 | 1.039 | 0.026 | 0.164 | 0.164 | 2.171 | 0.07 | 0.35 | 0.15 | 0.22 | 0.05 | 0.16 |
| 7 | 94545863 | rs4727304 | -1.355 | 0.577 | 0.786 | 1.293 | 0.021 | 0.026 | 0.026 | 0.292 | 0.51 | 0.7 | 0.57 | 0.59 | 0.56 | 0.67 |
| 7 | 94554764 | rs10953133 | -1.414 | 0.586 | 1.058 | 1.384 | 0.015 | 0.071 | 0.071 | 0.363 | 0.51 | 0.67 | 0.65 | 0.73 | 0.56 | 0.76 |
| 7 | 94565122 | rs1527676 | -1.394 | 0.579 | 1.1 | 1.362 | 0.013 | 0.054 | 0.054 | 0.322 | 0.51 | 0.65 | 0.56 | 0.73 | 0.56 | 0.75 |
| 7 | 94571332 | rs2204630 | -1.039 | 0.57 | 1.424 | 1.398 | -0.017 | -0.007 | -0.007 | 0 | 0.43 | 0.22 | 0.44 | 0.43 | 0.51 | 0.58 |
| 7 | 94573288 | rs6465448 | -1.64 | 0.554 | 1.375 | 1.366 | 0.006 | 0.081 | 0.081 | 0.446 | 0.53 | 0.81 | 0.57 | 0.73 | 0.57 | 0.76 |
| 7 | 94574220 | rs10499917 | -0.507 | 0.613 | 1.453 | 1.198 | -0.05 | -0.005 | -0.005 | 0.789 | 0.33 | 0.14 | 0.23 | 0.3 | 0.43 | 0.46 |
| 7 | 94600834 | rs854721 | 2.084 | 0.632 | 0.578 | 0.255 | -0.054 | 0.057 | 0.057 | 0 | 0.1 | 0.01 | 0.1 | 0 | 0.15 | 0.04 |
| 7 | 94601868 | rs13242203 | 0.772 | 0.577 | 0.682 | 0.293 | 0.001 | -0.002 | -0.002 | 0.511 | 0.11 | 0.1 | 0.22 | 0.16 | 0.08 | 0.12 |
| 7 | 94602096 | rs854722 | 0.736 | 0.69 | 0.669 | 0.295 | 0.003 | -0.02 | -0.02 | 0 | 0.58 | 0.62 | 0.57 | 0.18 | 0.51 | 0.29 |
| 7 | 94606271 | rs6950265 | 0.386 | 1.561 | 0.439 | -0.03 | -0.084 | -0.092 | -0.092 | 0 | 0.23 | 0.04 | 0.18 | 0.32 | 0.31 | 0.41 |
| 7 | 94615259 | rs854738 | 0.347 | 1.523 | 0.446 | -0.065 | -0.047 | -0.01 | -0.01 | 1.59 | 0.67 | 0.86 | 0.77 | 0.68 | 0.57 | 0.54 |
| 7 | 94623080 | rs854747 | 1.437 | 1.622 | 0.454 | -0.863 | -0.007 | 0.017 | 0.017 | 0.015 | 0.57 | 0.43 | 0.53 | 0.15 | 0.52 | 0.29 |
| 7 | 94626992 | rs854726 | 0.634 | 1.628 | 0.316 | -0.857 | -0.017 | 0.029 | 0.029 | 0.004 | 0.63 | 0.77 | 0.62 | 0.26 | 0.56 | 0.4 |
| 7 | 94633688 | rs854731 | 1.364 | 1.658 | 0.034 | -0.989 | -0.005 | 0.008 | 0.008 | 0.028 | 0.57 | 0.42 | 0.53 | 0.15 | 0.51 | 0.29 |
| 7 | 94633926 | rs17460316 | 1.977 | 1.725 | -0.414 | -1.057 | -0.039 | 0.059 | 0.059 | 2.087 | 0.1 | 0.01 | 0.04 | 0.01 | 0.11 | 0.28 |
| 7 | 94637458 | rs854734 | 2.225 | 1.569 | -0.191 | -0.928 | -0.012 | 0 | 0 | 0 | 0.1 | 0.07 | 0.11 | 0 | 0.15 | 0.04 |

| | | | | | | | | | | | | | | | | |
|---|---|---|---|---|---|---|---|---|---|---|---|---|---|---|---|---|
| 7 | 94646829 | rs10265388 | -0.972 | 1.141 | -0.031 | -0.518 | -0.005 | -0.009 | -0.009 | 0.639 | 0.9 | 0.94 | 0.96 | 1 | 0.89 | 0.95 |
| 7 | 94648239 | rs11981513 | 1.004 | 0.953 | -0.074 | -0.304 | 0.001 | -0.018 | -0.018 | 0 | 0.57 | 0.62 | 0.56 | 0.18 | 0.5 | 0.29 |
| 7 | 94653626 | rs13310108 | 0.327 | 0.515 | -0.418 | 0.453 | 0.003 | -0.026 | -0.026 | 0 | 0.1 | 0.02 | 0.21 | 0.15 | 0.08 | 0.12 |
| 7 | 94665282 | rs3801925 | 0.141 | 0.475 | -0.194 | 0.407 | -0.001 | 0 | 0 | 0 | 0.11 | 0.11 | 0.22 | 0.15 | 0.08 | 0.12 |
| 7 | 94685143 | rs6959219 | 1.215 | 0.085 | -0.135 | -0.62 | -0.002 | 0.046 | 0.046 | 1.474 | 0.47 | 0.63 | 0.4 | 0.12 | 0.46 | 0.25 |
| 7 | 94693970 | rs739504 | -0.023 | -0.451 | -0.298 | -0.138 | -0.043 | 0 | 0 | 0 | 0.65 | 0.85 | 0.77 | 0.68 | 0.58 | 0.52 |
| 7 | 94694136 | rs12532531 | 1.218 | -0.391 | -0.323 | -0.099 | 0.005 | -0.022 | -0.022 | 0 | 0.1 | 0.05 | 0.24 | 0.17 | 0.08 | 0.12 |
| 7 | 94696139 | rs6961415 | 0.91 | -0.416 | -0.269 | -0.305 | -0.006 | 0.126 | 0.126 | 0.326 | 0.53 | 0.73 | 0.44 | 0.12 | 0.49 | 0.27 |
| 7 | 94702805 | rs6465454 | 1.246 | -0.245 | -0.33 | -0.227 | -0.001 | 0.048 | 0.048 | 1.249 | 0.47 | 0.63 | 0.4 | 0.12 | 0.46 | 0.25 |
| 7 | 94722053 | rs6967937 | 1.31 | -0.227 | -0.503 | -0.217 | -0.001 | -0.033 | -0.033 | 0.005 | 0.46 | 0.53 | 0.39 | 0.11 | 0.46 | 0.25 |
| 7 | 94722785 | rs10485996 | 1.369 | 0.058 | -1.345 | -0.309 | -0.05 | -0.083 | -0.083 | 0 | 0.12 | 0.01 | 0.12 | 0.31 | 0.18 | 0.12 |
| 7 | 94728705 | rs6972772 | 1.83 | -0.283 | -1.01 | -0.468 | -0.042 | 0.159 | 0.159 | 2.153 | 0.39 | 0.16 | 0.3 | 0.03 | 0.41 | 0.13 |
| 7 | 94733521 | rs2374992 | 0.415 | 0.264 | -0.987 | -0.053 | -0.092 | -0.114 | -0.114 | 0 | 0.24 | 0.02 | 0.18 | 0.32 | 0.3 | 0.44 |
| 7 | 94733635 | rs10266293 | -0.221 | 0.131 | -0.092 | -0.091 | -0.009 | 0.007 | 0.007 | 0 | 0.06 | 0.15 | 0.07 | 0.09 | 0.05 | 0.12 |
| 7 | 94742832 | rs17305795 | 0.199 | 0.097 | -0.776 | -0.249 | -0.007 | -0.046 | -0.046 | 0 | 0.29 | 0.14 | 0.28 | 0.39 | 0.34 | 0.47 |
| 7 | 94747455 | rs17166665 | 0.395 | -0.181 | -0.506 | -0.292 | -0.006 | 0.005 | 0.005 | 0 | 0.14 | 0.16 | 0.25 | 0.16 | 0.11 | 0.11 |
| 7 | 94751895 | rs17305991 | -0.117 | -0.033 | -0.39 | -0.562 | -0.02 | -0.012 | -0.012 | 0 | 0.65 | 0.73 | 0.68 | 0.61 | 0.57 | 0.49 |
| 7 | 94754985 | rs9649194 | -0.118 | -0.032 | -0.403 | -0.537 | -0.023 | -0.011 | -0.011 | 0 | 0.64 | 0.75 | 0.67 | 0.61 | 0.54 | 0.49 |
| 7 | 94768272 | rs10953137 | -0.17 | -0.052 | -0.42 | -0.557 | -0.039 | -0.022 | -0.022 | 0 | 0.65 | 0.8 | 0.67 | 0.61 | 0.53 | 0.49 |
| 7 | 94786666 | rs4729176 | -1.32 | -0.91 | -0.496 | -0.735 | -0.01 | 0.004 | 0.004 | 0 | 0.28 | 0.7 | 0.43 | 0.59 | 0.19 | 0.35 |
| 7 | 94787641 | rs7802749 | 1.07 | -0.801 | 0.128 | -0.649 | 0.002 | 0.008 | 0.008 | 0.369 | 0.45 | 0.38 | 0.34 | 0.43 | 0.42 | 0.34 |
| 7 | 94788164 | rs1012367 | 0.503 | -0.818 | 0.163 | -0.644 | -0.005 | 0.005 | 0.005 | 0 | 0.15 | 0.15 | 0.26 | 0.17 | 0.12 | 0.14 |
| 7 | 94789681 | rs7777082 | 0.904 | -0.793 | 0.139 | -0.655 | 0.001 | 0.003 | 0.003 | 0.413 | 0.45 | 0.4 | 0.33 | 0.42 | 0.41 | 0.33 |
| 7 | 94802693 | rs1234476 | -0.478 | -0.685 | 0.013 | -0.632 | -0.008 | -0.004 | -0.004 | 2.018 | 0.65 | 0.7 | 0.66 | 0.63 | 0.55 | 0.51 |
| 7 | 94823371 | rs1527721 | 0.854 | -0.731 | -0.116 | -0.572 | 0.003 | -0.049 | -0.049 | 0 | 0.14 | 0 | 0.22 | 0.15 | 0.11 | 0.14 |
| 7 | 94831043 | rs10263800 | -2.535 | -0.97 | -0.652 | -1.053 | -0.001 | 0.088 | 0.088 | 0 | 0.86 | 0.99 | 0.9 | 1 | 0.88 | 0.98 |
| 7 | 94836537 | rs13229820 | 0.698 | -0.733 | -0.068 | -0.908 | -0.009 | -0.004 | -0.004 | 0 | 0.36 | 0.3 | 0.34 | 0.39 | 0.46 | 0.5 |
| 7 | 94839576 | rs6465459 | -2.816 | -0.825 | -0.307 | -1.002 | -0.008 | 0.047 | 0.047 | 0.426 | 0.92 | 0.99 | 0.95 | 1 | 0.91 | 0.98 |
| 7 | 94840916 | rs11767065 | -0.232 | -0.498 | 0.51 | -0.53 | -0.003 | -0.005 | -0.005 | 0 | 0.42 | 0.63 | 0.46 | 0.48 | 0.47 | 0.57 |
| 7 | 94850268 | rs6947420 | -0.15 | -0.464 | 0.508 | -0.524 | -0.001 | -0.002 | -0.002 | 0.261 | 0.4 | 0.6 | 0.42 | 0.46 | 0.47 | 0.56 |
| 7 | 94865281 | rs11766452 | -0.995 | -0.505 | 0.383 | -0.525 | -0.042 | 0.021 | 0.021 | 1.377 | 0.57 | 0.84 | 0.7 | 0.64 | 0.58 | 0.72 |
| 7 | 94871724 | rs2374983 | 0.399 | -0.523 | 0.607 | -0.513 | -0.028 | -0.015 | -0.015 | 1.203 | 0.36 | 0.24 | 0.34 | 0.39 | 0.46 | 0.49 |
| 7 | 94872226 | rs10499911 | 0.414 | -0.569 | 0.633 | -0.408 | -0.017 | 0.011 | 0.011 | 0 | 0.15 | 0.2 | 0.24 | 0.16 | 0.11 | 0.14 |
| 7 | 94884381 | rs16868732 | 0.532 | -0.492 | 0.411 | -0.393 | -0.008 | 0.004 | 0.004 | 0 | 0.14 | 0.15 | 0.24 | 0.16 | 0.11 | 0.14 |
| 7 | 94888158 | rs10953141 | -0.181 | -0.545 | 0.375 | -0.431 | -0.008 | 0 | 0 | 1.301 | 0.18 | 0.2 | 0.3 | 0.17 | 0.13 | 0.16 |
| 7 | 94898233 | rs854523 | -0.023 | -0.387 | 0.367 | -0.381 | -0.002 | -0.002 | -0.002 | 4.485 | 0.6 | 0.39 | 0.59 | 0.51 | 0.53 | 0.44 |
| 7 | 94898811 | rs854524 | -0.376 | -0.398 | 0.47 | -0.428 | -0.006 | -0.003 | -0.003 | 3.206 | 0.63 | 0.68 | 0.66 | 0.62 | 0.55 | 0.51 |
| 7 | 94900279 | rs854527 | 0.007 | -0.257 | -0.199 | -0.469 | -0.003 | -0.003 | -0.003 | 0.17 | 0.59 | 0.39 | 0.59 | 0.54 | 0.53 | 0.44 |
| 7 | 94900678 | rs705374 | -1.309 | 0.054 | -0.759 | -0.613 | -0.01 | 0 | 0 | 0.046 | 0.9 | 0.94 | 0.95 | 1 | 0.87 | 0.94 |
| 7 | 94900975 | rs705375 | 0.296 | -0.239 | -0.793 | -0.287 | -0.012 | 0.019 | 0.019 | 1.062 | 0.44 | 0.21 | 0.35 | 0.4 | 0.42 | 0.3 |
| 7 | 94902649 | rs854530 | 0.362 | -0.026 | -0.937 | -0.33 | -0.013 | -0.034 | -0.034 | 0 | 0.45 | 0.18 | 0.36 | 0.48 | 0.44 | 0.38 |
| 7 | 94903140 | rs16868737 | 1.175 | -0.342 | 0.102 | -0.175 | -0.033 | -0.035 | -0.035 | 0.649 | 0.07 | 0.25 | 0.02 | 0 | 0.01 | 0.01 |
| 7 | 94903420 | rs2240026 | -0.143 | -0.065 | 0.071 | 0.206 | -0.006 | 0.117 | 0.117 | 0 | 0.07 | 0.12 | 0.11 | 0.4 | 0.07 | 0.13 |
| 7 | 94903652 | rs2240027 | -0.765 | -0.05 | 0.29 | 0.163 | -0.007 | 0.184 | 0.184 | 0 | 0.08 | 0.18 | 0.12 | 0.41 | 0.07 | 0.15 |
| 7 | 94905790 | rs10156050 | 0.477 | -0.159 | 0.277 | 0.054 | -0.024 | 0.014 | 0.014 | 0.02 | 0.1 | 0.25 | 0.22 | 0.11 | 0.07 | 0.08 |
| 7 | 94907500 | rs12537967 | 0.477 | -0.125 | 0.277 | 0.054 | -0.021 | 0.014 | 0.014 | 0.02 | 0.1 | 0.25 | 0.22 | 0.11 | 0.07 | 0.08 |
| 7 | 94909495 | rs854534 | 0.785 | 0.228 | 0.309 | -0.456 | 0.025 | 0.025 | 0.025 | 0.749 | 0.48 | 0.26 | 0.34 | 0.44 | 0.42 | 0.31 |
| 7 | 94909899 | rs854536 | -0.462 | 0.787 | 0.511 | -0.411 | -0.014 | -0.011 | -0.011 | 0 | 0.67 | 0.75 | 0.69 | 0.67 | 0.59 | 0.51 |
| 7 | 94912404 | rs2285694 | 1.578 | 0.528 | 0.542 | -0.358 | -0.012 | 0.011 | 0.011 | 0 | 0.08 | 0.11 | 0.21 | 0.11 | 0.06 | 0.08 |
| 7 | 94920480 | rs854541 | -0.067 | 0.864 | 0.476 | -0.422 | -0.021 | -0.018 | -0.018 | 0.724 | 0.63 | 0.71 | 0.59 | 0.57 | 0.51 | 0.44 |
| 7 | 94920992 | rs11764692 | 1.722 | 0.885 | 0.243 | -0.333 | -0.014 | 0.075 | 0.075 | 0 | 0.13 | 0.01 | 0.05 | 0 | 0.12 | 0.28 |
| 7 | 94921491 | rs854542 | 0.028 | 1.255 | 0.724 | 0.119 | 0.025 | -0.007 | -0.007 | 0.537 | 0.87 | 0.81 | 0.86 | 0.88 | 0.8 | 0.86 |
| 7 | 94923856 | rs854547 | 1.671 | 1.081 | 0.746 | 0.046 | -0.002 | -0.003 | -0.003 | 1.403 | 0.63 | 0.32 | 0.48 | 0.35 | 0.62 | 0.59 |
| 7 | 94925820 | rs854548 | 0.627 | 1.211 | 0.94 | 0.198 | 0.043 | 0.018 | 0.018 | 0 | 0.79 | 0.6 | 0.64 | 0.77 | 0.75 | 0.79 |
| 7 | 94926321 | rs3917586 | 1.277 | 1.18 | 0.851 | 0.224 | 0.002 | 0.009 | 0.009 | 0 | 0.11 | 0.08 | 0.14 | 0.41 | 0.11 | 0.16 |
| 7 | 94927242 | rs854550 | -0.337 | 1.213 | 0.935 | 0.2 | 0.025 | -0.008 | -0.008 | 0 | 0.87 | 0.81 | 0.86 | 0.89 | 0.81 | 0.87 |
| 7 | 94927495 | rs3735590 | 0.694 | 1.16 | 1.038 | 0.191 | -0.023 | 0.035 | 0.035 | 0.241 | 0.11 | 0.35 | 0.23 | 0.11 | 0.07 | 0.08 |
| 7 | 94927677 | rs854551 | -0.19 | 1.173 | 1.074 | 0.189 | 0.056 | -0.012 | -0.012 | 0.221 | 0.87 | 0.76 | 0.85 | 0.89 | 0.81 | 0.87 |
| 7 | 94927708 | rs3917577 | 1.245 | 1.158 | 1.031 | 0.203 | 0.002 | 0.007 | 0.007 | 0 | 0.11 | 0.08 | 0.14 | 0.41 | 0.11 | 0.16 |
| 7 | 94927924 | rs854552 | 1.233 | 1.146 | 1.018 | 0.165 | 0.008 | -0.01 | -0.01 | 0.036 | 0.75 | 0.38 | 0.61 | 0.76 | 0.74 | 0.75 |
| 7 | 94928746 | rs854553 | 1.358 | 1.154 | 1.011 | 0.164 | 0.013 | -0.004 | -0.004 | 0.099 | 0.75 | 0.4 | 0.61 | 0.76 | 0.74 | 0.76 |
| 7 | 94929190 | rs3917572 | 0.635 | 1.024 | 0.945 | 0.043 | -0.024 | 0.034 | 0.034 | 0 | 0.1 | 0.23 | 0.22 | 0.13 | 0.07 | 0.12 |
| 7 | 94929439 | rs3917569 | 1.39 | 1.044 | 0.893 | 0.051 | -0.016 | 0.031 | 0.031 | 0 | 0.08 | 0.19 | 0.21 | 0.11 | 0.06 | 0.08 |
| 7 | 94930108 | rs2237580 | 1.212 | 1.063 | 0.727 | 0.09 | 0.002 | 0.007 | 0.007 | 0 | 0.11 | 0.08 | 0.14 | 0.41 | 0.11 | 0.18 |
| 7 | 94930391 | rs854555 | 1.715 | 1.113 | 0.92 | 0.032 | -0.002 | -0.029 | -0.029 | 0 | 0.63 | 0.26 | 0.48 | 0.39 | 0.64 | 0.58 |
| 7 | 94932904 | rs3917558 | 0.153 | 1.062 | 1.361 | 0.108 | -0.007 | 0.055 | 0.055 | 0.638 | 0.12 | 0.49 | 0.22 | 0.11 | 0.06 | 0.09 |
| 7 | 94933189 | rs3917556 | 1.178 | 1.096 | 1.131 | 0.128 | -0.014 | 0.021 | 0.021 | 0 | 0.09 | 0.18 | 0.2 | 0.11 | 0.06 | 0.09 |
| 7 | 94934455 | rs3917551 | 1.178 | 1.096 | 1.131 | 0.128 | -0.014 | 0.021 | 0.021 | 0 | 0.09 | 0.18 | 0.2 | 0.11 | 0.06 | 0.09 |
| 7 | 94934573 | rs3917550 | 0.602 | 1.276 | 1.046 | 0.12 | 0.026 | 0.003 | 0.003 | 0 | 0.07 | 0.12 | 0.13 | 0.11 | 0.12 | 0.1 |

| | | | | | | | | | | | | | | | | |
|---|---|---|---|---|---|---|---|---|---|---|---|---|---|---|---|---|
| 7 | 94935905 | rs3917548 | 1.109 | 1.235 | 1.069 | 0.117 | -0.015 | 0.021 | 0.021 | 0 | 0.09 | 0.2 | 0.21 | 0.11 | 0.06 | 0.08 |
| 7 | 94936129 | rs2269829 | -1.405 | 1.298 | 1.109 | 0.199 | 0.005 | 0.032 | 0.032 | 0.068 | 0.28 | 0.59 | 0.48 | 0.65 | 0.28 | 0.37 |
| 7 | 94936235 | rs3917545 | 0.643 | 1.313 | 1.114 | 0.23 | 0.026 | 0.008 | 0.008 | 0 | 0.07 | 0.12 | 0.13 | 0.11 | 0.12 | 0.1 |
| 7 | 94936692 | rs3917542 | -0.541 | 1.308 | 1.148 | 0.217 | 0.04 | 0.154 | 0.154 | 0 | 0.19 | 0.39 | 0.27 | 0.54 | 0.23 | 0.29 |
| 7 | 94937624 | rs3917541 | 1.152 | 1.347 | 1.154 | 0.218 | -0.012 | 0.017 | 0.017 | 0 | 0.09 | 0.2 | 0.21 | 0.11 | 0.06 | 0.08 |
| 7 | 94937893 | rs3917538 | -1.059 | 1.312 | 1.196 | 0.207 | 0.028 | 0.135 | 0.135 | 0 | 0.22 | 0.55 | 0.28 | 0.54 | 0.23 | 0.29 |
| 7 | 94938176 | rs2158155 | 1.374 | 1.308 | 1.191 | 0.204 | -0.014 | 0.014 | 0.014 | 0 | 0.09 | 0.18 | 0.2 | 0.11 | 0.06 | 0.08 |
| 7 | 94938786 | rs2299255 | 0.619 | 1.377 | 1.135 | 0.253 | 0.026 | 0.005 | 0.005 | 0 | 0.07 | 0.12 | 0.13 | 0.11 | 0.12 | 0.1 |
| 7 | 94939032 | rs2299256 | -1.092 | 1.372 | 1.141 | 0.255 | 0.029 | 0.135 | 0.135 | 0 | 0.22 | 0.55 | 0.28 | 0.54 | 0.23 | 0.29 |
| 7 | 94939830 | rs3917533 | -0.579 | 1.376 | 1.128 | 0.258 | 0.04 | 0.154 | 0.154 | 0 | 0.19 | 0.38 | 0.27 | 0.54 | 0.23 | 0.29 |
| 7 | 94940122 | rs3917531 | 1.035 | 1.329 | 1.114 | 0.267 | -0.003 | 0.155 | 0.155 | 0.03 | 0.12 | 0.26 | 0.15 | 0.43 | 0.11 | 0.18 |
| 7 | 94940240 | rs3917528 | 0.987 | 1.315 | 1.108 | 0.26 | -0.004 | 0.152 | 0.152 | 0.025 | 0.12 | 0.26 | 0.14 | 0.43 | 0.11 | 0.18 |
| 7 | 94940587 | rs2074354 | 1.037 | 1.322 | 1.11 | 0.263 | -0.003 | 0.155 | 0.155 | 0.03 | 0.12 | 0.26 | 0.14 | 0.43 | 0.11 | 0.18 |
| 7 | 94941415 | rs3917521 | -0.415 | 1.325 | 1.01 | 0.192 | 0.039 | 0.151 | 0.151 | 0 | 0.19 | 0.38 | 0.27 | 0.54 | 0.23 | 0.29 |
| 7 | 94942765 | rs2299257 | -1.829 | 1.915 | 0.46 | 0.043 | -0.004 | -0.003 | -0.003 | 0 | 0.35 | 0.63 | 0.53 | 0.67 | 0.4 | 0.43 |
| 7 | 94943397 | rs2299259 | -0.293 | 1.745 | 0.226 | 0.164 | -0.025 | -0.037 | -0.037 | 1.623 | 0.2 | 0.15 | 0.29 | 0.53 | 0.29 | 0.29 |
| 7 | 94945374 | rs854558 | -0.436 | 1.286 | -0.163 | -0.408 | -0.004 | 0.004 | 0.004 | 0.23 | 0.71 | 0.66 | 0.73 | 0.7 | 0.71 | 0.61 |
| 7 | 94945453 | rs3917503 | -1.18 | 1.36 | -0.028 | -0.35 | 0.015 | 0.021 | 0.021 | 0.155 | 0.32 | 0.51 | 0.52 | 0.67 | 0.36 | 0.43 |
| 7 | 94945659 | rs2301711 | 0.573 | 1.288 | 0.056 | -0.331 | -0.029 | 0.035 | 0.035 | 0.698 | 0.12 | 0.36 | 0.23 | 0.13 | 0.06 | 0.09 |
| 7 | 94947799 | rs2074351 | -0.183 | 1.454 | -0.12 | -0.292 | -0.012 | -0.018 | -0.018 | 0.875 | 0.21 | 0.17 | 0.31 | 0.55 | 0.3 | 0.34 |
| 7 | 94948028 | rs3917493 | 1.577 | 1.359 | -0.22 | -0.298 | -0.013 | 0.012 | 0.012 | 0 | 0.09 | 0.15 | 0.19 | 0.12 | 0.06 | 0.09 |
| 7 | 94948626 | rs2272365 | 1.383 | 1.616 | -0.599 | -0.295 | -0.053 | -0.09 | -0.09 | 1.125 | 0.12 | 0.01 | 0.17 | 0.42 | 0.18 | 0.19 |
| 7 | 94948784 | rs854567 | -0.719 | 1.27 | -0.341 | -1.023 | -0.013 | -0.093 | -0.093 | 0 | 0.79 | 0.67 | 0.9 | 1 | 0.81 | 0.73 |
| 7 | 94948841 | rs3917490 | -0.671 | 0.971 | -0.647 | -1.182 | 0 | -0.05 | -0.05 | 0 | 0.49 | 0.52 | 0.73 | 0.97 | 0.51 | 0.59 |
| 7 | 94949537 | rs2299260 | 0.207 | 1.037 | -0.297 | -0.188 | -0.009 | -0.013 | -0.013 | 0 | 0.19 | 0.33 | 0.11 | 0.11 | 0.18 | 0.27 |
| 7 | 94949663 | rs2299261 | -0.662 | 1.201 | -0.569 | -0.49 | -0.001 | -0.004 | -0.004 | 0.01 | 0.36 | 0.38 | 0.19 | 0.12 | 0.35 | 0.36 |
| 7 | 94949801 | rs854568 | 0.57 | 0.832 | -0.439 | 0.099 | -0.024 | 0.026 | 0.026 | 0.116 | 0.75 | 0.48 | 0.7 | 0.71 | 0.79 | 0.63 |
| 7 | 94949928 | rs2299262 | -0.688 | 0.859 | -0.78 | -0.143 | -0.008 | 0.014 | 0.014 | 0 | 0.33 | 0.23 | 0.48 | 0.72 | 0.42 | 0.43 |
| 7 | 94950055 | rs854569 | 0.77 | 0.303 | -0.155 | 0.473 | 0.032 | 0.007 | 0.007 | 0.493 | 0.7 | 0.23 | 0.67 | 0.72 | 0.78 | 0.6 |
| 7 | 94950177 | rs2237583 | -0.323 | 0.991 | -0.105 | 0.931 | -0.046 | -0.077 | -0.077 | 0 | 0.17 | 0.05 | 0.34 | 0.62 | 0.31 | 0.27 |
| 7 | 94950837 | rs2237584 | 1.711 | 0.749 | 0.032 | 1.624 | -0.009 | -0.007 | -0.007 | 0.608 | 0.06 | 0.01 | 0.19 | 0.25 | 0.08 | 0.13 |
| 7 | 94951569 | rs3917478 | 0.544 | 0.835 | 0.147 | 1.454 | -0.003 | -0.003 | -0.003 | 0 | 0.07 | 0.05 | 0.07 | 0.18 | 0.12 | 0.09 |
| 7 | 94953949 | rs705381 | 0.156 | 0.704 | 0.644 | 1.148 | 0.006 | -0.031 | -0.031 | 0.077 | 0.77 | 0.59 | 0.84 | 0.87 | 0.78 | 0.71 |
| 7 | 94954619 | rs854571 | 0.801 | 0.69 | 0.584 | 1.204 | -0.001 | -0.001 | -0.001 | 0 | 0.71 | 0.36 | 0.66 | 0.71 | 0.73 | 0.6 |
| 7 | 94954863 | rs854573 | 0.103 | 0.684 | 0.523 | 1.127 | 0.009 | -0.033 | -0.033 | 0 | 0.77 | 0.56 | 0.84 | 0.87 | 0.77 | 0.71 |
| 7 | 94955528 | rs757158 | -1.578 | 0.375 | 0.073 | 1.285 | -0.001 | -0.053 | -0.053 | 0 | 0.44 | 0.93 | 0.53 | 0.55 | 0.42 | 0.57 |
| 7 | 94958312 | rs17883318 | -0.862 | 0.232 | 0.929 | 0.883 | -0.023 | -0.033 | -0.033 | 0 | 0.31 | 0.61 | 0.26 | 0.15 | 0.25 | 0.41 |
| 7 | 94958628 | rs17166818 | -1.782 | 0.207 | 1.089 | 0.87 | 0.035 | 0.022 | 0.022 | 1.847 | 0.06 | 0.36 | 0.06 | 0.04 | 0.02 | 0.07 |
| 7 | 94960825 | rs13228784 | -0.263 | 0.215 | 1.051 | 0.876 | 0.001 | 0.001 | 0.001 | 0.015 | 0.25 | 0.25 | 0.23 | 0.13 | 0.24 | 0.39 |
| 7 | 94986960 | rs916864 | 0.443 | 0.392 | 0.868 | 0.815 | -0.05 | -0.078 | -0.078 | 0 | 0.14 | 0.03 | 0.38 | 0.6 | 0.23 | 0.25 |
| 7 | 94987727 | rs17886036 | -0.12 | 0.362 | 0.876 | 0.786 | 0.002 | 0.003 | 0.003 | 0.012 | 0.25 | 0.25 | 0.23 | 0.14 | 0.24 | 0.39 |
| 7 | 94989673 | rs2074352 | 0.452 | 0.363 | 0.875 | 0.786 | -0.05 | -0.078 | -0.078 | 0 | 0.14 | 0.03 | 0.38 | 0.6 | 0.23 | 0.25 |
| 7 | 94991632 | rs17884000 | 0.452 | 0.363 | 0.875 | 0.786 | -0.05 | -0.078 | -0.078 | 0.637 | 0.14 | 0.03 | 0.38 | 0.6 | 0.23 | 0.25 |
| 7 | 94992244 | rs10953143 | -1.257 | 0.362 | 0.875 | 0.786 | -0.013 | 0.047 | 0.047 | 0.37 | 0.4 | 0.29 | 0.61 | 0.74 | 0.48 | 0.64 |
| 7 | 94992468 | rs9641162 | 0.43 | 0.362 | 0.876 | 0.786 | -0.05 | -0.078 | -0.078 | 0.008 | 0.14 | 0.04 | 0.38 | 0.6 | 0.23 | 0.25 |
| 7 | 94992908 | rs2074353 | -1.246 | 0.362 | 0.876 | 0.786 | -0.014 | 0.047 | 0.047 | 0.345 | 0.39 | 0.28 | 0.61 | 0.74 | 0.48 | 0.64 |
| 7 | 94995519 | rs17166859 | -0.121 | 0.361 | 0.867 | 0.799 | 0.002 | 0.003 | 0.003 | 0.012 | 0.25 | 0.25 | 0.23 | 0.14 | 0.24 | 0.39 |
| 7 | 94996913 | rs3757708 | -1.72 | 0.314 | 0.865 | 0.797 | -0.001 | 0.018 | 0.018 | 0.423 | 0.45 | 0.64 | 0.64 | 0.76 | 0.49 | 0.67 |
| 7 | 94999438 | rs17883791 | -0.14 | 0.172 | 0.993 | 0.657 | 0.002 | 0.003 | 0.003 | 0.012 | 0.25 | 0.25 | 0.23 | 0.14 | 0.24 | 0.39 |
| 7 | 95000591 | rs11768074 | 1.456 | 0.168 | 0.876 | 0.644 | -0.031 | 0.002 | 0.002 | 1.56 | 0.11 | 0.01 | 0.09 | 0.06 | 0.14 | 0.24 |
| 7 | 95001132 | rs7807131 | -1.361 | 0.208 | 1.17 | 0.758 | 0.049 | 0.046 | 0.046 | 2.125 | 0.05 | 0.35 | 0.03 | 0.02 | 0.01 | 0.02 |
| 7 | 95019355 | rs6977389 | -1.039 | 0.206 | 1.067 | 0.715 | 0.001 | -0.033 | -0.033 | 1.795 | 0.44 | 0.54 | 0.66 | 0.76 | 0.49 | 0.69 |
| 7 | 95020061 | rs10487131 | 0.262 | 0.3 | 0.827 | 0.843 | -0.046 | -0.074 | -0.074 | 0 | 0.14 | 0.04 | 0.38 | 0.6 | 0.23 | 0.25 |
| 7 | 95021803 | rs740264 | 0.376 | 0.27 | 0.885 | 0.817 | 0.008 | 0.015 | 0.015 | 0.055 | 0.25 | 0.21 | 0.23 | 0.14 | 0.24 | 0.39 |
| 7 | 95022316 | rs17774169 | 0.243 | 0.405 | 0.521 | 0.919 | -0.05 | -0.078 | -0.078 | 0.941 | 0.14 | 0.03 | 0.38 | 0.6 | 0.23 | 0.25 |
| 7 | 95024046 | rs11970910 | -1.137 | 0.272 | 0.774 | 0.732 | -0.001 | -0.018 | -0.018 | 0 | 0.45 | 0.58 | 0.63 | 0.76 | 0.49 | 0.66 |
| 7 | 95025600 | rs13226149 | -0.035 | 0.02 | 0.857 | 0.618 | -0.014 | -0.023 | -0.023 | 0.377 | 0.28 | 0.4 | 0.25 | 0.17 | 0.26 | 0.41 |
| 7 | 95026229 | rs11764079 | -0.017 | 0.029 | 0.875 | 0.466 | -0.014 | -0.026 | -0.026 | 0 | 0.28 | 0.38 | 0.24 | 0.16 | 0.26 | 0.41 |
| 7 | 95026327 | rs11770903 | -0.017 | 0.029 | 0.875 | 0.466 | -0.014 | -0.026 | -0.026 | 0 | 0.28 | 0.39 | 0.24 | 0.16 | 0.26 | 0.41 |
| 7 | 95026408 | rs17882539 | -0.004 | 0.029 | 0.871 | 0.454 | -0.012 | -0.024 | -0.024 | 2.173 | 0.28 | 0.37 | 0.24 | 0.16 | 0.26 | 0.43 |
| 7 | 95029767 | rs11977702 | -0.176 | 0.016 | 1.022 | 0.455 | -0.027 | -0.045 | -0.045 | 0 | 0.29 | 0.49 | 0.24 | 0.16 | 0.25 | 0.4 |
| 7 | 95030632 | rs17879277 | 0.197 | 0.013 | 0.934 | 0.473 | -0.003 | -0.006 | -0.006 | 0 | 0.27 | 0.29 | 0.23 | 0.19 | 0.25 | 0.39 |
| 7 | 95030646 | rs17879517 | 0.157 | 0.013 | 0.961 | 0.473 | -0.012 | -0.024 | -0.024 | 0 | 0.27 | 0.35 | 0.23 | 0.19 | 0.25 | 0.39 |
| 7 | 95031164 | rs17880030 | 0.27 | 0.151 | 0.838 | 0.582 | -0.049 | -0.074 | -0.074 | 0 | 0.15 | 0.03 | 0.37 | 0.62 | 0.23 | 0.27 |
| 7 | 95031464 | rs17881071 | 0.126 | 0.124 | 0.976 | 0.57 | -0.033 | -0.053 | -0.053 | 0 | 0.15 | 0.06 | 0.37 | 0.62 | 0.23 | 0.27 |
| 7 | 95033416 | rs17166868 | 0.248 | 0.125 | 1.051 | 0.562 | -0.003 | -0.006 | -0.006 | 0 | 0.27 | 0.29 | 0.23 | 0.19 | 0.25 | 0.39 |
| 7 | 95033841 | rs7785846 | 0.222 | 0.124 | 1.065 | 0.562 | -0.01 | -0.02 | -0.02 | 0 | 0.27 | 0.33 | 0.23 | 0.19 | 0.25 | 0.39 |
| 7 | 95034001 | rs7786401 | 0.248 | 0.124 | 1.061 | 0.562 | -0.003 | -0.006 | -0.006 | 0.569 | 0.27 | 0.29 | 0.23 | 0.19 | 0.25 | 0.39 |
| 7 | 95035950 | rs10487133 | -0.72 | -0.007 | 1.274 | 0.398 | -0.008 | 0.006 | 0.006 | 0 | 0.89 | 0.84 | 0.91 | 0.83 | 0.88 | 0.94 |

| | | | | | | | | | | | | | | | | |
|---|---|---|---|---|---|---|---|---|---|---|---|---|---|---|---|---|
| 7 | 95037661 | rs6961624 | 0.245 | -0.001 | 1.276 | 0.497 | -0.01 | -0.02 | -0.02 | 0 | 0.27 | 0.33 | 0.23 | 0.19 | 0.25 | 0.39 |
| 7 | 95039445 | rs2286232 | 0.245 | -0.001 | 1.276 | 0.497 | -0.01 | -0.02 | -0.02 | 0.045 | 0.27 | 0.33 | 0.23 | 0.19 | 0.25 | 0.39 |
| 7 | 95041271 | rs17166875 | 0.245 | -0.009 | 1.294 | 0.5 | -0.01 | -0.02 | -0.02 | 0.008 | 0.27 | 0.33 | 0.23 | 0.19 | 0.25 | 0.39 |
| 7 | 95041542 | rs2068604 | 0.232 | -0.01 | 1.3 | 0.5 | -0.012 | -0.024 | -0.024 | 0 | 0.27 | 0.35 | 0.23 | 0.19 | 0.25 | 0.39 |
| 7 | 95043152 | rs12669182 | 0.1 | -0.167 | 1.201 | 0.42 | -0.048 | -0.082 | -0.082 | 0 | 0.17 | 0.03 | 0.39 | 0.62 | 0.24 | 0.27 |
| 7 | 95043523 | rs6973380 | 0.576 | -0.17 | 1.212 | 0.417 | 0 | 0.001 | 0.001 | 0 | 0.24 | 0.24 | 0.22 | 0.17 | 0.24 | 0.37 |
| 7 | 95043850 | rs1639 | 0.107 | -0.168 | 1.206 | 0.418 | -0.048 | -0.082 | -0.082 | 1.101 | 0.17 | 0.03 | 0.39 | 0.62 | 0.24 | 0.27 |
| 7 | 95044183 | rs3779492 | 0.287 | -0.171 | 1.218 | 0.416 | -0.01 | -0.02 | -0.02 | 0 | 0.27 | 0.33 | 0.23 | 0.19 | 0.25 | 0.39 |
| 7 | 95045307 | rs17876123 | 0.248 | -0.189 | 1.209 | 0.408 | -0.003 | -0.006 | -0.006 | 0 | 0.27 | 0.29 | 0.23 | 0.19 | 0.25 | 0.39 |
| 7 | 95046708 | rs2299264 | 0.284 | -0.135 | 1.234 | 0.432 | -0.01 | -0.02 | -0.02 | 1.169 | 0.27 | 0.33 | 0.23 | 0.19 | 0.25 | 0.39 |
| 7 | 95049073 | rs2299266 | 0.298 | -0.09 | 1.188 | 0.448 | -0.005 | -0.01 | -0.01 | 0 | 0.27 | 0.3 | 0.23 | 0.19 | 0.25 | 0.39 |
| 7 | 95049818 | rs2237585 | -1.088 | -0.103 | 1.415 | 0.44 | 0 | -0.043 | -0.043 | 0 | 0.44 | 0.52 | 0.64 | 0.81 | 0.49 | 0.66 |
| 7 | 95050946 | rs7801803 | 1.176 | -0.121 | 1.37 | 0.43 | 0 | 0.084 | 0.084 | 0.44 | 0.53 | 0.71 | 0.69 | 0.83 | 0.5 | 0.71 |
| 7 | 95051052 | rs2237586 | 0.552 | -0.053 | 0.914 | 0.476 | 0 | 0.001 | 0.001 | 0 | 0.24 | 0.24 | 0.22 | 0.17 | 0.24 | 0.37 |
| 7 | 95051998 | rs2158806 | 0.297 | -0.052 | 0.924 | 0.568 | -0.003 | -0.006 | -0.006 | 0 | 0.27 | 0.29 | 0.23 | 0.19 | 0.25 | 0.39 |
| 7 | 95052694 | rs17774346 | 0.012 | -0.012 | 0.9 | 0.589 | -0.048 | -0.084 | -0.084 | 0.015 | 0.17 | 0.03 | 0.39 | 0.62 | 0.24 | 0.27 |
| 7 | 95053404 | rs17876090 | 0.312 | -0.207 | 0.838 | 0.49 | 0.004 | 0.007 | 0.007 | 0 | 0.27 | 0.24 | 0.22 | 0.17 | 0.24 | 0.4 |
| 7 | 95054480 | rs11770901 | 0.624 | 0.066 | 0.961 | 0.615 | 0 | 0.001 | 0.001 | 0 | 0.25 | 0.24 | 0.22 | 0.17 | 0.24 | 0.37 |
| 7 | 95054977 | rs17876082 | -0.264 | 0.152 | 0.861 | 0.675 | -0.012 | -0.023 | -0.023 | 0 | 0.18 | 0.11 | 0.39 | 0.62 | 0.24 | 0.28 |
| 7 | 95055598 | rs11982486 | 1.537 | 0.085 | 0.686 | -0.289 | -0.012 | 0.044 | 0.044 | 0.146 | 0.33 | 0.25 | 0.16 | 0.01 | 0.3 | 0.23 |
| 7 | 95056939 | rs10259688 | 0.539 | 0.313 | 0.448 | 0.123 | 0 | 0.014 | 0.014 | 0 | 0.14 | 0.15 | 0.22 | 0.32 | 0.2 | 0.24 |
| 7 | 95057510 | rs12534274 | 0.578 | 0.153 | 0.342 | 0.204 | 0.003 | -0.014 | -0.014 | 0.381 | 0.29 | 0.26 | 0.39 | 0.47 | 0.26 | 0.39 |
| 7 | 95057959 | rs4729189 | -0.698 | 0.166 | 0.065 | 0.062 | 0.002 | 0.006 | 0.006 | 0 | 0.79 | 0.81 | 0.79 | 0.81 | 0.79 | 0.89 |
| 7 | 95060242 | rs7785039 | -0.749 | 0.195 | 0.052 | 0.052 | 0.001 | 0.006 | 0.006 | 0 | 0.79 | 0.82 | 0.8 | 0.81 | 0.78 | 0.89 |
| 7 | 95060313 | rs7802018 | 1.279 | 0.13 | 0.239 | -0.5 | -0.002 | -0.02 | -0.02 | 0 | 0.36 | 0.29 | 0.16 | 0.01 | 0.31 | 0.24 |
| 7 | 95061328 | rs10261470 | 1.571 | 0.086 | 0.555 | -0.282 | 0.001 | -0.04 | -0.04 | 0 | 0.15 | 0.23 | 0.12 | 0.05 | 0.13 | 0.02 |
| 7 | 95061921 | rs2299267 | 0.446 | 0.159 | 0.53 | 0.187 | -0.009 | -0.032 | -0.032 | 0.12 | 0.14 | 0.09 | 0.23 | 0.38 | 0.19 | 0.21 |
| 7 | 95063270 | rs740265 | 0.592 | 0.222 | 0.978 | 0.043 | 0.002 | 0.03 | 0.03 | 0 | 0.29 | 0.33 | 0.41 | 0.42 | 0.29 | 0.43 |
| 7 | 95063702 | rs2374993 | 1.486 | 0.289 | 0.868 | 0.048 | 0.005 | -0.043 | -0.043 | 0 | 0.15 | 0.29 | 0.13 | 0.06 | 0.13 | 0.03 |
| 7 | 95066406 | rs6946850 | -1.733 | 0.243 | 0.65 | 0.062 | 0.006 | -0.039 | -0.039 | 0 | 0.86 | 0.76 | 0.88 | 0.94 | 0.87 | 0.97 |
| 7 | 95066975 | rs3757707 | 0.474 | 0.187 | 0.496 | 0.102 | 0.001 | 0.01 | 0.01 | 0 | 0.29 | 0.31 | 0.46 | 0.43 | 0.31 | 0.47 |
| 7 | 95067006 | rs43037 | 0.92 | 0.174 | 0.328 | -0.468 | 0.006 | 0.107 | 0.107 | 0 | 0.41 | 0.22 | 0.19 | 0.14 | 0.37 | 0.29 |
| 7 | 95067311 | rs43038 | -1.28 | 0.365 | 0.495 | -0.149 | 0.001 | -0.049 | -0.049 | 0 | 0.84 | 0.68 | 0.85 | 0.94 | 0.87 | 0.97 |
| 7 | 95068684 | rs6978425 | 0.148 | 0.276 | 0.277 | 0.04 | 0 | 0.038 | 0.038 | 0 | 0.33 | 0.55 | 0.49 | 0.48 | 0.34 | 0.49 |
| 7 | 95069666 | rs12535571 | 0.559 | 0.238 | 0.625 | -0.023 | -0.001 | -0.004 | -0.004 | 1.497 | 0.27 | 0.25 | 0.36 | 0.47 | 0.33 | 0.44 |
| 7 | 95070838 | rs43041 | -0.841 | -0.282 | 0.255 | -0.697 | 0.013 | 0.033 | 0.033 | 0.002 | 0.76 | 0.82 | 0.83 | 0.95 | 0.83 | 0.91 |
| 7 | 95070850 | rs43042 | 1.407 | -0.202 | -0.059 | -0.847 | -0.056 | 0.131 | 0.131 | 0.051 | 0.31 | 0.07 | 0.15 | 0.13 | 0.31 | 0.18 |
| 7 | 95076247 | rs43048 | 1.486 | -0.158 | -0.416 | -0.795 | -0.059 | 0.125 | 0.125 | 0.093 | 0.3 | 0.07 | 0.15 | 0.13 | 0.31 | 0.18 |
| 7 | 95077501 | rs43050 | -0.666 | -0.304 | -0.072 | -0.728 | -0.031 | -0.112 | -0.112 | 6.322 | 0.73 | 0.47 | 0.8 | 0.95 | 0.83 | 0.91 |
| 7 | 95078175 | rs12530498 | 0.798 | -0.094 | -0.203 | -0.593 | -0.001 | 0.014 | 0.014 | 0 | 0.14 | 0.14 | 0.19 | 0.34 | 0.18 | 0.26 |
| 7 | 95078296 | rs7778623 | 0.895 | 0.103 | -0.426 | -0.654 | -0.003 | -0.023 | -0.023 | 0.382 | 0.33 | 0.25 | 0.39 | 0.47 | 0.35 | 0.43 |
| 7 | 95078950 | rs43052 | 0.448 | -0.104 | -0.513 | -0.219 | 0.005 | -0.007 | -0.007 | 0 | 0.62 | 0.68 | 0.64 | 0.61 | 0.64 | 0.65 |
| 7 | 95080342 | rs43057 | -1.477 | -0.694 | -0.611 | -0.737 | 0.001 | 0.039 | 0.039 | 0.716 | 0.85 | 0.95 | 0.88 | 0.95 | 0.87 | 0.93 |
| 7 | 95081804 | rs10240138 | -0.535 | -0.9 | -0.305 | -1.086 | 0.001 | 0.001 | 0.001 | 0.404 | 0.39 | 0.62 | 0.55 | 0.76 | 0.34 | 0.55 |
| 7 | 95082010 | rs10240398 | -0.992 | -0.955 | -0.407 | -1.258 | -0.068 | 0.18 | 0.18 | 0.47 | 0.6 | 0.84 | 0.68 | 0.95 | 0.55 | 0.61 |
| 7 | 95084622 | rs7809951 | 0.472 | -1.081 | 0.683 | -1.012 | -0.003 | -0.049 | -0.049 | 0 | 0.1 | 0.36 | 0.04 | 0 | 0.04 | 0.05 |
| 7 | 95085553 | rs10237960 | -0.038 | -0.839 | 0.413 | -0.842 | -0.002 | 0.007 | 0.007 | 0.011 | 0.27 | 0.22 | 0.39 | 0.74 | 0.3 | 0.44 |
| 7 | 95088489 | rs10228393 | -0.342 | -0.838 | 0.462 | -0.838 | 0 | -0.019 | -0.019 | 1.637 | 0.29 | 0.37 | 0.51 | 0.76 | 0.31 | 0.5 |
| 7 | 95096615 | rs1015882 | 0.105 | -0.764 | 0.235 | -0.299 | 0.008 | 0.025 | 0.025 | 0.116 | 0.18 | 0.19 | 0.34 | 0.4 | 0.25 | 0.29 |
| 7 | 95099157 | rs1557782 | 0.435 | -0.799 | 0.259 | -0.308 | -0.034 | -0.052 | -0.052 | 1.574 | 0.17 | 0.1 | 0.27 | 0.39 | 0.24 | 0.24 |
| 7 | 95102534 | rs43061 | 0.875 | -0.867 | 0.159 | -0.935 | -0.026 | 0.056 | 0.056 | 0.195 | 0.33 | 0.19 | 0.34 | 0.07 | 0.39 | 0.37 |
| 7 | 95110894 | rs43064 | 1.48 | -1.272 | 0.296 | -0.913 | -0.02 | -0.001 | -0.001 | 0 | 0.22 | 0.08 | 0.14 | 0.19 | 0.25 | 0.08 |
| 7 | 95112564 | rs10241004 | -0.571 | -1.299 | 0.809 | -0.972 | 0.006 | 0.015 | 0.015 | 0 | 0.44 | 0.41 | 0.56 | 0.76 | 0.35 | 0.57 |
| 7 | 95114704 | rs43067 | -1.179 | -1.165 | 0.641 | -0.899 | -0.052 | 0.123 | 0.123 | 0.264 | 0.68 | 0.86 | 0.72 | 0.95 | 0.6 | 0.66 |
| 7 | 95119261 | rs1476515 | -0.167 | -1.193 | 0.698 | -0.883 | -0.009 | -0.019 | -0.019 | 0 | 0.34 | 0.44 | 0.48 | 0.75 | 0.29 | 0.46 |
| 7 | 95119655 | rs3801918 | 1.181 | -1.113 | 0.435 | -0.712 | -0.012 | 0.069 | 0.069 | 0 | 0.09 | 0.13 | 0.17 | 0.35 | 0.04 | 0.19 |
| 7 | 95121019 | rs3801916 | -1.24 | -1.167 | 0.341 | -0.774 | -0.034 | 0.081 | 0.081 | 0.157 | 0.68 | 0.82 | 0.72 | 0.95 | 0.6 | 0.66 |
| 7 | 95127234 | rs12533957 | -0.184 | -1.207 | 0.436 | -0.758 | -0.037 | 0.066 | 0.066 | 0 | 0.18 | 0.36 | 0.21 | 0.36 | 0.08 | 0.25 |
| 7 | 95131217 | rs6973126 | 0.106 | -1.203 | 0.413 | -0.778 | -0.001 | 0.007 | 0.007 | 0 | 0.49 | 0.44 | 0.51 | 0.59 | 0.52 | 0.4 |
| 7 | 95131967 | rs17717907 | -0.216 | -0.9 | 0.111 | -0.552 | -0.026 | 0.046 | 0.046 | 0 | 0.17 | 0.26 | 0.2 | 0.36 | 0.08 | 0.25 |
| 7 | 95138755 | rs737902 | -1.709 | -0.655 | -0.86 | -1.073 | -0.053 | 0.082 | 0.082 | 1.985 | 0.82 | 0.95 | 0.8 | 0.99 | 0.71 | 0.86 |
| 7 | 95142280 | rs10225152 | 0.229 | -0.831 | -0.499 | -2.069 | 0 | 0.001 | 0.001 | 0 | 0.13 | 0.14 | 0.11 | 0.04 | 0.12 | 0.18 |
| 7 | 95151939 | rs2158047 | 1.435 | -0.407 | -1.721 | -1.907 | -0.048 | 0.047 | 0.047 | 0 | 0.17 | 0.04 | 0.2 | 0.06 | 0.2 | 0.14 |
| 7 | 95153462 | rs6979120 | -0.472 | -0.277 | -1.077 | -1.902 | 0.006 | 0.016 | 0.016 | 0.373 | 0.42 | 0.61 | 0.43 | 0.28 | 0.4 | 0.44 |
| 7 | 95157935 | rs10487135 | -0.027 | 0.255 | -1.535 | -1.558 | 0.017 | -0.002 | -0.002 | 0 | 0.05 | 0.09 | 0.08 | 0.04 | 0.09 | 0.14 |
| 7 | 95162010 | rs2375019 | -0.522 | 0.262 | -2.007 | -1.805 | -0.065 | 0.121 | 0.121 | 1.651 | 0.73 | 0.9 | 0.78 | 0.98 | 0.65 | 0.8 |
| 7 | 95162037 | rs3801909 | 0.578 | 0.614 | -1.946 | -1.392 | 0.001 | 0.04 | 0.04 | 0.419 | 0.34 | 0.17 | 0.39 | 0.74 | 0.3 | 0.44 |
| 7 | 95164907 | rs3757703 | 0.714 | 0.735 | -2.263 | -1.339 | -0.023 | 0.035 | 0.035 | 0.607 | 0.33 | 0.1 | 0.39 | 0.74 | 0.31 | 0.44 |
| 7 | 95167204 | rs6465467 | 0.517 | 0.918 | -1.56 | -1.329 | -0.065 | 0.283 | 0.283 | 1.716 | 0.53 | 0.12 | 0.5 | 0.88 | 0.51 | 0.52 |

| | | | | | | | | | | | | | | | | |
|---|---|---|---|---|---|---|---|---|---|---|---|---|---|---|---|---|
| 7 | 95167543 | rs7784619 | 0.521 | 0.924 | -1.657 | -1.326 | -0.076 | 0.297 | 0.297 | 1.817 | 0.53 | 0.1 | 0.5 | 0.88 | 0.51 | 0.52 |
| 7 | 95171652 | rs16868799 | -0.698 | 0.953 | -0.401 | -0.189 | 0.014 | -0.013 | -0.013 | 0 | 0.08 | 0.14 | 0.08 | 0.04 | 0.09 | 0.14 |
| 7 | 95180458 | rs4727315 | 0.435 | 0.764 | -0.073 | -1.129 | 0.001 | 0 | 0 | 0 | 0.55 | 0.45 | 0.33 | 0.86 | 0.46 | 0.45 |
| 7 | 95183373 | rs7802553 | 0.135 | 0.672 | 0.454 | -1.323 | -0.024 | 0.078 | 0.078 | 0 | 0.6 | 0.76 | 0.39 | 0.86 | 0.49 | 0.5 |
| 7 | 95183883 | rs7806705 | -0.135 | 0.71 | 0.408 | -1.482 | -0.053 | 0.063 | 0.063 | 0 | 0.66 | 0.82 | 0.42 | 0.86 | 0.52 | 0.52 |
| 7 | 95190753 | rs4269466 | 1.055 | 0.328 | 0.744 | -1.78 | 0.056 | 0.103 | 0.103 | 0 | 0.43 | 0.25 | 0.26 | 0.82 | 0.29 | 0.42 |
| 7 | 95198625 | rs11772378 | 1.047 | 0.304 | 1.262 | -1.791 | 0.056 | 0.104 | 0.104 | 0 | 0.43 | 0.25 | 0.25 | 0.83 | 0.3 | 0.42 |
| 7 | 95213059 | rs6930 | -0.34 | 0.447 | 1.559 | -1.777 | -0.029 | 0.058 | 0.058 | 0 | 0.64 | 0.76 | 0.39 | 0.87 | 0.49 | 0.5 |
| 7 | 95213702 | rs11531570 | 0.369 | 0.436 | 1.761 | -1.778 | -0.002 | 0.044 | 0.044 | 0 | 0.58 | 0.39 | 0.33 | 0.86 | 0.45 | 0.46 |
| 7 | 95213953 | rs11768781 | 0.37 | 0.441 | 1.774 | -1.78 | -0.003 | 0.041 | 0.041 | 0 | 0.58 | 0.4 | 0.33 | 0.86 | 0.46 | 0.46 |
| 7 | 95216247 | rs10247649 | 0.598 | 0.412 | 1.668 | -1.818 | 0.004 | 0.006 | 0.006 | 0 | 0.55 | 0.6 | 0.31 | 0.86 | 0.4 | 0.44 |
| 7 | 95216262 | rs4729201 | -0.475 | 0.456 | 1.456 | -1.896 | -0.03 | 0.031 | 0.031 | 0 | 0.67 | 0.76 | 0.39 | 0.94 | 0.49 | 0.67 |
| 7 | 95218438 | rs12668651 | 1.125 | 0.412 | 1.528 | -1.877 | 0.074 | 0.115 | 0.115 | 0 | 0.47 | 0.3 | 0.26 | 0.83 | 0.28 | 0.43 |
| 7 | 95220011 | rs17704516 | 0.475 | 0.31 | 1.206 | -1.741 | -0.002 | 0 | 0 | 0 | 0.07 | 0.07 | 0.04 | 0.04 | 0.11 | 0.01 |
| 7 | 95222819 | rs3779478 | -0.802 | 0.295 | 1.192 | -1.257 | 0.004 | -0.01 | -0.01 | 0 | 0.09 | 0.05 | 0.04 | 0.14 | 0.07 | 0.24 |
| 7 | 95225813 | rs10085637 | 0.596 | 0.185 | 1.541 | -1.509 | 0.018 | 0.029 | 0.029 | 0 | 0.52 | 0.54 | 0.29 | 0.81 | 0.35 | 0.46 |
| 7 | 95226733 | rs3807892 | 0.645 | 0.166 | 1.535 | -1.518 | 0 | -0.009 | -0.009 | 0 | 0.52 | 0.43 | 0.29 | 0.81 | 0.37 | 0.46 |
| 7 | 95229077 | rs10250330 | -0.705 | 0.136 | 1.721 | -1.465 | 0.009 | 0.133 | 0.133 | 0 | 0.08 | 0.28 | 0.05 | 0.13 | 0.05 | 0.22 |
| 7 | 95231456 | rs10259067 | -0.614 | 0.086 | 1.785 | -1.464 | -0.016 | 0.077 | 0.077 | 0 | 0.08 | 0.2 | 0.04 | 0.13 | 0.05 | 0.22 |
| 7 | 95236687 | rs2888979 | 0.692 | -0.213 | 1.405 | -1.776 | 0.01 | -0.015 | -0.015 | 0 | 0.47 | 0.38 | 0.27 | 0.9 | 0.26 | 0.63 |
| 7 | 95238104 | rs4379388 | -0.267 | -0.17 | 1.117 | -1.734 | -0.005 | 0.014 | 0.014 | 0 | 0.13 | 0.09 | 0.06 | 0.04 | 0.17 | 0.08 |
| 7 | 95238643 | rs17704630 | -0.8 | -0.039 | 1.203 | -1.76 | 0.004 | -0.028 | -0.028 | 0 | 0.16 | 0.21 | 0.08 | 0.04 | 0.19 | 0.09 |
| 7 | 95241819 | rs10238561 | 1.132 | -0.423 | 0.895 | -1.711 | 0.049 | 0.032 | 0.032 | 0 | 0.38 | 0.22 | 0.23 | 0.77 | 0.21 | 0.41 |
| 7 | 95248369 | rs12704811 | 0.966 | -0.443 | 0.373 | -1.638 | 0.049 | 0.023 | 0.023 | 0.539 | 0.39 | 0.03 | 0.22 | 0.71 | 0.22 | 0.4 |
| 7 | 95250693 | rs10264649 | -0.982 | -0.472 | 1.072 | -1.871 | -0.009 | -0.064 | -0.064 | 0 | 0.14 | 0.3 | 0.05 | 0.05 | 0.11 | 0.08 |
| 7 | 95250806 | rs970626 | 0.078 | 0.383 | 1.272 | -2.041 | 0.009 | -0.019 | -0.019 | 0 | 0.1 | 0.19 | 0.23 | 0.06 | 0.12 | 0.14 |
| 7 | 95253449 | rs12669414 | 0.361 | -0.108 | 1.339 | -2.238 | 0 | -0.05 | -0.05 | 0 | 0.51 | 0.5 | 0.49 | 0.89 | 0.37 | 0.74 |
| 7 | 95253612 | rs12704814 | 1.284 | 0.029 | 1.266 | -2.189 | 0.073 | 0.145 | 0.145 | 1.805 | 0.41 | 0.15 | 0.25 | 0.89 | 0.25 | 0.62 |
| 7 | 95260685 | rs1079916 | 0.276 | 0.212 | 1.379 | -1.867 | 0.011 | 0.037 | 0.037 | 0.752 | 0.54 | 0.38 | 0.51 | 0.92 | 0.38 | 0.76 |
| 7 | 95261859 | rs1485002 | -1.221 | 0.217 | 1.502 | -1.758 | 0.012 | -0.057 | -0.057 | 0 | 0.23 | 0.41 | 0.28 | 0.15 | 0.23 | 0.22 |
| 7 | 95262116 | rs1485003 | -0.32 | 0.261 | 1.455 | -1.738 | 0.007 | -0.081 | -0.081 | 0 | 0.65 | 0.61 | 0.55 | 0.99 | 0.49 | 0.82 |
| 7 | 95265377 | rs10275635 | -1.347 | -0.253 | 1.732 | -1.589 | -0.009 | -0.057 | -0.057 | 0.133 | 0.11 | 0.31 | 0.05 | 0 | 0.05 | 0.06 |
| 7 | 95270506 | rs1485004 | 0.705 | -0.74 | 1.259 | -1.388 | 0.045 | 0.061 | 0.061 | 0.009 | 0.44 | 0.25 | 0.26 | 0.61 | 0.29 | 0.54 |
| 7 | 95281099 | rs1633996 | 0.51 | -0.548 | 1.432 | -1.393 | 0.018 | 0.011 | 0.011 | 1.327 | 0.8 | 0.59 | 0.7 | 0.69 | 0.73 | 0.79 |
| 7 | 95282480 | rs1682624 | 0.744 | -0.615 | 1.284 | -1.464 | -0.054 | 0.076 | 0.076 | 0 | 0.29 | 0.17 | 0.4 | 0.08 | 0.42 | 0.19 |
| 7 | 95287743 | rs1682731 | 1.757 | -0.595 | 1.429 | -0.868 | 0.001 | 0.004 | 0.004 | 1.048 | 0.67 | 0.34 | 0.56 | 0.52 | 0.65 | 0.52 |
| 7 | 95291549 | rs854093 | 1.678 | -0.58 | 1.365 | -0.853 | 0.006 | 0.017 | 0.017 | 0.836 | 0.67 | 0.38 | 0.55 | 0.52 | 0.64 | 0.53 |
| 7 | 95294658 | rs12672152 | 0.974 | -0.587 | 1.482 | -0.831 | 0.012 | 0.046 | 0.046 | 0 | 0.07 | 0.13 | 0.11 | 0.16 | 0.06 | 0.23 |
| 7 | 95301853 | rs854083 | 1.197 | -0.68 | 1.219 | -0.774 | 0.005 | -0.01 | -0.01 | 0 | 0.74 | 0.5 | 0.66 | 0.68 | 0.71 | 0.76 |
| 7 | 95315537 | rs6979207 | -0.561 | -0.555 | 0.997 | -0.806 | 0.012 | -0.048 | -0.048 | 0 | 0.15 | 0.27 | 0.13 | 0.06 | 0.17 | 0.09 |
| 7 | 95318217 | rs1227518 | 0.451 | -0.404 | 0.594 | -0.696 | 0.007 | -0.019 | -0.019 | 0 | 0.09 | 0.11 | 0.09 | 0.05 | 0.13 | 0.05 |
| 7 | 95325141 | rs10487140 | 0.384 | -1.171 | -0.262 | -0.664 | 0.014 | -0.075 | -0.075 | 0 | 0.22 | 0.1 | 0.12 | 0.41 | 0.19 | 0.12 |
| 7 | 95325734 | rs854104 | -0.759 | -1.135 | -0.288 | -0.729 | -0.001 | -0.03 | -0.03 | 0.046 | 0.91 | 0.92 | 0.79 | 0.74 | 0.88 | 0.84 |
| 7 | 95340949 | rs2127185 | 0.557 | -1.113 | -0.092 | -1.032 | -0.039 | -0.115 | -0.115 | 1.944 | 0.3 | 0.47 | 0.17 | 0.07 | 0.2 | 0.25 |
| 7 | 95347221 | rs2618985 | -0.445 | 0.077 | -0.234 | -0.558 | -0.011 | -0.001 | -0.001 | 0 | 0.74 | 0.78 | 0.65 | 0.92 | 0.61 | 0.81 |
| 7 | 95348225 | rs6966287 | 0.072 | 0.1 | -0.626 | -0.083 | 0.009 | -0.032 | -0.032 | 0.704 | 0.23 | 0.13 | 0.15 | 0.38 | 0.2 | 0.12 |
| 7 | 95360479 | rs7797290 | 0.143 | 0.066 | -0.596 | -0.087 | 0.009 | -0.028 | -0.028 | 0.498 | 0.23 | 0.13 | 0.15 | 0.38 | 0.2 | 0.12 |
| 7 | 95368070 | rs2706859 | -0.425 | 0.48 | -0.84 | -0.408 | -0.062 | 0.077 | 0.077 | 0 | 0.74 | 0.88 | 0.64 | 0.9 | 0.6 | 0.84 |
| 7 | 95373302 | rs11973465 | -0.202 | 0.38 | -0.547 | -0.524 | 0.015 | -0.082 | -0.082 | 0 | 0.16 | 0.31 | 0.15 | 0.02 | 0.18 | 0.08 |
| 7 | 95373608 | rs2618995 | -0.202 | 0.38 | -0.547 | -0.518 | 0.015 | -0.074 | -0.074 | 0.02 | 0.16 | 0.31 | 0.15 | 0.02 | 0.18 | 0.08 |
| 7 | 95373688 | rs10265090 | -0.202 | 0.38 | -0.547 | -0.523 | 0.015 | -0.082 | -0.082 | 0 | 0.16 | 0.31 | 0.15 | 0.02 | 0.18 | 0.08 |
| 7 | 95377966 | rs7809206 | -0.831 | 0.459 | -0.63 | -0.555 | -0.04 | 0.039 | 0.039 | 0 | 0.78 | 0.86 | 0.64 | 0.9 | 0.63 | 0.87 |
| 7 | 95378808 | rs11980835 | 1.047 | 0.448 | -0.613 | -0.529 | 0.035 | -0.049 | -0.049 | 0.017 | 0.1 | 0.19 | 0.12 | 0.02 | 0.16 | 0.05 |
| 7 | 95384357 | rs10245223 | -0.809 | 0.609 | -1.548 | -0.457 | -0.067 | 0.065 | 0.065 | 0 | 0.78 | 0.9 | 0.65 | 0.9 | 0.64 | 0.87 |
| 7 | 95385180 | rs1382618 | 1.429 | 0.194 | -1.812 | -0.176 | -0.012 | 0.036 | 0.036 | 0 | 0.35 | 0.06 | 0.21 | 0.23 | 0.28 | 0.52 |
| 7 | 95395681 | rs3912439 | 0.425 | 0.271 | -1.094 | 0.167 | -0.006 | 0.064 | 0.064 | 0.243 | 0.1 | 0.18 | 0.22 | 0.34 | 0.1 | 0.17 |
| 7 | 95399229 | rs4729205 | 0.33 | 1.047 | -1.194 | -0.033 | -0.024 | 0.025 | 0.025 | 0 | 0.79 | 0.84 | 0.7 | 0.92 | 0.69 | 0.81 |
| 7 | 95401417 | rs7801111 | -0.618 | 1.242 | -1.436 | 0.13 | -0.037 | 0.026 | 0.026 | 0 | 0.86 | 0.94 | 0.74 | 0.92 | 0.75 | 0.85 |
| 7 | 95413253 | rs7798936 | 0.211 | 0.616 | -1.457 | 0.308 | -0.031 | 0.023 | 0.023 | 0.002 | 0.64 | 0.22 | 0.4 | 0.63 | 0.53 | 0.62 |
| 7 | 95419256 | rs1478984 | 0.444 | 0.352 | -1.473 | 0.279 | 0.016 | 0.059 | 0.059 | 0 | 0.43 | 0.18 | 0.35 | 0.26 | 0.37 | 0.54 |
| 7 | 95421030 | rs10265425 | 0.195 | 0.547 | -1.263 | 0.896 | -0.006 | -0.027 | -0.027 | 0 | 0.21 | 0.14 | 0.13 | 0.3 | 0.23 | 0.16 |
| 7 | 95421568 | rs1485001 | -0.093 | 0.454 | -0.688 | 1.039 | -0.005 | 0.04 | 0.04 | 3.663 | 0.16 | 0.23 | 0.37 | 0.32 | 0.13 | 0.27 |
| 7 | 95424457 | rs2706888 | -1.313 | 0.342 | -0.8 | 0.28 | -0.002 | 0.023 | 0.023 | 1.727 | 0.65 | 0.84 | 0.69 | 0.75 | 0.64 | 0.72 |
| 7 | 95431640 | rs2618971 | 0.709 | 0.567 | 0.226 | 0.39 | 0.02 | 0.027 | 0.027 | 0.039 | 0.07 | 0.2 | 0.08 | 0.1 | 0.08 | 0.14 |
| 7 | 95436435 | rs11767450 | 0.499 | 0.567 | 0.142 | 0.274 | 0.009 | -0.032 | -0.032 | 0 | 0.09 | 0.18 | 0.08 | 0.02 | 0.12 | 0.05 |
| 7 | 95441451 | rs17167184 | 1.087 | 0.68 | 0.143 | 0.367 | -0.005 | -0.006 | -0.006 | 0 | 0.06 | 0.07 | 0.08 | 0.1 | 0.08 | 0.14 |
| 7 | 95443892 | rs2690295 | -1.159 | 0.624 | 0.171 | 0.408 | 0.013 | 0.105 | 0.105 | 7.151 | 0.52 | 0.8 | 0.67 | 0.75 | 0.56 | 0.62 |
| 7 | 95446706 | rs2706880 | -0.724 | 0.726 | 0.269 | 0.415 | -0.013 | 0.023 | 0.023 | 6.647 | 0.45 | 0.71 | 0.58 | 0.64 | 0.49 | 0.47 |
| 7 | 95467621 | rs1585737 | -0.308 | 0.554 | 0.464 | 0.748 | 0 | 0.021 | 0.021 | 9.467 | 0.37 | 0.48 | 0.38 | 0.53 | 0.36 | 0.36 |

| | | | | | | | | | | | | | | | | |
|---|---|---|---|---|---|---|---|---|---|---|---|---|---|---|---|---|
| 7 | 95473550 | rs4729215 | 0.121 | -0.138 | 0.947 | 1.675 | 0.012 | 0.152 | 0.152 | 0 | 0.13 | 0.23 | 0.37 | 0.48 | 0.14 | 0.26 |
| 7 | 95484290 | rs2633939 | 0.867 | -0.446 | 0.84 | 1.607 | -0.048 | 0.089 | 0.089 | 12.326 | 0.75 | 0.48 | 0.61 | 0.38 | 0.85 | 0.63 |
| 7 | 95494080 | rs6957233 | -0.131 | -0.274 | 0.56 | 1.677 | 0.007 | 0.143 | 0.143 | 0 | 0.14 | 0.25 | 0.37 | 0.49 | 0.14 | 0.28 |
| 7 | 95500522 | rs1488503 | -0.651 | -0.442 | 0.859 | 1.749 | -0.026 | 0.071 | 0.071 | 9.766 | 0.21 | 0.36 | 0.12 | 0.48 | 0.11 | 0.35 |
| 7 | 95501480 | rs12535988 | 0.068 | -0.298 | 0.73 | 1.539 | 0.009 | 0.058 | 0.058 | 0 | 0.13 | 0.25 | 0.37 | 0.24 | 0.12 | 0.22 |
| 7 | 95505561 | rs995596 | 0.428 | -0.268 | 0.755 | 1.395 | 0.012 | 0.007 | 0.007 | 0 | 0.11 | 0.2 | 0.24 | 0.13 | 0.12 | 0.14 |
| 7 | 95507302 | rs10953165 | -0.222 | -0.266 | 0.629 | 1.096 | -0.007 | 0 | 0 | 0 | 0.16 | 0.12 | 0.3 | 0.02 | 0.22 | 0.16 |
| 7 | 95508212 | rs6465484 | -0.766 | -0.517 | 0.868 | 1.249 | -0.034 | 0.001 | 0.001 | 22.581 | 0.89 | 0.83 | 0.98 | 0.87 | 0.99 | 0.9 |
| 7 | 95509763 | rs17167209 | 0.486 | -0.537 | 1.207 | 1.482 | -0.002 | 0.027 | 0.027 | 0.123 | 0.11 | 0.12 | 0.09 | 0.18 | 0.08 | 0.25 |
| 7 | 95509806 | rs6967902 | -2.068 | -0.522 | 1.333 | 1.449 | -0.011 | 0.002 | 0.002 | 3.99 | 0.39 | 0.52 | 0.42 | 0.39 | 0.31 | 0.5 |
| 7 | 95512626 | rs10249566 | 0.813 | -0.37 | 1.089 | 1.232 | 0.007 | -0.023 | -0.023 | 0.196 | 0.08 | 0.11 | 0.22 | 0.02 | 0.09 | 0.11 |
| 7 | 95516569 | rs6465485 | 0.384 | -0.216 | 1.058 | 1.557 | -0.041 | 0.107 | 0.107 | 21.062 | 0.11 | 0.25 | 0.04 | 0.34 | 0.03 | 0.17 |
| 7 | 95529156 | rs984862 | 0.962 | -0.485 | 0.453 | 0.849 | -0.006 | 0.002 | 0.002 | 0.638 | 0.61 | 0.45 | 0.53 | 0.55 | 0.68 | 0.64 |
| 7 | 95529329 | rs10246503 | 0.172 | -0.546 | 0.332 | 0.684 | 0.007 | 0.051 | 0.051 | 0.969 | 0.2 | 0.09 | 0.13 | 0.02 | 0.18 | 0.12 |
| 7 | 95532370 | rs319299 | 1.776 | -0.574 | 0.3 | 0.77 | -0.01 | 0.085 | 0.085 | 0.018 | 0.47 | 0.25 | 0.38 | 0.27 | 0.56 | 0.38 |
| 7 | 95536269 | rs2188505 | 0.756 | -0.428 | 0.402 | 1.037 | 0.01 | 0.071 | 0.071 | 0 | 0.08 | 0.11 | 0.11 | 0.2 | 0.09 | 0.2 |
| 7 | 95543624 | rs319333 | 1.242 | -0.569 | 0.614 | 1.052 | -0.016 | 0.16 | 0.16 | 6.234 | 0.07 | 0.18 | 0.02 | 0.33 | 0.02 | 0.17 |
| 7 | 95544100 | rs319332 | -0.671 | -0.586 | 0.667 | 1.056 | -0.007 | 0.102 | 0.102 | 11.785 | 0.19 | 0.29 | 0.13 | 0.52 | 0.15 | 0.39 |
| 7 | 95544363 | rs319331 | 0.376 | -0.493 | 0.482 | 1.005 | -0.004 | 0.143 | 0.143 | 0.572 | 0.14 | 0.37 | 0.04 | 0.33 | 0.08 | 0.18 |
| 7 | 95546753 | rs6952893 | 0.791 | -0.536 | 0.044 | 0.839 | -0.001 | 0.059 | 0.059 | 0.659 | 0.16 | 0.04 | 0.08 | 0 | 0.15 | 0.07 |
| 7 | 95550100 | rs319327 | 0.178 | -0.198 | -0.463 | 0.371 | 0.008 | 0.043 | 0.043 | 0 | 0.24 | 0.45 | 0.12 | 0.34 | 0.23 | 0.22 |
| 7 | 95550340 | rs13227290 | 0.083 | -0.26 | -1.245 | 0.263 | 0.006 | -0.011 | -0.011 | 0.385 | 0.19 | 0.13 | 0.17 | 0.21 | 0.15 | 0.37 |
| 7 | 95553445 | rs319324 | -1.782 | -0.257 | -1.234 | 0.298 | -0.031 | 0.035 | 0.035 | 0 | 0.64 | 0.83 | 0.7 | 0.75 | 0.57 | 0.7 |
| 7 | 95554916 | rs10280608 | -0.162 | -0.027 | -1.179 | 0.332 | 0.004 | -0.011 | -0.011 | 3.648 | 0.19 | 0.13 | 0.17 | 0.21 | 0.15 | 0.33 |
| 7 | 95558358 | rs6950190 | 0.192 | 0.309 | -1.359 | -0.037 | -0.009 | 0.025 | 0.025 | 0.041 | 0.2 | 0.08 | 0.45 | 0.13 | 0.22 | 0.16 |
| 7 | 95559094 | rs2860011 | -0.406 | 0.157 | -1.294 | 0.022 | 0.003 | 0.032 | 0.032 | 0 | 0.32 | 0.21 | 0.2 | 0.22 | 0.31 | 0.27 |
| 7 | 95559291 | rs319312 | -0.337 | 0.75 | -1.269 | 0.058 | -0.017 | 0.042 | 0.042 | 0 | 0.76 | 0.88 | 0.86 | 0.86 | 0.77 | 0.8 |
| 7 | 95563391 | rs1488516 | 0.074 | 1.171 | -1.005 | 0.006 | -0.02 | 0.027 | 0.027 | 0 | 0.81 | 0.85 | 0.86 | 0.92 | 0.76 | 0.83 |
| 7 | 95565500 | rs319296 | 0.825 | 0.815 | -0.514 | -0.094 | -0.008 | -0.067 | -0.067 | 0.021 | 0.67 | 0.48 | 0.81 | 0.89 | 0.73 | 0.66 |
| 7 | 95577825 | rs319345 | 0.18 | 1.059 | -1.003 | 0.297 | -0.002 | 0.001 | 0.001 | 0.622 | 0.19 | 0.16 | 0.35 | 0.64 | 0.24 | 0.32 |
| 7 | 95594026 | rs12534478 | 1.063 | 1.158 | -1.212 | -0.57 | -0.022 | 0.088 | 0.088 | 2.843 | 0.47 | 0.2 | 0.4 | 0.22 | 0.48 | 0.31 |
| 7 | 95600368 | rs13231718 | 0.921 | 0.666 | -1.049 | -0.58 | -0.008 | 0.052 | 0.052 | 0 | 0.09 | 0 | 0.04 | 0 | 0.06 | 0.03 |
| 7 | 95609297 | rs17705297 | -0.045 | 0.842 | -0.884 | -0.701 | -0.001 | 0.023 | 0.023 | 0.132 | 0.16 | 0.06 | 0.09 | 0.03 | 0.17 | 0.13 |
| 7 | 95609957 | rs319322 | -0.435 | 0.044 | -0.171 | -0.682 | -0.031 | 0.056 | 0.056 | 0.809 | 0.22 | 0.45 | 0.32 | 0.64 | 0.17 | 0.37 |
| 7 | 95610634 | rs11770791 | 0.633 | 0.054 | -0.388 | -0.79 | 0.001 | 0.041 | 0.041 | 0 | 0.11 | 0.03 | 0.11 | 0 | 0.1 | 0.08 |
| 7 | 95614521 | rs17167270 | 0.197 | 0.063 | -0.075 | -0.858 | -0.006 | -0.048 | -0.048 | 0 | 0.17 | 0.17 | 0.11 | 0.02 | 0.18 | 0.13 |
| 7 | 95616309 | rs17705339 | 0.291 | 0.928 | -0.063 | -0.898 | 0.008 | -0.001 | -0.001 | 0 | 0.08 | 0.06 | 0.09 | 0.07 | 0.16 | 0.1 |
| 7 | 95619073 | rs2299268 | -1.013 | 0.45 | 0.444 | -0.846 | -0.015 | -0.014 | -0.014 | 0 | 0.39 | 0.52 | 0.47 | 0.67 | 0.38 | 0.53 |
| 7 | 95619377 | rs2299269 | -0.182 | 1.088 | 0.452 | -0.143 | -0.003 | 0.058 | 0.058 | 0 | 0.09 | 0.16 | 0.24 | 0.64 | 0.07 | 0.3 |
| 7 | 95624677 | rs2375048 | 0.449 | 0.94 | 0.508 | -0.489 | 0 | 0.001 | 0.001 | 1.597 | 0.56 | 0.47 | 0.52 | 0.63 | 0.5 | 0.47 |
| 7 | 95626133 | rs4729230 | -0.176 | 1.206 | 0.254 | -0.506 | -0.012 | 0.008 | 0.008 | 1.475 | 0.66 | 0.68 | 0.64 | 0.7 | 0.65 | 0.57 |
| 7 | 95627771 | rs1488514 | -0.627 | 1.592 | -0.152 | -0.416 | -0.044 | 0.012 | 0.012 | 1.34 | 0.72 | 0.83 | 0.67 | 0.76 | 0.68 | 0.66 |
| 7 | 95629078 | rs756860 | -0.654 | 1.269 | 0.208 | -0.161 | 0.019 | -0.005 | -0.005 | 0 | 0.08 | 0.13 | 0.12 | 0.06 | 0.14 | 0.1 |
| 7 | 95635148 | rs739513 | 0.309 | 0.933 | 0.139 | -0.346 | 0.016 | -0.033 | -0.033 | 0 | 0.09 | 0.1 | 0.09 | 0 | 0.13 | 0.01 |
| 7 | 95637097 | rs720780 | -0.564 | -0.17 | 0.199 | -0.447 | -0.021 | -0.017 | -0.017 | 0 | 0.4 | 0.47 | 0.23 | 0.28 | 0.3 | 0.3 |
| 7 | 95658291 | rs7782220 | 0.079 | -0.693 | 0.009 | -0.578 | 0.058 | -0.005 | -0.005 | 0 | 0.27 | 0.15 | 0.12 | 0.26 | 0.16 | 0.27 |
| 7 | 95659880 | rs2214097 | -0.144 | -0.525 | 0.591 | -0.552 | -0.015 | -0.003 | -0.003 | 0 | 0.31 | 0.36 | 0.16 | 0.27 | 0.2 | 0.35 |
| 7 | 95666462 | rs2299278 | -1.403 | -0.035 | 0.397 | 0.208 | -0.046 | 0.001 | 0.001 | 1.947 | 0.61 | 0.78 | 0.56 | 0.57 | 0.57 | 0.59 |
| 7 | 95666875 | rs2251779 | 0.088 | -0.195 | 0.873 | 0.23 | 0.019 | 0.073 | 0.073 | 0.054 | 0.19 | 0.27 | 0.21 | 0.31 | 0.24 | 0.08 |
| 7 | 95682275 | rs2299282 | 0.503 | -0.291 | 0.518 | 0.318 | 0.054 | 0.006 | 0.006 | 0 | 0.28 | 0.16 | 0.14 | 0.25 | 0.17 | 0.24 |
| 7 | 95685122 | rs2106513 | 0.092 | -0.078 | 1.386 | 0.495 | 0.038 | 0.116 | 0.116 | 0 | 0.18 | 0.35 | 0.21 | 0.31 | 0.24 | 0.07 |
| 7 | 95687885 | rs11983484 | -0.583 | -0.11 | 1.465 | 0.461 | 0.036 | 0.106 | 0.106 | 0.184 | 0.22 | 0.44 | 0.24 | 0.32 | 0.27 | 0.14 |
| 7 | 95697118 | rs2299284 | 0.088 | -0.051 | 1.187 | 0.505 | 0.03 | 0.098 | 0.098 | 0 | 0.18 | 0.31 | 0.21 | 0.31 | 0.24 | 0.07 |
| 7 | 95711278 | rs10228472 | -0.211 | -0.688 | 1.052 | 0.547 | -0.006 | -0.001 | -0.001 | 0 | 0.4 | 0.6 | 0.37 | 0.44 | 0.32 | 0.39 |
| 7 | 95712878 | rs2299288 | -0.09 | -0.771 | 0.817 | 0.578 | 0.001 | 0.016 | 0.016 | 0.016 | 0.07 | 0.05 | 0.06 | 0.23 | 0.04 | 0.14 |
| 7 | 95716383 | rs2299293 | 0.64 | -0.928 | 1.086 | 0.425 | 0.003 | -0.003 | -0.003 | 0 | 0.26 | 0.22 | 0.11 | 0.05 | 0.21 | 0.15 |
| 7 | 95716625 | rs42067 | 0.322 | -0.65 | 0.936 | 0.341 | 0.007 | 0.045 | 0.045 | 0 | 0.88 | 0.81 | 0.79 | 0.76 | 0.88 | 0.66 |
| 7 | 95718579 | rs42065 | 1.025 | -0.699 | 0.726 | 0.408 | -0.024 | 0.038 | 0.038 | 0 | 0.44 | 0.21 | 0.4 | 0.31 | 0.52 | 0.27 |
| 7 | 95725936 | rs916758 | -0.365 | -1.158 | 0.923 | 0.711 | -0.011 | 0.046 | 0.046 | 0 | 0.17 | 0.3 | 0.27 | 0.4 | 0.14 | 0.24 |
| 7 | 95727967 | rs6966540 | -0.886 | -1.682 | 0.577 | 0.235 | 0.001 | 0.002 | 0.002 | 0 | 0.59 | 0.62 | 0.47 | 0.89 | 0.38 | 0.65 |
| 7 | 95731159 | rs12670075 | -0.048 | -0.967 | 0.732 | 0.494 | 0.014 | 0.007 | 0.007 | 0.068 | 0.28 | 0.17 | 0.16 | 0.24 | 0.22 | 0.33 |
| 7 | 95740335 | rs10267710 | -1.221 | -1.018 | 0.769 | 0.405 | 0.048 | 0.007 | 0.007 | 0.075 | 0.48 | 0.72 | 0.41 | 0.63 | 0.37 | 0.62 |
| 7 | 95742412 | rs7808223 | 0.172 | -0.913 | 0.714 | 0.408 | -0.007 | -0.06 | -0.06 | 0.044 | 0.23 | 0.3 | 0.12 | 0.03 | 0.2 | 0.17 |
| 7 | 95743918 | rs12666041 | 0.297 | -0.86 | 0.39 | 0.444 | 0.001 | -0.011 | -0.011 | 0 | 0.23 | 0.19 | 0.11 | 0.03 | 0.2 | 0.17 |
| 7 | 95748323 | rs4729233 | -0.855 | -0.484 | 0.615 | 0.511 | 0.008 | 0.005 | 0.005 | 0.317 | 0.5 | 0.42 | 0.45 | 0.62 | 0.42 | 0.64 |
| 7 | 95755635 | rs10278888 | -0.924 | -0.428 | 0.576 | 0.508 | 0.023 | 0.006 | 0.006 | 0 | 0.46 | 0.34 | 0.43 | 0.62 | 0.4 | 0.57 |
| 7 | 95757895 | rs6465486 | -0.494 | -0.555 | 0.487 | 0.655 | 0.016 | -0.004 | -0.004 | 1.5 | 0.34 | 0.26 | 0.18 | 0.36 | 0.26 | 0.32 |
| 7 | 95758692 | rs9656035 | -1.326 | -0.588 | 1.085 | 0.587 | 0.019 | 0.008 | 0.008 | 0.066 | 0.5 | 0.64 | 0.41 | 0.62 | 0.39 | 0.57 |
| 7 | 95778402 | rs3779486 | -1.184 | -0.605 | 1.252 | 0.615 | 0.051 | 0.034 | 0.034 | 0.099 | 0.5 | 0.76 | 0.41 | 0.62 | 0.39 | 0.56 |

| | | | | | | | | | | | | | | | | |
|---|---|---|---|---|---|---|---|---|---|---|---|---|---|---|---|---|
| 7 | 95781748 | rs1007452 | -1.248 | -0.597 | 1.327 | 0.619 | -0.002 | -0.002 | -0.002 | 0 | 0.49 | 0.53 | 0.4 | 0.62 | 0.39 | 0.56 |
| 7 | 95787683 | rs6971996 | -0.976 | -0.542 | 1.001 | 0.632 | 0.028 | 0 | 0 | 0 | 0.46 | 0.29 | 0.39 | 0.61 | 0.38 | 0.52 |
| 7 | 95794801 | rs13241748 | -0.191 | -0.918 | 0.901 | 0.855 | -0.011 | -0.029 | -0.029 | 0 | 0.15 | 0.05 | 0.23 | 0.26 | 0.18 | 0.27 |
| 7 | 95800820 | rs2301629 | -1.214 | -1.212 | 0.966 | 0.708 | 0.041 | -0.006 | -0.006 | 0 | 0.44 | 0.25 | 0.34 | 0.61 | 0.35 | 0.54 |
| 7 | 95806578 | rs6959358 | -1.515 | -1.166 | 1.146 | 0.613 | 0.045 | 0.038 | 0.038 | 1.005 | 0.51 | 0.74 | 0.48 | 0.64 | 0.39 | 0.64 |
| 7 | 95817231 | rs1029391 | 1.254 | -1.274 | 1.474 | 0.679 | -0.002 | -0.001 | -0.001 | 0 | 0.53 | 0.49 | 0.64 | 0.37 | 0.65 | 0.43 |
| 7 | 95835862 | rs12667395 | 1.249 | -1.264 | 1.479 | 0.651 | -0.002 | -0.001 | -0.001 | 0 | 0.53 | 0.48 | 0.64 | 0.38 | 0.65 | 0.43 |
| 7 | 95853837 | rs6950250 | -0.236 | -1.267 | 1.063 | 0.408 | 0.027 | -0.022 | -0.022 | 0 | 0.28 | 0.2 | 0.11 | 0.35 | 0.17 | 0.27 |
| 7 | 95864108 | rs6957975 | 1.305 | -1.267 | 0.983 | 0.518 | 0.05 | -0.031 | -0.031 | 0 | 0.55 | 0.76 | 0.68 | 0.45 | 0.66 | 0.46 |
| 7 | 95900270 | rs35974282 | 1.365 | -1.669 | 1.265 | 0.624 | -0.001 | -0.001 | -0.001 | 0 | 0.53 | 0.46 | 0.67 | 0.44 | 0.69 | 0.44 |
| 7 | 95902397 | rs7779963 | 1.554 | -1.71 | 0.72 | 0.581 | 0.097 | 0.002 | 0.002 | 0.588 | 0.51 | 0.25 | 0.65 | 0.43 | 0.69 | 0.43 |
| 7 | 95931247 | rs10234418 | -0.657 | -1.781 | -0.233 | 0.618 | 0.021 | 0.023 | 0.023 | 0 | 0.8 | 0.89 | 0.94 | 0.96 | 0.9 | 0.87 |
| 7 | 95937098 | rs13245850 | -0.258 | -1.584 | -0.296 | 0.618 | -0.022 | -0.009 | -0.009 | 0 | 0.16 | 0.03 | 0.22 | 0.22 | 0.18 | 0.26 |
| 7 | 95947916 | rs11982522 | 0.721 | -1.23 | -0.084 | 0.599 | 0.032 | -0.008 | -0.008 | 0 | 0.55 | 0.63 | 0.68 | 0.45 | 0.7 | 0.47 |
| 7 | 95986043 | rs764453 | 0.495 | -0.235 | -0.282 | 0.24 | -0.042 | -0.046 | -0.046 | 1.258 | 0.89 | 0.99 | 0.82 | 0.79 | 0.85 | 0.74 |
| 7 | 96000818 | rs7789537 | 0.943 | -0.523 | 0.309 | -0.131 | -0.013 | -0.041 | -0.041 | 0 | 0.75 | 0.55 | 0.74 | 0.77 | 0.79 | 0.68 |
| 7 | 96013072 | rs2724028 | 0.741 | -1.184 | 0.223 | -0.498 | -0.043 | -0.042 | -0.042 | 0 | 0.66 | 0.5 | 0.72 | 0.73 | 0.8 | 0.52 |
| 7 | 96017011 | rs12536064 | 1.539 | -1.578 | 0.11 | 0.28 | 0.116 | 0.048 | 0.048 | 0.685 | 0.5 | 0.21 | 0.62 | 0.38 | 0.68 | 0.42 |
| 7 | 96020434 | rs6964209 | -0.861 | -1.125 | 0.625 | 0.277 | 0.04 | 0.002 | 0.002 | 0.102 | 0.08 | 0.44 | 0.05 | 0.01 | 0.03 | 0.08 |
| 7 | 96026380 | rs11760823 | -0.308 | -1.006 | 0.127 | 0.546 | -0.007 | -0.068 | -0.068 | 0.165 | 0.14 | 0.07 | 0.08 | 0.34 | 0.12 | 0.1 |
| 7 | 96029077 | rs6945303 | -0.308 | -1.006 | 0.127 | 0.546 | -0.007 | -0.068 | -0.068 | 0 | 0.14 | 0.06 | 0.08 | 0.34 | 0.12 | 0.1 |
| 7 | 96038377 | rs17147518 | 0.407 | -0.875 | -0.107 | -0.101 | -0.003 | -0.05 | -0.05 | 0 | 0.15 | 0.05 | 0.07 | 0.25 | 0.12 | 0.11 |
| 7 | 96039063 | rs12154661 | 0.034 | -0.494 | -0.228 | -0.655 | -0.013 | -0.059 | -0.059 | 0 | 0.34 | 0.57 | 0.22 | 0.06 | 0.29 | 0.27 |
| 7 | 96039267 | rs6971215 | 0.523 | -0.503 | 0.202 | -0.692 | -0.01 | -0.103 | -0.103 | 0 | 0.29 | 0.38 | 0.18 | 0.04 | 0.28 | 0.22 |
| 7 | 96046442 | rs2524983 | 0.339 | -0.008 | 0.065 | -0.594 | 0.001 | -0.009 | -0.009 | 1.373 | 0.77 | 0.76 | 0.78 | 0.59 | 0.8 | 0.54 |
| 7 | 96046527 | rs767206 | 1.313 | -0.008 | 0.044 | -0.614 | -0.012 | -0.013 | -0.013 | 0 | 0.35 | 0.27 | 0.54 | 0.54 | 0.42 | 0.3 |
| 7 | 96047597 | rs1963303 | 1.053 | -0.121 | 0.066 | -0.685 | 0.002 | -0.023 | -0.023 | 0 | 0.31 | 0.23 | 0.17 | 0.05 | 0.28 | 0.21 |
| 7 | 96048278 | rs9640640 | 1.006 | 0.307 | -0.416 | -0.358 | -0.024 | -0.042 | -0.042 | 2.115 | 0.35 | 0.21 | 0.52 | 0.53 | 0.42 | 0.31 |
| 7 | 96049191 | rs2524981 | -0.75 | 0.329 | -0.492 | -0.348 | -0.061 | -0.01 | -0.01 | 0 | 0.84 | 0.97 | 0.81 | 0.82 | 0.83 | 0.68 |
| 7 | 96049498 | rs1989968 | 0.672 | 0.368 | -0.37 | -0.382 | -0.005 | -0.019 | -0.019 | 0 | 0.41 | 0.4 | 0.24 | 0.28 | 0.39 | 0.29 |
| 7 | 96053137 | rs2214516 | -0.926 | 0.597 | -0.023 | -0.169 | -0.016 | -0.032 | -0.032 | 0 | 0.06 | 0.16 | 0.01 | 0 | 0.01 | 0.06 |
| 7 | 96065364 | rs2107284 | 0.828 | 0.764 | -0.234 | -0.129 | -0.001 | -0.018 | -0.018 | 0 | 0.77 | 0.63 | 0.76 | 0.82 | 0.81 | 0.61 |
| 7 | 96074122 | rs17167549 | -0.265 | 1.048 | -0.473 | -0.153 | 0.002 | 0.002 | 0.002 | 0 | 0.11 | 0.09 | 0.07 | 0.16 | 0.13 | 0.13 |
| 7 | 96076927 | rs2724041 | -0.59 | 1.073 | -1.368 | -1.001 | -0.033 | 0.154 | 0.154 | 0 | 0.6 | 0.81 | 0.64 | 0.88 | 0.58 | 0.59 |
| 7 | 96081911 | rs3094750 | 0.641 | 0.836 | -0.853 | -1.341 | -0.005 | 0.006 | 0.006 | 2.41 | 0.24 | 0.28 | 0.43 | 0.7 | 0.25 | 0.34 |
| 7 | 96093898 | rs3113040 | 0.628 | 0.83 | -0.856 | -1.539 | -0.005 | -0.004 | -0.004 | 1.447 | 0.24 | 0.28 | 0.43 | 0.75 | 0.25 | 0.35 |
| 7 | 96094480 | rs2724058 | 0.184 | 0.84 | -0.537 | -1.844 | -0.01 | -0.102 | -0.102 | 0 | 0.26 | 0.46 | 0.16 | 0.07 | 0.22 | 0.19 |
| 7 | 96096148 | rs3113041 | -0.189 | 0.793 | -0.283 | -1.812 | 0.001 | -0.051 | -0.051 | 0 | 0.36 | 0.37 | 0.5 | 0.91 | 0.38 | 0.47 |
| 7 | 96102582 | rs3113045 | -0.217 | 0.782 | -0.486 | -1.817 | 0.001 | -0.064 | -0.064 | 0 | 0.36 | 0.38 | 0.5 | 0.91 | 0.38 | 0.47 |
| 7 | 96119481 | rs10429035 | -0.976 | 0.171 | -0.177 | -1.65 | 0.011 | 0.03 | 0.03 | 0.42 | 0.35 | 0.73 | 0.26 | 0.19 | 0.27 | 0.31 |
| 7 | 96121426 | rs4507687 | 0.912 | 0.274 | -0.641 | -1.738 | -0.076 | 0.012 | 0.012 | 0 | 0.37 | 0.1 | 0.47 | 0.68 | 0.37 | 0.48 |
| 7 | 96125315 | rs4613908 | 0.985 | 0.261 | -0.679 | -1.766 | -0.098 | -0.007 | -0.007 | 0 | 0.37 | 0.07 | 0.47 | 0.68 | 0.36 | 0.47 |
| 7 | 96126368 | rs11760378 | 0.517 | 0.11 | 0.096 | -1.912 | 0.002 | -0.026 | -0.026 | 0.088 | 0.19 | 0.15 | 0.11 | 0.02 | 0.16 | 0.1 |
| 7 | 96130638 | rs10953178 | 0.23 | 0.143 | 0.389 | -1.781 | 0.008 | 0.039 | 0.039 | 0.162 | 0.56 | 0.32 | 0.58 | 0.7 | 0.52 | 0.58 |
| 7 | 96132827 | rs12674052 | 0.239 | 0.153 | 0.593 | -1.816 | -0.002 | -0.003 | -0.003 | 0 | 0.13 | 0.17 | 0.09 | 0.12 | 0.11 | 0.2 |
| 7 | 96137674 | rs10085588 | -1.157 | 0.043 | 0.457 | -1.921 | -0.001 | 0.036 | 0.036 | 2.494 | 0.72 | 0.82 | 0.74 | 0.88 | 0.64 | 0.79 |
| 7 | 96148254 | rs4367457 | 0.077 | 0.103 | 0.488 | -1.84 | -0.007 | -0.007 | -0.007 | 0.019 | 0.14 | 0.18 | 0.1 | 0.11 | 0.1 | 0.21 |
| 7 | 96150473 | rs6961784 | 0.716 | 0.093 | 0.705 | -1.93 | 0.002 | -0.009 | -0.009 | 0.27 | 0.39 | 0.44 | 0.47 | 0.75 | 0.37 | 0.48 |
| 7 | 96158820 | rs4490763 | -0.278 | -0.906 | 0.858 | -2.099 | 0.002 | -0.021 | -0.021 | 0 | 0.22 | 0.16 | 0.14 | 0.02 | 0.18 | 0.15 |
| 7 | 96162825 | rs7776689 | 0.809 | -1.094 | 1.029 | -2.055 | -0.022 | -0.034 | -0.034 | 0.084 | 0.74 | 0.49 | 0.72 | 0.82 | 0.78 | 0.74 |
| 7 | 96167634 | rs13239179 | 1.617 | -1.244 | 1.084 | -2.092 | 0.042 | 0.052 | 0.052 | 0.06 | 0.59 | 0.25 | 0.61 | 0.7 | 0.68 | 0.6 |
| 7 | 96168217 | rs4577892 | 1.01 | -1.126 | 0.722 | -2.416 | -0.001 | -0.064 | -0.064 | 0.11 | 0.29 | 0.3 | 0.23 | 0.01 | 0.26 | 0.21 |
| 7 | 96170054 | rs6963647 | -0.365 | -1.166 | 1.204 | -2.195 | 0.001 | -0.048 | -0.048 | 0.073 | 0.19 | 0.39 | 0.19 | 0.09 | 0.18 | 0.14 |
| 7 | 96173019 | rs7792972 | 1.364 | -1.04 | 0.979 | -2.103 | 0.004 | 0.005 | 0.005 | 0 | 0.6 | 0.4 | 0.63 | 0.7 | 0.68 | 0.6 |
| 7 | 96188993 | rs7784410 | 1.308 | -1.035 | 0.859 | -2.115 | 0.017 | 0.027 | 0.027 | 0.073 | 0.6 | 0.33 | 0.67 | 0.77 | 0.69 | 0.6 |
| 7 | 96190397 | rs13244919 | 1.528 | -1.066 | 0.606 | -2.115 | 0.063 | 0.085 | 0.085 | 0.816 | 0.57 | 0.15 | 0.61 | 0.7 | 0.68 | 0.59 |
| 7 | 96199201 | rs11769840 | 0.078 | -0.906 | 0.797 | -1.929 | 0.007 | -0.002 | -0.002 | 1.344 | 0.93 | 0.96 | 0.84 | 0.89 | 0.97 | 0.88 |
| 7 | 96208580 | rs6465527 | 0.037 | -0.906 | 0.794 | -1.929 | 0.01 | -0.003 | -0.003 | 0 | 0.93 | 0.97 | 0.84 | 0.89 | 0.97 | 0.88 |
| 7 | 96215832 | rs1880512 | 1.62 | -0.901 | 0.916 | -2.056 | 0.026 | 0.047 | 0.047 | 0.654 | 0.6 | 0.22 | 0.5 | 0.68 | 0.66 | 0.62 |
| 7 | 96218698 | rs6465529 | 1.009 | -0.894 | 1.716 | -2.047 | -0.039 | 0.024 | 0.024 | 2.346 | 0.9 | 0.57 | 0.81 | 0.89 | 0.97 | 0.87 |
| 7 | 96220622 | rs10953189 | -0.964 | -0.674 | 1.678 | -1.959 | 0.003 | -0.015 | -0.015 | 1.201 | 0.29 | 0.35 | 0.31 | 0.21 | 0.31 | 0.25 |
| 7 | 96228434 | rs1404508 | 0.069 | -0.71 | 1.894 | -2.007 | -0.009 | -0.021 | -0.021 | 0 | 0.84 | 0.68 | 0.86 | 0.88 | 0.86 | 0.88 |
| 7 | 96230631 | rs10240174 | 0.381 | -0.699 | 1.773 | -2.028 | 0.025 | -0.039 | -0.039 | 0 | 0.82 | 0.47 | 0.8 | 0.92 | 0.83 | 0.83 |
| 7 | 96230686 | rs10464592 | 1.568 | -0.656 | 1.134 | -2.135 | -0.067 | -0.002 | -0.002 | 0 | 0.32 | 0.11 | 0.33 | 0.67 | 0.43 | 0.33 |
| 7 | 96254519 | rs13233151 | 1.153 | -0.532 | 1.515 | -2.123 | 0 | 0.005 | 0.005 | 0 | 0.68 | 0.31 | 0.71 | 0.83 | 0.69 | 0.72 |
| 7 | 96259990 | rs1608739 | -0.335 | -0.334 | 1.698 | -2.008 | 0.041 | -0.045 | -0.045 | 3.447 | 0.91 | 0.65 | 0.92 | 1 | 0.93 | 0.99 |
| 7 | 96294270 | rs12537768 | -0.014 | -0.402 | 1.127 | -1.964 | -0.003 | -0.003 | -0.003 | 0.05 | 0.13 | 0.16 | 0.17 | 0.19 | 0.17 | 0.21 |
| 7 | 96309021 | rs2272225 | 0.098 | -0.758 | 1.168 | -2.069 | -0.007 | -0.002 | -0.002 | 0.003 | 0.87 | 0.89 | 0.83 | 0.89 | 0.93 | 0.83 |
| 7 | 96313582 | rs4729274 | -1.409 | -0.404 | 1.64 | -1.905 | 0.007 | -0.013 | -0.013 | 1.028 | 0.23 | 0.64 | 0.26 | 0.19 | 0.25 | 0.22 |

| | | | | | | | | | | | | | | | | |
|---|---|---|---|---|---|---|---|---|---|---|---|---|---|---|---|---|
| 7 | 96318575 | rs6968090 | -1.126 | -0.434 | 1.439 | -1.916 | -0.003 | -0.002 | -0.002 | 0 | 0.19 | 0.19 | 0.19 | 0.21 | 0.21 | 0.25 |
| 7 | 96334274 | rs12704871 | 1.66 | -1.058 | 1.024 | -2.069 | -0.154 | -0.06 | -0.06 | 0.034 | 0.29 | 0.02 | 0.36 | 0.68 | 0.41 | 0.32 |
| 7 | 96343642 | rs10257162 | -0.155 | -1.103 | 1.35 | -2.129 | 0.067 | 0.15 | 0.15 | 0 | 0.53 | 0.75 | 0.63 | 0.88 | 0.66 | 0.54 |
| 7 | 96343665 | rs10241838 | -0.13 | -1.099 | 1.369 | -2.127 | 0.053 | 0.104 | 0.104 | 0 | 0.53 | 0.7 | 0.63 | 0.87 | 0.66 | 0.54 |
| 7 | 96351843 | rs1404511 | -0.241 | -1.109 | 1.349 | -2.126 | 0.036 | 0.065 | 0.065 | 0 | 0.56 | 0.69 | 0.63 | 0.89 | 0.69 | 0.58 |
| 7 | 96357756 | rs10226466 | 0.567 | -1.014 | 1.583 | -2 | 0.088 | -0.016 | -0.016 | 0.396 | 0.08 | 0.41 | 0.09 | 0 | 0.07 | 0.01 |
| 7 | 96362344 | rs1464807 | 0.643 | -1.007 | 1.273 | -1.961 | -0.003 | -0.011 | -0.011 | 0 | 0.1 | 0.07 | 0.12 | 0.15 | 0.11 | 0.09 |
| 7 | 96362400 | rs10276660 | 0.866 | -0.969 | 1.452 | -1.95 | 0.056 | -0.027 | -0.027 | 0 | 0.08 | 0.28 | 0.07 | 0 | 0.07 | 0.01 |
| 7 | 96366342 | rs12704876 | 0.609 | -0.79 | 0.708 | -1.759 | -0.05 | 0.098 | 0.098 | 1.386 | 0.4 | 0.25 | 0.47 | 0.83 | 0.5 | 0.4 |
| 7 | 96371732 | rs1524927 | 1.342 | -0.951 | -0.028 | -1.422 | -0.062 | -0.028 | -0.028 | 0 | 0.29 | 0.14 | 0.36 | 0.63 | 0.42 | 0.31 |
| 7 | 96371760 | rs1474383 | 0.083 | -1.201 | 0.043 | -1.441 | 0.021 | 0.075 | 0.075 | 0 | 0.58 | 0.73 | 0.58 | 0.81 | 0.63 | 0.49 |
| 7 | 96377441 | rs11982457 | -0.701 | -1.157 | 0.641 | -0.917 | -0.068 | 0.078 | 0.078 | 0.132 | 0.14 | 0.41 | 0.22 | 0.23 | 0.05 | 0.24 |
| 7 | 96377488 | rs4729281 | -1.66 | -1.529 | -0.098 | -1.508 | 0.062 | 0.14 | 0.14 | 0.542 | 0.5 | 0.87 | 0.46 | 0.35 | 0.37 | 0.43 |
| 7 | 96377562 | rs12673272 | 0.416 | -1.747 | -0.126 | -1.517 | 0.024 | 0.038 | 0.038 | 0 | 0.15 | 0.03 | 0.08 | 0.11 | 0.1 | 0.1 |
| 7 | 96385092 | rs4729289 | -0.031 | -1.764 | -0.039 | -1.506 | -0.062 | 0.078 | 0.078 | 0.071 | 0.14 | 0.36 | 0.22 | 0.23 | 0.05 | 0.24 |
| 7 | 96396916 | rs12674116 | 0.391 | -1.63 | -0.662 | -1.554 | 0.023 | 0.043 | 0.043 | 0 | 0.15 | 0.03 | 0.08 | 0.11 | 0.1 | 0.1 |
| 7 | 96396971 | rs17167787 | 0.005 | -1.618 | -0.4 | -1.593 | 0.028 | -0.128 | -0.128 | 0.924 | 0.2 | 0.37 | 0.15 | 0 | 0.23 | 0.1 |
| 7 | 96399494 | rs6977057 | 1.148 | -1.638 | -0.413 | -1.532 | -0.035 | -0.006 | -0.006 | 0 | 0.67 | 0.45 | 0.57 | 0.65 | 0.78 | 0.58 |
| 7 | 96401045 | rs12375086 | 0.306 | -1.428 | -1.408 | -1.326 | 0.021 | 0.043 | 0.043 | 0 | 0.15 | 0.02 | 0.08 | 0.11 | 0.1 | 0.1 |
| 7 | 96407773 | rs6953983 | 0.534 | -1.393 | -1.363 | -1.342 | -0.01 | 0.057 | 0.057 | 0.442 | 0.64 | 0.15 | 0.51 | 0.76 | 0.65 | 0.59 |
| 7 | 96408925 | rs990902 | 1.017 | -1.059 | -1.431 | -1.097 | -0.058 | 0.079 | 0.079 | 0.596 | 0.47 | 0.05 | 0.42 | 0.65 | 0.56 | 0.48 |
| 7 | 96414506 | rs7786855 | 0.931 | -1.099 | -1.348 | -0.823 | -0.041 | -0.018 | -0.018 | 0 | 0.45 | 0.09 | 0.35 | 0.49 | 0.55 | 0.46 |
| 7 | 96419077 | rs2394502 | -0.115 | -1.311 | -0.691 | -0.833 | -0.024 | -0.025 | -0.025 | 0 | 0.89 | 0.86 | 0.97 | 0.96 | 0.96 | 0.9 |
| 7 | 96419248 | rs17167805 | 0.591 | -1.394 | -0.135 | -0.863 | -0.027 | -0.029 | -0.029 | 0.65 | 0.06 | 0.18 | 0.03 | 0 | 0.01 | 0 |
| 7 | 96425238 | rs17167814 | 0.269 | -1.447 | 0.208 | -0.897 | -0.014 | -0.022 | -0.022 | 0 | 0.07 | 0.28 | 0.03 | 0 | 0.01 | 0 |
| 7 | 96429251 | rs12704882 | 1.535 | -1.378 | -0.886 | -1.144 | -0.118 | -0.095 | -0.095 | 0 | 0.28 | 0.07 | 0.29 | 0.49 | 0.44 | 0.33 |
| 7 | 96431346 | rs10241355 | 0.991 | -1.297 | -0.027 | -1.336 | -0.006 | -0.054 | -0.054 | 0.067 | 0.65 | 0.41 | 0.82 | 0.92 | 0.75 | 0.81 |
| 7 | 96435891 | rs2922915 | 0.148 | -1.225 | 0.252 | -0.95 | -0.023 | -0.04 | -0.04 | 0 | 0.93 | 0.89 | 0.97 | 1 | 0.97 | 0.97 |
| 7 | 96440624 | rs9655758 | -0.228 | -1.877 | 0.393 | -0.548 | -0.033 | 0.065 | 0.065 | 0 | 0.09 | 0.12 | 0.17 | 0.35 | 0.02 | 0.28 |
| 7 | 96444600 | rs7805612 | 1.216 | -2.22 | 0.483 | -0.987 | -0.033 | -0.061 | -0.061 | 0.326 | 0.76 | 0.34 | 0.87 | 0.92 | 0.86 | 0.88 |
| 7 | 96446260 | rs767198 | 0.327 | -2.182 | 0.805 | -0.961 | -0.001 | -0.004 | -0.004 | 0.06 | 0.13 | 0.16 | 0.07 | 0.08 | 0.11 | 0.09 |
| 7 | 96457119 | rs2948276 | -1.325 | -2.221 | 0.83 | -1.101 | -0.027 | -0.066 | -0.066 | 1.127 | 0.26 | 0.63 | 0.16 | 0.11 | 0.17 | 0.12 |
| 7 | 96461649 | rs1947228 | 2.391 | -2.288 | 0.079 | -1.219 | -0.031 | -0.011 | -0.011 | 0.005 | 0.45 | 0.08 | 0.43 | 0.51 | 0.56 | 0.4 |
| 7 | 96475637 | rs7802312 | -0.685 | -2.251 | 0.254 | -1.198 | -0.003 | 0.007 | 0.007 | 0 | 0.25 | 0.29 | 0.4 | 0.37 | 0.24 | 0.46 |
| 7 | 96484741 | rs7785299 | 0.483 | -2.187 | 0.184 | -1.279 | -0.008 | 0.014 | 0.014 | 0 | 0.09 | 0.04 | 0.16 | 0.03 | 0.11 | 0.1 |
| 7 | 96484929 | rs7456227 | 1.197 | -2.338 | 0.493 | -1.369 | -0.024 | -0.061 | -0.061 | 0.192 | 0.69 | 0.43 | 0.72 | 0.9 | 0.75 | 0.79 |
| 7 | 96488196 | rs9656041 | -0.141 | -2.34 | 0.504 | -1.267 | 0 | 0.036 | 0.036 | 0 | 0.15 | 0.25 | 0.24 | 0.3 | 0.13 | 0.34 |
| 7 | 96493854 | rs1542250 | 0.046 | -2.287 | 0.656 | -1.203 | -0.003 | -0.009 | -0.009 | 0.088 | 0.12 | 0.1 | 0.17 | 0.03 | 0.12 | 0.1 |
| 7 | 96496562 | rs6979505 | -0.276 | -2.298 | 0.703 | -1.209 | -0.012 | -0.026 | -0.026 | 0 | 0.13 | 0.15 | 0.18 | 0.03 | 0.12 | 0.1 |
| 7 | 96496658 | rs2948244 | -1.281 | -2.232 | 0.581 | -1.226 | 0.05 | 0.045 | 0.045 | 0.055 | 0.44 | 0.86 | 0.51 | 0.4 | 0.36 | 0.52 |
| 7 | 96498913 | rs2948240 | -0.103 | -2.295 | 0.553 | -1.328 | 0.003 | 0.023 | 0.023 | 0 | 0.89 | 0.96 | 0.95 | 0.96 | 0.92 | 0.93 |
| 7 | 96503974 | rs1522529 | 1.29 | -2.801 | -0.076 | -1.802 | -0.011 | 0.036 | 0.036 | 0.032 | 0.39 | 0.35 | 0.59 | 0.84 | 0.53 | 0.58 |
| 7 | 96508334 | rs6957376 | 2.267 | -2.726 | -0.194 | -1.771 | -0.149 | -0.072 | -0.072 | 1.108 | 0.3 | 0.07 | 0.39 | 0.55 | 0.48 | 0.28 |
| 7 | 96525973 | rs10267527 | 0.82 | -2.733 | -0.011 | -1.775 | -0.001 | -0.009 | -0.009 | 0 | 0.43 | 0.45 | 0.61 | 0.84 | 0.55 | 0.58 |
| 7 | 96526030 | rs10267636 | 0.561 | -2.828 | -0.061 | -1.83 | -0.002 | -0.035 | -0.035 | 0 | 0.58 | 0.48 | 0.65 | 0.86 | 0.61 | 0.71 |
| 7 | 96530404 | rs10265665 | 0.13 | -2.758 | 0.07 | -1.686 | -0.009 | 0.121 | 0.121 | 0 | 0.09 | 0.26 | 0.2 | 0.29 | 0.05 | 0.26 |
| 7 | 96535299 | rs1522535 | -1.089 | -2.74 | 0.272 | -1.736 | 0.001 | -0.023 | -0.023 | 0.316 | 0.92 | 0.84 | 0.96 | 0.96 | 0.95 | 0.97 |
| 7 | 96540655 | rs1403185 | -0.517 | -2.867 | -0.017 | -1.855 | 0.027 | 0.049 | 0.049 | 0.973 | 0.48 | 0.34 | 0.32 | 0.12 | 0.37 | 0.38 |
| 7 | 96542982 | rs17168021 | -0.04 | -2.709 | -0.345 | -1.777 | 0.003 | 0.017 | 0.017 | 0.09 | 0.33 | 0.23 | 0.27 | 0.11 | 0.31 | 0.26 |
| 7 | 96554285 | rs1403183 | 1.207 | -1.884 | -0.478 | -1.309 | -0.066 | -0.016 | -0.016 | 0 | 0.36 | 0.19 | 0.45 | 0.55 | 0.52 | 0.32 |
| 7 | 96560984 | rs4727353 | -0.352 | -1.83 | -0.325 | -0.49 | -0.021 | 0.175 | 0.175 | 0 | 0.12 | 0.34 | 0.21 | 0.42 | 0.07 | 0.3 |
| 7 | 96561449 | rs4729322 | 0.03 | -1.54 | -0.279 | -0.277 | -0.01 | 0.207 | 0.207 | 0 | 0.08 | 0.23 | 0.19 | 0.41 | 0.04 | 0.29 |
| 7 | 96562031 | rs1207688 | -0.453 | -0.289 | -1.224 | -0.268 | -0.001 | -0.001 | -0.001 | 0 | 0.63 | 0.66 | 0.59 | 0.53 | 0.57 | 0.65 |
| 7 | 96562541 | rs17168056 | 0.811 | -0.097 | -1.975 | 0.073 | -0.02 | -0.053 | -0.053 | 0.625 | 0.14 | 0.08 | 0.25 | 0.36 | 0.18 | 0.31 |
| 7 | 96567080 | rs11976595 | 1.357 | 0.66 | -2.192 | 0.078 | -0.032 | -0.046 | -0.046 | 0.064 | 0.17 | 0.08 | 0.23 | 0.38 | 0.29 | 0.23 |
| 7 | 96569280 | rs13227263 | 0.421 | 0.317 | -2.284 | -0.627 | 0.021 | 0.213 | 0.213 | 3.074 | 0.47 | 0.18 | 0.36 | 0.16 | 0.41 | 0.35 |
| 7 | 96571373 | rs17656440 | 0.269 | 0.77 | -2.144 | 0.391 | -0.043 | -0.081 | -0.081 | 2.065 | 0.25 | 0.08 | 0.3 | 0.49 | 0.3 | 0.24 |
| 7 | 96572376 | rs2526684 | -0.053 | 0.46 | -1.722 | -0.186 | 0.017 | 0.118 | 0.118 | 0 | 0.49 | 0.68 | 0.42 | 0.2 | 0.45 | 0.47 |
| 7 | 96575827 | rs10486859 | -0.11 | 0.57 | -1.715 | 0.57 | -0.054 | -0.025 | -0.025 | 0.322 | 0.28 | 0.1 | 0.31 | 0.33 | 0.43 | 0.24 |
| 7 | 96578640 | rs1207692 | 0.015 | -0.677 | -1.124 | 0.704 | 0.001 | 0 | 0 | 0 | 0.67 | 0.67 | 0.52 | 0.79 | 0.75 | 0.76 |
| 7 | 96579339 | rs17598132 | 0.656 | -0.892 | -1.259 | 0.416 | 0.019 | 0.032 | 0.032 | 0 | 0.19 | 0.09 | 0.06 | 0 | 0.1 | 0.12 |
| 7 | 96593097 | rs12531148 | 0.77 | -0.757 | -1.517 | 0.821 | -0.072 | -0.081 | -0.081 | 0.317 | 0.39 | 0.07 | 0.53 | 0.54 | 0.58 | 0.44 |
| 7 | 96605368 | rs2214643 | -0.785 | -0.851 | -1.472 | 0.597 | 0.021 | 0.004 | 0.004 | 0 | 0.78 | 0.82 | 0.7 | 0.86 | 0.87 | 0.8 |
| 7 | 96608069 | rs2214644 | 0.19 | 0.327 | -1.539 | 0.496 | -0.053 | 0.004 | 0.004 | 0 | 0.39 | 0.09 | 0.4 | 0.44 | 0.57 | 0.44 |
| 7 | 96609059 | rs5012357 | -0.733 | 0.86 | -0.626 | 0.561 | 0.011 | -0.007 | -0.007 | 0 | 0.11 | 0.19 | 0.11 | 0.06 | 0.11 | 0.04 |
| 7 | 96615467 | rs1476604 | -1.056 | 0.364 | -0.695 | 0.619 | -0.008 | 0 | 0 | 0 | 0.17 | 0.24 | 0.26 | 0.16 | 0.17 | 0.17 |
| 7 | 96615819 | rs717483 | -0.519 | 0.935 | -1.773 | -0.305 | -0.079 | 0.126 | 0.126 | 6.608 | 0.29 | 0.06 | 0.23 | 0.06 | 0.36 | 0.31 |
| 7 | 96616012 | rs6956369 | -0.586 | -0.365 | -1.968 | -0.508 | -0.024 | -0.003 | -0.003 | 0 | 0.17 | 0.23 | 0.21 | 0.16 | 0.1 | 0.16 |
| 7 | 96622615 | rs11766968 | -0.196 | -0.138 | -2.183 | 0.573 | -0.02 | 0.034 | 0.034 | 0 | 0.11 | 0.03 | 0.13 | 0.03 | 0.14 | 0.16 |

| | | | | | | | | | | | | | | | | |
|---|---|---|---|---|---|---|---|---|---|---|---|---|---|---|---|---|
| 7 | 96622720 | rs2240293 | 0.813 | 0.031 | -3.15 | 0.356 | -0.04 | 0.04 | 0.04 | 2.655 | 0.49 | 0.06 | 0.36 | 0.36 | 0.51 | 0.5 |
| 7 | 96625589 | rs17657924 | 0.743 | 0.72 | -3.418 | 0.229 | -0.062 | 0.029 | 0.029 | 1.842 | 0.45 | 0.08 | 0.32 | 0.35 | 0.45 | 0.45 |
| 7 | 96626119 | rs6957108 | 0.355 | 0.567 | -1.738 | 0.387 | -0.004 | 0 | 0 | 0 | 0.2 | 0.31 | 0.15 | 0.17 | 0.18 | 0.12 |
| 7 | 96645329 | rs1207730 | 0.071 | 0.308 | -1.392 | 0.661 | 0.001 | 0 | 0 | 0.224 | 0.21 | 0.23 | 0.14 | 0.16 | 0.19 | 0.13 |
| 7 | 96649356 | rs1207731 | 0.303 | 0.305 | -1.383 | 0.648 | -0.002 | -0.001 | -0.001 | 0 | 0.19 | 0.16 | 0.14 | 0.16 | 0.19 | 0.12 |
| 7 | 96649522 | rs12533469 | -0.423 | 1.488 | -1.207 | 0.303 | -0.01 | -0.006 | -0.006 | 0 | 0.24 | 0.21 | 0.44 | 0.26 | 0.31 | 0.22 |
| 7 | 96651985 | rs9769385 | 0.959 | 1.184 | -0.892 | 0.117 | -0.001 | -0.008 | -0.008 | 0 | 0.06 | 0.09 | 0.05 | 0.03 | 0.06 | 0.11 |
| 7 | 96656572 | rs3757493 | 0.427 | 0.416 | -0.947 | -0.219 | 0 | -0.008 | -0.008 | 0.009 | 0.44 | 0.4 | 0.27 | 0.19 | 0.42 | 0.46 |
| 7 | 96661390 | rs17600042 | 0.521 | 0.624 | -1.603 | -0.055 | -0.002 | 0.017 | 0.017 | 0 | 0.09 | 0.02 | 0.06 | 0.03 | 0.07 | 0.12 |
| 7 | 96674424 | rs1207721 | 0.166 | 0.834 | -1.648 | 0.302 | -0.03 | 0.013 | 0.013 | 0 | 0.7 | 0.87 | 0.57 | 0.78 | 0.65 | 0.68 |
| 7 | 96679200 | rs1207725 | -0.187 | 0.979 | -1.583 | -0.146 | -0.017 | 0.043 | 0.043 | 0.089 | 0.76 | 0.85 | 0.74 | 0.9 | 0.7 | 0.75 |
| 7 | 96693047 | rs12670161 | -0.411 | 1.135 | -1.299 | 0.511 | 0.002 | 0.031 | 0.031 | 0 | 0.13 | 0.11 | 0.25 | 0.33 | 0.13 | 0.19 |
| 7 | 96694302 | rs11768581 | 1.408 | 0.882 | -1.618 | 0.29 | -0.07 | 0.108 | 0.108 | 0 | 0.32 | 0.02 | 0.18 | 0.14 | 0.34 | 0.33 |
| 7 | 96697572 | rs1526084 | -0.625 | 0.767 | -0.959 | 0.754 | 0 | 0.024 | 0.024 | 0 | 0.13 | 0.12 | 0.25 | 0.34 | 0.11 | 0.19 |
| 7 | 96715879 | rs12669100 | 0.842 | 0.801 | -0.977 | 0.436 | -0.019 | -0.034 | -0.034 | 0.08 | 0.32 | 0.14 | 0.32 | 0.43 | 0.34 | 0.35 |
| 7 | 96716864 | rs1917939 | 0.911 | 0.912 | -0.994 | 0.404 | -0.007 | -0.023 | -0.023 | 0 | 0.23 | 0.14 | 0.28 | 0.43 | 0.25 | 0.25 |
| 7 | 96719140 | rs2037719 | 0.543 | 0.813 | -1.203 | -0.123 | -0.009 | 0.057 | 0.057 | 0.292 | 0.24 | 0.12 | 0.26 | 0.1 | 0.29 | 0.29 |
| 7 | 96724452 | rs1343646 | 1.63 | 0.507 | -0.971 | 0.006 | -0.011 | 0.038 | 0.038 | 0.141 | 0.21 | 0.06 | 0.12 | 0.14 | 0.22 | 0.16 |
| 7 | 96724585 | rs1343645 | 0.306 | 0.456 | -0.836 | 0.143 | -0.002 | 0.006 | 0.006 | 0 | 0.14 | 0.12 | 0.26 | 0.34 | 0.12 | 0.25 |
| 7 | 96737576 | rs10499934 | 1.805 | 0.369 | -0.701 | 0.121 | 0 | 0.022 | 0.022 | 0 | 0.21 | 0.12 | 0.12 | 0.13 | 0.22 | 0.15 |
| 7 | 96738758 | rs10953201 | -1.361 | 0.41 | 0.024 | -0.006 | 0.005 | -0.031 | -0.031 | 0.057 | 0.65 | 0.78 | 0.69 | 0.59 | 0.68 | 0.71 |
| 7 | 96749012 | rs7794886 | 0.046 | 0.425 | 0.091 | 0.052 | 0 | 0.01 | 0.01 | 0 | 0.38 | 0.47 | 0.41 | 0.47 | 0.35 | 0.4 |
| 7 | 96757057 | rs4398831 | 0.905 | 0.481 | -0.027 | 0.072 | -0.022 | 0.055 | 0.055 | 0.328 | 0.25 | 0.12 | 0.28 | 0.12 | 0.33 | 0.28 |
| 7 | 96758671 | rs12056091 | -0.261 | 0.874 | 0.272 | 0.65 | -0.005 | -0.024 | -0.024 | 0 | 0.37 | 0.26 | 0.39 | 0.46 | 0.35 | 0.4 |
| 7 | 96786625 | rs12670377 | 0.878 | 0.726 | 0.287 | 0.832 | -0.001 | -0.029 | -0.029 | 0.34 | 0.24 | 0.16 | 0.3 | 0.4 | 0.24 | 0.26 |
| 7 | 96825451 | rs6465572 | -0.078 | 0.851 | 0.636 | 0.346 | 0.013 | -0.029 | -0.029 | 1.647 | 0.75 | 0.69 | 0.73 | 0.89 | 0.67 | 0.82 |
| 7 | 96827184 | rs6465573 | 0.36 | 0.649 | 1.331 | 1.158 | 0.012 | 0.051 | 0.051 | 0.216 | 0.71 | 0.44 | 0.56 | 0.55 | 0.66 | 0.59 |
| 7 | 96833078 | rs6967782 | 0.148 | 0.646 | 1.457 | 1.16 | 0.016 | 0.056 | 0.056 | 0 | 0.72 | 0.58 | 0.56 | 0.54 | 0.66 | 0.64 |
| 7 | 96836212 | rs1918523 | 0.649 | 0.524 | 0.756 | 0.988 | -0.007 | 0.024 | 0.024 | 0 | 0.12 | 0.02 | 0.16 | 0.05 | 0.1 | 0.12 |
| 7 | 96847312 | rs10273271 | 0.265 | 0.366 | 1.078 | 1.053 | 0.023 | 0.078 | 0.078 | 0.075 | 0.71 | 0.56 | 0.56 | 0.55 | 0.66 | 0.59 |
| 7 | 96854267 | rs12531233 | 0.422 | 0.263 | 0.555 | 0.996 | -0.009 | -0.044 | -0.044 | 0 | 0.69 | 0.24 | 0.54 | 0.54 | 0.66 | 0.59 |
| 7 | 96870082 | rs2394614 | 0.154 | 0.321 | 0.686 | 1.013 | -0.006 | 0.006 | 0.006 | 0 | 0.25 | 0.21 | 0.27 | 0.16 | 0.34 | 0.2 |
| 7 | 96878550 | rs705384 | -1.395 | 0.344 | -0.019 | 0.967 | 0.007 | 0.081 | 0.081 | 1.036 | 0.54 | 0.79 | 0.73 | 0.65 | 0.58 | 0.58 |
| 7 | 96880081 | rs7788665 | -0.033 | 0.447 | 0.691 | 1.011 | -0.014 | 0.018 | 0.018 | 0 | 0.25 | 0.19 | 0.26 | 0.15 | 0.33 | 0.2 |
| 7 | 96880443 | rs799609 | -0.768 | 0.441 | 0.518 | 1.059 | -0.009 | -0.036 | -0.036 | 0.08 | 0.31 | 0.76 | 0.43 | 0.5 | 0.34 | 0.38 |
| 7 | 96881176 | rs11767058 | -0.143 | 0.812 | -0.226 | 1.076 | -0.025 | 0.031 | 0.031 | 0 | 0.25 | 0.15 | 0.26 | 0.15 | 0.33 | 0.2 |
| 7 | 96896364 | rs799593 | -0.49 | 0 | 0.417 | 1.118 | -0.03 | 0.077 | 0.077 | 0 | 0.24 | 0.42 | 0.35 | 0.42 | 0.19 | 0.35 |
| 7 | 96902638 | rs799588 | -1.129 | 0.293 | 0.01 | 0.268 | 0.02 | 0.088 | 0.088 | 1.174 | 0.55 | 0.72 | 0.66 | 0.85 | 0.59 | 0.67 |
| 7 | 96911814 | rs10486857 | 0.313 | 0.207 | -0.234 | 0.303 | 0.007 | 0.047 | 0.047 | 0 | 0.21 | 0.11 | 0.18 | 0.09 | 0.18 | 0.19 |
| 7 | 96920447 | rs799581 | -1.005 | -0.047 | 0.326 | 0.676 | 0.003 | 0.003 | 0.003 | 1.024 | 0.54 | 0.56 | 0.61 | 0.7 | 0.59 | 0.68 |
| 7 | 96930837 | rs1362373 | 0.399 | 0.094 | 0.06 | 0.158 | -0.016 | -0.014 | -0.014 | 0.859 | 0.24 | 0.19 | 0.25 | 0.35 | 0.35 | 0.27 |
| 7 | 96937096 | rs1420172 | -1.29 | -0.266 | -0.035 | -0.109 | 0.004 | -0.003 | -0.003 | 0.323 | 0.57 | 0.4 | 0.52 | 0.46 | 0.58 | 0.54 |
| 7 | 96937807 | rs4729349 | 0.468 | -0.147 | 0.052 | -0.044 | 0.007 | 0.031 | 0.031 | 0.74 | 0.21 | 0.12 | 0.2 | 0.1 | 0.18 | 0.23 |
| 7 | 96946490 | rs7796570 | 0.26 | 0.497 | 0.017 | -0.072 | 0.004 | 0.009 | 0.009 | 0.437 | 0.25 | 0.28 | 0.25 | 0.36 | 0.29 | 0.25 |
| 7 | 96946643 | rs9692573 | -0.356 | 0.511 | -0.005 | -0.094 | -0.01 | -0.008 | -0.008 | 0 | 0.3 | 0.28 | 0.29 | 0.36 | 0.38 | 0.29 |
| 7 | 96950914 | rs11770839 | -0.216 | 0.42 | -0.139 | -0.15 | -0.013 | 0.047 | 0.047 | 0.038 | 0.24 | 0.13 | 0.24 | 0.1 | 0.27 | 0.22 |
| 7 | 96967600 | rs12539655 | 0.348 | 0.269 | 0.246 | -0.119 | -0.006 | -0.035 | -0.035 | 0 | 0.2 | 0.27 | 0.21 | 0.1 | 0.17 | 0.18 |
| 7 | 96968328 | rs2394625 | -0.011 | 0.303 | -0.18 | -0.196 | 0.005 | -0.011 | -0.011 | 0 | 0.63 | 0.64 | 0.72 | 0.79 | 0.57 | 0.66 |
| 7 | 96968676 | rs12704913 | -0.264 | 0.473 | -0.292 | -0.262 | 0.016 | -0.02 | -0.02 | 0 | 0.67 | 0.63 | 0.75 | 0.75 | 0.64 | 0.69 |
| 7 | 96968805 | rs12704914 | 0.253 | 0.467 | -0.299 | -0.268 | 0.002 | -0.008 | -0.008 | 0 | 0.62 | 0.63 | 0.71 | 0.76 | 0.55 | 0.64 |
| 7 | 96969163 | rs953687 | -0.184 | 0.458 | -0.15 | -0.143 | 0.011 | -0.014 | -0.014 | 0 | 0.67 | 0.65 | 0.75 | 0.76 | 0.64 | 0.67 |
| 7 | 96970705 | rs13225811 | 1.437 | 0.228 | -0.593 | 0.168 | 0.001 | -0.028 | -0.028 | 4.996 | 0.26 | 0.2 | 0.36 | 0.54 | 0.29 | 0.33 |
| 7 | 96972939 | rs13247934 | -0.383 | 0.22 | -0.626 | -0.054 | 0.011 | 0.009 | 0.009 | 0.222 | 0.35 | 0.26 | 0.37 | 0.26 | 0.34 | 0.35 |
| 7 | 96974137 | rs17168354 | -0.826 | 1.401 | -0.529 | -0.508 | -0.004 | -0.021 | -0.021 | 0 | 0.25 | 0.24 | 0.23 | 0.05 | 0.35 | 0.21 |
| 7 | 96981954 | rs10282426 | -0.632 | 0.882 | -0.332 | 0.222 | -0.014 | 0.087 | 0.087 | 0 | 0.27 | 0.45 | 0.19 | 0.45 | 0.18 | 0.39 |
| 7 | 96983589 | rs11983012 | -0.987 | 0.415 | -0.811 | -0.136 | -0.023 | 0.024 | 0.024 | 0 | 0.35 | 0.46 | 0.23 | 0.45 | 0.21 | 0.41 |
| 7 | 96983785 | rs11979492 | -0.477 | 0.368 | -0.853 | -0.158 | -0.043 | 0.053 | 0.053 | 0.009 | 0.28 | 0.44 | 0.17 | 0.4 | 0.14 | 0.34 |
| 7 | 96986462 | rs10280571 | 1.492 | 0.389 | -0.549 | 0.14 | -0.001 | 0.003 | 0.003 | 2.537 | 0.13 | 0.16 | 0.08 | 0.17 | 0.08 | 0.13 |
| 7 | 97003104 | rs2058667 | 0.516 | 0.278 | -0.44 | -0.289 | -0.008 | -0.031 | -0.031 | 0 | 0.2 | 0.28 | 0.23 | 0.13 | 0.22 | 0.24 |
| 7 | 97003503 | rs10435325 | 0.084 | 0.906 | 0.076 | 0.246 | 0.028 | 0.002 | 0.002 | 0 | 0.13 | 0.27 | 0.12 | 0.09 | 0.15 | 0.08 |
| 7 | 97006278 | rs12111926 | -0.86 | 0.991 | -0.143 | 0.212 | 0.005 | 0.015 | 0.015 | 1.642 | 0.33 | 0.52 | 0.28 | 0.37 | 0.31 | 0.37 |
| 7 | 97012300 | rs2394631 | -0.974 | 1.072 | -0.306 | 0.198 | -0.001 | -0.001 | -0.001 | 0 | 0.34 | 0.36 | 0.3 | 0.37 | 0.36 | 0.36 |
| 7 | 97012909 | rs10257214 | 0.019 | 1.017 | -0.215 | 0.266 | 0.004 | 0.012 | 0.012 | 0.871 | 0.27 | 0.34 | 0.23 | 0.32 | 0.28 | 0.3 |
| 7 | 97014330 | rs10264553 | -0.117 | 0.279 | -0.137 | 0.2 | -0.007 | 0.022 | 0.022 | 0 | 0.09 | 0.04 | 0.07 | 0.05 | 0.08 | 0.09 |
| 7 | 97015800 | rs1420182 | -0.744 | 0.2 | -0.121 | 0.45 | -0.007 | -0.007 | -0.007 | 0 | 0.41 | 0.51 | 0.49 | 0.6 | 0.4 | 0.37 |
| 7 | 97017795 | rs10255878 | -1.076 | 0.174 | 0.068 | 0.542 | -0.005 | 0.054 | 0.054 | 0.144 | 0.16 | 0.34 | 0.12 | 0.28 | 0.13 | 0.27 |
| 7 | 97018719 | rs6973117 | 1.829 | -0.125 | 0.263 | 0.243 | -0.011 | -0.059 | -0.059 | 0 | 0.11 | 0.23 | 0.07 | 0 | 0.11 | 0.08 |
| 7 | 97018885 | rs2216032 | -0.315 | -0.107 | -0.06 | 0.266 | -0.003 | 0.031 | 0.031 | 2.396 | 0.34 | 0.43 | 0.22 | 0.53 | 0.27 | 0.51 |
| 7 | 97019295 | rs10279061 | 0.245 | -0.094 | 0.009 | 0.287 | 0.027 | 0.218 | 0.218 | 1.811 | 0.06 | 0.29 | 0.05 | 0.23 | 0.05 | 0.18 |

| | | | | | | | | | | | | | | | | |
|---|---|---|---|---|---|---|---|---|---|---|---|---|---|---|---|---|
| 7 | 97020627 | rs1592472 | -0.261 | -0.114 | -0.101 | 0.378 | -0.015 | 0.026 | 0.026 | 0 | 0.52 | 0.15 | 0.4 | 0.56 | 0.5 | 0.44 |
| 7 | 97022110 | rs10275742 | 0.283 | -0.042 | 0.012 | 0.452 | 0.028 | 0.221 | 0.221 | 1.985 | 0.06 | 0.29 | 0.05 | 0.23 | 0.05 | 0.17 |
| 7 | 97022873 | rs9986738 | -0.995 | -0.082 | 0.018 | 0.401 | 0 | -0.01 | -0.01 | 0 | 0.6 | 0.59 | 0.47 | 0.8 | 0.55 | 0.63 |
| 7 | 97023012 | rs1420181 | -0.336 | -0.077 | -0.214 | 0.403 | -0.019 | 0.054 | 0.054 | 0 | 0.54 | 0.15 | 0.53 | 0.63 | 0.51 | 0.45 |
| 7 | 97023144 | rs1420180 | 1.202 | -0.088 | -0.27 | 0.443 | -0.038 | -0.117 | -0.117 | 0 | 0.25 | 0.03 | 0.36 | 0.32 | 0.27 | 0.11 |
| 7 | 97024877 | rs2216034 | 0.401 | -0.062 | -0.002 | 0.48 | 0.003 | 0.141 | 0.141 | 2.189 | 0.05 | 0.19 | 0.05 | 0.23 | 0.04 | 0.17 |
| 7 | 97028821 | rs10236201 | 0.131 | -0.044 | 0.026 | 0.487 | 0.027 | 0.218 | 0.218 | 1.811 | 0.06 | 0.29 | 0.05 | 0.23 | 0.05 | 0.18 |
| 7 | 97030294 | rs6970139 | 1.57 | 0.015 | -0.376 | 0.236 | -0.003 | -0.013 | -0.013 | 0 | 0.19 | 0.1 | 0.1 | 0.26 | 0.15 | 0.26 |
| 7 | 97030369 | rs6970178 | 0.325 | 0.02 | -0.392 | 0.202 | -0.006 | 0.002 | 0.002 | 3.777 | 0.29 | 0.13 | 0.17 | 0.31 | 0.23 | 0.35 |
| 7 | 97038975 | rs4729357 | -0.136 | 0.401 | -0.488 | -0.122 | -0.007 | -0.016 | -0.016 | 1.376 | 0.32 | 0.14 | 0.3 | 0.31 | 0.34 | 0.18 |
| 7 | 97041330 | rs2160259 | 0.844 | 0.453 | -0.547 | -0.092 | -0.002 | -0.034 | -0.034 | 0 | 0.25 | 0.13 | 0.24 | 0.26 | 0.27 | 0.11 |
| 7 | 97041696 | rs2110784 | -1.296 | 0.537 | -0.549 | -0.156 | -0.031 | 0.049 | 0.049 | 0 | 0.73 | 0.87 | 0.66 | 0.85 | 0.66 | 0.76 |
| 7 | 97043362 | rs10230834 | -0.601 | 0.127 | 0.252 | 0.66 | 0.006 | -0.041 | -0.041 | 0 | 0.1 | 0.21 | 0.08 | 0.05 | 0.08 | 0.09 |
| 7 | 97044705 | rs7794869 | -0.546 | 0.238 | -0.125 | 1.11 | 0.004 | -0.007 | -0.007 | 0.021 | 0.27 | 0.21 | 0.24 | 0.26 | 0.27 | 0.1 |
| 7 | 97061931 | rs10233010 | -0.863 | 0.315 | 0.173 | 1.201 | -0.005 | 0.015 | 0.015 | 1.803 | 0.33 | 0.55 | 0.54 | 0.41 | 0.34 | 0.28 |
| 7 | 97074956 | rs722065 | -0.732 | 0.333 | -0.375 | 1.239 | -0.003 | 0.011 | 0.011 | 0.479 | 0.29 | 0.33 | 0.52 | 0.39 | 0.32 | 0.26 |
| 7 | 97078281 | rs11763574 | -0.624 | 0.44 | -0.328 | 1.139 | -0.002 | 0.024 | 0.024 | 0.405 | 0.28 | 0.33 | 0.51 | 0.42 | 0.32 | 0.26 |
| 7 | 97087089 | rs12704920 | -0.91 | 0.503 | 0.141 | 1.246 | 0.02 | 0.108 | 0.108 | 0 | 0.18 | 0.29 | 0.35 | 0.53 | 0.2 | 0.26 |
| 7 | 97096681 | rs1003620 | -0.972 | 0.266 | 0.016 | 0.525 | -0.002 | -0.01 | -0.01 | 0.13 | 0.42 | 0.53 | 0.55 | 0.66 | 0.36 | 0.5 |
| 7 | 97098364 | rs11973256 | -0.91 | 0.349 | -0.567 | 0.563 | -0.001 | -0.009 | -0.009 | 0 | 0.42 | 0.38 | 0.53 | 0.66 | 0.36 | 0.51 |
| 7 | 97099739 | rs10485998 | -0.935 | -0.132 | -1.096 | 0.294 | 0.01 | 0.103 | 0.103 | 0.011 | 0.23 | 0.03 | 0.18 | 0.04 | 0.21 | 0.21 |
| 7 | 97103911 | rs7800682 | 1.169 | -0.293 | -0.836 | 0.769 | 0.038 | 0.032 | 0.032 | 0.584 | 0.58 | 0.2 | 0.45 | 0.35 | 0.62 | 0.45 |
| 7 | 97107457 | rs10234240 | 1.178 | -0.236 | -0.809 | 0.779 | 0.037 | 0.033 | 0.033 | 0.615 | 0.58 | 0.16 | 0.44 | 0.35 | 0.61 | 0.44 |
| 7 | 97114775 | rs17168696 | 0.266 | -0.11 | -0.661 | 1.587 | -0.005 | -0.038 | -0.038 | 0 | 0.08 | 0 | 0.16 | 0.28 | 0.1 | 0.11 |
| 7 | 97119556 | rs6976195 | 1.257 | -0.186 | -0.441 | 1.635 | 0.01 | -0.048 | -0.048 | 4.184 | 0.67 | 0.83 | 0.52 | 0.48 | 0.65 | 0.54 |
| 7 | 97122053 | rs11981782 | 1.244 | -0.249 | -0.479 | 1.644 | 0.012 | 0.002 | 0.002 | 0.99 | 0.63 | 0.8 | 0.49 | 0.37 | 0.61 | 0.44 |
| 7 | 97136298 | rs12666768 | 0.438 | 0.254 | -0.396 | 2.395 | -0.009 | -0.048 | -0.048 | 0.006 | 0.91 | 0.99 | 0.7 | 0.63 | 0.88 | 0.84 |
| 7 | 97148119 | rs17168717 | -0.814 | 0.541 | 0.015 | 1.79 | -0.022 | -0.031 | -0.031 | 0 | 0.06 | 0.01 | 0.06 | 0.13 | 0.08 | 0.11 |
| 7 | 97160590 | rs10256461 | -1.427 | 0.808 | 0.565 | 1.651 | 0.013 | 0.069 | 0.069 | 0 | 0.19 | 0.32 | 0.23 | 0.42 | 0.19 | 0.36 |
| 7 | 97180908 | rs6465597 | -1.365 | 0.435 | 0.837 | 1.296 | 0.005 | 0.031 | 0.031 | 0 | 0.08 | 0.28 | 0.04 | 0.07 | 0.06 | 0.12 |
| 7 | 97184423 | rs17252863 | -1.289 | 0.804 | 0.186 | 1.296 | -0.025 | -0.013 | -0.013 | 0 | 0.09 | 0.02 | 0.22 | 0.08 | 0.09 | 0.13 |
| 7 | 97192321 | rs10230081 | -1.472 | 0.262 | 0.588 | 0.931 | -0.004 | -0.023 | -0.023 | 0 | 0.37 | 0.45 | 0.54 | 0.82 | 0.32 | 0.64 |
| 7 | 97193864 | rs10278295 | -1.573 | 0.188 | 0.772 | 2.11 | -0.003 | 0.01 | 0.01 | 0.283 | 0.29 | 0.36 | 0.38 | 0.51 | 0.22 | 0.45 |
| 7 | 97195851 | rs17265767 | -2.038 | -0.47 | 0.935 | 1.557 | 0.035 | 0.121 | 0.121 | 0 | 0.48 | 0.78 | 0.61 | 0.77 | 0.38 | 0.65 |
| 7 | 97196411 | rs2394715 | 1.016 | -0.479 | 1.389 | 1.553 | -0.015 | -0.075 | -0.075 | 0.463 | 0.12 | 0.32 | 0.09 | 0 | 0.07 | 0.02 |
| 7 | 97196664 | rs6415328 | -1.972 | -0.389 | 1.252 | 1.557 | 0.028 | 0.116 | 0.116 | 0 | 0.48 | 0.77 | 0.6 | 0.77 | 0.38 | 0.65 |
| 7 | 97198136 | rs4729365 | -1.103 | -0.452 | 1.819 | 1.522 | -0.007 | 0.036 | 0.036 | 0.321 | 0.28 | 0.39 | 0.37 | 0.51 | 0.2 | 0.44 |
| 7 | 97207764 | rs7809014 | -2.121 | -0.653 | 2.244 | 1.903 | -0.001 | 0 | 0 | 0 | 0.39 | 0.63 | 0.51 | 0.6 | 0.31 | 0.54 |
| 7 | 97221135 | rs10225656 | -1.118 | -0.472 | 2.157 | 2.3 | 0.003 | 0.048 | 0.048 | 0.004 | 0.1 | 0.19 | 0.15 | 0.35 | 0.14 | 0.21 |
| 7 | 97228011 | rs17168832 | -2.051 | -0.628 | 2.275 | 2.188 | 0.027 | 0.082 | 0.082 | 0 | 0.05 | 0.31 | 0.03 | 0.08 | 0.04 | 0.05 |
| 7 | 97229778 | rs10486003 | 0.161 | -0.586 | 2.061 | 2.33 | -0.028 | -0.031 | -0.031 | 0 | 0.06 | 0.01 | 0.12 | 0.27 | 0.1 | 0.16 |
| 7 | 97231644 | rs4729366 | -0.04 | -0.123 | 1.601 | 1.858 | -0.027 | -0.072 | -0.072 | 0.962 | 0.23 | 0.07 | 0.24 | 0.38 | 0.26 | 0.28 |
| 7 | 97242254 | rs10486004 | -1.249 | -0.398 | 2.302 | 1.898 | 0.018 | 0.088 | 0.088 | 0 | 0.14 | 0.3 | 0.27 | 0.25 | 0.09 | 0.28 |
| 7 | 97249954 | rs1859033 | -2.082 | -0.456 | 2.142 | 1.857 | -0.01 | 0.086 | 0.086 | 0.102 | 0.19 | 0.56 | 0.3 | 0.33 | 0.12 | 0.37 |
| 7 | 97264565 | rs10242448 | -1.365 | -0.865 | 1.757 | 1.908 | -0.013 | 0.022 | 0.022 | 0 | 0.17 | 0.26 | 0.28 | 0.33 | 0.09 | 0.27 |
| 7 | 97265029 | rs17320229 | 1.056 | -0.896 | 1.661 | 1.528 | 0.001 | -0.016 | -0.016 | 0 | 0.11 | 0.11 | 0.08 | 0 | 0.09 | 0.02 |
| 7 | 97271508 | rs974060 | -2.265 | -1.023 | 1.916 | 1.966 | 0.018 | -0.012 | -0.012 | 0.783 | 0.37 | 0.72 | 0.51 | 0.56 | 0.29 | 0.51 |
| 7 | 97273056 | rs7809389 | -2.076 | -1.028 | 2.012 | 1.967 | -0.007 | 0.017 | 0.017 | 0.06 | 0.25 | 0.33 | 0.42 | 0.56 | 0.2 | 0.49 |
| 7 | 97281912 | rs1558896 | -2.339 | -1.081 | 1.932 | 1.907 | 0.069 | -0.045 | -0.045 | 0.817 | 0.4 | 0.84 | 0.52 | 0.57 | 0.3 | 0.56 |
| 7 | 97281981 | rs12537683 | -0.375 | -1.115 | 1.965 | 2.057 | -0.001 | 0.023 | 0.023 | 0.053 | 0.08 | 0.09 | 0.14 | 0.23 | 0.1 | 0.22 |
| 7 | 97283118 | rs2058893 | -1.04 | -1.089 | 1.929 | 2.028 | -0.012 | 0.023 | 0.023 | 0 | 0.17 | 0.25 | 0.28 | 0.34 | 0.1 | 0.28 |
| 7 | 97285625 | rs4729368 | -2.1 | -1.262 | 2.209 | 2.116 | -0.033 | 0.032 | 0.032 | 0 | 0.35 | 0.58 | 0.53 | 0.57 | 0.26 | 0.51 |
| 7 | 97298821 | rs10275909 | -1.944 | -1.2 | 2.121 | 2.165 | 0.036 | -0.04 | -0.04 | 0 | 0.31 | 0.74 | 0.31 | 0.57 | 0.22 | 0.52 |
| 7 | 97298855 | rs10246118 | -0.475 | -1.025 | 2.076 | 2.234 | -0.017 | 0.034 | 0.034 | 0 | 0.19 | 0.36 | 0.23 | 0.26 | 0.19 | 0.24 |
| 7 | 97303273 | rs2216369 | -0.257 | -1.039 | 1.397 | 2.368 | -0.015 | 0.059 | 0.059 | 0 | 0.08 | 0.17 | 0.08 | 0.29 | 0.03 | 0.19 |
| 7 | 97312449 | rs7796445 | -0.045 | -0.788 | 1.283 | 1.792 | -0.023 | 0.044 | 0.044 | 0 | 0.15 | 0.24 | 0.17 | 0.26 | 0.1 | 0.2 |
| 7 | 97318147 | rs759658 | 0.184 | -0.624 | 0.6 | 1.683 | -0.005 | -0.055 | -0.055 | 0 | 0.1 | 0.01 | 0.27 | 0.27 | 0.09 | 0.17 |
| 7 | 97324281 | rs10280384 | -0.151 | -0.894 | 1.591 | 1.675 | -0.054 | 0.088 | 0.088 | 0 | 0.15 | 0.47 | 0.14 | 0.21 | 0.06 | 0.24 |
| 7 | 97329406 | rs1588422 | 0.002 | -0.379 | 0.982 | 0.737 | 0.002 | -0.003 | -0.003 | 0.78 | 0.27 | 0.39 | 0.29 | 0.25 | 0.26 | 0.27 |
| 7 | 97332781 | rs2665460 | 0.472 | -0.098 | 0.782 | 0.154 | 0.013 | -0.046 | -0.046 | 0 | 0.27 | 0.31 | 0.2 | 0.07 | 0.34 | 0.27 |
| 7 | 97351623 | rs1229439 | 1.077 | -0.134 | 0.833 | 0.136 | 0.002 | -0.009 | -0.009 | 0 | 0.23 | 0.2 | 0.18 | 0.07 | 0.34 | 0.17 |
| 7 | 97352893 | rs1237992 | 0.461 | -0.219 | 1.05 | 0.238 | 0.023 | -0.052 | -0.052 | 0 | 0.27 | 0.33 | 0.24 | 0.08 | 0.33 | 0.25 |
| 7 | 97365931 | rs3779470 | 1.711 | -0.26 | 1.151 | 0.393 | -0.006 | -0.012 | -0.012 | 0.146 | 0.16 | 0.2 | 0.16 | 0.27 | 0.17 | 0.22 |
| 7 | 97366145 | rs7795107 | 2.295 | -0.31 | 1.148 | 0.337 | 0.012 | 0.034 | 0.034 | 0 | 0.09 | 0.03 | 0.06 | 0.01 | 0.06 | 0.01 |
| 7 | 97369391 | rs12532490 | 1.747 | -0.164 | 1.185 | 0.427 | -0.006 | -0.012 | -0.012 | 0.06 | 0.16 | 0.2 | 0.16 | 0.27 | 0.17 | 0.22 |
| 7 | 97369496 | rs12534885 | 1.767 | -0.162 | 1.165 | 0.427 | -0.009 | -0.017 | -0.017 | 0 | 0.16 | 0.18 | 0.15 | 0.27 | 0.17 | 0.22 |
| 7 | 97372593 | rs10247565 | -0.251 | 0.064 | 1.012 | 0.354 | -0.005 | 0.031 | 0.031 | 0 | 0.47 | 0.67 | 0.41 | 0.34 | 0.51 | 0.37 |
| 7 | 97379204 | rs6945396 | -0.633 | 0.114 | 0.708 | 0.297 | 0.003 | 0.047 | 0.047 | 0 | 0.49 | 0.73 | 0.43 | 0.35 | 0.53 | 0.43 |
| 7 | 97385344 | rs10486008 | 1.279 | 0.141 | 0.87 | 0.359 | 0.013 | 0.059 | 0.059 | 0 | 0.19 | 0.39 | 0.17 | 0.28 | 0.18 | 0.23 |

| | | | | | | | | | | | | | | | | |
|---|---|---|---|---|---|---|---|---|---|---|---|---|---|---|---|---|
| 7 | 97385511 | rs2017923 | 0.207 | 0.152 | 0.752 | 0.355 | -0.015 | 0.003 | 0.003 | 0 | 0.3 | 0.5 | 0.23 | 0.29 | 0.25 | 0.27 |
| 7 | 97386915 | rs7808126 | 1.052 | 0.167 | 0.757 | 0.121 | -0.022 | 0.039 | 0.039 | 0 | 0.21 | 0.1 | 0.18 | 0.07 | 0.29 | 0.19 |
| 7 | 97388270 | rs10254181 | -0.917 | 0.085 | 0.446 | 0.144 | 0.004 | -0.004 | -0.004 | 0.018 | 0.67 | 0.74 | 0.69 | 0.65 | 0.7 | 0.7 |
| 7 | 97389030 | rs10085387 | 1.041 | 0.355 | 0.43 | -0.004 | 0.011 | -0.024 | -0.024 | 2.884 | 0.2 | 0.21 | 0.19 | 0.08 | 0.28 | 0.15 |
| 7 | 97403141 | rs1510306 | 0.677 | -0.211 | 0.642 | -0.079 | -0.046 | -0.078 | -0.078 | 0 | 0.15 | 0.38 | 0.09 | 0.01 | 0.07 | 0.05 |
| 7 | 97404277 | rs11763921 | 0.902 | -0.233 | 0.731 | -0.089 | -0.039 | -0.071 | -0.071 | 0 | 0.14 | 0.38 | 0.09 | 0.01 | 0.07 | 0.01 |
| 7 | 97404380 | rs11767488 | 0.902 | -0.233 | 0.731 | -0.089 | -0.039 | -0.071 | -0.071 | 0.01 | 0.14 | 0.38 | 0.09 | 0.01 | 0.07 | 0.01 |
| 7 | 97406045 | rs1229569 | -1.495 | -0.217 | 0.319 | -0.276 | -0.038 | 0.044 | 0.044 | 0.011 | 0.8 | 0.91 | 0.83 | 0.92 | 0.74 | 0.87 |
| 7 | 97414110 | rs460 | -0.103 | -0.893 | 0.018 | -0.705 | -0.038 | -0.081 | -0.081 | 0 | 0.34 | 0.48 | 0.22 | 0.11 | 0.2 | 0.27 |
| 7 | 97422926 | rs1229542 | 0.828 | -0.515 | 0.108 | -0.452 | -0.013 | -0.037 | -0.037 | 1.73 | 0.17 | 0.48 | 0.13 | 0.09 | 0.13 | 0.04 |
| 7 | 97423394 | rs1229540 | -1.432 | -0.376 | 0.129 | -0.15 | -0.041 | 0.005 | 0.005 | 0 | 0.66 | 0.8 | 0.68 | 0.69 | 0.55 | 0.67 |
| 7 | 97425721 | rs1229568 | 0.967 | -0.229 | 0.17 | -0.046 | -0.009 | -0.033 | -0.033 | 1.157 | 0.17 | 0.48 | 0.13 | 0.09 | 0.13 | 0.04 |
| 7 | 97432533 | rs1229553 | 1.296 | -0.192 | 0.061 | -0.029 | -0.006 | -0.015 | -0.015 | 0.021 | 0.15 | 0.26 | 0.12 | 0.09 | 0.13 | 0.04 |
| 7 | 97446702 | rs2136704 | 1.106 | -0.172 | -0.258 | 0.321 | -0.033 | -0.062 | -0.062 | 0 | 0.18 | 0.06 | 0.31 | 0.37 | 0.22 | 0.31 |
| 7 | 97447728 | rs1912445 | 0.9 | -0.215 | -0.044 | 0.302 | -0.004 | -0.008 | -0.008 | 0.124 | 0.19 | 0.16 | 0.32 | 0.37 | 0.22 | 0.31 |
| 7 | 97454980 | rs2665441 | 0.087 | -0.295 | 0.03 | 0.19 | 0.003 | -0.014 | -0.014 | 0 | 0.38 | 0.26 | 0.45 | 0.46 | 0.36 | 0.52 |
| 7 | 97461115 | rs2665466 | -1.362 | -0.276 | 0.186 | 0.197 | -0.005 | 0.002 | 0.002 | 0 | 0.64 | 0.69 | 0.67 | 0.7 | 0.55 | 0.67 |
| 7 | 97464212 | rs17278667 | 1.664 | -0.158 | -0.168 | 0.403 | -0.01 | -0.038 | -0.038 | 0.257 | 0.15 | 0.06 | 0.27 | 0.29 | 0.16 | 0.28 |
| 7 | 97464809 | rs10269471 | 1.356 | -0.234 | 0.079 | 0.453 | 0 | -0.003 | -0.003 | 0 | 0.12 | 0.11 | 0.13 | 0.22 | 0.12 | 0.1 |
| 7 | 97465888 | rs12668997 | 1.406 | -0.303 | -0.041 | 0.239 | -0.006 | 0.058 | 0.058 | 2.385 | 0.18 | 0.03 | 0.12 | 0.1 | 0.14 | 0.21 |
| 7 | 97471538 | rs7800746 | 2.155 | -0.086 | -0.003 | 0.27 | -0.053 | -0.054 | -0.054 | 0 | 0.14 | 0.02 | 0.14 | 0.22 | 0.18 | 0.19 |
| 7 | 97476155 | rs7798362 | 0.142 | -0.394 | 0.607 | -0.007 | -0.006 | -0.011 | -0.011 | 0.217 | 0.24 | 0.3 | 0.18 | 0.19 | 0.2 | 0.29 |
| 7 | 97478318 | rs10486010 | 0.773 | -0.396 | 0.497 | -0.009 | -0.007 | -0.017 | -0.017 | 0 | 0.2 | 0.24 | 0.14 | 0.11 | 0.14 | 0.22 |
| 7 | 97480919 | rs10447748 | -0.334 | -0.44 | 0.532 | -0.047 | -0.012 | -0.021 | -0.021 | 1.393 | 0.35 | 0.44 | 0.24 | 0.2 | 0.27 | 0.3 |
| 7 | 97485402 | rs12113193 | 0.018 | -0.548 | 0.506 | -0.175 | -0.021 | -0.049 | -0.049 | 0 | 0.32 | 0.59 | 0.22 | 0.11 | 0.21 | 0.23 |
| 7 | 97488733 | rs7797354 | 0.364 | -0.388 | 0.132 | 0.006 | -0.009 | -0.019 | -0.019 | 0 | 0.32 | 0.29 | 0.33 | 0.53 | 0.36 | 0.36 |
| 7 | 97489701 | rs17169185 | 1.785 | -0.497 | 0.323 | -0.07 | -0.018 | -0.025 | -0.025 | 0 | 0.11 | 0.2 | 0.07 | 0.02 | 0.07 | 0.03 |
| 7 | 97489911 | rs13239311 | 1.174 | -0.364 | -0.09 | -0.044 | 0.014 | 0.052 | 0.052 | 4.077 | 0.18 | 0.07 | 0.11 | 0.1 | 0.13 | 0.22 |
| 7 | 97493340 | rs4727377 | 0.832 | -0.14 | -0.382 | -0.036 | -0.037 | -0.028 | -0.028 | 3.032 | 0.06 | 0.01 | 0.04 | 0.1 | 0.1 | 0.05 |
| 7 | 97494722 | rs7810919 | -0.899 | -0.228 | -0.636 | -0.071 | -0.137 | 0.015 | 0.015 | 0.187 | 0.73 | 0.96 | 0.78 | 0.84 | 0.62 | 0.84 |
| 7 | 97496179 | rs10263979 | 0.803 | -0.185 | -0.209 | -0.13 | -0.001 | -0.003 | -0.003 | 0 | 0.28 | 0.29 | 0.3 | 0.43 | 0.31 | 0.3 |
| 7 | 97499082 | rs3735557 | -0.006 | -0.117 | -0.167 | -0.09 | 0.005 | -0.003 | -0.003 | 0 | 0.47 | 0.41 | 0.41 | 0.53 | 0.43 | 0.51 |
| 7 | 97502737 | rs1974562 | -1.311 | 0.547 | -0.358 | 0.136 | -0.083 | 0.055 | 0.055 | 0 | 0.8 | 0.98 | 0.83 | 0.93 | 0.73 | 0.88 |
| 7 | 97503140 | rs3757674 | -0.091 | 0.654 | 0.657 | 0.61 | -0.002 | -0.011 | -0.011 | 1.064 | 0.28 | 0.27 | 0.3 | 0.42 | 0.3 | 0.28 |
| 7 | 97595184 | rs9692345 | 0.223 | 0.945 | 0.498 | 0.432 | 0.002 | 0.007 | 0.007 | 0 | 0.22 | 0.23 | 0.22 | 0.35 | 0.23 | 0.2 |
| 7 | 97595215 | rs11983172 | -0.256 | 0.559 | -0.333 | 0.108 | 0.003 | 0.128 | 0.128 | 0.127 | 0.24 | 0.06 | 0.16 | 0 | 0.23 | 0.2 |
| 7 | 97602656 | rs10228951 | -1.264 | 0.723 | -0.113 | 0.368 | 0.004 | -0.006 | -0.006 | 0.011 | 0.56 | 0.62 | 0.46 | 0.58 | 0.55 | 0.47 |
| 7 | 97611196 | rs10240723 | -0.606 | 0.937 | -0.391 | 0.342 | 0.002 | 0.003 | 0.003 | 0 | 0.32 | 0.3 | 0.28 | 0.58 | 0.36 | 0.35 |
| 7 | 97612024 | rs17169233 | -0.116 | 1.057 | -0.828 | 0.378 | -0.028 | -0.029 | -0.029 | 5.178 | 0.24 | 0.17 | 0.25 | 0.36 | 0.29 | 0.22 |
| 7 | 97620691 | rs13438327 | 0.461 | 0.78 | -0.779 | 0.397 | -0.003 | -0.027 | -0.027 | 4.42 | 0.15 | 0.19 | 0.09 | 0.01 | 0.09 | 0.15 |
| 7 | 97623394 | rs7799577 | 0.768 | 1.613 | -0.761 | 1.137 | -0.061 | -0.019 | -0.019 | 0 | 0.66 | 0.84 | 0.68 | 0.64 | 0.57 | 0.72 |
| 7 | 97626161 | rs1614702 | -0.723 | 1.087 | -0.408 | 1.358 | -0.004 | 0.003 | 0.003 | 1.298 | 0.29 | 0.62 | 0.27 | 0.37 | 0.25 | 0.28 |
| 7 | 97635841 | rs12672767 | -1.195 | 2.176 | -0.199 | 2.143 | 0.013 | 0.042 | 0.042 | 1.699 | 0.15 | 0.18 | 0.34 | 0.56 | 0.27 | 0.39 |
| 7 | 97644627 | rs1964235 | -0.878 | 2.479 | -0.669 | 1.897 | -0.022 | -0.015 | -0.015 | 0.404 | 0.13 | 0.05 | 0.15 | 0.13 | 0.14 | 0.16 |
| 7 | 97647191 | rs1799041 | -0.916 | 2.679 | -0.203 | 1.551 | 0.017 | -0.026 | -0.026 | 0 | 0.1 | 0.21 | 0.07 | 0.05 | 0.14 | 0.18 |
| 7 | 97653811 | rs740209 | -0.954 | 2.537 | 0.415 | 2.299 | 0.025 | 0.079 | 0.079 | 0.09 | 0.15 | 0.22 | 0.3 | 0.44 | 0.26 | 0.3 |
| 7 | 97676433 | rs1799053 | 1.02 | 1.95 | 0.355 | 2.224 | 0.002 | -0.004 | -0.004 | 0 | 0.86 | 0.88 | 0.79 | 0.83 | 0.89 | 0.84 |
| 7 | 97677267 | rs3857802 | 0.982 | 1.954 | 0.357 | 2.226 | 0.002 | -0.003 | -0.003 | 0 | 0.87 | 0.88 | 0.79 | 0.83 | 0.89 | 0.84 |
| 7 | 97684468 | rs12538837 | -1.778 | 2.353 | 0.727 | 3.215 | -0.011 | -0.007 | -0.007 | 0 | 0.15 | 0.12 | 0.21 | 0.44 | 0.27 | 0.26 |
| 7 | 97688936 | rs7777145 | -1.243 | 1.959 | 1.318 | 3.339 | -0.019 | 0.058 | 0.058 | 1.198 | 0.14 | 0.23 | 0.26 | 0.26 | 0.11 | 0.25 |
| 7 | 97691889 | rs1799049 | -0.977 | 1.901 | 1.324 | 3.253 | -0.023 | 0.006 | 0.006 | 0.837 | 0.1 | 0.15 | 0.1 | 0.09 | 0.06 | 0.17 |
| 7 | 97699292 | rs17169261 | -1.218 | 2.075 | 1.296 | 3.546 | -0.001 | 0.019 | 0.019 | 0 | 0.1 | 0.09 | 0.14 | 0.34 | 0.14 | 0.12 |
| 7 | 97731700 | rs10282336 | -1.039 | 1.998 | 1.416 | 3.4 | -0.023 | -0.005 | -0.005 | 0 | 0.1 | 0.14 | 0.1 | 0.08 | 0.05 | 0.17 |
| 7 | 97739317 | rs2394824 | -1.6 | 1.911 | 2.189 | 3.512 | -0.001 | -0.001 | -0.001 | 0 | 0.15 | 0.18 | 0.22 | 0.19 | 0.22 | 0.42 |
| 7 | 97741795 | rs10242903 | -2.979 | 1.722 | 2.984 | 3.899 | 0.032 | 0.054 | 0.054 | 3.081 | 0.3 | 0.54 | 0.35 | 0.41 | 0.36 | 0.56 |
| 7 | 97758335 | rs16869462 | -1.618 | 1.194 | 3.219 | 3.916 | -0.019 | 0.233 | 0.233 | 0.069 | 0.13 | 0.37 | 0.31 | 0.42 | 0.09 | 0.24 |
| 7 | 97764162 | rs10232153 | -0.975 | 1.175 | 3.208 | 3.757 | -0.039 | 0 | 0 | 0.373 | 0.11 | 0.31 | 0.11 | 0.08 | 0.05 | 0.18 |
| 7 | 97774300 | rs10275267 | -1.253 | 1.192 | 3.229 | 3.804 | -0.002 | 0 | 0 | 0 | 0.11 | 0.1 | 0.1 | 0.2 | 0.14 | 0.14 |
| 7 | 97781645 | rs10270280 | -0.919 | 1.177 | 3.269 | 3.774 | -0.037 | 0.003 | 0.003 | 0.389 | 0.1 | 0.3 | 0.11 | 0.08 | 0.05 | 0.18 |
| 7 | 97796093 | rs10248229 | -2.158 | 1.172 | 3.21 | 3.785 | 0.007 | 0.079 | 0.079 | 0.718 | 0.15 | 0.35 | 0.13 | 0.2 | 0.14 | 0.13 |
| 7 | 97835640 | rs2291748 | -1.874 | 1.218 | 3.414 | 3.949 | 0.001 | 0.039 | 0.039 | 0.139 | 0.13 | 0.2 | 0.21 | 0.21 | 0.14 | 0.2 |
| 7 | 97838142 | rs13221415 | -1.002 | 1.226 | 3.461 | 3.957 | -0.033 | 0.009 | 0.009 | 0.644 | 0.11 | 0.34 | 0.1 | 0.08 | 0.05 | 0.19 |
| 7 | 97839140 | rs17169462 | -1.381 | 1.247 | 2.921 | 3.751 | 0 | 0.014 | 0.014 | 0.212 | 0.12 | 0.12 | 0.21 | 0.21 | 0.14 | 0.2 |
| 7 | 97841169 | rs34228515 | -0.478 | 1.22 | 2.725 | 3.366 | 0.033 | 0.048 | 0.048 | 0.681 | 0.08 | 0.02 | 0.05 | 0 | 0.04 | 0.09 |
| 7 | 97842668 | rs754464 | -1.851 | 1.288 | 2.777 | 3.543 | -0.004 | 0.008 | 0.008 | 0.285 | 0.14 | 0.12 | 0.4 | 0.55 | 0.19 | 0.26 |
| 7 | 97844578 | rs3148 | -1.397 | 1.347 | 3.055 | 2.885 | -0.023 | -0.019 | -0.019 | 0.136 | 0.12 | 0.27 | 0.14 | 0.07 | 0.08 | 0.19 |
| 7 | 97853184 | rs3816943 | -1.198 | 1.575 | 3.209 | 3.214 | 0.091 | 0.134 | 0.134 | 0 | 0.1 | 0.24 | 0.29 | 0.26 | 0.19 | 0.11 |
| 7 | 97854393 | rs2291745 | -2.114 | 1.924 | 3.188 | 3.371 | -0.017 | -0.04 | -0.04 | 0 | 0.21 | 0.16 | 0.2 | 0.42 | 0.23 | 0.46 |
| 7 | 97863145 | rs2291750 | -2.11 | 1.881 | 3.333 | 3.351 | -0.005 | -0.004 | -0.004 | 0 | 0.21 | 0.2 | 0.19 | 0.43 | 0.23 | 0.45 |

| | | | | | | | | | | | | | | | | |
|---|---|---|---|---|---|---|---|---|---|---|---|---|---|---|---|---|
| 7 | 97866291 | rs3735258 | -1.509 | 1.907 | 3.384 | 3.292 | 0.087 | 0.166 | 0.166 | 0 | 0.12 | 0.18 | 0.32 | 0.45 | 0.26 | 0.28 |
| 7 | 97868033 | rs17169479 | -1.965 | 1.656 | 3.124 | 3.08 | 0 | -0.005 | -0.005 | 0 | 0.18 | 0.18 | 0.18 | 0.23 | 0.15 | 0.29 |
| 7 | 97921533 | rs1132971 | 2.391 | 0.631 | 2.798 | 2.264 | -0.016 | 0.073 | 0.073 | 0.536 | 0.43 | 0.22 | 0.35 | 0.27 | 0.48 | 0.16 |
| 7 | 97922851 | rs12728 | 2.443 | 0.617 | 2.785 | 2.257 | -0.04 | 0.09 | 0.09 | 0.649 | 0.43 | 0.14 | 0.35 | 0.27 | 0.48 | 0.16 |
| 7 | 97925240 | rs6970801 | 2.388 | 0.602 | 2.788 | 2.261 | -0.019 | 0.073 | 0.073 | 0.536 | 0.43 | 0.22 | 0.35 | 0.27 | 0.48 | 0.16 |
| 7 | 97925382 | rs6970990 | 2.437 | 0.602 | 2.787 | 2.261 | -0.039 | 0.089 | 0.089 | 0.589 | 0.43 | 0.14 | 0.35 | 0.27 | 0.48 | 0.16 |
| 7 | 97933601 | rs1045916 | 1.702 | 0.132 | 2.797 | 2.146 | 0.034 | 0.024 | 0.024 | 0 | 0.54 | 0.33 | 0.68 | 0.69 | 0.75 | 0.44 |
| 7 | 97938443 | rs1006169 | -0.863 | 0.381 | 2.812 | 2.255 | 0.064 | 0.168 | 0.168 | 0 | 0.12 | 0.18 | 0.35 | 0.44 | 0.25 | 0.28 |
| 7 | 97946902 | rs7797943 | 2.662 | 0.335 | 2.623 | 2.175 | -0.015 | 0.064 | 0.064 | 0.23 | 0.43 | 0.25 | 0.35 | 0.26 | 0.47 | 0.16 |
| 7 | 97957517 | rs7807555 | 2.756 | 0.337 | 2.439 | 2.176 | -0.04 | 0.087 | 0.087 | 0.338 | 0.43 | 0.14 | 0.34 | 0.26 | 0.48 | 0.16 |
| 7 | 97957622 | rs7807455 | 2.756 | 0.337 | 2.439 | 2.176 | -0.04 | 0.087 | 0.087 | 0.338 | 0.43 | 0.15 | 0.34 | 0.26 | 0.48 | 0.16 |
| 7 | 97996593 | rs4236537 | -1.159 | 0.392 | 2.625 | 2.258 | 0.058 | 0.16 | 0.16 | 0 | 0.14 | 0.21 | 0.35 | 0.45 | 0.25 | 0.29 |
| 7 | 98005398 | rs2107717 | 2.072 | 0.399 | 2.425 | 2.44 | 0.053 | 0.028 | 0.028 | 0.088 | 0.52 | 0.22 | 0.6 | 0.44 | 0.68 | 0.29 |
| 7 | 98005462 | rs2107718 | 0.24 | 0.407 | 2.419 | 2.443 | -0.005 | -0.005 | -0.005 | 0 | 0.08 | 0.06 | 0.25 | 0.17 | 0.2 | 0.13 |
| 7 | 98013107 | rs10953259 | 1.67 | 0.233 | 2.623 | 1.748 | 0 | -0.003 | -0.003 | 0 | 0.79 | 0.78 | 0.8 | 0.76 | 0.82 | 0.69 |
| 7 | 98013839 | rs12539761 | -0.552 | 0.432 | 2.797 | 1.776 | 0.003 | 0.039 | 0.039 | 0 | 0.12 | 0.09 | 0.35 | 0.44 | 0.25 | 0.28 |
| 7 | 98016530 | rs1635613 | -0.168 | 0.437 | 2.376 | 1.772 | 0.012 | 0.008 | 0.008 | 0.071 | 0.22 | 0.21 | 0.09 | 0.05 | 0.09 | 0.26 |
| 7 | 98021211 | rs9649213 | -1.484 | 0.429 | 2.391 | 1.785 | 0.001 | -0.008 | -0.008 | 0 | 0.36 | 0.61 | 0.45 | 0.5 | 0.34 | 0.53 |
| 7 | 98026236 | rs1688604 | -0.474 | -0.301 | 1.864 | 0.863 | 0.012 | 0.009 | 0.009 | 0.06 | 0.22 | 0.21 | 0.09 | 0.05 | 0.11 | 0.27 |
| 7 | 98039846 | rs1688605 | 1.745 | -0.232 | 2.048 | 1.047 | 0.011 | 0.05 | 0.05 | 0.094 | 0.85 | 0.61 | 0.86 | 0.77 | 0.9 | 0.76 |
| 7 | 98042527 | rs7783611 | 1.755 | -0.561 | 1.626 | 0.693 | 0.005 | 0.008 | 0.008 | 1.444 | 0.54 | 0.37 | 0.73 | 0.7 | 0.67 | 0.55 |
| 7 | 98051932 | rs2023971 | -1.352 | -0.465 | 1.27 | 0.742 | -0.008 | 0.006 | 0.006 | 0 | 0.1 | 0.1 | 0.08 | 0.06 | 0.1 | 0.12 |
| 7 | 98054504 | rs10240312 | 1.438 | -0.474 | 1.286 | 0.745 | 0.032 | -0.017 | -0.017 | 0 | 0.87 | 0.62 | 0.9 | 0.92 | 0.89 | 0.88 |
| 7 | 98092368 | rs903453 | 1.633 | -1.072 | 0.396 | 0.748 | -0.008 | -0.013 | -0.013 | 0 | 0.91 | 0.88 | 0.98 | 0.95 | 0.93 | 0.79 |
| 7 | 98097170 | rs705321 | -1.428 | -1.439 | 0.072 | 0.667 | 0.025 | 0.025 | 0.025 | 4.721 | 0.39 | 0.29 | 0.16 | 0.2 | 0.2 | 0.46 |
| 7 | 98106228 | rs17147641 | -1.058 | -1.138 | -0.469 | 0.665 | 0.062 | 0.079 | 0.079 | 3.602 | 0.33 | 0.18 | 0.12 | 0.16 | 0.17 | 0.36 |
| 7 | 98124854 | rs4144071 | -0.935 | -0.506 | -0.215 | 1.225 | -0.002 | -0.016 | -0.016 | 0 | 0.25 | 0.21 | 0.23 | 0.48 | 0.22 | 0.32 |
| 7 | 98125134 | rs4729451 | -0.813 | -1.072 | 0.498 | 1.234 | -0.018 | 0.195 | 0.195 | 0 | 0.11 | 0.25 | 0.25 | 0.39 | 0.09 | 0.15 |
| 7 | 98125908 | rs10272977 | -0.278 | -1.292 | 0.358 | 0.594 | -0.012 | -0.033 | -0.033 | 0.062 | 0.06 | 0.19 | 0.04 | 0 | 0.02 | 0.01 |
| 7 | 98128828 | rs2394868 | -0.115 | -1.173 | -0.174 | 0.797 | 0.002 | 0.058 | 0.058 | 12.591 | 0.2 | 0.1 | 0.14 | 0.06 | 0.16 | 0.12 |
| 7 | 98131499 | rs2625403 | -1.571 | -1.46 | 0.03 | 0.911 | -0.073 | 0.094 | 0.094 | 0 | 0.19 | 0.45 | 0.3 | 0.36 | 0.11 | 0.35 |
| 7 | 98143429 | rs2625400 | -0.768 | -0.862 | -1.201 | 0.701 | 0.009 | 0.012 | 0.012 | 5.817 | 0.37 | 0.66 | 0.35 | 0.73 | 0.26 | 0.5 |
| 7 | 98155105 | rs869029 | 0.732 | -0.391 | -2.383 | 0.363 | -0.072 | 0.047 | 0.047 | 0.099 | 0.42 | 0.08 | 0.57 | 0.25 | 0.58 | 0.29 |
| 7 | 98157318 | rs817764 | 0.665 | 0.392 | -2.076 | 0.826 | -0.073 | -0.096 | -0.096 | 0 | 0.78 | 0.94 | 0.75 | 0.37 | 0.72 | 0.5 |
| 7 | 98157619 | rs2711167 | 0.669 | 0.139 | -1.864 | 0.1 | -0.004 | 0.037 | 0.037 | 0.365 | 0.61 | 0.81 | 0.62 | 0.31 | 0.64 | 0.43 |
| 7 | 98162894 | rs705323 | 0.11 | 0.376 | -1.767 | -0.337 | 0.005 | 0.056 | 0.056 | 3.186 | 0.56 | 0.78 | 0.6 | 0.76 | 0.6 | 0.59 |
| 7 | 98163939 | rs817759 | 0.273 | 0.396 | -1.489 | -0.277 | 0 | 0.01 | 0.01 | 2.757 | 0.89 | 0.93 | 0.92 | 0.95 | 0.91 | 0.85 |
| 7 | 98164735 | rs817757 | -0.998 | -0.051 | -0.663 | -0.494 | -0.028 | -0.063 | -0.063 | 0 | 0.17 | 0.65 | 0.14 | 0.04 | 0.1 | 0.12 |
| 7 | 98183565 | rs705314 | -0.17 | 0.378 | -0.675 | -0.579 | -0.002 | 0 | 0 | 0.255 | 0.61 | 0.6 | 0.67 | 0.83 | 0.53 | 0.65 |
| 7 | 98187498 | rs17161409 | -0.177 | 0.118 | -0.922 | -0.105 | 0 | -0.006 | -0.006 | 0 | 0.16 | 0.12 | 0.12 | 0.25 | 0.13 | 0.27 |
| 7 | 98190906 | rs17161411 | 0.753 | -0.218 | -1.06 | -0.012 | -0.03 | -0.106 | -0.106 | 0.376 | 0.38 | 0.06 | 0.38 | 0.5 | 0.35 | 0.29 |
| 7 | 98192346 | rs2132044 | -0.232 | -0.223 | -0.869 | 0.047 | 0.002 | -0.01 | -0.01 | 0 | 0.16 | 0.12 | 0.11 | 0.25 | 0.13 | 0.28 |
| 7 | 98197405 | rs705330 | -1.101 | 0.276 | -0.106 | -0.144 | 0.042 | 0.013 | 0.013 | 0.759 | 0.11 | 0.6 | 0.27 | 0.1 | 0.09 | 0.15 |
| 7 | 98201136 | rs861006 | -1.521 | 0.173 | -0.613 | -0.253 | -0.089 | 0.077 | 0.077 | 0.251 | 0.71 | 0.95 | 0.83 | 0.89 | 0.64 | 0.69 |
| 7 | 98203886 | rs817749 | -1.516 | 0.099 | -0.607 | -0.268 | -0.09 | 0.08 | 0.08 | 0.212 | 0.71 | 0.95 | 0.83 | 0.89 | 0.64 | 0.71 |
| 7 | 98205072 | rs1494810 | 0.026 | 0.077 | -0.246 | -0.305 | 0.033 | -0.026 | -0.026 | 0.139 | 0.07 | 0.33 | 0.08 | 0 | 0.05 | 0.02 |
| 7 | 98205391 | rs12536662 | -0.117 | 0.042 | -0.761 | -0.28 | -0.001 | 0 | 0 | 0 | 0.09 | 0.08 | 0.09 | 0.06 | 0.11 | 0.14 |
| 7 | 98210558 | rs6465691 | 0.04 | 0.041 | -0.844 | -0.21 | 0.005 | 0.058 | 0.058 | 0.486 | 0.53 | 0.27 | 0.51 | 0.72 | 0.48 | 0.52 |
| 7 | 98225036 | rs817771 | -1.887 | -0.852 | -1.301 | -0.192 | 0.034 | -0.091 | -0.091 | 0 | 0.43 | 0.97 | 0.5 | 0.41 | 0.32 | 0.45 |
| 7 | 98226627 | rs705337 | -1.494 | -0.686 | -0.812 | -0.351 | 0.008 | 0.028 | 0.028 | 0.374 | 0.21 | 0.84 | 0.27 | 0.05 | 0.19 | 0.17 |
| 7 | 98230657 | rs817765 | -1.666 | -0.516 | -0.973 | 0.162 | 0.001 | -0.064 | -0.064 | 0 | 0.39 | 0.92 | 0.48 | 0.39 | 0.37 | 0.43 |
| 7 | 98238960 | rs13247122 | 0.668 | -0.366 | -1.265 | -0.089 | -0.023 | 0.053 | 0.053 | 0.138 | 0.09 | 0.01 | 0.07 | 0 | 0.1 | 0.05 |
| 7 | 98244565 | rs1681247 | 0.758 | -0.666 | -0.596 | 0.026 | -0.004 | -0.027 | -0.027 | 0 | 0.09 | 0.11 | 0.06 | 0 | 0.05 | 0.01 |
| 7 | 98257085 | rs705316 | -1.814 | -0.708 | -0.582 | 0.013 | -0.025 | -0.088 | -0.088 | 1.747 | 0.4 | 0.94 | 0.5 | 0.39 | 0.38 | 0.44 |
| 7 | 98258608 | rs2642895 | 0.052 | -0.701 | 0.176 | 0.094 | 0.001 | 0.058 | 0.058 | 9.251 | 0.11 | 0.25 | 0.12 | 0.22 | 0.08 | 0.11 |
| 7 | 98258736 | rs705315 | -0.028 | -0.689 | 0.275 | 0.083 | -0.012 | 0.067 | 0.067 | 9.547 | 0.12 | 0.28 | 0.1 | 0.21 | 0.07 | 0.11 |
| 7 | 98262328 | rs1115147 | 2.004 | -0.762 | 0.706 | 0.053 | 0.022 | -0.031 | -0.031 | 0.198 | 0.59 | 0.17 | 0.51 | 0.61 | 0.62 | 0.57 |
| 7 | 98269772 | rs10235866 | -0.746 | -0.811 | 0.867 | 0.004 | 0 | 0 | 0 | 0 | 0.19 | 0.15 | 0.28 | 0.18 | 0.2 | 0.24 |
| 7 | 98269815 | rs6979846 | 2.392 | -0.811 | 0.827 | 0.002 | 0.01 | -0.052 | -0.052 | 1.675 | 0.59 | 0.13 | 0.51 | 0.61 | 0.63 | 0.57 |
| 7 | 98273043 | rs17530700 | 0.479 | -0.719 | 0.83 | 0.018 | -0.018 | 0.002 | 0.002 | 0.163 | 0.1 | 0.02 | 0.1 | 0.06 | 0.14 | 0.16 |
| 7 | 98273346 | rs879187 | -1.054 | -0.729 | 1.224 | 0.018 | 0.014 | -0.044 | -0.044 | 0 | 0.18 | 0.36 | 0.18 | 0.06 | 0.23 | 0.24 |
| 7 | 98282745 | rs11760352 | 1.848 | -0.735 | 1.127 | 0.023 | -0.001 | -0.002 | -0.002 | 0 | 0.7 | 0.37 | 0.69 | 0.73 | 0.69 | 0.65 |
| 7 | 98282950 | rs13221723 | 1.854 | -0.725 | 1.137 | 0.028 | -0.001 | -0.001 | -0.001 | 0 | 0.7 | 0.33 | 0.69 | 0.73 | 0.69 | 0.65 |
| 7 | 98291653 | rs6964898 | 0.933 | -0.67 | 1.103 | 0.093 | 0.004 | 0.123 | 0.123 | 4.041 | 0.83 | 0.58 | 0.81 | 0.68 | 0.86 | 0.84 |
| 7 | 98293467 | rs4729469 | 0.479 | -0.719 | 1.122 | 0.108 | -0.019 | 0.103 | 0.103 | 4.521 | 0.86 | 0.7 | 0.84 | 0.68 | 0.91 | 0.86 |
| 7 | 98294265 | rs17440093 | 0.359 | -0.745 | 1.136 | 0.073 | -0.006 | 0.004 | 0.004 | 0.154 | 0.11 | 0.05 | 0.1 | 0.06 | 0.15 | 0.16 |
| 7 | 98296161 | rs10256945 | 1.381 | -0.766 | 1.19 | 0.089 | -0.006 | 0.048 | 0.048 | 4.641 | 0.74 | 0.64 | 0.74 | 0.62 | 0.76 | 0.7 |
| 7 | 98299157 | rs12704998 | -0.349 | -0.551 | 0.747 | -0.129 | -0.01 | 0.013 | 0.013 | 0 | 0.3 | 0.21 | 0.28 | 0.09 | 0.37 | 0.27 |
| 7 | 98301926 | rs4729472 | 0.264 | -0.65 | 0.768 | -0.165 | -0.023 | -0.053 | -0.053 | 1.235 | 0.12 | 0.29 | 0.18 | 0.01 | 0.1 | 0.1 |

| | | | | | | | | | | | | | | | | |
|---|---|---|---|---|---|---|---|---|---|---|---|---|---|---|---|---|
| 7 | 98304639 | rs6950545 | -1.231 | -0.677 | 0.739 | -0.199 | 0 | -0.031 | -0.031 | 0.121 | 0.42 | 0.57 | 0.46 | 0.09 | 0.45 | 0.37 |
| 7 | 98304764 | rs13223648 | 0.927 | -0.59 | 0.669 | -0.164 | -0.013 | 0.036 | 0.036 | 1.152 | 0.19 | 0.12 | 0.16 | 0.02 | 0.2 | 0.11 |
| 7 | 98311136 | rs10230224 | -0.458 | -0.636 | 0.964 | -0.049 | -0.014 | 0.046 | 0.046 | 6.929 | 0.89 | 0.77 | 0.89 | 0.8 | 0.94 | 0.9 |
| 7 | 98312921 | rs10238278 | -0.173 | -0.515 | 0.925 | -0.287 | 0.022 | -0.014 | -0.014 | 0.059 | 0.13 | 0.28 | 0.13 | 0.06 | 0.16 | 0.18 |
| 7 | 98316837 | rs1994410 | -0.08 | -0.509 | 0.909 | -0.284 | 0.01 | -0.01 | -0.01 | 0.815 | 0.13 | 0.18 | 0.11 | 0.06 | 0.16 | 0.18 |
| 7 | 98317100 | rs10262556 | 0.521 | -0.507 | 0.904 | -0.283 | 0.003 | -0.003 | -0.003 | 0 | 0.11 | 0.1 | 0.1 | 0.06 | 0.15 | 0.16 |
| 7 | 98319499 | rs4729475 | 1.338 | -0.786 | 0.786 | -0.391 | -0.001 | 0.009 | 0.009 | 0 | 0.51 | 0.39 | 0.52 | 0.6 | 0.52 | 0.58 |
| 7 | 98321862 | rs1038585 | 1.197 | -0.729 | 0.411 | -0.363 | 0.037 | 0.054 | 0.054 | 0.476 | 0.51 | 0.2 | 0.45 | 0.65 | 0.58 | 0.66 |
| 7 | 98326439 | rs980729 | 2.076 | -0.617 | -0.049 | -0.269 | -0.095 | -0.105 | -0.105 | 0.391 | 0.39 | 0.06 | 0.34 | 0.59 | 0.42 | 0.49 |
| 7 | 98330450 | rs7780073 | 0.068 | -0.734 | 0.105 | -0.242 | -0.001 | 0.001 | 0.001 | 0.314 | 0.68 | 0.68 | 0.68 | 0.66 | 0.69 | 0.77 |
| 7 | 98348297 | rs6465710 | -0.865 | -0.704 | -0.125 | -0.35 | -0.007 | -0.055 | -0.055 | 0 | 0.9 | 0.79 | 0.93 | 1 | 0.94 | 0.97 |
| 7 | 98358654 | rs12531849 | 1.849 | -0.754 | -0.432 | -0.181 | -0.088 | -0.09 | -0.09 | 0.459 | 0.39 | 0.1 | 0.36 | 0.56 | 0.43 | 0.49 |
| 7 | 98361817 | rs6972315 | -1.366 | -0.637 | -0.132 | 0.176 | 0.012 | 0.012 | 0.012 | 0 | 0.5 | 0.68 | 0.57 | 0.44 | 0.51 | 0.48 |
| 7 | 98366565 | rs11773403 | 0.134 | -0.614 | -0.501 | 0.803 | -0.007 | -0.061 | -0.061 | 0 | 0.22 | 0.12 | 0.32 | 0.39 | 0.24 | 0.22 |
| 7 | 98376458 | rs9986780 | -1.265 | -0.604 | 0.186 | 0.887 | -0.009 | 0.007 | 0.007 | 0.054 | 0.24 | 0.37 | 0.17 | 0.22 | 0.21 | 0.38 |
| 7 | 98399810 | rs4729491 | -1.317 | -1.152 | 0.083 | 0.648 | -0.005 | 0 | 0 | 0.201 | 0.31 | 0.71 | 0.25 | 0.2 | 0.17 | 0.14 |
| 7 | 98400450 | rs13233862 | -1.181 | -1.315 | 0.619 | 0.721 | -0.04 | -0.013 | -0.013 | 12.021 | 0.25 | 0.39 | 0.14 | 0.14 | 0.12 | 0.12 |
| 7 | 98400599 | rs12668051 | -0.76 | -1.496 | 0.585 | 0.739 | -0.016 | 0 | 0 | 0 | 0.21 | 0.27 | 0.17 | 0.19 | 0.11 | 0.1 |
| 7 | 98404438 | rs2029220 | -0.426 | -0.524 | 0.343 | 1.18 | 0.01 | 0.014 | 0.014 | 0.957 | 0.14 | 0.14 | 0.22 | 0.25 | 0.18 | 0.09 |
| 7 | 98405206 | rs4727416 | 1.7 | -1.238 | 0.481 | 1.159 | 0 | 0.001 | 0.001 | 0.14 | 0.47 | 0.49 | 0.38 | 0.52 | 0.55 | 0.41 |
| 7 | 98407758 | rs10245517 | -1.091 | -1.297 | 1.017 | 1.65 | 0.064 | 0.193 | 0.193 | 0.917 | 0.18 | 0.51 | 0.27 | 0.44 | 0.19 | 0.1 |
| 7 | 98411980 | rs10232758 | 1.634 | -1.264 | 0.673 | 1.503 | 0 | -0.001 | -0.001 | 0 | 0.51 | 0.52 | 0.43 | 0.52 | 0.54 | 0.42 |
| 7 | 98413656 | rs2394936 | 0.646 | -1.53 | -0.113 | 1.292 | 0.017 | -0.015 | -0.015 | 0 | 0.74 | 0.87 | 0.66 | 0.77 | 0.73 | 0.55 |
| 7 | 98434486 | rs4236538 | 1.181 | -1.169 | 0.195 | 1.328 | 0.008 | -0.024 | -0.024 | 0 | 0.55 | 0.29 | 0.48 | 0.53 | 0.62 | 0.4 |
| 7 | 98439544 | rs4526300 | -0.172 | -1.269 | 0.624 | 1.413 | -0.034 | -0.01 | -0.01 | 0 | 0.2 | 0.36 | 0.27 | 0.17 | 0.14 | 0.38 |
| 7 | 98439700 | rs4368910 | 1.639 | -1.12 | -0.114 | 1.453 | -0.007 | -0.03 | -0.03 | 0 | 0.48 | 0.17 | 0.45 | 0.54 | 0.58 | 0.38 |
| 7 | 98443953 | rs6968995 | -0.673 | -1.125 | -0.03 | 1.479 | -0.017 | 0.006 | 0.006 | 0 | 0.31 | 0.65 | 0.36 | 0.33 | 0.21 | 0.41 |
| 7 | 98444287 | rs1047035 | -1.173 | -0.62 | 0.009 | 1.344 | -0.016 | -0.019 | -0.019 | 0 | 0.23 | 0.12 | 0.22 | 0.29 | 0.23 | 0.22 |
| 7 | 98447353 | rs1203843 | 0.27 | -1.27 | 0.203 | 0.74 | 0.009 | 0.002 | 0.002 | 0 | 0.87 | 0.91 | 0.95 | 0.96 | 0.94 | 0.93 |
| 7 | 98451358 | rs7792235 | 1.668 | -1.027 | 0.788 | 0.856 | -0.005 | -0.016 | -0.016 | 0.016 | 0.81 | 0.73 | 0.82 | 0.88 | 0.86 | 0.9 |
| 7 | 98451597 | rs7795995 | 1.668 | -1.027 | 0.788 | 0.856 | -0.005 | -0.016 | -0.016 | 5.036 | 0.81 | 0.73 | 0.82 | 0.88 | 0.86 | 0.91 |
| 7 | 98460824 | rs10238224 | 2.332 | -1.162 | 1.279 | 0.77 | 0.008 | 0.013 | 0.013 | 0.325 | 0.77 | 0.22 | 0.76 | 0.88 | 0.85 | 0.9 |
| 7 | 98461062 | rs1548622 | 0.741 | -1.266 | 1.393 | 0.748 | 0.005 | 0.01 | 0.01 | 0 | 0.9 | 0.82 | 0.84 | 0.88 | 0.91 | 0.94 |
| 7 | 98462589 | rs6955886 | 0.726 | -1.263 | 1.439 | 1.041 | -0.002 | -0.005 | -0.005 | 0 | 0.9 | 0.91 | 0.84 | 0.76 | 0.91 | 0.94 |
| 7 | 98474128 | rs7457148 | -1.339 | -1.262 | 1.491 | 0.901 | 0.002 | 0.004 | 0.004 | 6.545 | 0.07 | 0.57 | 0.05 | 0 | 0.01 | 0 |
| 7 | 98478680 | rs6465726 | -2.215 | -1.264 | 2.087 | 0.9 | -0.026 | -0.086 | -0.086 | 0.069 | 0.13 | 0.5 | 0.06 | 0 | 0.05 | 0.04 |
| 7 | 98480722 | rs2299294 | 0.236 | -1.269 | 2.004 | 0.924 | -0.031 | -0.051 | -0.051 | 0.143 | 0.92 | 0.99 | 0.86 | 0.79 | 0.92 | 0.95 |
| 7 | 98481436 | rs2245385 | 0.25 | -1.517 | 1.357 | 0.96 | 0.02 | -0.02 | -0.02 | 0 | 0.37 | 0.17 | 0.28 | 0.4 | 0.34 | 0.52 |
| 7 | 98491261 | rs2249169 | 0.39 | -1.386 | 1.436 | 0.988 | 0.002 | 0.02 | 0.02 | 0 | 0.91 | 0.85 | 0.84 | 0.79 | 0.92 | 0.95 |
| 7 | 98522893 | rs6967096 | -1.435 | -1.38 | 1.517 | 0.676 | 0.015 | 0.006 | 0.006 | 4.157 | 0.07 | 0.52 | 0.05 | 0 | 0.01 | 0 |
| 7 | 98556962 | rs41633 | -0.076 | -1.303 | 1.241 | 0.698 | -0.034 | -0.05 | -0.05 | 0.01 | 0.08 | 0 | 0.14 | 0.21 | 0.08 | 0.05 |
| 7 | 98583791 | rs219802 | -1.224 | -1.362 | 1.277 | 0.556 | 0.005 | -0.003 | -0.003 | 0 | 0.07 | 0.57 | 0.05 | 0 | 0.01 | 0 |
| 7 | 98610595 | rs219816 | -2.642 | -1.38 | 1.096 | 0.58 | 0.006 | 0.004 | 0.004 | 0.424 | 0.23 | 0.77 | 0.25 | 0.22 | 0.15 | 0.1 |
| 7 | 98610684 | rs219815 | -2.605 | -1.376 | 1.09 | 0.579 | 0.009 | 0.006 | 0.006 | 0.464 | 0.23 | 0.77 | 0.22 | 0.19 | 0.13 | 0.09 |
| 7 | 98616657 | rs17720576 | -0.4 | -1.333 | 1.106 | 0.608 | 0 | 0.004 | 0.004 | 0 | 0.09 | 0.13 | 0.16 | 0.21 | 0.09 | 0.05 |
| 7 | 98624877 | rs1007007 | -0.4 | -1.334 | 1.205 | 0.603 | 0 | 0.003 | 0.003 | 0 | 0.09 | 0.13 | 0.16 | 0.19 | 0.09 | 0.05 |
| 7 | 98626536 | rs1007008 | 0.026 | -1.345 | 1.188 | 0.609 | -0.019 | -0.034 | -0.034 | 0.027 | 0.08 | 0.03 | 0.14 | 0.19 | 0.08 | 0.05 |
| 7 | 98627463 | rs219810 | -1.225 | -1.337 | 1.229 | 0.609 | -0.009 | -0.017 | -0.017 | 0 | 0.07 | 0.57 | 0.05 | 0 | 0.01 | 0 |
| 7 | 98644032 | rs219823 | -1.6 | -1.548 | 0.57 | 0.526 | -0.15 | -0.096 | -0.096 | 0.004 | 0.61 | 0.96 | 0.52 | 0.65 | 0.48 | 0.61 |
| 7 | 98645517 | rs219824 | -0.575 | -1.598 | 0.927 | 0.58 | 0.098 | 0.001 | 0.001 | 4.963 | 0.45 | 0.78 | 0.3 | 0.42 | 0.34 | 0.51 |
| 7 | 98647139 | rs219826 | -0.109 | -0.849 | 0.787 | 0.928 | 0.008 | -0.014 | -0.014 | 0 | 0.34 | 0.24 | 0.23 | 0.44 | 0.31 | 0.49 |
| 7 | 98676767 | rs925720 | -0.007 | -0.846 | 0.878 | 0.725 | 0.008 | -0.023 | -0.023 | 0.489 | 0.63 | 0.74 | 0.6 | 0.51 | 0.65 | 0.48 |
| 7 | 98689289 | rs10266176 | 1.561 | -0.474 | 1.843 | 0.654 | -0.011 | -0.003 | -0.003 | 0 | 0.81 | 0.22 | 0.78 | 0.8 | 0.89 | 0.93 |
| 7 | 98690685 | rs12705021 | 1.221 | -0.375 | 1.72 | 0.697 | -0.034 | 0.007 | 0.007 | 0.173 | 0.83 | 0.38 | 0.8 | 0.82 | 0.9 | 0.94 |
| 7 | 98705460 | rs2395015 | 1.22 | -0.258 | 0.498 | 0.723 | -0.029 | 0.005 | 0.005 | 0.166 | 0.82 | 0.34 | 0.79 | 0.81 | 0.89 | 0.93 |
| 7 | 98721885 | rs1986465 | 1.16 | 0.35 | 0.497 | 0.739 | 0.008 | 0.024 | 0.024 | 0 | 0.76 | 0.46 | 0.78 | 0.75 | 0.8 | 0.78 |
| 7 | 98724730 | rs12531875 | 0.048 | 0.75 | -0.388 | 0.285 | -0.05 | 0.044 | 0.044 | 0.359 | 0.2 | 0.06 | 0.17 | 0.11 | 0.22 | 0.36 |
| 7 | 98725348 | rs6465737 | 1.071 | 0.318 | 0.136 | -0.111 | 0.001 | 0 | 0 | 0.596 | 0.7 | 0.31 | 0.75 | 0.74 | 0.74 | 0.75 |
| 7 | 98732012 | rs10499947 | -0.373 | -0.028 | 0.855 | -0.044 | -0.018 | 0.092 | 0.092 | 0 | 0.15 | 0.36 | 0.14 | 0.23 | 0.1 | 0.16 |
| 7 | 98733337 | rs6952161 | 1.662 | -0.361 | 0.665 | -0.249 | 0.036 | 0.058 | 0.058 | 0 | 0.52 | 0.17 | 0.6 | 0.65 | 0.6 | 0.42 |
| 7 | 98739231 | rs7788750 | -0.706 | -0.446 | 1.778 | -0.275 | 0.009 | -0.038 | -0.038 | 2.7 | 0.09 | 0.32 | 0.05 | 0 | 0.06 | 0.04 |
| 7 | 98743596 | rs4729514 | 1.747 | -0.458 | 1.834 | -0.274 | -0.016 | 0 | 0 | 0.188 | 0.73 | 0.38 | 0.79 | 0.77 | 0.83 | 0.77 |
| 7 | 98750379 | rs2395022 | 0.257 | -0.428 | 2.27 | -0.28 | -0.011 | -0.044 | -0.044 | 0.264 | 0.92 | 0.85 | 0.95 | 1 | 0.94 | 0.96 |
| 7 | 98759117 | rs9297145 | 0.572 | -0.388 | 2.295 | -0.305 | -0.001 | -0.041 | -0.041 | 0.511 | 0.7 | 0.43 | 0.75 | 0.89 | 0.71 | 0.6 |
| 7 | 98766930 | rs6954584 | 0.365 | -0.4 | 1.861 | -0.306 | -0.015 | 0.034 | 0.034 | 0.423 | 0.21 | 0.14 | 0.18 | 0.11 | 0.23 | 0.36 |
| 7 | 98771462 | rs10499948 | -0.201 | -0.4 | 1.668 | -0.166 | 0.01 | -0.022 | -0.022 | 2.062 | 0.05 | 0.01 | 0.08 | 0.16 | 0.04 | 0.04 |
| 7 | 98772131 | rs10953281 | 1.529 | -0.377 | 1.919 | -0.16 | 0.019 | 0.026 | 0.026 | 0 | 0.53 | 0.35 | 0.61 | 0.65 | 0.6 | 0.41 |
| 7 | 98780395 | rs13243638 | -0.34 | -0.328 | 1.412 | -0.261 | -0.012 | -0.011 | -0.011 | 0.959 | 0.11 | 0.19 | 0.06 | 0.08 | 0.07 | 0.15 |
| 7 | 98780520 | rs4729521 | 1.545 | -0.346 | 0.707 | -0.224 | 0.027 | 0.067 | 0.067 | 0 | 0.51 | 0.14 | 0.6 | 0.67 | 0.6 | 0.41 |

| | | | | | | | | | | | | | | | | |
|---|---|---|---|---|---|---|---|---|---|---|---|---|---|---|---|---|
| 7 | 98788589 | rs7809188 | -0.503 | -0.451 | 1.088 | -0.09 | -0.007 | -0.008 | -0.008 | 0 | 0.08 | 0.1 | 0.08 | 0.07 | 0.07 | 0.19 |
| 7 | 98799772 | rs13224433 | -0.45 | -0.462 | 0.928 | -0.095 | 0.008 | 0.009 | 0.009 | 0.335 | 0.08 | 0.04 | 0.08 | 0.07 | 0.07 | 0.19 |
| 7 | 98799972 | rs12705034 | -0.546 | -0.466 | 0.928 | 0.097 | 0.003 | -0.005 | -0.005 | 0.012 | 0.08 | 0.07 | 0.09 | 0.11 | 0.07 | 0.2 |
| 7 | 98810612 | rs4727428 | 1.919 | -0.592 | 0.791 | 0.027 | 0.02 | -0.006 | -0.006 | 0 | 0.75 | 0.19 | 0.7 | 0.79 | 0.79 | 0.7 |
| 7 | 98830439 | rs4236541 | 1.892 | -0.636 | 0.372 | 0.014 | 0.048 | -0.002 | -0.002 | 0.532 | 0.58 | 0.03 | 0.61 | 0.73 | 0.7 | 0.5 |
| 7 | 98834127 | rs10274646 | 1.742 | -0.382 | 1.924 | 0.339 | -0.018 | 0.082 | 0.082 | 0.249 | 0.81 | 0.63 | 0.76 | 0.74 | 0.86 | 0.63 |
| 7 | 98836916 | rs4236542 | 2.232 | -0.462 | 2.246 | 0.335 | -0.027 | 0.019 | 0.019 | 0.647 | 0.77 | 0.37 | 0.73 | 0.74 | 0.85 | 0.61 |
| 7 | 98843919 | rs6465744 | 1.646 | -0.472 | 2.401 | 0.342 | -0.002 | 0.116 | 0.116 | 0 | 0.88 | 0.67 | 0.82 | 0.78 | 0.91 | 0.76 |
| 7 | 98846684 | rs4255064 | 1.246 | -0.451 | 2.466 | 0.337 | 0.014 | 0.091 | 0.091 | 0 | 0.9 | 0.71 | 0.83 | 0.84 | 0.91 | 0.91 |
| 7 | 98848399 | rs13438315 | -1.228 | -0.37 | 2.135 | 0.352 | -0.021 | -0.017 | -0.017 | 0 | 0.1 | 0.25 | 0.09 | 0.04 | 0.06 | 0.14 |
| 7 | 98858784 | rs12535654 | -1.942 | -0.479 | 2.205 | 0.404 | -0.028 | 0.061 | 0.061 | 1.391 | 0.15 | 0.42 | 0.22 | 0.19 | 0.11 | 0.16 |
| 7 | 98861915 | rs10281195 | -2.017 | -0.484 | 2.137 | 0.401 | -0.03 | 0.045 | 0.045 | 0.716 | 0.16 | 0.4 | 0.22 | 0.19 | 0.11 | 0.16 |
| 7 | 98880642 | rs6960805 | -1.677 | -0.234 | 2.461 | 0.496 | 0.026 | -0.01 | -0.01 | 0 | 0.08 | 0.29 | 0.1 | 0.02 | 0.06 | 0.1 |
| 7 | 98891438 | rs9969217 | -2.718 | -0.206 | 2.395 | 0.487 | 0.012 | 0.016 | 0.016 | 3.231 | 0.15 | 0.82 | 0.17 | 0.03 | 0.1 | 0.14 |
| 7 | 98931105 | rs2301889 | -2.484 | -0.136 | 2.987 | 1.3 | 0.012 | 0.299 | 0.299 | 0 | 0.06 | 0.22 | 0.15 | 0.35 | 0.06 | 0.3 |
| 7 | 98933250 | rs12705058 | 0.298 | -0.439 | 1.887 | 1.261 | -0.069 | -0.033 | -0.033 | 0 | 0.28 | 0.01 | 0.24 | 0.23 | 0.3 | 0.22 |
| 7 | 98953863 | rs17161692 | -1.98 | -0.076 | 2.697 | 1.59 | 0.015 | 0.296 | 0.296 | 0 | 0.05 | 0.2 | 0.15 | 0.35 | 0.06 | 0.3 |
| 7 | 98963340 | rs10268984 | -2.365 | -0.084 | 2.706 | 1.125 | -0.018 | -0.072 | -0.072 | 2.116 | 0.13 | 0.68 | 0.11 | 0 | 0.08 | 0.07 |
| 7 | 98974588 | rs6955490 | -3.166 | -0.145 | 2.599 | 1.133 | 0.022 | 0.048 | 0.048 | 3.16 | 0.16 | 0.89 | 0.18 | 0.02 | 0.12 | 0.13 |
| 7 | 99005506 | rs10279308 | -3.044 | -0.145 | 3.037 | 1.211 | -0.002 | -0.014 | -0.014 | 0 | 0.15 | 0.81 | 0.17 | 0.04 | 0.12 | 0.12 |
| 7 | 99005695 | rs6960810 | -2.424 | -0.12 | 3.024 | 1.218 | -0.023 | -0.077 | -0.077 | 0.003 | 0.13 | 0.64 | 0.11 | 0 | 0.08 | 0.08 |
| 7 | 99032399 | rs11973173 | -2.291 | 0.009 | 3.236 | 1.487 | 0.021 | -0.072 | -0.072 | 1.311 | 0.12 | 0.51 | 0.12 | 0 | 0.11 | 0.05 |
| 7 | 99053816 | rs12535424 | -1.368 | 0.012 | 3.478 | 1.465 | -0.001 | -0.004 | -0.004 | 0 | 0.08 | 0.05 | 0.07 | 0.01 | 0.1 | 0.06 |
| 7 | 99115677 | rs12533251 | -2.354 | 0.431 | 3.557 | 1.449 | 0.024 | -0.064 | -0.064 | 0 | 0.11 | 0.45 | 0.12 | 0.01 | 0.11 | 0.06 |
| 7 | 99150185 | rs34257469 | -2.354 | 0.475 | 3.398 | 1.471 | 0.024 | -0.064 | -0.064 | 0 | 0.11 | 0.45 | 0.12 | 0.01 | 0.11 | 0.06 |
| 7 | 99156446 | rs6947941 | -1.792 | 0.405 | 3.315 | 1.48 | 0.012 | -0.047 | -0.047 | 0.201 | 0.09 | 0.22 | 0.09 | 0.01 | 0.1 | 0.06 |
| 7 | 99156520 | rs6947826 | -1.858 | 0.405 | 3.369 | 1.48 | 0.019 | -0.046 | -0.046 | 0 | 0.09 | 0.28 | 0.09 | 0.01 | 0.1 | 0.06 |
| 7 | 99156934 | rs17161761 | -1.792 | 0.406 | 3.332 | 1.481 | 0.012 | -0.047 | -0.047 | 0.656 | 0.09 | 0.22 | 0.09 | 0.01 | 0.1 | 0.06 |
| 7 | 99236069 | rs34096782 | -0.331 | 0.288 | 3.121 | 1.501 | -0.001 | 0.027 | 0.027 | 0 | 0.05 | 0 | 0.03 | 0 | 0.03 | 0.03 |
| 7 | 99245013 | rs4646458 | -2.431 | 0.264 | 3.827 | 1.663 | 0.033 | 0.396 | 0.396 | 3.661 | 0.07 | 0.48 | 0.13 | 0.27 | 0.02 | 0.3 |
| 7 | 99260092 | rs1419745 | -2.431 | -0.09 | 3.89 | 1.689 | 0.033 | 0.396 | 0.396 | 3.661 | 0.07 | 0.48 | 0.13 | 0.26 | 0.02 | 0.3 |
| 7 | 99272310 | rs28365067 | -0.054 | -0.606 | 3.507 | 1.199 | 0.035 | 0.049 | 0.049 | 0.635 | 0.11 | 0.02 | 0.06 | 0.03 | 0.05 | 0.07 |
| 7 | 99277593 | rs28371764 | -0.176 | -0.556 | 3.295 | 1.177 | -0.001 | 0.027 | 0.027 | 0 | 0.05 | 0 | 0.03 | 0 | 0.03 | 0.03 |
| 7 | 99294472 | rs2687087 | 1.482 | -0.521 | 3.574 | 1.339 | -0.051 | 0.037 | 0.037 | 0.37 | 0.81 | 0.31 | 0.8 | 0.72 | 0.93 | 0.63 |
| 7 | 99294762 | rs1879859 | 1.482 | -0.521 | 3.574 | 1.339 | -0.051 | 0.037 | 0.037 | 0.37 | 0.81 | 0.31 | 0.8 | 0.71 | 0.93 | 0.63 |
| 7 | 99300758 | rs2687081 | 1.499 | -0.522 | 3.574 | 1.338 | -0.051 | 0.036 | 0.036 | 0.399 | 0.81 | 0.31 | 0.8 | 0.71 | 0.93 | 0.63 |
| 7 | 99302994 | rs10211 | 1.481 | -0.522 | 3.574 | 1.338 | -0.051 | 0.037 | 0.037 | 0.37 | 0.81 | 0.31 | 0.8 | 0.71 | 0.93 | 0.63 |
| 7 | 99306321 | rs1403196 | 1.483 | -0.525 | 3.568 | 1.337 | -0.051 | 0.037 | 0.037 | 0.39 | 0.81 | 0.31 | 0.8 | 0.71 | 0.93 | 0.63 |
| 7 | 99306566 | rs2687077 | 1.376 | -0.505 | 3.628 | 1.346 | -0.075 | 0.081 | 0.081 | 0.476 | 0.82 | 0.38 | 0.81 | 0.71 | 0.93 | 0.65 |
| 7 | 99307268 | rs2687076 | 1.393 | -0.505 | 3.628 | 1.346 | -0.075 | 0.08 | 0.08 | 0.437 | 0.82 | 0.38 | 0.81 | 0.71 | 0.93 | 0.65 |
| 7 | 99310181 | rs1357319 | 1.393 | -0.505 | 3.628 | 1.346 | -0.075 | 0.08 | 0.08 | 0.437 | 0.82 | 0.38 | 0.81 | 0.71 | 0.93 | 0.65 |
| 7 | 99311550 | rs2037498 | 1.393 | -0.505 | 3.628 | 1.346 | -0.075 | 0.08 | 0.08 | 0.437 | 0.82 | 0.38 | 0.81 | 0.71 | 0.93 | 0.65 |
| 7 | 99314986 | rs2687075 | 1.393 | -0.505 | 3.628 | 1.346 | -0.075 | 0.08 | 0.08 | 0.437 | 0.82 | 0.38 | 0.81 | 0.71 | 0.93 | 0.65 |
| 7 | 99318644 | rs2687145 | 1.393 | -0.439 | 3.635 | 1.375 | -0.075 | 0.08 | 0.08 | 0.437 | 0.82 | 0.38 | 0.81 | 0.71 | 0.93 | 0.65 |
| 7 | 99320570 | rs2687142 | 1.393 | -0.439 | 3.635 | 1.375 | -0.075 | 0.08 | 0.08 | 0.437 | 0.82 | 0.38 | 0.81 | 0.71 | 0.93 | 0.65 |
| 7 | 99325882 | rs2687136 | 1.393 | -0.439 | 3.635 | 1.375 | -0.075 | 0.08 | 0.08 | 0.437 | 0.82 | 0.38 | 0.81 | 0.71 | 0.93 | 0.65 |
| 7 | 99331042 | rs2687134 | 1.181 | -0.428 | 3.616 | 1.382 | -0.071 | 0.091 | 0.091 | 0.399 | 0.83 | 0.37 | 0.82 | 0.71 | 0.94 | 0.66 |
| 7 | 99331473 | rs4646461 | -0.954 | -0.417 | 3.043 | 1.387 | -0.037 | 0.056 | 0.056 | 0.031 | 0.14 | 0.22 | 0.13 | 0.29 | 0.05 | 0.34 |
| 7 | 99355975 | rs3735451 | -1.596 | -0.387 | 2.326 | 1.432 | 0.005 | -0.008 | -0.008 | 0 | 0.22 | 0.81 | 0.23 | 0.32 | 0.1 | 0.37 |
| 7 | 99365083 | rs4646437 | -1.656 | -0.359 | 0.7 | 0.814 | 0.017 | 0.013 | 0.013 | 0.261 | 0.23 | 0.82 | 0.15 | 0.16 | 0.09 | 0.38 |
| 7 | 99365719 | rs2246709 | -0.73 | 0.319 | 2.131 | 1.23 | 0.015 | 0.073 | 0.073 | 0.119 | 0.22 | 0.36 | 0.46 | 0.39 | 0.27 | 0.39 |
| 7 | 99419226 | rs667660 | -1.937 | -0.642 | 3.175 | 0.435 | -0.029 | -0.058 | -0.058 | 0 | 0.14 | 0.79 | 0.13 | 0.01 | 0.06 | 0.06 |
| 7 | 99426802 | rs585071 | -1.966 | -0.672 | 2.832 | 0.529 | -0.031 | -0.061 | -0.061 | 0 | 0.14 | 0.78 | 0.13 | 0.01 | 0.06 | 0.06 |
| 7 | 99429843 | rs651430 | -2.09 | 0.142 | 2.238 | 0.395 | -0.075 | -0.01 | -0.01 | 0 | 0.39 | 0.91 | 0.42 | 0.34 | 0.44 | 0.5 |
| 7 | 99436198 | rs800672 | -2.088 | 0.109 | 2.223 | 0.399 | -0.075 | -0.004 | -0.004 | 0.052 | 0.39 | 0.91 | 0.41 | 0.33 | 0.43 | 0.5 |
| 7 | 99438988 | rs688926 | -1.248 | 0.039 | 2.593 | 0.309 | -0.029 | -0.077 | -0.077 | 0 | 0.12 | 0.59 | 0.1 | 0 | 0.06 | 0.06 |
| 7 | 99440680 | rs660629 | -1.403 | 0.04 | 2.585 | 0.31 | -0.037 | -0.083 | -0.083 | 0.508 | 0.13 | 0.6 | 0.1 | 0 | 0.06 | 0.06 |
| 7 | 99441963 | rs501275 | -1.403 | 0.04 | 2.578 | 0.313 | -0.036 | -0.078 | -0.078 | 0.012 | 0.13 | 0.59 | 0.1 | 0 | 0.06 | 0.06 |
| 7 | 99444176 | rs549061 | -1.403 | 0.04 | 2.578 | 0.3 | -0.036 | -0.084 | -0.084 | 0 | 0.13 | 0.59 | 0.1 | 0 | 0.06 | 0.06 |
| 7 | 99444571 | rs545400 | -1.403 | 0.04 | 2.578 | 0.3 | -0.036 | -0.084 | -0.084 | 0 | 0.13 | 0.59 | 0.1 | 0 | 0.06 | 0.06 |
| 7 | 99446802 | rs679320 | -1.403 | 0.04 | 2.578 | 0.3 | -0.036 | -0.084 | -0.084 | 0 | 0.13 | 0.59 | 0.1 | 0 | 0.06 | 0.06 |
| 7 | 99447046 | rs678040 | -1.403 | 0.04 | 2.578 | 0.3 | -0.036 | -0.084 | -0.084 | 0 | 0.13 | 0.59 | 0.1 | 0 | 0.06 | 0.06 |
| 7 | 99447241 | rs800667 | -1.403 | 0.04 | 2.578 | 0.3 | -0.036 | -0.084 | -0.084 | 0.021 | 0.13 | 0.59 | 0.1 | 0 | 0.06 | 0.06 |
| 7 | 99448959 | rs10225908 | -1.403 | 0.04 | 2.578 | 0.3 | -0.036 | -0.084 | -0.084 | 0.553 | 0.13 | 0.59 | 0.1 | 0 | 0.06 | 0.06 |
| 7 | 99450700 | rs528144 | -1.403 | 0.04 | 2.578 | 0.3 | -0.036 | -0.084 | -0.084 | 0 | 0.13 | 0.59 | 0.1 | 0 | 0.06 | 0.06 |
| 7 | 99460458 | rs580123 | -0.809 | 1.255 | 2.916 | 0.146 | -0.016 | -0.055 | -0.055 | 0.981 | 0.1 | 0.32 | 0.04 | 0 | 0.05 | 0.03 |
| 7 | 99466558 | rs474229 | -1.756 | 0.748 | 2.036 | 0.02 | -0.019 | 0.12 | 0.12 | 0 | 0.49 | 0.93 | 0.46 | 0.29 | 0.54 | 0.57 |
| 7 | 99471072 | rs2099446 | 0.039 | 0.758 | 2.072 | 0.037 | -0.042 | -0.02 | -0.02 | 0 | 0.25 | 0.14 | 0.33 | 0.32 | 0.43 | 0.47 |
| 7 | 99472686 | rs35649099 | -0.267 | 0.743 | 2.086 | 0.055 | -0.009 | -0.057 | -0.057 | 0 | 0.1 | 0.39 | 0.06 | 0 | 0.06 | 0.03 |

| | | | | | | | | | | | | | | | | |
|---|---|---|---|---|---|---|---|---|---|---|---|---|---|---|---|---|
| 7 | 99481416 | rs2082744 | -1.973 | 0.713 | 1.497 | 0.039 | -0.063 | 0.024 | 0.024 | 0 | 0.48 | 0.94 | 0.47 | 0.33 | 0.53 | 0.57 |
| 7 | 99487071 | rs2572011 | 0.056 | 0.691 | 1.871 | -0.011 | 0.022 | 0.004 | 0.004 | 0.712 | 0.24 | 0.22 | 0.34 | 0.29 | 0.42 | 0.47 |
| 7 | 99489005 | rs2572009 | -2.056 | 0.694 | 1.811 | -0.008 | -0.063 | 0.079 | 0.079 | 0 | 0.48 | 0.94 | 0.47 | 0.3 | 0.53 | 0.57 |
| 7 | 99525241 | rs2527900 | 0.058 | 1.375 | 1.804 | -0.053 | -0.035 | -0.011 | -0.011 | 0 | 0.24 | 0.15 | 0.34 | 0.29 | 0.42 | 0.47 |
| 7 | 99526316 | rs11977827 | -0.068 | 1.426 | 2.18 | -0.112 | 0.028 | -0.033 | -0.033 | 2.578 | 0.09 | 0.37 | 0.07 | 0 | 0.07 | 0.03 |
| 7 | 99527977 | rs11974610 | -1.282 | 1.682 | 1.742 | -0.028 | 0.044 | -0.002 | -0.002 | 0 | 0.16 | 0.61 | 0.2 | 0.15 | 0.16 | 0.09 |
| 7 | 99529181 | rs2527899 | -2.25 | 1.667 | 1.569 | -0.058 | -0.069 | 0.077 | 0.077 | 0 | 0.48 | 0.94 | 0.46 | 0.3 | 0.53 | 0.56 |
| 7 | 99533111 | rs2525549 | -1.213 | 1.446 | 1.232 | -0.283 | -0.001 | 0.005 | 0.005 | 0 | 0.36 | 0.3 | 0.29 | 0.2 | 0.38 | 0.49 |
| 7 | 99533178 | rs2525550 | -0.404 | 1.419 | 1.087 | -0.288 | -0.009 | 0.017 | 0.017 | 0 | 0.31 | 0.2 | 0.26 | 0.2 | 0.34 | 0.46 |
| 7 | 99534080 | rs2527897 | -2.351 | 1.381 | 0.971 | -0.275 | -0.029 | 0.067 | 0.067 | 0 | 0.52 | 0.91 | 0.48 | 0.32 | 0.55 | 0.59 |
| 7 | 99535951 | rs2107351 | -2.355 | 1.362 | 0.963 | -0.284 | -0.03 | 0.066 | 0.066 | 0 | 0.52 | 0.91 | 0.48 | 0.32 | 0.55 | 0.59 |
| 7 | 99545309 | rs2525539 | 0.553 | 1.802 | 1.223 | -0.016 | -0.008 | -0.004 | -0.004 | 0 | 0.18 | 0.13 | 0.21 | 0.2 | 0.28 | 0.39 |
| 7 | 99557217 | rs6962405 | 2.294 | 1.288 | 0.63 | -0.264 | -0.024 | 0.064 | 0.064 | 0 | 0.51 | 0.1 | 0.49 | 0.66 | 0.44 | 0.41 |
| 7 | 99562258 | rs2404977 | 2.339 | 1.566 | 0.661 | -0.249 | 0.011 | 0.027 | 0.027 | 0 | 0.51 | 0.11 | 0.51 | 0.63 | 0.42 | 0.41 |
| 7 | 99567455 | rs2527923 | -1.308 | 1.754 | 0.967 | -0.214 | -0.013 | -0.001 | -0.001 | 0 | 0.33 | 0.69 | 0.4 | 0.34 | 0.47 | 0.46 |
| 7 | 99579032 | rs10953293 | 0.564 | 1.751 | 1.058 | -0.249 | 0.014 | 0.004 | 0.004 | 0 | 0.07 | 0.15 | 0.15 | 0.1 | 0.12 | 0.1 |
| 7 | 99580371 | rs6944766 | -1.032 | 1.801 | 0.972 | -0.273 | 0.024 | -0.036 | -0.036 | 0.082 | 0.16 | 0.59 | 0.21 | 0.1 | 0.17 | 0.12 |
| 7 | 99583286 | rs4729561 | -0.311 | 2.004 | 0.84 | -0.017 | 0.012 | 0.005 | 0.005 | 0 | 0.78 | 0.8 | 0.76 | 0.75 | 0.65 | 0.63 |
| 7 | 99593346 | rs4727443 | 2.132 | 1.566 | -0.045 | -0.247 | 0.012 | 0.041 | 0.041 | 0 | 0.49 | 0.11 | 0.48 | 0.64 | 0.4 | 0.38 |
| 7 | 99602191 | rs6967487 | 0.479 | 1.814 | 0.73 | -0.045 | 0.031 | -0.01 | -0.01 | 0.401 | 0.87 | 0.74 | 0.78 | 0.86 | 0.81 | 0.84 |
| 7 | 99605588 | rs11771331 | -0.385 | 1.874 | 0.829 | 0.875 | 0.059 | 0.107 | 0.107 | 0 | 0.78 | 0.69 | 0.72 | 0.56 | 0.66 | 0.6 |
| 7 | 99608557 | rs4729565 | -1.016 | 1.763 | 0.598 | 0.875 | -0.015 | -0.011 | -0.011 | 0.027 | 0.94 | 0.98 | 0.96 | 0.96 | 0.93 | 0.95 |
| 7 | 99614770 | rs35414630 | -0.399 | 2.181 | 0.824 | 0.838 | 0.013 | 0.001 | 0.001 | 0 | 0.1 | 0.17 | 0.18 | 0.15 | 0.17 | 0.12 |
| 7 | 99632982 | rs1133790 | 2.352 | 1.932 | 0.382 | 0.754 | 0.047 | -0.024 | -0.024 | 0 | 0.48 | 0.14 | 0.45 | 0.47 | 0.35 | 0.37 |
| 7 | 99637242 | rs4729567 | 1.193 | 2.054 | 0.692 | 0.566 | 0.007 | -0.05 | -0.05 | 0 | 0.66 | 0.39 | 0.64 | 0.85 | 0.61 | 0.71 |
| 7 | 99639445 | rs4729568 | 1.106 | 2.05 | 0.943 | 0.564 | 0.012 | -0.064 | -0.064 | 0.065 | 0.67 | 0.54 | 0.65 | 0.85 | 0.61 | 0.71 |
| 7 | 99654600 | rs11558475 | 2.133 | 2.045 | 0.875 | 0.627 | 0.074 | -0.046 | -0.046 | 0 | 0.51 | 0.12 | 0.4 | 0.45 | 0.33 | 0.37 |
| 7 | 99654689 | rs11558476 | 2.32 | 2.041 | 0.871 | 0.625 | 0.067 | -0.053 | -0.053 | 0.799 | 0.49 | 0.12 | 0.39 | 0.45 | 0.33 | 0.37 |
| 7 | 99655897 | rs10281368 | 1.688 | 2.064 | 1.017 | 0.634 | 0.03 | 0.066 | 0.066 | 0 | 0.15 | 0.16 | 0.23 | 0.41 | 0.28 | 0.34 |
| 7 | 99656927 | rs12705070 | 1.546 | 2.067 | 0.964 | 0.636 | 0.078 | 0.151 | 0.151 | 0.012 | 0.16 | 0.27 | 0.24 | 0.41 | 0.28 | 0.34 |
| 7 | 99663516 | rs12532238 | 2.287 | 2.062 | 0.776 | 0.621 | 0.068 | -0.053 | -0.053 | 2.33 | 0.49 | 0.12 | 0.39 | 0.44 | 0.33 | 0.37 |
| 7 | 99664817 | rs7778571 | 0.208 | 2.057 | 0.958 | 0.602 | 0 | -0.032 | -0.032 | 0 | 0.22 | 0.21 | 0.14 | 0.01 | 0.22 | 0.16 |
| 7 | 99668107 | rs6592 | 2.255 | 1.99 | 0.703 | 0.643 | 0.068 | -0.053 | -0.053 | 1.163 | 0.49 | 0.12 | 0.39 | 0.45 | 0.33 | 0.37 |
| 7 | 99669802 | rs11550034 | 1.372 | 1.995 | 0.833 | 0.659 | -0.016 | -0.014 | -0.014 | 0 | 0.07 | 0.04 | 0.15 | 0.14 | 0.12 | 0.12 |
| 7 | 99678622 | rs941290 | -0.819 | 1.928 | 0.857 | 0.676 | 0.025 | 0.001 | 0.001 | 0 | 0.89 | 0.77 | 0.8 | 0.86 | 0.83 | 0.88 |
| 7 | 99691740 | rs1527423 | 1.626 | 1.824 | 0.635 | 0.816 | -0.018 | 0.081 | 0.081 | 0 | 0.59 | 0.22 | 0.59 | 0.8 | 0.54 | 0.66 |
| 7 | 99692993 | rs2261360 | 2.366 | 1.843 | 0.474 | 0.902 | 0.082 | -0.047 | -0.047 | 0.001 | 0.44 | 0.06 | 0.35 | 0.41 | 0.26 | 0.31 |
| 7 | 99696797 | rs2070215 | 1.844 | 1.905 | 0.544 | 1.126 | -0.086 | -0.077 | -0.077 | 0.415 | 0.13 | 0.01 | 0.19 | 0.28 | 0.27 | 0.29 |
| 7 | 99703958 | rs2293479 | 1.358 | 1.987 | 0.484 | 0.951 | 0.028 | 0.055 | 0.055 | 0 | 0.17 | 0.17 | 0.24 | 0.48 | 0.29 | 0.36 |
| 7 | 99707712 | rs2272338 | 0.836 | 1.906 | 0.436 | 0.84 | 0.013 | -0.07 | -0.07 | 0 | 0.67 | 0.52 | 0.65 | 0.85 | 0.6 | 0.72 |
| 7 | 99711040 | rs4134903 | 1.537 | 1.975 | 0.277 | 0.879 | 0.025 | 0.055 | 0.055 | 0 | 0.16 | 0.15 | 0.23 | 0.4 | 0.28 | 0.35 |
| 7 | 99722266 | rs1880949 | 1.424 | 1.922 | 0.291 | 0.904 | 0.034 | 0.066 | 0.066 | 0.036 | 0.15 | 0.16 | 0.23 | 0.41 | 0.27 | 0.33 |
| 7 | 99736059 | rs11771139 | 1.42 | 1.922 | 0.279 | 0.818 | 0.037 | 0.084 | 0.084 | 0.031 | 0.15 | 0.16 | 0.28 | 0.44 | 0.28 | 0.33 |
| 7 | 99739362 | rs6975031 | -0.425 | 1.354 | 0.365 | 0.512 | 0.015 | -0.013 | -0.013 | 0.105 | 0.89 | 0.76 | 0.85 | 0.89 | 0.84 | 0.86 |
| 7 | 99747130 | rs12878 | 0.889 | 1.25 | 0.107 | 0.824 | 0.003 | -0.011 | -0.011 | 0 | 0.67 | 0.53 | 0.49 | 0.68 | 0.63 | 0.64 |
| 7 | 99751017 | rs3736591 | 0.938 | 1.318 | 0.098 | 0.762 | 0.003 | -0.011 | -0.011 | 0 | 0.67 | 0.53 | 0.47 | 0.68 | 0.61 | 0.64 |
| 7 | 99751281 | rs3736590 | 1.047 | 1.331 | 0.097 | 0.767 | 0.006 | -0.006 | -0.006 | 0.19 | 0.62 | 0.47 | 0.44 | 0.64 | 0.56 | 0.61 |
| 7 | 99757612 | rs3823646 | 1.244 | 1.228 | 0.204 | 0.719 | -0.001 | 0.001 | 0.001 | 0 | 0.6 | 0.33 | 0.42 | 0.64 | 0.56 | 0.61 |
| 7 | 99758136 | rs3800951 | 1.056 | 1.26 | 0.056 | 0.734 | 0.007 | -0.007 | -0.007 | 0 | 0.62 | 0.46 | 0.44 | 0.64 | 0.56 | 0.61 |
| 7 | 99771849 | rs3735242 | 1.433 | 1.071 | -0.505 | 0.403 | -0.033 | 0.029 | 0.029 | 0 | 0.46 | 0.11 | 0.59 | 0.68 | 0.48 | 0.41 |
| 7 | 99774122 | rs12705073 | 0.988 | 0.905 | -0.116 | 0.095 | -0.001 | 0.001 | 0.001 | 0 | 0.5 | 0.44 | 0.64 | 0.68 | 0.5 | 0.41 |
| 7 | 99796146 | rs3735241 | 1.232 | 0.932 | -0.106 | 0.079 | -0.003 | 0.009 | 0.009 | 0 | 0.47 | 0.37 | 0.63 | 0.68 | 0.5 | 0.41 |
| 7 | 99807473 | rs941287 | -1.284 | 0.948 | -0.106 | 0.079 | -0.002 | 0.009 | 0.009 | 0 | 0.53 | 0.63 | 0.37 | 0.32 | 0.51 | 0.59 |
| 7 | 99808097 | rs17249588 | 1.982 | 0.834 | -0.292 | 0.013 | -0.005 | 0.027 | 0.027 | 0 | 0.05 | 0 | 0.01 | 0 | 0.03 | 0.05 |
| 7 | 99818927 | rs705867 | -0.823 | 0.848 | -0.077 | -0.161 | 0 | 0 | 0 | 0 | 0.5 | 0.57 | 0.37 | 0.32 | 0.51 | 0.59 |
| 7 | 99939321 | rs11761253 | 0.812 | 0.575 | -0.41 | -0.419 | -0.004 | 0.032 | 0.032 | 0 | 0.17 | 0.1 | 0.1 | 0.03 | 0.18 | 0.11 |
| 7 | 99964661 | rs11769057 | 0.808 | 0.643 | -0.439 | -0.45 | -0.004 | 0.033 | 0.033 | 0 | 0.17 | 0.11 | 0.1 | 0.03 | 0.18 | 0.11 |
| 7 | 99965285 | rs705866 | 1.338 | 0.63 | -0.446 | -0.456 | -0.042 | 0.086 | 0.086 | 0 | 0.15 | 0.02 | 0.09 | 0.03 | 0.18 | 0.11 |
| 7 | 99970188 | rs6948685 | 0.777 | 0.651 | -0.444 | -0.455 | -0.006 | 0.033 | 0.033 | 0 | 0.17 | 0.11 | 0.1 | 0.03 | 0.18 | 0.11 |
| 7 | 99971313 | rs2405442 | -1.915 | 0.577 | -0.624 | -0.531 | -0.022 | -0.049 | -0.049 | 1.269 | 0.73 | 0.91 | 0.5 | 0.39 | 0.68 | 0.71 |
| 7 | 99971834 | rs1859788 | -1.937 | 0.569 | -0.616 | -0.534 | -0.021 | -0.046 | -0.046 | 0.082 | 0.73 | 0.9 | 0.5 | 0.39 | 0.68 | 0.71 |
| 7 | 99983135 | rs10085549 | 0.65 | 0.565 | -0.439 | -1.11 | 0 | -0.09 | -0.09 | 0.155 | 0.19 | 0.29 | 0.12 | 0.03 | 0.18 | 0.11 |
| 7 | 99998607 | rs11769886 | 1.362 | 0.594 | -0.412 | -1.066 | -0.046 | 0.09 | 0.09 | 0 | 0.16 | 0.03 | 0.11 | 0.03 | 0.19 | 0.11 |
| 7 | 100004446 | rs1476679 | -2.103 | 0.585 | -0.405 | -0.899 | -0.107 | -0.104 | -0.104 | 1.393 | 0.75 | 0.98 | 0.77 | 0.65 | 0.7 | 0.77 |
| 7 | 100004543 | rs1000215 | 0.473 | 0.661 | -0.314 | -0.905 | 0 | -0.093 | -0.093 | 0 | 0.19 | 0.3 | 0.13 | 0.03 | 0.19 | 0.11 |
| 7 | 100013402 | rs5015755 | 0.99 | 0.606 | 0.025 | -0.911 | 0.021 | 0.019 | 0.019 | 0 | 0.08 | 0.14 | 0.1 | 0.16 | 0.16 | 0.12 |
| 7 | 100013457 | rs5015756 | -1.019 | 0.431 | -0.015 | -1.047 | -0.002 | 0.008 | 0.008 | 0 | 0.6 | 0.68 | 0.38 | 0.26 | 0.57 | 0.56 |
| 7 | 100017454 | rs7783159 | 1.453 | -0.564 | 0.155 | -0.757 | -0.046 | 0.09 | 0.09 | 0 | 0.16 | 0.03 | 0.11 | 0.03 | 0.19 | 0.11 |
| 7 | 100026292 | rs3807475 | 1.072 | -0.537 | 0.175 | -0.74 | 0.024 | 0.023 | 0.023 | 0 | 0.08 | 0.14 | 0.09 | 0.17 | 0.16 | 0.12 |

| | | | | | | | | | | | | | | | | |
|---|---|---|---|---|---|---|---|---|---|---|---|---|---|---|---|---|
| 7 | 100031612 | rs1073 | 0.81 | -0.58 | -0.053 | -0.951 | -0.002 | 0.004 | 0.004 | 0 | 0.18 | 0.14 | 0.12 | 0.02 | 0.19 | 0.11 |
| 7 | 100032665 | rs4074838 | 0.796 | -0.58 | -0.053 | -0.951 | -0.002 | 0.004 | 0.004 | 0 | 0.18 | 0.13 | 0.12 | 0.02 | 0.19 | 0.11 |
| 7 | 100039815 | rs11761725 | 1.328 | -0.577 | -0.221 | -0.95 | -0.041 | 0.088 | 0.088 | 0 | 0.16 | 0.03 | 0.11 | 0.02 | 0.19 | 0.11 |
| 7 | 100039970 | rs11761784 | 0.505 | -0.579 | -0.231 | -0.951 | 0 | -0.093 | -0.093 | 0.013 | 0.19 | 0.3 | 0.13 | 0.02 | 0.19 | 0.11 |
| 7 | 100044754 | rs28578163 | 0.699 | -0.542 | -0.191 | -0.936 | 0.004 | -0.087 | -0.087 | 0 | 0.18 | 0.29 | 0.13 | 0.02 | 0.19 | 0.11 |
| 7 | 100053690 | rs2406255 | 1.471 | -0.538 | -0.255 | -0.934 | -0.047 | 0.086 | 0.086 | 0 | 0.15 | 0.03 | 0.11 | 0.02 | 0.19 | 0.11 |
| 7 | 100064719 | rs7806537 | -0.435 | -0.541 | -0.24 | -0.953 | 0.007 | 0.003 | 0.003 | 0 | 0.51 | 0.45 | 0.27 | 0.1 | 0.4 | 0.42 |
| 7 | 100070758 | rs4472444 | 0.199 | 0.37 | -0.064 | -0.506 | 0.005 | -0.095 | -0.095 | 0 | 0.18 | 0.3 | 0.13 | 0.02 | 0.19 | 0.11 |
| 7 | 100072031 | rs7809801 | 0.199 | 0.37 | -0.064 | -0.506 | 0.005 | -0.095 | -0.095 | 0.495 | 0.18 | 0.3 | 0.13 | 0.02 | 0.19 | 0.11 |
| 7 | 100073292 | rs11766752 | 0.164 | 0.368 | -0.065 | -0.507 | 0.004 | -0.095 | -0.095 | 6.326 | 0.19 | 0.3 | 0.13 | 0.02 | 0.2 | 0.11 |
| 7 | 100078754 | rs11760536 | 0.96 | 0.431 | -0.445 | -0.479 | -0.047 | 0.086 | 0.086 | 0 | 0.15 | 0.03 | 0.11 | 0.02 | 0.19 | 0.11 |
| 7 | 100081944 | rs11763511 | 0.42 | 0.529 | -0.391 | -0.434 | 0.002 | -0.077 | -0.077 | 0 | 0.18 | 0.27 | 0.13 | 0.02 | 0.19 | 0.11 |
| 7 | 100091795 | rs12539172 | -1.255 | 0.736 | -1.016 | -0.115 | -0.033 | -0.028 | -0.028 | 0.288 | 0.75 | 0.91 | 0.77 | 0.65 | 0.69 | 0.72 |
| 7 | 100110324 | rs11761624 | 0.017 | 0.05 | -0.659 | -0.59 | -0.005 | -0.068 | -0.068 | 0 | 0.26 | 0.29 | 0.16 | 0.03 | 0.27 | 0.14 |
| 7 | 100119643 | rs1962151 | 0.205 | 0.771 | -0.012 | 0.038 | -0.004 | -0.049 | -0.049 | 0.361 | 0.11 | 0.18 | 0.07 | 0.02 | 0.12 | 0.12 |
| 7 | 100125975 | rs11765024 | -0.201 | 0.777 | 0.112 | 0.331 | -0.011 | 0.033 | 0.033 | 0 | 0.11 | 0.24 | 0.07 | 0.14 | 0.09 | 0.13 |
| 7 | 100141861 | rs11977607 | 0.35 | 1.247 | -0.69 | 0.366 | -0.003 | -0.042 | -0.042 | 0 | 0.09 | 0.13 | 0.07 | 0.01 | 0.07 | 0.04 |
| 7 | 100156438 | rs11978096 | 0.513 | 1.274 | -1.043 | 0.378 | 0.001 | 0 | 0 | 0 | 0.08 | 0.06 | 0.07 | 0.01 | 0.07 | 0.04 |
| 7 | 100159567 | rs2734897 | -0.926 | 1.545 | -1.139 | 0.64 | -0.041 | 0.002 | 0.002 | 0 | 0.76 | 0.9 | 0.54 | 0.74 | 0.69 | 0.71 |
| 7 | 100160109 | rs11760595 | -0.048 | 1.463 | -0.544 | 0.957 | -0.02 | -0.038 | -0.038 | 0 | 0.07 | 0.01 | 0.06 | 0.14 | 0.1 | 0.11 |
| 7 | 100173481 | rs2075668 | -1.57 | 1.745 | -0.224 | 0.325 | 0.002 | -0.054 | -0.054 | 4.978 | 0.26 | 0.36 | 0.2 | 0.08 | 0.27 | 0.27 |
| 7 | 100180060 | rs34440977 | -0.555 | 1.197 | -0.558 | 0.622 | 0.001 | 0 | 0 | 0 | 0.08 | 0.06 | 0.07 | 0.01 | 0.07 | 0.03 |
| 7 | 100197871 | rs2272575 | -0.672 | 1.329 | 0.106 | 0.951 | 0.002 | 0.039 | 0.039 | 0 | 0.08 | 0.17 | 0.04 | 0.09 | 0.06 | 0.13 |
| 7 | 100198386 | rs11768465 | -1.06 | 1.402 | 0.554 | 0.29 | 0.022 | -0.122 | -0.122 | 0 | 0.18 | 0.38 | 0.21 | 0.02 | 0.21 | 0.21 |
| 7 | 100211556 | rs7812123 | -0.542 | 1.446 | 0.203 | 0.751 | -0.022 | 0 | 0 | 0 | 0.12 | 0.07 | 0.17 | 0.11 | 0.19 | 0.15 |
| 7 | 100212254 | rs12532878 | -0.806 | 1.418 | 0.517 | 0.745 | 0.002 | -0.001 | -0.001 | 0 | 0.13 | 0.15 | 0.17 | 0.11 | 0.19 | 0.17 |
| 7 | 100212754 | rs11764045 | -0.545 | 1.429 | 0.491 | 0.75 | -0.022 | -0.002 | -0.002 | 0 | 0.12 | 0.06 | 0.17 | 0.11 | 0.19 | 0.15 |
| 7 | 100214015 | rs7786877 | -1.449 | 1.015 | 0.312 | 0.323 | -0.001 | 0.008 | 0.008 | 0.012 | 0.22 | 0.19 | 0.22 | 0.11 | 0.26 | 0.2 |
| 7 | 100219167 | rs12673102 | 0.43 | 1.913 | 0.028 | 0.484 | -0.051 | 0.003 | 0.003 | 0 | 0.88 | 0.98 | 0.9 | 0.9 | 0.82 | 0.81 |
| 7 | 100225031 | rs2075674 | -0.521 | 1.997 | 0.178 | 0.499 | -0.016 | 0.003 | 0.003 | 0 | 0.13 | 0.09 | 0.17 | 0.12 | 0.18 | 0.11 |
| 7 | 100236593 | rs4727457 | 0.475 | 1.959 | 0.426 | 0.542 | -0.031 | 0.007 | 0.007 | 0 | 0.87 | 0.97 | 0.9 | 0.89 | 0.82 | 0.79 |
| 7 | 100240296 | rs2075672 | 1.815 | 2.125 | 0.681 | 0.515 | 0.019 | -0.016 | -0.016 | 0 | 0.72 | 0.66 | 0.71 | 0.77 | 0.62 | 0.67 |
| 7 | 100256228 | rs6952341 | -0.782 | 2.179 | 0.792 | 0.89 | 0.002 | 0.001 | 0.001 | 0 | 0.14 | 0.15 | 0.45 | 0.22 | 0.21 | 0.16 |
| 7 | 100271521 | rs221789 | 0.721 | 2.178 | 1.041 | 0.651 | 0.005 | -0.015 | -0.015 | 0 | 0.87 | 0.87 | 0.89 | 0.92 | 0.82 | 0.79 |
| 7 | 100276821 | rs4727459 | -0.362 | 2.416 | 1.158 | 0.666 | 0.014 | -0.001 | -0.001 | 0 | 0.13 | 0.19 | 0.18 | 0.13 | 0.23 | 0.13 |
| 7 | 100276994 | rs11520986 | -0.467 | 2.414 | 1.198 | 0.665 | 0.031 | 0 | 0 | 0 | 0.13 | 0.23 | 0.19 | 0.13 | 0.23 | 0.13 |
| 7 | 100281768 | rs221794 | 0.686 | 2.42 | 1.2 | 0.656 | 0.006 | -0.017 | -0.017 | 0.552 | 0.87 | 0.88 | 0.9 | 0.92 | 0.82 | 0.79 |
| 7 | 100282538 | rs11974395 | -0.353 | 2.43 | 1.207 | 0.657 | 0.014 | -0.001 | -0.001 | 0.456 | 0.13 | 0.19 | 0.18 | 0.13 | 0.23 | 0.13 |
| 7 | 100283261 | rs221795 | 1.145 | 2.429 | 1.208 | 0.675 | 0.088 | 0.004 | 0.004 | 0 | 0.82 | 0.66 | 0.75 | 0.79 | 0.65 | 0.71 |
| 7 | 100285888 | rs2272572 | -0.351 | 2.44 | 1.232 | 0.661 | 0.013 | -0.001 | -0.001 | 0 | 0.13 | 0.19 | 0.18 | 0.13 | 0.23 | 0.13 |
| 7 | 100292830 | rs221784 | 0.676 | 2.38 | 1.198 | 0.653 | 0.006 | -0.019 | -0.019 | 0 | 0.87 | 0.88 | 0.9 | 0.92 | 0.82 | 0.78 |
| 7 | 100293639 | rs11979818 | -0.185 | 2.386 | 1.194 | 0.746 | -0.031 | -0.011 | -0.011 | 0 | 0.12 | 0.08 | 0.17 | 0.14 | 0.23 | 0.13 |
| 7 | 100309544 | rs1734910 | 1.649 | 2.274 | 1.186 | 0.74 | 0.039 | -0.022 | -0.022 | 0 | 0.74 | 0.66 | 0.72 | 0.79 | 0.59 | 0.66 |
| 7 | 100313768 | rs1734908 | 1.658 | 2.271 | 1.237 | 0.739 | 0.047 | -0.026 | -0.026 | 0 | 0.74 | 0.63 | 0.71 | 0.79 | 0.59 | 0.66 |
| 7 | 100315517 | rs1734907 | 0.716 | 2.315 | 1.057 | 0.724 | 0.009 | -0.032 | -0.032 | 0 | 0.87 | 0.83 | 0.89 | 0.92 | 0.83 | 0.78 |
| 7 | 100321790 | rs4729606 | -0.379 | 1.888 | 0.847 | 0.862 | -0.029 | -0.01 | -0.01 | 0 | 0.13 | 0.1 | 0.18 | 0.14 | 0.24 | 0.13 |
| 7 | 100361675 | rs2293767 | -1.674 | 1.111 | 0.947 | 1.38 | -0.005 | -0.016 | -0.016 | 0 | 0.28 | 0.25 | 0.27 | 0.37 | 0.3 | 0.47 |
| 7 | 100364552 | rs7801875 | -0.298 | 1.005 | 0.971 | 1.396 | -0.019 | -0.023 | -0.023 | 0 | 0.11 | 0.19 | 0.08 | 0.1 | 0.06 | 0.25 |
| 7 | 100366284 | rs10247980 | -1.884 | 0.929 | 1.25 | 1.131 | 0.002 | 0.009 | 0.009 | 0 | 0.34 | 0.54 | 0.32 | 0.37 | 0.33 | 0.47 |
| 7 | 100371114 | rs314298 | 0.869 | 1.038 | 0.953 | 0.781 | -0.011 | 0.019 | 0.019 | 0.155 | 0.68 | 0.7 | 0.38 | 0.18 | 0.58 | 0.48 |
| 7 | 100373271 | rs540330 | -0.612 | 1.109 | 0.672 | 0.514 | -0.03 | 0.034 | 0.034 | 0.116 | 0.26 | 0.21 | 0.54 | 0.79 | 0.34 | 0.44 |
| 7 | 100386900 | rs191137 | -0.22 | 1.123 | 0.704 | 0.515 | -0.001 | -0.001 | -0.001 | 0.375 | 0.94 | 0.94 | 0.93 | 0.97 | 0.92 | 0.92 |
| 7 | 100410597 | rs2230585 | -0.457 | 0.992 | 0.636 | 0.649 | -0.051 | -0.05 | -0.05 | 0.836 | 0.27 | 0.16 | 0.29 | 0.31 | 0.37 | 0.29 |
| 7 | 100411278 | rs314359 | 0.661 | 0.854 | 1.209 | 0.548 | 0.028 | 0.031 | 0.031 | 0 | 0.68 | 0.46 | 0.67 | 0.63 | 0.63 | 0.56 |
| 7 | 100416250 | rs144173 | 0.698 | 0.846 | 1.238 | 0.544 | 0.022 | 0.025 | 0.025 | 0 | 0.67 | 0.41 | 0.67 | 0.64 | 0.63 | 0.56 |
| 7 | 100422156 | rs2250818 | 1.099 | 0.258 | -0.474 | -0.363 | 0.011 | -0.079 | -0.079 | 1.004 | 0.27 | 0.08 | 0.41 | 0.58 | 0.24 | 0.27 |
| 7 | 100423365 | rs314313 | 0.376 | 0.289 | -0.178 | -0.782 | 0.004 | -0.014 | -0.014 | 0 | 0.36 | 0.28 | 0.19 | 0.04 | 0.31 | 0.2 |
| 7 | 100446235 | rs7794532 | 0.173 | 0.383 | -0.938 | -0.535 | -0.034 | 0.097 | 0.097 | 0 | 0.39 | 0.22 | 0.3 | 0.18 | 0.44 | 0.35 |
| 7 | 100458795 | rs314378 | 0.094 | -0.03 | 0.148 | -0.676 | 0.003 | -0.035 | -0.035 | 0.045 | 0.56 | 0.42 | 0.65 | 0.81 | 0.52 | 0.57 |
| 7 | 100461565 | rs7784933 | 0.746 | -0.115 | -0.178 | 0.003 | -0.024 | -0.095 | -0.095 | 0 | 0.24 | 0.06 | 0.46 | 0.45 | 0.26 | 0.26 |
| 7 | 100466441 | rs2075756 | 0.746 | -0.115 | -0.178 | 0.003 | -0.024 | -0.095 | -0.095 | 0.388 | 0.24 | 0.06 | 0.46 | 0.45 | 0.26 | 0.26 |
| 7 | 100468103 | rs7800241 | 0.716 | -0.09 | -0.185 | -0.004 | -0.026 | -0.095 | -0.095 | 0 | 0.24 | 0.06 | 0.46 | 0.45 | 0.27 | 0.26 |
| 7 | 100471044 | rs6706 | 1.571 | -0.4 | -0.367 | -0.212 | -0.025 | 0.095 | 0.095 | 0 | 0.2 | 0.04 | 0.1 | 0.03 | 0.18 | 0.1 |
| 7 | 100471121 | rs12538287 | 1 | -0.401 | -0.368 | -0.21 | -0.023 | -0.095 | -0.095 | 0.448 | 0.24 | 0.06 | 0.46 | 0.45 | 0.27 | 0.26 |
| 7 | 100472380 | rs12705091 | 1.181 | -0.408 | -0.389 | -0.19 | -0.03 | -0.097 | -0.097 | 0.008 | 0.22 | 0.04 | 0.13 | 0.36 | 0.19 | 0.25 |
| 7 | 100473550 | rs12705092 | 1.587 | -0.411 | -0.389 | -0.269 | -0.026 | 0.094 | 0.094 | 0 | 0.2 | 0.04 | 0.1 | 0.03 | 0.18 | 0.1 |
| 7 | 100473729 | rs4729617 | -0.527 | -0.437 | -0.244 | -0.289 | 0.001 | -0.02 | -0.02 | 0 | 0.56 | 0.4 | 0.65 | 0.81 | 0.52 | 0.57 |
| 7 | 100474289 | rs12667888 | 1.598 | -0.433 | -0.255 | -0.286 | -0.026 | 0.094 | 0.094 | 0 | 0.2 | 0.04 | 0.1 | 0.03 | 0.18 | 0.1 |
| 7 | 100476110 | rs12669120 | 0.127 | -0.437 | -0.242 | -0.279 | -0.056 | 0.175 | 0.175 | 0.096 | 0.47 | 0.1 | 0.59 | 0.81 | 0.46 | 0.51 |

| | | | | | | | | | | | | | | | | |
|---|---|---|---|---|---|---|---|---|---|---|---|---|---|---|---|---|
| 7 | 100476229 | rs12666785 | -0.57 | -0.459 | -0.109 | -0.29 | -0.002 | 0.009 | 0.009 | 0 | 0.55 | 0.35 | 0.65 | 0.81 | 0.51 | 0.56 |
| 7 | 100477829 | rs12705094 | 0.141 | -0.455 | -0.121 | -0.288 | -0.056 | 0.174 | 0.174 | 0.092 | 0.46 | 0.1 | 0.59 | 0.81 | 0.46 | 0.51 |
| 7 | 100482026 | rs15624 | -0.521 | -0.452 | -0.119 | -0.284 | 0.003 | -0.033 | -0.033 | 0 | 0.56 | 0.41 | 0.65 | 0.81 | 0.52 | 0.57 |
| 7 | 100484381 | rs12705095 | 1.587 | -0.447 | -0.177 | -0.288 | -0.026 | 0.094 | 0.094 | 0 | 0.2 | 0.04 | 0.1 | 0.03 | 0.18 | 0.1 |
| 7 | 100484917 | rs6954151 | 0.146 | -0.501 | -0.172 | -0.35 | -0.056 | 0.174 | 0.174 | 0.097 | 0.46 | 0.1 | 0.59 | 0.81 | 0.46 | 0.52 |
| 7 | 100486035 | rs2720392 | 0.87 | -0.599 | -0.154 | -0.439 | -0.076 | 0.17 | 0.17 | 0.689 | 0.4 | 0.16 | 0.3 | 0.13 | 0.48 | 0.36 |
| 7 | 100486088 | rs3087504 | 0.096 | -0.416 | -0.114 | -0.435 | -0.056 | 0.175 | 0.175 | 0.085 | 0.47 | 0.1 | 0.59 | 0.81 | 0.46 | 0.52 |
| 7 | 100486110 | rs11171 | 1.228 | -0.419 | 0.028 | -0.436 | 0.013 | -0.03 | -0.03 | 2.828 | 0.08 | 0.21 | 0.07 | 0 | 0.05 | 0.04 |
| 7 | 100486656 | rs13241786 | -0.856 | -0.435 | 0.459 | -0.441 | -0.014 | 0.07 | 0.07 | 0 | 0.59 | 0.61 | 0.69 | 0.82 | 0.54 | 0.62 |
| 7 | 100487721 | rs17228616 | 1.071 | -0.44 | 0.19 | -0.622 | 0.013 | -0.032 | -0.032 | 0 | 0.08 | 0.25 | 0.06 | 0 | 0.05 | 0.04 |
| 7 | 100490077 | rs7636 | 0.916 | -0.34 | -0.063 | -0.617 | 0.009 | -0.037 | -0.037 | 0 | 0.09 | 0.29 | 0.07 | 0 | 0.05 | 0.04 |
| 7 | 100490797 | rs1799805 | 1.907 | -0.34 | -0.165 | -0.617 | -0.007 | 0.034 | 0.034 | 0 | 0.06 | 0 | 0.04 | 0 | 0.05 | 0.04 |
| 7 | 100492322 | rs3808356 | 1.144 | -0.344 | -0.173 | -0.557 | -0.028 | -0.096 | -0.096 | 0.781 | 0.23 | 0.05 | 0.46 | 0.44 | 0.27 | 0.25 |
| 7 | 100492568 | rs10953305 | 0.041 | -0.289 | -0.237 | -0.519 | -0.05 | 0.211 | 0.211 | 0.077 | 0.46 | 0.12 | 0.6 | 0.84 | 0.46 | 0.52 |
| 7 | 100493998 | rs3757869 | -0.738 | -0.287 | -0.242 | -0.528 | 0.003 | -0.048 | -0.048 | 0.563 | 0.57 | 0.46 | 0.67 | 0.84 | 0.51 | 0.58 |
| 7 | 100497131 | rs13245899 | 0.887 | 0.137 | -1.97 | -0.153 | -0.021 | -0.106 | -0.106 | 0 | 0.23 | 0.06 | 0.14 | 0.4 | 0.19 | 0.27 |
| 7 | 100498475 | rs3847063 | -0.059 | 0.327 | -1.811 | -1.039 | -0.05 | 0.215 | 0.215 | 0.077 | 0.46 | 0.12 | 0.6 | 0.84 | 0.46 | 0.52 |
| 7 | 100505792 | rs7803865 | -0.063 | 0.35 | -1.799 | -0.993 | -0.054 | 0.195 | 0.195 | 0.065 | 0.46 | 0.12 | 0.6 | 0.84 | 0.46 | 0.53 |
| 7 | 100516003 | rs10278546 | 0.395 | 0.514 | -1.23 | -0.641 | -0.003 | 0 | 0 | 0 | 0.19 | 0.17 | 0.1 | 0.01 | 0.2 | 0.12 |
| 7 | 100529261 | rs4729624 | -0.555 | 0.442 | -1.919 | -0.433 | -0.067 | 0.015 | 0.015 | 0.058 | 0.68 | 0.96 | 0.79 | 0.84 | 0.65 | 0.75 |
| 7 | 100532733 | rs4637750 | 0.462 | 0.301 | -1.421 | -0.313 | 0.001 | -0.003 | -0.003 | 0 | 0.3 | 0.37 | 0.51 | 0.34 | 0.29 | 0.39 |
| 7 | 100537743 | rs7787389 | -1.142 | 0.338 | -1.897 | -0.192 | -0.004 | 0.048 | 0.048 | 0.096 | 0.91 | 0.98 | 0.94 | 1 | 0.91 | 0.94 |
| 7 | 100542445 | rs6980219 | 1.122 | 0.505 | -1.843 | -0.025 | -0.061 | -0.039 | -0.039 | 0 | 0.23 | 0.01 | 0.16 | 0.15 | 0.27 | 0.19 |
| 7 | 100548261 | rs4341099 | 0.453 | 0.659 | -1.2 | -0.206 | -0.002 | -0.008 | -0.008 | 0 | 0.28 | 0.34 | 0.17 | 0.49 | 0.24 | 0.31 |
| 7 | 100548273 | rs4326315 | -0.623 | 0.362 | -0.826 | -0.082 | -0.003 | 0 | 0 | 1.108 | 0.54 | 0.61 | 0.37 | 0.53 | 0.53 | 0.61 |
| 7 | 100552017 | rs9691167 | -0.419 | 0.793 | -1.018 | 0.171 | 0 | -0.005 | -0.005 | 0 | 0.46 | 0.45 | 0.29 | 0.52 | 0.46 | 0.36 |
| 7 | 100606918 | rs4316092 | -0.028 | 0.983 | -1.1 | 0.419 | -0.005 | -0.013 | -0.013 | 0.012 | 0.71 | 0.73 | 0.84 | 0.61 | 0.76 | 0.72 |
| 7 | 100623849 | rs11765099 | 0.426 | 0.533 | -0.991 | 0.079 | 0 | -0.004 | -0.004 | 0 | 0.39 | 0.25 | 0.34 | 0.58 | 0.44 | 0.51 |
| 7 | 100626955 | rs13224928 | -0.642 | 0.676 | -0.962 | -0.169 | -0.027 | 0.004 | 0.004 | 0.535 | 0.79 | 0.87 | 0.82 | 0.8 | 0.73 | 0.71 |
| 7 | 100627938 | rs12530867 | 1.489 | 0.39 | -0.904 | -0.253 | 0.011 | -0.009 | -0.009 | 0.963 | 0.11 | 0.1 | 0.33 | 0.17 | 0.07 | 0.04 |
| 7 | 100628107 | rs4605982 | 0.24 | 0.366 | -0.44 | -0.276 | 0.005 | 0 | 0 | 0.311 | 0.67 | 0.6 | 0.47 | 0.63 | 0.66 | 0.67 |
| 7 | 100647920 | rs11772963 | 0.781 | 0.214 | -0.427 | -0.463 | -0.004 | 0.012 | 0.012 | 0 | 0.44 | 0.34 | 0.37 | 0.61 | 0.51 | 0.52 |
| 7 | 100648029 | rs10953315 | -0.773 | 0.239 | -0.382 | -0.455 | 0.006 | -0.002 | -0.002 | 0 | 0.78 | 0.77 | 0.81 | 0.8 | 0.74 | 0.73 |
| 7 | 100652181 | rs12538004 | 0.46 | 0.512 | -0.331 | -0.215 | 0 | -0.007 | -0.007 | 2.216 | 0.46 | 0.49 | 0.38 | 0.61 | 0.52 | 0.53 |
| 7 | 100666552 | rs13239478 | -0.468 | 0.251 | -0.021 | 0.11 | -0.033 | 0.014 | 0.014 | 0.015 | 0.2 | 0.38 | 0.11 | 0.23 | 0.12 | 0.29 |
| 7 | 100679809 | rs4992074 | -0.434 | 0.247 | -0.074 | 0.125 | -0.024 | 0.016 | 0.016 | 0 | 0.19 | 0.34 | 0.1 | 0.25 | 0.12 | 0.29 |
| 7 | 100692376 | rs6968731 | 0.467 | 0.503 | -0.782 | -0.487 | -0.01 | 0.068 | 0.068 | 0 | 0.19 | 0.08 | 0.16 | 0 | 0.22 | 0.09 |
| 7 | 100692680 | rs10246021 | -0.88 | 0.358 | -0.845 | -0.787 | -0.028 | 0.084 | 0.084 | 0 | 0.61 | 0.8 | 0.47 | 0.84 | 0.62 | 0.77 |
| 7 | 100693575 | rs7806980 | 1.033 | 0.096 | -0.271 | -0.653 | -0.003 | 0.027 | 0.027 | 0 | 0.24 | 0.3 | 0.2 | 0.58 | 0.29 | 0.37 |
| 7 | 100693849 | rs6978780 | 1.047 | 0.603 | -0.341 | -0.698 | -0.045 | 0.069 | 0.069 | 0 | 0.18 | 0.03 | 0.16 | 0.03 | 0.24 | 0.09 |
| 7 | 100700531 | rs6965671 | -0.864 | 0.064 | -0.018 | -0.553 | -0.017 | 0.093 | 0.093 | 0.076 | 0.67 | 0.87 | 0.54 | 0.84 | 0.73 | 0.79 |
| 7 | 100701610 | rs4729656 | -0.035 | -0.115 | 0.205 | -0.228 | -0.001 | 0.073 | 0.073 | 0 | 0.54 | 0.71 | 0.46 | 0.72 | 0.61 | 0.68 |
| 7 | 100707378 | rs13221286 | -1.058 | -0.104 | 0.466 | -0.347 | -0.018 | -0.031 | -0.031 | 0 | 0.27 | 0.58 | 0.19 | 0.15 | 0.23 | 0.15 |
| 7 | 100711147 | rs13228919 | 0.52 | -0.06 | 0.44 | -0.31 | -0.002 | -0.001 | -0.001 | 0 | 0.13 | 0.21 | 0.09 | 0.12 | 0.12 | 0.11 |
| 7 | 100713042 | rs7782171 | -0.832 | -0.084 | 0.227 | 0.11 | -0.006 | -0.039 | -0.039 | 0 | 0.66 | 0.85 | 0.53 | 0.6 | 0.73 | 0.6 |
| 7 | 100715933 | rs7797230 | 1.16 | -0.12 | 0.282 | 0.089 | -0.028 | -0.019 | -0.019 | 0 | 0.38 | 0.24 | 0.33 | 0.45 | 0.5 | 0.44 |
| 7 | 100721337 | rs10231529 | -0.861 | -0.439 | 0.459 | -0.042 | -0.04 | -0.047 | -0.047 | 1.575 | 0.27 | 0.47 | 0.16 | 0.15 | 0.2 | 0.15 |
| 7 | 100727412 | rs6968929 | -0.666 | -0.328 | 0.096 | 0.281 | 0 | -0.078 | -0.078 | 0.117 | 0.65 | 0.89 | 0.53 | 0.52 | 0.74 | 0.6 |
| 7 | 100731829 | rs6948536 | -0.795 | -0.312 | -0.084 | 0.271 | -0.039 | -0.11 | -0.11 | 0 | 0.66 | 0.96 | 0.54 | 0.55 | 0.75 | 0.61 |
| 7 | 100733565 | rs10900 | 0.008 | -0.319 | 0.257 | 0.205 | -0.034 | -0.079 | -0.079 | 0 | 0.13 | 0.34 | 0.08 | 0 | 0.07 | 0.03 |
| 7 | 100739682 | rs6942824 | 0.935 | -0.014 | 0.156 | 0.396 | -0.052 | -0.005 | -0.005 | 0.771 | 0.38 | 0.25 | 0.36 | 0.38 | 0.53 | 0.45 |
| 7 | 100740785 | rs12912 | 0.039 | -0.074 | 0.285 | 0.408 | -0.002 | -0.001 | -0.001 | 0 | 0.52 | 0.5 | 0.46 | 0.55 | 0.67 | 0.58 |
| 7 | 100746658 | rs10953322 | -0.226 | -0.083 | 0.6 | 0.489 | 0.013 | 0.048 | 0.048 | 0 | 0.14 | 0.29 | 0.11 | 0.16 | 0.14 | 0.13 |
| 7 | 100755709 | rs12667943 | 0.337 | 0.032 | 0.478 | 0.475 | 0.026 | 0.023 | 0.023 | 0 | 0.11 | 0.22 | 0.1 | 0.1 | 0.13 | 0.12 |
| 7 | 100759727 | rs6955537 | 0.156 | -0.648 | -0.02 | 0.224 | 0 | -0.004 | -0.004 | 0.043 | 0.3 | 0.32 | 0.44 | 0.17 | 0.22 | 0.19 |
| 7 | 100766603 | rs6950982 | 0.41 | -0.612 | -0.846 | 0.194 | -0.038 | 0.09 | 0.09 | 0 | 0.27 | 0.04 | 0.42 | 0.15 | 0.2 | 0.17 |
| 7 | 100769538 | rs2227631 | -0.821 | -0.412 | -0.571 | 0.629 | 0.005 | 0.034 | 0.034 | 0 | 0.54 | 0.75 | 0.63 | 0.61 | 0.41 | 0.51 |
| 7 | 100775686 | rs2227672 | 0.763 | 0.032 | -0.505 | 0.914 | -0.021 | -0.022 | -0.022 | 0 | 0.08 | 0.03 | 0.1 | 0.11 | 0.12 | 0.13 |
| 7 | 100776931 | rs2227684 | 1.099 | -0.194 | -0.284 | 0.707 | 0.017 | 0.015 | 0.015 | 0 | 0.38 | 0.49 | 0.29 | 0.46 | 0.45 | 0.4 |
| 7 | 100779244 | rs2227692 | 0.849 | -0.726 | -0.439 | 0.464 | 0.005 | -0.007 | -0.007 | 0 | 0.21 | 0.16 | 0.38 | 0.36 | 0.09 | 0.24 |
| 7 | 100781445 | rs7242 | 1.456 | -0.521 | -0.473 | 0.395 | 0.015 | 0.013 | 0.013 | 0 | 0.38 | 0.49 | 0.3 | 0.45 | 0.45 | 0.4 |
| 7 | 100781615 | rs1050813 | 0.671 | -0.413 | -0.747 | 0.174 | -0.08 | 0.114 | 0.114 | 0 | 0.18 | 0.01 | 0.11 | 0 | 0.21 | 0.11 |
| 7 | 100782460 | rs1050955 | 0.052 | -0.423 | -0.481 | 0.358 | 0.019 | -0.023 | -0.023 | 0 | 0.32 | 0.25 | 0.48 | 0.47 | 0.21 | 0.38 |
| 7 | 100783736 | rs13238709 | 0.916 | 0.238 | -0.302 | 0.706 | 0.011 | 0.01 | 0.01 | 0 | 0.38 | 0.46 | 0.3 | 0.45 | 0.46 | 0.4 |
| 7 | 100795046 | rs6944945 | -1.205 | 0.476 | 0.264 | 1.327 | -0.002 | 0.015 | 0.015 | 1.356 | 0.36 | 0.51 | 0.32 | 0.47 | 0.36 | 0.6 |
| 7 | 100798611 | rs2239537 | 1.004 | 0.68 | 0.453 | 1.613 | -0.017 | 0.061 | 0.061 | 0.355 | 0.7 | 0.52 | 0.77 | 0.55 | 0.74 | 0.53 |
| 7 | 100800131 | rs4727481 | -1.199 | 0.685 | 0.361 | 1.6 | -0.017 | -0.036 | -0.036 | 0 | 0.93 | 0.99 | 0.95 | 0.91 | 0.92 | 0.91 |
| 7 | 100800635 | rs2074686 | 1.856 | 0.798 | 0.4 | 1.565 | -0.003 | -0.002 | -0.002 | 0 | 0.58 | 0.36 | 0.62 | 0.44 | 0.53 | 0.34 |
| 7 | 100804140 | rs1048303 | 1.788 | 0.31 | 0.43 | 1.601 | 0 | -0.002 | -0.002 | 0 | 0.58 | 0.4 | 0.63 | 0.43 | 0.56 | 0.34 |

| | | | | | | | | | | | | | | | | |
|---|---|---|---|---|---|---|---|---|---|---|---|---|---|---|---|---|
| 7 | 100804430 | rs1048365 | -0.522 | 0.315 | 0.428 | 1.628 | -0.017 | 0.049 | 0.049 | 0.22 | 0.23 | 0.39 | 0.17 | 0.36 | 0.19 | 0.38 |
| 7 | 100805704 | rs9918635 | 1.009 | 0.328 | 0.406 | 1.642 | -0.015 | 0.062 | 0.062 | 0.664 | 0.71 | 0.59 | 0.79 | 0.52 | 0.74 | 0.55 |
| 7 | 100810983 | rs2898568 | 0.573 | 0.509 | 0.491 | 1.366 | -0.014 | -0.013 | -0.013 | 0 | 0.83 | 0.88 | 0.81 | 0.79 | 0.76 | 0.75 |
| 7 | 100814034 | rs1006507 | 1.854 | 0.475 | 0.529 | 1.496 | -0.007 | -0.047 | -0.047 | 0 | 0.66 | 0.24 | 0.66 | 0.47 | 0.61 | 0.51 |
| 7 | 100814078 | rs1859528 | 1.854 | 0.476 | 0.529 | 1.496 | -0.009 | -0.047 | -0.047 | 0 | 0.66 | 0.24 | 0.66 | 0.47 | 0.61 | 0.51 |
| 7 | 100814722 | rs10953326 | 0.095 | 0.498 | 0.527 | 1.517 | -0.007 | -0.001 | -0.001 | 2.564 | 0.11 | 0.07 | 0.15 | 0.13 | 0.17 | 0.18 |
| 7 | 100818846 | rs734688 | 1.695 | 0.502 | 0.679 | 1.513 | -0.003 | -0.017 | -0.017 | 0 | 0.66 | 0.3 | 0.66 | 0.47 | 0.61 | 0.51 |
| 7 | 100833153 | rs11971673 | -0.21 | 0.6 | 0.682 | 1.505 | 0.001 | 0 | 0 | 0 | 0.08 | 0.07 | 0.11 | 0.13 | 0.13 | 0.17 |
| 7 | 100835885 | rs7780766 | -0.126 | 0.597 | 0.918 | 1.54 | 0.001 | 0.099 | 0.099 | 2.552 | 0.11 | 0.34 | 0.11 | 0.23 | 0.09 | 0.19 |
| 7 | 100839401 | rs3735338 | -0.382 | 0.941 | -0.357 | 1.031 | -0.002 | -0.002 | -0.002 | 1.15 | 0.08 | 0.07 | 0.12 | 0.13 | 0.13 | 0.17 |
| 7 | 100850048 | rs4729671 | -0.677 | 0.21 | 0.037 | 0.89 | -0.046 | 0.012 | 0.012 | 0 | 0.2 | 0.46 | 0.12 | 0.19 | 0.11 | 0.26 |
| 7 | 100850659 | rs7802724 | -0.677 | 0.21 | 0.037 | 0.915 | -0.046 | 0.015 | 0.015 | 0 | 0.2 | 0.46 | 0.12 | 0.19 | 0.11 | 0.26 |
| 7 | 100853173 | rs1468358 | -1.203 | -0.076 | -0.35 | 1.252 | 0.007 | -0.005 | -0.005 | 0 | 0.39 | 0.6 | 0.28 | 0.49 | 0.31 | 0.5 |
| 7 | 100853231 | rs7777930 | 1.322 | 0.429 | -1.136 | 0.581 | -0.05 | 0.066 | 0.066 | 0 | 0.14 | 0.01 | 0.08 | 0.04 | 0.15 | 0.12 |
| 7 | 100861213 | rs17319250 | -0.313 | 0.168 | -0.699 | 0.679 | -0.006 | 0.012 | 0.012 | 0.06 | 0.26 | 0.23 | 0.22 | 0.17 | 0.25 | 0.32 |
| 7 | 100861452 | rs11769624 | -0.313 | 0.168 | -0.699 | 0.679 | -0.006 | 0.012 | 0.012 | 0.06 | 0.26 | 0.23 | 0.22 | 0.17 | 0.25 | 0.32 |
| 7 | 100865084 | rs10265206 | 0.683 | 0.227 | -0.346 | 0.628 | -0.008 | -0.026 | -0.026 | 0 | 0.06 | 0.12 | 0.03 | 0 | 0.04 | 0.06 |
| 7 | 100865847 | rs10953329 | -1.106 | 0.213 | -0.132 | 0.674 | -0.002 | -0.009 | -0.009 | 0 | 0.34 | 0.58 | 0.27 | 0.26 | 0.31 | 0.42 |
| 7 | 100873835 | rs10250685 | -0.949 | 0.271 | -0.317 | 0.578 | 0 | -0.021 | -0.021 | 0 | 0.33 | 0.44 | 0.26 | 0.17 | 0.3 | 0.41 |
| 7 | 100874354 | rs7778271 | 0.124 | 0.273 | -0.307 | 0.578 | -0.007 | -0.047 | -0.047 | 0 | 0.08 | 0.21 | 0.05 | 0 | 0.06 | 0.07 |
| 7 | 100881640 | rs3757460 | -0.971 | 0.3 | -0.251 | 0.604 | 0 | -0.022 | -0.022 | 4.257 | 0.33 | 0.45 | 0.26 | 0.17 | 0.3 | 0.41 |
| 7 | 100883973 | rs10241107 | -0.484 | 0.017 | -0.008 | 0.908 | -0.007 | -0.049 | -0.049 | 0.9 | 0.09 | 0.26 | 0.05 | 0 | 0.06 | 0.07 |
| 7 | 100894750 | rs6979122 | -0.368 | 0.04 | -0.092 | 0.918 | -0.007 | -0.046 | -0.046 | 0.038 | 0.08 | 0.22 | 0.05 | 0 | 0.06 | 0.07 |
| 7 | 100895480 | rs6942609 | 0.033 | 0.345 | -0.706 | 1.201 | -0.057 | -0.084 | -0.084 | 0.577 | 0.12 | 0.01 | 0.14 | 0.38 | 0.2 | 0.16 |
| 7 | 100928622 | rs4727484 | -0.047 | 0.044 | -0.419 | 0.359 | 0 | -0.002 | -0.002 | 0 | 0.65 | 0.66 | 0.77 | 0.41 | 0.65 | 0.7 |
| 7 | 100931873 | rs6961898 | 0.592 | 0.028 | -0.47 | 0.037 | -0.008 | 0.03 | 0.03 | 0 | 0.45 | 0.41 | 0.38 | 0.25 | 0.48 | 0.52 |
| 7 | 100938008 | rs10269821 | 0.243 | -0.539 | -0.144 | -0.252 | -0.021 | -0.096 | -0.096 | 3.131 | 0.26 | 0.31 | 0.09 | 0 | 0.18 | 0.16 |
| 7 | 100954700 | rs34532455 | 0.404 | -0.374 | -0.081 | -0.19 | -0.001 | -0.04 | -0.04 | 0 | 0.23 | 0.17 | 0.09 | 0 | 0.18 | 0.15 |
| 7 | 100958270 | rs11544858 | 0.404 | -0.374 | -0.081 | -0.19 | -0.001 | -0.04 | -0.04 | 0 | 0.23 | 0.17 | 0.09 | 0 | 0.18 | 0.15 |
| 7 | 100960954 | rs34169259 | 0.472 | -0.442 | -0.141 | -0.186 | -0.002 | -0.042 | -0.042 | 1.016 | 0.23 | 0.17 | 0.09 | 0 | 0.18 | 0.15 |
| 7 | 100964722 | rs28719520 | -0.264 | -0.372 | -0.148 | 0.193 | 0.005 | 0.036 | 0.036 | 0 | 0.08 | 0.12 | 0.33 | 0.25 | 0.1 | 0.09 |
| 7 | 100965601 | rs3807513 | -0.009 | -0.391 | 0.232 | 0.51 | 0.002 | 0.106 | 0.106 | 0 | 0.17 | 0.3 | 0.14 | 0.39 | 0.18 | 0.16 |
| 7 | 100968363 | rs7799285 | -1.771 | -0.632 | 0.057 | -0.058 | -0.104 | 0.093 | 0.093 | 0 | 0.56 | 0.95 | 0.6 | 0.71 | 0.49 | 0.48 |
| 7 | 100968533 | rs13236755 | 0.731 | -0.593 | 0.095 | -0.048 | -0.001 | -0.04 | -0.04 | 0 | 0.23 | 0.17 | 0.09 | 0 | 0.18 | 0.15 |
| 7 | 100979310 | rs6958889 | -1.622 | 0.036 | 0.62 | 0.35 | 0.01 | 0.03 | 0.03 | 0.034 | 0.27 | 0.56 | 0.49 | 0.64 | 0.29 | 0.25 |
| 7 | 100989201 | rs7798805 | -1.185 | 0.075 | 0.369 | 0.043 | 0.02 | 0.007 | 0.007 | 0 | 0.45 | 0.62 | 0.29 | 0.55 | 0.43 | 0.42 |
| 7 | 100995857 | rs4727485 | -0.224 | 0.195 | 0.132 | 0.027 | -0.008 | -0.039 | -0.039 | 1.545 | 0.29 | 0.4 | 0.19 | 0.13 | 0.28 | 0.21 |
| 7 | 101001016 | rs6943474 | -1.55 | 0.227 | -0.338 | 0.105 | -0.054 | 0.103 | 0.103 | 0 | 0.54 | 0.87 | 0.39 | 0.68 | 0.53 | 0.48 |
| 7 | 101020021 | rs10237610 | 0.132 | 0.354 | -0.089 | 0.408 | 0.015 | 0.165 | 0.165 | 0.058 | 0.84 | 0.69 | 0.85 | 0.51 | 0.8 | 0.82 |
| 7 | 101024309 | rs10268365 | -0.157 | 0.302 | -0.542 | 0.52 | -0.019 | 0.015 | 0.015 | 0 | 0.06 | 0.17 | 0.02 | 0.06 | 0.01 | 0.08 |
| 7 | 101029430 | rs6944691 | -1.795 | 0.348 | -0.646 | 0.415 | -0.002 | 0.005 | 0.005 | 0 | 0.16 | 0.26 | 0.12 | 0.19 | 0.15 | 0.15 |
| 7 | 101032269 | rs9640666 | 0.675 | 0.502 | -0.94 | 0.157 | -0.021 | 0.09 | 0.09 | 0 | 0.5 | 0.18 | 0.61 | 0.38 | 0.46 | 0.57 |
| 7 | 101051157 | rs4729704 | 0.94 | 0.446 | -1.118 | -0.017 | -0.045 | 0.036 | 0.036 | 0 | 0.46 | 0.1 | 0.62 | 0.42 | 0.49 | 0.55 |
| 7 | 101051502 | rs4729705 | 0.755 | 0.419 | -1.093 | -0.02 | -0.019 | 0.011 | 0.011 | 0 | 0.48 | 0.14 | 0.64 | 0.45 | 0.5 | 0.55 |
| 7 | 101063059 | rs10254310 | 0.3 | 0.521 | -0.378 | -0.166 | 0.089 | 0.351 | 0.351 | 0.105 | 0.89 | 0.65 | 0.88 | 0.51 | 0.85 | 0.8 |
| 7 | 101065028 | rs13229048 | 0.257 | 0.281 | -1.334 | -0.341 | -0.016 | 0.184 | 0.184 | 0 | 0.28 | 0.06 | 0.14 | 0 | 0.27 | 0.2 |
| 7 | 101067089 | rs4045 | -0.315 | 0.022 | -1.286 | -0.192 | -0.094 | -0.176 | -0.176 | 0 | 0.7 | 0.93 | 0.78 | 0.56 | 0.62 | 0.63 |
| 7 | 101070104 | rs4729707 | 0.936 | 0.231 | -0.935 | -0.113 | -0.011 | 0.013 | 0.013 | 0 | 0.48 | 0.18 | 0.65 | 0.45 | 0.5 | 0.53 |
| 7 | 101091470 | rs869127 | -0.941 | 0.033 | 0.128 | -0.493 | -0.027 | -0.033 | -0.033 | 0.122 | 0.11 | 0.67 | 0.07 | 0.06 | 0.04 | 0.03 |
| 7 | 101097366 | rs1476653 | -0.91 | 0.408 | -0.513 | -0.578 | -0.017 | -0.054 | -0.054 | 0 | 0.59 | 0.89 | 0.44 | 0.48 | 0.59 | 0.54 |
| 7 | 101102046 | rs11770740 | -1.095 | 0.665 | -0.083 | -0.815 | 0.005 | -0.025 | -0.025 | 0 | 0.14 | 0.22 | 0.1 | 0.04 | 0.16 | 0.09 |
| 7 | 101103457 | rs6973489 | -1.104 | 0.63 | -0.13 | -0.725 | -0.013 | 0.001 | 0.001 | 0 | 0.58 | 0.88 | 0.43 | 0.37 | 0.58 | 0.53 |
| 7 | 101105017 | rs7796259 | -0.012 | 0.85 | 0.006 | -0.724 | 0.01 | -0.006 | -0.006 | 0.22 | 0.07 | 0.12 | 0.04 | 0.04 | 0.09 | 0.07 |
| 7 | 101108299 | rs10229291 | -0.631 | 0.497 | 0.673 | -0.922 | -0.004 | 0.025 | 0.025 | 0.007 | 0.45 | 0.65 | 0.36 | 0.32 | 0.51 | 0.47 |
| 7 | 101109296 | rs2097907 | 0.143 | 0.675 | 0.969 | -0.979 | 0 | -0.073 | -0.073 | 0.114 | 0.35 | 0.28 | 0.21 | 0 | 0.38 | 0.24 |
| 7 | 101113729 | rs10953342 | -0.643 | 0.626 | 0.849 | -0.921 | -0.006 | 0.025 | 0.025 | 0.006 | 0.45 | 0.65 | 0.36 | 0.32 | 0.51 | 0.51 |
| 7 | 101115923 | rs1008062 | -0.126 | 0.637 | 0.578 | -0.941 | -0.016 | 0.046 | 0.046 | 0.014 | 0.4 | 0.27 | 0.24 | 0.04 | 0.46 | 0.31 |
| 7 | 101116218 | rs1008064 | -1.299 | 0.649 | 0.113 | -1.015 | -0.106 | -0.141 | -0.141 | 0 | 0.59 | 0.97 | 0.43 | 0.5 | 0.57 | 0.65 |
| 7 | 101127455 | rs13232646 | -0.981 | 0.397 | 0.453 | -2.161 | -0.034 | 0.156 | 0.156 | 0 | 0.56 | 0.78 | 0.33 | 0.21 | 0.5 | 0.45 |
| 7 | 101138286 | rs11766744 | -0.92 | 0.228 | 0.58 | -2.195 | 0.007 | -0.001 | -0.001 | 0 | 0.1 | 0.14 | 0.1 | 0.12 | 0.12 | 0.1 |
| 7 | 101138662 | rs6967458 | -0.458 | 0.014 | 1.024 | -2.266 | -0.031 | -0.046 | -0.046 | 0 | 0.1 | 0.3 | 0.04 | 0.03 | 0.03 | 0.04 |
| 7 | 101139424 | rs885501 | 0.217 | 0.179 | 0.811 | -2.39 | 0 | 0.129 | 0.129 | 0 | 0.35 | 0.13 | 0.2 | 0.04 | 0.37 | 0.28 |
| 7 | 101142190 | rs12538381 | -0.568 | 0.181 | 1.041 | -2.045 | 0.013 | 0.011 | 0.011 | 0.009 | 0.07 | 0.13 | 0.08 | 0.12 | 0.09 | 0.1 |
| 7 | 101145350 | rs1558011 | -2.39 | 0.106 | 1.081 | -2.158 | -0.012 | -0.024 | -0.024 | 0 | 0.23 | 0.7 | 0.16 | 0.13 | 0.14 | 0.19 |
| 7 | 101150073 | rs10250055 | -1.736 | 0.054 | 0.928 | -2.206 | -0.048 | -0.069 | -0.069 | 0 | 0.18 | 0.62 | 0.1 | 0.07 | 0.08 | 0.14 |
| 7 | 101154626 | rs7778986 | -2.484 | 0.064 | 0.745 | -2.206 | 0.022 | 0.027 | 0.027 | 0 | 0.27 | 0.76 | 0.18 | 0.17 | 0.16 | 0.25 |
| 7 | 101155321 | rs10241776 | 0.202 | 0.275 | 1.038 | -2.114 | 0.001 | -0.006 | -0.006 | 0 | 0.07 | 0.28 | 0.04 | 0.03 | 0.03 | 0.03 |
| 7 | 101158329 | rs6975738 | -0.02 | 0.486 | 0.579 | -2.425 | 0.001 | -0.05 | -0.05 | 0 | 0.39 | 0.36 | 0.24 | 0.03 | 0.39 | 0.28 |
| 7 | 101159870 | rs11770737 | -0.919 | -0.229 | 0.281 | -2.751 | 0.032 | 0.04 | 0.04 | 0 | 0.19 | 0.11 | 0.12 | 0.14 | 0.13 | 0.15 |

| | | | | | | | | | | | | | | | | |
|---|---|---|---|---|---|---|---|---|---|---|---|---|---|---|---|---|
| 7 | 101159913 | rs6947735 | -1.969 | 0.033 | 0.413 | -2.736 | -0.018 | 0.039 | 0.039 | 0 | 0.69 | 0.79 | 0.47 | 0.18 | 0.64 | 0.47 |
| 7 | 101161537 | rs1543885 | 0.188 | -0.209 | 0.349 | -2.74 | 0.007 | -0.039 | -0.039 | 0 | 0.41 | 0.36 | 0.24 | 0.03 | 0.37 | 0.28 |
| 7 | 101167569 | rs13236381 | -1.147 | -0.111 | 0.153 | -2.604 | -0.001 | 0.013 | 0.013 | 0 | 0.26 | 0.25 | 0.2 | 0.13 | 0.23 | 0.18 |
| 7 | 101171028 | rs10279545 | -1.239 | -0.209 | -0.12 | -2.618 | -0.026 | 0.033 | 0.033 | 0 | 0.63 | 0.72 | 0.4 | 0.1 | 0.53 | 0.37 |
| 7 | 101171175 | rs10264271 | -1.289 | -0.127 | 0.005 | -2.476 | -0.031 | -0.114 | -0.114 | 0 | 0.33 | 0.48 | 0.2 | 0.1 | 0.23 | 0.14 |
| 7 | 101174965 | rs13245946 | 0.194 | 0.22 | -0.804 | -2.14 | 0.005 | -0.001 | -0.001 | 0 | 0.07 | 0.07 | 0.09 | 0.08 | 0.14 | 0.12 |
| 7 | 101176140 | rs6949545 | -1.063 | -0.477 | -0.648 | -2.219 | -0.007 | -0.013 | -0.013 | 0 | 0.2 | 0.24 | 0.08 | 0.03 | 0.08 | 0.07 |
| 7 | 101176641 | rs6954348 | -1.078 | -0.477 | -0.648 | -2.22 | -0.007 | -0.012 | -0.012 | 0 | 0.2 | 0.24 | 0.08 | 0.03 | 0.08 | 0.07 |
| 7 | 101183108 | rs6465817 | -0.524 | -0.016 | -0.841 | -1.955 | -0.01 | 0.124 | 0.124 | 0 | 0.56 | 0.76 | 0.35 | 0.1 | 0.48 | 0.33 |
| 7 | 101190922 | rs17135617 | 0.009 | -0.33 | -1.199 | -1.589 | 0.047 | 0.018 | 0.018 | 0 | 0.11 | 0.05 | 0.04 | 0.05 | 0.03 | 0.04 |
| 7 | 101194424 | rs17135626 | -0.062 | 0.019 | -0.611 | -1.841 | 0.001 | -0.037 | -0.037 | 3.272 | 0.4 | 0.4 | 0.29 | 0.06 | 0.39 | 0.28 |
| 7 | 101200628 | rs11768650 | -0.342 | 0.178 | -0.964 | -1.912 | -0.001 | 0.045 | 0.045 | 0 | 0.17 | 0.1 | 0.13 | 0.03 | 0.2 | 0.29 |
| 7 | 101200727 | rs17393069 | 0.649 | -0.28 | -1.107 | -2.367 | 0.003 | -0.013 | -0.013 | 0 | 0.38 | 0.34 | 0.32 | 0.24 | 0.37 | 0.33 |
| 7 | 101201314 | rs10264081 | 0.622 | -0.441 | -1.006 | -2.844 | 0 | 0.004 | 0.004 | 0.05 | 0.2 | 0.2 | 0.35 | 0.71 | 0.18 | 0.32 |
| 7 | 101203066 | rs724574 | 0.412 | -0.534 | -0.947 | -2.641 | 0.004 | -0.022 | -0.022 | 0 | 0.58 | 0.55 | 0.68 | 0.97 | 0.56 | 0.66 |
| 7 | 101204556 | rs757824 | -0.94 | -0.468 | -0.647 | -2.495 | 0.006 | -0.027 | -0.027 | 0 | 0.94 | 0.88 | 0.97 | 1 | 0.95 | 0.98 |
| 7 | 101207510 | rs6955500 | 0.718 | -0.593 | -0.45 | -2.593 | -0.004 | -0.047 | -0.047 | 4.047 | 0.18 | 0.26 | 0.17 | 0 | 0.19 | 0.03 |
| 7 | 101208943 | rs4729731 | 0.817 | -0.661 | -0.433 | -2.749 | 0 | -0.007 | -0.007 | 0 | 0.51 | 0.49 | 0.62 | 0.94 | 0.45 | 0.57 |
| 7 | 101209445 | rs10275953 | 0.043 | -0.608 | -0.428 | -2.726 | 0 | -0.038 | -0.038 | 0.079 | 0.69 | 0.64 | 0.78 | 0.94 | 0.64 | 0.6 |
| 7 | 101210116 | rs6960162 | -0.099 | -1.169 | -0.698 | -3.02 | -0.02 | -0.059 | -0.059 | 0.679 | 0.62 | 0.53 | 0.71 | 0.87 | 0.68 | 0.87 |
| 7 | 101214296 | rs6465819 | -0.868 | -1.212 | -0.69 | -2.937 | -0.035 | -0.041 | -0.041 | 0 | 0.78 | 0.65 | 0.87 | 0.86 | 0.87 | 0.9 |
| 7 | 101220194 | rs2158723 | 0.831 | -0.984 | -0.813 | -2.801 | -0.001 | -0.005 | -0.005 | 0 | 0.35 | 0.36 | 0.52 | 0.82 | 0.37 | 0.5 |
| 7 | 101223428 | rs13228618 | 0.458 | -0.718 | -0.883 | -2.578 | -0.04 | 0.11 | 0.11 | 0 | 0.26 | 0.09 | 0.18 | 0.05 | 0.32 | 0.37 |
| 7 | 101224018 | rs11973739 | -0.788 | -0.959 | -0.774 | -2.652 | 0.001 | -0.011 | -0.011 | 0 | 0.54 | 0.57 | 0.68 | 0.72 | 0.58 | 0.67 |
| 7 | 101225827 | rs4729736 | 0.51 | -0.315 | -1.164 | -2.861 | -0.007 | 0.049 | 0.049 | 0 | 0.35 | 0.29 | 0.28 | 0.14 | 0.38 | 0.23 |
| 7 | 101226230 | rs7786803 | 1.578 | -0.282 | -1.289 | -2.636 | -0.031 | 0.019 | 0.019 | 0 | 0.18 | 0.1 | 0.14 | 0.13 | 0.2 | 0.17 |
| 7 | 101229969 | rs917182 | -0.312 | -0.162 | -1.245 | -2.714 | 0.005 | -0.006 | -0.006 | 0 | 0.88 | 0.92 | 0.94 | 1 | 0.93 | 0.95 |
| 7 | 101234771 | rs13240922 | 1.008 | -0.123 | -1.573 | -2.848 | -0.047 | 0.105 | 0.105 | 0 | 0.22 | 0.06 | 0.19 | 0.06 | 0.29 | 0.34 |
| 7 | 101238449 | rs4727499 | 1.546 | -0.028 | -1.403 | -2.581 | -0.007 | 0.007 | 0.007 | 0 | 0.18 | 0.21 | 0.14 | 0.16 | 0.21 | 0.21 |
| 7 | 101239926 | rs1468395 | 1.461 | 0.151 | -1.262 | -2.37 | -0.006 | 0.049 | 0.049 | 0 | 0.05 | 0.02 | 0.28 | 0.65 | 0.08 | 0.26 |
| 7 | 101241760 | rs7778019 | 1.526 | -0.186 | -1.284 | -2.46 | 0.012 | 0.044 | 0.044 | 0 | 0.16 | 0.08 | 0.13 | 0 | 0.18 | 0.05 |
| 7 | 101242973 | rs1981582 | 0.968 | 0.071 | -1.204 | -2.292 | -0.011 | 0.05 | 0.05 | 0 | 0.22 | 0.15 | 0.14 | 0.14 | 0.2 | 0.19 |
| 7 | 101247430 | rs11489598 | 0.63 | 0.244 | -0.696 | -2.304 | 0.008 | 0 | 0 | 0 | 0.26 | 0.3 | 0.44 | 0.7 | 0.32 | 0.53 |
| 7 | 101252365 | rs6952330 | -0.144 | 0.38 | -0.665 | -2.003 | -0.018 | 0.065 | 0.065 | 0 | 0.71 | 0.82 | 0.82 | 0.85 | 0.72 | 0.75 |
| 7 | 101256074 | rs7786466 | -0.695 | 0.568 | -0.514 | -1.663 | -0.029 | 0.039 | 0.039 | 0 | 0.93 | 1 | 0.96 | 1 | 0.92 | 0.94 |
| 7 | 101256833 | rs2240389 | 0.37 | 0.388 | -0.068 | -1.709 | 0 | -0.02 | -0.02 | 0 | 0.27 | 0.31 | 0.27 | 0.14 | 0.26 | 0.15 |
| 7 | 101259815 | rs740200 | 0.344 | 0.39 | 0.102 | -1.916 | 0 | -0.035 | -0.035 | 0 | 0.34 | 0.33 | 0.26 | 0.06 | 0.37 | 0.43 |
| 7 | 101267435 | rs876825 | 0.044 | 0.091 | 0.415 | -1.691 | -0.002 | 0.042 | 0.042 | 0 | 0.56 | 0.4 | 0.46 | 0.26 | 0.58 | 0.5 |
| 7 | 101277334 | rs848652 | 0.873 | -0.033 | 0.16 | -1.295 | -0.039 | 0.154 | 0.154 | 0 | 0.3 | 0.07 | 0.2 | 0.05 | 0.32 | 0.28 |
| 7 | 101280561 | rs10278829 | 1.109 | -0.046 | -0.337 | -1.247 | -0.016 | 0.01 | 0.01 | 0 | 0.06 | 0.09 | 0.03 | 0.11 | 0.02 | 0.01 |
| 7 | 101280652 | rs740202 | -0.586 | 0.082 | -0.535 | -1.406 | -0.035 | 0.131 | 0.131 | 0 | 0.63 | 0.79 | 0.71 | 0.94 | 0.57 | 0.59 |
| 7 | 101284930 | rs1620168 | -0.204 | 0.574 | -0.601 | -1.097 | 0.001 | 0.036 | 0.036 | 0 | 0.57 | 0.61 | 0.67 | 0.79 | 0.55 | 0.58 |
| 7 | 101286624 | rs11771949 | -0.021 | 0.677 | -1.092 | -1.14 | -0.008 | 0.071 | 0.071 | 0 | 0.32 | 0.15 | 0.22 | 0.09 | 0.32 | 0.28 |
| 7 | 101330734 | rs1722224 | -0.199 | 0.55 | -0.99 | -0.94 | 0.001 | 0.016 | 0.016 | 0.498 | 0.89 | 0.96 | 0.93 | 1 | 0.92 | 0.98 |
| 7 | 101331807 | rs1636443 | -0.388 | 0.537 | -0.956 | -0.946 | -0.001 | 0.011 | 0.011 | 0 | 0.9 | 0.97 | 0.93 | 1 | 0.92 | 0.98 |
| 7 | 101332332 | rs1636444 | 0.78 | 0.349 | -0.987 | -1.588 | 0.005 | 0.057 | 0.057 | 0 | 0.7 | 0.84 | 0.82 | 1 | 0.76 | 0.86 |
| 7 | 101340115 | rs11978556 | 0.002 | 0.283 | -0.13 | -1.592 | 0.001 | -0.127 | -0.127 | 0 | 0.2 | 0.41 | 0.14 | 0 | 0.18 | 0.12 |
| 7 | 101341246 | rs2529190 | -0.574 | 0.226 | -0.999 | -1.287 | 0.008 | -0.065 | -0.065 | 0.226 | 0.3 | 0.08 | 0.26 | 0.29 | 0.3 | 0.3 |
| 7 | 101350768 | rs2690897 | -0.385 | 0.587 | -0.515 | -1.216 | -0.006 | 0.025 | 0.025 | 1.958 | 0.21 | 0.29 | 0.22 | 0.25 | 0.2 | 0.16 |
| 7 | 101350790 | rs2529179 | -1.168 | 0.364 | -0.674 | -1.41 | -0.049 | 0.015 | 0.015 | 1.069 | 0.56 | 0.82 | 0.44 | 0.3 | 0.53 | 0.44 |
| 7 | 101356053 | rs2529243 | 0.309 | 0.592 | -0.798 | -1.745 | -0.007 | -0.005 | -0.005 | 0.844 | 0.27 | 0.28 | 0.29 | 0.25 | 0.35 | 0.28 |
| 7 | 101364599 | rs2529241 | -1.633 | 0.408 | -1.379 | -1.72 | -0.058 | -0.111 | -0.111 | 0 | 0.78 | 0.97 | 0.61 | 0.29 | 0.77 | 0.6 |
| 7 | 101372755 | rs1880345 | 0.195 | -0.162 | -0.618 | -2.052 | 0.007 | 0.019 | 0.019 | 0 | 0.31 | 0.25 | 0.2 | 0.08 | 0.27 | 0.27 |
| 7 | 101375291 | rs6465827 | -0.322 | 0.029 | -0.301 | -1.943 | -0.005 | -0.045 | -0.045 | 0 | 0.47 | 0.46 | 0.29 | 0.08 | 0.37 | 0.31 |
| 7 | 101381523 | rs11977362 | -0.748 | 0.553 | -0.162 | -1.652 | 0.007 | -0.055 | -0.055 | 1.082 | 0.36 | 0.4 | 0.24 | 0.05 | 0.4 | 0.3 |
| 7 | 101385552 | rs2690948 | 0.789 | 0.603 | -0.427 | -1.445 | -0.018 | -0.015 | -0.015 | 0.009 | 0.07 | 0 | 0.09 | 0.06 | 0.08 | 0.08 |
| 7 | 101387262 | rs2529192 | 0.287 | 0.605 | -0.426 | -1.219 | -0.01 | -0.039 | -0.039 | 0 | 0.09 | 0.03 | 0.12 | 0.23 | 0.1 | 0.17 |
| 7 | 101388981 | rs2970492 | -0.594 | 0.421 | -0.219 | -1.43 | -0.028 | -0.065 | -0.065 | 0 | 0.23 | 0.38 | 0.14 | 0.04 | 0.18 | 0.13 |
| 7 | 101392745 | rs2906725 | -0.594 | 0.421 | -0.219 | -1.43 | -0.028 | -0.065 | -0.065 | 0 | 0.23 | 0.38 | 0.13 | 0.04 | 0.18 | 0.13 |
| 7 | 101402732 | rs2690943 | -0.262 | 0.704 | -0.351 | -1.291 | -0.082 | 0.012 | 0.012 | 0 | 0.24 | 0.07 | 0.25 | 0.22 | 0.34 | 0.34 |
| 7 | 101406088 | rs2690939 | -0.437 | 0.592 | 0.375 | -1.323 | 0.018 | -0.015 | -0.015 | 3.324 | 0.26 | 0.33 | 0.27 | 0.22 | 0.34 | 0.34 |
| 7 | 101412077 | rs2906734 | -0.757 | -0.378 | 0.715 | -1.175 | -0.038 | -0.06 | -0.06 | 0 | 0.2 | 0.36 | 0.1 | 0.04 | 0.12 | 0.13 |
| 7 | 101425057 | rs2949326 | -0.455 | -0.199 | 0.454 | -1.105 | -0.015 | -0.028 | -0.028 | 0.901 | 0.17 | 0.24 | 0.08 | 0.04 | 0.12 | 0.13 |
| 7 | 101427497 | rs940482 | 0.07 | 0.004 | 0.457 | -1.107 | 0 | 0 | 0 | 0.452 | 0.11 | 0.11 | 0.16 | 0.13 | 0.15 | 0.18 |
| 7 | 101428464 | rs2949324 | -0.371 | 0.002 | 0.367 | -1.106 | -0.013 | -0.025 | -0.025 | 0 | 0.17 | 0.21 | 0.08 | 0.04 | 0.12 | 0.13 |
| 7 | 101429264 | rs2690926 | -1.282 | 0.021 | 0.356 | -1.08 | -0.015 | -0.047 | -0.047 | 2.468 | 0.3 | 0.46 | 0.25 | 0.17 | 0.27 | 0.31 |
| 7 | 101447844 | rs11974028 | -0.037 | -0.436 | 0.107 | -1.408 | -0.012 | -0.058 | -0.058 | 0 | 0.23 | 0.24 | 0.11 | 0 | 0.15 | 0.1 |
| 7 | 101454616 | rs2529232 | -0.683 | -0.013 | 0.199 | -0.118 | 0 | 0.042 | 0.042 | 0 | 0.17 | 0.25 | 0.27 | 0.34 | 0.17 | 0.27 |
| 7 | 101466998 | rs2906721 | 0.452 | 0.949 | 0.182 | -1.063 | 0.002 | -0.027 | -0.027 | 0.806 | 0.15 | 0.14 | 0.11 | 0.01 | 0.18 | 0.11 |

| | | | | | | | | | | | | | | | | |
|---|---|---|---|---|---|---|---|---|---|---|---|---|---|---|---|---|
| 7 | 101467854 | rs2970471 | -1.018 | 0.398 | 0.015 | -0.951 | -0.002 | -0.022 | -0.022 | 0.2 | 0.61 | 0.75 | 0.52 | 0.55 | 0.57 | 0.51 |
| 7 | 101482682 | rs2970463 | 0.47 | 0.74 | -0.098 | -0.912 | 0.038 | -0.085 | -0.085 | 0 | 0.17 | 0.24 | 0.12 | 0.01 | 0.19 | 0.11 |
| 7 | 101494907 | rs2970459 | -0.267 | -0.015 | -0.455 | -1.057 | 0 | -0.064 | -0.064 | 0 | 0.56 | 0.47 | 0.35 | 0.07 | 0.52 | 0.42 |
| 7 | 101497449 | rs13232623 | 1.062 | -0.034 | -1.097 | -0.544 | -0.01 | 0.06 | 0.06 | 0 | 0.05 | 0.02 | 0.18 | 0.46 | 0.06 | 0.11 |
| 7 | 101505732 | rs11773426 | 1.235 | -0.274 | -1.386 | -1.34 | 0.036 | 0.094 | 0.094 | 0 | 0.21 | 0.05 | 0.09 | 0 | 0.14 | 0.1 |
| 7 | 101513404 | rs10245008 | 0.433 | 0.07 | -1.38 | -1.27 | 0.003 | 0.103 | 0.103 | 0 | 0.38 | 0.24 | 0.58 | 0.88 | 0.4 | 0.47 |
| 7 | 101526248 | rs10270083 | -1.104 | 0.585 | -1.384 | -1.271 | -0.02 | 0.016 | 0.016 | 0 | 0.85 | 0.88 | 0.9 | 0.99 | 0.83 | 0.88 |
| 7 | 101528560 | rs4412307 | -0.858 | 0.546 | -1.402 | -1.288 | -0.022 | 0.025 | 0.025 | 0 | 0.83 | 0.88 | 0.9 | 0.99 | 0.83 | 0.88 |
| 7 | 101531781 | rs13238593 | 1.21 | 0.474 | -1.49 | -0.876 | -0.013 | 0.018 | 0.018 | 0 | 0.07 | 0 | 0.18 | 0.43 | 0.06 | 0.12 |
| 7 | 101532721 | rs11765661 | 1.347 | 0.324 | -1.758 | -1.741 | -0.073 | 0.13 | 0.13 | 0 | 0.25 | 0.02 | 0.16 | 0.04 | 0.3 | 0.13 |
| 7 | 101536726 | rs4644158 | 1.049 | 0.131 | -1.456 | -1.565 | -0.003 | 0.013 | 0.013 | 0 | 0.1 | 0.06 | 0.1 | 0.03 | 0.13 | 0.21 |
| 7 | 101545685 | rs6465836 | 0.243 | 0.089 | -0.992 | -1.595 | 0 | -0.037 | -0.037 | 0 | 0.43 | 0.57 | 0.31 | 0.1 | 0.4 | 0.38 |
| 7 | 101549007 | rs6465837 | -0.447 | 0.297 | -1.277 | -1.562 | -0.047 | 0.206 | 0.206 | 0 | 0.57 | 0.83 | 0.73 | 0.96 | 0.51 | 0.75 |
| 7 | 101556363 | rs11974778 | 0.504 | 0.301 | -0.815 | -1.26 | -0.014 | -0.102 | -0.102 | 0.375 | 0.33 | 0.48 | 0.2 | 0.11 | 0.27 | 0.18 |
| 7 | 101560734 | rs2906656 | 0.477 | 0.991 | -1.024 | -1.019 | -0.009 | 0.004 | 0.004 | 0.072 | 0.14 | 0.1 | 0.08 | 0.01 | 0.15 | 0.08 |
| 7 | 101562491 | rs2906652 | 0.027 | 0.844 | -1.168 | -0.488 | -0.003 | 0.092 | 0.092 | 0 | 0.1 | 0.1 | 0.2 | 0.48 | 0.08 | 0.15 |
| 7 | 101569302 | rs2906655 | -0.668 | 0.915 | -1.094 | -0.472 | -0.015 | -0.046 | -0.046 | 0.402 | 0.23 | 0.14 | 0.28 | 0.49 | 0.24 | 0.22 |
| 7 | 101577038 | rs2906649 | 0.591 | 1.049 | -1.104 | -0.921 | 0.001 | -0.017 | -0.017 | 0.301 | 0.68 | 0.24 | 0.57 | 0.59 | 0.7 | 0.5 |
| 7 | 101591561 | rs434564 | 0.631 | 1.249 | -0.976 | -0.866 | 0 | -0.008 | -0.008 | 0 | 0.68 | 0.29 | 0.57 | 0.59 | 0.7 | 0.52 |
| 7 | 101597464 | rs7810922 | -0.023 | 1.701 | -0.549 | -0.855 | 0.04 | -0.122 | -0.122 | 0 | 0.19 | 0.44 | 0.15 | 0.02 | 0.19 | 0.22 |
| 7 | 101604646 | rs407366 | 0.85 | 1.475 | -0.967 | -0.919 | -0.003 | 0.033 | 0.033 | 0 | 0.12 | 0.04 | 0.06 | 0.01 | 0.1 | 0.09 |
| 7 | 101606325 | rs2031085 | -0.292 | 1.565 | -0.518 | -0.49 | 0.018 | -0.111 | -0.111 | 0 | 0.19 | 0.51 | 0.16 | 0.02 | 0.17 | 0.21 |
| 7 | 101608080 | rs441601 | -0.47 | 1.45 | -0.994 | 0.214 | -0.02 | -0.017 | -0.017 | 0 | 0.89 | 0.91 | 0.73 | 0.62 | 0.85 | 0.73 |
| 7 | 101626640 | rs2410922 | 0.165 | 0.781 | -0.604 | -0.642 | -0.011 | -0.056 | -0.056 | 0 | 0.71 | 0.61 | 0.76 | 0.91 | 0.77 | 0.77 |
| 7 | 101626768 | rs439124 | 0.513 | 0.012 | -0.467 | -1.509 | -0.044 | 0 | 0 | 0 | 0.33 | 0.49 | 0.33 | 0.61 | 0.24 | 0.27 |
| 7 | 101630597 | rs12537526 | 0.54 | 0.936 | -0.628 | -1.826 | -0.008 | -0.003 | -0.003 | 0 | 0.16 | 0.09 | 0.12 | 0.02 | 0.2 | 0.21 |
| 7 | 101635335 | rs202161 | 1.034 | 1.103 | -0.477 | -1.805 | 0.004 | 0.055 | 0.055 | 0 | 0.12 | 0.15 | 0.34 | 0.33 | 0.12 | 0.3 |
| 7 | 101637246 | rs202159 | -0.831 | 1.122 | -0.721 | -2.181 | 0.002 | 0.132 | 0.132 | 0.023 | 0.68 | 0.83 | 0.82 | 0.99 | 0.7 | 0.83 |
| 7 | 101647143 | rs12668172 | 0.821 | 1.219 | -0.526 | -2.16 | 0.003 | 0.014 | 0.014 | 0 | 0.41 | 0.35 | 0.35 | 0.64 | 0.44 | 0.37 |
| 7 | 101649622 | rs10488043 | 0.982 | 1.187 | -0.505 | -2.182 | -0.002 | -0.013 | -0.013 | 0.012 | 0.13 | 0.08 | 0.11 | 0.02 | 0.16 | 0.17 |
| 7 | 101653744 | rs10259603 | 0.654 | 0.99 | -0.482 | -2.474 | 0.003 | -0.13 | -0.13 | 0 | 0.37 | 0.43 | 0.29 | 0.02 | 0.39 | 0.3 |
| 7 | 101667516 | rs11769397 | 1.335 | 0.697 | -0.992 | -2.581 | 0.014 | 0.037 | 0.037 | 0 | 0.29 | 0.2 | 0.21 | 0 | 0.3 | 0.16 |
| 7 | 101671721 | rs6465845 | -0.86 | 0.411 | -1.022 | -2.214 | -0.009 | 0.13 | 0.13 | 0.064 | 0.74 | 0.9 | 0.86 | 0.99 | 0.8 | 0.86 |
| 7 | 101671793 | rs12671456 | -0.104 | 0.494 | -0.665 | -1.751 | -0.001 | 0.007 | 0.007 | 0.092 | 0.19 | 0.2 | 0.19 | 0.61 | 0.19 | 0.24 |
| 7 | 101683563 | rs11971570 | -0.961 | -0.673 | -0.242 | -2.207 | -0.021 | -0.008 | -0.008 | 0 | 0.18 | 0.26 | 0.09 | 0.12 | 0.1 | 0.14 |
| 7 | 101693308 | rs10246310 | 0.07 | -0.186 | -0.198 | -2.506 | -0.003 | 0.021 | 0.021 | 0 | 0.19 | 0.1 | 0.14 | 0.02 | 0.16 | 0.17 |
| 7 | 101697980 | rs12112504 | 0.404 | -0.193 | 0.194 | -2.198 | 0.008 | -0.028 | -0.028 | 0 | 0.06 | 0.17 | 0.04 | 0 | 0.05 | 0.05 |
| 7 | 101722425 | rs380934 | -0.253 | -0.228 | -0.303 | -2.049 | -0.004 | -0.11 | -0.11 | 0 | 0.2 | 0.42 | 0.16 | 0.01 | 0.18 | 0.11 |
| 7 | 101731965 | rs446076 | -0.818 | -0.285 | -1.07 | -2.284 | 0.005 | 0.214 | 0.214 | 0.285 | 0.43 | 0.8 | 0.32 | 0.04 | 0.36 | 0.34 |
| 7 | 101736364 | rs16869529 | -0.067 | -0.559 | -0.498 | -2.182 | 0.004 | -0.029 | -0.029 | 0 | 0.06 | 0.16 | 0.04 | 0.01 | 0.04 | 0.03 |
| 7 | 101741571 | rs201460 | -0.178 | -1.559 | -0.363 | -3.074 | -0.034 | -0.118 | -0.118 | 1.457 | 0.44 | 0.53 | 0.2 | 0.03 | 0.25 | 0.19 |
| 7 | 101747575 | rs201480 | -1.57 | -1.009 | -0.563 | -3.057 | -0.04 | 0.047 | 0.047 | 0.82 | 0.72 | 0.82 | 0.54 | 0.09 | 0.57 | 0.55 |
| 7 | 101758584 | rs201501 | -0.259 | -1.368 | -0.341 | -3.245 | 0.002 | -0.093 | -0.093 | 0 | 0.55 | 0.47 | 0.36 | 0.05 | 0.47 | 0.44 |
| 7 | 101773105 | rs201525 | 0.6 | -1.403 | -0.409 | -3.218 | 0.004 | -0.021 | -0.021 | 0.159 | 0.33 | 0.27 | 0.23 | 0.03 | 0.19 | 0.22 |
| 7 | 101784718 | rs424028 | -0.885 | -1.08 | -0.344 | -2.966 | 0.003 | -0.088 | -0.088 | 0 | 0.39 | 0.56 | 0.31 | 0.06 | 0.38 | 0.33 |
| 7 | 101795994 | rs409419 | -0.478 | -1.153 | 0.002 | -2.634 | -0.046 | -0.111 | -0.111 | 0 | 0.2 | 0.5 | 0.11 | 0.04 | 0.11 | 0.11 |
| 7 | 101809851 | rs365836 | -0.468 | -1.379 | -0.266 | -2.801 | -0.021 | -0.063 | -0.063 | 0 | 0.36 | 0.5 | 0.29 | 0.07 | 0.24 | 0.23 |
| 7 | 101817795 | rs411030 | 0.726 | -1.263 | -0.166 | -2.734 | 0 | -0.007 | -0.007 | 0.236 | 0.06 | 0.08 | 0.05 | 0.01 | 0.05 | 0.04 |
| 7 | 101819315 | rs420437 | -0.364 | -1.329 | -0.227 | -2.971 | -0.001 | -0.062 | -0.062 | 0 | 0.44 | 0.43 | 0.31 | 0.05 | 0.39 | 0.33 |
| 7 | 101840903 | rs11760331 | 0.088 | -1.32 | 0.241 | -2.532 | 0.004 | 0.034 | 0.034 | 0 | 0.36 | 0.23 | 0.24 | 0.04 | 0.33 | 0.29 |
| 7 | 101846898 | rs6465852 | -1.019 | -1.086 | 0 | -2.119 | -0.009 | 0.134 | 0.134 | 0.322 | 0.55 | 0.77 | 0.35 | 0.13 | 0.44 | 0.42 |
| 7 | 101864836 | rs385417 | 0.621 | -1.07 | 0.141 | -2.331 | 0.014 | 0.004 | 0.004 | 0.083 | 0.51 | 0.41 | 0.7 | 0.93 | 0.61 | 0.62 |
| 7 | 101865210 | rs409224 | 0.481 | -1.021 | -0.238 | -2.309 | 0.013 | 0.013 | 0.013 | 0.064 | 0.53 | 0.56 | 0.71 | 0.93 | 0.61 | 0.62 |
| 7 | 101875641 | rs449806 | 1.018 | -1 | -0.259 | -2.258 | -0.002 | 0.118 | 0.118 | 0.228 | 0.45 | 0.28 | 0.65 | 0.93 | 0.56 | 0.59 |
| 7 | 101885814 | rs392606 | -0.387 | -1.278 | -0.373 | -2.316 | -0.057 | -0.11 | -0.11 | 0 | 0.74 | 0.6 | 0.91 | 0.97 | 0.88 | 0.84 |
| 7 | 101892833 | rs367634 | 0.707 | -0.704 | -0.346 | -1.964 | 0.007 | 0.124 | 0.124 | 0.438 | 0.46 | 0.28 | 0.66 | 0.91 | 0.58 | 0.6 |
| 7 | 101895864 | rs2734605 | 0.149 | -0.681 | -0.686 | -1.75 | 0.003 | -0.021 | -0.021 | 0 | 0.6 | 0.59 | 0.76 | 0.92 | 0.71 | 0.72 |
| 7 | 101901652 | rs1734729 | -0.267 | 0.237 | -0.875 | -0.137 | -0.016 | -0.055 | -0.055 | 0 | 0.21 | 0.16 | 0.25 | 0.5 | 0.23 | 0.29 |
| 7 | 101907824 | rs2694158 | -0.255 | -0.206 | -0.608 | -0.488 | 0.014 | 0.032 | 0.032 | 0 | 0.57 | 0.66 | 0.67 | 0.78 | 0.61 | 0.69 |
| 7 | 101909103 | rs2694159 | -0.575 | -0.501 | -0.119 | 0.097 | 0.01 | 0.037 | 0.037 | 0 | 0.44 | 0.64 | 0.38 | 0.21 | 0.37 | 0.43 |
| 7 | 101917521 | rs803064 | 0.34 | -0.644 | -0.447 | 0.429 | 0.009 | 0 | 0 | 0 | 0.5 | 0.62 | 0.51 | 0.5 | 0.54 | 0.6 |
| 7 | 103119863 | rs3823992 | -1.632 | 0.263 | 2.817 | 1.868 | 0.077 | 0.159 | 0.159 | 0 | 0.1 | 0.32 | 0.16 | 0.27 | 0.15 | 0.2 |
| 7 | 103151611 | rs2528858 | -1.645 | 0.184 | 2.857 | 1.798 | 0.015 | 0.079 | 0.079 | 0.652 | 0.08 | 0.24 | 0.04 | 0.17 | 0.05 | 0.09 |
| 7 | 103154037 | rs2711861 | -2.211 | -0.348 | 2.79 | 2.03 | -0.028 | 0.065 | 0.065 | 0 | 0.19 | 0.32 | 0.26 | 0.4 | 0.13 | 0.43 |
| 7 | 103157375 | rs2528865 | -2.352 | -0.314 | 2.833 | 2.181 | -0.003 | 0.308 | 0.308 | 0 | 0.15 | 0.41 | 0.24 | 0.55 | 0.12 | 0.33 |
| 7 | 103157510 | rs2528866 | -1.778 | -0.282 | 2.834 | 2.092 | 0.022 | 0.2 | 0.2 | 0 | 0.11 | 0.34 | 0.14 | 0.31 | 0.09 | 0.25 |
| 7 | 103162370 | rs10282605 | 1.054 | -0.631 | 2.599 | 1.929 | 0.003 | 0.053 | 0.053 | 0.087 | 0.81 | 0.58 | 0.83 | 0.75 | 0.82 | 0.84 |
| 7 | 103162847 | rs2075040 | 2.151 | -0.649 | 2.11 | 1.972 | 0.004 | -0.089 | -0.089 | 0.006 | 0.7 | 0.16 | 0.69 | 0.47 | 0.72 | 0.65 |
| 7 | 103167675 | rs2256670 | -2.297 | -0.623 | 2.175 | 1.891 | -0.027 | 0.028 | 0.028 | 0.001 | 0.2 | 0.62 | 0.19 | 0.31 | 0.12 | 0.36 |

| | | | | | | | | | | | | | | | |
|---|---|---|---|---|---|---|---|---|---|---|---|---|---|---|---|
| 7 | 103177947 | rs362710 | 1.791 | -0.318 | 1.348 | 1.618 | -0.016 | 0.041 | 0.041 | 0 | 0.71 | 0.5 | 0.63 | 0.37 | 0.73 | 0.6 |
| 7 | 103190197 | rs362777 | -1.326 | -0.436 | 1.073 | 1.447 | -0.035 | 0.011 | 0.011 | 0.889 | 0.14 | 0.28 | 0.13 | 0.14 | 0.11 | 0.19 |
| 7 | 103190840 | rs362784 | 0.364 | 0.86 | -0.642 | 1.107 | -0.005 | -0.005 | -0.005 | 0.152 | 0.85 | 0.85 | 0.82 | 0.77 | 0.75 | 0.79 |
| 7 | 103192532 | rs2535764 | -1.337 | 0.025 | -1.076 | 0.414 | 0.033 | 0.024 | 0.024 | 0.056 | 0.3 | 0.16 | 0.3 | 0.22 | 0.26 | 0.46 |
| 7 | 103192782 | rs362723 | -1.33 | 0.151 | -0.405 | 0.395 | -0.008 | -0.013 | -0.013 | 0 | 0.11 | 0.16 | 0.09 | 0.06 | 0.07 | 0.16 |
| 7 | 103201263 | rs362794 | -0.642 | 0.229 | -0.729 | -0.106 | 0.002 | 0.007 | 0.007 | 0.341 | 0.27 | 0.23 | 0.22 | 0.07 | 0.25 | 0.25 |
| 7 | 103207234 | rs362726 | -1.153 | -0.291 | -0.26 | 0.657 | 0.016 | -0.013 | -0.013 | 0 | 0.43 | 0.66 | 0.43 | 0.47 | 0.38 | 0.35 |
| 7 | 103208865 | rs362806 | -0.222 | -0.307 | 0.28 | 0.667 | 0 | 0.184 | 0.184 | 0 | 0.13 | 0.31 | 0.19 | 0.4 | 0.12 | 0.09 |
| 7 | 103209028 | rs362731 | -0.355 | -0.133 | -0.324 | 0.104 | 0.01 | 0.013 | 0.013 | 0.583 | 0.41 | 0.5 | 0.45 | 0.49 | 0.46 | 0.46 |
| 7 | 103210068 | rs362732 | 0.644 | -0.066 | -0.495 | 0.385 | -0.084 | 0.032 | 0.032 | 0.953 | 0.27 | 0.07 | 0.24 | 0.17 | 0.34 | 0.42 |
| 7 | 103220611 | rs2283021 | -0.059 | -0.206 | -0.571 | 0.275 | -0.008 | 0.083 | 0.083 | 0 | 0.17 | 0.31 | 0.22 | 0.33 | 0.16 | 0.1 |
| 7 | 103222306 | rs2237628 | -0.333 | -0.224 | -0.343 | 0.214 | 0 | 0 | 0 | 0.339 | 0.43 | 0.44 | 0.46 | 0.49 | 0.5 | 0.51 |
| 7 | 103226907 | rs2711870 | -1.127 | -0.091 | -0.538 | 0.618 | 0.007 | -0.015 | -0.015 | 0.164 | 0.7 | 0.83 | 0.69 | 0.66 | 0.73 | 0.73 |
| 7 | 103229035 | rs2245617 | -1.349 | -0.063 | -0.709 | 0.727 | -0.001 | 0.053 | 0.053 | 0.247 | 0.73 | 0.98 | 0.74 | 0.84 | 0.77 | 0.84 |
| 7 | 103231265 | rs82878 | 0.33 | -0.684 | -0.438 | 0.097 | 0.012 | -0.008 | -0.008 | 0 | 0.27 | 0.25 | 0.14 | 0.02 | 0.19 | 0.08 |
| 7 | 103231290 | rs123712 | -1.564 | -0.633 | -0.27 | 0.372 | 0.001 | -0.001 | -0.001 | 0 | 0.71 | 0.9 | 0.68 | 0.73 | 0.72 | 0.76 |
| 7 | 103233787 | rs362633 | -0.516 | -0.596 | 0.417 | 0.804 | -0.004 | -0.001 | -0.001 | 0 | 0.55 | 0.64 | 0.47 | 0.49 | 0.56 | 0.51 |
| 7 | 103235743 | rs2109698 | 0.67 | -0.568 | 0.44 | 0.842 | -0.006 | 0.009 | 0.009 | 0 | 0.11 | 0.08 | 0.12 | 0.07 | 0.11 | 0.21 |
| 7 | 103241465 | rs2159677 | -1.647 | -0.597 | 0.633 | 0.831 | -0.017 | -0.042 | -0.042 | 1.23 | 0.69 | 0.84 | 0.6 | 0.58 | 0.68 | 0.74 |
| 7 | 103241633 | rs362635 | -1.641 | -0.606 | 0.621 | 0.827 | -0.019 | -0.049 | -0.049 | 0 | 0.69 | 0.85 | 0.6 | 0.57 | 0.68 | 0.72 |
| 7 | 103255144 | rs2017179 | -0.542 | -0.734 | 0.975 | 0.95 | -0.021 | -0.034 | -0.034 | 0 | 0.37 | 0.54 | 0.26 | 0.19 | 0.24 | 0.28 |
| 7 | 103255195 | rs7783216 | -0.219 | -0.869 | 1.215 | 1.094 | -0.028 | -0.019 | -0.019 | 0.448 | 0.32 | 0.42 | 0.18 | 0.16 | 0.23 | 0.13 |
| 7 | 103259909 | rs977639 | 0.134 | -0.705 | 1.221 | 1.064 | 0.019 | 0.012 | 0.012 | 0 | 0.13 | 0.26 | 0.14 | 0.16 | 0.11 | 0.2 |
| 7 | 103262685 | rs12674323 | 0.878 | -0.611 | 0.878 | 1.069 | -0.013 | -0.037 | -0.037 | 0.109 | 0.1 | 0 | 0.1 | 0.16 | 0.07 | 0.19 |
| 7 | 103265213 | rs1122672 | 0.749 | -0.618 | 0.933 | 1.069 | -0.006 | -0.004 | -0.004 | 0.906 | 0.1 | 0.07 | 0.12 | 0.16 | 0.11 | 0.2 |
| 7 | 103265376 | rs997447 | 0.597 | -0.776 | 0.953 | 1.12 | 0.024 | 0.025 | 0.025 | 1.998 | 0.24 | 0.18 | 0.12 | 0.05 | 0.14 | 0.08 |
| 7 | 103277292 | rs362698 | -0.253 | -0.843 | 0.859 | 1.105 | -0.013 | -0.024 | -0.024 | 0 | 0.14 | 0.06 | 0.19 | 0.31 | 0.16 | 0.23 |
| 7 | 103278471 | rs10245257 | -2.016 | -0.872 | 0.851 | 1.151 | 0.004 | 0.006 | 0.006 | 0.088 | 0.69 | 0.84 | 0.64 | 0.73 | 0.7 | 0.74 |
| 7 | 103278744 | rs12539504 | 0.145 | -0.875 | 1.008 | 1.133 | -0.004 | -0.004 | -0.004 | 0 | 0.3 | 0.31 | 0.31 | 0.39 | 0.41 | 0.44 |
| 7 | 103283013 | rs720078 | 0.774 | -0.863 | 0.971 | 1.091 | 0.038 | 0.043 | 0.043 | 2.017 | 0.23 | 0.15 | 0.1 | 0.02 | 0.15 | 0.07 |
| 7 | 103291825 | rs6958081 | 0.068 | -0.773 | 1.041 | 1.183 | 0.021 | 0.024 | 0.024 | 0.05 | 0.29 | 0.38 | 0.31 | 0.39 | 0.39 | 0.42 |
| 7 | 103292487 | rs3919520 | -0.463 | -0.772 | 1.123 | 1.205 | 0.002 | 0.001 | 0.001 | 1.143 | 0.36 | 0.62 | 0.43 | 0.54 | 0.46 | 0.46 |
| 7 | 103300835 | rs2299355 | -1.295 | -0.905 | 1.273 | 1.091 | 0.002 | -0.008 | -0.008 | 0.646 | 0.63 | 0.67 | 0.59 | 0.6 | 0.63 | 0.7 |
| 7 | 103301455 | rs727531 | 0.792 | -0.786 | 1.276 | 1.122 | 0.028 | 0.005 | 0.005 | 0.147 | 0.23 | 0.3 | 0.26 | 0.26 | 0.32 | 0.38 |
| 7 | 103307425 | rs17154114 | -0.29 | -0.784 | 1.186 | 1.087 | -0.015 | -0.029 | -0.029 | 0.493 | 0.36 | 0.41 | 0.24 | 0.18 | 0.25 | 0.26 |
| 7 | 103309822 | rs2299356 | -0.774 | -0.749 | 1.319 | 1.145 | 0.002 | -0.004 | -0.004 | 0 | 0.57 | 0.52 | 0.49 | 0.44 | 0.57 | 0.65 |
| 7 | 103314344 | rs2237631 | -0.918 | -0.797 | 1.563 | 1.108 | -0.004 | 0.005 | 0.005 | 0.009 | 0.59 | 0.49 | 0.53 | 0.6 | 0.59 | 0.69 |
| 7 | 103318924 | rs2073559 | -0.487 | -0.864 | 1.406 | 1.136 | 0 | 0.014 | 0.014 | 0 | 0.49 | 0.65 | 0.46 | 0.45 | 0.51 | 0.49 |
| 7 | 103321872 | rs10487164 | 0.663 | -0.976 | 0.818 | 0.846 | 0.016 | 0.003 | 0.003 | 1.662 | 0.24 | 0.2 | 0.11 | 0.02 | 0.15 | 0.07 |
| 7 | 103325604 | rs12705140 | -0.251 | -0.392 | 0.807 | 0.875 | 0.044 | 0.073 | 0.073 | 1.239 | 0.35 | 0.21 | 0.2 | 0.16 | 0.25 | 0.27 |
| 7 | 103330211 | rs12538423 | 1.033 | -0.052 | 0.525 | 1.112 | -0.076 | -0.098 | -0.098 | 0.663 | 0.2 | 0.05 | 0.29 | 0.45 | 0.31 | 0.38 |
| 7 | 103330944 | rs727708 | -0.676 | -0.294 | 0.87 | 0.976 | -0.004 | 0.002 | 0.002 | 0 | 0.54 | 0.38 | 0.42 | 0.48 | 0.51 | 0.65 |
| 7 | 103333110 | rs10281066 | -1.414 | -0.422 | 1.133 | 0.999 | 0.035 | 0.186 | 0.186 | 0.097 | 0.13 | 0.41 | 0.19 | 0.28 | 0.14 | 0.09 |
| 7 | 103334488 | rs2106172 | -1.992 | -0.448 | 1.238 | 0.992 | 0.014 | 0.153 | 0.153 | 0.204 | 0.14 | 0.58 | 0.21 | 0.29 | 0.14 | 0.12 |
| 7 | 103334604 | rs12705148 | -0.14 | -0.47 | 1.164 | 0.957 | 0.06 | 0.109 | 0.109 | 0.964 | 0.35 | 0.15 | 0.19 | 0.16 | 0.25 | 0.27 |
| 7 | 103340098 | rs17154438 | -2.148 | -0.443 | 1.248 | 0.999 | 0.015 | 0.15 | 0.15 | 0.007 | 0.15 | 0.46 | 0.22 | 0.28 | 0.16 | 0.11 |
| 7 | 103346226 | rs7799028 | -0.876 | -0.493 | 1.039 | 0.963 | 0.039 | 0.008 | 0.008 | 0 | 0.39 | 0.74 | 0.31 | 0.32 | 0.28 | 0.19 |
| 7 | 103352577 | rs3819453 | -0.275 | -0.44 | 1.271 | 1.002 | 0.011 | 0.064 | 0.064 | 0 | 0.07 | 0.16 | 0.08 | 0.14 | 0.08 | 0.07 |
| 7 | 103353742 | rs10265101 | -0.863 | -0.458 | 1.199 | 0.949 | -0.024 | 0.031 | 0.031 | 0 | 0.56 | 0.78 | 0.53 | 0.62 | 0.55 | 0.55 |
| 7 | 103354223 | rs10281266 | 1.265 | -0.461 | 1.444 | 0.936 | 0.024 | 0.025 | 0.025 | 0.018 | 0.2 | 0.25 | 0.24 | 0.32 | 0.28 | 0.39 |
| 7 | 103356275 | rs12705150 | 0.575 | -0.507 | 1.333 | 0.866 | -0.016 | 0.01 | 0.01 | 0 | 0.15 | 0.1 | 0.14 | 0.14 | 0.17 | 0.21 |
| 7 | 103356296 | rs10247439 | -1.89 | -0.329 | 1.28 | 0.878 | -0.014 | 0.017 | 0.017 | 0.001 | 0.7 | 0.87 | 0.66 | 0.76 | 0.69 | 0.75 |
| 7 | 103358920 | rs2299371 | -0.828 | -0.362 | 1.311 | 0.883 | 0.023 | 0.13 | 0.13 | 0.039 | 0.1 | 0.24 | 0.16 | 0.29 | 0.12 | 0.08 |
| 7 | 103359720 | rs540058 | 0.95 | -0.403 | 1.29 | 0.849 | 0.04 | 0.045 | 0.045 | 1.815 | 0.23 | 0.13 | 0.1 | 0.02 | 0.14 | 0.06 |
| 7 | 103361285 | rs2299374 | 1.576 | -0.401 | 1.271 | 0.854 | -0.065 | -0.068 | -0.068 | 0 | 0.19 | 0.03 | 0.22 | 0.3 | 0.25 | 0.37 |
| 7 | 103362484 | rs496907 | -2.2 | -0.42 | 1.22 | 0.857 | -0.085 | -0.035 | -0.035 | 0 | 0.72 | 0.98 | 0.68 | 0.76 | 0.7 | 0.77 |
| 7 | 103365428 | rs476848 | 0.68 | -0.06 | 1.64 | 1.005 | 0.036 | 0.038 | 0.038 | 1.837 | 0.23 | 0.14 | 0.1 | 0.02 | 0.14 | 0.06 |
| 7 | 103365478 | rs17154829 | -1.762 | 0.023 | 1.777 | 1.085 | 0.051 | 0.239 | 0.239 | 0.282 | 0.12 | 0.42 | 0.18 | 0.29 | 0.12 | 0.09 |
| 7 | 103370682 | rs2299375 | 1.115 | 0.049 | 1.552 | 1.073 | 0.008 | 0.01 | 0.01 | 0 | 0.19 | 0.18 | 0.23 | 0.32 | 0.26 | 0.39 |
| 7 | 103371316 | rs580884 | -0.393 | -0.091 | 1.251 | 0.854 | -0.01 | 0.007 | 0.007 | 0 | 0.28 | 0.34 | 0.23 | 0.31 | 0.21 | 0.15 |
| 7 | 103373051 | rs2299379 | -0.446 | 0.428 | 1.195 | 0.843 | 0.006 | 0.015 | 0.015 | 0 | 0.47 | 0.33 | 0.46 | 0.43 | 0.56 | 0.55 |
| 7 | 103375920 | rs541768 | -0.268 | -0.052 | 1.073 | 0.692 | -0.014 | 0.039 | 0.039 | 0 | 0.46 | 0.3 | 0.43 | 0.37 | 0.52 | 0.45 |
| 7 | 103379897 | rs2237633 | 0.763 | -0.087 | 0.81 | 0.666 | -0.116 | -0.073 | -0.073 | 1.543 | 0.22 | 0.06 | 0.32 | 0.34 | 0.39 | 0.38 |
| 7 | 103380849 | rs471360 | 0.347 | -0.061 | 1.009 | 0.8 | -0.074 | -0.072 | -0.072 | 0 | 0.26 | 0.51 | 0.15 | 0.09 | 0.15 | 0.11 |
| 7 | 103389085 | rs528528 | -0.132 | 0.264 | 0.494 | 0.93 | 0 | -0.002 | -0.002 | 0 | 0.49 | 0.45 | 0.45 | 0.39 | 0.54 | 0.48 |
| 7 | 103391042 | rs3819466 | 0.032 | 0.486 | 0.215 | 1.067 | 0 | 0.057 | 0.057 | 0 | 0.05 | 0.11 | 0.11 | 0.19 | 0.05 | 0.04 |
| 7 | 103391409 | rs634500 | -0.666 | 0.462 | 0.121 | 1.047 | -0.033 | -0.044 | -0.044 | 0 | 0.59 | 0.85 | 0.58 | 0.57 | 0.59 | 0.56 |
| 7 | 103392761 | rs651097 | -0.326 | 0.538 | 0.085 | 0.631 | 0.002 | 0.035 | 0.035 | 0 | 0.53 | 0.65 | 0.47 | 0.39 | 0.55 | 0.52 |
| 7 | 103394799 | rs12705155 | -0.085 | 0.402 | -0.833 | 0.698 | 0 | 0.001 | 0.001 | 0 | 0.46 | 0.43 | 0.46 | 0.36 | 0.55 | 0.51 |

| | | | | | | | | | | | | | | | | |
|---|---|---|---|---|---|---|---|---|---|---|---|---|---|---|---|---|
| 7 | 103395352 | rs492063 | 0.023 | 0.416 | -0.819 | 0.66 | 0.005 | 0.009 | 0.009 | 0 | 0.51 | 0.43 | 0.48 | 0.36 | 0.59 | 0.53 |
| 7 | 103396604 | rs579030 | -0.118 | 0.516 | -1.088 | 0.744 | 0.006 | 0.009 | 0.009 | 0 | 0.41 | 0.4 | 0.3 | 0.3 | 0.38 | 0.26 |
| 7 | 103396663 | rs3819477 | 0.135 | -0.049 | -1.314 | 0.841 | 0.017 | -0.001 | -0.001 | 0 | 0.12 | 0.1 | 0.06 | 0.13 | 0.07 | 0.14 |
| 7 | 103396683 | rs650974 | -0.387 | 0.628 | -1.233 | 1.003 | 0.002 | 0.004 | 0.004 | 0 | 0.58 | 0.61 | 0.4 | 0.51 | 0.49 | 0.43 |
| 7 | 103397373 | rs475880 | 0.231 | 1.323 | -2.009 | 1.454 | -0.078 | -0.103 | -0.103 | 3.602 | 0.84 | 0.99 | 0.81 | 0.71 | 0.83 | 0.76 |
| 7 | 103397404 | rs694894 | 0.06 | 0.294 | -2.646 | 0.85 | 0.077 | 0.06 | 0.06 | 1.213 | 0.44 | 0.88 | 0.41 | 0.35 | 0.34 | 0.34 |
| 7 | 103397666 | rs499953 | 0.216 | 0.188 | -2.237 | 0.748 | 0.031 | -0.003 | -0.003 | 0.123 | 0.59 | 0.29 | 0.54 | 0.4 | 0.66 | 0.43 |
| 7 | 103398041 | rs661492 | -0.202 | -0.27 | -2.256 | 0.201 | 0.021 | 0.051 | 0.051 | 0 | 0.26 | 0.14 | 0.28 | 0.09 | 0.21 | 0.15 |
| 7 | 103398107 | rs661575 | -0.045 | 0.118 | -1.294 | 0.529 | 0.011 | 0.028 | 0.028 | 0 | 0.76 | 0.45 | 0.78 | 0.65 | 0.7 | 0.8 |
| 7 | 103398219 | rs2283029 | -0.502 | -0.507 | -1.076 | 0.067 | -0.006 | -0.009 | -0.009 | 0.013 | 0.24 | 0.27 | 0.13 | 0.11 | 0.17 | 0.12 |
| 7 | 103401172 | rs10487171 | -0.478 | 0.012 | -1.137 | 0.401 | -0.002 | -0.034 | -0.034 | 0.088 | 0.17 | 0.09 | 0.23 | 0.3 | 0.17 | 0.25 |
| 7 | 103403076 | rs1705107 | -0.44 | -0.293 | -1.467 | -0.015 | 0.072 | 0.029 | 0.029 | 1.835 | 0.5 | 0.87 | 0.52 | 0.43 | 0.36 | 0.37 |
| 7 | 103404815 | rs7341475 | 0.86 | -0.297 | -0.923 | -0.108 | 0.036 | 0.055 | 0.055 | 0.511 | 0.27 | 0.17 | 0.23 | 0.07 | 0.17 | 0.12 |
| 7 | 103417999 | rs2299381 | 1.156 | 0.646 | -0.21 | 0.104 | 0.028 | -0.004 | -0.004 | 0.387 | 0.48 | 0.32 | 0.4 | 0.41 | 0.41 | 0.5 |
| 7 | 103418983 | rs262342 | -0.425 | 1.203 | 0.257 | -0.008 | 0.051 | -0.018 | -0.018 | 0 | 0.84 | 0.72 | 0.77 | 0.86 | 0.75 | 0.88 |
| 7 | 103419656 | rs2299386 | 1.61 | 1.185 | 0.27 | -0.009 | 0.039 | -0.027 | -0.027 | 0 | 0.48 | 0.19 | 0.4 | 0.41 | 0.41 | 0.5 |
| 7 | 103420619 | rs10251365 | -0.84 | 1.1 | 0.553 | 0.053 | 0.002 | -0.002 | -0.002 | 0 | 0.35 | 0.36 | 0.35 | 0.43 | 0.33 | 0.37 |
| 7 | 103420755 | rs10280683 | -0.84 | 1.1 | 0.553 | 0.053 | 0.002 | -0.002 | -0.002 | 0 | 0.35 | 0.36 | 0.35 | 0.43 | 0.33 | 0.37 |
| 7 | 103421702 | rs1510846 | -0.503 | 1.108 | 0.516 | 0.002 | -0.006 | 0.007 | 0.007 | 0 | 0.31 | 0.38 | 0.33 | 0.45 | 0.28 | 0.36 |
| 7 | 103422656 | rs745541 | -0.243 | 1.258 | 0.629 | 0.012 | 0.08 | -0.011 | -0.011 | 0 | 0.83 | 0.72 | 0.75 | 0.84 | 0.74 | 0.87 |
| 7 | 103424598 | rs2237637 | -0.732 | 1.452 | 0.361 | 0.044 | 0 | -0.008 | -0.008 | 0 | 0.35 | 0.29 | 0.4 | 0.43 | 0.35 | 0.38 |
| 7 | 103435607 | rs16872603 | 0.554 | 1.328 | 0.243 | 0.029 | 0.007 | 0.008 | 0.008 | 0 | 0.17 | 0.13 | 0.1 | 0.13 | 0.16 | 0.14 |
| 7 | 103438842 | rs4729931 | 0.137 | 1.297 | 0.251 | 0.053 | -0.021 | -0.033 | -0.033 | 1.858 | 0.19 | 0.11 | 0.29 | 0.3 | 0.2 | 0.25 |
| 7 | 103438880 | rs4729932 | -0.439 | 1.323 | 0.374 | 0.05 | -0.003 | -0.003 | -0.003 | 0 | 0.24 | 0.24 | 0.32 | 0.33 | 0.22 | 0.26 |
| 7 | 103439114 | rs155333 | -0.211 | 1.487 | 0.245 | 0.06 | 0.06 | 0.048 | 0.048 | 0.172 | 0.86 | 0.72 | 0.81 | 0.74 | 0.75 | 0.77 |
| 7 | 103445791 | rs39329 | -0.319 | 1.433 | -0.106 | 0.098 | -0.022 | 0.008 | 0.008 | 0 | 0.26 | 0.19 | 0.26 | 0.24 | 0.36 | 0.24 |
| 7 | 103453057 | rs262375 | 0.471 | 1.178 | 0.338 | 0.179 | -0.004 | 0.038 | 0.038 | 0 | 0.77 | 0.59 | 0.68 | 0.65 | 0.81 | 0.73 |
| 7 | 103453099 | rs39334 | -0.095 | 0.675 | -0.281 | -0.266 | 0.043 | 0.028 | 0.028 | 4.217 | 0.49 | 0.13 | 0.42 | 0.57 | 0.65 | 0.62 |
| 7 | 103453569 | rs39335 | 1.796 | 0.688 | 0.056 | -0.275 | -0.026 | -0.062 | -0.062 | 0 | 0.24 | 0.32 | 0.24 | 0.05 | 0.15 | 0.11 |
| 7 | 103453953 | rs39336 | 0.766 | 1.182 | -0.11 | -0.304 | -0.025 | 0.011 | 0.011 | 0.333 | 0.13 | 0.07 | 0.18 | 0.09 | 0.26 | 0.09 |
| 7 | 103461273 | rs39341 | 1.569 | 1.196 | 0.159 | -0.257 | -0.004 | 0 | 0 | 0.037 | 0.09 | 0.07 | 0.16 | 0.09 | 0.21 | 0.08 |
| 7 | 103461909 | rs262359 | 0.997 | 1.13 | 0.303 | -0.321 | 0.009 | -0.057 | -0.057 | 0 | 0.34 | 0.44 | 0.41 | 0.17 | 0.36 | 0.19 |
| 7 | 103463998 | rs39343 | 1.009 | 0.976 | 0.166 | -0.303 | -0.006 | -0.001 | -0.001 | 0 | 0.39 | 0.42 | 0.45 | 0.38 | 0.25 | 0.35 |
| 7 | 103464318 | rs264375 | -0.366 | 1.189 | 0.331 | -0.323 | 0.004 | -0.065 | -0.065 | 0.081 | 0.88 | 0.75 | 0.87 | 0.98 | 0.86 | 0.97 |
| 7 | 103466268 | rs39346 | 1.183 | 1.574 | 0.071 | -0.166 | 0.033 | 0.007 | 0.007 | 0 | 0.09 | 0.11 | 0.15 | 0.12 | 0.18 | 0.08 |
| 7 | 103466498 | rs39347 | 0.276 | 1.164 | 0.093 | -0.091 | 0.01 | 0.028 | 0.028 | 0.345 | 0.54 | 0.33 | 0.51 | 0.39 | 0.42 | 0.5 |
| 7 | 103467134 | rs11977571 | 0.35 | 1.433 | 0.147 | -0.14 | 0.013 | -0.032 | -0.032 | 0 | 0.07 | 0.1 | 0.1 | 0 | 0.11 | 0.02 |
| 7 | 103470499 | rs39359 | -0.597 | 1.991 | 0.463 | -0.152 | 0.086 | -0.017 | -0.017 | 0 | 0.18 | 0.27 | 0.28 | 0.16 | 0.4 | 0.14 |
| 7 | 103479069 | rs39379 | -2.116 | 1.464 | 0.356 | -0.483 | 0.029 | 0.048 | 0.048 | 4.439 | 0.72 | 0.8 | 0.75 | 0.87 | 0.81 | 0.88 |
| 7 | 103485921 | rs39387 | -0.261 | 1.362 | 0.491 | 0.102 | 0.003 | -0.001 | -0.001 | 0 | 0.39 | 0.42 | 0.46 | 0.57 | 0.55 | 0.5 |
| 7 | 103489028 | rs10262803 | 0.393 | 1.316 | 0.551 | 0.025 | 0 | 0.012 | 0.012 | 0 | 0.16 | 0.21 | 0.19 | 0.29 | 0.1 | 0.24 |
| 7 | 103489729 | rs39395 | -1.162 | 1.357 | 0.6 | 0.007 | 0.006 | 0.014 | 0.014 | 0.307 | 0.36 | 0.61 | 0.49 | 0.52 | 0.5 | 0.38 |
| 7 | 103489992 | rs39399 | -0.258 | 1.388 | 0.658 | -0.054 | 0.114 | 0.023 | 0.023 | 0.118 | 0.19 | 0.33 | 0.3 | 0.23 | 0.4 | 0.14 |
| 7 | 103490420 | rs13223714 | 0.127 | 1.313 | 0.593 | -0.079 | 0.019 | 0.007 | 0.007 | 0.206 | 0.37 | 0.33 | 0.35 | 0.63 | 0.25 | 0.6 |
| 7 | 103492908 | rs1510858 | -0.517 | 1.28 | 0.692 | -0.111 | 0.004 | -0.002 | -0.002 | 0.39 | 0.32 | 0.28 | 0.3 | 0.35 | 0.26 | 0.38 |
| 7 | 103495393 | rs39401 | 0.598 | 1.592 | 0.782 | -0.2 | 0.081 | -0.066 | -0.066 | 0 | 0.13 | 0.23 | 0.18 | 0.04 | 0.29 | 0.12 |
| 7 | 103505675 | rs3914126 | -0.958 | 1.227 | 0.492 | -0.169 | 0.01 | -0.003 | -0.003 | 0 | 0.56 | 0.63 | 0.64 | 0.86 | 0.66 | 0.74 |
| 7 | 103507670 | rs9986961 | -0.355 | 1.48 | 0.616 | 0.253 | 0.08 | -0.015 | -0.015 | 0 | 0.19 | 0.28 | 0.27 | 0.16 | 0.4 | 0.16 |
| 7 | 103512401 | rs17133142 | 0.743 | 1.18 | 0.414 | 0.152 | 0.006 | 0.008 | 0.008 | 0 | 0.14 | 0.08 | 0.1 | 0.01 | 0.15 | 0.14 |
| 7 | 103519638 | rs2237640 | 1.316 | 1.003 | 0.393 | 0.198 | -0.01 | 0.036 | 0.036 | 0.029 | 0.17 | 0.17 | 0.16 | 0.46 | 0.12 | 0.31 |
| 7 | 103521696 | rs2237643 | -0.938 | 1.139 | 0.437 | 0.111 | 0.017 | 0.008 | 0.008 | 0.526 | 0.33 | 0.42 | 0.45 | 0.42 | 0.51 | 0.4 |
| 7 | 103522992 | rs3914124 | 0.841 | 1.05 | 0.036 | 0.068 | 0.003 | 0.002 | 0.002 | 0 | 0.14 | 0.1 | 0.1 | 0.01 | 0.15 | 0.14 |
| 7 | 103527369 | rs3914132 | -2.309 | 0.842 | 0.038 | 0.132 | 0.025 | 0.034 | 0.034 | 3.78 | 0.68 | 0.76 | 0.73 | 0.89 | 0.79 | 0.86 |
| 7 | 103527448 | rs9641319 | -0.631 | 1.735 | 0.114 | 0.729 | -0.022 | -0.026 | -0.026 | 0 | 0.23 | 0.2 | 0.32 | 0.4 | 0.29 | 0.35 |
| 7 | 103529040 | rs2299395 | 1.003 | 1.744 | 0.077 | 0.797 | -0.012 | -0.018 | -0.018 | 0.218 | 0.09 | 0.02 | 0.14 | 0.38 | 0.04 | 0.22 |
| 7 | 103535980 | rs3857816 | -0.703 | 1.367 | 0.178 | 0.077 | -0.029 | 0.094 | 0.094 | 3.635 | 0.39 | 0.23 | 0.46 | 0.76 | 0.41 | 0.66 |
| 7 | 103536824 | rs7811777 | -0.734 | 1.387 | 0.166 | -0.089 | -0.032 | 0.129 | 0.129 | 2.823 | 0.39 | 0.23 | 0.47 | 0.87 | 0.41 | 0.66 |
| 7 | 103538228 | rs17157125 | -0.166 | 0.463 | 0.809 | 0.55 | -0.011 | 0.1 | 0.1 | 0 | 0.16 | 0.32 | 0.23 | 0.3 | 0.08 | 0.23 |
| 7 | 103538333 | rs17157128 | 0.057 | 0.464 | 0.801 | 0.576 | 0.001 | -0.001 | -0.001 | 0.056 | 0.14 | 0.15 | 0.2 | 0.25 | 0.08 | 0.23 |
| 7 | 103538402 | rs3905915 | -0.043 | 0.315 | 0.577 | 0.599 | 0.008 | -0.029 | -0.029 | 0 | 0.26 | 0.2 | 0.18 | 0.42 | 0.26 | 0.38 |
| 7 | 103542060 | rs3898845 | -0.691 | 0.501 | 0.419 | 0.356 | 0 | -0.003 | -0.003 | 0 | 0.42 | 0.53 | 0.43 | 0.72 | 0.4 | 0.63 |
| 7 | 103545292 | rs4729939 | -0.506 | 1.163 | 0.135 | 0.074 | 0.019 | -0.022 | -0.022 | 0 | 0.19 | 0.24 | 0.29 | 0.05 | 0.39 | 0.19 |
| 7 | 103548787 | rs17157216 | 0.107 | 0.232 | 0.02 | 0.244 | -0.008 | 0.047 | 0.047 | 0 | 0.19 | 0.07 | 0.14 | 0.1 | 0.15 | 0.19 |
| 7 | 103549724 | rs802787 | -0.404 | -0.442 | 0.009 | 0.239 | 0.008 | 0.014 | 0.014 | 3.637 | 0.39 | 0.26 | 0.44 | 0.61 | 0.62 | 0.51 |
| 7 | 103555270 | rs7797395 | -0.746 | -0.512 | 0.57 | 0.276 | -0.035 | 0.001 | 0.001 | 0 | 0.27 | 0.47 | 0.3 | 0.3 | 0.17 | 0.34 |
| 7 | 103556774 | rs10245317 | -0.783 | -0.514 | 0.578 | 0.275 | -0.033 | 0 | 0 | 0.318 | 0.27 | 0.49 | 0.3 | 0.3 | 0.17 | 0.34 |
| 7 | 103560616 | rs802788 | -0.374 | -0.259 | 0.144 | 0.551 | -0.008 | -0.021 | -0.021 | 0.159 | 0.4 | 0.3 | 0.38 | 0.48 | 0.43 | 0.51 |
| 7 | 103561412 | rs802790 | -2.129 | -0.381 | 0.224 | 0.496 | 0.048 | 0.027 | 0.027 | 5.543 | 0.67 | 0.8 | 0.73 | 0.76 | 0.79 | 0.85 |
| 7 | 103561431 | rs16872993 | -0.287 | -0.286 | 0.344 | 0.524 | -0.018 | 0.145 | 0.145 | 1.173 | 0.1 | 0.4 | 0.15 | 0.19 | 0.02 | 0.12 |

| | | | | | | | | | | | | | | | | |
|---|---|---|---|---|---|---|---|---|---|---|---|---|---|---|---|---|
| 7 | 103562551 | rs10277802 | -0.294 | -0.356 | -0.174 | 0.491 | -0.029 | 0.198 | 0.198 | 1.619 | 0.1 | 0.49 | 0.16 | 0.23 | 0.02 | 0.12 |
| 7 | 103563145 | rs7792066 | -0.715 | -0.311 | -0.31 | 0.477 | 0.059 | 0.001 | 0.001 | 0 | 0.27 | 0.54 | 0.35 | 0.29 | 0.36 | 0.34 |
| 7 | 103570601 | rs6954835 | -0.938 | -0.051 | -0.945 | 0.066 | 0.014 | 0.074 | 0.074 | 1.441 | 0.49 | 0.7 | 0.53 | 0.71 | 0.46 | 0.63 |
| 7 | 103570673 | rs6954870 | -0.961 | -0.035 | -1.1 | 0.07 | 0.017 | 0.089 | 0.089 | 1.425 | 0.5 | 0.73 | 0.54 | 0.7 | 0.46 | 0.63 |
| 7 | 103571888 | rs6960627 | 0.664 | 0.678 | -2.031 | -0.476 | -0.123 | 0.05 | 0.05 | 0.085 | 0.16 | 0.01 | 0.18 | 0.06 | 0.34 | 0.23 |
| 7 | 103573436 | rs6950645 | 0.318 | 0.066 | -1.569 | -0.187 | -0.02 | 0.052 | 0.052 | 0 | 0.74 | 0.86 | 0.72 | 0.87 | 0.71 | 0.75 |
| 7 | 103575173 | rs3808019 | -0.445 | 0.508 | -1.825 | 0.299 | -0.018 | -0.011 | -0.011 | 0 | 0.87 | 1 | 0.89 | 0.9 | 0.92 | 0.83 |
| 7 | 103575689 | rs10266480 | -0.514 | 0.873 | -1.268 | 0.609 | -0.025 | 0.022 | 0.022 | 0.369 | 0.37 | 0.34 | 0.41 | 0.71 | 0.49 | 0.62 |
| 7 | 103580446 | rs17157475 | -1.178 | 0.401 | -0.931 | 1.046 | -0.001 | 0.008 | 0.008 | 0 | 0.12 | 0.17 | 0.21 | 0.2 | 0.12 | 0.22 |
| 7 | 103581626 | rs6965133 | 0.342 | -0.341 | -1.846 | 1.1 | -0.012 | -0.1 | -0.1 | 0 | 0.7 | 0.96 | 0.77 | 0.57 | 0.77 | 0.57 |
| 7 | 103581806 | rs4639449 | 0.974 | -0.853 | -1.457 | 0.653 | -0.004 | 0.011 | 0.011 | 0.046 | 0.36 | 0.38 | 0.38 | 0.43 | 0.39 | 0.27 |
| 7 | 103584701 | rs876425 | 0.442 | -0.42 | -1.24 | 0.549 | -0.03 | 0.029 | 0.029 | 0 | 0.23 | 0.15 | 0.32 | 0.06 | 0.34 | 0.21 |
| 7 | 103586190 | rs4236562 | 0.068 | -0.62 | -1.318 | 0.724 | 0.009 | -0.121 | -0.121 | 0 | 0.71 | 0.93 | 0.77 | 0.55 | 0.79 | 0.59 |
| 7 | 103590285 | rs4729942 | -0.219 | -0.758 | -0.698 | 0.942 | -0.009 | -0.008 | -0.008 | 0.269 | 0.74 | 0.79 | 0.6 | 0.67 | 0.69 | 0.64 |
| 7 | 103595652 | rs722277 | 0.963 | -1.249 | -0.492 | 0.737 | -0.006 | 0.112 | 0.112 | 0 | 0.42 | 0.24 | 0.38 | 0.14 | 0.47 | 0.2 |
| 7 | 103596176 | rs4727580 | 0.396 | -1.589 | 0.3 | 1.054 | -0.008 | 0.001 | 0.001 | 0 | 0.53 | 0.7 | 0.48 | 0.46 | 0.52 | 0.44 |
| 7 | 103597786 | rs10276794 | -0.723 | -1.642 | 0.359 | 1.218 | -0.036 | 0.047 | 0.047 | 0 | 0.29 | 0.56 | 0.2 | 0.43 | 0.2 | 0.39 |
| 7 | 103598553 | rs7794418 | 0.504 | -1.482 | 0.102 | 0.927 | 0.017 | -0.005 | -0.005 | 0 | 0.19 | 0.1 | 0.11 | 0.2 | 0.16 | 0.17 |
| 7 | 103598671 | rs6943822 | 1.154 | -1.474 | -0.028 | 0.583 | -0.008 | 0.036 | 0.036 | 0 | 0.35 | 0.24 | 0.34 | 0.24 | 0.43 | 0.12 |
| 7 | 103601185 | rs10246267 | -0.225 | -1.331 | 0.476 | 0.245 | -0.022 | -0.072 | -0.072 | 0.73 | 0.12 | 0.25 | 0.07 | 0.01 | 0.05 | 0.09 |
| 7 | 103601257 | rs7805541 | -0.485 | -1.887 | 0.634 | 0.505 | -0.036 | 0.005 | 0.005 | 0 | 0.3 | 0.56 | 0.12 | 0.29 | 0.15 | 0.28 |
| 7 | 103601382 | rs17313716 | 0.887 | -1.628 | 0.538 | 0.85 | -0.001 | -0.02 | -0.02 | 0 | 0.4 | 0.36 | 0.42 | 0.47 | 0.46 | 0.33 |
| 7 | 103602004 | rs961206 | 0.309 | -1.354 | 0.082 | 0.794 | -0.031 | -0.011 | -0.011 | 0 | 0.27 | 0.36 | 0.14 | 0.21 | 0.18 | 0.21 |
| 7 | 103603530 | rs6950391 | 0.471 | -1.475 | 0.013 | 0.815 | 0.001 | 0.006 | 0.006 | 0 | 0.49 | 0.6 | 0.48 | 0.42 | 0.53 | 0.43 |
| 7 | 103607121 | rs4483088 | -0.702 | -1.498 | -0.257 | 0.831 | -0.118 | -0.091 | -0.091 | 0.989 | 0.77 | 0.96 | 0.6 | 0.63 | 0.69 | 0.63 |
| 7 | 103609900 | rs10245730 | 0.473 | -1.463 | -0.113 | 0.632 | 0.017 | 0.005 | 0.005 | 0 | 0.26 | 0.2 | 0.1 | 0.2 | 0.16 | 0.2 |
| 7 | 103621132 | rs4460306 | 0.218 | -1.174 | -0.279 | 0.507 | 0.002 | -0.002 | -0.002 | 0 | 0.49 | 0.63 | 0.49 | 0.46 | 0.53 | 0.43 |
| 7 | 103625877 | rs4298437 | -0.097 | -0.714 | -0.691 | -0.051 | -0.008 | -0.006 | -0.006 | 0 | 0.7 | 0.78 | 0.71 | 0.95 | 0.66 | 0.63 |
| 7 | 103626322 | rs6952315 | 1.24 | -1.042 | -1.59 | -0.198 | -0.118 | 0.085 | 0.085 | 0 | 0.25 | 0.02 | 0.31 | 0.08 | 0.43 | 0.1 |
| 7 | 103628036 | rs17157903 | 0.393 | -1.502 | -1.249 | 0.423 | 0.041 | -0.025 | -0.025 | 0.114 | 0.23 | 0.1 | 0.1 | 0.29 | 0.15 | 0.19 |
| 7 | 103637906 | rs6955789 | 0.677 | -1.053 | -0.431 | 0.61 | 0.011 | -0.027 | -0.027 | 0.035 | 0.21 | 0.21 | 0.2 | 0.05 | 0.27 | 0.26 |
| 7 | 103639756 | rs6966089 | -1.34 | -1.083 | -0.138 | 0.455 | 0.001 | 0.022 | 0.022 | 0 | 0.67 | 0.72 | 0.67 | 0.84 | 0.58 | 0.88 |
| 7 | 103639821 | rs12533473 | 0.692 | -0.826 | -0.441 | 0.139 | 0.011 | -0.013 | -0.013 | 0 | 0.26 | 0.09 | 0.17 | 0.27 | 0.19 | 0.2 |
| 7 | 103640262 | rs12111672 | 0.714 | -0.52 | -0.322 | 0.03 | 0.025 | -0.047 | -0.047 | 0.005 | 0.21 | 0.24 | 0.21 | 0.05 | 0.27 | 0.26 |
| 7 | 103643666 | rs6465953 | -0.325 | -0.735 | -0.346 | 0.313 | -0.004 | -0.004 | -0.004 | 0 | 0.55 | 0.5 | 0.52 | 0.43 | 0.53 | 0.67 |
| 7 | 103646910 | rs10254364 | -1.038 | 0.013 | -1.573 | -0.094 | -0.128 | 0.067 | 0.067 | 0.284 | 0.59 | 0.97 | 0.62 | 0.72 | 0.53 | 0.64 |
| 7 | 103647122 | rs10224925 | -0.186 | -0.04 | -0.664 | -0.107 | -0.008 | 0.027 | 0.027 | 0 | 0.16 | 0.29 | 0.15 | 0.19 | 0.1 | 0.24 |
| 7 | 103650219 | rs17158144 | 0.21 | -0.065 | -0.643 | -0.091 | -0.007 | 0.051 | 0.051 | 0 | 0.09 | 0.16 | 0.05 | 0.19 | 0.05 | 0.19 |
| 7 | 103650246 | rs10260517 | 1.042 | -0.053 | -0.414 | -0.375 | 0.033 | -0.027 | -0.027 | 0 | 0.08 | 0.27 | 0.11 | 0 | 0.05 | 0.06 |
| 7 | 103658861 | rs10487183 | 0.421 | -0.031 | -0.969 | -0.41 | -0.06 | 0.115 | 0.115 | 0 | 0.22 | 0.02 | 0.15 | 0.04 | 0.26 | 0.23 |
| 7 | 103672960 | rs13233398 | -0.835 | 0.066 | -1.169 | -0.119 | -0.058 | 0.035 | 0.035 | 0 | 0.62 | 0.85 | 0.7 | 0.73 | 0.54 | 0.66 |
| 7 | 103679529 | rs1858812 | -0.17 | -0.356 | -0.55 | 0.541 | -0.045 | 0.245 | 0.245 | 0.036 | 0.15 | 0.55 | 0.39 | 0.44 | 0.07 | 0.18 |
| 7 | 103686753 | rs10953402 | 0.077 | -0.183 | 0.03 | 0.622 | -0.026 | 0.29 | 0.29 | 0 | 0.13 | 0.44 | 0.38 | 0.45 | 0.07 | 0.18 |
| 7 | 103690016 | rs10264307 | 0.128 | -0.442 | -0.483 | 0.279 | -0.036 | 0.001 | 0.001 | 0 | 0.08 | 0.18 | 0.03 | 0.05 | 0.01 | 0.2 |
| 7 | 103719248 | rs7780591 | -1.142 | -0.484 | -1.37 | -0.017 | -0.101 | 0.028 | 0.028 | 0 | 0.6 | 0.92 | 0.68 | 0.72 | 0.53 | 0.69 |
| 7 | 103734021 | rs6465964 | -0.248 | -0.619 | -1.262 | -0.062 | -0.022 | 0.058 | 0.058 | 0 | 0.48 | 0.88 | 0.63 | 0.67 | 0.48 | 0.44 |
| 7 | 103746862 | rs983960 | -0.606 | -1 | -0.838 | -0.391 | 0.004 | 0.011 | 0.011 | 0 | 0.34 | 0.3 | 0.21 | 0.22 | 0.26 | 0.38 |
| 7 | 103749889 | rs194824 | 0.268 | -1.022 | -0.632 | -0.478 | 0.003 | -0.018 | -0.018 | 0.707 | 0.28 | 0.75 | 0.5 | 0.5 | 0.27 | 0.32 |
| 7 | 103759385 | rs7808273 | 1.606 | -0.9 | -0.727 | -0.494 | -0.061 | -0.02 | -0.02 | 0 | 0.16 | 0.01 | 0.12 | 0.18 | 0.19 | 0.09 |
| 7 | 103762313 | rs194834 | -1.177 | -1.023 | -0.762 | -0.565 | -0.033 | 0.066 | 0.066 | 0 | 0.54 | 0.8 | 0.66 | 0.69 | 0.51 | 0.54 |
| 7 | 103765719 | rs194836 | 0.861 | -1.076 | -0.553 | -0.305 | 0.013 | 0.13 | 0.13 | 0 | 0.24 | 0.46 | 0.48 | 0.48 | 0.27 | 0.31 |
| 7 | 103766003 | rs194837 | -0.45 | -1.123 | -0.545 | -0.31 | 0 | 0.002 | 0.002 | 0 | 0.45 | 0.56 | 0.61 | 0.72 | 0.46 | 0.56 |
| 7 | 103770773 | rs194842 | 0.848 | -1.044 | -0.456 | -0.253 | 0.013 | 0.128 | 0.128 | 0 | 0.24 | 0.46 | 0.48 | 0.48 | 0.27 | 0.32 |
| 7 | 103771201 | rs194844 | -0.103 | 0.067 | -0.833 | -0.508 | 0.029 | 0.031 | 0.031 | 0 | 0.33 | 0.55 | 0.54 | 0.52 | 0.41 | 0.52 |
| 7 | 103779703 | rs11982003 | 1.585 | -0.258 | -0.739 | -0.439 | -0.032 | 0.068 | 0.068 | 0 | 0.19 | 0.07 | 0.14 | 0.05 | 0.24 | 0.23 |
| 7 | 103781065 | rs194855 | 0.276 | -0.196 | -0.831 | -0.306 | -0.008 | 0.055 | 0.055 | 0 | 0.29 | 0.56 | 0.49 | 0.52 | 0.28 | 0.47 |
| 7 | 103790469 | rs194868 | -1.324 | -0.21 | -1.114 | -0.411 | -0.099 | 0.053 | 0.053 | 0 | 0.58 | 0.91 | 0.66 | 0.73 | 0.5 | 0.66 |
| 7 | 103811808 | rs17326034 | 2.761 | -0.406 | -1.838 | -1.009 | -0.017 | 0.077 | 0.077 | 0 | 0.15 | 0.02 | 0.13 | 0 | 0.2 | 0.07 |
| 7 | 103811841 | rs3808008 | 0.411 | -0.622 | -1.537 | -0.805 | -0.013 | 0.105 | 0.105 | 0 | 0.51 | 0.12 | 0.37 | 0.28 | 0.53 | 0.44 |
| 7 | 103813499 | rs7804399 | 0.601 | -0.652 | -1.792 | -0.232 | -0.07 | -0.02 | -0.02 | 0 | 0.31 | 0.02 | 0.17 | 0.31 | 0.31 | 0.32 |
| 7 | 103827064 | rs11770973 | 2.721 | -0.629 | -1.406 | -0.117 | 0.006 | 0.07 | 0.07 | 0.419 | 0.12 | 0 | 0.03 | 0 | 0.06 | 0.05 |
| 7 | 103837333 | rs3808003 | -0.215 | -0.749 | -1.169 | -0.23 | -0.007 | -0.024 | -0.024 | 0 | 0.33 | 0.34 | 0.18 | 0.18 | 0.23 | 0.35 |
| 7 | 103838539 | rs7807600 | 1.134 | -0.679 | -1.16 | -0.232 | -0.011 | 0.021 | 0.021 | 0 | 0.2 | 0.05 | 0.12 | 0.18 | 0.18 | 0.24 |
| 7 | 103843369 | rs1035169 | 1.072 | -0.637 | -1.234 | -0.289 | -0.04 | 0.035 | 0.035 | 0 | 0.4 | 0.2 | 0.33 | 0.27 | 0.48 | 0.32 |
| 7 | 103844197 | rs10238218 | -0.083 | -0.554 | -1.233 | -0.249 | -0.006 | -0.022 | -0.022 | 0 | 0.33 | 0.33 | 0.27 | 0.18 | 0.23 | 0.35 |
| 7 | 103861035 | rs12537685 | 0.951 | -0.778 | -0.589 | 0.141 | 0.012 | 0.096 | 0.096 | 0 | 0.26 | 0.53 | 0.48 | 0.55 | 0.29 | 0.33 |
| 7 | 103864567 | rs2188284 | -0.21 | -0.842 | -0.984 | 0.087 | 0.016 | -0.041 | -0.041 | 0 | 0.38 | 0.82 | 0.55 | 0.55 | 0.34 | 0.43 |
| 7 | 103870718 | rs12535884 | 0.829 | -0.402 | -0.901 | 0.426 | 0.005 | 0.046 | 0.046 | 0 | 0.25 | 0.29 | 0.47 | 0.54 | 0.29 | 0.34 |
| 7 | 103871529 | rs6971455 | 1.006 | -0.945 | -1.374 | 0.722 | -0.036 | -0.058 | -0.058 | 0.926 | 0.18 | 0.1 | 0.21 | 0.31 | 0.25 | 0.21 |

| | | | | | | | | | | | | | | | | |
|---|---|---|---|---|---|---|---|---|---|---|---|---|---|---|---|---|
| 7 | 103872165 | rs193890 | -0.592 | -2.502 | -0.719 | 0.898 | -0.002 | -0.002 | -0.002 | 0 | 0.52 | 0.48 | 0.6 | 0.64 | 0.43 | 0.59 |
| 7 | 103878454 | rs4446676 | -0.3 | -2.902 | -0.925 | 0.898 | 0.012 | 0 | 0 | 0 | 0.3 | 0.14 | 0.21 | 0.27 | 0.27 | 0.29 |
| 7 | 103879276 | rs4727590 | 0.933 | -2.968 | -0.813 | 0.99 | -0.018 | 0.01 | 0.01 | 0 | 0.22 | 0.28 | 0.38 | 0.28 | 0.16 | 0.3 |
| 7 | 103880245 | rs990815 | -0.53 | -2.435 | -0.868 | 0.882 | -0.001 | 0 | 0 | 1.176 | 0.52 | 0.47 | 0.6 | 0.55 | 0.43 | 0.58 |
| 7 | 103893012 | rs10259086 | -0.725 | -2.537 | -1.427 | 0.428 | 0.074 | -0.041 | -0.041 | 0 | 0.43 | 0.79 | 0.59 | 0.57 | 0.32 | 0.37 |
| 7 | 103893163 | rs2385143 | -1.166 | -2.719 | -1.463 | 0.315 | 0.064 | -0.023 | -0.023 | 0.435 | 0.53 | 0.83 | 0.61 | 0.57 | 0.38 | 0.44 |
| 7 | 103895925 | rs4729985 | -0.322 | -2.803 | -1.323 | 0.203 | 0.029 | -0.014 | -0.014 | 0 | 0.34 | 0.7 | 0.48 | 0.43 | 0.22 | 0.17 |
| 7 | 103896585 | rs4729986 | 0.319 | -2.74 | -0.942 | 0.252 | -0.012 | 0.117 | 0.117 | 0.626 | 0.17 | 0.55 | 0.45 | 0.35 | 0.13 | 0.1 |
| 7 | 103904428 | rs2385147 | -1.25 | -2.668 | -1.341 | 0.372 | 0.046 | 0.063 | 0.063 | 0 | 0.53 | 0.87 | 0.57 | 0.68 | 0.39 | 0.44 |
| 7 | 103905649 | rs10275423 | 0.978 | -2.65 | -1.289 | 0.267 | 0.009 | 0.016 | 0.016 | 0 | 0.1 | 0.03 | 0.04 | 0 | 0.07 | 0.07 |
| 7 | 103908726 | rs10265838 | -0.636 | -2.582 | -1.113 | 0.449 | 0.045 | 0.031 | 0.031 | 0 | 0.43 | 0.68 | 0.46 | 0.35 | 0.29 | 0.24 |
| 7 | 103909059 | rs12056177 | 0.659 | -1.878 | -0.815 | 0.694 | -0.003 | 0.033 | 0.033 | 0.043 | 0.1 | 0.13 | 0.1 | 0.25 | 0.1 | 0.19 |
| 7 | 103924453 | rs41508 | -1.222 | -1.838 | -1.353 | 0.124 | -0.175 | 0.095 | 0.095 | 0 | 0.73 | 0.96 | 0.72 | 0.88 | 0.55 | 0.67 |
| 7 | 103931634 | rs41519 | -0.116 | -1.675 | -1.004 | 0.264 | 0.049 | 0.154 | 0.154 | 0 | 0.46 | 0.79 | 0.64 | 0.79 | 0.39 | 0.52 |
| 7 | 103936631 | rs7787013 | -0.056 | -1.41 | 0.14 | 0.653 | 0.001 | 0.001 | 0.001 | 0 | 0.48 | 0.48 | 0.23 | 0.33 | 0.37 | 0.45 |
| 7 | 103941535 | rs1858915 | 0.63 | -1.345 | 0.075 | 0.958 | -0.002 | 0.049 | 0.049 | 0.199 | 0.09 | 0.15 | 0.09 | 0.2 | 0.09 | 0.1 |
| 7 | 103942231 | rs9641326 | -0.455 | -1.382 | 0.455 | 0.925 | -0.002 | 0 | 0 | 0 | 0.56 | 0.51 | 0.3 | 0.52 | 0.47 | 0.55 |
| 7 | 103949760 | rs10268758 | 0.846 | -1.551 | -0.101 | 0.794 | 0.029 | 0.024 | 0.024 | 0 | 0.32 | 0.13 | 0.14 | 0.23 | 0.24 | 0.37 |
| 7 | 103950365 | rs10229129 | -0.103 | -1.467 | 0.076 | 0.86 | 0.044 | -0.005 | -0.005 | 0.38 | 0.46 | 0.28 | 0.24 | 0.44 | 0.37 | 0.5 |
| 7 | 103966977 | rs2106502 | -0.634 | -1.399 | 0.13 | 0.781 | 0.002 | 0.003 | 0.003 | 0 | 0.55 | 0.44 | 0.31 | 0.57 | 0.46 | 0.55 |
| 7 | 103967992 | rs11974897 | 0.49 | -1.373 | -0.123 | 0.881 | 0.043 | -0.046 | -0.046 | 0.095 | 0.37 | 0.16 | 0.17 | 0.42 | 0.28 | 0.39 |
| 7 | 103970510 | rs11770570 | 1.304 | -0.988 | 0.105 | 0.717 | -0.001 | -0.004 | -0.004 | 0.016 | 0.08 | 0.07 | 0.04 | 0 | 0.09 | 0.04 |
| 7 | 103971508 | rs1124994 | -0.168 | -1.093 | 0.535 | 1.019 | -0.002 | 0.323 | 0.323 | 0 | 0.13 | 0.42 | 0.36 | 0.47 | 0.12 | 0.15 |
| 7 | 103974305 | rs2891692 | -0.971 | -1.008 | -0.05 | 0.55 | -0.049 | 0.06 | 0.06 | 0 | 0.67 | 0.83 | 0.67 | 0.86 | 0.58 | 0.69 |
| 7 | 103975989 | rs2106503 | -0.491 | -1.124 | 0.059 | 0.549 | -0.025 | 0.148 | 0.148 | 0 | 0.58 | 0.81 | 0.63 | 0.86 | 0.49 | 0.64 |
| 7 | 103976833 | rs12673807 | 0.311 | -1.211 | 0.667 | 0.886 | -0.006 | 0.066 | 0.066 | 0 | 0.11 | 0.21 | 0.36 | 0.28 | 0.12 | 0.14 |
| 7 | 103977467 | rs6973111 | -0.058 | -1.584 | 0.836 | 0.592 | -0.023 | -0.088 | -0.088 | 0.01 | 0.13 | 0.4 | 0.05 | 0 | 0.08 | 0.08 |
| 7 | 103978882 | rs6979350 | -0.097 | -1.565 | 0.661 | 0.647 | 0.003 | 0.002 | 0.002 | 0 | 0.46 | 0.5 | 0.26 | 0.39 | 0.37 | 0.49 |
| 7 | 103987785 | rs7794181 | 0.483 | -1.091 | 0.684 | 0.874 | -0.001 | 0.003 | 0.003 | 0 | 0.34 | 0.29 | 0.22 | 0.4 | 0.31 | 0.41 |
| 7 | 103987924 | rs7777442 | -0.519 | -1.226 | 0.281 | 0.792 | 0.018 | 0.106 | 0.106 | 0 | 0.51 | 0.72 | 0.6 | 0.7 | 0.47 | 0.56 |
| 7 | 103990755 | rs6465981 | 1.291 | -1.22 | -0.074 | 0.937 | 0.001 | -0.035 | -0.035 | 0 | 0.25 | 0.05 | 0.13 | 0.25 | 0.21 | 0.32 |
| 7 | 103996031 | rs11977984 | 0.145 | -1.292 | -0.127 | 0.89 | -0.015 | -0.004 | -0.004 | 2.08 | 0.41 | 0.12 | 0.25 | 0.34 | 0.39 | 0.45 |
| 7 | 103996499 | rs6970753 | -0.204 | -1.366 | 0.253 | 1.021 | -0.021 | 0.066 | 0.066 | 0 | 0.17 | 0.3 | 0.37 | 0.36 | 0.16 | 0.16 |
| 7 | 103996931 | rs17331134 | 0.392 | -1.391 | 0.307 | 0.998 | -0.007 | 0.004 | 0.004 | 0 | 0.16 | 0.09 | 0.13 | 0.13 | 0.19 | 0.14 |
| 7 | 103997606 | rs1465197 | -1.133 | -1.449 | 0.38 | 1.031 | -0.078 | 0.15 | 0.15 | 0 | 0.61 | 0.82 | 0.65 | 0.87 | 0.56 | 0.63 |
| 7 | 103998714 | rs1003778 | 0.57 | -0.487 | 0.432 | 0.778 | -0.003 | 0.011 | 0.011 | 0 | 0.36 | 0.48 | 0.21 | 0.39 | 0.32 | 0.37 |
| 7 | 103999996 | rs17137016 | 1.214 | -0.863 | 0.457 | 0.818 | -0.027 | 0.063 | 0.063 | 0 | 0.27 | 0.4 | 0.15 | 0.39 | 0.21 | 0.31 |
| 7 | 104004677 | rs10487202 | 1.031 | -0.758 | 0.369 | 0.835 | -0.029 | -0.035 | -0.035 | 0 | 0.08 | 0 | 0.07 | 0.13 | 0.1 | 0.1 |
| 7 | 104006993 | rs7792560 | -0.751 | -0.199 | -0.053 | 0.587 | -0.001 | 0.007 | 0.007 | 0 | 0.33 | 0.35 | 0.21 | 0.48 | 0.3 | 0.24 |
| 7 | 104007886 | rs1557689 | 0.014 | -0.199 | -0.074 | 0.592 | -0.011 | -0.034 | -0.034 | 0 | 0.25 | 0.35 | 0.14 | 0.16 | 0.2 | 0.13 |
| 7 | 104008533 | rs4383914 | -0.835 | -0.179 | -0.249 | 0.582 | 0.001 | -0.003 | -0.003 | 0 | 0.41 | 0.39 | 0.26 | 0.47 | 0.39 | 0.29 |
| 7 | 104009435 | rs17137052 | 1.804 | 0.171 | -0.376 | 0.577 | 0 | -0.009 | -0.009 | 0 | 0.08 | 0.07 | 0.05 | 0 | 0.09 | 0.04 |
| 7 | 104010118 | rs6465983 | -0.389 | 0.224 | -0.285 | 0.865 | -0.041 | -0.006 | -0.006 | 0.159 | 0.68 | 0.81 | 0.41 | 0.68 | 0.59 | 0.6 |
| 7 | 104010267 | rs6465984 | -0.187 | 0.464 | 0.176 | 1.014 | -0.008 | 0.003 | 0.003 | 0 | 0.1 | 0.16 | 0.06 | 0.13 | 0.09 | 0.09 |
| 7 | 104017717 | rs6972034 | -0.366 | 0.84 | -0.139 | 1.384 | -0.034 | -0.03 | -0.03 | 0.449 | 0.94 | 0.99 | 0.92 | 0.79 | 0.89 | 0.86 |
| 7 | 104022201 | rs4730001 | -0.436 | 0.464 | 0.178 | 0.734 | -0.001 | 0 | 0 | 0.412 | 0.29 | 0.65 | 0.57 | 0.3 | 0.33 | 0.35 |
| 7 | 104023332 | rs1468144 | 0.01 | 0.429 | 0.846 | 0.718 | 0.002 | 0 | 0 | 0 | 0.26 | 0.28 | 0.56 | 0.3 | 0.33 | 0.35 |
| 7 | 104026849 | rs10953427 | -0.756 | 0.274 | 0.696 | 0.747 | -0.004 | -0.008 | -0.008 | 0 | 0.34 | 0.35 | 0.6 | 0.47 | 0.42 | 0.43 |
| 7 | 104026863 | rs10953428 | 0.081 | 0.293 | 0.766 | 0.756 | 0.002 | 0 | 0 | 0 | 0.26 | 0.28 | 0.56 | 0.3 | 0.33 | 0.35 |
| 7 | 104035816 | rs1989823 | -0.246 | 0.287 | 1.161 | 0.751 | 0.017 | 0.002 | 0.002 | 0.266 | 0.27 | 0.47 | 0.57 | 0.3 | 0.33 | 0.35 |
| 7 | 104042712 | rs6465985 | 0.069 | 0.332 | 1.051 | 0.772 | 0.001 | 0 | 0 | 0 | 0.26 | 0.27 | 0.56 | 0.3 | 0.33 | 0.35 |
| 7 | 104056266 | rs6978267 | -0.094 | -0.573 | 0.86 | 0.21 | 0 | 0.044 | 0.044 | 0 | 0.2 | 0.13 | 0.1 | 0.03 | 0.17 | 0.1 |
| 7 | 104060000 | rs17137518 | 0.058 | -0.692 | 1.102 | 0.38 | -0.001 | 0.009 | 0.009 | 0 | 0.08 | 0.13 | 0.07 | 0.1 | 0.09 | 0.08 |
| 7 | 104060973 | rs6976167 | 0.021 | -0.692 | 1.113 | 0.531 | -0.001 | 0.074 | 0.074 | 0 | 0.08 | 0.14 | 0.07 | 0.26 | 0.09 | 0.09 |
| 7 | 104067111 | rs10273435 | -0.048 | -0.903 | 1.106 | 0.221 | -0.045 | -0.042 | -0.042 | 0 | 0.21 | 0.45 | 0.36 | 0.14 | 0.17 | 0.25 |
| 7 | 104071382 | rs10255597 | 0.056 | -0.926 | 1.312 | 0.213 | -0.02 | 0.011 | 0.011 | 0.001 | 0.21 | 0.28 | 0.35 | 0.29 | 0.17 | 0.31 |
| 7 | 104076814 | rs6957611 | -1.174 | -1.038 | 1.381 | 0.049 | 0.028 | -0.001 | -0.001 | 0.177 | 0.32 | 0.73 | 0.46 | 0.37 | 0.26 | 0.38 |
| 7 | 104078323 | rs10262774 | 0.099 | -1.048 | 1.426 | 0.05 | -0.013 | -0.013 | -0.013 | 0 | 0.21 | 0.26 | 0.35 | 0.13 | 0.16 | 0.31 |
| 7 | 104078657 | rs2385168 | -1.214 | -1.038 | 1.152 | 0.051 | 0.053 | -0.008 | -0.008 | 0.371 | 0.32 | 0.79 | 0.46 | 0.37 | 0.26 | 0.38 |
| 7 | 104083719 | rs727927 | -0.441 | -0.929 | 1.54 | 0.093 | 0.021 | 0.135 | 0.135 | 0 | 0.11 | 0.47 | 0.11 | 0.23 | 0.1 | 0.07 |
| 7 | 104085146 | rs17137694 | 0.566 | -0.931 | 1.536 | -0.043 | -0.005 | -0.045 | -0.045 | 0 | 0.11 | 0.16 | 0.04 | 0 | 0.1 | 0.07 |
| 7 | 104088583 | rs10239251 | -0.084 | -0.83 | 1.371 | 0.216 | 0.002 | 0.005 | 0.005 | 0 | 0.2 | 0.17 | 0.13 | 0.12 | 0.16 | 0.1 |
| 7 | 104090417 | rs7806966 | -0.329 | -0.827 | 1.464 | 0.248 | -0.014 | 0.009 | 0.009 | 0 | 0.21 | 0.41 | 0.15 | 0.27 | 0.17 | 0.1 |
| 7 | 104091769 | rs11763095 | -0.047 | -0.805 | 1.119 | 0.234 | -0.003 | -0.011 | -0.011 | 0 | 0.2 | 0.23 | 0.13 | 0.12 | 0.16 | 0.1 |
| 7 | 104097453 | rs10268222 | -0.132 | -0.766 | 0.815 | 0.197 | -0.011 | 0.015 | 0.015 | 0 | 0.21 | 0.35 | 0.15 | 0.28 | 0.16 | 0.1 |
| 7 | 104100036 | rs1858780 | -0.13 | -0.768 | 0.814 | 0.175 | -0.01 | 0.013 | 0.013 | 0 | 0.21 | 0.36 | 0.15 | 0.28 | 0.16 | 0.1 |
| 7 | 104106026 | rs2385172 | 1.259 | -0.788 | 0.573 | 0.125 | -0.006 | -0.005 | -0.005 | 2.247 | 0.58 | 0.42 | 0.47 | 0.59 | 0.65 | 0.58 |
| 7 | 104107166 | rs6977743 | 0.148 | -0.831 | 0.037 | 0.048 | -0.004 | -0.007 | -0.007 | 0 | 0.8 | 0.77 | 0.62 | 0.87 | 0.81 | 0.69 |
| 7 | 104110492 | rs6949536 | 0.708 | -0.47 | -0.027 | 0.075 | -0.002 | 0.047 | 0.047 | 0.34 | 0.42 | 0.29 | 0.31 | 0.23 | 0.41 | 0.2 |

| | | | | | | | | | | | | | | | | |
|---|---|---|---|---|---|---|---|---|---|---|---|---|---|---|---|---|
| 7 | 104112216 | rs12705218 | 1.197 | -0.641 | -0.018 | -0.051 | -0.023 | 0.048 | 0.048 | 0 | 0.33 | 0.14 | 0.28 | 0.2 | 0.37 | 0.2 |
| 7 | 104113349 | rs6965697 | 0.301 | -0.957 | 0.469 | 0.011 | 0.015 | 0.051 | 0.051 | 1.164 | 0.49 | 0.62 | 0.36 | 0.28 | 0.44 | 0.22 |
| 7 | 104115294 | rs17137948 | 0.338 | -0.965 | 0.738 | 0.113 | 0.004 | 0.013 | 0.013 | 1.199 | 0.49 | 0.56 | 0.35 | 0.29 | 0.44 | 0.22 |
| 7 | 104117921 | rs12666599 | 0.312 | -1.008 | 0.172 | 0.465 | 0.001 | -0.001 | -0.001 | 0 | 0.17 | 0.15 | 0.18 | 0.2 | 0.14 | 0.27 |
| 7 | 104119381 | rs7776632 | 0.343 | -0.704 | -0.091 | 0.332 | -0.002 | -0.01 | -0.01 | 0 | 0.59 | 0.71 | 0.45 | 0.38 | 0.58 | 0.29 |
| 7 | 104123545 | rs11760238 | 1.24 | -0.698 | -0.081 | 0.335 | -0.027 | 0.116 | 0.116 | 0.068 | 0.45 | 0.18 | 0.33 | 0.2 | 0.47 | 0.3 |
| 7 | 104127901 | rs10234384 | -0.239 | -0.761 | 0.144 | 0.367 | -0.001 | -0.015 | -0.015 | 0 | 0.71 | 0.7 | 0.5 | 0.38 | 0.69 | 0.4 |
| 7 | 104128886 | rs2188172 | 1.287 | -0.782 | 0.11 | 0.22 | -0.04 | 0.132 | 0.132 | 0.465 | 0.44 | 0.15 | 0.32 | 0.21 | 0.47 | 0.31 |
| 7 | 104132489 | rs13234934 | 0.191 | -0.763 | 0.635 | 0.482 | 0.02 | 0.011 | 0.011 | 0 | 0.08 | 0.09 | 0.08 | 0.09 | 0.13 | 0.06 |
| 7 | 104133882 | rs10216100 | 0.41 | -0.881 | 0.76 | 0.419 | -0.052 | -0.074 | -0.074 | 2.738 | 0.16 | 0.39 | 0.07 | 0.03 | 0.06 | 0.02 |
| 7 | 104135160 | rs768974 | 1.489 | -0.841 | 0.446 | 0.726 | 0.02 | -0.002 | -0.002 | 0.223 | 0.55 | 0.28 | 0.45 | 0.4 | 0.55 | 0.52 |
| 7 | 104135332 | rs41015 | 0.011 | -0.364 | 0.408 | 1.05 | 0 | 0.003 | 0.003 | 0 | 0.79 | 0.77 | 0.6 | 0.54 | 0.75 | 0.63 |
| 7 | 104144632 | rs756669 | 0.923 | -0.201 | 0.464 | 0.381 | -0.025 | 0.083 | 0.083 | 0.644 | 0.4 | 0.2 | 0.28 | 0.21 | 0.36 | 0.17 |
| 7 | 104145122 | rs41035 | -0.845 | -0.027 | 0.172 | 0.619 | 0.03 | 0.073 | 0.073 | 0 | 0.1 | 0.21 | 0.1 | 0.22 | 0.14 | 0.1 |
| 7 | 104151867 | rs42209 | 0.074 | -0.045 | 0.125 | 0.601 | 0.005 | 0.009 | 0.009 | 1.456 | 0.52 | 0.6 | 0.39 | 0.43 | 0.5 | 0.26 |
| 7 | 104153691 | rs181974 | -0.922 | 0.099 | 0.402 | 0.616 | 0.049 | 0.181 | 0.181 | 0 | 0.06 | 0.21 | 0.06 | 0.21 | 0.06 | 0.08 |
| 7 | 104154069 | rs41055 | 0.557 | 0.206 | -0.374 | 0.418 | 0 | -0.001 | -0.001 | 0 | 0.4 | 0.45 | 0.29 | 0.36 | 0.36 | 0.21 |
| 7 | 104154850 | rs28253 | 0.999 | -0.025 | -0.501 | 0.625 | -0.032 | 0.044 | 0.044 | 0 | 0.4 | 0.25 | 0.38 | 0.25 | 0.4 | 0.27 |
| 7 | 104155283 | rs41056 | 0.136 | -0.196 | -0.618 | 0.67 | 0.023 | -0.015 | -0.015 | 0 | 0.7 | 0.9 | 0.66 | 0.7 | 0.7 | 0.58 |
| 7 | 104156150 | rs993262 | -1.073 | 0.515 | 0.43 | 0.656 | 0 | 0.034 | 0.034 | 0.106 | 0.09 | 0.29 | 0.06 | 0.11 | 0.08 | 0.03 |
| 7 | 104159886 | rs992692 | -1.394 | 0.641 | 0.286 | 0.518 | 0.009 | -0.001 | -0.001 | 0 | 0.09 | 0.2 | 0.1 | 0.13 | 0.12 | 0.09 |
| 7 | 104160202 | rs10243072 | -0.53 | 0.959 | 0.25 | 0.063 | -0.001 | -0.008 | -0.008 | 0 | 0.11 | 0.12 | 0.06 | 0.03 | 0.12 | 0.17 |
| 7 | 104164294 | rs1989781 | -0.076 | 1.043 | 0.337 | 0.167 | 0.018 | 0.026 | 0.026 | 0 | 0.62 | 0.53 | 0.61 | 0.49 | 0.61 | 0.48 |
| 7 | 104171423 | rs985409 | 0.736 | 1.172 | 0.389 | -0.053 | 0.053 | -0.019 | -0.019 | 0 | 0.47 | 0.27 | 0.45 | 0.47 | 0.44 | 0.29 |
| 7 | 104172232 | rs10255304 | -0.101 | 1.217 | 0.484 | -0.043 | 0.002 | -0.021 | -0.021 | 0 | 0.14 | 0.19 | 0.16 | 0.03 | 0.17 | 0.18 |
| 7 | 104181530 | rs6465994 | 0.627 | 1.033 | 0.886 | -0.174 | 0.058 | -0.025 | -0.025 | 0 | 0.44 | 0.25 | 0.38 | 0.47 | 0.4 | 0.28 |
| 7 | 104181707 | rs6465995 | -0.318 | 1.135 | 1.106 | -0.097 | 0.008 | 0.011 | 0.011 | 0 | 0.63 | 0.58 | 0.59 | 0.53 | 0.63 | 0.48 |
| 7 | 104192398 | rs10254402 | -0.836 | 0.406 | 1.13 | 0.052 | -0.015 | 0.056 | 0.056 | 0 | 0.13 | 0.45 | 0.05 | 0.15 | 0.06 | 0.05 |
| 7 | 104206297 | rs2193203 | 0.727 | 0.879 | 0.551 | 0.059 | -0.101 | -0.123 | -0.123 | 0 | 0.32 | 0.02 | 0.37 | 0.44 | 0.32 | 0.46 |
| 7 | 104213298 | rs7796437 | -0.725 | 1.112 | 1.01 | -0.131 | 0.022 | -0.021 | -0.021 | 0.839 | 0.19 | 0.24 | 0.22 | 0.11 | 0.27 | 0.23 |
| 7 | 104214487 | rs6969323 | -0.805 | 1.102 | 1.255 | -0.128 | 0.064 | -0.014 | -0.014 | 0 | 0.19 | 0.36 | 0.22 | 0.17 | 0.25 | 0.21 |
| 7 | 104215791 | rs4354244 | -0.701 | 1.142 | 0.929 | -0.122 | 0.026 | -0.006 | -0.006 | 0 | 0.19 | 0.23 | 0.22 | 0.17 | 0.27 | 0.22 |
| 7 | 104218575 | rs11761864 | -1.361 | 1.262 | 1.002 | -0.166 | 0.027 | -0.042 | -0.042 | 0 | 0.28 | 0.45 | 0.24 | 0.18 | 0.29 | 0.27 |
| 7 | 104220920 | rs6967399 | 0.241 | 0.79 | 0.963 | 0.049 | 0.001 | 0.038 | 0.038 | 0 | 0.92 | 0.8 | 0.96 | 0.87 | 0.95 | 0.94 |
| 7 | 104225738 | rs7805979 | -0.052 | -0.16 | 0.609 | 0.392 | -0.008 | 0.023 | 0.023 | 0 | 0.18 | 0.2 | 0.25 | 0.38 | 0.18 | 0.16 |
| 7 | 104227479 | rs9692289 | 1.063 | -0.23 | -0.22 | -0.325 | -0.079 | 0.119 | 0.119 | 0 | 0.22 | 0.02 | 0.18 | 0.08 | 0.24 | 0.11 |
| 7 | 104240200 | rs9641335 | 0.902 | -0.279 | -0.147 | -0.496 | 0.002 | 0.031 | 0.031 | 0 | 0.5 | 0.41 | 0.43 | 0.32 | 0.46 | 0.55 |
| 7 | 104245132 | rs17139096 | 1.322 | -0.12 | -0.22 | -0.487 | 0 | -0.001 | -0.001 | 0 | 0.11 | 0.09 | 0.16 | 0.13 | 0.15 | 0.05 |
| 7 | 104251618 | rs11764816 | 1.447 | -0.206 | -0.828 | -0.52 | -0.12 | -0.013 | -0.013 | 0.282 | 0.32 | 0.05 | 0.32 | 0.27 | 0.33 | 0.51 |
| 7 | 104257737 | rs4727605 | 0.328 | -0.471 | -0.651 | -0.613 | 0.003 | -0.005 | -0.005 | 0 | 0.65 | 0.6 | 0.67 | 0.71 | 0.63 | 0.72 |
| 7 | 104259720 | rs6948818 | 1.087 | -0.424 | -0.574 | -0.663 | -0.004 | 0.04 | 0.04 | 0 | 0.46 | 0.37 | 0.42 | 0.27 | 0.45 | 0.55 |
| 7 | 104263404 | rs1113933 | 0.946 | -0.382 | -0.841 | -0.706 | -0.076 | 0.137 | 0.137 | 0 | 0.25 | 0.06 | 0.2 | 0.04 | 0.29 | 0.21 |
| 7 | 104264665 | rs4730030 | 0.199 | -0.453 | -0.654 | -0.612 | -0.035 | 0.029 | 0.029 | 0 | 0.7 | 0.56 | 0.72 | 0.43 | 0.77 | 0.77 |
| 7 | 104265350 | rs4730031 | -1.414 | -0.332 | -0.747 | -0.694 | 0.006 | 0.029 | 0.029 | 0 | 0.66 | 0.75 | 0.74 | 0.8 | 0.74 | 0.53 |
| 7 | 104267189 | rs13222581 | 1.498 | -0.247 | -0.742 | -0.73 | -0.039 | 0.043 | 0.043 | 0 | 0.12 | 0.04 | 0.13 | 0.04 | 0.17 | 0.18 |
| 7 | 104268155 | rs1547977 | 0.794 | -0.263 | -0.573 | -0.695 | 0.028 | -0.111 | -0.111 | 0 | 0.23 | 0.4 | 0.2 | 0.08 | 0.24 | 0.13 |
| 7 | 104271093 | rs2891730 | -1.103 | -0.407 | -0.79 | -0.417 | 0.007 | -0.009 | -0.009 | 0 | 0.66 | 0.75 | 0.74 | 0.68 | 0.74 | 0.53 |
| 7 | 104272864 | rs10487220 | 2.008 | -0.392 | -0.973 | -0.482 | -0.048 | 0.051 | 0.051 | 0.037 | 0.28 | 0.01 | 0.23 | 0.14 | 0.24 | 0.43 |
| 7 | 104289858 | rs999378 | -0.14 | -0.246 | 0.052 | -0.617 | 0.014 | -0.055 | -0.055 | 0.386 | 0.35 | 0.4 | 0.33 | 0.14 | 0.4 | 0.29 |
| 7 | 104293093 | rs7779168 | 1.741 | -0.267 | 0.101 | -0.62 | -0.001 | -0.05 | -0.05 | 0 | 0.14 | 0.18 | 0.09 | 0 | 0.12 | 0.04 |
| 7 | 104295232 | rs10487222 | 0.985 | -0.362 | 0.096 | -0.282 | 0.03 | 0.006 | 0.006 | 0.107 | 0.43 | 0.26 | 0.48 | 0.61 | 0.4 | 0.61 |
| 7 | 104296144 | rs11766478 | 1.782 | -0.374 | 0.024 | -0.285 | 0 | 0.015 | 0.015 | 0 | 0.08 | 0.04 | 0.02 | 0.01 | 0.05 | 0.04 |
| 7 | 104297802 | rs11977439 | 1.334 | -0.387 | 0.152 | -0.214 | -0.004 | 0.036 | 0.036 | 0 | 0.09 | 0.15 | 0.03 | 0.14 | 0.05 | 0.06 |
| 7 | 104301597 | rs4265125 | -0.645 | -0.288 | 0.221 | -0.227 | 0.003 | -0.006 | -0.006 | 0.13 | 0.73 | 0.79 | 0.76 | 0.75 | 0.73 | 0.89 |
| 7 | 104304300 | rs12537797 | 0.646 | -0.305 | 0.269 | -0.411 | 0.013 | 0.044 | 0.044 | 0 | 0.52 | 0.37 | 0.51 | 0.76 | 0.44 | 0.7 |
| 7 | 104313331 | rs10280745 | 0.49 | -0.201 | 0.288 | -0.368 | -0.016 | 0.012 | 0.012 | 0 | 0.31 | 0.34 | 0.34 | 0.53 | 0.25 | 0.2 |
| 7 | 104314345 | rs17490551 | 1.956 | -0.297 | 0.528 | -0.342 | 0 | -0.002 | -0.002 | 0 | 0.07 | 0.05 | 0.03 | 0.01 | 0.05 | 0.01 |
| 7 | 104316059 | rs9648932 | 0.125 | -0.092 | 0.573 | -0.401 | 0.002 | 0.02 | 0.02 | 0 | 0.42 | 0.36 | 0.47 | 0.58 | 0.42 | 0.33 |
| 7 | 104316145 | rs11761950 | -0.069 | -0.02 | 0.093 | -0.397 | -0.009 | -0.004 | -0.004 | 0 | 0.16 | 0.09 | 0.13 | 0.2 | 0.16 | 0.11 |
| 7 | 104318103 | rs11767619 | 0.246 | 0.042 | 0.098 | -0.402 | 0.003 | 0.048 | 0.048 | 0 | 0.49 | 0.34 | 0.49 | 0.44 | 0.53 | 0.65 |
| 7 | 104319876 | rs10279375 | 0.19 | 0.07 | -0.169 | -0.389 | 0.002 | 0.013 | 0.013 | 0 | 0.5 | 0.41 | 0.49 | 0.44 | 0.53 | 0.66 |
| 7 | 104321056 | rs9886292 | 0.307 | 0.092 | -0.119 | -0.073 | -0.007 | -0.008 | -0.008 | 2.539 | 0.42 | 0.52 | 0.39 | 0.45 | 0.35 | 0.6 |
| 7 | 104326251 | rs12531723 | 0.048 | 0.135 | -0.177 | -0.09 | 0 | 0.018 | 0.018 | 0 | 0.5 | 0.42 | 0.59 | 0.57 | 0.52 | 0.35 |
| 7 | 104335994 | rs10249869 | 1.182 | 0.086 | -0.144 | -0.131 | -0.006 | 0.042 | 0.042 | 0 | 0.31 | 0.18 | 0.3 | 0.31 | 0.32 | 0.55 |
| 7 | 104337956 | rs7812002 | 0.05 | 0.126 | 0.728 | -0.278 | 0.043 | 0.054 | 0.054 | 0.044 | 0.89 | 0.66 | 0.9 | 0.89 | 0.88 | 0.91 |
| 7 | 104338193 | rs12705266 | -0.122 | 0.14 | 0.651 | -0.272 | 0.028 | 0.036 | 0.036 | 0.057 | 0.9 | 0.75 | 0.91 | 0.89 | 0.88 | 0.91 |
| 7 | 104347312 | rs6966224 | 0.12 | -0.174 | 0.413 | -0.272 | -0.006 | 0.121 | 0.121 | 0 | 0.48 | 0.33 | 0.4 | 0.21 | 0.49 | 0.61 |
| 7 | 104347834 | rs12705267 | 0.026 | -0.286 | 0.511 | -0.283 | -0.01 | 0.088 | 0.088 | 0 | 0.48 | 0.22 | 0.56 | 0.67 | 0.5 | 0.36 |
| 7 | 104351961 | rs3934669 | 0.499 | -0.119 | 0.171 | -0.417 | 0.028 | 0 | 0 | 0 | 0.36 | 0.16 | 0.43 | 0.62 | 0.34 | 0.22 |

| | | | | | | | | | | | | | | | | |
|---|---|---|---|---|---|---|---|---|---|---|---|---|---|---|---|---|
| 7 | 104356293 | rs4612276 | -0.508 | -0.257 | -0.024 | -0.163 | -0.004 | -0.03 | -0.03 | 0.557 | 0.35 | 0.5 | 0.44 | 0.7 | 0.35 | 0.34 |
| 7 | 104359226 | rs4338031 | 0.406 | -0.123 | -0.166 | -0.1 | 0.032 | 0.02 | 0.02 | 0 | 0.14 | 0.25 | 0.17 | 0.17 | 0.18 | 0.18 |
| 7 | 104362704 | rs4584085 | 1.163 | -0.317 | -0.335 | -0.028 | 0.029 | -0.06 | -0.06 | 0 | 0.18 | 0.08 | 0.25 | 0.54 | 0.16 | 0.17 |
| 7 | 104368522 | rs12672491 | 1.54 | -0.404 | -0.406 | -0.519 | 0 | 0.032 | 0.032 | 0 | 0.26 | 0.16 | 0.22 | 0.07 | 0.25 | 0.47 |
| 7 | 104376929 | rs4730039 | -0.199 | -0.436 | -0.314 | -0.669 | 0.003 | -0.065 | -0.065 | 0 | 0.59 | 0.55 | 0.66 | 0.89 | 0.56 | 0.8 |
| 7 | 104379606 | rs17568789 | 1.619 | -0.38 | -0.477 | -0.649 | 0 | -0.006 | -0.006 | 0 | 0.27 | 0.26 | 0.23 | 0.07 | 0.25 | 0.47 |
| 7 | 104380200 | rs6978452 | 0.251 | -0.232 | -1.177 | -0.447 | -0.01 | -0.023 | -0.023 | 3.231 | 0.3 | 0.18 | 0.39 | 0.64 | 0.31 | 0.28 |
| 7 | 104381939 | rs12705273 | 0.186 | -0.202 | -1.176 | -0.429 | -0.007 | 0.027 | 0.027 | 0 | 0.31 | 0.18 | 0.39 | 0.75 | 0.31 | 0.31 |
| 7 | 104399617 | rs17141284 | 1.548 | -0.303 | -1.096 | -0.12 | 0.016 | -0.025 | -0.025 | 0 | 0.16 | 0.08 | 0.23 | 0.59 | 0.13 | 0.16 |
| 7 | 104402709 | rs4132013 | 0.935 | 0.232 | -0.945 | -0.601 | 0.002 | 0.05 | 0.05 | 0 | 0.27 | 0.17 | 0.26 | 0.14 | 0.25 | 0.48 |
| 7 | 104411273 | rs13221774 | 0.439 | 0.702 | -0.964 | -0.311 | -0.066 | 0.017 | 0.017 | 0 | 0.11 | 0.02 | 0.16 | 0.05 | 0.19 | 0.14 |
| 7 | 104411976 | rs4236575 | -0.802 | 0.532 | -0.952 | 0.252 | 0 | -0.005 | -0.005 | 0 | 0.8 | 0.81 | 0.85 | 0.89 | 0.81 | 0.88 |
| 7 | 104413055 | rs12705275 | -0.88 | 0.523 | -0.893 | 0.247 | -0.001 | 0.013 | 0.013 | 0 | 0.81 | 0.89 | 0.85 | 0.89 | 0.81 | 0.88 |
| 7 | 104413768 | rs12540437 | 1.794 | 0.104 | -1.01 | 0.13 | 0.002 | -0.004 | -0.004 | 0 | 0.14 | 0.12 | 0.14 | 0.43 | 0.14 | 0.13 |
| 7 | 104414719 | rs13236549 | 0.392 | 0.162 | -0.868 | -0.22 | -0.001 | 0.002 | 0.002 | 0.228 | 0.26 | 0.27 | 0.19 | 0.45 | 0.23 | 0.2 |
| 7 | 104415415 | rs4530975 | 0.358 | 0.002 | -1.184 | -0.629 | -0.041 | 0.078 | 0.078 | 0.017 | 0.24 | 0.1 | 0.24 | 0.11 | 0.26 | 0.21 |
| 7 | 104416070 | rs12705276 | 2.349 | -0.175 | -0.918 | -0.462 | 0.003 | -0.005 | -0.005 | 0.693 | 0.15 | 0.12 | 0.14 | 0.44 | 0.14 | 0.13 |
| 7 | 104416645 | rs13310494 | -0.579 | -0.165 | -0.955 | -0.895 | -0.001 | 0.097 | 0.097 | 0 | 0.51 | 0.37 | 0.49 | 0.83 | 0.49 | 0.46 |
| 7 | 104417567 | rs12672558 | 1.834 | -0.252 | -0.417 | -0.979 | 0.002 | -0.051 | -0.051 | 0.182 | 0.27 | 0.32 | 0.34 | 0.06 | 0.31 | 0.43 |
| 7 | 104418428 | rs11771722 | 1.365 | -0.269 | -0.34 | -0.987 | 0.003 | -0.098 | -0.098 | 1.509 | 0.28 | 0.41 | 0.35 | 0.06 | 0.32 | 0.43 |
| 7 | 104420060 | rs4460308 | -0.307 | -0.365 | -0.504 | -0.83 | -0.011 | 0.061 | 0.061 | 0 | 0.37 | 0.31 | 0.3 | 0.11 | 0.35 | 0.29 |
| 7 | 104425811 | rs4283980 | -0.147 | -0.291 | -0.627 | -0.801 | 0.001 | -0.02 | -0.02 | 1.795 | 0.42 | 0.49 | 0.43 | 0.74 | 0.4 | 0.35 |
| 7 | 104430056 | rs4730049 | -0.298 | -0.087 | -0.368 | -0.576 | -0.003 | 0.028 | 0.028 | 0 | 0.52 | 0.47 | 0.5 | 0.83 | 0.5 | 0.45 |
| 7 | 104435213 | rs2470961 | 0.052 | 0.052 | 0.034 | -0.348 | 0.002 | 0 | 0 | 0.01 | 0.55 | 0.42 | 0.5 | 0.49 | 0.54 | 0.45 |
| 7 | 104435897 | rs13243476 | 1.581 | 0.117 | -0.365 | -0.291 | 0 | 0.009 | 0.009 | 0 | 0.15 | 0.15 | 0.15 | 0.45 | 0.14 | 0.14 |
| 7 | 104440625 | rs2470957 | -0.691 | 0.085 | -0.014 | -0.355 | -0.002 | -0.087 | -0.087 | 0.263 | 0.68 | 0.47 | 0.65 | 0.9 | 0.67 | 0.54 |
| 7 | 104441909 | rs10085776 | -0.112 | 0.235 | -0.671 | -0.293 | 0.012 | -0.006 | -0.006 | 0 | 0.54 | 0.25 | 0.54 | 0.45 | 0.54 | 0.42 |
| 7 | 104453866 | rs12667640 | 0.641 | 0.413 | -0.656 | -0.428 | -0.023 | 0.079 | 0.079 | 0.038 | 0.34 | 0.16 | 0.39 | 0.12 | 0.4 | 0.43 |
| 7 | 104453887 | rs10279083 | 1.17 | 0.518 | -0.824 | -0.203 | 0.001 | -0.039 | -0.039 | 0 | 0.15 | 0.13 | 0.21 | 0.33 | 0.12 | 0.16 |
| 7 | 104456874 | rs6975741 | 0.197 | 0.689 | -0.32 | -0.513 | 0 | -0.034 | -0.034 | 0 | 0.64 | 0.36 | 0.58 | 0.17 | 0.66 | 0.66 |
| 7 | 104460747 | rs10256308 | 1.538 | 0.457 | -0.188 | -0.238 | -0.001 | 0.032 | 0.032 | 0 | 0.15 | 0.18 | 0.16 | 0.44 | 0.15 | 0.14 |
| 7 | 104461230 | rs2465066 | 0.364 | 0.337 | -0.253 | -0.618 | 0.003 | 0.003 | 0.003 | 0 | 0.3 | 0.23 | 0.19 | 0.11 | 0.25 | 0.22 |
| 7 | 104466964 | rs4073894 | 1.085 | 0.546 | -0.678 | -0.441 | -0.01 | 0.049 | 0.049 | 0.716 | 0.17 | 0.11 | 0.11 | 0.05 | 0.19 | 0.1 |
| 7 | 104472647 | rs4382394 | 1.085 | 0.416 | -0.78 | -0.881 | 0.019 | 0.026 | 0.026 | 0.6 | 0.3 | 0.2 | 0.33 | 0.12 | 0.22 | 0.39 |
| 7 | 104482321 | rs4730052 | 1.202 | 0.396 | -0.809 | -0.681 | 0.021 | 0.022 | 0.022 | 0.353 | 0.3 | 0.19 | 0.33 | 0.17 | 0.22 | 0.42 |
| 7 | 104486707 | rs4073643 | 1.148 | 0.514 | -1.194 | -0.062 | -0.009 | -0.06 | -0.06 | 1.422 | 0.19 | 0.1 | 0.16 | 0.29 | 0.21 | 0.11 |
| 7 | 104497678 | rs12705284 | -0.363 | 0.705 | -0.431 | 0.048 | 0.018 | 0.005 | 0.005 | 0 | 0.47 | 0.33 | 0.48 | 0.52 | 0.56 | 0.46 |
| 7 | 104500549 | rs17706792 | 0.223 | 1.157 | -1.092 | -0.549 | -0.027 | 0.072 | 0.072 | 0 | 0.32 | 0.17 | 0.19 | 0.03 | 0.34 | 0.27 |
| 7 | 104524092 | rs11561931 | 0.729 | 1.028 | -0.909 | -0.264 | -0.031 | 0.123 | 0.123 | 0.007 | 0.4 | 0.17 | 0.35 | 0.11 | 0.44 | 0.44 |
| 7 | 104524466 | rs17173 | -0.382 | 1.014 | -0.969 | -0.437 | 0.007 | 0.198 | 0.198 | 0 | 0.58 | 0.84 | 0.67 | 0.96 | 0.62 | 0.61 |
| 7 | 104536482 | rs13232695 | -0.083 | 1.006 | -0.572 | 0.151 | 0.008 | -0.002 | -0.002 | 0 | 0.51 | 0.68 | 0.51 | 0.59 | 0.56 | 0.55 |
| 7 | 104536783 | rs2041830 | 0.342 | 1.049 | -0.665 | -0.083 | -0.024 | 0.153 | 0.153 | 0 | 0.45 | 0.21 | 0.4 | 0.12 | 0.54 | 0.44 |
| 7 | 104552636 | rs10953457 | -0.996 | 1.309 | 0.02 | 0.623 | 0.082 | 0.334 | 0.334 | 0.701 | 0.11 | 0.48 | 0.2 | 0.47 | 0.16 | 0.14 |
| 7 | 104568299 | rs7795893 | 0.487 | 0.958 | -0.499 | 0.256 | 0.005 | -0.003 | -0.003 | 0.466 | 0.45 | 0.29 | 0.31 | 0.52 | 0.38 | 0.5 |
| 7 | 104590780 | rs2470949 | 0.156 | 1.15 | -0.454 | 0.153 | -0.001 | -0.001 | -0.001 | 1.46 | 0.31 | 0.28 | 0.37 | 0.26 | 0.41 | 0.2 |
| 7 | 104592969 | rs2017598 | -1.845 | 1.197 | -0.529 | 0.033 | -0.035 | 0.045 | 0.045 | 0.02 | 0.88 | 0.96 | 0.82 | 1 | 0.85 | 0.88 |
| 7 | 104599208 | rs17777642 | -0.558 | 0.82 | 0.077 | -0.062 | -0.004 | 0.014 | 0.014 | 0 | 0.38 | 0.47 | 0.44 | 0.43 | 0.4 | 0.31 |
| 7 | 104605530 | rs7776707 | 0.056 | 0.841 | -0.04 | -0.167 | -0.001 | -0.002 | -0.002 | 0 | 0.29 | 0.24 | 0.34 | 0.25 | 0.34 | 0.19 |
| 7 | 104614664 | rs17639782 | 2.008 | 0.901 | -0.602 | -0.747 | -0.043 | 0.068 | 0.068 | 0.497 | 0.12 | 0.02 | 0.05 | 0 | 0.13 | 0.08 |
| 7 | 104626690 | rs888086 | -1.373 | 0.381 | 0.413 | -0.411 | -0.007 | 0.269 | 0.269 | 0 | 0.49 | 0.84 | 0.57 | 0.93 | 0.5 | 0.42 |
| 7 | 104632980 | rs6976394 | -1.283 | 0.368 | 0.426 | -0.417 | 0 | 0.275 | 0.275 | 0 | 0.5 | 0.84 | 0.57 | 0.93 | 0.5 | 0.42 |
| 7 | 104635597 | rs10274607 | -0.944 | 0.37 | 0.645 | -0.323 | -0.025 | 0.077 | 0.077 | 0.044 | 0.2 | 0.6 | 0.25 | 0.69 | 0.14 | 0.21 |
| 7 | 104636940 | rs6980244 | -1.508 | 0.374 | 0.62 | -0.334 | -0.004 | 0.274 | 0.274 | 0 | 0.49 | 0.84 | 0.57 | 0.93 | 0.5 | 0.42 |
| 7 | 104652671 | rs6950894 | 1.438 | 0.375 | 0.62 | -0.333 | -0.001 | 0.281 | 0.281 | 0 | 0.5 | 0.16 | 0.43 | 0.07 | 0.5 | 0.58 |
| 7 | 104653265 | rs2192932 | -0.158 | 0.407 | 0.661 | -0.309 | -0.004 | -0.001 | -0.001 | 0.401 | 0.71 | 0.76 | 0.67 | 0.76 | 0.64 | 0.8 |
| 7 | 104660672 | rs10224526 | -1.005 | -0.155 | 0.843 | -0.117 | -0.021 | 0.147 | 0.147 | 0.009 | 0.19 | 0.57 | 0.24 | 0.56 | 0.13 | 0.19 |
| 7 | 104717188 | rs2240454 | -0.67 | -0.175 | 1.269 | -0.128 | -0.01 | 0.142 | 0.142 | 0 | 0.17 | 0.39 | 0.23 | 0.56 | 0.13 | 0.19 |
| 7 | 104717517 | rs2240455 | -1.139 | -0.181 | 1.275 | -0.126 | -0.028 | 0.1 | 0.1 | 0.045 | 0.2 | 0.59 | 0.24 | 0.63 | 0.13 | 0.2 |
| 7 | 104750240 | rs17721388 | 0.774 | -0.242 | 1.374 | -0.045 | -0.007 | 0.049 | 0.049 | 0 | 0.08 | 0.18 | 0.13 | 0.18 | 0.06 | 0.12 |
| 7 | 104755514 | rs2385537 | 0.741 | -0.2 | 1.509 | -1.197 | 0.016 | 0.025 | 0.025 | 0 | 0.2 | 0.13 | 0.14 | 0.03 | 0.16 | 0.11 |
| 7 | 104755816 | rs1013016 | -0.279 | -0.142 | 1.594 | -1.134 | -0.006 | 0.067 | 0.067 | 0 | 0.84 | 0.75 | 0.78 | 0.43 | 0.87 | 0.79 |
| 7 | 104756344 | rs1143 | 0.315 | -0.403 | 1.438 | -1.361 | 0.001 | 0.004 | 0.004 | 0 | 0.71 | 0.67 | 0.69 | 0.44 | 0.68 | 0.72 |
| 7 | 104767295 | rs3815256 | 0.087 | -1.258 | 2.262 | -0.535 | -0.002 | 0.039 | 0.039 | 0 | 0.85 | 0.8 | 0.78 | 0.45 | 0.87 | 0.83 |
| 7 | 104772836 | rs1019040 | 0.517 | -1.255 | 2.306 | -0.533 | -0.01 | 0.148 | 0.148 | 0 | 0.83 | 0.6 | 0.77 | 0.44 | 0.87 | 0.8 |
| 7 | 104775968 | rs4730070 | -0.249 | -1.174 | 2.24 | -0.46 | -0.023 | -0.025 | -0.025 | 0 | 0.09 | 0.03 | 0.17 | 0.2 | 0.18 | 0.1 |
| 7 | 104781644 | rs12705300 | -0.513 | -1.172 | 2.201 | -0.555 | 0.031 | 0 | 0 | 0.306 | 0.07 | 0.18 | 0.08 | 0.07 | 0.09 | 0.03 |
| 7 | 104786147 | rs6466045 | 0.262 | -1.195 | 2.045 | -0.5 | -0.006 | 0.067 | 0.067 | 0 | 0.84 | 0.75 | 0.78 | 0.44 | 0.87 | 0.77 |
| 7 | 104793841 | rs2237606 | -0.008 | -1.193 | 1.844 | -0.498 | -0.001 | 0.023 | 0.023 | 0 | 0.85 | 0.84 | 0.79 | 0.44 | 0.87 | 0.78 |
| 7 | 104812129 | rs10266871 | -0.379 | -1.204 | 1.778 | -0.533 | -0.001 | -0.004 | -0.004 | 0.203 | 0.29 | 0.24 | 0.32 | 0.3 | 0.35 | 0.22 |

| | | | | | | | | | | | | | | | | |
|---|---|---|---|---|---|---|---|---|---|---|---|---|---|---|---|---|
| 7 | 104813097 | rs17640472 | 1.624 | -1.217 | 1.706 | -0.554 | 0.015 | 0.296 | 0.296 | 0 | 0.5 | 0.15 | 0.43 | 0.07 | 0.52 | 0.54 |
| 7 | 104828098 | rs3779210 | -0.342 | -1.274 | 1.723 | -0.512 | -0.01 | -0.018 | -0.018 | 0.301 | 0.29 | 0.14 | 0.31 | 0.3 | 0.34 | 0.22 |
| 7 | 104838050 | rs7796945 | 1.555 | -1.307 | 1.731 | -0.541 | 0.021 | 0.281 | 0.281 | 0 | 0.5 | 0.16 | 0.43 | 0.07 | 0.52 | 0.54 |
| 7 | 104845094 | rs41562 | -0.394 | -1.251 | 1.695 | -0.505 | -0.001 | -0.003 | -0.003 | 0 | 0.29 | 0.23 | 0.32 | 0.3 | 0.34 | 0.22 |
| 7 | 104870573 | rs2237615 | -0.459 | -1.231 | 1.62 | -0.488 | -0.005 | 0.054 | 0.054 | 0 | 0.16 | 0.21 | 0.22 | 0.57 | 0.13 | 0.23 |
| 7 | 104896470 | rs12538590 | 1.579 | -1.247 | 1.493 | -0.517 | 0.021 | 0.179 | 0.179 | 0 | 0.51 | 0.23 | 0.44 | 0.14 | 0.52 | 0.56 |
| 7 | 104948076 | rs10241415 | 0.323 | -1.398 | 1.048 | -0.427 | -0.006 | -0.01 | -0.01 | 0 | 0.71 | 0.81 | 0.69 | 0.7 | 0.66 | 0.78 |
| 7 | 104972800 | rs10228503 | -0.248 | -1.396 | 1.052 | -0.395 | -0.001 | 0.026 | 0.026 | 0 | 0.14 | 0.16 | 0.21 | 0.57 | 0.13 | 0.22 |
| 7 | 104995627 | rs12334245 | -0.385 | -1.404 | 0.987 | -0.472 | -0.005 | -0.021 | -0.021 | 0 | 0.29 | 0.14 | 0.3 | 0.3 | 0.34 | 0.22 |
| 7 | 105013788 | rs17270450 | 1.462 | -1.358 | 0.719 | -0.488 | 0.029 | 0.179 | 0.179 | 0 | 0.51 | 0.23 | 0.44 | 0.07 | 0.52 | 0.55 |
| 7 | 105035037 | rs4727622 | -0.604 | -1.337 | 0.439 | -0.197 | -0.006 | 0.077 | 0.077 | 0 | 0.16 | 0.23 | 0.22 | 0.58 | 0.14 | 0.24 |
| 7 | 105036690 | rs10267633 | -0.438 | -1.312 | 0.361 | -0.264 | -0.013 | 0.074 | 0.074 | 0 | 0.16 | 0.25 | 0.22 | 0.57 | 0.13 | 0.23 |
| 7 | 105059606 | rs4266584 | 0.963 | -1.041 | 0.834 | -0.217 | -0.004 | 0 | 0 | 0.411 | 0.66 | 0.37 | 0.67 | 0.67 | 0.66 | 0.8 |
| 7 | 105067229 | rs6945887 | 1.311 | -1.117 | 0.649 | -0.296 | 0.023 | 0.039 | 0.039 | 0 | 0.59 | 0.36 | 0.65 | 0.76 | 0.65 | 0.74 |
| 7 | 105067767 | rs6980024 | 1.36 | -1.11 | 0.595 | -0.281 | 0.028 | 0.038 | 0.038 | 0 | 0.58 | 0.33 | 0.65 | 0.69 | 0.65 | 0.72 |
| 7 | 105073788 | rs955056 | 1.373 | -1.077 | 0.606 | -0.27 | 0.027 | 0.037 | 0.037 | 0 | 0.58 | 0.33 | 0.65 | 0.69 | 0.65 | 0.72 |
| 7 | 105076977 | rs6970417 | -1.574 | -0.369 | 1.31 | 0.148 | -0.002 | -0.008 | -0.008 | 0 | 0.29 | 0.61 | 0.26 | 0.21 | 0.29 | 0.2 |
| 7 | 105088693 | rs6977560 | 0.877 | -0.462 | 1.174 | 0.222 | -0.017 | -0.013 | -0.013 | 0 | 0.75 | 0.43 | 0.8 | 0.78 | 0.81 | 0.7 |
| 7 | 105096661 | rs13437626 | -1.026 | -0.347 | 1.094 | 0.181 | 0.016 | 0.018 | 0.018 | 0.051 | 0.06 | 0.2 | 0.05 | 0.08 | 0.05 | 0.02 |
| 7 | 105110990 | rs2258851 | -1.201 | -0.717 | 0.954 | 0.17 | -0.051 | -0.022 | -0.022 | 0 | 0.21 | 0.47 | 0.15 | 0.2 | 0.13 | 0.21 |
| 7 | 105119308 | rs6466069 | -0.522 | -0.412 | 0.485 | -0.083 | -0.001 | -0.011 | -0.011 | 0 | 0.2 | 0.27 | 0.2 | 0.12 | 0.18 | 0.21 |
| 7 | 105123148 | rs818469 | -1.058 | -0.608 | 0.912 | -0.185 | -0.004 | -0.027 | -0.027 | 0.005 | 0.25 | 0.35 | 0.24 | 0.2 | 0.24 | 0.28 |
| 7 | 105127696 | rs6971072 | -0.384 | -0.586 | 0.73 | -0.245 | -0.001 | -0.006 | -0.006 | 0.438 | 0.17 | 0.22 | 0.2 | 0.12 | 0.17 | 0.2 |
| 7 | 105138947 | rs3735369 | -1.322 | -0.787 | 0.534 | -0.294 | 0.01 | 0.021 | 0.021 | 0.318 | 0.26 | 0.23 | 0.25 | 0.18 | 0.23 | 0.31 |
| 7 | 105141008 | rs3735368 | -0.123 | -0.788 | 0.529 | -0.296 | -0.006 | 0.015 | 0.015 | 0 | 0.16 | 0.12 | 0.19 | 0.12 | 0.17 | 0.2 |
| 7 | 105141152 | rs6957263 | -1.113 | -0.733 | 0.625 | -0.261 | 0.008 | 0.016 | 0.016 | 0.128 | 0.26 | 0.25 | 0.25 | 0.18 | 0.23 | 0.31 |
| 7 | 105144968 | rs3852332 | 0.193 | -0.576 | -0.495 | -0.802 | -0.046 | 0.08 | 0.08 | 0 | 0.19 | 0.01 | 0.11 | 0.04 | 0.2 | 0.23 |
| 7 | 105155930 | rs11764594 | -0.132 | -0.76 | -1.236 | -0.335 | -0.035 | -0.03 | -0.03 | 0.867 | 0.15 | 0.01 | 0.12 | 0.17 | 0.13 | 0.08 |
| 7 | 105161755 | rs711427 | -0.421 | -0.44 | -0.556 | -0.783 | -0.007 | -0.042 | -0.042 | 0.359 | 0.32 | 0.58 | 0.3 | 0.21 | 0.28 | 0.34 |
| 7 | 105167969 | rs711430 | -0.372 | -0.522 | -0.494 | -0.948 | -0.003 | -0.113 | -0.113 | 0 | 0.24 | 0.48 | 0.2 | 0.06 | 0.2 | 0.28 |
| 7 | 105170476 | rs1728648 | 0.057 | -0.515 | -0.462 | -1.056 | -0.005 | 0.007 | 0.007 | 0 | 0.87 | 0.8 | 0.81 | 0.79 | 0.87 | 0.85 |
| 7 | 105173029 | rs818621 | -0.143 | -0.512 | -0.408 | -1.052 | -0.003 | -0.113 | -0.113 | 0.079 | 0.24 | 0.47 | 0.21 | 0.06 | 0.2 | 0.28 |
| 7 | 105173974 | rs10273718 | 0.179 | -1.002 | -0.584 | -0.425 | 0.001 | -0.012 | -0.012 | 0 | 0.23 | 0.14 | 0.17 | 0.37 | 0.21 | 0.12 |
| 7 | 105174384 | rs6466070 | 0.285 | -0.792 | -1.011 | -0.775 | 0.003 | 0.071 | 0.071 | 0 | 0.43 | 0.23 | 0.42 | 0.72 | 0.42 | 0.51 |
| 7 | 105179176 | rs1355614 | 0.272 | -0.764 | -0.503 | -0.764 | -0.003 | -0.111 | -0.111 | 0.006 | 0.24 | 0.46 | 0.21 | 0.06 | 0.2 | 0.28 |
| 7 | 105193051 | rs11766216 | 2.156 | -0.685 | -0.721 | -0.681 | -0.053 | 0.106 | 0.106 | 0.352 | 0.21 | 0.03 | 0.18 | 0.02 | 0.25 | 0.1 |
| 7 | 105197175 | rs12538052 | 0.799 | -0.466 | -0.077 | 0.031 | -0.001 | 0.002 | 0.002 | 0 | 0.07 | 0.1 | 0.1 | 0.14 | 0.08 | 0.06 |
| 7 | 105205917 | rs864687 | -0.039 | -0.555 | 0.247 | 0.336 | 0.003 | 0.017 | 0.017 | 0.003 | 0.07 | 0.25 | 0.1 | 0.08 | 0.06 | 0.08 |
| 7 | 105215970 | rs4585682 | 0.645 | -0.343 | -0.306 | -0.022 | -0.001 | 0.001 | 0.001 | 0 | 0.07 | 0.1 | 0.1 | 0.15 | 0.08 | 0.06 |
| 7 | 105221097 | rs1706923 | -0.838 | -0.387 | -0.286 | -0.253 | -0.005 | -0.077 | -0.077 | 0 | 0.3 | 0.56 | 0.22 | 0.13 | 0.27 | 0.3 |
| 7 | 105226180 | rs1721492 | -1.232 | -0.357 | -0.218 | 0.032 | 0.002 | -0.012 | -0.012 | 0.888 | 0.29 | 0.5 | 0.27 | 0.26 | 0.28 | 0.31 |
| 7 | 105228825 | rs10260974 | 1.065 | -0.047 | -0.407 | 0.396 | 0.007 | -0.007 | -0.007 | 0.581 | 0.7 | 0.37 | 0.68 | 0.72 | 0.66 | 0.62 |
| 7 | 105235033 | rs7801900 | 0.391 | -0.414 | -0.344 | 1.37 | -0.002 | 0.04 | 0.04 | 0 | 0.05 | 0.08 | 0.1 | 0.2 | 0.06 | 0.07 |
| 7 | 105241647 | rs7809769 | 1.243 | -0.54 | 0.031 | 1.068 | -0.024 | 0.254 | 0.254 | 0 | 0.83 | 0.43 | 0.78 | 0.52 | 0.84 | 0.78 |
| 7 | 105246821 | rs12154239 | 0.973 | -0.488 | -1.046 | 0.747 | 0.001 | 0.087 | 0.087 | 2.586 | 0.48 | 0.29 | 0.45 | 0.24 | 0.48 | 0.49 |
| 7 | 105247470 | rs7777200 | 1.133 | 0.034 | -1.499 | 0.446 | -0.023 | -0.073 | -0.073 | 0.104 | 0.3 | 0.18 | 0.31 | 0.41 | 0.34 | 0.16 |
| 7 | 105249120 | rs6949597 | -0.521 | 0.227 | -0.905 | 0.638 | 0.014 | -0.008 | -0.008 | 0 | 0.67 | 0.62 | 0.69 | 0.69 | 0.66 | 0.73 |
| 7 | 105252718 | rs10435352 | 0.683 | 0.207 | -1.601 | 0.684 | -0.003 | 0.066 | 0.066 | 4.145 | 0.41 | 0.23 | 0.34 | 0.27 | 0.43 | 0.46 |
| 7 | 105252812 | rs16871759 | 0.189 | 0.165 | -1.52 | 1.44 | -0.004 | 0.059 | 0.059 | 0 | 0.05 | 0.05 | 0.14 | 0.36 | 0.07 | 0.06 |
| 7 | 105256374 | rs1706896 | 0.398 | 0.26 | -1.641 | 0.847 | 0.017 | 0.032 | 0.032 | 1.905 | 0.51 | 0.19 | 0.41 | 0.56 | 0.46 | 0.36 |
| 7 | 105273458 | rs1615197 | -0.679 | 0.356 | -0.841 | 1.065 | 0.017 | 0.096 | 0.096 | 1.901 | 0.55 | 0.72 | 0.73 | 0.79 | 0.62 | 0.71 |
| 7 | 105273720 | rs10242311 | 0.697 | 0.597 | -1.333 | 1.446 | 0.001 | -0.016 | -0.016 | 1.596 | 0.09 | 0.03 | 0.08 | 0.23 | 0.06 | 0.13 |
| 7 | 105276155 | rs4730097 | -0.416 | 0.527 | -0.883 | 1.236 | 0.005 | 0.018 | 0.018 | 0.616 | 0.47 | 0.41 | 0.65 | 0.58 | 0.57 | 0.63 |
| 7 | 105278201 | rs10266988 | 0.411 | 0.992 | -1.082 | 1.519 | -0.007 | -0.014 | -0.014 | 0 | 0.27 | 0.27 | 0.47 | 0.4 | 0.37 | 0.26 |
| 7 | 105291444 | rs562221 | 0.359 | 0.757 | -1.821 | 1.021 | -0.056 | -0.024 | -0.024 | 0 | 0.43 | 0.12 | 0.46 | 0.61 | 0.42 | 0.48 |
| 7 | 105297520 | rs471303 | -0.824 | 0.286 | -0.85 | 1.15 | -0.011 | 0.001 | 0.001 | 0 | 0.32 | 0.46 | 0.18 | 0.41 | 0.23 | 0.37 |
| 7 | 105299515 | rs590966 | 0.26 | 0.412 | -1.329 | 1.325 | -0.006 | -0.043 | -0.043 | 0 | 0.88 | 0.96 | 0.89 | 0.82 | 0.87 | 0.88 |
| 7 | 105306556 | rs7799245 | 1.047 | 0.223 | -0.957 | 0.888 | -0.002 | 0 | 0 | 0 | 0.56 | 0.54 | 0.61 | 0.46 | 0.62 | 0.46 |
| 7 | 105307121 | rs572003 | 0.842 | 0.458 | -0.625 | 0.635 | -0.013 | -0.059 | -0.059 | 0.175 | 0.81 | 0.72 | 0.92 | 0.98 | 0.84 | 0.89 |
| 7 | 105312942 | rs846926 | 0.697 | 1.019 | -1.009 | 1.175 | -0.021 | -0.037 | -0.037 | 0 | 0.9 | 0.97 | 0.91 | 0.76 | 0.86 | 0.89 |
| 7 | 105314571 | rs516507 | -0.764 | 0.639 | -0.623 | 1.12 | -0.012 | 0.048 | 0.048 | 1.648 | 0.27 | 0.55 | 0.29 | 0.37 | 0.22 | 0.49 |
| 7 | 105315630 | rs711437 | 0.309 | 0.83 | -0.905 | 0.943 | 0.001 | 0.006 | 0.006 | 0 | 0.21 | 0.22 | 0.29 | 0.28 | 0.23 | 0.17 |
| 7 | 105319649 | rs1790060 | 0.108 | 0.774 | -1.073 | 0.951 | -0.024 | -0.026 | -0.026 | 5.359 | 0.09 | 0.07 | 0.23 | 0.18 | 0.14 | 0.35 |
| 7 | 105321923 | rs11974376 | -0.836 | 0.781 | -1.118 | 0.837 | -0.01 | 0.002 | 0.002 | 0 | 0.05 | 0.04 | 0.06 | 0.06 | 0.09 | 0.1 |
| 7 | 105322023 | rs485832 | -0.259 | 1.154 | -1.404 | 0.325 | -0.009 | 0.029 | 0.029 | 0 | 0.79 | 0.86 | 0.85 | 0.87 | 0.77 | 0.93 |
| 7 | 105326131 | rs846931 | 1.164 | 1.319 | -1.28 | 0.357 | -0.001 | 0.002 | 0.002 | 0 | 0.19 | 0.16 | 0.26 | 0.27 | 0.18 | 0.15 |
| 7 | 105326172 | rs846933 | -0.487 | 1.208 | -0.798 | 0.276 | 0.011 | 0.033 | 0.033 | 0 | 0.33 | 0.42 | 0.55 | 0.5 | 0.39 | 0.6 |
| 7 | 105326263 | rs573015 | 0.38 | 1.338 | -0.978 | -0.018 | 0.004 | 0.003 | 0.003 | 0 | 0.58 | 0.52 | 0.67 | 0.5 | 0.57 | 0.65 |
| 7 | 105330067 | rs635267 | -0.39 | 1.272 | -1.089 | -0.092 | -0.007 | 0.044 | 0.044 | 0 | 0.71 | 0.6 | 0.79 | 0.63 | 0.75 | 0.71 |

| | | | | | | | | | | | | | | | | |
|---|---|---|---|---|---|---|---|---|---|---|---|---|---|---|---|---|
| 7 | 105331819 | rs10272208 | 0.714 | 1.171 | -1.08 | -0.084 | -0.001 | 0.009 | 0.009 | 0.019 | 0.53 | 0.32 | 0.6 | 0.45 | 0.51 | 0.53 |
| 7 | 105339137 | rs620806 | 0.432 | 0.953 | -0.98 | 0.023 | -0.016 | 0.012 | 0.012 | 0 | 0.25 | 0.31 | 0.26 | 0.31 | 0.18 | 0.23 |
| 7 | 105339909 | rs11760426 | 0.854 | 1.456 | -1.214 | -0.356 | -0.023 | -0.013 | -0.013 | 0.767 | 0.07 | 0.03 | 0.09 | 0.09 | 0.1 | 0.02 |
| 7 | 105341608 | rs567003 | -1.036 | 1.288 | -1.351 | -0.657 | 0 | 0.107 | 0.107 | 0 | 0.64 | 0.84 | 0.77 | 0.95 | 0.63 | 0.81 |
| 7 | 105341913 | rs495517 | -1.47 | 1.205 | -1.42 | -0.659 | 0.008 | 0.137 | 0.137 | 0.004 | 0.69 | 0.89 | 0.81 | 1 | 0.69 | 0.93 |
| 7 | 105345947 | rs17151401 | 0.538 | 1.461 | -0.98 | -0.106 | -0.001 | -0.001 | -0.001 | 0 | 0.07 | 0.07 | 0.1 | 0.09 | 0.1 | 0.02 |
| 7 | 105346117 | rs562850 | 0.016 | 1.379 | -0.721 | 0.156 | 0.006 | 0.127 | 0.127 | 0.142 | 0.23 | 0.46 | 0.17 | 0.39 | 0.19 | 0.31 |
| 7 | 105348730 | rs521572 | -0.668 | 1.482 | -0.617 | 0.097 | 0.031 | 0.042 | 0.042 | 0.093 | 0.12 | 0.27 | 0.22 | 0.21 | 0.16 | 0.17 |
| 7 | 105355368 | rs12668158 | 0.364 | 1.463 | -0.852 | 0.049 | -0.02 | -0.014 | -0.014 | 0 | 0.08 | 0.04 | 0.12 | 0.09 | 0.11 | 0.03 |
| 7 | 105356897 | rs614260 | -1.235 | 1.434 | -1.053 | 0.04 | -0.031 | -0.017 | -0.017 | 0.044 | 0.39 | 0.81 | 0.46 | 0.63 | 0.4 | 0.54 |
| 7 | 105360021 | rs538249 | -1.349 | 1.095 | -1.365 | -0.677 | -0.019 | 0.055 | 0.055 | 0 | 0.62 | 0.82 | 0.75 | 1 | 0.57 | 0.83 |
| 7 | 105361835 | rs505288 | -1.111 | 1.159 | -1.215 | -0.418 | 0 | 0.001 | 0.001 | 0 | 0.15 | 0.24 | 0.16 | 0.14 | 0.13 | 0.21 |
| 7 | 105364258 | rs669959 | -0.358 | 0.451 | -1.559 | -0.881 | -0.004 | 0 | 0 | 2.231 | 0.4 | 0.59 | 0.3 | 0.46 | 0.28 | 0.39 |
| 7 | 105373099 | rs668373 | -2.152 | 0.825 | -1.575 | -0.851 | -0.097 | 0.212 | 0.212 | 0 | 0.7 | 0.98 | 0.78 | 1 | 0.61 | 0.88 |
| 7 | 105373405 | rs567184 | 0.052 | 0.711 | -1.094 | -0.426 | -0.014 | 0.058 | 0.058 | 1.404 | 0.28 | 0.53 | 0.26 | 0.46 | 0.21 | 0.31 |
| 7 | 105376750 | rs672416 | 0.271 | 0.396 | -1.076 | -0.292 | -0.003 | -0.005 | -0.005 | 0 | 0.19 | 0.2 | 0.28 | 0.32 | 0.19 | 0.28 |
| 7 | 105377070 | rs484491 | -0.529 | 0.554 | -1.016 | -0.358 | 0.008 | 0.025 | 0.025 | 0 | 0.42 | 0.66 | 0.47 | 0.78 | 0.3 | 0.55 |
| 7 | 105383459 | rs651220 | -0.672 | 0.466 | -0.911 | -0.2 | 0.023 | 0.056 | 0.056 | 0.086 | 0.26 | 0.4 | 0.47 | 0.45 | 0.34 | 0.46 |
| 7 | 105383875 | rs577004 | -0.039 | 0.527 | -0.925 | -0.134 | -0.003 | -0.004 | -0.004 | 0 | 0.21 | 0.23 | 0.3 | 0.32 | 0.25 | 0.29 |
| 7 | 105384666 | rs516802 | 0.256 | 0.393 | -0.804 | -0.246 | -0.044 | 0.079 | 0.079 | 0 | 0.27 | 0.44 | 0.2 | 0.47 | 0.13 | 0.33 |
| 7 | 105384906 | rs12667632 | 0.2 | 0.496 | -0.893 | -0.227 | -0.011 | -0.016 | -0.016 | 0.406 | 0.2 | 0.19 | 0.29 | 0.33 | 0.26 | 0.29 |
| 7 | 105386461 | rs587741 | -1.143 | 0.291 | -1.177 | -0.865 | -0.052 | 0.161 | 0.161 | 0 | 0.59 | 0.85 | 0.68 | 0.91 | 0.49 | 0.79 |
| 7 | 105387724 | rs497552 | -0.539 | 0.469 | -1.101 | -0.778 | 0.003 | 0.048 | 0.048 | 0 | 0.47 | 0.63 | 0.49 | 0.79 | 0.38 | 0.62 |
| 7 | 105390559 | rs3601 | 1.226 | 0.761 | -1.172 | -0.879 | -0.028 | 0.053 | 0.053 | 0 | 0.26 | 0.15 | 0.21 | 0 | 0.35 | 0.21 |
| 7 | 105394489 | rs12535963 | -0.056 | 0.421 | -0.495 | -0.227 | 0.026 | 0.06 | 0.06 | 0 | 0.18 | 0.34 | 0.27 | 0.28 | 0.2 | 0.19 |
| 7 | 105394542 | rs11977943 | 1.159 | 0.175 | -1.105 | -0.313 | -0.012 | -0.002 | -0.002 | 0.015 | 0.12 | 0.15 | 0.18 | 0.14 | 0.12 | 0.21 |
| 7 | 105396333 | rs10262705 | 1.109 | 0.392 | -1.304 | -0.473 | -0.043 | 0.109 | 0.109 | 0 | 0.28 | 0.11 | 0.2 | 0 | 0.36 | 0.25 |
| 7 | 105396550 | rs1939549 | -0.703 | 0.642 | -0.267 | 0.595 | 0.04 | 0.101 | 0.101 | 0 | 0.19 | 0.48 | 0.28 | 0.3 | 0.2 | 0.19 |
| 7 | 105399126 | rs557657 | 0.187 | 0.572 | -1.556 | 0.655 | -0.067 | -0.096 | -0.096 | 0.303 | 0.65 | 0.91 | 0.69 | 0.42 | 0.6 | 0.61 |
| 7 | 105399253 | rs3999821 | 1.143 | 0.374 | -0.931 | 0.308 | 0.001 | -0.009 | -0.009 | 0.002 | 0.46 | 0.4 | 0.27 | 0.12 | 0.42 | 0.42 |
| 7 | 105400316 | rs7802434 | -0.216 | 0.839 | -0.747 | 0.987 | 0.001 | -0.012 | -0.012 | 0.048 | 0.06 | 0.13 | 0.05 | 0.02 | 0.05 | 0.03 |
| 7 | 105401015 | rs498280 | -1.139 | 0.981 | -0.876 | 0.888 | -0.041 | 0.043 | 0.043 | 0.025 | 0.89 | 0.97 | 0.89 | 0.97 | 0.81 | 0.96 |
| 7 | 105401049 | rs6953345 | -0.691 | 0.759 | -0.782 | 1.014 | 0.004 | -0.015 | -0.015 | 0 | 0.07 | 0.24 | 0.06 | 0.02 | 0.05 | 0.03 |
| 7 | 105406916 | rs514122 | -0.137 | 0.787 | -1.375 | 1.315 | -0.045 | -0.051 | -0.051 | 0 | 0.14 | 0.04 | 0.22 | 0.2 | 0.17 | 0.15 |
| 7 | 105407407 | rs17152089 | -0.527 | 0.433 | -1.267 | 1.681 | -0.022 | 0.037 | 0.037 | 0 | 0.12 | 0.19 | 0.13 | 0.25 | 0.04 | 0.22 |
| 7 | 105413922 | rs17152107 | -1.311 | 0.586 | -1.413 | 1.627 | -0.022 | 0.054 | 0.054 | 0 | 0.18 | 0.31 | 0.26 | 0.29 | 0.1 | 0.23 |
| 7 | 105416560 | rs2529369 | -1.688 | 0.704 | -1.472 | 1.953 | 0 | -0.001 | -0.001 | 0 | 0.34 | 0.4 | 0.47 | 0.66 | 0.3 | 0.4 |
| 7 | 105420269 | rs1204371 | -1.128 | 0.816 | -1.372 | 1.527 | 0.002 | 0.033 | 0.033 | 0 | 0.27 | 0.4 | 0.3 | 0.49 | 0.27 | 0.24 |
| 7 | 105421445 | rs4730105 | -0.023 | 1.21 | -2.339 | 0.97 | -0.053 | -0.046 | -0.046 | 0 | 0.12 | 0.03 | 0.21 | 0.19 | 0.18 | 0.16 |
| 7 | 105421568 | rs1204370 | -0.435 | 0.798 | -1.956 | 0.368 | -0.013 | 0.206 | 0.206 | 0 | 0.51 | 0.89 | 0.65 | 0.8 | 0.5 | 0.48 |
| 7 | 105431876 | rs1209199 | 0.34 | 0.655 | -1.241 | 0.626 | 0.003 | 0.024 | 0.024 | 0 | 0.51 | 0.55 | 0.62 | 0.78 | 0.53 | 0.33 |
| 7 | 105442523 | rs13231470 | 0.017 | 0.188 | -1.451 | 0.613 | -0.006 | 0.02 | 0.02 | 0 | 0.26 | 0.12 | 0.17 | 0.2 | 0.24 | 0.45 |
| 7 | 105448210 | rs11980210 | 0.399 | -0.428 | -0.928 | 0.914 | 0.009 | -0.017 | -0.017 | 0 | 0.29 | 0.26 | 0.24 | 0.45 | 0.19 | 0.18 |
| 7 | 105450966 | rs17152163 | -0.852 | -0.37 | -0.58 | 1.487 | -0.024 | 0.09 | 0.09 | 0 | 0.12 | 0.22 | 0.15 | 0.37 | 0.07 | 0.33 |
| 7 | 105453542 | rs9691488 | -0.848 | -0.046 | -0.366 | 1.004 | 0.004 | -0.002 | -0.002 | 0.058 | 0.3 | 0.35 | 0.4 | 0.66 | 0.3 | 0.51 |
| 7 | 105453745 | rs2012691 | -0.788 | -0.061 | -0.413 | 1.749 | 0.02 | 0.044 | 0.044 | 0.083 | 0.21 | 0.31 | 0.23 | 0.33 | 0.25 | 0.46 |
| 7 | 105456869 | rs740309 | -0.818 | -0.969 | -0.457 | 0.624 | 0.012 | -0.07 | -0.07 | 0 | 0.66 | 0.62 | 0.74 | 0.95 | 0.53 | 0.74 |
| 7 | 105462708 | rs10245474 | 1.157 | -1.212 | -0.151 | 0.683 | -0.004 | -0.002 | -0.002 | 0 | 0.35 | 0.33 | 0.28 | 0.25 | 0.23 | 0.24 |
| 7 | 105463975 | rs2158897 | 1.071 | -1.391 | -0.539 | 0.59 | -0.005 | -0.051 | -0.051 | 0 | 0.08 | 0 | 0.11 | 0.25 | 0.06 | 0.12 |
| 7 | 105472752 | rs997438 | 0.608 | -1.331 | -0.526 | 0.867 | -0.062 | -0.109 | -0.109 | 0.097 | 0.16 | 0.03 | 0.3 | 0.42 | 0.22 | 0.48 |
| 7 | 105482615 | rs212412 | 1.113 | -1.384 | -0.55 | 0.5 | 0.03 | -0.102 | -0.102 | 0 | 0.3 | 0.38 | 0.23 | 0.03 | 0.42 | 0.19 |
| 7 | 105483775 | rs17152217 | 1.645 | -1.459 | -0.664 | 0.597 | 0.011 | -0.059 | -0.059 | 0.175 | 0.1 | 0.01 | 0.11 | 0.22 | 0.04 | 0.02 |
| 7 | 105487655 | rs17152219 | -0.214 | -1.526 | -0.574 | 0.548 | 0.017 | -0.048 | -0.048 | 0 | 0.21 | 0.14 | 0.24 | 0.55 | 0.13 | 0.14 |
| 7 | 105487862 | rs212414 | -0.52 | -1.497 | -0.318 | 0.295 | 0.001 | -0.027 | -0.027 | 0 | 0.93 | 0.88 | 0.94 | 0.98 | 0.93 | 0.93 |
| 7 | 105487951 | rs4730112 | 0.606 | -1.484 | -0.488 | 0.349 | -0.066 | -0.097 | -0.097 | 0 | 0.15 | 0.03 | 0.24 | 0.28 | 0.22 | 0.46 |
| 7 | 105488330 | rs10487836 | 1.871 | -1.612 | -0.301 | -0.016 | 0.04 | 0.028 | 0.028 | 0 | 0.25 | 0.15 | 0.16 | 0 | 0.17 | 0.12 |
| 7 | 105494963 | rs212417 | 1.249 | -1.198 | 0.009 | 0.283 | 0.005 | 0.019 | 0.019 | 0 | 0.67 | 0.62 | 0.6 | 0.46 | 0.65 | 0.36 |
| 7 | 105503535 | rs1859814 | -0.053 | -0.696 | -0.114 | -0.045 | 0 | 0.008 | 0.008 | 0 | 0.56 | 0.38 | 0.57 | 0.89 | 0.49 | 0.72 |
| 7 | 105503900 | rs6947520 | -0.782 | -0.551 | -0.069 | 0.16 | 0.007 | 0.056 | 0.056 | 0 | 0.27 | 0.39 | 0.25 | 0.5 | 0.28 | 0.56 |
| 7 | 105506737 | rs17152230 | 1.111 | -1.151 | -0.365 | 0.541 | -0.023 | -0.04 | -0.04 | 0 | 0.07 | 0.01 | 0.11 | 0.27 | 0.09 | 0.1 |
| 7 | 105511660 | rs7777514 | 0.648 | -1.454 | -0.09 | 0.255 | -0.028 | 0.003 | 0.003 | 0 | 0.39 | 0.57 | 0.39 | 0.43 | 0.25 | 0.17 |
| 7 | 105517719 | rs212429 | -0.526 | -1.111 | -0.042 | -0.011 | -0.002 | -0.022 | -0.022 | 0 | 0.94 | 0.93 | 0.97 | 1 | 0.95 | 0.96 |
| 7 | 105520040 | rs212432 | -0.156 | -1.065 | -0.105 | 0.083 | 0.027 | 0.032 | 0.032 | 0 | 0.48 | 0.39 | 0.47 | 0.26 | 0.6 | 0.67 |
| 7 | 105522619 | rs212435 | 0.579 | -0.607 | -0.032 | -0.471 | 0.006 | -0.062 | -0.062 | 0.009 | 0.32 | 0.34 | 0.37 | 0.03 | 0.46 | 0.22 |
| 7 | 105528339 | rs212438 | 0.153 | 0.012 | 0.511 | 0.279 | 0.016 | -0.068 | -0.068 | 0.793 | 0.84 | 0.77 | 0.88 | 0.95 | 0.8 | 0.78 |
| 7 | 105533735 | rs11765906 | -0.129 | -0.598 | 0.214 | 0.727 | 0.001 | 0.026 | 0.026 | 0.884 | 0.11 | 0.12 | 0.2 | 0.39 | 0.06 | 0.05 |
| 7 | 105542580 | rs13438712 | 0.945 | -0.392 | 0.126 | -0.079 | 0.021 | 0.112 | 0.112 | 0 | 0.41 | 0.21 | 0.3 | 0.03 | 0.39 | 0.37 |
| 7 | 105545666 | rs9690395 | 0.388 | -0.451 | 0.796 | -0.002 | 0.006 | -0.017 | -0.017 | 0 | 0.07 | 0.26 | 0.1 | 0 | 0.04 | 0.03 |
| 7 | 105546918 | rs9656120 | -0.963 | -0.292 | 0.175 | 0.403 | -0.051 | -0.028 | -0.028 | 0.281 | 0.77 | 0.9 | 0.77 | 0.71 | 0.62 | 0.59 |

| | | | | | | | | | | | | | | | | |
|---|---|---|---|---|---|---|---|---|---|---|---|---|---|---|---|---|
| 7 | 105554522 | rs17152281 | 0.021 | -0.528 | -0.034 | 0.575 | 0.013 | 0.013 | 0.013 | 0 | 0.19 | 0.13 | 0.15 | 0.15 | 0.16 | 0.25 |
| 7 | 105555336 | rs2057945 | -1.017 | -0.327 | -0.194 | 0.671 | -0.071 | -0.046 | -0.046 | 0 | 0.78 | 0.92 | 0.78 | 0.69 | 0.62 | 0.62 |
| 7 | 105556832 | rs2240398 | 1.2 | -0.539 | 0.043 | 0.719 | 0.027 | 0.012 | 0.012 | 0.002 | 0.36 | 0.21 | 0.37 | 0.3 | 0.29 | 0.23 |
| 7 | 105560857 | rs10265194 | 0.268 | -0.476 | 0.556 | 1.313 | -0.007 | 0.083 | 0.083 | 2.158 | 0.05 | 0.08 | 0.15 | 0.23 | 0.03 | 0.01 |
| 7 | 105560892 | rs10487834 | -0.47 | -0.445 | 1.267 | 1.435 | 0.015 | 0.114 | 0.114 | 0 | 0.12 | 0.35 | 0.18 | 0.25 | 0.14 | 0.22 |
| 7 | 105561135 | rs234 | 0.545 | -0.198 | 0.703 | 0.946 | -0.005 | 0.015 | 0.015 | 0 | 0.58 | 0.69 | 0.64 | 0.68 | 0.5 | 0.52 |
| 7 | 105562645 | rs10487833 | -0.08 | -0.048 | 0.61 | 0.868 | -0.007 | -0.022 | -0.022 | 0 | 0.15 | 0.23 | 0.08 | 0.04 | 0.09 | 0.05 |
| 7 | 105562881 | rs10244533 | 0.927 | -0.05 | 0.592 | 0.885 | -0.001 | 0.008 | 0.008 | 0 | 0.57 | 0.59 | 0.62 | 0.67 | 0.49 | 0.5 |
| 7 | 105563314 | rs10248057 | -0.146 | 0.045 | 0.253 | 0.9 | -0.043 | -0.008 | -0.008 | 0 | 0.72 | 0.85 | 0.71 | 0.71 | 0.58 | 0.54 |
| 7 | 105563666 | rs3735367 | 0.809 | 0.166 | 0.327 | 0.375 | -0.004 | -0.023 | -0.023 | 0 | 0.44 | 0.44 | 0.41 | 0.13 | 0.35 | 0.26 |
| 7 | 105572183 | rs10755868 | -1.425 | -0.127 | 0.468 | 1.391 | -0.023 | 0.034 | 0.034 | 0 | 0.36 | 0.5 | 0.42 | 0.43 | 0.28 | 0.46 |
| 7 | 105574754 | rs12533809 | -0.872 | 0.385 | 0.387 | 1.234 | -0.009 | -0.022 | -0.022 | 0.853 | 0.1 | 0.2 | 0.11 | 0.05 | 0.08 | 0.22 |
| 7 | 105575202 | rs17152355 | -1.353 | 0.476 | 0.364 | 1.553 | 0.006 | 0.037 | 0.037 | 0 | 0.16 | 0.22 | 0.18 | 0.26 | 0.16 | 0.21 |
| 7 | 105576034 | rs11983066 | -0.937 | 0.577 | 0.033 | 1.432 | -0.001 | 0.002 | 0.002 | 0 | 0.07 | 0.1 | 0.06 | 0.05 | 0.05 | 0.02 |
| 7 | 105576245 | rs11983073 | -0.906 | 0.584 | -0.037 | 1.435 | -0.001 | 0.001 | 0.001 | 0 | 0.07 | 0.09 | 0.06 | 0.05 | 0.05 | 0.02 |
| 7 | 105576557 | rs193858 | -0.807 | 0.583 | -0.259 | 1.519 | -0.003 | -0.009 | -0.009 | 0.319 | 0.09 | 0.05 | 0.12 | 0.19 | 0.11 | 0.14 |
| 7 | 105577549 | rs2727763 | -1.531 | 0.518 | 0.265 | 2.153 | 0.017 | 0.259 | 0.259 | 0 | 0.09 | 0.3 | 0.17 | 0.38 | 0.09 | 0.25 |
| 7 | 105577762 | rs10262629 | -0.428 | 0.749 | -1.14 | 1.142 | -0.012 | 0.051 | 0.051 | 0 | 0.47 | 0.81 | 0.46 | 0.34 | 0.43 | 0.62 |
| 7 | 105579284 | rs193853 | -1.075 | 1.224 | -0.573 | 0.837 | 0.019 | 0.001 | 0.001 | 0 | 0.31 | 0.39 | 0.36 | 0.32 | 0.32 | 0.37 |
| 7 | 105581663 | rs17416073 | -0.749 | 1.243 | -0.671 | 0.202 | -0.009 | 0.009 | 0.009 | 0 | 0.1 | 0.05 | 0.16 | 0.04 | 0.12 | 0.17 |
| 7 | 105583256 | rs193843 | 0.186 | 1.115 | -0.499 | 0.283 | -0.009 | -0.019 | -0.019 | 0 | 0.1 | 0.15 | 0.05 | 0.1 | 0.08 | 0.16 |
| 7 | 105585666 | rs11763182 | 0.305 | 0.924 | -1.45 | -0.826 | -0.052 | 0.139 | 0.139 | 0 | 0.38 | 0.14 | 0.35 | 0.05 | 0.45 | 0.43 |
| 7 | 105586853 | rs193841 | -1.401 | 0.448 | -1.716 | -1.219 | 0.02 | 0.06 | 0.06 | 0 | 0.81 | 0.95 | 0.87 | 0.96 | 0.83 | 0.96 |
| 7 | 105586946 | rs2727744 | 0.041 | 0.319 | -1.405 | -0.887 | 0.001 | 0 | 0 | 0.184 | 0.6 | 0.41 | 0.69 | 0.71 | 0.68 | 0.73 |
| 7 | 105587100 | rs193840 | 0.459 | 0.482 | -1.124 | -0.672 | 0.012 | 0.017 | 0.017 | 1.303 | 0.59 | 0.37 | 0.53 | 0.43 | 0.66 | 0.7 |
| 7 | 105587321 | rs35496280 | 0.756 | 0.487 | -1.112 | -0.635 | -0.018 | 0.004 | 0.004 | 0 | 0.08 | 0.04 | 0.16 | 0.04 | 0.1 | 0.1 |
| 7 | 105587441 | rs408631 | 0.207 | 0.395 | -1.124 | -0.495 | 0.012 | 0.017 | 0.017 | 1.329 | 0.59 | 0.37 | 0.53 | 0.43 | 0.66 | 0.69 |
| 7 | 105592671 | rs2527853 | 0.276 | 0.382 | -1.332 | -0.332 | -0.001 | 0.026 | 0.026 | 0 | 0.13 | 0.12 | 0.17 | 0.49 | 0.08 | 0.18 |
| 7 | 105600568 | rs193814 | 0.606 | 0.498 | -1.448 | -0.867 | -0.001 | 0.11 | 0.11 | 0 | 0.48 | 0.3 | 0.37 | 0.26 | 0.49 | 0.58 |
| 7 | 105604722 | rs2727755 | -0.541 | 0.696 | -1.306 | -0.409 | -0.001 | -0.002 | -0.002 | 0 | 0.22 | 0.24 | 0.35 | 0.66 | 0.23 | 0.28 |
| 7 | 105605317 | rs193805 | -0.197 | 0.396 | -1.42 | 0.001 | 0.007 | -0.009 | -0.009 | 0 | 0.17 | 0.09 | 0.15 | 0.24 | 0.16 | 0.27 |
| 7 | 105619169 | rs41266 | 0.177 | 0.83 | -1.64 | 0.021 | -0.118 | -0.007 | -0.007 | 0 | 0.36 | 0.05 | 0.42 | 0.31 | 0.42 | 0.38 |
| 7 | 105619206 | rs41267 | 0.577 | 0.449 | -1.365 | 0.062 | -0.003 | 0 | 0 | 0 | 0.76 | 0.33 | 0.82 | 0.73 | 0.78 | 0.76 |
| 7 | 105619330 | rs6947411 | 0.634 | 0.849 | -1.443 | 0.014 | -0.02 | 0.006 | 0.006 | 0 | 0.07 | 0.02 | 0.16 | 0.05 | 0.07 | 0.21 |
| 7 | 105619565 | rs41268 | 0.659 | 0.76 | -0.828 | -0.017 | 0.053 | 0.005 | 0.005 | 0.114 | 0.84 | 0.53 | 0.74 | 0.85 | 0.84 | 0.71 |
| 7 | 105621512 | rs193795 | 0.311 | 0.497 | -0.465 | 0.1 | -0.014 | 0.003 | 0.003 | 0 | 0.48 | 0.28 | 0.56 | 0.49 | 0.44 | 0.61 |
| 7 | 105624196 | rs445736 | -0.447 | 0.634 | -0.517 | 0.159 | 0.001 | -0.009 | -0.009 | 0 | 0.61 | 0.5 | 0.64 | 0.65 | 0.58 | 0.76 |
| 7 | 105625564 | rs6980139 | -0.66 | 0.203 | -0.635 | -0.145 | -0.008 | 0.004 | 0.004 | 0 | 0.14 | 0.23 | 0.08 | 0.16 | 0.13 | 0.15 |
| 7 | 105630156 | rs375981 | -0.131 | 0.031 | -0.933 | -0.954 | 0.015 | 0.001 | 0.001 | 0 | 0.72 | 0.8 | 0.86 | 0.99 | 0.78 | 0.75 |
| 7 | 105630440 | rs3847076 | 0.375 | 0.399 | -1.262 | -1.395 | 0.015 | 0.054 | 0.054 | 0 | 0.39 | 0.26 | 0.28 | 0.19 | 0.36 | 0.44 |
| 7 | 105632516 | rs11764661 | -0.123 | 0.983 | -1.233 | -1.366 | 0.046 | -0.031 | -0.031 | 0.022 | 0.07 | 0.18 | 0.08 | 0 | 0.09 | 0.06 |
| 7 | 105634441 | rs2528883 | 0.147 | 1.183 | -1.171 | -0.699 | -0.023 | -0.045 | -0.045 | 0 | 0.27 | 0.23 | 0.33 | 0.37 | 0.34 | 0.35 |
| 7 | 105634589 | rs2526975 | -0.557 | 1.339 | -1.227 | -0.725 | -0.029 | -0.019 | -0.019 | 0 | 0.35 | 0.3 | 0.41 | 0.38 | 0.47 | 0.42 |
| 7 | 105636146 | rs2526977 | -0.801 | 1.121 | -1.112 | -0.852 | 0.018 | 0.118 | 0.118 | 0 | 0.79 | 0.56 | 0.67 | 0.58 | 0.81 | 0.75 |
| 7 | 105637007 | rs396600 | 0.422 | 1.466 | -0.762 | -1.661 | 0.002 | -0.031 | -0.031 | 0 | 0.18 | 0.21 | 0.13 | 0.01 | 0.24 | 0.17 |
| 7 | 105637532 | rs11981655 | -0.818 | 0.317 | -0.866 | -1.283 | 0.005 | -0.014 | -0.014 | 0 | 0.16 | 0.2 | 0.12 | 0.07 | 0.16 | 0.28 |
| 7 | 105639848 | rs7793669 | 0.885 | 0.167 | -1.388 | -1.227 | -0.002 | 0.021 | 0.021 | 0.878 | 0.23 | 0.02 | 0.29 | 0.19 | 0.26 | 0.12 |
| 7 | 105644137 | rs17288345 | 0.931 | -0.151 | -1.396 | -1.269 | -0.003 | 0.012 | 0.012 | 1.027 | 0.23 | 0.01 | 0.26 | 0.18 | 0.22 | 0.12 |
| 7 | 105645301 | rs10953488 | -0.309 | -0.424 | -1.037 | -1.417 | 0.063 | 0.065 | 0.065 | 2.167 | 0.39 | 0.28 | 0.32 | 0.24 | 0.27 | 0.31 |
| 7 | 105649492 | rs11770729 | 0.959 | -0.325 | -1.302 | -1.36 | 0.014 | 0.035 | 0.035 | 1.316 | 0.23 | 0.02 | 0.34 | 0.18 | 0.22 | 0.12 |
| 7 | 105651320 | rs7357197 | -0.672 | -0.142 | -0.765 | -1.497 | 0.004 | -0.085 | -0.085 | 0 | 0.26 | 0.43 | 0.2 | 0.08 | 0.23 | 0.28 |
| 7 | 105653095 | rs10953489 | -0.246 | 0.06 | -0.715 | -1.427 | 0.013 | -0.114 | -0.114 | 0.703 | 0.21 | 0.39 | 0.17 | 0.01 | 0.21 | 0.1 |
| 7 | 105653867 | rs3887998 | -0.131 | 0.286 | -0.706 | -0.921 | 0.005 | 0.006 | 0.006 | 2.06 | 0.48 | 0.58 | 0.44 | 0.24 | 0.41 | 0.39 |
| 7 | 105654035 | rs10257002 | -0.189 | 0.178 | -0.73 | -0.89 | 0.007 | 0.02 | 0.02 | 1.557 | 0.49 | 0.59 | 0.46 | 0.25 | 0.43 | 0.4 |
| 7 | 105662690 | rs10258293 | -0.621 | -0.661 | -0.924 | -1.026 | 0.023 | 0.02 | 0.02 | 0 | 0.29 | 0.22 | 0.13 | 0.08 | 0.2 | 0.22 |
| 7 | 105670992 | rs105347 | -0.73 | -0.186 | -0.792 | -1.173 | -0.001 | -0.01 | -0.01 | 0 | 0.63 | 0.57 | 0.4 | 0.19 | 0.6 | 0.52 |
| 7 | 105671267 | rs10270308 | 0.41 | -0.486 | -0.76 | 0.203 | 0.03 | 0.479 | 0.479 | 3.181 | 0.06 | 0.38 | 0.17 | 0.36 | 0.01 | 0.06 |
| 7 | 105674590 | rs381188 | -0.206 | -0.664 | -0.82 | -0.398 | 0.011 | 0.08 | 0.08 | 0.745 | 0.53 | 0.31 | 0.35 | 0.15 | 0.55 | 0.38 |
| 7 | 105677579 | rs17152490 | 0.413 | -0.646 | -0.957 | -0.195 | 0.044 | 0.03 | 0.03 | 0.089 | 0.22 | 0.1 | 0.12 | 0.16 | 0.18 | 0.1 |
| 7 | 105679551 | rs367311 | -0.048 | 0.014 | -0.514 | 0.082 | 0.019 | 0.017 | 0.017 | 0 | 0.38 | 0.54 | 0.4 | 0.5 | 0.38 | 0.3 |
| 7 | 105684875 | rs4730126 | 0.364 | 0.518 | -0.507 | 0.169 | -0.003 | 0.001 | 0.001 | 0 | 0.59 | 0.42 | 0.66 | 0.61 | 0.61 | 0.58 |
| 7 | 105687825 | rs7795448 | -0.204 | 0.839 | -0.394 | 0.023 | -0.023 | -0.009 | -0.009 | 0 | 0.68 | 0.87 | 0.72 | 0.66 | 0.68 | 0.67 |
| 7 | 105689148 | rs4730129 | 1.422 | 0.579 | -0.696 | 0.078 | -0.021 | -0.043 | -0.043 | 0.707 | 0.28 | 0.19 | 0.46 | 0.44 | 0.34 | 0.34 |
| 7 | 105692493 | rs7803117 | 0.518 | 0.316 | -0.766 | -0.105 | -0.021 | 0.027 | 0.027 | 0 | 0.5 | 0.18 | 0.58 | 0.59 | 0.51 | 0.53 |
| 7 | 105695500 | rs7782502 | 1.274 | 0.266 | -0.562 | -0.02 | -0.037 | -0.057 | -0.057 | 0 | 0.3 | 0.18 | 0.47 | 0.49 | 0.38 | 0.39 |
| 7 | 105700456 | rs7798028 | 0.788 | 0.31 | 0.101 | -0.058 | 0.016 | 0.046 | 0.046 | 0 | 0.34 | 0.58 | 0.51 | 0.48 | 0.4 | 0.38 |
| 7 | 105700868 | rs10487831 | -0.341 | -0.66 | 0.137 | -0.459 | 0.006 | 0.008 | 0.008 | 0 | 0.34 | 0.24 | 0.21 | 0.18 | 0.27 | 0.28 |
| 7 | 105709218 | rs7776827 | -0.855 | -0.799 | 0.222 | -0.547 | -0.026 | -0.03 | -0.03 | 0 | 0.19 | 0.33 | 0.11 | 0.1 | 0.13 | 0.16 |
| 7 | 105710737 | rs16872046 | 1.014 | -0.762 | -0.188 | -0.538 | 0.011 | -0.006 | -0.006 | 0 | 0.1 | 0.07 | 0.07 | 0.1 | 0.09 | 0.1 |

| | | | | | | | | | | | | | | | | |
|---|---|---|---|---|---|---|---|---|---|---|---|---|---|---|---|---|
| 7 | 105711325 | rs12705332 | 0.795 | -0.797 | -0.939 | -0.752 | -0.159 | -0.124 | -0.124 | 0 | 0.34 | 0.06 | 0.52 | 0.54 | 0.52 | 0.35 |
| 7 | 105714513 | rs6973699 | -0.809 | -1.342 | -0.708 | -1.003 | -0.014 | -0.002 | -0.002 | 0.013 | 0.43 | 0.46 | 0.25 | 0.35 | 0.28 | 0.34 |
| 7 | 105721017 | rs176470 | -0.038 | -1.399 | -1.292 | -1.063 | -0.044 | -0.063 | -0.063 | 0 | 0.7 | 0.61 | 0.86 | 0.9 | 0.8 | 0.86 |
| 7 | 105735042 | rs10953491 | 0.275 | -1.405 | -1.232 | -0.996 | -0.045 | 0.091 | 0.091 | 0 | 0.17 | 0.5 | 0.15 | 0.25 | 0.1 | 0.23 |
| 7 | 105735118 | rs6960866 | 1.622 | -1.409 | -1.42 | -1.076 | 0.017 | 0.029 | 0.029 | 1.174 | 0.3 | 0.31 | 0.49 | 0.52 | 0.42 | 0.33 |
| 7 | 105736971 | rs176489 | 1.308 | -1.345 | -1.355 | -0.965 | -0.089 | 0.098 | 0.098 | 0 | 0.27 | 0.06 | 0.23 | 0.13 | 0.27 | 0.31 |
| 7 | 105739546 | rs12540052 | 0.055 | -0.705 | -0.785 | -0.508 | -0.045 | 0.091 | 0.091 | 0 | 0.17 | 0.5 | 0.15 | 0.25 | 0.1 | 0.23 |
| 7 | 105741200 | rs6466083 | 0.156 | 0.171 | -0.54 | -0.272 | -0.004 | -0.004 | -0.004 | 0 | 0.37 | 0.32 | 0.54 | 0.52 | 0.51 | 0.34 |
| 7 | 105761160 | rs176512 | -0.03 | 0.022 | -0.294 | -0.239 | -0.009 | 0.016 | 0.016 | 1.177 | 0.91 | 0.96 | 0.95 | 1 | 0.89 | 0.97 |
| 7 | 105763426 | rs6466086 | -0.648 | -0.842 | 0.117 | -0.087 | -0.041 | 0.039 | 0.039 | 0.011 | 0.17 | 0.68 | 0.34 | 0.26 | 0.08 | 0.19 |
| 7 | 105766466 | rs4730136 | -1.325 | -0.835 | 0.056 | -0.176 | -0.003 | -0.001 | -0.001 | 0.322 | 0.25 | 0.74 | 0.4 | 0.27 | 0.17 | 0.26 |
| 7 | 105767172 | rs10274327 | -1.339 | -0.848 | 0.05 | -0.182 | -0.002 | -0.001 | -0.001 | 0.515 | 0.25 | 0.74 | 0.4 | 0.27 | 0.17 | 0.26 |
| 7 | 105773989 | rs6942777 | -1.339 | -0.866 | 0.373 | -0.19 | -0.002 | -0.001 | -0.001 | 0.054 | 0.25 | 0.73 | 0.4 | 0.26 | 0.17 | 0.26 |
| 7 | 105775564 | rs7785757 | -1.513 | -1.602 | -0.261 | -0.503 | 0.113 | 0.024 | 0.024 | 0.412 | 0.46 | 0.88 | 0.5 | 0.37 | 0.32 | 0.38 |
| 7 | 105776457 | rs10277426 | -1.544 | -1.06 | 0.274 | -0.273 | -0.001 | -0.001 | -0.001 | 0 | 0.25 | 0.74 | 0.4 | 0.26 | 0.17 | 0.26 |
| 7 | 105779739 | rs1558055 | -0.12 | -0.99 | 0.257 | -0.22 | 0.016 | -0.025 | -0.025 | 0.785 | 0.12 | 0.07 | 0.3 | 0.26 | 0.08 | 0.19 |
| 7 | 105782968 | rs7795101 | -1.215 | -0.965 | 0.841 | -0.209 | -0.047 | 0.022 | 0.022 | 0 | 0.23 | 0.53 | 0.38 | 0.26 | 0.17 | 0.26 |
| 7 | 105789603 | rs176516 | 0.479 | -0.289 | 0.285 | -0.312 | -0.031 | -0.059 | -0.059 | 0 | 0.2 | 0.1 | 0.38 | 0.39 | 0.28 | 0.27 |
| 7 | 105795338 | rs176523 | -1.656 | -0.112 | 0.544 | -0.745 | -0.002 | -0.011 | -0.011 | 0.017 | 0.65 | 0.72 | 0.64 | 0.45 | 0.62 | 0.53 |
| 7 | 105803764 | rs176557 | -0.334 | -0.112 | 0.507 | -0.825 | 0.013 | 0.115 | 0.115 | 4.631 | 0.41 | 0.22 | 0.24 | 0.06 | 0.36 | 0.21 |
| 7 | 105807547 | rs4730147 | 0.231 | 0.049 | 0.641 | -0.656 | 0.014 | 0.079 | 0.079 | 0.089 | 0.23 | 0.45 | 0.38 | 0.39 | 0.24 | 0.3 |
| 7 | 105814205 | rs176566 | 0.663 | 0.21 | 0.217 | -0.975 | -0.011 | 0.082 | 0.082 | 6.33 | 0.2 | 0.06 | 0.08 | 0 | 0.16 | 0.06 |
| 7 | 105816502 | rs176568 | 0.894 | 0.14 | 0.213 | -0.946 | -0.005 | 0.068 | 0.068 | 3.676 | 0.18 | 0.05 | 0.07 | 0 | 0.13 | 0.05 |
| 7 | 105817328 | rs4730148 | 0.138 | 0.039 | 0.323 | -0.933 | 0 | -0.004 | -0.004 | 1.32 | 0.19 | 0.15 | 0.16 | 0.06 | 0.2 | 0.15 |
| 7 | 105818666 | rs11765784 | 0.777 | 0.164 | 0.069 | -0.642 | 0 | 0.01 | 0.01 | 0 | 0.21 | 0.21 | 0.36 | 0.41 | 0.24 | 0.29 |
| 7 | 105820364 | rs176577 | 0.654 | -0.971 | -0.056 | -0.675 | 0.008 | 0.004 | 0.004 | 1.458 | 0.19 | 0.11 | 0.07 | 0 | 0.12 | 0.09 |
| 7 | 105827361 | rs176588 | -1.535 | -1.043 | -0.301 | -0.708 | -0.053 | -0.066 | -0.066 | 0 | 0.64 | 0.86 | 0.63 | 0.46 | 0.57 | 0.5 |
| 7 | 105831774 | rs6979063 | -0.697 | -1.031 | -0.319 | -0.581 | 0.005 | 0.001 | 0.001 | 0 | 0.44 | 0.65 | 0.55 | 0.46 | 0.44 | 0.41 |
| 7 | 105838305 | rs1029980 | -0.362 | -1.373 | -0.032 | -1.321 | -0.017 | -0.076 | -0.076 | 0 | 0.21 | 0.29 | 0.17 | 0.06 | 0.2 | 0.15 |
| 7 | 105840608 | rs10236515 | 0.41 | -1.155 | -0.272 | -1.058 | -0.013 | -0.081 | -0.081 | 0 | 0.59 | 0.55 | 0.75 | 0.94 | 0.68 | 0.8 |
| 7 | 105843081 | rs10245392 | -0.504 | -1.555 | -0.154 | -0.114 | -0.029 | 0.017 | 0.017 | 0 | 0.24 | 0.29 | 0.34 | 0.55 | 0.11 | 0.18 |
| 7 | 105858211 | rs3926927 | -0.236 | -1.502 | -0.028 | -0.041 | 0.016 | -0.022 | -0.022 | 0 | 0.21 | 0.2 | 0.27 | 0.4 | 0.11 | 0.11 |
| 7 | 105860188 | rs2192383 | -0.412 | -1.356 | 0.074 | -1.041 | 0.003 | -0.096 | -0.096 | 0 | 0.61 | 0.63 | 0.75 | 0.97 | 0.59 | 0.76 |
| 7 | 105863250 | rs2215816 | 0.13 | -1.259 | -0.127 | -1.175 | 0.01 | -0.068 | -0.068 | 0 | 0.66 | 0.55 | 0.75 | 0.85 | 0.65 | 0.69 |
| 7 | 105864236 | rs10953500 | 0.505 | -1.295 | -0.121 | -1.158 | 0.015 | -0.063 | -0.063 | 0.449 | 0.38 | 0.17 | 0.37 | 0.45 | 0.33 | 0.21 |
| 7 | 105865194 | rs7791378 | 0.207 | -1.29 | 0.004 | -1.182 | 0.017 | -0.015 | -0.015 | 0.008 | 0.41 | 0.37 | 0.4 | 0.45 | 0.35 | 0.23 |
| 7 | 105865405 | rs12535322 | 0.026 | -1.233 | 0.09 | -1.207 | 0.008 | -0.105 | -0.105 | 0 | 0.68 | 0.49 | 0.77 | 0.9 | 0.68 | 0.69 |
| 7 | 105879974 | rs7776987 | 0.29 | -1.11 | -0.167 | -1.115 | 0 | -0.009 | -0.009 | 0 | 0.62 | 0.4 | 0.73 | 0.88 | 0.66 | 0.66 |
| 7 | 105880915 | rs13236748 | 1.386 | -1.004 | -0.2 | -1.028 | -0.045 | -0.083 | -0.083 | 0.039 | 0.25 | 0.06 | 0.36 | 0.4 | 0.32 | 0.44 |
| 7 | 105884848 | rs13224516 | 1.386 | -0.988 | -0.237 | -0.964 | -0.052 | -0.064 | -0.064 | 2.372 | 0.25 | 0.05 | 0.36 | 0.34 | 0.32 | 0.44 |
| 7 | 105887559 | rs10487820 | 0.726 | -1.102 | -0.274 | -1.303 | 0.035 | 0.271 | 0.271 | 0 | 0.49 | 0.14 | 0.68 | 0.88 | 0.61 | 0.65 |
| 7 | 105896742 | rs3801270 | 0.671 | -1.104 | -0.275 | -1.286 | 0.057 | 0.28 | 0.28 | 0 | 0.49 | 0.14 | 0.7 | 0.87 | 0.63 | 0.65 |
| 7 | 105922523 | rs1319313 | -0.14 | -1.227 | 0.228 | -1.325 | 0 | -0.07 | -0.07 | 4.22 | 0.66 | 0.68 | 0.84 | 1 | 0.76 | 0.9 |
| 7 | 105924250 | rs3801266 | -0.294 | -1.012 | 0.04 | -1.223 | -0.058 | -0.093 | -0.093 | 0 | 0.29 | 0.52 | 0.19 | 0.13 | 0.18 | 0.26 |
| 7 | 105927752 | rs7789066 | 1.145 | -0.849 | -0.649 | -1.238 | 0.015 | -0.001 | -0.001 | 0 | 0.16 | 0.14 | 0.06 | 0 | 0.07 | 0.01 |
| 7 | 105928515 | rs11977021 | 1.231 | -0.798 | -0.817 | -1.057 | -0.05 | -0.133 | -0.133 | 0.285 | 0.23 | 0.07 | 0.31 | 0.47 | 0.28 | 0.27 |
| 7 | 105930388 | rs6971643 | -0.419 | 0.2 | -0.111 | -0.574 | 0.011 | -0.172 | -0.172 | 1.346 | 0.77 | 0.59 | 0.86 | 1 | 0.8 | 0.9 |
| 7 | 105931877 | rs12537455 | 0.211 | 0.539 | -0.658 | 0.619 | -0.048 | -0.09 | -0.09 | 0.427 | 0.23 | 0.06 | 0.32 | 0.4 | 0.31 | 0.35 |
| 7 | 105933203 | rs17152854 | -1 | 0.128 | 0.266 | 0.331 | 0.008 | -0.006 | -0.006 | 0 | 0.16 | 0.45 | 0.2 | 0.15 | 0.15 | 0.27 |
| 7 | 105951509 | rs997980 | 0.215 | -0.24 | -0.507 | 0.476 | 0.016 | -0.005 | -0.005 | 0.362 | 0.58 | 0.39 | 0.62 | 0.42 | 0.67 | 0.76 |
| 7 | 105951720 | rs2539847 | -0.445 | 0.394 | -0.17 | -0.394 | 0.014 | -0.04 | -0.04 | 0 | 0.3 | 0.55 | 0.22 | 0.23 | 0.34 | 0.5 |
| 7 | 105954740 | rs2704966 | 0.289 | 0.023 | -0.592 | -0.954 | -0.029 | 0.057 | 0.057 | 0.306 | 0.8 | 0.85 | 0.81 | 0.98 | 0.71 | 0.68 |
| 7 | 105969189 | rs10499961 | 1.038 | -1.564 | -1.269 | -1.121 | 0.065 | -0.083 | -0.083 | 0 | 0.26 | 0.12 | 0.37 | 0.56 | 0.13 | 0.08 |
| 7 | 105970988 | rs2106303 | -0.989 | -1.775 | -0.791 | -1.553 | 0.007 | -0.01 | -0.01 | 0 | 0.7 | 0.71 | 0.74 | 0.86 | 0.62 | 0.72 |
| 7 | 105971434 | rs2704958 | -0.361 | -1.86 | -0.426 | -1.584 | -0.007 | 0.021 | 0.021 | 0.521 | 0.38 | 0.38 | 0.29 | 0.19 | 0.43 | 0.58 |
| 7 | 105985078 | rs2014669 | 0.453 | -1.792 | -0.658 | -1.495 | 0.004 | -0.065 | -0.065 | 0 | 0.45 | 0.16 | 0.54 | 0.56 | 0.44 | 0.4 |
| 7 | 105990115 | rs4538930 | 0.051 | -1.762 | -0.469 | -1.426 | -0.063 | 0.075 | 0.075 | 0.199 | 0.6 | 0.22 | 0.64 | 0.78 | 0.55 | 0.64 |
| 7 | 105995670 | rs2040542 | -1.135 | -1.839 | -0.503 | -1.434 | -0.145 | 0.017 | 0.017 | 0.752 | 0.72 | 0.98 | 0.75 | 0.83 | 0.62 | 0.76 |
| 7 | 105997164 | rs2519672 | -0.255 | -1.811 | -0.053 | -1.344 | 0.026 | -0.027 | -0.027 | 0 | 0.63 | 0.57 | 0.66 | 0.76 | 0.55 | 0.7 |
| 7 | 105997946 | rs17356935 | 0.055 | -1.737 | -0.649 | -1.337 | -0.06 | 0.073 | 0.073 | 0.197 | 0.6 | 0.22 | 0.64 | 0.78 | 0.55 | 0.64 |
| 7 | 105999186 | rs11543784 | 0.486 | -2.02 | -0.589 | -1.169 | 0.094 | -0.115 | -0.115 | 0.333 | 0.32 | 0.08 | 0.39 | 0.6 | 0.19 | 0.14 |
| 7 | 106006593 | rs11526273 | 0.749 | -1.657 | -0.548 | -1.209 | 0.018 | -0.109 | -0.109 | 0 | 0.29 | 0.07 | 0.39 | 0.52 | 0.25 | 0.11 |
| 7 | 106016311 | rs2188278 | -0.45 | -1.898 | 0.027 | -1.668 | 0.009 | 0.121 | 0.121 | 0 | 0.43 | 0.09 | 0.24 | 0.24 | 0.38 | 0.51 |
| 7 | 106018358 | rs179869 | -0.75 | -2.095 | 0.57 | -1.743 | 0.025 | 0.037 | 0.037 | 0.134 | 0.5 | 0.61 | 0.28 | 0.24 | 0.39 | 0.59 |
| 7 | 106020924 | rs7779417 | 0.382 | -1.89 | 0.46 | -1.745 | 0.003 | -0.037 | -0.037 | 0 | 0.16 | 0.03 | 0.09 | 0.2 | 0.11 | 0.27 |
| 7 | 106021254 | rs179883 | -0.91 | -1.71 | 0.555 | -1.798 | 0.024 | 0.043 | 0.043 | 0.015 | 0.51 | 0.64 | 0.3 | 0.24 | 0.39 | 0.64 |
| 7 | 106021572 | rs120432 | 0.315 | -1.4 | 0.275 | -1.697 | -0.031 | -0.131 | -0.131 | 0 | 0.3 | 0.47 | 0.19 | 0.04 | 0.26 | 0.28 |
| 7 | 106023839 | rs179892 | -1.416 | -1.036 | 0.49 | -1.387 | -0.035 | -0.033 | -0.033 | 0.702 | 0.29 | 0.54 | 0.15 | 0.2 | 0.19 | 0.37 |
| 7 | 106024939 | rs17153016 | -0.35 | -0.488 | 0.645 | -1.082 | 0.009 | -0.008 | -0.008 | 0 | 0.2 | 0.1 | 0.12 | 0.2 | 0.15 | 0.3 |

| | | | | | | | | | | | | | | | | |
|---|---|---|---|---|---|---|---|---|---|---|---|---|---|---|---|---|
| 7 | 106039310 | rs179898 | -1.215 | -0.144 | 1.233 | -0.938 | 0.094 | 0.051 | 0.051 | 0.421 | 0.06 | 0.5 | 0.04 | 0 | 0.03 | 0.07 |
| 7 | 106052506 | rs4730173 | 0.31 | 0.052 | -0.662 | -0.858 | 0 | -0.042 | -0.042 | 0 | 0.62 | 0.37 | 0.78 | 0.94 | 0.68 | 0.65 |
| 7 | 106054927 | rs6966889 | -0.819 | 0.576 | -0.84 | 0.03 | 0.002 | 0.008 | 0.008 | 0 | 0.21 | 0.14 | 0.22 | 0.13 | 0.21 | 0.2 |
| 7 | 106055220 | rs2539828 | -1.423 | 0.593 | 0.196 | 1.041 | 0.01 | 0.254 | 0.254 | 0.32 | 0.18 | 0.45 | 0.37 | 0.39 | 0.19 | 0.29 |
| 7 | 106055504 | rs2893638 | -1.363 | 0.713 | 0.579 | 0.624 | 0.021 | 0.022 | 0.022 | 0.41 | 0.17 | 0.53 | 0.17 | 0.22 | 0.14 | 0.19 |
| 7 | 106062157 | rs11772805 | -0.697 | 0.162 | -0.2 | -0.065 | 0.017 | 0.043 | 0.043 | 0 | 0.15 | 0.06 | 0.05 | 0.04 | 0.12 | 0.07 |
| 7 | 106063668 | rs11981468 | -0.654 | 0.11 | -0.226 | -0.092 | 0.016 | 0.046 | 0.046 | 0 | 0.14 | 0.05 | 0.05 | 0.03 | 0.12 | 0.07 |
| 7 | 106064558 | rs2704976 | -0.902 | 0.182 | -1.414 | -0.671 | -0.112 | 0.06 | 0.06 | 0 | 0.67 | 0.92 | 0.57 | 0.81 | 0.59 | 0.74 |
| 7 | 106066658 | rs2704973 | 0.206 | 0.225 | -0.681 | -0.439 | 0.018 | 0.094 | 0.094 | 0 | 0.51 | 0.71 | 0.44 | 0.72 | 0.46 | 0.62 |
| 7 | 106068163 | rs9641363 | -0.065 | -0.089 | 0.355 | -0.711 | -0.011 | 0.048 | 0.048 | 0 | 0.55 | 0.33 | 0.49 | 0.8 | 0.53 | 0.68 |
| 7 | 106070167 | rs17153112 | -0.323 | -0.323 | 0.615 | -0.591 | 0.017 | -0.023 | -0.023 | 0 | 0.05 | 0.18 | 0.05 | 0 | 0.05 | 0.06 |
| 7 | 106083136 | rs2160062 | 0.646 | -0.038 | 0.221 | -0.74 | -0.019 | 0.027 | 0.027 | 0 | 0.05 | 0.01 | 0.04 | 0 | 0.06 | 0.01 |
| 7 | 106088278 | rs1024761 | 0.844 | -0.537 | 0.101 | -1.284 | -0.104 | 0.181 | 0.181 | 0.006 | 0.33 | 0.06 | 0.42 | 0.06 | 0.42 | 0.26 |
| 7 | 106089135 | rs987595 | -0.057 | -1.017 | 0.969 | -1.573 | 0 | -0.047 | -0.047 | 0 | 0.5 | 0.45 | 0.51 | 0.11 | 0.52 | 0.38 |
| 7 | 106089343 | rs2011125 | -0.955 | -2.034 | 0.076 | -1.952 | -0.062 | -0.03 | -0.03 | 0.87 | 0.75 | 0.89 | 0.63 | 0.27 | 0.64 | 0.51 |
| 7 | 106089978 | rs7798009 | 0.787 | -2.013 | 0.548 | -1.931 | 0.018 | 0.026 | 0.026 | 0 | 0.14 | 0.09 | 0.07 | 0.05 | 0.1 | 0.11 |
| 7 | 106102201 | rs7796310 | 0.35 | -2.041 | 0.886 | -1.931 | 0 | 0.018 | 0.018 | 0 | 0.53 | 0.61 | 0.53 | 0.26 | 0.53 | 0.39 |
| 7 | 106106606 | rs7785060 | 0.592 | -2.038 | 0.664 | -1.99 | 0.003 | 0.071 | 0.071 | 0 | 0.51 | 0.37 | 0.52 | 0.24 | 0.53 | 0.4 |
| 7 | 106107315 | rs10240320 | -0.701 | -2.044 | 0.33 | -1.893 | -0.069 | -0.036 | -0.036 | 0 | 0.73 | 0.89 | 0.63 | 0.28 | 0.63 | 0.49 |
| 7 | 106110799 | rs9986860 | 0.212 | -2.104 | 0.334 | -1.83 | 0.031 | 0.063 | 0.063 | 0 | 0.21 | 0.08 | 0.09 | 0.04 | 0.12 | 0.11 |
| 7 | 106117962 | rs6466117 | -0.088 | -2.043 | 0.702 | -1.73 | -0.033 | 0.006 | 0.006 | 0.13 | 0.17 | 0.25 | 0.08 | 0.18 | 0.09 | 0.15 |
| 7 | 106122462 | rs2392845 | 0.795 | -1.852 | 0.89 | -1.916 | 0 | -0.045 | -0.045 | 0.185 | 0.47 | 0.48 | 0.49 | 0.13 | 0.48 | 0.38 |
| 7 | 106125847 | rs17288359 | 0.292 | -1.843 | 0.882 | -1.879 | 0.018 | 0.002 | 0.002 | 0.037 | 0.21 | 0.15 | 0.09 | 0.15 | 0.09 | 0.12 |
| 7 | 106128329 | rs7809421 | -0.113 | -1.834 | 0.722 | -1.9 | 0.004 | -0.002 | -0.002 | 0 | 0.68 | 0.63 | 0.59 | 0.28 | 0.58 | 0.48 |
| 7 | 106129804 | rs2392847 | 1.092 | -1.757 | 0.877 | -2.106 | -0.026 | 0.137 | 0.137 | 0.037 | 0.45 | 0.23 | 0.48 | 0.13 | 0.48 | 0.37 |
| 7 | 106132346 | rs13438730 | 0.705 | -1.536 | 0.634 | -2.277 | -0.014 | -0.069 | -0.069 | 1.595 | 0.12 | 0.28 | 0.08 | 0 | 0.09 | 0.05 |
| 7 | 106147658 | rs17288506 | 2.083 | -1.159 | -0.006 | -2.317 | -0.041 | 0.107 | 0.107 | 1.23 | 0.35 | 0.12 | 0.39 | 0.15 | 0.37 | 0.31 |
| 7 | 106149219 | rs17153258 | -0.201 | -1.361 | 0.455 | -2.435 | -0.033 | -0.133 | -0.133 | 0 | 0.2 | 0.43 | 0.13 | 0 | 0.17 | 0.1 |
| 7 | 106150400 | rs17153267 | 1.111 | -1.401 | -0.199 | -2.282 | 0.001 | 0 | 0 | 2.238 | 0.17 | 0.17 | 0.09 | 0.11 | 0.08 | 0.1 |
| 7 | 106152687 | rs4140724 | -1.028 | -0.81 | -0.689 | -2.037 | -0.072 | -0.076 | -0.076 | 0 | 0.85 | 0.98 | 0.7 | 0.26 | 0.74 | 0.64 |
| 7 | 106156780 | rs6975426 | -0.149 | -1.002 | -0.49 | -2.057 | -0.002 | -0.011 | -0.011 | 0 | 0.36 | 0.68 | 0.23 | 0.11 | 0.24 | 0.18 |
| 7 | 106157161 | rs6958292 | 0.831 | -0.164 | -0.365 | -1.374 | -0.013 | 0.002 | 0.002 | 0 | 0.71 | 0.79 | 0.62 | 0.25 | 0.61 | 0.49 |
| 7 | 106164316 | rs17367541 | -0.051 | -0.327 | -0.468 | -1.615 | -0.002 | 0.001 | 0.001 | 0 | 0.14 | 0.1 | 0.09 | 0.01 | 0.16 | 0.14 |
| 7 | 106175920 | rs6965078 | 1.026 | -0.531 | -0.64 | -1.615 | -0.007 | 0.019 | 0.019 | 0 | 0.38 | 0.33 | 0.47 | 0.2 | 0.43 | 0.27 |
| 7 | 106178454 | rs6969980 | 0.367 | -0.796 | -0.484 | -1.6 | -0.001 | 0.03 | 0.03 | 0 | 0.49 | 0.61 | 0.53 | 0.23 | 0.48 | 0.3 |
| 7 | 106183083 | rs6466121 | 0.021 | -0.726 | -1 | -1.705 | -0.118 | -0.028 | -0.028 | 0.038 | 0.72 | 0.9 | 0.61 | 0.23 | 0.53 | 0.43 |
| 7 | 106184796 | rs17153276 | 0.497 | -0.747 | -0.852 | -1.675 | -0.02 | -0.008 | -0.008 | 0 | 0.12 | 0.25 | 0.06 | 0.05 | 0.07 | 0.04 |
| 7 | 106189195 | rs1859747 | 0.16 | 0.031 | -1.808 | -1.251 | -0.115 | -0.045 | -0.045 | 0.817 | 0.72 | 0.91 | 0.61 | 0.26 | 0.54 | 0.47 |
| 7 | 106194267 | rs6466122 | 0.594 | 0.251 | -1.529 | -1.124 | -0.003 | 0.016 | 0.016 | 0 | 0.42 | 0.38 | 0.47 | 0.19 | 0.42 | 0.33 |
| 7 | 106194724 | rs6951829 | 1.049 | 0.472 | -1.567 | -0.987 | -0.005 | 0.023 | 0.023 | 0 | 0.36 | 0.28 | 0.43 | 0.19 | 0.35 | 0.25 |
| 7 | 106206379 | rs12536313 | 0.649 | 1.567 | -1.579 | -0.115 | -0.081 | -0.036 | -0.036 | 0.753 | 0.75 | 0.9 | 0.66 | 0.23 | 0.61 | 0.47 |
| 7 | 106210542 | rs17290370 | 0.073 | 1.705 | -1.613 | -0.145 | -0.101 | -0.093 | -0.093 | 0.794 | 0.84 | 0.99 | 0.68 | 0.24 | 0.71 | 0.6 |
| 7 | 106210680 | rs6962878 | 0.058 | 0.005 | -1.454 | -1.203 | 0.046 | 0.05 | 0.05 | 0.009 | 0.43 | 0.64 | 0.23 | 0.07 | 0.24 | 0.27 |
| 7 | 106212359 | rs881358 | 0.041 | 0.141 | -1.373 | -0.805 | -0.01 | -0.005 | -0.005 | 0 | 0.16 | 0.15 | 0.36 | 0.17 | 0.25 | 0.21 |
| 7 | 106214257 | rs17369135 | -0.891 | -0.323 | -1.601 | -1.139 | -0.009 | 0.002 | 0.002 | 0.786 | 0.29 | 0.2 | 0.44 | 0.22 | 0.38 | 0.37 |
| 7 | 106216583 | rs2108237 | 0.174 | -0.364 | -1.277 | -1.133 | -0.02 | -0.01 | -0.01 | 0.525 | 0.17 | 0.14 | 0.37 | 0.17 | 0.29 | 0.27 |
| 7 | 106221448 | rs997803 | -0.221 | -0.594 | -0.754 | -1.218 | 0.008 | 0.025 | 0.025 | 0.076 | 0.19 | 0.32 | 0.36 | 0.21 | 0.26 | 0.29 |
| 7 | 106222328 | rs10258803 | -0.222 | -1.236 | -0.606 | -1.665 | -0.002 | -0.017 | -0.017 | 1.585 | 0.14 | 0.42 | 0.1 | 0.05 | 0.11 | 0.07 |
| 7 | 106229098 | rs11769222 | 1.025 | -1.165 | -1.177 | -1.637 | 0.012 | 0.054 | 0.054 | 0 | 0.11 | 0.02 | 0.07 | 0 | 0.09 | 0.06 |
| 7 | 106230778 | rs10486012 | 1.53 | -1.833 | -1.392 | -1.934 | 0.053 | 0.053 | 0.053 | 0 | 0.23 | 0.11 | 0.07 | 0.03 | 0.06 | 0.18 |
| 7 | 106234010 | rs41805 | -1.08 | -1.277 | -1.443 | -1.714 | -0.126 | -0.024 | -0.024 | 0.212 | 0.67 | 0.94 | 0.62 | 0.25 | 0.55 | 0.55 |
| 7 | 106241491 | rs41813 | 0.55 | -0.901 | -1.128 | -1.508 | -0.002 | -0.002 | -0.002 | 0.063 | 0.15 | 0.18 | 0.35 | 0.19 | 0.25 | 0.25 |
| 7 | 106243554 | rs17153336 | 0.236 | -0.85 | -1.155 | -1.775 | -0.003 | -0.017 | -0.017 | 0 | 0.42 | 0.38 | 0.15 | 0.09 | 0.2 | 0.28 |
| 7 | 106247144 | rs11764334 | 0.291 | -0.507 | -1.242 | -1.546 | -0.028 | -0.068 | -0.068 | 0.009 | 0.46 | 0.47 | 0.23 | 0.05 | 0.24 | 0.28 |
| 7 | 106251521 | rs41820 | 0.385 | 0.146 | -1.608 | -1.101 | -0.037 | -0.036 | -0.036 | 0 | 0.15 | 0.07 | 0.37 | 0.21 | 0.29 | 0.23 |
| 7 | 106251899 | rs41821 | 0.634 | 0.055 | -1.579 | -1.1 | -0.028 | -0.033 | -0.033 | 0.01 | 0.14 | 0.07 | 0.36 | 0.2 | 0.25 | 0.21 |
| 7 | 106257023 | rs41826 | -0.622 | 1.191 | -1.511 | 0.187 | -0.06 | -0.088 | -0.088 | 0.012 | 0.87 | 0.99 | 0.79 | 0.55 | 0.83 | 0.75 |
| 7 | 106257186 | rs41827 | -0.465 | 0.023 | -1.179 | -0.642 | -0.005 | 0.026 | 0.026 | 0 | 0.6 | 0.75 | 0.61 | 0.25 | 0.51 | 0.51 |
| 7 | 106259029 | rs17311264 | -0.711 | 0.186 | -1.17 | -0.494 | 0 | 0.003 | 0.003 | 0 | 0.22 | 0.17 | 0.16 | 0.05 | 0.21 | 0.13 |
| 7 | 106260226 | rs7795118 | -0.985 | 0.084 | -1.281 | -0.399 | 0 | 0.008 | 0.008 | 0 | 0.25 | 0.23 | 0.26 | 0.17 | 0.27 | 0.23 |
| 7 | 106260911 | rs41829 | 0.103 | 0.08 | -1.628 | 0.132 | -0.007 | -0.039 | -0.039 | 0 | 0.78 | 0.85 | 0.75 | 0.49 | 0.78 | 0.67 |
| 7 | 106266814 | rs41837 | 0.742 | -0.918 | -1.76 | -0.462 | 0.032 | -0.041 | -0.041 | 0 | 0.61 | 0.71 | 0.67 | 0.48 | 0.71 | 0.57 |
| 7 | 106267355 | rs41838 | -0.094 | -0.734 | -1.974 | -0.353 | -0.011 | -0.093 | -0.093 | 0.417 | 0.76 | 0.89 | 0.74 | 0.49 | 0.78 | 0.64 |
| 7 | 106275700 | rs2108239 | 0.646 | 0.219 | -2.443 | -0.598 | -0.037 | 0.004 | 0.004 | 0 | 0.26 | 0.12 | 0.26 | 0.24 | 0.38 | 0.29 |
| 7 | 106283830 | rs2057904 | -0.718 | 0.383 | -2.447 | -0.553 | -0.031 | 0.001 | 0.001 | 0 | 0.73 | 0.88 | 0.73 | 0.73 | 0.61 | 0.7 |
| 7 | 106287779 | rs12705380 | 0.706 | -0.453 | -2.914 | -1.272 | -0.056 | 0.053 | 0.053 | 0.248 | 0.38 | 0.14 | 0.55 | 0.73 | 0.4 | 0.6 |
| 7 | 106290520 | rs10225065 | -1.205 | -0.407 | -2.873 | -1.215 | -0.031 | 0.002 | 0.002 | 0 | 0.73 | 0.87 | 0.73 | 0.73 | 0.61 | 0.7 |
| 7 | 106291118 | rs17507636 | 1.224 | -0.021 | -2.788 | -0.773 | -0.019 | -0.005 | -0.005 | 0 | 0.25 | 0.09 | 0.35 | 0.24 | 0.26 | 0.26 |
| 7 | 106299404 | rs2190093 | 0.422 | 0.405 | -2.606 | -0.411 | -0.026 | -0.048 | -0.048 | 0 | 0.12 | 0.05 | 0.2 | 0.51 | 0.14 | 0.36 |

| | | | | | | | | | | | | | | | | |
|---|---|---|---|---|---|---|---|---|---|---|---|---|---|---|---|---|
| 7 | 106299792 | rs2286149 | 0.422 | 0.405 | -2.606 | -0.411 | -0.026 | -0.048 | -0.048 | 0 | 0.12 | 0.05 | 0.2 | 0.51 | 0.14 | 0.36 |
| 7 | 106302691 | rs342182 | 0.129 | 0.166 | -2.631 | -1.011 | -0.05 | 0.212 | 0.212 | 0.941 | 0.48 | 0.12 | 0.58 | 0.77 | 0.44 | 0.62 |
| 7 | 106306401 | rs342184 | 0.339 | 0.494 | -2.039 | -0.908 | -0.029 | 0.024 | 0.024 | 0 | 0.67 | 0.78 | 0.69 | 0.74 | 0.49 | 0.55 |
| 7 | 106309643 | rs342189 | 0.242 | 0.47 | -1.659 | -0.891 | 0.002 | -0.021 | -0.021 | 0.425 | 0.95 | 0.91 | 0.96 | 1 | 0.91 | 0.97 |
| 7 | 106315711 | rs342197 | 0.826 | 1.199 | -0.82 | -0.911 | 0.049 | -0.02 | -0.02 | 0.664 | 0.7 | 0.63 | 0.62 | 0.75 | 0.52 | 0.59 |
| 7 | 106322818 | rs6951084 | 0.742 | 1.125 | -0.707 | -0.99 | 0.008 | -0.005 | -0.005 | 0 | 0.69 | 0.68 | 0.62 | 0.76 | 0.5 | 0.6 |
| 7 | 106337250 | rs342240 | -0.268 | 1.199 | -0.654 | -0.999 | -0.002 | 0.001 | 0.001 | 0 | 0.25 | 0.18 | 0.33 | 0.24 | 0.41 | 0.37 |
| 7 | 106349163 | rs17153388 | 1.506 | 0.065 | -1.284 | -0.526 | -0.035 | -0.065 | -0.065 | 0 | 0.31 | 0.08 | 0.44 | 0.35 | 0.32 | 0.28 |
| 7 | 106356646 | rs10274553 | 0.098 | 0.032 | -0.955 | -0.509 | 0.04 | -0.028 | -0.028 | 0 | 0.68 | 0.59 | 0.62 | 0.76 | 0.5 | 0.6 |
| 7 | 106369026 | rs2107910 | 0.183 | -1.047 | -0.325 | -0.219 | -0.001 | -0.002 | -0.002 | 4.121 | 0.34 | 0.29 | 0.15 | 0.41 | 0.18 | 0.32 |
| 7 | 106377328 | rs342301 | -0.083 | -0.475 | -0.092 | -0.108 | -0.001 | -0.019 | -0.019 | 0 | 0.37 | 0.43 | 0.28 | 0.26 | 0.39 | 0.45 |
| 7 | 106388622 | rs11975816 | 0.695 | -0.575 | -0.784 | 0.11 | 0.012 | 0.047 | 0.047 | 0 | 0.49 | 0.4 | 0.62 | 0.3 | 0.52 | 0.26 |
| 7 | 106390179 | rs17153394 | 0.333 | -0.421 | -1.005 | 0.098 | 0.019 | -0.032 | -0.032 | 0.044 | 0.12 | 0.05 | 0.07 | 0.43 | 0.08 | 0.3 |
| 7 | 106390395 | rs6964378 | -1.991 | 0.09 | -0.911 | -0.183 | -0.002 | 0.004 | 0.004 | 0 | 0.85 | 0.89 | 0.91 | 0.9 | 0.84 | 0.86 |
| 7 | 106391804 | rs7803053 | 0.309 | 0.357 | -0.936 | -0.072 | -0.076 | 0.011 | 0.011 | 0 | 0.41 | 0.15 | 0.56 | 0.32 | 0.45 | 0.25 |
| 7 | 106398153 | rs10953517 | -0.126 | 0.371 | -1.472 | -0.044 | -0.024 | 0.079 | 0.079 | 0 | 0.46 | 0.15 | 0.6 | 0.28 | 0.51 | 0.28 |
| 7 | 106399168 | rs10235709 | -0.015 | 0.619 | -0.967 | 0.204 | -0.004 | 0.006 | 0.006 | 0 | 0.41 | 0.37 | 0.44 | 0.26 | 0.41 | 0.28 |
| 7 | 106399882 | rs10953518 | -1.329 | 0.753 | -1.057 | 0.367 | 0 | 0.019 | 0.019 | 0.256 | 0.11 | 0.06 | 0.14 | 0.11 | 0.11 | 0.22 |
| 7 | 106400535 | rs4517053 | -0.529 | 0.44 | -0.715 | 0.504 | -0.012 | 0.033 | 0.033 | 0 | 0.21 | 0.29 | 0.36 | 0.46 | 0.16 | 0.35 |
| 7 | 106401855 | rs6957683 | -0.844 | 0.835 | -0.62 | -0.348 | -0.002 | -0.003 | -0.003 | 0 | 0.41 | 0.61 | 0.61 | 0.56 | 0.44 | 0.53 |
| 7 | 106404514 | rs6973049 | -0.854 | 0.791 | -0.456 | -0.387 | -0.006 | -0.011 | -0.011 | 0 | 0.41 | 0.71 | 0.61 | 0.56 | 0.44 | 0.53 |
| 7 | 106407117 | rs1496140 | -0.854 | 0.826 | -0.223 | -0.507 | 0.023 | 0.07 | 0.07 | 0.209 | 0.28 | 0.43 | 0.54 | 0.55 | 0.36 | 0.43 |
| 7 | 106409897 | rs7787898 | -0.899 | 1.1 | -0.579 | -0.809 | 0.003 | -0.024 | -0.024 | 1.143 | 0.4 | 0.43 | 0.4 | 0.22 | 0.46 | 0.47 |
| 7 | 106410069 | rs7804859 | -0.288 | 1.115 | -0.574 | -0.798 | 0.032 | -0.084 | -0.084 | 3.178 | 0.17 | 0.37 | 0.25 | 0.03 | 0.24 | 0.17 |
| 7 | 106410811 | rs6959745 | 0.024 | 1.762 | -0.434 | -1.483 | -0.001 | -0.001 | -0.001 | 0 | 0.25 | 0.27 | 0.35 | 0.47 | 0.22 | 0.37 |
| 7 | 106411691 | rs17477143 | -0.588 | 1.404 | -0.385 | -1.618 | 0.078 | 0.005 | 0.005 | 0 | 0.14 | 0.37 | 0.24 | 0.08 | 0.23 | 0.12 |
| 7 | 106411858 | rs17477177 | -0.008 | 1.149 | -1.053 | -1.587 | -0.02 | 0.03 | 0.03 | 0 | 0.22 | 0.06 | 0.16 | 0.14 | 0.22 | 0.3 |
| 7 | 106422717 | rs2132462 | 0.525 | 1.377 | -0.821 | -1.678 | 0.001 | -0.001 | -0.001 | 0.516 | 0.29 | 0.31 | 0.27 | 0.24 | 0.39 | 0.32 |
| 7 | 106431237 | rs12535670 | -0.34 | 1.1 | -0.672 | -1.883 | -0.012 | 0.197 | 0.197 | 0.32 | 0.2 | 0.44 | 0.41 | 0.51 | 0.16 | 0.31 |
| 7 | 106431520 | rs7801016 | -0.656 | 0.99 | -0.7 | -2.035 | 0.059 | 0.143 | 0.143 | 1.747 | 0.52 | 0.76 | 0.72 | 0.81 | 0.64 | 0.64 |
| 7 | 106433166 | rs6974881 | -0.566 | 0.897 | -0.678 | -2.092 | 0.071 | 0.142 | 0.142 | 1.739 | 0.52 | 0.73 | 0.72 | 0.82 | 0.66 | 0.64 |
| 7 | 106433598 | rs10215920 | -0.248 | 0.792 | -0.573 | -2.108 | 0.024 | 0.041 | 0.041 | 0.849 | 0.47 | 0.38 | 0.62 | 0.81 | 0.65 | 0.58 |
| 7 | 106440032 | rs10268373 | 0.734 | 0.96 | -0.982 | -1.981 | -0.004 | -0.008 | -0.008 | 0 | 0.13 | 0.07 | 0.3 | 0.52 | 0.13 | 0.23 |
| 7 | 106440096 | rs6946226 | -0.455 | 0.725 | -1.157 | -2.139 | 0.003 | -0.002 | -0.002 | 0 | 0.34 | 0.3 | 0.46 | 0.66 | 0.29 | 0.48 |
| 7 | 106444790 | rs12113051 | -0.482 | 0.832 | -1.198 | -2.019 | -0.001 | 0 | 0 | 0 | 0.37 | 0.34 | 0.49 | 0.65 | 0.37 | 0.47 |
| 7 | 106446900 | rs4730197 | -0.019 | 0.531 | -1.181 | -1.27 | -0.005 | -0.031 | -0.031 | 0.649 | 0.13 | 0.07 | 0.3 | 0.53 | 0.13 | 0.23 |
| 7 | 106446949 | rs4730198 | -0.155 | 0.593 | -1.167 | -1.196 | -0.003 | 0.072 | 0.072 | 0.022 | 0.18 | 0.26 | 0.4 | 0.53 | 0.15 | 0.28 |
| 7 | 106449124 | rs916686 | 0.563 | 0.728 | -1.331 | -1.661 | 0.016 | 0.013 | 0.013 | 0 | 0.56 | 0.63 | 0.47 | 0.35 | 0.67 | 0.6 |
| 7 | 106453135 | rs10241504 | -0.179 | 1.262 | -1.982 | -1.256 | -0.031 | 0.188 | 0.188 | 1.786 | 0.66 | 0.99 | 0.84 | 0.94 | 0.71 | 0.75 |
| 7 | 106454351 | rs2028009 | -0.025 | 1.43 | -1.192 | -0.838 | 0.005 | -0.014 | -0.014 | 0 | 0.44 | 0.52 | 0.33 | 0.37 | 0.45 | 0.43 |
| 7 | 106459917 | rs6958639 | -0.661 | 1.535 | -1.433 | -1.063 | -0.017 | 0.014 | 0.014 | 0 | 0.31 | 0.18 | 0.26 | 0 | 0.32 | 0.14 |
| 7 | 106466998 | rs42154 | -0.712 | 1.382 | -1.855 | -1.017 | -0.021 | 0.12 | 0.12 | 0 | 0.32 | 0.16 | 0.22 | 0.11 | 0.31 | 0.22 |
| 7 | 106467455 | rs10274710 | -1.098 | 0.937 | -1.455 | 0.06 | -0.005 | 0.067 | 0.067 | 0.358 | 0.11 | 0.2 | 0.26 | 0.59 | 0.1 | 0.31 |
| 7 | 106475927 | rs42162 | 0.485 | 1.249 | -1.62 | 0.359 | -0.033 | -0.021 | -0.021 | 1.079 | 0.6 | 0.79 | 0.7 | 0.48 | 0.56 | 0.63 |
| 7 | 106480649 | rs42168 | 0.237 | 0.759 | -1.052 | -0.376 | -0.003 | -0.051 | -0.051 | 0 | 0.19 | 0.32 | 0.25 | 0.03 | 0.2 | 0.13 |
| 7 | 106482966 | rs4727663 | 0.617 | 0.441 | -0.376 | 0.033 | -0.007 | 0.01 | 0.01 | 0 | 0.4 | 0.44 | 0.46 | 0.46 | 0.37 | 0.5 |
| 7 | 106486974 | rs42172 | -0.435 | -0.046 | -0.814 | -0.639 | 0.001 | 0.003 | 0.003 | 0 | 0.41 | 0.36 | 0.6 | 0.33 | 0.43 | 0.41 |
| 7 | 106489559 | rs185087 | -0.087 | -0.369 | -0.065 | -1.022 | -0.016 | -0.074 | -0.074 | 0.314 | 0.28 | 0.42 | 0.26 | 0.07 | 0.25 | 0.17 |
| 7 | 106491633 | rs193740 | -1.102 | -0.676 | 0.203 | -1.04 | -0.024 | -0.02 | -0.02 | 0 | 0.67 | 0.72 | 0.71 | 0.51 | 0.61 | 0.63 |
| 7 | 106494318 | rs849376 | 0.201 | -0.868 | 0.263 | -0.934 | -0.009 | 0.133 | 0.133 | 0 | 0.09 | 0.17 | 0.2 | 0.32 | 0.08 | 0.08 |
| 7 | 106495809 | rs17153527 | 1.713 | -1.07 | 0.16 | -1.049 | 0.001 | -0.037 | -0.037 | 0.596 | 0.16 | 0.01 | 0.06 | 0.16 | 0.12 | 0.24 |
| 7 | 106496510 | rs849380 | 0.331 | -1.119 | 0.181 | -0.859 | 0.008 | -0.016 | -0.016 | 0.036 | 0.26 | 0.19 | 0.26 | 0.49 | 0.2 | 0.31 |
| 7 | 106499100 | rs849384 | -0.658 | -1.114 | -0.112 | -0.915 | -0.009 | 0.006 | 0.006 | 0 | 0.57 | 0.32 | 0.62 | 0.51 | 0.55 | 0.59 |
| 7 | 106500358 | rs757902 | 0.453 | -1.397 | 0.221 | -0.997 | -0.015 | -0.014 | -0.014 | 0.005 | 0.57 | 0.5 | 0.75 | 0.79 | 0.71 | 0.59 |
| 7 | 106500436 | rs757903 | -0.147 | -1.412 | 0.215 | -0.959 | -0.031 | -0.038 | -0.038 | 0 | 0.66 | 0.58 | 0.83 | 0.8 | 0.75 | 0.68 |
| 7 | 106505057 | rs4730204 | 0.384 | -1.424 | -0.127 | -0.948 | -0.068 | -0.075 | -0.075 | 0 | 0.25 | 0.08 | 0.43 | 0.31 | 0.32 | 0.27 |
| 7 | 106505478 | rs4727666 | 0.733 | -1.528 | -0.013 | -1.019 | -0.065 | -0.107 | -0.107 | 0 | 0.72 | 0.52 | 0.8 | 0.94 | 0.81 | 0.83 |
| 7 | 106508987 | rs849390 | -1.944 | -1.491 | 0.05 | -1 | -0.018 | -0.038 | -0.038 | 0 | 0.92 | 0.86 | 0.96 | 1 | 0.96 | 0.96 |
| 7 | 106518631 | rs6963859 | 0.361 | -1.194 | -0.037 | -1.066 | -0.072 | -0.058 | -0.058 | 0 | 0.23 | 0.08 | 0.43 | 0.29 | 0.31 | 0.27 |
| 7 | 106545228 | rs12536620 | 1.693 | -1.392 | 0.331 | 0.233 | 0.002 | -0.001 | -0.001 | 0 | 0.38 | 0.4 | 0.42 | 0.48 | 0.45 | 0.39 |
| 7 | 106546657 | rs12667819 | 1.74 | -1.421 | 0.394 | 0.243 | -0.002 | 0 | 0 | 0 | 0.38 | 0.38 | 0.42 | 0.47 | 0.45 | 0.44 |
| 7 | 106549902 | rs849412 | -0.222 | -1.433 | 0.413 | 0.202 | -0.029 | -0.051 | -0.051 | 0.935 | 0.18 | 0.34 | 0.17 | 0.06 | 0.13 | 0.13 |
| 7 | 106556805 | rs10226981 | -0.43 | -1.419 | 0.4 | 0.202 | 0.006 | -0.008 | -0.008 | 2.42 | 0.43 | 0.25 | 0.46 | 0.48 | 0.4 | 0.48 |
| 7 | 106565923 | rs849424 | 0.78 | -1.566 | 0.335 | 0.18 | 0.001 | -0.024 | -0.024 | 0.06 | 0.09 | 0.11 | 0.07 | 0.03 | 0.09 | 0.04 |
| 7 | 106568489 | rs17153616 | 1.928 | -1.565 | 0.236 | 0.221 | 0.01 | -0.042 | -0.042 | 0 | 0.17 | 0.01 | 0.05 | 0.16 | 0.11 | 0.19 |
| 7 | 106570828 | rs849428 | -1.415 | -1.453 | 0.535 | 0.122 | -0.021 | -0.041 | -0.041 | 0 | 0.91 | 0.84 | 0.96 | 1 | 0.95 | 0.96 |
| 7 | 106576858 | rs849430 | 0.383 | -1.354 | 0.309 | 0.188 | 0.012 | -0.039 | -0.039 | 0 | 0.1 | 0.2 | 0.08 | 0.03 | 0.09 | 0.04 |
| 7 | 106579678 | rs849434 | 0.379 | -1.565 | 0.369 | 0.341 | 0.006 | 0.009 | 0.009 | 0 | 0.1 | 0.21 | 0.09 | 0.07 | 0.1 | 0.05 |
| 7 | 106580081 | rs6466139 | 0.543 | -1.586 | 0.267 | 0.47 | -0.006 | 0.017 | 0.017 | 0 | 0.72 | 0.55 | 0.62 | 0.69 | 0.71 | 0.76 |

| | | | | | | | | | | | | | | | | |
|---|---|---|---|---|---|---|---|---|---|---|---|---|---|---|---|---|
| 7 | 106586936 | rs17153654 | 0.584 | -0.996 | -0.184 | 0.737 | -0.002 | -0.002 | -0.002 | 0.044 | 0.08 | 0.05 | 0.07 | 0.06 | 0.1 | 0.05 |
| 7 | 106587807 | rs6466140 | -1.311 | -1.263 | 0.068 | 0.592 | -0.008 | -0.045 | -0.045 | 0 | 0.33 | 0.46 | 0.3 | 0.1 | 0.33 | 0.28 |
| 7 | 106587964 | rs711440 | -1.515 | -1.529 | -0.233 | 0.537 | 0.033 | -0.001 | -0.001 | 0 | 0.76 | 0.99 | 0.92 | 0.87 | 0.84 | 0.77 |
| 7 | 106591092 | rs13438464 | 0.426 | -1.213 | 0.011 | 0.513 | 0.008 | -0.071 | -0.071 | 0.312 | 0.16 | 0.28 | 0.15 | 0.01 | 0.2 | 0.15 |
| 7 | 106591776 | rs17480945 | 1.206 | -1.304 | 0.051 | 0.599 | 0.01 | -0.007 | -0.007 | 0.265 | 0.4 | 0.45 | 0.42 | 0.48 | 0.46 | 0.44 |
| 7 | 106593608 | rs10263050 | 0.681 | -1.685 | 0.061 | 0.594 | 0.003 | -0.008 | -0.008 | 0 | 0.15 | 0.25 | 0.15 | 0.07 | 0.19 | 0.15 |
| 7 | 106601566 | rs11771974 | 0.213 | -1.819 | -0.122 | 0.533 | -0.002 | -0.024 | -0.024 | 1.637 | 0.19 | 0.33 | 0.17 | 0.07 | 0.21 | 0.15 |
| 7 | 106606478 | rs17315922 | 0.14 | -1.886 | -0.429 | 0.596 | 0.095 | 0.041 | 0.041 | 0 | 0.39 | 0.09 | 0.22 | 0.21 | 0.26 | 0.37 |
| 7 | 106609103 | rs953184 | 0.738 | -1.836 | -0.286 | 0.597 | 0.005 | -0.035 | -0.035 | 0 | 0.18 | 0.38 | 0.17 | 0.07 | 0.21 | 0.15 |
| 7 | 106612197 | rs6942461 | 1.112 | -1.679 | -0.392 | 0.766 | -0.001 | 0 | 0 | 0.358 | 0.38 | 0.4 | 0.41 | 0.46 | 0.5 | 0.43 |
| 7 | 106619192 | rs13246564 | 1.234 | -1.746 | -1.342 | 0.88 | 0.052 | -0.004 | -0.004 | 0 | 0.23 | 0.01 | 0.08 | 0.16 | 0.13 | 0.27 |
| 7 | 106625544 | rs6961257 | 0.693 | -0.827 | -0.554 | 1.371 | -0.017 | 0.064 | 0.064 | 0 | 0.08 | 0.14 | 0.25 | 0.27 | 0.04 | 0.09 |
| 7 | 106626928 | rs940938 | 0.27 | -0.713 | -0.717 | 0.7 | 0.006 | 0.018 | 0.018 | 0 | 0.76 | 0.82 | 0.78 | 0.92 | 0.77 | 0.77 |
| 7 | 106632113 | rs12530679 | -0.041 | 0.166 | -0.637 | 0.889 | -0.002 | -0.001 | -0.001 | 0 | 0.33 | 0.35 | 0.41 | 0.48 | 0.5 | 0.43 |
| 7 | 106634790 | rs7777867 | -0.027 | -0.429 | -0.202 | 1.234 | -0.034 | 0.061 | 0.061 | 0 | 0.1 | 0.22 | 0.25 | 0.18 | 0.04 | 0.05 |
| 7 | 106642123 | rs7786720 | -0.542 | -0.339 | -0.745 | 0.854 | 0.01 | -0.005 | -0.005 | 0.753 | 0.39 | 0.38 | 0.53 | 0.52 | 0.65 | 0.54 |
| 7 | 106656845 | rs711441 | 1.128 | 0.328 | -0.773 | 0.614 | -0.015 | 0.025 | 0.025 | 0 | 0.56 | 0.68 | 0.57 | 0.56 | 0.5 | 0.53 |
| 7 | 106660723 | rs12113322 | -0.58 | 0.122 | -0.069 | 0.791 | 0.026 | 0.079 | 0.079 | 1.39 | 0.09 | 0.42 | 0.05 | 0.05 | 0.04 | 0.04 |
| 7 | 106661150 | rs849309 | 0.215 | -0.018 | -0.439 | 0.828 | -0.109 | 0.019 | 0.019 | 0 | 0.69 | 0.85 | 0.62 | 0.69 | 0.57 | 0.71 |
| 7 | 106661176 | rs849310 | 1.248 | -0.1 | -0.32 | 0.821 | -0.043 | 0.042 | 0.042 | 0 | 0.56 | 0.78 | 0.57 | 0.58 | 0.49 | 0.51 |
| 7 | 106662083 | rs7798500 | -0.362 | -0.152 | -0.346 | 0.776 | 0.029 | 0.077 | 0.077 | 1.242 | 0.08 | 0.42 | 0.05 | 0.05 | 0.04 | 0.03 |
| 7 | 106663168 | rs849312 | -0.099 | 0.977 | -1.264 | 0.736 | -0.077 | -0.029 | -0.029 | 0.29 | 0.84 | 0.99 | 0.84 | 0.8 | 0.78 | 0.89 |
| 7 | 106663520 | rs12671537 | 1.944 | 0.503 | -1.199 | 1.144 | -0.067 | -0.002 | -0.002 | 0 | 0.41 | 0.02 | 0.38 | 0.29 | 0.38 | 0.45 |
| 7 | 106670288 | rs6946733 | 1.963 | 0.476 | -1.192 | 1.069 | -0.026 | -0.08 | -0.08 | 0 | 0.41 | 0.02 | 0.42 | 0.36 | 0.35 | 0.42 |
| 7 | 106672004 | rs7786279 | -0.478 | 0.935 | -0.654 | 1.105 | 0.013 | -0.028 | -0.028 | 0 | 0.05 | 0.18 | 0.03 | 0 | 0.04 | 0.03 |
| 7 | 106686448 | rs2395833 | 1.794 | 0.847 | -0.464 | 1.191 | 0.052 | -0.016 | -0.016 | 1.452 | 0.45 | 0.24 | 0.44 | 0.45 | 0.37 | 0.43 |
| 7 | 106687838 | rs1544582 | 0.111 | 1.023 | -0.733 | 1.171 | -0.059 | -0.019 | -0.019 | 0 | 0.75 | 0.88 | 0.63 | 0.63 | 0.56 | 0.7 |
| 7 | 106708765 | rs11766415 | 1.862 | 0.457 | -0.889 | 0.812 | -0.126 | -0.109 | -0.109 | 3.22 | 0.39 | 0.02 | 0.47 | 0.38 | 0.41 | 0.48 |
| 7 | 106711398 | rs2536501 | 0.288 | 1.669 | -0.109 | 1.148 | -0.128 | -0.057 | -0.057 | 0 | 0.78 | 0.98 | 0.68 | 0.69 | 0.67 | 0.7 |
| 7 | 106713815 | rs2536503 | -1.932 | 0.795 | 0.496 | 1.287 | -0.016 | 0.041 | 0.041 | 1.571 | 0.16 | 0.7 | 0.12 | 0.21 | 0.12 | 0.06 |
| 7 | 106713996 | rs17340479 | 0.018 | 0.703 | 0.359 | 1.257 | 0.016 | -0.001 | -0.001 | 0 | 0.13 | 0.01 | 0.05 | 0.08 | 0.09 | 0.15 |
| 7 | 106714165 | rs2536505 | -2.106 | 0.734 | 0.446 | 1.201 | -0.006 | 0.01 | 0.01 | 2.359 | 0.17 | 0.78 | 0.13 | 0.21 | 0.12 | 0.06 |
| 7 | 106719536 | rs2299419 | -1.245 | 0.726 | 1.033 | 1.209 | 0.015 | 0.217 | 0.217 | 0.274 | 0.09 | 0.32 | 0.08 | 0.21 | 0.07 | 0.04 |
| 7 | 106723467 | rs11767753 | -0.214 | 0.717 | 0.935 | 1.2 | -0.008 | -0.008 | -0.008 | 0 | 0.14 | 0.2 | 0.06 | 0.08 | 0.1 | 0.15 |
| 7 | 106723926 | rs2707374 | -1.402 | 0.674 | 0.967 | 1.205 | -0.003 | 0.205 | 0.205 | 0 | 0.1 | 0.38 | 0.08 | 0.21 | 0.07 | 0.04 |
| 7 | 106731934 | rs6466156 | 1.185 | 1.097 | 0.916 | 1.22 | 0.003 | 0.035 | 0.035 | 0 | 0.69 | 0.61 | 0.64 | 0.49 | 0.67 | 0.64 |
| 7 | 106733363 | rs1860805 | -0.219 | 1.127 | 1.066 | 1.336 | -0.01 | -0.01 | -0.01 | 0 | 0.14 | 0.18 | 0.06 | 0.08 | 0.1 | 0.15 |
| 7 | 106742048 | rs3735613 | 2.234 | 1.161 | 0.954 | 1.324 | -0.089 | -0.02 | -0.02 | 0 | 0.49 | 0.03 | 0.54 | 0.4 | 0.53 | 0.48 |
| 7 | 106768356 | rs9656135 | -1.071 | 1.252 | 1.247 | 1.465 | 0.038 | 0.217 | 0.217 | 0 | 0.08 | 0.37 | 0.07 | 0.18 | 0.07 | 0.03 |
| 7 | 106779840 | rs1860803 | 1.325 | 1.345 | 0.529 | 1.25 | -0.007 | -0.021 | -0.021 | 0.827 | 0.61 | 0.27 | 0.56 | 0.48 | 0.6 | 0.61 |
| 7 | 106792332 | rs11767623 | 0.723 | 1.037 | 0.852 | 1.133 | 0 | 0.013 | 0.013 | 0 | 0.81 | 0.9 | 0.89 | 0.92 | 0.81 | 0.81 |
| 7 | 106792554 | rs12705406 | -0.448 | 1.012 | 0.827 | 1.113 | -0.042 | 0.046 | 0.046 | 0 | 0.91 | 0.99 | 0.93 | 0.98 | 0.87 | 0.94 |
| 7 | 106796977 | rs257372 | -1.275 | 0.973 | 0.814 | 1.1 | 0.001 | 0.016 | 0.016 | 0 | 0.19 | 0.1 | 0.11 | 0.08 | 0.19 | 0.19 |
| 7 | 106801088 | rs257377 | -1.123 | 0.981 | 0.667 | 1.103 | -0.042 | 0.028 | 0.028 | 0 | 0.18 | 0.01 | 0.1 | 0.08 | 0.18 | 0.18 |
| 7 | 106808480 | rs257387 | -0.97 | 1.088 | 0.716 | 1.152 | -0.045 | 0.023 | 0.023 | 0 | 0.18 | 0.01 | 0.1 | 0.09 | 0.18 | 0.18 |
| 7 | 106826439 | rs7794965 | -1.07 | 1.146 | 0.749 | 1.195 | -0.012 | -0.014 | -0.014 | 0.36 | 0.18 | 0.06 | 0.12 | 0.16 | 0.2 | 0.18 |
| 7 | 106870746 | rs2301801 | -0.371 | 1.446 | 0.526 | 1.238 | -0.069 | -0.048 | -0.048 | 0 | 0.19 | 0.03 | 0.25 | 0.3 | 0.27 | 0.14 |
| 7 | 106875400 | rs2237664 | -1.214 | 1.274 | 1.3 | 1.416 | -0.018 | 0.141 | 0.141 | 0.254 | 0.1 | 0.36 | 0.09 | 0.17 | 0.06 | 0.11 |
| 7 | 106897237 | rs17349904 | -0.758 | 1.292 | 1.071 | 1.422 | -0.059 | -0.06 | -0.06 | 0 | 0.18 | 0.01 | 0.12 | 0.16 | 0.2 | 0.19 |
| 7 | 106900341 | rs10264687 | -2.674 | 1.146 | 1.07 | 1.396 | -0.113 | 0.053 | 0.053 | 0.232 | 0.53 | 0.97 | 0.52 | 0.64 | 0.54 | 0.45 |
| 7 | 106904218 | rs11766060 | 0.298 | 1.181 | 1.072 | 1.256 | 0.013 | -0.055 | -0.055 | 0.075 | 0.09 | 0 | 0.03 | 0.15 | 0.06 | 0.13 |
| 7 | 106944563 | rs1548524 | -0.387 | 1.233 | 1.139 | 1.272 | -0.011 | -0.012 | -0.012 | 0 | 0.19 | 0.06 | 0.12 | 0.16 | 0.2 | 0.19 |
| 7 | 106954721 | rs10241552 | -1.3 | 1.205 | 1.258 | 1.288 | -0.018 | 0.137 | 0.137 | 0.045 | 0.1 | 0.33 | 0.09 | 0.18 | 0.06 | 0.11 |
| 7 | 106973534 | rs982447 | -0.385 | 1.257 | 0.644 | 1.277 | -0.034 | -0.024 | -0.024 | 0 | 0.2 | 0.1 | 0.26 | 0.31 | 0.27 | 0.15 |
| 7 | 106977613 | rs7783181 | -0.614 | 1.324 | 0.769 | 1.228 | 0 | 0 | 0 | 0 | 0.2 | 0.22 | 0.14 | 0.16 | 0.2 | 0.19 |
| 7 | 106984288 | rs12538769 | -0.367 | 1.35 | 0.317 | 1.135 | -0.008 | 0.024 | 0.024 | 0 | 0.19 | 0.08 | 0.12 | 0.02 | 0.2 | 0.18 |
| 7 | 107002806 | rs2269970 | -2.136 | 1.326 | 0.098 | 1.199 | -0.121 | 0.033 | 0.033 | 0.171 | 0.52 | 0.98 | 0.51 | 0.64 | 0.54 | 0.42 |
| 7 | 107003177 | rs7804398 | -2.256 | 1.33 | 0.1 | 1.201 | -0.129 | 0.021 | 0.021 | 0.274 | 0.53 | 0.98 | 0.52 | 0.64 | 0.54 | 0.44 |
| 7 | 107029879 | rs2107763 | -2.169 | 1.522 | 0.185 | 1.259 | -0.12 | -0.025 | -0.025 | 0 | 0.51 | 0.98 | 0.49 | 0.6 | 0.54 | 0.41 |
| 7 | 107050221 | rs12537785 | -0.405 | 1.214 | 0.573 | 1.254 | -0.009 | 0.027 | 0.027 | 0 | 0.18 | 0.09 | 0.12 | 0.01 | 0.2 | 0.18 |
| 7 | 107066030 | rs6979121 | -2.243 | 1.257 | 0.473 | 1.423 | -0.099 | -0.092 | -0.092 | 0.045 | 0.46 | 0.98 | 0.69 | 0.5 | 0.5 | 0.37 |
| 7 | 107091321 | rs10276321 | -1.754 | 1.071 | 0.949 | 1.519 | -0.028 | -0.027 | -0.027 | 0.065 | 0.37 | 0.25 | 0.38 | 0.46 | 0.48 | 0.3 |
| 7 | 107096144 | rs4727681 | -1.311 | 1.054 | 1.095 | 1.516 | -0.006 | 0.001 | 0.001 | 0 | 0.25 | 0.22 | 0.3 | 0.32 | 0.32 | 0.16 |
| 7 | 107132241 | rs2712197 | -0.071 | 1.108 | 0.907 | 1.304 | -0.02 | -0.069 | -0.069 | 0.181 | 0.16 | 0.01 | 0.2 | 0.31 | 0.17 | 0.13 |
| 7 | 107135612 | rs1015411 | -1.32 | 1.057 | 1.006 | 1.369 | -0.012 | 0.001 | 0.001 | 0 | 0.24 | 0.19 | 0.3 | 0.32 | 0.32 | 0.15 |
| 7 | 107145564 | rs2712201 | -1.335 | 1.031 | 1.003 | 1.369 | -0.006 | 0.001 | 0.001 | 0 | 0.24 | 0.22 | 0.3 | 0.32 | 0.32 | 0.15 |
| 7 | 107184466 | rs10267907 | -1.331 | 1.032 | 0.994 | 1.37 | -0.011 | 0.001 | 0.001 | 0 | 0.24 | 0.19 | 0.3 | 0.32 | 0.32 | 0.15 |
| 7 | 107196723 | rs763386 | 0.28 | 1.042 | 0.95 | 1.389 | -0.051 | -0.065 | -0.065 | 0.633 | 0.13 | 0.01 | 0.09 | 0.15 | 0.16 | 0.15 |
| 7 | 107200319 | rs2253146 | -1.401 | 0.728 | 1.144 | 1.365 | -0.002 | 0 | 0 | 0 | 0.25 | 0.28 | 0.31 | 0.31 | 0.32 | 0.15 |

| | | | | | | | | | | | | | | | | |
|---|---|---|---|---|---|---|---|---|---|---|---|---|---|---|---|---|
| 7 | 107202502 | rs2237677 | -1.413 | 0.722 | 1.142 | 1.362 | -0.002 | 0 | 0 | 0 | 0.25 | 0.28 | 0.31 | 0.31 | 0.32 | 0.15 |
| 7 | 107210383 | rs11977187 | 1.163 | 0.734 | 0.904 | 1.448 | -0.021 | -0.017 | -0.017 | 0 | 0.76 | 0.84 | 0.7 | 0.64 | 0.67 | 0.84 |
| 7 | 107212966 | rs11761775 | 2.29 | 0.907 | 0.736 | 1.265 | -0.116 | 0.014 | 0.014 | 0.17 | 0.53 | 0.02 | 0.51 | 0.37 | 0.46 | 0.62 |
| 7 | 107216738 | rs10276604 | 1.009 | 1.267 | 1.045 | 1.546 | -0.024 | -0.009 | -0.009 | 0 | 0.76 | 0.89 | 0.71 | 0.68 | 0.68 | 0.85 |
| 7 | 107217020 | rs6957510 | -0.189 | 1.27 | 0.991 | 1.532 | 0.004 | 0.003 | 0.003 | 0 | 0.06 | 0.04 | 0.02 | 0 | 0.05 | 0.02 |
| 7 | 107218033 | rs711442 | 2.305 | 1.272 | 0.989 | 1.538 | -0.116 | 0.013 | 0.013 | 0.169 | 0.53 | 0.02 | 0.5 | 0.37 | 0.46 | 0.62 |
| 7 | 107226470 | rs4727684 | 1.036 | 1.39 | 1.05 | 1.581 | -0.039 | -0.02 | -0.02 | 0 | 0.77 | 0.92 | 0.71 | 0.68 | 0.68 | 0.85 |
| 7 | 107232369 | rs2107713 | 1.332 | 1.411 | 1.268 | 1.577 | 0.001 | -0.001 | -0.001 | 0 | 0.75 | 0.74 | 0.7 | 0.68 | 0.68 | 0.85 |
| 7 | 107243350 | rs10258385 | 1.04 | 1.36 | 1.084 | 1.588 | -0.039 | -0.021 | -0.021 | 0.671 | 0.77 | 0.92 | 0.71 | 0.68 | 0.68 | 0.85 |
| 7 | 107243995 | rs7785962 | 0.137 | 1.408 | 1.086 | 1.639 | -0.08 | -0.065 | -0.065 | 0 | 0.14 | 0.01 | 0.11 | 0.15 | 0.21 | 0.16 |
| 7 | 107249699 | rs10953542 | -1.449 | 0.646 | 2.293 | 2.29 | 0.056 | 0.326 | 0.326 | 2.37 | 0.05 | 0.4 | 0.06 | 0.17 | 0.01 | 0.07 |
| 7 | 107258121 | rs10273733 | 1.477 | 0.743 | 1.385 | 1.667 | -0.018 | -0.008 | -0.008 | 0 | 0.77 | 0.87 | 0.71 | 0.68 | 0.68 | 0.85 |
| 7 | 107260778 | rs2807 | 0.61 | 0.747 | 1.315 | 1.666 | -0.037 | -0.039 | -0.039 | 0.567 | 0.82 | 0.95 | 0.73 | 0.68 | 0.73 | 0.87 |
| 7 | 107260856 | rs2808 | 1.569 | 0.761 | 1.223 | 1.675 | 0 | 0 | 0 | 0.761 | 0.76 | 0.77 | 0.71 | 0.68 | 0.68 | 0.85 |
| 7 | 107294151 | rs2712230 | 0.785 | 0.925 | 0.302 | 0.909 | -0.005 | -0.012 | -0.012 | 0 | 0.8 | 0.8 | 0.76 | 0.47 | 0.73 | 0.8 |
| 7 | 107300340 | rs2712228 | 1.846 | 0.893 | -0.014 | 0.88 | -0.02 | -0.109 | -0.109 | 0 | 0.68 | 0.06 | 0.68 | 0.48 | 0.67 | 0.78 |
| 7 | 107303628 | rs2248465 | 0.995 | 1.339 | 0.325 | 1.156 | 0.013 | 0.059 | 0.059 | 0 | 0.79 | 0.65 | 0.7 | 0.48 | 0.7 | 0.78 |
| 7 | 107306421 | rs7799849 | -1.531 | 1.47 | 0.545 | 0.428 | 0.054 | -0.081 | -0.081 | 0 | 0.12 | 0.4 | 0.13 | 0 | 0.16 | 0.02 |
| 7 | 107307206 | rs2701688 | -1.26 | 2.104 | 0.103 | 0.743 | 0.013 | 0.002 | 0.002 | 0 | 0.22 | 0.25 | 0.22 | 0.24 | 0.28 | 0.23 |
| 7 | 107315124 | rs3817613 | -0.64 | 1.448 | -1.595 | 0.453 | -0.026 | -0.028 | -0.028 | 0 | 0.56 | 0.98 | 0.71 | 0.63 | 0.62 | 0.45 |
| 7 | 107317175 | rs10276013 | -0.553 | 1.699 | -1.374 | 0.731 | -0.037 | -0.05 | -0.05 | 0 | 0.38 | 0.79 | 0.44 | 0.49 | 0.43 | 0.32 |
| 7 | 107327852 | rs2712210 | 1.002 | 0.954 | -1.965 | 0.285 | -0.005 | -0.092 | -0.092 | 0.166 | 0.27 | 0.04 | 0.28 | 0.35 | 0.26 | 0.45 |
| 7 | 107328814 | rs1989837 | 0.163 | 0.873 | -1.293 | -0.126 | -0.009 | -0.056 | -0.056 | 0 | 0.11 | 0.19 | 0.03 | 0.01 | 0.06 | 0.06 |
| 7 | 107334026 | rs2237679 | 1.296 | 0.817 | -1.724 | -0.257 | -0.032 | -0.109 | -0.109 | 0.639 | 0.26 | 0.02 | 0.28 | 0.35 | 0.27 | 0.47 |
| 7 | 107334661 | rs2072065 | -0.563 | 1.389 | -1.256 | -0.935 | 0.003 | -0.008 | -0.008 | 0 | 0.42 | 0.47 | 0.47 | 0.64 | 0.44 | 0.39 |
| 7 | 107339911 | rs982915 | -0.738 | 1.228 | -0.819 | -1.028 | 0.002 | -0.067 | -0.067 | 0 | 0.7 | 0.66 | 0.76 | 0.99 | 0.71 | 0.86 |
| 7 | 107344033 | rs982663 | -0.584 | 1.171 | -0.531 | -0.999 | 0.002 | -0.006 | -0.006 | 0.724 | 0.42 | 0.45 | 0.47 | 0.64 | 0.44 | 0.39 |
| 7 | 107359384 | rs12667481 | 1.192 | 1.389 | -0.952 | -0.253 | -0.072 | -0.104 | -0.104 | 0 | 0.14 | 0.01 | 0.2 | 0.4 | 0.22 | 0.2 |
| 7 | 107361436 | rs4730253 | -0.431 | 1.172 | -0.6 | -0.317 | 0.003 | -0.145 | -0.145 | 1.055 | 0.67 | 0.45 | 0.75 | 0.98 | 0.7 | 0.85 |
| 7 | 107382352 | rs13224313 | 1.8 | 0.973 | -0.752 | -0.194 | 0.004 | -0.072 | -0.072 | 0 | 0.27 | 0.07 | 0.28 | 0.35 | 0.26 | 0.47 |
| 7 | 107383133 | rs12705420 | 1.008 | 1.247 | -0.969 | -0.262 | -0.047 | -0.065 | -0.065 | 0.011 | 0.18 | 0.01 | 0.3 | 0.24 | 0.21 | 0.15 |
| 7 | 107386810 | rs10274201 | -0.428 | 1.115 | -0.602 | -0.397 | -0.002 | 0 | 0 | 0 | 0.41 | 0.35 | 0.46 | 0.63 | 0.43 | 0.38 |
| 7 | 107408802 | rs41664 | -0.533 | 1.027 | -0.716 | -0.677 | -0.005 | -0.014 | -0.014 | 1.873 | 0.7 | 0.77 | 0.77 | 0.98 | 0.7 | 0.86 |
| 7 | 107410279 | rs10487275 | -0.052 | 0.818 | -0.459 | 0.207 | 0.001 | -0.003 | -0.003 | 0 | 0.26 | 0.26 | 0.32 | 0.29 | 0.25 | 0.16 |
| 7 | 107411716 | rs41668 | -0.397 | 0.597 | -0.344 | 0.147 | 0.017 | 0.021 | 0.021 | 0.19 | 0.79 | 0.79 | 0.65 | 0.72 | 0.72 | 0.62 |
| 7 | 107416602 | rs6973044 | -1.001 | -0.259 | 0.243 | -0.418 | -0.039 | -0.132 | -0.132 | 0.333 | 0.17 | 0.57 | 0.08 | 0 | 0.1 | 0.1 |
| 7 | 107420354 | rs10247487 | -0.618 | 0.017 | -0.449 | -0.039 | -0.004 | -0.009 | -0.009 | 1.876 | 0.36 | 0.38 | 0.21 | 0.31 | 0.26 | 0.27 |
| 7 | 107421072 | rs10263826 | -0.874 | 0.006 | -0.458 | -0.032 | -0.012 | -0.028 | -0.028 | 0 | 0.39 | 0.43 | 0.22 | 0.31 | 0.29 | 0.3 |
| 7 | 107423254 | rs3735605 | 0.137 | 0.86 | -0.722 | 0.447 | -0.02 | -0.013 | -0.013 | 1.162 | 0.14 | 0.06 | 0.14 | 0.28 | 0.18 | 0.14 |
| 7 | 107424320 | rs6466182 | -0.397 | 0.964 | -0.546 | 0.616 | -0.004 | -0.005 | -0.005 | 0 | 0.72 | 0.7 | 0.61 | 0.67 | 0.64 | 0.75 |
| 7 | 107433655 | rs6961051 | -0.157 | 0.853 | -0.091 | 0.555 | -0.001 | 0 | 0 | 0 | 0.22 | 0.13 | 0.35 | 0.2 | 0.23 | 0.3 |
| 7 | 107435187 | rs2237680 | -0.871 | 0.717 | 0.465 | 0.402 | 0.004 | -0.015 | -0.015 | 2.394 | 0.36 | 0.52 | 0.52 | 0.28 | 0.37 | 0.46 |
| 7 | 107435868 | rs17154444 | 0.409 | 0.744 | 0.35 | 0.657 | 0.002 | 0.01 | 0.01 | 0.091 | 0.06 | 0.06 | 0.06 | 0.09 | 0.05 | 0.09 |
| 7 | 107437048 | rs929393 | -0.799 | 0.783 | 0.533 | 0.55 | 0 | -0.002 | -0.002 | 0 | 0.37 | 0.58 | 0.52 | 0.29 | 0.38 | 0.46 |
| 7 | 107438305 | rs10248785 | -0.669 | 0.794 | 0.852 | 0.3 | 0.018 | -0.04 | -0.04 | 0 | 0.32 | 0.42 | 0.47 | 0.21 | 0.36 | 0.42 |
| 7 | 107440451 | rs2158837 | 0.435 | 0.823 | 0.372 | 0.32 | -0.012 | 0.018 | 0.018 | 0.006 | 0.19 | 0.05 | 0.32 | 0.15 | 0.2 | 0.28 |
| 7 | 107443871 | rs2301989 | -1.253 | 0.212 | 0.141 | 0 | 0.023 | 0.143 | 0.143 | 0.195 | 0.47 | 0.78 | 0.52 | 0.24 | 0.42 | 0.51 |
| 7 | 107446180 | rs6959729 | -0.72 | 0.263 | 0.294 | 0.321 | 0.009 | -0.01 | -0.01 | 0 | 0.37 | 0.41 | 0.48 | 0.35 | 0.37 | 0.48 |
| 7 | 107451045 | rs6466190 | -1.177 | 0.317 | -0.458 | 0.436 | 0.007 | 0.026 | 0.026 | 0.218 | 0.49 | 0.77 | 0.56 | 0.43 | 0.44 | 0.55 |
| 7 | 107453103 | rs2108225 | -1.204 | 0.341 | -0.581 | 0.446 | -0.006 | 0.019 | 0.019 | 0.355 | 0.49 | 0.84 | 0.56 | 0.42 | 0.44 | 0.56 |
| 7 | 107455282 | rs17408760 | -0.045 | 0.358 | -0.53 | 0.327 | 0.005 | 0.005 | 0.005 | 1.65 | 0.28 | 0.19 | 0.36 | 0.26 | 0.26 | 0.33 |
| 7 | 107456320 | rs6951457 | -1.403 | 0.463 | -1.056 | -0.317 | -0.071 | 0.145 | 0.145 | 0.442 | 0.72 | 0.94 | 0.79 | 0.96 | 0.68 | 0.86 |
| 7 | 107458561 | rs7804147 | -0.988 | 0.455 | -0.13 | 0.43 | -0.005 | -0.004 | -0.004 | 0 | 0.22 | 0.67 | 0.2 | 0.18 | 0.16 | 0.23 |
| 7 | 107462149 | rs10242548 | -0.053 | 1.269 | -0.038 | 0.976 | 0.083 | 0.134 | 0.134 | 0 | 0.14 | 0.3 | 0.33 | 0.48 | 0.22 | 0.3 |
| 7 | 107464932 | rs10241074 | -0.059 | 1.416 | 0.361 | 0.859 | 0.028 | 0.007 | 0.007 | 0.051 | 0.14 | 0.3 | 0.16 | 0.12 | 0.15 | 0.17 |
| 7 | 107465340 | rs2108227 | -0.796 | 0.851 | -0.115 | 0.632 | -0.001 | 0.018 | 0.018 | 0 | 0.28 | 0.31 | 0.31 | 0.45 | 0.25 | 0.31 |
| 7 | 107466818 | rs11762481 | 0.55 | 0.21 | -0.824 | 0.167 | -0.039 | -0.068 | -0.068 | 0 | 0.15 | 0.31 | 0.09 | 0.03 | 0.11 | 0.14 |
| 7 | 107472138 | rs4727688 | -1.838 | 0.141 | -2.145 | 0.007 | -0.059 | 0.235 | 0.235 | 0 | 0.62 | 0.95 | 0.74 | 0.94 | 0.61 | 0.78 |
| 7 | 107472872 | rs6466193 | -1.558 | 0.622 | -1.646 | 0.555 | -0.062 | -0.038 | -0.038 | 0 | 0.49 | 0.94 | 0.58 | 0.58 | 0.51 | 0.63 |
| 7 | 107473326 | rs714582 | 0.233 | 1.512 | -0.902 | 1.142 | -0.026 | -0.024 | -0.024 | 0 | 0.14 | 0.09 | 0.3 | 0.43 | 0.23 | 0.29 |
| 7 | 107475208 | rs12671393 | -0.808 | 0.954 | -0.672 | 0.883 | 0.004 | 0.003 | 0.003 | 0.017 | 0.34 | 0.62 | 0.5 | 0.57 | 0.39 | 0.49 |
| 7 | 107476546 | rs4730271 | -0.556 | 0.839 | -0.771 | 1.168 | 0 | 0.004 | 0.004 | 0.413 | 0.18 | 0.2 | 0.25 | 0.32 | 0.2 | 0.25 |
| 7 | 107479519 | rs4730273 | 0.511 | 0.744 | -0.985 | 1.082 | -0.014 | -0.041 | -0.041 | 1.748 | 0.26 | 0.16 | 0.3 | 0.39 | 0.27 | 0.24 |
| 7 | 107479719 | rs4730274 | -0.604 | 0.262 | -0.763 | 0.81 | -0.003 | 0.127 | 0.127 | 0.018 | 0.48 | 0.74 | 0.59 | 0.87 | 0.49 | 0.52 |
| 7 | 107483618 | rs6466199 | -0.495 | 0.249 | -0.758 | 0.945 | -0.022 | 0.02 | 0.02 | 1.453 | 0.31 | 0.4 | 0.3 | 0.43 | 0.29 | 0.41 |
| 7 | 107484437 | rs4730276 | 0.218 | 1.025 | -1.095 | 1.185 | -0.044 | -0.029 | -0.029 | 0 | 0.31 | 0.15 | 0.35 | 0.38 | 0.38 | 0.25 |
| 7 | 107494614 | rs6965857 | -0.49 | 0.624 | -0.842 | 1.38 | 0.003 | -0.01 | -0.01 | 0 | 0.18 | 0.14 | 0.22 | 0.34 | 0.19 | 0.23 |
| 7 | 107494658 | rs1017057 | -0.501 | 0.601 | -0.856 | 1.718 | 0.004 | -0.008 | -0.008 | 0.945 | 0.18 | 0.12 | 0.22 | 0.48 | 0.19 | 0.24 |
| 7 | 107495145 | rs2395932 | -0.881 | 0.493 | -0.6 | 1.449 | 0.002 | 0.065 | 0.065 | 0 | 0.57 | 0.71 | 0.63 | 0.8 | 0.61 | 0.55 |

| | | | | | | | | | | | | | | | | |
|---|---|---|---|---|---|---|---|---|---|---|---|---|---|---|---|---|
| 7 | 107497790 | rs10274614 | -0.111 | 0.558 | -0.63 | 1.601 | 0.004 | -0.01 | -0.01 | 0.322 | 0.18 | 0.13 | 0.22 | 0.46 | 0.2 | 0.24 |
| 7 | 107499761 | rs6964905 | -0.596 | 0.549 | -0.175 | 1.361 | 0.012 | 0.13 | 0.13 | 0 | 0.5 | 0.27 | 0.58 | 0.83 | 0.56 | 0.57 |
| 7 | 107503441 | rs4598195 | 0.347 | 0.422 | -0.222 | 1.243 | -0.055 | -0.015 | -0.015 | 0 | 0.34 | 0.15 | 0.38 | 0.35 | 0.4 | 0.31 |
| 7 | 107509273 | rs4348408 | -0.476 | 0.421 | -0.113 | 1.182 | 0.029 | 0.139 | 0.139 | 0 | 0.5 | 0.3 | 0.59 | 0.87 | 0.59 | 0.57 |
| 7 | 107525732 | rs6944829 | 0.275 | 0.39 | 0.887 | 1.264 | -0.052 | -0.015 | -0.015 | 0 | 0.34 | 0.17 | 0.38 | 0.34 | 0.39 | 0.3 |
| 7 | 107525846 | rs10238021 | -0.698 | 0.269 | 1.585 | 1.229 | -0.033 | 0.139 | 0.139 | 0.49 | 0.21 | 0.55 | 0.26 | 0.52 | 0.19 | 0.28 |
| 7 | 107531203 | rs34572011 | -0.559 | 0.294 | 1.66 | 1.368 | -0.036 | 0.186 | 0.186 | 0.365 | 0.21 | 0.5 | 0.26 | 0.52 | 0.19 | 0.27 |
| 7 | 107536417 | rs3801936 | 0.559 | 0.196 | 1.472 | 1.485 | 0.001 | -0.076 | -0.076 | 0 | 0.11 | 0.01 | 0.19 | 0.34 | 0.12 | 0.22 |
| 7 | 107545586 | rs17412104 | 0.55 | 0.186 | 1.506 | 0.902 | -0.106 | -0.052 | -0.052 | 1.885 | 0.33 | 0.07 | 0.37 | 0.34 | 0.39 | 0.3 |
| 7 | 107545799 | rs10263341 | -1.072 | 0.18 | 1.525 | 0.897 | -0.029 | 0.176 | 0.176 | 0 | 0.57 | 0.85 | 0.65 | 0.86 | 0.58 | 0.58 |
| 7 | 107546937 | rs10281520 | -1.216 | 0.169 | 1.576 | 0.918 | -0.024 | 0.066 | 0.066 | 1.343 | 0.23 | 0.65 | 0.27 | 0.52 | 0.19 | 0.27 |
| 7 | 107556623 | rs10271385 | -0.236 | 0.216 | 1.59 | 0.923 | -0.008 | 0.076 | 0.076 | 0.041 | 0.17 | 0.2 | 0.24 | 0.52 | 0.19 | 0.27 |
| 7 | 107560191 | rs4518 | -1.104 | 0.25 | 0.931 | 0.856 | -0.028 | 0.167 | 0.167 | 0 | 0.57 | 0.84 | 0.65 | 0.86 | 0.59 | 0.57 |
| 7 | 107563586 | rs3735602 | -1.045 | 0.476 | 0.95 | 0.749 | -0.025 | 0.172 | 0.172 | 0 | 0.57 | 0.84 | 0.64 | 0.86 | 0.58 | 0.61 |
| 7 | 107566430 | rs7806946 | -1.032 | 0.477 | 0.948 | 0.75 | -0.026 | 0.175 | 0.175 | 0 | 0.57 | 0.85 | 0.65 | 0.86 | 0.58 | 0.61 |
| 7 | 107575162 | rs2237686 | -0.019 | 0.546 | 1.135 | 0.826 | 0.004 | -0.004 | -0.004 | 0.463 | 0.38 | 0.54 | 0.39 | 0.33 | 0.38 | 0.3 |
| 7 | 107580839 | rs2158836 | -0.943 | 0.632 | 0.289 | 0.486 | 0.003 | 0.101 | 0.101 | 0 | 0.59 | 0.69 | 0.66 | 0.86 | 0.61 | 0.58 |
| 7 | 107585473 | rs2237687 | -0.342 | 1.146 | -0.489 | 0.271 | 0.002 | -0.028 | -0.028 | 0 | 0.45 | 0.5 | 0.6 | 0.82 | 0.5 | 0.51 |
| 7 | 107585844 | rs2237689 | 0.383 | 1.697 | -1.433 | 0.278 | -0.144 | -0.08 | -0.08 | 0.611 | 0.33 | 0.05 | 0.38 | 0.37 | 0.41 | 0.3 |
| 7 | 107586518 | rs757931 | -0.955 | 1.301 | -1.113 | -0.013 | 0.018 | 0.023 | 0.023 | 0 | 0.59 | 0.35 | 0.72 | 0.89 | 0.69 | 0.63 |
| 7 | 107591914 | rs2072208 | 0.166 | 1.523 | -1.566 | 0.341 | -0.116 | -0.087 | -0.087 | 0 | 0.31 | 0.06 | 0.37 | 0.37 | 0.38 | 0.35 |
| 7 | 107600558 | rs2528650 | -0.643 | 0.148 | -1.414 | -0.139 | 0.015 | 0.03 | 0.03 | 1.955 | 0.39 | 0.32 | 0.36 | 0.27 | 0.35 | 0.33 |
| 7 | 107601523 | rs7788778 | 0.078 | -0.007 | -1.388 | -0.07 | -0.045 | 0.002 | 0.002 | 1.011 | 0.29 | 0.06 | 0.31 | 0.26 | 0.3 | 0.29 |
| 7 | 107602624 | rs2528651 | -0.549 | -0.23 | -0.377 | 0.068 | -0.046 | -0.063 | -0.063 | 0.124 | 0.15 | 0.3 | 0.09 | 0.03 | 0.09 | 0.09 |
| 7 | 107604818 | rs1548638 | 0.005 | 0.333 | -0.752 | 0.327 | 0.001 | 0.002 | 0.002 | 0 | 0.53 | 0.52 | 0.65 | 0.71 | 0.59 | 0.62 |
| 7 | 107605412 | rs7789846 | 1.272 | 0.486 | -0.509 | 0.85 | -0.001 | 0.054 | 0.054 | 0 | 0.91 | 0.85 | 0.74 | 0.84 | 0.94 | 0.85 |
| 7 | 107608364 | rs13247770 | -0.031 | 0.535 | -1.349 | 0.192 | 0.003 | 0.104 | 0.104 | 0.128 | 0.48 | 0.19 | 0.47 | 0.3 | 0.47 | 0.39 |
| 7 | 107611650 | rs11766345 | 0.492 | 0.624 | -1.467 | 0.03 | -0.009 | 0.016 | 0.016 | 0 | 0.12 | 0.09 | 0.12 | 0.07 | 0.1 | 0.1 |
| 7 | 107612629 | rs1544474 | 0.338 | 0.185 | -0.708 | -0.142 | -0.033 | -0.004 | -0.004 | 0 | 0.33 | 0.18 | 0.38 | 0.68 | 0.43 | 0.51 |
| 7 | 107613379 | rs2701034 | -0.634 | 0.178 | -0.686 | -0.152 | 0.029 | 0.145 | 0.145 | 1.129 | 0.5 | 0.81 | 0.54 | 0.75 | 0.53 | 0.61 |
| 7 | 107615024 | rs737955 | -0.643 | 0.149 | -0.694 | -0.126 | 0.031 | 0.195 | 0.195 | 1.237 | 0.5 | 0.81 | 0.54 | 0.79 | 0.54 | 0.6 |
| 7 | 107616068 | rs11760290 | 0.69 | 0.224 | -0.674 | -0.067 | -0.004 | 0.008 | 0.008 | 0 | 0.12 | 0.11 | 0.12 | 0.07 | 0.1 | 0.09 |
| 7 | 107622150 | rs2237701 | -0.414 | 0.576 | -0.698 | -0.097 | -0.023 | 0.021 | 0.021 | 4.943 | 0.67 | 0.8 | 0.63 | 0.25 | 0.6 | 0.48 |
| 7 | 107622288 | rs2237702 | -0.403 | 0.575 | -0.698 | -0.102 | -0.023 | 0.023 | 0.023 | 5.118 | 0.66 | 0.8 | 0.62 | 0.25 | 0.6 | 0.48 |
| 7 | 107623542 | rs2701039 | -0.308 | 0.568 | -0.437 | 0.066 | -0.013 | 0.008 | 0.008 | 4.093 | 0.66 | 0.76 | 0.62 | 0.28 | 0.6 | 0.49 |
| 7 | 107626770 | rs25659 | 0.244 | 0.723 | -0.515 | -0.051 | -0.016 | 0.016 | 0.016 | 0 | 0.12 | 0.06 | 0.13 | 0.07 | 0.12 | 0.1 |
| 7 | 107638236 | rs10229305 | 0.139 | 0.286 | -0.095 | -0.557 | -0.008 | -0.022 | -0.022 | 0 | 0.41 | 0.51 | 0.43 | 0.23 | 0.4 | 0.36 |
| 7 | 107641616 | rs2249956 | -0.47 | 0.263 | -0.264 | -0.6 | -0.059 | 0.064 | 0.064 | 0.843 | 0.58 | 0.85 | 0.58 | 0.27 | 0.55 | 0.42 |
| 7 | 107643766 | rs2070917 | 0.082 | 0.259 | 0.077 | -0.619 | 0.011 | 0.106 | 0.106 | 0.485 | 0.5 | 0.31 | 0.52 | 0.23 | 0.49 | 0.39 |
| 7 | 107643977 | rs4320486 | 0.089 | 0.259 | 0.076 | -0.633 | 0.012 | 0.111 | 0.111 | 0.501 | 0.5 | 0.31 | 0.52 | 0.22 | 0.49 | 0.39 |
| 7 | 107644593 | rs3814473 | 0.064 | -0.057 | 0.429 | -0.255 | 0.005 | -0.007 | -0.007 | 0 | 0.07 | 0.11 | 0.16 | 0.05 | 0.1 | 0.08 |
| 7 | 107645133 | rs2395813 | 0.088 | -0.312 | -0.019 | -0.519 | 0.006 | 0.008 | 0.008 | 0 | 0.24 | 0.2 | 0.14 | 0.17 | 0.2 | 0.18 |
| 7 | 107645775 | rs3829896 | -0.363 | -0.081 | 0.018 | -0.534 | 0.006 | -0.002 | -0.002 | 0.242 | 0.94 | 0.92 | 0.83 | 0.95 | 0.9 | 0.93 |
| 7 | 107646766 | rs6943225 | -0.679 | -0.49 | -0.04 | -0.718 | 0.005 | 0.007 | 0.007 | 0 | 0.42 | 0.6 | 0.35 | 0.2 | 0.35 | 0.32 |
| 7 | 107649288 | rs1732164 | -0.594 | -0.728 | -0.775 | -0.436 | 0.084 | 0.079 | 0.079 | 0 | 0.43 | 0.81 | 0.23 | 0.24 | 0.28 | 0.32 |
| 7 | 107651115 | rs1732145 | -0.833 | -0.667 | -1.053 | -0.398 | 0.077 | 0.111 | 0.111 | 0 | 0.49 | 0.86 | 0.39 | 0.28 | 0.37 | 0.39 |
| 7 | 107652781 | rs6964934 | -0.311 | -0.637 | -0.184 | -0.25 | -0.004 | -0.005 | -0.005 | 0 | 0.25 | 0.27 | 0.13 | 0.17 | 0.18 | 0.25 |
| 7 | 107655604 | rs1636961 | -1.213 | -0.374 | 0.79 | -0.216 | 0.01 | -0.051 | -0.051 | 0.408 | 0.12 | 0.56 | 0.17 | 0.04 | 0.11 | 0.07 |
| 7 | 107656377 | rs983585 | -1.606 | -0.225 | 0.217 | 0.154 | 0 | 0 | 0 | 0 | 0.32 | 0.6 | 0.39 | 0.34 | 0.31 | 0.24 |
| 7 | 107661837 | rs1732146 | -0.969 | -0.607 | 0.77 | 0.132 | -0.02 | -0.055 | -0.055 | 0 | 0.12 | 0.66 | 0.09 | 0.03 | 0.08 | 0.05 |
| 7 | 107663059 | rs1613597 | -1.029 | -0.622 | 0.763 | 0.125 | -0.022 | -0.057 | -0.057 | 1.589 | 0.12 | 0.66 | 0.09 | 0.03 | 0.08 | 0.05 |
| 7 | 107664429 | rs1636944 | -0.99 | -0.615 | 0.766 | 0.129 | -0.02 | -0.064 | -0.064 | 0 | 0.12 | 0.66 | 0.09 | 0.02 | 0.08 | 0.05 |
| 7 | 107671407 | rs2528693 | -0.496 | -0.578 | 1.406 | 0.138 | 0.012 | -0.032 | -0.032 | 0 | 0.09 | 0.23 | 0.06 | 0.03 | 0.08 | 0.05 |
| 7 | 107673710 | rs1636956 | -0.751 | -0.58 | 1.563 | 0.138 | 0.02 | -0.031 | -0.031 | 0 | 0.1 | 0.34 | 0.06 | 0.03 | 0.08 | 0.05 |
| 7 | 107675223 | rs929395 | -1.136 | -0.565 | 0.48 | 0.111 | -0.044 | -0.019 | -0.019 | 0 | 0.85 | 0.99 | 0.86 | 0.87 | 0.82 | 0.87 |
| 7 | 107677984 | rs1627354 | -1.542 | -0.652 | 1.274 | 0.114 | -0.022 | -0.063 | -0.063 | 1.047 | 0.13 | 0.61 | 0.09 | 0.03 | 0.08 | 0.05 |
| 7 | 107678977 | rs2528699 | -1.201 | -0.743 | 1.043 | 0.229 | -0.003 | -0.005 | -0.005 | 0 | 0.29 | 0.67 | 0.26 | 0.24 | 0.25 | 0.16 |
| 7 | 107683100 | rs1636958 | -0.777 | -0.763 | 1.3 | -0.127 | -0.014 | -0.057 | -0.057 | 0 | 0.11 | 0.28 | 0.05 | 0.03 | 0.09 | 0.09 |
| 7 | 107685895 | rs1990158 | 0.211 | -0.682 | 0.976 | 0.328 | -0.034 | -0.116 | -0.116 | 0.23 | 0.3 | 0.05 | 0.4 | 0.52 | 0.35 | 0.28 |
| 7 | 107689844 | rs10260756 | -2.698 | -0.782 | 2.372 | 0.578 | -0.043 | -0.046 | -0.046 | 0 | 0.09 | 0.65 | 0.07 | 0 | 0.01 | 0.02 |
| 7 | 107692186 | rs17154848 | -1.658 | -0.779 | 2.556 | 0.578 | 0.035 | 0.031 | 0.031 | 0 | 0.05 | 0.3 | 0.02 | 0 | 0.01 | 0.02 |
| 7 | 107693740 | rs11487092 | -2.483 | -0.776 | 1.964 | 0.58 | -0.043 | -0.046 | -0.046 | 0 | 0.09 | 0.65 | 0.07 | 0 | 0.01 | 0.02 |
| 7 | 107710123 | rs11974886 | -1.189 | -0.477 | 1.667 | 0.309 | 0.024 | -0.045 | -0.045 | 0 | 0.1 | 0.35 | 0.09 | 0.01 | 0.1 | 0.08 |
| 7 | 107710599 | rs6973739 | 0.168 | -0.469 | 0.296 | 0.337 | 0.004 | -0.03 | -0.03 | 2.193 | 0.11 | 0.05 | 0.14 | 0.18 | 0.15 | 0.16 |
| 7 | 107715658 | rs6942816 | -1.81 | -0.145 | 0.942 | 0.227 | -0.023 | -0.07 | -0.07 | 0 | 0.14 | 0.64 | 0.13 | 0.01 | 0.1 | 0.08 |
| 7 | 107716767 | rs10226708 | -1.875 | -0.142 | 0.663 | 0.229 | -0.019 | -0.057 | -0.057 | 0.008 | 0.15 | 0.71 | 0.13 | 0.01 | 0.1 | 0.08 |
| 7 | 107720162 | rs9690688 | -1.448 | -0.088 | 1.036 | 0.273 | -0.001 | -0.069 | -0.069 | 0 | 0.12 | 0.48 | 0.09 | 0.01 | 0.08 | 0.07 |
| 7 | 107730749 | rs423686 | -0.495 | 0.331 | 0.426 | 0.571 | -0.002 | -0.001 | -0.001 | 0 | 0.35 | 0.62 | 0.58 | 0.62 | 0.43 | 0.42 |
| 7 | 107738836 | rs6974722 | 0.022 | 0.669 | 0.378 | 0.996 | -0.005 | 0 | 0 | 0.016 | 0.28 | 0.25 | 0.37 | 0.26 | 0.38 | 0.31 |

| | | | | | | | | | | | | | | | | |
|---|---|---|---|---|---|---|---|---|---|---|---|---|---|---|---|---|
| 7 | 107738948 | rs451252 | 0.033 | 0.675 | 0.381 | 0.999 | -0.005 | 0 | 0 | 0 | 0.28 | 0.25 | 0.37 | 0.26 | 0.38 | 0.31 |
| 7 | 107743664 | rs402113 | 0.041 | 0.706 | 0.362 | 0.966 | -0.006 | 0 | 0 | 0.008 | 0.28 | 0.23 | 0.37 | 0.26 | 0.38 | 0.32 |
| 7 | 107746253 | rs420753 | -0.163 | 0.611 | 0.356 | 1.292 | -0.01 | -0.018 | -0.018 | 0.052 | 0.31 | 0.23 | 0.4 | 0.4 | 0.39 | 0.39 |
| 7 | 107757843 | rs17155006 | -0.66 | 0.333 | -0.21 | 1.246 | -0.002 | -0.003 | -0.003 | 0 | 0.4 | 0.59 | 0.55 | 0.42 | 0.44 | 0.47 |
| 7 | 107764359 | rs12540374 | 1.194 | -0.459 | -1.969 | 0.765 | -0.047 | -0.122 | -0.122 | 0 | 0.27 | 0.01 | 0.18 | 0.38 | 0.24 | 0.25 |
| 7 | 107770624 | rs4730287 | 0.089 | 0.183 | -1.392 | 0.771 | -0.197 | -0.041 | -0.041 | 0 | 0.32 | 0.02 | 0.55 | 0.28 | 0.47 | 0.39 |
| 7 | 107771288 | rs10487840 | 0.006 | 0.254 | -0.842 | 1.004 | 0.008 | -0.028 | -0.028 | 0 | 0.05 | 0.01 | 0.04 | 0.16 | 0.02 | 0.11 |
| 7 | 107775581 | rs760355 | -0.238 | 0.284 | -0.645 | 0.34 | -0.012 | 0.011 | 0.011 | 0 | 0.39 | 0.33 | 0.54 | 0.35 | 0.51 | 0.35 |
| 7 | 107782190 | rs12669580 | -0.859 | 0.053 | 0.093 | 0.455 | -0.012 | 0.082 | 0.082 | 0.24 | 0.07 | 0.18 | 0.07 | 0.17 | 0.04 | 0.13 |
| 7 | 107782200 | rs382140 | 1.151 | -0.766 | 0.542 | -0.371 | -0.042 | -0.04 | -0.04 | 0 | 0.74 | 0.55 | 0.81 | 0.83 | 0.81 | 0.76 |
| 7 | 107782590 | rs405046 | 1.298 | -1.354 | 1.004 | -0.778 | -0.082 | -0.066 | -0.066 | 0.868 | 0.74 | 0.54 | 0.81 | 0.86 | 0.84 | 0.67 |
| 7 | 107782812 | rs7795932 | 1.166 | -1.492 | 1.648 | -0.859 | -0.06 | -0.064 | -0.064 | 0 | 0.77 | 0.59 | 0.85 | 0.9 | 0.87 | 0.75 |
| 7 | 107787327 | rs17155059 | 1.26 | -2.128 | 0.805 | -1.252 | 0.033 | 0.06 | 0.06 | 0.176 | 0.12 | 0.01 | 0.01 | 0.02 | 0.04 | 0.05 |
| 7 | 107789251 | rs439587 | 1.092 | -2.142 | 0.953 | -1.28 | -0.09 | -0.064 | -0.064 | 0.019 | 0.74 | 0.53 | 0.81 | 0.85 | 0.86 | 0.68 |
| 7 | 107789927 | rs449077 | -0.578 | -2.121 | 0.63 | -1.272 | 0.023 | 0.053 | 0.053 | 0 | 0.86 | 0.96 | 0.93 | 1 | 0.91 | 0.89 |
| 7 | 107797618 | rs409969 | 1.869 | -2.015 | 0.677 | -1.443 | 0.153 | 0.21 | 0.21 | 0.727 | 0.54 | 0.16 | 0.74 | 0.84 | 0.74 | 0.6 |
| 7 | 107800627 | rs11974486 | 1.189 | -1.053 | 0.16 | -1.726 | -0.099 | 0.011 | 0.011 | 5.93 | 0.13 | 0.01 | 0.23 | 0.07 | 0.36 | 0.19 |
| 7 | 107809412 | rs11767318 | 0.501 | -2.017 | 0.781 | -1.673 | -0.022 | -0.111 | -0.111 | 0.125 | 0.78 | 0.67 | 0.88 | 1 | 0.83 | 0.84 |
| 7 | 107809666 | rs11763780 | 1.833 | -2.226 | 0.755 | -1.722 | 0.106 | 0.135 | 0.135 | 0.599 | 0.64 | 0.08 | 0.75 | 0.85 | 0.78 | 0.64 |
| 7 | 107809927 | rs4730292 | 0.456 | -2.211 | 0.783 | -1.719 | -0.022 | -0.109 | -0.109 | 0 | 0.78 | 0.67 | 0.88 | 1 | 0.83 | 0.84 |
| 7 | 107811247 | rs383691 | 1.161 | -2.205 | 0.777 | -1.695 | -0.065 | -0.041 | -0.041 | 1.136 | 0.72 | 0.37 | 0.8 | 0.85 | 0.86 | 0.67 |
| 7 | 107811702 | rs2284280 | 1.892 | -2.285 | 0.448 | -1.762 | 0.179 | 0.247 | 0.247 | 1.007 | 0.53 | 0.08 | 0.73 | 0.84 | 0.74 | 0.6 |
| 7 | 107812100 | rs10487841 | 0.25 | -2.181 | 0.564 | -1.706 | 0.008 | -0.036 | -0.036 | 0.436 | 0.07 | 0.16 | 0.03 | 0 | 0.08 | 0.03 |
| 7 | 107815158 | rs11772445 | 1.728 | -2.227 | 0.43 | -1.725 | 0.107 | 0.132 | 0.132 | 0.943 | 0.64 | 0.08 | 0.75 | 0.85 | 0.78 | 0.64 |
| 7 | 107816299 | rs396136 | 0.891 | -2.26 | 0.847 | -1.727 | -0.039 | -0.046 | -0.046 | 0 | 0.66 | 0.57 | 0.83 | 0.88 | 0.84 | 0.71 |
| 7 | 107816651 | rs394921 | 0.161 | -2.134 | 0.908 | -1.779 | 0.024 | 0.002 | 0.002 | 1.571 | 0.73 | 0.76 | 0.89 | 0.99 | 0.87 | 0.84 |
| 7 | 107818851 | rs428459 | 0.601 | -2.103 | 0.978 | -1.778 | -0.07 | -0.12 | -0.12 | 0.646 | 0.69 | 0.49 | 0.84 | 0.94 | 0.85 | 0.76 |
| 7 | 107820911 | rs414077 | -0.695 | -2.036 | 0.798 | -1.773 | 0.009 | -0.007 | -0.007 | 0.019 | 0.86 | 0.88 | 0.92 | 1 | 0.91 | 0.88 |
| 7 | 107824678 | rs6970656 | 0.94 | -1.789 | 0.967 | -1.682 | 0.007 | -0.037 | -0.037 | 0 | 0.6 | 0.35 | 0.8 | 0.96 | 0.76 | 0.72 |
| 7 | 107831520 | rs6962066 | 0.935 | -1.811 | 0.946 | -1.674 | 0.009 | -0.025 | -0.025 | 0 | 0.6 | 0.37 | 0.8 | 0.95 | 0.76 | 0.71 |
| 7 | 107838464 | rs381318 | 1.031 | -1.531 | 1.266 | -1.551 | 0.013 | 0.007 | 0.007 | 0.074 | 0.58 | 0.39 | 0.77 | 0.88 | 0.74 | 0.65 |
| 7 | 107849908 | rs1269621 | 1.178 | -0.756 | 1.184 | -1.733 | 0.008 | 0.029 | 0.029 | 0 | 0.5 | 0.54 | 0.61 | 0.82 | 0.51 | 0.47 |
| 7 | 107862789 | rs17155196 | 0.756 | -0.659 | 1.183 | -1.716 | 0.03 | -0.039 | -0.039 | 0 | 0.15 | 0.23 | 0.2 | 0.06 | 0.28 | 0.2 |
| 7 | 107863395 | rs12534073 | 0.756 | -0.659 | 1.17 | -1.716 | 0.016 | -0.026 | -0.026 | 2.273 | 0.15 | 0.2 | 0.2 | 0.06 | 0.28 | 0.2 |
| 7 | 107867525 | rs926146 | -1.161 | -0.704 | 1.219 | -1.716 | 0.002 | -0.011 | -0.011 | 0 | 0.29 | 0.63 | 0.33 | 0.15 | 0.37 | 0.33 |
| 7 | 107868007 | rs17155207 | 0.742 | -0.744 | 1.235 | -1.652 | -0.013 | 0.021 | 0.021 | 0.054 | 0.06 | 0.14 | 0.07 | 0.09 | 0.03 | 0.11 |
| 7 | 107868331 | rs7784319 | 0.742 | -0.744 | 1.099 | -1.652 | -0.008 | 0.036 | 0.036 | 0.054 | 0.06 | 0.16 | 0.07 | 0.09 | 0.03 | 0.11 |
| 7 | 107872816 | rs2072546 | 2.086 | -0.976 | -0.011 | -1.388 | -0.024 | -0.127 | -0.127 | 0 | 0.16 | 0.01 | 0.27 | 0.55 | 0.12 | 0.2 |
| 7 | 107874906 | rs1269632 | -0.221 | -0.929 | 0.585 | -1.803 | 0.039 | 0.045 | 0.045 | 1.45 | 0.71 | 0.77 | 0.88 | 0.98 | 0.83 | 0.8 |
| 7 | 107880612 | rs1269634 | 1.6 | -0.914 | -0.166 | -1.815 | -0.076 | 0.173 | 0.173 | 0.01 | 0.43 | 0.08 | 0.55 | 0.83 | 0.47 | 0.48 |
| 7 | 107881205 | rs6970096 | -0.855 | -0.969 | -0.181 | -1.81 | 0.01 | -0.029 | -0.029 | 0 | 0.28 | 0.59 | 0.32 | 0.15 | 0.36 | 0.32 |
| 7 | 107883621 | rs1269637 | -0.504 | -0.904 | -0.878 | -1.955 | 0.043 | 0.164 | 0.164 | 2.046 | 0.73 | 0.96 | 0.9 | 0.98 | 0.83 | 0.8 |
| 7 | 107891529 | rs1269645 | 1.408 | -0.732 | -0.81 | -1.89 | -0.07 | 0.217 | 0.217 | 0.034 | 0.44 | 0.08 | 0.56 | 0.86 | 0.46 | 0.48 |
| 7 | 107893220 | rs2284284 | -0.582 | -0.424 | -0.33 | -1.574 | 0.009 | -0.037 | -0.037 | 0.332 | 0.23 | 0.66 | 0.3 | 0.13 | 0.31 | 0.29 |
| 7 | 107898586 | rs1269651 | 1.003 | -0.248 | -0.536 | -1.441 | 0.025 | 0.206 | 0.206 | 0 | 0.52 | 0.14 | 0.59 | 0.86 | 0.53 | 0.52 |
| 7 | 107911705 | rs1269662 | 0.726 | 0.589 | -0.15 | -0.889 | 0.031 | -0.078 | -0.078 | 0 | 0.76 | 0.62 | 0.72 | 0.95 | 0.63 | 0.7 |
| 7 | 107920667 | rs1269673 | 0.369 | 0.482 | -0.701 | -0.765 | 0.007 | 0.027 | 0.027 | 0 | 0.87 | 0.91 | 0.87 | 0.94 | 0.89 | 0.85 |
| 7 | 107921440 | rs7786371 | -0.263 | 0.492 | -0.283 | -0.894 | 0.047 | -0.062 | -0.062 | 0.065 | 0.18 | 0.41 | 0.21 | 0.05 | 0.26 | 0.18 |
| 7 | 107923131 | rs1269683 | 0.442 | 0.162 | -0.506 | -0.602 | 0.007 | 0.027 | 0.027 | 0 | 0.87 | 0.91 | 0.87 | 0.94 | 0.89 | 0.85 |
| 7 | 107929474 | rs1035204 | 0.197 | 0.214 | -1.043 | -0.293 | -0.013 | 0.005 | 0.005 | 0.271 | 0.43 | 0.28 | 0.46 | 0.36 | 0.56 | 0.39 |
| 7 | 107932781 | rs759548 | 0.208 | -0.028 | -0.747 | -0.481 | -0.007 | -0.044 | -0.044 | 0.801 | 0.88 | 0.77 | 0.93 | 1 | 0.9 | 0.98 |
| 7 | 107932852 | rs759549 | 0.987 | -0.677 | -1.127 | -0.678 | 0.018 | 0.021 | 0.021 | 0 | 0.77 | 0.76 | 0.86 | 0.92 | 0.85 | 0.84 |
| 7 | 107933191 | rs1990713 | 0.905 | -0.488 | -0.308 | -0.52 | 0 | -0.002 | -0.002 | 0.051 | 0.31 | 0.25 | 0.38 | 0.51 | 0.3 | 0.4 |
| 7 | 107935905 | rs2267881 | -0.65 | -0.069 | -0.444 | -0.741 | -0.006 | 0.036 | 0.036 | 0 | 0.66 | 0.7 | 0.66 | 0.89 | 0.6 | 0.78 |
| 7 | 107938376 | rs10241406 | 1.627 | 0.044 | 0.287 | -0.583 | 0.07 | -0.009 | -0.009 | 0 | 0.12 | 0.29 | 0.17 | 0.07 | 0.23 | 0.11 |
| 7 | 107940173 | rs4142284 | 0.722 | -0.31 | 0.119 | -0.166 | 0.016 | -0.112 | -0.112 | 0.185 | 0.41 | 0.09 | 0.36 | 0.59 | 0.28 | 0.49 |
| 7 | 107943818 | rs10262724 | 0.046 | -0.225 | 0.491 | -0.191 | 0.059 | 0.029 | 0.029 | 1.646 | 0.21 | 0.4 | 0.24 | 0.17 | 0.32 | 0.22 |
| 7 | 107946905 | rs2193241 | 0.669 | -0.318 | 0.523 | -0.246 | -0.02 | 0.009 | 0.009 | 0.934 | 0.69 | 0.26 | 0.67 | 0.83 | 0.59 | 0.76 |
| 7 | 107948721 | rs2267882 | 1.375 | -0.355 | 0.4 | -0.247 | -0.022 | 0.013 | 0.013 | 0 | 0.29 | 0.09 | 0.3 | 0.25 | 0.31 | 0.28 |
| 7 | 107950504 | rs17333297 | 1.799 | -0.403 | 0.197 | -0.131 | -0.01 | 0.034 | 0.034 | 0.483 | 0.11 | 0 | 0.03 | 0.02 | 0.07 | 0.08 |
| 7 | 107950821 | rs1548949 | -0.69 | -0.38 | 0.371 | -0.127 | -0.003 | -0.04 | -0.04 | 0.645 | 0.91 | 0.81 | 0.92 | 1 | 0.9 | 0.97 |
| 7 | 107952864 | rs17155421 | 0.925 | -0.451 | 0.255 | -0.133 | 0.003 | 0.009 | 0.009 | 0.811 | 0.15 | 0.09 | 0.12 | 0.12 | 0.13 | 0.16 |
| 7 | 107964618 | rs4727708 | -0.791 | -0.486 | 0.328 | -0.127 | 0 | -0.017 | -0.017 | 0 | 0.92 | 0.85 | 0.92 | 1 | 0.9 | 0.97 |
| 7 | 107965942 | rs2111201 | 0.376 | -0.492 | 0.305 | 0.191 | 0.042 | -0.06 | -0.06 | 0 | 0.42 | 0.17 | 0.37 | 0.56 | 0.28 | 0.49 |
| 7 | 107968803 | rs2267887 | 0.153 | -0.508 | 0.486 | 0.199 | 0.005 | 0 | 0 | 0 | 0.49 | 0.48 | 0.53 | 0.44 | 0.62 | 0.48 |
| 7 | 107974924 | rs2300026 | 0.403 | -0.435 | 0.447 | 0.16 | 0.008 | -0.029 | -0.029 | 0 | 0.64 | 0.46 | 0.66 | 0.81 | 0.6 | 0.73 |
| 7 | 107976561 | rs12539530 | 0.71 | -0.494 | 0.488 | 0.218 | 0.008 | -0.015 | -0.015 | 0 | 0.34 | 0.21 | 0.34 | 0.55 | 0.27 | 0.46 |
| 7 | 107984943 | rs2193243 | 1.067 | -0.471 | 0.384 | 0.253 | -0.001 | 0.031 | 0.031 | 0 | 0.07 | 0.1 | 0.06 | 0.11 | 0.06 | 0.1 |
| 7 | 107986372 | rs1861636 | 0.374 | -0.471 | 0.335 | 0.206 | 0.009 | -0.031 | -0.031 | 0.039 | 0.64 | 0.46 | 0.66 | 0.81 | 0.59 | 0.73 |

| | | | | | | | | | | | | | | | | |
|---|---|---|---|---|---|---|---|---|---|---|---|---|---|---|---|---|
| 7 | 107988922 | rs17155518 | 1.733 | -0.475 | 0.001 | 0.232 | -0.021 | -0.125 | -0.125 | 0.878 | 0.26 | 0.01 | 0.27 | 0.45 | 0.2 | 0.34 |
| 7 | 107992582 | rs10487851 | 1.534 | -0.5 | 0 | 0.174 | -0.044 | -0.023 | -0.023 | 0 | 0.29 | 0.09 | 0.3 | 0.26 | 0.33 | 0.28 |
| 7 | 107994117 | rs7804045 | 0.656 | -0.539 | 0.587 | 0.133 | -0.015 | 0.026 | 0.026 | 0 | 0.34 | 0.31 | 0.35 | 0.56 | 0.26 | 0.45 |
| 7 | 107997567 | rs13233068 | 0.497 | -0.526 | 0.287 | 0.078 | 0.011 | -0.025 | -0.025 | 0 | 0.25 | 0.37 | 0.2 | 0.09 | 0.3 | 0.17 |
| 7 | 108004957 | rs12538050 | 1.747 | -0.513 | -0.197 | 0.16 | 0.016 | -0.054 | -0.054 | 0 | 0.25 | 0.07 | 0.28 | 0.45 | 0.21 | 0.34 |
| 7 | 108006941 | rs2300033 | 1.168 | -0.492 | -0.298 | 0.134 | -0.049 | -0.052 | -0.052 | 0.969 | 0.32 | 0.09 | 0.36 | 0.35 | 0.34 | 0.36 |
| 7 | 108008952 | rs2300034 | -0.079 | -0.284 | -0.238 | 0.163 | -0.069 | 0.047 | 0.047 | 0.29 | 0.48 | 0.86 | 0.49 | 0.63 | 0.5 | 0.55 |
| 7 | 108010825 | rs17155592 | 1.437 | -0.736 | -0.086 | 0.06 | -0.005 | 0.007 | 0.007 | 0 | 0.1 | 0.09 | 0.06 | 0.11 | 0.06 | 0.11 |
| 7 | 108011268 | rs16872512 | 0.081 | -0.814 | 0.422 | 0.022 | -0.019 | 0.068 | 0.068 | 0.01 | 0.13 | 0.34 | 0.07 | 0.11 | 0.06 | 0.11 |
| 7 | 108011946 | rs16872515 | 0.202 | -0.804 | 0.426 | -0.065 | -0.015 | -0.062 | -0.062 | 0 | 0.13 | 0.34 | 0.07 | 0.03 | 0.06 | 0.1 |
| 7 | 108014337 | rs7783893 | 0.378 | -0.833 | 0.154 | -0.245 | 0.016 | 0.035 | 0.035 | 0 | 0.44 | 0.31 | 0.46 | 0.32 | 0.42 | 0.42 |
| 7 | 108015239 | rs2193248 | -0.898 | -0.439 | 0.056 | -0.24 | 0.049 | -0.042 | -0.042 | 0 | 0.81 | 0.61 | 0.76 | 0.93 | 0.68 | 0.84 |
| 7 | 108024132 | rs1990712 | 0.589 | -0.636 | -0.207 | -0.091 | -0.009 | 0.004 | 0.004 | 0 | 0.36 | 0.32 | 0.31 | 0.63 | 0.25 | 0.44 |
| 7 | 108025115 | rs16872525 | 0.193 | -0.226 | -0.946 | 0.181 | -0.004 | -0.087 | -0.087 | 0 | 0.25 | 0.07 | 0.23 | 0.35 | 0.18 | 0.33 |
| 7 | 108026243 | rs12333431 | 0.169 | -0.403 | -1.472 | 0.02 | -0.005 | -0.08 | -0.08 | 0 | 0.43 | 0.14 | 0.43 | 0.45 | 0.39 | 0.42 |
| 7 | 108026543 | rs722519 | 2.553 | -0.334 | -1.529 | 0 | -0.017 | 0.076 | 0.076 | 0 | 0.17 | 0.03 | 0.19 | 0.05 | 0.21 | 0.08 |
| 7 | 108034717 | rs11976021 | 1.782 | 0.09 | -1.077 | 0.433 | -0.09 | 0.042 | 0.042 | 2.501 | 0.14 | 0.01 | 0.17 | 0.06 | 0.26 | 0.14 |
| 7 | 108039065 | rs1859768 | -0.395 | 0.25 | -0.79 | 0.494 | 0.018 | -0.027 | -0.027 | 0 | 0.62 | 0.71 | 0.65 | 0.46 | 0.67 | 0.56 |
| 7 | 108043038 | rs1859776 | -0.538 | 0.138 | -0.85 | 0.608 | -0.004 | 0.005 | 0.005 | 0 | 0.89 | 0.86 | 0.9 | 0.89 | 0.9 | 0.93 |
| 7 | 108043641 | rs11983030 | 0.829 | 0.15 | -0.91 | 0.537 | 0.005 | -0.013 | -0.013 | 0.065 | 0.42 | 0.15 | 0.44 | 0.4 | 0.39 | 0.42 |
| 7 | 108044106 | rs17155687 | 2.274 | 0.131 | -0.978 | 0.639 | -0.005 | -0.028 | -0.028 | 0 | 0.08 | 0 | 0.06 | 0.09 | 0.04 | 0.09 |
| 7 | 108045033 | rs2300037 | 1.024 | 0.515 | -0.557 | 0.748 | 0.006 | -0.012 | -0.012 | 0.394 | 0.42 | 0.15 | 0.44 | 0.4 | 0.39 | 0.44 |
| 7 | 108049865 | rs2158846 | 1.017 | 0.566 | -0.311 | 0.757 | -0.017 | 0.02 | 0.02 | 0.195 | 0.66 | 0.2 | 0.68 | 0.76 | 0.6 | 0.78 |
| 7 | 108050027 | rs2300039 | 1.027 | 0.566 | -0.311 | 0.757 | -0.016 | 0.02 | 0.02 | 0.216 | 0.66 | 0.2 | 0.68 | 0.76 | 0.6 | 0.78 |
| 7 | 108055858 | rs7788503 | -0.841 | 0.373 | -0.159 | 0.448 | 0.004 | 0.02 | 0.02 | 0 | 0.94 | 0.95 | 0.96 | 1 | 0.95 | 0.98 |
| 7 | 108058574 | rs2057981 | 1.165 | 0.286 | 0.404 | 0.402 | -0.038 | -0.095 | -0.095 | 0 | 0.83 | 0.48 | 0.87 | 0.96 | 0.88 | 0.93 |
| 7 | 108058945 | rs4591957 | 1.165 | 0.286 | 0.407 | 0.402 | -0.038 | -0.095 | -0.095 | 0.043 | 0.83 | 0.48 | 0.88 | 0.96 | 0.88 | 0.93 |
| 7 | 108059247 | rs10236888 | 0.622 | 0.377 | 0.504 | 0.427 | -0.008 | -0.078 | -0.078 | 0 | 0.86 | 0.56 | 0.89 | 0.96 | 0.9 | 0.93 |
| 7 | 108063238 | rs2300045 | 0.324 | 0.575 | 0.125 | 0.557 | 0.008 | -0.004 | -0.004 | 0 | 0.41 | 0.55 | 0.44 | 0.43 | 0.46 | 0.49 |
| 7 | 108074766 | rs2284290 | 0.899 | 0.634 | -0.596 | 0.446 | 0.001 | -0.042 | -0.042 | 0 | 0.43 | 0.14 | 0.43 | 0.47 | 0.4 | 0.46 |
| 7 | 108075566 | rs10269351 | 1.3 | 0.774 | -0.765 | 0.373 | -0.013 | -0.101 | -0.101 | 2.798 | 0.26 | 0.09 | 0.25 | 0.41 | 0.23 | 0.35 |
| 7 | 108080565 | rs13221639 | -0.93 | 0.841 | -0.692 | 0.416 | 0.001 | -0.028 | -0.028 | 0 | 0.55 | 0.83 | 0.53 | 0.52 | 0.56 | 0.55 |
| 7 | 108083604 | rs1034826 | 0.159 | 1.138 | -0.154 | 0.401 | -0.024 | 0.029 | 0.029 | 0 | 0.34 | 0.47 | 0.31 | 0.45 | 0.26 | 0.36 |
| 7 | 108087080 | rs2284291 | -0.713 | 1.178 | -0.263 | 0.375 | 0.016 | -0.001 | -0.001 | 0 | 0.5 | 0.75 | 0.5 | 0.52 | 0.54 | 0.53 |
| 7 | 108089035 | rs13245242 | 1.232 | 1.491 | -0.604 | 0.561 | 0.01 | -0.014 | -0.014 | 0 | 0.16 | 0.23 | 0.19 | 0.07 | 0.27 | 0.14 |
| 7 | 108098223 | rs3763461 | 1.01 | 0.882 | -0.977 | 0.821 | 0.014 | -0.077 | -0.077 | 0.053 | 0.21 | 0.08 | 0.23 | 0.4 | 0.18 | 0.26 |
| 7 | 108102346 | rs726471 | -0.858 | 1.075 | -0.74 | 0.578 | 0.011 | -0.002 | -0.002 | 0 | 0.53 | 0.77 | 0.53 | 0.53 | 0.56 | 0.5 |
| 7 | 108105345 | rs10225561 | -0.343 | 1.262 | -0.008 | 0.326 | -0.003 | 0.004 | 0.004 | 0 | 0.34 | 0.33 | 0.32 | 0.47 | 0.29 | 0.38 |
| 7 | 108108759 | rs1859774 | -1.092 | 1.154 | 0.429 | 0.674 | 0.01 | 0.028 | 0.028 | 1.967 | 0.53 | 0.8 | 0.53 | 0.63 | 0.57 | 0.52 |
| 7 | 108108830 | rs1859775 | -1.164 | 1.164 | 0.477 | 0.687 | 0.008 | 0.021 | 0.021 | 1.628 | 0.53 | 0.79 | 0.53 | 0.62 | 0.57 | 0.52 |
| 7 | 108111356 | rs17134769 | 0.459 | 1.113 | 0.802 | 0.46 | 0 | -0.033 | -0.033 | 0.249 | 0.13 | 0.21 | 0.09 | 0.07 | 0.11 | 0.1 |
| 7 | 108111566 | rs6466238 | 1.105 | 1.084 | 1.061 | 0.475 | 0.056 | -0.076 | -0.076 | 0.495 | 0.19 | 0.38 | 0.2 | 0.07 | 0.28 | 0.14 |
| 7 | 108116481 | rs10249427 | 1.429 | 1.076 | 1.034 | 0.494 | 0.014 | -0.054 | -0.054 | 0 | 0.21 | 0.13 | 0.23 | 0.39 | 0.18 | 0.27 |
| 7 | 108117088 | rs10215206 | 1.458 | 1.075 | 1.003 | 0.493 | 0.012 | -0.075 | -0.075 | 0 | 0.21 | 0.08 | 0.23 | 0.39 | 0.18 | 0.25 |
| 7 | 108121463 | rs11981670 | 1.173 | 1.115 | 1.069 | 0.433 | 0.067 | -0.078 | -0.078 | 0.19 | 0.19 | 0.41 | 0.21 | 0.07 | 0.28 | 0.14 |
| 7 | 108129318 | rs10487859 | 1.302 | 1.12 | 1.067 | 0.437 | 0.053 | -0.068 | -0.068 | 1.324 | 0.18 | 0.35 | 0.2 | 0.07 | 0.28 | 0.14 |
| 7 | 108129884 | rs6964631 | 1.135 | 1.11 | 1.065 | 0.412 | 0.003 | -0.013 | -0.013 | 0 | 0.1 | 0.09 | 0.07 | 0.06 | 0.09 | 0.09 |
| 7 | 108145121 | rs40851 | -0.226 | 1.105 | 1.061 | 0.411 | 0.045 | -0.083 | -0.083 | 1.856 | 0.28 | 0.51 | 0.28 | 0.13 | 0.37 | 0.23 |
| 7 | 108156477 | rs10232596 | 1.381 | 1.001 | 0.723 | 0.547 | 0.013 | -0.072 | -0.072 | 0.013 | 0.21 | 0.08 | 0.23 | 0.4 | 0.18 | 0.25 |
| 7 | 108163237 | rs1014112 | 1.332 | 1.003 | 0.758 | 0.564 | 0.01 | -0.024 | -0.024 | 0.035 | 0.21 | 0.16 | 0.23 | 0.4 | 0.18 | 0.25 |
| 7 | 108164460 | rs40892 | -0.686 | 1.005 | 0.797 | 0.534 | -0.007 | 0.001 | 0.001 | 0.906 | 0.43 | 0.68 | 0.46 | 0.48 | 0.47 | 0.4 |
| 7 | 108164484 | rs10480679 | 1.252 | 0.602 | 0.227 | 0.686 | 0.01 | -0.026 | -0.026 | 0 | 0.21 | 0.16 | 0.23 | 0.39 | 0.18 | 0.25 |
| 7 | 108177962 | rs10225135 | 0.864 | 0.749 | -0.528 | 0.097 | 0.06 | 0.131 | 0.131 | 0.633 | 0.48 | 0.22 | 0.43 | 0.24 | 0.39 | 0.45 |
| 7 | 108191625 | rs40928 | 1.072 | -0.169 | -0.063 | 0.005 | 0.053 | -0.043 | -0.043 | 0 | 0.18 | 0.3 | 0.26 | 0.07 | 0.32 | 0.16 |
| 7 | 108202792 | rs40939 | 0.767 | -0.23 | -0.065 | -0.027 | 0.044 | -0.047 | -0.047 | 0 | 0.19 | 0.31 | 0.26 | 0.07 | 0.32 | 0.16 |
| 7 | 108204479 | rs40943 | -0.676 | -0.832 | -0.603 | 0.068 | -0.024 | 0.037 | 0.037 | 0.703 | 0.5 | 0.81 | 0.65 | 0.64 | 0.51 | 0.51 |
| 7 | 108209623 | rs2227272 | 0.239 | -0.964 | -0.647 | 0.079 | 0.062 | -0.101 | -0.101 | 0.043 | 0.3 | 0.09 | 0.36 | 0.56 | 0.2 | 0.42 |
| 7 | 108212353 | rs1043615 | -0.303 | -0.573 | 0.022 | 0.215 | -0.035 | 0.007 | 0.007 | 0.036 | 0.45 | 0.26 | 0.58 | 0.64 | 0.5 | 0.51 |
| 7 | 108215858 | rs4727713 | -0.038 | -0.379 | 0.024 | 0.659 | -0.011 | -0.023 | -0.023 | 0 | 0.08 | 0.03 | 0.1 | 0.23 | 0.08 | 0.11 |
| 7 | 108223326 | rs12532433 | 1.934 | -0.673 | 0.151 | -0.451 | 0 | 0.036 | 0.036 | 0 | 0.2 | 0.08 | 0.14 | 0.11 | 0.22 | 0.16 |
| 7 | 108239477 | rs11536567 | -0.973 | -0.732 | 0.986 | 0.027 | -0.008 | -0.01 | -0.01 | 0.098 | 0.35 | 0.25 | 0.31 | 0.41 | 0.35 | 0.28 |
| 7 | 108244234 | rs2215141 | 0.827 | -1.289 | 1.538 | -0.133 | -0.012 | 0.03 | 0.03 | 0 | 0.59 | 0.62 | 0.53 | 0.68 | 0.48 | 0.6 |
| 7 | 108253242 | rs726608 | 0.848 | -1.364 | 1.624 | -0.169 | -0.008 | 0.018 | 0.018 | 0 | 0.58 | 0.59 | 0.54 | 0.69 | 0.49 | 0.6 |
| 7 | 108260762 | rs1013307 | 1.044 | -1.178 | 1.922 | -0.2 | -0.053 | -0.041 | -0.041 | 0.065 | 0.79 | 0.5 | 0.91 | 0.8 | 0.88 | 0.88 |
| 7 | 108262714 | rs6978592 | 1.322 | -1.098 | 2.019 | -0.394 | 0.002 | 0.002 | 0.002 | 0 | 0.51 | 0.52 | 0.42 | 0.63 | 0.42 | 0.49 |
| 7 | 108265021 | rs7785225 | 0.696 | -1.008 | 1.56 | -0.497 | -0.019 | 0.03 | 0.03 | 0 | 0.61 | 0.73 | 0.49 | 0.73 | 0.49 | 0.54 |
| 7 | 108283606 | rs10480450 | -0.34 | -0.406 | 1.966 | -0.212 | -0.007 | 0.017 | 0.017 | 0 | 0.12 | 0.45 | 0.11 | 0.12 | 0.08 | 0.05 |
| 7 | 108320146 | rs1978100 | -1.565 | 0.149 | 1.281 | -0.403 | 0 | 0.046 | 0.046 | 0 | 0.46 | 0.61 | 0.54 | 0.19 | 0.51 | 0.46 |
| 7 | 108328266 | rs10228847 | -0.955 | 0.081 | 1.626 | -0.509 | -0.032 | -0.021 | -0.021 | 0 | 0.23 | 0.3 | 0.12 | 0.15 | 0.13 | 0.18 |

| | | | | | | | | | | | | | | | | |
|---|---|---|---|---|---|---|---|---|---|---|---|---|---|---|---|---|
| 7 | 108339862 | rs17156090 | -0.21 | 0.051 | 1.546 | -0.521 | 0.05 | 0.031 | 0.031 | 0 | 0.18 | 0 | 0.04 | 0.09 | 0.07 | 0.17 |
| 7 | 108342068 | rs6959354 | -0.414 | 0.096 | 1.662 | -0.531 | 0.024 | -0.044 | -0.044 | 4.811 | 0.23 | 0.28 | 0.41 | 0.04 | 0.38 | 0.28 |
| 7 | 108343278 | rs10253885 | -0.79 | 0.066 | 1.47 | -0.52 | 0.069 | -0.066 | -0.066 | 0.342 | 0.27 | 0.4 | 0.48 | 0.09 | 0.43 | 0.29 |
| 7 | 108348466 | rs1544346 | -1.447 | 0.06 | 1.482 | -0.518 | -0.004 | -0.004 | -0.004 | 0.403 | 0.42 | 0.36 | 0.45 | 0.13 | 0.45 | 0.45 |
| 7 | 108348490 | rs4321913 | -1.793 | 0.066 | 1.148 | -0.517 | 0 | 0.014 | 0.014 | 0 | 0.47 | 0.55 | 0.53 | 0.19 | 0.51 | 0.46 |
| 7 | 108348785 | rs12705489 | -0.257 | -0.052 | 1.013 | -0.695 | 0.059 | 0.041 | 0.041 | 0 | 0.18 | 0.01 | 0.04 | 0.09 | 0.07 | 0.17 |
| 7 | 108351652 | rs4730328 | -1.659 | 0.008 | 1.032 | -0.664 | 0.002 | -0.036 | -0.036 | 0 | 0.44 | 0.49 | 0.47 | 0.13 | 0.46 | 0.46 |
| 7 | 108355037 | rs17350594 | -0.907 | 1.164 | 1.268 | -0.275 | -0.006 | -0.012 | -0.012 | 0.038 | 0.23 | 0.22 | 0.4 | 0.03 | 0.37 | 0.27 |
| 7 | 108358839 | rs12539724 | -0.987 | 1.19 | 0.695 | -0.263 | 0.064 | -0.102 | -0.102 | 5.267 | 0.25 | 0.59 | 0.44 | 0.03 | 0.38 | 0.27 |
| 7 | 108377828 | rs2188598 | 0.898 | 0.239 | 0.536 | 0.176 | -0.009 | -0.003 | -0.003 | 0 | 0.79 | 0.31 | 0.72 | 0.86 | 0.88 | 0.91 |
| 7 | 108384142 | rs6977586 | 0.233 | 0.167 | 0.723 | 0.144 | 0.018 | -0.008 | -0.008 | 0 | 0.86 | 0.94 | 0.77 | 0.86 | 0.89 | 0.91 |
| 7 | 108385515 | rs6967009 | 1.688 | -0.048 | 0.892 | 0.155 | -0.006 | -0.001 | -0.001 | 0 | 0.74 | 0.32 | 0.71 | 0.8 | 0.84 | 0.88 |
| 7 | 108406238 | rs7795930 | -1.198 | 0.032 | 2.353 | 0.148 | 0.042 | 0.039 | 0.039 | 5.537 | 0.05 | 0.51 | 0.04 | 0 | 0.01 | 0 |
| 7 | 108410409 | rs12705500 | -0.401 | 1.149 | 2.318 | 0.965 | 0.003 | -0.004 | -0.004 | 0.036 | 0.1 | 0.1 | 0.24 | 0.2 | 0.25 | 0.27 |
| 7 | 108413876 | rs17156176 | -0.512 | 0.252 | 2.072 | 0.802 | -0.029 | -0.037 | -0.037 | 0 | 0.1 | 0 | 0.07 | 0.14 | 0.12 | 0.19 |
| 7 | 108415940 | rs6967523 | -0.195 | 0.132 | 2.177 | 0.829 | 0.014 | -0.021 | -0.021 | 0 | 0.86 | 0.97 | 0.77 | 0.84 | 0.89 | 0.92 |
| 7 | 108438172 | rs9784995 | -2.305 | 0.333 | 2.925 | 0.736 | -0.04 | -0.032 | -0.032 | 0 | 0.13 | 0.67 | 0.06 | 0.06 | 0.05 | 0.04 |
| 7 | 108455334 | rs4730333 | 2.136 | 0.332 | 2.841 | 0.735 | -0.033 | -0.027 | -0.027 | 0.38 | 0.87 | 0.29 | 0.93 | 0.94 | 0.95 | 0.96 |
| 7 | 108466719 | rs1568857 | 1.414 | 0.371 | 2.859 | 0.752 | -0.014 | -0.007 | -0.007 | 0 | 0.92 | 0.82 | 0.98 | 0.94 | 0.96 | 0.95 |
| 7 | 108471548 | rs1949853 | -0.963 | 0.414 | 2.856 | 0.751 | 0.003 | 0.001 | 0.001 | 0.057 | 0.06 | 0.04 | 0.02 | 0.06 | 0.04 | 0.05 |
| 7 | 108471968 | rs10953581 | 2.06 | 0.374 | 3.124 | 0.749 | -0.039 | -0.031 | -0.031 | 0 | 0.88 | 0.32 | 0.94 | 0.94 | 0.95 | 0.95 |
| 7 | 108490660 | rs10252772 | -2.13 | 0.368 | 3.165 | 0.74 | -0.032 | -0.027 | -0.027 | 0 | 0.12 | 0.57 | 0.09 | 0.06 | 0.06 | 0.05 |
| 7 | 108503008 | rs1525183 | 2.111 | 0.363 | 2.994 | 0.738 | -0.041 | -0.037 | -0.037 | 0 | 0.87 | 0.32 | 0.9 | 0.94 | 0.95 | 0.95 |
| 7 | 108506618 | rs9649314 | 0.003 | 0.458 | 2.138 | 0.862 | -0.015 | -0.036 | -0.036 | 0.23 | 0.06 | 0 | 0.03 | 0.13 | 0.05 | 0.11 |
| 7 | 108521842 | rs848405 | -2.268 | 0.671 | 2.406 | 0.664 | -0.024 | -0.049 | -0.049 | 0 | 0.15 | 0.63 | 0.11 | 0.07 | 0.09 | 0.06 |
| 7 | 108522844 | rs848403 | 1.175 | 0.329 | 1.446 | 0.609 | -0.009 | -0.006 | -0.006 | 0 | 0.62 | 0.46 | 0.76 | 0.61 | 0.71 | 0.55 |
| 7 | 108524003 | rs2280645 | -1.014 | 0.897 | 0.886 | 0.851 | -0.025 | -0.019 | -0.019 | 0.153 | 0.06 | 0.01 | 0.06 | 0.1 | 0.08 | 0.07 |
| 7 | 108531829 | rs848386 | 0.947 | 0.774 | 1.09 | 0.796 | -0.001 | -0.001 | -0.001 | 0 | 0.71 | 0.74 | 0.76 | 0.74 | 0.76 | 0.66 |
| 7 | 108534672 | rs12705521 | -0.189 | 1.641 | 0.336 | 0.899 | -0.101 | -0.047 | -0.047 | 0 | 0.13 | 0.01 | 0.4 | 0.18 | 0.3 | 0.08 |
| 7 | 108586855 | rs848321 | -1.861 | 0.727 | 1.64 | 0.801 | 0.075 | 0.112 | 0.112 | 0.87 | 0.07 | 0.41 | 0.1 | 0.14 | 0.07 | 0.07 |
| 7 | 108596014 | rs11768328 | -1.192 | 0.732 | 1.568 | 0.875 | 0.058 | 0.09 | 0.09 | 0 | 0.05 | 0.27 | 0.08 | 0.15 | 0.07 | 0.07 |
| 7 | 108606430 | rs1513921 | 1.273 | 0.632 | 1.58 | 0.797 | -0.014 | 0.01 | 0.01 | 0 | 0.68 | 0.41 | 0.69 | 0.61 | 0.71 | 0.6 |
| 7 | 108613292 | rs757152 | 0.832 | 0.54 | 1.641 | 0.76 | -0.026 | -0.012 | -0.012 | 0 | 0.72 | 0.54 | 0.76 | 0.75 | 0.78 | 0.71 |
| 7 | 108621943 | rs4727718 | 0.656 | 0.595 | 1.632 | 0.77 | -0.015 | -0.006 | -0.006 | 0 | 0.73 | 0.65 | 0.77 | 0.75 | 0.78 | 0.69 |
| 7 | 108624433 | rs1513922 | 0.777 | 0.658 | 1.636 | 0.77 | -0.018 | -0.009 | -0.009 | 0 | 0.72 | 0.6 | 0.76 | 0.75 | 0.78 | 0.7 |
| 7 | 108633265 | rs1513917 | 0.674 | 0.964 | 1.59 | 0.75 | -0.003 | -0.005 | -0.005 | 0 | 0.72 | 0.67 | 0.75 | 0.75 | 0.75 | 0.67 |
| 7 | 108641894 | rs1913409 | 1.126 | 0.952 | 1.515 | 0.649 | 0.002 | 0.003 | 0.003 | 0 | 0.68 | 0.61 | 0.68 | 0.61 | 0.68 | 0.62 |
| 7 | 108644659 | rs769306 | 1.102 | 0.952 | 1.503 | 0.649 | 0.002 | 0.003 | 0.003 | 0 | 0.68 | 0.63 | 0.68 | 0.61 | 0.68 | 0.63 |
| 7 | 108649872 | rs4727720 | 1.121 | 0.953 | 1.381 | 0.65 | 0.006 | 0.01 | 0.01 | 0 | 0.68 | 0.56 | 0.68 | 0.61 | 0.68 | 0.6 |
| 7 | 108657719 | rs7778952 | 1.127 | 0.962 | 1.256 | 0.653 | 0.004 | 0.007 | 0.007 | 0 | 0.68 | 0.57 | 0.68 | 0.62 | 0.68 | 0.6 |
| 7 | 108660736 | rs10280128 | 1.124 | 1.008 | 1.228 | 0.749 | 0.002 | 0.009 | 0.009 | 0 | 0.68 | 0.59 | 0.68 | 0.6 | 0.69 | 0.59 |
| 7 | 108692515 | rs11975647 | -0.606 | 0.736 | 0.811 | 0.608 | -0.029 | -0.039 | -0.039 | 0.081 | 0.26 | 0.05 | 0.24 | 0.27 | 0.26 | 0.29 |
| 7 | 108700623 | rs9969266 | 0.993 | 0.738 | 0.933 | 0.608 | -0.01 | -0.004 | -0.004 | 0 | 0.7 | 0.44 | 0.72 | 0.73 | 0.73 | 0.71 |
| 7 | 108713889 | rs1473892 | -0.68 | 0.845 | -0.101 | 0.699 | -0.009 | -0.02 | -0.02 | 0.045 | 0.26 | 0.1 | 0.23 | 0.28 | 0.26 | 0.3 |
| 7 | 108719134 | rs17457580 | -0.028 | 0.523 | -0.869 | 0.406 | -0.028 | 0.011 | 0.011 | 0 | 0.17 | 0.01 | 0.34 | 0.15 | 0.14 | 0.23 |
| 7 | 108720580 | rs11978924 | 0.82 | 0.185 | -0.426 | 0.434 | -0.001 | -0.001 | -0.001 | 0.067 | 0.62 | 0.4 | 0.66 | 0.37 | 0.68 | 0.57 |
| 7 | 108722532 | rs6944775 | 1.736 | 0.315 | -1.166 | 0.112 | -0.115 | 0.192 | 0.192 | 0.984 | 0.33 | 0.04 | 0.23 | 0.05 | 0.41 | 0.23 |
| 7 | 108729850 | rs7787600 | -0.625 | -0.151 | 0.226 | 1.19 | 0.009 | 0.088 | 0.088 | 0 | 0.12 | 0.21 | 0.08 | 0.22 | 0.13 | 0.15 |
| 7 | 108731147 | rs1513924 | 0.321 | -0.531 | 0.128 | 0.64 | 0.002 | -0.017 | -0.017 | 0 | 0.77 | 0.8 | 0.75 | 0.59 | 0.81 | 0.73 |
| 7 | 108756678 | rs10225252 | 1.356 | -0.87 | 0.977 | 0.284 | -0.028 | 0.177 | 0.177 | 0 | 0.83 | 0.56 | 0.64 | 0.43 | 0.87 | 0.71 |
| 7 | 108757561 | rs9768602 | -0.815 | -0.763 | 0.615 | 0.144 | 0.001 | -0.008 | -0.008 | 0 | 0.09 | 0.11 | 0.12 | 0 | 0.08 | 0.09 |
| 7 | 108760833 | rs4595060 | 1.76 | -1.179 | 1.329 | 0.132 | -0.048 | 0.153 | 0.153 | 0 | 0.82 | 0.62 | 0.82 | 0.47 | 0.89 | 0.72 |
| 7 | 108762225 | rs10953594 | 1.159 | -1.223 | 1.353 | 0.099 | -0.036 | 0.117 | 0.117 | 0 | 0.88 | 0.8 | 0.84 | 0.47 | 0.95 | 0.77 |
| 7 | 108771276 | rs11974628 | -1.091 | -1 | 1.643 | -0.253 | 0.002 | -0.058 | -0.058 | 0 | 0.1 | 0.23 | 0.05 | 0 | 0.09 | 0.07 |
| 7 | 108774899 | rs6968523 | -1.729 | -1.526 | 1.547 | -0.823 | -0.02 | -0.099 | -0.099 | 0.366 | 0.27 | 0.65 | 0.14 | 0 | 0.18 | 0.14 |
| 7 | 108790097 | rs6954523 | -0.494 | -1.268 | 1.876 | -0.659 | -0.033 | -0.094 | -0.094 | 0.535 | 0.15 | 0.33 | 0.08 | 0 | 0.1 | 0.08 |
| 7 | 108797466 | rs1917537 | -0.036 | -2.248 | 1.241 | -1.46 | -0.007 | -0.107 | -0.107 | 0 | 0.45 | 0.54 | 0.24 | 0 | 0.31 | 0.22 |
| 7 | 108798551 | rs1525035 | -0.219 | -2.275 | 1.314 | -1.471 | 0.061 | 0.038 | 0.038 | 0.091 | 0.47 | 0.66 | 0.25 | 0 | 0.31 | 0.22 |
| 7 | 108798860 | rs10231177 | 0.735 | -2.178 | 1.308 | -1.405 | -0.019 | -0.058 | -0.058 | 0.747 | 0.12 | 0.2 | 0.06 | 0 | 0.09 | 0.08 |
| 7 | 108801624 | rs6951925 | -0.095 | -1.456 | 0.987 | -1.371 | -0.003 | -0.003 | -0.003 | 0 | 0.83 | 0.86 | 0.72 | 0.21 | 0.78 | 0.65 |
| 7 | 108813468 | rs17458769 | 1.312 | -2.003 | 0.776 | -1.508 | 0.045 | 0.179 | 0.179 | 0.16 | 0.3 | 0.06 | 0.14 | 0 | 0.23 | 0.14 |
| 7 | 108818739 | rs6958440 | 1.259 | -1.999 | 0.94 | -1.506 | 0.035 | 0.038 | 0.038 | 0.378 | 0.31 | 0.19 | 0.15 | 0 | 0.22 | 0.14 |
| 7 | 108820800 | rs10256677 | -0.018 | -1.683 | 1.317 | -1.279 | -0.023 | -0.119 | -0.119 | 1.109 | 0.15 | 0.47 | 0.1 | 0 | 0.12 | 0.08 |
| 7 | 108821664 | rs10260614 | 0.682 | -1.581 | 1.053 | -1.228 | -0.013 | -0.058 | -0.058 | 0.014 | 0.13 | 0.2 | 0.06 | 0 | 0.11 | 0.08 |
| 7 | 108829829 | rs10953597 | 1.175 | -1.452 | 0.792 | -1.012 | -0.02 | -0.038 | -0.038 | 0.214 | 0.07 | 0.01 | 0.15 | 0.26 | 0.08 | 0.16 |
| 7 | 108839848 | rs10278149 | 0.844 | -1.466 | 1.035 | -1.803 | -0.007 | -0.041 | -0.041 | 0 | 0.12 | 0.15 | 0.06 | 0 | 0.11 | 0.08 |
| 7 | 108845262 | rs7781788 | 0.306 | -1.432 | 1.102 | -1.822 | -0.007 | -0.073 | -0.073 | 0 | 0.14 | 0.23 | 0.07 | 0 | 0.13 | 0.1 |
| 7 | 108845931 | rs1525021 | -0.284 | -1.712 | 1.045 | -2.123 | 0.05 | 0.057 | 0.057 | 0.043 | 0.52 | 0.65 | 0.29 | 0 | 0.38 | 0.25 |
| 7 | 108854537 | rs6952038 | 0.871 | -1.655 | 0.786 | -2.045 | 0.038 | 0.031 | 0.031 | 0.836 | 0.37 | 0.22 | 0.18 | 0 | 0.27 | 0.17 |

| | | | | | | | | | | | | | | | | |
|---|---|---|---|---|---|---|---|---|---|---|---|---|---|---|---|---|
| 7 | 108859003 | rs17459383 | 0.858 | -1.655 | 0.786 | -2.045 | 0.038 | 0.031 | 0.031 | 0.759 | 0.37 | 0.22 | 0.18 | 0 | 0.27 | 0.17 |
| 7 | 108862337 | rs10270197 | -0.062 | -1.591 | 0.685 | -1.793 | -0.013 | -0.118 | -0.118 | 0 | 0.17 | 0.42 | 0.11 | 0 | 0.13 | 0.1 |
| 7 | 108862937 | rs1464813 | -0.485 | -1.697 | 0.66 | -1.776 | 0 | -0.001 | -0.001 | 0.185 | 0.83 | 0.83 | 0.74 | 0.22 | 0.86 | 0.65 |
| 7 | 108864838 | rs10263650 | -0.242 | -1.418 | 0.672 | -1.749 | -0.015 | -0.125 | -0.125 | 0.069 | 0.17 | 0.42 | 0.11 | 0 | 0.13 | 0.1 |
| 7 | 108885892 | rs10953600 | -0.032 | -1.573 | 0.425 | -1.922 | 0.04 | -0.004 | -0.004 | 0 | 0.5 | 0.43 | 0.71 | 1 | 0.63 | 0.76 |
| 7 | 108901536 | rs10240609 | 0.733 | -1.376 | 0.625 | -1.944 | -0.007 | -0.06 | -0.06 | 2.115 | 0.12 | 0.21 | 0.08 | 0.01 | 0.12 | 0.08 |
| 7 | 108914605 | rs10243474 | 0.305 | -1.428 | 0.831 | -1.967 | -0.007 | -0.094 | -0.094 | 0.098 | 0.14 | 0.36 | 0.1 | 0 | 0.13 | 0.08 |
| 7 | 108919090 | rs4730374 | 0.083 | -1.655 | 1.085 | -2.158 | 0.043 | 0.007 | 0.007 | 0.077 | 0.5 | 0.42 | 0.72 | 0.99 | 0.61 | 0.75 |
| 7 | 108930349 | rs1541338 | 0.522 | -1.354 | 1.357 | -1.714 | 0 | 0 | 0 | 0 | 0.14 | 0.13 | 0.23 | 0.76 | 0.13 | 0.31 |
| 7 | 108936876 | rs10270894 | 0.21 | -0.899 | 1.607 | -2.01 | -0.012 | -0.08 | -0.08 | 0.011 | 0.13 | 0.29 | 0.09 | 0.01 | 0.12 | 0.08 |
| 7 | 108960029 | rs848896 | 0.647 | -0.799 | 0.726 | -1.971 | -0.003 | -0.028 | -0.028 | 0.2 | 0.11 | 0.1 | 0.06 | 0 | 0.11 | 0.1 |
| 7 | 108979199 | rs10277164 | -0.783 | -1.164 | 0.803 | -2.3 | -0.008 | -0.086 | -0.086 | 1.132 | 0.14 | 0.31 | 0.08 | 0 | 0.12 | 0.07 |
| 7 | 108982274 | rs711450 | -1.393 | -1.855 | -0.093 | -2.68 | 0.063 | 0.039 | 0.039 | 0 | 0.68 | 0.85 | 0.64 | 0.24 | 0.77 | 0.64 |
| 7 | 108987486 | rs848866 | -1.202 | -0.91 | 0.587 | -2.51 | -0.007 | -0.117 | -0.117 | 0.332 | 0.27 | 0.57 | 0.17 | 0.01 | 0.27 | 0.18 |
| 7 | 109027535 | rs1406298 | 0.9 | -0.564 | 0.839 | -2.184 | 0.073 | 0 | 0 | 0.064 | 0.92 | 0.52 | 0.93 | 1 | 0.95 | 0.96 |
| 7 | 109028297 | rs7794651 | 1.111 | -0.575 | 0.452 | -2.211 | 0.006 | -0.091 | -0.091 | 0 | 0.84 | 0.37 | 0.87 | 0.99 | 0.85 | 0.92 |
| 7 | 109052359 | rs10953607 | 0.681 | -0.605 | -0.513 | -2.729 | -0.092 | 0.069 | 0.069 | 0 | 0.36 | 0.13 | 0.4 | 0.22 | 0.44 | 0.28 |
| 7 | 109053967 | rs2396099 | 0.964 | -0.59 | 0.347 | -2.612 | 0.026 | -0.105 | -0.105 | 0.857 | 0.85 | 0.45 | 0.88 | 1 | 0.85 | 0.92 |
| 7 | 109064120 | rs12705587 | 0.52 | 0.856 | -0.359 | -2.907 | -0.08 | -0.009 | -0.009 | 0 | 0.12 | 0.01 | 0.21 | 0.12 | 0.23 | 0.09 |
| 7 | 109076184 | rs2091546 | 0.206 | 0.225 | 0.363 | -2.785 | 0 | -0.054 | -0.054 | 0.241 | 0.9 | 0.81 | 0.93 | 1 | 0.9 | 0.96 |
| 7 | 109081747 | rs11975565 | 0.793 | -0.033 | 0.329 | -2.824 | 0.063 | 0.036 | 0.036 | 0 | 0.32 | 0.17 | 0.37 | 0.77 | 0.23 | 0.52 |
| 7 | 109085999 | rs2396100 | 0.917 | -0.286 | 0.208 | -2.875 | 0.077 | 0.048 | 0.048 | 0 | 0.3 | 0.14 | 0.36 | 0.79 | 0.22 | 0.5 |
| 7 | 109098364 | rs13222164 | -0.353 | 1.06 | 1.01 | -1.969 | -0.024 | -0.007 | -0.007 | 0 | 0.11 | 0.06 | 0.09 | 0.08 | 0.18 | 0.11 |
| 7 | 109101870 | rs11763570 | -0.536 | 1.244 | 0.99 | -2.031 | -0.01 | 0.012 | 0.012 | 0 | 0.16 | 0.08 | 0.1 | 0.11 | 0.17 | 0.13 |
| 7 | 109102855 | rs11561993 | 2.354 | 1.198 | 1.146 | -2.259 | 0.019 | 0.11 | 0.11 | 0 | 0.52 | 0.26 | 0.53 | 0.78 | 0.48 | 0.61 |
| 7 | 109104920 | rs7794942 | -0.331 | 1.314 | 1.516 | -2.207 | 0.007 | -0.106 | -0.106 | 0 | 0.83 | 0.55 | 0.77 | 0.97 | 0.88 | 0.9 |
| 7 | 109109019 | rs7788553 | -2.027 | 1.089 | 1.46 | -2.146 | -0.001 | -0.001 | -0.001 | 0 | 0.36 | 0.35 | 0.23 | 0.19 | 0.4 | 0.27 |
| 7 | 109109099 | rs10499972 | -0.765 | 1.124 | 1.421 | -2.164 | -0.013 | 0.025 | 0.025 | 0 | 0.2 | 0.15 | 0.12 | 0.08 | 0.23 | 0.15 |
| 7 | 109115461 | rs11765085 | -0.289 | 1.124 | 1.415 | -2.16 | 0.001 | -0.004 | -0.004 | 1.049 | 0.15 | 0.13 | 0.11 | 0.11 | 0.17 | 0.13 |
| 7 | 109123917 | rs1880233 | 2.294 | 1.103 | 1.451 | -2.002 | -0.012 | 0.047 | 0.047 | 1.154 | 0.61 | 0.27 | 0.75 | 0.81 | 0.6 | 0.73 |
| 7 | 109130739 | rs6951122 | -0.481 | 1.261 | 0.16 | -1.859 | -0.003 | 0.004 | 0.004 | 0 | 0.16 | 0.12 | 0.11 | 0.11 | 0.17 | 0.13 |
| 7 | 109136112 | rs12705601 | -0.265 | 1.241 | 0.046 | -1.837 | -0.015 | 0.019 | 0.019 | 0 | 0.17 | 0.08 | 0.11 | 0.08 | 0.2 | 0.11 |
| 7 | 109142224 | rs12705602 | -0.247 | 1.252 | 0.052 | -1.836 | -0.013 | 0.018 | 0.018 | 0 | 0.17 | 0.08 | 0.11 | 0.08 | 0.2 | 0.1 |
| 7 | 109150237 | rs12113865 | -2.127 | 1.192 | 0.191 | -1.867 | 0.026 | 0.097 | 0.097 | 0 | 0.48 | 0.3 | 0.45 | 0.21 | 0.54 | 0.37 |
| 7 | 109153739 | rs6979859 | -0.73 | 1.244 | 0.044 | -1.787 | -0.02 | 0.017 | 0.017 | 0.029 | 0.17 | 0.1 | 0.14 | 0.11 | 0.22 | 0.17 |
| 7 | 109159007 | rs10953618 | -0.792 | 0.992 | -0.071 | -1.737 | -0.038 | 0.093 | 0.093 | 0 | 0.31 | 0.13 | 0.31 | 0.1 | 0.33 | 0.21 |
| 7 | 109161891 | rs12705610 | -1.065 | 1.058 | -0.083 | -1.733 | -0.024 | 0.102 | 0.102 | 0 | 0.35 | 0.18 | 0.33 | 0.1 | 0.36 | 0.23 |
| 7 | 109166664 | rs11770336 | -0.46 | 0.816 | 0.388 | -1.33 | -0.005 | 0.014 | 0.014 | 0 | 0.16 | 0.1 | 0.1 | 0.1 | 0.17 | 0.13 |
| 7 | 109168430 | rs12705612 | 0.074 | 0.718 | 0.276 | -1.381 | -0.002 | 0.003 | 0.003 | 0.029 | 0.12 | 0.09 | 0.07 | 0.08 | 0.15 | 0.09 |
| 7 | 109170014 | rs4730391 | -0.775 | 0.777 | 0.586 | -1.447 | -0.003 | 0.005 | 0.005 | 0 | 0.24 | 0.22 | 0.28 | 0.1 | 0.28 | 0.18 |
| 7 | 109174062 | rs12705614 | -0.016 | 0.512 | 0.394 | -1.389 | 0 | 0 | 0 | 0.05 | 0.13 | 0.13 | 0.07 | 0.08 | 0.15 | 0.07 |
| 7 | 109190835 | rs1357687 | 0.974 | 0.536 | 0.633 | -1.462 | 0.01 | -0.037 | -0.037 | 0 | 0.68 | 0.63 | 0.64 | 0.88 | 0.64 | 0.73 |
| 7 | 109204464 | rs1357683 | -0.205 | 0.754 | 0.138 | -1.321 | 0.009 | 0.069 | 0.069 | 0 | 0.14 | 0.03 | 0.04 | 0.01 | 0.11 | 0.11 |
| 7 | 109207197 | rs10499978 | 0.36 | 0.77 | 0.07 | -1.329 | -0.07 | -0.003 | -0.003 | 0 | 0.15 | 0.02 | 0.27 | 0.12 | 0.21 | 0.13 |
| 7 | 109214139 | rs1721447 | 1.611 | 0.71 | 0.549 | -1.332 | 0.001 | -0.021 | -0.021 | 0 | 0.54 | 0.52 | 0.55 | 0.78 | 0.48 | 0.61 |
| 7 | 109217197 | rs1524272 | 0.608 | 0.018 | 0.621 | -0.976 | -0.008 | -0.115 | -0.115 | 0 | 0.81 | 0.6 | 0.9 | 0.99 | 0.85 | 0.86 |
| 7 | 109222164 | rs954483 | -0.519 | 0.022 | -0.296 | -0.423 | -0.001 | -0.001 | -0.001 | 0 | 0.12 | 0.06 | 0.06 | 0.09 | 0.14 | 0.12 |
| 7 | 109230184 | rs2396135 | 1.08 | -0.403 | -0.443 | 0.247 | 0.042 | -0.02 | -0.02 | 0 | 0.56 | 0.24 | 0.6 | 0.47 | 0.64 | 0.53 |
| 7 | 109240787 | rs1464743 | 0.81 | -1.162 | 0.627 | 0.596 | -0.058 | 0.015 | 0.015 | 0.53 | 0.73 | 0.45 | 0.76 | 0.7 | 0.85 | 0.63 |
| 7 | 109251487 | rs6949142 | 0.115 | -1.135 | 1.189 | 0.616 | -0.033 | 0.248 | 0.248 | 1.788 | 0.89 | 0.55 | 0.83 | 0.71 | 0.95 | 0.76 |
| 7 | 109252043 | rs10255690 | 1.345 | -1.095 | 0.251 | 0.481 | 0.05 | -0.049 | -0.049 | 0 | 0.67 | 0.25 | 0.68 | 0.56 | 0.79 | 0.58 |
| 7 | 109262114 | rs10261639 | 0.446 | -0.74 | 0.152 | 0.083 | 0.001 | -0.009 | -0.009 | 0.365 | 0.25 | 0.3 | 0.21 | 0.16 | 0.25 | 0.22 |
| 7 | 109266627 | rs2063937 | 0.446 | -0.74 | 0.152 | 0.074 | 0.001 | -0.01 | -0.01 | 0 | 0.25 | 0.3 | 0.21 | 0.16 | 0.25 | 0.22 |
| 7 | 109267536 | rs10499980 | 1.13 | -1.104 | -0.758 | -0.156 | -0.096 | -0.101 | -0.101 | 0.46 | 0.4 | 0.08 | 0.49 | 0.53 | 0.52 | 0.35 |
| 7 | 109267610 | rs711459 | 0.02 | -1.228 | -0.351 | -0.352 | -0.014 | -0.084 | -0.084 | 0 | 0.23 | 0.28 | 0.11 | 0 | 0.17 | 0.23 |
| 7 | 109275509 | rs711467 | -0.313 | -1.195 | -0.493 | -0.156 | -0.037 | 0.075 | 0.075 | 1.138 | 0.88 | 0.77 | 0.83 | 0.69 | 0.94 | 0.79 |
| 7 | 109278577 | rs7784378 | 0.759 | -0.629 | -1.341 | -0.194 | -0.09 | -0.126 | -0.126 | 0 | 0.41 | 0.07 | 0.5 | 0.5 | 0.53 | 0.34 |
| 7 | 109282982 | rs11761734 | 0.014 | -0.596 | -1.157 | -0.159 | -0.006 | -0.05 | -0.05 | 2.803 | 0.2 | 0.21 | 0.1 | 0 | 0.14 | 0.25 |
| 7 | 109288483 | rs6976113 | 0.194 | 0.265 | -1.561 | 0.489 | -0.061 | -0.078 | -0.078 | 0 | 0.4 | 0.17 | 0.5 | 0.51 | 0.52 | 0.34 |
| 7 | 109297612 | rs820939 | -0.485 | 0.109 | -0.896 | 0.637 | -0.011 | 0.007 | 0.007 | 3.466 | 0.37 | 0.58 | 0.38 | 0.59 | 0.38 | 0.6 |
| 7 | 109308566 | rs820930 | -0.666 | 0.788 | -1.638 | 0.049 | -0.001 | -0.006 | -0.006 | 1.82 | 0.66 | 0.69 | 0.81 | 0.79 | 0.69 | 0.78 |
| 7 | 109314640 | rs4314580 | 0.419 | 0.761 | -1.701 | 0.044 | -0.107 | 0 | 0 | 0 | 0.3 | 0.09 | 0.48 | 0.21 | 0.43 | 0.19 |
| 7 | 109314773 | rs711481 | 0.108 | 0.301 | -1.694 | -0.107 | 0.004 | 0 | 0 | 0 | 0.55 | 0.63 | 0.42 | 0.62 | 0.45 | 0.62 |
| 7 | 109333611 | rs711499 | 0.09 | 0.228 | -1.695 | -0.114 | 0.009 | -0.002 | -0.002 | 0.186 | 0.55 | 0.67 | 0.42 | 0.62 | 0.44 | 0.63 |
| 7 | 109334594 | rs11767794 | 1.751 | 0.043 | -1.542 | -0.413 | -0.031 | 0.008 | 0.008 | 0.301 | 0.06 | 0.01 | 0.04 | 0.06 | 0.1 | 0.11 |
| 7 | 109345821 | rs990943 | 0.122 | -0.634 | -0.96 | -0.037 | 0.004 | 0.001 | 0.001 | 0 | 0.38 | 0.44 | 0.67 | 0.55 | 0.47 | 0.44 |
| 7 | 109349375 | rs1033242 | 1.415 | -0.627 | -1.033 | -0.015 | -0.082 | -0.028 | -0.028 | 0 | 0.22 | 0.09 | 0.44 | 0.21 | 0.37 | 0.15 |
| 7 | 109350011 | rs10275927 | 0.929 | -0.732 | -0.932 | -0.049 | -0.019 | 0.029 | 0.029 | 0 | 0.16 | 0.23 | 0.23 | 0.34 | 0.1 | 0.29 |
| 7 | 109355060 | rs28481921 | 0.563 | -0.748 | -0.897 | -0.075 | 0.005 | 0.019 | 0.019 | 1.962 | 0.38 | 0.31 | 0.2 | 0.27 | 0.34 | 0.33 |

| | | | | | | | | | | | | | | | | |
|---|---|---|---|---|---|---|---|---|---|---|---|---|---|---|---|---|
| 7 | 109359012 | rs12537146 | 0.868 | -0.722 | -0.774 | -0.065 | 0 | 0.004 | 0.004 | 0.011 | 0.16 | 0.19 | 0.24 | 0.35 | 0.13 | 0.34 |
| 7 | 109368769 | rs711513 | 0.326 | -0.7 | -0.788 | -0.229 | 0 | -0.001 | -0.001 | 0 | 0.4 | 0.38 | 0.2 | 0.27 | 0.35 | 0.34 |
| 7 | 109369391 | rs1980352 | -0.049 | -0.715 | -0.798 | -0.123 | -0.015 | 0.005 | 0.005 | 2.759 | 0.6 | 0.41 | 0.64 | 0.49 | 0.71 | 0.48 |
| 7 | 109377068 | rs711518 | 0.828 | -0.644 | -0.331 | -0.186 | 0.05 | 0.009 | 0.009 | 0 | 0.25 | 0.37 | 0.45 | 0.24 | 0.35 | 0.15 |
| 7 | 109383126 | rs1608655 | -0.098 | -1.01 | -0.351 | -0.183 | 0.016 | -0.049 | -0.049 | 0 | 0.26 | 0.21 | 0.29 | 0.46 | 0.24 | 0.39 |
| 7 | 109388584 | rs6974557 | -0.07 | -1.023 | -0.316 | -0.204 | 0.011 | -0.03 | -0.03 | 0 | 0.26 | 0.22 | 0.29 | 0.46 | 0.24 | 0.39 |
| 7 | 109400135 | rs711529 | -1.147 | -1.158 | -0.239 | -0.292 | -0.02 | -0.037 | -0.037 | 0 | 0.76 | 0.71 | 0.87 | 0.89 | 0.81 | 0.75 |
| 7 | 109407682 | rs1524466 | 0.394 | -1.109 | -0.214 | -0.297 | 0.007 | 0.019 | 0.019 | 0.079 | 0.52 | 0.4 | 0.28 | 0.31 | 0.46 | 0.47 |
| 7 | 109408487 | rs1608659 | 0.039 | -1.15 | -0.078 | -0.315 | -0.034 | 0.062 | 0.062 | 3.138 | 0.74 | 0.54 | 0.72 | 0.54 | 0.81 | 0.62 |
| 7 | 109409473 | rs10274990 | 1.297 | -1.059 | -0.092 | -0.244 | -0.006 | -0.004 | -0.004 | 0 | 0.23 | 0.19 | 0.44 | 0.24 | 0.35 | 0.15 |
| 7 | 109422023 | rs7797714 | 1.548 | -1.104 | -0.182 | -0.275 | 0.003 | 0.035 | 0.035 | 7.092 | 0.25 | 0.06 | 0.12 | 0.18 | 0.21 | 0.21 |
| 7 | 109424105 | rs3114815 | 0.55 | -0.926 | -0.021 | -0.173 | 0.009 | 0.015 | 0.015 | 0.045 | 0.49 | 0.41 | 0.26 | 0.3 | 0.41 | 0.48 |
| 7 | 109426960 | rs3114803 | -0.761 | -0.332 | 0.694 | 0.174 | 0.051 | 0.01 | 0.01 | 0 | 0.75 | 0.62 | 0.55 | 0.76 | 0.65 | 0.85 |
| 7 | 109455705 | rs3114767 | -0.569 | -0.412 | 0.629 | 0.144 | 0.085 | 0.015 | 0.015 | 0 | 0.73 | 0.48 | 0.54 | 0.76 | 0.65 | 0.86 |
| 7 | 109460110 | rs3114758 | -0.723 | -0.391 | 0.649 | 0.184 | 0.052 | 0.009 | 0.009 | 0 | 0.75 | 0.62 | 0.54 | 0.76 | 0.64 | 0.84 |
| 7 | 109484997 | rs3107968 | -0.141 | -0.523 | 0.554 | 0.143 | 0.039 | 0.018 | 0.018 | 7.183 | 0.5 | 0.32 | 0.41 | 0.59 | 0.44 | 0.65 |
| 7 | 109504136 | rs6949632 | 0.31 | 0.379 | 0.277 | 0.703 | -0.084 | -0.038 | -0.038 | 0 | 0.23 | 0.08 | 0.44 | 0.24 | 0.35 | 0.16 |
| 7 | 109510718 | rs6978512 | 0.227 | 0.454 | -0.09 | 0.673 | -0.13 | -0.051 | -0.051 | 0 | 0.21 | 0.05 | 0.42 | 0.24 | 0.37 | 0.18 |
| 7 | 109513917 | rs10953645 | 1.093 | 0.029 | 0.064 | 0.522 | 0.024 | 0.041 | 0.041 | 0.268 | 0.43 | 0.17 | 0.25 | 0.33 | 0.41 | 0.44 |
| 7 | 109542198 | rs4552825 | 1.021 | 0.438 | 1.07 | 0.706 | -0.008 | -0.018 | -0.018 | 2.014 | 0.95 | 0.76 | 0.97 | 1 | 0.99 | 0.94 |
| 7 | 109554669 | rs3094537 | -2.078 | 0.343 | 1.001 | 0.662 | -0.046 | -0.057 | -0.057 | 0 | 0.1 | 0.73 | 0.06 | 0.01 | 0.02 | 0.07 |
| 7 | 109568347 | rs3114807 | 0.594 | 1.158 | 0.296 | 1.057 | -0.143 | -0.098 | -0.098 | 0 | 0.79 | 0.98 | 0.63 | 0.73 | 0.64 | 0.83 |
| 7 | 109570757 | rs976399 | -1.298 | 1.213 | 0.972 | 1.185 | 0.056 | 0.009 | 0.009 | 0.442 | 0.27 | 0.43 | 0.42 | 0.25 | 0.35 | 0.17 |
| 7 | 109572509 | rs1115285 | -1.406 | 1.2 | 0.785 | 1.181 | 0.053 | 0.005 | 0.005 | 0.565 | 0.28 | 0.53 | 0.43 | 0.26 | 0.35 | 0.19 |
| 7 | 109590629 | rs1013745 | -1.762 | 0.249 | 0.804 | 1.522 | -0.017 | 0.038 | 0.038 | 0.703 | 0.22 | 0.68 | 0.27 | 0.41 | 0.13 | 0.36 |
| 7 | 109594847 | rs3114833 | -1.119 | 0.181 | 1.523 | 1.525 | -0.027 | 0.113 | 0.113 | 0 | 0.15 | 0.27 | 0.19 | 0.4 | 0.09 | 0.28 |
| 7 | 109596453 | rs2711380 | -1.675 | 0.159 | 1.762 | 1.532 | -0.046 | 0.133 | 0.133 | 0.251 | 0.18 | 0.61 | 0.22 | 0.41 | 0.09 | 0.28 |
| 7 | 109598161 | rs3114834 | -2.113 | 0.056 | 1.959 | 1.74 | -0.044 | 0.137 | 0.137 | 0.297 | 0.23 | 0.48 | 0.27 | 0.53 | 0.16 | 0.43 |
| 7 | 109600924 | rs2711378 | -1.66 | 0.03 | 1.457 | 1.812 | -0.053 | 0.149 | 0.149 | 0.129 | 0.2 | 0.53 | 0.22 | 0.41 | 0.11 | 0.28 |
| 7 | 109610387 | rs2523210 | 0.427 | 0.67 | 0.819 | 1.372 | -0.039 | 0.005 | 0.005 | 0.029 | 0.83 | 0.95 | 0.78 | 0.83 | 0.75 | 0.84 |
| 7 | 109634984 | rs2523234 | -1.186 | 0.481 | 0.735 | 0.722 | 0.013 | -0.007 | -0.007 | 0 | 0.05 | 0.24 | 0.02 | 0 | 0.02 | 0.02 |
| 7 | 109647829 | rs2711364 | 0.88 | 0.908 | 0.327 | 0.751 | 0.005 | -0.01 | -0.01 | 0.124 | 0.63 | 0.44 | 0.61 | 0.27 | 0.55 | 0.5 |
| 7 | 109649916 | rs2711368 | 0.626 | 0.767 | -0.261 | -0.125 | 0.027 | 0.174 | 0.174 | 0.822 | 0.49 | 0.21 | 0.45 | 0.15 | 0.43 | 0.3 |
| 7 | 109677042 | rs10251582 | -0.197 | 0.814 | 0.117 | 0.284 | 0 | 0 | 0 | 0 | 0.12 | 0.15 | 0.25 | 0.14 | 0.14 | 0.12 |
| 7 | 109703062 | rs2249789 | -1.274 | 0.865 | 0.189 | 0.273 | 0.059 | 0.11 | 0.11 | 1.295 | 0.54 | 0.74 | 0.62 | 0.82 | 0.64 | 0.77 |
| 7 | 109705384 | rs1525513 | -0.517 | 0.875 | 0.333 | 0.247 | -0.005 | 0.006 | 0.006 | 3.733 | 0.4 | 0.64 | 0.36 | 0.68 | 0.48 | 0.63 |
| 7 | 109711537 | rs7793344 | -0.067 | 0.616 | 0.695 | 0.936 | 0.077 | 0.088 | 0.088 | 1.08 | 0.29 | 0.48 | 0.27 | 0.5 | 0.41 | 0.47 |
| 7 | 109719099 | rs1723652 | 0.046 | 0.555 | 0.644 | 0.893 | 0.001 | 0 | 0 | 0 | 0.13 | 0.11 | 0.25 | 0.15 | 0.14 | 0.13 |
| 7 | 109720628 | rs12539392 | 0.444 | 0.566 | 0.616 | 0.898 | -0.026 | -0.028 | -0.028 | 0 | 0.11 | 0.01 | 0.24 | 0.15 | 0.14 | 0.13 |
| 7 | 109720986 | rs10245119 | -0.118 | 0.776 | 1.087 | 1.442 | 0.034 | -0.023 | -0.023 | 0 | 0.29 | 0.32 | 0.26 | 0.2 | 0.41 | 0.37 |
| 7 | 109722969 | rs13340481 | -0.024 | 0.783 | 0.946 | 1.488 | -0.039 | 0.048 | 0.048 | 0 | 0.08 | 0.16 | 0.06 | 0.17 | 0.02 | 0.13 |
| 7 | 109733975 | rs940770 | -0.937 | 0.795 | 1.256 | 1.534 | -0.01 | 0.013 | 0.013 | 3.944 | 0.41 | 0.69 | 0.36 | 0.66 | 0.48 | 0.63 |
| 7 | 109734425 | rs1525510 | 0.163 | 0.806 | 1.341 | 1.523 | 0.021 | -0.015 | -0.015 | 0 | 0.27 | 0.33 | 0.22 | 0.2 | 0.36 | 0.34 |
| 7 | 109736919 | rs1722034 | -0.754 | 0.808 | 1.245 | 1.525 | 0.018 | -0.001 | -0.001 | 0 | 0.38 | 0.51 | 0.32 | 0.38 | 0.43 | 0.5 |
| 7 | 109738166 | rs1722035 | -1.536 | 0.813 | 1.1 | 1.101 | 0.068 | 0.135 | 0.135 | 1.332 | 0.54 | 0.81 | 0.64 | 0.82 | 0.64 | 0.72 |
| 7 | 109742696 | rs17157405 | -1.528 | 0.769 | 1.459 | 1.229 | 0.024 | 0.22 | 0.22 | 0 | 0.14 | 0.34 | 0.13 | 0.46 | 0.12 | 0.3 |
| 7 | 109757243 | rs6948497 | -1.504 | 0.758 | 1.434 | 1.25 | 0.019 | 0.2 | 0.2 | 0 | 0.14 | 0.33 | 0.13 | 0.44 | 0.12 | 0.32 |
| 7 | 109774271 | rs1722047 | -1.346 | 0.498 | 0.616 | 0.861 | 0.035 | 0.076 | 0.076 | 1.883 | 0.59 | 0.77 | 0.65 | 0.8 | 0.64 | 0.8 |
| 7 | 109782499 | rs1722052 | -0.541 | 0.535 | 0.885 | 0.807 | -0.002 | -0.008 | -0.008 | 0.922 | 0.46 | 0.5 | 0.53 | 0.33 | 0.56 | 0.47 |
| 7 | 109783942 | rs10953652 | -0.55 | 0.576 | 0.837 | 0.826 | -0.005 | 0.088 | 0.088 | 0.048 | 0.15 | 0.23 | 0.13 | 0.44 | 0.11 | 0.34 |
| 7 | 109792895 | rs7804183 | -0.607 | 0.478 | 0.635 | 0.783 | 0.004 | -0.01 | -0.01 | 0.043 | 0.16 | 0.17 | 0.13 | 0.45 | 0.11 | 0.36 |
| 7 | 109793244 | rs1525500 | -0.093 | 0.788 | 0.734 | 0.701 | 0.008 | 0.029 | 0.029 | 0 | 0.21 | 0.29 | 0.32 | 0.36 | 0.24 | 0.26 |
| 7 | 109799611 | rs1625755 | -1.145 | 0.819 | 0.677 | 0.444 | 0.004 | -0.023 | -0.023 | 0 | 0.34 | 0.45 | 0.43 | 0.8 | 0.33 | 0.58 |
| 7 | 109802202 | rs10499987 | 0.291 | 0.882 | 0.823 | 0.079 | 0.006 | -0.068 | -0.068 | 0.746 | 0.23 | 0.29 | 0.2 | 0.01 | 0.29 | 0.18 |
| 7 | 109806673 | rs6980262 | 0.529 | 0.885 | 0.782 | 0.086 | -0.054 | 0.082 | 0.082 | 0.89 | 0.22 | 0.08 | 0.19 | 0.01 | 0.3 | 0.18 |
| 7 | 109809055 | rs7795982 | 0.227 | 0.901 | 0.725 | 0.12 | 0.019 | -0.114 | -0.114 | 0.038 | 0.24 | 0.34 | 0.21 | 0.01 | 0.29 | 0.18 |
| 7 | 109811300 | rs1723593 | -0.817 | 0.955 | 0.527 | 0.244 | 0 | -0.002 | -0.002 | 0 | 0.43 | 0.57 | 0.48 | 0.31 | 0.5 | 0.42 |
| 7 | 109814028 | rs13224529 | -0.102 | 0.857 | 0.223 | -0.119 | 0.009 | -0.077 | -0.077 | 0 | 0.23 | 0.29 | 0.2 | 0.01 | 0.28 | 0.18 |
| 7 | 109816138 | rs2254897 | 0.42 | -0.018 | -0.058 | 0.505 | -0.011 | -0.021 | -0.021 | 0 | 0.88 | 0.81 | 0.91 | 0.92 | 0.89 | 0.9 |
| 7 | 109816518 | rs940457 | -0.937 | 0.202 | -0.208 | 1.011 | -0.009 | 0.008 | 0.008 | 0.134 | 0.11 | 0.14 | 0.09 | 0.2 | 0.07 | 0.23 |
| 7 | 109819227 | rs6960899 | -1.201 | 0.568 | 0.084 | 1.543 | 0.01 | 0.109 | 0.109 | 0 | 0.19 | 0.36 | 0.24 | 0.5 | 0.21 | 0.24 |
| 7 | 109827761 | rs10249619 | -0.175 | 0.405 | -0.095 | 1.206 | 0.002 | -0.002 | -0.002 | 0 | 0.37 | 0.33 | 0.29 | 0.39 | 0.33 | 0.44 |
| 7 | 109829271 | rs11770896 | 0.692 | 0.701 | -0.1 | 0.76 | -0.018 | 0.018 | 0.018 | 2.199 | 0.47 | 0.27 | 0.54 | 0.53 | 0.52 | 0.37 |
| 7 | 109830780 | rs1723580 | -0.114 | 0.841 | 0.293 | 0.827 | 0.013 | -0.058 | -0.058 | 0 | 0.15 | 0.28 | 0.16 | 0.08 | 0.14 | 0.18 |
| 7 | 109842540 | rs1608837 | 0.572 | 0.448 | -0.387 | 0.755 | -0.052 | -0.028 | -0.028 | 0 | 0.4 | 0.26 | 0.56 | 0.53 | 0.54 | 0.39 |
| 7 | 109842734 | rs1608836 | -0.224 | 0.22 | -0.107 | 0.675 | 0.006 | 0.002 | 0.002 | 0.512 | 0.55 | 0.43 | 0.7 | 0.61 | 0.66 | 0.6 |
| 7 | 109848853 | rs17157477 | -0.245 | 0.265 | -0.954 | -0.01 | 0.017 | 0.057 | 0.057 | 0 | 0.25 | 0.14 | 0.21 | 0.1 | 0.21 | 0.33 |
| 7 | 109856125 | rs2396220 | 0.781 | 1.035 | -1.071 | -0.415 | -0.026 | 0.166 | 0.166 | 0.994 | 0.64 | 0.09 | 0.65 | 0.9 | 0.6 | 0.56 |
| 7 | 109862289 | rs4730423 | -0.35 | 0.947 | -1.403 | 0.033 | -0.081 | -0.02 | -0.02 | 0 | 0.29 | 0.11 | 0.39 | 0.32 | 0.44 | 0.4 |

| | | | | | | | | | | | | | | | | |
|---|---|---|---|---|---|---|---|---|---|---|---|---|---|---|---|---|
| 7 | 109862980 | rs10270509 | -1.065 | 0.857 | -0.95 | 0.03 | -0.04 | -0.066 | -0.066 | 0 | 0.11 | 0.63 | 0.08 | 0 | 0.05 | 0.13 |
| 7 | 109866785 | rs1989874 | 1.12 | 0.715 | -1.871 | -0.23 | 0 | 0.003 | 0.003 | 0.577 | 0.37 | 0.6 | 0.38 | 0.49 | 0.33 | 0.37 |
| 7 | 109866859 | rs1989875 | -0.013 | 0.66 | -2.665 | -0.386 | -0.099 | -0.104 | -0.104 | 1.884 | 0.72 | 0.95 | 0.62 | 0.58 | 0.59 | 0.72 |
| 7 | 109867062 | rs17157502 | 1.205 | 0.544 | -2.658 | -0.805 | 0.001 | 0.02 | 0.02 | 0 | 0.12 | 0.02 | 0.05 | 0.05 | 0.07 | 0.06 |
| 7 | 109871406 | rs9641438 | -1.209 | 0.767 | -2.546 | -0.641 | -0.051 | -0.065 | -0.065 | 0 | 0.76 | 0.98 | 0.72 | 0.69 | 0.77 | 0.61 |
| 7 | 109875729 | rs10258980 | 0.472 | -0.087 | -2.445 | -0.755 | 0.033 | -0.015 | -0.015 | 0.006 | 0.47 | 0.7 | 0.42 | 0.48 | 0.38 | 0.43 |
| 7 | 109877036 | rs13230336 | 1.118 | -0.045 | -2.438 | -1.105 | -0.071 | 0.09 | 0.09 | 0 | 0.25 | 0.1 | 0.28 | 0.1 | 0.37 | 0.34 |
| 7 | 109877875 | rs9640729 | 0.427 | -0.1 | -1.749 | -0.453 | 0.009 | -0.007 | -0.007 | 0.632 | 0.4 | 0.67 | 0.39 | 0.52 | 0.34 | 0.44 |
| 7 | 109881210 | rs2188814 | -0.983 | -0.368 | -1.792 | -0.551 | -0.078 | -0.085 | -0.085 | 0 | 0.15 | 0.61 | 0.06 | 0.01 | 0.04 | 0.21 |
| 7 | 109885059 | rs10259163 | 0.142 | 0.115 | -1.685 | -0.333 | 0.01 | -0.01 | -0.01 | 0 | 0.41 | 0.65 | 0.41 | 0.51 | 0.35 | 0.38 |
| 7 | 109888464 | rs12705672 | 0.965 | 0.774 | -2.1 | -0.089 | -0.06 | -0.125 | -0.125 | 0 | 0.28 | 0.05 | 0.33 | 0.51 | 0.3 | 0.23 |
| 7 | 109890965 | rs719530 | -0.288 | 0.945 | -1.658 | -0.06 | -0.009 | -0.019 | -0.019 | 0 | 0.57 | 0.69 | 0.66 | 0.53 | 0.55 | 0.47 |
| 7 | 109893076 | rs17417471 | 0.404 | 1.636 | -1.904 | -0.246 | -0.048 | 0.012 | 0.012 | 0 | 0.09 | 0.03 | 0.22 | 0.07 | 0.21 | 0.23 |
| 7 | 109893783 | rs4141364 | -0.096 | 0.478 | -1.481 | -0.164 | 0.008 | 0.103 | 0.103 | 0 | 0.51 | 0.27 | 0.55 | 0.75 | 0.5 | 0.59 |
| 7 | 109900921 | rs7796325 | -1.001 | 0.27 | -1.282 | 0.478 | -0.006 | 0.054 | 0.054 | 0.194 | 0.19 | 0.31 | 0.18 | 0.39 | 0.19 | 0.12 |
| 7 | 109910153 | rs7779508 | -0.595 | 0.201 | -1.229 | 0.51 | -0.007 | 0.082 | 0.082 | 0 | 0.15 | 0.29 | 0.15 | 0.39 | 0.13 | 0.13 |
| 7 | 109911695 | rs10487317 | -1.093 | -0.137 | -1.402 | 0.43 | 0.003 | -0.005 | -0.005 | 0.007 | 0.23 | 0.28 | 0.17 | 0.39 | 0.15 | 0.15 |
| 7 | 109924336 | rs7785319 | -0.32 | 0.206 | -1.804 | -0.151 | 0.021 | 0.08 | 0.08 | 0 | 0.52 | 0.76 | 0.46 | 0.36 | 0.54 | 0.44 |
| 7 | 109924415 | rs10953655 | -0.534 | 0.204 | -2.012 | -0.282 | 0.024 | 0.032 | 0.032 | 0.098 | 0.43 | 0.81 | 0.41 | 0.38 | 0.38 | 0.38 |
| 7 | 109931836 | rs6979705 | 0.914 | 0.219 | -1.601 | -0.2 | 0.001 | -0.007 | -0.007 | 0 | 0.06 | 0.04 | 0.03 | 0 | 0.04 | 0.02 |
| 7 | 109933856 | rs2396222 | -0.64 | -0.038 | -1.503 | -0.529 | 0.009 | 0.022 | 0.022 | 0 | 0.3 | 0.23 | 0.15 | 0.19 | 0.3 | 0.19 |
| 7 | 109938523 | rs17157600 | 0.712 | -0.058 | -1.436 | -0.525 | -0.022 | -0.038 | -0.038 | 0 | 0.07 | 0.14 | 0.05 | 0 | 0.04 | 0.02 |
| 7 | 109938926 | rs38755 | -0.464 | -0.014 | -1.189 | -0.204 | -0.006 | -0.002 | -0.002 | 0 | 0.62 | 0.63 | 0.41 | 0.58 | 0.54 | 0.62 |
| 7 | 109939167 | rs10272725 | -0.694 | 0.087 | -1.181 | -0.15 | 0.001 | -0.011 | -0.011 | 0.64 | 0.64 | 0.67 | 0.66 | 0.56 | 0.68 | 0.53 |
| 7 | 109939445 | rs38756 | 1.288 | 0.044 | -1.445 | -0.157 | -0.045 | -0.148 | -0.148 | 0 | 0.27 | 0.01 | 0.21 | 0.39 | 0.22 | 0.41 |
| 7 | 109943907 | rs10487320 | 0.496 | 0.281 | -1.116 | -0.084 | 0 | -0.001 | -0.001 | 0.681 | 0.36 | 0.37 | 0.51 | 0.37 | 0.38 | 0.36 |
| 7 | 109955092 | rs6466326 | -0.433 | 0.203 | -1.015 | -0.236 | -0.007 | -0.024 | -0.024 | 0 | 0.32 | 0.36 | 0.18 | 0.19 | 0.32 | 0.22 |
| 7 | 109987031 | rs1859027 | 0.155 | 0.666 | -1.067 | 0.113 | 0.019 | 0 | 0 | 0 | 0.19 | 0.32 | 0.13 | 0.18 | 0.25 | 0.16 |
| 7 | 110010335 | rs12536111 | 0.265 | 0.422 | -1.963 | 0.846 | -0.068 | -0.128 | -0.128 | 0 | 0.18 | 0.01 | 0.31 | 0.4 | 0.21 | 0.18 |
| 7 | 110018099 | rs17157641 | 0.62 | 0.202 | -1.862 | 0.363 | -0.001 | 0.007 | 0.007 | 0 | 0.21 | 0.08 | 0.1 | 0.15 | 0.16 | 0.27 |
| 7 | 110032926 | rs10224745 | -0.337 | 0.358 | -1.951 | 0.313 | -0.017 | -0.001 | -0.001 | 0.052 | 0.88 | 0.95 | 0.83 | 0.89 | 0.88 | 0.88 |
| 7 | 110036601 | rs10487324 | 0.855 | -0.18 | -2.094 | -0.012 | -0.036 | 0.051 | 0.051 | 0 | 0.36 | 0.17 | 0.31 | 0.24 | 0.47 | 0.32 |
| 7 | 110037249 | rs11764402 | 0.801 | -0.247 | -1.838 | -0.101 | 0.003 | 0.035 | 0.035 | 0 | 0.23 | 0.1 | 0.12 | 0.14 | 0.19 | 0.34 |
| 7 | 110043681 | rs13228785 | 0.028 | -0.381 | -0.935 | 0.087 | 0.003 | 0.007 | 0.007 | 0 | 0.45 | 0.45 | 0.48 | 0.63 | 0.38 | 0.61 |
| 7 | 110046716 | rs6974080 | -0.595 | -0.177 | -1.193 | 0.016 | -0.031 | -0.032 | -0.032 | 0.098 | 0.86 | 0.95 | 0.87 | 0.81 | 0.85 | 0.89 |
| 7 | 110054993 | rs211815 | 0.001 | -0.518 | -0.824 | -0.112 | 0.009 | -0.027 | -0.027 | 0.177 | 0.82 | 0.94 | 0.87 | 0.78 | 0.86 | 0.84 |
| 7 | 110074144 | rs6466334 | 0.277 | -0.75 | -0.363 | -0.312 | 0.003 | 0.006 | 0.006 | 0 | 0.5 | 0.55 | 0.49 | 0.63 | 0.39 | 0.59 |
| 7 | 110074276 | rs9656169 | 0.969 | -0.822 | -0.026 | -0.391 | -0.007 | 0.007 | 0.007 | 0 | 0.29 | 0.24 | 0.35 | 0.15 | 0.42 | 0.2 |
| 7 | 110075577 | rs211801 | 0.372 | -0.876 | -0.071 | -0.368 | 0 | -0.001 | -0.001 | 0.052 | 0.49 | 0.54 | 0.49 | 0.63 | 0.43 | 0.63 |
| 7 | 110085734 | rs12673482 | 0.795 | -0.832 | -0.379 | -0.533 | 0.006 | 0.041 | 0.041 | 0 | 0.46 | 0.36 | 0.46 | 0.67 | 0.4 | 0.61 |
| 7 | 110088623 | rs211809 | -0.212 | -0.49 | -0.699 | -0.327 | 0.002 | 0.041 | 0.041 | 0 | 0.83 | 0.93 | 0.87 | 0.94 | 0.85 | 0.89 |
| 7 | 110096274 | rs7795082 | -0.446 | -0.425 | -0.604 | -0.297 | 0.003 | 0.009 | 0.009 | 0 | 0.84 | 0.88 | 0.79 | 0.9 | 0.83 | 0.88 |
| 7 | 110097308 | rs1396176 | 0.879 | -0.252 | -0.461 | -0.338 | 0.031 | -0.054 | -0.054 | 0 | 0.27 | 0.31 | 0.28 | 0.11 | 0.38 | 0.18 |
| 7 | 110106697 | rs6953798 | 0.89 | -0.198 | -0.481 | -0.31 | 0.025 | -0.046 | -0.046 | 0 | 0.27 | 0.31 | 0.28 | 0.11 | 0.38 | 0.18 |
| 7 | 110108202 | rs407050 | 0.001 | -0.228 | -0.499 | -0.192 | 0.002 | -0.01 | -0.01 | 0.011 | 0.58 | 0.58 | 0.51 | 0.78 | 0.46 | 0.7 |
| 7 | 110109590 | rs2023788 | 0.362 | -0.573 | -0.862 | 0.291 | -0.027 | -0.102 | -0.102 | 0 | 0.17 | 0.04 | 0.3 | 0.59 | 0.18 | 0.22 |
| 7 | 110123697 | rs1508184 | 0.826 | -1.994 | 0.06 | -0.123 | 0.069 | -0.001 | -0.001 | 0 | 0.6 | 0.2 | 0.6 | 0.64 | 0.72 | 0.51 |
| 7 | 110125420 | rs2940378 | 0.278 | -2.234 | 0.501 | -0.136 | 0.01 | 0.004 | 0.004 | 0 | 0.71 | 0.65 | 0.64 | 0.76 | 0.75 | 0.6 |
| 7 | 110131520 | rs10277639 | 1.385 | -1.919 | 0.209 | -0.771 | -0.016 | 0.053 | 0.053 | 0 | 0.25 | 0.14 | 0.23 | 0.05 | 0.3 | 0.1 |
| 7 | 110132602 | rs6943600 | 1.086 | -1.946 | 0.359 | -0.677 | 0.002 | -0.02 | -0.02 | 4.162 | 0.28 | 0.33 | 0.24 | 0.15 | 0.3 | 0.14 |
| 7 | 110135183 | rs2940371 | 0.547 | -2.096 | 0.181 | -0.634 | 0.016 | 0.01 | 0.01 | 0.155 | 0.62 | 0.59 | 0.66 | 0.7 | 0.72 | 0.5 |
| 7 | 110138817 | rs12672609 | 1.284 | -1.871 | -0.01 | -0.531 | -0.169 | -0.184 | -0.184 | 0.16 | 0.33 | 0.02 | 0.4 | 0.53 | 0.42 | 0.32 |
| 7 | 110143899 | rs6974477 | 2.14 | -1.909 | 0.031 | -0.534 | 0.016 | -0.015 | -0.015 | 0 | 0.06 | 0.02 | 0.02 | 0.11 | 0.02 | 0.06 |
| 7 | 110144415 | rs7811230 | 1.245 | -1.961 | 0.17 | -0.63 | -0.009 | 0.017 | 0.017 | 0 | 0.28 | 0.22 | 0.26 | 0.07 | 0.33 | 0.14 |
| 7 | 110157546 | rs2966417 | 0.1 | -1.961 | 0.359 | -0.585 | -0.036 | -0.051 | -0.051 | 0 | 0.69 | 0.46 | 0.68 | 0.82 | 0.77 | 0.6 |
| 7 | 110158246 | rs1354991 | 0.895 | -1.925 | 0.13 | -0.545 | 0.062 | -0.009 | -0.009 | 0 | 0.61 | 0.24 | 0.66 | 0.59 | 0.76 | 0.47 |
| 7 | 110160252 | rs7799099 | -0.048 | -1.892 | -0.002 | -0.681 | -0.004 | 0.085 | 0.085 | 0.09 | 0.45 | 0.21 | 0.46 | 0.21 | 0.47 | 0.29 |
| 7 | 110160724 | rs4730438 | 0.026 | -1.703 | 0.193 | -0.53 | 0.024 | 0.028 | 0.028 | 0.959 | 0.51 | 0.32 | 0.65 | 0.7 | 0.64 | 0.55 |
| 7 | 110161348 | rs6975627 | 1.358 | -1.537 | 0.029 | -0.429 | -0.186 | -0.175 | -0.175 | 0.119 | 0.3 | 0.04 | 0.42 | 0.53 | 0.45 | 0.35 |
| 7 | 110164447 | rs6954042 | -0.654 | -1.997 | 0.465 | -0.812 | -0.051 | -0.027 | -0.027 | 0 | 0.81 | 0.56 | 0.9 | 0.84 | 0.91 | 0.68 |
| 7 | 110172048 | rs2966420 | -1.004 | -1.992 | 0.438 | -0.831 | -0.022 | -0.056 | -0.056 | 0 | 0.82 | 0.69 | 0.9 | 0.95 | 0.9 | 0.72 |
| 7 | 110172394 | rs4730439 | 0.363 | -2.085 | 0.379 | -0.783 | 0.006 | 0.001 | 0.001 | 0 | 0.18 | 0.07 | 0.22 | 0.15 | 0.17 | 0.16 |
| 7 | 110174041 | rs10260781 | 0.891 | -1.952 | 0.346 | -0.849 | 0.005 | 0.018 | 0.018 | 0 | 0.33 | 0.26 | 0.27 | 0.16 | 0.32 | 0.18 |
| 7 | 110176383 | rs10272579 | -0.921 | -1.911 | 0.374 | -0.857 | -0.045 | -0.101 | -0.101 | 0.064 | 0.82 | 0.59 | 0.9 | 0.95 | 0.9 | 0.71 |
| 7 | 110178548 | rs10808165 | 1.686 | -1.903 | -0.139 | -0.833 | -0.186 | -0.171 | -0.171 | 0.019 | 0.28 | 0.02 | 0.4 | 0.54 | 0.41 | 0.34 |
| 7 | 110178950 | rs2966422 | -1.057 | -1.881 | 0.074 | -0.831 | -0.007 | -0.03 | -0.03 | 0 | 0.82 | 0.75 | 0.91 | 0.95 | 0.9 | 0.71 |
| 7 | 110181022 | rs10238549 | 0.729 | -1.546 | -0.203 | -0.607 | 0.008 | 0.041 | 0.041 | 0 | 0.33 | 0.23 | 0.26 | 0.14 | 0.32 | 0.18 |
| 7 | 110192161 | rs4727740 | 0.394 | -1.466 | -0.184 | -0.605 | -0.008 | -0.005 | -0.005 | 0 | 0.61 | 0.4 | 0.65 | 0.66 | 0.72 | 0.51 |
| 7 | 110208243 | rs7800391 | 1.188 | -1.074 | -0.759 | -0.152 | -0.172 | -0.168 | -0.168 | 0.022 | 0.27 | 0.02 | 0.38 | 0.52 | 0.41 | 0.33 |

| | | | | | | | | | | | | | | | | |
|---|---|---|---|---|---|---|---|---|---|---|---|---|---|---|---|---|
| 7 | 110227761 | rs1734236 | -1.023 | -0.874 | -0.124 | -0.712 | -0.01 | 0.001 | 0.001 | 4.245 | 0.68 | 0.71 | 0.57 | 0.35 | 0.6 | 0.64 |
| 7 | 110232807 | rs7812132 | 1.263 | -1.067 | 0.103 | -0.604 | -0.026 | 0.051 | 0.051 | 0 | 0.06 | 0.2 | 0.02 | 0.09 | 0.01 | 0.05 |
| 7 | 110235056 | rs6953429 | 1.064 | -1.092 | 0.091 | -0.708 | -0.028 | 0.038 | 0.038 | 0 | 0.06 | 0.2 | 0.02 | 0.07 | 0.01 | 0.05 |
| 7 | 110236196 | rs1723744 | -0.675 | -0.99 | -0.457 | -1.16 | 0.014 | -0.036 | -0.036 | 0.043 | 0.64 | 0.5 | 0.55 | 0.27 | 0.6 | 0.59 |
| 7 | 110237043 | rs1723743 | 0.267 | -1.178 | -1.597 | -1.403 | -0.02 | 0.126 | 0.126 | 0 | 0.44 | 0.18 | 0.43 | 0.24 | 0.44 | 0.29 |
| 7 | 110240470 | rs7789139 | 0.323 | -1.362 | -1.383 | -1.516 | 0.019 | 0.031 | 0.031 | 0.008 | 0.21 | 0.14 | 0.12 | 0.03 | 0.16 | 0.33 |
| 7 | 110261886 | rs1404916 | 0.051 | -0.862 | -1.722 | -1.134 | -0.004 | 0.068 | 0.068 | 0 | 0.45 | 0.18 | 0.45 | 0.29 | 0.44 | 0.3 |
| 7 | 110272427 | rs1404915 | -0.839 | -0.488 | -1.28 | -1.648 | -0.056 | 0.213 | 0.213 | 1.121 | 0.57 | 0.82 | 0.43 | 0.11 | 0.51 | 0.49 |
| 7 | 110282027 | rs6949507 | 0.079 | -0.058 | -0.874 | -1.278 | -0.014 | 0.062 | 0.062 | 0 | 0.7 | 0.79 | 0.74 | 0.92 | 0.66 | 0.86 |
| 7 | 110287125 | rs17157870 | 0.081 | -0.321 | -0.716 | -1.107 | 0 | -0.005 | -0.005 | 1.081 | 0.06 | 0.06 | 0.02 | 0.01 | 0.04 | 0.06 |
| 7 | 110295142 | rs10230353 | -0.638 | -0.149 | 0.214 | -1.181 | 0.006 | -0.044 | -0.044 | 0 | 0.31 | 0.59 | 0.34 | 0.16 | 0.35 | 0.21 |
| 7 | 110302846 | rs1525675 | -0.582 | -0.23 | 0.307 | -1.314 | 0.034 | -0.063 | -0.063 | 0.144 | 0.27 | 0.51 | 0.25 | 0.13 | 0.34 | 0.17 |
| 7 | 110310858 | rs13247342 | -0.426 | -0.148 | -0.566 | -1.345 | -0.069 | 0.098 | 0.098 | 0 | 0.23 | 0.04 | 0.28 | 0.07 | 0.3 | 0.13 |
| 7 | 110320069 | rs4305834 | 1.483 | -0.491 | -1.266 | -1.509 | -0.154 | 0.082 | 0.082 | 7.84 | 0.43 | 0.03 | 0.51 | 0.73 | 0.52 | 0.44 |
| 7 | 110320244 | rs7807511 | 0.815 | -0.824 | -0.919 | -1.664 | -0.007 | -0.012 | -0.012 | 0 | 0.11 | 0.13 | 0.05 | 0.05 | 0.06 | 0.11 |
| 7 | 110320785 | rs10953663 | 1.798 | -0.824 | -1.146 | -1.664 | -0.167 | 0.053 | 0.053 | 7.93 | 0.42 | 0.03 | 0.51 | 0.71 | 0.51 | 0.42 |
| 7 | 110322162 | rs10499991 | -1.457 | -0.924 | -0.563 | -1.79 | -0.001 | 0.099 | 0.099 | 6.181 | 0.51 | 0.76 | 0.36 | 0.28 | 0.48 | 0.51 |
| 7 | 110322696 | rs7786068 | -0.263 | -0.923 | -0.424 | -1.78 | -0.019 | -0.022 | -0.022 | 0.22 | 0.26 | 0.39 | 0.15 | 0.16 | 0.19 | 0.36 |
| 7 | 110324741 | rs1464895 | -0.109 | -0.89 | -0.409 | -1.868 | -0.021 | -0.031 | -0.031 | 1.633 | 0.26 | 0.39 | 0.15 | 0.15 | 0.19 | 0.35 |
| 7 | 110324879 | rs4730447 | 1.826 | -0.817 | -0.809 | -1.822 | -0.156 | 0.084 | 0.084 | 8.361 | 0.43 | 0.03 | 0.51 | 0.73 | 0.5 | 0.42 |
| 7 | 110329655 | rs6961889 | -1.184 | -0.296 | -0.284 | -1.627 | 0 | -0.028 | -0.028 | 0.622 | 0.42 | 0.57 | 0.4 | 0.22 | 0.43 | 0.44 |
| 7 | 110338460 | rs10266246 | 0.051 | 0.042 | -0.917 | -1.34 | -0.001 | 0.037 | 0.037 | 0 | 0.79 | 0.86 | 0.86 | 0.89 | 0.79 | 0.63 |
| 7 | 110339035 | rs955158 | -0.304 | 0.224 | -0.866 | -1.336 | -0.037 | 0.036 | 0.036 | 0 | 0.18 | 0.09 | 0.26 | 0.08 | 0.27 | 0.1 |
| 7 | 110342924 | rs6965544 | 1.239 | -0.025 | -0.41 | -0.851 | -0.002 | 0.001 | 0.001 | 1.523 | 0.6 | 0.56 | 0.53 | 0.67 | 0.54 | 0.6 |
| 7 | 110343824 | rs7803158 | 1.819 | -0.454 | -0.854 | -1.159 | -0.095 | 0.02 | 0.02 | 5.63 | 0.44 | 0.09 | 0.44 | 0.61 | 0.47 | 0.49 |
| 7 | 110353450 | rs6957675 | 0.318 | -0.269 | -0.669 | -1.139 | -0.025 | 0.016 | 0.016 | 0.088 | 0.14 | 0.11 | 0.26 | 0.09 | 0.25 | 0.11 |
| 7 | 110355786 | rs1589883 | -0.265 | -0.283 | -0.592 | -1.211 | 0.002 | -0.005 | -0.005 | 0 | 0.13 | 0.12 | 0.06 | 0.03 | 0.12 | 0.09 |
| 7 | 110357889 | rs4730451 | 1.19 | -0.395 | -0.301 | -1.256 | -0.008 | -0.01 | -0.01 | 0.006 | 0.73 | 0.69 | 0.75 | 0.82 | 0.74 | 0.66 |
| 7 | 110358857 | rs17157920 | -0.319 | -0.239 | -0.299 | -1.187 | 0.001 | -0.008 | -0.008 | 0 | 0.13 | 0.12 | 0.06 | 0.03 | 0.12 | 0.1 |
| 7 | 110365212 | rs2030780 | 0.672 | -0.246 | -0.265 | -1.156 | -0.001 | -0.003 | -0.003 | 0.03 | 0.77 | 0.75 | 0.77 | 0.82 | 0.75 | 0.71 |
| 7 | 110365453 | rs2030779 | 0.826 | -0.247 | -0.289 | -1.138 | -0.003 | -0.005 | -0.005 | 0 | 0.76 | 0.72 | 0.77 | 0.8 | 0.75 | 0.68 |
| 7 | 110371763 | rs12155444 | 1.473 | 0.478 | -0.785 | -0.908 | -0.063 | -0.057 | -0.057 | 0 | 0.12 | 0.01 | 0.19 | 0.17 | 0.14 | 0.24 |
| 7 | 110376327 | rs2091309 | -1.298 | -0.185 | -0.296 | -1.35 | 0.01 | 0.06 | 0.06 | 1.591 | 0.39 | 0.7 | 0.37 | 0.21 | 0.37 | 0.24 |
| 7 | 110391589 | rs4376443 | 0.613 | -0.243 | 0.164 | -1.371 | 0.006 | -0.041 | -0.041 | 0 | 0.69 | 0.62 | 0.66 | 0.79 | 0.66 | 0.76 |
| 7 | 110400223 | rs17157942 | 2.04 | -0.266 | -0.478 | -1.469 | -0.082 | -0.007 | -0.007 | 0 | 0.39 | 0.04 | 0.31 | 0.66 | 0.37 | 0.36 |
| 7 | 110400604 | rs6954311 | -1.444 | -0.326 | -0.325 | -1.533 | -0.034 | 0.133 | 0.133 | 0.32 | 0.51 | 0.9 | 0.55 | 0.3 | 0.57 | 0.59 |
| 7 | 110405982 | rs13247446 | -1.263 | -0.376 | -0.096 | -1.609 | -0.005 | 0.025 | 0.025 | 0.73 | 0.33 | 0.28 | 0.33 | 0.11 | 0.38 | 0.23 |
| 7 | 110409361 | rs10499999 | -0.784 | -0.327 | -0.057 | -1.587 | -0.007 | 0.007 | 0.007 | 0.035 | 0.28 | 0.27 | 0.32 | 0.11 | 0.37 | 0.21 |
| 7 | 110418829 | rs12530971 | 0.106 | -0.392 | 0.139 | -1.591 | -0.01 | -0.023 | -0.023 | 0 | 0.21 | 0.41 | 0.21 | 0.11 | 0.2 | 0.33 |
| 7 | 110419154 | rs1528041 | -1.112 | -0.282 | 0.06 | -1.632 | -0.001 | 0.006 | 0.006 | 0 | 0.43 | 0.6 | 0.48 | 0.22 | 0.5 | 0.51 |
| 7 | 110420611 | rs6978082 | 0.211 | -0.316 | 0.047 | -1.635 | -0.022 | -0.043 | -0.043 | 0.19 | 0.85 | 0.65 | 0.91 | 0.89 | 0.91 | 0.9 |
| 7 | 110424487 | rs4730462 | 0.145 | -0.278 | -0.133 | -1.645 | -0.087 | 0.043 | 0.043 | 0.024 | 0.19 | 0.02 | 0.25 | 0.11 | 0.3 | 0.16 |
| 7 | 110429536 | rs1881164 | 2.416 | -0.222 | -0.086 | -1.58 | -0.09 | -0.031 | -0.031 | 0.129 | 0.37 | 0.03 | 0.3 | 0.65 | 0.36 | 0.34 |
| 7 | 110432334 | rs4730464 | -1.557 | -0.129 | 0.225 | -1.387 | -0.008 | -0.016 | -0.016 | 0.266 | 0.93 | 0.92 | 0.97 | 0.98 | 0.98 | 0.97 |
| 7 | 110435602 | rs1950004 | -1.594 | -0.004 | 0.441 | -1.401 | -0.001 | 0.011 | 0.011 | 0.559 | 0.47 | 0.56 | 0.49 | 0.23 | 0.51 | 0.54 |
| 7 | 110441855 | rs10500002 | 0.471 | -0.197 | 0.637 | -1.334 | -0.001 | 0.002 | 0.002 | 0 | 0.17 | 0.08 | 0.19 | 0.11 | 0.18 | 0.33 |
| 7 | 110443171 | rs1358434 | -0.542 | -0.134 | 0.5 | -1.353 | -0.011 | 0.027 | 0.027 | 0 | 0.23 | 0.16 | 0.27 | 0.12 | 0.28 | 0.19 |
| 7 | 110451383 | rs7778910 | -1.863 | -0.098 | 0.139 | -1.312 | -0.058 | 0.127 | 0.127 | 0.062 | 0.49 | 0.93 | 0.56 | 0.3 | 0.56 | 0.59 |
| 7 | 110452753 | rs1406054 | 0.566 | -0.185 | 0.5 | -1.117 | -0.014 | -0.057 | -0.057 | 0 | 0.84 | 0.64 | 0.9 | 0.9 | 0.88 | 0.9 |
| 7 | 110459927 | rs1569122 | -0.07 | -0.204 | 0.594 | -1.126 | -0.006 | -0.012 | -0.012 | 0 | 0.18 | 0.34 | 0.21 | 0.11 | 0.18 | 0.33 |
| 7 | 110468988 | rs1528034 | 1.844 | -0.071 | 0.294 | -1.102 | -0.02 | 0.147 | 0.147 | 0.085 | 0.5 | 0.16 | 0.48 | 0.74 | 0.48 | 0.43 |
| 7 | 110471452 | rs953143 | -0.525 | 0.044 | 0.276 | -1.029 | -0.012 | 0.013 | 0.013 | 0 | 0.21 | 0.15 | 0.24 | 0.16 | 0.26 | 0.18 |
| 7 | 110474808 | rs7806611 | -0.155 | 0.256 | 0.164 | -0.717 | 0.014 | -0.032 | -0.032 | 0 | 0.07 | 0.21 | 0.05 | 0 | 0.07 | 0.04 |
| 7 | 110475186 | rs12531989 | -1.161 | 0.053 | -0.049 | -0.638 | 0.007 | -0.014 | -0.014 | 0.339 | 0.35 | 0.45 | 0.42 | 0.27 | 0.41 | 0.45 |
| 7 | 110477073 | rs11767545 | 1.461 | -0.19 | -1.268 | -0.519 | -0.019 | 0.097 | 0.097 | 0.587 | 0.53 | 0.14 | 0.49 | 0.69 | 0.51 | 0.5 |
| 7 | 110477566 | rs10280436 | 0.887 | -1.557 | -0.929 | -0.463 | 0.038 | 0.026 | 0.026 | 0 | 0.72 | 0.87 | 0.76 | 0.84 | 0.78 | 0.61 |
| 7 | 110480458 | rs10233811 | 0.719 | -1.175 | -0.706 | -0.257 | -0.06 | -0.094 | -0.094 | 0 | 0.77 | 0.53 | 0.89 | 0.91 | 0.87 | 0.83 |
| 7 | 110482096 | rs12670438 | -0.162 | -0.202 | -1.412 | 0.436 | -0.005 | -0.018 | -0.018 | 5.234 | 0.05 | 0.01 | 0.13 | 0.13 | 0.06 | 0.28 |
| 7 | 110482753 | rs10500003 | -0.531 | -0.539 | -1.582 | 0.33 | 0.082 | 0.043 | 0.043 | 1.268 | 0.4 | 0.82 | 0.31 | 0.33 | 0.29 | 0.45 |
| 7 | 110483677 | rs799617 | -0.621 | 0.38 | -1.086 | 1.051 | 0.014 | 0.049 | 0.049 | 0.046 | 0.17 | 0.17 | 0.19 | 0.44 | 0.21 | 0.21 |
| 7 | 110484091 | rs714111 | 0.342 | -1.445 | -1.544 | -0.006 | 0.057 | 0.059 | 0.059 | 1.407 | 0.61 | 0.29 | 0.68 | 0.84 | 0.74 | 0.76 |
| 7 | 110485124 | rs2396282 | -0.455 | -1.608 | -1.147 | -0.939 | -0.002 | -0.034 | -0.034 | 0 | 0.54 | 0.52 | 0.46 | 0.81 | 0.47 | 0.7 |
| 7 | 110489975 | rs12537852 | 0.291 | -2.139 | -0.847 | -0.994 | -0.071 | -0.038 | -0.038 | 0 | 0.26 | 0.05 | 0.37 | 0.65 | 0.38 | 0.5 |
| 7 | 110494399 | rs6948090 | -0.787 | -2.13 | -0.314 | -1.092 | 0.001 | -0.011 | -0.011 | 0.032 | 0.52 | 0.55 | 0.45 | 0.81 | 0.44 | 0.7 |
| 7 | 110500979 | rs17158049 | 0.988 | -2.436 | -0.26 | -1.129 | -0.023 | -0.003 | -0.003 | 0 | 0.16 | 0.24 | 0.04 | 0.17 | 0.04 | 0.15 |
| 7 | 110509579 | rs17158079 | 0.607 | -2.679 | -0.139 | -1.406 | -0.016 | -0.023 | -0.023 | 0 | 0.07 | 0.1 | 0.05 | 0.05 | 0.04 | 0.1 |
| 7 | 110511038 | rs799624 | -1.393 | -2.714 | -0.125 | -1.408 | -0.069 | 0.136 | 0.136 | 0.144 | 0.62 | 0.89 | 0.53 | 0.85 | 0.51 | 0.79 |
| 7 | 110511679 | rs799625 | -1.851 | -3.051 | -0.316 | -1.757 | -0.098 | 0.065 | 0.065 | 0.012 | 0.71 | 0.93 | 0.57 | 0.85 | 0.59 | 0.83 |
| 7 | 110516563 | rs2190529 | -0.601 | -2.848 | 0.224 | -1.362 | -0.001 | -0.008 | -0.008 | 0.845 | 0.44 | 0.48 | 0.48 | 0.66 | 0.48 | 0.64 |

| | | | | | | | | | | | | | | | | |
|---|---|---|---|---|---|---|---|---|---|---|---|---|---|---|---|---|
| 7 | 110519790 | rs12705744 | 0.706 | -2.606 | -0.283 | -1.217 | -0.09 | -0.083 | -0.083 | 0 | 0.26 | 0.03 | 0.36 | 0.62 | 0.38 | 0.48 |
| 7 | 110543241 | rs258989 | -0.625 | -2.465 | 0.471 | -1.236 | 0.007 | 0.012 | 0.012 | 0.791 | 0.55 | 0.4 | 0.45 | 0.83 | 0.45 | 0.74 |
| 7 | 110546511 | rs258991 | -0.776 | -2.492 | 0.608 | -1.244 | 0.016 | 0.067 | 0.067 | 0.42 | 0.57 | 0.69 | 0.48 | 0.84 | 0.45 | 0.75 |
| 7 | 110548594 | rs258996 | -0.837 | -2.502 | 0.556 | -1.248 | 0.014 | 0.141 | 0.141 | 0.565 | 0.58 | 0.86 | 0.49 | 0.83 | 0.45 | 0.74 |
| 7 | 110559571 | rs259006 | -0.861 | -2.509 | 0.571 | -1.236 | 0.015 | 0.143 | 0.143 | 0.354 | 0.58 | 0.84 | 0.49 | 0.87 | 0.45 | 0.74 |
| 7 | 110562681 | rs115825 | -0.827 | -1.987 | 0.993 | -1.352 | 0.003 | -0.02 | -0.02 | 0 | 0.6 | 0.53 | 0.52 | 0.83 | 0.56 | 0.74 |
| 7 | 110572231 | rs259013 | -0.737 | -2.455 | 1.134 | -1.227 | 0.02 | 0.081 | 0.081 | 0.361 | 0.57 | 0.71 | 0.48 | 0.83 | 0.45 | 0.74 |
| 7 | 110577992 | rs6466365 | 0.076 | -2.86 | 1.301 | -1.265 | 0.022 | 0.036 | 0.036 | 0 | 0.6 | 0.81 | 0.59 | 0.33 | 0.59 | 0.37 |
| 7 | 110580701 | rs17158169 | 0.931 | -2.858 | 1.406 | -1.258 | -0.074 | -0.011 | -0.011 | 0 | 0.17 | 0.36 | 0.05 | 0.17 | 0.04 | 0.12 |
| 7 | 110582600 | rs10464637 | 0.806 | -2.829 | 1.311 | -1.265 | 0.02 | 0.082 | 0.082 | 0.324 | 0.43 | 0.29 | 0.49 | 0.16 | 0.55 | 0.26 |
| 7 | 110591124 | rs10252992 | 0.624 | -2.834 | 1.426 | -1.224 | -0.089 | 0.007 | 0.007 | 3.719 | 0.18 | 0.54 | 0.07 | 0.21 | 0.04 | 0.14 |
| 7 | 110602792 | rs7795011 | 0.096 | -2.883 | 1.132 | -1.276 | 0.021 | 0.034 | 0.034 | 0 | 0.6 | 0.8 | 0.59 | 0.33 | 0.59 | 0.38 |
| 7 | 110603427 | rs10225614 | 0.842 | -2.875 | 1.224 | -1.284 | 0.008 | 0.023 | 0.023 | 0.477 | 0.43 | 0.36 | 0.52 | 0.13 | 0.55 | 0.26 |
| 7 | 110611438 | rs2109863 | 0.073 | -2.821 | 1.133 | -1.252 | 0.016 | 0.023 | 0.023 | 0 | 0.6 | 0.73 | 0.58 | 0.34 | 0.58 | 0.39 |
| 7 | 110613685 | rs758953 | -0.257 | -2.977 | 0.933 | -1.294 | 0.007 | -0.006 | -0.006 | 0 | 0.66 | 0.86 | 0.63 | 0.39 | 0.62 | 0.45 |
| 7 | 110617845 | rs17528429 | 0.955 | -3.022 | 0.956 | -1.32 | 0.018 | 0.077 | 0.077 | 0.495 | 0.43 | 0.29 | 0.52 | 0.16 | 0.55 | 0.26 |
| 7 | 110631333 | rs16873249 | 0.291 | -2.995 | 0.833 | -1.272 | 0.015 | 0.006 | 0.006 | 1.567 | 0.21 | 0.22 | 0.08 | 0.23 | 0.07 | 0.22 |
| 7 | 110661337 | rs4730473 | -0.776 | -2.833 | 0.755 | -1.289 | -0.001 | -0.003 | -0.003 | 0.064 | 0.75 | 0.8 | 0.64 | 0.34 | 0.66 | 0.51 |
| 7 | 110678370 | rs17158277 | 0.425 | -2.885 | 0.323 | -1.287 | 0.055 | 0.018 | 0.018 | 3.843 | 0.23 | 0.12 | 0.06 | 0.17 | 0.08 | 0.17 |
| 7 | 110678928 | rs10487300 | -0.225 | -2.836 | 0.338 | -1.269 | 0.085 | 0.046 | 0.046 | 1.374 | 0.29 | 0.14 | 0.1 | 0.21 | 0.11 | 0.26 |
| 7 | 110681482 | rs6466371 | -0.767 | -2.759 | 0.253 | -1.249 | -0.005 | -0.02 | -0.02 | 0 | 0.75 | 0.84 | 0.65 | 0.36 | 0.67 | 0.51 |
| 7 | 110681641 | rs2191926 | 0.639 | -2.49 | 0.626 | -1.196 | 0.016 | 0.055 | 0.055 | 0.452 | 0.44 | 0.35 | 0.52 | 0.18 | 0.55 | 0.25 |
| 7 | 110691428 | rs10487302 | 0.991 | -3.053 | 1.077 | -0.952 | -0.044 | -0.025 | -0.025 | 0 | 0.73 | 0.39 | 0.87 | 0.83 | 0.9 | 0.8 |
| 7 | 110707029 | rs4727755 | 0.988 | -3.047 | 1.08 | -0.95 | -0.044 | -0.026 | -0.026 | 0 | 0.73 | 0.39 | 0.87 | 0.84 | 0.9 | 0.8 |
| 7 | 110713043 | rs7781676 | -0.561 | -3.048 | 1.233 | -0.899 | -0.063 | -0.029 | -0.029 | 0.287 | 0.21 | 0.29 | 0.09 | 0.16 | 0.07 | 0.18 |
| 7 | 110714407 | rs12669471 | -0.561 | -3.048 | 1.233 | -0.892 | -0.063 | -0.018 | -0.018 | 0 | 0.21 | 0.29 | 0.09 | 0.18 | 0.07 | 0.18 |
| 7 | 110718387 | rs7778044 | -0.826 | -2.929 | 1.329 | -0.949 | -0.022 | -0.017 | -0.017 | 0 | 0.22 | 0.23 | 0.12 | 0.15 | 0.11 | 0.2 |
| 7 | 110721800 | rs214897 | 0.69 | -2.663 | 1.252 | -0.951 | -0.041 | -0.043 | -0.043 | 0 | 0.78 | 0.72 | 0.88 | 0.88 | 0.89 | 0.8 |
| 7 | 110722482 | rs757219 | 1.515 | -2.75 | 1.466 | -0.993 | 0.021 | 0.016 | 0.016 | 0.551 | 0.71 | 0.25 | 0.82 | 0.8 | 0.86 | 0.78 |
| 7 | 110724876 | rs214896 | 0.673 | -2.708 | 1.421 | -0.992 | -0.044 | -0.03 | -0.03 | 0.741 | 0.78 | 0.71 | 0.88 | 0.84 | 0.89 | 0.8 |
| 7 | 110733717 | rs2107147 | 1.409 | -2.376 | 1.478 | -1.092 | -0.015 | -0.031 | -0.031 | 0 | 0.71 | 0.38 | 0.83 | 0.87 | 0.82 | 0.78 |
| 7 | 110735577 | rs10270683 | 1.37 | -2.382 | 1.533 | -1.087 | 0.013 | 0.013 | 0.013 | 0.458 | 0.7 | 0.35 | 0.83 | 0.84 | 0.82 | 0.77 |
| 7 | 110737149 | rs2301733 | 1.163 | -2.481 | 0.709 | -1.166 | 0.019 | 0.039 | 0.039 | 0.229 | 0.43 | 0.22 | 0.48 | 0.28 | 0.54 | 0.3 |
| 7 | 110746608 | rs214884 | -0.11 | -2.262 | 1.128 | -1.296 | 0.001 | -0.003 | -0.003 | 0 | 0.83 | 0.85 | 0.92 | 0.93 | 0.91 | 0.87 |
| 7 | 110762899 | rs9942557 | 1.788 | -2.242 | 0.812 | -1.501 | 0.003 | 0.008 | 0.008 | 0 | 0.07 | 0.04 | 0.02 | 0 | 0.06 | 0.01 |
| 7 | 110763563 | rs214865 | 0.818 | -2.233 | 1.21 | -1.461 | 0.035 | 0.012 | 0.012 | 0 | 0.06 | 0.3 | 0.04 | 0 | 0.03 | 0.01 |
| 7 | 110778571 | rs10487304 | 1.382 | -2.378 | 1.018 | -1.545 | 0.017 | 0.014 | 0.014 | 0.177 | 0.7 | 0.26 | 0.82 | 0.89 | 0.83 | 0.78 |
| 7 | 110780471 | rs4730479 | 1.098 | -2.411 | 1.029 | -1.538 | 0.005 | 0.001 | 0.001 | 0.161 | 0.72 | 0.27 | 0.85 | 0.9 | 0.88 | 0.81 |
| 7 | 110811863 | rs4730481 | 1.11 | -2.359 | 1.064 | -1.535 | -0.03 | -0.041 | -0.041 | 0 | 0.72 | 0.38 | 0.86 | 0.9 | 0.87 | 0.8 |
| 7 | 110812463 | rs10953682 | 0.719 | -2.406 | 1.24 | -1.542 | -0.096 | -0.1 | -0.1 | 0 | 0.76 | 0.56 | 0.9 | 0.9 | 0.9 | 0.82 |
| 7 | 110813808 | rs2189325 | 0.623 | -2.494 | 1.281 | -1.573 | -0.097 | -0.097 | -0.097 | 0 | 0.79 | 0.47 | 0.9 | 0.93 | 0.92 | 0.88 |
| 7 | 110814695 | rs7789480 | -0.001 | -2.521 | 1.253 | -1.536 | 0.013 | 0.008 | 0.008 | 1.825 | 0.82 | 0.86 | 0.92 | 0.86 | 0.91 | 0.87 |
| 7 | 110820818 | rs10953683 | 1.348 | -2.486 | 1.22 | -1.598 | -0.014 | -0.033 | -0.033 | 0 | 0.7 | 0.38 | 0.85 | 0.9 | 0.83 | 0.8 |
| 7 | 110820901 | rs10257931 | 0.634 | -2.364 | 1.226 | -1.541 | 0.018 | -0.035 | -0.035 | 0.533 | 0.07 | 0.18 | 0.05 | 0 | 0.08 | 0.02 |
| 7 | 110825467 | rs10487306 | 1.464 | -2.46 | 1.04 | -1.584 | 0.005 | 0.009 | 0.009 | 0 | 0.07 | 0.04 | 0.02 | 0 | 0.06 | 0.01 |
| 7 | 110827173 | rs769130 | 0.618 | -2.277 | 1.222 | -1.54 | -0.044 | -0.044 | -0.044 | 0 | 0.75 | 0.66 | 0.87 | 0.86 | 0.87 | 0.8 |
| 7 | 110827830 | rs6979841 | 1.692 | -2.263 | 1.022 | -1.521 | 0.008 | 0.008 | 0.008 | 0 | 0.67 | 0.36 | 0.82 | 0.83 | 0.79 | 0.77 |
| 7 | 110836294 | rs17441325 | 1 | -2.093 | 0.547 | -1.408 | -0.007 | -0.021 | -0.021 | 0 | 0.06 | 0.01 | 0.04 | 0.08 | 0.06 | 0.08 |
| 7 | 110840236 | rs214454 | 1.009 | -1.938 | 0.532 | -1.045 | 0.005 | 0.004 | 0.004 | 0.484 | 0.41 | 0.43 | 0.47 | 0.42 | 0.49 | 0.26 |
| 7 | 110843795 | rs214455 | 0.384 | -1.987 | 0.568 | -1.059 | 0.017 | -0.002 | -0.002 | 0.335 | 0.49 | 0.6 | 0.53 | 0.43 | 0.59 | 0.28 |
| 7 | 110851553 | rs214460 | -0.285 | -2.071 | 0.462 | -1.088 | 0.031 | -0.071 | -0.071 | 0 | 0.58 | 0.87 | 0.59 | 0.47 | 0.63 | 0.36 |
| 7 | 110854261 | rs214463 | -0.057 | -2.079 | 0.548 | -1.091 | 0.033 | -0.016 | -0.016 | 0.156 | 0.57 | 0.71 | 0.57 | 0.47 | 0.64 | 0.37 |
| 7 | 110858268 | rs214467 | -0.1 | -2.138 | 0.528 | -1.111 | 0.048 | -0.04 | -0.04 | 0 | 0.57 | 0.81 | 0.58 | 0.43 | 0.63 | 0.36 |
| 7 | 110861883 | rs214468 | 1.063 | -2.142 | 0.631 | -1.113 | 0 | 0 | 0 | 0.349 | 0.42 | 0.37 | 0.46 | 0.43 | 0.49 | 0.26 |
| 7 | 110864299 | rs214469 | -0.221 | -2.139 | 0.479 | -1.102 | 0.042 | -0.045 | -0.045 | 0 | 0.58 | 0.85 | 0.59 | 0.47 | 0.64 | 0.36 |
| 7 | 110867535 | rs214475 | -0.004 | -2.056 | 0.478 | -1.065 | 0.036 | -0.01 | -0.01 | 0 | 0.56 | 0.71 | 0.58 | 0.47 | 0.63 | 0.36 |
| 7 | 110879251 | rs37715 | 1.109 | -2.026 | 0.272 | -1.048 | -0.018 | -0.013 | -0.013 | 6.331 | 0.41 | 0.27 | 0.46 | 0.39 | 0.49 | 0.26 |
| 7 | 110879776 | rs37717 | 1.14 | -2.025 | 0.277 | -1.047 | -0.017 | -0.012 | -0.012 | 0 | 0.41 | 0.27 | 0.46 | 0.39 | 0.49 | 0.26 |
| 7 | 110892226 | rs7779264 | 1.542 | -1.973 | 0.186 | -1.019 | -0.005 | 0.014 | 0.014 | 0 | 0.07 | 0.01 | 0.04 | 0.04 | 0.05 | 0.08 |
| 7 | 110895375 | rs17539223 | 1.542 | -1.973 | 0.186 | -1.019 | -0.005 | 0.014 | 0.014 | 0 | 0.07 | 0.01 | 0.04 | 0.04 | 0.05 | 0.09 |
| 7 | 110898122 | rs10487307 | 1.542 | -1.973 | 0.186 | -1.019 | -0.005 | 0.014 | 0.014 | 0 | 0.07 | 0.01 | 0.04 | 0.04 | 0.05 | 0.09 |
| 7 | 110899634 | rs2109298 | 2.025 | -2.302 | -0.04 | -1.178 | -0.056 | -0.128 | -0.128 | 0.138 | 0.2 | 0.02 | 0.33 | 0.46 | 0.26 | 0.47 |
| 7 | 110915339 | rs37743 | -0.943 | -1.618 | 0.342 | -0.841 | -0.009 | -0.009 | -0.009 | 0 | 0.59 | 0.64 | 0.53 | 0.58 | 0.51 | 0.74 |
| 7 | 110918507 | rs37746 | -0.993 | -1.634 | 0.368 | -0.855 | -0.013 | -0.016 | -0.016 | 0 | 0.6 | 0.7 | 0.54 | 0.58 | 0.52 | 0.74 |
| 7 | 110921994 | rs37752 | -0.954 | -1.606 | 0.365 | -0.789 | -0.005 | -0.004 | -0.004 | 8.157 | 0.59 | 0.61 | 0.53 | 0.6 | 0.51 | 0.74 |
| 7 | 110924289 | rs10487308 | 1.393 | -1.593 | 0.304 | -0.803 | -0.005 | 0.014 | 0.014 | 0 | 0.07 | 0.01 | 0.04 | 0.04 | 0.05 | 0.08 |
| 7 | 110931475 | rs37757 | -0.945 | -1.627 | 0.316 | -0.811 | -0.009 | -0.009 | -0.009 | 0 | 0.59 | 0.64 | 0.53 | 0.6 | 0.51 | 0.74 |
| 7 | 110935979 | rs11534045 | 0.52 | -1.605 | 0.335 | -0.85 | 0.037 | -0.031 | -0.031 | 0 | 0.42 | 0.19 | 0.41 | 0.52 | 0.36 | 0.63 |
| 7 | 110939653 | rs253376 | -0.986 | -1.67 | 0.425 | -0.884 | 0.003 | 0.003 | 0.003 | 0.228 | 0.59 | 0.56 | 0.53 | 0.56 | 0.51 | 0.74 |

| | | | | | | | | | | | | | | | | |
|---|---|---|---|---|---|---|---|---|---|---|---|---|---|---|---|---|
| 7 | 110949566 | rs6962665 | 0.614 | -1.48 | 0.363 | -0.795 | 0.035 | -0.045 | -0.045 | 0 | 0.4 | 0.19 | 0.39 | 0.53 | 0.34 | 0.62 |
| 7 | 110971919 | rs37637 | -0.943 | -1.309 | 0.539 | -0.822 | 0.002 | -0.098 | -0.098 | 0 | 0.17 | 0.53 | 0.12 | 0.03 | 0.14 | 0.1 |
| 7 | 110977344 | rs37642 | -2.434 | -0.029 | 0.157 | -1.013 | -0.039 | 0.035 | 0.035 | 0 | 0.93 | 1 | 0.94 | 1 | 0.81 | 0.97 |
| 7 | 110983575 | rs37646 | -0.616 | -0.767 | 0.489 | -0.882 | 0.021 | -0.055 | -0.055 | 0 | 0.13 | 0.52 | 0.12 | 0.03 | 0.1 | 0.09 |
| 7 | 110986897 | rs17443228 | 1.193 | -0.756 | 0.495 | -0.876 | 0.001 | 0.003 | 0.003 | 0 | 0.07 | 0.05 | 0.04 | 0.03 | 0.05 | 0.08 |
| 7 | 110990633 | rs37651 | -0.911 | -0.913 | 0.423 | -0.877 | -0.02 | -0.007 | -0.007 | 0 | 0.6 | 0.74 | 0.55 | 0.64 | 0.52 | 0.74 |
| 7 | 111014240 | rs37662 | -0.927 | -0.845 | 0.692 | -0.884 | 0.006 | -0.101 | -0.101 | 0 | 0.17 | 0.53 | 0.13 | 0.03 | 0.16 | 0.1 |
| 7 | 111025533 | rs6955272 | 0.287 | -0.896 | 0.707 | -0.853 | 0 | -0.023 | -0.023 | 0 | 0.09 | 0.27 | 0.05 | 0.03 | 0.05 | 0.08 |
| 7 | 111025948 | rs12532143 | 0.418 | -0.97 | 0.387 | -0.772 | 0.036 | -0.028 | -0.028 | 0 | 0.43 | 0.23 | 0.44 | 0.6 | 0.38 | 0.64 |
| 7 | 111031094 | rs37674 | 1.367 | -0.81 | 0.472 | -0.832 | 0.005 | -0.035 | -0.035 | 1.797 | 0.08 | 0.16 | 0.06 | 0 | 0.09 | 0.02 |
| 7 | 111044377 | rs39725 | -0.889 | -0.856 | 0.619 | -0.859 | -0.001 | -0.102 | -0.102 | 0 | 0.17 | 0.52 | 0.12 | 0.03 | 0.14 | 0.1 |
| 7 | 111053822 | rs10267546 | 0.708 | -0.789 | 0.679 | -0.847 | 0.022 | -0.045 | -0.045 | 1.374 | 0.09 | 0.33 | 0.06 | 0 | 0.09 | 0.01 |
| 7 | 111069946 | rs38751 | -0.93 | -0.898 | 0.603 | -0.831 | -0.004 | -0.002 | -0.002 | 0 | 0.6 | 0.67 | 0.6 | 0.64 | 0.52 | 0.74 |
| 7 | 111080241 | rs10215967 | 1.681 | -0.821 | 0.468 | -0.78 | 0.003 | -0.028 | -0.028 | 0 | 0.07 | 0.15 | 0.05 | 0 | 0.09 | 0.01 |
| 7 | 111092478 | rs10282292 | -0.467 | -0.681 | 0.296 | -0.618 | 0.02 | -0.026 | -0.026 | 0 | 0.56 | 0.84 | 0.56 | 0.39 | 0.61 | 0.36 |
| 7 | 111108260 | rs10258236 | 0.286 | -0.699 | 0.504 | -0.737 | 0.043 | -0.052 | -0.052 | 0 | 0.1 | 0.52 | 0.08 | 0 | 0.09 | 0.01 |
| 7 | 111112853 | rs12671676 | -1.84 | -1.036 | 0.138 | -0.911 | -0.052 | -0.135 | -0.135 | 0 | 0.73 | 0.97 | 0.64 | 0.53 | 0.71 | 0.52 |
| 7 | 111122474 | rs1558078 | 0.297 | -0.649 | 0.694 | -0.816 | 0.03 | -0.048 | -0.048 | 0.471 | 0.1 | 0.42 | 0.07 | 0 | 0.09 | 0.01 |
| 7 | 111132643 | rs10251897 | 1.365 | -0.633 | 0.719 | -0.809 | 0.007 | -0.041 | -0.041 | 0 | 0.08 | 0.21 | 0.06 | 0 | 0.09 | 0.01 |
| 7 | 111151253 | rs6466378 | -0.787 | -0.634 | 0.45 | -0.829 | -0.005 | -0.104 | -0.104 | 0 | 0.18 | 0.53 | 0.12 | 0.03 | 0.14 | 0.09 |
| 7 | 111158861 | rs2529488 | -0.785 | -0.64 | 0.446 | -0.832 | -0.005 | -0.104 | -0.104 | 0 | 0.18 | 0.52 | 0.12 | 0.03 | 0.14 | 0.09 |
| 7 | 111159896 | rs2529489 | -0.825 | -0.679 | 0.372 | -0.833 | 0.024 | -0.048 | -0.048 | 0.158 | 0.57 | 0.84 | 0.56 | 0.41 | 0.61 | 0.36 |
| 7 | 111166165 | rs4730488 | 0.475 | -0.722 | 0.41 | -0.831 | -0.02 | -0.019 | -0.019 | 0 | 0.41 | 0.27 | 0.44 | 0.44 | 0.48 | 0.27 |
| 7 | 111180544 | rs2613590 | -0.887 | -0.651 | 0.294 | -0.842 | 0.024 | -0.059 | -0.059 | 3.971 | 0.58 | 0.84 | 0.56 | 0.46 | 0.61 | 0.35 |
| 7 | 111183610 | rs2613588 | -1.009 | -0.371 | 0.661 | -0.768 | -0.022 | -0.092 | -0.092 | 0 | 0.18 | 0.42 | 0.11 | 0.04 | 0.13 | 0.09 |
| 7 | 111199771 | rs12705779 | 0.597 | -0.403 | 0.624 | -0.722 | -0.003 | -0.099 | -0.099 | 3.727 | 0.83 | 0.49 | 0.88 | 0.97 | 0.86 | 0.91 |
| 7 | 111228173 | rs17158652 | 0.817 | -0.239 | 0.532 | -0.716 | 0 | -0.019 | -0.019 | 0 | 0.09 | 0.14 | 0.07 | 0 | 0.11 | 0.02 |
| 7 | 111228448 | rs2528691 | -0.34 | -0.415 | 0.395 | -0.831 | -0.012 | -0.008 | -0.008 | 1.217 | 0.53 | 0.73 | 0.58 | 0.57 | 0.52 | 0.72 |
| 7 | 111228993 | rs2969506 | -0.697 | -0.472 | 0.42 | -0.882 | 0.003 | -0.104 | -0.104 | 0 | 0.18 | 0.46 | 0.13 | 0.03 | 0.16 | 0.1 |
| 7 | 111230244 | rs2613596 | -0.517 | -0.634 | -0.104 | -1.02 | -0.157 | -0.168 | -0.168 | 0.022 | 0.65 | 0.98 | 0.69 | 0.61 | 0.58 | 0.75 |
| 7 | 111246951 | rs2613611 | -0.933 | -0.197 | 0.433 | -0.613 | 0.01 | -0.11 | -0.11 | 0 | 0.18 | 0.5 | 0.13 | 0.03 | 0.16 | 0.1 |
| 7 | 111251449 | rs2528665 | 0.082 | 0.204 | 0.085 | -0.501 | -0.001 | -0.016 | -0.016 | 0 | 0.67 | 0.77 | 0.73 | 0.63 | 0.68 | 0.82 |
| 7 | 111265262 | rs2969502 | -0.616 | 0.632 | 0.164 | -0.847 | -0.01 | -0.074 | -0.074 | 0 | 0.15 | 0.29 | 0.1 | 0 | 0.13 | 0.02 |
| 7 | 111275653 | rs10253118 | -1.248 | 0.718 | 0.229 | -0.734 | -0.004 | -0.07 | -0.07 | 0 | 0.18 | 0.35 | 0.11 | 0.04 | 0.15 | 0.09 |
| 7 | 111278445 | rs11978935 | -0.353 | 0.596 | -0.188 | -0.569 | 0.004 | 0.007 | 0.007 | 0 | 0.1 | 0.08 | 0.04 | 0.04 | 0.05 | 0.08 |
| 7 | 111285540 | rs1859744 | 1.068 | 0.475 | -0.086 | -0.505 | 0.005 | 0.011 | 0.011 | 1.064 | 0.49 | 0.25 | 0.61 | 0.57 | 0.51 | 0.67 |
| 7 | 111287980 | rs2528679 | 0.488 | 0.414 | 0.546 | -0.605 | -0.018 | -0.024 | -0.024 | 0.112 | 0.61 | 0.74 | 0.68 | 0.58 | 0.61 | 0.67 |
| 7 | 111287998 | rs2528680 | 0.584 | 0.397 | 1.006 | -0.612 | 0.006 | 0.009 | 0.009 | 0.838 | 0.6 | 0.54 | 0.66 | 0.58 | 0.61 | 0.67 |
| 7 | 111297991 | rs2528689 | 0.347 | 0.504 | 0.621 | -0.546 | -0.022 | -0.038 | -0.038 | 0.137 | 0.64 | 0.79 | 0.68 | 0.58 | 0.61 | 0.69 |
| 7 | 111309224 | rs2396412 | -0.438 | 0.718 | 0.902 | -0.319 | 0.033 | 0.065 | 0.065 | 0 | 0.77 | 0.68 | 0.72 | 0.49 | 0.67 | 0.83 |
| 7 | 111317076 | rs6957983 | -0.264 | 0.611 | 0.875 | -0.439 | 0.016 | 0.046 | 0.046 | 0 | 0.77 | 0.71 | 0.73 | 0.53 | 0.68 | 0.84 |
| 7 | 111317219 | rs6978071 | 1.336 | 0.494 | 0.75 | -0.619 | -0.005 | -0.009 | -0.009 | 0 | 0.6 | 0.32 | 0.64 | 0.53 | 0.56 | 0.82 |
| 7 | 111319190 | rs1833105 | -0.446 | 0.621 | 0.691 | -0.464 | -0.024 | 0.108 | 0.108 | 1.704 | 0.28 | 0.46 | 0.27 | 0.47 | 0.22 | 0.14 |
| 7 | 111322755 | rs17158713 | 0.441 | 0.251 | 0.572 | -0.682 | 0.003 | 0.019 | 0.019 | 0 | 0.06 | 0.15 | 0.12 | 0.07 | 0.05 | 0.11 |
| 7 | 111323799 | rs10263444 | 1.192 | 0.27 | 0.522 | -0.74 | -0.001 | -0.028 | -0.028 | 0.684 | 0.07 | 0.13 | 0.03 | 0 | 0.05 | 0.01 |
| 7 | 111329336 | rs10236334 | -0.994 | 0.342 | 0.603 | -0.784 | 0.003 | -0.05 | -0.05 | 0 | 0.15 | 0.41 | 0.17 | 0.06 | 0.12 | 0.13 |
| 7 | 111333872 | rs6466385 | -0.498 | 0.582 | 0.573 | -0.502 | -0.011 | -0.019 | -0.019 | 0 | 0.79 | 0.81 | 0.8 | 0.52 | 0.69 | 0.88 |
| 7 | 111339228 | rs1582274 | -1.382 | 0.291 | 0.463 | -0.424 | 0.011 | -0.014 | -0.014 | 0 | 0.1 | 0.34 | 0.07 | 0.04 | 0.08 | 0.11 |
| 7 | 111353361 | rs17158744 | -1.262 | 0.312 | 0.416 | -0.412 | 0.013 | -0.003 | -0.003 | 0 | 0.09 | 0.27 | 0.14 | 0.06 | 0.08 | 0.11 |
| 7 | 111362942 | rs957143 | 1.609 | 0.342 | -0.105 | -1.243 | 0.028 | -0.025 | -0.025 | 2.048 | 0.39 | 0.21 | 0.45 | 0.49 | 0.34 | 0.67 |
| 7 | 111365687 | rs2057974 | 0.718 | 0.19 | -0.1 | -1.659 | 0.014 | 0.014 | 0.014 | 0 | 0.49 | 0.4 | 0.54 | 0.4 | 0.38 | 0.66 |
| 7 | 111365743 | rs9641464 | -0.36 | 0.193 | -0.123 | -1.651 | 0.006 | 0.01 | 0.01 | 0 | 0.1 | 0.08 | 0.06 | 0.03 | 0.07 | 0.09 |
| 7 | 111367623 | rs3801790 | 1.5 | 0.221 | -0.105 | -1.627 | 0.009 | 0.003 | 0.003 | 0.157 | 0.41 | 0.33 | 0.5 | 0.42 | 0.33 | 0.68 |
| 7 | 111379582 | rs2074115 | -1.69 | 0.119 | -0.238 | -1.809 | 0.004 | 0.001 | 0.001 | 0 | 0.62 | 0.59 | 0.5 | 0.59 | 0.68 | 0.4 |
| 7 | 111385025 | rs3801786 | -1.376 | 0.059 | -0.492 | -1.963 | 0.015 | 0.017 | 0.017 | 0 | 0.54 | 0.56 | 0.52 | 0.61 | 0.65 | 0.34 |
| 7 | 111386690 | rs3735534 | -1.341 | 0.064 | -0.517 | -1.962 | 0.006 | 0.007 | 0.007 | 0 | 0.53 | 0.53 | 0.52 | 0.6 | 0.64 | 0.34 |
| 7 | 111388472 | rs11979621 | -1.057 | 0.045 | -0.546 | -1.979 | -0.003 | -0.003 | -0.003 | 0 | 0.52 | 0.48 | 0.48 | 0.59 | 0.64 | 0.33 |
| 7 | 111390462 | rs3801783 | 2.659 | 0.098 | -0.536 | -1.974 | 0.019 | 0 | 0 | 0 | 0.31 | 0.21 | 0.36 | 0.35 | 0.26 | 0.52 |
| 7 | 111392415 | rs6979050 | -0.337 | 0.65 | -0.334 | -1.942 | 0.013 | -0.105 | -0.105 | 0 | 0.27 | 0.34 | 0.23 | 0.03 | 0.29 | 0.22 |
| 7 | 111395390 | rs2074117 | 0.773 | 0.658 | -0.366 | -1.94 | -0.053 | -0.072 | -0.072 | 0 | 0.19 | 0.07 | 0.23 | 0.56 | 0.31 | 0.12 |
| 7 | 111395987 | rs2074118 | -2.042 | 0.796 | -0.215 | -1.859 | -0.01 | -0.007 | -0.007 | 6.105 | 0.62 | 0.57 | 0.54 | 0.62 | 0.69 | 0.4 |
| 7 | 111396632 | rs13232211 | -0.323 | 0.525 | -0.294 | -1.829 | 0.014 | -0.078 | -0.078 | 0.13 | 0.26 | 0.34 | 0.22 | 0.04 | 0.28 | 0.21 |
| 7 | 111396880 | rs10252167 | 1.124 | 0.195 | -0.445 | -1.711 | 0.039 | 0.048 | 0.048 | 0 | 0.51 | 0.4 | 0.47 | 0.37 | 0.35 | 0.64 |
| 7 | 111399633 | rs6968514 | 0.075 | 0.22 | -0.582 | -1.488 | -0.062 | -0.092 | -0.092 | 0.273 | 0.25 | 0.09 | 0.27 | 0.56 | 0.34 | 0.12 |
| 7 | 111399977 | rs2074119 | 1.644 | -0.318 | -0.287 | -1.198 | 0.027 | 0.004 | 0.004 | 0 | 0.36 | 0.23 | 0.36 | 0.36 | 0.29 | 0.54 |
| 7 | 111402117 | rs10235868 | 1.279 | 0.382 | 0.356 | -0.687 | 0.041 | 0.045 | 0.045 | 0 | 0.48 | 0.29 | 0.37 | 0.37 | 0.37 | 0.64 |
| 7 | 111403763 | rs4730499 | 0.446 | 0.01 | 1.328 | -0.47 | 0.062 | -0.101 | -0.101 | 0.251 | 0.84 | 0.61 | 0.83 | 0.96 | 0.83 | 0.83 |
| 7 | 111420124 | rs17158813 | -0.422 | -1.002 | 1.343 | 0.167 | 0.023 | 0.02 | 0.02 | 0.164 | 0.07 | 0.24 | 0.03 | 0.02 | 0.03 | 0.05 |
| 7 | 111426041 | rs3757646 | -0.385 | -0.994 | 1.138 | 0.413 | 0.005 | 0.048 | 0.048 | 0 | 0.12 | 0.15 | 0.26 | 0.51 | 0.17 | 0.07 |

| | | | | | | | | | | | | | | | | |
|---|---|---|---|---|---|---|---|---|---|---|---|---|---|---|---|---|
| 7 | 111426682 | rs3757645 | -1.024 | -1.004 | 1.194 | 0.412 | -0.022 | -0.013 | -0.013 | 0 | 0.09 | 0.64 | 0.06 | 0.04 | 0.04 | 0.07 |
| 7 | 111435243 | rs757032 | -1.607 | -1.053 | -0.123 | 0.376 | 0.003 | -0.034 | -0.034 | 0 | 0.23 | 0.16 | 0.28 | 0.52 | 0.21 | 0.07 |
| 7 | 111438198 | rs7790671 | -2.029 | -0.521 | -0.157 | 0.725 | 0.013 | -0.043 | -0.043 | 0.086 | 0.33 | 0.87 | 0.37 | 0.54 | 0.26 | 0.18 |
| 7 | 111439939 | rs2078987 | -1.667 | -0.271 | 0.067 | 1.096 | -0.009 | 0.042 | 0.042 | 0 | 0.24 | 0.39 | 0.3 | 0.52 | 0.2 | 0.14 |
| 7 | 111440600 | rs17158825 | -1.329 | 0.45 | -0.52 | 1.352 | -0.018 | -0.117 | -0.117 | 0 | 0.24 | 0.07 | 0.29 | 0.57 | 0.23 | 0.08 |
| 7 | 111473040 | rs10238664 | -1.571 | 0.519 | 1.365 | 1.596 | 0 | -0.059 | -0.059 | 0 | 0.23 | 0.86 | 0.21 | 0.32 | 0.21 | 0.16 |
| 7 | 111487026 | rs10275038 | -1.656 | 0.511 | 1.317 | 1.724 | -0.017 | -0.031 | -0.031 | 0 | 0.24 | 0.91 | 0.21 | 0.23 | 0.21 | 0.18 |
| 7 | 111497826 | rs1476517 | -1.581 | 1.462 | 1.886 | 1.505 | 0.058 | 0.034 | 0.034 | 0 | 0.25 | 0.35 | 0.29 | 0.33 | 0.33 | 0.14 |
| 7 | 111500553 | rs10085411 | -0.869 | 1.229 | 1.998 | 1.333 | -0.02 | -0.016 | -0.016 | 0 | 0.08 | 0.22 | 0.06 | 0.03 | 0.08 | 0.14 |
| 7 | 111509224 | rs2074134 | -0.915 | 1.411 | 2.432 | 1.387 | 0.026 | 0.008 | 0.008 | 0 | 0.14 | 0.13 | 0.17 | 0.14 | 0.2 | 0.11 |
| 7 | 111511138 | rs17158930 | -1.283 | 1.349 | 3.026 | 1.35 | 0.091 | 0.025 | 0.025 | 0 | 0.16 | 0.43 | 0.2 | 0.14 | 0.2 | 0.13 |
| 7 | 111515077 | rs17451013 | -0.904 | 1.419 | 2.999 | 1.25 | -0.021 | 0.015 | 0.015 | 0 | 0.13 | 0.09 | 0.16 | 0.07 | 0.21 | 0.11 |
| 7 | 111518413 | rs757036 | -2.363 | 1.338 | 3.163 | 1.301 | -0.046 | 0.005 | 0.005 | 0.664 | 0.35 | 0.76 | 0.34 | 0.29 | 0.4 | 0.29 |
| 7 | 111523264 | rs12539182 | -0.452 | 1.326 | 3.582 | 1.316 | 0.108 | 0.164 | 0.164 | 0 | 0.11 | 0.35 | 0.12 | 0.21 | 0.12 | 0.03 |
| 7 | 111524959 | rs6466389 | -2.608 | 1.33 | 3.05 | 1.321 | -0.096 | -0.012 | -0.012 | 0 | 0.37 | 0.86 | 0.36 | 0.3 | 0.41 | 0.3 |
| 7 | 111527560 | rs10255299 | -1.233 | 0.737 | 3.214 | 1.056 | -0.026 | -0.051 | -0.051 | 0.083 | 0.1 | 0.22 | 0.06 | 0.01 | 0.08 | 0.15 |
| 7 | 111533520 | rs10236850 | -1.079 | 0.774 | 3.105 | 1.072 | -0.014 | -0.033 | -0.033 | 0 | 0.09 | 0.13 | 0.06 | 0.01 | 0.08 | 0.15 |
| 7 | 111541692 | rs12154775 | -1.451 | 1.104 | 3.07 | 1.273 | 0.086 | -0.007 | -0.007 | 0 | 0.15 | 0.35 | 0.17 | 0.1 | 0.2 | 0.11 |
| 7 | 111550387 | rs4607530 | -1.754 | 1.202 | 2.81 | 0.828 | 0.048 | -0.08 | -0.08 | 0 | 0.19 | 0.4 | 0.14 | 0.05 | 0.23 | 0.19 |
| 7 | 111558797 | rs6954980 | -1.995 | 1.032 | 2.785 | 1.3 | 0.037 | 0.08 | 0.08 | 0 | 0.23 | 0.38 | 0.18 | 0.35 | 0.28 | 0.21 |
| 7 | 111567391 | rs17159016 | -1.825 | 0.773 | 2.71 | 1.51 | 0.046 | 0.242 | 0.242 | 1.092 | 0.11 | 0.37 | 0.18 | 0.38 | 0.13 | 0.09 |
| 7 | 111577642 | rs7811376 | -1.589 | 0.486 | 2.723 | 1.498 | 0.018 | 0.248 | 0.248 | 0.344 | 0.1 | 0.34 | 0.11 | 0.35 | 0.07 | 0.08 |
| 7 | 111589392 | rs13229792 | -1.274 | 0.351 | 2.617 | 1.165 | 0.027 | -0.043 | -0.043 | 0.124 | 0.09 | 0.21 | 0.1 | 0.02 | 0.12 | 0.1 |
| 7 | 111590391 | rs6944619 | -1.88 | -0.567 | 2.28 | 0.805 | -0.04 | 0.017 | 0.017 | 1.989 | 0.31 | 0.4 | 0.15 | 0.38 | 0.17 | 0.21 |
| 7 | 111601473 | rs17159057 | -1.994 | -0.509 | 1.633 | 0.783 | -0.018 | 0.014 | 0.014 | 0.842 | 0.32 | 0.51 | 0.17 | 0.39 | 0.21 | 0.22 |
| 7 | 111617978 | rs3852333 | -1.005 | -0.514 | 1.17 | 0.775 | 0.004 | 0.005 | 0.005 | 2.193 | 0.2 | 0.11 | 0.07 | 0.03 | 0.14 | 0.12 |
| 7 | 111619752 | rs6466393 | -1.197 | -0.367 | 0.799 | 0.892 | -0.021 | 0.031 | 0.031 | 0 | 0.3 | 0.52 | 0.16 | 0.4 | 0.21 | 0.22 |
| 7 | 111627007 | rs6951506 | -1.12 | -0.582 | -0.461 | 0.783 | 0.057 | -0.037 | -0.037 | 0 | 0.43 | 0.87 | 0.26 | 0.48 | 0.32 | 0.31 |
| 7 | 111628355 | rs183724 | 1.137 | -0.272 | 0.132 | 0.58 | -0.009 | -0.035 | -0.035 | 0 | 0.94 | 0.88 | 0.96 | 1 | 0.93 | 0.96 |
| 7 | 111629366 | rs2074129 | -0.723 | 1.174 | 0.303 | 1.34 | 0.105 | 0.196 | 0.196 | 0 | 0.19 | 0.49 | 0.29 | 0.44 | 0.33 | 0.25 |
| 7 | 111630024 | rs29465 | -1.008 | 1.41 | 0.03 | 1.032 | 0.073 | 0.102 | 0.102 | 0 | 0.24 | 0.52 | 0.46 | 0.44 | 0.4 | 0.3 |
| 7 | 111633529 | rs173860 | -0.311 | 0.741 | -1.102 | 0.68 | -0.032 | 0.136 | 0.136 | 0.488 | 0.66 | 0.88 | 0.76 | 0.87 | 0.62 | 0.74 |
| 7 | 111634898 | rs10240134 | 1.268 | 0.085 | -1.071 | 0.621 | 0.051 | 0.04 | 0.04 | 0.269 | 0.33 | 0.23 | 0.19 | 0.25 | 0.18 | 0.32 |
| 7 | 111636575 | rs10248098 | 0.939 | 0.124 | -0.751 | 0.632 | 0.011 | -0.001 | -0.001 | 0.899 | 0.38 | 0.36 | 0.31 | 0.41 | 0.22 | 0.45 |
| 7 | 111637820 | rs39774 | -0.06 | 1.071 | -0.268 | 1.009 | 0.092 | 0.146 | 0.146 | 0 | 0.23 | 0.43 | 0.44 | 0.42 | 0.33 | 0.33 |
| 7 | 111651812 | rs1018915 | 0.231 | 0.668 | 0.034 | 1.064 | 0.016 | 0.025 | 0.025 | 1.461 | 0.32 | 0.44 | 0.41 | 0.36 | 0.43 | 0.38 |
| 7 | 111654794 | rs6960113 | 0.201 | 0.451 | -0.337 | 0.433 | -0.033 | 0.078 | 0.078 | 0.451 | 0.33 | 0.19 | 0.27 | 0.14 | 0.44 | 0.31 |
| 7 | 111658510 | rs680701 | -0.172 | 0.77 | -0.978 | 0.735 | -0.136 | -0.079 | -0.079 | 1.183 | 0.78 | 0.98 | 0.63 | 0.79 | 0.65 | 0.78 |
| 7 | 111661836 | rs2522217 | 1.023 | -0.022 | -0.561 | 0.709 | 0.037 | -0.057 | -0.057 | 0.683 | 0.34 | 0.16 | 0.34 | 0.51 | 0.22 | 0.4 |
| 7 | 111664992 | rs254869 | 0.319 | -0.145 | -0.321 | 0.374 | 0.056 | 0.032 | 0.032 | 1.374 | 0.44 | 0.69 | 0.43 | 0.65 | 0.24 | 0.46 |
| 7 | 111665703 | rs9649349 | -0.019 | 0.762 | -0.017 | -0.146 | 0.008 | -0.07 | -0.07 | 0 | 0.27 | 0.28 | 0.2 | 0.01 | 0.39 | 0.18 |
| 7 | 111666943 | rs6466396 | -0.866 | 1.081 | -0.387 | 0.493 | -0.076 | -0.071 | -0.071 | 3.564 | 0.34 | 0.85 | 0.44 | 0.42 | 0.44 | 0.34 |
| 7 | 111707742 | rs11973245 | -1.012 | 0.881 | 0.581 | 0.728 | 0.063 | 0.421 | 0.421 | 0.244 | 0.1 | 0.38 | 0.23 | 0.41 | 0.07 | 0.15 |
| 7 | 111718012 | rs6959338 | -0.815 | 1.175 | -0.752 | 0.098 | -0.088 | -0.084 | -0.084 | 2.026 | 0.39 | 0.86 | 0.43 | 0.44 | 0.43 | 0.36 |
| 7 | 111722814 | rs2894634 | 0.195 | 1.178 | -0.244 | 0.22 | -0.013 | -0.013 | -0.013 | 0.432 | 0.27 | 0.27 | 0.36 | 0.43 | 0.39 | 0.35 |
| 7 | 111730967 | rs2729538 | 0.618 | -0.09 | -1.092 | -0.449 | 0.02 | 0.041 | 0.041 | 0 | 0.37 | 0.18 | 0.24 | 0.31 | 0.29 | 0.28 |
| 7 | 111732480 | rs10273453 | 0.268 | -0.235 | -0.574 | -0.846 | 0.007 | -0.132 | -0.132 | 0 | 0.34 | 0.48 | 0.22 | 0.01 | 0.41 | 0.21 |
| 7 | 111732556 | rs40143 | 0.021 | -0.499 | -0.652 | -0.743 | -0.008 | -0.022 | -0.022 | 0 | 0.63 | 0.55 | 0.39 | 0.3 | 0.66 | 0.47 |
| 7 | 111734491 | rs17455646 | 0.275 | -0.641 | -0.807 | -1.306 | 0.011 | 0.021 | 0.021 | 0 | 0.41 | 0.32 | 0.21 | 0.02 | 0.42 | 0.27 |
| 7 | 111734612 | rs2522220 | -0.274 | -0.686 | -0.577 | -0.743 | 0.015 | -0.025 | -0.025 | 0.013 | 0.32 | 0.48 | 0.3 | 0.71 | 0.29 | 0.34 |
| 7 | 111739842 | rs194590 | 1.435 | -0.476 | -0.573 | -0.083 | 0.009 | 0.007 | 0.007 | 0.962 | 0.3 | 0.32 | 0.3 | 0.35 | 0.45 | 0.39 |
| 7 | 111740653 | rs2106913 | 0.032 | -0.615 | -0.37 | 0.011 | -0.009 | 0.008 | 0.008 | 0.022 | 0.2 | 0.23 | 0.15 | 0.29 | 0.16 | 0.17 |
| 7 | 111740689 | rs259307 | 1.426 | -0.967 | -0.61 | 0.203 | 0.013 | 0.01 | 0.01 | 0.557 | 0.29 | 0.31 | 0.3 | 0.35 | 0.45 | 0.39 |
| 7 | 111741870 | rs259312 | 1.726 | -0.916 | -0.579 | -0.105 | 0.004 | -0.052 | -0.052 | 0 | 0.27 | 0.26 | 0.21 | 0.01 | 0.39 | 0.25 |
| 7 | 111745058 | rs2158127 | -0.417 | -1.223 | -0.66 | 0.171 | 0.001 | 0.025 | 0.025 | 0.586 | 0.3 | 0.45 | 0.24 | 0.45 | 0.33 | 0.23 |
| 7 | 111749409 | rs2729549 | 0.109 | -1.293 | -0.656 | 0.182 | 0.05 | 0.049 | 0.049 | 1.469 | 0.35 | 0.22 | 0.44 | 0.18 | 0.19 | 0.34 |
| 7 | 111749447 | rs849014 | 1.164 | -1.156 | -1.433 | 0.435 | -0.113 | -0.098 | -0.098 | 0 | 0.32 | 0.07 | 0.24 | 0.38 | 0.47 | 0.4 |
| 7 | 111750606 | rs453 | -0.084 | -0.479 | -0.672 | 1.114 | 0.003 | -0.015 | -0.015 | 0 | 0.7 | 0.83 | 0.73 | 0.68 | 0.68 | 0.8 |
| 7 | 111750634 | rs454 | 0.882 | -0.484 | -0.908 | 1.051 | 0.075 | -0.028 | -0.028 | 0 | 0.49 | 0.19 | 0.61 | 0.49 | 0.61 | 0.67 |
| 7 | 111751157 | rs849015 | 0.952 | -0.171 | -0.876 | 1.056 | -0.024 | 0.058 | 0.058 | 0.507 | 0.49 | 0.14 | 0.58 | 0.36 | 0.55 | 0.58 |
| 7 | 111756578 | rs11981541 | -0.414 | -0.017 | -0.295 | 1.358 | 0.007 | 0.036 | 0.036 | 0 | 0.3 | 0.51 | 0.34 | 0.56 | 0.34 | 0.25 |
| 7 | 111759584 | rs6974167 | -0.323 | -0.148 | -0.572 | 0.684 | 0.001 | 0.021 | 0.021 | 0 | 0.28 | 0.38 | 0.28 | 0.36 | 0.31 | 0.21 |
| 7 | 111771254 | rs40144 | 0.581 | -0.603 | -1.469 | 0.049 | -0.159 | 0.196 | 0.196 | 0.825 | 0.41 | 0.02 | 0.52 | 0.14 | 0.49 | 0.53 |
| 7 | 111772043 | rs35846 | 0.037 | -0.517 | -0.503 | 0.704 | -0.012 | 0.093 | 0.093 | 1.39 | 0.8 | 0.38 | 0.83 | 0.57 | 0.86 | 0.81 |
| 7 | 111773164 | rs42876 | 0.246 | -0.5 | -0.273 | 0.527 | -0.011 | 0.087 | 0.087 | 0.902 | 0.77 | 0.49 | 0.81 | 0.62 | 0.82 | 0.81 |
| 7 | 111773284 | rs35845 | 0.883 | -0.774 | -0.834 | -0.128 | -0.106 | 0.088 | 0.088 | 3.207 | 0.32 | 0.09 | 0.32 | 0.15 | 0.45 | 0.47 |
| 7 | 111778586 | rs13232973 | 0.567 | -1.272 | -0.46 | -0.222 | -0.021 | -0.04 | -0.04 | 0 | 0.19 | 0.2 | 0.23 | 0.01 | 0.05 | 0.11 |
| 7 | 111782891 | rs6466401 | 0.63 | 0.016 | -0.246 | 0.574 | 0.011 | -0.001 | -0.001 | 0 | 0.45 | 0.35 | 0.48 | 0.49 | 0.38 | 0.42 |
| 7 | 111783764 | rs2729536 | -0.219 | 0.502 | -0.882 | 0.526 | -0.012 | 0.04 | 0.04 | 1.68 | 0.83 | 0.78 | 0.8 | 0.73 | 0.87 | 0.96 |

| | | | | | | | | | | | | | | | | |
|---|---|---|---|---|---|---|---|---|---|---|---|---|---|---|---|---|
| 7 | 111783871 | rs7786289 | 0.086 | 0.602 | -0.716 | 0.553 | 0.013 | 0.001 | 0.001 | 0.364 | 0.5 | 0.65 | 0.54 | 0.54 | 0.41 | 0.54 |
| 7 | 111785581 | rs2560640 | -0.978 | 1.723 | -0.47 | 0.362 | 0.059 | -0.021 | -0.021 | 2.307 | 0.15 | 0.31 | 0.16 | 0.08 | 0.27 | 0.09 |
| 7 | 111785993 | rs2560639 | -0.553 | 1.147 | -1.635 | 0.253 | -0.041 | -0.007 | -0.007 | 0 | 0.24 | 0.02 | 0.16 | 0.12 | 0.25 | 0.21 |
| 7 | 111786013 | rs10266737 | 0.225 | 0.625 | -1.621 | 0.501 | 0.019 | -0.045 | -0.045 | 0.173 | 0.33 | 0.18 | 0.21 | 0.45 | 0.29 | 0.49 |
| 7 | 111789767 | rs10260793 | 0.603 | 0.706 | -1.045 | 0.259 | 0.011 | 0.018 | 0.018 | 0 | 0.52 | 0.38 | 0.36 | 0.59 | 0.5 | 0.69 |
| 7 | 111795949 | rs2214432 | -0.06 | 1.225 | -0.597 | -0.119 | 0.008 | -0.005 | -0.005 | 0 | 0.76 | 0.75 | 0.54 | 0.85 | 0.67 | 0.86 |
| 7 | 111796377 | rs2217262 | 0.293 | 0.934 | -0.817 | 0.005 | 0 | -0.003 | -0.003 | 1.446 | 0.08 | 0.07 | 0.15 | 0.07 | 0.08 | 0.15 |
| 7 | 111797422 | rs2189286 | 0.403 | 0.935 | -0.134 | -0.095 | 0.033 | -0.001 | -0.001 | 0 | 0.64 | 0.54 | 0.46 | 0.7 | 0.53 | 0.71 |
| 7 | 111803700 | rs2523016 | -1.089 | 0.25 | 0.157 | -0.219 | -0.047 | -0.09 | -0.09 | 0 | 0.15 | 0.37 | 0.08 | 0.01 | 0.05 | 0.03 |
| 7 | 111807776 | rs2523017 | -0.802 | 0.942 | -0.117 | 0.111 | 0.007 | -0.001 | -0.001 | 0.057 | 0.14 | 0.17 | 0.12 | 0.08 | 0.16 | 0.15 |
| 7 | 111809140 | rs10953703 | 1.046 | 1.37 | 0.063 | 0.18 | 0.015 | 0.008 | 0.008 | 3.15 | 0.92 | 0.86 | 0.7 | 0.91 | 0.9 | 0.93 |
| 7 | 111809195 | rs10953704 | 1.604 | 1.344 | 0.108 | -0.308 | 0.002 | -0.011 | -0.011 | 0 | 0.75 | 0.29 | 0.5 | 0.87 | 0.75 | 0.77 |
| 7 | 111810894 | rs10487347 | 0.248 | 1.44 | -0.262 | -0.186 | -0.02 | -0.049 | -0.049 | 0 | 0.12 | 0.01 | 0.07 | 0.09 | 0.11 | 0.13 |
| 7 | 111811004 | rs10487348 | 1.168 | 1.28 | -1.449 | -0.888 | -0.109 | -0.152 | -0.152 | 1.952 | 0.23 | 0.01 | 0.18 | 0.62 | 0.31 | 0.36 |
| 7 | 111817242 | rs2729548 | 0.091 | 0.507 | -0.654 | -1.588 | -0.023 | -0.096 | -0.096 | 2.308 | 0.14 | 0.37 | 0.2 | 0 | 0.05 | 0.06 |
| 7 | 111817732 | rs2729547 | -1.315 | 0.55 | -0.369 | -1.694 | 0.02 | 0.046 | 0.046 | 0.863 | 0.38 | 0.73 | 0.48 | 0.13 | 0.29 | 0.34 |
| 7 | 111818296 | rs12705815 | 1.366 | 0.592 | -0.54 | -1.72 | 0.01 | 0.124 | 0.124 | 0 | 0.35 | 0.09 | 0.2 | 0.14 | 0.34 | 0.27 |
| 7 | 111820090 | rs2729544 | -1.127 | 0.157 | -0.552 | -1.672 | -0.062 | -0.04 | -0.04 | 0 | 0.71 | 0.88 | 0.68 | 0.26 | 0.63 | 0.61 |
| 7 | 111820439 | rs10428921 | -0.254 | -0.116 | 0.491 | -1.599 | -0.027 | -0.02 | -0.02 | 0 | 0.24 | 0.33 | 0.12 | 0.09 | 0.15 | 0.16 |
| 7 | 111829456 | rs6971023 | 0.178 | 0.206 | 0.192 | -1.499 | -0.003 | 0.008 | 0.008 | 0 | 0.1 | 0.07 | 0.16 | 0.02 | 0.1 | 0.17 |
| 7 | 111832301 | rs6949234 | 0.273 | -0.073 | 0.317 | -1.498 | 0.004 | 0.002 | 0.002 | 0 | 0.09 | 0.07 | 0.13 | 0.02 | 0.05 | 0.15 |
| 7 | 111837561 | rs17159351 | -0.994 | -0.185 | 0.732 | -1.469 | -0.019 | -0.021 | -0.021 | 0.553 | 0.25 | 0.47 | 0.26 | 0.17 | 0.19 | 0.32 |
| 7 | 111838904 | rs12154685 | 1.372 | 0.079 | 0.609 | -1.558 | 0.003 | 0.008 | 0.008 | 0.753 | 0.67 | 0.25 | 0.61 | 0.69 | 0.74 | 0.61 |
| 7 | 111838957 | rs12154681 | 1.525 | -0.041 | 0.04 | -1.884 | -0.075 | -0.15 | -0.15 | 0.209 | 0.22 | 0.01 | 0.33 | 0.6 | 0.27 | 0.31 |
| 7 | 111839373 | rs12705816 | 0.843 | 0.231 | 0.517 | -1.675 | 0.004 | 0.009 | 0.009 | 4.311 | 0.76 | 0.37 | 0.74 | 0.71 | 0.79 | 0.76 |
| 7 | 111845773 | rs3807884 | 1.359 | -0.13 | 0.133 | -1.651 | -0.036 | -0.052 | -0.052 | 0 | 0.11 | 0.01 | 0.07 | 0.14 | 0.12 | 0.1 |
| 7 | 111852682 | rs960348 | 1.167 | -0.189 | 0.43 | -1.687 | 0.004 | -0.001 | -0.001 | 0.685 | 0.71 | 0.26 | 0.62 | 0.7 | 0.77 | 0.65 |
| 7 | 111868610 | rs12534171 | 1.963 | -0.088 | -0.315 | -2.097 | -0.073 | 0.07 | 0.07 | 0.109 | 0.12 | 0.02 | 0.16 | 0 | 0.28 | 0.05 |
| 7 | 111872577 | rs11768793 | 0.957 | -0.467 | 0.019 | -1.836 | -0.006 | -0.015 | -0.015 | 0 | 0.14 | 0.09 | 0.09 | 0.14 | 0.13 | 0.12 |
| 7 | 111877893 | rs11768608 | 0.98 | -0.467 | 0.03 | -1.836 | -0.005 | -0.014 | -0.014 | 0 | 0.14 | 0.09 | 0.09 | 0.14 | 0.13 | 0.12 |
| 7 | 111884988 | rs10953708 | 0.969 | -0.464 | 0.031 | -1.835 | -0.005 | -0.014 | -0.014 | 0 | 0.14 | 0.1 | 0.1 | 0.14 | 0.14 | 0.12 |
| 7 | 111910388 | rs10272188 | 0.969 | -0.464 | 0.109 | -1.816 | -0.002 | -0.003 | -0.003 | 0 | 0.14 | 0.12 | 0.1 | 0.13 | 0.13 | 0.12 |
| 7 | 111915984 | rs4730520 | 0.468 | -0.491 | 0.501 | -1.845 | 0.005 | -0.002 | -0.002 | 0 | 0.16 | 0.32 | 0.12 | 0.13 | 0.14 | 0.15 |
| 7 | 111955948 | rs11769219 | 0.844 | -0.481 | 0.491 | -1.842 | 0 | -0.005 | -0.005 | 0 | 0.15 | 0.11 | 0.1 | 0.14 | 0.14 | 0.14 |
| 7 | 111958382 | rs4727766 | -1.258 | -0.599 | 0.417 | -1.881 | 0.013 | -0.002 | -0.002 | 0 | 0.3 | 0.8 | 0.39 | 0.29 | 0.23 | 0.35 |
| 7 | 111960100 | rs7802828 | -1.534 | -1.145 | 0.028 | -2.185 | 0.066 | 0.074 | 0.074 | 0 | 0.41 | 0.85 | 0.42 | 0.27 | 0.27 | 0.39 |
| 7 | 111967040 | rs10272976 | -0.95 | -1.047 | 0.266 | -2.118 | 0.011 | 0.014 | 0.014 | 0.218 | 0.28 | 0.76 | 0.28 | 0.16 | 0.2 | 0.32 |
| 7 | 111972769 | rs2301725 | -0.086 | -1.014 | 0.644 | -2.103 | -0.012 | -0.019 | -0.019 | 0 | 0.2 | 0.56 | 0.14 | 0.12 | 0.15 | 0.17 |
| 7 | 111975558 | rs2107117 | -1.288 | -1.158 | 0.498 | -2.165 | 0.038 | 0.054 | 0.054 | 0.637 | 0.36 | 0.74 | 0.31 | 0.14 | 0.22 | 0.36 |
| 7 | 111976546 | rs10276369 | 1.149 | -1.169 | 0.539 | -2.152 | 0.007 | 0.01 | 0.01 | 0.015 | 0.08 | 0.05 | 0.13 | 0.02 | 0.04 | 0.15 |
| 7 | 111977110 | rs10247540 | 0.608 | -1.196 | 0.832 | -2.161 | -0.019 | -0.062 | -0.062 | 1.151 | 0.11 | 0.36 | 0.06 | 0 | 0.04 | 0.05 |
| 7 | 111981007 | rs11539696 | 1.786 | -1.872 | -0.026 | -2.542 | 0.024 | 0.03 | 0.03 | 0 | 0.12 | 0.04 | 0.03 | 0.02 | 0.07 | 0.05 |
| 7 | 111983071 | rs1047550 | 1.42 | -1.446 | 0.201 | -2.313 | 0.004 | 0.008 | 0.008 | 0.11 | 0.07 | 0.05 | 0.13 | 0.02 | 0.04 | 0.15 |
| 7 | 111986673 | rs2249157 | -1.043 | -1.443 | 0.254 | -2.33 | 0.022 | 0.026 | 0.026 | 1.026 | 0.36 | 0.69 | 0.3 | 0.16 | 0.22 | 0.35 |
| 7 | 111986803 | rs2249151 | -1.059 | -1.44 | 0.256 | -2.329 | 0.022 | 0.027 | 0.027 | 1.115 | 0.36 | 0.69 | 0.3 | 0.16 | 0.22 | 0.35 |
| 7 | 111999497 | rs7789649 | -0.82 | -1.707 | 0.193 | -2.641 | -0.003 | -0.01 | -0.01 | 0 | 0.6 | 0.49 | 0.45 | 0.32 | 0.38 | 0.51 |
| 7 | 112002804 | rs17492961 | 1.412 | -1.86 | 0.134 | -2.697 | 0.029 | 0.036 | 0.036 | 4.711 | 0.27 | 0.18 | 0.19 | 0.16 | 0.17 | 0.16 |
| 7 | 112005565 | rs2708600 | -0.955 | -1.76 | 0.328 | -2.644 | -0.005 | -0.011 | -0.011 | 0 | 0.6 | 0.54 | 0.46 | 0.32 | 0.38 | 0.48 |
| 7 | 112009331 | rs2708603 | 1.574 | -1.762 | 0.035 | -2.652 | -0.001 | 0 | 0 | 0.392 | 0.14 | 0.15 | 0.08 | 0.14 | 0.13 | 0.13 |
| 7 | 112012838 | rs6960498 | 0.464 | -1.753 | -0.192 | -2.636 | 0.044 | 0.068 | 0.068 | 2.099 | 0.36 | 0.23 | 0.33 | 0.18 | 0.23 | 0.34 |
| 7 | 112016581 | rs10435366 | 1.739 | -0.867 | -0.448 | -2.667 | -0.057 | -0.056 | -0.056 | 0 | 0.24 | 0.08 | 0.34 | 0.67 | 0.32 | 0.4 |
| 7 | 112044527 | rs2520480 | -0.874 | -0.988 | -0.121 | -1.995 | -0.017 | -0.033 | -0.033 | 0 | 0.58 | 0.86 | 0.48 | 0.4 | 0.4 | 0.62 |
| 7 | 112046687 | rs10275406 | 1.67 | -0.879 | -0.268 | -1.959 | -0.002 | -0.017 | -0.017 | 0 | 0.06 | 0.11 | 0.04 | 0.02 | 0.05 | 0.15 |
| 7 | 112047345 | rs2520479 | -0.691 | -0.956 | -0.007 | -1.982 | -0.002 | -0.001 | -0.001 | 0 | 0.55 | 0.55 | 0.44 | 0.4 | 0.39 | 0.59 |
| 7 | 112050314 | rs2188534 | 0.389 | -1.012 | -0.313 | -1.898 | 0.002 | 0 | 0 | 0 | 0.2 | 0.18 | 0.21 | 0.16 | 0.17 | 0.28 |
| 7 | 112062683 | rs7780160 | 0.975 | -1.016 | -0.77 | -1.877 | 0.03 | 0.016 | 0.016 | 4.247 | 0.29 | 0.17 | 0.21 | 0.23 | 0.2 | 0.23 |
| 7 | 112063642 | rs2520482 | 0.073 | -0.87 | -1.029 | -1.588 | -0.008 | 0 | 0 | 0 | 0.24 | 0.26 | 0.21 | 0.19 | 0.18 | 0.27 |
| 7 | 112064376 | rs9640740 | 1.134 | -0.532 | -1.283 | -1.597 | -0.11 | -0.154 | -0.154 | 2.47 | 0.29 | 0.02 | 0.36 | 0.6 | 0.32 | 0.35 |
| 7 | 112071795 | rs3109114 | -0.985 | -1.349 | -0.594 | -1.049 | 0.005 | -0.011 | -0.011 | 0 | 0.84 | 0.88 | 0.91 | 0.8 | 0.87 | 0.86 |
| 7 | 112076358 | rs3109117 | 0.414 | -1.231 | -0.276 | -1.293 | -0.007 | -0.027 | -0.027 | 0 | 0.08 | 0.12 | 0.13 | 0.02 | 0.06 | 0.14 |
| 7 | 112082922 | rs3109107 | -0.679 | -1.434 | -0.432 | -1.382 | -0.015 | -0.023 | -0.023 | 0 | 0.4 | 0.56 | 0.35 | 0.21 | 0.25 | 0.41 |
| 7 | 112083308 | rs17402962 | -0.03 | -0.315 | -0.575 | -0.833 | 0.006 | 0.02 | 0.02 | 0 | 0.08 | 0.04 | 0.12 | 0.02 | 0.06 | 0.14 |
| 7 | 112086893 | rs10155986 | -0.138 | -0.359 | -0.343 | -0.809 | -0.022 | -0.007 | -0.007 | 0 | 0.59 | 0.51 | 0.7 | 0.62 | 0.7 | 0.58 |
| 7 | 112087033 | rs10155882 | -0.009 | -0.029 | -0.362 | -0.743 | 0.002 | 0.001 | 0.001 | 0.111 | 0.52 | 0.47 | 0.66 | 0.55 | 0.61 | 0.5 |
| 7 | 112089060 | rs6963343 | -0.778 | -0.112 | 0.089 | -0.963 | 0.019 | 0.013 | 0.013 | 0 | 0.16 | 0.33 | 0.21 | 0.15 | 0.12 | 0.35 |
| 7 | 112090360 | rs17403123 | 0.505 | 0.397 | 0.102 | -1.426 | -0.004 | -0.006 | -0.006 | 0 | 0.39 | 0.32 | 0.26 | 0 | 0.45 | 0.19 |
| 7 | 112094085 | rs11768354 | 0.339 | 0.366 | 0.361 | -1.423 | -0.001 | -0.124 | -0.124 | 0 | 0.41 | 0.45 | 0.28 | 0 | 0.45 | 0.22 |
| 7 | 112094743 | rs6956741 | 0.779 | 0.373 | 0.233 | -1.409 | 0.017 | 0.011 | 0.011 | 0 | 0.1 | 0.04 | 0.14 | 0.08 | 0.06 | 0.23 |
| 7 | 112095782 | rs6944364 | -0.222 | 0.168 | 0.196 | -1.671 | 0.008 | -0.033 | -0.033 | 0.674 | 0.39 | 0.48 | 0.42 | 0.16 | 0.5 | 0.44 |

| | | | | | | | | | | | | | | | | |
|---|---|---|---|---|---|---|---|---|---|---|---|---|---|---|---|---|
| 7 | 112097121 | rs6968084 | -0.294 | 0.308 | 0.219 | -1.62 | 0.01 | 0.007 | 0.007 | 0 | 0.14 | 0.23 | 0.2 | 0.16 | 0.13 | 0.33 |
| 7 | 112098521 | rs12155248 | 1.335 | 0.308 | -0.102 | -1.624 | -0.007 | -0.034 | -0.034 | 0.922 | 0.26 | 0.27 | 0.22 | 0 | 0.37 | 0.11 |
| 7 | 112102355 | rs2253962 | 1.213 | 0.846 | -0.384 | -1.185 | -0.002 | -0.003 | -0.003 | 0 | 0.33 | 0.33 | 0.16 | 0.19 | 0.25 | 0.2 |
| 7 | 112103055 | rs2529588 | 1.42 | 0.843 | -0.413 | -1.186 | 0.035 | 0.042 | 0.042 | 0.806 | 0.32 | 0.19 | 0.14 | 0.19 | 0.25 | 0.17 |
| 7 | 112105754 | rs3807213 | -0.422 | 1.006 | -0.788 | -1.124 | 0.009 | 0.048 | 0.048 | 0.025 | 0.54 | 0.62 | 0.46 | 0.16 | 0.58 | 0.52 |
| 7 | 112107887 | rs1024568 | -0.395 | 1.029 | -0.768 | -1.114 | -0.013 | -0.01 | -0.01 | 0.265 | 0.13 | 0.08 | 0.18 | 0.16 | 0.13 | 0.33 |
| 7 | 112109331 | rs2708607 | -0.795 | 1.038 | -0.756 | -1.111 | -0.002 | 0.014 | 0.014 | 0.521 | 0.87 | 0.91 | 0.62 | 0.35 | 0.83 | 0.72 |
| 7 | 112112279 | rs2074796 | 1.088 | 0.228 | -0.841 | -0.958 | -0.008 | -0.01 | -0.01 | 0 | 0.33 | 0.36 | 0.16 | 0.19 | 0.25 | 0.2 |
| 7 | 112119702 | rs11773335 | -0.442 | 1.059 | -0.953 | 0.006 | 0.017 | 0.01 | 0.01 | 0 | 0.1 | 0.04 | 0.15 | 0.08 | 0.06 | 0.23 |
| 7 | 112122640 | rs3128383 | 1.031 | 1.137 | -0.877 | -0.246 | -0.007 | -0.018 | -0.018 | 0 | 0.77 | 0.87 | 0.47 | 0.27 | 0.77 | 0.49 |
| 7 | 112128978 | rs3095031 | 0.326 | 1.588 | -0.795 | -0.553 | -0.022 | -0.024 | -0.024 | 0 | 0.88 | 0.92 | 0.65 | 0.24 | 0.83 | 0.73 |
| 7 | 112129333 | rs3128376 | -0.135 | 1.105 | -1.115 | -2.149 | 0.004 | 0.089 | 0.089 | 0 | 0.41 | 0.67 | 0.43 | 0.04 | 0.43 | 0.25 |
| 7 | 112130128 | rs10225870 | -0.479 | 0.754 | -0.274 | -2.625 | -0.026 | -0.074 | -0.074 | 0.648 | 0.15 | 0.37 | 0.21 | 0.03 | 0.09 | 0.13 |
| 7 | 112130494 | rs12531501 | -0.479 | 0.754 | -0.274 | -2.625 | -0.026 | -0.074 | -0.074 | 0 | 0.15 | 0.37 | 0.21 | 0.03 | 0.09 | 0.13 |
| 7 | 112140430 | rs11772829 | 0.593 | 1.066 | -0.884 | -2.825 | -0.043 | 0.128 | 0.128 | 0 | 0.25 | 0.03 | 0.18 | 0.04 | 0.3 | 0.32 |
| 7 | 112152119 | rs10273549 | -1.28 | 1.109 | -0.11 | -2.859 | -0.004 | -0.035 | -0.035 | 0 | 0.11 | 0.14 | 0.03 | 0 | 0.07 | 0.1 |
| 7 | 112153368 | rs2396552 | -0.046 | 1.046 | 0.239 | -2.998 | 0.002 | -0.019 | -0.019 | 2.659 | 0.35 | 0.33 | 0.32 | 0.07 | 0.35 | 0.33 |
| 7 | 112155176 | rs2966473 | 0.665 | 0.782 | 0.147 | -3.24 | 0.003 | 0.051 | 0.051 | 0.082 | 0.47 | 0.34 | 0.44 | 0.92 | 0.43 | 0.49 |
| 7 | 112158484 | rs16873403 | 0.246 | 0.765 | -0.029 | -3.25 | -0.01 | -0.035 | -0.035 | 0.247 | 0.12 | 0.14 | 0.04 | 0.01 | 0.09 | 0.08 |
| 7 | 112159178 | rs5009270 | 0.737 | 0.9 | -0.272 | -3.342 | 0.001 | -0.042 | -0.042 | 0 | 0.45 | 0.53 | 0.24 | 0.09 | 0.34 | 0.31 |
| 7 | 112162658 | rs2190591 | -0.823 | 0.897 | -0.266 | -3.355 | -0.063 | 0.116 | 0.116 | 0.856 | 0.71 | 0.87 | 0.65 | 0.96 | 0.62 | 0.66 |
| 7 | 112171249 | rs3095043 | -1.112 | 0.857 | 0.318 | -3.022 | -0.015 | -0.138 | -0.138 | 0.619 | 0.2 | 0.42 | 0.11 | 0.01 | 0.18 | 0.09 |
| 7 | 112194084 | rs3111450 | -0.604 | 0.829 | 0.295 | -3.015 | -0.058 | -0.113 | -0.113 | 0 | 0.17 | 0.36 | 0.05 | 0.01 | 0.1 | 0.06 |
| 7 | 112198998 | rs3111446 | -0.979 | 0.843 | 0.245 | -3.005 | -0.018 | -0.135 | -0.135 | 2.588 | 0.21 | 0.41 | 0.11 | 0.01 | 0.18 | 0.1 |
| 7 | 112211290 | rs2079530 | 0.294 | 1.172 | -0.144 | -2.765 | 0.002 | 0.002 | 0.002 | 0 | 0.21 | 0.25 | 0.32 | 0.76 | 0.23 | 0.31 |
| 7 | 112212543 | rs3111438 | -1.224 | 1.202 | -0.351 | -2.985 | -0.021 | 0.054 | 0.054 | 0 | 0.42 | 0.68 | 0.55 | 0.77 | 0.46 | 0.47 |
| 7 | 112212780 | rs3128399 | -0.92 | 1.201 | -0.328 | -2.971 | -0.003 | 0.022 | 0.022 | 0 | 0.37 | 0.66 | 0.51 | 0.77 | 0.39 | 0.42 |
| 7 | 112224898 | rs3111434 | -0.021 | 0.87 | -0.898 | -3.187 | 0.008 | 0.244 | 0.244 | 0 | 0.53 | 0.79 | 0.29 | 0.08 | 0.42 | 0.31 |
| 7 | 112225536 | rs1234352 | 0.672 | 0.899 | -1.091 | -3.131 | -0.008 | -0.025 | -0.025 | 0.381 | 0.21 | 0.1 | 0.38 | 0.76 | 0.21 | 0.37 |
| 7 | 112230510 | rs7802221 | 1.887 | 0.705 | -1.134 | -3.182 | 0.06 | 0.142 | 0.142 | 1.567 | 0.28 | 0.04 | 0.12 | 0.06 | 0.18 | 0.13 |
| 7 | 112235033 | rs2188547 | -0.85 | 1.174 | -0.978 | -2.903 | -0.021 | -0.061 | -0.061 | 0 | 0.8 | 0.95 | 0.74 | 0.25 | 0.81 | 0.64 |
| 7 | 112247187 | rs43083 | -0.788 | 1.106 | -0.923 | -2.902 | 0.003 | -0.028 | -0.028 | 0 | 0.8 | 0.89 | 0.64 | 0.25 | 0.81 | 0.65 |
| 7 | 112250044 | rs43089 | 0.635 | 1.306 | -0.705 | -2.593 | -0.128 | 0.002 | 0.002 | 0.942 | 0.26 | 0.04 | 0.34 | 0.17 | 0.39 | 0.33 |
| 7 | 112257009 | rs4730556 | 0.179 | 1.344 | -0.571 | -2.399 | -0.033 | -0.027 | -0.027 | 2.709 | 0.05 | 0 | 0.16 | 0.17 | 0.09 | 0.23 |
| 7 | 112261226 | rs2966478 | -0.037 | 0.505 | -0.075 | -2.645 | 0.011 | -0.035 | -0.035 | 0 | 0.64 | 0.44 | 0.55 | 0.84 | 0.47 | 0.68 |
| 7 | 112262301 | rs1234380 | -0.069 | 0.417 | -0.55 | -2.749 | -0.13 | 0.1 | 0.1 | 1.737 | 0.63 | 0.94 | 0.62 | 0.83 | 0.5 | 0.59 |
| 7 | 112276681 | rs2520570 | 0.717 | 0.159 | -0.221 | -2.727 | 0.001 | -0.015 | -0.015 | 0.349 | 0.84 | 0.84 | 0.92 | 1 | 0.84 | 0.9 |
| 7 | 112278724 | rs1634112 | 0.663 | 0.164 | 0.006 | -2.814 | -0.001 | -0.013 | -0.013 | 0 | 0.54 | 0.51 | 0.52 | 0.83 | 0.39 | 0.54 |
| 7 | 112278813 | rs1634113 | 1.14 | 0.157 | 0.124 | -2.708 | -0.012 | -0.132 | -0.132 | 7.075 | 0.81 | 0.55 | 0.88 | 0.99 | 0.84 | 0.89 |
| 7 | 112278964 | rs1796507 | 0.773 | 0.047 | -0.326 | -2.837 | -0.054 | -0.064 | -0.064 | 0 | 0.21 | 0.03 | 0.34 | 0.76 | 0.21 | 0.34 |
| 7 | 112290310 | rs2966479 | 0.156 | -0.086 | 0.116 | -2.853 | -0.025 | -0.034 | -0.034 | 2.014 | 0.66 | 0.52 | 0.61 | 0.22 | 0.74 | 0.56 |
| 7 | 112295934 | rs1989835 | 1.012 | -0.142 | -0.098 | -3.013 | 0.038 | 0.04 | 0.04 | 0 | 0.34 | 0.22 | 0.18 | 0.06 | 0.21 | 0.17 |
| 7 | 112295954 | rs2952690 | 0.879 | 0.087 | -0.464 | -2.954 | -0.172 | 0.019 | 0.019 | 0.873 | 0.27 | 0.03 | 0.36 | 0.16 | 0.46 | 0.35 |
| 7 | 112308895 | rs1922899 | -0.782 | 0.095 | -0.324 | -2.833 | -0.024 | -0.03 | -0.03 | 0 | 0.74 | 0.97 | 0.66 | 0.23 | 0.76 | 0.61 |
| 7 | 112314910 | rs2952683 | -0.021 | 0.159 | 0.162 | -2.84 | -0.002 | 0.008 | 0.008 | 0 | 0.67 | 0.67 | 0.58 | 0.23 | 0.65 | 0.56 |
| 7 | 112318992 | rs2028028 | -0.304 | -0.329 | -0.002 | -2.898 | 0.103 | 0.361 | 0.361 | 0.822 | 0.49 | 0.93 | 0.31 | 0.07 | 0.31 | 0.36 |
| 7 | 112321309 | rs10269975 | 1.334 | -0.475 | 0.085 | -3.005 | 0.087 | 0.126 | 0.126 | 0 | 0.29 | 0.08 | 0.16 | 0.06 | 0.17 | 0.14 |
| 7 | 112321368 | rs10226544 | -0.421 | -0.436 | 0.054 | -2.993 | 0.103 | 0.361 | 0.361 | 0.822 | 0.49 | 0.93 | 0.31 | 0.07 | 0.31 | 0.36 |
| 7 | 112321378 | rs10270000 | 0.589 | -0.529 | 0.742 | -2.959 | -0.023 | -0.048 | -0.048 | 0 | 0.37 | 0.52 | 0.2 | 0.07 | 0.18 | 0.19 |
| 7 | 112373996 | rs1614000 | 0.336 | -0.511 | 0.75 | -2.959 | 0.126 | 0.152 | 0.152 | 0.244 | 0.41 | 0.75 | 0.22 | 0.07 | 0.2 | 0.23 |
| 7 | 112386091 | rs1227158 | -0.371 | -0.452 | 0.715 | -2.921 | 0.126 | 0.307 | 0.307 | 0.496 | 0.49 | 0.87 | 0.3 | 0.08 | 0.32 | 0.39 |
| 7 | 112400060 | rs10271752 | -0.061 | -0.452 | 1.045 | -2.924 | 0.078 | 0.119 | 0.119 | 0.163 | 0.46 | 0.66 | 0.27 | 0.08 | 0.29 | 0.37 |
| 7 | 112411379 | rs1227171 | -0.331 | -0.452 | 0.92 | -2.926 | 0.121 | 0.28 | 0.28 | 0.105 | 0.48 | 0.83 | 0.29 | 0.08 | 0.31 | 0.39 |
| 7 | 112420072 | rs7806781 | 0.496 | -0.485 | 1.534 | -2.923 | 0.027 | 0.013 | 0.013 | 0 | 0.38 | 0.29 | 0.2 | 0.07 | 0.22 | 0.25 |
| 7 | 112434003 | rs10238492 | 1.247 | -0.438 | 1.25 | -2.897 | 0.098 | 0.156 | 0.156 | 0 | 0.3 | 0.05 | 0.15 | 0.06 | 0.17 | 0.15 |
| 7 | 112443541 | rs2396570 | 1.186 | -0.438 | 1.327 | -2.894 | 0.061 | 0.074 | 0.074 | 0 | 0.3 | 0.15 | 0.17 | 0.06 | 0.18 | 0.15 |
| 7 | 112464841 | rs821785 | -0.226 | -0.405 | 1.052 | -2.926 | 0.106 | 0.238 | 0.238 | 0.386 | 0.47 | 0.81 | 0.29 | 0.08 | 0.31 | 0.38 |
| 7 | 112469673 | rs7781871 | 0.792 | -0.413 | 1.388 | -2.921 | -0.067 | -0.118 | -0.118 | 0 | 0.35 | 0.4 | 0.19 | 0.07 | 0.19 | 0.22 |
| 7 | 112469778 | rs821783 | -0.201 | -0.318 | 1.354 | -2.915 | 0.092 | 0.23 | 0.23 | 0.311 | 0.46 | 0.81 | 0.29 | 0.08 | 0.31 | 0.39 |
| 7 | 112499924 | rs6948575 | -0.107 | -0.473 | 1.115 | -2.886 | 0.082 | 0.133 | 0.133 | 0.354 | 0.46 | 0.71 | 0.27 | 0.12 | 0.29 | 0.36 |
| 7 | 112516977 | rs6973324 | 0.825 | -0.206 | 0.899 | -3.031 | -0.095 | 0.03 | 0.03 | 1.035 | 0.28 | 0.1 | 0.42 | 0.17 | 0.46 | 0.35 |
| 7 | 112517826 | rs660743 | -0.082 | -0.239 | 0.924 | -3.103 | 0.111 | 0.219 | 0.219 | 0.383 | 0.54 | 0.2 | 0.72 | 0.92 | 0.71 | 0.64 |
| 7 | 112538378 | rs11762737 | 1.338 | -0.264 | 0.822 | -3.092 | 0.021 | 0.007 | 0.007 | 0 | 0.3 | 0.2 | 0.13 | 0.06 | 0.17 | 0.11 |
| 7 | 112547374 | rs498377 | 1.051 | -0.582 | 0.789 | -2.982 | -0.041 | -0.046 | -0.046 | 0.27 | 0.25 | 0.03 | 0.29 | 0.76 | 0.23 | 0.29 |
| 7 | 112553037 | rs1530756 | 0.59 | -0.599 | 0.887 | -2.957 | -0.054 | -0.1 | -0.1 | 2.949 | 0.35 | 0.37 | 0.15 | 0.07 | 0.18 | 0.19 |
| 7 | 112565907 | rs17484145 | 0.861 | -0.504 | 0.473 | -2.92 | -0.013 | 0.009 | 0.009 | 0 | 0.06 | 0.03 | 0.06 | 0 | 0.09 | 0.14 |
| 7 | 112574233 | rs12539085 | 0.12 | -0.53 | 0.42 | -2.911 | 0.1 | 0.296 | 0.296 | 0.933 | 0.53 | 0.12 | 0.7 | 0.92 | 0.69 | 0.63 |
| 7 | 112587413 | rs1992988 | 0.927 | -0.475 | -0.523 | -2.656 | -0.028 | -0.056 | -0.056 | 0 | 0.31 | 0.29 | 0.14 | 0.06 | 0.16 | 0.11 |
| 7 | 112587963 | rs2056798 | 0.682 | 0.013 | -1.615 | -2.419 | -0.121 | 0.035 | 0.035 | 0.925 | 0.28 | 0.08 | 0.42 | 0.16 | 0.46 | 0.38 |

| | | | | | | | | | | | | | | | | |
|---|---|---|---|---|---|---|---|---|---|---|---|---|---|---|---|---|
| 7 | 112597791 | rs7806102 | 0.475 | -0.594 | -0.836 | -2.365 | 0.066 | 0.142 | 0.142 | 0 | 0.28 | 0.09 | 0.14 | 0.06 | 0.19 | 0.13 |
| 7 | 112614215 | rs10246678 | -0.248 | -0.052 | -0.489 | -2.11 | 0.035 | 0.063 | 0.063 | 0.098 | 0.44 | 0.67 | 0.29 | 0.08 | 0.32 | 0.34 |
| 7 | 112625841 | rs7797689 | 0.163 | -0.043 | -0.93 | -1.965 | -0.07 | 0.08 | 0.08 | 0.124 | 0.42 | 0.07 | 0.64 | 0.76 | 0.57 | 0.33 |
| 7 | 112628373 | rs10229603 | 0.449 | -0.741 | -0.865 | -2.381 | 0.015 | 0.019 | 0.019 | 0 | 0.37 | 0.35 | 0.22 | 0.21 | 0.3 | 0.45 |
| 7 | 112631636 | rs1569277 | 0.334 | -0.82 | -1.314 | -2.772 | 0.039 | 0.135 | 0.135 | 0 | 0.41 | 0.16 | 0.25 | 0.19 | 0.36 | 0.46 |
| 7 | 112633232 | rs10261051 | 0.676 | -0.421 | -0.334 | -2.827 | 0.005 | -0.015 | -0.015 | 0 | 0.37 | 0.35 | 0.62 | 0.8 | 0.55 | 0.37 |
| 7 | 112652854 | rs7801538 | -1.19 | -0.257 | 0.945 | -2.888 | -0.037 | -0.168 | -0.168 | 5.317 | 0.3 | 0.52 | 0.19 | 0 | 0.18 | 0.19 |
| 7 | 112666609 | rs6957202 | -0.291 | -0.314 | 1.05 | -2.889 | -0.014 | -0.069 | -0.069 | 0 | 0.19 | 0.28 | 0.1 | 0 | 0.09 | 0.18 |
| 7 | 112673539 | rs13236467 | 0.656 | -0.225 | 1.011 | -2.831 | 0.002 | 0.014 | 0.014 | 4.28 | 0.1 | 0.05 | 0.07 | 0 | 0.09 | 0.02 |
| 7 | 112675601 | rs2014707 | 0.695 | -0.239 | 1.092 | -2.841 | 0.023 | -0.094 | -0.094 | 0 | 0.83 | 0.48 | 0.86 | 0.97 | 0.82 | 0.9 |
| 7 | 112694930 | rs1830469 | 0.116 | -0.297 | 0.902 | -2.773 | -0.008 | -0.073 | -0.073 | 2.361 | 0.87 | 0.54 | 0.9 | 0.99 | 0.91 | 0.98 |
| 7 | 112695484 | rs10808170 | 1.22 | -0.248 | 0.243 | -2.771 | 0.019 | 0.092 | 0.092 | 3.859 | 0.64 | 0.19 | 0.77 | 0.95 | 0.74 | 0.73 |
| 7 | 112697690 | rs11972227 | 0.146 | -0.121 | 0.309 | -2.079 | 0 | -0.053 | -0.053 | 0.119 | 0.09 | 0.26 | 0.06 | 0 | 0.08 | 0.02 |
| 7 | 112709272 | rs1608882 | 0.395 | -0.296 | -0.075 | -2.238 | -0.006 | -0.03 | -0.03 | 0.084 | 0.81 | 0.75 | 0.88 | 0.96 | 0.83 | 0.8 |
| 7 | 112716152 | rs13221267 | -0.013 | -0.782 | -0.573 | -3.029 | 0.004 | -0.06 | -0.06 | 0 | 0.5 | 0.48 | 0.74 | 0.95 | 0.61 | 0.69 |
| 7 | 112718961 | rs2255442 | 0.406 | 0.165 | -0.567 | -2.962 | -0.004 | -0.074 | -0.074 | 0.374 | 0.63 | 0.63 | 0.44 | 0.06 | 0.64 | 0.48 |
| 7 | 112723093 | rs1581688 | 0.566 | 0.338 | -0.836 | -3.084 | 0.007 | 0.037 | 0.037 | 0.453 | 0.42 | 0.28 | 0.24 | 0.04 | 0.36 | 0.41 |
| 7 | 112724417 | rs2256044 | 0.133 | -0.1 | -0.888 | -2.906 | 0.027 | 0.056 | 0.056 | 0 | 0.84 | 0.94 | 0.93 | 0.97 | 0.88 | 0.83 |
| 7 | 112728053 | rs2140913 | 1.146 | 0.114 | -0.63 | -3.014 | 0 | 0.002 | 0.002 | 0 | 0.8 | 0.78 | 0.9 | 0.94 | 0.84 | 0.67 |
| 7 | 112734225 | rs2245985 | 1.048 | 0.121 | -0.61 | -2.874 | -0.006 | 0.035 | 0.035 | 0 | 0.81 | 0.88 | 0.9 | 0.92 | 0.84 | 0.67 |
| 7 | 112737222 | rs9656200 | -1.428 | 0.132 | -0.607 | -2.87 | 0 | 0.006 | 0.006 | 0 | 0.2 | 0.22 | 0.1 | 0.08 | 0.16 | 0.33 |
| 7 | 112750891 | rs2253486 | -0.433 | 0.576 | -0.547 | -2.961 | 0.004 | -0.007 | -0.007 | 0 | 0.74 | 0.75 | 0.46 | 0.14 | 0.62 | 0.64 |
| 7 | 112751685 | rs1599836 | 1.784 | 0.151 | -0.411 | -3.255 | 0.008 | 0.142 | 0.142 | 1.146 | 0.43 | 0.19 | 0.29 | 0.06 | 0.41 | 0.3 |
| 7 | 112752594 | rs2253640 | -2.007 | -0.056 | 0.007 | -2.762 | -0.021 | -0.096 | -0.096 | 0 | 0.31 | 0.57 | 0.18 | 0.08 | 0.21 | 0.35 |
| 7 | 112755548 | rs2256872 | -1.289 | -0.004 | 0.105 | -2.787 | -0.005 | -0.042 | -0.042 | 0 | 0.15 | 0.32 | 0.1 | 0.04 | 0.12 | 0.18 |
| 7 | 112757945 | rs2272261 | -0.396 | -0.004 | 0.101 | -2.808 | 0.006 | 0.024 | 0.024 | 0 | 0.15 | 0.09 | 0.07 | 0.04 | 0.09 | 0.17 |
| 7 | 112761420 | rs2177800 | -2.009 | 0.041 | 0.041 | -2.778 | -0.02 | -0.097 | -0.097 | 0 | 0.31 | 0.5 | 0.18 | 0.08 | 0.21 | 0.35 |
| 7 | 112766082 | rs2246845 | -1.936 | -0.246 | 0.096 | -2.721 | -0.029 | -0.108 | -0.108 | 0 | 0.28 | 0.48 | 0.15 | 0.04 | 0.17 | 0.26 |
| 7 | 112771116 | rs10241676 | -0.72 | -0.158 | 0.158 | -2.492 | -0.002 | -0.006 | -0.006 | 0 | 0.08 | 0.1 | 0.03 | 0.04 | 0.04 | 0.15 |
| 7 | 112778922 | rs12672405 | -0.587 | -0.14 | 0.146 | -2.484 | 0 | -0.001 | -0.001 | 0 | 0.07 | 0.07 | 0.03 | 0.04 | 0.04 | 0.15 |
| 7 | 112786875 | rs12670645 | 0.149 | -0.091 | 0.209 | -2.435 | -0.002 | -0.005 | -0.005 | 0 | 0.07 | 0.09 | 0.03 | 0.04 | 0.04 | 0.14 |
| 7 | 112791331 | rs10225200 | -1.737 | -0.357 | 0.515 | -2.639 | -0.019 | -0.072 | -0.072 | 0.37 | 0.19 | 0.37 | 0.1 | 0.04 | 0.12 | 0.22 |
| 7 | 112802475 | rs10278248 | -2.196 | -0.443 | 0.538 | -2.676 | -0.036 | -0.101 | -0.101 | 0 | 0.23 | 0.51 | 0.13 | 0.05 | 0.13 | 0.23 |
| 7 | 112804274 | rs10254201 | -0.952 | -0.207 | 0.617 | -2.448 | -0.008 | -0.018 | -0.018 | 0 | 0.1 | 0.24 | 0.05 | 0.05 | 0.06 | 0.16 |
| 7 | 112854661 | rs1917763 | -0.369 | -0.141 | 0.487 | -2.381 | -0.002 | 0.003 | 0.003 | 0.774 | 0.4 | 0.34 | 0.61 | 0.75 | 0.55 | 0.56 |
| 7 | 112862269 | rs7797620 | -0.604 | -0.279 | 0.617 | -2.605 | 0.004 | 0.005 | 0.005 | 1.175 | 0.46 | 0.37 | 0.69 | 0.89 | 0.59 | 0.66 |
| 7 | 112862789 | rs12705851 | 1.196 | -0.134 | 0.878 | -2.431 | 0.051 | 0.063 | 0.063 | 2.464 | 0.93 | 0.58 | 0.95 | 0.95 | 0.96 | 0.88 |
| 7 | 112870469 | rs4481522 | 0.906 | -0.127 | 0.837 | -2.428 | 0.035 | 0.047 | 0.047 | 1.534 | 0.93 | 0.66 | 0.95 | 0.95 | 0.96 | 0.88 |
| 7 | 112873045 | rs6956889 | 1.67 | -0.238 | 0.601 | -2.477 | 0 | -0.009 | -0.009 | 0.083 | 0.06 | 0.04 | 0.02 | 0 | 0.02 | 0.03 |
| 7 | 112879751 | rs11971013 | 1.599 | -0.496 | 0.504 | -2.412 | 0.002 | 0.017 | 0.017 | 0.618 | 0.39 | 0.31 | 0.24 | 0.08 | 0.36 | 0.21 |
| 7 | 112889512 | rs4464895 | 0.107 | -0.373 | 0.672 | -2.314 | -0.025 | -0.031 | -0.031 | 0 | 0.88 | 0.68 | 0.93 | 0.95 | 0.94 | 0.83 |
| 7 | 112896363 | rs10273065 | -0.764 | -0.171 | 0.995 | -2.114 | 0.003 | 0.067 | 0.067 | 0.09 | 0.49 | 0.39 | 0.56 | 0.81 | 0.54 | 0.61 |
| 7 | 112908269 | rs7789056 | -0.705 | -0.109 | 0.839 | -2.086 | 0 | -0.012 | -0.012 | 0.07 | 0.51 | 0.53 | 0.58 | 0.81 | 0.54 | 0.61 |
| 7 | 112908788 | rs6466440 | -0.077 | 0.132 | 0.89 | -1.978 | -0.006 | -0.023 | -0.023 | 0 | 0.87 | 0.78 | 0.81 | 0.95 | 0.9 | 0.8 |
| 7 | 112920080 | rs2140636 | 2.158 | -0.01 | 0.769 | -2.042 | -0.032 | 0.149 | 0.149 | 5.327 | 0.37 | 0.08 | 0.22 | 0.14 | 0.37 | 0.19 |
| 7 | 112922865 | rs6962650 | 0.311 | 0.144 | 1.217 | -1.603 | -0.004 | -0.031 | -0.031 | 0 | 0.86 | 0.71 | 0.8 | 0.95 | 0.9 | 0.8 |
| 7 | 112934872 | rs1722305 | -1.101 | 0.695 | 1.051 | -1.532 | 0.001 | -0.014 | -0.014 | 0 | 0.45 | 0.51 | 0.61 | 0.8 | 0.51 | 0.59 |
| 7 | 112947322 | rs1701822 | 0.074 | 0.72 | 1.064 | -1.493 | 0.015 | -0.013 | -0.013 | 0 | 0.87 | 0.67 | 0.8 | 0.94 | 0.89 | 0.81 |
| 7 | 112960975 | rs1608654 | -0.049 | 0.801 | 0.853 | -1.509 | 0.006 | 0.01 | 0.01 | 0 | 0.17 | 0.11 | 0.18 | 0.16 | 0.14 | 0.27 |
| 7 | 112971214 | rs1723472 | -0.028 | 0.802 | 0.929 | -1.263 | -0.001 | -0.007 | -0.007 | 2.55 | 0.87 | 0.81 | 0.81 | 0.94 | 0.89 | 0.8 |
| 7 | 112978243 | rs1524450 | -0.083 | 0.896 | 1.027 | -1.294 | -0.002 | -0.001 | -0.001 | 0.42 | 0.31 | 0.28 | 0.43 | 0.63 | 0.41 | 0.3 |
| 7 | 112980076 | rs1701850 | -0.178 | 0.888 | 1.066 | -1.301 | 0 | 0.001 | 0.001 | 0 | 0.22 | 0.21 | 0.21 | 0.17 | 0.18 | 0.33 |
| 7 | 112986019 | rs10248657 | 0.64 | 0.895 | 1.007 | -1.318 | -0.008 | -0.018 | -0.018 | 0.142 | 0.74 | 0.61 | 0.63 | 0.76 | 0.79 | 0.52 |
| 7 | 112987650 | rs4476935 | -0.122 | 0.976 | 0.589 | -1.262 | 0 | 0 | 0 | 0 | 0.31 | 0.3 | 0.43 | 0.61 | 0.41 | 0.29 |
| 7 | 112994137 | rs7805547 | 1.8 | 0.867 | 0.446 | -1.104 | 0.018 | 0.132 | 0.132 | 4.455 | 0.42 | 0.19 | 0.2 | 0.15 | 0.38 | 0.22 |
| 7 | 112994204 | rs2692198 | 1.365 | 0.97 | 0.37 | -1.065 | 0.001 | -0.001 | -0.001 | 0.807 | 0.09 | 0.07 | 0.13 | 0.16 | 0.08 | 0.12 |
| 7 | 112998089 | rs9648969 | 0.452 | 1.194 | 0.428 | -0.834 | -0.002 | -0.001 | -0.001 | 0 | 0.22 | 0.21 | 0.39 | 0.62 | 0.33 | 0.24 |
| 7 | 113017399 | rs7784560 | 1.186 | 1.136 | 0.182 | -0.995 | 0.02 | 0.097 | 0.097 | 3.554 | 0.43 | 0.25 | 0.2 | 0.15 | 0.39 | 0.24 |
| 7 | 113020044 | rs11977351 | 0.293 | 1.208 | 0.011 | -0.981 | 0.002 | 0.008 | 0.008 | 1.022 | 0.31 | 0.25 | 0.25 | 0.19 | 0.26 | 0.37 |
| 7 | 113029219 | rs1524443 | 0.481 | 1.123 | -0.287 | -0.721 | 0.014 | 0.052 | 0.052 | 3.238 | 0.43 | 0.33 | 0.2 | 0.15 | 0.39 | 0.25 |
| 7 | 113040467 | rs1701827 | -0.171 | 1.117 | -0.13 | 0.394 | 0.026 | 0.054 | 0.054 | 0.024 | 0.16 | 0.25 | 0.35 | 0.61 | 0.24 | 0.2 |
| 7 | 113057370 | rs25456 | -0.808 | -0.1 | -0.114 | -0.614 | 0.004 | 0.064 | 0.064 | 0 | 0.52 | 0.63 | 0.62 | 0.92 | 0.54 | 0.68 |
| 7 | 113065006 | rs2030808 | -0.797 | -0.078 | -0.079 | -0.616 | 0.008 | 0.064 | 0.064 | 0 | 0.51 | 0.6 | 0.64 | 0.92 | 0.54 | 0.67 |
| 7 | 113072070 | rs12705868 | -0.61 | -0.3 | 0.224 | -0.573 | 0.001 | -0.001 | -0.001 | 0.234 | 0.32 | 0.35 | 0.39 | 0.73 | 0.39 | 0.35 |
| 7 | 113101159 | rs10500033 | 1.039 | -0.34 | 0.345 | -0.604 | 0 | 0.005 | 0.005 | 0 | 0.12 | 0.12 | 0.22 | 0.18 | 0.12 | 0.19 |
| 7 | 113105607 | rs4286878 | 1.058 | -0.342 | 0.345 | -0.605 | 0.001 | 0.006 | 0.006 | 0 | 0.12 | 0.12 | 0.22 | 0.18 | 0.12 | 0.19 |
| 7 | 113122401 | rs7808348 | -0.296 | -0.503 | 0.549 | -0.52 | -0.099 | 0.138 | 0.138 | 0 | 0.27 | 0.04 | 0.19 | 0.05 | 0.36 | 0.2 |
| 7 | 113130984 | rs3922816 | -1.265 | -0.551 | 0.563 | -0.644 | -0.003 | 0.012 | 0.012 | 0 | 0.45 | 0.59 | 0.61 | 0.91 | 0.51 | 0.54 |
| 7 | 113141308 | rs953023 | -1.253 | -0.516 | 1.208 | 0.036 | 0.024 | 0.02 | 0.02 | 0 | 0.33 | 0.51 | 0.37 | 0.63 | 0.38 | 0.34 |

| | | | | | | | | | | | | | | | | |
|---|---|---|---|---|---|---|---|---|---|---|---|---|---|---|---|---|
| 7 | 113148087 | rs6950364 | -1.792 | -0.637 | 1.516 | 0.167 | -0.006 | -0.038 | -0.038 | 0 | 0.09 | 0.32 | 0.05 | 0.01 | 0.03 | 0.15 |
| 7 | 113181591 | rs10228955 | -1.927 | -0.654 | 1.52 | 0.166 | 0.001 | -0.03 | -0.03 | 5.524 | 0.09 | 0.41 | 0.06 | 0.01 | 0.03 | 0.15 |
| 7 | 113207209 | rs6973592 | -1.403 | -0.642 | 1.415 | 0.172 | -0.014 | -0.035 | -0.035 | 0 | 0.08 | 0.29 | 0.04 | 0.01 | 0.03 | 0.14 |
| 7 | 113214707 | rs2396653 | -2.135 | -0.693 | 1.32 | 0.229 | -0.034 | 0.071 | 0.071 | 0 | 0.24 | 0.6 | 0.22 | 0.59 | 0.14 | 0.35 |
| 7 | 113222518 | rs2964930 | -0.716 | -0.673 | 1.332 | 0.241 | -0.006 | 0.056 | 0.056 | 0 | 0.15 | 0.17 | 0.16 | 0.58 | 0.11 | 0.19 |
| 7 | 113237912 | rs2456955 | -1.198 | -0.588 | 1.25 | 0.258 | -0.015 | 0.253 | 0.253 | 0 | 0.17 | 0.5 | 0.2 | 0.58 | 0.12 | 0.21 |
| 7 | 113254761 | rs2520297 | -0.792 | -0.658 | 1.017 | 0.258 | -0.013 | 0.178 | 0.178 | 0 | 0.16 | 0.31 | 0.18 | 0.55 | 0.12 | 0.21 |
| 7 | 113274372 | rs10278962 | -1.707 | -0.539 | 0.625 | -0.611 | -0.009 | -0.035 | -0.035 | 0 | 0.08 | 0.3 | 0.05 | 0.01 | 0.03 | 0.13 |
| 7 | 113291922 | rs6977215 | -1.951 | -0.576 | 0.122 | -0.772 | 0.008 | 0.227 | 0.227 | 3.154 | 0.45 | 0.9 | 0.31 | 0.14 | 0.38 | 0.31 |
| 7 | 113300330 | rs7808000 | -0.116 | -0.393 | 0.431 | -0.771 | 0.013 | -0.039 | -0.039 | 0 | 0.11 | 0.24 | 0.07 | 0.04 | 0.11 | 0.03 |
| 7 | 113302262 | rs10487510 | -0.116 | -0.393 | 0.431 | -0.771 | 0.013 | -0.039 | -0.039 | 0 | 0.11 | 0.24 | 0.07 | 0.04 | 0.11 | 0.03 |
| 7 | 113315926 | rs10244620 | -2.006 | -0.497 | 0.13 | -0.822 | 0 | 0.032 | 0.032 | 0.545 | 0.46 | 0.64 | 0.43 | 0.3 | 0.45 | 0.37 |
| 7 | 113322649 | rs9784924 | -0.905 | 0.286 | -0.256 | -0.354 | -0.045 | 0.051 | 0.051 | 0 | 0.2 | 0.34 | 0.23 | 0.25 | 0.1 | 0.28 |
| 7 | 113327166 | rs2396676 | -1.522 | 0.39 | -0.566 | -0.285 | 0.076 | -0.008 | -0.008 | 0.143 | 0.38 | 0.9 | 0.35 | 0.29 | 0.23 | 0.31 |
| 7 | 113333114 | rs17136593 | -0.878 | 0.689 | -0.009 | -0.448 | 0.011 | -0.033 | -0.033 | 0 | 0.21 | 0.27 | 0.15 | 0.09 | 0.23 | 0.23 |
| 7 | 113339175 | rs2188422 | -1.155 | 0.655 | -0.282 | -0.031 | -0.004 | -0.009 | -0.009 | 1.228 | 0.27 | 0.65 | 0.21 | 0.19 | 0.24 | 0.29 |
| 7 | 113347199 | rs10228266 | -1.081 | 1.542 | -0.112 | 0.26 | 0.035 | 0.018 | 0.018 | 0 | 0.23 | 0.31 | 0.38 | 0.34 | 0.35 | 0.4 |
| 7 | 113347411 | rs7779574 | 0.162 | 0.614 | -0.652 | -0.593 | -0.04 | 0.004 | 0.004 | 2.591 | 0.67 | 0.78 | 0.59 | 0.34 | 0.59 | 0.65 |
| 7 | 113353572 | rs1548395 | 0.744 | 1.055 | 0.157 | -0.612 | 0.056 | 0.047 | 0.047 | 0 | 0.84 | 0.7 | 0.6 | 0.7 | 0.76 | 0.74 |
| 7 | 113364319 | rs2968837 | -0.469 | 0.71 | 0.89 | -1.04 | -0.009 | -0.085 | -0.085 | 0 | 0.17 | 0.26 | 0.1 | 0.01 | 0.14 | 0.04 |
| 7 | 113368805 | rs2106441 | -0.276 | -0.069 | 0.815 | -1.368 | -0.057 | 0.066 | 0.066 | 0 | 0.17 | 0.36 | 0.19 | 0.69 | 0.1 | 0.28 |
| 7 | 113392228 | rs2968844 | 0.572 | 0.393 | 1.484 | -1.411 | 0.042 | 0.026 | 0.026 | 0 | 0.83 | 0.72 | 0.59 | 0.7 | 0.74 | 0.73 |
| 7 | 113436542 | rs10240330 | 0.495 | 0.342 | 1.439 | -1.424 | 0.004 | -0.005 | -0.005 | 0 | 0.12 | 0.1 | 0.05 | 0.01 | 0.09 | 0.02 |
| 7 | 113439063 | rs17136650 | 0.075 | 0.237 | 2.02 | -1.409 | -0.017 | 0.019 | 0.019 | 0.005 | 0.14 | 0.18 | 0.17 | 0.69 | 0.1 | 0.29 |
| 7 | 113439283 | rs2023691 | 0.098 | 0.334 | 1.96 | -1.427 | 0.041 | 0.033 | 0.033 | 0 | 0.85 | 0.76 | 0.61 | 0.71 | 0.76 | 0.74 |
| 7 | 113442839 | rs1119660 | -0.09 | 0.287 | 1.974 | -1.419 | 0.017 | 0.026 | 0.026 | 0 | 0.12 | 0.17 | 0.37 | 0.29 | 0.18 | 0.25 |
| 7 | 113446199 | rs721996 | -1.403 | 0.213 | 1.971 | -1.429 | -0.005 | 0 | 0 | 0 | 0.31 | 0.66 | 0.27 | 0.7 | 0.21 | 0.32 |
| 7 | 113472031 | rs12333960 | 1.544 | -0.684 | 1.504 | -1.993 | 0.005 | -0.079 | -0.079 | 0.067 | 0.5 | 0.44 | 0.34 | 0.01 | 0.56 | 0.37 |
| 7 | 113493772 | rs4490752 | 1.439 | -0.661 | 1.567 | -1.708 | -0.009 | 0.043 | 0.043 | 1.333 | 0.59 | 0.23 | 0.5 | 0.65 | 0.61 | 0.72 |
| 7 | 113503703 | rs10247621 | 1.27 | -0.689 | 1.91 | -1.685 | 0.008 | -0.019 | -0.019 | 0.391 | 0.61 | 0.45 | 0.52 | 0.64 | 0.61 | 0.71 |
| 7 | 113516227 | rs7789198 | 0.817 | -0.7 | 2.02 | -1.703 | 0.004 | -0.003 | -0.003 | 0 | 0.67 | 0.6 | 0.56 | 0.65 | 0.65 | 0.78 |
| 7 | 113516616 | rs2968845 | 0.628 | -0.725 | 2.096 | -1.714 | -0.013 | 0.004 | 0.004 | 0.927 | 0.69 | 0.71 | 0.57 | 0.65 | 0.66 | 0.78 |
| 7 | 113520262 | rs11973095 | -0.76 | -0.721 | 2.078 | -1.712 | -0.011 | 0.004 | 0.004 | 0.996 | 0.31 | 0.29 | 0.43 | 0.35 | 0.34 | 0.22 |
| 7 | 113528961 | rs10228303 | 1.036 | -0.733 | 2.071 | -1.718 | 0.013 | -0.019 | -0.019 | 0 | 0.65 | 0.43 | 0.55 | 0.65 | 0.65 | 0.77 |
| 7 | 113547890 | rs7792710 | 0.991 | -0.694 | 1.763 | -1.708 | 0.024 | -0.013 | -0.013 | 0 | 0.66 | 0.53 | 0.54 | 0.64 | 0.62 | 0.75 |
| 7 | 113568382 | rs13242366 | -0.64 | -0.569 | 0.914 | -1.768 | -0.024 | 0.012 | 0.012 | 0.019 | 0.21 | 0.25 | 0.23 | 0.65 | 0.15 | 0.43 |
| 7 | 113570129 | rs9641507 | -0.598 | -0.571 | 0.931 | -1.775 | -0.01 | 0.003 | 0.003 | 0.114 | 0.2 | 0.21 | 0.22 | 0.65 | 0.15 | 0.43 |
| 7 | 113585089 | rs13223841 | 0.832 | -0.683 | 0.625 | -1.975 | -0.011 | 0.022 | 0.022 | 3.711 | 0.63 | 0.3 | 0.52 | 0.65 | 0.62 | 0.75 |
| 7 | 113600387 | rs4342528 | 0.832 | -0.683 | 0.601 | -1.975 | -0.012 | 0.023 | 0.023 | 3.711 | 0.63 | 0.3 | 0.52 | 0.65 | 0.62 | 0.75 |
| 7 | 113631742 | rs1493949 | -0.468 | -0.721 | 0.564 | -1.909 | -0.026 | 0.021 | 0.021 | 0.189 | 0.18 | 0.25 | 0.22 | 0.63 | 0.12 | 0.42 |
| 7 | 113636951 | rs10953750 | 0.446 | -0.193 | 0.713 | -1.57 | -0.053 | -0.05 | -0.05 | 0 | 0.08 | 0.01 | 0.34 | 0.35 | 0.18 | 0.2 |
| 7 | 113664061 | rs13222524 | 0.925 | -0.312 | 0.298 | -1.44 | -0.005 | 0.017 | 0.017 | 2.758 | 0.61 | 0.34 | 0.52 | 0.64 | 0.6 | 0.74 |
| 7 | 113735848 | rs13228816 | -1.073 | -0.494 | -0.056 | 0.022 | 0.012 | -0.027 | -0.027 | 0.007 | 0.11 | 0.06 | 0.18 | 0.57 | 0.1 | 0.22 |
| 7 | 113737983 | rs1826607 | -0.311 | -0.521 | -0.105 | 0.028 | -0.007 | 0.005 | 0.005 | 0 | 0.06 | 0 | 0.17 | 0.57 | 0.07 | 0.19 |
| 7 | 113745701 | rs7795683 | -2.213 | -0.911 | 0.563 | 0.099 | -0.052 | 0.121 | 0.121 | 0 | 0.21 | 0.59 | 0.18 | 0.5 | 0.12 | 0.25 |
| 7 | 113751991 | rs1868757 | -1.311 | -1.331 | -0.756 | -0.296 | 0.027 | 0.009 | 0.009 | 0 | 0.46 | 0.66 | 0.31 | 0.51 | 0.4 | 0.38 |
| 7 | 113752297 | rs13243235 | 0.361 | -1.055 | -1.02 | -0.355 | -0.029 | 0.152 | 0.152 | 0 | 0.25 | 0.04 | 0.13 | 0.02 | 0.28 | 0.13 |
| 7 | 113767770 | rs16869678 | -0.719 | -0.64 | 0.144 | 0.792 | -0.017 | 0.233 | 0.233 | 0.104 | 0.11 | 0.3 | 0.13 | 0.39 | 0.07 | 0.22 |
| 7 | 113767895 | rs6948447 | -0.775 | -0.919 | 0.317 | 0.645 | -0.014 | -0.021 | -0.021 | 0.011 | 0.05 | 0.08 | 0.01 | 0 | 0.01 | 0.03 |
| 7 | 113769768 | rs1376243 | 2.095 | -1.048 | 1.013 | 0.588 | -0.06 | 0.158 | 0.158 | 0 | 0.82 | 0.45 | 0.84 | 0.61 | 0.92 | 0.75 |
| 7 | 113781291 | rs1450836 | 0.913 | -0.986 | 1.159 | 0.637 | -0.017 | 0.352 | 0.352 | 0.398 | 0.88 | 0.55 | 0.85 | 0.61 | 0.93 | 0.78 |
| 7 | 113792523 | rs2244419 | 0.328 | -0.975 | 1.289 | 0.682 | -0.017 | 0.17 | 0.17 | 0.11 | 0.89 | 0.74 | 0.87 | 0.63 | 0.93 | 0.78 |
| 7 | 113796122 | rs4730626 | -0.201 | 0.395 | 0.813 | 0.868 | -0.002 | 0.023 | 0.023 | 0 | 0.12 | 0.14 | 0.29 | 0.41 | 0.19 | 0.24 |
| 7 | 113837389 | rs7778560 | -0.606 | 1.173 | 0.761 | 0.83 | 0.139 | -0.018 | -0.018 | 0.011 | 0.08 | 0.19 | 0.1 | 0.05 | 0.2 | 0.04 |
| 7 | 113842998 | rs1830282 | -0.136 | 0.955 | 0.769 | 1.376 | -0.017 | 0.254 | 0.254 | 0.187 | 0.89 | 0.67 | 0.94 | 0.65 | 0.93 | 0.77 |
| 7 | 113845251 | rs1528090 | 2.223 | 1.336 | 0.543 | 1.368 | -0.014 | -0.016 | -0.016 | 0 | 0.77 | 0.28 | 0.81 | 0.6 | 0.69 | 0.71 |
| 7 | 113849929 | rs12538008 | 0.871 | 1.233 | 0.474 | 1.214 | -0.001 | 0.012 | 0.012 | 0.081 | 0.06 | 0.06 | 0.1 | 0.11 | 0.02 | 0.12 |
| 7 | 113853089 | rs717233 | -1.635 | 1.37 | 0.794 | 1.012 | 0.23 | -0.044 | -0.044 | 0 | 0.12 | 0.33 | 0.13 | 0.05 | 0.24 | 0.06 |
| 7 | 113858363 | rs12535428 | 1.752 | 0.953 | -0.39 | 0.127 | -0.058 | 0.162 | 0.162 | 0 | 0.26 | 0.02 | 0.17 | 0.05 | 0.33 | 0.15 |
| 7 | 113863384 | rs1450832 | 0.433 | 0.384 | -0.409 | -0.177 | 0.12 | -0.121 | -0.121 | 0 | 0.2 | 0.49 | 0.16 | 0.03 | 0.27 | 0.08 |
| 7 | 113898450 | rs1194328 | 0.957 | 0.101 | -0.881 | 0.266 | -0.035 | -0.047 | -0.047 | 0 | 0.7 | 0.19 | 0.79 | 0.58 | 0.68 | 0.68 |
| 7 | 113904061 | rs11771168 | 0.791 | -0.502 | -1.507 | -0.118 | 0.024 | 0.028 | 0.028 | 0 | 0.34 | 0.12 | 0.4 | 0.23 | 0.25 | 0.43 |
| 7 | 113917539 | rs1209254 | -0.089 | -0.339 | -0.776 | 0.232 | 0.137 | 0.167 | 0.167 | 2.333 | 0.82 | 0.45 | 0.84 | 0.59 | 0.69 | 0.69 |
| 7 | 113924219 | rs12112572 | 2.774 | -0.397 | -0.775 | 0.211 | -0.036 | -0.041 | -0.041 | 0 | 0.1 | 0.13 | 0.03 | 0 | 0.01 | 0 |
| 7 | 113945495 | rs12705958 | 1.507 | -0.34 | -1.003 | 0.234 | -0.007 | -0.055 | -0.055 | 0 | 0.09 | 0.03 | 0.21 | 0.3 | 0.1 | 0.09 |
| 7 | 113956930 | rs1527152 | 0.787 | -0.088 | -0.359 | 0.398 | -0.016 | -0.022 | -0.022 | 0 | 0.7 | 0.24 | 0.8 | 0.6 | 0.68 | 0.69 |
| 7 | 113963334 | rs1358278 | -0.078 | -0.545 | -0.262 | 0.531 | 0.053 | 0.042 | 0.042 | 0.537 | 0.27 | 0.62 | 0.37 | 0.5 | 0.42 | 0.37 |
| 7 | 113973098 | rs2694933 | -0.502 | -0.12 | 0.009 | 0.159 | -0.003 | 0 | 0 | 0.164 | 0.66 | 0.38 | 0.61 | 0.83 | 0.61 | 0.74 |
| 7 | 113993281 | rs1916986 | 0.904 | -0.485 | -0.233 | 0.425 | 0.045 | -0.052 | -0.052 | 0.994 | 0.17 | 0.1 | 0.23 | 0.35 | 0.09 | 0.22 |

| | | | | | | | | | | | | | | | | |
|---|---|---|---|---|---|---|---|---|---|---|---|---|---|---|---|---|
| 7 | 113997624 | rs2690833 | -0.607 | -0.381 | 0 | 0.111 | 0.01 | -0.019 | -0.019 | 0 | 0.66 | 0.4 | 0.62 | 0.83 | 0.61 | 0.74 |
| 7 | 114004477 | rs7785799 | 0.419 | -0.222 | 0.163 | 0.027 | 0.018 | -0.064 | -0.064 | 0.1 | 0.37 | 0.52 | 0.41 | 0.14 | 0.49 | 0.24 |
| 7 | 114007270 | rs2140619 | 0.294 | -0.242 | -0.122 | -0.024 | 0.016 | 0.177 | 0.177 | 0 | 0.46 | 0.21 | 0.5 | 0.86 | 0.45 | 0.63 |
| 7 | 114009931 | rs2694943 | 0.23 | -0.239 | -0.115 | -0.022 | 0.02 | 0.154 | 0.154 | 0.102 | 0.47 | 0.25 | 0.5 | 0.86 | 0.45 | 0.63 |
| 7 | 114014574 | rs10249531 | 0.923 | -0.191 | 0.206 | -0.318 | 0.038 | -0.121 | -0.121 | 0.134 | 0.31 | 0.44 | 0.27 | 0.07 | 0.36 | 0.2 |
| 7 | 114015707 | rs12705960 | 0.2 | -0.331 | 0.24 | -0.118 | -0.012 | 0.089 | 0.089 | 0.255 | 0.5 | 0.2 | 0.44 | 0.59 | 0.56 | 0.55 |
| 7 | 114016025 | rs1830198 | -0.709 | -0.339 | -0.155 | -0.092 | -0.072 | 0.068 | 0.068 | 0.001 | 0.81 | 0.64 | 0.71 | 0.66 | 0.92 | 0.75 |
| 7 | 114027269 | rs1852470 | -0.19 | -0.102 | -0.708 | -0.224 | -0.056 | 0.334 | 0.334 | 0.653 | 0.52 | 0.13 | 0.61 | 0.94 | 0.57 | 0.69 |
| 7 | 114072443 | rs10255943 | -1.56 | 0.234 | -0.508 | 0.222 | -0.061 | 0.074 | 0.074 | 0 | 0.73 | 0.98 | 0.84 | 0.91 | 0.7 | 0.84 |
| 7 | 114091753 | rs10262192 | 0.779 | -1.305 | 0.5 | 0.706 | -0.004 | 0.001 | 0.001 | 0 | 0.39 | 0.36 | 0.47 | 0.4 | 0.43 | 0.44 |
| 7 | 114091844 | rs10262103 | -0.292 | -1.225 | 0.112 | 0.791 | -0.016 | 0.015 | 0.015 | 0 | 0.6 | 0.81 | 0.61 | 0.41 | 0.51 | 0.59 |
| 7 | 114109349 | rs727644 | -0.306 | -1.262 | -0.044 | 0.629 | -0.022 | -0.001 | -0.001 | 0 | 0.59 | 0.83 | 0.6 | 0.44 | 0.52 | 0.59 |
| 7 | 114110568 | rs4727799 | -1.194 | -1.232 | -0.206 | 0.574 | -0.088 | 0.158 | 0.158 | 0 | 0.68 | 0.98 | 0.77 | 0.95 | 0.61 | 0.85 |
| 7 | 114159242 | rs17372842 | 0.882 | -1.283 | -0.142 | 0.565 | 0.042 | 0.04 | 0.04 | 0 | 0.16 | 0.06 | 0.07 | 0 | 0.07 | 0.14 |
| 7 | 114162740 | rs6980093 | -0.685 | -1.255 | -0.143 | 0.696 | -0.049 | -0.069 | -0.069 | 0.058 | 0.63 | 0.88 | 0.69 | 0.57 | 0.53 | 0.72 |
| 7 | 114175727 | rs7788341 | -0.634 | -1.398 | 0.446 | 0.723 | -0.002 | 0.133 | 0.133 | 0 | 0.07 | 0.29 | 0.1 | 0.15 | 0.02 | 0.13 |
| 7 | 114190643 | rs7784315 | -0.71 | -1.742 | 0.507 | -0.328 | -0.072 | -0.026 | -0.026 | 1.255 | 0.24 | 0.35 | 0.17 | 0.15 | 0.09 | 0.27 |
| 7 | 114203949 | rs12532920 | -0.768 | -1.832 | 0.272 | -0.332 | -0.098 | 0.011 | 0.011 | 0.153 | 0.24 | 0.47 | 0.19 | 0.23 | 0.1 | 0.27 |
| 7 | 114211447 | rs10269986 | 0.668 | -1.838 | 0.142 | -0.518 | 0.004 | 0.029 | 0.029 | 0 | 0.43 | 0.38 | 0.59 | 0.75 | 0.53 | 0.6 |
| 7 | 114214354 | rs12671286 | -0.602 | -1.765 | 0.191 | -0.449 | 0.015 | 0.299 | 0.299 | 0.46 | 0.08 | 0.4 | 0.12 | 0.23 | 0.02 | 0.13 |
| 7 | 114223723 | rs1229761 | -0.805 | -1.648 | -0.663 | -0.534 | -0.021 | 0.115 | 0.115 | 0 | 0.67 | 0.85 | 0.78 | 0.98 | 0.62 | 0.87 |
| 7 | 114248851 | rs12705966 | 0.843 | -1.632 | -0.955 | -0.496 | -0.094 | 0.195 | 0.195 | 0 | 0.31 | 0.02 | 0.21 | 0.02 | 0.37 | 0.13 |
| 7 | 114254407 | rs10230087 | 1.185 | -1.859 | 0.432 | 0.505 | -0.011 | 0 | 0 | 0.223 | 0.74 | 0.3 | 0.79 | 0.77 | 0.9 | 0.73 |
| 7 | 114270077 | rs10227893 | -0.702 | -1.319 | -0.241 | -0.056 | -0.046 | -0.082 | -0.082 | 0 | 0.19 | 0.3 | 0.1 | 0 | 0.07 | 0.14 |
| 7 | 114293859 | rs11765385 | -1.122 | -0.748 | -0.555 | 0.924 | 0.008 | 0.003 | 0.003 | 0 | 0.11 | 0.08 | 0.11 | 0.35 | 0.07 | 0.27 |
| 7 | 114296104 | rs6966051 | -0.128 | -1.412 | 0.171 | 0.397 | -0.03 | -0.067 | -0.067 | 0 | 0.11 | 0.3 | 0.07 | 0 | 0.05 | 0 |
| 7 | 114298708 | rs17213159 | 0.423 | -1.145 | -0.611 | -0.019 | -0.029 | 0.054 | 0.054 | 0 | 0.1 | 0.01 | 0.07 | 0 | 0.13 | 0.02 |
| 7 | 114303189 | rs1563408 | -0.228 | -0.644 | -0.737 | -0.065 | -0.023 | -0.024 | -0.024 | 0.36 | 0.26 | 0.21 | 0.38 | 0.48 | 0.43 | 0.43 |
| 7 | 114304969 | rs2396765 | 0.192 | -0.514 | -0.715 | -0.205 | -0.016 | 0.007 | 0.007 | 0.714 | 0.22 | 0.18 | 0.29 | 0.13 | 0.41 | 0.27 |
| 7 | 114310053 | rs12705971 | 0.352 | -0.531 | -0.675 | -0.237 | -0.069 | 0.013 | 0.013 | 1.713 | 0.2 | 0.09 | 0.29 | 0.13 | 0.41 | 0.27 |
| 7 | 114337652 | rs10261050 | -0.591 | -0.613 | -0.286 | -0.244 | 0.015 | 0.019 | 0.019 | 0 | 0.36 | 0.48 | 0.44 | 0.48 | 0.44 | 0.42 |
| 7 | 114338312 | rs11769707 | -1.041 | -0.998 | -0.278 | -0.476 | -0.011 | -0.004 | -0.004 | 0 | 0.46 | 0.68 | 0.47 | 0.48 | 0.48 | 0.51 |
| 7 | 114347154 | rs1378766 | 0.821 | -0.867 | -0.509 | -0.665 | -0.03 | 0.03 | 0.03 | 0 | 0.07 | 0.01 | 0.07 | 0 | 0.13 | 0.02 |
| 7 | 114349036 | rs605421 | -0.421 | -1.206 | -0.044 | -0.635 | -0.005 | -0.05 | -0.05 | 0.11 | 0.91 | 0.8 | 0.96 | 1 | 0.95 | 0.9 |
| 7 | 114352126 | rs651902 | 1.038 | -2.321 | -0.855 | -1.025 | -0.045 | 0.122 | 0.122 | 0 | 0.37 | 0.19 | 0.32 | 0.11 | 0.42 | 0.25 |
| 7 | 114352862 | rs776472 | 0.399 | -2.601 | -1.058 | -1.199 | -0.027 | -0.053 | -0.053 | 0 | 0.62 | 0.19 | 0.53 | 0.52 | 0.58 | 0.64 |
| 7 | 114355002 | rs12705988 | -0.297 | -2.21 | -0.835 | -0.991 | 0.01 | 0.016 | 0.016 | 0 | 0.35 | 0.59 | 0.35 | 0.48 | 0.42 | 0.4 |
| 7 | 114355097 | rs1378769 | 0.765 | -2.352 | -0.81 | -0.935 | -0.068 | 0.23 | 0.23 | 2.802 | 0.15 | 0.47 | 0.16 | 0.36 | 0.06 | 0.14 |
| 7 | 114357543 | rs1869839 | 0.469 | -2.332 | -0.835 | -0.945 | -0.084 | 0.011 | 0.011 | 1.946 | 0.2 | 0.06 | 0.18 | 0.12 | 0.36 | 0.26 |
| 7 | 114361551 | rs6970540 | 0.687 | -2.354 | -0.805 | -0.929 | -0.07 | 0.224 | 0.224 | 2.998 | 0.15 | 0.51 | 0.16 | 0.36 | 0.06 | 0.14 |
| 7 | 114363566 | rs1378770 | 0.277 | -2.331 | -1.026 | -0.958 | -0.056 | 0.02 | 0.02 | 0.895 | 0.21 | 0.12 | 0.19 | 0.12 | 0.37 | 0.26 |
| 7 | 114375216 | rs10269071 | 0.653 | -2.464 | -0.467 | -0.934 | -0.069 | 0.196 | 0.196 | 3.241 | 0.15 | 0.42 | 0.15 | 0.35 | 0.06 | 0.14 |
| 7 | 114378032 | rs666187 | -0.628 | -2.211 | -0.494 | -0.921 | 0.005 | 0.013 | 0.013 | 0 | 0.36 | 0.61 | 0.34 | 0.48 | 0.4 | 0.39 |
| 7 | 114383452 | rs681914 | 0.281 | -1.89 | -0.517 | -1.007 | -0.025 | 0.011 | 0.011 | 0.581 | 0.2 | 0.16 | 0.18 | 0.12 | 0.34 | 0.25 |
| 7 | 114398325 | rs4727803 | -0.525 | -2.279 | -0.117 | -0.856 | 0.011 | -0.054 | -0.054 | 0 | 0.9 | 0.64 | 0.94 | 1 | 0.94 | 0.89 |
| 7 | 114405945 | rs1448345 | 0.904 | -2.734 | -0.402 | -0.745 | 0.171 | 0.064 | 0.064 | 2.429 | 0.57 | 0.15 | 0.6 | 0.25 | 0.79 | 0.52 |
| 7 | 114411742 | rs17137252 | -0.266 | -2.659 | -0.374 | -0.602 | -0.074 | -0.002 | -0.002 | 0.208 | 0.32 | 0.45 | 0.34 | 0.73 | 0.15 | 0.38 |
| 7 | 114421923 | rs1448346 | 0.423 | -2.532 | 0.033 | -0.386 | -0.051 | 0.015 | 0.015 | 0.099 | 0.71 | 0.41 | 0.73 | 0.58 | 0.84 | 0.67 |
| 7 | 114425799 | rs12667130 | 0.423 | -2.532 | 0.033 | -0.386 | -0.051 | 0.015 | 0.015 | 0.099 | 0.71 | 0.41 | 0.73 | 0.58 | 0.84 | 0.67 |
| 7 | 114430569 | rs1448347 | -0.616 | -2.252 | 0.433 | -0.328 | 0.025 | -0.036 | -0.036 | 0.001 | 0.91 | 0.68 | 0.95 | 1 | 0.95 | 0.9 |
| 7 | 114436126 | rs4366023 | 1.119 | -2.127 | -0.055 | -0.606 | -0.048 | -0.066 | -0.066 | 0.217 | 0.55 | 0.12 | 0.61 | 0.56 | 0.58 | 0.49 |
| 7 | 114436380 | rs1375093 | -0.759 | -2.053 | 0.532 | -0.314 | 0.013 | -0.041 | -0.041 | 0 | 0.91 | 0.74 | 0.95 | 1 | 0.95 | 0.9 |
| 7 | 114454337 | rs1011449 | 0.622 | -2.124 | 0.4 | -0.136 | -0.04 | 0.111 | 0.111 | 0 | 0.84 | 0.38 | 0.84 | 0.67 | 0.93 | 0.77 |
| 7 | 114465993 | rs6944734 | -0.173 | -2.157 | 0.755 | -0.349 | -0.004 | -0.066 | -0.066 | 0.25 | 0.89 | 0.6 | 0.94 | 1 | 0.95 | 0.9 |
| 7 | 114493274 | rs6466499 | 0.637 | -1.556 | 0.633 | -0.063 | 0.002 | 0.058 | 0.058 | 1.319 | 0.8 | 0.62 | 0.86 | 0.72 | 0.82 | 0.75 |
| 7 | 114499932 | rs2188307 | 1.419 | -1.977 | 0.246 | -0.62 | 0.027 | -0.003 | -0.003 | 1.843 | 0.53 | 0.25 | 0.7 | 0.56 | 0.62 | 0.6 |
| 7 | 114506265 | rs4730650 | 0.197 | -1.849 | 0.371 | -0.522 | -0.038 | 0.044 | 0.044 | 0 | 0.85 | 0.5 | 0.92 | 0.84 | 0.93 | 0.79 |
| 7 | 114514125 | rs4265116 | 0.913 | -1.853 | 0.314 | -0.505 | -0.017 | -0.02 | -0.02 | 0 | 0.09 | 0.04 | 0.07 | 0.28 | 0.13 | 0.14 |
| 7 | 114517317 | rs12706000 | -1.13 | -2.033 | 0.608 | -0.506 | -0.003 | -0.049 | -0.049 | 0 | 0.91 | 0.8 | 0.95 | 1 | 0.95 | 0.9 |
| 7 | 114549625 | rs7802139 | -0.205 | -1.997 | 0.46 | -0.479 | -0.013 | 0.056 | 0.056 | 0.367 | 0.05 | 0.1 | 0.01 | 0.16 | 0.02 | 0.11 |
| 7 | 114551039 | rs10232096 | 0.534 | -2.23 | 0.354 | -0.633 | -0.005 | -0.007 | -0.007 | 0.007 | 0.14 | 0.07 | 0.1 | 0 | 0.16 | 0.02 |
| 7 | 114560827 | rs12706002 | 0.814 | -2.794 | -0.295 | -0.826 | -0.019 | 0.011 | 0.011 | 0.156 | 0.27 | 0.27 | 0.38 | 0.5 | 0.18 | 0.41 |
| 7 | 114569809 | rs7777885 | 0.14 | -2 | 0.159 | -0.897 | -0.035 | 0.034 | 0.034 | 0.013 | 0.85 | 0.48 | 0.92 | 0.84 | 0.93 | 0.82 |
| 7 | 114582191 | rs4615508 | 0.257 | -1.737 | -0.328 | -0.833 | -0.025 | 0.06 | 0.06 | 0 | 0.15 | 0.06 | 0.1 | 0 | 0.16 | 0.03 |
| 7 | 114590228 | rs10247983 | -0.078 | -1.659 | 0.283 | -0.413 | -0.006 | 0.074 | 0.074 | 0 | 0.88 | 0.57 | 0.92 | 0.84 | 0.94 | 0.83 |
| 7 | 114592049 | rs4727809 | 0.724 | -0.182 | -0.815 | -1.322 | -0.013 | 0.01 | 0.01 | 3.704 | 0.39 | 0.34 | 0.37 | 0.03 | 0.46 | 0.25 |
| 7 | 114592233 | rs7800698 | -0.168 | -0.282 | -1.422 | -1.341 | -0.002 | 0.017 | 0.017 | 0 | 0.16 | 0.09 | 0.1 | 0.1 | 0.16 | 0.15 |
| 7 | 114592267 | rs4730659 | 0.754 | 0.123 | -1.95 | -1.542 | -0.095 | 0.191 | 0.191 | 0 | 0.36 | 0.05 | 0.34 | 0.06 | 0.38 | 0.21 |
| 7 | 114601094 | rs7783637 | 0.908 | -0.561 | -1.047 | -0.962 | 0.007 | 0.002 | 0.002 | 0 | 0.1 | 0.05 | 0.05 | 0 | 0.08 | 0.09 |

| | | | | | | | | | | | | | | | | |
|---|---|---|---|---|---|---|---|---|---|---|---|---|---|---|---|---|
| 7 | 114603410 | rs6974433 | 0.022 | -0.838 | -0.868 | -0.705 | 0 | -0.001 | -0.001 | 0 | 0.49 | 0.44 | 0.34 | 0.45 | 0.52 | 0.32 |
| 7 | 114606579 | rs10273599 | 0.691 | -0.697 | -0.834 | -0.619 | -0.003 | -0.001 | -0.001 | 0.879 | 0.39 | 0.34 | 0.29 | 0.45 | 0.44 | 0.23 |
| 7 | 114612109 | rs2594452 | 0.058 | -0.7 | -1.036 | -0.395 | -0.012 | -0.016 | -0.016 | 0 | 0.42 | 0.29 | 0.54 | 0.49 | 0.42 | 0.57 |
| 7 | 114613275 | rs12538651 | 1.169 | -0.859 | -1.342 | -0.043 | -0.005 | -0.07 | -0.07 | 0.405 | 0.11 | 0 | 0.23 | 0.37 | 0.1 | 0.2 |
| 7 | 114613838 | rs4730660 | 1.685 | -0.79 | -1.135 | 0.056 | 0.021 | -0.049 | -0.049 | 0 | 0.13 | 0.05 | 0.07 | 0.45 | 0.09 | 0.14 |
| 7 | 114629285 | rs7784447 | 0.479 | -0.519 | -0.882 | -0.068 | -0.017 | 0.045 | 0.045 | 0.022 | 0.3 | 0.24 | 0.29 | 0.06 | 0.29 | 0.26 |
| 7 | 114640826 | rs2709517 | 0.604 | -0.641 | -0.8 | -0.028 | -0.002 | -0.005 | -0.005 | 0 | 0.45 | 0.45 | 0.32 | 0.44 | 0.45 | 0.28 |
| 7 | 114648223 | rs2594442 | 0.142 | -0.718 | -0.72 | 0.153 | -0.03 | 0.042 | 0.042 | 0.096 | 0.75 | 0.6 | 0.67 | 0.64 | 0.82 | 0.7 |
| 7 | 114648253 | rs2709501 | -0.904 | -0.606 | -0.544 | -0.433 | -0.033 | -0.089 | -0.089 | 0.159 | 0.87 | 0.62 | 0.92 | 1 | 0.95 | 0.9 |
| 7 | 114651031 | rs13227167 | 1.115 | -0.079 | -0.816 | -0.02 | -0.092 | 0.056 | 0.056 | 0 | 0.19 | 0.03 | 0.22 | 0.05 | 0.25 | 0.17 |
| 7 | 114658486 | rs13237006 | -0.393 | -0.467 | -0.686 | 0.387 | 0.009 | -0.052 | -0.052 | 0 | 0.21 | 0.12 | 0.14 | 0.33 | 0.2 | 0.36 |
| 7 | 114663499 | rs998903 | 0.86 | -0.456 | -0.751 | 0.414 | -0.027 | -0.072 | -0.072 | 0 | 0.11 | 0 | 0.09 | 0.26 | 0.11 | 0.18 |
| 7 | 114664779 | rs1123065 | 1.235 | -0.387 | -0.623 | 0.178 | -0.048 | 0.049 | 0.049 | 0 | 0.19 | 0.05 | 0.22 | 0.05 | 0.24 | 0.17 |
| 7 | 114672410 | rs4281059 | -1.086 | -0.724 | -0.367 | 0.046 | -0.031 | -0.037 | -0.037 | 0 | 0.93 | 0.87 | 0.97 | 1 | 0.99 | 0.95 |
| 7 | 114672720 | rs7791565 | 1.567 | -1.025 | -0.322 | -0.286 | -0.012 | -0.019 | -0.019 | 1.393 | 0.37 | 0.32 | 0.28 | 0.01 | 0.42 | 0.19 |
| 7 | 114676724 | rs2520600 | 0.126 | -1.05 | -0.325 | 0.045 | -0.01 | 0.01 | 0.01 | 0 | 0.6 | 0.55 | 0.56 | 0.49 | 0.67 | 0.62 |
| 7 | 114678739 | rs2709494 | 0.389 | -1.11 | -0.257 | 0.118 | -0.101 | 0.138 | 0.138 | 0 | 0.21 | 0.42 | 0.22 | 0.46 | 0.09 | 0.21 |
| 7 | 114719858 | rs425956 | -0.848 | -0.898 | -0.227 | -0.022 | -0.019 | -0.014 | -0.014 | 0 | 0.4 | 0.39 | 0.35 | 0.78 | 0.28 | 0.57 |
| 7 | 114722787 | rs641459 | -1.328 | -0.728 | -0.216 | 0.004 | -0.028 | 0.046 | 0.046 | 0 | 0.62 | 0.7 | 0.58 | 0.83 | 0.54 | 0.74 |
| 7 | 114732621 | rs2520575 | -0.83 | -0.866 | -0.313 | 0.066 | -0.026 | -0.017 | -0.017 | 0 | 0.32 | 0.45 | 0.33 | 0.8 | 0.27 | 0.56 |
| 7 | 114735229 | rs2428194 | 0.263 | -0.731 | -0.506 | 0.255 | -0.036 | 0.052 | 0.052 | 0 | 0.23 | 0.32 | 0.23 | 0.47 | 0.18 | 0.34 |
| 7 | 114735547 | rs371004 | -1.136 | -0.851 | -0.548 | 0.126 | 0.039 | 0.055 | 0.055 | 0 | 0.46 | 0.62 | 0.46 | 0.84 | 0.34 | 0.66 |
| 7 | 114736595 | rs7795179 | -0.093 | -0.84 | -0.538 | 0.155 | -0.007 | 0.009 | 0.009 | 0 | 0.23 | 0.25 | 0.14 | 0.3 | 0.15 | 0.26 |
| 7 | 114739166 | rs1476479 | -0.067 | -0.584 | -0.256 | 0.35 | -0.034 | 0.083 | 0.083 | 0 | 0.24 | 0.34 | 0.31 | 0.54 | 0.19 | 0.4 |
| 7 | 114750436 | rs12706009 | -1.103 | -0.114 | -0.711 | -0.162 | 0 | -0.017 | -0.017 | 0 | 0.67 | 0.65 | 0.66 | 0.91 | 0.57 | 0.84 |
| 7 | 114754867 | rs12706011 | 0.351 | -0.223 | -0.689 | 0.483 | 0.009 | -0.015 | -0.015 | 0 | 0.75 | 0.77 | 0.75 | 0.56 | 0.78 | 0.66 |
| 7 | 114763318 | rs4613914 | 0.913 | -0.461 | -0.538 | 0.677 | -0.013 | 0.027 | 0.027 | 0.047 | 0.13 | 0.16 | 0.17 | 0.38 | 0.06 | 0.15 |
| 7 | 114769689 | rs10279639 | 0.57 | -0.562 | -0.453 | 0.376 | -0.023 | -0.027 | -0.027 | 0 | 0.07 | 0.08 | 0.02 | 0 | 0.01 | 0.05 |
| 7 | 114772802 | rs13229627 | -0.736 | -0.43 | 0 | 0.601 | -0.018 | 0.094 | 0.094 | 0 | 0.21 | 0.36 | 0.14 | 0.35 | 0.17 | 0.22 |
| 7 | 114778640 | rs10249920 | 1.416 | -0.228 | -0.31 | 0.137 | -0.018 | 0.025 | 0.025 | 0 | 0.34 | 0.31 | 0.4 | 0.15 | 0.42 | 0.21 |
| 7 | 114784028 | rs2460481 | 0.437 | -0.339 | -0.417 | 0.083 | 0.006 | -0.011 | -0.011 | 0 | 0.23 | 0.22 | 0.25 | 0.44 | 0.18 | 0.38 |
| 7 | 114784608 | rs11773798 | 0.838 | -0.134 | -0.3 | 0.382 | -0.002 | 0.001 | 0.001 | 0.021 | 0.57 | 0.57 | 0.58 | 0.52 | 0.59 | 0.53 |
| 7 | 114792582 | rs10215709 | 0.026 | -0.157 | -0.376 | 0.357 | 0.007 | -0.021 | -0.021 | 0 | 0.81 | 0.88 | 0.88 | 0.65 | 0.85 | 0.78 |
| 7 | 114793930 | rs17293743 | 1.04 | -0.164 | -0.428 | 0.352 | -0.067 | 0.044 | 0.044 | 0.264 | 0.21 | 0.01 | 0.2 | 0.06 | 0.23 | 0.2 |
| 7 | 114798962 | rs2251136 | 0.348 | -0.155 | -0.271 | 0.393 | -0.025 | 0.088 | 0.088 | 0 | 0.8 | 0.7 | 0.86 | 0.65 | 0.84 | 0.78 |
| 7 | 114799909 | rs2464884 | 0.649 | -0.153 | -0.329 | 0.394 | -0.013 | 0.141 | 0.141 | 0 | 0.78 | 0.62 | 0.78 | 0.58 | 0.82 | 0.74 |
| 7 | 114839127 | rs11974449 | -0.408 | -0.085 | -1.122 | 0.186 | -0.001 | -0.063 | -0.063 | 0 | 0.39 | 0.13 | 0.32 | 0.45 | 0.37 | 0.42 |
| 7 | 114860847 | rs17137685 | 0.123 | 0.097 | -1.39 | 0.207 | -0.077 | -0.153 | -0.153 | 0 | 0.3 | 0.02 | 0.26 | 0.44 | 0.32 | 0.33 |
| 7 | 114862948 | rs10279558 | 0.61 | 0.172 | -0.94 | -0.02 | 0.014 | -0.015 | -0.015 | 0 | 0.64 | 0.32 | 0.61 | 0.51 | 0.76 | 0.46 |
| 7 | 114870466 | rs11765534 | 0.202 | 0.517 | -0.478 | -0.226 | 0.019 | 0.015 | 0.015 | 0 | 0.87 | 0.65 | 0.81 | 0.88 | 0.87 | 0.69 |
| 7 | 114883390 | rs7782721 | 0.327 | 1.013 | -0.586 | -0.93 | 0.049 | -0.067 | -0.067 | 1.175 | 0.15 | 0.38 | 0.15 | 0.04 | 0.18 | 0.11 |
| 7 | 114897575 | rs9942713 | -0.403 | 0.468 | -1.095 | -1.089 | 0.006 | -0.06 | -0.06 | 0.069 | 0.42 | 0.5 | 0.38 | 0.11 | 0.44 | 0.33 |
| 7 | 114900684 | rs2188538 | -0.414 | 0.464 | -1.102 | -1.091 | 0.005 | -0.056 | -0.056 | 0.741 | 0.42 | 0.5 | 0.38 | 0.1 | 0.44 | 0.33 |
| 7 | 114918175 | rs7786940 | -0.526 | 0.504 | -1.484 | -0.514 | 0.039 | -0.028 | -0.028 | 0 | 0.57 | 0.83 | 0.64 | 0.47 | 0.6 | 0.57 |
| 7 | 114929930 | rs2396812 | 0.38 | 0.73 | -1.022 | -1.01 | 0 | -0.008 | -0.008 | 0 | 0.77 | 0.72 | 0.83 | 0.89 | 0.8 | 0.77 |
| 7 | 114934232 | rs2214108 | 0.561 | 0.824 | -1.061 | -1.145 | -0.02 | 0.053 | 0.053 | 0 | 0.59 | 0.64 | 0.66 | 0.85 | 0.51 | 0.62 |
| 7 | 114945308 | rs2023702 | 0.014 | 0.32 | -0.729 | -0.72 | -0.001 | -0.004 | -0.004 | 0.257 | 0.1 | 0.07 | 0.03 | 0 | 0.07 | 0.07 |
| 7 | 114947770 | rs34984260 | 0.794 | 0.049 | -0.786 | -0.045 | 0.022 | -0.057 | -0.057 | 0 | 0.15 | 0.07 | 0.23 | 0.39 | 0.11 | 0.19 |
| 7 | 114948351 | rs10270358 | 1.096 | 0.139 | -0.091 | -0.367 | -0.015 | 0.04 | 0.04 | 0 | 0.66 | 0.56 | 0.6 | 0.51 | 0.68 | 0.63 |
| 7 | 114961423 | rs6466523 | -1.682 | 0.037 | 0.243 | -1.057 | 0 | -0.068 | -0.068 | 0 | 0.24 | 0.63 | 0.19 | 0.04 | 0.21 | 0.17 |
| 7 | 114964446 | rs2188540 | 0.075 | 0.329 | 0.452 | -0.967 | 0.002 | -0.048 | -0.048 | 0 | 0.84 | 0.53 | 0.89 | 0.94 | 0.88 | 0.84 |
| 7 | 114971787 | rs6978038 | -0.777 | 0.432 | 0.472 | -1.002 | 0.048 | -0.112 | -0.112 | 0.435 | 0.21 | 0.38 | 0.2 | 0.04 | 0.24 | 0.14 |
| 7 | 114982393 | rs11768031 | -0.756 | 0.314 | 0.355 | -0.744 | -0.004 | 0.014 | 0.014 | 0 | 0.29 | 0.4 | 0.41 | 0.32 | 0.29 | 0.28 |
| 7 | 115026740 | rs10267100 | 1.469 | 0.364 | 0.707 | -0.788 | 0.014 | -0.053 | -0.053 | 0.252 | 0.7 | 0.65 | 0.76 | 0.9 | 0.69 | 0.72 |
| 7 | 115031575 | rs9641538 | 1.716 | 0.406 | 0.569 | -0.752 | -0.001 | 0.014 | 0.014 | 0 | 0.53 | 0.41 | 0.51 | 0.62 | 0.57 | 0.53 |
| 7 | 115047851 | rs13246549 | 0.976 | 0.46 | 0.363 | -0.748 | 0.004 | -0.027 | -0.027 | 0.769 | 0.08 | 0.08 | 0.03 | 0 | 0.07 | 0.04 |
| 7 | 115050501 | rs10500042 | 0.67 | 0.476 | 0.3 | -0.737 | -0.043 | 0.037 | 0.037 | 0.191 | 0.1 | 0.02 | 0.15 | 0.04 | 0.17 | 0.1 |
| 7 | 115052164 | rs7797141 | 1.23 | 0.361 | 0.424 | -0.669 | -0.008 | 0.002 | 0.002 | 0 | 0.65 | 0.61 | 0.67 | 0.66 | 0.74 | 0.63 |
| 7 | 115082872 | rs9641540 | -0.963 | 0.29 | -0.11 | -0.452 | -0.016 | 0.024 | 0.024 | 0 | 0.82 | 0.77 | 0.76 | 0.7 | 0.88 | 0.81 |
| 7 | 115089527 | rs4644177 | -0.951 | 0.291 | -0.149 | -0.452 | -0.021 | 0.031 | 0.031 | 0 | 0.82 | 0.76 | 0.76 | 0.7 | 0.88 | 0.81 |
| 7 | 115094152 | rs11531592 | 1.862 | 0.305 | -0.547 | -0.661 | -0.037 | 0.025 | 0.025 | 0.009 | 0.47 | 0.17 | 0.46 | 0.61 | 0.53 | 0.51 |
| 7 | 115099020 | rs11514944 | -0.595 | 0.971 | -0.538 | -0.887 | -0.04 | 0.055 | 0.055 | 0 | 0.21 | 0.08 | 0.2 | 0.09 | 0.25 | 0.24 |
| 7 | 115110126 | rs10260121 | -0.534 | 0.646 | 0.034 | -0.561 | 0.02 | -0.091 | -0.091 | 0 | 0.14 | 0.38 | 0.08 | 0 | 0.11 | 0.06 |
| 7 | 115132030 | rs1918524 | 0.094 | 0.622 | -1.288 | -0.129 | -0.008 | -0.047 | -0.047 | 0.522 | 0.43 | 0.16 | 0.55 | 0.51 | 0.44 | 0.49 |
| 7 | 115133095 | rs16869824 | 0.689 | 0.571 | -0.946 | -0.06 | 0.014 | -0.117 | -0.117 | 0 | 0.3 | 0.08 | 0.37 | 0.45 | 0.28 | 0.32 |
| 7 | 115144978 | rs9640766 | 0.885 | 0.486 | -0.201 | -0.191 | 0 | -0.004 | -0.004 | 0 | 0.28 | 0.26 | 0.38 | 0.43 | 0.27 | 0.33 |
| 7 | 115148127 | rs10215218 | 0.847 | 0.457 | 0.094 | -0.204 | -0.004 | 0.02 | 0.02 | 0.79 | 0.29 | 0.34 | 0.39 | 0.43 | 0.27 | 0.33 |
| 7 | 115154793 | rs7783313 | -0.259 | 0.515 | 0.31 | -0.341 | -0.001 | 0.002 | 0.002 | 0 | 0.25 | 0.25 | 0.18 | 0.29 | 0.21 | 0.14 |
| 7 | 115155837 | rs6955743 | -0.33 | 0.513 | 0.246 | -0.351 | 0 | -0.007 | -0.007 | 0 | 0.53 | 0.5 | 0.57 | 0.73 | 0.48 | 0.47 |

| | | | | | | | | | | | | | | | | |
|---|---|---|---|---|---|---|---|---|---|---|---|---|---|---|---|---|
| 7 | 115169187 | rs11983531 | -0.613 | 0.424 | -0.515 | -0.421 | 0.006 | -0.006 | -0.006 | 0.108 | 0.3 | 0.18 | 0.2 | 0.29 | 0.26 | 0.23 |
| 7 | 115175707 | rs6967569 | 0.906 | 0.932 | -0.457 | -0.177 | -0.113 | 0.029 | 0.029 | 4.294 | 0.38 | 0.06 | 0.38 | 0.28 | 0.47 | 0.45 |
| 7 | 115181720 | rs2037783 | 1.078 | 0.639 | -0.374 | -0.182 | -0.041 | -0.097 | -0.097 | 0.863 | 0.28 | 0.09 | 0.39 | 0.43 | 0.31 | 0.34 |
| 7 | 115201385 | rs13227701 | 0.922 | 0.875 | -0.2 | -0.098 | -0.09 | 0.033 | 0.033 | 3.405 | 0.4 | 0.08 | 0.33 | 0.28 | 0.48 | 0.48 |
| 7 | 115202534 | rs2896159 | 0.356 | 1.343 | 0.616 | -0.108 | 0.005 | 0.013 | 0.013 | 0.201 | 0.29 | 0.3 | 0.5 | 0.39 | 0.34 | 0.29 |
| 7 | 115216261 | rs6955690 | -0.999 | 0.928 | 0.941 | -0.263 | 0.11 | -0.105 | -0.105 | 0 | 0.15 | 0.36 | 0.15 | 0 | 0.24 | 0.09 |
| 7 | 115255384 | rs4730685 | 0.884 | -0.189 | 1.19 | 0.126 | -0.043 | 0.086 | 0.086 | 0.418 | 0.42 | 0.15 | 0.3 | 0.27 | 0.44 | 0.42 |
| 7 | 115257756 | rs1986847 | -1.176 | -0.16 | 1.113 | 0.218 | -0.018 | -0.068 | -0.068 | 0 | 0.12 | 0.56 | 0.06 | 0 | 0.04 | 0.09 |
| 7 | 115277575 | rs10953787 | -0.273 | -0.101 | 0.596 | 0.525 | -0.006 | 0.035 | 0.035 | 0 | 0.84 | 0.82 | 0.82 | 0.64 | 0.87 | 0.88 |
| 7 | 115283613 | rs10251693 | -0.41 | -0.086 | 0 | 0.536 | -0.004 | -0.091 | -0.091 | 0.206 | 0.85 | 0.98 | 0.85 | 0.64 | 0.89 | 0.9 |
| 7 | 115295298 | rs13227817 | 0.603 | 0.012 | 0.054 | 0.492 | -0.043 | -0.05 | -0.05 | 0 | 0.28 | 0.12 | 0.43 | 0.37 | 0.37 | 0.36 |
| 7 | 115300183 | rs12706043 | 1.318 | -0.622 | 1.124 | 0.819 | -0.011 | 0.021 | 0.021 | 0 | 0.75 | 0.68 | 0.78 | 0.64 | 0.83 | 0.79 |
| 7 | 115311330 | rs10249355 | 1.519 | -0.524 | 0.906 | 0.852 | -0.027 | 0.043 | 0.043 | 0 | 0.74 | 0.48 | 0.76 | 0.64 | 0.83 | 0.79 |
| 7 | 115317894 | rs10268441 | -0.419 | -0.01 | -0.225 | 1.11 | 0.003 | -0.007 | -0.007 | 0 | 0.15 | 0.13 | 0.18 | 0.36 | 0.13 | 0.12 |
| 7 | 115320308 | rs4073131 | -1.21 | 0.254 | 0.03 | 0.763 | 0 | -0.017 | -0.017 | 0 | 0.05 | 0.07 | 0.03 | 0 | 0.04 | 0.02 |
| 7 | 115323654 | rs10264239 | -1.384 | 0.038 | 0.452 | 1.491 | 0.014 | 0.386 | 0.386 | 0 | 0.06 | 0.3 | 0.05 | 0.4 | 0.06 | 0.08 |
| 7 | 115327840 | rs10267413 | -2.345 | -0.519 | 0.812 | 1.182 | -0.015 | 0.039 | 0.039 | 1.482 | 0.15 | 0.77 | 0.11 | 0.4 | 0.09 | 0.06 |
| 7 | 115347652 | rs9641549 | -1.2 | -0.494 | 0.821 | 1.181 | -0.001 | 0.378 | 0.378 | 0.056 | 0.06 | 0.3 | 0.05 | 0.4 | 0.05 | 0.04 |
| 7 | 115358201 | rs10281558 | -1.63 | -0.56 | 0.845 | 1.128 | 0.001 | -0.047 | -0.047 | 0 | 0.1 | 0.52 | 0.05 | 0 | 0.04 | 0.02 |
| 7 | 115392863 | rs4327800 | -0.183 | -0.456 | -0.637 | 0.451 | 0.027 | 0.101 | 0.101 | 0 | 0.52 | 0.78 | 0.49 | 0.3 | 0.59 | 0.49 |
| 7 | 115412350 | rs12706046 | 0.155 | -0.662 | -0.878 | -0.014 | -0.055 | -0.023 | -0.023 | 0 | 0.26 | 0.04 | 0.44 | 0.25 | 0.3 | 0.25 |
| 7 | 115413714 | rs7810586 | -1.255 | -0.799 | 0.184 | 0.007 | 0 | -0.051 | -0.051 | 0 | 0.1 | 0.55 | 0.09 | 0 | 0.07 | 0.02 |
| 7 | 115428104 | rs10250832 | -0.839 | -0.698 | 0.197 | 0.082 | 0.045 | 0.005 | 0.005 | 1.573 | 0.07 | 0.51 | 0.08 | 0 | 0.06 | 0.01 |
| 7 | 115437853 | rs4412312 | 0.49 | -0.285 | -1.351 | -0.052 | -0.18 | 0.029 | 0.029 | 0.105 | 0.38 | 0.02 | 0.43 | 0.22 | 0.48 | 0.45 |
| 7 | 115439353 | rs6958700 | 0.388 | -0.952 | -1.884 | -0.049 | -0.073 | 0 | 0 | 0 | 0.28 | 0.02 | 0.34 | 0.18 | 0.28 | 0.3 |
| 7 | 115443656 | rs7785169 | -0.217 | -0.95 | -0.636 | -0.2 | -0.007 | -0.007 | -0.007 | 0.478 | 0.45 | 0.49 | 0.45 | 0.64 | 0.38 | 0.49 |
| 7 | 115447435 | rs11979121 | -0.361 | -0.757 | -0.434 | -0.333 | -0.015 | 0.02 | 0.02 | 0 | 0.06 | 0.02 | 0.03 | 0 | 0.06 | 0.03 |
| 7 | 115447643 | rs4472446 | -0.834 | -1.004 | -0.272 | -0.228 | -0.005 | -0.001 | -0.001 | 0 | 0.62 | 0.61 | 0.56 | 0.61 | 0.51 | 0.66 |
| 7 | 115465720 | rs7807451 | -0.076 | -0.821 | 0.01 | -0.56 | -0.016 | -0.076 | -0.076 | 0.036 | 0.16 | 0.25 | 0.08 | 0 | 0.12 | 0.23 |
| 7 | 115468751 | rs12531777 | 1.473 | -0.949 | -0.17 | -0.557 | -0.143 | -0.074 | -0.074 | 1.278 | 0.31 | 0.03 | 0.39 | 0.27 | 0.44 | 0.27 |
| 7 | 115479274 | rs2896168 | -0.395 | -0.89 | 0.158 | -0.367 | -0.001 | -0.004 | -0.004 | 0.098 | 0.95 | 0.9 | 0.97 | 0.92 | 0.97 | 0.97 |
| 7 | 115480789 | rs2402009 | 0.319 | -0.997 | 0.383 | -0.436 | 0.018 | 0.006 | 0.006 | 0 | 0.4 | 0.26 | 0.45 | 0.64 | 0.35 | 0.44 |
| 7 | 115485311 | rs7801942 | 0.01 | -0.969 | 0.474 | -0.448 | -0.015 | -0.073 | -0.073 | 0 | 0.16 | 0.25 | 0.08 | 0 | 0.12 | 0.23 |
| 7 | 115490279 | rs2896169 | -0.135 | -0.608 | 0.284 | 0.005 | 0.016 | 0.006 | 0.006 | 0 | 0.4 | 0.25 | 0.45 | 0.65 | 0.35 | 0.44 |
| 7 | 115498079 | rs6948338 | -0.133 | -0.582 | -0.151 | 0.033 | 0.01 | 0.005 | 0.005 | 0 | 0.39 | 0.32 | 0.46 | 0.65 | 0.35 | 0.44 |
| 7 | 115508663 | rs6960629 | 0.038 | -0.377 | -0.382 | 0.107 | -0.001 | -0.005 | -0.005 | 0 | 0.95 | 0.89 | 0.96 | 0.92 | 0.97 | 0.97 |
| 7 | 115511488 | rs10246828 | -0.21 | -0.529 | -0.761 | 0.189 | 0.012 | -0.002 | -0.002 | 1.513 | 0.39 | 0.25 | 0.45 | 0.64 | 0.35 | 0.43 |
| 7 | 115513145 | rs13233933 | -0.818 | -0.269 | -0.358 | 0.155 | 0.002 | -0.042 | -0.042 | 0.217 | 0.27 | 0.33 | 0.19 | 0.14 | 0.25 | 0.38 |
| 7 | 115513397 | rs12667263 | 0.771 | 0.199 | -0.657 | 0.483 | -0.018 | -0.008 | -0.008 | 0 | 0.58 | 0.36 | 0.45 | 0.54 | 0.51 | 0.65 |
| 7 | 115513497 | rs6976503 | -1.045 | 0.543 | -0.264 | 1.133 | 0 | 0.027 | 0.027 | 0 | 0.35 | 0.54 | 0.61 | 0.52 | 0.38 | 0.36 |
| 7 | 115513634 | rs6976817 | -1.41 | -0.109 | -0.414 | 1.328 | -0.014 | 0.046 | 0.046 | 1.329 | 0.2 | 0.72 | 0.35 | 0.45 | 0.17 | 0.17 |
| 7 | 115518049 | rs10279797 | 0.411 | -0.054 | -0.362 | 0.623 | -0.005 | -0.004 | -0.004 | 0 | 0.67 | 0.56 | 0.62 | 0.68 | 0.68 | 0.69 |
| 7 | 115527544 | rs7792371 | 0.383 | -0.164 | -0.135 | 0.416 | 0.009 | -0.039 | -0.039 | 0 | 0.12 | 0.12 | 0.11 | 0 | 0.2 | 0.06 |
| 7 | 115534399 | rs6967997 | -0.382 | -0.207 | 0.065 | 0.441 | 0.011 | -0.018 | -0.018 | 0 | 0.8 | 0.83 | 0.74 | 0.68 | 0.87 | 0.76 |
| 7 | 115544763 | rs10246289 | -0.348 | -0.231 | 0.054 | 0.431 | 0.012 | -0.019 | -0.019 | 0 | 0.8 | 0.83 | 0.74 | 0.68 | 0.87 | 0.76 |
| 7 | 115546486 | rs2401996 | 1.126 | -0.193 | 0.141 | 0.398 | -0.005 | -0.007 | -0.007 | 0 | 0.66 | 0.55 | 0.62 | 0.68 | 0.67 | 0.7 |
| 7 | 115551123 | rs4532544 | -0.383 | -0.2 | 0.068 | 0.403 | 0.011 | -0.018 | -0.018 | 0 | 0.8 | 0.83 | 0.74 | 0.68 | 0.87 | 0.76 |
| 7 | 115553705 | rs1316857 | 0.903 | -0.219 | 0.129 | 0.394 | 0 | 0 | 0 | 0 | 0.68 | 0.79 | 0.64 | 0.68 | 0.68 | 0.7 |
| 7 | 115566788 | rs10264545 | 1.167 | -1.284 | -0.132 | -0.098 | 0.058 | -0.007 | -0.007 | 0 | 0.37 | 0.29 | 0.3 | 0.42 | 0.22 | 0.29 |
| 7 | 115574930 | rs2896167 | -1.149 | -1.205 | -0.182 | 0.025 | 0.011 | -0.018 | -0.018 | 0 | 0.8 | 0.83 | 0.74 | 0.68 | 0.87 | 0.76 |
| 7 | 115578104 | rs17241507 | 1.457 | -1.231 | -0.199 | 0.008 | -0.056 | 0.066 | 0.066 | 1.125 | 0.11 | 0.01 | 0.1 | 0 | 0.2 | 0.06 |
| 7 | 115578124 | rs10229962 | -1.16 | -1.232 | -0.199 | 0.008 | 0.011 | -0.018 | -0.018 | 0.096 | 0.8 | 0.82 | 0.74 | 0.68 | 0.87 | 0.76 |
| 7 | 115578441 | rs17241584 | 1.449 | -1.214 | -0.198 | -0.002 | -0.056 | 0.066 | 0.066 | 1.125 | 0.11 | 0.01 | 0.1 | 0 | 0.2 | 0.06 |
| 7 | 115585871 | rs12154579 | -1.244 | -1.261 | -0.183 | 0.103 | 0.009 | -0.013 | -0.013 | 0.021 | 0.81 | 0.82 | 0.74 | 0.7 | 0.88 | 0.85 |
| 7 | 115600956 | rs10281257 | 0.239 | -2.142 | -0.508 | -0.405 | 0.052 | 0.062 | 0.062 | 1.131 | 0.52 | 0.72 | 0.46 | 0.3 | 0.61 | 0.58 |
| 7 | 115608355 | rs10248479 | 1.309 | -2.272 | -0.618 | -0.534 | 0.012 | 0.029 | 0.029 | 0 | 0.17 | 0.14 | 0.1 | 0.04 | 0.13 | 0.14 |
| 7 | 115619298 | rs11767053 | 2.19 | -2.083 | -0.982 | -0.619 | -0.146 | 0.015 | 0.015 | 0.056 | 0.22 | 0.04 | 0.24 | 0.13 | 0.37 | 0.3 |
| 7 | 115622561 | rs13222870 | 1.287 | -2.132 | -0.118 | -0.537 | 0.003 | -0.001 | -0.001 | 0.011 | 0.33 | 0.33 | 0.35 | 0.26 | 0.48 | 0.44 |
| 7 | 115628138 | rs6978353 | 1.287 | -2.132 | -0.146 | -0.537 | 0.003 | -0.001 | -0.001 | 0 | 0.33 | 0.33 | 0.35 | 0.26 | 0.48 | 0.44 |
| 7 | 115638026 | rs10238185 | -0.357 | -1.883 | -0.24 | -0.183 | -0.005 | -0.008 | -0.008 | 0 | 0.69 | 0.81 | 0.65 | 0.68 | 0.69 | 0.7 |
| 7 | 115653998 | rs17138238 | 1.771 | -2.004 | -0.102 | -0.028 | 0.001 | -0.003 | -0.003 | 0 | 0.17 | 0.14 | 0.19 | 0.38 | 0.08 | 0.11 |
| 7 | 115655686 | rs6946453 | -0.029 | -2.082 | -0.004 | -0.081 | -0.054 | -0.011 | -0.011 | 0 | 0.62 | 0.49 | 0.54 | 0.3 | 0.8 | 0.65 |
| 7 | 115655978 | rs1554630 | -1.006 | -1.556 | -0.113 | -0.044 | -0.012 | 0.017 | 0.017 | 0 | 0.8 | 0.75 | 0.74 | 0.68 | 0.87 | 0.76 |
| 7 | 115665688 | rs10258989 | 0.476 | -1.057 | -0.227 | -0.15 | -0.024 | 0.027 | 0.027 | 1.681 | 0.13 | 0.1 | 0.13 | 0 | 0.22 | 0.08 |
| 7 | 115665933 | rs6972100 | 0.964 | -1.445 | 0.109 | 0.087 | -0.033 | 0.171 | 0.171 | 2.165 | 0.12 | 0.24 | 0.18 | 0.39 | 0.07 | 0.14 |
| 7 | 115667227 | rs7793323 | -0.483 | -1.373 | -0.311 | -0.273 | 0.04 | 0.074 | 0.074 | 0.601 | 0.48 | 0.28 | 0.55 | 0.75 | 0.38 | 0.43 |
| 7 | 115671435 | rs11767151 | 0.192 | -1.446 | -0.345 | -0.191 | 0 | -0.006 | -0.006 | 0 | 0.2 | 0.18 | 0.28 | 0.39 | 0.24 | 0.2 |
| 7 | 115678646 | rs1113295 | -1.256 | -1.521 | -0.163 | -0.317 | -0.008 | 0.013 | 0.013 | 0 | 0.59 | 0.71 | 0.62 | 0.75 | 0.56 | 0.52 |
| 7 | 115680998 | rs7781921 | 0.686 | -1.314 | -0.565 | -0.192 | -0.003 | -0.024 | -0.024 | 0.816 | 0.24 | 0.14 | 0.23 | 0.32 | 0.21 | 0.22 |

| | | | | | | | | | | | | | | | | |
|---|---|---|---|---|---|---|---|---|---|---|---|---|---|---|---|---|
| 7 | 115699678 | rs4419752 | 0.188 | -1.281 | -0.503 | -0.188 | -0.001 | 0.024 | 0.024 | 0 | 0.2 | 0.22 | 0.29 | 0.39 | 0.25 | 0.2 |
| 7 | 115705799 | rs1917059 | -0.124 | -1.093 | -0.91 | -0.193 | -0.062 | 0.002 | 0.002 | 0 | 0.33 | 0.12 | 0.32 | 0.31 | 0.4 | 0.3 |
| 7 | 115713980 | rs12669287 | -1.294 | -1.425 | -0.616 | -0.219 | -0.064 | 0.041 | 0.041 | 0 | 0.66 | 0.85 | 0.68 | 0.8 | 0.6 | 0.58 |
| 7 | 115715094 | rs10953804 | 0.82 | -1.565 | -0.184 | -0.065 | -0.009 | 0.041 | 0.041 | 0 | 0.16 | 0.17 | 0.23 | 0.37 | 0.15 | 0.18 |
| 7 | 115723698 | rs7804495 | 1.638 | -1.764 | 0.266 | -0.334 | -0.017 | 0.011 | 0.011 | 0 | 0.11 | 0.25 | 0.08 | 0.12 | 0.07 | 0.06 |
| 7 | 115728072 | rs2402030 | 1.812 | -1.765 | 0.272 | -0.335 | -0.004 | 0.002 | 0.002 | 0 | 0.1 | 0.14 | 0.07 | 0.12 | 0.07 | 0.06 |
| 7 | 115742657 | rs9691405 | -0.865 | -1.934 | 0.408 | -0.13 | 0.021 | 0.046 | 0.046 | 0 | 0.47 | 0.71 | 0.48 | 0.71 | 0.44 | 0.46 |
| 7 | 115744180 | rs10269713 | 1.56 | -1.999 | 0.386 | -0.189 | 0.009 | 0.01 | 0.01 | 0 | 0.17 | 0.08 | 0.1 | 0.01 | 0.16 | 0.12 |
| 7 | 115747757 | rs12374699 | 1.078 | -2.074 | 0.486 | -0.175 | -0.009 | 0.016 | 0.016 | 0 | 0.16 | 0.15 | 0.13 | 0.28 | 0.15 | 0.17 |
| 7 | 115751974 | rs7785501 | -0.172 | -2.07 | 0.471 | -0.171 | -0.008 | 0.031 | 0.031 | 0 | 0.32 | 0.61 | 0.36 | 0.43 | 0.28 | 0.29 |
| 7 | 115757442 | rs10246006 | -1.729 | -2.004 | 0.291 | -0.149 | -0.062 | -0.008 | -0.008 | 0 | 0.65 | 0.84 | 0.59 | 0.72 | 0.6 | 0.58 |
| 7 | 115769311 | rs9641558 | -0.494 | -1.714 | 0.614 | 0.184 | -0.002 | 0.024 | 0.024 | 0 | 0.36 | 0.44 | 0.39 | 0.6 | 0.36 | 0.39 |
| 7 | 115771982 | rs7780647 | 1.353 | -1.675 | 0.388 | 0.174 | -0.01 | -0.022 | -0.022 | 0.091 | 0.17 | 0.1 | 0.22 | 0.32 | 0.18 | 0.21 |
| 7 | 115795139 | rs10224339 | -1.013 | -1.71 | 0.604 | 0.145 | 0.006 | 0.013 | 0.013 | 0 | 0.46 | 0.64 | 0.48 | 0.71 | 0.44 | 0.46 |
| 7 | 115796254 | rs17138365 | 0.009 | -1.698 | 0.691 | -0.044 | -0.019 | 0.031 | 0.031 | 0.034 | 0.2 | 0.4 | 0.18 | 0.28 | 0.18 | 0.19 |
| 7 | 115801846 | rs9649390 | -0.76 | -1.719 | 0.608 | -0.046 | -0.029 | 0.022 | 0.022 | 0 | 0.3 | 0.48 | 0.26 | 0.39 | 0.25 | 0.25 |
| 7 | 115806524 | rs11773521 | -0.189 | -1.618 | 0.282 | 0.077 | 0 | -0.002 | -0.002 | 0.655 | 0.33 | 0.25 | 0.35 | 0.69 | 0.34 | 0.38 |
| 7 | 115807797 | rs11764493 | 1.33 | -1.476 | 0.175 | 0.177 | -0.003 | 0.001 | 0.001 | 0 | 0.11 | 0.16 | 0.08 | 0.11 | 0.07 | 0.07 |
| 7 | 115817269 | rs12668723 | 0.983 | -1.459 | 0.201 | 0.204 | 0.003 | 0.038 | 0.038 | 0 | 0.18 | 0.2 | 0.25 | 0.42 | 0.2 | 0.2 |
| 7 | 115819785 | rs10270061 | 1.337 | -1.539 | 0.295 | 0.108 | -0.002 | 0 | 0 | 0 | 0.11 | 0.14 | 0.09 | 0.11 | 0.1 | 0.08 |
| 7 | 115823384 | rs6968419 | -0.856 | -1.694 | 0.414 | 0.005 | -0.007 | -0.027 | -0.027 | 0 | 0.45 | 0.51 | 0.45 | 0.8 | 0.42 | 0.46 |
| 7 | 115830736 | rs10227194 | 1.402 | -1.695 | 0.414 | -0.051 | -0.001 | 0 | 0 | 0 | 0.11 | 0.14 | 0.09 | 0.11 | 0.1 | 0.08 |
| 7 | 115831260 | rs11764793 | -0.926 | -1.764 | 0.65 | 0.063 | 0.012 | 0.036 | 0.036 | 0 | 0.45 | 0.62 | 0.46 | 0.8 | 0.42 | 0.44 |
| 7 | 115835771 | rs10225986 | 1.057 | -1.729 | 0.565 | 0.083 | -0.011 | 0.003 | 0.003 | 0 | 0.12 | 0.2 | 0.11 | 0.11 | 0.11 | 0.08 |
| 7 | 115838228 | rs10250473 | 0.956 | -1.731 | 0.527 | 0.082 | -0.013 | 0.004 | 0.004 | 0 | 0.13 | 0.24 | 0.11 | 0.11 | 0.11 | 0.08 |
| 7 | 115839253 | rs6946058 | -0.14 | -1.68 | 0.169 | 0.129 | 0.002 | -0.004 | -0.004 | 0 | 0.32 | 0.27 | 0.34 | 0.69 | 0.31 | 0.37 |
| 7 | 115844871 | rs12668487 | 1.047 | -0.766 | -0.539 | -0.375 | -0.068 | -0.011 | -0.011 | 0.294 | 0.41 | 0.15 | 0.57 | 0.59 | 0.47 | 0.46 |
| 7 | 115846757 | rs10279056 | 0.641 | -0.786 | -0.395 | -0.456 | 0.017 | 0.098 | 0.098 | 0 | 0.49 | 0.25 | 0.65 | 0.7 | 0.54 | 0.51 |
| 7 | 115847524 | rs10808180 | 1.087 | -0.623 | -0.065 | -0.675 | -0.014 | 0.014 | 0.014 | 0 | 0.41 | 0.3 | 0.52 | 0.68 | 0.46 | 0.45 |
| 7 | 115851487 | rs13223583 | 0.439 | -0.717 | 0.028 | -0.768 | -0.005 | -0.046 | -0.046 | 3.012 | 0.56 | 0.47 | 0.67 | 0.85 | 0.58 | 0.61 |
| 7 | 115856602 | rs1528246 | -2.038 | -0.957 | 0.453 | -0.555 | -0.025 | 0.001 | 0.001 | 0.005 | 0.68 | 0.71 | 0.6 | 0.69 | 0.64 | 0.57 |
| 7 | 115883585 | rs10256347 | 0.588 | -1.029 | 0.398 | -0.607 | 0.006 | -0.041 | -0.041 | 0 | 0.1 | 0.13 | 0.09 | 0 | 0.14 | 0.04 |
| 7 | 115893362 | rs3807979 | 1.524 | -1.046 | 0.415 | -0.578 | 0.019 | 0.13 | 0.13 | 0 | 0.17 | 0.33 | 0.22 | 0.4 | 0.18 | 0.17 |
| 7 | 115895718 | rs6966770 | 0.084 | -1.065 | 0.405 | -0.613 | -0.011 | 0.043 | 0.043 | 0 | 0.32 | 0.41 | 0.36 | 0.57 | 0.3 | 0.34 |
| 7 | 115897287 | rs2402060 | 1.34 | -1.069 | 0.413 | -0.611 | 0 | -0.001 | -0.001 | 0 | 0.15 | 0.13 | 0.15 | 0.16 | 0.12 | 0.17 |
| 7 | 115897884 | rs2896181 | 0.118 | -1.044 | 0.37 | -0.676 | -0.011 | 0.043 | 0.043 | 0 | 0.32 | 0.41 | 0.36 | 0.57 | 0.3 | 0.34 |
| 7 | 115898710 | rs9777 | 0.982 | -1.048 | 0.264 | -0.658 | 0.031 | 0.04 | 0.04 | 0 | 0.26 | 0.14 | 0.17 | 0.12 | 0.2 | 0.2 |
| 7 | 115900262 | rs10242959 | -0.629 | -1.017 | 0.333 | -0.629 | 0 | 0.001 | 0.001 | 0 | 0.41 | 0.49 | 0.45 | 0.56 | 0.44 | 0.38 |
| 7 | 115904091 | rs11766908 | 0.955 | -0.956 | 0.201 | -0.591 | 0.031 | 0.037 | 0.037 | 0 | 0.26 | 0.14 | 0.17 | 0.13 | 0.2 | 0.2 |
| 7 | 115906373 | rs1918912 | 0.088 | -0.862 | 0.261 | -0.539 | -0.007 | 0.043 | 0.043 | 0 | 0.33 | 0.45 | 0.37 | 0.57 | 0.3 | 0.34 |
| 7 | 115910811 | rs10266135 | -0.164 | -0.861 | 0.254 | -0.48 | 0.01 | 0.048 | 0.048 | 0 | 0.28 | 0.38 | 0.31 | 0.4 | 0.32 | 0.23 |
| 7 | 115911312 | rs10269717 | -0.164 | -0.861 | 0.254 | -0.48 | 0.01 | 0.048 | 0.048 | 0 | 0.28 | 0.38 | 0.31 | 0.4 | 0.32 | 0.23 |
| 7 | 115920902 | rs10953818 | 1.372 | -0.852 | 0.107 | -0.447 | 0.009 | 0.093 | 0.093 | 0 | 0.16 | 0.26 | 0.22 | 0.4 | 0.18 | 0.19 |
| 7 | 115924557 | rs17248419 | 0.692 | -1.45 | -0.04 | -0.496 | 0.025 | 0.032 | 0.032 | 0 | 0.26 | 0.14 | 0.17 | 0.12 | 0.2 | 0.17 |
| 7 | 115943514 | rs2204679 | -0.459 | -1.46 | 0.075 | -0.622 | -0.029 | -0.034 | -0.034 | 0 | 0.39 | 0.4 | 0.28 | 0.24 | 0.27 | 0.31 |
| 7 | 115945391 | rs6952197 | 0.358 | -1.483 | -0.016 | -0.65 | -0.003 | -0.029 | -0.029 | 0.429 | 0.3 | 0.27 | 0.17 | 0.07 | 0.23 | 0.19 |
| 7 | 115949014 | rs6466573 | -0.302 | -1.316 | 0.011 | -0.57 | -0.008 | 0.002 | 0.002 | 0 | 0.44 | 0.43 | 0.37 | 0.48 | 0.37 | 0.38 |
| 7 | 115951746 | rs17308500 | -1.966 | -1.204 | -0.222 | -0.758 | -0.022 | -0.01 | -0.01 | 0.02 | 0.67 | 0.68 | 0.57 | 0.64 | 0.6 | 0.59 |
| 7 | 115953518 | rs10953820 | -1.102 | -0.684 | 0.109 | -1.114 | 0.002 | 0.002 | 0.002 | 0 | 0.5 | 0.49 | 0.43 | 0.42 | 0.47 | 0.41 |
| 7 | 115970622 | rs12154695 | -0.595 | -0.626 | -0.103 | -0.828 | -0.025 | -0.109 | -0.109 | 1.839 | 0.39 | 0.43 | 0.22 | 0.07 | 0.3 | 0.26 |
| 7 | 115970664 | rs1528243 | 0.894 | -0.209 | -0.433 | -0.19 | -0.001 | 0.002 | 0.002 | 0 | 0.13 | 0.1 | 0.19 | 0.35 | 0.14 | 0.19 |
| 7 | 115976844 | rs12540030 | 0.894 | -0.209 | -0.433 | -0.203 | -0.001 | 0.002 | 0.002 | 0 | 0.13 | 0.1 | 0.19 | 0.35 | 0.14 | 0.18 |
| 7 | 115979468 | rs7806219 | -0.379 | -0.373 | -0.142 | -1.158 | -0.001 | -0.027 | -0.027 | 0 | 0.14 | 0.13 | 0.08 | 0 | 0.09 | 0.07 |
| 7 | 115992733 | rs17138624 | -0.235 | -0.343 | -0.208 | -1.134 | 0.003 | -0.017 | -0.017 | 0 | 0.13 | 0.09 | 0.06 | 0 | 0.07 | 0.07 |
| 7 | 115992849 | rs2402061 | 1.398 | -0.837 | -0.191 | -1.833 | 0.085 | 0.161 | 0.161 | 0.45 | 0.5 | 0.32 | 0.75 | 0.93 | 0.68 | 0.69 |
| 7 | 115996856 | rs760444 | -0.517 | -0.336 | -0.485 | -1.599 | -0.005 | 0 | 0 | 0 | 0.19 | 0.09 | 0.2 | 0.16 | 0.18 | 0.22 |
| 7 | 115997905 | rs1633722 | 0.019 | -0.56 | -0.034 | -1.764 | 0.018 | 0.024 | 0.024 | 0.03 | 0.71 | 0.81 | 0.86 | 0.93 | 0.83 | 0.78 |
| 7 | 116003445 | rs10235210 | 1.802 | -0.581 | 0.498 | -1.747 | 0.004 | 0.06 | 0.06 | 0 | 0.44 | 0.31 | 0.61 | 0.76 | 0.58 | 0.56 |
| 7 | 116006105 | rs926201 | -0.342 | -0.085 | 0.908 | -1.206 | -0.004 | -0.025 | -0.025 | 2.732 | 0.88 | 0.82 | 0.93 | 0.94 | 0.9 | 0.9 |
| 7 | 116018179 | rs1728723 | 1.354 | 0.015 | 1.057 | -1.184 | 0.003 | -0.014 | -0.014 | 0.373 | 0.7 | 0.61 | 0.73 | 0.75 | 0.73 | 0.7 |
| 7 | 116019562 | rs1633716 | -0.323 | -0.082 | 0.626 | -1.166 | -0.01 | -0.022 | -0.022 | 0 | 0.87 | 0.82 | 0.94 | 0.94 | 0.91 | 0.88 |
| 7 | 116028923 | rs2049902 | 1.681 | 0.19 | 0.099 | -1.17 | -0.004 | 0.047 | 0.047 | 0.091 | 0.62 | 0.24 | 0.67 | 0.75 | 0.63 | 0.67 |
| 7 | 116036160 | rs1633731 | -1.661 | -0.058 | 0.162 | -1.48 | 0.033 | 0.094 | 0.094 | 1.021 | 0.39 | 0.7 | 0.23 | 0.06 | 0.29 | 0.27 |
| 7 | 116043100 | rs10252986 | -0.377 | -0.136 | 0.728 | -1.548 | -0.011 | -0.078 | -0.078 | 0 | 0.18 | 0.23 | 0.09 | 0 | 0.1 | 0.14 |
| 7 | 116055528 | rs17138675 | 0.247 | -0.164 | 1.091 | -1.565 | -0.017 | -0.033 | -0.033 | 1.728 | 0.14 | 0.19 | 0.08 | 0.06 | 0.09 | 0.12 |
| 7 | 116067883 | rs4540346 | 1.702 | 0.139 | 1.196 | -1.511 | 0.009 | 0.029 | 0.029 | 0.783 | 0.61 | 0.38 | 0.76 | 0.93 | 0.67 | 0.73 |
| 7 | 116072835 | rs6963408 | -0.457 | 0.134 | 1.212 | -1.508 | -0.013 | -0.082 | -0.082 | 0 | 0.19 | 0.25 | 0.09 | 0 | 0.1 | 0.13 |
| 7 | 116073567 | rs10464649 | 1.901 | 0.132 | 1.124 | -1.502 | 0.006 | 0.126 | 0.126 | 1.481 | 0.55 | 0.23 | 0.63 | 0.78 | 0.59 | 0.62 |
| 7 | 116081121 | rs4730738 | 0.385 | 0.132 | 1.112 | -1.404 | 0.006 | -0.067 | -0.067 | 0 | 0.8 | 0.7 | 0.85 | 0.93 | 0.78 | 0.86 |

| | | | | | | | | | | | | | | | | |
|---|---|---|---|---|---|---|---|---|---|---|---|---|---|---|---|---|
| 7 | 116118330 | rs926197 | 1.907 | 0.262 | 1.099 | -1.504 | -0.004 | 0.073 | 0.073 | 0.383 | 0.52 | 0.38 | 0.58 | 0.78 | 0.53 | 0.58 |
| 7 | 116121575 | rs879211 | -0.314 | 0.26 | 1.069 | -1.538 | 0.002 | 0.001 | 0.001 | 0.617 | 0.21 | 0.13 | 0.13 | 0.01 | 0.15 | 0.17 |
| 7 | 116122845 | rs6975771 | -0.524 | 0.312 | 1.112 | -1.434 | 0.002 | -0.032 | -0.032 | 0 | 0.84 | 0.78 | 0.89 | 0.93 | 0.86 | 0.87 |
| 7 | 116134179 | rs17515508 | 0.215 | 0.33 | 0.945 | -1.409 | 0.004 | -0.021 | -0.021 | 0.206 | 0.05 | 0.08 | 0.05 | 0 | 0.09 | 0.01 |
| 7 | 116135253 | rs17138756 | -0.549 | 0.313 | 1.04 | -1.417 | 0.014 | -0.037 | -0.037 | 0 | 0.07 | 0.19 | 0.05 | 0 | 0.1 | 0.01 |
| 7 | 116137892 | rs4730742 | -0.462 | 0.276 | 1.054 | -1.525 | 0.003 | -0.004 | -0.004 | 0.521 | 0.22 | 0.15 | 0.14 | 0.01 | 0.16 | 0.18 |
| 7 | 116138666 | rs2191500 | 1.259 | 0.274 | 1.127 | -1.546 | -0.009 | -0.092 | -0.092 | 0.76 | 0.62 | 0.59 | 0.75 | 0.92 | 0.7 | 0.7 |
| 7 | 116147401 | rs10278782 | -0.542 | 0.242 | 1.105 | -1.482 | 0 | -0.03 | -0.03 | 0 | 0.23 | 0.19 | 0.15 | 0.01 | 0.16 | 0.18 |
| 7 | 116148370 | rs1052990 | -1.655 | 0.243 | 1.065 | -1.458 | 0.004 | -0.04 | -0.04 | 0.33 | 0.34 | 0.42 | 0.32 | 0.16 | 0.34 | 0.3 |
| 7 | 116151155 | rs6466579 | 1.945 | 0.232 | 1.067 | -1.466 | -0.003 | 0.1 | 0.1 | 0.367 | 0.5 | 0.37 | 0.57 | 0.78 | 0.52 | 0.57 |
| 7 | 116151338 | rs10281637 | -1.268 | 0.253 | 1.103 | -1.591 | -0.001 | -0.129 | -0.129 | 0 | 0.29 | 0.4 | 0.21 | 0.01 | 0.26 | 0.19 |
| 7 | 116165463 | rs2742125 | -0.257 | 0.158 | 0.888 | -1.513 | -0.011 | -0.049 | -0.049 | 0 | 0.83 | 0.69 | 0.89 | 0.93 | 0.86 | 0.87 |
| 7 | 116167208 | rs926198 | 1.346 | 0.194 | 0.661 | -1.564 | -0.001 | -0.038 | -0.038 | 0.27 | 0.6 | 0.5 | 0.75 | 0.92 | 0.7 | 0.7 |
| 7 | 116175711 | rs10256914 | -0.916 | 0.147 | 0.188 | -1.567 | 0.001 | -0.011 | -0.011 | 0 | 0.28 | 0.21 | 0.17 | 0.01 | 0.24 | 0.2 |
| 7 | 116182054 | rs959173 | -0.088 | -0.046 | 0.617 | -1.293 | 0.005 | -0.056 | -0.056 | 0 | 0.83 | 0.57 | 0.88 | 0.93 | 0.85 | 0.87 |
| 7 | 116186241 | rs3807989 | 1.96 | -0.009 | 0.355 | -1.124 | -0.002 | 0.056 | 0.056 | 0 | 0.58 | 0.3 | 0.64 | 0.73 | 0.61 | 0.65 |
| 7 | 116190693 | rs3815412 | -1.647 | 0.051 | 0.06 | -1.069 | 0.001 | -0.016 | -0.016 | 0.03 | 0.26 | 0.4 | 0.25 | 0.2 | 0.25 | 0.23 |
| 7 | 116198001 | rs6466587 | -1.368 | 0.052 | 0.012 | -1.08 | -0.006 | -0.071 | -0.071 | 0 | 0.22 | 0.32 | 0.13 | 0.01 | 0.16 | 0.14 |
| 7 | 116199797 | rs8713 | -1.368 | 0.052 | 0.013 | -1.08 | -0.006 | -0.074 | -0.074 | 0.355 | 0.22 | 0.32 | 0.12 | 0.01 | 0.16 | 0.14 |
| 7 | 116200587 | rs1049337 | 1.315 | -0.162 | -0.843 | -0.728 | -0.023 | -0.13 | -0.13 | 0.63 | 0.31 | 0.04 | 0.32 | 0.57 | 0.31 | 0.31 |
| 7 | 116217029 | rs7795356 | 0.421 | -0.324 | 0.8 | -0.056 | -0.003 | -0.058 | -0.058 | 0 | 0.79 | 0.74 | 0.87 | 0.93 | 0.82 | 0.82 |
| 7 | 116217657 | rs2109517 | 1.578 | -0.476 | 0.921 | 0.178 | 0.013 | 0.012 | 0.012 | 0 | 0.61 | 0.3 | 0.58 | 0.32 | 0.68 | 0.53 |
| 7 | 116218086 | rs2191502 | -0.799 | -0.397 | 0.884 | 0.546 | -0.021 | 0.128 | 0.128 | 0.151 | 0.17 | 0.4 | 0.28 | 0.61 | 0.14 | 0.29 |
| 7 | 116226422 | rs6978354 | 0.682 | -0.296 | 0.095 | 0.128 | 0 | 0 | 0 | 0 | 0.61 | 0.44 | 0.47 | 0.26 | 0.68 | 0.44 |
| 7 | 116233338 | rs10227421 | -1.26 | -0.344 | 0.595 | 0.815 | -0.019 | 0.012 | 0.012 | 2.516 | 0.19 | 0.69 | 0.13 | 0.23 | 0.11 | 0.21 |
| 7 | 116241322 | rs10808181 | -0.76 | -0.337 | 0.966 | 0.813 | -0.03 | 0.094 | 0.094 | 1.19 | 0.16 | 0.44 | 0.1 | 0.23 | 0.08 | 0.17 |
| 7 | 116266792 | rs6965534 | -1.394 | -0.34 | 0.678 | 0.773 | -0.017 | 0.009 | 0.009 | 2.291 | 0.19 | 0.7 | 0.13 | 0.22 | 0.11 | 0.22 |
| 7 | 116285715 | rs1002399 | 0.182 | -0.153 | 0.091 | 0.838 | -0.017 | -0.014 | -0.014 | 0 | 0.89 | 0.95 | 0.83 | 0.88 | 0.85 | 0.84 |
| 7 | 116290966 | rs10280998 | -0.453 | -0.192 | 0.252 | 0.888 | -0.024 | 0.064 | 0.064 | 1.193 | 0.16 | 0.35 | 0.1 | 0.2 | 0.08 | 0.21 |
| 7 | 116296896 | rs437 | 1.635 | -0.178 | -0.056 | 0.879 | 0.012 | -0.012 | -0.012 | 0.081 | 0.69 | 0.2 | 0.7 | 0.69 | 0.74 | 0.6 |
| 7 | 116314868 | rs11767567 | -0.545 | -0.117 | -0.035 | 1.086 | -0.021 | 0.079 | 0.079 | 0.543 | 0.15 | 0.36 | 0.1 | 0.2 | 0.08 | 0.26 |
| 7 | 116317877 | rs40239 | 0.25 | 0.082 | -1.141 | 1.078 | 0.006 | -0.026 | -0.026 | 0.055 | 0.85 | 0.95 | 0.82 | 0.81 | 0.88 | 0.86 |
| 7 | 116319002 | rs714180 | 0.312 | 0.467 | -2.067 | 1.031 | -0.091 | -0.1 | -0.1 | 0 | 0.59 | 0.91 | 0.37 | 0.57 | 0.52 | 0.61 |
| 7 | 116319926 | rs38841 | 0.266 | 0.837 | -2.084 | 1.081 | -0.177 | -0.08 | -0.08 | 0 | 0.27 | 0.02 | 0.46 | 0.25 | 0.39 | 0.39 |
| 7 | 116322791 | rs9641562 | -1.09 | 0.631 | -0.77 | 0.993 | 0.05 | 0.07 | 0.07 | 0.174 | 0.07 | 0.38 | 0.12 | 0.09 | 0.06 | 0.1 |
| 7 | 116332197 | rs7798983 | -0.7 | -0.245 | -1.546 | 0.905 | 0.007 | -0.003 | -0.003 | 1.113 | 0.34 | 0.35 | 0.26 | 0.39 | 0.3 | 0.35 |
| 7 | 116336880 | rs2237708 | -0.445 | -0.611 | -1.699 | 0.74 | 0 | -0.013 | -0.013 | 1.33 | 0.23 | 0.18 | 0.23 | 0.33 | 0.26 | 0.24 |
| 7 | 116338658 | rs2237712 | -0.582 | -0.497 | -1.645 | 0.75 | -0.007 | -0.005 | -0.005 | 0.366 | 0.1 | 0.17 | 0.03 | 0.05 | 0.04 | 0.11 |
| 7 | 116346603 | rs10243024 | -1.315 | -0.71 | -0.505 | 0.438 | -0.031 | -0.014 | -0.014 | 0 | 0.29 | 0.5 | 0.2 | 0.3 | 0.26 | 0.29 |
| 7 | 116358044 | rs38855 | 1.496 | -0.794 | -1.198 | 0.246 | -0.052 | 0.013 | 0.013 | 0 | 0.43 | 0.15 | 0.56 | 0.57 | 0.48 | 0.58 |
| 7 | 116370287 | rs38858 | 1.501 | -0.827 | -0.588 | 0.231 | -0.05 | 0.013 | 0.013 | 0 | 0.43 | 0.15 | 0.55 | 0.58 | 0.48 | 0.58 |
| 7 | 116371475 | rs7804778 | 0.11 | -0.31 | -0.458 | 0.242 | 0.01 | 0.007 | 0.007 | 0 | 0.09 | 0.1 | 0.05 | 0.02 | 0.03 | 0.04 |
| 7 | 116379094 | rs38859 | 1.944 | 0.109 | -0.733 | 0.279 | -0.143 | -0.136 | -0.136 | 0.158 | 0.4 | 0.03 | 0.46 | 0.47 | 0.44 | 0.5 |
| 7 | 116397572 | rs13223756 | 0.291 | 0.063 | -0.522 | 0.254 | -0.002 | -0.016 | -0.016 | 0 | 0.17 | 0.11 | 0.13 | 0.27 | 0.2 | 0.23 |
| 7 | 116399131 | rs10215153 | 0.172 | 0.041 | -0.192 | 0.193 | 0.003 | -0.065 | -0.065 | 1.103 | 0.28 | 0.4 | 0.25 | 0.11 | 0.28 | 0.13 |
| 7 | 116428573 | rs2402118 | -2.584 | 0.826 | -0.421 | 0.252 | -0.07 | 0.206 | 0.206 | 1.105 | 0.64 | 0.98 | 0.69 | 0.92 | 0.63 | 0.79 |
| 7 | 116431427 | rs193686 | -0.41 | 0.814 | -0.337 | 0.271 | 0.001 | -0.057 | -0.057 | 0.062 | 0.72 | 0.6 | 0.75 | 0.88 | 0.72 | 0.87 |
| 7 | 116435123 | rs13239139 | 0.288 | 0.823 | -0.374 | 0.299 | -0.002 | -0.009 | -0.009 | 0 | 0.17 | 0.15 | 0.13 | 0.28 | 0.2 | 0.24 |
| 7 | 116435768 | rs41736 | 2.018 | 0.815 | -0.443 | 0.29 | -0.138 | -0.143 | -0.143 | 0 | 0.4 | 0.03 | 0.46 | 0.47 | 0.44 | 0.49 |
| 7 | 116436022 | rs2023748 | 2.009 | 0.848 | -0.434 | 0.299 | -0.137 | -0.142 | -0.142 | 0 | 0.41 | 0.03 | 0.47 | 0.47 | 0.44 | 0.49 |
| 7 | 116436097 | rs41737 | 2 | 0.847 | -0.434 | 0.299 | -0.132 | -0.137 | -0.137 | 0 | 0.41 | 0.04 | 0.47 | 0.47 | 0.44 | 0.49 |
| 7 | 116437206 | rs41738 | 1.99 | 0.847 | -0.434 | 0.298 | -0.136 | -0.141 | -0.141 | 1.562 | 0.41 | 0.04 | 0.47 | 0.47 | 0.44 | 0.49 |
| 7 | 116437504 | rs41739 | 2.01 | 0.847 | -0.434 | 0.298 | -0.137 | -0.142 | -0.142 | 0.185 | 0.41 | 0.03 | 0.47 | 0.47 | 0.44 | 0.49 |
| 7 | 116437606 | rs1621 | 0.349 | 0.831 | -0.452 | 0.27 | -0.012 | -0.08 | -0.08 | 0.003 | 0.64 | 0.5 | 0.7 | 0.86 | 0.68 | 0.85 |
| 7 | 116438418 | rs6566 | 1.935 | 0.868 | -0.66 | 0.297 | -0.131 | -0.137 | -0.137 | 3.115 | 0.41 | 0.04 | 0.47 | 0.47 | 0.44 | 0.49 |
| 7 | 116456018 | rs41756 | 1.448 | 1.309 | -0.685 | 0.216 | -0.039 | -0.039 | -0.039 | 1.718 | 0.46 | 0.1 | 0.4 | 0.51 | 0.43 | 0.5 |
| 7 | 116456466 | rs41758 | 0.462 | 1.334 | -0.569 | 0.22 | -0.002 | 0.01 | 0.01 | 0 | 0.65 | 0.35 | 0.52 | 0.8 | 0.64 | 0.75 |
| 7 | 116457242 | rs2402120 | 0.25 | 1.322 | -0.587 | 0.235 | -0.002 | -0.012 | -0.012 | 0.328 | 0.17 | 0.13 | 0.11 | 0.29 | 0.2 | 0.25 |
| 7 | 116475257 | rs6977473 | 0.494 | 1.316 | -0.624 | 0.412 | -0.003 | -0.013 | -0.013 | 0 | 0.07 | 0.07 | 0.06 | 0.03 | 0.03 | 0.04 |
| 7 | 116481603 | rs41779 | -0.161 | 1.321 | -0.531 | 0.395 | 0.008 | -0.074 | -0.074 | 0.236 | 0.7 | 0.58 | 0.62 | 0.91 | 0.67 | 0.85 |
| 7 | 116482219 | rs41780 | 2.358 | 1.332 | -0.688 | 0.458 | -0.089 | -0.126 | -0.126 | 0 | 0.39 | 0.04 | 0.35 | 0.49 | 0.39 | 0.46 |
| 7 | 116489938 | rs438 | 0.589 | 1.214 | -0.428 | 0.379 | -0.002 | -0.075 | -0.075 | 0.056 | 0.62 | 0.51 | 0.56 | 0.88 | 0.63 | 0.75 |
| 7 | 116528240 | rs4808 | -0.687 | 1.399 | 0.555 | 0.822 | -0.011 | 0.039 | 0.039 | 0 | 0.23 | 0.39 | 0.31 | 0.39 | 0.23 | 0.26 |
| 7 | 116534747 | rs10953830 | -0.7 | 1.398 | 0.516 | 0.823 | -0.012 | 0.046 | 0.046 | 0 | 0.23 | 0.4 | 0.31 | 0.4 | 0.24 | 0.26 |
| 7 | 116534820 | rs964449 | 1.711 | 1.45 | -0.098 | 0.815 | -0.146 | -0.121 | -0.121 | 0 | 0.44 | 0.03 | 0.37 | 0.45 | 0.45 | 0.56 |
| 7 | 116558716 | rs1049618 | 1.715 | 1.494 | -0.103 | 0.81 | -0.142 | -0.121 | -0.121 | 0 | 0.44 | 0.03 | 0.37 | 0.45 | 0.44 | 0.56 |
| 7 | 116562931 | rs1978105 | -0.894 | 1.711 | 0.18 | 0.721 | -0.031 | 0.073 | 0.073 | 0 | 0.83 | 0.96 | 0.84 | 1 | 0.8 | 0.92 |
| 7 | 116567649 | rs9649395 | 0.284 | 1.648 | 0.314 | 0.875 | -0.001 | 0.003 | 0.003 | 0 | 0.7 | 0.73 | 0.71 | 0.85 | 0.69 | 0.85 |
| 7 | 116569219 | rs7804651 | -1.227 | 1.741 | -0.145 | 1.321 | -0.004 | 0.001 | 0.001 | 0.128 | 0.27 | 0.26 | 0.45 | 0.77 | 0.3 | 0.6 |

| | | | | | | | | | | | | | | | | |
|---|---|---|---|---|---|---|---|---|---|---|---|---|---|---|---|---|
| 7 | 116570027 | rs12706123 | -0.498 | 1.805 | -0.372 | 1.533 | -0.046 | -0.058 | -0.058 | 0 | 0.2 | 0.02 | 0.21 | 0.3 | 0.25 | 0.25 |
| 7 | 116571673 | rs10953831 | -0.844 | 1.701 | -0.315 | 0.823 | -0.043 | -0.057 | -0.057 | 0 | 0.24 | 0.13 | 0.38 | 0.66 | 0.31 | 0.52 |
| 7 | 116573864 | rs7802605 | -0.591 | 0.829 | 0.295 | 0.582 | -0.026 | -0.046 | -0.046 | 0 | 0.28 | 0.51 | 0.24 | 0.12 | 0.23 | 0.12 |
| 7 | 116577124 | rs7782376 | -1.421 | 0.638 | 0.396 | 1.119 | -0.016 | 0.051 | 0.051 | 0.06 | 0.25 | 0.59 | 0.27 | 0.53 | 0.19 | 0.35 |
| 7 | 116577942 | rs10244937 | -1.165 | 0.526 | 0.698 | 0.46 | -0.023 | 0.011 | 0.011 | 0.062 | 0.25 | 0.6 | 0.22 | 0.29 | 0.17 | 0.18 |
| 7 | 116635048 | rs12706135 | 1.19 | 1.065 | -0.157 | 0.369 | -0.062 | -0.028 | -0.028 | 0 | 0.16 | 0.01 | 0.1 | 0.12 | 0.23 | 0.26 |
| 7 | 116660803 | rs7788320 | 1.532 | 0.873 | 0.01 | 0.448 | 0.011 | 0.002 | 0.002 | 0 | 0.56 | 0.3 | 0.6 | 0.46 | 0.57 | 0.52 |
| 7 | 116663029 | rs11764134 | 1.192 | 0.89 | 0.048 | 0.455 | -0.061 | -0.027 | -0.027 | 0.022 | 0.16 | 0.01 | 0.11 | 0.12 | 0.23 | 0.26 |
| 7 | 116672764 | rs4489259 | 0.083 | 0.875 | 0.079 | 0.496 | -0.008 | 0.055 | 0.055 | 2.151 | 0.87 | 0.39 | 0.88 | 0.86 | 0.9 | 0.94 |
| 7 | 116684434 | rs2428290 | -0.705 | 0.851 | 0.066 | 0.491 | 0 | 0.005 | 0.005 | 0 | 0.29 | 0.62 | 0.22 | 0.26 | 0.33 | 0.32 |
| 7 | 116701357 | rs13222576 | 1.249 | 0.883 | 0.084 | 0.556 | -0.039 | -0.028 | -0.028 | 0 | 0.16 | 0.01 | 0.09 | 0.12 | 0.2 | 0.27 |
| 7 | 116705218 | rs10260368 | -0.278 | 0.859 | 0.129 | 0.55 | 0.021 | 0.066 | 0.066 | 3.038 | 0.13 | 0.61 | 0.13 | 0.14 | 0.13 | 0.05 |
| 7 | 116708026 | rs7785492 | 0.554 | 1.259 | -0.255 | 0.375 | -0.003 | -0.006 | -0.006 | 0 | 0.23 | 0.23 | 0.43 | 0.44 | 0.25 | 0.36 |
| 7 | 116709570 | rs6961817 | -0.696 | 1.038 | 0.225 | 1.195 | 0.008 | -0.021 | -0.021 | 0 | 0.15 | 0.07 | 0.17 | 0.27 | 0.12 | 0.17 |
| 7 | 116716800 | rs13308921 | 0.557 | 1.051 | 0.236 | 1.207 | -0.039 | -0.028 | -0.028 | 0 | 0.16 | 0.01 | 0.09 | 0.12 | 0.2 | 0.27 |
| 7 | 116754688 | rs3808222 | 0.232 | -0.029 | -0.415 | 0.633 | -0.053 | -0.005 | -0.005 | 0.635 | 0.22 | 0.01 | 0.13 | 0.12 | 0.26 | 0.27 |
| 7 | 116754862 | rs739557 | -0.899 | -0.048 | 0.809 | 0.918 | -0.015 | 0.007 | 0.007 | 0 | 0.12 | 0 | 0.11 | 0.12 | 0.08 | 0.12 |
| 7 | 116758761 | rs3808221 | -1.946 | 0.437 | 1.306 | 1.167 | 0.02 | 0.235 | 0.235 | 2.292 | 0.08 | 0.42 | 0.12 | 0.18 | 0.04 | 0.08 |
| 7 | 116780499 | rs10224415 | -2.051 | 0.618 | 1.203 | 0.53 | 0.063 | -0.009 | -0.009 | 0 | 0.08 | 0.41 | 0.05 | 0 | 0.05 | 0.06 |
| 7 | 116782226 | rs17139390 | -0.745 | -0.058 | 0.98 | 0.977 | -0.019 | 0.019 | 0.019 | 0 | 0.1 | 0.11 | 0.09 | 0.19 | 0.03 | 0.11 |
| 7 | 116790493 | rs10487358 | -1.907 | -0.006 | 1.448 | 0.378 | -0.035 | -0.014 | -0.014 | 0.932 | 0.25 | 0.52 | 0.23 | 0.19 | 0.17 | 0.14 |
| 7 | 116799466 | rs2402157 | -0.821 | -0.018 | 1.654 | 0.381 | -0.053 | 0.086 | 0.086 | 0 | 0.12 | 0.29 | 0.1 | 0.19 | 0.03 | 0.11 |
| 7 | 116804930 | rs10258677 | -1.489 | -0.401 | 0.763 | -0.308 | 0.002 | 0 | 0 | 0 | 0.5 | 0.44 | 0.39 | 0.24 | 0.41 | 0.33 |
| 7 | 116811037 | rs193562 | -1.587 | -0.197 | 1.205 | 0.037 | -0.031 | -0.012 | -0.012 | 0 | 0.25 | 0.5 | 0.22 | 0.2 | 0.19 | 0.16 |
| 7 | 116812784 | rs193568 | -0.527 | -0.17 | 1.264 | 0.091 | -0.05 | 0.133 | 0.133 | 0 | 0.12 | 0.33 | 0.1 | 0.21 | 0.03 | 0.11 |
| 7 | 116812974 | rs13229745 | -0.393 | -0.14 | 1.179 | 0.052 | 0.001 | -0.034 | -0.034 | 0 | 0.14 | 0.15 | 0.13 | 0 | 0.17 | 0.05 |
| 7 | 116819510 | rs38869 | -0.852 | -0.324 | 1.145 | 0.346 | -0.052 | 0.128 | 0.128 | 0.684 | 0.12 | 0.4 | 0.11 | 0.2 | 0.03 | 0.11 |
| 7 | 116834863 | rs38892 | 0.859 | 1.129 | 0.891 | 0.546 | 0.022 | -0.017 | -0.017 | 0.067 | 0.64 | 0.45 | 0.55 | 0.69 | 0.54 | 0.72 |
| 7 | 116838311 | rs193573 | 0.865 | 1.307 | 1.193 | 0.404 | 0.012 | -0.098 | -0.098 | 0 | 0.72 | 0.51 | 0.76 | 0.93 | 0.65 | 0.79 |
| 7 | 116839950 | rs193579 | 1.13 | 1.216 | 1.245 | 0.431 | -0.001 | 0.041 | 0.041 | 0.01 | 0.69 | 0.23 | 0.72 | 0.89 | 0.64 | 0.8 |
| 7 | 116840492 | rs193580 | 0.842 | 1.239 | 1.432 | 0.441 | 0.001 | -0.027 | -0.027 | 0.005 | 0.72 | 0.67 | 0.77 | 0.93 | 0.65 | 0.8 |
| 7 | 116864331 | rs6942674 | -0.141 | 1.318 | 0.09 | 1.118 | -0.016 | -0.056 | -0.056 | 0.944 | 0.11 | 0.04 | 0.28 | 0.49 | 0.09 | 0.36 |
| 7 | 116866811 | rs6954357 | -1.17 | 1.321 | 0.363 | 0.418 | -0.001 | 0.006 | 0.006 | 0 | 0.3 | 0.7 | 0.27 | 0.1 | 0.36 | 0.19 |
| 7 | 116867295 | rs7788285 | -0.887 | 1.599 | -0.625 | 0.416 | -0.01 | 0.026 | 0.026 | 0 | 0.26 | 0.19 | 0.22 | 0.06 | 0.35 | 0.18 |
| 7 | 116878314 | rs38895 | -1.055 | 1.577 | -1.135 | 1 | -0.029 | -0.099 | -0.099 | 0 | 0.3 | 0.14 | 0.39 | 0.54 | 0.29 | 0.35 |
| 7 | 116879224 | rs38896 | -1.04 | 1.792 | -0.762 | 1.036 | 0.007 | 0.004 | 0.004 | 0 | 0.34 | 0.35 | 0.38 | 0.37 | 0.37 | 0.39 |
| 7 | 116879607 | rs10247427 | -0.525 | 2.103 | -0.631 | 0.985 | 0 | -0.001 | -0.001 | 0.013 | 0.21 | 0.18 | 0.28 | 0.25 | 0.18 | 0.3 |
| 7 | 116889750 | rs11763514 | -0.52 | 2.029 | -0.347 | 0.947 | 0.001 | -0.002 | -0.002 | 0 | 0.21 | 0.18 | 0.27 | 0.29 | 0.16 | 0.3 |
| 7 | 116890886 | rs38903 | -0.851 | 2.08 | -0.231 | 0.698 | -0.027 | 0.035 | 0.035 | 0 | 0.16 | 0.1 | 0.13 | 0.01 | 0.22 | 0.13 |
| 7 | 116892846 | rs38904 | -2.042 | 2.185 | 0.591 | 0.584 | -0.02 | 0.033 | 0.033 | 0.04 | 0.43 | 0.69 | 0.47 | 0.31 | 0.44 | 0.47 |
| 7 | 116894626 | rs7776871 | -0.369 | 2.277 | 0.848 | 0.931 | 0.003 | 0.134 | 0.134 | 0 | 0.06 | 0.1 | 0.16 | 0.43 | 0.06 | 0.28 |
| 7 | 116895588 | rs38913 | -1.517 | 2.273 | 0.824 | 0.903 | 0.021 | -0.068 | -0.068 | 4.058 | 0.2 | 0.23 | 0.18 | 0.01 | 0.28 | 0.15 |
| 7 | 116897490 | rs12669592 | -0.449 | 2.018 | 0.562 | 1.077 | -0.002 | 0.09 | 0.09 | 0 | 0.06 | 0.07 | 0.15 | 0.43 | 0.06 | 0.28 |
| 7 | 116916228 | rs887575 | 1.721 | 2.08 | 0.641 | 0.791 | -0.005 | 0.011 | 0.011 | 0 | 0.56 | 0.61 | 0.43 | 0.22 | 0.55 | 0.29 |
| 7 | 116917118 | rs4730775 | 1.882 | 2.062 | 0.702 | 0.738 | 0.011 | 0.1 | 0.1 | 0.233 | 0.51 | 0.31 | 0.37 | 0.2 | 0.49 | 0.27 |
| 7 | 116917427 | rs2024233 | 1.699 | 2.187 | 0.728 | 1.02 | 0.001 | 0.005 | 0.005 | 0 | 0.73 | 0.69 | 0.69 | 0.52 | 0.69 | 0.56 |
| 7 | 116918085 | rs2228946 | -1.73 | 1.819 | 0.781 | 0.419 | -0.002 | -0.11 | -0.11 | 0 | 0.22 | 0.34 | 0.15 | 0.01 | 0.21 | 0.1 |
| 7 | 116918792 | rs733153 | -1.997 | 1.854 | 0.308 | 1.164 | -0.001 | 0.044 | 0.044 | 1.006 | 0.13 | 0.22 | 0.27 | 0.52 | 0.09 | 0.34 |
| 7 | 116919978 | rs2896218 | 1.383 | 1.972 | 0.359 | 0.826 | 0.005 | -0.001 | -0.001 | 0 | 0.66 | 0.63 | 0.65 | 0.7 | 0.57 | 0.64 |
| 7 | 116920970 | rs916725 | 1.165 | 2.04 | 0.418 | 0.811 | -0.001 | -0.001 | -0.001 | 0 | 0.68 | 0.68 | 0.67 | 0.73 | 0.59 | 0.65 |
| 7 | 116928371 | rs4727847 | 2.142 | 2.185 | 0.514 | 0.845 | -0.002 | -0.025 | -0.025 | 0 | 0.57 | 0.48 | 0.39 | 0.33 | 0.5 | 0.29 |
| 7 | 116929065 | rs10487362 | -1.033 | 1.949 | 0.487 | 0.467 | -0.019 | -0.026 | -0.026 | 2.041 | 0.1 | 0.05 | 0.28 | 0.44 | 0.07 | 0.37 |
| 7 | 116935478 | rs17139625 | -1.229 | 1.943 | 0.597 | 0.463 | 0.005 | 0.124 | 0.124 | 0.123 | 0.11 | 0.18 | 0.3 | 0.43 | 0.07 | 0.37 |
| 7 | 116938020 | rs2301633 | -2.155 | 1.794 | 0.785 | 0.379 | 0.008 | 0.183 | 0.183 | 0.409 | 0.17 | 0.39 | 0.33 | 0.45 | 0.13 | 0.41 |
| 7 | 116940048 | rs10227271 | -2.081 | 1.799 | 0.718 | 0.368 | 0.006 | 0.164 | 0.164 | 0.327 | 0.16 | 0.35 | 0.33 | 0.47 | 0.12 | 0.41 |
| 7 | 116943135 | rs6947329 | 1.701 | 1.501 | 0.153 | -0.369 | 0.007 | 0.003 | 0.003 | 0 | 0.48 | 0.48 | 0.4 | 0.59 | 0.39 | 0.37 |
| 7 | 116954785 | rs39312 | -1.764 | 1.775 | 0.156 | -0.737 | 0.006 | -0.001 | -0.001 | 0.014 | 0.4 | 0.51 | 0.45 | 0.39 | 0.39 | 0.47 |
| 7 | 116988685 | rs916727 | -1.582 | 2.126 | 0.171 | -0.858 | -0.003 | -0.157 | -0.157 | 0 | 0.33 | 0.48 | 0.21 | 0 | 0.32 | 0.19 |
| 7 | 116998176 | rs10248393 | -2.238 | 1.856 | 0.046 | -0.827 | -0.004 | -0.006 | -0.006 | 0 | 0.4 | 0.33 | 0.44 | 0.4 | 0.38 | 0.47 |
| 7 | 117024820 | rs1029396 | -0.299 | 1.774 | -0.011 | -0.765 | -0.007 | 0.028 | 0.028 | 0.005 | 0.09 | 0.08 | 0.25 | 0.42 | 0.08 | 0.36 |
| 7 | 117040117 | rs2188554 | -0.723 | 1.751 | -0.044 | -0.975 | 0.002 | -0.055 | -0.055 | 0 | 0.2 | 0.24 | 0.12 | 0 | 0.22 | 0.07 |
| 7 | 117045256 | rs1119800 | -1.923 | 1.671 | -0.775 | -1.457 | -0.039 | 0.061 | 0.061 | 0.005 | 0.89 | 1 | 0.95 | 1 | 0.88 | 1 |
| 7 | 117052915 | rs994996 | -2.679 | 1.513 | -0.303 | -1.298 | -0.078 | -0.041 | -0.041 | 5.535 | 0.44 | 0.9 | 0.51 | 0.39 | 0.41 | 0.51 |
| 7 | 117058220 | rs10487364 | 0.375 | 1.49 | -0.235 | -1.277 | 0.006 | 0.031 | 0.031 | 3.816 | 0.11 | 0.03 | 0.07 | 0 | 0.09 | 0.04 |
| 7 | 117059654 | rs10264450 | -1.602 | 1.532 | -0.116 | -1.316 | -0.001 | -0.012 | -0.012 | 0.112 | 0.31 | 0.24 | 0.19 | 0 | 0.31 | 0.11 |
| 7 | 117077110 | rs10953846 | -2.128 | 1.474 | 0.101 | -1.108 | -0.005 | -0.003 | -0.003 | 0.112 | 0.41 | 0.37 | 0.45 | 0.4 | 0.38 | 0.47 |
| 7 | 117100046 | rs13222797 | 2.389 | 1.49 | 0.299 | -1.093 | 0.005 | -0.003 | -0.003 | 0 | 0.57 | 0.49 | 0.53 | 0.59 | 0.56 | 0.49 |
| 7 | 117105755 | rs43029 | -2.572 | 1.37 | -0.071 | -1.21 | -0.088 | -0.035 | -0.035 | 0 | 0.46 | 0.9 | 0.51 | 0.41 | 0.44 | 0.51 |
| 7 | 117128942 | rs756665 | -2.575 | 1.378 | -0.067 | -1.391 | -0.088 | 0 | 0 | 0 | 0.46 | 0.9 | 0.51 | 0.39 | 0.44 | 0.5 |

| | | | | | | | | | | | | | | | | |
|---|---|---|---|---|---|---|---|---|---|---|---|---|---|---|---|---|
| 7 | 117140095 | rs10487368 | -1.326 | 1.475 | 0.048 | -1.397 | -0.001 | -0.003 | -0.003 | 1.808 | 0.21 | 0.22 | 0.34 | 0.38 | 0.2 | 0.43 |
| 7 | 117147547 | rs2237723 | -1.341 | 1.478 | 0.048 | -1.397 | -0.002 | -0.004 | -0.004 | 0 | 0.21 | 0.22 | 0.34 | 0.38 | 0.2 | 0.43 |
| 7 | 117164439 | rs213938 | -1.348 | 1.446 | 0.149 | -1.428 | 0.008 | -0.097 | -0.097 | 0 | 0.25 | 0.64 | 0.17 | 0 | 0.24 | 0.07 |
| 7 | 117176738 | rs1800503 | 0.387 | 1.482 | 0.056 | -1.352 | 0.006 | 0.008 | 0.008 | 4.709 | 0.11 | 0.05 | 0.07 | 0 | 0.09 | 0.04 |
| 7 | 117178754 | rs17449197 | 0.3 | 1.567 | -0.004 | -1.367 | -0.03 | 0.073 | 0.073 | 0 | 0.12 | 0.01 | 0.07 | 0 | 0.15 | 0.04 |
| 7 | 117181509 | rs4148703 | -0.332 | 1.459 | 0.061 | -1.289 | -0.006 | 0.022 | 0.022 | 0.008 | 0.09 | 0.08 | 0.25 | 0.38 | 0.08 | 0.36 |
| 7 | 117190703 | rs213945 | -0.681 | 1.513 | 0.151 | -1.346 | -0.03 | 0.039 | 0.039 | 1.669 | 0.2 | 0.13 | 0.12 | 0 | 0.23 | 0.07 |
| 7 | 117199533 | rs213950 | -2.71 | 1.449 | 0.064 | -1.374 | -0.114 | -0.02 | -0.02 | 0 | 0.46 | 0.93 | 0.51 | 0.39 | 0.44 | 0.5 |
| 7 | 117200899 | rs10487372 | -0.206 | 1.482 | 0.222 | -1.372 | -0.002 | -0.038 | -0.038 | 0 | 0.12 | 0.15 | 0.09 | 0 | 0.13 | 0.08 |
| 7 | 117207860 | rs718829 | -2.224 | 1.379 | 0.417 | -1.329 | -0.011 | 0.001 | 0.001 | 0 | 0.42 | 0.36 | 0.46 | 0.39 | 0.43 | 0.5 |
| 7 | 117214089 | rs3808185 | -1.054 | 1.484 | 0.52 | -1.251 | 0.038 | 0.191 | 0.191 | 0.589 | 0.11 | 0.29 | 0.28 | 0.39 | 0.11 | 0.39 |
| 7 | 117214125 | rs3808184 | -1.901 | 1.407 | 0.471 | -1.288 | 0.005 | 0.003 | 0.003 | 2.103 | 0.31 | 0.37 | 0.39 | 0.39 | 0.34 | 0.45 |
| 7 | 117225159 | rs6949974 | -2.373 | 1.401 | 0.396 | -1.287 | -0.026 | -0.019 | -0.019 | 0 | 0.34 | 0.74 | 0.43 | 0.39 | 0.33 | 0.44 |
| 7 | 117229167 | rs213963 | -2.62 | 1.371 | 0.201 | -1.324 | -0.097 | -0.022 | -0.022 | 0 | 0.44 | 0.91 | 0.52 | 0.39 | 0.43 | 0.5 |
| 7 | 117231852 | rs11978434 | -1.018 | 1.259 | 0.424 | -1.702 | -0.033 | 0.037 | 0.037 | 1.676 | 0.21 | 0.15 | 0.14 | 0 | 0.26 | 0.1 |
| 7 | 117246315 | rs2299445 | -1.007 | 0.78 | 1.335 | -1.478 | 0.016 | 0.126 | 0.126 | 0.947 | 0.11 | 0.28 | 0.19 | 0.29 | 0.08 | 0.21 |
| 7 | 117252874 | rs213980 | -0.883 | 0.785 | 1.208 | -1.673 | -0.008 | 0.007 | 0.007 | 0.47 | 0.21 | 0.18 | 0.13 | 0.01 | 0.22 | 0.19 |
| 7 | 117256712 | rs17451754 | 0.314 | 0.818 | 0.955 | -1.694 | -0.044 | 0.076 | 0.076 | 0 | 0.13 | 0.01 | 0.11 | 0 | 0.16 | 0.04 |
| 7 | 117261293 | rs213987 | -2.148 | 0.738 | 1.097 | -1.663 | -0.04 | 0.024 | 0.024 | 0.889 | 0.37 | 0.86 | 0.38 | 0.3 | 0.33 | 0.44 |
| 7 | 117282644 | rs1800130 | -0.618 | 0.747 | 1.164 | -1.641 | -0.01 | -0.033 | -0.033 | 0 | 0.07 | 0.18 | 0.03 | 0 | 0.04 | 0.05 |
| 7 | 117296630 | rs11974545 | -0.456 | 0.795 | 1.163 | -1.64 | -0.034 | 0.089 | 0.089 | 0.774 | 0.2 | 0.08 | 0.12 | 0.01 | 0.21 | 0.19 |
| 7 | 117307108 | rs1800136 | -0.882 | 0.747 | 1.169 | -1.632 | -0.001 | -0.027 | -0.027 | 0 | 0.23 | 0.25 | 0.15 | 0.01 | 0.25 | 0.23 |
| 7 | 117308413 | rs1042180 | -0.897 | 0.759 | 0.96 | -1.627 | -0.001 | -0.03 | -0.03 | 0 | 0.23 | 0.25 | 0.15 | 0.02 | 0.25 | 0.23 |
| 7 | 117312350 | rs11764070 | -0.904 | 0.766 | 0.934 | -1.624 | -0.002 | -0.024 | -0.024 | 0 | 0.23 | 0.25 | 0.15 | 0.01 | 0.25 | 0.23 |
| 7 | 117314915 | rs13237279 | -0.21 | 0.77 | 0.92 | -1.519 | 0 | 0.025 | 0.025 | 0 | 0.09 | 0.09 | 0.17 | 0.29 | 0.08 | 0.21 |
| 7 | 117324019 | rs6466618 | -1.258 | 0.677 | 0.684 | -1.703 | -0.006 | 0.044 | 0.044 | 0.02 | 0.33 | 0.24 | 0.2 | 0.01 | 0.34 | 0.27 |
| 7 | 117324588 | rs17452984 | -1.236 | 0.678 | 0.689 | -1.702 | -0.006 | 0.039 | 0.039 | 0.02 | 0.33 | 0.23 | 0.2 | 0.01 | 0.34 | 0.27 |
| 7 | 117337484 | rs1208526 | -2.041 | 0.645 | 0.448 | -1.646 | -0.039 | 0.09 | 0.09 | 0.236 | 0.48 | 0.91 | 0.43 | 0.32 | 0.42 | 0.59 |
| 7 | 117347082 | rs11768272 | -0.756 | 0.659 | 0.909 | -1.739 | 0.002 | -0.045 | -0.045 | 0.179 | 0.24 | 0.25 | 0.16 | 0.01 | 0.28 | 0.22 |
| 7 | 117352536 | rs4730797 | -0.754 | 0.721 | 0.916 | -1.739 | 0.003 | -0.048 | -0.048 | 3.06 | 0.24 | 0.25 | 0.16 | 0.01 | 0.29 | 0.22 |
| 7 | 117354866 | rs3735645 | -2.171 | 0.383 | 0.564 | -1.813 | -0.034 | 0.061 | 0.061 | 0.357 | 0.52 | 0.89 | 0.43 | 0.35 | 0.43 | 0.59 |
| 7 | 117364998 | rs6973257 | -1.342 | 0.41 | 1.118 | -2.259 | 0.003 | -0.074 | -0.074 | 0 | 0.27 | 0.66 | 0.19 | 0.01 | 0.29 | 0.24 |
| 7 | 117372998 | rs4730799 | -1.161 | 0.48 | 1.227 | -2.249 | 0.026 | -0.157 | -0.157 | 3.575 | 0.26 | 0.5 | 0.18 | 0.01 | 0.28 | 0.26 |
| 7 | 117378356 | rs739798 | -1.792 | 0.356 | 1.026 | -2.309 | -0.015 | 0.216 | 0.216 | 0 | 0.39 | 0.88 | 0.26 | 0.01 | 0.35 | 0.28 |
| 7 | 117381393 | rs10278481 | -0.775 | 0.385 | 1.177 | -2.288 | 0.032 | -0.112 | -0.112 | 1.033 | 0.23 | 0.31 | 0.16 | 0.01 | 0.28 | 0.22 |
| 7 | 117387108 | rs10953849 | 0.388 | 0.36 | 1.216 | -2.191 | -0.03 | -0.068 | -0.068 | 0 | 0.12 | 0.25 | 0.07 | 0 | 0.06 | 0.03 |
| 7 | 117389972 | rs10247163 | -0.46 | 0.481 | 1.171 | -2.237 | 0.002 | -0.04 | -0.04 | 0 | 0.21 | 0.18 | 0.14 | 0.02 | 0.26 | 0.21 |
| 7 | 117393516 | rs11974874 | -0.293 | 0.463 | 1.198 | -2.232 | -0.046 | -0.095 | -0.095 | 0 | 0.15 | 0.64 | 0.1 | 0 | 0.07 | 0.03 |
| 7 | 117401398 | rs10258815 | -1.559 | 0.488 | 0.925 | -2.226 | -0.024 | 0.177 | 0.177 | 0 | 0.37 | 0.88 | 0.26 | 0.02 | 0.34 | 0.26 |
| 7 | 117406238 | rs10254610 | -1.743 | 0.494 | 0.911 | -2.227 | -0.031 | 0.223 | 0.223 | 0 | 0.39 | 0.92 | 0.27 | 0.02 | 0.35 | 0.29 |
| 7 | 117411796 | rs10249282 | -0.658 | 0.6 | 1.374 | -1.955 | -0.017 | -0.111 | -0.111 | 0 | 0.17 | 0.44 | 0.12 | 0 | 0.12 | 0.09 |
| 7 | 117414080 | rs10235887 | -0.883 | 0.609 | 0.822 | -1.95 | -0.024 | -0.123 | -0.123 | 0.852 | 0.18 | 0.63 | 0.13 | 0 | 0.12 | 0.09 |
| 7 | 117416677 | rs6977929 | -0.74 | 0.632 | -0.078 | -1.94 | -0.011 | -0.063 | -0.063 | 0 | 0.18 | 0.79 | 0.13 | 0 | 0.12 | 0.06 |
| 7 | 117445664 | rs929668 | 0.139 | 0.384 | -1.056 | -1.896 | -0.008 | 0.043 | 0.043 | 1.403 | 0.75 | 0.84 | 0.87 | 0.97 | 0.75 | 0.81 |
| 7 | 117449844 | rs10226236 | 0.741 | 0.505 | -1.441 | -1.911 | 0.004 | 0.264 | 0.264 | 0.554 | 0.52 | 0.15 | 0.71 | 0.97 | 0.59 | 0.67 |
| 7 | 117451124 | rs6973399 | 0.613 | 0.919 | -1.139 | -0.848 | -0.045 | 0.057 | 0.057 | 0 | 0.55 | 0.21 | 0.64 | 0.68 | 0.57 | 0.4 |
| 7 | 117453157 | rs1477113 | 0.385 | 0.664 | -0.694 | -0.701 | 0.003 | 0.022 | 0.022 | 0 | 0.51 | 0.49 | 0.61 | 0.72 | 0.57 | 0.47 |
| 7 | 117456439 | rs9969163 | 0.133 | 0.684 | -0.431 | -0.799 | 0.011 | 0.041 | 0.041 | 0 | 0.56 | 0.68 | 0.65 | 0.72 | 0.64 | 0.53 |
| 7 | 117460254 | rs10487377 | 0.3 | 0.559 | -0.633 | -0.62 | 0.002 | 0.011 | 0.011 | 0 | 0.51 | 0.58 | 0.62 | 0.72 | 0.57 | 0.48 |
| 7 | 117471549 | rs13224934 | 0.331 | 0.618 | -1.02 | -0.596 | 0.004 | 0.029 | 0.029 | 0 | 0.5 | 0.48 | 0.61 | 0.71 | 0.57 | 0.48 |
| 7 | 117478028 | rs2193261 | -0.03 | 0.566 | -1.84 | 0.167 | -0.066 | 0.005 | 0.005 | 0.606 | 0.58 | 0.93 | 0.68 | 0.64 | 0.58 | 0.51 |
| 7 | 117486312 | rs17140446 | -0.436 | 0.465 | -1.513 | 0.645 | -0.001 | 0.001 | 0.001 | 0 | 0.35 | 0.42 | 0.52 | 0.56 | 0.32 | 0.49 |
| 7 | 117490334 | rs2024492 | -0.232 | 0.397 | -1.815 | 0.585 | -0.053 | 0.031 | 0.031 | 0.403 | 0.55 | 0.93 | 0.67 | 0.65 | 0.57 | 0.5 |
| 7 | 117493510 | rs2193264 | -0.281 | -0.519 | -1.307 | 0.762 | -0.012 | -0.001 | -0.001 | 0 | 0.17 | 0.23 | 0.09 | 0.15 | 0.08 | 0.34 |
| 7 | 117496319 | rs7776980 | 1.16 | -0.404 | -1.037 | 0.354 | -0.002 | 0.017 | 0.017 | 0.015 | 0.31 | 0.22 | 0.2 | 0.17 | 0.32 | 0.35 |
| 7 | 117497811 | rs2402267 | -0.866 | -0.406 | -1.763 | 0.493 | -0.127 | -0.025 | -0.025 | 0.232 | 0.74 | 0.97 | 0.65 | 0.76 | 0.64 | 0.92 |
| 7 | 117498461 | rs10245031 | -0.866 | -0.406 | -1.763 | 0.493 | -0.127 | -0.025 | -0.025 | 0.071 | 0.74 | 0.97 | 0.65 | 0.75 | 0.64 | 0.92 |
| 7 | 117502574 | rs7807677 | 0.421 | -0.468 | -1.474 | 0.685 | 0.001 | 0.011 | 0.011 | 0.012 | 0.53 | 0.66 | 0.4 | 0.41 | 0.55 | 0.49 |
| 7 | 117517293 | rs10953856 | 0.042 | -0.625 | -1.214 | 0.905 | -0.002 | 0.006 | 0.006 | 2.368 | 0.2 | 0.23 | 0.32 | 0.4 | 0.1 | 0.43 |
| 7 | 117533230 | rs1966438 | -0.686 | -0.459 | -0.929 | 0.777 | 0.006 | 0.003 | 0.003 | 2.345 | 0.41 | 0.59 | 0.52 | 0.64 | 0.33 | 0.57 |
| 7 | 117544180 | rs17488728 | 0.509 | -0.208 | -0.662 | 0.765 | 0.004 | 0.022 | 0.022 | 0 | 0.44 | 0.33 | 0.33 | 0.35 | 0.47 | 0.41 |
| 7 | 117544391 | rs17488749 | 0.926 | -0.264 | -0.757 | 0.605 | -0.004 | 0.008 | 0.008 | 0 | 0.11 | 0.07 | 0.1 | 0 | 0.15 | 0.04 |
| 7 | 117572304 | rs10272923 | 0.302 | -0.327 | -0.562 | 0.59 | 0.006 | 0.024 | 0.024 | 0 | 0.45 | 0.35 | 0.34 | 0.35 | 0.47 | 0.41 |
| 7 | 117576675 | rs6466637 | -0.578 | -0.241 | -0.291 | 0.35 | 0.004 | -0.003 | -0.003 | 0.064 | 0.77 | 0.79 | 0.77 | 0.81 | 0.72 | 0.89 |
| 7 | 117580053 | rs1024434 | 0.964 | -0.213 | -0.68 | 0.372 | 0.002 | 0.027 | 0.027 | 0 | 0.44 | 0.31 | 0.33 | 0.35 | 0.47 | 0.41 |
| 7 | 117585779 | rs7782815 | 1.936 | -0.201 | -0.926 | 0.223 | -0.016 | 0.107 | 0.107 | 0.393 | 0.31 | 0.07 | 0.19 | 0.12 | 0.3 | 0.35 |
| 7 | 117592956 | rs2188834 | -0.933 | -0.121 | -0.596 | 0.424 | -0.082 | -0.04 | -0.04 | 0.217 | 0.82 | 0.98 | 0.79 | 0.81 | 0.76 | 0.94 |
| 7 | 117599514 | rs916784 | 1.151 | -0.644 | -0.953 | 0.05 | -0.044 | 0.163 | 0.163 | 0.984 | 0.39 | 0.1 | 0.26 | 0.1 | 0.4 | 0.4 |
| 7 | 117617360 | rs1859031 | -0.877 | -0.326 | -0.882 | 0.992 | -0.026 | -0.068 | -0.068 | 2.447 | 0.89 | 0.99 | 0.69 | 0.74 | 0.87 | 0.83 |

| | | | | | | | | | | | | | | | | |
|---|---|---|---|---|---|---|---|---|---|---|---|---|---|---|---|---|
| 7 | 117621877 | rs7777391 | 0.539 | -0.2 | -0.354 | 0.485 | 0.001 | 0.019 | 0.019 | 0.363 | 0.59 | 0.61 | 0.4 | 0.19 | 0.63 | 0.48 |
| 7 | 117629898 | rs10487381 | 0.624 | -0.283 | -0.64 | 0.459 | -0.027 | 0.011 | 0.011 | 0 | 0.16 | 0.05 | 0.11 | 0.11 | 0.22 | 0.06 |
| 7 | 117631978 | rs1859032 | 0.319 | -0.267 | -0.32 | 0.57 | -0.01 | -0.044 | -0.044 | 0.178 | 0.64 | 0.83 | 0.46 | 0.55 | 0.64 | 0.57 |
| 7 | 117642252 | rs989996 | 1.933 | -0.374 | 0.389 | 0.542 | -0.002 | 0.028 | 0.028 | 0 | 0.47 | 0.36 | 0.32 | 0.09 | 0.43 | 0.51 |
| 7 | 117643268 | rs2074136 | 0.164 | -0.379 | 0.289 | 0.853 | 0.049 | 0.457 | 0.457 | 0.296 | 0.05 | 0.34 | 0.07 | 0.35 | 0.01 | 0.07 |
| 7 | 117656762 | rs13226962 | 0.504 | -0.226 | 0.104 | 0.674 | 0.009 | -0.006 | -0.006 | 0.565 | 0.15 | 0.18 | 0.11 | 0.12 | 0.22 | 0.06 |
| 7 | 117667791 | rs2402296 | 0.309 | -0.244 | -0.229 | 0.716 | -0.001 | 0.009 | 0.009 | 0.197 | 0.73 | 0.77 | 0.53 | 0.56 | 0.74 | 0.72 |
| 7 | 117669962 | rs7456706 | 0.652 | -0.243 | -0.192 | 0.773 | -0.01 | 0.003 | 0.003 | 0 | 0.14 | 0.08 | 0.11 | 0.12 | 0.22 | 0.06 |
| 7 | 117670654 | rs1319470 | 1.107 | -0.261 | -0.319 | 0.786 | 0.001 | -0.005 | -0.005 | 0 | 0.66 | 0.75 | 0.5 | 0.56 | 0.68 | 0.64 |
| 7 | 117679958 | rs2214226 | 0.428 | -0.339 | -0.378 | 0.814 | -0.072 | 0.003 | 0.003 | 0 | 0.77 | 0.97 | 0.81 | 0.81 | 0.67 | 0.83 |
| 7 | 117686907 | rs12673201 | 0.509 | -0.646 | 0.184 | 1.123 | -0.032 | 0.088 | 0.088 | 0.212 | 0.14 | 0.24 | 0.33 | 0.29 | 0.1 | 0.2 |
| 7 | 117704706 | rs12537079 | 2.048 | -0.826 | 0.106 | 0.856 | 0.037 | 0.26 | 0.26 | 2.263 | 0.5 | 0.14 | 0.34 | 0.11 | 0.53 | 0.52 |
| 7 | 117705257 | rs39745 | 1.366 | -0.772 | 0.137 | 0.893 | -0.021 | -0.003 | -0.003 | 0 | 0.11 | 0.07 | 0.1 | 0.12 | 0.19 | 0.07 |
| 7 | 117707028 | rs39746 | -1.275 | -0.861 | 0.492 | 0.984 | -0.009 | 0.035 | 0.035 | 0.168 | 0.32 | 0.38 | 0.48 | 0.5 | 0.33 | 0.33 |
| 7 | 117707293 | rs38818 | 1.789 | -1.375 | 0.991 | 1.245 | -0.003 | -0.003 | -0.003 | 0 | 0.62 | 0.37 | 0.45 | 0.22 | 0.73 | 0.59 |
| 7 | 117708747 | rs38819 | 1.086 | -1.274 | 0.726 | 1.318 | 0.002 | -0.004 | -0.004 | 0 | 0.78 | 0.73 | 0.8 | 0.52 | 0.83 | 0.79 |
| 7 | 117713573 | rs38825 | -0.017 | -0.934 | 0.127 | 1.434 | 0.004 | -0.022 | -0.022 | 0 | 0.85 | 0.86 | 0.84 | 0.6 | 0.86 | 0.85 |
| 7 | 117713915 | rs17140751 | -0.077 | -0.983 | 0.236 | 1.113 | -0.03 | 0.075 | 0.075 | 0.192 | 0.14 | 0.23 | 0.34 | 0.29 | 0.1 | 0.16 |
| 7 | 117721635 | rs10235327 | 0.61 | -0.435 | -0.323 | 0.15 | -0.002 | 0 | 0 | 0 | 0.37 | 0.32 | 0.33 | 0.17 | 0.46 | 0.49 |
| 7 | 117726480 | rs10272234 | 0.295 | -0.514 | -0.123 | 0.52 | 0 | -0.001 | -0.001 | 0 | 0.58 | 0.46 | 0.38 | 0.45 | 0.58 | 0.63 |
| 7 | 117739400 | rs6466643 | 0.696 | -0.625 | -0.153 | 0.418 | -0.019 | 0.166 | 0.166 | 7.454 | 0.52 | 0.13 | 0.32 | 0.22 | 0.56 | 0.56 |
| 7 | 117744660 | rs6953304 | 0.69 | -0.646 | -0.183 | 0.389 | -0.02 | 0.169 | 0.169 | 6.688 | 0.52 | 0.12 | 0.31 | 0.22 | 0.56 | 0.56 |
| 7 | 117781277 | rs4730826 | -1.105 | -0.636 | -0.011 | 0.695 | 0.003 | 0.068 | 0.068 | 0 | 0.09 | 0.24 | 0.17 | 0.24 | 0.06 | 0.08 |
| 7 | 117783250 | rs7811055 | -0.013 | -0.618 | -0.073 | 0.731 | -0.003 | -0.012 | -0.012 | 1.052 | 0.06 | 0.03 | 0.12 | 0.18 | 0.05 | 0.05 |
| 7 | 117800733 | rs12112697 | 0.025 | -0.739 | -0.226 | 0.418 | -0.001 | -0.002 | -0.002 | 0 | 0.65 | 0.37 | 0.49 | 0.46 | 0.63 | 0.66 |
| 7 | 117823289 | rs3757800 | 0.524 | -0.978 | 0.426 | 0.918 | -0.031 | 0.264 | 0.264 | 0.086 | 0.11 | 0.39 | 0.17 | 0.28 | 0.04 | 0.2 |
| 7 | 117827631 | rs2215628 | 1.092 | -1.105 | 0.229 | 0.862 | -0.026 | 0.06 | 0.06 | 1.704 | 0.57 | 0.19 | 0.41 | 0.29 | 0.57 | 0.57 |
| 7 | 117828710 | rs7792264 | 0.287 | -0.956 | 0.043 | 0.823 | -0.001 | 0 | 0 | 0 | 0.06 | 0.1 | 0.12 | 0.1 | 0.05 | 0.05 |
| 7 | 117832138 | rs1061731 | -0.669 | -0.962 | 0.047 | 0.834 | -0.013 | -0.004 | -0.004 | 0 | 0.77 | 0.77 | 0.74 | 0.75 | 0.67 | 0.85 |
| 7 | 117832457 | rs3808175 | 0.311 | -0.953 | 0.046 | 0.822 | -0.001 | 0 | 0 | 0 | 0.06 | 0.08 | 0.12 | 0.1 | 0.05 | 0.05 |
| 7 | 117835284 | rs17580223 | 1.749 | -1.071 | -0.397 | 0.423 | -0.026 | 0.051 | 0.051 | 7.456 | 0.09 | 0.01 | 0.07 | 0 | 0.09 | 0.01 |
| 7 | 117848384 | rs10487390 | -0.795 | -1.028 | -0.137 | 0.675 | -0.017 | -0.005 | -0.005 | 0 | 0.77 | 0.77 | 0.75 | 0.75 | 0.67 | 0.85 |
| 7 | 117856291 | rs2058418 | 0.098 | -1.064 | -0.111 | 0.695 | 0 | 0 | 0 | 0.686 | 0.69 | 0.63 | 0.58 | 0.58 | 0.6 | 0.77 |
| 7 | 117869066 | rs11981239 | 0.494 | -1.258 | 0.092 | 0.927 | -0.039 | 0.281 | 0.281 | 0.377 | 0.12 | 0.44 | 0.18 | 0.29 | 0.04 | 0.2 |
| 7 | 117874308 | rs10267938 | 0.631 | -1.222 | 0.006 | 0.897 | -0.007 | -0.002 | -0.002 | 0.895 | 0.24 | 0.25 | 0.25 | 0.25 | 0.34 | 0.15 |
| 7 | 117903588 | rs17141162 | -0.459 | -1.236 | 0.003 | 0.895 | -0.002 | 0.013 | 0.013 | 0 | 0.08 | 0.15 | 0.16 | 0.17 | 0.06 | 0.08 |
| 7 | 117907104 | rs41914 | 0.604 | -1.229 | -0.175 | 0.856 | -0.007 | -0.002 | -0.002 | 0 | 0.24 | 0.25 | 0.25 | 0.25 | 0.34 | 0.15 |
| 7 | 117917719 | rs41923 | -0.951 | -1.232 | -0.343 | 0.725 | -0.067 | 0.07 | 0.07 | 2.904 | 0.44 | 0.91 | 0.59 | 0.71 | 0.44 | 0.42 |
| 7 | 117960973 | rs41943 | -1.017 | -1.16 | -0.923 | 0.544 | -0.136 | 0.174 | 0.174 | 8.005 | 0.56 | 0.97 | 0.7 | 0.83 | 0.52 | 0.52 |
| 7 | 117965390 | rs41953 | 0.085 | -0.536 | 0.318 | 1.151 | -0.07 | -0.038 | -0.038 | 0.638 | 0.25 | 0.1 | 0.25 | 0.31 | 0.36 | 0.17 |
| 7 | 117972361 | rs42488 | -0.396 | -0.542 | 0.361 | 1.199 | -0.062 | -0.043 | -0.043 | 0 | 0.29 | 0.19 | 0.37 | 0.42 | 0.4 | 0.25 |
| 7 | 118041733 | rs6959273 | -0.372 | -0.36 | -0.054 | 1.257 | -0.099 | -0.054 | -0.054 | 0 | 0.76 | 0.95 | 0.76 | 0.71 | 0.64 | 0.82 |
| 7 | 118046168 | rs917861 | 0.309 | -0.649 | 0.496 | 1.415 | -0.03 | 0.279 | 0.279 | 0.384 | 0.11 | 0.42 | 0.18 | 0.29 | 0.04 | 0.2 |
| 7 | 118062404 | rs4730864 | 0.239 | -0.648 | 0.258 | 1.4 | -0.054 | -0.033 | -0.033 | 0.606 | 0.71 | 0.8 | 0.63 | 0.59 | 0.6 | 0.75 |
| 7 | 118069861 | rs7782386 | 0.239 | -0.65 | 0.249 | 1.398 | -0.056 | -0.035 | -0.035 | 0 | 0.71 | 0.8 | 0.63 | 0.59 | 0.6 | 0.75 |
| 7 | 118070496 | rs4730865 | 0.238 | -0.656 | 0.243 | 1.395 | -0.063 | -0.039 | -0.039 | 2.853 | 0.71 | 0.82 | 0.63 | 0.59 | 0.6 | 0.75 |
| 7 | 118099945 | rs881802 | -0.362 | -0.64 | 0.258 | 1.402 | -0.076 | -0.037 | -0.037 | 0 | 0.76 | 0.88 | 0.76 | 0.71 | 0.64 | 0.82 |
| 7 | 118106436 | rs4727871 | 0.469 | -0.665 | 0.708 | 1.387 | -0.039 | 0.14 | 0.14 | 0 | 0.09 | 0.21 | 0.16 | 0.28 | 0.04 | 0.2 |
| 7 | 118119143 | rs39457 | 0.326 | -0.323 | 0.219 | 1.274 | -0.001 | 0.005 | 0.005 | 0 | 0.95 | 0.92 | 0.86 | 0.88 | 0.96 | 0.92 |
| 7 | 118151455 | rs39489 | 0.04 | -0.281 | 0.346 | 1.33 | -0.04 | -0.018 | -0.018 | 0 | 0.26 | 0.16 | 0.26 | 0.31 | 0.36 | 0.19 |
| 7 | 118156957 | rs1029405 | 0.495 | -0.303 | 0.466 | 1.38 | -0.004 | -0.003 | -0.003 | 0 | 0.68 | 0.65 | 0.59 | 0.56 | 0.6 | 0.72 |
| 7 | 118160303 | rs1010181 | -0.05 | -0.326 | 0.345 | 1.379 | -0.089 | -0.074 | -0.074 | 0 | 0.27 | 0.08 | 0.28 | 0.41 | 0.35 | 0.25 |
| 7 | 118170515 | rs9641597 | 0.045 | -0.311 | 0.373 | 1.335 | 0.006 | -0.007 | -0.007 | 0.056 | 0.95 | 0.85 | 0.88 | 0.97 | 0.96 | 0.98 |
| 7 | 118190093 | rs4615491 | 0.712 | -0.337 | 0.51 | 1.333 | 0.027 | 0.019 | 0.019 | 0.029 | 0.63 | 0.49 | 0.56 | 0.51 | 0.56 | 0.71 |
| 7 | 118203255 | rs11984235 | 0.252 | -0.611 | 0.355 | 1.339 | -0.02 | 0.088 | 0.088 | 1.453 | 0.19 | 0.41 | 0.25 | 0.27 | 0.15 | 0.21 |
| 7 | 118216736 | rs1510498 | 1.288 | -0.567 | -0.121 | 1.335 | -0.007 | 0.096 | 0.096 | 0.268 | 0.46 | 0.08 | 0.33 | 0.29 | 0.45 | 0.51 |
| 7 | 118222185 | rs1510495 | 0.003 | -0.61 | 0.012 | 1.355 | -0.028 | 0.035 | 0.035 | 3.34 | 0.22 | 0.62 | 0.26 | 0.27 | 0.15 | 0.21 |
| 7 | 118222312 | rs1355313 | 1.09 | -0.516 | -0.41 | 1.14 | -0.04 | 0.074 | 0.074 | 0.828 | 0.45 | 0.03 | 0.33 | 0.29 | 0.45 | 0.51 |
| 7 | 118225898 | rs778880 | 1.259 | -0.584 | -0.449 | 1.147 | -0.035 | 0.055 | 0.055 | 0.299 | 0.37 | 0.1 | 0.33 | 0.26 | 0.42 | 0.33 |
| 7 | 118228470 | rs1588447 | 1.387 | -0.102 | 0.04 | 1.41 | -0.085 | 0.151 | 0.151 | 0.577 | 0.46 | 0.07 | 0.36 | 0.24 | 0.52 | 0.37 |
| 7 | 118229822 | rs304525 | 1.025 | -0.092 | 0.538 | 1.525 | -0.008 | 0.05 | 0.05 | 0 | 0.52 | 0.33 | 0.44 | 0.37 | 0.54 | 0.45 |
| 7 | 118252557 | rs7781794 | 1.227 | -0.19 | 0.786 | 1.492 | 0 | 0.112 | 0.112 | 0.103 | 0.5 | 0.3 | 0.41 | 0.25 | 0.54 | 0.39 |
| 7 | 118263380 | rs13237572 | 1.041 | -0.216 | 0.992 | 1.504 | -0.001 | 0.029 | 0.029 | 0.003 | 0.52 | 0.46 | 0.43 | 0.26 | 0.54 | 0.4 |
| 7 | 118267904 | rs6954612 | -0.172 | -0.133 | 1.169 | 1.65 | -0.005 | 0.025 | 0.025 | 2.149 | 0.23 | 0.29 | 0.35 | 0.31 | 0.21 | 0.2 |
| 7 | 118274228 | rs960885 | 0.738 | 0.084 | 0.93 | 1.191 | 0.001 | 0.08 | 0.08 | 0.086 | 0.93 | 0.75 | 0.9 | 0.84 | 0.97 | 0.89 |
| 7 | 118275496 | rs7800988 | -0.615 | 0.103 | 0.91 | 1.202 | -0.005 | 0.028 | 0.028 | 0.015 | 0.46 | 0.64 | 0.53 | 0.62 | 0.46 | 0.52 |
| 7 | 118291975 | rs6968432 | -0.552 | -0.005 | 0.916 | 1.209 | 0.002 | -0.009 | -0.009 | 0.026 | 0.44 | 0.47 | 0.5 | 0.62 | 0.45 | 0.52 |
| 7 | 118297844 | rs9641609 | 1.036 | 0.049 | 0.471 | 1.448 | -0.098 | 0.064 | 0.064 | 0.397 | 0.46 | 0.05 | 0.39 | 0.3 | 0.52 | 0.39 |
| 7 | 118300390 | rs925351 | 1.039 | 0.089 | 0.498 | 1.062 | -0.091 | 0.159 | 0.159 | 0.266 | 0.46 | 0.06 | 0.38 | 0.22 | 0.52 | 0.38 |

| | | | | | | | | | | | | | | | | |
|---|---|---|---|---|---|---|---|---|---|---|---|---|---|---|---|---|
| 7 | 118301283 | rs4345498 | 0.576 | -0.011 | 1.007 | 1.104 | -0.014 | 0.044 | 0.044 | 0.858 | 0.85 | 0.63 | 0.79 | 0.77 | 0.9 | 0.86 |
| 7 | 118302687 | rs12532763 | -0.085 | 0.003 | 0.99 | 1.094 | 0.004 | 0.008 | 0.008 | 0.104 | 0.37 | 0.41 | 0.4 | 0.55 | 0.37 | 0.48 |
| 7 | 118316310 | rs6978807 | 0.982 | -0.743 | 0.63 | 0.896 | 0.006 | 0.201 | 0.201 | 1.177 | 0.49 | 0.06 | 0.46 | 0.2 | 0.59 | 0.41 |
| 7 | 118322312 | rs6972870 | 1.001 | -0.756 | 0.726 | 0.896 | 0.005 | 0.199 | 0.199 | 1.088 | 0.49 | 0.05 | 0.46 | 0.2 | 0.58 | 0.4 |
| 7 | 118333259 | rs778862 | -0.231 | -0.623 | 0.731 | 0.936 | -0.006 | -0.014 | -0.014 | 0.023 | 0.12 | 0.26 | 0.09 | 0.09 | 0.09 | 0.06 |
| 7 | 118338267 | rs1523795 | 0.083 | -0.646 | 0.208 | 0.962 | 0.004 | -0.016 | -0.016 | 0 | 0.33 | 0.25 | 0.36 | 0.54 | 0.3 | 0.45 |
| 7 | 118341243 | rs1033064 | 0.922 | -0.556 | -0.44 | 1.003 | 0.009 | 0.168 | 0.168 | 0.702 | 0.49 | 0.05 | 0.46 | 0.25 | 0.59 | 0.41 |
| 7 | 118345178 | rs10265433 | -0.529 | -0.229 | 0.028 | 1.252 | -0.015 | 0.022 | 0.022 | 0.013 | 0.35 | 0.46 | 0.37 | 0.54 | 0.3 | 0.45 |
| 7 | 118355075 | rs6967747 | 0.546 | 0.085 | -0.303 | 1.57 | -0.015 | -0.075 | -0.075 | 0.01 | 0.75 | 0.83 | 0.7 | 0.5 | 0.77 | 0.84 |
| 7 | 118356108 | rs12531671 | -0.861 | 0.058 | 0.764 | 1.291 | 0.075 | -0.006 | -0.006 | 0 | 0.07 | 0.27 | 0.06 | 0 | 0.09 | 0.03 |
| 7 | 118358453 | rs6466675 | 0.725 | -0.079 | 0.972 | 1.248 | -0.013 | -0.039 | -0.039 | 0 | 0.71 | 0.76 | 0.66 | 0.5 | 0.68 | 0.84 |
| 7 | 118367111 | rs1523794 | -1.334 | -0.335 | 0.828 | 1.263 | -0.012 | 0.012 | 0.012 | 1.239 | 0.16 | 0.73 | 0.21 | 0.21 | 0.13 | 0.13 |
| 7 | 118372845 | rs1916706 | -0.273 | -0.339 | 1.379 | 1.263 | 0.021 | 0.116 | 0.116 | 0 | 0.1 | 0.3 | 0.16 | 0.21 | 0.12 | 0.11 |
| 7 | 118377700 | rs1523810 | -0.864 | -0.337 | 1.443 | 1.264 | 0.015 | 0.125 | 0.125 | 0.159 | 0.12 | 0.42 | 0.17 | 0.21 | 0.12 | 0.11 |
| 7 | 118394453 | rs1916717 | -1.118 | -0.346 | 1.621 | 1.178 | 0.05 | 0.041 | 0.041 | 1.191 | 0.06 | 0.37 | 0.04 | 0 | 0.01 | 0.02 |
| 7 | 118405195 | rs1403861 | 1.419 | -0.527 | 1.448 | 1.132 | 0.019 | 0.074 | 0.074 | 0.421 | 0.56 | 0.15 | 0.48 | 0.29 | 0.63 | 0.7 |
| 7 | 118417803 | rs13233937 | 1.676 | -0.53 | 1.363 | 1.109 | -0.056 | 0.098 | 0.098 | 1.376 | 0.51 | 0.04 | 0.44 | 0.29 | 0.54 | 0.7 |
| 7 | 118422475 | rs722593 | 0.895 | -0.653 | 1.471 | 1.15 | -0.001 | -0.004 | -0.004 | 0 | 0.61 | 0.56 | 0.52 | 0.29 | 0.64 | 0.72 |
| 7 | 118429175 | rs10241665 | 0.041 | -0.669 | 1.319 | 1.497 | -0.024 | -0.077 | -0.077 | 0 | 0.25 | 0.08 | 0.3 | 0.4 | 0.22 | 0.11 |
| 7 | 118458376 | rs2699729 | 0.542 | -0.578 | 1.548 | 1.439 | 0.005 | 0.068 | 0.068 | 0 | 0.87 | 0.79 | 0.83 | 0.69 | 0.85 | 0.81 |
| 7 | 118473755 | rs1919912 | 1.742 | -0.569 | 1.469 | 1.511 | -0.018 | 0.106 | 0.106 | 1.089 | 0.51 | 0.07 | 0.46 | 0.28 | 0.61 | 0.66 |
| 7 | 118489031 | rs2699750 | -0.562 | -0.532 | 1.541 | 1.509 | -0.011 | 0.033 | 0.033 | 0.902 | 0.3 | 0.54 | 0.33 | 0.43 | 0.22 | 0.13 |
| 7 | 118489424 | rs6973556 | -0.102 | -0.478 | 0.897 | 1.535 | -0.018 | -0.069 | -0.069 | 0 | 0.25 | 0.13 | 0.29 | 0.43 | 0.21 | 0.11 |
| 7 | 118489794 | rs12056021 | -0.102 | -0.478 | 0.783 | 1.535 | -0.032 | -0.094 | -0.094 | 0 | 0.25 | 0.09 | 0.29 | 0.43 | 0.21 | 0.11 |
| 7 | 118515013 | rs7798483 | 1.642 | -0.517 | 0.546 | 1.384 | -0.054 | 0.057 | 0.057 | 0.595 | 0.48 | 0.07 | 0.49 | 0.33 | 0.59 | 0.66 |
| 7 | 118518479 | rs1554904 | 1.644 | -0.54 | 0.545 | 1.374 | -0.055 | 0.057 | 0.057 | 0.909 | 0.48 | 0.07 | 0.49 | 0.34 | 0.59 | 0.66 |
| 7 | 118534383 | rs2590634 | -0.736 | -0.421 | 1.183 | 1.589 | -0.013 | 0.004 | 0.004 | 1.961 | 0.32 | 0.52 | 0.31 | 0.33 | 0.22 | 0.13 |
| 7 | 118569946 | rs17141871 | -0.272 | -0.097 | 1.248 | 1.685 | 0.015 | 0.06 | 0.06 | 0.094 | 0.07 | 0.08 | 0.12 | 0.23 | 0.12 | 0.08 |
| 7 | 118570281 | rs9641615 | -1.333 | -0.337 | 1.785 | 1.443 | -0.003 | 0.004 | 0.004 | 0.615 | 0.46 | 0.54 | 0.46 | 0.59 | 0.47 | 0.24 |
| 7 | 118570452 | rs2402428 | 1.116 | -0.452 | 1.851 | 1.409 | -0.019 | -0.028 | -0.028 | 0 | 0.85 | 0.66 | 0.9 | 0.92 | 0.93 | 0.95 |
| 7 | 118570487 | rs10500071 | 1.824 | -0.462 | 1.789 | 1.403 | -0.084 | 0.048 | 0.048 | 1.362 | 0.45 | 0.08 | 0.47 | 0.33 | 0.5 | 0.73 |
| 7 | 118571724 | rs4370456 | -0.45 | -0.441 | 1.895 | 1.431 | -0.002 | 0.007 | 0.007 | 1.798 | 0.33 | 0.4 | 0.31 | 0.36 | 0.31 | 0.14 |
| 7 | 118572158 | rs2402431 | 2.08 | -0.473 | 1.886 | 1.342 | 0.002 | 0.002 | 0.002 | 1.118 | 0.69 | 0.4 | 0.7 | 0.69 | 0.68 | 0.85 |
| 7 | 118572753 | rs2896261 | 1.58 | -0.479 | 1.607 | 1.161 | 0.026 | 0.011 | 0.011 | 0.413 | 0.53 | 0.24 | 0.59 | 0.56 | 0.62 | 0.81 |
| 7 | 118612923 | rs12672097 | 0.232 | -0.426 | 1.58 | 1.296 | 0 | -0.006 | -0.006 | 0.725 | 0.22 | 0.16 | 0.22 | 0.36 | 0.18 | 0.12 |
| 7 | 118615427 | rs10242193 | 0.949 | -0.25 | 1.819 | 1.025 | -0.009 | -0.017 | -0.017 | 0 | 0.87 | 0.77 | 0.91 | 0.92 | 0.93 | 0.95 |
| 7 | 118634585 | rs7782688 | 1.672 | -0.073 | 1.91 | 1.279 | 0.025 | 0.049 | 0.049 | 1.122 | 0.78 | 0.42 | 0.76 | 0.67 | 0.77 | 0.86 |
| 7 | 118635459 | rs7787447 | -0.128 | -0.07 | 2.16 | 1.286 | 0.017 | 0.039 | 0.039 | 0.335 | 0.08 | 0.07 | 0.13 | 0.23 | 0.16 | 0.09 |
| 7 | 118636817 | rs6466686 | 2.268 | -0.095 | 2.098 | 1.11 | 0.003 | -0.001 | -0.001 | 0.048 | 0.68 | 0.34 | 0.7 | 0.67 | 0.65 | 0.83 |
| 7 | 118645137 | rs4730907 | -1.619 | -0.072 | 2.246 | 1.108 | 0.013 | 0.012 | 0.012 | 0.013 | 0.19 | 0.21 | 0.19 | 0.23 | 0.22 | 0.11 |
| 7 | 118669625 | rs4615492 | 0.104 | -0.562 | 1.918 | 1.236 | 0.011 | -0.028 | -0.028 | 0.274 | 0.24 | 0.19 | 0.26 | 0.42 | 0.15 | 0.11 |
| 7 | 118701195 | rs4730911 | 0.009 | -0.188 | 1.957 | 1.232 | 0.021 | 0.027 | 0.027 | 0.32 | 0.08 | 0.06 | 0.13 | 0.23 | 0.17 | 0.09 |
| 7 | 118742974 | rs10230384 | -0.849 | -0.368 | 2.199 | 0.716 | 0.024 | -0.052 | -0.052 | 0 | 0.13 | 0.49 | 0.09 | 0.05 | 0.13 | 0.04 |
| 7 | 118769366 | rs10275972 | -1.069 | -0.369 | 2.049 | 0.718 | 0.004 | -0.057 | -0.057 | 0 | 0.14 | 0.61 | 0.1 | 0.05 | 0.13 | 0.04 |
| 7 | 118777401 | rs10271820 | 1.582 | -0.423 | 1.799 | 0.757 | 0 | 0.012 | 0.012 | 1.307 | 0.46 | 0.47 | 0.47 | 0.27 | 0.5 | 0.71 |
| 7 | 118792218 | rs2402455 | 1.624 | -0.455 | 1.743 | 0.862 | -0.001 | -0.01 | -0.01 | 0.097 | 0.45 | 0.51 | 0.47 | 0.32 | 0.5 | 0.71 |
| 7 | 118796603 | rs10248244 | 1.436 | -0.482 | 1.729 | 0.845 | 0 | -0.011 | -0.011 | 0 | 0.45 | 0.52 | 0.47 | 0.32 | 0.5 | 0.71 |
| 7 | 118801966 | rs4449727 | 1.032 | -0.52 | 1.747 | 0.804 | 0 | 0.008 | 0.008 | 0 | 0.56 | 0.63 | 0.53 | 0.32 | 0.62 | 0.74 |
| 7 | 118850537 | rs6964033 | 0.809 | -0.481 | 1.805 | 0.791 | -0.004 | 0.013 | 0.013 | 0 | 0.58 | 0.67 | 0.53 | 0.33 | 0.62 | 0.74 |
| 7 | 118868426 | rs1489119 | 1.854 | -0.482 | 1.212 | 0.788 | -0.12 | 0.037 | 0.037 | 3.247 | 0.42 | 0.03 | 0.43 | 0.27 | 0.49 | 0.71 |
| 7 | 118873720 | rs12114016 | -0.216 | -0.451 | 1.584 | 0.783 | -0.001 | -0.01 | -0.01 | 0.078 | 0.11 | 0.08 | 0.06 | 0 | 0.12 | 0.03 |
| 7 | 118883867 | rs2896279 | -0.241 | -0.444 | 1.613 | 0.785 | -0.001 | -0.023 | -0.023 | 0 | 0.11 | 0.1 | 0.06 | 0 | 0.12 | 0.02 |
| 7 | 118920503 | rs10272061 | -1.311 | -0.42 | 1.193 | 0.735 | -0.015 | -0.103 | -0.103 | 0 | 0.16 | 0.61 | 0.1 | 0.01 | 0.13 | 0.03 |
| 7 | 118978523 | rs11975286 | 1.32 | -0.346 | 0.872 | 1.088 | -0.027 | 0.088 | 0.088 | 0.15 | 0.51 | 0.12 | 0.46 | 0.33 | 0.59 | 0.73 |
| 7 | 119000212 | rs170099 | 2.06 | -0.037 | 1.436 | 1.113 | 0.139 | 0.197 | 0.197 | 1.317 | 0.94 | 0.48 | 0.89 | 0.9 | 0.97 | 0.96 |
| 7 | 119006416 | rs1682503 | 2.06 | -0.037 | 1.436 | 1.113 | 0.139 | 0.197 | 0.197 | 1.317 | 0.94 | 0.48 | 0.89 | 0.9 | 0.97 | 0.96 |
| 7 | 119154756 | rs7785413 | 0.125 | -0.902 | 1.455 | 1.317 | 0.013 | -0.02 | -0.02 | 0 | 0.73 | 0.78 | 0.85 | 0.52 | 0.87 | 0.89 |
| 7 | 119161743 | rs4730922 | -1.372 | -0.463 | 1.504 | 1.213 | 0.039 | 0.011 | 0.011 | 0 | 0.19 | 0.59 | 0.18 | 0.23 | 0.25 | 0.13 |
| 7 | 119177845 | rs7455060 | -0.907 | -0.48 | 1.73 | 1.144 | 0.054 | -0.015 | -0.015 | 0.285 | 0.08 | 0.41 | 0.08 | 0.01 | 0.05 | 0.03 |
| 7 | 119189729 | rs4236610 | -1.861 | -0.476 | 1.706 | 1.151 | 0.022 | 0.003 | 0.003 | 0 | 0.2 | 0.67 | 0.2 | 0.23 | 0.25 | 0.13 |
| 7 | 119198764 | rs10279834 | -0.765 | -0.482 | 1.893 | 1.143 | 0.053 | -0.013 | -0.013 | 0 | 0.08 | 0.38 | 0.08 | 0.01 | 0.05 | 0.03 |
| 7 | 119201366 | rs12537239 | -0.37 | -0.417 | 1.77 | 1.209 | 0.044 | 0.043 | 0.043 | 0 | 0.11 | 0.23 | 0.11 | 0.22 | 0.2 | 0.1 |
| 7 | 119206096 | rs10228248 | -1.106 | -0.452 | 1.634 | 0.862 | 0.033 | -0.066 | -0.066 | 0.057 | 0.14 | 0.66 | 0.13 | 0.07 | 0.18 | 0.12 |
| 7 | 119270386 | rs10227401 | -1.102 | -0.452 | 1.913 | 0.862 | 0.053 | -0.072 | -0.072 | 0 | 0.14 | 0.57 | 0.13 | 0.07 | 0.18 | 0.12 |
| 7 | 119277137 | rs4130671 | 2.195 | -1.026 | 0.843 | 0.6 | -0.103 | -0.077 | -0.077 | 0 | 0.47 | 0.03 | 0.58 | 0.46 | 0.51 | 0.71 |
| 7 | 119301786 | rs11532787 | 0.573 | -1.065 | 0.957 | 1.051 | 0.01 | -0.071 | -0.071 | 0 | 0.15 | 0.04 | 0.09 | 0.3 | 0.1 | 0.07 |
| 7 | 119313222 | rs11978304 | 1.861 | -1.145 | 0.547 | 0.897 | -0.064 | -0.155 | -0.155 | 1.215 | 0.59 | 0.04 | 0.7 | 0.5 | 0.65 | 0.75 |
| 7 | 119326227 | rs10245797 | -0.305 | -1.036 | 1.321 | 0.661 | 0.082 | -0.061 | -0.061 | 0 | 0.12 | 0.39 | 0.12 | 0.06 | 0.19 | 0.13 |
| 7 | 119329075 | rs4449718 | 0.728 | -1.056 | 1.481 | 0.658 | 0.008 | -0.054 | -0.054 | 0 | 0.9 | 0.54 | 0.93 | 0.99 | 0.96 | 0.97 |

| | | | | | | | | | | | | | | | | |
|---|---|---|---|---|---|---|---|---|---|---|---|---|---|---|---|---|
| 7 | 119383120 | rs11978043 | 1.622 | -0.562 | 1.408 | 0.478 | -0.011 | 0.064 | 0.064 | 1.402 | 0.77 | 0.16 | 0.81 | 0.93 | 0.77 | 0.84 |
| 7 | 119388127 | rs2402489 | 0.002 | -0.559 | 1.663 | 0.48 | -0.006 | -0.045 | -0.045 | 0.225 | 0.92 | 0.8 | 0.96 | 1 | 0.96 | 0.98 |
| 7 | 119393615 | rs10500075 | 0.016 | -0.563 | 1.797 | 0.477 | 0.039 | -0.05 | -0.05 | 0.709 | 0.12 | 0.23 | 0.12 | 0.06 | 0.19 | 0.12 |
| 7 | 119399411 | rs12706258 | 1.091 | -0.511 | 1.805 | 0.453 | 0.04 | -0.122 | -0.122 | 0 | 0.8 | 0.55 | 0.84 | 0.94 | 0.77 | 0.86 |
| 7 | 119401720 | rs6973035 | 0.248 | -0.489 | 1.681 | 0.473 | 0.006 | -0.016 | -0.016 | 2.119 | 0.11 | 0.12 | 0.11 | 0.06 | 0.19 | 0.12 |
| 7 | 119408979 | rs17142320 | -0.665 | -0.484 | 1.634 | 0.472 | 0.065 | -0.08 | -0.08 | 1.26 | 0.14 | 0.52 | 0.14 | 0.07 | 0.19 | 0.13 |
| 7 | 119428982 | rs12536246 | 1.55 | -0.486 | 0.999 | 0.455 | -0.018 | 0.097 | 0.097 | 2.474 | 0.77 | 0.11 | 0.81 | 0.93 | 0.77 | 0.84 |
| 7 | 119456162 | rs4730939 | 1.362 | -0.479 | 1.091 | 0.457 | -0.001 | -0.02 | -0.02 | 0 | 0.78 | 0.26 | 0.81 | 0.93 | 0.77 | 0.85 |
| 7 | 119463832 | rs10085666 | 0.768 | -0.499 | 1.083 | 0.471 | -0.029 | -0.068 | -0.068 | 0.028 | 0.88 | 0.31 | 0.91 | 0.99 | 0.96 | 0.96 |
| 7 | 119469564 | rs12540390 | 0.185 | -0.458 | 0.398 | 0.34 | -0.008 | -0.076 | -0.076 | 0.401 | 0.19 | 0.05 | 0.12 | 0.3 | 0.15 | 0.08 |
| 7 | 119493570 | rs846427 | -1.985 | -0.216 | 0.364 | -0.106 | -0.078 | -0.108 | -0.108 | 0.653 | 0.56 | 0.97 | 0.47 | 0.54 | 0.57 | 0.32 |
| 7 | 119502677 | rs846433 | -0.838 | -0.065 | 1.248 | -0.69 | 0.002 | -0.059 | -0.059 | 0.504 | 0.11 | 0.51 | 0.07 | 0.01 | 0.04 | 0.03 |
| 7 | 119537370 | rs11764046 | -0.378 | 0.039 | 0.766 | -0.71 | 0.054 | -0.054 | -0.054 | 2.275 | 0.12 | 0.32 | 0.12 | 0.06 | 0.2 | 0.13 |
| 7 | 119564205 | rs10254361 | -1.801 | 0.006 | 0.198 | -0.751 | -0.005 | 0.051 | 0.051 | 1.801 | 0.23 | 0.78 | 0.19 | 0.07 | 0.24 | 0.16 |
| 7 | 119565056 | rs4730945 | -0.254 | 0.374 | 0.116 | -0.485 | 0.002 | -0.013 | -0.013 | 0 | 0.11 | 0.1 | 0.1 | 0.06 | 0.19 | 0.12 |
| 7 | 119572391 | rs10231641 | -1.21 | 0.037 | -0.725 | -1.002 | -0.034 | 0.138 | 0.138 | 1.981 | 0.79 | 0.98 | 0.85 | 1 | 0.76 | 0.85 |
| 7 | 119575680 | rs12112212 | -1.066 | 0.316 | 0.542 | -0.079 | -0.004 | 0.046 | 0.046 | 1.712 | 0.23 | 0.27 | 0.25 | 0.52 | 0.19 | 0.11 |
| 7 | 119577292 | rs12670725 | -1.066 | 0.316 | 0.542 | -0.079 | -0.004 | 0.046 | 0.046 | 1.712 | 0.23 | 0.27 | 0.25 | 0.52 | 0.19 | 0.11 |
| 7 | 119579473 | rs13234712 | -1.562 | 0.387 | 0.574 | -0.075 | -0.001 | 0 | 0 | 0 | 0.38 | 0.63 | 0.39 | 0.53 | 0.35 | 0.14 |
| 7 | 119605536 | rs10281213 | -1.398 | 0.118 | 1.785 | -0.682 | -0.017 | 0.023 | 0.023 | 1.98 | 0.09 | 0.54 | 0.07 | 0.07 | 0.01 | 0.05 |
| 7 | 119605900 | rs2190311 | -1.125 | 0.15 | 1.69 | -0.83 | 0.001 | -0.015 | -0.015 | 0 | 0.19 | 0.19 | 0.18 | 0.12 | 0.18 | 0.05 |
| 7 | 119659037 | rs3898475 | -2.757 | -0.008 | 1.662 | -0.825 | 0.063 | 0.061 | 0.061 | 5.054 | 0.3 | 0.9 | 0.28 | 0.25 | 0.17 | 0.12 |
| 7 | 119672864 | rs17520243 | 1.1 | 0.166 | 1.298 | -1.221 | -0.056 | 0.072 | 0.072 | 0.374 | 0.24 | 0 | 0.14 | 0.06 | 0.23 | 0.46 |
| 7 | 119695355 | rs6973461 | -3.447 | 0.811 | 2.762 | -0.292 | 0.041 | 0.079 | 0.079 | 5.569 | 0.31 | 0.88 | 0.28 | 0.22 | 0.22 | 0.12 |
| 7 | 119705787 | rs9655832 | -2.143 | 0.95 | 3.284 | -0.219 | -0.052 | -0.017 | -0.017 | 0 | 0.1 | 0.64 | 0.08 | 0.07 | 0.03 | 0.06 |
| 7 | 119706135 | rs10280627 | -1.726 | 0.956 | 3.485 | -0.218 | -0.004 | 0.024 | 0.024 | 2.16 | 0.08 | 0.56 | 0.06 | 0.05 | 0.03 | 0.05 |
| 7 | 119735196 | rs10226287 | -3.413 | 0.973 | 3.015 | -0.251 | 0.029 | 0.078 | 0.078 | 6.387 | 0.3 | 0.9 | 0.28 | 0.21 | 0.22 | 0.11 |
| 7 | 119761318 | rs757708 | -2.781 | 0.8 | 3.218 | -0.235 | -0.027 | -0.064 | -0.064 | 0 | 0.28 | 0.52 | 0.26 | 0.16 | 0.22 | 0.1 |
| 7 | 119765939 | rs7804857 | -3.307 | 1.271 | 2.925 | -0.004 | 0.015 | 0.102 | 0.102 | 0.875 | 0.29 | 0.86 | 0.28 | 0.16 | 0.27 | 0.1 |
| 7 | 119772461 | rs10258425 | -2.649 | 1.71 | 3.754 | 0.198 | 0.024 | -0.067 | -0.067 | 0 | 0.2 | 0.41 | 0.21 | 0.12 | 0.23 | 0.07 |
| 7 | 119809612 | rs7777091 | -2.37 | 1.95 | 3.592 | 0.243 | 0.04 | -0.06 | -0.06 | 0.373 | 0.16 | 0.38 | 0.18 | 0.11 | 0.19 | 0.07 |
| 7 | 119829589 | rs10224700 | -2.889 | 1.831 | 3.475 | 0.28 | -0.001 | -0.014 | -0.014 | 2.154 | 0.21 | 0.73 | 0.21 | 0.14 | 0.19 | 0.09 |
| 7 | 119862301 | rs2190183 | -3.242 | 1.764 | 4.05 | 0.258 | 0.001 | 0.052 | 0.052 | 2.159 | 0.23 | 0.85 | 0.23 | 0.14 | 0.23 | 0.1 |
| 7 | 119888754 | rs6960301 | -2.081 | 1.74 | 4.098 | 0.256 | 0.003 | -0.023 | -0.023 | 0 | 0.14 | 0.17 | 0.16 | 0.1 | 0.15 | 0.05 |
| 7 | 119912849 | rs3814463 | -2.889 | 1.798 | 4.454 | 0.281 | 0.002 | -0.031 | -0.031 | 0 | 0.18 | 0.73 | 0.22 | 0.14 | 0.19 | 0.09 |
| 7 | 119917158 | rs4730954 | 2.74 | 1.722 | 4.527 | 0.402 | -0.005 | -0.024 | -0.024 | 0 | 0.83 | 0.3 | 0.81 | 0.86 | 0.85 | 0.92 |
| 7 | 119944579 | rs12706276 | 3.09 | 1.723 | 4.512 | 0.4 | 0.001 | 0.008 | 0.008 | 2.229 | 0.81 | 0.2 | 0.79 | 0.86 | 0.81 | 0.91 |
| 7 | 119950420 | rs34850246 | -1.463 | 1.724 | 4.233 | 0.401 | 0.008 | -0.015 | -0.015 | 0 | 0.1 | 0.15 | 0.13 | 0.1 | 0.11 | 0.04 |
| 7 | 119955585 | rs2402526 | 3.411 | 1.876 | 4.298 | 0.712 | 0.004 | 0.034 | 0.034 | 3.69 | 0.81 | 0.15 | 0.78 | 0.86 | 0.81 | 0.91 |
| 7 | 120000704 | rs12706279 | 3.026 | 1.868 | 4.309 | 0.769 | -0.002 | -0.014 | -0.014 | 0 | 0.82 | 0.27 | 0.79 | 0.85 | 0.81 | 0.91 |
| 7 | 120003915 | rs7778213 | -1.541 | 1.898 | 4.332 | 0.782 | 0.005 | -0.011 | -0.011 | 0 | 0.09 | 0.11 | 0.13 | 0.06 | 0.1 | 0.03 |
| 7 | 120010469 | rs6943092 | 2.816 | 1.879 | 4.307 | 0.733 | 0.133 | 0.114 | 0.114 | 0.048 | 0.94 | 0.48 | 0.93 | 0.92 | 0.91 | 0.94 |
| 7 | 120017847 | rs10953910 | -0.822 | 1.829 | 3.196 | 0.739 | 0.009 | -0.011 | -0.011 | 0.039 | 0.07 | 0.09 | 0.12 | 0.04 | 0.09 | 0.03 |
| 7 | 120041216 | rs11769050 | -0.822 | 1.829 | 3.22 | 0.739 | 0.014 | -0.014 | -0.014 | 0 | 0.07 | 0.09 | 0.12 | 0.04 | 0.09 | 0.03 |
| 7 | 120073573 | rs6980067 | -2.609 | 1.733 | 3.204 | 1.441 | -0.002 | 0.258 | 0.258 | 1.21 | 0.16 | 0.52 | 0.35 | 0.57 | 0.13 | 0.09 |
| 7 | 120109606 | rs10276390 | -2.785 | 1.826 | 2.855 | 1.328 | 0.007 | 0.245 | 0.245 | 2.229 | 0.16 | 0.55 | 0.33 | 0.51 | 0.15 | 0.08 |
| 7 | 120110248 | rs10277228 | -1.852 | 1.841 | 2.715 | 1.294 | 0.049 | -0.027 | -0.027 | 0 | 0.09 | 0.3 | 0.15 | 0.04 | 0.12 | 0.04 |
| 7 | 120122040 | rs12531730 | -1.616 | 1.692 | 2.471 | 1.333 | -0.001 | 0.061 | 0.061 | 0.279 | 0.11 | 0.13 | 0.25 | 0.51 | 0.09 | 0.08 |
| 7 | 120136191 | rs723908 | -1.572 | 1.302 | 2.283 | 1.386 | 0.003 | 0.025 | 0.025 | 0.026 | 0.11 | 0.1 | 0.25 | 0.51 | 0.09 | 0.08 |
| 7 | 120143507 | rs1609181 | -3.263 | 1.374 | 2.885 | 1.551 | 0.138 | 0.083 | 0.083 | 0.365 | 0.05 | 0.42 | 0.07 | 0.03 | 0.06 | 0.04 |
| 7 | 120156307 | rs802333 | -1.833 | 1.38 | 2.664 | 1.59 | 0.017 | 0.378 | 0.378 | 0.369 | 0.07 | 0.31 | 0.19 | 0.51 | 0.04 | 0.07 |
| 7 | 120167316 | rs10488301 | -0.921 | 1.392 | 2.135 | 1.066 | -0.058 | 0.152 | 0.152 | 0.465 | 0.24 | 0.02 | 0.19 | 0 | 0.29 | 0.14 |
| 7 | 120194784 | rs10085878 | 1.045 | 1.064 | 3.008 | 2.055 | -0.015 | 0.221 | 0.221 | 0.018 | 0.95 | 0.87 | 0.85 | 0.52 | 0.98 | 0.93 |
| 7 | 120206385 | rs12537274 | 2.233 | 1.008 | 3.104 | 2.024 | -0.013 | 0.125 | 0.125 | 0 | 0.87 | 0.75 | 0.76 | 0.52 | 0.89 | 0.85 |
| 7 | 120224300 | rs4730964 | -0.928 | 1.053 | 2.534 | 1.929 | 0.001 | -0.018 | -0.018 | 0.088 | 0.06 | 0.09 | 0.09 | 0 | 0.09 | 0.07 |
| 7 | 120243576 | rs1527650 | -1.754 | 1.057 | 2.396 | 1.931 | 0.022 | -0.007 | -0.007 | 0 | 0.09 | 0.35 | 0.12 | 0.05 | 0.09 | 0.09 |
| 7 | 120259518 | rs7791822 | -2.064 | 1.089 | 1.717 | 1.968 | 0.002 | 0.23 | 0.23 | 0.652 | 0.18 | 0.53 | 0.31 | 0.52 | 0.18 | 0.18 |
| 7 | 120268303 | rs2402539 | -1.737 | 0.773 | 1.43 | 2.001 | -0.017 | 0.33 | 0.33 | 0.023 | 0.14 | 0.47 | 0.26 | 0.51 | 0.12 | 0.17 |
| 7 | 120273226 | rs4730967 | -2.205 | 1.354 | 1.524 | 2.031 | 0.003 | 0.061 | 0.061 | 4.188 | 0.24 | 0.67 | 0.41 | 0.51 | 0.27 | 0.23 |
| 7 | 120305149 | rs728115 | -1.947 | 1.342 | 1.943 | 2.293 | -0.001 | 0.093 | 0.093 | 0.276 | 0.07 | 0.09 | 0.23 | 0.41 | 0.1 | 0.09 |
| 7 | 120327284 | rs2896298 | -2.532 | 1.104 | 2.065 | 2.303 | -0.006 | 0.074 | 0.074 | 0.034 | 0.1 | 0.13 | 0.24 | 0.39 | 0.1 | 0.09 |
| 7 | 120335338 | rs4727914 | -3.232 | 1.319 | 2.41 | 2.284 | -0.002 | 0.058 | 0.058 | 1.904 | 0.27 | 0.62 | 0.4 | 0.49 | 0.27 | 0.27 |
| 7 | 120339674 | rs6944490 | -1.835 | 1.379 | 2.82 | 2.197 | 0.038 | -0.058 | -0.058 | 3.684 | 0.12 | 0.28 | 0.13 | 0.01 | 0.16 | 0.12 |
| 7 | 120359727 | rs11983106 | -2.068 | 1.418 | 2.636 | 2.298 | 0.003 | 0.001 | 0.001 | 0 | 0.05 | 0.08 | 0.09 | 0.05 | 0.08 | 0.1 |
| 7 | 120364028 | rs12538990 | -2.903 | 1.353 | 2.799 | 2.3 | 0.013 | 0.118 | 0.118 | 0.021 | 0.2 | 0.46 | 0.3 | 0.44 | 0.2 | 0.15 |
| 7 | 120372127 | rs734317 | -2.436 | 1.447 | 2.58 | 2.039 | -0.003 | 0.002 | 0.002 | 0.041 | 0.06 | 0.07 | 0.08 | 0.04 | 0.06 | 0.09 |
| 7 | 120405335 | rs10239799 | -3.383 | 1.155 | 3.629 | 2.443 | 0.005 | -0.009 | -0.009 | 3.224 | 0.23 | 0.77 | 0.32 | 0.42 | 0.2 | 0.22 |
| 7 | 120428607 | rs41622 | 2.409 | 1.478 | 3.776 | 2.223 | 0.039 | -0.043 | -0.043 | 0 | 0.86 | 0.76 | 0.85 | 0.96 | 0.77 | 0.92 |
| 7 | 120428799 | rs41623 | 2.375 | 1.494 | 3.81 | 2.223 | 0.032 | -0.035 | -0.035 | 0 | 0.86 | 0.79 | 0.86 | 0.96 | 0.76 | 0.92 |

| | | | | | | | | | | | | | | | | |
|---|---|---|---|---|---|---|---|---|---|---|---|---|---|---|---|---|
| 7 | 120458978 | rs17142976 | -2.532 | 1.437 | 3.807 | 2.211 | 0.037 | -0.044 | -0.044 | 0 | 0.14 | 0.25 | 0.15 | 0.03 | 0.22 | 0.08 |
| 7 | 120483178 | rs3779386 | -3.04 | 1.45 | 4.1 | 2.48 | -0.021 | 0.175 | 0.175 | 0 | 0.1 | 0.28 | 0.19 | 0.34 | 0.06 | 0.15 |
| 7 | 120507617 | rs17132731 | -2.023 | 1.678 | 3.788 | 2.348 | 0.037 | -0.033 | -0.033 | 0 | 0.12 | 0.21 | 0.11 | 0.03 | 0.21 | 0.07 |
| 7 | 120519615 | rs2158324 | -1.625 | 1.625 | 3.727 | 2.478 | -0.001 | 0.056 | 0.056 | 0 | 0.06 | 0.05 | 0.18 | 0.3 | 0.03 | 0.13 |
| 7 | 120558675 | rs2525708 | -3.748 | 1.585 | 4.102 | 2.46 | -0.043 | 0.231 | 0.231 | 0.041 | 0.12 | 0.57 | 0.24 | 0.34 | 0.05 | 0.15 |
| 7 | 120559538 | rs2525709 | -3.974 | 1.672 | 4.055 | 2.399 | 0 | -0.019 | -0.019 | 0.06 | 0.25 | 0.76 | 0.34 | 0.37 | 0.26 | 0.22 |
| 7 | 120577688 | rs6975372 | -2.149 | 1.643 | 4.218 | 2.385 | -0.021 | 0.137 | 0.137 | 0 | 0.07 | 0.16 | 0.2 | 0.32 | 0.03 | 0.14 |
| 7 | 120579637 | rs12706298 | 1.167 | 1.607 | 2.495 | 1.471 | -0.09 | 0.085 | 0.085 | 7.36 | 0.32 | 0.01 | 0.2 | 0.15 | 0.29 | 0.26 |
| 7 | 120587502 | rs2097806 | -3.373 | 1.646 | 3.115 | 1.712 | 0.001 | -0.03 | -0.03 | 0.002 | 0.26 | 0.77 | 0.37 | 0.41 | 0.28 | 0.25 |
| 7 | 120591936 | rs6964344 | -2.967 | 1.387 | 3.177 | 1.679 | -0.041 | 0.224 | 0.224 | 0.043 | 0.12 | 0.56 | 0.24 | 0.33 | 0.05 | 0.14 |
| 7 | 120611163 | rs2074577 | -2.796 | 1.149 | 2.365 | 1.401 | 0.018 | -0.021 | -0.021 | 2.828 | 0.3 | 0.78 | 0.35 | 0.35 | 0.27 | 0.24 |
| 7 | 120616010 | rs13881 | -1.877 | 1.23 | 2.835 | 1.106 | 0.04 | -0.081 | -0.081 | 0 | 0.18 | 0.49 | 0.14 | 0.05 | 0.22 | 0.1 |
| 7 | 120629481 | rs2525740 | -1.558 | 1.273 | 2.807 | 1.13 | 0.004 | -0.018 | -0.018 | 0 | 0.17 | 0.25 | 0.11 | 0.07 | 0.22 | 0.11 |
| 7 | 120639617 | rs41700 | 0.606 | 1.874 | 2.626 | 1.284 | -0.004 | 0 | 0 | 0 | 0.84 | 0.87 | 0.64 | 0.6 | 0.77 | 0.8 |
| 7 | 120640410 | rs41701 | 0.697 | 1.875 | 2.603 | 1.285 | 0.002 | 0.019 | 0.019 | 0 | 0.83 | 0.82 | 0.63 | 0.59 | 0.77 | 0.8 |
| 7 | 120646520 | rs2215789 | 1.137 | 1.881 | 2.497 | 1.258 | 0.005 | -0.018 | -0.018 | 0.037 | 0.83 | 0.75 | 0.89 | 0.92 | 0.78 | 0.89 |
| 7 | 120664459 | rs1981711 | 2.698 | 1.794 | 2.351 | 1.265 | -0.053 | -0.061 | -0.061 | 0 | 0.62 | 0.15 | 0.48 | 0.49 | 0.54 | 0.68 |
| 7 | 120666122 | rs7808535 | 0.72 | 1.706 | 2.379 | 1.274 | 0 | -0.01 | -0.01 | 0.285 | 0.85 | 0.8 | 0.9 | 0.94 | 0.8 | 0.9 |
| 7 | 120667666 | rs7782014 | -1.171 | 1.622 | 1.881 | 1.263 | 0.001 | 0.015 | 0.015 | 0.011 | 0.19 | 0.2 | 0.38 | 0.42 | 0.25 | 0.22 |
| 7 | 120671034 | rs6944789 | 2.094 | 1.648 | 1.142 | 1.215 | -0.027 | -0.038 | -0.038 | 0 | 0.62 | 0.23 | 0.49 | 0.49 | 0.55 | 0.67 |
| 7 | 120683751 | rs4730988 | 1.592 | 0.876 | 1.314 | 0.861 | -0.009 | -0.006 | -0.006 | 0 | 0.74 | 0.36 | 0.83 | 0.76 | 0.76 | 0.75 |
| 7 | 120705246 | rs2538492 | 0.968 | 0.968 | 1.55 | 0.644 | 0.029 | -0.015 | -0.015 | 0 | 0.92 | 0.75 | 0.94 | 0.96 | 0.92 | 0.79 |
| 7 | 120706892 | rs2192289 | 0.911 | 0.971 | 1.557 | 0.646 | 0.016 | -0.018 | -0.018 | 0.418 | 0.92 | 0.79 | 0.94 | 0.96 | 0.92 | 0.79 |
| 7 | 120711855 | rs10261386 | 0.421 | 0.892 | 0.691 | 0.114 | -0.01 | -0.009 | -0.009 | 0 | 0.4 | 0.23 | 0.57 | 0.61 | 0.41 | 0.33 |
| 7 | 120714700 | rs10251692 | -0.021 | 0.33 | 0.933 | -0.289 | 0.005 | 0.011 | 0.011 | 1.373 | 0.49 | 0.43 | 0.62 | 0.61 | 0.46 | 0.36 |
| 7 | 120719961 | rs17143147 | 0.695 | 0.201 | 0.588 | -0.275 | -0.035 | -0.007 | -0.007 | 0 | 0.41 | 0.24 | 0.63 | 0.61 | 0.48 | 0.37 |
| 7 | 120720819 | rs7790569 | -0.039 | 0.346 | 0.566 | -0.884 | 0.005 | -0.006 | -0.006 | 0 | 0.93 | 0.84 | 0.94 | 0.96 | 0.91 | 0.79 |
| 7 | 120724454 | rs2908572 | -0.039 | 0.346 | 0.575 | -0.884 | 0.006 | -0.007 | -0.007 | 0 | 0.93 | 0.85 | 0.94 | 0.96 | 0.91 | 0.79 |
| 7 | 120726926 | rs2968349 | -0.291 | -0.003 | 1.205 | -1.018 | 0.004 | 0.001 | 0.001 | 0 | 0.53 | 0.44 | 0.29 | 0.12 | 0.43 | 0.41 |
| 7 | 120735062 | rs1917113 | 0.57 | 0.002 | 1.098 | -1.147 | 0.034 | 0.08 | 0.08 | 0.504 | 0.39 | 0.21 | 0.21 | 0.12 | 0.32 | 0.34 |
| 7 | 120737058 | rs12673968 | 0.474 | 0.043 | 0.999 | -1.129 | 0.024 | 0.046 | 0.046 | 0.319 | 0.4 | 0.26 | 0.22 | 0.12 | 0.32 | 0.34 |
| 7 | 120763212 | rs798947 | -0.475 | 0.391 | 1.58 | -0.852 | 0.015 | -0.006 | -0.006 | 0 | 0.94 | 0.86 | 0.94 | 0.96 | 0.9 | 0.8 |
| 7 | 120765954 | rs798949 | 0.762 | 0.141 | 1.453 | -1.069 | -0.026 | 0.094 | 0.094 | 0 | 0.42 | 0.25 | 0.62 | 0.86 | 0.48 | 0.43 |
| 7 | 120778814 | rs10248011 | 0.834 | 0.129 | 1.43 | -1.074 | -0.037 | 0.111 | 0.111 | 0.041 | 0.42 | 0.21 | 0.62 | 0.86 | 0.48 | 0.44 |
| 7 | 120785064 | rs13245690 | 0.206 | 0.137 | 1.428 | -1.069 | 0.055 | 0.136 | 0.136 | 0 | 0.45 | 0.22 | 0.26 | 0.11 | 0.36 | 0.35 |
| 7 | 120800837 | rs1357755 | 0.724 | 0.156 | 1.189 | -1.033 | -0.025 | 0.095 | 0.095 | 0.037 | 0.43 | 0.25 | 0.62 | 0.85 | 0.48 | 0.44 |
| 7 | 120819121 | rs4730998 | 0.75 | 0.214 | 1.142 | -0.927 | -0.013 | 0.057 | 0.057 | 0 | 0.42 | 0.29 | 0.48 | 0.81 | 0.46 | 0.42 |
| 7 | 120822386 | rs7798060 | 0.061 | 0.113 | 0.605 | -0.873 | 0.038 | 0.102 | 0.102 | 0 | 0.46 | 0.28 | 0.28 | 0.12 | 0.38 | 0.35 |
| 7 | 120845747 | rs13229354 | 1.002 | 0.237 | 0.345 | -0.808 | -0.005 | -0.005 | -0.005 | 0.039 | 0.05 | 0.05 | 0.18 | 0.07 | 0.1 | 0.2 |
| 7 | 120852193 | rs1357756 | 0.189 | -0.325 | 0.375 | -0.958 | 0.018 | 0.025 | 0.025 | 0 | 0.52 | 0.55 | 0.7 | 0.88 | 0.62 | 0.65 |
| 7 | 120869464 | rs17284876 | -0.017 | -0.33 | -0.192 | -0.86 | -0.035 | 0.037 | 0.037 | 0.097 | 0.14 | 0.01 | 0.11 | 0.07 | 0.17 | 0.09 |
| 7 | 120872121 | rs17357115 | 1.249 | -1.249 | 0.099 | -1.406 | 0.017 | 0.225 | 0.225 | 0.167 | 0.45 | 0.11 | 0.64 | 0.87 | 0.54 | 0.62 |
| 7 | 120872569 | rs12672898 | -0.878 | -1.371 | 0.899 | -1.48 | 0.02 | 0.038 | 0.038 | 0 | 0.51 | 0.57 | 0.32 | 0.13 | 0.46 | 0.37 |
| 7 | 120873078 | rs1534016 | -0.256 | -1.483 | 1.08 | -1.555 | 0.044 | 0.112 | 0.112 | 0 | 0.42 | 0.21 | 0.26 | 0.13 | 0.38 | 0.34 |
| 7 | 120880573 | rs10480747 | -0.128 | -1.499 | 0.848 | -1.53 | 0.024 | 0.113 | 0.113 | 0 | 0.37 | 0.17 | 0.23 | 0.13 | 0.36 | 0.32 |
| 7 | 120883911 | rs7805374 | -0.971 | -1.446 | 0.846 | -1.485 | -0.004 | -0.035 | -0.035 | 0 | 0.47 | 0.48 | 0.3 | 0.13 | 0.44 | 0.35 |
| 7 | 120895302 | rs11770502 | -0.896 | -1.445 | 0.692 | -1.456 | -0.004 | -0.03 | -0.03 | 0 | 0.46 | 0.47 | 0.3 | 0.14 | 0.44 | 0.36 |
| 7 | 120897650 | rs12706333 | -1.083 | -1.682 | 0.568 | -1.601 | 0.001 | -0.017 | -0.017 | 0 | 0.49 | 0.47 | 0.3 | 0.14 | 0.44 | 0.36 |
| 7 | 120904659 | rs4731007 | -1.009 | -1.564 | 0.604 | -1.455 | -0.001 | -0.02 | -0.02 | 0.278 | 0.48 | 0.47 | 0.3 | 0.14 | 0.44 | 0.39 |
| 7 | 120908082 | rs2536150 | 0.206 | -1.476 | 0.47 | -1.226 | -0.022 | -0.046 | -0.046 | 0 | 0.25 | 0.34 | 0.13 | 0.07 | 0.19 | 0.26 |
| 7 | 120908822 | rs2536149 | 0.221 | -1.572 | 0.031 | -1.265 | 0.011 | 0.012 | 0.012 | 0.285 | 0.18 | 0.16 | 0.09 | 0.07 | 0.11 | 0.27 |
| 7 | 120912918 | rs1860753 | 0.396 | -1.504 | -0.36 | -1.351 | -0.023 | 0.049 | 0.049 | 0 | 0.17 | 0.04 | 0.21 | 0.09 | 0.2 | 0.12 |
| 7 | 120914373 | rs17508690 | 0.591 | -1.502 | -0.359 | -1.35 | -0.026 | 0.043 | 0.043 | 0 | 0.16 | 0.03 | 0.2 | 0.09 | 0.2 | 0.12 |
| 7 | 120915268 | rs798907 | -1.732 | -1.708 | -0.367 | -1.476 | 0.052 | 0.199 | 0.199 | 0.815 | 0.45 | 0.91 | 0.38 | 0.15 | 0.38 | 0.39 |
| 7 | 120918780 | rs798905 | -1.423 | -1.561 | -0.323 | -1.417 | 0.015 | 0.104 | 0.104 | 2.337 | 0.29 | 0.88 | 0.21 | 0.07 | 0.25 | 0.27 |
| 7 | 120919228 | rs1541509 | -0.375 | -1.133 | -0.492 | -1.521 | -0.037 | 0.094 | 0.094 | 0.014 | 0.27 | 0.09 | 0.28 | 0.1 | 0.31 | 0.15 |
| 7 | 120919495 | rs798903 | -1.567 | -1.123 | 0.059 | -1.512 | 0 | -0.023 | -0.023 | 0 | 0.49 | 0.55 | 0.42 | 0.15 | 0.49 | 0.41 |
| 7 | 120920406 | rs798901 | -2.171 | -1.197 | 0.04 | -0.364 | -0.168 | -0.151 | -0.151 | 0.22 | 0.59 | 0.98 | 0.55 | 0.43 | 0.52 | 0.64 |
| 7 | 120920827 | rs12534075 | -1.513 | -1.21 | 0.957 | -0.161 | 0.002 | 0 | 0 | 0 | 0.4 | 0.39 | 0.27 | 0.33 | 0.35 | 0.44 |
| 7 | 120927706 | rs17509082 | 0.41 | -0.983 | 1.182 | 0.013 | -0.003 | 0.02 | 0.02 | 0 | 0.17 | 0.12 | 0.12 | 0.08 | 0.19 | 0.19 |
| 7 | 120936006 | rs10241888 | -0.606 | -0.829 | 1.229 | 0.181 | -0.01 | -0.002 | -0.002 | 0 | 0.36 | 0.43 | 0.27 | 0.33 | 0.33 | 0.42 |
| 7 | 120940436 | rs2707521 | -1.163 | -0.94 | 1.306 | 0.281 | 0.057 | -0.046 | -0.046 | 0 | 0.43 | 0.91 | 0.36 | 0.45 | 0.35 | 0.58 |
| 7 | 120941964 | rs10282303 | -0.848 | -0.789 | 1.202 | 0.388 | -0.001 | -0.001 | -0.001 | 0 | 0.41 | 0.58 | 0.33 | 0.45 | 0.34 | 0.58 |
| 7 | 120952582 | rs10231005 | -0.836 | -0.654 | 0.848 | 0.34 | -0.001 | -0.001 | -0.001 | 0.93 | 0.41 | 0.58 | 0.33 | 0.45 | 0.34 | 0.59 |
| 7 | 120962164 | rs2908007 | -1.246 | -0.665 | -0.029 | 0.31 | 0.035 | -0.041 | -0.041 | 0.017 | 0.49 | 0.95 | 0.41 | 0.54 | 0.39 | 0.64 |
| 7 | 120963965 | rs3757552 | 1.072 | -0.721 | -0.027 | 0.467 | -0.025 | -0.09 | -0.09 | 0.608 | 0.12 | 0.01 | 0.16 | 0.36 | 0.11 | 0.2 |
| 7 | 120964405 | rs2707483 | 0.005 | -1.048 | 0.781 | 0.279 | 0.056 | 0.028 | 0.028 | 0 | 0.05 | 0.27 | 0.04 | 0 | 0.03 | 0.01 |
| 7 | 120966790 | rs3779381 | -0.266 | -1.148 | 0.564 | 0.261 | -0.015 | -0.05 | -0.05 | 0 | 0.29 | 0.4 | 0.22 | 0.12 | 0.26 | 0.26 |
| 7 | 120975115 | rs2707473 | -0.541 | -1.125 | 0.408 | 0.087 | -0.005 | -0.07 | -0.07 | 0 | 0.19 | 0.5 | 0.18 | 0.05 | 0.18 | 0.21 |

| | | | | | | | | | | | | | | | | |
|---|---|---|---|---|---|---|---|---|---|---|---|---|---|---|---|---|
| 7 | 120976677 | rs2707471 | 0.239 | -1.115 | 0.233 | 0.086 | 0.026 | -0.109 | -0.109 | 0.602 | 0.15 | 0.39 | 0.16 | 0 | 0.16 | 0.04 |
| 7 | 120977543 | rs3801385 | 1.148 | -1.112 | 0.082 | 0.203 | -0.003 | -0.092 | -0.092 | 0 | 0.13 | 0 | 0.12 | 0.37 | 0.09 | 0.2 |
| 7 | 120979089 | rs2707466 | -1.658 | -0.399 | 0.77 | -0.553 | 0.014 | 0.221 | 0.221 | 0.574 | 0.48 | 0.9 | 0.41 | 0.17 | 0.44 | 0.44 |
| 7 | 120986360 | rs2707460 | -0.805 | -0.326 | 1.749 | 0.078 | 0.029 | -0.112 | -0.112 | 0.118 | 0.15 | 0.4 | 0.18 | 0 | 0.18 | 0.04 |
| 7 | 120988757 | rs2536184 | -1.133 | -0.357 | 1.706 | 0.074 | -0.009 | -0.083 | -0.083 | 0 | 0.2 | 0.46 | 0.19 | 0.05 | 0.19 | 0.21 |
| 7 | 120991489 | rs11768996 | -0.798 | -0.251 | 1.683 | 0.121 | -0.004 | 0.05 | 0.05 | 0 | 0.18 | 0.09 | 0.16 | 0.05 | 0.19 | 0.21 |
| 7 | 120998784 | rs2707507 | -0.973 | -0.23 | 1.757 | 0.131 | -0.009 | -0.085 | -0.085 | 0 | 0.2 | 0.46 | 0.19 | 0.05 | 0.19 | 0.21 |
| 7 | 121054213 | rs6950058 | -1.299 | 0.004 | 1.473 | 1.051 | 0.085 | 0.414 | 0.414 | 1.149 | 0.07 | 0.41 | 0.17 | 0.27 | 0.02 | 0.13 |
| 7 | 121072904 | rs2697189 | 1.38 | 0.073 | -0.442 | 0.916 | 0.013 | -0.018 | -0.018 | 0 | 0.69 | 0.23 | 0.65 | 0.63 | 0.75 | 0.56 |
| 7 | 121073357 | rs2697188 | 1.38 | 0.073 | -0.442 | 0.916 | 0.013 | -0.018 | -0.018 | 0 | 0.69 | 0.23 | 0.66 | 0.63 | 0.75 | 0.56 |
| 7 | 121082818 | rs12706348 | 1.342 | 0.128 | -0.411 | 0.942 | 0.004 | -0.006 | -0.006 | 0 | 0.7 | 0.27 | 0.66 | 0.63 | 0.75 | 0.56 |
| 7 | 121090453 | rs11521035 | -0.659 | 0.853 | -0.883 | 1.351 | -0.062 | -0.003 | -0.003 | 0 | 0.15 | 0.02 | 0.12 | 0.1 | 0.21 | 0.29 |
| 7 | 121090531 | rs11521036 | -0.935 | 0.857 | -0.837 | 1.303 | -0.031 | -0.078 | -0.078 | 0.035 | 0.17 | 0.06 | 0.24 | 0.34 | 0.21 | 0.32 |
| 7 | 121098085 | rs2178185 | -1.012 | 0.892 | -0.857 | 0.618 | -0.026 | 0.027 | 0.027 | 0 | 0.18 | 0.07 | 0.14 | 0.11 | 0.25 | 0.31 |
| 7 | 121100885 | rs7806789 | -0.726 | 0.76 | -0.499 | 0.51 | 0.01 | -0.001 | -0.001 | 0 | 0.5 | 0.46 | 0.45 | 0.45 | 0.62 | 0.38 |
| 7 | 121103232 | rs2178192 | -0.808 | 1.445 | 0.137 | -0.143 | 0.005 | -0.024 | -0.024 | 1.357 | 0.28 | 0.31 | 0.21 | 0.09 | 0.35 | 0.2 |
| 7 | 121108696 | rs6466790 | -0.956 | 1.539 | 0.621 | -0.195 | 0.021 | -0.052 | -0.052 | 0 | 0.32 | 0.46 | 0.23 | 0.12 | 0.42 | 0.23 |
| 7 | 121123423 | rs4731018 | 1.305 | 0.892 | 0.768 | 0.265 | 0.052 | 0.01 | 0.01 | 0.604 | 0.95 | 0.72 | 0.96 | 1 | 0.95 | 0.98 |
| 7 | 121130580 | rs2402574 | -1.081 | 0.783 | 0.345 | 0.507 | -0.003 | 0.007 | 0.007 | 0.014 | 0.09 | 0.15 | 0.07 | 0.12 | 0.09 | 0.1 |
| 7 | 121130887 | rs10240090 | -0.039 | 0.865 | -0.138 | 0.097 | -0.026 | 0.02 | 0.02 | 0 | 0.36 | 0.26 | 0.38 | 0.28 | 0.43 | 0.29 |
| 7 | 121132213 | rs6954048 | 0.983 | 0.873 | 0.282 | -0.177 | 0.007 | -0.009 | -0.009 | 0 | 0.39 | 0.32 | 0.33 | 0.51 | 0.35 | 0.39 |
| 7 | 121148166 | rs7455326 | 0.105 | 0.957 | 0.538 | -0.15 | -0.001 | 0.008 | 0.008 | 0.458 | 0.09 | 0.1 | 0.07 | 0.14 | 0.1 | 0.07 |
| 7 | 121151589 | rs1983496 | 0.441 | 0.892 | 0.507 | -0.613 | 0.002 | 0.002 | 0.002 | 0 | 0.49 | 0.44 | 0.47 | 0.51 | 0.47 | 0.56 |
| 7 | 121152479 | rs1406194 | 0.441 | 0.892 | 0.507 | -0.613 | 0.002 | 0.002 | 0.002 | 0 | 0.49 | 0.44 | 0.47 | 0.51 | 0.47 | 0.56 |
| 7 | 121160264 | rs1881374 | 1.597 | 0.794 | 0.296 | -0.655 | 0.007 | -0.023 | -0.023 | 0 | 0.36 | 0.28 | 0.33 | 0.51 | 0.34 | 0.39 |
| 7 | 121162276 | rs6953922 | 1.752 | 0.745 | 0.496 | -0.677 | -0.011 | -0.085 | -0.085 | 2.109 | 0.35 | 0.12 | 0.32 | 0.51 | 0.34 | 0.39 |
| 7 | 121165809 | rs1528353 | -0.451 | 0.491 | 0.997 | -0.674 | 0.003 | -0.001 | -0.001 | 0 | 0.66 | 0.65 | 0.56 | 0.66 | 0.58 | 0.64 |
| 7 | 121167875 | rs1528351 | -0.451 | 0.491 | 1.009 | -0.674 | 0.002 | -0.001 | -0.001 | 0 | 0.66 | 0.66 | 0.56 | 0.66 | 0.58 | 0.64 |
| 7 | 121169049 | rs7795140 | 1.357 | 0.759 | 0.603 | -0.588 | -0.001 | -0.021 | -0.021 | 0 | 0.37 | 0.29 | 0.36 | 0.5 | 0.38 | 0.39 |
| 7 | 121178271 | rs11770273 | 0.068 | 0.679 | 0.548 | -0.603 | -0.007 | 0.038 | 0.038 | 0 | 0.13 | 0.04 | 0.09 | 0.01 | 0.13 | 0.17 |
| 7 | 121202525 | rs12538176 | -1.166 | 0.368 | 0.649 | -0.557 | 0 | -0.001 | -0.001 | 0 | 0.6 | 0.57 | 0.58 | 0.44 | 0.59 | 0.57 |
| 7 | 121217473 | rs5000116 | -0.6 | 0.166 | 1 | -0.133 | 0.001 | 0.001 | 0.001 | 0 | 0.46 | 0.5 | 0.47 | 0.44 | 0.42 | 0.39 |
| 7 | 121229994 | rs10234711 | -0.078 | 0.874 | 1.038 | -0.319 | 0.009 | -0.036 | -0.036 | 0 | 0.27 | 0.33 | 0.32 | 0.12 | 0.32 | 0.27 |
| 7 | 121230079 | rs10250499 | -0.725 | 0.838 | 1.274 | -0.226 | 0.02 | -0.02 | -0.02 | 0 | 0.33 | 0.54 | 0.4 | 0.27 | 0.38 | 0.33 |
| 7 | 121237869 | rs6955079 | -0.953 | 0.783 | 1.217 | -0.25 | -0.016 | 0.024 | 0.024 | 1.155 | 0.35 | 0.75 | 0.43 | 0.27 | 0.38 | 0.33 |
| 7 | 121247936 | rs10488051 | 0.006 | 0.718 | 0.62 | -0.222 | -0.002 | -0.001 | -0.001 | 0 | 0.24 | 0.22 | 0.31 | 0.12 | 0.29 | 0.24 |
| 7 | 121249593 | rs724813 | -0.588 | 0.671 | -0.273 | -0.123 | -0.011 | 0.015 | 0.015 | 2.611 | 0.33 | 0.73 | 0.41 | 0.3 | 0.37 | 0.31 |
| 7 | 121269679 | rs988022 | -0.595 | 0.595 | -1.372 | -0.097 | -0.043 | 0.044 | 0.044 | 0 | 0.54 | 0.89 | 0.55 | 0.39 | 0.56 | 0.54 |
| 7 | 121270857 | rs10258169 | -0.36 | -0.021 | -1.168 | -0.51 | -0.011 | -0.026 | -0.026 | 0 | 0.13 | 0.15 | 0.06 | 0 | 0.04 | 0.05 |
| 7 | 121270977 | rs13225566 | -0.129 | 0.214 | -0.339 | -0.516 | -0.001 | 0.022 | 0.022 | 3.45 | 0.17 | 0.28 | 0.16 | 0.28 | 0.16 | 0.22 |
| 7 | 121275229 | rs10253361 | -1.593 | 0.335 | -0.007 | -0.457 | -0.017 | 0.046 | 0.046 | 0 | 0.55 | 0.83 | 0.55 | 0.39 | 0.56 | 0.54 |
| 7 | 121275477 | rs10227063 | 0.441 | 0.349 | -0.019 | -0.455 | 0.026 | -0.047 | -0.047 | 0 | 0.27 | 0.38 | 0.33 | 0.11 | 0.36 | 0.27 |
| 7 | 121277383 | rs17143591 | -1.126 | 0.309 | -0.037 | -0.439 | -0.027 | -0.017 | -0.017 | 0 | 0.28 | 0.44 | 0.21 | 0.28 | 0.2 | 0.27 |
| 7 | 121277853 | rs17527997 | 0.821 | 0.601 | 0.072 | -0.266 | -0.026 | 0.018 | 0.018 | 2.277 | 0.2 | 0.1 | 0.29 | 0.11 | 0.34 | 0.25 |
| 7 | 121279041 | rs10240185 | -0.524 | 0.412 | 0.115 | -0.283 | 0.018 | -0.001 | -0.001 | 0 | 0.35 | 0.4 | 0.44 | 0.39 | 0.49 | 0.48 |
| 7 | 121282782 | rs2057876 | -0.472 | 1.519 | 0.258 | 0.348 | 0.001 | -0.01 | -0.01 | 0 | 0.46 | 0.49 | 0.45 | 0.31 | 0.49 | 0.4 |
| 7 | 121282913 | rs10243011 | -0.645 | 1.841 | 0.091 | 0.402 | -0.002 | -0.003 | -0.003 | 0 | 0.11 | 0.12 | 0.13 | 0.14 | 0.17 | 0.11 |
| 7 | 121283078 | rs4731030 | -0.235 | 1.855 | 0.409 | 0.909 | -0.013 | -0.007 | -0.007 | 0 | 0.43 | 0.5 | 0.43 | 0.63 | 0.4 | 0.31 |
| 7 | 121298820 | rs7812162 | -0.752 | 2.156 | 0.481 | 0.899 | 0.009 | -0.009 | -0.009 | 0 | 0.38 | 0.42 | 0.38 | 0.67 | 0.42 | 0.46 |
| 7 | 121299469 | rs11770667 | -0.788 | 2.075 | 0.538 | 0.8 | -0.01 | -0.032 | -0.032 | 1.561 | 0.07 | 0.11 | 0.04 | 0.01 | 0.04 | 0.04 |
| 7 | 121300485 | rs10228122 | -0.158 | 2.086 | 0.529 | 0.813 | -0.007 | -0.002 | -0.002 | 0 | 0.31 | 0.25 | 0.33 | 0.67 | 0.38 | 0.42 |
| 7 | 121323059 | rs11982968 | -1.683 | 2.15 | 0.607 | 0.649 | 0.053 | -0.051 | -0.051 | 0 | 0.09 | 0.3 | 0.08 | 0.01 | 0.09 | 0.06 |
| 7 | 121336037 | rs11763157 | -0.085 | 2.192 | -0.073 | 0.504 | 0.001 | -0.043 | -0.043 | 0 | 0.32 | 0.51 | 0.27 | 0.2 | 0.32 | 0.43 |
| 7 | 121339238 | rs6959329 | -1.461 | 2.078 | -0.008 | 0.599 | 0.088 | -0.015 | -0.015 | 0 | 0.07 | 0.3 | 0.08 | 0 | 0.08 | 0.03 |
| 7 | 121340319 | rs996635 | -1.241 | 2.188 | -0.431 | 0.681 | 0.018 | 0.129 | 0.129 | 2.785 | 0.63 | 0.95 | 0.63 | 0.85 | 0.7 | 0.86 |
| 7 | 121356943 | rs6466804 | -1.116 | 2.011 | -0.346 | 0.872 | 0.006 | -0.076 | -0.076 | 0.68 | 0.62 | 0.89 | 0.6 | 0.49 | 0.64 | 0.83 |
| 7 | 121373997 | rs11983078 | -1.405 | 1.768 | -0.167 | 0.436 | 0.054 | -0.027 | -0.027 | 0 | 0.06 | 0.21 | 0.07 | 0 | 0.07 | 0.03 |
| 7 | 121379616 | rs2041650 | -0.828 | 1.374 | -0.445 | 0.576 | -0.001 | 0.006 | 0.006 | 0.221 | 0.49 | 0.52 | 0.5 | 0.4 | 0.52 | 0.62 |
| 7 | 121388419 | rs17143719 | 0.533 | 0.961 | -0.713 | 0.622 | 0.007 | 0 | 0 | 0 | 0.23 | 0.21 | 0.17 | 0.27 | 0.24 | 0.38 |
| 7 | 121391810 | rs17438627 | 0.399 | 1.12 | -0.974 | 0.608 | -0.069 | -0.001 | -0.001 | 0 | 0.18 | 0.01 | 0.23 | 0.12 | 0.21 | 0.19 |
| 7 | 121391916 | rs10953942 | -0.662 | 1.063 | -0.88 | 0.592 | 0.007 | 0.021 | 0.021 | 0.081 | 0.46 | 0.41 | 0.44 | 0.39 | 0.45 | 0.57 |
| 7 | 121398174 | rs1476926 | -0.444 | 0.727 | -0.145 | 0.967 | -0.024 | -0.013 | -0.013 | 0.378 | 0.29 | 0.45 | 0.21 | 0.27 | 0.24 | 0.39 |
| 7 | 121406624 | rs6972380 | -0.881 | 1.149 | -0.988 | 1.112 | -0.013 | 0.096 | 0.096 | 0.333 | 0.51 | 0.9 | 0.64 | 0.67 | 0.52 | 0.68 |
| 7 | 121407003 | rs2192300 | -0.923 | 1.448 | -0.379 | 1.568 | 0.012 | 0.031 | 0.031 | 0 | 0.09 | 0.17 | 0.24 | 0.18 | 0.08 | 0.13 |
| 7 | 121427405 | rs6466807 | -0.558 | 1.079 | 0.773 | 1.395 | 0.001 | 0.002 | 0.002 | 0.294 | 0.39 | 0.6 | 0.39 | 0.5 | 0.44 | 0.54 |
| 7 | 121438897 | rs740962 | -1.311 | 1.021 | 1.407 | 1.47 | 0.001 | 0.009 | 0.009 | 0 | 0.5 | 0.54 | 0.53 | 0.6 | 0.49 | 0.67 |
| 7 | 121439019 | rs11983105 | -1.201 | 1.018 | 1.379 | 1.473 | 0.003 | 0.02 | 0.02 | 0 | 0.51 | 0.6 | 0.53 | 0.61 | 0.49 | 0.69 |
| 7 | 121442599 | rs17143823 | -0.428 | 1.464 | 1.513 | 1.903 | 0.054 | 0.055 | 0.055 | 0 | 0.07 | 0.14 | 0.1 | 0.2 | 0.15 | 0.07 |
| 7 | 121449928 | rs4731040 | -0.498 | 1.247 | 1.651 | 1.889 | 0.001 | -0.002 | -0.002 | 0 | 0.31 | 0.65 | 0.29 | 0.28 | 0.33 | 0.42 |

| | | | | | | | | | | | | | | | | |
|---|---|---|---|---|---|---|---|---|---|---|---|---|---|---|---|---|
| 7 | 121453246 | rs10225335 | -0.244 | 1.269 | 1.451 | 1.878 | 0.003 | -0.001 | -0.001 | 0.053 | 0.27 | 0.3 | 0.25 | 0.28 | 0.33 | 0.42 |
| 7 | 121467697 | rs7784550 | 0.646 | 1.183 | 1.373 | 1.928 | 0.014 | 0.029 | 0.029 | 0 | 0.21 | 0.14 | 0.15 | 0.13 | 0.19 | 0.34 |
| 7 | 121482703 | rs7798109 | -1.822 | 1.432 | 1.589 | 2.185 | -0.011 | 0.083 | 0.083 | 0 | 0.23 | 0.53 | 0.32 | 0.32 | 0.16 | 0.26 |
| 7 | 121486702 | rs4731041 | -0.15 | 1.474 | 1.546 | 2.255 | -0.001 | -0.001 | -0.001 | 0 | 0.28 | 0.29 | 0.24 | 0.3 | 0.31 | 0.42 |
| 7 | 121488137 | rs7797305 | -1.678 | 1.463 | 1.382 | 2.233 | -0.035 | 0.1 | 0.1 | 0.509 | 0.5 | 0.88 | 0.57 | 0.62 | 0.47 | 0.67 |
| 7 | 121495548 | rs12535129 | -0.144 | 1.538 | 1.351 | 2.234 | 0.063 | 0.074 | 0.074 | 0 | 0.06 | 0.17 | 0.08 | 0.18 | 0.12 | 0.05 |
| 7 | 121505847 | rs6965735 | -0.604 | 1.121 | 1.138 | 2.292 | -0.02 | -0.085 | -0.085 | 0.945 | 0.14 | 0.01 | 0.23 | 0.3 | 0.12 | 0.21 |
| 7 | 121512949 | rs740964 | -1.79 | 1.142 | 1.092 | 2.466 | -0.054 | 0.14 | 0.14 | 0.635 | 0.52 | 0.94 | 0.62 | 0.66 | 0.45 | 0.66 |
| 7 | 121513561 | rs740965 | -0.628 | 1.158 | 1.12 | 2.464 | -0.019 | -0.086 | -0.086 | 0 | 0.14 | 0.01 | 0.23 | 0.3 | 0.12 | 0.21 |
| 7 | 121524600 | rs2402593 | -0.481 | 1.141 | 1.537 | 2.442 | -0.023 | 0 | 0 | 0 | 0.32 | 0.51 | 0.22 | 0.3 | 0.24 | 0.39 |
| 7 | 121527480 | rs1007522 | -1.667 | 1.265 | 1.526 | 2.309 | -0.016 | 0.105 | 0.105 | 0.373 | 0.51 | 0.85 | 0.54 | 0.62 | 0.46 | 0.64 |
| 7 | 121530086 | rs1916888 | -1.718 | 1.277 | 1.686 | 2.315 | 0.005 | 0.062 | 0.062 | 0 | 0.49 | 0.69 | 0.53 | 0.62 | 0.45 | 0.64 |
| 7 | 121545143 | rs12669706 | -0.82 | 1.292 | 1.322 | 2.436 | -0.007 | -0.042 | -0.042 | 0 | 0.15 | 0.07 | 0.27 | 0.29 | 0.18 | 0.24 |
| 7 | 121554791 | rs6946211 | -0.92 | 1.106 | 1.321 | 2.387 | 0.002 | -0.003 | -0.003 | 0 | 0.16 | 0.11 | 0.26 | 0.32 | 0.15 | 0.24 |
| 7 | 121567350 | rs1206504 | -2.248 | 0.789 | 0.581 | 1.761 | -0.118 | 0.031 | 0.031 | 0.702 | 0.64 | 0.96 | 0.65 | 0.67 | 0.55 | 0.73 |
| 7 | 121586237 | rs960930 | -0.954 | 0.593 | 0.873 | 1.892 | -0.002 | 0.014 | 0.014 | 0 | 0.12 | 0.2 | 0.08 | 0.18 | 0.11 | 0.04 |
| 7 | 121587980 | rs6466808 | -0.595 | 0.546 | 0.994 | 1.91 | 0.007 | 0 | 0 | 0 | 0.39 | 0.4 | 0.27 | 0.34 | 0.33 | 0.46 |
| 7 | 121590463 | rs1196488 | -1.689 | 0.549 | 1.23 | 2.113 | -0.024 | 0.019 | 0.019 | 0.678 | 0.56 | 0.71 | 0.57 | 0.55 | 0.5 | 0.67 |
| 7 | 121595105 | rs1196485 | -0.593 | 0.554 | 1.312 | 2.104 | 0.02 | 0.047 | 0.047 | 0.189 | 0.16 | 0.25 | 0.29 | 0.22 | 0.17 | 0.21 |
| 7 | 121596593 | rs1196481 | -1.98 | 0.523 | 1.206 | 2.082 | -0.04 | 0.047 | 0.047 | 0.033 | 0.6 | 0.78 | 0.58 | 0.64 | 0.52 | 0.73 |
| 7 | 121601907 | rs6970897 | 0.431 | 0.649 | 1.305 | 2.044 | 0.007 | 0.014 | 0.014 | 0 | 0.1 | 0.03 | 0.06 | 0 | 0.08 | 0.08 |
| 7 | 121604718 | rs10278079 | -0.258 | 0.631 | 1.248 | 2.109 | 0 | 0 | 0 | 0.876 | 0.35 | 0.4 | 0.25 | 0.3 | 0.3 | 0.45 |
| 7 | 121605726 | rs2690271 | -1.342 | 0.648 | 1.24 | 2.14 | -0.008 | 0.037 | 0.037 | 2.204 | 0.51 | 0.7 | 0.55 | 0.52 | 0.48 | 0.66 |
| 7 | 121608449 | rs1196510 | -1.099 | 0.605 | 1.251 | 2.182 | 0.006 | 0.011 | 0.011 | 0.564 | 0.46 | 0.62 | 0.51 | 0.51 | 0.42 | 0.61 |
| 7 | 121611833 | rs1209633 | -0.613 | 0.699 | 1.165 | 2.117 | 0.016 | 0.031 | 0.031 | 0.074 | 0.16 | 0.22 | 0.29 | 0.21 | 0.18 | 0.21 |
| 7 | 121617017 | rs3817483 | 0.762 | 0.503 | 0.479 | 1.982 | 0.002 | -0.012 | -0.012 | 0.019 | 0.22 | 0.01 | 0.13 | 0.13 | 0.15 | 0.37 |
| 7 | 121628842 | rs1196505 | 1.758 | 0.724 | 1.232 | 2.21 | 0.048 | 0.236 | 0.236 | 0 | 0.88 | 0.62 | 0.83 | 0.67 | 0.87 | 0.9 |
| 7 | 121629854 | rs1196502 | 1.031 | 0.657 | 1.115 | 1.944 | 0.011 | 0.094 | 0.094 | 0 | 0.92 | 0.76 | 0.9 | 0.84 | 0.96 | 0.93 |
| 7 | 121637265 | rs2693657 | -0.759 | 0.549 | 0.833 | 1.6 | -0.008 | -0.001 | -0.001 | 0 | 0.43 | 0.47 | 0.45 | 0.38 | 0.37 | 0.6 |
| 7 | 121647107 | rs1196473 | 1.589 | 0.558 | 0.9 | 1.876 | 0.031 | 0.196 | 0.196 | 0 | 0.87 | 0.63 | 0.9 | 0.71 | 0.9 | 0.91 |
| 7 | 121649223 | rs1196475 | -0.593 | 0.427 | 0.704 | 1.738 | -0.006 | 0.003 | 0.003 | 0.168 | 0.39 | 0.54 | 0.47 | 0.36 | 0.37 | 0.57 |
| 7 | 121653398 | rs1147504 | -1.404 | 0.473 | 0.454 | 1.664 | -0.089 | 0.112 | 0.112 | 0.832 | 0.53 | 0.93 | 0.57 | 0.65 | 0.47 | 0.67 |
| 7 | 121676924 | rs1147492 | -1.8 | 1.005 | 0.414 | 1.33 | -0.115 | 0.003 | 0.003 | 0.022 | 0.59 | 0.95 | 0.58 | 0.61 | 0.51 | 0.68 |
| 7 | 121677000 | rs1147491 | -1.281 | 1.144 | 0.846 | 1.384 | 0.107 | 0.254 | 0.254 | 0 | 0.09 | 0.34 | 0.1 | 0.25 | 0.1 | 0.1 |
| 7 | 121679133 | rs1147489 | -1.319 | 0.939 | 0.863 | 1.356 | 0.01 | -0.031 | -0.031 | 0.018 | 0.24 | 0.5 | 0.31 | 0.17 | 0.24 | 0.25 |
| 7 | 121684718 | rs1147487 | -1.321 | 0.68 | 0.914 | 1.468 | 0.02 | 0.018 | 0.018 | 0 | 0.49 | 0.62 | 0.49 | 0.35 | 0.41 | 0.6 |
| 7 | 121698827 | rs1206381 | -1.1 | 0.669 | 0.91 | 1.464 | 0.028 | 0.017 | 0.017 | 0 | 0.47 | 0.65 | 0.49 | 0.35 | 0.39 | 0.59 |
| 7 | 121712619 | rs1206378 | -0.384 | 0.587 | 1.234 | 1.374 | -0.024 | -0.049 | -0.049 | 0 | 0.1 | 0.2 | 0.06 | 0 | 0.06 | 0.07 |
| 7 | 121714653 | rs12217 | -1.768 | 0.631 | 1.092 | 1.469 | -0.115 | 0.023 | 0.023 | 0 | 0.56 | 0.95 | 0.56 | 0.61 | 0.48 | 0.63 |
| 7 | 121717282 | rs1206367 | -1.768 | 0.63 | 1.092 | 1.469 | -0.119 | 0.023 | 0.023 | 0 | 0.56 | 0.95 | 0.56 | 0.61 | 0.48 | 0.63 |
| 7 | 121726812 | rs7787099 | 0.801 | 0.49 | 1.15 | 1.459 | 0.024 | -0.002 | -0.002 | 0 | 0.23 | 0.13 | 0.18 | 0.19 | 0.16 | 0.33 |
| 7 | 121728068 | rs1206376 | -2.056 | 0.482 | 1.073 | 1.455 | -0.119 | 0.017 | 0.017 | 0 | 0.57 | 0.95 | 0.56 | 0.61 | 0.48 | 0.64 |
| 7 | 121733901 | rs1206362 | -0.208 | 0.482 | 1.147 | 1.451 | -0.025 | -0.051 | -0.051 | 0 | 0.09 | 0.2 | 0.06 | 0 | 0.06 | 0.07 |
| 7 | 121737773 | rs1206361 | -0.285 | 0.481 | 1.14 | 1.451 | -0.031 | -0.056 | -0.056 | 0.235 | 0.1 | 0.23 | 0.07 | 0 | 0.06 | 0.07 |
| 7 | 121750775 | rs11771234 | 0.769 | 0.437 | 1.124 | 1.457 | 0.038 | -0.01 | -0.01 | 0 | 0.23 | 0.09 | 0.17 | 0.19 | 0.16 | 0.33 |
| 7 | 121754695 | rs4140745 | 0.155 | 0.554 | 1.105 | 1.468 | -0.009 | 0.049 | 0.049 | 0 | 0.3 | 0.36 | 0.26 | 0.44 | 0.25 | 0.39 |
| 7 | 121757026 | rs279676 | -0.495 | 0.353 | 1.235 | 1.142 | -0.027 | -0.056 | -0.056 | 0 | 0.1 | 0.23 | 0.07 | 0 | 0.06 | 0.07 |
| 7 | 121774649 | rs10488107 | -0.707 | 0.57 | 0.968 | 1.314 | 0.013 | 0.008 | 0.008 | 0 | 0.17 | 0.35 | 0.23 | 0.17 | 0.16 | 0.18 |
| 7 | 121787676 | rs2109333 | -0.551 | 0.458 | 1 | 1.308 | 0.005 | 0.003 | 0.003 | 0 | 0.16 | 0.31 | 0.23 | 0.16 | 0.16 | 0.22 |
| 7 | 121790934 | rs279686 | -1.624 | 0.278 | 1.116 | 1.292 | -0.02 | -0.009 | -0.009 | 0.487 | 0.28 | 0.58 | 0.3 | 0.26 | 0.23 | 0.31 |
| 7 | 121797265 | rs279698 | -1.661 | 0.449 | 0.409 | 1.201 | -0.053 | 0.038 | 0.038 | 0 | 0.56 | 0.93 | 0.53 | 0.62 | 0.45 | 0.6 |
| 7 | 121802068 | rs17144133 | 2.11 | 0.864 | -0.018 | 1.041 | -0.116 | -0.042 | -0.042 | 0 | 0.37 | 0.04 | 0.41 | 0.38 | 0.43 | 0.39 |
| 7 | 121818068 | rs2120365 | 1.504 | 0.266 | 0.184 | 1.139 | -0.044 | 0.039 | 0.039 | 0 | 0.44 | 0.09 | 0.47 | 0.38 | 0.54 | 0.41 |
| 7 | 121822246 | rs2083860 | 1.488 | 0.263 | 0.206 | 1.138 | -0.037 | 0.045 | 0.045 | 0.041 | 0.44 | 0.1 | 0.47 | 0.38 | 0.54 | 0.41 |
| 7 | 121833023 | rs12706389 | 0.062 | 0.259 | 0.513 | 0.917 | -0.01 | -0.031 | -0.031 | 0.865 | 0.1 | 0.1 | 0.05 | 0 | 0.08 | 0.05 |
| 7 | 121853616 | rs2134057 | 0.062 | 0.264 | 0.528 | 0.907 | -0.011 | -0.035 | -0.035 | 0 | 0.1 | 0.1 | 0.05 | 0 | 0.08 | 0.05 |
| 7 | 121857984 | rs537162 | -0.882 | 0.272 | 0.646 | 1.078 | -0.014 | 0.032 | 0.032 | 0 | 0.14 | 0.31 | 0.11 | 0.17 | 0.13 | 0.07 |
| 7 | 121878287 | rs7802078 | 1.45 | 0.199 | 0.46 | 1.118 | 0.003 | 0.002 | 0.002 | 0 | 0.45 | 0.24 | 0.49 | 0.49 | 0.55 | 0.44 |
| 7 | 121885449 | rs284394 | 0.117 | 0.19 | 0.532 | 1.102 | -0.017 | -0.046 | -0.046 | 0 | 0.11 | 0.14 | 0.06 | 0 | 0.08 | 0.05 |
| 7 | 121888291 | rs10953953 | -0.018 | 0.123 | 0.534 | 1.203 | -0.013 | -0.027 | -0.027 | 0 | 0.16 | 0.09 | 0.24 | 0.24 | 0.18 | 0.25 |
| 7 | 121890781 | rs284391 | -0.259 | 0.206 | 0.447 | 1.29 | -0.04 | 0.021 | 0.021 | 0 | 0.72 | 0.5 | 0.79 | 0.73 | 0.82 | 0.74 |
| 7 | 121896770 | rs284387 | -0.591 | 0.199 | 0.462 | 1.03 | -0.011 | -0.011 | -0.011 | 0 | 0.75 | 0.74 | 0.82 | 0.9 | 0.84 | 0.76 |
| 7 | 121897109 | rs284386 | -1.004 | 0.335 | 0.172 | 1.077 | -0.007 | -0.004 | -0.004 | 0 | 0.27 | 0.38 | 0.31 | 0.25 | 0.27 | 0.29 |
| 7 | 121911013 | rs6946696 | 0.647 | -0.164 | -0.311 | 0.614 | 0.015 | 0.009 | 0.009 | 0 | 0.21 | 0.15 | 0.15 | 0.11 | 0.14 | 0.17 |
| 7 | 121913741 | rs11765191 | 1.336 | 0.862 | -0.178 | 0.753 | -0.032 | -0.041 | -0.041 | 0 | 0.58 | 0.22 | 0.55 | 0.53 | 0.54 | 0.57 |
| 7 | 121926270 | rs1604328 | 1.277 | 0.977 | -0.195 | 0.503 | 0.01 | 0.001 | 0.001 | 0.814 | 0.6 | 0.47 | 0.57 | 0.63 | 0.54 | 0.59 |
| 7 | 121927932 | rs4329221 | 0.838 | 1.541 | -0.59 | 1.09 | 0.007 | 0.011 | 0.011 | 2.319 | 0.76 | 0.72 | 0.67 | 0.63 | 0.67 | 0.67 |
| 7 | 121934519 | rs284345 | 0.313 | 0.723 | -1.17 | 0.76 | 0.011 | -0.023 | -0.023 | 0.312 | 0.6 | 0.27 | 0.52 | 0.41 | 0.58 | 0.55 |
| 7 | 121934655 | rs284346 | -0.384 | -0.36 | -0.883 | 0.642 | -0.043 | 0.02 | 0.02 | 0 | 0.27 | 0.45 | 0.33 | 0.3 | 0.19 | 0.23 |

| | | | | | | | | | | | | | | | | |
|---|---|---|---|---|---|---|---|---|---|---|---|---|---|---|---|---|
| 7 | 121937298 | rs11762051 | 0.028 | 0.545 | -1.452 | 0.433 | -0.066 | -0.158 | -0.158 | 0.799 | 0.2 | 0.01 | 0.32 | 0.54 | 0.18 | 0.26 |
| 7 | 121939771 | rs5007345 | -0.008 | 0.568 | -1.315 | 0.443 | -0.014 | -0.092 | -0.092 | 0.097 | 0.22 | 0.06 | 0.33 | 0.54 | 0.18 | 0.26 |
| 7 | 121944527 | rs11761299 | -0.486 | 0.323 | -0.929 | 0.137 | -0.001 | 0 | 0 | 0 | 0.59 | 0.59 | 0.61 | 0.6 | 0.69 | 0.63 |
| 7 | 121947456 | rs11767283 | -0.178 | 0.999 | -1.224 | 0.315 | -0.015 | -0.071 | -0.071 | 0.615 | 0.24 | 0.1 | 0.34 | 0.55 | 0.25 | 0.3 |
| 7 | 121949318 | rs1443753 | -0.588 | 0.621 | -0.843 | -0.244 | 0.038 | 0 | 0 | 1.317 | 0.47 | 0.34 | 0.54 | 0.55 | 0.66 | 0.52 |
| 7 | 121950026 | rs1867335 | -0.808 | 0.364 | -0.088 | -0.341 | 0.003 | 0 | 0 | 0.713 | 0.49 | 0.46 | 0.57 | 0.53 | 0.66 | 0.52 |
| 7 | 121950965 | rs7779018 | 0.279 | 0.92 | -0.066 | -0.458 | -0.04 | 0.015 | 0.015 | 0 | 0.22 | 0.16 | 0.19 | 0 | 0.41 | 0.21 |
| 7 | 121958195 | rs13241158 | 0.75 | 0.264 | -0.77 | -0.8 | 0.012 | 0.007 | 0.007 | 0 | 0.25 | 0.11 | 0.24 | 0.2 | 0.25 | 0.26 |
| 7 | 121959746 | rs12706405 | 0.745 | 0.259 | -0.779 | -0.807 | 0.009 | 0.006 | 0.006 | 0 | 0.25 | 0.11 | 0.24 | 0.2 | 0.26 | 0.26 |
| 7 | 121964349 | rs1899689 | 0.42 | 1.099 | -0.426 | -0.67 | -0.005 | -0.022 | -0.022 | 0.82 | 0.44 | 0.27 | 0.44 | 0.47 | 0.38 | 0.41 |
| 7 | 121967298 | rs11972628 | 0.204 | 1.006 | -0.041 | -0.676 | 0.018 | 0.187 | 0.187 | 0 | 0.11 | 0.36 | 0.18 | 0.33 | 0.12 | 0.13 |
| 7 | 121970118 | rs920802 | 0.211 | 1.207 | -0.293 | -0.955 | -0.002 | 0.037 | 0.037 | 0 | 0.39 | 0.29 | 0.36 | 0.23 | 0.41 | 0.4 |
| 7 | 121987158 | rs798682 | -1.055 | 1.43 | -0.251 | -1.012 | -0.034 | 0.053 | 0.053 | 0 | 0.79 | 0.92 | 0.87 | 1 | 0.73 | 0.8 |
| 7 | 121988404 | rs798681 | 0.615 | 1.338 | -0.156 | -0.881 | -0.011 | 0.028 | 0.028 | 0 | 0.33 | 0.39 | 0.41 | 0.52 | 0.35 | 0.4 |
| 7 | 121990181 | rs798678 | 1.279 | 1.294 | 0.337 | -0.839 | -0.001 | 0 | 0 | 0.043 | 0.28 | 0.24 | 0.29 | 0.28 | 0.26 | 0.3 |
| 7 | 121992989 | rs702421 | -1.432 | 1.127 | -0.283 | -1.072 | -0.049 | -0.143 | -0.143 | 0 | 0.71 | 0.98 | 0.67 | 0.63 | 0.76 | 0.73 |
| 7 | 121999274 | rs10487223 | 1.129 | 0.999 | 0.055 | -1.183 | -0.005 | 0 | 0 | 0 | 0.12 | 0.13 | 0.08 | 0.03 | 0.14 | 0.14 |
| 7 | 121999971 | rs7785641 | 0.339 | 1.011 | 0.154 | -1.218 | 0.02 | -0.14 | -0.14 | 0 | 0.23 | 0.41 | 0.15 | 0.01 | 0.26 | 0.17 |
| 7 | 122003326 | rs7778318 | 0.851 | 1.075 | -0.249 | -1.212 | -0.048 | 0.089 | 0.089 | 0 | 0.2 | 0.06 | 0.12 | 0.01 | 0.26 | 0.16 |
| 7 | 122003582 | rs6944990 | -0.415 | 1.046 | -0.277 | -1.154 | 0.02 | -0.132 | -0.132 | 0.009 | 0.34 | 0.57 | 0.23 | 0.03 | 0.4 | 0.31 |
| 7 | 122007319 | rs6965749 | 0.651 | 1.237 | -0.189 | -0.738 | 0 | 0 | 0 | 0 | 0.33 | 0.32 | 0.45 | 0.58 | 0.38 | 0.42 |
| 7 | 122011039 | rs17463319 | -0.659 | 1.38 | -0.273 | -0.826 | 0.001 | -0.057 | -0.057 | 0 | 0.23 | 0.28 | 0.14 | 0 | 0.26 | 0.17 |
| 7 | 122030296 | rs10268511 | 0.305 | 0.869 | -0.453 | 0.215 | -0.011 | 0.015 | 0.015 | 0 | 0.37 | 0.57 | 0.42 | 0.43 | 0.31 | 0.3 |
| 7 | 122031424 | rs4731056 | -0.241 | 1.031 | -0.537 | 0.062 | -0.006 | 0.054 | 0.054 | 0 | 0.53 | 0.7 | 0.61 | 0.65 | 0.5 | 0.59 |
| 7 | 122048354 | rs7785453 | -0.548 | 1.201 | -0.377 | -0.195 | -0.018 | -0.001 | -0.001 | 0.249 | 0.69 | 0.79 | 0.67 | 0.64 | 0.63 | 0.63 |
| 7 | 122052524 | rs3807875 | 0.399 | 1.11 | -0.17 | -0.225 | 0 | -0.001 | -0.001 | 0 | 0.43 | 0.54 | 0.48 | 0.42 | 0.4 | 0.34 |
| 7 | 122057923 | rs3823855 | 1.079 | 0.918 | 0.02 | -0.075 | 0.007 | -0.039 | -0.039 | 2.592 | 0.33 | 0.16 | 0.37 | 0.35 | 0.29 | 0.28 |
| 7 | 122071191 | rs11972946 | 0.499 | 0.959 | 0.132 | -0.088 | -0.009 | 0.017 | 0.017 | 0 | 0.39 | 0.45 | 0.44 | 0.42 | 0.37 | 0.3 |
| 7 | 122078414 | rs2074589 | 0.967 | 1.108 | 0.047 | -0.095 | 0.018 | -0.057 | -0.057 | 0.265 | 0.35 | 0.13 | 0.38 | 0.39 | 0.3 | 0.3 |
| 7 | 122080536 | rs3757534 | 1.384 | 1.029 | 0.117 | -0.084 | 0.002 | -0.006 | -0.006 | 0 | 0.09 | 0.07 | 0.09 | 0.13 | 0.08 | 0.23 |
| 7 | 122088595 | rs4727941 | 0.506 | 1.012 | 0.477 | -0.224 | -0.003 | 0.004 | 0.004 | 0 | 0.4 | 0.53 | 0.46 | 0.42 | 0.38 | 0.31 |
| 7 | 122091852 | rs6956211 | 1.918 | 1.486 | -0.035 | -0.56 | -0.059 | 0.07 | 0.07 | 0.141 | 0.13 | 0.02 | 0.08 | 0 | 0.19 | 0.04 |
| 7 | 122106514 | rs6959220 | 0.608 | 0.757 | 0.339 | -0.348 | -0.013 | 0.006 | 0.006 | 0 | 0.07 | 0.19 | 0.04 | 0.04 | 0.03 | 0.04 |
| 7 | 122139167 | rs7778137 | 0.291 | 0.787 | 0.552 | -0.4 | 0.005 | -0.004 | -0.004 | 0 | 0.4 | 0.57 | 0.46 | 0.38 | 0.38 | 0.29 |
| 7 | 122148998 | rs1159907 | -1.747 | 0.686 | 0.222 | -0.619 | -0.021 | 0.089 | 0.089 | 0.082 | 0.85 | 0.99 | 0.92 | 1 | 0.85 | 0.97 |
| 7 | 122149225 | rs10256973 | -0.584 | 0.555 | 0.683 | -0.276 | 0.021 | -0.011 | -0.011 | 0 | 0.1 | 0.35 | 0.11 | 0.04 | 0.1 | 0.06 |
| 7 | 122157469 | rs10242162 | -0.513 | 0.663 | 0.053 | -0.202 | 0.003 | 0.034 | 0.034 | 0 | 0.49 | 0.71 | 0.55 | 0.42 | 0.5 | 0.5 |
| 7 | 122157895 | rs11764665 | 0.929 | 0.655 | -0.413 | -0.474 | -0.045 | 0.175 | 0.175 | 0 | 0.25 | 0.02 | 0.12 | 0 | 0.22 | 0.16 |
| 7 | 122185149 | rs4727942 | -0.372 | 0.016 | 0.392 | -0.223 | 0.037 | 0.158 | 0.158 | 0.896 | 0.17 | 0.41 | 0.38 | 0.42 | 0.25 | 0.31 |
| 7 | 122186718 | rs10238396 | -0.851 | 0.177 | 0.298 | -0.552 | 0.009 | 0.041 | 0.041 | 0.177 | 0.51 | 0.78 | 0.57 | 0.43 | 0.55 | 0.52 |
| 7 | 122186974 | rs740812 | 0.175 | 0.175 | 0.409 | -0.54 | -0.038 | 0.009 | 0.009 | 0 | 0.36 | 0.26 | 0.42 | 0.34 | 0.45 | 0.44 |
| 7 | 122195452 | rs6964649 | -0.34 | 0.182 | 0.257 | -0.889 | -0.001 | -0.045 | -0.045 | 0.763 | 0.41 | 0.42 | 0.27 | 0.09 | 0.38 | 0.26 |
| 7 | 122197763 | rs929506 | -2.444 | 0.291 | -0.175 | -1.109 | -0.025 | 0.05 | 0.05 | 0 | 0.88 | 0.97 | 0.95 | 1 | 0.87 | 0.97 |
| 7 | 122202472 | rs6466821 | 0.83 | 0.208 | 0.161 | -0.739 | -0.092 | 0.18 | 0.18 | 0 | 0.26 | 0.02 | 0.13 | 0 | 0.28 | 0.18 |
| 7 | 122202593 | rs2429582 | -1.189 | 0.027 | 0.372 | -0.294 | -0.004 | -0.002 | -0.002 | 0 | 0.62 | 0.85 | 0.68 | 0.68 | 0.65 | 0.76 |
| 7 | 122219899 | rs10266303 | -1.304 | 0.308 | 0.96 | -0.326 | -0.03 | -0.03 | -0.03 | 0.107 | 0.15 | 0.37 | 0.13 | 0.08 | 0.11 | 0.08 |
| 7 | 122238571 | rs4363139 | -0.235 | 0.448 | 0.616 | -0.266 | 0.014 | 0 | 0 | 0 | 0.08 | 0.23 | 0.1 | 0.05 | 0.08 | 0.03 |
| 7 | 122247051 | rs4604372 | -0.812 | 0.436 | 0.299 | -0.304 | -0.021 | -0.031 | -0.031 | 0 | 0.17 | 0.25 | 0.13 | 0.09 | 0.12 | 0.11 |
| 7 | 122264411 | rs961628 | 0.745 | 0.839 | -0.068 | -0.241 | 0.002 | -0.021 | -0.021 | 0 | 0.58 | 0.46 | 0.53 | 0.72 | 0.54 | 0.66 |
| 7 | 122286939 | rs2471194 | 0.277 | 1.213 | 0.084 | -0.192 | 0.111 | 0.141 | 0.141 | 0 | 0.91 | 0.65 | 0.75 | 0.78 | 0.83 | 0.9 |
| 7 | 122287279 | rs2471193 | 0.894 | 1.162 | -0.008 | -0.279 | 0.015 | -0.031 | -0.031 | 0 | 0.62 | 0.51 | 0.56 | 0.75 | 0.53 | 0.72 |
| 7 | 122290999 | rs7797968 | 0.101 | 1.091 | 0.201 | -0.155 | 0.061 | 0.081 | 0.081 | 0 | 0.08 | 0.17 | 0.23 | 0.22 | 0.17 | 0.1 |
| 7 | 122303321 | rs2251761 | 0.873 | 0.955 | 0.105 | -0.286 | 0.009 | 0.014 | 0.014 | 0 | 0.8 | 0.65 | 0.66 | 0.76 | 0.73 | 0.79 |
| 7 | 122307692 | rs2430031 | 0.655 | 1.736 | 0.371 | -0.03 | 0.067 | -0.039 | -0.039 | 0 | 0.71 | 0.5 | 0.59 | 0.79 | 0.52 | 0.79 |
| 7 | 122316592 | rs1968147 | 0.039 | 1.813 | 0.143 | 0.001 | 0.041 | 0.002 | 0.002 | 0 | 0.07 | 0.12 | 0.09 | 0.08 | 0.17 | 0.1 |
| 7 | 122316632 | rs7777183 | 1.299 | 1.775 | 0.182 | 0.062 | 0.052 | 0.001 | 0.001 | 0.107 | 0.82 | 0.49 | 0.73 | 0.82 | 0.71 | 0.71 |
| 7 | 122317278 | rs4140960 | -0.312 | 1.269 | -0.349 | 0.145 | 0.009 | 0.028 | 0.028 | 0.112 | 0.08 | 0.22 | 0.12 | 0.1 | 0.07 | 0.18 |
| 7 | 122328750 | rs12668965 | 0.777 | 1.235 | -1.703 | 0.397 | 0 | -0.091 | -0.091 | 1.138 | 0.22 | 0.02 | 0.19 | 0.28 | 0.18 | 0.22 |
| 7 | 122330642 | rs9640810 | -0.007 | 1.947 | -0.894 | 0.006 | -0.001 | -0.001 | -0.001 | 0.187 | 0.07 | 0.06 | 0.1 | 0.1 | 0.07 | 0.18 |
| 7 | 122335352 | rs12154838 | 0.138 | 1.981 | -1.051 | -0.009 | -0.092 | 0.036 | 0.036 | 0 | 0.18 | 0.02 | 0.25 | 0.09 | 0.26 | 0.1 |
| 7 | 122367349 | rs2159824 | 0.725 | 1.02 | -0.302 | 0.456 | 0.024 | 0.021 | 0.021 | 0.109 | 0.91 | 0.82 | 0.85 | 0.9 | 0.87 | 0.8 |
| 7 | 122371707 | rs9969220 | -0.047 | 0.936 | -1.369 | 0.258 | 0.001 | 0.041 | 0.041 | 0.726 | 0.41 | 0.18 | 0.35 | 0.68 | 0.37 | 0.57 |
| 7 | 122375615 | rs7782606 | 0.215 | 0.612 | -0.824 | -0.512 | -0.011 | -0.027 | -0.027 | 0 | 0.33 | 0.36 | 0.26 | 0.11 | 0.27 | 0.2 |
| 7 | 122376318 | rs929569 | 0.441 | 1.32 | -0.926 | -0.647 | 0.027 | 0.079 | 0.079 | 0.221 | 0.56 | 0.23 | 0.6 | 0.72 | 0.58 | 0.6 |
| 7 | 122392897 | rs37908 | -0.433 | 1.312 | -0.986 | -0.615 | 0.048 | 0.068 | 0.068 | 0.543 | 0.43 | 0.77 | 0.36 | 0.28 | 0.36 | 0.39 |
| 7 | 122415644 | rs13227529 | 0.571 | 1.329 | -0.274 | -0.675 | 0.035 | 0.048 | 0.048 | 0 | 0.31 | 0.21 | 0.23 | 0.11 | 0.22 | 0.18 |
| 7 | 122424605 | rs12706448 | 0.41 | 1.611 | -0.343 | 0.054 | 0.03 | 0.017 | 0.017 | 0 | 0.33 | 0.23 | 0.29 | 0.25 | 0.24 | 0.37 |
| 7 | 122447312 | rs10277410 | 0.466 | 1.624 | 0.05 | -0.142 | 0.001 | 0.002 | 0.002 | 0 | 0.09 | 0.08 | 0.11 | 0.17 | 0.15 | 0.21 |
| 7 | 122492340 | rs13240237 | 0.586 | 1.611 | 0.2 | -0.155 | 0.005 | 0.001 | 0.001 | 0 | 0.31 | 0.28 | 0.23 | 0.11 | 0.22 | 0.18 |

| | | | | | | | | | | | | | | | | |
|---|---|---|---|---|---|---|---|---|---|---|---|---|---|---|---|---|
| 7 | 122493144 | rs2501448 | 0.009 | 1.752 | 0.275 | -0.007 | 0.072 | 0.084 | 0.084 | 0 | 0.11 | 0.33 | 0.13 | 0.17 | 0.15 | 0.21 |
| 7 | 122495375 | rs3919490 | 0.981 | 1.818 | -0.727 | 0.143 | -0.084 | -0.134 | -0.134 | 0 | 0.2 | 0.03 | 0.24 | 0.53 | 0.27 | 0.43 |
| 7 | 122522740 | rs2501440 | -0.111 | 1.306 | -0.401 | -0.134 | -0.003 | -0.025 | -0.025 | 3.968 | 0.3 | 0.42 | 0.25 | 0.26 | 0.33 | 0.36 |
| 7 | 122525329 | rs981321 | 0.765 | 0.801 | -0.43 | -0.052 | -0.012 | -0.009 | -0.009 | 0.421 | 0.29 | 0.35 | 0.2 | 0.14 | 0.13 | 0.15 |
| 7 | 122534888 | rs10260099 | 0.178 | 0.683 | -0.791 | 0.075 | -0.009 | -0.017 | -0.017 | 0 | 0.32 | 0.26 | 0.35 | 0.61 | 0.39 | 0.45 |
| 7 | 122542284 | rs2428770 | 0.177 | 0.422 | -0.833 | -0.127 | -0.001 | -0.002 | -0.002 | 0.984 | 0.41 | 0.54 | 0.48 | 0.31 | 0.48 | 0.41 |
| 7 | 122542645 | rs2428769 | -0.59 | 0.424 | -0.919 | -0.089 | -0.027 | -0.04 | -0.04 | 1.831 | 0.68 | 0.81 | 0.65 | 0.4 | 0.6 | 0.55 |
| 7 | 122542790 | rs1006091 | 0.963 | 0.141 | -0.923 | -0.029 | 0.03 | 0.032 | 0.032 | 0 | 0.27 | 0.2 | 0.17 | 0.09 | 0.13 | 0.14 |
| 7 | 122547130 | rs6952251 | -0.435 | -0.326 | -1.03 | -0.255 | 0.004 | 0.03 | 0.03 | 0 | 0.52 | 0.71 | 0.56 | 0.43 | 0.47 | 0.46 |
| 7 | 122566809 | rs6945932 | 0.166 | -0.391 | -0.605 | -0.406 | -0.008 | -0.016 | -0.016 | 0.359 | 0.4 | 0.5 | 0.46 | 0.27 | 0.29 | 0.24 |
| 7 | 122567652 | rs9632696 | -0.826 | -0.022 | -0.944 | -0.115 | -0.09 | -0.132 | -0.132 | 0 | 0.68 | 0.95 | 0.69 | 0.46 | 0.54 | 0.53 |
| 7 | 122571553 | rs886621 | 0.554 | 0.024 | -0.14 | -0.196 | 0.008 | 0.046 | 0.046 | 0 | 0.34 | 0.21 | 0.39 | 0.16 | 0.29 | 0.19 |
| 7 | 122591543 | rs10269123 | 0.433 | 0.027 | -0.124 | -0.195 | -0.009 | -0.032 | -0.032 | 0 | 0.36 | 0.35 | 0.42 | 0.16 | 0.29 | 0.19 |
| 7 | 122595412 | rs10263466 | 1.714 | -0.088 | -0.351 | -0.28 | -0.043 | 0.068 | 0.068 | 0 | 0.19 | 0.03 | 0.27 | 0.06 | 0.2 | 0.06 |
| 7 | 122596254 | rs6466845 | 0.667 | 0.027 | -0.074 | -0.158 | 0.006 | 0.063 | 0.063 | 0 | 0.33 | 0.15 | 0.38 | 0.16 | 0.29 | 0.19 |
| 7 | 122601667 | rs17145186 | 0.092 | 0.137 | 0.091 | -0.06 | 0.007 | 0.004 | 0.004 | 0.067 | 0.13 | 0.21 | 0.11 | 0.14 | 0.18 | 0.2 |
| 7 | 122611508 | rs10249362 | 0.496 | 0.041 | 0.137 | -0.104 | 0.005 | 0.02 | 0.02 | 0 | 0.37 | 0.31 | 0.42 | 0.19 | 0.31 | 0.19 |
| 7 | 122614719 | rs6965704 | -0.014 | 0.045 | -0.101 | -0.079 | 0.001 | -0.002 | -0.002 | 0 | 0.37 | 0.43 | 0.33 | 0.33 | 0.37 | 0.47 |
| 7 | 122627618 | rs7811926 | 0.775 | 0.062 | -0.703 | -0.081 | -0.001 | 0.063 | 0.063 | 0 | 0.33 | 0.15 | 0.39 | 0.19 | 0.31 | 0.19 |
| 7 | 122635024 | rs860170 | -1.363 | 0.046 | -0.819 | -0.074 | -0.052 | -0.155 | -0.155 | 0 | 0.75 | 0.95 | 0.76 | 0.52 | 0.69 | 0.66 |
| 7 | 122636413 | rs702424 | -1.292 | 0.062 | -0.79 | -0.067 | -0.029 | -0.129 | -0.129 | 0.667 | 0.74 | 0.91 | 0.74 | 0.52 | 0.68 | 0.66 |
| 7 | 122637976 | rs9691345 | 0.785 | 0.101 | -0.678 | -0.166 | -0.004 | 0.079 | 0.079 | 0 | 0.33 | 0.11 | 0.39 | 0.16 | 0.29 | 0.19 |
| 7 | 122648394 | rs4319013 | -0.055 | 0.183 | -0.45 | 0.054 | -0.028 | -0.003 | -0.003 | 0 | 0.39 | 0.22 | 0.37 | 0.62 | 0.5 | 0.52 |
| 7 | 122659026 | rs11767634 | -0.684 | 0.188 | -0.192 | 0.004 | -0.001 | -0.007 | -0.007 | 0.299 | 0.71 | 0.34 | 0.8 | 0.81 | 0.78 | 0.7 |
| 7 | 122659240 | rs11770924 | 0.878 | 0.125 | -0.502 | -0.07 | 0 | 0.064 | 0.064 | 0 | 0.32 | 0.1 | 0.43 | 0.19 | 0.27 | 0.19 |
| 7 | 122668487 | rs2056633 | 0.764 | 0.16 | -0.044 | -0.02 | 0 | -0.007 | -0.007 | 0 | 0.28 | 0.38 | 0.18 | 0.19 | 0.22 | 0.3 |
| 7 | 122670718 | rs13221696 | 1.206 | 0.158 | -0.387 | 0.216 | -0.058 | -0.152 | -0.152 | 0 | 0.25 | 0.04 | 0.25 | 0.46 | 0.32 | 0.33 |
| 7 | 122673115 | rs4727959 | -0.718 | 0.182 | -0.01 | 0.099 | 0.001 | -0.02 | -0.02 | 0 | 0.71 | 0.44 | 0.81 | 0.81 | 0.78 | 0.7 |
| 7 | 122674826 | rs11972945 | 0.566 | 0.051 | -0.029 | 0.471 | -0.001 | -0.001 | -0.001 | 1.695 | 0.28 | 0.38 | 0.21 | 0.3 | 0.2 | 0.35 |
| 7 | 122679919 | rs13221935 | 1.486 | 0.553 | -0.577 | 0.828 | -0.06 | -0.125 | -0.125 | 0.356 | 0.2 | 0.02 | 0.17 | 0.31 | 0.29 | 0.27 |
| 7 | 122680146 | rs17370453 | 0.452 | 0.599 | -0.37 | 0.72 | -0.014 | -0.008 | -0.008 | 0 | 0.1 | 0.06 | 0.12 | 0.14 | 0.22 | 0.17 |
| 7 | 122691242 | rs10249143 | 0.388 | 0.354 | -0.394 | 0.678 | -0.043 | 0.01 | 0.01 | 0 | 0.57 | 0.21 | 0.61 | 0.56 | 0.58 | 0.48 |
| 7 | 122696223 | rs1404838 | 0.87 | 0.259 | 0.405 | 0.998 | -0.002 | 0.031 | 0.031 | 0 | 0.51 | 0.44 | 0.55 | 0.37 | 0.43 | 0.48 |
| 7 | 122708549 | rs4288315 | 0.69 | 0.165 | 0.367 | 1.097 | -0.007 | -0.001 | -0.001 | 0.065 | 0.15 | 0.19 | 0.13 | 0.12 | 0.06 | 0.11 |
| 7 | 122708700 | rs4527807 | -0.165 | 0.201 | 0.333 | 1.303 | 0.022 | -0.066 | -0.066 | 0.311 | 0.35 | 0.22 | 0.3 | 0.44 | 0.33 | 0.37 |
| 7 | 122745653 | rs10500089 | 1.154 | 0.245 | 0.439 | 1.298 | -0.054 | -0.11 | -0.11 | 0.974 | 0.21 | 0.03 | 0.18 | 0.34 | 0.3 | 0.25 |
| 7 | 122753122 | rs12706485 | 1.154 | 0.245 | 0.439 | 1.298 | -0.054 | -0.11 | -0.11 | 0.301 | 0.21 | 0.03 | 0.18 | 0.34 | 0.3 | 0.25 |
| 7 | 122758190 | rs12706486 | 1.154 | 0.237 | 0.574 | 1.294 | -0.054 | -0.11 | -0.11 | 0 | 0.21 | 0.03 | 0.18 | 0.34 | 0.3 | 0.25 |
| 7 | 122759876 | rs4352781 | -0.766 | 0.196 | 0.559 | 1.187 | 0.003 | -0.002 | -0.002 | 0 | 0.32 | 0.31 | 0.28 | 0.31 | 0.3 | 0.27 |
| 7 | 122759980 | rs4727963 | 1.101 | 0.198 | 0.617 | 1.188 | -0.003 | 0.019 | 0.019 | 0 | 0.46 | 0.46 | 0.53 | 0.35 | 0.4 | 0.48 |
| 7 | 122762539 | rs10255262 | 1.274 | 0.199 | 0.62 | 1.189 | -0.021 | -0.063 | -0.063 | 0 | 0.21 | 0.07 | 0.18 | 0.34 | 0.3 | 0.25 |
| 7 | 122763787 | rs12706487 | 1.326 | 0.199 | 0.615 | 1.198 | -0.054 | -0.112 | -0.112 | 0.117 | 0.21 | 0.03 | 0.18 | 0.34 | 0.3 | 0.25 |
| 7 | 122766270 | rs10487980 | -0.638 | 0.195 | 0.632 | 1.196 | 0.004 | -0.01 | -0.01 | 1.19 | 0.31 | 0.25 | 0.27 | 0.31 | 0.3 | 0.27 |
| 7 | 122766538 | rs10487979 | -0.708 | 0.197 | 0.619 | 1.176 | 0.003 | -0.002 | -0.002 | 0 | 0.31 | 0.29 | 0.27 | 0.31 | 0.3 | 0.27 |
| 7 | 122766907 | rs2204292 | -1.129 | 0.197 | 0.621 | 1.176 | -0.003 | 0.02 | 0.02 | 0 | 0.54 | 0.54 | 0.47 | 0.65 | 0.6 | 0.52 |
| 7 | 122767805 | rs12538634 | -0.689 | 0.212 | 0.607 | 1.187 | 0.003 | -0.003 | -0.003 | 0.017 | 0.31 | 0.29 | 0.28 | 0.31 | 0.3 | 0.27 |
| 7 | 122768213 | rs12706489 | 1.2 | 0.243 | 0.341 | 1.227 | -0.02 | -0.064 | -0.064 | 0 | 0.21 | 0.07 | 0.18 | 0.34 | 0.3 | 0.25 |
| 7 | 122769628 | rs13229376 | 1.216 | 0.458 | 0.426 | 0.989 | -0.019 | -0.061 | -0.061 | 0 | 0.21 | 0.07 | 0.18 | 0.34 | 0.3 | 0.25 |
| 7 | 122770085 | rs13244086 | 1.263 | 0.464 | 0.309 | 0.992 | -0.054 | -0.112 | -0.112 | 1.947 | 0.21 | 0.03 | 0.18 | 0.34 | 0.3 | 0.25 |
| 7 | 122773032 | rs12706494 | 1.402 | 0.472 | 0.328 | 0.991 | -0.022 | -0.068 | -0.068 | 2.581 | 0.2 | 0.06 | 0.17 | 0.34 | 0.28 | 0.23 |
| 7 | 122774603 | rs2470983 | -0.103 | 0.563 | 0.549 | 0.944 | -0.025 | 0.007 | 0.007 | 2.049 | 0.24 | 0.58 | 0.22 | 0.21 | 0.16 | 0.19 |
| 7 | 122774729 | rs2470984 | -0.746 | 0.648 | 0.691 | 0.856 | -0.008 | -0.003 | -0.003 | 0 | 0.33 | 0.55 | 0.3 | 0.28 | 0.3 | 0.27 |
| 7 | 122777190 | rs6466855 | -0.63 | 0.646 | 0.729 | 0.855 | -0.003 | -0.001 | -0.001 | 0 | 0.32 | 0.42 | 0.29 | 0.28 | 0.3 | 0.27 |
| 7 | 122780027 | rs6964272 | -0.576 | 0.635 | 0.739 | 0.857 | -0.004 | 0 | 0 | 0 | 0.32 | 0.42 | 0.28 | 0.28 | 0.29 | 0.27 |
| 7 | 122781859 | rs12706496 | 1.47 | 0.64 | 0.718 | 0.864 | -0.067 | -0.124 | -0.124 | 0.097 | 0.2 | 0.03 | 0.18 | 0.34 | 0.29 | 0.24 |
| 7 | 122783013 | rs10236058 | 1.437 | 0.64 | 0.729 | 0.864 | -0.024 | -0.067 | -0.067 | 0 | 0.2 | 0.06 | 0.18 | 0.34 | 0.3 | 0.24 |
| 7 | 122783281 | rs10251971 | -0.592 | 0.638 | 0.735 | 0.862 | -0.004 | -0.001 | -0.001 | 0 | 0.32 | 0.42 | 0.28 | 0.28 | 0.29 | 0.27 |
| 7 | 122785428 | rs2470975 | -1.106 | 0.642 | 0.703 | 0.855 | -0.006 | 0.043 | 0.043 | 0 | 0.54 | 0.64 | 0.48 | 0.65 | 0.6 | 0.52 |
| 7 | 122785918 | rs12666250 | 1.399 | 0.698 | 0.54 | 0.892 | -0.024 | -0.067 | -0.067 | 0.057 | 0.2 | 0.06 | 0.18 | 0.34 | 0.3 | 0.24 |
| 7 | 122790109 | rs10258700 | 1.415 | 0.599 | 0.564 | 0.912 | -0.035 | -0.084 | -0.084 | 3.314 | 0.2 | 0.04 | 0.18 | 0.34 | 0.3 | 0.24 |
| 7 | 122791856 | rs13245959 | 1.432 | 0.596 | 0.595 | 0.911 | -0.067 | -0.124 | -0.124 | 0.568 | 0.2 | 0.03 | 0.18 | 0.34 | 0.29 | 0.24 |
| 7 | 122793182 | rs10953973 | -0.959 | 0.562 | 0.62 | 0.891 | 0 | -0.003 | -0.003 | 0 | 0.53 | 0.53 | 0.46 | 0.62 | 0.58 | 0.51 |
| 7 | 122798636 | rs2462152 | -1.031 | 0.549 | 0.604 | 0.884 | -0.01 | 0.045 | 0.045 | 0 | 0.55 | 0.67 | 0.49 | 0.65 | 0.6 | 0.52 |
| 7 | 122799019 | rs12671045 | 1.363 | 0.523 | 0.544 | 0.915 | -0.065 | -0.124 | -0.124 | 0.238 | 0.21 | 0.03 | 0.18 | 0.34 | 0.29 | 0.24 |
| 7 | 122799737 | rs10262500 | -0.972 | 0.558 | 0.563 | 0.931 | 0 | -0.006 | -0.006 | 0.087 | 0.53 | 0.53 | 0.46 | 0.63 | 0.58 | 0.51 |
| 7 | 122799767 | rs12537296 | -0.566 | 0.554 | 0.483 | 0.926 | 0.001 | 0 | 0 | 0 | 0.32 | 0.38 | 0.28 | 0.28 | 0.29 | 0.27 |
| 7 | 122802824 | rs10231621 | 1.338 | 0.597 | 0.394 | 0.956 | -0.034 | -0.085 | -0.085 | 0 | 0.2 | 0.05 | 0.18 | 0.34 | 0.3 | 0.24 |
| 7 | 122805355 | rs10280430 | -0.761 | 0.676 | 0.652 | 0.908 | -0.006 | -0.002 | -0.002 | 0 | 0.33 | 0.45 | 0.28 | 0.28 | 0.29 | 0.27 |
| 7 | 122807680 | rs1343905 | 0.718 | 1.458 | -0.479 | 1.261 | -0.001 | -0.001 | -0.001 | 0.706 | 0.22 | 0.22 | 0.19 | 0.32 | 0.3 | 0.23 |

| | | | | | | | | | | | | | | | | |
|---|---|---|---|---|---|---|---|---|---|---|---|---|---|---|---|---|
| 7 | 122807917 | rs6953295 | 0.01 | 1.486 | -0.598 | 1.24 | -0.044 | -0.03 | -0.03 | 0.735 | 0.15 | 0.06 | 0.13 | 0.16 | 0.21 | 0.15 |
| 7 | 122808528 | rs11976455 | -0.102 | 1.493 | -0.595 | 1.245 | -0.044 | -0.032 | -0.032 | 0 | 0.15 | 0.06 | 0.13 | 0.17 | 0.21 | 0.15 |
| 7 | 122809234 | rs2140516 | 0.234 | 1.405 | -0.165 | 1.09 | 0 | 0.001 | 0.001 | 0 | 0.24 | 0.24 | 0.2 | 0.35 | 0.34 | 0.24 |
| 7 | 122809564 | rs2204291 | 0.532 | 1.378 | -0.146 | 0.666 | 0.006 | -0.005 | -0.005 | 0 | 0.21 | 0.24 | 0.18 | 0.16 | 0.31 | 0.22 |
| 7 | 122812281 | rs17445780 | 0.716 | 0.736 | 0.099 | 0.396 | 0.023 | 0.018 | 0.018 | 0.452 | 0.43 | 0.2 | 0.43 | 0.38 | 0.38 | 0.42 |
| 7 | 122825956 | rs10253693 | 0.281 | 0.731 | 0.299 | 0.377 | 0.018 | 0.023 | 0.023 | 0.079 | 0.47 | 0.59 | 0.48 | 0.4 | 0.42 | 0.43 |
| 7 | 122828502 | rs10265711 | 0.277 | 0.728 | 0.214 | 0.365 | 0.017 | 0.023 | 0.023 | 0.059 | 0.47 | 0.58 | 0.48 | 0.4 | 0.42 | 0.43 |
| 7 | 122835155 | rs7780966 | 0.532 | 0.703 | 0.362 | 0.347 | 0.009 | 0.009 | 0.009 | 0.002 | 0.46 | 0.51 | 0.46 | 0.38 | 0.38 | 0.43 |
| 7 | 122835742 | rs2204294 | 0.823 | 0.613 | 0.797 | 0.3 | 0.02 | 0.017 | 0.017 | 0.254 | 0.42 | 0.27 | 0.44 | 0.37 | 0.38 | 0.4 |
| 7 | 122838633 | rs1880179 | 0.811 | 0.592 | 0.78 | 0.29 | 0.022 | 0.016 | 0.016 | 0.667 | 0.42 | 0.16 | 0.43 | 0.38 | 0.38 | 0.42 |
| 7 | 122841575 | rs3824026 | 0.877 | 0.601 | 0.807 | 0.282 | 0.02 | 0.016 | 0.016 | 0.346 | 0.42 | 0.23 | 0.44 | 0.37 | 0.38 | 0.42 |
| 7 | 122842029 | rs3824025 | 0.204 | 0.919 | 0.824 | 0.178 | -0.066 | -0.08 | -0.08 | 0 | 0.35 | 0.14 | 0.39 | 0.53 | 0.44 | 0.43 |
| 7 | 122842075 | rs17544071 | 1.226 | 0.934 | 0.839 | 0.243 | 0.013 | 0.007 | 0.007 | 0.627 | 0.41 | 0.15 | 0.43 | 0.37 | 0.37 | 0.42 |
| 7 | 122845289 | rs10487975 | 1.055 | 0.93 | 0.82 | 0.242 | 0.025 | 0.019 | 0.019 | 0.439 | 0.42 | 0.2 | 0.43 | 0.37 | 0.38 | 0.42 |
| 7 | 122846211 | rs7810239 | 1.105 | 0.942 | 0.798 | 0.247 | 0.019 | 0.016 | 0.016 | 0.25 | 0.42 | 0.27 | 0.44 | 0.37 | 0.38 | 0.42 |
| 7 | 122846527 | rs2222514 | 0.79 | 0.973 | 0.57 | 0.263 | 0.009 | 0.009 | 0.009 | 0.002 | 0.46 | 0.51 | 0.46 | 0.37 | 0.38 | 0.42 |
| 7 | 122858776 | rs7776703 | 1.034 | 0.921 | 0.31 | 0.111 | 0.002 | 0.007 | 0.007 | 0.954 | 0.44 | 0.15 | 0.43 | 0.38 | 0.39 | 0.41 |
| 7 | 122864924 | rs4731086 | 1.19 | 0.885 | 0.294 | 0.095 | -0.006 | -0.001 | -0.001 | 0.539 | 0.42 | 0.17 | 0.43 | 0.38 | 0.39 | 0.4 |
| 7 | 122865373 | rs4731088 | 1.19 | 0.885 | 0.294 | 0.095 | -0.006 | -0.001 | -0.001 | 0.086 | 0.42 | 0.17 | 0.43 | 0.38 | 0.39 | 0.4 |
| 7 | 122869067 | rs17447609 | 1.193 | 1.388 | 0.099 | 0.147 | -0.035 | -0.002 | -0.002 | 0 | 0.09 | 0.01 | 0.12 | 0.07 | 0.15 | 0.03 |
| 7 | 122874252 | rs12670549 | 0.587 | 1.07 | -0.009 | 0.321 | -0.002 | -0.018 | -0.018 | 0 | 0.15 | 0.09 | 0.16 | 0.32 | 0.16 | 0.18 |
| 7 | 122877637 | rs12533726 | 0.488 | 0.987 | -0.072 | 0.271 | -0.05 | 0.014 | 0.014 | 1.944 | 0.22 | 0.02 | 0.23 | 0.18 | 0.27 | 0.27 |
| 7 | 122878459 | rs7807163 | 0.758 | 0.924 | 0.247 | 0.267 | 0.01 | 0.02 | 0.02 | 0.256 | 0.5 | 0.55 | 0.48 | 0.41 | 0.43 | 0.43 |
| 7 | 122885312 | rs6963870 | -1.151 | 1.164 | 0.295 | 0.205 | 0 | 0 | 0 | 0.326 | 0.55 | 0.45 | 0.54 | 0.62 | 0.64 | 0.61 |
| 7 | 122894149 | rs6961894 | -1.632 | 0.591 | -0.03 | -0.274 | -0.011 | -0.06 | -0.06 | 0 | 0.83 | 0.7 | 0.92 | 0.97 | 0.88 | 0.92 |
| 7 | 122895308 | rs10953976 | 1.481 | 0.502 | -0.146 | -0.229 | 0.008 | -0.01 | -0.01 | 1.171 | 0.29 | 0.22 | 0.39 | 0.35 | 0.26 | 0.33 |
| 7 | 122896288 | rs7786287 | 1.224 | 0.916 | -0.423 | -0.195 | -0.024 | -0.003 | -0.003 | 0 | 0.19 | 0.08 | 0.22 | 0.21 | 0.24 | 0.23 |
| 7 | 122903158 | rs10953977 | -0.263 | 0.654 | -0.405 | -0.077 | 0.028 | 0.015 | 0.015 | 0.684 | 0.47 | 0.22 | 0.5 | 0.58 | 0.59 | 0.57 |
| 7 | 122914243 | rs4142046 | 1.515 | 0.583 | -0.095 | -0.066 | 0.024 | -0.02 | -0.02 | 5.385 | 0.31 | 0.24 | 0.38 | 0.36 | 0.26 | 0.34 |
| 7 | 122915565 | rs10808198 | 0.05 | 1.17 | -0.289 | -0.01 | 0.017 | 0.037 | 0.037 | 0 | 0.1 | 0.1 | 0.18 | 0.32 | 0.23 | 0.2 |
| 7 | 122917060 | rs4731094 | -0.436 | 0.432 | -0.209 | -0.149 | 0.064 | 0.032 | 0.032 | 1.351 | 0.49 | 0.21 | 0.51 | 0.58 | 0.61 | 0.58 |
| 7 | 122918068 | rs12671529 | 1.017 | 0.28 | 0.152 | 0.088 | -0.017 | 0.007 | 0.007 | 0 | 0.33 | 0.42 | 0.4 | 0.38 | 0.26 | 0.34 |
| 7 | 122927649 | rs7805826 | 1.158 | 0.463 | -0.037 | 0.148 | 0.021 | -0.026 | -0.026 | 0 | 0.29 | 0.25 | 0.37 | 0.34 | 0.25 | 0.31 |
| 7 | 122932940 | rs7795788 | -0.227 | 0.469 | -0.081 | -0.084 | 0.043 | 0.013 | 0.013 | 1.087 | 0.47 | 0.21 | 0.46 | 0.6 | 0.59 | 0.61 |
| 7 | 122935503 | rs12538914 | 0.895 | 0.718 | -0.246 | -0.113 | -0.037 | -0.043 | -0.043 | 0 | 0.24 | 0.11 | 0.25 | 0.37 | 0.32 | 0.33 |
| 7 | 122935640 | rs12706507 | 1.687 | 0.574 | -0.022 | -0.202 | -0.009 | 0.008 | 0.008 | 0 | 0.3 | 0.39 | 0.4 | 0.37 | 0.26 | 0.3 |
| 7 | 122950090 | rs4731100 | 0.686 | 0.754 | -0.346 | -0.103 | -0.005 | -0.006 | -0.006 | 0 | 0.26 | 0.18 | 0.26 | 0.37 | 0.32 | 0.33 |
| 7 | 122955347 | rs11984044 | -0.058 | 0.734 | -0.52 | -0.175 | -0.016 | 0.004 | 0.004 | 0 | 0.23 | 0.14 | 0.2 | 0.24 | 0.26 | 0.28 |
| 7 | 122968560 | rs11977834 | 1.071 | 1.037 | -0.793 | -0.047 | -0.033 | -0.034 | -0.034 | 0.696 | 0.06 | 0.01 | 0.11 | 0.19 | 0.18 | 0.15 |
| 7 | 122978928 | rs2191800 | -1.05 | 0.772 | -0.488 | -0.174 | -0.008 | -0.037 | -0.037 | 0.413 | 0.79 | 0.74 | 0.86 | 0.95 | 0.84 | 0.91 |
| 7 | 122986940 | rs4731102 | 0.238 | 0.946 | -0.811 | 0.044 | 0.003 | 0.023 | 0.023 | 0 | 0.55 | 0.44 | 0.65 | 0.71 | 0.58 | 0.65 |
| 7 | 122990979 | rs2191801 | 0.031 | 1.503 | -1.108 | 0.388 | -0.026 | -0.024 | -0.024 | 0.365 | 0.24 | 0.14 | 0.2 | 0.35 | 0.31 | 0.34 |
| 7 | 122992488 | rs7778004 | -0.084 | 1.687 | -1.321 | 0.374 | -0.07 | -0.037 | -0.037 | 0.025 | 0.26 | 0.09 | 0.23 | 0.36 | 0.39 | 0.37 |
| 7 | 122992937 | rs4731104 | 0.255 | 1.389 | -1.758 | 0.264 | -0.063 | -0.021 | -0.021 | 0 | 0.63 | 0.89 | 0.76 | 0.64 | 0.57 | 0.62 |
| 7 | 122993794 | rs11769355 | -0.553 | 0.904 | -0.948 | 0.216 | 0 | 0 | 0 | 0 | 0.55 | 0.49 | 0.47 | 0.44 | 0.57 | 0.69 |
| 7 | 123001083 | rs7809779 | 0.122 | 0.841 | -0.747 | 0.257 | 0.003 | 0.023 | 0.023 | 0 | 0.12 | 0.2 | 0.21 | 0.2 | 0.14 | 0.23 |
| 7 | 123010773 | rs13247719 | 1.346 | 1.006 | -1.04 | 0.033 | -0.036 | -0.011 | -0.011 | 0 | 0.18 | 0.08 | 0.13 | 0.16 | 0.21 | 0.19 |
| 7 | 123014728 | rs17132820 | 0.101 | 0.832 | -0.711 | 0.281 | 0 | 0.006 | 0.006 | 0 | 0.12 | 0.12 | 0.21 | 0.21 | 0.14 | 0.24 |
| 7 | 123017883 | rs12536968 | 0.242 | 1.023 | -0.439 | 0.234 | 0.003 | 0.049 | 0.049 | 0.004 | 0.45 | 0.34 | 0.24 | 0.25 | 0.37 | 0.46 |
| 7 | 123037157 | rs1860722 | -0.436 | 1.074 | 0.47 | 0.307 | 0.023 | 0.04 | 0.04 | 0 | 0.13 | 0.27 | 0.22 | 0.21 | 0.16 | 0.32 |
| 7 | 123037887 | rs10278912 | 0.267 | 1.111 | 0.275 | 0.323 | 0.013 | 0.022 | 0.022 | 0 | 0.1 | 0.14 | 0.22 | 0.21 | 0.15 | 0.2 |
| 7 | 123051342 | rs805801 | -0.718 | 0.875 | 0.249 | 0.149 | -0.035 | -0.045 | -0.045 | 0 | 0.61 | 0.82 | 0.5 | 0.45 | 0.54 | 0.77 |
| 7 | 123051352 | rs6967772 | -0.464 | 0.842 | 0.597 | 0.204 | 0.018 | 0.032 | 0.032 | 0 | 0.13 | 0.3 | 0.22 | 0.21 | 0.16 | 0.3 |
| 7 | 123066710 | rs10244905 | 0.557 | 1.051 | 0.395 | 0.288 | 0.018 | 0.023 | 0.023 | 0 | 0.09 | 0.12 | 0.2 | 0.2 | 0.15 | 0.18 |
| 7 | 123076048 | rs1554716 | 0.051 | 0.76 | 0.409 | 0.114 | 0.013 | 0.051 | 0.051 | 0.083 | 0.5 | 0.42 | 0.68 | 0.69 | 0.61 | 0.53 |
| 7 | 123097194 | rs1525627 | -0.661 | 1.057 | 0.806 | 0.837 | 0.042 | 0.092 | 0.092 | 0 | 0.21 | 0.39 | 0.28 | 0.39 | 0.3 | 0.29 |
| 7 | 123100220 | rs4727971 | -1.498 | 1.059 | 0.727 | 0.66 | -0.001 | -0.001 | -0.001 | 0 | 0.8 | 0.84 | 0.83 | 0.79 | 0.82 | 0.78 |
| 7 | 123128500 | rs7807724 | -0.745 | 1.395 | 0.925 | 0.554 | 0.045 | 0.038 | 0.038 | 0.021 | 0.23 | 0.36 | 0.3 | 0.37 | 0.38 | 0.23 |
| 7 | 123129020 | rs7791660 | -1.484 | 1.303 | 0.966 | 0.536 | 0.002 | 0.007 | 0.007 | 0 | 0.8 | 0.81 | 0.83 | 0.77 | 0.82 | 0.77 |
| 7 | 123144234 | rs10280991 | -0.294 | 1.351 | 1.064 | 0.546 | 0.013 | 0.028 | 0.028 | 0 | 0.2 | 0.22 | 0.26 | 0.37 | 0.29 | 0.21 |
| 7 | 123150236 | rs4276609 | 1.435 | 0.966 | 0.787 | 0.109 | -0.007 | -0.044 | -0.044 | 0 | 0.33 | 0.16 | 0.43 | 0.34 | 0.3 | 0.23 |
| 7 | 123150816 | rs10250681 | 0.231 | 0.938 | 0.894 | 0.096 | -0.018 | -0.027 | -0.027 | 0 | 0.19 | 0.1 | 0.25 | 0.37 | 0.28 | 0.21 |
| 7 | 123152019 | rs10255061 | -0.01 | 0.957 | 0.833 | 0.104 | 0.013 | 0.028 | 0.028 | 0 | 0.2 | 0.22 | 0.26 | 0.37 | 0.29 | 0.21 |
| 7 | 123152035 | rs10270705 | -0.447 | 0.95 | 0.843 | 0.1 | 0.04 | 0.035 | 0.035 | 0.013 | 0.23 | 0.34 | 0.3 | 0.37 | 0.37 | 0.23 |
| 7 | 123155952 | rs12535069 | -1.873 | 0.956 | 0.879 | 0.09 | 0.005 | 0.01 | 0.01 | 0 | 0.8 | 0.77 | 0.82 | 0.77 | 0.82 | 0.77 |
| 7 | 123162177 | rs2037697 | 1.376 | 1 | 0.8 | 0.226 | -0.007 | 0.009 | 0.009 | 1.537 | 0.33 | 0.16 | 0.4 | 0.27 | 0.3 | 0.22 |
| 7 | 123174364 | rs10081283 | -0.652 | 1.013 | 0.962 | 0.358 | 0.063 | 0.056 | 0.056 | 0.002 | 0.23 | 0.44 | 0.3 | 0.37 | 0.38 | 0.28 |
| 7 | 123174842 | rs10428991 | -1.809 | 0.968 | 0.867 | 0.371 | 0 | 0.005 | 0.005 | 0 | 0.81 | 0.83 | 0.81 | 0.7 | 0.81 | 0.76 |
| 7 | 123179787 | rs4147635 | -0.344 | 1.051 | 0.953 | 0.249 | 0.02 | 0.04 | 0.04 | 0 | 0.2 | 0.27 | 0.27 | 0.37 | 0.29 | 0.27 |

| | | | | | | | | | | | | | | | | |
|---|---|---|---|---|---|---|---|---|---|---|---|---|---|---|---|---|
| 7 | 123191974 | rs12666974 | 0.149 | 1.044 | 0.695 | 0.246 | -0.035 | -0.048 | -0.048 | 0 | 0.17 | 0.07 | 0.25 | 0.37 | 0.29 | 0.27 |
| 7 | 123198709 | rs2037696 | 0.242 | 1.053 | 0.8 | 0.25 | 0.045 | 0.087 | 0.087 | 0 | 0.79 | 0.57 | 0.72 | 0.63 | 0.71 | 0.74 |
| 7 | 123200547 | rs12706537 | 0.627 | 1.05 | 0.766 | 0.244 | 0.058 | 0.051 | 0.051 | 0.012 | 0.77 | 0.5 | 0.69 | 0.63 | 0.62 | 0.73 |
| 7 | 123207609 | rs12538107 | 0.639 | 0.572 | 0.417 | -0.052 | 0.007 | -0.001 | -0.001 | 0.706 | 0.25 | 0.23 | 0.1 | 0.06 | 0.14 | 0.25 |
| 7 | 123210624 | rs940800 | 1.099 | 1.107 | 0.429 | 0.323 | 0.001 | 0.008 | 0.008 | 0 | 0.08 | 0.07 | 0.19 | 0.15 | 0.13 | 0.12 |
| 7 | 123212712 | rs12671023 | -0.658 | 0.948 | 0.549 | 0.209 | -0.003 | -0.01 | -0.01 | 0 | 0.46 | 0.5 | 0.39 | 0.37 | 0.5 | 0.42 |
| 7 | 123214835 | rs805779 | -1.532 | 0.844 | 0.298 | 0.216 | -0.02 | 0.013 | 0.013 | 0.1 | 0.72 | 0.81 | 0.61 | 0.75 | 0.71 | 0.75 |
| 7 | 123218522 | rs10231717 | -0.111 | 0.858 | 0.349 | 0.203 | -0.012 | -0.01 | -0.01 | 0.789 | 0.41 | 0.45 | 0.34 | 0.38 | 0.4 | 0.4 |
| 7 | 123219972 | rs805785 | 0.998 | 0.881 | 0.402 | 0.278 | -0.001 | 0.013 | 0.013 | 0 | 0.26 | 0.23 | 0.21 | 0.38 | 0.21 | 0.33 |
| 7 | 123243261 | rs9649463 | 0.377 | 0.875 | 0.393 | 0.285 | 0.017 | 0.03 | 0.03 | 0 | 0.17 | 0.2 | 0.26 | 0.31 | 0.27 | 0.16 |
| 7 | 123245010 | rs7798658 | -0.578 | 0.812 | 0.336 | 0.226 | 0.007 | 0.047 | 0.047 | 0 | 0.47 | 0.6 | 0.51 | 0.69 | 0.57 | 0.51 |
| 7 | 123248738 | rs35947027 | 0.578 | 0.752 | 0.299 | 0.2 | 0.002 | 0.011 | 0.011 | 0 | 0.29 | 0.3 | 0.24 | 0.37 | 0.29 | 0.35 |
| 7 | 123251748 | rs7797819 | 0.916 | 0.515 | -0.114 | -0.234 | 0.032 | 0.045 | 0.045 | 0 | 0.25 | 0.17 | 0.1 | 0.09 | 0.15 | 0.2 |
| 7 | 123253494 | rs6966714 | 0.005 | 0.76 | -0.101 | -0.069 | -0.001 | -0.001 | -0.001 | 0 | 0.42 | 0.35 | 0.47 | 0.7 | 0.48 | 0.49 |
| 7 | 123256427 | rs6962756 | -1.476 | 1.052 | -0.427 | -0.157 | -0.005 | 0.004 | 0.004 | 1.305 | 0.77 | 0.81 | 0.64 | 0.79 | 0.77 | 0.83 |
| 7 | 123271192 | rs12669165 | 0.744 | 1.233 | -0.476 | -0.027 | -0.037 | -0.025 | -0.025 | 0.999 | 0.09 | 0.01 | 0.08 | 0.13 | 0.14 | 0.04 |
| 7 | 123285578 | rs17146203 | 0.006 | 1.709 | 0.066 | 0.166 | 0.13 | 0.14 | 0.14 | 0 | 0.05 | 0.25 | 0.21 | 0.16 | 0.12 | 0.06 |
| 7 | 123295964 | rs7779099 | -1.17 | 1.556 | -0.371 | -0.077 | -0.01 | 0.037 | 0.037 | 0.193 | 0.71 | 0.83 | 0.74 | 0.84 | 0.71 | 0.85 |
| 7 | 123296320 | rs12538876 | 0.515 | 1.183 | -0.21 | -0.262 | 0.014 | 0.024 | 0.024 | 0 | 0.51 | 0.32 | 0.43 | 0.56 | 0.44 | 0.71 |
| 7 | 123301940 | rs7809453 | 0.386 | 1.147 | -0.273 | -0.278 | 0.004 | 0.007 | 0.007 | 0 | 0.52 | 0.4 | 0.43 | 0.56 | 0.44 | 0.71 |
| 7 | 123303477 | rs2286831 | 0.395 | 1.147 | -0.27 | -0.278 | 0.004 | 0.008 | 0.008 | 0 | 0.52 | 0.39 | 0.43 | 0.56 | 0.44 | 0.71 |
| 7 | 123305444 | rs12706544 | 1.451 | 1.066 | -0.477 | -0.25 | 0.014 | -0.029 | -0.029 | 0 | 0.25 | 0.09 | 0.24 | 0.26 | 0.25 | 0.34 |
| 7 | 123312847 | rs4731117 | 1.264 | 1.443 | -0.336 | -0.07 | -0.035 | -0.026 | -0.026 | 0 | 0.08 | 0.02 | 0.08 | 0.13 | 0.14 | 0.04 |
| 7 | 123314446 | rs10487806 | 0.526 | 1.463 | -0.319 | -0.057 | 0.017 | 0.031 | 0.031 | 0 | 0.51 | 0.29 | 0.43 | 0.56 | 0.44 | 0.71 |
| 7 | 123336804 | rs10270793 | -0.223 | 1.484 | -0.369 | -0.039 | 0.047 | 0.036 | 0.036 | 0 | 0.19 | 0.41 | 0.31 | 0.28 | 0.27 | 0.13 |
| 7 | 123337873 | rs1006764 | 0.283 | 1.527 | -0.758 | 0.022 | 0.004 | 0.007 | 0.007 | 0 | 0.52 | 0.4 | 0.43 | 0.56 | 0.44 | 0.71 |
| 7 | 123395983 | rs608447 | -0.101 | 1.002 | -0.976 | 0.198 | 0.01 | 0.017 | 0.017 | 0 | 0.49 | 0.64 | 0.57 | 0.44 | 0.56 | 0.29 |
| 7 | 123407275 | rs623201 | -1.019 | 1.025 | -1.404 | 0.387 | -0.059 | -0.086 | -0.086 | 0.375 | 0.81 | 0.98 | 0.84 | 0.7 | 0.81 | 0.74 |
| 7 | 123408842 | rs9886090 | 0.31 | 1.121 | -0.774 | 0.21 | 0.001 | -0.002 | -0.002 | 0 | 0.08 | 0.08 | 0.07 | 0.13 | 0.1 | 0.04 |
| 7 | 123441559 | rs10260159 | -0.231 | 0.936 | -0.565 | 0.432 | -0.005 | 0.017 | 0.017 | 0 | 0.09 | 0.15 | 0.15 | 0.17 | 0.1 | 0.05 |
| 7 | 123442156 | rs633590 | 0.143 | 1.018 | -0.73 | -0.022 | 0 | 0.027 | 0.027 | 0 | 0.39 | 0.33 | 0.26 | 0.2 | 0.39 | 0.21 |
| 7 | 123446388 | rs13240375 | 0.827 | 0.854 | -1.132 | 0.042 | -0.016 | 0.02 | 0.02 | 0.046 | 0.32 | 0.14 | 0.35 | 0.25 | 0.34 | 0.39 |
| 7 | 123450154 | rs12672990 | 0.42 | 0.809 | -0.732 | 0.353 | 0 | 0 | 0 | 1.29 | 0.15 | 0.12 | 0.08 | 0.16 | 0.15 | 0.07 |
| 7 | 123454976 | rs11764394 | 0.704 | 0.8 | -0.733 | 0.274 | -0.005 | 0.001 | 0.001 | 1.861 | 0.13 | 0.06 | 0.07 | 0.14 | 0.14 | 0.07 |
| 7 | 123463159 | rs2109352 | -0.523 | 0.774 | -0.394 | 0.485 | 0.002 | -0.003 | -0.003 | 1.263 | 0.82 | 0.84 | 0.82 | 0.65 | 0.85 | 0.72 |
| 7 | 123465342 | rs693998 | -0.437 | 0.791 | 0.017 | 0.519 | 0.039 | 0.039 | 0.039 | 0.599 | 0.2 | 0.39 | 0.3 | 0.26 | 0.24 | 0.12 |
| 7 | 123466528 | rs17624713 | 1.833 | 0.738 | -0.543 | 0.198 | -0.001 | -0.003 | -0.003 | 0.179 | 0.05 | 0.06 | 0.02 | 0 | 0.04 | 0.04 |
| 7 | 123467419 | rs1506639 | 1.162 | 0.753 | -0.653 | 0.273 | 0.002 | 0.01 | 0.01 | 0.108 | 0.56 | 0.33 | 0.5 | 0.4 | 0.58 | 0.55 |
| 7 | 123488985 | rs10953985 | -0.267 | 0.752 | -0.712 | 0.26 | -0.001 | -0.003 | -0.003 | 0 | 0.15 | 0.15 | 0.08 | 0.14 | 0.15 | 0.07 |
| 7 | 123498400 | rs10276917 | -0.267 | 0.752 | -0.712 | 0.26 | -0.001 | -0.003 | -0.003 | 2.715 | 0.15 | 0.15 | 0.08 | 0.14 | 0.15 | 0.07 |
| 7 | 123504320 | rs17703063 | 0.157 | 0.356 | -1.338 | -0.49 | 0.019 | 0.031 | 0.031 | 0 | 0.2 | 0.1 | 0.09 | 0 | 0.16 | 0.15 |
| 7 | 123505744 | rs36929 | -0.361 | 0.238 | -0.957 | -0.082 | -0.001 | 0.078 | 0.078 | 0 | 0.83 | 0.78 | 0.66 | 0.55 | 0.81 | 0.71 |
| 7 | 123506615 | rs12706554 | 0.644 | 0.579 | -1.099 | -0.131 | 0.005 | 0.032 | 0.032 | 0 | 0.2 | 0.1 | 0.1 | 0 | 0.18 | 0.15 |
| 7 | 123508309 | rs1034955 | 0.511 | 0.522 | -0.927 | -0.136 | 0.001 | -0.027 | -0.027 | 0 | 0.21 | 0.17 | 0.1 | 0 | 0.18 | 0.15 |
| 7 | 123510483 | rs36930 | 2.119 | 0.665 | -1.191 | -0.092 | 0.007 | 0.042 | 0.042 | 2.33 | 0.28 | 0.17 | 0.16 | 0.23 | 0.24 | 0.21 |
| 7 | 123514896 | rs6949082 | 0.944 | 0.515 | -1.027 | -0.105 | 0.005 | 0.004 | 0.004 | 0 | 0.2 | 0.13 | 0.1 | 0 | 0.18 | 0.15 |
| 7 | 123526959 | rs250641 | -1.113 | 0.398 | -0.967 | -0.12 | 0.002 | 0.065 | 0.065 | 0 | 0.92 | 0.93 | 0.85 | 0.66 | 0.87 | 0.8 |
| 7 | 123542668 | rs2402676 | -1.71 | 0.346 | -0.713 | -0.498 | -0.027 | 0.002 | 0.002 | 0.131 | 0.59 | 0.7 | 0.54 | 0.56 | 0.53 | 0.58 |
| 7 | 123550573 | rs6466886 | 0.447 | 0.385 | -0.443 | -0.484 | 0 | -0.039 | -0.039 | 0 | 0.21 | 0.19 | 0.1 | 0 | 0.18 | 0.15 |
| 7 | 123552968 | rs13221454 | 0.711 | 0.47 | -0.714 | -0.446 | -0.002 | 0.06 | 0.06 | 0 | 0.19 | 0.04 | 0.09 | 0 | 0.18 | 0.15 |
| 7 | 123578101 | rs9986737 | 0.135 | 0.483 | -0.662 | -0.321 | -0.003 | -0.004 | -0.004 | 0 | 0.26 | 0.22 | 0.4 | 0.3 | 0.29 | 0.24 |
| 7 | 123580016 | rs2189684 | -2.23 | 0.28 | -0.767 | -0.473 | 0.002 | 0.027 | 0.027 | 0.613 | 0.73 | 0.82 | 0.84 | 0.8 | 0.75 | 0.8 |
| 7 | 123582328 | rs1014439 | 0.169 | 0.16 | -0.629 | -0.435 | 0.003 | 0.065 | 0.065 | 0 | 0.2 | 0.3 | 0.31 | 0.42 | 0.21 | 0.3 |
| 7 | 123584796 | rs10269135 | 1.096 | 0.219 | -0.452 | -0.583 | 0 | -0.025 | -0.025 | 0 | 0.16 | 0.11 | 0.09 | 0 | 0.13 | 0.13 |
| 7 | 123588444 | rs3808168 | -1.08 | 0.213 | -0.488 | -0.658 | 0.007 | 0.051 | 0.051 | 0.931 | 0.49 | 0.65 | 0.54 | 0.37 | 0.53 | 0.49 |
| 7 | 123588893 | rs4727988 | 0.948 | 0.19 | -0.475 | -0.691 | 0.003 | 0.007 | 0.007 | 0 | 0.16 | 0.08 | 0.08 | 0 | 0.14 | 0.14 |
| 7 | 123592946 | rs7804904 | -1.41 | 0.054 | -0.532 | -0.76 | -0.006 | 0.036 | 0.036 | 1.362 | 0.55 | 0.75 | 0.56 | 0.37 | 0.55 | 0.52 |
| 7 | 123594206 | rs2285996 | 0.985 | 0.124 | -0.481 | -0.705 | -0.01 | -0.011 | -0.011 | 0 | 0.19 | 0.13 | 0.38 | 0.3 | 0.28 | 0.23 |
| 7 | 123597121 | rs17146516 | 0.888 | 0.172 | -0.703 | -0.548 | -0.038 | -0.114 | -0.114 | 0 | 0.16 | 0.01 | 0.09 | 0.38 | 0.16 | 0.22 |
| 7 | 123598503 | rs2253692 | -0.809 | 0.318 | -0.091 | -0.722 | -0.008 | -0.001 | -0.001 | 0.015 | 0.36 | 0.29 | 0.45 | 0.36 | 0.41 | 0.38 |
| 7 | 123607158 | rs12670939 | 0.615 | 0.437 | -0.107 | -0.643 | -0.01 | -0.011 | -0.011 | 0 | 0.19 | 0.13 | 0.38 | 0.3 | 0.28 | 0.22 |
| 7 | 123607276 | rs6953544 | -1.384 | 0.118 | 0.373 | -0.696 | -0.016 | -0.084 | -0.084 | 1.021 | 0.36 | 0.56 | 0.17 | 0.07 | 0.27 | 0.29 |
| 7 | 123608302 | rs13229421 | -1.384 | 0.129 | 0.373 | -0.598 | -0.015 | -0.042 | -0.042 | 0.385 | 0.36 | 0.56 | 0.22 | 0.15 | 0.28 | 0.29 |
| 7 | 123610077 | rs12706565 | -0.266 | 0.025 | 0.412 | -1.038 | -0.009 | -0.123 | -0.123 | 1.534 | 0.17 | 0.4 | 0.1 | 0 | 0.14 | 0.14 |
| 7 | 123610348 | rs11975640 | -1.667 | 0.099 | 0.385 | -1.023 | -0.01 | -0.093 | -0.093 | 0 | 0.36 | 0.56 | 0.18 | 0.05 | 0.29 | 0.3 |
| 7 | 123613318 | rs17146532 | -1.113 | 0.378 | 0.448 | -0.591 | -0.015 | -0.06 | -0.06 | 0.912 | 0.2 | 0.31 | 0.1 | 0.05 | 0.15 | 0.16 |
| 7 | 123624149 | rs11973429 | -1.023 | 0.354 | 0.325 | -0.563 | -0.005 | -0.025 | -0.025 | 0.049 | 0.19 | 0.21 | 0.08 | 0.05 | 0.14 | 0.16 |
| 7 | 123633233 | rs12668777 | -0.521 | 0.516 | 0.358 | -0.501 | 0.007 | -0.037 | -0.037 | 0 | 0.11 | 0.22 | 0.07 | 0.04 | 0.1 | 0.13 |
| 7 | 123662321 | rs11764422 | -0.728 | 0.391 | 0.46 | -0.614 | -0.012 | -0.105 | -0.105 | 2.286 | 0.21 | 0.29 | 0.09 | 0 | 0.13 | 0.12 |

| | | | | | | | | | | | | | | | | |
|---|---|---|---|---|---|---|---|---|---|---|---|---|---|---|---|---|
| 7 | 123673952 | rs11771905 | -0.731 | 0.359 | 0.437 | -0.616 | -0.013 | -0.099 | -0.099 | 0.296 | 0.21 | 0.29 | 0.09 | 0 | 0.13 | 0.12 |
| 7 | 123679743 | rs10255200 | -1.628 | 0.565 | 0.428 | -0.463 | -0.005 | 0.021 | 0.021 | 1.093 | 0.45 | 0.55 | 0.5 | 0.29 | 0.48 | 0.43 |
| 7 | 123681237 | rs11771846 | -0.896 | 0.37 | 0.49 | -0.592 | 0.016 | 0.025 | 0.025 | 0 | 0.28 | 0.19 | 0.11 | 0.04 | 0.18 | 0.21 |
| 7 | 123692725 | rs10271762 | -1.817 | 0.484 | 0.535 | -0.513 | 0.004 | 0.039 | 0.039 | 0.743 | 0.49 | 0.54 | 0.51 | 0.29 | 0.51 | 0.49 |
| 7 | 123700519 | rs2052094 | -0.012 | 0.514 | 0.662 | -0.522 | 0.059 | 0.017 | 0.017 | 0.06 | 0.21 | 0.35 | 0.4 | 0.25 | 0.32 | 0.27 |
| 7 | 123701345 | rs13224429 | -1.816 | 0.507 | 0.614 | -0.533 | -0.001 | 0.035 | 0.035 | 0.329 | 0.48 | 0.54 | 0.5 | 0.29 | 0.5 | 0.49 |
| 7 | 123706922 | rs10242895 | 0.428 | 0.447 | 0.542 | -0.598 | 0 | -0.005 | -0.005 | 3.724 | 0.15 | 0.11 | 0.06 | 0 | 0.11 | 0.11 |
| 7 | 123711663 | rs10487808 | -0.923 | 0.449 | 0.553 | -0.597 | 0.019 | 0.045 | 0.045 | 0 | 0.27 | 0.15 | 0.1 | 0.04 | 0.18 | 0.21 |
| 7 | 123712562 | rs1860486 | -0.239 | 0.74 | 0.904 | -0.236 | 0.066 | 0.015 | 0.015 | 0.048 | 0.21 | 0.36 | 0.4 | 0.25 | 0.32 | 0.27 |
| 7 | 123719096 | rs10255489 | -0.703 | 0.825 | 0.892 | -0.255 | -0.013 | -0.027 | -0.027 | 0 | 0.16 | 0.19 | 0.07 | 0.04 | 0.1 | 0.11 |
| 7 | 123719407 | rs10258541 | 0.435 | 0.811 | 0.895 | -0.278 | -0.037 | -0.05 | -0.05 | 0 | 0.11 | 0.15 | 0.03 | 0 | 0.02 | 0.03 |
| 7 | 123721563 | rs9918670 | 0.059 | 0.916 | 0.926 | -0.234 | 0.04 | 0.018 | 0.018 | 0.309 | 0.18 | 0.24 | 0.37 | 0.26 | 0.29 | 0.28 |
| 7 | 123732469 | rs10270484 | -0.539 | 0.747 | 0.608 | -0.248 | -0.014 | -0.031 | -0.031 | 0 | 0.16 | 0.19 | 0.07 | 0.04 | 0.1 | 0.11 |
| 7 | 123751137 | rs6951821 | 0.008 | 0.931 | 0.591 | -0.196 | 0.091 | 0.034 | 0.034 | 0.107 | 0.18 | 0.31 | 0.38 | 0.25 | 0.31 | 0.28 |
| 7 | 123757883 | rs2215348 | -1.252 | 0.846 | 0.375 | -0.175 | 0.004 | -0.008 | -0.008 | 0 | 0.34 | 0.54 | 0.45 | 0.29 | 0.39 | 0.38 |
| 7 | 123768153 | rs2191280 | -1.467 | 0.845 | 0.052 | 0.187 | 0.001 | 0.007 | 0.007 | 0.017 | 0.47 | 0.41 | 0.67 | 0.65 | 0.55 | 0.59 |
| 7 | 123786924 | rs7788439 | 0.459 | 0.703 | -0.341 | -0.305 | -0.02 | 0.066 | 0.066 | 0 | 0.21 | 0.1 | 0.12 | 0 | 0.19 | 0.11 |
| 7 | 123795935 | rs10216192 | 0.18 | 0.634 | -0.261 | -0.2 | 0 | -0.01 | -0.01 | 0 | 0.27 | 0.33 | 0.15 | 0.07 | 0.23 | 0.26 |
| 7 | 123797641 | rs6947191 | 0.589 | 0.654 | -0.639 | 0.311 | -0.025 | -0.056 | -0.056 | 0 | 0.16 | 0.04 | 0.26 | 0.29 | 0.19 | 0.22 |
| 7 | 123809048 | rs12154449 | -1.209 | 0.518 | -0.576 | 0.273 | -0.029 | 0.065 | 0.065 | 0.309 | 0.44 | 0.2 | 0.61 | 0.68 | 0.48 | 0.61 |
| 7 | 123819155 | rs11560338 | 0.821 | 0.633 | -0.486 | 0.619 | -0.069 | -0.103 | -0.103 | 0 | 0.15 | 0.01 | 0.29 | 0.3 | 0.21 | 0.27 |
| 7 | 123836711 | rs1540352 | -0.985 | 0.606 | -0.269 | 0.294 | -0.009 | 0.006 | 0.006 | 0 | 0.36 | 0.28 | 0.6 | 0.67 | 0.43 | 0.58 |
| 7 | 123845784 | rs6466898 | -2.208 | 0.567 | -0.118 | 0.303 | 0.006 | 0.033 | 0.033 | 0.041 | 0.64 | 0.77 | 0.76 | 0.77 | 0.65 | 0.84 |
| 7 | 123847391 | rs4727992 | 0.215 | 0.657 | -0.494 | 0.336 | -0.003 | -0.035 | -0.035 | 0 | 0.27 | 0.39 | 0.16 | 0.09 | 0.22 | 0.26 |
| 7 | 123850419 | rs2159570 | -0.595 | 0.585 | -0.706 | 0.545 | 0 | -0.001 | -0.001 | 0 | 0.47 | 0.55 | 0.45 | 0.45 | 0.43 | 0.57 |
| 7 | 123854695 | rs1990393 | -0.307 | 0.738 | -0.907 | 1.298 | -0.031 | -0.067 | -0.067 | 0 | 0.17 | 0.09 | 0.3 | 0.31 | 0.22 | 0.26 |
| 7 | 123855507 | rs2215426 | -0.418 | 0.739 | -0.899 | 1.299 | -0.02 | -0.048 | -0.048 | 0.406 | 0.17 | 0.14 | 0.3 | 0.31 | 0.22 | 0.26 |
| 7 | 123858915 | rs2098144 | -0.092 | 0.697 | -0.814 | 1.307 | -0.017 | -0.039 | -0.039 | 0 | 0.12 | 0.08 | 0.11 | 0.26 | 0.15 | 0.22 |
| 7 | 123860214 | rs6466899 | 0.714 | 0.219 | -0.429 | 1.06 | -0.002 | 0.013 | 0.013 | 0.429 | 0.74 | 0.75 | 0.65 | 0.54 | 0.78 | 0.54 |
| 7 | 123863237 | rs2191468 | -0.982 | -0.149 | -0.862 | 0.885 | 0.027 | -0.036 | -0.036 | 0.104 | 0.28 | 0.12 | 0.35 | 0.35 | 0.21 | 0.25 |
| 7 | 123863638 | rs13245041 | 0.009 | -0.039 | -0.201 | 0.82 | -0.003 | 0.021 | 0.021 | 2.56 | 0.85 | 0.61 | 0.89 | 0.85 | 0.87 | 0.72 |
| 7 | 123864147 | rs4644181 | 1.648 | -0.302 | 0.019 | 0.767 | 0.07 | 0.001 | 0.001 | 0.33 | 0.53 | 0.21 | 0.5 | 0.53 | 0.65 | 0.46 |
| 7 | 123864185 | rs4573205 | -0.083 | -0.19 | 0.183 | 0.872 | -0.002 | 0.019 | 0.019 | 2.471 | 0.85 | 0.6 | 0.89 | 0.85 | 0.87 | 0.72 |
| 7 | 123865926 | rs17146827 | 1.493 | 0.226 | -0.5 | 0.61 | -0.027 | -0.014 | -0.014 | 0.739 | 0.45 | 0.24 | 0.44 | 0.53 | 0.52 | 0.41 |
| 7 | 123869567 | rs10487814 | -0.28 | 0.042 | -0.349 | 0.594 | 0.002 | 0.005 | 0.005 | 0 | 0.07 | 0.05 | 0.05 | 0.04 | 0.06 | 0.07 |
| 7 | 123870082 | rs12534491 | 1.544 | -0.272 | 0.022 | 0.983 | -0.007 | -0.002 | -0.002 | 2.739 | 0.6 | 0.42 | 0.62 | 0.55 | 0.66 | 0.46 |
| 7 | 123875796 | rs4361708 | -0.399 | -0.332 | 0.136 | 0.988 | 0.002 | -0.008 | -0.008 | 3.098 | 0.17 | 0.12 | 0.22 | 0.3 | 0.16 | 0.2 |
| 7 | 123878512 | rs2007127 | 0.647 | -0.341 | 0.248 | 0.965 | -0.015 | -0.038 | -0.038 | 0 | 0.79 | 0.62 | 0.85 | 0.85 | 0.82 | 0.66 |
| 7 | 123883427 | rs6972553 | 1.251 | -0.066 | 0.148 | 1.073 | -0.011 | 0.042 | 0.042 | 1.706 | 0.69 | 0.53 | 0.67 | 0.53 | 0.74 | 0.54 |
| 7 | 123892162 | rs1019197 | 1.66 | -0.013 | 0.302 | 1.054 | -0.003 | -0.009 | -0.009 | 0.251 | 0.59 | 0.51 | 0.51 | 0.28 | 0.64 | 0.43 |
| 7 | 123895098 | rs7799219 | -0.322 | 0.002 | 0.251 | 1.066 | 0.002 | -0.018 | -0.018 | 0 | 0.17 | 0.09 | 0.22 | 0.3 | 0.16 | 0.19 |
| 7 | 123898418 | rs11978415 | 0.328 | -0.021 | 0.125 | 1.025 | -0.032 | 0.082 | 0.082 | 0 | 0.13 | 0 | 0.06 | 0 | 0.14 | 0.07 |
| 7 | 123911882 | rs1860577 | 0.766 | -0.018 | 0.299 | 1.059 | -0.007 | 0.052 | 0.052 | 1.064 | 0.76 | 0.6 | 0.73 | 0.58 | 0.8 | 0.62 |
| 7 | 123914885 | rs917657 | 2.015 | -0.071 | 0.364 | 0.975 | -0.001 | -0.007 | -0.007 | 0 | 0.52 | 0.41 | 0.46 | 0.23 | 0.59 | 0.35 |
| 7 | 123916425 | rs10808205 | 1.998 | -0.092 | 0.401 | 0.985 | -0.001 | -0.007 | -0.007 | 0 | 0.52 | 0.41 | 0.46 | 0.23 | 0.59 | 0.35 |
| 7 | 123918641 | rs13233306 | -0.172 | -0.058 | 0.44 | 0.779 | -0.005 | -0.03 | -0.03 | 0 | 0.08 | 0.15 | 0.06 | 0.05 | 0.05 | 0.08 |
| 7 | 123922971 | rs12668123 | 0.724 | -0.087 | 0.368 | 1.15 | -0.002 | 0.016 | 0.016 | 1.349 | 0.77 | 0.6 | 0.71 | 0.71 | 0.78 | 0.63 |
| 7 | 123930549 | rs4304277 | -0.874 | -0.082 | 0.379 | 1.153 | -0.002 | 0.016 | 0.016 | 1.277 | 0.23 | 0.4 | 0.28 | 0.29 | 0.22 | 0.37 |
| 7 | 123936486 | rs12534665 | 1.973 | -0.077 | 0.363 | 1.171 | 0.01 | 0.009 | 0.009 | 0.379 | 0.52 | 0.35 | 0.45 | 0.36 | 0.6 | 0.36 |
| 7 | 123951585 | rs7784625 | -0.636 | -0.144 | 0.395 | 1.003 | 0.003 | -0.004 | -0.004 | 0.162 | 0.1 | 0.13 | 0.09 | 0.16 | 0.06 | 0.1 |
| 7 | 123952475 | rs995996 | -0.251 | -0.025 | 0.348 | 0.844 | 0.015 | 0.004 | 0.004 | 0.619 | 0.86 | 0.74 | 0.8 | 0.87 | 0.82 | 0.73 |
| 7 | 123958189 | rs11765479 | -1.272 | -0.199 | -0.118 | 0.797 | -0.009 | -0.003 | -0.003 | 0 | 0.17 | 0.2 | 0.14 | 0.18 | 0.1 | 0.19 |
| 7 | 123966517 | rs2051906 | 0.254 | -0.128 | 0.142 | 0.693 | 0.009 | -0.025 | -0.025 | 0 | 0.83 | 0.44 | 0.76 | 0.88 | 0.81 | 0.74 |
| 7 | 123967269 | rs6976230 | 0.254 | -0.128 | 0.148 | 0.693 | 0.009 | -0.025 | -0.025 | 0.118 | 0.83 | 0.44 | 0.76 | 0.88 | 0.81 | 0.74 |
| 7 | 123972862 | rs12539157 | 1.132 | -0.183 | 0.049 | 0.688 | -0.014 | -0.031 | -0.031 | 0 | 0.76 | 0.33 | 0.71 | 0.84 | 0.78 | 0.65 |
| 7 | 123977335 | rs6949182 | 1.687 | -0.221 | 0.053 | 0.698 | 0.015 | 0.017 | 0.017 | 1.078 | 0.67 | 0.19 | 0.62 | 0.71 | 0.71 | 0.55 |
| 7 | 123994631 | rs12535600 | -0.102 | -0.134 | -0.009 | 0.722 | -0.002 | 0.009 | 0.009 | 0 | 0.08 | 0.13 | 0.1 | 0.13 | 0.06 | 0.11 |
| 7 | 124008463 | rs10215793 | -0.84 | -0.102 | 0.161 | 0.672 | -0.002 | -0.048 | -0.048 | 0 | 0.21 | 0.54 | 0.26 | 0.16 | 0.21 | 0.33 |
| 7 | 124010036 | rs10487263 | 1.923 | 0.041 | -0.697 | 0.202 | 0.01 | 0.082 | 0.082 | 3.049 | 0.34 | 0.08 | 0.18 | 0.14 | 0.31 | 0.11 |
| 7 | 124027861 | rs972172 | 1.368 | -0.169 | -0.12 | 0.55 | -0.003 | -0.005 | -0.005 | 0 | 0.67 | 0.32 | 0.63 | 0.71 | 0.69 | 0.54 |
| 7 | 124029567 | rs10238997 | -0.773 | -0.15 | -0.07 | 0.55 | -0.012 | -0.064 | -0.064 | 0.088 | 0.25 | 0.54 | 0.28 | 0.15 | 0.24 | 0.35 |
| 7 | 124035800 | rs975033 | 0.855 | -0.163 | 0.006 | 0.551 | -0.013 | -0.003 | -0.003 | 0 | 0.86 | 0.76 | 0.86 | 0.84 | 0.9 | 0.82 |
| 7 | 124035929 | rs10226954 | -0.675 | -0.187 | -0.011 | 1.044 | -0.001 | 0.093 | 0.093 | 0.19 | 0.22 | 0.5 | 0.38 | 0.39 | 0.23 | 0.4 |
| 7 | 124037765 | rs2190117 | -0.174 | -0.183 | -0.017 | 1.027 | 0.012 | -0.041 | -0.041 | 0.052 | 0.19 | 0.46 | 0.23 | 0.13 | 0.21 | 0.28 |
| 7 | 124042325 | rs2040933 | 1.082 | -0.371 | -0.221 | 1.135 | -0.006 | 0.026 | 0.026 | 1.196 | 0.83 | 0.79 | 0.73 | 0.58 | 0.87 | 0.7 |
| 7 | 124066585 | rs721949 | 0.442 | -0.395 | -0.169 | 1.124 | -0.014 | 0.001 | 0.001 | 0 | 0.89 | 0.75 | 0.88 | 0.87 | 0.92 | 0.91 |
| 7 | 124067250 | rs6974521 | 1.67 | -0.414 | -0.339 | 1.011 | 0.052 | -0.057 | -0.057 | 0 | 0.61 | 0.14 | 0.47 | 0.45 | 0.68 | 0.45 |
| 7 | 124068753 | rs1468517 | 0.322 | 0.013 | -0.132 | 0.93 | -0.006 | -0.02 | -0.02 | 0 | 0.79 | 0.49 | 0.75 | 0.85 | 0.77 | 0.74 |
| 7 | 124069511 | rs7784948 | -0.867 | -0.104 | -0.188 | 1.139 | -0.006 | 0.039 | 0.039 | 0 | 0.13 | 0.16 | 0.25 | 0.38 | 0.1 | 0.22 |

| | | | | | | | | | | | | | | | | |
|---|---|---|---|---|---|---|---|---|---|---|---|---|---|---|---|---|
| 7 | 124069721 | rs13238690 | 0.656 | -0.785 | -0.614 | 0.673 | 0.031 | 0.03 | 0.03 | 0.637 | 0.16 | 0.07 | 0.07 | 0.04 | 0.05 | 0.07 |
| 7 | 124069748 | rs2159183 | -0.593 | -0.596 | -0.559 | 0.794 | -0.006 | 0.04 | 0.04 | 0 | 0.13 | 0.16 | 0.25 | 0.38 | 0.1 | 0.22 |
| 7 | 124075494 | rs720547 | 0.127 | -0.468 | -0.564 | 0.532 | -0.007 | -0.031 | -0.031 | 0 | 0.77 | 0.46 | 0.73 | 0.85 | 0.76 | 0.73 |
| 7 | 124078501 | rs17147142 | -0.835 | -1.131 | -1.133 | 0.607 | -0.007 | 0 | 0 | 0 | 0.29 | 0.32 | 0.19 | 0.34 | 0.14 | 0.31 |
| 7 | 124078609 | rs13239910 | 2.183 | -0.293 | -1.428 | 0.335 | -0.108 | 0.181 | 0.181 | 0.496 | 0.35 | 0.08 | 0.22 | 0 | 0.45 | 0.1 |
| 7 | 124081532 | rs6466925 | 0.678 | -0.279 | -0.783 | 0.882 | -0.024 | 0.031 | 0.031 | 1.183 | 0.7 | 0.46 | 0.58 | 0.42 | 0.73 | 0.47 |
| 7 | 124084779 | rs11983138 | -0.319 | -0.273 | -0.884 | 0.909 | -0.008 | -0.016 | -0.016 | 0.047 | 0.11 | 0.07 | 0.2 | 0.3 | 0.19 | 0.32 |
| 7 | 124085853 | rs10435284 | -0.525 | -0.299 | -0.712 | 0.812 | -0.051 | -0.096 | -0.096 | 0 | 0.83 | 0.56 | 0.92 | 0.98 | 0.93 | 0.91 |
| 7 | 124096727 | rs6973149 | -0.438 | -0.058 | -0.899 | 0.889 | -0.038 | -0.036 | -0.036 | 0 | 0.15 | 0.09 | 0.29 | 0.28 | 0.25 | 0.29 |
| 7 | 124109931 | rs11762379 | -0.552 | -0.076 | -0.782 | 0.881 | -0.002 | -0.002 | -0.002 | 0 | 0.16 | 0.19 | 0.3 | 0.28 | 0.25 | 0.29 |
| 7 | 124112155 | rs17132871 | 0.439 | -0.257 | -0.711 | 0.909 | -0.077 | 0.164 | 0.164 | 0.989 | 0.77 | 0.54 | 0.77 | 0.56 | 0.91 | 0.65 |
| 7 | 124117874 | rs11761806 | 0.439 | -0.257 | -0.711 | 0.909 | -0.077 | 0.164 | 0.164 | 0.989 | 0.77 | 0.54 | 0.77 | 0.56 | 0.91 | 0.65 |
| 7 | 124126038 | rs2079644 | 1.731 | -0.226 | -0.709 | 0.898 | 0.005 | 0.009 | 0.009 | 1.891 | 0.62 | 0.36 | 0.47 | 0.27 | 0.66 | 0.37 |
| 7 | 124126280 | rs7811975 | 0.542 | -0.188 | -0.761 | 0.917 | -0.013 | -0.022 | -0.022 | 0 | 0.08 | 0.04 | 0.1 | 0.15 | 0.07 | 0.1 |
| 7 | 124134350 | rs35138317 | -0.467 | -0.241 | -0.701 | 0.954 | -0.001 | 0.016 | 0.016 | 0.546 | 0.19 | 0.22 | 0.44 | 0.54 | 0.27 | 0.41 |
| 7 | 124138187 | rs2058136 | 0.359 | -0.001 | -0.96 | 0.439 | -0.003 | -0.014 | -0.014 | 0 | 0.26 | 0.14 | 0.24 | 0.27 | 0.23 | 0.27 |
| 7 | 124151281 | rs4728000 | -0.289 | -0.122 | -0.568 | -0.586 | -0.054 | -0.101 | -0.101 | 0.011 | 0.83 | 0.56 | 0.92 | 0.98 | 0.93 | 0.91 |
| 7 | 124153392 | rs4731184 | 1.187 | 0.473 | -1.104 | -0.308 | -0.008 | -0.024 | -0.024 | 0 | 0.37 | 0.32 | 0.24 | 0 | 0.43 | 0.1 |
| 7 | 124155970 | rs1544521 | 0.745 | 0.349 | -1.258 | 0.752 | 0.007 | -0.04 | -0.04 | 0.123 | 0.77 | 0.8 | 0.65 | 0.39 | 0.8 | 0.64 |
| 7 | 124164127 | rs2190921 | 0.511 | 0.221 | -1.036 | 0.148 | 0.014 | 0.021 | 0.021 | 0.124 | 0.61 | 0.7 | 0.65 | 0.67 | 0.6 | 0.53 |
| 7 | 124165999 | rs10225755 | 0.486 | 0.217 | -1.072 | 0.016 | 0.017 | 0.033 | 0.033 | 0.49 | 0.61 | 0.76 | 0.64 | 0.68 | 0.59 | 0.53 |
| 7 | 124172307 | rs17147231 | 1.204 | 0.168 | -0.513 | -0.671 | 0.002 | -0.082 | -0.082 | 0 | 0.22 | 0.23 | 0.14 | 0.02 | 0.21 | 0.23 |
| 7 | 124173789 | rs17394006 | 0.141 | -0.377 | -0.76 | -0.842 | 0.026 | 0.161 | 0.161 | 0.15 | 0.39 | 0.14 | 0.3 | 0.19 | 0.39 | 0.39 |
| 7 | 124177591 | rs12535612 | 1.195 | -0.069 | -0.75 | -0.672 | -0.005 | -0.003 | -0.003 | 0 | 0.12 | 0.1 | 0.22 | 0.18 | 0.18 | 0.08 |
| 7 | 124182414 | rs1181804 | 0.346 | -0.224 | -0.534 | -0.759 | 0.027 | 0.095 | 0.095 | 0.144 | 0.39 | 0.25 | 0.31 | 0.19 | 0.38 | 0.39 |
| 7 | 124189490 | rs1404407 | 0.721 | 0.009 | 0.091 | -0.638 | -0.018 | -0.041 | -0.041 | 0 | 0.28 | 0.43 | 0.22 | 0.15 | 0.22 | 0.33 |
| 7 | 124189726 | rs12706612 | 0.992 | -0.053 | -0.114 | -0.737 | -0.001 | 0.001 | 0.001 | 2.676 | 0.2 | 0.18 | 0.23 | 0.18 | 0.18 | 0.11 |
| 7 | 124192333 | rs1357794 | -1.71 | -0.053 | 0.022 | -0.799 | 0 | -0.029 | -0.029 | 1.38 | 0.78 | 0.82 | 0.87 | 0.95 | 0.79 | 0.76 |
| 7 | 124196444 | rs1917353 | 0.032 | 0.122 | -0.188 | -0.546 | -0.008 | 0.012 | 0.012 | 2.841 | 0.4 | 0.37 | 0.48 | 0.66 | 0.4 | 0.3 |
| 7 | 124207343 | rs2402741 | 0.788 | 0.14 | -0.339 | -0.467 | 0 | -0.001 | -0.001 | 0 | 0.2 | 0.2 | 0.23 | 0.2 | 0.18 | 0.11 |
| 7 | 124212465 | rs6952573 | -0.309 | 0.226 | -0.212 | -0.684 | -0.01 | 0.011 | 0.011 | 2.045 | 0.61 | 0.69 | 0.53 | 0.33 | 0.61 | 0.7 |
| 7 | 124222007 | rs1568814 | -0.988 | 0.337 | -0.872 | -0.612 | -0.116 | -0.092 | -0.092 | 4.589 | 0.72 | 0.96 | 0.57 | 0.32 | 0.65 | 0.68 |
| 7 | 124234030 | rs1524770 | 0.028 | 0.022 | -0.227 | -0.511 | -0.022 | -0.01 | -0.01 | 0.159 | 0.32 | 0.47 | 0.32 | 0.27 | 0.29 | 0.39 |
| 7 | 124237138 | rs10954042 | -0.131 | -0.001 | -0.07 | -0.715 | 0.007 | 0.026 | 0.026 | 0.25 | 0.51 | 0.54 | 0.44 | 0.23 | 0.49 | 0.55 |
| 7 | 124237196 | rs1524756 | 1.269 | -0.251 | -0.587 | -1.023 | -0.024 | -0.016 | -0.016 | 0 | 0.16 | 0.02 | 0.23 | 0.18 | 0.19 | 0.21 |
| 7 | 124240099 | rs9649007 | -0.633 | -0.228 | -0.237 | -0.8 | -0.016 | 0.054 | 0.054 | 0.442 | 0.56 | 0.81 | 0.48 | 0.32 | 0.51 | 0.63 |
| 7 | 124240284 | rs1524760 | -0.898 | -0.046 | -0.526 | -0.715 | -0.016 | 0.013 | 0.013 | 0 | 0.75 | 0.87 | 0.72 | 0.77 | 0.71 | 0.84 |
| 7 | 124240452 | rs13227017 | 1.818 | -0.173 | -0.679 | -0.907 | -0.005 | 0.062 | 0.062 | 0 | 0.21 | 0.08 | 0.14 | 0.05 | 0.21 | 0.25 |
| 7 | 124270732 | rs2049437 | 0.812 | -0.201 | -0.545 | -0.858 | 0.008 | 0.041 | 0.041 | 0.215 | 0.27 | 0.17 | 0.28 | 0.2 | 0.27 | 0.3 |
| 7 | 124270979 | rs4728003 | 1.325 | -0.21 | -0.311 | -0.865 | -0.003 | -0.032 | -0.032 | 0 | 0.24 | 0.27 | 0.15 | 0.06 | 0.21 | 0.25 |
| 7 | 124272939 | rs12670293 | 2.03 | 0.015 | -0.476 | -0.667 | -0.08 | -0.157 | -0.157 | 0.051 | 0.19 | 0.01 | 0.24 | 0.43 | 0.21 | 0.2 |
| 7 | 124278363 | rs1917362 | -0.227 | -0.061 | -0.138 | -0.505 | 0.014 | 0.094 | 0.094 | 0.45 | 0.49 | 0.32 | 0.42 | 0.25 | 0.49 | 0.56 |
| 7 | 124284298 | rs1917365 | 0.701 | -0.148 | -0.287 | -0.459 | -0.001 | 0.014 | 0.014 | 0.149 | 0.22 | 0.17 | 0.28 | 0.2 | 0.27 | 0.31 |
| 7 | 124298853 | rs7795796 | 0.245 | -0.072 | 0.071 | -0.429 | 0.016 | 0.157 | 0.157 | 0.393 | 0.15 | 0.45 | 0.26 | 0.31 | 0.16 | 0.18 |
| 7 | 124300762 | rs12673211 | 1.342 | -0.298 | -0.69 | -0.56 | -0.024 | 0.114 | 0.114 | 0.361 | 0.26 | 0.03 | 0.13 | 0.07 | 0.22 | 0.26 |
| 7 | 124322409 | rs10954043 | 1.188 | -0.054 | -0.686 | -0.449 | -0.087 | -0.107 | -0.107 | 0 | 0.35 | 0.08 | 0.29 | 0.4 | 0.35 | 0.24 |
| 7 | 124322510 | rs7808448 | -0.822 | -0.226 | -0.203 | -0.398 | 0.011 | -0.016 | -0.016 | 1.935 | 0.4 | 0.65 | 0.56 | 0.53 | 0.42 | 0.52 |
| 7 | 124322993 | rs10262687 | 0.518 | -0.176 | -0.524 | -0.358 | -0.004 | -0.011 | -0.011 | 0 | 0.23 | 0.29 | 0.28 | 0.22 | 0.26 | 0.33 |
| 7 | 124323241 | rs11496161 | 0.706 | -0.105 | -0.53 | -0.324 | 0 | 0.009 | 0.009 | 0.044 | 0.22 | 0.19 | 0.27 | 0.22 | 0.26 | 0.33 |
| 7 | 124326717 | rs7777180 | -0.593 | 0.111 | -0.928 | -0.263 | 0.02 | -0.02 | -0.02 | 0 | 0.42 | 0.73 | 0.57 | 0.53 | 0.43 | 0.52 |
| 7 | 124337929 | rs12669312 | 0.91 | 0.42 | -0.828 | -0.357 | -0.011 | 0.032 | 0.032 | 0.634 | 0.21 | 0.11 | 0.14 | 0.1 | 0.22 | 0.25 |
| 7 | 124347899 | rs1100506 | 0.084 | 0.386 | -0.81 | -0.225 | 0 | 0.043 | 0.043 | 0 | 0.52 | 0.72 | 0.56 | 0.61 | 0.52 | 0.58 |
| 7 | 124348319 | rs11981435 | 0.773 | 0.373 | -0.786 | -0.237 | -0.003 | 0.002 | 0.002 | 0 | 0.22 | 0.12 | 0.28 | 0.2 | 0.24 | 0.3 |
| 7 | 124349606 | rs10046543 | 0.812 | 0.417 | -0.561 | -0.178 | 0.008 | 0.055 | 0.055 | 0.233 | 0.3 | 0.49 | 0.27 | 0.4 | 0.27 | 0.27 |
| 7 | 124360711 | rs702426 | -1.67 | 0.368 | -1.215 | -0.289 | -0.004 | -0.041 | -0.041 | 0 | 0.83 | 0.95 | 0.76 | 0.68 | 0.83 | 0.9 |
| 7 | 124363797 | rs13239353 | 0.695 | 0.708 | -0.812 | -0.162 | 0 | -0.004 | -0.004 | 0 | 0.3 | 0.32 | 0.21 | 0.13 | 0.3 | 0.36 |
| 7 | 124366328 | rs17147389 | 1.633 | 0.763 | -1.276 | 0.131 | -0.069 | -0.119 | -0.119 | 0 | 0.2 | 0.02 | 0.19 | 0.33 | 0.21 | 0.16 |
| 7 | 124367812 | rs1319514 | -0.054 | 0.452 | -0.908 | -0.075 | 0.005 | 0 | 0 | 0 | 0.71 | 0.78 | 0.68 | 0.66 | 0.74 | 0.83 |
| 7 | 124378805 | rs717240 | 0.26 | 0.548 | -0.754 | -0.541 | -0.003 | -0.079 | -0.079 | 0 | 0.65 | 0.44 | 0.82 | 1 | 0.74 | 0.83 |
| 7 | 124379392 | rs10464672 | 1.877 | 1.017 | -1.097 | -0.329 | -0.059 | -0.077 | -0.077 | 0.411 | 0.21 | 0.06 | 0.15 | 0.34 | 0.24 | 0.16 |
| 7 | 124381635 | rs1008290 | 0.473 | 0.948 | -1.355 | -0.383 | 0.018 | 0.109 | 0.109 | 0 | 0.49 | 0.34 | 0.54 | 0.81 | 0.53 | 0.54 |
| 7 | 124381689 | rs1544452 | 0.834 | 0.934 | -1.341 | -0.372 | 0.001 | -0.012 | -0.012 | 0 | 0.28 | 0.27 | 0.4 | 0.47 | 0.3 | 0.38 |
| 7 | 124382714 | rs6466946 | 0.095 | 0.812 | -1.408 | -0.433 | 0.032 | 0.071 | 0.071 | 0 | 0.6 | 0.31 | 0.79 | 0.89 | 0.7 | 0.78 |
| 7 | 124383948 | rs17147421 | 0.975 | 0.877 | -1.377 | -0.278 | -0.02 | -0.054 | -0.054 | 1.067 | 0.3 | 0.16 | 0.19 | 0.45 | 0.29 | 0.21 |
| 7 | 124384821 | rs2190099 | 0.907 | 0.847 | -1.311 | -0.142 | -0.059 | 0.058 | 0.058 | 0 | 0.25 | 0.02 | 0.19 | 0.1 | 0.27 | 0.35 |
| 7 | 124387871 | rs2284215 | 0.134 | 0.833 | -1.166 | -0.089 | -0.011 | -0.063 | -0.063 | 0 | 0.33 | 0.17 | 0.4 | 0.57 | 0.35 | 0.42 |
| 7 | 124388204 | rs2284214 | -0.359 | -0.004 | -0.411 | -0.468 | 0 | -0.001 | -0.001 | 0 | 0.46 | 0.41 | 0.47 | 0.55 | 0.51 | 0.51 |
| 7 | 124397026 | rs2267707 | 0.471 | -0.244 | -0.435 | -0.666 | -0.033 | -0.044 | -0.044 | 0 | 0.29 | 0.21 | 0.41 | 0.5 | 0.41 | 0.4 |
| 7 | 124400705 | rs3807544 | -0.081 | -0.179 | -0.005 | -0.55 | 0.008 | 0.006 | 0.006 | 0.183 | 0.42 | 0.46 | 0.47 | 0.5 | 0.49 | 0.49 |

| | | | | | | | | | | | | | | | | |
|---|---|---|---|---|---|---|---|---|---|---|---|---|---|---|---|---|
| 7 | 124410577 | rs4141075 | -0.321 | -0.139 | -0.018 | -0.643 | 0.004 | 0.007 | 0.007 | 0 | 0.67 | 0.71 | 0.67 | 0.81 | 0.74 | 0.65 |
| 7 | 124425618 | rs17833562 | 1.699 | -0.221 | -0.44 | -0.655 | -0.026 | 0.025 | 0.025 | 0 | 0.13 | 0.01 | 0.09 | 0.06 | 0.12 | 0.22 |
| 7 | 124434069 | rs716991 | -0.116 | -0.875 | 0.449 | -0.143 | 0.001 | 0.008 | 0.008 | 0 | 0.67 | 0.77 | 0.66 | 0.8 | 0.71 | 0.62 |
| 7 | 124436063 | rs7777947 | 0.014 | -0.939 | 0.566 | -0.238 | -0.017 | -0.002 | -0.002 | 0.692 | 0.33 | 0.48 | 0.22 | 0.31 | 0.27 | 0.23 |
| 7 | 124462081 | rs727505 | 0.978 | -0.866 | 0.405 | -0.294 | -0.012 | -0.046 | -0.046 | 0 | 0.24 | 0.13 | 0.18 | 0.31 | 0.26 | 0.19 |
| 7 | 124462661 | rs17246404 | 1.311 | -0.883 | 0.347 | -0.302 | -0.067 | -0.111 | -0.111 | 0 | 0.22 | 0.01 | 0.17 | 0.31 | 0.25 | 0.2 |
| 7 | 124512647 | rs11770818 | -0.117 | -0.966 | 0.708 | -0.347 | -0.004 | -0.018 | -0.018 | 0 | 0.34 | 0.33 | 0.35 | 0.2 | 0.28 | 0.38 |
| 7 | 124513770 | rs4463363 | -1.339 | -0.995 | 0.846 | -0.362 | -0.003 | -0.004 | -0.004 | 0.028 | 0.65 | 0.64 | 0.55 | 0.51 | 0.56 | 0.61 |
| 7 | 124520308 | rs10464529 | 1.206 | -0.992 | 0.817 | -0.357 | -0.037 | -0.08 | -0.08 | 0 | 0.23 | 0.08 | 0.18 | 0.31 | 0.25 | 0.2 |
| 7 | 124526739 | rs4360236 | 0.448 | -1.047 | 0.716 | -0.417 | -0.052 | 0.016 | 0.016 | 0 | 0.2 | 0.31 | 0.26 | 0.2 | 0.1 | 0.31 |
| 7 | 124550133 | rs12706627 | -0.39 | -1.023 | 0.732 | -0.402 | -0.004 | -0.018 | -0.018 | 0 | 0.34 | 0.33 | 0.35 | 0.2 | 0.28 | 0.38 |
| 7 | 124558339 | rs962863 | -1.322 | -0.905 | 0.625 | -0.383 | -0.012 | -0.015 | -0.015 | 0 | 0.66 | 0.72 | 0.56 | 0.51 | 0.56 | 0.61 |
| 7 | 124569382 | rs2291207 | -0.098 | -1.135 | 0.865 | -0.585 | 0.001 | 0.004 | 0.004 | 0 | 0.33 | 0.24 | 0.35 | 0.2 | 0.3 | 0.38 |
| 7 | 124578931 | rs10252443 | -1.446 | -1.014 | 0.768 | -0.538 | -0.074 | 0.115 | 0.115 | 0 | 0.68 | 0.91 | 0.75 | 0.92 | 0.58 | 0.65 |
| 7 | 124584834 | rs12112909 | -1.25 | -1.096 | 0.71 | -0.666 | -0.004 | -0.002 | -0.002 | 0 | 0.67 | 0.7 | 0.57 | 0.57 | 0.56 | 0.61 |
| 7 | 124590871 | rs6961441 | -1.212 | -1.096 | 0.713 | -0.666 | 0.007 | 0.002 | 0.002 | 0 | 0.66 | 0.61 | 0.56 | 0.57 | 0.56 | 0.61 |
| 7 | 124619404 | rs12539580 | -1.038 | -1.112 | 0.752 | -0.673 | 0.019 | 0.003 | 0.003 | 0 | 0.64 | 0.52 | 0.55 | 0.57 | 0.56 | 0.61 |
| 7 | 124638034 | rs7802007 | -1.214 | -1.112 | 0.763 | -0.674 | 0.008 | 0.002 | 0.002 | 0 | 0.66 | 0.61 | 0.56 | 0.57 | 0.56 | 0.62 |
| 7 | 124656377 | rs6957525 | -1.042 | -1.104 | 0.707 | -0.67 | 0.019 | 0.003 | 0.003 | 0 | 0.64 | 0.52 | 0.55 | 0.57 | 0.56 | 0.62 |
| 7 | 124667796 | rs1481337 | -1.217 | -1.054 | 0.613 | -0.662 | -0.002 | -0.001 | -0.001 | 0 | 0.66 | 0.65 | 0.57 | 0.57 | 0.58 | 0.62 |
| 7 | 124674939 | rs1871773 | -1.238 | -1.053 | 0.574 | -0.661 | -0.011 | -0.005 | -0.005 | 0 | 0.67 | 0.73 | 0.57 | 0.57 | 0.58 | 0.62 |
| 7 | 124722617 | rs628365 | 0.857 | -2.178 | 0.848 | -0.583 | 0.008 | -0.003 | -0.003 | 0 | 0.39 | 0.44 | 0.45 | 0.42 | 0.48 | 0.41 |
| 7 | 124767295 | rs664062 | 0.134 | -2.239 | 1.007 | -0.508 | 0.031 | 0.022 | 0.022 | 0 | 0.68 | 0.81 | 0.67 | 0.77 | 0.78 | 0.62 |
| 7 | 124767997 | rs11534064 | 0.596 | -2.313 | 0.874 | -0.481 | -0.001 | -0.002 | -0.002 | 0 | 0.28 | 0.29 | 0.22 | 0.35 | 0.32 | 0.19 |
| 7 | 124798169 | rs669064 | 0.141 | -2.618 | 1.014 | -0.511 | -0.005 | -0.002 | -0.002 | 0 | 0.57 | 0.61 | 0.73 | 0.64 | 0.65 | 0.74 |
| 7 | 124810399 | rs2968865 | 1.308 | -2.656 | 1.093 | -0.536 | -0.024 | -0.015 | -0.015 | 0 | 0.35 | 0.24 | 0.44 | 0.41 | 0.51 | 0.4 |
| 7 | 124813749 | rs792615 | -0.927 | -2.88 | 1.206 | -0.552 | -0.031 | -0.057 | -0.057 | 0 | 0.84 | 0.86 | 0.94 | 1 | 0.93 | 0.93 |
| 7 | 124824917 | rs792639 | -0.595 | -2.878 | 1.184 | -0.578 | -0.072 | -0.113 | -0.113 | 1.134 | 0.82 | 0.72 | 0.93 | 1 | 0.93 | 0.93 |
| 7 | 124827236 | rs4731236 | 1.106 | -2.903 | 1.028 | -0.571 | -0.05 | -0.132 | -0.132 | 0.693 | 0.25 | 0.01 | 0.19 | 0.35 | 0.28 | 0.2 |
| 7 | 124828243 | rs715937 | 1.04 | -2.668 | 1.244 | -0.486 | 0.088 | 0.06 | 0.06 | 0 | 0.61 | 0.26 | 0.64 | 0.76 | 0.8 | 0.6 |
| 7 | 124855591 | rs12534920 | -0.048 | -2.517 | 0.636 | -0.398 | -0.029 | 0.016 | 0.016 | 0 | 0.2 | 0.33 | 0.26 | 0.24 | 0.13 | 0.32 |
| 7 | 124883581 | rs11770950 | 0.089 | -2.372 | 0.464 | -0.358 | -0.043 | -0.072 | -0.072 | 0.533 | 0.18 | 0.27 | 0.07 | 0 | 0.06 | 0.07 |
| 7 | 124890172 | rs1108984 | 0.19 | -1.219 | 0.954 | 0.244 | 0.001 | 0.012 | 0.012 | 0 | 0.16 | 0.19 | 0.28 | 0.26 | 0.17 | 0.35 |
| 7 | 124890364 | rs1108985 | 0.17 | -1.402 | 0.875 | 0.233 | 0.009 | -0.027 | -0.027 | 0.188 | 0.28 | 0.21 | 0.22 | 0.35 | 0.3 | 0.18 |
| 7 | 124893729 | rs12706645 | -0.588 | -1.45 | 0.908 | 0.193 | -0.014 | 0.003 | 0.003 | 0 | 0.38 | 0.49 | 0.28 | 0.36 | 0.35 | 0.22 |
| 7 | 124897780 | rs6957121 | -1.45 | -1.353 | -0.237 | 0.169 | 0.001 | 0.03 | 0.03 | 0.339 | 0.55 | 0.78 | 0.55 | 0.61 | 0.51 | 0.57 |
| 7 | 124905060 | rs11771079 | 0.395 | -0.822 | -1.024 | 0.589 | -0.003 | -0.01 | -0.01 | 0.01 | 0.15 | 0.08 | 0.26 | 0.26 | 0.16 | 0.35 |
| 7 | 124910833 | rs10279685 | -0.866 | -0.739 | -0.85 | 0.625 | 0.02 | -0.031 | -0.031 | 0 | 0.32 | 0.23 | 0.28 | 0.36 | 0.32 | 0.22 |
| 7 | 124921028 | rs2162587 | -0.95 | -0.725 | -0.913 | 0.635 | 0.012 | -0.015 | -0.015 | 0 | 0.33 | 0.32 | 0.29 | 0.36 | 0.34 | 0.22 |
| 7 | 124926770 | rs4728020 | -0.4 | -0.198 | -1.563 | 1.067 | -0.045 | -0.145 | -0.145 | 0 | 0.23 | 0.02 | 0.23 | 0.37 | 0.28 | 0.16 |
| 7 | 124930633 | rs2114748 | -0.698 | -0.485 | -0.693 | 1.079 | -0.026 | 0.034 | 0.034 | 0.538 | 0.34 | 0.53 | 0.46 | 0.55 | 0.28 | 0.41 |
| 7 | 124938998 | rs12674092 | -0.479 | -0.355 | -1.065 | 1.542 | -0.02 | -0.104 | -0.104 | 3.734 | 0.13 | 0.01 | 0.13 | 0.41 | 0.13 | 0.18 |
| 7 | 124941244 | rs1427585 | -1.419 | -0.485 | -0.253 | 1.324 | -0.023 | 0.034 | 0.034 | 0 | 0.33 | 0.5 | 0.32 | 0.55 | 0.3 | 0.34 |
| 7 | 124956491 | rs9692521 | -0.799 | 0.185 | 0.067 | 1.056 | 0.019 | -0.011 | -0.011 | 0 | 0.17 | 0.23 | 0.21 | 0.14 | 0.22 | 0.17 |
| 7 | 124958806 | rs6973477 | -0.724 | 0.062 | 0.149 | 1.041 | -0.015 | 0.002 | 0.002 | 0 | 0.17 | 0.12 | 0.2 | 0.15 | 0.22 | 0.21 |
| 7 | 124977736 | rs7795867 | -0.59 | 0.066 | 0.125 | 1.043 | -0.036 | -0.003 | -0.003 | 0 | 0.16 | 0.07 | 0.19 | 0.15 | 0.22 | 0.21 |
| 7 | 124979488 | rs17148196 | -1.781 | 0.152 | 0.683 | 0.976 | 0.018 | -0.006 | -0.006 | 0 | 0.3 | 0.48 | 0.29 | 0.29 | 0.33 | 0.5 |
| 7 | 124983020 | rs6970401 | -0.543 | -0.009 | 0.735 | 0.948 | -0.006 | 0.004 | 0.004 | 0 | 0.17 | 0.16 | 0.18 | 0.15 | 0.18 | 0.2 |
| 7 | 124989897 | rs17148220 | 0.241 | -0.108 | 0.388 | 1.108 | -0.019 | -0.121 | -0.121 | 0 | 0.16 | 0.01 | 0.11 | 0.39 | 0.13 | 0.25 |
| 7 | 124990498 | rs6965898 | -1.134 | -0.106 | 0.678 | 1.002 | 0.008 | -0.072 | -0.072 | 0 | 0.21 | 0.4 | 0.21 | 0.1 | 0.2 | 0.25 |
| 7 | 124991885 | rs1427575 | -1.404 | 0.008 | 0.569 | 1.086 | 0 | -0.002 | -0.002 | 2.596 | 0.24 | 0.32 | 0.23 | 0.18 | 0.24 | 0.36 |
| 7 | 124992232 | rs10500107 | 0.24 | 0.038 | 0.496 | 1.082 | -0.001 | -0.003 | -0.003 | 0 | 0.07 | 0.09 | 0.04 | 0.09 | 0.07 | 0.17 |
| 7 | 125000368 | rs11768153 | 1.773 | -0.091 | 1.188 | 1.016 | 0.002 | 0.005 | 0.005 | 1.087 | 0.69 | 0.25 | 0.71 | 0.72 | 0.73 | 0.58 |
| 7 | 125000684 | rs1593756 | 0.329 | -0.282 | 1.507 | 1.079 | -0.007 | -0.042 | -0.042 | 0 | 0.83 | 0.75 | 0.82 | 0.91 | 0.84 | 0.75 |
| 7 | 125001305 | rs1365938 | 1.549 | -0.277 | 1.546 | 1.083 | -0.013 | 0.004 | 0.004 | 0.517 | 0.73 | 0.44 | 0.76 | 0.72 | 0.78 | 0.58 |
| 7 | 125011270 | rs4728022 | 0.453 | -0.317 | 1.567 | 0.976 | -0.013 | -0.055 | -0.055 | 0 | 0.82 | 0.62 | 0.82 | 0.91 | 0.86 | 0.76 |
| 7 | 125017393 | rs6967299 | 1.175 | -0.303 | 1.554 | 1.071 | -0.012 | -0.021 | -0.021 | 0 | 0.76 | 0.62 | 0.79 | 0.83 | 0.8 | 0.58 |
| 7 | 125021576 | rs7784640 | 2.076 | -0.346 | 1.431 | 1.19 | 0.037 | 0.042 | 0.042 | 0 | 0.56 | 0.24 | 0.65 | 0.34 | 0.67 | 0.34 |
| 7 | 125027977 | rs10954071 | 1.087 | -0.299 | 1.495 | 1.043 | -0.02 | -0.03 | -0.03 | 0 | 0.75 | 0.54 | 0.79 | 0.83 | 0.8 | 0.58 |
| 7 | 125028870 | rs7804579 | 1.099 | -0.302 | 1.504 | 1.042 | -0.019 | -0.029 | -0.029 | 0.003 | 0.75 | 0.53 | 0.79 | 0.83 | 0.8 | 0.59 |
| 7 | 125031758 | rs7783813 | 1.096 | -0.298 | 1.465 | 1.044 | -0.018 | -0.028 | -0.028 | 0.05 | 0.75 | 0.52 | 0.78 | 0.82 | 0.8 | 0.59 |
| 7 | 125050529 | rs12154752 | 1.253 | -0.08 | 0.291 | 0.06 | 0 | 0.034 | 0.034 | 0 | 0.29 | 0.17 | 0.23 | 0.01 | 0.28 | 0.23 |
| 7 | 125052182 | rs6945564 | -0.881 | -0.111 | 0.621 | 0.24 | -0.015 | 0.014 | 0.014 | 0.246 | 0.25 | 0.47 | 0.22 | 0.27 | 0.2 | 0.42 |
| 7 | 125061480 | rs1834819 | 0.308 | -0.105 | 0.562 | 0.253 | -0.002 | 0.01 | 0.01 | 0 | 0.08 | 0.17 | 0.04 | 0.09 | 0.06 | 0.17 |
| 7 | 125068580 | rs10500112 | 1.445 | -0.153 | 0.324 | 0.333 | -0.13 | -0.097 | -0.097 | 0.012 | 0.27 | 0.02 | 0.41 | 0.34 | 0.39 | 0.11 |
| 7 | 125070806 | rs7456179 | 0.125 | -0.64 | 1.016 | 0.864 | 0 | 0.018 | 0.018 | 0 | 0.15 | 0.07 | 0.15 | 0.09 | 0.14 | 0.24 |
| 7 | 125072969 | rs1358148 | -1.289 | -0.643 | 1.096 | 0.867 | -0.013 | 0.011 | 0.011 | 0.126 | 0.25 | 0.44 | 0.22 | 0.27 | 0.2 | 0.42 |
| 7 | 125073353 | rs6956995 | -0.242 | -0.482 | 0.907 | 1.02 | 0 | -0.001 | -0.001 | 0.006 | 0.08 | 0.12 | 0.04 | 0.09 | 0.06 | 0.18 |
| 7 | 125075396 | rs17132917 | -0.162 | -0.48 | 0.893 | 1.021 | 0 | -0.002 | -0.002 | 0 | 0.08 | 0.11 | 0.04 | 0.09 | 0.07 | 0.18 |

| | | | | | | | | | | | | | | | | |
|---|---|---|---|---|---|---|---|---|---|---|---|---|---|---|---|---|
| 7 | 125088151 | rs1526413 | 1.173 | -0.506 | 0.849 | 0.992 | 0.006 | -0.009 | -0.009 | 7.445 | 0.71 | 0.26 | 0.79 | 0.43 | 0.8 | 0.58 |
| 7 | 125104792 | rs10267468 | -1.401 | -0.408 | 1.343 | 1.317 | -0.023 | 0.098 | 0.098 | 0 | 0.24 | 0.58 | 0.18 | 0.56 | 0.17 | 0.31 |
| 7 | 125105696 | rs17148452 | -1.337 | -0.413 | 1.534 | 1.314 | -0.022 | 0.118 | 0.118 | 0 | 0.23 | 0.45 | 0.18 | 0.56 | 0.17 | 0.31 |
| 7 | 125111326 | rs6962166 | 0.015 | -0.394 | 1.286 | 1.309 | -0.041 | -0.097 | -0.097 | 0.957 | 0.14 | 0.01 | 0.12 | 0.38 | 0.14 | 0.22 |
| 7 | 125116814 | rs6960986 | -2.631 | -0.236 | 1.507 | 0.891 | -0.033 | 0.053 | 0.053 | 1.233 | 0.1 | 0.57 | 0.06 | 0.09 | 0.02 | 0.09 |
| 7 | 125124029 | rs2049459 | -0.714 | -0.116 | 1.414 | 1.384 | 0.006 | 0.117 | 0.117 | 0 | 0.14 | 0.26 | 0.15 | 0.41 | 0.14 | 0.24 |
| 7 | 125130235 | rs7791922 | 2.485 | -0.225 | 1.748 | 1.045 | -0.016 | 0.167 | 0.167 | 1.885 | 0.9 | 0.49 | 0.91 | 0.84 | 0.97 | 0.89 |
| 7 | 125134765 | rs12534594 | 0.454 | -0.195 | 1.339 | 1.266 | -0.039 | -0.066 | -0.066 | 0 | 0.1 | 0.01 | 0.09 | 0.36 | 0.12 | 0.22 |
| 7 | 125136777 | rs6467012 | 1.405 | -0.267 | 1.967 | 1.269 | -0.009 | 0.154 | 0.154 | 0 | 0.81 | 0.53 | 0.82 | 0.43 | 0.85 | 0.65 |
| 7 | 125149625 | rs1534101 | -0.348 | -0.272 | 1.962 | 1.267 | -0.003 | 0.019 | 0.019 | 0 | 0.13 | 0.12 | 0.13 | 0.49 | 0.14 | 0.31 |
| 7 | 125156846 | rs11762899 | 1.489 | -0.33 | 2.38 | 1.246 | -0.012 | 0.148 | 0.148 | 0.025 | 0.81 | 0.42 | 0.81 | 0.44 | 0.84 | 0.65 |
| 7 | 125165453 | rs1464995 | 0.486 | -0.087 | 1.376 | 0.166 | -0.031 | -0.005 | -0.005 | 0.42 | 0.32 | 0.23 | 0.37 | 0.02 | 0.48 | 0.1 |
| 7 | 125169947 | rs6963279 | 0.628 | 0.581 | 1.311 | 0.129 | 0.034 | -0.047 | -0.047 | 0 | 0.09 | 0.16 | 0.15 | 0.01 | 0.15 | 0.04 |
| 7 | 125186170 | rs3915766 | -0.111 | 0.494 | 1.064 | 0.2 | 0.003 | 0.094 | 0.094 | 0 | 0.86 | 0.75 | 0.84 | 0.44 | 0.86 | 0.69 |
| 7 | 125196970 | rs10954076 | 0.678 | 0.044 | 0.96 | 0.436 | 0.025 | -0.054 | -0.054 | 3.701 | 0.47 | 0.05 | 0.43 | 0.42 | 0.36 | 0.55 |
| 7 | 125203939 | rs792416 | -0.832 | 0.301 | 1.166 | 0.114 | -0.07 | -0.019 | -0.019 | 0 | 0.67 | 0.9 | 0.72 | 0.64 | 0.58 | 0.67 |
| 7 | 125224602 | rs3857841 | -0.27 | 0.284 | 1.536 | 0.117 | -0.002 | 0 | 0 | 0 | 0.62 | 0.38 | 0.65 | 0.6 | 0.56 | 0.67 |
| 7 | 125231398 | rs2524886 | -0.978 | 0.303 | 1.423 | 0.111 | -0.094 | -0.039 | -0.039 | 0 | 0.69 | 0.93 | 0.75 | 0.64 | 0.59 | 0.7 |
| 7 | 125232739 | rs6952116 | 0.52 | 0.321 | 1.465 | 0.06 | 0.022 | 0.059 | 0.059 | 0 | 0.51 | 0.15 | 0.48 | 0.6 | 0.4 | 0.61 |
| 7 | 125239058 | rs3847096 | -0.419 | 0.26 | 1.669 | 0.229 | 0.011 | -0.006 | -0.006 | 0.48 | 0.6 | 0.53 | 0.66 | 0.6 | 0.52 | 0.65 |
| 7 | 125240393 | rs4731254 | -0.511 | 0.31 | 1.673 | 0.172 | 0.003 | -0.007 | -0.007 | 0 | 0.6 | 0.59 | 0.72 | 0.66 | 0.53 | 0.69 |
| 7 | 125257092 | rs6958258 | -0.459 | -0.253 | 1.558 | 0.183 | -0.018 | 0.025 | 0.025 | 0.243 | 0.57 | 0.74 | 0.59 | 0.62 | 0.47 | 0.64 |
| 7 | 125265983 | rs10954078 | -0.47 | -0.248 | 1.546 | 0.185 | -0.019 | 0.024 | 0.024 | 0.348 | 0.58 | 0.74 | 0.58 | 0.62 | 0.47 | 0.64 |
| 7 | 125267702 | rs2189417 | -1.03 | -0.298 | 1.448 | 0.178 | -0.107 | 0.026 | 0.026 | 2.26 | 0.63 | 0.9 | 0.66 | 0.66 | 0.5 | 0.69 |
| 7 | 125278195 | rs10278107 | 0.171 | -0.327 | 1.532 | 0.144 | -0.003 | 0.004 | 0.004 | 1.361 | 0.44 | 0.53 | 0.43 | 0.39 | 0.52 | 0.36 |
| 7 | 125284039 | rs6467022 | 0.096 | -0.431 | 1.296 | -0.018 | 0 | -0.058 | -0.058 | 0 | 0.87 | 0.77 | 0.84 | 1 | 0.87 | 0.96 |
| 7 | 125287479 | rs2040724 | 0.1 | -0.4 | 1.31 | 0.139 | 0.028 | -0.066 | -0.066 | 0 | 0.89 | 0.63 | 0.83 | 1 | 0.9 | 0.97 |
| 7 | 125289360 | rs1419617 | 1.052 | -0.529 | 0.523 | 0.458 | -0.046 | -0.081 | -0.081 | 0.105 | 0.12 | 0.01 | 0.21 | 0.38 | 0.12 | 0.24 |
| 7 | 125321117 | rs1419708 | 0.076 | -0.251 | 1.025 | -0.19 | 0.053 | 0.059 | 0.059 | 0.359 | 0.46 | 0.73 | 0.47 | 0.66 | 0.31 | 0.6 |
| 7 | 125330439 | rs7809559 | -0.575 | -0.226 | 1.003 | -0.171 | 0.088 | 0.115 | 0.115 | 2.12 | 0.05 | 0.47 | 0.1 | 0.03 | 0.01 | 0 |
| 7 | 125339318 | rs402608 | 0.288 | 0.009 | -0.203 | -0.181 | -0.005 | -0.009 | -0.009 | 0 | 0.13 | 0.09 | 0.17 | 0.01 | 0.16 | 0.07 |
| 7 | 125372123 | rs12706681 | 0.705 | 0.274 | -1.533 | 0.101 | 0.028 | -0.051 | -0.051 | 0 | 0.37 | 0.16 | 0.34 | 0.55 | 0.28 | 0.55 |
| 7 | 125375744 | rs2107098 | -0.087 | 1.076 | -1.318 | 0.284 | -0.059 | 0.044 | 0.044 | 0.573 | 0.4 | 0.15 | 0.34 | 0.34 | 0.54 | 0.34 |
| 7 | 125405273 | rs1362048 | -0.006 | 1.952 | -1.366 | 0.401 | -0.069 | -0.017 | -0.017 | 0 | 0.33 | 0.1 | 0.31 | 0.37 | 0.48 | 0.34 |
| 7 | 125405545 | rs4731262 | -0.819 | 0.843 | -2.29 | 0.325 | 0.034 | -0.13 | -0.13 | 0 | 0.72 | 0.97 | 0.81 | 0.61 | 0.84 | 0.65 |
| 7 | 125407937 | rs6974130 | 0.432 | 0.935 | -2.136 | 0.204 | -0.021 | -0.003 | -0.003 | 0.232 | 0.38 | 0.24 | 0.4 | 0.4 | 0.58 | 0.41 |
| 7 | 125408170 | rs1419605 | -0.416 | 0.663 | -2.013 | -0.042 | 0.04 | -0.027 | -0.027 | 0.971 | 0.59 | 0.85 | 0.63 | 0.41 | 0.7 | 0.48 |
| 7 | 125425608 | rs7801590 | 0.59 | 0.394 | -0.635 | -0.178 | 0 | -0.005 | -0.005 | 0 | 0.4 | 0.4 | 0.36 | 0.37 | 0.6 | 0.42 |
| 7 | 125438880 | rs7805133 | -0.134 | 0.24 | -0.692 | -0.242 | 0.058 | 0.03 | 0.03 | 0 | 0.55 | 0.74 | 0.56 | 0.38 | 0.7 | 0.48 |
| 7 | 125446062 | rs10266914 | -0.189 | 0.361 | -0.709 | -0.24 | 0.044 | 0.027 | 0.027 | 0 | 0.56 | 0.78 | 0.55 | 0.39 | 0.68 | 0.49 |
| 7 | 125448866 | rs10954086 | 0.962 | 0.282 | -0.318 | -0.302 | -0.005 | -0.02 | -0.02 | 0 | 0.06 | 0.1 | 0.12 | 0 | 0.04 | 0.02 |
| 7 | 125449666 | rs2107086 | -0.62 | 0.249 | -0.459 | -0.308 | 0.022 | -0.029 | -0.029 | 0 | 0.6 | 0.9 | 0.61 | 0.41 | 0.7 | 0.5 |
| 7 | 125453254 | rs4728029 | 1.307 | 0.355 | -0.293 | -0.332 | -0.068 | -0.024 | -0.024 | 0.564 | 0.33 | 0.06 | 0.29 | 0.37 | 0.49 | 0.36 |
| 7 | 125461881 | rs12706688 | 0.563 | 0.333 | 0.55 | -0.344 | -0.005 | 0.007 | 0.007 | 0 | 0.43 | 0.4 | 0.34 | 0.37 | 0.53 | 0.39 |
| 7 | 125463717 | rs6963314 | -1.052 | 0.37 | 0.763 | -0.255 | 0.072 | -0.047 | -0.047 | 0 | 0.14 | 0.48 | 0.26 | 0.03 | 0.15 | 0.11 |
| 7 | 125467854 | rs725092 | 0.128 | 0.641 | 0.53 | -0.296 | -0.002 | 0.005 | 0.005 | 0 | 0.44 | 0.43 | 0.34 | 0.37 | 0.52 | 0.39 |
| 7 | 125468407 | rs10272687 | -0.291 | 0.626 | 0.693 | -0.177 | -0.014 | -0.065 | -0.065 | 2.027 | 0.12 | 0.28 | 0.06 | 0 | 0.06 | 0.03 |
| 7 | 125473932 | rs11764707 | 1.451 | 0.633 | 0.639 | -0.142 | 0.004 | -0.032 | -0.032 | 0 | 0.06 | 0.13 | 0.03 | 0 | 0.04 | 0.03 |
| 7 | 125479578 | rs1362043 | 0.571 | 0.429 | 0.775 | -0.257 | -0.015 | -0.036 | -0.036 | 0 | 0.08 | 0.2 | 0.15 | 0.01 | 0.05 | 0.03 |
| 7 | 125481946 | rs2107093 | -0.56 | 0.598 | 0.931 | -0.218 | 0.014 | -0.001 | -0.001 | 0 | 0.59 | 0.8 | 0.57 | 0.39 | 0.68 | 0.45 |
| 7 | 125484796 | rs10281994 | -1.023 | 0.804 | 1.077 | -0.225 | 0.037 | -0.053 | -0.053 | 0 | 0.09 | 0.26 | 0.07 | 0 | 0.13 | 0.05 |
| 7 | 125488428 | rs6467054 | -0.973 | 0.797 | 1.195 | 0.134 | 0.044 | -0.098 | -0.098 | 0.045 | 0.15 | 0.38 | 0.1 | 0 | 0.17 | 0.08 |
| 7 | 125490642 | rs4275146 | 1.637 | 0.76 | 1.114 | 0.181 | 0.006 | -0.031 | -0.031 | 0.783 | 0.06 | 0.12 | 0.03 | 0 | 0.04 | 0.03 |
| 7 | 125492934 | rs6948932 | 0.622 | 0.713 | 1.025 | 0.164 | -0.015 | -0.036 | -0.036 | 0.592 | 0.08 | 0.2 | 0.15 | 0.01 | 0.05 | 0.03 |
| 7 | 125494850 | rs2107094 | 0.049 | 0.735 | 1.016 | 0.231 | 0.015 | 0.019 | 0.019 | 0 | 0.51 | 0.59 | 0.42 | 0.38 | 0.63 | 0.42 |
| 7 | 125508794 | rs9649018 | 0.689 | 0.814 | 1.029 | 0.248 | -0.002 | 0.003 | 0.003 | 0 | 0.43 | 0.33 | 0.35 | 0.38 | 0.5 | 0.37 |
| 7 | 125515354 | rs7781587 | -1.167 | 0.842 | 0.702 | 0.316 | 0.013 | -0.046 | -0.046 | 0 | 0.28 | 0.46 | 0.29 | 0.16 | 0.29 | 0.25 |
| 7 | 125521995 | rs721118 | -0.202 | 0.927 | 0.601 | 0.319 | -0.019 | 0.022 | 0.022 | 0 | 0.59 | 0.73 | 0.44 | 0.67 | 0.6 | 0.56 |
| 7 | 125525026 | rs11973696 | -1.078 | 0.937 | 0.69 | 0.31 | 0.017 | -0.025 | -0.025 | 0 | 0.68 | 0.87 | 0.51 | 0.67 | 0.75 | 0.63 |
| 7 | 125530879 | rs12154397 | 0.012 | 0.994 | 0.712 | 0.334 | 0.028 | -0.04 | -0.04 | 0.259 | 0.09 | 0.15 | 0.07 | 0 | 0.15 | 0.07 |
| 7 | 125543552 | rs1362034 | 1.265 | 0.898 | 0.414 | 0.453 | -0.069 | -0.108 | -0.108 | 0.372 | 0.34 | 0.04 | 0.28 | 0.41 | 0.43 | 0.34 |
| 7 | 125543863 | rs1419584 | -0.621 | 0.752 | 0.495 | 0.423 | 0.012 | 0.033 | 0.033 | 0.006 | 0.62 | 0.88 | 0.49 | 0.7 | 0.69 | 0.61 |
| 7 | 125566215 | rs1419557 | 0.781 | 1.238 | 0.815 | 0.534 | 0.007 | -0.005 | -0.005 | 0 | 0.38 | 0.26 | 0.3 | 0.42 | 0.38 | 0.36 |
| 7 | 125566293 | rs1419560 | 1.362 | 1.355 | 0.642 | 0.489 | -0.01 | -0.069 | -0.069 | 0.009 | 0.31 | 0.04 | 0.25 | 0.37 | 0.35 | 0.33 |
| 7 | 125569353 | rs7807653 | 0.093 | 1.512 | 0.971 | 0.526 | 0.018 | 0.007 | 0.007 | 0 | 0.81 | 0.73 | 0.85 | 0.74 | 0.72 | 0.76 |
| 7 | 125569441 | rs1419555 | 0.66 | 1.55 | 1.201 | 0.533 | -0.013 | 0.006 | 0.006 | 0 | 0.38 | 0.41 | 0.31 | 0.42 | 0.38 | 0.36 |
| 7 | 125576498 | rs9641778 | 1.065 | 1.564 | 1.092 | 0.432 | -0.047 | -0.075 | -0.075 | 0.964 | 0.12 | 0.01 | 0.08 | 0.27 | 0.13 | 0.14 |
| 7 | 125576985 | rs11767202 | 1.688 | 1.51 | 1.324 | 0.41 | -0.001 | -0.021 | -0.021 | 0 | 0.06 | 0.09 | 0.03 | 0 | 0.04 | 0.04 |
| 7 | 125582500 | rs13243887 | 0.978 | 1.463 | 1.374 | 0.385 | -0.004 | -0.021 | -0.021 | 0.112 | 0.36 | 0.13 | 0.3 | 0.38 | 0.41 | 0.35 |

| | | | | | | | | | | | | | | | | |
|---|---|---|---|---|---|---|---|---|---|---|---|---|---|---|---|---|
| 7 | 125582748 | rs1833075 | -0.151 | 1.541 | 1.449 | 0.392 | 0.04 | 0.024 | 0.024 | 0 | 0.81 | 0.7 | 0.86 | 0.73 | 0.73 | 0.81 |
| 7 | 125587606 | rs1034651 | 0.469 | 1.054 | 1.431 | 0.456 | -0.001 | 0 | 0 | 0.074 | 0.38 | 0.37 | 0.32 | 0.37 | 0.4 | 0.35 |
| 7 | 125604657 | rs10245403 | 0.186 | 1.088 | 1.388 | 0.65 | -0.001 | -0.003 | -0.003 | 0 | 0.37 | 0.39 | 0.32 | 0.35 | 0.4 | 0.38 |
| 7 | 125614933 | rs1362026 | 0.829 | 1.101 | 1.309 | 0.681 | -0.01 | -0.019 | -0.019 | 0 | 0.29 | 0.12 | 0.25 | 0.35 | 0.34 | 0.37 |
| 7 | 125619396 | rs11978171 | 0.861 | 1.284 | 1.35 | 0.674 | 0.011 | 0.037 | 0.037 | 0 | 0.43 | 0.19 | 0.53 | 0.34 | 0.37 | 0.43 |
| 7 | 125623142 | rs4731279 | -0.134 | 1.306 | 1.398 | 0.694 | 0.011 | 0.016 | 0.016 | 0 | 0.82 | 0.72 | 0.86 | 0.72 | 0.77 | 0.84 |
| 7 | 125623199 | rs10239988 | 0.56 | 1.19 | 1.455 | 0.712 | -0.007 | 0.026 | 0.026 | 0 | 0.15 | 0.22 | 0.1 | 0.28 | 0.12 | 0.11 |
| 7 | 125644656 | rs1419540 | -0.017 | 1.185 | 1.246 | 0.588 | 0.001 | -0.006 | -0.006 | 0.003 | 0.54 | 0.46 | 0.62 | 0.39 | 0.47 | 0.49 |
| 7 | 125648081 | rs17149165 | -0.143 | 1.258 | 1.365 | 0.65 | 0.023 | 0.002 | 0.002 | 0 | 0.12 | 0.32 | 0.1 | 0.07 | 0.1 | 0.06 |
| 7 | 125664894 | rs6965646 | -0.495 | 1.192 | 1.055 | 0.681 | 0.045 | 0.03 | 0.03 | 0.104 | 0.14 | 0.48 | 0.13 | 0.08 | 0.11 | 0.06 |
| 7 | 125667555 | rs17149212 | -0.474 | 1.196 | 1.061 | 0.683 | 0.047 | 0.032 | 0.032 | 0.118 | 0.13 | 0.48 | 0.14 | 0.08 | 0.11 | 0.06 |
| 7 | 125671593 | rs10487432 | 1.169 | 1.109 | 0.121 | 0.7 | -0.021 | -0.033 | -0.033 | 0 | 0.26 | 0.03 | 0.19 | 0.3 | 0.29 | 0.35 |
| 7 | 125705705 | rs10248306 | 0.167 | 1.439 | 0.991 | 0.588 | -0.001 | 0.005 | 0.005 | 0 | 0.38 | 0.32 | 0.31 | 0.35 | 0.4 | 0.42 |
| 7 | 125707685 | rs2079055 | 0.917 | 1.293 | 1.058 | 0.664 | 0.007 | 0.056 | 0.056 | 0 | 0.41 | 0.18 | 0.54 | 0.31 | 0.38 | 0.43 |
| 7 | 125709760 | rs6979551 | 0.375 | 1.28 | 1.027 | 0.707 | -0.001 | -0.001 | -0.001 | 0 | 0.38 | 0.32 | 0.31 | 0.38 | 0.4 | 0.42 |
| 7 | 125716635 | rs10228474 | -0.575 | 1.278 | 0.936 | 0.811 | -0.007 | 0.022 | 0.022 | 0 | 0.54 | 0.6 | 0.42 | 0.65 | 0.51 | 0.54 |
| 7 | 125716866 | rs10272238 | 0.538 | 1.252 | 1.005 | 0.758 | -0.011 | 0.039 | 0.039 | 0 | 0.15 | 0.25 | 0.1 | 0.28 | 0.12 | 0.11 |
| 7 | 125719132 | rs8180794 | -0.868 | 1.101 | 0.461 | 0.616 | -0.012 | -0.002 | -0.002 | 0 | 0.68 | 0.75 | 0.74 | 0.67 | 0.59 | 0.59 |
| 7 | 125725649 | rs7457541 | 0.485 | 1.151 | 0.897 | 0.851 | 0.013 | 0.061 | 0.061 | 0 | 0.78 | 0.55 | 0.86 | 0.69 | 0.8 | 0.83 |
| 7 | 125727151 | rs11975021 | 1.111 | 0.949 | 0.697 | 0.648 | -0.068 | -0.003 | -0.003 | 0 | 0.29 | 0.07 | 0.47 | 0.28 | 0.34 | 0.37 |
| 7 | 125730694 | rs10228582 | -0.049 | 1.408 | 1.136 | 1.056 | -0.04 | 0.083 | 0.083 | 0 | 0.17 | 0.35 | 0.09 | 0.31 | 0.09 | 0.13 |
| 7 | 125733306 | rs7778354 | -0.049 | 1.408 | 1.136 | 1.056 | -0.04 | 0.083 | 0.083 | 0 | 0.17 | 0.35 | 0.09 | 0.31 | 0.09 | 0.13 |
| 7 | 125733885 | rs6958702 | -0.049 | 1.408 | 1.136 | 1.056 | -0.04 | 0.083 | 0.083 | 0 | 0.17 | 0.35 | 0.09 | 0.31 | 0.09 | 0.13 |
| 7 | 125735050 | rs10954099 | 0.819 | 1.417 | 1.14 | 1.022 | -0.004 | 0.041 | 0.041 | 0 | 0.4 | 0.26 | 0.55 | 0.29 | 0.4 | 0.39 |
| 7 | 125763255 | rs10237506 | -0.086 | 0.869 | 1.038 | 1.116 | -0.04 | 0.101 | 0.101 | 0 | 0.17 | 0.36 | 0.09 | 0.33 | 0.09 | 0.13 |
| 7 | 125772120 | rs3857846 | 0.781 | 0.929 | 0.364 | 0.995 | -0.011 | 0.074 | 0.074 | 0 | 0.4 | 0.18 | 0.53 | 0.26 | 0.39 | 0.39 |
| 7 | 125773460 | rs9655857 | 0.281 | 1.339 | 0.516 | 0.869 | -0.01 | 0 | 0 | 0 | 0.57 | 0.64 | 0.65 | 0.59 | 0.49 | 0.52 |
| 7 | 125776626 | rs626762 | -2.022 | 1.719 | 1.298 | 1.443 | 0.003 | 0.037 | 0.037 | 4.101 | 0.18 | 0.69 | 0.2 | 0.3 | 0.21 | 0.25 |
| 7 | 125794295 | rs507803 | -1.109 | 1.772 | 1.979 | 1.501 | 0.014 | 0.076 | 0.076 | 0 | 0.1 | 0.21 | 0.13 | 0.27 | 0.14 | 0.23 |
| 7 | 125803961 | rs510319 | -2.899 | 1.691 | 2.014 | 1.502 | 0 | -0.011 | -0.011 | 0 | 0.2 | 0.8 | 0.21 | 0.31 | 0.22 | 0.26 |
| 7 | 125816381 | rs630935 | -2.082 | 1.573 | 2.28 | 1.607 | -0.01 | 0.083 | 0.083 | 4.822 | 0.15 | 0.66 | 0.18 | 0.31 | 0.15 | 0.24 |
| 7 | 125827089 | rs668717 | -1.192 | 1.809 | 1.579 | 1.547 | -0.001 | 0.008 | 0.008 | 0 | 0.09 | 0.09 | 0.17 | 0.27 | 0.16 | 0.22 |
| 7 | 125834329 | rs672097 | -1.85 | 2.228 | 1.996 | 1.56 | 0.061 | 0.1 | 0.1 | 0 | 0.12 | 0.33 | 0.17 | 0.28 | 0.17 | 0.27 |
| 7 | 125846665 | rs11983395 | -0.365 | 2.371 | 1.638 | 1.184 | -0.004 | 0 | 0 | 0 | 0.06 | 0.05 | 0.07 | 0 | 0.11 | 0.07 |
| 7 | 125848263 | rs471173 | -1.406 | 2.87 | 1.942 | 1.394 | 0.051 | 0.052 | 0.052 | 0.022 | 0.08 | 0.12 | 0.15 | 0.18 | 0.15 | 0.11 |
| 7 | 125853688 | rs727594 | -1.848 | 3.012 | 1.998 | 1.754 | 0.012 | 0.006 | 0.006 | 0 | 0.22 | 0.36 | 0.19 | 0.22 | 0.27 | 0.25 |
| 7 | 125867303 | rs1419503 | 0.276 | 3.313 | 1.939 | 1.474 | 0.005 | -0.009 | -0.009 | 0 | 0.12 | 0.17 | 0.25 | 0.05 | 0.15 | 0.11 |
| 7 | 125869122 | rs2687481 | -2.442 | 3.172 | 2.371 | 1.434 | 0.086 | 0.048 | 0.048 | 0.907 | 0.18 | 0.4 | 0.2 | 0.24 | 0.22 | 0.16 |
| 7 | 125878744 | rs2687473 | -0.993 | 3.188 | 2.394 | 1.437 | 0.038 | 0.017 | 0.017 | 0 | 0.09 | 0.09 | 0.15 | 0.16 | 0.16 | 0.13 |
| 7 | 125892242 | rs1673366 | -1.278 | 3.19 | 2.424 | 1.44 | 0.097 | 0.052 | 0.052 | 0.129 | 0.1 | 0.19 | 0.17 | 0.16 | 0.16 | 0.13 |
| 7 | 125901145 | rs13241603 | 0.637 | 3.154 | 2.32 | 1.417 | 0.005 | -0.009 | -0.009 | 0 | 0.11 | 0.17 | 0.25 | 0.05 | 0.14 | 0.1 |
| 7 | 125903591 | rs7776473 | -1.094 | 3.091 | 2.12 | 1.423 | -0.01 | 0 | 0 | 1.901 | 0.08 | 0.21 | 0.04 | 0.08 | 0.06 | 0.03 |
| 7 | 125908665 | rs10256624 | -1.206 | 3.147 | 2.07 | 1.438 | -0.004 | -0.003 | -0.003 | 0 | 0.21 | 0.22 | 0.18 | 0.22 | 0.27 | 0.25 |
| 7 | 125915063 | rs337304 | -2.549 | 3.058 | 1.896 | 1.45 | 0.015 | -0.018 | -0.018 | 0 | 0.43 | 0.72 | 0.61 | 0.45 | 0.57 | 0.48 |
| 7 | 125922203 | rs337317 | -2.552 | 3.056 | 1.895 | 1.449 | 0.015 | -0.018 | -0.018 | 0 | 0.43 | 0.72 | 0.61 | 0.46 | 0.57 | 0.48 |
| 7 | 125931382 | rs17149532 | -0.229 | 3.087 | 1.331 | 1.46 | -0.022 | -0.041 | -0.041 | 0.632 | 0.13 | 0.06 | 0.11 | 0.22 | 0.16 | 0.18 |
| 7 | 125945936 | rs6952074 | 0.18 | 2.971 | 1.614 | 1.385 | 0.016 | -0.028 | -0.028 | 0 | 0.12 | 0.24 | 0.26 | 0.05 | 0.14 | 0.1 |
| 7 | 125946683 | rs336673 | 1.519 | 2.933 | 1.025 | 1.324 | -0.053 | -0.038 | -0.038 | 6.58 | 0.74 | 0.87 | 0.72 | 0.62 | 0.62 | 0.67 |
| 7 | 125961043 | rs485 | 2.674 | 2.939 | 1.065 | 1.564 | 0.094 | 0.049 | 0.049 | 0.232 | 0.5 | 0.14 | 0.4 | 0.39 | 0.39 | 0.44 |
| 7 | 125972744 | rs917451 | 0.488 | 2.899 | 1.035 | 1.389 | -0.003 | 0.003 | 0.003 | 0 | 0.11 | 0.1 | 0.21 | 0.05 | 0.14 | 0.1 |
| 7 | 125975251 | rs496 | 0.591 | 3.205 | 1.001 | 1.529 | 0.027 | 0.012 | 0.012 | 0 | 0.92 | 0.93 | 0.85 | 0.84 | 0.84 | 0.86 |
| 7 | 125990516 | rs335915 | 0.38 | 3.245 | 0.915 | 1.672 | -0.026 | -0.026 | -0.026 | 0 | 0.88 | 0.93 | 0.87 | 0.77 | 0.79 | 0.81 |
| 7 | 125992885 | rs10954109 | 2.632 | 2.816 | 0.804 | 1.589 | 0.058 | 0.035 | 0.035 | 0.17 | 0.44 | 0.15 | 0.38 | 0.37 | 0.36 | 0.36 |
| 7 | 126006996 | rs17149630 | -0.472 | 2.863 | 0.782 | 1.469 | -0.003 | -0.006 | -0.006 | 0 | 0.13 | 0.06 | 0.14 | 0.2 | 0.15 | 0.15 |
| 7 | 126022653 | rs6976095 | -1.318 | 2.821 | 1.424 | 1.445 | 0.004 | -0.004 | -0.004 | 0 | 0.27 | 0.33 | 0.36 | 0.28 | 0.29 | 0.3 |
| 7 | 126037381 | rs12539616 | 2.968 | 2.733 | 1.889 | 1.464 | 0.029 | 0.002 | 0.002 | 0.476 | 0.47 | 0.34 | 0.42 | 0.45 | 0.37 | 0.42 |
| 7 | 126040117 | rs887025 | 1.917 | 2.773 | 1.977 | 1.476 | 0.029 | 0 | 0 | 0 | 0.73 | 0.61 | 0.77 | 0.75 | 0.65 | 0.72 |
| 7 | 126042050 | rs2188191 | 2.451 | 2.751 | 1.902 | 1.475 | 0.001 | 0.002 | 0.002 | 0 | 0.62 | 0.62 | 0.63 | 0.53 | 0.49 | 0.58 |
| 7 | 126045700 | rs10271617 | 0.074 | 2.667 | 1.624 | 1.588 | -0.024 | -0.023 | -0.023 | 0.378 | 0.88 | 0.92 | 0.88 | 0.8 | 0.8 | 0.82 |
| 7 | 126045783 | rs10241921 | 0.074 | 2.667 | 1.624 | 1.572 | -0.024 | -0.02 | -0.02 | 0 | 0.88 | 0.92 | 0.88 | 0.82 | 0.8 | 0.82 |
| 7 | 126055187 | rs12666584 | 0.952 | 2.555 | 1.647 | 1.51 | -0.033 | -0.002 | -0.002 | 0 | 0.76 | 0.83 | 0.68 | 0.74 | 0.67 | 0.68 |
| 7 | 126060738 | rs2106197 | 2.211 | 2.519 | 2.03 | 1.515 | 0.015 | -0.011 | -0.011 | 0 | 0.62 | 0.46 | 0.56 | 0.66 | 0.51 | 0.53 |
| 7 | 126063696 | rs1016973 | 2.072 | 2.508 | 2.09 | 1.537 | 0.05 | 0.017 | 0.017 | 0 | 0.73 | 0.55 | 0.76 | 0.72 | 0.65 | 0.66 |
| 7 | 126066990 | rs10226718 | -0.915 | 2.502 | 2.067 | 1.499 | -0.009 | 0.038 | 0.038 | 0 | 0.22 | 0.15 | 0.27 | 0.08 | 0.24 | 0.2 |
| 7 | 126071308 | rs9918568 | 1.906 | 2.909 | 2.103 | 1.494 | 0.035 | 0.044 | 0.044 | 0 | 0.71 | 0.66 | 0.7 | 0.49 | 0.6 | 0.61 |
| 7 | 126072066 | rs10487449 | -0.656 | 2.932 | 2.034 | 1.489 | -0.016 | -0.019 | -0.019 | 0.673 | 0.15 | 0.07 | 0.16 | 0.26 | 0.2 | 0.16 |
| 7 | 126073820 | rs1419458 | 2.345 | 2.976 | 2.003 | 1.498 | 0.021 | 0.015 | 0.015 | 0.635 | 0.54 | 0.36 | 0.46 | 0.38 | 0.42 | 0.43 |
| 7 | 126074431 | rs1419459 | -0.162 | 2.975 | 2.006 | 1.498 | -0.027 | -0.022 | -0.022 | 0 | 0.88 | 0.93 | 0.88 | 0.82 | 0.8 | 0.81 |
| 7 | 126077198 | rs10276102 | 0.07 | 2.978 | 2.002 | 1.499 | -0.018 | -0.014 | -0.014 | 0.992 | 0.87 | 0.91 | 0.88 | 0.82 | 0.8 | 0.81 |

| | | | | | | | | | | | | | | | | |
|---|---|---|---|---|---|---|---|---|---|---|---|---|---|---|---|---|
| 7 | 126079144 | rs712723 | 2.253 | 2.862 | 1.911 | 1.432 | 0.02 | 0.015 | 0.015 | 0.568 | 0.54 | 0.36 | 0.46 | 0.38 | 0.42 | 0.43 |
| 7 | 126081733 | rs2283059 | 2.927 | 2.817 | 1.67 | 1.451 | 0.087 | -0.002 | -0.002 | 0 | 0.44 | 0.13 | 0.37 | 0.38 | 0.32 | 0.37 |
| 7 | 126090509 | rs13247338 | 0.857 | 2.816 | 1.625 | 1.359 | -0.011 | 0.037 | 0.037 | 0 | 0.13 | 0.03 | 0.2 | 0.08 | 0.13 | 0.14 |
| 7 | 126090571 | rs17679839 | 2.016 | 2.888 | 1.98 | 1.385 | -0.001 | 0.003 | 0.003 | 0.133 | 0.58 | 0.46 | 0.5 | 0.43 | 0.47 | 0.47 |
| 7 | 126091804 | rs17149799 | -0.597 | 2.957 | 1.938 | 1.422 | -0.034 | -0.032 | -0.032 | 0 | 0.14 | 0.06 | 0.16 | 0.23 | 0.2 | 0.14 |
| 7 | 126093581 | rs11978873 | 1.569 | 2.736 | 2.119 | 1.369 | 0.006 | 0.008 | 0.008 | 0.12 | 0.73 | 0.68 | 0.71 | 0.51 | 0.6 | 0.61 |
| 7 | 126099842 | rs13235082 | 0.674 | 2.655 | 1.896 | 1.348 | 0 | 0.029 | 0.029 | 0 | 0.14 | 0.09 | 0.2 | 0.08 | 0.13 | 0.14 |
| 7 | 126101126 | rs2283061 | 1.938 | 2.701 | 2.006 | 1.376 | 0.07 | 0.085 | 0.085 | 0.22 | 0.7 | 0.46 | 0.69 | 0.51 | 0.6 | 0.61 |
| 7 | 126101191 | rs2283062 | 2.983 | 2.497 | 1.535 | 1.382 | 0.081 | -0.031 | -0.031 | 0 | 0.43 | 0.09 | 0.38 | 0.41 | 0.32 | 0.39 |
| 7 | 126107001 | rs6467082 | 1.744 | 2.93 | 1.656 | 1.787 | -0.022 | -0.025 | -0.025 | 0 | 0.68 | 0.77 | 0.57 | 0.53 | 0.51 | 0.63 |
| 7 | 126112702 | rs2108849 | 1.66 | 2.831 | 1.591 | 1.79 | -0.029 | -0.024 | -0.024 | 1.362 | 0.7 | 0.76 | 0.61 | 0.51 | 0.55 | 0.66 |
| 7 | 126113335 | rs7796391 | 2.548 | 2.667 | 1.751 | 1.745 | 0.013 | 0.012 | 0.012 | 0 | 0.61 | 0.5 | 0.56 | 0.42 | 0.49 | 0.57 |
| 7 | 126122354 | rs1582258 | -0.912 | 2.935 | 1.689 | 1.572 | -0.071 | -0.01 | -0.01 | 1.833 | 0.28 | 0.12 | 0.38 | 0.3 | 0.38 | 0.3 |
| 7 | 126125053 | rs2283064 | 1.962 | 2.81 | 1.817 | 1.589 | 0.043 | -0.006 | -0.006 | 0 | 0.53 | 0.7 | 0.4 | 0.41 | 0.41 | 0.48 |
| 7 | 126145179 | rs6943531 | -1.475 | 2.693 | 2.424 | 1.617 | 0.007 | -0.008 | -0.008 | 0 | 0.35 | 0.4 | 0.46 | 0.32 | 0.43 | 0.34 |
| 7 | 126149319 | rs1419472 | 0.834 | 2.743 | 2.403 | 1.642 | -0.009 | -0.009 | -0.009 | 0 | 0.83 | 0.88 | 0.78 | 0.71 | 0.79 | 0.76 |
| 7 | 126150163 | rs728226 | 2.761 | 2.747 | 2.601 | 1.634 | 0.065 | -0.016 | -0.016 | 0 | 0.44 | 0.14 | 0.3 | 0.38 | 0.36 | 0.42 |
| 7 | 126153376 | rs2237734 | -0.82 | 2.862 | 2.363 | 1.624 | -0.044 | -0.024 | -0.024 | 0.343 | 0.13 | 0.06 | 0.19 | 0.2 | 0.28 | 0.14 |
| 7 | 126154236 | rs2283065 | 2.864 | 2.757 | 2.349 | 2.006 | 0.023 | 0.048 | 0.048 | 0.051 | 0.44 | 0.02 | 0.26 | 0.26 | 0.36 | 0.4 |
| 7 | 126160548 | rs13246399 | -1.28 | 2.853 | 2.422 | 1.406 | -0.025 | 0.011 | 0.011 | 0 | 0.32 | 0.25 | 0.44 | 0.32 | 0.43 | 0.34 |
| 7 | 126162920 | rs1362178 | 0.632 | 2.831 | 2.395 | 1.435 | -0.025 | -0.008 | -0.008 | 0 | 0.85 | 0.87 | 0.78 | 0.81 | 0.79 | 0.82 |
| 7 | 126167037 | rs2237739 | 2.786 | 2.6 | 2.413 | 1.264 | 0.051 | -0.001 | -0.001 | 0 | 0.45 | 0.23 | 0.3 | 0.38 | 0.36 | 0.42 |
| 7 | 126168766 | rs2299456 | 0.21 | 2.614 | 2.407 | 1.258 | 0.004 | 0.013 | 0.013 | 0 | 0.18 | 0.16 | 0.22 | 0.1 | 0.14 | 0.18 |
| 7 | 126170622 | rs6949287 | 2.6 | 2.585 | 2.202 | 1.172 | 0.007 | 0 | 0 | 0 | 0.46 | 0.42 | 0.31 | 0.39 | 0.36 | 0.42 |
| 7 | 126183236 | rs1990247 | 0.218 | 2.411 | 1.834 | 0.695 | 0.001 | -0.008 | -0.008 | 0 | 0.18 | 0.18 | 0.21 | 0.03 | 0.14 | 0.18 |
| 7 | 126186307 | rs7806558 | 1.565 | 2.334 | 1.787 | 0.663 | 0.002 | -0.042 | -0.042 | 0 | 0.6 | 0.45 | 0.53 | 0.86 | 0.65 | 0.59 |
| 7 | 126197081 | rs2299465 | 2.417 | 2.18 | 1.706 | 0.746 | 0.055 | -0.047 | -0.047 | 0.125 | 0.43 | 0.15 | 0.3 | 0.43 | 0.38 | 0.42 |
| 7 | 126221861 | rs12536122 | 2.393 | 2.223 | 1.942 | 0.753 | 0.05 | -0.047 | -0.047 | 0 | 0.44 | 0.11 | 0.3 | 0.43 | 0.38 | 0.43 |
| 7 | 126225566 | rs7804379 | 2.408 | 2.207 | 1.941 | 0.752 | 0.057 | -0.048 | -0.048 | 0.39 | 0.44 | 0.11 | 0.29 | 0.43 | 0.38 | 0.43 |
| 7 | 126228289 | rs10254233 | 0.283 | 2.392 | 2.109 | 0.589 | 0.012 | -0.088 | -0.088 | 2.073 | 0.76 | 0.58 | 0.66 | 0.95 | 0.79 | 0.76 |
| 7 | 126265711 | rs17684282 | -0.439 | 2.606 | 2.046 | 0.533 | -0.037 | 0.042 | 0.042 | 0.04 | 0.11 | 0.05 | 0.08 | 0 | 0.19 | 0.06 |
| 7 | 126269475 | rs7795337 | 0.188 | 2.645 | 2.268 | 0.498 | 0.005 | -0.057 | -0.057 | 0 | 0.77 | 0.65 | 0.66 | 0.95 | 0.8 | 0.76 |
| 7 | 126284782 | rs1361985 | -0.305 | 2.608 | 2.172 | 0.495 | -0.054 | 0.07 | 0.07 | 1.447 | 0.8 | 0.93 | 0.7 | 0.95 | 0.8 | 0.8 |
| 7 | 126288566 | rs11767915 | -0.409 | 2.688 | 2.1 | 0.603 | -0.049 | -0.051 | -0.051 | 0 | 0.13 | 0.02 | 0.1 | 0.18 | 0.16 | 0.21 |
| 7 | 126295919 | rs7792723 | 1.962 | 2.599 | 2.582 | 0.742 | -0.017 | 0.015 | 0.015 | 0.246 | 0.59 | 0.31 | 0.42 | 0.62 | 0.54 | 0.64 |
| 7 | 126296428 | rs12706740 | -0.813 | 2.57 | 2.768 | 0.803 | -0.001 | 0.022 | 0.022 | 1.888 | 0.26 | 0.42 | 0.46 | 0.38 | 0.23 | 0.29 |
| 7 | 126297078 | rs4236623 | -2.243 | 2.68 | 2.468 | 0.722 | -0.052 | -0.042 | -0.042 | 1.124 | 0.44 | 0.85 | 0.61 | 0.56 | 0.43 | 0.5 |
| 7 | 126299008 | rs734524 | -2.341 | 2.527 | 2.616 | 0.642 | 0.034 | -0.028 | -0.028 | 0 | 0.3 | 0.5 | 0.25 | 0.2 | 0.38 | 0.29 |
| 7 | 126305659 | rs7779831 | -2.715 | 2.475 | 2.481 | 0.641 | 0.048 | -0.049 | -0.049 | 0.584 | 0.55 | 0.89 | 0.64 | 0.56 | 0.62 | 0.57 |
| 7 | 126308312 | rs13243112 | -1.735 | 2.534 | 2.477 | 0.686 | -0.008 | 0.149 | 0.149 | 2.285 | 0.09 | 0.25 | 0.17 | 0.33 | 0.05 | 0.08 |
| 7 | 126312844 | rs1074728 | -2.053 | 2.421 | 2.19 | 0.534 | -0.004 | -0.012 | -0.012 | 0 | 0.47 | 0.57 | 0.49 | 0.23 | 0.57 | 0.48 |
| 7 | 126324030 | rs4731323 | 0.257 | 2.32 | 1.915 | 0.502 | -0.007 | 0.007 | 0.007 | 0 | 0.2 | 0.2 | 0.24 | 0.04 | 0.22 | 0.2 |
| 7 | 126327399 | rs1008907 | -2.79 | 2.355 | 1.837 | 0.515 | 0.013 | -0.067 | -0.067 | 0 | 0.56 | 0.91 | 0.62 | 0.57 | 0.61 | 0.57 |
| 7 | 126328230 | rs17613247 | -0.503 | 2.399 | 1.725 | 0.476 | -0.039 | 0.054 | 0.054 | 0 | 0.1 | 0.04 | 0.08 | 0 | 0.18 | 0.06 |
| 7 | 126331987 | rs3808154 | 0.354 | 2.414 | 1.77 | 0.516 | -0.012 | 0.021 | 0.021 | 0 | 0.19 | 0.15 | 0.22 | 0.03 | 0.21 | 0.2 |
| 7 | 126357927 | rs2299498 | -1.837 | 2.329 | 1.774 | 0.759 | 0.002 | 0.016 | 0.016 | 0.026 | 0.18 | 0.24 | 0.25 | 0.37 | 0.24 | 0.2 |
| 7 | 126368292 | rs10256873 | -2.701 | 2.399 | 1.758 | 0.777 | -0.005 | -0.009 | -0.009 | 0 | 0.35 | 0.71 | 0.35 | 0.37 | 0.39 | 0.37 |
| 7 | 126372269 | rs1345211 | 0.652 | 2.258 | 2.116 | 0.736 | 0.019 | 0.026 | 0.026 | 0 | 0.87 | 0.76 | 0.86 | 0.84 | 0.84 | 0.83 |
| 7 | 126376133 | rs2299505 | -0.767 | 2.327 | 2.071 | 0.098 | -0.034 | 0.13 | 0.13 | 0.447 | 0.36 | 0.2 | 0.31 | 0.07 | 0.4 | 0.33 |
| 7 | 126378143 | rs10487458 | 0.974 | 2.521 | 2.388 | 0.257 | 0.034 | 0.145 | 0.145 | 0.029 | 0.83 | 0.67 | 0.75 | 0.5 | 0.79 | 0.77 |
| 7 | 126378868 | rs10250459 | -0.951 | 2.315 | 2.148 | 0.254 | -0.053 | 0.155 | 0.155 | 0 | 0.4 | 0.21 | 0.36 | 0.11 | 0.49 | 0.36 |
| 7 | 126380847 | rs2157977 | -1.089 | 2.319 | 2.09 | 0.202 | -0.062 | 0.168 | 0.168 | 0 | 0.4 | 0.2 | 0.36 | 0.07 | 0.49 | 0.35 |
| 7 | 126381249 | rs2237768 | 1.912 | 2.317 | 2.04 | 0.422 | -0.01 | -0.021 | -0.021 | 0 | 0.45 | 0.03 | 0.39 | 0.34 | 0.39 | 0.43 |
| 7 | 126393139 | rs1946116 | 0.705 | 2.349 | 2.3 | 0.392 | 0.024 | 0.299 | 0.299 | 0 | 0.88 | 0.63 | 0.79 | 0.49 | 0.88 | 0.79 |
| 7 | 126396962 | rs10954135 | 1.957 | 2.374 | 2.15 | 0.378 | 0.039 | -0.022 | -0.022 | 0 | 0.46 | 0.19 | 0.41 | 0.41 | 0.39 | 0.44 |
| 7 | 126399506 | rs11769491 | -1.057 | 2.278 | 2.127 | 0.147 | -0.084 | 0.205 | 0.205 | 0 | 0.39 | 0.16 | 0.35 | 0.05 | 0.49 | 0.31 |
| 7 | 126403956 | rs11768413 | -1.043 | 2.267 | 2.133 | 0.156 | -0.085 | 0.191 | 0.191 | 0 | 0.39 | 0.16 | 0.35 | 0.07 | 0.47 | 0.39 |
| 7 | 126409064 | rs2299514 | 0.345 | 2.498 | 1.999 | -0.102 | 0.006 | 0.081 | 0.081 | 0 | 0.87 | 0.81 | 0.79 | 0.49 | 0.84 | 0.78 |
| 7 | 126415127 | rs2299517 | -0.85 | 2.437 | 1.805 | -0.107 | -0.098 | 0.194 | 0.194 | 0 | 0.38 | 0.13 | 0.35 | 0.07 | 0.47 | 0.37 |
| 7 | 126425745 | rs10954138 | 1.885 | 2.387 | 1.834 | -0.136 | 0.051 | -0.04 | -0.04 | 0 | 0.43 | 0.18 | 0.4 | 0.41 | 0.35 | 0.41 |
| 7 | 126429517 | rs6951643 | -1.15 | 2.345 | 1.798 | -0.451 | 0.003 | -0.054 | -0.054 | 0.544 | 0.42 | 0.57 | 0.38 | 0.07 | 0.48 | 0.37 |
| 7 | 126447564 | rs4731330 | -0.907 | 2.41 | 0.924 | -0.452 | -0.07 | 0.157 | 0.157 | 0 | 0.38 | 0.17 | 0.35 | 0.06 | 0.47 | 0.32 |
| 7 | 126462415 | rs1557646 | -0.717 | 2.385 | 0.943 | -0.508 | -0.059 | 0.116 | 0.116 | 0.009 | 0.36 | 0.21 | 0.37 | 0.07 | 0.49 | 0.36 |
| 7 | 126464966 | rs2283075 | -1.496 | 2.197 | 1.356 | -0.347 | 0.016 | 0.003 | 0.003 | 0 | 0.51 | 0.46 | 0.57 | 0.59 | 0.65 | 0.58 |
| 7 | 126472747 | rs2157752 | -1.792 | 2.133 | 1.069 | -0.376 | 0.03 | 0.017 | 0.017 | 0 | 0.54 | 0.68 | 0.59 | 0.65 | 0.65 | 0.59 |
| 7 | 126479218 | rs929175 | 0.32 | 2.925 | 0.538 | -0.291 | -0.112 | 0.099 | 0.099 | 0.955 | 0.22 | 0.03 | 0.28 | 0.05 | 0.36 | 0.32 |
| 7 | 126487481 | rs1002744 | -1.097 | 2.839 | 0.811 | -0.218 | 0.015 | 0.228 | 0.228 | 0.449 | 0.17 | 0.45 | 0.22 | 0.58 | 0.16 | 0.22 |
| 7 | 126493619 | rs2237773 | -0.175 | 2.653 | 0.344 | -0.172 | 0.003 | 0.06 | 0.06 | 0.064 | 0.13 | 0.16 | 0.2 | 0.52 | 0.15 | 0.22 |
| 7 | 126503680 | rs1204518 | -0.674 | 2.453 | -0.157 | -0.42 | -0.008 | 0.167 | 0.167 | 0.505 | 0.19 | 0.55 | 0.22 | 0.58 | 0.16 | 0.18 |

| | | | | | | | | | | | | | | | | |
|---|---|---|---|---|---|---|---|---|---|---|---|---|---|---|---|---|
| 7 | 126512514 | rs1204526 | -1.814 | 2.401 | -0.36 | -0.626 | 0.039 | 0.03 | 0.03 | 0 | 0.54 | 0.74 | 0.59 | 0.65 | 0.63 | 0.54 |
| 7 | 126513625 | rs7809518 | -1.18 | 2.689 | -0.804 | -1.322 | -0.047 | 0.082 | 0.082 | 0.217 | 0.35 | 0.23 | 0.37 | 0.07 | 0.48 | 0.39 |
| 7 | 126514557 | rs4731334 | -1.184 | 2.056 | -0.477 | -0.533 | -0.007 | 0.01 | 0.01 | 0 | 0.1 | 0.1 | 0.08 | 0.02 | 0.11 | 0.03 |
| 7 | 126519669 | rs2051681 | -1.65 | 2.315 | -0.358 | -0.666 | 0.002 | -0.021 | -0.021 | 0 | 0.15 | 0.17 | 0.16 | 0.04 | 0.22 | 0.2 |
| 7 | 126535600 | rs12669064 | -0.847 | 2.902 | 0.433 | -0.276 | 0.067 | 0.136 | 0.136 | 0 | 0.12 | 0.28 | 0.18 | 0.28 | 0.18 | 0.17 |
| 7 | 126539911 | rs916614 | 0.742 | 3.832 | 0.168 | -0.485 | -0.049 | 0.037 | 0.037 | 0 | 0.92 | 0.99 | 0.78 | 0.98 | 0.8 | 0.88 |
| 7 | 126563643 | rs3808143 | -0.372 | 3.928 | 0.29 | -0.415 | -0.018 | -0.025 | -0.025 | 0 | 0.09 | 0.05 | 0.14 | 0.2 | 0.14 | 0.15 |
| 7 | 126582814 | rs1419471 | -0.721 | 3.313 | 0.194 | -1.262 | 0 | -0.001 | -0.001 | 0 | 0.32 | 0.27 | 0.27 | 0.3 | 0.35 | 0.33 |
| 7 | 126583521 | rs10232276 | 0.139 | 3.303 | 0.287 | -1.273 | -0.004 | -0.075 | -0.075 | 0.471 | 0.24 | 0.32 | 0.15 | 0.1 | 0.21 | 0.18 |
| 7 | 126584795 | rs1204556 | 1.856 | 2.892 | 0.521 | -1.397 | -0.003 | -0.024 | -0.024 | 0 | 0.77 | 0.74 | 0.7 | 0.97 | 0.69 | 0.78 |
| 7 | 126593467 | rs17863168 | -0.215 | 2.877 | 0.514 | -1.315 | -0.005 | -0.001 | -0.001 | 0.021 | 0.09 | 0.12 | 0.07 | 0.01 | 0.1 | 0.09 |
| 7 | 126602285 | rs1204579 | 1.694 | 2.6 | 0.636 | -1.318 | 0.032 | 0.028 | 0.028 | 0 | 0.49 | 0.58 | 0.44 | 0.68 | 0.34 | 0.45 |
| 7 | 126613163 | rs1155597 | 1.686 | 2.613 | 0.573 | -1.316 | -0.004 | -0.002 | -0.002 | 0 | 0.5 | 0.52 | 0.44 | 0.68 | 0.34 | 0.45 |
| 7 | 126628880 | rs1419438 | -0.998 | 2.275 | 0.521 | -1.236 | 0.002 | -0.004 | -0.004 | 0 | 0.34 | 0.3 | 0.28 | 0.29 | 0.38 | 0.34 |
| 7 | 126634489 | rs1582255 | -0.939 | 1.837 | 0.052 | -1.4 | -0.001 | 0.002 | 0.002 | 0 | 0.37 | 0.27 | 0.3 | 0.3 | 0.38 | 0.36 |
| 7 | 126639525 | rs10260443 | -0.86 | 1.924 | -0.406 | -1.422 | 0.002 | -0.004 | -0.004 | 0 | 0.41 | 0.44 | 0.43 | 0.26 | 0.44 | 0.43 |
| 7 | 126640633 | rs4728060 | -1.934 | 0.981 | -0.793 | -1.898 | 0.001 | 0.003 | 0.003 | 0 | 0.63 | 0.9 | 0.56 | 0.32 | 0.59 | 0.56 |
| 7 | 126642455 | rs2237777 | -0.602 | 1.472 | -0.835 | -1.711 | -0.001 | -0.062 | -0.062 | 1.206 | 0.38 | 0.54 | 0.33 | 0.09 | 0.34 | 0.36 |
| 7 | 126642603 | rs929174 | 0.011 | 1.565 | -1.279 | -0.909 | -0.03 | -0.048 | -0.048 | 0.587 | 0.09 | 0.03 | 0.12 | 0.21 | 0.13 | 0.11 |
| 7 | 126644787 | rs2402848 | -0.372 | 1.058 | -0.893 | -0.575 | -0.001 | 0.014 | 0.014 | 0 | 0.15 | 0.14 | 0.14 | 0.25 | 0.15 | 0.11 |
| 7 | 126645463 | rs11563410 | -0.19 | 1.164 | -1.245 | -0.897 | -0.013 | 0.05 | 0.05 | 1.731 | 0.2 | 0.06 | 0.13 | 0.04 | 0.19 | 0.2 |
| 7 | 126645953 | rs2283088 | -0.611 | 1.053 | -1.48 | -0.317 | 0.023 | 0.001 | 0.001 | 0 | 0.48 | 0.77 | 0.45 | 0.55 | 0.41 | 0.57 |
| 7 | 126648289 | rs10242944 | -0.518 | 1.286 | -1.193 | -0.646 | 0.001 | -0.021 | -0.021 | 0.152 | 0.09 | 0.13 | 0.07 | 0 | 0.1 | 0.16 |
| 7 | 126653699 | rs12673312 | -0.29 | 1.29 | -1.517 | -0.734 | -0.019 | 0.001 | 0.001 | 0 | 0.38 | 0.27 | 0.37 | 0.33 | 0.39 | 0.35 |
| 7 | 126655460 | rs11563361 | 0.178 | 1.094 | -1.734 | -0.775 | -0.066 | 0.102 | 0.102 | 0 | 0.28 | 0.06 | 0.18 | 0.09 | 0.34 | 0.19 |
| 7 | 126657545 | rs1018854 | -1.639 | 0.899 | -0.692 | -0.696 | 0 | 0 | 0 | 0 | 0.49 | 0.55 | 0.45 | 0.34 | 0.49 | 0.51 |
| 7 | 126664152 | rs978874 | -1.089 | 0.763 | -1.066 | -0.503 | -0.021 | -0.047 | -0.047 | 0 | 0.72 | 0.86 | 0.42 | 0.28 | 0.68 | 0.55 |
| 7 | 126665052 | rs2299525 | 1.251 | 0.739 | -1.026 | -0.534 | -0.03 | -0.03 | -0.03 | 0 | 0.33 | 0.43 | 0.16 | 0.19 | 0.24 | 0.2 |
| 7 | 126674192 | rs2237780 | 1.903 | 0.526 | -0.654 | -0.714 | -0.003 | -0.001 | -0.001 | 0 | 0.25 | 0.23 | 0.1 | 0.19 | 0.2 | 0.16 |
| 7 | 126676366 | rs7778308 | -0.023 | 0.557 | -0.931 | -0.735 | 0.024 | 0.083 | 0.083 | 0 | 0.43 | 0.67 | 0.23 | 0.19 | 0.34 | 0.36 |
| 7 | 126681865 | rs2237782 | -1.874 | 0.532 | -1.047 | -0.677 | -0.015 | -0.098 | -0.098 | 0.183 | 0.77 | 0.91 | 0.49 | 0.41 | 0.72 | 0.69 |
| 7 | 126694082 | rs11563766 | 0.81 | 0.603 | -0.999 | -0.742 | -0.003 | 0.018 | 0.018 | 0 | 0.06 | 0.12 | 0.08 | 0.13 | 0.05 | 0.14 |
| 7 | 126708411 | rs2402851 | 0.276 | 0.76 | -0.81 | -0.621 | 0.002 | 0.024 | 0.024 | 0.101 | 0.58 | 0.25 | 0.34 | 0.33 | 0.57 | 0.48 |
| 7 | 126708532 | rs2402852 | -1.871 | 0.793 | -0.805 | -0.61 | -0.005 | -0.068 | -0.068 | 0 | 0.77 | 0.88 | 0.48 | 0.41 | 0.72 | 0.69 |
| 7 | 126712118 | rs1156653 | 0.371 | 0.765 | -0.776 | -0.547 | -0.028 | 0.035 | 0.035 | 0 | 0.34 | 0.23 | 0.26 | 0.22 | 0.38 | 0.33 |
| 7 | 126713350 | rs1006728 | 0.901 | 0.708 | -0.696 | -0.961 | -0.021 | -0.021 | -0.021 | 3.451 | 0.32 | 0.49 | 0.16 | 0.19 | 0.24 | 0.2 |
| 7 | 126716624 | rs916610 | 0.72 | 0.855 | -0.719 | -0.96 | -0.01 | 0.015 | 0.015 | 0 | 0.29 | 0.23 | 0.19 | 0.09 | 0.34 | 0.19 |
| 7 | 126717992 | rs2237785 | -0.324 | 0.826 | -0.647 | -0.836 | -0.003 | 0.012 | 0.012 | 0 | 0.62 | 0.69 | 0.36 | 0.28 | 0.59 | 0.4 |
| 7 | 126734794 | rs10954144 | 1.677 | 0.693 | -0.543 | -0.899 | 0.005 | -0.006 | -0.006 | 0 | 0.24 | 0.07 | 0.1 | 0.19 | 0.2 | 0.16 |
| 7 | 126737888 | rs7794734 | -0.492 | 0.639 | -0.554 | -0.992 | 0 | -0.028 | -0.028 | 0.049 | 0.19 | 0.23 | 0.18 | 0.04 | 0.17 | 0.29 |
| 7 | 126742861 | rs17864129 | 0.695 | 0.689 | -0.755 | -0.981 | -0.038 | 0.071 | 0.071 | 0.033 | 0.12 | 0.01 | 0.1 | 0 | 0.13 | 0.09 |
| 7 | 126745550 | rs6975798 | -0.62 | 0.61 | -0.658 | -1.033 | 0.036 | 0.106 | 0.106 | 0 | 0.48 | 0.75 | 0.3 | 0.24 | 0.38 | 0.47 |
| 7 | 126747051 | rs1557644 | 1.131 | 0.757 | -0.814 | -1.076 | -0.07 | 0.079 | 0.079 | 0 | 0.26 | 0.05 | 0.15 | 0.09 | 0.3 | 0.21 |
| 7 | 126748966 | rs2237790 | 1.737 | 0.637 | -0.667 | -1.013 | 0 | 0 | 0 | 0 | 0.24 | 0.07 | 0.1 | 0.2 | 0.21 | 0.17 |
| 7 | 126753700 | rs13239559 | 0.784 | 0.931 | -1.107 | -1.132 | -0.076 | 0.12 | 0.12 | 0.006 | 0.3 | 0.06 | 0.16 | 0.07 | 0.33 | 0.22 |
| 7 | 126762244 | rs11563302 | 0.418 | 0.892 | -0.943 | -1.122 | 0.005 | -0.042 | -0.042 | 0 | 0.1 | 0.19 | 0.09 | 0 | 0.09 | 0.16 |
| 7 | 126775198 | rs2040502 | -1.453 | 0.904 | -1.237 | -1.008 | -0.009 | -0.045 | -0.045 | 0 | 0.74 | 0.86 | 0.42 | 0.3 | 0.67 | 0.67 |
| 7 | 126776219 | rs1419424 | 0.098 | 0.897 | -1.08 | -0.999 | 0.025 | 0.071 | 0.071 | 0 | 0.4 | 0.73 | 0.22 | 0.2 | 0.31 | 0.33 |
| 7 | 126789945 | rs12538500 | 0.495 | 1.489 | -1.322 | -1.305 | -0.047 | 0.051 | 0.051 | 0.613 | 0.26 | 0.11 | 0.16 | 0.09 | 0.32 | 0.23 |
| 7 | 126792164 | rs2237795 | -1.059 | 0.85 | -1.221 | -1.72 | 0 | -0.009 | -0.009 | 0 | 0.76 | 0.79 | 0.42 | 0.28 | 0.68 | 0.63 |
| 7 | 126792213 | rs2237796 | 2.018 | 0.681 | -1.476 | -2.168 | 0.01 | 0.087 | 0.087 | 0.003 | 0.28 | 0.08 | 0.09 | 0.1 | 0.25 | 0.21 |
| 7 | 126792834 | rs11266838 | 0.777 | 0.725 | -1.52 | -2.294 | -0.019 | 0.044 | 0.044 | 0 | 0.25 | 0.15 | 0.15 | 0.08 | 0.25 | 0.14 |
| 7 | 126795617 | rs11563701 | -0.691 | 0.748 | -1.503 | -2.182 | 0.001 | -0.046 | -0.046 | 0 | 0.22 | 0.54 | 0.17 | 0.1 | 0.17 | 0.28 |
| 7 | 126796034 | rs11563405 | 1.225 | 0.772 | -2.156 | -2.122 | -0.025 | 0.069 | 0.069 | 0 | 0.12 | 0.01 | 0.07 | 0 | 0.1 | 0.09 |
| 7 | 126803474 | rs17863228 | 0.207 | 0.727 | -1.6 | -1.801 | 0.009 | 0.052 | 0.052 | 0 | 0.49 | 0.58 | 0.26 | 0.19 | 0.42 | 0.49 |
| 7 | 126811206 | rs17862309 | 0.381 | 0.426 | -1.624 | -1.879 | 0.014 | 0.009 | 0.009 | 0 | 0.06 | 0.2 | 0.05 | 0.03 | 0.03 | 0.12 |
| 7 | 126827871 | rs2299546 | 0.06 | 0.503 | -2.352 | -1.893 | -0.05 | 0.069 | 0.069 | 0 | 0.24 | 0.07 | 0.15 | 0.08 | 0.25 | 0.14 |
| 7 | 126845237 | rs2237798 | 0.443 | -0.722 | -2.304 | -1.977 | -0.01 | 0.145 | 0.145 | 0.263 | 0.31 | 0.01 | 0.11 | 0.08 | 0.26 | 0.21 |
| 7 | 126847016 | rs886003 | 0.601 | -0.505 | -2.779 | -2.147 | -0.073 | 0.021 | 0.021 | 0 | 0.18 | 0.02 | 0.11 | 0.1 | 0.22 | 0.17 |
| 7 | 126847044 | rs886004 | -1.409 | -0.806 | -2.639 | -2.303 | -0.029 | 0.01 | 0.01 | 2.4 | 0.69 | 0.83 | 0.39 | 0.25 | 0.63 | 0.53 |
| 7 | 126849879 | rs2237803 | -1.483 | -0.844 | -2.933 | -2.312 | -0.082 | -0.019 | -0.019 | 0 | 0.71 | 0.91 | 0.4 | 0.26 | 0.63 | 0.55 |
| 7 | 126854653 | rs17862325 | -0.057 | -0.763 | -2.7 | -2.201 | -0.016 | 0.07 | 0.07 | 0 | 0.27 | 0.11 | 0.15 | 0.1 | 0.29 | 0.18 |
| 7 | 126856229 | rs13232763 | 1.206 | -1.114 | -2.18 | -2.178 | -0.021 | 0.068 | 0.068 | 0.077 | 0.11 | 0.01 | 0.07 | 0 | 0.11 | 0.01 |
| 7 | 126857240 | rs2299555 | -0.166 | -1.449 | -1.846 | -1.745 | 0.021 | 0.124 | 0.124 | 0 | 0.35 | 0.08 | 0.15 | 0.13 | 0.29 | 0.24 |
| 7 | 126863196 | rs17863247 | 0.296 | -1.417 | -1.518 | -1.539 | 0.05 | 0.088 | 0.088 | 0 | 0.27 | 0.06 | 0.1 | 0.09 | 0.2 | 0.21 |
| 7 | 126866037 | rs10487472 | 0.064 | -0.74 | -0.345 | -1.003 | -0.009 | -0.01 | -0.01 | 0 | 0.06 | 0.16 | 0.04 | 0.02 | 0.03 | 0.08 |
| 7 | 126867288 | rs2106149 | -0.754 | -0.637 | -1.276 | -1.361 | -0.167 | -0.025 | -0.025 | 0 | 0.64 | 0.95 | 0.44 | 0.29 | 0.47 | 0.47 |
| 7 | 126867650 | rs2237807 | -0.04 | -0.85 | -0.351 | -1.663 | 0 | -0.026 | -0.026 | 0 | 0.45 | 0.4 | 0.23 | 0.11 | 0.31 | 0.3 |
| 7 | 126870612 | rs7806403 | -0.432 | -0.41 | 0.227 | -1.277 | 0.016 | -0.02 | -0.02 | 0 | 0.07 | 0.17 | 0.08 | 0.01 | 0.05 | 0.03 |

| | | | | | | | | | | | | | | | | |
|---|---|---|---|---|---|---|---|---|---|---|---|---|---|---|---|---|
| 7 | 126873928 | rs11563675 | 0.447 | -0.684 | -0.055 | -1.873 | -0.004 | 0.065 | 0.065 | 0 | 0.18 | 0.07 | 0.13 | 0.03 | 0.15 | 0.05 |
| 7 | 126880627 | rs11563673 | -1.442 | -0.993 | 0.571 | -2.104 | -0.007 | -0.06 | -0.06 | 0 | 0.35 | 0.41 | 0.29 | 0.06 | 0.26 | 0.2 |
| 7 | 126881029 | rs3779537 | -0.431 | -0.983 | 0.579 | -2.112 | 0.025 | -0.003 | -0.003 | 0 | 0.08 | 0.18 | 0.1 | 0.02 | 0.07 | 0.11 |
| 7 | 126881216 | rs13246388 | -0.188 | -1.25 | 0.563 | -2.082 | 0.003 | 0.016 | 0.016 | 0 | 0.23 | 0.18 | 0.18 | 0.04 | 0.19 | 0.08 |
| 7 | 126885911 | rs7776557 | -1.243 | -1.301 | 0.599 | -2.113 | 0.001 | -0.014 | -0.014 | 0 | 0.33 | 0.33 | 0.27 | 0.06 | 0.25 | 0.15 |
| 7 | 126889071 | rs17863255 | 0.364 | -1.299 | 0.595 | -2.11 | 0.006 | 0.053 | 0.053 | 0 | 0.19 | 0.1 | 0.13 | 0.03 | 0.15 | 0.06 |
| 7 | 126890706 | rs10270765 | -1.469 | -1.422 | 1.172 | -2.163 | -0.008 | -0.045 | -0.045 | 0 | 0.36 | 0.58 | 0.3 | 0.06 | 0.25 | 0.15 |
| 7 | 126891392 | rs1592376 | 0.037 | -1.391 | 0.972 | -2.149 | 0.001 | 0.001 | 0.001 | 0 | 0.22 | 0.2 | 0.17 | 0.05 | 0.18 | 0.08 |
| 7 | 126891451 | rs1592377 | -1.447 | -1.403 | 0.982 | -2.156 | -0.014 | -0.066 | -0.066 | 0.366 | 0.34 | 0.42 | 0.29 | 0.06 | 0.25 | 0.15 |
| 7 | 126891893 | rs1582259 | -0.964 | -1.412 | 0.979 | -2.152 | -0.021 | -0.074 | -0.074 | 0.008 | 0.31 | 0.4 | 0.25 | 0.05 | 0.21 | 0.1 |
| 7 | 126896545 | rs17866960 | -1.108 | -1.444 | 0.952 | -2.179 | -0.006 | -0.017 | -0.017 | 0 | 0.08 | 0.12 | 0.04 | 0.01 | 0.04 | 0.05 |
| 7 | 126896796 | rs17862334 | -1.096 | -1.403 | 1.084 | -2.136 | -0.003 | -0.027 | -0.027 | 0 | 0.3 | 0.35 | 0.21 | 0.08 | 0.22 | 0.13 |
| 7 | 126899880 | rs17875093 | -0.881 | -1.416 | 1.056 | -2.106 | -0.014 | -0.011 | -0.011 | 0 | 0.1 | 0.14 | 0.1 | 0.04 | 0.06 | 0.04 |
| 7 | 126902211 | rs17864174 | 0.601 | -1.412 | 1.065 | -2.103 | -0.003 | -0.013 | -0.013 | 0 | 0.05 | 0.16 | 0.08 | 0 | 0.03 | 0.02 |
| 7 | 126904423 | rs10225587 | -1.464 | -1.405 | 1.002 | -2.144 | -0.019 | -0.074 | -0.074 | 0.316 | 0.36 | 0.54 | 0.31 | 0.08 | 0.25 | 0.15 |
| 7 | 126914820 | rs17862340 | -1.026 | -1.258 | 0.418 | -2.067 | 0.001 | 0.004 | 0.004 | 0 | 0.09 | 0.09 | 0.06 | 0.05 | 0.07 | 0.07 |
| 7 | 126919592 | rs10487475 | 0.595 | -1.112 | 0.282 | -1.97 | -0.006 | 0.069 | 0.069 | 0 | 0.18 | 0.07 | 0.13 | 0.03 | 0.15 | 0.05 |
| 7 | 126922129 | rs10245698 | 0.43 | -1.084 | 0.606 | -1.871 | 0.008 | -0.007 | -0.007 | 0 | 0.05 | 0.18 | 0.08 | 0 | 0.03 | 0.02 |
| 7 | 126922628 | rs10278805 | 0.283 | -1.662 | -0.108 | -2.222 | -0.005 | -0.012 | -0.012 | 0.517 | 0.37 | 0.56 | 0.23 | 0.17 | 0.24 | 0.24 |
| 7 | 126924753 | rs1419391 | -1.089 | -0.895 | -0.894 | -1.933 | -0.194 | -0.056 | -0.056 | 0.29 | 0.72 | 0.97 | 0.5 | 0.27 | 0.51 | 0.39 |
| 7 | 126926891 | rs916598 | -0.831 | -0.879 | -0.854 | -2.294 | -0.171 | 0.051 | 0.051 | 2.844 | 0.68 | 0.93 | 0.43 | 0.17 | 0.51 | 0.34 |
| 7 | 126932903 | rs11563399 | 0.609 | -0.984 | -0.584 | -2.41 | 0.002 | -0.018 | -0.018 | 0 | 0.19 | 0.15 | 0.12 | 0.01 | 0.15 | 0.06 |
| 7 | 126932944 | rs10248665 | 0.32 | -1.078 | -0.469 | -2.323 | 0.051 | 0.073 | 0.073 | 0.061 | 0.45 | 0.61 | 0.25 | 0.09 | 0.32 | 0.25 |
| 7 | 126937952 | rs10247414 | 0.32 | -1.057 | -0.453 | -2.323 | 0.048 | 0.069 | 0.069 | 0.061 | 0.45 | 0.61 | 0.26 | 0.09 | 0.32 | 0.25 |
| 7 | 126942291 | rs11563658 | -0.015 | -0.942 | -0.65 | -2.282 | 0.069 | 0.148 | 0.148 | 0.054 | 0.49 | 0.74 | 0.3 | 0.16 | 0.37 | 0.29 |
| 7 | 126943420 | rs4731347 | 0.416 | -0.618 | -0.309 | -2.254 | -0.015 | -0.09 | -0.09 | 0.128 | 0.71 | 0.61 | 0.82 | 0.96 | 0.82 | 0.91 |
| 7 | 126962488 | rs6957751 | -0.127 | -0.168 | -0.917 | -2.314 | -0.053 | 0.098 | 0.098 | 0 | 0.78 | 0.97 | 0.84 | 0.93 | 0.77 | 0.89 |
| 7 | 126963445 | rs17864190 | 0.408 | 0.056 | -0.727 | -2.472 | -0.001 | -0.017 | -0.017 | 0 | 0.1 | 0.13 | 0.04 | 0.03 | 0.07 | 0.08 |
| 7 | 126970005 | rs3944151 | -0.578 | -0.086 | -0.183 | -3.092 | 0.012 | 0.071 | 0.071 | 0.678 | 0.64 | 0.84 | 0.81 | 0.94 | 0.75 | 0.83 |
| 7 | 127001657 | rs11563465 | 1.448 | -0.119 | -0.562 | -3.049 | -0.012 | 0.048 | 0.048 | 0 | 0.15 | 0.02 | 0.1 | 0.06 | 0.16 | 0.08 |
| 7 | 127006928 | rs11563463 | 1.29 | 0.423 | -0.845 | -3.245 | -0.045 | -0.022 | -0.022 | 0 | 0.21 | 0.11 | 0.38 | 0.75 | 0.34 | 0.44 |
| 7 | 127009515 | rs1361944 | -0.797 | 0.373 | -0.776 | -3.283 | 0.019 | -0.043 | -0.043 | 0.868 | 0.91 | 0.89 | 0.93 | 1 | 0.93 | 0.97 |
| 7 | 127017001 | rs7796747 | 1.373 | 0.569 | -0.649 | -3.098 | 0.046 | 0.072 | 0.072 | 1.547 | 0.28 | 0.1 | 0.13 | 0.1 | 0.18 | 0.15 |
| 7 | 127040012 | rs11563460 | 1.437 | 0.313 | -0.869 | -3.286 | 0.016 | 0.115 | 0.115 | 1.692 | 0.28 | 0.01 | 0.11 | 0.06 | 0.18 | 0.14 |
| 7 | 127040067 | rs1419415 | -1.156 | 0.433 | -0.799 | -3.283 | -0.015 | 0.011 | 0.011 | 0 | 0.92 | 0.96 | 0.93 | 1 | 0.93 | 0.97 |
| 7 | 127047641 | rs1557639 | 1.709 | -0.036 | -0.689 | -3.257 | 0.001 | 0.003 | 0.003 | 0 | 0.15 | 0.08 | 0.1 | 0.06 | 0.16 | 0.08 |
| 7 | 127053001 | rs11563459 | 1.208 | -0.086 | -0.437 | -3.338 | -0.005 | -0.028 | -0.028 | 0 | 0.13 | 0.19 | 0.25 | 0.04 | 0.14 | 0.18 |
| 7 | 127053246 | rs11563624 | 1.683 | -0.146 | -0.517 | -3.398 | 0.016 | 0.115 | 0.115 | 1.562 | 0.28 | 0.01 | 0.11 | 0.06 | 0.18 | 0.14 |
| 7 | 127055564 | rs1361948 | 1.239 | 0.219 | 0.504 | -2.946 | -0.009 | -0.045 | -0.045 | 0 | 0.16 | 0.2 | 0.11 | 0.06 | 0.16 | 0.08 |
| 7 | 127056775 | rs17865261 | 0.608 | 0.491 | 0.304 | -2.628 | -0.005 | -0.028 | -0.028 | 0 | 0.13 | 0.19 | 0.25 | 0.04 | 0.14 | 0.18 |
| 7 | 127058129 | rs7801218 | -0.304 | 0.524 | 0.086 | -2.624 | 0.004 | -0.041 | -0.041 | 0.158 | 0.18 | 0.53 | 0.31 | 0.09 | 0.19 | 0.24 |
| 7 | 127059236 | rs11563620 | 1.169 | 0.573 | -1.139 | -2.618 | -0.125 | -0.016 | -0.016 | 0 | 0.26 | 0.02 | 0.36 | 0.79 | 0.38 | 0.51 |
| 7 | 127060796 | rs17867852 | 0.156 | 0.244 | -1.313 | -3.133 | -0.009 | 0.112 | 0.112 | 1.086 | 0.65 | 0.18 | 0.7 | 0.89 | 0.69 | 0.76 |
| 7 | 127072159 | rs7809156 | -0.509 | 0.302 | -0.717 | -3.186 | 0.103 | -0.002 | -0.002 | 0.859 | 0.89 | 0.58 | 0.88 | 0.95 | 0.89 | 0.89 |
| 7 | 127074138 | rs10487483 | 0.536 | 0.088 | -1.099 | -3.188 | 0.028 | 0.202 | 0.202 | 0 | 0.39 | 0.13 | 0.33 | 0.08 | 0.31 | 0.25 |
| 7 | 127077326 | rs10250993 | 0.584 | 0.119 | -0.584 | -2.827 | -0.013 | -0.055 | -0.055 | 0 | 0.22 | 0.3 | 0.15 | 0.06 | 0.2 | 0.13 |
| 7 | 127079816 | rs17713208 | 1.54 | 0.201 | -1.086 | -2.614 | 0 | -0.008 | -0.008 | 0 | 0.07 | 0.1 | 0.21 | 0.04 | 0.1 | 0.11 |
| 7 | 127080377 | rs10247993 | -0.37 | 0.161 | -0.952 | -2.661 | 0.009 | 0.176 | 0.176 | 0 | 0.54 | 0.77 | 0.59 | 0.92 | 0.62 | 0.72 |
| 7 | 127082497 | rs4731365 | 0.675 | 0.496 | -1.289 | -2.074 | -0.114 | -0.046 | -0.046 | 0.062 | 0.29 | 0.03 | 0.35 | 0.74 | 0.39 | 0.42 |
| 7 | 127084563 | rs17133014 | 0.481 | 0.106 | -1.08 | -2.32 | 0.004 | 0.008 | 0.008 | 0 | 0.27 | 0.23 | 0.14 | 0.12 | 0.22 | 0.19 |
| 7 | 127105414 | rs10487484 | 1.66 | 0.01 | -1.091 | -2.367 | 0.009 | 0.022 | 0.022 | 0 | 0.17 | 0.1 | 0.09 | 0.05 | 0.15 | 0.14 |
| 7 | 127108363 | rs1419407 | -0.031 | 0.074 | -1.051 | -2.386 | -0.003 | 0.009 | 0.009 | 0 | 0.31 | 0.29 | 0.23 | 0.2 | 0.31 | 0.29 |
| 7 | 127143881 | rs7809567 | -0.986 | -0.009 | -0.507 | -2.454 | -0.01 | -0.105 | -0.105 | 0 | 0.65 | 0.59 | 0.62 | 0.95 | 0.71 | 0.72 |
| 7 | 127145723 | rs6949899 | 0.274 | 0.17 | -0.607 | -2.2 | -0.001 | -0.01 | -0.01 | 0 | 0.25 | 0.33 | 0.21 | 0.13 | 0.25 | 0.25 |
| 7 | 127147205 | rs17714284 | 0.288 | 0.702 | -0.633 | -1.753 | 0.023 | 0.021 | 0.021 | 0 | 0.06 | 0.13 | 0.1 | 0.09 | 0.1 | 0.1 |
| 7 | 127147763 | rs6978712 | 0.353 | 0.691 | -0.65 | -1.75 | 0.02 | 0.02 | 0.02 | 0 | 0.06 | 0.13 | 0.1 | 0.09 | 0.09 | 0.1 |
| 7 | 127148632 | rs12668127 | 0.647 | 0.609 | -1.009 | -1.718 | -0.115 | -0.058 | -0.058 | 0.293 | 0.28 | 0.02 | 0.35 | 0.73 | 0.39 | 0.43 |
| 7 | 127154042 | rs7778682 | -0.875 | 0.43 | -0.015 | -1.913 | 0.042 | 0.001 | 0.001 | 0.024 | 0.15 | 0.42 | 0.16 | 0.09 | 0.13 | 0.15 |
| 7 | 127154121 | rs7778716 | -0.231 | 0.402 | -0.019 | -1.947 | 0.008 | -0.049 | -0.049 | 0 | 0.1 | 0.4 | 0.07 | 0 | 0.04 | 0.03 |
| 7 | 127155539 | rs10487490 | -0.762 | 0.148 | -0.07 | -2.402 | 0.027 | 0.111 | 0.111 | 0 | 0.56 | 0.73 | 0.55 | 0.88 | 0.59 | 0.61 |
| 7 | 127156445 | rs1107491 | 0.649 | 0.126 | 0.151 | -2.377 | 0.028 | -0.028 | -0.028 | 1.5 | 0.08 | 0.31 | 0.06 | 0 | 0.04 | 0.03 |
| 7 | 127158623 | rs10258162 | 0.233 | 0.1 | 0.313 | -2.38 | -0.018 | 0.057 | 0.057 | 0 | 0.41 | 0.26 | 0.39 | 0.79 | 0.46 | 0.51 |
| 7 | 127175236 | rs4551267 | 0.405 | 0.061 | 0.021 | -2.393 | -0.07 | 0.091 | 0.091 | 0 | 0.38 | 0.14 | 0.37 | 0.79 | 0.45 | 0.51 |
| 7 | 127178162 | rs4731367 | -0.737 | 0.023 | 0.009 | -2.402 | 0.032 | 0.138 | 0.138 | 0 | 0.55 | 0.74 | 0.55 | 0.89 | 0.59 | 0.66 |
| 7 | 127185250 | rs17151006 | 0.444 | 0.262 | -0.003 | -2.259 | 0.01 | 0.022 | 0.022 | 0 | 0.18 | 0.1 | 0.1 | 0.11 | 0.12 | 0.18 |
| 7 | 127190237 | rs2896394 | -1.335 | -0.355 | -0.245 | -2.78 | 0.017 | 0.105 | 0.105 | 0 | 0.68 | 0.83 | 0.64 | 0.93 | 0.74 | 0.73 |
| 7 | 127195238 | rs1419429 | -0.771 | -0.057 | -0.067 | -2.658 | 0.04 | 0.174 | 0.174 | 0 | 0.49 | 0.73 | 0.52 | 0.88 | 0.56 | 0.62 |
| 7 | 127195294 | rs1419430 | 0.184 | -0.137 | 0.119 | -2.613 | 0.003 | -0.012 | -0.012 | 0 | 0.37 | 0.39 | 0.38 | 0.8 | 0.44 | 0.49 |
| 7 | 127203830 | rs6977152 | 0.069 | -0.144 | 0.226 | -2.608 | 0.006 | -0.06 | -0.06 | 0 | 0.11 | 0.43 | 0.08 | 0 | 0.05 | 0.03 |

| | | | | | | | | | | | | | | | |
|---|---|---|---|---|---|---|---|---|---|---|---|---|---|---|---|
| 7 | 127204844 | rs10281056 | -0.544 | -0.143 | -0.284 | -2.629 | -0.029 | 0.126 | 0.126 | 0.004 | 0.46 | 0.74 | 0.45 | 0.8 | 0.48 | 0.52 |
| 7 | 127206134 | rs1557643 | 0.218 | 0.251 | -0.679 | -2.401 | -0.04 | 0.046 | 0.046 | 0 | 0.34 | 0.2 | 0.35 | 0.8 | 0.44 | 0.49 |
| 7 | 127219417 | rs10232298 | -0.882 | 0.287 | -1.093 | -2.353 | -0.037 | 0.209 | 0.209 | 0 | 0.67 | 0.93 | 0.61 | 0.95 | 0.68 | 0.72 |
| 7 | 127220827 | rs989099 | 1.887 | 0.731 | -1.015 | -2.125 | -0.027 | -0.058 | -0.058 | 0.785 | 0.16 | 0.06 | 0.22 | 0.7 | 0.21 | 0.27 |
| 7 | 127220918 | rs989100 | 2.036 | 0.724 | -1.05 | -2.128 | -0.061 | -0.096 | -0.096 | 0.007 | 0.15 | 0 | 0.22 | 0.7 | 0.21 | 0.27 |
| 7 | 127224286 | rs3735642 | 1.647 | 0.688 | -0.573 | -2.143 | 0.002 | 0.016 | 0.016 | 0 | 0.17 | 0.15 | 0.22 | 0.7 | 0.21 | 0.27 |
| 7 | 127241980 | rs3824006 | 1.482 | 0.853 | -0.144 | -2.219 | -0.032 | -0.02 | -0.02 | 0 | 0.07 | 0.01 | 0.24 | 0.1 | 0.11 | 0.18 |
| 7 | 127246903 | rs10229583 | 0.431 | 0.607 | 0.118 | -2.498 | 0 | -0.004 | -0.004 | 0 | 0.24 | 0.23 | 0.19 | 0.15 | 0.24 | 0.34 |
| 7 | 127247022 | rs12536698 | 2.066 | 0.464 | -0.377 | -2.633 | -0.004 | 0.058 | 0.058 | 0.574 | 0.16 | 0.03 | 0.1 | 0.04 | 0.14 | 0.07 |
| 7 | 127250907 | rs712700 | -1.817 | 0.602 | -0.027 | -2.492 | 0.011 | 0.064 | 0.064 | 0 | 0.81 | 0.72 | 0.76 | 0.36 | 0.78 | 0.74 |
| 7 | 127251188 | rs712701 | -1.851 | 0.602 | -0.023 | -2.492 | 0.011 | 0.064 | 0.064 | 0 | 0.81 | 0.73 | 0.76 | 0.36 | 0.78 | 0.74 |
| 7 | 127251792 | rs327518 | 0.81 | 0.541 | -0.174 | -2.552 | -0.075 | 0.193 | 0.193 | 0.007 | 0.36 | 0.06 | 0.41 | 0.1 | 0.39 | 0.28 |
| 7 | 127257894 | rs327510 | -0.967 | 0.509 | 0.126 | -2.131 | 0.003 | 0.054 | 0.054 | 0 | 0.77 | 0.52 | 0.71 | 0.31 | 0.77 | 0.7 |
| 7 | 127283170 | rs11977915 | 0.901 | 0.64 | -0.226 | -2.036 | -0.008 | 0.001 | 0.001 | 0.124 | 0.08 | 0.07 | 0.26 | 0.09 | 0.12 | 0.19 |
| 7 | 127283699 | rs806180 | -0.865 | 0.49 | -0.492 | -2.377 | 0.033 | 0.095 | 0.095 | 0.061 | 0.47 | 0.87 | 0.55 | 0.31 | 0.37 | 0.59 |
| 7 | 127288424 | rs806176 | -0.241 | 0.392 | -0.252 | -2.286 | 0.03 | 0.045 | 0.045 | 0.459 | 0.39 | 0.76 | 0.29 | 0.22 | 0.26 | 0.4 |
| 7 | 127297002 | rs806172 | -1.148 | 0.437 | -0.725 | -2.25 | -0.089 | -0.063 | -0.063 | 0 | 0.72 | 0.97 | 0.71 | 0.31 | 0.66 | 0.62 |
| 7 | 127311863 | rs2582717 | -1.948 | 0.465 | -0.563 | -2.17 | -0.069 | -0.096 | -0.096 | 0.438 | 0.79 | 0.98 | 0.74 | 0.29 | 0.73 | 0.66 |
| 7 | 127339415 | rs741599 | 0.642 | 0.433 | -0.052 | -2.185 | -0.002 | -0.009 | -0.009 | 0 | 0.33 | 0.26 | 0.43 | 0.08 | 0.4 | 0.22 |
| 7 | 127357644 | rs712719 | 1.302 | 0.336 | -0.202 | -2.302 | -0.057 | -0.041 | -0.041 | 1.059 | 0.28 | 0.06 | 0.29 | 0.71 | 0.34 | 0.38 |
| 7 | 127364960 | rs3808095 | 0.676 | 0.421 | -0.115 | -2.233 | -0.002 | -0.009 | -0.009 | 0 | 0.33 | 0.26 | 0.43 | 0.08 | 0.4 | 0.22 |
| 7 | 127412379 | rs1440972 | 0.9 | 0.254 | -0.094 | -2.279 | 0.008 | -0.002 | -0.002 | 1.129 | 0.16 | 0.37 | 0.17 | 0.11 | 0.15 | 0.25 |
| 7 | 127428382 | rs6467148 | -0.237 | 0.308 | -0.22 | -2.327 | -0.015 | -0.035 | -0.035 | 0 | 0.35 | 0.46 | 0.23 | 0.17 | 0.24 | 0.39 |
| 7 | 127430825 | rs6467152 | 0.886 | 0.34 | -0.178 | -2.325 | 0.009 | -0.006 | -0.006 | 0.712 | 0.17 | 0.4 | 0.17 | 0.11 | 0.15 | 0.25 |
| 7 | 127454026 | rs737242 | 2.021 | 0.317 | -0.422 | -2.295 | 0.011 | 0.054 | 0.054 | 0.682 | 0.15 | 0.03 | 0.05 | 0.04 | 0.08 | 0.07 |
| 7 | 127464386 | rs325446 | -0.95 | 0.423 | -0.247 | -2.327 | -0.022 | 0.065 | 0.065 | 0 | 0.68 | 0.84 | 0.57 | 0.92 | 0.6 | 0.79 |
| 7 | 127506080 | rs17151466 | 1.545 | 0.594 | -0.205 | -2.4 | 0 | 0.006 | 0.006 | 0 | 0.25 | 0.15 | 0.17 | 0 | 0.28 | 0.03 |
| 7 | 127507991 | rs10260037 | -1.575 | 0.681 | -0.325 | -2.398 | -0.03 | -0.013 | -0.013 | 0 | 0.92 | 0.98 | 0.75 | 0.92 | 0.88 | 0.81 |
| 7 | 127511705 | rs2896395 | 0.197 | 0.629 | -0.134 | -2.468 | 0.013 | -0.014 | -0.014 | 0 | 0.73 | 0.55 | 0.57 | 0.81 | 0.72 | 0.5 |
| 7 | 127526038 | rs3808088 | 1.303 | 0.609 | -0.4 | -2.399 | -0.077 | -0.051 | -0.051 | 0 | 0.3 | 0.04 | 0.29 | 0.71 | 0.35 | 0.39 |
| 7 | 127553442 | rs12375023 | 1.568 | 0.507 | -0.216 | -2.404 | -0.007 | 0.054 | 0.054 | 0 | 0.24 | 0.09 | 0.17 | 0 | 0.28 | 0.03 |
| 7 | 127561187 | rs3808083 | 0.9 | 0.409 | 0.062 | -2.365 | 0.007 | -0.007 | -0.007 | 0.117 | 0.16 | 0.39 | 0.17 | 0.11 | 0.15 | 0.24 |
| 7 | 127562736 | rs3808082 | 0.604 | 0.377 | -0.036 | -2.493 | -0.002 | -0.015 | -0.015 | 0.028 | 0.58 | 0.53 | 0.52 | 0.77 | 0.63 | 0.43 |
| 7 | 127575665 | rs3808080 | 1.393 | 0.38 | 0.044 | -2.449 | 0 | 0 | 0 | 0.041 | 0.14 | 0.13 | 0.14 | 0.11 | 0.15 | 0.24 |
| 7 | 127596408 | rs3757763 | 1.434 | 0.432 | -0.379 | -2.508 | -0.099 | -0.06 | -0.06 | 0.08 | 0.31 | 0.03 | 0.3 | 0.71 | 0.37 | 0.4 |
| 7 | 127636958 | rs17676986 | 0.838 | 0.501 | -0.145 | -2.505 | -0.023 | -0.001 | -0.001 | 1.092 | 0.1 | 0.04 | 0.26 | 0.08 | 0.14 | 0.27 |
| 7 | 127638703 | rs1362897 | 1.448 | 0.507 | -0.145 | -2.503 | -0.021 | -0.006 | -0.006 | 0 | 0.08 | 0.03 | 0.25 | 0.08 | 0.13 | 0.2 |
| 7 | 127639414 | rs1345509 | -0.564 | 0.432 | -0.024 | -2.574 | -0.002 | -0.017 | -0.017 | 0 | 0.42 | 0.46 | 0.48 | 0.23 | 0.37 | 0.53 |
| 7 | 127645852 | rs17151653 | 1.646 | 0.446 | 0.01 | -2.565 | -0.002 | 0.038 | 0.038 | 0 | 0.24 | 0.14 | 0.17 | 0 | 0.26 | 0.03 |
| 7 | 127660288 | rs6968751 | 1.032 | 0.422 | 0.119 | -2.571 | 0.012 | -0.004 | -0.004 | 0 | 0.16 | 0.35 | 0.16 | 0.11 | 0.15 | 0.19 |
| 7 | 127660763 | rs6467157 | 1.468 | 0.42 | -0.005 | -2.699 | -0.005 | 0.015 | 0.015 | 0 | 0.25 | 0.16 | 0.19 | 0 | 0.28 | 0.04 |
| 7 | 127666089 | rs3823995 | -0.659 | 0.294 | -0.039 | -2.622 | -0.003 | -0.021 | -0.021 | 0.915 | 0.42 | 0.47 | 0.47 | 0.24 | 0.36 | 0.52 |
| 7 | 127667516 | rs6944446 | -0.739 | 0.845 | -0.358 | -2.97 | 0.018 | 0.12 | 0.12 | 0.104 | 0.53 | 0.79 | 0.61 | 0.21 | 0.53 | 0.57 |
| 7 | 127668071 | rs3808058 | 1.161 | -0.218 | -0.184 | -2.474 | -0.018 | -0.003 | -0.003 | 1.739 | 0.08 | 0.04 | 0.25 | 0.08 | 0.13 | 0.19 |
| 7 | 127675108 | rs13240628 | 1.642 | -0.339 | -0.166 | -2.424 | 0.014 | 0.059 | 0.059 | 1.488 | 0.17 | 0.03 | 0.05 | 0.04 | 0.08 | 0.07 |
| 7 | 127693537 | rs10250463 | 1.115 | -0.225 | 0.226 | -2.386 | 0.003 | 0.011 | 0.011 | 0 | 0.14 | 0.28 | 0.15 | 0.11 | 0.14 | 0.21 |
| 7 | 127698403 | rs6949446 | 0.412 | -0.24 | -0.175 | -2.418 | -0.014 | -0.005 | -0.005 | 0.539 | 0.08 | 0.2 | 0.06 | 0.08 | 0.04 | 0.03 |
| 7 | 127703544 | rs13223326 | 1.829 | -0.318 | -0.467 | -2.445 | 0.007 | 0.08 | 0.08 | 1.299 | 0.16 | 0.03 | 0.05 | 0 | 0.08 | 0.07 |
| 7 | 127705174 | rs3808055 | -0.981 | -0.279 | -0.307 | -2.27 | 0.033 | 0.073 | 0.073 | 0.986 | 0.5 | 0.8 | 0.53 | 0.33 | 0.39 | 0.58 |
| 7 | 127707175 | rs3779522 | -1.752 | -0.294 | -0.485 | -2.291 | -0.082 | -0.088 | -0.088 | 0 | 0.73 | 0.96 | 0.71 | 0.33 | 0.66 | 0.62 |
| 7 | 127712244 | rs322834 | -0.415 | -0.306 | -0.286 | -2.297 | -0.006 | -0.019 | -0.019 | 0 | 0.41 | 0.51 | 0.46 | 0.25 | 0.35 | 0.51 |
| 7 | 127721507 | rs322825 | -1.58 | -0.049 | -0.732 | -2.112 | -0.083 | -0.089 | -0.089 | 0.071 | 0.73 | 0.96 | 0.71 | 0.33 | 0.66 | 0.62 |
| 7 | 127731618 | rs322820 | -1.523 | -0.049 | -0.734 | -2.086 | -0.06 | -0.068 | -0.068 | 0 | 0.73 | 0.92 | 0.7 | 0.33 | 0.66 | 0.62 |
| 7 | 127733822 | rs17151739 | 1.245 | -0.101 | -0.575 | -2.239 | -0.001 | -0.003 | -0.003 | 0 | 0.25 | 0.21 | 0.19 | 0 | 0.29 | 0.06 |
| 7 | 127737045 | rs322818 | 0.866 | -0.257 | -1.153 | -2.268 | 0.005 | 0.034 | 0.034 | 0 | 0.17 | 0.44 | 0.34 | 0.15 | 0.17 | 0.24 |
| 7 | 127739666 | rs6953698 | -0.191 | -0.371 | -1.183 | -2.272 | -0.016 | -0.041 | -0.041 | 0 | 0.32 | 0.45 | 0.19 | 0.18 | 0.2 | 0.32 |
| 7 | 127743022 | rs12534062 | 0.936 | -0.269 | -0.917 | -1.81 | -0.065 | 0.162 | 0.162 | 0 | 0.24 | 0.03 | 0.17 | 0 | 0.3 | 0.06 |
| 7 | 127744787 | rs322812 | -0.332 | -0.306 | -0.73 | -1.747 | 0.002 | 0.008 | 0.008 | 0 | 0.43 | 0.58 | 0.52 | 0.15 | 0.46 | 0.35 |
| 7 | 127771601 | rs322764 | -1.402 | -0.416 | -1.229 | -1.29 | -0.125 | -0.045 | -0.045 | 0 | 0.58 | 0.95 | 0.66 | 0.36 | 0.55 | 0.6 |
| 7 | 127772339 | rs322762 | -0.121 | -0.488 | -0.928 | -1.339 | -0.011 | -0.047 | -0.047 | 0 | 0.42 | 0.47 | 0.5 | 0.16 | 0.42 | 0.3 |
| 7 | 127773388 | rs322760 | -0.445 | -0.476 | -0.958 | -1.334 | 0.031 | 0.093 | 0.093 | 0 | 0.44 | 0.7 | 0.52 | 0.16 | 0.42 | 0.34 |
| 7 | 127778512 | rs4731401 | 0.061 | -0.286 | -0.856 | -1.187 | 0.029 | 0.044 | 0.044 | 0.207 | 0.11 | 0.26 | 0.13 | 0.18 | 0.13 | 0.25 |
| 7 | 127785381 | rs322738 | -0.002 | -0.614 | -0.878 | -1.242 | -0.069 | -0.021 | -0.021 | 0 | 0.22 | 0.47 | 0.34 | 0.15 | 0.13 | 0.24 |
| 7 | 127788296 | rs322731 | -0.961 | -0.296 | -0.985 | -1.072 | -0.022 | 0.009 | 0.009 | 1.62 | 0.57 | 0.78 | 0.64 | 0.38 | 0.56 | 0.56 |
| 7 | 127793648 | rs1421309 | -0.355 | 0.35 | -0.84 | -0.5 | 0.002 | 0 | 0 | 0 | 0.14 | 0.18 | 0.14 | 0.2 | 0.23 | 0.29 |
| 7 | 127793861 | rs322728 | 0.194 | -0.19 | -0.407 | -0.515 | -0.048 | 0.041 | 0.041 | 0 | 0.17 | 0.31 | 0.36 | 0.17 | 0.09 | 0.23 |
| 7 | 127795804 | rs10447854 | 0.893 | -0.235 | -0.941 | -0.865 | -0.027 | 0.093 | 0.093 | 0 | 0.21 | 0.1 | 0.17 | 0 | 0.27 | 0.06 |
| 7 | 127799143 | rs10954169 | 0.663 | -0.596 | -0.871 | -0.691 | 0.079 | 0.138 | 0.138 | 0 | 0.53 | 0.28 | 0.47 | 0.83 | 0.64 | 0.66 |
| 7 | 127801942 | rs4728089 | -0.518 | -0.547 | -0.546 | -0.64 | -0.067 | 0.011 | 0.011 | 0 | 0.82 | 0.64 | 0.67 | 0.86 | 0.92 | 0.73 |

| | | | | | | | | | | | | | | | |
|---|---|---|---|---|---|---|---|---|---|---|---|---|---|---|---|
| 7 | 127806005 | rs12155383 | 0.59 | -0.314 | -0.501 | -0.64 | 0 | -0.017 | -0.017 | 2.099 | 0.25 | 0.23 | 0.18 | 0.02 | 0.27 | 0.06 |
| 7 | 127814357 | rs1017607 | 0.155 | -0.028 | -0.821 | -0.337 | -0.062 | 0.062 | 0.062 | 0 | 0.36 | 0.21 | 0.31 | 0.15 | 0.5 | 0.26 |
| 7 | 127828450 | rs12706827 | 0.05 | -0.319 | -0.611 | -0.042 | -0.065 | -0.016 | -0.016 | 1.527 | 0.81 | 0.67 | 0.67 | 0.91 | 0.92 | 0.74 |
| 7 | 127830114 | rs4728090 | 0.883 | 0.1 | -1.901 | 0.003 | -0.001 | -0.042 | -0.042 | 1.511 | 0.61 | 0.06 | 0.49 | 0.71 | 0.69 | 0.56 |
| 7 | 127831580 | rs896183 | 1.16 | 0.594 | -2.015 | -0.107 | -0.106 | -0.135 | -0.135 | 1.384 | 0.36 | 0.04 | 0.34 | 0.62 | 0.41 | 0.41 |
| 7 | 127833995 | rs6976221 | -0.005 | 0.158 | -0.976 | -1.167 | -0.001 | -0.046 | -0.046 | 0 | 0.31 | 0.33 | 0.18 | 0.04 | 0.27 | 0.17 |
| 7 | 127841020 | rs6956123 | 0.163 | 0.251 | -0.785 | -0.826 | 0.004 | 0.008 | 0.008 | 0 | 0.85 | 0.88 | 0.73 | 0.95 | 0.87 | 0.8 |
| 7 | 127842526 | rs6467165 | -1.444 | 0.628 | 0.23 | -0.594 | 0.07 | 0.066 | 0.066 | 1.23 | 0.09 | 0.36 | 0.08 | 0.15 | 0.11 | 0.14 |
| 7 | 127847013 | rs11772985 | 0.93 | 0.845 | -0.751 | -1.069 | -0.047 | 0.002 | 0.002 | 0.263 | 0.34 | 0.14 | 0.32 | 0.64 | 0.38 | 0.3 |
| 7 | 127850795 | rs4731419 | 0.155 | 0.571 | -0.157 | -1.437 | 0.022 | 0.048 | 0.048 | 1.642 | 0.52 | 0.64 | 0.5 | 0.78 | 0.6 | 0.68 |
| 7 | 127851628 | rs7778167 | 0.304 | -0.1 | -0.103 | -1.796 | 0.005 | 0.006 | 0.006 | 0 | 0.31 | 0.24 | 0.19 | 0.13 | 0.27 | 0.23 |
| 7 | 127852065 | rs11981584 | 0.574 | -0.135 | 0.104 | -1.7 | 0 | -0.019 | -0.019 | 0.042 | 0.8 | 0.73 | 0.67 | 0.91 | 0.81 | 0.8 |
| 7 | 127856276 | rs6979832 | 0.856 | 0.056 | -0.043 | -1.711 | 0 | 0.005 | 0.005 | 0 | 0.5 | 0.5 | 0.47 | 0.72 | 0.51 | 0.47 |
| 7 | 127857326 | rs6947095 | -1.156 | 0.048 | -0.262 | -1.771 | -0.025 | 0 | 0 | 0 | 0.88 | 0.91 | 0.69 | 0.92 | 0.81 | 0.86 |
| 7 | 127857891 | rs2167289 | 1.686 | -0.281 | -0.481 | -1.9 | -0.078 | -0.01 | -0.01 | 0 | 0.35 | 0.13 | 0.38 | 0.66 | 0.42 | 0.27 |
| 7 | 127862802 | rs791595 | 0.27 | 0.575 | 0.053 | -1.531 | -0.05 | -0.087 | -0.087 | 0 | 0.68 | 0.57 | 0.87 | 0.88 | 0.82 | 0.79 |
| 7 | 127864005 | rs10954172 | -0.159 | 1.054 | -0.225 | -1.185 | -0.085 | -0.025 | -0.025 | 0.101 | 0.18 | 0.07 | 0.4 | 0.16 | 0.31 | 0.34 |
| 7 | 127869691 | rs2021808 | 0.904 | -0.111 | 0.849 | -1.246 | 0.021 | -0.01 | -0.01 | 0 | 0.92 | 0.4 | 0.91 | 0.94 | 0.93 | 0.92 |
| 7 | 127878155 | rs10487506 | 1.369 | -0.135 | 0.199 | -1.542 | -0.092 | 0.115 | 0.115 | 2.651 | 0.44 | 0.06 | 0.42 | 0.73 | 0.44 | 0.51 |
| 7 | 127881349 | rs2167270 | -0.064 | -0.204 | 0.839 | -1.608 | 0.002 | -0.036 | -0.036 | 0 | 0.36 | 0.44 | 0.41 | 0.2 | 0.37 | 0.28 |
| 7 | 127881851 | rs2278815 | 1.176 | -0.2 | 0.448 | -1.608 | -0.027 | 0.158 | 0.158 | 1.809 | 0.52 | 0.06 | 0.5 | 0.73 | 0.55 | 0.58 |
| 7 | 127887139 | rs12706832 | 1.114 | -0.233 | 0.602 | -1.573 | -0.003 | 0.119 | 0.119 | 1.049 | 0.52 | 0.15 | 0.5 | 0.72 | 0.54 | 0.54 |
| 7 | 127890062 | rs11763517 | -0.425 | -0.284 | 0.879 | -1.665 | -0.061 | 0.069 | 0.069 | 0 | 0.39 | 0.16 | 0.48 | 0.21 | 0.51 | 0.34 |
| 7 | 127891087 | rs11760956 | -0.19 | -0.143 | 0.944 | -1.162 | -0.035 | 0.04 | 0.04 | 0 | 0.34 | 0.15 | 0.39 | 0.21 | 0.38 | 0.32 |
| 7 | 127895216 | rs17151922 | -0.02 | -0.024 | 1.213 | -1.106 | 0.003 | -0.006 | -0.006 | 0 | 0.07 | 0.31 | 0.04 | 0 | 0.01 | 0.04 |
| 7 | 127899129 | rs2060715 | 0.812 | 0.112 | -1.054 | -0.977 | -0.048 | 0.111 | 0.111 | 0.215 | 0.53 | 0.12 | 0.48 | 0.71 | 0.53 | 0.53 |
| 7 | 127903788 | rs1376268 | -0.082 | 0.817 | -0.967 | -0.679 | -0.053 | 0.044 | 0.044 | 0.195 | 0.31 | 0.11 | 0.4 | 0.19 | 0.38 | 0.3 |
| 7 | 127907319 | rs10954176 | 0.213 | 0.547 | -1.169 | -0.725 | 0.004 | 0.145 | 0.145 | 0 | 0.52 | 0.24 | 0.43 | 0.81 | 0.54 | 0.56 |
| 7 | 127910047 | rs1116656 | -0.044 | 0.296 | -0.496 | -0.524 | 0.003 | 0.057 | 0.057 | 0 | 0.52 | 0.36 | 0.47 | 0.69 | 0.53 | 0.48 |
| 7 | 127911408 | rs12537998 | 0.805 | 0.258 | -1.116 | -0.849 | -0.109 | -0.062 | -0.062 | 0 | 0.28 | 0.03 | 0.46 | 0.69 | 0.35 | 0.32 |
| 7 | 127913274 | rs10231132 | 0.294 | 0.383 | -0.68 | -1.31 | 0.006 | 0.035 | 0.035 | 1.372 | 0.75 | 0.87 | 0.84 | 0.92 | 0.8 | 0.78 |
| 7 | 127915376 | rs1466146 | 0.345 | 0.394 | -0.551 | -1.285 | -0.001 | -0.002 | -0.002 | 0 | 0.18 | 0.16 | 0.12 | 0.08 | 0.13 | 0.17 |
| 7 | 127916208 | rs6979784 | -0.408 | 0.421 | -0.556 | -1.291 | 0.005 | 0.021 | 0.021 | 1.086 | 0.24 | 0.17 | 0.16 | 0.08 | 0.19 | 0.2 |
| 7 | 127923207 | rs12532565 | 0.473 | 0.396 | -0.619 | -1.288 | 0.013 | 0.02 | 0.02 | 0.108 | 0.17 | 0.12 | 0.12 | 0.08 | 0.13 | 0.18 |
| 7 | 127923778 | rs7811892 | 0.566 | 0.087 | -0.166 | -1.275 | -0.012 | 0.074 | 0.074 | 0 | 0.41 | 0.23 | 0.34 | 0.25 | 0.45 | 0.47 |
| 7 | 127927963 | rs17151967 | 0.253 | 0.565 | 0.779 | -0.642 | -0.019 | -0.027 | -0.027 | 0 | 0.05 | 0.15 | 0.03 | 0 | 0.01 | 0 |
| 7 | 127929066 | rs12534540 | 0.801 | 0.569 | 0.767 | -0.654 | -0.09 | -0.062 | -0.062 | 1.948 | 0.27 | 0.07 | 0.46 | 0.67 | 0.34 | 0.31 |
| 7 | 127931324 | rs2122628 | 1.169 | 0.243 | 1.373 | -0.6 | 0 | -0.016 | -0.016 | 0.489 | 0.92 | 0.59 | 0.96 | 1 | 0.99 | 1 |
| 7 | 127939461 | rs7778230 | -0.195 | 0.304 | 1.012 | -0.573 | 0.002 | -0.009 | -0.009 | 0 | 0.06 | 0.24 | 0.03 | 0 | 0.01 | 0 |
| 7 | 127941334 | rs6954653 | 0.737 | 0.339 | 0.22 | -0.983 | 0.034 | 0.105 | 0.105 | 0 | 0.35 | 0.14 | 0.26 | 0.15 | 0.31 | 0.36 |
| 7 | 127943131 | rs6964936 | 0.876 | 0.541 | 0.222 | -0.902 | 0.033 | 0.118 | 0.118 | 0 | 0.35 | 0.1 | 0.25 | 0.15 | 0.31 | 0.36 |
| 7 | 127946173 | rs3750035 | 0.019 | 1.043 | 0.162 | -0.745 | -0.024 | -0.004 | -0.004 | 0.286 | 0.22 | 0.14 | 0.18 | 0.24 | 0.29 | 0.27 |
| 7 | 127950725 | rs12850 | 1.055 | 1.189 | 0.328 | -0.728 | -0.072 | -0.071 | -0.071 | 0.004 | 0.28 | 0.09 | 0.47 | 0.6 | 0.35 | 0.31 |
| 7 | 127953243 | rs7804796 | -0.064 | 0.892 | 0.527 | -0.826 | 0.036 | -0.045 | -0.045 | 0 | 0.09 | 0.29 | 0.05 | 0 | 0.06 | 0.07 |
| 7 | 127957602 | rs2290225 | 0.181 | 1.021 | -0.535 | -0.808 | -0.049 | 0.116 | 0.116 | 0.378 | 0.42 | 0.22 | 0.56 | 0.79 | 0.48 | 0.48 |
| 7 | 127962835 | rs12536858 | 0.679 | 1.141 | -1.064 | -0.774 | 0 | 0 | 0 | 0 | 0.09 | 0.08 | 0.1 | 0.06 | 0.15 | 0.1 |
| 7 | 127966626 | rs12706849 | 0.99 | 1.156 | -1.473 | -0.598 | -0.049 | -0.073 | -0.073 | 0 | 0.27 | 0.09 | 0.45 | 0.6 | 0.28 | 0.3 |
| 7 | 127967998 | rs723627 | 0.312 | 0.668 | -0.854 | -0.928 | 0.024 | -0.035 | -0.035 | 0 | 0.07 | 0.17 | 0.05 | 0 | 0.05 | 0.06 |
| 7 | 127976595 | rs359652 | -0.476 | 0.518 | -1.721 | -0.74 | -0.081 | 0.19 | 0.19 | 1.587 | 0.57 | 0.84 | 0.44 | 0.17 | 0.51 | 0.51 |
| 7 | 127984275 | rs3763398 | -0.283 | -0.11 | -1.201 | -0.106 | 0.04 | 0.107 | 0.107 | 0 | 0.33 | 0.12 | 0.22 | 0.13 | 0.25 | 0.35 |
| 7 | 127989672 | rs359664 | 1.339 | 0.375 | -0.141 | 1.565 | 0.009 | 0.297 | 0.297 | 0 | 0.91 | 0.72 | 0.77 | 0.49 | 0.92 | 0.72 |
| 7 | 128000091 | rs359642 | 1.323 | 0.182 | -0.165 | 1.627 | -0.01 | 0.098 | 0.098 | 0 | 0.79 | 0.64 | 0.78 | 0.39 | 0.83 | 0.78 |
| 7 | 128002433 | rs359648 | 1.358 | -0.174 | -0.46 | 1.22 | -0.002 | -0.025 | -0.025 | 0.474 | 0.7 | 0.25 | 0.68 | 0.41 | 0.69 | 0.71 |
| 7 | 128006395 | rs983828 | -1.57 | -0.162 | 0.208 | 1.163 | 0.018 | 0.078 | 0.078 | 2.126 | 0.22 | 0.32 | 0.25 | 0.54 | 0.28 | 0.2 |
| 7 | 128020100 | rs880084 | 1.038 | -0.285 | -1.049 | -0.266 | -0.052 | 0.203 | 0.203 | 0 | 0.43 | 0.15 | 0.31 | 0.05 | 0.47 | 0.24 |
| 7 | 128023522 | rs1077322 | -0.836 | 0.115 | -1.5 | -0.119 | 0.003 | -0.025 | -0.025 | 1.848 | 0.62 | 0.72 | 0.48 | 0.61 | 0.68 | 0.59 |
| 7 | 128024834 | rs4728106 | 1.151 | 0.22 | -1.129 | -0.23 | -0.002 | -0.031 | -0.031 | 0 | 0.18 | 0.13 | 0.14 | 0 | 0.23 | 0.1 |
| 7 | 128024863 | rs885343 | -0.865 | 0.092 | -0.723 | -0.238 | 0.005 | -0.017 | -0.017 | 0.428 | 0.62 | 0.69 | 0.47 | 0.62 | 0.68 | 0.58 |
| 7 | 128027391 | rs6944392 | 1.366 | 0.747 | -0.82 | -0.398 | 0.004 | -0.049 | -0.049 | 1.018 | 0.13 | 0.14 | 0.08 | 0 | 0.14 | 0.02 |
| 7 | 128031298 | rs714510 | 0.111 | 0.525 | -0.886 | 0.127 | -0.004 | 0.001 | 0.001 | 0.977 | 0.2 | 0.22 | 0.13 | 0.2 | 0.16 | 0.18 |
| 7 | 128040699 | rs2278294 | -0.538 | 0.189 | -0.28 | 0.444 | -0.004 | 0.027 | 0.027 | 0.031 | 0.34 | 0.43 | 0.38 | 0.57 | 0.35 | 0.46 |
| 7 | 128041339 | rs4731448 | -2.442 | 0.245 | -0.231 | 0.09 | 0.001 | -0.011 | -0.011 | 0 | 0.73 | 0.73 | 0.81 | 0.95 | 0.76 | 0.85 |
| 7 | 128044162 | rs2288548 | -2.26 | 0.268 | -0.199 | 0.077 | 0.005 | 0.003 | 0.003 | 0 | 0.71 | 0.73 | 0.8 | 0.95 | 0.75 | 0.85 |
| 7 | 128047757 | rs11769333 | 2.418 | 0.166 | -0.156 | 0.134 | 0.007 | -0.007 | -0.007 | 0.48 | 0.1 | 0.05 | 0.05 | 0.12 | 0.09 | 0.13 |
| 7 | 128048194 | rs11770116 | -1.835 | 0.255 | -0.169 | 0.113 | -0.008 | 0.035 | 0.035 | 0 | 0.66 | 0.72 | 0.75 | 0.95 | 0.66 | 0.77 |
| 7 | 128054208 | rs13221784 | 0.222 | 0.189 | -0.091 | 0.202 | 0.003 | 0.009 | 0.009 | 0.231 | 0.27 | 0.31 | 0.31 | 0.39 | 0.33 | 0.3 |
| 7 | 128059295 | rs11982673 | -0.109 | 0.164 | -0.106 | 0.187 | 0.001 | 0.002 | 0.002 | 0 | 0.31 | 0.34 | 0.33 | 0.39 | 0.37 | 0.37 |
| 7 | 128069662 | rs1035971 | -1.988 | 0.183 | -0.073 | 0.239 | -0.004 | 0 | 0 | 0 | 0.67 | 0.76 | 0.57 | 0.72 | 0.71 | 0.76 |
| 7 | 128073326 | rs12540762 | 1.35 | 0.23 | -0.068 | 0.298 | -0.016 | 0.002 | 0.002 | 1.706 | 0.15 | 0.04 | 0.1 | 0.12 | 0.18 | 0.21 |

| | | | | | | | | | | | | | | | | |
|---|---|---|---|---|---|---|---|---|---|---|---|---|---|---|---|---|
| 7 | 128083454 | rs4566975 | 1.254 | 0.289 | 0.268 | 0.292 | 0.004 | 0.118 | 0.118 | 0.156 | 0.06 | 0.08 | 0.16 | 0.35 | 0.08 | 0.16 |
| 7 | 128088973 | rs11765126 | -0.338 | 0.21 | 0.064 | 0.273 | 0.003 | 0.012 | 0.012 | 0.074 | 0.28 | 0.35 | 0.31 | 0.38 | 0.33 | 0.31 |
| 7 | 128098071 | rs1048531 | 1.355 | 0.285 | -0.304 | 0.105 | -0.019 | 0.011 | 0.011 | 2.092 | 0.14 | 0.05 | 0.11 | 0.1 | 0.18 | 0.21 |
| 7 | 128101609 | rs11765023 | -0.628 | 0.045 | 0.107 | 0.228 | 0.019 | 0.039 | 0.039 | 0 | 0.24 | 0.42 | 0.28 | 0.37 | 0.29 | 0.26 |
| 7 | 128112709 | rs12530672 | 0.919 | 0.114 | -0.145 | 0.114 | 0 | -0.01 | -0.01 | 0 | 0.11 | 0.11 | 0.06 | 0.03 | 0.13 | 0.07 |
| 7 | 128130546 | rs34742251 | -0.037 | -0.009 | -0.227 | 0.212 | 0.003 | -0.008 | -0.008 | 0 | 0.17 | 0.13 | 0.11 | 0.23 | 0.16 | 0.23 |
| 7 | 128147119 | rs7806863 | 0.963 | 0.684 | -0.235 | 0.073 | 0.024 | 0.033 | 0.033 | 0 | 0.46 | 0.41 | 0.31 | 0.28 | 0.38 | 0.37 |
| 7 | 128149789 | rs12535956 | 1.222 | 0.964 | -0.567 | -0.01 | -0.052 | 0.047 | 0.047 | 0.679 | 0.19 | 0.04 | 0.14 | 0.07 | 0.24 | 0.26 |
| 7 | 128150653 | rs4371935 | -0.356 | 0.62 | -0.432 | -0.258 | -0.02 | -0.014 | -0.014 | 0 | 0.62 | 0.74 | 0.54 | 0.64 | 0.57 | 0.62 |
| 7 | 128155563 | rs13224784 | -0.106 | 0.623 | -0.535 | -0.429 | -0.004 | -0.05 | -0.05 | 0 | 0.16 | 0.22 | 0.1 | 0.03 | 0.16 | 0.14 |
| 7 | 128169884 | rs3802004 | 1.994 | 0.759 | -0.928 | -0.38 | -0.04 | -0.026 | -0.026 | 0.669 | 0.15 | 0.01 | 0.09 | 0.1 | 0.15 | 0.17 |
| 7 | 128169979 | rs3802003 | 2.03 | -0.042 | -1.232 | -0.89 | 0.004 | 0.095 | 0.095 | 0.226 | 0.2 | 0.04 | 0.1 | 0.03 | 0.17 | 0.07 |
| 7 | 128307573 | rs10954205 | 0.939 | 0.25 | -0.666 | -0.737 | 0.062 | -0.008 | -0.008 | 0.032 | 0.09 | 0.3 | 0.07 | 0.03 | 0.12 | 0.08 |
| 7 | 128308813 | rs7811083 | 0.939 | 0.25 | -0.666 | -0.737 | 0.062 | -0.008 | -0.008 | 0.025 | 0.09 | 0.3 | 0.07 | 0.03 | 0.12 | 0.08 |
| 7 | 128312022 | rs6979575 | -0.748 | 0.144 | -0.599 | -0.715 | 0.013 | 0.022 | 0.022 | 0.009 | 0.36 | 0.47 | 0.42 | 0.59 | 0.45 | 0.5 |
| 7 | 128312176 | rs17169024 | 0.796 | 0.166 | -0.709 | -0.699 | -0.004 | -0.009 | -0.009 | 0 | 0.24 | 0.18 | 0.31 | 0.56 | 0.29 | 0.36 |
| 7 | 128312450 | rs11769769 | 0.464 | 0.146 | -0.645 | -0.782 | 0.016 | -0.011 | -0.011 | 1.465 | 0.43 | 0.25 | 0.43 | 0.36 | 0.37 | 0.44 |
| 7 | 128317658 | rs6467210 | 0.663 | 0.361 | -0.607 | -0.474 | -0.004 | -0.009 | -0.009 | 0 | 0.24 | 0.18 | 0.31 | 0.56 | 0.29 | 0.36 |
| 7 | 128323034 | rs1109552 | 0.663 | 0.361 | -0.607 | -0.474 | -0.004 | -0.009 | -0.009 | 0 | 0.24 | 0.18 | 0.31 | 0.56 | 0.29 | 0.36 |
| 7 | 128348405 | rs7790478 | 0.22 | 0.352 | -0.337 | -0.532 | 0.03 | -0.046 | -0.046 | 0.001 | 0.13 | 0.28 | 0.11 | 0.03 | 0.16 | 0.14 |
| 7 | 128353840 | rs6946321 | -0.131 | 0.477 | -0.622 | -0.263 | 0.001 | 0.008 | 0.008 | 0 | 0.19 | 0.18 | 0.47 | 0.62 | 0.25 | 0.33 |
| 7 | 128354937 | rs3807125 | 0.464 | 0.502 | -0.61 | -0.252 | 0.011 | 0.04 | 0.04 | 0 | 0.16 | 0.18 | 0.44 | 0.62 | 0.21 | 0.27 |
| 7 | 128357300 | rs339092 | -1.76 | 0.324 | -0.678 | -0.493 | 0.002 | 0.038 | 0.038 | 0.227 | 0.69 | 0.79 | 0.75 | 0.78 | 0.68 | 0.74 |
| 7 | 128358839 | rs17162583 | 1.033 | 0.331 | -0.661 | -0.35 | -0.005 | 0 | 0 | 0 | 0.09 | 0.09 | 0.25 | 0.24 | 0.09 | 0.17 |
| 7 | 128358891 | rs745229 | 2.003 | 0.111 | -0.72 | -0.53 | -0.002 | 0.003 | 0.003 | 0 | 0.23 | 0.22 | 0.16 | 0.1 | 0.24 | 0.27 |
| 7 | 128359133 | rs6949056 | 0.682 | 0.105 | -0.619 | -0.459 | 0.007 | -0.014 | -0.014 | 0.12 | 0.27 | 0.16 | 0.27 | 0.41 | 0.24 | 0.22 |
| 7 | 128360997 | rs339078 | -1.014 | 0.118 | -0.705 | -0.71 | -0.001 | 0 | 0 | 6.662 | 0.45 | 0.51 | 0.59 | 0.67 | 0.45 | 0.47 |
| 7 | 128362426 | rs339075 | -1.904 | 0.28 | -0.824 | -0.741 | -0.002 | -0.035 | -0.035 | 0 | 0.74 | 0.71 | 0.82 | 0.95 | 0.79 | 0.88 |
| 7 | 128363287 | rs6971091 | 1.943 | 0.414 | -0.711 | -0.346 | -0.032 | 0.021 | 0.021 | 0.282 | 0.2 | 0.06 | 0.16 | 0.11 | 0.23 | 0.27 |
| 7 | 128364080 | rs339072 | -1.341 | 0.532 | -0.695 | -0.386 | -0.017 | 0.072 | 0.072 | 0.054 | 0.61 | 0.75 | 0.69 | 0.78 | 0.6 | 0.71 |
| 7 | 128364895 | rs6952563 | 0.992 | 0.415 | -0.414 | -0.025 | -0.039 | 0.068 | 0.068 | 0.056 | 0.23 | 0.31 | 0.22 | 0.43 | 0.17 | 0.19 |
| 7 | 128365787 | rs339069 | 0.402 | 0.853 | -0.505 | 0.169 | 0.005 | 0.026 | 0.026 | 0 | 0.12 | 0.16 | 0.26 | 0.25 | 0.16 | 0.19 |
| 7 | 128366485 | rs731770 | 1.317 | 0.547 | -0.756 | 0.011 | -0.012 | 0.051 | 0.051 | 0 | 0.38 | 0.22 | 0.3 | 0.25 | 0.42 | 0.34 |
| 7 | 128368205 | rs6951397 | 0.584 | 1.419 | -0.428 | 0.687 | -0.058 | 0.037 | 0.037 | 1.021 | 0.25 | 0.11 | 0.23 | 0.13 | 0.34 | 0.3 |
| 7 | 128369156 | rs12531956 | 1.224 | 0.92 | -0.295 | 0.982 | -0.039 | 0.006 | 0.006 | 0.581 | 0.19 | 0.06 | 0.18 | 0.13 | 0.24 | 0.27 |
| 7 | 128371298 | rs339065 | -0.634 | 0.455 | 0.501 | 0.712 | 0.01 | 0.08 | 0.08 | 0.017 | 0.13 | 0.3 | 0.26 | 0.19 | 0.1 | 0.17 |
| 7 | 128373534 | rs4731512 | -0.062 | 0.589 | -0.497 | 0.902 | -0.012 | 0.006 | 0.006 | 0 | 0.07 | 0.01 | 0.22 | 0.34 | 0.06 | 0.12 |
| 7 | 128373828 | rs339061 | -0.746 | 0.543 | -0.675 | -0.447 | -0.009 | 0.019 | 0.019 | 0 | 0.6 | 0.7 | 0.62 | 0.69 | 0.56 | 0.45 |
| 7 | 128376662 | rs7780294 | -0.35 | 0.961 | -0.547 | -0.147 | -0.008 | -0.03 | -0.03 | 0 | 0.09 | 0.03 | 0.29 | 0.62 | 0.09 | 0.19 |
| 7 | 128376900 | rs339057 | -0.515 | 0.748 | -0.574 | -0.367 | 0.014 | 0.025 | 0.025 | 0 | 0.55 | 0.65 | 0.62 | 0.69 | 0.6 | 0.41 |
| 7 | 128379209 | rs7795071 | 0.067 | 1.281 | -0.487 | -1.025 | -0.01 | 0.009 | 0.009 | 0.002 | 0.24 | 0.14 | 0.15 | 0.01 | 0.26 | 0.13 |
| 7 | 128382511 | rs339054 | -0.73 | 1.591 | -0.236 | -0.943 | -0.027 | 0.082 | 0.082 | 0.002 | 0.41 | 0.75 | 0.38 | 0.25 | 0.5 | 0.32 |
| 7 | 128389554 | rs1006023 | 0.056 | 1.819 | -0.042 | -1.167 | -0.021 | 0.148 | 0.148 | 0 | 0.33 | 0.11 | 0.22 | 0.07 | 0.37 | 0.15 |
| 7 | 128393950 | rs109829 | -1.7 | 1.415 | -0.686 | -1.779 | -0.028 | 0.188 | 0.188 | 0.565 | 0.69 | 0.95 | 0.81 | 0.94 | 0.72 | 0.72 |
| 7 | 128409225 | rs1043550 | -0.227 | 1.507 | 0.062 | -1.101 | -0.039 | 0.152 | 0.152 | 0 | 0.33 | 0.11 | 0.24 | 0.07 | 0.42 | 0.16 |
| 7 | 128410012 | rs1043595 | 1.183 | 1.684 | -0.026 | -1.017 | -0.068 | 0.048 | 0.048 | 0 | 0.14 | 0.02 | 0.15 | 0.07 | 0.28 | 0.06 |
| 7 | 128416587 | rs339038 | 0.802 | 1.474 | 0.321 | -1.058 | 0.025 | -0.051 | -0.051 | 0 | 0.7 | 0.46 | 0.77 | 0.81 | 0.63 | 0.78 |
| 7 | 128422986 | rs339088 | -0.441 | 1.27 | -0.299 | -0.955 | 0 | 0.086 | 0.086 | 0 | 0.35 | 0.2 | 0.22 | 0.05 | 0.39 | 0.15 |
| 7 | 128423763 | rs339085 | -0.447 | 1.291 | -0.284 | -0.953 | 0.001 | 0.005 | 0.005 | 0 | 0.36 | 0.29 | 0.22 | 0.05 | 0.39 | 0.15 |
| 7 | 128434900 | rs3736279 | -0.641 | 1.585 | -0.018 | 0.14 | 0.011 | 0.009 | 0.009 | 0 | 0.12 | 0.15 | 0.13 | 0.2 | 0.15 | 0.13 |
| 7 | 128445370 | rs1565629 | -0.665 | 1.51 | 0.043 | 0.129 | 0.014 | 0.014 | 0.014 | 0 | 0.11 | 0.19 | 0.12 | 0.2 | 0.14 | 0.09 |
| 7 | 128449387 | rs1565628 | 0.63 | 1.416 | -0.559 | 0.181 | -0.025 | 0.101 | 0.101 | 4.837 | 0.39 | 0.06 | 0.36 | 0.21 | 0.36 | 0.25 |
| 7 | 128449405 | rs2307037 | -0.435 | 1.529 | -0.428 | 0.055 | 0.007 | 0.019 | 0.019 | 0 | 0.12 | 0.21 | 0.12 | 0.23 | 0.11 | 0.12 |
| 7 | 128450532 | rs7799186 | 0.519 | 1.378 | 0.065 | 0.448 | -0.026 | 0.081 | 0.081 | 0.042 | 0.4 | 0.21 | 0.35 | 0.21 | 0.48 | 0.46 |
| 7 | 128451004 | rs17165191 | -0.84 | 1.029 | 0.887 | 0.846 | 0.035 | 0.038 | 0.038 | 0 | 0.12 | 0.23 | 0.13 | 0.23 | 0.14 | 0.14 |
| 7 | 128458649 | rs4731517 | -1.264 | 0.915 | 0.878 | 0.815 | 0.013 | 0.035 | 0.035 | 0 | 0.13 | 0.28 | 0.13 | 0.24 | 0.11 | 0.2 |
| 7 | 128461720 | rs3734974 | -0.716 | 0.983 | 0.648 | 0.84 | 0.006 | 0.015 | 0.015 | 0 | 0.11 | 0.19 | 0.11 | 0.2 | 0.11 | 0.09 |
| 7 | 128470838 | rs3734972 | -0.059 | 1.021 | 0.599 | 0.88 | -0.002 | -0.002 | -0.002 | 0 | 0.09 | 0.09 | 0.09 | 0.2 | 0.08 | 0.08 |
| 7 | 128477472 | rs2291560 | -0.465 | 1.169 | 0.665 | 0.71 | -0.001 | -0.001 | -0.001 | 1.353 | 0.1 | 0.12 | 0.06 | 0.12 | 0.11 | 0.17 |
| 7 | 128477547 | rs2291561 | -0.465 | 1.169 | 0.665 | 0.71 | -0.001 | -0.001 | -0.001 | 0 | 0.1 | 0.12 | 0.06 | 0.12 | 0.11 | 0.17 |
| 7 | 128478103 | rs2291562 | -0.699 | 1.073 | 1.395 | 1.05 | 0.055 | 0.421 | 0.421 | 0.34 | 0.06 | 0.33 | 0.15 | 0.32 | 0.03 | 0.09 |
| 7 | 128483227 | rs2291567 | -1.69 | 1.093 | 1.334 | 1.016 | 0.048 | 0.258 | 0.258 | 0 | 0.16 | 0.54 | 0.22 | 0.49 | 0.16 | 0.3 |
| 7 | 128487893 | rs2291568 | -1.619 | 1.041 | 1.321 | 1.039 | 0.048 | 0.457 | 0.457 | 0.29 | 0.08 | 0.42 | 0.16 | 0.38 | 0.04 | 0.21 |
| 7 | 128495338 | rs3816885 | -1.619 | 1.041 | 1.321 | 1.048 | 0.048 | 0.451 | 0.451 | 0.29 | 0.08 | 0.42 | 0.16 | 0.38 | 0.04 | 0.21 |
| 7 | 128500139 | rs902842 | -1.699 | 0.994 | 1.327 | 0.802 | 0.061 | 0.022 | 0.022 | 0.257 | 0.07 | 0.3 | 0.04 | 0.06 | 0.04 | 0.16 |
| 7 | 128503543 | rs1472147 | -2.121 | 0.899 | 1.159 | 0.737 | 0.013 | 0.078 | 0.078 | 0.371 | 0.22 | 0.67 | 0.25 | 0.49 | 0.2 | 0.34 |
| 7 | 128511731 | rs2279668 | -0.734 | 0.444 | 1.089 | 0.666 | 0 | 0.293 | 0.293 | 0.283 | 0.06 | 0.22 | 0.19 | 0.34 | 0.02 | 0.1 |
| 7 | 128515846 | rs7797420 | -1.619 | 0.449 | 1.015 | 0.631 | -0.002 | 0.25 | 0.25 | 1.587 | 0.11 | 0.57 | 0.22 | 0.34 | 0.06 | 0.2 |
| 7 | 128521240 | rs9656375 | -1.183 | 0.107 | 0.471 | -0.352 | -0.011 | 0.055 | 0.055 | 1.6 | 0.28 | 0.55 | 0.29 | 0.53 | 0.2 | 0.37 |

| | | | | | | | | | | | | | | | | |
|---|---|---|---|---|---|---|---|---|---|---|---|---|---|---|---|---|
| 7 | 128521431 | rs10954211 | -0.049 | 0.224 | 0.072 | -0.339 | 0.005 | -0.03 | -0.03 | 0.154 | 0.62 | 0.86 | 0.56 | 0.55 | 0.63 | 0.6 |
| 7 | 128530276 | rs4731523 | -0.051 | 0.108 | 0.31 | -0.443 | -0.008 | -0.035 | -0.035 | 2.151 | 0.29 | 0.34 | 0.34 | 0.15 | 0.25 | 0.36 |
| 7 | 128548463 | rs7786945 | -0.208 | 0.247 | 0.311 | 0.16 | -0.011 | 0.092 | 0.092 | 0.152 | 0.23 | 0.55 | 0.18 | 0.54 | 0.17 | 0.29 |
| 7 | 128551740 | rs1495461 | -0.432 | 0.409 | -0.239 | -0.308 | -0.005 | -0.037 | -0.037 | 0 | 0.34 | 0.38 | 0.28 | 0.13 | 0.32 | 0.27 |
| 7 | 128554093 | rs10247518 | -0.62 | 0.352 | -0.241 | 0.178 | -0.011 | -0.021 | -0.021 | 0.062 | 0.05 | 0.2 | 0.02 | 0 | 0.01 | 0.03 |
| 7 | 128557370 | rs6955705 | -0.249 | 0.581 | -0.718 | 0.555 | 0 | 0.009 | 0.009 | 0 | 0.21 | 0.19 | 0.28 | 0.61 | 0.17 | 0.28 |
| 7 | 128557406 | rs4731528 | -0.563 | 0.8 | -1.655 | -0.141 | -0.034 | 0.11 | 0.11 | 0 | 0.56 | 0.86 | 0.63 | 0.74 | 0.5 | 0.65 |
| 7 | 128560761 | rs960633 | 0.009 | 0.867 | -1.712 | -0.48 | -0.062 | 0.098 | 0.098 | 0.008 | 0.27 | 0.07 | 0.22 | 0.1 | 0.33 | 0.19 |
| 7 | 128568960 | rs729302 | 0.079 | 0.774 | -2.265 | -0.17 | -0.007 | -0.035 | -0.035 | 1.288 | 0.31 | 0.18 | 0.19 | 0.32 | 0.33 | 0.3 |
| 7 | 128573967 | rs4728142 | 0.679 | 0.974 | -2.056 | -0.865 | 0.006 | 0.103 | 0.103 | 0 | 0.42 | 0.24 | 0.31 | 0.1 | 0.45 | 0.39 |
| 7 | 128577304 | rs3757385 | 0.106 | 0.837 | -1.786 | -0.721 | 0.014 | -0.012 | -0.012 | 0 | 0.63 | 0.49 | 0.42 | 0.33 | 0.64 | 0.62 |
| 7 | 128579420 | rs752637 | 0.101 | 0.742 | -1.7 | -0.769 | 0.017 | -0.015 | -0.015 | 0 | 0.63 | 0.51 | 0.42 | 0.34 | 0.64 | 0.61 |
| 7 | 128580680 | rs3807306 | 0.866 | 1.035 | -1.707 | -0.866 | -0.005 | 0.126 | 0.126 | 0.172 | 0.49 | 0.3 | 0.34 | 0.15 | 0.51 | 0.45 |
| 7 | 128581835 | rs11761199 | 1.37 | 1.257 | -2.007 | -1.162 | -0.03 | 0.001 | 0.001 | 0 | 0.43 | 0.12 | 0.34 | 0.32 | 0.47 | 0.46 |
| 7 | 128585104 | rs1874328 | 0.987 | 0.97 | -2.302 | -1.416 | -0.004 | 0.003 | 0.003 | 0 | 0.37 | 0.46 | 0.32 | 0.36 | 0.42 | 0.38 |
| 7 | 128594183 | rs10488631 | 1.79 | 0.93 | -1.641 | -1.45 | -0.017 | 0.081 | 0.081 | 0 | 0.13 | 0.01 | 0.07 | 0 | 0.1 | 0.14 |
| 7 | 128594725 | rs2280714 | -0.499 | 0.98 | -1.671 | -1.492 | 0.011 | 0.021 | 0.021 | 0.925 | 0.67 | 0.58 | 0.47 | 0.54 | 0.69 | 0.7 |
| 7 | 128597587 | rs10236569 | -0.53 | 0.98 | -1.673 | -1.491 | 0.011 | 0.022 | 0.022 | 0.987 | 0.67 | 0.58 | 0.47 | 0.54 | 0.69 | 0.7 |
| 7 | 128605460 | rs6957529 | 2.066 | 0.957 | -1.693 | -1.499 | 0.004 | 0.003 | 0.003 | 0 | 0.11 | 0.05 | 0.05 | 0.08 | 0.11 | 0.11 |
| 7 | 128607384 | rs8043 | 0.687 | 1.305 | -0.965 | -1.243 | -0.001 | 0 | 0 | 0 | 0.48 | 0.51 | 0.37 | 0.44 | 0.53 | 0.49 |
| 7 | 128625788 | rs1154330 | 1.395 | 1.318 | -0.737 | -1.345 | 0.002 | 0.022 | 0.022 | 4.998 | 0.19 | 0.15 | 0.11 | 0.01 | 0.16 | 0.12 |
| 7 | 128644533 | rs3807304 | 1.776 | 1.351 | -0.762 | -1.351 | 0.024 | 0.114 | 0.114 | 0 | 0.11 | 0.15 | 0.39 | 0.46 | 0.14 | 0.15 |
| 7 | 128668510 | rs10239340 | 0.264 | 1.43 | -0.869 | -1.263 | 0.009 | 0.013 | 0.013 | 0.948 | 0.62 | 0.57 | 0.44 | 0.45 | 0.63 | 0.63 |
| 7 | 128672645 | rs9649520 | 0.737 | 1.383 | -0.881 | -1.277 | -0.001 | 0 | 0 | 0 | 0.48 | 0.51 | 0.36 | 0.44 | 0.53 | 0.49 |
| 7 | 128683547 | rs10279821 | -0.264 | 1.384 | -0.857 | -1.239 | 0.014 | 0.026 | 0.026 | 1.342 | 0.68 | 0.58 | 0.47 | 0.54 | 0.69 | 0.7 |
| 7 | 128689413 | rs2167273 | 1.399 | 1.322 | -1.004 | -1.266 | -0.011 | -0.058 | -0.058 | 4.087 | 0.4 | 0.14 | 0.32 | 0.4 | 0.48 | 0.45 |
| 7 | 128692657 | rs6960994 | -0.366 | 1.72 | -0.706 | -0.832 | 0.05 | 0.162 | 0.162 | 0 | 0.8 | 0.66 | 0.54 | 0.45 | 0.79 | 0.75 |
| 7 | 128701331 | rs13246321 | 0.938 | 1.04 | -0.834 | -1.222 | -0.017 | 0.081 | 0.081 | 0 | 0.13 | 0.01 | 0.07 | 0 | 0.1 | 0.13 |
| 7 | 128711699 | rs7783840 | 0.016 | 1.207 | -0.007 | -1.065 | 0.01 | -0.059 | -0.059 | 0 | 0.22 | 0.33 | 0.15 | 0.07 | 0.22 | 0.17 |
| 7 | 128721724 | rs17425212 | 0.102 | 1.14 | 0.148 | -0.68 | 0.005 | 0.054 | 0.054 | 0 | 0.15 | 0.07 | 0.1 | 0 | 0.13 | 0.23 |
| 7 | 128722514 | rs17340646 | -0.898 | 1.182 | 0.352 | -0.771 | 0.001 | -0.002 | -0.002 | 0 | 0.23 | 0.19 | 0.16 | 0.01 | 0.21 | 0.2 |
| 7 | 128725550 | rs6959557 | -1.721 | 1.214 | 0.639 | -0.082 | 0.006 | 0.015 | 0.015 | 0.369 | 0.34 | 0.58 | 0.42 | 0.49 | 0.34 | 0.42 |
| 7 | 128732013 | rs7791525 | -1.473 | 1.286 | 0.229 | -0.179 | -0.006 | 0.014 | 0.014 | 1.01 | 0.44 | 0.58 | 0.56 | 0.36 | 0.56 | 0.53 |
| 7 | 128736014 | rs13225818 | -0.387 | 1.175 | 0.179 | -0.161 | 0.003 | 0.003 | 0.003 | 0 | 0.2 | 0.19 | 0.11 | 0.17 | 0.15 | 0.26 |
| 7 | 128738544 | rs4728152 | 1.708 | 1.28 | -0.067 | -0.063 | 0.019 | 0.002 | 0.002 | 2.432 | 0.51 | 0.28 | 0.42 | 0.48 | 0.41 | 0.36 |
| 7 | 128741913 | rs4731549 | 1.606 | 1.002 | -0.144 | 0.169 | -0.008 | 0.023 | 0.023 | 0.208 | 0.59 | 0.39 | 0.69 | 0.81 | 0.55 | 0.58 |
| 7 | 128742164 | rs7779637 | 1.601 | 1.009 | -0.141 | 0.172 | -0.008 | 0.023 | 0.023 | 0.189 | 0.59 | 0.39 | 0.69 | 0.81 | 0.55 | 0.58 |
| 7 | 128743124 | rs1077483 | -1.369 | 1.232 | 0.687 | 0.903 | 0.087 | 0.109 | 0.109 | 0.284 | 0.12 | 0.5 | 0.11 | 0.17 | 0.12 | 0.15 |
| 7 | 128755306 | rs17425565 | -0.241 | 1.247 | -0.643 | 0.345 | -0.009 | 0.012 | 0.012 | 0 | 0.15 | 0.09 | 0.14 | 0.02 | 0.23 | 0.14 |
| 7 | 128773770 | rs11975493 | -1.383 | 0.828 | -0.395 | 1.052 | -0.001 | 0.017 | 0.017 | 0 | 0.06 | 0.1 | 0.05 | 0.2 | 0.06 | 0.12 |
| 7 | 128782249 | rs6971037 | -0.353 | 0.483 | -0.315 | 0.189 | -0.004 | -0.048 | -0.048 | 0.004 | 0.17 | 0.18 | 0.11 | 0 | 0.13 | 0.05 |
| 7 | 128785008 | rs13243618 | 1.16 | 0.631 | -0.079 | 0.51 | -0.003 | -0.015 | -0.015 | 0.383 | 0.68 | 0.61 | 0.77 | 0.79 | 0.74 | 0.71 |
| 7 | 128787548 | rs3993434 | 1.063 | 0.326 | 0.326 | 0.975 | -0.01 | -0.018 | -0.018 | 0.169 | 0.64 | 0.49 | 0.6 | 0.34 | 0.68 | 0.5 |
| 7 | 128788002 | rs719335 | 0.962 | 0.142 | 0.312 | 1.493 | 0.012 | -0.026 | -0.026 | 1.866 | 0.7 | 0.75 | 0.73 | 0.48 | 0.75 | 0.58 |
| 7 | 128789884 | rs10216028 | 1.004 | 0.01 | 0.512 | 1.484 | 0.005 | -0.008 | -0.008 | 0 | 0.71 | 0.71 | 0.73 | 0.47 | 0.77 | 0.61 |
| 7 | 128802936 | rs919551 | 0.926 | 0.215 | 0.403 | 1.167 | 0 | 0 | 0 | 0.246 | 0.79 | 0.76 | 0.78 | 0.81 | 0.8 | 0.74 |
| 7 | 128804041 | rs7798321 | -1.022 | 0.211 | 0.411 | 1.165 | 0.002 | 0.001 | 0.001 | 0.574 | 0.21 | 0.2 | 0.22 | 0.19 | 0.2 | 0.26 |
| 7 | 128804218 | rs2566892 | 0.931 | 0.213 | 0.396 | 1.167 | 0 | 0 | 0 | 0 | 0.79 | 0.76 | 0.78 | 0.81 | 0.8 | 0.74 |
| 7 | 128817136 | rs4386907 | -1.123 | 0.446 | 0.372 | 0.961 | 0.001 | -0.006 | -0.006 | 0 | 0.2 | 0.23 | 0.23 | 0.15 | 0.24 | 0.24 |
| 7 | 128838659 | rs2718107 | 0.717 | 0.606 | 0.042 | 0.947 | 0.006 | 0.03 | 0.03 | 0.021 | 0.55 | 0.33 | 0.49 | 0.4 | 0.53 | 0.49 |
| 7 | 128843169 | rs2566871 | 0.811 | 0.936 | 0.032 | 0.699 | -0.001 | 0.014 | 0.014 | 0 | 0.52 | 0.46 | 0.52 | 0.42 | 0.49 | 0.44 |
| 7 | 128845702 | rs2075778 | 0.559 | 1.042 | 0.674 | 1.066 | 0.058 | 0.062 | 0.062 | 0 | 0.85 | 0.67 | 0.79 | 0.76 | 0.82 | 0.77 |
| 7 | 128853057 | rs9607 | 0.934 | 0.979 | 0.846 | 1 | 0.057 | 0.062 | 0.062 | 0 | 0.82 | 0.53 | 0.77 | 0.76 | 0.82 | 0.76 |
| 7 | 128853123 | rs1061280 | -0.792 | 1.009 | 0.362 | 1.068 | 0.003 | 0.007 | 0.007 | 0 | 0.14 | 0.17 | 0.17 | 0.24 | 0.15 | 0.23 |
| 7 | 128853308 | rs1061285 | -0.77 | 1.013 | 0.053 | 1.07 | -0.008 | -0.013 | -0.013 | 0 | 0.14 | 0.12 | 0.17 | 0.24 | 0.15 | 0.23 |
| 7 | 128866389 | rs3734969 | -0.379 | 0.528 | 0.033 | 1.007 | -0.003 | 0.005 | 0.005 | 0 | 0.3 | 0.28 | 0.28 | 0.38 | 0.31 | 0.26 |
| 7 | 128932749 | rs4110091 | -0.076 | 0.841 | -0.972 | 0.175 | -0.043 | 0.013 | 0.013 | 0 | 0.18 | 0.02 | 0.14 | 0.17 | 0.18 | 0.27 |
| 7 | 128954595 | rs2718098 | 0.041 | 0.814 | -0.298 | -0.036 | -0.005 | -0.041 | -0.041 | 0.988 | 0.87 | 0.74 | 0.94 | 0.97 | 0.92 | 0.89 |
| 7 | 128957954 | rs1526897 | 1.499 | 0.667 | -1.071 | -0.805 | -0.075 | 0.134 | 0.134 | 0 | 0.25 | 0.04 | 0.18 | 0.01 | 0.3 | 0.14 |
| 7 | 128959427 | rs7803052 | 0.373 | 0.184 | -0.817 | -0.434 | 0.032 | 0.019 | 0.019 | 0.049 | 0.34 | 0.21 | 0.33 | 0.33 | 0.24 | 0.41 |
| 7 | 129006202 | rs6966595 | 0.25 | 0.236 | -1.002 | -0.321 | -0.024 | -0.015 | -0.015 | 0 | 0.36 | 0.49 | 0.38 | 0.36 | 0.26 | 0.41 |
| 7 | 129015511 | rs1000676 | 1.164 | 1.232 | -1.427 | -0.567 | -0.106 | -0.11 | -0.11 | 0.172 | 0.27 | 0.03 | 0.37 | 0.61 | 0.39 | 0.35 |
| 7 | 129015894 | rs1559486 | 0.476 | 1.155 | -1.257 | -0.614 | -0.012 | -0.008 | -0.008 | 0 | 0.35 | 0.38 | 0.36 | 0.36 | 0.25 | 0.41 |
| 7 | 129022102 | rs10954230 | 1.092 | 1.209 | -1.23 | -0.564 | -0.079 | -0.081 | -0.081 | 0 | 0.27 | 0.06 | 0.38 | 0.6 | 0.39 | 0.35 |
| 7 | 129023597 | rs7787531 | -0.716 | 1.295 | -1.092 | -0.609 | -0.003 | -0.043 | -0.043 | 1.316 | 0.87 | 0.74 | 0.95 | 0.97 | 0.93 | 0.89 |
| 7 | 129029093 | rs10954231 | 0.289 | 1.257 | -1.029 | -0.619 | 0.009 | -0.016 | -0.016 | 0.47 | 0.58 | 0.49 | 0.52 | 0.37 | 0.51 | 0.53 |
| 7 | 129040018 | rs17552411 | 1.451 | 1.295 | -1.135 | -0.67 | -0.038 | 0.109 | 0.109 | 0 | 0.23 | 0.06 | 0.16 | 0.01 | 0.26 | 0.12 |
| 7 | 129045986 | rs10224011 | 0.094 | 1.198 | -1.056 | -0.713 | -0.012 | -0.006 | -0.006 | 0 | 0.9 | 0.59 | 0.95 | 0.97 | 0.97 | 0.95 |
| 7 | 129047368 | rs17552570 | 1.544 | 1.278 | -1.016 | -0.695 | -0.018 | 0.056 | 0.056 | 0 | 0.22 | 0.09 | 0.17 | 0.01 | 0.26 | 0.12 |

| | | | | | | | | | | | | | | | | |
|---|---|---|---|---|---|---|---|---|---|---|---|---|---|---|---|---|
| 7 | 129065477 | rs822038 | 0.651 | 1.229 | -1.033 | -0.597 | -0.018 | -0.006 | -0.006 | 0.194 | 0.35 | 0.4 | 0.36 | 0.37 | 0.25 | 0.41 |
| 7 | 129067053 | rs4728164 | 1.22 | 1.364 | -1.165 | -0.573 | -0.102 | -0.122 | -0.122 | 0 | 0.27 | 0.03 | 0.38 | 0.59 | 0.39 | 0.35 |
| 7 | 129068978 | rs1665105 | 0.391 | 1.505 | -1.33 | -0.564 | 0.011 | 0.015 | 0.015 | 0 | 0.48 | 0.51 | 0.48 | 0.38 | 0.44 | 0.43 |
| 7 | 129072024 | rs6971551 | 0.839 | 1.154 | -0.991 | -0.62 | -0.019 | 0.044 | 0.044 | 0 | 0.2 | 0.09 | 0.14 | 0.01 | 0.24 | 0.1 |
| 7 | 129077531 | rs587499 | 0.872 | 1.21 | -0.939 | -0.533 | 0 | -0.026 | -0.026 | 0 | 0.6 | 0.55 | 0.76 | 0.95 | 0.63 | 0.72 |
| 7 | 129094195 | rs581829 | -0.255 | 1.06 | -0.845 | -0.423 | -0.007 | -0.033 | -0.033 | 0 | 0.87 | 0.75 | 0.95 | 0.97 | 0.92 | 0.87 |
| 7 | 129096534 | rs753336 | 1.495 | 1.205 | -1.09 | -0.296 | -0.058 | -0.086 | -0.086 | 0.89 | 0.29 | 0.07 | 0.4 | 0.6 | 0.39 | 0.38 |
| 7 | 129098439 | rs691124 | 1.08 | 1.2 | -1.019 | -1.124 | -0.014 | 0.002 | 0.002 | 1.049 | 0.31 | 0.44 | 0.35 | 0.36 | 0.25 | 0.34 |
| 7 | 129108676 | rs12667232 | 1.497 | 1.244 | -1.299 | -1.122 | -0.058 | -0.086 | -0.086 | 0 | 0.29 | 0.07 | 0.4 | 0.6 | 0.39 | 0.38 |
| 7 | 129129740 | rs13223151 | 0.654 | 1.542 | -1.085 | -0.902 | -0.09 | 0.141 | 0.141 | 0.656 | 0.23 | 0.01 | 0.16 | 0.02 | 0.27 | 0.15 |
| 7 | 129132554 | rs692597 | 0.359 | 0.599 | -0.436 | -0.356 | -0.008 | 0.006 | 0.006 | 0.837 | 0.31 | 0.41 | 0.36 | 0.36 | 0.27 | 0.34 |
| 7 | 129146415 | rs657664 | 0.493 | 0.644 | 0.143 | -0.438 | -0.024 | -0.11 | -0.11 | 0.525 | 0.65 | 0.55 | 0.74 | 0.96 | 0.74 | 0.72 |
| 7 | 129147485 | rs11771549 | 0.234 | 0.733 | -0.695 | -0.301 | -0.006 | -0.061 | -0.061 | 0 | 0.63 | 0.5 | 0.72 | 0.94 | 0.71 | 0.71 |
| 7 | 129148058 | rs11761878 | 0.456 | 0.718 | -0.987 | -0.03 | 0.014 | 0.033 | 0.033 | 0.022 | 0.58 | 0.37 | 0.56 | 0.77 | 0.68 | 0.57 |
| 7 | 129157420 | rs12539828 | -0.596 | 1.321 | -1.092 | 0.38 | -0.027 | -0.035 | -0.035 | 0 | 0.15 | 0.09 | 0.31 | 0.78 | 0.2 | 0.37 |
| 7 | 129162363 | rs634484 | 0.976 | 1.014 | -0.341 | 0.058 | 0.017 | -0.037 | -0.037 | 0 | 0.93 | 0.73 | 0.91 | 1 | 0.93 | 0.96 |
| 7 | 129165838 | rs691740 | 0.252 | 1.134 | -0.227 | -0.62 | 0.007 | -0.043 | -0.043 | 0 | 0.21 | 0.29 | 0.24 | 0.09 | 0.25 | 0.17 |
| 7 | 129169483 | rs4574773 | 0.619 | 0.621 | -1.022 | -0.979 | 0.047 | 0.141 | 0.141 | 2.648 | 0.51 | 0.26 | 0.37 | 0.3 | 0.5 | 0.37 |
| 7 | 129170723 | rs4731588 | -0.019 | 0.883 | -0.532 | -0.91 | 0.029 | -0.011 | -0.011 | 0 | 0.65 | 0.52 | 0.52 | 0.37 | 0.6 | 0.46 |
| 7 | 129171989 | rs712735 | -0.132 | 0.906 | -0.577 | -0.893 | 0.018 | -0.009 | -0.009 | 0 | 0.37 | 0.47 | 0.49 | 0.59 | 0.43 | 0.55 |
| 7 | 129176106 | rs725473 | 0.294 | 0.159 | -1.379 | -1.238 | 0.014 | 0.014 | 0.014 | 0 | 0.29 | 0.24 | 0.33 | 0.26 | 0.3 | 0.27 |
| 7 | 129179844 | rs1248465 | -0.209 | 0.534 | -1.067 | -1.121 | 0 | -0.002 | -0.002 | 0 | 0.39 | 0.53 | 0.55 | 0.75 | 0.45 | 0.55 |
| 7 | 129186854 | rs2896417 | 0.385 | 0.492 | -0.987 | -0.478 | 0.002 | -0.003 | -0.003 | 0.298 | 0.08 | 0.04 | 0.11 | 0.25 | 0.09 | 0.14 |
| 7 | 129187430 | rs2598179 | 0.957 | 0.59 | -0.783 | -0.699 | 0.001 | 0.015 | 0.015 | 0 | 0.75 | 0.75 | 0.74 | 0.58 | 0.72 | 0.73 |
| 7 | 129192039 | rs12667965 | -0.192 | 0.581 | -0.875 | -0.775 | -0.023 | -0.001 | -0.001 | 1.211 | 0.16 | 0.03 | 0.14 | 0.17 | 0.19 | 0.17 |
| 7 | 129194092 | rs659495 | 0.016 | 0.575 | -0.666 | -0.739 | 0.001 | 0.061 | 0.061 | 0 | 0.1 | 0.17 | 0.12 | 0.25 | 0.11 | 0.15 |
| 7 | 129196392 | rs12671559 | -0.13 | 0.6 | -0.764 | -0.761 | -0.025 | -0.003 | -0.003 | 0 | 0.16 | 0.03 | 0.14 | 0.17 | 0.18 | 0.16 |
| 7 | 129196616 | rs2044455 | 0.292 | 0.432 | -0.813 | -1.615 | 0.028 | 0.027 | 0.027 | 0.039 | 0.28 | 0.43 | 0.44 | 0.57 | 0.27 | 0.43 |
| 7 | 129196930 | rs595444 | -1.407 | -0.389 | -0.699 | -1.71 | -0.015 | -0.004 | -0.004 | 0 | 0.39 | 0.47 | 0.34 | 0.42 | 0.38 | 0.33 |
| 7 | 129197408 | rs691184 | -0.613 | -0.455 | -0.594 | -1.763 | -0.014 | -0.034 | -0.034 | 0.121 | 0.89 | 0.88 | 0.94 | 1 | 0.91 | 0.99 |
| 7 | 129199328 | rs10245560 | 1.139 | -0.293 | -0.629 | -1.857 | 0.017 | 0.105 | 0.105 | 0 | 0.41 | 0.21 | 0.28 | 0.01 | 0.45 | 0.21 |
| 7 | 129199487 | rs11765308 | -0.475 | -0.124 | -0.518 | -1.616 | 0 | -0.026 | -0.026 | 0 | 0.45 | 0.52 | 0.59 | 0.74 | 0.44 | 0.62 |
| 7 | 129210535 | rs692489 | 1.427 | 0.074 | -0.716 | -1.716 | -0.008 | 0.143 | 0.143 | 0 | 0.45 | 0.23 | 0.37 | 0.05 | 0.5 | 0.3 |
| 7 | 129214174 | rs754386 | 0.519 | 0.167 | -0.266 | -1.665 | -0.004 | -0.009 | -0.009 | 0 | 0.18 | 0.19 | 0.1 | 0.13 | 0.13 | 0.16 |
| 7 | 129214387 | rs754385 | -1.828 | 0.136 | -0.221 | -1.542 | -0.006 | 0.122 | 0.122 | 0 | 0.55 | 0.79 | 0.61 | 0.86 | 0.5 | 0.64 |
| 7 | 129219112 | rs6948627 | 1.575 | 0.362 | -0.063 | -1.537 | 0.005 | 0.014 | 0.014 | 0 | 0.52 | 0.47 | 0.56 | 0.66 | 0.57 | 0.63 |
| 7 | 129227770 | rs7788769 | -1.386 | 0.447 | -0.386 | -1.443 | 0.016 | -0.02 | -0.02 | 0.057 | 0.9 | 0.84 | 0.88 | 0.96 | 0.9 | 0.92 |
| 7 | 129232720 | rs13233824 | 0.57 | 0.353 | -0.659 | -1.471 | 0 | -0.004 | -0.004 | 0 | 0.15 | 0.12 | 0.19 | 0.17 | 0.16 | 0.12 |
| 7 | 129242076 | rs12154955 | 1.465 | 0.497 | -0.564 | -1.378 | -0.072 | 0.12 | 0.12 | 0.893 | 0.55 | 0.17 | 0.48 | 0.27 | 0.61 | 0.42 |
| 7 | 129250536 | rs1562833 | -1.267 | 0.45 | -0.382 | -1.198 | 0.012 | -0.009 | -0.009 | 0 | 0.9 | 0.81 | 0.88 | 0.95 | 0.9 | 0.92 |
| 7 | 129257508 | rs10245818 | 1.058 | 0.491 | 0.019 | -0.876 | 0.011 | 0.066 | 0.066 | 0 | 0.14 | 0.22 | 0.25 | 0.48 | 0.16 | 0.34 |
| 7 | 129259743 | rs10225103 | -0.148 | 0.364 | -0.029 | -0.947 | -0.04 | -0.048 | -0.048 | 0 | 0.22 | 0.52 | 0.13 | 0.14 | 0.13 | 0.16 |
| 7 | 129265143 | rs4469397 | 0.154 | 0.526 | 0.142 | -1.031 | -0.021 | -0.033 | -0.033 | 0 | 0.21 | 0.27 | 0.16 | 0.14 | 0.17 | 0.17 |
| 7 | 129286408 | rs6969441 | -1.606 | 0.449 | 0.003 | -0.972 | -0.001 | 0.023 | 0.023 | 0.008 | 0.39 | 0.76 | 0.42 | 0.66 | 0.33 | 0.49 |
| 7 | 129286436 | rs6949152 | -0.539 | 0.354 | 0.248 | -0.958 | -0.046 | -0.039 | -0.039 | 0 | 0.24 | 0.44 | 0.16 | 0.19 | 0.17 | 0.18 |
| 7 | 129304991 | rs6467257 | -1.314 | 0.524 | 0.169 | -0.905 | 0.01 | -0.014 | -0.014 | 0.044 | 0.89 | 0.83 | 0.87 | 0.95 | 0.89 | 0.87 |
| 7 | 129312570 | rs10276606 | 1.392 | 0.919 | 0.123 | -0.849 | 0.006 | 0.073 | 0.073 | 0 | 0.11 | 0.16 | 0.24 | 0.48 | 0.14 | 0.3 |
| 7 | 129321191 | rs6467259 | 0.624 | 0.619 | 0.121 | -1.114 | -0.005 | -0.005 | -0.005 | 0 | 0.69 | 0.47 | 0.64 | 0.71 | 0.69 | 0.75 |
| 7 | 129321647 | rs2402968 | -1.514 | 0.647 | 0.131 | -1.132 | 0.027 | 0.007 | 0.007 | 0 | 0.93 | 0.83 | 0.88 | 0.95 | 0.92 | 0.87 |
| 7 | 129329261 | rs2402974 | 1.403 | 0.63 | -0.001 | -1.132 | -0.04 | -0.049 | -0.049 | 0.017 | 0.11 | 0.01 | 0.17 | 0.18 | 0.15 | 0.11 |
| 7 | 129332946 | rs1984823 | 1.887 | 0.554 | -0.048 | -1.26 | 0.037 | 0.154 | 0.154 | 0.933 | 0.49 | 0.25 | 0.34 | 0.16 | 0.48 | 0.27 |
| 7 | 129333905 | rs9656383 | -0.478 | 0.578 | -0.351 | -1.232 | 0.018 | 0.022 | 0.022 | 0 | 0.31 | 0.43 | 0.36 | 0.61 | 0.29 | 0.49 |
| 7 | 129342404 | rs2402976 | 1.747 | 0.509 | -0.398 | -1.188 | -0.011 | 0.049 | 0.049 | 1.715 | 0.56 | 0.32 | 0.42 | 0.24 | 0.53 | 0.44 |
| 7 | 129343967 | rs11773023 | 2.551 | 0.518 | -0.352 | -1.177 | 0.001 | 0.177 | 0.177 | 2.522 | 0.39 | 0.16 | 0.3 | 0.1 | 0.41 | 0.26 |
| 7 | 129350170 | rs3736626 | 2.551 | 0.518 | -0.352 | -1.177 | 0.001 | 0.177 | 0.177 | 2.522 | 0.39 | 0.16 | 0.3 | 0.1 | 0.41 | 0.26 |
| 7 | 129356936 | rs17161218 | 1.348 | 0.555 | -0.465 | -1.095 | -0.012 | -0.011 | -0.011 | 0 | 0.11 | 0.12 | 0.08 | 0.11 | 0.07 | 0.09 |
| 7 | 129360201 | rs3800602 | 1.653 | 0.548 | -0.722 | -1.094 | 0.005 | 0 | 0 | 0 | 0.09 | 0 | 0.03 | 0.06 | 0.07 | 0.02 |
| 7 | 129362665 | rs11766629 | 1.557 | 0.584 | -0.689 | -1.071 | -0.002 | -0.006 | -0.006 | 0 | 0.1 | 0 | 0.07 | 0.09 | 0.07 | 0.1 |
| 7 | 129368612 | rs10239118 | -0.576 | 0.692 | -0.482 | -1.013 | -0.005 | -0.002 | -0.002 | 0.496 | 0.29 | 0.31 | 0.32 | 0.32 | 0.28 | 0.29 |
| 7 | 129380243 | rs7796553 | 1.28 | 0.877 | -0.471 | -0.978 | 0.005 | 0.051 | 0.051 | 0 | 0.13 | 0.18 | 0.21 | 0.48 | 0.15 | 0.31 |
| 7 | 129385136 | rs7794909 | -1.384 | 0.681 | -0.507 | -0.917 | 0.01 | 0 | 0 | 4.497 | 0.92 | 0.91 | 0.88 | 0.95 | 0.91 | 0.87 |
| 7 | 129387380 | rs10244147 | -1.542 | 0.366 | -0.665 | -0.972 | 0.009 | 0.006 | 0.006 | 0 | 0.4 | 0.32 | 0.36 | 0.38 | 0.35 | 0.31 |
| 7 | 129393341 | rs10500120 | -1.134 | 0.712 | -0.364 | -0.762 | -0.005 | -0.002 | -0.002 | 0.105 | 0.29 | 0.31 | 0.32 | 0.33 | 0.3 | 0.29 |
| 7 | 129394491 | rs13438321 | -0.338 | 0.584 | -0.367 | -0.702 | -0.011 | -0.02 | -0.02 | 0 | 0.2 | 0.25 | 0.14 | 0.14 | 0.14 | 0.19 |
| 7 | 129396342 | rs13241028 | 0.128 | 0.908 | -1.034 | -0.581 | -0.033 | -0.018 | -0.018 | 0.684 | 0.15 | 0.04 | 0.19 | 0.19 | 0.19 | 0.11 |
| 7 | 129405681 | rs2402961 | -0.813 | 0.719 | -0.684 | -0.56 | 0.007 | -0.004 | -0.004 | 0 | 0.91 | 0.91 | 0.88 | 0.95 | 0.9 | 0.8 |
| 7 | 129405807 | rs2402959 | -0.85 | 0.486 | -0.632 | -0.752 | -0.001 | 0.001 | 0.001 | 0 | 0.28 | 0.25 | 0.21 | 0.32 | 0.28 | 0.21 |
| 7 | 129407261 | rs6467264 | -1.336 | 0.325 | -0.982 | -1.213 | -0.005 | 0.227 | 0.227 | 0 | 0.5 | 0.87 | 0.53 | 0.87 | 0.43 | 0.51 |
| 7 | 129407564 | rs6965643 | -0.527 | 0.894 | -0.316 | -0.747 | 0.017 | 0.013 | 0.013 | 0 | 0.24 | 0.36 | 0.38 | 0.78 | 0.23 | 0.41 |

| | | | | | | | | | | | | | | | | |
|---|---|---|---|---|---|---|---|---|---|---|---|---|---|---|---|---|
| 7 | 129413188 | rs4626538 | 0.72 | 0.253 | -0.508 | -1.233 | -0.001 | -0.022 | -0.022 | 0 | 0.55 | 0.59 | 0.42 | 0.06 | 0.51 | 0.3 |
| 7 | 129417730 | rs2693727 | 0.132 | -0.017 | 0.684 | -0.601 | 0.021 | 0 | 0 | 1.937 | 0.91 | 0.69 | 0.91 | 0.97 | 0.91 | 0.98 |
| 7 | 129425910 | rs10224663 | -1.198 | -0.147 | 0.33 | -0.618 | -0.001 | 0.008 | 0.008 | 0 | 0.52 | 0.54 | 0.47 | 0.35 | 0.46 | 0.49 |
| 7 | 129427201 | rs4731630 | -1.073 | -0.222 | 0.295 | -0.557 | 0.002 | 0.013 | 0.013 | 0 | 0.48 | 0.39 | 0.42 | 0.36 | 0.43 | 0.46 |
| 7 | 129430994 | rs2693737 | -0.52 | 0.019 | 0.74 | -0.226 | 0.009 | -0.009 | -0.009 | 0 | 0.93 | 0.88 | 0.94 | 0.97 | 0.91 | 0.98 |
| 7 | 129431812 | rs12706902 | 1.281 | -0.084 | 0.761 | -0.354 | 0 | -0.017 | -0.017 | 0 | 0.64 | 0.62 | 0.7 | 0.77 | 0.66 | 0.64 |
| 7 | 129433814 | rs11764547 | 0.166 | 0.106 | 0.572 | -0.971 | -0.005 | -0.027 | -0.027 | 0 | 0.11 | 0.12 | 0.07 | 0 | 0.06 | 0.04 |
| 7 | 129435124 | rs868056 | 1.603 | 0.29 | 0.387 | -0.823 | -0.012 | 0.014 | 0.014 | 0.048 | 0.4 | 0.34 | 0.51 | 0.63 | 0.43 | 0.42 |
| 7 | 129435191 | rs868055 | 0.227 | 0.399 | 0.555 | -0.345 | 0.025 | -0.008 | -0.008 | 0 | 0.93 | 0.84 | 0.93 | 0.97 | 0.91 | 0.98 |
| 7 | 129444951 | rs4728178 | -1.058 | 0.154 | 0.438 | -0.525 | -0.001 | -0.056 | -0.056 | 0.146 | 0.26 | 0.35 | 0.19 | 0.14 | 0.25 | 0.35 |
| 7 | 129450855 | rs10225672 | -0.912 | -0.341 | 0.419 | -0.577 | -0.008 | -0.007 | -0.007 | 0 | 0.27 | 0.28 | 0.25 | 0.21 | 0.22 | 0.27 |
| 7 | 129454847 | rs2727455 | 2.048 | -0.262 | 0.054 | -0.658 | 0.005 | 0.062 | 0.062 | 0 | 0.48 | 0.16 | 0.52 | 0.67 | 0.54 | 0.47 |
| 7 | 129459578 | rs4298432 | -1.393 | -0.202 | 0.438 | -0.748 | -0.001 | -0.008 | -0.008 | 0 | 0.28 | 0.33 | 0.25 | 0.12 | 0.28 | 0.25 |
| 7 | 129464791 | rs13238927 | 0.095 | -0.303 | 0.682 | -0.922 | 0.003 | -0.059 | -0.059 | 0.938 | 0.1 | 0.21 | 0.07 | 0 | 0.08 | 0.11 |
| 7 | 129473739 | rs1046691 | -0.783 | 0.002 | 0.109 | -0.654 | -0.009 | -0.039 | -0.039 | 0 | 0.19 | 0.28 | 0.18 | 0.02 | 0.15 | 0.09 |
| 7 | 129501926 | rs17558874 | -0.476 | 0.049 | 0.104 | -0.576 | -0.013 | -0.011 | -0.011 | 0 | 0.1 | 0.09 | 0.04 | 0.04 | 0.05 | 0.14 |
| 7 | 129505141 | rs3807120 | -1.157 | 0.183 | -0.058 | -0.327 | -0.031 | -0.037 | -0.037 | 0.328 | 0.26 | 0.09 | 0.33 | 0.3 | 0.29 | 0.28 |
| 7 | 129525832 | rs1471876 | 1.103 | 0.241 | 0.116 | -0.53 | -0.018 | -0.112 | -0.112 | 0 | 0.72 | 0.51 | 0.74 | 0.94 | 0.74 | 0.88 |
| 7 | 129569997 | rs1725077 | 1.864 | 0.258 | 0.055 | -0.451 | -0.011 | -0.01 | -0.01 | 0 | 0.6 | 0.45 | 0.6 | 0.66 | 0.64 | 0.59 |
| 7 | 129579325 | rs2727507 | 1.629 | 0.158 | 0.081 | -0.501 | -0.028 | -0.08 | -0.08 | 0.129 | 0.65 | 0.46 | 0.72 | 0.9 | 0.71 | 0.76 |
| 7 | 129586354 | rs10215972 | -1.378 | 0.781 | -0.105 | -0.294 | 0.002 | -0.001 | -0.001 | 0.257 | 0.3 | 0.33 | 0.37 | 0.3 | 0.37 | 0.31 |
| 7 | 129591773 | rs1723728 | 1.171 | 0.753 | -0.378 | -0.588 | -0.006 | 0.028 | 0.028 | 0 | 0.83 | 0.91 | 0.86 | 0.97 | 0.78 | 0.9 |
| 7 | 129593339 | rs6963861 | 1.388 | 0.917 | 0.122 | -0.744 | 0.014 | -0.061 | -0.061 | 0 | 0.69 | 0.52 | 0.73 | 0.78 | 0.67 | 0.74 |
| 7 | 129598336 | rs6980213 | 1.213 | 1.183 | 0.153 | -0.607 | 0.007 | -0.001 | -0.001 | 0 | 0.83 | 0.81 | 0.81 | 0.78 | 0.74 | 0.85 |
| 7 | 129603129 | rs2464893 | -0.928 | 1.131 | 0.354 | -0.637 | 0.03 | 0.004 | 0.004 | 0 | 0.09 | 0.15 | 0.13 | 0.16 | 0.15 | 0.11 |
| 7 | 129607187 | rs2708635 | 0.019 | 1.211 | 0.647 | -0.674 | 0.02 | -0.004 | -0.004 | 0 | 0.95 | 0.88 | 0.94 | 0.97 | 0.92 | 0.98 |
| 7 | 129609330 | rs2692437 | -0.199 | 0.714 | -0.187 | -1.079 | 0.016 | -0.021 | -0.021 | 0 | 0.61 | 0.5 | 0.36 | 0.35 | 0.45 | 0.47 |
| 7 | 129609681 | rs7805545 | -0.307 | 0.701 | -0.002 | -1.085 | 0.002 | -0.061 | -0.061 | 3.142 | 0.19 | 0.23 | 0.11 | 0 | 0.16 | 0.16 |
| 7 | 129615937 | rs6943541 | 1.01 | 0.821 | -0.095 | -0.929 | 0.001 | 0 | 0 | 0.028 | 0.85 | 0.82 | 0.77 | 0.79 | 0.78 | 0.88 |
| 7 | 129617357 | rs2177773 | -0.622 | 0.71 | 0.216 | -1.234 | 0 | -0.146 | -0.146 | 2.067 | 0.25 | 0.42 | 0.15 | 0 | 0.19 | 0.19 |
| 7 | 129617727 | rs10808241 | 1.167 | 0.638 | 0.153 | -1.156 | -0.004 | 0.016 | 0.016 | 0 | 0.65 | 0.32 | 0.65 | 0.78 | 0.62 | 0.68 |
| 7 | 129634289 | rs12533403 | 0.734 | 0.642 | -0.548 | -1.255 | -0.024 | 0.087 | 0.087 | 0 | 0.46 | 0.15 | 0.53 | 0.66 | 0.5 | 0.54 |
| 7 | 129637694 | rs4731641 | 0.763 | 0.535 | -0.643 | -1.26 | -0.05 | 0.084 | 0.084 | 0 | 0.46 | 0.11 | 0.54 | 0.66 | 0.54 | 0.56 |
| 7 | 129649530 | rs7787207 | -0.054 | -0.272 | 0.511 | -1.19 | -0.018 | 0.024 | 0.024 | 0 | 0.14 | 0.17 | 0.11 | 0.31 | 0.07 | 0.16 |
| 7 | 129649928 | rs11769085 | 0.109 | -0.148 | 0.352 | -1.703 | -0.001 | -0.046 | -0.046 | 0 | 0.25 | 0.29 | 0.16 | 0 | 0.23 | 0.22 |
| 7 | 129653231 | rs12113749 | 0.288 | -0.183 | 0.528 | -1.689 | 0 | -0.048 | -0.048 | 0.515 | 0.23 | 0.24 | 0.16 | 0.01 | 0.22 | 0.21 |
| 7 | 129655876 | rs10265116 | 0.288 | -0.183 | 0.528 | -1.689 | 0 | -0.048 | -0.048 | 0 | 0.23 | 0.24 | 0.16 | 0.01 | 0.22 | 0.21 |
| 7 | 129657882 | rs2242487 | 1.159 | -0.198 | 0.222 | -1.675 | 0.031 | 0.057 | 0.057 | 0 | 0.55 | 0.33 | 0.68 | 0.7 | 0.65 | 0.61 |
| 7 | 129663496 | rs11556924 | 1.677 | -0.162 | -1.021 | -2.04 | -0.115 | 0.101 | 0.101 | 0 | 0.21 | 0.03 | 0.21 | 0.05 | 0.38 | 0.17 |
| 7 | 129664312 | rs1464890 | 1.001 | -0.424 | -0.328 | -1.543 | 0.023 | 0.135 | 0.135 | 0 | 0.51 | 0.1 | 0.56 | 0.67 | 0.61 | 0.58 |
| 7 | 129668931 | rs11974220 | 1.117 | -0.048 | 0.012 | -1.574 | 0 | -0.008 | -0.008 | 0 | 0.06 | 0.05 | 0.04 | 0 | 0.07 | 0.06 |
| 7 | 129672975 | rs12706910 | -0.107 | -0.093 | 0.288 | -1.452 | -0.004 | -0.152 | -0.152 | 0 | 0.73 | 0.52 | 0.79 | 0.98 | 0.75 | 0.79 |
| 7 | 129698762 | rs13245505 | -0.731 | -0.082 | -0.315 | -0.645 | -0.018 | -0.008 | -0.008 | 0 | 0.34 | 0.47 | 0.29 | 0.31 | 0.28 | 0.3 |
| 7 | 129702226 | rs6943859 | -0.058 | -0.057 | -0.565 | -0.646 | 0 | -0.011 | -0.011 | 0 | 0.09 | 0.1 | 0.06 | 0 | 0.07 | 0.07 |
| 7 | 129710136 | rs10277874 | -0.635 | -0.674 | -1.425 | -1.231 | -0.059 | 0.064 | 0.064 | 0 | 0.59 | 0.84 | 0.46 | 0.28 | 0.47 | 0.49 |
| 7 | 129712524 | rs10241456 | 0.464 | -0.568 | -1.526 | -0.405 | -0.031 | -0.048 | -0.048 | 1.289 | 0.34 | 0.17 | 0.35 | 0.44 | 0.43 | 0.28 |
| 7 | 129725858 | rs10279425 | 0.546 | -0.395 | -1.523 | -0.422 | -0.037 | -0.048 | -0.048 | 1.68 | 0.33 | 0.13 | 0.36 | 0.42 | 0.44 | 0.3 |
| 7 | 129726745 | rs6961836 | -0.243 | -1.024 | -1.18 | -0.67 | -0.016 | 0.018 | 0.018 | 0 | 0.79 | 0.7 | 0.76 | 0.73 | 0.88 | 0.79 |
| 7 | 129734173 | rs6945486 | 0.354 | -1.032 | -1.142 | -0.741 | -0.008 | 0.042 | 0.042 | 0.021 | 0.46 | 0.63 | 0.41 | 0.31 | 0.44 | 0.49 |
| 7 | 129735494 | rs10279986 | 0.628 | -1.345 | -1.293 | -0.811 | 0.001 | -0.032 | -0.032 | 0 | 0.2 | 0.19 | 0.08 | 0 | 0.1 | 0.05 |
| 7 | 129744700 | rs6467280 | -0.119 | -1.075 | -1.124 | -0.566 | -0.008 | 0.008 | 0.008 | 0 | 0.8 | 0.75 | 0.76 | 0.73 | 0.88 | 0.76 |
| 7 | 129749447 | rs4731659 | 1.367 | -0.998 | -1.204 | -0.626 | 0.002 | 0.03 | 0.03 | 0 | 0.1 | 0.14 | 0.21 | 0.3 | 0.12 | 0.23 |
| 7 | 129756523 | rs17641278 | 2.255 | -1.638 | -1.7 | -1.166 | 0.026 | 0.098 | 0.098 | 0 | 0.16 | 0.03 | 0.04 | 0 | 0.07 | 0.12 |
| 7 | 129756758 | rs4731660 | 1.557 | -1.41 | -0.897 | -0.982 | 0.024 | 0.118 | 0.118 | 0 | 0.1 | 0.24 | 0.21 | 0.3 | 0.12 | 0.23 |
| 7 | 129761083 | rs4324873 | 1.408 | -1.344 | -1.335 | -1.067 | -0.044 | -0.067 | -0.067 | 5.003 | 0.31 | 0.12 | 0.35 | 0.43 | 0.43 | 0.27 |
| 7 | 129771117 | rs12534379 | 0.548 | -1.477 | -0.95 | -1.126 | 0 | 0.002 | 0.002 | 0 | 0.42 | 0.42 | 0.42 | 0.46 | 0.5 | 0.28 |
| 7 | 129772788 | rs3800562 | 1.458 | -0.941 | -0.856 | -0.746 | 0.001 | 0.016 | 0.016 | 0 | 0.09 | 0.11 | 0.2 | 0.28 | 0.11 | 0.23 |
| 7 | 129774714 | rs6821 | 1.124 | -1.099 | -1.037 | -1.271 | -0.023 | 0.027 | 0.027 | 0 | 0.25 | 0.16 | 0.15 | 0.01 | 0.28 | 0.17 |
| 7 | 129775950 | rs6952837 | 0.349 | -1.362 | -0.69 | -0.674 | -0.006 | 0.13 | 0.13 | 0 | 0.77 | 0.58 | 0.6 | 0.47 | 0.83 | 0.55 |
| 7 | 129782377 | rs4731662 | 0.198 | -0.913 | -1.185 | -0.552 | 0.004 | 0.011 | 0.011 | 0 | 0.26 | 0.24 | 0.19 | 0.22 | 0.19 | 0.3 |
| 7 | 129786406 | rs10264186 | 1.062 | -1.272 | -1.22 | -0.1 | 0.011 | -0.014 | -0.014 | 0 | 0.13 | 0.08 | 0.1 | 0.37 | 0.07 | 0.2 |
| 7 | 129787300 | rs4363128 | 0.299 | -0.52 | -1.723 | -0.552 | 0.008 | -0.041 | -0.041 | 0 | 0.29 | 0.09 | 0.29 | 0.33 | 0.31 | 0.25 |
| 7 | 129788624 | rs6971899 | 0.261 | 0.026 | -1.13 | -1.504 | -0.003 | -0.091 | -0.091 | 0 | 0.38 | 0.36 | 0.23 | 0.03 | 0.32 | 0.24 |
| 7 | 129799845 | rs10241132 | 0.554 | -0.756 | -0.67 | -1.516 | -0.038 | -0.123 | -0.123 | 0 | 0.2 | 0.38 | 0.13 | 0.02 | 0.11 | 0.12 |
| 7 | 129802128 | rs4443601 | -0.193 | -0.917 | -1.213 | -1.417 | 0.045 | -0.05 | -0.05 | 0 | 0.43 | 0.15 | 0.44 | 0.49 | 0.34 | 0.53 |
| 7 | 129803086 | rs11770708 | 1.622 | -0.177 | -0.61 | -1.462 | -0.076 | 0.127 | 0.127 | 0.488 | 0.21 | 0.04 | 0.12 | 0.01 | 0.25 | 0.11 |
| 7 | 129811276 | rs4520105 | 0.415 | -0.231 | -0.499 | -1.412 | -0.015 | 0.087 | 0.087 | 0 | 0.23 | 0.34 | 0.32 | 0.41 | 0.18 | 0.37 |
| 7 | 129813738 | rs3823482 | 1.042 | -0.156 | -0.813 | -1.561 | -0.009 | -0.027 | -0.027 | 0.047 | 0.21 | 0.13 | 0.27 | 0.55 | 0.26 | 0.34 |
| 7 | 129817504 | rs1881257 | 1.218 | -0.189 | -0.816 | -1.578 | -0.016 | -0.043 | -0.043 | 0 | 0.2 | 0.1 | 0.26 | 0.55 | 0.25 | 0.33 |

| | | | | | | | | | | | | | | | | |
|---|---|---|---|---|---|---|---|---|---|---|---|---|---|---|---|---|
| 7 | 129825215 | rs4731668 | 1.811 | -0.427 | -0.653 | -1.68 | -0.026 | 0.05 | 0.05 | 0.603 | 0.21 | 0.09 | 0.11 | 0.01 | 0.23 | 0.11 |
| 7 | 129827755 | rs2633387 | 0.884 | -0.308 | -0.646 | -1.464 | -0.013 | -0.03 | -0.03 | 0 | 0.21 | 0.13 | 0.3 | 0.58 | 0.28 | 0.35 |
| 7 | 129840285 | rs17164727 | 0.802 | -0.39 | -0.891 | -1.343 | -0.002 | -0.093 | -0.093 | 1.737 | 0.2 | 0.08 | 0.3 | 0.4 | 0.18 | 0.37 |
| 7 | 129842229 | rs17164729 | 1.071 | -0.238 | -0.819 | -1.249 | 0.017 | 0.009 | 0.009 | 0 | 0.17 | 0.13 | 0.06 | 0 | 0.08 | 0.12 |
| 7 | 129847093 | rs1994467 | 0.409 | 0.124 | -0.638 | -1.054 | 0.033 | 0.113 | 0.113 | 0.021 | 0.23 | 0.38 | 0.33 | 0.58 | 0.29 | 0.36 |
| 7 | 129847238 | rs1994468 | 0.496 | 0.203 | -0.849 | -1.176 | -0.005 | 0.036 | 0.036 | 0 | 0.22 | 0.26 | 0.31 | 0.4 | 0.18 | 0.37 |
| 7 | 129854753 | rs2087242 | 0.784 | 0.523 | -0.858 | -0.968 | 0.007 | 0.032 | 0.032 | 0 | 0.2 | 0.19 | 0.31 | 0.58 | 0.28 | 0.35 |
| 7 | 129855837 | rs4728190 | 1.137 | 0.488 | -0.951 | -1.175 | 0.006 | -0.1 | -0.1 | 0 | 0.24 | 0.36 | 0.15 | 0.01 | 0.24 | 0.12 |
| 7 | 129856473 | rs7808330 | 1.137 | 0.488 | -0.951 | -1.175 | 0.006 | -0.1 | -0.1 | 0 | 0.24 | 0.36 | 0.15 | 0.01 | 0.24 | 0.12 |
| 7 | 129857820 | rs10808242 | -0.545 | 0.77 | -1.235 | -0.436 | 0.007 | 0.032 | 0.032 | 0 | 0.8 | 0.81 | 0.69 | 0.42 | 0.72 | 0.65 |
| 7 | 129873928 | rs6467289 | -0.223 | 0.775 | -1.664 | -0.542 | 0.004 | -0.022 | -0.022 | 0 | 0.65 | 0.81 | 0.64 | 0.36 | 0.68 | 0.6 |
| 7 | 129881673 | rs10233902 | -0.605 | 0.549 | -1.249 | -0.299 | 0.015 | -0.047 | -0.047 | 0 | 0.29 | 0.39 | 0.31 | 0.15 | 0.35 | 0.31 |
| 7 | 129906147 | rs2171493 | 0.539 | 0.384 | -0.837 | -0.125 | 0.001 | -0.037 | -0.037 | 0.459 | 0.17 | 0.16 | 0.18 | 0.01 | 0.27 | 0.06 |
| 7 | 129912602 | rs2242337 | -0.307 | 0.362 | -0.788 | -0.034 | 0.042 | -0.06 | -0.06 | 0 | 0.22 | 0.39 | 0.24 | 0.14 | 0.32 | 0.2 |
| 7 | 129916030 | rs9641869 | -0.15 | -0.69 | -0.732 | 0.945 | -0.034 | 0.05 | 0.05 | 0 | 0.24 | 0.31 | 0.29 | 0.56 | 0.17 | 0.36 |
| 7 | 129923765 | rs2178158 | -0.041 | -0.888 | -1.568 | 0.983 | 0.013 | -0.017 | -0.017 | 0 | 0.18 | 0.14 | 0.24 | 0.44 | 0.11 | 0.22 |
| 7 | 129927778 | rs6467293 | -0.571 | 0.226 | -1.407 | 0.376 | 0.02 | 0.015 | 0.015 | 0 | 0.22 | 0.16 | 0.17 | 0.23 | 0.14 | 0.29 |
| 7 | 129931402 | rs901799 | -1.187 | 1.253 | -1.177 | 0.468 | -0.01 | 0.001 | 0.001 | 0 | 0.12 | 0.08 | 0.12 | 0.14 | 0.17 | 0.15 |
| 7 | 129933481 | rs10954267 | 0.207 | 0.843 | -0.929 | 0.554 | 0.001 | 0.012 | 0.012 | 0 | 0.63 | 0.54 | 0.65 | 0.58 | 0.61 | 0.64 |
| 7 | 129934219 | rs3807344 | 0.482 | 0.813 | -1.074 | 0.065 | -0.001 | -0.007 | -0.007 | 0 | 0.35 | 0.33 | 0.48 | 0.51 | 0.36 | 0.32 |
| 7 | 129944685 | rs3735052 | 0.658 | 0.304 | -0.723 | -0.152 | -0.014 | -0.024 | -0.024 | 0 | 0.25 | 0.2 | 0.26 | 0.35 | 0.33 | 0.35 |
| 7 | 129944994 | rs1038627 | -0.317 | 0.001 | -0.094 | -0.41 | 0.011 | 0.008 | 0.008 | 0 | 0.37 | 0.48 | 0.35 | 0.36 | 0.42 | 0.38 |
| 7 | 129946939 | rs6953458 | -0.157 | -0.061 | 0.152 | -0.254 | -0.002 | -0.003 | -0.003 | 0 | 0.34 | 0.33 | 0.33 | 0.36 | 0.38 | 0.36 |
| 7 | 129950740 | rs2171492 | -0.012 | -0.026 | 0.279 | -0.313 | -0.015 | -0.04 | -0.04 | 0 | 0.33 | 0.21 | 0.32 | 0.37 | 0.36 | 0.36 |
| 7 | 129951108 | rs729167 | 0.734 | -0.031 | 0.235 | -0.295 | -0.019 | -0.05 | -0.05 | 3.594 | 0.25 | 0.17 | 0.25 | 0.36 | 0.3 | 0.35 |
| 7 | 129954274 | rs12534255 | 1.06 | -0.076 | 0.234 | -0.336 | -0.013 | 0.019 | 0.019 | 0 | 0.07 | 0 | 0.05 | 0.01 | 0.06 | 0.02 |
| 7 | 129958098 | rs1488009 | -0.072 | -0.096 | 0.751 | -0.359 | 0.017 | -0.045 | -0.045 | 1.696 | 0.9 | 0.78 | 0.92 | 0.99 | 0.91 | 0.97 |
| 7 | 129958167 | rs3800775 | 0.06 | -0.166 | 0.793 | -0.369 | 0.001 | 0.007 | 0.007 | 0 | 0.66 | 0.63 | 0.68 | 0.62 | 0.63 | 0.65 |
| 7 | 129962414 | rs2306848 | 0.808 | -0.188 | 0.765 | -0.371 | -0.002 | -0.011 | -0.011 | 0 | 0.08 | 0.06 | 0.06 | 0.01 | 0.06 | 0.02 |
| 7 | 129963356 | rs3778858 | 0.784 | -1.096 | 0.613 | -1.281 | -0.011 | -0.016 | -0.016 | 0 | 0.33 | 0.31 | 0.49 | 0.52 | 0.39 | 0.39 |
| 7 | 129963481 | rs12539755 | 0.513 | -0.722 | 0.635 | -0.978 | 0.003 | -0.025 | -0.025 | 0 | 0.1 | 0.12 | 0.07 | 0.01 | 0.1 | 0.03 |
| 7 | 129973050 | rs7793804 | 1.213 | -0.754 | 0.39 | -0.95 | -0.013 | -0.038 | -0.038 | 0.055 | 0.26 | 0.22 | 0.25 | 0.37 | 0.31 | 0.37 |
| 7 | 129973093 | rs4731675 | 0.579 | -1.07 | 0.504 | -1.084 | -0.001 | 0.017 | 0.017 | 0 | 0.51 | 0.35 | 0.47 | 0.5 | 0.49 | 0.53 |
| 7 | 129976506 | rs6948984 | 0.156 | -0.957 | 0.17 | -1.049 | -0.005 | -0.035 | -0.035 | 0 | 0.37 | 0.29 | 0.35 | 0.46 | 0.41 | 0.45 |
| 7 | 129978082 | rs11765940 | 0.486 | -1.188 | -0.163 | -1.37 | 0.032 | 0.049 | 0.049 | 0.219 | 0.22 | 0.08 | 0.13 | 0.09 | 0.15 | 0.08 |
| 7 | 129979092 | rs7797072 | 1.532 | -1.06 | -0.119 | -1.235 | -0.013 | -0.011 | -0.011 | 0.218 | 0.15 | 0.15 | 0.22 | 0.37 | 0.26 | 0.37 |
| 7 | 129979363 | rs7797371 | 1.413 | -1.053 | -0.101 | -1.236 | -0.016 | -0.012 | -0.012 | 0 | 0.16 | 0.16 | 0.22 | 0.37 | 0.29 | 0.38 |
| 7 | 129983735 | rs11764240 | 0.922 | -1.127 | -0.326 | -1.257 | 0.006 | 0.004 | 0.004 | 0 | 0.12 | 0.06 | 0.09 | 0.09 | 0.11 | 0.08 |
| 7 | 129984574 | rs1532044 | 0.112 | -1.054 | 0.041 | -1.237 | 0 | 0.006 | 0.006 | 0 | 0.62 | 0.64 | 0.65 | 0.53 | 0.59 | 0.55 |
| 7 | 129984686 | rs1532046 | 0.084 | -1.055 | 0.048 | -1.238 | -0.001 | 0.001 | 0.001 | 0 | 0.62 | 0.65 | 0.65 | 0.53 | 0.59 | 0.55 |
| 7 | 129984990 | rs10226437 | 0.46 | -1.089 | 0.052 | -1.236 | -0.002 | 0 | 0 | 0 | 0.54 | 0.61 | 0.58 | 0.52 | 0.49 | 0.52 |
| 7 | 129986961 | rs10234851 | -0.254 | -0.62 | 0.16 | -0.956 | 0.008 | 0.032 | 0.032 | 0 | 0.78 | 0.73 | 0.72 | 0.58 | 0.68 | 0.61 |
| 7 | 129987100 | rs4731681 | 0.758 | -0.411 | 0.197 | -1.205 | -0.007 | -0.081 | -0.081 | 0 | 0.2 | 0.28 | 0.09 | 0.02 | 0.16 | 0.12 |
| 7 | 129989878 | rs7808823 | 0.499 | -0.24 | 0.254 | -1.164 | -0.002 | -0.002 | -0.002 | 0 | 0.58 | 0.45 | 0.63 | 0.57 | 0.52 | 0.49 |
| 7 | 129990103 | rs10247573 | 0.552 | -0.222 | 0.244 | -1.166 | -0.001 | -0.006 | -0.006 | 0 | 0.57 | 0.42 | 0.64 | 0.55 | 0.57 | 0.5 |
| 7 | 129990192 | rs10247483 | -0.189 | -0.2 | 0.248 | -1.151 | 0.002 | 0.01 | 0.01 | 0 | 0.76 | 0.72 | 0.72 | 0.56 | 0.66 | 0.61 |
| 7 | 129990763 | rs4560737 | -0.198 | -0.2 | 0.246 | -1.057 | 0.002 | 0.008 | 0.008 | 0 | 0.76 | 0.73 | 0.73 | 0.61 | 0.66 | 0.62 |
| 7 | 129991322 | rs888397 | 0.874 | -0.214 | -0.14 | -1.232 | -0.021 | -0.116 | -0.116 | 0 | 0.19 | 0.29 | 0.09 | 0.01 | 0.14 | 0.12 |
| 7 | 129992509 | rs11760909 | 0.617 | -0.097 | -0.288 | -1.215 | -0.004 | -0.003 | -0.003 | 1.077 | 0.57 | 0.42 | 0.6 | 0.53 | 0.53 | 0.5 |
| 7 | 130000319 | rs17164837 | 0.799 | -0.105 | 0.01 | -1.354 | -0.006 | -0.064 | -0.064 | 0 | 0.19 | 0.25 | 0.07 | 0 | 0.13 | 0.11 |
| 7 | 130002317 | rs1544705 | -0.167 | -0.12 | 0.043 | -1.337 | -0.008 | -0.083 | -0.083 | 0.243 | 0.77 | 0.37 | 0.88 | 0.98 | 0.83 | 0.88 |
| 7 | 130003487 | rs17164851 | 0.602 | -0.019 | -0.195 | -1.347 | 0.001 | -0.034 | -0.034 | 0.055 | 0.21 | 0.2 | 0.08 | 0 | 0.16 | 0.13 |
| 7 | 130007381 | rs11761888 | -0.024 | -0.091 | -0.081 | -1.373 | 0 | -0.115 | -0.115 | 4.457 | 0.24 | 0.43 | 0.12 | 0.03 | 0.2 | 0.14 |
| 7 | 130008065 | rs11772922 | -0.04 | -0.091 | -0.081 | -1.383 | 0 | -0.143 | -0.143 | 0.274 | 0.24 | 0.42 | 0.12 | 0 | 0.2 | 0.14 |
| 7 | 130013024 | rs10954268 | 0.439 | 0.323 | -0.616 | -1.102 | 0.029 | 0.03 | 0.03 | 0 | 0.47 | 0.3 | 0.54 | 0.42 | 0.4 | 0.28 |
| 7 | 130013638 | rs1465370 | 0.379 | 0.741 | -0.504 | -0.739 | -0.015 | -0.009 | -0.009 | 0 | 0.42 | 0.47 | 0.53 | 0.41 | 0.36 | 0.3 |
| 7 | 130017309 | rs5011386 | -0.297 | 0.99 | -0.495 | -0.608 | -0.002 | -0.001 | -0.001 | 0 | 0.54 | 0.57 | 0.62 | 0.62 | 0.51 | 0.49 |
| 7 | 130017940 | rs4731690 | -0.349 | 1.039 | -0.497 | -0.61 | -0.001 | -0.001 | -0.001 | 0.009 | 0.55 | 0.57 | 0.62 | 0.62 | 0.54 | 0.5 |
| 7 | 130018863 | rs10954269 | 1.052 | 0.634 | -0.697 | -0.652 | -0.029 | -0.027 | -0.027 | 1.559 | 0.08 | 0.03 | 0.07 | 0.18 | 0.11 | 0.19 |
| 7 | 130019491 | rs13226219 | 1.11 | 0.534 | -0.689 | -0.643 | -0.033 | -0.027 | -0.027 | 0 | 0.08 | 0.03 | 0.07 | 0.18 | 0.12 | 0.19 |
| 7 | 130021331 | rs2239604 | 0.274 | 0.384 | -0.651 | -0.813 | -0.013 | -0.008 | -0.008 | 0 | 0.43 | 0.49 | 0.45 | 0.41 | 0.37 | 0.29 |
| 7 | 130022041 | rs968404 | 0.553 | 0.343 | -0.48 | -1.19 | 0.011 | 0.038 | 0.038 | 0 | 0.2 | 0.13 | 0.07 | 0.01 | 0.14 | 0.12 |
| 7 | 130023539 | rs12706927 | 1.04 | 0.398 | -0.479 | -0.96 | 0.006 | 0.043 | 0.043 | 0 | 0.16 | 0.1 | 0.06 | 0.01 | 0.11 | 0.1 |
| 7 | 130033630 | rs1990790 | -0.441 | 0.778 | -1.161 | -0.529 | -0.005 | 0 | 0 | 0 | 0.57 | 0.65 | 0.51 | 0.51 | 0.54 | 0.52 |
| 7 | 130034695 | rs7793239 | 0.007 | 0.402 | -1.26 | -0.797 | 0.006 | 0.009 | 0.009 | 0 | 0.3 | 0.29 | 0.26 | 0.19 | 0.22 | 0.2 |
| 7 | 130036801 | rs17133175 | 0.826 | 0.633 | -1.215 | -0.88 | 0.008 | 0.038 | 0.038 | 0 | 0.24 | 0.14 | 0.11 | 0.05 | 0.19 | 0.21 |
| 7 | 130047408 | rs11972262 | -0.064 | 1.196 | -1.1 | -0.386 | -0.013 | -0.007 | -0.007 | 4.416 | 0.15 | 0.13 | 0.15 | 0.18 | 0.18 | 0.2 |
| 7 | 130052404 | rs11973495 | -0.093 | 1.001 | -1.069 | -0.384 | 0.003 | 0 | 0 | 0 | 0.16 | 0.22 | 0.17 | 0.2 | 0.21 | 0.2 |
| 7 | 130065429 | rs10260920 | 0.026 | 0.764 | -1.007 | -0.015 | 0.002 | -0.005 | -0.005 | 0 | 0.6 | 0.58 | 0.49 | 0.4 | 0.52 | 0.51 |

| | | | | | | | | | | | | | | | | |
|---|---|---|---|---|---|---|---|---|---|---|---|---|---|---|---|---|
| 7 | 130079834 | rs17164973 | -0.093 | 0.526 | -1.409 | -0.253 | -0.025 | 0.031 | 0.031 | 0 | 0.37 | 0.21 | 0.54 | 0.66 | 0.39 | 0.58 |
| 7 | 130080812 | rs2287371 | -0.054 | 0.431 | -1.068 | -0.286 | -0.013 | 0.016 | 0.016 | 0 | 0.37 | 0.29 | 0.54 | 0.67 | 0.39 | 0.58 |
| 7 | 130101373 | rs17164981 | 0.457 | -0.905 | -1.383 | -0.804 | 0.076 | 0.155 | 0.155 | 0.086 | 0.35 | 0.09 | 0.12 | 0.11 | 0.24 | 0.24 |
| 7 | 130113070 | rs13234660 | 0.526 | -1.559 | -1.309 | -1.344 | 0.012 | 0.011 | 0.011 | 0.202 | 0.67 | 0.28 | 0.69 | 0.66 | 0.74 | 0.75 |
| 7 | 130117211 | rs17164992 | 0.241 | -1.962 | -1.602 | -1.848 | 0.002 | -0.025 | -0.025 | 0.065 | 0.16 | 0.15 | 0.07 | 0 | 0.11 | 0.12 |
| 7 | 130118002 | rs12672192 | 0.876 | -2.152 | -1.419 | -1.898 | 0.041 | 0.083 | 0.083 | 0 | 0.58 | 0.26 | 0.66 | 0.77 | 0.67 | 0.65 |
| 7 | 130118545 | rs12706936 | 0.26 | -2.129 | -1.386 | -1.881 | 0.001 | 0 | 0 | 0 | 0.67 | 0.33 | 0.7 | 0.77 | 0.75 | 0.74 |
| 7 | 130123625 | rs1421140 | 1.046 | -1.822 | -1.378 | -1.695 | 0.037 | 0.219 | 0.219 | 0 | 0.5 | 0.06 | 0.62 | 0.77 | 0.63 | 0.62 |
| 7 | 130140766 | rs765205 | 1.377 | -1.769 | -1.576 | -1.244 | -0.059 | -0.03 | -0.03 | 0 | 0.33 | 0.16 | 0.54 | 0.61 | 0.55 | 0.51 |
| 7 | 130145589 | rs10863 | -0.462 | -1.978 | -0.545 | -1.344 | -0.007 | 0.003 | 0.003 | 0.412 | 0.72 | 0.73 | 0.76 | 0.72 | 0.85 | 0.75 |
| 7 | 130148817 | rs1047456 | 0.265 | -2.07 | -0.384 | -1.435 | 0.011 | 0.011 | 0.011 | 0 | 0.34 | 0.32 | 0.16 | 0.11 | 0.27 | 0.24 |
| 7 | 130148903 | rs12706941 | 1.289 | -1.987 | -0.631 | -1.387 | -0.083 | -0.046 | -0.046 | 0 | 0.35 | 0.12 | 0.56 | 0.61 | 0.57 | 0.51 |
| 7 | 130151694 | rs10128 | 1.194 | -1.92 | -0.295 | -1.299 | 0.024 | -0.017 | -0.017 | 0 | 0.2 | 0.08 | 0.2 | 0.21 | 0.1 | 0.16 |
| 7 | 130313744 | rs6467310 | -0.564 | -1.9 | 0.234 | -1.298 | 0.012 | 0.007 | 0.007 | 0.035 | 0.92 | 0.83 | 0.96 | 0.94 | 0.95 | 0.9 |
| 7 | 130329325 | rs17562397 | 1.724 | -2.001 | 0.047 | -1.378 | -0.135 | -0.09 | -0.09 | 0 | 0.31 | 0.06 | 0.52 | 0.61 | 0.54 | 0.5 |
| 7 | 130330493 | rs1814 | 0.785 | -2.024 | 0.161 | -1.368 | 0.021 | 0.055 | 0.055 | 0 | 0.27 | 0.13 | 0.13 | 0.12 | 0.24 | 0.21 |
| 7 | 130344936 | rs3847101 | -0.104 | -2.053 | 0.237 | -1.162 | -0.004 | 0.001 | 0.001 | 0.176 | 0.7 | 0.72 | 0.75 | 0.7 | 0.85 | 0.74 |
| 7 | 130349166 | rs10263009 | -0.116 | -2.035 | 0.245 | -1.257 | -0.004 | 0 | 0 | 0.241 | 0.7 | 0.71 | 0.75 | 0.72 | 0.85 | 0.74 |
| 7 | 130351884 | rs3804620 | 0.571 | -1.747 | -0.025 | -1.221 | 0.021 | 0.054 | 0.054 | 0 | 0.27 | 0.13 | 0.13 | 0.12 | 0.24 | 0.21 |
| 7 | 130369644 | rs4731699 | 0.231 | -1.548 | -0.119 | -1.273 | -0.034 | 0.108 | 0.108 | 0 | 0.33 | 0.06 | 0.19 | 0.2 | 0.3 | 0.35 |
| 7 | 130370267 | rs17165066 | -0.35 | -1.75 | 0.481 | -1.404 | -0.024 | -0.074 | -0.074 | 0 | 0.12 | 0.32 | 0.08 | 0 | 0.07 | 0.05 |
| 7 | 130379944 | rs12539726 | 0.315 | -1.769 | 0.189 | -1.4 | -0.034 | 0.113 | 0.113 | 0 | 0.33 | 0.06 | 0.19 | 0.2 | 0.3 | 0.35 |
| 7 | 130388801 | rs17165072 | -0.19 | -1.642 | 0.411 | -1.353 | -0.019 | -0.055 | -0.055 | 0.046 | 0.11 | 0.17 | 0.06 | 0 | 0.06 | 0.04 |
| 7 | 130391504 | rs12154966 | 1.231 | -1.926 | 0.454 | -1.343 | -0.029 | 0.008 | 0.008 | 0 | 0.36 | 0.28 | 0.62 | 0.63 | 0.57 | 0.5 |
| 7 | 130399691 | rs4728198 | 0.331 | -1.996 | 0.589 | -1.331 | -0.003 | 0 | 0 | 0 | 0.71 | 0.39 | 0.82 | 0.82 | 0.87 | 0.86 |
| 7 | 130400698 | rs3847104 | -0.197 | -1.973 | 0.826 | -1.321 | -0.045 | -0.018 | -0.018 | 0 | 0.75 | 0.62 | 0.86 | 0.83 | 0.88 | 0.87 |
| 7 | 130404776 | rs4067228 | 0.795 | -1.611 | 0.775 | -0.939 | -0.004 | -0.002 | -0.002 | 0.004 | 0.41 | 0.58 | 0.65 | 0.53 | 0.56 | 0.5 |
| 7 | 130406287 | rs7804963 | 0.25 | -1.373 | 1.024 | -1.003 | -0.041 | -0.003 | -0.003 | 0.425 | 0.25 | 0.48 | 0.26 | 0.19 | 0.14 | 0.17 |
| 7 | 130407589 | rs10266864 | -0.104 | -1.47 | 1.13 | -0.992 | -0.002 | -0.001 | -0.001 | 0 | 0.27 | 0.67 | 0.28 | 0.19 | 0.14 | 0.17 |
| 7 | 130408380 | rs12665998 | 0.339 | -1.348 | 1.015 | -0.91 | 0.002 | 0.004 | 0.004 | 0 | 0.54 | 0.42 | 0.72 | 0.71 | 0.62 | 0.6 |
| 7 | 130408959 | rs13238028 | 1.069 | -1.294 | 0.942 | -0.877 | 0.012 | 0.009 | 0.009 | 0.022 | 0.37 | 0.4 | 0.52 | 0.51 | 0.51 | 0.42 |
| 7 | 130410981 | rs13232691 | 0.172 | -1.177 | 0.761 | -0.728 | -0.002 | -0.011 | -0.011 | 0 | 0.61 | 0.54 | 0.74 | 0.74 | 0.66 | 0.63 |
| 7 | 130413311 | rs3909552 | 0.158 | -1.123 | 0.785 | -0.703 | -0.002 | -0.012 | -0.012 | 0 | 0.61 | 0.54 | 0.74 | 0.74 | 0.65 | 0.63 |
| 7 | 130415672 | rs290796 | -0.464 | -1.265 | 0.562 | -0.174 | 0.001 | -0.002 | -0.002 | 3.445 | 0.79 | 0.78 | 0.78 | 0.79 | 0.9 | 0.82 |
| 7 | 130416394 | rs290794 | -0.788 | -0.586 | 0.56 | -0.199 | -0.01 | -0.013 | -0.013 | 0 | 0.25 | 0.39 | 0.2 | 0.26 | 0.2 | 0.2 |
| 7 | 130437476 | rs7810507 | 0.217 | -0.217 | 0.012 | 0.513 | -0.01 | -0.027 | -0.027 | 0 | 0.22 | 0.17 | 0.3 | 0.39 | 0.28 | 0.36 |
| 7 | 130438005 | rs287624 | -0.367 | -0.4 | 0.087 | 0.284 | 0.005 | 0.004 | 0.004 | 4.525 | 0.37 | 0.36 | 0.3 | 0.28 | 0.24 | 0.25 |
| 7 | 130438214 | rs13234407 | 1.379 | -0.292 | -0.449 | 0.307 | -0.027 | 0.022 | 0.022 | 0.038 | 0.38 | 0.24 | 0.38 | 0.31 | 0.45 | 0.38 |
| 7 | 130445981 | rs1364422 | 0.646 | -0.001 | -1.369 | 0.34 | -0.006 | -0.015 | -0.015 | 0.233 | 0.22 | 0.18 | 0.3 | 0.41 | 0.28 | 0.36 |
| 7 | 130453382 | rs1596972 | -0.422 | 0.014 | -1.994 | -0.25 | 0.014 | 0.077 | 0.077 | 0 | 0.52 | 0.79 | 0.58 | 0.64 | 0.47 | 0.56 |
| 7 | 130458874 | rs7795357 | -0.185 | 0.135 | -1.953 | 0.086 | -0.005 | -0.012 | -0.012 | 0.041 | 0.13 | 0.15 | 0.05 | 0.01 | 0.06 | 0.08 |
| 7 | 130466854 | rs972283 | -0.398 | 1.173 | -2.055 | 0.554 | -0.087 | 0.01 | 0.01 | 0.198 | 0.64 | 0.88 | 0.64 | 0.68 | 0.56 | 0.63 |
| 7 | 130469097 | rs4731706 | -1.172 | 1.193 | -1.457 | 0.748 | -0.027 | -0.03 | -0.03 | 0 | 0.09 | 0.03 | 0.2 | 0.18 | 0.09 | 0.09 |
| 7 | 130476833 | rs10954285 | -0.779 | 1.016 | -1.095 | 0.77 | 0.001 | 0.013 | 0.013 | 0.473 | 0.22 | 0.27 | 0.24 | 0.35 | 0.27 | 0.3 |
| 7 | 130480244 | rs12673358 | -0.758 | 1.019 | -1.091 | 0.771 | 0.001 | 0.013 | 0.013 | 0.608 | 0.22 | 0.25 | 0.23 | 0.35 | 0.27 | 0.3 |
| 7 | 130489853 | rs6979302 | 0.028 | 0.554 | -1.869 | -0.193 | 0.018 | 0.122 | 0.122 | 0.115 | 0.25 | 0.07 | 0.1 | 0.03 | 0.18 | 0.19 |
| 7 | 130492122 | rs11763341 | -0.579 | 0.665 | -1.298 | -0.001 | 0.008 | 0.012 | 0.012 | 0.83 | 0.13 | 0.2 | 0.09 | 0.18 | 0.17 | 0.15 |
| 7 | 130493672 | rs6954253 | -0.919 | 0.534 | -1.174 | -0.06 | 0.005 | 0.01 | 0.01 | 1.989 | 0.14 | 0.27 | 0.1 | 0.18 | 0.17 | 0.16 |
| 7 | 130505073 | rs4269476 | -0.262 | 0.816 | -2.07 | 0.211 | 0 | -0.004 | -0.004 | 0 | 0.68 | 0.32 | 0.65 | 0.6 | 0.72 | 0.67 |
| 7 | 130513816 | rs11974458 | 0.345 | 0.213 | -2.455 | 0.034 | 0.021 | -0.028 | -0.028 | 0 | 0.24 | 0.12 | 0.27 | 0.29 | 0.21 | 0.19 |
| 7 | 130523468 | rs207204 | 0.584 | 0.987 | -2.968 | -0.091 | -0.014 | -0.002 | -0.002 | 0.014 | 0.52 | 0.25 | 0.39 | 0.54 | 0.5 | 0.57 |
| 7 | 130535477 | rs207210 | 0.071 | 1.051 | -2.076 | 0.252 | 0.005 | 0.007 | 0.007 | 0.652 | 0.3 | 0.25 | 0.45 | 0.45 | 0.4 | 0.44 |
| 7 | 130536392 | rs11982376 | 0.009 | 1.04 | -1.764 | 0.158 | 0.01 | -0.021 | -0.021 | 0 | 0.05 | 0.09 | 0.09 | 0 | 0.13 | 0.02 |
| 7 | 130536491 | rs4728200 | 1.106 | 0.881 | -1.488 | 0.363 | -0.001 | 0.009 | 0.009 | 0 | 0.37 | 0.25 | 0.24 | 0.34 | 0.37 | 0.28 |
| 7 | 130550812 | rs4731714 | -0.197 | 0.523 | -0.855 | 0.752 | 0.032 | -0.01 | -0.01 | 0 | 0.24 | 0.3 | 0.38 | 0.18 | 0.37 | 0.22 |
| 7 | 130552007 | rs11766402 | 1.092 | 0.196 | -0.994 | 0.708 | -0.005 | -0.004 | -0.004 | 0 | 0.12 | 0.05 | 0.07 | 0.16 | 0.18 | 0.14 |
| 7 | 130553262 | rs6949271 | -0.813 | -0.012 | -0.652 | 0.515 | 0.05 | -0.027 | -0.027 | 0 | 0.3 | 0.45 | 0.44 | 0.24 | 0.42 | 0.22 |
| 7 | 130558179 | rs7349991 | 0.609 | 0.182 | -0.65 | 0.638 | 0.047 | 0.014 | 0.014 | 0.585 | 0.18 | 0.28 | 0.33 | 0.19 | 0.3 | 0.17 |
| 7 | 130562824 | rs7781163 | 0.527 | 0.023 | -0.595 | 0.694 | 0.005 | 0.002 | 0.002 | 0.799 | 0.25 | 0.25 | 0.23 | 0.21 | 0.13 | 0.25 |
| 7 | 130566594 | rs2293746 | 0.974 | -0.043 | -0.543 | 0.713 | 0.03 | 0.013 | 0.013 | 0 | 0.17 | 0.25 | 0.28 | 0.18 | 0.21 | 0.17 |
| 7 | 130566755 | rs731730 | 0.769 | -0.088 | -0.597 | 0.902 | -0.014 | -0.001 | -0.001 | 0 | 0.21 | 0.28 | 0.21 | 0.22 | 0.1 | 0.19 |
| 7 | 130566795 | rs731729 | 0.769 | -0.088 | -0.597 | 0.901 | -0.014 | 0 | 0 | 1.06 | 0.21 | 0.28 | 0.21 | 0.22 | 0.1 | 0.19 |
| 7 | 130576178 | rs157920 | 0.118 | -0.177 | -0.967 | 0.853 | -0.092 | -0.065 | -0.065 | 0 | 0.41 | 0.07 | 0.33 | 0.55 | 0.5 | 0.49 |
| 7 | 130576263 | rs157921 | -0.252 | -0.31 | -0.593 | 0.856 | -0.031 | -0.031 | -0.031 | 0 | 0.71 | 0.44 | 0.66 | 0.75 | 0.76 | 0.78 |
| 7 | 130579162 | rs7781295 | 1.169 | 0.013 | -1.994 | 0.956 | 0.026 | -0.008 | -0.008 | 0 | 0.2 | 0.11 | 0.23 | 0.2 | 0.12 | 0.2 |
| 7 | 130580165 | rs157925 | -0.14 | 0.077 | -1.994 | 1.055 | -0.054 | -0.03 | -0.03 | 0 | 0.43 | 0.09 | 0.37 | 0.54 | 0.52 | 0.5 |
| 7 | 130580343 | rs205755 | 0.589 | 0.37 | -1.787 | 1.179 | 0.022 | -0.009 | -0.009 | 0.015 | 0.61 | 0.8 | 0.67 | 0.61 | 0.66 | 0.62 |
| 7 | 130581076 | rs205756 | -0.474 | 0.355 | -1.796 | 1.338 | -0.033 | -0.028 | -0.028 | 0 | 0.31 | 0.15 | 0.25 | 0.37 | 0.37 | 0.34 |
| 7 | 130582204 | rs4731717 | 0.188 | 0.266 | -1.366 | 0.995 | -0.056 | -0.036 | -0.036 | 0.004 | 0.11 | 0.01 | 0.05 | 0.12 | 0.17 | 0.13 |

| | | | | | | | | | | | | | | | | |
|---|---|---|---|---|---|---|---|---|---|---|---|---|---|---|---|---|
| 7 | 130582724 | rs157931 | 0.117 | 0.085 | -1.158 | 0.797 | -0.004 | -0.052 | -0.052 | 0 | 0.3 | 0.15 | 0.29 | 0.42 | 0.33 | 0.34 |
| 7 | 130583944 | rs3750176 | 0.35 | -0.011 | -0.774 | 0.858 | 0 | -0.006 | -0.006 | 0 | 0.41 | 0.54 | 0.4 | 0.33 | 0.37 | 0.32 |
| 7 | 130585553 | rs157935 | -0.437 | -0.006 | -0.689 | 1.262 | 0.001 | -0.029 | -0.029 | 1.615 | 0.3 | 0.2 | 0.31 | 0.44 | 0.33 | 0.36 |
| 7 | 130586612 | rs157939 | 1.394 | 0.109 | -0.03 | 1.239 | -0.01 | 0.143 | 0.143 | 0 | 0.92 | 0.76 | 0.9 | 0.73 | 0.94 | 0.88 |
| 7 | 130589791 | rs205761 | 0.269 | 0.007 | -0.697 | 0.063 | 0.027 | 0.038 | 0.038 | 0 | 0.48 | 0.22 | 0.43 | 0.37 | 0.57 | 0.63 |
| 7 | 130593685 | rs285522 | 1.097 | 0.523 | 0.404 | -0.149 | 0.098 | -0.042 | -0.042 | 0.018 | 0.9 | 0.51 | 0.86 | 1 | 0.88 | 0.97 |
| 7 | 130595047 | rs205763 | 1.053 | 0.41 | 0.273 | -0.18 | 0.007 | -0.052 | -0.052 | 0 | 0.81 | 0.43 | 0.83 | 0.88 | 0.81 | 0.75 |
| 7 | 130598236 | rs205764 | 1.483 | 0.41 | -0.078 | -0.263 | -0.001 | -0.01 | -0.01 | 0 | 0.72 | 0.31 | 0.8 | 0.85 | 0.75 | 0.71 |
| 7 | 130598454 | rs547311 | -1.129 | 0.63 | 0.166 | -0.065 | 0.007 | -0.047 | -0.047 | 0 | 0.2 | 0.61 | 0.19 | 0.14 | 0.2 | 0.23 |
| 7 | 130616236 | rs205716 | 0.711 | 0.362 | 0.465 | -0.416 | -0.014 | -0.13 | -0.13 | 0 | 0.81 | 0.62 | 0.88 | 1 | 0.84 | 0.88 |
| 7 | 130620191 | rs157905 | -0.573 | 0.308 | 0.744 | -0.459 | -0.008 | -0.044 | -0.044 | 0 | 0.13 | 0.16 | 0.09 | 0 | 0.09 | 0.07 |
| 7 | 130621451 | rs205723 | 0.995 | 0.711 | 0.271 | -0.362 | 0.008 | 0.012 | 0.012 | 0 | 0.5 | 0.5 | 0.5 | 0.34 | 0.37 | 0.34 |
| 7 | 130622564 | rs17165295 | 0.043 | 1.448 | -0.094 | 0.338 | -0.016 | -0.039 | -0.039 | 0 | 0.19 | 0.06 | 0.16 | 0.32 | 0.21 | 0.26 |
| 7 | 130623437 | rs205725 | -0.58 | 1.431 | -0.052 | 0.304 | 0.008 | 0.039 | 0.039 | 0.837 | 0.87 | 0.95 | 0.92 | 1 | 0.9 | 0.93 |
| 7 | 130627014 | rs12533596 | -0.688 | 1.725 | 0.174 | 0.383 | -0.005 | 0.012 | 0.012 | 0 | 0.23 | 0.3 | 0.18 | 0.31 | 0.23 | 0.27 |
| 7 | 130629493 | rs2293748 | -1.591 | 1.748 | 0.44 | 0.663 | 0.003 | 0.002 | 0.002 | 0 | 0.39 | 0.37 | 0.43 | 0.48 | 0.56 | 0.6 |
| 7 | 130629901 | rs3823556 | -1.543 | 1.732 | 0.433 | 0.89 | 0.006 | 0.001 | 0.001 | 0 | 0.39 | 0.38 | 0.42 | 0.58 | 0.54 | 0.61 |
| 7 | 130641577 | rs205774 | 1.266 | 1.881 | 0.338 | 0.885 | -0.022 | -0.001 | -0.001 | 0 | 0.69 | 0.75 | 0.64 | 0.72 | 0.54 | 0.59 |
| 7 | 130651215 | rs205781 | 1.355 | 1.34 | -0.281 | 0.801 | -0.02 | -0.001 | -0.001 | 0 | 0.52 | 0.1 | 0.43 | 0.44 | 0.56 | 0.43 |
| 7 | 130653527 | rs7798603 | -0.663 | 1.184 | -0.096 | 0.634 | -0.004 | 0.035 | 0.035 | 0.311 | 0.27 | 0.61 | 0.42 | 0.45 | 0.26 | 0.41 |
| 7 | 130653616 | rs10228040 | -1.076 | 1.053 | -0.2 | 0.652 | -0.003 | -0.035 | -0.035 | 0.008 | 0.33 | 0.75 | 0.45 | 0.45 | 0.33 | 0.46 |
| 7 | 130655734 | rs205730 | 0.368 | 1.014 | 0.07 | 0.561 | -0.021 | -0.043 | -0.043 | 0.014 | 0.75 | 0.5 | 0.65 | 0.85 | 0.83 | 0.7 |
| 7 | 130664881 | rs205743 | 1.077 | 0.849 | 0.067 | 0.558 | 0.002 | 0.006 | 0.006 | 0 | 0.53 | 0.45 | 0.45 | 0.43 | 0.55 | 0.38 |
| 7 | 130665254 | rs205744 | 1.29 | 0.806 | -0.298 | 0.538 | -0.012 | 0.016 | 0.016 | 0 | 0.51 | 0.13 | 0.43 | 0.43 | 0.54 | 0.38 |
| 7 | 130666105 | rs205750 | 1.306 | 0.81 | -0.296 | 0.532 | -0.011 | 0.017 | 0.017 | 0 | 0.51 | 0.13 | 0.43 | 0.43 | 0.54 | 0.39 |
| 7 | 130667258 | rs4141794 | -0.86 | 0.69 | 0.269 | 0.932 | -0.001 | 0 | 0 | 0.164 | 0.32 | 0.25 | 0.44 | 0.28 | 0.36 | 0.37 |
| 7 | 130667323 | rs205752 | 1.393 | 0.557 | -0.013 | 0.752 | 0.019 | 0.006 | 0.006 | 0 | 0.53 | 0.18 | 0.44 | 0.46 | 0.57 | 0.43 |
| 7 | 130668383 | rs6973318 | -0.267 | -0.049 | -0.326 | 0.033 | 0.009 | 0.007 | 0.007 | 0 | 0.67 | 0.54 | 0.51 | 0.37 | 0.61 | 0.67 |
| 7 | 130668586 | rs17688449 | -0.366 | 0.725 | -0.222 | 0.113 | 0.044 | 0.012 | 0.012 | 0 | 0.18 | 0.29 | 0.23 | 0.2 | 0.26 | 0.28 |
| 7 | 130668618 | rs6957511 | -1.355 | 0.078 | -0.849 | -0.106 | 0.011 | 0.025 | 0.025 | 0.113 | 0.5 | 0.53 | 0.47 | 0.36 | 0.57 | 0.65 |
| 7 | 130673153 | rs6467328 | -0.595 | -0.421 | -0.698 | -0.467 | -0.014 | -0.033 | -0.033 | 0 | 0.31 | 0.56 | 0.23 | 0.09 | 0.27 | 0.31 |
| 7 | 130678154 | rs6978148 | 0.931 | -0.383 | -1.018 | -0.308 | 0.005 | 0.034 | 0.034 | 0 | 0.67 | 0.75 | 0.66 | 0.77 | 0.66 | 0.55 |
| 7 | 130680521 | rs6971499 | 0.423 | -0.681 | -0.867 | -0.547 | 0.004 | 0.007 | 0.007 | 0.032 | 0.15 | 0.14 | 0.1 | 0.03 | 0.13 | 0.19 |
| 7 | 130683644 | rs6467332 | 0.176 | -0.953 | -0.952 | -0.724 | -0.017 | -0.035 | -0.035 | 0 | 0.84 | 0.77 | 0.9 | 0.97 | 0.91 | 0.92 |
| 7 | 130688225 | rs10488383 | 0.214 | -0.866 | -0.366 | -0.556 | 0.066 | 0.094 | 0.094 | 0 | 0.06 | 0.23 | 0.03 | 0.07 | 0.05 | 0.06 |
| 7 | 130688577 | rs983173 | 1.87 | -1.328 | -0.364 | -0.656 | 0.025 | 0.051 | 0.051 | 0 | 0.52 | 0.31 | 0.57 | 0.74 | 0.58 | 0.41 |
| 7 | 130691182 | rs10237038 | 0.223 | -1.765 | -0.286 | -0.189 | 0.006 | 0.02 | 0.02 | 0.086 | 0.15 | 0.15 | 0.1 | 0.04 | 0.13 | 0.21 |
| 7 | 130693823 | rs1588770 | -0.328 | -2.007 | 0.413 | 0.136 | -0.006 | 0.023 | 0.023 | 0.011 | 0.13 | 0.27 | 0.15 | 0.2 | 0.09 | 0.29 |
| 7 | 130696720 | rs1588769 | 1.795 | -2.205 | 0.458 | -0.037 | -0.044 | -0.029 | -0.029 | 0 | 0.71 | 0.45 | 0.74 | 0.78 | 0.82 | 0.62 |
| 7 | 130704375 | rs765965 | -1.122 | -2.129 | -0.083 | 0.393 | 0.007 | -0.031 | -0.031 | 0 | 0.2 | 0.11 | 0.19 | 0.29 | 0.14 | 0.4 |
| 7 | 130706995 | rs10259462 | 0.79 | -2.271 | 0.277 | 0.319 | -0.044 | 0.014 | 0.014 | 0.011 | 0.83 | 0.61 | 0.73 | 0.84 | 0.92 | 0.85 |
| 7 | 130711775 | rs17738298 | -0.805 | -1.58 | 0.107 | 0.341 | -0.007 | -0.02 | -0.02 | 0.468 | 0.16 | 0.06 | 0.19 | 0.18 | 0.12 | 0.41 |
| 7 | 130713621 | rs11980527 | -1.397 | -1.605 | 0.901 | 0.114 | 0.13 | 0.087 | 0.087 | 0 | 0.05 | 0.39 | 0.05 | 0 | 0.04 | 0.01 |
| 7 | 130715096 | rs4480053 | -1.921 | -1.622 | 0.924 | 0.106 | 0.093 | 0.053 | 0.053 | 0 | 0.06 | 0.47 | 0.05 | 0 | 0.04 | 0.01 |
| 7 | 130717723 | rs350650 | -2.001 | -1.839 | 0.661 | 0.075 | -0.006 | -0.007 | -0.007 | 0 | 0.17 | 0.79 | 0.11 | 0.05 | 0.09 | 0.09 |
| 7 | 130720805 | rs10265693 | -2.182 | -1.563 | 0.824 | 0.105 | 0.033 | -0.05 | -0.05 | 0 | 0.11 | 0.55 | 0.11 | 0.01 | 0.11 | 0.11 |
| 7 | 130727436 | rs11978185 | -1.278 | -1.729 | 0.871 | 0.199 | 0.062 | 0.015 | 0.015 | 0 | 0.07 | 0.39 | 0.06 | 0.01 | 0.06 | 0.01 |
| 7 | 130742066 | rs7803075 | 1.26 | -2.526 | -0.382 | -0.027 | 0.1 | 0.253 | 0.253 | 0 | 0.49 | 0.04 | 0.43 | 0.11 | 0.69 | 0.39 |
| 7 | 130743218 | rs2117495 | 1.323 | -2.728 | 0.935 | 0.686 | 0.002 | -0.002 | -0.002 | 0 | 0.59 | 0.37 | 0.59 | 0.49 | 0.74 | 0.69 |
| 7 | 130751832 | rs13242887 | -0.864 | -3.049 | 1.067 | 0.802 | 0.027 | -0.063 | -0.063 | 0 | 0.26 | 0.16 | 0.15 | 0.44 | 0.23 | 0.23 |
| 7 | 130754944 | rs4731743 | 1.703 | -2.932 | 1.036 | 0.762 | 0.004 | -0.002 | -0.002 | 0.154 | 0.59 | 0.37 | 0.6 | 0.49 | 0.74 | 0.69 |
| 7 | 130764475 | rs7801167 | -1.638 | -2.922 | 1.082 | 0.752 | -0.005 | 0 | 0 | 0 | 0.37 | 0.37 | 0.37 | 0.51 | 0.26 | 0.31 |
| 7 | 130766797 | rs6467345 | 0.812 | -2.2 | 1.205 | 0.554 | -0.006 | -0.031 | -0.031 | 0.012 | 0.91 | 0.78 | 0.96 | 0.97 | 0.95 | 0.92 |
| 7 | 130767116 | rs6467346 | 1.724 | -2.44 | 0.89 | 0.601 | 0.027 | -0.012 | -0.012 | 0.537 | 0.62 | 0.26 | 0.59 | 0.45 | 0.75 | 0.68 |
| 7 | 130773240 | rs17165404 | -0.783 | -2.747 | 0.229 | 0.811 | 0.016 | 0.002 | 0.002 | 0 | 0.1 | 0.06 | 0.06 | 0.45 | 0.07 | 0.15 |
| 7 | 130775266 | rs2398733 | 0.329 | -2.77 | 0.427 | 0.723 | -0.005 | -0.073 | -0.073 | 1.016 | 0.87 | 0.74 | 0.92 | 1 | 0.88 | 0.94 |
| 7 | 130783121 | rs6974804 | 1.364 | -2.975 | 0.055 | 0.639 | -0.075 | -0.055 | -0.055 | 0 | 0.69 | 0.48 | 0.63 | 0.88 | 0.85 | 0.86 |
| 7 | 130784051 | rs7812207 | -0.527 | -2.604 | -0.318 | 0.766 | 0.002 | -0.012 | -0.012 | 0 | 0.19 | 0.14 | 0.13 | 0.45 | 0.2 | 0.21 |
| 7 | 130787448 | rs7791880 | -0.829 | -2.408 | -0.296 | 0.701 | 0.007 | 0.031 | 0.031 | 1.22 | 0.13 | 0.03 | 0.07 | 0.04 | 0.1 | 0.05 |
| 7 | 130788445 | rs10954302 | 1.932 | -2.378 | -0.281 | 0.719 | -0.053 | -0.037 | -0.037 | 0 | 0.7 | 0.58 | 0.64 | 0.88 | 0.85 | 0.86 |
| 7 | 130791653 | rs3800567 | -1.033 | -1.762 | -1.106 | 1.066 | 0.015 | -0.099 | -0.099 | 0.084 | 0.16 | 0.01 | 0.09 | 0.3 | 0.11 | 0.16 |
| 7 | 130796652 | rs10253511 | -1.133 | -1.137 | -0.762 | 0.401 | -0.011 | -0.003 | -0.003 | 0 | 0.17 | 0.22 | 0.31 | 0.07 | 0.08 | 0.1 |
| 7 | 130800451 | rs12669991 | 0.096 | -0.18 | -1.132 | 0.463 | 0 | 0.025 | 0.025 | 0 | 0.51 | 0.37 | 0.52 | 0.37 | 0.51 | 0.32 |
| 7 | 130808100 | rs6467350 | 0.253 | -0.226 | -1.435 | -0.095 | -0.004 | 0.044 | 0.044 | 0.052 | 0.68 | 0.75 | 0.7 | 0.87 | 0.43 | 0.85 |
| 7 | 130810098 | rs892706 | 0.689 | -0.672 | -1.391 | -0.178 | 0.018 | 0.036 | 0.036 | 0 | 0.61 | 0.67 | 0.65 | 0.81 | 0.37 | 0.78 |
| 7 | 130810971 | rs4445168 | -0.041 | -0.899 | -1.45 | -0.171 | -0.003 | -0.001 | -0.001 | 0.297 | 0.07 | 0.07 | 0.16 | 0.06 | 0.06 | 0.14 |
| 7 | 130813097 | rs6974649 | -0.185 | -0.66 | -1.468 | -0.166 | -0.009 | 0.059 | 0.059 | 0 | 0.32 | 0.21 | 0.3 | 0.13 | 0.57 | 0.15 |
| 7 | 130819459 | rs12706962 | -0.851 | -0.31 | -1.461 | 0.399 | -0.006 | -0.013 | -0.013 | 0 | 0.38 | 0.33 | 0.64 | 0.42 | 0.59 | 0.39 |
| 7 | 130819754 | rs9690297 | -0.584 | -0.289 | -1.255 | 0.31 | 0 | -0.002 | -0.002 | 1.761 | 0.13 | 0.12 | 0.08 | 0.1 | 0.13 | 0.1 |

| | | | | | | | | | | | | | | | | |
|---|---|---|---|---|---|---|---|---|---|---|---|---|---|---|---|---|
| 7 | 130823696 | rs4731756 | -0.209 | -0.16 | -1.652 | -0.222 | -0.094 | 0.079 | 0.079 | 0.785 | 0.21 | 0.08 | 0.24 | 0.01 | 0.47 | 0.15 |
| 7 | 130824877 | rs2398758 | -1.275 | -0.636 | -1.057 | -0.426 | -0.003 | 0.008 | 0.008 | 0.83 | 0.35 | 0.31 | 0.45 | 0.08 | 0.6 | 0.37 |
| 7 | 130828889 | rs10246495 | -0.666 | -0.916 | -0.684 | -0.465 | -0.019 | -0.079 | -0.079 | 0.041 | 0.18 | 0.38 | 0.25 | 0.07 | 0.15 | 0.23 |
| 7 | 130829836 | rs11977045 | 1.391 | -2.801 | -1.827 | -0.626 | -0.013 | -0.174 | -0.174 | 0.892 | 0.24 | 0.02 | 0.36 | 0.52 | 0.2 | 0.2 |
| 7 | 130832989 | rs12533615 | 0.084 | -3.085 | -1.227 | -0.876 | -0.009 | 0.018 | 0.018 | 0.869 | 0.32 | 0.31 | 0.32 | 0.02 | 0.57 | 0.22 |
| 7 | 130833874 | rs12534322 | -0.554 | -3.279 | -1.141 | -0.952 | 0 | -0.024 | -0.024 | 0.149 | 0.39 | 0.44 | 0.49 | 0.1 | 0.65 | 0.43 |
| 7 | 130834659 | rs1439794 | 2.045 | -3.355 | -1.199 | -0.873 | -0.009 | -0.154 | -0.154 | 1.054 | 0.23 | 0.04 | 0.36 | 0.53 | 0.2 | 0.19 |
| 7 | 130834716 | rs7792497 | 1.848 | -3.753 | -1.332 | -0.997 | 0.091 | -0.033 | -0.033 | 0.001 | 0.39 | 0.07 | 0.39 | 0.69 | 0.23 | 0.47 |
| 7 | 130835547 | rs12112721 | 1.58 | -3.124 | -1.055 | -1.05 | -0.004 | -0.006 | -0.006 | 0 | 0.07 | 0.1 | 0.17 | 0.07 | 0.08 | 0.2 |
| 7 | 130838759 | rs7795830 | 2.226 | -2.938 | -1.035 | -1.029 | -0.003 | -0.025 | -0.025 | 0 | 0.08 | 0.1 | 0.04 | 0 | 0.06 | 0.03 |
| 7 | 130839735 | rs4731759 | 0.006 | -2.759 | -1.077 | -0.945 | -0.002 | -0.018 | -0.018 | 0 | 0.32 | 0.34 | 0.32 | 0.03 | 0.57 | 0.22 |
| 7 | 130848109 | rs6963416 | 0.689 | -2.707 | -1.091 | -0.846 | -0.002 | 0.011 | 0.011 | 0 | 0.25 | 0.25 | 0.39 | 0.58 | 0.23 | 0.26 |
| 7 | 130850903 | rs2580813 | -0.453 | -2.441 | -1.051 | -1.062 | -0.003 | -0.009 | -0.009 | 0 | 0.36 | 0.45 | 0.59 | 0.65 | 0.36 | 0.45 |
| 7 | 130854036 | rs16873924 | 1.822 | -2.404 | -1.196 | -0.886 | -0.06 | -0.144 | -0.144 | 0 | 0.17 | 0.01 | 0.32 | 0.49 | 0.18 | 0.17 |
| 7 | 130854175 | rs12530892 | 1.56 | -2.337 | -1.082 | -1.808 | -0.099 | 0.092 | 0.092 | 0 | 0.3 | 0.07 | 0.26 | 0.16 | 0.42 | 0.32 |
| 7 | 130854668 | rs2043924 | -0.189 | -2.216 | -0.197 | -1.691 | 0.009 | -0.012 | -0.012 | 0 | 0.93 | 0.69 | 0.95 | 0.98 | 0.96 | 0.97 |
| 7 | 130855220 | rs10259386 | 1.738 | -2.168 | -0.588 | -1.638 | -0.001 | -0.02 | -0.02 | 0.017 | 0.13 | 0.11 | 0.06 | 0 | 0.12 | 0.04 |
| 7 | 130867682 | rs326226 | 0.284 | -1.244 | -0.711 | -1.543 | 0.016 | 0.009 | 0.009 | 1.054 | 0.48 | 0.4 | 0.35 | 0.41 | 0.3 | 0.47 |
| 7 | 130868175 | rs2117496 | 1.519 | -0.995 | -0.705 | -1.418 | -0.002 | 0.001 | 0.001 | 0 | 0.07 | 0.07 | 0.18 | 0.07 | 0.1 | 0.18 |
| 7 | 130871361 | rs7798605 | 0.825 | -0.958 | -0.507 | -1.572 | -0.003 | -0.009 | -0.009 | 2.673 | 0.33 | 0.26 | 0.43 | 0.57 | 0.34 | 0.42 |
| 7 | 130872884 | rs326231 | 2.514 | -0.402 | -0.073 | -1.599 | 0.029 | -0.021 | -0.021 | 0 | 0.07 | 0.3 | 0.06 | 0 | 0.06 | 0.02 |
| 7 | 130873953 | rs326232 | -0.919 | -0.341 | -0.239 | -1.731 | 0.026 | -0.05 | -0.05 | 0 | 0.81 | 0.76 | 0.78 | 0.98 | 0.64 | 0.9 |
| 7 | 130875552 | rs326234 | 0.972 | -0.589 | -0.122 | -1.603 | 0.017 | -0.072 | -0.072 | 0.368 | 0.15 | 0.39 | 0.26 | 0.04 | 0.16 | 0.17 |
| 7 | 130886083 | rs2895224 | -0.618 | -0.84 | -0.382 | -1.644 | 0 | -0.001 | -0.001 | 0.004 | 0.58 | 0.61 | 0.59 | 0.47 | 0.44 | 0.67 |
| 7 | 130887581 | rs6467365 | -0.05 | -0.992 | -0.322 | -1.769 | -0.017 | -0.001 | -0.001 | 0 | 0.43 | 0.48 | 0.51 | 0.47 | 0.32 | 0.62 |
| 7 | 130887687 | rs6467366 | 0.374 | -1.342 | -0.419 | -1.94 | -0.015 | -0.076 | -0.076 | 0.915 | 0.24 | 0.35 | 0.27 | 0.06 | 0.21 | 0.22 |
| 7 | 130887794 | rs10250721 | 2.163 | -1.306 | -0.479 | -1.902 | 0 | -0.024 | -0.024 | 0 | 0.15 | 0.14 | 0.08 | 0 | 0.12 | 0.04 |
| 7 | 130888388 | rs17165483 | 2.095 | -1.154 | -0.491 | -1.677 | 0.023 | -0.035 | -0.035 | 0 | 0.19 | 0.13 | 0.25 | 0.42 | 0.12 | 0.4 |
| 7 | 130889128 | rs2029223 | 0.274 | -1.041 | -0.564 | -1.548 | 0.012 | 0 | 0 | 0 | 0.23 | 0.13 | 0.26 | 0.22 | 0.2 | 0.32 |
| 7 | 130890470 | rs1863011 | -0.238 | -0.652 | -0.46 | -1.304 | 0.023 | 0.012 | 0.012 | 0 | 0.51 | 0.6 | 0.43 | 0.48 | 0.4 | 0.64 |
| 7 | 130896696 | rs7777467 | 1.314 | -0.508 | -0.457 | -1.548 | 0 | -0.001 | -0.001 | 0 | 0.24 | 0.26 | 0.23 | 0.66 | 0.22 | 0.35 |
| 7 | 130897018 | rs10954306 | 1.659 | 0.342 | -0.58 | -1.768 | -0.02 | 0.037 | 0.037 | 0.5 | 0.18 | 0.08 | 0.15 | 0.02 | 0.22 | 0.07 |
| 7 | 130898325 | rs10270974 | 2.941 | 0.304 | -0.511 | -1.796 | 0.002 | -0.033 | -0.033 | 0 | 0.08 | 0.1 | 0.04 | 0 | 0.08 | 0.01 |
| 7 | 130898544 | rs10954308 | 0.26 | 0.301 | -0.273 | -1.796 | -0.001 | -0.001 | -0.001 | 0 | 0.42 | 0.46 | 0.38 | 0.68 | 0.44 | 0.42 |
| 7 | 130901516 | rs7798625 | -0.214 | 0.021 | -0.396 | -2.117 | -0.015 | -0.117 | -0.117 | 0.399 | 0.66 | 0.59 | 0.85 | 0.98 | 0.75 | 0.87 |
| 7 | 130901669 | rs10488388 | 1.567 | -0.081 | -0.723 | -1.983 | 0.017 | 0.017 | 0.017 | 0 | 0.16 | 0.09 | 0.06 | 0.02 | 0.1 | 0.09 |
| 7 | 130903178 | rs1979567 | 1.001 | 0.587 | -0.552 | -1.622 | -0.004 | -0.001 | -0.001 | 0 | 0.25 | 0.18 | 0.48 | 0.3 | 0.31 | 0.44 |
| 7 | 130903368 | rs10240502 | 0.225 | 0.185 | -0.487 | -1.354 | -0.011 | 0.022 | 0.022 | 0 | 0.42 | 0.35 | 0.62 | 0.32 | 0.53 | 0.5 |
| 7 | 130905076 | rs2129557 | -0.456 | 0.236 | -0.301 | -0.682 | -0.001 | -0.043 | -0.043 | 1.477 | 0.9 | 0.85 | 0.96 | 1 | 0.93 | 0.97 |
| 7 | 130907924 | rs11982958 | 0.64 | -0.227 | -0.362 | -0.446 | 0.013 | 0.034 | 0.034 | 0 | 0.72 | 0.62 | 0.76 | 0.56 | 0.66 | 0.59 |
| 7 | 130908112 | rs17814135 | 0.404 | 0.239 | -0.704 | -0.536 | -0.021 | -0.025 | -0.025 | 0 | 0.09 | 0.03 | 0.14 | 0.26 | 0.18 | 0.27 |
| 7 | 130913725 | rs9640862 | 0.644 | -0.528 | -0.868 | -1.104 | 0.003 | 0.009 | 0.009 | 0 | 0.46 | 0.37 | 0.52 | 0.55 | 0.51 | 0.54 |
| 7 | 130916954 | rs10954310 | 0.136 | -1.121 | -0.761 | -1.469 | 0.053 | 0.064 | 0.064 | 2.069 | 0.52 | 0.39 | 0.75 | 0.79 | 0.68 | 0.85 |
| 7 | 130917360 | rs10276927 | 2.248 | -1.229 | -0.446 | -1.581 | 0.012 | -0.051 | -0.051 | 2.113 | 0.09 | 0.16 | 0.05 | 0 | 0.09 | 0.01 |
| 7 | 130919350 | rs7800081 | -0.134 | -1.277 | -0.498 | -1.592 | -0.017 | -0.011 | -0.011 | 0.93 | 0.61 | 0.58 | 0.81 | 0.79 | 0.77 | 0.86 |
| 7 | 130922209 | rs715810 | 0.811 | -0.937 | -0.706 | -1.188 | -0.018 | 0.02 | 0.02 | 0 | 0.17 | 0.22 | 0.09 | 0.19 | 0.13 | 0.06 |
| 7 | 130923681 | rs13235516 | 0.677 | 0.052 | -1.185 | -1.274 | -0.028 | -0.068 | -0.068 | 0 | 0.24 | 0.11 | 0.47 | 0.43 | 0.25 | 0.38 |
| 7 | 130926304 | rs12671602 | 0.62 | -0.246 | -1.012 | -1.48 | -0.017 | -0.04 | -0.04 | 0 | 0.27 | 0.16 | 0.52 | 0.5 | 0.29 | 0.34 |
| 7 | 130927457 | rs1965692 | 0.471 | -0.685 | -0.606 | -2.792 | 0.004 | -0.002 | -0.002 | 0 | 0.61 | 0.35 | 0.71 | 0.93 | 0.62 | 0.8 |
| 7 | 130928101 | rs7782739 | 0.465 | 0.646 | -0.719 | -2.208 | 0.002 | -0.01 | -0.01 | 0 | 0.13 | 0.16 | 0.12 | 0.05 | 0.23 | 0.11 |
| 7 | 130928188 | rs17814449 | 0.499 | 0.664 | -0.838 | -2.2 | -0.01 | 0.006 | 0.006 | 0 | 0.13 | 0.12 | 0.11 | 0.04 | 0.23 | 0.08 |
| 7 | 130929680 | rs10249099 | 1.048 | 0.479 | -0.939 | -1.897 | -0.006 | 0.007 | 0.007 | 0 | 0.08 | 0.03 | 0.04 | 0 | 0.09 | 0.01 |
| 7 | 130932541 | rs4728213 | 0.257 | 0.262 | -0.743 | -1.955 | 0 | -0.055 | -0.055 | 0.099 | 0.56 | 0.47 | 0.74 | 0.92 | 0.59 | 0.65 |
| 7 | 130937474 | rs4731774 | -0.197 | -0.22 | -0.559 | -1.151 | 0.003 | 0.002 | 0.002 | 0 | 0.62 | 0.55 | 0.27 | 0.36 | 0.4 | 0.57 |
| 7 | 130937760 | rs17165508 | 0.36 | -1.026 | -1.024 | -1.632 | 0.02 | 0.003 | 0.003 | 0 | 0.27 | 0.2 | 0.12 | 0 | 0.17 | 0.14 |
| 7 | 130938612 | rs6978995 | 0.702 | -0.127 | -1.169 | -1.11 | -0.001 | -0.018 | -0.018 | 0.154 | 0.09 | 0.11 | 0.05 | 0 | 0.08 | 0.01 |
| 7 | 130943749 | rs11973084 | 0.161 | -0.375 | -1.015 | -1.022 | 0.011 | 0.008 | 0.008 | 0 | 0.4 | 0.31 | 0.69 | 0.64 | 0.63 | 0.34 |
| 7 | 130944529 | rs4731775 | 0.113 | -0.358 | -1.008 | -1.008 | 0.013 | 0.009 | 0.009 | 0 | 0.41 | 0.32 | 0.69 | 0.64 | 0.63 | 0.34 |
| 7 | 130948666 | rs11971101 | -0.489 | 0.19 | -0.603 | -0.699 | 0.015 | -0.043 | -0.043 | 0 | 0.08 | 0.16 | 0.07 | 0 | 0.1 | 0.05 |
| 7 | 130949902 | rs12534288 | -0.322 | 0.149 | -1.23 | -0.582 | -0.003 | -0.031 | -0.031 | 0.585 | 0.22 | 0.14 | 0.11 | 0.34 | 0.17 | 0.43 |
| 7 | 130950421 | rs4731777 | 0.654 | -0.023 | -1.772 | -0.855 | -0.07 | 0.018 | 0.018 | 0.496 | 0.41 | 0.18 | 0.7 | 0.63 | 0.56 | 0.33 |
| 7 | 130953394 | rs10236300 | -0.321 | -0.614 | -1.403 | -0.687 | 0.038 | -0.006 | -0.006 | 0.018 | 0.22 | 0.15 | 0.09 | 0.28 | 0.12 | 0.37 |
| 7 | 130957256 | rs6943595 | -0.579 | -0.402 | -0.81 | -1.065 | 0.016 | 0.043 | 0.043 | 0.177 | 0.52 | 0.62 | 0.26 | 0.28 | 0.38 | 0.59 |
| 7 | 130958828 | rs17165530 | -1.087 | -0.873 | 0.281 | -0.563 | 0 | 0.083 | 0.083 | 0 | 0.14 | 0.48 | 0.09 | 0.21 | 0.1 | 0.14 |
| 7 | 130960827 | rs4728215 | -0.801 | -1.024 | -0.015 | -0.792 | 0 | 0 | 0 | 0.003 | 0.38 | 0.62 | 0.23 | 0.38 | 0.28 | 0.4 |
| 7 | 130963771 | rs2129561 | 1.332 | -1.538 | -0.67 | -1.204 | -0.072 | 0.003 | 0.003 | 0 | 0.44 | 0.07 | 0.64 | 0.62 | 0.55 | 0.46 |
| 7 | 130963871 | rs2171356 | -0.537 | -1.28 | -0.337 | -1.117 | -0.006 | -0.024 | -0.024 | 0 | 0.91 | 0.91 | 0.94 | 0.99 | 0.92 | 0.92 |
| 7 | 130973720 | rs13237471 | 0.124 | -1.29 | 0.294 | -0.989 | 0.052 | 0.07 | 0.07 | 0 | 0.13 | 0.28 | 0.12 | 0.22 | 0.18 | 0.16 |
| 7 | 130979676 | rs2060722 | -0.39 | -1.734 | -0.146 | -1.302 | 0.052 | 0.065 | 0.065 | 0 | 0.78 | 0.91 | 0.91 | 0.97 | 0.89 | 0.78 |

| | | | | | | | | | | | | | | | | |
|---|---|---|---|---|---|---|---|---|---|---|---|---|---|---|---|---|
| 7 | 130983641 | rs6467369 | -0.418 | -1.807 | 0.146 | -1.357 | 0.015 | -0.006 | -0.006 | 0 | 0.79 | 0.86 | 0.9 | 0.99 | 0.89 | 0.86 |
| 7 | 130983818 | rs7807337 | 1.222 | -2.405 | -0.631 | -1.497 | -0.041 | -0.025 | -0.025 | 0 | 0.38 | 0.2 | 0.65 | 0.57 | 0.55 | 0.45 |
| 7 | 130985792 | rs7781339 | 0.333 | -2.225 | -0.269 | -1.543 | 0.025 | 0.041 | 0.041 | 0 | 0.14 | 0.39 | 0.1 | 0.16 | 0.15 | 0.2 |
| 7 | 130987321 | rs17165557 | 2.037 | -2.42 | -0.753 | -1.63 | 0.009 | -0.001 | -0.001 | 0 | 0.12 | 0.02 | 0.06 | 0.08 | 0.07 | 0.09 |
| 7 | 130989335 | rs4728218 | 1.627 | -2.117 | -0.842 | -1.682 | -0.042 | -0.024 | -0.024 | 0 | 0.35 | 0.2 | 0.62 | 0.58 | 0.46 | 0.43 |
| 7 | 130998106 | rs10268409 | -0.178 | -2.311 | -0.788 | -1.662 | 0.025 | 0.066 | 0.066 | 0 | 0.7 | 0.89 | 0.83 | 0.82 | 0.76 | 0.74 |
| 7 | 131010943 | rs3823483 | 1.448 | -2.3 | -0.686 | -1.388 | -0.013 | -0.007 | -0.007 | 0 | 0.36 | 0.34 | 0.48 | 0.56 | 0.48 | 0.42 |
| 7 | 131021956 | rs13238544 | 1.521 | -1.183 | -0.559 | -1.09 | -0.021 | 0.06 | 0.06 | 0 | 0.27 | 0.02 | 0.32 | 0.14 | 0.23 | 0.22 |
| 7 | 131027214 | rs17742759 | 1.373 | -1.145 | -0.512 | -1.073 | -0.01 | 0.068 | 0.068 | 0 | 0.28 | 0.04 | 0.32 | 0.14 | 0.23 | 0.22 |
| 7 | 131029996 | rs12539822 | 1.156 | -0.418 | -0.359 | -0.705 | 0.074 | 0.11 | 0.11 | 0.495 | 0.18 | 0.32 | 0.36 | 0.51 | 0.34 | 0.3 |
| 7 | 131071701 | rs3735030 | 2.236 | -0.657 | -0.242 | -0.694 | -0.004 | 0.026 | 0.026 | 0 | 0.07 | 0.1 | 0.03 | 0.15 | 0.06 | 0.1 |
| 7 | 131080047 | rs3847106 | -2.39 | -0.831 | -0.182 | -0.822 | -0.002 | -0.016 | -0.016 | 0 | 0.94 | 0.96 | 0.95 | 0.99 | 0.93 | 0.99 |
| 7 | 131100319 | rs17165599 | 2.232 | -0.753 | -0.085 | -0.736 | -0.006 | 0.039 | 0.039 | 0 | 0.07 | 0.11 | 0.04 | 0.16 | 0.06 | 0.11 |
| 7 | 131126531 | rs10259329 | 1.536 | -1.061 | 0.255 | -0.963 | -0.032 | -0.016 | -0.016 | 0 | 0.2 | 0.24 | 0.08 | 0.17 | 0.1 | 0.21 |
| 7 | 131132340 | rs13228664 | 1.466 | -0.576 | 0.288 | -0.861 | 0.046 | 0.046 | 0.046 | 0.019 | 0.17 | 0.19 | 0.39 | 0.52 | 0.41 | 0.31 |
| 7 | 131179797 | rs1043893 | 0.463 | -0.898 | 0.149 | -1.056 | 0.015 | 0.028 | 0.028 | 0 | 0.55 | 0.32 | 0.47 | 0.31 | 0.41 | 0.43 |
| 7 | 131180918 | rs15956 | -0.853 | -0.626 | 0.299 | -0.841 | 0.096 | 0.091 | 0.091 | 0.872 | 0.79 | 0.66 | 0.6 | 0.47 | 0.59 | 0.66 |
| 7 | 131183218 | rs3847112 | -1.08 | -0.656 | 0.161 | -0.811 | 0.08 | 0.097 | 0.097 | 0.414 | 0.79 | 0.66 | 0.62 | 0.48 | 0.65 | 0.66 |
| 7 | 131185003 | rs3735034 | 2.473 | -0.821 | 0.009 | -0.666 | 0.005 | -0.01 | -0.01 | 0.124 | 0.06 | 0.02 | 0.03 | 0.16 | 0.06 | 0.1 |
| 7 | 131185394 | rs8232 | 2.179 | -0.82 | 0.062 | -0.668 | -0.005 | 0.034 | 0.034 | 0 | 0.06 | 0.09 | 0.04 | 0.16 | 0.06 | 0.11 |
| 7 | 131186424 | rs1045045 | -1.17 | -1.058 | -0.068 | -0.826 | -0.032 | -0.017 | -0.017 | 0 | 0.78 | 0.74 | 0.9 | 0.82 | 0.89 | 0.78 |
| 7 | 131190696 | rs6651125 | 0.939 | -0.473 | -0.248 | -0.716 | 0 | 0.002 | 0.002 | 0 | 0.38 | 0.37 | 0.42 | 0.3 | 0.39 | 0.32 |
| 7 | 131192475 | rs3800688 | 1.5 | -0.013 | -0.435 | -0.151 | 0.028 | 0.044 | 0.044 | 0 | 0.13 | 0.15 | 0.32 | 0.45 | 0.31 | 0.2 |
| 7 | 131192500 | rs1733858 | 0.52 | -0.649 | -0.595 | -0.233 | 0.003 | 0 | 0 | 0 | 0.47 | 0.48 | 0.69 | 0.52 | 0.58 | 0.53 |
| 7 | 131193485 | rs1733859 | -0.359 | -0.853 | -0.247 | -0.938 | -0.013 | -0.102 | -0.102 | 0 | 0.75 | 0.71 | 0.84 | 0.99 | 0.81 | 0.84 |
| 7 | 131193695 | rs3212299 | 0.826 | -0.905 | -0.266 | -0.928 | -0.01 | 0.177 | 0.177 | 0.287 | 0.09 | 0.39 | 0.05 | 0.2 | 0.06 | 0.1 |
| 7 | 131193739 | rs3212298 | 2.423 | -0.875 | -0.901 | -0.9 | 0.007 | -0.011 | -0.011 | 1.071 | 0.05 | 0.02 | 0.02 | 0.2 | 0.06 | 0.1 |
| 7 | 131195712 | rs12670788 | 0.894 | -0.874 | -1.084 | -1.064 | -0.142 | -0.087 | -0.087 | 0 | 0.33 | 0.04 | 0.39 | 0.65 | 0.43 | 0.42 |
| 7 | 131195959 | rs3735035 | 1.039 | -0.872 | -1.084 | -1.062 | -0.143 | -0.088 | -0.088 | 0 | 0.33 | 0.04 | 0.39 | 0.65 | 0.44 | 0.41 |
| 7 | 131196062 | rs17744792 | 0.928 | -1.032 | -0.891 | -1.084 | -0.003 | 0.058 | 0.058 | 0.011 | 0.2 | 0.06 | 0.14 | 0.01 | 0.18 | 0.14 |
| 7 | 131198028 | rs1733866 | -0.67 | -0.908 | -0.703 | -1.064 | -0.006 | -0.126 | -0.126 | 0 | 0.78 | 0.61 | 0.86 | 1 | 0.81 | 0.83 |
| 7 | 131198175 | rs4731799 | 0.929 | -0.847 | -1.131 | -0.972 | -0.125 | -0.089 | -0.089 | 0 | 0.32 | 0.04 | 0.38 | 0.66 | 0.41 | 0.41 |
| 7 | 131200639 | rs1013368 | 0.528 | -0.6 | -0.941 | -1.252 | -0.019 | 0.094 | 0.094 | 0 | 0.39 | 0.2 | 0.45 | 0.15 | 0.41 | 0.33 |
| 7 | 131201947 | rs11768640 | 1.264 | -0.427 | -1.297 | -0.998 | -0.008 | 0.074 | 0.074 | 0 | 0.28 | 0.12 | 0.35 | 0.14 | 0.26 | 0.3 |
| 7 | 131203424 | rs10264841 | -1.127 | -0.676 | -1.108 | -0.917 | -0.019 | -0.084 | -0.084 | 0 | 0.83 | 0.73 | 0.91 | 1 | 0.89 | 0.86 |
| 7 | 131204362 | rs11763812 | 1.178 | -0.534 | -1.647 | -0.871 | 0.008 | 0.071 | 0.071 | 0 | 0.28 | 0.15 | 0.45 | 0.14 | 0.25 | 0.28 |
| 7 | 131212431 | rs1625957 | -0.808 | -0.473 | -1.02 | -0.511 | -0.012 | -0.07 | -0.07 | 0 | 0.83 | 0.78 | 0.91 | 1 | 0.88 | 0.85 |
| 7 | 131212459 | rs9649034 | 0.972 | -0.662 | -1.708 | -0.287 | 0.022 | -0.066 | -0.066 | 0.35 | 0.56 | 0.08 | 0.71 | 0.61 | 0.64 | 0.62 |
| 7 | 131218957 | rs1643256 | -0.579 | -0.769 | -0.716 | -0.719 | -0.042 | -0.076 | -0.076 | 0 | 0.85 | 0.8 | 0.95 | 1 | 0.95 | 0.86 |
| 7 | 131221194 | rs1619267 | -0.363 | 0.138 | -0.818 | -0.352 | 0.006 | 0 | 0 | 0.559 | 0.35 | 0.44 | 0.37 | 0.67 | 0.54 | 0.43 |
| 7 | 131222808 | rs10273876 | 0.09 | 0.309 | -1.128 | -0.495 | 0.011 | 0.008 | 0.008 | 0 | 0.39 | 0.47 | 0.36 | 0.55 | 0.45 | 0.32 |
| 7 | 131222935 | rs2398793 | -0.268 | 0.823 | -1.883 | -1.393 | -0.072 | 0.071 | 0.071 | 0.137 | 0.34 | 0.16 | 0.63 | 0.18 | 0.49 | 0.4 |
| 7 | 131223177 | rs6953894 | -0.13 | 0.552 | -1.465 | -0.866 | 0.008 | 0.065 | 0.065 | 0 | 0.6 | 0.72 | 0.5 | 0.83 | 0.63 | 0.69 |
| 7 | 131231032 | rs1465387 | -0.094 | 0.174 | -1.746 | -0.449 | -0.032 | 0.068 | 0.068 | 0.942 | 0.43 | 0.26 | 0.49 | 0.69 | 0.54 | 0.41 |
| 7 | 131233690 | rs2398795 | -0.229 | -0.727 | -1.472 | -0.664 | 0.029 | 0.045 | 0.045 | 0 | 0.22 | 0.12 | 0.09 | 0.07 | 0.19 | 0.22 |
| 7 | 131236868 | rs11763366 | -0.347 | -0.951 | -1.703 | -0.779 | 0.029 | 0.093 | 0.093 | 0 | 0.24 | 0.02 | 0.09 | 0.07 | 0.2 | 0.23 |
| 7 | 131239009 | rs1643257 | -0.468 | -1.186 | -0.61 | -0.323 | -0.018 | 0.002 | 0.002 | 0 | 0.34 | 0.61 | 0.49 | 0.32 | 0.26 | 0.46 |
| 7 | 131239744 | rs12667410 | 0.09 | -0.751 | -0.721 | -0.338 | -0.003 | -0.023 | -0.023 | 0 | 0.24 | 0.16 | 0.44 | 0.32 | 0.26 | 0.36 |
| 7 | 131245155 | rs1733877 | -0.733 | -1.057 | -0.603 | -0.42 | 0.016 | 0.011 | 0.011 | 0.104 | 0.52 | 0.63 | 0.6 | 0.4 | 0.46 | 0.68 |
| 7 | 131247420 | rs876340 | 0.436 | -0.868 | -0.619 | -0.643 | -0.003 | -0.035 | -0.035 | 0 | 0.14 | 0.12 | 0.07 | 0 | 0.1 | 0.09 |
| 7 | 131253456 | rs4728221 | 0.158 | -0.601 | -0.65 | 0.095 | -0.002 | -0.005 | -0.005 | 0.321 | 0.42 | 0.33 | 0.52 | 0.4 | 0.52 | 0.22 |
| 7 | 131257288 | rs17165717 | 0.126 | -0.807 | -0.66 | 0.235 | 0.006 | -0.007 | -0.007 | 0 | 0.08 | 0.05 | 0.04 | 0.16 | 0.03 | 0.26 |
| 7 | 131258203 | rs1643275 | -0.621 | -0.858 | -0.51 | 0.203 | -0.037 | 0.026 | 0.026 | 0 | 0.12 | 0.3 | 0.06 | 0.16 | 0.04 | 0.29 |
| 7 | 131266059 | rs11760907 | -0.133 | -0.356 | -1.414 | 0.041 | 0.026 | 0.07 | 0.07 | 0 | 0.5 | 0.19 | 0.55 | 0.58 | 0.54 | 0.51 |
| 7 | 131268207 | rs6961544 | -0.003 | -0.34 | -1.419 | 0.057 | 0.022 | 0.069 | 0.069 | 0 | 0.5 | 0.21 | 0.55 | 0.58 | 0.53 | 0.51 |
| 7 | 131271113 | rs1643287 | 0.856 | -0.128 | -0.912 | 0.113 | 0.005 | 0.017 | 0.017 | 0 | 0.5 | 0.47 | 0.29 | 0.24 | 0.48 | 0.47 |
| 7 | 131271820 | rs9690213 | -0.087 | 0.543 | -1.189 | 0.58 | -0.004 | -0.014 | -0.014 | 0 | 0.86 | 0.9 | 0.82 | 0.73 | 0.83 | 0.85 |
| 7 | 131281224 | rs10155893 | 0.175 | 0.21 | -1.084 | 0.326 | 0.006 | 0.013 | 0.013 | 0 | 0.5 | 0.56 | 0.5 | 0.39 | 0.52 | 0.42 |
| 7 | 131281811 | rs4336534 | -0.894 | 0.293 | -1.638 | 0.419 | 0.001 | -0.003 | -0.003 | 0.166 | 0.92 | 0.93 | 0.84 | 0.81 | 0.98 | 0.84 |
| 7 | 131287849 | rs1862083 | 0.229 | 0.464 | -1.797 | 0.277 | -0.05 | -0.042 | -0.042 | 0 | 0.39 | 0.12 | 0.32 | 0.41 | 0.46 | 0.41 |
| 7 | 131287990 | rs12534221 | 0.363 | 0.341 | -1.596 | 0.448 | -0.027 | -0.042 | -0.042 | 0 | 0.1 | 0.04 | 0.17 | 0.27 | 0.16 | 0.15 |
| 7 | 131291677 | rs7780168 | 0.746 | -0.629 | -1.185 | -0.103 | -0.014 | -0.048 | -0.048 | 0 | 0.26 | 0.24 | 0.12 | 0.08 | 0.21 | 0.21 |
| 7 | 131295022 | rs888604 | 0.001 | 0.065 | -0.725 | 0.384 | 0.001 | -0.007 | -0.007 | 0.3 | 0.5 | 0.43 | 0.4 | 0.23 | 0.4 | 0.51 |
| 7 | 131295088 | rs888605 | 0.388 | 0.206 | -1.023 | 0.323 | 0.004 | -0.012 | -0.012 | 0 | 0.39 | 0.3 | 0.18 | 0.08 | 0.31 | 0.3 |
| 7 | 131300258 | rs2909668 | 0.27 | 0.062 | -1.388 | 0.909 | -0.006 | -0.006 | -0.006 | 5.041 | 0.53 | 0.57 | 0.44 | 0.48 | 0.49 | 0.45 |
| 7 | 131301614 | rs17165751 | -0.83 | -0.418 | -1.434 | 1.085 | -0.008 | -0.038 | -0.038 | 0.06 | 0.11 | 0.01 | 0.2 | 0.16 | 0.09 | 0.21 |
| 7 | 131304940 | rs746017 | 0.692 | -1.062 | -2.133 | 1.035 | 0.072 | 0.056 | 0.056 | 0 | 0.49 | 0.04 | 0.5 | 0.29 | 0.69 | 0.47 |
| 7 | 131307658 | rs1559019 | 0.561 | -1.917 | -1.764 | 1.598 | 0.034 | -0.002 | -0.002 | 0.563 | 0.71 | 0.21 | 0.64 | 0.71 | 0.84 | 0.69 |
| 7 | 131315092 | rs929731 | 1.47 | -1.665 | -1.987 | 1.666 | 0.034 | 0.163 | 0.163 | 0.018 | 0.48 | 0.03 | 0.49 | 0.18 | 0.66 | 0.31 |

| | | | | | | | | | | | | | | | | |
|---|---|---|---|---|---|---|---|---|---|---|---|---|---|---|---|---|
| 7 | 131317195 | rs12706980 | 1.292 | -1.959 | -0.089 | 1.963 | -0.024 | 0.01 | 0.01 | 0 | 0.72 | 0.4 | 0.64 | 0.31 | 0.81 | 0.69 |
| 7 | 131317893 | rs1862084 | -1.075 | -2.031 | 0.999 | 2.005 | -0.1 | 0.113 | 0.113 | 0 | 0.23 | 0.5 | 0.12 | 0.46 | 0.09 | 0.22 |
| 7 | 131320832 | rs10272488 | 1.337 | -1.6 | 1.705 | 2.087 | 0.051 | 0.23 | 0.23 | 1.955 | 0.9 | 0.59 | 0.73 | 0.75 | 0.9 | 0.89 |
| 7 | 131327818 | rs918339 | 1.531 | -1.637 | 1.528 | 2.08 | 0.023 | 0.166 | 0.166 | 0.019 | 0.87 | 0.61 | 0.72 | 0.75 | 0.89 | 0.84 |
| 7 | 131334001 | rs2909678 | 1.326 | -2.01 | 1.081 | 2.196 | -0.057 | 0.104 | 0.104 | 0 | 0.75 | 0.47 | 0.67 | 0.38 | 0.87 | 0.66 |
| 7 | 131334612 | rs2909679 | -1.463 | -2.277 | 0.277 | 1.911 | 0.004 | -0.006 | -0.006 | 0 | 0.23 | 0.8 | 0.14 | 0.38 | 0.06 | 0.2 |
| 7 | 131336523 | rs2909680 | 1.286 | -2.017 | -0.125 | 1.854 | 0.061 | -0.044 | -0.044 | 0 | 0.67 | 0.23 | 0.56 | 0.48 | 0.86 | 0.58 |
| 7 | 131356066 | rs11978264 | -0.985 | -2.024 | 0.282 | 1.687 | -0.046 | 0.009 | 0.009 | 3.286 | 0.13 | 0.17 | 0.1 | 0.11 | 0.06 | 0.06 |
| 7 | 131356509 | rs11971513 | 1.416 | -2.386 | 1.338 | 2.692 | 0.056 | 0.427 | 0.427 | 3.68 | 0.94 | 0.47 | 0.79 | 0.63 | 0.99 | 0.98 |
| 7 | 131358281 | rs4078096 | 1.843 | -1.953 | 0.589 | 2.429 | 0.027 | -0.064 | -0.064 | 0 | 0.63 | 0.16 | 0.41 | 0.41 | 0.7 | 0.7 |
| 7 | 131360928 | rs11765576 | 2.27 | -1.128 | 0.433 | 2.284 | -0.021 | -0.05 | -0.05 | 0.045 | 0.6 | 0.14 | 0.37 | 0.39 | 0.62 | 0.62 |
| 7 | 131363167 | rs10215358 | 0.906 | -0.543 | -0.409 | 0.687 | -0.002 | -0.029 | -0.029 | 0 | 0.61 | 0.57 | 0.55 | 0.13 | 0.62 | 0.47 |
| 7 | 131365443 | rs7795093 | 1.814 | -0.451 | -0.069 | 0.668 | 0.008 | 0.001 | 0.001 | 2.62 | 0.4 | 0.26 | 0.24 | 0.01 | 0.39 | 0.24 |
| 7 | 131370039 | rs11761231 | 0.352 | -0.095 | -0.527 | 1.101 | -0.038 | -0.016 | -0.016 | 3.26 | 0.75 | 0.8 | 0.51 | 0.24 | 0.64 | 0.55 |
| 7 | 131371921 | rs4073301 | -0.314 | -0.686 | -1.143 | 0.996 | 0.027 | 0.01 | 0.01 | 4.742 | 0.17 | 0 | 0.1 | 0.07 | 0.11 | 0.12 |
| 7 | 131372051 | rs4076265 | 1.019 | -1.349 | -1.765 | 0.556 | -0.001 | 0.026 | 0.026 | 1.259 | 0.26 | 0.11 | 0.13 | 0.18 | 0.23 | 0.2 |
| 7 | 131374946 | rs11771003 | 0.707 | -1.781 | -2.472 | 0.775 | 0.051 | 0.036 | 0.036 | 0 | 0.52 | 0.04 | 0.53 | 0.33 | 0.67 | 0.64 |
| 7 | 131380008 | rs4076505 | -0.209 | -1.024 | -1.846 | 0.979 | 0.003 | -0.012 | -0.012 | 0 | 0.83 | 0.93 | 0.87 | 0.84 | 0.83 | 0.79 |
| 7 | 131381498 | rs10266232 | -0.115 | -0.88 | -0.748 | 1.259 | -0.001 | 0.114 | 0.114 | 0.011 | 0.17 | 0.35 | 0.28 | 0.4 | 0.15 | 0.11 |
| 7 | 131381906 | rs12669428 | 0.435 | -0.495 | -0.923 | 0.902 | 0.011 | -0.03 | -0.03 | 0 | 0.46 | 0.26 | 0.36 | 0.49 | 0.55 | 0.47 |
| 7 | 131383556 | rs10282190 | -1.271 | 0.297 | -0.614 | 0.642 | -0.021 | -0.053 | -0.053 | 0 | 0.42 | 0.91 | 0.43 | 0.37 | 0.36 | 0.41 |
| 7 | 131385220 | rs10231383 | 0.188 | 0.286 | -0.157 | 0.618 | -0.003 | 0.008 | 0.008 | 0.207 | 0.15 | 0.17 | 0.12 | 0.23 | 0.08 | 0.14 |
| 7 | 131387145 | rs11978012 | -1.707 | 0.367 | 0.112 | 0.603 | 0.01 | -0.021 | -0.021 | 0 | 0.43 | 0.84 | 0.44 | 0.38 | 0.36 | 0.41 |
| 7 | 131389655 | rs10251246 | -0.895 | 0.649 | 0.459 | 0.529 | 0 | -0.01 | -0.01 | 0 | 0.27 | 0.66 | 0.32 | 0.14 | 0.28 | 0.27 |
| 7 | 131397286 | rs11772080 | -0.762 | 0.699 | 0.468 | 0.551 | 0.005 | -0.034 | -0.034 | 0 | 0.26 | 0.36 | 0.29 | 0.14 | 0.28 | 0.26 |
| 7 | 131407473 | rs12540278 | -0.76 | 1.3 | 0.322 | 0.775 | -0.035 | 0 | 0 | 0 | 0.24 | 0.19 | 0.37 | 0.19 | 0.38 | 0.37 |
| 7 | 131407670 | rs2971772 | -0.572 | 0.555 | 0.64 | 1.121 | -0.008 | 0.104 | 0.104 | 0.041 | 0.15 | 0.65 | 0.13 | 0.34 | 0.13 | 0.1 |
| 7 | 131407940 | rs2971769 | -1.313 | 0.573 | 0.474 | 0.915 | -0.017 | -0.023 | -0.023 | 0.017 | 0.33 | 0.7 | 0.47 | 0.47 | 0.38 | 0.4 |
| 7 | 131412636 | rs2333890 | -1.609 | 0.404 | 1.147 | 0.848 | 0.024 | 0.413 | 0.413 | 3.292 | 0.07 | 0.46 | 0.05 | 0.29 | 0.01 | 0.04 |
| 7 | 131414457 | rs2971760 | -1.498 | 0.413 | 0.533 | 0.618 | -0.045 | -0.018 | -0.018 | 0 | 0.64 | 0.94 | 0.43 | 0.62 | 0.62 | 0.72 |
| 7 | 131429246 | rs17165866 | 0.828 | 0.54 | -0.605 | -0.034 | 0 | 0.009 | 0.009 | 0.02 | 0.1 | 0.07 | 0.08 | 0 | 0.09 | 0.09 |
| 7 | 131438872 | rs2971753 | -0.885 | 0.581 | -0.684 | -0.028 | -0.057 | 0.055 | 0.055 | 2.2 | 0.54 | 0.91 | 0.36 | 0.61 | 0.53 | 0.63 |
| 7 | 131453525 | rs1106683 | 0.364 | 1.04 | 0.133 | 0.151 | 0.038 | 0.01 | 0.01 | 0 | 0.09 | 0.17 | 0.07 | 0.09 | 0.15 | 0.12 |
| 7 | 131454000 | rs2333894 | 0.565 | 0.861 | -0.152 | 0.093 | -0.021 | -0.008 | -0.008 | 0.12 | 0.45 | 0.27 | 0.63 | 0.52 | 0.53 | 0.41 |
| 7 | 131454208 | rs888539 | 0.603 | 0.907 | -0.164 | 0.106 | 0.038 | 0.01 | 0.01 | 0 | 0.09 | 0.17 | 0.07 | 0.09 | 0.15 | 0.12 |
| 7 | 131454275 | rs1076497 | 0.728 | 0.739 | -0.111 | 0.062 | 0.013 | -0.039 | -0.039 | 0.322 | 0.31 | 0.17 | 0.16 | 0.4 | 0.27 | 0.2 |
| 7 | 131455573 | rs1477229 | 0.492 | 1.143 | -0.153 | 0.253 | 0 | -0.037 | -0.037 | 0 | 0.76 | 0.61 | 0.81 | 0.91 | 0.8 | 0.59 |
| 7 | 131459782 | rs1421361 | 0.364 | 0.865 | -0.868 | 0.413 | 0.011 | 0.003 | 0.003 | 0 | 0.52 | 0.38 | 0.7 | 0.51 | 0.62 | 0.46 |
| 7 | 131463902 | rs1073259 | -0.443 | 1.287 | -0.761 | 0.394 | -0.005 | 0.004 | 0.004 | 0 | 0.13 | 0.06 | 0.09 | 0.07 | 0.12 | 0.29 |
| 7 | 131465908 | rs11773898 | -0.64 | 1.134 | -0.548 | 0.512 | 0.014 | 0.002 | 0.002 | 0 | 0.08 | 0.1 | 0.07 | 0.09 | 0.15 | 0.12 |
| 7 | 131467857 | rs12706991 | -0.394 | 0.413 | -0.82 | 0.204 | 0.008 | 0.015 | 0.015 | 0 | 0.35 | 0.3 | 0.2 | 0.23 | 0.27 | 0.39 |
| 7 | 131468631 | rs6980306 | 0.49 | 0.469 | -1.002 | -0.079 | -0.007 | -0.013 | -0.013 | 1.479 | 0.43 | 0.28 | 0.63 | 0.47 | 0.47 | 0.28 |
| 7 | 131473147 | rs13227914 | 0.021 | -0.165 | -0.009 | -0.578 | -0.028 | -0.102 | -0.102 | 0.203 | 0.16 | 0.3 | 0.07 | 0.01 | 0.08 | 0.09 |
| 7 | 131483386 | rs10954336 | 0.545 | 0.013 | 0.673 | -0.524 | 0.008 | -0.009 | -0.009 | 0.006 | 0.6 | 0.38 | 0.78 | 0.45 | 0.71 | 0.55 |
| 7 | 131491505 | rs741535 | -0.95 | -0.123 | 0.674 | -0.729 | 0.003 | 0.011 | 0.011 | 0 | 0.49 | 0.4 | 0.32 | 0.57 | 0.5 | 0.72 |
| 7 | 131491623 | rs741536 | 1.68 | -0.722 | -0.122 | -1.386 | -0.093 | -0.135 | -0.135 | 0 | 0.31 | 0.05 | 0.56 | 0.43 | 0.37 | 0.21 |
| 7 | 131492300 | rs17165941 | 1.204 | -0.782 | -0.073 | -1.848 | 0.01 | 0.013 | 0.013 | 0 | 0.12 | 0.09 | 0.07 | 0 | 0.1 | 0.06 |
| 7 | 131496807 | rs10272905 | 0.243 | -0.728 | 0.024 | -1.457 | 0.014 | 0.007 | 0.007 | 0 | 0.33 | 0.21 | 0.19 | 0.71 | 0.26 | 0.46 |
| 7 | 131497995 | rs4728236 | -0.842 | -0.693 | 0.167 | -1.438 | -0.019 | 0.084 | 0.084 | 0 | 0.54 | 0.72 | 0.34 | 0.72 | 0.5 | 0.74 |
| 7 | 131505116 | rs1122732 | 0.428 | -0.263 | -0.068 | -1.377 | -0.045 | 0.063 | 0.063 | 0 | 0.18 | 0.07 | 0.24 | 0.01 | 0.27 | 0.33 |
| 7 | 131511877 | rs11976075 | 1.251 | -0.816 | -0.586 | -1.644 | -0.089 | -0.018 | -0.018 | 0 | 0.36 | 0.06 | 0.48 | 0.29 | 0.38 | 0.16 |
| 7 | 131517925 | rs1004592 | 0.058 | -0.817 | -0.178 | -1.633 | 0.012 | 0.014 | 0.014 | 0 | 0.39 | 0.34 | 0.23 | 0.71 | 0.34 | 0.51 |
| 7 | 131520482 | rs12706996 | 0.703 | -0.876 | -0.236 | -1.717 | 0.008 | -0.002 | -0.002 | 0 | 0.3 | 0.25 | 0.18 | 0.7 | 0.25 | 0.45 |
| 7 | 131522786 | rs10954340 | -0.347 | -0.613 | 0.044 | -2.191 | 0.002 | 0.063 | 0.063 | 0 | 0.21 | 0.13 | 0.12 | 0.03 | 0.18 | 0.08 |
| 7 | 131522963 | rs4731820 | 1.564 | -0.588 | 0.093 | -2.263 | -0.015 | 0.195 | 0.195 | 0 | 0.53 | 0.23 | 0.58 | 0.95 | 0.53 | 0.58 |
| 7 | 131523692 | rs1425087 | 0.305 | -0.353 | 0.632 | -2.111 | 0.026 | -0.007 | -0.007 | 0 | 0.93 | 0.67 | 0.96 | 1 | 0.97 | 0.98 |
| 7 | 131526745 | rs11771563 | 0.669 | -0.333 | 0.661 | -2.117 | -0.024 | 0.025 | 0.025 | 0 | 0.17 | 0.11 | 0.24 | 0.01 | 0.25 | 0.32 |
| 7 | 131526762 | rs4731821 | 0.39 | -0.636 | 1.173 | -1.589 | 0.038 | 0.003 | 0.003 | 0 | 0.94 | 0.67 | 0.96 | 1 | 0.97 | 0.98 |
| 7 | 131529852 | rs17165969 | 0.176 | -0.618 | 1.002 | -1.577 | 0.001 | -0.001 | -0.001 | 0 | 0.09 | 0.09 | 0.04 | 0 | 0.08 | 0.05 |
| 7 | 131532931 | rs3885260 | 0.257 | -0.721 | 1.194 | -1.577 | -0.024 | -0.102 | -0.102 | 0 | 0.83 | 0.55 | 0.9 | 0.97 | 0.9 | 0.96 |
| 7 | 131533816 | rs2333896 | 1.161 | -0.54 | 1.126 | -1.673 | -0.002 | -0.094 | -0.094 | 0 | 0.75 | 0.45 | 0.71 | 0.96 | 0.76 | 0.67 |
| 7 | 131539363 | rs11975170 | -0.845 | -1.062 | 1.193 | -1.631 | -0.003 | -0.051 | -0.051 | 0 | 0.21 | 0.24 | 0.25 | 0.04 | 0.21 | 0.33 |
| 7 | 131546516 | rs10954341 | -1.519 | -1.072 | 1.052 | -1.65 | -0.006 | -0.081 | -0.081 | 0.743 | 0.29 | 0.37 | 0.29 | 0.04 | 0.29 | 0.36 |
| 7 | 131555001 | rs731203 | 0.144 | -1.018 | 1 | -1.614 | 0.015 | -0.032 | -0.032 | 0.06 | 0.08 | 0.13 | 0.19 | 0.01 | 0.14 | 0.3 |
| 7 | 131555959 | rs7796957 | 0.318 | -1.197 | 1.305 | -1.458 | 0.016 | -0.051 | -0.051 | 0 | 0.9 | 0.59 | 0.95 | 1 | 0.92 | 0.97 |
| 7 | 131564200 | rs888536 | 0.86 | -1.255 | 1.277 | -1.51 | -0.058 | -0.122 | -0.122 | 0 | 0.8 | 0.53 | 0.89 | 0.97 | 0.91 | 0.95 |
| 7 | 131570695 | rs10085839 | 1.599 | -1.271 | 1.213 | -1.54 | 0.001 | -0.03 | -0.03 | 0 | 0.71 | 0.33 | 0.82 | 0.96 | 0.77 | 0.71 |
| 7 | 131571695 | rs7799455 | 1.812 | -1.184 | 0.978 | -1.506 | -0.021 | -0.108 | -0.108 | 0 | 0.68 | 0.41 | 0.8 | 0.96 | 0.74 | 0.69 |
| 7 | 131572510 | rs9641911 | -0.464 | -1.137 | 0.862 | -1.468 | 0.016 | 0.02 | 0.02 | 0 | 0.17 | 0.13 | 0.09 | 0.03 | 0.09 | 0.05 |

| | | | | | | | | | | | | | | | | |
|---|---|---|---|---|---|---|---|---|---|---|---|---|---|---|---|---|
| 7 | 131574321 | rs10488343 | -0.234 | -0.859 | 0.846 | -1.254 | 0.038 | -0.02 | -0.02 | 0.91 | 0.05 | 0.17 | 0.06 | 0.01 | 0.09 | 0.22 |
| 7 | 131590226 | rs1991190 | 0.131 | -1.01 | 0.401 | -1.106 | -0.001 | -0.015 | -0.015 | 0 | 0.08 | 0.11 | 0.04 | 0 | 0.08 | 0.03 |
| 7 | 131597409 | rs12666394 | 0.446 | -0.987 | 0.343 | -0.889 | -0.042 | -0.1 | -0.1 | 0 | 0.81 | 0.67 | 0.91 | 0.97 | 0.91 | 0.95 |
| 7 | 131597501 | rs2216982 | 0.484 | -0.982 | 0.345 | -0.887 | -0.043 | -0.101 | -0.101 | 0 | 0.8 | 0.67 | 0.91 | 0.97 | 0.91 | 0.95 |
| 7 | 131613053 | rs12540215 | 0.354 | -0.305 | 0.168 | -0.36 | -0.015 | -0.023 | -0.023 | 0 | 0.05 | 0.17 | 0.02 | 0 | 0.01 | 0.02 |
| 7 | 131613277 | rs11983666 | -0.601 | -0.62 | 0.406 | -0.512 | -0.008 | -0.014 | -0.014 | 0.436 | 0.05 | 0.21 | 0.02 | 0 | 0.02 | 0.04 |
| 7 | 131613763 | rs1582678 | -0.082 | -0.861 | -0.438 | -0.202 | 0.01 | -0.029 | -0.029 | 0 | 0.59 | 0.76 | 0.77 | 0.53 | 0.63 | 0.56 |
| 7 | 131613963 | rs1593306 | 0.307 | -0.975 | -0.778 | -0.55 | -0.007 | -0.076 | -0.076 | 3.333 | 0.87 | 0.55 | 0.93 | 1 | 0.93 | 0.94 |
| 7 | 131619847 | rs1593304 | 0.801 | -0.926 | -2.2 | -0.421 | 0.056 | 0.072 | 0.072 | 1.95 | 0.68 | 0.07 | 0.79 | 0.84 | 0.82 | 0.67 |
| 7 | 131625606 | rs4731831 | 0.766 | -0.063 | -1.918 | 0.084 | -0.149 | 0.04 | 0.04 | 0 | 0.33 | 0.04 | 0.56 | 0.25 | 0.45 | 0.19 |
| 7 | 131628225 | rs7806149 | -0.288 | -0.276 | -1.157 | 0.122 | -0.001 | 0.004 | 0.004 | 0 | 0.23 | 0.19 | 0.18 | 0.16 | 0.25 | 0.14 |
| 7 | 131628692 | rs11773206 | 0.941 | -0.829 | -1.904 | 0.598 | 0.04 | -0.014 | -0.014 | 0.157 | 0.49 | 0.08 | 0.68 | 0.67 | 0.6 | 0.68 |
| 7 | 131631982 | rs12707008 | 0.865 | -0.717 | -1.423 | 0.282 | 0.071 | 0.113 | 0.113 | 0.153 | 0.5 | 0.2 | 0.7 | 0.76 | 0.61 | 0.47 |
| 7 | 131632308 | rs7800092 | -0.331 | -0.388 | -0.855 | 1.101 | 0.025 | -0.019 | -0.019 | 0 | 0.17 | 0.12 | 0.07 | 0.16 | 0.09 | 0.15 |
| 7 | 131634654 | rs6943990 | 0.885 | 0.353 | -0.284 | 1.531 | 0.025 | 0.109 | 0.109 | 0.02 | 0.92 | 0.87 | 0.9 | 0.74 | 0.88 | 0.74 |
| 7 | 131634774 | rs10488345 | -0.259 | -0.157 | -0.187 | 0.409 | 0.015 | 0.012 | 0.012 | 0 | 0.48 | 0.45 | 0.49 | 0.87 | 0.36 | 0.74 |
| 7 | 131644243 | rs10224200 | -1.058 | 0.454 | 0.254 | 1.356 | 0.009 | 0.026 | 0.026 | 0 | 0.15 | 0.2 | 0.14 | 0.39 | 0.21 | 0.23 |
| 7 | 131647414 | rs726284 | -1.38 | 0.474 | 0.411 | 1.325 | 0.025 | 0.08 | 0.08 | 0.226 | 0.19 | 0.34 | 0.16 | 0.42 | 0.23 | 0.39 |
| 7 | 131679290 | rs1514055 | -1.009 | 0.257 | 0.554 | 1.077 | 0.014 | 0.088 | 0.088 | 0.966 | 0.07 | 0.28 | 0.04 | 0.16 | 0.08 | 0.1 |
| 7 | 131685534 | rs10954350 | 0.063 | -0.09 | -0.112 | 0.312 | 0.008 | 0.056 | 0.056 | 0 | 0.83 | 0.92 | 0.89 | 0.99 | 0.86 | 0.92 |
| 7 | 131686224 | rs10231527 | -0.15 | -0.894 | 0.047 | 0.488 | -0.018 | 0.019 | 0.019 | 0 | 0.17 | 0.25 | 0.11 | 0.18 | 0.14 | 0.28 |
| 7 | 131688552 | rs11767564 | -0.413 | -0.852 | 0 | 0.476 | 0.008 | 0.055 | 0.055 | 0 | 0.83 | 0.92 | 0.89 | 0.99 | 0.86 | 0.92 |
| 7 | 131702716 | rs1514061 | -0.69 | -0.782 | 0.145 | 1.05 | 0.005 | 0.064 | 0.064 | 0.103 | 0.09 | 0.19 | 0.19 | 0.32 | 0.14 | 0.24 |
| 7 | 131705144 | rs6943360 | -0.324 | -1.021 | 0.78 | 0.894 | -0.012 | -0.065 | -0.065 | 1.111 | 0.82 | 0.77 | 0.87 | 0.98 | 0.85 | 0.91 |
| 7 | 131708955 | rs10253104 | 1.016 | -1.59 | -0.011 | 0.193 | 0.016 | 0.035 | 0.035 | 0 | 0.21 | 0.13 | 0.13 | 0.06 | 0.15 | 0.12 |
| 7 | 131710854 | rs9641920 | 0.559 | -1.531 | 0.135 | 0.226 | -0.031 | -0.037 | -0.037 | 0 | 0.65 | 0.55 | 0.77 | 0.8 | 0.72 | 0.62 |
| 7 | 131713689 | rs6944714 | 0.881 | -1.734 | 0.152 | 0.182 | -0.001 | -0.002 | -0.002 | 0.81 | 0.51 | 0.46 | 0.54 | 0.63 | 0.55 | 0.59 |
| 7 | 131714470 | rs10282350 | 1.005 | -1.794 | 0.008 | 0.122 | 0.011 | 0.048 | 0.048 | 0 | 0.21 | 0.12 | 0.14 | 0.06 | 0.18 | 0.11 |
| 7 | 131714663 | rs10954351 | 1.913 | -1.648 | 0.071 | 0.151 | -0.037 | -0.026 | -0.026 | 0 | 0.34 | 0.21 | 0.45 | 0.45 | 0.44 | 0.28 |
| 7 | 131720635 | rs996278 | 0.054 | -1.823 | -0.177 | 0.072 | 0.005 | 0.066 | 0.066 | 0.152 | 0.63 | 0.8 | 0.63 | 0.78 | 0.64 | 0.66 |
| 7 | 131726132 | rs4731839 | 0.884 | -1.708 | 0.138 | 0.254 | -0.011 | 0.014 | 0.014 | 0.624 | 0.11 | 0.2 | 0.08 | 0.15 | 0.09 | 0.08 |
| 7 | 131730526 | rs879404 | -0.91 | -1.78 | -0.061 | 0.24 | 0.021 | -0.011 | -0.011 | 0 | 0.3 | 0.74 | 0.2 | 0.34 | 0.19 | 0.39 |
| 7 | 131730844 | rs6467419 | -0.838 | -1.655 | -0.333 | 0.249 | -0.034 | -0.023 | -0.023 | 0 | 0.84 | 0.98 | 0.77 | 0.83 | 0.82 | 0.78 |
| 7 | 131736352 | rs7349978 | 0.271 | -1.56 | 0.156 | 0.459 | -0.031 | 0.019 | 0.019 | 0 | 0.17 | 0.29 | 0.09 | 0.18 | 0.11 | 0.31 |
| 7 | 131749933 | rs6977162 | 0.884 | -1.696 | 0.321 | 0.44 | -0.038 | 0.073 | 0.073 | 0 | 0.09 | 0.22 | 0.05 | 0.18 | 0.03 | 0.27 |
| 7 | 131750195 | rs10954355 | 1.255 | -1.743 | 0.181 | 0.365 | 0.039 | -0.055 | -0.055 | 2.395 | 0.66 | 0.23 | 0.65 | 0.52 | 0.74 | 0.47 |
| 7 | 131761018 | rs4731841 | -1.361 | -1.795 | 0.523 | 0.302 | -0.024 | 0.02 | 0.02 | 0.047 | 0.34 | 0.57 | 0.33 | 0.45 | 0.26 | 0.54 |
| 7 | 131770198 | rs4039585 | 1.359 | -1.599 | 0.581 | 0.07 | 0.032 | 0.013 | 0.013 | 0.006 | 0.55 | 0.34 | 0.58 | 0.63 | 0.63 | 0.39 |
| 7 | 131770215 | rs6963716 | -0.475 | -1.599 | 0.325 | 0.07 | -0.021 | 0.028 | 0.028 | 0 | 0.23 | 0.33 | 0.25 | 0.31 | 0.18 | 0.47 |
| 7 | 131771336 | rs2175859 | 0.267 | -1.272 | -0.097 | -0.148 | 0.003 | 0.012 | 0.012 | 0 | 0.21 | 0.14 | 0.15 | 0.06 | 0.19 | 0.14 |
| 7 | 131775747 | rs1355883 | 0.798 | -0.951 | -0.223 | -0.041 | -0.001 | -0.006 | -0.006 | 1.226 | 0.72 | 0.47 | 0.74 | 0.71 | 0.71 | 0.62 |
| 7 | 131780255 | rs13224052 | 1.268 | -1.539 | -0.754 | -0.426 | 0.023 | 0.071 | 0.071 | 0.42 | 0.59 | 0.13 | 0.68 | 0.71 | 0.66 | 0.55 |
| 7 | 131783062 | rs1983285 | 0.004 | -1.518 | -0.343 | -0.37 | 0.016 | -0.024 | -0.024 | 0 | 0.12 | 0.32 | 0.08 | 0.07 | 0.12 | 0.07 |
| 7 | 131783961 | rs13245659 | 0.724 | -1.522 | -0.767 | -0.371 | 0.002 | 0.012 | 0.012 | 0 | 0.14 | 0.03 | 0.1 | 0.09 | 0.1 | 0.21 |
| 7 | 131789045 | rs3099913 | -1.526 | -1.245 | -0.913 | -0.502 | -0.019 | 0.074 | 0.074 | 0.207 | 0.42 | 0.89 | 0.34 | 0.29 | 0.39 | 0.45 |
| 7 | 131791194 | rs204291 | -1.562 | -1.35 | -2.077 | -0.212 | -0.172 | 0.029 | 0.029 | 0.399 | 0.6 | 0.97 | 0.56 | 0.75 | 0.51 | 0.67 |
| 7 | 131794686 | rs281909 | 0.619 | -1.077 | -1.11 | -0.374 | 0.051 | -0.005 | -0.005 | 0 | 0.1 | 0.29 | 0.07 | 0.06 | 0.13 | 0.04 |
| 7 | 131795474 | rs204292 | -0.414 | -1.123 | -1.144 | -0.386 | -0.005 | -0.041 | -0.041 | 1.7 | 0.24 | 0.33 | 0.17 | 0.14 | 0.22 | 0.25 |
| 7 | 131797860 | rs281913 | -1.341 | -0.91 | -1.816 | -0.437 | -0.022 | 0.036 | 0.036 | 1.978 | 0.49 | 0.94 | 0.32 | 0.38 | 0.38 | 0.49 |
| 7 | 131802832 | rs281916 | -0.393 | -1.114 | -0.306 | -0.545 | 0.03 | -0.096 | -0.096 | 0.013 | 0.14 | 0.63 | 0.1 | 0 | 0.16 | 0.07 |
| 7 | 131808274 | rs281893 | -0.474 | -1.259 | -0.808 | -0.613 | 0.001 | -0.002 | -0.002 | 0 | 0.84 | 0.91 | 0.92 | 0.93 | 0.87 | 0.94 |
| 7 | 131813407 | rs2278816 | 0.675 | -1.602 | -0.688 | -0.696 | -0.04 | -0.073 | -0.073 | 0.031 | 0.69 | 0.48 | 0.6 | 0.83 | 0.77 | 0.69 |
| 7 | 131813462 | rs11772555 | 1.206 | -1.59 | -1.112 | -0.736 | 0.071 | 0.183 | 0.183 | 0.015 | 0.61 | 0.08 | 0.55 | 0.83 | 0.72 | 0.64 |
| 7 | 131815118 | rs10954359 | 1.181 | -1.569 | -1.102 | -0.727 | 0.068 | 0.179 | 0.179 | 0.06 | 0.61 | 0.08 | 0.55 | 0.83 | 0.72 | 0.64 |
| 7 | 131815220 | rs10954360 | 1.181 | -1.569 | -1.102 | -0.727 | 0.068 | 0.179 | 0.179 | 0.06 | 0.61 | 0.08 | 0.55 | 0.83 | 0.72 | 0.64 |
| 7 | 131815310 | rs2671103 | -0.808 | -1.426 | -0.98 | -0.639 | -0.006 | -0.024 | -0.024 | 0 | 0.86 | 0.86 | 0.92 | 0.93 | 0.87 | 0.94 |
| 7 | 131816243 | rs4731850 | 0.354 | -1.394 | -1.017 | -0.624 | -0.012 | -0.028 | -0.028 | 0.156 | 0.73 | 0.68 | 0.63 | 0.83 | 0.77 | 0.72 |
| 7 | 131822719 | rs281878 | -2.141 | -1.108 | -1.53 | -0.136 | -0.157 | 0.022 | 0.022 | 0.701 | 0.69 | 0.98 | 0.77 | 0.84 | 0.63 | 0.74 |
| 7 | 131824908 | rs6954317 | -1.162 | -1.084 | -1.106 | -0.4 | 0.074 | 0.075 | 0.075 | 1.15 | 0.3 | 0.91 | 0.4 | 0.16 | 0.21 | 0.35 |
| 7 | 131828336 | rs4728251 | 1.722 | -1.072 | -1.214 | -0.382 | -0.022 | 0.115 | 0.115 | 0.153 | 0.53 | 0.06 | 0.45 | 0.74 | 0.56 | 0.52 |
| 7 | 131829128 | rs1399090 | -0.562 | -2.328 | -0.505 | -0.293 | 0.036 | 0.035 | 0.035 | 2.136 | 0.16 | 0.87 | 0.1 | 0.06 | 0.07 | 0.08 |
| 7 | 131832158 | rs1562835 | -0.428 | -2.278 | -0.906 | -0.27 | 0.008 | 0.002 | 0.002 | 0 | 0.86 | 0.97 | 0.68 | 0.89 | 0.88 | 0.76 |
| 7 | 131834858 | rs2598202 | -0.656 | -2.081 | -0.777 | -0.184 | 0.036 | 0.034 | 0.034 | 2.263 | 0.16 | 0.87 | 0.1 | 0.06 | 0.07 | 0.08 |
| 7 | 131845112 | rs9649039 | 1.225 | -1.927 | -0.902 | -0.133 | 0.107 | 0.127 | 0.127 | 0.918 | 0.6 | 0.07 | 0.53 | 0.81 | 0.73 | 0.53 |
| 7 | 131849670 | rs896169 | 0.231 | -1.476 | -0.428 | 0.139 | 0.008 | -0.014 | -0.014 | 0 | 0.7 | 0.76 | 0.57 | 0.61 | 0.73 | 0.62 |
| 7 | 131852910 | rs11763166 | 1.429 | -1.548 | -0.704 | 0.101 | 0.033 | -0.013 | -0.013 | 0 | 0.46 | 0.17 | 0.43 | 0.55 | 0.56 | 0.44 |
| 7 | 131853601 | rs12386622 | -1.227 | -1.227 | 0.138 | -0.121 | -0.055 | -0.13 | -0.13 | 0 | 0.25 | 0.59 | 0.14 | 0.04 | 0.17 | 0.19 |
| 7 | 131855165 | rs10954365 | -1.213 | -0.907 | 0.251 | 0.091 | -0.014 | -0.022 | -0.022 | 0 | 0.37 | 0.6 | 0.45 | 0.16 | 0.27 | 0.48 |
| 7 | 131855368 | rs10954366 | -1.017 | -0.956 | 0.41 | 0.358 | -0.046 | -0.045 | -0.045 | 0 | 0.32 | 0.57 | 0.42 | 0.16 | 0.19 | 0.46 |

| | | | | | | | | | | | | | | | | |
|---|---|---|---|---|---|---|---|---|---|---|---|---|---|---|---|---|
| 7 | 131856536 | rs9641933 | -0.947 | -0.803 | -0.06 | 0.399 | -0.027 | -0.034 | -0.034 | 0.617 | 0.3 | 0.4 | 0.39 | 0.14 | 0.18 | 0.41 |
| 7 | 131857841 | rs7802114 | -1.233 | -0.046 | 0.368 | 1.505 | 0.056 | 0.319 | 0.319 | 2.173 | 0.06 | 0.32 | 0.34 | 0.25 | 0.04 | 0.18 |
| 7 | 131859077 | rs10277421 | -0.355 | 0.731 | -0.991 | 1.437 | -0.131 | -0.094 | -0.094 | 0 | 0.77 | 0.98 | 0.72 | 0.69 | 0.61 | 0.89 |
| 7 | 131859395 | rs9649040 | -0.001 | -0.403 | -1.491 | 1.236 | 0.04 | -0.023 | -0.023 | 0 | 0.21 | 0.08 | 0.1 | 0.2 | 0.12 | 0.33 |
| 7 | 131862530 | rs3734992 | 0.794 | -0.463 | -1.759 | 0.347 | -0.009 | 0.095 | 0.095 | 1.187 | 0.29 | 0.13 | 0.21 | 0.13 | 0.26 | 0.21 |
| 7 | 131864591 | rs3734989 | -1.445 | -0.242 | -1.713 | 0.742 | -0.127 | -0.125 | -0.125 | 0.064 | 0.77 | 0.98 | 0.73 | 0.63 | 0.61 | 0.88 |
| 7 | 131865092 | rs896170 | 0.945 | -0.509 | -2.101 | 0.232 | -0.093 | 0.132 | 0.132 | 0.004 | 0.29 | 0.04 | 0.18 | 0.12 | 0.29 | 0.23 |
| 7 | 131865683 | rs7809724 | 0.313 | -0.644 | -1.316 | 0.263 | -0.021 | -0.048 | -0.048 | 0.005 | 0.1 | 0.18 | 0.06 | 0 | 0.06 | 0.03 |
| 7 | 131866817 | rs3734985 | 0.61 | -0.915 | -0.459 | 0.557 | -0.024 | 0.078 | 0.078 | 0 | 0.15 | 0.21 | 0.1 | 0.33 | 0.09 | 0.36 |
| 7 | 131867842 | rs6968634 | -1.841 | -0.742 | -0.677 | 0.51 | -0.137 | -0.123 | -0.123 | 0 | 0.76 | 0.98 | 0.72 | 0.65 | 0.61 | 0.86 |
| 7 | 131870845 | rs1020508 | -0.018 | -0.813 | 0.388 | 0.381 | 0.014 | 0.021 | 0.021 | 0 | 0.15 | 0.4 | 0.33 | 0.16 | 0.11 | 0.19 |
| 7 | 131875371 | rs7810362 | -1.579 | -0.894 | -0.294 | 0.41 | 0.072 | -0.005 | -0.005 | 0 | 0.49 | 0.91 | 0.54 | 0.55 | 0.36 | 0.67 |
| 7 | 131875684 | rs7778553 | -1.239 | -0.7 | -0.102 | 0.301 | 0.032 | 0.079 | 0.079 | 0.384 | 0.34 | 0.84 | 0.45 | 0.21 | 0.26 | 0.31 |
| 7 | 131876234 | rs7779444 | -0.657 | -0.435 | 0.011 | 0.107 | -0.001 | -0.126 | -0.126 | 1.235 | 0.24 | 0.4 | 0.16 | 0.02 | 0.22 | 0.26 |
| 7 | 131879262 | rs10954367 | -0.95 | -0.166 | -0.224 | 0.481 | 0.007 | 0.005 | 0.005 | 0 | 0.69 | 0.67 | 0.66 | 0.6 | 0.57 | 0.85 |
| 7 | 131881620 | rs1450885 | -1.046 | -0.307 | -0.217 | 0.534 | 0.002 | 0.002 | 0.002 | 0 | 0.39 | 0.62 | 0.62 | 0.61 | 0.37 | 0.67 |
| 7 | 131886428 | rs9649574 | 0.532 | -0.478 | -0.224 | 0.735 | 0.004 | -0.044 | -0.044 | 0 | 0.15 | 0.07 | 0.21 | 0.44 | 0.15 | 0.38 |
| 7 | 131886839 | rs2198051 | 0.759 | -0.221 | -0.418 | 0.529 | 0.003 | -0.008 | -0.008 | 0.009 | 0.38 | 0.36 | 0.19 | 0.18 | 0.27 | 0.21 |
| 7 | 131887029 | rs930686 | -1.716 | 0.35 | -0.679 | 0.795 | -0.117 | -0.051 | -0.051 | 0 | 0.77 | 0.98 | 0.82 | 0.79 | 0.66 | 0.88 |
| 7 | 131891191 | rs4731855 | -1.732 | 0.629 | -0.774 | 0.786 | -0.11 | -0.04 | -0.04 | 0 | 0.78 | 0.98 | 0.84 | 0.81 | 0.68 | 0.88 |
| 7 | 131893917 | rs17215844 | 0.239 | 0.393 | -0.141 | 0.796 | 0.016 | -0.048 | -0.048 | 0 | 0.11 | 0.18 | 0.11 | 0.01 | 0.12 | 0.14 |
| 7 | 131899254 | rs13227818 | -1.404 | 0.248 | 0.016 | 0.611 | 0.003 | 0.037 | 0.037 | 0.016 | 0.43 | 0.7 | 0.68 | 0.64 | 0.44 | 0.65 |
| 7 | 131901200 | rs2060724 | -1.036 | 0.311 | 0.178 | 0.677 | 0 | 0.025 | 0.025 | 0.42 | 0.32 | 0.57 | 0.37 | 0.47 | 0.31 | 0.51 |
| 7 | 131901251 | rs17216066 | -0.382 | 0.628 | -0.055 | 0.308 | 0.016 | -0.06 | -0.06 | 0 | 0.15 | 0.2 | 0.15 | 0.01 | 0.18 | 0.18 |
| 7 | 131907045 | rs1450879 | -1.019 | 1.041 | -0.106 | 0.794 | 0.008 | 0.002 | 0.002 | 0 | 0.72 | 0.72 | 0.66 | 0.7 | 0.56 | 0.81 |
| 7 | 131908468 | rs2290232 | 0.294 | 0.658 | -0.209 | 0.682 | -0.001 | 0 | 0 | 0 | 0.53 | 0.54 | 0.28 | 0.58 | 0.36 | 0.62 |
| 7 | 131908954 | rs977267 | -0.06 | 0.693 | -0.182 | 0.698 | -0.001 | 0.004 | 0.004 | 0 | 0.59 | 0.57 | 0.33 | 0.58 | 0.41 | 0.64 |
| 7 | 131911739 | rs1026192 | 0.15 | 0.732 | -0.27 | 0.627 | 0.003 | -0.025 | -0.025 | 0.753 | 0.13 | 0.12 | 0.08 | 0.01 | 0.14 | 0.13 |
| 7 | 131913645 | rs1026194 | -1.452 | 1.036 | -0.179 | 0.828 | -0.068 | -0.057 | -0.057 | 0 | 0.75 | 0.92 | 0.71 | 0.7 | 0.63 | 0.87 |
| 7 | 131921239 | rs1991118 | -1.487 | 1.029 | -0.207 | 0.804 | -0.03 | -0.025 | -0.025 | 0.817 | 0.76 | 0.89 | 0.72 | 0.7 | 0.63 | 0.87 |
| 7 | 131922332 | rs10231824 | 0.515 | 0.759 | 0.005 | 0.668 | 0.023 | 0.001 | 0.001 | 0 | 0.48 | 0.4 | 0.24 | 0.58 | 0.33 | 0.58 |
| 7 | 131924557 | rs1364513 | -0.506 | 0.797 | -0.249 | 0.243 | -0.002 | -0.023 | -0.023 | 0 | 0.23 | 0.23 | 0.19 | 0.01 | 0.22 | 0.18 |
| 7 | 131934509 | rs17805585 | -1.004 | 0.506 | -0.963 | 0.463 | -0.018 | -0.03 | -0.03 | 0 | 0.29 | 0.11 | 0.21 | 0.31 | 0.27 | 0.41 |
| 7 | 131934762 | rs7792482 | 0.119 | 0.305 | -0.573 | -0.024 | 0.045 | 0.069 | 0.069 | 0 | 0.51 | 0.65 | 0.49 | 0.74 | 0.34 | 0.62 |
| 7 | 131937820 | rs11762143 | 0.089 | 0.515 | -0.655 | 0 | 0.027 | 0.071 | 0.071 | 0 | 0.51 | 0.6 | 0.54 | 0.74 | 0.4 | 0.68 |
| 7 | 131939267 | rs10237001 | -0.398 | 1.087 | -0.909 | -0.55 | -0.005 | -0.009 | -0.009 | 0 | 0.22 | 0.24 | 0.18 | 0.01 | 0.24 | 0.16 |
| 7 | 131939805 | rs11760515 | 0.76 | 0.328 | -1.609 | -1.067 | 0.012 | 0.063 | 0.063 | 1.036 | 0.2 | 0.07 | 0.09 | 0 | 0.15 | 0.07 |
| 7 | 131942569 | rs10271349 | -1.863 | 0.129 | -1.97 | -1.018 | -0.11 | -0.075 | -0.075 | 0 | 0.72 | 0.98 | 0.73 | 0.69 | 0.64 | 0.79 |
| 7 | 131942949 | rs10274622 | 1.179 | 0.224 | -1.826 | -1.183 | -0.005 | 0.003 | 0.003 | 0 | 0.15 | 0.12 | 0.12 | 0.01 | 0.16 | 0.12 |
| 7 | 131942999 | rs11772044 | 0.996 | 0.056 | -1.667 | -1.037 | 0.016 | 0.061 | 0.061 | 1.513 | 0.19 | 0.07 | 0.09 | 0 | 0.15 | 0.05 |
| 7 | 131944273 | rs10954368 | -0.382 | 0.251 | -1.389 | -0.803 | -0.007 | 0.062 | 0.062 | 0 | 0.54 | 0.7 | 0.6 | 0.69 | 0.47 | 0.67 |
| 7 | 131947532 | rs4731857 | 0.38 | -0.2 | -0.619 | -0.229 | -0.039 | 0.172 | 0.172 | 0.541 | 0.17 | 0.36 | 0.34 | 0.46 | 0.1 | 0.47 |
| 7 | 131947817 | rs7776974 | 0.331 | -0.112 | -0.907 | -0.488 | 0 | -0.001 | -0.001 | 0 | 0.42 | 0.57 | 0.31 | 0.21 | 0.4 | 0.2 |
| 7 | 131950297 | rs717007 | 1.231 | -0.083 | -0.852 | -0.2 | -0.021 | 0.06 | 0.06 | 0 | 0.15 | 0.26 | 0.09 | 0.37 | 0.09 | 0.44 |
| 7 | 131951844 | rs1364512 | -0.556 | 0.167 | -1.11 | -0.196 | -0.141 | 0.004 | 0.004 | 0.008 | 0.59 | 0.94 | 0.66 | 0.69 | 0.51 | 0.67 |
| 7 | 131955001 | rs1424593 | -0.566 | 0.174 | -1.117 | -0.226 | -0.157 | -0.021 | -0.021 | 0.13 | 0.59 | 0.96 | 0.66 | 0.68 | 0.51 | 0.67 |
| 7 | 131964396 | rs7778725 | -0.831 | -0.24 | -0.917 | -0.426 | -0.132 | -0.06 | -0.06 | 0.771 | 0.61 | 0.93 | 0.73 | 0.63 | 0.51 | 0.69 |
| 7 | 131978162 | rs10954371 | 1.247 | -0.432 | -0.567 | -0.574 | -0.005 | 0 | 0 | 0 | 0.28 | 0.29 | 0.45 | 0.25 | 0.27 | 0.21 |
| 7 | 131980522 | rs6964570 | 0.347 | -0.565 | 0.366 | -0.316 | -0.007 | 0.006 | 0.006 | 0 | 0.08 | 0.05 | 0.06 | 0 | 0.09 | 0.02 |
| 7 | 131980776 | rs4731860 | 0.495 | -0.653 | 0.576 | -0.225 | -0.023 | 0.085 | 0.085 | 0.004 | 0.15 | 0.3 | 0.17 | 0.38 | 0.1 | 0.44 |
| 7 | 131983930 | rs6957015 | -0.084 | -0.508 | 0.214 | -0.294 | 0.002 | -0.058 | -0.058 | 0 | 0.16 | 0.21 | 0.1 | 0 | 0.15 | 0.05 |
| 7 | 131986419 | rs1424596 | -1.537 | -0.416 | -0.275 | -0.179 | 0.017 | -0.006 | -0.006 | 0.402 | 0.49 | 0.71 | 0.64 | 0.54 | 0.42 | 0.66 |
| 7 | 131995999 | rs3906454 | -1.538 | -0.256 | 0.12 | -0.779 | 0 | -0.018 | -0.018 | 0 | 0.29 | 0.34 | 0.43 | 0.16 | 0.28 | 0.23 |
| 7 | 131996949 | rs13231950 | -0.796 | -0.312 | 0.304 | -0.584 | -0.009 | -0.004 | -0.004 | 1.182 | 0.21 | 0.08 | 0.37 | 0.16 | 0.2 | 0.21 |
| 7 | 131997701 | rs16874167 | -1.593 | -0.307 | 0.491 | -0.562 | 0.001 | -0.009 | -0.009 | 0 | 0.25 | 0.3 | 0.42 | 0.16 | 0.25 | 0.22 |
| 7 | 131997968 | rs17166328 | -1.661 | -0.809 | 0.747 | -0.565 | -0.01 | -0.016 | -0.016 | 0 | 0.25 | 0.42 | 0.41 | 0.2 | 0.21 | 0.24 |
| 7 | 131998202 | rs17166330 | -1.706 | -1.381 | 1.001 | -0.659 | 0.003 | -0.04 | -0.04 | 0.18 | 0.26 | 0.39 | 0.39 | 0.17 | 0.24 | 0.21 |
| 7 | 131999807 | rs11762270 | -1.637 | -1.559 | 1.273 | -0.575 | -0.047 | -0.033 | -0.033 | 0 | 0.27 | 0.46 | 0.36 | 0.19 | 0.16 | 0.21 |
| 7 | 132001341 | rs11763352 | -1.779 | -1.554 | 1.148 | -0.571 | -0.018 | -0.017 | -0.017 | 0 | 0.31 | 0.58 | 0.39 | 0.19 | 0.19 | 0.23 |
| 7 | 132002753 | rs17166338 | -0.992 | -1.496 | 1.016 | -0.522 | -0.031 | 0.012 | 0.012 | 0.542 | 0.18 | 0.3 | 0.32 | 0.19 | 0.1 | 0.19 |
| 7 | 132007379 | rs2042513 | 1.557 | -1.518 | 0.371 | -1.173 | 0.042 | 0.229 | 0.229 | 0 | 0.51 | 0.19 | 0.46 | 0.81 | 0.6 | 0.67 |
| 7 | 132009356 | rs1582639 | -0.463 | -1.258 | 0.421 | -1.02 | -0.002 | 0.018 | 0.018 | 0 | 0.79 | 0.88 | 0.84 | 0.82 | 0.78 | 0.82 |
| 7 | 132012358 | rs17166347 | 0.024 | -1.051 | 0.247 | -0.97 | -0.013 | -0.039 | -0.039 | 0.794 | 0.23 | 0.37 | 0.17 | 0.17 | 0.21 | 0.18 |
| 7 | 132014616 | rs732490 | -1.649 | -1.08 | -0.233 | -0.87 | 0.028 | 0.142 | 0.142 | 0 | 0.51 | 0.86 | 0.58 | 0.3 | 0.41 | 0.43 |
| 7 | 132016733 | rs17166360 | 0.604 | -1.171 | -0.158 | -0.966 | 0 | -0.014 | -0.014 | 0 | 0.1 | 0.07 | 0.28 | 0.01 | 0.08 | 0.1 |
| 7 | 132017158 | rs12707034 | -1.704 | -1.167 | -0.137 | -0.973 | 0.043 | 0.225 | 0.225 | 0 | 0.49 | 0.83 | 0.55 | 0.23 | 0.41 | 0.36 |
| 7 | 132017849 | rs10263402 | -1.623 | -1.164 | -0.122 | -0.967 | 0.041 | 0.209 | 0.209 | 0 | 0.49 | 0.81 | 0.55 | 0.22 | 0.41 | 0.35 |
| 7 | 132019979 | rs10272059 | 0.184 | -1.013 | -0.254 | -0.846 | -0.003 | -0.032 | -0.032 | 0 | 0.14 | 0.12 | 0.29 | 0.01 | 0.09 | 0.11 |
| 7 | 132023921 | rs1364504 | -0.914 | -1.124 | -0.62 | -0.838 | 0.025 | 0.064 | 0.064 | 0 | 0.37 | 0.78 | 0.25 | 0.24 | 0.31 | 0.23 |

| | | | | | | | | | | | | | | | | |
|---|---|---|---|---|---|---|---|---|---|---|---|---|---|---|---|---|
| 7 | 132025936 | rs10231544 | -0.402 | -0.637 | -0.66 | -0.464 | 0.026 | 0.043 | 0.043 | 0 | 0.4 | 0.66 | 0.49 | 0.26 | 0.31 | 0.31 |
| 7 | 132030716 | rs6467431 | -0.767 | -0.598 | -0.784 | -0.332 | -0.025 | -0.052 | -0.052 | 1.7 | 0.17 | 0.37 | 0.11 | 0.04 | 0.11 | 0.13 |
| 7 | 132032495 | rs6959579 | -0.51 | 0.309 | -1.749 | 0.052 | -0.047 | 0.139 | 0.139 | 0 | 0.56 | 0.86 | 0.57 | 0.25 | 0.49 | 0.43 |
| 7 | 132042198 | rs1863015 | -0.642 | -0.035 | -1.166 | 0.603 | 0.043 | 0.064 | 0.064 | 0 | 0.51 | 0.84 | 0.52 | 0.32 | 0.4 | 0.48 |
| 7 | 132056155 | rs10231676 | -1.143 | -0.105 | -0.097 | 0.854 | 0.014 | -0.036 | -0.036 | 0 | 0.37 | 0.81 | 0.52 | 0.39 | 0.32 | 0.45 |
| 7 | 132063153 | rs10954373 | -0.933 | -0.261 | 1.087 | 0.801 | -0.027 | 0.053 | 0.053 | 0.101 | 0.28 | 0.46 | 0.44 | 0.4 | 0.23 | 0.38 |
| 7 | 132063522 | rs7807933 | -1.5 | -0.225 | 1.933 | 0.728 | 0.067 | -0.039 | -0.039 | 0.415 | 0.08 | 0.21 | 0.27 | 0.01 | 0.12 | 0.05 |
| 7 | 132064886 | rs10269601 | -1.71 | -0.233 | 1.896 | 0.706 | 0.074 | -0.064 | -0.064 | 0 | 0.11 | 0.31 | 0.3 | 0.01 | 0.13 | 0.06 |
| 7 | 132069727 | rs4731867 | 0.69 | -0.257 | 1.508 | 0.745 | -0.006 | 0.018 | 0.018 | 0.042 | 0.75 | 0.69 | 0.81 | 0.62 | 0.8 | 0.66 |
| 7 | 132084779 | rs10250019 | -2.156 | -0.231 | 2.009 | 0.393 | 0.036 | -0.093 | -0.093 | 0 | 0.13 | 0.42 | 0.31 | 0.01 | 0.12 | 0.05 |
| 7 | 132093935 | rs2341824 | -2.019 | -0.257 | 2.014 | 0.468 | 0.077 | 0.003 | 0.003 | 0.99 | 0.07 | 0.45 | 0.27 | 0 | 0.05 | 0.04 |
| 7 | 132094453 | rs2341825 | -2.481 | -0.542 | 2.071 | 0.316 | -0.045 | -0.09 | -0.09 | 0.014 | 0.15 | 0.66 | 0.32 | 0 | 0.06 | 0.12 |
| 7 | 132096426 | rs1004674 | -1.61 | -0.275 | 2.05 | 0.455 | 0.04 | -0.016 | -0.016 | 0.57 | 0.06 | 0.28 | 0.26 | 0 | 0.05 | 0.04 |
| 7 | 132097016 | rs1000058 | -0.385 | -0.61 | 1.063 | 0.644 | 0.016 | -0.012 | -0.012 | 0 | 0.12 | 0.02 | 0.09 | 0.09 | 0.07 | 0.1 |
| 7 | 132103982 | rs1347229 | -0.585 | -0.328 | 1.07 | 1.175 | -0.019 | -0.025 | -0.025 | 0.033 | 0.05 | 0 | 0.13 | 0.22 | 0.07 | 0.13 |
| 7 | 132115060 | rs2241728 | -0.539 | -0.693 | 1.037 | 1.382 | -0.015 | -0.083 | -0.083 | 0 | 0.19 | 0.06 | 0.15 | 0.36 | 0.19 | 0.31 |
| 7 | 132126343 | rs277492 | 1.508 | -0.482 | 1.839 | 1.442 | 0.017 | -0.041 | -0.041 | 0 | 0.92 | 0.78 | 0.75 | 1 | 0.94 | 0.89 |
| 7 | 132129571 | rs10488374 | 2.192 | -0.493 | 1.819 | 1.407 | -0.009 | 0.016 | 0.016 | 1.888 | 0.82 | 0.27 | 0.65 | 0.71 | 0.88 | 0.78 |
| 7 | 132137171 | rs17166480 | -0.443 | -0.595 | 1.397 | 1.38 | 0.001 | -0.094 | -0.094 | 0.045 | 0.28 | 0.08 | 0.42 | 0.36 | 0.26 | 0.35 |
| 7 | 132138906 | rs156940 | -2.181 | -0.33 | 2.633 | 1.809 | 0.046 | 0.252 | 0.252 | 0.052 | 0.08 | 0.3 | 0.06 | 0.31 | 0.07 | 0.19 |
| 7 | 132150710 | rs156931 | 1.945 | -0.377 | 2.091 | 1.734 | -0.016 | 0.042 | 0.042 | 0.872 | 0.82 | 0.35 | 0.67 | 0.69 | 0.89 | 0.8 |
| 7 | 132166973 | rs4731871 | -1.774 | -0.333 | 1.271 | 1.791 | -0.013 | 0.127 | 0.127 | 0 | 0.16 | 0.41 | 0.32 | 0.31 | 0.1 | 0.19 |
| 7 | 132172359 | rs156927 | 1.098 | -0.145 | 0.009 | 1.753 | -0.018 | 0.05 | 0.05 | 0 | 0.81 | 0.38 | 0.82 | 0.69 | 0.86 | 0.85 |
| 7 | 132176719 | rs2075421 | -1.325 | 0.345 | -0.599 | 1.587 | 0 | 0.052 | 0.052 | 0 | 0.06 | 0.1 | 0.16 | 0.3 | 0.07 | 0.11 |
| 7 | 132187403 | rs3847115 | 0.034 | 0.775 | -1.728 | 0.395 | -0.132 | 0.098 | 0.098 | 0 | 0.29 | 0.02 | 0.2 | 0.1 | 0.35 | 0.26 |
| 7 | 132192949 | rs156962 | -0.154 | 0.215 | -1.131 | 0.458 | 0.009 | -0.017 | -0.017 | 0 | 0.35 | 0.52 | 0.36 | 0.27 | 0.39 | 0.38 |
| 7 | 132196251 | rs156966 | -0.09 | 0.184 | -0.834 | 0.147 | 0.001 | -0.004 | -0.004 | 0 | 0.3 | 0.65 | 0.33 | 0.27 | 0.33 | 0.38 |
| 7 | 132196753 | rs4728258 | 1 | -0.434 | -1.185 | 0.67 | -0.074 | -0.096 | -0.096 | 0 | 0.44 | 0.1 | 0.29 | 0.49 | 0.44 | 0.53 |
| 7 | 132197234 | rs156968 | -1.328 | -0.749 | -1.057 | 0.605 | -0.019 | 0.021 | 0.021 | 0 | 0.9 | 0.91 | 0.77 | 0.77 | 0.97 | 0.95 |
| 7 | 132197554 | rs156969 | 0.638 | -0.98 | -0.121 | 0.464 | 0.012 | 0.001 | 0.001 | 0.425 | 0.26 | 0.32 | 0.29 | 0.27 | 0.32 | 0.37 |
| 7 | 132200868 | rs156974 | -0.065 | -1.055 | 0.033 | 0.473 | 0 | -0.011 | -0.011 | 0 | 0.32 | 0.58 | 0.32 | 0.27 | 0.33 | 0.34 |
| 7 | 132208640 | rs6974251 | -1.412 | -1.021 | -0.15 | 0.498 | -0.005 | 0.004 | 0.004 | 0 | 0.9 | 0.93 | 0.98 | 0.77 | 0.97 | 0.95 |
| 7 | 132217201 | rs17166546 | 1.221 | -0.897 | -0.228 | 0.504 | -0.12 | -0.069 | -0.069 | 0 | 0.48 | 0.06 | 0.54 | 0.52 | 0.5 | 0.57 |
| 7 | 132217786 | rs10260766 | 0.537 | -0.849 | 0.29 | 0.511 | 0.013 | -0.002 | -0.002 | 0 | 0.26 | 0.32 | 0.3 | 0.26 | 0.32 | 0.34 |
| 7 | 132226627 | rs7789101 | 0.833 | -1.04 | 0.951 | 0.377 | -0.021 | -0.013 | -0.013 | 0 | 0.74 | 0.34 | 0.64 | 0.78 | 0.83 | 0.91 |
| 7 | 132237410 | rs4728260 | 1.652 | -1.176 | 1.619 | 0.178 | -0.012 | -0.038 | -0.038 | 0 | 0.91 | 0.62 | 0.97 | 1 | 0.97 | 0.95 |
| 7 | 132241580 | rs963046 | 1.236 | -0.744 | 1.469 | 0.584 | -0.002 | -0.004 | -0.004 | 0.016 | 0.57 | 0.34 | 0.46 | 0.49 | 0.57 | 0.66 |
| 7 | 132254917 | rs1108116 | 0.297 | -0.944 | 1.012 | 0.57 | 0.021 | -0.026 | -0.026 | 0 | 0.08 | 0.03 | 0.06 | 0.4 | 0.04 | 0.28 |
| 7 | 132263595 | rs2288275 | 0.707 | -0.938 | 1.291 | 0.589 | 0.008 | 0.007 | 0.007 | 0 | 0.62 | 0.45 | 0.47 | 0.47 | 0.57 | 0.68 |
| 7 | 132275850 | rs7806862 | -0.505 | -0.818 | 0.941 | 0.782 | -0.02 | 0.007 | 0.007 | 0 | 0.29 | 0.21 | 0.19 | 0.29 | 0.34 | 0.27 |
| 7 | 132281079 | rs1426495 | 1.471 | -1.026 | 1.372 | 0.641 | -0.024 | 0.033 | 0.033 | 0.031 | 0.77 | 0.45 | 0.59 | 0.73 | 0.81 | 0.9 |
| 7 | 132294607 | rs1991313 | 1.489 | -0.934 | 1.34 | 0.638 | -0.023 | 0.046 | 0.046 | 0.321 | 0.76 | 0.49 | 0.61 | 0.73 | 0.81 | 0.9 |
| 7 | 132309592 | rs6957896 | 1.786 | -0.589 | 0.789 | 0.399 | -0.032 | 0.175 | 0.175 | 0 | 0.44 | 0.22 | 0.36 | 0.1 | 0.48 | 0.35 |
| 7 | 132311000 | rs1499300 | -1.047 | -0.456 | 1.228 | 0.565 | -0.011 | 0.064 | 0.064 | 1.124 | 0.19 | 0.35 | 0.37 | 0.27 | 0.17 | 0.07 |
| 7 | 132316317 | rs17429020 | 0.223 | -0.41 | 1.11 | 0.465 | 0.008 | -0.04 | -0.04 | 0.571 | 0.11 | 0.11 | 0.11 | 0 | 0.15 | 0.04 |
| 7 | 132317536 | rs10256188 | 2.219 | -0.552 | 0.747 | 0.278 | -0.021 | 0.126 | 0.126 | 0 | 0.4 | 0.23 | 0.24 | 0.1 | 0.42 | 0.25 |
| 7 | 132321422 | rs10273006 | 0.572 | -0.43 | 0.782 | 0.367 | -0.009 | 0.002 | 0.002 | 0.159 | 0.6 | 0.7 | 0.61 | 0.37 | 0.57 | 0.33 |
| 7 | 132322563 | rs6966705 | 1.975 | -0.292 | 0.587 | 0.215 | -0.011 | 0.082 | 0.082 | 0 | 0.4 | 0.29 | 0.24 | 0.1 | 0.42 | 0.26 |
| 7 | 132329246 | rs6965827 | 0.061 | -0.02 | 0.332 | 0.581 | 0.004 | -0.005 | -0.005 | 0 | 0.7 | 0.69 | 0.69 | 0.74 | 0.67 | 0.69 |
| 7 | 132335088 | rs13224271 | 1.944 | -0.098 | 0.354 | 0.825 | -0.027 | 0.02 | 0.02 | 0 | 0.45 | 0.15 | 0.5 | 0.38 | 0.45 | 0.3 |
| 7 | 132336527 | rs6975497 | -1.379 | -0.09 | 0.51 | 0.754 | -0.032 | 0.082 | 0.082 | 5.842 | 0.25 | 0.46 | 0.19 | 0.38 | 0.22 | 0.39 |
| 7 | 132354348 | rs924368 | 1.925 | -0.068 | 0.195 | 0.648 | -0.023 | 0.04 | 0.04 | 0 | 0.44 | 0.15 | 0.28 | 0.34 | 0.45 | 0.29 |
| 7 | 132354682 | rs17166672 | 0.144 | -0.102 | 0.3 | 1.044 | 0.015 | 0.003 | 0.003 | 0 | 0.09 | 0.05 | 0.08 | 0.37 | 0.05 | 0.31 |
| 7 | 132357208 | rs924369 | 0.167 | 0.007 | 0.34 | 0.866 | -0.004 | 0.002 | 0.002 | 0.196 | 0.71 | 0.75 | 0.69 | 0.71 | 0.66 | 0.68 |
| 7 | 132367230 | rs728469 | 0.836 | 0.436 | -0.73 | 0.525 | 0.03 | 0.066 | 0.066 | 0 | 0.52 | 0.32 | 0.35 | 0.31 | 0.59 | 0.34 |
| 7 | 132369580 | rs604943 | 0.962 | 1.169 | -0.742 | 0.692 | -0.004 | -0.004 | -0.004 | 0 | 0.3 | 0.22 | 0.42 | 0.33 | 0.3 | 0.23 |
| 7 | 132371110 | rs562761 | 0.983 | 1.181 | -1.115 | 0.702 | -0.044 | -0.045 | -0.045 | 0 | 0.28 | 0.06 | 0.4 | 0.32 | 0.28 | 0.23 |
| 7 | 132372798 | rs677153 | 0.376 | 1.187 | -1.433 | 0.612 | -0.018 | 0.058 | 0.058 | 0.115 | 0.55 | 0.14 | 0.41 | 0.67 | 0.57 | 0.64 |
| 7 | 132373204 | rs7790859 | -1.286 | 1.03 | -0.57 | 0.46 | -0.006 | 0.059 | 0.059 | 0.01 | 0.5 | 0.79 | 0.38 | 0.35 | 0.5 | 0.54 |
| 7 | 132373271 | rs10277610 | -1.467 | 1.103 | -0.216 | 0.438 | -0.001 | 0.04 | 0.04 | 0.653 | 0.51 | 0.72 | 0.36 | 0.34 | 0.5 | 0.56 |
| 7 | 132377487 | rs548221 | 0.591 | 0.752 | 0.556 | 0.323 | -0.033 | 0.021 | 0.021 | 0 | 0.73 | 0.44 | 0.77 | 0.67 | 0.82 | 0.79 |
| 7 | 132378413 | rs10239652 | -1.06 | 0.896 | 0.354 | 0.203 | -0.004 | -0.021 | -0.021 | 0 | 0.24 | 0.22 | 0.16 | 0.02 | 0.34 | 0.33 |
| 7 | 132379644 | rs1424598 | -1.039 | 0.968 | 0.6 | 0.236 | -0.009 | -0.004 | -0.004 | 0 | 0.24 | 0.19 | 0.16 | 0.02 | 0.34 | 0.33 |
| 7 | 132383800 | rs4449730 | -1.127 | 0.95 | 0.781 | 0.338 | -0.1 | -0.025 | -0.025 | 0 | 0.36 | 0.16 | 0.32 | 0.35 | 0.5 | 0.34 |
| 7 | 132387111 | rs10488668 | 0.012 | 0.646 | 0.861 | 0.398 | 0.005 | -0.043 | -0.043 | 0.05 | 0.21 | 0.15 | 0.18 | 0.33 | 0.18 | 0.27 |
| 7 | 132387461 | rs688257 | 1.752 | 0.59 | 1.215 | 0.637 | 0 | 0 | 0 | 0 | 0.55 | 0.52 | 0.67 | 0.45 | 0.53 | 0.39 |
| 7 | 132398788 | rs631653 | -2.133 | 0.698 | 1.174 | 0.61 | 0.02 | 0.003 | 0.003 | 0 | 0.5 | 0.74 | 0.36 | 0.55 | 0.44 | 0.61 |
| 7 | 132403581 | rs10954386 | -0.755 | 0.673 | 1.138 | 0.469 | -0.009 | -0.002 | -0.002 | 0 | 0.21 | 0.16 | 0.14 | 0.02 | 0.28 | 0.29 |
| 7 | 132404942 | rs10226516 | -1.468 | 0.377 | 1.234 | 0.692 | -0.001 | 0.004 | 0.004 | 0 | 0.48 | 0.56 | 0.35 | 0.36 | 0.47 | 0.56 |

| | | | | | | | | | | | | | | | | |
|---|---|---|---|---|---|---|---|---|---|---|---|---|---|---|---|---|
| 7 | 132405108 | rs10226545 | -1.275 | 0.385 | 1.175 | 0.696 | -0.002 | 0.001 | 0.001 | 0 | 0.44 | 0.58 | 0.34 | 0.37 | 0.47 | 0.56 |
| 7 | 132407635 | rs672421 | 0.711 | 0.32 | 1.315 | 0.812 | 0.034 | -0.042 | -0.042 | 19.543 | 0.67 | 0.56 | 0.72 | 0.79 | 0.6 | 0.81 |
| 7 | 132407741 | rs504719 | 0.116 | 0.335 | 1.502 | 0.796 | 0.029 | -0.014 | -0.014 | 0.246 | 0.73 | 0.66 | 0.74 | 0.78 | 0.61 | 0.82 |
| 7 | 132408676 | rs687405 | 1.5 | 0.43 | 1.334 | 0.942 | -0.006 | -0.017 | -0.017 | 0.054 | 0.53 | 0.24 | 0.65 | 0.47 | 0.54 | 0.41 |
| 7 | 132409504 | rs17166729 | -0.852 | 0.343 | 1.316 | 0.929 | -0.024 | 0.002 | 0.002 | 0 | 0.12 | 0.2 | 0.05 | 0.09 | 0.05 | 0.21 |
| 7 | 132411901 | rs10233448 | -1.089 | 0.236 | 1.209 | 0.799 | -0.02 | -0.013 | -0.013 | 0 | 0.17 | 0.21 | 0.06 | 0.09 | 0.08 | 0.22 |
| 7 | 132413024 | rs640339 | 1.936 | 0.252 | 1.19 | 0.798 | -0.04 | -0.033 | -0.033 | 0 | 0.47 | 0.18 | 0.63 | 0.47 | 0.51 | 0.4 |
| 7 | 132422248 | rs9656412 | -0.299 | 0.297 | 1.203 | 0.676 | -0.018 | -0.045 | -0.045 | 0.121 | 0.11 | 0.15 | 0.03 | 0 | 0.04 | 0.02 |
| 7 | 132426603 | rs2342807 | -1.399 | 0.3 | 1.251 | 0.703 | -0.002 | 0.026 | 0.026 | 0 | 0.46 | 0.39 | 0.28 | 0.32 | 0.47 | 0.54 |
| 7 | 132430093 | rs12707053 | -1.576 | 0.379 | 1.23 | 0.741 | -0.001 | -0.004 | -0.004 | 0 | 0.48 | 0.49 | 0.28 | 0.31 | 0.47 | 0.55 |
| 7 | 132430348 | rs13224732 | -0.402 | 0.42 | 0.771 | 0.662 | -0.001 | -0.002 | -0.002 | 0.234 | 0.31 | 0.33 | 0.23 | 0.21 | 0.39 | 0.32 |
| 7 | 132433567 | rs11772710 | -1.385 | 0.59 | -0.117 | 0.77 | -0.011 | -0.014 | -0.014 | 0 | 0.52 | 0.76 | 0.37 | 0.51 | 0.49 | 0.6 |
| 7 | 132441207 | rs7795475 | -1.242 | 0.278 | 0.005 | 0.647 | -0.005 | 0.056 | 0.056 | 0 | 0.48 | 0.79 | 0.37 | 0.41 | 0.46 | 0.37 |
| 7 | 132445685 | rs7793248 | -1.036 | 0.664 | 0.322 | 1.294 | 0.007 | 0.021 | 0.021 | 0 | 0.34 | 0.56 | 0.54 | 0.49 | 0.36 | 0.33 |
| 7 | 132455518 | rs17166752 | -1.197 | -0.023 | 0.32 | 1.122 | -0.018 | 0.05 | 0.05 | 0 | 0.07 | 0.24 | 0.05 | 0.1 | 0.02 | 0.13 |
| 7 | 132469174 | rs1318460 | 0.301 | -0.07 | -0.731 | 0.971 | 0.01 | -0.017 | -0.017 | 0.013 | 0.37 | 0.31 | 0.25 | 0.39 | 0.3 | 0.28 |
| 7 | 132475678 | rs7786571 | -0.206 | 0.421 | -0.956 | 0.792 | -0.037 | 0.007 | 0.007 | 0 | 0.75 | 0.86 | 0.79 | 0.79 | 0.65 | 0.72 |
| 7 | 132476411 | rs10259095 | 0.734 | 0.071 | -0.974 | 0.465 | -0.008 | 0.004 | 0.004 | 0 | 0.09 | 0.07 | 0.06 | 0 | 0.15 | 0.06 |
| 7 | 132483188 | rs10236673 | 0.496 | -0.057 | -0.846 | 0.381 | -0.001 | -0.022 | -0.022 | 0 | 0.11 | 0.13 | 0.06 | 0 | 0.14 | 0.06 |
| 7 | 132483938 | rs6947991 | 0.793 | -0.634 | -0.317 | 0.083 | 0.04 | 0.084 | 0.084 | 0 | 0.49 | 0.36 | 0.51 | 0.72 | 0.36 | 0.48 |
| 7 | 132489250 | rs4728265 | 0.844 | -1.194 | -0.399 | 0.092 | 0.006 | 0.016 | 0.016 | 0 | 0.48 | 0.26 | 0.4 | 0.4 | 0.43 | 0.4 |
| 7 | 132495299 | rs10435414 | 1.818 | -1.175 | -0.441 | 0.104 | -0.037 | -0.11 | -0.11 | 0 | 0.35 | 0.07 | 0.23 | 0.4 | 0.28 | 0.32 |
| 7 | 132503145 | rs6975749 | 0.94 | -1.318 | -0.425 | 0.038 | 0.026 | 0.083 | 0.083 | 0 | 0.47 | 0.31 | 0.5 | 0.72 | 0.37 | 0.47 |
| 7 | 132505658 | rs4731898 | 0.681 | -1.349 | -0.058 | -0.065 | -0.026 | -0.025 | -0.025 | 0 | 0.11 | 0.23 | 0.08 | 0.06 | 0.09 | 0.16 |
| 7 | 132521183 | rs4989636 | 0.642 | -1.357 | 0.036 | -0.005 | -0.003 | 0.002 | 0.002 | 0 | 0.09 | 0.14 | 0.03 | 0.11 | 0.08 | 0.1 |
| 7 | 132521753 | rs4731904 | 1.092 | -1.385 | -0.011 | 0.13 | -0.005 | 0.02 | 0.02 | 0 | 0.47 | 0.21 | 0.39 | 0.38 | 0.44 | 0.39 |
| 7 | 132522698 | rs6967294 | 1.458 | -1.478 | 0.037 | 0.181 | -0.011 | -0.002 | -0.002 | 0.21 | 0.4 | 0.46 | 0.25 | 0.38 | 0.31 | 0.33 |
| 7 | 132523462 | rs3807325 | 1.678 | -1.494 | -0.016 | 0.17 | 0.002 | -0.001 | -0.001 | 0.033 | 0.38 | 0.29 | 0.25 | 0.38 | 0.3 | 0.33 |
| 7 | 132525147 | rs6467447 | -1.033 | -1.607 | -0.029 | 0.155 | -0.041 | -0.002 | -0.002 | 0 | 0.75 | 0.86 | 0.77 | 0.76 | 0.63 | 0.72 |
| 7 | 132525243 | rs4731905 | -0.014 | -1.622 | -0.03 | 0.142 | 0.002 | -0.001 | -0.001 | 0 | 0.22 | 0.25 | 0.24 | 0.07 | 0.24 | 0.24 |
| 7 | 132526350 | rs4731907 | 0.544 | -1.591 | -0.013 | 0.163 | 0.01 | 0.005 | 0.005 | 0 | 0.59 | 0.5 | 0.49 | 0.45 | 0.54 | 0.56 |
| 7 | 132527075 | rs9649043 | 0.383 | -2.693 | -0.51 | 0.01 | -0.002 | 0.003 | 0.003 | 0 | 0.45 | 0.39 | 0.42 | 0.39 | 0.48 | 0.49 |
| 7 | 132532731 | rs2032949 | 0.098 | -2.401 | -0.503 | -0.013 | 0.009 | 0.005 | 0.005 | 0 | 0.58 | 0.5 | 0.48 | 0.45 | 0.54 | 0.57 |
| 7 | 132536559 | rs10808266 | 0.227 | -2.277 | -0.432 | -0.002 | 0.006 | 0.01 | 0.01 | 0 | 0.22 | 0.24 | 0.25 | 0.09 | 0.24 | 0.24 |
| 7 | 132552767 | rs10488210 | -0.632 | -2.315 | -0.409 | 0.038 | -0.056 | 0.011 | 0.011 | 0.242 | 0.29 | 0.55 | 0.5 | 0.3 | 0.25 | 0.29 |
| 7 | 132553318 | rs10488211 | 1.058 | -2.185 | -0.594 | -0.085 | -0.037 | 0.009 | 0.009 | 0 | 0.27 | 0.17 | 0.23 | 0.22 | 0.37 | 0.29 |
| 7 | 132555476 | rs4728273 | 0.247 | -2.077 | -0.481 | 0.071 | 0.01 | 0.031 | 0.031 | 0 | 0.21 | 0.16 | 0.24 | 0.09 | 0.24 | 0.24 |
| 7 | 132568255 | rs11982852 | 0.392 | -2.028 | -0.495 | 0.086 | 0.011 | 0.037 | 0.037 | 0 | 0.2 | 0.13 | 0.23 | 0.09 | 0.23 | 0.25 |
| 7 | 132573366 | rs10245279 | -0.222 | -2.087 | -0.381 | 0.286 | -0.013 | 0.209 | 0.209 | 0 | 0.05 | 0.19 | 0.26 | 0.31 | 0.03 | 0.14 |
| 7 | 132579694 | rs1468974 | -1.269 | -2.308 | -0.526 | 0.076 | 0 | 0 | 0 | 0 | 0.68 | 0.75 | 0.79 | 0.63 | 0.7 | 0.67 |
| 7 | 132611681 | rs2042066 | -1.095 | -1.648 | -0.643 | -0.072 | -0.006 | -0.002 | -0.002 | 0 | 0.65 | 0.75 | 0.75 | 0.64 | 0.63 | 0.63 |
| 7 | 132616793 | rs6962471 | 0.687 | -2.058 | -0.671 | -0.066 | 0.01 | 0.046 | 0.046 | 0 | 0.2 | 0.11 | 0.22 | 0.09 | 0.23 | 0.24 |
| 7 | 132624847 | rs10233020 | 0.526 | -2.118 | -0.592 | -0.054 | -0.021 | 0.013 | 0.013 | 0 | 0.37 | 0.25 | 0.29 | 0.34 | 0.44 | 0.29 |
| 7 | 132644903 | rs6951454 | -0.008 | -2.011 | -0.48 | -0.046 | -0.011 | -0.031 | -0.031 | 0 | 0.24 | 0.36 | 0.24 | 0.09 | 0.23 | 0.3 |
| 7 | 132651302 | rs2042067 | -0.089 | -2.071 | -0.334 | 0.087 | -0.003 | -0.005 | -0.005 | 0 | 0.43 | 0.39 | 0.55 | 0.55 | 0.46 | 0.36 |
| 7 | 132712250 | rs13226173 | -0.282 | -1.323 | -0.516 | 0.301 | 0.004 | 0.005 | 0.005 | 0 | 0.29 | 0.35 | 0.49 | 0.39 | 0.38 | 0.22 |
| 7 | 132716502 | rs7808471 | 0.866 | -1.278 | -0.929 | 0.033 | 0.047 | 0.116 | 0.116 | 0 | 0.37 | 0.16 | 0.3 | 0.11 | 0.31 | 0.29 |
| 7 | 132732992 | rs13239083 | -0.531 | -1.26 | -0.823 | 0.042 | 0.006 | 0.018 | 0.018 | 0 | 0.35 | 0.64 | 0.51 | 0.55 | 0.39 | 0.38 |
| 7 | 132741757 | rs12531471 | 1.242 | -1.333 | -0.843 | 0.001 | -0.11 | -0.104 | -0.104 | 0.805 | 0.27 | 0.03 | 0.18 | 0.34 | 0.3 | 0.34 |
| 7 | 132741896 | rs6953845 | -1.002 | -0.831 | -0.006 | 0.53 | 0.006 | 0.018 | 0.018 | 0 | 0.35 | 0.64 | 0.51 | 0.55 | 0.39 | 0.38 |
| 7 | 132774918 | rs17339165 | 0.799 | -0.686 | -0.19 | 0.624 | 0.005 | -0.037 | -0.037 | 0 | 0.64 | 0.2 | 0.49 | 0.45 | 0.63 | 0.6 |
| 7 | 132786126 | rs10274483 | 0.66 | 0.45 | -0.395 | 0.729 | 0.038 | 0.047 | 0.047 | 0.569 | 0.76 | 0.59 | 0.78 | 0.63 | 0.66 | 0.65 |
| 7 | 132800527 | rs10046599 | 0.391 | 0.796 | -1.047 | 0.864 | 0.021 | 0.021 | 0.021 | 0.215 | 0.76 | 0.72 | 0.76 | 0.64 | 0.66 | 0.62 |
| 7 | 132805104 | rs2343323 | 0.242 | 1.449 | -1.792 | 1.119 | -0.048 | -0.083 | -0.083 | 0.473 | 0.21 | 0.04 | 0.17 | 0.3 | 0.23 | 0.3 |
| 7 | 132805848 | rs17166902 | -0.593 | 1.853 | -2.024 | 1.132 | -0.091 | -0.099 | -0.099 | 0.183 | 0.22 | 0.01 | 0.35 | 0.29 | 0.26 | 0.39 |
| 7 | 132813492 | rs4728280 | -0.072 | 1.813 | -2.115 | 0.805 | -0.069 | -0.083 | -0.083 | 0 | 0.36 | 0.13 | 0.55 | 0.47 | 0.45 | 0.35 |
| 7 | 132813934 | rs4038363 | -0.169 | 1.58 | -2.156 | 0.398 | -0.038 | 0.048 | 0.048 | 0.953 | 0.09 | 0.01 | 0.09 | 0.01 | 0.13 | 0.17 |
| 7 | 132817561 | rs10249856 | 0.057 | 0.708 | -1.073 | 0.106 | -0.003 | -0.006 | -0.006 | 0 | 0.92 | 0.43 | 0.96 | 1 | 1 | 1 |
| 7 | 132820026 | rs2880602 | -0.874 | 0.338 | -0.601 | -0.149 | -0.009 | -0.053 | -0.053 | 0.187 | 0.82 | 0.65 | 0.87 | 0.92 | 0.82 | 0.9 |
| 7 | 132822204 | rs7459152 | 0.896 | 0.207 | -0.521 | -0.073 | 0.054 | -0.064 | -0.064 | 0 | 0.32 | 0.18 | 0.1 | 0.49 | 0.19 | 0.4 |
| 7 | 132822216 | rs7456888 | 0.224 | 0.349 | -0.593 | -0.122 | 0.012 | -0.085 | -0.085 | 0 | 0.73 | 0.55 | 0.78 | 0.92 | 0.69 | 0.76 |
| 7 | 132822540 | rs2343319 | -1.436 | 0.433 | -0.645 | -0.043 | -0.001 | -0.016 | -0.016 | 0.003 | 0.85 | 0.82 | 0.89 | 0.93 | 0.82 | 0.92 |
| 7 | 132839865 | rs4142542 | 0.654 | 0.503 | -0.58 | -0.088 | -0.047 | 0.039 | 0.039 | 0 | 0.35 | 0.51 | 0.14 | 0.48 | 0.21 | 0.41 |
| 7 | 132849038 | rs6969448 | 0.46 | 0.624 | -0.342 | 0.095 | -0.06 | -0.031 | -0.031 | 0 | 0.39 | 0.2 | 0.65 | 0.45 | 0.47 | 0.35 |
| 7 | 132850309 | rs6975382 | -1.143 | 0.572 | -0.32 | 0.082 | 0.007 | -0.056 | -0.056 | 0 | 0.84 | 0.72 | 0.87 | 0.94 | 0.82 | 0.93 |
| 7 | 132865353 | rs6954960 | -1.037 | 0.472 | -0.325 | 0.09 | 0.003 | -0.068 | -0.068 | 4.939 | 0.83 | 0.68 | 0.87 | 0.94 | 0.81 | 0.92 |
| 7 | 132866718 | rs6944686 | -0.132 | 0.404 | -0.3 | 0.383 | -0.011 | 0.004 | 0.004 | 0 | 0.44 | 0.5 | 0.23 | 0.49 | 0.34 | 0.57 |
| 7 | 132879799 | rs12533456 | -0.205 | 0.431 | -0.644 | 0.414 | 0.005 | -0.003 | -0.003 | 0.178 | 0.45 | 0.57 | 0.24 | 0.48 | 0.34 | 0.58 |
| 7 | 132888811 | rs7800830 | -0.203 | 0.588 | -1.274 | 0.508 | -0.009 | -0.046 | -0.046 | 0 | 0.4 | 0.2 | 0.34 | 0.54 | 0.37 | 0.48 |

| | | | | | | | | | | | | | | | | |
|---|---|---|---|---|---|---|---|---|---|---|---|---|---|---|---|---|
| 7 | 132891647 | rs6955404 | -0.393 | -0.118 | -1.943 | 0.315 | -0.13 | -0.056 | -0.056 | 0 | 0.56 | 0.94 | 0.38 | 0.58 | 0.52 | 0.63 |
| 7 | 132899833 | rs4272312 | 0.432 | -0.72 | -0.777 | 0.354 | -0.009 | 0.081 | 0.081 | 1.51 | 0.24 | 0.5 | 0.19 | 0.35 | 0.24 | 0.31 |
| 7 | 132900540 | rs6962059 | 0.177 | -0.765 | -0.625 | 0.186 | -0.012 | 0.029 | 0.029 | 3.386 | 0.28 | 0.67 | 0.2 | 0.52 | 0.25 | 0.38 |
| 7 | 132907651 | rs11979455 | 0.177 | -0.765 | -0.625 | 0.186 | -0.012 | 0.029 | 0.029 | 3.386 | 0.28 | 0.67 | 0.2 | 0.52 | 0.25 | 0.38 |
| 7 | 132927070 | rs7778404 | 0.039 | -1.146 | -0.205 | -0.253 | -0.023 | -0.003 | -0.003 | 0.111 | 0.11 | 0.44 | 0.07 | 0.05 | 0.05 | 0.03 |
| 7 | 132927272 | rs2880925 | 0.274 | -1.228 | -0.792 | -0.536 | -0.003 | 0.04 | 0.04 | 0 | 0.16 | 0.07 | 0.1 | 0 | 0.16 | 0.11 |
| 7 | 132927399 | rs11770426 | 0.546 | -1.169 | -0.763 | -0.506 | -0.007 | 0.032 | 0.032 | 0 | 0.15 | 0.06 | 0.1 | 0 | 0.16 | 0.11 |
| 7 | 132932589 | rs6467484 | -0.486 | -1.511 | -0.267 | -0.101 | -0.077 | 0.088 | 0.088 | 2.376 | 0.16 | 0.55 | 0.09 | 0.22 | 0.04 | 0.19 |
| 7 | 132938858 | rs6962538 | -0.898 | -1.553 | -0.645 | -0.394 | 0.023 | 0.015 | 0.015 | 0.216 | 0.37 | 0.68 | 0.24 | 0.23 | 0.24 | 0.32 |
| 7 | 132945754 | rs10215367 | -1.365 | -0.68 | -1.531 | -0.35 | -0.136 | -0.187 | -0.187 | 0.043 | 0.7 | 0.97 | 0.45 | 0.48 | 0.59 | 0.72 |
| 7 | 132956182 | rs10263196 | 0.924 | -0.722 | -1.263 | -0.46 | -0.029 | 0.022 | 0.022 | 0 | 0.31 | 0.23 | 0.2 | 0.25 | 0.36 | 0.39 |
| 7 | 132960079 | rs12112133 | -0.59 | -1.233 | -0.719 | -0.475 | -0.026 | -0.01 | -0.01 | 0 | 0.36 | 0.58 | 0.21 | 0.23 | 0.22 | 0.32 |
| 7 | 132971305 | rs11770490 | -0.156 | -1.65 | -1.008 | -0.912 | 0.03 | 0.135 | 0.135 | 0 | 0.25 | 0.05 | 0.12 | 0 | 0.2 | 0.15 |
| 7 | 132973522 | rs1317057 | -0.617 | -1.303 | -0.516 | -0.622 | -0.053 | -0.04 | -0.04 | 0.279 | 0.36 | 0.5 | 0.21 | 0.17 | 0.22 | 0.31 |
| 7 | 133019325 | rs7798267 | -0.718 | -1.303 | -0.468 | -0.622 | 0.005 | 0.006 | 0.006 | 0.408 | 0.36 | 0.63 | 0.22 | 0.18 | 0.23 | 0.32 |
| 7 | 133045944 | rs1149560 | 0.603 | -1.027 | -0.672 | -0.617 | -0.005 | -0.006 | -0.006 | 0.075 | 0.64 | 0.39 | 0.78 | 0.82 | 0.75 | 0.68 |
| 7 | 133053154 | rs12534801 | -0.336 | -1.201 | -0.34 | -0.648 | -0.005 | -0.006 | -0.006 | 0 | 0.3 | 0.29 | 0.18 | 0.13 | 0.22 | 0.3 |
| 7 | 133062665 | rs1149555 | -2.192 | -0.566 | -0.747 | -0.631 | -0.013 | 0.032 | 0.032 | 0 | 0.85 | 0.91 | 0.9 | 0.93 | 0.81 | 0.92 |
| 7 | 133074480 | rs1149552 | -0.062 | -0.576 | -1.257 | -0.691 | -0.059 | -0.135 | -0.135 | 0 | 0.32 | 0.07 | 0.55 | 0.5 | 0.34 | 0.26 |
| 7 | 133103048 | rs4731953 | 1.157 | -0.629 | -1.032 | -0.718 | -0.075 | -0.137 | -0.137 | 0.232 | 0.16 | 0.02 | 0.5 | 0.5 | 0.25 | 0.17 |
| 7 | 133105815 | rs12540212 | 1.059 | -0.78 | -0.608 | -0.751 | -0.003 | 0.011 | 0.011 | 0 | 0.12 | 0.21 | 0.11 | 0.12 | 0.13 | 0.24 |
| 7 | 133113977 | rs12707089 | 2.109 | -0.657 | -1.33 | -1.111 | -0.076 | 0.038 | 0.038 | 0 | 0.19 | 0.01 | 0.11 | 0.07 | 0.24 | 0.25 |
| 7 | 133118553 | rs2059376 | -1.932 | -0.703 | -0.792 | -1.167 | -0.003 | -0.077 | -0.077 | 0.107 | 0.8 | 0.65 | 0.87 | 0.94 | 0.81 | 0.92 |
| 7 | 133123294 | rs9649045 | -2.929 | -0.689 | -0.799 | -1.145 | -0.026 | 0.043 | 0.043 | 0 | 0.85 | 0.93 | 0.89 | 0.94 | 0.81 | 0.93 |
| 7 | 133130178 | rs6956293 | -1.713 | -0.603 | -0.792 | -1.201 | 0.01 | -0.079 | -0.079 | 0.021 | 0.78 | 0.65 | 0.86 | 0.95 | 0.78 | 0.91 |
| 7 | 133136872 | rs12538505 | -0.564 | -0.821 | -0.618 | -1.188 | -0.002 | 0.01 | 0.01 | 0 | 0.45 | 0.45 | 0.67 | 0.62 | 0.48 | 0.51 |
| 7 | 133136919 | rs12530513 | 1.527 | -0.849 | -0.573 | -1.181 | 0.003 | 0.045 | 0.045 | 0 | 0.12 | 0.32 | 0.11 | 0.12 | 0.13 | 0.23 |
| 7 | 133137644 | rs10488165 | 0.33 | -0.82 | -0.644 | -1.177 | -0.059 | -0.134 | -0.134 | 0 | 0.32 | 0.07 | 0.56 | 0.5 | 0.36 | 0.28 |
| 7 | 133146663 | rs7785217 | -0.36 | -0.866 | -0.57 | -1.095 | 0.001 | 0.008 | 0.008 | 0 | 0.44 | 0.44 | 0.66 | 0.62 | 0.47 | 0.51 |
| 7 | 133159583 | rs1424583 | 0.599 | -0.922 | -0.57 | -1.076 | -0.007 | 0.011 | 0.011 | 0 | 0.36 | 0.21 | 0.21 | 0.32 | 0.34 | 0.4 |
| 7 | 133185749 | rs12707091 | 1.077 | -0.881 | -0.505 | -1.071 | -0.033 | 0.019 | 0.019 | 0 | 0.33 | 0.12 | 0.21 | 0.28 | 0.34 | 0.4 |
| 7 | 133191127 | rs11975951 | -1.553 | -0.671 | -0.412 | -0.863 | -0.006 | -0.087 | -0.087 | 0.465 | 0.79 | 0.63 | 0.87 | 0.94 | 0.8 | 0.92 |
| 7 | 133193425 | rs17656452 | 2.543 | -0.652 | -0.543 | -0.854 | -0.035 | 0.056 | 0.056 | 0 | 0.15 | 0.04 | 0.1 | 0.06 | 0.19 | 0.07 |
| 7 | 133223017 | rs17167135 | 1.077 | -0.697 | -0.706 | -0.856 | -0.069 | -0.159 | -0.159 | 0.02 | 0.2 | 0.02 | 0.49 | 0.52 | 0.25 | 0.18 |
| 7 | 133225181 | rs9640874 | 1.861 | -0.589 | -0.138 | -0.83 | -0.01 | 0.012 | 0.012 | 0 | 0.13 | 0.05 | 0.07 | 0.09 | 0.1 | 0.12 |
| 7 | 133234255 | rs6947831 | -0.002 | -0.601 | -0.005 | -0.885 | -0.001 | -0.004 | -0.004 | 0 | 0.61 | 0.42 | 0.76 | 0.72 | 0.7 | 0.75 |
| 7 | 133235334 | rs17167141 | 1.762 | -0.527 | -0.065 | -0.824 | -0.009 | -0.021 | -0.021 | 0.115 | 0.13 | 0.05 | 0.08 | 0.19 | 0.1 | 0.14 |
| 7 | 133243248 | rs6967776 | 1.409 | -0.459 | 0.011 | -0.814 | 0.001 | -0.089 | -0.089 | 0.087 | 0.21 | 0.37 | 0.14 | 0.06 | 0.19 | 0.08 |
| 7 | 133269783 | rs2432636 | -1.293 | -0.544 | 0.074 | -0.621 | -0.001 | 0.001 | 0.001 | 0 | 0.84 | 0.83 | 0.9 | 0.79 | 0.9 | 0.84 |
| 7 | 133280339 | rs11526285 | 1.044 | -0.545 | 0.052 | -0.621 | -0.001 | 0.002 | 0.002 | 0 | 0.17 | 0.16 | 0.1 | 0.21 | 0.1 | 0.16 |
| 7 | 133304255 | rs10239061 | 0.808 | -0.544 | 0.091 | -0.527 | -0.007 | 0.018 | 0.018 | 0 | 0.18 | 0.27 | 0.11 | 0.24 | 0.11 | 0.17 |
| 7 | 133305729 | rs6956399 | 1.485 | -0.359 | -0.112 | -0.658 | 0.001 | -0.035 | -0.035 | 0 | 0.19 | 0.24 | 0.13 | 0.07 | 0.2 | 0.07 |
| 7 | 133330853 | rs17167187 | 1.307 | -0.425 | -0.041 | -0.558 | -0.001 | 0.009 | 0.009 | 0 | 0.16 | 0.16 | 0.09 | 0.23 | 0.1 | 0.17 |
| 7 | 133337635 | rs2345941 | 0.597 | -0.485 | -0.317 | -0.609 | -0.039 | -0.031 | -0.031 | 0 | 0.35 | 0.21 | 0.62 | 0.63 | 0.45 | 0.26 |
| 7 | 133338271 | rs7800006 | -0.1 | -0.493 | -0.042 | -0.532 | 0 | -0.003 | -0.003 | 0.475 | 0.5 | 0.46 | 0.27 | 0.21 | 0.45 | 0.56 |
| 7 | 133361412 | rs2345944 | 0.219 | -0.495 | -0.126 | -0.564 | -0.001 | -0.015 | -0.015 | 0.309 | 0.48 | 0.5 | 0.28 | 0.15 | 0.46 | 0.54 |
| 7 | 133388056 | rs10954428 | 0.89 | -0.778 | -0.217 | -0.758 | 0.002 | -0.006 | -0.006 | 0 | 0.33 | 0.41 | 0.23 | 0.27 | 0.36 | 0.47 |
| 7 | 133409403 | rs10274328 | -1.54 | -0.804 | -0.232 | -0.777 | -0.023 | -0.06 | -0.06 | 0.121 | 0.85 | 0.8 | 0.94 | 1 | 0.91 | 0.92 |
| 7 | 133409869 | rs1362736 | -0.972 | -0.816 | -0.208 | -0.782 | -0.042 | -0.088 | -0.088 | 0 | 0.82 | 0.73 | 0.93 | 1 | 0.91 | 0.91 |
| 7 | 133424668 | rs17167240 | 1.304 | -0.551 | -0.627 | -0.393 | -0.064 | -0.131 | -0.131 | 0 | 0.17 | 0.02 | 0.5 | 0.46 | 0.24 | 0.2 |
| 7 | 133431038 | rs10247014 | 0.806 | -0.544 | -0.546 | -0.38 | -0.088 | -0.117 | -0.117 | 2.865 | 0.33 | 0.12 | 0.61 | 0.55 | 0.43 | 0.27 |
| 7 | 133440786 | rs6963221 | -0.108 | -0.58 | -0.342 | -0.418 | 0.009 | 0.016 | 0.016 | 0 | 0.48 | 0.54 | 0.29 | 0.3 | 0.46 | 0.56 |
| 7 | 133466346 | rs13222377 | 1.945 | -0.275 | -0.936 | -0.444 | -0.054 | -0.061 | -0.061 | 0 | 0.19 | 0.01 | 0.12 | 0.2 | 0.23 | 0.27 |
| 7 | 133469109 | rs17167267 | 1.281 | -0.306 | -0.389 | -0.546 | -0.014 | -0.01 | -0.01 | 0.108 | 0.11 | 0.22 | 0.07 | 0.07 | 0.08 | 0.18 |
| 7 | 133476680 | rs1005342 | -1.174 | -0.019 | -0.578 | -0.479 | -0.008 | -0.048 | -0.048 | 1.447 | 0.84 | 0.81 | 0.93 | 1 | 0.88 | 0.91 |
| 7 | 133480664 | rs2542264 | 2.284 | 0.083 | -0.787 | -0.576 | -0.075 | 0.042 | 0.042 | 0 | 0.16 | 0.01 | 0.12 | 0.1 | 0.21 | 0.07 |
| 7 | 133491109 | rs2551009 | -0.935 | 0.023 | -0.605 | -0.535 | -0.022 | -0.075 | -0.075 | 0 | 0.82 | 0.74 | 0.92 | 1 | 0.88 | 0.91 |
| 7 | 133492717 | rs17167289 | 1.133 | 0.044 | -0.647 | -0.525 | -0.001 | -0.031 | -0.031 | 0 | 0.16 | 0.13 | 0.07 | 0 | 0.12 | 0.09 |
| 7 | 133502387 | rs3757406 | 0.96 | 0.126 | -0.839 | -0.343 | -0.069 | -0.143 | -0.143 | 0.044 | 0.18 | 0.02 | 0.5 | 0.47 | 0.25 | 0.2 |
| 7 | 133506955 | rs13243553 | 0.272 | -0.005 | -0.459 | -0.332 | 0.001 | -0.004 | -0.004 | 0 | 0.45 | 0.54 | 0.27 | 0.43 | 0.41 | 0.63 |
| 7 | 133529648 | rs17167306 | 0.395 | -0.04 | -0.371 | -0.457 | -0.022 | -0.083 | -0.083 | 0.029 | 0.19 | 0.29 | 0.1 | 0 | 0.13 | 0.1 |
| 7 | 133555720 | rs17167309 | 0.871 | 0.045 | -0.584 | -0.419 | -0.001 | -0.032 | -0.032 | 8.63 | 0.16 | 0.2 | 0.08 | 0 | 0.13 | 0.09 |
| 7 | 133569556 | rs17167332 | 1.214 | 0.055 | -0.589 | -0.415 | 0.007 | -0.002 | -0.002 | 0 | 0.15 | 0.09 | 0.07 | 0 | 0.13 | 0.09 |
| 7 | 133573039 | rs10246164 | -0.886 | 0.093 | -0.879 | -0.374 | -0.147 | -0.157 | -0.157 | 0 | 0.66 | 0.96 | 0.38 | 0.45 | 0.54 | 0.74 |
| 7 | 133580447 | rs2042456 | 0.067 | 0.052 | -0.778 | -0.372 | 0.027 | 0.001 | 0.001 | 0 | 0.49 | 0.73 | 0.32 | 0.46 | 0.42 | 0.64 |
| 7 | 133589846 | rs10279261 | -1.079 | 0.374 | -0.945 | -0.397 | -0.105 | -0.163 | -0.163 | 0 | 0.68 | 0.97 | 0.43 | 0.46 | 0.63 | 0.75 |
| 7 | 133601862 | rs10226267 | -0.319 | 0.233 | -0.435 | 0.023 | 0.027 | 0 | 0 | 0 | 0.49 | 0.73 | 0.32 | 0.46 | 0.42 | 0.65 |
| 7 | 133606631 | rs1030268 | 0.597 | 0.017 | -0.057 | 0.042 | -0.009 | 0 | 0 | 0.409 | 0.31 | 0.4 | 0.21 | 0.3 | 0.3 | 0.46 |
| 7 | 133626334 | rs11773892 | 1.563 | 0.303 | -0.98 | -0.018 | -0.093 | 0.081 | 0.081 | 0.077 | 0.31 | 0.01 | 0.18 | 0.11 | 0.33 | 0.19 |

| | | | | | | | | | | | | | | | | |
|---|---|---|---|---|---|---|---|---|---|---|---|---|---|---|---|---|
| 7 | 133627436 | rs4266574 | 0.761 | 0.34 | -1.111 | 0.051 | -0.056 | -0.12 | -0.12 | 0 | 0.18 | 0.02 | 0.5 | 0.44 | 0.24 | 0.19 |
| 7 | 133633152 | rs10259686 | -0.022 | 0.37 | -0.759 | 0.023 | 0.011 | -0.009 | -0.009 | 0 | 0.53 | 0.27 | 0.7 | 0.53 | 0.59 | 0.41 |
| 7 | 133638952 | rs2001618 | -1.445 | 0.366 | -0.078 | -0.054 | 0.032 | 0.047 | 0.047 | 0 | 0.85 | 0.65 | 0.9 | 0.83 | 0.88 | 0.81 |
| 7 | 133643848 | rs2971971 | -1.342 | 0.206 | -0.697 | -0.221 | -0.009 | 0.083 | 0.083 | 0 | 0.68 | 0.83 | 0.81 | 0.91 | 0.67 | 0.81 |
| 7 | 133648625 | rs2860599 | -1.218 | 0.199 | -0.693 | -0.224 | -0.003 | 0.072 | 0.072 | 0 | 0.67 | 0.79 | 0.81 | 0.91 | 0.67 | 0.79 |
| 7 | 133653884 | rs2971957 | -1.416 | 0.225 | -0.814 | -0.203 | -0.047 | 0.156 | 0.156 | 0.022 | 0.68 | 0.93 | 0.82 | 0.91 | 0.67 | 0.8 |
| 7 | 133657458 | rs2953626 | -1.416 | 0.229 | -0.823 | -0.201 | -0.027 | 0.126 | 0.126 | 0.024 | 0.68 | 0.89 | 0.82 | 0.91 | 0.67 | 0.8 |
| 7 | 133680601 | rs3823572 | -0.327 | 0.23 | -0.95 | 0.049 | -0.008 | -0.009 | -0.009 | 0 | 0.49 | 0.75 | 0.33 | 0.45 | 0.49 | 0.6 |
| 7 | 133681101 | rs7792010 | 0.624 | 0.223 | -0.555 | 0.107 | -0.012 | -0.01 | -0.01 | 0.9 | 0.31 | 0.43 | 0.22 | 0.26 | 0.33 | 0.41 |
| 7 | 133684991 | rs7785128 | -1.182 | 0.153 | -1.212 | 0.279 | -0.043 | 0.167 | 0.167 | 0 | 0.66 | 0.9 | 0.8 | 0.91 | 0.67 | 0.78 |
| 7 | 133692667 | rs11769734 | 1.378 | 0.157 | -1.204 | 0.34 | 0 | 0.003 | 0.003 | 0 | 0.07 | 0.06 | 0.05 | 0.08 | 0.1 | 0.14 |
| 7 | 133693897 | rs10488419 | 0.913 | 0.194 | -1.441 | 0.352 | -0.008 | -0.104 | -0.104 | 0.504 | 0.15 | 0.02 | 0.2 | 0.43 | 0.15 | 0.17 |
| 7 | 133697429 | rs4596578 | -0.104 | 0.196 | -1.392 | 0.331 | 0.007 | 0.001 | 0.001 | 0 | 0.5 | 0.71 | 0.56 | 0.47 | 0.52 | 0.61 |
| 7 | 133702217 | rs6966335 | 0.061 | 0.2 | -1.414 | 0.33 | -0.008 | -0.003 | -0.003 | 0 | 0.49 | 0.72 | 0.33 | 0.47 | 0.49 | 0.6 |
| 7 | 133705907 | rs2346442 | 0.061 | 0.2 | -1.414 | 0.33 | -0.008 | -0.003 | -0.003 | 0 | 0.49 | 0.72 | 0.33 | 0.47 | 0.49 | 0.6 |
| 7 | 133709787 | rs6952869 | 0.378 | 0.213 | -1.353 | 0.407 | 0.005 | -0.022 | -0.022 | 2.568 | 0.17 | 0.15 | 0.24 | 0.44 | 0.18 | 0.19 |
| 7 | 133715210 | rs2346444 | 0.544 | 0.211 | -1.382 | 0.414 | 0.011 | -0.06 | -0.06 | 0 | 0.16 | 0.08 | 0.23 | 0.43 | 0.15 | 0.18 |
| 7 | 133721697 | rs6978383 | 0.035 | 0.243 | -1.401 | 0.321 | -0.009 | -0.006 | -0.006 | 0.364 | 0.49 | 0.72 | 0.33 | 0.47 | 0.49 | 0.6 |
| 7 | 133727497 | rs6950383 | -0.093 | 0.272 | -1.483 | 0.352 | -0.011 | -0.003 | -0.003 | 0 | 0.5 | 0.74 | 0.56 | 0.48 | 0.5 | 0.6 |
| 7 | 133730639 | rs4728307 | 0.237 | 0.271 | -1.096 | 1.081 | -0.021 | -0.108 | -0.108 | 0 | 0.15 | 0.03 | 0.21 | 0.44 | 0.18 | 0.18 |
| 7 | 133735533 | rs9656434 | -0.277 | 0.305 | -0.869 | 1.127 | 0 | 0.018 | 0.018 | 0 | 0.18 | 0.2 | 0.48 | 0.52 | 0.19 | 0.19 |
| 7 | 133737030 | rs6955114 | -0.671 | 0.098 | -0.685 | 1.008 | -0.017 | 0.006 | 0.006 | 0 | 0.41 | 0.47 | 0.63 | 0.6 | 0.38 | 0.35 |
| 7 | 133738113 | rs10249912 | 0.254 | -0.087 | -0.601 | 1.206 | -0.037 | -0.02 | -0.02 | 3.782 | 0.23 | 0.32 | 0.18 | 0.17 | 0.17 | 0.12 |
| 7 | 133742010 | rs6467515 | -1.247 | 0.085 | -1.394 | 0.826 | -0.098 | 0.056 | 0.056 | 0.793 | 0.68 | 0.93 | 0.79 | 0.81 | 0.61 | 0.55 |
| 7 | 133743393 | rs1986692 | -0.051 | -0.033 | -1.21 | 0.95 | 0.025 | -0.026 | -0.026 | 0.114 | 0.51 | 0.8 | 0.35 | 0.44 | 0.41 | 0.33 |
| 7 | 133748235 | rs7457999 | 0.897 | -0.127 | -0.784 | 0.831 | 0.007 | -0.003 | -0.003 | 0 | 0.27 | 0.47 | 0.4 | 0.21 | 0.24 | 0.22 |
| 7 | 133749164 | rs6953296 | -0.432 | -0.391 | -0.736 | 0.772 | -0.023 | 0.049 | 0.049 | 0 | 0.63 | 0.71 | 0.76 | 0.76 | 0.6 | 0.53 |
| 7 | 133749964 | rs3735047 | -0.006 | -0.478 | -0.58 | 0.551 | -0.003 | -0.015 | -0.015 | 1.457 | 0.52 | 0.53 | 0.46 | 0.73 | 0.56 | 0.56 |
| 7 | 133755346 | rs12539923 | -0.959 | -0.327 | -1.095 | 0.58 | -0.023 | 0.092 | 0.092 | 0 | 0.7 | 0.86 | 0.8 | 0.88 | 0.7 | 0.66 |
| 7 | 133760551 | rs7357258 | 0.29 | -0.403 | -0.846 | 0.656 | -0.072 | -0.01 | -0.01 | 0.007 | 0.22 | 0.49 | 0.16 | 0.17 | 0.14 | 0.12 |
| 7 | 133768460 | rs6954364 | -0.852 | 0.43 | -0.83 | 0.92 | -0.095 | 0.109 | 0.109 | 0.008 | 0.73 | 0.98 | 0.83 | 0.91 | 0.71 | 0.67 |
| 7 | 133774841 | rs11762999 | -0.829 | 0.44 | -0.826 | 0.943 | -0.095 | 0.105 | 0.105 | 0.01 | 0.73 | 0.98 | 0.83 | 0.91 | 0.71 | 0.67 |
| 7 | 133810818 | rs12707136 | 0.719 | 0.155 | -0.488 | 1.284 | -0.002 | 0.003 | 0.003 | 0 | 0.55 | 0.43 | 0.66 | 0.48 | 0.56 | 0.48 |
| 7 | 133815391 | rs11971451 | -0.459 | -0.093 | -1.041 | 1.362 | 0 | -0.002 | -0.002 | 0 | 0.15 | 0.17 | 0.16 | 0.31 | 0.08 | 0.17 |
| 7 | 133821619 | rs13246630 | 0.619 | -0.068 | -1.029 | 1.391 | 0.025 | -0.005 | -0.005 | 0 | 0.58 | 0.71 | 0.66 | 0.49 | 0.64 | 0.49 |
| 7 | 133825945 | rs442352 | 0.415 | 0.196 | -1.249 | 1.344 | -0.104 | 0.016 | 0.016 | 0.049 | 0.28 | 0.02 | 0.2 | 0.2 | 0.34 | 0.39 |
| 7 | 133826448 | rs691130 | -0.687 | -0.058 | -1.103 | 1.512 | 0.004 | -0.042 | -0.042 | 0.114 | 0.4 | 0.17 | 0.33 | 0.5 | 0.37 | 0.52 |
| 7 | 133827457 | rs7782175 | 0.63 | -0.067 | -1.014 | 1.415 | 0.016 | -0.019 | -0.019 | 0 | 0.6 | 0.74 | 0.67 | 0.5 | 0.63 | 0.48 |
| 7 | 133829889 | rs511411 | 0.678 | -0.073 | -1.063 | 1.402 | -0.079 | 0.007 | 0.007 | 0.565 | 0.24 | 0.02 | 0.17 | 0.19 | 0.29 | 0.36 |
| 7 | 133835731 | rs725392 | -0.902 | -0.4 | -0.224 | 1.736 | 0.005 | -0.006 | -0.006 | 0 | 0.14 | 0.15 | 0.16 | 0.32 | 0.08 | 0.16 |
| 7 | 133843638 | rs423455 | 0.897 | -0.353 | -0.218 | 1.718 | -0.013 | 0.014 | 0.014 | 0 | 0.84 | 0.77 | 0.84 | 0.67 | 0.91 | 0.84 |
| 7 | 133848257 | rs17167553 | -0.926 | -0.371 | -0.335 | 1.763 | 0.005 | -0.006 | -0.006 | 0 | 0.14 | 0.16 | 0.16 | 0.32 | 0.08 | 0.16 |
| 7 | 133874831 | rs11487152 | -0.519 | -0.242 | -0.335 | 1.566 | 0.01 | -0.011 | -0.011 | 0 | 0.12 | 0.09 | 0.36 | 0.38 | 0.07 | 0.14 |
| 7 | 133890232 | rs10488426 | -0.025 | -0.13 | -0.457 | 1.605 | -0.002 | -0.059 | -0.059 | 0 | 0.1 | 0.01 | 0.13 | 0.38 | 0.07 | 0.12 |
| 7 | 133908453 | rs17167646 | 0.001 | -0.125 | -0.168 | 1.675 | -0.015 | 0.028 | 0.028 | 0.119 | 0.06 | 0.13 | 0.04 | 0.14 | 0.01 | 0.08 |
| 7 | 133915226 | rs7799320 | 0.485 | -0.122 | -0.206 | 1.597 | -0.002 | -0.009 | -0.009 | 0 | 0.05 | 0.07 | 0.01 | 0 | 0.01 | 0.04 |
| 7 | 133915760 | rs1222430 | -1.488 | -0.305 | 0.297 | 1.723 | -0.042 | 0.112 | 0.112 | 0 | 0.18 | 0.34 | 0.2 | 0.54 | 0.08 | 0.18 |
| 7 | 133919032 | rs1421477 | 0.514 | -0.325 | 0.758 | 1.412 | 0.006 | 0.092 | 0.092 | 0 | 0.94 | 0.84 | 0.93 | 0.82 | 0.93 | 0.88 |
| 7 | 133920443 | rs2544165 | -0.14 | 0.018 | 0.564 | 0.856 | -0.001 | -0.009 | -0.009 | 0 | 0.48 | 0.47 | 0.56 | 0.21 | 0.45 | 0.58 |
| 7 | 133930361 | rs1993058 | 0.818 | 0.004 | 0.467 | 0.832 | -0.112 | 0.173 | 0.173 | 0 | 0.36 | 0.05 | 0.5 | 0.11 | 0.44 | 0.51 |
| 7 | 133932334 | rs2598264 | 0.565 | 0.006 | 0.776 | 0.991 | 0.005 | 0.158 | 0.158 | 0 | 0.06 | 0.25 | 0.05 | 0.14 | 0.01 | 0.06 |
| 7 | 133932915 | rs2544176 | 0.561 | 0.006 | 0.776 | 0.991 | 0.005 | 0.158 | 0.158 | 0 | 0.06 | 0.25 | 0.05 | 0.14 | 0.01 | 0.06 |
| 7 | 133944902 | rs2598276 | 0.517 | 0.005 | 0.778 | 0.991 | 0.001 | 0.149 | 0.149 | 0.016 | 0.06 | 0.24 | 0.05 | 0.14 | 0.01 | 0.06 |
| 7 | 133958114 | rs1421480 | -0.109 | -0.014 | 0.882 | 0.857 | 0.001 | 0.015 | 0.015 | 0 | 0.47 | 0.57 | 0.58 | 0.25 | 0.47 | 0.58 |
| 7 | 133974519 | rs8134 | -0.132 | -0.061 | 0.923 | 0.794 | 0.007 | 0.037 | 0.037 | 0 | 0.48 | 0.59 | 0.58 | 0.27 | 0.47 | 0.55 |
| 7 | 133975183 | rs7486 | -0.127 | -0.061 | 0.923 | 0.794 | 0.006 | 0.037 | 0.037 | 0 | 0.47 | 0.59 | 0.58 | 0.27 | 0.47 | 0.55 |
| 7 | 133976029 | rs2598286 | -0.097 | -0.063 | 0.946 | 0.793 | 0.004 | 0.021 | 0.021 | 0 | 0.47 | 0.58 | 0.57 | 0.27 | 0.47 | 0.55 |
| 7 | 133978419 | rs1035328 | 0.145 | -0.099 | 0.955 | 0.772 | 0 | -0.007 | -0.007 | 0 | 0.43 | 0.43 | 0.55 | 0.27 | 0.45 | 0.55 |
| 7 | 133979368 | rs2241335 | -0.081 | -0.098 | 1.011 | 0.773 | -0.001 | -0.006 | -0.006 | 0.068 | 0.47 | 0.52 | 0.57 | 0.27 | 0.46 | 0.55 |
| 7 | 133979578 | rs2241336 | -0.081 | -0.098 | 1.011 | 0.773 | -0.001 | -0.006 | -0.006 | 0 | 0.47 | 0.52 | 0.57 | 0.27 | 0.46 | 0.55 |
| 7 | 133979795 | rs1421484 | -0.149 | -0.091 | 0.725 | 0.778 | 0.006 | 0.037 | 0.037 | 0 | 0.47 | 0.6 | 0.58 | 0.27 | 0.47 | 0.55 |
| 7 | 133981227 | rs1862052 | 0.817 | -0.065 | 0.387 | 0.781 | -0.104 | 0.171 | 0.171 | 0 | 0.36 | 0.05 | 0.49 | 0.12 | 0.44 | 0.51 |
| 7 | 133985896 | rs1901204 | 0.087 | -0.072 | 0.402 | 0.783 | 0 | -0.006 | -0.006 | 0 | 0.44 | 0.44 | 0.56 | 0.27 | 0.45 | 0.55 |
| 7 | 133988069 | rs12669902 | 0.978 | -0.119 | 0.289 | 1.009 | 0.01 | 0.003 | 0.003 | 0 | 0.08 | 0 | 0.11 | 0.47 | 0.04 | 0.14 |
| 7 | 133993062 | rs11766132 | 0.645 | 0.122 | 0.45 | 0.627 | -0.055 | 0.157 | 0.157 | 0 | 0.37 | 0.09 | 0.5 | 0.12 | 0.44 | 0.51 |
| 7 | 133994652 | rs722658 | -0.439 | -0.006 | 0.556 | 0.72 | -0.001 | -0.001 | -0.001 | 0.275 | 0.55 | 0.55 | 0.67 | 0.41 | 0.59 | 0.64 |
| 7 | 134003557 | rs1643031 | -0.225 | -0.039 | 0.979 | 1.064 | 0.017 | 0.277 | 0.277 | 2.02 | 0.07 | 0.33 | 0.06 | 0.22 | 0.01 | 0.03 |
| 7 | 134013464 | rs1643042 | -1.305 | -0.201 | 0.997 | 0.98 | -0.029 | 0.232 | 0.232 | 7.048 | 0.11 | 0.46 | 0.07 | 0.22 | 0.02 | 0.02 |
| 7 | 134013902 | rs1643043 | -0.442 | -0.107 | 0.395 | 0.311 | -0.013 | 0.12 | 0.12 | 0 | 0.55 | 0.76 | 0.7 | 0.84 | 0.49 | 0.67 |

| | | | | | | | | | | | | | | | | |
|---|---|---|---|---|---|---|---|---|---|---|---|---|---|---|---|---|
| 7 | 134019192 | rs13235296 | 0.664 | 0.096 | 0.252 | 0.531 | -0.006 | 0.114 | 0.114 | 0 | 0.36 | 0.15 | 0.49 | 0.15 | 0.39 | 0.47 |
| 7 | 134021784 | rs12666682 | 0.58 | 0.243 | 0.213 | 0.492 | -0.017 | 0.119 | 0.119 | 0 | 0.37 | 0.15 | 0.5 | 0.15 | 0.41 | 0.47 |
| 7 | 134022714 | rs1978411 | 0.578 | 0.185 | 0.287 | 0.627 | 0.002 | 0.127 | 0.127 | 0 | 0.08 | 0.11 | 0.13 | 0.47 | 0.08 | 0.17 |
| 7 | 134025825 | rs6955248 | 0.016 | 0.129 | 0.343 | 0.13 | 0 | 0.019 | 0.019 | 0.246 | 0.45 | 0.28 | 0.65 | 0.62 | 0.47 | 0.64 |
| 7 | 134026821 | rs1364423 | 0.346 | 0.238 | 0.617 | 0.27 | 0.029 | 0.292 | 0.292 | 3.049 | 0.06 | 0.32 | 0.04 | 0.21 | 0.01 | 0.02 |
| 7 | 134030156 | rs6977112 | 0.007 | 0.234 | -0.19 | -0.073 | -0.035 | 0.028 | 0.028 | 0.013 | 0.44 | 0.18 | 0.64 | 0.62 | 0.47 | 0.61 |
| 7 | 134031799 | rs1362985 | -0.674 | -0.008 | -0.305 | -0.661 | -0.02 | 0.043 | 0.043 | 0.065 | 0.66 | 0.81 | 0.78 | 0.81 | 0.58 | 0.62 |
| 7 | 134031847 | rs1364424 | 0.628 | -0.465 | -0.577 | -1.266 | -0.002 | -0.001 | -0.001 | 0.13 | 0.23 | 0.3 | 0.15 | 0.21 | 0.22 | 0.22 |
| 7 | 134033934 | rs1646663 | -0.02 | -0.582 | -0.775 | -1.174 | 0 | 0 | 0 | 0 | 0.37 | 0.66 | 0.25 | 0.21 | 0.33 | 0.25 |
| 7 | 134035080 | rs1646667 | -0.109 | -0.488 | -0.875 | -1.275 | 0.002 | 0.007 | 0.007 | 0.363 | 0.39 | 0.67 | 0.25 | 0.21 | 0.35 | 0.26 |
| 7 | 134036313 | rs12667801 | 0.407 | -0.172 | -1.133 | -0.929 | 0.004 | -0.04 | -0.04 | 0 | 0.28 | 0.14 | 0.53 | 0.6 | 0.23 | 0.36 |
| 7 | 134039630 | rs1646691 | -0.218 | 0.493 | -0.392 | -1.344 | 0.012 | -0.128 | -0.128 | 0 | 0.3 | 0.42 | 0.21 | 0.02 | 0.35 | 0.25 |
| 7 | 134040308 | rs1643067 | -0.384 | 0.516 | -0.503 | -1.42 | 0.003 | -0.038 | -0.038 | 0 | 0.23 | 0.23 | 0.19 | 0.03 | 0.3 | 0.17 |
| 7 | 134040681 | rs4728317 | 0.478 | 0.05 | -0.738 | -1.636 | -0.002 | -0.008 | -0.008 | 0 | 0.43 | 0.55 | 0.53 | 0.69 | 0.41 | 0.64 |
| 7 | 134042887 | rs6971727 | 0.498 | -0.109 | -0.637 | -1.676 | -0.005 | -0.006 | -0.006 | 0.037 | 0.43 | 0.57 | 0.49 | 0.69 | 0.34 | 0.63 |
| 7 | 134043175 | rs2161001 | 0.51 | -0.138 | -0.583 | -1.689 | 0.001 | 0 | 0 | 0 | 0.42 | 0.58 | 0.49 | 0.69 | 0.34 | 0.64 |
| 7 | 134043986 | rs1646696 | -0.443 | 0.083 | -0.577 | -1.365 | -0.003 | -0.001 | -0.001 | 0 | 0.71 | 0.7 | 0.84 | 0.71 | 0.75 | 0.83 |
| 7 | 134057403 | rs1643094 | -0.252 | 0.022 | -0.457 | -0.915 | 0.018 | -0.015 | -0.015 | 0 | 0.34 | 0.41 | 0.37 | 0.19 | 0.42 | 0.35 |
| 7 | 134061252 | rs1643097 | -0.878 | -0.422 | -0.436 | -1.037 | 0.012 | 0.006 | 0.006 | 0 | 0.7 | 0.78 | 0.83 | 0.71 | 0.76 | 0.84 |
| 7 | 134064285 | rs1646726 | -0.072 | -0.458 | -0.252 | -1.071 | 0.02 | -0.015 | -0.015 | 0 | 0.34 | 0.4 | 0.37 | 0.19 | 0.42 | 0.35 |
| 7 | 134067297 | rs1424437 | 1.187 | -0.689 | -0.294 | -1.172 | 0.002 | -0.004 | -0.004 | 0 | 0.35 | 0.33 | 0.46 | 0.52 | 0.33 | 0.5 |
| 7 | 134081571 | rs880458 | 0.702 | -0.092 | 0.003 | -1.506 | 0.055 | -0.091 | -0.091 | 1.071 | 0.18 | 0.29 | 0.15 | 0.01 | 0.26 | 0.04 |
| 7 | 134083383 | rs11534074 | -0.571 | -0.667 | 0.119 | -1.577 | -0.026 | -0.025 | -0.025 | 0.126 | 0.63 | 0.5 | 0.78 | 0.77 | 0.72 | 0.68 |
| 7 | 134083411 | rs1646744 | 0.494 | -0.679 | 0.2 | -1.583 | -0.011 | -0.011 | -0.011 | 0 | 0.37 | 0.45 | 0.21 | 0.23 | 0.28 | 0.33 |
| 7 | 134083535 | rs6951767 | 0.167 | -0.67 | 0.007 | -1.573 | -0.001 | -0.005 | -0.005 | 2.3 | 0.49 | 0.49 | 0.59 | 0.74 | 0.56 | 0.51 |
| 7 | 134085621 | rs4732021 | 0.249 | -0.79 | -0.293 | -1.575 | 0.001 | 0.009 | 0.009 | 0 | 0.46 | 0.43 | 0.58 | 0.87 | 0.54 | 0.52 |
| 7 | 134090200 | rs17167825 | 0.492 | -0.587 | -0.259 | -1.075 | 0.063 | 0.105 | 0.105 | 0.111 | 0.21 | 0.4 | 0.19 | 0.38 | 0.27 | 0.2 |
| 7 | 134091434 | rs741606 | 1.095 | -0.732 | -0.349 | -1.259 | -0.037 | 0.065 | 0.065 | 0.014 | 0.14 | 0.03 | 0.17 | 0.03 | 0.16 | 0.15 |
| 7 | 134092974 | rs6975650 | -1.168 | -1.122 | -0.365 | -1.451 | -0.031 | -0.109 | -0.109 | 0.192 | 0.63 | 0.55 | 0.79 | 0.92 | 0.73 | 0.76 |
| 7 | 134093813 | rs17773014 | 0.421 | -0.987 | -0.505 | -1.349 | -0.012 | 0.014 | 0.014 | 0 | 0.41 | 0.34 | 0.55 | 0.65 | 0.52 | 0.42 |
| 7 | 134094025 | rs1646747 | 1.328 | -1.029 | -0.33 | -1.356 | 0.032 | 0.093 | 0.093 | 0.012 | 0.34 | 0.19 | 0.17 | 0.08 | 0.27 | 0.24 |
| 7 | 134095375 | rs12672751 | 0.567 | -0.646 | -0.101 | -1.021 | 0.073 | 0.122 | 0.122 | 0.253 | 0.21 | 0.43 | 0.19 | 0.38 | 0.27 | 0.14 |
| 7 | 134100241 | rs949283 | -0.944 | -0.889 | -0.263 | -1.1 | 0.01 | -0.006 | -0.006 | 0 | 0.71 | 0.84 | 0.56 | 0.31 | 0.69 | 0.49 |
| 7 | 134100446 | rs1791003 | -0.966 | -0.892 | -0.264 | -1.117 | 0.01 | -0.004 | -0.004 | 0.458 | 0.71 | 0.84 | 0.56 | 0.28 | 0.7 | 0.49 |
| 7 | 134100951 | rs7795758 | 0.904 | -0.842 | -0.311 | -1.163 | 0.046 | 0.005 | 0.005 | 0.104 | 0.19 | 0.32 | 0.18 | 0.16 | 0.27 | 0.11 |
| 7 | 134103235 | rs706157 | 0.724 | -0.908 | -0.618 | -1.227 | 0.017 | 0.207 | 0.207 | 0 | 0.47 | 0.2 | 0.35 | 0.11 | 0.43 | 0.38 |
| 7 | 134103298 | rs3800756 | 1.413 | -0.76 | -0.676 | -0.974 | -0.046 | -0.161 | -0.161 | 0 | 0.23 | 0.03 | 0.39 | 0.49 | 0.28 | 0.32 |
| 7 | 134109303 | rs759852 | 0.035 | -0.467 | -0.236 | -0.886 | 0.058 | -0.033 | -0.033 | 0 | 0.23 | 0.45 | 0.18 | 0.13 | 0.26 | 0.1 |
| 7 | 134110166 | rs706151 | 1.198 | -0.902 | -1.464 | -1.118 | -0.058 | -0.115 | -0.115 | 0.029 | 0.47 | 0.04 | 0.53 | 0.52 | 0.49 | 0.4 |
| 7 | 134122440 | rs13225343 | -0.134 | -0.128 | -0.798 | -1.579 | -0.013 | 0.001 | 0.001 | 0 | 0.21 | 0.21 | 0.24 | 0.03 | 0.29 | 0.3 |
| 7 | 134123831 | rs1708416 | -1.363 | -0.449 | -0.874 | -1.802 | 0.029 | 0.046 | 0.046 | 0 | 0.65 | 0.79 | 0.83 | 0.93 | 0.76 | 0.86 |
| 7 | 134124005 | rs1708414 | -1.225 | -0.232 | -0.664 | -1.496 | 0.003 | -0.012 | -0.012 | 0 | 0.77 | 0.76 | 0.83 | 0.91 | 0.77 | 0.91 |
| 7 | 134125384 | rs969309 | -1.073 | -0.304 | -0.102 | -0.683 | -0.017 | 0.044 | 0.044 | 0 | 0.24 | 0.41 | 0.25 | 0.3 | 0.2 | 0.46 |
| 7 | 134125918 | rs2734665 | 0.192 | 0.031 | -0.713 | -0.376 | -0.001 | -0.002 | -0.002 | 0 | 0.82 | 0.81 | 0.81 | 0.8 | 0.82 | 0.72 |
| 7 | 134126134 | rs2701020 | 0.8 | 0.069 | -1.016 | -0.385 | -0.006 | -0.06 | -0.06 | 0 | 0.21 | 0.08 | 0.41 | 0.64 | 0.28 | 0.2 |
| 7 | 134126580 | rs2734663 | -0.106 | 0.106 | -1.18 | -0.159 | -0.038 | -0.084 | -0.084 | 0.019 | 0.31 | 0.07 | 0.54 | 0.64 | 0.39 | 0.35 |
| 7 | 134135751 | rs2734649 | -0.728 | -0.465 | -0.502 | -0.734 | 0.014 | -0.008 | -0.008 | 0 | 0.53 | 0.53 | 0.79 | 0.92 | 0.7 | 0.75 |
| 7 | 134137705 | rs3896278 | -0.093 | 0.139 | -0.25 | -0.38 | -0.03 | 0.03 | 0.03 | 0.269 | 0.3 | 0.21 | 0.32 | 0.18 | 0.38 | 0.33 |
| 7 | 134144485 | rs918825 | -0.109 | -0.129 | 0.313 | 0.276 | -0.016 | 0.112 | 0.112 | 0 | 0.07 | 0.16 | 0.04 | 0.22 | 0.04 | 0.07 |
| 7 | 134171722 | rs7788953 | 1.16 | -0.026 | -0.194 | 0.639 | -0.069 | -0.098 | -0.098 | 0.017 | 0.15 | 0.01 | 0.42 | 0.36 | 0.29 | 0.21 |
| 7 | 134175017 | rs782535 | -0.515 | -0.223 | 0.415 | 0.665 | 0.006 | 0.239 | 0.239 | 0.015 | 0.08 | 0.43 | 0.07 | 0.22 | 0.04 | 0.07 |
| 7 | 134178291 | rs782527 | -0.869 | -0.313 | 0.144 | 0.453 | -0.034 | -0.004 | -0.004 | 0 | 0.42 | 0.79 | 0.41 | 0.58 | 0.42 | 0.42 |
| 7 | 134179315 | rs7789596 | 0.526 | -0.252 | -0.154 | 0.308 | 0.007 | -0.005 | -0.005 | 0 | 0.15 | 0.13 | 0.07 | 0.19 | 0.1 | 0.26 |
| 7 | 134188002 | rs782544 | -0.968 | -0.349 | -0.363 | 0.421 | -0.074 | 0.007 | 0.007 | 0.826 | 0.47 | 0.88 | 0.42 | 0.58 | 0.45 | 0.56 |
| 7 | 134195690 | rs782513 | -0.533 | -0.297 | -0.298 | 0.32 | -0.017 | -0.002 | -0.002 | 0 | 0.4 | 0.71 | 0.38 | 0.39 | 0.41 | 0.38 |
| 7 | 134196661 | rs6953277 | 0.35 | -0.683 | -0.033 | 0.204 | 0.01 | -0.014 | -0.014 | 0 | 0.31 | 0.31 | 0.32 | 0.17 | 0.37 | 0.3 |
| 7 | 134202349 | rs10247315 | 0.408 | -0.674 | -0.042 | 0.207 | -0.01 | 0.008 | 0.008 | 0.449 | 0.31 | 0.26 | 0.32 | 0.17 | 0.37 | 0.3 |
| 7 | 134209317 | rs1424450 | 0.202 | -0.646 | 0.126 | 0.216 | 0.034 | -0.046 | -0.046 | 2.442 | 0.32 | 0.39 | 0.33 | 0.17 | 0.36 | 0.3 |
| 7 | 134212038 | rs10236737 | 0.225 | -0.724 | 0.045 | 0.209 | 0.014 | -0.021 | -0.021 | 0.724 | 0.32 | 0.34 | 0.32 | 0.18 | 0.37 | 0.3 |
| 7 | 134212611 | rs2037004 | 0.219 | -0.72 | 0.047 | 0.211 | 0.014 | -0.021 | -0.021 | 1.34 | 0.32 | 0.33 | 0.32 | 0.18 | 0.37 | 0.31 |
| 7 | 134214862 | rs1708429 | -0.111 | -0.569 | -0.548 | 0.276 | 0.013 | 0.004 | 0.004 | 1.551 | 0.38 | 0.58 | 0.37 | 0.41 | 0.41 | 0.38 |
| 7 | 134215403 | rs1722883 | -0.431 | -0.633 | -1.1 | 0.404 | -0.022 | 0.013 | 0.013 | 1.487 | 0.45 | 0.78 | 0.39 | 0.41 | 0.45 | 0.51 |
| 7 | 134217729 | rs3735041 | 0.379 | -0.513 | -1.212 | 0.624 | -0.059 | -0.095 | -0.095 | 1.381 | 0.14 | 0.01 | 0.35 | 0.36 | 0.28 | 0.21 |
| 7 | 134227299 | rs4732037 | -0.436 | -0.768 | -0.525 | -0.097 | -0.001 | 0.001 | 0.001 | 1.175 | 0.42 | 0.58 | 0.47 | 0.31 | 0.48 | 0.42 |
| 7 | 134232615 | rs889877 | -0.627 | -0.838 | -0.363 | 0.153 | 0.018 | 0 | 0 | 0 | 0.48 | 0.71 | 0.51 | 0.53 | 0.5 | 0.45 |
| 7 | 134252639 | rs17775910 | -0.211 | -1.002 | -0.318 | -0.177 | 0.017 | -0.065 | -0.065 | 1.059 | 0.39 | 0.46 | 0.36 | 0.15 | 0.44 | 0.25 |
| 7 | 134254029 | rs3792574 | -0.708 | -1.001 | -0.394 | -0.134 | 0.011 | -0.005 | -0.005 | 0 | 0.47 | 0.62 | 0.5 | 0.48 | 0.5 | 0.44 |
| 7 | 134262441 | rs10241998 | -0.206 | -0.703 | -0.461 | -0.008 | 0 | -0.017 | -0.017 | 0 | 0.36 | 0.56 | 0.31 | 0.15 | 0.34 | 0.24 |
| 7 | 134263249 | rs10488429 | 0.729 | -0.695 | -0.466 | 0.023 | -0.013 | 0.018 | 0.018 | 0 | 0.06 | 0.01 | 0.13 | 0.33 | 0.06 | 0.18 |

| | | | | | | | | | | | | | | | | |
|---|---|---|---|---|---|---|---|---|---|---|---|---|---|---|---|---|
| 7 | 134263676 | rs6955887 | -0.685 | -0.72 | -0.471 | 0.013 | -0.007 | -0.014 | -0.014 | 0.579 | 0.41 | 0.62 | 0.44 | 0.47 | 0.4 | 0.43 |
| 7 | 134264286 | rs6467538 | -0.25 | -0.68 | -0.396 | -0.785 | 0.001 | -0.018 | -0.018 | 0.854 | 0.36 | 0.56 | 0.31 | 0.14 | 0.34 | 0.25 |
| 7 | 134269909 | rs706193 | -0.471 | -0.722 | -0.428 | -0.899 | -0.002 | 0.022 | 0.022 | 0.245 | 0.42 | 0.68 | 0.35 | 0.35 | 0.36 | 0.35 |
| 7 | 134288060 | rs12666996 | 0.198 | -0.728 | -0.514 | -0.964 | -0.045 | -0.012 | -0.012 | 0 | 0.3 | 0.15 | 0.29 | 0.32 | 0.32 | 0.25 |
| 7 | 134288628 | rs1961330 | -0.961 | -0.974 | -0.447 | -1.501 | 0.001 | -0.039 | -0.039 | 0.033 | 0.7 | 0.77 | 0.81 | 0.98 | 0.73 | 0.87 |
| 7 | 134290869 | rs6951927 | -0.405 | -1.18 | -0.588 | -1.356 | -0.001 | 0 | 0 | 0 | 0.43 | 0.6 | 0.35 | 0.38 | 0.4 | 0.48 |
| 7 | 134296821 | rs7788959 | 0.308 | -0.812 | -1.184 | -1.124 | -0.014 | 0 | 0 | 0 | 0.07 | 0.17 | 0.02 | 0.04 | 0.05 | 0.12 |
| 7 | 134298890 | rs6966285 | 0.805 | -0.74 | -1.259 | -1.183 | -0.038 | -0.074 | -0.074 | 0 | 0.23 | 0.11 | 0.45 | 0.61 | 0.33 | 0.31 |
| 7 | 134301397 | rs10263716 | 0.59 | -0.84 | -1.282 | -1.234 | -0.054 | -0.03 | -0.03 | 0 | 0.29 | 0.12 | 0.3 | 0.32 | 0.32 | 0.24 |
| 7 | 134303336 | rs11766218 | -0.56 | -0.482 | -1.329 | -0.117 | -0.006 | -0.01 | -0.01 | 0 | 0.1 | 0.17 | 0.05 | 0.04 | 0.08 | 0.15 |
| 7 | 134305426 | rs4142278 | -0.184 | -0.097 | -1.652 | 0.214 | -0.042 | -0.084 | -0.084 | 1.629 | 0.23 | 0.09 | 0.44 | 0.61 | 0.32 | 0.3 |
| 7 | 134308254 | rs964546 | -0.873 | -1.006 | -1.078 | -0.085 | -0.058 | -0.023 | -0.023 | 0 | 0.28 | 0.44 | 0.28 | 0.2 | 0.17 | 0.24 |
| 7 | 134314404 | rs2032943 | 0.092 | -1.111 | -0.999 | -0.159 | 0.002 | 0.002 | 0.002 | 0.729 | 0.15 | 0.11 | 0.21 | 0.07 | 0.13 | 0.1 |
| 7 | 134325411 | rs12672388 | -1.328 | -0.671 | -1.853 | -0.809 | -0.15 | 0.178 | 0.178 | 0 | 0.7 | 0.97 | 0.78 | 0.96 | 0.62 | 0.85 |
| 7 | 134328370 | rs6962033 | -0.317 | -0.624 | -1.562 | -0.658 | -0.006 | 0.16 | 0.16 | 0 | 0.54 | 0.76 | 0.57 | 0.89 | 0.49 | 0.75 |
| 7 | 134330759 | rs3757401 | 0.434 | -0.695 | -1.542 | -0.446 | 0 | -0.005 | -0.005 | 0 | 0.42 | 0.54 | 0.5 | 0.66 | 0.45 | 0.61 |
| 7 | 134343327 | rs3800691 | -0.081 | -0.596 | -1.395 | -0.165 | -0.039 | 0.055 | 0.055 | 0 | 0.11 | 0.23 | 0.07 | 0.23 | 0.04 | 0.15 |
| 7 | 134349078 | rs6979233 | 0.295 | -0.479 | -1.549 | -0.365 | 0 | -0.002 | -0.002 | 0 | 0.42 | 0.55 | 0.5 | 0.66 | 0.45 | 0.61 |
| 7 | 134351368 | rs17200009 | -0.112 | -0.762 | -1.729 | -0.562 | 0.001 | -0.001 | -0.001 | 3 | 0.21 | 0.22 | 0.26 | 0.21 | 0.15 | 0.15 |
| 7 | 134355631 | rs3823563 | -0.759 | -0.746 | -1.735 | -0.576 | -0.027 | 0.064 | 0.064 | 0 | 0.64 | 0.76 | 0.76 | 0.87 | 0.6 | 0.76 |
| 7 | 134355857 | rs7783456 | -0.327 | -0.499 | -1.525 | -0.201 | 0.01 | 0.118 | 0.118 | 0 | 0.49 | 0.75 | 0.52 | 0.75 | 0.47 | 0.7 |
| 7 | 134362448 | rs889876 | -0.058 | -0.229 | -1.395 | -0.068 | 0 | 0.011 | 0.011 | 0 | 0.43 | 0.61 | 0.49 | 0.66 | 0.43 | 0.61 |
| 7 | 134364997 | rs17200700 | 0.52 | -0.111 | -1.692 | 0.047 | -0.053 | 0.043 | 0.043 | 0 | 0.19 | 0.01 | 0.31 | 0.03 | 0.2 | 0.18 |
| 7 | 134373997 | rs4728334 | -0.393 | -0.225 | -1.466 | 0.964 | 0.013 | -0.03 | -0.03 | 0.026 | 0.36 | 0.74 | 0.45 | 0.46 | 0.3 | 0.37 |
| 7 | 134380776 | rs2347694 | 0.033 | -0.054 | -0.615 | 0.76 | 0.001 | 0.005 | 0.005 | 0 | 0.31 | 0.28 | 0.35 | 0.16 | 0.25 | 0.28 |
| 7 | 134382784 | rs4524734 | -0.311 | 0.475 | -0.576 | 0.641 | -0.002 | 0 | 0 | 0 | 0.35 | 0.33 | 0.39 | 0.15 | 0.32 | 0.29 |
| 7 | 134388255 | rs2347698 | -0.485 | 0.35 | 0.021 | 1.056 | -0.001 | 0 | 0 | 0 | 0.38 | 0.58 | 0.44 | 0.3 | 0.32 | 0.3 |
| 7 | 134388544 | rs6973339 | -0.189 | 0.357 | 0.065 | 0.905 | 0.009 | -0.051 | -0.051 | 0 | 0.36 | 0.45 | 0.4 | 0.15 | 0.32 | 0.29 |
| 7 | 134389334 | rs4269477 | -0.633 | 0.343 | -0.849 | 0.985 | -0.052 | -0.026 | -0.026 | 0 | 0.57 | 0.84 | 0.53 | 0.56 | 0.46 | 0.7 |
| 7 | 134391526 | rs12539076 | 0.51 | -0.711 | -1.042 | 0.033 | 0.027 | 0.019 | 0.019 | 0 | 0.18 | 0.02 | 0.1 | 0.12 | 0.16 | 0.19 |
| 7 | 134391555 | rs2347700 | 0.148 | -0.56 | -0.908 | -0.331 | -0.008 | -0.003 | -0.003 | 0 | 0.28 | 0.22 | 0.45 | 0.66 | 0.31 | 0.54 |
| 7 | 134391874 | rs10488449 | -0.018 | -0.666 | -0.278 | -0.676 | -0.008 | -0.024 | -0.024 | 0 | 0.28 | 0.38 | 0.16 | 0.23 | 0.23 | 0.19 |
| 7 | 134393422 | rs2347704 | -0.955 | -1.052 | 0.283 | -0.331 | -0.014 | -0.012 | -0.012 | 0.029 | 0.38 | 0.61 | 0.26 | 0.36 | 0.34 | 0.39 |
| 7 | 134395461 | rs6944407 | -0.697 | -1.052 | 0.287 | -0.368 | -0.02 | -0.013 | -0.013 | 0 | 0.21 | 0.6 | 0.15 | 0.26 | 0.18 | 0.27 |
| 7 | 134397507 | rs10275303 | 0.377 | -1.067 | 0.428 | -0.37 | 0.003 | 0.003 | 0.003 | 0 | 0.21 | 0.2 | 0.12 | 0.12 | 0.16 | 0.24 |
| 7 | 134398634 | rs6960688 | 0.436 | -1.026 | 0.367 | -0.254 | 0.011 | 0.004 | 0.004 | 0 | 0.2 | 0.17 | 0.13 | 0.17 | 0.17 | 0.25 |
| 7 | 134403741 | rs2347709 | -1.656 | -0.89 | 0.136 | -0.499 | -0.075 | 0.15 | 0.15 | 0 | 0.66 | 0.92 | 0.8 | 0.99 | 0.65 | 0.88 |
| 7 | 134405310 | rs10488451 | -0.986 | -0.765 | 0.426 | -0.279 | 0.014 | 0.007 | 0.007 | 0 | 0.4 | 0.7 | 0.27 | 0.4 | 0.34 | 0.4 |
| 7 | 134417742 | rs10229310 | 0.528 | -0.65 | 0.502 | -0.285 | 0 | -0.001 | -0.001 | 0 | 0.28 | 0.26 | 0.53 | 0.6 | 0.31 | 0.46 |
| 7 | 134417820 | rs10258722 | -1.062 | -0.792 | 1.024 | -0.387 | -0.002 | -0.005 | -0.005 | 0 | 0.57 | 0.58 | 0.76 | 0.9 | 0.63 | 0.76 |
| 7 | 134419894 | rs7778776 | -1.28 | -0.803 | 1.108 | -0.392 | -0.02 | 0.095 | 0.095 | 0 | 0.59 | 0.75 | 0.77 | 0.9 | 0.63 | 0.77 |
| 7 | 134422732 | rs10155941 | -0.182 | -0.721 | 0.802 | -0.348 | 0.007 | 0.054 | 0.054 | 0 | 0.43 | 0.27 | 0.63 | 0.67 | 0.47 | 0.6 |
| 7 | 134425170 | rs10808276 | 0.09 | -0.951 | 0.647 | -0.45 | -0.012 | 0.025 | 0.025 | 0.618 | 0.37 | 0.3 | 0.61 | 0.8 | 0.46 | 0.6 |
| 7 | 134430108 | rs1993204 | -0.64 | -0.978 | 0.722 | -0.49 | 0 | 0.005 | 0.005 | 0 | 0.54 | 0.44 | 0.74 | 0.87 | 0.63 | 0.73 |
| 7 | 134430732 | rs17792160 | -1.071 | -1.034 | 0.879 | -0.513 | 0 | -0.032 | -0.032 | 0.235 | 0.58 | 0.6 | 0.76 | 0.91 | 0.63 | 0.77 |
| 7 | 134430805 | rs17168014 | 1.154 | -1.093 | 0.717 | -0.428 | -0.002 | -0.013 | -0.013 | 1.548 | 0.15 | 0.01 | 0.11 | 0.13 | 0.16 | 0.13 |
| 7 | 134433852 | rs2084909 | -0.718 | -1.146 | 0.825 | -0.405 | 0.002 | 0.031 | 0.031 | 0 | 0.54 | 0.41 | 0.73 | 0.85 | 0.63 | 0.71 |
| 7 | 134435912 | rs6973790 | -0.011 | -1.069 | 0.859 | -0.37 | -0.03 | -0.02 | -0.02 | 0 | 0.21 | 0.35 | 0.13 | 0.16 | 0.16 | 0.16 |
| 7 | 134436060 | rs6973845 | -0.514 | -1.148 | 0.976 | -0.409 | -0.056 | 0.013 | 0.013 | 0.036 | 0.26 | 0.45 | 0.16 | 0.27 | 0.17 | 0.27 |
| 7 | 134438524 | rs10488456 | 0.192 | -1.099 | 0.772 | -0.406 | -0.008 | -0.003 | -0.003 | 0 | 0.2 | 0.24 | 0.12 | 0.16 | 0.16 | 0.16 |
| 7 | 134441703 | rs10282404 | -0.864 | -0.945 | 0.845 | -0.391 | 0 | -0.061 | -0.061 | 0 | 0.58 | 0.58 | 0.75 | 0.96 | 0.63 | 0.85 |
| 7 | 134441990 | rs10279045 | -1.498 | -0.888 | 0.648 | -0.544 | -0.03 | 0.071 | 0.071 | 0 | 0.64 | 0.82 | 0.77 | 0.99 | 0.64 | 0.88 |
| 7 | 134447630 | rs17133290 | 0.578 | -0.747 | 0.473 | -0.41 | -0.001 | -0.001 | -0.001 | 0.097 | 0.16 | 0.19 | 0.12 | 0.14 | 0.17 | 0.12 |
| 7 | 134447642 | rs17133294 | 0.118 | -0.615 | 0.973 | 0.206 | -0.001 | 0.098 | 0.098 | 0.166 | 0.06 | 0.24 | 0.03 | 0.11 | 0.01 | 0.1 |
| 7 | 134449218 | rs10954469 | -0.484 | -0.411 | -0.508 | 0.266 | -0.007 | 0.041 | 0.041 | 0 | 0.4 | 0.28 | 0.61 | 0.74 | 0.46 | 0.65 |
| 7 | 134456444 | rs17168029 | -0.082 | -0.465 | -1.232 | 0.66 | -0.115 | -0.092 | -0.092 | 0 | 0.32 | 0.12 | 0.53 | 0.51 | 0.46 | 0.5 |
| 7 | 134457538 | rs11975997 | -1.416 | -0.658 | -1.177 | 0.585 | -0.02 | 0.155 | 0.155 | 0 | 0.54 | 0.77 | 0.67 | 0.88 | 0.53 | 0.78 |
| 7 | 134462521 | rs3807333 | 0.192 | 1.041 | -1.434 | 1.127 | -0.009 | 0.007 | 0.007 | 0.117 | 0.06 | 0.07 | 0.02 | 0.14 | 0.01 | 0.09 |
| 7 | 134463822 | rs3807337 | -0.922 | 1.647 | -1.47 | 1.2 | -0.119 | -0.065 | -0.065 | 0 | 0.34 | 0.11 | 0.54 | 0.51 | 0.48 | 0.5 |
| 7 | 134464523 | rs17168032 | -1.589 | 1.295 | -0.452 | 0.886 | -0.012 | -0.012 | -0.012 | 0 | 0.07 | 0.06 | 0.08 | 0.06 | 0.14 | 0.09 |
| 7 | 134472116 | rs3800707 | -1.468 | 1.475 | -0.487 | 0.757 | 0.003 | 0.002 | 0.002 | 0 | 0.15 | 0.14 | 0.26 | 0.24 | 0.23 | 0.2 |
| 7 | 134476735 | rs9641994 | -0.737 | 1.448 | -0.997 | 0.829 | -0.007 | -0.025 | -0.025 | 0 | 0.1 | 0.01 | 0.07 | 0.09 | 0.1 | 0.06 |
| 7 | 134478668 | rs3778844 | 0.132 | 1.976 | -2.001 | 0.325 | -0.136 | -0.135 | -0.135 | 0.217 | 0.2 | 0.02 | 0.33 | 0.61 | 0.32 | 0.22 |
| 7 | 134488289 | rs13225311 | -0.724 | 2.102 | -1.882 | 0.142 | -0.018 | -0.018 | -0.018 | 0 | 0.11 | 0.07 | 0.11 | 0.21 | 0.2 | 0.19 |
| 7 | 134489431 | rs6467559 | -1.621 | 0.925 | -1.254 | -0.555 | 0.054 | 0.059 | 0.059 | 0 | 0.51 | 0.64 | 0.58 | 0.93 | 0.68 | 0.51 |
| 7 | 134493909 | rs3778847 | -1.726 | 0.872 | -0.723 | -0.586 | 0.044 | 0.028 | 0.028 | 0 | 0.26 | 0.54 | 0.23 | 0.31 | 0.35 | 0.29 |
| 7 | 134494688 | rs2009372 | -1.595 | 0.303 | -0.706 | -0.608 | 0.053 | 0.056 | 0.056 | 0 | 0.51 | 0.63 | 0.58 | 0.93 | 0.68 | 0.51 |
| 7 | 134497231 | rs3304 | -1.601 | 0.516 | -0.401 | -0.536 | 0.001 | -0.017 | -0.017 | 0 | 0.21 | 0.16 | 0.1 | 0.3 | 0.18 | 0.27 |
| 7 | 134503166 | rs9642002 | -0.354 | 0.489 | 0.232 | -0.554 | -0.01 | 0.008 | 0.008 | 0 | 0.11 | 0.02 | 0.06 | 0.07 | 0.1 | 0.07 |

| | | | | | | | | | | | | | | | | |
|---|---|---|---|---|---|---|---|---|---|---|---|---|---|---|---|---|
| 7 | 134503247 | rs1869261 | -1.741 | 0.445 | 0.701 | -0.626 | 0.029 | 0.009 | 0.009 | 0 | 0.49 | 0.59 | 0.55 | 0.95 | 0.63 | 0.51 |
| 7 | 134504019 | rs7794696 | -0.024 | 0.589 | 0.651 | -0.522 | 0.027 | 0.013 | 0.013 | 0 | 0.27 | 0.32 | 0.42 | 0.63 | 0.43 | 0.23 |
| 7 | 134505463 | rs7800283 | 0 | -0.82 | -0.171 | -1.447 | -0.008 | 0.016 | 0.016 | 0 | 0.4 | 0.33 | 0.48 | 0.68 | 0.52 | 0.32 |
| 7 | 134508066 | rs1452913 | 0.549 | -0.291 | -0.283 | -1.233 | -0.044 | -0.028 | -0.028 | 0 | 0.24 | 0.14 | 0.39 | 0.62 | 0.4 | 0.22 |
| 7 | 134510505 | rs3800717 | -0.736 | -1.062 | -0.461 | -1.364 | 0.01 | -0.018 | -0.018 | 0 | 0.18 | 0.1 | 0.09 | 0.23 | 0.15 | 0.39 |
| 7 | 134512591 | rs2075464 | -1.173 | -1.473 | -0.136 | -1.621 | 0.032 | 0.028 | 0.028 | 0.046 | 0.48 | 0.63 | 0.3 | 0.37 | 0.36 | 0.53 |
| 7 | 134514731 | rs6954307 | -0.701 | -1.546 | 0.158 | -1.676 | 0.007 | -0.055 | -0.055 | 0 | 0.11 | 0.63 | 0.1 | 0.01 | 0.11 | 0.06 |
| 7 | 134519965 | rs7777356 | 1.738 | -1.612 | -0.186 | -1.706 | 0.001 | -0.024 | -0.024 | 0 | 0.16 | 0.13 | 0.07 | 0.01 | 0.07 | 0.05 |
| 7 | 134526525 | rs10255828 | -0.723 | -1.503 | -0.027 | -1.652 | 0.008 | -0.009 | -0.009 | 0 | 0.16 | 0.45 | 0.14 | 0.12 | 0.15 | 0.14 |
| 7 | 134528383 | rs2290363 | -1.347 | -1.577 | -0.526 | -1.685 | 0.025 | 0.045 | 0.045 | 0.579 | 0.53 | 0.85 | 0.34 | 0.37 | 0.42 | 0.57 |
| 7 | 134529143 | rs17801306 | 0.421 | -1.559 | -0.462 | -1.673 | -0.004 | -0.005 | -0.005 | 0 | 0.09 | 0.07 | 0.07 | 0.12 | 0.09 | 0.11 |
| 7 | 134530573 | rs17207352 | -0.879 | -1.615 | -0.463 | -1.713 | 0.075 | 0.089 | 0.089 | 0.47 | 0.44 | 0.78 | 0.26 | 0.25 | 0.33 | 0.46 |
| 7 | 134530915 | rs3778850 | -0.897 | -1.614 | -0.462 | -1.712 | 0.076 | 0.09 | 0.09 | 0.443 | 0.44 | 0.78 | 0.26 | 0.25 | 0.33 | 0.46 |
| 7 | 134531076 | rs3778851 | 0.15 | -1.434 | -0.468 | -1.579 | 0.014 | 0.069 | 0.069 | 0.003 | 0.11 | 0.21 | 0.05 | 0.2 | 0.12 | 0.2 |
| 7 | 134532702 | rs3800730 | -1.019 | -1.321 | -0.956 | -1.536 | 0.031 | 0.063 | 0.063 | 0.664 | 0.53 | 0.85 | 0.33 | 0.36 | 0.42 | 0.57 |
| 7 | 134540385 | rs7789417 | -1.596 | -1.374 | -1.013 | -1.835 | 0.017 | 0.043 | 0.043 | 0 | 0.76 | 0.88 | 0.73 | 0.99 | 0.81 | 0.77 |
| 7 | 134546119 | rs1377297 | -0.731 | -0.953 | -0.97 | -2.08 | 0.072 | 0.176 | 0.176 | 0 | 0.49 | 0.73 | 0.59 | 0.94 | 0.66 | 0.48 |
| 7 | 134546567 | rs17168108 | 0.137 | -1.022 | -0.973 | -2.025 | -0.003 | -0.019 | -0.019 | 0.002 | 0.11 | 0.14 | 0.07 | 0.04 | 0.07 | 0.21 |
| 7 | 134551474 | rs11974887 | -0.67 | -1.073 | -0.542 | -2.047 | -0.003 | -0.017 | -0.017 | 0 | 0.47 | 0.49 | 0.59 | 0.77 | 0.57 | 0.51 |
| 7 | 134552146 | rs4732064 | -1.542 | -1.153 | -1.274 | -2.239 | 0.058 | 0.19 | 0.19 | 0.974 | 0.63 | 0.89 | 0.67 | 0.97 | 0.74 | 0.72 |
| 7 | 134555102 | rs2290362 | -0.218 | -0.507 | -0.921 | -2.096 | -0.024 | -0.054 | -0.054 | 0 | 0.16 | 0.36 | 0.09 | 0.07 | 0.12 | 0.08 |
| 7 | 134556318 | rs7781307 | -0.293 | -0.473 | -0.879 | -2.088 | -0.002 | -0.078 | -0.078 | 0 | 0.2 | 0.37 | 0.14 | 0.07 | 0.19 | 0.13 |
| 7 | 134561241 | rs2218989 | -1.573 | -0.487 | -1.165 | -2.033 | 0 | -0.001 | -0.001 | 0 | 0.4 | 0.6 | 0.24 | 0.31 | 0.38 | 0.51 |
| 7 | 134561819 | rs10488465 | 1.284 | -0.825 | -1.833 | -1.955 | -0.097 | -0.126 | -0.126 | 0 | 0.2 | 0.02 | 0.39 | 0.63 | 0.31 | 0.18 |
| 7 | 134562744 | rs10488466 | 0.167 | -1.004 | -2.006 | -2.261 | -0.121 | 0.09 | 0.09 | 0 | 0.31 | 0.06 | 0.46 | 0.85 | 0.38 | 0.46 |
| 7 | 134569177 | rs4732066 | -1.68 | -1.229 | -1.713 | -2.311 | 0.029 | 0.18 | 0.18 | 0 | 0.61 | 0.85 | 0.66 | 0.93 | 0.69 | 0.68 |
| 7 | 134577965 | rs17168120 | -0.024 | -0.952 | -0.898 | -1.931 | -0.009 | 0 | 0 | 0.233 | 0.12 | 0.26 | 0.09 | 0.1 | 0.08 | 0.26 |
| 7 | 134581864 | rs1542309 | -0.321 | -1.168 | -1.008 | -2.076 | -0.002 | -0.046 | -0.046 | 0 | 0.61 | 0.63 | 0.61 | 0.9 | 0.64 | 0.68 |
| 7 | 134583666 | rs7776948 | -1.268 | -1.119 | -0.576 | -1.831 | -0.016 | -0.111 | -0.111 | 0 | 0.76 | 0.68 | 0.7 | 0.99 | 0.78 | 0.79 |
| 7 | 134584374 | rs4732067 | 1.197 | -0.831 | -0.797 | -0.217 | 0.009 | -0.007 | -0.007 | 0.74 | 0.17 | 0.16 | 0.11 | 0.28 | 0.09 | 0.13 |
| 7 | 134616524 | rs12667528 | 0.127 | 0.008 | -0.487 | -0.994 | -0.014 | 0.022 | 0.022 | 0 | 0.4 | 0.32 | 0.35 | 0.86 | 0.48 | 0.37 |
| 7 | 134618710 | rs6973420 | 0.143 | -0.052 | -0.209 | -1.032 | 0.003 | -0.042 | -0.042 | 0 | 0.41 | 0.44 | 0.37 | 0.86 | 0.48 | 0.37 |
| 7 | 134627147 | rs11975285 | 0.393 | -0.454 | -0.075 | -1.206 | -0.02 | -0.066 | -0.066 | 0 | 0.31 | 0.65 | 0.45 | 0.08 | 0.17 | 0.4 |
| 7 | 134630642 | rs3800752 | 0.034 | 0.646 | -0.273 | -0.338 | 0.047 | 0.065 | 0.065 | 0 | 0.15 | 0.2 | 0.22 | 0.29 | 0.3 | 0.15 |
| 7 | 134635040 | rs7797991 | 1.396 | 0.265 | -0.948 | -0.397 | 0 | 0.109 | 0.109 | 0.021 | 0.23 | 0.04 | 0.41 | 0.07 | 0.16 | 0.37 |
| 7 | 134637542 | rs6976629 | -0.683 | 0.569 | -0.763 | -0.306 | 0.002 | 0.035 | 0.035 | 0 | 0.49 | 0.75 | 0.7 | 0.39 | 0.57 | 0.58 |
| 7 | 134650544 | rs3823571 | 0.657 | 1.335 | -0.091 | 0.32 | 0.109 | 0.149 | 0.149 | 0 | 0.8 | 0.56 | 0.73 | 0.48 | 0.64 | 0.71 |
| 7 | 134651624 | rs17168134 | 0.991 | 1.176 | -0.686 | -0.229 | 0.012 | 0.084 | 0.084 | 0 | 0.27 | 0.15 | 0.42 | 0.07 | 0.22 | 0.39 |
| 7 | 134652029 | rs17168135 | 0.454 | 1.759 | -0.353 | 0.391 | 0 | 0.023 | 0.023 | 0 | 0.06 | 0.04 | 0.02 | 0.14 | 0.06 | 0.15 |
| 7 | 134652247 | rs2244307 | 0.624 | 1.754 | -0.085 | 0.379 | 0.01 | 0.015 | 0.015 | 0.061 | 0.82 | 0.83 | 0.76 | 0.49 | 0.63 | 0.72 |
| 7 | 134653930 | rs11204 | -0.634 | 1.642 | -0.259 | 0.298 | -0.017 | -0.005 | -0.005 | 0.495 | 0.89 | 0.93 | 0.78 | 0.64 | 0.7 | 0.86 |
| 7 | 134671030 | rs17168163 | 0.474 | 0.44 | -0.958 | -0.565 | 0.022 | 0.037 | 0.037 | 0 | 0.37 | 0.29 | 0.18 | 0.18 | 0.27 | 0.19 |
| 7 | 134678463 | rs12540686 | 1.341 | 0.624 | -1.825 | -0.643 | -0.062 | 0.036 | 0.036 | 2.1 | 0.31 | 0.03 | 0.37 | 0.28 | 0.3 | 0.44 |
| 7 | 134678523 | rs2252826 | -0.541 | 1.092 | -1.345 | -0.114 | -0.033 | -0.039 | -0.039 | 0.077 | 0.85 | 0.91 | 0.68 | 0.66 | 0.72 | 0.85 |
| 7 | 134696740 | rs4329203 | 0.66 | 0.804 | -1.247 | -1.531 | -0.014 | -0.032 | -0.032 | 0 | 0.11 | 0.06 | 0.09 | 0.19 | 0.14 | 0.18 |
| 7 | 134708675 | rs4366014 | 0.467 | 0.271 | -1.219 | -2.227 | 0.012 | 0.059 | 0.059 | 0 | 0.51 | 0.34 | 0.41 | 0.04 | 0.42 | 0.56 |
| 7 | 134729244 | rs10265256 | -0.056 | 0.397 | -1.023 | -2.229 | -0.005 | 0.009 | 0.009 | 0 | 0.58 | 0.41 | 0.48 | 0.23 | 0.48 | 0.7 |
| 7 | 134738607 | rs7776519 | -0.053 | 0.428 | -1.032 | -2.231 | -0.005 | 0.01 | 0.01 | 0 | 0.58 | 0.39 | 0.48 | 0.23 | 0.48 | 0.7 |
| 7 | 134746551 | rs7809191 | -0.046 | 0.432 | -1.031 | -2.229 | -0.005 | 0.011 | 0.011 | 0 | 0.58 | 0.39 | 0.48 | 0.23 | 0.48 | 0.7 |
| 7 | 134765148 | rs10215144 | -0.046 | 0.432 | -1.031 | -2.229 | -0.005 | 0.011 | 0.011 | 0 | 0.58 | 0.39 | 0.48 | 0.23 | 0.48 | 0.7 |
| 7 | 134772484 | rs10239305 | -0.048 | 0.453 | -1.013 | -2.2 | -0.005 | 0.01 | 0.01 | 0.002 | 0.58 | 0.39 | 0.48 | 0.23 | 0.48 | 0.7 |
| 7 | 134782352 | rs11771999 | -0.06 | 0.467 | -1.008 | -2.194 | -0.005 | 0.01 | 0.01 | 0 | 0.58 | 0.39 | 0.48 | 0.23 | 0.48 | 0.7 |
| 7 | 134786349 | rs1962522 | -0.068 | 0.284 | -1.255 | -2.271 | -0.005 | 0.011 | 0.011 | 0 | 0.58 | 0.39 | 0.48 | 0.23 | 0.48 | 0.7 |
| 7 | 134794015 | rs10227500 | -0.2 | 0.242 | -1.284 | -2.294 | -0.005 | 0.006 | 0.006 | 0 | 0.59 | 0.39 | 0.48 | 0.23 | 0.48 | 0.7 |
| 7 | 134814411 | rs4728353 | 0.937 | 0.288 | -1.331 | -2.19 | -0.043 | -0.052 | -0.052 | 0 | 0.15 | 0.05 | 0.36 | 0.58 | 0.3 | 0.17 |
| 7 | 134827792 | rs12707217 | 2.114 | 0.499 | -1.372 | -2.252 | -0.002 | 0.05 | 0.05 | 0 | 0.08 | 0.05 | 0.33 | 0.58 | 0.24 | 0.14 |
| 7 | 134831843 | rs10238516 | -0.068 | -0.185 | -1.34 | -2.119 | -0.005 | 0.01 | 0.01 | 0 | 0.58 | 0.39 | 0.48 | 0.23 | 0.48 | 0.7 |
| 7 | 134833870 | rs3800594 | 1.758 | -0.105 | -1.619 | -2.105 | 0 | 0 | 0 | 0 | 0.07 | 0.05 | 0.07 | 0.19 | 0.06 | 0.13 |
| 7 | 134834499 | rs11543821 | 1.179 | 0.091 | -1.406 | -1.909 | -0.021 | -0.024 | -0.024 | 0 | 0.16 | 0.13 | 0.37 | 0.58 | 0.29 | 0.17 |
| 7 | 134835350 | rs7795497 | 0.151 | -0.001 | -1.305 | -2.045 | -0.019 | 0.026 | 0.026 | 0.071 | 0.39 | 0.32 | 0.49 | 0.76 | 0.51 | 0.29 |
| 7 | 134853888 | rs11761021 | 0.001 | -0.515 | -1.582 | -2.018 | -0.007 | 0.167 | 0.167 | 0 | 0.35 | 0.13 | 0.31 | 0.04 | 0.27 | 0.49 |
| 7 | 134868824 | rs10457 | -0.548 | -0.899 | -1.034 | -1.837 | -0.018 | -0.054 | -0.054 | 0 | 0.71 | 0.5 | 0.84 | 0.82 | 0.71 | 0.81 |
| 7 | 134870201 | rs8345 | -0.548 | -0.87 | -1.034 | -1.806 | -0.015 | -0.047 | -0.047 | 0 | 0.71 | 0.5 | 0.84 | 0.81 | 0.71 | 0.81 |
| 7 | 134871775 | rs292557 | -0.548 | -0.939 | -1.034 | -1.828 | -0.018 | -0.054 | -0.054 | 1.303 | 0.71 | 0.5 | 0.84 | 0.82 | 0.71 | 0.81 |
| 7 | 134879830 | rs292575 | -0.527 | -0.956 | -1.042 | -1.634 | -0.018 | 0.053 | 0.053 | 0 | 0.71 | 0.5 | 0.8 | 0.61 | 0.71 | 0.76 |
| 7 | 134882592 | rs292581 | 1.297 | -0.773 | -1.177 | -1.889 | -0.006 | -0.002 | -0.002 | 0 | 0.15 | 0.11 | 0.38 | 0.55 | 0.35 | 0.18 |
| 7 | 134883991 | rs17807215 | 1.315 | -0.459 | -0.808 | -1.51 | -0.001 | -0.01 | -0.01 | 0 | 0.11 | 0.08 | 0.06 | 0 | 0.17 | 0.07 |
| 7 | 134884526 | rs10266018 | -0.058 | -0.678 | -0.891 | -1.476 | -0.032 | 0.198 | 0.198 | 0 | 0.33 | 0.09 | 0.33 | 0.04 | 0.27 | 0.46 |
| 7 | 134889141 | rs292592 | -0.43 | -0.407 | -0.669 | -1.228 | -0.019 | 0.052 | 0.052 | 0 | 0.71 | 0.5 | 0.8 | 0.61 | 0.71 | 0.75 |

| | | | | | | | | | | | | | | | | |
|---|---|---|---|---|---|---|---|---|---|---|---|---|---|---|---|---|
| 7 | 134889264 | rs17233604 | 1.482 | -0.437 | -0.759 | -1.316 | -0.053 | 0.043 | 0.043 | 0.379 | 0.1 | 0.01 | 0.23 | 0.03 | 0.15 | 0.29 |
| 7 | 134891308 | rs292594 | -0.376 | -0.467 | -0.552 | -1.126 | -0.019 | 0.052 | 0.052 | 0 | 0.71 | 0.5 | 0.8 | 0.61 | 0.71 | 0.75 |
| 7 | 134892508 | rs292597 | 0.072 | -0.577 | -0.548 | -1.188 | -0.013 | 0.01 | 0.01 | 0 | 0.65 | 0.43 | 0.76 | 0.61 | 0.68 | 0.75 |
| 7 | 134913049 | rs292624 | -0.287 | -0.375 | -0.449 | -0.797 | -0.009 | 0.029 | 0.029 | 0 | 0.72 | 0.43 | 0.8 | 0.62 | 0.72 | 0.76 |
| 7 | 134918608 | rs12665884 | 0.401 | -0.121 | -1.106 | -0.752 | 0 | 0.012 | 0.012 | 0 | 0.24 | 0.2 | 0.13 | 0.18 | 0.26 | 0.17 |
| 7 | 134923406 | rs10281202 | -0.259 | -0.092 | -0.937 | -0.683 | 0 | 0.107 | 0.107 | 0.337 | 0.48 | 0.28 | 0.41 | 0.21 | 0.47 | 0.57 |
| 7 | 134939767 | rs17168332 | -0.239 | -0.453 | -0.178 | -0.385 | -0.009 | -0.066 | -0.066 | 0 | 0.43 | 0.44 | 0.19 | 0.07 | 0.33 | 0.31 |
| 7 | 134941141 | rs7806817 | 0.009 | -0.443 | -0.738 | -0.398 | 0.038 | 0.1 | 0.1 | 0 | 0.4 | 0.21 | 0.17 | 0.06 | 0.33 | 0.31 |
| 7 | 134941877 | rs6951100 | -0.401 | -0.478 | -0.534 | -0.212 | 0.037 | 0.015 | 0.015 | 0 | 0.46 | 0.66 | 0.24 | 0.38 | 0.33 | 0.37 |
| 7 | 134942937 | rs17808352 | -0.132 | -0.045 | -1.556 | -0.756 | 0.041 | 0.095 | 0.095 | 0 | 0.41 | 0.24 | 0.17 | 0.06 | 0.32 | 0.31 |
| 7 | 134944039 | rs10280844 | -0.128 | 0.846 | -1.407 | 0.211 | -0.009 | 0.016 | 0.016 | 1.148 | 0.46 | 0.24 | 0.66 | 0.63 | 0.53 | 0.5 |
| 7 | 134944544 | rs889929 | -0.865 | 1.867 | -1.113 | 1.164 | -0.043 | -0.048 | -0.048 | 0 | 0.18 | 0.11 | 0.44 | 0.49 | 0.31 | 0.22 |
| 7 | 134945250 | rs12666242 | -0.608 | 1.823 | -0.497 | 1.057 | 0.018 | 0.046 | 0.046 | 0 | 0.14 | 0.15 | 0.39 | 0.52 | 0.25 | 0.22 |
| 7 | 134945413 | rs292637 | -0.03 | 1.205 | -0.393 | 0.882 | -0.005 | -0.003 | -0.003 | 0 | 0.1 | 0.08 | 0.04 | 0.04 | 0.06 | 0.13 |
| 7 | 134946179 | rs292639 | 0.32 | 0.457 | -0.41 | 0.206 | -0.01 | -0.01 | -0.01 | 0 | 0.58 | 0.71 | 0.41 | 0.51 | 0.5 | 0.6 |
| 7 | 134946554 | rs292640 | 0.447 | 0.239 | 0.144 | 0.272 | 0 | 0.001 | 0.001 | 0 | 0.54 | 0.46 | 0.32 | 0.23 | 0.48 | 0.56 |
| 7 | 134947623 | rs1019621 | 1.011 | 0.225 | 0.11 | 0.239 | 0.006 | 0.052 | 0.052 | 0 | 0.44 | 0.4 | 0.28 | 0.2 | 0.42 | 0.42 |
| 7 | 134954478 | rs51511 | -0.518 | 0.148 | -0.047 | 0.32 | -0.05 | 0.179 | 0.179 | 0 | 0.14 | 0.29 | 0.13 | 0.32 | 0.08 | 0.21 |
| 7 | 134960640 | rs17168349 | 0.985 | 0.368 | -0.301 | -0.235 | 0.012 | 0.063 | 0.063 | 0 | 0.43 | 0.33 | 0.27 | 0.16 | 0.4 | 0.42 |
| 7 | 134965705 | rs292672 | -0.322 | 0.43 | -0.659 | -0.308 | -0.02 | -0.084 | -0.084 | 0 | 0.39 | 0.39 | 0.26 | 0.05 | 0.33 | 0.32 |
| 7 | 134965728 | rs17235308 | 0.993 | 0.53 | -1.445 | -0.135 | -0.013 | 0.213 | 0.213 | 0 | 0.41 | 0.13 | 0.25 | 0.17 | 0.39 | 0.38 |
| 7 | 134965925 | rs292673 | -0.097 | -0.308 | -0.858 | 0.212 | -0.019 | -0.008 | -0.008 | 0 | 0.1 | 0.13 | 0.05 | 0.04 | 0.07 | 0.14 |
| 7 | 134970088 | rs4732112 | 0.398 | -0.334 | -1.497 | 0.335 | 0.118 | 0.008 | 0.008 | 0.268 | 0.53 | 0.17 | 0.64 | 0.69 | 0.66 | 0.61 |
| 7 | 134972126 | rs10954491 | 0.538 | 0.09 | -1.778 | 0.051 | 0.021 | 0.155 | 0.155 | 0 | 0.49 | 0.17 | 0.34 | 0.23 | 0.48 | 0.5 |
| 7 | 134972291 | rs17168358 | 0.252 | -0.106 | -1.77 | -0.357 | 0.014 | 0.006 | 0.006 | 0 | 0.16 | 0.1 | 0.09 | 0 | 0.1 | 0.16 |
| 7 | 134972466 | rs7805912 | -0.21 | 0.556 | -1.161 | 0.45 | -0.023 | -0.026 | -0.026 | 0 | 0.2 | 0.17 | 0.46 | 0.7 | 0.26 | 0.23 |
| 7 | 134986157 | rs10954493 | -0.083 | 0.515 | -0.374 | 0.97 | -0.06 | -0.056 | -0.056 | 0.325 | 0.19 | 0.1 | 0.38 | 0.39 | 0.25 | 0.2 |
| 7 | 134988039 | rs292703 | -1.237 | 0.452 | 0.301 | 0.927 | -0.008 | 0 | 0 | 0.057 | 0.29 | 0.34 | 0.54 | 0.71 | 0.37 | 0.3 |
| 7 | 134988792 | rs292706 | -1.166 | 0.41 | 0.45 | 0.906 | 0.036 | -0.002 | -0.002 | 1.087 | 0.3 | 0.45 | 0.54 | 0.71 | 0.38 | 0.3 |
| 7 | 134988899 | rs292707 | -0.009 | 0.474 | 0.16 | 0.594 | 0.009 | -0.004 | -0.004 | 0 | 0.1 | 0.2 | 0.09 | 0.04 | 0.11 | 0.06 |
| 7 | 134989301 | rs292708 | 0.125 | 0.494 | -0.204 | 0.674 | 0.013 | 0.011 | 0.011 | 0.025 | 0.79 | 0.76 | 0.61 | 0.61 | 0.74 | 0.8 |
| 7 | 134994297 | rs17235985 | 0.983 | -0.251 | -1.411 | -0.771 | 0.054 | 0.209 | 0.209 | 0 | 0.42 | 0.08 | 0.22 | 0.11 | 0.33 | 0.4 |
| 7 | 134995116 | rs17809576 | -0.172 | 0.673 | -1.327 | -0.388 | -0.076 | 0.005 | 0.005 | 0 | 0.17 | 0.02 | 0.16 | 0.13 | 0.26 | 0.19 |
| 7 | 134998517 | rs10954494 | 0.261 | 0.952 | -1.121 | -0.078 | -0.002 | 0.003 | 0.003 | 0.698 | 0.16 | 0.14 | 0.31 | 0.57 | 0.25 | 0.24 |
| 7 | 134998591 | rs17236136 | 0.419 | 0.945 | -1.33 | -0.281 | -0.035 | 0.054 | 0.054 | 0 | 0.15 | 0.02 | 0.13 | 0.1 | 0.18 | 0.16 |
| 7 | 134998730 | rs1863000 | 0.287 | 1.097 | -1.105 | -0.072 | -0.002 | 0.004 | 0.004 | 0.192 | 0.16 | 0.14 | 0.45 | 0.57 | 0.25 | 0.25 |
| 7 | 135001908 | rs13240667 | -0.703 | 0.772 | -0.795 | -0.386 | 0.001 | -0.006 | -0.006 | 0 | 0.22 | 0.2 | 0.34 | 0.13 | 0.31 | 0.24 |
| 7 | 135003377 | rs13230718 | 0.558 | 0.543 | -0.143 | -0.448 | 0.003 | 0.023 | 0.023 | 0 | 0.11 | 0.08 | 0.09 | 0.09 | 0.14 | 0.1 |
| 7 | 135003596 | rs12154454 | -0.182 | 0.171 | -0.695 | -1.302 | 0.002 | 0.024 | 0.024 | 0 | 0.45 | 0.38 | 0.65 | 0.83 | 0.5 | 0.42 |
| 7 | 135005120 | rs9642017 | 0.84 | 0.269 | -0.813 | -1.147 | -0.019 | -0.018 | -0.018 | 0 | 0.18 | 0.15 | 0.49 | 0.58 | 0.29 | 0.25 |
| 7 | 135007412 | rs2059364 | -1.607 | 0.095 | -0.949 | -1.35 | -0.024 | 0.016 | 0.016 | 0 | 0.89 | 0.98 | 0.92 | 0.97 | 0.84 | 0.93 |
| 7 | 135013131 | rs6467580 | -0.761 | 0.036 | -0.969 | -1.354 | -0.007 | -0.027 | -0.027 | 0 | 0.56 | 0.48 | 0.74 | 0.86 | 0.63 | 0.51 |
| 7 | 135014808 | rs11762135 | 1.407 | -0.012 | -1.004 | -1.395 | 0.033 | 0.144 | 0.144 | 0 | 0.23 | 0.03 | 0.13 | 0 | 0.16 | 0.12 |
| 7 | 135015407 | rs2881947 | -1.295 | -0.011 | -0.954 | -1.401 | 0.043 | -0.026 | -0.026 | 0 | 0.88 | 0.69 | 0.9 | 0.97 | 0.84 | 0.93 |
| 7 | 135015546 | rs12532231 | 1.899 | -0.009 | -1.007 | -1.387 | 0.036 | 0.05 | 0.05 | 0 | 0.31 | 0.2 | 0.16 | 0.11 | 0.21 | 0.42 |
| 7 | 135016454 | rs7777931 | 0.567 | 0.118 | -0.911 | -1.244 | 0.027 | 0.023 | 0.023 | 0 | 0.21 | 0.22 | 0.51 | 0.57 | 0.32 | 0.29 |
| 7 | 135017035 | rs4732119 | -0.046 | -0.163 | -0.874 | -1.104 | 0.001 | 0.025 | 0.025 | 0 | 0.45 | 0.39 | 0.65 | 0.76 | 0.49 | 0.42 |
| 7 | 135018924 | rs4075517 | 0.755 | 0.517 | -0.277 | -0.446 | 0.008 | 0.08 | 0.08 | 0.184 | 0.1 | 0.15 | 0.3 | 0.54 | 0.13 | 0.16 |
| 7 | 135020123 | rs7798021 | 0.174 | 0.534 | -0.351 | -0.434 | 0.055 | 0.223 | 0.223 | 0.226 | 0.13 | 0.33 | 0.33 | 0.54 | 0.15 | 0.19 |
| 7 | 135021399 | rs7803365 | -0.904 | 0.472 | -0.452 | -0.505 | 0.014 | 0.059 | 0.059 | 0.119 | 0.25 | 0.43 | 0.42 | 0.63 | 0.29 | 0.29 |
| 7 | 135021898 | rs11767423 | 1.052 | 0.238 | -0.725 | -0.704 | -0.022 | 0.018 | 0.018 | 0 | 0.63 | 0.25 | 0.48 | 0.34 | 0.55 | 0.64 |
| 7 | 135023914 | rs11772685 | 1.052 | 0.238 | -0.725 | -0.706 | -0.022 | 0.017 | 0.017 | 0 | 0.63 | 0.26 | 0.48 | 0.34 | 0.55 | 0.64 |
| 7 | 135026353 | rs6951296 | 0.092 | 0.098 | -0.567 | -0.902 | -0.009 | -0.079 | -0.079 | 0 | 0.16 | 0.39 | 0.11 | 0.09 | 0.15 | 0.1 |
| 7 | 135031468 | rs6958241 | 0.084 | -0.402 | -1.026 | -1.144 | 0.019 | 0.018 | 0.018 | 0 | 0.6 | 0.64 | 0.77 | 0.71 | 0.69 | 0.8 |
| 7 | 135031484 | rs6978334 | -1.045 | -0.341 | -1.497 | -1.009 | 0.084 | 0.019 | 0.019 | 0 | 0.74 | 0.91 | 0.86 | 0.8 | 0.85 | 0.88 |
| 7 | 135034378 | rs1599874 | 0.086 | 0.711 | -1.145 | -0.12 | -0.007 | -0.005 | -0.005 | 0 | 0.19 | 0.17 | 0.3 | 0.43 | 0.26 | 0.21 |
| 7 | 135038388 | rs889933 | 0.212 | 0.817 | -1.045 | -0.454 | -0.017 | 0.014 | 0.014 | 0 | 0.34 | 0.23 | 0.41 | 0.27 | 0.38 | 0.56 |
| 7 | 135043388 | rs2348526 | 0.166 | 1.027 | -0.813 | -0.387 | 0.003 | -0.009 | -0.009 | 0 | 0.31 | 0.32 | 0.38 | 0.24 | 0.34 | 0.38 |
| 7 | 135045786 | rs6962887 | -0.746 | 1.332 | -0.922 | -0.24 | 0.008 | 0.101 | 0.101 | 0 | 0.35 | 0.18 | 0.2 | 0.18 | 0.32 | 0.29 |
| 7 | 135046552 | rs1003226 | 0.29 | 1.1 | -0.166 | -0.36 | 0.004 | -0.029 | -0.029 | 0.369 | 0.92 | 0.67 | 0.94 | 1 | 0.96 | 0.96 |
| 7 | 135047964 | rs3750067 | 0.353 | 1.052 | -0.423 | -0.471 | -0.023 | -0.037 | -0.037 | 0 | 0.1 | 0.03 | 0.07 | 0.16 | 0.11 | 0.14 |
| 7 | 135053638 | rs1863004 | 1.327 | 0.947 | -0.242 | -0.591 | 0 | 0.016 | 0.016 | 0 | 0.75 | 0.79 | 0.83 | 0.82 | 0.72 | 0.77 |
| 7 | 135054371 | rs3828955 | 0.355 | 1.543 | -0.329 | -0.47 | -0.007 | 0.009 | 0.009 | 0 | 0.14 | 0.11 | 0.32 | 0.52 | 0.25 | 0.18 |
| 7 | 135061329 | rs3812274 | 1.957 | 1.471 | -0.321 | -0.544 | 0.006 | -0.014 | -0.014 | 0 | 0.6 | 0.54 | 0.49 | 0.3 | 0.48 | 0.59 |
| 7 | 135062056 | rs7455447 | 0.856 | 1.426 | -0.458 | -0.613 | 0.019 | 0.054 | 0.054 | 0 | 0.14 | 0.06 | 0.07 | 0 | 0.08 | 0.09 |
| 7 | 135072141 | rs3812278 | 0.509 | 1.499 | -0.458 | -0.583 | -0.018 | -0.009 | -0.009 | 0.382 | 0.13 | 0.09 | 0.32 | 0.52 | 0.24 | 0.2 |
| 7 | 135073047 | rs11772832 | 2.387 | 1.359 | -0.022 | -0.623 | 0.012 | 0.058 | 0.058 | 0 | 0.43 | 0.31 | 0.42 | 0.25 | 0.4 | 0.5 |
| 7 | 135082387 | rs7810911 | -0.782 | 1.27 | 0.004 | -0.581 | -0.011 | -0.031 | -0.031 | 2.608 | 0.94 | 0.88 | 0.96 | 1 | 0.96 | 0.96 |
| 7 | 135094763 | rs10279987 | 0.482 | 1.322 | -0.027 | -0.533 | -0.019 | -0.045 | -0.045 | 0 | 0.16 | 0.19 | 0.08 | 0.05 | 0.09 | 0.1 |

| | | | | | | | | | | | | | | | | |
|---|---|---|---|---|---|---|---|---|---|---|---|---|---|---|---|---|
| 7 | 135096934 | rs3812284 | 1.435 | 1.473 | -0.15 | -0.425 | -0.037 | 0.019 | 0.019 | 0 | 0.08 | 0.01 | 0.05 | 0.01 | 0.13 | 0.06 |
| 7 | 135103660 | rs2696879 | -0.404 | 1.752 | -0.049 | -0.403 | 0.019 | 0.036 | 0.036 | 0 | 0.17 | 0.21 | 0.38 | 0.59 | 0.3 | 0.21 |
| 7 | 135105533 | rs3812287 | -0.299 | 1.676 | -0.296 | -0.316 | -0.066 | -0.027 | -0.027 | 0 | 0.18 | 0.03 | 0.12 | 0.17 | 0.23 | 0.18 |
| 7 | 135119417 | rs2551774 | -2.343 | 1.376 | -0.226 | -0.466 | 0.009 | 0.055 | 0.055 | 0 | 0.58 | 0.7 | 0.58 | 0.75 | 0.6 | 0.5 |
| 7 | 135165275 | rs10250306 | 0.524 | 1.312 | -0.456 | -0.46 | 0.007 | 0.004 | 0.004 | 0 | 0.16 | 0.13 | 0.07 | 0.05 | 0.09 | 0.1 |
| 7 | 135199511 | rs2348775 | -2.242 | 1.469 | -0.396 | -0.123 | 0.003 | 0.022 | 0.022 | 0 | 0.57 | 0.66 | 0.71 | 0.75 | 0.59 | 0.55 |
| 7 | 135211446 | rs10265128 | 0.578 | 1.555 | -0.748 | -0.436 | -0.003 | 0.083 | 0.083 | 0 | 0.16 | 0.04 | 0.08 | 0 | 0.12 | 0.1 |
| 7 | 135213352 | rs16874692 | -0.776 | 1.757 | -0.665 | -0.629 | 0 | -0.052 | -0.052 | 0 | 0.19 | 0.24 | 0.13 | 0.01 | 0.22 | 0.2 |
| 7 | 135214060 | rs17403842 | 0.268 | 1.833 | -0.71 | -0.215 | -0.009 | -0.029 | -0.029 | 0.047 | 0.11 | 0.05 | 0.07 | 0.17 | 0.11 | 0.15 |
| 7 | 135221210 | rs7799709 | -0.364 | 1.82 | -0.567 | -0.282 | 0.006 | 0.015 | 0.015 | 0 | 0.19 | 0.15 | 0.12 | 0.06 | 0.16 | 0.24 |
| 7 | 135223938 | rs17404165 | -0.922 | 1.953 | -0.598 | -0.108 | 0.04 | 0.071 | 0.071 | 0 | 0.34 | 0.12 | 0.16 | 0.2 | 0.24 | 0.3 |
| 7 | 135227513 | rs11768238 | 2.038 | 2.194 | -0.209 | -0.006 | 0.009 | 0.016 | 0.016 | 0 | 0.41 | 0.31 | 0.21 | 0.2 | 0.33 | 0.34 |
| 7 | 135228262 | rs3112379 | 0.573 | 3.707 | 0.634 | 1.307 | 0.025 | 0.071 | 0.071 | 0 | 0.85 | 0.82 | 0.52 | 0.46 | 0.75 | 0.8 |
| 7 | 135231872 | rs3112383 | 0.592 | 3.705 | 0.633 | 1.306 | 0.025 | 0.069 | 0.069 | 0 | 0.84 | 0.82 | 0.52 | 0.46 | 0.75 | 0.8 |
| 7 | 135240951 | rs6961391 | -0.158 | 3.642 | 0.57 | 1.305 | 0.005 | -0.027 | -0.027 | 0.179 | 0.08 | 0.03 | 0.07 | 0.25 | 0.07 | 0.17 |
| 7 | 135241045 | rs13311326 | -0.848 | 3.628 | 0.774 | 1.289 | 0.015 | 0.049 | 0.049 | 0 | 0.15 | 0.18 | 0.48 | 0.54 | 0.25 | 0.19 |
| 7 | 135247752 | rs6467595 | 0.779 | 3.463 | 0.879 | 1.307 | 0.043 | 0.148 | 0.148 | 0 | 0.84 | 0.7 | 0.53 | 0.48 | 0.78 | 0.73 |
| 7 | 135254692 | rs11768962 | -0.123 | 3.632 | 0.667 | 1.183 | 0.032 | 0.157 | 0.157 | 0 | 0.07 | 0.1 | 0.41 | 0.46 | 0.16 | 0.13 |
| 7 | 135271762 | rs7782920 | -0.902 | 4.01 | 0.447 | 0.923 | -0.005 | -0.005 | -0.005 | 0 | 0.11 | 0.14 | 0.14 | 0.23 | 0.22 | 0.18 |
| 7 | 135302785 | rs12536760 | -0.757 | 3.045 | 0.573 | 0.656 | -0.011 | -0.009 | -0.009 | 0.037 | 0.1 | 0.15 | 0.06 | 0.07 | 0.06 | 0.19 |
| 7 | 135311120 | rs7810260 | -2.319 | 3.448 | 0.473 | 0.371 | -0.002 | 0.022 | 0.022 | 0 | 0.43 | 0.43 | 0.36 | 0.26 | 0.51 | 0.49 |
| 7 | 135312572 | rs4073627 | -0.06 | 3.6 | 0.157 | 0.419 | -0.02 | -0.017 | -0.017 | 0 | 0.08 | 0.05 | 0.13 | 0.18 | 0.14 | 0.17 |
| 7 | 135318779 | rs7806221 | -1.243 | 3.62 | 0.147 | 0.389 | -0.081 | 0.05 | 0.05 | 0.446 | 0.24 | 0.1 | 0.26 | 0.19 | 0.38 | 0.28 |
| 7 | 135328110 | rs10260691 | 2.591 | 3.434 | 0.357 | 0.643 | 0.091 | 0.116 | 0.116 | 0 | 0.49 | 0.21 | 0.26 | 0.26 | 0.36 | 0.38 |
| 7 | 135329690 | rs12540688 | -1.151 | 2.877 | 0.536 | 0.812 | 0.004 | 0.003 | 0.003 | 0 | 0.19 | 0.24 | 0.2 | 0.25 | 0.2 | 0.36 |
| 7 | 135336264 | rs12531468 | -1.023 | 2.888 | 0.541 | 0.693 | 0.006 | -0.004 | -0.004 | 0 | 0.18 | 0.24 | 0.19 | 0.2 | 0.19 | 0.33 |
| 7 | 135342501 | rs3884652 | -0.369 | 2.527 | 0.643 | 0.988 | -0.01 | 0.048 | 0.048 | 0 | 0.12 | 0.03 | 0.07 | 0 | 0.1 | 0.06 |
| 7 | 135364526 | rs3110789 | -1.747 | 2.533 | 0.891 | 1.22 | 0.023 | 0.108 | 0.108 | 0 | 0.24 | 0.43 | 0.46 | 0.55 | 0.26 | 0.26 |
| 7 | 135376893 | rs3112335 | -1.025 | 2.639 | 1.386 | 1.607 | 0.092 | 0.28 | 0.28 | 0 | 0.11 | 0.3 | 0.39 | 0.44 | 0.18 | 0.22 |
| 7 | 135387495 | rs3112346 | 0.636 | 2.53 | 0.94 | 0.998 | -0.03 | 0.064 | 0.064 | 0.542 | 0.69 | 0.78 | 0.71 | 0.89 | 0.61 | 0.72 |
| 7 | 135392322 | rs3112353 | 1.024 | 2.476 | 1.02 | 0.931 | -0.004 | -0.004 | -0.004 | 0.664 | 0.63 | 0.65 | 0.66 | 0.88 | 0.56 | 0.68 |
| 7 | 135392876 | rs3110800 | 1.017 | 2.474 | 1.019 | 0.93 | -0.004 | -0.004 | -0.004 | 0.532 | 0.63 | 0.64 | 0.66 | 0.88 | 0.56 | 0.68 |
| 7 | 135398094 | rs4236657 | 1.742 | 2.274 | 1.067 | 1.014 | 0.06 | 0.034 | 0.034 | 2.534 | 0.49 | 0.22 | 0.26 | 0.38 | 0.37 | 0.39 |
| 7 | 135400731 | rs3112358 | -0.493 | 1.619 | 1.228 | 1.714 | 0.018 | 0.104 | 0.104 | 0 | 0.1 | 0.21 | 0.37 | 0.24 | 0.13 | 0.21 |
| 7 | 135404841 | rs3112362 | -1.725 | 1.725 | 1.375 | 1.805 | 0.032 | 0.141 | 0.141 | 0 | 0.18 | 0.36 | 0.45 | 0.33 | 0.23 | 0.34 |
| 7 | 135405904 | rs6948713 | 2.107 | 1.614 | 1.343 | 1.689 | 0.041 | 0.109 | 0.109 | 3.565 | 0.48 | 0.2 | 0.25 | 0.3 | 0.4 | 0.28 |
| 7 | 135406176 | rs4596594 | 1.769 | 1.63 | 1.465 | 1.722 | -0.001 | -0.001 | -0.001 | 0.301 | 0.55 | 0.51 | 0.32 | 0.44 | 0.46 | 0.47 |
| 7 | 135411152 | rs3110817 | 0.415 | 1.618 | 1.405 | 1.691 | -0.042 | 0.01 | 0.01 | 1.62 | 0.71 | 0.86 | 0.73 | 0.7 | 0.64 | 0.71 |
| 7 | 135412273 | rs3112369 | 0.849 | 1.64 | 1.43 | 1.691 | -0.031 | 0.025 | 0.025 | 1.931 | 0.66 | 0.82 | 0.69 | 0.69 | 0.59 | 0.67 |
| 7 | 135412568 | rs3800769 | 1.743 | 1.641 | 1.484 | 1.714 | 0 | 0 | 0 | 2.878 | 0.53 | 0.46 | 0.3 | 0.45 | 0.45 | 0.44 |
| 7 | 135412747 | rs3112370 | -0.429 | 1.403 | 1.24 | 1.718 | 0.015 | 0.099 | 0.099 | 0 | 0.1 | 0.22 | 0.37 | 0.24 | 0.13 | 0.21 |
| 7 | 135420018 | rs4732139 | 1.644 | 1.387 | 0.911 | 1.584 | -0.003 | -0.007 | -0.007 | 0.324 | 0.56 | 0.48 | 0.32 | 0.39 | 0.43 | 0.4 |
| 7 | 135428793 | rs4430037 | 1.265 | 1.042 | 0.847 | 1.186 | -0.019 | 0 | 0 | 0.026 | 0.62 | 0.36 | 0.64 | 0.62 | 0.53 | 0.6 |
| 7 | 135432657 | rs10954502 | 1.905 | 1.108 | 0.917 | 1.225 | 0.023 | 0.019 | 0.019 | 1.849 | 0.49 | 0.31 | 0.55 | 0.38 | 0.39 | 0.29 |
| 7 | 135439915 | rs10257539 | 0.63 | 1.009 | 0.813 | 0.999 | -0.058 | -0.008 | -0.008 | 0 | 0.72 | 0.92 | 0.75 | 0.72 | 0.65 | 0.71 |
| 7 | 135441131 | rs7808197 | 1.443 | 1.697 | 1.304 | 1.226 | 0.03 | 0.024 | 0.024 | 1.519 | 0.66 | 0.47 | 0.68 | 0.55 | 0.55 | 0.53 |
| 7 | 135455224 | rs6467606 | 0.495 | 1.13 | 1.229 | 0.604 | 0.009 | 0.006 | 0.006 | 0 | 0.75 | 0.68 | 0.8 | 0.7 | 0.72 | 0.73 |
| 7 | 135456092 | rs6961992 | 1.304 | 1.265 | 0.924 | 0.693 | 0.009 | 0.006 | 0.006 | 0.519 | 0.66 | 0.44 | 0.71 | 0.62 | 0.63 | 0.62 |
| 7 | 135459752 | rs6467607 | -0.968 | 0.902 | 0.874 | 0.613 | 0.01 | 0.014 | 0.014 | 1.068 | 0.09 | 0.28 | 0.08 | 0.07 | 0.1 | 0.11 |
| 7 | 135466041 | rs10279370 | -1.252 | 0.852 | 0.579 | 0.536 | -0.002 | -0.02 | -0.02 | 0 | 0.16 | 0.66 | 0.16 | 0.09 | 0.16 | 0.17 |
| 7 | 135466740 | rs7357254 | 0.543 | 0.791 | 0.077 | 0.577 | -0.005 | 0.009 | 0.009 | 0.002 | 0.71 | 0.63 | 0.82 | 0.7 | 0.77 | 0.75 |
| 7 | 135469462 | rs6977424 | 0.395 | 1.106 | -0.423 | -0.038 | -0.004 | 0.019 | 0.019 | 0 | 0.59 | 0.68 | 0.61 | 0.8 | 0.48 | 0.57 |
| 7 | 135472073 | rs4130689 | 0.448 | 1.04 | -0.611 | -0.162 | 0.024 | 0.04 | 0.04 | 0.281 | 0.52 | 0.69 | 0.46 | 0.67 | 0.39 | 0.51 |
| 7 | 135478701 | rs4732145 | 0.168 | 1.568 | -0.128 | 0.16 | -0.083 | 0.034 | 0.034 | 0.014 | 0.61 | 0.88 | 0.72 | 0.7 | 0.53 | 0.58 |
| 7 | 135478903 | rs4732146 | -0.396 | 2.26 | -0.457 | -0.056 | -0.036 | -0.029 | -0.029 | 0.424 | 0.88 | 0.97 | 0.9 | 0.83 | 0.82 | 0.7 |
| 7 | 135480428 | rs11761160 | 1.328 | 1.848 | -0.229 | 0.137 | 0.027 | -0.012 | -0.012 | 0 | 0.44 | 0.35 | 0.4 | 0.53 | 0.33 | 0.42 |
| 7 | 135482945 | rs4602829 | -0.449 | 2.013 | -0.427 | 0.113 | -0.038 | -0.029 | -0.029 | 0 | 0.87 | 0.97 | 0.87 | 0.83 | 0.8 | 0.69 |
| 7 | 135483051 | rs10233321 | 0.486 | 2.116 | -0.599 | -0.08 | -0.099 | 0.115 | 0.115 | 0.268 | 0.59 | 0.9 | 0.72 | 0.77 | 0.5 | 0.57 |
| 7 | 135483229 | rs11983139 | 0.122 | 2.126 | -0.05 | 0.163 | 0.056 | 0.056 | 0.056 | 0 | 0.08 | 0.19 | 0.19 | 0.1 | 0.11 | 0.08 |
| 7 | 135483806 | rs4260843 | 0.427 | 2.052 | 0.169 | 0.064 | -0.005 | -0.003 | -0.003 | 0 | 0.79 | 0.8 | 0.85 | 0.77 | 0.77 | 0.65 |
| 7 | 135484024 | rs4551283 | -1.238 | 1.574 | -0.092 | 0.405 | -0.006 | 0.013 | 0.013 | 0.141 | 0.63 | 0.58 | 0.55 | 0.45 | 0.67 | 0.58 |
| 7 | 135484792 | rs9656461 | -0.377 | 1.352 | 0.088 | 0.386 | 0 | 0.002 | 0.002 | 0 | 0.44 | 0.55 | 0.44 | 0.26 | 0.46 | 0.26 |
| 7 | 135485052 | rs9656462 | 0.792 | 1.356 | -0.088 | 0.108 | 0.008 | -0.004 | -0.004 | 0 | 0.24 | 0.36 | 0.31 | 0.21 | 0.3 | 0.16 |
| 7 | 135487308 | rs11762744 | 0.488 | 1.336 | -0.214 | 0.082 | -0.005 | 0.07 | 0.07 | 0 | 0.19 | 0.06 | 0.12 | 0.04 | 0.16 | 0.1 |
| 7 | 135487851 | rs4732148 | 1.802 | 1.232 | -0.178 | 0.088 | 0.008 | 0.022 | 0.022 | 0 | 0.17 | 0.12 | 0.09 | 0.09 | 0.19 | 0.08 |
| 7 | 135488335 | rs10954507 | -1.001 | 1.325 | -0.154 | 0.357 | -0.001 | 0.005 | 0.005 | 0 | 0.44 | 0.57 | 0.47 | 0.34 | 0.48 | 0.27 |
| 7 | 135488621 | rs6946494 | 1.607 | 1.258 | -0.37 | 0.427 | 0.008 | 0.022 | 0.022 | 0 | 0.18 | 0.12 | 0.09 | 0.1 | 0.19 | 0.08 |
| 7 | 135491262 | rs1443759 | 0.531 | 1.315 | -0.153 | 0.781 | -0.002 | -0.002 | -0.002 | 0.042 | 0.59 | 0.42 | 0.67 | 0.46 | 0.56 | 0.54 |
| 7 | 135494535 | rs2920486 | -1.094 | 1.446 | -0.286 | 0.814 | -0.002 | 0.017 | 0.017 | 0 | 0.78 | 0.68 | 0.8 | 0.73 | 0.76 | 0.89 |

| | | | | | | | | | | | | | | | | |
|---|---|---|---|---|---|---|---|---|---|---|---|---|---|---|---|---|
| 7 | 135495586 | rs2920487 | -1.106 | 1.457 | -0.282 | 0.269 | -0.001 | -0.023 | -0.023 | 0 | 0.78 | 0.68 | 0.81 | 0.86 | 0.76 | 0.9 |
| 7 | 135508122 | rs17810062 | 0.621 | 1.297 | -0.784 | 0.289 | -0.015 | -0.018 | -0.018 | 0 | 0.12 | 0.07 | 0.09 | 0.18 | 0.16 | 0.28 |
| 7 | 135508956 | rs17168469 | -0.267 | 0.548 | -0.849 | -0.017 | 0 | 0.048 | 0.048 | 0.25 | 0.23 | 0.11 | 0.26 | 0.07 | 0.22 | 0.18 |
| 7 | 135512610 | rs4636148 | 1.12 | 0.852 | -0.712 | 0.387 | -0.024 | -0.014 | -0.014 | 0 | 0.2 | 0.14 | 0.3 | 0.23 | 0.3 | 0.14 |
| 7 | 135512761 | rs6956576 | -0.904 | 0.343 | -0.301 | 0.322 | 0.001 | 0.002 | 0.002 | 2.59 | 0.51 | 0.52 | 0.46 | 0.42 | 0.46 | 0.33 |
| 7 | 135514121 | rs6962270 | 1.235 | 0.164 | -0.596 | 0.318 | -0.049 | 0.003 | 0.003 | 0.882 | 0.42 | 0.2 | 0.55 | 0.39 | 0.49 | 0.41 |
| 7 | 135514146 | rs6962277 | 0.542 | -0.044 | -0.484 | 0.335 | -0.007 | -0.001 | -0.001 | 0.024 | 0.71 | 0.36 | 0.83 | 0.81 | 0.87 | 0.85 |
| 7 | 135519460 | rs17168475 | 1.271 | 0.547 | 0.049 | 0.668 | -0.002 | 0.007 | 0.007 | 0 | 0.09 | 0.09 | 0.03 | 0.15 | 0.06 | 0.07 |
| 7 | 135521869 | rs4732153 | 1.207 | 0.437 | -0.003 | 0.638 | 0.007 | -0.008 | -0.008 | 5.154 | 0.67 | 0.32 | 0.79 | 0.6 | 0.77 | 0.51 |
| 7 | 135522547 | rs4333513 | 1.196 | 0.462 | -0.002 | 0.638 | 0.006 | -0.007 | -0.007 | 0.01 | 0.67 | 0.35 | 0.79 | 0.6 | 0.77 | 0.51 |
| 7 | 135524678 | rs17133353 | -0.154 | 0.463 | 0.163 | 0.632 | -0.007 | 0.002 | 0.002 | 0 | 0.19 | 0.2 | 0.1 | 0.18 | 0.11 | 0.14 |
| 7 | 135526123 | rs13244788 | -0.152 | 0.461 | 0.162 | 0.551 | -0.007 | -0.001 | -0.001 | 1.26 | 0.19 | 0.2 | 0.1 | 0.16 | 0.11 | 0.14 |
| 7 | 135527643 | rs9918682 | 0.406 | 0.559 | 0.329 | 0.56 | -0.003 | 0.007 | 0.007 | 0.306 | 0.61 | 0.48 | 0.67 | 0.59 | 0.64 | 0.75 |
| 7 | 135529279 | rs10262451 | 1.841 | 0.663 | 0.334 | 0.654 | -0.023 | -0.024 | -0.024 | 0 | 0.18 | 0.1 | 0.21 | 0.23 | 0.25 | 0.11 |
| 7 | 135535762 | rs1579783 | -0.027 | 0.61 | 0.429 | 0.647 | -0.013 | 0.003 | 0.003 | 0.205 | 0.77 | 0.68 | 0.84 | 0.76 | 0.83 | 0.59 |
| 7 | 135539396 | rs13247578 | 1.115 | 0.612 | 0.64 | 0.652 | 0.001 | 0.037 | 0.037 | 0 | 0.09 | 0.23 | 0.04 | 0.12 | 0.06 | 0.07 |
| 7 | 135543143 | rs1000433 | -0.206 | 0.599 | 0.241 | 0.578 | 0 | 0 | 0 | 0 | 0.79 | 0.8 | 0.88 | 0.8 | 0.81 | 0.62 |
| 7 | 135547247 | rs4732156 | 1.952 | 0.635 | 0.124 | 0.599 | -0.03 | -0.031 | -0.031 | 2.401 | 0.18 | 0.09 | 0.21 | 0.23 | 0.25 | 0.11 |
| 7 | 135547365 | rs17168496 | 1.129 | 0.671 | 0.122 | 0.633 | -0.041 | -0.056 | -0.056 | 0 | 0.14 | 0.02 | 0.1 | 0.2 | 0.14 | 0.35 |
| 7 | 135553732 | rs1562028 | 1.513 | 0.613 | 0.185 | 0.644 | -0.009 | 0.042 | 0.042 | 1.517 | 0.45 | 0.21 | 0.56 | 0.38 | 0.47 | 0.37 |
| 7 | 135555255 | rs2348910 | 0.514 | 0.685 | 0.199 | 0.652 | 0 | 0.009 | 0.009 | 0 | 0.61 | 0.49 | 0.68 | 0.58 | 0.61 | 0.72 |
| 7 | 135556323 | rs994678 | 0.624 | 0.489 | 0.317 | 0.539 | -0.028 | -0.012 | -0.012 | 0 | 0.79 | 0.6 | 0.89 | 0.84 | 0.86 | 0.82 |
| 7 | 135561276 | rs978738 | 0 | 0.498 | 0.271 | 0.511 | -0.014 | -0.01 | -0.01 | 7.758 | 0.88 | 0.7 | 0.92 | 0.94 | 0.92 | 0.89 |
| 7 | 135562501 | rs894795 | 0.417 | 0.623 | 0.143 | 0.548 | -0.005 | -0.005 | -0.005 | 0 | 0.56 | 0.64 | 0.62 | 0.53 | 0.53 | 0.45 |
| 7 | 135565457 | rs7787315 | 0.786 | 0.66 | 0.049 | 0.572 | -0.002 | -0.002 | -0.002 | 0 | 0.47 | 0.49 | 0.49 | 0.42 | 0.45 | 0.39 |
| 7 | 135569910 | rs10085478 | 0.313 | 0.707 | -0.134 | 0.62 | -0.008 | -0.01 | -0.01 | 0 | 0.55 | 0.39 | 0.51 | 0.5 | 0.51 | 0.46 |
| 7 | 135573240 | rs4732158 | 0.768 | 0.851 | -0.803 | 0.752 | 0.006 | 0.042 | 0.042 | 0.034 | 0.62 | 0.27 | 0.77 | 0.76 | 0.64 | 0.69 |
| 7 | 135577509 | rs17816783 | 0.892 | 0.917 | -1.443 | 0.549 | -0.035 | 0.086 | 0.086 | 1.477 | 0.19 | 0.04 | 0.12 | 0.03 | 0.22 | 0.2 |
| 7 | 135577585 | rs1982292 | -0.134 | 0.961 | -1.025 | 0.915 | 0.069 | 0.076 | 0.076 | 0 | 0.9 | 0.72 | 0.89 | 0.85 | 0.91 | 0.94 |
| 7 | 135579485 | rs12674250 | 0.267 | 0.401 | -2.02 | 0.788 | 0.007 | -0.001 | -0.001 | 0.065 | 0.05 | 0.03 | 0.05 | 0.03 | 0.03 | 0.07 |
| 7 | 135583537 | rs7789182 | 0.979 | 0.609 | -2.068 | 0.769 | -0.033 | -0.011 | -0.011 | 0.034 | 0.39 | 0.11 | 0.5 | 0.37 | 0.41 | 0.25 |
| 7 | 135586482 | rs10229676 | -0.789 | 0.458 | -1.101 | 1.113 | 0.012 | -0.015 | -0.015 | 0 | 0.2 | 0.14 | 0.19 | 0.27 | 0.14 | 0.4 |
| 7 | 135590031 | rs10275042 | 0.525 | 0.938 | -0.74 | 1.08 | 0.001 | 0.002 | 0.002 | 0.433 | 0.55 | 0.54 | 0.4 | 0.5 | 0.52 | 0.42 |
| 7 | 135594129 | rs1373234 | 1.686 | 1.375 | -0.891 | 1.384 | -0.012 | 0.044 | 0.044 | 0 | 0.4 | 0.1 | 0.49 | 0.34 | 0.39 | 0.3 |
| 7 | 135600091 | rs12667644 | 1.919 | 1.543 | 0.299 | 2.007 | -0.011 | 0.065 | 0.065 | 0 | 0.54 | 0.28 | 0.62 | 0.38 | 0.52 | 0.51 |
| 7 | 135603088 | rs10155818 | -2.52 | 2.5 | 1.179 | 2.404 | 0.036 | -0.01 | -0.01 | 0.621 | 0.36 | 0.5 | 0.35 | 0.61 | 0.48 | 0.51 |
| 7 | 135608912 | rs7794499 | -1.192 | 2.66 | 1.135 | 2.753 | -0.043 | -0.04 | -0.04 | 3.163 | 0.08 | 0.02 | 0.08 | 0.19 | 0.16 | 0.12 |
| 7 | 135617507 | rs4732162 | -2.916 | 2.88 | 1.464 | 3.107 | 0.04 | -0.012 | -0.012 | 0 | 0.34 | 0.49 | 0.34 | 0.61 | 0.46 | 0.5 |
| 7 | 135620639 | rs11772267 | -1.356 | 2.715 | 1.79 | 3.219 | -0.022 | -0.023 | -0.023 | 0 | 0.11 | 0.04 | 0.14 | 0.16 | 0.15 | 0.08 |
| 7 | 135632137 | rs13246297 | -3.42 | 2.805 | 1.922 | 3.17 | 0.031 | -0.01 | -0.01 | 0.381 | 0.36 | 0.55 | 0.35 | 0.61 | 0.45 | 0.44 |
| 7 | 135632351 | rs999250 | -1.195 | 2.847 | 1.822 | 2.906 | -0.002 | -0.001 | -0.001 | 0 | 0.09 | 0.11 | 0.03 | 0.07 | 0.07 | 0.16 |
| 7 | 135637264 | rs12531496 | -2.238 | 2.777 | 1.805 | 2.934 | 0.043 | 0.055 | 0.055 | 0 | 0.16 | 0.25 | 0.21 | 0.28 | 0.28 | 0.18 |
| 7 | 135641580 | rs1443762 | -0.236 | 2.657 | 1.818 | 2.838 | -0.008 | -0.005 | -0.005 | 0 | 0.06 | 0.03 | 0.09 | 0.08 | 0.09 | 0.04 |
| 7 | 135675538 | rs4728373 | -3.602 | 2.653 | 1.827 | 2.898 | 0.035 | 0.049 | 0.049 | 0 | 0.28 | 0.46 | 0.25 | 0.47 | 0.36 | 0.4 |
| 7 | 135678299 | rs11772302 | -1.688 | 2.829 | 1.863 | 3.095 | -0.001 | 0 | 0 | 0 | 0.15 | 0.13 | 0.12 | 0.15 | 0.16 | 0.19 |
| 7 | 135678346 | rs4728374 | -1.696 | 2.854 | 1.84 | 3.103 | 0.018 | 0.078 | 0.078 | 0 | 0.11 | 0.12 | 0.14 | 0.4 | 0.21 | 0.2 |
| 7 | 135682419 | rs1994464 | -2.427 | 2.736 | 1.892 | 2.662 | 0.035 | 0.02 | 0.02 | 0 | 0.27 | 0.33 | 0.25 | 0.55 | 0.37 | 0.39 |
| 7 | 135688926 | rs6467623 | -2.249 | 3.062 | 2.245 | 3.042 | 0.047 | 0.075 | 0.075 | 0 | 0.19 | 0.3 | 0.12 | 0.33 | 0.26 | 0.31 |
| 7 | 135691400 | rs10278297 | -2.621 | 2.865 | 2.323 | 3.081 | 0.074 | 0.188 | 0.188 | 0 | 0.19 | 0.57 | 0.22 | 0.45 | 0.3 | 0.23 |
| 7 | 135699857 | rs7793221 | -1.962 | 2.728 | 2.433 | 2.864 | 0.024 | 0.081 | 0.081 | 0 | 0.17 | 0.39 | 0.1 | 0.26 | 0.16 | 0.22 |
| 7 | 135704136 | rs7804115 | -0.67 | 2.744 | 2.06 | 2.87 | -0.022 | -0.027 | -0.027 | 0 | 0.1 | 0.02 | 0.05 | 0.15 | 0.12 | 0.11 |
| 7 | 135704269 | rs7804029 | -1.901 | 2.832 | 2.11 | 2.931 | -0.008 | -0.007 | -0.007 | 0 | 0.2 | 0.17 | 0.32 | 0.27 | 0.27 | 0.22 |
| 7 | 135704751 | rs10262219 | -1.374 | 2.82 | 2.179 | 2.9 | 0.053 | 0.139 | 0.139 | 0 | 0.06 | 0.39 | 0.05 | 0.1 | 0.04 | 0.11 |
| 7 | 135705059 | rs1552476 | -0.941 | 2.851 | 1.296 | 2.674 | 0.018 | 0.007 | 0.007 | 0 | 0.1 | 0.17 | 0.28 | 0.12 | 0.15 | 0.11 |
| 7 | 135709862 | rs7798284 | -0.973 | 2.854 | 1.259 | 2.684 | 0.019 | 0.062 | 0.062 | 0 | 0.16 | 0.28 | 0.09 | 0.26 | 0.16 | 0.22 |
| 7 | 135710436 | rs7798858 | -2.19 | 2.356 | 0.659 | 2.324 | 0.061 | 0.108 | 0.108 | 0 | 0.13 | 0.23 | 0.35 | 0.32 | 0.25 | 0.13 |
| 7 | 135711388 | rs1488002 | -1.483 | 1.315 | 0.131 | 1.264 | 0.002 | 0 | 0 | 0 | 0.1 | 0.11 | 0.27 | 0.12 | 0.15 | 0.11 |
| 7 | 135718444 | rs7776674 | -1.396 | 0.457 | 0.315 | 1.203 | -0.01 | 0.014 | 0.014 | 0 | 0.1 | 0.14 | 0.07 | 0.15 | 0.07 | 0.06 |
| 7 | 135718968 | rs901796 | -2.42 | 0.378 | 1.284 | 1.078 | -0.012 | 0.036 | 0.036 | 0.166 | 0.15 | 0.56 | 0.11 | 0.16 | 0.1 | 0.15 |
| 7 | 135720387 | rs1290737 | -2.057 | 0.26 | 1.364 | 1.085 | -0.025 | 0.067 | 0.067 | 0 | 0.13 | 0.44 | 0.1 | 0.16 | 0.07 | 0.11 |
| 7 | 135731036 | rs2129800 | 0.063 | -0.024 | 0.351 | 1.171 | -0.03 | -0.116 | -0.116 | 0 | 0.29 | 0.03 | 0.36 | 0.45 | 0.24 | 0.38 |
| 7 | 135736414 | rs322213 | -2.408 | 0.042 | 1.347 | 1.002 | -0.026 | 0.032 | 0.032 | 0.484 | 0.14 | 0.41 | 0.1 | 0.16 | 0.09 | 0.16 |
| 7 | 135737781 | rs322211 | -2.207 | 0.087 | 1.074 | 1 | -0.022 | 0.018 | 0.018 | 0.496 | 0.14 | 0.33 | 0.09 | 0.15 | 0.09 | 0.09 |
| 7 | 135742442 | rs7781896 | -1.786 | 0.35 | 0.793 | 1.069 | 0.002 | 0.014 | 0.014 | 0.043 | 0.08 | 0.16 | 0.07 | 0.11 | 0.06 | 0.07 |
| 7 | 135752140 | rs10246095 | -1.908 | -0.344 | 1.322 | 0.657 | -0.065 | -0.054 | -0.054 | 0 | 0.13 | 0.52 | 0.07 | 0.05 | 0.06 | 0.07 |
| 7 | 135754469 | rs17168576 | -2.21 | -0.641 | 1.449 | 1.024 | -0.033 | 0.01 | 0.01 | 0.202 | 0.14 | 0.24 | 0.07 | 0.13 | 0.06 | 0.08 |
| 7 | 135762274 | rs10229838 | -2.405 | -0.515 | 2.079 | 1.195 | -0.017 | 0.081 | 0.081 | 1.78 | 0.15 | 0.33 | 0.07 | 0.25 | 0.1 | 0.13 |
| 7 | 135768477 | rs7782458 | -2.522 | 0.054 | 2.125 | 1.93 | 0.04 | 0.149 | 0.149 | 0 | 0.11 | 0.24 | 0.07 | 0.27 | 0.15 | 0.24 |
| 7 | 135772668 | rs11766224 | -1.643 | 0.057 | 2.13 | 1.911 | 0.071 | 0.193 | 0.193 | 0.811 | 0.06 | 0.22 | 0.05 | 0.15 | 0.1 | 0.14 |

| | | | | | | | | | | | | | | | | |
|---|---|---|---|---|---|---|---|---|---|---|---|---|---|---|---|---|
| 7 | 135774180 | rs4728378 | -2.715 | 0.032 | 2.009 | 1.769 | 0.008 | 0.201 | 0.201 | 1.568 | 0.2 | 0.46 | 0.12 | 0.4 | 0.2 | 0.28 |
| 7 | 135774345 | rs1488003 | -2.127 | 0.024 | 2.023 | 1.648 | 0.01 | 0.216 | 0.216 | 1.605 | 0.15 | 0.42 | 0.1 | 0.28 | 0.15 | 0.17 |
| 7 | 135776776 | rs6467636 | -0.028 | 0.523 | 0.503 | 0.793 | -0.024 | -0.039 | -0.039 | 0 | 0.75 | 0.84 | 0.48 | 0.29 | 0.67 | 0.7 |
| 7 | 135785380 | rs9801253 | 0.034 | 0.306 | -0.175 | -1.814 | -0.035 | 0.056 | 0.056 | 0 | 0.27 | 0.17 | 0.23 | 0.05 | 0.35 | 0.21 |
| 7 | 135788599 | rs12535285 | 0.226 | 0.256 | -0.168 | -1.835 | -0.062 | 0.091 | 0.091 | 0 | 0.25 | 0.09 | 0.22 | 0.05 | 0.35 | 0.21 |
| 7 | 135800804 | rs12112159 | -0.414 | -0.56 | 0.052 | -2.742 | 0.001 | -0.052 | -0.052 | 0 | 0.5 | 0.55 | 0.37 | 0.06 | 0.49 | 0.48 |
| 7 | 135801215 | rs17241838 | 0.427 | -0.86 | -0.176 | -2.921 | -0.054 | 0.103 | 0.103 | 0.429 | 0.32 | 0.16 | 0.25 | 0.05 | 0.39 | 0.24 |
| 7 | 135802554 | rs6959619 | -1.003 | -0.995 | 0.359 | -2.704 | -0.029 | -0.106 | -0.106 | 0 | 0.26 | 0.65 | 0.18 | 0.01 | 0.14 | 0.28 |
| 7 | 135804051 | rs17168588 | -0.423 | -0.922 | 0.379 | -2.607 | -0.054 | -0.147 | -0.147 | 0 | 0.2 | 0.54 | 0.14 | 0.01 | 0.1 | 0.24 |
| 7 | 135806402 | rs10281536 | 0.059 | -0.954 | -0.289 | -3.325 | 0.001 | -0.043 | -0.043 | 0 | 0.25 | 0.21 | 0.14 | 0 | 0.18 | 0.17 |
| 7 | 135806732 | rs12707271 | -0.39 | -0.3 | -0.359 | -3.62 | 0 | -0.043 | -0.043 | 0 | 0.22 | 0.18 | 0.15 | 0.01 | 0.26 | 0.09 |
| 7 | 135807663 | rs10270418 | -1.05 | -0.484 | -0.681 | -3.778 | 0 | -0.042 | -0.042 | 0.048 | 0.31 | 0.71 | 0.21 | 0.01 | 0.31 | 0.13 |
| 7 | 135813034 | rs6951864 | -0.903 | -0.755 | -0.459 | -3.937 | 0.002 | -0.139 | -0.139 | 0 | 0.29 | 0.54 | 0.19 | 0.01 | 0.31 | 0.13 |
| 7 | 135815196 | rs2129801 | -1.418 | -0.693 | -0.902 | -3.989 | -0.12 | 0.381 | 0.381 | 0 | 0.48 | 0.95 | 0.35 | 0.05 | 0.47 | 0.4 |
| 7 | 135815329 | rs2171491 | 1.224 | -0.455 | -0.657 | -3.869 | -0.015 | 0.02 | 0.02 | 1.634 | 0.14 | 0.09 | 0.11 | 0 | 0.15 | 0.16 |
| 7 | 135821703 | rs12534779 | -2.003 | -0.697 | -1.017 | -4.363 | -0.081 | 0.233 | 0.233 | 0 | 0.62 | 0.94 | 0.39 | 0.11 | 0.56 | 0.48 |
| 7 | 135831413 | rs10267117 | 0.143 | -0.915 | -0.321 | -4.383 | -0.03 | -0.088 | -0.088 | 0.441 | 0.19 | 0.33 | 0.07 | 0.02 | 0.1 | 0.09 |
| 7 | 135834934 | rs12707272 | -1.615 | -0.841 | -0.482 | -4.438 | -0.014 | 0.232 | 0.232 | 0.043 | 0.47 | 0.77 | 0.27 | 0.08 | 0.43 | 0.34 |
| 7 | 135838528 | rs1002292 | -0.212 | -0.733 | -0.457 | -4.445 | 0.004 | -0.056 | -0.056 | 0 | 0.24 | 0.23 | 0.18 | 0.01 | 0.32 | 0.14 |
| 7 | 135838892 | rs834753 | -1.511 | -0.727 | -0.619 | -4.41 | -0.008 | 0.172 | 0.172 | 0.047 | 0.47 | 0.76 | 0.28 | 0.06 | 0.43 | 0.34 |
| 7 | 135840925 | rs12531124 | -0.716 | -1.007 | -0.503 | -4.059 | -0.007 | -0.09 | -0.09 | 0 | 0.33 | 0.37 | 0.18 | 0.01 | 0.23 | 0.22 |
| 7 | 135842250 | rs834764 | 0.833 | -0.782 | -1.063 | -3.753 | 0 | -0.018 | -0.018 | 2.593 | 0.16 | 0.15 | 0.12 | 0.01 | 0.14 | 0.16 |
| 7 | 135845461 | rs834767 | -1.566 | -0.455 | -1.729 | -4.121 | -0.103 | 0.257 | 0.257 | 0 | 0.63 | 0.93 | 0.53 | 0.04 | 0.54 | 0.58 |
| 7 | 135848396 | rs834771 | 0.548 | -0.351 | -1.039 | -3.74 | 0.007 | 0.008 | 0.008 | 0.041 | 0.19 | 0.16 | 0.11 | 0.02 | 0.15 | 0.19 |
| 7 | 135849744 | rs4113948 | 0.465 | -0.249 | -0.454 | -3.71 | 0.006 | -0.055 | -0.055 | 0 | 0.1 | 0.25 | 0.06 | 0 | 0.07 | 0.03 |
| 7 | 135854002 | rs834780 | -0.078 | -0.267 | -0.476 | -4 | -0.015 | -0.102 | -0.102 | 0 | 0.26 | 0.58 | 0.17 | 0.02 | 0.21 | 0.19 |
| 7 | 135854856 | rs834781 | -1.158 | -0.224 | -0.796 | -4.313 | -0.002 | 0.138 | 0.138 | 0 | 0.43 | 0.74 | 0.29 | 0.04 | 0.41 | 0.39 |
| 7 | 135854941 | rs741632 | 0.721 | -0.214 | -0.881 | -4.294 | -0.016 | 0.022 | 0.022 | 0 | 0.17 | 0.1 | 0.1 | 0.02 | 0.2 | 0.19 |
| 7 | 135856359 | rs12534419 | 0.886 | -0.359 | -1.041 | -4.403 | 0.008 | 0.005 | 0.005 | 0.271 | 0.13 | 0.1 | 0.04 | 0 | 0.06 | 0.06 |
| 7 | 135859750 | rs706582 | -2.146 | -0.318 | -1.116 | -4.484 | -0.06 | 0.233 | 0.233 | 0 | 0.59 | 0.86 | 0.34 | 0.04 | 0.5 | 0.47 |
| 7 | 135860469 | rs10278510 | 1.52 | -0.351 | -0.834 | -4.353 | -0.01 | -0.09 | -0.09 | 0 | 0.15 | 0.28 | 0.11 | 0 | 0.11 | 0.13 |
| 7 | 135861304 | rs834787 | -1.545 | -0.106 | -1.903 | -4.141 | -0.071 | 0.274 | 0.274 | 0 | 0.55 | 0.85 | 0.33 | 0.04 | 0.47 | 0.45 |
| 7 | 135862877 | rs12532042 | 0.739 | -0.108 | -1.846 | -4.091 | 0.007 | 0.004 | 0.004 | 0.364 | 0.12 | 0.1 | 0.04 | 0 | 0.06 | 0.06 |
| 7 | 135864820 | rs834795 | 0.031 | 0.271 | -1.933 | -3.757 | 0.032 | 0.056 | 0.056 | 0 | 0.38 | 0.7 | 0.22 | 0.02 | 0.27 | 0.25 |
| 7 | 135865186 | rs17243303 | -0.678 | 0.741 | -1.006 | -2.456 | -0.007 | -0.001 | -0.001 | 0 | 0.17 | 0.15 | 0.11 | 0.02 | 0.2 | 0.2 |
| 7 | 135870953 | rs834800 | -0.133 | 0.374 | -1.067 | -1.989 | -0.003 | 0.09 | 0.09 | 0 | 0.47 | 0.7 | 0.58 | 0.17 | 0.55 | 0.46 |
| 7 | 135873681 | rs4732179 | 0.063 | 0.104 | -0.126 | -2.209 | -0.001 | -0.009 | -0.009 | 0 | 0.41 | 0.45 | 0.36 | 0.14 | 0.51 | 0.39 |
| 7 | 135879132 | rs1121038 | 0.312 | -0.412 | 0.342 | -2.195 | -0.076 | 0.024 | 0.024 | 0 | 0.31 | 0.16 | 0.32 | 0.19 | 0.49 | 0.34 |
| 7 | 135880446 | rs1097147 | -1.305 | -0.997 | 0.467 | -2.525 | -0.003 | 0.013 | 0.013 | 0 | 0.62 | 0.72 | 0.63 | 0.26 | 0.64 | 0.67 |
| 7 | 135883871 | rs1097151 | -0.047 | -0.973 | 0.487 | -2.523 | 0.006 | -0.018 | -0.018 | 1.162 | 0.4 | 0.6 | 0.37 | 0.19 | 0.5 | 0.38 |
| 7 | 135884571 | rs834811 | -0.826 | -0.971 | 0.476 | -2.534 | 0.004 | 0.035 | 0.035 | 0 | 0.55 | 0.68 | 0.42 | 0.23 | 0.59 | 0.59 |
| 7 | 135889329 | rs2162332 | 0.479 | -0.891 | 0.328 | -2.495 | 0.014 | 0.021 | 0.021 | 0.303 | 0.15 | 0.09 | 0.05 | 0.05 | 0.08 | 0.2 |
| 7 | 135889681 | rs7779031 | 0.637 | -0.905 | 0.292 | -2.461 | -0.073 | 0.023 | 0.023 | 0 | 0.3 | 0.15 | 0.31 | 0.19 | 0.48 | 0.34 |
| 7 | 135896926 | rs834822 | -0.958 | -0.495 | 0.277 | -2.697 | -0.012 | -0.066 | -0.066 | 0 | 0.29 | 0.39 | 0.37 | 0.09 | 0.25 | 0.33 |
| 7 | 135897003 | rs10278072 | 0.231 | -0.593 | 0.298 | -2.753 | 0.001 | -0.001 | -0.001 | 0 | 0.31 | 0.34 | 0.24 | 0.2 | 0.38 | 0.34 |
| 7 | 135901545 | rs2349456 | 1.485 | -0.512 | 0.224 | -2.744 | -0.001 | 0.022 | 0.022 | 0 | 0.4 | 0.26 | 0.39 | 0.72 | 0.37 | 0.33 |
| 7 | 135904371 | rs1346208 | 0.26 | -0.623 | 0.233 | -2.697 | 0.003 | -0.055 | -0.055 | 0.921 | 0.6 | 0.47 | 0.71 | 0.81 | 0.57 | 0.5 |
| 7 | 135904504 | rs1346209 | 0.56 | -0.67 | 0.197 | -2.712 | 0 | 0.003 | 0.003 | 0 | 0.54 | 0.43 | 0.51 | 0.73 | 0.52 | 0.4 |
| 7 | 135904832 | rs4732187 | 1.444 | -0.852 | 0.001 | -3.038 | -0.023 | 0.001 | 0.001 | 0.358 | 0.32 | 0.17 | 0.4 | 0.72 | 0.36 | 0.26 |
| 7 | 135905859 | rs1365358 | 1.694 | -0.811 | -0.109 | -3.111 | -0.02 | 0.057 | 0.057 | 0.147 | 0.17 | 0.06 | 0.07 | 0 | 0.16 | 0.17 |
| 7 | 135906817 | rs1426479 | 1.074 | -0.823 | -0.128 | -3.068 | -0.021 | 0.022 | 0.022 | 0 | 0.23 | 0.16 | 0.22 | 0.13 | 0.35 | 0.26 |
| 7 | 135915023 | rs2067624 | -0.094 | -0.835 | -0.074 | -3.059 | -0.02 | -0.006 | -0.006 | 0 | 0.21 | 0.26 | 0.26 | 0.15 | 0.11 | 0.28 |
| 7 | 135920630 | rs1426483 | -0.982 | -0.437 | 0.092 | -2.845 | 0.009 | -0.013 | -0.013 | 0.233 | 0.77 | 0.72 | 0.78 | 0.87 | 0.65 | 0.72 |
| 7 | 135925039 | rs975789 | 1.549 | -0.575 | 0.011 | -2.821 | 0.001 | 0.001 | 0.001 | 0.188 | 0.09 | 0.07 | 0.03 | 0.07 | 0.05 | 0.16 |
| 7 | 135938900 | rs7789574 | 1.081 | -0.46 | -0.006 | -2.872 | -0.007 | 0.052 | 0.052 | 0 | 0.46 | 0.29 | 0.39 | 0.71 | 0.42 | 0.42 |
| 7 | 135945310 | rs10464717 | 1.006 | -0.424 | 0.035 | -2.856 | -0.004 | 0.054 | 0.054 | 0 | 0.47 | 0.3 | 0.39 | 0.71 | 0.42 | 0.43 |
| 7 | 135950203 | rs730694 | -0.35 | -0.364 | 0.078 | -2.86 | -0.003 | 0.001 | 0.001 | 0 | 0.69 | 0.68 | 0.7 | 0.86 | 0.55 | 0.71 |
| 7 | 135953018 | rs11526287 | 0.709 | -0.352 | 0.355 | -2.582 | -0.024 | 0.011 | 0.011 | 0 | 0.15 | 0.25 | 0.25 | 0.15 | 0.09 | 0.26 |
| 7 | 135957494 | rs12707290 | 1.505 | -0.304 | 0.273 | -2.578 | -0.003 | -0.039 | -0.039 | 0.041 | 0.07 | 0.14 | 0.06 | 0 | 0.04 | 0.02 |
| 7 | 135958755 | rs11767724 | 1.505 | -0.304 | 0.27 | -2.578 | -0.004 | -0.038 | -0.038 | 0 | 0.07 | 0.14 | 0.06 | 0 | 0.04 | 0.02 |
| 7 | 135959346 | rs13238853 | -0.574 | -0.339 | 0.215 | -2.683 | 0.007 | 0.016 | 0.016 | 0 | 0.49 | 0.54 | 0.46 | 0.16 | 0.39 | 0.5 |
| 7 | 135960417 | rs2349458 | 0.992 | -0.418 | 0.159 | -2.602 | 0.004 | -0.003 | -0.003 | 1.053 | 0.14 | 0.11 | 0.23 | 0.14 | 0.09 | 0.26 |
| 7 | 135960581 | rs6975667 | 1.494 | -0.277 | -0.181 | -2.568 | -0.005 | -0.013 | -0.013 | 0.188 | 0.28 | 0.17 | 0.31 | 0.71 | 0.26 | 0.23 |
| 7 | 135960799 | rs7811360 | 0.999 | 0.114 | -0.278 | -2.293 | 0 | -0.012 | -0.012 | 0.099 | 0.09 | 0.08 | 0.06 | 0 | 0.09 | 0.06 |
| 7 | 135967572 | rs9801179 | 1.273 | 0.032 | -0.206 | -2.076 | 0 | 0.008 | 0.008 | 1.202 | 0.35 | 0.28 | 0.37 | 0.71 | 0.31 | 0.28 |
| 7 | 135967904 | rs4141426 | 0.901 | 0.802 | -0.539 | -2.025 | 0.002 | -0.013 | -0.013 | 1.421 | 0.5 | 0.54 | 0.63 | 0.86 | 0.4 | 0.54 |
| 7 | 135972132 | rs10275033 | -0.56 | 0.266 | -1.181 | -1.758 | -0.003 | 0.029 | 0.029 | 0 | 0.26 | 0.14 | 0.15 | 0.01 | 0.25 | 0.22 |
| 7 | 135985050 | rs2217092 | 0.209 | 0.102 | -0.888 | -1.145 | 0.004 | 0.03 | 0.03 | 0 | 0.49 | 0.64 | 0.63 | 0.69 | 0.53 | 0.76 |
| 7 | 136018322 | rs4487700 | 0.831 | -0.059 | -0.668 | -1.076 | -0.037 | -0.045 | -0.045 | 1.202 | 0.33 | 0.14 | 0.51 | 0.61 | 0.41 | 0.58 |

| | | | | | | | | | | | | | | | | |
|---|---|---|---|---|---|---|---|---|---|---|---|---|---|---|---|---|
| 7 | 136020256 | rs4579471 | -0.108 | -0.028 | -0.72 | -1.33 | -0.001 | 0 | 0 | 0 | 0.72 | 0.7 | 0.78 | 0.76 | 0.68 | 0.88 |
| 7 | 136028899 | rs11760528 | -0.099 | -0.666 | -1.237 | -2.173 | -0.018 | 0.06 | 0.06 | 0 | 0.2 | 0.02 | 0.16 | 0.05 | 0.18 | 0.06 |
| 7 | 136029472 | rs6966878 | -1.635 | -0.566 | -1.494 | -2.205 | -0.124 | -0.013 | -0.013 | 0 | 0.64 | 0.97 | 0.53 | 0.32 | 0.54 | 0.47 |
| 7 | 136038492 | rs10488413 | -0.51 | -0.85 | -0.784 | -2.37 | 0.037 | 0.097 | 0.097 | 0 | 0.44 | 0.6 | 0.34 | 0.13 | 0.33 | 0.26 |
| 7 | 136053854 | rs2882248 | -1.226 | -0.396 | -0.17 | -2.187 | 0.031 | -0.006 | -0.006 | 1.825 | 0.19 | 0.47 | 0.22 | 0.19 | 0.23 | 0.2 |
| 7 | 136056458 | rs10242359 | -0.351 | -0.754 | 0.173 | -2.166 | 0.007 | 0.038 | 0.038 | 0.191 | 0.46 | 0.31 | 0.34 | 0.13 | 0.39 | 0.28 |
| 7 | 136057432 | rs10246453 | -1.425 | -0.131 | 0.792 | -1.579 | 0.004 | 0.068 | 0.068 | 2.701 | 0.5 | 0.72 | 0.43 | 0.34 | 0.45 | 0.45 |
| 7 | 136057530 | rs10276178 | -0.834 | -0.128 | 0.742 | -1.577 | -0.001 | 0.016 | 0.016 | 0.563 | 0.13 | 0.37 | 0.15 | 0.19 | 0.14 | 0.18 |
| 7 | 136057883 | rs4732202 | -1.182 | -0.178 | 0.687 | -1.518 | 0.002 | 0.032 | 0.032 | 1.531 | 0.49 | 0.67 | 0.43 | 0.34 | 0.45 | 0.44 |
| 7 | 136067679 | rs10275448 | -0.921 | -0.24 | 0.905 | -1.175 | 0.003 | -0.023 | -0.023 | 0 | 0.39 | 0.53 | 0.32 | 0.3 | 0.39 | 0.4 |
| 7 | 136074356 | rs7796512 | -0.236 | -0.264 | 0.859 | -1.181 | 0.002 | -0.037 | -0.037 | 0 | 0.32 | 0.31 | 0.24 | 0.15 | 0.29 | 0.29 |
| 7 | 136078041 | rs11981211 | -0.809 | -0.143 | 0.171 | -1.168 | -0.006 | 0.054 | 0.054 | 0.636 | 0.41 | 0.8 | 0.35 | 0.3 | 0.39 | 0.41 |
| 7 | 136106307 | rs10488416 | -1.015 | 0.495 | -0.575 | 0.057 | -0.001 | 0.004 | 0.004 | 0 | 0.08 | 0.12 | 0.06 | 0.19 | 0.1 | 0.11 |
| 7 | 136120119 | rs1806430 | -0.937 | 0.234 | -0.471 | -0.674 | -0.003 | -0.035 | -0.035 | 0 | 0.14 | 0.16 | 0.1 | 0 | 0.13 | 0.12 |
| 7 | 136124848 | rs10215946 | -0.657 | -0.369 | 0.369 | -1.207 | -0.015 | -0.061 | -0.061 | 0.039 | 0.12 | 0.34 | 0.06 | 0.02 | 0.09 | 0.13 |
| 7 | 136131088 | rs12707307 | 0.882 | -0.155 | -0.254 | -2.173 | -0.007 | 0.107 | 0.107 | 0 | 0.35 | 0.19 | 0.26 | 0.09 | 0.3 | 0.24 |
| 7 | 136134819 | rs4732209 | 0.129 | -0.217 | 0.132 | -2.271 | -0.016 | -0.105 | -0.105 | 0.03 | 0.81 | 0.64 | 0.86 | 0.96 | 0.84 | 0.73 |
| 7 | 136142209 | rs4421308 | 0.228 | -0.287 | 0.358 | -2.302 | -0.02 | -0.126 | -0.126 | 0.052 | 0.81 | 0.57 | 0.86 | 0.96 | 0.84 | 0.73 |
| 7 | 136146536 | rs10250997 | 0.392 | -0.28 | 0.806 | -2.338 | -0.009 | -0.055 | -0.055 | 0.064 | 0.12 | 0.29 | 0.06 | 0.02 | 0.09 | 0.12 |
| 7 | 136150869 | rs11761139 | 0.297 | -0.263 | 0.746 | -2.275 | -0.012 | -0.066 | -0.066 | 0 | 0.15 | 0.22 | 0.1 | 0 | 0.13 | 0.11 |
| 7 | 136166189 | rs1946330 | -0.78 | -0.369 | 0.726 | -2.313 | -0.002 | 0.033 | 0.033 | 0 | 0.62 | 0.74 | 0.71 | 0.9 | 0.66 | 0.55 |
| 7 | 136173789 | rs1364742 | -1.141 | -0.404 | 0.706 | -2.331 | -0.023 | 0.14 | 0.14 | 0 | 0.66 | 0.89 | 0.74 | 0.91 | 0.69 | 0.75 |
| 7 | 136179342 | rs10258191 | -0.796 | -0.55 | 0.885 | -2.48 | -0.002 | -0.071 | -0.071 | 0 | 0.64 | 0.61 | 0.71 | 0.89 | 0.68 | 0.56 |
| 7 | 136186762 | rs10954532 | 1.197 | -0.517 | 0.585 | -2.447 | -0.018 | 0.119 | 0.119 | 0 | 0.31 | 0.1 | 0.24 | 0.1 | 0.28 | 0.24 |
| 7 | 136188728 | rs10258427 | 1.139 | -0.556 | 0.574 | -2.466 | -0.007 | 0.078 | 0.078 | 0 | 0.31 | 0.16 | 0.24 | 0.11 | 0.28 | 0.24 |
| 7 | 136200127 | rs6467670 | 0.669 | -0.569 | 0.725 | -2.438 | -0.006 | -0.049 | -0.049 | 1.388 | 0.14 | 0.17 | 0.12 | 0 | 0.13 | 0.11 |
| 7 | 136205020 | rs11767346 | 0.665 | -0.702 | 0.596 | -2.553 | -0.012 | 0.016 | 0.016 | 0 | 0.38 | 0.3 | 0.53 | 0.86 | 0.47 | 0.3 |
| 7 | 136205736 | rs1473220 | -0.386 | -0.726 | 0.693 | -2.573 | -0.004 | -0.011 | -0.011 | 0 | 0.58 | 0.61 | 0.44 | 0.13 | 0.5 | 0.48 |
| 7 | 136228650 | rs10277960 | -0.964 | -0.836 | 0.772 | -2.627 | 0 | 0.035 | 0.035 | 0 | 0.65 | 0.73 | 0.73 | 0.9 | 0.69 | 0.54 |
| 7 | 136231030 | rs1946332 | 0.715 | -0.864 | 0.812 | -2.657 | -0.027 | 0.063 | 0.063 | 0 | 0.38 | 0.24 | 0.52 | 0.87 | 0.47 | 0.31 |
| 7 | 136254878 | rs36088642 | 1.391 | -0.862 | 0.651 | -2.636 | -0.034 | 0.142 | 0.142 | 0 | 0.31 | 0.07 | 0.24 | 0.09 | 0.28 | 0.25 |
| 7 | 136272383 | rs10271343 | -0.637 | -1.064 | 0.675 | -2.734 | -0.033 | -0.142 | -0.142 | 0 | 0.26 | 0.47 | 0.2 | 0.02 | 0.22 | 0.24 |
| 7 | 136273464 | rs10954538 | -0.948 | -1.166 | 0.616 | -2.728 | 0.004 | 0.125 | 0.125 | 0 | 0.64 | 0.81 | 0.73 | 0.9 | 0.69 | 0.54 |
| 7 | 136283910 | rs17168728 | -0.211 | -1.218 | 0.644 | -2.721 | -0.016 | -0.133 | -0.133 | 0 | 0.19 | 0.44 | 0.17 | 0.01 | 0.17 | 0.14 |
| 7 | 136308528 | rs11763296 | 0.157 | -1.375 | 0.47 | -2.85 | 0 | 0.041 | 0.041 | 0 | 0.47 | 0.36 | 0.56 | 0.89 | 0.52 | 0.4 |
| 7 | 136312729 | rs10247745 | -0.173 | -1.294 | 0.489 | -2.816 | -0.016 | -0.144 | -0.144 | 0.119 | 0.19 | 0.47 | 0.17 | 0.01 | 0.17 | 0.15 |
| 7 | 136319271 | rs2350176 | 1.107 | -1.315 | 0.436 | -2.827 | -0.035 | 0.057 | 0.057 | 0 | 0.36 | 0.24 | 0.53 | 0.87 | 0.47 | 0.29 |
| 7 | 136326080 | rs7778878 | 0.493 | -1.41 | 0.484 | -2.765 | 0.007 | 0.012 | 0.012 | 0 | 0.08 | 0.05 | 0.02 | 0.02 | 0.05 | 0.1 |
| 7 | 136327805 | rs11982756 | -0.006 | -1.396 | 0.414 | -2.871 | 0.037 | 0.121 | 0.121 | 0 | 0.49 | 0.32 | 0.61 | 0.89 | 0.62 | 0.41 |
| 7 | 136333854 | rs7459286 | 0.659 | -1.389 | 0.399 | -2.868 | -0.01 | 0.094 | 0.094 | 0 | 0.42 | 0.25 | 0.59 | 0.87 | 0.56 | 0.31 |
| 7 | 136334971 | rs10239526 | -1.014 | -1.442 | 0.441 | -2.896 | 0.006 | 0.098 | 0.098 | 0 | 0.63 | 0.8 | 0.73 | 0.89 | 0.69 | 0.54 |
| 7 | 136335761 | rs4282513 | 0.859 | -1.461 | 0.36 | -2.89 | -0.053 | -0.095 | -0.095 | 0 | 0.81 | 0.4 | 0.85 | 0.97 | 0.9 | 0.67 |
| 7 | 136339058 | rs10435421 | 0.932 | -1.453 | 0.111 | -2.85 | -0.133 | 0.175 | 0.175 | 0 | 0.39 | 0.03 | 0.57 | 0.86 | 0.56 | 0.31 |
| 7 | 136339927 | rs10244704 | -0.763 | -1.381 | 0.318 | -2.782 | -0.066 | -0.132 | -0.132 | 0 | 0.21 | 0.45 | 0.13 | 0.03 | 0.13 | 0.22 |
| 7 | 136340237 | rs13227663 | 1.075 | -1.176 | -0.213 | -2.671 | -0.016 | 0.133 | 0.133 | 0 | 0.33 | 0.13 | 0.24 | 0.09 | 0.28 | 0.26 |
| 7 | 136341233 | rs4518582 | 0.508 | 0.313 | -0.834 | -1.611 | -0.097 | -0.132 | -0.132 | 0 | 0.2 | 0.04 | 0.43 | 0.52 | 0.34 | 0.19 |
| 7 | 136348038 | rs4321934 | -1.564 | 0.051 | -1.381 | -2.709 | -0.007 | 0.122 | 0.122 | 0.089 | 0.64 | 0.89 | 0.82 | 0.81 | 0.71 | 0.52 |
| 7 | 136348356 | rs10447873 | 0.282 | 0.458 | -0.885 | -2.922 | 0.01 | -0.058 | -0.058 | 1.47 | 0.11 | 0.24 | 0.11 | 0 | 0.14 | 0.05 |
| 7 | 136350740 | rs10235715 | 0.225 | 0.328 | -0.401 | -3.073 | -0.01 | -0.075 | -0.075 | 0.64 | 0.12 | 0.43 | 0.07 | 0 | 0.06 | 0.03 |
| 7 | 136364995 | rs9649615 | 0.255 | 0.267 | -0.584 | -3.236 | -0.057 | 0.04 | 0.04 | 0 | 0.28 | 0.12 | 0.61 | 0.79 | 0.42 | 0.31 |
| 7 | 136370812 | rs10257276 | -2.08 | 0.128 | -0.513 | -3.21 | 0.001 | 0.11 | 0.11 | 0.06 | 0.64 | 0.87 | 0.82 | 0.8 | 0.71 | 0.52 |
| 7 | 136372424 | rs10269811 | 0.242 | 0.108 | -0.109 | -2.952 | -0.015 | -0.071 | -0.071 | 0 | 0.11 | 0.34 | 0.07 | 0 | 0.06 | 0.03 |
| 7 | 136375979 | rs10954541 | 0.412 | 0.364 | -0.385 | -3.132 | -0.07 | -0.014 | -0.014 | 0 | 0.17 | 0.05 | 0.53 | 0.79 | 0.32 | 0.28 |
| 7 | 136380996 | rs7790740 | -0.884 | 0.29 | -0.344 | -3.151 | -0.052 | -0.16 | -0.16 | 0 | 0.22 | 0.49 | 0.11 | 0 | 0.14 | 0.11 |
| 7 | 136389040 | rs10259281 | -1.669 | 0.333 | -0.498 | -3.141 | -0.007 | 0.083 | 0.083 | 0.153 | 0.44 | 0.71 | 0.65 | 0.8 | 0.48 | 0.43 |
| 7 | 136398712 | rs10275735 | -1.267 | 0.327 | -0.47 | -3.15 | -0.032 | -0.098 | -0.098 | 0.997 | 0.26 | 0.64 | 0.12 | 0.02 | 0.15 | 0.15 |
| 7 | 136399178 | rs10249472 | -1.652 | 0.349 | -0.505 | -3.138 | -0.009 | 0.087 | 0.087 | 0.073 | 0.44 | 0.72 | 0.66 | 0.8 | 0.48 | 0.43 |
| 7 | 136419814 | rs7809111 | -0.46 | 0.308 | -0.314 | -3.095 | -0.029 | -0.089 | -0.089 | 2.949 | 0.14 | 0.5 | 0.07 | 0.02 | 0.06 | 0.06 |
| 7 | 136421981 | rs10249508 | -1.445 | 0.301 | -0.453 | -3.108 | -0.018 | -0.072 | -0.072 | 0 | 0.25 | 0.68 | 0.12 | 0.02 | 0.14 | 0.14 |
| 7 | 136433776 | rs4728404 | 1.91 | 0.212 | -0.656 | -3.283 | -0.007 | 0.136 | 0.136 | 0 | 0.51 | 0.26 | 0.37 | 0.2 | 0.48 | 0.56 |
| 7 | 136442955 | rs7809426 | 1.551 | -0.211 | -0.043 | -3.008 | -0.025 | -0.102 | -0.102 | 0.057 | 0.72 | 0.38 | 0.86 | 0.98 | 0.82 | 0.83 |
| 7 | 136452082 | rs10954546 | 0.245 | -0.217 | 0.152 | -2.983 | -0.012 | -0.104 | -0.104 | 0 | 0.86 | 0.57 | 0.9 | 1 | 0.91 | 0.95 |
| 7 | 136508139 | rs7810141 | 1.24 | -0.162 | -0.123 | -2.947 | -0.031 | -0.114 | -0.114 | 0 | 0.73 | 0.41 | 0.87 | 0.98 | 0.82 | 0.82 |
| 7 | 136524926 | rs12539762 | -0.764 | 0.023 | -0.345 | -2.443 | -0.003 | -0.101 | -0.101 | 0 | 0.29 | 0.34 | 0.18 | 0.02 | 0.27 | 0.15 |
| 7 | 136526263 | rs7811987 | -0.842 | -0.339 | -0.53 | -2.518 | -0.001 | -0.079 | -0.079 | 0 | 0.41 | 0.4 | 0.23 | 0.02 | 0.34 | 0.21 |
| 7 | 136530327 | rs1424385 | 1.687 | -0.3 | -0.517 | -2.424 | 0.009 | 0.027 | 0.027 | 0 | 0.12 | 0.05 | 0.05 | 0 | 0.07 | 0.06 |
| 7 | 136533997 | rs1424557 | -0.641 | -0.394 | -0.412 | -2.344 | -0.01 | 0.009 | 0.009 | 0 | 0.91 | 0.85 | 0.88 | 0.86 | 0.91 | 0.75 |
| 7 | 136534657 | rs9656474 | 1.137 | -0.347 | -0.46 | -2.456 | 0.008 | -0.07 | -0.07 | 0.211 | 0.11 | 0.27 | 0.05 | 0 | 0.09 | 0.04 |
| 7 | 136536427 | rs6967616 | -0.639 | -0.373 | -0.494 | -2.525 | -0.005 | -0.111 | -0.111 | 0 | 0.38 | 0.41 | 0.23 | 0.02 | 0.33 | 0.18 |

| | | | | | | | | | | | | | | | | |
|---|---|---|---|---|---|---|---|---|---|---|---|---|---|---|---|---|
| 7 | 136537218 | rs1364487 | 1.649 | -0.282 | -0.652 | -2.49 | 0.008 | 0.048 | 0.048 | 0 | 0.12 | 0.04 | 0.06 | 0.01 | 0.09 | 0.06 |
| 7 | 136537510 | rs1424554 | -0.307 | 0.051 | -0.384 | -2.141 | 0.002 | -0.117 | -0.117 | 0.094 | 0.25 | 0.33 | 0.17 | 0.01 | 0.24 | 0.12 |
| 7 | 136542202 | rs10244320 | 0.898 | 0.056 | -0.382 | -2.101 | 0.006 | -0.066 | -0.066 | 0 | 0.11 | 0.27 | 0.05 | 0 | 0.09 | 0.04 |
| 7 | 136546916 | rs6467694 | 0.282 | 0.202 | -0.368 | -1.775 | -0.025 | 0.003 | 0.003 | 0 | 0.87 | 0.77 | 0.86 | 0.85 | 0.91 | 0.71 |
| 7 | 136549033 | rs1424574 | 0.53 | 0.994 | -0.804 | -1.71 | -0.007 | 0.018 | 0.018 | 0.357 | 0.14 | 0.06 | 0.12 | 0.01 | 0.16 | 0.09 |
| 7 | 136558158 | rs3735028 | 0.525 | 0.916 | -0.983 | -1.54 | -0.001 | -0.002 | -0.002 | 0 | 0.06 | 0.04 | 0.12 | 0.12 | 0.06 | 0.08 |
| 7 | 136560248 | rs10954565 | 0.013 | 0.955 | -1.075 | -1.513 | -0.034 | 0.09 | 0.09 | 0.005 | 0.17 | 0.05 | 0.15 | 0.01 | 0.22 | 0.1 |
| 7 | 136563813 | rs11771119 | 0.339 | 0.882 | -0.957 | -1.436 | -0.007 | 0.036 | 0.036 | 0.039 | 0.15 | 0.07 | 0.13 | 0.01 | 0.18 | 0.08 |
| 7 | 136566533 | rs2113550 | -0.17 | 0.951 | -0.789 | -1.356 | 0.001 | -0.044 | -0.044 | 0 | 0.35 | 0.41 | 0.29 | 0.12 | 0.34 | 0.21 |
| 7 | 136569416 | rs1424569 | -0.524 | 0.909 | -0.851 | -1.327 | 0 | -0.028 | -0.028 | 1.718 | 0.51 | 0.53 | 0.42 | 0.13 | 0.51 | 0.28 |
| 7 | 136570312 | rs17494846 | -0.382 | 1.302 | -1.329 | -1.445 | -0.054 | 0.125 | 0.125 | 0 | 0.22 | 0.05 | 0.16 | 0.01 | 0.3 | 0.1 |
| 7 | 136571707 | rs10242108 | -0.221 | 0.513 | -0.868 | -0.828 | -0.001 | 0 | 0 | 0 | 0.54 | 0.56 | 0.41 | 0.48 | 0.61 | 0.5 |
| 7 | 136575199 | rs4475425 | -0.151 | 0.676 | -0.876 | -1.165 | 0.007 | 0.066 | 0.066 | 0 | 0.25 | 0.41 | 0.21 | 0.44 | 0.27 | 0.35 |
| 7 | 136577349 | rs889934 | 0.729 | 0.89 | -0.66 | -1.297 | 0.068 | -0.038 | -0.038 | 0.08 | 0.08 | 0.28 | 0.06 | 0 | 0.11 | 0.03 |
| 7 | 136579008 | rs12537962 | 0.892 | 0.811 | -0.514 | -1.371 | -0.006 | 0.026 | 0.026 | 1.718 | 0.29 | 0.2 | 0.19 | 0.03 | 0.35 | 0.13 |
| 7 | 136582393 | rs10228048 | -0.523 | 0.789 | -0.51 | -1.334 | 0.007 | 0.073 | 0.073 | 0.023 | 0.25 | 0.43 | 0.21 | 0.44 | 0.27 | 0.35 |
| 7 | 136584075 | rs12535371 | 0.977 | 0.8 | -0.539 | -1.335 | -0.019 | 0.088 | 0.088 | 1.569 | 0.28 | 0.15 | 0.19 | 0.04 | 0.34 | 0.13 |
| 7 | 136588109 | rs6963819 | -1.17 | 0.669 | -0.335 | -1.618 | -0.001 | -0.029 | -0.029 | 0 | 0.63 | 0.8 | 0.46 | 0.47 | 0.69 | 0.56 |
| 7 | 136588827 | rs1364403 | -1.254 | 0.807 | -0.424 | -0.945 | 0.017 | 0 | 0 | 0 | 0.65 | 0.8 | 0.74 | 0.75 | 0.75 | 0.69 |
| 7 | 136596457 | rs7810473 | 0.497 | 0.814 | -0.189 | -1.043 | 0.013 | -0.102 | -0.102 | 0 | 0.36 | 0.45 | 0.26 | 0.04 | 0.46 | 0.16 |
| 7 | 136597265 | rs10267949 | 1.228 | 0.234 | -0.19 | -0.876 | 0.008 | -0.033 | -0.033 | 0 | 0.06 | 0.17 | 0.05 | 0 | 0.05 | 0.04 |
| 7 | 136605458 | rs10488600 | 0.736 | 0.161 | -0.305 | -0.794 | 0.004 | -0.054 | -0.054 | 0 | 0.14 | 0.06 | 0.14 | 0.4 | 0.15 | 0.33 |
| 7 | 136608159 | rs1364405 | 1.343 | 0.422 | -0.466 | -0.964 | -0.126 | 0.136 | 0.136 | 3.276 | 0.27 | 0.02 | 0.21 | 0.04 | 0.36 | 0.12 |
| 7 | 136609925 | rs1364407 | -0.868 | 0.299 | -0.218 | -0.976 | 0.029 | -0.009 | -0.009 | 0 | 0.61 | 0.76 | 0.47 | 0.43 | 0.71 | 0.51 |
| 7 | 136612314 | rs10256854 | 1.274 | 0.379 | -0.086 | -0.931 | 0.004 | -0.032 | -0.032 | 0 | 0.06 | 0.14 | 0.04 | 0 | 0.05 | 0.04 |
| 7 | 136620152 | rs7806357 | -0.976 | 0.371 | -0.175 | -0.94 | 0.001 | 0.004 | 0.004 | 0.659 | 0.32 | 0.64 | 0.26 | 0.4 | 0.34 | 0.35 |
| 7 | 136624320 | rs7800170 | -0.136 | 0.393 | -0.191 | -1.1 | 0.001 | 0.012 | 0.012 | 2.077 | 0.44 | 0.65 | 0.33 | 0.04 | 0.55 | 0.19 |
| 7 | 136624659 | rs10271552 | 0.367 | 0.457 | 0.093 | -0.871 | 0.076 | -0.053 | -0.053 | 0 | 0.1 | 0.35 | 0.07 | 0 | 0.12 | 0.03 |
| 7 | 136630812 | rs10225215 | -0.463 | 0.311 | -0.428 | -0.591 | -0.002 | -0.014 | -0.014 | 0 | 0.32 | 0.72 | 0.26 | 0.46 | 0.33 | 0.34 |
| 7 | 136631703 | rs1455858 | -0.514 | 0.11 | -0.053 | -0.213 | 0.014 | -0.054 | -0.054 | 0 | 0.63 | 0.83 | 0.48 | 0.48 | 0.71 | 0.5 |
| 7 | 136635001 | rs1378646 | -0.166 | 0.092 | -0.041 | -0.223 | 0.022 | -0.028 | -0.028 | 2.059 | 0.61 | 0.77 | 0.47 | 0.48 | 0.7 | 0.47 |
| 7 | 136639941 | rs1455857 | -0.513 | 0.186 | -0.02 | -0.165 | 0.015 | -0.025 | -0.025 | 0 | 0.64 | 0.83 | 0.48 | 0.45 | 0.72 | 0.51 |
| 7 | 136643694 | rs1824024 | -0.22 | 0.249 | -0.021 | -0.153 | 0.006 | 0 | 0 | 0 | 0.61 | 0.66 | 0.46 | 0.42 | 0.71 | 0.49 |
| 7 | 136652138 | rs324588 | 0.784 | 0.527 | 0.513 | 0.367 | 0.089 | 0.245 | 0.245 | 1.006 | 0.95 | 0.72 | 0.74 | 0.7 | 0.94 | 0.82 |
| 7 | 136655323 | rs324594 | 0.122 | -0.106 | 0.31 | 0.069 | 0.024 | -0.018 | -0.018 | 0 | 0.33 | 0.21 | 0.3 | 0.32 | 0.21 | 0.29 |
| 7 | 136662741 | rs2113545 | -0.054 | -0.078 | 0.29 | -0.233 | 0.001 | 0 | 0 | 0 | 0.64 | 0.39 | 0.44 | 0.39 | 0.73 | 0.57 |
| 7 | 136671341 | rs12707339 | 0.081 | 0.346 | 0.131 | -0.453 | 0.011 | -0.059 | -0.059 | 6.075 | 0.15 | 0.21 | 0.11 | 0 | 0.18 | 0.07 |
| 7 | 136676570 | rs2350786 | -0.229 | 0.052 | -0.13 | 0.003 | 0.006 | 0.056 | 0.056 | 0.125 | 0.73 | 0.55 | 0.47 | 0.41 | 0.77 | 0.62 |
| 7 | 136682188 | rs17414604 | 1.211 | 0.41 | -1.045 | -0.696 | -0.08 | 0.105 | 0.105 | 0.64 | 0.25 | 0.09 | 0.23 | 0.06 | 0.36 | 0.16 |
| 7 | 136687403 | rs420817 | 0.28 | -0.144 | -0.43 | -0.841 | -0.002 | -0.031 | -0.031 | 0.121 | 0.43 | 0.49 | 0.32 | 0.06 | 0.52 | 0.21 |
| 7 | 136688996 | rs324640 | 0.089 | -0.041 | -0.891 | -0.772 | -0.007 | 0.125 | 0.125 | 1.331 | 0.45 | 0.67 | 0.34 | 0.08 | 0.55 | 0.21 |
| 7 | 136690503 | rs10488602 | 0.791 | -0.259 | -1.463 | 0.136 | -0.024 | -0.179 | -0.179 | 0 | 0.23 | 0.01 | 0.24 | 0.42 | 0.16 | 0.26 |
| 7 | 136703376 | rs6967953 | -0.152 | -0.033 | -0.87 | -0.45 | 0.042 | 0.125 | 0.125 | 0.063 | 0.54 | 0.75 | 0.39 | 0.11 | 0.63 | 0.26 |
| 7 | 136707227 | rs17506733 | 1.16 | 0.344 | -0.439 | -0.825 | -0.015 | 0.008 | 0.008 | 0 | 0.31 | 0.24 | 0.23 | 0.01 | 0.38 | 0.15 |
| 7 | 136709168 | rs11762896 | 0.744 | 0.318 | -0.455 | -0.509 | -0.017 | 0.026 | 0.026 | 0 | 0.34 | 0.24 | 0.25 | 0.01 | 0.41 | 0.16 |
| 7 | 136712058 | rs17168896 | 0.763 | 0.228 | -0.024 | -0.527 | 0.017 | -0.057 | -0.057 | 1.455 | 0.11 | 0.18 | 0.08 | 0 | 0.15 | 0.06 |
| 7 | 136715254 | rs7779506 | 1.008 | 0.389 | -0.142 | -0.391 | -0.049 | 0.072 | 0.072 | 0 | 0.32 | 0.17 | 0.26 | 0.01 | 0.42 | 0.18 |
| 7 | 136716653 | rs10954569 | 0.693 | 0.381 | -0.133 | -0.33 | -0.046 | 0.083 | 0.083 | 0 | 0.36 | 0.2 | 0.27 | 0.09 | 0.45 | 0.21 |
| 7 | 136727451 | rs324666 | 0.357 | 0.327 | 0.205 | 0.198 | 0.012 | 0.064 | 0.064 | 0 | 0.89 | 0.7 | 0.88 | 0.74 | 0.91 | 0.78 |
| 7 | 136732759 | rs324669 | 1.041 | 0.456 | -0.48 | 0.19 | -0.025 | 0.048 | 0.048 | 0 | 0.45 | 0.2 | 0.37 | 0.3 | 0.49 | 0.42 |
| 7 | 136733228 | rs324670 | 0.585 | 0.323 | 0.217 | 0.396 | 0.013 | 0.111 | 0.111 | 0.229 | 0.89 | 0.66 | 0.91 | 0.71 | 0.91 | 0.8 |
| 7 | 136744954 | rs11977730 | 0.768 | 0.464 | 0.467 | 0.416 | 0.051 | 0.031 | 0.031 | 1.334 | 0.07 | 0.21 | 0.07 | 0.09 | 0.08 | 0.04 |
| 7 | 136752785 | rs10954570 | -0.585 | 0.168 | 0.236 | 0.311 | -0.021 | 0.026 | 0.026 | 0 | 0.32 | 0.39 | 0.45 | 0.47 | 0.24 | 0.3 |
| 7 | 136761961 | rs324568 | 1.519 | 0.211 | 0.046 | 0.225 | 0.006 | 0.067 | 0.067 | 0.234 | 0.52 | 0.21 | 0.4 | 0.3 | 0.56 | 0.46 |
| 7 | 136765572 | rs324572 | -0.221 | 0.13 | 0.413 | 0.395 | 0.04 | 0.159 | 0.159 | 0.373 | 0.95 | 0.77 | 0.93 | 0.77 | 0.95 | 0.82 |
| 7 | 136772275 | rs1345934 | -1.057 | 0.009 | 0.453 | 0.326 | 0.007 | -0.011 | -0.011 | 0 | 0.33 | 0.34 | 0.47 | 0.55 | 0.26 | 0.48 |
| 7 | 136779028 | rs1834168 | -1.364 | 0.071 | -0.075 | 0.264 | 0 | 0 | 0 | 0 | 0.38 | 0.64 | 0.54 | 0.7 | 0.33 | 0.52 |
| 7 | 136782819 | rs11761344 | 0.38 | -0.262 | -0.41 | 0.336 | 0.042 | -0.038 | -0.038 | 0 | 0.21 | 0.11 | 0.37 | 0.31 | 0.15 | 0.25 |
| 7 | 136793350 | rs7792052 | -1.475 | -0.232 | -0.378 | 0.29 | 0.025 | 0.016 | 0.016 | 0.335 | 0.42 | 0.7 | 0.53 | 0.7 | 0.31 | 0.51 |
| 7 | 136794060 | rs12674354 | 0.266 | -0.226 | -0.717 | 0.4 | 0.028 | -0.032 | -0.032 | 0 | 0.19 | 0.11 | 0.35 | 0.31 | 0.13 | 0.24 |
| 7 | 136794884 | rs16874907 | -0.888 | -0.19 | -0.59 | 0.438 | 0.025 | -0.003 | -0.003 | 0 | 0.29 | 0.21 | 0.41 | 0.34 | 0.21 | 0.28 |
| 7 | 136797182 | rs2113886 | -0.596 | -0.089 | -1.157 | -0.209 | 0.019 | 0.008 | 0.008 | 0 | 0.66 | 0.88 | 0.8 | 0.72 | 0.73 | 0.69 |
| 7 | 136798166 | rs1035688 | -1.254 | 0.126 | -0.333 | 0.211 | -0.009 | 0.001 | 0.001 | 0.858 | 0.4 | 0.82 | 0.56 | 0.66 | 0.37 | 0.5 |
| 7 | 136798427 | rs1964318 | -1.296 | 0.142 | -0.282 | 0.225 | -0.006 | -0.015 | -0.015 | 0 | 0.38 | 0.75 | 0.55 | 0.61 | 0.37 | 0.48 |
| 7 | 136804676 | rs1477525 | -1.416 | 0.089 | 0.049 | 0.166 | 0.021 | 0.022 | 0.022 | 0 | 0.44 | 0.65 | 0.54 | 0.68 | 0.36 | 0.52 |
| 7 | 136804753 | rs1364716 | -0.414 | -0.004 | 0.655 | 0.166 | 0.002 | -0.041 | -0.041 | 0.031 | 0.18 | 0.32 | 0.1 | 0.08 | 0.17 | 0.09 |
| 7 | 136811372 | rs4732242 | 1.893 | 0.135 | 0.639 | 0.151 | 0.018 | 0.101 | 0.101 | 0 | 0.5 | 0.16 | 0.39 | 0.32 | 0.56 | 0.46 |
| 7 | 136817769 | rs4090714 | -1.535 | -0.084 | 0.677 | 0.183 | 0.021 | 0.022 | 0.022 | 0 | 0.43 | 0.66 | 0.54 | 0.68 | 0.36 | 0.52 |
| 7 | 136838997 | rs832997 | -1.495 | -0.091 | 0.806 | 0.179 | 0.019 | 0.019 | 0.019 | 0 | 0.43 | 0.66 | 0.54 | 0.68 | 0.36 | 0.52 |

| | | | | | | | | | | | | | | | | |
|---|---|---|---|---|---|---|---|---|---|---|---|---|---|---|---|---|
| 7 | 136849386 | rs832960 | -1.457 | -0.057 | 0.855 | 0.177 | 0.012 | 0.012 | 0.012 | 0 | 0.42 | 0.63 | 0.54 | 0.68 | 0.36 | 0.52 |
| 7 | 136855232 | rs706553 | 0.98 | 0.011 | 1.16 | 0.048 | 0.016 | -0.065 | -0.065 | 0 | 0.74 | 0.63 | 0.87 | 0.95 | 0.73 | 0.87 |
| 7 | 136872377 | rs833018 | -0.33 | -0.001 | 1.235 | 0.236 | -0.012 | 0.023 | 0.023 | 0 | 0.24 | 0.27 | 0.27 | 0.6 | 0.19 | 0.42 |
| 7 | 136874568 | rs833016 | -0.582 | -0.087 | 1.189 | 0.208 | -0.032 | 0.059 | 0.059 | 0 | 0.24 | 0.37 | 0.28 | 0.64 | 0.19 | 0.38 |
| 7 | 136880693 | rs11980852 | -0.207 | -0.039 | 0.951 | 0.087 | 0.007 | -0.04 | -0.04 | 0 | 0.18 | 0.27 | 0.09 | 0.04 | 0.18 | 0.08 |
| 7 | 136882925 | rs6467700 | 2.105 | -0.116 | 0.705 | 0.549 | -0.051 | 0.06 | 0.06 | 0 | 0.49 | 0.12 | 0.54 | 0.33 | 0.56 | 0.48 |
| 7 | 136888610 | rs832984 | -1.546 | -0.246 | 0.86 | 0.616 | -0.003 | -0.003 | -0.003 | 0.03 | 0.43 | 0.56 | 0.4 | 0.67 | 0.37 | 0.52 |
| 7 | 136891459 | rs4732247 | 1.858 | -0.231 | 0.422 | 0.639 | -0.007 | 0.064 | 0.064 | 0 | 0.52 | 0.14 | 0.58 | 0.33 | 0.58 | 0.51 |
| 7 | 136894484 | rs11773575 | 0.493 | -0.53 | 0.486 | 0.766 | 0.017 | -0.08 | -0.08 | 0 | 0.15 | 0.01 | 0.2 | 0.28 | 0.12 | 0.26 |
| 7 | 136904386 | rs1579354 | -0.206 | -0.567 | 0.557 | 0.781 | 0.012 | -0.026 | -0.026 | 0.128 | 0.19 | 0.14 | 0.25 | 0.6 | 0.17 | 0.43 |
| 7 | 136913223 | rs10256915 | -1.547 | -0.665 | 0.575 | 0.6 | 0.039 | 0.04 | 0.04 | 0 | 0.41 | 0.74 | 0.4 | 0.69 | 0.35 | 0.49 |
| 7 | 136914815 | rs10954577 | -0.018 | -0.671 | 0.624 | 0.611 | 0.009 | -0.02 | -0.02 | 0 | 0.19 | 0.14 | 0.25 | 0.61 | 0.17 | 0.4 |
| 7 | 136915059 | rs17168990 | -0.209 | -0.656 | 0.969 | 0.595 | 0.001 | -0.049 | -0.049 | 0 | 0.18 | 0.29 | 0.09 | 0.04 | 0.17 | 0.07 |
| 7 | 136915408 | rs1451528 | -2.041 | -0.629 | 0.462 | 0.586 | 0.022 | 0.108 | 0.108 | 0 | 0.51 | 0.84 | 0.42 | 0.69 | 0.42 | 0.58 |
| 7 | 136916976 | rs10488606 | -0.79 | -0.534 | 0.739 | 0.635 | -0.006 | -0.076 | -0.076 | 0 | 0.21 | 0.57 | 0.11 | 0.05 | 0.18 | 0.1 |
| 7 | 136921062 | rs322295 | -1.787 | -0.414 | 0.524 | 0.633 | 0.005 | 0.016 | 0.016 | 0 | 0.49 | 0.58 | 0.46 | 0.67 | 0.45 | 0.54 |
| 7 | 136921270 | rs322294 | -1.729 | -0.395 | 0.068 | 0.646 | 0.012 | 0.086 | 0.086 | 0 | 0.51 | 0.77 | 0.46 | 0.66 | 0.45 | 0.56 |
| 7 | 136923730 | rs891586 | 0.055 | -0.785 | -0.08 | 0.276 | -0.01 | -0.089 | -0.089 | 0.015 | 0.14 | 0.02 | 0.23 | 0.31 | 0.12 | 0.28 |
| 7 | 136925471 | rs864428 | -2.058 | -0.867 | -0.319 | -0.062 | 0.023 | 0.021 | 0.021 | 0 | 0.51 | 0.85 | 0.41 | 0.38 | 0.42 | 0.43 |
| 7 | 136926419 | rs1864609 | -0.599 | -0.764 | 0.12 | -0.323 | -0.001 | -0.141 | -0.141 | 0 | 0.18 | 0.46 | 0.09 | 0 | 0.15 | 0.04 |
| 7 | 136931552 | rs322302 | 0.9 | -0.718 | 0.026 | -0.262 | -0.001 | -0.031 | -0.031 | 0 | 0.1 | 0.1 | 0.09 | 0.01 | 0.11 | 0.09 |
| 7 | 136933042 | rs11763248 | 0.468 | -0.761 | -0.066 | -0.201 | -0.017 | -0.086 | -0.086 | 1.289 | 0.13 | 0.01 | 0.2 | 0.27 | 0.12 | 0.26 |
| 7 | 136935459 | rs322297 | -0.194 | -0.737 | 0.142 | -0.238 | -0.005 | -0.055 | -0.055 | 0 | 0.15 | 0.19 | 0.1 | 0.01 | 0.13 | 0.09 |
| 7 | 136940560 | rs322323 | -1.785 | -0.711 | -0.222 | -0.119 | 0.008 | 0.007 | 0.007 | 0 | 0.47 | 0.55 | 0.4 | 0.35 | 0.4 | 0.43 |
| 7 | 136940800 | rs10435425 | -0.37 | -0.746 | -0.405 | -0.096 | 0.02 | -0.05 | -0.05 | 0.107 | 0.21 | 0.07 | 0.22 | 0.27 | 0.15 | 0.28 |
| 7 | 136946672 | rs322313 | -1.573 | -0.484 | -0.322 | -0.083 | 0.01 | 0.009 | 0.009 | 0 | 0.47 | 0.57 | 0.4 | 0.36 | 0.4 | 0.43 |
| 7 | 136951940 | rs11764598 | 0.068 | 0.06 | -0.555 | -0.128 | -0.024 | -0.068 | -0.068 | 0 | 0.13 | 0.01 | 0.08 | 0.19 | 0.13 | 0.23 |
| 7 | 136954899 | rs322236 | -1.292 | -0.013 | -0.103 | -0.61 | 0.008 | 0.033 | 0.033 | 0 | 0.31 | 0.77 | 0.21 | 0.07 | 0.27 | 0.15 |
| 7 | 136956607 | rs322239 | 0.085 | -0.192 | 0.471 | -0.478 | -0.003 | -0.078 | -0.078 | 0.492 | 0.86 | 0.72 | 0.9 | 0.98 | 0.89 | 0.97 |
| 7 | 136956990 | rs322240 | 1.03 | -0.252 | 0.197 | -0.207 | -0.003 | -0.001 | -0.001 | 0 | 0.66 | 0.39 | 0.76 | 0.69 | 0.72 | 0.65 |
| 7 | 136958339 | rs322243 | 0.332 | 0.131 | -0.481 | -0.667 | -0.002 | -0.012 | -0.012 | 0 | 0.86 | 0.87 | 0.93 | 1 | 0.86 | 0.95 |
| 7 | 136960167 | rs322246 | 0.947 | 0.046 | -0.579 | -0.437 | 0.007 | 0.04 | 0.04 | 0 | 0.54 | 0.36 | 0.7 | 0.75 | 0.59 | 0.56 |
| 7 | 136966278 | rs1880790 | -0.403 | 0.261 | -1.247 | -0.258 | 0.049 | 0.003 | 0.003 | 0 | 0.53 | 0.3 | 0.45 | 0.48 | 0.62 | 0.64 |
| 7 | 136966331 | rs6970530 | 0.134 | 0.018 | -1.086 | -0.407 | -0.014 | -0.037 | -0.037 | 0 | 0.09 | 0.17 | 0.05 | 0 | 0.04 | 0.06 |
| 7 | 136968621 | rs17169033 | 0.566 | 0.592 | -1.56 | -0.558 | -0.037 | -0.039 | -0.039 | 0 | 0.14 | 0.08 | 0.15 | 0.3 | 0.23 | 0.25 |
| 7 | 136969917 | rs7786318 | 0.827 | 0.564 | -1.821 | -0.872 | -0.021 | 0.08 | 0.08 | 0.294 | 0.23 | 0.1 | 0.14 | 0 | 0.25 | 0.16 |
| 7 | 136971101 | rs3959914 | -1.372 | 0.191 | -1.528 | -0.606 | 0.003 | -0.003 | -0.003 | 0 | 0.61 | 0.46 | 0.47 | 0.48 | 0.66 | 0.7 |
| 7 | 136974025 | rs17169047 | 0.247 | 0.578 | -2.055 | -0.367 | -0.036 | -0.053 | -0.053 | 1.775 | 0.16 | 0.01 | 0.16 | 0.18 | 0.14 | 0.24 |
| 7 | 136980174 | rs919581 | 0.166 | 0.569 | -2.058 | -0.358 | -0.034 | -0.054 | -0.054 | 0 | 0.16 | 0.01 | 0.16 | 0.19 | 0.15 | 0.24 |
| 7 | 136985504 | rs6977749 | -0.284 | 0.896 | -1.505 | -0.416 | -0.054 | 0.038 | 0.038 | 0 | 0.37 | 0.21 | 0.28 | 0.3 | 0.47 | 0.4 |
| 7 | 136989304 | rs13245564 | -0.314 | 0.852 | -1.466 | -0.262 | -0.047 | 0.022 | 0.022 | 0 | 0.36 | 0.21 | 0.28 | 0.31 | 0.46 | 0.46 |
| 7 | 136989606 | rs13245911 | -0.843 | 0.298 | -1.099 | -0.25 | 0.006 | 0.001 | 0.001 | 0 | 0.54 | 0.4 | 0.36 | 0.47 | 0.56 | 0.69 |
| 7 | 136990665 | rs1731954 | 1.278 | -0.219 | -1.733 | -0.309 | -0.064 | -0.13 | -0.13 | 2.678 | 0.33 | 0.07 | 0.59 | 0.46 | 0.41 | 0.29 |
| 7 | 136990721 | rs12666148 | 0.408 | -0.042 | -1.715 | -0.322 | -0.008 | -0.008 | -0.008 | 0.828 | 0.13 | 0.13 | 0.12 | 0.28 | 0.19 | 0.33 |
| 7 | 136994013 | rs322345 | 0.105 | -0.487 | -1.543 | -0.339 | -0.013 | 0.003 | 0.003 | 3.514 | 0.79 | 0.3 | 0.89 | 0.75 | 0.91 | 0.83 |
| 7 | 137003404 | rs322335 | -0.859 | -0.544 | -0.767 | -0.377 | -0.011 | 0.014 | 0.014 | 0.821 | 0.84 | 0.78 | 0.94 | 0.79 | 0.91 | 0.83 |
| 7 | 137004109 | rs322334 | 1.633 | -0.344 | -1.56 | -0.494 | -0.051 | -0.099 | -0.099 | 0 | 0.34 | 0.09 | 0.6 | 0.47 | 0.42 | 0.29 |
| 7 | 137005808 | rs2242045 | -0.296 | -0.033 | -1.444 | -0.505 | -0.047 | 0.105 | 0.105 | 0 | 0.45 | 0.2 | 0.29 | 0.29 | 0.49 | 0.53 |
| 7 | 137011207 | rs10268599 | 0.077 | -0.014 | -1.305 | -0.502 | -0.01 | -0.019 | -0.019 | 0 | 0.21 | 0.17 | 0.15 | 0.35 | 0.25 | 0.31 |
| 7 | 137016297 | rs1431093 | -0.409 | -0.458 | -1.292 | -0.486 | -0.011 | -0.014 | -0.014 | 0 | 0.39 | 0.31 | 0.29 | 0.44 | 0.43 | 0.46 |
| 7 | 137021176 | rs13240024 | 0.664 | -0.781 | -1.679 | -0.338 | -0.009 | 0.001 | 0.001 | 0 | 0.22 | 0.01 | 0.09 | 0.17 | 0.13 | 0.18 |
| 7 | 137038092 | rs320682 | -0.253 | -0.679 | -1.273 | -0.254 | -0.012 | -0.013 | -0.013 | 0.334 | 0.38 | 0.29 | 0.41 | 0.42 | 0.44 | 0.46 |
| 7 | 137040948 | rs320686 | -0.287 | -0.789 | -0.661 | -0.526 | 0.001 | -0.002 | -0.002 | 0.152 | 0.44 | 0.52 | 0.46 | 0.37 | 0.46 | 0.36 |
| 7 | 137043484 | rs320692 | -0.165 | -0.803 | -0.615 | -0.531 | 0.002 | -0.004 | -0.004 | 0 | 0.43 | 0.44 | 0.45 | 0.37 | 0.46 | 0.35 |
| 7 | 137051175 | rs320704 | -1.816 | -0.846 | -0.731 | -0.395 | -0.016 | -0.008 | -0.008 | 8.753 | 0.7 | 0.78 | 0.63 | 0.69 | 0.65 | 0.67 |
| 7 | 137053658 | rs161338 | 0.952 | -0.818 | -0.758 | -0.419 | -0.042 | -0.021 | -0.021 | 0 | 0.27 | 0.12 | 0.22 | 0.3 | 0.34 | 0.35 |
| 7 | 137055525 | rs161339 | 1.267 | -0.613 | -0.478 | -0.015 | 0.002 | 0.051 | 0.051 | 0 | 0.08 | 0.07 | 0.08 | 0.29 | 0.08 | 0.19 |
| 7 | 137059416 | rs2599358 | 0.246 | -1.158 | -0.224 | -0.062 | -0.052 | 0.02 | 0.02 | 0.011 | 0.73 | 0.56 | 0.82 | 0.7 | 0.82 | 0.7 |
| 7 | 137068963 | rs2351136 | 0.168 | -0.904 | -0.593 | 0.147 | -0.04 | 0.048 | 0.048 | 0 | 0.23 | 0.4 | 0.18 | 0.3 | 0.16 | 0.29 |
| 7 | 137072096 | rs4728408 | 0.977 | -0.608 | -1.445 | 0.182 | -0.028 | -0.001 | -0.001 | 1.884 | 0.3 | 0.2 | 0.37 | 0.29 | 0.36 | 0.34 |
| 7 | 137072364 | rs728054 | 1.134 | -0.571 | -1.587 | 0.193 | -0.037 | -0.009 | -0.009 | 0 | 0.28 | 0.14 | 0.37 | 0.28 | 0.36 | 0.34 |
| 7 | 137072931 | rs2278829 | 1.245 | -0.571 | -1.879 | 0.193 | -0.095 | -0.051 | -0.051 | 1.486 | 0.28 | 0.06 | 0.36 | 0.29 | 0.36 | 0.34 |
| 7 | 137073259 | rs13230189 | 1.18 | -0.574 | -1.876 | 0.192 | -0.069 | -0.031 | -0.031 | 0 | 0.28 | 0.08 | 0.36 | 0.29 | 0.36 | 0.3 |
| 7 | 137074540 | rs7779548 | 1.172 | -0.48 | -1.961 | 0.23 | -0.104 | -0.061 | -0.061 | 0.028 | 0.28 | 0.05 | 0.36 | 0.29 | 0.36 | 0.34 |
| 7 | 137074761 | rs1376608 | 1.172 | -0.502 | -1.961 | 0.262 | -0.093 | -0.068 | -0.068 | 0 | 0.28 | 0.06 | 0.36 | 0.32 | 0.36 | 0.34 |
| 7 | 137074839 | rs3735026 | 1.172 | -0.502 | -2.032 | 0.262 | -0.107 | -0.08 | -0.08 | 0.156 | 0.28 | 0.05 | 0.36 | 0.32 | 0.36 | 0.34 |
| 7 | 137080244 | rs7792568 | -0.338 | -1.107 | -0.839 | 0.462 | 0.008 | 0.001 | 0.001 | 0 | 0.2 | 0.07 | 0.1 | 0.22 | 0.15 | 0.18 |
| 7 | 137081223 | rs1451526 | -1.113 | -1.091 | -0.658 | 0.441 | -0.001 | 0 | 0 | 0 | 0.3 | 0.28 | 0.19 | 0.3 | 0.2 | 0.33 |
| 7 | 137086826 | rs11760855 | -0.353 | -0.995 | -0.832 | 0.401 | 0.016 | 0.019 | 0.019 | 0 | 0.23 | 0.11 | 0.09 | 0.19 | 0.16 | 0.19 |

| | | | | | | | | | | | | | | | | |
|---|---|---|---|---|---|---|---|---|---|---|---|---|---|---|---|---|
| 7 | 137090425 | rs10224080 | 0.719 | -0.908 | -0.676 | 0.125 | 0.003 | 0.014 | 0.014 | 0 | 0.17 | 0.14 | 0.11 | 0.09 | 0.19 | 0.09 |
| 7 | 137090469 | rs4728409 | 1.506 | -1.017 | -0.839 | 0.733 | -0.074 | -0.085 | -0.085 | 0 | 0.26 | 0.04 | 0.34 | 0.28 | 0.28 | 0.3 |
| 7 | 137090935 | rs1863037 | -0.063 | -0.795 | 0.313 | 0.985 | 0.01 | 0.009 | 0.009 | 0 | 0.76 | 0.76 | 0.8 | 0.71 | 0.68 | 0.73 |
| 7 | 137093065 | rs2278102 | -0.387 | -1.042 | 0.986 | 0.514 | -0.01 | -0.006 | -0.006 | 0 | 0.42 | 0.52 | 0.43 | 0.45 | 0.37 | 0.42 |
| 7 | 137097262 | rs4732254 | 0.31 | -0.992 | 1.545 | 0.951 | 0.001 | -0.022 | -0.022 | 0.052 | 0.41 | 0.12 | 0.49 | 0.65 | 0.35 | 0.48 |
| 7 | 137098995 | rs4732255 | -0.04 | -0.984 | 1.822 | 0.798 | 0.04 | 0.037 | 0.037 | 0 | 0.46 | 0.17 | 0.53 | 0.66 | 0.37 | 0.52 |
| 7 | 137118980 | rs10234672 | -0.39 | -1.041 | 2.68 | 0.773 | 0.001 | -0.004 | -0.004 | 0.261 | 0.5 | 0.49 | 0.61 | 0.77 | 0.38 | 0.53 |
| 7 | 137121760 | rs10244101 | -0.285 | -1.025 | 2.569 | 0.785 | 0.003 | 0 | 0 | 0 | 0.51 | 0.53 | 0.61 | 0.76 | 0.36 | 0.52 |
| 7 | 137124915 | rs1731913 | -0.424 | -1.028 | 2.605 | 0.783 | 0.035 | 0.079 | 0.079 | 0 | 0.52 | 0.7 | 0.64 | 0.77 | 0.38 | 0.58 |
| 7 | 137128390 | rs10488609 | -0.099 | -0.948 | 1.726 | 0.673 | 0.044 | 0.035 | 0.035 | 0 | 0.47 | 0.21 | 0.53 | 0.66 | 0.36 | 0.57 |
| 7 | 137128830 | rs1918837 | -0.099 | -0.948 | 1.726 | 0.673 | 0.044 | 0.035 | 0.035 | 0 | 0.47 | 0.21 | 0.53 | 0.66 | 0.36 | 0.57 |
| 7 | 137131235 | rs12155075 | 0.452 | -1.147 | 1.685 | 0.646 | -0.011 | -0.019 | -0.019 | 0.573 | 0.39 | 0.09 | 0.49 | 0.65 | 0.37 | 0.48 |
| 7 | 137131588 | rs1918839 | 1.41 | -0.784 | 1.883 | 0.802 | 0.068 | 0.375 | 0.375 | 1.095 | 0.92 | 0.57 | 0.83 | 0.71 | 0.9 | 0.93 |
| 7 | 137132243 | rs12707359 | 0.265 | -0.662 | 1.037 | 0.387 | -0.006 | 0.023 | 0.023 | 0 | 0.44 | 0.34 | 0.3 | 0.06 | 0.52 | 0.36 |
| 7 | 137139529 | rs975965 | 1.771 | -0.852 | 1.984 | 2.019 | 0.009 | 0.219 | 0.219 | 0 | 0.92 | 0.74 | 0.86 | 0.73 | 0.92 | 0.92 |
| 7 | 137140515 | rs1406089 | 2.212 | -1.279 | 2.208 | 2.077 | -0.012 | 0.014 | 0.014 | 0.251 | 0.76 | 0.32 | 0.71 | 0.63 | 0.86 | 0.67 |
| 7 | 137146452 | rs6942907 | -1.987 | -1.41 | 2.913 | 1.624 | -0.038 | -0.084 | -0.084 | 0 | 0.13 | 0.38 | 0.05 | 0 | 0.04 | 0.06 |
| 7 | 137151096 | rs1528098 | -2.482 | -1.461 | 2.957 | 1.712 | -0.056 | 0.049 | 0.049 | 0.237 | 0.14 | 0.5 | 0.12 | 0.12 | 0.04 | 0.07 |
| 7 | 137156237 | rs6467704 | -2.317 | -1.465 | 3.223 | 1.617 | -0.025 | 0.013 | 0.013 | 0 | 0.16 | 0.46 | 0.23 | 0.11 | 0.09 | 0.14 |
| 7 | 137170734 | rs6971080 | -1.89 | -1.464 | 2.823 | 1.384 | -0.008 | -0.056 | -0.056 | 0 | 0.13 | 0.21 | 0.1 | 0 | 0.09 | 0.12 |
| 7 | 137175198 | rs10480523 | -2.382 | -1.461 | 2.462 | 1.385 | -0.037 | -0.114 | -0.114 | 0.352 | 0.18 | 0.63 | 0.12 | 0 | 0.09 | 0.13 |
| 7 | 137182446 | rs4732260 | 2.331 | -1.48 | 1.794 | 1.729 | 0.044 | -0.056 | -0.056 | 0 | 0.7 | 0.18 | 0.62 | 0.55 | 0.83 | 0.66 |
| 7 | 137187798 | rs7781490 | -2.426 | -1.382 | 2.59 | 1.397 | -0.054 | 0.036 | 0.036 | 1.14 | 0.14 | 0.57 | 0.19 | 0.11 | 0.04 | 0.23 |
| 7 | 137203122 | rs10261618 | -1.32 | -1.351 | 2.789 | 1.394 | -0.027 | -0.012 | -0.012 | 0 | 0.07 | 0.19 | 0.03 | 0.02 | 0.01 | 0.11 |
| 7 | 137208391 | rs6964837 | -1.757 | -1.341 | 2.962 | 1.446 | -0.03 | 0.101 | 0.101 | 0.075 | 0.08 | 0.26 | 0.15 | 0.12 | 0.02 | 0.11 |
| 7 | 137213552 | rs10252668 | -0.905 | -1.316 | 2.872 | 1.429 | -0.027 | -0.013 | -0.013 | 0 | 0.06 | 0.12 | 0.01 | 0.02 | 0.01 | 0.11 |
| 7 | 137227544 | rs834063 | -2.596 | -1.803 | 3.108 | 0.815 | -0.048 | -0.116 | -0.116 | 0 | 0.18 | 0.36 | 0.09 | 0 | 0.07 | 0.19 |
| 7 | 137229997 | rs834059 | -3.112 | -1.818 | 3.062 | 0.872 | -0.053 | -0.019 | -0.019 | 0 | 0.25 | 0.58 | 0.24 | 0.13 | 0.09 | 0.3 |
| 7 | 137246531 | rs1629172 | -1.687 | -1.758 | 2.967 | 0.83 | -0.019 | 0.139 | 0.139 | 0.144 | 0.08 | 0.35 | 0.16 | 0.11 | 0.02 | 0.11 |
| 7 | 137253784 | rs7784757 | -2.22 | -1.749 | 2.806 | 0.295 | -0.019 | -0.061 | -0.061 | 0 | 0.15 | 0.2 | 0.07 | 0 | 0.07 | 0.18 |
| 7 | 137263481 | rs834100 | -2.127 | -1.665 | 2.705 | 0.339 | -0.037 | -0.035 | -0.035 | 0 | 0.1 | 0.28 | 0.03 | 0.02 | 0.02 | 0.15 |
| 7 | 137271543 | rs10245413 | -1.111 | -1.608 | 2.403 | 0.401 | -0.033 | -0.02 | -0.02 | 0.869 | 0.07 | 0.14 | 0.02 | 0.02 | 0.02 | 0.11 |
| 7 | 137272621 | rs7456122 | -2.1 | -1.671 | 2.054 | 0.297 | -0.022 | -0.069 | -0.069 | 0.175 | 0.16 | 0.22 | 0.08 | 0 | 0.07 | 0.15 |
| 7 | 137283212 | rs706567 | -2.621 | -1.487 | 2.076 | 0.475 | -0.083 | -0.111 | -0.111 | 3.01 | 0.18 | 0.4 | 0.08 | 0.03 | 0.06 | 0.22 |
| 7 | 137284359 | rs6949094 | -1.893 | -0.912 | 1.276 | 0.631 | -0.014 | -0.052 | -0.052 | 0 | 0.1 | 0.18 | 0.06 | 0 | 0.04 | 0.11 |
| 7 | 137298910 | rs706578 | -0.594 | -0.737 | -0.36 | 0.745 | -0.076 | -0.079 | -0.079 | 0 | 0.52 | 0.97 | 0.42 | 0.54 | 0.44 | 0.6 |
| 7 | 137315924 | rs11489681 | -1.736 | -1.224 | 1.647 | 0.407 | 0.025 | 0.022 | 0.022 | 6.744 | 0.06 | 0.34 | 0.03 | 0 | 0 | 0.02 |
| 7 | 137319214 | rs11982810 | -2.48 | -1.427 | 2.537 | 0.47 | -0.042 | -0.013 | -0.013 | 0 | 0.1 | 0.58 | 0.06 | 0.04 | 0.01 | 0.06 |
| 7 | 137335279 | rs6946901 | -1.799 | -1.386 | 3.297 | 0.422 | 0.073 | 0.075 | 0.075 | 0.011 | 0.06 | 0.38 | 0.04 | 0.03 | 0.01 | 0.07 |
| 7 | 137351769 | rs3800621 | -2.802 | -1.455 | 3.233 | 0.391 | -0.043 | -0.042 | -0.042 | 0 | 0.11 | 0.68 | 0.06 | 0.03 | 0.01 | 0.07 |
| 7 | 137356651 | rs270904 | -0.497 | -1.361 | 2.868 | 0.728 | 0.001 | -0.002 | -0.002 | 0 | 0.38 | 0.41 | 0.42 | 0.41 | 0.3 | 0.45 |
| 7 | 137366467 | rs270896 | -1.017 | -1.193 | 2.231 | 0.766 | 0.074 | -0.039 | -0.039 | 2.011 | 0.46 | 0.93 | 0.48 | 0.41 | 0.31 | 0.52 |
| 7 | 137380919 | rs6976725 | -0.877 | -1.26 | 2.411 | 0.756 | 0.05 | -0.008 | -0.008 | 0 | 0.43 | 0.72 | 0.46 | 0.44 | 0.31 | 0.55 |
| 7 | 137399820 | rs12056089 | -0.446 | -0.941 | 2.029 | 0.528 | 0.004 | -0.006 | -0.006 | 0.162 | 0.39 | 0.3 | 0.47 | 0.45 | 0.33 | 0.5 |
| 7 | 137401641 | rs3800631 | -0.99 | -1.156 | 1.695 | 0.548 | 0.004 | 0.021 | 0.021 | 0 | 0.35 | 0.68 | 0.27 | 0.14 | 0.3 | 0.28 |
| 7 | 137402718 | rs3800636 | -1.366 | -1.848 | 1.127 | 0.14 | 0.083 | 0.015 | 0.015 | 0.176 | 0.5 | 0.79 | 0.51 | 0.45 | 0.32 | 0.49 |
| 7 | 137420389 | rs3800642 | -2.136 | -1.966 | 1.671 | 0.401 | -0.057 | 0.072 | 0.072 | 0.414 | 0.13 | 0.66 | 0.23 | 0.22 | 0.06 | 0.25 |
| 7 | 137439592 | rs3800654 | -0.43 | -1.722 | 0.161 | -0.704 | 0.036 | 0.129 | 0.129 | 0 | 0.4 | 0.19 | 0.32 | 0.24 | 0.27 | 0.33 |
| 7 | 137446778 | rs3778811 | -0.665 | -1.662 | 1.385 | -0.587 | -0.005 | -0.03 | -0.03 | 0.062 | 0.44 | 0.59 | 0.37 | 0.24 | 0.31 | 0.34 |
| 7 | 137447398 | rs3778812 | -0.933 | -1.702 | 1.287 | -0.674 | 0.017 | 0.035 | 0.035 | 0.21 | 0.48 | 0.71 | 0.42 | 0.35 | 0.35 | 0.38 |
| 7 | 137461581 | rs7785642 | -1.168 | -1.946 | 0.385 | -0.804 | 0.011 | 0.091 | 0.091 | 2.52 | 0.51 | 0.91 | 0.37 | 0.39 | 0.38 | 0.5 |
| 7 | 137467829 | rs2351857 | -1.233 | -1.476 | 0.319 | -0.854 | -0.07 | 0.061 | 0.061 | 0.859 | 0.53 | 0.92 | 0.42 | 0.4 | 0.46 | 0.5 |
| 7 | 137477912 | rs1559490 | -0.173 | -1.627 | 1.129 | -0.97 | -0.006 | 0.02 | 0.02 | 0.545 | 0.37 | 0.34 | 0.25 | 0.12 | 0.35 | 0.34 |
| 7 | 137479043 | rs3778818 | -1.561 | -1.61 | 1.479 | -0.851 | -0.018 | 0.107 | 0.107 | 0 | 0.15 | 0.49 | 0.16 | 0.29 | 0.11 | 0.15 |
| 7 | 137483664 | rs13243717 | -0.576 | -1.344 | 1.747 | -1.191 | -0.003 | 0.014 | 0.014 | 0.112 | 0.39 | 0.4 | 0.26 | 0.13 | 0.35 | 0.34 |
| 7 | 137493265 | rs10267736 | -0.65 | -1.411 | 1.556 | -1.221 | -0.004 | 0.044 | 0.044 | 0 | 0.48 | 0.32 | 0.4 | 0.07 | 0.51 | 0.4 |
| 7 | 137496871 | rs6966474 | -0.688 | -1.406 | 1.632 | -1.213 | 0 | -0.04 | -0.04 | 0 | 0.51 | 0.48 | 0.42 | 0.07 | 0.52 | 0.4 |
| 7 | 137506041 | rs889869 | -1.021 | -1.341 | 0.871 | -1.163 | -0.018 | 0.123 | 0.123 | 0 | 0.54 | 0.74 | 0.46 | 0.15 | 0.52 | 0.43 |
| 7 | 137514791 | rs7796821 | -2.369 | -1.329 | 1.495 | -0.503 | -0.026 | 0.003 | 0.003 | 5.749 | 0.12 | 0.76 | 0.12 | 0.13 | 0.06 | 0.05 |
| 7 | 137516329 | rs10260637 | -2.135 | -1.296 | 1.772 | -0.526 | -0.034 | 0.013 | 0.013 | 3.244 | 0.11 | 0.7 | 0.11 | 0.11 | 0.05 | 0.04 |
| 7 | 137529578 | rs6978230 | -2.382 | -1.306 | 1.459 | -0.591 | -0.001 | 0.001 | 0.001 | 4.414 | 0.12 | 0.82 | 0.12 | 0.05 | 0.06 | 0.04 |
| 7 | 137534627 | rs1125727 | 0.974 | -1.629 | 1.377 | -1.156 | 0.048 | 0.085 | 0.085 | 0 | 0.5 | 0.4 | 0.67 | 0.9 | 0.63 | 0.65 |
| 7 | 137548919 | rs4732272 | 0.179 | -1.714 | 1.704 | -1.193 | 0.028 | 0.084 | 0.084 | 0 | 0.6 | 0.76 | 0.74 | 0.9 | 0.69 | 0.69 |
| 7 | 137553110 | rs1020961 | 0.684 | -1.648 | 2.026 | -0.808 | 0.027 | 0.056 | 0.056 | 0 | 0.56 | 0.67 | 0.7 | 0.93 | 0.64 | 0.65 |
| 7 | 137556148 | rs10250405 | -1.988 | -1.364 | 2.035 | -0.495 | 0.017 | 0.211 | 0.211 | 3.33 | 0.07 | 0.55 | 0.07 | 0.21 | 0.01 | 0.1 |
| 7 | 137559455 | rs3735016 | -1.988 | -1.364 | 1.799 | -0.495 | 0.032 | 0.244 | 0.244 | 3.33 | 0.07 | 0.51 | 0.06 | 0.21 | 0.01 | 0.1 |
| 7 | 137562504 | rs6959012 | -1.24 | -1.312 | 1.218 | -0.472 | 0.062 | 0.287 | 0.287 | 0.857 | 0.05 | 0.39 | 0.05 | 0.21 | 0.01 | 0.1 |
| 7 | 137562834 | rs3735020 | -1.486 | -1.319 | 1.086 | -0.473 | 0.053 | 0.279 | 0.279 | 1.698 | 0.06 | 0.45 | 0.06 | 0.21 | 0.01 | 0.1 |
| 7 | 137563434 | rs3735021 | 0.884 | -1.446 | -0.184 | -0.115 | 0.101 | -0.075 | -0.075 | 0.446 | 0.48 | 0.09 | 0.48 | 0.48 | 0.64 | 0.55 |

| | | | | | | | | | | | | | | | | |
|---|---|---|---|---|---|---|---|---|---|---|---|---|---|---|---|---|
| 7 | 137571931 | rs6963146 | -0.824 | -0.543 | -0.385 | -0.563 | 0.03 | 0.05 | 0.05 | 0.077 | 0.35 | 0.82 | 0.25 | 0.33 | 0.24 | 0.35 |
| 7 | 137580565 | rs273938 | 0.527 | -0.446 | 0.163 | -0.687 | -0.001 | 0.015 | 0.015 | 0 | 0.66 | 0.7 | 0.75 | 0.77 | 0.71 | 0.65 |
| 7 | 137584781 | rs1677499 | 0.468 | -0.337 | -0.333 | -0.696 | -0.002 | -0.031 | -0.031 | 0 | 0.65 | 0.5 | 0.73 | 0.69 | 0.71 | 0.58 |
| 7 | 137590369 | rs273968 | 0.066 | -0.027 | -1.639 | -1.028 | -0.127 | 0.143 | 0.143 | 0 | 0.64 | 0.97 | 0.68 | 0.86 | 0.61 | 0.61 |
| 7 | 137590808 | rs273967 | 0.907 | -0.146 | -1.508 | -1.048 | -0.024 | 0.212 | 0.212 | 0.03 | 0.53 | 0.11 | 0.61 | 0.78 | 0.59 | 0.55 |
| 7 | 137595253 | rs273962 | 1.033 | -0.502 | -1.644 | -0.603 | -0.027 | -0.019 | -0.019 | 0 | 0.45 | 0.06 | 0.42 | 0.61 | 0.59 | 0.53 |
| 7 | 137600690 | rs273957 | 0.035 | -0.069 | -1.449 | -1.348 | -0.104 | 0.195 | 0.195 | 0 | 0.66 | 0.97 | 0.69 | 0.94 | 0.63 | 0.7 |
| 7 | 137605669 | rs273950 | 0.524 | 0.011 | -1.058 | -1.288 | 0.028 | 0.123 | 0.123 | 0 | 0.58 | 0.23 | 0.63 | 0.92 | 0.64 | 0.69 |
| 7 | 137611566 | rs273945 | 0.782 | 0.062 | -1.468 | -1.201 | -0.052 | 0.227 | 0.227 | 0.074 | 0.52 | 0.05 | 0.6 | 0.83 | 0.59 | 0.64 |
| 7 | 137612571 | rs273972 | 0.284 | 0.316 | -1.464 | -1.461 | -0.043 | 0.238 | 0.238 | 0 | 0.6 | 0.92 | 0.67 | 0.94 | 0.62 | 0.64 |
| 7 | 137618152 | rs273978 | 0.042 | -0.055 | -1.588 | -1.48 | -0.033 | 0.125 | 0.125 | 0.122 | 0.63 | 0.93 | 0.68 | 0.82 | 0.63 | 0.63 |
| 7 | 137618171 | rs10224250 | -1.064 | 0.224 | -0.77 | -1.641 | 0.035 | -0.037 | -0.037 | 0.529 | 0.08 | 0.24 | 0.25 | 0.02 | 0.11 | 0.07 |
| 7 | 137618312 | rs273979 | 0.479 | 0.194 | -0.366 | -1.681 | 0.01 | 0.092 | 0.092 | 0.163 | 0.56 | 0.29 | 0.61 | 0.8 | 0.6 | 0.61 |
| 7 | 137620895 | rs112417 | 0.864 | 0.203 | -0.127 | -1.669 | -0.003 | 0.07 | 0.07 | 0.098 | 0.51 | 0.34 | 0.39 | 0.77 | 0.51 | 0.54 |
| 7 | 137629146 | rs273995 | 1.069 | 0.487 | -0.642 | -1.536 | -0.155 | 0.157 | 0.157 | 0 | 0.44 | 0.05 | 0.36 | 0.77 | 0.49 | 0.52 |
| 7 | 137639026 | rs10267160 | -1.037 | 0.168 | 0.148 | -1.267 | 0.026 | -0.033 | -0.033 | 0 | 0.08 | 0.25 | 0.26 | 0.03 | 0.1 | 0.08 |
| 7 | 137646711 | rs274007 | 0.911 | 0.178 | -0.169 | -1.148 | 0.009 | 0.172 | 0.172 | 0 | 0.49 | 0.13 | 0.38 | 0.74 | 0.56 | 0.55 |
| 7 | 137647262 | rs274009 | -0.109 | 0.011 | 0.248 | -1.105 | 0.041 | 0.052 | 0.052 | 0 | 0.63 | 0.74 | 0.73 | 0.83 | 0.76 | 0.65 |
| 7 | 137650044 | rs274012 | 0.16 | 0.18 | 0.275 | -1.013 | 0.079 | 0.114 | 0.114 | 0 | 0.57 | 0.8 | 0.72 | 0.82 | 0.72 | 0.6 |
| 7 | 137652152 | rs889870 | -2.294 | 0.956 | 1.526 | -0.299 | 0.165 | 0.082 | 0.082 | 0.264 | 0.09 | 0.47 | 0.31 | 0.08 | 0.13 | 0.07 |
| 7 | 137652622 | rs1871214 | -1.403 | 0.662 | 0.715 | -0.341 | 0.117 | 0.064 | 0.064 | 0.278 | 0.05 | 0.38 | 0.04 | 0 | 0.03 | 0.01 |
| 7 | 137660460 | rs6943342 | -1.068 | 0.534 | -0.654 | -0.195 | 0.005 | 0.006 | 0.006 | 0 | 0.28 | 0.73 | 0.38 | 0.23 | 0.26 | 0.18 |
| 7 | 137661023 | rs17276467 | 0.008 | 0.486 | -1.599 | -0.305 | 0.009 | 0.037 | 0.037 | 0 | 0.36 | 0.06 | 0.31 | 0.33 | 0.29 | 0.51 |
| 7 | 137661589 | rs6948572 | 1.402 | 1.232 | -0.234 | -0.456 | 0.013 | -0.111 | -0.111 | 0 | 0.81 | 0.6 | 0.82 | 0.98 | 0.79 | 0.93 |
| 7 | 137663477 | rs13232030 | -0.075 | 1.696 | -0.654 | -0.026 | -0.087 | -0.028 | -0.028 | 0 | 0.15 | 0.01 | 0.13 | 0.14 | 0.25 | 0.25 |
| 7 | 137664771 | rs6948022 | -1.467 | 0.979 | 0.204 | 0.05 | 0.048 | -0.046 | -0.046 | 0.013 | 0.09 | 0.31 | 0.08 | 0 | 0.11 | 0.02 |
| 7 | 137669111 | rs12707374 | 2.058 | 1.098 | 0.457 | 0.615 | 0.006 | 0.009 | 0.009 | 0.83 | 0.82 | 0.29 | 0.76 | 0.76 | 0.77 | 0.9 |
| 7 | 137669251 | rs10260498 | 1.519 | 0.749 | 0.618 | 0.452 | 0.043 | 0.036 | 0.036 | 0 | 0.9 | 0.78 | 0.82 | 0.88 | 0.84 | 0.94 |
| 7 | 137671487 | rs10954592 | 1.522 | -0.122 | 0.685 | 0.366 | -0.039 | -0.015 | -0.015 | 0.177 | 0.83 | 0.36 | 0.85 | 0.88 | 0.91 | 0.93 |
| 7 | 137671540 | rs11770714 | 1.489 | -0.116 | 0.775 | 0.369 | -0.041 | -0.014 | -0.014 | 0 | 0.83 | 0.38 | 0.86 | 0.88 | 0.91 | 0.94 |
| 7 | 137672006 | rs1076517 | 0.184 | 0.08 | -0.342 | 0.349 | -0.032 | 0.009 | 0.009 | 0.054 | 0.15 | 0.01 | 0.11 | 0.08 | 0.15 | 0.17 |
| 7 | 137675576 | rs6467728 | 0.866 | -0.031 | 0.133 | 0.028 | -0.007 | -0.025 | -0.025 | 0 | 0.67 | 0.64 | 0.81 | 0.88 | 0.75 | 0.79 |
| 7 | 137679345 | rs4732278 | 1.444 | -0.104 | 1.241 | 1.345 | -0.015 | 0.07 | 0.07 | 3.578 | 0.8 | 0.38 | 0.59 | 0.62 | 0.85 | 0.84 |
| 7 | 137682885 | rs7783688 | 1.323 | -0.035 | 1.75 | 1.327 | 0.071 | 0.219 | 0.219 | 0.062 | 0.9 | 0.6 | 0.69 | 0.74 | 0.9 | 0.93 |
| 7 | 137687989 | rs1424376 | 0.869 | -0.415 | 0.994 | 1.229 | 0.042 | 0.081 | 0.081 | 2.02 | 0.5 | 0.29 | 0.45 | 0.24 | 0.59 | 0.52 |
| 7 | 137689901 | rs10271444 | -0.621 | -0.31 | 1.341 | 1.736 | 0.076 | 0.223 | 0.223 | 0 | 0.05 | 0.24 | 0.05 | 0.22 | 0.06 | 0.09 |
| 7 | 137694764 | rs10954596 | 0.881 | -0.507 | 1.356 | 1.787 | 0.018 | 0.035 | 0.035 | 1.355 | 0.52 | 0.4 | 0.45 | 0.25 | 0.59 | 0.57 |
| 7 | 137695178 | rs10243728 | 0.534 | -0.592 | 0.893 | 1.683 | 0.035 | -0.038 | -0.038 | 0 | 0.58 | 0.76 | 0.51 | 0.46 | 0.66 | 0.67 |
| 7 | 137716835 | rs4732283 | 0.506 | -0.758 | 0.686 | 1.855 | 0.032 | -0.037 | -0.037 | 0.215 | 0.59 | 0.74 | 0.55 | 0.48 | 0.66 | 0.66 |
| 7 | 137731418 | rs6947017 | 0.119 | -0.799 | 0.312 | 1.808 | -0.022 | -0.023 | -0.023 | 0 | 0.93 | 0.97 | 0.74 | 0.82 | 0.89 | 0.97 |
| 7 | 137732209 | rs998767 | 0.717 | -0.692 | 0.279 | 1.678 | 0.023 | -0.044 | -0.044 | 0.016 | 0.6 | 0.8 | 0.54 | 0.48 | 0.66 | 0.7 |
| 7 | 137737546 | rs6467729 | 0.901 | -0.968 | 0.518 | 1.517 | -0.011 | 0.018 | 0.018 | 0 | 0.67 | 0.58 | 0.59 | 0.49 | 0.73 | 0.74 |
| 7 | 137737600 | rs6467730 | 0.526 | -0.909 | -0.247 | 1.548 | -0.003 | -0.111 | -0.111 | 0.372 | 0.7 | 0.92 | 0.61 | 0.49 | 0.73 | 0.74 |
| 7 | 137743167 | rs12112517 | 1.097 | -0.251 | -0.02 | 1.086 | 0.003 | 0.033 | 0.033 | 0 | 0.5 | 0.39 | 0.44 | 0.31 | 0.53 | 0.51 |
| 7 | 137743236 | rs10245239 | 0.28 | -0.138 | -0.033 | 1.505 | 0.002 | -0.029 | -0.029 | 0 | 0.73 | 0.82 | 0.65 | 0.64 | 0.73 | 0.77 |
| 7 | 137753091 | rs1025967 | 0.573 | -0.173 | -0.14 | 1.42 | 0.002 | -0.064 | -0.064 | 0 | 0.7 | 0.86 | 0.64 | 0.64 | 0.72 | 0.76 |
| 7 | 137753520 | rs10265064 | 1.466 | -0.693 | -0.101 | 1.229 | -0.001 | -0.004 | -0.004 | 0 | 0.52 | 0.45 | 0.52 | 0.44 | 0.56 | 0.57 |
| 7 | 137753675 | rs7812287 | 0.355 | -0.704 | -0.122 | 1.203 | -0.021 | -0.032 | -0.032 | 0.17 | 0.1 | 0.05 | 0.07 | 0.2 | 0.12 | 0.17 |
| 7 | 137755143 | rs17277937 | 2.023 | -1.203 | -1.113 | 0.794 | -0.144 | -0.028 | -0.028 | 0.023 | 0.23 | 0.02 | 0.32 | 0.21 | 0.36 | 0.26 |
| 7 | 137759000 | rs3757393 | 0.232 | -0.684 | -0.002 | 1.158 | 0.008 | -0.031 | -0.031 | 0 | 0.7 | 0.82 | 0.69 | 0.68 | 0.72 | 0.76 |
| 7 | 137764147 | rs6945659 | 0.221 | -0.684 | 0.04 | 1.146 | 0.006 | -0.035 | -0.035 | 0 | 0.71 | 0.83 | 0.64 | 0.64 | 0.72 | 0.75 |
| 7 | 137767051 | rs6943498 | 1.533 | -0.627 | -0.02 | 1.076 | -0.001 | -0.005 | -0.005 | 0 | 0.53 | 0.45 | 0.51 | 0.44 | 0.56 | 0.57 |
| 7 | 137773291 | rs2120846 | 0.217 | -0.627 | -0.101 | 0.922 | -0.01 | -0.097 | -0.097 | 0.132 | 0.71 | 0.91 | 0.65 | 0.65 | 0.72 | 0.76 |
| 7 | 137776481 | rs6943935 | 0.712 | -0.523 | -0.019 | 0.805 | -0.002 | -0.025 | -0.025 | 0 | 0.05 | 0.08 | 0.02 | 0 | 0.04 | 0.02 |
| 7 | 137786315 | rs6467735 | -0.623 | -0.594 | -0.159 | 0.945 | -0.029 | -0.047 | -0.047 | 0 | 0.88 | 0.98 | 0.72 | 0.88 | 0.87 | 0.97 |
| 7 | 137794388 | rs3805362 | -0.614 | -0.506 | 0.013 | 0.836 | 0.001 | -0.001 | -0.001 | 0 | 0.17 | 0.19 | 0.11 | 0.18 | 0.16 | 0.19 |
| 7 | 137800665 | rs2035736 | 1.623 | -0.67 | -0.169 | 0.902 | 0.002 | -0.003 | -0.003 | 0 | 0.55 | 0.37 | 0.53 | 0.56 | 0.6 | 0.57 |
| 7 | 137801444 | rs1872930 | 0.384 | -0.797 | -0.15 | 0.809 | 0.018 | -0.035 | -0.035 | 0.267 | 0.74 | 0.88 | 0.66 | 0.74 | 0.78 | 0.77 |
| 7 | 137802825 | rs17169522 | -1.557 | -0.447 | 0.764 | 0.544 | 0.077 | 0.049 | 0.049 | 2.365 | 0.06 | 0.45 | 0.05 | 0 | 0.02 | 0 |
| 7 | 137808839 | rs17169524 | 0.027 | -0.08 | 0.045 | 0.88 | -0.007 | -0.035 | -0.035 | 0 | 0.08 | 0.01 | 0.09 | 0.17 | 0.08 | 0.15 |
| 7 | 137809469 | rs17169525 | 0.104 | -0.073 | 0.049 | 0.883 | -0.008 | -0.034 | -0.034 | 0 | 0.07 | 0.01 | 0.09 | 0.17 | 0.08 | 0.15 |
| 7 | 137811868 | rs2440799 | 0.292 | -0.355 | 0.052 | -0.091 | 0.004 | 0.025 | 0.025 | 0 | 0.28 | 0.19 | 0.13 | 0.1 | 0.24 | 0.21 |
| 7 | 137812369 | rs2709984 | -0.401 | -0.096 | 0.298 | 0.633 | -0.04 | -0.044 | -0.044 | 0.187 | 0.93 | 0.99 | 0.76 | 0.88 | 0.86 | 0.97 |
| 7 | 137812605 | rs2465911 | 0.759 | -0.151 | 1.134 | 0.565 | 0.053 | 0.055 | 0.055 | 0 | 0.89 | 0.67 | 0.72 | 0.88 | 0.84 | 0.96 |
| 7 | 137815855 | rs17169527 | -1.47 | -0.289 | 1.13 | 0.45 | 0.058 | 0.032 | 0.032 | 2.74 | 0.06 | 0.38 | 0.04 | 0 | 0.02 | 0 |
| 7 | 137819998 | rs2440804 | 1.162 | -0.555 | 0.331 | 0.591 | -0.002 | 0 | 0 | 0.122 | 0.63 | 0.34 | 0.59 | 0.61 | 0.72 | 0.77 |
| 7 | 137821588 | rs2465072 | -0.196 | -0.105 | 0.407 | 0.751 | 0.003 | 0.005 | 0.005 | 0 | 0.91 | 0.91 | 0.75 | 0.87 | 0.85 | 0.96 |
| 7 | 137835785 | rs2451233 | 1.143 | -0.343 | 0.282 | 0.617 | -0.005 | 0.018 | 0.018 | 0.513 | 0.71 | 0.31 | 0.63 | 0.68 | 0.73 | 0.74 |
| 7 | 137842872 | rs10240712 | -0.219 | -0.496 | -0.086 | 0.498 | -0.003 | 0.012 | 0.012 | 0 | 0.12 | 0.12 | 0.07 | 0.16 | 0.1 | 0.24 |

| | | | | | | | | | | | | | | | | |
|---|---|---|---|---|---|---|---|---|---|---|---|---|---|---|---|---|
| 7 | 137846601 | rs6969282 | 0.931 | -0.819 | -0.637 | 0.079 | 0.01 | 0.035 | 0.035 | 0.188 | 0.11 | 0.01 | 0.05 | 0.04 | 0.08 | 0.03 |
| 7 | 137848071 | rs6961222 | 0.031 | -0.778 | -0.933 | -0.826 | 0.025 | 0.083 | 0.083 | 0.276 | 0.54 | 0.73 | 0.29 | 0.23 | 0.42 | 0.46 |
| 7 | 137848078 | rs6979366 | 0.592 | -1.458 | -0.223 | -1.24 | -0.03 | -0.094 | -0.094 | 0.155 | 0.18 | 0.28 | 0.13 | 0.04 | 0.13 | 0.1 |
| 7 | 137848981 | rs10238886 | -0.891 | -0.077 | 0.456 | -0.814 | 0.035 | -0.043 | -0.043 | 0 | 0.13 | 0.35 | 0.09 | 0.03 | 0.17 | 0.11 |
| 7 | 137851317 | rs2450847 | 0.625 | -0.18 | 0.067 | -0.675 | -0.005 | 0.004 | 0.004 | 0 | 0.63 | 0.7 | 0.43 | 0.28 | 0.54 | 0.45 |
| 7 | 137852311 | rs2633358 | 0.524 | -0.012 | 0.041 | -0.832 | -0.001 | -0.001 | -0.001 | 0 | 0.43 | 0.55 | 0.25 | 0.32 | 0.36 | 0.46 |
| 7 | 137854038 | rs2465075 | 0.448 | 0.16 | -0.031 | -1.337 | 0 | 0.073 | 0.073 | 1.367 | 0.48 | 0.36 | 0.45 | 0.74 | 0.43 | 0.76 |
| 7 | 137854699 | rs6945615 | 0.456 | 0.126 | -0.203 | -1.275 | 0.016 | 0.063 | 0.063 | 0.629 | 0.31 | 0.21 | 0.2 | 0.11 | 0.28 | 0.17 |
| 7 | 137856017 | rs10232331 | -0.083 | 0.429 | -0.069 | -0.912 | 0 | 0.005 | 0.005 | 0 | 0.2 | 0.26 | 0.33 | 0.68 | 0.19 | 0.47 |
| 7 | 137858766 | rs10255295 | 0.368 | -0.033 | -0.379 | -0.964 | 0.011 | -0.001 | -0.001 | 0.013 | 0.16 | 0.1 | 0.11 | 0 | 0.12 | 0.06 |
| 7 | 137873527 | rs6964417 | -0.042 | 0.664 | 0.306 | -0.263 | -0.001 | 0 | 0 | 0 | 0.52 | 0.45 | 0.47 | 0.32 | 0.45 | 0.4 |
| 7 | 137879848 | rs1552583 | -0.392 | 0.659 | 0.407 | 0.183 | -0.022 | 0.106 | 0.106 | 0.597 | 0.68 | 0.86 | 0.82 | 0.93 | 0.68 | 0.74 |
| 7 | 137885181 | rs1906889 | 0.441 | 0.788 | 0.629 | 0.705 | 0.006 | 0.068 | 0.068 | 0 | 0.07 | 0.08 | 0.21 | 0.33 | 0.1 | 0.29 |
| 7 | 137889941 | rs10268501 | 0.055 | 0.758 | 0.439 | 0.593 | -0.001 | -0.011 | -0.011 | 0 | 0.64 | 0.62 | 0.6 | 0.74 | 0.66 | 0.74 |
| 7 | 137894519 | rs12538071 | -0.847 | 0.826 | 1.011 | 1.393 | 0.003 | 0.053 | 0.053 | 0 | 0.08 | 0.25 | 0.24 | 0.13 | 0.05 | 0.03 |
| 7 | 137896908 | rs6955437 | -0.85 | 1.016 | 0.537 | 1.352 | 0.05 | 0.301 | 0.301 | 0 | 0.1 | 0.37 | 0.22 | 0.38 | 0.1 | 0.29 |
| 7 | 137917364 | rs2352564 | -0.735 | 1.236 | -0.156 | 0.286 | 0.009 | -0.028 | -0.028 | 0.02 | 0.24 | 0.32 | 0.17 | 0.14 | 0.3 | 0.19 |
| 7 | 137920895 | rs10272694 | -0.604 | 0.404 | -0.346 | -0.256 | -0.022 | -0.038 | -0.038 | 0 | 0.15 | 0.25 | 0.27 | 0.07 | 0.1 | 0.05 |
| 7 | 137925419 | rs2431049 | 0.528 | 0.471 | -1.698 | -0.47 | -0.093 | 0.02 | 0.02 | 1.216 | 0.42 | 0.04 | 0.48 | 0.6 | 0.4 | 0.6 |
| 7 | 137926501 | rs2465904 | 0.698 | 0.576 | -2.102 | -0.559 | -0.03 | 0.107 | 0.107 | 1.574 | 0.3 | 0.01 | 0.14 | 0.11 | 0.24 | 0.3 |
| 7 | 137930112 | rs2465908 | 0.076 | 0.606 | -2.17 | -1.066 | -0.001 | -0.03 | -0.03 | 0.052 | 0.83 | 0.84 | 0.83 | 1 | 0.77 | 0.94 |
| 7 | 137930588 | rs2465909 | 0.55 | 0.215 | -2.633 | -1.231 | -0.094 | 0.01 | 0.01 | 2.199 | 0.39 | 0.09 | 0.49 | 0.56 | 0.44 | 0.59 |
| 7 | 137931633 | rs10237044 | 0.304 | 0.294 | -2.639 | -2.196 | 0.002 | 0.083 | 0.083 | 0 | 0.36 | 0.24 | 0.28 | 0.07 | 0.33 | 0.26 |
| 7 | 137932264 | rs10954608 | -0.029 | 0.222 | -2.141 | -2.047 | 0.021 | 0.029 | 0.029 | 0 | 0.44 | 0.35 | 0.56 | 0.88 | 0.34 | 0.57 |
| 7 | 137934757 | rs4099093 | 0.607 | 0.742 | -2.043 | -1.299 | -0.015 | 0.014 | 0.014 | 0 | 0.13 | 0.1 | 0.14 | 0.08 | 0.21 | 0.16 |
| 7 | 137939326 | rs2352479 | -0.566 | 0.334 | -2.176 | -1.259 | 0.014 | 0.107 | 0.107 | 0 | 0.49 | 0.79 | 0.48 | 0.29 | 0.47 | 0.33 |
| 7 | 137942208 | rs2253200 | -0.404 | 0.529 | -1.455 | -1.317 | 0.024 | -0.02 | -0.02 | 0 | 0.41 | 0.21 | 0.36 | 0.45 | 0.39 | 0.4 |
| 7 | 137946207 | rs12670058 | 0.084 | 0.502 | -0.809 | -2.16 | -0.032 | 0.016 | 0.016 | 0 | 0.29 | 0.16 | 0.32 | 0.67 | 0.35 | 0.27 |
| 7 | 137947218 | rs4732302 | -0.871 | 0.344 | -0.629 | -2.141 | 0.002 | -0.046 | -0.046 | 0.672 | 0.64 | 0.47 | 0.54 | 0.79 | 0.65 | 0.61 |
| 7 | 137956892 | rs4732304 | -0.137 | 0.421 | -0.499 | -1.807 | 0.009 | -0.027 | -0.027 | 0.731 | 0.37 | 0.42 | 0.4 | 0.7 | 0.43 | 0.28 |
| 7 | 137962186 | rs11763343 | -0.19 | -0.64 | -1.044 | -1.71 | 0.022 | 0.052 | 0.052 | 0.009 | 0.27 | 0.16 | 0.17 | 0.09 | 0.2 | 0.23 |
| 7 | 137972605 | rs10246131 | -0.259 | 0.043 | -0.627 | -1.494 | -0.012 | -0.062 | -0.062 | 0 | 0.34 | 0.5 | 0.46 | 0.78 | 0.32 | 0.32 |
| 7 | 137972636 | rs17361616 | -0.159 | 0.186 | -1.217 | -1.43 | 0 | -0.005 | -0.005 | 0 | 0.33 | 0.32 | 0.45 | 0.78 | 0.32 | 0.3 |
| 7 | 137973629 | rs6467756 | -0.031 | 0.305 | -1.524 | -1.005 | 0.007 | 0.107 | 0.107 | 0 | 0.73 | 0.91 | 0.85 | 0.93 | 0.76 | 0.79 |
| 7 | 137976868 | rs357365 | -0.442 | 0.298 | -0.779 | -0.298 | -0.008 | -0.018 | -0.018 | 0 | 0.27 | 0.36 | 0.44 | 0.72 | 0.26 | 0.28 |
| 7 | 137982771 | rs357372 | -0.306 | -0.007 | -1.579 | 0.196 | 0.005 | -0.079 | -0.079 | 0 | 0.21 | 0.03 | 0.17 | 0.32 | 0.19 | 0.4 |
| 7 | 137983003 | rs357373 | 0.241 | 0.343 | -2.467 | -0.458 | -0.076 | -0.121 | -0.121 | 0 | 0.76 | 0.91 | 0.53 | 0.42 | 0.66 | 0.76 |
| 7 | 137985801 | rs1421487 | -1.004 | -0.521 | -1.646 | -0.619 | -0.019 | -0.02 | -0.02 | 0 | 0.23 | 0.49 | 0.23 | 0.26 | 0.17 | 0.24 |
| 7 | 138005671 | rs357382 | 1.264 | -0.583 | -1.375 | -0.701 | 0 | 0 | 0 | 0 | 0.63 | 0.36 | 0.49 | 0.32 | 0.64 | 0.59 |
| 7 | 138012161 | rs380650 | 0.846 | -0.612 | -1.596 | -0.744 | 0.005 | -0.093 | -0.093 | 0.095 | 0.21 | 0.02 | 0.19 | 0.31 | 0.2 | 0.42 |
| 7 | 138013414 | rs11769851 | 1.58 | -0.58 | -1.426 | -0.792 | -0.024 | 0.06 | 0.06 | 0 | 0.41 | 0.27 | 0.3 | 0.01 | 0.44 | 0.2 |
| 7 | 138033787 | rs357420 | -0.475 | -0.38 | -1.339 | -0.496 | -0.022 | -0.06 | -0.06 | 0 | 0.86 | 0.93 | 0.72 | 0.53 | 0.81 | 0.84 |
| 7 | 138037255 | rs10262321 | -0.991 | -0.591 | -1.164 | -0.627 | -0.011 | -0.036 | -0.036 | 0 | 0.18 | 0.59 | 0.19 | 0.15 | 0.14 | 0.13 |
| 7 | 138041329 | rs357405 | -0.427 | -0.351 | -1.751 | -0.333 | -0.045 | -0.099 | -0.099 | 0.084 | 0.86 | 0.98 | 0.71 | 0.56 | 0.8 | 0.86 |
| 7 | 138042100 | rs357403 | 0.754 | -0.487 | -1.843 | -0.61 | -0.065 | 0.031 | 0.031 | 0.062 | 0.74 | 0.94 | 0.61 | 0.84 | 0.7 | 0.77 |
| 7 | 138043043 | rs357401 | 0.688 | -0.544 | -1.646 | -0.904 | 0.001 | 0.009 | 0.009 | 0 | 0.71 | 0.73 | 0.72 | 0.94 | 0.73 | 0.72 |
| 7 | 138044576 | rs357399 | 0.78 | -0.556 | -1.982 | -0.698 | 0.005 | 0.03 | 0.03 | 0.01 | 0.63 | 0.74 | 0.58 | 0.74 | 0.67 | 0.64 |
| 7 | 138066098 | rs434084 | 0.295 | -0.128 | -2.066 | -0.335 | -0.062 | -0.068 | -0.068 | 1.016 | 0.22 | 0.02 | 0.26 | 0.3 | 0.27 | 0.36 |
| 7 | 138067176 | rs427508 | -0.495 | -0.303 | -2.048 | -0.443 | 0.015 | 0.045 | 0.045 | 0.057 | 0.85 | 0.93 | 0.77 | 0.98 | 0.87 | 0.86 |
| 7 | 138071567 | rs475811 | -0.624 | -0.224 | -2.066 | -0.434 | 0.013 | 0.038 | 0.038 | 0.046 | 0.86 | 0.92 | 0.77 | 0.98 | 0.87 | 0.86 |
| 7 | 138073611 | rs370042 | -0.542 | -0.226 | -2.16 | -0.533 | -0.02 | -0.02 | -0.02 | 0.442 | 0.47 | 0.66 | 0.5 | 0.51 | 0.46 | 0.64 |
| 7 | 138074052 | rs403760 | 0.401 | -0.17 | -2.103 | -0.801 | 0.01 | 0.023 | 0.023 | 0 | 0.61 | 0.39 | 0.68 | 0.78 | 0.69 | 0.58 |
| 7 | 138077393 | rs11762037 | 0.236 | 0.252 | -1.968 | -0.628 | 0.001 | -0.014 | -0.014 | 0 | 0.27 | 0.47 | 0.25 | 0.23 | 0.21 | 0.29 |
| 7 | 138079079 | rs13232937 | 0.285 | 0.758 | -2.612 | -0.023 | 0.043 | 0.087 | 0.087 | 0 | 0.44 | 0.72 | 0.33 | 0.23 | 0.33 | 0.43 |
| 7 | 138081425 | rs7791577 | 0.376 | 0.795 | -2.806 | -0.007 | 0.046 | 0.091 | 0.091 | 0 | 0.44 | 0.73 | 0.33 | 0.23 | 0.33 | 0.42 |
| 7 | 138082575 | rs7796720 | 0.737 | 1.143 | -2.245 | 0.582 | -0.046 | -0.054 | -0.054 | 0.413 | 0.78 | 0.97 | 0.75 | 0.65 | 0.73 | 0.64 |
| 7 | 138083138 | rs4473955 | 0.564 | 1.142 | -2.232 | 0.619 | -0.031 | -0.052 | -0.052 | 0.157 | 0.82 | 0.94 | 0.79 | 0.65 | 0.77 | 0.63 |
| 7 | 138086495 | rs357379 | -0.959 | 1.673 | -2.122 | 0.776 | -0.089 | -0.088 | -0.088 | 0.067 | 0.22 | 0.03 | 0.25 | 0.38 | 0.3 | 0.32 |
| 7 | 138095400 | rs4732314 | 1.02 | 1.548 | -2.015 | 0.632 | -0.051 | -0.063 | -0.063 | 0 | 0.77 | 0.94 | 0.77 | 0.6 | 0.73 | 0.65 |
| 7 | 138098694 | rs2352570 | 1.111 | 1.072 | -1.683 | -0.617 | -0.022 | -0.137 | -0.137 | 0.632 | 0.35 | 0.46 | 0.23 | 0.03 | 0.31 | 0.15 |
| 7 | 138111275 | rs2353348 | 1.002 | 0.851 | -1.719 | -0.16 | -0.012 | -0.056 | -0.056 | 0 | 0.75 | 0.88 | 0.78 | 0.53 | 0.73 | 0.56 |
| 7 | 138125434 | rs7808971 | -0.144 | 0.763 | -1.391 | -0.221 | -0.004 | -0.001 | -0.001 | 0 | 0.07 | 0.08 | 0.03 | 0.03 | 0.04 | 0.09 |
| 7 | 138143811 | rs13231375 | -0.701 | 1.239 | -1.562 | -0.498 | -0.009 | 0.024 | 0.024 | 0 | 0.18 | 0.1 | 0.12 | 0.01 | 0.17 | 0.16 |
| 7 | 138144257 | rs7798391 | -1.177 | 0.939 | -1.282 | -0.494 | 0 | 0.001 | 0.001 | 0 | 0.47 | 0.56 | 0.54 | 0.55 | 0.46 | 0.57 |
| 7 | 138157079 | rs749546 | 0.188 | 1.227 | -0.987 | -0.304 | -0.004 | 0.007 | 0.007 | 0 | 0.17 | 0.1 | 0.4 | 0.49 | 0.28 | 0.26 |
| 7 | 138178222 | rs6467778 | -0.266 | 1.161 | -1.075 | -0.486 | -0.003 | -0.054 | -0.054 | 0 | 0.81 | 0.86 | 0.78 | 0.52 | 0.8 | 0.66 |
| 7 | 138188932 | rs10488642 | 2.179 | 1.122 | -1.101 | -0.749 | -0.009 | -0.148 | -0.148 | 0 | 0.37 | 0.43 | 0.26 | 0 | 0.36 | 0.13 |
| 7 | 138222987 | rs3807155 | -0.725 | 1.152 | -1.026 | -0.59 | 0.002 | 0.008 | 0.008 | 0 | 0.28 | 0.33 | 0.47 | 0.51 | 0.31 | 0.37 |
| 7 | 138238844 | rs7811152 | 1.151 | 0.963 | -0.997 | -0.885 | -0.004 | -0.038 | -0.038 | 0 | 0.53 | 0.51 | 0.32 | 0.01 | 0.5 | 0.28 |

| | | | | | | | | | | | | | | | | |
|---|---|---|---|---|---|---|---|---|---|---|---|---|---|---|---|---|
| 7 | 138281355 | rs1659832 | -0.335 | 1.328 | -1.537 | -0.126 | -0.025 | -0.031 | -0.031 | 0.024 | 0.83 | 0.93 | 0.62 | 0.38 | 0.74 | 0.73 |
| 7 | 138287097 | rs1673214 | -0.942 | 1.001 | -1.651 | 0.205 | -0.033 | -0.072 | -0.072 | 0 | 0.28 | 0.16 | 0.45 | 0.54 | 0.34 | 0.35 |
| 7 | 138287452 | rs13231090 | -0.332 | 1.005 | -1.588 | 0.202 | -0.014 | 0.058 | 0.058 | 0 | 0.17 | 0.05 | 0.07 | 0 | 0.15 | 0.16 |
| 7 | 138292828 | rs1673218 | -0.919 | 0.229 | -1.498 | 0.333 | 0.001 | -0.002 | -0.002 | 0 | 0.31 | 0.31 | 0.32 | 0.59 | 0.27 | 0.43 |
| 7 | 138297714 | rs1673223 | 0.732 | 0.366 | -1.684 | 0.246 | -0.054 | -0.086 | -0.086 | 0 | 0.16 | 0.01 | 0.22 | 0.28 | 0.2 | 0.25 |
| 7 | 138298688 | rs1673224 | 0.994 | 0.621 | -1.58 | 0.154 | 0.057 | 0.136 | 0.136 | 0.613 | 0.5 | 0.75 | 0.34 | 0.2 | 0.38 | 0.33 |
| 7 | 138299511 | rs1659821 | -1.124 | 0.61 | -1.576 | -0.08 | 0.057 | 0.132 | 0.132 | 0.454 | 0.51 | 0.27 | 0.69 | 0.99 | 0.62 | 0.79 |
| 7 | 138303027 | rs1673227 | -0.573 | 0.566 | -1.592 | 0.066 | 0.005 | -0.041 | -0.041 | 0 | 0.34 | 0.19 | 0.35 | 0.58 | 0.31 | 0.53 |
| 7 | 138304579 | rs753223 | -0.701 | 0.934 | -0.579 | 0.979 | -0.021 | -0.05 | -0.05 | 0 | 0.11 | 0.19 | 0.07 | 0 | 0.04 | 0.05 |
| 7 | 138305782 | rs3734945 | -1.352 | 1.083 | -0.6 | 1.08 | -0.008 | -0.014 | -0.014 | 0 | 0.28 | 0.25 | 0.42 | 0.39 | 0.35 | 0.31 |
| 7 | 138309274 | rs17837468 | -1.07 | -0.073 | -0.477 | 0.895 | -0.004 | -0.001 | -0.001 | 0 | 0.23 | 0.25 | 0.2 | 0.25 | 0.1 | 0.31 |
| 7 | 138314452 | rs6467784 | 1.154 | 0.394 | -0.506 | 1.196 | -0.005 | -0.029 | -0.029 | 0 | 0.83 | 0.89 | 0.85 | 0.74 | 0.84 | 0.78 |
| 7 | 138314465 | rs1659809 | -1.827 | 0.261 | 0.136 | 1.12 | 0.038 | 0.198 | 0.198 | 1.832 | 0.08 | 0.37 | 0.11 | 0.26 | 0.06 | 0.03 |
| 7 | 138314555 | rs6467785 | 1.419 | 0.302 | 0.085 | 1.155 | -0.007 | 0.016 | 0.016 | 0 | 0.8 | 0.65 | 0.83 | 0.74 | 0.84 | 0.78 |
| 7 | 138323516 | rs1631216 | 1.242 | 0.821 | 0.336 | 0.987 | 0.022 | -0.008 | -0.008 | 0 | 0.84 | 0.71 | 0.85 | 0.85 | 0.82 | 0.84 |
| 7 | 138326540 | rs2919140 | 1.641 | 0.059 | 1.212 | 1.115 | -0.006 | 0.016 | 0.016 | 0.896 | 0.83 | 0.42 | 0.85 | 0.8 | 0.87 | 0.85 |
| 7 | 138335246 | rs6956225 | -1.753 | 0.074 | 1.673 | 1.495 | 0.021 | -0.005 | -0.005 | 0 | 0.16 | 0.32 | 0.2 | 0.16 | 0.14 | 0.16 |
| 7 | 138336608 | rs1659794 | -2.444 | 0.1 | 1.678 | 1.795 | -0.007 | 0.032 | 0.032 | 2.175 | 0.27 | 0.6 | 0.26 | 0.38 | 0.2 | 0.34 |
| 7 | 138343936 | rs11762109 | 0.546 | -0.879 | 0.292 | 1.231 | -0.024 | 0.062 | 0.062 | 0.064 | 0.36 | 0.1 | 0.3 | 0.17 | 0.33 | 0.19 |
| 7 | 138346011 | rs11770967 | -0.807 | -0.831 | 0.418 | 1.143 | -0.014 | -0.008 | -0.008 | 0.15 | 0.59 | 0.69 | 0.45 | 0.54 | 0.43 | 0.48 |
| 7 | 138358456 | rs17160449 | -0.62 | -1.257 | 0.908 | 1.072 | 0 | -0.001 | -0.001 | 0 | 0.54 | 0.61 | 0.45 | 0.52 | 0.4 | 0.49 |
| 7 | 138363896 | rs12535253 | 0.437 | -2.005 | 1.106 | 0.818 | -0.001 | -0.001 | -0.001 | 0.366 | 0.37 | 0.35 | 0.16 | 0.18 | 0.25 | 0.17 |
| 7 | 138364652 | rs17160459 | 0.754 | -2.14 | 0.464 | 0.801 | 0.049 | 0.033 | 0.033 | 1.939 | 0.31 | 0.14 | 0.1 | 0.23 | 0.19 | 0.16 |
| 7 | 138366052 | rs7804639 | -0.231 | -1.341 | 0.739 | 1.045 | 0.004 | -0.005 | -0.005 | 0 | 0.58 | 0.73 | 0.34 | 0.56 | 0.4 | 0.46 |
| 7 | 138367358 | rs10250086 | -0.307 | -1.284 | 0.029 | 1.071 | -0.016 | -0.047 | -0.047 | 0.571 | 0.6 | 0.89 | 0.35 | 0.56 | 0.41 | 0.46 |
| 7 | 138376184 | rs296913 | -0.752 | -1.128 | 0.21 | 0.619 | 0.029 | 0.029 | 0.029 | 0 | 0.51 | 0.68 | 0.59 | 0.55 | 0.4 | 0.47 |
| 7 | 138377805 | rs296917 | -0.884 | -1.469 | 0.359 | -0.169 | 0.034 | 0.083 | 0.083 | 0 | 0.29 | 0.14 | 0.27 | 0.14 | 0.25 | 0.2 |
| 7 | 138378905 | rs11769043 | -0.228 | -1.149 | 0.886 | -0.165 | 0.023 | -0.011 | -0.011 | 0.59 | 0.06 | 0.24 | 0.04 | 0 | 0.04 | 0.03 |
| 7 | 138380477 | rs6944016 | -0.278 | -1.259 | 0.304 | -0.223 | 0.039 | 0.002 | 0.002 | 0.001 | 0.13 | 0.04 | 0.09 | 0.14 | 0.05 | 0.15 |
| 7 | 138382677 | rs400800 | -1.253 | -1.144 | 0.585 | -0.188 | -0.026 | -0.056 | -0.056 | 0 | 0.31 | 0.41 | 0.29 | 0.14 | 0.24 | 0.2 |
| 7 | 138387779 | rs6957669 | -1.254 | -1.034 | 0.395 | -0.253 | -0.012 | -0.045 | -0.045 | 0.235 | 0.27 | 0.32 | 0.17 | 0.04 | 0.21 | 0.21 |
| 7 | 138396125 | rs12707395 | -0.794 | -0.385 | 0.438 | 0.035 | 0.005 | -0.029 | -0.029 | 0 | 0.35 | 0.53 | 0.52 | 0.39 | 0.39 | 0.28 |
| 7 | 138396282 | rs3778703 | -0.614 | -0.508 | 0.532 | 0.491 | -0.04 | 0.02 | 0.02 | 0 | 0.14 | 0.25 | 0.07 | 0.14 | 0.08 | 0.21 |
| 7 | 138396349 | rs3778701 | -0.248 | -0.5 | 0.319 | 0.462 | 0.006 | 0.005 | 0.005 | 0 | 0.13 | 0.12 | 0.04 | 0.04 | 0.07 | 0.16 |
| 7 | 138397297 | rs6977910 | -1.472 | -0.47 | 0.141 | 0.489 | 0.036 | 0.023 | 0.023 | 0 | 0.48 | 0.79 | 0.58 | 0.51 | 0.47 | 0.47 |
| 7 | 138397399 | rs7810105 | -0.995 | -0.441 | 0.536 | 0.626 | -0.049 | 0.031 | 0.031 | 0 | 0.15 | 0.33 | 0.07 | 0.14 | 0.08 | 0.21 |
| 7 | 138400796 | rs999229 | -0.548 | 0.44 | -0.195 | 0.763 | 0.001 | 0.001 | 0.001 | 0 | 0.29 | 0.32 | 0.26 | 0.38 | 0.37 | 0.28 |
| 7 | 138400858 | rs999228 | -0.885 | 0.229 | -0.28 | 0.403 | -0.001 | 0.002 | 0.002 | 0 | 0.49 | 0.66 | 0.36 | 0.52 | 0.49 | 0.45 |
| 7 | 138401167 | rs6943456 | 0.071 | 0.493 | -0.266 | 0.377 | -0.012 | -0.028 | -0.028 | 0 | 0.2 | 0.35 | 0.11 | 0.14 | 0.12 | 0.17 |
| 7 | 138406368 | rs3823501 | -0.116 | 1.448 | -1.106 | 0.424 | -0.036 | -0.006 | -0.006 | 0 | 0.77 | 0.88 | 0.77 | 0.78 | 0.71 | 0.82 |
| 7 | 138406404 | rs3734942 | 0.505 | 1.328 | -1.026 | 0.585 | 0.001 | 0 | 0 | 0 | 0.15 | 0.14 | 0.17 | 0.25 | 0.24 | 0.09 |
| 7 | 138411425 | rs3800569 | 0.075 | 1.051 | -1.103 | 0.204 | -0.024 | -0.008 | -0.008 | 0 | 0.79 | 0.86 | 0.78 | 0.78 | 0.73 | 0.83 |
| 7 | 138413753 | rs3734941 | 0.367 | 1.154 | -0.969 | 0.165 | -0.019 | -0.001 | -0.001 | 0 | 0.77 | 0.8 | 0.75 | 0.76 | 0.67 | 0.85 |
| 7 | 138417718 | rs3807154 | 0.841 | 0.6 | -0.576 | -0.508 | -0.003 | -0.068 | -0.068 | 0 | 0.66 | 0.42 | 0.78 | 0.85 | 0.72 | 0.72 |
| 7 | 138417791 | rs3807153 | -0.464 | 0.248 | -0.399 | -0.523 | -0.022 | -0.016 | -0.016 | 0 | 0.15 | 0.21 | 0.03 | 0.05 | 0.03 | 0.15 |
| 7 | 138418204 | rs1344381 | 0.704 | 0.129 | -0.535 | -0.752 | 0.005 | 0.023 | 0.023 | 0.054 | 0.63 | 0.66 | 0.77 | 0.84 | 0.71 | 0.7 |
| 7 | 138418910 | rs1026435 | 0.752 | 0.117 | -0.453 | -0.745 | 0.001 | -0.001 | -0.001 | 0 | 0.63 | 0.61 | 0.77 | 0.84 | 0.71 | 0.7 |
| 7 | 138419085 | rs1037692 | 0.534 | 0.158 | -0.746 | -0.976 | 0.005 | 0.114 | 0.114 | 0.003 | 0.65 | 0.78 | 0.81 | 0.95 | 0.72 | 0.71 |
| 7 | 138424023 | rs3734939 | 0.257 | 0.185 | -0.517 | -1.309 | 0 | 0.083 | 0.083 | 0.649 | 0.5 | 0.62 | 0.69 | 0.92 | 0.55 | 0.58 |
| 7 | 138429851 | rs6467797 | -0.276 | 0.591 | -0.543 | -0.404 | 0.005 | 0.075 | 0.075 | 0.406 | 0.21 | 0.49 | 0.12 | 0.32 | 0.19 | 0.13 |
| 7 | 138430954 | rs1562853 | 0.017 | 0.416 | -0.974 | -1.219 | -0.002 | 0.103 | 0.103 | 1.281 | 0.52 | 0.71 | 0.7 | 0.91 | 0.57 | 0.57 |
| 7 | 138432983 | rs430262 | -0.132 | 0.278 | -0.806 | -1.626 | -0.012 | 0.06 | 0.06 | 0 | 0.45 | 0.64 | 0.53 | 0.22 | 0.48 | 0.39 |
| 7 | 138433506 | rs7781292 | 0.788 | 0.491 | -0.449 | -1.693 | -0.006 | 0.006 | 0.006 | 0 | 0.18 | 0.14 | 0.27 | 0.07 | 0.24 | 0.19 |
| 7 | 138434485 | rs366193 | -0.789 | 0.614 | -0.995 | -1.076 | -0.09 | -0.093 | -0.093 | 0.708 | 0.63 | 0.93 | 0.62 | 0.57 | 0.58 | 0.48 |
| 7 | 138436480 | rs381821 | 0.347 | 0.452 | -0.517 | -1.169 | -0.004 | 0.051 | 0.051 | 0.32 | 0.22 | 0.27 | 0.13 | 0.49 | 0.2 | 0.11 |
| 7 | 138436498 | rs17442029 | 1.054 | 0.496 | -0.812 | -1.187 | 0.012 | 0.036 | 0.036 | 0 | 0.25 | 0.17 | 0.22 | 0.02 | 0.22 | 0.16 |
| 7 | 138438848 | rs411467 | -0.292 | 0.587 | -0.596 | -1.083 | -0.002 | -0.005 | -0.005 | 0.229 | 0.35 | 0.62 | 0.39 | 0.54 | 0.36 | 0.33 |
| 7 | 138440260 | rs3807147 | 2.625 | 0.943 | -1.206 | -0.837 | -0.041 | 0.015 | 0.015 | 0 | 0.1 | 0.03 | 0.23 | 0.05 | 0.15 | 0.18 |
| 7 | 138440607 | rs296911 | 1.961 | 0.902 | -1.144 | -0.814 | 0.004 | -0.001 | -0.001 | 0 | 0.12 | 0.11 | 0.28 | 0.19 | 0.22 | 0.21 |
| 7 | 138441124 | rs296910 | -0.45 | 0.625 | -1.234 | -0.717 | -0.037 | -0.045 | -0.045 | 0.339 | 0.61 | 0.85 | 0.62 | 0.56 | 0.58 | 0.51 |
| 7 | 138446998 | rs3823499 | 1.07 | 0.52 | -0.711 | -0.369 | 0.001 | 0.044 | 0.044 | 0 | 0.1 | 0.13 | 0.06 | 0.38 | 0.09 | 0.09 |
| 7 | 138447395 | rs3734936 | 0.924 | 0.081 | -1.003 | -0.402 | 0.005 | 0.005 | 0.005 | 0 | 0.24 | 0.16 | 0.15 | 0.18 | 0.16 | 0.28 |
| 7 | 138449419 | rs10241449 | -1.141 | 0.555 | -1.3 | -0.495 | -0.03 | -0.034 | -0.034 | 0 | 0.83 | 0.88 | 0.59 | 0.64 | 0.69 | 0.77 |
| 7 | 138450098 | rs4732330 | -0.16 | -0.504 | -0.786 | -0.098 | -0.041 | 0.023 | 0.023 | 0 | 0.34 | 0.39 | 0.22 | 0.56 | 0.25 | 0.37 |
| 7 | 138455108 | rs7799249 | 0.419 | 0.016 | -0.825 | 0.066 | -0.004 | -0.026 | -0.026 | 0.177 | 0.7 | 0.38 | 0.61 | 0.26 | 0.73 | 0.67 |
| 7 | 138455988 | rs10258719 | 0.121 | 0.641 | -1.008 | 0.654 | 0.002 | 0.016 | 0.016 | 0 | 0.74 | 0.74 | 0.56 | 0.53 | 0.74 | 0.78 |
| 7 | 138463568 | rs6979424 | 0.32 | 0.389 | -1.764 | 0.404 | -0.112 | -0.095 | -0.095 | 0 | 0.32 | 0.07 | 0.53 | 0.48 | 0.43 | 0.44 |
| 7 | 138463724 | rs6979610 | -0.379 | -0.379 | -1.553 | -0.595 | 0.007 | 0 | 0 | 0 | 0.71 | 0.29 | 0.8 | 0.94 | 0.75 | 0.79 |
| 7 | 138466720 | rs4732334 | 0.485 | 0.147 | -0.975 | -0.187 | -0.009 | -0.017 | -0.017 | 0.05 | 0.33 | 0.34 | 0.57 | 0.5 | 0.38 | 0.53 |

| | | | | | | | | | | | | | | | | |
|---|---|---|---|---|---|---|---|---|---|---|---|---|---|---|---|---|
| 7 | 138467140 | rs7791537 | -0.753 | -0.436 | -0.868 | -0.038 | 0.047 | 0.056 | 0.056 | 0.003 | 0.66 | 0.87 | 0.79 | 0.76 | 0.72 | 0.74 |
| 7 | 138472215 | rs6467802 | -0.367 | -0.39 | -0.369 | -0.032 | 0.008 | 0.004 | 0.004 | 0.929 | 0.51 | 0.66 | 0.66 | 0.54 | 0.52 | 0.62 |
| 7 | 138479309 | rs3778689 | 0.251 | 0.141 | -0.195 | 0.316 | 0.004 | 0.007 | 0.007 | 3.305 | 0.36 | 0.44 | 0.59 | 0.5 | 0.43 | 0.62 |
| 7 | 138481857 | rs3823493 | -0.53 | -0.054 | 0.496 | 0.326 | -0.005 | -0.019 | -0.019 | 0.021 | 0.1 | 0.23 | 0.06 | 0.04 | 0.06 | 0.08 |
| 7 | 138483581 | rs3807328 | 0.279 | 0.09 | 0.554 | 0.413 | 0 | 0.001 | 0.001 | 0.361 | 0.54 | 0.49 | 0.58 | 0.52 | 0.48 | 0.44 |
| 7 | 138486536 | rs6969815 | -0.242 | 0.186 | 0.611 | 0.482 | -0.014 | -0.001 | -0.001 | 4.604 | 0.66 | 0.74 | 0.66 | 0.69 | 0.63 | 0.6 |
| 7 | 138488116 | rs896189 | 0.285 | 0.263 | 0.699 | 0.564 | 0.001 | -0.004 | -0.004 | 0 | 0.09 | 0.11 | 0.07 | 0.04 | 0.12 | 0.05 |
| 7 | 138491500 | rs6976426 | -1.01 | 0.018 | 1.061 | 0.523 | -0.015 | 0.027 | 0.027 | 0 | 0.16 | 0.51 | 0.12 | 0.15 | 0.14 | 0.1 |
| 7 | 138494176 | rs10435432 | -0.121 | -0.108 | 0.586 | 0.363 | -0.016 | 0.028 | 0.028 | 0 | 0.59 | 0.74 | 0.64 | 0.7 | 0.6 | 0.58 |
| 7 | 138494588 | rs10435433 | 0.758 | -0.038 | 0.66 | 0.374 | -0.001 | -0.007 | -0.007 | 3.783 | 0.44 | 0.36 | 0.53 | 0.55 | 0.47 | 0.5 |
| 7 | 138498772 | rs6948137 | -0.247 | 0.126 | 0.554 | 0.277 | -0.001 | 0.003 | 0.003 | 0 | 0.09 | 0.08 | 0.08 | 0.04 | 0.12 | 0.06 |
| 7 | 138502417 | rs6965588 | -0.388 | 0.131 | 0.575 | 0.279 | 0 | -0.011 | -0.011 | 0 | 0.1 | 0.11 | 0.09 | 0.04 | 0.12 | 0.06 |
| 7 | 138518056 | rs896193 | -0.386 | 0.804 | -0.765 | 0.003 | -0.107 | 0.036 | 0.036 | 1.21 | 0.79 | 0.99 | 0.73 | 0.89 | 0.73 | 0.69 |
| 7 | 138518783 | rs878992 | 0.002 | 0.158 | -0.119 | 0.375 | -0.041 | 0.096 | 0.096 | 0 | 0.2 | 0.4 | 0.14 | 0.34 | 0.15 | 0.17 |
| 7 | 138523590 | rs11973431 | 0.792 | 0.507 | 0.173 | -0.509 | -0.004 | -0.001 | -0.001 | 0.196 | 0.6 | 0.65 | 0.6 | 0.86 | 0.54 | 0.54 |
| 7 | 138525605 | rs12668833 | -0.005 | 0.446 | 0.152 | -0.539 | 0.007 | -0.035 | -0.035 | 0 | 0.72 | 0.69 | 0.7 | 0.9 | 0.7 | 0.69 |
| 7 | 138532408 | rs7781030 | -0.048 | 0.637 | -0.003 | -0.518 | -0.004 | 0.003 | 0.003 | 0 | 0.12 | 0.09 | 0.11 | 0.05 | 0.18 | 0.15 |
| 7 | 138534762 | rs10243651 | -0.012 | 0.49 | 0.176 | -0.486 | 0.005 | -0.029 | -0.029 | 0 | 0.72 | 0.7 | 0.68 | 0.88 | 0.71 | 0.68 |
| 7 | 138538228 | rs10227043 | -0.689 | 0.432 | 0.097 | -0.155 | 0.018 | -0.033 | -0.033 | 0 | 0.13 | 0.23 | 0.12 | 0.05 | 0.17 | 0.15 |
| 7 | 138541488 | rs6467814 | -0.864 | 0.112 | -0.578 | 0.186 | -0.01 | -0.046 | -0.046 | 0.196 | 0.37 | 0.83 | 0.38 | 0.38 | 0.33 | 0.33 |
| 7 | 138550994 | rs10229557 | -0.846 | 0.096 | -0.518 | 0.18 | -0.006 | -0.034 | -0.034 | 0 | 0.36 | 0.77 | 0.36 | 0.37 | 0.33 | 0.31 |
| 7 | 138561260 | rs10278730 | -0.288 | 0.085 | -0.148 | -0.099 | 0.005 | 0.006 | 0.006 | 0.257 | 0.58 | 0.62 | 0.28 | 0.38 | 0.43 | 0.41 |
| 7 | 138562547 | rs9918611 | -0.595 | 0.629 | 0.139 | -0.015 | -0.014 | 0.122 | 0.122 | 0 | 0.15 | 0.39 | 0.07 | 0.23 | 0.08 | 0.04 |
| 7 | 138566810 | rs6973895 | -1.269 | 0.911 | -0.325 | -0.034 | -0.084 | -0.127 | -0.127 | 0 | 0.63 | 0.94 | 0.44 | 0.52 | 0.6 | 0.53 |
| 7 | 138567110 | rs6974263 | -0.852 | -0.024 | 0.855 | 0.012 | 0.046 | 0.058 | 0.058 | 0 | 0.5 | 0.63 | 0.28 | 0.31 | 0.41 | 0.39 |
| 7 | 138568430 | rs6942581 | -0.12 | 0.092 | 0.735 | 0.31 | 0.002 | 0.049 | 0.049 | 0 | 0.07 | 0.08 | 0.04 | 0.22 | 0.07 | 0.05 |
| 7 | 138573471 | rs6944423 | -0.752 | -0.04 | 0.908 | -0.206 | 0.01 | 0.232 | 0.232 | 0.186 | 0.12 | 0.41 | 0.09 | 0.25 | 0.07 | 0.05 |
| 7 | 138574993 | rs6467815 | -0.651 | 0.188 | -0.518 | -0.335 | 0.04 | 0.064 | 0.064 | 0 | 0.51 | 0.76 | 0.27 | 0.33 | 0.42 | 0.43 |
| 7 | 138577911 | rs10270572 | -1.179 | -0.035 | -0.21 | -0.466 | 0.004 | 0.001 | 0.001 | 0.901 | 0.3 | 0.72 | 0.2 | 0.23 | 0.23 | 0.12 |
| 7 | 138580906 | rs6944822 | -0.782 | 0.071 | 0.129 | -0.331 | -0.007 | 0.037 | 0.037 | 0.099 | 0.09 | 0.12 | 0.04 | 0.17 | 0.05 | 0.05 |
| 7 | 138583840 | rs7802841 | 0.592 | -0.184 | 0.008 | -1.451 | 0.011 | 0.069 | 0.069 | 0 | 0.63 | 0.75 | 0.82 | 0.9 | 0.65 | 0.68 |
| 7 | 138588953 | rs6979681 | -1.141 | -0.286 | 0.443 | -1.525 | 0.046 | 0.132 | 0.132 | 0 | 0.45 | 0.74 | 0.23 | 0.1 | 0.36 | 0.31 |
| 7 | 138590990 | rs10954638 | 0.105 | -0.31 | 0.197 | -1.626 | -0.005 | 0.121 | 0.121 | 0 | 0.25 | 0.05 | 0.1 | 0.05 | 0.22 | 0.24 |
| 7 | 138596001 | rs16873582 | -1.649 | 0.004 | 0.739 | -1.587 | -0.005 | -0.004 | -0.004 | 0.95 | 0.1 | 0.14 | 0.05 | 0.06 | 0.07 | 0.06 |
| 7 | 138596064 | rs6969074 | -1.952 | 0.336 | 0.686 | -1.81 | -0.06 | 0.138 | 0.138 | 0 | 0.56 | 0.87 | 0.31 | 0.19 | 0.47 | 0.43 |
| 7 | 138597036 | rs3735013 | -0.793 | 0.332 | 1.013 | -1.815 | 0 | -0.03 | -0.03 | 0 | 0.39 | 0.62 | 0.21 | 0.06 | 0.33 | 0.26 |
| 7 | 138597434 | rs17160610 | -2.159 | 0.334 | 1.002 | -1.803 | 0.006 | -0.014 | -0.014 | 1.503 | 0.18 | 0.3 | 0.12 | 0.13 | 0.14 | 0.18 |
| 7 | 138600299 | rs2774965 | -1.838 | 0.538 | 0.686 | -1.718 | -0.063 | 0.154 | 0.154 | 0 | 0.54 | 0.9 | 0.29 | 0.19 | 0.42 | 0.44 |
| 7 | 138601572 | rs2150283 | -0.43 | 0.532 | 0.647 | -1.778 | -0.003 | 0.018 | 0.018 | 0.583 | 0.08 | 0.03 | 0.02 | 0 | 0.07 | 0.15 |
| 7 | 138601706 | rs2354336 | -0.43 | 0.532 | 0.647 | -1.778 | -0.003 | 0.018 | 0.018 | 0.583 | 0.08 | 0.03 | 0.02 | 0 | 0.07 | 0.15 |
| 7 | 138602417 | rs2774960 | -1.487 | 0.517 | 1.135 | -1.794 | 0.037 | 0.098 | 0.098 | 0 | 0.44 | 0.72 | 0.25 | 0.2 | 0.34 | 0.29 |
| 7 | 138602523 | rs2774962 | -2.076 | 0.54 | 0.868 | -1.754 | -0.076 | 0.133 | 0.133 | 0 | 0.54 | 0.94 | 0.29 | 0.2 | 0.42 | 0.44 |
| 7 | 138615546 | rs6942786 | 0.09 | 0.456 | 1.602 | -1.553 | -0.01 | -0.036 | -0.036 | 0 | 0.24 | 0.36 | 0.13 | 0.11 | 0.2 | 0.12 |
| 7 | 138615831 | rs6963169 | -2.269 | 0.456 | 1.384 | -1.604 | -0.033 | 0.147 | 0.147 | 0.124 | 0.5 | 0.94 | 0.31 | 0.24 | 0.39 | 0.46 |
| 7 | 138617682 | rs17160621 | -1.146 | 0.459 | 1.799 | -1.562 | 0.006 | -0.068 | -0.068 | 0 | 0.13 | 0.22 | 0.05 | 0 | 0.11 | 0.21 |
| 7 | 138619660 | rs17160622 | -1.658 | 0.428 | 1.995 | -1.549 | -0.034 | 0.019 | 0.019 | 0.359 | 0.12 | 0.29 | 0.09 | 0.13 | 0.06 | 0.11 |
| 7 | 138620876 | rs17160623 | -2.512 | 0.464 | 1.747 | -1.555 | -0.01 | -0.058 | -0.058 | 0 | 0.25 | 0.5 | 0.18 | 0.14 | 0.19 | 0.33 |
| 7 | 138622572 | rs2718136 | -1.53 | 0.711 | 1.185 | -1.593 | 0 | 0.11 | 0.11 | 0 | 0.39 | 0.8 | 0.23 | 0.11 | 0.34 | 0.35 |
| 7 | 138626175 | rs2774961 | -0.089 | 0.428 | 1.413 | -1.441 | -0.022 | -0.051 | -0.051 | 0.02 | 0.26 | 0.52 | 0.13 | 0.13 | 0.2 | 0.15 |
| 7 | 138633905 | rs2774969 | -1.975 | 0.401 | 0.846 | -1.383 | 0.027 | 0.102 | 0.102 | 0.024 | 0.48 | 0.96 | 0.24 | 0.28 | 0.34 | 0.46 |
| 7 | 138636475 | rs9656509 | -0.848 | 0.374 | 1.236 | -1.404 | -0.004 | -0.04 | -0.04 | 1.707 | 0.11 | 0.12 | 0.03 | 0 | 0.08 | 0.2 |
| 7 | 138638846 | rs661079 | -1.419 | 0.392 | 1.252 | -1.428 | 0.039 | 0.114 | 0.114 | 0 | 0.41 | 0.89 | 0.19 | 0.24 | 0.29 | 0.38 |
| 7 | 138639832 | rs540649 | -0.164 | 0.587 | 1.608 | -1.435 | -0.013 | -0.022 | -0.022 | 1.502 | 0.27 | 0.53 | 0.14 | 0.24 | 0.23 | 0.18 |
| 7 | 138640631 | rs569048 | -0.074 | 0.643 | 1.744 | -1.404 | -0.008 | -0.015 | -0.015 | 0 | 0.26 | 0.4 | 0.13 | 0.24 | 0.23 | 0.18 |
| 7 | 138641054 | rs583749 | -1.383 | 0.652 | 1.194 | -1.403 | 0.03 | 0.101 | 0.101 | 0 | 0.4 | 0.83 | 0.18 | 0.24 | 0.31 | 0.39 |
| 7 | 138642602 | rs552946 | -1.406 | 0.666 | 1.206 | -1.41 | 0.017 | 0.082 | 0.082 | 0 | 0.41 | 0.78 | 0.19 | 0.24 | 0.33 | 0.42 |
| 7 | 138644441 | rs501492 | -2.144 | 0.676 | 1.045 | -1.412 | -0.003 | 0.168 | 0.168 | 0 | 0.49 | 0.92 | 0.27 | 0.25 | 0.38 | 0.47 |
| 7 | 138647227 | rs1463914 | -2.327 | 0.675 | 1.232 | -1.435 | -0.015 | -0.08 | -0.08 | 0.476 | 0.22 | 0.37 | 0.13 | 0.04 | 0.16 | 0.29 |
| 7 | 138648537 | rs512927 | -1.936 | 0.808 | 0.608 | -1.359 | -0.009 | 0.124 | 0.124 | 0 | 0.48 | 0.92 | 0.27 | 0.27 | 0.38 | 0.47 |
| 7 | 138660337 | rs6952395 | -1.492 | 1.323 | 1.194 | -1.147 | 0.017 | -0.014 | -0.014 | 0 | 0.11 | 0.18 | 0.05 | 0.1 | 0.13 | 0.2 |
| 7 | 138664132 | rs1888349 | -1.429 | 2.108 | 1.17 | -1.162 | 0.042 | -0.013 | -0.013 | 0.947 | 0.11 | 0.16 | 0.09 | 0.1 | 0.19 | 0.2 |
| 7 | 138667788 | rs6977460 | -2.566 | 1.873 | 1.227 | -1.269 | 0.015 | -0.047 | -0.047 | 1.315 | 0.25 | 0.63 | 0.17 | 0.11 | 0.24 | 0.29 |
| 7 | 138673567 | rs4732343 | -2.134 | 1.881 | 0.962 | -0.958 | 0.089 | -0.06 | -0.06 | 0.347 | 0.16 | 0.38 | 0.13 | 0.08 | 0.22 | 0.22 |
| 7 | 138692417 | rs1604912 | -0.604 | 0.984 | 0.266 | -1.151 | -0.01 | 0.006 | 0.006 | 0 | 0.22 | 0.19 | 0.18 | 0.07 | 0.3 | 0.22 |
| 7 | 138693945 | rs7789244 | -0.442 | -0.474 | -0.011 | -1.428 | -0.003 | 0.001 | 0.001 | 0 | 0.39 | 0.57 | 0.18 | 0.24 | 0.21 | 0.35 |
| 7 | 138698825 | rs6945033 | -0.846 | -0.053 | 1.193 | -0.826 | -0.001 | 0.006 | 0.006 | 0.019 | 0.54 | 0.45 | 0.34 | 0.32 | 0.51 | 0.56 |
| 7 | 138704135 | rs10252810 | -1.262 | -0.141 | 1.286 | -0.822 | -0.04 | 0.026 | 0.026 | 0 | 0.59 | 0.74 | 0.38 | 0.32 | 0.53 | 0.58 |
| 7 | 138712700 | rs10085553 | -0.564 | -0.416 | 1.512 | -0.929 | 0.002 | 0.038 | 0.038 | 0 | 0.5 | 0.55 | 0.36 | 0.22 | 0.51 | 0.4 |
| 7 | 138715731 | rs10280239 | -0.44 | -0.428 | 1.711 | -0.893 | -0.003 | 0.023 | 0.023 | 0 | 0.47 | 0.38 | 0.33 | 0.22 | 0.5 | 0.4 |

| | | | | | | | | | | | | | | | | |
|---|---|---|---|---|---|---|---|---|---|---|---|---|---|---|---|---|
| 7 | 138719231 | rs7810237 | -1.217 | -0.39 | 1.491 | -0.753 | -0.032 | 0.021 | 0.021 | 0 | 0.58 | 0.78 | 0.36 | 0.34 | 0.53 | 0.59 |
| 7 | 138729795 | rs10271373 | -0.707 | -0.525 | 1.745 | -0.673 | 0.004 | 0.019 | 0.019 | 0 | 0.49 | 0.47 | 0.36 | 0.34 | 0.51 | 0.43 |
| 7 | 138730025 | rs1044551 | -0.711 | -0.556 | 1.751 | -0.668 | 0.007 | 0.019 | 0.019 | 0 | 0.49 | 0.47 | 0.35 | 0.34 | 0.49 | 0.42 |
| 7 | 138730361 | rs7803490 | -1.445 | -0.799 | 1.855 | -0.757 | -0.015 | 0.006 | 0.006 | 0 | 0.6 | 0.71 | 0.38 | 0.36 | 0.51 | 0.58 |
| 7 | 138731343 | rs3187141 | -1.272 | -0.747 | 1.663 | -0.748 | -0.034 | -0.032 | -0.032 | 0 | 0.1 | 0.19 | 0.02 | 0.03 | 0.02 | 0.16 |
| 7 | 138732107 | rs5022943 | -0.597 | -0.277 | 1.155 | -0.492 | 0.015 | 0.062 | 0.062 | 0 | 0.48 | 0.27 | 0.35 | 0.33 | 0.49 | 0.43 |
| 7 | 138732457 | rs3735008 | -0.843 | -0.205 | 0.891 | -0.46 | 0.007 | 0.018 | 0.018 | 0 | 0.49 | 0.48 | 0.35 | 0.33 | 0.49 | 0.43 |
| 7 | 138735512 | rs10273974 | -0.804 | -0.185 | 0.831 | -0.444 | 0.003 | 0.009 | 0.009 | 0 | 0.51 | 0.48 | 0.35 | 0.36 | 0.5 | 0.58 |
| 7 | 138742775 | rs17133444 | -0.585 | -1.049 | 0.463 | 0.249 | 0.057 | 0.04 | 0.04 | 0 | 0.24 | 0.06 | 0.11 | 0.16 | 0.18 | 0.2 |
| 7 | 138742981 | rs17133451 | -0.881 | -1.054 | 0.721 | 0.417 | 0.006 | -0.002 | -0.002 | 0 | 0.28 | 0.24 | 0.15 | 0.32 | 0.17 | 0.24 |
| 7 | 138746752 | rs7779972 | -1.677 | -0.013 | 1.499 | 1.317 | 0.021 | 0.012 | 0.012 | 0 | 0.3 | 0.52 | 0.51 | 0.66 | 0.29 | 0.26 |
| 7 | 138758992 | rs12672266 | -1.522 | -0.395 | 2.132 | 1.817 | -0.013 | 0.035 | 0.035 | 0.342 | 0.27 | 0.63 | 0.46 | 0.4 | 0.18 | 0.16 |
| 7 | 138763654 | rs2236423 | -1.342 | -0.331 | 2.247 | 1.806 | -0.021 | 0.123 | 0.123 | 0.006 | 0.26 | 0.49 | 0.44 | 0.4 | 0.18 | 0.21 |
| 7 | 138764233 | rs2236426 | -0.786 | -0.324 | 2.118 | 1.832 | -0.013 | 0.061 | 0.061 | 0 | 0.21 | 0.22 | 0.34 | 0.38 | 0.12 | 0.13 |
| 7 | 138770415 | rs9690859 | -2.15 | -0.318 | 1.658 | 1.622 | 0.042 | 0.117 | 0.117 | 0 | 0.19 | 0.53 | 0.28 | 0.4 | 0.2 | 0.42 |
| 7 | 138771905 | rs12535022 | -2.158 | -0.32 | 1.658 | 1.622 | 0.042 | 0.117 | 0.117 | 0 | 0.19 | 0.53 | 0.28 | 0.4 | 0.2 | 0.42 |
| 7 | 138773011 | rs6467822 | -1.406 | -0.631 | 1.697 | 1.642 | 0.03 | 0.144 | 0.144 | 0 | 0.16 | 0.44 | 0.2 | 0.4 | 0.14 | 0.38 |
| 7 | 138778604 | rs7801123 | -1.237 | -0.605 | 1.515 | 1.659 | -0.006 | -0.021 | -0.021 | 0.247 | 0.14 | 0.16 | 0.18 | 0.43 | 0.16 | 0.4 |
| 7 | 138781422 | rs4728461 | -1.71 | -0.584 | 1.613 | 1.649 | 0.024 | 0.098 | 0.098 | 0 | 0.17 | 0.36 | 0.23 | 0.44 | 0.2 | 0.42 |
| 7 | 138792539 | rs4732354 | -1.482 | -0.56 | 1.323 | 1.652 | 0.012 | 0.06 | 0.06 | 0 | 0.17 | 0.3 | 0.23 | 0.45 | 0.2 | 0.41 |
| 7 | 138793127 | rs4732356 | -1.794 | -0.706 | 1.241 | 1.675 | 0.011 | 0.085 | 0.085 | 0 | 0.18 | 0.35 | 0.44 | 0.45 | 0.2 | 0.42 |
| 7 | 138794618 | rs7811342 | -2.228 | -1.347 | 1.274 | 1.809 | -0.03 | 0.216 | 0.216 | 0.581 | 0.15 | 0.59 | 0.13 | 0.45 | 0.1 | 0.31 |
| 7 | 138794879 | rs7792792 | -1.73 | -1.669 | 0 | 1.689 | 0.069 | 0.046 | 0.046 | 1.422 | 0.4 | 0.89 | 0.47 | 0.72 | 0.24 | 0.45 |
| 7 | 138797842 | rs12056143 | -2.42 | -0.982 | 0.568 | 1.89 | 0.009 | 0.064 | 0.064 | 0.047 | 0.16 | 0.34 | 0.15 | 0.47 | 0.2 | 0.37 |
| 7 | 138799596 | rs2134955 | -2.426 | -0.973 | 0.54 | 1.895 | 0.008 | 0.058 | 0.058 | 0.04 | 0.16 | 0.33 | 0.15 | 0.47 | 0.2 | 0.37 |
| 7 | 138800342 | rs2174362 | -1.666 | -1.13 | 0.165 | 1.57 | -0.002 | -0.002 | -0.002 | 1.912 | 0.09 | 0.12 | 0.06 | 0.15 | 0.04 | 0.21 |
| 7 | 138816931 | rs7778825 | -1.272 | -1.885 | 0.004 | 1.695 | 0.067 | -0.048 | -0.048 | 0 | 0.33 | 0.8 | 0.39 | 0.61 | 0.15 | 0.29 |
| 7 | 138817240 | rs7798474 | -1.731 | -1.952 | 0.656 | 1.689 | 0.039 | 0.003 | 0.003 | 0.138 | 0.38 | 0.71 | 0.43 | 0.7 | 0.25 | 0.48 |
| 7 | 138817539 | rs10215341 | -2.288 | -1.888 | 1.141 | 2.14 | -0.042 | 0.016 | 0.016 | 2.017 | 0.24 | 0.69 | 0.35 | 0.37 | 0.08 | 0.34 |
| 7 | 138821602 | rs7804077 | -2.385 | -1.871 | 1.695 | 2.063 | -0.025 | 0.005 | 0.005 | 1.488 | 0.24 | 0.69 | 0.34 | 0.37 | 0.08 | 0.33 |
| 7 | 138827235 | rs7803251 | -2.385 | -1.871 | 1.702 | 2.06 | -0.028 | 0.007 | 0.007 | 1.488 | 0.24 | 0.69 | 0.34 | 0.38 | 0.08 | 0.33 |
| 7 | 138831707 | rs10248924 | -2.385 | -1.871 | 1.718 | 2.06 | -0.03 | 0.009 | 0.009 | 1.488 | 0.24 | 0.69 | 0.34 | 0.38 | 0.08 | 0.33 |
| 7 | 138846477 | rs6944116 | -1.288 | -1.898 | 2.103 | 2.116 | -0.069 | 0.159 | 0.159 | 0.798 | 0.14 | 0.37 | 0.28 | 0.26 | 0.04 | 0.12 |
| 7 | 138853500 | rs17133455 | -2.451 | -1.862 | 1.501 | 2.034 | -0.014 | -0.002 | -0.002 | 0 | 0.24 | 0.73 | 0.34 | 0.37 | 0.08 | 0.33 |
| 7 | 138853727 | rs10085848 | -2.459 | -1.855 | 1.504 | 2.037 | -0.014 | -0.003 | -0.003 | 0.013 | 0.24 | 0.73 | 0.34 | 0.37 | 0.08 | 0.33 |
| 7 | 138886066 | rs28549275 | -1.615 | -1.406 | 1.321 | 1.658 | 0.032 | -0.007 | -0.007 | 0 | 0.36 | 0.74 | 0.48 | 0.36 | 0.27 | 0.27 |
| 7 | 138918990 | rs10245229 | -0.96 | -1.304 | 1.751 | 1.5 | 0.006 | 0.021 | 0.021 | 0 | 0.06 | 0.2 | 0.27 | 0.09 | 0.04 | 0.06 |
| 7 | 138922666 | rs11762145 | 0.602 | -1.343 | 1.538 | 1.381 | -0.007 | 0.041 | 0.041 | 0 | 0.12 | 0.03 | 0.12 | 0 | 0.18 | 0.03 |
| 7 | 138929093 | rs2355149 | -2.507 | -1.484 | 1.829 | 1.381 | -0.041 | -0.033 | -0.033 | 0.146 | 0.18 | 0.66 | 0.35 | 0.13 | 0.09 | 0.19 |
| 7 | 138941823 | rs6467832 | -2.598 | -1.506 | 1.77 | 1.455 | -0.034 | -0.02 | -0.02 | 0 | 0.19 | 0.71 | 0.35 | 0.14 | 0.09 | 0.19 |
| 7 | 138952700 | rs17160838 | 0.106 | -0.539 | 0.936 | 1.637 | 0.004 | 0.002 | 0.002 | 1.642 | 0.09 | 0.08 | 0.19 | 0.27 | 0.19 | 0.15 |
| 7 | 138963364 | rs7796255 | -2.079 | -0.707 | 0.565 | 1.189 | -0.113 | 0.033 | 0.033 | 0.349 | 0.48 | 0.94 | 0.7 | 0.69 | 0.53 | 0.43 |
| 7 | 138977206 | rs10954644 | -1.678 | -0.788 | 1.459 | 1.062 | 0.006 | 0.046 | 0.046 | 0.091 | 0.08 | 0.3 | 0.28 | 0.1 | 0.04 | 0.06 |
| 7 | 138997907 | rs6953376 | -0.727 | -0.691 | 0.397 | 0.841 | 0.005 | 0.014 | 0.014 | 0 | 0.18 | 0.13 | 0.39 | 0.11 | 0.22 | 0.09 |
| 7 | 139003602 | rs10260850 | -1.376 | -0.967 | 1.316 | 0.685 | -0.03 | -0.071 | -0.071 | 0 | 0.22 | 0.43 | 0.41 | 0.11 | 0.23 | 0.1 |
| 7 | 139007755 | rs2355150 | -1.889 | -1.285 | 1.413 | 0.574 | 0.001 | -0.003 | -0.003 | 0 | 0.3 | 0.7 | 0.46 | 0.13 | 0.27 | 0.21 |
| 7 | 139025094 | rs11765421 | -2.024 | -1.652 | 0.967 | 0.773 | 0.064 | 0.052 | 0.052 | 1.813 | 0.33 | 0.88 | 0.27 | 0.2 | 0.28 | 0.26 |
| 7 | 139034207 | rs4732368 | -0.633 | -1.731 | 1.333 | 0.734 | 0.014 | -0.001 | -0.001 | 0 | 0.19 | 0.12 | 0.17 | 0.19 | 0.21 | 0.17 |
| 7 | 139034806 | rs6962186 | -2.258 | -1.739 | 1.239 | 0.731 | 0.063 | 0.053 | 0.053 | 1.666 | 0.33 | 0.88 | 0.27 | 0.2 | 0.28 | 0.26 |
| 7 | 139036058 | rs28477480 | -0.867 | -1.793 | 1.65 | 0.729 | -0.028 | -0.034 | -0.034 | 0 | 0.2 | 0.34 | 0.15 | 0.11 | 0.2 | 0.12 |
| 7 | 139036598 | rs10273527 | -2.412 | -2.006 | 1.295 | 1.61 | 0.104 | -0.077 | -0.077 | 0 | 0.39 | 0.87 | 0.27 | 0.42 | 0.29 | 0.3 |
| 7 | 139054107 | rs6949890 | -0.367 | -0.849 | 0.875 | 1.645 | -0.034 | -0.044 | -0.044 | 1.848 | 0.16 | 0.08 | 0.25 | 0.52 | 0.27 | 0.26 |
| 7 | 139055752 | rs17160882 | -2.266 | -1.571 | 1.683 | 1.339 | 0 | -0.006 | -0.006 | 0 | 0.06 | 0.23 | 0.06 | 0.01 | 0.04 | 0.02 |
| 7 | 139058803 | rs11772771 | -1.24 | -1.553 | 1.714 | 1.336 | -0.028 | -0.032 | -0.032 | 0.044 | 0.26 | 0.46 | 0.25 | 0.19 | 0.26 | 0.24 |
| 7 | 139122814 | rs4732379 | -1.507 | -1.486 | -0.094 | 1.207 | 0.05 | 0.037 | 0.037 | 0.634 | 0.32 | 0.81 | 0.28 | 0.21 | 0.28 | 0.27 |
| 7 | 139136670 | rs4732381 | -1.32 | -1.362 | -0.33 | 1.331 | 0.027 | -0.013 | -0.013 | 0 | 0.32 | 0.78 | 0.32 | 0.35 | 0.27 | 0.31 |
| 7 | 139138551 | rs10954649 | -1.147 | -1.073 | -0.592 | 0.911 | 0.03 | 0.016 | 0.016 | 0 | 0.41 | 0.76 | 0.36 | 0.33 | 0.36 | 0.33 |
| 7 | 139141670 | rs12112028 | -0.701 | -1.356 | -0.683 | 0.578 | 0.017 | 0.033 | 0.033 | 0 | 0.21 | 0.13 | 0.14 | 0.05 | 0.16 | 0.17 |
| 7 | 139142117 | rs10441417 | -1.765 | -2.381 | -0.872 | 0.744 | 0.079 | -0.011 | -0.011 | 0 | 0.4 | 0.78 | 0.29 | 0.4 | 0.23 | 0.32 |
| 7 | 139143798 | rs17160922 | -0.564 | -2.136 | -0.685 | 0.958 | 0.014 | -0.012 | -0.012 | 0 | 0.09 | 0.05 | 0.09 | 0.33 | 0.05 | 0.11 |
| 7 | 139151335 | rs10250296 | -0.431 | -1.987 | -0.51 | 0.767 | -0.003 | -0.002 | -0.002 | 0 | 0.11 | 0.12 | 0.08 | 0.06 | 0.09 | 0.08 |
| 7 | 139163463 | rs7779375 | -1.358 | -1.867 | -1.68 | 0.483 | -0.058 | 0.012 | 0.012 | 1.307 | 0.67 | 0.84 | 0.48 | 0.68 | 0.52 | 0.49 |
| 7 | 139163618 | rs6945850 | -0.67 | -1.596 | -0.896 | 0.338 | 0.028 | 0.02 | 0.02 | 0.075 | 0.47 | 0.7 | 0.36 | 0.42 | 0.36 | 0.33 |
| 7 | 139168616 | rs1006482 | -0.051 | -1.451 | -0.135 | 0.26 | -0.002 | -0.051 | -0.051 | 0.063 | 0.14 | 0.18 | 0.07 | 0 | 0.11 | 0.1 |
| 7 | 139174266 | rs7804582 | -0.717 | -1.245 | -0.093 | 0.84 | -0.001 | 0.034 | 0.034 | 0 | 0.12 | 0.19 | 0.1 | 0.26 | 0.11 | 0.2 |
| 7 | 139174424 | rs2256682 | 0.212 | -1.144 | 0.206 | 0.208 | 0.001 | -0.036 | -0.036 | 0 | 0.8 | 0.81 | 0.81 | 0.98 | 0.79 | 0.88 |
| 7 | 139174872 | rs758260 | 1.075 | -2.211 | 0.155 | 0.509 | -0.031 | 0.03 | 0.03 | 0 | 0.66 | 0.49 | 0.71 | 0.44 | 0.74 | 0.69 |
| 7 | 139176219 | rs2530733 | 0.969 | -1.829 | -0.187 | 0.713 | 0.027 | -0.011 | -0.011 | 0 | 0.61 | 0.31 | 0.69 | 0.5 | 0.73 | 0.72 |
| 7 | 139176277 | rs2718233 | 1.492 | -1.79 | -1.137 | 0.516 | -0.106 | -0.105 | -0.105 | 0 | 0.41 | 0.03 | 0.56 | 0.41 | 0.55 | 0.47 |

| | | | | | | | | | | | | | | | | |
|---|---|---|---|---|---|---|---|---|---|---|---|---|---|---|---|---|
| 7 | 139180317 | rs17160954 | 0.085 | -1.509 | -0.671 | 0.16 | 0.002 | 0.004 | 0.004 | 0 | 0.18 | 0.17 | 0.1 | 0.09 | 0.16 | 0.25 |
| 7 | 139180806 | rs2530730 | 0.292 | -1.529 | -0.697 | 0.389 | -0.023 | -0.022 | -0.022 | 1.676 | 0.63 | 0.56 | 0.71 | 0.74 | 0.69 | 0.65 |
| 7 | 139184786 | rs17160967 | 0.893 | -0.674 | -1.714 | 0.354 | -0.058 | -0.06 | -0.06 | 1.593 | 0.4 | 0.15 | 0.48 | 0.42 | 0.47 | 0.43 |
| 7 | 139185749 | rs7385378 | 0.826 | -0.886 | -1.61 | 0.407 | -0.05 | -0.093 | -0.093 | 0 | 0.41 | 0.1 | 0.5 | 0.43 | 0.53 | 0.45 |
| 7 | 139188055 | rs12374796 | -0.703 | -0.918 | -1.268 | 0.098 | 0.039 | 0.048 | 0.048 | 4.142 | 0.23 | 0.15 | 0.1 | 0.05 | 0.16 | 0.1 |
| 7 | 139188909 | rs10240456 | -0.68 | -0.896 | -1.258 | 0.108 | 0.038 | 0.047 | 0.047 | 4.342 | 0.23 | 0.15 | 0.1 | 0.05 | 0.16 | 0.1 |
| 7 | 139197147 | rs10435440 | -0.493 | 0.015 | -0.852 | 0.709 | -0.002 | 0.017 | 0.017 | 0.076 | 0.37 | 0.5 | 0.38 | 0.42 | 0.33 | 0.21 |
| 7 | 139209767 | rs12665981 | -0.401 | 0.043 | -1.247 | 0.553 | 0 | -0.001 | -0.001 | 0 | 0.31 | 0.73 | 0.37 | 0.29 | 0.3 | 0.16 |
| 7 | 139230464 | rs12672312 | -0.101 | -0.195 | -1.585 | 0.503 | 0.007 | -0.018 | -0.018 | 0 | 0.24 | 0.16 | 0.18 | 0.55 | 0.19 | 0.39 |
| 7 | 139231393 | rs7799361 | -0.624 | -0.64 | -1.579 | -0.788 | -0.016 | -0.075 | -0.075 | 0.776 | 0.29 | 0.68 | 0.14 | 0.01 | 0.2 | 0.14 |
| 7 | 139240471 | rs4732386 | -0.115 | 0.226 | -2.195 | -0.662 | -0.105 | 0.066 | 0.066 | 0 | 0.67 | 0.96 | 0.52 | 0.78 | 0.61 | 0.69 |
| 7 | 139263879 | rs11974162 | -0.589 | -0.207 | -0.333 | -0.72 | -0.016 | -0.021 | -0.021 | 0 | 0.05 | 0.16 | 0.04 | 0 | 0.02 | 0.01 |
| 7 | 139275081 | rs4732389 | -1.204 | -0.638 | 0.31 | -1.267 | -0.049 | -0.127 | -0.127 | 0.692 | 0.23 | 0.54 | 0.13 | 0.02 | 0.18 | 0.12 |
| 7 | 139285490 | rs3735197 | -0.49 | -0.637 | 0.655 | -1.253 | -0.009 | -0.011 | -0.011 | 0 | 0.05 | 0.18 | 0.04 | 0.01 | 0.02 | 0.02 |
| 7 | 139294931 | rs2030712 | -0.944 | -0.686 | 0.389 | -1.816 | 0.027 | 0.115 | 0.115 | 0 | 0.5 | 0.66 | 0.26 | 0.08 | 0.44 | 0.33 |
| 7 | 139297145 | rs10266114 | -0.136 | -0.791 | 0.539 | -1.568 | 0 | -0.023 | -0.023 | 0 | 0.16 | 0.11 | 0.08 | 0.01 | 0.16 | 0.11 |
| 7 | 139302846 | rs7788362 | -0.744 | -0.739 | 0.402 | -1.576 | -0.05 | -0.127 | -0.127 | 1.592 | 0.19 | 0.44 | 0.09 | 0.01 | 0.13 | 0.11 |
| 7 | 139307411 | rs7782233 | -0.928 | -0.623 | 0.25 | -1.569 | -0.029 | -0.118 | -0.118 | 1.595 | 0.2 | 0.37 | 0.11 | 0.01 | 0.17 | 0.11 |
| 7 | 139307835 | rs12539357 | -0.602 | -0.61 | 0.276 | -1.501 | -0.031 | -0.108 | -0.108 | 0.156 | 0.18 | 0.36 | 0.09 | 0.01 | 0.13 | 0.1 |
| 7 | 139310477 | rs10256326 | -0.664 | -0.594 | 0.264 | -1.5 | -0.031 | -0.119 | -0.119 | 1.067 | 0.18 | 0.42 | 0.09 | 0.01 | 0.13 | 0.1 |
| 7 | 139326552 | rs10954650 | -0.226 | 0 | -0.06 | -1.26 | 0.004 | -0.086 | -0.086 | 0.532 | 0.38 | 0.52 | 0.36 | 0.1 | 0.36 | 0.28 |
| 7 | 139333338 | rs10954651 | 0.167 | -0.42 | -0.56 | -0.496 | 0.001 | 0.038 | 0.038 | 0.256 | 0.36 | 0.21 | 0.33 | 0.23 | 0.34 | 0.36 |
| 7 | 139336576 | rs7781224 | -0.095 | -0.842 | -0.288 | 0.425 | -0.002 | -0.056 | -0.056 | 0.301 | 0.83 | 0.88 | 0.67 | 0.51 | 0.84 | 0.7 |
| 7 | 139338133 | rs11761839 | 1.225 | -1.107 | 0.069 | 0.122 | -0.017 | 0.005 | 0.005 | 0.627 | 0.56 | 0.59 | 0.53 | 0.45 | 0.66 | 0.45 |
| 7 | 139343366 | rs9969246 | 0.026 | -1.144 | -0.351 | 0.077 | -0.001 | 0.003 | 0.003 | 0 | 0.73 | 0.76 | 0.6 | 0.46 | 0.8 | 0.52 |
| 7 | 139343779 | rs10954652 | 0.543 | -0.549 | -0.937 | 0.389 | 0.026 | -0.006 | -0.006 | 0 | 0.55 | 0.47 | 0.52 | 0.46 | 0.66 | 0.45 |
| 7 | 139348866 | rs12668118 | -0.513 | -0.677 | -0.691 | 0.627 | 0.014 | -0.003 | -0.003 | 0 | 0.41 | 0.54 | 0.49 | 0.57 | 0.29 | 0.54 |
| 7 | 139352954 | rs10954653 | -0.752 | 0.225 | -0.679 | 0.856 | -0.001 | 0.002 | 0.002 | 0 | 0.07 | 0.1 | 0.12 | 0.15 | 0.06 | 0.17 |
| 7 | 139357580 | rs9719152 | -0.045 | 0.243 | -1.122 | 0.017 | -0.01 | 0.06 | 0.06 | 0 | 0.35 | 0.23 | 0.51 | 0.15 | 0.56 | 0.22 |
| 7 | 139359730 | rs10262234 | 0.128 | -0.663 | -0.437 | 0.337 | 0.003 | 0.046 | 0.046 | 0 | 0.8 | 0.5 | 0.77 | 0.77 | 0.81 | 0.78 |
| 7 | 139360198 | rs9691662 | 0.186 | -1.195 | -0.28 | 0.209 | 0.006 | 0.011 | 0.011 | 0 | 0.42 | 0.37 | 0.57 | 0.2 | 0.61 | 0.3 |
| 7 | 139366112 | rs9690885 | -1.297 | -1.042 | 0.478 | -0.055 | -0.008 | -0.058 | -0.058 | 0 | 0.22 | 0.61 | 0.18 | 0.07 | 0.21 | 0.13 |
| 7 | 139372966 | rs9690295 | 1.005 | -0.907 | 0.064 | -0.387 | -0.112 | 0.072 | 0.072 | 0 | 0.26 | 0.1 | 0.28 | 0.14 | 0.45 | 0.18 |
| 7 | 139409674 | rs10952346 | -1.997 | -0.865 | 1.367 | 0.213 | -0.003 | 0.148 | 0.148 | 0.786 | 0.1 | 0.52 | 0.3 | 0.16 | 0.05 | 0.05 |
| 7 | 139415723 | rs10224408 | -0.678 | -0.87 | 1.364 | 0.378 | 0.005 | 0.001 | 0.001 | 0 | 0.11 | 0.11 | 0.17 | 0.11 | 0.16 | 0.07 |
| 7 | 139421180 | rs10230692 | -0.678 | -0.87 | 1.364 | 0.41 | 0.005 | 0.002 | 0.002 | 0 | 0.11 | 0.11 | 0.17 | 0.11 | 0.16 | 0.07 |
| 7 | 139434215 | rs12673068 | -2.692 | -0.93 | 1.617 | 0.39 | 0 | -0.003 | -0.003 | 0 | 0.24 | 0.8 | 0.49 | 0.27 | 0.21 | 0.12 |
| 7 | 139434903 | rs6605557 | -1.047 | -0.902 | 1.714 | 0.411 | -0.004 | 0.053 | 0.053 | 0 | 0.05 | 0.07 | 0.26 | 0.16 | 0.04 | 0.05 |
| 7 | 139448995 | rs12674180 | -1.116 | -0.905 | 1.713 | 0.389 | -0.004 | 0.046 | 0.046 | 0 | 0.06 | 0.08 | 0.26 | 0.16 | 0.05 | 0.05 |
| 7 | 139454165 | rs6464214 | -2.521 | -0.901 | 1.651 | 0.27 | 0.001 | -0.005 | -0.005 | 0 | 0.24 | 0.8 | 0.49 | 0.26 | 0.21 | 0.13 |
| 7 | 139462249 | rs2058265 | -2.126 | -0.896 | 1.749 | 0.282 | 0.01 | 0.066 | 0.066 | 0.315 | 0.21 | 0.62 | 0.47 | 0.26 | 0.21 | 0.12 |
| 7 | 139486394 | rs10244528 | -2.11 | -0.765 | 1.982 | 0.141 | 0.087 | 0.083 | 0.083 | 7.884 | 0.05 | 0.49 | 0.04 | 0 | 0 | 0 |
| 7 | 139496215 | rs6958644 | -0.423 | -0.579 | 1.302 | -0.541 | 0.035 | -0.097 | -0.097 | 0 | 0.69 | 0.55 | 0.72 | 0.95 | 0.53 | 0.79 |
| 7 | 139502073 | rs10257313 | -0.721 | 0.266 | 1.216 | -0.558 | -0.06 | 0.068 | 0.068 | 0.122 | 0.73 | 0.89 | 0.75 | 0.95 | 0.56 | 0.79 |
| 7 | 139514352 | rs6960593 | 0.691 | 0.493 | 1.153 | 0.112 | 0.026 | 0.005 | 0.005 | 0 | 0.54 | 0.19 | 0.42 | 0.58 | 0.42 | 0.74 |
| 7 | 139517307 | rs10487662 | 0.186 | 0.588 | 1.211 | 0.187 | -0.019 | 0.01 | 0.01 | 0 | 0.08 | 0.03 | 0.06 | 0.04 | 0.14 | 0.04 |
| 7 | 139519064 | rs10277434 | 0.914 | 0.523 | 1.272 | 0.164 | 0.021 | 0.003 | 0.003 | 0 | 0.55 | 0.24 | 0.42 | 0.58 | 0.4 | 0.73 |
| 7 | 139524992 | rs6969517 | 0.504 | 0.709 | 1.161 | -0.098 | 0 | -0.028 | -0.028 | 0 | 0.59 | 0.54 | 0.67 | 0.89 | 0.43 | 0.75 |
| 7 | 139525212 | rs12667080 | 1.04 | 0.696 | 0.762 | 0.175 | -0.035 | -0.033 | -0.033 | 2.705 | 0.06 | 0 | 0.05 | 0.12 | 0.12 | 0.12 |
| 7 | 139525978 | rs10253277 | -0.471 | 0.707 | 0.902 | -0.009 | -0.113 | 0.159 | 0.159 | 0 | 0.7 | 0.95 | 0.76 | 0.93 | 0.56 | 0.79 |
| 7 | 139528444 | rs10256282 | -0.484 | 0.723 | 0.887 | -0.009 | -0.128 | 0.164 | 0.164 | 0 | 0.7 | 0.96 | 0.76 | 0.93 | 0.56 | 0.79 |
| 7 | 139528662 | rs4590360 | -0.471 | 0.706 | 0.887 | -0.009 | -0.125 | 0.165 | 0.165 | 0 | 0.7 | 0.95 | 0.76 | 0.93 | 0.56 | 0.79 |
| 7 | 139530089 | rs764746 | -0.467 | 0.693 | 0.882 | -0.015 | -0.125 | 0.166 | 0.166 | 0 | 0.7 | 0.95 | 0.76 | 0.93 | 0.56 | 0.79 |
| 7 | 139538522 | rs12534299 | -0.467 | 0.734 | 0.878 | -0.018 | -0.123 | 0.166 | 0.166 | 0 | 0.7 | 0.95 | 0.76 | 0.93 | 0.56 | 0.79 |
| 7 | 139539105 | rs8192803 | 0.804 | 0.653 | 1.14 | -0.063 | -0.001 | -0.054 | -0.054 | 0 | 0.56 | 0.46 | 0.65 | 0.9 | 0.41 | 0.72 |
| 7 | 139539218 | rs2267679 | -0.499 | 0.662 | 1.104 | -0.057 | -0.123 | 0.166 | 0.166 | 0 | 0.7 | 0.95 | 0.76 | 0.93 | 0.56 | 0.79 |
| 7 | 139540293 | rs2267680 | 0.441 | 0.696 | 1.318 | -0.036 | -0.012 | 0.08 | 0.08 | 0 | 0.59 | 0.8 | 0.67 | 0.9 | 0.41 | 0.72 |
| 7 | 139540601 | rs2107900 | -0.273 | 0.694 | 1.299 | -0.043 | -0.02 | 0.023 | 0.023 | 0 | 0.68 | 0.81 | 0.73 | 0.93 | 0.55 | 0.77 |
| 7 | 139540935 | rs2267681 | -0.783 | 0.687 | 1.223 | -0.029 | -0.002 | -0.064 | -0.064 | 6.401 | 0.43 | 0.46 | 0.65 | 0.9 | 0.41 | 0.71 |
| 7 | 139542268 | rs6952056 | -0.034 | 0.465 | 0.915 | 0.037 | -0.062 | 0.112 | 0.112 | 0 | 0.66 | 0.87 | 0.72 | 0.93 | 0.52 | 0.76 |
| 7 | 139543904 | rs4725857 | -0.016 | 0.469 | 0.917 | 0.076 | -0.054 | 0.097 | 0.097 | 0 | 0.66 | 0.84 | 0.71 | 0.89 | 0.52 | 0.76 |
| 7 | 139544674 | rs7810415 | 0.195 | 0.128 | 0.705 | -0.049 | 0.002 | -0.042 | -0.042 | 0 | 0.6 | 0.54 | 0.68 | 0.89 | 0.49 | 0.75 |
| 7 | 139546846 | rs41708 | 1.144 | -0.052 | 1.142 | 0.567 | -0.027 | 0.107 | 0.107 | 0 | 0.88 | 0.68 | 0.72 | 0.79 | 0.94 | 0.93 |
| 7 | 139548136 | rs727359 | -1.316 | 0.925 | 1.554 | 0.725 | 0.094 | 0.183 | 0.183 | 0.546 | 0.15 | 0.56 | 0.47 | 0.3 | 0.16 | 0.1 |
| 7 | 139548604 | rs1859569 | -1.069 | 0.932 | 1.599 | 0.743 | 0.11 | 0.208 | 0.208 | 0 | 0.13 | 0.46 | 0.46 | 0.3 | 0.16 | 0.1 |
| 7 | 139549172 | rs2284198 | -1.245 | 1.191 | 1.594 | 0.816 | 0.13 | 0.195 | 0.195 | 0 | 0.14 | 0.47 | 0.46 | 0.3 | 0.18 | 0.1 |
| 7 | 139549385 | rs41707 | 1.632 | 0.027 | 0.932 | 0.604 | -0.014 | -0.023 | -0.023 | 0 | 0.5 | 0.19 | 0.34 | 0.52 | 0.56 | 0.36 |
| 7 | 139550679 | rs2214757 | -0.56 | 0.458 | 1.148 | 0.828 | 0.112 | 0.205 | 0.205 | 0 | 0.13 | 0.45 | 0.46 | 0.3 | 0.16 | 0.1 |
| 7 | 139550843 | rs9801544 | -0.722 | 0.461 | 1.144 | 0.826 | 0.119 | 0.198 | 0.198 | 0 | 0.14 | 0.47 | 0.46 | 0.3 | 0.17 | 0.12 |

| | | | | | | | | | | | | | | | | |
|---|---|---|---|---|---|---|---|---|---|---|---|---|---|---|---|---|
| 7 | 139552588 | rs3779134 | 2.078 | 0.442 | 0.972 | 0.409 | -0.031 | -0.026 | -0.026 | 2.2 | 0.05 | 0 | 0.04 | 0.06 | 0.09 | 0.11 |
| 7 | 139553090 | rs12532529 | -0.33 | 0.514 | 1.126 | 0.44 | 0.104 | 0.088 | 0.088 | 0 | 0.12 | 0.45 | 0.2 | 0.15 | 0.14 | 0.07 |
| 7 | 139558522 | rs194149 | 1.005 | 0.105 | 0.929 | 0.358 | 0.009 | 0 | 0 | 0 | 0.61 | 0.37 | 0.68 | 0.62 | 0.71 | 0.4 |
| 7 | 139563081 | rs12536965 | 0.598 | 0.317 | 0.928 | 0.315 | 0.005 | -0.004 | -0.004 | 0 | 0.11 | 0.15 | 0.19 | 0.06 | 0.15 | 0.06 |
| 7 | 139564806 | rs8192810 | 0.493 | 0.313 | 0.922 | 0.313 | 0.008 | -0.007 | -0.007 | 2.392 | 0.11 | 0.17 | 0.19 | 0.06 | 0.15 | 0.06 |
| 7 | 139583360 | rs7778515 | -1.29 | 0 | 1.31 | 0.527 | -0.003 | 0.014 | 0.014 | 0 | 0.46 | 0.6 | 0.47 | 0.38 | 0.43 | 0.61 |
| 7 | 139583829 | rs2299887 | -1.297 | 0.001 | 1.31 | 0.527 | -0.003 | 0.014 | 0.014 | 0 | 0.46 | 0.6 | 0.47 | 0.38 | 0.43 | 0.61 |
| 7 | 139584056 | rs2299888 | -1.277 | -0.021 | 1.412 | 0.517 | 0.003 | -0.011 | -0.011 | 0 | 0.45 | 0.46 | 0.45 | 0.38 | 0.43 | 0.61 |
| 7 | 139584261 | rs2299889 | -1.277 | -0.021 | 1.412 | 0.517 | 0.003 | -0.012 | -0.012 | 0 | 0.45 | 0.46 | 0.45 | 0.38 | 0.43 | 0.61 |
| 7 | 139584505 | rs2299890 | -1.27 | -0.023 | 1.41 | 0.516 | 0.002 | -0.009 | -0.009 | 0 | 0.45 | 0.46 | 0.45 | 0.38 | 0.43 | 0.61 |
| 7 | 139584796 | rs8192817 | -1.34 | -0.022 | 1.396 | 0.516 | -0.002 | 0.014 | 0.014 | 0 | 0.46 | 0.6 | 0.47 | 0.38 | 0.43 | 0.61 |
| 7 | 139586614 | rs1015571 | -1.224 | -0.057 | 1.211 | 0.519 | 0.002 | -0.004 | -0.004 | 0 | 0.44 | 0.43 | 0.45 | 0.38 | 0.42 | 0.58 |
| 7 | 139586807 | rs1015570 | -1.219 | -0.051 | 1.212 | 0.522 | 0 | -0.002 | -0.002 | 0 | 0.44 | 0.42 | 0.44 | 0.38 | 0.42 | 0.58 |
| 7 | 139587509 | rs2267685 | -1.349 | -0.06 | 1.164 | 0.519 | -0.002 | 0.015 | 0.015 | 0 | 0.47 | 0.6 | 0.47 | 0.38 | 0.43 | 0.61 |
| 7 | 139588176 | rs3801153 | 0.858 | 0.111 | 0.995 | 0.293 | -0.016 | 0.008 | 0.008 | 0 | 0.11 | 0.07 | 0.17 | 0.07 | 0.15 | 0.06 |
| 7 | 139588629 | rs4726420 | 0.822 | 0.113 | 1.034 | 0.293 | 0 | 0.001 | 0.001 | 0 | 0.11 | 0.1 | 0.17 | 0.07 | 0.15 | 0.06 |
| 7 | 139589797 | rs2284200 | 1.882 | 0.046 | 0.903 | 0.268 | -0.028 | -0.008 | -0.008 | 1.785 | 0.07 | 0.01 | 0.05 | 0.06 | 0.1 | 0.11 |
| 7 | 139590723 | rs6464432 | -1.223 | -0.01 | 0.996 | 0.318 | -0.001 | 0.001 | 0.001 | 0 | 0.41 | 0.68 | 0.45 | 0.39 | 0.34 | 0.51 |
| 7 | 139591731 | rs6464435 | -1.242 | -0.012 | 1.1 | 0.317 | 0.003 | 0.001 | 0.001 | 0 | 0.39 | 0.49 | 0.42 | 0.39 | 0.34 | 0.53 |
| 7 | 139591860 | rs7810727 | -1.242 | -0.012 | 1.1 | 0.317 | 0.003 | 0.001 | 0.001 | 0 | 0.39 | 0.49 | 0.42 | 0.39 | 0.34 | 0.53 |
| 7 | 139591947 | rs7810857 | -1.151 | -0.011 | 1.106 | 0.316 | 0.003 | 0.003 | 0.003 | 0 | 0.38 | 0.48 | 0.42 | 0.39 | 0.34 | 0.52 |
| 7 | 139593026 | rs4726434 | -1.756 | 0.027 | 0.813 | 0.13 | 0.002 | -0.008 | -0.008 | 1.209 | 0.56 | 0.69 | 0.53 | 0.45 | 0.5 | 0.66 |
| 7 | 139596859 | rs2267691 | 1.793 | 0.035 | 0.737 | 0.109 | -0.009 | -0.001 | -0.001 | 0 | 0.07 | 0.08 | 0.05 | 0.06 | 0.09 | 0.11 |
| 7 | 139596953 | rs2267692 | -1.171 | -0.015 | 0.854 | 0.102 | -0.002 | -0.006 | -0.006 | 7.373 | 0.47 | 0.46 | 0.45 | 0.38 | 0.39 | 0.52 |
| 7 | 139597298 | rs11981804 | -1.15 | 0.007 | 0.863 | 0.094 | -0.002 | -0.006 | -0.006 | 0.016 | 0.46 | 0.46 | 0.45 | 0.38 | 0.39 | 0.52 |
| 7 | 139597880 | rs2267696 | -1.234 | -0.004 | 0.861 | 0.095 | 0 | 0.001 | 0.001 | 0.037 | 0.47 | 0.5 | 0.46 | 0.39 | 0.4 | 0.55 |
| 7 | 139607500 | rs11973494 | -0.511 | -0.178 | 0.702 | 0.085 | -0.025 | -0.006 | -0.006 | 2.215 | 0.36 | 0.38 | 0.28 | 0.32 | 0.24 | 0.46 |
| 7 | 139608697 | rs6977672 | -0.356 | -0.149 | 0.706 | 0.098 | 0.001 | 0 | 0 | 0.357 | 0.35 | 0.27 | 0.27 | 0.32 | 0.24 | 0.46 |
| 7 | 139612333 | rs2284203 | -1.693 | 0.11 | 0.037 | 0.259 | -0.033 | -0.067 | -0.067 | 0 | 0.58 | 0.86 | 0.54 | 0.5 | 0.5 | 0.67 |
| 7 | 139613072 | rs6464448 | -0.253 | 0.671 | 0.853 | 0.029 | 0.038 | -0.013 | -0.013 | 0 | 0.13 | 0.31 | 0.19 | 0.1 | 0.15 | 0.06 |
| 7 | 139613430 | rs3819733 | 0.592 | 0.696 | 0.634 | 0.04 | -0.054 | -0.01 | -0.01 | 0 | 0.1 | 0.02 | 0.17 | 0.1 | 0.15 | 0.07 |
| 7 | 139613566 | rs8192827 | -2.528 | 0.177 | 1.122 | 0.289 | 0.08 | 0.136 | 0.136 | 0.531 | 0.05 | 0.43 | 0.15 | 0.1 | 0.02 | 0.07 |
| 7 | 139618876 | rs6952982 | -0.83 | -0.167 | 1.197 | 0.013 | 0.029 | 0.053 | 0.053 | 0 | 0.1 | 0.43 | 0.12 | 0.15 | 0.06 | 0.17 |
| 7 | 139620219 | rs2284204 | -0.902 | -0.214 | 1.207 | -0.008 | 0.017 | 0.041 | 0.041 | 0.064 | 0.1 | 0.44 | 0.12 | 0.15 | 0.06 | 0.16 |
| 7 | 139620388 | rs2284205 | -1.569 | -0.51 | 1.008 | -0.302 | -0.037 | -0.09 | -0.09 | 0 | 0.26 | 0.56 | 0.19 | 0.15 | 0.18 | 0.24 |
| 7 | 139621001 | rs6464450 | -1.871 | -0.339 | 1.053 | -0.169 | 0.004 | 0.019 | 0.019 | 0.063 | 0.13 | 0.53 | 0.13 | 0.15 | 0.08 | 0.2 |
| 7 | 139621137 | rs6464451 | -1.088 | -0.317 | 0.995 | -0.169 | 0.04 | 0.047 | 0.047 | 0 | 0.08 | 0.37 | 0.09 | 0.1 | 0.05 | 0.07 |
| 7 | 139623170 | rs4726457 | 1.329 | -0.573 | -0.016 | -0.342 | 0.048 | 0.083 | 0.083 | 0.019 | 0.56 | 0.19 | 0.6 | 0.69 | 0.67 | 0.5 |
| 7 | 139631494 | rs193961 | 0.694 | -0.468 | -0.111 | -0.146 | -0.008 | -0.002 | -0.002 | 0.11 | 0.94 | 0.83 | 0.95 | 0.94 | 0.97 | 0.87 |
| 7 | 139635469 | rs41732 | 0.285 | -0.467 | -0.326 | -0.166 | -0.015 | -0.006 | -0.006 | 0 | 0.95 | 0.9 | 0.96 | 0.94 | 0.99 | 0.92 |
| 7 | 139640567 | rs4725560 | -0.015 | -0.31 | -0.926 | -0.203 | 0.005 | 0.002 | 0.002 | 0 | 0.77 | 0.91 | 0.85 | 0.76 | 0.76 | 0.74 |
| 7 | 139640896 | rs4726473 | 0.171 | -0.32 | -0.907 | -0.207 | 0.007 | 0.005 | 0.005 | 0 | 0.76 | 0.79 | 0.84 | 0.76 | 0.76 | 0.73 |
| 7 | 139641247 | rs2299894 | 1.292 | -0.696 | -0.987 | -0.523 | 0.016 | 0.024 | 0.024 | 0 | 0.59 | 0.56 | 0.78 | 0.76 | 0.64 | 0.61 |
| 7 | 139641589 | rs2107901 | 1.301 | -0.7 | -0.989 | -0.524 | 0.016 | 0.025 | 0.025 | 0 | 0.59 | 0.55 | 0.78 | 0.76 | 0.64 | 0.61 |
| 7 | 139643890 | rs2284210 | 1.279 | -0.635 | -1.139 | -0.513 | 0.044 | 0.086 | 0.086 | 0 | 0.59 | 0.74 | 0.79 | 0.76 | 0.64 | 0.61 |
| 7 | 139644600 | rs757760 | -0.147 | -0.622 | -1.162 | -0.501 | -0.025 | -0.022 | -0.022 | 0.632 | 0.76 | 0.97 | 0.86 | 0.76 | 0.76 | 0.72 |
| 7 | 139646209 | rs4140996 | -0.016 | -0.588 | -1.087 | -0.478 | 0.006 | 0.008 | 0.008 | 0 | 0.75 | 0.79 | 0.84 | 0.76 | 0.76 | 0.73 |
| 7 | 139646469 | rs1544451 | 0.011 | -0.589 | -1.086 | -0.478 | 0.006 | 0.007 | 0.007 | 0 | 0.74 | 0.78 | 0.84 | 0.76 | 0.76 | 0.72 |
| 7 | 139646514 | rs1034791 | -0.086 | -0.581 | -1.115 | -0.475 | -0.028 | -0.021 | -0.021 | 0.049 | 0.76 | 0.97 | 0.86 | 0.76 | 0.76 | 0.72 |
| 7 | 139648902 | rs886697 | -0.074 | -0.586 | -1.126 | -0.473 | -0.024 | -0.02 | -0.02 | 6.252 | 0.76 | 0.97 | 0.86 | 0.76 | 0.76 | 0.73 |
| 7 | 139648942 | rs4726477 | -0.074 | -0.586 | -1.126 | -0.473 | -0.024 | -0.02 | -0.02 | 0 | 0.76 | 0.97 | 0.86 | 0.76 | 0.76 | 0.72 |
| 7 | 139649210 | rs4629769 | 1.324 | -0.631 | -1.096 | -0.51 | 0.028 | 0.049 | 0.049 | 0 | 0.58 | 0.63 | 0.79 | 0.76 | 0.64 | 0.61 |
| 7 | 139649561 | rs2190087 | -0.033 | -0.548 | -1.345 | -0.457 | -0.022 | -0.019 | -0.019 | 0 | 0.75 | 0.97 | 0.86 | 0.76 | 0.76 | 0.72 |
| 7 | 139649697 | rs757759 | 0.003 | -0.552 | -1.346 | -0.448 | -0.015 | -0.015 | -0.015 | 0 | 0.75 | 0.97 | 0.86 | 0.76 | 0.76 | 0.72 |
| 7 | 139650762 | rs1557968 | 0.971 | -0.776 | -1.296 | -0.35 | 0.024 | 0.079 | 0.079 | 0 | 0.63 | 0.78 | 0.8 | 0.79 | 0.67 | 0.62 |
| 7 | 139653612 | rs1003816 | 0.274 | -1.429 | -0.821 | 0.245 | -0.01 | 0.008 | 0.008 | 0 | 0.88 | 0.84 | 0.91 | 0.76 | 0.89 | 0.83 |
| 7 | 139654710 | rs917147 | 0.16 | -1.423 | -0.85 | 0.247 | 0.004 | -0.005 | -0.005 | 0.791 | 0.88 | 0.92 | 0.92 | 0.76 | 0.89 | 0.82 |
| 7 | 139656609 | rs8192835 | 1.388 | -1.584 | -0.83 | 0.248 | 0.007 | 0.004 | 0.004 | 0 | 0.68 | 0.64 | 0.83 | 0.71 | 0.75 | 0.71 |
| 7 | 139657221 | rs3823717 | 0.974 | -1.39 | -0.792 | 0.026 | 0.066 | -0.059 | -0.059 | 0 | 0.29 | 0.39 | 0.26 | 0.12 | 0.37 | 0.33 |
| 7 | 139659141 | rs41728 | 1.137 | -2.013 | -0.513 | 0.464 | -0.02 | 0.001 | 0.001 | 0 | 0.83 | 0.79 | 0.87 | 0.79 | 0.88 | 0.81 |
| 7 | 139662586 | rs41726 | 1.049 | -1.912 | -0.761 | 0.297 | -0.001 | -0.001 | -0.001 | 0 | 0.83 | 0.85 | 0.88 | 0.79 | 0.88 | 0.82 |
| 7 | 139662904 | rs41724 | 0.936 | -1.893 | -0.75 | 0.309 | -0.001 | -0.001 | -0.001 | 0 | 0.84 | 0.86 | 0.88 | 0.79 | 0.89 | 0.82 |
| 7 | 139664598 | rs17181314 | -0.628 | -1.681 | -0.603 | 0.043 | 0.008 | -0.006 | -0.006 | 0.126 | 0.15 | 0.16 | 0.06 | 0 | 0.11 | 0.08 |
| 7 | 139664675 | rs41723 | 1.462 | -1.732 | -0.565 | 0.049 | -0.018 | -0.029 | -0.029 | 0 | 0.75 | 0.61 | 0.83 | 0.87 | 0.82 | 0.79 |
| 7 | 139666054 | rs17181356 | -0.352 | -1.7 | -0.766 | 0.041 | 0.009 | 0.063 | 0.063 | 0 | 0.14 | 0.05 | 0.06 | 0 | 0.12 | 0.08 |
| 7 | 139666331 | rs41720 | 0.41 | -1.617 | -0.678 | 0.14 | -0.007 | 0 | 0 | 0 | 0.91 | 0.85 | 0.9 | 0.88 | 0.94 | 0.88 |
| 7 | 139668438 | rs41718 | 1.047 | -1.608 | -0.662 | 0.036 | -0.012 | -0.022 | -0.022 | 0 | 0.75 | 0.62 | 0.82 | 0.87 | 0.81 | 0.79 |
| 7 | 139670417 | rs41717 | 0.694 | -2.311 | -1.561 | 0.348 | 0.119 | 0.014 | 0.014 | 0 | 0.52 | 0.7 | 0.59 | 0.57 | 0.68 | 0.59 |
| 7 | 139670494 | rs42334 | 0.877 | -1.722 | -0.97 | 0.453 | -0.029 | -0.015 | -0.015 | 0.039 | 0.82 | 0.74 | 0.86 | 0.86 | 0.88 | 0.81 |

| | | | | | | | | | | | | | | | | |
|---|---|---|---|---|---|---|---|---|---|---|---|---|---|---|---|---|
| 7 | 139672617 | rs10260531 | 0.107 | -1.501 | -0.913 | 0.491 | -0.025 | -0.04 | -0.04 | 0 | 0.08 | 0.16 | 0.04 | 0 | 0.05 | 0.06 |
| 7 | 139673336 | rs41716 | -0.002 | -1.542 | -0.97 | 0.52 | 0.002 | -0.028 | -0.028 | 0.006 | 0.94 | 0.99 | 0.92 | 0.86 | 0.96 | 0.87 |
| 7 | 139674671 | rs41715 | 1.088 | -2.023 | -1.188 | 0.495 | 0.04 | -0.011 | -0.011 | 1.412 | 0.55 | 0.61 | 0.61 | 0.52 | 0.68 | 0.46 |
| 7 | 139675510 | rs740150 | -0.036 | -1.912 | -1.302 | 0.567 | 0.022 | -0.02 | -0.02 | 0 | 0.31 | 0.26 | 0.27 | 0.32 | 0.23 | 0.34 |
| 7 | 139676790 | rs2284211 | 0.129 | -1.904 | -1.367 | 0.581 | 0.036 | -0.041 | -0.041 | 0 | 0.28 | 0.17 | 0.27 | 0.32 | 0.2 | 0.36 |
| 7 | 139678960 | rs41712 | -0.15 | -1.542 | -1.224 | 0.509 | 0.005 | -0.011 | -0.011 | 0.098 | 0.94 | 0.97 | 0.92 | 0.84 | 0.96 | 0.87 |
| 7 | 139679245 | rs11971428 | -1.078 | -1.272 | -1.301 | 0.566 | 0 | -0.03 | -0.03 | 0 | 0.58 | 0.58 | 0.61 | 0.78 | 0.46 | 0.62 |
| 7 | 139680431 | rs193950 | 0.01 | -0.979 | -1.507 | 0.852 | 0.002 | -0.03 | -0.03 | 1.617 | 0.94 | 0.99 | 0.92 | 0.83 | 0.96 | 0.87 |
| 7 | 139682412 | rs8192846 | -0.075 | -1.33 | -1.674 | 0.796 | 0.049 | -0.039 | -0.039 | 0 | 0.31 | 0.16 | 0.26 | 0.31 | 0.22 | 0.36 |
| 7 | 139682643 | rs193947 | 0.91 | -1.214 | -1.445 | 0.403 | 0 | 0.005 | 0.005 | 0 | 0.42 | 0.48 | 0.39 | 0.23 | 0.54 | 0.38 |
| 7 | 139682681 | rs193946 | 0.917 | -1.214 | -1.445 | 0.403 | 0 | 0.005 | 0.005 | 0 | 0.42 | 0.48 | 0.39 | 0.22 | 0.54 | 0.38 |
| 7 | 139683674 | rs10487666 | -0.478 | -1.303 | -1.28 | 0.614 | -0.015 | -0.005 | -0.005 | 0 | 0.12 | 0.18 | 0.08 | 0.15 | 0.08 | 0.17 |
| 7 | 139684043 | rs10487667 | -0.389 | -0.682 | -1.405 | 0.797 | 0.019 | -0.011 | -0.011 | 0 | 0.34 | 0.21 | 0.34 | 0.3 | 0.28 | 0.45 |
| 7 | 139685695 | rs3801150 | -0.437 | -0.448 | -1.052 | 0.4 | 0.002 | 0.002 | 0.002 | 0 | 0.44 | 0.39 | 0.57 | 0.69 | 0.34 | 0.49 |
| 7 | 139687783 | rs2267703 | -0.432 | 0.784 | -0.715 | 0.524 | 0.039 | -0.002 | -0.002 | 0 | 0.24 | 0.35 | 0.59 | 0.78 | 0.32 | 0.23 |
| 7 | 139688004 | rs4726499 | -0.523 | 1.149 | -1.371 | -0.188 | -0.018 | 0.062 | 0.062 | 0 | 0.82 | 0.92 | 0.86 | 0.98 | 0.77 | 0.85 |
| 7 | 139689377 | rs7785584 | 0.202 | 1.162 | -1.42 | -0.502 | -0.009 | 0.097 | 0.097 | 0 | 0.66 | 0.78 | 0.82 | 0.9 | 0.68 | 0.65 |
| 7 | 139691838 | rs8192858 | -0.633 | 1.744 | -0.473 | -0.257 | 0.04 | -0.012 | -0.012 | 0 | 0.13 | 0.2 | 0.25 | 0.1 | 0.17 | 0.11 |
| 7 | 139692481 | rs2267704 | -0.235 | 1.444 | -0.176 | 0.259 | 0.002 | 0.095 | 0.095 | 0 | 0.21 | 0.37 | 0.2 | 0.42 | 0.26 | 0.28 |
| 7 | 139694500 | rs3801148 | 1.468 | 1.339 | -0.966 | -0.375 | -0.096 | -0.106 | -0.106 | 0.609 | 0.53 | 0.07 | 0.49 | 0.46 | 0.51 | 0.56 |
| 7 | 139694661 | rs8192859 | 1.411 | 1.054 | -1.308 | -0.902 | -0.066 | 0.117 | 0.117 | 0 | 0.19 | 0.04 | 0.11 | 0 | 0.19 | 0.18 |
| 7 | 139695144 | rs7795996 | -1.298 | 0.744 | -0.109 | -0.152 | 0.018 | -0.017 | -0.017 | 0 | 0.36 | 0.75 | 0.26 | 0.43 | 0.33 | 0.34 |
| 7 | 139695316 | rs2284212 | -0.749 | 0.676 | 0.551 | -0.2 | -0.008 | 0.066 | 0.066 | 0.009 | 0.27 | 0.55 | 0.23 | 0.43 | 0.29 | 0.33 |
| 7 | 139695594 | rs2284213 | -1.502 | 0.679 | 0.429 | -0.08 | 0.013 | -0.012 | -0.012 | 1.905 | 0.36 | 0.73 | 0.25 | 0.44 | 0.33 | 0.34 |
| 7 | 139696248 | rs2267705 | -1.708 | 0.789 | 0.427 | -0.053 | 0.007 | -0.008 | -0.008 | 0.023 | 0.35 | 0.7 | 0.24 | 0.42 | 0.33 | 0.34 |
| 7 | 139696268 | rs2267706 | -0.313 | 1.036 | 0.946 | 0.062 | 0.002 | 0.084 | 0.084 | 0 | 0.19 | 0.3 | 0.17 | 0.41 | 0.24 | 0.27 |
| 7 | 139701692 | rs17161334 | -0.702 | 0.797 | 1.311 | 0.024 | -0.02 | -0.051 | -0.051 | 0.148 | 0.09 | 0.24 | 0.05 | 0 | 0.04 | 0.01 |
| 7 | 139704563 | rs11761051 | -0.401 | 0.895 | 1.31 | 0.375 | 0.013 | 0.078 | 0.078 | 0 | 0.17 | 0.29 | 0.17 | 0.38 | 0.24 | 0.24 |
| 7 | 139705609 | rs17161335 | -0.673 | 0.766 | 1.199 | 0.305 | -0.02 | -0.052 | -0.052 | 1.899 | 0.09 | 0.24 | 0.05 | 0 | 0.04 | 0.01 |
| 7 | 139705822 | rs17161337 | -0.743 | 0.76 | 1.165 | 0.303 | -0.015 | -0.05 | -0.05 | 0 | 0.09 | 0.27 | 0.05 | 0 | 0.04 | 0.01 |
| 7 | 139707140 | rs740204 | 1.761 | 0.851 | 0.539 | 0.403 | 0.001 | -0.01 | -0.01 | 0 | 0.66 | 0.23 | 0.72 | 0.63 | 0.63 | 0.71 |
| 7 | 139713109 | rs13227152 | -0.497 | 0.214 | 0.621 | 0.054 | 0.005 | -0.005 | -0.005 | 0.722 | 0.09 | 0.04 | 0.03 | 0 | 0.04 | 0.11 |
| 7 | 139717178 | rs757835 | -1.398 | 0.616 | 1.219 | 0.639 | 0.05 | 0.18 | 0.18 | 0.365 | 0.14 | 0.34 | 0.37 | 0.54 | 0.2 | 0.17 |
| 7 | 139717200 | rs757834 | -1.385 | 0.596 | 1.223 | 0.63 | 0.048 | 0.176 | 0.176 | 0.239 | 0.15 | 0.34 | 0.37 | 0.54 | 0.2 | 0.17 |
| 7 | 139719184 | rs2072190 | -2.642 | 0.454 | 0.469 | 0.265 | -0.027 | -0.083 | -0.083 | 0 | 0.44 | 0.88 | 0.53 | 0.52 | 0.38 | 0.35 |
| 7 | 139723596 | rs1044718 | 1.038 | 0.424 | 0.466 | 0.117 | -0.017 | 0.051 | 0.051 | 0 | 0.1 | 0.01 | 0.07 | 0 | 0.08 | 0.02 |
| 7 | 139723742 | rs1044703 | 1.081 | 0.434 | 0.464 | 0.116 | -0.015 | 0.05 | 0.05 | 0 | 0.1 | 0.01 | 0.06 | 0 | 0.08 | 0.01 |
| 7 | 139724169 | rs13760 | -2.828 | 0.421 | 0.491 | 0.145 | -0.02 | -0.07 | -0.07 | 0 | 0.44 | 0.88 | 0.53 | 0.53 | 0.38 | 0.35 |
| 7 | 139724358 | rs2269996 | -0.617 | 0.444 | 1.024 | 0.172 | 0.062 | 0.224 | 0.224 | 0 | 0.13 | 0.33 | 0.36 | 0.53 | 0.2 | 0.16 |
| 7 | 139724742 | rs3735351 | 0.791 | 0.512 | 0.522 | 0.227 | -0.022 | 0.011 | 0.011 | 0 | 0.05 | 0.02 | 0.06 | 0.04 | 0.11 | 0.03 |
| 7 | 139726207 | rs2286198 | -1.53 | 0.496 | 0.679 | 0.279 | 0.084 | 0.322 | 0.322 | 0 | 0.15 | 0.45 | 0.39 | 0.5 | 0.21 | 0.17 |
| 7 | 139728584 | rs2286195 | -1.803 | 0.5 | 0.653 | 0.242 | 0.067 | 0.264 | 0.264 | 0 | 0.17 | 0.45 | 0.39 | 0.53 | 0.21 | 0.18 |
| 7 | 139736834 | rs12703513 | 0.785 | 0.495 | 0.567 | 0.124 | -0.022 | 0.011 | 0.011 | 0 | 0.05 | 0.02 | 0.06 | 0.03 | 0.11 | 0.03 |
| 7 | 139738367 | rs13244572 | -0.201 | 0.322 | 0.73 | 0.101 | -0.017 | -0.043 | -0.043 | 0 | 0.09 | 0.1 | 0.04 | 0 | 0.05 | 0.13 |
| 7 | 139741766 | rs10267006 | -1.582 | 0.336 | 0.812 | 0.061 | -0.015 | -0.1 | -0.1 | 0 | 0.25 | 0.29 | 0.14 | 0 | 0.17 | 0.17 |
| 7 | 139744802 | rs17161378 | -1.697 | 0.26 | 0.763 | 0.224 | -0.048 | -0.116 | -0.116 | 0.105 | 0.17 | 0.41 | 0.08 | 0 | 0.09 | 0.15 |
| 7 | 139757136 | rs10237377 | -1.328 | 0.857 | 0.13 | 0.639 | -0.006 | -0.03 | -0.03 | 0 | 0.35 | 0.77 | 0.49 | 0.53 | 0.34 | 0.23 |
| 7 | 139763994 | rs10808024 | -0.886 | 0.483 | 0.349 | 0.756 | 0.011 | 0.046 | 0.046 | 2.539 | 0.27 | 0.36 | 0.47 | 0.52 | 0.29 | 0.31 |
| 7 | 139770813 | rs7803170 | -1.189 | 0.287 | -0.136 | 0.341 | 0.002 | -0.014 | -0.014 | 1.597 | 0.39 | 0.57 | 0.52 | 0.67 | 0.36 | 0.44 |
| 7 | 139772295 | rs12538215 | -0.987 | -0.576 | -0.026 | 0.497 | -0.009 | 0.007 | 0.007 | 0 | 0.18 | 0.17 | 0.42 | 0.23 | 0.11 | 0.19 |
| 7 | 139774014 | rs12539669 | 0.583 | -1.157 | -0.146 | -0.38 | 0.041 | 0.009 | 0.009 | 0 | 0.43 | 0.32 | 0.45 | 0.4 | 0.31 | 0.47 |
| 7 | 139776384 | rs13227771 | -0.585 | -0.972 | 0.074 | -0.611 | -0.018 | -0.058 | -0.058 | 0 | 0.24 | 0.46 | 0.2 | 0.16 | 0.22 | 0.12 |
| 7 | 139794009 | rs12375063 | 0.586 | -1.637 | -0.391 | -0.942 | -0.017 | 0.142 | 0.142 | 0.905 | 0.33 | 0.04 | 0.19 | 0.1 | 0.29 | 0.28 |
| 7 | 139801777 | rs1062277 | -0.177 | -1.604 | -0.3 | -0.933 | 0.017 | 0.032 | 0.032 | 0.465 | 0.45 | 0.42 | 0.28 | 0.25 | 0.34 | 0.35 |
| 7 | 139831585 | rs13222813 | 0.417 | -1.483 | -0.226 | -0.893 | 0.01 | 0.036 | 0.036 | 0 | 0.35 | 0.29 | 0.22 | 0.1 | 0.29 | 0.28 |
| 7 | 139837061 | rs11983076 | 0.071 | -1.445 | -0.511 | -0.865 | 0.001 | -0.008 | -0.008 | 0 | 0.39 | 0.36 | 0.26 | 0.1 | 0.37 | 0.28 |
| 7 | 139839774 | rs12056023 | 1.393 | -1.118 | -0.629 | -0.672 | -0.032 | 0.034 | 0.034 | 0 | 0.2 | 0.34 | 0.32 | 0.31 | 0.11 | 0.25 |
| 7 | 139846749 | rs10237170 | 1.384 | -1.111 | -0.626 | -0.669 | -0.032 | 0.033 | 0.033 | 0 | 0.2 | 0.34 | 0.32 | 0.31 | 0.11 | 0.25 |
| 7 | 139855216 | rs17192222 | 0.906 | -0.152 | -0.961 | -0.321 | -0.018 | 0.055 | 0.055 | 0.352 | 0.1 | 0.01 | 0.06 | 0 | 0.09 | 0.04 |
| 7 | 139884494 | rs1005261 | -0.889 | 0.167 | -0.657 | 0.053 | -0.165 | -0.169 | -0.169 | 0 | 0.79 | 0.98 | 0.68 | 0.55 | 0.58 | 0.67 |
| 7 | 139902694 | rs10279097 | -0.575 | 0.2 | -0.311 | -0.011 | -0.013 | -0.012 | -0.012 | 0 | 0.75 | 0.83 | 0.66 | 0.55 | 0.56 | 0.67 |
| 7 | 139904130 | rs4626515 | -1.031 | 0.556 | 0.487 | -0.652 | 0.067 | -0.026 | -0.026 | 0.226 | 0.09 | 0.51 | 0.11 | 0 | 0.1 | 0.01 |
| 7 | 139913599 | rs2363101 | -0.867 | 0.461 | 0.164 | -0.788 | 0.022 | 0.155 | 0.155 | 0 | 0.5 | 0.76 | 0.36 | 0.24 | 0.47 | 0.36 |
| 7 | 139918895 | rs4725655 | 0.16 | 0.267 | -0.28 | -0.489 | 0.004 | 0.029 | 0.029 | 0 | 0.31 | 0.16 | 0.18 | 0.02 | 0.28 | 0.27 |
| 7 | 139920973 | rs13223890 | -0.744 | 0.372 | -0.22 | -0.443 | -0.004 | -0.017 | -0.017 | 0 | 0.43 | 0.4 | 0.26 | 0.24 | 0.36 | 0.35 |
| 7 | 139924127 | rs6969577 | -0.289 | 0.507 | -0.512 | 0.308 | -0.005 | 0.019 | 0.019 | 1.724 | 0.11 | 0.14 | 0.08 | 0.22 | 0.09 | 0.07 |
| 7 | 139935860 | rs6956556 | -0.409 | 0.707 | -0.534 | -0.573 | 0 | -0.032 | -0.032 | 0.066 | 0.48 | 0.43 | 0.31 | 0.07 | 0.49 | 0.36 |
| 7 | 139936877 | rs2471281 | 0.62 | 2.091 | -0.955 | 0.037 | -0.06 | -0.015 | -0.015 | 0 | 0.74 | 0.85 | 0.74 | 0.7 | 0.64 | 0.77 |
| 7 | 139937237 | rs2471279 | 0.605 | 2.191 | -0.713 | 0.182 | -0.014 | -0.01 | -0.01 | 0 | 0.81 | 0.87 | 0.84 | 0.76 | 0.73 | 0.81 |

| | | | | | | | | | | | | | | | | |
|---|---|---|---|---|---|---|---|---|---|---|---|---|---|---|---|---|
| 7 | 139937791 | rs12703639 | 1.201 | 1.68 | -0.938 | 0.088 | 0.104 | -0.018 | -0.018 | 0.179 | 0.48 | 0.15 | 0.48 | 0.53 | 0.3 | 0.5 |
| 7 | 139939203 | rs17161443 | 1.289 | 1.532 | -0.972 | 0.152 | 0.085 | -0.023 | -0.023 | 2.56 | 0.46 | 0.15 | 0.38 | 0.49 | 0.32 | 0.52 |
| 7 | 139942304 | rs11977670 | 1.167 | 0.924 | -0.608 | 0.116 | 0.024 | 0.017 | 0.017 | 0.237 | 0.49 | 0.25 | 0.54 | 0.47 | 0.43 | 0.51 |
| 7 | 139942889 | rs2253729 | 0.124 | 1.162 | -0.945 | 0.195 | -0.06 | -0.119 | -0.119 | 0 | 0.77 | 0.94 | 0.74 | 0.58 | 0.69 | 0.75 |
| 7 | 139944473 | rs17161447 | -0.85 | 0.888 | -0.128 | 0.148 | 0.012 | -0.063 | -0.063 | 0.056 | 0.16 | 0.55 | 0.12 | 0.09 | 0.13 | 0.18 |
| 7 | 139949176 | rs7799803 | -0.775 | 0.934 | -0.068 | 0.12 | 0.025 | -0.08 | -0.08 | 1.292 | 0.14 | 0.51 | 0.11 | 0.02 | 0.12 | 0.15 |
| 7 | 139954640 | rs12703654 | 0.865 | 0.887 | -0.726 | 0.318 | 0.011 | 0.021 | 0.021 | 0.049 | 0.5 | 0.2 | 0.53 | 0.46 | 0.45 | 0.49 |
| 7 | 139956452 | rs7777438 | -0.933 | 0.636 | -0.245 | 0.092 | -0.002 | -0.035 | -0.035 | 0 | 0.1 | 0.16 | 0.06 | 0.03 | 0.07 | 0.17 |
| 7 | 139958544 | rs2429216 | 1.259 | 0.46 | -0.772 | 0.457 | -0.012 | 0.006 | 0.006 | 0.19 | 0.46 | 0.26 | 0.64 | 0.59 | 0.5 | 0.46 |
| 7 | 139959782 | rs11971015 | 1.175 | 0.494 | -0.199 | 0.44 | 0 | -0.038 | -0.038 | 0 | 0.68 | 0.5 | 0.75 | 0.77 | 0.67 | 0.58 |
| 7 | 139960630 | rs2429213 | 0.71 | 0.553 | -0.143 | 0.643 | 0.007 | 0.023 | 0.023 | 0.628 | 0.78 | 0.48 | 0.82 | 0.76 | 0.79 | 0.65 |
| 7 | 139963396 | rs6947787 | -1.283 | 0.085 | 0.21 | 0.381 | -0.008 | -0.031 | -0.031 | 0.139 | 0.14 | 0.41 | 0.1 | 0.08 | 0.1 | 0.11 |
| 7 | 139964350 | rs1034864 | -0.979 | -0.209 | -0.411 | 0.26 | 0.034 | 0.006 | 0.006 | 0 | 0.49 | 0.65 | 0.59 | 0.48 | 0.41 | 0.42 |
| 7 | 139964597 | rs12154965 | 0.006 | -0.501 | -0.792 | -0.309 | 0.03 | 0.069 | 0.069 | 1.894 | 0.25 | 0.1 | 0.13 | 0.08 | 0.22 | 0.15 |
| 7 | 139969051 | rs2471312 | -0.4 | 0.29 | -0.824 | -0.453 | -0.017 | 0.057 | 0.057 | 0 | 0.56 | 0.64 | 0.7 | 0.71 | 0.49 | 0.62 |
| 7 | 139975028 | rs2471282 | 0.534 | 0.335 | -1.021 | -0.536 | -0.02 | 0.015 | 0.015 | 0 | 0.06 | 0 | 0.02 | 0.03 | 0.08 | 0.09 |
| 7 | 139975457 | rs764891 | -0.395 | 0.113 | -0.747 | -0.234 | -0.013 | 0.057 | 0.057 | 0 | 0.29 | 0.49 | 0.5 | 0.45 | 0.26 | 0.23 |
| 7 | 139976963 | rs6948262 | 0.277 | 0 | -0.505 | -0.161 | -0.015 | -0.074 | -0.074 | 0 | 0.21 | 0.4 | 0.18 | 0.11 | 0.19 | 0.12 |
| 7 | 139976992 | rs6960670 | 0.787 | 0.787 | -0.665 | -0.146 | -0.008 | 0.011 | 0.011 | 0 | 0.58 | 0.62 | 0.42 | 0.41 | 0.56 | 0.46 |
| 7 | 139979843 | rs11769504 | 0.871 | 0.655 | -0.882 | -0.015 | -0.018 | -0.057 | -0.057 | 0 | 0.19 | 0.02 | 0.16 | 0.23 | 0.17 | 0.3 |
| 7 | 139983172 | rs17194049 | 0.565 | 0.641 | -0.767 | -0.028 | 0.003 | -0.01 | -0.01 | 0.043 | 0.21 | 0.15 | 0.16 | 0.22 | 0.17 | 0.3 |
| 7 | 139985962 | rs2429215 | 0.429 | 0.669 | -0.185 | -0.902 | -0.009 | -0.028 | -0.028 | 0 | 0.43 | 0.53 | 0.29 | 0.23 | 0.36 | 0.3 |
| 7 | 139986455 | rs2429211 | 0.578 | 0.479 | 0.233 | -0.999 | -0.012 | -0.031 | -0.031 | 0 | 0.41 | 0.49 | 0.26 | 0.26 | 0.34 | 0.33 |
| 7 | 139986527 | rs12532353 | 0.977 | 0.312 | -0.127 | -1.111 | 0.034 | 0.066 | 0.066 | 0 | 0.37 | 0.19 | 0.21 | 0.26 | 0.28 | 0.28 |
| 7 | 139987072 | rs2471318 | -0.156 | 0.329 | -0.208 | -1.035 | 0.053 | 0.079 | 0.079 | 0.089 | 0.47 | 0.18 | 0.63 | 0.62 | 0.61 | 0.65 |
| 7 | 139987531 | rs11763495 | 0.201 | 0.731 | -0.21 | -0.941 | -0.015 | -0.011 | -0.011 | 0 | 0.4 | 0.15 | 0.53 | 0.53 | 0.39 | 0.57 |
| 7 | 139992324 | rs4236486 | 0.904 | 0.945 | -0.064 | -0.993 | -0.002 | 0.021 | 0.021 | 0 | 0.41 | 0.35 | 0.28 | 0.28 | 0.4 | 0.33 |
| 7 | 139992387 | rs4236488 | -0.315 | 0.923 | -0.059 | -0.931 | -0.023 | -0.019 | -0.019 | 0 | 0.86 | 0.63 | 0.89 | 0.9 | 0.9 | 0.94 |
| 7 | 139992942 | rs12703712 | -0.456 | 1.467 | -0.832 | -0.699 | -0.016 | 0.035 | 0.035 | 0 | 0.44 | 0.24 | 0.6 | 0.62 | 0.49 | 0.61 |
| 7 | 139995647 | rs4275141 | 0.319 | 1.341 | -1.23 | -0.647 | -0.019 | 0.015 | 0.015 | 1.73 | 0.54 | 0.8 | 0.4 | 0.46 | 0.54 | 0.39 |
| 7 | 140001939 | rs12538711 | 0.002 | 1.09 | -1.319 | -0.513 | -0.079 | -0.031 | -0.031 | 0 | 0.38 | 0.13 | 0.6 | 0.54 | 0.46 | 0.54 |
| 7 | 140009332 | rs7792748 | 0.417 | 1.256 | -1.436 | -0.535 | -0.02 | -0.065 | -0.065 | 0 | 0.28 | 0.11 | 0.49 | 0.58 | 0.28 | 0.48 |
| 7 | 140009905 | rs4725675 | 0.493 | 1.455 | -1.479 | -0.829 | -0.026 | 0.039 | 0.039 | 0 | 0.83 | 0.87 | 0.87 | 0.96 | 0.77 | 0.87 |
| 7 | 140010108 | rs11764865 | 0.168 | 1.368 | -1.483 | -0.828 | -0.002 | 0.019 | 0.019 | 0 | 0.88 | 0.89 | 0.93 | 0.96 | 0.84 | 0.89 |
| 7 | 140012638 | rs3924239 | 0.462 | 1.001 | -1.128 | -1.273 | -0.001 | -0.002 | -0.002 | 0 | 0.36 | 0.37 | 0.25 | 0.26 | 0.39 | 0.36 |
| 7 | 140013037 | rs4349904 | -0.557 | 0.961 | -1.299 | -1.131 | 0.005 | 0.012 | 0.012 | 0 | 0.9 | 0.91 | 0.95 | 0.96 | 0.89 | 0.9 |
| 7 | 140014556 | rs11772239 | 0.704 | 0.973 | -1.414 | -1.219 | -0.077 | -0.069 | -0.069 | 0 | 0.32 | 0.13 | 0.54 | 0.58 | 0.36 | 0.5 |
| 7 | 140015104 | rs4416746 | -0.347 | 0.735 | -1.268 | -1.163 | 0.005 | 0.011 | 0.011 | 0 | 0.9 | 0.91 | 0.95 | 0.96 | 0.9 | 0.9 |
| 7 | 140020787 | rs6956771 | 0.331 | 0.673 | -0.999 | -1.487 | 0.009 | -0.065 | -0.065 | 0 | 0.3 | 0.41 | 0.26 | 0.1 | 0.32 | 0.13 |
| 7 | 140024052 | rs12703744 | 0.952 | 0.681 | -1.364 | -1.518 | -0.004 | 0.085 | 0.085 | 0.082 | 0.52 | 0.34 | 0.64 | 0.83 | 0.51 | 0.75 |
| 7 | 140025370 | rs12533908 | 1.638 | 0.332 | -1.123 | -1.352 | 0.008 | 0.017 | 0.017 | 0.95 | 0.17 | 0.05 | 0.13 | 0.1 | 0.15 | 0.04 |
| 7 | 140026436 | rs4373438 | -0.42 | 0.393 | -1.102 | -1.277 | 0.003 | 0.003 | 0.003 | 0 | 0.64 | 0.62 | 0.79 | 0.7 | 0.67 | 0.61 |
| 7 | 140030555 | rs6947292 | 0.441 | 0.905 | -0.406 | -1.112 | -0.003 | -0.07 | -0.07 | 0 | 0.85 | 0.79 | 0.91 | 0.97 | 0.85 | 0.9 |
| 7 | 140033631 | rs558 | 0.862 | 0.685 | -1.015 | -1.139 | 0.012 | 0.008 | 0.008 | 0 | 0.38 | 0.28 | 0.22 | 0.01 | 0.35 | 0.14 |
| 7 | 140038907 | rs9640226 | 0.13 | 0.949 | -1.483 | -0.383 | -0.077 | 0.12 | 0.12 | 0.153 | 0.61 | 0.93 | 0.41 | 0.17 | 0.58 | 0.3 |
| 7 | 140039914 | rs6964148 | 0.177 | 0.943 | -1.445 | -0.386 | -0.06 | 0.12 | 0.12 | 0.01 | 0.61 | 0.91 | 0.41 | 0.17 | 0.58 | 0.3 |
| 7 | 140042556 | rs6464696 | -0.24 | 1.156 | -1.744 | 0.445 | -0.077 | -0.14 | -0.14 | 0.522 | 0.69 | 0.96 | 0.64 | 0.46 | 0.66 | 0.42 |
| 7 | 140047210 | rs12703767 | -0.108 | 1.19 | -0.693 | 1.077 | 0 | 0.048 | 0.048 | 0.023 | 0.08 | 0.1 | 0.24 | 0.3 | 0.08 | 0.14 |
| 7 | 140047236 | rs6959565 | -0.688 | 1.445 | 0.094 | 0.834 | 0.102 | 0.111 | 0.111 | 0 | 0.08 | 0.3 | 0.1 | 0.15 | 0.13 | 0.09 |
| 7 | 140048890 | rs6964909 | -0.603 | 0.796 | -0.028 | 0.412 | 0.003 | 0.002 | 0.002 | 0 | 0.6 | 0.61 | 0.77 | 0.8 | 0.65 | 0.66 |
| 7 | 140050018 | rs12703771 | 0.403 | 1 | 0.042 | 0.736 | -0.003 | -0.002 | -0.002 | 0 | 0.39 | 0.3 | 0.37 | 0.42 | 0.46 | 0.39 |
| 7 | 140067576 | rs12703792 | 0.56 | 0.968 | -0.011 | 0.74 | -0.051 | -0.046 | -0.046 | 0.397 | 0.38 | 0.12 | 0.35 | 0.42 | 0.46 | 0.38 |
| 7 | 140070378 | rs12703799 | -0.425 | 0.902 | 0.615 | 0.718 | 0.001 | -0.001 | -0.001 | 0.228 | 0.55 | 0.51 | 0.45 | 0.52 | 0.57 | 0.59 |
| 7 | 140071315 | rs7777468 | -0.571 | 0.88 | 0.67 | 0.636 | 0.028 | -0.004 | -0.004 | 0 | 0.13 | 0.29 | 0.09 | 0.1 | 0.1 | 0.2 |
| 7 | 140082363 | rs6968618 | -1.368 | 0.841 | 0.567 | 0.818 | 0.005 | 0.017 | 0.017 | 0 | 0.67 | 0.73 | 0.8 | 0.81 | 0.68 | 0.83 |
| 7 | 140092314 | rs6978505 | 0.655 | 0.656 | 0.605 | 1.187 | -0.005 | 0.007 | 0.007 | 0 | 0.39 | 0.3 | 0.34 | 0.36 | 0.43 | 0.27 |
| 7 | 140096157 | rs9640465 | 0.107 | 0.758 | 0.521 | 1.331 | -0.002 | -0.003 | -0.003 | 0.356 | 0.89 | 0.9 | 0.92 | 0.84 | 0.9 | 0.79 |
| 7 | 140096203 | rs9640467 | 2.278 | 0.803 | 0.704 | 1.38 | -0.008 | 0.064 | 0.064 | 1.86 | 0.72 | 0.46 | 0.56 | 0.54 | 0.81 | 0.38 |
| 7 | 140097554 | rs12539339 | -1.518 | 0.702 | 0.615 | 1.563 | -0.002 | 0.011 | 0.011 | 0 | 0.14 | 0.17 | 0.33 | 0.29 | 0.1 | 0.37 |
| 7 | 140098605 | rs4407796 | -0.266 | 0.618 | 0.44 | 0.92 | 0.008 | 0.011 | 0.011 | 0 | 0.49 | 0.61 | 0.66 | 0.61 | 0.45 | 0.56 |
| 7 | 140100890 | rs6954902 | -0.025 | 0.81 | 0.316 | 0.549 | 0.006 | 0.019 | 0.019 | 0 | 0.47 | 0.56 | 0.65 | 0.73 | 0.41 | 0.52 |
| 7 | 140106504 | rs13237949 | -0.013 | 0.683 | -0.105 | 0.601 | 0.011 | 0.044 | 0.044 | 0 | 0.49 | 0.41 | 0.64 | 0.73 | 0.42 | 0.47 |
| 7 | 140108096 | rs10235144 | -0.313 | 0.786 | -0.512 | 0.414 | -0.002 | 0.003 | 0.003 | 0 | 0.61 | 0.66 | 0.74 | 0.81 | 0.53 | 0.65 |
| 7 | 140117411 | rs2001942 | -0.493 | 0.068 | -0.748 | 0.856 | 0.011 | -0.013 | -0.013 | 0 | 0.14 | 0.09 | 0.21 | 0.21 | 0.1 | 0.17 |
| 7 | 140120546 | rs11971702 | 0.15 | 0.032 | -0.607 | 0.749 | -0.005 | -0.001 | -0.001 | 0.128 | 0.44 | 0.48 | 0.58 | 0.56 | 0.45 | 0.43 |
| 7 | 140122481 | rs4574768 | -0.422 | -0.295 | -0.558 | 0.673 | -0.041 | 0.071 | 0.071 | 0 | 0.59 | 0.78 | 0.67 | 0.74 | 0.55 | 0.57 |
| 7 | 140125750 | rs2948386 | 0.868 | 0.173 | -1.068 | 0.36 | -0.079 | 0.078 | 0.078 | 0.112 | 0.2 | 0.04 | 0.12 | 0.07 | 0.24 | 0.3 |
| 7 | 140126507 | rs11765962 | 0.55 | -0.307 | -0.202 | 0.488 | -0.031 | -0.006 | -0.006 | 0 | 0.16 | 0.27 | 0.1 | 0.15 | 0.1 | 0.13 |
| 7 | 140129507 | rs17161539 | 1.231 | -0.186 | -0.277 | 0.548 | -0.019 | -0.029 | -0.029 | 0.005 | 0.05 | 0.01 | 0.05 | 0.09 | 0.06 | 0.1 |

| | | | | | | | | | | | | | | | | |
|---|---|---|---|---|---|---|---|---|---|---|---|---|---|---|---|---|
| 7 | 140129626 | rs1007435 | 1.632 | -0.504 | -0.111 | 0.861 | -0.011 | 0.048 | 0.048 | 0.964 | 0.13 | 0.24 | 0.23 | 0.27 | 0.08 | 0.2 |
| 7 | 140132266 | rs1976851 | 1.933 | -0.521 | -0.098 | 0.867 | 0.002 | -0.006 | -0.006 | 0 | 0.11 | 0.11 | 0.22 | 0.27 | 0.08 | 0.2 |
| 7 | 140132891 | rs9648835 | -0.228 | -0.604 | -0.127 | 0.791 | -0.048 | 0.045 | 0.045 | 0 | 0.3 | 0.49 | 0.33 | 0.42 | 0.19 | 0.34 |
| 7 | 140133730 | rs7786260 | -0.055 | -0.653 | -0.191 | 0.794 | -0.027 | 0.027 | 0.027 | 0 | 0.29 | 0.64 | 0.32 | 0.45 | 0.17 | 0.33 |
| 7 | 140145558 | rs6464773 | 0.249 | -0.707 | 0.262 | 0.381 | -0.052 | -0.001 | -0.001 | 0 | 0.21 | 0.55 | 0.12 | 0.18 | 0.11 | 0.15 |
| 7 | 140145753 | rs12155423 | 1.952 | -0.507 | -0.321 | 0.327 | -0.056 | -0.003 | -0.003 | 0.238 | 0.2 | 0.02 | 0.16 | 0.17 | 0.31 | 0.1 |
| 7 | 140152722 | rs11760895 | 0.233 | -0.891 | -0.096 | -0.185 | -0.065 | -0.007 | -0.007 | 0.24 | 0.22 | 0.59 | 0.12 | 0.18 | 0.11 | 0.15 |
| 7 | 140155296 | rs2272096 | -0.031 | -0.892 | -0.348 | -0.186 | 0.019 | 0.006 | 0.006 | 0.593 | 0.24 | 0.74 | 0.13 | 0.18 | 0.11 | 0.15 |
| 7 | 140155800 | rs17621014 | 0.939 | -0.334 | -0.406 | -0.358 | -0.016 | -0.024 | -0.024 | 0.015 | 0.09 | 0.01 | 0.06 | 0.08 | 0.08 | 0.11 |
| 7 | 140169141 | rs7776957 | -1.082 | 0.023 | -0.777 | -0.436 | -0.033 | 0.018 | 0.018 | 0 | 0.87 | 0.93 | 0.84 | 0.89 | 0.85 | 0.96 |
| 7 | 140176596 | rs17621391 | 0.743 | -0.413 | -0.306 | -0.327 | -0.041 | 0.083 | 0.083 | 0 | 0.21 | 0.02 | 0.16 | 0.09 | 0.29 | 0.11 |
| 7 | 140179500 | rs7811509 | -1.525 | -0.368 | 0.926 | 0.32 | 0.058 | 0.091 | 0.091 | 2.749 | 0.07 | 0.38 | 0.03 | 0.03 | 0.02 | 0.04 |
| 7 | 140182402 | rs7785094 | -1.009 | -0.623 | 0.562 | 0.157 | -0.069 | -0.06 | -0.06 | 0.415 | 0.24 | 0.5 | 0.15 | 0.13 | 0.15 | 0.14 |
| 7 | 140185397 | rs4726826 | -1.102 | 0.233 | 0.206 | 0.717 | -0.018 | -0.036 | -0.036 | 0.024 | 0.29 | 0.22 | 0.38 | 0.42 | 0.31 | 0.38 |
| 7 | 140187074 | rs10253030 | -0.251 | 0.063 | -0.214 | 0.123 | -0.044 | -0.066 | -0.066 | 1.613 | 0.33 | 0.19 | 0.28 | 0.47 | 0.39 | 0.34 |
| 7 | 140195463 | rs12534545 | -0.238 | 0.089 | -0.516 | -0.077 | 0.008 | 0.021 | 0.021 | 0.154 | 0.22 | 0.15 | 0.24 | 0.05 | 0.16 | 0.15 |
| 7 | 140198159 | rs7781853 | 0.712 | -0.013 | -0.398 | 0.361 | 0.003 | 0.01 | 0.01 | 0 | 0.38 | 0.3 | 0.46 | 0.38 | 0.43 | 0.4 |
| 7 | 140209941 | rs6945832 | -0.582 | 1.032 | -0.369 | 0.617 | 0.028 | 0.05 | 0.05 | 0 | 0.32 | 0.42 | 0.32 | 0.42 | 0.43 | 0.3 |
| 7 | 140213880 | rs4726876 | -0.623 | 0.972 | -0.866 | 0.584 | -0.016 | -0.021 | -0.021 | 1.02 | 0.37 | 0.28 | 0.32 | 0.46 | 0.46 | 0.36 |
| 7 | 140218464 | rs6852 | 0.266 | 0.533 | -1.56 | -0.34 | -0.023 | 0.141 | 0.141 | 0.046 | 0.46 | 0.18 | 0.46 | 0.77 | 0.53 | 0.58 |
| 7 | 140219044 | rs6464818 | -0.517 | 0.082 | -0.373 | -0.887 | -0.003 | -0.079 | -0.079 | 0 | 0.12 | 0.44 | 0.07 | 0.01 | 0.08 | 0.08 |
| 7 | 140224024 | rs6948786 | -0.565 | -0.033 | 0.71 | -1.02 | 0 | -0.018 | -0.018 | 0.006 | 0.47 | 0.5 | 0.47 | 0.21 | 0.43 | 0.39 |
| 7 | 140227293 | rs6464833 | -1.032 | -0.058 | 1.019 | -1.06 | 0.032 | -0.066 | -0.066 | 0.187 | 0.11 | 0.47 | 0.08 | 0.01 | 0.08 | 0.09 |
| 7 | 140228118 | rs6975257 | 0.194 | -0.153 | 0.614 | -1.437 | 0.008 | 0.017 | 0.017 | 0.053 | 0.25 | 0.2 | 0.23 | 0.12 | 0.2 | 0.25 |
| 7 | 140228192 | rs6963099 | -0.589 | -0.068 | 0.719 | -1.348 | -0.006 | 0.093 | 0.093 | 0 | 0.41 | 0.22 | 0.43 | 0.19 | 0.4 | 0.32 |
| 7 | 140230269 | rs10280493 | -0.861 | -0.069 | 1.292 | -1.381 | 0.032 | -0.053 | -0.053 | 0 | 0.1 | 0.43 | 0.07 | 0 | 0.07 | 0.09 |
| 7 | 140231130 | rs10237118 | -0.001 | -0.025 | 1.085 | -1.361 | 0.012 | -0.039 | -0.039 | 0 | 0.07 | 0.16 | 0.05 | 0 | 0.07 | 0.07 |
| 7 | 140232243 | rs4726889 | -0.987 | -0.505 | 1.055 | -1.343 | 0.011 | 0.042 | 0.042 | 0 | 0.5 | 0.62 | 0.62 | 0.87 | 0.47 | 0.71 |
| 7 | 140233585 | rs10254777 | -0.288 | -0.438 | 1.449 | -1.572 | -0.018 | -0.064 | -0.064 | 0.052 | 0.35 | 0.48 | 0.29 | 0.14 | 0.31 | 0.31 |
| 7 | 140239809 | rs10808050 | -0.814 | -0.765 | 1.692 | -1.707 | -0.001 | -0.023 | -0.023 | 0 | 0.43 | 0.46 | 0.47 | 0.15 | 0.43 | 0.35 |
| 7 | 140240862 | rs1109939 | -1.058 | -0.819 | 1.592 | -1.731 | 0.021 | 0.17 | 0.17 | 0 | 0.47 | 0.78 | 0.49 | 0.16 | 0.43 | 0.38 |
| 7 | 140244560 | rs2293177 | 0.02 | -0.682 | 1.44 | -1.717 | 0 | 0.007 | 0.007 | 0 | 0.32 | 0.28 | 0.27 | 0.14 | 0.32 | 0.29 |
| 7 | 140254890 | rs4571657 | 0.144 | -1.198 | 0.905 | -1.854 | 0.019 | 0.044 | 0.044 | 0 | 0.88 | 0.94 | 0.82 | 0.99 | 0.91 | 0.94 |
| 7 | 140265434 | rs685182 | -0.995 | -1.471 | 1.057 | -2.069 | 0.067 | 0.144 | 0.144 | 0 | 0.51 | 0.73 | 0.49 | 0.15 | 0.4 | 0.39 |
| 7 | 140278758 | rs269216 | -0.793 | -1.586 | 1.233 | -2.15 | -0.003 | -0.028 | -0.028 | 0.926 | 0.47 | 0.46 | 0.4 | 0.15 | 0.39 | 0.37 |
| 7 | 140316236 | rs10952779 | 0.216 | -1.487 | 0.725 | -2.302 | -0.01 | -0.038 | -0.038 | 0.048 | 0.33 | 0.31 | 0.27 | 0.16 | 0.28 | 0.33 |
| 7 | 140317120 | rs7776899 | -0.748 | -1.634 | 0.524 | -2.195 | -0.056 | -0.066 | -0.066 | 0 | 0.11 | 0.45 | 0.05 | 0 | 0.02 | 0.02 |
| 7 | 140330281 | rs269264 | -0.912 | -0.659 | -1.605 | -2.196 | -0.159 | 0.118 | 0.118 | 0 | 0.73 | 0.98 | 0.76 | 0.93 | 0.58 | 0.82 |
| 7 | 140351925 | rs10441215 | -0.686 | -1.583 | -0.428 | -1.481 | -0.048 | -0.097 | -0.097 | 0 | 0.15 | 0.61 | 0.1 | 0.01 | 0.07 | 0.04 |
| 7 | 140358461 | rs2930328 | -1.108 | -1.776 | -0.683 | -1.806 | -0.029 | -0.06 | -0.06 | 0 | 0.19 | 0.71 | 0.11 | 0.01 | 0.08 | 0.06 |
| 7 | 140369114 | rs4727011 | -1.351 | -2.319 | -0.693 | -1.76 | -0.036 | -0.029 | -0.029 | 0 | 0.28 | 0.65 | 0.21 | 0.11 | 0.11 | 0.27 |
| 7 | 140372299 | rs269267 | -1.247 | -2.274 | -1.368 | -1.042 | -0.049 | -0.106 | -0.106 | 0.627 | 0.87 | 0.99 | 0.79 | 0.48 | 0.83 | 0.76 |
| 7 | 140372579 | rs6464937 | -1.262 | -2.303 | -0.997 | -1.432 | -0.002 | 0.001 | 0.001 | 2.802 | 0.28 | 0.72 | 0.21 | 0.11 | 0.11 | 0.27 |
| 7 | 140374049 | rs1140034 | -1.282 | -2.308 | -0.999 | -1.434 | -0.001 | 0.001 | 0.001 | 2.661 | 0.28 | 0.72 | 0.21 | 0.11 | 0.11 | 0.27 |
| 7 | 140382156 | rs2968553 | -1.236 | -2.161 | -1.159 | -1.523 | 0.023 | 0.023 | 0.023 | 1.449 | 0.31 | 0.74 | 0.25 | 0.11 | 0.16 | 0.27 |
| 7 | 140390656 | rs12375245 | 0.446 | -2.195 | -0.521 | -1.535 | -0.015 | 0.004 | 0.004 | 0 | 0.12 | 0.19 | 0.16 | 0.1 | 0.08 | 0.21 |
| 7 | 140394587 | rs1046515 | 0.519 | -2.19 | -0.518 | -1.533 | -0.014 | 0.005 | 0.005 | 0 | 0.12 | 0.19 | 0.16 | 0.1 | 0.08 | 0.21 |
| 7 | 140397792 | rs1267673 | -1.971 | -2.504 | -1.391 | -1.86 | 0.133 | 0.201 | 0.201 | 0.353 | 0.5 | 0.93 | 0.35 | 0.19 | 0.28 | 0.4 |
| 7 | 140405706 | rs2364397 | -0.859 | -2.287 | -1.11 | -1.688 | 0.02 | 0.02 | 0.02 | 1.533 | 0.3 | 0.74 | 0.25 | 0.11 | 0.16 | 0.27 |
| 7 | 140415661 | rs2930325 | 1.421 | -1.743 | -1.188 | -1.468 | -0.002 | 0.054 | 0.054 | 0 | 0.1 | 0.01 | 0.04 | 0.01 | 0.07 | 0.06 |
| 7 | 140432893 | rs11771946 | 0.601 | -1.791 | -1.094 | -1.465 | 0.01 | 0.014 | 0.014 | 0 | 0.1 | 0.06 | 0.04 | 0 | 0.07 | 0.05 |
| 7 | 140473446 | rs1267638 | 0.513 | -2.288 | -0.203 | -1.607 | -0.023 | 0.001 | 0.001 | 0 | 0.13 | 0.21 | 0.07 | 0.1 | 0.08 | 0.21 |
| 7 | 140499107 | rs10487888 | 2.219 | -2.212 | -0.725 | -1.867 | -0.184 | 0.053 | 0.053 | 0.537 | 0.33 | 0.03 | 0.31 | 0.16 | 0.54 | 0.15 |
| 7 | 140505189 | rs1476868 | 0.979 | -2.597 | -0.172 | -1.904 | 0.024 | 0.011 | 0.011 | 0 | 0.14 | 0.01 | 0.04 | 0.08 | 0.07 | 0.11 |
| 7 | 140544165 | rs1267601 | 0.743 | -2.608 | -0.152 | -1.907 | -0.022 | 0.003 | 0.003 | 0 | 0.12 | 0.21 | 0.07 | 0.1 | 0.08 | 0.21 |
| 7 | 140556167 | rs1267621 | -1.066 | -2.612 | -0.197 | -1.919 | -0.04 | -0.017 | -0.017 | 0.168 | 0.3 | 0.39 | 0.12 | 0.18 | 0.15 | 0.33 |
| 7 | 140563901 | rs17695707 | 0.965 | -2.563 | -0.265 | -1.877 | 0.029 | 0.018 | 0.018 | 0 | 0.14 | 0.03 | 0.04 | 0.08 | 0.07 | 0.12 |
| 7 | 140585871 | rs6954652 | -1.402 | -2.671 | -0.162 | -1.928 | 0.063 | 0.045 | 0.045 | 0.469 | 0.37 | 0.75 | 0.16 | 0.18 | 0.15 | 0.33 |
| 7 | 140592227 | rs17623382 | 2.614 | -2.592 | -0.539 | -2.264 | -0.028 | 0.056 | 0.056 | 1.206 | 0.1 | 0.01 | 0.08 | 0 | 0.12 | 0.01 |
| 7 | 140603652 | rs4726020 | -2.355 | -3.014 | -0.022 | -1.775 | 0.166 | 0.191 | 0.191 | 0.564 | 0.48 | 0.93 | 0.32 | 0.18 | 0.24 | 0.4 |
| 7 | 140625102 | rs7810757 | -0.822 | -2.63 | 0.353 | -1.611 | -0.07 | -0.067 | -0.067 | 0.032 | 0.22 | 0.36 | 0.08 | 0.08 | 0.08 | 0.12 |
| 7 | 140650632 | rs801088 | -1.784 | -2.599 | 0.481 | -1.619 | -0.057 | -0.145 | -0.145 | 0 | 0.2 | 0.5 | 0.19 | 0.01 | 0.09 | 0.08 |
| 7 | 140651523 | rs801089 | -1.769 | -2.584 | 0.291 | -1.612 | 0.009 | -0.004 | -0.004 | 0 | 0.35 | 0.64 | 0.24 | 0.09 | 0.16 | 0.19 |
| 7 | 140658283 | rs2533307 | -0.832 | -1.756 | 0.051 | -1.257 | -0.004 | -0.052 | -0.052 | 1.595 | 0.11 | 0.18 | 0.16 | 0 | 0.08 | 0.06 |
| 7 | 140690296 | rs246744 | 0.395 | -1.541 | 0.018 | -1.181 | 0.008 | -0.038 | -0.038 | 0 | 0.9 | 0.88 | 0.84 | 1 | 0.86 | 0.92 |
| 7 | 140690655 | rs9886312 | 1.753 | -2.119 | 0.32 | -1.366 | 0.006 | 0.007 | 0.007 | 0.035 | 0.61 | 0.36 | 0.7 | 0.82 | 0.73 | 0.58 |
| 7 | 140695812 | rs38723 | 0.216 | -2.128 | 0.251 | -1.375 | 0.029 | 0.026 | 0.026 | 0 | 0.15 | 0.05 | 0.05 | 0.09 | 0.06 | 0.1 |
| 7 | 140696374 | rs10952364 | 1.449 | -2.132 | 0.394 | -1.378 | -0.03 | -0.096 | -0.096 | 0 | 0.71 | 0.55 | 0.85 | 0.9 | 0.79 | 0.81 |
| 7 | 140710848 | rs38739 | 0.248 | -2.109 | 0.209 | -1.322 | 0.032 | 0.028 | 0.028 | 0 | 0.15 | 0.04 | 0.05 | 0.09 | 0.07 | 0.1 |

| | | | | | | | | | | | | | | | | |
|---|---|---|---|---|---|---|---|---|---|---|---|---|---|---|---|---|
| 7 | 140712618 | rs13236704 | 1.523 | -2.144 | 0.38 | -1.389 | -0.027 | -0.094 | -0.094 | 0.73 | 0.71 | 0.55 | 0.86 | 0.9 | 0.79 | 0.81 |
| 7 | 140714283 | rs38741 | 0.377 | -2.07 | 0.447 | -1.355 | 0.015 | -0.053 | -0.053 | 2.216 | 0.89 | 0.83 | 0.93 | 1 | 0.86 | 0.92 |
| 7 | 140754785 | rs11764061 | 0.122 | -2.026 | 0.508 | -1.281 | 0.059 | -0.026 | -0.026 | 0 | 0.05 | 0.13 | 0.05 | 0 | 0.12 | 0.05 |
| 7 | 140758393 | rs4726225 | 1.65 | -2.341 | 0.521 | -1.427 | -0.039 | -0.103 | -0.103 | 2.284 | 0.68 | 0.54 | 0.85 | 0.91 | 0.79 | 0.8 |
| 7 | 140761247 | rs471817 | 0.974 | -2.329 | 0.453 | -1.404 | -0.017 | -0.021 | -0.021 | 0 | 0.74 | 0.7 | 0.91 | 0.91 | 0.91 | 0.86 |
| 7 | 140769459 | rs528957 | 0.974 | -2.329 | 0.453 | -1.403 | -0.017 | -0.023 | -0.023 | 0 | 0.74 | 0.7 | 0.91 | 0.92 | 0.91 | 0.86 |
| 7 | 140775825 | rs498933 | 1.962 | -2.494 | 0.4 | -1.356 | -0.012 | -0.017 | -0.017 | 0 | 0.59 | 0.46 | 0.8 | 0.78 | 0.73 | 0.63 |
| 7 | 140779740 | rs557962 | 0.109 | -2.479 | 0.294 | -1.33 | 0.043 | 0.052 | 0.052 | 0 | 0.81 | 0.95 | 0.94 | 0.91 | 0.92 | 0.86 |
| 7 | 140791820 | rs4726268 | 0.015 | -2.444 | 0.3 | -1.281 | -0.006 | -0.002 | -0.002 | 0 | 0.86 | 0.91 | 0.92 | 1 | 0.83 | 0.88 |
| 7 | 140799480 | rs7805034 | -1.721 | -1.92 | 0.377 | -1.056 | -0.059 | -0.02 | -0.02 | 0 | 0.28 | 0.52 | 0.14 | 0.23 | 0.16 | 0.27 |
| 7 | 140800690 | rs10277943 | 0.357 | -1.902 | 0.318 | -1.039 | -0.021 | 0.021 | 0.021 | 0 | 0.1 | 0.21 | 0.05 | 0.14 | 0.06 | 0.19 |
| 7 | 140803520 | rs6464330 | -1.451 | -1.756 | 0.069 | -0.988 | -0.02 | -0.032 | -0.032 | 0 | 0.33 | 0.49 | 0.18 | 0.22 | 0.27 | 0.33 |
| 7 | 140812939 | rs10215085 | -1.202 | -1.256 | 0.043 | -0.479 | 0.029 | -0.002 | -0.002 | 0 | 0.14 | 0.37 | 0.23 | 0.17 | 0.15 | 0.26 |
| 7 | 140813017 | rs4725499 | 0.811 | -1.398 | -1.326 | -0.378 | -0.174 | 0.009 | 0.009 | 0 | 0.3 | 0.03 | 0.39 | 0.16 | 0.48 | 0.17 |
| 7 | 140813472 | rs17161914 | 0.322 | -1.695 | -0.9 | -0.094 | 0.013 | -0.001 | -0.001 | 0.055 | 0.14 | 0.06 | 0.04 | 0.13 | 0.06 | 0.21 |
| 7 | 140815840 | rs17161916 | -0.788 | -1.885 | -1 | -0.058 | 0.009 | 0.006 | 0.006 | 0 | 0.29 | 0.16 | 0.26 | 0.31 | 0.22 | 0.47 |
| 7 | 140823118 | rs2270214 | -1.039 | -2.287 | -0.599 | -0.129 | -0.019 | 0.001 | 0.001 | 0 | 0.36 | 0.45 | 0.28 | 0.41 | 0.23 | 0.56 |
| 7 | 140831341 | rs10242784 | -1.62 | -2.006 | -0.866 | -0.247 | 0 | 0.034 | 0.034 | 0.084 | 0.94 | 1 | 0.97 | 0.99 | 0.96 | 0.98 |
| 7 | 140835959 | rs10232772 | 1.24 | -2.041 | -1.039 | -0.504 | -0.127 | 0.112 | 0.112 | 0.002 | 0.28 | 0.07 | 0.39 | 0.06 | 0.47 | 0.11 |
| 7 | 140837624 | rs10952458 | 0.24 | -2.24 | -0.212 | 0.198 | -0.005 | 0.001 | 0.001 | 0 | 0.85 | 0.88 | 0.76 | 0.83 | 0.82 | 0.74 |
| 7 | 140855094 | rs4726342 | 1.194 | -2.423 | -0.012 | 0.429 | 0.021 | -0.02 | -0.02 | 0 | 0.65 | 0.83 | 0.71 | 0.53 | 0.74 | 0.41 |
| 7 | 140860574 | rs6976212 | 1.053 | -2.409 | -0.114 | 0.426 | 0.016 | -0.026 | -0.026 | 0 | 0.66 | 0.87 | 0.71 | 0.55 | 0.74 | 0.41 |
| 7 | 140865752 | rs1005281 | 1.276 | -2.589 | -0.058 | 0.386 | -0.007 | -0.001 | -0.001 | 0 | 0.61 | 0.59 | 0.69 | 0.54 | 0.75 | 0.4 |
| 7 | 140870388 | rs11768129 | 0.122 | -2.26 | -0.153 | 0.616 | -0.013 | -0.011 | -0.011 | 0 | 0.08 | 0.05 | 0.19 | 0.17 | 0.13 | 0.25 |
| 7 | 140903256 | rs10952496 | 0.883 | -2.127 | -0.116 | 0.673 | 0.003 | -0.002 | -0.002 | 0 | 0.6 | 0.63 | 0.68 | 0.49 | 0.71 | 0.4 |
| 7 | 140906152 | rs2041535 | 0.937 | -2.131 | -0.287 | 0.674 | -0.021 | 0.004 | 0.004 | 0 | 0.6 | 0.56 | 0.69 | 0.49 | 0.72 | 0.39 |
| 7 | 140913710 | rs7789917 | 0.902 | -2.226 | -0.286 | 0.632 | -0.015 | 0.002 | 0.002 | 0 | 0.56 | 0.48 | 0.65 | 0.42 | 0.71 | 0.36 |
| 7 | 140914659 | rs7790291 | 0.577 | -1.844 | 0.245 | 0.261 | -0.027 | -0.013 | -0.013 | 0 | 0.7 | 0.5 | 0.72 | 0.72 | 0.78 | 0.46 |
| 7 | 140917556 | rs6959814 | 0.727 | -1.46 | -0.295 | -0.115 | 0.028 | 0.035 | 0.035 | 0 | 0.48 | 0.37 | 0.36 | 0.71 | 0.39 | 0.36 |
| 7 | 140918672 | rs6979775 | 0.167 | -0.442 | 0.158 | 0.155 | -0.002 | 0.024 | 0.024 | 0 | 0.84 | 0.72 | 0.75 | 0.81 | 0.86 | 0.7 |
| 7 | 140918819 | rs1005461 | 0.098 | -0.433 | 0.172 | -0.049 | -0.001 | 0.018 | 0.018 | 0 | 0.84 | 0.74 | 0.75 | 0.82 | 0.86 | 0.71 |
| 7 | 140923414 | rs11978917 | -0.031 | -0.031 | -0.108 | 0.007 | -0.021 | 0.015 | 0.015 | 1.555 | 0.25 | 0.16 | 0.35 | 0.14 | 0.39 | 0.16 |
| 7 | 140923630 | rs4637730 | 0.785 | -0.111 | -0.047 | 0.033 | 0.001 | 0.01 | 0.01 | 1.955 | 0.62 | 0.77 | 0.4 | 0.68 | 0.43 | 0.58 |
| 7 | 140930833 | rs13236601 | -0.147 | -0.057 | -0.249 | 0.073 | 0.002 | -0.004 | -0.004 | 0 | 0.05 | 0.03 | 0.12 | 0.15 | 0.04 | 0.08 |
| 7 | 140936088 | rs1024722 | -0.411 | -0.021 | -0.275 | 0.067 | -0.007 | -0.017 | -0.017 | 0 | 0.87 | 0.93 | 0.76 | 0.81 | 0.84 | 0.74 |
| 7 | 140936468 | rs1016665 | -0.296 | -0.017 | -0.276 | 0.025 | -0.013 | -0.017 | -0.017 | 0 | 0.87 | 0.93 | 0.76 | 0.81 | 0.82 | 0.74 |
| 7 | 140937078 | rs4726432 | 1.116 | 0.012 | 0.026 | -0.069 | 0.008 | 0.01 | 0.01 | 0.587 | 0.56 | 0.38 | 0.37 | 0.65 | 0.43 | 0.54 |
| 7 | 140946310 | rs17162012 | -0.575 | -0.221 | -0.033 | -0.067 | -0.007 | 0.013 | 0.013 | 0 | 0.07 | 0.11 | 0.12 | 0.15 | 0.04 | 0.08 |
| 7 | 140954613 | rs4726433 | 0.109 | -0.372 | -0.001 | -0.219 | 0.004 | -0.107 | -0.107 | 0 | 0.84 | 0.64 | 0.88 | 0.99 | 0.88 | 0.84 |
| 7 | 140955139 | rs10256379 | 1.098 | -0.033 | -0.068 | -0.168 | 0.001 | 0.002 | 0.002 | 1.141 | 0.57 | 0.42 | 0.38 | 0.65 | 0.46 | 0.57 |
| 7 | 140964847 | rs4726441 | 2.078 | -0.578 | -0.627 | -0.229 | -0.034 | -0.175 | -0.175 | 0 | 0.38 | 0.02 | 0.3 | 0.51 | 0.39 | 0.47 |
| 7 | 140970382 | rs216996 | -0.198 | 0.374 | 0.171 | -0.326 | -0.109 | 0.013 | 0.013 | 1.466 | 0.75 | 0.85 | 0.63 | 0.8 | 0.58 | 0.83 |
| 7 | 140973781 | rs216997 | -0.727 | 0.137 | 0.679 | -0.45 | -0.017 | -0.103 | -0.103 | 0.412 | 0.16 | 0.34 | 0.11 | 0.01 | 0.1 | 0.17 |
| 7 | 140979634 | rs217002 | -0.705 | 0.134 | 0.607 | -0.429 | -0.017 | -0.087 | -0.087 | 2.574 | 0.15 | 0.29 | 0.11 | 0.01 | 0.1 | 0.17 |
| 7 | 140979971 | rs10276017 | -0.773 | 0.127 | 0.198 | -0.35 | -0.006 | -0.027 | -0.027 | 1.278 | 0.1 | 0.11 | 0.04 | 0 | 0.04 | 0.05 |
| 7 | 140980634 | rs217003 | 1.36 | 0.326 | 0.328 | -0.189 | -0.002 | -0.003 | -0.003 | 0 | 0.88 | 0.82 | 0.88 | 0.85 | 0.91 | 0.87 |
| 7 | 140981282 | rs217004 | 1.589 | 0.21 | 0.135 | -0.372 | 0.003 | -0.013 | -0.013 | 0 | 0.86 | 0.91 | 0.94 | 1 | 0.91 | 0.94 |
| 7 | 140983999 | rs217007 | 0.132 | 0.071 | 0.571 | -0.516 | -0.021 | -0.058 | -0.058 | 0 | 0.1 | 0.26 | 0.09 | 0 | 0.04 | 0.14 |
| 7 | 140984375 | rs11973211 | -0.384 | 0.578 | 0.401 | -0.454 | -0.036 | -0.022 | -0.022 | 0.69 | 0.27 | 0.19 | 0.48 | 0.36 | 0.42 | 0.25 |
| 7 | 140988611 | rs217013 | -1.031 | 0.822 | 0.424 | -0.482 | 0.018 | 0.003 | 0.003 | 1.368 | 0.37 | 0.52 | 0.57 | 0.37 | 0.46 | 0.42 |
| 7 | 140991143 | rs217020 | -0.03 | 0.687 | 0.44 | -0.44 | -0.03 | -0.035 | -0.035 | 0 | 0.12 | 0.26 | 0.1 | 0.05 | 0.06 | 0.17 |
| 7 | 140996913 | rs7790976 | 1.442 | 0.668 | 0.31 | -0.382 | 0.027 | 0.015 | 0.015 | 0.644 | 0.52 | 0.39 | 0.37 | 0.62 | 0.44 | 0.51 |
| 7 | 141004310 | rs6967493 | -1.822 | 0.331 | 0.324 | -0.498 | -0.001 | -0.007 | -0.007 | 0 | 0.15 | 0.17 | 0.08 | 0.11 | 0.12 | 0.2 |
| 7 | 141006004 | rs217034 | 1.16 | 1.135 | 0.396 | -0.563 | -0.026 | 0.03 | 0.03 | 0.077 | 0.71 | 0.75 | 0.8 | 0.88 | 0.65 | 0.75 |
| 7 | 141007010 | rs2057915 | 0.858 | 1.164 | 0.274 | -0.55 | -0.108 | 0.091 | 0.091 | 0.271 | 0.73 | 0.93 | 0.82 | 0.88 | 0.65 | 0.77 |
| 7 | 141007717 | rs7458320 | -1.313 | 1.266 | 0.46 | -0.596 | -0.002 | -0.006 | -0.006 | 0 | 0.34 | 0.59 | 0.44 | 0.27 | 0.32 | 0.41 |
| 7 | 141008038 | rs17489724 | 1.02 | 1.311 | 0.386 | -0.571 | -0.051 | 0.063 | 0.063 | 0.209 | 0.13 | 0.03 | 0.11 | 0.01 | 0.2 | 0.05 |
| 7 | 141008377 | rs2158680 | -1.228 | 1.318 | 0.569 | -0.561 | 0.008 | -0.012 | -0.012 | 0 | 0.34 | 0.39 | 0.4 | 0.17 | 0.37 | 0.36 |
| 7 | 141012015 | rs7797098 | 1.158 | 1.22 | 0.509 | -0.463 | -0.042 | 0.012 | 0.012 | 0 | 0.86 | 0.96 | 0.93 | 0.89 | 0.85 | 0.82 |
| 7 | 141012313 | rs2040929 | 1.813 | 1.144 | 0.655 | -0.482 | -0.001 | 0.012 | 0.012 | 0 | 0.68 | 0.48 | 0.61 | 0.64 | 0.75 | 0.5 |
| 7 | 141012923 | rs17162075 | 0.378 | -0.038 | 0.726 | -0.364 | -0.013 | 0.077 | 0.077 | 0 | 0.13 | 0.33 | 0.26 | 0.16 | 0.09 | 0.22 |
| 7 | 141017055 | rs10254214 | 0.032 | 0.123 | -0.017 | -0.491 | -0.042 | 0.065 | 0.065 | 0 | 0.19 | 0.07 | 0.13 | 0.02 | 0.24 | 0.13 |
| 7 | 141017643 | rs2366081 | -1.333 | 0.038 | 0.158 | -0.273 | -0.023 | 0.035 | 0.035 | 0 | 0.27 | 0.62 | 0.37 | 0.4 | 0.17 | 0.43 |
| 7 | 141020747 | rs1557973 | -1.778 | 0.135 | 0.143 | -0.256 | 0.024 | 0.017 | 0.017 | 0 | 0.5 | 0.76 | 0.5 | 0.43 | 0.4 | 0.56 |
| 7 | 141021265 | rs7807179 | -1.258 | 0.07 | 0.178 | -0.214 | -0.009 | 0.006 | 0.006 | 0 | 0.3 | 0.66 | 0.37 | 0.41 | 0.17 | 0.42 |
| 7 | 141024444 | rs6980372 | -1.591 | 0.301 | 0.207 | -0.113 | -0.021 | 0.064 | 0.064 | 0.176 | 0.14 | 0.27 | 0.1 | 0.27 | 0.07 | 0.22 |
| 7 | 141025713 | rs10258837 | -1.545 | 0.339 | 0.238 | -0.142 | 0.001 | -0.01 | -0.01 | 0 | 0.34 | 0.37 | 0.24 | 0.29 | 0.31 | 0.35 |
| 7 | 141030552 | rs6975836 | -1.3 | 0.338 | 0.454 | 0.039 | 0.015 | 0.037 | 0.037 | 1.826 | 0.06 | 0.26 | 0.05 | 0.09 | 0.02 | 0.1 |
| 7 | 141040182 | rs917158 | -1.339 | 0.507 | -0.329 | -0.095 | -0.014 | 0.003 | 0.003 | 0.143 | 0.53 | 0.78 | 0.53 | 0.44 | 0.45 | 0.58 |

| | | | | | | | | | | | | | | | | |
|---|---|---|---|---|---|---|---|---|---|---|---|---|---|---|---|---|
| 7 | 141047961 | rs17490790 | -0.324 | 1.289 | -0.263 | -0.191 | -0.002 | 0.001 | 0.001 | 0.191 | 0.15 | 0.12 | 0.19 | 0.08 | 0.2 | 0.14 |
| 7 | 141072134 | rs1859575 | -2.112 | 1.255 | -0.029 | -0.392 | 0.01 | -0.033 | -0.033 | 1.3 | 0.33 | 0.41 | 0.29 | 0.23 | 0.36 | 0.39 |
| 7 | 141084969 | rs10264996 | -1.341 | 0.719 | 0.246 | -0.606 | 0.006 | -0.02 | -0.02 | 0.009 | 0.07 | 0.11 | 0.08 | 0.03 | 0.07 | 0.13 |
| 7 | 141085514 | rs10215677 | -1.934 | 0.678 | 0.227 | -0.664 | -0.003 | -0.106 | -0.106 | 0 | 0.27 | 0.4 | 0.23 | 0.06 | 0.25 | 0.26 |
| 7 | 141085654 | rs10258537 | -1.923 | 0.678 | 0.227 | -0.664 | -0.003 | -0.106 | -0.106 | 0.112 | 0.27 | 0.4 | 0.23 | 0.06 | 0.25 | 0.22 |
| 7 | 141087803 | rs10277430 | -0.174 | 0.721 | -0.223 | -0.644 | -0.013 | 0.053 | 0.053 | 0 | 0.14 | 0.06 | 0.12 | 0.02 | 0.17 | 0.06 |
| 7 | 141088498 | rs10259042 | -1.286 | 0.74 | -0.224 | -0.633 | 0.007 | -0.02 | -0.02 | 0 | 0.07 | 0.13 | 0.08 | 0.03 | 0.07 | 0.13 |
| 7 | 141091477 | rs10261257 | -1.831 | 0.739 | -0.213 | -0.581 | -0.002 | -0.056 | -0.056 | 0.054 | 0.26 | 0.32 | 0.23 | 0.08 | 0.25 | 0.25 |
| 7 | 141093612 | rs10266594 | -2.659 | 0.551 | 0.04 | -0.109 | 0 | -0.028 | -0.028 | 0.593 | 0.11 | 0.19 | 0.1 | 0.07 | 0.09 | 0.15 |
| 7 | 141101902 | rs11983421 | -1.612 | 0.491 | -0.212 | -0.085 | -0.005 | -0.027 | -0.027 | 0 | 0.06 | 0.18 | 0.03 | 0 | 0.02 | 0.03 |
| 7 | 141106552 | rs11764687 | -1.955 | 0.655 | -0.432 | 0.062 | -0.002 | -0.011 | -0.011 | 0 | 0.08 | 0.11 | 0.09 | 0.06 | 0.06 | 0.12 |
| 7 | 141109797 | rs2077023 | -1.235 | 0.746 | -0.68 | 0.128 | -0.016 | -0.002 | -0.002 | 0.094 | 0.05 | 0 | 0.07 | 0.06 | 0.05 | 0.11 |
| 7 | 141110794 | rs1181724 | -0.694 | 0.97 | -0.67 | -0.428 | -0.001 | -0.004 | -0.004 | 0 | 0.24 | 0.18 | 0.15 | 0.15 | 0.2 | 0.14 |
| 7 | 141110923 | rs35122961 | -0.458 | 0.505 | -0.97 | -0.063 | 0.026 | 0.005 | 0.005 | 0 | 0.46 | 0.31 | 0.39 | 0.44 | 0.36 | 0.46 |
| 7 | 141116780 | rs9969247 | 0.084 | 0.871 | -1.368 | 0.055 | -0.075 | -0.06 | -0.06 | 0 | 0.23 | 0.05 | 0.21 | 0.34 | 0.33 | 0.23 |
| 7 | 141117744 | rs1558211 | -0.171 | 0.666 | -0.894 | -0.021 | -0.014 | -0.01 | -0.01 | 1.719 | 0.26 | 0.19 | 0.22 | 0.34 | 0.33 | 0.24 |
| 7 | 141119143 | rs11760722 | 0.463 | 0.513 | -0.644 | -0.197 | -0.008 | 0.023 | 0.023 | 0 | 0.29 | 0.34 | 0.46 | 0.38 | 0.29 | 0.37 |
| 7 | 141119428 | rs1181728 | -0.554 | 0.465 | -0.522 | -0.314 | -0.009 | 0.08 | 0.08 | 0.028 | 0.57 | 0.82 | 0.71 | 0.71 | 0.62 | 0.61 |
| 7 | 141124286 | rs1181729 | 0.341 | -0.207 | 0.167 | -0.134 | -0.004 | -0.008 | -0.008 | 0 | 0.22 | 0.3 | 0.15 | 0.18 | 0.21 | 0.15 |
| 7 | 141125596 | rs1181731 | -1.145 | -0.535 | 0.092 | -0.283 | -0.003 | -0.003 | -0.003 | 0 | 0.64 | 0.67 | 0.44 | 0.59 | 0.52 | 0.73 |
| 7 | 141130465 | rs736871 | 0.409 | -0.659 | -0.175 | -0.19 | -0.006 | 0.004 | 0.004 | 0 | 0.3 | 0.35 | 0.25 | 0.38 | 0.25 | 0.49 |
| 7 | 141130890 | rs10952500 | 1.193 | -0.541 | -0.478 | -0.114 | 0.019 | 0.026 | 0.026 | 0 | 0.08 | 0.02 | 0.03 | 0 | 0.04 | 0.06 |
| 7 | 141130932 | rs1181736 | 0.233 | -0.189 | -0.157 | 0.155 | -0.012 | -0.01 | -0.01 | 0 | 0.24 | 0.35 | 0.18 | 0.2 | 0.22 | 0.16 |
| 7 | 141132843 | rs1181739 | -0.226 | 0.581 | -0.571 | 0.465 | -0.065 | -0.065 | -0.065 | 0.469 | 0.64 | 0.82 | 0.45 | 0.58 | 0.51 | 0.71 |
| 7 | 141142060 | rs1181752 | -1.168 | 0.002 | 1.027 | 0.783 | -0.007 | 0.005 | 0.005 | 0.328 | 0.29 | 0.65 | 0.27 | 0.61 | 0.2 | 0.54 |
| 7 | 141142132 | rs1181753 | -0.846 | -0.102 | 1.267 | 0.8 | -0.014 | 0.037 | 0.037 | 0 | 0.24 | 0.34 | 0.24 | 0.59 | 0.18 | 0.52 |
| 7 | 141147241 | rs1181763 | -0.736 | 0.273 | 1.321 | 1.207 | 0.001 | -0.011 | -0.011 | 0.02 | 0.26 | 0.18 | 0.2 | 0.32 | 0.23 | 0.44 |
| 7 | 141149608 | rs1181768 | -0.388 | 1.205 | 1.537 | 1.365 | -0.004 | -0.005 | -0.005 | 0 | 0.83 | 0.89 | 0.7 | 0.72 | 0.79 | 0.87 |
| 7 | 141151394 | rs1181770 | 0.115 | 1.051 | 1.569 | 1.399 | 0.007 | 0.019 | 0.019 | 0 | 0.8 | 0.69 | 0.69 | 0.7 | 0.79 | 0.86 |
| 7 | 141152680 | rs1181775 | 0.453 | 1.056 | 1.208 | 1.403 | 0.006 | 0.031 | 0.031 | 0 | 0.77 | 0.57 | 0.68 | 0.7 | 0.79 | 0.86 |
| 7 | 141153209 | rs1181777 | 0.349 | 1.109 | 0.98 | 1.424 | 0.007 | 0.029 | 0.029 | 0 | 0.78 | 0.56 | 0.68 | 0.7 | 0.79 | 0.86 |
| 7 | 141172474 | rs10280774 | 0.812 | 0.84 | 0.649 | 1.89 | 0 | 0.017 | 0.017 | 0 | 0.87 | 0.84 | 0.84 | 0.64 | 0.86 | 0.57 |
| 7 | 141194823 | rs10224311 | -1.437 | 0.216 | 1.103 | 0.357 | 0.065 | 0.043 | 0.043 | 4.485 | 0.06 | 0.48 | 0.04 | 0.01 | 0.02 | 0.01 |
| 7 | 141211816 | rs1108250 | 0.498 | 0.256 | 0.091 | -0.389 | 0.004 | 0.066 | 0.066 | 0 | 0.46 | 0.36 | 0.3 | 0.09 | 0.5 | 0.27 |
| 7 | 141224952 | rs2366254 | -0.65 | 0.236 | 0.549 | 0.467 | 0.008 | 0.001 | 0.001 | 0.375 | 0.37 | 0.62 | 0.61 | 0.4 | 0.51 | 0.23 |
| 7 | 141241540 | rs1569085 | -0.663 | -0.019 | 0.804 | 0.677 | 0.008 | -0.007 | -0.007 | 0 | 0.36 | 0.43 | 0.56 | 0.6 | 0.47 | 0.51 |
| 7 | 141248475 | rs3757470 | -0.3 | -0.073 | 0.848 | 0.881 | -0.063 | -0.061 | -0.061 | 0.04 | 0.3 | 0.14 | 0.52 | 0.52 | 0.46 | 0.35 |
| 7 | 141287652 | rs4725559 | -0.247 | -0.083 | 0.926 | 0.862 | -0.063 | -0.063 | -0.063 | 0 | 0.29 | 0.14 | 0.52 | 0.53 | 0.46 | 0.36 |
| 7 | 141316834 | rs11773340 | -0.356 | -0.091 | 1.042 | 0.861 | -0.048 | -0.045 | -0.045 | 3.579 | 0.3 | 0.19 | 0.53 | 0.53 | 0.46 | 0.36 |
| 7 | 141329651 | rs12703398 | -0.669 | -0.09 | 1.095 | 0.862 | -0.005 | -0.003 | -0.003 | 0.489 | 0.34 | 0.35 | 0.56 | 0.58 | 0.48 | 0.51 |
| 7 | 141360585 | rs3800989 | -0.728 | 0.023 | 1.11 | 0.846 | 0.02 | 0.006 | 0.006 | 0.347 | 0.36 | 0.47 | 0.57 | 0.57 | 0.49 | 0.51 |
| 7 | 141363842 | rs12534759 | -0.397 | -0.437 | 0.328 | 0.742 | -0.001 | 0.018 | 0.018 | 0 | 0.11 | 0.08 | 0.14 | 0.33 | 0.07 | 0.21 |
| 7 | 141364712 | rs2286150 | -0.947 | -0.412 | 0.237 | 0.733 | 0.009 | -0.012 | -0.012 | 0 | 0.14 | 0.09 | 0.16 | 0.37 | 0.09 | 0.36 |
| 7 | 141389640 | rs7791469 | -1.113 | -0.354 | 0.094 | 0.55 | -0.011 | 0.057 | 0.057 | 0 | 0.45 | 0.29 | 0.55 | 0.74 | 0.51 | 0.59 |
| 7 | 141398707 | rs10261515 | 0.864 | -0.603 | 0.157 | 0.424 | 0 | 0.063 | 0.063 | 0 | 0.55 | 0.72 | 0.45 | 0.25 | 0.49 | 0.43 |
| 7 | 141398942 | rs13231650 | 0.087 | -0.608 | 0.201 | 0.665 | -0.048 | -0.056 | -0.056 | 0.056 | 0.71 | 0.84 | 0.48 | 0.49 | 0.55 | 0.74 |
| 7 | 141407716 | rs6963959 | 0.132 | -0.629 | 0.121 | 0.682 | -0.053 | -0.065 | -0.065 | 0 | 0.69 | 0.88 | 0.58 | 0.49 | 0.55 | 0.74 |
| 7 | 141413051 | rs768055 | 1.13 | -1.081 | 0.263 | -0.155 | 0 | -0.001 | -0.001 | 0 | 0.47 | 0.53 | 0.25 | 0.24 | 0.41 | 0.39 |
| 7 | 141421031 | rs6947065 | 1.025 | -1.263 | 0.233 | 0.251 | -0.003 | 0.068 | 0.068 | 0 | 0.55 | 0.74 | 0.45 | 0.26 | 0.51 | 0.45 |
| 7 | 141450588 | rs17162459 | 0.246 | -1.582 | 1.024 | 0.817 | 0.004 | -0.003 | -0.003 | 0 | 0.08 | 0.06 | 0.03 | 0.21 | 0.04 | 0.13 |
| 7 | 141456192 | rs12540335 | -1.068 | -1.599 | 0.875 | 0.591 | -0.004 | 0.075 | 0.075 | 0 | 0.46 | 0.17 | 0.52 | 0.67 | 0.46 | 0.43 |
| 7 | 141463914 | rs765007 | -1.383 | -1.602 | 0.91 | 0.589 | -0.001 | -0.006 | -0.006 | 1.349 | 0.52 | 0.5 | 0.6 | 0.75 | 0.47 | 0.6 |
| 7 | 141464765 | rs2270009 | -1.026 | -1.602 | 0.823 | 0.59 | -0.005 | 0.074 | 0.074 | 0 | 0.46 | 0.17 | 0.52 | 0.67 | 0.46 | 0.43 |
| 7 | 141478800 | rs2234002 | -1.283 | -1.596 | 0.811 | 0.621 | 0.006 | 0.127 | 0.127 | 0 | 0.49 | 0.19 | 0.57 | 0.75 | 0.47 | 0.6 |
| 7 | 141483508 | rs12539227 | -1.025 | -1.595 | 0.811 | 0.62 | -0.005 | 0.074 | 0.074 | 0 | 0.46 | 0.17 | 0.52 | 0.67 | 0.47 | 0.43 |
| 7 | 141484094 | rs2190245 | -1.025 | -1.595 | 0.811 | 0.62 | -0.005 | 0.074 | 0.074 | 0 | 0.46 | 0.17 | 0.52 | 0.67 | 0.47 | 0.43 |
| 7 | 141490107 | rs2234012 | -1.219 | -1.649 | 0.915 | 0.556 | 0.01 | 0.101 | 0.101 | 0 | 0.48 | 0.21 | 0.56 | 0.7 | 0.47 | 0.43 |
| 7 | 141490238 | rs2227264 | -1.112 | -1.373 | 0.99 | 0.602 | 0.005 | 0.093 | 0.093 | 0 | 0.47 | 0.21 | 0.56 | 0.7 | 0.47 | 0.43 |
| 7 | 141495604 | rs1859646 | -1.315 | -1.345 | 0.998 | 0.613 | 0.008 | 0.062 | 0.062 | 0 | 0.51 | 0.35 | 0.59 | 0.75 | 0.47 | 0.6 |
| 7 | 141532187 | rs10464444 | -0.475 | -1.022 | 0.728 | 0.665 | -0.013 | -0.054 | -0.054 | 0 | 0.29 | 0.13 | 0.39 | 0.46 | 0.29 | 0.3 |
| 7 | 141532934 | rs12703410 | -1.576 | -1.035 | 0.765 | 0.679 | 0.01 | 0.116 | 0.116 | 0 | 0.48 | 0.23 | 0.57 | 0.76 | 0.46 | 0.58 |
| 7 | 141535363 | rs11765852 | -0.417 | -1.034 | 0.745 | 0.683 | 0.02 | -0.035 | -0.035 | 0.747 | 0.17 | 0.07 | 0.13 | 0.22 | 0.16 | 0.18 |
| 7 | 141537541 | rs3800990 | -0.502 | -1.033 | 0.744 | 0.684 | 0.022 | -0.028 | -0.028 | 0 | 0.18 | 0.08 | 0.14 | 0.22 | 0.16 | 0.18 |
| 7 | 141537735 | rs12669721 | -1.67 | -1.039 | 0.801 | 0.679 | 0.012 | 0.076 | 0.076 | 0 | 0.49 | 0.35 | 0.58 | 0.76 | 0.46 | 0.58 |
| 7 | 141537968 | rs11765575 | -0.534 | -1.014 | 0.823 | 0.706 | -0.002 | -0.012 | -0.012 | 0.206 | 0.31 | 0.26 | 0.44 | 0.53 | 0.3 | 0.4 |
| 7 | 141543098 | rs9640357 | -0.534 | -1.011 | 0.807 | 0.702 | -0.002 | -0.014 | -0.014 | 0 | 0.31 | 0.27 | 0.4 | 0.5 | 0.3 | 0.4 |
| 7 | 141546847 | rs12703413 | -0.355 | -1.01 | 0.741 | 0.704 | -0.049 | -0.122 | -0.122 | 0.129 | 0.28 | 0.05 | 0.38 | 0.46 | 0.29 | 0.3 |
| 7 | 141563970 | rs1433594 | 1.479 | -0.977 | 0.817 | 0.673 | 0.008 | 0.038 | 0.038 | 0 | 0.52 | 0.62 | 0.47 | 0.32 | 0.55 | 0.53 |
| 7 | 141569606 | rs6959360 | -0.419 | -0.97 | 0.768 | 0.696 | -0.017 | -0.06 | -0.06 | 0 | 0.3 | 0.13 | 0.43 | 0.53 | 0.3 | 0.43 |

| | | | | | | | | | | | | | | | | |
|---|---|---|---|---|---|---|---|---|---|---|---|---|---|---|---|---|
| 7 | 141581325 | rs1285961 | 1.436 | -0.824 | 0.753 | 0.574 | 0.001 | -0.002 | -0.002 | 0 | 0.55 | 0.48 | 0.54 | 0.25 | 0.64 | 0.43 |
| 7 | 141589060 | rs1285958 | 0.213 | -0.907 | 0.494 | 0.755 | 0.018 | -0.043 | -0.043 | 0 | 0.83 | 0.95 | 0.87 | 0.77 | 0.84 | 0.83 |
| 7 | 141605084 | rs4726479 | 1.321 | -0.967 | 0.495 | 0.717 | 0.026 | -0.024 | -0.024 | 0 | 0.76 | 0.84 | 0.79 | 0.69 | 0.81 | 0.63 |
| 7 | 141607385 | rs2471744 | 0.336 | -0.973 | 0.484 | 0.717 | 0.019 | -0.024 | -0.024 | 3.105 | 0.82 | 0.86 | 0.87 | 0.76 | 0.84 | 0.84 |
| 7 | 141612621 | rs11767947 | 0.247 | -0.99 | 0.439 | 0.691 | -0.062 | -0.13 | -0.13 | 2.002 | 0.23 | 0.04 | 0.36 | 0.45 | 0.28 | 0.24 |
| 7 | 141612717 | rs6976028 | -0.453 | -0.99 | 0.465 | 0.693 | -0.029 | -0.084 | -0.084 | 0.033 | 0.29 | 0.07 | 0.39 | 0.45 | 0.29 | 0.31 |
| 7 | 141616506 | rs745162 | 1.564 | -0.969 | 0.523 | 0.674 | 0.012 | 0.075 | 0.075 | 0 | 0.51 | 0.66 | 0.42 | 0.24 | 0.53 | 0.39 |
| 7 | 141627149 | rs1285933 | 1.604 | -0.969 | 0.513 | 0.606 | 0.014 | 0.082 | 0.082 | 0 | 0.51 | 0.65 | 0.42 | 0.23 | 0.53 | 0.39 |
| 7 | 141628704 | rs11770855 | -0.141 | -0.7 | 0.817 | 0.934 | -0.065 | -0.142 | -0.142 | 1.005 | 0.23 | 0.04 | 0.35 | 0.46 | 0.28 | 0.24 |
| 7 | 141630267 | rs12539499 | -0.163 | -0.7 | 0.831 | 0.934 | -0.042 | -0.107 | -0.107 | 0.417 | 0.23 | 0.05 | 0.35 | 0.46 | 0.28 | 0.24 |
| 7 | 141636563 | rs1285950 | 1.243 | -0.768 | 1.143 | 0.97 | -0.005 | 0.021 | 0.021 | 0.904 | 0.65 | 0.49 | 0.52 | 0.47 | 0.68 | 0.56 |
| 7 | 141639589 | rs1285946 | 1.64 | -0.663 | 0.673 | 0.626 | -0.019 | 0.008 | 0.008 | 0 | 0.9 | 0.68 | 0.9 | 0.93 | 0.97 | 0.79 |
| 7 | 141653637 | rs7794708 | 0.738 | -0.561 | 0.227 | 0.845 | -0.003 | -0.004 | -0.004 | 0 | 0.7 | 0.79 | 0.68 | 0.69 | 0.69 | 0.7 |
| 7 | 141654892 | rs2570407 | 0.282 | -0.464 | -0.019 | 0.937 | -0.012 | -0.024 | -0.024 | 0.047 | 0.76 | 0.85 | 0.71 | 0.69 | 0.71 | 0.76 |
| 7 | 141656487 | rs11762634 | -0.241 | -0.416 | -0.378 | 0.856 | -0.059 | -0.09 | -0.09 | 4.227 | 0.23 | 0.04 | 0.32 | 0.34 | 0.29 | 0.24 |
| 7 | 141672604 | rs10246939 | -0.482 | 0.29 | 0.277 | 0.209 | 0 | -0.009 | -0.009 | 0 | 0.42 | 0.48 | 0.69 | 0.68 | 0.46 | 0.36 |
| 7 | 141672705 | rs1726866 | 0.542 | 0.268 | -0.033 | 0.21 | 0.003 | 0.032 | 0.032 | 0.034 | 0.56 | 0.33 | 0.29 | 0.32 | 0.54 | 0.64 |
| 7 | 141686694 | rs4726484 | -0.853 | 1.607 | -0.791 | 0.086 | -0.098 | 0.038 | 0.038 | 0.133 | 0.19 | 0.05 | 0.19 | 0.08 | 0.31 | 0.24 |
| 7 | 141687327 | rs10225228 | -1.107 | 0.394 | 0.269 | 0.029 | -0.001 | -0.032 | -0.032 | 0 | 0.08 | 0.29 | 0.06 | 0 | 0.03 | 0.04 |
| 7 | 141692808 | rs6947481 | -0.808 | -0.247 | -0.035 | 0.428 | -0.045 | 0.044 | 0.044 | 0 | 0.25 | 0.43 | 0.25 | 0.31 | 0.14 | 0.23 |
| 7 | 141701356 | rs1285938 | -0.106 | 0.027 | -0.838 | -0.021 | -0.019 | 0.115 | 0.115 | 0 | 0.48 | 0.79 | 0.51 | 0.82 | 0.51 | 0.54 |
| 7 | 141705540 | rs1285965 | 0.572 | -0.453 | -0.069 | -0.013 | -0.035 | 0.008 | 0.008 | 0 | 0.37 | 0.25 | 0.45 | 0.68 | 0.5 | 0.46 |
| 7 | 141711308 | rs10248676 | 0.53 | -0.617 | 0.717 | 0.008 | 0.015 | -0.001 | -0.001 | 0 | 0.36 | 0.38 | 0.46 | 0.67 | 0.49 | 0.45 |
| 7 | 141712467 | rs11764581 | 0.853 | -0.17 | 0.229 | 0.035 | -0.03 | -0.022 | -0.022 | 0 | 0.07 | 0.03 | 0.16 | 0.18 | 0.12 | 0.09 |
| 7 | 141724799 | rs10270832 | 0.143 | -0.329 | 0.568 | -0.039 | -0.014 | 0.008 | 0.008 | 1.888 | 0.06 | 0.15 | 0.02 | 0.12 | 0.01 | 0.01 |
| 7 | 141724852 | rs10255702 | 0.089 | -0.33 | 0.567 | -0.043 | -0.014 | 0.006 | 0.006 | 2.112 | 0.06 | 0.15 | 0.02 | 0.12 | 0.01 | 0.01 |
| 7 | 141725973 | rs2960748 | 0.528 | -0.294 | 0.849 | 0.068 | 0.008 | 0.035 | 0.035 | 0 | 0.08 | 0.12 | 0.11 | 0.18 | 0.11 | 0.09 |
| 7 | 141727526 | rs2272330 | 0.074 | -0.649 | 0.738 | -0.183 | -0.014 | 0.007 | 0.007 | 1.888 | 0.06 | 0.15 | 0.02 | 0.12 | 0.01 | 0.01 |
| 7 | 141728209 | rs10225000 | -0.681 | -0.679 | 0.969 | -0.138 | -0.02 | 0.035 | 0.035 | 6.452 | 0.07 | 0.27 | 0.04 | 0.12 | 0.01 | 0.01 |
| 7 | 141728622 | rs12540186 | 0.181 | -1.824 | 0.472 | -0.801 | 0 | -0.016 | -0.016 | 0 | 0.19 | 0.17 | 0.26 | 0.01 | 0.22 | 0.19 |
| 7 | 141728635 | rs4607527 | -0.225 | -3.04 | 0.376 | -0.335 | -0.009 | -0.004 | -0.004 | 0 | 0.37 | 0.4 | 0.23 | 0.26 | 0.19 | 0.33 |
| 7 | 141729628 | rs6956284 | 0.326 | -2.744 | 0.117 | -0.227 | -0.001 | -0.005 | -0.005 | 0.029 | 0.55 | 0.45 | 0.56 | 0.8 | 0.59 | 0.62 |
| 7 | 141731207 | rs11761379 | 0.607 | -2.669 | -0.22 | -0.334 | 0.027 | 0.05 | 0.05 | 0 | 0.27 | 0.05 | 0.15 | 0.18 | 0.2 | 0.18 |
| 7 | 141731340 | rs4374898 | 1.122 | -2.975 | -0.031 | -0.222 | -0.011 | -0.015 | -0.015 | 0 | 0.32 | 0.26 | 0.42 | 0.6 | 0.4 | 0.49 |
| 7 | 141733848 | rs4416750 | -0.848 | -2.907 | -0.341 | -0.189 | -0.128 | -0.004 | -0.004 | 0 | 0.73 | 0.94 | 0.65 | 0.79 | 0.63 | 0.7 |
| 7 | 141735114 | rs7811738 | 0.027 | -3.83 | 0.692 | 0.082 | 0.037 | 0.019 | 0.019 | 0 | 0.28 | 0.08 | 0.2 | 0.2 | 0.21 | 0.18 |
| 7 | 141736798 | rs4132774 | -0.463 | -3.237 | 0.367 | 0.369 | -0.025 | -0.023 | -0.023 | 0.089 | 0.41 | 0.49 | 0.26 | 0.2 | 0.24 | 0.22 |
| 7 | 141761438 | rs7810984 | -0.789 | -1.841 | 0.499 | 0.9 | -0.055 | 0.222 | 0.222 | 0.081 | 0.15 | 0.37 | 0.06 | 0.47 | 0.05 | 0.37 |
| 7 | 141761754 | rs11762139 | -0.458 | -1.06 | -0.069 | 0.656 | 0.011 | 0.004 | 0.004 | 0 | 0.54 | 0.56 | 0.28 | 0.6 | 0.37 | 0.59 |
| 7 | 141774290 | rs4281037 | -1.513 | -0.336 | 0.195 | 0.681 | -0.032 | 0.005 | 0.005 | 0 | 0.09 | 0.26 | 0.04 | 0.04 | 0.04 | 0.13 |
| 7 | 141774390 | rs3922664 | -1.543 | -0.335 | 0.196 | 0.682 | -0.034 | 0.002 | 0.002 | 0 | 0.1 | 0.26 | 0.04 | 0.04 | 0.04 | 0.13 |
| 7 | 141808573 | rs4428591 | 1.325 | 0.141 | 1.519 | 0.927 | 0.039 | 0.068 | 0.068 | 0.301 | 0.91 | 0.62 | 0.91 | 0.9 | 0.93 | 0.93 |
| 7 | 141813014 | rs4593441 | -0.71 | -0.646 | 1.214 | -0.101 | -0.034 | -0.104 | -0.104 | 0 | 0.23 | 0.42 | 0.3 | 0.06 | 0.15 | 0.35 |
| 7 | 141816117 | rs6951410 | -1.208 | -0.552 | 0.249 | 0.132 | 0.02 | 0.054 | 0.054 | 1.162 | 0.46 | 0.82 | 0.55 | 0.37 | 0.4 | 0.45 |
| 7 | 141817070 | rs7788775 | -0.795 | -0.833 | 0.942 | 0.022 | -0.009 | 0 | 0 | 0.056 | 0.3 | 0.36 | 0.23 | 0.36 | 0.23 | 0.2 |
| 7 | 141819278 | rs10248368 | 0.45 | -0.788 | 0.802 | 0.032 | -0.011 | -0.06 | -0.06 | 0.125 | 0.17 | 0.07 | 0.16 | 0.36 | 0.18 | 0.14 |
| 7 | 141823702 | rs4628182 | 0.112 | -0.842 | 0.866 | 0.02 | -0.002 | 0.013 | 0.013 | 0 | 0.11 | 0.08 | 0.26 | 0.04 | 0.1 | 0.19 |
| 7 | 141825321 | rs7790470 | -1.323 | -0.774 | 0.864 | 0.065 | 0.056 | 0.035 | 0.035 | 1.625 | 0.44 | 0.83 | 0.46 | 0.41 | 0.34 | 0.45 |
| 7 | 141835869 | rs6944985 | -1.043 | -0.777 | 0.936 | 0.087 | -0.005 | -0.003 | -0.003 | 1.04 | 0.42 | 0.54 | 0.42 | 0.41 | 0.34 | 0.45 |
| 7 | 141845717 | rs10215277 | -1.072 | -1.45 | 0.873 | 0.364 | -0.023 | 0.032 | 0.032 | 0 | 0.25 | 0.39 | 0.3 | 0.41 | 0.25 | 0.38 |
| 7 | 141845864 | rs13224443 | -0.157 | -1.386 | 0.83 | 0.452 | 0.001 | 0.018 | 0.018 | 0 | 0.13 | 0.2 | 0.27 | 0.25 | 0.18 | 0.15 |
| 7 | 141847782 | rs6954522 | -0.158 | -0.911 | 0.017 | -0.394 | -0.023 | 0.05 | 0.05 | 0.012 | 0.43 | 0.33 | 0.59 | 0.35 | 0.49 | 0.49 |
| 7 | 141850305 | rs7780248 | 0.344 | -0.476 | -0.749 | 0.237 | -0.033 | -0.044 | -0.044 | 0 | 0.24 | 0.13 | 0.25 | 0.38 | 0.29 | 0.26 |
| 7 | 141850772 | rs13223215 | -0.808 | -0.722 | -0.342 | -0.067 | -0.008 | -0.027 | -0.027 | 0 | 0.89 | 0.88 | 0.88 | 0.97 | 0.89 | 0.92 |
| 7 | 141851610 | rs10270056 | -0.199 | 0.185 | 0.096 | 0.395 | -0.017 | 0.001 | 0.001 | 0 | 0.35 | 0.26 | 0.3 | 0.64 | 0.38 | 0.44 |
| 7 | 141852521 | rs6464466 | -0.023 | 0.016 | 0.005 | 0.125 | 0.019 | -0.066 | -0.066 | 0 | 0.7 | 0.62 | 0.58 | 0.92 | 0.6 | 0.66 |
| 7 | 141854659 | rs11770705 | 0.426 | -0.034 | -0.5 | 0.102 | 0.023 | -0.05 | -0.05 | 0 | 0.22 | 0.31 | 0.32 | 0.05 | 0.28 | 0.26 |
| 7 | 141855193 | rs10224453 | 1.986 | -0.102 | -0.472 | 0.123 | 0.043 | 0.02 | 0.02 | 0 | 0.3 | 0.17 | 0.26 | 0.28 | 0.23 | 0.21 |
| 7 | 141858688 | rs5024722 | 0.185 | -0.054 | -0.423 | 0.194 | -0.025 | 0.042 | 0.042 | 0.311 | 0.76 | 0.62 | 0.82 | 0.72 | 0.78 | 0.74 |
| 7 | 141860400 | rs12703439 | 1.941 | 0.099 | -1.028 | 0.164 | 0.043 | 0.019 | 0.019 | 0 | 0.3 | 0.17 | 0.26 | 0.28 | 0.23 | 0.21 |
| 7 | 141861235 | rs11771612 | 0.297 | 0.552 | -0.821 | -0.068 | 0.02 | -0.034 | -0.034 | 0 | 0.19 | 0.25 | 0.31 | 0.05 | 0.28 | 0.26 |
| 7 | 141861251 | rs11772365 | -0.192 | 0.547 | -0.787 | 0.007 | 0.033 | -0.019 | -0.019 | 0 | 0.23 | 0.33 | 0.32 | 0.13 | 0.33 | 0.28 |
| 7 | 141861458 | rs7808303 | 0.809 | 0.53 | -1.263 | -0.396 | 0.001 | -0.052 | -0.052 | 2.684 | 0.1 | 0 | 0.03 | 0.15 | 0.06 | 0.15 |
| 7 | 141861469 | rs11761774 | 0.102 | 0.793 | -1.039 | -0.491 | 0.02 | -0.034 | -0.034 | 0 | 0.19 | 0.25 | 0.31 | 0.05 | 0.28 | 0.26 |
| 7 | 141862840 | rs11526220 | -0.263 | -0.02 | -1.156 | 0.208 | 0.015 | 0.01 | 0.01 | 0.929 | 0.27 | 0.21 | 0.19 | 0.21 | 0.22 | 0.29 |
| 7 | 141872874 | rs9986802 | -1.014 | 0.285 | -1.877 | 0.516 | 0.011 | 0 | 0 | 0.009 | 0.49 | 0.8 | 0.5 | 0.59 | 0.43 | 0.63 |
| 7 | 141875267 | rs12532545 | -0.48 | 0.975 | -1.334 | 0.799 | -0.013 | -0.029 | -0.029 | 0 | 0.19 | 0.11 | 0.27 | 0.36 | 0.19 | 0.31 |
| 7 | 141888080 | rs10226440 | -0.278 | 0.49 | -1.33 | 0.001 | 0.016 | 0.059 | 0.059 | 0.291 | 0.37 | 0.68 | 0.27 | 0.15 | 0.31 | 0.22 |
| 7 | 141897595 | rs3110843 | -0.293 | 0.613 | -1.435 | 0.401 | 0.006 | -0.035 | -0.035 | 0 | 0.7 | 0.8 | 0.78 | 0.66 | 0.72 | 0.63 |

| | | | | | | | | | | | | | | | | |
|---|---|---|---|---|---|---|---|---|---|---|---|---|---|---|---|---|
| 7 | 141898023 | rs4260813 | 0.412 | 0.295 | -1.721 | 0.17 | -0.031 | -0.018 | -0.018 | 0.181 | 0.4 | 0.26 | 0.56 | 0.5 | 0.45 | 0.44 |
| 7 | 141898680 | rs2961065 | -0.438 | 0.352 | -0.946 | 0.187 | 0.002 | -0.007 | -0.007 | 3.53 | 0.69 | 0.76 | 0.77 | 0.67 | 0.72 | 0.64 |
| 7 | 141909140 | rs6977351 | 0.576 | 0.421 | 0.031 | -0.153 | 0 | -0.035 | -0.035 | 0 | 0.22 | 0.2 | 0.16 | 0 | 0.21 | 0.16 |
| 7 | 141937889 | rs6464473 | 0.08 | 0.004 | 0.436 | -0.679 | -0.015 | -0.134 | -0.134 | 0.065 | 0.63 | 0.54 | 0.73 | 0.98 | 0.69 | 0.74 |
| 7 | 141938235 | rs6973013 | -0.548 | 0.846 | 0.276 | -0.203 | 0.025 | 0.012 | 0.012 | 0 | 0.18 | 0.22 | 0.3 | 0.24 | 0.28 | 0.26 |
| 7 | 141941327 | rs7456100 | -0.636 | 0.81 | 0.302 | -0.148 | 0.011 | 0 | 0 | 0 | 0.23 | 0.25 | 0.19 | 0.23 | 0.31 | 0.3 |
| 7 | 141942895 | rs5012030 | 1.002 | -0.007 | 0.559 | -0.236 | 0 | 0.005 | 0.005 | 0 | 0.62 | 0.39 | 0.57 | 0.67 | 0.6 | 0.6 |
| 7 | 141943262 | rs4329193 | 1.052 | -0.191 | 0.399 | -0.494 | -0.015 | -0.005 | -0.005 | 0 | 0.34 | 0.12 | 0.34 | 0.66 | 0.31 | 0.37 |
| 7 | 141943552 | rs6944336 | 1.095 | -0.194 | 0.382 | -0.495 | -0.019 | -0.009 | -0.009 | 0 | 0.34 | 0.1 | 0.34 | 0.66 | 0.31 | 0.37 |
| 7 | 141943578 | rs17689193 | 1.113 | -0.198 | 0.44 | -0.506 | -0.001 | -0.007 | -0.007 | 0.365 | 0.28 | 0.25 | 0.22 | 0.01 | 0.29 | 0.23 |
| 7 | 141944150 | rs4256513 | 1.102 | -0.23 | 0.413 | -0.501 | -0.019 | -0.009 | -0.009 | 0 | 0.34 | 0.1 | 0.34 | 0.66 | 0.31 | 0.37 |
| 7 | 141945160 | rs6464476 | 0.542 | -0.508 | 0.708 | -0.425 | -0.037 | -0.021 | -0.021 | 0 | 0.14 | 0.26 | 0.22 | 0.09 | 0.1 | 0.1 |
| 7 | 141947082 | rs916011 | 0.802 | -0.482 | 0.675 | -0.373 | -0.002 | 0.007 | 0.007 | 0 | 0.63 | 0.37 | 0.59 | 0.7 | 0.6 | 0.62 |
| 7 | 141947398 | rs6966761 | 0.264 | -0.509 | 0.693 | -0.342 | -0.044 | -0.003 | -0.003 | 0 | 0.14 | 0.28 | 0.24 | 0.12 | 0.1 | 0.11 |
| 7 | 141948152 | rs12703448 | 0.13 | -0.444 | 0.498 | -0.493 | -0.008 | -0.022 | -0.022 | 0 | 0.47 | 0.47 | 0.59 | 0.79 | 0.35 | 0.47 |
| 7 | 141948871 | rs2960763 | 1.198 | -0.391 | 0.369 | -0.253 | 0.01 | -0.047 | -0.047 | 0 | 0.32 | 0.08 | 0.33 | 0.65 | 0.25 | 0.34 |
| 7 | 141950745 | rs7801392 | 1.36 | -0.433 | 0.316 | 0.15 | -0.03 | -0.175 | -0.175 | 0 | 0.3 | 0.02 | 0.33 | 0.51 | 0.25 | 0.33 |
| 7 | 141951780 | rs6963035 | 0.341 | -0.512 | 0.453 | 0.125 | -0.051 | 0.023 | 0.023 | 0 | 0.15 | 0.34 | 0.26 | 0.14 | 0.1 | 0.13 |
| 7 | 141951998 | rs1052410 | 0.341 | -0.512 | 0.453 | 0.125 | -0.051 | 0.023 | 0.023 | 0 | 0.15 | 0.34 | 0.26 | 0.14 | 0.1 | 0.13 |
| 7 | 141952023 | rs1052408 | 0.341 | -0.512 | 0.453 | 0.125 | -0.051 | 0.023 | 0.023 | 0 | 0.15 | 0.34 | 0.26 | 0.14 | 0.1 | 0.13 |
| 7 | 141952110 | rs1052406 | 1.18 | -0.501 | 0.512 | 0.075 | -0.015 | 0.012 | 0.012 | 0 | 0.3 | 0.21 | 0.23 | 0.01 | 0.35 | 0.24 |
| 7 | 141955128 | rs1894317 | 0.368 | -0.504 | 0.489 | 0.081 | -0.051 | 0.026 | 0.026 | 0 | 0.15 | 0.34 | 0.26 | 0.14 | 0.1 | 0.13 |
| 7 | 141957504 | rs7786497 | 1.318 | -0.496 | 0.429 | 0.083 | 0.001 | -0.136 | -0.136 | 0.112 | 0.31 | 0.06 | 0.33 | 0.51 | 0.25 | 0.33 |
| 7 | 141968747 | rs2960773 | 1.446 | -0.507 | 0.422 | 0.115 | -0.03 | -0.175 | -0.175 | 0 | 0.3 | 0.02 | 0.33 | 0.51 | 0.25 | 0.33 |
| 7 | 141968974 | rs4726513 | 0.549 | -0.509 | 0.458 | 0.121 | -0.042 | 0.024 | 0.024 | 0 | 0.14 | 0.38 | 0.24 | 0.11 | 0.1 | 0.1 |
| 7 | 141969003 | rs9640366 | 0.034 | -0.504 | 0.46 | 0.112 | 0.017 | 0.011 | 0.011 | 0 | 0.23 | 0.26 | 0.17 | 0.33 | 0.3 | 0.29 |
| 7 | 141969572 | rs2001011 | 1.327 | -0.501 | 0.441 | 0.113 | 0.001 | -0.136 | -0.136 | 0 | 0.31 | 0.06 | 0.33 | 0.52 | 0.25 | 0.33 |
| 7 | 141969592 | rs2001012 | 1.327 | -0.501 | 0.441 | 0.113 | 0.001 | -0.136 | -0.136 | 0 | 0.31 | 0.06 | 0.33 | 0.52 | 0.25 | 0.33 |
| 7 | 141971825 | rs6976829 | 0.308 | -0.484 | 0.46 | 0.013 | -0.055 | 0.041 | 0.041 | 0 | 0.15 | 0.4 | 0.26 | 0.14 | 0.1 | 0.13 |
| 7 | 141972370 | rs980385 | 0.106 | -0.472 | 0.541 | 0.01 | 0.001 | -0.001 | -0.001 | 0.951 | 0.22 | 0.2 | 0.17 | 0.21 | 0.3 | 0.29 |
| 7 | 141974617 | rs10251247 | 0.274 | -0.566 | 0.224 | 0.121 | -0.055 | 0.053 | 0.053 | 0 | 0.15 | 0.41 | 0.26 | 0.15 | 0.1 | 0.14 |
| 7 | 141977608 | rs28838390 | 0.274 | -0.566 | 0.224 | 0.121 | -0.055 | 0.053 | 0.053 | 0 | 0.15 | 0.41 | 0.26 | 0.15 | 0.1 | 0.14 |
| 7 | 141983565 | rs1018771 | -0.206 | -0.542 | 0.137 | 0.021 | 0.005 | 0.001 | 0.001 | 0 | 0.52 | 0.45 | 0.4 | 0.21 | 0.65 | 0.53 |
| 7 | 142007325 | rs7357130 | 0.254 | -0.455 | 0.342 | -0.102 | 0.015 | 0.113 | 0.113 | 0 | 0.53 | 0.25 | 0.52 | 0.86 | 0.56 | 0.67 |
| 7 | 142010982 | rs12112030 | 0.122 | -0.681 | 0.139 | 0.078 | -0.046 | 0.072 | 0.072 | 0 | 0.16 | 0.32 | 0.26 | 0.27 | 0.11 | 0.13 |
| 7 | 142014213 | rs965274 | -0.076 | -0.678 | 0.111 | 0.079 | -0.055 | 0.081 | 0.081 | 0 | 0.17 | 0.41 | 0.26 | 0.27 | 0.11 | 0.15 |
| 7 | 142014445 | rs10246910 | 0.38 | -0.08 | 0.007 | 0.042 | -0.007 | 0.094 | 0.094 | 0 | 0.52 | 0.26 | 0.48 | 0.72 | 0.47 | 0.63 |
| 7 | 142016272 | rs1014957 | -0.311 | -0.069 | -0.069 | 0.047 | -0.003 | 0.026 | 0.026 | 0 | 0.47 | 0.59 | 0.51 | 0.28 | 0.53 | 0.37 |
| 7 | 142022250 | rs7803851 | 0.279 | -0.565 | 0.32 | 0.324 | -0.055 | -0.006 | -0.006 | 0 | 0.83 | 0.53 | 0.73 | 0.87 | 0.89 | 0.85 |
| 7 | 142022633 | rs17162991 | -0.291 | -0.485 | 0.102 | 0.36 | -0.042 | -0.002 | -0.002 | 0.063 | 0.16 | 0.36 | 0.26 | 0.13 | 0.11 | 0.15 |
| 7 | 142023396 | rs17162994 | 0.138 | -0.228 | -0.843 | 0.576 | 0.002 | 0.002 | 0.002 | 0.044 | 0.5 | 0.46 | 0.64 | 0.62 | 0.41 | 0.58 |
| 7 | 142025955 | rs2960760 | -0.095 | -0.353 | -0.897 | 0.509 | -0.013 | -0.032 | -0.032 | 0 | 0.35 | 0.53 | 0.38 | 0.21 | 0.27 | 0.39 |
| 7 | 142030318 | rs12674023 | 0.885 | -0.935 | -1.629 | 0.736 | 0.053 | -0.049 | -0.049 | 0 | 0.26 | 0.11 | 0.22 | 0.43 | 0.15 | 0.29 |
| 7 | 142032983 | rs2156936 | 0.134 | 0.358 | -1.151 | 0.892 | -0.011 | -0.002 | -0.002 | 0 | 0.63 | 0.69 | 0.62 | 0.63 | 0.42 | 0.68 |
| 7 | 142040894 | rs1155458 | -0.379 | 1.64 | -1.48 | 0.941 | -0.153 | -0.069 | -0.069 | 0 | 0.21 | 0.03 | 0.32 | 0.28 | 0.39 | 0.19 |
| 7 | 142041961 | rs2855836 | -1.254 | 1.337 | -0.752 | 1.025 | -0.003 | -0.003 | -0.003 | 0 | 0.32 | 0.33 | 0.41 | 0.43 | 0.56 | 0.31 |
| 7 | 142044060 | rs2011381 | -0.291 | 1.534 | -0.937 | 1.009 | -0.156 | -0.092 | -0.092 | 0 | 0.22 | 0.04 | 0.33 | 0.36 | 0.44 | 0.19 |
| 7 | 142099013 | rs10277702 | -1.113 | 0.511 | 0.161 | 0.329 | -0.022 | -0.044 | -0.044 | 0 | 0.2 | 0.53 | 0.2 | 0.11 | 0.15 | 0.27 |
| 7 | 142122115 | rs6958838 | -1.507 | 0.658 | -0.22 | -0.019 | -0.006 | 0.076 | 0.076 | 0 | 0.42 | 0.74 | 0.5 | 0.24 | 0.46 | 0.4 |
| 7 | 142137119 | rs2040369 | -0.733 | 0.759 | -0.162 | -0.075 | -0.028 | 0.017 | 0.017 | 0.023 | 0.24 | 0.15 | 0.37 | 0.16 | 0.38 | 0.25 |
| 7 | 142141507 | rs7793482 | -0.903 | 0.429 | 0.602 | 0.113 | -0.051 | -0.059 | -0.059 | 0 | 0.17 | 0.4 | 0.1 | 0.08 | 0.07 | 0.19 |
| 7 | 142146497 | rs7778566 | 0.837 | 0.344 | 0.54 | 0.034 | -0.069 | -0.091 | -0.091 | 0.032 | 0.82 | 0.55 | 0.89 | 0.94 | 0.94 | 0.81 |
| 7 | 142152369 | rs2011310 | -1.361 | 0.753 | -0.01 | 0.124 | -0.001 | 0 | 0 | 0.514 | 0.24 | 0.22 | 0.42 | 0.21 | 0.3 | 0.35 |
| 7 | 142162715 | rs10487531 | 1.366 | 0.975 | -1.009 | 0.224 | -0.032 | -0.115 | -0.115 | 1.487 | 0.43 | 0.05 | 0.34 | 0.52 | 0.4 | 0.44 |
| 7 | 142163046 | rs1987475 | 1.129 | 0.807 | -0.353 | -0.036 | -0.001 | -0.001 | -0.001 | 0 | 0.73 | 0.73 | 0.57 | 0.79 | 0.7 | 0.61 |
| 7 | 142166888 | rs2011726 | -1.044 | 1.177 | 0.143 | 0.326 | 0.008 | 0.009 | 0.009 | 0.868 | 0.21 | 0.2 | 0.42 | 0.29 | 0.3 | 0.34 |
| 7 | 142175816 | rs2734153 | -1.346 | 1.377 | 0.326 | 0.152 | 0.021 | -0.001 | -0.001 | 0 | 0.23 | 0.28 | 0.43 | 0.27 | 0.31 | 0.35 |
| 7 | 142177451 | rs1882723 | -0.44 | 1.429 | 0.259 | 0.147 | -0.002 | -0.002 | -0.002 | 1.084 | 0.15 | 0.14 | 0.37 | 0.22 | 0.21 | 0.23 |
| 7 | 142182298 | rs6979469 | 1.401 | 1.331 | 0.121 | -0.001 | 0.03 | -0.025 | -0.025 | 0 | 0.91 | 0.76 | 0.93 | 0.95 | 0.91 | 0.87 |
| 7 | 142190774 | rs17232 | 0.094 | 1.129 | -0.686 | -0.32 | -0.035 | 0.059 | 0.059 | 0 | 0.67 | 0.77 | 0.68 | 0.82 | 0.58 | 0.79 |
| 7 | 142191647 | rs10276363 | -0.685 | 1.22 | -0.251 | -0.087 | 0.017 | 0.017 | 0.017 | 0 | 0.13 | 0.18 | 0.26 | 0.16 | 0.16 | 0.24 |
| 7 | 142192931 | rs10271793 | 1.429 | 0.279 | -0.026 | -0.029 | -0.004 | -0.001 | -0.001 | 1.086 | 0.45 | 0.44 | 0.36 | 0.57 | 0.32 | 0.41 |
| 7 | 142194668 | rs10260565 | 0.711 | 0.278 | 0.02 | -0.025 | -0.001 | -0.001 | -0.001 | 0 | 0.54 | 0.53 | 0.41 | 0.65 | 0.42 | 0.58 |
| 7 | 142196434 | rs2040366 | -1.171 | 0.419 | -0.04 | 0.083 | 0.025 | 0.007 | 0.007 | 0.015 | 0.51 | 0.46 | 0.64 | 0.45 | 0.67 | 0.59 |
| 7 | 142198547 | rs10216024 | 0.696 | 0.42 | -0.233 | 0.068 | 0.004 | 0.053 | 0.053 | 0 | 0.79 | 0.62 | 0.68 | 0.73 | 0.77 | 0.58 |
| 7 | 142198611 | rs6976636 | 1.167 | 0.407 | -0.241 | 0.061 | -0.003 | -0.001 | -0.001 | 0 | 0.49 | 0.48 | 0.36 | 0.53 | 0.32 | 0.41 |
| 7 | 142200371 | rs11768792 | 0.035 | 0.751 | -0.771 | -0.252 | -0.12 | 0.071 | 0.071 | 0 | 0.3 | 0.14 | 0.32 | 0.2 | 0.46 | 0.18 |
| 7 | 142210014 | rs4726526 | -0.447 | -0.262 | -0.139 | -0.581 | -0.015 | -0.043 | -0.043 | 0 | 0.84 | 0.61 | 0.89 | 0.93 | 0.9 | 0.98 |
| 7 | 142217780 | rs2213192 | 0.118 | -0.127 | -0.527 | -0.891 | -0.033 | 0.053 | 0.053 | 0 | 0.14 | 0.05 | 0.07 | 0.02 | 0.17 | 0.22 |

| | | | | | | | | | | | | | | | | |
|---|---|---|---|---|---|---|---|---|---|---|---|---|---|---|---|---|
| 7 | 142222364 | rs6961143 | 0.965 | -0.217 | -0.432 | -0.931 | 0 | -0.087 | -0.087 | 0.002 | 0.71 | 0.5 | 0.81 | 0.91 | 0.73 | 0.76 |
| 7 | 142222582 | rs975494 | 1.648 | -1.001 | -1.132 | -0.838 | 0.079 | -0.131 | -0.131 | 0 | 0.31 | 0.03 | 0.22 | 0.59 | 0.16 | 0.28 |
| 7 | 142224458 | rs6950275 | 0.607 | 0.221 | 0.082 | -0.734 | -0.007 | -0.073 | -0.073 | 0 | 0.13 | 0.25 | 0.09 | 0 | 0.09 | 0.02 |
| 7 | 142231701 | rs17248 | 0.554 | 0.502 | -0.033 | -0.888 | 0 | -0.002 | -0.002 | 1.328 | 0.63 | 0.62 | 0.58 | 0.71 | 0.4 | 0.61 |
| 7 | 142235861 | rs2367191 | 0.783 | 0.792 | -0.056 | -0.855 | -0.092 | 0.015 | 0.015 | 0 | 0.21 | 0.11 | 0.33 | 0.2 | 0.44 | 0.17 |
| 7 | 142236708 | rs11770471 | 0.823 | 0.787 | -0.065 | -0.857 | -0.096 | 0.013 | 0.013 | 0 | 0.21 | 0.08 | 0.32 | 0.2 | 0.44 | 0.17 |
| 7 | 142237142 | rs2734118 | 0.277 | 0.737 | -0.061 | -0.841 | -0.03 | 0.047 | 0.047 | 0 | 0.13 | 0.05 | 0.07 | 0.02 | 0.16 | 0.16 |
| 7 | 142237788 | rs2855918 | 1.278 | 0.547 | 0.011 | -0.858 | 0.047 | 0.046 | 0.046 | 0 | 0.5 | 0.36 | 0.49 | 0.71 | 0.31 | 0.65 |
| 7 | 142239119 | rs11768398 | -0.426 | 0.824 | 0 | -0.894 | -0.016 | 0.022 | 0.022 | 0 | 0.34 | 0.28 | 0.39 | 0.2 | 0.51 | 0.19 |
| 7 | 142239946 | rs17209 | 0.058 | 0.877 | 0.107 | -0.922 | 0.023 | -0.046 | -0.046 | 0 | 0.14 | 0.25 | 0.12 | 0.02 | 0.19 | 0.17 |
| 7 | 142244415 | rs1573618 | -0.872 | 0.924 | -0.067 | -0.788 | -0.038 | -0.013 | -0.013 | 0 | 0.78 | 0.84 | 0.68 | 0.71 | 0.59 | 0.83 |
| 7 | 142245627 | rs17304 | -0.791 | 0.889 | -0.103 | -0.813 | 0.002 | -0.042 | -0.042 | 0 | 0.27 | 0.32 | 0.18 | 0.02 | 0.29 | 0.19 |
| 7 | 142247347 | rs361359 | -0.8 | 0.891 | -0.102 | -0.812 | 0.001 | -0.042 | -0.042 | 0.612 | 0.27 | 0.32 | 0.18 | 0.02 | 0.29 | 0.19 |
| 7 | 142251132 | rs361436 | -0.231 | 0.906 | -0.273 | -0.703 | -0.012 | 0.015 | 0.015 | 0 | 0.86 | 0.87 | 0.9 | 0.98 | 0.81 | 0.83 |
| 7 | 142258330 | rs2855896 | 1.173 | 0.65 | -0.219 | -0.722 | 0.048 | 0.042 | 0.042 | 0 | 0.5 | 0.37 | 0.49 | 0.69 | 0.31 | 0.64 |
| 7 | 142272619 | rs2855882 | 0.01 | 1.29 | -0.522 | -0.936 | -0.058 | 0.059 | 0.059 | 0 | 0.14 | 0.05 | 0.11 | 0.02 | 0.24 | 0.17 |
| 7 | 142273037 | rs361489 | -0.687 | 1.327 | 0.23 | -0.593 | 0.035 | -0.033 | -0.033 | 0 | 0.34 | 0.42 | 0.39 | 0.22 | 0.46 | 0.19 |
| 7 | 142274257 | rs6464489 | 0.265 | 1.521 | 0.076 | -0.496 | 0.023 | -0.002 | -0.002 | 0.315 | 0.22 | 0.23 | 0.32 | 0.22 | 0.38 | 0.17 |
| 7 | 142275743 | rs949425 | -0.113 | 0.642 | 0.237 | -0.768 | 0.001 | -0.018 | -0.018 | 0 | 0.12 | 0.11 | 0.08 | 0 | 0.1 | 0.02 |
| 7 | 142328771 | rs6974518 | 2.02 | 0.742 | 0.976 | -0.568 | -0.017 | 0.034 | 0.034 | 5.768 | 0.85 | 0.33 | 0.7 | 0.76 | 0.89 | 0.93 |
| 7 | 142330328 | rs7384887 | -0.845 | 0.826 | 1.038 | -0.534 | 0.004 | 0.034 | 0.034 | 0 | 0.08 | 0.11 | 0.23 | 0.23 | 0.11 | 0.07 |
| 7 | 142344964 | rs2285027 | 1.508 | 0.753 | 1.807 | -0.544 | 0.03 | 0.166 | 0.166 | 0.101 | 0.9 | 0.66 | 0.75 | 0.76 | 0.91 | 0.93 |
| 7 | 142347046 | rs6943492 | 0.918 | 0.916 | 0.942 | -0.987 | -0.017 | 0.074 | 0.074 | 0.185 | 0.47 | 0.16 | 0.45 | 0.69 | 0.55 | 0.66 |
| 7 | 142356859 | rs2078176 | -0.883 | 0.924 | 0.953 | -0.971 | 0.009 | 0.087 | 0.087 | 0.089 | 0.52 | 0.75 | 0.55 | 0.31 | 0.45 | 0.34 |
| 7 | 142369835 | rs6975391 | 0.165 | 0.908 | 1.143 | -1.077 | -0.013 | -0.078 | -0.078 | 0 | 0.37 | 0.55 | 0.28 | 0.06 | 0.29 | 0.2 |
| 7 | 142372872 | rs10260647 | -1.987 | 0.929 | 1.115 | -0.897 | 0.005 | 0.034 | 0.034 | 0.181 | 0.16 | 0.22 | 0.27 | 0.26 | 0.16 | 0.15 |
| 7 | 142377014 | rs17837474 | -1.419 | 0.833 | 1.24 | -0.795 | 0.004 | 0.103 | 0.103 | 0.215 | 0.1 | 0.22 | 0.23 | 0.27 | 0.09 | 0.07 |
| 7 | 142384755 | rs2156966 | -0.806 | 0.698 | 1.1 | -0.892 | 0.015 | 0.04 | 0.04 | 0 | 0.5 | 0.59 | 0.52 | 0.33 | 0.44 | 0.34 |
| 7 | 142386873 | rs731521 | -0.38 | 0.652 | 0.961 | -0.871 | -0.002 | -0.039 | -0.039 | 0 | 0.07 | 0.02 | 0.21 | 0.22 | 0.06 | 0.07 |
| 7 | 142389219 | rs17837475 | -0.111 | 0.736 | 0.999 | -1.028 | -0.006 | -0.029 | -0.029 | 0 | 0.07 | 0.02 | 0.21 | 0.22 | 0.06 | 0.07 |
| 7 | 142389366 | rs6968949 | -0.926 | 0.754 | 1.095 | -1.073 | 0.011 | 0.032 | 0.032 | 0 | 0.49 | 0.6 | 0.52 | 0.33 | 0.44 | 0.33 |
| 7 | 142401786 | rs6946770 | -0.935 | 0.881 | 1.219 | -0.999 | 0.013 | 0.041 | 0.041 | 0 | 0.49 | 0.66 | 0.53 | 0.33 | 0.43 | 0.34 |
| 7 | 142418236 | rs4726568 | 0.877 | 0.882 | 1.112 | -1.197 | -0.015 | 0.103 | 0.103 | 0.734 | 0.5 | 0.16 | 0.47 | 0.69 | 0.55 | 0.67 |
| 7 | 142423289 | rs17251 | -0.252 | 0.867 | 1.299 | -1.417 | 0 | -0.029 | -0.029 | 0.242 | 0.4 | 0.4 | 0.29 | 0.12 | 0.39 | 0.24 |
| 7 | 142424782 | rs4726571 | -1.095 | 0.907 | 0.754 | -1.389 | 0.007 | 0.097 | 0.097 | 0.237 | 0.49 | 0.74 | 0.52 | 0.3 | 0.45 | 0.33 |
| 7 | 142424843 | rs4726572 | -0.17 | 0.47 | 0.34 | -1.249 | 0 | 0.02 | 0.02 | 0 | 0.06 | 0.09 | 0.21 | 0.18 | 0.06 | 0.09 |
| 7 | 142434960 | rs6959895 | -0.426 | 1.204 | 0.17 | -1.934 | -0.027 | 0.138 | 0.138 | 2.702 | 0.43 | 0.14 | 0.48 | 0.18 | 0.57 | 0.28 |
| 7 | 142456928 | rs10273639 | -0.666 | 1.327 | 0.89 | -1.802 | 0.001 | 0.008 | 0.008 | 0.306 | 0.44 | 0.39 | 0.53 | 0.22 | 0.59 | 0.28 |
| 7 | 142487836 | rs2734222 | -0.649 | 1.155 | 1.444 | -1.85 | 0.004 | -0.02 | -0.02 | 0.218 | 0.43 | 0.5 | 0.54 | 0.23 | 0.54 | 0.3 |
| 7 | 142488688 | rs2734224 | -0.42 | 1.189 | 1.566 | -1.833 | -0.017 | 0.029 | 0.029 | 0 | 0.4 | 0.33 | 0.52 | 0.23 | 0.54 | 0.27 |
| 7 | 142492130 | rs2367486 | -0.612 | 1.177 | 1.733 | -1.845 | 0.002 | -0.01 | -0.01 | 0 | 0.42 | 0.45 | 0.52 | 0.23 | 0.54 | 0.27 |
| 7 | 142493129 | rs3134902 | -0.43 | 1.198 | 1.353 | -1.836 | -0.048 | 0.055 | 0.055 | 0 | 0.39 | 0.24 | 0.5 | 0.23 | 0.54 | 0.27 |
| 7 | 142496293 | rs12539089 | -0.599 | 1.202 | 1.086 | -1.806 | 0.001 | -0.007 | -0.007 | 0.123 | 0.43 | 0.44 | 0.5 | 0.23 | 0.51 | 0.21 |
| 7 | 142498109 | rs1800907 | -0.944 | 1.318 | 0.748 | -1.729 | -0.036 | 0.056 | 0.056 | 0 | 0.43 | 0.78 | 0.55 | 0.26 | 0.51 | 0.22 |
| 7 | 142500409 | rs11327 | -1.386 | 1.644 | 1.402 | -2.104 | -0.017 | 0.019 | 0.019 | 0 | 0.35 | 0.3 | 0.3 | 0.18 | 0.44 | 0.16 |
| 7 | 142505154 | rs3134906 | -1.953 | 1.351 | 1.466 | -2.209 | 0.016 | 0.13 | 0.13 | 0.109 | 0.45 | 0.79 | 0.41 | 0.21 | 0.55 | 0.26 |
| 7 | 142508376 | rs4281045 | -0.775 | 0.024 | 1.517 | -2.268 | -0.089 | 0.058 | 0.058 | 0 | 0.25 | 0.05 | 0.22 | 0.11 | 0.28 | 0.14 |
| 7 | 142510174 | rs17270 | -0.777 | 0.023 | 1.517 | -2.268 | -0.089 | 0.058 | 0.058 | 0 | 0.25 | 0.05 | 0.22 | 0.11 | 0.28 | 0.14 |
| 7 | 142512803 | rs7805607 | -2.325 | 0.024 | 2.054 | -2.28 | -0.006 | -0.061 | -0.061 | 0 | 0.13 | 0.4 | 0.08 | 0.03 | 0.11 | 0.14 |
| 7 | 142515811 | rs17163745 | -2.079 | 0.009 | 2.129 | -2.284 | 0.004 | -0.058 | -0.058 | 0 | 0.39 | 0.52 | 0.31 | 0.14 | 0.39 | 0.28 |
| 7 | 142525934 | rs2049819 | -0.974 | -0.048 | 1.987 | -2.177 | -0.002 | 0.002 | 0.002 | 0 | 0.05 | 0.05 | 0.04 | 0.03 | 0.09 | 0.05 |
| 7 | 142531590 | rs17722379 | -0.666 | -0.095 | 1.981 | -2.301 | 0.004 | 0.015 | 0.015 | 0 | 0.22 | 0.14 | 0.12 | 0.04 | 0.17 | 0.09 |
| 7 | 142536951 | rs12537777 | 0.019 | 0.218 | 1.796 | -2.138 | -0.036 | -0.013 | -0.013 | 0 | 0.08 | 0.03 | 0.12 | 0.07 | 0.13 | 0.06 |
| 7 | 142545218 | rs2063993 | -1.485 | 0.042 | 2.114 | -2.055 | 0.008 | 0.078 | 0.078 | 0.22 | 0.42 | 0.71 | 0.33 | 0.15 | 0.4 | 0.33 |
| 7 | 142552373 | rs7789029 | -2.387 | 0.03 | 3.01 | -1.342 | -0.032 | -0.036 | -0.036 | 0 | 0.09 | 0.38 | 0.05 | 0.01 | 0.03 | 0.11 |
| 7 | 142562736 | rs6947538 | -2.831 | 0.233 | 3.345 | -1.285 | -0.027 | -0.06 | -0.06 | 0.006 | 0.15 | 0.69 | 0.13 | 0.04 | 0.1 | 0.12 |
| 7 | 142563253 | rs8177146 | -1.773 | 0.23 | 3.515 | -1.284 | -0.024 | -0.018 | -0.018 | 0 | 0.05 | 0.13 | 0.04 | 0.01 | 0.03 | 0.11 |
| 7 | 142565665 | rs8177155 | -0.902 | 0.287 | 3.679 | -1.261 | 0.014 | -0.02 | -0.02 | 0 | 0.06 | 0.21 | 0.04 | 0 | 0.03 | 0 |
| 7 | 142568878 | rs4987687 | -2.495 | 0.287 | 3.5 | -1.262 | 0.006 | -0.053 | -0.053 | 0.285 | 0.11 | 0.6 | 0.11 | 0.03 | 0.08 | 0.01 |
| 7 | 142568934 | rs4987686 | -2.15 | 0.292 | 3.572 | -1.261 | 0.05 | -0.021 | -0.021 | 0 | 0.09 | 0.48 | 0.09 | 0.03 | 0.07 | 0.01 |
| 7 | 142569596 | rs4987682 | -2.421 | 0.312 | 3.444 | -1.252 | 0.01 | -0.05 | -0.05 | 0.053 | 0.1 | 0.6 | 0.11 | 0.03 | 0.08 | 0.01 |
| 7 | 142572649 | rs4987668 | -2.346 | 0.31 | 3.427 | -1.253 | 0.015 | -0.047 | -0.047 | 0.135 | 0.1 | 0.57 | 0.1 | 0.03 | 0.08 | 0.01 |
| 7 | 142572908 | rs4987667 | -2.363 | 0.311 | 3.427 | -1.252 | 0.014 | -0.048 | -0.048 | 0 | 0.1 | 0.58 | 0.1 | 0.03 | 0.08 | 0.01 |
| 7 | 142573263 | rs4987665 | -2.363 | 0.311 | 3.427 | -1.252 | 0.014 | -0.048 | -0.048 | 0 | 0.1 | 0.58 | 0.1 | 0.03 | 0.08 | 0.01 |
| 7 | 142605221 | rs4252512 | -0.391 | 0.253 | 3.175 | -1.236 | -0.001 | -0.026 | -0.026 | 0 | 0.05 | 0.13 | 0.03 | 0 | 0.02 | 0 |
| 7 | 142605336 | rs4252511 | -3.204 | 0.2 | 3.126 | -1.278 | -0.018 | -0.042 | -0.042 | 2.216 | 0.16 | 0.71 | 0.12 | 0.04 | 0.1 | 0.17 |
| 7 | 142617301 | rs4252460 | -2.963 | 0.205 | 3.281 | -1.276 | -0.024 | -0.073 | -0.073 | 0 | 0.15 | 0.57 | 0.1 | 0.05 | 0.11 | 0.17 |
| 7 | 142619784 | rs4252448 | -3.008 | 0.19 | 3.264 | -1.282 | -0.031 | -0.07 | -0.07 | 0 | 0.16 | 0.61 | 0.11 | 0.05 | 0.11 | 0.17 |
| 7 | 142622714 | rs4252435 | -2.972 | 0.21 | 3.255 | -1.274 | -0.03 | -0.07 | -0.07 | 0.233 | 0.16 | 0.61 | 0.11 | 0.05 | 0.11 | 0.17 |

| | | | | | | | | | | | | | | | | |
|---|---|---|---|---|---|---|---|---|---|---|---|---|---|---|---|---|
| 7 | 142624333 | rs4252424 | -0.268 | 0.258 | 2.938 | -1.513 | -0.058 | 0.066 | 0.066 | 0 | 0.17 | 0.02 | 0.13 | 0.04 | 0.2 | 0.11 |
| 7 | 142625417 | rs4252416 | -2.065 | 0.337 | 3.039 | -1.472 | 0.008 | 0.047 | 0.047 | 0.02 | 0.33 | 0.73 | 0.26 | 0.09 | 0.3 | 0.28 |
| 7 | 142627138 | rs4252402 | -1.375 | 0.372 | 2.84 | -1.456 | 0.003 | 0.008 | 0.008 | 0 | 0.27 | 0.2 | 0.18 | 0.05 | 0.25 | 0.27 |
| 7 | 142629375 | rs4252381 | -2.948 | 0.244 | 3.633 | -0.793 | -0.017 | -0.063 | -0.063 | 0 | 0.13 | 0.67 | 0.1 | 0.04 | 0.08 | 0.01 |
| 7 | 142630534 | rs4252372 | -0.842 | 0.247 | 3.573 | -0.785 | 0.007 | -0.02 | -0.02 | 0 | 0.05 | 0.14 | 0.03 | 0 | 0.02 | 0 |
| 7 | 142634920 | rs12333618 | -0.59 | 0.257 | 3.31 | -0.78 | -0.006 | -0.028 | -0.028 | 0 | 0.05 | 0.1 | 0.03 | 0 | 0.02 | 0 |
| 7 | 142636599 | rs11979330 | -2.15 | 0.285 | 2.997 | -0.783 | -0.014 | -0.063 | -0.063 | 0.524 | 0.13 | 0.67 | 0.1 | 0.04 | 0.09 | 0.01 |
| 7 | 142657775 | rs8175963 | -1.785 | 0.254 | 1.261 | -0.695 | 0.023 | -0.056 | -0.056 | 1.423 | 0.12 | 0.47 | 0.08 | 0.04 | 0.06 | 0.01 |
| 7 | 142661695 | rs8175950 | -1.292 | -0.581 | 1.495 | -0.993 | -0.051 | -0.096 | -0.096 | 0 | 0.14 | 0.39 | 0.08 | 0 | 0.06 | 0.01 |
| 7 | 142686688 | rs10265609 | -1.264 | -0.673 | 1.797 | -1.029 | -0.041 | -0.072 | -0.072 | 0 | 0.14 | 0.28 | 0.06 | 0 | 0.06 | 0.01 |
| 7 | 142689472 | rs6464536 | -0.703 | -1.669 | 2.179 | -1.523 | -0.035 | -0.037 | -0.037 | 0 | 0.16 | 0.21 | 0.08 | 0.04 | 0.07 | 0.13 |
| 7 | 142700712 | rs10256437 | -2.137 | -1.752 | 2.561 | -1.562 | 0.013 | 0.005 | 0.005 | 0.106 | 0.31 | 0.7 | 0.16 | 0.04 | 0.13 | 0.14 |
| 7 | 142713375 | rs2367618 | -0.475 | -1.737 | 2.626 | -1.539 | -0.005 | -0.02 | -0.02 | 0 | 0.12 | 0.1 | 0.05 | 0 | 0.06 | 0.01 |
| 7 | 142718902 | rs10277182 | -2.211 | -1.791 | 2.716 | -1.569 | 0.021 | 0.014 | 0.014 | 0.255 | 0.31 | 0.69 | 0.16 | 0.04 | 0.14 | 0.14 |
| 7 | 142721361 | rs1980316 | -2.154 | -1.793 | 3.218 | -1.57 | -0.085 | -0.116 | -0.116 | 0.515 | 0.3 | 0.45 | 0.12 | 0.04 | 0.13 | 0.14 |
| 7 | 142724062 | rs9885986 | -0.766 | -1.872 | 3.175 | -1.609 | -0.021 | -0.019 | -0.019 | 3.464 | 0.14 | 0.14 | 0.05 | 0.04 | 0.06 | 0.12 |
| 7 | 142727560 | rs6960104 | -2.32 | -1.862 | 2.755 | -1.604 | 0.013 | 0.005 | 0.005 | 0.256 | 0.31 | 0.69 | 0.16 | 0.04 | 0.13 | 0.14 |
| 7 | 142747291 | rs4395807 | -0.929 | -1.509 | 2.831 | -1.6 | -0.021 | -0.036 | -0.036 | 0 | 0.14 | 0.19 | 0.07 | 0.03 | 0.08 | 0.09 |
| 7 | 142748205 | rs7801166 | -2.083 | -1.559 | 2.914 | -1.628 | -0.072 | -0.12 | -0.12 | 0 | 0.28 | 0.43 | 0.13 | 0.03 | 0.15 | 0.1 |
| 7 | 142749737 | rs17164103 | -1.061 | -1.555 | 2.944 | -1.626 | -0.028 | -0.053 | -0.053 | 0 | 0.15 | 0.32 | 0.08 | 0.03 | 0.08 | 0.09 |
| 7 | 142750147 | rs10245778 | -0.852 | -1.56 | 3.06 | -1.628 | -0.005 | -0.009 | -0.009 | 0 | 0.14 | 0.12 | 0.07 | 0.03 | 0.08 | 0.09 |
| 7 | 142750834 | rs1506404 | -2.163 | -2.91 | 2.224 | -2.329 | 0.093 | 0.099 | 0.099 | 0.105 | 0.43 | 0.69 | 0.2 | 0.04 | 0.2 | 0.18 |
| 7 | 142750951 | rs1506403 | -2.163 | -2.91 | 2.224 | -2.329 | 0.093 | 0.099 | 0.099 | 0.105 | 0.43 | 0.69 | 0.2 | 0.04 | 0.2 | 0.18 |
| 7 | 142792658 | rs10230444 | -0.158 | -2.773 | 2.215 | -2.285 | -0.027 | -0.057 | -0.057 | 0 | 0.13 | 0.21 | 0.06 | 0 | 0.07 | 0.01 |
| 7 | 142816927 | rs1994497 | -1.661 | -2.869 | 2.22 | -2.332 | -0.043 | -0.07 | -0.07 | 0 | 0.28 | 0.54 | 0.14 | 0.04 | 0.14 | 0.09 |
| 7 | 142822210 | rs10243198 | -0.778 | -3 | 2.229 | -2.359 | -0.04 | -0.054 | -0.054 | 2.33 | 0.16 | 0.38 | 0.09 | 0.04 | 0.07 | 0.08 |
| 7 | 142827954 | rs315284 | -2.178 | -3.088 | 2.07 | -2.405 | 0.091 | 0.098 | 0.098 | 0.051 | 0.43 | 0.69 | 0.19 | 0.04 | 0.2 | 0.18 |
| 7 | 142830623 | rs315283 | -0.1 | -2.912 | 1.701 | -2.278 | 0.011 | -0.001 | -0.001 | 0.036 | 0.15 | 0.13 | 0.05 | 0 | 0.06 | 0.09 |
| 7 | 142835239 | rs9986765 | -0.728 | -2.705 | 1.74 | -2.179 | -0.03 | -0.04 | -0.04 | 0 | 0.14 | 0.27 | 0.07 | 0.04 | 0.07 | 0.08 |
| 7 | 142839168 | rs12703516 | -0.944 | -2.017 | 0.657 | -1.153 | -0.048 | -0.06 | -0.06 | 0.772 | 0.9 | 0.98 | 0.67 | 0.34 | 0.82 | 0.78 |
| 7 | 142840947 | rs10277938 | 0.049 | -2.481 | -0.048 | -2.167 | 0.008 | 0.012 | 0.012 | 0 | 0.26 | 0.16 | 0.13 | 0.04 | 0.19 | 0.19 |
| 7 | 142852814 | rs1386942 | 0.558 | -2.653 | -0.176 | -2.223 | 0.024 | 0.016 | 0.016 | 0.621 | 0.13 | 0.08 | 0.05 | 0 | 0.08 | 0.02 |
| 7 | 142869453 | rs315294 | -1.676 | -2.594 | -0.29 | -1.379 | -0.028 | -0.015 | -0.015 | 0 | 0.94 | 0.99 | 0.94 | 0.69 | 0.93 | 0.86 |
| 7 | 142872381 | rs4452723 | -0.584 | -3.072 | -0.215 | -2.086 | 0 | -0.04 | -0.04 | 0.07 | 0.27 | 0.27 | 0.11 | 0 | 0.13 | 0.07 |
| 7 | 142877619 | rs11976378 | 1.052 | -3.06 | -0.394 | -2.078 | 0.037 | 0.044 | 0.044 | 7.415 | 0.12 | 0.04 | 0.05 | 0 | 0.05 | 0.06 |
| 7 | 142881540 | rs4726600 | 1.568 | -1.487 | -0.824 | -2.383 | -0.049 | 0.021 | 0.021 | 0 | 0.09 | 0.03 | 0.12 | 0.01 | 0.25 | 0.08 |
| 7 | 142899393 | rs11983741 | 0.642 | -2.242 | -0.506 | -2.208 | 0.031 | 0.033 | 0.033 | 1.079 | 0.13 | 0.07 | 0.05 | 0 | 0.08 | 0.02 |
| 7 | 142913569 | rs7780478 | 0.389 | -2.056 | -0.3 | -2.093 | -0.031 | -0.061 | -0.061 | 4.41 | 0.14 | 0.24 | 0.07 | 0 | 0.05 | 0.06 |
| 7 | 142916844 | rs17164163 | 0.88 | -2.077 | -0.366 | -2.103 | 0.037 | 0.047 | 0.047 | 7.519 | 0.13 | 0.04 | 0.04 | 0 | 0.05 | 0.06 |
| 7 | 142919731 | rs10260248 | 0.746 | -2.378 | -0.424 | -2.23 | -0.013 | -0.019 | -0.019 | 0 | 0.1 | 0.14 | 0.07 | 0.02 | 0.06 | 0.1 |
| 7 | 142936243 | rs10256011 | -0.122 | -2.294 | -0.672 | -2.27 | 0.021 | 0.022 | 0.022 | 0 | 0.39 | 0.22 | 0.19 | 0.04 | 0.26 | 0.22 |
| 7 | 142942447 | rs12703520 | 1.647 | -2.103 | -1.048 | -2.328 | -0.043 | 0.037 | 0.037 | 0 | 0.07 | 0.02 | 0.09 | 0.02 | 0.15 | 0.17 |
| 7 | 142954903 | rs6464540 | -0.156 | -2.683 | -0.779 | -2.103 | 0.021 | 0.022 | 0.022 | 0 | 0.39 | 0.22 | 0.19 | 0.04 | 0.26 | 0.22 |
| 7 | 143009487 | rs2140904 | 0.208 | -2.781 | -0.447 | -1.76 | -0.032 | -0.06 | -0.06 | 1.545 | 0.14 | 0.24 | 0.06 | 0 | 0.05 | 0.05 |
| 7 | 143014244 | rs10500136 | -0.759 | -2.212 | -0.235 | -1.46 | -0.097 | -0.121 | -0.121 | 0 | 0.25 | 0.44 | 0.11 | 0.03 | 0.1 | 0.16 |
| 7 | 143020649 | rs2272256 | -0.326 | -1.172 | -0.603 | -0.854 | 0.007 | 0.024 | 0.024 | 0.052 | 0.29 | 0.26 | 0.18 | 0.13 | 0.23 | 0.16 |
| 7 | 143020780 | rs2272255 | -0.942 | -1.842 | -0.411 | -1.324 | 0.055 | 0.119 | 0.119 | 0 | 0.52 | 0.71 | 0.29 | 0.16 | 0.34 | 0.32 |
| 7 | 143021110 | rs12667732 | 0.162 | -1.398 | -0.05 | -0.951 | -0.057 | -0.095 | -0.095 | 0.035 | 0.2 | 0.37 | 0.1 | 0.03 | 0.11 | 0.16 |
| 7 | 143023994 | rs940867 | 0.605 | -1.033 | -0.319 | -0.39 | 0.026 | 0.028 | 0.028 | 0.18 | 0.51 | 0.41 | 0.69 | 0.64 | 0.65 | 0.48 |
| 7 | 143024733 | rs4568530 | 0.411 | -0.523 | 0.45 | 0.333 | 0.033 | 0.137 | 0.137 | 0 | 0.92 | 0.68 | 0.94 | 0.83 | 0.92 | 0.85 |
| 7 | 143025152 | rs4327779 | 0.53 | -0.45 | 0.278 | 0.404 | 0.023 | 0.122 | 0.122 | 0 | 0.91 | 0.73 | 0.94 | 0.78 | 0.9 | 0.85 |
| 7 | 143025985 | rs4725613 | -0.018 | 0.052 | -0.47 | 0.049 | -0.001 | 0.036 | 0.036 | 0 | 0.47 | 0.37 | 0.32 | 0.2 | 0.52 | 0.33 |
| 7 | 143027055 | rs6954291 | -0.564 | -0.074 | -0.219 | 0.559 | -0.022 | 0.04 | 0.04 | 0.478 | 0.24 | 0.55 | 0.28 | 0.28 | 0.22 | 0.27 |
| 7 | 143028536 | rs2242493 | -0.314 | -0.003 | -0.747 | 0.06 | -0.017 | -0.037 | -0.037 | 0 | 0.56 | 0.5 | 0.7 | 0.83 | 0.67 | 0.57 |
| 7 | 143029983 | rs7794560 | 0.81 | 1.494 | -1.297 | 0.113 | -0.054 | 0.001 | 0.001 | 0 | 0.11 | 0.03 | 0.13 | 0.09 | 0.2 | 0.04 |
| 7 | 143031527 | rs6947788 | -1.055 | 1.035 | -1.307 | -0.119 | -0.002 | 0 | 0 | 0 | 0.87 | 0.95 | 0.9 | 0.88 | 0.86 | 0.85 |
| 7 | 143036318 | rs2272253 | 0.252 | 1.194 | -1.036 | 0.109 | 0.003 | 0.013 | 0.013 | 0 | 0.47 | 0.59 | 0.31 | 0.28 | 0.41 | 0.4 |
| 7 | 143042837 | rs2272251 | 0.39 | 1.225 | -1.059 | 0.136 | -0.004 | 0.001 | 0.001 | 0 | 0.4 | 0.36 | 0.59 | 0.61 | 0.46 | 0.44 |
| 7 | 143044381 | rs4726610 | 0.632 | 1.267 | -0.694 | 0.117 | 0.005 | 0.002 | 0.002 | 0 | 0.15 | 0.19 | 0.16 | 0.23 | 0.22 | 0.14 |
| 7 | 143045402 | rs10249881 | 0.07 | 1.242 | -0.683 | 0.322 | -0.003 | -0.017 | -0.017 | 0 | 0.46 | 0.49 | 0.29 | 0.28 | 0.41 | 0.41 |
| 7 | 143049965 | rs7806322 | 0.063 | 1.563 | -0.61 | 1.145 | 0.025 | 0.106 | 0.106 | 0 | 0.86 | 0.74 | 0.74 | 0.68 | 0.8 | 0.7 |
| 7 | 143049973 | rs7802536 | -0.138 | 1.874 | -1.019 | 0.825 | 0.011 | 0.011 | 0.011 | 0 | 0.14 | 0.18 | 0.14 | 0.26 | 0.21 | 0.15 |
| 7 | 143050370 | rs11771082 | 0.651 | 0.576 | -1.659 | 0.48 | -0.006 | 0.026 | 0.026 | 0.014 | 0.66 | 0.22 | 0.69 | 0.73 | 0.67 | 0.75 |
| 7 | 143054309 | rs4236482 | -0.64 | 0.19 | -1.325 | 0.637 | 0.009 | -0.017 | -0.017 | 0 | 0.22 | 0.11 | 0.24 | 0.26 | 0.21 | 0.15 |
| 7 | 143060291 | rs4726612 | 0.134 | 0.422 | -1.132 | 0.398 | 0.001 | 0.027 | 0.027 | 0 | 0.77 | 0.66 | 0.62 | 0.47 | 0.77 | 0.81 |
| 7 | 143066020 | rs10233736 | 0.483 | 0.355 | -1.23 | -0.531 | 0.006 | 0.017 | 0.017 | 0 | 0.5 | 0.58 | 0.39 | 0.18 | 0.54 | 0.54 |
| 7 | 143077036 | rs1135411 | -1.034 | 0.226 | -1.427 | 0.093 | 0.006 | 0.007 | 0.007 | 0 | 0.13 | 0.07 | 0.14 | 0.09 | 0.09 | 0.09 |
| 7 | 144241058 | rs17169957 | -1.973 | -0.363 | 2.388 | -0.663 | -0.02 | -0.089 | -0.089 | 0 | 0.15 | 0.65 | 0.1 | 0 | 0.09 | 0.05 |
| 7 | 144274735 | rs7792518 | -1.129 | 0.116 | 0.626 | -0.455 | 0.023 | -0.088 | -0.088 | 0 | 0.13 | 0.5 | 0.08 | 0 | 0.09 | 0.05 |

| | | | | | | | | | | | | | | | | |
|---|---|---|---|---|---|---|---|---|---|---|---|---|---|---|---|---|
| 7 | 144277770 | rs7804157 | 0.53 | -0.327 | 0.167 | -0.134 | -0.002 | 0 | 0 | 0.275 | 0.61 | 0.37 | 0.62 | 0.61 | 0.67 | 0.64 |
| 7 | 144277865 | rs9640426 | 0.04 | 0.458 | -0.404 | 0.153 | -0.039 | -0.04 | -0.04 | 1.538 | 0.1 | 0.04 | 0.23 | 0.42 | 0.17 | 0.19 |
| 7 | 144287580 | rs17170200 | -0.881 | -0.566 | 0.662 | -0.182 | 0.048 | 0.332 | 0.332 | 0.93 | 0.12 | 0.51 | 0.24 | 0.41 | 0.13 | 0.14 |
| 7 | 144295970 | rs12703621 | -0.048 | -0.773 | 0.01 | -0.607 | -0.018 | 0.095 | 0.095 | 0 | 0.79 | 0.95 | 0.87 | 0.97 | 0.81 | 0.85 |
| 7 | 144298474 | rs17170231 | -1.364 | -0.728 | 1.035 | 0.123 | 0.053 | 0.275 | 0.275 | 1.337 | 0.13 | 0.53 | 0.26 | 0.41 | 0.16 | 0.15 |
| 7 | 144301347 | rs10487637 | -1.43 | -0.761 | 1.211 | -0.166 | 0.003 | -0.041 | -0.041 | 0 | 0.09 | 0.29 | 0.06 | 0 | 0.05 | 0.01 |
| 7 | 144303474 | rs13245975 | -0.752 | -0.629 | 1.211 | 0.066 | 0.032 | 0.176 | 0.176 | 0 | 0.09 | 0.23 | 0.21 | 0.38 | 0.15 | 0.21 |
| 7 | 144308410 | rs10231525 | -1.997 | -1.804 | 1.483 | 0.099 | -0.053 | 0.123 | 0.123 | 1.131 | 0.17 | 0.61 | 0.25 | 0.37 | 0.1 | 0.19 |
| 7 | 144318049 | rs11981245 | -1.257 | -1.452 | 1.382 | -0.793 | -0.057 | -0.104 | -0.104 | 0.059 | 0.24 | 0.55 | 0.11 | 0 | 0.08 | 0.04 |
| 7 | 144358244 | rs17170428 | -1.096 | -0.687 | 1.19 | 0.608 | -0.006 | 0.025 | 0.025 | 0 | 0.07 | 0.08 | 0.13 | 0.24 | 0.03 | 0.22 |
| 7 | 144358948 | rs10269062 | -0.681 | -1.885 | 1.255 | -0.107 | -0.092 | -0.128 | -0.128 | 0 | 0.19 | 0.45 | 0.08 | 0 | 0.06 | 0.03 |
| 7 | 144368758 | rs1476612 | -0.441 | -1.81 | 0.53 | -0.274 | 0.014 | -0.014 | -0.014 | 3.823 | 0.42 | 0.74 | 0.35 | 0.45 | 0.34 | 0.42 |
| 7 | 144372803 | rs11978358 | 0.698 | -2.007 | 0.693 | -0.969 | 0.019 | -0.063 | -0.063 | 0 | 0.1 | 0.24 | 0.07 | 0 | 0.12 | 0.03 |
| 7 | 144377107 | rs7801499 | -0.425 | -2.107 | 0.575 | -1.014 | -0.019 | -0.098 | -0.098 | 1.063 | 0.15 | 0.6 | 0.11 | 0 | 0.12 | 0.03 |
| 7 | 144379429 | rs6956805 | -1.133 | -2.281 | 0.453 | -1.093 | 0.018 | -0.018 | -0.018 | 0 | 0.31 | 0.68 | 0.16 | 0 | 0.17 | 0.05 |
| 7 | 144392407 | rs6979644 | -1.498 | -2.397 | 0.372 | -0.941 | -0.068 | -0.062 | -0.062 | 0 | 0.65 | 0.82 | 0.46 | 0.45 | 0.52 | 0.47 |
| 7 | 144406318 | rs6954373 | 0.15 | -2.349 | 0.564 | -0.955 | -0.056 | -0.084 | -0.084 | 0.886 | 0.21 | 0.61 | 0.1 | 0 | 0.06 | 0.03 |
| 7 | 144408433 | rs228578 | -1.6 | -2.573 | -0.133 | -1.166 | -0.09 | -0.04 | -0.04 | 0 | 0.81 | 0.96 | 0.81 | 0.81 | 0.71 | 0.79 |
| 7 | 144419877 | rs228582 | -0.118 | -2.446 | -0.342 | -0.998 | -0.081 | -0.079 | -0.079 | 0.019 | 0.54 | 0.87 | 0.64 | 0.53 | 0.52 | 0.53 |
| 7 | 144425718 | rs6976400 | 0.803 | -2.368 | -0.154 | -0.967 | -0.065 | -0.11 | -0.11 | 0 | 0.18 | 0.36 | 0.08 | 0 | 0.06 | 0.03 |
| 7 | 144429379 | rs228587 | 1.098 | -1.8 | -0.448 | -0.909 | -0.057 | -0.09 | -0.09 | 0 | 0.31 | 0.16 | 0.51 | 0.53 | 0.43 | 0.44 |
| 7 | 144441187 | rs1547941 | 0.395 | -1.084 | -0.709 | -0.937 | -0.01 | -0.013 | -0.013 | 0 | 0.38 | 0.3 | 0.45 | 0.48 | 0.41 | 0.44 |
| 7 | 144452100 | rs17170937 | -0.211 | -1.391 | -0.603 | -0.954 | -0.023 | 0.103 | 0.103 | 0.209 | 0.18 | 0.41 | 0.24 | 0.32 | 0.14 | 0.3 |
| 7 | 144461670 | rs4725657 | -0.517 | -1.352 | -0.777 | -1.28 | -0.002 | 0.048 | 0.048 | 0.489 | 0.54 | 0.66 | 0.67 | 0.8 | 0.55 | 0.73 |
| 7 | 144461855 | rs4725658 | -0.517 | -1.352 | -0.777 | -1.28 | -0.002 | 0.048 | 0.048 | 0.489 | 0.54 | 0.65 | 0.67 | 0.8 | 0.55 | 0.73 |
| 7 | 144463792 | rs17170990 | 0.497 | -1.371 | -0.351 | -1.143 | -0.007 | -0.033 | -0.033 | 0 | 0.12 | 0.23 | 0.15 | 0.05 | 0.1 | 0.1 |
| 7 | 144465912 | rs1731117 | -0.764 | -1.045 | -1.284 | -1.406 | -0.098 | 0.057 | 0.057 | 0.873 | 0.63 | 0.91 | 0.43 | 0.19 | 0.51 | 0.3 |
| 7 | 144472276 | rs1731108 | -0.094 | -1.011 | -1.202 | -1.574 | 0.009 | 0.023 | 0.023 | 0.902 | 0.36 | 0.29 | 0.2 | 0.16 | 0.29 | 0.17 |
| 7 | 144472632 | rs1320825 | -0.094 | -1.011 | -1.202 | -1.574 | 0.009 | 0.023 | 0.023 | 0.902 | 0.36 | 0.29 | 0.2 | 0.16 | 0.29 | 0.17 |
| 7 | 144478662 | rs1635354 | 0.417 | -0.985 | -1.314 | -1.556 | -0.009 | -0.085 | -0.085 | 0 | 0.27 | 0.59 | 0.22 | 0.05 | 0.22 | 0.11 |
| 7 | 144480549 | rs1635357 | -0.315 | -0.464 | -1.168 | -1.157 | -0.018 | 0.047 | 0.047 | 1.429 | 0.58 | 0.73 | 0.42 | 0.19 | 0.5 | 0.23 |
| 7 | 144480830 | rs17171050 | 0.507 | 0.056 | -1.585 | -0.934 | -0.022 | -0.055 | -0.055 | 0.265 | 0.12 | 0.02 | 0.09 | 0.15 | 0.12 | 0.08 |
| 7 | 144497672 | rs2699471 | -0.84 | 0.31 | -1.428 | 0.249 | -0.116 | -0.128 | -0.128 | 0.825 | 0.7 | 0.95 | 0.53 | 0.39 | 0.54 | 0.48 |
| 7 | 144507976 | rs7455032 | 0.116 | -0.333 | -1.4 | 0.103 | -0.069 | 0.076 | 0.076 | 3.374 | 0.57 | 0.89 | 0.4 | 0.22 | 0.44 | 0.24 |
| 7 | 144513549 | rs2371642 | -0.539 | -0.241 | -1.39 | 0.36 | -0.142 | -0.139 | -0.139 | 0.183 | 0.66 | 0.9 | 0.52 | 0.46 | 0.51 | 0.55 |
| 7 | 144524224 | rs6962350 | -0.147 | -0.521 | -1 | 0.16 | -0.031 | 0.063 | 0.063 | 5.151 | 0.59 | 0.8 | 0.45 | 0.28 | 0.46 | 0.44 |
| 7 | 144532104 | rs10261806 | -0.042 | -0.577 | -1.001 | 0.148 | -0.025 | 0.079 | 0.079 | 3.061 | 0.57 | 0.84 | 0.45 | 0.28 | 0.44 | 0.43 |
| 7 | 144543167 | rs1881849 | -0.028 | -0.672 | -0.414 | 0.073 | -0.014 | 0.086 | 0.086 | 3.572 | 0.57 | 0.8 | 0.45 | 0.28 | 0.44 | 0.43 |
| 7 | 144566296 | rs6958341 | 0.641 | -0.772 | -0.49 | -0.12 | 0.039 | 0.104 | 0.104 | 1.536 | 0.43 | 0.7 | 0.3 | 0.05 | 0.31 | 0.14 |
| 7 | 144571200 | rs1881847 | -1.272 | -0.732 | -0.533 | 0.075 | -0.001 | 0.004 | 0.004 | 0 | 0.24 | 0.27 | 0.23 | 0.4 | 0.2 | 0.43 |
| 7 | 144574232 | rs2371645 | 0.189 | -0.609 | -0.883 | 0.107 | -0.015 | -0.045 | -0.045 | 0.386 | 0.08 | 0.01 | 0.11 | 0.23 | 0.09 | 0.29 |
| 7 | 144582157 | rs11761362 | -1.012 | -0.565 | -0.881 | 0.085 | -0.189 | -0.185 | -0.185 | 5.48 | 0.67 | 0.96 | 0.52 | 0.45 | 0.51 | 0.56 |
| 7 | 144603425 | rs2692390 | 0.773 | -0.614 | -0.568 | 0.059 | -0.103 | -0.099 | -0.099 | 0 | 0.34 | 0.12 | 0.49 | 0.55 | 0.49 | 0.44 |
| 7 | 144612152 | rs10231646 | 0.76 | -0.883 | -0.013 | -0.07 | 0.011 | 0.015 | 0.015 | 1.02 | 0.39 | 0.27 | 0.26 | 0.06 | 0.3 | 0.14 |
| 7 | 144621695 | rs2699467 | -0.159 | -0.795 | -0.071 | 0.124 | -0.002 | 0.123 | 0.123 | 0.528 | 0.87 | 0.54 | 0.91 | 0.84 | 0.92 | 0.86 |
| 7 | 144622959 | rs2699473 | -0.351 | -0.364 | -0.794 | 0.267 | 0.04 | 0.107 | 0.107 | 2.248 | 0.49 | 0.22 | 0.65 | 0.78 | 0.62 | 0.72 |
| 7 | 144633224 | rs2692394 | 0.518 | -0.215 | -1.348 | 0.449 | -0.177 | -0.183 | -0.183 | 0 | 0.36 | 0.05 | 0.52 | 0.54 | 0.52 | 0.5 |
| 7 | 144638953 | rs10500143 | -0.397 | -0.442 | -0.543 | 0.554 | -0.02 | 0.039 | 0.039 | 0 | 0.12 | 0.17 | 0.11 | 0.32 | 0.08 | 0.26 |
| 7 | 144639494 | rs1525463 | 0.522 | -0.888 | -1.095 | 0.329 | -0.013 | 0.069 | 0.069 | 0 | 0.44 | 0.22 | 0.58 | 0.29 | 0.44 | 0.35 |
| 7 | 144649240 | rs6464631 | 0.869 | -0.69 | -0.688 | 0.305 | -0.01 | 0.022 | 0.022 | 0 | 0.18 | 0.27 | 0.08 | 0.2 | 0.11 | 0.15 |
| 7 | 144652109 | rs10500145 | 1.296 | -0.417 | -0.82 | -0.047 | -0.003 | -0.002 | -0.002 | 0 | 0.14 | 0.11 | 0.06 | 0 | 0.1 | 0.03 |
| 7 | 144653582 | rs6980235 | 0.822 | -0.283 | -0.567 | 0.521 | -0.082 | 0.002 | 0.002 | 0.493 | 0.23 | 0.07 | 0.22 | 0.18 | 0.38 | 0.24 |
| 7 | 144654042 | rs4397312 | 0.261 | 0.033 | -0.564 | 0.695 | 0.004 | 0.015 | 0.015 | 0 | 0.61 | 0.48 | 0.66 | 0.51 | 0.56 | 0.5 |
| 7 | 144654138 | rs11767967 | -0.142 | -0.002 | -0.863 | 1.178 | -0.001 | -0.005 | -0.005 | 0 | 0.05 | 0 | 0.06 | 0.32 | 0.04 | 0.25 |
| 7 | 144656932 | rs10500146 | -0.398 | -0.001 | -0.843 | 1.179 | 0.006 | 0.015 | 0.015 | 0 | 0.06 | 0.03 | 0.07 | 0.32 | 0.04 | 0.25 |
| 7 | 144672571 | rs1554699 | 0.385 | -0.114 | -0.85 | 0.693 | 0.012 | -0.012 | -0.012 | 0 | 0.56 | 0.39 | 0.51 | 0.51 | 0.63 | 0.45 |
| 7 | 144673883 | rs12703633 | 0.813 | 0.552 | -1.618 | 0.664 | -0.072 | -0.088 | -0.088 | 0.166 | 0.16 | 0.02 | 0.12 | 0.25 | 0.18 | 0.15 |
| 7 | 144676336 | rs12673350 | 0.274 | 0.209 | -1.193 | 0.499 | -0.013 | -0.077 | -0.077 | 0.658 | 0.39 | 0.16 | 0.45 | 0.5 | 0.37 | 0.55 |
| 7 | 144679309 | rs10258763 | 1.136 | -0.131 | -0.966 | 0.219 | -0.012 | 0.024 | 0.024 | 0.076 | 0.31 | 0.25 | 0.37 | 0.25 | 0.42 | 0.27 |
| 7 | 144684050 | rs1525459 | -0.08 | -0.46 | -0.643 | 0.163 | -0.048 | 0.009 | 0.009 | 2.293 | 0.75 | 0.55 | 0.86 | 0.72 | 0.87 | 0.74 |
| 7 | 144687913 | rs6464634 | -1.044 | -0.269 | -0.16 | -0.009 | -0.001 | -0.025 | -0.025 | 0 | 0.92 | 0.7 | 0.96 | 1 | 0.98 | 0.99 |
| 7 | 144688140 | rs7797520 | 0.157 | 0.066 | -0.565 | 0.16 | 0.006 | 0.029 | 0.029 | 0 | 0.26 | 0.31 | 0.2 | 0.53 | 0.24 | 0.42 |
| 7 | 144695212 | rs10250889 | 0.541 | 0.433 | -0.857 | 0.147 | -0.017 | 0.028 | 0.028 | 0.058 | 0.31 | 0.19 | 0.37 | 0.26 | 0.42 | 0.27 |
| 7 | 144702577 | rs10251934 | 0.097 | 0.261 | -0.588 | 0.216 | 0.001 | 0.002 | 0.002 | 0.005 | 0.35 | 0.33 | 0.19 | 0.23 | 0.32 | 0.21 |
| 7 | 144703987 | rs17394323 | -0.079 | 0.161 | -0.678 | 0.026 | -0.027 | -0.016 | -0.016 | 0 | 0.35 | 0.27 | 0.51 | 0.59 | 0.47 | 0.47 |
| 7 | 144706092 | rs12671406 | 0.339 | 0.794 | -0.749 | -0.103 | -0.003 | -0.01 | -0.01 | 0 | 0.28 | 0.22 | 0.44 | 0.59 | 0.34 | 0.4 |
| 7 | 144713191 | rs11526228 | 0.387 | 0.703 | -0.396 | -0.189 | -0.037 | 0.053 | 0.053 | 0 | 0.41 | 0.28 | 0.26 | 0.25 | 0.45 | 0.27 |
| 7 | 144720781 | rs7799590 | -1.068 | 0.691 | -0.331 | -0.195 | -0.037 | -0.055 | -0.055 | 0.45 | 0.7 | 0.5 | 0.7 | 0.84 | 0.79 | 0.68 |
| 7 | 144721441 | rs7781772 | 0.495 | 0.664 | -0.63 | -0.325 | -0.05 | 0.04 | 0.04 | 0 | 0.38 | 0.23 | 0.24 | 0.25 | 0.42 | 0.26 |

| | | | | | | | | | | | | | | | | |
|---|---|---|---|---|---|---|---|---|---|---|---|---|---|---|---|---|
| 7 | 144735177 | rs10808030 | 0.767 | 0.42 | -0.473 | -0.034 | -0.061 | -0.132 | -0.132 | 1.467 | 0.25 | 0.05 | 0.41 | 0.49 | 0.34 | 0.38 |
| 7 | 144743405 | rs883839 | 0.104 | 0.53 | -0.405 | -0.114 | 0.015 | -0.02 | -0.02 | 0.157 | 0.08 | 0.11 | 0.08 | 0.01 | 0.14 | 0.07 |
| 7 | 144744047 | rs885367 | 0.753 | 0.251 | -0.407 | -0.045 | -0.055 | -0.123 | -0.123 | 0.534 | 0.25 | 0.05 | 0.41 | 0.48 | 0.34 | 0.39 |
| 7 | 144747898 | rs6464644 | 0.45 | 0.055 | 0.328 | -0.197 | -0.007 | -0.016 | -0.016 | 0.186 | 0.37 | 0.57 | 0.22 | 0.23 | 0.32 | 0.2 |
| 7 | 144757945 | rs954072 | -0.269 | 0.241 | 0.324 | -0.191 | 0.002 | 0.013 | 0.013 | 0 | 0.45 | 0.61 | 0.28 | 0.24 | 0.41 | 0.26 |
| 7 | 144789291 | rs7804647 | 0.22 | 0.367 | 0.109 | -0.106 | -0.01 | 0.012 | 0.012 | 0.254 | 0.07 | 0.07 | 0.05 | 0.01 | 0.09 | 0.06 |
| 7 | 144789432 | rs17728337 | 0.888 | 0.292 | 0.069 | -0.167 | 0.005 | 0.011 | 0.011 | 0 | 0.33 | 0.22 | 0.19 | 0.22 | 0.3 | 0.2 |
| 7 | 144795359 | rs17795722 | 0.02 | 0.37 | -0.081 | -0.116 | -0.01 | 0.011 | 0.011 | 0.209 | 0.07 | 0.07 | 0.06 | 0.01 | 0.09 | 0.05 |
| 7 | 144798554 | rs7789134 | -0.249 | 0.119 | -0.046 | -0.022 | 0.003 | 0.034 | 0.034 | 0 | 0.58 | 0.69 | 0.62 | 0.75 | 0.59 | 0.5 |
| 7 | 144801464 | rs9640440 | 0.31 | 0.191 | -0.411 | -0.014 | -0.041 | 0.071 | 0.071 | 0 | 0.44 | 0.22 | 0.51 | 0.68 | 0.49 | 0.51 |
| 7 | 144801924 | rs17173004 | 1.351 | 0.221 | -0.455 | 0.07 | -0.021 | -0.007 | -0.007 | 0 | 0.13 | 0.04 | 0.1 | 0.17 | 0.13 | 0.06 |
| 7 | 144810436 | rs4726739 | 0.703 | 0.447 | -0.483 | 0.376 | -0.002 | 0.037 | 0.037 | 0.013 | 0.12 | 0.14 | 0.27 | 0.45 | 0.2 | 0.35 |
| 7 | 144810487 | rs13328363 | -0.392 | 0.291 | -0.049 | 0.221 | -0.019 | -0.038 | -0.038 | 0 | 0.07 | 0.2 | 0.04 | 0 | 0.05 | 0.03 |
| 7 | 144810953 | rs509853 | 1.51 | 0.253 | -0.73 | 0.145 | -0.046 | 0.028 | 0.028 | 0.314 | 0.28 | 0.03 | 0.27 | 0.15 | 0.24 | 0.37 |
| 7 | 144811326 | rs605627 | 0.855 | 0.261 | -0.378 | 0.196 | 0.001 | 0.006 | 0.006 | 0 | 0.35 | 0.29 | 0.34 | 0.15 | 0.33 | 0.4 |
| 7 | 144822060 | rs17796261 | 1.408 | 0.295 | -1.033 | 0.219 | -0.085 | -0.003 | -0.003 | 4.362 | 0.31 | 0.04 | 0.24 | 0.22 | 0.31 | 0.17 |
| 7 | 144824230 | rs642782 | 0.701 | 0.478 | -0.487 | 0.467 | -0.001 | 0.037 | 0.037 | 0 | 0.34 | 0.21 | 0.33 | 0.15 | 0.33 | 0.4 |
| 7 | 144825976 | rs1997089 | 0.269 | 0.713 | -0.582 | 1.099 | -0.007 | 0.01 | 0.01 | 0.033 | 0.12 | 0.09 | 0.27 | 0.42 | 0.18 | 0.34 |
| 7 | 144828398 | rs564697 | 1.173 | 0.589 | -0.876 | 0.855 | -0.063 | -0.058 | -0.058 | 0 | 0.21 | 0.04 | 0.26 | 0.19 | 0.26 | 0.38 |
| 7 | 144828813 | rs10228724 | 0.417 | 0.16 | -0.808 | 0.576 | -0.004 | 0.011 | 0.011 | 0 | 0.55 | 0.61 | 0.46 | 0.42 | 0.53 | 0.29 |
| 7 | 144830345 | rs7781379 | -0.329 | -0.266 | -0.755 | 0.454 | -0.001 | -0.003 | -0.003 | 0 | 0.48 | 0.53 | 0.47 | 0.64 | 0.45 | 0.48 |
| 7 | 144832263 | rs10487525 | 1.533 | -0.025 | -0.956 | 0.201 | -0.056 | -0.029 | -0.029 | 0 | 0.23 | 0.05 | 0.2 | 0.22 | 0.25 | 0.13 |
| 7 | 144840057 | rs1525734 | -1.302 | 0.058 | -0.872 | 0.351 | -0.02 | 0.007 | 0.007 | 0 | 0.73 | 0.87 | 0.66 | 0.81 | 0.73 | 0.59 |
| 7 | 144840724 | rs725849 | -0.386 | 0.027 | -0.696 | 0.326 | 0.008 | 0.021 | 0.021 | 3.766 | 0.37 | 0.47 | 0.41 | 0.57 | 0.45 | 0.45 |
| 7 | 144841531 | rs10487524 | 0.146 | 0.336 | -0.904 | 0.613 | -0.027 | -0.018 | -0.018 | 1.772 | 0.1 | 0.05 | 0.15 | 0.47 | 0.18 | 0.33 |
| 7 | 144842546 | rs4726744 | -1.034 | 0.081 | -0.656 | 0.044 | -0.001 | 0.006 | 0.006 | 0 | 0.72 | 0.73 | 0.65 | 0.81 | 0.73 | 0.59 |
| 7 | 144855345 | rs2040979 | -0.612 | -0.296 | -0.332 | -0.277 | -0.008 | -0.029 | -0.029 | 0 | 0.4 | 0.51 | 0.3 | 0.13 | 0.31 | 0.15 |
| 7 | 144865200 | rs10238114 | 0.397 | -0.031 | 0.115 | -0.153 | -0.007 | -0.029 | -0.029 | 0 | 0.18 | 0.29 | 0.19 | 0.09 | 0.16 | 0.09 |
| 7 | 144866710 | rs16884210 | 0.721 | -0.026 | -0.226 | 0.194 | 0.003 | 0.035 | 0.035 | 3.635 | 0.31 | 0.09 | 0.2 | 0.22 | 0.26 | 0.15 |
| 7 | 144868611 | rs6953302 | -0.797 | 0.144 | 0.302 | 0.604 | 0.01 | 0.121 | 0.121 | 0 | 0.15 | 0.28 | 0.15 | 0.44 | 0.19 | 0.32 |
| 7 | 144872496 | rs17173023 | 0.145 | 0.254 | 0.149 | 0.595 | 0.012 | 0.105 | 0.105 | 0 | 0.08 | 0.15 | 0.11 | 0.44 | 0.15 | 0.3 |
| 7 | 144872969 | rs17173024 | 0.412 | -0.15 | 0.176 | 0.274 | 0.013 | 0.012 | 0.012 | 0.25 | 0.36 | 0.29 | 0.22 | 0.23 | 0.27 | 0.17 |
| 7 | 144874336 | rs6947029 | 0.159 | -0.163 | 0.305 | 0.262 | -0.028 | -0.027 | -0.027 | 0.723 | 0.38 | 0.44 | 0.24 | 0.23 | 0.27 | 0.18 |
| 7 | 144875551 | rs9640443 | 0.497 | -0.036 | -0.375 | 0.305 | 0.026 | 0.03 | 0.03 | 1.314 | 0.34 | 0.18 | 0.22 | 0.23 | 0.26 | 0.17 |
| 7 | 144880214 | rs7800213 | 0.205 | -0.005 | -0.17 | -0.034 | 0 | 0 | 0 | 0.191 | 0.39 | 0.35 | 0.31 | 0.67 | 0.39 | 0.52 |
| 7 | 144880643 | rs7804383 | -0.43 | 0.157 | -0.131 | -0.188 | 0.002 | -0.011 | -0.011 | 0.05 | 0.54 | 0.51 | 0.43 | 0.7 | 0.51 | 0.56 |
| 7 | 144883188 | rs4726747 | -0.527 | -0.011 | 0.111 | -0.478 | -0.029 | -0.053 | -0.053 | 0.776 | 0.11 | 0.56 | 0.09 | 0.01 | 0.05 | 0.02 |
| 7 | 144886066 | rs10232519 | -1.109 | 0.506 | -0.481 | -0.314 | -0.003 | -0.001 | -0.001 | 0.039 | 0.74 | 0.84 | 0.67 | 0.75 | 0.72 | 0.62 |
| 7 | 144891618 | rs13221722 | -0.633 | 0.622 | -0.668 | -0.248 | -0.018 | 0.036 | 0.036 | 0 | 0.24 | 0.13 | 0.3 | 0.12 | 0.26 | 0.09 |
| 7 | 144892086 | rs12531488 | -0.638 | 0.635 | -0.651 | -0.242 | -0.005 | 0.016 | 0.016 | 0 | 0.26 | 0.22 | 0.31 | 0.12 | 0.26 | 0.09 |
| 7 | 144893242 | rs7788595 | -0.456 | 0.626 | -0.863 | 0.041 | -0.016 | 0.032 | 0.032 | 0.545 | 0.13 | 0.05 | 0.13 | 0.02 | 0.12 | 0.02 |
| 7 | 144906316 | rs10272139 | 0.554 | 1.095 | -0.624 | -0.027 | 0 | -0.018 | -0.018 | 0 | 0.89 | 0.85 | 0.9 | 0.98 | 0.89 | 0.95 |
| 7 | 144906538 | rs8180879 | -0.039 | 1.155 | -0.751 | 0.496 | -0.001 | 0.014 | 0.014 | 0 | 0.11 | 0.06 | 0.09 | 0.51 | 0.1 | 0.28 |
| 7 | 144906844 | rs7791708 | 1.9 | 1.165 | -0.615 | 0.384 | -0.003 | -0.004 | -0.004 | 0 | 0.72 | 0.68 | 0.72 | 0.46 | 0.7 | 0.66 |
| 7 | 144909144 | rs12673241 | -0.824 | 1.372 | -0.755 | 0.371 | -0.027 | -0.036 | -0.036 | 0 | 0.16 | 0.11 | 0.2 | 0.53 | 0.24 | 0.3 |
| 7 | 144916029 | rs10281160 | 0.195 | 1.334 | -0.726 | 0.33 | -0.001 | 0 | 0 | 0.524 | 0.9 | 0.88 | 0.91 | 0.98 | 0.89 | 0.96 |
| 7 | 144940098 | rs4725670 | 1.403 | 1.135 | -0.289 | 0.348 | 0.021 | 0.057 | 0.057 | 0 | 0.75 | 0.64 | 0.73 | 0.45 | 0.7 | 0.67 |
| 7 | 144945903 | rs7810304 | 0.908 | 1.033 | -0.759 | -0.119 | -0.003 | -0.015 | -0.015 | 0 | 0.36 | 0.36 | 0.33 | 0.15 | 0.32 | 0.29 |
| 7 | 144948010 | rs7777430 | 0.981 | 1.078 | -0.864 | -0.11 | 0.023 | 0.065 | 0.065 | 0 | 0.74 | 0.6 | 0.72 | 0.45 | 0.7 | 0.67 |
| 7 | 144948360 | rs4725672 | 0.762 | 0.526 | -1.615 | 0.077 | -0.062 | -0.04 | -0.04 | 0 | 0.37 | 0.03 | 0.38 | 0.3 | 0.39 | 0.37 |
| 7 | 144955976 | rs3919487 | 0.888 | 0.628 | -1.551 | 0.06 | 0.012 | 0.025 | 0.025 | 0 | 0.36 | 0.21 | 0.31 | 0.28 | 0.36 | 0.37 |
| 7 | 144958662 | rs4726752 | -0.4 | 0.716 | -1.216 | 0.655 | 0.005 | 0 | 0 | 1.033 | 0.08 | 0.08 | 0.06 | 0 | 0.07 | 0.01 |
| 7 | 144959531 | rs957452 | -0.592 | 0.963 | -1.099 | 1.043 | -0.008 | 0.024 | 0.024 | 0.174 | 0.1 | 0.05 | 0.16 | 0.53 | 0.12 | 0.27 |
| 7 | 144971709 | rs2190073 | -0.491 | 0.468 | -1.478 | -0.077 | 0.02 | 0.088 | 0.088 | 0 | 0.29 | 0.12 | 0.19 | 0.12 | 0.27 | 0.17 |
| 7 | 144976210 | rs1118226 | -0.291 | 0.621 | -2.168 | -0.773 | -0.007 | 0.041 | 0.041 | 0 | 0.67 | 0.28 | 0.66 | 0.92 | 0.68 | 0.79 |
| 7 | 144979610 | rs1074131 | 0.853 | 0.573 | -2.189 | -0.801 | -0.046 | 0.117 | 0.117 | 0.051 | 0.39 | 0.16 | 0.47 | 0.8 | 0.41 | 0.61 |
| 7 | 144986475 | rs10231714 | -0.278 | 0.579 | -2.186 | -0.732 | -0.007 | 0.04 | 0.04 | 0 | 0.68 | 0.28 | 0.66 | 0.92 | 0.68 | 0.78 |
| 7 | 144986643 | rs10280702 | 0.478 | 0.615 | -2.217 | -0.763 | 0.001 | 0.111 | 0.111 | 0.071 | 0.28 | 0.09 | 0.19 | 0.12 | 0.27 | 0.18 |
| 7 | 144986806 | rs1859566 | 1.27 | 0.437 | -2.52 | 0.157 | -0.066 | -0.105 | -0.105 | 0 | 0.27 | 0.05 | 0.28 | 0.33 | 0.29 | 0.34 |
| 7 | 144990171 | rs12673814 | 0.541 | 0.193 | -2.34 | -0.621 | -0.048 | 0.103 | 0.103 | 0.047 | 0.39 | 0.16 | 0.47 | 0.8 | 0.41 | 0.61 |
| 7 | 144990352 | rs7806648 | 0.524 | -0.488 | -1.046 | -0.94 | -0.029 | -0.017 | -0.017 | 0 | 0.09 | 0.39 | 0.04 | 0 | 0.01 | 0 |
| 7 | 144991291 | rs2023995 | 0.176 | -0.422 | -0.874 | -0.995 | 0.002 | 0.001 | 0.001 | 0 | 0.47 | 0.58 | 0.51 | 0.81 | 0.41 | 0.62 |
| 7 | 145000958 | rs7779216 | -1.21 | 0.262 | -1.001 | -1.198 | -0.013 | 0.011 | 0.011 | 0 | 0.94 | 0.97 | 0.93 | 0.99 | 0.87 | 0.97 |
| 7 | 145005608 | rs2190083 | -0.169 | 0.136 | -0.741 | -1.319 | 0.018 | -0.018 | -0.018 | 0 | 0.79 | 0.8 | 0.71 | 0.92 | 0.69 | 0.79 |
| 7 | 145018727 | rs996342 | 0.98 | -0.001 | -0.348 | -1.405 | -0.013 | 0.086 | 0.086 | 0.024 | 0.4 | 0.27 | 0.47 | 0.81 | 0.41 | 0.62 |
| 7 | 145024241 | rs11770321 | 0.976 | -0.008 | -0.368 | -1.427 | -0.003 | -0.019 | -0.019 | 0.027 | 0.15 | 0.11 | 0.11 | 0 | 0.18 | 0.16 |
| 7 | 145024376 | rs2107890 | 0.274 | -0.015 | -0.421 | -1.43 | -0.004 | -0.052 | -0.052 | 0.26 | 0.56 | 0.46 | 0.69 | 0.87 | 0.59 | 0.79 |
| 7 | 145024780 | rs7792751 | -0.096 | -0.047 | -0.499 | -1.434 | -0.005 | 0.069 | 0.069 | 0.316 | 0.44 | 0.28 | 0.42 | 0.18 | 0.45 | 0.35 |
| 7 | 145034721 | rs4726759 | 0.981 | -0.06 | -0.512 | -1.436 | 0.006 | 0.116 | 0.116 | 0.077 | 0.28 | 0.09 | 0.19 | 0.12 | 0.27 | 0.18 |

| | | | | | | | | | | | | | | | | |
|---|---|---|---|---|---|---|---|---|---|---|---|---|---|---|---|---|
| 7 | 145037728 | rs6958143 | -0.091 | -0.101 | -0.497 | -1.431 | -0.005 | 0.069 | 0.069 | 0.348 | 0.44 | 0.28 | 0.42 | 0.18 | 0.45 | 0.36 |
| 7 | 145045961 | rs10252914 | 0.994 | -0.007 | -0.58 | -1.432 | 0.014 | 0.112 | 0.112 | 0.133 | 0.27 | 0.09 | 0.18 | 0.12 | 0.26 | 0.17 |
| 7 | 145047897 | rs17173069 | 1.16 | -0.025 | -0.712 | -1.529 | 0.032 | 0.079 | 0.079 | 0.064 | 0.23 | 0.03 | 0.13 | 0.12 | 0.18 | 0.16 |
| 7 | 145048236 | rs193463 | -0.056 | 0.062 | -0.091 | -1.482 | 0.006 | -0.094 | -0.094 | 0 | 0.86 | 0.66 | 0.9 | 0.99 | 0.86 | 0.97 |
| 7 | 145048455 | rs10264696 | 0.538 | 0.178 | -0.221 | -1.33 | 0.027 | -0.027 | -0.027 | 0 | 0.17 | 0.24 | 0.24 | 0.07 | 0.19 | 0.18 |
| 7 | 145055797 | rs2023551 | -0.043 | 0.02 | -0.34 | -1.372 | 0.014 | 0.062 | 0.062 | 0.899 | 0.41 | 0.25 | 0.35 | 0.2 | 0.36 | 0.32 |
| 7 | 145068124 | rs41675 | 1.151 | -0.068 | -0.153 | -1.232 | -0.012 | 0.017 | 0.017 | 0.059 | 0.35 | 0.3 | 0.47 | 0.74 | 0.39 | 0.6 |
| 7 | 145069168 | rs10233618 | -0.423 | 0.169 | 0.761 | -0.981 | 0.053 | -0.06 | -0.06 | 0.915 | 0.22 | 0.38 | 0.29 | 0.08 | 0.28 | 0.18 |
| 7 | 145070430 | rs768152 | -0.371 | 0.149 | 0.752 | -0.919 | 0.064 | -0.041 | -0.041 | 0 | 0.21 | 0.38 | 0.29 | 0.1 | 0.27 | 0.17 |
| 7 | 145071087 | rs10257092 | -0.308 | 0.144 | 0.926 | -0.818 | 0.046 | -0.025 | -0.025 | 0 | 0.21 | 0.32 | 0.3 | 0.11 | 0.28 | 0.16 |
| 7 | 145071775 | rs10952617 | 0.264 | 0.15 | 0.932 | -0.993 | 0.001 | -0.027 | -0.027 | 0.035 | 0.11 | 0.09 | 0.08 | 0.01 | 0.12 | 0.04 |
| 7 | 145081764 | rs1100687 | -0.228 | 0.19 | 0.95 | -0.959 | 0.028 | -0.019 | -0.019 | 0 | 0.19 | 0.29 | 0.28 | 0.1 | 0.25 | 0.16 |
| 7 | 145092128 | rs17169669 | -0.017 | 0.202 | 0.712 | -0.983 | 0 | -0.019 | -0.019 | 0 | 0.12 | 0.09 | 0.09 | 0.01 | 0.14 | 0.05 |
| 7 | 145092324 | rs850345 | -0.027 | 0.268 | 0.596 | -0.832 | 0.004 | -0.002 | -0.002 | 0.56 | 0.18 | 0.17 | 0.27 | 0.1 | 0.25 | 0.16 |
| 7 | 145105169 | rs1406039 | 0.172 | 0.254 | 0.39 | -0.95 | -0.039 | 0.04 | 0.04 | 1.391 | 0.16 | 0.1 | 0.16 | 0 | 0.24 | 0.14 |
| 7 | 145120099 | rs1528023 | -0.416 | 0.048 | 0.551 | -1.042 | -0.006 | -0.021 | -0.021 | 0 | 0.2 | 0.21 | 0.17 | 0 | 0.24 | 0.14 |
| 7 | 145121559 | rs850323 | -1.556 | -0.031 | 0.742 | -1.146 | 0.014 | -0.059 | -0.059 | 0.157 | 0.35 | 0.54 | 0.38 | 0.1 | 0.37 | 0.2 |
| 7 | 145123344 | rs9640453 | 0.607 | -0.412 | 0.192 | -1.368 | 0.014 | 0.012 | 0.012 | 0 | 0.34 | 0.3 | 0.15 | 0.22 | 0.2 | 0.19 |
| 7 | 145130880 | rs850398 | -0.104 | -0.137 | 0.559 | -1.075 | 0 | 0.005 | 0.005 | 0.017 | 0.53 | 0.48 | 0.43 | 0.3 | 0.46 | 0.33 |
| 7 | 145130995 | rs10236257 | 0.127 | -0.112 | 0.361 | -1.084 | 0.013 | 0.09 | 0.09 | 0.083 | 0.51 | 0.36 | 0.3 | 0.22 | 0.45 | 0.3 |
| 7 | 145133150 | rs850390 | -0.403 | -0.19 | 0.451 | -0.85 | 0.016 | -0.01 | -0.01 | 0 | 0.15 | 0.36 | 0.26 | 0.1 | 0.14 | 0.06 |
| 7 | 145141776 | rs850377 | -0.267 | -0.257 | -0.075 | -1.11 | 0.002 | -0.004 | -0.004 | 6.183 | 0.44 | 0.61 | 0.25 | 0.18 | 0.36 | 0.25 |
| 7 | 145145326 | rs850365 | 0.029 | 0.163 | 0.231 | -1.102 | 0.012 | -0.036 | -0.036 | 0 | 0.16 | 0.25 | 0.24 | 0.05 | 0.19 | 0.11 |
| 7 | 145149739 | rs850360 | -1.227 | -0.095 | 0.111 | -1.027 | 0 | -0.004 | -0.004 | 0.361 | 0.54 | 0.48 | 0.58 | 0.38 | 0.52 | 0.38 |
| 7 | 145154459 | rs10269546 | 0.001 | 0.049 | -0.156 | -0.903 | -0.029 | -0.049 | -0.049 | 0 | 0.23 | 0.16 | 0.4 | 0.34 | 0.29 | 0.22 |
| 7 | 145158233 | rs4725676 | -0.069 | -0.387 | 0.855 | -1.288 | -0.034 | -0.128 | -0.128 | 0 | 0.79 | 0.64 | 0.9 | 1 | 0.89 | 0.91 |
| 7 | 145160477 | rs13228818 | -0.001 | -0.282 | 0.404 | -1.256 | 0.009 | -0.003 | -0.003 | 0 | 0.19 | 0.15 | 0.08 | 0 | 0.11 | 0.09 |
| 7 | 145167006 | rs1523955 | 0.073 | -0.25 | 0.183 | -1.266 | 0.005 | -0.027 | -0.027 | 0 | 0.1 | 0.13 | 0.07 | 0.04 | 0.12 | 0.05 |
| 7 | 145169598 | rs6974092 | 0.506 | -0.237 | -0.131 | -1.197 | -0.022 | -0.123 | -0.123 | 0 | 0.69 | 0.56 | 0.83 | 0.96 | 0.77 | 0.86 |
| 7 | 145171914 | rs10808033 | 0.213 | -0.118 | -0.559 | -1.001 | -0.01 | -0.069 | -0.069 | 1.04 | 0.61 | 0.47 | 0.81 | 0.87 | 0.68 | 0.79 |
| 7 | 145175716 | rs6947759 | 0.451 | 0.274 | -1.02 | 0.008 | -0.038 | -0.071 | -0.071 | 0 | 0.21 | 0.1 | 0.27 | 0.34 | 0.24 | 0.34 |
| 7 | 145182425 | rs2372052 | -0.605 | -0.23 | -0.77 | 0.016 | 0 | 0.001 | 0.001 | 0 | 0.46 | 0.48 | 0.42 | 0.56 | 0.42 | 0.53 |
| 7 | 145182535 | rs2372053 | 0.465 | 0.229 | -0.92 | -0.063 | -0.033 | -0.059 | -0.059 | 0 | 0.2 | 0.11 | 0.28 | 0.34 | 0.24 | 0.33 |
| 7 | 145190638 | rs12532655 | 0.629 | 0.287 | -1.003 | -0.05 | -0.067 | -0.084 | -0.084 | 0 | 0.19 | 0.07 | 0.28 | 0.33 | 0.26 | 0.33 |
| 7 | 145193259 | rs10441206 | -0.6 | -0.097 | -0.002 | -0.12 | 0.006 | 0.058 | 0.058 | 0.084 | 0.06 | 0.33 | 0.04 | 0.07 | 0.01 | 0.02 |
| 7 | 145206498 | rs6952800 | -0.705 | -0.547 | -0.253 | -0.512 | -0.002 | 0 | 0 | 0 | 0.45 | 0.58 | 0.43 | 0.46 | 0.41 | 0.54 |
| 7 | 145208265 | rs7804968 | 0.611 | -0.653 | -0.355 | -0.567 | 0.035 | 0.063 | 0.063 | 0 | 0.17 | 0.02 | 0.09 | 0.05 | 0.13 | 0.18 |
| 7 | 145209049 | rs10280300 | -0.065 | -0.409 | 0.002 | -0.414 | 0.013 | 0.071 | 0.071 | 0.454 | 0.27 | 0.53 | 0.33 | 0.4 | 0.27 | 0.36 |
| 7 | 145223795 | rs17475339 | 1.089 | 0.077 | -0.684 | -1.265 | -0.061 | 0.097 | 0.097 | 0 | 0.23 | 0.05 | 0.15 | 0.07 | 0.28 | 0.16 |
| 7 | 145225338 | rs17169739 | 0.754 | 0.059 | 0.071 | -0.734 | -0.004 | 0.094 | 0.094 | 0 | 0.11 | 0.16 | 0.26 | 0.39 | 0.11 | 0.15 |
| 7 | 145231464 | rs7780152 | -0.408 | 0.147 | 0.09 | -0.719 | -0.001 | -0.002 | -0.002 | 0 | 0.46 | 0.4 | 0.46 | 0.48 | 0.45 | 0.57 |
| 7 | 145236490 | rs6952652 | 0.735 | 0.226 | -0.004 | -0.747 | -0.01 | -0.03 | -0.03 | 0 | 0.24 | 0.17 | 0.32 | 0.42 | 0.28 | 0.36 |
| 7 | 145286533 | rs10500154 | -0.032 | 0.1 | 0.044 | -0.755 | -0.008 | -0.055 | -0.055 | 0.076 | 0.22 | 0.26 | 0.15 | 0.05 | 0.17 | 0.2 |
| 7 | 145301236 | rs850449 | 0.326 | 0.28 | 0.017 | -0.722 | 0.004 | -0.003 | -0.003 | 0.882 | 0.54 | 0.63 | 0.49 | 0.53 | 0.56 | 0.42 |
| 7 | 145315057 | rs850491 | -0.842 | 0.132 | -0.162 | -0.564 | -0.025 | -0.092 | -0.092 | 0 | 0.75 | 0.91 | 0.64 | 0.58 | 0.73 | 0.58 |
| 7 | 145324078 | rs850476 | -0.448 | 0.005 | 0.42 | -0.612 | -0.006 | 0.033 | 0.033 | 0 | 0.72 | 0.62 | 0.62 | 0.58 | 0.73 | 0.58 |
| 7 | 145327959 | rs11761986 | 0.64 | -0.131 | 0.224 | -0.697 | 0.006 | 0.08 | 0.08 | 0.195 | 0.17 | 0.02 | 0.08 | 0.04 | 0.13 | 0.18 |
| 7 | 145333229 | rs850462 | -0.734 | -0.062 | 0.407 | -0.693 | 0 | 0.009 | 0.009 | 0 | 0.73 | 0.71 | 0.61 | 0.58 | 0.72 | 0.59 |
| 7 | 145346374 | rs17169790 | 0.581 | -0.052 | 0.673 | -0.675 | 0.012 | 0.022 | 0.022 | 0.144 | 0.17 | 0.09 | 0.09 | 0.05 | 0.13 | 0.19 |
| 7 | 145353563 | rs1990347 | -0.389 | -0.099 | 0.615 | -0.694 | 0.002 | -0.052 | -0.052 | 0.441 | 0.68 | 0.88 | 0.62 | 0.58 | 0.7 | 0.57 |
| 7 | 145363652 | rs12703731 | 0.232 | 0.147 | 0.472 | -0.536 | 0.004 | 0 | 0 | 2.282 | 0.54 | 0.46 | 0.52 | 0.53 | 0.59 | 0.4 |
| 7 | 145380791 | rs1523730 | 0.066 | 1.261 | -0.548 | -0.145 | -0.022 | -0.008 | -0.008 | 0.082 | 0.11 | 0.09 | 0.12 | 0.16 | 0.22 | 0.1 |
| 7 | 145382259 | rs850570 | -0.568 | 0.87 | -0.554 | -0.2 | -0.006 | -0.032 | -0.032 | 0 | 0.74 | 0.82 | 0.63 | 0.58 | 0.72 | 0.61 |
| 7 | 145395771 | rs850546 | 0.749 | 0.284 | -0.618 | -1.201 | 0.026 | 0.004 | 0.004 | 0 | 0.35 | 0.26 | 0.17 | 0 | 0.25 | 0.22 |
| 7 | 145412108 | rs6948622 | -0.71 | 1.237 | -0.328 | -0.243 | 0 | -0.005 | -0.005 | 0 | 0.29 | 0.33 | 0.21 | 0.23 | 0.33 | 0.2 |
| 7 | 145418387 | rs1860482 | -0.913 | 1.105 | -0.114 | 0.234 | 0.004 | 0.013 | 0.013 | 0 | 0.3 | 0.41 | 0.21 | 0.34 | 0.28 | 0.37 |
| 7 | 145422736 | rs1523727 | -0.463 | 0.76 | -0.191 | -0.086 | -0.003 | -0.035 | -0.035 | 0 | 0.08 | 0.11 | 0.06 | 0 | 0.06 | 0.03 |
| 7 | 145436630 | rs11975765 | -1.161 | 0.208 | 0.072 | -0.362 | -0.065 | -0.097 | -0.097 | 0 | 0.28 | 0.45 | 0.15 | 0.1 | 0.14 | 0.24 |
| 7 | 145454215 | rs956885 | -1.231 | -0.03 | 0.508 | -0.478 | 0.023 | 0.024 | 0.024 | 0 | 0.3 | 0.7 | 0.18 | 0.1 | 0.15 | 0.23 |
| 7 | 145469247 | rs6946400 | -1.236 | 1.303 | 0.546 | -0.382 | -0.015 | -0.029 | -0.029 | 0.568 | 0.29 | 0.6 | 0.17 | 0.1 | 0.14 | 0.23 |
| 7 | 145482898 | rs10249365 | -1.231 | 1.294 | 0.615 | -0.386 | -0.006 | -0.016 | -0.016 | 0 | 0.29 | 0.6 | 0.17 | 0.1 | 0.14 | 0.23 |
| 7 | 145549570 | rs4726773 | -0.052 | 1.845 | 0.739 | -0.54 | -0.031 | 0 | 0 | 0 | 0.05 | 0 | 0.04 | 0.06 | 0.09 | 0.02 |
| 7 | 145580411 | rs4601231 | -0.424 | 1.547 | 0.909 | -0.689 | -0.053 | -0.08 | -0.08 | 0 | 0.13 | 0.24 | 0.06 | 0 | 0.04 | 0.02 |
| 7 | 145581082 | rs7809901 | 1.112 | 1.23 | -0.271 | -1.104 | -0.006 | 0.094 | 0.094 | 0 | 0.41 | 0.26 | 0.33 | 0.19 | 0.39 | 0.26 |
| 7 | 145600035 | rs12703746 | -0.614 | 1.258 | -0.435 | -0.882 | -0.011 | 0.007 | 0.007 | 0 | 0.06 | 0.06 | 0.03 | 0.06 | 0.08 | 0.03 |
| 7 | 145605216 | rs12670972 | -0.616 | 1.088 | -0.426 | -0.907 | -0.001 | -0.008 | -0.008 | 0 | 0.21 | 0.25 | 0.12 | 0.17 | 0.16 | 0.24 |
| 7 | 145616446 | rs7803306 | -1.003 | 0.933 | -0.398 | -0.976 | -0.019 | -0.034 | -0.034 | 0.018 | 0.28 | 0.35 | 0.16 | 0.12 | 0.15 | 0.24 |
| 7 | 145626980 | rs4314573 | 0.493 | 0.728 | -1.199 | -1.081 | -0.097 | -0.072 | -0.072 | 0.324 | 0.33 | 0.05 | 0.58 | 0.61 | 0.44 | 0.51 |
| 7 | 145661661 | rs13238234 | -1.323 | 0.032 | -0.511 | -1.338 | -0.024 | -0.059 | -0.059 | 0 | 0.33 | 0.53 | 0.2 | 0.17 | 0.2 | 0.24 |

| | | | | | | | | | | | | | | | | |
|---|---|---|---|---|---|---|---|---|---|---|---|---|---|---|---|---|
| 7 | 145670723 | rs6973802 | -1.457 | 0.076 | -0.442 | -1.287 | -0.024 | -0.059 | -0.059 | 0 | 0.33 | 0.53 | 0.19 | 0.17 | 0.21 | 0.23 |
| 7 | 145671873 | rs4591940 | 0.903 | 0.308 | -1.139 | -1.003 | -0.01 | -0.004 | -0.004 | 0 | 0.17 | 0.04 | 0.3 | 0.15 | 0.16 | 0.16 |
| 7 | 145680148 | rs6952571 | 0.567 | 0.146 | -0.641 | -1.004 | -0.005 | -0.004 | -0.004 | 0 | 0.11 | 0.15 | 0.06 | 0.1 | 0.08 | 0.18 |
| 7 | 145681191 | rs11977340 | -0.925 | 0.118 | -0.633 | -1.02 | 0.004 | 0.003 | 0.003 | 0 | 0.22 | 0.2 | 0.11 | 0.1 | 0.12 | 0.21 |
| 7 | 145683160 | rs866992 | 0.254 | 0.169 | -0.659 | -0.981 | 0.029 | 0.028 | 0.028 | 0 | 0.11 | 0.05 | 0.05 | 0 | 0.04 | 0.02 |
| 7 | 145700081 | rs6968866 | -1.181 | 0.194 | -0.6 | -0.996 | -0.002 | -0.019 | -0.019 | 0 | 0.17 | 0.24 | 0.09 | 0.07 | 0.1 | 0.03 |
| 7 | 145701716 | rs17169914 | -0.688 | 0.266 | -0.631 | -0.962 | 0.027 | -0.006 | -0.006 | 0 | 0.06 | 0.19 | 0.04 | 0.07 | 0.06 | 0.01 |
| 7 | 145709165 | rs11972913 | -0.828 | 0.222 | -0.643 | -0.984 | 0.003 | 0.002 | 0.002 | 0 | 0.22 | 0.19 | 0.11 | 0.1 | 0.12 | 0.2 |
| 7 | 145717550 | rs6965855 | -0.59 | 0.896 | -0.418 | -0.549 | 0.011 | -0.012 | -0.012 | 0.127 | 0.18 | 0.33 | 0.09 | 0.17 | 0.13 | 0.2 |
| 7 | 145719949 | rs6943324 | 0.223 | 1.028 | -0.731 | -0.933 | -0.001 | -0.01 | -0.01 | 0.106 | 0.59 | 0.59 | 0.71 | 0.81 | 0.62 | 0.68 |
| 7 | 145721952 | rs10272217 | -0.279 | 0.973 | -1.762 | -1.092 | 0.018 | 0.01 | 0.01 | 0 | 0.66 | 0.58 | 0.35 | 0.43 | 0.57 | 0.56 |
| 7 | 145725923 | rs11772501 | 1.539 | 0.699 | -1.854 | -1.557 | -0.001 | -0.006 | -0.006 | 0.016 | 0.16 | 0.16 | 0.1 | 0.21 | 0.17 | 0.26 |
| 7 | 145728574 | rs4725687 | -1.757 | 0.783 | -1.493 | -1.619 | 0.043 | -0.054 | -0.054 | 0.315 | 0.86 | 0.75 | 0.83 | 0.96 | 0.78 | 0.89 |
| 7 | 145728709 | rs4726778 | 0.607 | 0.693 | -1.92 | -1.616 | -0.013 | 0.07 | 0.07 | 0 | 0.36 | 0.2 | 0.57 | 0.74 | 0.38 | 0.61 |
| 7 | 145730478 | rs17400895 | 1.569 | -0.016 | -1.758 | -1.453 | -0.008 | 0.065 | 0.065 | 0.195 | 0.16 | 0.05 | 0.1 | 0 | 0.16 | 0.05 |
| 7 | 145732003 | rs6979491 | 0.625 | 0.026 | -1.482 | -1.272 | 0.031 | 0.076 | 0.076 | 1.344 | 0.28 | 0.07 | 0.11 | 0.16 | 0.18 | 0.21 |
| 7 | 145746333 | rs1554690 | -0.481 | 0.131 | -1.391 | -1.234 | 0.001 | 0.001 | 0.001 | 0.606 | 0.49 | 0.53 | 0.26 | 0.46 | 0.4 | 0.51 |
| 7 | 145750217 | rs1404803 | -0.807 | 0.016 | -1.516 | -1.041 | 0.01 | 0.094 | 0.094 | 0.272 | 0.44 | 0.83 | 0.27 | 0.29 | 0.41 | 0.38 |
| 7 | 145768088 | rs10277243 | -0.85 | -0.196 | -1.128 | -0.975 | 0.04 | 0.084 | 0.084 | 0.041 | 0.4 | 0.8 | 0.26 | 0.26 | 0.34 | 0.28 |
| 7 | 145768267 | rs1917651 | 0.086 | -0.237 | -0.278 | -1.03 | -0.031 | -0.127 | -0.127 | 0 | 0.21 | 0.38 | 0.15 | 0.02 | 0.17 | 0.05 |
| 7 | 145770180 | rs10256906 | -0.611 | 0.19 | -0.263 | -0.699 | 0.006 | 0.02 | 0.02 | 0 | 0.84 | 0.79 | 0.84 | 0.73 | 0.82 | 0.75 |
| 7 | 145773704 | rs2204465 | 0.677 | 0.083 | -0.104 | -1.023 | -0.001 | -0.007 | -0.007 | 0 | 0.62 | 0.44 | 0.69 | 0.71 | 0.64 | 0.69 |
| 7 | 145773791 | rs2204466 | 1.126 | -0.091 | -1.023 | -0.961 | -0.024 | -0.057 | -0.057 | 0.109 | 0.28 | 0.14 | 0.49 | 0.52 | 0.32 | 0.49 |
| 7 | 145777665 | rs1404807 | 0.516 | -1.692 | -1.088 | -1.42 | -0.007 | 0.086 | 0.086 | 0.159 | 0.28 | 0.07 | 0.11 | 0.14 | 0.19 | 0.27 |
| 7 | 145780820 | rs2056629 | 0.269 | -1.97 | -0.621 | -1.129 | -0.01 | -0.005 | -0.005 | 0 | 0.68 | 0.46 | 0.74 | 0.7 | 0.73 | 0.78 |
| 7 | 145781803 | rs1404810 | 0.958 | -1.916 | -0.53 | -1.111 | -0.008 | 0 | 0 | 0 | 0.41 | 0.36 | 0.63 | 0.56 | 0.54 | 0.51 |
| 7 | 145782848 | rs9655686 | -0.808 | -1.943 | -0.507 | -1.047 | -0.004 | 0.014 | 0.014 | 0 | 0.57 | 0.38 | 0.29 | 0.39 | 0.46 | 0.5 |
| 7 | 145794331 | rs10247330 | 0.813 | -1.857 | -0.452 | -0.964 | -0.027 | -0.039 | -0.039 | 1.194 | 0.14 | 0.16 | 0.07 | 0.04 | 0.07 | 0.04 |
| 7 | 145795152 | rs7797682 | -0.78 | -1.403 | -0.818 | -0.721 | -0.018 | -0.031 | -0.031 | 1.892 | 0.85 | 0.91 | 0.83 | 0.74 | 0.79 | 0.85 |
| 7 | 145796382 | rs6953776 | 1.11 | -1.375 | -0.792 | -0.755 | -0.015 | -0.014 | -0.014 | 0.26 | 0.13 | 0.09 | 0.1 | 0.19 | 0.19 | 0.13 |
| 7 | 145796758 | rs6975019 | 0.468 | -1.584 | -0.663 | -1.387 | 0.07 | 0.096 | 0.096 | 1.044 | 0.48 | 0.3 | 0.71 | 0.72 | 0.7 | 0.6 |
| 7 | 145803445 | rs7795836 | -0.412 | -1.768 | -0.01 | -1.857 | 0.009 | 0.035 | 0.035 | 0 | 0.5 | 0.59 | 0.27 | 0.22 | 0.42 | 0.46 |
| 7 | 145807447 | rs2692359 | -1.056 | -1.813 | -0.147 | -1.87 | -0.025 | -0.032 | -0.032 | 0 | 0.86 | 0.94 | 0.72 | 0.72 | 0.78 | 0.8 |
| 7 | 145808436 | rs12703766 | 1.031 | -2.008 | -0.371 | -2.027 | 0.01 | 0.145 | 0.145 | 0.374 | 0.23 | 0.03 | 0.09 | 0.01 | 0.12 | 0.14 |
| 7 | 145809668 | rs17480644 | 0.551 | -1.927 | -0.162 | -1.949 | 0.014 | 0.014 | 0.014 | 0 | 0.08 | 0.06 | 0.04 | 0.05 | 0.04 | 0.09 |
| 7 | 145813854 | rs2462603 | 0.987 | -1.972 | 0.054 | -2.122 | 0 | -0.002 | -0.002 | 0 | 0.44 | 0.42 | 0.51 | 0.62 | 0.54 | 0.5 |
| 7 | 145825216 | rs7797553 | -0.43 | -1.646 | 0.452 | -1.863 | -0.014 | -0.059 | -0.059 | 0 | 0.42 | 0.54 | 0.19 | 0.09 | 0.23 | 0.27 |
| 7 | 145880266 | rs13230938 | -0.835 | -1.54 | 0.487 | -1.978 | 0.017 | 0.003 | 0.003 | 0.462 | 0.95 | 0.82 | 0.97 | 0.98 | 0.97 | 0.95 |
| 7 | 145882001 | rs4612252 | 1.603 | -1.66 | 0.239 | -2.078 | -0.101 | 0.042 | 0.042 | 0.368 | 0.39 | 0.12 | 0.5 | 0.63 | 0.52 | 0.49 |
| 7 | 145883566 | rs1464961 | -0.436 | -1.742 | 0.792 | -2.024 | 0.019 | 0.01 | 0.01 | 1.198 | 0.4 | 0.35 | 0.17 | 0.07 | 0.23 | 0.26 |
| 7 | 145913594 | rs17577839 | 0.744 | -1.736 | 0.531 | -1.992 | 0.014 | 0.051 | 0.051 | 0 | 0.16 | 0.08 | 0.05 | 0.06 | 0.09 | 0.11 |
| 7 | 145917253 | rs802531 | -0.998 | -1.799 | 0.594 | -2.041 | 0.016 | 0.01 | 0.01 | 0.168 | 0.78 | 0.83 | 0.89 | 0.99 | 0.86 | 0.85 |
| 7 | 145917377 | rs802532 | 0.564 | -1.672 | 0.533 | -1.979 | -0.017 | -0.083 | -0.083 | 0 | 0.58 | 0.48 | 0.82 | 0.93 | 0.76 | 0.74 |
| 7 | 145924072 | rs11761213 | 0.371 | -0.73 | 1.07 | -0.853 | -0.063 | -0.073 | -0.073 | 2.9 | 0.12 | 0.01 | 0.29 | 0.3 | 0.22 | 0.19 |
| 7 | 145928429 | rs1526079 | 0.091 | -0.911 | 1.118 | -0.964 | 0.015 | 0.045 | 0.045 | 0.018 | 0.16 | 0.07 | 0.06 | 0.06 | 0.09 | 0.14 |
| 7 | 145933013 | rs11765641 | 0.374 | -0.461 | 0.693 | -0.911 | -0.064 | -0.071 | -0.071 | 0 | 0.12 | 0.01 | 0.29 | 0.3 | 0.23 | 0.19 |
| 7 | 145936334 | rs802556 | -1.646 | -1.028 | 1.496 | -1.228 | -0.021 | -0.051 | -0.051 | 0.447 | 0.28 | 0.33 | 0.11 | 0.1 | 0.13 | 0.28 |
| 7 | 145951642 | rs802524 | -2.474 | -0.86 | 1.77 | -0.861 | -0.004 | -0.007 | -0.007 | 0 | 0.19 | 0.74 | 0.1 | 0.05 | 0.04 | 0.11 |
| 7 | 145952946 | rs2430327 | -2.107 | -0.838 | 1.849 | -0.851 | -0.068 | -0.075 | -0.075 | 0.519 | 0.17 | 0.56 | 0.08 | 0.05 | 0.04 | 0.1 |
| 7 | 145962186 | rs802571 | -1.861 | -0.815 | 1.287 | -0.96 | -0.018 | -0.028 | -0.028 | 0 | 0.15 | 0.77 | 0.09 | 0.03 | 0.03 | 0.09 |
| 7 | 145975963 | rs1405109 | 1.773 | -0.433 | 0.998 | -0.949 | -0.06 | 0.02 | 0.02 | 1.08 | 0.56 | 0.04 | 0.45 | 0.6 | 0.63 | 0.69 |
| 7 | 145991131 | rs7810054 | -0.199 | -0.688 | 1.127 | -0.971 | -0.024 | -0.033 | -0.033 | 3.303 | 0.06 | 0.15 | 0.03 | 0 | 0.02 | 0.02 |
| 7 | 145993544 | rs10276255 | 0.042 | -0.656 | 0.715 | -0.959 | -0.011 | -0.026 | -0.026 | 0 | 0.07 | 0.09 | 0.02 | 0 | 0.02 | 0.03 |
| 7 | 145998111 | rs17170033 | 0.11 | -0.423 | -0.798 | -0.858 | -0.043 | -0.019 | -0.019 | 0 | 0.23 | 0.09 | 0.42 | 0.34 | 0.31 | 0.2 |
| 7 | 146020936 | rs6464749 | -1.494 | -0.842 | -0.497 | -0.742 | 0.048 | 0.048 | 0.048 | 4.484 | 0.21 | 0.84 | 0.11 | 0.05 | 0.05 | 0.11 |
| 7 | 146032524 | rs7788551 | 0.183 | -0.007 | -0.639 | -0.506 | -0.033 | -0.022 | -0.022 | 9.988 | 0.25 | 0.09 | 0.42 | 0.31 | 0.28 | 0.16 |
| 7 | 146035708 | rs17170039 | -0.687 | -0.3 | 0.004 | -0.689 | -0.011 | -0.046 | -0.046 | 1.984 | 0.08 | 0.18 | 0.04 | 0 | 0.04 | 0.05 |
| 7 | 146036942 | rs7808440 | -1.993 | -0.608 | 0.686 | -0.832 | -0.052 | -0.092 | -0.092 | 1.324 | 0.2 | 0.59 | 0.11 | 0.03 | 0.09 | 0.12 |
| 7 | 146043176 | rs17503262 | 0.843 | -0.438 | 0.017 | -1.056 | -0.043 | 0.011 | 0.011 | 0.635 | 0.12 | 0.01 | 0.17 | 0.05 | 0.18 | 0.16 |
| 7 | 146046444 | rs965003 | -2.275 | -0.611 | 0.748 | -0.832 | -0.014 | -0.036 | -0.036 | 0 | 0.22 | 0.72 | 0.12 | 0.03 | 0.09 | 0.13 |
| 7 | 146053416 | rs10226754 | -0.671 | -0.4 | 0.978 | -0.783 | 0.001 | 0.012 | 0.012 | 0 | 0.33 | 0.27 | 0.46 | 0.32 | 0.32 | 0.22 |
| 7 | 146054350 | rs12703801 | -0.671 | -0.383 | 1.019 | -0.783 | 0 | 0.011 | 0.011 | 0 | 0.33 | 0.28 | 0.46 | 0.32 | 0.32 | 0.22 |
| 7 | 146056420 | rs1860681 | -1.729 | -0.571 | 1.457 | -0.817 | -0.031 | -0.046 | -0.046 | 0.016 | 0.11 | 0.52 | 0.08 | 0.03 | 0.03 | 0.06 |
| 7 | 146058754 | rs10225925 | -0.478 | -0.549 | 1.337 | -0.814 | -0.015 | -0.034 | -0.034 | 1.19 | 0.08 | 0.12 | 0.03 | 0 | 0.03 | 0.05 |
| 7 | 146060451 | rs4437559 | -2.385 | -0.607 | 1.551 | -0.832 | -0.057 | -0.09 | -0.09 | 0.062 | 0.2 | 0.61 | 0.12 | 0.03 | 0.08 | 0.12 |
| 7 | 146065117 | rs6464757 | -0.708 | -0.449 | 1.969 | -0.795 | -0.004 | -0.015 | -0.015 | 0 | 0.35 | 0.41 | 0.48 | 0.32 | 0.32 | 0.22 |
| 7 | 146065457 | rs17170068 | -0.245 | -0.058 | 2.108 | -0.788 | 0.015 | -0.002 | -0.002 | 0.208 | 0.3 | 0.38 | 0.5 | 0.36 | 0.34 | 0.21 |
| 7 | 146074359 | rs17170073 | -1.408 | -0.076 | 2.099 | -0.827 | -0.024 | -0.056 | -0.056 | 0 | 0.11 | 0.22 | 0.05 | 0 | 0.05 | 0.06 |
| 7 | 146098510 | rs1724527 | -1.938 | -0.064 | 2.336 | -0.836 | -0.003 | -0.073 | -0.073 | 0.468 | 0.12 | 0.38 | 0.08 | 0 | 0.08 | 0.05 |

| | | | | | | | | | | | | | | | | |
|---|---|---|---|---|---|---|---|---|---|---|---|---|---|---|---|---|
| 7 | 146098829 | rs1639489 | -1.824 | -0.063 | 2.221 | -0.836 | -0.013 | -0.067 | -0.067 | 0 | 0.12 | 0.26 | 0.06 | 0 | 0.08 | 0.05 |
| 7 | 146104454 | rs802196 | -0.658 | -0.006 | 1.932 | -0.723 | 0.002 | 0.042 | 0.042 | 0.286 | 0.34 | 0.83 | 0.51 | 0.29 | 0.3 | 0.22 |
| 7 | 146112428 | rs6978796 | -0.191 | 0.118 | 1.881 | -0.698 | -0.002 | -0.005 | -0.005 | 1.198 | 0.31 | 0.66 | 0.49 | 0.29 | 0.28 | 0.2 |
| 7 | 146121539 | rs2021559 | -0.191 | 0.118 | 1.862 | -0.678 | -0.001 | -0.005 | -0.005 | 0.727 | 0.31 | 0.66 | 0.49 | 0.28 | 0.28 | 0.21 |
| 7 | 146168418 | rs17170100 | -2.52 | 0.286 | 2.206 | -0.347 | 0.003 | -0.089 | -0.089 | 0 | 0.14 | 0.52 | 0.1 | 0.03 | 0.13 | 0.1 |
| 7 | 146175409 | rs10279409 | -2.308 | 0.382 | 2.215 | -0.337 | 0.03 | -0.077 | -0.077 | 0 | 0.13 | 0.5 | 0.11 | 0.03 | 0.15 | 0.1 |
| 7 | 146189057 | rs700276 | 0.331 | 0.605 | 1.416 | -0.196 | -0.004 | 0.002 | 0.002 | 0 | 0.54 | 0.46 | 0.62 | 0.44 | 0.52 | 0.5 |
| 7 | 146189136 | rs700277 | 0.44 | 0.582 | 1.108 | -0.178 | -0.007 | 0.016 | 0.016 | 0 | 0.53 | 0.33 | 0.59 | 0.42 | 0.51 | 0.46 |
| 7 | 146190783 | rs696773 | -0.345 | 0.803 | 0.914 | -0.065 | -0.01 | -0.027 | -0.027 | 2.035 | 0.65 | 0.8 | 0.7 | 0.43 | 0.63 | 0.54 |
| 7 | 146192877 | rs700281 | 0.482 | 0.637 | 0.893 | -0.079 | -0.01 | 0.037 | 0.037 | 0 | 0.52 | 0.33 | 0.59 | 0.39 | 0.51 | 0.46 |
| 7 | 146204005 | rs1089448 | 0.137 | 0.483 | 1.352 | 0.328 | 0.002 | -0.045 | -0.045 | 2.479 | 0.26 | 0.17 | 0.22 | 0.4 | 0.27 | 0.33 |
| 7 | 146204352 | rs1089447 | -0.918 | 0.402 | 1.62 | 0.208 | 0.012 | -0.025 | -0.025 | 0 | 0.39 | 0.75 | 0.31 | 0.47 | 0.36 | 0.39 |
| 7 | 146239545 | rs2693406 | -1.566 | 0.5 | 1.803 | 0.048 | 0.068 | -0.001 | -0.001 | 2.624 | 0.08 | 0.36 | 0.07 | 0.05 | 0.07 | 0.06 |
| 7 | 146239908 | rs2693405 | -0.38 | 0.639 | 1.067 | 0.05 | -0.004 | -0.013 | -0.013 | 0 | 0.84 | 0.88 | 0.63 | 0.7 | 0.85 | 0.92 |
| 7 | 146245753 | rs2693394 | -0.409 | 0.631 | 1.057 | 0.044 | -0.008 | -0.025 | -0.025 | 0 | 0.84 | 0.89 | 0.64 | 0.7 | 0.85 | 0.91 |
| 7 | 146254508 | rs12154459 | 0.448 | 0.443 | 0.95 | -0.072 | -0.02 | -0.038 | -0.038 | 0 | 0.19 | 0.11 | 0.2 | 0.47 | 0.29 | 0.3 |
| 7 | 146266525 | rs10249436 | 0.02 | 0.18 | 0.684 | -0.247 | 0 | -0.038 | -0.038 | 0 | 0.74 | 0.92 | 0.62 | 0.67 | 0.81 | 0.89 |
| 7 | 146281412 | rs17587202 | 2.271 | 0.059 | 0.126 | -0.732 | -0.056 | 0.138 | 0.138 | 0 | 0.27 | 0.02 | 0.21 | 0.05 | 0.3 | 0.24 |
| 7 | 146292088 | rs454612 | -0.381 | -0.169 | 1.533 | 0.069 | 0.027 | 0.131 | 0.131 | 0.122 | 0.24 | 0.45 | 0.23 | 0.45 | 0.3 | 0.3 |
| 7 | 146297650 | rs347180 | 0.397 | -0.161 | 1.419 | -0.031 | -0.068 | 0.074 | 0.074 | 0 | 0.68 | 0.89 | 0.47 | 0.21 | 0.61 | 0.65 |
| 7 | 146317074 | rs347223 | 0.447 | -0.119 | 1.228 | -0.043 | -0.055 | 0.071 | 0.071 | 0 | 0.67 | 0.85 | 0.47 | 0.2 | 0.61 | 0.66 |
| 7 | 146323187 | rs347220 | 1.639 | -0.34 | 1.311 | -0.087 | -0.006 | 0.045 | 0.045 | 0 | 0.55 | 0.39 | 0.35 | 0.19 | 0.49 | 0.56 |
| 7 | 146325412 | rs724216 | 1.431 | -0.299 | 0.869 | -0.074 | 0.004 | 0.096 | 0.096 | 0 | 0.5 | 0.37 | 0.35 | 0.16 | 0.54 | 0.58 |
| 7 | 146332198 | rs12536200 | 1.134 | -0.178 | 0.948 | 0.175 | -0.068 | -0.12 | -0.12 | 0.048 | 0.15 | 0.01 | 0.16 | 0.46 | 0.22 | 0.26 |
| 7 | 146332953 | rs766094 | -1.281 | 0.192 | 1.282 | 0.261 | 0.029 | -0.048 | -0.048 | 0.728 | 0.09 | 0.32 | 0.07 | 0.01 | 0.09 | 0.05 |
| 7 | 146350629 | rs344488 | -0.235 | 0.059 | 1.075 | 0.682 | -0.022 | -0.047 | -0.047 | 0 | 0.83 | 0.93 | 0.63 | 0.67 | 0.84 | 0.91 |
| 7 | 146354025 | rs714584 | 1.182 | 0.036 | 1.037 | 0.548 | -0.001 | 0.088 | 0.088 | 0 | 0.48 | 0.35 | 0.34 | 0.19 | 0.53 | 0.59 |
| 7 | 146355594 | rs7810856 | 0.501 | 0.33 | 1.687 | 1.167 | 0.024 | 0.097 | 0.097 | 0.2 | 0.83 | 0.75 | 0.62 | 0.62 | 0.84 | 0.91 |
| 7 | 146360917 | rs10952650 | 2.04 | 0.089 | 1.193 | 0.662 | 0.011 | -0.017 | -0.017 | 0.282 | 0.69 | 0.18 | 0.45 | 0.61 | 0.77 | 0.85 |
| 7 | 146367793 | rs717171 | 0.741 | 0.102 | 1.514 | 0.716 | -0.008 | -0.009 | -0.009 | 0 | 0.81 | 0.83 | 0.57 | 0.72 | 0.83 | 0.91 |
| 7 | 146373024 | rs1353235 | -1.898 | 0.07 | 2.156 | 0.707 | 0.02 | 0.098 | 0.098 | 1.44 | 0.08 | 0.5 | 0.07 | 0.11 | 0.03 | 0.02 |
| 7 | 146374523 | rs173168 | -1.898 | 0.07 | 2.156 | 0.707 | 0.02 | 0.098 | 0.098 | 1.44 | 0.08 | 0.5 | 0.07 | 0.11 | 0.03 | 0.02 |
| 7 | 146374576 | rs6959595 | 0.844 | 0.348 | 2.031 | 0.817 | -0.008 | -0.009 | -0.009 | 0.254 | 0.81 | 0.83 | 0.57 | 0.72 | 0.83 | 0.92 |
| 7 | 146377256 | rs7802801 | 0.844 | 0.348 | 2.031 | 0.817 | -0.008 | -0.009 | -0.009 | 0 | 0.81 | 0.83 | 0.57 | 0.72 | 0.83 | 0.92 |
| 7 | 146389259 | rs2642483 | 0.603 | 0.252 | 2.241 | 1.018 | -0.007 | -0.014 | -0.014 | 0.895 | 0.89 | 0.9 | 0.61 | 0.73 | 0.89 | 0.96 |
| 7 | 146392116 | rs2131930 | -1.933 | 0.183 | 2.304 | 0.994 | 0.051 | 0.104 | 0.104 | 0 | 0.08 | 0.42 | 0.1 | 0.1 | 0.05 | 0.05 |
| 7 | 146397210 | rs2642504 | 1.281 | 0.209 | 1.943 | 0.995 | 0.007 | 0.037 | 0.037 | 0.056 | 0.87 | 0.73 | 0.6 | 0.72 | 0.88 | 0.93 |
| 7 | 146406379 | rs2533096 | -3.032 | -0.373 | 2.194 | 0.575 | 0.019 | 0.01 | 0.01 | 3.889 | 0.17 | 0.78 | 0.13 | 0.13 | 0.07 | 0.08 |
| 7 | 146406407 | rs6951437 | -2.583 | -0.334 | 2.481 | 0.584 | 0.052 | 0.032 | 0.032 | 0.344 | 0.06 | 0.35 | 0.06 | 0.01 | 0.02 | 0.02 |
| 7 | 146413497 | rs344470 | -3.444 | -0.008 | 2.346 | 0.528 | 0.012 | 0.019 | 0.019 | 3.965 | 0.18 | 0.82 | 0.18 | 0.13 | 0.14 | 0.09 |
| 7 | 146416662 | rs7777225 | -2.916 | 0.053 | 2.465 | 0.523 | 0.124 | 0.012 | 0.012 | 0.691 | 0.06 | 0.5 | 0.11 | 0 | 0.09 | 0.02 |
| 7 | 146419326 | rs344468 | -2.817 | -0.093 | 2.091 | 0.571 | -0.039 | -0.068 | -0.068 | 3.127 | 0.12 | 0.32 | 0.08 | 0.01 | 0.05 | 0.06 |
| 7 | 146420571 | rs344465 | -2.001 | -0.013 | 1.386 | 0.686 | -0.003 | 0.001 | 0.001 | 0.062 | 0.08 | 0.1 | 0.05 | 0.12 | 0.06 | 0.06 |
| 7 | 146431210 | rs1472258 | -0.944 | -0.492 | 1.6 | 0.452 | -0.007 | -0.002 | -0.002 | 0 | 0.15 | 0.26 | 0.23 | 0.15 | 0.14 | 0.07 |
| 7 | 146439869 | rs7790238 | -1.452 | -0.668 | 1.078 | 1.084 | 0.057 | -0.045 | -0.045 | 0 | 0.41 | 0.85 | 0.38 | 0.55 | 0.34 | 0.43 |
| 7 | 146454421 | rs986062 | -1.401 | -0.996 | 1.741 | 1.151 | 0.026 | -0.014 | -0.014 | 0.078 | 0.39 | 0.72 | 0.23 | 0.43 | 0.31 | 0.33 |
| 7 | 146461144 | rs10230326 | -2.126 | -1.021 | 2.229 | 1.129 | 0.03 | -0.035 | -0.035 | 0.06 | 0.11 | 0.47 | 0.08 | 0.03 | 0.09 | 0.03 |
| 7 | 146464971 | rs11977396 | -2.879 | -1.051 | 2.491 | 1.184 | -0.01 | -0.004 | -0.004 | 0.187 | 0.16 | 0.72 | 0.14 | 0.15 | 0.14 | 0.06 |
| 7 | 146477026 | rs2372765 | -0.432 | -1.104 | 2.137 | 1.276 | 0.048 | -0.052 | -0.052 | 0 | 0.25 | 0.1 | 0.11 | 0.32 | 0.18 | 0.3 |
| 7 | 146481597 | rs10241577 | -1.697 | -1.086 | 1.899 | 1.275 | 0.08 | -0.028 | -0.028 | 0 | 0.41 | 0.82 | 0.25 | 0.44 | 0.32 | 0.35 |
| 7 | 146486101 | rs10239703 | -1.737 | -1.196 | 1.726 | 1.25 | 0.087 | -0.038 | -0.038 | 0 | 0.42 | 0.86 | 0.26 | 0.44 | 0.32 | 0.35 |
| 7 | 146487961 | rs6968847 | -2.567 | -0.777 | 2.078 | 1.364 | 0.05 | 0.054 | 0.054 | 0 | 0.09 | 0.28 | 0.09 | 0.12 | 0.13 | 0.05 |
| 7 | 146497318 | rs10952662 | -1.142 | -1.444 | 1.063 | 0.918 | 0.16 | 0.081 | 0.081 | 0.238 | 0.48 | 0.82 | 0.22 | 0.33 | 0.27 | 0.38 |
| 7 | 146501346 | rs10239811 | -2.015 | -1.535 | 1.204 | 0.845 | -0.044 | -0.038 | -0.038 | 0 | 0.11 | 0.73 | 0.07 | 0.03 | 0.01 | 0.02 |
| 7 | 146516749 | rs7806453 | -2.002 | -1.533 | 0.953 | 0.846 | -0.05 | -0.045 | -0.045 | 0.007 | 0.11 | 0.71 | 0.06 | 0.03 | 0.01 | 0.02 |
| 7 | 146527939 | rs6942840 | -1.532 | -1.524 | 0.769 | 0.841 | -0.016 | -0.02 | -0.02 | 0.108 | 0.08 | 0.54 | 0.05 | 0 | 0.01 | 0.01 |
| 7 | 146528042 | rs6464774 | -1.257 | -1.523 | -0.567 | 0.974 | 0.128 | -0.074 | -0.074 | 0.169 | 0.51 | 0.96 | 0.27 | 0.47 | 0.3 | 0.37 |
| 7 | 146532528 | rs10282480 | -0.408 | -1.015 | -0.272 | 1.044 | 0.069 | -0.036 | -0.036 | 0 | 0.37 | 0.08 | 0.22 | 0.33 | 0.26 | 0.34 |
| 7 | 146544492 | rs9801492 | 0.216 | -0.383 | -0.233 | 0.618 | 0.001 | 0.043 | 0.043 | 0 | 0.25 | 0.14 | 0.34 | 0.15 | 0.21 | 0.19 |
| 7 | 146547048 | rs6955130 | 0.562 | 0.457 | 0.561 | 0.709 | 0.059 | 0.026 | 0.026 | 0 | 0.18 | 0.35 | 0.27 | 0.2 | 0.28 | 0.17 |
| 7 | 146550239 | rs7780475 | 0.349 | 0.453 | 0.572 | 0.784 | 0.09 | 0.071 | 0.071 | 0 | 0.19 | 0.5 | 0.29 | 0.24 | 0.29 | 0.17 |
| 7 | 146553618 | rs9648688 | -0.134 | 0.781 | 0.151 | 1.007 | 0.002 | 0 | 0 | 0 | 0.37 | 0.6 | 0.52 | 0.32 | 0.37 | 0.29 |
| 7 | 146559474 | rs4367451 | -1.307 | 0.507 | -0.299 | 0.855 | 0.007 | -0.008 | -0.008 | 0.236 | 0.68 | 0.96 | 0.84 | 0.71 | 0.74 | 0.58 |
| 7 | 146569738 | rs4480030 | 0.027 | 1.457 | 0.106 | 1.068 | 0.003 | 0.004 | 0.004 | 0 | 0.32 | 0.36 | 0.45 | 0.33 | 0.33 | 0.25 |
| 7 | 146569874 | rs4537227 | 0.64 | 1.417 | -0.633 | 1.089 | 0.004 | 0.005 | 0.005 | 0 | 0.26 | 0.18 | 0.35 | 0.21 | 0.21 | 0.2 |
| 7 | 146571183 | rs12703845 | 0.486 | 1.533 | -0.699 | 1.157 | -0.004 | -0.003 | -0.003 | 0 | 0.17 | 0.15 | 0.26 | 0.24 | 0.29 | 0.17 |
| 7 | 146575177 | rs9648840 | -0.227 | 0.693 | -0.993 | 0.893 | 0.073 | 0.025 | 0.025 | 0 | 0.55 | 0.89 | 0.78 | 0.56 | 0.65 | 0.45 |
| 7 | 146575682 | rs10230479 | -0.646 | 1.314 | 0.255 | 0.848 | 0.111 | 0.12 | 0.12 | 0 | 0.08 | 0.31 | 0.1 | 0.12 | 0.12 | 0.05 |
| 7 | 146578167 | rs6963027 | -1.291 | 1.019 | 0.364 | 0.228 | 0.024 | -0.081 | -0.081 | 0 | 0.13 | 0.42 | 0.1 | 0.03 | 0.11 | 0.09 |

| | | | | | | | | | | | | | | | | |
|---|---|---|---|---|---|---|---|---|---|---|---|---|---|---|---|---|
| 7 | 146580713 | rs7341537 | -0.527 | 1.041 | -0.503 | 0.283 | 0.002 | 0.004 | 0.004 | 0 | 0.47 | 0.36 | 0.34 | 0.44 | 0.44 | 0.38 |
| 7 | 146582565 | rs12703846 | -0.684 | 1.113 | -1.607 | -0.925 | -0.09 | 0.213 | 0.213 | 0 | 0.37 | 0.1 | 0.23 | 0.02 | 0.44 | 0.38 |
| 7 | 146590038 | rs7810174 | -1.54 | 0.831 | -1.933 | -0.641 | 0.024 | 0.047 | 0.047 | 0 | 0.61 | 0.89 | 0.56 | 0.69 | 0.65 | 0.68 |
| 7 | 146591888 | rs10216156 | -1.18 | 0.798 | -1.644 | -0.687 | 0.026 | 0.043 | 0.043 | 0 | 0.57 | 0.74 | 0.53 | 0.67 | 0.61 | 0.63 |
| 7 | 146596362 | rs11982298 | -0.984 | 0.722 | -1.727 | -1.001 | -0.002 | 0 | 0 | 0 | 0.38 | 0.36 | 0.2 | 0.04 | 0.4 | 0.4 |
| 7 | 146597770 | rs12703848 | -0.202 | 0.588 | -2.034 | -1.071 | -0.094 | 0.151 | 0.151 | 0 | 0.27 | 0.06 | 0.17 | 0.04 | 0.36 | 0.31 |
| 7 | 146618729 | rs4467861 | -2.39 | 0.567 | -1.945 | -0.855 | -0.038 | 0.017 | 0.017 | 0 | 0.63 | 0.96 | 0.57 | 0.7 | 0.64 | 0.68 |
| 7 | 146655312 | rs10251619 | -0.196 | 0.224 | -1.031 | -0.821 | -0.022 | -0.046 | -0.046 | 0 | 0.08 | 0.17 | 0.02 | 0 | 0.03 | 0.08 |
| 7 | 146660924 | rs10081175 | -0.56 | 0.259 | -1.041 | -0.808 | -0.007 | -0.03 | -0.03 | 0.02 | 0.12 | 0.17 | 0.04 | 0.03 | 0.08 | 0.12 |
| 7 | 146673884 | rs10243212 | -1.681 | 0.276 | -1.203 | -0.973 | -0.001 | -0.011 | -0.011 | 0 | 0.38 | 0.35 | 0.2 | 0.05 | 0.4 | 0.41 |
| 7 | 146677310 | rs12703872 | -0.444 | 0.53 | -1.623 | -0.646 | -0.095 | 0.143 | 0.143 | 0.09 | 0.23 | 0.02 | 0.15 | 0.02 | 0.32 | 0.28 |
| 7 | 146682211 | rs12670868 | -0.915 | 0.362 | -0.779 | 0.313 | 0 | -0.003 | -0.003 | 0.125 | 0.26 | 0.21 | 0.29 | 0.23 | 0.3 | 0.37 |
| 7 | 146683899 | rs4368897 | 0.529 | 0.667 | -1.272 | 0.979 | -0.047 | -0.068 | -0.068 | 0 | 0.1 | 0.01 | 0.11 | 0.24 | 0.14 | 0.08 |
| 7 | 146687912 | rs10270807 | -0.742 | 0.538 | -1.033 | 0.643 | -0.011 | 0.007 | 0.007 | 0 | 0.25 | 0.12 | 0.28 | 0.21 | 0.3 | 0.34 |
| 7 | 146688750 | rs10215441 | -0.459 | 0.275 | -1.061 | 0.65 | -0.018 | 0.025 | 0.025 | 0 | 0.08 | 0.07 | 0.15 | 0.15 | 0.03 | 0.07 |
| 7 | 146690926 | rs7800935 | 1.232 | 0.195 | -0.583 | 0.183 | 0.006 | -0.115 | -0.115 | 0 | 0.66 | 0.53 | 0.78 | 0.94 | 0.63 | 0.65 |
| 7 | 146691220 | rs6945085 | -0.48 | 0.24 | -0.559 | 0.212 | -0.017 | 0.031 | 0.031 | 0 | 0.93 | 0.98 | 0.94 | 1 | 0.91 | 0.94 |
| 7 | 146692561 | rs10280931 | -1.142 | -0.051 | -0.198 | 0.386 | -0.014 | -0.019 | -0.019 | 0.275 | 0.31 | 0.41 | 0.32 | 0.23 | 0.3 | 0.35 |
| 7 | 146694441 | rs10248780 | -2.051 | -0.401 | -0.494 | 0.777 | -0.075 | 0.073 | 0.073 | 0 | 0.54 | 0.92 | 0.49 | 0.71 | 0.49 | 0.69 |
| 7 | 146695674 | rs1608631 | -0.537 | -0.505 | 0.04 | 0.897 | 0.005 | 0.023 | 0.023 | 0 | 0.31 | 0.4 | 0.28 | 0.5 | 0.36 | 0.44 |
| 7 | 146706689 | rs1524341 | -2.432 | -0.627 | -0.113 | 0.829 | -0.095 | 0.052 | 0.052 | 0.166 | 0.59 | 0.92 | 0.5 | 0.72 | 0.54 | 0.74 |
| 7 | 146707835 | rs4598172 | -1.713 | -0.613 | 0.136 | 0.865 | -0.001 | 0 | 0 | 0.117 | 0.48 | 0.74 | 0.35 | 0.48 | 0.51 | 0.63 |
| 7 | 146707932 | rs1608629 | -1.058 | -0.61 | 0.176 | 0.825 | 0.001 | 0.002 | 0.002 | 0 | 0.38 | 0.43 | 0.29 | 0.47 | 0.44 | 0.54 |
| 7 | 146711789 | rs1404156 | -0.752 | -0.651 | 0.253 | 0.821 | 0.003 | 0.02 | 0.02 | 0 | 0.33 | 0.6 | 0.28 | 0.48 | 0.36 | 0.44 |
| 7 | 146712903 | rs6953708 | 0.738 | -0.545 | 0.243 | 0.799 | -0.005 | -0.023 | -0.023 | 0.214 | 0.06 | 0.2 | 0.03 | 0 | 0.02 | 0.03 |
| 7 | 146720738 | rs2109305 | -0.687 | -0.724 | 0.179 | 0.869 | -0.047 | 0.071 | 0.071 | 0 | 0.12 | 0.22 | 0.17 | 0.2 | 0.04 | 0.11 |
| 7 | 146731365 | rs2372808 | -2.38 | -0.657 | -0.19 | 0.858 | -0.09 | 0.017 | 0.017 | 0.185 | 0.59 | 0.93 | 0.5 | 0.67 | 0.54 | 0.73 |
| 7 | 146731727 | rs2098114 | -1.627 | -0.639 | 0.012 | 0.848 | -0.001 | 0 | 0 | 0.555 | 0.48 | 0.51 | 0.34 | 0.53 | 0.51 | 0.64 |
| 7 | 146740577 | rs7812091 | -0.545 | -0.482 | -0.156 | 0.772 | 0.001 | 0.006 | 0.006 | 0 | 0.31 | 0.37 | 0.28 | 0.5 | 0.36 | 0.43 |
| 7 | 146743606 | rs2159472 | -1.496 | -1.085 | 0.036 | 0.569 | -0.049 | 0.021 | 0.021 | 0 | 0.28 | 0.47 | 0.23 | 0.32 | 0.17 | 0.28 |
| 7 | 146748970 | rs17222184 | 0.445 | -0.576 | -0.01 | 0.26 | -0.006 | 0.007 | 0.007 | 0 | 0.09 | 0.05 | 0.04 | 0 | 0.09 | 0.1 |
| 7 | 146750254 | rs6980283 | -0.059 | -0.526 | -0.044 | 0.485 | -0.021 | 0.095 | 0.095 | 0.862 | 0.18 | 0.44 | 0.25 | 0.31 | 0.13 | 0.28 |
| 7 | 146751635 | rs10266239 | 0.251 | -0.577 | -0.189 | 0.351 | 0.022 | 0.003 | 0.003 | 0 | 0.27 | 0.13 | 0.24 | 0.27 | 0.21 | 0.22 |
| 7 | 146751925 | rs10266622 | -0.494 | -0.485 | -0.285 | -0.083 | 0.002 | 0.001 | 0.001 | 0 | 0.46 | 0.6 | 0.49 | 0.59 | 0.34 | 0.49 |
| 7 | 146759484 | rs6973868 | -1.14 | -0.045 | -0.999 | 0.124 | -0.137 | -0.079 | -0.079 | 0 | 0.61 | 0.97 | 0.57 | 0.63 | 0.44 | 0.65 |
| 7 | 146767404 | rs740809 | 0.138 | 0 | 0.464 | -0.204 | -0.001 | 0.003 | 0.003 | 0 | 0.71 | 0.72 | 0.7 | 0.72 | 0.73 | 0.6 |
| 7 | 146768776 | rs7777098 | -0.279 | 0.263 | 0.338 | -0.09 | 0.023 | 0 | 0 | 0.505 | 0.06 | 0.36 | 0.06 | 0 | 0.02 | 0.01 |
| 7 | 146780248 | rs10261458 | 0.453 | 0.384 | -0.37 | 0.018 | 0.043 | 0.073 | 0.073 | 0.206 | 0.58 | 0.17 | 0.57 | 0.69 | 0.69 | 0.55 |
| 7 | 146796125 | rs10273726 | -0.893 | 1.381 | 0.145 | 0.298 | -0.022 | 0.034 | 0.034 | 4.176 | 0.1 | 0.56 | 0.08 | 0.12 | 0.04 | 0.21 |
| 7 | 146800106 | rs6962089 | -1.709 | 0.567 | -0.723 | 0.88 | -0.034 | -0.101 | -0.101 | 0 | 0.4 | 0.89 | 0.45 | 0.47 | 0.37 | 0.44 |
| 7 | 146814569 | rs4726818 | 1.407 | 0.083 | -0.491 | 0.646 | -0.036 | -0.021 | -0.021 | 0 | 0.59 | 0.17 | 0.46 | 0.63 | 0.59 | 0.63 |
| 7 | 146816475 | rs6968536 | 1.027 | 0.032 | 0.621 | 0.654 | 0.009 | 0.006 | 0.006 | 0 | 0.66 | 0.58 | 0.51 | 0.66 | 0.63 | 0.69 |
| 7 | 146819356 | rs7806761 | -0.17 | -0.153 | 0.674 | 0.677 | -0.025 | 0.051 | 0.051 | 0.011 | 0.81 | 0.74 | 0.75 | 0.65 | 0.89 | 0.83 |
| 7 | 146831849 | rs9640492 | 1.278 | -0.402 | 0.318 | 0.242 | 0.011 | -0.006 | -0.006 | 0 | 0.47 | 0.25 | 0.39 | 0.55 | 0.44 | 0.56 |
| 7 | 146842776 | rs924146 | 0.839 | -0.402 | 0.891 | 0.161 | 0.006 | 0.009 | 0.009 | 0 | 0.11 | 0.06 | 0.07 | 0 | 0.11 | 0.03 |
| 7 | 146854213 | rs7791332 | 1.096 | -0.443 | 0.861 | 0.152 | 0.007 | 0.007 | 0.007 | 0 | 0.5 | 0.34 | 0.41 | 0.56 | 0.47 | 0.6 |
| 7 | 146856881 | rs7807033 | 0.322 | 0.018 | 0.708 | 0.094 | 0.006 | 0.05 | 0.05 | 0 | 0.13 | 0.05 | 0.07 | 0 | 0.13 | 0.1 |
| 7 | 146867339 | rs10952674 | 1.316 | 0.014 | 0.771 | 0.133 | 0.031 | -0.011 | -0.011 | 0 | 0.47 | 0.2 | 0.37 | 0.56 | 0.41 | 0.53 |
| 7 | 146868611 | rs4725704 | 0.969 | 0.015 | 0.664 | 0.078 | 0 | 0.044 | 0.044 | 0 | 0.52 | 0.25 | 0.41 | 0.67 | 0.48 | 0.63 |
| 7 | 146876502 | rs7457112 | -0.257 | 0.011 | 0.642 | 0.05 | 0.001 | -0.042 | -0.042 | 0 | 0.16 | 0.17 | 0.11 | 0.02 | 0.18 | 0.17 |
| 7 | 146879634 | rs4559158 | 1.214 | -0.125 | 0.601 | 0.081 | 0.001 | 0.031 | 0.031 | 0.109 | 0.48 | 0.2 | 0.4 | 0.62 | 0.47 | 0.55 |
| 7 | 146879704 | rs4282491 | 1.846 | -0.822 | -0.363 | -0.166 | -0.096 | -0.019 | -0.019 | 0 | 0.28 | 0.02 | 0.21 | 0.2 | 0.33 | 0.22 |
| 7 | 146884426 | rs1908765 | -0.251 | -0.587 | 0.106 | -0.529 | 0.009 | 0.003 | 0.003 | 0 | 0.65 | 0.41 | 0.49 | 0.65 | 0.61 | 0.73 |
| 7 | 146897211 | rs10085697 | 0.906 | -0.711 | -0.018 | -0.551 | -0.003 | 0.021 | 0.021 | 0.161 | 0.47 | 0.15 | 0.38 | 0.63 | 0.44 | 0.55 |
| 7 | 146897403 | rs10273775 | -0.386 | -0.556 | 0.016 | -0.585 | 0.006 | 0.002 | 0.002 | 0 | 0.65 | 0.41 | 0.48 | 0.64 | 0.59 | 0.72 |
| 7 | 146901009 | rs6966651 | 0.146 | -0.352 | -0.375 | -1.127 | 0.012 | 0.038 | 0.038 | 0 | 0.14 | 0.09 | 0.07 | 0 | 0.12 | 0.1 |
| 7 | 146910565 | rs1390720 | 0.086 | -0.558 | 0.075 | -1.206 | 0.014 | -0.067 | -0.067 | 0 | 0.35 | 0.44 | 0.38 | 0.06 | 0.42 | 0.31 |
| 7 | 146913540 | rs1603450 | 0.193 | -0.856 | -0.329 | -1.292 | -0.004 | -0.001 | -0.001 | 0.072 | 0.27 | 0.27 | 0.35 | 0.73 | 0.18 | 0.39 |
| 7 | 146923861 | rs1496547 | -0.906 | -0.661 | -0.735 | -1.234 | -0.007 | 0.059 | 0.059 | 0 | 0.68 | 0.88 | 0.78 | 0.79 | 0.66 | 0.76 |
| 7 | 146934096 | rs6943628 | 0.988 | -0.775 | -0.848 | -1.276 | 0.003 | -0.004 | -0.004 | 0 | 0.09 | 0.07 | 0.05 | 0 | 0.09 | 0.05 |
| 7 | 146935930 | rs1390723 | -1.455 | -0.633 | -0.911 | -1.139 | -0.063 | -0.019 | -0.019 | 0 | 0.76 | 0.98 | 0.82 | 0.8 | 0.73 | 0.8 |
| 7 | 146947702 | rs985419 | 0.595 | 0.368 | -0.909 | -1.908 | 0.003 | -0.009 | -0.009 | 0 | 0.28 | 0.24 | 0.21 | 0.03 | 0.23 | 0.18 |
| 7 | 146952081 | rs4725709 | -0.334 | -0.086 | -0.601 | -2.063 | 0.035 | 0.065 | 0.065 | 0 | 0.46 | 0.64 | 0.51 | 0.81 | 0.34 | 0.53 |
| 7 | 146954397 | rs9769600 | -1.416 | 0.045 | -0.641 | -2.26 | -0.033 | 0.036 | 0.036 | 0 | 0.63 | 0.74 | 0.62 | 0.76 | 0.52 | 0.67 |
| 7 | 146955688 | rs17170378 | 0.75 | -0.13 | -0.318 | -2.22 | -0.028 | 0.038 | 0.038 | 0 | 0.18 | 0.27 | 0.29 | 0.73 | 0.11 | 0.36 |
| 7 | 146958195 | rs12112963 | 0.643 | 0.068 | -0.754 | -2.27 | 0.003 | 0.001 | 0.001 | 0 | 0.14 | 0.09 | 0.08 | 0 | 0.14 | 0.1 |
| 7 | 146959088 | rs2132591 | -0.86 | 0.223 | -0.582 | -2.003 | -0.023 | 0.066 | 0.066 | 0 | 0.6 | 0.7 | 0.59 | 0.83 | 0.49 | 0.64 |
| 7 | 146968295 | rs10279700 | 0.517 | 0.472 | -0.115 | -1.585 | -0.003 | 0.005 | 0.005 | 0 | 0.17 | 0.18 | 0.3 | 0.77 | 0.13 | 0.35 |
| 7 | 146969399 | rs1472361 | 0.353 | 1.014 | -0.203 | -1.685 | 0.025 | -0.045 | -0.045 | 0.141 | 0.14 | 0.18 | 0.18 | 0 | 0.26 | 0.13 |

| | | | | | | | | | | | | | | | | |
|---|---|---|---|---|---|---|---|---|---|---|---|---|---|---|---|---|
| 7 | 146970047 | rs4637733 | -2.136 | 0.622 | -0.604 | -1.841 | -0.054 | -0.03 | -0.03 | 0 | 0.72 | 0.97 | 0.75 | 0.81 | 0.72 | 0.75 |
| 7 | 146971312 | rs10230132 | 0.582 | 0.34 | -0.193 | -2.041 | 0.004 | -0.028 | -0.028 | 0.03 | 0.32 | 0.27 | 0.22 | 0.03 | 0.28 | 0.24 |
| 7 | 146974832 | rs12703890 | 0.438 | 0.589 | -0.038 | -1.793 | 0.005 | -0.031 | -0.031 | 0.018 | 0.09 | 0.11 | 0.04 | 0 | 0.09 | 0.11 |
| 7 | 146975480 | rs10268597 | -1.393 | 0.775 | 0.089 | -1.778 | 0.004 | -0.007 | -0.007 | 0 | 0.38 | 0.59 | 0.48 | 0.73 | 0.4 | 0.47 |
| 7 | 146980556 | rs12531764 | -1.013 | 0.718 | 0.034 | -1.876 | 0.012 | -0.023 | -0.023 | 0 | 0.35 | 0.52 | 0.46 | 0.78 | 0.36 | 0.44 |
| 7 | 146984325 | rs12111644 | 1.013 | 0.42 | 0.124 | -1.71 | 0.004 | -0.034 | -0.034 | 0 | 0.06 | 0.13 | 0.02 | 0 | 0.04 | 0.06 |
| 7 | 146985808 | rs10248220 | 0.664 | 0.447 | -0.007 | -1.684 | -0.015 | 0.022 | 0.022 | 0 | 0.16 | 0.21 | 0.29 | 0.78 | 0.11 | 0.33 |
| 7 | 147001975 | rs2620441 | -0.951 | 0.452 | -0.089 | -1.598 | -0.026 | 0.022 | 0.022 | 0.461 | 0.28 | 0.53 | 0.33 | 0.72 | 0.19 | 0.42 |
| 7 | 147006000 | rs12154311 | 1.114 | 0.485 | -0.188 | -1.904 | 0.008 | 0.027 | 0.027 | 0 | 0.27 | 0.16 | 0.22 | 0.02 | 0.26 | 0.19 |
| 7 | 147008093 | rs1525209 | -0.405 | 1.064 | -0.177 | -1.765 | -0.002 | 0.046 | 0.046 | 0.194 | 0.55 | 0.68 | 0.55 | 0.74 | 0.46 | 0.61 |
| 7 | 147011388 | rs10500174 | 1.139 | 0.902 | -0.263 | -1.743 | -0.002 | -0.026 | -0.026 | 0 | 0.06 | 0.11 | 0.02 | 0.01 | 0.04 | 0.05 |
| 7 | 147013993 | rs2620452 | -0.112 | 1.032 | -0.361 | -1.632 | 0.021 | 0.035 | 0.035 | 0.327 | 0.52 | 0.6 | 0.52 | 0.74 | 0.41 | 0.56 |
| 7 | 147021164 | rs10500175 | 0.495 | 1.287 | -1.128 | -1.348 | -0.075 | 0.077 | 0.077 | 0 | 0.13 | 0.03 | 0.16 | 0.01 | 0.24 | 0.11 |
| 7 | 147022896 | rs2692136 | -1.147 | 0.726 | -1.123 | -1.066 | -0.046 | -0.09 | -0.09 | 0 | 0.71 | 0.96 | 0.69 | 0.71 | 0.7 | 0.71 |
| 7 | 147023537 | rs2692133 | 0.173 | 0.694 | -0.425 | -0.896 | -0.016 | -0.018 | -0.018 | 0 | 0.38 | 0.45 | 0.35 | 0.37 | 0.34 | 0.27 |
| 7 | 147023864 | rs2620472 | 0.213 | 0.651 | -0.064 | -0.906 | -0.003 | -0.004 | -0.004 | 0 | 0.37 | 0.55 | 0.37 | 0.37 | 0.34 | 0.24 |
| 7 | 147025943 | rs1479842 | -0.669 | 0.855 | -0.073 | -1.128 | 0.013 | 0.064 | 0.064 | 0 | 0.52 | 0.62 | 0.52 | 0.38 | 0.57 | 0.35 |
| 7 | 147025978 | rs13224673 | 1.121 | 0.853 | -0.025 | -1.136 | -0.003 | -0.016 | -0.016 | 0 | 0.14 | 0.12 | 0.15 | 0.01 | 0.22 | 0.11 |
| 7 | 147027426 | rs826790 | 0.345 | 0.865 | -0.029 | -1.12 | -0.007 | -0.009 | -0.009 | 0.117 | 0.38 | 0.51 | 0.37 | 0.37 | 0.35 | 0.24 |
| 7 | 147027457 | rs826791 | -1.851 | 0.775 | -0.338 | -1.066 | -0.041 | -0.049 | -0.049 | 0 | 0.72 | 0.96 | 0.81 | 0.77 | 0.75 | 0.72 |
| 7 | 147027789 | rs10500176 | 0.956 | 0.828 | -0.205 | -1.008 | -0.001 | -0.011 | -0.011 | 0 | 0.19 | 0.18 | 0.24 | 0.36 | 0.18 | 0.39 |
| 7 | 147028591 | rs826792 | 1.366 | 0.763 | -0.227 | -1.016 | -0.015 | -0.002 | -0.002 | 0.379 | 0.06 | 0.02 | 0.13 | 0.32 | 0.1 | 0.07 |
| 7 | 147029028 | rs6973542 | -0.157 | 0.799 | -0.057 | -0.887 | -0.006 | -0.004 | -0.004 | 0.491 | 0.33 | 0.32 | 0.4 | 0.37 | 0.4 | 0.5 |
| 7 | 147035067 | rs826802 | 0.442 | 0.709 | -0.013 | -0.915 | 0.005 | 0.006 | 0.006 | 0.166 | 0.38 | 0.63 | 0.37 | 0.37 | 0.34 | 0.24 |
| 7 | 147039722 | rs6945022 | -0.392 | 0.806 | 0.038 | -0.881 | -0.006 | -0.004 | -0.004 | 0.02 | 0.34 | 0.31 | 0.4 | 0.37 | 0.4 | 0.5 |
| 7 | 147041579 | rs17497254 | 0.257 | 0.933 | 0.045 | -0.963 | -0.014 | 0.003 | 0.003 | 0 | 0.2 | 0.13 | 0.17 | 0.02 | 0.27 | 0.16 |
| 7 | 147042196 | rs10276770 | 1.027 | 0.628 | 0.264 | -1.136 | 0.001 | -0.072 | -0.072 | 1.96 | 0.17 | 0.38 | 0.16 | 0.05 | 0.16 | 0.08 |
| 7 | 147054730 | rs10500179 | 0.329 | 0.628 | 0.473 | -1.138 | 0.038 | -0.104 | -0.104 | 0.472 | 0.21 | 0.32 | 0.19 | 0.02 | 0.26 | 0.16 |
| 7 | 147058991 | rs826824 | -0.855 | 0.523 | 0.369 | -1.145 | 0.007 | 0.034 | 0.034 | 0 | 0.58 | 0.76 | 0.64 | 0.69 | 0.57 | 0.64 |
| 7 | 147061654 | rs10268483 | 1.611 | 0.422 | 0.43 | -1.126 | 0.001 | -0.016 | -0.016 | 0 | 0.14 | 0.19 | 0.14 | 0.05 | 0.17 | 0.08 |
| 7 | 147067719 | rs826809 | -0.127 | 0.515 | 0.097 | -1.097 | 0.014 | 0.058 | 0.058 | 0 | 0.5 | 0.65 | 0.58 | 0.75 | 0.47 | 0.57 |
| 7 | 147073363 | rs10250570 | -0.492 | 0.53 | 0.093 | -1.149 | 0.024 | -0.118 | -0.118 | 0 | 0.35 | 0.51 | 0.34 | 0.07 | 0.43 | 0.24 |
| 7 | 147073554 | rs10500181 | 1.747 | 0.224 | 0.194 | -0.996 | 0 | 0.02 | 0.02 | 0 | 0.15 | 0.18 | 0.23 | 0.35 | 0.14 | 0.33 |
| 7 | 147074235 | rs1525221 | 0.344 | 0.235 | 0.128 | -1.026 | -0.012 | -0.103 | -0.103 | 1.375 | 0.22 | 0.43 | 0.18 | 0.06 | 0.21 | 0.14 |
| 7 | 147074441 | rs10500184 | 1.012 | 0.283 | 0.204 | -0.971 | -0.007 | 0.124 | 0.124 | 0 | 0.19 | 0.37 | 0.25 | 0.35 | 0.14 | 0.33 |
| 7 | 147078658 | rs1404709 | 1.218 | 0.303 | 0.203 | -0.978 | 0.006 | -0.069 | -0.069 | 0.137 | 0.16 | 0.41 | 0.16 | 0.05 | 0.17 | 0.08 |
| 7 | 147086045 | rs700303 | 0.17 | 0.121 | 0.132 | -0.856 | -0.011 | -0.003 | -0.003 | 0.024 | 0.37 | 0.5 | 0.43 | 0.67 | 0.3 | 0.49 |
| 7 | 147098651 | rs12531630 | 1.557 | 0.232 | -0.158 | -0.761 | -0.001 | 0.044 | 0.044 | 0 | 0.16 | 0.23 | 0.23 | 0.34 | 0.14 | 0.33 |
| 7 | 147099227 | rs826660 | 1.3 | 0.253 | -0.317 | -0.794 | 0.01 | -0.03 | -0.03 | 1.522 | 0.19 | 0.07 | 0.18 | 0.33 | 0.16 | 0.17 |
| 7 | 147117454 | rs10251377 | 0.278 | 0.476 | -0.092 | -0.911 | 0.009 | -0.087 | -0.087 | 1.501 | 0.22 | 0.46 | 0.18 | 0.07 | 0.23 | 0.14 |
| 7 | 147129593 | rs826645 | 0.47 | 0.681 | -0.282 | -0.959 | -0.052 | 0.019 | 0.019 | 0 | 0.32 | 0.12 | 0.34 | 0.34 | 0.38 | 0.28 |
| 7 | 147131270 | rs4621717 | 1.243 | 0.598 | -0.177 | -0.895 | -0.001 | 0.012 | 0.012 | 0.142 | 0.2 | 0.22 | 0.25 | 0.35 | 0.18 | 0.37 |
| 7 | 147144359 | rs12539907 | 1.261 | 0.597 | -0.231 | -0.895 | -0.001 | 0.008 | 0.008 | 0.128 | 0.2 | 0.2 | 0.25 | 0.35 | 0.18 | 0.37 |
| 7 | 147148420 | rs1528532 | 1.747 | 0.655 | -0.38 | -0.919 | 0.003 | -0.033 | -0.033 | 0 | 0.17 | 0.07 | 0.18 | 0.33 | 0.16 | 0.16 |
| 7 | 147149788 | rs13221396 | 1.271 | 0.746 | -0.377 | -0.978 | -0.042 | 0.042 | 0.042 | 0 | 0.14 | 0.05 | 0.16 | 0.01 | 0.23 | 0.12 |
| 7 | 147155060 | rs4726860 | -1.595 | 0.555 | -0.355 | -0.96 | 0.033 | -0.003 | -0.003 | 0 | 0.7 | 0.86 | 0.76 | 0.75 | 0.75 | 0.75 |
| 7 | 147157357 | rs2022225 | -1.574 | 0.526 | -0.364 | -0.93 | 0.023 | -0.001 | -0.001 | 0 | 0.71 | 0.89 | 0.73 | 0.76 | 0.74 | 0.75 |
| 7 | 147157377 | rs7790550 | 1.033 | 0.525 | -0.153 | -0.993 | 0.018 | -0.041 | -0.041 | 3.587 | 0.17 | 0.4 | 0.19 | 0.08 | 0.19 | 0.09 |
| 7 | 147157421 | rs2022226 | -0.389 | 0.536 | -0.211 | -0.975 | -0.002 | -0.003 | -0.003 | 0 | 0.53 | 0.49 | 0.54 | 0.68 | 0.55 | 0.67 |
| 7 | 147168231 | rs6464810 | -1.631 | 0.621 | -0.87 | -0.947 | 0.03 | 0.008 | 0.008 | 0 | 0.7 | 0.9 | 0.8 | 0.78 | 0.75 | 0.75 |
| 7 | 147168314 | rs7783425 | -1.631 | 0.621 | -0.87 | -0.947 | 0.03 | 0.008 | 0.008 | 0 | 0.7 | 0.9 | 0.8 | 0.78 | 0.75 | 0.75 |
| 7 | 147168331 | rs7783548 | -1.651 | 0.606 | -0.903 | -0.954 | -0.01 | -0.036 | -0.036 | 0 | 0.71 | 0.96 | 0.81 | 0.78 | 0.75 | 0.75 |
| 7 | 147169167 | rs10952684 | -1.459 | 0.611 | -0.895 | -0.952 | 0.032 | 0.011 | 0.011 | 0 | 0.7 | 0.87 | 0.8 | 0.78 | 0.75 | 0.75 |
| 7 | 147175188 | rs10265626 | 1.795 | 0.632 | -0.895 | -0.898 | 0.004 | -0.029 | -0.029 | 1.562 | 0.17 | 0.08 | 0.18 | 0.3 | 0.15 | 0.17 |
| 7 | 147176188 | rs4725734 | 0.905 | 0.595 | -0.824 | -0.918 | 0.011 | -0.039 | -0.039 | 0.142 | 0.18 | 0.53 | 0.21 | 0.12 | 0.19 | 0.08 |
| 7 | 147177691 | rs1528523 | -0.088 | 0.676 | -1.027 | -0.921 | -0.001 | -0.047 | -0.047 | 0 | 0.38 | 0.75 | 0.52 | 0.47 | 0.36 | 0.42 |
| 7 | 147183399 | rs1468370 | -1.114 | 0.505 | -0.876 | -0.662 | 0.001 | 0.033 | 0.033 | 0 | 0.71 | 0.95 | 0.81 | 0.84 | 0.74 | 0.72 |
| 7 | 147190372 | rs6973129 | 0.749 | 0.536 | -0.37 | -0.762 | -0.005 | 0.008 | 0.008 | 0.156 | 0.19 | 0.25 | 0.24 | 0.31 | 0.16 | 0.2 |
| 7 | 147191337 | rs1015932 | -1.035 | 0.359 | -0.761 | -0.926 | -0.001 | -0.029 | -0.029 | 0 | 0.68 | 0.75 | 0.6 | 0.59 | 0.7 | 0.55 |
| 7 | 147192393 | rs1528510 | 1.371 | 0.452 | -1.112 | -0.901 | 0.02 | -0.015 | -0.015 | 0 | 0.17 | 0.01 | 0.1 | 0.12 | 0.13 | 0.14 |
| 7 | 147194880 | rs16883528 | 1.353 | 0.453 | -1.11 | -0.901 | 0.03 | -0.004 | -0.004 | 0 | 0.17 | 0.02 | 0.1 | 0.12 | 0.13 | 0.14 |
| 7 | 147200383 | rs10266867 | -0.138 | -0.171 | -0.807 | -0.825 | -0.001 | -0.001 | -0.001 | 0.215 | 0.39 | 0.39 | 0.35 | 0.43 | 0.3 | 0.34 |
| 7 | 147200576 | rs7792855 | 0.426 | 0.004 | -0.819 | -0.872 | 0.033 | -0.049 | -0.049 | 2.837 | 0.29 | 0.36 | 0.24 | 0.15 | 0.4 | 0.21 |
| 7 | 147201446 | rs7781317 | -0.09 | -0.055 | -1.254 | -0.826 | -0.003 | 0.025 | 0.025 | 0 | 0.47 | 0.33 | 0.34 | 0.27 | 0.54 | 0.35 |
| 7 | 147202658 | rs11762071 | 0.084 | 0.665 | -2.02 | -0.738 | 0.001 | -0.008 | -0.008 | 0 | 0.33 | 0.63 | 0.3 | 0.18 | 0.4 | 0.21 |
| 7 | 147203197 | rs971818 | -0.684 | 0.699 | -2.092 | -0.694 | -0.006 | -0.053 | -0.053 | 1.022 | 0.69 | 0.78 | 0.6 | 0.58 | 0.7 | 0.55 |
| 7 | 147203767 | rs7808274 | -0.561 | 1.218 | -1.289 | -0.876 | 0.004 | -0.012 | -0.012 | 0 | 0.32 | 0.62 | 0.3 | 0.18 | 0.4 | 0.21 |
| 7 | 147205421 | rs1541415 | 0.592 | 1.233 | -2.074 | -1.1 | -0.092 | 0.183 | 0.183 | 2.56 | 0.7 | 0.98 | 0.83 | 0.96 | 0.65 | 0.82 |
| 7 | 147206439 | rs10226360 | 0.478 | 1.122 | -1.611 | -0.488 | -0.003 | 0.02 | 0.02 | 0 | 0.49 | 0.24 | 0.44 | 0.56 | 0.51 | 0.66 |

| | | | | | | | | | | | | | | | | |
|---|---|---|---|---|---|---|---|---|---|---|---|---|---|---|---|---|
| 7 | 147209008 | rs10282028 | 0.651 | 1.304 | -0.851 | -0.061 | 0.002 | 0.005 | 0.005 | 0 | 0.69 | 0.36 | 0.69 | 0.63 | 0.69 | 0.76 |
| 7 | 147209601 | rs1919188 | 0.474 | 1.633 | -1.021 | 0.106 | 0.001 | 0.002 | 0.002 | 0 | 0.73 | 0.72 | 0.72 | 0.63 | 0.7 | 0.76 |
| 7 | 147223640 | rs17225888 | -0.91 | 1.652 | -1.131 | 0.269 | -0.014 | -0.021 | -0.021 | 0.052 | 0.28 | 0.18 | 0.28 | 0.34 | 0.27 | 0.19 |
| 7 | 147226358 | rs851814 | -0.672 | 1.614 | -0.796 | 0.138 | 0 | -0.002 | -0.002 | 2.194 | 0.46 | 0.49 | 0.45 | 0.59 | 0.48 | 0.51 |
| 7 | 147226891 | rs6464816 | -0.326 | 1.759 | -0.999 | 0.26 | -0.021 | -0.054 | -0.054 | 0 | 0.16 | 0.08 | 0.19 | 0.37 | 0.15 | 0.18 |
| 7 | 147234727 | rs12703926 | 0.316 | 1.578 | -1.309 | 0.029 | -0.052 | -0.045 | -0.045 | 0 | 0.22 | 0.05 | 0.15 | 0.24 | 0.24 | 0.29 |
| 7 | 147235384 | rs10487690 | -0.089 | 1.696 | -1.036 | 0.175 | 0.007 | 0.039 | 0.039 | 0 | 0.12 | 0.15 | 0.19 | 0.4 | 0.14 | 0.18 |
| 7 | 147236636 | rs17170583 | -1.083 | 1.639 | -0.815 | 0.136 | 0.012 | 0.038 | 0.038 | 0 | 0.18 | 0.24 | 0.23 | 0.41 | 0.19 | 0.2 |
| 7 | 147236757 | rs17170584 | 0.214 | 1.174 | -0.425 | -0.276 | -0.007 | -0.052 | -0.052 | 0 | 0.21 | 0.23 | 0.3 | 0.02 | 0.2 | 0.14 |
| 7 | 147242624 | rs851821 | -0.732 | 1.166 | 0.019 | -0.355 | 0 | -0.001 | -0.001 | 0 | 0.42 | 0.46 | 0.41 | 0.63 | 0.46 | 0.48 |
| 7 | 147247139 | rs12666634 | 0.537 | 1.157 | -0.118 | -0.509 | -0.009 | -0.002 | -0.002 | 0 | 0.12 | 0.09 | 0.18 | 0.4 | 0.13 | 0.17 |
| 7 | 147257915 | rs6464824 | -1.645 | 0.536 | 0.228 | -0.705 | -0.001 | -0.015 | -0.015 | 0 | 0.65 | 0.84 | 0.73 | 0.66 | 0.67 | 0.63 |
| 7 | 147258432 | rs6946710 | -0.542 | 0.721 | 0.359 | -0.626 | 0.003 | -0.002 | -0.002 | 0.133 | 0.42 | 0.5 | 0.41 | 0.64 | 0.44 | 0.49 |
| 7 | 147259779 | rs2240376 | -0.401 | 0.765 | 0.394 | -0.607 | -0.004 | 0 | 0 | 0 | 0.4 | 0.41 | 0.4 | 0.64 | 0.44 | 0.49 |
| 7 | 147268460 | rs2190004 | -0.511 | 0.769 | 0.406 | -0.605 | -0.002 | 0 | 0 | 0 | 0.4 | 0.42 | 0.39 | 0.64 | 0.43 | 0.48 |
| 7 | 147277640 | rs5003024 | -1.193 | 0.589 | 0.177 | -0.682 | 0.005 | 0.073 | 0.073 | 0 | 0.59 | 0.85 | 0.7 | 0.74 | 0.63 | 0.6 |
| 7 | 147281247 | rs995891 | -1.184 | 0.588 | 0.176 | -0.683 | 0.006 | 0.074 | 0.074 | 0 | 0.59 | 0.84 | 0.7 | 0.74 | 0.63 | 0.6 |
| 7 | 147284801 | rs6954225 | -1.228 | 0.604 | 0.176 | -0.679 | 0.001 | 0.074 | 0.074 | 0 | 0.6 | 0.86 | 0.71 | 0.74 | 0.63 | 0.6 |
| 7 | 147292345 | rs1013872 | -0.343 | 0.903 | 0.489 | -0.579 | 0.027 | 0.074 | 0.074 | 0 | 0.18 | 0.32 | 0.25 | 0.38 | 0.21 | 0.22 |
| 7 | 147292995 | rs6464830 | -1.432 | 0.891 | 0.39 | -0.547 | 0.004 | 0.048 | 0.048 | 0 | 0.6 | 0.81 | 0.7 | 0.74 | 0.63 | 0.6 |
| 7 | 147294703 | rs10259411 | -1.461 | 0.889 | 0.322 | -0.548 | -0.003 | 0.072 | 0.072 | 0 | 0.6 | 0.88 | 0.71 | 0.74 | 0.63 | 0.6 |
| 7 | 147296158 | rs7794693 | 1.285 | 0.849 | 0.15 | -0.547 | -0.032 | -0.113 | -0.113 | 0 | 0.2 | 0.05 | 0.11 | 0.34 | 0.2 | 0.24 |
| 7 | 147302816 | rs11982531 | 0.033 | 0.86 | 0.459 | -0.477 | 0.007 | 0.015 | 0.015 | 0 | 0.41 | 0.34 | 0.42 | 0.31 | 0.42 | 0.38 |
| 7 | 147305180 | rs11773848 | -1.152 | 0.455 | 0.306 | -0.639 | -0.005 | 0.02 | 0.02 | 0 | 0.63 | 0.81 | 0.7 | 0.69 | 0.63 | 0.61 |
| 7 | 147307694 | rs12703951 | 0.18 | 0.493 | 0.046 | -0.626 | -0.012 | -0.045 | -0.045 | 0 | 0.18 | 0.07 | 0.22 | 0.37 | 0.18 | 0.22 |
| 7 | 147310299 | rs10244661 | -1.06 | 0.534 | -0.096 | -0.607 | -0.001 | 0.002 | 0.002 | 0 | 0.62 | 0.71 | 0.68 | 0.69 | 0.63 | 0.61 |
| 7 | 147311353 | rs11975946 | -0.049 | 0.257 | -0.402 | -0.472 | 0.002 | 0.008 | 0.008 | 0 | 0.4 | 0.35 | 0.42 | 0.33 | 0.42 | 0.39 |
| 7 | 147313809 | rs727716 | 0.303 | -0.172 | -0.742 | -1.102 | -0.021 | 0.04 | 0.04 | 0 | 0.43 | 0.22 | 0.49 | 0.62 | 0.45 | 0.62 |
| 7 | 147317086 | rs10251488 | -1.069 | -1.443 | 0.386 | -1.723 | -0.024 | -0.123 | -0.123 | 0.41 | 0.28 | 0.54 | 0.36 | 0.01 | 0.22 | 0.11 |
| 7 | 147320252 | rs10274432 | -1.537 | -1.539 | 0.716 | -1.708 | 0.041 | 0.043 | 0.043 | 1.145 | 0.05 | 0.31 | 0.19 | 0.01 | 0 | 0 |
| 7 | 147333930 | rs1860135 | -1.038 | -1.466 | -0.01 | -1.712 | -0.011 | -0.09 | -0.09 | 0.004 | 0.25 | 0.34 | 0.34 | 0.01 | 0.22 | 0.11 |
| 7 | 147333967 | rs4272288 | -0.667 | -1.424 | -0.031 | -1.694 | 0 | -0.037 | -0.037 | 0 | 0.23 | 0.21 | 0.16 | 0 | 0.22 | 0.11 |
| 7 | 147335023 | rs10243377 | -0.868 | -1.405 | -0.118 | -1.744 | -0.002 | -0.008 | -0.008 | 0 | 0.08 | 0.14 | 0.23 | 0.03 | 0.08 | 0.09 |
| 7 | 147337742 | rs7783626 | -1.108 | -1.391 | -0.056 | -1.677 | -0.01 | -0.099 | -0.099 | 0.45 | 0.31 | 0.45 | 0.4 | 0.04 | 0.3 | 0.21 |
| 7 | 147344718 | rs10259590 | -1.64 | -1.719 | -0.291 | -0.982 | -0.035 | -0.032 | -0.032 | 0.035 | 0.14 | 0.16 | 0.19 | 0.01 | 0 | 0 |
| 7 | 147351071 | rs10244236 | -1.154 | -1.602 | 0.025 | -0.885 | -0.017 | -0.015 | -0.015 | 0 | 0.3 | 0.63 | 0.14 | 0.11 | 0.15 | 0.14 |
| 7 | 147357473 | rs17170632 | -0.717 | -1.623 | -0.4 | -1.235 | -0.004 | 0.029 | 0.029 | 0 | 0.13 | 0.06 | 0.05 | 0.05 | 0.09 | 0.15 |
| 7 | 147359961 | rs6947112 | -0.508 | -1.863 | 0.496 | -1.388 | -0.042 | -0.095 | -0.095 | 0 | 0.17 | 0.33 | 0.07 | 0.03 | 0.09 | 0.11 |
| 7 | 147360817 | rs17327575 | -0.937 | -1.745 | -0.094 | -1.161 | 0.002 | 0.005 | 0.005 | 0 | 0.16 | 0.07 | 0.08 | 0.1 | 0.13 | 0.15 |
| 7 | 147362512 | rs12672896 | -1.576 | -2.069 | -0.373 | -1.591 | 0.032 | 0.07 | 0.07 | 0.187 | 0.28 | 0.12 | 0.12 | 0.1 | 0.18 | 0.15 |
| 7 | 147366954 | rs851727 | -1.185 | -2.044 | 0.314 | -1.793 | -0.054 | -0.129 | -0.129 | 0 | 0.2 | 0.49 | 0.29 | 0 | 0.07 | 0.04 |
| 7 | 147369658 | rs4383899 | -0.047 | -1.941 | 0.125 | -1.742 | 0.014 | 0.011 | 0.011 | 1.956 | 0.14 | 0.09 | 0.25 | 0 | 0.05 | 0.01 |
| 7 | 147370955 | rs700315 | -2.073 | -2.038 | 0.031 | -1.879 | 0.096 | 0.228 | 0.228 | 0.18 | 0.48 | 0.82 | 0.42 | 0.08 | 0.28 | 0.29 |
| 7 | 147398856 | rs851723 | -1.387 | -0.953 | -0.076 | -1.956 | 0 | 0.133 | 0.133 | 0 | 0.45 | 0.75 | 0.48 | 0.12 | 0.36 | 0.4 |
| 7 | 147405082 | rs851721 | -0.007 | -0.787 | 0.119 | -1.876 | 0.043 | -0.088 | -0.088 | 0.047 | 0.23 | 0.32 | 0.17 | 0.07 | 0.27 | 0.25 |
| 7 | 147406262 | rs731565 | 0.171 | -0.758 | -0.328 | -1.865 | -0.009 | -0.044 | -0.044 | 1.67 | 0.14 | 0.2 | 0.26 | 0 | 0.05 | 0.01 |
| 7 | 147407010 | rs700317 | -1.407 | -0.507 | -0.467 | -1.712 | -0.041 | 0.168 | 0.168 | 0 | 0.57 | 0.83 | 0.53 | 0.13 | 0.46 | 0.41 |
| 7 | 147414399 | rs2373177 | 0.731 | 0.455 | 0.082 | -0.556 | 0.039 | -0.026 | -0.026 | 0 | 0.74 | 0.57 | 0.74 | 0.75 | 0.72 | 0.61 |
| 7 | 147425466 | rs12056208 | -0.555 | 0.183 | -0.3 | -0.279 | -0.001 | 0 | 0 | 0 | 0.07 | 0.1 | 0.09 | 0.08 | 0.05 | 0.1 |
| 7 | 147426374 | rs10225143 | -0.992 | -1.047 | -0.464 | -1.477 | 0.095 | 0.163 | 0.163 | 0 | 0.5 | 0.71 | 0.35 | 0.11 | 0.3 | 0.41 |
| 7 | 147436087 | rs4392814 | 0.157 | -1.363 | 0.359 | -1.669 | 0.04 | 0.037 | 0.037 | 0 | 0.62 | 0.77 | 0.87 | 1 | 0.74 | 0.67 |
| 7 | 147444123 | rs10500192 | -0.077 | -1.432 | 0.374 | -1.688 | 0.054 | 0.22 | 0.22 | 0 | 0.37 | 0.09 | 0.25 | 0.04 | 0.26 | 0.33 |
| 7 | 147449063 | rs7791979 | -0.867 | -1.512 | 0.819 | -1.714 | 0.068 | 0.094 | 0.094 | 0 | 0.44 | 0.64 | 0.32 | 0.08 | 0.27 | 0.36 |
| 7 | 147454562 | rs7785603 | -1.305 | -1.509 | 0.863 | -1.712 | -0.013 | 0.006 | 0.006 | 0.406 | 0.13 | 0.46 | 0.23 | 0.11 | 0.05 | 0.1 |
| 7 | 147456946 | rs851654 | -0.654 | -1.437 | 0.802 | -1.688 | 0.004 | -0.029 | -0.029 | 0 | 0.43 | 0.55 | 0.31 | 0.08 | 0.27 | 0.35 |
| 7 | 147460586 | rs851650 | -1.244 | -1.04 | 0.399 | -0.832 | 0.042 | 0.088 | 0.088 | 0 | 0.54 | 0.75 | 0.46 | 0.27 | 0.38 | 0.46 |
| 7 | 147469901 | rs700319 | -0.838 | -1.139 | 0.331 | -0.877 | 0.128 | 0.119 | 0.119 | 0 | 0.47 | 0.77 | 0.36 | 0.24 | 0.3 | 0.39 |
| 7 | 147471710 | rs851674 | -1.112 | -1.067 | 0.356 | -0.799 | 0.086 | 0.097 | 0.097 | 0 | 0.53 | 0.78 | 0.39 | 0.27 | 0.34 | 0.46 |
| 7 | 147488371 | rs851697 | -1.489 | -0.904 | -0.056 | -0.652 | 0.084 | 0.093 | 0.093 | 0 | 0.49 | 0.81 | 0.46 | 0.3 | 0.38 | 0.47 |
| 7 | 147494803 | rs851685 | -0.532 | -0.907 | 0.601 | -0.879 | -0.043 | -0.137 | -0.137 | 0 | 0.33 | 0.49 | 0.29 | 0.04 | 0.25 | 0.29 |
| 7 | 147495973 | rs10271363 | -0.537 | -0.75 | 0.771 | -0.801 | 0.03 | 0.166 | 0.166 | 0 | 0.07 | 0.21 | 0.14 | 0.19 | 0.1 | 0.05 |
| 7 | 147505893 | rs1404733 | -0.36 | -0.827 | 0.679 | -0.958 | -0.003 | 0.006 | 0.006 | 0 | 0.1 | 0.09 | 0.06 | 0.07 | 0.07 | 0.22 |
| 7 | 147511889 | rs851663 | -1.076 | -0.933 | 0.81 | -0.928 | 0.037 | 0.06 | 0.06 | 0 | 0.4 | 0.69 | 0.42 | 0.17 | 0.34 | 0.34 |
| 7 | 147512619 | rs7785839 | -0.581 | -0.847 | 0.66 | -0.854 | 0.009 | 0.079 | 0.079 | 0 | 0.07 | 0.22 | 0.13 | 0.12 | 0.08 | 0.04 |
| 7 | 147513184 | rs851661 | -1.574 | -0.983 | 0.059 | -0.932 | 0.053 | 0.177 | 0.177 | 0.556 | 0.47 | 0.89 | 0.32 | 0.21 | 0.36 | 0.44 |
| 7 | 147520674 | rs1723430 | 1.46 | -0.161 | -0.145 | -0.317 | 0.011 | 0.025 | 0.025 | 0 | 0.42 | 0.28 | 0.34 | 0.42 | 0.35 | 0.36 |
| 7 | 147522124 | rs851716 | 1.46 | -0.161 | -0.145 | -0.288 | 0.011 | 0.023 | 0.023 | 0 | 0.42 | 0.28 | 0.34 | 0.41 | 0.35 | 0.36 |
| 7 | 147526360 | rs13238480 | 0.02 | -0.394 | -0.255 | -0.445 | 0.026 | 0.028 | 0.028 | 0.065 | 0.25 | 0.15 | 0.11 | 0.17 | 0.23 | 0.34 |
| 7 | 147527038 | rs17825525 | 0.242 | -0.438 | -0.264 | -0.475 | 0.005 | 0.02 | 0.02 | 0 | 0.11 | 0.03 | 0.19 | 0.07 | 0.1 | 0.04 |

| | | | | | | | | | | | | | | | | |
|---|---|---|---|---|---|---|---|---|---|---|---|---|---|---|---|---|
| 7 | 147528731 | rs851712 | 0.35 | -0.405 | 0.288 | -0.098 | -0.015 | 0.012 | 0.012 | 0 | 0.61 | 0.67 | 0.62 | 0.73 | 0.54 | 0.52 |
| 7 | 147533267 | rs851708 | 2.736 | -0.811 | -0.014 | 0.07 | 0.011 | 0.027 | 0.027 | 0.001 | 0.3 | 0.19 | 0.22 | 0.25 | 0.24 | 0.25 |
| 7 | 147538107 | rs851705 | 1.483 | -0.707 | 0.042 | 0.109 | -0.005 | 0.002 | 0.002 | 0 | 0.44 | 0.48 | 0.37 | 0.53 | 0.36 | 0.43 |
| 7 | 147549713 | rs2011815 | -0.867 | -0.305 | 0.029 | 0.355 | 0.004 | -0.009 | -0.009 | 0 | 0.16 | 0.13 | 0.24 | 0.2 | 0.18 | 0.08 |
| 7 | 147558680 | rs1013335 | 0.498 | -0.317 | 0.123 | 0.321 | 0.013 | 0.005 | 0.005 | 0.006 | 0.68 | 0.66 | 0.47 | 0.68 | 0.59 | 0.76 |
| 7 | 147566163 | rs4725747 | 0.369 | 0.047 | 0.214 | 0.141 | 0.02 | 0.035 | 0.035 | 0 | 0.71 | 0.66 | 0.49 | 0.59 | 0.63 | 0.72 |
| 7 | 147574390 | rs2710102 | 0.542 | 0.216 | 0.271 | 0.334 | -0.02 | -0.002 | -0.002 | 0 | 0.58 | 0.7 | 0.6 | 0.63 | 0.51 | 0.46 |
| 7 | 147574509 | rs2538994 | -1.082 | 0.183 | 0.245 | 0.32 | -0.09 | -0.103 | -0.103 | 0.132 | 0.82 | 0.97 | 0.71 | 0.65 | 0.7 | 0.75 |
| 7 | 147575112 | rs759178 | 0.515 | 0.144 | 0.212 | 0.35 | -0.02 | -0.002 | -0.002 | 0 | 0.58 | 0.7 | 0.6 | 0.63 | 0.51 | 0.46 |
| 7 | 147575230 | rs759177 | 0.515 | 0.144 | 0.212 | 0.35 | -0.02 | -0.002 | -0.002 | 0 | 0.58 | 0.7 | 0.6 | 0.63 | 0.51 | 0.46 |
| 7 | 147579619 | rs2538991 | 0.602 | 0.153 | 0.488 | 0.353 | 0.009 | -0.001 | -0.001 | 0 | 0.57 | 0.57 | 0.59 | 0.63 | 0.51 | 0.45 |
| 7 | 147581743 | rs11771110 | 0.22 | 0.3 | 0.493 | 0.272 | 0.007 | 0.023 | 0.023 | 0 | 0.12 | 0.2 | 0.09 | 0.17 | 0.13 | 0.09 |
| 7 | 147581989 | rs17236183 | 2.023 | 0.248 | 0.472 | 0.239 | 0.007 | 0.079 | 0.079 | 0.856 | 0.39 | 0.07 | 0.41 | 0.27 | 0.35 | 0.31 |
| 7 | 147585819 | rs2538976 | 0.869 | 0.288 | -0.123 | 0.24 | -0.002 | 0.002 | 0.002 | 0.143 | 0.55 | 0.46 | 0.53 | 0.43 | 0.5 | 0.44 |
| 7 | 147591839 | rs2538973 | -0.101 | 0.376 | 0.067 | 0.512 | -0.005 | -0.016 | -0.016 | 0 | 0.33 | 0.41 | 0.18 | 0.17 | 0.28 | 0.27 |
| 7 | 147595362 | rs2710109 | -0.164 | 0.401 | 0.079 | 0.472 | -0.004 | -0.007 | -0.007 | 0.016 | 0.34 | 0.42 | 0.32 | 0.21 | 0.28 | 0.29 |
| 7 | 147595540 | rs4725752 | 1.985 | 0.485 | -0.048 | 0.239 | 0.032 | 0.09 | 0.09 | 1.404 | 0.37 | 0.07 | 0.26 | 0.24 | 0.3 | 0.28 |
| 7 | 147597251 | rs4431524 | 1.543 | 0.466 | -0.028 | 0.248 | 0.036 | 0.07 | 0.07 | 0.209 | 0.4 | 0.27 | 0.29 | 0.24 | 0.32 | 0.33 |
| 7 | 147598261 | rs2538966 | 1.474 | 0.471 | -0.025 | 0.26 | 0.036 | 0.069 | 0.069 | 0.184 | 0.4 | 0.28 | 0.29 | 0.24 | 0.32 | 0.33 |
| 7 | 147598395 | rs7794222 | -0.238 | 0.537 | 0.057 | 0.351 | 0.002 | -0.005 | -0.005 | 0 | 0.35 | 0.33 | 0.36 | 0.4 | 0.32 | 0.27 |
| 7 | 147607585 | rs10238572 | 0.845 | 0.673 | -0.11 | 0.51 | 0.011 | 0.009 | 0.009 | 0.242 | 0.69 | 0.38 | 0.46 | 0.62 | 0.57 | 0.54 |
| 7 | 147613506 | rs13438769 | -1.222 | 0.406 | 0.085 | -0.062 | -0.001 | -0.068 | -0.068 | 0 | 0.13 | 0.24 | 0.09 | 0 | 0.11 | 0.08 |
| 7 | 147613758 | rs2538959 | -0.019 | 0.276 | -0.596 | -0.277 | -0.002 | 0.073 | 0.073 | 0 | 0.67 | 0.9 | 0.85 | 0.8 | 0.7 | 0.62 |
| 7 | 147614835 | rs7794874 | 1.617 | 0.589 | -0.635 | -0.157 | 0.051 | 0.089 | 0.089 | 0 | 0.4 | 0.23 | 0.23 | 0.15 | 0.27 | 0.32 |
| 7 | 147619772 | rs2539000 | 0.546 | 0.49 | -0.452 | -0.23 | -0.034 | 0.15 | 0.15 | 0 | 0.57 | 0.89 | 0.78 | 0.8 | 0.58 | 0.54 |
| 7 | 147619904 | rs2538999 | 0.524 | 0.511 | -0.443 | -0.22 | -0.036 | 0.147 | 0.147 | 0 | 0.58 | 0.89 | 0.78 | 0.8 | 0.58 | 0.54 |
| 7 | 147629487 | rs2710091 | 1.669 | 0.819 | 0.078 | -0.087 | 0 | 0 | 0 | 0 | 0.78 | 0.31 | 0.81 | 0.74 | 0.83 | 0.75 |
| 7 | 147641383 | rs2707579 | 0.256 | 0.93 | 0.485 | -0.11 | -0.006 | -0.027 | -0.027 | 1.601 | 0.85 | 0.73 | 0.87 | 0.92 | 0.87 | 0.93 |
| 7 | 147643746 | rs1922886 | 0.011 | 0.758 | 0.157 | -0.398 | 0.003 | -0.012 | -0.012 | 0.008 | 0.26 | 0.23 | 0.12 | 0.02 | 0.23 | 0.32 |
| 7 | 147643987 | rs1922884 | 0.774 | 1.001 | 0.001 | -0.212 | -0.071 | -0.064 | -0.064 | 0 | 0.1 | 0.02 | 0.42 | 0.56 | 0.27 | 0.21 |
| 7 | 147647187 | rs1882687 | 0.161 | 0.788 | 0.063 | -0.293 | 0.003 | -0.012 | -0.012 | 0 | 0.26 | 0.23 | 0.12 | 0.02 | 0.23 | 0.31 |
| 7 | 147654425 | rs2249958 | 0.033 | 0.967 | 0.629 | -0.055 | 0 | -0.039 | -0.039 | 0 | 0.84 | 0.59 | 0.85 | 0.92 | 0.87 | 0.92 |
| 7 | 147655743 | rs2710084 | 1.779 | 0.911 | 0.535 | -0.063 | 0.034 | -0.019 | -0.019 | 0 | 0.75 | 0.15 | 0.78 | 0.74 | 0.83 | 0.9 |
| 7 | 147661356 | rs10229688 | -1.282 | 1.011 | 0.709 | 0.025 | 0.021 | 0.141 | 0.141 | 0.596 | 0.09 | 0.33 | 0.07 | 0.18 | 0.07 | 0.04 |
| 7 | 147666964 | rs13225815 | -1.526 | 1.026 | 0.841 | 0.035 | 0.012 | 0.137 | 0.137 | 0.366 | 0.09 | 0.34 | 0.07 | 0.18 | 0.06 | 0.04 |
| 7 | 147669194 | rs12670399 | 0.404 | 1.233 | 0.653 | 0.057 | -0.048 | -0.041 | -0.041 | 0 | 0.11 | 0.03 | 0.42 | 0.56 | 0.28 | 0.22 |
| 7 | 147674559 | rs10243164 | -0.957 | 0.901 | 0.859 | 0.015 | 0.019 | 0.128 | 0.128 | 0.007 | 0.08 | 0.24 | 0.06 | 0.18 | 0.07 | 0.04 |
| 7 | 147674978 | rs10240503 | -2.765 | 0.933 | 0.957 | -0.022 | 0.02 | 0.049 | 0.049 | 1.508 | 0.15 | 0.53 | 0.1 | 0.2 | 0.12 | 0.2 |
| 7 | 147681603 | rs10269250 | -0.939 | 0.869 | 0.774 | -0.003 | 0.017 | 0.118 | 0.118 | 0.01 | 0.08 | 0.23 | 0.06 | 0.18 | 0.06 | 0.04 |
| 7 | 147687165 | rs2710079 | -1.55 | 0.819 | 0.522 | -0.031 | 0 | 0.036 | 0.036 | 0 | 0.26 | 0.48 | 0.26 | 0.34 | 0.23 | 0.26 |
| 7 | 147689625 | rs6958434 | -0.791 | 0.908 | 0.599 | 0.131 | -0.006 | 0.038 | 0.038 | 0 | 0.21 | 0.37 | 0.24 | 0.32 | 0.18 | 0.1 |
| 7 | 147695550 | rs17170777 | -0.492 | 0.849 | 0.47 | -0.006 | 0.007 | 0.008 | 0.008 | 0 | 0.35 | 0.33 | 0.23 | 0.08 | 0.27 | 0.22 |
| 7 | 147698173 | rs10249855 | -1.53 | 1.022 | -0.187 | 0.061 | 0.016 | -0.01 | -0.01 | 0 | 0.53 | 0.84 | 0.65 | 0.57 | 0.58 | 0.46 |
| 7 | 147702130 | rs6942912 | -1.725 | 0.994 | -0.171 | -0.015 | 0.026 | 0.154 | 0.154 | 0 | 0.56 | 0.79 | 0.74 | 0.81 | 0.63 | 0.48 |
| 7 | 147714911 | rs12539260 | 0.199 | 1.088 | -0.238 | 0.082 | -0.053 | -0.055 | -0.055 | 0 | 0.13 | 0.03 | 0.37 | 0.5 | 0.29 | 0.23 |
| 7 | 147715702 | rs4725759 | -0.789 | 1.063 | -0.025 | 0.061 | -0.011 | -0.007 | -0.007 | 0 | 0.4 | 0.54 | 0.34 | 0.33 | 0.33 | 0.24 |
| 7 | 147720679 | rs10952724 | 0.487 | 1.284 | -0.174 | 0.137 | -0.077 | -0.077 | -0.077 | 2.357 | 0.11 | 0.02 | 0.36 | 0.5 | 0.26 | 0.22 |
| 7 | 147722994 | rs993715 | -2.019 | 0.979 | -0.2 | -0.036 | 0.039 | 0.127 | 0.127 | 0.192 | 0.61 | 0.86 | 0.75 | 0.86 | 0.68 | 0.66 |
| 7 | 147724273 | rs12703990 | -0.32 | 1.163 | -1.596 | 0.164 | -0.036 | -0.024 | -0.024 | 0 | 0.16 | 0.07 | 0.45 | 0.76 | 0.33 | 0.28 |
| 7 | 147725036 | rs7800075 | -0.211 | 1.134 | -1.694 | 0.285 | -0.001 | 0.039 | 0.039 | 0 | 0.07 | 0.05 | 0.1 | 0.29 | 0.09 | 0.18 |
| 7 | 147725495 | rs2710157 | -2.017 | 1.09 | -1.759 | 0.015 | -0.002 | 0.107 | 0.107 | 0.151 | 0.65 | 0.97 | 0.78 | 0.86 | 0.7 | 0.68 |
| 7 | 147749825 | rs16882145 | -0.442 | 1.647 | -0.625 | 0.988 | -0.052 | -0.053 | -0.053 | 0.054 | 0.13 | 0.03 | 0.42 | 0.49 | 0.29 | 0.23 |
| 7 | 147753561 | rs7797894 | -0.693 | 0.352 | -0.176 | 0.722 | 0.014 | 0.004 | 0.004 | 0 | 0.48 | 0.63 | 0.7 | 0.6 | 0.58 | 0.47 |
| 7 | 147754273 | rs10271435 | -1.55 | -0.068 | 0.424 | 0.758 | -0.026 | -0.016 | -0.016 | 0 | 0.13 | 0.23 | 0.07 | 0.07 | 0.07 | 0.18 |
| 7 | 147754816 | rs10485845 | -1.726 | 0.589 | 0.69 | 1.352 | 0.038 | 0.09 | 0.09 | 0 | 0.14 | 0.23 | 0.34 | 0.65 | 0.2 | 0.26 |
| 7 | 147755266 | rs6971184 | -1.626 | -0.008 | 1.007 | 1.279 | -0.025 | 0.007 | 0.007 | 0 | 0.23 | 0.3 | 0.34 | 0.24 | 0.16 | 0.41 |
| 7 | 147795145 | rs13246816 | -2.617 | 0.408 | 1.703 | 1.87 | 0.037 | 0.231 | 0.231 | 0 | 0.14 | 0.34 | 0.2 | 0.47 | 0.15 | 0.22 |
| 7 | 147798729 | rs10235838 | -1.899 | 0.124 | 1.756 | 1.753 | 0.02 | 0.001 | 0.001 | 3.226 | 0.07 | 0.33 | 0.04 | 0 | 0.02 | 0.02 |
| 7 | 147803861 | rs7798592 | -2.202 | 0.119 | 0.877 | 1.83 | -0.052 | 0.119 | 0.119 | 4.081 | 0.13 | 0.6 | 0.13 | 0.27 | 0.04 | 0.08 |
| 7 | 147810841 | rs4725763 | -2.254 | 0.102 | 0.669 | 1.959 | -0.012 | 0.046 | 0.046 | 0.234 | 0.21 | 0.54 | 0.28 | 0.67 | 0.19 | 0.23 |
| 7 | 147812192 | rs17170840 | -1.024 | 0.618 | -0.207 | 1.896 | -0.044 | -0.079 | -0.079 | 0.601 | 0.11 | 0.02 | 0.16 | 0.4 | 0.14 | 0.15 |
| 7 | 147812591 | rs10256700 | -1.469 | 0.787 | 0.176 | 0.933 | 0.023 | 0.006 | 0.006 | 0 | 0.07 | 0.12 | 0.19 | 0.07 | 0.11 | 0.08 |
| 7 | 147821033 | rs12533593 | -1.487 | 0.29 | 0.369 | 0.584 | -0.011 | -0.043 | -0.043 | 0 | 0.08 | 0.2 | 0.03 | 0 | 0.05 | 0.01 |
| 7 | 147829431 | rs6967121 | -2.137 | -0.502 | 0.993 | 1.155 | -0.056 | 0.028 | 0.028 | 0 | 0.2 | 0.41 | 0.09 | 0.24 | 0.1 | 0.14 |
| 7 | 147839308 | rs2906308 | -0.093 | 0.013 | 0.673 | 1.115 | 0.019 | 0.025 | 0.025 | 0 | 0.25 | 0.33 | 0.34 | 0.35 | 0.26 | 0.3 |
| 7 | 147852552 | rs2906302 | -1.528 | -0.133 | 0.68 | 1.118 | 0.004 | 0.011 | 0.011 | 0 | 0.44 | 0.63 | 0.42 | 0.6 | 0.38 | 0.44 |
| 7 | 147854767 | rs2906300 | -0.892 | -0.208 | 0.736 | 1.179 | 0.009 | 0.002 | 0.002 | 0 | 0.33 | 0.44 | 0.37 | 0.34 | 0.31 | 0.34 |
| 7 | 147855874 | rs10264662 | -0.703 | -0.118 | 0.306 | 1.206 | 0.001 | 0.002 | 0.002 | 0 | 0.12 | 0.11 | 0.1 | 0.26 | 0.14 | 0.14 |
| 7 | 147862900 | rs2906294 | -1.696 | -0.194 | 0.498 | 0.963 | 0.005 | -0.005 | -0.005 | 0 | 0.34 | 0.67 | 0.27 | 0.57 | 0.23 | 0.4 |

| | | | | | | | | | | | | | | | | |
|---|---|---|---|---|---|---|---|---|---|---|---|---|---|---|---|---|
| 7 | 147865869 | rs2972110 | -1.062 | -0.476 | -0.124 | 0.435 | 0.025 | 0.034 | 0.034 | 0 | 0.43 | 0.67 | 0.38 | 0.32 | 0.3 | 0.36 |
| 7 | 147868281 | rs6973203 | -0.603 | -0.515 | 0.104 | 0.285 | -0.008 | -0.037 | -0.037 | 0 | 0.38 | 0.41 | 0.27 | 0.19 | 0.34 | 0.26 |
| 7 | 147870422 | rs2972112 | -1.356 | -0.163 | -0.368 | 0.378 | -0.048 | 0.065 | 0.065 | 0 | 0.61 | 0.78 | 0.59 | 0.75 | 0.47 | 0.67 |
| 7 | 147870469 | rs10255956 | -0.775 | 0.023 | -0.294 | 0.904 | -0.001 | 0.025 | 0.025 | 0 | 0.18 | 0.23 | 0.35 | 0.51 | 0.17 | 0.44 |
| 7 | 147872505 | rs1013738 | -0.711 | 0.611 | -0.769 | 0.528 | -0.071 | -0.017 | -0.017 | 0.125 | 0.74 | 0.9 | 0.67 | 0.72 | 0.59 | 0.78 |
| 7 | 147878619 | rs2906283 | -0.375 | -0.096 | -0.325 | 0.585 | -0.005 | -0.007 | -0.007 | 0.023 | 0.27 | 0.37 | 0.24 | 0.25 | 0.18 | 0.25 |
| 7 | 147879557 | rs2972106 | -1.055 | 0.209 | -0.371 | 0.814 | 0 | 0 | 0 | 0 | 0.71 | 0.62 | 0.67 | 0.68 | 0.68 | 0.7 |
| 7 | 147881158 | rs2972102 | -1.348 | 0.153 | -0.134 | 0.852 | -0.008 | 0.024 | 0.024 | 0 | 0.73 | 0.84 | 0.7 | 0.84 | 0.68 | 0.76 |
| 7 | 147883608 | rs17170876 | 0.76 | 0.035 | -0.329 | 0.932 | -0.041 | -0.052 | -0.052 | 5.054 | 0.4 | 0.19 | 0.38 | 0.42 | 0.48 | 0.44 |
| 7 | 147884662 | rs2972100 | -0.363 | -0.541 | -0.146 | 0.912 | -0.003 | -0.001 | -0.001 | 0 | 0.27 | 0.31 | 0.26 | 0.26 | 0.18 | 0.27 |
| 7 | 147885534 | rs13230925 | 0.816 | -0.543 | -0.665 | 1.156 | -0.031 | -0.006 | -0.006 | 0.016 | 0.19 | 0.01 | 0.08 | 0.13 | 0.2 | 0.22 |
| 7 | 147898538 | rs6956460 | 0.803 | -0.353 | -0.174 | 1.322 | -0.005 | 0.013 | 0.013 | 0 | 0.1 | 0.13 | 0.14 | 0.23 | 0.07 | 0.18 |
| 7 | 147899223 | rs4726925 | 0.098 | -0.157 | -0.058 | 1.375 | 0.005 | 0.002 | 0.002 | 0 | 0.8 | 0.73 | 0.84 | 0.81 | 0.79 | 0.79 |
| 7 | 147900046 | rs10245185 | 0.784 | -0.103 | 0.305 | 1.365 | -0.043 | 0.064 | 0.064 | 0 | 0.43 | 0.17 | 0.53 | 0.29 | 0.52 | 0.33 |
| 7 | 147900962 | rs4382390 | -0.537 | -0.089 | 0.275 | 1.413 | 0.034 | 0.033 | 0.033 | 0 | 0.49 | 0.58 | 0.4 | 0.67 | 0.4 | 0.63 |
| 7 | 147900985 | rs4486103 | -0.198 | -0.032 | 0.197 | 1.318 | 0.009 | -0.004 | -0.004 | 0 | 0.46 | 0.38 | 0.34 | 0.51 | 0.39 | 0.6 |
| 7 | 147902079 | rs4421277 | 0.109 | -0.032 | 0.172 | 1.324 | 0.026 | 0.058 | 0.058 | 0 | 0.82 | 0.55 | 0.77 | 0.76 | 0.84 | 0.78 |
| 7 | 147902886 | rs6944384 | 0.221 | -0.021 | -0.236 | 0.862 | 0.004 | 0.003 | 0.003 | 0 | 0.36 | 0.3 | 0.21 | 0.31 | 0.32 | 0.42 |
| 7 | 147905440 | rs10254641 | 0.433 | 0.539 | -0.595 | 0.427 | -0.04 | 0.108 | 0.108 | 0 | 0.45 | 0.19 | 0.53 | 0.24 | 0.51 | 0.35 |
| 7 | 147905623 | rs10254881 | 0.778 | 0.71 | 0.59 | 0.643 | 0.064 | -0.011 | -0.011 | 0 | 0.89 | 0.63 | 0.9 | 0.97 | 0.92 | 0.95 |
| 7 | 147908658 | rs4520084 | -0.598 | 0.629 | -0.105 | 0.8 | 0.009 | 0.009 | 0.009 | 0 | 0.47 | 0.43 | 0.53 | 0.61 | 0.54 | 0.53 |
| 7 | 147908694 | rs4513897 | 0.067 | 1.124 | -0.538 | 0.733 | -0.001 | 0.061 | 0.061 | 0 | 0.18 | 0.04 | 0.22 | 0.09 | 0.16 | 0.16 |
| 7 | 147910432 | rs4345484 | 0.688 | 0.958 | 0.55 | 1.398 | 0.026 | 0.114 | 0.114 | 0.145 | 0.93 | 0.72 | 0.95 | 0.91 | 0.96 | 0.92 |
| 7 | 147910970 | rs3923876 | -1.494 | 0.779 | 0.54 | 1.346 | 0.013 | 0.017 | 0.017 | 0 | 0.38 | 0.53 | 0.36 | 0.54 | 0.41 | 0.43 |
| 7 | 147914885 | rs12333931 | 0.645 | 0.577 | -0.286 | 2.086 | -0.009 | 0.033 | 0.033 | 0 | 0.78 | 0.65 | 0.83 | 0.62 | 0.82 | 0.82 |
| 7 | 147918464 | rs6947963 | -0.391 | 0.736 | -0.564 | 1.808 | -0.034 | -0.043 | -0.043 | 0 | 0.1 | 0.06 | 0.17 | 0.23 | 0.15 | 0.2 |
| 7 | 147920324 | rs2373347 | -1.02 | 0.487 | -0.242 | 1.213 | -0.007 | 0.002 | 0.002 | 0 | 0.23 | 0.31 | 0.15 | 0.31 | 0.19 | 0.19 |
| 7 | 147925492 | rs10263021 | 0.87 | 0.81 | -0.685 | 1.272 | -0.001 | 0 | 0 | 0.01 | 0.7 | 0.7 | 0.63 | 0.5 | 0.71 | 0.62 |
| 7 | 147931329 | rs6952506 | -0.227 | 0.371 | -1.541 | 0.787 | 0.027 | -0.003 | -0.003 | 0 | 0.4 | 0.17 | 0.53 | 0.62 | 0.39 | 0.61 |
| 7 | 147933546 | rs4725770 | -0.331 | -0.184 | -1.731 | 0.299 | 0.012 | 0.075 | 0.075 | 0.481 | 0.33 | 0.2 | 0.31 | 0.11 | 0.34 | 0.26 |
| 7 | 147933618 | rs1881726 | -0.771 | 0.835 | -1.316 | -0.155 | 0.118 | 0.052 | 0.052 | 0.049 | 0.84 | 0.51 | 0.82 | 0.83 | 0.78 | 0.94 |
| 7 | 147936803 | rs1534702 | -0.355 | 0.717 | -0.957 | -0.401 | 0.082 | -0.081 | -0.081 | 0 | 0.81 | 0.52 | 0.82 | 0.95 | 0.78 | 0.94 |
| 7 | 147937001 | rs4726931 | -0.657 | 0.708 | -0.175 | -0.434 | 0.038 | -0.037 | -0.037 | 0.001 | 0.84 | 0.72 | 0.85 | 0.95 | 0.8 | 0.94 |
| 7 | 147937551 | rs12673040 | 1.04 | 0.296 | -0.269 | 0.081 | 0.01 | -0.021 | -0.021 | 0.248 | 0.12 | 0.05 | 0.23 | 0.55 | 0.11 | 0.35 |
| 7 | 147938567 | rs11772596 | 0.909 | 0.23 | -0.298 | 0.061 | 0.011 | -0.027 | -0.027 | 1.766 | 0.13 | 0.05 | 0.23 | 0.55 | 0.11 | 0.35 |
| 7 | 147938808 | rs10952737 | 1.021 | 0.23 | -0.278 | -0.571 | 0.002 | 0.069 | 0.069 | 0.019 | 0.37 | 0.23 | 0.27 | 0.2 | 0.34 | 0.34 |
| 7 | 147942578 | rs6945371 | 1.263 | -0.373 | -0.381 | -1.135 | -0.001 | -0.005 | -0.005 | 0.681 | 0.2 | 0.19 | 0.13 | 0.09 | 0.17 | 0.15 |
| 7 | 147951195 | rs17170924 | 1.047 | -0.742 | -0.552 | -1.294 | -0.007 | -0.013 | -0.013 | 0 | 0.2 | 0.18 | 0.12 | 0.07 | 0.12 | 0.13 |
| 7 | 147953655 | rs1406596 | -0.567 | -0.253 | -0.367 | -1.506 | 0.08 | 0.013 | 0.013 | 0 | 0.79 | 0.61 | 0.77 | 0.8 | 0.72 | 0.86 |
| 7 | 147954909 | rs4726937 | 2.305 | -0.381 | -0.834 | -1.801 | -0.043 | 0.012 | 0.012 | 0 | 0.12 | 0.03 | 0.22 | 0.08 | 0.21 | 0.08 |
| 7 | 147954976 | rs4726938 | -0.737 | -0.167 | -0.623 | -1.547 | 0.089 | 0.044 | 0.044 | 0 | 0.79 | 0.59 | 0.77 | 0.72 | 0.73 | 0.86 |
| 7 | 147955394 | rs17433476 | 1.321 | -0.762 | -1.043 | -1.945 | -0.008 | -0.108 | -0.108 | 0.001 | 0.22 | 0.05 | 0.3 | 0.55 | 0.23 | 0.52 |
| 7 | 147956916 | rs11980200 | 0.744 | -0.995 | -0.984 | -2.1 | 0.004 | 0.016 | 0.016 | 0 | 0.39 | 0.34 | 0.38 | 0.22 | 0.37 | 0.22 |
| 7 | 147957456 | rs1881722 | 0.007 | -1 | -0.939 | -2.102 | 0 | -0.002 | -0.002 | 0 | 0.62 | 0.41 | 0.67 | 0.77 | 0.59 | 0.74 |
| 7 | 147957814 | rs12538984 | 0.864 | -1.069 | -0.893 | -2.111 | -0.015 | -0.055 | -0.055 | 0 | 0.17 | 0.22 | 0.1 | 0.02 | 0.13 | 0.11 |
| 7 | 147958134 | rs10952739 | 1.003 | -1.031 | -0.903 | -2.057 | 0.002 | 0.014 | 0.014 | 0 | 0.36 | 0.32 | 0.36 | 0.15 | 0.36 | 0.23 |
| 7 | 147958844 | rs12670839 | 0.868 | -0.59 | -0.54 | -2.149 | 0 | -0.004 | -0.004 | 0.392 | 0.26 | 0.19 | 0.31 | 0.53 | 0.25 | 0.55 |
| 7 | 147958867 | rs4501499 | 0.248 | -0.693 | -0.335 | -2.025 | 0.018 | -0.017 | -0.017 | 0 | 0.68 | 0.54 | 0.7 | 0.78 | 0.64 | 0.81 |
| 7 | 147959204 | rs11982930 | 0.934 | -0.496 | -0.212 | -1.875 | -0.03 | -0.055 | -0.055 | 0 | 0.28 | 0.36 | 0.15 | 0.16 | 0.18 | 0.2 |
| 7 | 147959930 | rs2141512 | -0.603 | 0.16 | 0.256 | -1.481 | -0.002 | -0.002 | -0.002 | 0 | 0.83 | 0.79 | 0.81 | 0.8 | 0.79 | 0.89 |
| 7 | 147960310 | rs2888553 | -0.307 | 0.281 | 0.685 | -1.576 | 0.021 | -0.041 | -0.041 | 0 | 0.08 | 0.29 | 0.06 | 0.02 | 0.06 | 0.04 |
| 7 | 147960469 | rs17170929 | 0.01 | 0.295 | 0.492 | -1.599 | 0.001 | -0.041 | -0.041 | 1.413 | 0.08 | 0.21 | 0.04 | 0.02 | 0.05 | 0.06 |
| 7 | 147961673 | rs17170932 | 1.406 | 0.021 | -0.065 | -1.531 | 0.044 | 0.047 | 0.047 | 0 | 0.31 | 0.16 | 0.15 | 0.16 | 0.2 | 0.2 |
| 7 | 147964016 | rs3801975 | -0.824 | 0.201 | -0.048 | -1.519 | -0.01 | -0.007 | -0.007 | 0.905 | 0.82 | 0.78 | 0.81 | 0.77 | 0.77 | 0.88 |
| 7 | 147964377 | rs13240892 | 1.052 | 0.178 | -0.015 | -1.631 | -0.007 | 0.004 | 0.004 | 0 | 0.66 | 0.29 | 0.69 | 0.74 | 0.62 | 0.78 |
| 7 | 147966812 | rs10952742 | 1.189 | 0.115 | -0.075 | -1.65 | -0.048 | -0.093 | -0.093 | 0.154 | 0.33 | 0.06 | 0.51 | 0.58 | 0.39 | 0.59 |
| 7 | 147967593 | rs10464461 | 0.997 | 0.111 | -0.008 | -1.639 | 0.005 | -0.004 | -0.004 | 0 | 0.66 | 0.33 | 0.69 | 0.74 | 0.61 | 0.78 |
| 7 | 147968301 | rs12539824 | 0.885 | 0.139 | -0.081 | -1.528 | -0.043 | -0.071 | -0.071 | 0 | 0.34 | 0.09 | 0.53 | 0.58 | 0.41 | 0.59 |
| 7 | 147973957 | rs17170936 | -0.643 | 0.377 | 0.229 | -1.377 | 0.019 | -0.122 | -0.122 | 0 | 0.16 | 0.44 | 0.12 | 0.03 | 0.16 | 0.1 |
| 7 | 147976770 | rs7809486 | 1.19 | 0.361 | -0.063 | -1.339 | 0 | -0.021 | -0.021 | 0.582 | 0.33 | 0.2 | 0.53 | 0.58 | 0.39 | 0.55 |
| 7 | 147977039 | rs6964680 | -0.117 | 0.415 | 0.337 | -1.413 | 0.027 | -0.078 | -0.078 | 0 | 0.12 | 0.35 | 0.09 | 0 | 0.13 | 0.06 |
| 7 | 147980926 | rs10952748 | 0.043 | 0.328 | 0.181 | -1.374 | 0.054 | 0.008 | 0.008 | 0 | 0.75 | 0.57 | 0.76 | 0.74 | 0.73 | 0.8 |
| 7 | 147988263 | rs7779924 | 1.034 | 0.162 | -0.448 | -1.331 | -0.004 | -0.003 | -0.003 | 0 | 0.33 | 0.28 | 0.56 | 0.63 | 0.41 | 0.55 |
| 7 | 147989522 | rs17434745 | 1.178 | 0.168 | -0.935 | -1.028 | 0.064 | 0.095 | 0.095 | 0 | 0.29 | 0.1 | 0.17 | 0.13 | 0.18 | 0.21 |
| 7 | 147990737 | rs10225169 | 0 | 0.938 | -0.24 | -0.445 | -0.007 | -0.035 | -0.035 | 0.224 | 0.07 | 0.14 | 0.04 | 0.03 | 0.04 | 0.08 |
| 7 | 147991656 | rs9640521 | 1.04 | 1.036 | -0.267 | -0.452 | 0.045 | -0.015 | -0.015 | 0 | 0.71 | 0.47 | 0.77 | 0.76 | 0.67 | 0.81 |
| 7 | 147992571 | rs6968094 | 0.377 | 1.479 | -0.733 | -0.071 | 0.05 | 0.013 | 0.013 | 0 | 0.79 | 0.65 | 0.82 | 0.79 | 0.72 | 0.88 |
| 7 | 147995434 | rs4562230 | 0.627 | 0.992 | -1.491 | -0.244 | -0.037 | 0.057 | 0.057 | 0 | 0.27 | 0.1 | 0.25 | 0.16 | 0.29 | 0.18 |
| 7 | 147995460 | rs13244714 | 0.159 | 1.126 | -1.282 | 0.143 | -0.052 | -0.027 | -0.027 | 0 | 0.14 | 0.03 | 0.08 | 0.13 | 0.15 | 0.07 |

| | | | | | | | | | | | | | | | | |
|---|---|---|---|---|---|---|---|---|---|---|---|---|---|---|---|---|
| 7 | 148000692 | rs1110038 | 0.604 | 0.494 | -0.71 | 0.25 | 0.001 | 0.005 | 0.005 | 0 | 0.66 | 0.29 | 0.73 | 0.76 | 0.68 | 0.53 |
| 7 | 148001291 | rs1110144 | -0.629 | 0.291 | -0.434 | 0.633 | -0.002 | -0.013 | -0.013 | 1.071 | 0.32 | 0.6 | 0.26 | 0.24 | 0.32 | 0.47 |
| 7 | 148002581 | rs6956705 | -0.131 | 0.156 | 0.115 | 0.358 | -0.031 | -0.112 | -0.112 | 0 | 0.16 | 0.38 | 0.08 | 0 | 0.11 | 0.08 |
| 7 | 148013853 | rs11771834 | 1.151 | 0.101 | -0.523 | 0.081 | 0.013 | 0.137 | 0.137 | 0 | 0.33 | 0.11 | 0.21 | 0.15 | 0.27 | 0.24 |
| 7 | 148015957 | rs6946212 | 1.072 | 0.016 | -0.503 | 0.244 | -0.03 | -0.05 | -0.05 | 0 | 0.29 | 0.15 | 0.35 | 0.49 | 0.37 | 0.29 |
| 7 | 148020020 | rs10264551 | -0.256 | -0.036 | -0.3 | 0.216 | 0.008 | 0.046 | 0.046 | 0.011 | 0.2 | 0.29 | 0.34 | 0.37 | 0.25 | 0.41 |
| 7 | 148022695 | rs10273912 | 0.452 | 0.107 | -0.394 | -0.256 | -0.01 | 0.048 | 0.048 | 0 | 0.46 | 0.31 | 0.39 | 0.4 | 0.47 | 0.61 |
| 7 | 148023570 | rs12704008 | 0.937 | 0.392 | -0.665 | -0.006 | -0.027 | 0.033 | 0.033 | 0 | 0.29 | 0.15 | 0.34 | 0.15 | 0.36 | 0.27 |
| 7 | 148024202 | rs1918296 | 0.072 | 0.402 | -1.157 | -0.032 | -0.008 | -0.016 | -0.016 | 0 | 0.81 | 0.85 | 0.83 | 0.77 | 0.78 | 0.6 |
| 7 | 148030828 | rs7779793 | -0.776 | 0.502 | -0.981 | -0.109 | 0.009 | 0.014 | 0.014 | 0.06 | 0.62 | 0.65 | 0.74 | 0.87 | 0.68 | 0.75 |
| 7 | 148033848 | rs1177935 | 0.248 | 1.099 | -1.197 | 0.501 | -0.058 | 0.007 | 0.007 | 0 | 0.28 | 0.08 | 0.33 | 0.23 | 0.34 | 0.26 |
| 7 | 148034688 | rs886969 | 0.779 | 0.894 | -0.043 | 0.702 | 0.007 | 0.03 | 0.03 | 0 | 0.8 | 0.61 | 0.8 | 0.77 | 0.8 | 0.6 |
| 7 | 148035192 | rs10277969 | -1.138 | 1.04 | -0.312 | 0.656 | 0.003 | 0.004 | 0.004 | 0 | 0.24 | 0.28 | 0.28 | 0.24 | 0.25 | 0.43 |
| 7 | 148036191 | rs17170956 | -1.233 | 0.844 | -0.18 | 0.898 | -0.004 | 0.051 | 0.051 | 0.33 | 0.08 | 0.15 | 0.14 | 0.27 | 0.1 | 0.29 |
| 7 | 148036293 | rs7792210 | -0.134 | 0.543 | -0.523 | -0.363 | -0.003 | -0.003 | -0.003 | 0.849 | 0.66 | 0.68 | 0.8 | 0.96 | 0.65 | 0.77 |
| 7 | 148036303 | rs7792214 | -0.946 | 0.511 | -0.04 | -0.689 | -0.003 | 0.078 | 0.078 | 1.696 | 0.19 | 0.46 | 0.22 | 0.31 | 0.22 | 0.38 |
| 7 | 148036377 | rs7809658 | -1.172 | 0.284 | -0.137 | -0.617 | -0.027 | 0.036 | 0.036 | 0.165 | 0.22 | 0.46 | 0.16 | 0.27 | 0.22 | 0.39 |
| 7 | 148039047 | rs9648853 | 0.799 | -0.051 | -0.379 | -0.969 | -0.002 | 0.053 | 0.053 | 0 | 0.45 | 0.23 | 0.58 | 0.67 | 0.4 | 0.39 |
| 7 | 148040248 | rs1177924 | 0.342 | -0.172 | -0.251 | -1.077 | -0.005 | 0.022 | 0.022 | 0 | 0.37 | 0.29 | 0.21 | 0.06 | 0.39 | 0.24 |
| 7 | 148044472 | rs10278502 | -0.741 | -0.225 | -0.252 | -1.062 | 0.001 | 0.069 | 0.069 | 0.401 | 0.17 | 0.31 | 0.13 | 0.3 | 0.2 | 0.38 |
| 7 | 148044517 | rs4487670 | 1.159 | 0.229 | -1.07 | -0.798 | -0.043 | -0.056 | -0.056 | 0 | 0.1 | 0.02 | 0.29 | 0.5 | 0.15 | 0.1 |
| 7 | 148046130 | rs2011699 | 1.128 | -0.026 | -0.957 | -1.119 | 0.009 | 0.067 | 0.067 | 0 | 0.33 | 0.16 | 0.33 | 0.14 | 0.24 | 0.26 |
| 7 | 148049715 | rs4573177 | -0.579 | 0.127 | -0.579 | -1.04 | -0.008 | -0.002 | -0.002 | 0 | 0.63 | 0.66 | 0.78 | 0.94 | 0.58 | 0.76 |
| 7 | 148052603 | rs1177952 | 0.72 | 0.045 | -0.718 | -1.061 | -0.037 | 0.089 | 0.089 | 0 | 0.33 | 0.17 | 0.18 | 0.06 | 0.39 | 0.23 |
| 7 | 148055591 | rs1177951 | 0.532 | 0.033 | -0.505 | -1.079 | -0.006 | -0.003 | -0.003 | 0 | 0.36 | 0.32 | 0.2 | 0.06 | 0.39 | 0.23 |
| 7 | 148058265 | rs10952755 | 1.488 | -0.091 | -0.859 | -1.354 | 0.025 | 0.02 | 0.02 | 0.018 | 0.23 | 0.05 | 0.24 | 0.15 | 0.14 | 0.28 |
| 7 | 148069482 | rs1637857 | 0.378 | 0.213 | -0.261 | -1.307 | -0.011 | 0.009 | 0.009 | 0 | 0.34 | 0.27 | 0.19 | 0.06 | 0.39 | 0.23 |
| 7 | 148080647 | rs1637841 | 1.22 | 0.083 | 0.008 | -0.822 | -0.003 | 0.077 | 0.077 | 0.691 | 0.81 | 0.48 | 0.83 | 0.71 | 0.8 | 0.61 |
| 7 | 148084632 | rs4726964 | -0.714 | 0.34 | -0.15 | -1.136 | 0.001 | 0.065 | 0.065 | 1.461 | 0.45 | 0.67 | 0.25 | 0.06 | 0.42 | 0.29 |
| 7 | 148087337 | rs4726968 | -1.163 | 0.614 | -0.26 | -1.179 | 0.021 | 0.06 | 0.06 | 0.086 | 0.49 | 0.65 | 0.27 | 0.32 | 0.45 | 0.6 |
| 7 | 148092100 | rs886972 | -0.503 | 0.718 | -0.181 | -1.033 | -0.006 | -0.011 | -0.011 | 0.021 | 0.12 | 0.12 | 0.13 | 0.27 | 0.18 | 0.32 |
| 7 | 148092983 | rs10242598 | -0.081 | 0.641 | -0.096 | -1.133 | -0.016 | -0.103 | -0.103 | 0.114 | 0.34 | 0.5 | 0.15 | 0.05 | 0.27 | 0.28 |
| 7 | 148094425 | rs10273784 | -0.485 | 0.299 | 0.03 | -1.008 | 0.029 | -0.023 | -0.023 | 0 | 0.08 | 0.35 | 0.05 | 0 | 0.05 | 0.07 |
| 7 | 148096608 | rs17170987 | -0.115 | 0.32 | -0.086 | -0.831 | -0.014 | -0.021 | -0.021 | 0 | 0.11 | 0.07 | 0.12 | 0.32 | 0.18 | 0.34 |
| 7 | 148100402 | rs1034865 | -1.449 | 0.304 | -0.018 | -0.81 | -0.001 | 0.061 | 0.061 | 0.93 | 0.21 | 0.52 | 0.17 | 0.32 | 0.23 | 0.4 |
| 7 | 148100862 | rs10225245 | 0.487 | 0.369 | -0.147 | -0.906 | -0.02 | 0.061 | 0.061 | 0 | 0.3 | 0.16 | 0.16 | 0.05 | 0.32 | 0.21 |
| 7 | 148103405 | rs10265613 | 0.491 | 0.129 | -0.213 | -0.748 | -0.036 | 0.091 | 0.091 | 0 | 0.3 | 0.14 | 0.16 | 0.07 | 0.33 | 0.23 |
| 7 | 148104909 | rs4725782 | -0.947 | 0.078 | -0.212 | -0.61 | 0.014 | 0.037 | 0.037 | 0 | 0.51 | 0.68 | 0.33 | 0.39 | 0.56 | 0.63 |
| 7 | 148106490 | rs9648691 | -1.136 | 0.121 | -0.218 | -0.59 | 0.007 | 0.028 | 0.028 | 0 | 0.53 | 0.71 | 0.34 | 0.39 | 0.57 | 0.63 |
| 7 | 148112987 | rs987456 | 1.108 | 0.157 | -0.055 | -0.412 | -0.016 | 0.058 | 0.058 | 0 | 0.78 | 0.49 | 0.83 | 0.69 | 0.79 | 0.6 |
| 7 | 148114265 | rs3194 | 0.814 | -0.067 | -0.348 | -0.135 | -0.084 | 0.038 | 0.038 | 0.101 | 0.45 | 0.06 | 0.64 | 0.65 | 0.44 | 0.36 |
| 7 | 148114865 | rs1062072 | 0.189 | -0.069 | 0 | -0.26 | -0.001 | 0.004 | 0.004 | 0 | 0.53 | 0.52 | 0.7 | 0.63 | 0.46 | 0.43 |
| 7 | 148115852 | rs17170999 | -0.862 | -0.039 | -0.259 | -0.242 | 0.046 | 0.025 | 0.025 | 1.021 | 0.07 | 0.4 | 0.05 | 0 | 0.01 | 0.08 |
| 7 | 148116450 | rs2530311 | 0.232 | -0.04 | -0.272 | -0.245 | -0.001 | 0 | 0 | 0 | 0.52 | 0.46 | 0.69 | 0.63 | 0.45 | 0.43 |
| 7 | 148117547 | rs10280038 | -1.676 | -0.213 | 0.984 | 0.141 | 0.046 | 0.025 | 0.025 | 1.021 | 0.07 | 0.4 | 0.05 | 0 | 0.01 | 0.08 |
| 7 | 148119698 | rs10229693 | -1.336 | -0.058 | 0.478 | 0.183 | 0.042 | -0.005 | -0.005 | 0 | 0.06 | 0.3 | 0.07 | 0 | 0.04 | 0.08 |
| 7 | 148130329 | rs6963244 | -0.829 | -0.121 | -0.236 | 0.711 | -0.001 | 0.038 | 0.038 | 1.778 | 0.3 | 0.49 | 0.37 | 0.51 | 0.3 | 0.28 |
| 7 | 148139557 | rs17616515 | 0.582 | -0.511 | -1.499 | -0.318 | -0.129 | 0.203 | 0.203 | 1.643 | 0.38 | 0.09 | 0.41 | 0.11 | 0.44 | 0.33 |
| 7 | 148143416 | rs7780778 | -1.074 | -1.573 | -0.28 | 0.656 | -0.009 | 0.186 | 0.186 | 0.095 | 0.1 | 0.33 | 0.13 | 0.21 | 0.04 | 0.08 |
| 7 | 148153437 | rs2530293 | 0.062 | -2.228 | -1.03 | 0.777 | 0.057 | -0.037 | -0.037 | 0 | 0.68 | 0.8 | 0.75 | 0.62 | 0.78 | 0.7 |
| 7 | 148154833 | rs735787 | 0.409 | -1.842 | -0.711 | -0.008 | 0.041 | 0.037 | 0.037 | 0 | 0.5 | 0.59 | 0.67 | 0.33 | 0.64 | 0.37 |
| 7 | 148155186 | rs17532635 | 0.115 | -1.959 | -0.78 | 0.035 | 0.074 | -0.04 | -0.04 | 0 | 0.39 | 0.1 | 0.27 | 0.6 | 0.32 | 0.57 |
| 7 | 148158904 | rs17532809 | 0.086 | -1.952 | -0.764 | 0.006 | 0.087 | 0.027 | 0.027 | 0 | 0.4 | 0.14 | 0.28 | 0.65 | 0.32 | 0.57 |
| 7 | 148163784 | rs6970519 | -0.549 | -1.674 | -0.529 | 0.158 | -0.001 | 0.049 | 0.049 | 1.867 | 0.08 | 0.29 | 0.06 | 0.07 | 0.04 | 0.05 |
| 7 | 148176490 | rs11767412 | 0.407 | -1.364 | -0.674 | -0.133 | 0.03 | 0.035 | 0.035 | 0 | 0.51 | 0.56 | 0.67 | 0.26 | 0.64 | 0.37 |
| 7 | 148179678 | rs10952761 | 0.228 | -0.69 | -1.303 | -0.343 | 0.086 | 0.175 | 0.175 | 0 | 0.53 | 0.31 | 0.59 | 0.14 | 0.64 | 0.34 |
| 7 | 148188150 | rs2538483 | -0.564 | -0.056 | -1.431 | -0.209 | 0.022 | 0.071 | 0.071 | 0 | 0.4 | 0.79 | 0.45 | 0.26 | 0.36 | 0.3 |
| 7 | 148188533 | rs6464907 | -0.545 | 0.429 | -1.189 | -0.3 | -0.008 | 0.052 | 0.052 | 0 | 0.4 | 0.71 | 0.46 | 0.26 | 0.41 | 0.3 |
| 7 | 148195384 | rs749609 | -0.07 | 0.493 | -1.607 | -0.32 | -0.13 | 0.011 | 0.011 | 0.057 | 0.27 | 0.02 | 0.38 | 0.21 | 0.37 | 0.19 |
| 7 | 148204948 | rs2717799 | -0.693 | 0.496 | -1.008 | -0.504 | -0.022 | 0.06 | 0.06 | 0 | 0.41 | 0.22 | 0.46 | 0.26 | 0.44 | 0.29 |
| 7 | 148211140 | rs2717803 | -0.281 | 0.845 | -1.417 | -0.362 | -0.075 | 0.034 | 0.034 | 0 | 0.78 | 0.92 | 0.69 | 0.78 | 0.68 | 0.81 |
| 7 | 148211886 | rs2717804 | 0.265 | 0.705 | -0.587 | 0.366 | -0.001 | -0.001 | -0.001 | 0 | 0.32 | 0.31 | 0.43 | 0.29 | 0.37 | 0.21 |
| 7 | 148213724 | rs2717808 | -0.388 | 0.68 | 0.21 | 0.371 | 0.001 | -0.003 | -0.003 | 0 | 0.76 | 0.77 | 0.68 | 0.78 | 0.67 | 0.81 |
| 7 | 148218371 | rs2538474 | -0.507 | 0.634 | 0.187 | 0.372 | -0.073 | 0.068 | 0.068 | 0 | 0.77 | 0.89 | 0.69 | 0.81 | 0.67 | 0.81 |
| 7 | 148218875 | rs2538473 | 0.143 | 0.582 | 0.934 | 0.435 | 0.043 | -0.072 | -0.072 | 0 | 0.72 | 0.55 | 0.63 | 0.79 | 0.65 | 0.74 |
| 7 | 148225528 | rs759149 | 1.194 | 0.53 | 0.787 | 0.553 | 0.075 | -0.069 | -0.069 | 0 | 0.4 | 0.18 | 0.2 | 0.45 | 0.27 | 0.54 |
| 7 | 148226973 | rs11769463 | 0.524 | 0.443 | 0.825 | 0.498 | -0.015 | -0.016 | -0.016 | 2.416 | 0.31 | 0.21 | 0.42 | 0.33 | 0.38 | 0.22 |
| 7 | 148228005 | rs4726984 | 0.686 | 0.451 | 0.931 | 0.564 | -0.06 | 0.005 | 0.005 | 0 | 0.28 | 0.07 | 0.41 | 0.22 | 0.37 | 0.19 |
| 7 | 148233208 | rs2717772 | -0.411 | 0.505 | 0.955 | 0.563 | -0.015 | 0.003 | 0.003 | 0 | 0.74 | 0.76 | 0.68 | 0.69 | 0.67 | 0.78 |

| | | | | | | | | | | | | | | | | |
|---|---|---|---|---|---|---|---|---|---|---|---|---|---|---|---|---|
| 7 | 148239179 | rs6957883 | -0.092 | 0.386 | 0.738 | 0.55 | -0.024 | 0.003 | 0.003 | 0 | 0.71 | 0.76 | 0.65 | 0.64 | 0.63 | 0.72 |
| 7 | 148245573 | rs10280054 | -0.112 | 0 | 0.143 | 0.65 | -0.037 | -0.025 | -0.025 | 0 | 0.64 | 0.77 | 0.62 | 0.34 | 0.56 | 0.49 |
| 7 | 148245802 | rs10280425 | 1.206 | 0.091 | -0.191 | 0.61 | 0.048 | 0.053 | 0.053 | 2.223 | 0.32 | 0.14 | 0.16 | 0.15 | 0.21 | 0.29 |
| 7 | 148246393 | rs10253719 | 1.17 | 0.102 | -0.189 | 0.614 | 0.049 | 0.053 | 0.053 | 1.913 | 0.32 | 0.16 | 0.16 | 0.15 | 0.21 | 0.29 |
| 7 | 148251285 | rs12704034 | 0.612 | 0.476 | -0.236 | 0.508 | -0.064 | 0.011 | 0.011 | 0 | 0.27 | 0.07 | 0.4 | 0.2 | 0.36 | 0.2 |
| 7 | 148253738 | rs6971115 | 0.019 | 0.083 | -0.07 | 0.564 | -0.02 | 0.01 | 0.01 | 0 | 0.57 | 0.69 | 0.57 | 0.34 | 0.55 | 0.48 |
| 7 | 148254031 | rs12704035 | -0.306 | 0.608 | 0.453 | 0.688 | 0.038 | -0.004 | -0.004 | 0 | 0.26 | 0.44 | 0.41 | 0.21 | 0.33 | 0.14 |
| 7 | 148258161 | rs12704036 | -0.082 | 0.799 | -0.549 | 0.807 | -0.014 | -0.003 | -0.003 | 0 | 0.23 | 0.16 | 0.37 | 0.21 | 0.31 | 0.13 |
| 7 | 148265389 | rs12704038 | -0.473 | 0.963 | -0.582 | 1.325 | -0.042 | -0.035 | -0.035 | 0.036 | 0.08 | 0.01 | 0.1 | 0.33 | 0.11 | 0.11 |
| 7 | 148265863 | rs2014450 | 0.521 | -0.059 | -0.844 | 0.087 | 0.012 | 0.05 | 0.05 | 0 | 0.27 | 0.12 | 0.16 | 0.12 | 0.22 | 0.12 |
| 7 | 148266044 | rs7777242 | 0.403 | 0.027 | -0.722 | 0.06 | -0.006 | 0.044 | 0.044 | 0 | 0.3 | 0.17 | 0.19 | 0.12 | 0.3 | 0.13 |
| 7 | 148268652 | rs10242631 | 0.102 | -0.331 | -0.187 | -0.095 | -0.011 | 0.04 | 0.04 | 0 | 0.2 | 0.32 | 0.24 | 0.3 | 0.17 | 0.5 |
| 7 | 148269912 | rs10233671 | -0.032 | -0.637 | -0.033 | -0.293 | -0.008 | -0.005 | -0.005 | 0 | 0.13 | 0.18 | 0.18 | 0.08 | 0.08 | 0.13 |
| 7 | 148270503 | rs6978741 | -0.13 | -0.672 | 0.104 | -0.346 | -0.004 | -0.029 | -0.029 | 0 | 0.44 | 0.49 | 0.39 | 0.2 | 0.38 | 0.26 |
| 7 | 148272945 | rs17171051 | -0.892 | -0.826 | 0.041 | -0.083 | -0.038 | 0.003 | 0.003 | 0 | 0.3 | 0.44 | 0.28 | 0.28 | 0.17 | 0.46 |
| 7 | 148273685 | rs10253271 | -0.302 | -0.711 | -0.169 | -0.091 | -0.017 | -0.007 | -0.007 | 0 | 0.23 | 0.25 | 0.12 | 0.2 | 0.13 | 0.36 |
| 7 | 148274786 | rs6948227 | -1.159 | -0.33 | -0.754 | 0.002 | -0.034 | -0.039 | -0.039 | 0 | 0.7 | 0.8 | 0.52 | 0.43 | 0.56 | 0.72 |
| 7 | 148277923 | rs1859912 | -1.504 | -1.398 | -0.306 | -0.028 | -0.013 | -0.009 | -0.009 | 0.132 | 0.38 | 0.41 | 0.17 | 0.23 | 0.21 | 0.52 |
| 7 | 148280705 | rs10260314 | -1.119 | -0.593 | -0.285 | 0.602 | -0.015 | 0.002 | 0.002 | 0 | 0.22 | 0.3 | 0.12 | 0.23 | 0.14 | 0.44 |
| 7 | 148287005 | rs17171055 | -0.794 | -0.425 | -0.73 | 0.929 | 0.016 | -0.025 | -0.025 | 0 | 0.12 | 0.05 | 0.07 | 0.2 | 0.08 | 0.28 |
| 7 | 148296304 | rs6947121 | -1.265 | -0.739 | -0.765 | 0.875 | 0.018 | -0.02 | -0.02 | 0.163 | 0.35 | 0.75 | 0.22 | 0.38 | 0.25 | 0.45 |
| 7 | 148305986 | rs4726986 | -0.613 | -0.66 | -0.152 | 0.677 | 0.016 | 0.025 | 0.025 | 0.924 | 0.5 | 0.65 | 0.6 | 0.67 | 0.56 | 0.7 |
| 7 | 148306949 | rs11971205 | -0.119 | -0.859 | 0.009 | 0.612 | -0.059 | -0.08 | -0.08 | 0 | 0.37 | 0.13 | 0.41 | 0.55 | 0.51 | 0.56 |
| 7 | 148308179 | rs4263667 | -0.636 | -0.868 | 0.275 | 0.659 | 0 | 0.003 | 0.003 | 0.335 | 0.44 | 0.53 | 0.45 | 0.56 | 0.53 | 0.57 |
| 7 | 148310654 | rs11979232 | -1.028 | -0.738 | 0.546 | 0.694 | 0.002 | -0.002 | -0.002 | 0.09 | 0.45 | 0.66 | 0.6 | 0.58 | 0.52 | 0.67 |
| 7 | 148312520 | rs6962292 | -1.015 | -0.759 | 0.552 | 0.699 | 0.001 | -0.002 | -0.002 | 0 | 0.45 | 0.66 | 0.6 | 0.58 | 0.52 | 0.67 |
| 7 | 148321180 | rs1024577 | -0.157 | -0.506 | 0.495 | 0.732 | -0.036 | -0.032 | -0.032 | 0.172 | 0.31 | 0.26 | 0.4 | 0.51 | 0.43 | 0.53 |
| 7 | 148325326 | rs10267756 | -0.462 | -0.544 | 0.64 | 0.456 | 0.006 | 0.007 | 0.007 | 0.925 | 0.37 | 0.42 | 0.58 | 0.56 | 0.47 | 0.62 |
| 7 | 148326035 | rs4726988 | -0.292 | -0.602 | 0.837 | 0.469 | -0.008 | -0.005 | -0.005 | 1.114 | 0.37 | 0.32 | 0.49 | 0.51 | 0.5 | 0.51 |
| 7 | 148327260 | rs713590 | -1.012 | -0.945 | 0.795 | 0.583 | -0.004 | 0.058 | 0.058 | 0.565 | 0.07 | 0.38 | 0.14 | 0.05 | 0.01 | 0.11 |
| 7 | 148330977 | rs11983635 | -1.22 | -0.772 | 0.539 | 0.849 | -0.019 | 0.113 | 0.113 | 0 | 0.09 | 0.17 | 0.21 | 0.35 | 0.04 | 0.28 |
| 7 | 148331040 | rs7793987 | -0.322 | -0.765 | 0.039 | 0.617 | 0.019 | 0.002 | 0.002 | 0 | 0.5 | 0.66 | 0.59 | 0.55 | 0.56 | 0.62 |
| 7 | 148340864 | rs11773375 | 0.305 | -0.377 | -0.439 | 0.618 | 0.052 | 0.199 | 0.199 | 0.003 | 0.53 | 0.16 | 0.48 | 0.19 | 0.6 | 0.42 |
| 7 | 148347171 | rs7794085 | -0.623 | -0.701 | -0.504 | 0.99 | -0.014 | 0.221 | 0.221 | 0.348 | 0.06 | 0.34 | 0.11 | 0.28 | 0.01 | 0.13 |
| 7 | 148355456 | rs2041074 | -0.059 | -0.045 | -1.271 | 0.008 | 0.032 | 0.115 | 0.115 | 0.123 | 0.54 | 0.29 | 0.43 | 0.24 | 0.54 | 0.39 |
| 7 | 148357580 | rs10952773 | 0.122 | 0.195 | -1.859 | 0.107 | 0.03 | 0.183 | 0.183 | 0.01 | 0.5 | 0.11 | 0.42 | 0.24 | 0.54 | 0.38 |
| 7 | 148361277 | rs7812293 | -0.011 | 0.601 | -1.115 | 0.391 | 0.002 | -0.001 | -0.001 | 0 | 0.62 | 0.58 | 0.54 | 0.61 | 0.48 | 0.65 |
| 7 | 148362546 | rs9784937 | 0.116 | -0.357 | -0.846 | -0.138 | 0.01 | 0.106 | 0.106 | 0 | 0.52 | 0.73 | 0.54 | 0.9 | 0.44 | 0.66 |
| 7 | 148368016 | rs6961761 | -0.071 | -0.708 | -0.092 | -0.377 | 0.001 | 0.028 | 0.028 | 0.064 | 0.54 | 0.66 | 0.56 | 0.97 | 0.45 | 0.68 |
| 7 | 148368954 | rs6966980 | 0.061 | -1.14 | -0.443 | 0.214 | 0.001 | 0 | 0 | 0 | 0.36 | 0.61 | 0.41 | 0.63 | 0.26 | 0.3 |
| 7 | 148385725 | rs10257756 | -0.284 | -1.134 | 0.045 | 0.232 | 0.005 | 0.024 | 0.024 | 0.036 | 0.51 | 0.52 | 0.46 | 0.65 | 0.43 | 0.62 |
| 7 | 148394595 | rs122571 | -0.604 | -0.995 | -0.264 | -0.13 | 0.01 | 0.103 | 0.103 | 0 | 0.49 | 0.77 | 0.52 | 0.64 | 0.45 | 0.69 |
| 7 | 148400466 | rs10235843 | -0.154 | -1.075 | -0.082 | -0.099 | 0.001 | -0.001 | -0.001 | 0 | 0.05 | 0.02 | 0.06 | 0.04 | 0.03 | 0.02 |
| 7 | 148404379 | rs243538 | -0.003 | -1.269 | -0.291 | -0.039 | -0.045 | -0.046 | -0.046 | 0 | 0.24 | 0.1 | 0.21 | 0.3 | 0.33 | 0.43 |
| 7 | 148409941 | rs243536 | -0.984 | -1.563 | -0.037 | -0.437 | 0 | 0.017 | 0.017 | 0 | 0.39 | 0.36 | 0.32 | 0.3 | 0.42 | 0.52 |
| 7 | 148419066 | rs10488072 | -0.103 | -1.657 | -0.343 | -0.213 | 0.001 | -0.001 | -0.001 | 0 | 0.09 | 0.12 | 0.08 | 0.08 | 0.09 | 0.18 |
| 7 | 148419484 | rs243523 | -0.063 | -1.659 | -0.444 | -0.186 | -0.086 | -0.017 | -0.017 | 0.885 | 0.27 | 0.02 | 0.21 | 0.23 | 0.34 | 0.34 |
| 7 | 148422581 | rs11760399 | 0.795 | -1.622 | -0.404 | -0.164 | 0.009 | 0.07 | 0.07 | 0 | 0.12 | 0 | 0.07 | 0 | 0.08 | 0.09 |
| 7 | 148423624 | rs243519 | -1.176 | -1.856 | 0.016 | -0.261 | 0.009 | 0.02 | 0.02 | 0 | 0.42 | 0.61 | 0.35 | 0.31 | 0.42 | 0.51 |
| 7 | 148444355 | rs11764570 | 1.004 | -1.695 | 0.376 | -0.353 | -0.015 | -0.045 | -0.045 | 0.367 | 0.09 | 0.14 | 0.05 | 0 | 0.06 | 0.05 |
| 7 | 148445808 | rs243492 | -0.362 | -1.553 | 0.027 | -0.718 | 0.012 | -0.028 | -0.028 | 0 | 0.6 | 0.5 | 0.6 | 0.82 | 0.52 | 0.63 |
| 7 | 148448211 | rs243487 | 0.249 | -1.183 | 0.016 | -0.497 | -0.082 | -0.083 | -0.083 | 3.777 | 0.23 | 0.05 | 0.19 | 0.31 | 0.32 | 0.37 |
| 7 | 148450521 | rs243482 | -0.745 | -0.976 | 0.287 | -0.291 | -0.001 | 0.001 | 0.001 | 0 | 0.54 | 0.57 | 0.52 | 0.55 | 0.52 | 0.61 |
| 7 | 148452905 | rs243480 | 0.584 | -0.936 | 0.044 | -0.273 | -0.09 | -0.088 | -0.088 | 0 | 0.22 | 0.04 | 0.19 | 0.31 | 0.32 | 0.37 |
| 7 | 148456154 | rs243477 | -1.998 | -0.839 | 0.038 | -0.32 | -0.166 | -0.137 | -0.137 | 0 | 0.68 | 0.98 | 0.65 | 0.57 | 0.54 | 0.67 |
| 7 | 148458644 | rs11773084 | 0.528 | -0.854 | 0.235 | -0.36 | -0.003 | 0.006 | 0.006 | 0 | 0.47 | 0.36 | 0.51 | 0.45 | 0.53 | 0.4 |
| 7 | 148460536 | rs10246773 | 0.994 | -0.752 | 0.091 | -0.325 | 0.001 | -0.001 | -0.001 | 0 | 0.09 | 0.05 | 0.13 | 0.03 | 0.04 | 0.04 |
| 7 | 148463860 | rs1014095 | -1.021 | -0.765 | 0.067 | -0.32 | -0.073 | -0.093 | -0.093 | 0 | 0.78 | 0.96 | 0.85 | 0.7 | 0.73 | 0.65 |
| 7 | 148467447 | rs3807446 | 0.979 | -0.806 | 0.123 | -0.316 | 0.001 | 0 | 0 | 0.094 | 0.09 | 0.05 | 0.14 | 0.03 | 0.05 | 0.04 |
| 7 | 148480114 | rs1558420 | -0.114 | -0.812 | 0.3 | -0.325 | -0.017 | -0.012 | -0.012 | 0 | 0.69 | 0.84 | 0.81 | 0.7 | 0.66 | 0.6 |
| 7 | 148480990 | rs10271133 | -0.375 | -1.067 | 0.288 | -0.472 | 0.058 | 0.041 | 0.041 | 0.464 | 0.36 | 0.82 | 0.46 | 0.28 | 0.21 | 0.27 |
| 7 | 148484978 | rs7778273 | 1.617 | -0.832 | 0.451 | -0.399 | -0.011 | -0.044 | -0.044 | 0 | 0.1 | 0.14 | 0.05 | 0 | 0.07 | 0.05 |
| 7 | 148501703 | rs10252308 | 1.976 | -0.733 | 0.306 | -0.534 | -0.066 | -0.051 | -0.051 | 0 | 0.32 | 0.06 | 0.3 | 0.37 | 0.4 | 0.33 |
| 7 | 148501758 | rs9987055 | 0.451 | -0.499 | 0.578 | -0.157 | -0.01 | -0.002 | -0.002 | 0 | 0.67 | 0.71 | 0.73 | 0.7 | 0.6 | 0.59 |
| 7 | 148503377 | rs757624 | 0.877 | -0.577 | 0.54 | -0.175 | -0.033 | 0.021 | 0.021 | 0 | 0.2 | 0.32 | 0.27 | 0.28 | 0.13 | 0.2 |
| 7 | 148506363 | rs10268879 | 0.53 | -0.522 | 0.154 | -0.243 | -0.024 | -0.084 | -0.084 | 0 | 0.13 | 0.29 | 0.08 | 0 | 0.07 | 0.06 |
| 7 | 148517304 | rs17171118 | 1.314 | -0.238 | -0.648 | -0.017 | 0.021 | -0.041 | -0.041 | 0.01 | 0.17 | 0.06 | 0.22 | 0.28 | 0.12 | 0.2 |
| 7 | 148524752 | rs3757441 | -0.995 | 0.096 | -0.643 | 0.088 | -0.04 | -0.034 | -0.034 | 0 | 0.83 | 0.92 | 0.84 | 0.77 | 0.74 | 0.76 |
| 7 | 148562935 | rs12670401 | 0.477 | 0.183 | -0.261 | 0.227 | -0.014 | 0.014 | 0.014 | 0 | 0.45 | 0.3 | 0.46 | 0.43 | 0.47 | 0.4 |

| | | | | | | | | | | | | | | | | |
|---|---|---|---|---|---|---|---|---|---|---|---|---|---|---|---|---|
| 7 | 148578753 | rs2177567 | 0.432 | 0.323 | -0.481 | 0.278 | -0.002 | 0.003 | 0.003 | 0 | 0.46 | 0.48 | 0.47 | 0.42 | 0.47 | 0.4 |
| 7 | 148589869 | rs11984309 | -0.006 | 0.065 | -0.696 | 0.273 | 0.004 | -0.004 | -0.004 | 0 | 0.21 | 0.2 | 0.36 | 0.28 | 0.15 | 0.27 |
| 7 | 148600310 | rs7790642 | 0.185 | 0.785 | -0.62 | 0.075 | 0.002 | 0.002 | 0.002 | 0 | 0.45 | 0.4 | 0.31 | 0.37 | 0.4 | 0.42 |
| 7 | 148610111 | rs10952786 | -0.13 | 1.242 | -0.61 | -0.28 | 0.002 | -0.033 | -0.033 | 0 | 0.17 | 0.18 | 0.28 | 0.01 | 0.2 | 0.21 |
| 7 | 148615643 | rs7804456 | -0.352 | 1.205 | -0.595 | -0.262 | 0.018 | -0.08 | -0.08 | 0.003 | 0.21 | 0.36 | 0.41 | 0.06 | 0.22 | 0.25 |
| 7 | 148620581 | rs12531078 | -0.014 | 0.56 | -0.934 | 0.142 | -0.008 | 0.001 | 0.001 | 0 | 0.57 | 0.58 | 0.61 | 0.46 | 0.63 | 0.4 |
| 7 | 148629759 | rs822531 | 0.092 | 0.762 | -1.535 | -0.218 | -0.072 | 0.018 | 0.018 | 0 | 0.81 | 0.99 | 0.78 | 0.85 | 0.79 | 0.89 |
| 7 | 148635117 | rs9986856 | 0.713 | 0.399 | -1.943 | -0.416 | -0.042 | -0.018 | -0.018 | 0 | 0.35 | 0.16 | 0.42 | 0.61 | 0.42 | 0.42 |
| 7 | 148636850 | rs822533 | 0.601 | 0.23 | -1.564 | -0.5 | 0.002 | 0.024 | 0.024 | 0 | 0.06 | 0.09 | 0.06 | 0.11 | 0.09 | 0.1 |
| 7 | 148637765 | rs12667281 | 0.766 | -0.528 | -0.764 | -0.981 | 0 | -0.007 | -0.007 | 0 | 0.56 | 0.48 | 0.56 | 0.7 | 0.5 | 0.69 |
| 7 | 148638223 | rs822532 | 0.42 | -0.593 | -0.173 | -1.004 | 0.019 | 0.069 | 0.069 | 0 | 0.07 | 0.2 | 0.07 | 0.12 | 0.09 | 0.1 |
| 7 | 148638572 | rs2035216 | 0.742 | -0.684 | 0.172 | -0.989 | 0.004 | -0.021 | -0.021 | 0.731 | 0.56 | 0.53 | 0.56 | 0.67 | 0.5 | 0.71 |
| 7 | 148640449 | rs4727001 | 0.148 | -0.577 | 0.036 | -0.931 | -0.002 | 0.027 | 0.027 | 0.566 | 0.68 | 0.84 | 0.7 | 0.74 | 0.68 | 0.82 |
| 7 | 148645341 | rs4727003 | 0.297 | -0.935 | 0.483 | -0.739 | -0.027 | -0.017 | -0.017 | 0 | 0.18 | 0.23 | 0.13 | 0.07 | 0.07 | 0.26 |
| 7 | 148650818 | rs822553 | -1.642 | -0.557 | 1.064 | -0.737 | 0.035 | 0.01 | 0.01 | 0.097 | 0.19 | 0.38 | 0.18 | 0.19 | 0.28 | 0.2 |
| 7 | 148652565 | rs1676910 | -1.096 | -0.791 | 1.235 | -0.542 | 0.014 | 0.04 | 0.04 | 0 | 0.12 | 0.27 | 0.1 | 0.13 | 0.14 | 0.13 |
| 7 | 148656280 | rs1676904 | -1.197 | -0.774 | 1.419 | -0.643 | 0.008 | -0.001 | -0.001 | 0 | 0.17 | 0.45 | 0.13 | 0.14 | 0.16 | 0.17 |
| 7 | 148661502 | rs922411 | -1.21 | -1.014 | 1.297 | -0.766 | -0.03 | -0.095 | -0.095 | 0.361 | 0.14 | 0.45 | 0.08 | 0 | 0.05 | 0.11 |
| 7 | 148672346 | rs822546 | -0.683 | -0.907 | 1.458 | -0.546 | -0.004 | 0.011 | 0.011 | 0 | 0.14 | 0.23 | 0.2 | 0.15 | 0.14 | 0.16 |
| 7 | 148673365 | rs10952787 | 1.138 | -0.832 | 0.826 | -0.578 | 0.012 | 0.036 | 0.036 | 0 | 0.57 | 0.18 | 0.59 | 0.63 | 0.58 | 0.71 |
| 7 | 148680989 | rs2709331 | -0.055 | -0.715 | 0.868 | -0.651 | -0.001 | 0.035 | 0.035 | 0 | 0.26 | 0.18 | 0.17 | 0.1 | 0.28 | 0.12 |
| 7 | 148694939 | rs716817 | 1.615 | -1.115 | 1.213 | 0.131 | 0.046 | -0.022 | -0.022 | 0.002 | 0.56 | 0.2 | 0.61 | 0.51 | 0.65 | 0.66 |
| 7 | 148694971 | rs716818 | -0.224 | -1.089 | 1.472 | 0.064 | -0.002 | 0.072 | 0.072 | 0 | 0.88 | 0.85 | 0.79 | 0.61 | 0.87 | 0.81 |
| 7 | 148698474 | rs10236808 | -1.163 | -1.153 | 2.017 | -0.104 | -0.038 | -0.133 | -0.133 | 0 | 0.31 | 0.6 | 0.17 | 0.06 | 0.22 | 0.16 |
| 7 | 148700542 | rs1052549 | -1.183 | -1.149 | 2.013 | -0.108 | -0.036 | -0.133 | -0.133 | 0 | 0.31 | 0.6 | 0.17 | 0.06 | 0.23 | 0.16 |
| 7 | 148705490 | rs10272564 | -1.224 | -1.159 | 1.986 | -0.113 | -0.029 | -0.122 | -0.122 | 0 | 0.3 | 0.56 | 0.17 | 0.06 | 0.23 | 0.16 |
| 7 | 148711590 | rs10269104 | -0.627 | -1.119 | 1.286 | -0.006 | 0 | -0.043 | -0.043 | 0 | 0.25 | 0.3 | 0.15 | 0.01 | 0.22 | 0.15 |
| 7 | 148719656 | rs1551927 | 1.057 | -1.271 | 0.651 | 0.03 | 0.022 | 0.005 | 0.005 | 0.189 | 0.69 | 0.26 | 0.81 | 0.71 | 0.84 | 0.85 |
| 7 | 148737377 | rs899130 | 0.177 | -0.97 | 0.397 | -0.299 | -0.003 | -0.036 | -0.036 | 0 | 0.85 | 0.86 | 0.89 | 1 | 0.88 | 0.95 |
| 7 | 148738938 | rs10952789 | 2.016 | -1.233 | -0.149 | 0.086 | 0.051 | -0.004 | -0.004 | 0 | 0.61 | 0.1 | 0.68 | 0.66 | 0.68 | 0.84 |
| 7 | 148744033 | rs7789362 | 1.527 | -1.083 | 0.746 | 0.196 | -0.023 | 0.003 | 0.003 | 0 | 0.71 | 0.38 | 0.76 | 0.71 | 0.76 | 0.87 |
| 7 | 148746224 | rs6946304 | -1.704 | -0.567 | 1.081 | 0.45 | -0.002 | -0.013 | -0.013 | 0.066 | 0.17 | 0.36 | 0.13 | 0.14 | 0.2 | 0.15 |
| 7 | 148760850 | rs3801978 | 1.465 | -0.474 | -0.086 | -0.828 | -0.001 | 0.175 | 0.175 | 0 | 0.63 | 0.12 | 0.7 | 0.9 | 0.66 | 0.85 |
| 7 | 148779400 | rs1405123 | 0.491 | 0.047 | 0.787 | -0.978 | -0.003 | -0.031 | -0.031 | 0 | 0.79 | 0.75 | 0.82 | 0.95 | 0.8 | 0.94 |
| 7 | 148796253 | rs12666337 | 1.894 | 0.111 | 0.889 | -0.987 | -0.019 | 0.137 | 0.137 | 0 | 0.64 | 0.15 | 0.67 | 0.91 | 0.62 | 0.84 |
| 7 | 148799125 | rs11524138 | 2.011 | 0.066 | 0.875 | -0.993 | -0.024 | 0.168 | 0.168 | 0 | 0.63 | 0.13 | 0.67 | 0.91 | 0.61 | 0.84 |
| 7 | 148799572 | rs7796282 | 0.045 | 0.022 | 1.351 | -1.034 | 0.001 | -0.056 | -0.056 | 0 | 0.83 | 0.76 | 0.86 | 1 | 0.84 | 0.97 |
| 7 | 148801091 | rs11525060 | 2.093 | 0.005 | 2.112 | -1.039 | 0 | -0.02 | -0.02 | 0.186 | 0.67 | 0.35 | 0.69 | 0.91 | 0.62 | 0.84 |
| 7 | 148801658 | rs1137432 | 2.173 | 0.038 | 1.993 | -1.05 | -0.02 | 0.141 | 0.141 | 0 | 0.64 | 0.15 | 0.66 | 0.91 | 0.62 | 0.84 |
| 7 | 148806658 | rs10265213 | -0.66 | 0.041 | 2.147 | -1.051 | 0.092 | -0.002 | -0.002 | 0 | 0.07 | 0.22 | 0.08 | 0.04 | 0.14 | 0.12 |
| 7 | 148819519 | rs7794500 | 2.272 | 0.014 | 2.151 | -1.067 | -0.001 | -0.01 | -0.01 | 0 | 0.66 | 0.33 | 0.68 | 0.9 | 0.62 | 0.81 |
| 7 | 148822062 | rs4727017 | 2.378 | 0.011 | 2.042 | -1.068 | -0.029 | 0.171 | 0.171 | 0 | 0.63 | 0.12 | 0.66 | 0.9 | 0.61 | 0.81 |
| 7 | 148842558 | rs1202437 | -0.363 | 0.022 | 2.197 | -1.067 | 0.001 | -0.07 | -0.07 | 0 | 0.17 | 0.28 | 0.14 | 0 | 0.16 | 0.03 |
| 7 | 148851213 | rs917124 | 1.501 | 0.033 | 2.187 | -1.072 | 0.014 | -0.097 | -0.097 | 0.792 | 0.73 | 0.39 | 0.76 | 0.95 | 0.7 | 0.85 |
| 7 | 148864652 | rs1202449 | 0.178 | 0.025 | 2.013 | -1.067 | 0.008 | 0.018 | 0.018 | 0 | 0.15 | 0.12 | 0.08 | 0 | 0.1 | 0.02 |
| 7 | 148870108 | rs6961605 | 2.367 | -0.005 | 1.907 | -0.891 | -0.028 | 0.137 | 0.137 | 0 | 0.63 | 0.12 | 0.66 | 0.88 | 0.61 | 0.8 |
| 7 | 148874162 | rs2158632 | -0.563 | 0.063 | 1.816 | -0.895 | -0.004 | -0.001 | -0.001 | 0 | 0.08 | 0.08 | 0.08 | 0.05 | 0.09 | 0.04 |
| 7 | 148877605 | rs12704069 | -0.645 | 0.064 | 1.822 | -0.895 | -0.005 | -0.001 | -0.001 | 0 | 0.08 | 0.09 | 0.08 | 0.05 | 0.09 | 0.04 |
| 7 | 148880035 | rs1202462 | 0.667 | 0.234 | 1.95 | -0.914 | 0.088 | -0.003 | -0.003 | 0 | 0.93 | 0.78 | 0.91 | 0.95 | 0.86 | 0.88 |
| 7 | 148887137 | rs1202468 | -0.413 | 0.234 | 1.606 | -0.949 | -0.001 | -0.084 | -0.084 | 0.016 | 0.17 | 0.31 | 0.14 | 0 | 0.15 | 0.02 |
| 7 | 148898389 | rs1202407 | 1.378 | 0.107 | 1.646 | -0.975 | 0.086 | -0.032 | -0.032 | 1.619 | 0.9 | 0.66 | 0.91 | 0.96 | 0.86 | 0.9 |
| 7 | 148908562 | rs2888600 | 2.315 | -0.051 | 0.901 | -1.113 | -0.013 | 0.195 | 0.195 | 0.008 | 0.59 | 0.11 | 0.66 | 0.87 | 0.59 | 0.81 |
| 7 | 148913645 | rs1202389 | 0.502 | -0.061 | 0.994 | -1.142 | 0.013 | 0.007 | 0.007 | 0 | 0.14 | 0.12 | 0.06 | 0 | 0.06 | 0.03 |
| 7 | 148914122 | rs1202390 | 2.171 | -0.067 | 1.182 | -1.148 | -0.001 | -0.017 | -0.017 | 1.259 | 0.74 | 0.35 | 0.79 | 0.87 | 0.72 | 0.84 |
| 7 | 148914740 | rs1202392 | 0.109 | -0.433 | 1.16 | -1.064 | -0.001 | -0.033 | -0.033 | 0.038 | 0.15 | 0.2 | 0.13 | 0 | 0.13 | 0.03 |
| 7 | 148916244 | rs11762610 | -0.608 | -0.379 | 1.288 | -1.023 | 0.127 | 0.059 | 0.059 | 0 | 0.05 | 0.33 | 0.06 | 0.04 | 0.07 | 0.05 |
| 7 | 148916627 | rs1202397 | -1.57 | -0.518 | 1.136 | -1.053 | -0.002 | -0.019 | -0.019 | 0 | 0.31 | 0.35 | 0.23 | 0.09 | 0.27 | 0.09 |
| 7 | 148922666 | rs11976830 | 0.314 | -0.646 | 0.354 | -1.567 | -0.001 | -0.027 | -0.027 | 0.019 | 0.15 | 0.15 | 0.09 | 0 | 0.12 | 0.14 |
| 7 | 148926605 | rs1724329 | -0.548 | -0.524 | -0.059 | -1.51 | 0.003 | -0.019 | -0.019 | 0.017 | 0.41 | 0.52 | 0.5 | 0.26 | 0.37 | 0.36 |
| 7 | 148931522 | rs2693319 | 0.012 | -0.447 | -0.413 | -1.586 | -0.024 | 0.008 | 0.008 | 0 | 0.29 | 0.16 | 0.42 | 0.23 | 0.25 | 0.29 |
| 7 | 148933942 | rs7779475 | -0.444 | -0.717 | -0.464 | -1.914 | -0.008 | -0.067 | -0.067 | 0 | 0.27 | 0.32 | 0.15 | 0.01 | 0.25 | 0.2 |
| 7 | 148937136 | rs1202427 | -1.187 | -0.909 | -0.554 | -1.39 | 0.001 | -0.001 | -0.001 | 0 | 0.54 | 0.59 | 0.62 | 0.42 | 0.51 | 0.48 |
| 7 | 148940873 | rs10241873 | -0.822 | -1.258 | -0.442 | -1.364 | 0 | 0 | 0 | 0 | 0.42 | 0.59 | 0.53 | 0.46 | 0.34 | 0.35 |
| 7 | 148947279 | rs11769362 | 1.782 | -1.277 | -0.607 | -1.414 | -0.029 | 0.025 | 0.025 | 0.103 | 0.08 | 0 | 0.04 | 0.04 | 0.1 | 0.07 |
| 7 | 148950023 | rs10487889 | 1.93 | -1.192 | -0.571 | -1.488 | -0.013 | 0.069 | 0.069 | 0 | 0.12 | 0.02 | 0.06 | 0 | 0.14 | 0.13 |
| 7 | 148951992 | rs1053298 | -0.615 | -1.181 | -0.492 | -1.475 | -0.008 | 0.012 | 0.012 | 0 | 0.38 | 0.52 | 0.53 | 0.46 | 0.33 | 0.32 |
| 7 | 148952312 | rs3087882 | 0.444 | -0.916 | -0.685 | -1.444 | 0.016 | -0.004 | -0.004 | 0.02 | 0.28 | 0.16 | 0.36 | 0.25 | 0.18 | 0.29 |
| 7 | 148964513 | rs12704090 | -2.475 | -0.888 | -0.833 | -1.485 | -0.094 | -0.106 | -0.106 | 0 | 0.68 | 0.92 | 0.7 | 0.48 | 0.6 | 0.57 |
| 7 | 148976137 | rs7788681 | 0.524 | -0.874 | -1.014 | -1.334 | -0.003 | -0.026 | -0.026 | 0 | 0.07 | 0.13 | 0.04 | 0 | 0.09 | 0.02 |

| | | | | | | | | | | | | | | | | |
|---|---|---|---|---|---|---|---|---|---|---|---|---|---|---|---|---|
| 7 | 148979649 | rs3735315 | -1.362 | -0.878 | -0.996 | -1.343 | -0.001 | -0.001 | -0.001 | 0 | 0.56 | 0.59 | 0.6 | 0.46 | 0.48 | 0.49 |
| 7 | 148990933 | rs6977581 | -0.464 | -0.919 | -1.278 | -1.305 | -0.017 | -0.053 | -0.053 | 0.107 | 0.11 | 0.21 | 0.06 | 0 | 0.11 | 0.05 |
| 7 | 148991712 | rs13236978 | -2.657 | -0.789 | -2.031 | -1.026 | -0.145 | -0.15 | -0.15 | 0.309 | 0.74 | 0.98 | 0.72 | 0.58 | 0.63 | 0.61 |
| 7 | 148994241 | rs1044678 | -0.437 | -0.802 | -1.084 | -0.611 | 0.014 | -0.035 | -0.035 | 0 | 0.29 | 0.22 | 0.4 | 0.5 | 0.18 | 0.29 |
| 7 | 148998960 | rs6974847 | -0.771 | -0.769 | -1.194 | -0.866 | 0.019 | 0.033 | 0.033 | 0.011 | 0.27 | 0.2 | 0.19 | 0 | 0.23 | 0.08 |
| 7 | 149005429 | rs17171195 | 1.216 | -0.679 | -1.322 | -0.712 | -0.038 | -0.07 | -0.07 | 0 | 0.11 | 0.01 | 0.27 | 0.26 | 0.1 | 0.28 |
| 7 | 149007531 | rs4727044 | -1.246 | -0.734 | -1.138 | -0.739 | -0.01 | 0.002 | 0.002 | 0 | 0.43 | 0.51 | 0.52 | 0.47 | 0.3 | 0.39 |
| 7 | 149011410 | rs13231525 | 0.866 | -0.652 | -1.127 | -0.725 | 0 | -0.007 | -0.007 | 0 | 0.06 | 0.06 | 0.03 | 0 | 0.08 | 0 |
| 7 | 149011836 | rs4727046 | -0.934 | -0.661 | -1.144 | -0.794 | 0.009 | 0.046 | 0.046 | 0 | 0.42 | 0.34 | 0.49 | 0.26 | 0.37 | 0.45 |
| 7 | 149014262 | rs6959353 | -0.935 | -0.715 | -0.831 | -0.359 | 0.01 | 0.03 | 0.03 | 0.002 | 0.42 | 0.38 | 0.48 | 0.26 | 0.36 | 0.35 |
| 7 | 149015369 | rs757795 | -0.568 | -0.239 | -0.065 | 0.493 | -0.005 | 0.006 | 0.006 | 0 | 0.25 | 0.37 | 0.33 | 0.26 | 0.23 | 0.31 |
| 7 | 149015443 | rs10229125 | -1.279 | 0.065 | -0.256 | 0.915 | -0.029 | 0.081 | 0.081 | 0 | 0.06 | 0.13 | 0.08 | 0.21 | 0.03 | 0.05 |
| 7 | 149017788 | rs11761128 | -0.628 | 0.658 | -0.502 | 0.541 | 0.003 | -0.002 | -0.002 | 0.872 | 0.3 | 0.44 | 0.41 | 0.24 | 0.28 | 0.47 |
| 7 | 149018308 | rs11761219 | 0.277 | 0.321 | -1.435 | 0.247 | -0.021 | 0.078 | 0.078 | 0 | 0.46 | 0.14 | 0.46 | 0.39 | 0.55 | 0.5 |
| 7 | 149018975 | rs886714 | -0.315 | 0.431 | -0.57 | -0.029 | 0.03 | -0.054 | -0.054 | 0 | 0.09 | 0.17 | 0.12 | 0 | 0.14 | 0.03 |
| 7 | 149019712 | rs4725805 | -0.084 | 0.44 | -0.602 | -0.026 | 0.015 | -0.041 | -0.041 | 0.034 | 0.09 | 0.1 | 0.12 | 0 | 0.14 | 0.03 |
| 7 | 149026686 | rs1473127 | 0.392 | -0.465 | -0.523 | -0.007 | -0.056 | 0.012 | 0.012 | 0.021 | 0.78 | 0.38 | 0.85 | 0.73 | 0.86 | 0.88 |
| 7 | 149028452 | rs757796 | 0.396 | -0.584 | -0.253 | -0.079 | -0.011 | 0.053 | 0.053 | 0 | 0.12 | 0.22 | 0.06 | 0.26 | 0.11 | 0.11 |
| 7 | 149028791 | rs736548 | 0.864 | -0.395 | -0.941 | -0.018 | 0.018 | 0.02 | 0.02 | 1.135 | 0.68 | 0.24 | 0.73 | 0.73 | 0.72 | 0.85 |
| 7 | 149029106 | rs11762571 | 1.396 | -0.3 | -2.238 | -0.295 | -0.158 | 0.029 | 0.029 | 0.184 | 0.39 | 0.03 | 0.35 | 0.32 | 0.47 | 0.48 |
| 7 | 149043045 | rs7779412 | 0.365 | 0.018 | -1.809 | 0.051 | -0.049 | -0.068 | -0.068 | 7.828 | 0.23 | 0.02 | 0.34 | 0.22 | 0.21 | 0.33 |
| 7 | 149053677 | rs10464475 | 0.502 | 0.107 | -1.685 | 0.132 | -0.054 | -0.094 | -0.094 | 0 | 0.2 | 0.04 | 0.19 | 0.23 | 0.22 | 0.3 |
| 7 | 149054004 | rs765951 | 1.626 | -0.192 | -2.193 | -0.201 | -0.106 | -0.005 | -0.005 | 0 | 0.36 | 0.03 | 0.45 | 0.33 | 0.39 | 0.45 |
| 7 | 149060494 | rs2462608 | -0.2 | -0.471 | -1.76 | -0.217 | -0.006 | -0.008 | -0.008 | 0 | 0.79 | 0.66 | 0.84 | 0.79 | 0.82 | 0.87 |
| 7 | 149062209 | rs1525524 | -0.382 | -0.377 | -1.441 | -0.19 | 0.002 | 0.03 | 0.03 | 0 | 0.23 | 0.46 | 0.21 | 0.24 | 0.21 | 0.13 |
| 7 | 149064937 | rs1636375 | 0.255 | -0.477 | -1.421 | -0.241 | 0.016 | -0.104 | -0.104 | 0.178 | 0.16 | 0.54 | 0.16 | 0 | 0.15 | 0.05 |
| 7 | 149067575 | rs1608849 | -0.811 | -0.645 | -1.52 | -0.271 | -0.003 | 0 | 0 | 0.21 | 0.27 | 0.64 | 0.2 | 0.24 | 0.21 | 0.14 |
| 7 | 149072406 | rs1404862 | 0.294 | -0.661 | -1.606 | -0.237 | 0 | -0.006 | -0.006 | 0 | 0.6 | 0.65 | 0.64 | 0.58 | 0.6 | 0.58 |
| 7 | 149074314 | rs1636396 | 0.1 | -1.101 | -1.081 | -0.435 | 0.001 | 0.002 | 0.002 | 0 | 0.8 | 0.77 | 0.84 | 0.79 | 0.82 | 0.88 |
| 7 | 149075131 | rs1525538 | -1.459 | -1.291 | -1.019 | -0.439 | -0.011 | -0.003 | -0.003 | 0.85 | 0.3 | 0.51 | 0.2 | 0.24 | 0.23 | 0.15 |
| 7 | 149090233 | rs1636407 | -0.44 | -0.857 | -0.694 | -0.212 | 0.005 | -0.104 | -0.104 | 0 | 0.16 | 0.54 | 0.17 | 0 | 0.15 | 0.05 |
| 7 | 149091240 | rs1723692 | 0.781 | -0.704 | -0.805 | -0.097 | 0.013 | -0.024 | -0.024 | 0 | 0.7 | 0.72 | 0.81 | 0.54 | 0.76 | 0.79 |
| 7 | 149091469 | rs1636408 | -0.554 | 0.046 | -0.118 | -0.492 | 0.05 | -0.104 | -0.104 | 0 | 0.14 | 0.38 | 0.16 | 0 | 0.15 | 0.06 |
| 7 | 149093007 | rs1076283 | -0.865 | -0.215 | -0.5 | -0.009 | 0.003 | 0.014 | 0.014 | 0 | 0.14 | 0.25 | 0.1 | 0.21 | 0.14 | 0.12 |
| 7 | 149094701 | rs1636409 | 1.024 | -0.574 | -1.308 | -0.23 | 0.009 | -0.003 | -0.003 | 0.219 | 0.77 | 0.96 | 0.75 | 0.76 | 0.8 | 0.78 |
| 7 | 149095024 | rs1636410 | -0.68 | -1.117 | -0.429 | -0.655 | -0.067 | -0.067 | -0.067 | 0 | 0.21 | 0.41 | 0.1 | 0.11 | 0.07 | 0.17 |
| 7 | 149097526 | rs6947354 | 1.11 | -0.583 | -0.54 | -0.753 | -0.006 | -0.017 | -0.017 | 0 | 0.65 | 0.6 | 0.68 | 0.76 | 0.67 | 0.79 |
| 7 | 149099314 | rs13231611 | -0.927 | -0.081 | -0.478 | -0.54 | 0.007 | 0.001 | 0.001 | 0 | 0.14 | 0.2 | 0.18 | 0.15 | 0.16 | 0.15 |
| 7 | 149102171 | rs886712 | -0.208 | 0.222 | -0.829 | -0.334 | 0.007 | 0.054 | 0.054 | 1.333 | 0.46 | 0.7 | 0.43 | 0.66 | 0.51 | 0.67 |
| 7 | 149118535 | rs6965442 | -0.604 | 1.146 | -0.876 | -0.26 | -0.009 | -0.002 | -0.002 | 0 | 0.14 | 0.11 | 0.15 | 0.15 | 0.19 | 0.16 |
| 7 | 149127365 | rs6970608 | 0.279 | 0.736 | -1.046 | -0.751 | -0.019 | 0.041 | 0.041 | 0.097 | 0.67 | 0.81 | 0.72 | 0.88 | 0.61 | 0.76 |
| 7 | 149133916 | rs11762125 | -0.298 | 0.358 | -1.016 | -0.539 | -0.024 | 0.052 | 0.052 | 0 | 0.11 | 0.02 | 0.05 | 0 | 0.1 | 0.08 |
| 7 | 149145139 | rs6973105 | -0.304 | -0.57 | -0.389 | 0.395 | -0.042 | 0.071 | 0.071 | 0 | 0.22 | 0.33 | 0.23 | 0.52 | 0.11 | 0.25 |
| 7 | 149148319 | rs4727071 | 0.125 | 0.081 | -0.056 | -0.268 | -0.021 | 0.067 | 0.067 | 0 | 0.58 | 0.76 | 0.68 | 0.81 | 0.56 | 0.73 |
| 7 | 149152770 | rs3735319 | 0.143 | 0.028 | 0.356 | -0.309 | 0.002 | -0.017 | -0.017 | 0 | 0.58 | 0.55 | 0.65 | 0.82 | 0.56 | 0.76 |
| 7 | 149152906 | rs3735318 | 0.261 | -0.075 | 0.676 | -0.317 | -0.008 | 0.063 | 0.063 | 0 | 0.54 | 0.39 | 0.62 | 0.81 | 0.54 | 0.74 |
| 7 | 149158435 | rs3735317 | -0.128 | -0.006 | 0.299 | -0.463 | -0.019 | 0.094 | 0.094 | 0 | 0.6 | 0.82 | 0.69 | 0.88 | 0.57 | 0.74 |
| 7 | 149170236 | rs1046140 | 0.49 | -0.437 | 0.319 | -0.707 | 0.01 | 0.072 | 0.072 | 0 | 0.29 | 0.14 | 0.16 | 0.11 | 0.26 | 0.22 |
| 7 | 149173752 | rs10266573 | -0.812 | -0.468 | 0.434 | -0.568 | -0.011 | 0.058 | 0.058 | 0.009 | 0.64 | 0.83 | 0.72 | 0.83 | 0.62 | 0.74 |
| 7 | 149184850 | rs6971236 | 0.726 | -0.388 | 0.747 | -0.508 | 0.009 | 0.031 | 0.031 | 0 | 0.3 | 0.21 | 0.17 | 0.12 | 0.26 | 0.23 |
| 7 | 149191027 | rs13236631 | -0.599 | -0.483 | 0.772 | -0.541 | 0 | 0.035 | 0.035 | 0 | 0.61 | 0.73 | 0.71 | 0.83 | 0.62 | 0.73 |
| 7 | 149197030 | rs709067 | -0.324 | -0.547 | 0.952 | 0.34 | 0.003 | 0.009 | 0.009 | 0.021 | 0.73 | 0.68 | 0.51 | 0.49 | 0.65 | 0.61 |
| 7 | 149201258 | rs6464978 | -1.275 | -0.565 | 1.114 | 0.411 | 0.015 | 0.324 | 0.324 | 1.21 | 0.06 | 0.33 | 0.08 | 0.22 | 0.01 | 0.1 |
| 7 | 149204548 | rs855912 | 0.956 | -0.741 | 0.944 | 0.251 | -0.021 | 0.074 | 0.074 | 0.084 | 0.35 | 0.15 | 0.28 | 0.16 | 0.38 | 0.27 |
| 7 | 149267567 | rs4727090 | -0.105 | -0.474 | 0.828 | 0.556 | 0.009 | 0.01 | 0.01 | 0.222 | 0.76 | 0.74 | 0.76 | 0.69 | 0.67 | 0.67 |
| 7 | 149277045 | rs2373493 | 0.306 | -0.28 | 0.851 | 0.626 | 0.013 | 0.142 | 0.142 | 0 | 0.09 | 0.23 | 0.32 | 0.37 | 0.07 | 0.09 |
| 7 | 149277121 | rs12536745 | 0.844 | -0.577 | 0.856 | 0.611 | -0.025 | 0.075 | 0.075 | 0.204 | 0.35 | 0.15 | 0.27 | 0.16 | 0.37 | 0.27 |
| 7 | 149286695 | rs4255049 | 0.772 | -0.573 | 0.882 | 0.643 | -0.007 | 0.175 | 0.175 | 0.435 | 0.08 | 0.23 | 0.31 | 0.37 | 0.04 | 0.08 |
| 7 | 149297104 | rs354042 | 1.229 | -0.595 | 0.553 | 0.538 | 0.02 | 0.081 | 0.081 | 0 | 0.31 | 0.15 | 0.15 | 0.12 | 0.26 | 0.24 |
| 7 | 149298552 | rs354044 | -0.784 | -0.408 | 0.308 | 0.549 | -0.102 | 0.003 | 0.003 | 0.581 | 0.8 | 0.97 | 0.82 | 0.87 | 0.68 | 0.7 |
| 7 | 149304067 | rs10808056 | 0.739 | -0.312 | 0.366 | 0.158 | -0.026 | 0.073 | 0.073 | 0.312 | 0.34 | 0.15 | 0.27 | 0.16 | 0.37 | 0.27 |
| 7 | 149327723 | rs354072 | -0.244 | -0.135 | 0.539 | 0.325 | 0.007 | 0.014 | 0.014 | 0.291 | 0.77 | 0.74 | 0.75 | 0.66 | 0.67 | 0.6 |
| 7 | 149331027 | rs11773063 | 0.503 | -0.079 | -0.047 | 0.269 | -0.014 | 0.06 | 0.06 | 0 | 0.34 | 0.16 | 0.26 | 0.19 | 0.35 | 0.25 |
| 7 | 149334212 | rs354082 | 0.126 | 0.129 | -0.016 | 0.633 | -0.006 | 0.091 | 0.091 | 0 | 0.1 | 0.22 | 0.29 | 0.32 | 0.06 | 0.09 |
| 7 | 149344332 | rs12234594 | 0.608 | 0.169 | -0.12 | -0.407 | 0.032 | -0.025 | -0.025 | 0.954 | 0.22 | 0.32 | 0.23 | 0.16 | 0.27 | 0.17 |
| 7 | 149345681 | rs10250289 | -1.062 | 0.02 | -0.667 | -0.514 | 0.019 | 0.044 | 0.044 | 0 | 0.67 | 0.76 | 0.82 | 0.85 | 0.73 | 0.78 |
| 7 | 149361282 | rs731489 | 0.118 | 0.928 | -0.252 | 0.013 | 0.054 | -0.039 | -0.039 | 1.507 | 0.23 | 0.4 | 0.25 | 0.16 | 0.28 | 0.17 |
| 7 | 149363363 | rs855736 | 0.378 | 0.282 | -0.758 | -0.26 | 0.044 | 0.074 | 0.074 | 0 | 0.49 | 0.37 | 0.24 | 0.19 | 0.35 | 0.41 |
| 7 | 149363857 | rs12536519 | 0.296 | 1.423 | -0.888 | -0.291 | -0.024 | 0.024 | 0.024 | 0 | 0.16 | 0.07 | 0.14 | 0.05 | 0.24 | 0.1 |

| | | | | | | | | | | | | | | | | |
|---|---|---|---|---|---|---|---|---|---|---|---|---|---|---|---|---|
| 7 | 149364008 | rs855737 | 0.297 | 0.667 | -1.405 | -0.167 | 0.003 | 0 | 0 | 0 | 0.45 | 0.26 | 0.45 | 0.58 | 0.44 | 0.46 |
| 7 | 149364246 | rs6953356 | -0.764 | -0.462 | -0.556 | -1.724 | -0.023 | -0.124 | -0.124 | 0 | 0.24 | 0.4 | 0.17 | 0.01 | 0.18 | 0.23 |
| 7 | 149365696 | rs11982494 | -0.687 | -0.688 | -0.786 | -1.697 | -0.009 | 0.105 | 0.105 | 2.76 | 0.54 | 0.3 | 0.32 | 0.26 | 0.55 | 0.5 |
| 7 | 149367686 | rs10487554 | 0.542 | -0.693 | -1.144 | -1.858 | -0.09 | 0.027 | 0.027 | 0.341 | 0.19 | 0.02 | 0.1 | 0.1 | 0.26 | 0.21 |
| 7 | 149367798 | rs855740 | 1.866 | -1.006 | -0.686 | -1.612 | 0 | -0.004 | -0.004 | 1.437 | 0.28 | 0.26 | 0.21 | 0.15 | 0.26 | 0.2 |
| 7 | 149381716 | rs1101865 | 1.767 | -1.013 | -0.675 | -1.451 | 0 | 0 | 0 | 0 | 0.29 | 0.28 | 0.24 | 0.31 | 0.26 | 0.21 |
| 7 | 149384913 | rs17809023 | 1.256 | -1.115 | -0.971 | -1.652 | -0.034 | 0.051 | 0.051 | 0 | 0.21 | 0.01 | 0.15 | 0.13 | 0.22 | 0.17 |
| 7 | 149385122 | rs2888583 | 0.258 | -1.531 | -0.294 | -1.067 | -0.078 | 0.039 | 0.039 | 0.208 | 0.19 | 0.04 | 0.11 | 0.1 | 0.26 | 0.21 |
| 7 | 149394446 | rs6974082 | 1.017 | -1.604 | -0.042 | -0.906 | -0.005 | -0.017 | -0.017 | 0.01 | 0.13 | 0.17 | 0.06 | 0.04 | 0.07 | 0.14 |
| 7 | 149401731 | rs10952836 | -0.213 | -1.51 | 0.076 | -0.863 | 0.028 | 0.026 | 0.026 | 1.575 | 0.36 | 0.34 | 0.16 | 0.25 | 0.21 | 0.24 |
| 7 | 149403770 | rs6974985 | -0.476 | -1.545 | 0.216 | -0.879 | -0.003 | -0.001 | -0.001 | 0.715 | 0.39 | 0.66 | 0.19 | 0.25 | 0.21 | 0.25 |
| 7 | 149410638 | rs855749 | 2.122 | -1.384 | 0.132 | -1.274 | -0.063 | 0.081 | 0.081 | 0 | 0.34 | 0.13 | 0.3 | 0.17 | 0.39 | 0.35 |
| 7 | 149421210 | rs740136 | -0.038 | -0.765 | -0.243 | -1.067 | -0.049 | -0.07 | -0.07 | 0 | 0.78 | 0.88 | 0.53 | 0.54 | 0.62 | 0.64 |
| 7 | 149421773 | rs7791608 | 0.045 | -1.048 | -0.107 | -0.882 | -0.054 | -0.064 | -0.064 | 0 | 0.28 | 0.57 | 0.15 | 0.15 | 0.15 | 0.15 |
| 7 | 149422718 | rs855754 | 0.274 | -0.076 | -0.794 | -0.42 | -0.055 | -0.084 | -0.084 | 1.396 | 0.79 | 0.91 | 0.53 | 0.54 | 0.63 | 0.59 |
| 7 | 149427728 | rs6966081 | -0.625 | -1.65 | -0.037 | -0.167 | 0.024 | 0.024 | 0.024 | 1.451 | 0.36 | 0.34 | 0.16 | 0.21 | 0.21 | 0.24 |
| 7 | 149437827 | rs1439104 | -0.877 | -0.814 | 0.169 | 0.148 | -0.011 | 0.001 | 0.001 | 0 | 0.36 | 0.43 | 0.51 | 0.65 | 0.3 | 0.43 |
| 7 | 149438267 | rs11763878 | -0.763 | -1.046 | 0.091 | 0.264 | -0.057 | -0.008 | -0.008 | 2.494 | 0.32 | 0.46 | 0.18 | 0.36 | 0.21 | 0.22 |
| 7 | 149444248 | rs886659 | 0.11 | -0.79 | -0.778 | 0.287 | 0.009 | 0.013 | 0.013 | 0 | 0.11 | 0.08 | 0.06 | 0.02 | 0.08 | 0.1 |
| 7 | 149445585 | rs929344 | -0.613 | -0.288 | -0.603 | 0.645 | 0.028 | 0.09 | 0.09 | 0 | 0.13 | 0.23 | 0.46 | 0.6 | 0.18 | 0.24 |
| 7 | 149445992 | rs11769599 | 0.411 | -0.234 | -0.911 | 0.553 | 0.011 | 0.007 | 0.007 | 0.283 | 0.16 | 0.01 | 0.05 | 0.18 | 0.12 | 0.13 |
| 7 | 149446246 | rs10235552 | -0.319 | 0.398 | -0.3 | 1.057 | 0.033 | 0.217 | 0.217 | 0.886 | 0.07 | 0.16 | 0.38 | 0.44 | 0.11 | 0.22 |
| 7 | 149450379 | rs7785602 | -0.414 | -0.127 | -0.88 | -0.168 | 0.013 | 0.012 | 0.012 | 0.003 | 0.36 | 0.73 | 0.62 | 0.65 | 0.32 | 0.32 |
| 7 | 149450584 | rs855653 | -0.166 | 0.181 | -0.98 | -1.19 | 0.001 | -0.033 | -0.033 | 0.122 | 0.87 | 0.86 | 0.92 | 1 | 0.85 | 0.98 |
| 7 | 149450663 | rs12674335 | -0.336 | 0.157 | -1.017 | -1.197 | 0.029 | 0.067 | 0.067 | 2.372 | 0.43 | 0.67 | 0.3 | 0.25 | 0.37 | 0.34 |
| 7 | 149451480 | rs855654 | 0.212 | 0.516 | -0.833 | -0.926 | -0.018 | 0.017 | 0.017 | 1.225 | 0.7 | 0.82 | 0.8 | 0.83 | 0.66 | 0.59 |
| 7 | 149456359 | rs855656 | -1.123 | -0.345 | -0.17 | -1.13 | -0.053 | -0.109 | -0.109 | 0.194 | 0.27 | 0.56 | 0.14 | 0.07 | 0.16 | 0.1 |
| 7 | 149458128 | rs855663 | -0.808 | -0.344 | -0.078 | -1.153 | -0.032 | -0.112 | -0.112 | 0.525 | 0.2 | 0.47 | 0.09 | 0.02 | 0.11 | 0.02 |
| 7 | 149464433 | rs855673 | -1.485 | -0.29 | -0.041 | -1.16 | -0.032 | -0.083 | -0.083 | 0.158 | 0.29 | 0.61 | 0.14 | 0.08 | 0.17 | 0.09 |
| 7 | 149465674 | rs855675 | -0.555 | 0.28 | -0.92 | -0.961 | -0.061 | 0.022 | 0.022 | 1.756 | 0.78 | 0.99 | 0.8 | 0.9 | 0.73 | 0.66 |
| 7 | 149472312 | rs855676 | -1.328 | 0.163 | 0.528 | -0.559 | -0.018 | -0.037 | -0.037 | 2.566 | 0.19 | 0.44 | 0.12 | 0.1 | 0.16 | 0.06 |
| 7 | 149472508 | rs709063 | 0.097 | 0.503 | 0.005 | -0.048 | -0.049 | -0.045 | -0.045 | 0.904 | 0.91 | 0.99 | 0.92 | 0.79 | 0.86 | 0.93 |
| 7 | 149478321 | rs855679 | -2.106 | -0.39 | 0.943 | -0.372 | -0.018 | -0.028 | -0.028 | 0 | 0.15 | 0.47 | 0.09 | 0.1 | 0.09 | 0.17 |
| 7 | 149483373 | rs855686 | -2.318 | -0.557 | 0.352 | -0.361 | -0.03 | -0.042 | -0.042 | 0.484 | 0.17 | 0.39 | 0.11 | 0.1 | 0.1 | 0.12 |
| 7 | 149485059 | rs709060 | -1.873 | -0.366 | -0.182 | -0.274 | 0.001 | 0.001 | 0.001 | 0 | 0.12 | 0.14 | 0.09 | 0.09 | 0.1 | 0.08 |
| 7 | 149486246 | rs855690 | -1.589 | -0.255 | 0.055 | -0.039 | -0.008 | 0.001 | 0.001 | 0 | 0.08 | 0.13 | 0.06 | 0.1 | 0.07 | 0.07 |
| 7 | 149489491 | rs1076277 | -0.847 | -0.23 | -0.078 | -0.025 | -0.001 | -0.018 | -0.018 | 0.006 | 0.06 | 0.01 | 0.05 | 0.1 | 0.07 | 0.05 |
| 7 | 149492218 | rs1635807 | -1.364 | -0.221 | -0.068 | -0.022 | -0.004 | 0 | 0 | 0 | 0.08 | 0.1 | 0.06 | 0.1 | 0.07 | 0.1 |
| 7 | 149499412 | rs2074691 | -1.523 | -0.801 | 0.039 | -0.373 | -0.029 | -0.028 | -0.028 | 0 | 0.21 | 0.31 | 0.09 | 0.1 | 0.11 | 0.16 |
| 7 | 149499469 | rs2074692 | -1.784 | -0.973 | 0.674 | -0.455 | -0.073 | -0.069 | -0.069 | 0 | 0.2 | 0.56 | 0.1 | 0.1 | 0.09 | 0.12 |
| 7 | 149500992 | rs4725830 | -0.056 | -0.592 | -0.047 | -0.512 | 0.005 | 0.046 | 0.046 | 0 | 0.48 | 0.64 | 0.42 | 0.29 | 0.45 | 0.38 |
| 7 | 149503843 | rs10260959 | -0.455 | -1.441 | -0.361 | -0.521 | 0.049 | 0.013 | 0.013 | 1.187 | 0.33 | 0.8 | 0.27 | 0.2 | 0.16 | 0.25 |
| 7 | 149511985 | rs10952230 | -0.728 | -0.259 | -0.711 | 0.537 | -0.05 | -0.052 | -0.052 | 0 | 0.92 | 1 | 0.82 | 0.72 | 0.83 | 0.94 |
| 7 | 149512177 | rs11979187 | 0.394 | -1.205 | 0.235 | 0.256 | -0.005 | 0 | 0 | 2.551 | 0.23 | 0.27 | 0.23 | 0.17 | 0.16 | 0.22 |
| 7 | 149512674 | rs2074694 | 0.432 | -1.031 | 0.186 | 0.566 | 0.008 | 0.023 | 0.023 | 0 | 0.09 | 0.11 | 0.14 | 0.24 | 0.14 | 0.08 |
| 7 | 149513780 | rs2074688 | -0.95 | -1.408 | 0.043 | 0.341 | 0.081 | -0.015 | -0.015 | 0 | 0.45 | 0.87 | 0.43 | 0.42 | 0.32 | 0.35 |
| 7 | 149513846 | rs2072169 | -0.175 | -1.365 | 0.887 | 0.472 | 0.005 | -0.006 | -0.006 | 0 | 0.35 | 0.67 | 0.39 | 0.43 | 0.3 | 0.31 |
| 7 | 149522366 | rs2074697 | -0.893 | -1.308 | 0.632 | 0.482 | 0.066 | -0.042 | -0.042 | 0 | 0.43 | 0.85 | 0.39 | 0.43 | 0.31 | 0.34 |
| 7 | 149528262 | rs10261977 | 0.895 | -1.312 | 1.467 | 0.542 | 0.007 | -0.004 | -0.004 | 0 | 0.22 | 0.19 | 0.19 | 0.17 | 0.15 | 0.22 |
| 7 | 149530894 | rs6960187 | -0.021 | -1.055 | 1.41 | 0.477 | -0.007 | -0.004 | -0.004 | 0 | 0.32 | 0.58 | 0.31 | 0.18 | 0.2 | 0.26 |
| 7 | 149536214 | rs7782781 | -0.63 | -0.89 | 1.242 | 0.52 | -0.015 | 0.004 | 0.004 | 0 | 0.41 | 0.42 | 0.42 | 0.42 | 0.35 | 0.35 |
| 7 | 149542524 | rs10239759 | -1.457 | -1.072 | 1.274 | 0.601 | -0.012 | 0.02 | 0.02 | 0 | 0.17 | 0.37 | 0.16 | 0.24 | 0.15 | 0.14 |
| 7 | 149547703 | rs2074699 | -0.48 | -1.105 | 0.683 | 1.131 | 0.021 | -0.02 | -0.02 | 0.053 | 0.41 | 0.22 | 0.35 | 0.51 | 0.38 | 0.42 |
| 7 | 149551993 | rs4141214 | 0.642 | -0.963 | 0.444 | 1.04 | -0.001 | -0.059 | -0.059 | 0.802 | 0.28 | 0.08 | 0.22 | 0.26 | 0.27 | 0.31 |
| 7 | 149556518 | rs4725833 | -0.269 | -0.553 | 0.001 | 0.85 | -0.009 | -0.011 | -0.011 | 0 | 0.6 | 0.8 | 0.45 | 0.33 | 0.58 | 0.37 |
| 7 | 149560348 | rs10270385 | 0.078 | -0.652 | 0.767 | 0.945 | -0.001 | -0.002 | -0.002 | 3.231 | 0.56 | 0.52 | 0.49 | 0.33 | 0.57 | 0.33 |
| 7 | 149563894 | rs3823698 | 1.231 | -0.55 | 0.426 | 1.433 | -0.02 | -0.021 | -0.021 | 0 | 0.07 | 0.01 | 0.11 | 0.35 | 0.09 | 0.1 |
| 7 | 149564037 | rs11544265 | 0.919 | -0.494 | 0.792 | 0.599 | -0.015 | 0.014 | 0.014 | 0.456 | 0.23 | 0.15 | 0.21 | 0.19 | 0.27 | 0.1 |
| 7 | 149565489 | rs3735334 | -0.499 | -0.612 | 1.111 | 0.563 | 0.002 | -0.007 | -0.007 | 0 | 0.6 | 0.72 | 0.52 | 0.38 | 0.59 | 0.39 |
| 7 | 149567233 | rs13243007 | 0.093 | -0.476 | 0.406 | 0.535 | 0.025 | 0.072 | 0.072 | 1.6 | 0.52 | 0.24 | 0.47 | 0.32 | 0.57 | 0.34 |
| 7 | 149568903 | rs10952234 | 0.057 | -0.476 | 0.419 | 0.535 | 0.023 | 0.065 | 0.065 | 1.723 | 0.53 | 0.28 | 0.47 | 0.32 | 0.57 | 0.34 |
| 7 | 149577591 | rs1133480 | 0.052 | -0.452 | 0.32 | 0.548 | 0.022 | 0.059 | 0.059 | 1.434 | 0.53 | 0.31 | 0.48 | 0.32 | 0.57 | 0.34 |
| 7 | 149582906 | rs12703055 | 0.569 | -0.313 | 0.085 | 0.432 | -0.016 | 0.013 | 0.013 | 0.253 | 0.24 | 0.18 | 0.23 | 0.2 | 0.28 | 0.1 |
| 7 | 149583170 | rs10807976 | -0.583 | -0.378 | -0.129 | 0.554 | -0.002 | -0.022 | -0.022 | 0.078 | 0.61 | 0.77 | 0.53 | 0.4 | 0.6 | 0.42 |
| 7 | 149583591 | rs12540296 | -0.118 | -0.346 | -0.137 | 0.52 | 0.014 | -0.007 | -0.007 | 3.02 | 0.57 | 0.74 | 0.54 | 0.39 | 0.6 | 0.43 |
| 7 | 149585379 | rs6954118 | -1.189 | -0.148 | -0.204 | 0.12 | -0.008 | -0.03 | -0.03 | 0.159 | 0.71 | 0.68 | 0.73 | 0.89 | 0.73 | 0.73 |
| 7 | 149585639 | rs10952235 | 0.782 | 0.395 | -0.601 | 0.395 | -0.096 | -0.041 | -0.041 | 0 | 0.16 | 0.02 | 0.13 | 0.21 | 0.24 | 0.09 |
| 7 | 149596182 | rs7793500 | 0.367 | -0.378 | -0.021 | 0.559 | -0.006 | 0.008 | 0.008 | 0.003 | 0.22 | 0.1 | 0.21 | 0.16 | 0.2 | 0.43 |
| 7 | 149599046 | rs10952238 | 1.257 | -0.63 | 0.892 | 0.562 | 0.007 | -0.002 | -0.002 | 0 | 0.93 | 0.66 | 0.96 | 1 | 0.99 | 1 |

| | | | | | | | | | | | | | | | | |
|---|---|---|---|---|---|---|---|---|---|---|---|---|---|---|---|---|
| 7 | 149630727 | rs11770881 | 0.255 | -0.408 | -0.016 | 0.635 | -0.006 | 0.087 | 0.087 | 0 | 0.48 | 0.19 | 0.47 | 0.29 | 0.52 | 0.42 |
| 7 | 149639889 | rs13438249 | 0.005 | -0.582 | 0.643 | 1.255 | -0.039 | -0.021 | -0.021 | 0.293 | 0.24 | 0.59 | 0.19 | 0.22 | 0.11 | 0.15 |
| 7 | 149642019 | rs1872176 | -0.173 | 0.249 | 1.262 | 1.33 | -0.002 | 0.015 | 0.015 | 0 | 0.3 | 0.19 | 0.28 | 0.2 | 0.34 | 0.4 |
| 7 | 149678982 | rs11981739 | -1.59 | -0.313 | 1.573 | 1.332 | 0.032 | 0.289 | 0.289 | 3.768 | 0.09 | 0.43 | 0.1 | 0.26 | 0.04 | 0.06 |
| 7 | 149706925 | rs3864474 | -0.116 | 0.255 | 0.438 | 0.693 | -0.027 | 0.061 | 0.061 | 0.066 | 0.66 | 0.75 | 0.58 | 0.88 | 0.55 | 0.62 |
| 7 | 149707006 | rs3912373 | 0.721 | -0.008 | 0.297 | 0.961 | -0.001 | -0.004 | -0.004 | 0 | 0.42 | 0.3 | 0.33 | 0.46 | 0.38 | 0.36 |
| 7 | 149711187 | rs6944359 | -0.312 | 0.498 | 0.223 | 1.069 | -0.003 | 0.204 | 0.204 | 2.438 | 0.05 | 0.23 | 0.07 | 0.24 | 0.02 | 0.02 |
| 7 | 149711224 | rs2450027 | -0.358 | 0.909 | -0.668 | 0.163 | -0.077 | 0.086 | 0.086 | 0.289 | 0.33 | 0.17 | 0.42 | 0.17 | 0.45 | 0.39 |
| 7 | 149756192 | rs6958741 | -1.164 | 0.967 | 0.088 | 0.646 | 0.006 | 0.012 | 0.012 | 0.595 | 0.11 | 0.15 | 0.09 | 0.14 | 0.15 | 0.26 |
| 7 | 149787501 | rs2373674 | 0.037 | 1.289 | 0.033 | 0.508 | -0.008 | 0.025 | 0.025 | 0.437 | 0.58 | 0.31 | 0.63 | 0.79 | 0.58 | 0.69 |
| 7 | 149806076 | rs17173853 | -1.394 | 1.412 | 0.583 | 0.936 | 0.078 | 0.07 | 0.07 | 0.75 | 0.06 | 0.23 | 0.08 | 0.12 | 0.13 | 0.09 |
| 7 | 149866114 | rs6464052 | 0.767 | 1.309 | -0.186 | 0.565 | -0.012 | 0.007 | 0.007 | 0 | 0.78 | 0.84 | 0.46 | 0.84 | 0.67 | 0.8 |
| 7 | 149900486 | rs11975282 | -1.611 | 0.089 | 0.766 | 0.793 | -0.045 | 0.048 | 0.048 | 0 | 0.18 | 0.52 | 0.17 | 0.22 | 0.1 | 0.16 |
| 7 | 149925431 | rs7787192 | -0.824 | 0.1 | 0.666 | 0.885 | -0.004 | -0.03 | -0.03 | 0 | 0.05 | 0 | 0.05 | 0.12 | 0.06 | 0.08 |
| 7 | 149934267 | rs4591941 | -1.835 | 0.026 | 1.561 | 0.867 | -0.009 | 0.088 | 0.088 | 0 | 0.1 | 0.43 | 0.09 | 0.12 | 0.06 | 0.08 |
| 7 | 149937532 | rs6943821 | -1.503 | 0.037 | 1.782 | 0.899 | -0.011 | 0.062 | 0.062 | 0 | 0.08 | 0.27 | 0.07 | 0.12 | 0.06 | 0.08 |
| 7 | 149937855 | rs10273192 | -0.21 | -0.095 | 0.838 | 0.766 | -0.005 | 0.004 | 0.004 | 0 | 0.25 | 0.16 | 0.27 | 0.17 | 0.25 | 0.26 |
| 7 | 149948084 | rs10488347 | -1.9 | -0.564 | 1.401 | 0.75 | -0.021 | 0.098 | 0.098 | 0 | 0.14 | 0.49 | 0.15 | 0.19 | 0.09 | 0.13 |
| 7 | 149977731 | rs10262634 | -1.133 | -1.771 | 1.025 | 0.275 | -0.101 | -0.031 | -0.031 | 0 | 0.25 | 0.41 | 0.09 | 0.2 | 0.1 | 0.37 |
| 7 | 149983201 | rs2286450 | -1.25 | -1.817 | 1.281 | 0.255 | -0.092 | -0.029 | -0.029 | 0.094 | 0.26 | 0.53 | 0.1 | 0.2 | 0.1 | 0.35 |
| 7 | 150017408 | rs2108854 | -0.333 | -1.741 | 0.918 | 0.223 | 0.003 | -0.013 | -0.013 | 0 | 0.26 | 0.22 | 0.18 | 0.4 | 0.27 | 0.28 |
| 7 | 150020295 | rs998920 | -0.973 | -1.391 | 0.036 | -0.64 | -0.065 | 0.102 | 0.102 | 7.314 | 0.84 | 0.99 | 0.87 | 1 | 0.8 | 0.97 |
| 7 | 150025915 | rs11764936 | 0.379 | -1.544 | 0.382 | -0.42 | 0 | -0.001 | -0.001 | 0.489 | 0.22 | 0.22 | 0.15 | 0.29 | 0.24 | 0.29 |
| 7 | 150025966 | rs10952250 | -0.305 | -1.676 | 0.575 | -0.478 | -0.024 | 0.044 | 0.044 | 0 | 0.28 | 0.45 | 0.18 | 0.4 | 0.26 | 0.31 |
| 7 | 150028958 | rs3735172 | -0.391 | -1.751 | 0.542 | -0.482 | -0.028 | 0.038 | 0.038 | 0.121 | 0.29 | 0.44 | 0.18 | 0.41 | 0.26 | 0.33 |
| 7 | 150034184 | rs3735170 | 0.178 | -1.673 | 0.468 | -0.462 | -0.005 | 0.002 | 0.002 | 0.018 | 0.24 | 0.26 | 0.16 | 0.29 | 0.25 | 0.31 |
| 7 | 150036664 | rs17173608 | 0.238 | -2.005 | 0.344 | -0.619 | -0.033 | -0.04 | -0.04 | 0.22 | 0.13 | 0.2 | 0.05 | 0.06 | 0.05 | 0.14 |
| 7 | 150039555 | rs3735167 | 0.67 | -2.132 | 0.033 | -0.624 | -0.056 | -0.063 | -0.063 | 0.832 | 0.22 | 0.02 | 0.14 | 0.28 | 0.26 | 0.3 |
| 7 | 150041610 | rs10244748 | -0.637 | -2.186 | 0.128 | -0.645 | 0.004 | 0 | 0 | 0 | 0.16 | 0.15 | 0.07 | 0.2 | 0.08 | 0.15 |
| 7 | 150045302 | rs10282458 | 0.215 | -2.166 | 0.373 | -0.614 | 0.001 | -0.001 | -0.001 | 0 | 0.25 | 0.26 | 0.16 | 0.29 | 0.25 | 0.34 |
| 7 | 150050427 | rs10259796 | 0.316 | -2.324 | 0.055 | -0.657 | -0.016 | -0.018 | -0.018 | 0 | 0.25 | 0.09 | 0.15 | 0.28 | 0.27 | 0.39 |
| 7 | 150060163 | rs10263613 | -0.956 | -2.16 | 0.416 | -0.706 | -0.02 | -0.009 | -0.009 | 0 | 0.06 | 0.26 | 0.04 | 0.04 | 0.02 | 0.06 |
| 7 | 150067445 | rs4725336 | -1.303 | -1.818 | -0.148 | -0.818 | -0.05 | 0.056 | 0.056 | 0.39 | 0.9 | 0.99 | 0.89 | 1 | 0.84 | 0.97 |
| 7 | 150068933 | rs6722 | -0.697 | -2.75 | 0.196 | -0.921 | -0.014 | -0.003 | -0.003 | 0 | 0.43 | 0.49 | 0.22 | 0.47 | 0.32 | 0.5 |
| 7 | 150070630 | rs1051760 | -0.826 | -2.752 | 0.13 | -0.922 | 0.007 | 0.003 | 0.003 | 0.346 | 0.44 | 0.55 | 0.22 | 0.47 | 0.32 | 0.5 |
| 7 | 150074994 | rs6971465 | -0.44 | -2.375 | 0.348 | -0.598 | 0.005 | -0.003 | -0.003 | 0.04 | 0.37 | 0.6 | 0.26 | 0.46 | 0.31 | 0.45 |
| 7 | 150079459 | rs11981330 | -0.068 | -2.393 | 0.145 | -0.633 | 0.008 | -0.026 | -0.026 | 0 | 0.36 | 0.2 | 0.27 | 0.47 | 0.34 | 0.46 |
| 7 | 150081485 | rs7795021 | -1.434 | -2.248 | 0.888 | -0.473 | -0.017 | 0.002 | 0.002 | 0.249 | 0.11 | 0.38 | 0.08 | 0.06 | 0.07 | 0.09 |
| 7 | 150084460 | rs7810240 | 0.664 | -2.042 | 0.041 | -0.28 | 0.009 | 0.003 | 0.003 | 1.871 | 0.36 | 0.39 | 0.56 | 0.31 | 0.46 | 0.41 |
| 7 | 150092031 | rs1880316 | 0.617 | -1.957 | -0.102 | -0.245 | 0.007 | 0.003 | 0.003 | 1.429 | 0.37 | 0.42 | 0.56 | 0.31 | 0.45 | 0.4 |
| 7 | 150095271 | rs7393 | -0.137 | -1.448 | -0.714 | 0.004 | -0.007 | -0.001 | -0.001 | 1.168 | 0.43 | 0.28 | 0.32 | 0.47 | 0.44 | 0.4 |
| 7 | 150103137 | rs3800784 | -0.003 | -1.273 | -1.511 | 0.019 | -0.153 | -0.157 | -0.157 | 0 | 0.4 | 0.02 | 0.26 | 0.57 | 0.43 | 0.38 |
| 7 | 150115183 | rs4725896 | 0.331 | -0.429 | -0.747 | 0.295 | 0.001 | -0.002 | -0.002 | 0 | 0.7 | 0.35 | 0.63 | 0.61 | 0.75 | 0.62 |
| 7 | 150131997 | rs12531132 | 0.525 | -0.04 | -0.917 | -0.293 | -0.029 | 0.077 | 0.077 | 0 | 0.37 | 0.21 | 0.38 | 0.23 | 0.38 | 0.36 |
| 7 | 150135758 | rs1916931 | -0.341 | 0.061 | -0.723 | 0.061 | 0.015 | -0.017 | -0.017 | 0 | 0.34 | 0.28 | 0.36 | 0.38 | 0.3 | 0.43 |
| 7 | 150136323 | rs7777181 | -0.643 | -0.028 | -0.5 | -0.443 | 0 | 0.001 | 0.001 | 0 | 0.59 | 0.47 | 0.6 | 0.78 | 0.66 | 0.67 |
| 7 | 150140762 | rs7806921 | -0.376 | 0.07 | -0.233 | -0.349 | -0.009 | 0.056 | 0.056 | 0 | 0.46 | 0.63 | 0.54 | 0.78 | 0.55 | 0.55 |
| 7 | 150141809 | rs1104881 | 0.05 | 0.442 | 0.264 | 0.135 | 0.056 | 0.1 | 0.1 | 0 | 0.22 | 0.46 | 0.24 | 0.38 | 0.32 | 0.18 |
| 7 | 150146181 | rs13222973 | 0.101 | 0.131 | -0.123 | -0.104 | 0.005 | 0 | 0 | 0 | 0.08 | 0.04 | 0.03 | 0 | 0.06 | 0.06 |
| 7 | 150152520 | rs4146035 | 0.484 | -0.026 | 0.037 | -0.032 | 0 | -0.004 | -0.004 | 1.332 | 0.32 | 0.26 | 0.22 | 0.41 | 0.35 | 0.27 |
| 7 | 150153813 | rs7810209 | -0.084 | 0.388 | 0.197 | -0.266 | 0.005 | 0.001 | 0.001 | 0 | 0.46 | 0.41 | 0.41 | 0.44 | 0.58 | 0.39 |
| 7 | 150160458 | rs11769211 | 0.96 | 0.389 | 0.124 | -0.118 | -0.022 | -0.07 | -0.07 | 0.086 | 0.3 | 0.14 | 0.22 | 0.42 | 0.37 | 0.22 |
| 7 | 150163837 | rs2140595 | 0.7 | 0.443 | 0.291 | -0.094 | -0.001 | 0.004 | 0.004 | 0 | 0.33 | 0.31 | 0.23 | 0.42 | 0.37 | 0.25 |
| 7 | 150167583 | rs6969414 | 0.684 | 0.422 | 0.282 | -0.104 | -0.001 | 0.003 | 0.003 | 0 | 0.33 | 0.31 | 0.23 | 0.42 | 0.37 | 0.26 |
| 7 | 150170776 | rs6947973 | 0.782 | 0.444 | 0.42 | -0.094 | 0 | 0.018 | 0.018 | 0 | 0.32 | 0.31 | 0.23 | 0.42 | 0.37 | 0.25 |
| 7 | 150176933 | rs7776885 | 0.15 | 0.034 | 0.267 | 0.005 | -0.021 | -0.002 | -0.002 | 0.019 | 0.09 | 0.13 | 0.09 | 0.11 | 0.04 | 0.1 |
| 7 | 150176946 | rs7776515 | -0.283 | 0.102 | 0.199 | -0.17 | 0.004 | -0.026 | -0.026 | 0 | 0.64 | 0.69 | 0.55 | 0.54 | 0.69 | 0.54 |
| 7 | 150177524 | rs2091143 | 0.562 | -0.216 | 0.406 | 0.153 | -0.002 | -0.001 | -0.001 | 0 | 0.46 | 0.44 | 0.35 | 0.48 | 0.45 | 0.35 |
| 7 | 150182338 | rs10266589 | -0.555 | -0.59 | 0.474 | 0.041 | -0.05 | -0.058 | -0.058 | 0 | 0.16 | 0.34 | 0.07 | 0.09 | 0.08 | 0.16 |
| 7 | 150183718 | rs10226213 | 0.149 | -0.098 | -0.018 | 0.596 | -0.002 | -0.002 | -0.002 | 5.069 | 0.25 | 0.3 | 0.41 | 0.34 | 0.31 | 0.24 |
| 7 | 150184564 | rs7789612 | 0.09 | -0.149 | 0.049 | 0.576 | -0.032 | -0.024 | -0.024 | 0.56 | 0.11 | 0.22 | 0.05 | 0.08 | 0.05 | 0.14 |
| 7 | 150187245 | rs2222524 | 0 | 0.41 | -0.373 | 0.915 | 0.007 | 0.009 | 0.009 | 0 | 0.25 | 0.35 | 0.42 | 0.34 | 0.31 | 0.27 |
| 7 | 150200185 | rs7796182 | 0.183 | 0.39 | -0.657 | 1.01 | -0.018 | -0.05 | -0.05 | 1.207 | 0.78 | 0.95 | 0.84 | 0.76 | 0.82 | 0.65 |
| 7 | 150206362 | rs4272282 | 0.836 | -0.107 | -0.021 | 0.666 | 0.005 | 0.024 | 0.024 | 0 | 0.54 | 0.28 | 0.42 | 0.25 | 0.58 | 0.5 |
| 7 | 150208112 | rs4342497 | 0.731 | -0.247 | 0.451 | 0.67 | 0 | -0.004 | -0.004 | 0 | 0.6 | 0.53 | 0.43 | 0.31 | 0.58 | 0.54 |
| 7 | 150211193 | rs6974067 | 0.657 | 0.091 | 0.221 | 0.304 | 0.016 | -0.062 | -0.062 | 0 | 0.19 | 0.18 | 0.13 | 0.02 | 0.2 | 0.12 |
| 7 | 150212654 | rs4725906 | 0.378 | 0.301 | 0.091 | 0.583 | -0.013 | -0.012 | -0.012 | 0 | 0.09 | 0.05 | 0.11 | 0.18 | 0.13 | 0.1 |
| 7 | 150213019 | rs10270302 | 0.637 | 0.256 | 0.19 | 0.524 | 0.027 | -0.087 | -0.087 | 0 | 0.19 | 0.22 | 0.14 | 0.02 | 0.21 | 0.12 |
| 7 | 150213314 | rs7784050 | -0.52 | 0.122 | 0.363 | 0.961 | -0.001 | 0.008 | 0.008 | 0.155 | 0.77 | 0.8 | 0.64 | 0.84 | 0.77 | 0.77 |

| | | | | | | | | | | | | | | | | |
|---|---|---|---|---|---|---|---|---|---|---|---|---|---|---|---|---|
| 7 | 150217089 | rs7800829 | -0.947 | -0.034 | 0.311 | 1.168 | -0.047 | 0.241 | 0.241 | 2.143 | 0.14 | 0.49 | 0.11 | 0.38 | 0.07 | 0.17 |
| 7 | 150217248 | rs3735081 | -2.252 | 0.094 | -0.276 | 1.166 | -0.059 | -0.027 | -0.027 | 0 | 0.67 | 0.97 | 0.73 | 0.74 | 0.63 | 0.76 |
| 7 | 150217309 | rs3735080 | 0.712 | 0.077 | -0.052 | 1.117 | -0.066 | -0.021 | -0.021 | 0.053 | 0.19 | 0.01 | 0.33 | 0.16 | 0.22 | 0.22 |
| 7 | 150217335 | rs3735079 | 0.055 | 0.146 | 0.136 | 1.042 | -0.017 | -0.016 | -0.016 | 0 | 0.09 | 0.04 | 0.11 | 0.19 | 0.13 | 0.1 |
| 7 | 150235664 | rs1522597 | -0.542 | 0.2 | 0.523 | 1.146 | -0.003 | -0.004 | -0.004 | 0 | 0.46 | 0.55 | 0.54 | 0.49 | 0.45 | 0.45 |
| 7 | 150235783 | rs1522596 | -0.507 | 0.194 | 0.67 | 1.143 | 0 | -0.001 | -0.001 | 0 | 0.46 | 0.46 | 0.53 | 0.49 | 0.45 | 0.45 |
| 7 | 150238170 | rs2090954 | -0.572 | 0.193 | 0.622 | 1.142 | -0.002 | -0.004 | -0.004 | 0.115 | 0.46 | 0.55 | 0.54 | 0.49 | 0.45 | 0.45 |
| 7 | 150239113 | rs1568629 | -1.425 | 0.183 | 0.591 | 1.097 | 0.006 | 0.03 | 0.03 | 1.706 | 0.17 | 0.39 | 0.17 | 0.27 | 0.15 | 0.3 |
| 7 | 150239676 | rs1568627 | -0.967 | 0.196 | 0.641 | 1.103 | 0.01 | 0.027 | 0.027 | 0.73 | 0.15 | 0.32 | 0.15 | 0.27 | 0.15 | 0.17 |
| 7 | 150251708 | rs11768988 | 0.145 | 0.232 | 0.398 | 1.106 | -0.033 | -0.03 | -0.03 | 0 | 0.1 | 0.02 | 0.14 | 0.22 | 0.15 | 0.13 |
| 7 | 150253913 | rs1403222 | -1.736 | 0.163 | 0.428 | 1.066 | 0.01 | 0.052 | 0.052 | 0.198 | 0.61 | 0.83 | 0.69 | 0.75 | 0.6 | 0.61 |
| 7 | 150257768 | rs939958 | -1.824 | 0.166 | 0.394 | 1.054 | -0.001 | 0.053 | 0.053 | 0.247 | 0.62 | 0.88 | 0.7 | 0.75 | 0.6 | 0.61 |
| 7 | 150260388 | rs6464080 | -2.01 | 0.162 | 0.358 | 1.052 | -0.036 | 0.03 | 0.03 | 0.529 | 0.63 | 0.93 | 0.71 | 0.75 | 0.6 | 0.76 |
| 7 | 150261328 | rs13231883 | 0.116 | 0.248 | 0.543 | 1.097 | 0.003 | 0.002 | 0.002 | 0 | 0.1 | 0.11 | 0.15 | 0.22 | 0.15 | 0.13 |
| 7 | 150262436 | rs6951484 | -0.475 | 0.205 | 0.877 | 1.16 | -0.017 | -0.004 | -0.004 | 0.498 | 0.31 | 0.21 | 0.49 | 0.41 | 0.38 | 0.35 |
| 7 | 150263323 | rs2177189 | -0.433 | 0.654 | 0.923 | 1.263 | -0.045 | -0.03 | -0.03 | 0 | 0.12 | 0.02 | 0.17 | 0.22 | 0.18 | 0.15 |
| 7 | 150263761 | rs2177188 | -1.246 | 0.388 | 1.278 | 1.311 | 0.029 | 0.057 | 0.057 | 1.108 | 0.16 | 0.46 | 0.17 | 0.27 | 0.16 | 0.17 |
| 7 | 150264603 | rs6464084 | -0.795 | 0.392 | 1.288 | 1.308 | 0.001 | 0.051 | 0.051 | 3.16 | 0.06 | 0.34 | 0.02 | 0.05 | 0 | 0.04 |
| 7 | 150266835 | rs11769703 | 0.672 | 0.335 | 1.215 | 1.275 | -0.01 | 0.003 | 0.003 | 0.657 | 0.19 | 0.09 | 0.34 | 0.2 | 0.22 | 0.23 |
| 7 | 150268098 | rs9640279 | 0.218 | 0.317 | 1.205 | 1.303 | -0.001 | 0.028 | 0.028 | 0.334 | 0.25 | 0.31 | 0.18 | 0.29 | 0.22 | 0.22 |
| 7 | 150269542 | rs2293172 | 0.032 | 0.381 | 1.1 | 0.982 | -0.014 | -0.013 | -0.013 | 0 | 0.1 | 0.07 | 0.14 | 0.22 | 0.16 | 0.12 |
| 7 | 150279532 | rs11974030 | -0.867 | 0.31 | 1.377 | 0.939 | -0.003 | 0.027 | 0.027 | 2.878 | 0.06 | 0.31 | 0.02 | 0.04 | 0 | 0.03 |
| 7 | 150281362 | rs1960024 | -1.156 | 0.298 | 1.411 | 1.228 | -0.005 | -0.025 | -0.025 | 0 | 0.41 | 0.73 | 0.34 | 0.52 | 0.38 | 0.37 |
| 7 | 150284200 | rs6950421 | -0.105 | 0.68 | 1.533 | 1.533 | 0.002 | -0.005 | -0.005 | 0 | 0.22 | 0.33 | 0.36 | 0.22 | 0.22 | 0.25 |
| 7 | 150292545 | rs6969250 | -0.365 | 0.68 | 1.459 | 1.534 | -0.001 | -0.012 | -0.012 | 0.13 | 0.24 | 0.46 | 0.37 | 0.22 | 0.22 | 0.25 |
| 7 | 150292701 | rs10240209 | -0.053 | 0.986 | 1.509 | 1.736 | -0.002 | 0.016 | 0.016 | 0.494 | 0.25 | 0.3 | 0.17 | 0.26 | 0.22 | 0.22 |
| 7 | 150300735 | rs11984138 | -0.033 | 0.988 | 1.509 | 1.737 | -0.001 | 0.018 | 0.018 | 0.563 | 0.24 | 0.3 | 0.17 | 0.26 | 0.22 | 0.22 |
| 7 | 150317472 | rs10248553 | 0.287 | 0.995 | 1.791 | 1.724 | 0 | 0.002 | 0.002 | 0.402 | 0.23 | 0.2 | 0.17 | 0.25 | 0.23 | 0.16 |
| 7 | 150319290 | rs6952669 | -1.629 | 1.015 | 1.615 | 1.639 | -0.075 | -0.044 | -0.044 | 0 | 0.53 | 0.93 | 0.62 | 0.52 | 0.51 | 0.65 |
| 7 | 150321507 | rs10280903 | -1.093 | 1.041 | 1.844 | 1.662 | 0 | 0 | 0 | 0 | 0.46 | 0.41 | 0.57 | 0.49 | 0.51 | 0.62 |
| 7 | 150323647 | rs3735086 | 1.034 | 1.051 | 1.835 | 1.664 | -0.017 | -0.005 | -0.005 | 0.743 | 0.15 | 0.06 | 0.31 | 0.19 | 0.19 | 0.21 |
| 7 | 150323805 | rs3735085 | -0.24 | 1.054 | 1.841 | 1.665 | 0 | 0.005 | 0.005 | 2.2 | 0.3 | 0.29 | 0.25 | 0.29 | 0.31 | 0.27 |
| 7 | 150324157 | rs11762727 | 0.348 | 1.115 | 1.784 | 1.666 | -0.023 | -0.022 | -0.022 | 0 | 0.07 | 0.02 | 0.13 | 0.21 | 0.15 | 0.11 |
| 7 | 150324570 | rs10235151 | -0.367 | 0.823 | 1.782 | 1.657 | 0 | 0.004 | 0.004 | 2.152 | 0.3 | 0.28 | 0.25 | 0.3 | 0.31 | 0.27 |
| 7 | 150326819 | rs17542143 | 1.017 | 0.802 | 1.559 | 1.507 | -0.017 | -0.005 | -0.005 | 0 | 0.15 | 0.07 | 0.31 | 0.19 | 0.19 | 0.21 |
| 7 | 150327024 | rs3735082 | -0.791 | 0.841 | 1.514 | 1.531 | -0.004 | -0.006 | -0.006 | 0.014 | 0.37 | 0.32 | 0.37 | 0.5 | 0.44 | 0.39 |
| 7 | 150328326 | rs4725925 | -1.78 | 0.751 | 0.854 | 1.399 | 0.051 | 0.062 | 0.062 | 2.019 | 0.58 | 0.78 | 0.72 | 0.73 | 0.65 | 0.62 |
| 7 | 150343304 | rs6948827 | -0.845 | 0.811 | 1.022 | 1.356 | -0.004 | -0.001 | -0.001 | 1.698 | 0.35 | 0.59 | 0.28 | 0.31 | 0.34 | 0.44 |
| 7 | 150345692 | rs11766040 | -0.834 | 0.811 | 1.022 | 1.356 | -0.004 | -0.001 | -0.001 | 0 | 0.35 | 0.59 | 0.28 | 0.31 | 0.34 | 0.44 |
| 7 | 150345987 | rs1879868 | -0.834 | 0.811 | 1.022 | 1.356 | -0.004 | -0.001 | -0.001 | 0 | 0.35 | 0.59 | 0.28 | 0.31 | 0.34 | 0.44 |
| 7 | 150349937 | rs10266069 | -1.24 | 0.894 | 1.007 | 1.39 | 0.001 | -0.005 | -0.005 | 0.011 | 0.5 | 0.67 | 0.59 | 0.5 | 0.51 | 0.64 |
| 7 | 150350975 | rs10256482 | -1.046 | 0.88 | 1.228 | 1.39 | -0.001 | -0.004 | -0.004 | 0 | 0.44 | 0.55 | 0.52 | 0.48 | 0.4 | 0.54 |
| 7 | 150352887 | rs2140144 | -1.884 | 0.935 | 1.254 | 1.405 | 0.042 | 0.027 | 0.027 | 2.423 | 0.58 | 0.74 | 0.72 | 0.68 | 0.65 | 0.74 |
| 7 | 150353227 | rs6970218 | -1.023 | 0.864 | 1.251 | 1.392 | -0.001 | -0.003 | -0.003 | 0 | 0.44 | 0.55 | 0.52 | 0.48 | 0.4 | 0.54 |
| 7 | 150363735 | rs16884586 | -0.951 | 0.679 | 1.219 | 1.438 | 0.003 | 0.001 | 0.001 | 0 | 0.14 | 0.21 | 0.15 | 0.18 | 0.2 | 0.2 |
| 7 | 150369447 | rs10275417 | 0.13 | 0.561 | 1.185 | 1.45 | -0.009 | 0.026 | 0.026 | 0.721 | 0.26 | 0.41 | 0.19 | 0.29 | 0.23 | 0.17 |
| 7 | 150372452 | rs3936752 | -1.674 | 0.535 | 0.612 | 1.329 | 0.04 | 0.012 | 0.012 | 2.315 | 0.58 | 0.78 | 0.7 | 0.66 | 0.65 | 0.74 |
| 7 | 150381031 | rs1234358 | 1.58 | 0.539 | 0.573 | 1.255 | 0.04 | 0.026 | 0.026 | 2.18 | 0.42 | 0.22 | 0.28 | 0.32 | 0.35 | 0.26 |
| 7 | 150385900 | rs1234354 | 1.755 | 0.55 | 0.404 | 1.148 | 0.027 | -0.01 | -0.01 | 0 | 0.4 | 0.13 | 0.26 | 0.33 | 0.36 | 0.22 |
| 7 | 150389593 | rs1860871 | 0.48 | 0.757 | 0.399 | 1.23 | -0.014 | -0.015 | -0.015 | 0 | 0.06 | 0.03 | 0.08 | 0.22 | 0.09 | 0.09 |
| 7 | 150389856 | rs2075078 | -1.522 | 0.688 | 1.102 | 1.117 | 0.019 | 0.05 | 0.05 | 0.629 | 0.17 | 0.44 | 0.19 | 0.24 | 0.2 | 0.23 |
| 7 | 150390504 | rs1055240 | -1.262 | 0.713 | 0.956 | 1.166 | 0.001 | -0.006 | -0.006 | 1.191 | 0.41 | 0.65 | 0.4 | 0.51 | 0.42 | 0.54 |
| 7 | 150396134 | rs4725356 | -0.782 | 0.783 | 0.694 | 1.069 | -0.011 | 0 | 0 | 2.175 | 0.34 | 0.49 | 0.31 | 0.29 | 0.33 | 0.29 |
| 7 | 150397117 | rs6951867 | -0.917 | 1.045 | -0.122 | 1.091 | -0.091 | -0.028 | -0.028 | 0 | 0.85 | 0.99 | 0.65 | 0.84 | 0.78 | 0.8 |
| 7 | 150412100 | rs1016193 | -0.126 | 0.501 | 0.342 | 1.094 | 0.001 | -0.008 | -0.008 | 0 | 0.24 | 0.17 | 0.18 | 0.26 | 0.22 | 0.18 |
| 7 | 150412317 | rs10488372 | -0.784 | 0.552 | 0.627 | 1.235 | 0 | 0.005 | 0.005 | 1.256 | 0.31 | 0.29 | 0.27 | 0.48 | 0.32 | 0.27 |
| 7 | 150414572 | rs6464094 | -1.681 | 0.322 | 0.399 | 1.096 | -0.018 | -0.085 | -0.085 | 0.023 | 0.45 | 0.92 | 0.44 | 0.52 | 0.42 | 0.57 |
| 7 | 150418045 | rs2286900 | 0.137 | 0.613 | 0.659 | 0.92 | 0.011 | 0.046 | 0.046 | 0 | 0.07 | 0.12 | 0.09 | 0.21 | 0.09 | 0.09 |
| 7 | 150420704 | rs3735078 | -0.185 | 0.603 | 0.771 | 0.916 | 0.014 | 0.058 | 0.058 | 0.001 | 0.07 | 0.17 | 0.1 | 0.21 | 0.09 | 0.09 |
| 7 | 150424509 | rs6978377 | -2.071 | 0.549 | 1.045 | 0.96 | -0.009 | 0.017 | 0.017 | 1.355 | 0.2 | 0.7 | 0.24 | 0.31 | 0.2 | 0.4 |
| 7 | 150425351 | rs6945478 | 0.112 | 0.655 | 0.777 | 0.989 | 0.011 | 0.046 | 0.046 | 0 | 0.07 | 0.12 | 0.09 | 0.21 | 0.09 | 0.09 |
| 7 | 150428109 | rs11772925 | 0.71 | 0.581 | 0.609 | 0.832 | -0.022 | 0.042 | 0.042 | 0.079 | 0.15 | 0.05 | 0.1 | 0 | 0.16 | 0.06 |
| 7 | 150428885 | rs9690530 | -0.877 | 0.516 | 1.176 | 1.019 | 0.044 | 0.127 | 0.127 | 1.085 | 0.09 | 0.38 | 0.11 | 0.21 | 0.09 | 0.09 |
| 7 | 150430711 | rs1860870 | -1.563 | 0.377 | 0.605 | 0.834 | 0.003 | 0.026 | 0.026 | 2.771 | 0.42 | 0.8 | 0.4 | 0.37 | 0.43 | 0.53 |
| 7 | 150434158 | rs3807383 | -0.507 | 0.362 | 1.178 | 0.779 | -0.006 | -0.108 | -0.108 | 4.968 | 0.29 | 0.41 | 0.26 | 0.08 | 0.29 | 0.23 |
| 7 | 150434472 | rs6965571 | 0.434 | 0.347 | 1.258 | 0.738 | -0.006 | -0.089 | -0.089 | 1.04 | 0.2 | 0.22 | 0.12 | 0.01 | 0.17 | 0.11 |
| 7 | 150437087 | rs11973400 | 0.357 | 0.35 | 1.232 | 0.739 | -0.008 | -0.106 | -0.106 | 0 | 0.21 | 0.26 | 0.12 | 0.01 | 0.17 | 0.11 |
| 7 | 150438558 | rs9657886 | -0.62 | 0.378 | 1.217 | 0.789 | 0.005 | -0.004 | -0.004 | 0 | 0.29 | 0.32 | 0.24 | 0.28 | 0.31 | 0.23 |
| 7 | 150439500 | rs759011 | -1.072 | 0.424 | 0.889 | 0.811 | -0.004 | -0.008 | -0.008 | 0.197 | 0.33 | 0.57 | 0.27 | 0.3 | 0.31 | 0.41 |

| | | | | | | | | | | | | | | | | |
|---|---|---|---|---|---|---|---|---|---|---|---|---|---|---|---|---|
| 7 | 150440043 | rs1046355 | -0.94 | 0.448 | 0.886 | 0.822 | -0.004 | -0.009 | -0.009 | 0.553 | 0.32 | 0.54 | 0.26 | 0.3 | 0.31 | 0.26 |
| 7 | 150441163 | rs9657901 | -0.097 | 0.36 | 0.322 | 1.102 | 0.013 | 0.011 | 0.011 | 0 | 0.08 | 0.2 | 0.14 | 0.08 | 0.13 | 0.11 |
| 7 | 150447027 | rs28505524 | -0.245 | 0.361 | -0.103 | 1.155 | 0.02 | 0.076 | 0.076 | 0 | 0.09 | 0.15 | 0.13 | 0.26 | 0.14 | 0.12 |
| 7 | 150447194 | rs28418764 | -0.601 | 0.3 | -0.129 | 1.128 | 0.012 | 0.058 | 0.058 | 0 | 0.1 | 0.15 | 0.13 | 0.26 | 0.14 | 0.12 |
| 7 | 150447249 | rs11767311 | 1.456 | 0.095 | -0.922 | 0.839 | -0.063 | -0.057 | -0.057 | 0 | 0.36 | 0.01 | 0.2 | 0.33 | 0.34 | 0.24 |
| 7 | 150455295 | rs10266177 | -0.605 | -0.341 | -0.297 | 0.917 | 0.008 | -0.001 | -0.001 | 1.049 | 0.42 | 0.54 | 0.64 | 0.64 | 0.44 | 0.39 |
| 7 | 150456598 | rs4725360 | 0.98 | -0.57 | -0.602 | 0.97 | 0.012 | -0.022 | -0.022 | 1.489 | 0.4 | 0.27 | 0.34 | 0.45 | 0.37 | 0.4 |
| 7 | 150462394 | rs10265670 | 1.022 | -0.552 | -0.594 | 0.937 | 0.01 | -0.02 | -0.02 | 0.367 | 0.4 | 0.27 | 0.34 | 0.46 | 0.37 | 0.4 |
| 7 | 150464171 | rs7803700 | -0.363 | -0.554 | -1.029 | 0.888 | -0.01 | -0.048 | -0.048 | 0 | 0.29 | 0.48 | 0.3 | 0.15 | 0.25 | 0.3 |
| 7 | 150467723 | rs4725363 | 1.131 | -0.603 | -1.071 | 0.878 | -0.008 | 0.009 | 0.009 | 0.112 | 0.41 | 0.43 | 0.35 | 0.46 | 0.37 | 0.41 |
| 7 | 150476888 | rs7806458 | -0.287 | -0.407 | -1.59 | 0.755 | -0.078 | -0.036 | -0.036 | 0.512 | 0.7 | 0.92 | 0.65 | 0.61 | 0.63 | 0.71 |
| 7 | 150477384 | rs17173596 | 0.963 | -0.09 | -1.363 | 0.97 | -0.063 | -0.02 | -0.02 | 0.014 | 0.36 | 0.01 | 0.21 | 0.27 | 0.36 | 0.4 |
| 7 | 150483512 | rs12669824 | 0.433 | 0.071 | -0.751 | 0.529 | 0 | 0.001 | 0.001 | 1.009 | 0.51 | 0.43 | 0.51 | 0.52 | 0.47 | 0.53 |
| 7 | 150485401 | rs10281675 | 0.123 | 0.847 | -1.223 | 0.564 | -0.105 | -0.045 | -0.045 | 0 | 0.81 | 0.99 | 0.7 | 0.78 | 0.71 | 0.78 |
| 7 | 150486169 | rs4326309 | -0.051 | 0.85 | -0.724 | 0.604 | 0 | 0 | 0 | 0 | 0.2 | 0.14 | 0.32 | 0.24 | 0.18 | 0.18 |
| 7 | 150488468 | rs2302481 | 0.663 | 0.123 | -0.465 | 0.21 | 0.013 | -0.005 | -0.005 | 0 | 0.11 | 0.19 | 0.11 | 0.09 | 0.17 | 0.07 |
| 7 | 150488616 | rs2302479 | -0.284 | -0.027 | -0.397 | 0.235 | -0.071 | -0.011 | -0.011 | 0.234 | 0.66 | 0.81 | 0.51 | 0.63 | 0.59 | 0.61 |
| 7 | 150490130 | rs10215721 | 0.667 | 0.491 | -0.63 | 0.405 | 0.013 | -0.006 | -0.006 | 0 | 0.11 | 0.17 | 0.12 | 0.08 | 0.17 | 0.07 |
| 7 | 150490238 | rs17256042 | 0.958 | 0.437 | -0.847 | 0.258 | -0.011 | 0.035 | 0.035 | 0 | 0.07 | 0.01 | 0.03 | 0 | 0.06 | 0.01 |
| 7 | 150490376 | rs2072443 | 0.584 | 0.398 | -0.342 | 0.508 | 0.037 | 0.008 | 0.008 | 0 | 0.48 | 0.29 | 0.34 | 0.53 | 0.42 | 0.48 |
| 7 | 150491084 | rs3173833 | 0.253 | 0.349 | -0.56 | 0.525 | 0.008 | -0.007 | -0.007 | 0 | 0.59 | 0.5 | 0.46 | 0.61 | 0.58 | 0.55 |
| 7 | 150496104 | rs10230736 | -0.63 | -0.385 | -0.935 | 0.831 | 0.014 | -0.009 | -0.009 | 0 | 0.26 | 0.22 | 0.31 | 0.34 | 0.21 | 0.29 |
| 7 | 150496668 | rs11972506 | -0.213 | -1.303 | -1.753 | 0.399 | 0.041 | 0.068 | 0.068 | 0 | 0.43 | 0.19 | 0.54 | 0.74 | 0.36 | 0.61 |
| 7 | 150500455 | rs741067 | -0.2 | -0.302 | -2.151 | -0.342 | -0.013 | 0.004 | 0.004 | 0 | 0.76 | 0.79 | 0.65 | 0.76 | 0.68 | 0.79 |
| 7 | 150501245 | rs741061 | 0.224 | -0.837 | -1.36 | -0.958 | 0.004 | 0.049 | 0.049 | 0 | 0.47 | 0.35 | 0.35 | 0.08 | 0.53 | 0.4 |
| 7 | 150501516 | rs9088 | -0.583 | -0.551 | -1.382 | -1.008 | -0.013 | 0.004 | 0.004 | 0 | 0.76 | 0.79 | 0.64 | 0.76 | 0.68 | 0.79 |
| 7 | 150503079 | rs11772167 | -1.576 | -0.028 | -1.509 | -1.119 | -0.062 | 0.057 | 0.057 | 0 | 0.86 | 0.98 | 0.88 | 0.93 | 0.8 | 0.86 |
| 7 | 150510915 | rs2888674 | -0.206 | -0.878 | -1.109 | -1.621 | -0.015 | 0.057 | 0.057 | 0.059 | 0.62 | 0.78 | 0.52 | 0.73 | 0.46 | 0.66 |
| 7 | 150512100 | rs11982695 | 1.368 | -1.202 | -1.077 | -1.566 | 0.024 | -0.031 | -0.031 | 0.915 | 0.26 | 0.15 | 0.29 | 0.63 | 0.15 | 0.38 |
| 7 | 150512172 | rs10231759 | 0.161 | -1.523 | -0.599 | -1.737 | 0.042 | 0.043 | 0.043 | 0 | 0.45 | 0.67 | 0.39 | 0.69 | 0.29 | 0.55 |
| 7 | 150517680 | rs10231216 | 0.219 | -1.697 | -0.346 | -1.809 | -0.031 | -0.055 | -0.055 | 0.873 | 0.26 | 0.42 | 0.16 | 0.08 | 0.2 | 0.23 |
| 7 | 150518393 | rs10278673 | 0.881 | -1.499 | -0.064 | -1.5 | 0.024 | -0.008 | -0.008 | 0.113 | 0.37 | 0.31 | 0.32 | 0.54 | 0.21 | 0.47 |
| 7 | 150518895 | rs10235739 | -0.216 | -1.55 | 0.03 | -1.553 | -0.008 | 0.019 | 0.019 | 0 | 0.13 | 0.19 | 0.23 | 0.27 | 0.12 | 0.12 |
| 7 | 150521657 | rs917807 | -1.06 | -0.92 | -0.003 | -1.76 | 0.017 | -0.031 | -0.031 | 0 | 0.87 | 0.84 | 0.85 | 0.96 | 0.79 | 0.87 |
| 7 | 150522565 | rs13437864 | 0.723 | -0.954 | 0.034 | -1.803 | -0.032 | 0.063 | 0.063 | 0 | 0.36 | 0.25 | 0.3 | 0.11 | 0.46 | 0.27 |
| 7 | 150523061 | rs10952288 | 0.734 | -0.954 | 0.027 | -1.803 | -0.041 | 0.084 | 0.084 | 0 | 0.36 | 0.21 | 0.3 | 0.11 | 0.46 | 0.27 |
| 7 | 150523344 | rs11767582 | 1.552 | -0.932 | -0.068 | -1.822 | -0.053 | 0.066 | 0.066 | 0.456 | 0.21 | 0.11 | 0.18 | 0.07 | 0.3 | 0.16 |
| 7 | 150525410 | rs6980179 | 0.971 | -0.972 | 0.307 | -1.697 | -0.012 | -0.001 | -0.001 | 0 | 0.37 | 0.43 | 0.33 | 0.57 | 0.21 | 0.47 |
| 7 | 150529363 | rs12540772 | 0.575 | -0.943 | 0.16 | -1.704 | -0.035 | -0.019 | -0.019 | 0 | 0.41 | 0.55 | 0.36 | 0.64 | 0.22 | 0.48 |
| 7 | 150529449 | rs17173637 | 1.521 | -0.901 | 0.138 | -1.708 | 0.001 | -0.001 | -0.001 | 0 | 0.07 | 0.06 | 0.06 | 0.01 | 0.09 | 0.06 |
| 7 | 150533523 | rs4725956 | -1.334 | -0.846 | 0.039 | -1.698 | 0.007 | -0.013 | -0.013 | 0.562 | 0.88 | 0.91 | 0.86 | 0.96 | 0.79 | 0.87 |
| 7 | 150533765 | rs10952291 | 0.494 | -0.887 | 0.027 | -1.426 | -0.022 | 0.015 | 0.015 | 0 | 0.39 | 0.28 | 0.44 | 0.36 | 0.48 | 0.33 |
| 7 | 150534076 | rs10236614 | 0.462 | -0.904 | 0.036 | -1.398 | -0.03 | -0.003 | -0.003 | 0 | 0.42 | 0.53 | 0.35 | 0.56 | 0.22 | 0.47 |
| 7 | 150536172 | rs4725372 | -0.641 | -0.406 | 0.304 | -1.062 | -0.005 | 0.003 | 0.003 | 0 | 0.87 | 0.93 | 0.86 | 0.95 | 0.78 | 0.85 |
| 7 | 150536587 | rs4725960 | 0.06 | -0.434 | 0.352 | -1.046 | 0.018 | 0.01 | 0.01 | 0 | 0.83 | 0.77 | 0.7 | 0.78 | 0.76 | 0.8 |
| 7 | 150538501 | rs759008 | 0.832 | -0.473 | 0.339 | -1.06 | 0.018 | -0.008 | -0.008 | 0 | 0.74 | 0.62 | 0.63 | 0.77 | 0.67 | 0.75 |
| 7 | 150542515 | rs6977081 | 0.014 | -0.501 | 0.019 | -1.141 | 0.005 | 0.005 | 0.005 | 0.425 | 0.41 | 0.61 | 0.52 | 0.66 | 0.3 | 0.52 |
| 7 | 150542646 | rs6977381 | -0.814 | -0.518 | -0.327 | -1.175 | -0.055 | 0.132 | 0.132 | 1.907 | 0.57 | 0.88 | 0.69 | 0.76 | 0.52 | 0.67 |
| 7 | 150543138 | rs887588 | -1.411 | 0.047 | -0.2 | -0.885 | -0.084 | 0.04 | 0.04 | 8.41 | 0.74 | 0.93 | 0.82 | 0.8 | 0.7 | 0.79 |
| 7 | 150548433 | rs10273373 | -1.054 | 0.263 | 1.1 | -0.524 | 0.034 | -0.038 | -0.038 | 0.427 | 0.21 | 0.51 | 0.22 | 0.12 | 0.25 | 0.16 |
| 7 | 150548972 | rs2052129 | -0.243 | 0.438 | 1.129 | -0.469 | 0.048 | -0.009 | -0.009 | 0 | 0.15 | 0.23 | 0.19 | 0.12 | 0.25 | 0.16 |
| 7 | 150553605 | rs10156191 | -1.27 | 0.288 | 0.953 | -0.453 | 0.008 | -0.044 | -0.044 | 2.226 | 0.25 | 0.56 | 0.25 | 0.12 | 0.27 | 0.27 |
| 7 | 150555547 | rs2301257 | 0.472 | -0.217 | 0.911 | -0.075 | -0.023 | 0.074 | 0.074 | 0 | 0.23 | 0.35 | 0.31 | 0.45 | 0.15 | 0.38 |
| 7 | 150555915 | rs10893 | -0.551 | -0.161 | 1.068 | -0.157 | -0.006 | 0.019 | 0.019 | 0.326 | 0.35 | 0.56 | 0.4 | 0.48 | 0.27 | 0.53 |
| 7 | 150556002 | rs1049748 | -0.238 | -0.279 | 0.721 | -0.44 | -0.001 | -0.001 | -0.001 | 0 | 0.44 | 0.59 | 0.46 | 0.56 | 0.39 | 0.54 |
| 7 | 150557578 | rs11771771 | 1.358 | -0.009 | 0.38 | -0.481 | -0.042 | 0.006 | 0.006 | 0.326 | 0.07 | 0 | 0.09 | 0.04 | 0.13 | 0.07 |
| 7 | 150558366 | rs12539 | 1.024 | -0.081 | 0.458 | -0.469 | 0 | -0.001 | -0.001 | 0 | 0.21 | 0.22 | 0.15 | 0.1 | 0.23 | 0.16 |
| 7 | 150558399 | rs2071517 | 0.974 | -0.201 | 0.57 | -0.476 | -0.013 | 0.046 | 0.046 | 0 | 0.22 | 0.29 | 0.31 | 0.46 | 0.15 | 0.38 |
| 7 | 150558611 | rs4725970 | 0.952 | -0.032 | -0.206 | -0.282 | 0.001 | 0.003 | 0.003 | 0 | 0.21 | 0.21 | 0.15 | 0.1 | 0.23 | 0.16 |
| 7 | 150560322 | rs11978239 | 0.019 | -0.035 | -0.176 | -0.278 | -0.003 | 0.007 | 0.007 | 0.566 | 0.35 | 0.58 | 0.4 | 0.48 | 0.27 | 0.53 |
| 7 | 150562451 | rs4725377 | -0.447 | 0.061 | -0.742 | -0.264 | -0.034 | 0.018 | 0.018 | 4.293 | 0.56 | 0.81 | 0.55 | 0.58 | 0.5 | 0.69 |
| 7 | 150562812 | rs4725971 | -0.869 | 0.122 | -0.94 | -0.234 | -0.061 | -0.009 | -0.009 | 6.585 | 0.57 | 0.84 | 0.55 | 0.57 | 0.5 | 0.69 |
| 7 | 150563964 | rs2215564 | 1.014 | 0.33 | -0.896 | -0.239 | -0.044 | 0.007 | 0.007 | 0.466 | 0.07 | 0 | 0.1 | 0.05 | 0.13 | 0.07 |
| 7 | 150564361 | rs10250599 | -0.039 | -0.088 | -0.738 | -0.228 | 0.024 | -0.026 | -0.026 | 0.006 | 0.38 | 0.65 | 0.39 | 0.5 | 0.26 | 0.54 |
| 7 | 150564418 | rs10250704 | 0.878 | -0.073 | -0.764 | -0.262 | -0.002 | 0.031 | 0.031 | 0 | 0.2 | 0.16 | 0.16 | 0.1 | 0.24 | 0.16 |
| 7 | 150565220 | rs10254616 | -0.722 | 0.172 | -0.805 | -0.165 | -0.043 | 0.015 | 0.015 | 4.356 | 0.58 | 0.8 | 0.55 | 0.6 | 0.5 | 0.7 |
| 7 | 150572570 | rs12670489 | -0.26 | 0.419 | -0.56 | -0.166 | -0.008 | 0.012 | 0.012 | 1.367 | 0.13 | 0.09 | 0.12 | 0.05 | 0.15 | 0.12 |
| 7 | 150585880 | rs12674178 | -0.875 | 0.043 | 0.108 | -0.257 | 0.011 | -0.042 | -0.042 | 0.478 | 0.41 | 0.52 | 0.37 | 0.23 | 0.47 | 0.39 |
| 7 | 150586697 | rs12530900 | -0.64 | 0.097 | 0.277 | -0.301 | 0.002 | -0.025 | -0.025 | 0 | 0.36 | 0.41 | 0.32 | 0.23 | 0.4 | 0.36 |

| | | | | | | | | | | | | | | | | |
|---|---|---|---|---|---|---|---|---|---|---|---|---|---|---|---|---|
| 7 | 150588800 | rs10277774 | -1.632 | -0.377 | 0.699 | -0.134 | -0.004 | 0.016 | 0.016 | 11.629 | 0.63 | 0.72 | 0.65 | 0.67 | 0.63 | 0.81 |
| 7 | 150592741 | rs11762854 | 0.953 | -0.237 | 0.711 | 0.03 | -0.007 | 0.004 | 0.004 | 0.207 | 0.08 | 0.04 | 0.1 | 0.05 | 0.12 | 0.06 |
| 7 | 150594658 | rs12703105 | 0.55 | -0.302 | 0.713 | 0.127 | -0.009 | 0.029 | 0.029 | 0.384 | 0.23 | 0.23 | 0.29 | 0.48 | 0.16 | 0.45 |
| 7 | 150595644 | rs17173651 | -0.511 | -0.033 | 0.982 | 0.016 | 0.039 | -0.07 | -0.07 | 0 | 0.15 | 0.37 | 0.16 | 0.05 | 0.15 | 0.12 |
| 7 | 150596929 | rs11761679 | -0.394 | 0.003 | 0.406 | 0.237 | 0.004 | -0.01 | -0.01 | 0 | 0.14 | 0.17 | 0.14 | 0.06 | 0.16 | 0.1 |
| 7 | 150598492 | rs740954 | -0.083 | -0.279 | 0.345 | 0.904 | -0.002 | -0.006 | -0.006 | 0 | 0.43 | 0.43 | 0.44 | 0.4 | 0.42 | 0.63 |
| 7 | 150598659 | rs17173656 | -0.254 | -0.404 | 0.011 | 1.507 | 0.023 | -0.029 | -0.029 | 0 | 0.15 | 0.16 | 0.22 | 0.32 | 0.12 | 0.13 |
| 7 | 150599205 | rs10216051 | -1.565 | -0.632 | 0.02 | 1.032 | 0.009 | 0.025 | 0.025 | 0.718 | 0.39 | 0.7 | 0.6 | 0.71 | 0.35 | 0.35 |
| 7 | 150599259 | rs17173657 | 0.9 | -0.479 | -0.297 | 1.419 | 0.019 | -0.039 | -0.039 | 0 | 0.09 | 0.01 | 0.21 | 0.11 | 0.08 | 0.1 |
| 7 | 150600845 | rs9640171 | -0.024 | -0.585 | -0.286 | 1.142 | -0.001 | -0.003 | -0.003 | 1.194 | 0.28 | 0.26 | 0.33 | 0.35 | 0.29 | 0.33 |
| 7 | 150606288 | rs2968858 | -1.048 | -0.621 | 0.086 | 1.258 | 0 | 0.005 | 0.005 | 0.017 | 0.47 | 0.47 | 0.67 | 0.59 | 0.5 | 0.49 |
| 7 | 150611161 | rs17173658 | 0.796 | -0.752 | -0.228 | 1.557 | 0.037 | -0.061 | -0.061 | 0.028 | 0.09 | 0.01 | 0.2 | 0.23 | 0.06 | 0.1 |
| 7 | 150614712 | rs7781870 | 0.435 | -0.486 | 0.431 | 1.436 | 0.038 | -0.003 | -0.003 | 0.277 | 0.08 | 0.17 | 0.09 | 0.07 | 0.11 | 0.05 |
| 7 | 150614858 | rs7782274 | -1.298 | -0.755 | 0.956 | 1.633 | 0.018 | 0.052 | 0.052 | 0 | 0.17 | 0.5 | 0.15 | 0.24 | 0.15 | 0.09 |
| 7 | 150620701 | rs6972137 | 0.139 | -0.772 | 0.288 | 1.4 | 0 | -0.045 | -0.045 | 0 | 0.18 | 0.11 | 0.12 | 0.37 | 0.15 | 0.22 |
| 7 | 150634448 | rs758887 | -0.714 | -0.86 | 0.589 | 0.982 | 0.01 | -0.008 | -0.008 | 0 | 0.29 | 0.27 | 0.33 | 0.59 | 0.21 | 0.33 |
| 7 | 150637863 | rs4725982 | -0.714 | -0.858 | 0.589 | 0.982 | 0.01 | -0.008 | -0.008 | 0.054 | 0.29 | 0.27 | 0.33 | 0.58 | 0.2 | 0.33 |
| 7 | 150639871 | rs2968853 | -1.898 | -0.676 | 0.027 | 0.181 | -0.025 | 0.102 | 0.102 | 2.053 | 0.65 | 0.88 | 0.65 | 0.88 | 0.62 | 0.7 |
| 7 | 150640285 | rs1547958 | 1.248 | -0.622 | -0.077 | 0.205 | -0.075 | 0.092 | 0.092 | 0 | 0.19 | 0.01 | 0.17 | 0.05 | 0.25 | 0.25 |
| 7 | 150645534 | rs1805123 | 1.31 | -0.62 | -0.251 | 0.212 | -0.079 | 0.088 | 0.088 | 0 | 0.19 | 0.01 | 0.17 | 0.05 | 0.25 | 0.25 |
| 7 | 150645682 | rs2968860 | -1.304 | -0.608 | 0.008 | 0.326 | -0.017 | 0.084 | 0.084 | 1.639 | 0.6 | 0.76 | 0.61 | 0.88 | 0.56 | 0.69 |
| 7 | 150667210 | rs3807375 | -1.184 | -0.763 | -0.052 | 0.475 | 0.027 | 0.071 | 0.071 | 0.508 | 0.42 | 0.76 | 0.53 | 0.76 | 0.34 | 0.39 |
| 7 | 150668070 | rs10236214 | -0.961 | -0.369 | -0.818 | -0.198 | -0.044 | 0.052 | 0.052 | 0 | 0.73 | 0.85 | 0.76 | 0.95 | 0.6 | 0.74 |
| 7 | 150669057 | rs3807372 | -0.02 | -0.786 | -0.369 | 0.475 | -0.024 | 0.097 | 0.097 | 0.406 | 0.21 | 0.3 | 0.32 | 0.48 | 0.14 | 0.21 |
| 7 | 150671437 | rs748693 | 1.577 | -0.589 | -1.025 | 0.17 | -0.036 | 0.068 | 0.068 | 0.281 | 0.32 | 0.1 | 0.33 | 0.14 | 0.32 | 0.33 |
| 7 | 150673314 | rs3807370 | -1.178 | -0.45 | -1.075 | 0.246 | -0.113 | 0.075 | 0.075 | 0 | 0.74 | 0.94 | 0.76 | 0.88 | 0.58 | 0.75 |
| 7 | 150678080 | rs2373885 | -0.869 | -0.72 | -0.6 | 0.199 | -0.035 | -0.014 | -0.014 | 0 | 0.37 | 0.53 | 0.39 | 0.74 | 0.22 | 0.41 |
| 7 | 150681210 | rs2373961 | 1.678 | -0.67 | -1.083 | 0.19 | -0.024 | 0.05 | 0.05 | 0.375 | 0.32 | 0.16 | 0.33 | 0.13 | 0.35 | 0.3 |
| 7 | 150683083 | rs10247107 | -0.838 | 0.022 | -1.134 | 0.285 | -0.088 | 0.076 | 0.076 | 0 | 0.73 | 0.93 | 0.76 | 0.88 | 0.58 | 0.72 |
| 7 | 150687687 | rs11771443 | -0.246 | -0.198 | -0.799 | 0.709 | 0.022 | -0.066 | -0.066 | 0 | 0.2 | 0.08 | 0.31 | 0.44 | 0.15 | 0.21 |
| 7 | 150699877 | rs2853792 | 0.333 | 0.259 | -0.89 | 0.197 | -0.042 | -0.017 | -0.017 | 0.063 | 0.67 | 0.86 | 0.73 | 0.69 | 0.57 | 0.72 |
| 7 | 150700946 | rs3918227 | 1.442 | 0.432 | -0.934 | 0.319 | -0.032 | -0.023 | -0.023 | 0.229 | 0.06 | 0 | 0.06 | 0.08 | 0.09 | 0.06 |
| 7 | 150702219 | rs3918184 | 1.145 | -0.024 | -0.543 | 0.129 | 0 | -0.002 | -0.002 | 0 | 0.4 | 0.37 | 0.23 | 0.29 | 0.31 | 0.31 |
| 7 | 150703242 | rs2853795 | 1.126 | -0.016 | -0.716 | 0.269 | -0.058 | -0.079 | -0.079 | 0 | 0.12 | 0.03 | 0.41 | 0.3 | 0.18 | 0.22 |
| 7 | 150703915 | rs2853796 | 0.765 | -0.141 | -0.509 | -0.088 | 0 | 0.003 | 0.003 | 0.528 | 0.52 | 0.43 | 0.65 | 0.59 | 0.49 | 0.53 |
| 7 | 150704843 | rs891511 | -0.304 | -0.912 | 0.076 | -0.183 | -0.005 | 0 | 0 | 0 | 0.41 | 0.58 | 0.54 | 0.58 | 0.34 | 0.56 |
| 7 | 150705842 | rs3918196 | 0.886 | -1.044 | 0.122 | 0.281 | 0.024 | -0.02 | -0.02 | 0.196 | 0.12 | 0.05 | 0.06 | 0.16 | 0.08 | 0.1 |
| 7 | 150706383 | rs753482 | -0.233 | -0.192 | 0.79 | 0.077 | 0.01 | -0.034 | -0.034 | 0.014 | 0.82 | 0.72 | 0.84 | 0.95 | 0.75 | 0.86 |
| 7 | 150706915 | rs743506 | 0.884 | 0.093 | 0.774 | 0.34 | -0.003 | -0.02 | -0.02 | 0.075 | 0.75 | 0.53 | 0.82 | 0.8 | 0.71 | 0.82 |
| 7 | 150707488 | rs743507 | 0.435 | 0.211 | 0.277 | 0.394 | 0.004 | -0.002 | -0.002 | 1.689 | 0.79 | 0.73 | 0.83 | 0.8 | 0.71 | 0.83 |
| 7 | 150708089 | rs891512 | -0.759 | 0.513 | -0.28 | 0.467 | -0.069 | 0.011 | 0.011 | 3.954 | 0.86 | 0.99 | 0.86 | 0.95 | 0.75 | 0.85 |
| 7 | 150708302 | rs1808593 | 0.115 | 0.505 | -0.109 | 0.469 | 0.004 | -0.003 | -0.003 | 0.021 | 0.79 | 0.73 | 0.83 | 0.8 | 0.72 | 0.83 |
| 7 | 150709571 | rs7830 | -0.325 | 0.853 | -0.647 | 0.666 | -0.039 | -0.048 | -0.048 | 0 | 0.29 | 0.19 | 0.52 | 0.41 | 0.35 | 0.44 |
| 7 | 150713636 | rs3800787 | -0.4 | 0.67 | -1.106 | 0.507 | -0.106 | -0.092 | -0.092 | 0 | 0.35 | 0.08 | 0.51 | 0.42 | 0.39 | 0.45 |
| 7 | 150714812 | rs2373929 | -0.273 | 1.578 | -0.576 | 0.617 | -0.019 | -0.017 | -0.017 | 0 | 0.24 | 0.19 | 0.57 | 0.44 | 0.44 | 0.23 |
| 7 | 150717153 | rs6464119 | -0.409 | 0.586 | -0.714 | 0.517 | 0.001 | -0.005 | -0.005 | 0 | 0.79 | 0.89 | 0.83 | 0.77 | 0.75 | 0.72 |
| 7 | 150723265 | rs3763486 | 0.429 | 0.886 | -0.103 | 0.703 | 0.001 | 0.064 | 0.064 | 0 | 0.18 | 0.23 | 0.42 | 0.46 | 0.18 | 0.15 |
| 7 | 150725722 | rs2303922 | 0.204 | 0.024 | 0.161 | -0.038 | -0.004 | -0.009 | -0.009 | 0.359 | 0.4 | 0.57 | 0.27 | 0.3 | 0.31 | 0.36 |
| 7 | 150726920 | rs6944935 | 0.536 | -0.022 | 0.481 | -0.11 | 0 | 0.001 | 0.001 | 0 | 0.36 | 0.33 | 0.21 | 0.28 | 0.25 | 0.34 |
| 7 | 150729212 | rs6947821 | 0.087 | -0.014 | 0.459 | -0.109 | -0.004 | -0.009 | -0.009 | 0 | 0.4 | 0.57 | 0.27 | 0.3 | 0.31 | 0.36 |
| 7 | 150730220 | rs10952299 | 1.286 | -0.245 | 0.488 | -0.226 | 0 | 0.056 | 0.056 | 2.76 | 0.25 | 0.09 | 0.12 | 0.01 | 0.18 | 0.24 |
| 7 | 150732463 | rs2303925 | 1.155 | -0.034 | 0.417 | 0.335 | 0 | 0.034 | 0.034 | 0 | 0.17 | 0.22 | 0.41 | 0.34 | 0.19 | 0.19 |
| 7 | 150736382 | rs2373926 | 0.194 | -0.081 | 0.453 | 0.302 | 0.019 | 0.106 | 0.106 | 0.321 | 0.22 | 0.46 | 0.48 | 0.36 | 0.25 | 0.18 |
| 7 | 150736965 | rs4148848 | 0.967 | -0.128 | 0.378 | 0.359 | -0.017 | 0.029 | 0.029 | 0 | 0.15 | 0.2 | 0.1 | 0.26 | 0.11 | 0.12 |
| 7 | 150738624 | rs4148850 | 1.025 | 0.013 | -0.073 | 0.44 | -0.001 | 0.029 | 0.029 | 0 | 0.18 | 0.23 | 0.41 | 0.34 | 0.2 | 0.19 |
| 7 | 150742055 | rs4148852 | 1.057 | 0.076 | -0.609 | 0.421 | -0.001 | 0.025 | 0.025 | 0 | 0.17 | 0.21 | 0.41 | 0.34 | 0.2 | 0.19 |
| 7 | 150742700 | rs891506 | -0.678 | 0.062 | -0.851 | 0.284 | -0.049 | 0.022 | 0.022 | 0 | 0.65 | 0.88 | 0.75 | 0.7 | 0.59 | 0.62 |
| 7 | 150742985 | rs4148853 | 1.264 | 0.081 | -0.78 | 0.317 | -0.001 | 0.014 | 0.014 | 0 | 0.17 | 0.17 | 0.4 | 0.34 | 0.19 | 0.19 |
| 7 | 150743820 | rs4148854 | -0.489 | 0.042 | -1.07 | -0.035 | -0.049 | 0.022 | 0.022 | 0 | 0.65 | 0.88 | 0.75 | 0.7 | 0.59 | 0.62 |
| 7 | 150744922 | rs11770520 | 0.918 | -0.161 | -0.861 | -0.034 | -0.01 | 0.105 | 0.105 | 2.731 | 0.24 | 0.06 | 0.12 | 0.02 | 0.18 | 0.24 |
| 7 | 150747746 | rs2288645 | -0.764 | -0.201 | -1.145 | 0.051 | 0.007 | -0.021 | -0.021 | 0 | 0.81 | 0.94 | 0.89 | 0.77 | 0.83 | 0.78 |
| 7 | 150747937 | rs11977275 | 0.953 | -0.117 | -0.914 | -0.222 | 0.001 | 0.019 | 0.019 | 1.722 | 0.25 | 0.16 | 0.13 | 0.01 | 0.18 | 0.24 |
| 7 | 150751590 | rs2069459 | 0.475 | -0.035 | -0.367 | 0.19 | 0.008 | 0.007 | 0.007 | 0 | 0.35 | 0.46 | 0.29 | 0.38 | 0.4 | 0.32 |
| 7 | 150752608 | rs2069456 | 0.713 | -0.099 | -0.135 | 0.415 | 0.002 | 0.019 | 0.019 | 0.03 | 0.19 | 0.21 | 0.47 | 0.32 | 0.25 | 0.21 |
| 7 | 150755173 | rs2069443 | -0.027 | -0.544 | -0.015 | 0.319 | -0.014 | 0.013 | 0.013 | 0.876 | 0.28 | 0.38 | 0.48 | 0.3 | 0.26 | 0.27 |
| 7 | 150755839 | rs1549760 | 0.793 | -0.268 | -0.105 | 0.406 | 0.004 | 0.018 | 0.018 | 0.061 | 0.19 | 0.24 | 0.46 | 0.29 | 0.24 | 0.21 |
| 7 | 150762871 | rs3793336 | -0.749 | 0.019 | 0.038 | 0.178 | -0.002 | 0.003 | 0.003 | 0 | 0.88 | 0.93 | 0.9 | 0.94 | 0.84 | 0.82 |
| 7 | 150765236 | rs12703112 | 1.154 | -0.161 | -0.259 | -0.223 | -0.001 | 0.08 | 0.08 | 1.666 | 0.24 | 0.06 | 0.12 | 0.01 | 0.18 | 0.28 |
| 7 | 150766799 | rs2303933 | 0.39 | -0.503 | 0.579 | -0.019 | 0 | -0.001 | -0.001 | 0 | 0.56 | 0.53 | 0.71 | 0.72 | 0.61 | 0.49 |

| | | | | | | | | | | | | | | | | |
|---|---|---|---|---|---|---|---|---|---|---|---|---|---|---|---|---|
| 7 | 150768786 | rs2303937 | 0.253 | -0.938 | 0.747 | 0.296 | 0.017 | 0.022 | 0.022 | 0.027 | 0.5 | 0.34 | 0.57 | 0.42 | 0.43 | 0.57 |
| 7 | 150768849 | rs11542749 | 0.353 | -0.951 | 0.761 | 0.332 | -0.016 | -0.026 | -0.026 | 2.213 | 0.06 | 0.2 | 0.04 | 0 | 0.03 | 0.01 |
| 7 | 150770832 | rs11765015 | 1.172 | -0.898 | 0.517 | 0.394 | 0.009 | -0.044 | -0.044 | 0.2 | 0.26 | 0.19 | 0.45 | 0.38 | 0.25 | 0.28 |
| 7 | 150775306 | rs2303942 | -0.172 | -0.755 | 0.869 | 0.308 | -0.004 | 0 | 0 | 0.213 | 0.57 | 0.68 | 0.62 | 0.41 | 0.46 | 0.58 |
| 7 | 150782071 | rs10271154 | 1.043 | -0.716 | 0.501 | 0.376 | 0.013 | -0.043 | -0.043 | 0 | 0.27 | 0.17 | 0.46 | 0.35 | 0.25 | 0.29 |
| 7 | 150782617 | rs4725388 | 0.334 | -0.837 | 0.945 | 0.43 | -0.035 | 0.032 | 0.032 | 0.652 | 0.35 | 0.54 | 0.53 | 0.55 | 0.28 | 0.3 |
| 7 | 150785321 | rs6464126 | 0.177 | -0.842 | 0.879 | 0.339 | 0.004 | -0.004 | -0.004 | 0 | 0.36 | 0.63 | 0.54 | 0.57 | 0.29 | 0.31 |
| 7 | 150785753 | rs11769150 | 1.176 | -0.72 | 0.45 | 0.048 | 0 | 0.001 | 0.001 | 0 | 0.23 | 0.22 | 0.13 | 0.02 | 0.17 | 0.29 |
| 7 | 150787046 | rs10270107 | 0.907 | -0.893 | 0.562 | 0.28 | 0.011 | -0.016 | -0.016 | 0 | 0.28 | 0.23 | 0.46 | 0.36 | 0.26 | 0.3 |
| 7 | 150800739 | rs10273337 | -0.848 | -0.737 | 0.383 | -0.174 | -0.01 | 0.016 | 0.016 | 0 | 0.75 | 0.8 | 0.84 | 0.98 | 0.76 | 0.69 |
| 7 | 150801238 | rs4725991 | -0.622 | -0.798 | 0.766 | 0.175 | 0.01 | 0.234 | 0.234 | 5.308 | 0.08 | 0.39 | 0.08 | 0.22 | 0.04 | 0.01 |
| 7 | 150808109 | rs891509 | 0.574 | -0.608 | -0.069 | 0.041 | 0.013 | -0.005 | -0.005 | 0 | 0.39 | 0.32 | 0.55 | 0.61 | 0.38 | 0.41 |
| 7 | 150809164 | rs6951528 | -0.393 | -0.456 | -0.344 | -0.044 | -0.002 | -0.021 | -0.021 | 0 | 0.68 | 0.4 | 0.77 | 0.75 | 0.71 | 0.68 |
| 7 | 150809280 | rs4725392 | 0.245 | -0.159 | -0.781 | 0.112 | -0.029 | 0.045 | 0.045 | 0.348 | 0.28 | 0.11 | 0.22 | 0.16 | 0.34 | 0.27 |
| 7 | 150822192 | rs729712 | 0.025 | -0.119 | -0.955 | 0.167 | -0.024 | -0.019 | -0.019 | 1.749 | 0.34 | 0.13 | 0.34 | 0.33 | 0.35 | 0.29 |
| 7 | 150825087 | rs6970983 | -0.676 | 0.02 | -0.693 | 0.108 | -0.002 | -0.002 | -0.002 | 0 | 0.75 | 0.79 | 0.82 | 0.69 | 0.73 | 0.68 |
| 7 | 150825492 | rs2288651 | 0.852 | -0.14 | -0.667 | -0.133 | 0.009 | -0.014 | -0.014 | 1.771 | 0.36 | 0.29 | 0.44 | 0.4 | 0.35 | 0.37 |
| 7 | 150829458 | rs1966602 | -0.777 | 0.232 | -0.823 | -0.14 | -0.125 | 0.164 | 0.164 | 0 | 0.73 | 0.98 | 0.86 | 0.98 | 0.68 | 0.56 |
| 7 | 150830015 | rs6968877 | -0.07 | 0.336 | 0.227 | 0.336 | 0.007 | 0.06 | 0.06 | 0 | 0.32 | 0.45 | 0.37 | 0.51 | 0.32 | 0.2 |
| 7 | 150837529 | rs10230113 | 0.698 | 0.245 | -0.066 | 0.217 | 0.017 | -0.006 | -0.006 | 0 | 0.39 | 0.25 | 0.44 | 0.39 | 0.34 | 0.31 |
| 7 | 150844687 | rs4725999 | -0.98 | 0.739 | 1.314 | 1.145 | 0.005 | 0.014 | 0.014 | 0 | 0.34 | 0.57 | 0.39 | 0.6 | 0.37 | 0.24 |
| 7 | 150847749 | rs1001760 | -0.987 | 0.738 | 0.953 | 1.147 | -0.004 | -0.012 | -0.012 | 0.905 | 0.36 | 0.73 | 0.41 | 0.61 | 0.37 | 0.25 |
| 7 | 150849164 | rs6464132 | -0.998 | 0.741 | 0.946 | 1.149 | -0.005 | -0.012 | -0.012 | 0 | 0.36 | 0.74 | 0.41 | 0.61 | 0.37 | 0.25 |
| 7 | 150851566 | rs4725394 | -0.381 | 0.842 | 1.01 | 1.219 | 0.022 | 0.023 | 0.023 | 0 | 0.25 | 0.43 | 0.32 | 0.35 | 0.28 | 0.18 |
| 7 | 150869437 | rs7782650 | -0.605 | 0.886 | 0.628 | 1.246 | 0.026 | -0.009 | -0.009 | 0 | 0.24 | 0.55 | 0.35 | 0.19 | 0.29 | 0.17 |
| 7 | 150872363 | rs1469550 | 0.656 | 0.841 | 0.48 | 1.216 | -0.016 | 0.015 | 0.015 | 0.462 | 0.17 | 0.11 | 0.26 | 0.08 | 0.22 | 0.14 |
| 7 | 150873046 | rs310597 | -1.066 | 0.869 | 0.012 | 1.413 | -0.01 | -0.01 | -0.01 | 0 | 0.36 | 0.74 | 0.42 | 0.35 | 0.38 | 0.25 |
| 7 | 150882626 | rs2487150 | -0.904 | 0.892 | 0.186 | 1.19 | -0.001 | -0.019 | -0.019 | 0 | 0.36 | 0.76 | 0.45 | 0.59 | 0.37 | 0.26 |
| 7 | 150883761 | rs2257090 | -0.269 | 1.357 | -0.141 | 0.722 | 0.001 | -0.018 | -0.018 | 1.873 | 0.26 | 0.3 | 0.29 | 0.09 | 0.26 | 0.19 |
| 7 | 150888788 | rs2288649 | 0.018 | 1.818 | -0.992 | 1.038 | 0.008 | 0.025 | 0.025 | 0 | 0.83 | 0.8 | 0.64 | 0.67 | 0.77 | 0.77 |
| 7 | 150890241 | rs2487151 | -0.545 | 1.449 | -1.323 | 0.487 | -0.017 | 0.063 | 0.063 | 0.047 | 0.24 | 0.16 | 0.27 | 0.09 | 0.24 | 0.19 |
| 7 | 150892599 | rs310589 | -0.262 | 1.699 | -1.856 | 0.425 | -0.145 | 0.068 | 0.068 | 0.306 | 0.3 | 0.05 | 0.36 | 0.14 | 0.41 | 0.37 |
| 7 | 150892950 | rs310588 | -0.19 | 1.138 | -1.248 | 0.097 | 0.004 | 0.008 | 0.008 | 0 | 0.53 | 0.49 | 0.54 | 0.29 | 0.62 | 0.43 |
| 7 | 150893629 | rs310586 | 0.012 | 0.892 | -1.19 | -0.815 | 0.002 | 0.062 | 0.062 | 0.552 | 0.55 | 0.32 | 0.37 | 0.15 | 0.53 | 0.48 |
| 7 | 150901609 | rs310583 | -0.196 | 0.661 | -1.158 | -0.714 | -0.021 | 0.093 | 0.093 | 0 | 0.59 | 0.78 | 0.5 | 0.79 | 0.47 | 0.67 |
| 7 | 150905444 | rs2608291 | -0.131 | 0.871 | -0.732 | -0.776 | -0.013 | 0.013 | 0.013 | 0 | 0.6 | 0.71 | 0.44 | 0.26 | 0.45 | 0.57 |
| 7 | 150907194 | rs6967282 | -1.094 | 0.651 | -0.237 | -0.928 | 0.013 | -0.031 | -0.031 | 0 | 0.13 | 0.18 | 0.19 | 0.02 | 0.18 | 0.13 |
| 7 | 150912786 | rs7786151 | -0.618 | 0.664 | -0.119 | -0.881 | 0.018 | -0.019 | -0.019 | 0 | 0.07 | 0.16 | 0.12 | 0.02 | 0.08 | 0.05 |
| 7 | 150916228 | rs6949587 | 0.672 | 0.509 | -0.489 | -0.895 | -0.025 | -0.01 | -0.01 | 0.122 | 0.33 | 0.21 | 0.41 | 0.72 | 0.46 | 0.36 |
| 7 | 150927763 | rs2792432 | -0.305 | 0.029 | -0.902 | -1.142 | -0.001 | -0.015 | -0.015 | 0 | 0.17 | 0.15 | 0.22 | 0.14 | 0.16 | 0.11 |
| 7 | 150928545 | rs4726009 | 0.849 | 0.074 | -0.842 | -1.121 | -0.02 | -0.005 | -0.005 | 0 | 0.34 | 0.25 | 0.42 | 0.72 | 0.46 | 0.36 |
| 7 | 150940657 | rs122815 | -0.587 | 0.001 | -1.533 | -1.139 | 0.004 | -0.039 | -0.039 | 0 | 0.91 | 0.97 | 0.85 | 0.87 | 0.93 | 0.97 |
| 7 | 150940867 | rs219229 | 0.238 | 0.241 | -1.186 | -2.094 | 0.032 | 0.124 | 0.124 | 0 | 0.41 | 0.18 | 0.23 | 0.12 | 0.31 | 0.48 |
| 7 | 150951819 | rs3789823 | -0.029 | 0.282 | -1.052 | -1.955 | 0.006 | 0.149 | 0.149 | 0 | 0.55 | 0.8 | 0.65 | 0.85 | 0.6 | 0.51 |
| 7 | 150955765 | rs219235 | -0.317 | 0.19 | -1.161 | -1.926 | 0 | -0.041 | -0.041 | 0.14 | 0.24 | 0.22 | 0.21 | 0.01 | 0.19 | 0.13 |
| 7 | 150956627 | rs219236 | -0.339 | 0.786 | -1.567 | -1.245 | -0.023 | -0.038 | -0.038 | 0 | 0.86 | 0.97 | 0.88 | 0.85 | 0.81 | 0.78 |
| 7 | 150959826 | rs10487756 | 0.949 | 1.264 | -1.282 | -1.959 | -0.03 | 0.026 | 0.026 | 0 | 0.16 | 0.13 | 0.19 | 0.02 | 0.28 | 0.11 |
| 7 | 150963758 | rs11973301 | 0.204 | 0.088 | -1.053 | -2.049 | 0.012 | 0.072 | 0.072 | 0 | 0.61 | 0.75 | 0.5 | 0.16 | 0.59 | 0.56 |
| 7 | 150971430 | rs3789818 | 0.136 | -0.582 | -0.675 | -2.037 | -0.012 | -0.051 | -0.051 | 1.529 | 0.63 | 0.42 | 0.56 | 0.16 | 0.67 | 0.54 |
| 7 | 150972613 | rs2530457 | -0.678 | -1.123 | -0.768 | -2.073 | -0.031 | -0.127 | -0.127 | 0 | 0.32 | 0.54 | 0.24 | 0.03 | 0.17 | 0.14 |
| 7 | 150991795 | rs219249 | 0.535 | -0.439 | -1.255 | -1.969 | -0.037 | 0.175 | 0.175 | 0 | 0.45 | 0.74 | 0.43 | 0.14 | 0.47 | 0.42 |
| 7 | 150993833 | rs219261 | -0.469 | -0.816 | -1.568 | -2.309 | 0.008 | 0.074 | 0.074 | 0.484 | 0.71 | 0.85 | 0.63 | 0.16 | 0.72 | 0.54 |
| 7 | 150995567 | rs7804424 | -0.298 | -0.418 | -1.276 | -2.049 | -0.009 | 0.094 | 0.094 | 1.335 | 0.26 | 0.11 | 0.2 | 0.02 | 0.24 | 0.12 |
| 7 | 151005056 | rs369410 | 0.944 | -0.562 | -0.787 | -1.685 | -0.003 | 0.019 | 0.019 | 0 | 0.47 | 0.4 | 0.44 | 0.21 | 0.52 | 0.38 |
| 7 | 151006529 | rs6964641 | -0.695 | -0.618 | -0.678 | -0.937 | 0.002 | -0.011 | -0.011 | 0.015 | 0.29 | 0.48 | 0.35 | 0.27 | 0.26 | 0.19 |
| 7 | 151009517 | rs378140 | 0.78 | -0.847 | -0.513 | -1.116 | 0.022 | -0.011 | -0.011 | 0 | 0.56 | 0.68 | 0.64 | 0.48 | 0.65 | 0.49 |
| 7 | 151013263 | rs432519 | -0.163 | -0.59 | -0.678 | -1.038 | 0.016 | 0.027 | 0.027 | 2.097 | 0.8 | 0.88 | 0.86 | 0.99 | 0.85 | 0.9 |
| 7 | 151014615 | rs7811116 | 0.278 | -0.101 | -0.529 | -0.358 | -0.012 | -0.107 | -0.107 | 0 | 0.2 | 0.07 | 0.2 | 0.51 | 0.2 | 0.41 |
| 7 | 151016648 | rs2975199 | -0.053 | -0.521 | -0.448 | -0.313 | -0.039 | 0.088 | 0.088 | 0.505 | 0.25 | 0.38 | 0.2 | 0.51 | 0.16 | 0.4 |
| 7 | 151024986 | rs17705456 | 0.057 | 0.573 | -1.041 | -0.364 | -0.039 | 0.019 | 0.019 | 0 | 0.06 | 0.01 | 0.06 | 0.04 | 0.11 | 0.06 |
| 7 | 151031507 | rs7807594 | -0.609 | 0.044 | -0.859 | -0.313 | 0 | 0.003 | 0.003 | 0 | 0.24 | 0.31 | 0.14 | 0.25 | 0.23 | 0.26 |
| 7 | 151033628 | rs2487161 | -0.609 | -0.953 | -0.776 | -0.204 | -0.043 | 0.021 | 0.021 | 0.228 | 0.28 | 0.56 | 0.5 | 0.65 | 0.16 | 0.36 |
| 7 | 151037121 | rs2487149 | -0.427 | -0.458 | -1.139 | 0.131 | 0.004 | -0.005 | -0.005 | 0 | 0.18 | 0.21 | 0.26 | 0.35 | 0.14 | 0.19 |
| 7 | 151040394 | rs2063995 | -0.335 | 0.559 | -1.702 | -1.156 | -0.072 | 0.189 | 0.189 | 0.73 | 0.61 | 0.89 | 0.75 | 0.96 | 0.56 | 0.58 |
| 7 | 151043272 | rs7805834 | -0.525 | -0.17 | -0.607 | -0.866 | -0.03 | -0.027 | -0.027 | 0 | 0.17 | 0.29 | 0.07 | 0.11 | 0.11 | 0.16 |
| 7 | 151043700 | rs438559 | -0.536 | -0.168 | -0.428 | -1.868 | -0.001 | -0.062 | -0.062 | 0.154 | 0.74 | 0.71 | 0.77 | 0.98 | 0.78 | 0.71 |
| 7 | 151049337 | rs394259 | -0.824 | -0.134 | -0.315 | -1.933 | 0.003 | 0.083 | 0.083 | 0 | 0.78 | 0.93 | 0.88 | 0.98 | 0.83 | 0.76 |
| 7 | 151050158 | rs367547 | 0.623 | -0.329 | -0.158 | -2.024 | -0.022 | -0.074 | -0.074 | 0 | 0.6 | 0.57 | 0.8 | 0.88 | 0.72 | 0.6 |
| 7 | 151052147 | rs434705 | 0.598 | -0.398 | -0.103 | -2.055 | -0.016 | -0.058 | -0.058 | 0 | 0.59 | 0.51 | 0.79 | 0.87 | 0.72 | 0.6 |

| | | | | | | | | | | | | | | | | |
|---|---|---|---|---|---|---|---|---|---|---|---|---|---|---|---|---|
| 7 | 151053043 | rs446886 | 0.762 | -0.4 | -0.137 | -2.05 | -0.002 | -0.029 | -0.029 | 0.882 | 0.58 | 0.48 | 0.71 | 0.87 | 0.67 | 0.55 |
| 7 | 151059941 | rs10275580 | 0.184 | -0.661 | -0.07 | -2.022 | -0.051 | -0.026 | -0.026 | 0 | 0.16 | 0.3 | 0.06 | 0.11 | 0.07 | 0.15 |
| 7 | 151060074 | rs446079 | 0.405 | -0.826 | 0.206 | -2.121 | 0.028 | 0.052 | 0.052 | 1.444 | 0.6 | 0.38 | 0.72 | 0.87 | 0.72 | 0.56 |
| 7 | 151062348 | rs405281 | -0.055 | -0.193 | -0.282 | -1.26 | -0.068 | -0.08 | -0.08 | 0 | 0.25 | 0.15 | 0.34 | 0.41 | 0.4 | 0.26 |
| 7 | 151064422 | rs367567 | 0.018 | -2.029 | -0.417 | -1.457 | 0.009 | -0.042 | -0.042 | 0 | 0.39 | 0.25 | 0.47 | 0.47 | 0.63 | 0.3 |
| 7 | 151064872 | rs874607 | -0.627 | -2.051 | 0.119 | -1.182 | -0.001 | 0.253 | 0.253 | 0 | 0.13 | 0.41 | 0.29 | 0.41 | 0.09 | 0.16 |
| 7 | 151066541 | rs13234689 | 0.668 | -1.863 | -0.178 | -1.293 | -0.038 | -0.052 | -0.052 | 0 | 0.3 | 0.22 | 0.42 | 0.45 | 0.54 | 0.24 |
| 7 | 151068262 | rs6957858 | -2.047 | -2.095 | -0.137 | -1.45 | 0.008 | 0.007 | 0.007 | 0.097 | 0.78 | 0.83 | 0.89 | 0.99 | 0.84 | 0.7 |
| 7 | 151069296 | rs6963048 | 0.167 | -1.922 | -0.341 | -1.304 | -0.015 | -0.037 | -0.037 | 0 | 0.09 | 0.02 | 0.26 | 0.38 | 0.09 | 0.12 |
| 7 | 151069411 | rs2108726 | -0.62 | -2.285 | -0.26 | -1.515 | 0.039 | 0.073 | 0.073 | 0.219 | 0.53 | 0.39 | 0.73 | 0.87 | 0.68 | 0.46 |
| 7 | 151070531 | rs12668261 | 0.694 | -1.833 | -0.519 | -1.341 | -0.044 | -0.058 | -0.058 | 0 | 0.3 | 0.22 | 0.43 | 0.45 | 0.54 | 0.24 |
| 7 | 151074296 | rs2159158 | 0.533 | -1.809 | -0.681 | -1.33 | -0.078 | -0.083 | -0.083 | 0.242 | 0.3 | 0.12 | 0.4 | 0.46 | 0.52 | 0.24 |
| 7 | 151074479 | rs6952675 | -0.061 | -2.142 | -0.396 | -1.47 | -0.016 | 0.014 | 0.014 | 0 | 0.31 | 0.38 | 0.42 | 0.48 | 0.23 | 0.34 |
| 7 | 151076574 | rs7795959 | -0.48 | -2.146 | -1.204 | -1.464 | 0.035 | 0.02 | 0.02 | 0 | 0.52 | 0.29 | 0.47 | 0.51 | 0.62 | 0.44 |
| 7 | 151078218 | rs2286418 | 1.286 | -1.505 | -1.321 | -1.215 | 0.043 | 0.026 | 0.026 | 0.024 | 0.13 | 0.01 | 0.04 | 0.05 | 0.06 | 0.11 |
| 7 | 151079871 | rs758250 | -0.854 | -1.125 | -0.976 | -1.139 | -0.01 | -0.095 | -0.095 | 0 | 0.63 | 0.58 | 0.83 | 0.98 | 0.73 | 0.6 |
| 7 | 151086797 | rs1558164 | -0.06 | -0.133 | -1.469 | 0.014 | -0.079 | -0.091 | -0.091 | 2.015 | 0.29 | 0.12 | 0.48 | 0.43 | 0.54 | 0.23 |
| 7 | 151088648 | rs2240627 | -0.482 | -0.195 | -1.374 | 0.414 | 0.028 | -0.07 | -0.07 | 0.85 | 0.22 | 0.14 | 0.26 | 0.45 | 0.17 | 0.23 |
| 7 | 151091890 | rs11760463 | -0.397 | 0.275 | -1.555 | 0.796 | -0.143 | -0.086 | -0.086 | 0 | 0.23 | 0.08 | 0.37 | 0.34 | 0.49 | 0.27 |
| 7 | 151092775 | rs3815332 | -0.049 | -0.475 | -0.996 | 0.64 | -0.043 | 0.047 | 0.047 | 0 | 0.3 | 0.37 | 0.26 | 0.5 | 0.18 | 0.28 |
| 7 | 151094048 | rs11761588 | -0.398 | -0.448 | -0.639 | 0.038 | 0.011 | 0.012 | 0.012 | 0.48 | 0.56 | 0.58 | 0.74 | 0.82 | 0.7 | 0.56 |
| 7 | 151098249 | rs2019202 | -0.196 | -0.006 | -0.47 | 0.271 | 0.001 | 0.001 | 0.001 | 0 | 0.5 | 0.51 | 0.64 | 0.68 | 0.61 | 0.43 |
| 7 | 151099634 | rs1004340 | -0.673 | 0.052 | -0.642 | 0.037 | -0.001 | -0.002 | -0.002 | 0 | 0.45 | 0.62 | 0.33 | 0.33 | 0.44 | 0.26 |
| 7 | 151101253 | rs877155 | -0.53 | -0.008 | -0.498 | -0.536 | -0.007 | -0.077 | -0.077 | 0.973 | 0.26 | 0.33 | 0.2 | 0.02 | 0.27 | 0.13 |
| 7 | 151106066 | rs12533730 | -0.043 | -0.462 | -0.744 | -0.049 | 0.039 | -0.022 | -0.022 | 0 | 0.41 | 0.26 | 0.24 | 0.51 | 0.27 | 0.43 |
| 7 | 151119265 | rs10231083 | -0.02 | -0.552 | -0.436 | -0.15 | 0.026 | -0.03 | -0.03 | 0 | 0.12 | 0.24 | 0.13 | 0.05 | 0.16 | 0.07 |
| 7 | 151121676 | rs1558299 | 0.398 | -0.914 | -0.678 | -0.217 | 0.088 | -0.097 | -0.097 | 0 | 0.3 | 0.1 | 0.12 | 0.49 | 0.11 | 0.38 |
| 7 | 151122503 | rs7806340 | -0.468 | -0.858 | -0.426 | -0.277 | -0.015 | -0.003 | -0.003 | 0.005 | 0.47 | 0.56 | 0.27 | 0.55 | 0.28 | 0.45 |
| 7 | 151123529 | rs7792547 | -0.293 | 0.017 | -0.8 | -0.576 | 0.003 | 0 | 0 | 0 | 0.59 | 0.44 | 0.39 | 0.58 | 0.52 | 0.57 |
| 7 | 151125123 | rs10227136 | -0.34 | -0.042 | -0.798 | -0.448 | 0.035 | 0.042 | 0.042 | 0.501 | 0.53 | 0.73 | 0.3 | 0.62 | 0.36 | 0.54 |
| 7 | 151126822 | rs7808369 | 0.179 | 0.109 | -0.447 | -0.023 | 0.011 | 0.006 | 0.006 | 0 | 0.12 | 0.12 | 0.17 | 0.15 | 0.19 | 0.07 |
| 7 | 151126851 | rs11768968 | 1.411 | 0.643 | -0.714 | -0.197 | -0.004 | 0.004 | 0.004 | 0 | 0.09 | 0.07 | 0.06 | 0.04 | 0.1 | 0.15 |
| 7 | 151131811 | rs10252902 | -0.938 | -0.607 | -0.488 | -0.363 | 0.003 | 0.004 | 0.004 | 1.81 | 0.33 | 0.69 | 0.25 | 0.24 | 0.26 | 0.21 |
| 7 | 151132665 | rs2075002 | 1.334 | -0.711 | -0.752 | -0.067 | 0.049 | -0.079 | -0.079 | 0 | 0.21 | 0.06 | 0.04 | 0.42 | 0.1 | 0.37 |
| 7 | 151133271 | rs2075001 | 1.307 | -0.731 | -0.761 | -0.042 | 0.049 | -0.081 | -0.081 | 0 | 0.21 | 0.06 | 0.04 | 0.43 | 0.1 | 0.37 |
| 7 | 151133792 | rs2075000 | 1.099 | -0.779 | -0.34 | 0.087 | 0.014 | -0.02 | -0.02 | 0 | 0.22 | 0.15 | 0.04 | 0.45 | 0.1 | 0.37 |
| 7 | 151135503 | rs12539860 | -0.797 | -0.489 | -0.296 | 0.005 | 0.047 | 0.141 | 0.141 | 2.23 | 0.49 | 0.72 | 0.34 | 0.81 | 0.36 | 0.59 |
| 7 | 151140431 | rs6464142 | 0.049 | -0.406 | 0.172 | 0.27 | -0.01 | 0.019 | 0.019 | 0.51 | 0.33 | 0.41 | 0.21 | 0.6 | 0.28 | 0.43 |
| 7 | 151140792 | rs6464144 | -0.008 | -0.409 | 0.18 | 0.181 | -0.011 | 0.001 | 0.001 | 1.042 | 0.33 | 0.43 | 0.22 | 0.66 | 0.28 | 0.45 |
| 7 | 151147122 | rs7784512 | -0.082 | 0.238 | -0.233 | 0.375 | 0.002 | 0.015 | 0.015 | 0 | 0.52 | 0.46 | 0.54 | 0.67 | 0.55 | 0.62 |
| 7 | 151152665 | rs9640291 | -0.044 | 0.27 | 0.327 | 0.271 | 0 | -0.003 | -0.003 | 0 | 0.48 | 0.5 | 0.51 | 0.65 | 0.51 | 0.61 |
| 7 | 151160602 | rs12669535 | 0.932 | 0.12 | 0.239 | 0.185 | -0.015 | 0.011 | 0.011 | 0 | 0.09 | 0.18 | 0.1 | 0.11 | 0.04 | 0.03 |
| 7 | 151171662 | rs6972955 | 0.443 | -0.28 | 0.338 | 0.67 | 0.003 | 0.002 | 0.002 | 0 | 0.38 | 0.42 | 0.44 | 0.53 | 0.48 | 0.58 |
| 7 | 151172878 | rs12671654 | 1.163 | -0.014 | -0.399 | 0.45 | -0.068 | -0.069 | -0.069 | 1.737 | 0.14 | 0.01 | 0.12 | 0.21 | 0.2 | 0.17 |
| 7 | 151177134 | rs2299962 | -0.141 | -0.525 | -0.349 | 0.13 | -0.001 | 0.01 | 0.01 | 0.11 | 0.6 | 0.72 | 0.59 | 0.65 | 0.59 | 0.69 |
| 7 | 151184985 | rs3753151 | 0.493 | -0.51 | -0.087 | 0.415 | -0.001 | -0.01 | -0.01 | 0.008 | 0.58 | 0.65 | 0.57 | 0.54 | 0.58 | 0.66 |
| 7 | 151185656 | rs13224450 | -0.206 | -0.348 | 0.063 | 0.104 | 0 | -0.003 | -0.003 | 0.779 | 0.78 | 0.75 | 0.78 | 0.87 | 0.8 | 0.87 |
| 7 | 151215950 | rs6943752 | 0.447 | -0.291 | -0.056 | 0.418 | 0 | 0.002 | 0.002 | 0 | 0.51 | 0.6 | 0.51 | 0.57 | 0.53 | 0.63 |
| 7 | 151221936 | rs2058495 | 0.58 | 0.15 | -0.739 | 0.227 | -0.029 | -0.028 | -0.028 | 1.67 | 0.14 | 0.08 | 0.12 | 0.21 | 0.21 | 0.18 |
| 7 | 151225904 | rs4549696 | 0.759 | -1.14 | -0.769 | -0.621 | 0.072 | -0.015 | -0.015 | 0 | 0.56 | 0.08 | 0.54 | 0.58 | 0.7 | 0.65 |
| 7 | 151226122 | rs6951571 | -0.566 | -0.778 | -0.251 | -1.013 | -0.007 | -0.077 | -0.077 | 0.166 | 0.73 | 0.67 | 0.77 | 0.99 | 0.77 | 0.86 |
| 7 | 151243232 | rs12703141 | -0.218 | -0.838 | -0.127 | -0.972 | -0.019 | -0.144 | -0.144 | 2.023 | 0.7 | 0.48 | 0.75 | 0.99 | 0.78 | 0.85 |
| 7 | 151247714 | rs6963512 | 0.441 | -0.72 | -0.264 | -0.587 | -0.003 | 0.026 | 0.026 | 0 | 0.12 | 0.18 | 0.17 | 0.26 | 0.09 | 0.21 |
| 7 | 151250404 | rs12703143 | 0.85 | -1.13 | -0.385 | -0.669 | 0.028 | 0.018 | 0.018 | 0 | 0.56 | 0.17 | 0.52 | 0.6 | 0.67 | 0.64 |
| 7 | 151251745 | rs2041635 | -0.636 | -1.118 | 0.095 | -0.777 | -0.011 | -0.131 | -0.131 | 0 | 0.72 | 0.49 | 0.76 | 0.99 | 0.78 | 0.85 |
| 7 | 151253765 | rs1051956 | 0.665 | -1.157 | -0.03 | -0.739 | 0.019 | 0.061 | 0.061 | 0 | 0.59 | 0.24 | 0.57 | 0.73 | 0.68 | 0.64 |
| 7 | 151254175 | rs8961 | 0.526 | -1.113 | -0.053 | -0.78 | 0.015 | 0.05 | 0.05 | 0 | 0.6 | 0.25 | 0.57 | 0.73 | 0.68 | 0.64 |
| 7 | 151255160 | rs5017429 | -0.934 | -0.893 | 0.178 | -0.751 | -0.001 | -0.063 | -0.063 | 0 | 0.76 | 0.74 | 0.78 | 0.99 | 0.78 | 0.85 |
| 7 | 151260097 | rs11982878 | -0.852 | -0.901 | -0.286 | -0.874 | -0.007 | 0.013 | 0.013 | 2.856 | 0.32 | 0.61 | 0.38 | 0.42 | 0.28 | 0.29 |
| 7 | 151265714 | rs2241053 | -0.753 | -0.806 | -0.337 | -0.831 | -0.006 | 0.021 | 0.021 | 2.703 | 0.31 | 0.6 | 0.38 | 0.42 | 0.28 | 0.29 |
| 7 | 151268920 | rs17173197 | 0.059 | -1.027 | -0.659 | -1.142 | -0.016 | -0.034 | -0.034 | 0 | 0.31 | 0.21 | 0.4 | 0.47 | 0.41 | 0.38 |
| 7 | 151272464 | rs7791529 | -1.523 | -0.971 | -0.347 | -0.887 | -0.01 | -0.028 | -0.028 | 0.089 | 0.2 | 0.56 | 0.22 | 0.17 | 0.17 | 0.07 |
| 7 | 151275806 | rs1029946 | -0.288 | -0.904 | -0.264 | -0.661 | -0.035 | 0.051 | 0.051 | 0 | 0.22 | 0.38 | 0.25 | 0.41 | 0.11 | 0.24 |
| 7 | 151280892 | rs6953318 | 0.459 | -0.741 | -0.391 | -0.635 | -0.014 | 0.01 | 0.01 | 0 | 0.19 | 0.25 | 0.19 | 0.26 | 0.11 | 0.23 |
| 7 | 151283664 | rs2538039 | -0.128 | -0.721 | -0.587 | -0.876 | -0.002 | 0.023 | 0.023 | 0 | 0.57 | 0.41 | 0.58 | 0.82 | 0.58 | 0.77 |
| 7 | 151283992 | rs2538038 | 0.905 | -0.719 | -0.767 | -0.852 | -0.144 | -0.122 | -0.122 | 0 | 0.38 | 0.02 | 0.38 | 0.56 | 0.47 | 0.55 |
| 7 | 151285976 | rs12164128 | 1.376 | -0.711 | -0.222 | -0.258 | -0.006 | 0.047 | 0.047 | 0 | 0.14 | 0.19 | 0.19 | 0.31 | 0.11 | 0.23 |
| 7 | 151286730 | rs4726058 | 0.398 | -0.707 | -0.234 | -0.607 | -0.033 | 0.032 | 0.032 | 0 | 0.19 | 0.39 | 0.2 | 0.26 | 0.11 | 0.23 |
| 7 | 151306444 | rs2727567 | 0.249 | -0.579 | -1.045 | -0.739 | -0.028 | -0.047 | -0.047 | 0 | 0.47 | 0.04 | 0.49 | 0.56 | 0.63 | 0.59 |

| | | | | | | | | | | | | | | | | |
|---|---|---|---|---|---|---|---|---|---|---|---|---|---|---|---|---|
| 7 | 151314525 | rs2538045 | -0.666 | -0.534 | -0.576 | -0.708 | -0.006 | -0.019 | -0.019 | 0 | 0.59 | 0.51 | 0.6 | 0.72 | 0.66 | 0.62 |
| 7 | 151317512 | rs2538034 | -0.476 | -0.483 | -0.734 | -0.69 | 0.006 | 0.013 | 0.013 | 0 | 0.57 | 0.42 | 0.59 | 0.73 | 0.66 | 0.62 |
| 7 | 151321161 | rs2727565 | 1.062 | -0.206 | -1.015 | -0.716 | -0.047 | -0.176 | -0.176 | 0 | 0.23 | 0.01 | 0.27 | 0.46 | 0.27 | 0.31 |
| 7 | 151324155 | rs870441 | 0.767 | 0.403 | -0.271 | -1.022 | 0.036 | -0.033 | -0.033 | 0 | 0.07 | 0.19 | 0.1 | 0.01 | 0.13 | 0.05 |
| 7 | 151328218 | rs4726070 | -1.085 | 0.43 | -0.397 | -1.218 | 0.014 | 0.077 | 0.077 | 0 | 0.57 | 0.81 | 0.54 | 0.25 | 0.58 | 0.38 |
| 7 | 151330390 | rs6964957 | -0.953 | 0.188 | 0.12 | -1.403 | -0.001 | -0.044 | -0.044 | 0.426 | 0.38 | 0.66 | 0.36 | 0.03 | 0.37 | 0.17 |
| 7 | 151335260 | rs4726075 | -1.056 | 0.2 | 0.314 | -0.396 | 0.038 | -0.048 | -0.048 | 0 | 0.1 | 0.27 | 0.1 | 0.01 | 0.14 | 0.05 |
| 7 | 151350262 | rs4726081 | -0.262 | -0.038 | -0.596 | -0.143 | -0.003 | -0.006 | -0.006 | 0 | 0.34 | 0.35 | 0.36 | 0.2 | 0.34 | 0.27 |
| 7 | 151354400 | rs1860735 | -1.404 | 0.155 | -0.975 | -0.139 | -0.033 | -0.032 | -0.032 | 0 | 0.9 | 1 | 0.97 | 0.84 | 0.89 | 0.89 |
| 7 | 151356274 | rs2727529 | 0.171 | 0.377 | -0.76 | -0.213 | 0.008 | 0.015 | 0.015 | 0 | 0.57 | 0.66 | 0.61 | 0.63 | 0.56 | 0.62 |
| 7 | 151359434 | rs953221 | 0.687 | 0.412 | -1.193 | -0.529 | -0.098 | 0.08 | 0.08 | 0.043 | 0.25 | 0.05 | 0.38 | 0.1 | 0.37 | 0.2 |
| 7 | 151375542 | rs2727541 | 0.45 | -0.351 | -0.164 | -0.08 | -0.001 | -0.001 | -0.001 | 0 | 0.4 | 0.39 | 0.45 | 0.28 | 0.45 | 0.36 |
| 7 | 151378505 | rs6978142 | -0.126 | -0.995 | 0.144 | 0.125 | -0.003 | -0.003 | -0.003 | 0 | 0.44 | 0.42 | 0.36 | 0.35 | 0.38 | 0.35 |
| 7 | 151381200 | rs2536090 | 0.368 | -0.41 | 0.532 | -0.22 | -0.017 | 0.048 | 0.048 | 0 | 0.41 | 0.29 | 0.47 | 0.28 | 0.49 | 0.36 |
| 7 | 151384503 | rs4726084 | 0.057 | -0.382 | 0.365 | -0.103 | 0.004 | -0.001 | -0.001 | 1.072 | 0.24 | 0.18 | 0.21 | 0.23 | 0.21 | 0.15 |
| 7 | 151387197 | rs10277859 | 0.015 | -0.445 | 0.519 | -1.286 | -0.018 | -0.103 | -0.103 | 0 | 0.26 | 0.37 | 0.15 | 0.01 | 0.21 | 0.07 |
| 7 | 151387520 | rs6980380 | 0.185 | -0.419 | -0.199 | -1.268 | 0.008 | 0 | 0 | 5.185 | 0.24 | 0.19 | 0.13 | 0.01 | 0.2 | 0.07 |
| 7 | 151387531 | rs2536085 | 0.106 | -0.111 | -0.102 | -0.4 | 0.01 | 0.017 | 0.017 | 0 | 0.65 | 0.75 | 0.63 | 0.31 | 0.69 | 0.42 |
| 7 | 151391653 | rs2727551 | -1.706 | 0.197 | 0.696 | 0.312 | -0.011 | 0.137 | 0.137 | 1.053 | 0.13 | 0.38 | 0.21 | 0.26 | 0.07 | 0.2 |
| 7 | 151391821 | rs2536083 | -1.694 | 0.255 | 0.088 | 0.306 | 0.008 | 0.045 | 0.045 | 0 | 0.06 | 0.17 | 0.06 | 0.17 | 0.07 | 0.09 |
| 7 | 151392009 | rs7778356 | -0.923 | 0.585 | -0.379 | 0.028 | 0.003 | -0.003 | -0.003 | 0.264 | 0.21 | 0.24 | 0.24 | 0.18 | 0.28 | 0.14 |
| 7 | 151393484 | rs2727552 | -1.423 | 0.387 | -1.286 | -0.106 | -0.103 | 0.016 | 0.016 | 1.874 | 0.44 | 0.97 | 0.54 | 0.34 | 0.47 | 0.36 |
| 7 | 151395349 | rs7780804 | -0.766 | 0.687 | -0.379 | -0.081 | 0.019 | -0.009 | -0.009 | 0.949 | 0.2 | 0.29 | 0.22 | 0.17 | 0.27 | 0.17 |
| 7 | 151395637 | rs9648723 | -1.097 | 0.227 | 0.113 | -0.092 | -0.014 | -0.018 | -0.018 | 0.818 | 0.22 | 0.67 | 0.31 | 0.17 | 0.19 | 0.17 |
| 7 | 151396253 | rs6952398 | -1.123 | -0.441 | 0.153 | -0.535 | 0.007 | 0.009 | 0.009 | 0 | 0.41 | 0.62 | 0.41 | 0.15 | 0.38 | 0.3 |
| 7 | 151397343 | rs6947064 | -0.902 | 0.095 | 0.384 | -0.115 | 0.007 | -0.004 | -0.004 | 0 | 0.27 | 0.36 | 0.28 | 0.21 | 0.38 | 0.2 |
| 7 | 151400498 | rs10235949 | -1.098 | -0.418 | 0.728 | 0.133 | -0.023 | 0.005 | 0.005 | 0.139 | 0.1 | 0.28 | 0.17 | 0.08 | 0.03 | 0.12 |
| 7 | 151402500 | rs2727564 | -1.047 | -0.775 | -0.542 | 0.269 | -0.128 | -0.119 | -0.119 | 0.019 | 0.61 | 0.91 | 0.54 | 0.55 | 0.51 | 0.65 |
| 7 | 151402790 | rs2374270 | -0.714 | -0.767 | 0.471 | 0.332 | 0.001 | -0.01 | -0.01 | 0 | 0.35 | 0.36 | 0.35 | 0.26 | 0.37 | 0.3 |
| 7 | 151403089 | rs4725423 | -1.186 | -1.167 | 1.079 | 0.061 | -0.074 | -0.015 | -0.015 | 0.039 | 0.18 | 0.39 | 0.23 | 0.17 | 0.06 | 0.17 |
| 7 | 151405818 | rs10480299 | 0.402 | -0.935 | 1.005 | -0.05 | 0.038 | -0.109 | -0.109 | 0.05 | 0.2 | 0.32 | 0.17 | 0 | 0.29 | 0.07 |
| 7 | 151409480 | rs4725424 | -0.727 | -1.993 | 0.156 | 0.711 | 0.033 | 0.031 | 0.031 | 0 | 0.42 | 0.3 | 0.43 | 0.68 | 0.25 | 0.58 |
| 7 | 151415181 | rs2536076 | -0.021 | -1.511 | -0.356 | 1.379 | -0.027 | -0.169 | -0.169 | 0 | 0.22 | 0.01 | 0.24 | 0.49 | 0.17 | 0.31 |
| 7 | 151415554 | rs10952317 | -1.313 | -1.097 | 0.642 | 0.754 | -0.016 | 0.053 | 0.053 | 3.313 | 0.08 | 0.31 | 0.1 | 0.14 | 0.02 | 0.06 |
| 7 | 151419602 | rs2536075 | -0.174 | -1.133 | 0.008 | 0.854 | 0.025 | 0.001 | 0.001 | 0.244 | 0.32 | 0.2 | 0.19 | 0.26 | 0.21 | 0.43 |
| 7 | 151422947 | rs2536069 | -0.429 | -1.26 | 0.644 | 0.796 | -0.015 | -0.003 | -0.003 | 0 | 0.34 | 0.35 | 0.21 | 0.26 | 0.21 | 0.43 |
| 7 | 151423345 | rs2727548 | -0.996 | -0.493 | -0.311 | 0.068 | -0.092 | 0.197 | 0.197 | 0.011 | 0.6 | 0.87 | 0.68 | 0.88 | 0.49 | 0.81 |
| 7 | 151425449 | rs2536067 | -1.102 | -0.451 | -0.283 | 0.031 | -0.082 | 0.173 | 0.173 | 0.547 | 0.62 | 0.85 | 0.68 | 0.88 | 0.5 | 0.84 |
| 7 | 151426105 | rs2536066 | 0.386 | -0.821 | 0.53 | 0.202 | -0.017 | -0.001 | -0.001 | 0.347 | 0.34 | 0.35 | 0.19 | 0.27 | 0.18 | 0.43 |
| 7 | 151430709 | rs4726088 | 0.274 | -0.028 | 0.56 | 0.32 | 0.011 | 0.018 | 0.018 | 0 | 0.28 | 0.34 | 0.49 | 0.53 | 0.4 | 0.42 |
| 7 | 151431466 | rs1108722 | -1.436 | -0.171 | -0.156 | 0.079 | -0.059 | 0.155 | 0.155 | 0.063 | 0.66 | 0.88 | 0.77 | 0.89 | 0.59 | 0.82 |
| 7 | 151432574 | rs1104897 | 0.59 | -0.38 | 0.152 | 0.845 | 0.042 | -0.024 | -0.024 | 0 | 0.34 | 0.23 | 0.33 | 0.49 | 0.16 | 0.43 |
| 7 | 151436352 | rs885273 | -1.233 | -0.074 | -0.138 | 0.567 | -0.056 | 0.172 | 0.172 | 0.128 | 0.66 | 0.88 | 0.78 | 0.92 | 0.59 | 0.8 |
| 7 | 151437958 | rs6464167 | -1.844 | -0.229 | 0.888 | 1.163 | 0.034 | 0.317 | 0.317 | 3.018 | 0.07 | 0.47 | 0.13 | 0.18 | 0.01 | 0.04 |
| 7 | 151440162 | rs10278273 | 1.386 | 0.122 | -0.126 | 0.794 | 0.001 | 0.01 | 0.01 | 0 | 0.13 | 0.08 | 0.08 | 0.12 | 0.13 | 0.14 |
| 7 | 151440364 | rs1001116 | 0.529 | 0.887 | -0.393 | 0.941 | -0.027 | -0.02 | -0.02 | 0 | 0.22 | 0.13 | 0.2 | 0.3 | 0.3 | 0.31 |
| 7 | 151442744 | rs11771330 | -0.577 | 0.772 | -0.189 | 0.83 | 0.007 | -0.001 | -0.001 | 0 | 0.39 | 0.58 | 0.54 | 0.63 | 0.44 | 0.45 |
| 7 | 151444775 | rs7788160 | -0.743 | 0.565 | -1.117 | 0.693 | -0.046 | -0.09 | -0.09 | 0.306 | 0.43 | 0.93 | 0.48 | 0.52 | 0.42 | 0.46 |
| 7 | 151445557 | rs17173258 | -0.123 | -0.785 | -0.919 | -0.659 | 0 | -0.004 | -0.004 | 0.033 | 0.22 | 0.24 | 0.34 | 0.19 | 0.24 | 0.28 |
| 7 | 151446080 | rs6960717 | 0.248 | -1.233 | -1.272 | -1.034 | 0.045 | -0.068 | -0.068 | 0 | 0.27 | 0.13 | 0.3 | 0.35 | 0.17 | 0.36 |
| 7 | 151449553 | rs6970522 | -0.352 | -1.146 | -0.913 | -2.116 | 0.001 | 0.046 | 0.046 | 0.378 | 0.5 | 0.45 | 0.57 | 0.81 | 0.47 | 0.75 |
| 7 | 151452031 | rs7778077 | -0.351 | -0.784 | -1.045 | -2.716 | 0 | -0.05 | -0.05 | 0 | 0.57 | 0.45 | 0.74 | 0.95 | 0.56 | 0.85 |
| 7 | 151452628 | rs7801616 | -0.663 | -0.11 | -0.974 | -2.237 | 0.003 | -0.07 | -0.07 | 0 | 0.61 | 0.62 | 0.73 | 0.95 | 0.53 | 0.88 |
| 7 | 151456082 | rs6965771 | 0.642 | 0.593 | -1.311 | -1.33 | -0.063 | 0.005 | 0.005 | 0 | 0.16 | 0.05 | 0.3 | 0.13 | 0.26 | 0.23 |
| 7 | 151462537 | rs7780008 | -0.19 | -0.033 | -0.709 | -1.793 | -0.012 | 0.074 | 0.074 | 0 | 0.64 | 0.78 | 0.76 | 0.94 | 0.61 | 0.87 |
| 7 | 151463164 | rs7800069 | 0.664 | 0.393 | -0.781 | -1.574 | -0.012 | 0.07 | 0.07 | 0 | 0.42 | 0.28 | 0.39 | 0.73 | 0.45 | 0.57 |
| 7 | 151466187 | rs3934597 | -0.117 | 0.616 | -1.06 | -1.324 | -0.002 | -0.052 | -0.052 | 0 | 0.69 | 0.74 | 0.78 | 0.99 | 0.65 | 0.9 |
| 7 | 151471431 | rs6965834 | -0.63 | 0.107 | -1.111 | -0.408 | 0.005 | 0.005 | 0.005 | 0 | 0.28 | 0.18 | 0.42 | 0.27 | 0.24 | 0.45 |
| 7 | 151473179 | rs4725431 | -1.17 | -0.627 | -0.805 | -0.678 | 0.003 | 0.004 | 0.004 | 0 | 0.21 | 0.21 | 0.3 | 0.2 | 0.17 | 0.26 |
| 7 | 151473418 | rs4725432 | -0.926 | -1.315 | -0.344 | -0.886 | -0.027 | 0.021 | 0.021 | 0 | 0.16 | 0.31 | 0.37 | 0.2 | 0.13 | 0.28 |
| 7 | 151473511 | rs7792110 | -1.31 | -1.911 | -0.768 | -1.18 | 0.139 | 0.109 | 0.109 | 0.391 | 0.46 | 0.87 | 0.54 | 0.32 | 0.29 | 0.49 |
| 7 | 151473813 | rs7809589 | -0.539 | -1.823 | -0.343 | -1.137 | -0.04 | -0.016 | -0.016 | 0.014 | 0.24 | 0.39 | 0.42 | 0.25 | 0.19 | 0.33 |
| 7 | 151473958 | rs7809503 | -0.781 | -1.367 | -1.242 | -1.021 | 0.138 | 0.104 | 0.104 | 2.324 | 0.45 | 0.86 | 0.55 | 0.31 | 0.29 | 0.44 |
| 7 | 151474607 | rs4418266 | 0.445 | 0.357 | -0.669 | -1.078 | -0.008 | 0.028 | 0.028 | 0 | 0.69 | 0.75 | 0.79 | 0.92 | 0.67 | 0.84 |
| 7 | 151484157 | rs4425665 | 1.109 | 0.267 | 0.177 | -0.664 | 0.019 | -0.074 | -0.074 | 0 | 0.88 | 0.59 | 0.92 | 1 | 0.9 | 0.98 |
| 7 | 151491136 | rs7789699 | 0.187 | 0.592 | -0.743 | -0.208 | 0.007 | 0.034 | 0.034 | 0.754 | 0.53 | 0.75 | 0.49 | 0.38 | 0.54 | 0.32 |
| 7 | 151496826 | rs4726096 | 0.253 | 0.835 | -1.148 | 0.197 | -0.015 | -0.025 | -0.025 | 0.014 | 0.6 | 0.72 | 0.55 | 0.44 | 0.52 | 0.38 |
| 7 | 151498404 | rs4128396 | 1.194 | 1.114 | -1.787 | -0.046 | -0.091 | -0.003 | -0.003 | 0.478 | 0.17 | 0.03 | 0.13 | 0.12 | 0.22 | 0.07 |
| 7 | 151500379 | rs4725434 | 0.223 | 0.541 | -1.537 | -0.041 | -0.021 | -0.041 | -0.041 | 0 | 0.28 | 0.22 | 0.4 | 0.38 | 0.29 | 0.12 |

| | | | | | | | | | | | | | | | | |
|---|---|---|---|---|---|---|---|---|---|---|---|---|---|---|---|---|
| 7 | 151502302 | rs9648724 | 0.436 | 0.587 | -1.84 | 0.193 | -0.007 | -0.005 | -0.005 | 0 | 0.22 | 0.14 | 0.18 | 0.23 | 0.26 | 0.25 |
| 7 | 151503301 | rs4442045 | -0.817 | 0.556 | -1.763 | 0.245 | -0.007 | -0.019 | -0.019 | 0.101 | 0.83 | 0.9 | 0.85 | 0.69 | 0.82 | 0.68 |
| 7 | 151504498 | rs12703159 | 0.299 | -0.123 | -1.864 | -0.356 | 0.004 | 0.019 | 0.019 | 0 | 0.19 | 0.12 | 0.21 | 0.1 | 0.19 | 0.33 |
| 7 | 151504499 | rs10257529 | 0.425 | 0.035 | -1.333 | -0.55 | -0.001 | 0.004 | 0.004 | 0 | 0.56 | 0.48 | 0.34 | 0.31 | 0.53 | 0.4 |
| 7 | 151508257 | rs12539356 | 0.956 | 0.316 | -1.614 | -0.365 | -0.055 | 0.045 | 0.045 | 0 | 0.24 | 0.08 | 0.16 | 0.11 | 0.23 | 0.14 |
| 7 | 151510737 | rs10952318 | -0.235 | -0.296 | -0.867 | -0.387 | -0.013 | -0.036 | -0.036 | 0.083 | 0.38 | 0.39 | 0.23 | 0.13 | 0.29 | 0.35 |
| 7 | 151512177 | rs6954429 | 0.661 | -0.039 | -0.463 | 0.592 | -0.005 | 0.043 | 0.043 | 0 | 0.82 | 0.78 | 0.75 | 0.46 | 0.84 | 0.77 |
| 7 | 151512849 | rs11768925 | 1.396 | 0.139 | -1.034 | -0.299 | -0.035 | 0.149 | 0.149 | 2.146 | 0.3 | 0.08 | 0.22 | 0 | 0.27 | 0.1 |
| 7 | 151513282 | rs4726101 | 0.185 | -0.113 | -0.49 | -0.139 | -0.008 | 0.035 | 0.035 | 0 | 0.47 | 0.36 | 0.44 | 0.35 | 0.52 | 0.64 |
| 7 | 151513634 | rs1881633 | -0.885 | 0.005 | -1.125 | 0.33 | -0.003 | 0.047 | 0.047 | 0 | 0.9 | 0.89 | 0.92 | 0.59 | 0.92 | 0.92 |
| 7 | 151516076 | rs6975614 | 1.65 | 0.209 | -1.391 | 0.093 | -0.081 | 0.163 | 0.163 | 1.69 | 0.28 | 0.07 | 0.22 | 0 | 0.29 | 0.1 |
| 7 | 151516341 | rs1881632 | 1.215 | 0.265 | -1.496 | 0.032 | -0.021 | 0.005 | 0.005 | 0.361 | 0.11 | 0.06 | 0.16 | 0.11 | 0.18 | 0.29 |
| 7 | 151517142 | rs1881628 | 1.155 | 0.11 | -1 | 0.249 | -0.041 | 0.036 | 0.036 | 0 | 0.76 | 0.56 | 0.7 | 0.33 | 0.82 | 0.67 |
| 7 | 151517839 | rs13309585 | 0.077 | 0.518 | -0.692 | -0.227 | -0.001 | -0.006 | -0.006 | 0 | 0.58 | 0.54 | 0.57 | 0.75 | 0.62 | 0.66 |
| 7 | 151519128 | rs11770376 | -0.087 | 0.71 | -1.801 | -0.22 | -0.011 | 0.061 | 0.061 | 2.68 | 0.41 | 0.15 | 0.41 | 0.23 | 0.37 | 0.28 |
| 7 | 151520931 | rs7805942 | -0.651 | 0.596 | -1.51 | 0.147 | 0 | 0.013 | 0.013 | 0 | 0.11 | 0.09 | 0.19 | 0.24 | 0.1 | 0.18 |
| 7 | 151525466 | rs11773373 | 0.715 | 0.972 | -1.434 | -0.772 | -0.065 | 0.165 | 0.165 | 1.323 | 0.28 | 0.06 | 0.2 | 0 | 0.27 | 0.08 |
| 7 | 151526091 | rs11760502 | 0.582 | 0.96 | -1.308 | -0.777 | -0.029 | 0.102 | 0.102 | 1.131 | 0.29 | 0.12 | 0.21 | 0 | 0.27 | 0.08 |
| 7 | 151530671 | rs13238117 | -0.166 | 1.101 | -1.138 | -0.215 | -0.02 | 0.002 | 0.002 | 0 | 0.23 | 0.16 | 0.22 | 0.26 | 0.33 | 0.24 |
| 7 | 151530738 | rs11764602 | 1.133 | 1.218 | -1.288 | -0.715 | -0.034 | 0.109 | 0.109 | 7.215 | 0.29 | 0.1 | 0.2 | 0 | 0.3 | 0.07 |
| 7 | 151531019 | rs11765309 | 1.092 | 1.19 | -1.301 | -0.728 | -0.033 | 0.112 | 0.112 | 8.407 | 0.29 | 0.11 | 0.2 | 0 | 0.3 | 0.08 |
| 7 | 151531083 | rs1881624 | 0.574 | 0.551 | -0.573 | -0.539 | 0.006 | 0.054 | 0.054 | 1.704 | 0.54 | 0.39 | 0.4 | 0.11 | 0.52 | 0.47 |
| 7 | 151533285 | rs1881623 | -1.141 | 0.395 | -0.347 | 0.674 | 0.001 | 0.152 | 0.152 | 0 | 0.07 | 0.13 | 0.2 | 0.43 | 0.06 | 0.14 |
| 7 | 151540659 | rs10261999 | 0.315 | 0.454 | 0.02 | 0.164 | -0.003 | 0.032 | 0.032 | 2.522 | 0.49 | 0.37 | 0.4 | 0.33 | 0.49 | 0.36 |
| 7 | 151544991 | rs11772236 | -0.159 | 0.447 | -0.039 | -0.085 | 0.004 | 0.003 | 0.003 | 0.004 | 0.4 | 0.35 | 0.4 | 0.4 | 0.38 | 0.34 |
| 7 | 151546589 | rs7795096 | 0.294 | 0.541 | -0.329 | -0.01 | -0.002 | 0.039 | 0.039 | 2.605 | 0.53 | 0.66 | 0.6 | 0.71 | 0.51 | 0.37 |
| 7 | 151551793 | rs3813852 | 0.877 | 0.409 | -1.009 | 0.091 | 0.038 | 0.083 | 0.083 | 0 | 0.23 | 0.04 | 0.16 | 0.11 | 0.18 | 0.2 |
| 7 | 151556824 | rs11971243 | 0.48 | 0.698 | -0.599 | 0.431 | -0.009 | -0.046 | -0.046 | 0 | 0.14 | 0.05 | 0.23 | 0.28 | 0.17 | 0.13 |
| 7 | 151557843 | rs7789674 | 0.477 | 1.159 | -0.144 | 0.452 | 0.035 | -0.045 | -0.045 | 1.132 | 0.48 | 0.16 | 0.45 | 0.56 | 0.46 | 0.49 |
| 7 | 151558997 | rs9640300 | 0.328 | 1.334 | 0.052 | 0.5 | -0.004 | -0.016 | -0.016 | 0 | 0.54 | 0.31 | 0.41 | 0.57 | 0.5 | 0.54 |
| 7 | 151559168 | rs6961971 | 0.244 | 1.677 | -0.322 | -0.652 | 0.007 | -0.075 | -0.075 | 0.016 | 0.6 | 0.6 | 0.7 | 0.93 | 0.54 | 0.71 |
| 7 | 151561573 | rs9640302 | 0.848 | 1.68 | 0.406 | 0.294 | 0.06 | -0.061 | -0.061 | 0 | 0.89 | 0.71 | 0.88 | 0.98 | 0.86 | 0.79 |
| 7 | 151575404 | rs4078431 | -0.919 | 2.083 | -0.04 | 1.186 | -0.002 | 0.005 | 0.005 | 2.617 | 0.2 | 0.2 | 0.48 | 0.49 | 0.34 | 0.37 |
| 7 | 151576111 | rs2031026 | -1.367 | 2.736 | 0.276 | 0.954 | 0.033 | 0.036 | 0.036 | 1.273 | 0.24 | 0.32 | 0.53 | 0.49 | 0.41 | 0.37 |
| 7 | 151580648 | rs12540735 | -0.61 | 1.903 | 0.281 | 0.308 | 0 | 0.016 | 0.016 | 0 | 0.41 | 0.35 | 0.52 | 0.31 | 0.42 | 0.41 |
| 7 | 151581224 | rs1881643 | -0.862 | 1.799 | 0.829 | 0.223 | -0.002 | 0.002 | 0.002 | 0 | 0.45 | 0.58 | 0.59 | 0.31 | 0.5 | 0.42 |
| 7 | 151584802 | rs6950348 | -0.864 | 1.4 | 0.768 | 0.464 | 0.048 | 0.019 | 0.019 | 0.348 | 0.28 | 0.49 | 0.46 | 0.37 | 0.32 | 0.35 |
| 7 | 151588043 | rs10215172 | -1.075 | 0.656 | 0.479 | 0.23 | -0.016 | 0.06 | 0.06 | 0 | 0.52 | 0.75 | 0.62 | 0.41 | 0.5 | 0.46 |
| 7 | 151590285 | rs6464182 | -0.726 | 0.413 | 1.186 | 0.502 | 0.019 | 0.019 | 0.019 | 0.802 | 0.25 | 0.42 | 0.34 | 0.32 | 0.25 | 0.34 |
| 7 | 151593217 | rs10952328 | 0.126 | -0.135 | 0.483 | 0.512 | 0.013 | 0.005 | 0.005 | 0 | 0.37 | 0.23 | 0.26 | 0.31 | 0.29 | 0.44 |
| 7 | 151594907 | rs7788961 | 0.262 | -0.644 | 0.674 | 0.263 | 0.014 | -0.009 | -0.009 | 0.511 | 0.65 | 0.59 | 0.7 | 0.71 | 0.6 | 0.58 |
| 7 | 151596307 | rs2374271 | 0.445 | -1.308 | 0.798 | 0.607 | -0.055 | 0.09 | 0.09 | 0 | 0.76 | 0.62 | 0.6 | 0.6 | 0.84 | 0.67 |
| 7 | 151598556 | rs2374273 | -0.764 | -1.146 | 0.784 | 0.523 | -0.004 | -0.014 | -0.014 | 0 | 0.56 | 0.5 | 0.68 | 0.65 | 0.57 | 0.73 |
| 7 | 151598925 | rs4272280 | 1.021 | -1.14 | 0.356 | 0.503 | -0.012 | 0.039 | 0.039 | 1.873 | 0.41 | 0.19 | 0.29 | 0.34 | 0.43 | 0.27 |
| 7 | 151599503 | rs1534665 | -0.03 | -0.888 | 0.436 | 0.651 | -0.024 | -0.01 | -0.01 | 0 | 0.23 | 0.16 | 0.23 | 0.28 | 0.31 | 0.28 |
| 7 | 151606466 | rs17490405 | 0.553 | -1.019 | -0.026 | 0.331 | -0.028 | 0.045 | 0.045 | 0 | 0.41 | 0.24 | 0.31 | 0.26 | 0.49 | 0.3 |
| 7 | 151608112 | rs13238381 | -0.96 | -1.31 | 0.531 | 0.722 | -0.01 | -0.023 | -0.023 | 0.001 | 0.09 | 0.13 | 0.05 | 0.02 | 0.07 | 0.15 |
| 7 | 151611931 | rs1919520 | 0.354 | -1.489 | 0.291 | 0.68 | -0.036 | 0.07 | 0.07 | 0 | 0.12 | 0.01 | 0.07 | 0 | 0.11 | 0.1 |
| 7 | 151612836 | rs10952331 | -0.144 | -1.66 | 0.477 | 0.981 | -0.013 | 0.023 | 0.023 | 0 | 0.17 | 0.15 | 0.34 | 0.36 | 0.11 | 0.35 |
| 7 | 151613736 | rs13231246 | 0.067 | -1.045 | 0.024 | 0.502 | -0.001 | -0.023 | -0.023 | 0.028 | 0.13 | 0.1 | 0.08 | 0 | 0.11 | 0.1 |
| 7 | 151617897 | rs12703181 | -0.374 | -0.91 | -0.221 | 1.05 | 0.003 | 0.003 | 0.003 | 0 | 0.06 | 0.05 | 0.04 | 0.02 | 0.05 | 0.14 |
| 7 | 151625056 | rs4726114 | 0.64 | -0.62 | -0.108 | 1.494 | -0.021 | 0.07 | 0.07 | 0 | 0.83 | 0.72 | 0.65 | 0.63 | 0.88 | 0.6 |
| 7 | 151629636 | rs7805925 | -0.719 | -0.437 | -0.324 | 1.021 | -0.005 | -0.002 | -0.002 | 1.133 | 0.3 | 0.35 | 0.47 | 0.26 | 0.31 | 0.46 |
| 7 | 151633265 | rs10807990 | 0.848 | -0.396 | -0.238 | 1.452 | -0.01 | 0.012 | 0.012 | 0 | 0.68 | 0.61 | 0.6 | 0.65 | 0.67 | 0.61 |
| 7 | 151634721 | rs10264919 | -0.469 | -0.38 | -0.255 | 1.289 | -0.006 | -0.003 | -0.003 | 2.637 | 0.08 | 0.06 | 0.11 | 0 | 0.15 | 0.16 |
| 7 | 151636309 | rs10952333 | 1.11 | -0.209 | -0.732 | 1.143 | -0.002 | -0.015 | -0.015 | 0.112 | 0.33 | 0.23 | 0.34 | 0.37 | 0.33 | 0.3 |
| 7 | 151639562 | rs10271879 | 0.377 | -0.416 | -0.46 | 0.933 | -0.008 | 0.006 | 0.006 | 1.458 | 0.13 | 0.17 | 0.07 | 0.13 | 0.08 | 0.28 |
| 7 | 151640476 | rs12703185 | 1.078 | -0.356 | -0.394 | 0.887 | -0.011 | 0.01 | 0.01 | 0 | 0.23 | 0.24 | 0.22 | 0.15 | 0.29 | 0.16 |
| 7 | 151641278 | rs17569415 | 0.875 | -0.372 | -0.375 | 0.879 | -0.005 | 0.003 | 0.003 | 0.008 | 0.25 | 0.26 | 0.22 | 0.15 | 0.29 | 0.16 |
| 7 | 151642570 | rs11981199 | -0.259 | -0.684 | -0.465 | 0.814 | -0.013 | 0 | 0 | 0 | 0.21 | 0.3 | 0.16 | 0.18 | 0.19 | 0.41 |
| 7 | 151642607 | rs2374313 | 0.74 | -0.791 | -0.308 | 0.765 | -0.001 | 0.038 | 0.038 | 0 | 0.52 | 0.38 | 0.61 | 0.66 | 0.51 | 0.43 |
| 7 | 151644043 | rs2374315 | -0.153 | -0.795 | -0.46 | 0.895 | -0.009 | 0.052 | 0.052 | 1.275 | 0.44 | 0.34 | 0.46 | 0.3 | 0.45 | 0.35 |
| 7 | 151645934 | rs10259821 | -0.755 | -0.945 | -0.639 | 0.779 | -0.006 | -0.001 | -0.001 | 0 | 0.43 | 0.56 | 0.42 | 0.42 | 0.35 | 0.67 |
| 7 | 151648954 | rs7794520 | -0.38 | -0.518 | -0.914 | 0.776 | -0.066 | -0.008 | -0.008 | 0 | 0.72 | 0.89 | 0.75 | 0.73 | 0.64 | 0.86 |
| 7 | 151650180 | rs11763061 | 0.26 | -0.965 | -1.162 | 0.431 | 0.003 | 0.052 | 0.052 | 0.323 | 0.2 | 0.01 | 0.25 | 0.1 | 0.16 | 0.24 |
| 7 | 151650240 | rs11766254 | 0.838 | -1.201 | -1.185 | 0.573 | 0.006 | 0.23 | 0.23 | 0.521 | 0.51 | 0.11 | 0.48 | 0.22 | 0.49 | 0.45 |
| 7 | 151651690 | rs1881648 | -0.265 | -0.104 | -0.232 | 1.2 | 0.016 | 0.029 | 0.029 | 0 | 0.79 | 0.7 | 0.76 | 0.69 | 0.77 | 0.76 |
| 7 | 151652711 | rs7797169 | 0.759 | 0.009 | 0.349 | 1.308 | 0.022 | 0.022 | 0.022 | 0.028 | 0.7 | 0.34 | 0.58 | 0.62 | 0.67 | 0.68 |
| 7 | 151653797 | rs12667405 | -0.287 | -0.121 | 0.167 | 1.191 | -0.022 | 0.016 | 0.016 | 0 | 0.19 | 0.32 | 0.37 | 0.24 | 0.15 | 0.26 |

| | | | | | | | | | | | | | | | | |
|---|---|---|---|---|---|---|---|---|---|---|---|---|---|---|---|---|
| 7 | 151654972 | rs13240848 | -0.06 | -0.149 | 0.06 | 1.236 | -0.036 | -0.029 | -0.029 | 0.707 | 0.75 | 0.84 | 0.72 | 0.61 | 0.67 | 0.69 |
| 7 | 151656464 | rs13221211 | -0.805 | -0.259 | 0.288 | 1.286 | -0.029 | 0.043 | 0.043 | 0 | 0.23 | 0.45 | 0.41 | 0.29 | 0.19 | 0.35 |
| 7 | 151657351 | rs6953624 | -0.475 | -0.299 | 0.268 | 1.279 | -0.038 | 0.034 | 0.034 | 0 | 0.2 | 0.42 | 0.38 | 0.25 | 0.15 | 0.25 |
| 7 | 151660102 | rs1534664 | 0.64 | 0.067 | 0.165 | 1.045 | -0.001 | -0.001 | -0.001 | 0.63 | 0.59 | 0.4 | 0.39 | 0.36 | 0.61 | 0.49 |
| 7 | 151660298 | rs11978256 | 1.036 | 0.109 | 0.151 | 1.039 | 0 | 0.011 | 0.011 | 0 | 0.53 | 0.32 | 0.32 | 0.36 | 0.51 | 0.43 |
| 7 | 151660361 | rs11974761 | 0.714 | -0.053 | 0.221 | 1.13 | -0.001 | -0.001 | -0.001 | 0 | 0.59 | 0.39 | 0.39 | 0.36 | 0.61 | 0.48 |
| 7 | 151662184 | rs10278073 | 0.18 | 0.242 | 0.041 | 0.764 | 0.005 | -0.003 | -0.003 | 0 | 0.76 | 0.76 | 0.78 | 0.81 | 0.72 | 0.82 |
| 7 | 151666018 | rs10243234 | 1.183 | 0.013 | 0.038 | 0.865 | 0.006 | 0.013 | 0.013 | 0 | 0.52 | 0.27 | 0.49 | 0.54 | 0.49 | 0.5 |
| 7 | 151668435 | rs7799325 | 0.257 | 0.433 | 0.581 | 0.738 | 0.008 | -0.019 | -0.019 | 0.297 | 0.69 | 0.47 | 0.76 | 0.76 | 0.72 | 0.82 |
| 7 | 151675777 | rs4726124 | 1.299 | 0.427 | 0.453 | 1.141 | 0.034 | 0.077 | 0.077 | 0.084 | 0.49 | 0.17 | 0.48 | 0.36 | 0.47 | 0.44 |
| 7 | 151676152 | rs7780792 | 1.277 | 0.401 | 1 | 1.128 | 0.019 | 0.035 | 0.035 | 0 | 0.51 | 0.35 | 0.5 | 0.36 | 0.47 | 0.44 |
| 7 | 151677501 | rs7799395 | -0.497 | 0.433 | 1.136 | 1.122 | 0.002 | 0.004 | 0.004 | 0 | 0.83 | 0.79 | 0.77 | 0.78 | 0.79 | 0.71 |
| 7 | 151677673 | rs11770365 | 0.085 | 0.339 | 1.15 | 1.08 | -0.002 | 0.001 | 0.001 | 0.905 | 0.71 | 0.75 | 0.78 | 0.76 | 0.71 | 0.81 |
| 7 | 151678416 | rs9648731 | -1.067 | -0.127 | 1.418 | 1.029 | -0.011 | 0.032 | 0.032 | 0.409 | 0.27 | 0.43 | 0.29 | 0.39 | 0.21 | 0.35 |
| 7 | 151680072 | rs6960270 | -0.224 | -0.848 | 0.946 | 0.921 | -0.015 | -0.003 | -0.003 | 0 | 0.78 | 0.74 | 0.86 | 0.84 | 0.84 | 0.76 |
| 7 | 151681844 | rs4726126 | 0.323 | -0.418 | 1.423 | 1.012 | -0.014 | 0.155 | 0.155 | 0 | 0.78 | 0.54 | 0.59 | 0.53 | 0.81 | 0.68 |
| 7 | 151682382 | rs1919522 | 0.107 | -0.38 | 1.158 | 0.988 | -0.007 | -0.001 | -0.001 | 0 | 0.19 | 0.26 | 0.24 | 0.2 | 0.17 | 0.24 |
| 7 | 151684773 | rs10245251 | 1.158 | -0.186 | 1.452 | 0.909 | 0.083 | 0.379 | 0.379 | 2.293 | 0.94 | 0.63 | 0.79 | 0.74 | 0.95 | 0.92 |
| 7 | 151693502 | rs11766982 | -0.48 | -0.303 | 0.931 | 0.376 | -0.001 | -0.002 | -0.002 | 0 | 0.41 | 0.47 | 0.53 | 0.68 | 0.36 | 0.56 |
| 7 | 151694401 | rs10235029 | -0.395 | -0.326 | 0.448 | 0.605 | -0.007 | 0.206 | 0.206 | 0.783 | 0.09 | 0.26 | 0.25 | 0.34 | 0.06 | 0.11 |
| 7 | 151694438 | rs4591942 | 1.346 | 0.189 | -0.476 | -0.092 | 0 | 0.029 | 0.029 | 0 | 0.29 | 0.21 | 0.18 | 0.16 | 0.28 | 0.22 |
| 7 | 151694486 | rs4236435 | 0.085 | 0.803 | -0.442 | -0.049 | -0.037 | 0.051 | 0.051 | 0.198 | 0.72 | 0.85 | 0.72 | 0.88 | 0.63 | 0.79 |
| 7 | 151697032 | rs11765654 | 0.481 | 0.602 | -0.324 | 0.045 | -0.006 | -0.009 | -0.009 | 0 | 0.61 | 0.67 | 0.65 | 0.56 | 0.6 | 0.61 |
| 7 | 151698373 | rs10251856 | -0.313 | 0.65 | 0.213 | -0.596 | 0.013 | -0.041 | -0.041 | 0.971 | 0.39 | 0.51 | 0.3 | 0.22 | 0.43 | 0.15 |
| 7 | 151703287 | rs4726130 | -0.325 | 0.712 | 0.256 | -0.527 | 0.008 | -0.023 | -0.023 | 1.491 | 0.37 | 0.44 | 0.29 | 0.21 | 0.43 | 0.14 |
| 7 | 151705546 | rs10229388 | -0.007 | 0.734 | 0.038 | -0.412 | -0.109 | 0.055 | 0.055 | 0.23 | 0.3 | 0.09 | 0.23 | 0.22 | 0.37 | 0.13 |
| 7 | 151713518 | rs9655613 | -0.32 | 0.391 | 1.237 | -0.439 | -0.003 | 0.001 | 0.001 | 0.72 | 0.35 | 0.64 | 0.27 | 0.23 | 0.38 | 0.15 |
| 7 | 151716449 | rs7808913 | -0.287 | 0.381 | 1.481 | -0.447 | 0.019 | -0.035 | -0.035 | 0 | 0.36 | 0.55 | 0.27 | 0.22 | 0.4 | 0.2 |
| 7 | 151733340 | rs10267245 | -1.366 | 0.101 | 2.651 | 0.244 | 0.038 | 0.188 | 0.188 | 0 | 0.06 | 0.26 | 0.09 | 0.19 | 0.02 | 0.13 |
| 7 | 151755007 | rs10277069 | -2.24 | 0.049 | 2.788 | 0.256 | 0.04 | 0.228 | 0.228 | 0.478 | 0.08 | 0.47 | 0.11 | 0.19 | 0.02 | 0.13 |
| 7 | 151772124 | rs6944002 | -2.484 | 0.046 | 2.796 | 0.254 | 0.011 | 0.175 | 0.175 | 1.196 | 0.08 | 0.5 | 0.12 | 0.19 | 0.02 | 0.13 |
| 7 | 151776212 | rs10231910 | -2.699 | 0.027 | 2.831 | 0.245 | -0.003 | 0.144 | 0.144 | 1.845 | 0.09 | 0.53 | 0.12 | 0.19 | 0.02 | 0.13 |
| 7 | 151800653 | rs6464203 | -2.632 | 0.034 | 2.76 | 0.161 | -0.001 | 0.131 | 0.131 | 1.639 | 0.09 | 0.53 | 0.12 | 0.17 | 0.02 | 0.13 |
| 7 | 151819022 | rs3778921 | -2.694 | 0.026 | 2.697 | 0.184 | -0.003 | 0.136 | 0.136 | 1.629 | 0.09 | 0.54 | 0.12 | 0.18 | 0.02 | 0.12 |
| 7 | 151826268 | rs3735160 | -0.877 | 0.03 | 2.169 | 0.196 | -0.007 | 0.035 | 0.035 | 0 | 0.05 | 0.1 | 0.03 | 0.13 | 0.02 | 0.12 |
| 7 | 151826337 | rs3735158 | -1.761 | 0.013 | 2.229 | 0.156 | 0.012 | 0.084 | 0.084 | 0 | 0.07 | 0.23 | 0.06 | 0.14 | 0.01 | 0.11 |
| 7 | 151830873 | rs6955966 | -1.386 | 0.218 | 1.791 | 0.172 | 0.055 | 0.214 | 0.214 | 1.054 | 0.06 | 0.35 | 0.11 | 0.16 | 0.01 | 0.1 |
| 7 | 151833692 | rs9969322 | -0.987 | 0.193 | 1.758 | 0.134 | 0.036 | 0.172 | 0.172 | 0 | 0.05 | 0.25 | 0.1 | 0.16 | 0.01 | 0.1 |
| 7 | 151840461 | rs6942743 | -0.987 | 0.194 | 1.662 | 0.135 | 0.016 | 0.135 | 0.135 | 0 | 0.05 | 0.21 | 0.09 | 0.16 | 0.01 | 0.1 |
| 7 | 151847070 | rs3800836 | -0.845 | 0.148 | 1.051 | -0.051 | -0.026 | -0.033 | -0.033 | 5.845 | 0.58 | 0.86 | 0.5 | 0.46 | 0.47 | 0.4 |
| 7 | 151873853 | rs6464211 | -1.595 | -0.543 | 1.265 | 0.312 | 0.037 | -0.007 | -0.007 | 0.849 | 0.28 | 0.75 | 0.24 | 0.3 | 0.17 | 0.28 |
| 7 | 151874498 | rs10252263 | -1.219 | 0.15 | 1.234 | 0.569 | 0.054 | 0.206 | 0.206 | 0.898 | 0.06 | 0.34 | 0.11 | 0.16 | 0.01 | 0.09 |
| 7 | 151888002 | rs3800833 | -0.216 | 0.12 | 1.088 | 0.438 | 0 | -0.004 | -0.004 | 0 | 0.08 | 0.09 | 0.06 | 0.15 | 0.09 | 0.08 |
| 7 | 151955297 | rs6464215 | -1.198 | 0.044 | 1.676 | 0.432 | 0.048 | 0.101 | 0.101 | 0.464 | 0.06 | 0.34 | 0.1 | 0.07 | 0.01 | 0.09 |
| 7 | 152003627 | rs12533279 | -0.998 | -0.599 | 2.152 | 0.236 | -0.024 | 0.004 | 0.004 | 0 | 0.22 | 0.28 | 0.13 | 0.24 | 0.16 | 0.21 |
| 7 | 152009647 | rs7787666 | -1.203 | -0.61 | 2.347 | 0.23 | 0.055 | 0.112 | 0.112 | 0.906 | 0.07 | 0.47 | 0.11 | 0.07 | 0.01 | 0.09 |
| 7 | 152036934 | rs7799662 | -0.898 | -0.608 | 2.417 | 0.231 | 0.066 | 0.123 | 0.123 | 0.397 | 0.06 | 0.39 | 0.11 | 0.07 | 0.01 | 0.09 |
| 7 | 152051190 | rs7781309 | -1.797 | -0.654 | 1.749 | 0.168 | 0.043 | -0.006 | -0.006 | 0 | 0.29 | 0.76 | 0.24 | 0.31 | 0.17 | 0.3 |
| 7 | 152052595 | rs4570064 | 0.851 | 0.035 | 0.832 | -0.305 | -0.095 | 0.063 | 0.063 | 0.837 | 0.3 | 0.02 | 0.27 | 0.16 | 0.34 | 0.16 |
| 7 | 152162209 | rs3087586 | 0.32 | 0.162 | 0.708 | -0.289 | -0.001 | -0.006 | -0.006 | 0 | 0.36 | 0.42 | 0.27 | 0.21 | 0.35 | 0.24 |
| 7 | 152178955 | rs10235893 | -0.325 | 0.768 | 0.459 | 0.406 | 0 | -0.002 | -0.002 | 0 | 0.63 | 0.55 | 0.63 | 0.63 | 0.69 | 0.56 |
| 7 | 152189795 | rs12669177 | -0.802 | 0.9 | 0.028 | -0.157 | 0.006 | 0.063 | 0.063 | 0.448 | 0.21 | 0.64 | 0.31 | 0.41 | 0.2 | 0.22 |
| 7 | 152192121 | rs1107394 | -0.38 | 1.007 | -0.254 | -0.381 | -0.02 | -0.025 | -0.025 | 0.327 | 0.67 | 0.84 | 0.62 | 0.62 | 0.67 | 0.61 |
| 7 | 152199898 | rs4726162 | 0.253 | 0.682 | 0.351 | -1.531 | -0.002 | -0.055 | -0.055 | 0.312 | 0.56 | 0.45 | 0.67 | 0.89 | 0.57 | 0.72 |
| 7 | 152207309 | rs10260295 | -1.365 | 0.45 | 0.766 | -1.218 | 0.027 | 0.011 | 0.011 | 1.281 | 0.07 | 0.41 | 0.05 | 0 | 0.01 | 0 |
| 7 | 152215574 | rs6944891 | -0.185 | 0.611 | -0.159 | -1.498 | -0.083 | 0.125 | 0.125 | 0 | 0.66 | 0.9 | 0.74 | 0.91 | 0.58 | 0.63 |
| 7 | 152220077 | rs10248875 | 0.236 | 0.741 | 0.035 | -1.455 | 0.002 | -0.048 | -0.048 | 0 | 0.58 | 0.56 | 0.7 | 0.91 | 0.58 | 0.63 |
| 7 | 152227620 | rs6952647 | -0.029 | 0.806 | -0.015 | -1.429 | 0.001 | 0.013 | 0.013 | 0 | 0.52 | 0.57 | 0.61 | 0.84 | 0.5 | 0.68 |
| 7 | 152231124 | rs12532839 | 0.255 | 0.748 | -0.384 | -1.611 | 0.003 | 0.005 | 0.005 | 0 | 0.32 | 0.28 | 0.53 | 0.75 | 0.28 | 0.49 |
| 7 | 152242235 | rs6945852 | -0.413 | 0.985 | -0.521 | -1.675 | 0 | -0.025 | -0.025 | 0.415 | 0.56 | 0.55 | 0.69 | 0.84 | 0.55 | 0.66 |
| 7 | 152248829 | rs7784546 | -0.375 | 1.328 | -0.737 | -1.08 | -0.004 | 0.001 | 0.001 | 0 | 0.14 | 0.13 | 0.1 | 0.08 | 0.19 | 0.15 |
| 7 | 152255494 | rs10156134 | 0.772 | 1.357 | -1.045 | -1.289 | -0.013 | 0.035 | 0.035 | 0 | 0.79 | 0.82 | 0.83 | 0.87 | 0.72 | 0.7 |
| 7 | 152255539 | rs10155833 | -0.227 | 1.352 | -0.792 | -1.094 | 0.048 | 0.014 | 0.014 | 0 | 0.3 | 0.45 | 0.37 | 0.37 | 0.38 | 0.3 |
| 7 | 152256468 | rs9640309 | -0.746 | 1.229 | -1.132 | -0.705 | 0.011 | 0.006 | 0.006 | 0 | 0.45 | 0.35 | 0.38 | 0.63 | 0.43 | 0.48 |
| 7 | 152261892 | rs6965541 | 1.148 | 0.694 | -1.424 | -0.691 | -0.008 | -0.004 | -0.004 | 0 | 0.36 | 0.48 | 0.48 | 0.34 | 0.34 | 0.24 |
| 7 | 152262508 | rs36094861 | 1.443 | 0.969 | -1.654 | -0.405 | -0.041 | -0.068 | -0.068 | 0.657 | 0.13 | 0.03 | 0.13 | 0.33 | 0.18 | 0.18 |
| 7 | 152269523 | rs10248434 | 0.126 | 0.597 | -1.27 | -0.666 | 0.01 | -0.023 | -0.023 | 0 | 0.31 | 0.27 | 0.31 | 0.63 | 0.3 | 0.39 |
| 7 | 152272282 | rs4726177 | -1.016 | 0.989 | -1.102 | -0.437 | -0.033 | 0.037 | 0.037 | 0.143 | 0.83 | 0.89 | 0.87 | 0.96 | 0.78 | 0.72 |
| 7 | 152272347 | rs4726178 | 1.576 | 0.172 | -1.04 | -0.429 | -0.021 | 0.05 | 0.05 | 0 | 0.21 | 0.32 | 0.3 | 0.29 | 0.18 | 0.19 |

| | | | | | | | | | | | | | | | | |
|---|---|---|---|---|---|---|---|---|---|---|---|---|---|---|---|---|
| 7 | 152273471 | rs4480029 | -0.641 | 0.133 | -1.175 | -0.152 | 0.013 | 0.011 | 0.011 | 0 | 0.45 | 0.37 | 0.39 | 0.63 | 0.44 | 0.48 |
| 7 | 152274493 | rs10435212 | 0.218 | 0.659 | -1.069 | 0.149 | -0.02 | -0.05 | -0.05 | 0 | 0.16 | 0.08 | 0.14 | 0.33 | 0.2 | 0.19 |
| 7 | 152277531 | rs7789371 | -1.106 | 0.49 | -0.101 | -0.345 | 0.042 | -0.097 | -0.097 | 0 | 0.14 | 0.39 | 0.11 | 0 | 0.15 | 0.09 |
| 7 | 152285741 | rs13237261 | -0.331 | 0.599 | -1.088 | 0.417 | -0.08 | -0.039 | -0.039 | 0 | 0.68 | 0.85 | 0.65 | 0.62 | 0.52 | 0.66 |
| 7 | 152292180 | rs6464250 | -0.448 | 0.18 | -0.657 | 0.252 | 0.002 | 0.005 | 0.005 | 0 | 0.63 | 0.57 | 0.43 | 0.46 | 0.54 | 0.58 |
| 7 | 152297038 | rs10239037 | -0.081 | -0.131 | -1.163 | 0.293 | -0.001 | -0.017 | -0.017 | 0 | 0.36 | 0.22 | 0.22 | 0.38 | 0.31 | 0.33 |
| 7 | 152304317 | rs6967921 | -0.218 | 0.116 | -1.202 | 0.207 | 0 | 0.012 | 0.012 | 0 | 0.1 | 0.06 | 0.04 | 0.06 | 0.07 | 0.12 |
| 7 | 152311405 | rs11762192 | 0.518 | -0.04 | -1.841 | 0.345 | -0.1 | -0.099 | -0.099 | 0 | 0.37 | 0.09 | 0.31 | 0.53 | 0.47 | 0.34 |
| 7 | 152318718 | rs6948868 | 0.233 | -0.697 | -1.29 | -0.01 | 0.009 | 0.012 | 0.012 | 0 | 0.62 | 0.32 | 0.47 | 0.77 | 0.69 | 0.58 |
| 7 | 152323520 | rs2010091 | 0.083 | -0.914 | -0.463 | -0.231 | -0.017 | -0.061 | -0.061 | 0 | 0.68 | 0.43 | 0.54 | 0.83 | 0.72 | 0.7 |
| 7 | 152323962 | rs2310928 | 0.74 | -0.804 | -0.864 | 0.013 | -0.007 | -0.005 | -0.005 | 0.224 | 0.24 | 0.11 | 0.15 | 0.21 | 0.25 | 0.29 |
| 7 | 152330745 | rs4726191 | 0.001 | -1.235 | -0.645 | -0.252 | -0.016 | 0.019 | 0.019 | 0.817 | 0.08 | 0.05 | 0.06 | 0 | 0.08 | 0.05 |
| 7 | 152332052 | rs10952368 | 1.202 | -1.665 | -1.162 | -0.076 | -0.085 | -0.052 | -0.052 | 0 | 0.38 | 0.12 | 0.35 | 0.44 | 0.49 | 0.33 |
| 7 | 152333982 | rs757052 | -1.18 | -1.657 | -0.939 | -0.001 | 0.004 | -0.004 | -0.004 | 0 | 0.89 | 0.91 | 0.92 | 0.85 | 0.95 | 0.95 |
| 7 | 152351316 | rs3218504 | 0.569 | -1.892 | -0.188 | 0.096 | -0.032 | 0.039 | 0.039 | 0.223 | 0.09 | 0.23 | 0.05 | 0.14 | 0.03 | 0.09 |
| 7 | 152360184 | rs3218458 | -0.101 | -1.812 | -0.206 | 0.146 | -0.029 | 0.024 | 0.024 | 0 | 0.07 | 0.12 | 0.25 | 0.15 | 0.05 | 0.11 |
| 7 | 152374163 | rs3218373 | -0.171 | -1.866 | -0.176 | 0.133 | -0.043 | 0.056 | 0.056 | 0 | 0.14 | 0.24 | 0.32 | 0.31 | 0.08 | 0.16 |
| 7 | 152375188 | rs2040639 | -1.191 | -1.604 | -0.466 | -0.024 | -0.081 | -0.022 | -0.022 | 0 | 0.61 | 0.88 | 0.64 | 0.6 | 0.51 | 0.67 |
| 7 | 152380451 | rs6970449 | -0.912 | -0.785 | 0.112 | -0.127 | 0.008 | 0.034 | 0.034 | 0 | 0.45 | 0.7 | 0.52 | 0.3 | 0.43 | 0.51 |
| 7 | 152381150 | rs6464268 | -0.779 | -0.004 | 0.669 | 0.458 | 0.012 | 0.064 | 0.064 | 0.472 | 0.13 | 0.48 | 0.32 | 0.16 | 0.12 | 0.14 |
| 7 | 152381266 | rs12534240 | 0.701 | 0.929 | -0.019 | 0.199 | -0.045 | -0.016 | -0.016 | 0.985 | 0.17 | 0.04 | 0.12 | 0.14 | 0.2 | 0.22 |
| 7 | 152391765 | rs7796764 | 0.761 | 1.214 | 0.762 | 0.388 | 0.016 | 0.019 | 0.019 | 0.523 | 0.75 | 0.71 | 0.65 | 0.49 | 0.63 | 0.72 |
| 7 | 152394108 | rs13236792 | -0.284 | 1.486 | 0.136 | 0.4 | 0.057 | 0.054 | 0.054 | 1.491 | 0.06 | 0.11 | 0.11 | 0.13 | 0.12 | 0.1 |
| 7 | 152397077 | rs4609118 | -0.274 | 1.137 | -0.511 | 0.404 | -0.014 | -0.018 | -0.018 | 0 | 0.83 | 0.86 | 0.74 | 0.62 | 0.75 | 0.82 |
| 7 | 152406533 | rs10253827 | -0.949 | 1.654 | -0.951 | -0.343 | 0.019 | 0.002 | 0.002 | 0 | 0.13 | 0.18 | 0.14 | 0.13 | 0.18 | 0.15 |
| 7 | 152408081 | rs12539937 | -0.088 | 0.952 | -1.289 | -0.78 | -0.007 | 0.125 | 0.125 | 0 | 0.42 | 0.25 | 0.31 | 0.03 | 0.44 | 0.42 |
| 7 | 152420498 | rs7809850 | 0.63 | 0.883 | -0.604 | 0.004 | 0.003 | 0.024 | 0.024 | 0 | 0.68 | 0.4 | 0.48 | 0.68 | 0.65 | 0.53 |
| 7 | 152430770 | rs4601228 | 0.363 | 0.275 | -0.334 | -0.476 | -0.001 | 0.008 | 0.008 | 0 | 0.56 | 0.59 | 0.38 | 0.3 | 0.6 | 0.46 |
| 7 | 152431642 | rs4072875 | -0.288 | -0.131 | -0.71 | -0.788 | 0.006 | 0 | 0 | 0.862 | 0.34 | 0.29 | 0.19 | 0.38 | 0.22 | 0.33 |
| 7 | 152431851 | rs4476919 | 0.713 | -0.378 | -0.203 | -0.853 | 0.008 | 0.059 | 0.059 | 1.316 | 0.7 | 0.51 | 0.43 | 0.5 | 0.67 | 0.62 |
| 7 | 152433972 | rs4496879 | 0.832 | -0.517 | -0.53 | -0.959 | 0.009 | -0.019 | -0.019 | 0 | 0.65 | 0.8 | 0.49 | 0.66 | 0.69 | 0.66 |
| 7 | 152440382 | rs9969169 | 0.637 | -0.581 | -0.178 | -1.651 | 0.003 | -0.007 | -0.007 | 0.064 | 0.26 | 0.2 | 0.5 | 0.71 | 0.21 | 0.35 |
| 7 | 152457803 | rs7809363 | 1.121 | -0.682 | 0.211 | -1.715 | -0.003 | -0.001 | -0.001 | 0 | 0.41 | 0.41 | 0.27 | 0.15 | 0.48 | 0.29 |
| 7 | 152496656 | rs4726207 | 1.338 | 0.523 | 0.757 | -0.957 | 0.004 | -0.002 | -0.002 | 0.095 | 0.7 | 0.34 | 0.75 | 0.75 | 0.65 | 0.69 |
| 7 | 152529878 | rs12535713 | 2.078 | 0.569 | 0.352 | -0.928 | 0.033 | 0.051 | 0.051 | 0 | 0.55 | 0.23 | 0.69 | 0.71 | 0.59 | 0.58 |
| 7 | 152540119 | rs10276445 | -0.338 | 0.486 | 0.121 | -1.029 | -0.032 | -0.008 | -0.008 | 0 | 0.85 | 0.92 | 0.84 | 0.86 | 0.75 | 0.76 |
| 7 | 152544637 | rs940262 | -0.415 | 0.509 | 0.118 | -0.985 | -0.037 | -0.023 | -0.023 | 0 | 0.85 | 0.9 | 0.84 | 0.8 | 0.75 | 0.73 |
| 7 | 152545351 | rs940261 | 2.039 | 0.452 | 0.197 | -0.996 | 0.036 | 0.045 | 0.045 | 0 | 0.55 | 0.25 | 0.7 | 0.67 | 0.6 | 0.55 |
| 7 | 152550499 | rs4726215 | -0.101 | 0.558 | 0.373 | -0.75 | -0.02 | -0.009 | -0.009 | 0 | 0.84 | 0.85 | 0.83 | 0.8 | 0.73 | 0.74 |
| 7 | 152561613 | rs7792217 | 1.684 | 0.143 | -0.029 | -0.833 | -0.034 | 0.097 | 0.097 | 0 | 0.51 | 0.17 | 0.61 | 0.67 | 0.5 | 0.52 |
| 7 | 152567213 | rs1966134 | 1.402 | 0.072 | 0.678 | -0.181 | 0.002 | 0 | 0 | 0.529 | 0.68 | 0.39 | 0.72 | 0.67 | 0.64 | 0.62 |
| 7 | 152573935 | rs1880124 | -0.713 | 0.2 | 0.102 | -0.13 | 0.031 | 0 | 0 | 0 | 0.22 | 0.24 | 0.24 | 0.2 | 0.34 | 0.36 |
| 7 | 152581108 | rs7799736 | 0.931 | -1.399 | -0.112 | -0.247 | -0.052 | -0.015 | -0.015 | 0 | 0.78 | 0.35 | 0.91 | 0.86 | 0.93 | 0.79 |
| 7 | 152600817 | rs2535969 | -1.267 | -0.759 | 0.433 | -0.332 | -0.023 | -0.051 | -0.051 | 0.032 | 0.19 | 0.42 | 0.07 | 0.09 | 0.11 | 0.15 |
| 7 | 152601631 | rs2535967 | -1.369 | -0.85 | 0.884 | -0.372 | -0.027 | -0.055 | -0.055 | 0 | 0.19 | 0.45 | 0.09 | 0.09 | 0.11 | 0.16 |
| 7 | 152604342 | rs1523641 | -0.765 | -0.471 | 1.268 | -0.393 | 0.012 | -0.004 | -0.004 | 0.054 | 0.12 | 0.22 | 0.09 | 0.05 | 0.13 | 0.07 |
| 7 | 152604776 | rs10241971 | 0.021 | -0.085 | 0.715 | -1.08 | 0.01 | -0.071 | -0.071 | 0.523 | 0.15 | 0.2 | 0.14 | 0 | 0.18 | 0.04 |
| 7 | 152605672 | rs10275845 | -0.009 | -0.063 | 0.622 | -1.072 | 0.002 | -0.05 | -0.05 | 0.009 | 0.15 | 0.13 | 0.13 | 0 | 0.17 | 0.04 |
| 7 | 152607732 | rs6950216 | -1.186 | -0.554 | 0.758 | -1.194 | 0.003 | -0.027 | -0.027 | 0.038 | 0.42 | 0.5 | 0.3 | 0.15 | 0.43 | 0.27 |
| 7 | 152620372 | rs2535957 | -0.595 | -1.257 | 0.378 | -1.201 | 0.004 | 0.01 | 0.01 | 0.001 | 0.39 | 0.62 | 0.31 | 0.24 | 0.34 | 0.24 |
| 7 | 152626132 | rs1860711 | -1.089 | -1.053 | -0.261 | -1.393 | -0.04 | 0.065 | 0.065 | 0 | 0.55 | 0.86 | 0.49 | 0.27 | 0.42 | 0.38 |
| 7 | 152631037 | rs940270 | -1.27 | -1.364 | 0.095 | -1.691 | -0.037 | -0.106 | -0.106 | 0.941 | 0.27 | 0.53 | 0.2 | 0.04 | 0.16 | 0.21 |
| 7 | 152633164 | rs6952411 | 0.282 | -1.024 | -0.011 | -1.47 | -0.001 | 0.002 | 0.002 | 0 | 0.27 | 0.25 | 0.2 | 0.21 | 0.26 | 0.15 |
| 7 | 152636385 | rs7804787 | 0.256 | -1.026 | -0.027 | -1.35 | -0.002 | -0.006 | -0.006 | 1.006 | 0.27 | 0.25 | 0.21 | 0.26 | 0.28 | 0.13 |
| 7 | 152636629 | rs1523638 | -1.132 | -0.379 | -0.034 | -1.118 | -0.002 | -0.064 | -0.064 | 1.506 | 0.14 | 0.26 | 0.11 | 0.01 | 0.08 | 0.1 |
| 7 | 152647687 | rs2311367 | -0.53 | -0.695 | -1.098 | -0.816 | -0.026 | 0.003 | 0.003 | 0 | 0.55 | 0.86 | 0.39 | 0.35 | 0.42 | 0.32 |
| 7 | 152648941 | rs4726229 | -1.092 | 0.019 | -0.829 | -0.773 | -0.001 | 0 | 0 | 0 | 0.14 | 0.12 | 0.08 | 0.14 | 0.12 | 0.1 |
| 7 | 152653503 | rs10278177 | -0.368 | -0.266 | -0.68 | -1.235 | 0.026 | 0.035 | 0.035 | 0.334 | 0.51 | 0.62 | 0.4 | 0.12 | 0.36 | 0.27 |
| 7 | 152655921 | rs12532611 | 0.62 | 0.263 | 0.042 | -1.49 | 0.056 | -0.147 | -0.147 | 0.202 | 0.81 | 0.5 | 0.86 | 1 | 0.78 | 0.89 |
| 7 | 152660734 | rs10952379 | -0.61 | -0.845 | -0.089 | -0.794 | 0.02 | 0.002 | 0.002 | 0 | 0.38 | 0.63 | 0.29 | 0.37 | 0.28 | 0.28 |
| 7 | 152661626 | rs11761769 | -0.85 | -0.766 | -0.409 | -0.923 | 0.056 | 0.044 | 0.044 | 0 | 0.46 | 0.73 | 0.47 | 0.36 | 0.36 | 0.3 |
| 7 | 152664190 | rs6968914 | -0.887 | -0.905 | 0.37 | -0.707 | 0.009 | -0.001 | -0.001 | 0.34 | 0.4 | 0.56 | 0.43 | 0.36 | 0.31 | 0.29 |
| 7 | 152664481 | rs10265883 | -1.009 | -1.181 | 0.464 | -0.624 | 0.013 | -0.003 | -0.003 | 0 | 0.38 | 0.61 | 0.39 | 0.36 | 0.28 | 0.28 |
| 7 | 152668222 | rs10282370 | -0.729 | -0.837 | 0.701 | -0.305 | -0.04 | 0.049 | 0.049 | 0.386 | 0.28 | 0.5 | 0.35 | 0.35 | 0.22 | 0.25 |
| 7 | 152677202 | rs12674417 | -0.679 | -0.078 | 0.385 | -0.712 | -0.001 | -0.003 | -0.003 | 0 | 0.18 | 0.17 | 0.1 | 0.1 | 0.18 | 0.07 |
| 7 | 152682512 | rs6974677 | -1.824 | -0.779 | 0.903 | -0.849 | 0.005 | 0.003 | 0.003 | 1.1 | 0.34 | 0.65 | 0.34 | 0.25 | 0.26 | 0.15 |
| 7 | 152683529 | rs12534924 | -0.749 | -0.899 | 1.393 | -0.645 | -0.015 | 0.11 | 0.11 | 1.359 | 0.1 | 0.47 | 0.21 | 0.15 | 0.04 | 0.03 |
| 7 | 152686657 | rs6978760 | -1.293 | -0.927 | 1.267 | -0.757 | -0.02 | 0.051 | 0.051 | 0 | 0.09 | 0.28 | 0.07 | 0.09 | 0.07 | 0.05 |
| 7 | 152691200 | rs7785760 | -2.382 | -0.919 | 1.29 | -0.808 | -0.071 | 0.02 | 0.02 | 0 | 0.21 | 0.52 | 0.28 | 0.19 | 0.14 | 0.12 |

| | | | | | | | | | | | | | | | | |
|---|---|---|---|---|---|---|---|---|---|---|---|---|---|---|---|---|
| 7 | 152695859 | rs6464295 | -0.667 | -0.877 | 1.283 | -0.808 | -0.019 | 0.029 | 0.029 | 0 | 0.08 | 0.16 | 0.06 | 0.09 | 0.07 | 0.05 |
| 7 | 152699457 | rs12668490 | -2.306 | -0.83 | 1.291 | -0.781 | -0.064 | 0.048 | 0.048 | 0 | 0.21 | 0.47 | 0.29 | 0.27 | 0.16 | 0.13 |
| 7 | 152700215 | rs11486922 | -2.187 | -0.67 | 1.05 | -0.795 | -0.043 | 0.042 | 0.042 | 0 | 0.19 | 0.41 | 0.28 | 0.24 | 0.15 | 0.11 |
| 7 | 152700399 | rs12531982 | -1.197 | -0.7 | 1.152 | -0.829 | -0.031 | 0.021 | 0.021 | 0 | 0.12 | 0.24 | 0.21 | 0.17 | 0.06 | 0.06 |
| 7 | 152700965 | rs495658 | 0.525 | -0.746 | 0.476 | -1.326 | -0.008 | -0.117 | -0.117 | 0 | 0.66 | 0.49 | 0.85 | 0.96 | 0.72 | 0.88 |
| 7 | 152701498 | rs11763048 | 0.858 | -0.53 | -0.26 | -1.191 | 0.036 | 0.104 | 0.104 | 0.904 | 0.57 | 0.26 | 0.66 | 0.79 | 0.67 | 0.85 |
| 7 | 152703396 | rs520556 | 1.012 | -0.486 | -0.101 | -1.314 | 0.04 | 0.058 | 0.058 | 0.122 | 0.67 | 0.21 | 0.83 | 0.95 | 0.8 | 0.82 |
| 7 | 152704536 | rs10807999 | 1.333 | 0.059 | 0.985 | -1.084 | 0.024 | -0.064 | -0.064 | 0.386 | 0.89 | 0.6 | 0.93 | 1 | 0.91 | 0.98 |
| 7 | 152704794 | rs12533514 | -0.698 | 0.219 | 0.99 | -1.417 | 0.012 | -0.037 | -0.037 | 0 | 0.21 | 0.37 | 0.16 | 0.12 | 0.2 | 0.12 |
| 7 | 152706440 | rs10234783 | -1.259 | 0.019 | 0.131 | -2.091 | 0.014 | 0.203 | 0.203 | 4.329 | 0.46 | 0.81 | 0.29 | 0.1 | 0.39 | 0.37 |
| 7 | 152707641 | rs7800857 | 0.959 | -0.024 | 0.589 | -2.24 | 0.011 | -0.061 | -0.061 | 0 | 0.62 | 0.53 | 0.72 | 0.9 | 0.57 | 0.66 |
| 7 | 152708016 | rs13223696 | 1.578 | 0.133 | -0.112 | -2.11 | -0.089 | 0.272 | 0.272 | 2.106 | 0.48 | 0.08 | 0.66 | 0.9 | 0.52 | 0.61 |
| 7 | 152708415 | rs10227803 | 0.194 | 0.554 | 0.671 | -1.536 | 0.001 | -0.026 | -0.026 | 0 | 0.12 | 0.15 | 0.09 | 0.02 | 0.14 | 0.2 |
| 7 | 152709937 | rs7792747 | -0.898 | 0.595 | 1.033 | -0.621 | 0.039 | 0.03 | 0.03 | 2.275 | 0.2 | 0.52 | 0.3 | 0.23 | 0.23 | 0.16 |
| 7 | 152711442 | rs6951424 | -1.002 | 0.403 | 0.346 | -0.537 | -0.013 | -0.001 | -0.001 | 0 | 0.26 | 0.53 | 0.25 | 0.24 | 0.23 | 0.16 |
| 7 | 152718942 | rs567778 | -0.934 | 0.626 | -0.504 | -0.598 | -0.027 | 0.051 | 0.051 | 0.742 | 0.18 | 0.07 | 0.1 | 0.03 | 0.22 | 0.25 |
| 7 | 152719593 | rs11977276 | -0.861 | -0.237 | -0.08 | 0.089 | -0.027 | 0.02 | 0.02 | 0 | 0.32 | 0.54 | 0.35 | 0.4 | 0.23 | 0.36 |
| 7 | 152719887 | rs4726238 | -1.266 | 0.098 | -0.046 | 0.013 | 0.001 | -0.001 | -0.001 | 4.756 | 0.44 | 0.43 | 0.37 | 0.51 | 0.38 | 0.44 |
| 7 | 152726381 | rs6464297 | -1.198 | -0.368 | -0.557 | -0.023 | 0.019 | -0.145 | -0.145 | 0.253 | 0.4 | 0.95 | 0.47 | 0.48 | 0.29 | 0.39 |
| 7 | 152728354 | rs7798285 | -0.57 | -0.497 | 0.168 | -0.247 | -0.006 | -0.073 | -0.073 | 0 | 0.14 | 0.27 | 0.08 | 0.01 | 0.12 | 0.11 |
| 7 | 152728801 | rs7799215 | -0.302 | -0.568 | 0.119 | 0.112 | -0.014 | 0.145 | 0.145 | 0.01 | 0.09 | 0.16 | 0.21 | 0.46 | 0.05 | 0.18 |
| 7 | 152728896 | rs10276233 | -1.452 | -0.509 | 0.385 | -0.364 | 0.004 | -0.044 | -0.044 | 0 | 0.29 | 0.72 | 0.24 | 0.01 | 0.2 | 0.16 |
| 7 | 152735972 | rs7798882 | -1.118 | -0.263 | 0.667 | -0.362 | -0.009 | -0.147 | -0.147 | 0 | 0.2 | 0.55 | 0.18 | 0.01 | 0.19 | 0.12 |
| 7 | 152736800 | rs1525041 | -0.75 | -0.241 | 0.912 | -0.355 | -0.014 | -0.112 | -0.112 | 0.071 | 0.17 | 0.39 | 0.13 | 0 | 0.15 | 0.12 |
| 7 | 152737256 | rs6464300 | -1.981 | -0.464 | 0.681 | -0.125 | 0.025 | -0.047 | -0.047 | 0 | 0.37 | 0.79 | 0.49 | 0.48 | 0.31 | 0.36 |
| 7 | 152737733 | rs954359 | -1.112 | -0.185 | 0.783 | 0.262 | -0.009 | 0.187 | 0.187 | 0.47 | 0.19 | 0.4 | 0.36 | 0.48 | 0.16 | 0.24 |
| 7 | 152741593 | rs11773000 | -0.386 | 0.533 | 0.101 | 0.263 | -0.001 | 0 | 0 | 0 | 0.13 | 0.15 | 0.09 | 0.11 | 0.11 | 0.16 |
| 7 | 152745810 | rs4725479 | 1.374 | 1.401 | -0.288 | 0.322 | -0.022 | -0.024 | -0.024 | 0 | 0.68 | 0.76 | 0.63 | 0.42 | 0.62 | 0.59 |
| 7 | 152745936 | rs4725480 | 1.392 | 1.252 | -0.756 | 0.256 | 0.009 | -0.033 | -0.033 | 0 | 0.25 | 0.13 | 0.25 | 0.28 | 0.25 | 0.31 |
| 7 | 152751486 | rs2037929 | -0.045 | 1.478 | -0.671 | 0.306 | -0.021 | -0.04 | -0.04 | 0 | 0.83 | 0.92 | 0.73 | 0.55 | 0.76 | 0.75 |
| 7 | 152753455 | rs2311857 | -0.255 | 1.463 | -0.748 | 0.224 | -0.045 | -0.085 | -0.085 | 0 | 0.84 | 0.96 | 0.73 | 0.54 | 0.76 | 0.74 |
| 7 | 152756180 | rs1076578 | 1.545 | 1.184 | -0.809 | 0.219 | -0.034 | -0.086 | -0.086 | 0 | 0.24 | 0.02 | 0.25 | 0.28 | 0.25 | 0.31 |
| 7 | 152762916 | rs926102 | 1.215 | 1.078 | -0.014 | 0.155 | 0.023 | 0.153 | 0.153 | 0 | 0.77 | 0.49 | 0.69 | 0.53 | 0.76 | 0.75 |
| 7 | 152765284 | rs2205168 | -0.169 | 0.865 | -0.043 | 0.224 | -0.013 | -0.029 | -0.029 | 0.008 | 0.84 | 0.91 | 0.74 | 0.54 | 0.8 | 0.75 |
| 7 | 152767561 | rs6959411 | 0.749 | 1.157 | 0.225 | 0.188 | 0 | 0 | 0 | 0 | 0.3 | 0.31 | 0.28 | 0.28 | 0.32 | 0.32 |
| 7 | 152770019 | rs1404591 | -0.054 | 1.106 | 0.421 | 0.084 | 0.001 | 0.001 | 0.001 | 0.096 | 0.44 | 0.41 | 0.35 | 0.38 | 0.38 | 0.49 |
| 7 | 152770512 | rs7790428 | 0.351 | 1.055 | -0.261 | 0.136 | -0.005 | 0 | 0 | 0.582 | 0.15 | 0.25 | 0.13 | 0.13 | 0.11 | 0.27 |
| 7 | 152771892 | rs510588 | -1.095 | 1.246 | -0.988 | 0.113 | 0.001 | 0.113 | 0.113 | 0 | 0.92 | 0.9 | 0.79 | 0.52 | 0.92 | 0.83 |
| 7 | 152773086 | rs542535 | -0.414 | 1.278 | -1.043 | 0.048 | 0.001 | 0.001 | 0.001 | 0.389 | 0.49 | 0.53 | 0.4 | 0.4 | 0.47 | 0.55 |
| 7 | 152774552 | rs626190 | 0.327 | 1.586 | -1.076 | 0.198 | -0.001 | -0.012 | -0.012 | 0 | 0.07 | 0.05 | 0.06 | 0 | 0.12 | 0.06 |
| 7 | 152775442 | rs640368 | 0.327 | 1.586 | -1.076 | 0.198 | -0.001 | -0.012 | -0.012 | 0 | 0.07 | 0.05 | 0.07 | 0 | 0.12 | 0.06 |
| 7 | 152780524 | rs12112082 | 0.998 | 1.39 | -1.321 | 0.428 | -0.001 | 0.001 | 0.001 | 0 | 0.63 | 0.35 | 0.56 | 0.31 | 0.65 | 0.48 |
| 7 | 152782900 | rs11764857 | -0.77 | 1.488 | -2.2 | 0.846 | -0.045 | -0.022 | -0.022 | 0.043 | 0.23 | 0.04 | 0.26 | 0.19 | 0.26 | 0.31 |
| 7 | 152783129 | rs486390 | 0.078 | 1.441 | -1.856 | 1.749 | -0.032 | -0.019 | -0.019 | 0.058 | 0.92 | 1 | 0.79 | 0.58 | 0.92 | 0.82 |
| 7 | 152784173 | rs2710243 | 0.248 | 1.271 | -2.407 | 0.681 | -0.027 | 0.06 | 0.06 | 0.437 | 0.56 | 0.86 | 0.64 | 0.73 | 0.55 | 0.52 |
| 7 | 152784233 | rs7779529 | -0.433 | 0.939 | -2.256 | 0.562 | -0.032 | -0.051 | -0.051 | 0.378 | 0.13 | 0.01 | 0.19 | 0.16 | 0.15 | 0.15 |
| 7 | 152784551 | rs2710244 | -0.218 | 0.657 | -1.682 | -0.485 | 0.015 | 0.038 | 0.038 | 0 | 0.44 | 0.29 | 0.25 | 0 | 0.39 | 0.25 |
| 7 | 152786529 | rs7790094 | -1.255 | 0.376 | -0.239 | 1.117 | 0.047 | 0.256 | 0.256 | 0 | 0.06 | 0.25 | 0.08 | 0.26 | 0.06 | 0.2 |
| 7 | 152793253 | rs12703241 | -0.915 | 0.183 | -0.376 | 0.861 | 0 | 0 | 0 | 0.849 | 0.1 | 0.1 | 0.16 | 0.11 | 0.05 | 0.12 |
| 7 | 152799577 | rs6951874 | -1.472 | 0.26 | 0.484 | 1.268 | 0.131 | 0.348 | 0.348 | 0 | 0.07 | 0.37 | 0.11 | 0.27 | 0.09 | 0.19 |
| 7 | 152800787 | rs629598 | -1.796 | -0.547 | 0.55 | 0.916 | 0 | -0.001 | -0.001 | 0 | 0.24 | 0.7 | 0.3 | 0.37 | 0.14 | 0.35 |
| 7 | 152801121 | rs17520767 | 1.278 | -0.72 | -0.179 | 0.48 | -0.079 | 0.163 | 0.163 | 2.707 | 0.4 | 0.09 | 0.25 | 0.11 | 0.48 | 0.36 |
| 7 | 152801568 | rs2710236 | -0.999 | -0.459 | 0.364 | 0.693 | 0.048 | 0.285 | 0.285 | 0.181 | 0.08 | 0.56 | 0.12 | 0.26 | 0.07 | 0.2 |
| 7 | 152801831 | rs2710234 | -0.987 | -0.458 | 0.364 | 0.694 | 0.05 | 0.288 | 0.288 | 0.135 | 0.08 | 0.55 | 0.12 | 0.26 | 0.07 | 0.2 |
| 7 | 152802220 | rs760226 | -0.377 | -0.497 | 0.097 | 0.579 | -0.025 | -0.083 | -0.083 | 0 | 0.68 | 0.8 | 0.56 | 0.49 | 0.62 | 0.72 |
| 7 | 152805455 | rs594243 | -0.416 | -0.602 | 0.073 | 0.598 | -0.019 | -0.097 | -0.097 | 1.043 | 0.69 | 0.84 | 0.57 | 0.5 | 0.66 | 0.72 |
| 7 | 152811263 | rs590733 | 0.168 | -0.75 | 0.121 | 0.523 | -0.006 | -0.007 | -0.007 | 0 | 0.14 | 0.13 | 0.17 | 0.12 | 0.06 | 0.11 |
| 7 | 152822713 | rs7793882 | -0.998 | -0.76 | 0.852 | 0.501 | 0.005 | 0.04 | 0.04 | 2.169 | 0.51 | 0.78 | 0.68 | 0.64 | 0.46 | 0.46 |
| 7 | 152823941 | rs2311855 | -0.698 | -0.814 | 0.819 | 0.862 | 0.01 | 0.051 | 0.051 | 0.305 | 0.45 | 0.35 | 0.49 | 0.26 | 0.4 | 0.31 |
| 7 | 152824231 | rs940650 | 0.018 | -0.687 | 0.767 | 0.987 | -0.005 | -0.016 | -0.016 | 0 | 0.14 | 0.13 | 0.17 | 0.1 | 0.06 | 0.11 |
| 7 | 152826374 | rs4397307 | 0.688 | -0.619 | 0.784 | 1.226 | -0.006 | 0.16 | 0.16 | 0 | 0.94 | 0.83 | 0.93 | 0.75 | 0.97 | 0.79 |
| 7 | 152844101 | rs7810943 | -0.428 | -0.962 | 0.729 | 1.04 | 0.011 | 0.042 | 0.042 | 0 | 0.46 | 0.33 | 0.49 | 0.26 | 0.4 | 0.27 |
| 7 | 152845911 | rs715851 | 1.148 | -0.944 | 0.093 | 1.176 | -0.15 | 0.174 | 0.174 | 0.606 | 0.43 | 0.02 | 0.27 | 0.14 | 0.51 | 0.38 |
| 7 | 152847328 | rs11976958 | -0.727 | -0.947 | 0.126 | 1.232 | 0.008 | 0.03 | 0.03 | 0.317 | 0.51 | 0.81 | 0.66 | 0.62 | 0.45 | 0.41 |
| 7 | 152848982 | rs12333356 | -0.505 | -0.914 | 0.158 | 1.251 | -0.005 | 0.062 | 0.062 | 0 | 0.06 | 0.09 | 0.06 | 0.24 | 0.03 | 0.21 |
| 7 | 152853444 | rs960600 | -0.743 | -0.94 | 0.06 | 1.228 | -0.005 | 0.009 | 0.009 | 0.314 | 0.51 | 0.85 | 0.66 | 0.61 | 0.45 | 0.41 |
| 7 | 152853534 | rs4346923 | 1.01 | -0.984 | 0.839 | 1.765 | -0.011 | 0.097 | 0.097 | 0 | 0.94 | 0.87 | 0.93 | 0.76 | 0.97 | 0.79 |
| 7 | 152855875 | rs7792156 | -1.077 | -1.343 | 0.684 | 1.632 | -0.014 | 0.024 | 0.024 | 1.376 | 0.23 | 0.61 | 0.33 | 0.47 | 0.11 | 0.24 |
| 7 | 152859520 | rs10278457 | -2.234 | -0.715 | 1.133 | 1.484 | -0.002 | 0.103 | 0.103 | 0.157 | 0.11 | 0.42 | 0.18 | 0.22 | 0.09 | 0.21 |

| | | | | | | | | | | | | | | | | |
|---|---|---|---|---|---|---|---|---|---|---|---|---|---|---|---|---|
| 7 | 152867025 | rs10259781 | -1.3 | -0.459 | 0.658 | 1.05 | 0.012 | -0.018 | -0.018 | 0 | 0.05 | 0.2 | 0.06 | 0 | 0.07 | 0.06 |
| 7 | 152873210 | rs4313080 | 0.185 | -0.676 | 0.568 | 0.928 | -0.011 | -0.024 | -0.024 | 0 | 0.7 | 0.56 | 0.62 | 0.71 | 0.71 | 0.78 |
| 7 | 152876977 | rs10246727 | 0.408 | -0.924 | 0.795 | 1.535 | -0.015 | -0.029 | -0.029 | 0.001 | 0.68 | 0.47 | 0.61 | 0.71 | 0.71 | 0.78 |
| 7 | 152878333 | rs4285401 | 0.897 | -0.312 | -0.107 | 1.724 | -0.026 | 0.093 | 0.093 | 0 | 0.52 | 0.1 | 0.34 | 0.26 | 0.54 | 0.46 |
| 7 | 152884999 | rs9648748 | 1.149 | -0.504 | -0.445 | 1.11 | -0.156 | 0.184 | 0.184 | 0.514 | 0.4 | 0.04 | 0.24 | 0.09 | 0.46 | 0.33 |
| 7 | 152885590 | rs4999662 | -0.491 | -0.569 | 0.727 | 1.619 | -0.017 | -0.05 | -0.05 | 0 | 0.34 | 0.48 | 0.39 | 0.27 | 0.31 | 0.26 |
| 7 | 152887205 | rs12533454 | 0.592 | -0.568 | 0.989 | 1.859 | 0.029 | -0.042 | -0.042 | 0 | 0.63 | 0.23 | 0.52 | 0.4 | 0.68 | 0.63 |
| 7 | 152888917 | rs2312600 | 0.559 | -0.576 | 1.114 | 1.887 | 0.002 | 0.001 | 0.001 | 0.019 | 0.65 | 0.33 | 0.59 | 0.61 | 0.69 | 0.73 |
| 7 | 152898575 | rs10952397 | -2.314 | -0.353 | 1.235 | 1.683 | 0.047 | 0.27 | 0.27 | 0 | 0.05 | 0.26 | 0.1 | 0.21 | 0.04 | 0.11 |
| 7 | 152905314 | rs12533677 | -0.269 | -0.432 | 1.396 | 1.611 | -0.01 | -0.027 | -0.027 | 0 | 0.32 | 0.42 | 0.37 | 0.31 | 0.3 | 0.25 |
| 7 | 152907497 | rs13245385 | 0.886 | -0.453 | 1.359 | 1.655 | 0.033 | -0.055 | -0.055 | 0 | 0.61 | 0.16 | 0.51 | 0.49 | 0.67 | 0.64 |
| 7 | 152914739 | rs4439038 | -0.609 | -0.349 | 1.233 | 1.542 | -0.007 | -0.039 | -0.039 | 0 | 0.34 | 0.5 | 0.4 | 0.31 | 0.34 | 0.31 |
| 7 | 152916616 | rs4262255 | 2.17 | -0.171 | 1.94 | 1.802 | 0.139 | 0.418 | 0.418 | 0.32 | 0.93 | 0.59 | 0.86 | 0.75 | 0.93 | 0.86 |
| 7 | 152924893 | rs2710201 | -2.396 | -0.265 | 2.035 | 1.661 | 0.038 | 0.035 | 0.035 | 1.607 | 0.09 | 0.62 | 0.17 | 0.06 | 0.07 | 0.09 |
| 7 | 152934288 | rs2710194 | 1.016 | 0.609 | 0.94 | 1.461 | -0.043 | -0.027 | -0.027 | 0.019 | 0.6 | 0.16 | 0.53 | 0.55 | 0.6 | 0.57 |
| 7 | 152942680 | rs1882828 | 1.045 | 0.72 | 1.071 | 1.517 | -0.047 | -0.06 | -0.06 | 0 | 0.59 | 0.16 | 0.53 | 0.47 | 0.59 | 0.57 |
| 7 | 152945148 | rs2735769 | 0.358 | 0.4 | 0.764 | 1.444 | 0.005 | -0.049 | -0.049 | 1.075 | 0.68 | 0.8 | 0.73 | 0.53 | 0.71 | 0.66 |
| 7 | 152945615 | rs968916 | 0.703 | 0.421 | 0.94 | 1.456 | 0.003 | 0.018 | 0.018 | 0.437 | 0.63 | 0.49 | 0.66 | 0.5 | 0.65 | 0.62 |
| 7 | 152959100 | rs4726267 | 1.078 | 0.399 | -0.491 | 1.014 | -0.005 | 0.021 | 0.021 | 2.192 | 0.69 | 0.2 | 0.56 | 0.76 | 0.7 | 0.65 |
| 7 | 152960298 | rs1404556 | -0.521 | -0.364 | -1.191 | 0.346 | 0.01 | 0.077 | 0.077 | 0 | 0.37 | 0.22 | 0.26 | 0.18 | 0.32 | 0.31 |
| 7 | 152963895 | rs1404557 | -0.638 | -0.551 | -1.955 | 0.364 | 0.029 | 0.027 | 0.027 | 1.584 | 0.54 | 0.71 | 0.52 | 0.31 | 0.41 | 0.56 |
| 7 | 152964089 | rs1735075 | -0.343 | -0.621 | -1.927 | 0.395 | 0.04 | 0.053 | 0.053 | 0.807 | 0.5 | 0.66 | 0.44 | 0.29 | 0.4 | 0.46 |
| 7 | 152964855 | rs1735078 | -0.073 | -0.073 | -2.575 | 0.239 | -0.036 | 0.167 | 0.167 | 0 | 0.41 | 0.06 | 0.29 | 0.12 | 0.38 | 0.34 |
| 7 | 152991697 | rs1735086 | 0.544 | 0.402 | -2.219 | 0.758 | 0.044 | 0.039 | 0.039 | 0 | 0.52 | 0.75 | 0.57 | 0.62 | 0.58 | 0.4 |
| 7 | 152995701 | rs17174014 | 0.15 | 1.162 | -2.009 | 1.354 | 0.004 | 0.039 | 0.039 | 0 | 0.07 | 0.09 | 0.15 | 0.26 | 0.12 | 0.1 |
| 7 | 152996005 | rs1735091 | -0.102 | 0.639 | -2.255 | 0.383 | -0.022 | 0.021 | 0.021 | 0 | 0.89 | 0.94 | 0.92 | 0.88 | 0.85 | 0.85 |
| 7 | 152996101 | rs16884836 | 0.006 | -0.027 | -2.114 | 0.447 | 0.037 | -0.006 | -0.006 | 0.03 | 0.18 | 0.03 | 0.09 | 0.11 | 0.14 | 0.22 |
| 7 | 152997887 | rs1735093 | 0.172 | 0.578 | -1.61 | 0.154 | -0.014 | 0.028 | 0.028 | 0 | 0.42 | 0.35 | 0.57 | 0.66 | 0.46 | 0.48 |
| 7 | 152998473 | rs1722928 | -0.052 | 0.402 | -1.741 | -0.117 | -0.011 | 0.026 | 0.026 | 0.054 | 0.71 | 0.83 | 0.74 | 0.78 | 0.69 | 0.71 |
| 7 | 152999066 | rs1735094 | 0.847 | 0.115 | -2.072 | -0.164 | -0.075 | 0.08 | 0.08 | 1.021 | 0.41 | 0.12 | 0.43 | 0.25 | 0.49 | 0.4 |
| 7 | 153001234 | rs588958 | 0.557 | 0.149 | -1.087 | -0.761 | -0.001 | -0.043 | -0.043 | 0.308 | 0.63 | 0.63 | 0.76 | 0.86 | 0.66 | 0.73 |
| 7 | 153005933 | rs401246 | -1.041 | 0.27 | -1.419 | -0.615 | -0.033 | 0.004 | 0.004 | 0 | 0.92 | 1 | 0.97 | 0.94 | 0.89 | 0.91 |
| 7 | 153006925 | rs7341501 | 1.137 | 0.124 | -1.311 | -0.405 | 0.004 | -0.012 | -0.012 | 0 | 0.17 | 0.13 | 0.3 | 0.27 | 0.16 | 0.2 |
| 7 | 153017006 | rs396672 | 0.473 | 0.055 | -0.958 | -0.362 | 0.006 | 0.013 | 0.013 | 0 | 0.53 | 0.4 | 0.52 | 0.55 | 0.58 | 0.5 |
| 7 | 153020742 | rs6946476 | 2.006 | 0.171 | -0.922 | -0.343 | 0.001 | -0.014 | -0.014 | 0 | 0.05 | 0.05 | 0.04 | 0.02 | 0.05 | 0.06 |
| 7 | 153022374 | rs422283 | 1.426 | 0.263 | -1.378 | -0.51 | -0.077 | 0.078 | 0.078 | 0.517 | 0.24 | 0.06 | 0.21 | 0.09 | 0.34 | 0.22 |
| 7 | 153022606 | rs2312325 | 0.472 | -0.149 | -0.405 | 0.084 | -0.007 | 0.039 | 0.039 | 0 | 0.22 | 0.31 | 0.23 | 0.51 | 0.16 | 0.31 |
| 7 | 153025110 | rs440704 | 0.6 | 0.027 | -0.873 | -0.051 | 0.005 | 0.07 | 0.07 | 0.052 | 0.51 | 0.25 | 0.58 | 0.36 | 0.54 | 0.46 |
| 7 | 153027200 | rs1076582 | 0.435 | 0.102 | -1.012 | -0.046 | -0.003 | -0.007 | -0.007 | 0 | 0.23 | 0.16 | 0.35 | 0.31 | 0.23 | 0.25 |
| 7 | 153027485 | rs383389 | 1.278 | 0 | -0.922 | -0.476 | -0.032 | 0.115 | 0.115 | 0.14 | 0.33 | 0.1 | 0.27 | 0.1 | 0.37 | 0.25 |
| 7 | 153027587 | rs741836 | 0.583 | 0.015 | -0.959 | 0.069 | -0.003 | -0.035 | -0.035 | 0.242 | 0.23 | 0.16 | 0.37 | 0.43 | 0.23 | 0.26 |
| 7 | 153027652 | rs84034 | 0.073 | 0.048 | 0.14 | 0.595 | 0.002 | 0.025 | 0.025 | 0.928 | 0.72 | 0.68 | 0.74 | 0.66 | 0.7 | 0.71 |
| 7 | 153027843 | rs84035 | 1.131 | 0.051 | 0.117 | 0.593 | 0.012 | -0.02 | -0.02 | 0 | 0.56 | 0.25 | 0.63 | 0.53 | 0.6 | 0.53 |
| 7 | 153028287 | rs932252 | -0.701 | 0.049 | -0.092 | 0.546 | 0.004 | -0.029 | -0.029 | 0 | 0.35 | 0.44 | 0.55 | 0.74 | 0.37 | 0.47 |
| 7 | 153030102 | rs10487555 | -0.918 | 0.042 | 0.016 | 0.78 | -0.002 | 0.078 | 0.078 | 0 | 0.3 | 0.54 | 0.42 | 0.54 | 0.28 | 0.44 |
| 7 | 153033766 | rs4726277 | -1.166 | 0.149 | 0.03 | 0.79 | -0.001 | 0.035 | 0.035 | 0.195 | 0.32 | 0.49 | 0.38 | 0.49 | 0.31 | 0.32 |
| 7 | 153034805 | rs2872000 | 0.185 | 0.244 | -0.506 | 0.911 | -0.037 | -0.029 | -0.029 | 0 | 0.08 | 0 | 0.14 | 0.09 | 0.1 | 0.09 |
| 7 | 153036049 | rs2053640 | -1.22 | 0.099 | -0.749 | 0.627 | -0.069 | 0.193 | 0.193 | 4.014 | 0.54 | 0.85 | 0.67 | 0.82 | 0.54 | 0.62 |
| 7 | 153039060 | rs12667691 | -0.449 | -0.129 | 0.318 | 0.956 | 0.008 | 0.075 | 0.075 | 0 | 0.12 | 0.2 | 0.19 | 0.4 | 0.15 | 0.16 |
| 7 | 153040137 | rs6464326 | -0.559 | -0.349 | 0.193 | 0.136 | -0.026 | -0.061 | -0.061 | 0.45 | 0.12 | 0.2 | 0.05 | 0 | 0.06 | 0.15 |
| 7 | 153045913 | rs6464327 | -1.002 | -0.436 | 0.389 | 0.248 | -0.034 | -0.031 | -0.031 | 0 | 0.17 | 0.28 | 0.16 | 0.14 | 0.12 | 0.25 |
| 7 | 153046117 | rs1589836 | -0.375 | -0.305 | -0.069 | 0.15 | 0.019 | 0.005 | 0.005 | 0.018 | 0.52 | 0.36 | 0.48 | 0.53 | 0.47 | 0.58 |
| 7 | 153046345 | rs2312322 | -0.979 | 0.222 | 0.43 | 0.226 | 0.031 | 0.06 | 0.06 | 0 | 0.11 | 0.23 | 0.22 | 0.24 | 0.17 | 0.15 |
| 7 | 153050837 | rs4726278 | 0.443 | 0.11 | 0.307 | 0.205 | 0.013 | -0.051 | -0.051 | 0 | 0.4 | 0.14 | 0.39 | 0.54 | 0.39 | 0.44 |
| 7 | 153051073 | rs1596605 | -0.523 | 0.243 | 0.33 | -0.092 | -0.003 | 0.001 | 0.001 | 0 | 0.58 | 0.61 | 0.58 | 0.68 | 0.56 | 0.62 |
| 7 | 153054702 | rs11767432 | 1.495 | 0.133 | -0.346 | 0.17 | 0.021 | -0.102 | -0.102 | 0 | 0.26 | 0.02 | 0.15 | 0.29 | 0.22 | 0.25 |
| 7 | 153057390 | rs1373517 | -0.55 | 0.271 | 0.016 | 0.289 | -0.012 | 0.029 | 0.029 | 0 | 0.56 | 0.71 | 0.57 | 0.62 | 0.57 | 0.55 |
| 7 | 153061413 | rs960840 | 0.028 | 0.396 | -0.028 | 0.386 | 0.002 | -0.002 | -0.002 | 0 | 0.45 | 0.43 | 0.35 | 0.39 | 0.43 | 0.4 |
| 7 | 153069111 | rs7793529 | -1.352 | 0.652 | -0.537 | 0.573 | -0.059 | 0.004 | 0.004 | 0.915 | 0.7 | 0.91 | 0.74 | 0.77 | 0.67 | 0.76 |
| 7 | 153071114 | rs11761744 | 0.278 | 0.856 | -0.264 | 1.002 | 0.011 | -0.012 | -0.012 | 0 | 0.31 | 0.21 | 0.2 | 0.29 | 0.29 | 0.26 |
| 7 | 153078544 | rs10952413 | 0.276 | 0.746 | -0.252 | 0.984 | 0.017 | -0.015 | -0.015 | 0 | 0.31 | 0.2 | 0.19 | 0.29 | 0.29 | 0.25 |
| 7 | 153079426 | rs6951736 | -0.857 | 1.091 | 0.159 | 1.085 | 0.013 | 0.023 | 0.023 | 0 | 0.09 | 0.15 | 0.22 | 0.24 | 0.15 | 0.13 |
| 7 | 153081698 | rs6944993 | -1.555 | 1.01 | 0.402 | 1.054 | 0.057 | 0.116 | 0.116 | 0.843 | 0.17 | 0.51 | 0.36 | 0.34 | 0.22 | 0.2 |
| 7 | 153084208 | rs1822707 | 0.736 | 0.664 | -0.191 | 1.062 | 0.002 | -0.018 | -0.018 | 0 | 0.79 | 0.86 | 0.79 | 0.63 | 0.76 | 0.75 |
| 7 | 153085940 | rs982112 | 0.946 | 0.39 | -0.149 | 0.961 | 0.004 | -0.009 | -0.009 | 0 | 0.77 | 0.82 | 0.79 | 0.63 | 0.77 | 0.76 |
| 7 | 153088772 | rs10271744 | 1.118 | 0.764 | -0.291 | 0.822 | -0.004 | -0.015 | -0.015 | 0 | 0.76 | 0.86 | 0.75 | 0.63 | 0.72 | 0.74 |
| 7 | 153089510 | rs10808003 | 0.709 | 0.833 | -0.275 | 0.848 | -0.004 | -0.018 | -0.018 | 0 | 0.79 | 0.86 | 0.79 | 0.62 | 0.75 | 0.73 |
| 7 | 153093112 | rs1867488 | 0.52 | 0.894 | -0.317 | 0.923 | 0 | 0.029 | 0.029 | 0 | 0.05 | 0.05 | 0.13 | 0.22 | 0.09 | 0.09 |
| 7 | 153094739 | rs10245542 | 0.84 | 0.374 | -0.386 | 0.464 | -0.002 | -0.015 | -0.015 | 0 | 0.08 | 0.09 | 0.04 | 0 | 0.04 | 0.04 |

| | | | | | | | | | | | | | | | | |
|---|---|---|---|---|---|---|---|---|---|---|---|---|---|---|---|---|
| 7 | 153096195 | rs9640332 | 0.767 | 0.962 | -0.897 | 0.543 | -0.026 | -0.029 | -0.029 | 0.769 | 0.11 | 0.02 | 0.13 | 0.13 | 0.15 | 0.13 |
| 7 | 153101151 | rs10952417 | 1.091 | 0.445 | -1.91 | 0.595 | 0.007 | -0.057 | -0.057 | 0.044 | 0.19 | 0.02 | 0.27 | 0.21 | 0.16 | 0.27 |
| 7 | 153101349 | rs10216124 | 0.354 | 0.506 | -1.255 | 0.523 | 0.006 | -0.05 | -0.05 | 0.096 | 0.31 | 0.1 | 0.4 | 0.34 | 0.3 | 0.41 |
| 7 | 153109508 | rs13223314 | 1.041 | -0.044 | -1.429 | 0.186 | -0.073 | 0.035 | 0.035 | 0.053 | 0.42 | 0.12 | 0.34 | 0.29 | 0.45 | 0.35 |
| 7 | 153120067 | rs1444458 | 1.58 | 0.005 | -0.917 | 0.216 | -0.136 | -0.041 | -0.041 | 0.667 | 0.34 | 0.04 | 0.3 | 0.29 | 0.41 | 0.31 |
| 7 | 153134549 | rs7805637 | 0.131 | 0.597 | -0.525 | 0.045 | -0.044 | -0.006 | -0.006 | 8.922 | 0.68 | 0.87 | 0.71 | 0.67 | 0.63 | 0.72 |
| 7 | 153134888 | rs2872008 | 0.131 | 0.597 | -0.525 | 0.045 | -0.044 | -0.006 | -0.006 | 0.078 | 0.68 | 0.87 | 0.71 | 0.67 | 0.63 | 0.72 |
| 7 | 153136207 | rs2312340 | 1.877 | 0.541 | 0.052 | 0.018 | -0.002 | -0.002 | -0.002 | 0.089 | 0.28 | 0.21 | 0.26 | 0.31 | 0.33 | 0.3 |
| 7 | 153137459 | rs880798 | 1.829 | 0.524 | 0.049 | 0.015 | -0.002 | -0.002 | -0.002 | 0.698 | 0.28 | 0.21 | 0.26 | 0.31 | 0.33 | 0.3 |
| 7 | 153144983 | rs13230671 | 0.673 | -0.005 | 0.661 | 0.359 | -0.019 | 0.061 | 0.061 | 0 | 0.47 | 0.75 | 0.39 | 0.31 | 0.45 | 0.34 |
| 7 | 153145278 | rs13233777 | 1.337 | 0.063 | 0.702 | 0.389 | -0.065 | 0.022 | 0.022 | 0.061 | 0.39 | 0.1 | 0.33 | 0.31 | 0.44 | 0.32 |
| 7 | 153147715 | rs12703305 | 1.064 | 0.06 | 0.945 | 0.388 | -0.014 | 0.013 | 0.013 | 0 | 0.41 | 0.29 | 0.35 | 0.31 | 0.44 | 0.32 |
| 7 | 153150189 | rs1444461 | -0.245 | 0.607 | 1.086 | 0.438 | -0.025 | -0.033 | -0.033 | 0 | 0.79 | 0.87 | 0.78 | 0.67 | 0.71 | 0.75 |
| 7 | 153154107 | rs11773882 | 0.804 | 0.233 | 1.077 | 0.443 | -0.014 | 0.005 | 0.005 | 0.995 | 0.71 | 0.33 | 0.74 | 0.67 | 0.74 | 0.71 |
| 7 | 153154926 | rs10952420 | 0.402 | 0.302 | 0.768 | 0.531 | 0.006 | -0.032 | -0.032 | 0 | 0.31 | 0.11 | 0.41 | 0.35 | 0.3 | 0.39 |
| 7 | 153155162 | rs7782578 | 0.402 | 0.302 | 0.769 | 0.525 | 0.006 | -0.033 | -0.033 | 0 | 0.31 | 0.11 | 0.41 | 0.36 | 0.3 | 0.39 |
| 7 | 153164787 | rs4131584 | 0.336 | 0.256 | 0.893 | 0.64 | 0.015 | -0.042 | -0.042 | 0 | 0.32 | 0.15 | 0.4 | 0.39 | 0.3 | 0.39 |
| 7 | 153167347 | rs7785341 | 0.843 | 0.246 | 1.099 | 0.535 | 0.001 | -0.012 | -0.012 | 0.366 | 0.46 | 0.44 | 0.37 | 0.28 | 0.45 | 0.33 |
| 7 | 153169452 | rs13245532 | 1.154 | 0.219 | 0.899 | 0.542 | -0.036 | 0.055 | 0.055 | 0 | 0.41 | 0.16 | 0.33 | 0.28 | 0.45 | 0.33 |
| 7 | 153175815 | rs4431510 | -0.613 | 0.353 | 0.942 | 0.567 | 0.005 | 0.01 | 0.01 | 0 | 0.85 | 0.85 | 0.91 | 0.92 | 0.86 | 0.82 |
| 7 | 153176705 | rs4629773 | -0.613 | 0.358 | 0.944 | 0.569 | 0.005 | 0.01 | 0.01 | 0 | 0.85 | 0.85 | 0.91 | 0.92 | 0.86 | 0.81 |
| 7 | 153192052 | rs4563798 | 0.991 | 0.28 | 0.752 | 0.64 | -0.032 | 0.096 | 0.096 | 0 | 0.43 | 0.12 | 0.34 | 0.26 | 0.46 | 0.38 |
| 7 | 153194160 | rs6464341 | 0.081 | 0.552 | 0.563 | 0.832 | 0.015 | -0.05 | -0.05 | 0 | 0.59 | 0.8 | 0.51 | 0.53 | 0.58 | 0.52 |
| 7 | 153194173 | rs10243460 | 0.124 | 0.444 | 0.984 | 0.977 | 0.046 | 0.167 | 0.167 | 0 | 0.09 | 0.31 | 0.14 | 0.26 | 0.12 | 0.1 |
| 7 | 153197219 | rs4726301 | 0.223 | 0.239 | 0.129 | 0.177 | -0.002 | 0.111 | 0.111 | 0.017 | 0.45 | 0.25 | 0.49 | 0.16 | 0.4 | 0.42 |
| 7 | 153200474 | rs3110879 | -1.352 | 0.248 | -0.104 | 0.499 | -0.007 | -0.017 | -0.017 | 0 | 0.87 | 0.92 | 0.88 | 0.77 | 0.82 | 0.8 |
| 7 | 153201572 | rs3111266 | 0.02 | 0.15 | 0.02 | 0.462 | -0.002 | 0.003 | 0.003 | 0 | 0.7 | 0.71 | 0.73 | 0.63 | 0.69 | 0.66 |
| 7 | 153205490 | rs3110887 | 0.012 | 0.162 | 0.026 | 0.468 | -0.002 | 0.003 | 0.003 | 0 | 0.7 | 0.71 | 0.73 | 0.63 | 0.69 | 0.66 |
| 7 | 153210132 | rs3110893 | 0.315 | 0.547 | 0.217 | 0.748 | 0.018 | 0.078 | 0.078 | 0.278 | 0.27 | 0.41 | 0.23 | 0.47 | 0.29 | 0.24 |
| 7 | 153214451 | rs3111275 | -0.972 | 0.483 | -0.172 | 0.619 | -0.003 | -0.01 | -0.01 | 0 | 0.86 | 0.9 | 0.88 | 0.74 | 0.82 | 0.82 |
| 7 | 153215127 | rs3111276 | -0.009 | 0.31 | -0.041 | 0.394 | 0.011 | 0.028 | 0.028 | 0 | 0.38 | 0.57 | 0.4 | 0.48 | 0.42 | 0.32 |
| 7 | 153216346 | rs4626512 | 0.738 | 0.431 | -0.218 | 0.456 | -0.007 | -0.013 | -0.013 | 0.189 | 0.26 | 0.25 | 0.21 | 0.44 | 0.31 | 0.24 |
| 7 | 153220960 | rs10264981 | 0.567 | 0.491 | -0.901 | 0.127 | 0.004 | 0.035 | 0.035 | 0 | 0.4 | 0.25 | 0.35 | 0.26 | 0.34 | 0.48 |
| 7 | 153231010 | rs11766646 | 0.427 | 0.642 | -0.251 | 0.579 | -0.024 | -0.021 | -0.021 | 0.195 | 0.14 | 0.23 | 0.09 | 0.08 | 0.11 | 0.16 |
| 7 | 153231422 | rs10952429 | -0.534 | 0.565 | -0.573 | 0.522 | -0.014 | 0.023 | 0.023 | 0 | 0.3 | 0.33 | 0.28 | 0.43 | 0.29 | 0.34 |
| 7 | 153237813 | rs3110871 | 0.616 | 0.852 | -0.584 | 0.435 | 0.044 | 0.03 | 0.03 | 0 | 0.17 | 0.25 | 0.19 | 0.25 | 0.25 | 0.25 |
| 7 | 153238468 | rs7778208 | -0.219 | 0.467 | -0.601 | -0.108 | 0.018 | 0.022 | 0.022 | 0 | 0.67 | 0.59 | 0.66 | 0.45 | 0.58 | 0.55 |
| 7 | 153240425 | rs4255050 | -0.554 | -0.178 | 0.227 | 0.618 | -0.014 | 0.136 | 0.136 | 0.002 | 0.15 | 0.37 | 0.25 | 0.35 | 0.12 | 0.08 |
| 7 | 153241355 | rs6960275 | -0.596 | -0.236 | -0.192 | 0.462 | -0.048 | 0.028 | 0.028 | 0 | 0.3 | 0.48 | 0.38 | 0.36 | 0.22 | 0.22 |
| 7 | 153242365 | rs13241381 | -0.751 | 0.278 | -0.449 | -0.447 | -0.001 | -0.031 | -0.031 | 0 | 0.28 | 0.3 | 0.19 | 0.07 | 0.28 | 0.15 |
| 7 | 153249729 | rs2953087 | 0.354 | 0.208 | -0.907 | -0.516 | 0.038 | 0.109 | 0.109 | 0.204 | 0.51 | 0.23 | 0.68 | 0.68 | 0.6 | 0.53 |
| 7 | 153254763 | rs10282321 | -0.04 | -0.424 | -0.295 | -0.805 | -0.042 | -0.051 | -0.051 | 0.742 | 0.13 | 0.5 | 0.08 | 0.06 | 0.05 | 0.22 |
| 7 | 153263477 | rs13227282 | 0.429 | -0.419 | -0.508 | -0.944 | -0.019 | 0.068 | 0.068 | 0 | 0.26 | 0.11 | 0.14 | 0.13 | 0.25 | 0.18 |
| 7 | 153263779 | rs3110853 | -0.509 | -0.434 | -0.366 | -0.889 | -0.005 | 0.011 | 0.011 | 0 | 0.9 | 0.81 | 0.92 | 0.86 | 0.91 | 0.93 |
| 7 | 153266855 | rs7806552 | 0.04 | -0.464 | 0.296 | -0.924 | -0.003 | 0.014 | 0.014 | 0.141 | 0.07 | 0.34 | 0.04 | 0.06 | 0.01 | 0.16 |
| 7 | 153271877 | rs11771429 | 1.497 | 0.149 | -0.676 | -0.336 | -0.063 | -0.086 | -0.086 | 0 | 0.13 | 0.01 | 0.2 | 0.31 | 0.18 | 0.19 |
| 7 | 153278953 | rs7777758 | -0.65 | -0.11 | -0.617 | -1.239 | 0.028 | 0.183 | 0.183 | 0 | 0.46 | 0.8 | 0.33 | 0.18 | 0.41 | 0.44 |
| 7 | 153281003 | rs10247334 | -0.822 | -0.175 | -0.582 | -1.31 | -0.021 | -0.026 | -0.026 | 0 | 0.19 | 0.65 | 0.14 | 0.13 | 0.13 | 0.25 |
| 7 | 153281321 | rs7810934 | 0.837 | -0.132 | -0.647 | -1.244 | -0.007 | 0.08 | 0.08 | 0 | 0.27 | 0.14 | 0.18 | 0.05 | 0.28 | 0.19 |
| 7 | 153282382 | rs4725502 | 0.544 | 0.023 | -0.375 | -1.038 | -0.002 | 0.003 | 0.003 | 0 | 0.1 | 0.18 | 0.07 | 0.12 | 0.08 | 0.07 |
| 7 | 153285674 | rs13223818 | 0.808 | 0 | -0.541 | -1.368 | -0.005 | 0.059 | 0.059 | 0 | 0.24 | 0.14 | 0.16 | 0.05 | 0.24 | 0.19 |
| 7 | 153285755 | rs6942954 | 0.101 | 0.629 | -0.524 | -0.882 | -0.015 | 0.028 | 0.028 | 0 | 0.39 | 0.28 | 0.53 | 0.74 | 0.43 | 0.5 |
| 7 | 153287728 | rs7796025 | 0.702 | 0.503 | -1.081 | -0.86 | 0.015 | 0.142 | 0.142 | 0 | 0.45 | 0.26 | 0.27 | 0.09 | 0.43 | 0.29 |
| 7 | 153287960 | rs9648757 | 0.041 | 1.12 | -1.656 | -0.463 | -0.021 | -0.015 | -0.015 | 0 | 0.15 | 0.09 | 0.19 | 0.18 | 0.21 | 0.14 |
| 7 | 153290520 | rs4726308 | -0.27 | 0.741 | -1.411 | -0.014 | -0.019 | 0.039 | 0.039 | 0 | 0.36 | 0.2 | 0.52 | 0.75 | 0.38 | 0.46 |
| 7 | 153293199 | rs4266558 | 0.843 | 1.363 | -0.633 | 0.81 | -0.015 | -0.021 | -0.021 | 0 | 0.73 | 0.75 | 0.7 | 0.58 | 0.64 | 0.72 |
| 7 | 153294306 | rs4584067 | -0.652 | 0.804 | 0.114 | 0.806 | 0.003 | 0.144 | 0.144 | 0 | 0.21 | 0.37 | 0.4 | 0.48 | 0.18 | 0.32 |
| 7 | 153296249 | rs10250553 | 0.994 | 0.366 | 0.071 | 0.492 | -0.001 | 0.013 | 0.013 | 0 | 0.65 | 0.53 | 0.68 | 0.53 | 0.63 | 0.64 |
| 7 | 153301445 | rs1986703 | 0.036 | 0.459 | -0.375 | 0.661 | -0.01 | -0.026 | -0.026 | 0 | 0.16 | 0.04 | 0.14 | 0.18 | 0.17 | 0.15 |
| 7 | 153303583 | rs4292607 | 2.454 | 0.367 | -0.351 | 0.459 | -0.038 | 0.194 | 0.194 | 0 | 0.41 | 0.14 | 0.3 | 0.09 | 0.44 | 0.32 |
| 7 | 153304575 | rs6949962 | 0.661 | 0.437 | -0.183 | 0.49 | -0.001 | 0.016 | 0.016 | 0 | 0.74 | 0.3 | 0.63 | 0.51 | 0.73 | 0.65 |
| 7 | 153305945 | rs11763266 | 2.541 | 0.439 | -0.187 | 0.47 | -0.039 | 0.193 | 0.193 | 0 | 0.41 | 0.14 | 0.3 | 0.09 | 0.44 | 0.32 |
| 7 | 153308177 | rs13243682 | -1.428 | 0.487 | 0.023 | 0.657 | 0.006 | -0.06 | -0.06 | 0 | 0.32 | 0.16 | 0.33 | 0.42 | 0.3 | 0.33 |
| 7 | 153312599 | rs7797575 | 1.366 | 0.536 | 0.356 | 0.539 | 0.001 | -0.026 | -0.026 | 0 | 0.92 | 0.32 | 0.92 | 0.98 | 0.92 | 0.96 |
| 7 | 153320044 | rs7806949 | 1.243 | 0.533 | 0.298 | 0.594 | 0.002 | 0.003 | 0.003 | 0 | 0.61 | 0.61 | 0.56 | 0.35 | 0.63 | 0.53 |
| 7 | 153320323 | rs12539866 | -1.036 | 0.592 | 0.305 | 0.592 | -0.005 | -0.011 | -0.011 | 0 | 0.35 | 0.28 | 0.42 | 0.63 | 0.36 | 0.45 |
| 7 | 153325658 | rs4074106 | 2.059 | 0.62 | 0.119 | 0.502 | -0.025 | 0.17 | 0.17 | 0 | 0.42 | 0.14 | 0.3 | 0.1 | 0.44 | 0.34 |
| 7 | 153331061 | rs4595033 | 1.533 | 0.789 | 0.327 | 0.743 | 0.012 | 0.038 | 0.038 | 0 | 0.57 | 0.16 | 0.51 | 0.34 | 0.57 | 0.49 |
| 7 | 153338113 | rs4726318 | -1.264 | 1.238 | 1.305 | 1.205 | 0.007 | 0.036 | 0.036 | 0 | 0.27 | 0.33 | 0.33 | 0.4 | 0.32 | 0.28 |

| | | | | | | | | | | | | | | | | |
|---|---|---|---|---|---|---|---|---|---|---|---|---|---|---|---|---|
| 7 | 153338339 | rs4726319 | 0.793 | 1.701 | 1.625 | 1.883 | -0.003 | 0.027 | 0.027 | 0 | 0.89 | 0.89 | 0.7 | 0.56 | 0.88 | 0.77 |
| 7 | 153341750 | rs6979534 | 1.173 | 1.505 | 2.404 | 1.183 | 0.099 | 0.092 | 0.092 | 0 | 0.87 | 0.71 | 0.8 | 0.82 | 0.79 | 0.81 |
| 7 | 153344736 | rs10231537 | -1.088 | 1.334 | 2.321 | 0.894 | 0.005 | -0.005 | -0.005 | 0 | 0.23 | 0.31 | 0.17 | 0.18 | 0.27 | 0.23 |
| 7 | 153346402 | rs6947489 | -0.76 | 1.442 | 2.315 | 0.772 | 0.023 | -0.054 | -0.054 | 0 | 0.2 | 0.36 | 0.16 | 0.08 | 0.25 | 0.15 |
| 7 | 153350882 | rs4410821 | 1.968 | 1.339 | 1.589 | 0.799 | -0.009 | 0.003 | 0.003 | 0 | 0.66 | 0.37 | 0.67 | 0.74 | 0.6 | 0.61 |
| 7 | 153350972 | rs4410822 | 1.309 | 1.021 | 1.507 | 1.229 | 0.006 | 0.008 | 0.008 | 0 | 0.89 | 0.91 | 0.83 | 0.83 | 0.83 | 0.76 |
| 7 | 153356432 | rs4621714 | -1.877 | 0.875 | 1.666 | 1.308 | 0.006 | 0.054 | 0.054 | 0.117 | 0.14 | 0.65 | 0.2 | 0.17 | 0.14 | 0.26 |
| 7 | 153361179 | rs11980518 | 1.397 | 0.942 | 1.253 | 1.268 | -0.001 | 0.077 | 0.077 | 0.381 | 0.84 | 0.31 | 0.79 | 0.73 | 0.85 | 0.69 |
| 7 | 153363132 | rs13438300 | -1.734 | 0.791 | 1.692 | 0.567 | 0.019 | 0.012 | 0.012 | 0 | 0.07 | 0.3 | 0.05 | 0.03 | 0.05 | 0.06 |
| 7 | 153367707 | rs13221692 | -1.924 | 0.848 | 1.846 | 0.408 | 0.009 | -0.026 | -0.026 | 0.307 | 0.09 | 0.24 | 0.06 | 0.02 | 0.1 | 0.08 |
| 7 | 153368459 | rs10228200 | -1.816 | 0.794 | 2.215 | 0.395 | 0.039 | 0.011 | 0.011 | 0 | 0.08 | 0.33 | 0.06 | 0.02 | 0.06 | 0.06 |
| 7 | 153374988 | rs6969143 | -2.4 | 0.639 | 2.425 | 0.346 | -0.013 | -0.037 | -0.037 | 0.129 | 0.11 | 0.57 | 0.09 | 0.02 | 0.06 | 0.06 |
| 7 | 153378283 | rs7794175 | -2.783 | 0.642 | 2.044 | 0.347 | -0.028 | -0.039 | -0.039 | 0 | 0.13 | 0.69 | 0.1 | 0.03 | 0.07 | 0.08 |
| 7 | 153393602 | rs6955628 | -2.663 | 0.629 | 2.06 | 0.341 | -0.028 | -0.039 | -0.039 | 0 | 0.13 | 0.71 | 0.09 | 0.05 | 0.07 | 0.08 |
| 7 | 153397733 | rs7801720 | -1.211 | 1.098 | 1.238 | 0.302 | -0.003 | 0.033 | 0.033 | 3.837 | 0.38 | 0.76 | 0.58 | 0.7 | 0.39 | 0.31 |
| 7 | 153399165 | rs1558301 | -1.211 | 1.1 | 1.232 | 0.303 | -0.003 | 0.031 | 0.031 | 3.853 | 0.38 | 0.72 | 0.58 | 0.7 | 0.39 | 0.31 |
| 7 | 153435344 | rs2109283 | -0.245 | 1.198 | 1.641 | 0.366 | 0.005 | 0.008 | 0.008 | 2.42 | 0.27 | 0.26 | 0.51 | 0.68 | 0.32 | 0.26 |
| 7 | 153437424 | rs2862148 | -0.348 | 1.29 | 1.77 | 0.369 | 0.018 | 0.018 | 0.018 | 2.673 | 0.28 | 0.4 | 0.52 | 0.69 | 0.33 | 0.28 |
| 7 | 153461606 | rs6962199 | -0.276 | 1.215 | 0.92 | 0.331 | 0.005 | -0.003 | -0.003 | 0 | 0.38 | 0.37 | 0.59 | 0.65 | 0.45 | 0.32 |
| 7 | 153462806 | rs2533148 | 0.3 | 1.286 | 0.358 | -0.134 | -0.004 | 0.042 | 0.042 | 3.244 | 0.52 | 0.43 | 0.36 | 0.24 | 0.47 | 0.58 |
| 7 | 153478281 | rs980266 | 0.8 | 1.245 | 0.286 | -0.187 | -0.003 | 0.122 | 0.122 | 5.302 | 0.46 | 0.2 | 0.3 | 0.22 | 0.41 | 0.52 |
| 7 | 153485627 | rs10268402 | 0.398 | 1.306 | 0.043 | -0.033 | -0.001 | 0.018 | 0.018 | 2.895 | 0.53 | 0.46 | 0.35 | 0.22 | 0.46 | 0.58 |
| 7 | 153515648 | rs2622190 | -1.383 | 1.954 | 0.045 | 0.393 | 0.024 | 0 | 0 | 0 | 0.09 | 0.16 | 0.09 | 0.11 | 0.12 | 0.03 |
| 7 | 153518025 | rs2622254 | 0.813 | 1.635 | 0.189 | 0.702 | -0.014 | 0.017 | 0.017 | 0 | 0.79 | 0.66 | 0.86 | 0.77 | 0.85 | 0.78 |
| 7 | 153522140 | rs2538421 | -0.707 | 2.215 | 0.65 | 0.907 | 0.036 | 0.034 | 0.034 | 0 | 0.17 | 0.34 | 0.23 | 0.23 | 0.23 | 0.25 |
| 7 | 153522175 | rs2538420 | -0.097 | 1.793 | -0.61 | 0.415 | -0.042 | -0.038 | -0.038 | 0 | 0.53 | 0.86 | 0.63 | 0.58 | 0.56 | 0.64 |
| 7 | 153522368 | rs2622262 | -0.902 | 1.755 | -0.617 | 0.372 | -0.004 | -0.044 | -0.044 | 0 | 0.85 | 0.95 | 0.9 | 0.79 | 0.87 | 0.79 |
| 7 | 153524142 | rs6973414 | 1.209 | 1.994 | -0.542 | 0.263 | -0.004 | -0.007 | -0.007 | 0 | 0.08 | 0.05 | 0.11 | 0.01 | 0.11 | 0.08 |
| 7 | 153524514 | rs1544612 | -0.916 | 1.879 | -0.546 | 0.372 | -0.004 | -0.045 | -0.045 | 0.368 | 0.85 | 0.95 | 0.9 | 0.79 | 0.87 | 0.79 |
| 7 | 153524878 | rs2538416 | -0.325 | 2.047 | -0.366 | 0.312 | 0.006 | 0.008 | 0.008 | 0.155 | 0.5 | 0.44 | 0.49 | 0.43 | 0.54 | 0.4 |
| 7 | 153527304 | rs2622244 | -0.207 | 1.963 | 0.038 | 0.281 | 0 | -0.009 | -0.009 | 0 | 0.45 | 0.46 | 0.45 | 0.35 | 0.47 | 0.36 |
| 7 | 153528964 | rs10264523 | 0.292 | 1.532 | 0.209 | 0.218 | 0.006 | 0.065 | 0.065 | 0 | 0.25 | 0.33 | 0.42 | 0.53 | 0.3 | 0.4 |
| 7 | 153530887 | rs2215781 | -0.935 | 1.613 | 0.391 | -0.14 | 0.024 | -0.037 | -0.037 | 2.683 | 0.25 | 0.38 | 0.22 | 0.14 | 0.29 | 0.21 |
| 7 | 153531897 | rs7783202 | 0.004 | 0.815 | 0.128 | -0.212 | -0.013 | -0.051 | -0.051 | 0.063 | 0.12 | 0.17 | 0.05 | 0 | 0.08 | 0.09 |
| 7 | 153533128 | rs10241157 | 0.161 | 0.989 | -0.122 | -0.733 | 0.009 | -0.042 | -0.042 | 1.168 | 0.63 | 0.57 | 0.66 | 0.84 | 0.62 | 0.82 |
| 7 | 153534165 | rs1029956 | 0.04 | 0.699 | -0.879 | 0.026 | 0.005 | -0.007 | -0.007 | 0 | 0.48 | 0.23 | 0.6 | 0.44 | 0.59 | 0.36 |
| 7 | 153535315 | rs7783446 | -0.338 | 0.769 | -0.353 | -0.157 | 0.013 | -0.013 | -0.013 | 0.253 | 0.86 | 0.77 | 0.87 | 0.89 | 0.85 | 0.9 |
| 7 | 153538274 | rs2622128 | -0.373 | 0.978 | -0.92 | -0.163 | -0.076 | -0.019 | -0.019 | 0.035 | 0.83 | 0.99 | 0.83 | 0.86 | 0.78 | 0.89 |
| 7 | 153549221 | rs2192272 | 0.607 | 0.941 | -0.516 | -0.229 | 0 | -0.015 | -0.015 | 0.423 | 0.35 | 0.32 | 0.27 | 0.17 | 0.33 | 0.32 |
| 7 | 153549296 | rs2192271 | 1.389 | 0.801 | -0.703 | -0.289 | 0.01 | 0.023 | 0.023 | 0 | 0.25 | 0.13 | 0.16 | 0.17 | 0.2 | 0.24 |
| 7 | 153553871 | rs17173801 | 0.477 | 0.874 | -0.575 | 0.077 | 0 | 0.013 | 0.013 | 0 | 0.1 | 0.13 | 0.14 | 0.25 | 0.08 | 0.29 |
| 7 | 153555351 | rs6464369 | 0.619 | 0.543 | -0.691 | 0.471 | 0 | 0.005 | 0.005 | 0.66 | 0.62 | 0.36 | 0.66 | 0.7 | 0.62 | 0.72 |
| 7 | 153555808 | rs13246647 | 1.778 | 0.619 | -0.8 | 0.549 | -0.067 | -0.048 | -0.048 | 0.828 | 0.29 | 0.06 | 0.37 | 0.29 | 0.37 | 0.23 |
| 7 | 153555866 | rs12670107 | 0.851 | 0.653 | -0.693 | 0.63 | 0.005 | 0.066 | 0.066 | 0 | 0.05 | 0.13 | 0.12 | 0.22 | 0.05 | 0.23 |
| 7 | 153558949 | rs12671878 | 1.461 | 0.643 | -0.851 | 0.544 | 0.007 | 0.009 | 0.009 | 0 | 0.26 | 0.17 | 0.17 | 0.2 | 0.2 | 0.24 |
| 7 | 153559059 | rs17173803 | 0.812 | 0.905 | -0.768 | 0.905 | -0.01 | 0.011 | 0.011 | 0 | 0.05 | 0.02 | 0.1 | 0.27 | 0.06 | 0.23 |
| 7 | 153560296 | rs2538415 | -0.318 | 0.632 | -0.528 | 0.465 | 0.02 | 0.033 | 0.033 | 0 | 0.56 | 0.7 | 0.7 | 0.68 | 0.64 | 0.64 |
| 7 | 153561830 | rs12539925 | 0.608 | 0.756 | -0.66 | 0.661 | 0.003 | 0.003 | 0.003 | 0.605 | 0.15 | 0.17 | 0.14 | 0.16 | 0.17 | 0.17 |
| 7 | 153580769 | rs38987 | -0.643 | 0.69 | -0.764 | 0.719 | -0.005 | -0.019 | -0.019 | 0 | 0.65 | 0.74 | 0.5 | 0.5 | 0.58 | 0.54 |
| 7 | 153582136 | rs38982 | -0.471 | 0.653 | -0.78 | 0.703 | -0.002 | -0.022 | -0.022 | 0 | 0.63 | 0.74 | 0.5 | 0.5 | 0.58 | 0.53 |
| 7 | 153586831 | rs740576 | 1.171 | 0.865 | -0.805 | 0.709 | -0.001 | -0.003 | -0.003 | 0 | 0.23 | 0.22 | 0.2 | 0.21 | 0.25 | 0.17 |
| 7 | 153587503 | rs929455 | 0.741 | 1.169 | -0.834 | 0.607 | -0.003 | -0.003 | -0.003 | 0 | 0.17 | 0.17 | 0.22 | 0.2 | 0.19 | 0.17 |
| 7 | 153591994 | rs38979 | 0.088 | 1.135 | -0.893 | 0.595 | 0.007 | 0.008 | 0.008 | 0 | 0.31 | 0.42 | 0.36 | 0.3 | 0.32 | 0.33 |
| 7 | 153593372 | rs38978 | -0.957 | 1.253 | -0.935 | 0.551 | 0.001 | 0 | 0 | 0.864 | 0.81 | 0.78 | 0.69 | 0.79 | 0.79 | 0.81 |
| 7 | 153596033 | rs10245002 | 1.923 | 0.943 | -1.117 | 0.565 | -0.003 | 0.007 | 0.007 | 0.008 | 0.23 | 0.24 | 0.21 | 0.35 | 0.26 | 0.35 |
| 7 | 153597256 | rs38974 | 0.85 | 1.082 | -1.59 | 0.397 | -0.059 | -0.06 | -0.06 | 0 | 0.18 | 0.02 | 0.21 | 0.17 | 0.21 | 0.13 |
| 7 | 153601721 | rs764723 | -1.155 | 1.506 | -1.2 | 0.305 | -0.008 | -0.006 | -0.006 | 0 | 0.83 | 0.82 | 0.77 | 0.79 | 0.79 | 0.84 |
| 7 | 153607630 | rs10487688 | 1.028 | 1.191 | -1.154 | -0.041 | 0.022 | 0.061 | 0.061 | 0.322 | 0.32 | 0.14 | 0.27 | 0.18 | 0.25 | 0.16 |
| 7 | 153608121 | rs12673076 | -0.955 | 1.221 | -0.891 | 0.322 | -0.005 | -0.002 | -0.002 | 0 | 0.83 | 0.81 | 0.84 | 0.81 | 0.81 | 0.84 |
| 7 | 153608618 | rs10952453 | 0.102 | 1.092 | -0.796 | -0.093 | 0 | 0.002 | 0.002 | 0 | 0.51 | 0.55 | 0.5 | 0.34 | 0.47 | 0.34 |
| 7 | 153612143 | rs10260433 | -1.155 | 1.269 | -1.013 | -0.838 | -0.029 | 0.04 | 0.04 | 0.413 | 0.89 | 0.98 | 0.9 | 1 | 0.85 | 0.88 |
| 7 | 153615099 | rs1981632 | -0.11 | 0.434 | -0.711 | -1.279 | -0.001 | -0.051 | -0.051 | 0 | 0.32 | 0.59 | 0.28 | 0.08 | 0.3 | 0.18 |
| 7 | 153616857 | rs10255973 | 0.609 | -0.196 | -0.111 | -1.055 | -0.003 | 0.022 | 0.022 | 0 | 0.33 | 0.24 | 0.4 | 0.21 | 0.29 | 0.22 |
| 7 | 153617952 | rs6464371 | -0.618 | -0.074 | 0.405 | -1.041 | 0.012 | -0.028 | -0.028 | 0 | 0.79 | 0.75 | 0.8 | 0.9 | 0.74 | 0.81 |
| 7 | 153618948 | rs879629 | 0.334 | -0.276 | 0.422 | -1.133 | -0.001 | -0.007 | -0.007 | 0 | 0.41 | 0.44 | 0.45 | 0.22 | 0.35 | 0.26 |
| 7 | 153622306 | rs2190997 | -0.242 | -0.262 | 0.421 | -1.211 | -0.007 | 0.034 | 0.034 | 0.251 | 0.58 | 0.76 | 0.61 | 0.31 | 0.54 | 0.4 |
| 7 | 153623251 | rs10155813 | 0.796 | -0.203 | 0.345 | -1.24 | -0.005 | -0.035 | -0.035 | 0 | 0.33 | 0.1 | 0.3 | 0.65 | 0.35 | 0.55 |
| 7 | 153627177 | rs17338902 | 1.028 | -0.753 | 0.548 | -1.225 | -0.009 | 0.014 | 0.014 | 0.412 | 0.33 | 0.21 | 0.41 | 0.21 | 0.31 | 0.21 |
| 7 | 153631648 | rs758316 | -0.483 | -0.807 | 0.692 | -1.244 | 0.009 | -0.01 | -0.01 | 0 | 0.61 | 0.79 | 0.64 | 0.4 | 0.59 | 0.45 |

| | | | | | | | | | | | | | | | | |
|---|---|---|---|---|---|---|---|---|---|---|---|---|---|---|---|---|
| 7 | 153631739 | rs4726348 | 1.243 | -0.85 | 0.814 | -1.263 | -0.009 | -0.001 | -0.001 | 0 | 0.3 | 0.32 | 0.4 | 0.26 | 0.3 | 0.2 |
| 7 | 153634958 | rs9691820 | 1.933 | -0.946 | 0.547 | -1.297 | -0.007 | -0.007 | -0.007 | 0 | 0.22 | 0.17 | 0.31 | 0.26 | 0.25 | 0.17 |
| 7 | 153637120 | rs6955332 | 0.282 | -1.004 | 0.611 | -1.356 | 0.004 | 0.012 | 0.012 | 0 | 0.25 | 0.2 | 0.19 | 0.11 | 0.24 | 0.19 |
| 7 | 153637760 | rs7784947 | -0.867 | -1.018 | 0.368 | -1.33 | -0.007 | -0.031 | -0.031 | 0 | 0.64 | 0.86 | 0.66 | 0.41 | 0.6 | 0.46 |
| 7 | 153638248 | rs6973850 | 0.717 | -0.919 | 0.655 | -1.102 | -0.002 | -0.038 | -0.038 | 0 | 0.13 | 0.26 | 0.14 | 0.04 | 0.1 | 0.06 |
| 7 | 153640474 | rs10487687 | -0.449 | -0.645 | 0.572 | -0.896 | -0.003 | -0.06 | -0.06 | 0 | 0.38 | 0.47 | 0.32 | 0.14 | 0.35 | 0.28 |
| 7 | 153643429 | rs11975562 | 0.952 | -0.512 | 0.248 | -0.31 | -0.015 | -0.005 | -0.005 | 0 | 0.28 | 0.2 | 0.33 | 0.25 | 0.3 | 0.19 |
| 7 | 153645299 | rs10480321 | 1.094 | 0.014 | 0.813 | 0.083 | -0.006 | 0.094 | 0.094 | 0 | 0.52 | 0.31 | 0.65 | 0.82 | 0.51 | 0.67 |
| 7 | 153645852 | rs1860218 | -0.27 | 0.391 | -0.019 | 0.047 | -0.007 | -0.022 | -0.022 | 0 | 0.78 | 0.85 | 0.83 | 1 | 0.7 | 0.86 |
| 7 | 153647231 | rs11974308 | 0.357 | 0.525 | -0.625 | 0.147 | 0.009 | 0.019 | 0.019 | 0.179 | 0.16 | 0.11 | 0.11 | 0.1 | 0.14 | 0.16 |
| 7 | 153652188 | rs10229774 | 0.937 | 0.804 | -0.766 | 0.327 | -0.001 | -0.001 | -0.001 | 0 | 0.47 | 0.47 | 0.58 | 0.4 | 0.44 | 0.28 |
| 7 | 153654045 | rs7803838 | -0.719 | 0.105 | -0.419 | 1.397 | -0.005 | 0.006 | 0.006 | 0.159 | 0.32 | 0.61 | 0.28 | 0.6 | 0.27 | 0.59 |
| 7 | 153670448 | rs39156 | -1.054 | 0.533 | -0.525 | 1.662 | 0.002 | 0.009 | 0.009 | 0.006 | 0.1 | 0.09 | 0.13 | 0.22 | 0.06 | 0.23 |
| 7 | 153696268 | rs6464378 | -1.149 | 1.242 | 0.406 | 1.145 | 0.15 | 0.037 | 0.037 | 0.467 | 0.07 | 0.31 | 0.07 | 0.02 | 0.1 | 0.03 |
| 7 | 153697418 | rs4546569 | -0.474 | 2.234 | -0.521 | 1.738 | -0.032 | -0.041 | -0.041 | 0.519 | 0.18 | 0.13 | 0.33 | 0.56 | 0.31 | 0.24 |
| 7 | 153707385 | rs17134265 | -0.904 | 1.152 | -0.645 | 1.747 | 0.002 | 0.003 | 0.003 | 0 | 0.13 | 0.09 | 0.04 | 0.35 | 0.09 | 0.12 |
| 7 | 153709408 | rs10085865 | -0.472 | -0.138 | -0.67 | 0.786 | -0.038 | -0.004 | -0.004 | 0 | 0.28 | 0.63 | 0.23 | 0.26 | 0.19 | 0.33 |
| 7 | 153734323 | rs6464381 | -0.431 | 0.053 | -1.578 | 0.36 | 0.014 | 0.035 | 0.035 | 0 | 0.27 | 0.12 | 0.26 | 0.2 | 0.26 | 0.24 |
| 7 | 153743749 | rs3864483 | -0.185 | 0.29 | -1.869 | -0.061 | 0.012 | 0.114 | 0.114 | 0 | 0.41 | 0.19 | 0.35 | 0.22 | 0.45 | 0.28 |
| 7 | 153747353 | rs1476818 | -0.502 | 1.037 | -1.92 | 0.144 | -0.012 | -0.002 | -0.002 | 0 | 0.69 | 0.91 | 0.56 | 0.69 | 0.71 | 0.61 |
| 7 | 153768982 | rs11766679 | -0.408 | 1.48 | -2.098 | -0.255 | -0.075 | 0.046 | 0.046 | 0 | 0.17 | 0.02 | 0.3 | 0.09 | 0.26 | 0.21 |
| 7 | 153780244 | rs7357097 | -0.273 | 1.076 | -1.701 | 0.037 | -0.01 | 0 | 0 | 0.245 | 0.65 | 0.74 | 0.6 | 0.65 | 0.56 | 0.67 |
| 7 | 153789777 | rs4725525 | -0.151 | 0.326 | -1.445 | -0.146 | -0.004 | 0.002 | 0.002 | 0 | 0.6 | 0.7 | 0.58 | 0.64 | 0.5 | 0.61 |
| 7 | 153811655 | rs3115192 | 0.153 | 0.192 | -1.249 | -0.39 | -0.018 | 0.026 | 0.026 | 0 | 0.81 | 0.92 | 0.71 | 0.87 | 0.79 | 0.79 |
| 7 | 153865198 | rs4131646 | 0.534 | 0.012 | -1.237 | -0.752 | -0.011 | 0.06 | 0.06 | 0.184 | 0.81 | 0.93 | 0.81 | 1 | 0.8 | 0.74 |
| 7 | 153865752 | rs7790046 | -0.794 | -0.83 | -0.697 | -0.925 | -0.023 | -0.024 | -0.024 | 0 | 0.16 | 0.71 | 0.12 | 0.07 | 0.09 | 0.09 |
| 7 | 153871963 | rs7794112 | 0.882 | -0.93 | -0.309 | -0.98 | 0.001 | 0.014 | 0.014 | 0.465 | 0.77 | 0.82 | 0.78 | 0.94 | 0.79 | 0.75 |
| 7 | 153872322 | rs12532924 | 1.553 | -0.848 | -0.051 | -0.544 | 0.016 | 0.001 | 0.001 | 0 | 0.53 | 0.31 | 0.5 | 0.59 | 0.55 | 0.52 |
| 7 | 153875187 | rs12154291 | -0.956 | -0.906 | 1.137 | -0.198 | 0.001 | 0.009 | 0.009 | 0 | 0.26 | 0.27 | 0.29 | 0.34 | 0.32 | 0.2 |
| 7 | 153884957 | rs11975478 | -2.181 | -1.158 | 2.049 | 0.31 | 0.008 | 0.183 | 0.183 | 0 | 0.18 | 0.41 | 0.19 | 0.35 | 0.17 | 0.24 |
| 7 | 153886615 | rs10251943 | -2.27 | -1.139 | 2.121 | 0.318 | 0.004 | 0.175 | 0.175 | 0 | 0.18 | 0.49 | 0.2 | 0.35 | 0.16 | 0.23 |
| 7 | 153888535 | rs4507681 | -0.772 | -1.344 | 1.451 | 0.146 | 0.03 | 0.083 | 0.083 | 0 | 0.16 | 0.02 | 0.05 | 0 | 0.1 | 0.1 |
| 7 | 153893160 | rs7811481 | -2.23 | -1.326 | 1.819 | 0.354 | 0.013 | -0.018 | -0.018 | 1.123 | 0.31 | 0.73 | 0.26 | 0.36 | 0.23 | 0.33 |
| 7 | 153893727 | rs6945869 | -1.61 | -1.243 | 2.401 | 0.274 | 0.006 | 0.046 | 0.046 | 1.357 | 0.07 | 0.22 | 0.05 | 0.07 | 0.05 | 0.07 |
| 7 | 153895189 | rs4074568 | 1.397 | -1.389 | 2.334 | 0.379 | -0.044 | 0.129 | 0.129 | 0 | 0.78 | 0.55 | 0.79 | 0.62 | 0.84 | 0.79 |
| 7 | 153895742 | rs4074817 | 1.71 | -1.367 | 2.085 | 0.34 | 0.031 | -0.046 | -0.046 | 1.729 | 0.75 | 0.19 | 0.75 | 0.63 | 0.82 | 0.77 |
| 7 | 153897575 | rs12703320 | 1.842 | -1.35 | 2.065 | 0.279 | 0.008 | -0.01 | -0.01 | 0 | 0.75 | 0.25 | 0.75 | 0.65 | 0.79 | 0.78 |
| 7 | 153903590 | rs4380850 | 2.353 | -1.328 | 1.647 | 0.161 | 0.07 | -0.012 | -0.012 | 3.155 | 0.62 | 0.19 | 0.69 | 0.65 | 0.72 | 0.63 |
| 7 | 153907603 | rs12703326 | 0.526 | -1.22 | 1.403 | 0.334 | 0.003 | -0.008 | -0.008 | 0 | 0.83 | 0.84 | 0.81 | 0.73 | 0.84 | 0.85 |
| 7 | 153908837 | rs4302768 | 1.144 | -1.204 | 1.431 | 0.34 | -0.007 | 0.024 | 0.024 | 0 | 0.8 | 0.74 | 0.79 | 0.66 | 0.81 | 0.79 |
| 7 | 153911539 | rs9791911 | 1.609 | -1.215 | 1.061 | 0.297 | 0.004 | 0 | 0 | 0 | 0.7 | 0.72 | 0.76 | 0.72 | 0.79 | 0.72 |
| 7 | 153911616 | rs11505645 | -0.334 | -1.078 | 0.941 | 0.166 | -0.006 | -0.023 | -0.023 | 3.59 | 0.13 | 0.11 | 0.05 | 0 | 0.05 | 0.09 |
| 7 | 153913467 | rs4726386 | 1.354 | -0.229 | 1.164 | 0.727 | -0.01 | 0.017 | 0.017 | 0 | 0.76 | 0.31 | 0.75 | 0.66 | 0.78 | 0.78 |
| 7 | 153946551 | rs9640345 | -0.163 | 0.935 | -1.265 | 0.188 | -0.103 | 0.002 | 0.002 | 0.749 | 0.28 | 0.03 | 0.23 | 0.27 | 0.34 | 0.29 |
| 7 | 153947649 | rs10235250 | -1.058 | 0.357 | -2.478 | 0.157 | -0.031 | -0.172 | -0.172 | 0 | 0.69 | 0.98 | 0.65 | 0.51 | 0.72 | 0.59 |
| 7 | 153950342 | rs4129040 | 0.034 | -0.019 | -1.997 | -0.237 | 0.008 | -0.012 | -0.012 | 0.076 | 0.21 | 0.24 | 0.22 | 0.12 | 0.28 | 0.05 |
| 7 | 153951624 | rs10267219 | -0.663 | 0.011 | -2.191 | -0.223 | -0.093 | -0.107 | -0.107 | 0.737 | 0.58 | 0.94 | 0.55 | 0.51 | 0.54 | 0.55 |
| 7 | 153959855 | rs7384911 | -0.067 | 0.466 | -1.684 | -0.163 | -0.077 | 0.05 | 0.05 | 0 | 0.38 | 0.18 | 0.3 | 0.3 | 0.43 | 0.3 |
| 7 | 153967174 | rs10244110 | -0.037 | 0.415 | -1.241 | -0.156 | -0.087 | 0.044 | 0.044 | 0 | 0.62 | 0.85 | 0.7 | 0.71 | 0.57 | 0.7 |
| 7 | 153967280 | rs10274025 | 0.14 | 0.231 | -0.542 | -0.213 | -0.046 | -0.061 | -0.061 | 0.265 | 0.18 | 0.3 | 0.1 | 0.06 | 0.09 | 0.15 |
| 7 | 153970329 | rs4078461 | 0.793 | 0.979 | -0.497 | 0.328 | -0.044 | -0.068 | -0.068 | 0 | 0.55 | 0.1 | 0.51 | 0.48 | 0.51 | 0.52 |
| 7 | 153973881 | rs10952471 | -0.318 | 1.192 | -0.594 | 0.067 | -0.018 | 0.026 | 0.026 | 0 | 0.94 | 0.97 | 0.95 | 1 | 0.91 | 0.98 |
| 7 | 153975178 | rs12703336 | 0.315 | 1.183 | -0.278 | 0.099 | 0.065 | -0.003 | -0.003 | 0 | 0.91 | 0.75 | 0.83 | 0.92 | 0.86 | 0.87 |
| 7 | 153976598 | rs10230969 | -0.307 | 0.67 | -0.916 | 0.264 | -0.032 | 0.003 | 0.003 | 0 | 0.29 | 0.15 | 0.23 | 0.3 | 0.32 | 0.28 |
| 7 | 153981261 | rs4327776 | -0.719 | 0.22 | -0.472 | 0.493 | -0.003 | 0.006 | 0.006 | 0.045 | 0.34 | 0.66 | 0.49 | 0.52 | 0.32 | 0.41 |
| 7 | 153987075 | rs9655634 | -0.244 | 0.183 | -0.612 | 0.643 | 0.003 | -0.05 | -0.05 | 0 | 0.27 | 0.13 | 0.23 | 0.31 | 0.23 | 0.29 |
| 7 | 153988834 | rs6955612 | -0.714 | -0.036 | -0.31 | 0.259 | 0.01 | -0.001 | -0.001 | 0.191 | 0.4 | 0.33 | 0.31 | 0.32 | 0.31 | 0.37 |
| 7 | 153992610 | rs4278085 | -1.037 | -0.15 | -0.521 | 0.279 | 0.031 | -0.014 | -0.014 | 0.612 | 0.46 | 0.76 | 0.56 | 0.42 | 0.37 | 0.45 |
| 7 | 154001207 | rs4726403 | -0.978 | -0.441 | -0.691 | 0.362 | 0.036 | -0.025 | -0.025 | 0 | 0.47 | 0.84 | 0.56 | 0.41 | 0.37 | 0.51 |
| 7 | 154001559 | rs10231911 | -0.111 | -0.275 | -0.514 | -0.295 | -0.003 | 0.008 | 0.008 | 0 | 0.47 | 0.58 | 0.47 | 0.7 | 0.51 | 0.47 |
| 7 | 154007342 | rs6944113 | -1.37 | -0.888 | 0.178 | -0.456 | -0.055 | 0.015 | 0.015 | 0 | 0.23 | 0.47 | 0.2 | 0.19 | 0.14 | 0.19 |
| 7 | 154008127 | rs9648765 | -0.173 | -0.094 | -0.29 | -0.994 | -0.02 | 0.02 | 0.02 | 0 | 0.19 | 0.13 | 0.19 | 0.09 | 0.29 | 0.17 |
| 7 | 154009325 | rs6464401 | -2.052 | -0.251 | -0.54 | -1.34 | -0.014 | 0.003 | 0.003 | 0.24 | 0.57 | 0.77 | 0.57 | 0.36 | 0.56 | 0.53 |
| 7 | 154009637 | rs7805488 | -0.795 | 0.029 | 0.353 | -1.195 | -0.047 | 0.046 | 0.046 | 0 | 0.2 | 0.45 | 0.19 | 0.24 | 0.12 | 0.24 |
| 7 | 154009697 | rs7805372 | -1.37 | 0 | 0.302 | -1.159 | -0.037 | -0.002 | -0.002 | 0.529 | 0.31 | 0.49 | 0.23 | 0.25 | 0.2 | 0.29 |
| 7 | 154011984 | rs7780487 | 1.785 | 0.695 | 0.483 | -1.071 | 0.004 | 0.011 | 0.011 | 0.382 | 0.54 | 0.22 | 0.47 | 0.66 | 0.56 | 0.55 |
| 7 | 154012916 | rs1468756 | 1.415 | 0.896 | 1.059 | -1.009 | 0.029 | 0.041 | 0.041 | 0.055 | 0.69 | 0.46 | 0.61 | 0.67 | 0.62 | 0.65 |
| 7 | 154019581 | rs2024365 | 0.781 | 0.991 | 0.549 | -1.18 | -0.053 | -0.047 | -0.047 | 0 | 0.8 | 0.91 | 0.71 | 0.78 | 0.7 | 0.72 |
| 7 | 154019767 | rs10278563 | -1.167 | 0.104 | 1.15 | -1.347 | -0.051 | -0.127 | -0.127 | 0.419 | 0.21 | 0.38 | 0.07 | 0 | 0.12 | 0.14 |

| | | | | | | | | | | | | | | | | |
|---|---|---|---|---|---|---|---|---|---|---|---|---|---|---|---|---|
| 7 | 154023564 | rs11773122 | -0.675 | 0.015 | 1.253 | -1.102 | -0.034 | -0.06 | -0.06 | 0 | 0.2 | 0.3 | 0.09 | 0.04 | 0.13 | 0.26 |
| 7 | 154023803 | rs1035097 | -0.318 | 0.345 | 1.076 | -1.049 | -0.042 | -0.024 | -0.024 | 0 | 0.93 | 0.99 | 0.88 | 0.94 | 0.85 | 0.91 |
| 7 | 154025175 | rs11765037 | -0.689 | 0.246 | 1.224 | -1.044 | -0.033 | -0.069 | -0.069 | 0 | 0.2 | 0.33 | 0.1 | 0.04 | 0.14 | 0.21 |
| 7 | 154026312 | rs11765968 | 2.395 | 0.197 | 1.166 | -1.201 | -0.027 | 0.035 | 0.035 | 2.909 | 0.44 | 0.19 | 0.52 | 0.68 | 0.5 | 0.43 |
| 7 | 154026438 | rs6959816 | -0.736 | 0.305 | 1.344 | -1.162 | -0.041 | -0.076 | -0.076 | 0 | 0.23 | 0.43 | 0.15 | 0.04 | 0.15 | 0.25 |
| 7 | 154028052 | rs12531152 | 1.451 | 0.831 | 1.425 | -0.664 | 0.007 | -0.002 | -0.002 | 0.201 | 0.7 | 0.65 | 0.66 | 0.74 | 0.65 | 0.69 |
| 7 | 154032974 | rs2024366 | 1.538 | 0.846 | 1.559 | -0.612 | 0.012 | -0.001 | -0.001 | 0 | 0.71 | 0.63 | 0.73 | 0.73 | 0.66 | 0.71 |
| 7 | 154033152 | rs6964613 | -1.241 | 0.768 | 1.773 | -0.707 | 0.003 | 0.006 | 0.006 | 0.942 | 0.05 | 0.28 | 0.02 | 0.01 | 0 | 0.04 |
| 7 | 154034329 | rs887926 | 0.739 | 1.048 | 1.484 | -0.716 | -0.014 | 0.006 | 0.006 | 0.024 | 0.89 | 0.91 | 0.86 | 0.94 | 0.78 | 0.87 |
| 7 | 154037662 | rs6464410 | -1.732 | 0.273 | 2.023 | -0.628 | 0.015 | 0.031 | 0.031 | 0.045 | 0.11 | 0.5 | 0.09 | 0.09 | 0.07 | 0.14 |
| 7 | 154040152 | rs759264 | -0.646 | 0.275 | 2.123 | -0.627 | 0.038 | 0.056 | 0.056 | 0 | 0.07 | 0.25 | 0.07 | 0.08 | 0.07 | 0.11 |
| 7 | 154047636 | rs13229985 | 0.207 | 0.181 | 0.984 | -0.723 | -0.043 | -0.029 | -0.029 | 0 | 0.15 | 0.01 | 0.1 | 0.14 | 0.13 | 0.16 |
| 7 | 154055798 | rs7807092 | 0.042 | 0.227 | 1.043 | -0.734 | 0.002 | 0 | 0 | 0 | 0.92 | 0.91 | 0.87 | 0.94 | 0.84 | 0.88 |
| 7 | 154057993 | rs9918610 | 0.041 | 0.076 | 1.248 | -0.703 | 0.022 | 0.027 | 0.027 | 0 | 0.06 | 0.15 | 0.06 | 0.06 | 0.06 | 0.11 |
| 7 | 154058474 | rs12536378 | 2.072 | 0.036 | 0.589 | -0.777 | -0.069 | 0.039 | 0.039 | 0.067 | 0.58 | 0.11 | 0.62 | 0.71 | 0.59 | 0.51 |
| 7 | 154058755 | rs10228095 | -2.461 | -0.353 | 1.427 | -0.593 | -0.004 | -0.005 | -0.005 | 0 | 0.16 | 0.77 | 0.12 | 0.07 | 0.09 | 0.16 |
| 7 | 154064225 | rs17174375 | -1.404 | -0.24 | 1.504 | -0.569 | 0.005 | -0.016 | -0.016 | 0.462 | 0.07 | 0.27 | 0.02 | 0.01 | 0.02 | 0.05 |
| 7 | 154067543 | rs4725537 | -1.746 | -0.187 | 0.283 | -0.548 | -0.014 | -0.022 | -0.022 | 0 | 0.15 | 0.71 | 0.11 | 0.08 | 0.09 | 0.13 |
| 7 | 154070594 | rs12703349 | 0.966 | -0.349 | 0.002 | -0.392 | 0.002 | 0.002 | 0.002 | 3.983 | 0.68 | 0.35 | 0.72 | 0.78 | 0.74 | 0.68 |
| 7 | 154072020 | rs6977820 | 1.217 | -0.602 | -0.404 | -0.551 | 0.049 | 0.066 | 0.066 | 2.732 | 0.6 | 0.18 | 0.75 | 0.76 | 0.69 | 0.69 |
| 7 | 154077301 | rs7776495 | 1.622 | -0.564 | -0.786 | -0.689 | -0.033 | 0.028 | 0.028 | 1.022 | 0.46 | 0.12 | 0.64 | 0.64 | 0.57 | 0.52 |
| 7 | 154079635 | rs7804017 | -0.652 | -0.569 | -0.004 | -0.7 | 0.066 | 0.095 | 0.095 | 0.085 | 0.08 | 0.3 | 0.06 | 0.12 | 0.09 | 0.11 |
| 7 | 154080404 | rs7808582 | 0.266 | -0.673 | -0.132 | -0.733 | -0.008 | 0.022 | 0.022 | 0 | 0.16 | 0.06 | 0.14 | 0.12 | 0.14 | 0.18 |
| 7 | 154083325 | rs6464419 | 0.032 | -0.784 | 0.245 | -0.823 | 0.027 | 0.031 | 0.031 | 0 | 0.75 | 0.83 | 0.88 | 0.99 | 0.83 | 0.86 |
| 7 | 154083710 | rs756438 | 1.206 | -0.688 | -0.309 | -0.793 | 0.008 | 0.013 | 0.013 | 0 | 0.54 | 0.41 | 0.71 | 0.76 | 0.66 | 0.66 |
| 7 | 154084658 | rs10256276 | 1.415 | -0.007 | 0.365 | -0.131 | -0.009 | -0.018 | -0.018 | 0 | 0.7 | 0.42 | 0.78 | 0.76 | 0.77 | 0.79 |
| 7 | 154090156 | rs1292312 | 1.608 | 0.013 | 0.386 | -0.022 | 0.001 | -0.007 | -0.007 | 0 | 0.8 | 0.23 | 0.84 | 0.74 | 0.84 | 0.85 |
| 7 | 154090508 | rs17174400 | 0.25 | 0.013 | 0.513 | -0.022 | 0.04 | 0.067 | 0.067 | 0 | 0.05 | 0.12 | 0.05 | 0.12 | 0.08 | 0.11 |
| 7 | 154097732 | rs6955904 | 1.336 | -0.005 | 0.529 | 0.028 | -0.009 | -0.007 | -0.007 | 0 | 0.7 | 0.41 | 0.77 | 0.73 | 0.77 | 0.78 |
| 7 | 154109958 | rs1491839 | 2.091 | -0.015 | 0.366 | 0.112 | -0.015 | -0.067 | -0.067 | 0 | 0.61 | 0.11 | 0.7 | 0.58 | 0.68 | 0.67 |
| 7 | 154111242 | rs286852 | -0.277 | 0.039 | 0.403 | 0.135 | -0.002 | 0.012 | 0.012 | 0 | 0.18 | 0.14 | 0.14 | 0.14 | 0.16 | 0.18 |
| 7 | 154111855 | rs286855 | 1.457 | 0.056 | 0.95 | 0.446 | -0.024 | 0.054 | 0.054 | 4.233 | 0.86 | 0.48 | 0.89 | 0.84 | 0.92 | 0.96 |
| 7 | 154113518 | rs1491840 | -1.754 | 0.195 | 0.86 | 0.513 | 0.06 | 0.245 | 0.245 | 0 | 0.11 | 0.4 | 0.1 | 0.27 | 0.1 | 0.14 |
| 7 | 154122060 | rs10273396 | -1.158 | 0.016 | -0.245 | 0.236 | -0.03 | 0.017 | 0.017 | 5.172 | 0.15 | 0.67 | 0.12 | 0.15 | 0.08 | 0.04 |
| 7 | 154129140 | rs6979626 | -0.11 | 0.954 | -1.519 | 0.026 | -0.142 | -0.026 | -0.026 | 1.43 | 0.27 | 0.05 | 0.36 | 0.74 | 0.4 | 0.26 |
| 7 | 154131148 | rs4726417 | 0.617 | 0.182 | -0.75 | -0.407 | 0.015 | -0.017 | -0.017 | 0 | 0.75 | 0.65 | 0.76 | 0.81 | 0.73 | 0.77 |
| 7 | 154133742 | rs923712 | 0.096 | 0.311 | -1.022 | -0.321 | -0.058 | -0.042 | -0.042 | 0.05 | 0.87 | 0.98 | 0.84 | 0.82 | 0.82 | 0.79 |
| 7 | 154133785 | rs923713 | 0.668 | -0.94 | -0.709 | -1.021 | 0.002 | 0.006 | 0.006 | 0 | 0.66 | 0.67 | 0.73 | 0.79 | 0.68 | 0.6 |
| 7 | 154136555 | rs6965651 | -0.606 | -1.035 | 0.182 | 0.425 | 0.012 | 0.314 | 0.314 | 0.013 | 0.11 | 0.29 | 0.21 | 0.37 | 0.08 | 0.15 |
| 7 | 154137194 | rs11981812 | -0.087 | -1.751 | -0.291 | -0.675 | -0.003 | -0.04 | -0.04 | 0.91 | 0.28 | 0.28 | 0.15 | 0.03 | 0.2 | 0.29 |
| 7 | 154142551 | rs7783212 | -0.339 | -1.536 | 0.295 | -0.166 | -0.025 | 0.068 | 0.068 | 0.18 | 0.15 | 0.43 | 0.11 | 0.21 | 0.11 | 0.08 |
| 7 | 154145593 | rs2337377 | -0.787 | -2.011 | -0.612 | -0.263 | -0.025 | 0.004 | 0.004 | 0.238 | 0.58 | 0.94 | 0.38 | 0.37 | 0.47 | 0.49 |
| 7 | 154149688 | rs6978845 | -0.641 | -2.278 | -0.374 | 0.131 | -0.033 | 0.035 | 0.035 | 0 | 0.27 | 0.51 | 0.41 | 0.41 | 0.19 | 0.35 |
| 7 | 154150550 | rs12703354 | -0.632 | -1.777 | -0.32 | 0.74 | 0.008 | -0.013 | -0.013 | 1.23 | 0.22 | 0.18 | 0.35 | 0.46 | 0.13 | 0.4 |
| 7 | 154152834 | rs6974964 | -0.923 | -1.939 | -0.553 | 0.185 | -0.017 | -0.077 | -0.077 | 0 | 0.59 | 0.49 | 0.71 | 0.9 | 0.62 | 0.78 |
| 7 | 154156519 | rs7791484 | 0.911 | -1.519 | -0.692 | 0.544 | -0.102 | -0.028 | -0.028 | 0 | 0.3 | 0.1 | 0.3 | 0.3 | 0.42 | 0.27 |
| 7 | 154157438 | rs6966462 | -0.443 | -1.025 | 0.069 | 1.447 | -0.01 | 0.084 | 0.084 | 0 | 0.16 | 0.26 | 0.36 | 0.44 | 0.13 | 0.3 |
| 7 | 154157519 | rs13246835 | -0.488 | -0.094 | -0.18 | 0.687 | 0.002 | -0.02 | -0.02 | 0 | 0.08 | 0.03 | 0.04 | 0.14 | 0.06 | 0.13 |
| 7 | 154161235 | rs10268315 | -1.379 | 0.132 | 0.257 | 0.356 | 0.024 | 0.014 | 0.014 | 0.778 | 0.16 | 0.51 | 0.13 | 0.12 | 0.15 | 0.18 |
| 7 | 154165496 | rs4726422 | 0.02 | 0.309 | -0.433 | 0.033 | -0.009 | 0.026 | 0.026 | 0 | 0.45 | 0.7 | 0.63 | 0.67 | 0.42 | 0.58 |
| 7 | 154167651 | rs9640200 | -0.544 | 0.176 | -0.518 | -0.651 | 0.017 | 0.069 | 0.069 | 0 | 0.59 | 0.75 | 0.72 | 0.94 | 0.59 | 0.81 |
| 7 | 154169390 | rs4725545 | -0.61 | 0.55 | 0.041 | -0.283 | -0.007 | -0.04 | -0.04 | 0 | 0.23 | 0.39 | 0.33 | 0.17 | 0.22 | 0.16 |
| 7 | 154172154 | rs3750042 | 0.451 | 0.234 | -0.142 | -0.571 | 0.003 | 0 | 0 | 0.255 | 0.36 | 0.35 | 0.25 | 0.09 | 0.33 | 0.32 |
| 7 | 154172823 | rs13230019 | 0.244 | 0.343 | -0.619 | 0.045 | -0.004 | 0.026 | 0.026 | 0 | 0.46 | 0.19 | 0.43 | 0.46 | 0.51 | 0.41 |
| 7 | 154174813 | rs10278985 | 0.025 | 0.044 | -0.53 | -0.545 | -0.007 | -0.038 | -0.038 | 0 | 0.43 | 0.52 | 0.29 | 0.1 | 0.36 | 0.44 |
| 7 | 154174868 | rs10235672 | 0.002 | 0.217 | -0.543 | -0.545 | -0.005 | -0.034 | -0.034 | 1.709 | 0.43 | 0.53 | 0.29 | 0.1 | 0.37 | 0.43 |
| 7 | 154175955 | rs12535735 | 0.078 | 0.008 | -0.604 | -0.626 | -0.003 | -0.039 | -0.039 | 0 | 0.45 | 0.52 | 0.29 | 0.08 | 0.36 | 0.43 |
| 7 | 154177426 | rs6959127 | 0.241 | 0.661 | -1.064 | -0.084 | 0.019 | 0.079 | 0.079 | 0.386 | 0.51 | 0.32 | 0.3 | 0.25 | 0.45 | 0.26 |
| 7 | 154186526 | rs10263063 | 0.067 | 0.973 | -1.088 | 0.491 | 0.001 | 0.013 | 0.013 | 0 | 0.49 | 0.43 | 0.59 | 0.33 | 0.47 | 0.49 |
| 7 | 154194246 | rs10274160 | -0.223 | 0.919 | -1.083 | 0.809 | -0.01 | 0.001 | 0.001 | 0 | 0.59 | 0.43 | 0.74 | 0.66 | 0.63 | 0.68 |
| 7 | 154196001 | rs744684 | -0.888 | 1.379 | -0.394 | 1.18 | 0.056 | 0.13 | 0.13 | 0.892 | 0.16 | 0.36 | 0.2 | 0.36 | 0.17 | 0.22 |
| 7 | 154196985 | rs11771157 | 0.482 | 1.143 | -0.557 | 0.812 | 0.001 | 0.005 | 0.005 | 0 | 0.59 | 0.35 | 0.35 | 0.64 | 0.4 | 0.55 |
| 7 | 154197382 | rs12668503 | -1.309 | 1.111 | 0.106 | 1.864 | 0.037 | 0.315 | 0.315 | 0.027 | 0.06 | 0.25 | 0.06 | 0.35 | 0.07 | 0.23 |
| 7 | 154197672 | rs2087720 | -0.213 | 1.047 | -0.309 | 0.701 | -0.002 | 0.006 | 0.006 | 0 | 0.55 | 0.52 | 0.52 | 0.73 | 0.51 | 0.61 |
| 7 | 154201075 | rs12703361 | -0.09 | 1.004 | -0.619 | 0.106 | -0.001 | -0.004 | -0.004 | 0 | 0.59 | 0.46 | 0.51 | 0.3 | 0.56 | 0.48 |
| 7 | 154201314 | rs6955044 | 0.042 | 1.485 | -0.669 | 0.054 | -0.001 | -0.001 | -0.001 | 0 | 0.57 | 0.45 | 0.5 | 0.29 | 0.51 | 0.44 |
| 7 | 154201817 | rs10245134 | 0.461 | 1.201 | -0.993 | -0.422 | 0 | -0.001 | -0.001 | 0.007 | 0.28 | 0.25 | 0.16 | 0.11 | 0.28 | 0.2 |
| 7 | 154201853 | rs13226031 | 0.699 | 1.26 | -0.996 | -0.425 | -0.004 | 0.046 | 0.046 | 0 | 0.31 | 0.2 | 0.34 | 0.18 | 0.27 | 0.27 |
| 7 | 154202183 | rs10232716 | 0.462 | 1.228 | -0.998 | -0.422 | 0 | -0.001 | -0.001 | 0 | 0.28 | 0.25 | 0.16 | 0.11 | 0.28 | 0.19 |

| | | | | | | | | | | | | | | | | |
|---|---|---|---|---|---|---|---|---|---|---|---|---|---|---|---|---|
| 7 | 154203428 | rs12703362 | 0.761 | 1.216 | -1.009 | -0.442 | 0.005 | 0.047 | 0.047 | 0 | 0.3 | 0.18 | 0.23 | 0.18 | 0.25 | 0.25 |
| 7 | 154203454 | rs4725547 | -0.598 | 1.252 | -1.029 | -0.464 | -0.001 | 0.002 | 0.002 | 0 | 0.58 | 0.43 | 0.4 | 0.29 | 0.54 | 0.45 |
| 7 | 154205291 | rs11763727 | 0.687 | 1.143 | -1.082 | -0.473 | 0.004 | 0.05 | 0.05 | 0 | 0.3 | 0.18 | 0.35 | 0.19 | 0.26 | 0.25 |
| 7 | 154205894 | rs10239794 | -0.635 | 1.159 | -0.822 | -0.395 | 0 | -0.013 | -0.013 | 0 | 0.59 | 0.49 | 0.52 | 0.3 | 0.57 | 0.48 |
| 7 | 154208824 | rs13232525 | 0.45 | 0.869 | -1.181 | -0.717 | -0.028 | 0.06 | 0.06 | 0.009 | 0.18 | 0.08 | 0.28 | 0.07 | 0.17 | 0.16 |
| 7 | 154209858 | rs12668377 | -0.159 | 1.008 | -1.13 | -1.097 | -0.017 | 0.035 | 0.035 | 0 | 0.37 | 0.36 | 0.29 | 0.14 | 0.39 | 0.35 |
| 7 | 154210798 | rs10260404 | -0.086 | 1.011 | -1.055 | -1.107 | -0.009 | 0.021 | 0.021 | 0 | 0.37 | 0.35 | 0.28 | 0.15 | 0.38 | 0.35 |
| 7 | 154213814 | rs6464429 | -0.894 | 0.823 | -1.076 | -1.198 | 0.002 | 0.023 | 0.023 | 0 | 0.46 | 0.44 | 0.33 | 0.13 | 0.41 | 0.35 |
| 7 | 154216037 | rs11772385 | 2.37 | 0.595 | -1.087 | -1.196 | 0.017 | 0.007 | 0.007 | 1.062 | 0.13 | 0.06 | 0.07 | 0.1 | 0.09 | 0.08 |
| 7 | 154216346 | rs1388543 | 1.645 | 0.61 | -1.079 | -1.243 | -0.005 | 0.006 | 0.006 | 0.018 | 0.18 | 0.15 | 0.11 | 0.06 | 0.19 | 0.17 |
| 7 | 154217955 | rs10952491 | -0.95 | 0.677 | -1.135 | -1.261 | -0.003 | 0.03 | 0.03 | 0 | 0.47 | 0.44 | 0.34 | 0.13 | 0.45 | 0.34 |
| 7 | 154218976 | rs7785513 | 0.976 | 0.629 | -1.078 | -1.199 | -0.008 | 0.04 | 0.04 | 3.336 | 0.3 | 0.15 | 0.18 | 0.17 | 0.29 | 0.25 |
| 7 | 154219524 | rs6976924 | 0.914 | 0.392 | -0.743 | -0.966 | 0.007 | -0.064 | -0.064 | 0 | 0.11 | 0.24 | 0.08 | 0 | 0.1 | 0.03 |
| 7 | 154224041 | rs1552734 | -1.071 | 0.416 | -0.715 | -1.034 | 0.001 | 0.005 | 0.005 | 0 | 0.49 | 0.56 | 0.37 | 0.21 | 0.51 | 0.4 |
| 7 | 154225119 | rs10464419 | 1.837 | -0.314 | -0.53 | -0.538 | -0.024 | -0.025 | -0.025 | 0.997 | 0.23 | 0.11 | 0.46 | 0.37 | 0.31 | 0.3 |
| 7 | 154225365 | rs2055017 | 0.643 | -0.183 | -0.38 | -0.932 | 0.038 | -0.044 | -0.044 | 0 | 0.22 | 0.36 | 0.17 | 0.08 | 0.3 | 0.17 |
| 7 | 154225655 | rs7803828 | -0.547 | -0.171 | -0.475 | -0.99 | 0.002 | -0.011 | -0.011 | 0 | 0.4 | 0.44 | 0.27 | 0.22 | 0.43 | 0.35 |
| 7 | 154232976 | rs11975187 | -0.555 | -0.135 | -0.464 | -0.932 | -0.004 | -0.017 | -0.017 | 0 | 0.4 | 0.53 | 0.25 | 0.22 | 0.4 | 0.34 |
| 7 | 154233833 | rs10276300 | 0.101 | -0.259 | -0.279 | -0.839 | -0.062 | -0.021 | -0.021 | 0 | 0.21 | 0.46 | 0.14 | 0.13 | 0.13 | 0.2 |
| 7 | 154246235 | rs11971454 | 1.842 | 0.503 | -0.068 | -0.698 | 0.075 | -0.013 | -0.013 | 0 | 0.14 | 0.36 | 0.11 | 0.07 | 0.19 | 0.1 |
| 7 | 154248268 | rs902743 | -0.361 | 0.056 | -0.286 | -0.881 | 0.008 | 0.014 | 0.014 | 0 | 0.5 | 0.56 | 0.3 | 0.22 | 0.44 | 0.33 |
| 7 | 154248407 | rs902742 | -0.368 | 0.053 | -0.354 | -0.882 | 0.018 | 0.035 | 0.035 | 0 | 0.5 | 0.59 | 0.3 | 0.22 | 0.44 | 0.33 |
| 7 | 154249412 | rs902739 | -0.749 | -0.099 | -0.347 | -0.876 | 0 | 0.027 | 0.027 | 0 | 0.52 | 0.6 | 0.32 | 0.23 | 0.49 | 0.36 |
| 7 | 154250592 | rs6962496 | -0.581 | -0.64 | -0.591 | -0.37 | 0.031 | 0.029 | 0.029 | 0 | 0.36 | 0.32 | 0.2 | 0.15 | 0.25 | 0.21 |
| 7 | 154254861 | rs2337393 | 0.52 | -0.157 | -0.126 | 0.445 | 0.037 | 0.035 | 0.035 | 0.003 | 0.16 | 0.31 | 0.45 | 0.25 | 0.25 | 0.27 |
| 7 | 154255814 | rs11764641 | -0.506 | -0.284 | -0.262 | 0.705 | -0.004 | 0.044 | 0.044 | 0.029 | 0.19 | 0.27 | 0.22 | 0.55 | 0.17 | 0.31 |
| 7 | 154257107 | rs2878957 | 0.429 | -0.339 | 0.289 | 0.507 | 0.009 | 0 | 0 | 0.406 | 0.15 | 0.17 | 0.46 | 0.2 | 0.29 | 0.31 |
| 7 | 154261961 | rs10952495 | -1.023 | -1.083 | 0.513 | 0.879 | -0.01 | 0.241 | 0.241 | 0.429 | 0.13 | 0.36 | 0.19 | 0.55 | 0.1 | 0.26 |
| 7 | 154262562 | rs12672881 | 0.584 | -0.938 | -0.529 | 0.297 | 0.027 | -0.033 | -0.033 | 0.846 | 0.06 | 0 | 0.01 | 0.12 | 0.02 | 0.12 |
| 7 | 154371838 | rs11769619 | 1.262 | -0.838 | -0.093 | 0.008 | 0.02 | 0.019 | 0.019 | 0 | 0.61 | 0.33 | 0.68 | 0.29 | 0.72 | 0.54 |
| 7 | 154374448 | rs7458507 | 0.878 | -1.063 | 0.118 | 0.235 | -0.029 | 0.028 | 0.028 | 0 | 0.74 | 0.36 | 0.81 | 0.42 | 0.84 | 0.61 |
| 7 | 154377155 | rs7795467 | 1.247 | -0.876 | -0.088 | -0.273 | 0.04 | 0.046 | 0.046 | 0.352 | 0.57 | 0.32 | 0.64 | 0.29 | 0.71 | 0.51 |
| 7 | 154378331 | rs7800754 | -0.792 | -0.432 | 0.756 | -0.306 | 0.03 | 0.023 | 0.023 | 0 | 0.05 | 0.15 | 0.06 | 0.07 | 0.08 | 0.07 |
| 7 | 154385773 | rs6964189 | -0.9 | -0.618 | 0.134 | -0.556 | -0.013 | 0.047 | 0.047 | 0 | 0.55 | 0.73 | 0.64 | 0.68 | 0.52 | 0.7 |
| 7 | 154386019 | rs6952728 | -0.612 | 0.172 | 0.351 | -0.257 | 0.044 | -0.07 | -0.07 | 0 | 0.11 | 0.44 | 0.23 | 0.01 | 0.1 | 0.07 |
| 7 | 154404012 | rs6597439 | -0.147 | 0.406 | -0.363 | 0.236 | 0.009 | -0.002 | -0.002 | 0.009 | 0.05 | 0.07 | 0.05 | 0.03 | 0.07 | 0.06 |
| 7 | 154406965 | rs747561 | -0.239 | -0.001 | -0.649 | 0.356 | -0.032 | -0.005 | -0.005 | 1.672 | 0.41 | 0.2 | 0.54 | 0.64 | 0.44 | 0.63 |
| 7 | 154407807 | rs749120 | -0.155 | 0.39 | -0.146 | 0.685 | 0.007 | -0.026 | -0.026 | 0 | 0.42 | 0.21 | 0.46 | 0.62 | 0.43 | 0.55 |
| 7 | 154412918 | rs6597443 | 0.193 | 0.896 | 0.43 | 0.472 | 0.022 | -0.021 | -0.021 | 0 | 0.62 | 0.46 | 0.55 | 0.67 | 0.57 | 0.78 |
| 7 | 154413406 | rs6597446 | -0.321 | -0.89 | 0.369 | 0.875 | -0.037 | 0.018 | 0.018 | 0.094 | 0.34 | 0.42 | 0.35 | 0.48 | 0.17 | 0.29 |
| 7 | 154414010 | rs2046748 | 0.87 | -1.024 | 0.56 | 1.493 | -0.021 | 0.11 | 0.11 | 0 | 0.82 | 0.73 | 0.83 | 0.44 | 0.9 | 0.64 |
| 7 | 154414361 | rs6980443 | -0.179 | -0.747 | -0.362 | 0.474 | 0.01 | 0.057 | 0.057 | 0 | 0.52 | 0.71 | 0.61 | 0.35 | 0.54 | 0.39 |
| 7 | 154419301 | rs11976061 | 0.07 | 0.613 | -0.145 | 0.06 | 0.004 | -0.021 | -0.021 | 0.965 | 0.38 | 0.57 | 0.32 | 0.25 | 0.39 | 0.25 |
| 7 | 154420513 | rs12154602 | -1.03 | 0.682 | -0.067 | 0.101 | 0.02 | -0.009 | -0.009 | 0.578 | 0.87 | 0.84 | 0.7 | 0.9 | 0.81 | 0.83 |
| 7 | 154423082 | rs9642649 | 1.678 | 0.888 | 0.2 | -0.177 | 0 | -0.019 | -0.019 | 0 | 0.13 | 0.16 | 0.08 | 0.05 | 0.13 | 0.05 |
| 7 | 154427998 | rs1532090 | -0.158 | 0.872 | -0.033 | -0.267 | 0.027 | -0.006 | -0.006 | 0 | 0.2 | 0.31 | 0.18 | 0.2 | 0.25 | 0.19 |
| 7 | 154428808 | rs7800130 | 0.045 | 0.719 | 0.139 | -0.269 | 0.005 | -0.014 | -0.014 | 0 | 0.78 | 0.78 | 0.67 | 0.89 | 0.73 | 0.77 |
| 7 | 154428941 | rs7800390 | 0.045 | 0.719 | 0.139 | -0.269 | 0.005 | -0.014 | -0.014 | 0 | 0.78 | 0.78 | 0.67 | 0.89 | 0.73 | 0.77 |
| 7 | 154429195 | rs11243338 | 0.829 | 0.585 | -0.032 | 0.485 | 0.02 | 0.024 | 0.024 | 1.698 | 0.7 | 0.45 | 0.54 | 0.58 | 0.67 | 0.59 |
| 7 | 154429560 | rs11243339 | 0.393 | 0.545 | -0.031 | 0.251 | 0.035 | -0.088 | -0.088 | 0 | 0.75 | 0.51 | 0.64 | 0.89 | 0.72 | 0.76 |
| 7 | 154431180 | rs1979601 | 0.37 | 0.529 | 0.026 | 0.252 | 0.032 | -0.082 | -0.082 | 4.393 | 0.75 | 0.53 | 0.63 | 0.89 | 0.73 | 0.76 |
| 7 | 154431550 | rs1979600 | 1.117 | 0.226 | -0.589 | 0.218 | 0.065 | 0.015 | 0.015 | 1.166 | 0.42 | 0.2 | 0.28 | 0.33 | 0.36 | 0.39 |
| 7 | 154432282 | rs11767661 | -0.635 | 0.368 | -0.47 | -0.601 | -0.002 | -0.032 | -0.032 | 0 | 0.42 | 0.48 | 0.35 | 0.26 | 0.39 | 0.29 |
| 7 | 154432760 | rs10276974 | -0.254 | -0.127 | -0.256 | -0.749 | -0.019 | -0.023 | -0.023 | 0 | 0.32 | 0.37 | 0.22 | 0.2 | 0.19 | 0.22 |
| 7 | 154433651 | rs2130029 | 0.437 | -0.004 | -0.253 | -0.859 | -0.004 | -0.031 | -0.031 | 0 | 0.62 | 0.48 | 0.68 | 0.8 | 0.64 | 0.71 |
| 7 | 154435656 | rs10262749 | 0.354 | -0.138 | -0.179 | -0.819 | -0.025 | 0.015 | 0.015 | 0 | 0.14 | 0.24 | 0.14 | 0.18 | 0.09 | 0.16 |
| 7 | 154435853 | rs10272326 | 0.354 | -0.138 | -0.179 | -0.819 | -0.025 | 0.015 | 0.015 | 0 | 0.14 | 0.24 | 0.14 | 0.18 | 0.09 | 0.16 |
| 7 | 154436619 | rs1387183 | 1.73 | 0.009 | -0.587 | -1.006 | -0.052 | 0.037 | 0.037 | 0 | 0.21 | 0.08 | 0.28 | 0.13 | 0.27 | 0.19 |
| 7 | 154437256 | rs7787205 | 0.855 | 0.075 | -0.285 | -0.756 | -0.001 | 0 | 0 | 0 | 0.39 | 0.37 | 0.39 | 0.48 | 0.36 | 0.45 |
| 7 | 154439597 | rs1387180 | -0.145 | 0.095 | -0.174 | -0.581 | 0 | -0.004 | -0.004 | 0 | 0.25 | 0.26 | 0.23 | 0.19 | 0.26 | 0.26 |
| 7 | 154440685 | rs2316233 | -0.108 | -0.118 | -0.142 | -0.763 | -0.037 | -0.002 | -0.002 | 0 | 0.18 | 0.32 | 0.15 | 0.18 | 0.11 | 0.16 |
| 7 | 154441210 | rs10262218 | 0.93 | -0.111 | -0.137 | -0.769 | -0.011 | 0 | 0 | 0 | 0.13 | 0.16 | 0.12 | 0.14 | 0.08 | 0.16 |
| 7 | 154443285 | rs936995 | -0.66 | -0.224 | -0.044 | -0.958 | -0.001 | -0.071 | -0.071 | 0 | 0.7 | 0.46 | 0.78 | 0.8 | 0.71 | 0.82 |
| 7 | 154443713 | rs6962118 | -2.347 | -0.245 | 0.047 | -0.965 | -0.018 | -0.081 | -0.081 | 0 | 0.85 | 0.73 | 0.92 | 0.99 | 0.89 | 0.93 |
| 7 | 154444205 | rs6597490 | -1.714 | -0.102 | -0.074 | -1.027 | -0.048 | -0.033 | -0.033 | 1.576 | 0.81 | 0.48 | 0.87 | 0.8 | 0.88 | 0.92 |
| 7 | 154444461 | rs7787632 | 1.16 | -0.032 | -0.348 | -1.012 | 0.034 | 0.126 | 0.126 | 0 | 0.5 | 0.21 | 0.63 | 0.63 | 0.59 | 0.62 |
| 7 | 154458446 | rs6947173 | 0.522 | -0.187 | -0.212 | -0.849 | 0.018 | 0.046 | 0.046 | 0 | 0.57 | 0.28 | 0.64 | 0.63 | 0.63 | 0.67 |
| 7 | 154461112 | rs3807218 | -0.445 | -0.262 | -0.036 | -0.779 | -0.038 | 0.024 | 0.024 | 0 | 0.85 | 0.62 | 0.84 | 0.84 | 0.91 | 0.85 |
| 7 | 154462071 | rs3807219 | -0.406 | -0.386 | -0.876 | -0.611 | -0.016 | 0.012 | 0.012 | 0 | 0.31 | 0.41 | 0.37 | 0.68 | 0.24 | 0.37 |

| | | | | | | | | | | | | | | | | |
|---|---|---|---|---|---|---|---|---|---|---|---|---|---|---|---|---|
| 7 | 154466191 | rs3807222 | 1.046 | 0.002 | -1.661 | -0.926 | -0.159 | 0.091 | 0.091 | 0.274 | 0.3 | 0.04 | 0.35 | 0.13 | 0.43 | 0.31 |
| 7 | 154468976 | rs916514 | 2.232 | -0.121 | -1.448 | -0.798 | -0.006 | -0.034 | -0.034 | 0 | 0.16 | 0.18 | 0.09 | 0.02 | 0.11 | 0.06 |
| 7 | 154470392 | rs10247867 | 0.971 | 0.551 | -1.81 | -0.183 | -0.016 | 0.006 | 0.006 | 0 | 0.12 | 0.13 | 0.13 | 0.16 | 0.08 | 0.14 |
| 7 | 154471035 | rs13227354 | 1.505 | 1.6 | -2.012 | -0.067 | -0.051 | 0.058 | 0.058 | 0 | 0.11 | 0.02 | 0.09 | 0 | 0.17 | 0.1 |
| 7 | 154472155 | rs7793584 | -0.006 | 0.896 | -1.362 | 0.56 | -0.001 | 0.009 | 0.009 | 0 | 0.17 | 0.22 | 0.22 | 0.32 | 0.15 | 0.17 |
| 7 | 154473064 | rs13223942 | 0.07 | 1.11 | -1.317 | -0.236 | -0.093 | 0.021 | 0.021 | 0 | 0.21 | 0.02 | 0.27 | 0.13 | 0.27 | 0.21 |
| 7 | 154476131 | rs7792530 | -0.295 | 0.764 | -0.568 | -0.411 | 0 | -0.003 | -0.003 | 1.807 | 0.43 | 0.46 | 0.59 | 0.57 | 0.41 | 0.5 |
| 7 | 154476409 | rs3823518 | -1.248 | 0.462 | -0.019 | -0.239 | 0 | 0.002 | 0.002 | 0.003 | 0.4 | 0.4 | 0.38 | 0.44 | 0.4 | 0.4 |
| 7 | 154476491 | rs7807148 | -0.598 | 0.696 | -0.191 | -0.194 | 0.009 | 0.002 | 0.002 | 0 | 0.13 | 0.18 | 0.1 | 0.22 | 0.16 | 0.18 |
| 7 | 154476554 | rs3807230 | -0.777 | 0.551 | -0.093 | -0.181 | -0.008 | -0.002 | -0.002 | 0 | 0.28 | 0.25 | 0.23 | 0.33 | 0.31 | 0.31 |
| 7 | 154477378 | rs3807235 | 0.422 | 0.287 | 0.444 | -0.12 | -0.005 | -0.039 | -0.039 | 0 | 0.87 | 0.76 | 0.82 | 0.92 | 0.88 | 0.92 |
| 7 | 154482366 | rs3823519 | 0.961 | -0.001 | 1.27 | 0.232 | -0.022 | 0.013 | 0.013 | 0 | 0.81 | 0.43 | 0.75 | 0.74 | 0.86 | 0.85 |
| 7 | 154485710 | rs6973074 | 0.522 | 0.01 | 1.587 | 0.245 | -0.002 | -0.032 | -0.032 | 0.036 | 0.86 | 0.79 | 0.9 | 0.92 | 0.87 | 0.86 |
| 7 | 154492321 | rs4960593 | 0.918 | -0.036 | 1.059 | 0.401 | -0.028 | 0.031 | 0.031 | 0 | 0.8 | 0.44 | 0.75 | 0.69 | 0.86 | 0.83 |
| 7 | 154498135 | rs10261505 | 0.783 | -0.021 | 0.335 | 0.027 | 0.003 | -0.029 | -0.029 | 0.29 | 0.41 | 0.18 | 0.32 | 0.5 | 0.4 | 0.47 |
| 7 | 154501870 | rs880728 | -0.487 | -0.015 | 0.927 | 0.497 | -0.021 | -0.067 | -0.067 | 0.007 | 0.16 | 0.29 | 0.07 | 0.04 | 0.09 | 0.15 |
| 7 | 154502635 | rs753450 | 0.578 | 0.059 | -0.087 | 0.506 | -0.028 | 0.026 | 0.026 | 0 | 0.8 | 0.45 | 0.76 | 0.7 | 0.86 | 0.83 |
| 7 | 154507441 | rs10273747 | 0.065 | 0.46 | -1.109 | 0.41 | -0.008 | 0 | 0 | 0 | 0.44 | 0.32 | 0.38 | 0.5 | 0.57 | 0.54 |
| 7 | 154509324 | rs12666280 | 0.033 | 0.71 | -2.297 | 0.46 | -0.06 | -0.03 | -0.03 | 0 | 0.49 | 0.06 | 0.55 | 0.46 | 0.51 | 0.56 |
| 7 | 154511163 | rs3823523 | -0.638 | 1.138 | -1.779 | 0.091 | 0.007 | -0.01 | -0.01 | 1.204 | 0.35 | 0.56 | 0.32 | 0.29 | 0.42 | 0.28 |
| 7 | 154512110 | rs729269 | 0.991 | 0.534 | -2.311 | -0.035 | 0.005 | 0.001 | 0.001 | 0 | 0.2 | 0.1 | 0.26 | 0.11 | 0.2 | 0.16 |
| 7 | 154515442 | rs4960603 | 1.204 | 1.105 | -2.596 | 0.097 | -0.067 | -0.057 | -0.057 | 0.181 | 0.31 | 0.08 | 0.33 | 0.38 | 0.34 | 0.43 |
| 7 | 154516381 | rs10224371 | -0.24 | 0.775 | -1.445 | -0.127 | 0.009 | -0.05 | -0.05 | 0 | 0.16 | 0.37 | 0.17 | 0.09 | 0.16 | 0.12 |
| 7 | 154521737 | rs6976945 | -0.244 | 0.735 | -1.088 | -0.145 | 0.01 | -0.046 | -0.046 | 0 | 0.16 | 0.36 | 0.17 | 0.1 | 0.15 | 0.12 |
| 7 | 154522644 | rs547894 | 1.692 | 0.511 | -1.723 | -0.405 | -0.036 | -0.098 | -0.098 | 0.281 | 0.19 | 0.02 | 0.13 | 0.34 | 0.21 | 0.28 |
| 7 | 154523979 | rs505245 | -0.686 | 0.607 | -1.174 | -0.215 | -0.021 | -0.023 | -0.023 | 0 | 0.75 | 0.8 | 0.77 | 0.59 | 0.65 | 0.77 |
| 7 | 154526318 | rs478414 | 0.382 | 0.482 | -1.018 | -0.393 | -0.007 | 0.006 | 0.006 | 0.002 | 0.59 | 0.43 | 0.61 | 0.62 | 0.5 | 0.67 |
| 7 | 154528018 | rs515906 | 0.602 | 0.544 | -1.005 | -0.691 | 0.015 | -0.005 | -0.005 | 0 | 0.37 | 0.18 | 0.33 | 0.39 | 0.3 | 0.52 |
| 7 | 154539863 | rs579864 | -1.046 | 0.89 | -0.371 | -0.494 | -0.001 | 0.001 | 0.001 | 0 | 0.67 | 0.67 | 0.52 | 0.43 | 0.63 | 0.68 |
| 7 | 154540471 | rs12666918 | 0.902 | 0.894 | -1.075 | -0.636 | -0.082 | -0.086 | -0.086 | 0 | 0.34 | 0.03 | 0.32 | 0.39 | 0.29 | 0.45 |
| 7 | 154544544 | rs6974316 | -0.848 | 1.446 | 0.346 | -0.588 | 0.004 | -0.013 | -0.013 | 0 | 0.19 | 0.27 | 0.11 | 0.1 | 0.19 | 0.19 |
| 7 | 154545224 | rs10278751 | -1.164 | 1.661 | 0.195 | -0.473 | 0.014 | -0.034 | -0.034 | 0 | 0.21 | 0.62 | 0.15 | 0.11 | 0.22 | 0.21 |
| 7 | 154549403 | rs12112562 | -1.27 | 1.545 | -0.58 | -0.221 | 0 | 0.001 | 0.001 | 0.254 | 0.13 | 0.09 | 0.08 | 0 | 0.15 | 0.05 |
| 7 | 154551171 | rs961232 | -1.014 | 1.768 | -0.524 | 0.016 | 0.009 | -0.014 | -0.014 | 0 | 0.23 | 0.25 | 0.18 | 0.13 | 0.32 | 0.25 |
| 7 | 154552343 | rs12532804 | 0.284 | 0.837 | -0.896 | 0.321 | 0.023 | 0.014 | 0.014 | 0.099 | 0.46 | 0.46 | 0.44 | 0.34 | 0.34 | 0.37 |
| 7 | 154553149 | rs6955735 | -0.009 | 1.887 | -1.099 | 0.438 | -0.049 | 0.001 | 0.001 | 0 | 0.13 | 0.04 | 0.25 | 0.09 | 0.26 | 0.18 |
| 7 | 154553766 | rs6597423 | -0.319 | 0.922 | -0.879 | 0.493 | 0.012 | -0.012 | -0.012 | 0.024 | 0.45 | 0.39 | 0.51 | 0.41 | 0.62 | 0.57 |
| 7 | 154563276 | rs11763741 | -0.222 | 1.272 | -1.255 | 0.781 | 0.005 | 0.002 | 0.002 | 0 | 0.18 | 0.23 | 0.19 | 0.24 | 0.2 | 0.25 |
| 7 | 154566409 | rs6943853 | -0.138 | -0.158 | -1.428 | 0.47 | 0.04 | 0.023 | 0.023 | 0.208 | 0.61 | 0.69 | 0.66 | 0.64 | 0.74 | 0.77 |
| 7 | 154587081 | rs3807269 | 0.605 | 0.73 | -1.751 | 0.661 | -0.063 | -0.053 | -0.053 | 0 | 0.8 | 0.97 | 0.75 | 0.76 | 0.78 | 0.72 |
| 7 | 154587793 | rs10236495 | 0.277 | 0.441 | -1.574 | 0.554 | 0.014 | 0.031 | 0.031 | 0 | 0.45 | 0.6 | 0.49 | 0.27 | 0.37 | 0.47 |
| 7 | 154596517 | rs2280659 | 0.42 | 0.515 | -1.73 | 0.291 | -0.022 | -0.071 | -0.071 | 0 | 0.36 | 0.14 | 0.44 | 0.49 | 0.42 | 0.32 |
| 7 | 154599918 | rs10264427 | 0.403 | -0.169 | -1.071 | 0.197 | -0.007 | 0.009 | 0.009 | 0 | 0.81 | 0.77 | 0.79 | 0.74 | 0.83 | 0.74 |
| 7 | 154608793 | rs6957712 | 0.495 | -0.189 | -0.702 | 0.188 | -0.014 | 0.022 | 0.022 | 0 | 0.81 | 0.71 | 0.78 | 0.74 | 0.83 | 0.74 |
| 7 | 154616612 | rs4960621 | 0.787 | -0.607 | -0.713 | -0.108 | 0.047 | 0.009 | 0.009 | 0.735 | 0.62 | 0.22 | 0.65 | 0.66 | 0.74 | 0.61 |
| 7 | 154620262 | rs9649875 | 0.28 | -0.595 | -0.642 | -0.119 | 0.009 | -0.009 | -0.009 | 0 | 0.79 | 0.84 | 0.8 | 0.75 | 0.83 | 0.76 |
| 7 | 154621545 | rs2030791 | 0.681 | -0.715 | -0.021 | -0.119 | -0.037 | 0.018 | 0.018 | 0.034 | 0.77 | 0.54 | 0.78 | 0.72 | 0.83 | 0.75 |
| 7 | 154622647 | rs7807857 | 0.469 | -0.749 | -0.098 | -0.683 | -0.011 | -0.051 | -0.051 | 0 | 0.78 | 0.72 | 0.81 | 0.99 | 0.83 | 0.77 |
| 7 | 154623992 | rs4265142 | 0.941 | -0.572 | -0.768 | -0.32 | -0.121 | -0.086 | -0.086 | 0 | 0.42 | 0.07 | 0.49 | 0.61 | 0.51 | 0.4 |
| 7 | 154627209 | rs1997093 | 0.997 | -0.52 | 0.402 | -0.457 | -0.01 | 0.01 | 0.01 | 0 | 0.12 | 0.19 | 0.09 | 0.14 | 0.07 | 0.14 |
| 7 | 154632627 | rs13241033 | 0.329 | -0.584 | 0.53 | -0.528 | -0.003 | -0.002 | -0.002 | 0 | 0.15 | 0.21 | 0.11 | 0.13 | 0.12 | 0.2 |
| 7 | 154636784 | rs11243322 | 0.362 | -0.798 | 0.268 | -0.849 | 0.003 | 0.031 | 0.031 | 0 | 0.74 | 0.84 | 0.77 | 0.86 | 0.81 | 0.7 |
| 7 | 154643569 | rs6597432 | -0.277 | -0.85 | 1.077 | -0.85 | -0.017 | -0.028 | -0.028 | 0 | 0.21 | 0.36 | 0.14 | 0.15 | 0.15 | 0.22 |
| 7 | 154643901 | rs6948431 | -0.33 | -0.882 | 1.063 | -0.864 | -0.018 | -0.03 | -0.03 | 0.341 | 0.22 | 0.36 | 0.14 | 0.15 | 0.15 | 0.22 |
| 7 | 154645406 | rs6962284 | 0.269 | -0.958 | 0.907 | -1.317 | -0.001 | -0.108 | -0.108 | 0 | 0.66 | 0.56 | 0.62 | 0.97 | 0.7 | 0.64 |
| 7 | 154645576 | rs3778732 | 0.192 | -1.003 | 0.568 | -1.241 | 0 | 0 | 0 | 0 | 0.2 | 0.17 | 0.13 | 0.15 | 0.15 | 0.21 |
| 7 | 154647433 | rs11760986 | 1.115 | -0.754 | 0.236 | -1.043 | -0.126 | 0.112 | 0.112 | 0 | 0.39 | 0.07 | 0.46 | 0.8 | 0.53 | 0.4 |
| 7 | 154647453 | rs11243323 | 0.722 | -0.501 | 0.639 | -0.867 | 0 | -0.014 | -0.014 | 0.526 | 0.47 | 0.48 | 0.51 | 0.8 | 0.55 | 0.4 |
| 7 | 154648330 | rs3823527 | -0.358 | -0.329 | 0.391 | -0.373 | -0.002 | 0.025 | 0.025 | 0 | 0.25 | 0.19 | 0.21 | 0.12 | 0.22 | 0.26 |
| 7 | 154652279 | rs878742 | 0.252 | -0.094 | 0.451 | 0.78 | -0.003 | 0.079 | 0.079 | 0.408 | 0.11 | 0.15 | 0.25 | 0.46 | 0.13 | 0.18 |
| 7 | 154652608 | rs4960626 | 0.394 | -0.098 | 0.587 | 0.654 | -0.011 | 0.126 | 0.126 | 0.343 | 0.1 | 0.18 | 0.1 | 0.47 | 0.11 | 0.14 |
| 7 | 154656737 | rs868880 | -1.352 | -0.396 | 0.787 | 0.557 | -0.035 | 0.194 | 0.194 | 0.021 | 0.19 | 0.53 | 0.37 | 0.54 | 0.16 | 0.22 |
| 7 | 154657703 | rs4960629 | -1.495 | -0.335 | 1.073 | 0.581 | 0.073 | 0.097 | 0.097 | 0.038 | 0.07 | 0.35 | 0.24 | 0.06 | 0.05 | 0.07 |
| 7 | 154658504 | rs3823529 | 0.574 | -0.302 | 0.342 | 0.658 | 0.007 | 0.009 | 0.009 | 0.342 | 0.09 | 0.06 | 0.1 | 0.48 | 0.11 | 0.13 |
| 7 | 154663549 | rs11768385 | -1.069 | 0.349 | -0.289 | -0.325 | -0.03 | -0.035 | -0.035 | 0 | 0.46 | 0.79 | 0.6 | 0.56 | 0.53 | 0.43 |
| 7 | 154665152 | rs3807285 | 0.507 | 0.234 | -0.068 | -0.349 | -0.001 | 0.046 | 0.046 | 0 | 0.13 | 0.15 | 0.14 | 0.53 | 0.15 | 0.2 |
| 7 | 154667628 | rs2293353 | -1.167 | 0.351 | -0.083 | -0.359 | -0.031 | -0.025 | -0.025 | 0 | 0.46 | 0.77 | 0.59 | 0.56 | 0.52 | 0.43 |
| 7 | 154667907 | rs10269751 | -0.91 | -0.266 | 0.449 | 0.013 | 0.014 | 0.01 | 0.01 | 0.401 | 0.08 | 0.39 | 0.24 | 0.01 | 0.02 | 0.04 |
| 7 | 154669251 | rs3823531 | -0.225 | 0.192 | -0.416 | 0.246 | -0.005 | 0.076 | 0.076 | 0 | 0.13 | 0.17 | 0.14 | 0.54 | 0.15 | 0.19 |

| | | | | | | | | | | | | | | | | |
|---|---|---|---|---|---|---|---|---|---|---|---|---|---|---|---|---|
| 7 | 154671880 | rs3807294 | 0.602 | 0.223 | -1.301 | -0.366 | -0.056 | -0.049 | -0.049 | 0 | 0.37 | 0.2 | 0.26 | 0.36 | 0.35 | 0.42 |
| 7 | 154671923 | rs3807295 | -0.3 | -1.109 | -0.833 | -0.805 | -0.028 | 0.02 | 0.02 | 0 | 0.31 | 0.5 | 0.5 | 0.62 | 0.29 | 0.34 |
| 7 | 154672646 | rs3817522 | -0.266 | -1.659 | -0.222 | -0.919 | -0.075 | -0.073 | -0.073 | 0 | 0.26 | 0.49 | 0.4 | 0.1 | 0.19 | 0.12 |
| 7 | 154672839 | rs3817517 | -0.872 | -1.811 | -0.317 | -0.783 | -0.056 | 0.017 | 0.017 | 0 | 0.37 | 0.5 | 0.49 | 0.56 | 0.29 | 0.27 |
| 7 | 154673469 | rs3800577 | 1.676 | -1.218 | -0.486 | -1.144 | -0.04 | -0.017 | -0.017 | 0 | 0.38 | 0.24 | 0.26 | 0.37 | 0.38 | 0.44 |
| 7 | 154673686 | rs4960633 | -0.588 | -1.409 | -0.435 | -0.827 | -0.024 | 0.008 | 0.008 | 0.094 | 0.32 | 0.36 | 0.27 | 0.46 | 0.22 | 0.23 |
| 7 | 154674654 | rs7783483 | -0.931 | -1.407 | -0.436 | -0.852 | -0.015 | 0.005 | 0.005 | 0 | 0.36 | 0.39 | 0.28 | 0.51 | 0.28 | 0.25 |
| 7 | 154681383 | rs2280656 | -0.454 | -1.319 | -0.421 | -0.75 | -0.049 | 0.248 | 0.248 | 0.379 | 0.18 | 0.5 | 0.34 | 0.51 | 0.13 | 0.19 |
| 7 | 154681835 | rs7782222 | -0.936 | -1.317 | -0.428 | -0.753 | -0.045 | 0.154 | 0.154 | 0.012 | 0.22 | 0.53 | 0.36 | 0.56 | 0.19 | 0.2 |
| 7 | 154682141 | rs3807296 | 0.637 | -1.241 | -0.409 | -0.711 | 0.003 | 0.023 | 0.023 | 0 | 0.13 | 0.15 | 0.11 | 0.47 | 0.11 | 0.16 |
| 7 | 154684153 | rs3734960 | -0.859 | -1.294 | -0.591 | -0.516 | -0.052 | -0.031 | -0.031 | 0 | 0.35 | 0.52 | 0.46 | 0.27 | 0.26 | 0.23 |
| 7 | 154684971 | rs1047053 | -0.579 | -1.488 | -1.57 | -1.056 | -0.04 | -0.08 | -0.08 | 0 | 0.77 | 0.98 | 0.83 | 0.69 | 0.81 | 0.75 |
| 7 | 154685805 | rs1047064 | -0.613 | -1.05 | -0.952 | -0.937 | -0.004 | -0.006 | -0.006 | 0.062 | 0.34 | 0.32 | 0.44 | 0.21 | 0.27 | 0.26 |
| 7 | 154689120 | rs3863238 | 0.82 | -0.802 | -1.175 | -0.875 | -0.034 | -0.012 | -0.012 | 0 | 0.12 | 0.01 | 0.25 | 0.14 | 0.16 | 0.16 |
| 7 | 154689868 | rs4286904 | -0.976 | -0.785 | -1.087 | -0.915 | 0.01 | 0.019 | 0.019 | 0.063 | 0.44 | 0.39 | 0.52 | 0.24 | 0.42 | 0.31 |
| 7 | 154693614 | rs4294147 | 1.222 | -1.116 | -1.179 | -0.582 | 0.006 | 0.007 | 0.007 | 0 | 0.48 | 0.53 | 0.61 | 0.6 | 0.57 | 0.61 |
| 7 | 154694387 | rs2002865 | 0.065 | -0.94 | -1.837 | -0.316 | -0.038 | -0.063 | -0.063 | 0 | 0.8 | 0.94 | 0.82 | 0.71 | 0.8 | 0.76 |
| 7 | 154695553 | rs4960563 | 0.41 | -0.676 | -1.675 | -0.175 | 0.047 | 0.03 | 0.03 | 0.252 | 0.64 | 0.91 | 0.71 | 0.71 | 0.73 | 0.68 |
| 7 | 154701338 | rs882467 | -0.198 | -0.388 | -0.968 | -0.208 | -0.003 | -0.021 | -0.021 | 0.233 | 0.16 | 0.16 | 0.13 | 0.04 | 0.13 | 0.16 |
| 7 | 154701353 | rs882466 | -0.255 | -0.455 | -0.819 | -0.171 | 0.011 | -0.01 | -0.01 | 0 | 0.41 | 0.74 | 0.59 | 0.49 | 0.42 | 0.42 |
| 7 | 154705781 | rs11768600 | -0.002 | -0.454 | -0.234 | -0.132 | -0.008 | -0.047 | -0.047 | 0 | 0.17 | 0.27 | 0.13 | 0.04 | 0.13 | 0.16 |
| 7 | 154708716 | rs11771552 | 0.184 | -0.466 | -0.282 | -0.103 | -0.001 | -0.011 | -0.011 | 0 | 0.17 | 0.15 | 0.15 | 0.05 | 0.19 | 0.2 |
| 7 | 154720259 | rs11543936 | -0.664 | -0.537 | 0.129 | -0.162 | 0.005 | 0.015 | 0.015 | 0 | 0.46 | 0.43 | 0.49 | 0.26 | 0.45 | 0.48 |
| 7 | 154727008 | rs2316539 | -1.244 | -0.152 | 0.151 | 0.273 | -0.002 | -0.002 | -0.002 | 0 | 0.63 | 0.72 | 0.76 | 0.6 | 0.63 | 0.63 |
| 7 | 154734575 | rs751840 | 0.599 | -0.285 | -0.522 | 0.186 | -0.092 | 0.051 | 0.051 | 0 | 0.27 | 0.02 | 0.23 | 0.18 | 0.33 | 0.26 |
| 7 | 154735950 | rs930436 | 0.693 | -0.5 | -0.615 | 0.259 | -0.06 | 0.039 | 0.039 | 0.532 | 0.26 | 0.02 | 0.29 | 0.16 | 0.29 | 0.19 |
| 7 | 154736077 | rs930437 | -1.095 | -0.537 | -0.128 | 0.208 | -0.002 | -0.003 | -0.003 | 0.3 | 0.59 | 0.39 | 0.71 | 0.6 | 0.63 | 0.6 |
| 7 | 154751774 | rs2272174 | -0.746 | -0.542 | -0.028 | 0.229 | 0 | 0.039 | 0.039 | 0.165 | 0.57 | 0.23 | 0.62 | 0.39 | 0.62 | 0.57 |
| 7 | 154753854 | rs2293261 | -1.11 | -0.54 | -0.029 | 0.232 | -0.002 | -0.002 | -0.002 | 0 | 0.6 | 0.39 | 0.71 | 0.61 | 0.63 | 0.6 |
| 7 | 154765672 | rs11243329 | 0.482 | -0.367 | -0.019 | 0.306 | -0.031 | 0.011 | 0.011 | 0.259 | 0.26 | 0.07 | 0.36 | 0.21 | 0.29 | 0.3 |
| 7 | 154782114 | rs306283 | -0.052 | -0.332 | 0.049 | 0.284 | 0.001 | -0.011 | -0.011 | 0.011 | 0.29 | 0.28 | 0.45 | 0.42 | 0.3 | 0.32 |
| 7 | 154789678 | rs306276 | -0.404 | -0.415 | 0.087 | 0.447 | 0.018 | 0.035 | 0.035 | 0.006 | 0.53 | 0.26 | 0.62 | 0.44 | 0.61 | 0.55 |
| 7 | 154796579 | rs431085 | -0.286 | -0.374 | -0.458 | 0.358 | -0.001 | 0.029 | 0.029 | 0.101 | 0.52 | 0.15 | 0.57 | 0.44 | 0.6 | 0.54 |
| 7 | 154812516 | rs414315 | 0.039 | -0.314 | -0.453 | 0.316 | -0.028 | 0.014 | 0.014 | 0.361 | 0.31 | 0.2 | 0.43 | 0.24 | 0.34 | 0.34 |
| 7 | 154812950 | rs12539028 | -0.287 | -0.396 | -0.17 | 0.479 | -0.002 | -0.021 | -0.021 | 0 | 0.33 | 0.29 | 0.52 | 0.39 | 0.35 | 0.37 |
| 7 | 154815796 | rs11760608 | 0.722 | -0.369 | 0.463 | 0.492 | -0.001 | 0.003 | 0.003 | 0 | 0.29 | 0.23 | 0.24 | 0.24 | 0.32 | 0.31 |
| 7 | 154820318 | rs11765456 | -1.12 | -0.434 | 0.59 | 0.42 | 0 | 0.001 | 0.001 | 0 | 0.64 | 0.64 | 0.79 | 0.63 | 0.67 | 0.68 |
| 7 | 154822949 | rs6972688 | 0.301 | -0.443 | 0.614 | 0.465 | -0.007 | 0.009 | 0.009 | 0 | 0.33 | 0.35 | 0.35 | 0.4 | 0.33 | 0.34 |
| 7 | 154823361 | rs1946916 | 0.003 | -0.246 | 0.355 | 0.411 | -0.037 | 0.028 | 0.028 | 0.114 | 0.33 | 0.19 | 0.43 | 0.23 | 0.35 | 0.34 |
| 7 | 154832111 | rs1074939 | 0.05 | 0.102 | 0.96 | 1.169 | 0.085 | 0.51 | 0.51 | 1.963 | 0.95 | 0.64 | 0.92 | 0.65 | 0.95 | 0.89 |
| 7 | 154836010 | rs1583830 | 0.36 | 0.107 | 0.182 | 0.87 | 0.011 | 0.058 | 0.058 | 0.659 | 0.61 | 0.25 | 0.56 | 0.25 | 0.62 | 0.55 |
| 7 | 154846527 | rs2698497 | 1.283 | 0.187 | -0.252 | 0.488 | -0.032 | 0.083 | 0.083 | 0.17 | 0.2 | 0.02 | 0.12 | 0.06 | 0.19 | 0.17 |
| 7 | 154848855 | rs6597451 | 0.149 | 0.17 | 0.28 | 0.941 | -0.003 | 0.017 | 0.017 | 0 | 0.29 | 0.21 | 0.26 | 0.25 | 0.32 | 0.35 |
| 7 | 154855641 | rs1561602 | 0.351 | 0.221 | -0.176 | 0.969 | -0.049 | 0.034 | 0.034 | 0 | 0.27 | 0.07 | 0.25 | 0.25 | 0.32 | 0.34 |
| 7 | 154862434 | rs3734966 | -1.285 | 0.418 | 0.38 | 1.004 | 0.011 | 0.008 | 0.008 | 0.018 | 0.11 | 0.27 | 0.18 | 0.11 | 0.08 | 0.12 |
| 7 | 154863467 | rs2241859 | 0.621 | 0.598 | -0.369 | 0.757 | -0.093 | -0.017 | -0.017 | 1.798 | 0.26 | 0.02 | 0.24 | 0.26 | 0.32 | 0.31 |
| 7 | 154876386 | rs1946915 | 1.285 | 0.343 | 1.107 | 0.912 | 0.027 | 0.124 | 0.124 | 1.044 | 0.91 | 0.5 | 0.83 | 0.84 | 0.95 | 0.89 |
| 7 | 154876689 | rs1631327 | -0.916 | 0.375 | 0.999 | 0.884 | -0.087 | -0.018 | -0.018 | 0 | 0.74 | 0.98 | 0.76 | 0.74 | 0.68 | 0.72 |
| 7 | 154876724 | rs2581831 | -1.4 | 0.374 | 1.009 | 0.886 | 0.027 | 0.124 | 0.124 | 1.044 | 0.09 | 0.5 | 0.17 | 0.16 | 0.05 | 0.11 |
| 7 | 154876991 | rs1730209 | -0.901 | 0.377 | 0.843 | 0.884 | -0.087 | -0.018 | -0.018 | 0 | 0.74 | 0.98 | 0.76 | 0.74 | 0.68 | 0.7 |
| 7 | 154877352 | rs1371818 | -1.651 | 0.406 | 0.96 | 0.96 | -0.017 | 0.043 | 0.043 | 2.655 | 0.11 | 0.67 | 0.18 | 0.16 | 0.05 | 0.11 |
| 7 | 154880169 | rs1657273 | 1.023 | 0.543 | 0.498 | 0.733 | -0.018 | -0.038 | -0.038 | 0 | 0.26 | 0.18 | 0.31 | 0.48 | 0.29 | 0.35 |
| 7 | 154880341 | rs893113 | 0.225 | 0.373 | 0.419 | 0.66 | 0.006 | 0.026 | 0.026 | 0 | 0.58 | 0.25 | 0.55 | 0.58 | 0.6 | 0.57 |
| 7 | 154882572 | rs1657275 | -0.363 | 0.391 | 0.297 | 0.668 | -0.042 | -0.017 | -0.017 | 0 | 0.64 | 0.89 | 0.63 | 0.58 | 0.61 | 0.61 |
| 7 | 154885237 | rs1436818 | -0.284 | 0.386 | 0.583 | 0.666 | -0.028 | -0.005 | -0.005 | 0.079 | 0.63 | 0.88 | 0.63 | 0.58 | 0.61 | 0.61 |
| 7 | 154885525 | rs1657280 | -0.363 | 0.387 | 0.58 | 0.666 | -0.044 | -0.018 | -0.018 | 0.134 | 0.64 | 0.9 | 0.63 | 0.58 | 0.61 | 0.61 |
| 7 | 154886833 | rs1730136 | -0.373 | 0.424 | -0.018 | 0.687 | -0.049 | -0.022 | -0.022 | 0 | 0.64 | 0.91 | 0.63 | 0.58 | 0.61 | 0.61 |
| 7 | 154887865 | rs1730138 | -0.322 | 0.425 | -0.013 | 0.688 | -0.023 | -0.003 | -0.003 | 0.68 | 0.63 | 0.86 | 0.62 | 0.58 | 0.61 | 0.61 |
| 7 | 154889184 | rs1730186 | -0.244 | 0.459 | -0.007 | 0.692 | -0.015 | 0.008 | 0.008 | 0 | 0.63 | 0.83 | 0.62 | 0.59 | 0.61 | 0.63 |
| 7 | 154889716 | rs1730185 | -0.233 | 0.459 | -0.019 | 0.692 | -0.012 | 0.008 | 0.008 | 0 | 0.63 | 0.8 | 0.62 | 0.59 | 0.61 | 0.63 |
| 7 | 154890285 | rs1317905 | -0.89 | 0.54 | -0.678 | 0.663 | -0.097 | 0.012 | 0.012 | 0 | 0.69 | 0.98 | 0.75 | 0.75 | 0.65 | 0.7 |
| 7 | 154895108 | rs6974689 | 0.701 | 0.649 | -0.761 | 0.639 | -0.047 | 0.124 | 0.124 | 0 | 0.31 | 0.02 | 0.26 | 0.1 | 0.32 | 0.26 |
| 7 | 154898033 | rs1657265 | 1.326 | 0.753 | -0.641 | 0.952 | -0.016 | -0.049 | -0.049 | 0 | 0.23 | 0.15 | 0.17 | 0.33 | 0.24 | 0.27 |
| 7 | 154898078 | rs1619015 | -0.355 | 0.739 | -0.255 | 0.82 | 0 | 0.007 | 0.007 | 0 | 0.66 | 0.71 | 0.71 | 0.75 | 0.66 | 0.7 |
| 7 | 154899391 | rs2951007 | 0.027 | 0.771 | -0.623 | 0.855 | 0 | -0.028 | -0.028 | 0 | 0.61 | 0.56 | 0.66 | 0.74 | 0.61 | 0.63 |
| 7 | 154899824 | rs4960681 | 0.107 | 0.843 | -0.72 | 0.861 | 0.004 | -0.021 | -0.021 | 0 | 0.35 | 0.38 | 0.29 | 0.29 | 0.36 | 0.31 |
| 7 | 154900857 | rs2969540 | -0.893 | 1.312 | -0.935 | 1.501 | 0.056 | 0.287 | 0.287 | 3.048 | 0.1 | 0.42 | 0.24 | 0.33 | 0.09 | 0.2 |
| 7 | 154901217 | rs1730146 | -0.038 | 1.077 | -2.521 | 0.8 | -0.118 | -0.063 | -0.063 | 0 | 0.44 | 0.95 | 0.51 | 0.61 | 0.41 | 0.54 |
| 7 | 154903792 | rs1730148 | -0.173 | 1.149 | -2.441 | 0.242 | -0.061 | -0.002 | -0.002 | 0 | 0.22 | 0.03 | 0.26 | 0.16 | 0.27 | 0.22 |

| | | | | | | | | | | | | | | | | |
|---|---|---|---|---|---|---|---|---|---|---|---|---|---|---|---|---|
| 7 | 154904070 | rs1730149 | 0.037 | 0.954 | -1.887 | 0.762 | -0.028 | -0.027 | -0.027 | 0.817 | 0.11 | 0.04 | 0.16 | 0.14 | 0.17 | 0.11 |
| 7 | 154905252 | rs4352798 | 0.265 | 0.186 | -1.629 | 0.387 | 0.021 | 0.159 | 0.159 | 0.015 | 0.31 | 0.07 | 0.16 | 0.05 | 0.27 | 0.23 |
| 7 | 154905319 | rs4355727 | 1.185 | 0.837 | -0.417 | 1.493 | 0.018 | 0.071 | 0.071 | 0 | 0.94 | 0.85 | 0.87 | 0.8 | 0.94 | 0.87 |
| 7 | 154914851 | rs6597458 | 0.658 | 0.322 | -0.481 | 1.274 | -0.019 | 0.031 | 0.031 | 0 | 0.67 | 0.54 | 0.58 | 0.47 | 0.66 | 0.63 |
| 7 | 154918806 | rs6597459 | -0.362 | -0.377 | -0.85 | 0.929 | 0.045 | 0.031 | 0.031 | 0 | 0.55 | 0.36 | 0.63 | 0.66 | 0.78 | 0.74 |
| 7 | 154925087 | rs1881690 | 1.188 | -0.73 | 0.192 | 1.727 | -0.026 | 0.183 | 0.183 | 0.227 | 0.89 | 0.7 | 0.82 | 0.66 | 0.92 | 0.78 |
| 7 | 154932794 | rs4960686 | 1.485 | -0.985 | 0.634 | 1.566 | -0.036 | 0.165 | 0.165 | 0.42 | 0.84 | 0.6 | 0.77 | 0.58 | 0.86 | 0.86 |
| 7 | 154934501 | rs4146256 | -0.7 | -1.793 | -0.208 | 0.999 | 0.001 | 0.015 | 0.015 | 0.002 | 0.73 | 0.78 | 0.71 | 0.89 | 0.81 | 0.78 |
| 7 | 154935141 | rs9642653 | 0.929 | -1.075 | -1.081 | 0.669 | -0.084 | -0.001 | -0.001 | 0 | 0.27 | 0.08 | 0.27 | 0.24 | 0.4 | 0.29 |
| 7 | 154935982 | rs2951003 | -0.588 | -1.857 | -0.407 | 0.539 | -0.017 | -0.004 | -0.004 | 0.021 | 0.6 | 0.54 | 0.68 | 0.71 | 0.78 | 0.78 |
| 7 | 154936196 | rs7800040 | -0.224 | -1.925 | -0.585 | 0.375 | 0.028 | 0.017 | 0.017 | 0.106 | 0.28 | 0.2 | 0.29 | 0.16 | 0.16 | 0.36 |
| 7 | 154937034 | rs1615832 | -0.208 | -1.132 | -0.632 | 0.255 | 0.005 | 0.038 | 0.038 | 0 | 0.41 | 0.31 | 0.48 | 0.24 | 0.36 | 0.47 |
| 7 | 154938294 | rs1657249 | -0.91 | -1.3 | -0.593 | 0.25 | -0.008 | -0.004 | -0.004 | 0 | 0.64 | 0.45 | 0.72 | 0.36 | 0.7 | 0.7 |
| 7 | 154938659 | rs4246095 | -1.814 | -1.342 | -0.257 | 0.46 | -0.005 | -0.004 | -0.004 | 0 | 0.72 | 0.68 | 0.76 | 0.78 | 0.78 | 0.82 |
| 7 | 154938965 | rs1436810 | 0.867 | -1.419 | -0.251 | 0.574 | 0.024 | -0.002 | -0.002 | 0 | 0.23 | 0.14 | 0.2 | 0.23 | 0.11 | 0.14 |
| 7 | 154941814 | rs1529083 | 0.162 | -1.279 | -0.26 | 0.735 | -0.021 | 0.184 | 0.184 | 0 | 0.13 | 0.28 | 0.09 | 0.42 | 0.14 | 0.14 |
| 7 | 154945146 | rs4613951 | -1.787 | -1.411 | -0.102 | 0.783 | 0.003 | 0.003 | 0.003 | 0 | 0.7 | 0.75 | 0.73 | 0.78 | 0.77 | 0.82 |
| 7 | 154946061 | rs1730202 | 0.711 | -1.362 | -0.07 | 0.883 | -0.019 | 0.265 | 0.265 | 0.358 | 0.08 | 0.22 | 0.05 | 0.4 | 0.08 | 0.13 |
| 7 | 154946276 | rs7795216 | 0.914 | -1.207 | -0.244 | 0.928 | 0.025 | -0.01 | -0.01 | 0 | 0.23 | 0.08 | 0.22 | 0.21 | 0.15 | 0.15 |
| 7 | 154947071 | rs7800026 | -1.694 | -1.193 | -0.497 | 0.785 | 0.001 | 0.001 | 0.001 | 0 | 0.69 | 0.7 | 0.73 | 0.78 | 0.77 | 0.82 |
| 7 | 154948500 | rs2969550 | 1.501 | -1.196 | -0.721 | 0.761 | -0.087 | 0.072 | 0.072 | 0 | 0.16 | 0.01 | 0.25 | 0.04 | 0.25 | 0.33 |
| 7 | 154952461 | rs1657263 | -0.308 | -1.215 | -0.659 | 0.757 | 0.001 | 0.004 | 0.004 | 0 | 0.5 | 0.51 | 0.46 | 0.74 | 0.5 | 0.49 |
| 7 | 154955663 | rs1730175 | 1.503 | -1.097 | -1.14 | 0.71 | -0.087 | 0.072 | 0.072 | 0 | 0.16 | 0.01 | 0.25 | 0.04 | 0.25 | 0.33 |
| 7 | 154957101 | rs12531753 | -0.233 | -0.891 | -0.801 | 1.074 | 0 | -0.003 | -0.003 | 0 | 0.44 | 0.45 | 0.43 | 0.44 | 0.44 | 0.37 |
| 7 | 154959070 | rs1657293 | -0.68 | -1.525 | -0.596 | 0.966 | -0.037 | 0.016 | 0.016 | 0 | 0.33 | 0.42 | 0.23 | 0.61 | 0.22 | 0.3 |
| 7 | 154959808 | rs10242760 | 0.035 | -0.007 | -0.636 | 0.924 | -0.003 | -0.032 | -0.032 | 0.039 | 0.39 | 0.24 | 0.33 | 0.44 | 0.36 | 0.28 |
| 7 | 154963787 | rs1657298 | -1.134 | -0.401 | -0.039 | 1.164 | -0.024 | 0.079 | 0.079 | 1.036 | 0.1 | 0.21 | 0.07 | 0.31 | 0.07 | 0.12 |
| 7 | 154965379 | rs930339 | 0.024 | -0.181 | -0.24 | 0.026 | 0.007 | -0.032 | -0.032 | 0.704 | 0.43 | 0.54 | 0.45 | 0.27 | 0.49 | 0.48 |
| 7 | 154965392 | rs12536841 | 0.686 | -0.568 | -0.104 | 0.232 | -0.049 | 0.012 | 0.012 | 0 | 0.83 | 0.69 | 0.86 | 0.8 | 0.91 | 0.9 |
| 7 | 154966534 | rs10246702 | 0.742 | -0.636 | -0.097 | 0.264 | -0.061 | 0.022 | 0.022 | 0 | 0.83 | 0.68 | 0.87 | 0.8 | 0.92 | 0.86 |
| 7 | 154971868 | rs7784910 | 0.689 | -0.022 | -0.818 | -0.176 | -0.017 | 0.049 | 0.049 | 0.226 | 0.75 | 0.83 | 0.74 | 0.88 | 0.7 | 0.74 |
| 7 | 154972924 | rs1657302 | -0.354 | -0.208 | -0.526 | -0.101 | -0.018 | 0 | 0 | 0 | 0.44 | 0.35 | 0.47 | 0.52 | 0.5 | 0.57 |
| 7 | 154973603 | rs1730214 | 0.327 | -0.338 | -0.386 | -0.124 | 0.006 | -0.001 | -0.001 | 0 | 0.3 | 0.22 | 0.28 | 0.35 | 0.27 | 0.4 |
| 7 | 154975445 | rs2951012 | 0.887 | -0.708 | 0.217 | -0.068 | -0.001 | -0.003 | -0.003 | 0 | 0.43 | 0.38 | 0.37 | 0.45 | 0.44 | 0.35 |
| 7 | 154975497 | rs12719660 | 1.182 | -0.636 | -0.064 | -0.484 | -0.006 | -0.074 | -0.074 | 0.362 | 0.3 | 0.3 | 0.18 | 0.04 | 0.25 | 0.14 |
| 7 | 154975758 | rs3111876 | 0.453 | -1.311 | -0.301 | -0.838 | 0.01 | -0.018 | -0.018 | 0 | 0.68 | 0.73 | 0.69 | 0.6 | 0.76 | 0.71 |
| 7 | 154977756 | rs7783824 | 0.398 | -0.968 | 0.049 | -0.798 | -0.01 | -0.134 | -0.134 | 0 | 0.31 | 0.41 | 0.27 | 0.04 | 0.28 | 0.16 |
| 7 | 154977941 | rs7808464 | 0.398 | -0.968 | 0.049 | -0.798 | -0.01 | -0.134 | -0.134 | 0 | 0.31 | 0.41 | 0.27 | 0.04 | 0.28 | 0.16 |
| 7 | 154978874 | rs6597472 | 0.097 | -0.991 | 0.002 | -0.762 | 0.001 | -0.012 | -0.012 | 0 | 0.09 | 0.1 | 0.09 | 0 | 0.04 | 0.05 |
| 7 | 154980168 | rs7790169 | -0.451 | -0.94 | -0.07 | -0.725 | 0.005 | 0.073 | 0.073 | 0 | 0.48 | 0.65 | 0.51 | 0.25 | 0.53 | 0.37 |
| 7 | 154983942 | rs1657307 | 0.841 | -0.794 | -0.052 | -0.574 | 0.05 | 0.052 | 0.052 | 2.958 | 0.41 | 0.23 | 0.29 | 0.34 | 0.29 | 0.42 |
| 7 | 154987620 | rs11243342 | 0.703 | -0.343 | 0.037 | -0.712 | -0.005 | -0.001 | -0.001 | 0 | 0.14 | 0.12 | 0.18 | 0.16 | 0.23 | 0.18 |
| 7 | 154988042 | rs11525892 | 0.598 | -0.38 | 0.021 | -0.728 | -0.008 | -0.001 | -0.001 | 0.825 | 0.14 | 0.12 | 0.18 | 0.16 | 0.23 | 0.18 |
| 7 | 154998951 | rs4960580 | -0.601 | -0.663 | -0.211 | -0.716 | 0.006 | 0.073 | 0.073 | 0.388 | 0.47 | 0.66 | 0.51 | 0.26 | 0.55 | 0.39 |
| 7 | 155003983 | rs722465 | -0.11 | -0.109 | -0.484 | -0.569 | 0.028 | -0.043 | -0.043 | 2.545 | 0.36 | 0.42 | 0.35 | 0.22 | 0.49 | 0.3 |
| 7 | 155009761 | rs12719690 | -0.183 | -0.082 | -1.603 | -0.489 | -0.065 | 0.111 | 0.111 | 0.003 | 0.4 | 0.11 | 0.4 | 0.23 | 0.56 | 0.36 |
| 7 | 155012482 | rs11765868 | 0.249 | 0.005 | -1.328 | -0.315 | -0.053 | 0.123 | 0.123 | 0.119 | 0.3 | 0.07 | 0.22 | 0.08 | 0.33 | 0.16 |
| 7 | 155014996 | rs1124174 | -0.901 | -0.003 | -0.174 | 0.956 | -0.014 | 0.208 | 0.208 | 0.022 | 0.12 | 0.24 | 0.21 | 0.42 | 0.12 | 0.2 |
| 7 | 155019546 | rs12666834 | -1.029 | 0.392 | -0.337 | 0.394 | -0.001 | -0.009 | -0.009 | 0 | 0.2 | 0.26 | 0.34 | 0.82 | 0.2 | 0.38 |
| 7 | 155021487 | rs4599765 | 0.134 | 0.79 | -0.935 | 0.086 | -0.054 | -0.008 | -0.008 | 0 | 0.13 | 0.03 | 0.14 | 0.13 | 0.19 | 0.2 |
| 7 | 155021663 | rs4434576 | 0.071 | 0.764 | -0.947 | 0.221 | -0.054 | -0.032 | -0.032 | 0 | 0.13 | 0.03 | 0.15 | 0.15 | 0.2 | 0.21 |
| 7 | 155023320 | rs4246100 | -1.561 | 0.592 | -0.271 | -0.262 | -0.001 | -0.078 | -0.078 | 1.001 | 0.4 | 0.49 | 0.52 | 0.95 | 0.4 | 0.61 |
| 7 | 155024830 | rs10279638 | 0.365 | 0.685 | -0.438 | 0.67 | 0.007 | 0.186 | 0.186 | 0 | 0.07 | 0.11 | 0.13 | 0.41 | 0.1 | 0.16 |
| 7 | 155025772 | rs6962739 | -0.98 | 0.947 | -0.43 | 0.513 | 0 | 0.005 | 0.005 | 0 | 0.24 | 0.26 | 0.29 | 0.54 | 0.32 | 0.4 |
| 7 | 155036231 | rs4960710 | 0.071 | 0.497 | -0.821 | -0.785 | 0.004 | -0.106 | -0.106 | 0 | 0.4 | 0.51 | 0.26 | 0.01 | 0.41 | 0.22 |
| 7 | 155039544 | rs6967312 | 0.537 | 0.183 | -1.354 | -0.35 | -0.017 | 0.037 | 0.037 | 0 | 0.35 | 0.21 | 0.46 | 0.23 | 0.45 | 0.41 |
| 7 | 155046542 | rs6961125 | 0.348 | 0.192 | -1.338 | -0.258 | -0.007 | 0.04 | 0.04 | 0 | 0.41 | 0.25 | 0.49 | 0.27 | 0.48 | 0.47 |
| 7 | 155046789 | rs6969791 | -0.328 | 0.103 | -1.789 | -0.298 | -0.039 | 0.088 | 0.088 | 0.018 | 0.64 | 0.87 | 0.69 | 0.73 | 0.64 | 0.76 |
| 7 | 155050159 | rs6959431 | -0.225 | 0.461 | -1.425 | -0.144 | -0.021 | -0.008 | -0.008 | 0 | 0.7 | 0.73 | 0.68 | 0.64 | 0.63 | 0.74 |
| 7 | 155052274 | rs12719705 | 0.559 | 0.425 | -1.84 | -0.294 | -0.058 | 0.172 | 0.172 | 1.043 | 0.44 | 0.06 | 0.46 | 0.22 | 0.49 | 0.39 |
| 7 | 155056302 | rs6963411 | 0.273 | 0.585 | -1.094 | -0.201 | 0.001 | -0.01 | -0.01 | 0.413 | 0.28 | 0.3 | 0.21 | 0.17 | 0.31 | 0.23 |
| 7 | 155057929 | rs10254344 | -0.008 | 0.619 | -0.734 | -0.167 | 0.001 | 0.003 | 0.003 | 0 | 0.46 | 0.54 | 0.38 | 0.62 | 0.41 | 0.55 |
| 7 | 155057997 | rs4506162 | 0.465 | 0.922 | -0.54 | 0.194 | 0.004 | 0.013 | 0.013 | 0 | 0.88 | 0.85 | 0.91 | 0.82 | 0.9 | 0.85 |
| 7 | 155059150 | rs3923644 | 0.085 | 0.606 | -0.611 | -0.466 | -0.007 | -0.047 | -0.047 | 0 | 0.61 | 0.53 | 0.54 | 0.82 | 0.66 | 0.64 |
| 7 | 155059374 | rs4540368 | -0.202 | 1.084 | -0.953 | -0.366 | -0.006 | 0.046 | 0.046 | 0 | 0.45 | 0.65 | 0.39 | 0.32 | 0.5 | 0.44 |
| 7 | 155060730 | rs3923736 | 0.135 | 0.973 | -0.917 | -0.225 | 0.009 | -0.026 | -0.026 | 0 | 0.36 | 0.46 | 0.34 | 0.24 | 0.42 | 0.26 |
| 7 | 155064705 | rs4401804 | -0.788 | 1.04 | -1.317 | -0.36 | 0.019 | -0.033 | -0.033 | 0 | 0.07 | 0.12 | 0.04 | 0 | 0.08 | 0.05 |
| 7 | 155066386 | rs10901042 | -0.084 | 0.51 | -1.674 | -0.072 | 0 | 0.141 | 0.141 | 0 | 0.46 | 0.19 | 0.42 | 0.23 | 0.34 | 0.45 |
| 7 | 155082893 | rs9692071 | -0.404 | 0.047 | -0.458 | 1.026 | 0.006 | -0.01 | -0.01 | 0 | 0.84 | 0.88 | 0.85 | 0.74 | 0.9 | 0.72 |

| | | | | | | | | | | | | | | | | |
|---|---|---|---|---|---|---|---|---|---|---|---|---|---|---|---|---|
| 7 | 155085716 | rs9690040 | 0.195 | 0.058 | -0.573 | 0.507 | 0.003 | -0.003 | -0.003 | 0 | 0.44 | 0.59 | 0.32 | 0.46 | 0.41 | 0.26 |
| 7 | 155087486 | rs9690258 | 0.901 | 0.002 | -0.527 | -0.328 | 0.006 | 0.154 | 0.154 | 0.7 | 0.47 | 0.25 | 0.56 | 0.01 | 0.51 | 0.2 |
| 7 | 155088065 | rs10271719 | 0.696 | -0.055 | -0.691 | 0.001 | -0.003 | 0.011 | 0.011 | 0 | 0.16 | 0.15 | 0.11 | 0.33 | 0.15 | 0.24 |
| 7 | 155091406 | rs9767875 | 0.854 | -0.033 | -0.649 | 0.005 | -0.007 | 0.017 | 0.017 | 0 | 0.16 | 0.15 | 0.12 | 0.33 | 0.15 | 0.24 |
| 7 | 155099855 | rs9769506 | 0.896 | 0.031 | -0.386 | -0.09 | -0.002 | -0.105 | -0.105 | 1.492 | 0.49 | 0.55 | 0.58 | 0.01 | 0.52 | 0.2 |
| 7 | 155103781 | rs13223383 | 0.26 | -0.196 | -0.538 | 0.123 | 0.005 | 0.003 | 0.003 | 0.549 | 0.67 | 0.73 | 0.72 | 0.34 | 0.71 | 0.46 |
| 7 | 155104949 | rs9328528 | 0.671 | 0.109 | -0.847 | -0.4 | 0.002 | -0.071 | -0.071 | 0 | 0.48 | 0.51 | 0.59 | 0.01 | 0.53 | 0.21 |
| 7 | 155105274 | rs35353544 | 0.207 | 0.57 | -1.273 | 0.942 | -0.015 | -0.053 | -0.053 | 0 | 0.89 | 0.98 | 0.92 | 0.74 | 0.86 | 0.88 |
| 7 | 155124428 | rs6969303 | -0.396 | 0.694 | -0.939 | 0.526 | -0.004 | 0.007 | 0.007 | 1.292 | 0.42 | 0.39 | 0.51 | 0.31 | 0.43 | 0.5 |
| 7 | 155134684 | rs6973712 | 0.372 | 0.929 | -0.863 | 0.472 | 0 | -0.001 | -0.001 | 0 | 0.21 | 0.21 | 0.37 | 0.16 | 0.25 | 0.19 |
| 7 | 155136357 | rs6966802 | 0.082 | 0.907 | -0.965 | 0.733 | 0 | -0.005 | -0.005 | 0 | 0.18 | 0.17 | 0.13 | 0.16 | 0.17 | 0.3 |
| 7 | 155146549 | rs7806718 | -0.245 | 0.63 | -1.214 | 0.448 | -0.026 | -0.008 | -0.008 | 0 | 0.43 | 0.27 | 0.36 | 0.33 | 0.47 | 0.49 |
| 7 | 155148805 | rs7456406 | -0.427 | 0.395 | -1.068 | 0.454 | 0.003 | -0.003 | -0.003 | 0 | 0.8 | 0.81 | 0.84 | 0.67 | 0.86 | 0.58 |
| 7 | 155149037 | rs12719966 | 0.338 | 0.331 | -0.981 | 0.211 | 0.004 | -0.001 | -0.001 | 0 | 0.5 | 0.6 | 0.63 | 0.59 | 0.48 | 0.44 |
| 7 | 155153490 | rs7798126 | 0.268 | 0.314 | -0.65 | -0.087 | -0.018 | -0.042 | -0.042 | 0 | 0.61 | 0.51 | 0.73 | 0.24 | 0.69 | 0.42 |
| 7 | 155162399 | rs1861968 | 0.145 | 0.563 | -0.619 | 0.324 | -0.001 | -0.003 | -0.003 | 0 | 0.28 | 0.29 | 0.29 | 0.5 | 0.35 | 0.28 |
| 7 | 155164799 | rs7780353 | -0.183 | 0.376 | -0.526 | 0.373 | -0.001 | -0.001 | -0.001 | 0 | 0.54 | 0.6 | 0.41 | 0.53 | 0.55 | 0.38 |
| 7 | 155169143 | rs1990808 | 0.064 | 0.438 | -0.711 | -0.316 | -0.005 | -0.076 | -0.076 | 0.039 | 0.25 | 0.36 | 0.41 | 0.1 | 0.23 | 0.14 |
| 7 | 155169830 | rs1421318 | -0.349 | 0.672 | -1.244 | 0.502 | 0.003 | -0.008 | -0.008 | 0.3 | 0.83 | 0.9 | 0.85 | 0.83 | 0.84 | 0.75 |
| 7 | 155175544 | rs4716442 | -0.454 | 0.918 | -1.365 | 0.922 | -0.028 | -0.083 | -0.083 | 0 | 0.16 | 0.04 | 0.18 | 0.7 | 0.19 | 0.36 |
| 7 | 155182326 | rs2058950 | -0.649 | 0.851 | -0.607 | 0.602 | -0.006 | -0.023 | -0.023 | 0.43 | 0.26 | 0.52 | 0.32 | 0.21 | 0.2 | 0.27 |
| 7 | 155183950 | rs12698090 | -1.076 | 1.84 | -0.563 | 1.315 | -0.022 | -0.021 | -0.021 | 4.115 | 0.27 | 0.22 | 0.42 | 0.55 | 0.38 | 0.43 |
| 7 | 155184004 | rs12334121 | -0.271 | 1.183 | -0.811 | 0.397 | 0 | -0.018 | -0.018 | 0 | 0.82 | 0.81 | 0.87 | 0.98 | 0.81 | 0.91 |
| 7 | 155184295 | rs11760227 | -0.271 | 1.391 | -0.824 | 0.765 | -0.015 | -0.017 | -0.017 | 0 | 0.21 | 0.16 | 0.37 | 0.51 | 0.33 | 0.42 |
| 7 | 155184309 | rs11769883 | -1.221 | 1.287 | -0.545 | 0.487 | -0.01 | -0.004 | -0.004 | 0.158 | 0.34 | 0.31 | 0.46 | 0.6 | 0.42 | 0.49 |
| 7 | 155185261 | rs7781545 | -0.133 | 1.583 | -0.577 | 0.707 | 0.036 | 0.059 | 0.059 | 0.041 | 0.14 | 0.29 | 0.15 | 0.23 | 0.21 | 0.18 |
| 7 | 155195631 | rs4716542 | -0.548 | 0.542 | -1.117 | -0.227 | 0.042 | 0.032 | 0.032 | 0.352 | 0.47 | 0.75 | 0.51 | 0.56 | 0.61 | 0.5 |
| 7 | 155195759 | rs4716545 | -0.303 | 0.6 | -0.6 | -0.253 | -0.001 | 0 | 0 | 0 | 0.43 | 0.6 | 0.45 | 0.54 | 0.53 | 0.47 |
| 7 | 155196066 | rs6965086 | -0.302 | -0.001 | -1.471 | 0.062 | 0.037 | -0.069 | -0.069 | 0.785 | 0.84 | 0.99 | 0.77 | 0.73 | 0.91 | 0.73 |
| 7 | 155197610 | rs12112950 | -0.366 | 0.379 | -1.093 | -0.175 | 0.014 | -0.018 | -0.018 | 1.485 | 0.06 | 0.19 | 0.08 | 0.02 | 0.08 | 0.01 |
| 7 | 155206518 | rs4716915 | 0.823 | 0.226 | -1.787 | -0.082 | 0.017 | -0.026 | -0.026 | 0.02 | 0.66 | 0.78 | 0.65 | 0.42 | 0.68 | 0.52 |
| 7 | 155206748 | rs4716916 | -0.466 | 0.361 | -1.547 | 0.388 | -0.012 | 0.077 | 0.077 | 0 | 0.08 | 0.15 | 0.09 | 0.27 | 0.08 | 0.11 |
| 7 | 155209171 | rs4621673 | 0.735 | 0.871 | -1.691 | 0.046 | -0.006 | -0.007 | -0.007 | 0 | 0.3 | 0.26 | 0.33 | 0.43 | 0.37 | 0.33 |
| 7 | 155212392 | rs10264222 | -0.102 | 0.967 | -1.111 | 0.53 | -0.014 | 0.112 | 0.112 | 0 | 0.16 | 0.35 | 0.2 | 0.26 | 0.1 | 0.32 |
| 7 | 155213088 | rs6978591 | -0.39 | 0.969 | -1.434 | 0.186 | -0.026 | 0.18 | 0.18 | 0 | 0.19 | 0.41 | 0.25 | 0.5 | 0.12 | 0.38 |
| 7 | 155213800 | rs10279206 | -0.779 | 0.992 | -1.475 | 0.167 | -0.028 | 0.105 | 0.105 | 0 | 0.23 | 0.5 | 0.28 | 0.53 | 0.17 | 0.43 |
| 7 | 155218662 | rs749538 | -0.853 | 1.171 | -1.833 | 0.02 | -0.01 | 0.022 | 0.022 | 0.091 | 0.33 | 0.5 | 0.44 | 0.55 | 0.33 | 0.52 |
| 7 | 155219731 | rs12534280 | 0.787 | 1.09 | -2.393 | 0.019 | -0.018 | -0.065 | -0.065 | 1.162 | 0.25 | 0.09 | 0.22 | 0.33 | 0.24 | 0.32 |
| 7 | 155220964 | rs6459971 | -0.558 | 1.49 | -2.17 | 0.563 | 0.007 | -0.005 | -0.005 | 0 | 0.68 | 0.74 | 0.68 | 0.62 | 0.73 | 0.71 |
| 7 | 155221617 | rs12531084 | -1.043 | 1.914 | -2.057 | 0.601 | -0.045 | -0.039 | -0.039 | 2.443 | 0.23 | 0.05 | 0.3 | 0.27 | 0.3 | 0.27 |
| 7 | 155224288 | rs11764795 | -0.915 | 2.138 | -1.839 | 0.805 | -0.059 | -0.065 | -0.065 | 0.011 | 0.16 | 0.04 | 0.28 | 0.27 | 0.29 | 0.28 |
| 7 | 155224950 | rs4716964 | 0.109 | 1.552 | -1.909 | -0.502 | 0.001 | 0.104 | 0.104 | 0 | 0.45 | 0.74 | 0.35 | 0.1 | 0.44 | 0.36 |
| 7 | 155225289 | rs11773296 | 0.697 | 1.399 | -2.31 | -0.53 | -0.029 | 0.055 | 0.055 | 5.934 | 0.25 | 0.03 | 0.15 | 0.16 | 0.22 | 0.13 |
| 7 | 155226760 | rs17837543 | 0.1 | 0.365 | -1.733 | -0.368 | -0.02 | 0.02 | 0.02 | 0 | 0.27 | 0.32 | 0.24 | 0.54 | 0.14 | 0.38 |
| 7 | 155227977 | rs10949796 | 1.124 | -0.166 | -1.134 | -0.902 | 0.005 | 0.014 | 0.014 | 1.001 | 0.33 | 0.26 | 0.21 | 0.17 | 0.29 | 0.15 |
| 7 | 155228360 | rs11768588 | 0.514 | -0.056 | -1.404 | -1.224 | -0.124 | -0.006 | -0.006 | 0.284 | 0.24 | 0.07 | 0.46 | 0.18 | 0.47 | 0.37 |
| 7 | 155229162 | rs7777187 | -0.831 | 0.105 | -1.149 | -1.27 | -0.004 | -0.03 | -0.03 | 0 | 0.86 | 0.75 | 0.9 | 0.91 | 0.87 | 0.92 |
| 7 | 155229920 | rs12538270 | 0.369 | -0.055 | -1.126 | -1.276 | -0.008 | -0.003 | -0.003 | 0.126 | 0.59 | 0.44 | 0.68 | 0.37 | 0.77 | 0.54 |
| 7 | 155230541 | rs6948976 | -0.973 | 0.087 | -0.884 | -1.212 | -0.012 | -0.061 | -0.061 | 0 | 0.87 | 0.81 | 0.93 | 1 | 0.91 | 0.94 |
| 7 | 155231076 | rs732398 | -0.428 | 0.122 | -0.99 | -1.21 | 0.023 | 0.059 | 0.059 | 0.94 | 0.51 | 0.37 | 0.69 | 0.72 | 0.61 | 0.75 |
| 7 | 155232287 | rs10949799 | 0.665 | -0.276 | -0.694 | -1.457 | 0 | 0 | 0 | 0 | 0.57 | 0.43 | 0.67 | 0.36 | 0.73 | 0.55 |
| 7 | 155234875 | rs730109 | 0.24 | 0.505 | -1.142 | -1.194 | -0.146 | -0.03 | -0.03 | 0 | 0.25 | 0.03 | 0.42 | 0.18 | 0.4 | 0.41 |
| 7 | 155235978 | rs1477216 | -0.791 | 0.278 | -0.71 | -1.508 | 0.025 | 0.107 | 0.107 | 1.645 | 0.56 | 0.73 | 0.72 | 0.82 | 0.62 | 0.79 |
| 7 | 155241247 | rs4717002 | -0.473 | 0.611 | -0.261 | -1.179 | -0.002 | -0.012 | -0.012 | 0 | 0.33 | 0.44 | 0.38 | 0.75 | 0.32 | 0.52 |
| 7 | 155246892 | rs2885339 | 0.09 | 1.263 | -0.41 | -1.346 | -0.004 | 0.002 | 0.002 | 0.05 | 0.88 | 0.9 | 0.76 | 0.88 | 0.86 | 0.89 |
| 7 | 155248865 | rs10949805 | -0.825 | 1.071 | -0.249 | -1.706 | 0.011 | -0.054 | -0.054 | 0.33 | 0.34 | 0.54 | 0.39 | 0.79 | 0.39 | 0.5 |
| 7 | 155254145 | rs1861973 | 1.165 | 0.68 | 0.057 | -1.492 | -0.001 | -0.001 | -0.001 | 1.734 | 0.75 | 0.71 | 0.77 | 0.95 | 0.71 | 0.67 |
| 7 | 155257894 | rs1861958 | 0.471 | 0.556 | 0.29 | -1.8 | -0.014 | -0.028 | -0.028 | 0 | 0.17 | 0.25 | 0.11 | 0.11 | 0.11 | 0.19 |
| 7 | 155258641 | rs10243118 | -0.282 | 0.461 | 0.426 | -1.888 | -0.014 | -0.046 | -0.046 | 0 | 0.23 | 0.36 | 0.3 | 0.11 | 0.14 | 0.25 |
| 7 | 155259391 | rs10237786 | 2.035 | 0.854 | 0.169 | -1.653 | -0.01 | 0.011 | 0.011 | 7.213 | 0.55 | 0.38 | 0.52 | 0.64 | 0.5 | 0.46 |
| 7 | 155259640 | rs10247302 | 1.007 | 0.583 | 0.295 | -1.677 | -0.001 | -0.008 | -0.008 | 0.003 | 0.75 | 0.69 | 0.77 | 0.94 | 0.7 | 0.69 |
| 7 | 155261445 | rs4716599 | 1.483 | 0.979 | -0.248 | -1.565 | 0.001 | -0.016 | -0.016 | 0.04 | 0.7 | 0.34 | 0.54 | 0.84 | 0.69 | 0.59 |
| 7 | 155264377 | rs10251163 | 0.892 | 0.258 | -0.781 | -1.172 | 0.077 | 0.167 | 0.167 | 0 | 0.52 | 0.24 | 0.64 | 0.8 | 0.61 | 0.53 |
| 7 | 155269627 | rs4716609 | 0.995 | 0.359 | -0.731 | -0.836 | 0.006 | 0.011 | 0.011 | 0 | 0.8 | 0.91 | 0.82 | 0.84 | 0.76 | 0.74 |
| 7 | 155271041 | rs12697964 | -0.996 | 0.717 | -0.678 | -0.847 | -0.026 | 0.02 | 0.02 | 0 | 0.15 | 0.07 | 0.16 | 0.09 | 0.24 | 0.14 |
| 7 | 155271116 | rs1895092 | 0.409 | 0.472 | -0.114 | -0.819 | 0.011 | -0.063 | -0.063 | 0 | 0.9 | 0.75 | 0.91 | 1 | 0.89 | 0.97 |
| 7 | 155271875 | rs11771253 | 1.044 | 0.852 | -1.39 | -0.596 | -0.077 | -0.054 | -0.054 | 0 | 0.39 | 0.12 | 0.56 | 0.6 | 0.45 | 0.43 |
| 7 | 155275406 | rs11772113 | 1.106 | 0.873 | -1.383 | -0.688 | -0.056 | -0.043 | -0.043 | 0 | 0.38 | 0.13 | 0.54 | 0.59 | 0.44 | 0.38 |
| 7 | 155276541 | rs12538214 | 0.495 | 0.895 | -1.38 | -0.862 | -0.044 | -0.045 | -0.045 | 1.641 | 0.12 | 0.03 | 0.14 | 0.21 | 0.23 | 0.15 |

| | | | | | | | | | | | | | | | | |
|---|---|---|---|---|---|---|---|---|---|---|---|---|---|---|---|---|
| 7 | 155277026 | rs6942469 | 0.347 | 0.635 | -0.885 | -0.934 | -0.031 | -0.129 | -0.129 | 0.12 | 0.27 | 0.57 | 0.17 | 0.01 | 0.14 | 0.17 |
| 7 | 155279366 | rs2160924 | -0.925 | 0.63 | -1.222 | -0.787 | -0.091 | -0.029 | -0.029 | 0 | 0.59 | 0.88 | 0.45 | 0.38 | 0.52 | 0.53 |
| 7 | 155280693 | rs10488639 | -0.284 | 0.513 | -0.069 | -0.446 | -0.049 | -0.157 | -0.157 | 0.014 | 0.24 | 0.45 | 0.15 | 0.01 | 0.12 | 0.16 |
| 7 | 155285135 | rs1861960 | 0.447 | 1.734 | -0.12 | 0.699 | 0.025 | 0.068 | 0.068 | 0.139 | 0.84 | 0.7 | 0.71 | 0.35 | 0.77 | 0.72 |
| 7 | 155285212 | rs17580083 | -1.036 | 1.179 | -0.358 | 0.163 | 0.009 | -0.017 | -0.017 | 0.159 | 0.12 | 0.18 | 0.15 | 0.09 | 0.18 | 0.18 |
| 7 | 155292020 | rs722445 | -0.869 | 0.69 | -0.715 | 0.092 | 0.009 | 0.006 | 0.006 | 0.194 | 0.16 | 0.16 | 0.21 | 0.14 | 0.16 | 0.28 |
| 7 | 155295273 | rs4716653 | -0.651 | 0.451 | -0.73 | 0.204 | 0.057 | -0.01 | -0.01 | 0 | 0.49 | 0.81 | 0.63 | 0.49 | 0.46 | 0.58 |
| 7 | 155301781 | rs6947243 | 0.22 | 1.117 | -0.242 | -0.077 | -0.002 | -0.002 | -0.002 | 0.98 | 0.24 | 0.19 | 0.24 | 0.35 | 0.29 | 0.27 |
| 7 | 155303512 | rs10266572 | -0.197 | 0.547 | -0.301 | -0.895 | -0.03 | -0.097 | -0.097 | 0 | 0.35 | 0.45 | 0.23 | 0.15 | 0.24 | 0.18 |
| 7 | 155303563 | rs10272570 | -0.194 | 0.542 | -0.428 | -0.952 | -0.018 | -0.066 | -0.066 | 0 | 0.43 | 0.48 | 0.32 | 0.22 | 0.3 | 0.3 |
| 7 | 155303698 | rs2098521 | 1.095 | 0.696 | -0.805 | -0.8 | 0.014 | -0.041 | -0.041 | 0.106 | 0.24 | 0.18 | 0.37 | 0.33 | 0.27 | 0.23 |
| 7 | 155305104 | rs6459707 | 0.014 | 1.029 | -0.635 | -0.632 | 0 | 0 | 0 | 0 | 0.65 | 0.68 | 0.69 | 0.54 | 0.55 | 0.52 |
| 7 | 155306750 | rs1421321 | 0.148 | 1.379 | -1.351 | -0.598 | -0.049 | -0.09 | -0.09 | 0 | 0.31 | 0.12 | 0.24 | 0.43 | 0.3 | 0.45 |
| 7 | 155308238 | rs10233924 | -0.86 | 1.517 | -1.015 | -0.3 | -0.025 | -0.023 | -0.023 | 0 | 0.76 | 0.82 | 0.61 | 0.7 | 0.58 | 0.79 |
| 7 | 155312325 | rs7786816 | -0.297 | 1.226 | -1.342 | -0.408 | -0.011 | -0.008 | -0.008 | 0 | 0.68 | 0.65 | 0.57 | 0.54 | 0.55 | 0.67 |
| 7 | 155317658 | rs7801315 | 0.911 | 1.158 | -2.024 | -0.347 | -0.063 | -0.119 | -0.119 | 0.495 | 0.25 | 0.04 | 0.33 | 0.29 | 0.33 | 0.19 |
| 7 | 155317825 | rs6968788 | 0.214 | 0.825 | -1.804 | -0.333 | 0.006 | 0.008 | 0.008 | 0 | 0.1 | 0.04 | 0.08 | 0.09 | 0.08 | 0.19 |
| 7 | 155320784 | rs2111846 | 0.047 | 0.751 | -1.534 | -0.394 | 0 | -0.001 | -0.001 | 0.385 | 0.47 | 0.48 | 0.48 | 0.4 | 0.51 | 0.48 |
| 7 | 155328331 | rs12538189 | 0.098 | 0.646 | -2.479 | -0.063 | -0.088 | -0.034 | -0.034 | 1.588 | 0.29 | 0.02 | 0.2 | 0.27 | 0.24 | 0.17 |
| 7 | 155329203 | rs4568502 | 0.604 | 0.477 | -2.64 | -0.215 | -0.03 | 0.106 | 0.106 | 0 | 0.51 | 0.12 | 0.58 | 0.66 | 0.52 | 0.56 |
| 7 | 155332127 | rs17837556 | 0.272 | 0.36 | -2.542 | 0.101 | 0.044 | -0.064 | -0.064 | 3.761 | 0.26 | 0.09 | 0.25 | 0.39 | 0.18 | 0.35 |
| 7 | 155336563 | rs889954 | 0.902 | 0.773 | -1.747 | 0.473 | -0.01 | -0.009 | -0.009 | 0.335 | 0.59 | 0.58 | 0.65 | 0.47 | 0.54 | 0.53 |
| 7 | 155336638 | rs6975765 | -0.07 | 0.796 | -1.869 | 0.094 | -0.034 | 0.051 | 0.051 | 0 | 0.2 | 0.13 | 0.09 | 0.1 | 0.17 | 0.09 |
| 7 | 155343673 | rs956406 | 0.455 | -0.049 | -1.243 | 0.347 | -0.017 | 0.029 | 0.029 | 0.244 | 0.69 | 0.57 | 0.66 | 0.42 | 0.74 | 0.58 |
| 7 | 155355522 | rs10264499 | -0.439 | -0.104 | -1.553 | 0.531 | 0.023 | -0.016 | -0.016 | 0 | 0.29 | 0.19 | 0.2 | 0.37 | 0.22 | 0.33 |
| 7 | 155356960 | rs6966832 | 0.684 | 0.728 | -1.989 | 0.474 | -0.017 | -0.045 | -0.045 | 0.986 | 0.17 | 0.04 | 0.16 | 0.21 | 0.18 | 0.12 |
| 7 | 155358802 | rs10277542 | -0.14 | 1.563 | -2.019 | 0.28 | -0.035 | 0 | 0 | 0.531 | 0.8 | 0.83 | 0.59 | 0.79 | 0.62 | 0.76 |
| 7 | 155360511 | rs11772335 | 0.205 | 0.546 | -1.963 | 0.262 | -0.041 | -0.072 | -0.072 | 0.319 | 0.2 | 0.02 | 0.19 | 0.2 | 0.21 | 0.16 |
| 7 | 155374152 | rs2113583 | -0.039 | 0.208 | -1.558 | 0.347 | -0.025 | -0.105 | -0.105 | 1.383 | 0.75 | 0.88 | 0.64 | 0.51 | 0.72 | 0.66 |
| 7 | 155374700 | rs6459794 | 0.231 | -0.127 | -1.221 | -0.049 | -0.011 | -0.027 | -0.027 | 0 | 0.21 | 0.24 | 0.1 | 0.09 | 0.14 | 0.15 |
| 7 | 155377430 | rs6953751 | 0.375 | 0.399 | -0.609 | 0.508 | -0.001 | 0.032 | 0.032 | 0 | 0.7 | 0.58 | 0.69 | 0.54 | 0.71 | 0.63 |
| 7 | 155382621 | rs7785787 | -0.386 | 0.197 | -0.433 | -0.256 | -0.009 | -0.046 | -0.046 | 0 | 0.41 | 0.5 | 0.27 | 0.18 | 0.32 | 0.34 |
| 7 | 155386972 | rs4716763 | -0.355 | -0.022 | 0.316 | -0.69 | 0.001 | -0.062 | -0.062 | 0 | 0.29 | 0.47 | 0.2 | 0.14 | 0.27 | 0.3 |
| 7 | 155395948 | rs9654697 | -0.966 | -0.7 | 0.252 | -0.848 | 0.021 | 0.07 | 0.07 | 0 | 0.46 | 0.7 | 0.34 | 0.23 | 0.37 | 0.42 |
| 7 | 155396914 | rs889952 | -0.829 | -0.777 | 0.158 | -0.591 | 0.003 | 0.041 | 0.041 | 0 | 0.42 | 0.7 | 0.35 | 0.31 | 0.4 | 0.42 |
| 7 | 155398784 | rs6944909 | -0.133 | -0.982 | 0.115 | -0.853 | 0.016 | -0.013 | -0.013 | 0 | 0.16 | 0.11 | 0.14 | 0.16 | 0.14 | 0.23 |
| 7 | 155402033 | rs6959315 | -0.334 | -0.793 | 0.721 | -0.715 | 0.011 | 0.036 | 0.036 | 0 | 0.35 | 0.25 | 0.25 | 0.19 | 0.32 | 0.35 |
| 7 | 155403168 | rs28694796 | -1.951 | -0.633 | 1.675 | -0.728 | 0.037 | -0.018 | -0.018 | 1.474 | 0.09 | 0.32 | 0.1 | 0.06 | 0.06 | 0.07 |
| 7 | 155403802 | rs6965868 | -0.653 | -0.624 | 0.877 | -0.479 | -0.003 | -0.037 | -0.037 | 0.316 | 0.87 | 0.94 | 0.66 | 0.81 | 0.87 | 0.88 |
| 7 | 155404870 | rs889949 | 0.559 | -0.627 | 1.133 | -0.448 | 0 | -0.002 | -0.002 | 0 | 0.8 | 0.81 | 0.57 | 0.73 | 0.82 | 0.81 |
| 7 | 155416363 | rs6946081 | 1.085 | -0.51 | 1.141 | -0.881 | 0.002 | 0 | 0 | 0.502 | 0.66 | 0.34 | 0.48 | 0.69 | 0.74 | 0.79 |
| 7 | 155428591 | rs7778376 | -0.981 | -0.5 | 1.529 | -0.902 | 0.027 | -0.015 | -0.015 | 0 | 0.07 | 0.32 | 0.07 | 0.02 | 0.04 | 0.02 |
| 7 | 155429505 | rs12673528 | 0.499 | -0.552 | 1.521 | -0.925 | 0.009 | -0.007 | -0.007 | 0 | 0.72 | 0.79 | 0.57 | 0.71 | 0.77 | 0.8 |
| 7 | 155436997 | rs2241444 | 1.171 | -0.551 | 1.67 | -0.927 | -0.015 | -0.005 | -0.005 | 0 | 0.66 | 0.5 | 0.48 | 0.69 | 0.73 | 0.78 |
| 7 | 155442850 | rs6459848 | 1.191 | -0.755 | 1.577 | -0.929 | -0.007 | -0.002 | -0.002 | 1.055 | 0.66 | 0.45 | 0.49 | 0.69 | 0.75 | 0.79 |
| 7 | 155445053 | rs984543 | -0.179 | -0.825 | 1.533 | -0.972 | 0.01 | -0.076 | -0.076 | 0.587 | 0.81 | 0.94 | 0.62 | 0.71 | 0.84 | 0.84 |
| 7 | 155450991 | rs10246957 | 0.433 | -0.759 | 1.56 | -0.937 | 0.029 | -0.048 | -0.048 | 0 | 0.73 | 0.93 | 0.59 | 0.71 | 0.79 | 0.81 |
| 7 | 155453131 | rs6954802 | 1.19 | -0.723 | 1.516 | -0.923 | -0.017 | -0.003 | -0.003 | 0 | 0.66 | 0.47 | 0.49 | 0.69 | 0.75 | 0.79 |
| 7 | 155453280 | rs6955098 | 1.009 | -0.723 | 1.56 | -0.923 | -0.019 | 0 | 0 | 0 | 0.67 | 0.59 | 0.51 | 0.69 | 0.75 | 0.79 |
| 7 | 155470821 | rs2113586 | 0.549 | -0.707 | 1.553 | -0.909 | -0.002 | 0 | 0 | 0.482 | 0.72 | 0.74 | 0.57 | 0.71 | 0.79 | 0.81 |
| 7 | 155507042 | rs10227012 | -1.58 | -0.732 | 1.427 | -0.803 | 0.03 | 0 | 0 | 0.614 | 0.33 | 0.72 | 0.28 | 0.31 | 0.18 | 0.18 |
| 7 | 155507387 | rs4716900 | -1.26 | -0.719 | 1.481 | -0.807 | -0.02 | 0.005 | 0.005 | 0.207 | 0.29 | 0.36 | 0.24 | 0.31 | 0.18 | 0.18 |
| 7 | 155523922 | rs4501473 | -1.262 | -0.755 | 1.396 | -0.804 | -0.035 | 0.006 | 0.006 | 0.194 | 0.31 | 0.49 | 0.27 | 0.31 | 0.19 | 0.18 |
| 7 | 155531080 | rs3735576 | -1.564 | -0.744 | 0.719 | -0.799 | 0.03 | -0.001 | -0.001 | 0 | 0.33 | 0.72 | 0.28 | 0.31 | 0.19 | 0.18 |
| 7 | 155544740 | rs6961346 | -1.338 | -0.461 | 0.137 | -0.672 | -0.029 | 0.01 | 0.01 | 0.19 | 0.29 | 0.38 | 0.25 | 0.31 | 0.18 | 0.18 |
| 7 | 155546241 | rs13222019 | -0.82 | -0.458 | -0.176 | -0.67 | 0.012 | -0.009 | -0.009 | 0.282 | 0.25 | 0.19 | 0.18 | 0.29 | 0.15 | 0.16 |
| 7 | 155555930 | rs11769826 | -1.282 | -0.461 | -0.21 | -0.665 | -0.022 | 0.007 | 0.007 | 0.267 | 0.29 | 0.36 | 0.24 | 0.31 | 0.19 | 0.17 |
| 7 | 155557706 | rs11771062 | -1.282 | -0.461 | -0.21 | -0.665 | -0.022 | 0.007 | 0.007 | 0.267 | 0.29 | 0.36 | 0.24 | 0.31 | 0.19 | 0.18 |
| 7 | 155563015 | rs11761850 | -0.303 | 0.293 | -0.721 | -1.273 | 0.038 | -0.016 | -0.016 | 0.819 | 0.05 | 0.16 | 0.08 | 0.02 | 0.06 | 0.03 |
| 7 | 155569983 | rs2007922 | -0.308 | 0.28 | -1.817 | -1.277 | -0.02 | 0.085 | 0.085 | 0 | 0.61 | 0.81 | 0.52 | 0.79 | 0.57 | 0.58 |
| 7 | 155572107 | rs17550812 | -0.653 | 0.573 | -1.19 | -0.783 | 0.012 | 0.015 | 0.015 | 0 | 0.08 | 0.03 | 0.03 | 0 | 0.05 | 0.03 |
| 7 | 155577275 | rs2158813 | 0.799 | 1.142 | -1.359 | -0.78 | -0.035 | 0.079 | 0.079 | 0 | 0.75 | 0.95 | 0.85 | 0.91 | 0.71 | 0.9 |
| 7 | 155592357 | rs2239966 | 0.776 | 0.934 | -0.672 | -0.458 | 0.008 | -0.037 | -0.037 | 0.168 | 0.66 | 0.48 | 0.7 | 0.73 | 0.61 | 0.72 |
| 7 | 155600417 | rs1233556 | 0.752 | 0.171 | -0.895 | -0.986 | 0.004 | -0.027 | -0.027 | 0 | 0.74 | 0.72 | 0.89 | 0.98 | 0.82 | 0.94 |
| 7 | 155604941 | rs9333594 | -0.509 | -0.142 | -0.899 | -0.962 | 0.003 | -0.007 | -0.007 | 0.005 | 0.12 | 0.13 | 0.05 | 0.02 | 0.08 | 0.03 |
| 7 | 155605744 | rs872723 | -0.364 | 0.264 | -0.652 | -1.174 | 0.01 | -0.009 | -0.009 | 0 | 0.09 | 0.13 | 0.14 | 0.08 | 0.2 | 0.12 |
| 7 | 155606672 | rs288746 | 0.864 | 0.456 | -0.846 | -1.518 | 0.012 | 0.021 | 0.021 | 2.633 | 0.77 | 0.83 | 0.88 | 0.98 | 0.84 | 0.93 |
| 7 | 155609971 | rs288747 | 0.288 | -0.652 | -1.485 | -2.034 | 0.027 | 0.041 | 0.041 | 0.646 | 0.26 | 0.13 | 0.22 | 0.19 | 0.19 | 0.31 |
| 7 | 155615627 | rs172310 | 0.489 | -0.081 | -1.674 | -2.213 | -0.006 | 0.06 | 0.06 | 0.413 | 0.7 | 0.81 | 0.74 | 0.99 | 0.67 | 0.89 |

| | | | | | | | | | | | | | | | | |
|---|---|---|---|---|---|---|---|---|---|---|---|---|---|---|---|---|
| 7 | 155619733 | rs167020 | 0.224 | -0.137 | -1.075 | -1.478 | -0.009 | 0.09 | 0.09 | 1.225 | 0.73 | 0.86 | 0.75 | 0.99 | 0.7 | 0.91 |
| 7 | 155619980 | rs1233563 | -0.248 | -0.649 | -0.069 | -1.571 | -0.026 | -0.037 | -0.037 | 0 | 0.34 | 0.43 | 0.27 | 0.22 | 0.23 | 0.37 |
| 7 | 155624226 | rs288756 | 0.654 | -0.642 | 0.068 | -0.895 | -0.001 | 0.01 | 0.01 | 0.48 | 0.42 | 0.61 | 0.57 | 0.67 | 0.53 | 0.67 |
| 7 | 155625512 | rs721347 | 2.028 | -0.218 | -0.087 | -0.685 | -0.071 | -0.07 | -0.07 | 0.017 | 0.23 | 0.08 | 0.34 | 0.41 | 0.31 | 0.34 |
| 7 | 155626047 | rs288757 | 1.452 | -0.18 | 0.218 | -0.697 | -0.024 | -0.027 | -0.027 | 0 | 0.28 | 0.24 | 0.39 | 0.5 | 0.37 | 0.43 |
| 7 | 155629975 | rs1151790 | 0.096 | -0.363 | 0.339 | -0.824 | -0.029 | -0.024 | -0.024 | 0 | 0.33 | 0.4 | 0.28 | 0.19 | 0.22 | 0.42 |
| 7 | 155631538 | rs7804368 | 1.21 | -0.321 | 0.028 | -0.803 | -0.004 | -0.004 | -0.004 | 0 | 0.06 | 0.04 | 0.04 | 0 | 0.09 | 0.08 |
| 7 | 155632315 | rs916733 | 1.767 | -0.414 | -0.234 | -0.89 | -0.079 | -0.087 | -0.087 | 0 | 0.28 | 0.08 | 0.38 | 0.43 | 0.36 | 0.35 |
| 7 | 155633481 | rs288761 | 0.433 | -0.817 | -0.003 | -0.87 | -0.002 | 0.004 | 0.004 | 0 | 0.63 | 0.65 | 0.69 | 0.7 | 0.61 | 0.84 |
| 7 | 155638278 | rs1675547 | -0.697 | -1.639 | -0.249 | -0.541 | 0.038 | -0.097 | -0.097 | 0 | 0.78 | 0.92 | 0.81 | 0.53 | 0.88 | 0.61 |
| 7 | 155641885 | rs1225382 | -0.923 | -2.131 | 0.176 | -1.138 | 0.007 | 0.01 | 0.01 | 0 | 0.38 | 0.33 | 0.27 | 0.32 | 0.36 | 0.23 |
| 7 | 155643326 | rs1225387 | -0.176 | -2.155 | 0.069 | -1.269 | 0.011 | 0.04 | 0.04 | 1.819 | 0.25 | 0.16 | 0.21 | 0.01 | 0.27 | 0.1 |
| 7 | 155648007 | rs10249283 | 0.932 | -2.384 | 0.372 | -1.4 | -0.026 | -0.06 | -0.06 | 0.178 | 0.21 | 0.21 | 0.17 | 0.04 | 0.08 | 0.25 |
| 7 | 155649488 | rs1731835 | -0.155 | -2.486 | 0.478 | -1.356 | 0.013 | 0.028 | 0.028 | 1.138 | 0.26 | 0.2 | 0.22 | 0.01 | 0.27 | 0.1 |
| 7 | 155650354 | rs1731837 | -0.983 | -2.532 | 0.677 | -1.259 | 0.03 | 0.042 | 0.042 | 0.084 | 0.52 | 0.61 | 0.45 | 0.14 | 0.38 | 0.42 |
| 7 | 155650687 | rs1675569 | -0.664 | -2.535 | 0.694 | -1.262 | -0.003 | -0.029 | -0.029 | 0 | 0.32 | 0.38 | 0.28 | 0.11 | 0.33 | 0.21 |
| 7 | 155651102 | rs1731839 | -0.569 | -1.508 | 0.795 | -0.609 | 0.044 | -0.007 | -0.007 | 0 | 0.56 | 0.83 | 0.48 | 0.47 | 0.39 | 0.48 |
| 7 | 155655522 | rs1731847 | 1.104 | -1.118 | 0.747 | -1.282 | -0.023 | 0.04 | 0.04 | 0 | 0.39 | 0.25 | 0.52 | 0.63 | 0.54 | 0.43 |
| 7 | 155656603 | rs10261156 | 0.451 | -0.909 | 1.284 | -1.559 | -0.002 | -0.028 | -0.028 | 0 | 0.58 | 0.41 | 0.65 | 0.87 | 0.75 | 0.61 |
| 7 | 155657469 | rs10247896 | 0.901 | -0.867 | 1.138 | -1.554 | 0.068 | 0.147 | 0.147 | 0.06 | 0.51 | 0.26 | 0.57 | 0.86 | 0.63 | 0.6 |
| 7 | 155657480 | rs11770631 | -1.264 | -0.634 | 0.926 | -1.412 | 0.03 | -0.041 | -0.041 | 0.361 | 0.13 | 0.37 | 0.15 | 0.07 | 0.15 | 0.13 |
| 7 | 155657834 | rs11768525 | -0.244 | -0.445 | 0.45 | -1.182 | 0.013 | 0.007 | 0.007 | 0 | 0.06 | 0.2 | 0.06 | 0.07 | 0.05 | 0.11 |
| 7 | 155659348 | rs6459952 | 0.342 | -0.429 | -0.018 | -1.19 | -0.006 | -0.059 | -0.059 | 0.581 | 0.58 | 0.57 | 0.64 | 0.93 | 0.69 | 0.71 |
| 7 | 155661335 | rs10281731 | 1.17 | 0.381 | 0.538 | -0.594 | 0.015 | -0.087 | -0.087 | 0 | 0.86 | 0.72 | 0.89 | 1 | 0.85 | 0.96 |
| 7 | 155662910 | rs11766361 | 0.106 | 0.36 | -0.053 | -0.53 | 0.021 | -0.005 | -0.005 | 0 | 0.34 | 0.17 | 0.28 | 0.36 | 0.28 | 0.3 |
| 7 | 155664686 | rs6971211 | -0.247 | 0.862 | 0.586 | -0.502 | -0.002 | 0.001 | 0.001 | 0 | 0.42 | 0.37 | 0.36 | 0.47 | 0.4 | 0.4 |
| 7 | 155665339 | rs11767572 | -0.97 | 0.846 | 0.817 | -0.562 | 0 | -0.003 | -0.003 | 0 | 0.53 | 0.51 | 0.43 | 0.47 | 0.49 | 0.43 |
| 7 | 155665835 | rs13438255 | -1.06 | 0.62 | 0.936 | -0.552 | 0 | 0 | 0 | 0 | 0.54 | 0.54 | 0.46 | 0.47 | 0.54 | 0.42 |
| 7 | 155666929 | rs10267147 | -0.207 | 0.691 | 0.926 | -0.549 | 0.074 | -0.038 | -0.038 | 0 | 0.08 | 0.19 | 0.08 | 0.01 | 0.12 | 0.09 |
| 7 | 155669500 | rs11772416 | 0.272 | 0.668 | 0.77 | -0.512 | 0.002 | 0.005 | 0.005 | 0 | 0.35 | 0.2 | 0.31 | 0.37 | 0.33 | 0.3 |
| 7 | 155670997 | rs898695 | 0.44 | 0.822 | 1.035 | -0.376 | 0.023 | -0.017 | -0.017 | 0 | 0.95 | 0.82 | 0.97 | 1 | 0.97 | 0.99 |
| 7 | 155671277 | rs754469 | -0.566 | 0.517 | 1.013 | -0.476 | 0 | 0.001 | 0.001 | 0 | 0.49 | 0.48 | 0.43 | 0.49 | 0.52 | 0.45 |
| 7 | 155672649 | rs898696 | -0.038 | 0.697 | 0.47 | -0.206 | 0 | -0.002 | -0.002 | 1.43 | 0.55 | 0.63 | 0.63 | 0.44 | 0.57 | 0.56 |
| 7 | 155672944 | rs1106697 | -1.069 | 0.842 | 0.549 | -0.118 | 0.023 | -0.049 | -0.049 | 1.139 | 0.09 | 0.17 | 0.07 | 0 | 0.11 | 0.03 |
| 7 | 155676753 | rs7777688 | -1.16 | 0.626 | 0.801 | -0.031 | 0.022 | -0.018 | -0.018 | 0.677 | 0.11 | 0.36 | 0.09 | 0.04 | 0.08 | 0.08 |
| 7 | 155677439 | rs1461208 | -1.05 | 1.19 | 0.607 | 0.457 | 0.005 | 0.011 | 0.011 | 0 | 0.3 | 0.4 | 0.25 | 0.35 | 0.31 | 0.26 |
| 7 | 155677943 | rs7786445 | -0.15 | 0.856 | -0.065 | 0.073 | 0 | 0.014 | 0.014 | 0 | 0.78 | 0.82 | 0.85 | 0.85 | 0.79 | 0.67 |
| 7 | 155681732 | rs12698335 | 0.743 | 1.016 | 0.377 | -0.273 | 0.001 | -0.094 | -0.094 | 0.221 | 0.77 | 0.49 | 0.72 | 0.88 | 0.76 | 0.87 |
| 7 | 155682156 | rs7780318 | 0.919 | 0.813 | -0.525 | 0.385 | 0.001 | -0.02 | -0.02 | 0 | 0.23 | 0.12 | 0.39 | 0.34 | 0.22 | 0.3 |
| 7 | 155690938 | rs1531381 | 1.252 | 0.567 | -0.662 | 0.427 | -0.006 | -0.033 | -0.033 | 0.232 | 0.19 | 0.11 | 0.3 | 0.33 | 0.18 | 0.28 |
| 7 | 155693166 | rs2018408 | 0.669 | 0.758 | -0.258 | -0.067 | 0 | 0 | 0 | 0 | 0.35 | 0.38 | 0.3 | 0.31 | 0.33 | 0.46 |
| 7 | 155693269 | rs3857649 | 0.787 | 0.678 | -0.133 | -0.458 | 0.001 | -0.006 | -0.006 | 0 | 0.28 | 0.32 | 0.2 | 0.26 | 0.32 | 0.19 |
| 7 | 155693481 | rs1824552 | -0.566 | 0.821 | -0.192 | -0.53 | -0.002 | 0.007 | 0.007 | 0.282 | 0.45 | 0.36 | 0.33 | 0.35 | 0.51 | 0.24 |
| 7 | 155695674 | rs6950784 | 0.076 | 1.019 | -0.15 | -0.355 | 0.004 | 0.011 | 0.011 | 0 | 0.5 | 0.45 | 0.55 | 0.65 | 0.51 | 0.46 |
| 7 | 155698022 | rs6459958 | 1.512 | 1.032 | -0.569 | -0.077 | -0.025 | -0.051 | -0.051 | 0 | 0.19 | 0.09 | 0.26 | 0.36 | 0.18 | 0.27 |
| 7 | 155698169 | rs6459960 | 1.495 | 1.026 | -0.572 | -0.08 | -0.024 | -0.051 | -0.051 | 0.095 | 0.19 | 0.09 | 0.26 | 0.36 | 0.18 | 0.27 |
| 7 | 155698255 | rs6459961 | -0.68 | 1.125 | -0.044 | -0.169 | 0.03 | 0.024 | 0.024 | 0.034 | 0.74 | 0.65 | 0.53 | 0.64 | 0.66 | 0.61 |
| 7 | 155698742 | rs10262718 | 0.558 | -0.101 | -0.354 | -0.493 | 0.045 | -0.014 | -0.014 | 0.981 | 0.28 | 0.19 | 0.19 | 0.26 | 0.19 | 0.4 |
| 7 | 155700996 | rs1870375 | -0.633 | 0.488 | 1.381 | -0.551 | 0.053 | -0.07 | -0.07 | 0.449 | 0.2 | 0.4 | 0.3 | 0.08 | 0.27 | 0.13 |
| 7 | 155701296 | rs4716923 | 0.487 | -0.117 | 1.654 | -0.306 | 0.028 | -0.05 | -0.05 | 1.245 | 0.67 | 0.52 | 0.57 | 0.82 | 0.56 | 0.65 |
| 7 | 155715081 | rs6459968 | -0.876 | -1.175 | 1.772 | 0.215 | -0.005 | -0.001 | -0.001 | 0 | 0.42 | 0.47 | 0.44 | 0.4 | 0.4 | 0.42 |
| 7 | 155716520 | rs4716928 | -0.042 | -1.23 | 1.706 | 0.253 | -0.037 | 0.038 | 0.038 | 0 | 0.88 | 0.84 | 0.85 | 0.8 | 0.94 | 0.76 |
| 7 | 155718241 | rs4716930 | 1.578 | -1.405 | 1.713 | 0.047 | 0.013 | 0.005 | 0.005 | 0 | 0.69 | 0.31 | 0.72 | 0.76 | 0.8 | 0.62 |
| 7 | 155720971 | rs10949787 | 0.282 | -1.44 | 2.312 | 0.026 | 0.029 | -0.018 | -0.018 | 0 | 0.89 | 0.52 | 0.9 | 0.96 | 0.89 | 0.9 |
| 7 | 155721324 | rs11764820 | 1.748 | -1.455 | 2.325 | 0.025 | 0.011 | 0.004 | 0.004 | 0 | 0.7 | 0.32 | 0.72 | 0.76 | 0.8 | 0.62 |
| 7 | 155721386 | rs11769663 | 1.748 | -1.455 | 2.325 | 0.025 | 0.011 | 0.004 | 0.004 | 0 | 0.7 | 0.32 | 0.72 | 0.76 | 0.8 | 0.62 |
| 7 | 155722231 | rs6971588 | -0.856 | -1.436 | 2.153 | 0.04 | -0.032 | 0.004 | 0.004 | 0.964 | 0.19 | 0.2 | 0.18 | 0.2 | 0.09 | 0.28 |
| 7 | 155728192 | rs4716934 | 1.514 | -1.421 | 2.002 | -0.017 | 0.007 | -0.003 | -0.003 | 0 | 0.77 | 0.28 | 0.84 | 0.96 | 0.85 | 0.86 |
| 7 | 155730519 | rs6964974 | 0.862 | -1.378 | 1.863 | 0.077 | -0.001 | 0.006 | 0.006 | 0 | 0.59 | 0.4 | 0.63 | 0.72 | 0.54 | 0.72 |
| 7 | 155732792 | rs10263106 | 0.401 | -1.191 | 1.716 | 0.208 | -0.05 | 0.004 | 0.004 | 0.795 | 0.8 | 0.73 | 0.81 | 0.8 | 0.91 | 0.72 |
| 7 | 155733949 | rs4716941 | 0.65 | -0.881 | 1.736 | 0.161 | -0.001 | 0.002 | 0.002 | 0 | 0.55 | 0.44 | 0.54 | 0.52 | 0.51 | 0.47 |
| 7 | 155734665 | rs4716944 | 0.838 | -0.765 | 1.253 | 0.22 | 0.008 | -0.005 | -0.005 | 0 | 0.52 | 0.31 | 0.51 | 0.52 | 0.5 | 0.47 |
| 7 | 155737659 | rs10225087 | 0.246 | -0.508 | 1.153 | 0.078 | -0.015 | -0.02 | -0.02 | 0.26 | 0.2 | 0.25 | 0.08 | 0.13 | 0.1 | 0.06 |
| 7 | 155741872 | rs6945743 | 0.024 | 0.48 | 0.72 | 0.376 | 0.038 | 0.001 | 0.001 | 0 | 0.23 | 0.39 | 0.28 | 0.23 | 0.35 | 0.2 |
| 7 | 155747907 | rs7810074 | -0.228 | 0.086 | 0.233 | 0.582 | -0.014 | 0.009 | 0.009 | 3.924 | 0.61 | 0.69 | 0.63 | 0.72 | 0.56 | 0.78 |
| 7 | 155755336 | rs7807731 | -0.335 | -0.146 | -0.036 | 0.77 | 0.003 | 0.005 | 0.005 | 2.117 | 0.55 | 0.58 | 0.55 | 0.62 | 0.58 | 0.48 |
| 7 | 155755826 | rs2007015 | 0.644 | -0.339 | 0.29 | 1.104 | -0.035 | 0.123 | 0.123 | 0.044 | 0.81 | 0.62 | 0.79 | 0.6 | 0.85 | 0.72 |
| 7 | 155759926 | rs4716572 | -0.141 | -0.399 | -0.415 | 0.862 | 0.036 | -0.012 | -0.012 | 0 | 0.42 | 0.22 | 0.39 | 0.59 | 0.34 | 0.48 |
| 7 | 155760353 | rs6459974 | -0.86 | -0.043 | -0.174 | 1.219 | 0.003 | 0.038 | 0.038 | 0 | 0.24 | 0.33 | 0.15 | 0.39 | 0.27 | 0.22 |

| | | | | | | | | | | | | | | | | |
|---|---|---|---|---|---|---|---|---|---|---|---|---|---|---|---|---|
| 7 | 155761671 | rs12113238 | 0.147 | 0.398 | -0.088 | 1.103 | 0.005 | -0.006 | -0.006 | 0 | 0.63 | 0.55 | 0.56 | 0.6 | 0.57 | 0.54 |
| 7 | 155761773 | rs6459976 | -0.868 | 0.543 | 0.322 | 1.573 | 0.012 | 0.148 | 0.148 | 0.062 | 0.09 | 0.3 | 0.19 | 0.3 | 0.09 | 0.14 |
| 7 | 155765548 | rs12534938 | -0.638 | 0.656 | -0.271 | 1.559 | 0 | 0.038 | 0.038 | 0.337 | 0.08 | 0.13 | 0.17 | 0.33 | 0.08 | 0.13 |
| 7 | 155766830 | rs4716573 | 0.138 | 0.95 | -0.582 | 1.277 | -0.031 | -0.032 | -0.032 | 0 | 0.05 | 0.01 | 0.14 | 0.12 | 0.1 | 0.18 |
| 7 | 155767477 | rs3801899 | -0.39 | 0.458 | -0.732 | 0.919 | 0.013 | 0.12 | 0.12 | 0.813 | 0.52 | 0.8 | 0.57 | 0.73 | 0.46 | 0.46 |
| 7 | 155767806 | rs13224793 | 0.323 | 0.343 | -0.331 | 0.614 | -0.002 | -0.027 | -0.027 | 0 | 0.05 | 0.11 | 0.06 | 0 | 0.03 | 0.03 |
| 7 | 155768061 | rs1458806 | 0.353 | 0.278 | -0.499 | 0.786 | 0.017 | 0.023 | 0.023 | 0.096 | 0.53 | 0.22 | 0.48 | 0.42 | 0.58 | 0.54 |
| 7 | 155770280 | rs1531284 | 1.103 | -0.017 | 0.456 | 1.563 | -0.02 | 0.089 | 0.089 | 0 | 0.84 | 0.69 | 0.8 | 0.61 | 0.91 | 0.8 |
| 7 | 155771394 | rs10434949 | -0.339 | -0.017 | 0.364 | 1.581 | 0 | 0.158 | 0.158 | 0.044 | 0.08 | 0.19 | 0.15 | 0.39 | 0.06 | 0.17 |
| 7 | 155773435 | rs11980508 | 0.788 | 0.104 | 0.06 | 1.394 | 0.017 | 0.059 | 0.059 | 0 | 0.52 | 0.31 | 0.42 | 0.25 | 0.56 | 0.46 |
| 7 | 155777228 | rs11769678 | 1.03 | 0.078 | 0.473 | 1.849 | -0.027 | 0.159 | 0.159 | 0 | 0.85 | 0.59 | 0.79 | 0.61 | 0.92 | 0.81 |
| 7 | 155791386 | rs7799059 | -1.122 | 0.361 | 0.579 | 2.076 | 0.034 | 0.299 | 0.299 | 0 | 0.07 | 0.29 | 0.16 | 0.3 | 0.05 | 0.1 |
| 7 | 155791773 | rs7784908 | -0.353 | 0.212 | 0.144 | 1.147 | 0.006 | -0.014 | -0.014 | 0.625 | 0.35 | 0.29 | 0.29 | 0.56 | 0.34 | 0.4 |
| 7 | 155791799 | rs4716963 | 0.7 | 0.66 | -1.077 | -0.221 | -0.057 | 0.143 | 0.143 | 0 | 0.3 | 0.05 | 0.37 | 0.08 | 0.32 | 0.21 |
| 7 | 155793895 | rs7776456 | -0.316 | 0.734 | -0.435 | -0.141 | -0.012 | 0.037 | 0.037 | 0 | 0.74 | 0.82 | 0.83 | 0.93 | 0.7 | 0.69 |
| 7 | 155797686 | rs6459977 | -0.23 | 1.198 | -0.556 | -0.127 | -0.011 | 0.012 | 0.012 | 0 | 0.83 | 0.87 | 0.86 | 0.94 | 0.73 | 0.81 |
| 7 | 155800508 | rs2363923 | -0.167 | 0.802 | -0.418 | 0.109 | -0.002 | 0 | 0 | 0.926 | 0.69 | 0.7 | 0.78 | 0.86 | 0.67 | 0.75 |
| 7 | 155809123 | rs2305944 | -0.235 | 0.383 | -0.056 | 0.759 | 0.007 | 0.004 | 0.004 | 0 | 0.5 | 0.46 | 0.34 | 0.48 | 0.46 | 0.47 |
| 7 | 155810088 | rs1458804 | 0.751 | 0.5 | -0.333 | 0.739 | -0.022 | 0.018 | 0.018 | 0 | 0.26 | 0.17 | 0.4 | 0.23 | 0.25 | 0.3 |
| 7 | 155810671 | rs10949794 | 0.35 | 0.387 | -0.048 | 0.851 | -0.012 | -0.001 | -0.001 | 0.703 | 0.46 | 0.33 | 0.35 | 0.48 | 0.48 | 0.43 |
| 7 | 155811043 | rs6962792 | -0.076 | -0.152 | 0.701 | 0.926 | -0.014 | -0.006 | -0.006 | 0 | 0.35 | 0.44 | 0.46 | 0.33 | 0.24 | 0.38 |
| 7 | 155819065 | rs7778072 | -1.546 | 0.251 | 1.819 | 2.037 | 0.089 | 0.326 | 0.326 | 2.322 | 0.06 | 0.43 | 0.05 | 0.22 | 0.03 | 0.05 |
| 7 | 155819301 | rs7788600 | -1.377 | 0.401 | 1.772 | 2.348 | 0.02 | 0.241 | 0.241 | 0 | 0.17 | 0.41 | 0.13 | 0.51 | 0.18 | 0.2 |
| 7 | 155820825 | rs4716580 | 1.451 | 0.646 | 1.602 | 2.281 | 0.025 | 0.176 | 0.176 | 0 | 0.81 | 0.44 | 0.83 | 0.44 | 0.79 | 0.8 |
| 7 | 155822088 | rs10235613 | 0.87 | 0.7 | 1.396 | 2.023 | 0.046 | 0.123 | 0.123 | 0 | 0.85 | 0.62 | 0.87 | 0.71 | 0.81 | 0.85 |
| 7 | 155825162 | rs11768024 | -0.766 | 0.761 | 1.414 | 1.992 | 0.045 | 0.135 | 0.135 | 0.082 | 0.1 | 0.27 | 0.09 | 0.28 | 0.14 | 0.29 |
| 7 | 155826004 | rs6963951 | -0.425 | 0.468 | 0.688 | 1.708 | -0.014 | 0.016 | 0.016 | 0 | 0.2 | 0.26 | 0.11 | 0.27 | 0.13 | 0.11 |
| 7 | 155831189 | rs11773897 | 0.325 | 0.584 | -0.349 | 1.415 | -0.029 | -0.016 | -0.016 | 0 | 0.43 | 0.22 | 0.5 | 0.44 | 0.5 | 0.45 |
| 7 | 155831817 | rs6969015 | 1.119 | 0.571 | -1.111 | 0.688 | -0.014 | 0.03 | 0.03 | 0.656 | 0.3 | 0.11 | 0.33 | 0.23 | 0.31 | 0.16 |
| 7 | 155836562 | rs12673011 | 0.851 | 0.38 | -1.558 | 0.258 | -0.044 | 0.067 | 0.067 | 0 | 0.34 | 0.09 | 0.44 | 0.17 | 0.41 | 0.4 |
| 7 | 155839287 | rs6952416 | 0.41 | 0.366 | -1.318 | 0.231 | 0.005 | 0.064 | 0.064 | 0.145 | 0.31 | 0.16 | 0.29 | 0.12 | 0.28 | 0.16 |
| 7 | 155851624 | rs4294053 | 0.278 | 0.859 | -1.438 | 0.796 | -0.025 | -0.003 | -0.003 | 0.92 | 0.31 | 0.08 | 0.4 | 0.24 | 0.34 | 0.43 |
| 7 | 155853315 | rs11763938 | 0.58 | 0.261 | -1.114 | 0.701 | 0.05 | 0.111 | 0.111 | 0.194 | 0.46 | 0.23 | 0.31 | 0.29 | 0.36 | 0.37 |
| 7 | 155860445 | rs10275088 | -0.561 | -0.575 | -0.881 | 1.076 | 0.081 | 0.04 | 0.04 | 0.534 | 0.4 | 0.81 | 0.5 | 0.67 | 0.3 | 0.62 |
| 7 | 155863009 | rs10248131 | -0.526 | -0.631 | -0.49 | 1.692 | 0.049 | -0.039 | -0.039 | 3.509 | 0.38 | 0.73 | 0.46 | 0.46 | 0.29 | 0.61 |
| 7 | 155868777 | rs6963817 | -0.48 | -0.585 | 1.71 | 1.835 | -0.014 | -0.001 | -0.001 | 0 | 0.37 | 0.52 | 0.42 | 0.38 | 0.3 | 0.62 |
| 7 | 155870431 | rs715363 | -0.515 | -0.589 | 1.718 | 1.834 | 0.005 | 0 | 0 | 0 | 0.37 | 0.58 | 0.42 | 0.38 | 0.3 | 0.62 |
| 7 | 155879851 | rs10228290 | -0.161 | -0.629 | 1.927 | 1.832 | 0.008 | -0.004 | -0.004 | 0 | 0.32 | 0.25 | 0.38 | 0.37 | 0.28 | 0.52 |
| 7 | 155884468 | rs4716985 | -0.809 | -0.611 | 2.004 | 1.769 | 0.051 | 0.014 | 0.014 | 0 | 0.38 | 0.78 | 0.48 | 0.65 | 0.31 | 0.61 |
| 7 | 155887659 | rs6459990 | -1.404 | -0.593 | 2.068 | 1.767 | 0.08 | 0.053 | 0.053 | 0 | 0.06 | 0.37 | 0.04 | 0.01 | 0.02 | 0.08 |
| 7 | 155888203 | rs2178860 | -0.436 | -0.578 | 1.589 | 1.84 | 0.007 | -0.001 | -0.001 | 0 | 0.34 | 0.61 | 0.45 | 0.64 | 0.29 | 0.6 |
| 7 | 155897958 | rs10264945 | -0.376 | -0.887 | 1.312 | 1.63 | -0.019 | 0.007 | 0.007 | 1.314 | 0.35 | 0.54 | 0.46 | 0.55 | 0.26 | 0.55 |
| 7 | 155914367 | rs6979899 | -2.276 | -0.43 | 2.111 | 1.848 | 0.099 | 0.45 | 0.45 | 0.622 | 0.05 | 0.54 | 0.12 | 0.26 | 0.04 | 0.08 |
| 7 | 155923616 | rs741828 | -0.584 | -0.934 | 1.275 | 1.152 | -0.015 | -0.005 | -0.005 | 0.161 | 0.41 | 0.52 | 0.43 | 0.4 | 0.29 | 0.51 |
| 7 | 155928944 | rs10251592 | -0.299 | -1.206 | 1.581 | 0.935 | -0.011 | -0.054 | -0.054 | 0.241 | 0.21 | 0.33 | 0.09 | 0.08 | 0.17 | 0.18 |
| 7 | 155929196 | rs6955791 | -0.328 | -1.212 | 1.724 | 0.933 | -0.011 | -0.062 | -0.062 | 0 | 0.21 | 0.38 | 0.09 | 0.08 | 0.17 | 0.18 |
| 7 | 155929913 | rs4716589 | -0.409 | -0.71 | 1.575 | 0.786 | 0.018 | -0.057 | -0.057 | 0 | 0.28 | 0.37 | 0.18 | 0.08 | 0.32 | 0.22 |
| 7 | 155930663 | rs6969259 | -1.201 | -0.869 | 1.727 | 1.067 | 0.001 | 0.029 | 0.029 | 1.156 | 0.52 | 0.65 | 0.61 | 0.62 | 0.48 | 0.58 |
| 7 | 155931223 | rs6966416 | -1.032 | -0.974 | 1.449 | 1.036 | -0.007 | 0.002 | 0.002 | 0 | 0.57 | 0.59 | 0.62 | 0.6 | 0.51 | 0.57 |
| 7 | 155931361 | rs2178638 | -0.369 | -0.953 | 1.732 | 0.88 | -0.04 | 0.008 | 0.008 | 0.38 | 0.36 | 0.45 | 0.47 | 0.62 | 0.23 | 0.44 |
| 7 | 155932549 | rs17829622 | 0.43 | -0.918 | 1.628 | 0.386 | 0.006 | -0.049 | -0.049 | 0.811 | 0.21 | 0.19 | 0.14 | 0 | 0.29 | 0.15 |
| 7 | 155933093 | rs10267433 | -1.099 | -0.884 | 1.445 | 0.504 | -0.014 | 0.003 | 0.003 | 0 | 0.56 | 0.61 | 0.61 | 0.46 | 0.5 | 0.58 |
| 7 | 155935568 | rs11770182 | -0.025 | -1.461 | 1.16 | 1.039 | 0.031 | -0.016 | -0.016 | 3.274 | 0.27 | 0.2 | 0.42 | 0.37 | 0.18 | 0.31 |
| 7 | 155938607 | rs11760861 | -0.577 | -1.637 | 0.908 | 0.991 | -0.014 | 0.002 | 0.002 | 0.076 | 0.35 | 0.38 | 0.48 | 0.45 | 0.2 | 0.41 |
| 7 | 155942186 | rs11771793 | -0.159 | -1.212 | 0.379 | 1.18 | 0.033 | -0.012 | -0.012 | 0 | 0.28 | 0.22 | 0.31 | 0.35 | 0.16 | 0.31 |
| 7 | 155946592 | rs7793353 | -0.362 | -0.945 | 0.275 | 1.195 | -0.009 | 0 | 0 | 0 | 0.4 | 0.58 | 0.36 | 0.46 | 0.22 | 0.47 |
| 7 | 155952990 | rs4716999 | -0.248 | 0.533 | 0.424 | 1.714 | 0.052 | 0.071 | 0.071 | 0 | 0.2 | 0.36 | 0.28 | 0.38 | 0.3 | 0.15 |
| 7 | 155959222 | rs741829 | -0.663 | -0.747 | 0.502 | 1.781 | -0.073 | 0.105 | 0.105 | 1.081 | 0.28 | 0.48 | 0.37 | 0.54 | 0.17 | 0.47 |
| 7 | 155965622 | rs4717004 | 0.674 | -0.68 | 0.364 | 1.699 | -0.061 | 0.076 | 0.076 | 0.302 | 0.7 | 0.53 | 0.56 | 0.43 | 0.81 | 0.48 |
| 7 | 155971365 | rs7798069 | -1.589 | -0.202 | 0.852 | 2.208 | 0.022 | 0.351 | 0.351 | 0.619 | 0.11 | 0.51 | 0.22 | 0.36 | 0.07 | 0.2 |
| 7 | 155973540 | rs1181908 | 0.31 | -0.031 | -0.913 | 1.507 | -0.081 | 0.039 | 0.039 | 0.474 | 0.42 | 0.05 | 0.23 | 0.29 | 0.4 | 0.48 |
| 7 | 155974701 | rs6961071 | -0.097 | 0.624 | 0.249 | 1.972 | 0.032 | 0.011 | 0.011 | 0.419 | 0.5 | 0.31 | 0.39 | 0.43 | 0.56 | 0.46 |
| 7 | 155974712 | rs12113582 | 0.356 | 0.873 | 0.396 | 1.885 | 0.012 | 0.018 | 0.018 | 0 | 0.09 | 0.11 | 0.16 | 0.17 | 0.16 | 0.05 |
| 7 | 155975007 | rs12113716 | 0.175 | 0.835 | 0.221 | 1.704 | -0.003 | -0.003 | -0.003 | 0 | 0.13 | 0.11 | 0.17 | 0.17 | 0.21 | 0.08 |
| 7 | 155977986 | rs1181954 | -0.453 | 0.462 | 0.065 | 1.012 | 0.002 | -0.007 | -0.007 | 0.807 | 0.64 | 0.35 | 0.48 | 0.53 | 0.61 | 0.66 |
| 7 | 155978138 | rs13232734 | 0.442 | 0.438 | -0.115 | 0.526 | -0.042 | 0.008 | 0.008 | 0 | 0.16 | 0.01 | 0.27 | 0.12 | 0.18 | 0.26 |
| 7 | 155979858 | rs1024158 | -0.661 | 0.752 | 0.086 | 0.402 | 0.015 | -0.003 | -0.003 | 0.478 | 0.45 | 0.61 | 0.56 | 0.44 | 0.42 | 0.55 |
| 7 | 155980396 | rs1014851 | 0.902 | 0.636 | 0.318 | 0.959 | -0.048 | -0.077 | -0.077 | 1.024 | 0.37 | 0.09 | 0.29 | 0.47 | 0.35 | 0.33 |
| 7 | 155989669 | rs10243418 | -0.592 | 0.598 | 0.819 | 0.98 | 0.049 | 0.245 | 0.245 | 0.016 | 0.16 | 0.37 | 0.4 | 0.4 | 0.21 | 0.26 |

| | | | | | | | | | | | | | | | | |
|---|---|---|---|---|---|---|---|---|---|---|---|---|---|---|---|---|
| 7 | 155990813 | rs737681 | -0.36 | 0.256 | 0.816 | 0.699 | -0.002 | -0.044 | -0.044 | 0.025 | 0.57 | 0.5 | 0.7 | 0.86 | 0.61 | 0.59 |
| 7 | 155995102 | rs6460004 | -0.766 | 0.133 | 1.346 | 1.041 | 0.043 | 0.126 | 0.126 | 0.8 | 0.07 | 0.39 | 0.08 | 0.1 | 0.05 | 0.06 |
| 7 | 155997521 | rs1181930 | -1.515 | 0.271 | 1.08 | 1.005 | -0.007 | -0.007 | -0.007 | 0 | 0.24 | 0.19 | 0.51 | 0.62 | 0.34 | 0.47 |
| 7 | 155999146 | rs9638161 | -0.451 | 0.305 | 1.066 | 0.986 | 0.005 | 0.013 | 0.013 | 0.12 | 0.16 | 0.13 | 0.46 | 0.62 | 0.27 | 0.39 |
| 7 | 156002990 | rs1005229 | -0.922 | 0.129 | 1.087 | 0.821 | -0.014 | 0.003 | 0.003 | 0.009 | 0.64 | 0.55 | 0.74 | 0.66 | 0.7 | 0.78 |
| 7 | 156003168 | rs7802594 | 1.23 | 0.115 | 1.049 | 0.706 | -0.011 | 0.162 | 0.162 | 0.176 | 0.37 | 0.15 | 0.21 | 0.04 | 0.36 | 0.17 |
| 7 | 156005470 | rs1181942 | -1.065 | 0.559 | 1.144 | 1.211 | -0.007 | -0.006 | -0.006 | 0 | 0.15 | 0.12 | 0.15 | 0.23 | 0.15 | 0.22 |
| 7 | 156009288 | rs1181912 | -1.573 | 0.54 | 1.314 | 1.368 | -0.008 | -0.002 | -0.002 | 0 | 0.28 | 0.21 | 0.46 | 0.31 | 0.35 | 0.5 |
| 7 | 156010077 | rs13223801 | -0.918 | 0.486 | 1.53 | 1.384 | 0.021 | 0.099 | 0.099 | 1.453 | 0.08 | 0.37 | 0.08 | 0.11 | 0.05 | 0.05 |
| 7 | 156011617 | rs1181911 | -0.24 | 0.487 | 1.553 | 1.458 | -0.009 | 0.038 | 0.038 | 0 | 0.12 | 0.16 | 0.14 | 0.3 | 0.09 | 0.11 |
| 7 | 156011879 | rs4717021 | 1.401 | 0.463 | 1.507 | 1.349 | -0.014 | 0.095 | 0.095 | 0.067 | 0.33 | 0.18 | 0.2 | 0.04 | 0.34 | 0.17 |
| 7 | 156012230 | rs10281910 | -0.11 | 0.42 | 1.509 | 1.411 | 0.013 | 0.078 | 0.078 | 0.198 | 0.06 | 0.24 | 0.07 | 0.11 | 0.05 | 0.05 |
| 7 | 156012707 | rs1181936 | -1.743 | 0.437 | 1.488 | 0.988 | -0.011 | 0.076 | 0.076 | 0 | 0.25 | 0.51 | 0.31 | 0.62 | 0.22 | 0.31 |
| 7 | 156013230 | rs11769720 | 0.078 | 0.443 | 1.39 | 0.989 | 0.006 | 0 | 0 | 1.425 | 0.11 | 0.09 | 0.3 | 0.1 | 0.18 | 0.24 |
| 7 | 156015269 | rs7779135 | 0.436 | 0.38 | 1.327 | 0.929 | -0.007 | 0.076 | 0.076 | 0.773 | 0.44 | 0.29 | 0.51 | 0.14 | 0.52 | 0.41 |
| 7 | 156015899 | rs7783638 | 1.257 | -0.25 | 1.788 | 1.231 | 0.015 | 0.01 | 0.01 | 0.206 | 0.3 | 0.21 | 0.19 | 0.23 | 0.25 | 0.15 |
| 7 | 156016226 | rs1922086 | 0.41 | -0.247 | 1.774 | 1.242 | 0.065 | 0.071 | 0.071 | 0.323 | 0.49 | 0.29 | 0.61 | 0.36 | 0.62 | 0.56 |
| 7 | 156019500 | rs11761957 | 1.032 | -0.081 | 1.352 | 1.122 | 0.013 | 0.009 | 0.009 | 0.114 | 0.3 | 0.21 | 0.19 | 0.24 | 0.26 | 0.16 |
| 7 | 156019596 | rs1689205 | 0.314 | -0.162 | 1.273 | 1.132 | 0.005 | -0.004 | -0.004 | 0 | 0.09 | 0.06 | 0.05 | 0 | 0.07 | 0.08 |
| 7 | 156020486 | rs1493290 | 0.678 | -0.033 | 1.297 | 1.219 | 0 | -0.001 | -0.001 | 0 | 0.46 | 0.48 | 0.45 | 0.67 | 0.48 | 0.38 |
| 7 | 156025396 | rs1175446 | 0.305 | -0.123 | 1.268 | 1.3 | 0.083 | 0.054 | 0.054 | 0.054 | 0.52 | 0.75 | 0.59 | 0.62 | 0.62 | 0.56 |
| 7 | 156025546 | rs4717026 | 0.392 | -0.068 | 0.844 | 1.331 | 0.009 | 0.008 | 0.008 | 0.42 | 0.49 | 0.44 | 0.53 | 0.62 | 0.61 | 0.55 |
| 7 | 156028556 | rs1873884 | -0.567 | 0.33 | 0.712 | 1.783 | -0.022 | 0.031 | 0.031 | 0 | 0.69 | 0.62 | 0.71 | 0.47 | 0.73 | 0.81 |
| 7 | 156029667 | rs1175449 | 0.74 | 0.552 | 0.127 | 1.031 | 0.001 | 0.003 | 0.003 | 0 | 0.53 | 0.45 | 0.63 | 0.57 | 0.52 | 0.44 |
| 7 | 156032479 | rs11766938 | 0.004 | 0.749 | -0.459 | 0.945 | -0.03 | -0.012 | -0.012 | 0 | 0.09 | 0.04 | 0.12 | 0.08 | 0.16 | 0.22 |
| 7 | 156033299 | rs1157520 | -1.03 | 0.761 | -0.457 | 0.959 | -0.012 | -0.009 | -0.009 | 0 | 0.15 | 0.13 | 0.15 | 0.19 | 0.21 | 0.29 |
| 7 | 156037171 | rs1845249 | 0.035 | 0.708 | -0.636 | 0.842 | 0.003 | -0.01 | -0.01 | 0.669 | 0.06 | 0.04 | 0.03 | 0.11 | 0.05 | 0.07 |
| 7 | 156037378 | rs1689199 | 0.274 | 0.742 | -0.66 | 0.825 | 0 | -0.003 | -0.003 | 0.095 | 0.54 | 0.51 | 0.64 | 0.59 | 0.53 | 0.52 |
| 7 | 156040788 | rs1882458 | -1.089 | 1.203 | -0.021 | 0.961 | 0.046 | 0.011 | 0.011 | 0.05 | 0.17 | 0.39 | 0.22 | 0.21 | 0.22 | 0.31 |
| 7 | 156041056 | rs1733657 | 0.856 | 0.364 | -0.164 | 1.304 | -0.014 | 0.085 | 0.085 | 0 | 0.82 | 0.76 | 0.58 | 0.54 | 0.86 | 0.75 |
| 7 | 156043964 | rs7780536 | 0.575 | 0.559 | -0.903 | -0.189 | 0.024 | 0.24 | 0.24 | 0 | 0.48 | 0.19 | 0.46 | 0.07 | 0.58 | 0.47 |
| 7 | 156046163 | rs12673654 | 0.483 | 0.302 | -0.905 | 0.334 | 0.004 | 0.028 | 0.028 | 0.107 | 0.13 | 0.2 | 0.12 | 0.29 | 0.14 | 0.13 |
| 7 | 156046295 | rs12698388 | 0.455 | 0.3 | -0.903 | 0.333 | 0.003 | 0.024 | 0.024 | 0.116 | 0.13 | 0.2 | 0.12 | 0.29 | 0.14 | 0.13 |
| 7 | 156047259 | rs7789165 | 0.031 | 0.812 | -0.342 | 0.117 | -0.032 | -0.086 | -0.086 | 0.916 | 0.8 | 0.55 | 0.84 | 0.92 | 0.88 | 0.78 |
| 7 | 156048322 | rs9638077 | 0.01 | 0.835 | -0.339 | 0.192 | -0.022 | 0.062 | 0.062 | 0 | 0.19 | 0.24 | 0.26 | 0.52 | 0.16 | 0.17 |
| 7 | 156050809 | rs6957969 | -0.679 | 0.883 | -0.191 | -0.018 | 0.013 | -0.098 | -0.098 | 0 | 0.35 | 0.51 | 0.42 | 0.89 | 0.4 | 0.4 |
| 7 | 156053063 | rs11973944 | 0.597 | 0.808 | 0.139 | -0.077 | -0.012 | -0.023 | -0.023 | 0 | 0.1 | 0.12 | 0.03 | 0.04 | 0.05 | 0.11 |
| 7 | 156056343 | rs12386611 | 1.727 | 0.895 | 0.121 | -0.098 | -0.073 | 0.322 | 0.322 | 0.007 | 0.45 | 0.09 | 0.42 | 0.03 | 0.48 | 0.38 |
| 7 | 156056480 | rs9638080 | 0.548 | 1.022 | 0.198 | 0.111 | 0.02 | 0.071 | 0.071 | 0 | 0.84 | 0.65 | 0.88 | 0.68 | 0.83 | 0.71 |
| 7 | 156057327 | rs11977766 | 0.321 | 0.708 | -0.261 | -0.669 | -0.011 | -0.022 | -0.022 | 0.074 | 0.11 | 0.12 | 0.04 | 0.04 | 0.05 | 0.12 |
| 7 | 156062171 | rs9654739 | 0.434 | 0.844 | -0.48 | -0.544 | 0.001 | 0.024 | 0.024 | 0 | 0.54 | 0.38 | 0.54 | 0.08 | 0.55 | 0.5 |
| 7 | 156063460 | rs4717028 | 0.653 | 0.679 | -0.502 | -0.438 | 0 | 0.081 | 0.081 | 0 | 0.48 | 0.31 | 0.51 | 0.07 | 0.49 | 0.41 |
| 7 | 156067200 | rs11769116 | -1.104 | 0.63 | -0.236 | 0.212 | 0.001 | 0.006 | 0.006 | 0.014 | 0.06 | 0.15 | 0.07 | 0.06 | 0.06 | 0.09 |
| 7 | 156067956 | rs4716596 | 0.267 | -0.061 | -1.286 | -1.119 | 0.071 | 0.162 | 0.162 | 0.07 | 0.47 | 0.23 | 0.27 | 0.15 | 0.37 | 0.36 |
| 7 | 156068882 | rs4129489 | -0.239 | -0.36 | -1.078 | -0.616 | 0.01 | -0.008 | -0.008 | 0 | 0.77 | 0.87 | 0.89 | 0.69 | 0.87 | 0.75 |
| 7 | 156069045 | rs4129487 | 0.978 | 0.979 | -0.885 | -0.641 | 0.035 | 0.077 | 0.077 | 0.044 | 0.56 | 0.3 | 0.5 | 0.19 | 0.63 | 0.55 |
| 7 | 156078799 | rs6460015 | -0.564 | 0.667 | -0.955 | -0.73 | -0.02 | -0.041 | -0.041 | 0 | 0.91 | 0.89 | 0.97 | 1 | 0.95 | 0.97 |
| 7 | 156088843 | rs4479988 | 0.144 | 0.646 | -0.294 | -0.432 | 0.015 | 0.017 | 0.017 | 0 | 0.09 | 0.2 | 0.12 | 0.08 | 0.11 | 0.09 |
| 7 | 156094690 | rs13241845 | 0.892 | 0.733 | -0.331 | -0.467 | -0.052 | 0.008 | 0.008 | 1.219 | 0.09 | 0.01 | 0.23 | 0.04 | 0.26 | 0.18 |
| 7 | 156096029 | rs13224994 | -0.903 | 0.701 | -0.218 | -0.477 | 0.035 | -0.002 | -0.002 | 0 | 0.18 | 0.25 | 0.35 | 0.12 | 0.38 | 0.27 |
| 7 | 156099821 | rs6965591 | -1.583 | 0.422 | -0.184 | -0.332 | 0.008 | -0.015 | -0.015 | 0 | 0.29 | 0.72 | 0.41 | 0.52 | 0.28 | 0.22 |
| 7 | 156099856 | rs6969411 | -0.033 | 0.466 | 0.362 | -0.24 | 0 | 0.135 | 0.135 | 0 | 0.12 | 0.21 | 0.31 | 0.49 | 0.15 | 0.15 |
| 7 | 156102061 | rs4435995 | -0.602 | 0.423 | 0.271 | -0.279 | -0.025 | 0.037 | 0.037 | 0.672 | 0.23 | 0.42 | 0.17 | 0.35 | 0.15 | 0.31 |
| 7 | 156111133 | rs4717035 | 0.873 | 0.408 | -0.289 | -0.046 | 0.002 | -0.057 | -0.057 | 0.374 | 0.12 | 0 | 0.03 | 0.16 | 0.07 | 0.2 |
| 7 | 156113163 | rs10281068 | -0.144 | 0.438 | 0.356 | 0.509 | 0.006 | 0.203 | 0.203 | 0.282 | 0.13 | 0.37 | 0.23 | 0.39 | 0.14 | 0.1 |
| 7 | 156114595 | rs6976733 | -0.194 | 0.435 | 0.412 | 0.508 | 0.008 | 0.219 | 0.219 | 0.305 | 0.13 | 0.38 | 0.23 | 0.39 | 0.14 | 0.1 |
| 7 | 156117114 | rs4424129 | -0.974 | -0.714 | -0.607 | -0.318 | 0.042 | 0.047 | 0.047 | 0 | 0.53 | 0.75 | 0.74 | 0.69 | 0.59 | 0.63 |
| 7 | 156117638 | rs6977899 | 0.446 | -0.239 | -0.243 | -0.068 | 0.008 | 0.218 | 0.218 | 0.282 | 0.13 | 0.38 | 0.24 | 0.39 | 0.14 | 0.1 |
| 7 | 156120820 | rs4324820 | -0.007 | 0.108 | -0.801 | -0.303 | -0.001 | 0.013 | 0.013 | 0 | 0.87 | 0.79 | 0.89 | 0.86 | 0.86 | 0.86 |
| 7 | 156122267 | rs4446618 | 1.07 | 0.431 | -1.353 | -0.492 | 0.032 | 0.051 | 0.051 | 0 | 0.2 | 0.06 | 0.14 | 0.04 | 0.12 | 0.25 |
| 7 | 156128006 | rs9638177 | 0.557 | 1.437 | -0.92 | 0.223 | -0.059 | -0.02 | -0.02 | 0 | 0.09 | 0.01 | 0.28 | 0.13 | 0.26 | 0.21 |
| 7 | 156135838 | rs10252968 | 0.306 | -0.828 | -0.689 | 0.302 | 0.037 | -0.006 | -0.006 | 0 | 0.79 | 0.88 | 0.92 | 0.83 | 0.91 | 0.78 |
| 7 | 156142059 | rs9647707 | 1.236 | -0.764 | -0.04 | 0.916 | -0.034 | 0.053 | 0.053 | 0.026 | 0.68 | 0.55 | 0.71 | 0.51 | 0.76 | 0.58 |
| 7 | 156142309 | rs4717043 | 0.089 | -0.643 | -0.475 | 0.364 | -0.002 | 0.013 | 0.013 | 0 | 0.32 | 0.43 | 0.42 | 0.4 | 0.31 | 0.38 |
| 7 | 156142454 | rs4716598 | 0.975 | -0.586 | -0.382 | 0.464 | -0.037 | 0.053 | 0.053 | 0.144 | 0.68 | 0.51 | 0.7 | 0.51 | 0.76 | 0.58 |
| 7 | 156148608 | rs6945016 | -0.184 | -0.114 | -0.346 | 0.594 | 0.004 | 0.158 | 0.158 | 0.01 | 0.1 | 0.27 | 0.19 | 0.31 | 0.11 | 0.09 |
| 7 | 156151697 | rs6950584 | 1.5 | -0.206 | -0.899 | -0.017 | -0.105 | 0.193 | 0.193 | 0.052 | 0.37 | 0.08 | 0.28 | 0.1 | 0.44 | 0.25 |
| 7 | 156155083 | rs4555974 | 0.461 | -1.34 | -0.343 | 1.107 | -0.028 | 0.033 | 0.033 | 0 | 0.8 | 0.67 | 0.88 | 0.67 | 0.9 | 0.78 |
| 7 | 156157753 | rs10258969 | 2 | -1.643 | -0.481 | 0.9 | 0.042 | 0.159 | 0.159 | 1.35 | 0.49 | 0.19 | 0.48 | 0.21 | 0.57 | 0.53 |

| | | | | | | | | | | | | | | | | |
|---|---|---|---|---|---|---|---|---|---|---|---|---|---|---|---|---|
| 7 | 156158751 | rs4716600 | 0.237 | -1.721 | -0.554 | 1.135 | 0.008 | -0.051 | -0.051 | 0.267 | 0.17 | 0.01 | 0.06 | 0.2 | 0.09 | 0.22 |
| 7 | 156159448 | rs10249494 | 1.155 | -1.821 | -0.076 | 1.234 | -0.009 | 0.013 | 0.013 | 0 | 0.63 | 0.52 | 0.7 | 0.54 | 0.71 | 0.66 |
| 7 | 156159585 | rs12669717 | -0.889 | -1.821 | -0.1 | 1.243 | -0.025 | 0.145 | 0.145 | 0.257 | 0.14 | 0.34 | 0.22 | 0.33 | 0.14 | 0.13 |
| 7 | 156160671 | rs4717047 | 1.112 | -1.851 | -0.077 | 1.194 | -0.001 | 0.003 | 0.003 | 0 | 0.65 | 0.58 | 0.71 | 0.54 | 0.71 | 0.66 |
| 7 | 156161905 | rs11978423 | -1.017 | -1.767 | -0.208 | 1.239 | -0.028 | 0.143 | 0.143 | 0.559 | 0.15 | 0.35 | 0.22 | 0.33 | 0.14 | 0.12 |
| 7 | 156165232 | rs12535876 | 2.863 | -1.505 | -0.666 | 0.83 | -0.082 | 0.121 | 0.121 | 0.715 | 0.35 | 0.05 | 0.36 | 0.13 | 0.4 | 0.28 |
| 7 | 156169927 | rs3912927 | -1.15 | -1.169 | -0.07 | 1.258 | 0.002 | 0.102 | 0.102 | 0 | 0.27 | 0.51 | 0.36 | 0.51 | 0.31 | 0.36 |
| 7 | 156171735 | rs11763840 | 1.42 | -1.362 | 0.126 | 1.431 | -0.041 | -0.011 | -0.011 | 0.043 | 0.7 | 0.46 | 0.66 | 0.24 | 0.78 | 0.68 |
| 7 | 156172079 | rs6953059 | -1.061 | -1.013 | 0.549 | 1.256 | 0.009 | -0.032 | -0.032 | 0 | 0.3 | 0.35 | 0.26 | 0.1 | 0.35 | 0.35 |
| 7 | 156173632 | rs10235979 | 0.962 | -1.166 | 0.327 | 1.363 | -0.038 | 0.03 | 0.03 | 0 | 0.76 | 0.6 | 0.83 | 0.68 | 0.84 | 0.79 |
| 7 | 156178305 | rs4716603 | 0.281 | -1.068 | -0.524 | 1.371 | -0.003 | -0.045 | -0.045 | 1.64 | 0.14 | 0.01 | 0.05 | 0.19 | 0.09 | 0.16 |
| 7 | 156184748 | rs10246481 | 1.885 | -0.973 | -0.4 | 1.33 | -0.002 | -0.048 | -0.048 | 0.278 | 0.44 | 0.14 | 0.5 | 0.47 | 0.44 | 0.39 |
| 7 | 156187901 | rs11769797 | 0.568 | -0.853 | -0.319 | 1.317 | -0.006 | 0.002 | 0.002 | 0 | 0.11 | 0.1 | 0.11 | 0.08 | 0.16 | 0.22 |
| 7 | 156194185 | rs7808581 | 1.488 | -0.856 | -0.456 | 1.304 | -0.03 | -0.049 | -0.049 | 0 | 0.48 | 0.12 | 0.53 | 0.48 | 0.5 | 0.39 |
| 7 | 156207092 | rs1917035 | -0.77 | -0.804 | 0.228 | 1.69 | -0.015 | 0.108 | 0.108 | 0 | 0.1 | 0.2 | 0.18 | 0.32 | 0.1 | 0.11 |
| 7 | 156208176 | rs1880342 | 1.574 | -1.032 | 0.324 | 1.61 | -0.002 | 0.132 | 0.132 | 0.797 | 0.51 | 0.19 | 0.41 | 0.26 | 0.51 | 0.47 |
| 7 | 156208807 | rs2365275 | -0.542 | -0.898 | 0.388 | 1.755 | -0.015 | 0.109 | 0.109 | 0 | 0.09 | 0.19 | 0.18 | 0.31 | 0.09 | 0.1 |
| 7 | 156220710 | rs4716606 | -0.697 | -0.869 | 0.413 | 1.759 | -0.008 | 0.114 | 0.114 | 0 | 0.1 | 0.23 | 0.2 | 0.33 | 0.1 | 0.11 |
| 7 | 156226401 | rs10949589 | -0.233 | -0.846 | 0.105 | 1.752 | -0.04 | -0.079 | -0.079 | 0.058 | 0.16 | 0.06 | 0.28 | 0.31 | 0.18 | 0.17 |
| 7 | 156227374 | rs6954868 | 0.487 | -0.836 | 0.762 | 1.859 | 0 | 0 | 0 | 0 | 0.69 | 0.35 | 0.68 | 0.56 | 0.68 | 0.64 |
| 7 | 156228281 | rs1917030 | 0.516 | -0.836 | 0.763 | 1.86 | 0 | -0.001 | -0.001 | 0.34 | 0.69 | 0.35 | 0.68 | 0.56 | 0.68 | 0.64 |
| 7 | 156231105 | rs6961453 | 1.142 | -0.591 | 0.04 | 1.275 | -0.008 | 0.07 | 0.07 | 1.524 | 0.48 | 0.29 | 0.38 | 0.27 | 0.47 | 0.42 |
| 7 | 156236883 | rs2365281 | 0.029 | -0.264 | 0.14 | 1.434 | -0.003 | 0.005 | 0.005 | 0 | 0.07 | 0.09 | 0.14 | 0.16 | 0.05 | 0.11 |
| 7 | 156238371 | rs2140634 | 0.417 | -0.225 | 0.421 | 1.588 | 0.003 | 0.014 | 0.014 | 0 | 0.78 | 0.72 | 0.8 | 0.73 | 0.73 | 0.73 |
| 7 | 156239128 | rs1534002 | 1.052 | -0.559 | 0.819 | 1.709 | -0.017 | 0.142 | 0.142 | 0 | 0.87 | 0.63 | 0.78 | 0.67 | 0.89 | 0.82 |
| 7 | 156243562 | rs4716610 | 1.166 | -0.467 | 0.229 | 0.915 | -0.002 | 0.01 | 0.01 | 0.152 | 0.4 | 0.34 | 0.38 | 0.22 | 0.38 | 0.36 |
| 7 | 156249769 | rs10270285 | 1.72 | -0.957 | 0.865 | 1.935 | -0.015 | 0.017 | 0.017 | 1.114 | 0.75 | 0.34 | 0.78 | 0.47 | 0.88 | 0.81 |
| 7 | 156250444 | rs7778219 | 1.745 | -0.975 | 1.416 | 1.909 | -0.006 | 0.048 | 0.048 | 1.402 | 0.86 | 0.39 | 0.84 | 0.79 | 0.89 | 0.85 |
| 7 | 156251505 | rs2140631 | 1.7 | -1.322 | 1.597 | 1.835 | -0.055 | 0.05 | 0.05 | 0.595 | 0.73 | 0.58 | 0.81 | 0.65 | 0.85 | 0.83 |
| 7 | 156252939 | rs868822 | 2.617 | -1.327 | 1.53 | 1.819 | 0.071 | -0.012 | -0.012 | 0 | 0.61 | 0.19 | 0.67 | 0.44 | 0.74 | 0.68 |
| 7 | 156253492 | rs2280716 | -1.916 | -1.392 | 1.66 | 2.315 | 0.017 | -0.013 | -0.013 | 0 | 0.27 | 0.22 | 0.27 | 0.33 | 0.19 | 0.25 |
| 7 | 156254799 | rs7795670 | -2.302 | -1.23 | 1.99 | 2.288 | -0.035 | 0.025 | 0.025 | 0 | 0.27 | 0.39 | 0.26 | 0.56 | 0.19 | 0.27 |
| 7 | 156255115 | rs6459678 | 2.5 | -1.112 | 1.801 | 2.518 | 0.034 | 0.222 | 0.222 | 1.228 | 0.92 | 0.59 | 0.9 | 0.77 | 0.92 | 0.9 |
| 7 | 156255343 | rs6967289 | 2.215 | -1.11 | 1.765 | 2.523 | 0.024 | 0.234 | 0.234 | 1.662 | 0.93 | 0.63 | 0.91 | 0.77 | 0.93 | 0.9 |
| 7 | 156261602 | rs11766070 | 1.947 | -1.094 | 1.363 | 2.318 | -0.043 | 0.026 | 0.026 | 0.661 | 0.67 | 0.48 | 0.74 | 0.63 | 0.74 | 0.72 |
| 7 | 156266712 | rs1405809 | -1.93 | -0.747 | 0.48 | 1.77 | 0.013 | -0.004 | -0.004 | 0.265 | 0.36 | 0.66 | 0.32 | 0.58 | 0.26 | 0.3 |
| 7 | 156268109 | rs6957217 | -1.587 | -0.433 | 0.75 | 2.032 | -0.009 | 0.012 | 0.012 | 0.296 | 0.28 | 0.64 | 0.26 | 0.58 | 0.19 | 0.25 |
| 7 | 156274754 | rs6459682 | -1.748 | -0.257 | 1.245 | 2.25 | -0.024 | 0.224 | 0.224 | 1.248 | 0.19 | 0.43 | 0.16 | 0.49 | 0.16 | 0.23 |
| 7 | 156284040 | rs12669392 | 0.686 | 0.839 | 1.259 | 1.265 | 0.017 | -0.052 | -0.052 | 0.213 | 0.74 | 0.67 | 0.64 | 0.9 | 0.64 | 0.8 |
| 7 | 156296060 | rs2177559 | 1.849 | 0.848 | 1.64 | 2.471 | 0.013 | 0.233 | 0.233 | 2.291 | 0.88 | 0.59 | 0.79 | 0.6 | 0.85 | 0.72 |
| 7 | 156298516 | rs10235614 | 1.221 | 0.735 | 0.697 | 1.827 | 0.069 | 0.097 | 0.097 | 0 | 0.48 | 0.37 | 0.25 | 0.19 | 0.33 | 0.43 |
| 7 | 156299271 | rs1527575 | 0.052 | 2.046 | 0.18 | 2.278 | -0.025 | -0.025 | -0.025 | 0 | 0.8 | 0.9 | 0.66 | 0.38 | 0.61 | 0.76 |
| 7 | 156305215 | rs6459686 | 1.121 | 2.138 | 0.589 | 2.365 | 0.003 | -0.005 | -0.005 | 0.321 | 0.7 | 0.73 | 0.63 | 0.77 | 0.7 | 0.54 |
| 7 | 156307861 | rs12668409 | -0.083 | 2.262 | 0.091 | 2.333 | -0.039 | -0.04 | -0.04 | 0 | 0.18 | 0.08 | 0.33 | 0.63 | 0.36 | 0.23 |
| 7 | 156308688 | rs2177558 | -1.005 | 2.192 | 0.304 | 2.341 | -0.01 | 0.018 | 0.018 | 0.196 | 0.24 | 0.17 | 0.3 | 0.08 | 0.26 | 0.31 |
| 7 | 156321315 | rs6972602 | -1.167 | 2.383 | 0.026 | 2.477 | -0.005 | 0.002 | 0.002 | 1.426 | 0.24 | 0.28 | 0.31 | 0.07 | 0.27 | 0.31 |
| 7 | 156325320 | rs11771276 | -0.237 | 3.085 | -0.69 | 2.682 | -0.02 | -0.012 | -0.012 | 0 | 0.14 | 0.07 | 0.33 | 0.5 | 0.34 | 0.21 |
| 7 | 156327884 | rs1543989 | 0.041 | 2.928 | -0.662 | 2.642 | -0.013 | -0.011 | -0.011 | 0.014 | 0.13 | 0.07 | 0.12 | 0.21 | 0.26 | 0.12 |
| 7 | 156332575 | rs1534012 | -0.162 | 2.931 | -0.68 | 2.354 | 0.02 | -0.004 | -0.004 | 0 | 0.15 | 0.15 | 0.14 | 0.17 | 0.25 | 0.14 |
| 7 | 156334560 | rs6459690 | 0.719 | 2.639 | -1.093 | 2.814 | -0.036 | -0.037 | -0.037 | 0.187 | 0.88 | 0.95 | 0.77 | 0.7 | 0.77 | 0.78 |
| 7 | 156343858 | rs12534057 | -0.489 | 2.487 | -1.125 | 2.618 | -0.023 | -0.063 | -0.063 | 0 | 0.12 | 0.01 | 0.26 | 0.29 | 0.14 | 0.17 |
| 7 | 156347201 | rs9918628 | 0.854 | 2.386 | -0.673 | 2.357 | -0.022 | -0.015 | -0.015 | 1.171 | 0.88 | 0.93 | 0.77 | 0.69 | 0.77 | 0.8 |
| 7 | 156351099 | rs6459692 | -1.215 | 0.925 | -0.071 | 1.914 | -0.036 | 0.052 | 0.052 | 3.751 | 0.17 | 0.62 | 0.27 | 0.22 | 0.12 | 0.19 |
| 7 | 156352337 | rs13236262 | -0.215 | 0.726 | -0.819 | 1.538 | -0.001 | 0 | 0 | 0.746 | 0.37 | 0.4 | 0.3 | 0.49 | 0.38 | 0.44 |
| 7 | 156353029 | rs849084 | 0.416 | 1.16 | -0.922 | 1.715 | -0.001 | 0 | 0 | 0.672 | 0.54 | 0.54 | 0.57 | 0.44 | 0.51 | 0.57 |
| 7 | 156353250 | rs10244562 | 0.406 | 0.34 | -1.031 | 1.521 | -0.069 | 0.029 | 0.029 | 1.774 | 0.17 | 0.39 | 0.1 | 0.13 | 0.05 | 0.09 |
| 7 | 156354315 | rs849082 | 0.689 | 0.216 | -1.062 | 1.643 | -0.032 | 0.098 | 0.098 | 0 | 0.73 | 0.56 | 0.74 | 0.62 | 0.84 | 0.72 |
| 7 | 156354359 | rs849081 | 0.441 | 0.22 | -1.06 | 1.656 | -0.024 | 0.037 | 0.037 | 0.045 | 0.75 | 0.59 | 0.86 | 0.73 | 0.84 | 0.8 |
| 7 | 156354448 | rs849080 | 0.767 | 0.27 | -1.182 | 1.51 | -0.015 | 0.044 | 0.044 | 0.001 | 0.72 | 0.44 | 0.71 | 0.53 | 0.82 | 0.6 |
| 7 | 156359656 | rs6964104 | 0.087 | 0.159 | -1.139 | 1.349 | -0.082 | 0.089 | 0.089 | 0.538 | 0.19 | 0.49 | 0.22 | 0.23 | 0.06 | 0.18 |
| 7 | 156362735 | rs849095 | 1.117 | 0.488 | -1.55 | 0.91 | -0.109 | 0.193 | 0.193 | 0 | 0.34 | 0.02 | 0.43 | 0.13 | 0.43 | 0.31 |
| 7 | 156368530 | rs4716636 | 1.028 | 0.03 | -1.18 | 1.552 | 0.041 | -0.009 | -0.009 | 0.24 | 0.64 | 0.23 | 0.68 | 0.46 | 0.79 | 0.57 |
| 7 | 156371216 | rs2695582 | 0.772 | 0.013 | -0.66 | 1.237 | -0.012 | 0.007 | 0.007 | 0.561 | 0.56 | 0.51 | 0.65 | 0.56 | 0.67 | 0.65 |
| 7 | 156373401 | rs2695573 | 0.573 | 0.053 | -0.91 | 0.805 | -0.029 | 0.058 | 0.058 | 0 | 0.6 | 0.71 | 0.64 | 0.8 | 0.57 | 0.63 |
| 7 | 156374358 | rs6960855 | -0.194 | 0.08 | -1.051 | 0.774 | -0.086 | 0.083 | 0.083 | 0 | 0.71 | 0.94 | 0.86 | 0.87 | 0.67 | 0.72 |
| 7 | 156376362 | rs849063 | -0.077 | 0.024 | -1.071 | 0.723 | -0.081 | 0.034 | 0.034 | 0 | 0.69 | 0.89 | 0.73 | 0.8 | 0.65 | 0.65 |
| 7 | 156382758 | rs17837523 | 0.047 | -0.232 | -0.18 | 1.118 | -0.004 | 0.066 | 0.066 | 0 | 0.1 | 0.22 | 0.08 | 0.19 | 0.11 | 0.13 |
| 7 | 156383237 | rs878715 | 0.759 | -0.424 | -0.165 | 1.554 | -0.002 | 0.009 | 0.009 | 0 | 0.49 | 0.55 | 0.52 | 0.46 | 0.54 | 0.53 |
| 7 | 156395495 | rs2365502 | 0.404 | -0.395 | -0.119 | 1.807 | -0.03 | 0.179 | 0.179 | 0 | 0.1 | 0.18 | 0.16 | 0.45 | 0.05 | 0.17 |

| | | | | | | | | | | | | | | | | |
|---|---|---|---|---|---|---|---|---|---|---|---|---|---|---|---|---|
| 7 | 156396471 | rs3778929 | -0.529 | 0.474 | -1.047 | 0.588 | -0.02 | 0.018 | 0.018 | 0 | 0.35 | 0.26 | 0.23 | 0.3 | 0.34 | 0.37 |
| 7 | 156396540 | rs3778928 | 1.044 | 0.836 | -0.953 | 0.754 | -0.031 | -0.013 | -0.013 | 0 | 0.09 | 0.02 | 0.06 | 0.08 | 0.07 | 0.12 |
| 7 | 156398403 | rs740539 | 1.135 | 0.917 | -0.524 | 0.895 | -0.007 | 0.043 | 0.043 | 0.386 | 0.42 | 0.3 | 0.42 | 0.29 | 0.44 | 0.45 |
| 7 | 156404062 | rs10266933 | -0.151 | 0.842 | -0.961 | 1.217 | 0.003 | -0.077 | -0.077 | 0 | 0.7 | 0.87 | 0.63 | 0.51 | 0.72 | 0.66 |
| 7 | 156405131 | rs2365504 | 0.93 | 0.815 | -0.605 | 0.904 | -0.002 | 0.001 | 0.001 | 0 | 0.16 | 0.16 | 0.12 | 0.11 | 0.18 | 0.15 |
| 7 | 156405911 | rs7801996 | -0.987 | 0.177 | -0.046 | 1.299 | -0.054 | 0.123 | 0.123 | 0.946 | 0.22 | 0.39 | 0.18 | 0.43 | 0.1 | 0.18 |
| 7 | 156407224 | rs737982 | -1.056 | 0.843 | -0.661 | 0.97 | -0.044 | 0.049 | 0.049 | 0 | 0.81 | 0.96 | 0.87 | 0.9 | 0.77 | 0.8 |
| 7 | 156410334 | rs740541 | -1.181 | 0.624 | -0.033 | 1.508 | 0.004 | 0.113 | 0.113 | 0.204 | 0.14 | 0.58 | 0.14 | 0.25 | 0.11 | 0.11 |
| 7 | 156413196 | rs849072 | 0.647 | 0.533 | -0.566 | 1.177 | -0.003 | 0 | 0 | 0.182 | 0.62 | 0.62 | 0.73 | 0.62 | 0.67 | 0.66 |
| 7 | 156413721 | rs6459694 | 0.084 | 0.79 | -0.577 | 1.181 | 0.049 | 0.004 | 0.004 | 0 | 0.26 | 0.5 | 0.24 | 0.26 | 0.29 | 0.18 |
| 7 | 156419984 | rs849087 | 1.315 | 0.783 | -1.342 | 0.827 | 0.025 | 0.003 | 0.003 | 0 | 0.39 | 0.17 | 0.51 | 0.36 | 0.39 | 0.48 |
| 7 | 156420066 | rs849086 | -0.305 | 0.525 | -1.3 | 0.93 | -0.01 | 0.034 | 0.034 | 0 | 0.73 | 0.84 | 0.81 | 0.84 | 0.71 | 0.7 |
| 7 | 156426292 | rs13237363 | -0.523 | 0.646 | -0.792 | 1.366 | 0.001 | 0.011 | 0.011 | 0 | 0.32 | 0.63 | 0.3 | 0.48 | 0.29 | 0.21 |
| 7 | 156428211 | rs1016061 | -0.918 | 0.686 | -0.945 | 1.183 | -0.009 | -0.001 | -0.001 | 0 | 0.37 | 0.67 | 0.49 | 0.6 | 0.36 | 0.35 |
| 7 | 156431554 | rs3735189 | 0.517 | 0.554 | -1.055 | 1.711 | 0.002 | 0.012 | 0.012 | 0 | 0.09 | 0.06 | 0.14 | 0.28 | 0.05 | 0.11 |
| 7 | 156431731 | rs7805578 | -0.668 | 0.333 | -0.595 | 1.868 | -0.032 | 0.175 | 0.175 | 0 | 0.13 | 0.29 | 0.16 | 0.42 | 0.06 | 0.14 |
| 7 | 156438462 | rs10243532 | -0.016 | 0.587 | -0.87 | 1.509 | 0.079 | 0.068 | 0.068 | 0 | 0.25 | 0.53 | 0.41 | 0.35 | 0.34 | 0.25 |
| 7 | 156442035 | rs4716437 | 0.076 | 0.669 | -0.884 | 1.524 | 0.063 | 0.049 | 0.049 | 0 | 0.24 | 0.52 | 0.37 | 0.31 | 0.3 | 0.19 |
| 7 | 156463980 | rs2365509 | -0.441 | -0.23 | -1.915 | 0.684 | -0.064 | 0.056 | 0.056 | 0.303 | 0.59 | 0.87 | 0.64 | 0.7 | 0.49 | 0.61 |
| 7 | 156464214 | rs7790357 | 0.81 | -0.362 | -1.9 | 0.703 | 0.048 | -0.003 | -0.003 | 0 | 0.33 | 0.14 | 0.25 | 0.29 | 0.2 | 0.34 |
| 7 | 156467241 | rs7805523 | -0.375 | -0.147 | -2.018 | 0.717 | -0.045 | 0.074 | 0.074 | 0.042 | 0.58 | 0.84 | 0.64 | 0.72 | 0.49 | 0.61 |
| 7 | 156469096 | rs2302149 | -0.509 | 0.178 | -1.56 | 0.611 | 0.023 | 0.002 | 0.002 | 0 | 0.37 | 0.53 | 0.45 | 0.41 | 0.42 | 0.34 |
| 7 | 156469179 | rs2302146 | -0.227 | 0.464 | -1.032 | 1.106 | 0.072 | 0.046 | 0.046 | 0 | 0.28 | 0.53 | 0.42 | 0.41 | 0.36 | 0.26 |
| 7 | 156470258 | rs4716439 | -0.193 | 0.588 | -1.16 | 1.128 | 0.084 | 0.057 | 0.057 | 0 | 0.27 | 0.5 | 0.41 | 0.41 | 0.35 | 0.26 |
| 7 | 156471867 | rs2240642 | 0.181 | -0.518 | -1.679 | 1.01 | 0.06 | -0.028 | -0.028 | 0 | 0.19 | 0.04 | 0.06 | 0.17 | 0.09 | 0.25 |
| 7 | 156474683 | rs17837686 | -1.233 | 0.535 | -0.424 | 1.484 | 0.094 | 0.182 | 0.182 | 0 | 0.06 | 0.28 | 0.11 | 0.16 | 0.09 | 0.17 |
| 7 | 156476389 | rs3487 | -0.13 | 1.107 | -1.379 | 1.597 | -0.082 | -0.133 | -0.133 | 0 | 0.2 | 0.02 | 0.15 | 0.43 | 0.23 | 0.21 |
| 7 | 156477064 | rs10234525 | -1.366 | 1.16 | -0.966 | 1.539 | 0.083 | 0.119 | 0.119 | 0 | 0.09 | 0.33 | 0.15 | 0.15 | 0.11 | 0.18 |
| 7 | 156484201 | rs4716650 | 0.346 | 1.144 | -1.341 | 0.768 | -0.015 | -0.027 | -0.027 | 0 | 0.38 | 0.52 | 0.42 | 0.26 | 0.34 | 0.36 |
| 7 | 156497533 | rs12671098 | 1.9 | 0.894 | -1.642 | 0.407 | -0.032 | 0.011 | 0.011 | 0.429 | 0.12 | 0.01 | 0.14 | 0.08 | 0.16 | 0.22 |
| 7 | 156497668 | rs10228997 | -1.259 | 1.155 | -1.318 | 0.483 | -0.006 | 0.064 | 0.064 | 0.118 | 0.66 | 0.87 | 0.7 | 0.83 | 0.65 | 0.75 |
| 7 | 156507281 | rs4716655 | -0.507 | 1.135 | -1.302 | 0.507 | 0.001 | -0.008 | -0.008 | 0 | 0.58 | 0.55 | 0.57 | 0.68 | 0.57 | 0.57 |
| 7 | 156516951 | rs7793618 | -0.858 | 1.311 | -1.195 | 0.52 | 0.064 | 0.102 | 0.102 | 0 | 0.08 | 0.28 | 0.11 | 0.15 | 0.11 | 0.17 |
| 7 | 156518007 | rs2072468 | -0.085 | 1.504 | -1.383 | 0.823 | -0.004 | 0.001 | 0.001 | 0 | 0.45 | 0.52 | 0.49 | 0.51 | 0.41 | 0.45 |
| 7 | 156521894 | rs10265924 | -0.214 | 1.481 | -1.127 | 0.605 | 0.029 | 0.079 | 0.079 | 0 | 0.06 | 0.16 | 0.09 | 0.15 | 0.09 | 0.17 |
| 7 | 156524010 | rs10260986 | -1.35 | 1.451 | -1.161 | 0.58 | -0.041 | 0.074 | 0.074 | 0.557 | 0.66 | 0.95 | 0.72 | 0.84 | 0.66 | 0.75 |
| 7 | 156525513 | rs10265774 | -2.1 | 1.72 | -0.398 | 1.054 | 0.087 | 0.078 | 0.078 | 0 | 0.1 | 0.28 | 0.17 | 0.15 | 0.16 | 0.18 |
| 7 | 156537630 | rs6956930 | -0.653 | 1.677 | -0.406 | 1.049 | 0.022 | 0.051 | 0.051 | 0 | 0.06 | 0.12 | 0.09 | 0.14 | 0.09 | 0.17 |
| 7 | 156555475 | rs11977054 | -2.383 | 1.628 | -0.265 | 1.093 | 0.059 | 0.075 | 0.075 | 0 | 0.12 | 0.32 | 0.18 | 0.19 | 0.16 | 0.2 |
| 7 | 156562304 | rs7778704 | -2.355 | 1.627 | -0.059 | 1.086 | 0.102 | 0.124 | 0.124 | 0.018 | 0.13 | 0.43 | 0.19 | 0.19 | 0.16 | 0.2 |
| 7 | 156577354 | rs7788421 | 0.212 | 1.483 | -0.52 | 1.032 | -0.015 | -0.005 | -0.005 | 0 | 0.08 | 0.03 | 0.04 | 0.07 | 0.1 | 0.06 |
| 7 | 156586675 | rs1017137 | 0.553 | 2.319 | -0.655 | 1.248 | -0.033 | -0.038 | -0.038 | 2.388 | 0.15 | 0.1 | 0.34 | 0.4 | 0.23 | 0.19 |
| 7 | 156593270 | rs6945929 | -1.209 | 2.403 | -0.449 | 0.971 | 0.072 | 0.051 | 0.051 | 0 | 0.28 | 0.41 | 0.55 | 0.59 | 0.42 | 0.41 |
| 7 | 156599974 | rs9638092 | 0.382 | 2.104 | -0.085 | 0.831 | 0 | 0 | 0 | 0 | 0.12 | 0.11 | 0.04 | 0.07 | 0.07 | 0.07 |
| 7 | 156605489 | rs1861119 | -0.682 | 2.505 | 0.014 | 1.142 | 0.034 | 0.053 | 0.053 | 0 | 0.23 | 0.3 | 0.45 | 0.54 | 0.34 | 0.28 |
| 7 | 156632237 | rs2192510 | -0.986 | 2.459 | -0.197 | 1.112 | -0.03 | -0.056 | -0.056 | 2.526 | 0.91 | 0.99 | 0.9 | 0.77 | 0.89 | 0.94 |
| 7 | 156635651 | rs11770690 | 1.357 | 2.467 | 0.321 | 1.211 | -0.05 | -0.066 | -0.066 | 0 | 0.1 | 0.01 | 0.2 | 0.28 | 0.12 | 0.13 |
| 7 | 156643794 | rs11766294 | -0.86 | 2.364 | 0.389 | 1.111 | 0.035 | 0.004 | 0.004 | 0 | 0.05 | 0.12 | 0.1 | 0.06 | 0.09 | 0.13 |
| 7 | 156655570 | rs2192509 | -0.86 | 2.396 | 0.425 | 1.115 | 0.063 | 0.09 | 0.09 | 0 | 0.23 | 0.37 | 0.46 | 0.54 | 0.34 | 0.28 |
| 7 | 156664783 | rs6968944 | -1.211 | 2.283 | 0.254 | 1.029 | 0.069 | 0.065 | 0.065 | 0.212 | 0.25 | 0.39 | 0.49 | 0.46 | 0.35 | 0.34 |
| 7 | 156679621 | rs11764953 | -0.238 | 2.236 | -0.104 | 1.073 | 0.006 | 0.01 | 0.01 | 0 | 0.19 | 0.2 | 0.38 | 0.4 | 0.27 | 0.21 |
| 7 | 156698469 | rs870917 | -1.17 | 2.132 | -0.129 | 0.993 | 0.016 | 0.018 | 0.018 | 0 | 0.29 | 0.39 | 0.52 | 0.58 | 0.36 | 0.42 |
| 7 | 156698770 | rs870916 | -0.323 | 2.268 | -0.085 | 1.098 | 0.008 | 0.026 | 0.026 | 0 | 0.19 | 0.26 | 0.35 | 0.46 | 0.22 | 0.24 |
| 7 | 156701918 | rs11760288 | -0.312 | 2.262 | -0.087 | 1.095 | 0.008 | 0.026 | 0.026 | 0 | 0.19 | 0.26 | 0.35 | 0.46 | 0.22 | 0.24 |
| 7 | 156708467 | rs2969123 | -1.493 | 1.533 | -0.002 | 0.426 | 0.025 | 0.033 | 0.033 | 0.108 | 0.71 | 0.79 | 0.86 | 0.98 | 0.78 | 0.82 |
| 7 | 156711661 | rs2969124 | 0.37 | 1.63 | 0.064 | 0.5 | -0.014 | -0.002 | -0.002 | 0 | 0.43 | 0.42 | 0.32 | 0.4 | 0.38 | 0.4 |
| 7 | 156722649 | rs11980132 | -0.23 | 1.76 | 0.376 | 0.682 | -0.002 | 0.001 | 0.001 | 0.531 | 0.2 | 0.23 | 0.37 | 0.47 | 0.25 | 0.35 |
| 7 | 156726012 | rs1541506 | 1.197 | 1.264 | 0.118 | 0.283 | 0.018 | 0.063 | 0.063 | 0 | 0.25 | 0.09 | 0.19 | 0.09 | 0.25 | 0.3 |
| 7 | 156728668 | rs2969114 | -0.704 | 1.482 | 0.492 | 0.505 | 0.001 | -0.013 | -0.013 | 0.399 | 0.69 | 0.69 | 0.78 | 0.92 | 0.75 | 0.7 |
| 7 | 156732238 | rs4716445 | -0.226 | 1.412 | 0.475 | 0.612 | 0 | -0.002 | -0.002 | 0 | 0.61 | 0.56 | 0.65 | 0.7 | 0.69 | 0.69 |
| 7 | 156736705 | rs4716661 | -0.877 | 1.288 | 0.774 | 0.604 | -0.003 | -0.049 | -0.049 | 2.231 | 0.74 | 0.67 | 0.81 | 0.93 | 0.77 | 0.82 |
| 7 | 156748041 | rs12670999 | -1.048 | 0.807 | 0.681 | 0.972 | 0.002 | -0.002 | -0.002 | 0 | 0.76 | 0.75 | 0.85 | 0.74 | 0.8 | 0.79 |
| 7 | 156750695 | rs887614 | 0.461 | 0.827 | 0.886 | 1.042 | 0 | -0.007 | -0.007 | 0 | 0.54 | 0.42 | 0.51 | 0.27 | 0.56 | 0.44 |
| 7 | 156751230 | rs887611 | 0.379 | 0.847 | 0.88 | 1.052 | -0.003 | 0.008 | 0.008 | 0.051 | 0.54 | 0.53 | 0.52 | 0.27 | 0.56 | 0.44 |
| 7 | 156754981 | rs3750131 | 0.143 | 1 | 0.953 | 1.211 | 0.019 | 0.016 | 0.016 | 1.07 | 0.17 | 0.27 | 0.28 | 0.29 | 0.2 | 0.28 |
| 7 | 156757453 | rs10274639 | -1.278 | 0.903 | 1.011 | 1.231 | 0.011 | -0.013 | -0.013 | 1.79 | 0.77 | 0.84 | 0.86 | 0.74 | 0.81 | 0.8 |
| 7 | 156757513 | rs10256184 | -1.038 | 0.922 | 1.013 | 1.248 | 0.006 | -0.005 | -0.005 | 0 | 0.75 | 0.81 | 0.82 | 0.73 | 0.76 | 0.67 |
| 7 | 156761818 | rs2302445 | 0.452 | 1.001 | 0.942 | 1.272 | 0.002 | -0.016 | -0.016 | 0 | 0.56 | 0.43 | 0.52 | 0.28 | 0.57 | 0.46 |
| 7 | 156762248 | rs12919 | 0.284 | 1.009 | 0.862 | 1.277 | -0.004 | 0.01 | 0.01 | 0.337 | 0.57 | 0.56 | 0.54 | 0.28 | 0.57 | 0.46 |

| | | | | | | | | | | | | | | | | |
|---|---|---|---|---|---|---|---|---|---|---|---|---|---|---|---|---|
| 7 | 156762446 | rs3087905 | 0.418 | 1.03 | 0.671 | 1.287 | 0.002 | -0.012 | -0.012 | 0.975 | 0.56 | 0.49 | 0.53 | 0.28 | 0.57 | 0.46 |
| 7 | 156765043 | rs7803794 | 0.411 | 1.015 | 0.578 | 1.207 | 0.002 | -0.012 | -0.012 | 0.358 | 0.56 | 0.49 | 0.53 | 0.28 | 0.57 | 0.46 |
| 7 | 156772542 | rs6963005 | 0.162 | 1.456 | 1.023 | 1.635 | 0.02 | 0.029 | 0.029 | 0.54 | 0.14 | 0.2 | 0.24 | 0.28 | 0.15 | 0.15 |
| 7 | 156788065 | rs4716674 | 1.021 | 0.65 | 0.138 | 0.661 | -0.018 | 0.212 | 0.212 | 0.025 | 0.43 | 0.07 | 0.26 | 0.15 | 0.42 | 0.38 |
| 7 | 156799081 | rs2270210 | -1.049 | -0.088 | 1.581 | 1.271 | -0.035 | 0.002 | 0.002 | 0.278 | 0.16 | 0.56 | 0.18 | 0.11 | 0.09 | 0.22 |
| 7 | 156801023 | rs3793333 | -1.501 | -0.064 | 1.647 | 1.444 | -0.023 | 0.035 | 0.035 | 0 | 0.19 | 0.69 | 0.21 | 0.28 | 0.11 | 0.26 |
| 7 | 156804510 | rs10262191 | -0.922 | -0.03 | 1.481 | 1.365 | -0.036 | 0.007 | 0.007 | 0.468 | 0.16 | 0.45 | 0.18 | 0.11 | 0.09 | 0.21 |
| 7 | 156811242 | rs9918633 | -0.636 | 0.055 | -0.059 | 0.932 | -0.023 | -0.002 | -0.002 | 0 | 0.16 | 0.62 | 0.17 | 0.16 | 0.09 | 0.16 |
| 7 | 156815971 | rs4716447 | 0.531 | 0.411 | -0.526 | 1.383 | 0.006 | -0.017 | -0.017 | 0 | 0.72 | 0.78 | 0.57 | 0.43 | 0.74 | 0.64 |
| 7 | 156826585 | rs13244987 | -1.119 | 0.374 | -0.319 | 2.166 | 0 | 0.05 | 0.05 | 0 | 0.05 | 0.04 | 0.04 | 0.27 | 0.05 | 0.1 |
| 7 | 156829579 | rs13438768 | -2.454 | 0.198 | 0.908 | 2.535 | 0.057 | 0.306 | 0.306 | 0 | 0.05 | 0.32 | 0.15 | 0.23 | 0.03 | 0.06 |
| 7 | 156834995 | rs4278056 | -1.484 | 0.418 | 0.91 | 2.079 | 0.007 | 0.03 | 0.03 | 0 | 0.2 | 0.62 | 0.25 | 0.26 | 0.2 | 0.24 |
| 7 | 156840049 | rs12698021 | -1.717 | 0.85 | 1.072 | 1.863 | 0.021 | 0.042 | 0.042 | 0 | 0.08 | 0.19 | 0.16 | 0.13 | 0.1 | 0.09 |
| 7 | 156847704 | rs746508 | -2.12 | 0.55 | 1.122 | 2.054 | 0.009 | 0.075 | 0.075 | 0 | 0.21 | 0.54 | 0.36 | 0.36 | 0.24 | 0.34 |
| 7 | 156848531 | rs13234547 | -0.57 | 0.507 | 0.602 | 1.742 | 0 | 0.005 | 0.005 | 0.036 | 0.06 | 0.05 | 0.15 | 0.15 | 0.08 | 0.07 |
| 7 | 156852081 | rs6978657 | -0.543 | 0.368 | 0.088 | 1.097 | 0 | -0.007 | -0.007 | 0 | 0.54 | 0.47 | 0.57 | 0.66 | 0.51 | 0.55 |
| 7 | 156858239 | rs11768251 | -0.598 | 0.323 | -0.439 | 0.607 | -0.032 | -0.079 | -0.079 | 0.032 | 0.34 | 0.12 | 0.46 | 0.45 | 0.36 | 0.34 |
| 7 | 156867134 | rs7799010 | -1.787 | 0.064 | -0.979 | 0.333 | -0.118 | -0.125 | -0.125 | 0.202 | 0.6 | 0.97 | 0.7 | 0.55 | 0.54 | 0.61 |
| 7 | 156869431 | rs6953995 | -0.824 | 0.047 | -0.053 | 0.26 | 0.011 | -0.003 | -0.003 | 0 | 0.12 | 0.44 | 0.17 | 0.09 | 0.09 | 0.16 |
| 7 | 156876217 | rs6651030 | -0.574 | 0.291 | -0.598 | 0.344 | -0.021 | -0.069 | -0.069 | 0 | 0.33 | 0.17 | 0.54 | 0.51 | 0.37 | 0.33 |
| 7 | 156880398 | rs6966038 | -0.939 | 0.143 | -0.384 | 0.059 | -0.01 | -0.094 | -0.094 | 0 | 0.23 | 0.56 | 0.15 | 0.05 | 0.16 | 0.22 |
| 7 | 156882526 | rs10272338 | -0.665 | -0.177 | -0.747 | 0.147 | -0.015 | -0.084 | -0.084 | 0 | 0.36 | 0.09 | 0.44 | 0.41 | 0.33 | 0.28 |
| 7 | 156883839 | rs7805113 | -0.932 | -0.118 | -0.368 | 0.363 | -0.002 | 0 | 0 | 0 | 0.3 | 0.35 | 0.3 | 0.27 | 0.25 | 0.24 |
| 7 | 156885979 | rs12534197 | -1.265 | -0.207 | -0.471 | -0.049 | 0.005 | 0.009 | 0.009 | 0.036 | 0.37 | 0.62 | 0.33 | 0.2 | 0.3 | 0.36 |
| 7 | 156897590 | rs6943214 | -0.927 | 0.032 | -0.169 | 0.242 | 0.005 | -0.011 | -0.011 | 0.007 | 0.31 | 0.29 | 0.4 | 0.41 | 0.26 | 0.28 |
| 7 | 156904943 | rs6955655 | -1.132 | 0.248 | 0.248 | 0.287 | 0.002 | -0.006 | -0.006 | 0 | 0.3 | 0.64 | 0.43 | 0.46 | 0.26 | 0.28 |
| 7 | 156910420 | rs7802473 | -1.667 | 0.188 | 0.842 | 0.216 | 0.086 | 0.114 | 0.114 | 1.399 | 0.05 | 0.48 | 0.07 | 0.07 | 0.02 | 0.03 |
| 7 | 156923507 | rs7791545 | -1.095 | 0.585 | -0.015 | -0.037 | 0.018 | 0.016 | 0.016 | 0 | 0.45 | 0.82 | 0.46 | 0.44 | 0.38 | 0.42 |
| 7 | 156952133 | rs733483 | -1.743 | 0.905 | 1.204 | 0.189 | 0.066 | 0.032 | 0.032 | 0.085 | 0.08 | 0.48 | 0.11 | 0.06 | 0.08 | 0.07 |
| 7 | 156962874 | rs6956046 | -1.388 | 0.608 | 0.744 | -0.057 | 0.023 | -0.013 | -0.013 | 0 | 0.42 | 0.83 | 0.39 | 0.45 | 0.36 | 0.37 |
| 7 | 156983426 | rs17718923 | 0.692 | 0.859 | 0.599 | -0.193 | -0.022 | 0.027 | 0.027 | 0 | 0.05 | 0.01 | 0.12 | 0 | 0.11 | 0.04 |
| 7 | 157000176 | rs2301914 | -1.405 | 0.568 | 0.806 | -0.001 | 0.02 | -0.013 | -0.013 | 0.358 | 0.42 | 0.83 | 0.39 | 0.45 | 0.36 | 0.37 |
| 7 | 157005863 | rs6979947 | -1.044 | 0.589 | 0.91 | 0.04 | 0.023 | -0.015 | -0.015 | 3.094 | 0.36 | 0.79 | 0.34 | 0.42 | 0.3 | 0.3 |
| 7 | 157007665 | rs17837729 | -1.591 | 0.657 | 1.005 | -0.031 | 0.017 | -0.047 | -0.047 | 0 | 0.12 | 0.67 | 0.13 | 0.06 | 0.12 | 0.1 |
| 7 | 157027108 | rs6459743 | -0.678 | 0.714 | 0.644 | -0.005 | 0.008 | -0.009 | -0.009 | 0.219 | 0.3 | 0.71 | 0.28 | 0.45 | 0.19 | 0.3 |
| 7 | 157031407 | rs1182397 | 1.004 | 0.685 | 0.643 | -0.047 | 0.017 | -0.04 | -0.04 | 0.015 | 0.88 | 0.85 | 0.88 | 0.99 | 0.83 | 0.92 |
| 7 | 157045557 | rs929354 | 1.369 | 0.77 | 0.188 | 0.063 | -0.002 | -0.055 | -0.055 | 0 | 0.58 | 0.12 | 0.6 | 0.53 | 0.63 | 0.6 |
| 7 | 157058363 | rs1182414 | 1.2 | 0.916 | 0.755 | 0.003 | 0.005 | 0.004 | 0.004 | 0 | 0.6 | 0.41 | 0.64 | 0.62 | 0.64 | 0.67 |
| 7 | 157061642 | rs7807 | 0.512 | 0.653 | 0.316 | 0.275 | 0 | -0.002 | -0.002 | 0 | 0.71 | 0.72 | 0.74 | 0.62 | 0.77 | 0.73 |
| 7 | 157079586 | rs1182419 | -0.458 | 1.36 | 0.253 | 0.398 | -0.002 | 0.003 | 0.003 | 0 | 0.35 | 0.39 | 0.45 | 0.49 | 0.38 | 0.41 |
| 7 | 157082607 | rs12533310 | 1.135 | 1.462 | -0.367 | 0.146 | -0.103 | -0.113 | -0.113 | 0.018 | 0.48 | 0.04 | 0.38 | 0.5 | 0.44 | 0.5 |
| 7 | 157087825 | rs757854 | -0.861 | 1.323 | 0.683 | 0.301 | 0.013 | -0.003 | -0.003 | 0 | 0.45 | 0.65 | 0.55 | 0.51 | 0.46 | 0.48 |
| 7 | 157090296 | rs2527866 | -0.331 | 1.228 | 0.561 | 0.4 | -0.003 | 0.004 | 0.004 | 0 | 0.34 | 0.39 | 0.46 | 0.5 | 0.36 | 0.39 |
| 7 | 157094415 | rs1182369 | -0.623 | 1.314 | 0.528 | 0.423 | 0.001 | -0.003 | -0.003 | 1.613 | 0.39 | 0.39 | 0.47 | 0.52 | 0.41 | 0.44 |
| 7 | 157103038 | rs17837752 | -1.386 | 1.224 | 0.717 | 0.814 | 0.005 | 0.014 | 0.014 | 0 | 0.08 | 0.18 | 0.11 | 0.12 | 0.1 | 0.04 |
| 7 | 157103093 | rs6942806 | -2.588 | 1.406 | 1.098 | 1.372 | 0.003 | 0.029 | 0.029 | 0.029 | 0.16 | 0.69 | 0.18 | 0.26 | 0.17 | 0.08 |
| 7 | 157110720 | rs10253147 | 0.056 | 1.45 | 0.523 | 0.919 | 0.004 | 0.019 | 0.019 | 0 | 0.44 | 0.67 | 0.36 | 0.38 | 0.38 | 0.44 |
| 7 | 157111112 | rs10228030 | 0.292 | 1.101 | 0.836 | 0.572 | 0.021 | -0.02 | -0.02 | 0.384 | 0.14 | 0.5 | 0.1 | 0.06 | 0.12 | 0.02 |
| 7 | 157112604 | rs6459751 | -1.097 | 1.157 | 0.427 | 1.121 | -0.065 | 0.03 | 0.03 | 0 | 0.56 | 0.89 | 0.46 | 0.64 | 0.48 | 0.5 |
| 7 | 157115908 | rs7779677 | 0.941 | 0.84 | 0.577 | 1.149 | 0.009 | 0.027 | 0.027 | 0.887 | 0.26 | 0.11 | 0.23 | 0.24 | 0.21 | 0.37 |
| 7 | 157116750 | rs6947158 | -0.053 | 0.991 | 1.187 | 1.282 | 0.003 | -0.001 | -0.001 | 0 | 0.43 | 0.6 | 0.33 | 0.58 | 0.32 | 0.44 |
| 7 | 157122526 | rs4716702 | -1.516 | 1.311 | 0.783 | 0.957 | 0.07 | -0.036 | -0.036 | 0 | 0.09 | 0.5 | 0.09 | 0 | 0.08 | 0.03 |
| 7 | 157134356 | rs13224364 | 0.157 | 1.542 | 0.52 | 1.173 | 0.004 | 0.021 | 0.021 | 0 | 0.53 | 0.43 | 0.6 | 0.74 | 0.57 | 0.57 |
| 7 | 157140090 | rs9647660 | 1.003 | 1.732 | 0.085 | 1.237 | -0.068 | 0.046 | 0.046 | 0 | 0.34 | 0.05 | 0.46 | 0.22 | 0.37 | 0.37 |
| 7 | 157141237 | rs4716468 | -0.473 | 1.632 | 0.319 | 1.222 | 0.074 | -0.006 | -0.006 | 0 | 0.07 | 0.4 | 0.07 | 0 | 0.06 | 0.03 |
| 7 | 157145431 | rs12668448 | 0.061 | 1.79 | -0.301 | 1.27 | 0.012 | 0.043 | 0.043 | 0 | 0.49 | 0.15 | 0.56 | 0.42 | 0.55 | 0.52 |
| 7 | 157151234 | rs3802100 | -0.253 | 1.253 | -0.154 | 1.363 | 0.007 | 0.035 | 0.035 | 0 | 0.53 | 0.42 | 0.6 | 0.75 | 0.57 | 0.56 |
| 7 | 157160586 | rs4716707 | -0.955 | 1.28 | -0.194 | 1.136 | -0.005 | 0.026 | 0.026 | 0 | 0.61 | 0.87 | 0.68 | 0.69 | 0.63 | 0.58 |
| 7 | 157168469 | rs6459770 | -0.423 | 1.522 | -0.588 | 1.465 | 0.006 | 0.035 | 0.035 | 0 | 0.52 | 0.32 | 0.58 | 0.63 | 0.56 | 0.55 |
| 7 | 157175511 | rs3802088 | -0.023 | 1.493 | -0.693 | 1.256 | -0.051 | 0.045 | 0.045 | 0 | 0.47 | 0.13 | 0.53 | 0.39 | 0.51 | 0.5 |
| 7 | 157179909 | rs4716712 | -0.447 | 1.486 | -0.745 | 1.203 | 0.012 | 0.044 | 0.044 | 0 | 0.52 | 0.35 | 0.59 | 0.69 | 0.57 | 0.56 |
| 7 | 157186627 | rs917738 | -0.558 | 1.313 | -1.629 | 0.492 | -0.095 | 0.069 | 0.069 | 0 | 0.64 | 0.95 | 0.54 | 0.79 | 0.59 | 0.63 |
| 7 | 157193887 | rs10440911 | -0.021 | 1.21 | -1.128 | 0.674 | 0.016 | 0.065 | 0.065 | 0 | 0.51 | 0.81 | 0.46 | 0.62 | 0.43 | 0.5 |
| 7 | 157198031 | rs1009328 | 0.32 | 0.855 | -0.773 | 0.419 | -0.004 | 0.007 | 0.007 | 0 | 0.63 | 0.71 | 0.75 | 0.73 | 0.58 | 0.75 |
| 7 | 157198266 | rs6459771 | 0.635 | 0.714 | -0.449 | 0.498 | -0.046 | 0.027 | 0.027 | 0 | 0.58 | 0.23 | 0.69 | 0.69 | 0.55 | 0.7 |
| 7 | 157200711 | rs758929 | 0.498 | 0.827 | -0.419 | 0.389 | -0.018 | 0.015 | 0.015 | 0 | 0.62 | 0.22 | 0.73 | 0.7 | 0.63 | 0.7 |
| 7 | 157201648 | rs10264615 | 0.22 | 0.803 | -0.322 | 0.355 | 0 | -0.001 | -0.001 | 0 | 0.66 | 0.69 | 0.77 | 0.74 | 0.64 | 0.72 |
| 7 | 157205618 | rs1005435 | 0.572 | 0.5 | 0.583 | 0.224 | -0.04 | 0.018 | 0.018 | 0 | 0.6 | 0.2 | 0.71 | 0.71 | 0.6 | 0.68 |
| 7 | 157208617 | rs1009135 | 0.461 | 0.177 | 0.619 | 0.069 | 0.003 | -0.008 | -0.008 | 0 | 0.58 | 0.52 | 0.73 | 0.7 | 0.58 | 0.67 |

| | | | | | | | | | | | | | | | | |
|---|---|---|---|---|---|---|---|---|---|---|---|---|---|---|---|---|
| 7 | 157212045 | rs10252297 | 0.345 | 0.067 | 1.015 | 0.028 | 0.003 | -0.005 | -0.005 | 0 | 0.6 | 0.58 | 0.74 | 0.71 | 0.59 | 0.63 |
| 7 | 157212152 | rs7791873 | -0.126 | -0.821 | 0.441 | 0.484 | -0.033 | 0.025 | 0.025 | 0 | 0.23 | 0.36 | 0.21 | 0.3 | 0.15 | 0.16 |
| 7 | 157212935 | rs7455999 | 0.242 | -0.641 | 0.452 | 0.568 | -0.035 | 0.056 | 0.056 | 0 | 0.75 | 0.55 | 0.77 | 0.66 | 0.82 | 0.77 |
| 7 | 157213644 | rs1024716 | -0.44 | -0.878 | 0.247 | 0.877 | 0.012 | -0.02 | -0.02 | 0.108 | 0.12 | 0.04 | 0.11 | 0.21 | 0.1 | 0.1 |
| 7 | 157213719 | rs2109827 | 0.88 | -0.901 | 0.34 | 0.873 | -0.042 | 0.066 | 0.066 | 0 | 0.74 | 0.49 | 0.77 | 0.63 | 0.8 | 0.77 |
| 7 | 157214782 | rs6957319 | 0.896 | -0.844 | 0.302 | 0.899 | -0.032 | 0.017 | 0.017 | 0.189 | 0.13 | 0.33 | 0.09 | 0.13 | 0.08 | 0.05 |
| 7 | 157215026 | rs6975648 | -0.778 | -0.848 | 0.298 | 0.906 | -0.022 | 0.037 | 0.037 | 0.27 | 0.86 | 0.65 | 0.89 | 0.83 | 0.9 | 0.87 |
| 7 | 157216093 | rs2286842 | 1.696 | -1.05 | -0.062 | 0.715 | -0.115 | -0.066 | -0.066 | 0 | 0.31 | 0.02 | 0.5 | 0.37 | 0.38 | 0.46 |
| 7 | 157216837 | rs13307078 | 0.43 | -1.386 | 0.13 | 0.334 | -0.001 | 0.001 | 0.001 | 0 | 0.4 | 0.36 | 0.25 | 0.25 | 0.39 | 0.29 |
| 7 | 157217055 | rs10257910 | 0.962 | -1.033 | 0.003 | 0.571 | -0.024 | 0.014 | 0.014 | 0.048 | 0.12 | 0.22 | 0.09 | 0.13 | 0.08 | 0.05 |
| 7 | 157219256 | rs10266536 | -0.32 | -1.307 | -0.636 | 0.37 | -0.006 | -0.017 | -0.017 | 0 | 0.44 | 0.87 | 0.29 | 0.36 | 0.38 | 0.39 |
| 7 | 157221158 | rs6955908 | -1.065 | -0.706 | -0.282 | 0.865 | -0.142 | -0.137 | -0.137 | 0 | 0.68 | 0.98 | 0.48 | 0.53 | 0.6 | 0.47 |
| 7 | 157224794 | rs6978435 | -0.234 | -0.801 | 0 | 0.78 | -0.041 | -0.034 | -0.034 | 0.541 | 0.6 | 0.79 | 0.38 | 0.4 | 0.53 | 0.35 |
| 7 | 157226739 | rs1024718 | 0.086 | -0.831 | 0.384 | 0.751 | -0.011 | -0.006 | -0.006 | 0 | 0.56 | 0.69 | 0.37 | 0.4 | 0.52 | 0.35 |
| 7 | 157230236 | rs7804019 | -0.064 | -0.876 | 0.316 | 0.752 | -0.026 | -0.018 | -0.018 | 0 | 0.57 | 0.76 | 0.37 | 0.4 | 0.53 | 0.35 |
| 7 | 157233868 | rs7777477 | -1.094 | -1.065 | 0.89 | 0.533 | -0.03 | -0.027 | -0.027 | 0 | 0.24 | 0.4 | 0.14 | 0.16 | 0.16 | 0.15 |
| 7 | 157237369 | rs2366558 | 0.751 | -1.05 | 0.704 | 0.533 | -0.019 | 0.01 | 0.01 | 0.236 | 0.12 | 0.25 | 0.1 | 0.12 | 0.08 | 0.06 |
| 7 | 157237427 | rs2366559 | -1.337 | -1.069 | 0.696 | 0.576 | -0.038 | -0.006 | -0.006 | 0.032 | 0.26 | 0.58 | 0.23 | 0.25 | 0.18 | 0.21 |
| 7 | 157250133 | rs11981578 | 0.856 | -1.16 | 0.5 | 0.567 | 0.002 | 0.011 | 0.011 | 0.734 | 0.41 | 0.34 | 0.24 | 0.1 | 0.41 | 0.23 |
| 7 | 157250749 | rs6965466 | 0.877 | -1.173 | 0.494 | 0.592 | 0.001 | -0.005 | -0.005 | 0.036 | 0.41 | 0.34 | 0.24 | 0.06 | 0.41 | 0.23 |
| 7 | 157250854 | rs4716739 | -1.022 | -1.163 | 0.544 | 0.693 | -0.029 | 0.018 | 0.018 | 0 | 0.24 | 0.43 | 0.21 | 0.29 | 0.18 | 0.21 |
| 7 | 157252849 | rs917740 | -0.985 | -0.854 | -0.29 | 0.692 | -0.067 | -0.065 | -0.065 | 1.004 | 0.68 | 0.87 | 0.47 | 0.55 | 0.6 | 0.48 |
| 7 | 157253683 | rs17837764 | -0.386 | -0.529 | -0.226 | 0.758 | 0 | 0.008 | 0.008 | 0 | 0.11 | 0.04 | 0.1 | 0.09 | 0.09 | 0.09 |
| 7 | 157258109 | rs10237947 | 0.307 | -0.367 | -0.387 | 0.599 | 0.023 | 0.019 | 0.019 | 0 | 0.52 | 0.76 | 0.46 | 0.53 | 0.48 | 0.43 |
| 7 | 157263532 | rs7785035 | 0.891 | -1.066 | 0.02 | 0.147 | -0.05 | -0.053 | -0.053 | 0 | 0.72 | 0.44 | 0.88 | 0.85 | 0.85 | 0.81 |
| 7 | 157270499 | rs917741 | 0.283 | -0.909 | -0.253 | 0.076 | 0.001 | -0.016 | -0.016 | 0.153 | 0.83 | 0.84 | 0.92 | 0.98 | 0.9 | 0.85 |
| 7 | 157276327 | rs9768425 | 0.484 | -0.324 | -0.331 | 0.529 | 0.008 | -0.005 | -0.005 | 0 | 0.56 | 0.38 | 0.61 | 0.54 | 0.65 | 0.45 |
| 7 | 157276616 | rs9768449 | -0.301 | -0.581 | -0.242 | 0.944 | -0.017 | 0.155 | 0.155 | 0 | 0.1 | 0.23 | 0.18 | 0.38 | 0.08 | 0.1 |
| 7 | 157278626 | rs1076434 | -0.21 | -0.085 | -0.426 | 0.713 | -0.002 | -0.002 | -0.002 | 0 | 0.68 | 0.67 | 0.6 | 0.62 | 0.61 | 0.5 |
| 7 | 157278764 | rs1076435 | 0.797 | 0.115 | -1.031 | 0.517 | -0.007 | 0.108 | 0.108 | 0 | 0.48 | 0.25 | 0.4 | 0.22 | 0.49 | 0.4 |
| 7 | 157279376 | rs221213 | 0.047 | 1.108 | -1.398 | 0.934 | -0.132 | -0.096 | -0.096 | 0 | 0.24 | 0.01 | 0.33 | 0.29 | 0.32 | 0.29 |
| 7 | 157284101 | rs221219 | -0.224 | 1.198 | -1.224 | 1.082 | -0.103 | -0.068 | -0.068 | 6.518 | 0.22 | 0.04 | 0.34 | 0.27 | 0.36 | 0.31 |
| 7 | 157289534 | rs11980462 | 1.038 | 0.792 | -1.087 | 0.736 | -0.004 | 0.062 | 0.062 | 4.053 | 0.45 | 0.27 | 0.37 | 0.25 | 0.42 | 0.39 |
| 7 | 157289684 | rs887626 | -0.553 | 0.577 | -0.663 | 1.256 | -0.011 | 0.04 | 0.04 | 0.425 | 0.1 | 0.14 | 0.11 | 0.34 | 0.06 | 0.09 |
| 7 | 157298250 | rs10237006 | 0.538 | 0.864 | -0.03 | 1.407 | 0.001 | -0.001 | -0.001 | 0 | 0.53 | 0.51 | 0.51 | 0.55 | 0.49 | 0.44 |
| 7 | 157298637 | rs10240671 | 0.531 | 1.106 | 0.027 | 1.361 | -0.006 | 0.001 | 0.001 | 0 | 0.56 | 0.51 | 0.57 | 0.55 | 0.56 | 0.46 |
| 7 | 157299223 | rs11764996 | 1.108 | 1.068 | -0.077 | 1.294 | -0.007 | 0.097 | 0.097 | 2.031 | 0.45 | 0.26 | 0.29 | 0.19 | 0.42 | 0.32 |
| 7 | 157308110 | rs4716752 | 0.442 | 0.832 | -0.014 | 1.561 | 0 | -0.001 | -0.001 | 0.041 | 0.55 | 0.42 | 0.56 | 0.56 | 0.53 | 0.46 |
| 7 | 157308335 | rs740993 | 0.44 | 0.562 | -0.267 | 1.593 | -0.121 | -0.091 | -0.091 | 0 | 0.25 | 0.01 | 0.28 | 0.29 | 0.34 | 0.37 |
| 7 | 157308348 | rs740994 | 0.431 | 0.904 | 0.125 | 1.098 | -0.001 | 0.006 | 0.006 | 0.116 | 0.54 | 0.39 | 0.4 | 0.4 | 0.48 | 0.39 |
| 7 | 157309078 | rs1468649 | 0.414 | 0.909 | 0.127 | 1.1 | -0.001 | 0.005 | 0.005 | 0.134 | 0.55 | 0.39 | 0.4 | 0.4 | 0.48 | 0.39 |
| 7 | 157314762 | rs10263303 | 0.398 | 1.275 | 0.465 | 0.754 | -0.001 | 0.001 | 0.001 | 2.638 | 0.54 | 0.52 | 0.41 | 0.18 | 0.5 | 0.31 |
| 7 | 157316657 | rs11766985 | 0.557 | 1.234 | 0.591 | 0.717 | -0.001 | 0.032 | 0.032 | 2.612 | 0.52 | 0.54 | 0.36 | 0.15 | 0.48 | 0.27 |
| 7 | 157321633 | rs221232 | -0.389 | 1.222 | 0.373 | 0.843 | 0.005 | 0.019 | 0.019 | 0.688 | 0.49 | 0.43 | 0.55 | 0.65 | 0.48 | 0.64 |
| 7 | 157323039 | rs11768450 | 0.761 | 1.156 | 0.44 | 0.673 | -0.002 | 0.021 | 0.021 | 2.708 | 0.43 | 0.31 | 0.27 | 0.04 | 0.39 | 0.21 |
| 7 | 157330266 | rs221250 | 1.489 | 1.222 | 0.503 | 0.838 | 0.05 | 0.201 | 0.201 | 0.282 | 0.94 | 0.78 | 0.83 | 0.68 | 0.89 | 0.85 |
| 7 | 157334598 | rs221257 | -0.329 | 1.159 | 0.346 | 0.754 | -0.001 | -0.002 | -0.002 | 2.594 | 0.48 | 0.5 | 0.54 | 0.64 | 0.48 | 0.64 |
| 7 | 157338580 | rs4716755 | -1.263 | 1.138 | 0.31 | 0.618 | -0.001 | -0.026 | -0.026 | 0 | 0.12 | 0.14 | 0.09 | 0 | 0.12 | 0.14 |
| 7 | 157340076 | rs221272 | -0.009 | 1.018 | 0.408 | 0.639 | 0.001 | 0.012 | 0.012 | 0.305 | 0.48 | 0.45 | 0.6 | 0.68 | 0.48 | 0.67 |
| 7 | 157350699 | rs221281 | -0.037 | 0.982 | 0.443 | 0.65 | 0 | 0.03 | 0.03 | 0.491 | 0.48 | 0.37 | 0.56 | 0.67 | 0.48 | 0.68 |
| 7 | 157359289 | rs221293 | 0.104 | 1.003 | 0.443 | 0.634 | 0 | 0.025 | 0.025 | 0.586 | 0.46 | 0.37 | 0.55 | 0.67 | 0.48 | 0.68 |
| 7 | 157364630 | rs221298 | -0.127 | 0.921 | 0.494 | 0.611 | 0.002 | 0.014 | 0.014 | 0.072 | 0.49 | 0.44 | 0.56 | 0.68 | 0.47 | 0.68 |
| 7 | 157365106 | rs6459797 | 0.514 | 0.932 | 0.547 | 0.424 | 0 | -0.031 | -0.031 | 0.085 | 0.44 | 0.44 | 0.3 | 0.14 | 0.41 | 0.23 |
| 7 | 157368550 | rs7791552 | 0.117 | 1.071 | 0.527 | 0.503 | -0.001 | -0.009 | -0.009 | 0 | 0.51 | 0.51 | 0.34 | 0.14 | 0.48 | 0.27 |
| 7 | 157371513 | rs221305 | 1.647 | 0.921 | 1.066 | 1.179 | 0.03 | 0.05 | 0.05 | 2.142 | 0.87 | 0.68 | 0.76 | 0.81 | 0.82 | 0.83 |
| 7 | 157387441 | rs10274279 | -1.838 | 0.91 | 1.071 | 1.005 | 0.013 | -0.081 | -0.081 | 0.112 | 0.13 | 0.34 | 0.11 | 0 | 0.12 | 0.12 |
| 7 | 157400389 | rs10229026 | -1.915 | 0.855 | 0.944 | 1.393 | 0.037 | 0.24 | 0.24 | 2.426 | 0.13 | 0.48 | 0.22 | 0.44 | 0.15 | 0.17 |
| 7 | 157408733 | rs2008341 | 0.126 | 0.719 | 0.441 | 1.139 | 0.007 | -0.004 | -0.004 | 0 | 0.52 | 0.61 | 0.46 | 0.53 | 0.52 | 0.39 |
| 7 | 157409757 | rs7784733 | -2.161 | 0.494 | 1.065 | 1.572 | 0.037 | 0.146 | 0.146 | 5.354 | 0.09 | 0.41 | 0.14 | 0.18 | 0.08 | 0.08 |
| 7 | 157412024 | rs10249676 | -1.099 | 0.433 | 0.6 | 1.074 | -0.002 | -0.034 | -0.034 | 0 | 0.06 | 0.18 | 0.06 | 0 | 0.05 | 0.04 |
| 7 | 157415516 | rs1263560 | -0.786 | 0.312 | 0.546 | 1.423 | -0.017 | 0.021 | 0.021 | 0 | 0.28 | 0.41 | 0.37 | 0.32 | 0.26 | 0.42 |
| 7 | 157419695 | rs1263563 | -0.847 | 0.242 | 0.576 | 1.404 | -0.022 | 0.02 | 0.02 | 0.187 | 0.29 | 0.42 | 0.35 | 0.33 | 0.26 | 0.41 |
| 7 | 157421108 | rs1263542 | -0.823 | 0.39 | 0.653 | 1.494 | -0.01 | 0.03 | 0.03 | 0.225 | 0.27 | 0.52 | 0.47 | 0.59 | 0.27 | 0.45 |
| 7 | 157426990 | rs6970138 | -1.248 | 0.642 | 0.562 | 1.371 | 0.005 | -0.003 | -0.003 | 0 | 0.39 | 0.53 | 0.49 | 0.36 | 0.44 | 0.47 |
| 7 | 157427927 | rs17837786 | -0.618 | 0.693 | 1.407 | 1.215 | 0.038 | -0.018 | -0.018 | 0 | 0.06 | 0.24 | 0.03 | 0 | 0.04 | 0.03 |
| 7 | 157432193 | rs13239831 | 0.012 | 0.706 | 1.531 | 1.242 | -0.005 | 0.004 | 0.004 | 0 | 0.17 | 0.12 | 0.23 | 0.07 | 0.24 | 0.17 |
| 7 | 157435350 | rs1263573 | 0.834 | 0.39 | 1.335 | 1.384 | -0.001 | 0.001 | 0.001 | 0.499 | 0.68 | 0.71 | 0.68 | 0.71 | 0.64 | 0.57 |
| 7 | 157435421 | rs17837789 | 0.152 | 0.356 | 1.373 | 1.433 | 0.001 | -0.017 | -0.017 | 0.727 | 0.16 | 0.1 | 0.22 | 0.3 | 0.15 | 0.28 |
| 7 | 157437584 | rs1638020 | 1.915 | 0.294 | 1.398 | 1.395 | -0.007 | 0.018 | 0.018 | 2.526 | 0.45 | 0.24 | 0.42 | 0.41 | 0.45 | 0.27 |

| | | | | | | | | | | | | | | | | |
|---|---|---|---|---|---|---|---|---|---|---|---|---|---|---|---|---|
| 7 | 157438037 | rs1638021 | -0.406 | -0.013 | 1.411 | 1.269 | -0.008 | 0.012 | 0.012 | 0.761 | 0.84 | 0.8 | 0.91 | 0.78 | 0.88 | 0.74 |
| 7 | 157452854 | rs12333884 | 2.195 | -0.228 | 0.956 | 0.754 | -0.042 | 0.095 | 0.095 | 0.024 | 0.36 | 0.19 | 0.34 | 0.15 | 0.44 | 0.21 |
| 7 | 157455459 | rs1270639 | -0.998 | -0.556 | 1.272 | 0.708 | -0.005 | -0.023 | -0.023 | 0 | 0.84 | 0.85 | 0.92 | 0.97 | 0.88 | 0.84 |
| 7 | 157455933 | rs1263566 | 0.038 | -0.56 | 1.391 | 0.675 | -0.013 | -0.072 | -0.072 | 0 | 0.79 | 0.72 | 0.87 | 0.97 | 0.84 | 0.82 |
| 7 | 157458872 | rs11764539 | 0.03 | -0.543 | 1.269 | 1.129 | -0.002 | -0.002 | -0.002 | 0 | 0.21 | 0.2 | 0.32 | 0.24 | 0.24 | 0.19 |
| 7 | 157458997 | rs6979750 | -0.149 | -0.529 | 1.031 | 1.062 | -0.008 | 0.031 | 0.031 | 0 | 0.1 | 0.15 | 0.07 | 0.21 | 0.04 | 0.15 |
| 7 | 157461886 | rs1079060 | 1.261 | -0.518 | 0.938 | 1.055 | 0.007 | 0.018 | 0.018 | 0 | 0.49 | 0.46 | 0.47 | 0.44 | 0.52 | 0.51 |
| 7 | 157462425 | rs755703 | 0.71 | -0.496 | 0.703 | 1.161 | 0.003 | -0.002 | -0.002 | 1.402 | 0.61 | 0.66 | 0.55 | 0.49 | 0.63 | 0.57 |
| 7 | 157464422 | rs17837793 | -0.039 | -0.6 | 0.596 | 1.273 | -0.003 | 0.029 | 0.029 | 0 | 0.09 | 0.09 | 0.06 | 0.24 | 0.05 | 0.18 |
| 7 | 157464857 | rs12698098 | 0.14 | -0.612 | 0.57 | 0.877 | -0.013 | 0.029 | 0.029 | 0 | 0.2 | 0.11 | 0.22 | 0.1 | 0.23 | 0.15 |
| 7 | 157465014 | rs17731604 | 2.036 | -0.601 | 0.652 | 0.674 | -0.035 | 0.104 | 0.104 | 0.434 | 0.4 | 0.23 | 0.36 | 0.13 | 0.46 | 0.21 |
| 7 | 157465367 | rs1263546 | -0.529 | -1.055 | 0.821 | 1.084 | -0.03 | -0.001 | -0.001 | 0 | 0.84 | 0.68 | 0.83 | 0.86 | 0.89 | 0.81 |
| 7 | 157469223 | rs6459801 | 0.037 | -0.869 | 0.574 | 0.863 | 0.001 | -0.052 | -0.052 | 0.559 | 0.13 | 0.19 | 0.08 | 0 | 0.12 | 0.08 |
| 7 | 157470212 | rs17837795 | 0.387 | -1.052 | 0.689 | 1.345 | -0.007 | 0.002 | 0.002 | 0 | 0.08 | 0.07 | 0.11 | 0.12 | 0.02 | 0.14 |
| 7 | 157470412 | rs1263550 | -0.603 | -1.16 | 0.879 | 1.451 | -0.026 | 0.05 | 0.05 | 2.96 | 0.83 | 0.74 | 0.92 | 0.7 | 0.88 | 0.72 |
| 7 | 157473506 | rs1922739 | 0.074 | -1.21 | 0.739 | 1.33 | -0.003 | 0.009 | 0.009 | 4.172 | 0.8 | 0.75 | 0.82 | 0.43 | 0.84 | 0.51 |
| 7 | 157480939 | rs10250971 | -0.914 | -1.105 | 0.357 | 0.865 | -0.004 | 0.018 | 0.018 | 0 | 0.33 | 0.22 | 0.39 | 0.34 | 0.32 | 0.35 |
| 7 | 157485165 | rs1638006 | 0.123 | -1.071 | 0.256 | 1.031 | -0.022 | -0.033 | -0.033 | 0.365 | 0.15 | 0.11 | 0.22 | 0.33 | 0.23 | 0.15 |
| 7 | 157487013 | rs6962998 | -1.608 | -1.375 | 0.488 | 0.915 | 0 | 0 | 0 | 0.23 | 0.44 | 0.44 | 0.51 | 0.45 | 0.35 | 0.5 |
| 7 | 157493836 | rs1796236 | 0.078 | -1.505 | 0.492 | 0.93 | -0.004 | 0.003 | 0.003 | 0 | 0.15 | 0.19 | 0.21 | 0.18 | 0.13 | 0.09 |
| 7 | 157495354 | rs1242787 | -1.264 | -1.57 | 0.484 | 0.921 | -0.037 | 0.017 | 0.017 | 0 | 0.25 | 0.39 | 0.33 | 0.28 | 0.16 | 0.25 |
| 7 | 157495783 | rs1242786 | -0.949 | -1.432 | 0.135 | 0.979 | -0.003 | 0.013 | 0.013 | 2.192 | 0.85 | 0.84 | 0.91 | 0.69 | 0.88 | 0.73 |
| 7 | 157496629 | rs12672702 | 0.168 | -0.825 | -0.597 | 0.174 | -0.023 | 0.041 | 0.041 | 0 | 0.21 | 0.06 | 0.23 | 0.1 | 0.22 | 0.24 |
| 7 | 157502916 | rs1796258 | -0.44 | -0.826 | -0.423 | 0.193 | 0.001 | 0 | 0 | 0 | 0.2 | 0.19 | 0.22 | 0.18 | 0.15 | 0.12 |
| 7 | 157508424 | rs2952635 | -1.023 | -0.566 | -0.735 | 0.083 | -0.026 | 0.049 | 0.049 | 0 | 0.85 | 0.98 | 0.87 | 0.92 | 0.82 | 0.89 |
| 7 | 157510294 | rs6459805 | 1.129 | -0.783 | -0.255 | 0.191 | 0 | 0.003 | 0.003 | 0 | 0.46 | 0.49 | 0.41 | 0.35 | 0.43 | 0.44 |
| 7 | 157512750 | rs9654699 | -0.084 | -0.65 | -0.538 | -0.279 | -0.024 | -0.028 | -0.028 | 0 | 0.77 | 0.92 | 0.66 | 0.73 | 0.73 | 0.81 |
| 7 | 157514189 | rs10256772 | -0.062 | -0.658 | -0.43 | -0.284 | -0.007 | -0.01 | -0.01 | 0 | 0.77 | 0.86 | 0.66 | 0.73 | 0.73 | 0.81 |
| 7 | 157514269 | rs933423 | 0.839 | -0.735 | -0.227 | -0.322 | 0.001 | 0.069 | 0.069 | 4.146 | 0.15 | 0.17 | 0.14 | 0.47 | 0.14 | 0.32 |
| 7 | 157515275 | rs2109302 | 0.52 | -0.685 | -0.006 | -0.995 | 0.001 | 0.012 | 0.012 | 1.896 | 0.59 | 0.64 | 0.48 | 0.11 | 0.58 | 0.31 |
| 7 | 157516956 | rs10253909 | 0.436 | -0.589 | -0.115 | -0.949 | -0.001 | 0.012 | 0.012 | 1.173 | 0.61 | 0.65 | 0.52 | 0.22 | 0.58 | 0.49 |
| 7 | 157531174 | rs867923 | 0.133 | -0.327 | -0.314 | -0.822 | 0.002 | 0.02 | 0.02 | 0 | 0.48 | 0.42 | 0.46 | 0.29 | 0.55 | 0.41 |
| 7 | 157531809 | rs11976357 | 0.677 | -0.656 | 0.052 | -0.645 | -0.003 | -0.035 | -0.035 | 0 | 0.07 | 0.15 | 0.02 | 0 | 0.03 | 0.01 |
| 7 | 157532126 | rs1242775 | -0.321 | -0.374 | -0.418 | -0.592 | 0.001 | -0.011 | -0.011 | 0 | 0.67 | 0.63 | 0.61 | 0.69 | 0.65 | 0.54 |
| 7 | 157534105 | rs10949662 | 0.327 | 0.322 | -0.276 | 0.102 | 0.001 | -0.01 | -0.01 | 0 | 0.25 | 0.26 | 0.26 | 0.17 | 0.33 | 0.19 |
| 7 | 157538552 | rs1638742 | -0.939 | -0.344 | -0.083 | 0.415 | 0 | 0.001 | 0.001 | 0 | 0.15 | 0.15 | 0.26 | 0.17 | 0.1 | 0.12 |
| 7 | 157543338 | rs2707941 | -0.939 | -0.344 | -0.083 | 0.415 | 0 | 0.001 | 0.001 | 0 | 0.15 | 0.15 | 0.26 | 0.17 | 0.1 | 0.12 |
| 7 | 157544760 | rs7781481 | -0.387 | -0.284 | 0.019 | 0.201 | -0.001 | -0.013 | -0.013 | 0 | 0.09 | 0.07 | 0.06 | 0 | 0.07 | 0.1 |
| 7 | 157546894 | rs887417 | -1.315 | -1.085 | 0.657 | -0.008 | -0.05 | -0.054 | -0.054 | 0 | 0.22 | 0.57 | 0.15 | 0.12 | 0.12 | 0.12 |
| 7 | 157547384 | rs10272636 | -1.13 | -1.401 | 0.974 | -0.075 | -0.077 | -0.016 | -0.016 | 0 | 0.17 | 0.51 | 0.13 | 0.12 | 0.07 | 0.15 |
| 7 | 157548925 | rs1242782 | -0.494 | -1.243 | 0.612 | 0.063 | 0.001 | -0.002 | -0.002 | 0 | 0.12 | 0.11 | 0.23 | 0.2 | 0.08 | 0.04 |
| 7 | 157549838 | rs1242780 | -2.116 | -1.415 | 0.941 | -0.126 | -0.008 | 0.003 | 0.003 | 0 | 0.31 | 0.32 | 0.37 | 0.31 | 0.21 | 0.3 |
| 7 | 157550645 | rs11984338 | -0.992 | -1.135 | 0.937 | -0.365 | 0 | -0.026 | -0.026 | 0 | 0.14 | 0.15 | 0.08 | 0.01 | 0.13 | 0.13 |
| 7 | 157551358 | rs2952649 | -0.866 | -1.261 | 0.291 | -0.278 | 0.025 | -0.045 | -0.045 | 0.191 | 0.64 | 0.8 | 0.65 | 0.5 | 0.67 | 0.34 |
| 7 | 157560763 | rs1242778 | -1.711 | -0.905 | 0.912 | -0.526 | 0.017 | -0.002 | -0.002 | 0.002 | 0.33 | 0.7 | 0.38 | 0.33 | 0.22 | 0.3 |
| 7 | 157563249 | rs1638751 | -0.925 | -0.82 | 0.912 | -0.228 | -0.06 | 0.064 | 0.064 | 0 | 0.16 | 0.36 | 0.24 | 0.26 | 0.1 | 0.07 |
| 7 | 157570455 | rs1362218 | -0.743 | -0.256 | 0.74 | -0.989 | -0.03 | -0.061 | -0.061 | 0 | 0.2 | 0.38 | 0.12 | 0.04 | 0.12 | 0.24 |
| 7 | 157571598 | rs7807362 | -1.071 | -0.357 | 0.692 | -1.048 | -0.036 | -0.066 | -0.066 | 0 | 0.23 | 0.39 | 0.12 | 0.06 | 0.13 | 0.25 |
| 7 | 157583166 | rs4716494 | 0.675 | 0.646 | 0.732 | -1.239 | 0.037 | -0.071 | -0.071 | 0 | 0.74 | 0.62 | 0.63 | 0.95 | 0.59 | 0.75 |
| 7 | 157600780 | rs13222859 | -1.266 | -0.374 | 0.659 | -1.051 | -0.024 | -0.163 | -0.163 | 0.064 | 0.21 | 0.48 | 0.13 | 0 | 0.17 | 0.05 |
| 7 | 157605468 | rs870020 | -1.202 | -0.924 | 0.473 | -0.55 | -0.027 | -0.023 | -0.023 | 0 | 0.4 | 0.55 | 0.17 | 0.21 | 0.2 | 0.37 |
| 7 | 157605815 | rs2159469 | -1.531 | -0.487 | 1.004 | 0.132 | -0.046 | 0.062 | 0.062 | 0 | 0.18 | 0.51 | 0.13 | 0.21 | 0.09 | 0.32 |
| 7 | 157625051 | rs9654707 | -1.556 | -0.191 | 0.79 | 0.435 | -0.012 | 0.165 | 0.165 | 0 | 0.1 | 0.39 | 0.1 | 0.21 | 0.05 | 0.25 |
| 7 | 157634922 | rs7777198 | 0.66 | 0.67 | -0.297 | 0.572 | -0.021 | -0.02 | -0.02 | 2.18 | 0.66 | 0.76 | 0.55 | 0.4 | 0.58 | 0.6 |
| 7 | 157637962 | rs1865462 | 0.058 | 0.429 | -0.488 | 0.107 | 0.002 | 0.008 | 0.008 | 0 | 0.08 | 0.08 | 0.05 | 0 | 0.07 | 0.08 |
| 7 | 157641669 | rs1529835 | 0.499 | 0.086 | -1.095 | -0.393 | -0.008 | 0.118 | 0.118 | 0.083 | 0.43 | 0.18 | 0.52 | 0.81 | 0.55 | 0.47 |
| 7 | 157642767 | rs3952723 | -0.2 | -0.301 | -0.479 | -0.644 | 0.016 | -0.053 | -0.053 | 0 | 0.79 | 0.68 | 0.73 | 0.95 | 0.74 | 0.86 |
| 7 | 157643516 | rs1529839 | 0.725 | -0.634 | -0.457 | -0.749 | -0.004 | -0.076 | -0.076 | 0 | 0.65 | 0.42 | 0.63 | 0.89 | 0.65 | 0.67 |
| 7 | 157644152 | rs1000137 | 0.569 | 0.2 | -0.614 | 0.074 | -0.002 | -0.002 | -0.002 | 0 | 0.48 | 0.4 | 0.47 | 0.55 | 0.48 | 0.55 |
| 7 | 157646874 | rs4716507 | 1.105 | 0.473 | -0.84 | 0.135 | -0.007 | -0.008 | -0.008 | 0 | 0.37 | 0.3 | 0.39 | 0.51 | 0.42 | 0.41 |
| 7 | 157649261 | rs4716793 | -0.084 | 0.138 | -1.403 | -0.16 | 0.039 | 0.081 | 0.081 | 0 | 0.27 | 0.09 | 0.16 | 0.15 | 0.18 | 0.31 |
| 7 | 157650340 | rs12667537 | 0.467 | 0.657 | -0.64 | -0.237 | -0.002 | -0.024 | -0.024 | 0 | 0.54 | 0.45 | 0.64 | 0.85 | 0.55 | 0.62 |
| 7 | 157650469 | rs11769878 | -0.938 | 0.793 | -0.466 | 0.516 | -0.021 | 0.019 | 0.019 | 0.915 | 0.33 | 0.47 | 0.25 | 0.36 | 0.28 | 0.3 |
| 7 | 157657666 | rs13239010 | -1.142 | 1.678 | -0.64 | 0.503 | 0.001 | -0.002 | -0.002 | 0 | 0.16 | 0.19 | 0.21 | 0.1 | 0.22 | 0.12 |
| 7 | 157658696 | rs7788108 | -0.918 | 1.525 | -0.739 | 1.067 | -0.001 | 0.008 | 0.008 | 0 | 0.22 | 0.72 | 0.15 | 0.38 | 0.23 | 0.3 |
| 7 | 157674708 | rs4645466 | 0.536 | 1.979 | -1.079 | 0.446 | -0.037 | 0.051 | 0.051 | 0 | 0.87 | 0.97 | 0.88 | 1 | 0.83 | 0.83 |
| 7 | 157682613 | rs6952172 | -0.396 | 2.701 | -0.742 | 1.259 | 0.037 | 0.102 | 0.102 | 0 | 0.05 | 0.11 | 0.25 | 0.2 | 0.12 | 0.12 |
| 7 | 157688363 | rs4716801 | 1.442 | 2.852 | -0.408 | 1.102 | -0.009 | -0.005 | -0.005 | 0 | 0.61 | 0.35 | 0.5 | 0.57 | 0.53 | 0.6 |
| 7 | 157689081 | rs7802866 | 0.258 | 2.822 | 0.377 | 0.933 | 0.041 | -0.084 | -0.084 | 0 | 0.85 | 0.74 | 0.86 | 1 | 0.81 | 0.82 |

| | | | | | | | | | | | | | | | | |
|---|---|---|---|---|---|---|---|---|---|---|---|---|---|---|---|---|
| 7 | 157692304 | rs10265651 | -0.681 | 2.961 | 0.035 | 1.343 | -0.021 | -0.008 | -0.008 | 0.014 | 0.08 | 0.05 | 0.33 | 0.25 | 0.17 | 0.13 |
| 7 | 157693378 | rs6963993 | -1.284 | 2.101 | -0.678 | 0.761 | 0.03 | 0.039 | 0.039 | 0 | 0.64 | 0.79 | 0.78 | 0.8 | 0.73 | 0.51 |
| 7 | 157695943 | rs10949670 | 0.495 | 2.013 | -0.847 | 0.836 | -0.038 | 0.053 | 0.053 | 0 | 0.36 | 0.11 | 0.26 | 0.3 | 0.37 | 0.17 |
| 7 | 157710728 | rs10245118 | 0.002 | 1.336 | -0.5 | 0.533 | -0.016 | 0.022 | 0.022 | 0 | 0.14 | 0.07 | 0.11 | 0 | 0.18 | 0.17 |
| 7 | 157713213 | rs10253175 | -0.743 | 0.576 | -0.195 | 0.266 | -0.033 | 0.052 | 0.052 | 1.777 | 0.7 | 0.46 | 0.78 | 0.54 | 0.78 | 0.54 |
| 7 | 157714638 | rs4716807 | 0.258 | 0.362 | -0.386 | 0.071 | 0.011 | 0.003 | 0.003 | 0.166 | 0.53 | 0.41 | 0.66 | 0.54 | 0.58 | 0.36 |
| 7 | 157718481 | rs4716810 | -1.342 | 0.523 | -1.039 | 0.128 | 0.02 | -0.039 | -0.039 | 0.394 | 0.74 | 0.87 | 0.82 | 0.63 | 0.78 | 0.54 |
| 7 | 157720553 | rs12674025 | 0.152 | 0.556 | -0.99 | 0.181 | 0.011 | 0.032 | 0.032 | 0.056 | 0.56 | 0.78 | 0.7 | 0.63 | 0.58 | 0.37 |
| 7 | 157728792 | rs11772354 | -0.855 | 0.176 | -1.287 | 0.061 | 0.033 | 0.035 | 0.035 | 2.293 | 0.71 | 0.89 | 0.82 | 0.79 | 0.79 | 0.58 |
| 7 | 157732404 | rs10226480 | -1.606 | -0.035 | -1.402 | 0.068 | -0.024 | -0.026 | -0.026 | 0 | 0.9 | 1 | 0.86 | 0.9 | 0.88 | 0.75 |
| 7 | 157734314 | rs920023 | -0.102 | -0.162 | -1.511 | -0.264 | 0.01 | 0 | 0 | 0 | 0.35 | 0.69 | 0.38 | 0.64 | 0.25 | 0.31 |
| 7 | 157735966 | rs12670619 | -0.597 | 0.036 | -1.422 | -0.542 | -0.005 | 0.043 | 0.043 | 0 | 0.54 | 0.7 | 0.55 | 0.66 | 0.45 | 0.49 |
| 7 | 157736733 | rs13241820 | 0.891 | 0.02 | -1.611 | -0.638 | -0.098 | 0.063 | 0.063 | 0 | 0.34 | 0.06 | 0.27 | 0.22 | 0.42 | 0.2 |
| 7 | 157736936 | rs10249811 | -0.422 | 0.048 | -1.582 | -0.622 | -0.001 | 0.051 | 0.051 | 0 | 0.53 | 0.7 | 0.55 | 0.66 | 0.44 | 0.49 |
| 7 | 157743364 | rs2117516 | 1.045 | -0.045 | -1.273 | -0.562 | 0 | -0.024 | -0.024 | 0 | 0.14 | 0.13 | 0.09 | 0.02 | 0.14 | 0.17 |
| 7 | 157743465 | rs2241749 | 0.479 | 0.121 | -1.054 | 0.045 | 0.004 | 0.346 | 0.346 | 0.243 | 0.15 | 0.48 | 0.35 | 0.47 | 0.12 | 0.19 |
| 7 | 157744057 | rs1982126 | 1.283 | 0.66 | -1.49 | -0.45 | -0.088 | 0.085 | 0.085 | 0 | 0.21 | 0.02 | 0.22 | 0.08 | 0.31 | 0.12 |
| 7 | 157746669 | rs9654710 | 0.074 | -0.072 | -0.83 | 0.028 | 0.005 | 0.006 | 0.006 | 0 | 0.52 | 0.54 | 0.66 | 0.84 | 0.61 | 0.48 |
| 7 | 157747326 | rs4716828 | 0.195 | -0.047 | -0.833 | 0.528 | 0.009 | 0.009 | 0.009 | 0 | 0.51 | 0.52 | 0.66 | 0.59 | 0.61 | 0.4 |
| 7 | 157750173 | rs11773070 | -1.484 | -0.181 | -1.014 | 0.488 | -0.01 | -0.005 | -0.005 | 0.387 | 0.86 | 0.69 | 0.82 | 0.89 | 0.88 | 0.74 |
| 7 | 157754482 | rs7801805 | 0.287 | -0.062 | -1.665 | 0.185 | 0 | -0.04 | -0.04 | 0.612 | 0.35 | 0.29 | 0.17 | 0.05 | 0.25 | 0.27 |
| 7 | 157754963 | rs2164219 | 0.242 | 0.883 | -2.4 | 0.826 | -0.142 | 0.127 | 0.127 | 0 | 0.39 | 0.05 | 0.31 | 0.23 | 0.51 | 0.23 |
| 7 | 157755225 | rs2164220 | -0.193 | 1.304 | -0.969 | 1.695 | -0.028 | -0.034 | -0.034 | 0 | 0.09 | 0.02 | 0.31 | 0.36 | 0.11 | 0.16 |
| 7 | 157758481 | rs6956867 | -0.018 | 1.241 | -0.492 | 1.174 | -0.005 | -0.009 | -0.009 | 0 | 0.31 | 0.31 | 0.51 | 0.4 | 0.36 | 0.33 |
| 7 | 157761075 | rs4716515 | -0.044 | 1.079 | -0.14 | 0.235 | 0 | -0.011 | -0.011 | 0 | 0.54 | 0.5 | 0.59 | 0.69 | 0.52 | 0.64 |
| 7 | 157764649 | rs4236185 | 0.269 | 0.961 | -0.202 | -0.018 | 0.008 | 0.012 | 0.012 | 0 | 0.48 | 0.58 | 0.55 | 0.52 | 0.45 | 0.45 |
| 7 | 157768397 | rs4716858 | 0.283 | 1.079 | -0.316 | 0.001 | 0 | 0 | 0 | 0 | 0.51 | 0.49 | 0.56 | 0.53 | 0.5 | 0.48 |
| 7 | 157769055 | rs2043496 | -0.071 | 0.856 | -1.123 | -0.521 | -0.004 | 0.055 | 0.055 | 0.085 | 0.74 | 0.9 | 0.77 | 0.84 | 0.74 | 0.71 |
| 7 | 157777861 | rs4716866 | 0.111 | 0.279 | -0.766 | -0.527 | 0.002 | -0.001 | -0.001 | 1.265 | 0.38 | 0.33 | 0.35 | 0.58 | 0.32 | 0.25 |
| 7 | 157782708 | rs920027 | -0.136 | 0.375 | -0.994 | -0.864 | 0.012 | 0.035 | 0.035 | 0.039 | 0.78 | 0.91 | 0.83 | 0.84 | 0.81 | 0.69 |
| 7 | 157791029 | rs11984274 | 0.341 | 0.222 | -0.204 | -1.635 | -0.021 | -0.097 | -0.097 | 0.186 | 0.6 | 0.54 | 0.72 | 0.87 | 0.71 | 0.72 |
| 7 | 157799412 | rs1865464 | 0.037 | -0.111 | 0.68 | -1.581 | -0.027 | -0.076 | -0.076 | 0 | 0.73 | 0.67 | 0.8 | 0.87 | 0.85 | 0.8 |
| 7 | 157802672 | rs1001099 | -0.055 | -0.273 | 0.886 | -1.62 | -0.015 | -0.043 | -0.043 | 0 | 0.75 | 0.75 | 0.81 | 0.87 | 0.87 | 0.78 |
| 7 | 157803711 | rs12698121 | 0.75 | 0.337 | 0.84 | -1.732 | 0.001 | -0.002 | -0.002 | 0 | 0.63 | 0.38 | 0.54 | 0.8 | 0.68 | 0.68 |
| 7 | 157816130 | rs4716880 | 0.779 | 0.186 | 0.896 | -1.556 | 0.014 | 0.038 | 0.038 | 2.97 | 0.63 | 0.3 | 0.52 | 0.75 | 0.69 | 0.67 |
| 7 | 157817047 | rs4716881 | 0.81 | 0.24 | 0.884 | -1.554 | 0.015 | 0.046 | 0.046 | 2.886 | 0.63 | 0.29 | 0.51 | 0.75 | 0.68 | 0.67 |
| 7 | 157819325 | rs11772608 | -0.222 | 0.203 | 1.013 | -1.528 | -0.057 | -0.06 | -0.06 | 1.753 | 0.75 | 0.66 | 0.81 | 0.81 | 0.88 | 0.77 |
| 7 | 157823810 | rs4716535 | 0.553 | -0.056 | 1.032 | -1.461 | -0.032 | -0.034 | -0.034 | 0.555 | 0.68 | 0.53 | 0.62 | 0.75 | 0.76 | 0.73 |
| 7 | 157825009 | rs9918708 | -0.023 | -0.25 | 0.751 | -1.39 | -0.052 | -0.044 | -0.044 | 0 | 0.76 | 0.68 | 0.81 | 0.81 | 0.9 | 0.77 |
| 7 | 157827547 | rs6945759 | -1.131 | 0.23 | -0.367 | -1.526 | 0.007 | 0.029 | 0.029 | 0 | 0.63 | 0.86 | 0.61 | 0.31 | 0.66 | 0.51 |
| 7 | 157830676 | rs11760246 | -0.103 | 1.125 | 0.611 | -1.6 | 0.005 | -0.017 | -0.017 | 0.621 | 0.26 | 0.26 | 0.22 | 0.06 | 0.38 | 0.2 |
| 7 | 157834728 | rs6459830 | -0.875 | 0.576 | 1.252 | -1.606 | 0.012 | -0.06 | -0.06 | 0 | 0.37 | 0.51 | 0.42 | 0.12 | 0.56 | 0.27 |
| 7 | 157835260 | rs10949693 | -0.478 | 0.557 | 1.279 | -1.634 | -0.012 | 0.012 | 0.012 | 0 | 0.32 | 0.3 | 0.37 | 0.12 | 0.53 | 0.23 |
| 7 | 157837357 | rs7783724 | -1.242 | 0.525 | 1.118 | -1.722 | -0.001 | -0.05 | -0.05 | 0 | 0.39 | 0.45 | 0.47 | 0.13 | 0.57 | 0.29 |
| 7 | 157838234 | rs10234599 | -0.688 | 0.748 | 0.946 | -1.559 | -0.01 | 0.008 | 0.008 | 0 | 0.33 | 0.3 | 0.38 | 0.12 | 0.53 | 0.22 |
| 7 | 157842635 | rs10949694 | -1.048 | 0.854 | 0.745 | -1.515 | -0.011 | 0.032 | 0.032 | 0 | 0.38 | 0.68 | 0.47 | 0.15 | 0.56 | 0.26 |
| 7 | 157844387 | rs10256092 | -0.824 | 1.022 | 0.524 | -1.387 | 0.04 | -0.058 | -0.058 | 0 | 0.34 | 0.48 | 0.43 | 0.14 | 0.52 | 0.22 |
| 7 | 157848770 | rs10254965 | 0.013 | 0.719 | 0.233 | -0.721 | -0.011 | -0.015 | -0.015 | 0 | 0.4 | 0.57 | 0.29 | 0.32 | 0.29 | 0.45 |
| 7 | 157854787 | rs10276323 | -0.882 | 1.023 | -0.114 | -0.863 | -0.013 | 0.016 | 0.016 | 0.906 | 0.59 | 0.7 | 0.55 | 0.39 | 0.55 | 0.61 |
| 7 | 157860945 | rs10949695 | 0.581 | 0.83 | -0.018 | -0.678 | 0.009 | 0.048 | 0.048 | 0.893 | 0.34 | 0.16 | 0.22 | 0.22 | 0.27 | 0.4 |
| 7 | 157866856 | rs6961856 | 0.005 | 1.126 | -0.149 | -0.614 | 0.004 | 0.012 | 0.012 | 0.284 | 0.45 | 0.61 | 0.3 | 0.32 | 0.37 | 0.55 |
| 7 | 157870786 | rs6978278 | 1.028 | 1.589 | 0.225 | 0.337 | 0.074 | 0.036 | 0.036 | 0 | 0.78 | 0.58 | 0.62 | 0.74 | 0.62 | 0.79 |
| 7 | 157873975 | rs3800863 | 0.873 | 1.667 | 0.295 | 0.259 | 0.077 | -0.077 | -0.077 | 0.273 | 0.8 | 0.58 | 0.62 | 0.92 | 0.67 | 0.85 |
| 7 | 157875083 | rs4909143 | 1.56 | 1.33 | 0.076 | 0.196 | 0.023 | -0.056 | -0.056 | 0.073 | 0.72 | 0.55 | 0.61 | 0.84 | 0.65 | 0.7 |
| 7 | 157875779 | rs2335487 | 1.458 | 1.38 | 0.506 | 0.361 | 0.036 | 0.018 | 0.018 | 0.06 | 0.75 | 0.47 | 0.6 | 0.67 | 0.62 | 0.78 |
| 7 | 157875903 | rs2335486 | 0.75 | 1.167 | 0.756 | 0.379 | 0.082 | -0.021 | -0.021 | 0 | 0.92 | 0.68 | 0.85 | 1 | 0.91 | 0.97 |
| 7 | 157880135 | rs7780849 | 0.671 | 1.112 | 0.78 | 0.419 | 0.062 | 0.023 | 0.023 | 0 | 0.92 | 0.68 | 0.84 | 0.9 | 0.92 | 0.92 |
| 7 | 157903478 | rs3800861 | 0.806 | 0.638 | 0.551 | 0.553 | -0.01 | -0.002 | -0.002 | 0 | 0.67 | 0.64 | 0.71 | 0.64 | 0.67 | 0.56 |
| 7 | 157903504 | rs3800860 | 1.056 | 1.014 | -0.237 | 0.286 | -0.06 | 0.097 | 0.097 | 1.779 | 0.34 | 0.1 | 0.25 | 0.14 | 0.35 | 0.28 |
| 7 | 157918314 | rs4909256 | -0.908 | 0.765 | -0.214 | 0.181 | -0.069 | 0.015 | 0.015 | 1.151 | 0.63 | 0.86 | 0.73 | 0.69 | 0.59 | 0.69 |
| 7 | 157918582 | rs4909094 | 0.786 | 0.659 | 0.078 | 0.538 | 0.001 | 0.008 | 0.008 | 0 | 0.3 | 0.3 | 0.43 | 0.39 | 0.29 | 0.22 |
| 7 | 157925600 | rs2335851 | -1.13 | 0.588 | -0.005 | 0.477 | -0.072 | 0.01 | 0.01 | 0.644 | 0.64 | 0.88 | 0.73 | 0.69 | 0.59 | 0.69 |
| 7 | 157928710 | rs4475429 | 0.664 | 0.814 | 0.306 | 1.025 | 0.002 | 0.011 | 0.011 | 0 | 0.3 | 0.36 | 0.43 | 0.36 | 0.29 | 0.22 |
| 7 | 157930785 | rs10434933 | 1.117 | 0.81 | 0.528 | 0.932 | -0.005 | -0.052 | -0.052 | 0.599 | 0.25 | 0.14 | 0.31 | 0.44 | 0.23 | 0.19 |
| 7 | 157931119 | rs1130500 | 0.42 | 0.817 | 0.576 | 0.854 | 0.004 | 0.052 | 0.052 | 0 | 0.31 | 0.45 | 0.44 | 0.47 | 0.29 | 0.24 |
| 7 | 157931144 | rs1130499 | 0.42 | 0.817 | 0.574 | 0.854 | 0.003 | 0.044 | 0.044 | 0 | 0.31 | 0.44 | 0.44 | 0.47 | 0.29 | 0.24 |
| 7 | 157934557 | rs6459842 | -0.113 | 0.839 | 0.448 | 0.733 | -0.027 | 0.005 | 0.005 | 0 | 0.38 | 0.48 | 0.36 | 0.36 | 0.32 | 0.37 |
| 7 | 157954008 | rs7801641 | -0.763 | 0.422 | 0.244 | 0.539 | -0.044 | -0.09 | -0.09 | 0 | 0.24 | 0.57 | 0.17 | 0.06 | 0.13 | 0.16 |
| 7 | 157954904 | rs4909289 | 0.593 | 0.506 | -0.172 | 0.411 | -0.005 | 0.005 | 0.005 | 0.279 | 0.8 | 0.77 | 0.8 | 0.82 | 0.79 | 0.76 |

| | | | | | | | | | | | | | | | | |
|---|---|---|---|---|---|---|---|---|---|---|---|---|---|---|---|---|
| 7 | 157959879 | rs1130497 | 1.156 | 1.042 | -0.618 | 0.599 | -0.048 | -0.021 | -0.021 | 0 | 0.44 | 0.14 | 0.42 | 0.41 | 0.45 | 0.36 |
| 7 | 157959895 | rs1130496 | 1.156 | 1.042 | -0.641 | 0.599 | -0.05 | -0.022 | -0.022 | 0 | 0.44 | 0.14 | 0.42 | 0.41 | 0.45 | 0.36 |
| 7 | 157976267 | rs4909119 | 0.941 | 0.736 | -0.399 | 0.323 | -0.007 | -0.018 | -0.018 | 0 | 0.64 | 0.41 | 0.74 | 0.75 | 0.73 | 0.62 |
| 7 | 157981977 | rs6459851 | 0.263 | 0.791 | -0.859 | 0.457 | -0.005 | 0.037 | 0.037 | 0.992 | 0.91 | 0.99 | 0.95 | 0.94 | 0.94 | 0.91 |
| 7 | 157982756 | rs6966759 | -0.294 | 0.855 | -0.861 | 0.507 | 0.002 | 0.004 | 0.004 | 0 | 0.52 | 0.65 | 0.67 | 0.47 | 0.59 | 0.55 |
| 7 | 157983353 | rs10231253 | 0.789 | 0.765 | -0.593 | 0.566 | -0.014 | 0.011 | 0.011 | 0 | 0.23 | 0.36 | 0.23 | 0.25 | 0.17 | 0.23 |
| 7 | 157992083 | rs2335832 | 1.64 | 1.001 | -1.423 | 0.637 | -0.062 | -0.042 | -0.042 | 0 | 0.25 | 0.01 | 0.32 | 0.19 | 0.25 | 0.22 |
| 7 | 158015301 | rs10215289 | 0.284 | 0.07 | -0.672 | 0.59 | -0.004 | -0.001 | -0.001 | 0 | 0.31 | 0.65 | 0.29 | 0.3 | 0.19 | 0.28 |
| 7 | 158015560 | rs6944416 | -0.585 | 0.262 | -0.95 | 0.536 | 0.018 | 0.049 | 0.049 | 0 | 0.45 | 0.75 | 0.44 | 0.37 | 0.39 | 0.41 |
| 7 | 158019437 | rs10256080 | 0.481 | 0.282 | -0.966 | -0.018 | -0.043 | 0.077 | 0.077 | 0 | 0.29 | 0.43 | 0.27 | 0.48 | 0.19 | 0.3 |
| 7 | 158020147 | rs6970487 | 0.043 | 0.53 | -1.606 | 0.197 | 0.023 | -0.025 | -0.025 | 1.258 | 0.46 | 0.78 | 0.44 | 0.54 | 0.4 | 0.45 |
| 7 | 158022103 | rs2335848 | 0.013 | 0.5 | -1.64 | 0.201 | 0.022 | -0.025 | -0.025 | 0.048 | 0.46 | 0.8 | 0.45 | 0.54 | 0.4 | 0.45 |
| 7 | 158028901 | rs4909140 | 0.097 | 1.307 | -0.959 | 0.094 | -0.04 | 0.017 | 0.017 | 0 | 0.37 | 0.22 | 0.47 | 0.29 | 0.5 | 0.41 |
| 7 | 158031109 | rs6963236 | 0.845 | 1.035 | -0.133 | -0.279 | -0.003 | -0.074 | -0.074 | 2.15 | 0.78 | 0.71 | 0.81 | 0.96 | 0.83 | 0.86 |
| 7 | 158031556 | rs3752372 | -1.183 | 1.155 | 0.583 | 0.111 | 0.037 | 0.022 | 0.022 | 0.129 | 0.14 | 0.58 | 0.19 | 0.22 | 0.14 | 0.07 |
| 7 | 158031749 | rs7777448 | -0.171 | 1.595 | 0.553 | -0.423 | 0.026 | -0.125 | -0.125 | 0 | 0.27 | 0.49 | 0.26 | 0.06 | 0.36 | 0.32 |
| 7 | 158035784 | rs1986588 | 0.865 | 0.351 | 0.444 | 0.104 | -0.005 | 0.002 | 0.002 | 0 | 0.17 | 0.16 | 0.25 | 0.2 | 0.09 | 0.21 |
| 7 | 158038989 | rs10235911 | -0.12 | 0.505 | 0.222 | 0.343 | 0 | 0 | 0 | 0 | 0.44 | 0.49 | 0.49 | 0.5 | 0.6 | 0.58 |
| 7 | 158039697 | rs7791230 | 0.76 | 0.558 | -0.214 | 0.433 | -0.015 | 0.038 | 0.038 | 0 | 0.18 | 0.25 | 0.31 | 0.32 | 0.13 | 0.23 |
| 7 | 158043004 | rs10949703 | -0.484 | 0.879 | -0.625 | 0.092 | 0.014 | 0.077 | 0.077 | 1.387 | 0.64 | 0.88 | 0.81 | 0.82 | 0.74 | 0.81 |
| 7 | 158050239 | rs12698186 | -0.517 | 1.121 | 0.147 | 0.617 | 0.028 | 0.052 | 0.052 | 0 | 0.11 | 0.23 | 0.2 | 0.27 | 0.12 | 0.09 |
| 7 | 158050807 | rs12698189 | -0.256 | 1.132 | 0.154 | 0.625 | 0.016 | 0.014 | 0.014 | 0.097 | 0.09 | 0.18 | 0.11 | 0.14 | 0.09 | 0.06 |
| 7 | 158051502 | rs11773209 | -0.281 | 1.039 | 0.035 | 0.462 | 0.001 | -0.009 | -0.009 | 0 | 0.42 | 0.52 | 0.29 | 0.33 | 0.39 | 0.2 |
| 7 | 158053180 | rs6976603 | -0.492 | 0.998 | -0.063 | 0.362 | -0.008 | 0.003 | 0.003 | 0.021 | 0.45 | 0.67 | 0.38 | 0.46 | 0.42 | 0.24 |
| 7 | 158064729 | rs7796032 | -0.363 | 0.585 | -0.005 | -0.007 | -0.01 | -0.028 | -0.028 | 0 | 0.31 | 0.42 | 0.25 | 0.21 | 0.26 | 0.13 |
| 7 | 158071511 | rs7795387 | -0.97 | 0.308 | 0.124 | -0.031 | -0.017 | -0.007 | -0.007 | 0.024 | 0.27 | 0.48 | 0.28 | 0.24 | 0.25 | 0.09 |
| 7 | 158074146 | rs1982450 | -0.587 | 0.184 | -0.446 | 0.331 | 0.023 | -0.069 | -0.069 | 0.044 | 0.29 | 0.13 | 0.43 | 0.4 | 0.26 | 0.27 |
| 7 | 158075769 | rs11973244 | 1.01 | -0.085 | -0.191 | 0.194 | 0.003 | 0 | 0 | 0 | 0.54 | 0.47 | 0.38 | 0.44 | 0.58 | 0.6 |
| 7 | 158103844 | rs4909062 | -0.8 | 1.026 | -0.44 | 0.383 | -0.038 | -0.052 | -0.052 | 0 | 0.39 | 0.22 | 0.55 | 0.5 | 0.48 | 0.38 |
| 7 | 158113390 | rs896779 | -0.664 | -0.07 | 0.129 | -0.183 | 0.003 | 0.005 | 0.005 | 0 | 0.46 | 0.45 | 0.41 | 0.3 | 0.4 | 0.37 |
| 7 | 158115424 | rs10266663 | 0.735 | -0.245 | 0.343 | -0.18 | 0.034 | 0.092 | 0.092 | 0 | 0.93 | 0.65 | 0.87 | 0.86 | 0.97 | 0.94 |
| 7 | 158116911 | rs10434944 | 0.817 | -0.225 | 0.778 | -0.198 | 0.006 | 0.005 | 0.005 | 0 | 0.51 | 0.59 | 0.47 | 0.56 | 0.58 | 0.57 |
| 7 | 158122287 | rs10156158 | -0.669 | -0.139 | 0.452 | -0.297 | -0.002 | -0.004 | -0.004 | 0 | 0.45 | 0.48 | 0.43 | 0.3 | 0.39 | 0.38 |
| 7 | 158136204 | rs12698214 | -0.292 | -0.517 | -0.272 | -0.224 | 0.027 | 0.03 | 0.03 | 0.01 | 0.32 | 0.13 | 0.16 | 0.2 | 0.27 | 0.13 |
| 7 | 158145606 | rs12698219 | -0.376 | 0.161 | -0.248 | -0.419 | -0.013 | -0.022 | -0.022 | 0 | 0.11 | 0.02 | 0.14 | 0.12 | 0.09 | 0.24 |
| 7 | 158155259 | rs6459868 | 0.674 | -0.071 | 0.424 | -0.619 | -0.001 | 0.001 | 0.001 | 0 | 0.94 | 0.88 | 0.86 | 0.86 | 0.95 | 0.91 |
| 7 | 158161501 | rs6942394 | 1.037 | -0.402 | 0.211 | -0.944 | 0 | -0.007 | -0.007 | 0 | 0.72 | 0.68 | 0.73 | 0.84 | 0.76 | 0.82 |
| 7 | 158164771 | rs7810678 | 0.562 | -0.197 | 0.537 | -0.696 | 0.003 | -0.095 | -0.095 | 0.662 | 0.8 | 0.67 | 0.87 | 0.99 | 0.81 | 0.92 |
| 7 | 158166170 | rs10245268 | -1.661 | -0.806 | 0.909 | -0.277 | -0.051 | -0.01 | -0.01 | 1.453 | 0.17 | 0.34 | 0.27 | 0.21 | 0.11 | 0.08 |
| 7 | 158170316 | rs7803342 | 1.778 | -0.675 | 0.836 | -0.53 | -0.012 | -0.05 | -0.05 | 0 | 0.9 | 0.75 | 0.86 | 0.99 | 0.91 | 0.9 |
| 7 | 158172229 | rs7806066 | 1.841 | -0.836 | 1.028 | -0.499 | -0.031 | -0.04 | -0.04 | 0.266 | 0.82 | 0.74 | 0.9 | 0.93 | 0.93 | 0.89 |
| 7 | 158176488 | rs1453096 | 1.702 | -0.798 | 1.106 | -0.441 | -0.043 | -0.071 | -0.071 | 0 | 0.81 | 0.5 | 0.87 | 0.93 | 0.89 | 0.88 |
| 7 | 158182029 | rs896762 | 0.316 | -0.414 | -0.19 | -0.463 | -0.028 | 0.036 | 0.036 | 0.007 | 0.39 | 0.17 | 0.58 | 0.75 | 0.45 | 0.55 |
| 7 | 158194935 | rs1563066 | 0.147 | -0.374 | 0.041 | -0.495 | -0.012 | 0.038 | 0.038 | 0 | 0.42 | 0.26 | 0.63 | 0.75 | 0.48 | 0.61 |
| 7 | 158196003 | rs10949725 | 0.606 | -0.581 | 0.67 | -0.634 | 0.002 | -0.014 | -0.014 | 0 | 0.43 | 0.42 | 0.29 | 0.16 | 0.46 | 0.32 |
| 7 | 158209394 | rs2123498 | -0.035 | -0.558 | 0.566 | -0.553 | -0.007 | -0.014 | -0.014 | 0 | 0.94 | 0.91 | 0.97 | 0.96 | 0.96 | 0.93 |
| 7 | 158215549 | rs1377373 | 0.041 | -0.392 | 0.426 | -0.542 | -0.002 | 0.042 | 0.042 | 0 | 0.53 | 0.61 | 0.33 | 0.21 | 0.49 | 0.33 |
| 7 | 158223106 | rs7786808 | -0.19 | 0.004 | 0.136 | -0.063 | -0.002 | 0.012 | 0.012 | 0 | 0.59 | 0.68 | 0.43 | 0.38 | 0.56 | 0.6 |
| 7 | 158230595 | rs10247026 | 0.099 | -0.171 | -0.226 | 0.063 | -0.015 | 0.003 | 0.003 | 0 | 0.42 | 0.15 | 0.55 | 0.6 | 0.43 | 0.41 |
| 7 | 158235670 | rs10260982 | -0.73 | 0.249 | -0.131 | -0.068 | 0.001 | -0.001 | -0.001 | 0 | 0.08 | 0.13 | 0.07 | 0.08 | 0.08 | 0.05 |
| 7 | 158240776 | rs11773729 | -0.102 | -0.105 | -0.736 | 0.214 | 0.019 | 0.001 | 0.001 | 0 | 0.26 | 0.18 | 0.34 | 0.27 | 0.23 | 0.25 |
| 7 | 158245174 | rs1670377 | 0.027 | 0.107 | -0.41 | 0.117 | -0.001 | -0.003 | -0.003 | 2.108 | 0.52 | 0.49 | 0.53 | 0.39 | 0.54 | 0.34 |
| 7 | 158251657 | rs1733161 | -0.07 | 0.076 | -0.353 | 0.126 | 0.006 | 0.015 | 0.015 | 0 | 0.53 | 0.59 | 0.54 | 0.43 | 0.54 | 0.42 |
| 7 | 158259728 | rs1733148 | -1.369 | 0 | -0.542 | 0.537 | 0.007 | -0.028 | -0.028 | 0 | 0.81 | 0.88 | 0.85 | 0.8 | 0.82 | 0.72 |
| 7 | 158260724 | rs1670357 | -0.221 | -0.09 | -0.333 | 0.449 | -0.001 | 0.005 | 0.005 | 0.374 | 0.72 | 0.59 | 0.78 | 0.73 | 0.73 | 0.63 |
| 7 | 158265425 | rs1670350 | -0.257 | -0.057 | -0.859 | 0.405 | 0 | -0.001 | -0.001 | 0 | 0.72 | 0.65 | 0.78 | 0.73 | 0.72 | 0.61 |
| 7 | 158268596 | rs1630862 | 0.492 | -0.024 | -1.022 | 0.348 | 0 | 0.004 | 0.004 | 0 | 0.45 | 0.47 | 0.41 | 0.31 | 0.44 | 0.29 |
| 7 | 158271022 | rs1670340 | -0.534 | 0.015 | -1.063 | 0.26 | 0 | 0.022 | 0.022 | 0 | 0.51 | 0.49 | 0.56 | 0.67 | 0.52 | 0.68 |
| 7 | 158271139 | rs1670339 | 1.323 | -0.119 | -1.191 | 0.484 | 0.011 | -0.029 | -0.029 | 0 | 0.18 | 0.09 | 0.13 | 0.19 | 0.16 | 0.26 |
| 7 | 158276926 | rs7810884 | 0.528 | -0.078 | -1.152 | 0.494 | 0.013 | 0.007 | 0.007 | 0.205 | 0.29 | 0.21 | 0.2 | 0.22 | 0.26 | 0.34 |
| 7 | 158279412 | rs5016019 | 0.246 | 0.13 | -1.089 | 0.692 | -0.008 | -0.106 | -0.106 | 0 | 0.22 | 0.06 | 0.32 | 0.43 | 0.23 | 0.3 |
| 7 | 158299094 | rs10256091 | -0.14 | 0.114 | -0.916 | 0.731 | -0.004 | 0.023 | 0.023 | 0 | 0.25 | 0.36 | 0.34 | 0.45 | 0.23 | 0.33 |
| 7 | 158304646 | rs2091718 | 0.182 | 0.19 | -1.172 | 0.696 | -0.012 | -0.099 | -0.099 | 0.086 | 0.22 | 0.06 | 0.31 | 0.41 | 0.23 | 0.29 |
| 7 | 158322716 | rs10081235 | -0.307 | -0.069 | -1.295 | 0.386 | 0.007 | 0.071 | 0.071 | 0 | 0.5 | 0.37 | 0.53 | 0.68 | 0.49 | 0.71 |
| 7 | 158350293 | rs2335160 | -0.29 | 0.392 | -0.834 | 0.935 | -0.01 | -0.06 | -0.06 | 0.339 | 0.19 | 0.11 | 0.32 | 0.42 | 0.22 | 0.29 |
| 7 | 158376086 | rs10275533 | -0.562 | 0.481 | -0.814 | 0.945 | -0.002 | -0.074 | -0.074 | 0.07 | 0.23 | 0.12 | 0.39 | 0.45 | 0.23 | 0.3 |
| 7 | 158389294 | rs1882247 | 0.254 | 0.639 | -0.142 | 1.23 | -0.004 | 0.081 | 0.081 | 0 | 0.07 | 0.11 | 0.24 | 0.36 | 0.08 | 0.22 |
| 7 | 158400183 | rs10262207 | 0.577 | 1.423 | 0.676 | 0.55 | 0.068 | -0.086 | -0.086 | 0 | 0.8 | 0.68 | 0.79 | 0.96 | 0.72 | 0.9 |
| 7 | 158412279 | rs11983468 | 0.925 | 0.603 | 1.103 | 0.419 | 0 | -0.031 | -0.031 | 0.003 | 0.72 | 0.72 | 0.81 | 0.96 | 0.7 | 0.83 |

| | | | | | | | | | | | | | | | | |
|---|---|---|---|---|---|---|---|---|---|---|---|---|---|---|---|---|
| 7 | 158416203 | rs11763040 | 0.853 | 0.584 | 1.239 | 0.456 | -0.001 | -0.016 | -0.016 | 0 | 0.73 | 0.73 | 0.81 | 0.92 | 0.71 | 0.83 |
| 7 | 158424127 | rs10772 | 1.452 | 0.585 | 1.451 | 0.789 | 0.001 | 0.006 | 0.006 | 2.065 | 0.65 | 0.34 | 0.54 | 0.45 | 0.62 | 0.59 |
| 7 | 158427496 | rs6972008 | 1.685 | 0.493 | 1.647 | 0.827 | 0.004 | 0.007 | 0.007 | 0.654 | 0.63 | 0.43 | 0.55 | 0.56 | 0.62 | 0.59 |
| 7 | 158430907 | rs877279 | 1.936 | 0.363 | 1.772 | 0.799 | 0.001 | -0.004 | -0.004 | 0 | 0.6 | 0.35 | 0.61 | 0.45 | 0.62 | 0.58 |
| 7 | 158433732 | rs4909245 | 1.917 | 0.391 | 1.795 | 0.795 | 0 | -0.001 | -0.001 | 0 | 0.61 | 0.4 | 0.62 | 0.45 | 0.63 | 0.58 |
| 7 | 158433916 | rs12670677 | -0.508 | 0.312 | 1.922 | 0.786 | 0.007 | 0.01 | 0.01 | 0 | 0.11 | 0.07 | 0.18 | 0.4 | 0.09 | 0.22 |
| 7 | 158436547 | rs2290395 | -1.067 | 0.316 | 1.878 | 0.788 | -0.026 | 0.174 | 0.174 | 0.054 | 0.12 | 0.3 | 0.2 | 0.4 | 0.1 | 0.22 |
| 7 | 158438186 | rs2290393 | 2.022 | 0.29 | 1.869 | 0.773 | -0.001 | 0.002 | 0.002 | 0.664 | 0.61 | 0.43 | 0.62 | 0.45 | 0.63 | 0.58 |
| 7 | 158439800 | rs6970571 | -0.903 | 0.403 | 2.069 | 0.807 | 0.015 | -0.005 | -0.005 | 0 | 0.93 | 0.93 | 0.96 | 0.97 | 0.89 | 0.97 |
| 7 | 158444634 | rs1869362 | -0.697 | 0.311 | 2.02 | 0.849 | -0.018 | 0.21 | 0.21 | 0 | 0.11 | 0.3 | 0.2 | 0.4 | 0.09 | 0.2 |
| 7 | 158468437 | rs1530967 | -1.506 | 0.488 | 1.742 | 0.82 | -0.021 | 0.021 | 0.021 | 0.404 | 0.32 | 0.45 | 0.34 | 0.41 | 0.27 | 0.38 |
| 7 | 158474325 | rs10266006 | -0.677 | 0.538 | 1.779 | 0.813 | 0.014 | -0.005 | -0.005 | 0.262 | 0.93 | 0.92 | 0.95 | 0.95 | 0.89 | 0.94 |
| 7 | 158476148 | rs6969248 | -0.353 | 0.532 | 1.766 | 0.854 | 0.028 | 0.055 | 0.055 | 0 | 0.92 | 0.85 | 0.95 | 0.85 | 0.89 | 0.93 |
| 7 | 158489040 | rs13242357 | 1.878 | 0.48 | 1.603 | 0.844 | 0 | -0.001 | -0.001 | 0.855 | 0.61 | 0.4 | 0.61 | 0.44 | 0.62 | 0.55 |
| 7 | 158498994 | rs4621754 | -0.627 | 0.335 | 1.4 | 0.796 | 0.029 | 0.087 | 0.087 | 0.153 | 0.1 | 0.29 | 0.07 | 0.15 | 0.1 | 0.07 |
| 7 | 158511628 | rs10949741 | 1.849 | 0.392 | 0.904 | 0.728 | 0.003 | 0.01 | 0.01 | 1.804 | 0.57 | 0.25 | 0.53 | 0.38 | 0.59 | 0.51 |
| 7 | 158515660 | rs12698277 | 1.848 | 0.447 | 0.873 | 0.687 | 0.006 | 0.004 | 0.004 | 1.885 | 0.57 | 0.18 | 0.55 | 0.37 | 0.59 | 0.51 |
| 7 | 158519615 | rs1869363 | -1.568 | 0.194 | 1.776 | 0.985 | -0.015 | 0.176 | 0.176 | 1.441 | 0.2 | 0.44 | 0.3 | 0.49 | 0.21 | 0.29 |
| 7 | 158524763 | rs3763406 | -1.922 | -0.076 | 2.184 | 0.905 | -0.045 | 0.127 | 0.127 | 2.884 | 0.22 | 0.44 | 0.26 | 0.6 | 0.17 | 0.28 |
| 7 | 158524830 | rs3763409 | -1.922 | -0.076 | 2.189 | 0.905 | -0.045 | 0.127 | 0.127 | 2.884 | 0.22 | 0.44 | 0.26 | 0.6 | 0.17 | 0.28 |
| 7 | 158525086 | rs3763412 | -1.922 | -0.076 | 2.189 | 0.905 | -0.045 | 0.127 | 0.127 | 2.884 | 0.22 | 0.44 | 0.26 | 0.6 | 0.17 | 0.28 |
| 7 | 158526185 | rs712192 | -2.221 | -0.106 | 2.416 | 0.892 | -0.035 | 0.062 | 0.062 | 3.872 | 0.23 | 0.61 | 0.28 | 0.6 | 0.17 | 0.28 |
| 7 | 158526354 | rs1061739 | -1.918 | -0.098 | 2.36 | 0.896 | -0.045 | 0.127 | 0.127 | 2.884 | 0.22 | 0.44 | 0.26 | 0.6 | 0.17 | 0.28 |
| 7 | 158526455 | rs1061735 | -1.918 | -0.098 | 2.36 | 0.896 | -0.045 | 0.127 | 0.127 | 2.884 | 0.22 | 0.44 | 0.26 | 0.6 | 0.17 | 0.28 |
| 7 | 158526495 | rs12670487 | -1.691 | -0.133 | 2.643 | 0.943 | -0.068 | 0.205 | 0.205 | 2.355 | 0.15 | 0.33 | 0.22 | 0.55 | 0.07 | 0.22 |
| 7 | 158526661 | rs1061733 | 0.454 | -0.106 | 2.592 | 0.944 | 0.009 | -0.004 | -0.004 | 0 | 0.07 | 0.11 | 0.05 | 0.05 | 0.1 | 0.06 |
| 7 | 158526880 | rs3750051 | -1.022 | -0.128 | 2.614 | 0.986 | -0.058 | 0.311 | 0.311 | 1.825 | 0.13 | 0.37 | 0.21 | 0.55 | 0.07 | 0.21 |
| 7 | 158527605 | rs3763414 | -2.42 | -0.207 | 2.738 | 0.981 | -0.029 | 0.267 | 0.267 | 0.006 | 0.17 | 0.48 | 0.21 | 0.5 | 0.15 | 0.25 |
| 7 | 158527734 | rs896862 | 1.074 | -0.986 | 1.882 | 0.37 | -0.021 | 0.07 | 0.07 | 0.002 | 0.78 | 0.66 | 0.78 | 0.36 | 0.82 | 0.68 |
| 7 | 158527838 | rs3816458 | -1.782 | -1.02 | 1.595 | 0.353 | -0.062 | 0.173 | 0.173 | 2.374 | 0.2 | 0.46 | 0.17 | 0.6 | 0.14 | 0.28 |
| 7 | 158527900 | rs3816460 | -1.48 | -1.171 | 1.658 | 0.463 | -0.047 | 0.177 | 0.177 | 1.764 | 0.2 | 0.51 | 0.19 | 0.59 | 0.16 | 0.26 |
| 7 | 158528233 | rs3816462 | -1.307 | -0.986 | 1.638 | 0.506 | -0.044 | 0.144 | 0.144 | 2.197 | 0.21 | 0.44 | 0.18 | 0.6 | 0.17 | 0.29 |
| 7 | 158529704 | rs6960873 | -1.125 | -0.905 | 1.596 | 0.533 | -0.035 | 0.128 | 0.128 | 3.007 | 0.2 | 0.4 | 0.18 | 0.58 | 0.17 | 0.28 |
| 7 | 158532706 | rs11761517 | 2.098 | -1.254 | 1.183 | 0.289 | -0.125 | 0.114 | 0.114 | 1.781 | 0.39 | 0.06 | 0.36 | 0.22 | 0.47 | 0.2 |
| 7 | 158536267 | rs2305473 | -1.015 | -1.177 | 1.4 | 0.418 | -0.035 | 0.125 | 0.125 | 3.007 | 0.2 | 0.4 | 0.18 | 0.58 | 0.17 | 0.27 |
| 7 | 158536345 | rs2305475 | -1.015 | -1.177 | 1.4 | 0.418 | -0.035 | 0.125 | 0.125 | 3.007 | 0.2 | 0.4 | 0.18 | 0.58 | 0.17 | 0.28 |
| 7 | 158543734 | rs4548125 | 0.284 | -1.164 | 1.27 | 0.511 | 0 | 0.033 | 0.033 | 0 | 0.1 | 0.15 | 0.05 | 0.22 | 0.11 | 0.07 |
| 7 | 158546543 | rs710422 | 2.014 | -1.256 | 1.422 | 0.48 | -0.008 | 0.002 | 0.002 | 2.888 | 0.7 | 0.3 | 0.72 | 0.3 | 0.77 | 0.65 |
| 7 | 158552398 | rs896860 | 0.02 | -1.226 | 1.432 | 0.484 | -0.018 | 0.102 | 0.102 | 0.701 | 0.85 | 0.76 | 0.82 | 0.53 | 0.89 | 0.72 |
| 7 | 158553291 | rs959489 | 0.233 | -1.23 | 1.449 | 0.482 | -0.026 | 0.154 | 0.154 | 0.924 | 0.84 | 0.67 | 0.81 | 0.53 | 0.89 | 0.72 |
| 7 | 158560621 | rs2602542 | 1.693 | -1.228 | 1.441 | 0.491 | -0.028 | 0.075 | 0.075 | 2.408 | 0.73 | 0.47 | 0.75 | 0.42 | 0.78 | 0.66 |
| 7 | 158566367 | rs842449 | 0.14 | -1.207 | 1.159 | 0.518 | -0.023 | 0.134 | 0.134 | 1.097 | 0.84 | 0.73 | 0.81 | 0.53 | 0.89 | 0.72 |
| 7 | 158572273 | rs11971753 | 0.628 | -1.206 | 0.995 | 0.522 | -0.032 | 0.083 | 0.083 | 1.874 | 0.12 | 0.18 | 0.12 | 0.43 | 0.07 | 0.21 |
| 7 | 158577140 | rs842454 | 2.147 | -1.291 | 0.404 | -0.115 | 0.077 | 0.035 | 0.035 | 1.807 | 0.55 | 0.08 | 0.66 | 0.3 | 0.66 | 0.56 |
| 7 | 158584933 | rs7783880 | -0.281 | -1.015 | 1.283 | 0.715 | 0.006 | 0.002 | 0.002 | 0 | 0.1 | 0.2 | 0.06 | 0.11 | 0.11 | 0.06 |
| 7 | 158589340 | rs6459904 | -0.267 | -1.01 | 1.267 | 0.737 | 0.006 | 0.014 | 0.014 | 0 | 0.1 | 0.2 | 0.06 | 0.15 | 0.1 | 0.06 |
| 7 | 158604515 | rs9638144 | -0.389 | -0.853 | 1.253 | 0.753 | -0.051 | 0.203 | 0.203 | 2.005 | 0.13 | 0.28 | 0.13 | 0.55 | 0.07 | 0.22 |
| 7 | 158618491 | rs1153999 | 1.941 | -0.674 | 0.742 | 0.517 | 0.029 | 0.027 | 0.027 | 1.874 | 0.57 | 0.31 | 0.69 | 0.3 | 0.66 | 0.55 |
| 7 | 158626029 | rs2602565 | 1.002 | -0.681 | 0.586 | 0.97 | -0.036 | 0.139 | 0.139 | 2.533 | 0.8 | 0.58 | 0.79 | 0.41 | 0.86 | 0.7 |
| 7 | 158649005 | rs2788474 | -1.319 | -0.423 | 0.2 | 0.711 | 0.013 | 0.008 | 0.008 | 0.641 | 0.44 | 0.59 | 0.37 | 0.7 | 0.36 | 0.46 |
| 7 | 158651087 | rs2657381 | -0.861 | -0.603 | 0.522 | 0.59 | 0.003 | -0.027 | -0.027 | 0.875 | 0.21 | 0.23 | 0.15 | 0.09 | 0.23 | 0.21 |
| 7 | 158665748 | rs1188974 | -0.641 | -0.6 | 0.566 | 0.558 | -0.007 | 0.039 | 0.039 | 0 | 0.18 | 0.07 | 0.1 | 0.07 | 0.19 | 0.12 |
| 7 | 158668575 | rs710421 | -2.056 | -0.818 | 0.9 | 0.63 | 0.024 | 0.019 | 0.019 | 0.759 | 0.44 | 0.63 | 0.38 | 0.7 | 0.36 | 0.46 |
| 7 | 158672619 | rs2788478 | -2.11 | -0.718 | 1.022 | 0.614 | 0.043 | 0.064 | 0.064 | 0.897 | 0.46 | 0.74 | 0.43 | 0.72 | 0.38 | 0.5 |
| 7 | 158684152 | rs12672574 | 0.041 | -0.794 | 1.371 | 1.035 | -0.012 | 0.019 | 0.019 | 9.192 | 0.21 | 0.24 | 0.21 | 0.48 | 0.12 | 0.22 |
| 7 | 158687998 | rs1188960 | -0.585 | -0.789 | 1.292 | 0.969 | -0.006 | 0.05 | 0.05 | 0 | 0.18 | 0.06 | 0.09 | 0.06 | 0.19 | 0.15 |
| 7 | 158704054 | rs3815215 | -0.622 | -0.808 | 1.542 | 1.066 | -0.037 | 0.072 | 0.072 | 8.226 | 0.24 | 0.37 | 0.27 | 0.51 | 0.16 | 0.26 |
| 7 | 158710642 | rs10275341 | -1.009 | 0.04 | 1.515 | 0.815 | 0.11 | -0.004 | -0.004 | 0.576 | 0.09 | 0.3 | 0.13 | 0.04 | 0.17 | 0.07 |
| 7 | 158710879 | rs9654726 | -0.76 | 0.057 | 1.5 | 0.816 | 0.1 | -0.003 | -0.003 | 0 | 0.08 | 0.27 | 0.12 | 0.04 | 0.17 | 0.07 |
| 7 | 158711584 | rs10279428 | -0.76 | 0.042 | 1.5 | 0.816 | 0.104 | -0.003 | -0.003 | 0.598 | 0.08 | 0.27 | 0.12 | 0.04 | 0.17 | 0.07 |
| 7 | 158715264 | rs6459913 | -1.144 | -0.085 | 1.429 | 1.069 | -0.009 | 0.051 | 0.051 | 5.477 | 0.29 | 0.53 | 0.35 | 0.62 | 0.29 | 0.32 |
| 7 | 158726462 | rs6965685 | 1.436 | -0.171 | 1.135 | 0.957 | -0.001 | -0.001 | -0.001 | 0 | 0.61 | 0.6 | 0.7 | 0.5 | 0.7 | 0.64 |
| 7 | 158726655 | rs2657330 | 2.269 | -0.129 | 1.21 | 0.906 | 0.002 | 0.05 | 0.05 | 0 | 0.5 | 0.24 | 0.53 | 0.34 | 0.51 | 0.49 |
| 7 | 158727910 | rs12154715 | 0.018 | -0.093 | 1.002 | 0.9 | 0.094 | -0.017 | -0.017 | 0.45 | 0.07 | 0.2 | 0.11 | 0.01 | 0.17 | 0.08 |
| 7 | 158738611 | rs2527201 | 1.635 | -0.32 | 0.311 | 0.968 | 0.001 | -0.001 | -0.001 | 3.52 | 0.6 | 0.6 | 0.7 | 0.36 | 0.7 | 0.61 |
| 7 | 158738849 | rs2527202 | 0.832 | 0.17 | 0.201 | 0.744 | -0.002 | 0.05 | 0.05 | 0 | 0.82 | 0.92 | 0.92 | 0.95 | 0.82 | 0.85 |
| 7 | 158754581 | rs2527214 | -1.929 | 0.049 | 0.984 | 0.432 | -0.003 | -0.014 | -0.014 | 0 | 0.13 | 0.16 | 0.06 | 0.07 | 0.1 | 0.12 |
| 7 | 158755257 | rs10949749 | -1.038 | 0.247 | 1.198 | 0.582 | 0.037 | 0.008 | 0.008 | 0 | 0.07 | 0.18 | 0.11 | 0.1 | 0.14 | 0.09 |
| 7 | 158759451 | rs2730268 | 1.248 | -0.178 | 1.858 | 0.8 | -0.021 | 0.04 | 0.04 | 0 | 0.89 | 0.69 | 0.89 | 0.89 | 0.93 | 0.91 |

| | | | | | | | | | | | | | | | | |
|---|---|---|---|---|---|---|---|---|---|---|---|---|---|---|---|---|
| 7 | 158775172 | rs262152 | 1.18 | -0.222 | 1.66 | 0.783 | -0.012 | -0.009 | -0.009 | 0.261 | 0.87 | 0.82 | 0.91 | 0.83 | 0.86 | 0.78 |
| 8 | 266953 | rs17080309 | -0.418 | -1.319 | -1.112 | -1.833 | -0.034 | -0.023 | -0.023 | 0 | 0.28 | 0.6 | 0.21 | 0.14 | 0.11 | 0.33 |
| 8 | 280645 | rs7018431 | -0.679 | -2.062 | -0.964 | -1.122 | -0.018 | -0.003 | -0.003 | 0 | 0.16 | 0.2 | 0.07 | 0.16 | 0.1 | 0.18 |
| 8 | 281022 | rs11997640 | -1.556 | -2.211 | -0.828 | -1.198 | 0.093 | 0.08 | 0.08 | 0.296 | 0.43 | 0.79 | 0.27 | 0.26 | 0.25 | 0.51 |
| 8 | 281572 | rs6990287 | -1.584 | -2.268 | -0.921 | -1.356 | 0.131 | 0.105 | 0.105 | 0 | 0.45 | 0.84 | 0.27 | 0.29 | 0.23 | 0.49 |
| 8 | 282497 | rs918323 | -1.11 | -2.006 | 0.654 | -1.007 | -0.018 | -0.001 | -0.001 | 0 | 0.15 | 0.35 | 0.09 | 0.14 | 0.09 | 0.18 |
| 8 | 283603 | rs11136555 | -1.955 | -2.187 | 0.241 | -0.874 | 0.071 | -0.025 | -0.025 | 0 | 0.42 | 0.93 | 0.3 | 0.36 | 0.26 | 0.5 |
| 8 | 285124 | rs17063980 | -0.899 | -2.295 | 1.285 | -0.773 | -0.024 | 0.02 | 0.02 | 0.68 | 0.13 | 0.41 | 0.07 | 0.16 | 0.07 | 0.13 |
| 8 | 286045 | rs17063999 | -1.232 | -2.238 | 1.304 | -0.824 | -0.041 | -0.021 | -0.021 | 0 | 0.17 | 0.39 | 0.09 | 0.14 | 0.09 | 0.18 |
| 8 | 286941 | rs759775 | -0.2 | -2.288 | 1.141 | -0.838 | 0.018 | -0.023 | -0.023 | 0 | 0.18 | 0.05 | 0.14 | 0.16 | 0.13 | 0.28 |
| 8 | 287831 | rs11774820 | -1.448 | -2.301 | 1.578 | -0.874 | -0.055 | -0.042 | -0.042 | 0.021 | 0.2 | 0.54 | 0.12 | 0.14 | 0.11 | 0.2 |
| 8 | 287897 | rs918325 | 1.763 | -2.442 | 1.238 | -0.908 | 0.112 | 0.052 | 0.052 | 0.535 | 0.55 | 0.21 | 0.72 | 0.69 | 0.74 | 0.46 |
| 8 | 294150 | rs6993101 | -1.682 | -2.052 | 1.543 | -0.713 | -0.05 | -0.054 | -0.054 | 0 | 0.22 | 0.62 | 0.12 | 0.14 | 0.12 | 0.26 |
| 8 | 296848 | rs17812491 | -0.402 | -2.034 | 1.119 | -0.711 | 0.03 | -0.012 | -0.012 | 0 | 0.19 | 0.09 | 0.14 | 0.16 | 0.12 | 0.3 |
| 8 | 297118 | rs12546771 | -0.827 | -2.15 | 1.48 | -0.663 | -0.011 | -0.054 | -0.054 | 0 | 0.11 | 0.19 | 0.04 | 0.01 | 0.07 | 0.12 |
| 8 | 298447 | rs4330668 | -1.78 | -2.277 | 1.666 | -0.675 | -0.051 | -0.008 | -0.008 | 0 | 0.36 | 0.42 | 0.2 | 0.3 | 0.18 | 0.52 |
| 8 | 300058 | rs11781813 | -0.768 | -1.799 | 1.294 | -0.566 | -0.021 | 0.023 | 0.023 | 0 | 0.08 | 0.21 | 0.05 | 0.13 | 0.04 | 0.09 |
| 8 | 304569 | rs11991825 | -1.792 | -1.787 | 0.656 | -0.589 | -0.033 | -0.04 | -0.04 | 0 | 0.22 | 0.61 | 0.13 | 0.14 | 0.13 | 0.21 |
| 8 | 307738 | rs11775114 | -1.759 | -1.785 | 0.329 | -0.133 | -0.024 | -0.004 | -0.004 | 0 | 0.22 | 0.67 | 0.11 | 0.17 | 0.11 | 0.21 |
| 8 | 311009 | rs7816059 | -1.379 | -1.802 | 0.241 | -0.289 | 0.016 | 0.007 | 0.007 | 0.008 | 0.37 | 0.62 | 0.23 | 0.29 | 0.2 | 0.46 |
| 8 | 314765 | rs6558696 | -1.155 | -1.279 | 0.172 | -0.185 | 0.038 | 0.004 | 0.004 | 0.101 | 0.29 | 0.75 | 0.22 | 0.25 | 0.15 | 0.37 |
| 8 | 330328 | rs7461388 | -1.54 | -0.972 | 1.237 | -0.123 | 0.028 | 0.03 | 0.03 | 0 | 0.05 | 0.21 | 0.04 | 0.09 | 0.04 | 0.06 |
| 8 | 332249 | rs17064450 | -1.595 | -0.979 | 1.408 | -0.127 | 0.035 | 0.037 | 0.037 | 0 | 0.05 | 0.23 | 0.04 | 0.09 | 0.04 | 0.06 |
| 8 | 334288 | rs1979081 | -1.11 | -1.378 | 1.338 | -0.379 | -0.043 | -0.01 | -0.01 | 0.01 | 0.24 | 0.41 | 0.19 | 0.15 | 0.16 | 0.16 |
| 8 | 335646 | rs1550948 | -0.77 | -1.379 | 0.808 | -0.335 | 0.006 | -0.019 | -0.019 | 0 | 0.2 | 0.11 | 0.18 | 0.24 | 0.18 | 0.19 |
| 8 | 336284 | rs12547490 | 1.361 | -0.908 | -0.617 | 0.078 | -0.134 | 0.011 | 0.011 | 4.062 | 0.42 | 0.06 | 0.45 | 0.31 | 0.5 | 0.38 |
| 8 | 340067 | rs7825229 | -0.848 | -1.036 | 0.176 | -0.082 | -0.021 | -0.033 | -0.033 | 0.379 | 0.27 | 0.4 | 0.25 | 0.14 | 0.22 | 0.25 |
| 8 | 342004 | rs935561 | -0.185 | -1.168 | -0.303 | -0.257 | 0 | 0.03 | 0.03 | 0.019 | 0.11 | 0.06 | 0.06 | 0 | 0.09 | 0.08 |
| 8 | 349748 | rs11774596 | 0.665 | -0.851 | -0.251 | -0.154 | -0.007 | -0.02 | -0.02 | 0.26 | 0.12 | 0.04 | 0.11 | 0.1 | 0.12 | 0.14 |
| 8 | 351546 | rs6988801 | -1.369 | -0.93 | -1.043 | -0.141 | -0.012 | -0.008 | -0.008 | 0.863 | 0.56 | 0.8 | 0.44 | 0.55 | 0.45 | 0.55 |
| 8 | 360081 | rs13264582 | 1.058 | -0.535 | -0.822 | -0.18 | -0.003 | -0.029 | -0.029 | 0 | 0.12 | 0.03 | 0.12 | 0.09 | 0.11 | 0.14 |
| 8 | 365478 | rs17812876 | 0.787 | -0.449 | -0.862 | -0.213 | -0.007 | -0.026 | -0.026 | 1.07 | 0.13 | 0.03 | 0.15 | 0.1 | 0.12 | 0.18 |
| 8 | 376084 | rs17665382 | -0.711 | -0.261 | -0.424 | -0.086 | 0.004 | 0.004 | 0.004 | 0 | 0.08 | 0.17 | 0.07 | 0.1 | 0.08 | 0.07 |
| 8 | 400600 | rs17064974 | -1.121 | -0.33 | 0.281 | -0.386 | -0.044 | -0.052 | -0.052 | 0.258 | 0.17 | 0.51 | 0.13 | 0.09 | 0.12 | 0.1 |
| 8 | 400916 | rs13340594 | -0.836 | -0.192 | -0.341 | -0.353 | -0.03 | 0.016 | 0.016 | 1.302 | 0.29 | 0.53 | 0.22 | 0.4 | 0.22 | 0.27 |
| 8 | 401493 | rs6651443 | -0.479 | -0.093 | -0.114 | -0.378 | -0.004 | 0.008 | 0.008 | 0.669 | 0.26 | 0.32 | 0.19 | 0.4 | 0.25 | 0.26 |
| 8 | 401635 | rs10090308 | 0.914 | -0.291 | -0.344 | -0.729 | -0.004 | -0.004 | -0.004 | 0 | 0.21 | 0.21 | 0.2 | 0.1 | 0.14 | 0.17 |
| 8 | 402692 | rs1530662 | -1.949 | -0.131 | -0.366 | -0.722 | -0.04 | -0.054 | -0.054 | 0 | 0.56 | 0.86 | 0.46 | 0.5 | 0.46 | 0.48 |
| 8 | 403304 | rs3735925 | 1.116 | -0.117 | -0.319 | -0.623 | -0.006 | -0.039 | -0.039 | 0.049 | 0.15 | 0.09 | 0.13 | 0.41 | 0.15 | 0.24 |
| 8 | 406195 | rs10092971 | -1.387 | 0.324 | 0.05 | -0.704 | 0.042 | 0.082 | 0.082 | 0 | 0.43 | 0.73 | 0.34 | 0.16 | 0.32 | 0.31 |
| 8 | 406455 | rs17064981 | 0.692 | 0.051 | 0.148 | -0.869 | -0.005 | -0.066 | -0.066 | 0 | 0.11 | 0.26 | 0.07 | 0 | 0.09 | 0.02 |
| 8 | 406963 | rs2278765 | 0.339 | 0.141 | 0.173 | -0.755 | -0.018 | -0.02 | -0.02 | 0 | 0.24 | 0.29 | 0.21 | 0.09 | 0.15 | 0.19 |
| 8 | 410640 | rs10088894 | 0.069 | -0.025 | -0.075 | -0.936 | -0.006 | -0.01 | -0.01 | 0 | 0.3 | 0.33 | 0.22 | 0.11 | 0.19 | 0.24 |
| 8 | 411865 | rs12546599 | 0.718 | 0.049 | -0.125 | -0.72 | 0.047 | -0.027 | -0.027 | 0 | 0.06 | 0.2 | 0.05 | 0 | 0.07 | 0.02 |
| 8 | 415401 | rs10503146 | 0.716 | -0.015 | -0.573 | -0.734 | 0.03 | 0.02 | 0.02 | 0 | 0.2 | 0.08 | 0.14 | 0.1 | 0.12 | 0.17 |
| 8 | 416697 | rs10109251 | -1.964 | 0.516 | -0.942 | -0.724 | -0.077 | 0.007 | 0.007 | 0.692 | 0.58 | 0.85 | 0.47 | 0.35 | 0.46 | 0.47 |
| 8 | 420276 | rs17665621 | -1.675 | -0.082 | -0.923 | -0.987 | 0.049 | 0.167 | 0.167 | 0 | 0.49 | 0.82 | 0.49 | 0.18 | 0.4 | 0.38 |
| 8 | 421858 | rs7013349 | -1.44 | 0.09 | -0.876 | -0.843 | 0.03 | 0.023 | 0.023 | 0.003 | 0.39 | 0.81 | 0.41 | 0.4 | 0.36 | 0.33 |
| 8 | 423541 | rs11775200 | -0.859 | 0.049 | -0.268 | -0.888 | -0.014 | -0.007 | -0.007 | 0.558 | 0.34 | 0.49 | 0.38 | 0.4 | 0.34 | 0.33 |
| 8 | 423694 | rs11774384 | -1.523 | 0.006 | -0.354 | -0.953 | 0.001 | 0.004 | 0.004 | 0 | 0.42 | 0.61 | 0.43 | 0.4 | 0.4 | 0.37 |
| 8 | 425541 | rs11776266 | 0.315 | -0.021 | -0.136 | -0.969 | 0.001 | -0.042 | -0.042 | 0 | 0.15 | 0.18 | 0.21 | 0 | 0.18 | 0.09 |
| 8 | 426479 | rs1128051 | -1.54 | -0.241 | -0.538 | -0.943 | 0.038 | 0.004 | 0.004 | 2.882 | 0.51 | 0.63 | 0.46 | 0.49 | 0.38 | 0.48 |
| 8 | 427140 | rs2100160 | 1.282 | -0.229 | -0.81 | -0.846 | -0.001 | -0.018 | -0.018 | 0.322 | 0.18 | 0.01 | 0.15 | 0.1 | 0.14 | 0.17 |
| 8 | 427913 | rs7864 | -1.977 | 0.007 | -0.789 | -0.949 | -0.062 | -0.046 | -0.046 | 0.011 | 0.6 | 0.8 | 0.6 | 0.48 | 0.54 | 0.56 |
| 8 | 431161 | rs17813254 | 1.125 | -0.344 | -0.276 | -0.913 | 0.022 | -0.002 | -0.002 | 0 | 0.14 | 0.08 | 0.13 | 0.1 | 0.09 | 0.18 |
| 8 | 431501 | rs3735918 | 1.008 | -0.33 | -0.151 | -0.921 | 0.01 | 0.001 | 0.001 | 0 | 0.15 | 0.13 | 0.14 | 0.1 | 0.11 | 0.18 |
| 8 | 432146 | rs7001183 | -1.189 | -0.352 | -0.055 | -1.05 | -0.018 | -0.064 | -0.064 | 0 | 0.38 | 0.49 | 0.39 | 0.11 | 0.32 | 0.28 |
| 8 | 432585 | rs6994164 | -0.8 | -0.045 | -0.105 | -0.745 | -0.011 | -0.042 | -0.042 | 0.177 | 0.3 | 0.44 | 0.36 | 0.1 | 0.28 | 0.26 |
| 8 | 436230 | rs17065022 | -1.424 | 0.299 | -0.614 | -0.451 | 0.004 | 0.008 | 0.008 | 0.524 | 0.33 | 0.68 | 0.4 | 0.2 | 0.29 | 0.34 |
| 8 | 437030 | rs4735847 | -1.094 | 0.349 | -0.648 | -0.222 | 0.002 | 0.011 | 0.011 | 0.344 | 0.3 | 0.7 | 0.4 | 0.1 | 0.29 | 0.28 |
| 8 | 438673 | rs6983189 | -0.983 | -0.277 | -1.228 | 0.186 | 0.063 | -0.013 | -0.013 | 0.091 | 0.45 | 0.69 | 0.36 | 0.47 | 0.32 | 0.48 |
| 8 | 438821 | rs10098619 | 0.386 | 0.145 | -1.785 | 0.232 | 0.018 | -0.077 | -0.077 | 3.726 | 0.26 | 0.09 | 0.2 | 0.37 | 0.23 | 0.33 |
| 8 | 440424 | rs13429 | 0.67 | 0.692 | -1.238 | 0.55 | -0.001 | -0.007 | -0.007 | 0 | 0.47 | 0.46 | 0.41 | 0.37 | 0.48 | 0.43 |
| 8 | 440678 | rs1043720 | -0.631 | 0.589 | -1.361 | 0.613 | -0.005 | -0.101 | -0.101 | 0 | 0.68 | 0.91 | 0.73 | 0.48 | 0.68 | 0.64 |
| 8 | 441457 | rs1064416 | 0.424 | 0.718 | -1.044 | 0.474 | 0.013 | 0.047 | 0.047 | 0.009 | 0.54 | 0.73 | 0.47 | 0.37 | 0.55 | 0.46 |
| 8 | 441776 | rs2278764 | 1.809 | 0.55 | -1.019 | 0.715 | 0.025 | -0.032 | -0.032 | 0 | 0.16 | 0.07 | 0.3 | 0.3 | 0.15 | 0.19 |
| 8 | 442079 | rs1760 | 0.069 | 0.817 | -0.732 | 0.536 | 0.003 | -0.094 | -0.094 | 0.066 | 0.16 | 0.33 | 0.13 | 0 | 0.15 | 0.05 |
| 8 | 442755 | rs17813338 | 1.153 | 0.743 | -0.248 | 0.726 | -0.003 | -0.013 | -0.013 | 0.064 | 0.09 | 0.07 | 0.12 | 0.12 | 0.09 | 0.17 |
| 8 | 444485 | rs6999653 | -0.03 | 0.6 | 0.057 | 0.636 | -0.016 | -0.012 | -0.012 | 0 | 0.13 | 0.17 | 0.06 | 0.08 | 0.08 | 0.11 |

| | | | | | | | | | | | | | | | | |
|---|---|---|---|---|---|---|---|---|---|---|---|---|---|---|---|---|
| 8 | 445791 | rs11781389 | 0.509 | 0.567 | 0.046 | 0.684 | -0.036 | -0.017 | -0.017 | 0 | 0.26 | 0.39 | 0.21 | 0.28 | 0.2 | 0.17 |
| 8 | 446359 | rs17738513 | 0.156 | 0.719 | -0.295 | 0.714 | 0 | 0 | 0 | 0 | 0.1 | 0.08 | 0.06 | 0.09 | 0.07 | 0.11 |
| 8 | 448295 | rs10110988 | 1.107 | 0.743 | -0.398 | 0.74 | 0.009 | -0.005 | -0.005 | 0 | 0.2 | 0.18 | 0.18 | 0.29 | 0.17 | 0.21 |
| 8 | 449706 | rs7014579 | -1.112 | 0.869 | -0.595 | 0.869 | 0.009 | -0.033 | -0.033 | 0 | 0.81 | 0.89 | 0.89 | 0.75 | 0.83 | 0.82 |
| 8 | 450369 | rs6558856 | -1.093 | 1.425 | -0.68 | 1.325 | -0.004 | -0.022 | -0.022 | 0.295 | 0.52 | 0.82 | 0.6 | 0.5 | 0.46 | 0.54 |
| 8 | 453317 | rs4735851 | -0.892 | 1.021 | -0.742 | 0.529 | -0.001 | -0.04 | -0.04 | 1.598 | 0.78 | 0.87 | 0.88 | 0.75 | 0.79 | 0.8 |
| 8 | 455360 | rs2242 | 2.072 | 1.491 | -0.283 | 0.335 | 0.006 | -0.012 | -0.012 | 0 | 0.27 | 0.29 | 0.26 | 0.26 | 0.3 | 0.22 |
| 8 | 457459 | rs7821637 | 1.103 | 0.693 | 0.008 | 0.626 | -0.015 | 0.026 | 0.026 | 0 | 0.39 | 0.3 | 0.46 | 0.29 | 0.47 | 0.37 |
| 8 | 460350 | rs7002094 | 1.484 | 0.625 | 0.263 | 0.743 | -0.011 | 0.014 | 0.014 | 0 | 0.35 | 0.29 | 0.4 | 0.27 | 0.41 | 0.33 |
| 8 | 464571 | rs6990438 | -0.852 | -0.983 | 0.242 | 0.695 | -0.001 | 0 | 0 | 0 | 0.19 | 0.19 | 0.19 | 0.16 | 0.16 | 0.27 |
| 8 | 464784 | rs1345197 | 0.332 | -1.056 | 0.566 | 0.63 | -0.005 | -0.003 | -0.003 | 0 | 0.54 | 0.53 | 0.65 | 0.44 | 0.59 | 0.61 |
| 8 | 466262 | rs13266503 | 1.223 | -1.459 | 0.75 | 1.005 | 0.048 | 0.092 | 0.092 | 0.006 | 0.47 | 0.22 | 0.53 | 0.38 | 0.58 | 0.48 |
| 8 | 469429 | rs1448087 | 0.878 | -1.263 | 1.008 | 1.086 | -0.003 | -0.005 | -0.005 | 0 | 0.55 | 0.54 | 0.64 | 0.36 | 0.59 | 0.54 |
| 8 | 479107 | rs7008696 | 0.156 | -1.178 | 0.59 | 1.242 | -0.003 | 0.003 | 0.003 | 1.284 | 0.66 | 0.66 | 0.72 | 0.45 | 0.68 | 0.66 |
| 8 | 481767 | rs17813452 | 0.957 | -0.965 | -0.339 | 1.352 | 0.002 | -0.017 | -0.017 | 0.391 | 0.16 | 0.07 | 0.19 | 0.29 | 0.16 | 0.17 |
| 8 | 486020 | rs7837485 | -0.141 | -0.985 | -0.189 | 1.339 | -0.009 | 0.018 | 0.018 | 0 | 0.06 | 0.17 | 0.04 | 0.08 | 0.04 | 0.09 |
| 8 | 488249 | rs12541726 | 1.191 | -0.968 | -0.347 | 1.35 | 0.008 | -0.008 | -0.008 | 0 | 0.15 | 0.09 | 0.13 | 0.28 | 0.11 | 0.13 |
| 8 | 491497 | rs2084415 | 0.144 | -0.846 | -0.501 | 1.087 | 0.003 | -0.012 | -0.012 | 0.295 | 0.24 | 0.21 | 0.32 | 0.38 | 0.22 | 0.34 |
| 8 | 491735 | rs2084414 | 0.222 | -0.676 | -0.896 | 1.102 | 0.004 | -0.007 | -0.007 | 0 | 0.72 | 0.76 | 0.8 | 0.72 | 0.74 | 0.77 |
| 8 | 496764 | rs1669693 | 0.543 | -0.235 | -0.808 | 1.514 | -0.002 | -0.049 | -0.049 | 0 | 0.66 | 0.84 | 0.64 | 0.55 | 0.69 | 0.54 |
| 8 | 504401 | rs1703940 | -1.232 | 0.284 | -0.923 | 1.236 | -0.043 | -0.069 | -0.069 | 0.24 | 0.88 | 1 | 0.92 | 0.82 | 0.84 | 0.92 |
| 8 | 505001 | rs1018224 | 0.208 | 0.26 | -0.511 | 0.47 | 0.005 | 0.083 | 0.083 | 0 | 0.56 | 0.75 | 0.57 | 0.28 | 0.57 | 0.4 |
| 8 | 506821 | rs233697 | -0.143 | 0.404 | -0.405 | 0.709 | 0.009 | 0.027 | 0.027 | 0 | 0.59 | 0.72 | 0.6 | 0.37 | 0.62 | 0.47 |
| 8 | 507994 | rs184596 | 0.262 | 0.688 | -0.474 | 0.541 | 0 | -0.012 | -0.012 | 0.266 | 0.48 | 0.47 | 0.53 | 0.27 | 0.53 | 0.36 |
| 8 | 516479 | rs722782 | 1.247 | 1.051 | -0.065 | 1.954 | 0.008 | 0.176 | 0.176 | 0 | 0.9 | 0.75 | 0.85 | 0.46 | 0.91 | 0.8 |
| 8 | 517596 | rs2166842 | -0.773 | 1.219 | 0.409 | 1.966 | 0.014 | 0.079 | 0.079 | 0 | 0.17 | 0.36 | 0.24 | 0.31 | 0.16 | 0.14 |
| 8 | 519919 | rs734944 | 1.13 | 1.215 | 0.453 | 1.997 | 0.004 | 0.043 | 0.043 | 0 | 0.73 | 0.63 | 0.62 | 0.43 | 0.75 | 0.8 |
| 8 | 520044 | rs896523 | 0.205 | 1.205 | 0.179 | 2.109 | 0.001 | 0.005 | 0.005 | 0 | 0.91 | 0.93 | 0.89 | 0.93 | 0.91 | 0.95 |
| 8 | 521306 | rs12682073 | -0.135 | 1.424 | 0.363 | 2.158 | -0.003 | 0.031 | 0.031 | 0 | 0.13 | 0.12 | 0.23 | 0.5 | 0.15 | 0.14 |
| 8 | 527220 | rs3930083 | -0.124 | 1.391 | 0.292 | 2.172 | -0.009 | -0.011 | -0.011 | 0.085 | 0.13 | 0.09 | 0.23 | 0.48 | 0.13 | 0.19 |
| 8 | 531083 | rs11136864 | 1.187 | 0.801 | -0.573 | 1.409 | 0.049 | 0.064 | 0.064 | 0 | 0.29 | 0.08 | 0.17 | 0.12 | 0.23 | 0.24 |
| 8 | 531421 | rs11136866 | 0.222 | 1.207 | 0.158 | 1.33 | 0.008 | 0.005 | 0.005 | 0 | 0.43 | 0.3 | 0.44 | 0.6 | 0.42 | 0.43 |
| 8 | 533699 | rs11136871 | -0.248 | 1.496 | 0.401 | 0.603 | -0.001 | -0.026 | -0.026 | 0 | 0.64 | 0.64 | 0.71 | 0.93 | 0.62 | 0.67 |
| 8 | 540949 | rs1869075 | -0.714 | 1.4 | 0.1 | 0.43 | -0.074 | 0.16 | 0.16 | 0.771 | 0.72 | 0.94 | 0.82 | 0.99 | 0.7 | 0.72 |
| 8 | 541111 | rs1669684 | 0.543 | 1.149 | 0.539 | 0.753 | -0.003 | 0.005 | 0.005 | 0 | 0.2 | 0.24 | 0.27 | 0.31 | 0.18 | 0.23 |
| 8 | 541617 | rs896529 | 0.489 | 1.152 | 0.572 | 0.748 | 0.005 | -0.022 | -0.022 | 2.348 | 0.19 | 0.14 | 0.27 | 0.31 | 0.18 | 0.23 |
| 8 | 548856 | rs4735886 | 1.63 | 1.02 | 0.811 | 0.582 | 0.014 | 0.014 | 0.014 | 0 | 0.31 | 0.25 | 0.17 | 0.11 | 0.23 | 0.22 |
| 8 | 549870 | rs876412 | 0.672 | 1.346 | 0.83 | 0.973 | 0.04 | 0.181 | 0.181 | 0 | 0.14 | 0.22 | 0.19 | 0.52 | 0.18 | 0.19 |
| 8 | 549908 | rs906892 | 0.302 | 1.433 | 0.743 | 0.905 | 0.01 | -0.012 | -0.012 | 0 | 0.2 | 0.16 | 0.33 | 0.3 | 0.15 | 0.25 |
| 8 | 551687 | rs17813827 | -0.281 | 1.683 | 1.128 | 1.037 | 0.076 | 0.207 | 0.207 | 0 | 0.19 | 0.42 | 0.31 | 0.57 | 0.24 | 0.24 |
| 8 | 557126 | rs7014189 | 0.508 | 1.688 | 0.853 | 0.86 | 0.029 | 0.119 | 0.119 | 0 | 0.16 | 0.21 | 0.21 | 0.51 | 0.2 | 0.2 |
| 8 | 559071 | rs11136912 | 1.304 | 1.706 | 0.166 | 0.412 | 0.032 | 0.123 | 0.123 | 0 | 0.36 | 0.05 | 0.22 | 0.12 | 0.29 | 0.29 |
| 8 | 563371 | rs3736436 | -0.103 | 1.815 | 0.483 | 0.553 | 0.048 | 0.101 | 0.101 | 0.022 | 0.22 | 0.39 | 0.34 | 0.59 | 0.28 | 0.25 |
| 8 | 564000 | rs1669699 | -0.777 | 2.242 | 0.06 | 0.365 | -0.027 | 0.028 | 0.028 | 0 | 0.76 | 0.84 | 0.85 | 0.99 | 0.72 | 0.75 |
| 8 | 565959 | rs11781454 | 0.155 | 1.339 | 0.07 | 0.732 | -0.001 | 0 | 0 | 0 | 0.19 | 0.2 | 0.32 | 0.3 | 0.15 | 0.23 |
| 8 | 570463 | rs336419 | 0.548 | 0.989 | -0.111 | 0.716 | -0.004 | 0.01 | 0.01 | 0 | 0.12 | 0.14 | 0.26 | 0.27 | 0.08 | 0.16 |
| 8 | 572628 | rs336426 | 0.343 | 0.979 | 0.006 | 0.727 | -0.016 | 0.044 | 0.044 | 0 | 0.13 | 0.22 | 0.26 | 0.27 | 0.08 | 0.16 |
| 8 | 573646 | rs12716634 | 1.407 | 0.853 | -0.434 | 0.528 | 0.067 | 0.073 | 0.073 | 0 | 0.32 | 0.13 | 0.17 | 0.12 | 0.24 | 0.28 |
| 8 | 576209 | rs336430 | 0.035 | 1.13 | -0.496 | 0.617 | -0.001 | 0.004 | 0.004 | 0 | 0.14 | 0.18 | 0.29 | 0.27 | 0.11 | 0.17 |
| 8 | 576324 | rs336431 | 0.501 | 1.042 | -0.635 | 0.332 | 0.086 | 0.032 | 0.032 | 0 | 0.47 | 0.2 | 0.46 | 0.39 | 0.36 | 0.46 |
| 8 | 577101 | rs336433 | -0.571 | 1.37 | -0.88 | 0.572 | -0.157 | 0.036 | 0.036 | 0.077 | 0.69 | 0.95 | 0.75 | 0.84 | 0.56 | 0.67 |
| 8 | 578742 | rs12115088 | -0.156 | 1.448 | -0.318 | 0.211 | 0.012 | 0.018 | 0.018 | 0.185 | 0.4 | 0.41 | 0.41 | 0.54 | 0.5 | 0.48 |
| 8 | 587113 | rs89975 | 0.055 | 1.043 | -0.143 | 0.416 | -0.012 | 0.107 | 0.107 | 0 | 0.13 | 0.27 | 0.28 | 0.34 | 0.09 | 0.16 |
| 8 | 593041 | rs3735933 | -0.088 | 0.97 | -0.193 | 0.267 | -0.009 | 0.008 | 0.008 | 1.954 | 0.42 | 0.41 | 0.45 | 0.54 | 0.58 | 0.51 |
| 8 | 593641 | rs1669715 | -1.12 | 0.994 | -0.16 | 0.334 | 0 | 0 | 0 | 0 | 0.92 | 0.9 | 0.9 | 0.94 | 0.91 | 0.95 |
| 8 | 593707 | rs1669716 | -0.696 | 1.022 | -0.158 | 0.329 | -0.001 | 0.002 | 0.002 | 0 | 0.89 | 0.88 | 0.87 | 0.94 | 0.89 | 0.94 |
| 8 | 600729 | rs4735895 | 0.762 | 1.015 | -0.145 | 0.341 | -0.017 | 0.089 | 0.089 | 0.15 | 0.73 | 0.42 | 0.58 | 0.59 | 0.79 | 0.78 |
| 8 | 601043 | rs1703911 | 0.363 | 0.835 | -0.227 | 0.241 | 0.019 | 0.007 | 0.007 | 0.06 | 0.49 | 0.48 | 0.46 | 0.4 | 0.33 | 0.46 |
| 8 | 603256 | rs2335877 | 1.443 | 0.839 | -0.457 | 0.131 | 0.076 | 0.196 | 0.196 | 0.719 | 0.33 | 0.04 | 0.17 | 0.05 | 0.25 | 0.28 |
| 8 | 604680 | rs17751994 | 0.122 | 0.888 | -0.175 | 0.333 | -0.003 | 0.138 | 0.138 | 0 | 0.13 | 0.31 | 0.28 | 0.34 | 0.09 | 0.18 |
| 8 | 606117 | rs1669681 | 0.304 | 0.956 | -0.227 | 0.25 | 0.009 | 0.005 | 0.005 | 0.006 | 0.5 | 0.52 | 0.47 | 0.38 | 0.35 | 0.47 |
| 8 | 606356 | rs13252487 | 0.705 | 0.916 | -0.271 | 0.41 | -0.001 | -0.01 | -0.01 | 1.732 | 0.09 | 0.05 | 0.18 | 0.25 | 0.07 | 0.09 |
| 8 | 606696 | rs1669733 | -1.293 | 1.243 | -0.34 | 0.201 | -0.017 | 0.001 | 0.001 | 0.162 | 0.92 | 0.98 | 0.92 | 0.95 | 0.91 | 0.96 |
| 8 | 607251 | rs10105657 | 1.457 | 0.99 | -0.367 | 0.061 | 0.056 | 0.215 | 0.215 | 0.46 | 0.34 | 0.04 | 0.17 | 0.04 | 0.26 | 0.29 |
| 8 | 608018 | rs6993769 | 0.067 | 1.287 | -0.234 | 0.165 | 0.01 | 0.015 | 0.015 | 2.109 | 0.39 | 0.44 | 0.41 | 0.55 | 0.52 | 0.48 |
| 8 | 612758 | rs9314595 | 0.969 | 1.461 | -0.461 | 0.08 | -0.144 | 0.095 | 0.095 | 2.08 | 0.24 | 0.04 | 0.24 | 0.07 | 0.41 | 0.3 |
| 8 | 613620 | rs1703927 | -1.19 | 1.338 | -0.332 | 0.197 | -0.003 | -0.001 | -0.001 | 1.282 | 0.91 | 0.91 | 0.9 | 0.93 | 0.9 | 0.96 |
| 8 | 615117 | rs3824148 | 0.021 | 1.23 | -0.564 | 0.227 | -0.011 | 0.116 | 0.116 | 0.084 | 0.22 | 0.49 | 0.21 | 0.47 | 0.15 | 0.18 |
| 8 | 616646 | rs2280001 | -0.025 | 1.062 | -0.661 | 0.396 | -0.013 | 0.12 | 0.12 | 0 | 0.09 | 0.17 | 0.26 | 0.35 | 0.06 | 0.15 |

| | | | | | | | | | | | | | | | | |
|---|---|---|---|---|---|---|---|---|---|---|---|---|---|---|---|---|
| 8 | 617625 | rs17752041 | -0.05 | 1.072 | -0.723 | 0.384 | -0.009 | 0.077 | 0.077 | 0 | 0.1 | 0.14 | 0.26 | 0.35 | 0.06 | 0.16 |
| 8 | 617885 | rs876356 | 0.231 | 1.626 | -0.959 | -0.113 | -0.012 | -0.004 | -0.004 | 0 | 0.06 | 0.07 | 0.11 | 0.06 | 0.13 | 0.05 |
| 8 | 633684 | rs11137016 | 1.429 | 1.03 | -0.955 | -0.78 | 0.046 | 0.075 | 0.075 | 0 | 0.31 | 0.18 | 0.16 | 0.05 | 0.24 | 0.28 |
| 8 | 636251 | rs336437 | -0.807 | 1.235 | -1.062 | -0.472 | -0.002 | 0.005 | 0.005 | 0.196 | 0.91 | 0.92 | 0.9 | 0.94 | 0.9 | 0.95 |
| 8 | 642842 | rs3735917 | 0.779 | 0.959 | -1.151 | -0.789 | 0.071 | 0.172 | 0.172 | 0 | 0.39 | 0.16 | 0.19 | 0.07 | 0.3 | 0.3 |
| 8 | 644824 | rs7012226 | 0.75 | 1.182 | -0.58 | -0.359 | 0.014 | 0.189 | 0.189 | 0 | 0.15 | 0.31 | 0.17 | 0.46 | 0.12 | 0.18 |
| 8 | 649190 | rs12156406 | 1.033 | 1.609 | -1.135 | -0.704 | -0.082 | 0.092 | 0.092 | 1.18 | 0.2 | 0.05 | 0.15 | 0.01 | 0.29 | 0.26 |
| 8 | 655142 | rs7004577 | 0.854 | 1.158 | -0.56 | -0.651 | 0.033 | 0.001 | 0.001 | 0 | 0.46 | 0.32 | 0.33 | 0.51 | 0.37 | 0.44 |
| 8 | 657650 | rs1616976 | -0.242 | 0.824 | -0.905 | -0.862 | 0.02 | 0.005 | 0.005 | 0.118 | 0.45 | 0.36 | 0.55 | 0.41 | 0.56 | 0.48 |
| 8 | 665238 | rs11137063 | -0.167 | 0.863 | -0.862 | -0.846 | -0.003 | -0.002 | -0.002 | 0 | 0.22 | 0.26 | 0.38 | 0.39 | 0.25 | 0.24 |
| 8 | 665858 | rs1869074 | 1.301 | 1.278 | -1.162 | -0.973 | -0.04 | 0.107 | 0.107 | 1.395 | 0.21 | 0.04 | 0.15 | 0.02 | 0.21 | 0.22 |
| 8 | 673561 | rs10094564 | -0.059 | 0.78 | -0.739 | -1.001 | -0.001 | 0.004 | 0.004 | 0 | 0.51 | 0.53 | 0.45 | 0.57 | 0.46 | 0.47 |
| 8 | 676191 | rs12549425 | 1.335 | 0.613 | -1.02 | -1.2 | 0.022 | 0.039 | 0.039 | 0 | 0.27 | 0.16 | 0.15 | 0.05 | 0.23 | 0.25 |
| 8 | 677305 | rs4735916 | 0.935 | 0.751 | -0.891 | -0.804 | -0.005 | 0.07 | 0.07 | 0.019 | 0.12 | 0.18 | 0.17 | 0.47 | 0.09 | 0.16 |
| 8 | 679799 | rs11784674 | 0.76 | 0.826 | -1.307 | -1.16 | -0.007 | 0.06 | 0.06 | 1.291 | 0.55 | 0.29 | 0.4 | 0.13 | 0.51 | 0.53 |
| 8 | 698978 | rs7010117 | -0.381 | 0.727 | -2.045 | -0.697 | 0.007 | -0.046 | -0.046 | 0 | 0.58 | 0.82 | 0.64 | 0.49 | 0.63 | 0.58 |
| 8 | 706008 | rs10100132 | 1.371 | 0.566 | -2.197 | -0.855 | -0.027 | 0.111 | 0.111 | 0 | 0.2 | 0.01 | 0.16 | 0.06 | 0.18 | 0.24 |
| 8 | 707884 | rs13274614 | -0.19 | 0.665 | -1.892 | -0.547 | 0.044 | -0.014 | -0.014 | 0 | 0.52 | 0.7 | 0.45 | 0.45 | 0.62 | 0.43 |
| 8 | 714383 | rs6986946 | 0.297 | 1.162 | -1.878 | -0.173 | -0.015 | 0.041 | 0.041 | 0 | 0.06 | 0.03 | 0.09 | 0.32 | 0.1 | 0.13 |
| 8 | 714648 | rs7825497 | -0.237 | 1.155 | -1.575 | -1.1 | -0.005 | -0.014 | -0.014 | 0.171 | 0.78 | 0.63 | 0.68 | 0.86 | 0.83 | 0.78 |
| 8 | 719043 | rs12386927 | 1.273 | 1.102 | -1.694 | -1.186 | -0.036 | 0.08 | 0.08 | 0.307 | 0.26 | 0.09 | 0.2 | 0.02 | 0.33 | 0.2 |
| 8 | 720491 | rs6559181 | 0.227 | 1 | -1.107 | -0.49 | 0.002 | -0.012 | -0.012 | 0.023 | 0.77 | 0.64 | 0.66 | 0.84 | 0.79 | 0.72 |
| 8 | 722942 | rs4735919 | -0.045 | 1.308 | -1.545 | -0.302 | -0.025 | -0.009 | -0.009 | 0.987 | 0.17 | 0.07 | 0.46 | 0.19 | 0.23 | 0.27 |
| 8 | 737663 | rs10094226 | -0.152 | 0.751 | -0.927 | -0.092 | -0.014 | 0.054 | 0.054 | 0 | 0.32 | 0.44 | 0.18 | 0.53 | 0.24 | 0.31 |
| 8 | 737782 | rs6995315 | 1.258 | 0.319 | -1.296 | -0.645 | 0.013 | -0.007 | -0.007 | 0.039 | 0.18 | 0.19 | 0.08 | 0.22 | 0.13 | 0.1 |
| 8 | 742490 | rs11137104 | -0.422 | 0.56 | -1.171 | -1.097 | -0.004 | 0.027 | 0.027 | 0 | 0.46 | 0.56 | 0.61 | 0.2 | 0.46 | 0.46 |
| 8 | 749998 | rs12675507 | 0.297 | 0.575 | -1.387 | -1.093 | -0.04 | -0.008 | -0.008 | 0 | 0.18 | 0.05 | 0.47 | 0.2 | 0.25 | 0.33 |
| 8 | 750278 | rs7009367 | 0.997 | 0.563 | -1.335 | -1.216 | -0.014 | -0.164 | -0.164 | 0.239 | 0.28 | 0.49 | 0.14 | 0 | 0.2 | 0.14 |
| 8 | 751914 | rs4735934 | 0.29 | 0.674 | -1.476 | -1.113 | -0.04 | -0.006 | -0.006 | 0 | 0.18 | 0.05 | 0.48 | 0.2 | 0.26 | 0.33 |
| 8 | 752709 | rs10108289 | 1.972 | 0.853 | -1.599 | -1.369 | -0.037 | 0.082 | 0.082 | 0 | 0.22 | 0.08 | 0.17 | 0.02 | 0.27 | 0.23 |
| 8 | 755494 | rs13262566 | 0.667 | 0.93 | -1.334 | -0.81 | -0.005 | -0.013 | -0.013 | 0 | 0.13 | 0.05 | 0.14 | 0.53 | 0.15 | 0.22 |
| 8 | 758066 | rs9314632 | 1.225 | 0.757 | -0.922 | -1.207 | 0.001 | -0.155 | -0.155 | 2.051 | 0.22 | 0.38 | 0.12 | 0 | 0.17 | 0.12 |
| 8 | 769589 | rs12680347 | 0.788 | 1.079 | -0.948 | -0.894 | -0.019 | -0.019 | -0.019 | 0 | 0.12 | 0.06 | 0.44 | 0.22 | 0.2 | 0.32 |
| 8 | 773986 | rs6982473 | 0.809 | 0.89 | -0.802 | -1.321 | 0.019 | -0.159 | -0.159 | 0 | 0.28 | 0.47 | 0.16 | 0 | 0.26 | 0.15 |
| 8 | 779714 | rs2055516 | 0.84 | 1.984 | -0.099 | -1.333 | 0.013 | -0.176 | -0.176 | 0 | 0.28 | 0.44 | 0.16 | 0 | 0.25 | 0.15 |
| 8 | 782845 | rs1393703 | 0.628 | 0.684 | -0.85 | -1.191 | 0.031 | 0 | 0 | 0 | 0.28 | 0.19 | 0.17 | 0.27 | 0.22 | 0.2 |
| 8 | 783099 | rs1393701 | -0.574 | 0.654 | -0.995 | -1.188 | 0.016 | -0.05 | -0.05 | 0 | 0.19 | 0.1 | 0.25 | 0.56 | 0.19 | 0.39 |
| 8 | 783664 | rs10106538 | 0.04 | 0.606 | -0.979 | -1.198 | 0.042 | 0.022 | 0.022 | 0.063 | 0.63 | 0.69 | 0.81 | 0.73 | 0.73 | 0.75 |
| 8 | 785085 | rs1553408 | 0.71 | 1.016 | -0.919 | -0.885 | 0.009 | -0.037 | -0.037 | 0 | 0.12 | 0.05 | 0.12 | 0.52 | 0.11 | 0.22 |
| 8 | 799059 | rs2055515 | -0.455 | 1.344 | -0.93 | -0.856 | -0.041 | -0.046 | -0.046 | 0 | 0.12 | 0.03 | 0.42 | 0.22 | 0.19 | 0.22 |
| 8 | 799383 | rs9721194 | -1.564 | 1.393 | -0.509 | -1.228 | 0.023 | -0.134 | -0.134 | 0 | 0.75 | 0.61 | 0.84 | 0.98 | 0.75 | 0.79 |
| 8 | 799627 | rs7845440 | -0.354 | 1.105 | -0.233 | -1.472 | 0.006 | 0.029 | 0.029 | 0.122 | 0.46 | 0.56 | 0.64 | 0.22 | 0.59 | 0.5 |
| 8 | 801584 | rs17739674 | 0.107 | 0.882 | -0.737 | -1.462 | -0.002 | 0 | 0 | 0 | 0.34 | 0.34 | 0.52 | 0.2 | 0.42 | 0.39 |
| 8 | 801912 | rs17752292 | 1.059 | 1.065 | -1.005 | -0.625 | 0.011 | -0.096 | -0.096 | 0.26 | 0.17 | 0.04 | 0.13 | 0.51 | 0.13 | 0.23 |
| 8 | 809054 | rs2174098 | 1.746 | 0.637 | -0.915 | -1.513 | 0.008 | -0.006 | -0.006 | 1.91 | 0.18 | 0.18 | 0.07 | 0.22 | 0.12 | 0.06 |
| 8 | 809079 | rs2336409 | 0.763 | 0.656 | -0.788 | -1.691 | -0.002 | -0.002 | -0.002 | 0 | 0.39 | 0.34 | 0.29 | 0.55 | 0.39 | 0.44 |
| 8 | 809489 | rs3860879 | 0.619 | 0.785 | -1.089 | -1.732 | -0.036 | 0.09 | 0.09 | 0 | 0.29 | 0.12 | 0.28 | 0.05 | 0.35 | 0.34 |
| 8 | 809497 | rs3860880 | -0.562 | 0.828 | -1.091 | -1.699 | 0.002 | -0.018 | -0.018 | 0 | 0.42 | 0.44 | 0.64 | 0.22 | 0.48 | 0.5 |
| 8 | 809859 | rs12680727 | 0.863 | 0.868 | -1.123 | -1.724 | 0 | 0.029 | 0.029 | 0 | 0.08 | 0.03 | 0.12 | 0.02 | 0.08 | 0.11 |
| 8 | 813942 | rs2174097 | 1.566 | 0.735 | -1.118 | -1.777 | -0.012 | 0.006 | 0.006 | 0 | 0.19 | 0.14 | 0.17 | 0.02 | 0.27 | 0.23 |
| 8 | 814529 | rs7823711 | 1.157 | 0.627 | -1.101 | -1.61 | -0.003 | -0.015 | -0.015 | 0 | 0.4 | 0.54 | 0.23 | 0.33 | 0.37 | 0.26 |
| 8 | 817587 | rs881430 | 0.166 | 0.992 | -1.211 | -1.516 | -0.028 | 0.035 | 0.035 | 0 | 0.1 | 0.03 | 0.17 | 0.03 | 0.17 | 0.13 |
| 8 | 817847 | rs1105954 | -0.172 | 0.764 | -1.162 | -1.575 | -0.022 | 0.035 | 0.035 | 0 | 0.37 | 0.27 | 0.36 | 0.05 | 0.49 | 0.38 |
| 8 | 823449 | rs982352 | -0.587 | 0.974 | -1.192 | -1.237 | 0 | 0.004 | 0.004 | 0 | 0.61 | 0.63 | 0.76 | 0.55 | 0.67 | 0.57 |
| 8 | 823897 | rs4735946 | -0.811 | 1.034 | -1.524 | -0.775 | 0.019 | -0.073 | -0.073 | 0 | 0.83 | 0.94 | 0.9 | 0.55 | 0.9 | 0.77 |
| 8 | 827824 | rs1875664 | -0.219 | 1.133 | -1.551 | -0.852 | -0.002 | -0.045 | -0.045 | 0 | 0.8 | 0.88 | 0.61 | 0.38 | 0.84 | 0.63 |
| 8 | 828266 | rs11777864 | -0.525 | 1.316 | -1.288 | -0.927 | -0.016 | 0.034 | 0.034 | 0 | 0.13 | 0.03 | 0.18 | 0.02 | 0.19 | 0.13 |
| 8 | 847144 | rs4735825 | -0.676 | 1.476 | -1.238 | -0.939 | -0.025 | 0.023 | 0.023 | 0 | 0.31 | 0.25 | 0.65 | 0.22 | 0.5 | 0.46 |
| 8 | 849254 | rs11774254 | -1.158 | 1.433 | -1.237 | -0.755 | -0.006 | -0.001 | -0.001 | 0 | 0.14 | 0.13 | 0.45 | 0.19 | 0.22 | 0.25 |
| 8 | 853544 | rs1466919 | -0.135 | 1.428 | -1.266 | -0.595 | -0.007 | -0.005 | -0.005 | 0.108 | 0.05 | 0.04 | 0.32 | 0.18 | 0.11 | 0.14 |
| 8 | 856874 | rs885484 | 1.019 | 1.022 | -1.399 | -0.768 | 0.014 | -0.027 | -0.027 | 0 | 0.2 | 0.16 | 0.08 | 0.31 | 0.15 | 0.09 |
| 8 | 857459 | rs17740061 | 1.235 | 1.377 | -1.534 | -0.913 | -0.069 | 0.074 | 0.074 | 0.436 | 0.17 | 0.03 | 0.17 | 0.02 | 0.28 | 0.21 |
| 8 | 859982 | rs955980 | -0.426 | 1.422 | -1.064 | -0.664 | 0.01 | 0.019 | 0.019 | 0 | 0.06 | 0.07 | 0.32 | 0.18 | 0.1 | 0.15 |
| 8 | 860464 | rs17065459 | -1.321 | 1.378 | -0.935 | -0.801 | 0.014 | 0.005 | 0.005 | 0 | 0.14 | 0.23 | 0.47 | 0.2 | 0.21 | 0.23 |
| 8 | 860907 | rs13257920 | -1.04 | 1.155 | -0.743 | -1.016 | -0.002 | 0.021 | 0.021 | 0 | 0.64 | 0.64 | 0.78 | 0.53 | 0.71 | 0.59 |
| 8 | 864408 | rs17667487 | -0.813 | 1.185 | -0.701 | -0.53 | -0.002 | 0 | 0 | 0.719 | 0.07 | 0.06 | 0.32 | 0.17 | 0.1 | 0.15 |
| 8 | 866207 | rs17740145 | -1.063 | 1.433 | -0.696 | -0.684 | 0.022 | -0.003 | -0.003 | 0 | 0.18 | 0.28 | 0.45 | 0.2 | 0.3 | 0.3 |
| 8 | 870224 | rs11137111 | 0.845 | 0.915 | -1.098 | -0.713 | 0.038 | -0.019 | -0.019 | 0 | 0.25 | 0.18 | 0.11 | 0.29 | 0.16 | 0.09 |
| 8 | 872081 | rs4735959 | -1.34 | 1.684 | -0.819 | -0.223 | -0.007 | -0.065 | -0.065 | 0 | 0.88 | 0.99 | 0.93 | 0.55 | 0.92 | 0.86 |

| | | | | | | | | | | | | | | | | |
|---|---|---|---|---|---|---|---|---|---|---|---|---|---|---|---|---|
| 8 | 877433 | rs7822510 | -0.679 | 2.067 | 0.226 | -0.038 | 0.024 | -0.044 | -0.044 | 0 | 0.14 | 0.23 | 0.13 | 0.02 | 0.21 | 0.16 |
| 8 | 887909 | rs11777063 | -2.006 | 2.145 | 0.374 | -0.069 | 0.064 | -0.011 | -0.011 | 0.216 | 0.23 | 0.39 | 0.59 | 0.24 | 0.4 | 0.38 |
| 8 | 893453 | rs11137112 | 0.771 | 1.997 | 0.318 | -0.102 | 0.01 | -0.034 | -0.034 | 0.338 | 0.17 | 0.15 | 0.09 | 0.33 | 0.16 | 0.1 |
| 8 | 894193 | rs11777133 | 0.025 | 2.614 | 0.859 | -0.106 | -0.042 | 0.036 | 0.036 | 1.939 | 0.1 | 0.02 | 0.11 | 0.02 | 0.17 | 0.13 |
| 8 | 899726 | rs11782142 | -2.299 | 2.566 | 1.023 | -0.083 | 0.104 | -0.009 | -0.009 | 0 | 0.23 | 0.41 | 0.61 | 0.23 | 0.43 | 0.39 |
| 8 | 902064 | rs17667613 | -0.921 | 2.376 | 0.966 | -0.148 | 0.057 | -0.066 | -0.066 | 0.05 | 0.14 | 0.3 | 0.15 | 0.02 | 0.21 | 0.16 |
| 8 | 913042 | rs2701918 | -1.375 | 2.18 | 1.182 | -0.082 | 0.062 | -0.02 | -0.02 | 0 | 0.2 | 0.53 | 0.47 | 0.19 | 0.3 | 0.28 |
| 8 | 914780 | rs1553410 | -1.505 | 2.366 | 0.668 | -0.033 | 0.054 | -0.005 | -0.005 | 0 | 0.22 | 0.3 | 0.6 | 0.23 | 0.41 | 0.39 |
| 8 | 926941 | rs924696 | -1.869 | 2.245 | 0.719 | -0.037 | 0.069 | -0.02 | -0.02 | 0 | 0.25 | 0.39 | 0.62 | 0.24 | 0.42 | 0.37 |
| 8 | 927722 | rs13270728 | -1.794 | 2.218 | 0.427 | -0.027 | 0.036 | -0.01 | -0.01 | 0 | 0.24 | 0.3 | 0.6 | 0.24 | 0.41 | 0.36 |
| 8 | 930451 | rs10099591 | -1.81 | 2.2 | 0.396 | -0.106 | 0.037 | -0.016 | -0.016 | 4.018 | 0.25 | 0.32 | 0.61 | 0.23 | 0.41 | 0.36 |
| 8 | 931580 | rs7003378 | -1.162 | 2.267 | 0.421 | -0.088 | 0.043 | -0.013 | -0.013 | 0 | 0.19 | 0.28 | 0.47 | 0.17 | 0.32 | 0.25 |
| 8 | 933961 | rs7018425 | -0.93 | 1.614 | 0.057 | 0.019 | 0.004 | -0.009 | -0.009 | 0 | 0.06 | 0.11 | 0.15 | 0.05 | 0.1 | 0.1 |
| 8 | 951785 | rs4236863 | -1.202 | 2.074 | -0.115 | -0.194 | 0.007 | -0.014 | -0.014 | 0 | 0.27 | 0.28 | 0.39 | 0.17 | 0.38 | 0.34 |
| 8 | 952089 | rs4735965 | -1.48 | 2.247 | -0.113 | -0.205 | 0.044 | 0.009 | 0.009 | 0.179 | 0.12 | 0.25 | 0.29 | 0.15 | 0.18 | 0.22 |
| 8 | 955147 | rs6983944 | 0.946 | 2.012 | -0.054 | -0.069 | 0.009 | 0.006 | 0.006 | 0 | 0.44 | 0.42 | 0.39 | 0.38 | 0.37 | 0.3 |
| 8 | 959338 | rs4415346 | -0.7 | 2.051 | 0.522 | -0.018 | -0.001 | -0.029 | -0.029 | 0 | 0.71 | 0.81 | 0.79 | 0.55 | 0.75 | 0.63 |
| 8 | 968281 | rs6559202 | -0.118 | 1.948 | 1.1 | -0.004 | 0.001 | 0.002 | 0.002 | 0.007 | 0.64 | 0.64 | 0.67 | 0.52 | 0.59 | 0.55 |
| 8 | 973607 | rs6559205 | 1.034 | 1.75 | 0.885 | 0.051 | 0.043 | -0.028 | -0.028 | 0 | 0.45 | 0.17 | 0.53 | 0.5 | 0.39 | 0.42 |
| 8 | 974472 | rs10867028 | 0.061 | 2.425 | 0.79 | 0.479 | -0.023 | -0.028 | -0.028 | 0 | 0.85 | 0.84 | 0.8 | 0.47 | 0.75 | 0.73 |
| 8 | 980635 | rs7460543 | -0.376 | 2.05 | 1.142 | 0.203 | 0.019 | -0.046 | -0.046 | 0 | 0.21 | 0.4 | 0.22 | 0.15 | 0.21 | 0.27 |
| 8 | 986301 | rs4735969 | -0.474 | 2.022 | 1.277 | 0.395 | 0.036 | -0.015 | -0.015 | 0 | 0.2 | 0.42 | 0.22 | 0.16 | 0.24 | 0.3 |
| 8 | 994175 | rs2600514 | 1.277 | 1.93 | 0.933 | 0.462 | 0.045 | 0.119 | 0.119 | 0 | 0.44 | 0.17 | 0.32 | 0.3 | 0.35 | 0.29 |
| 8 | 996634 | rs4735971 | -1.162 | 1.949 | 0.942 | 0.626 | 0 | 0.186 | 0.186 | 1.278 | 0.94 | 0.9 | 0.88 | 0.57 | 0.95 | 0.93 |
| 8 | 998618 | rs4735972 | 0.713 | 1.848 | 1.052 | 0.441 | 0.011 | 0.011 | 0.011 | 0 | 0.52 | 0.61 | 0.47 | 0.45 | 0.4 | 0.46 |
| 8 | 1001246 | rs2701910 | 1.483 | 1.867 | 0.878 | 0.281 | 0.029 | 0.069 | 0.069 | 0.025 | 0.43 | 0.14 | 0.32 | 0.37 | 0.35 | 0.28 |
| 8 | 1003292 | rs2701907 | 0.886 | 1.901 | 1.187 | 0.274 | -0.002 | -0.001 | -0.001 | 0.015 | 0.49 | 0.55 | 0.45 | 0.51 | 0.38 | 0.42 |
| 8 | 1008232 | rs10503156 | 0.705 | 2.189 | 0.82 | 0.239 | -0.056 | 0.058 | 0.058 | 0 | 0.13 | 0.02 | 0.11 | 0.02 | 0.22 | 0.14 |
| 8 | 1009313 | rs2123060 | 0.959 | 2.049 | 1.128 | 0.252 | 0.035 | 0.017 | 0.017 | 2.401 | 0.51 | 0.65 | 0.44 | 0.52 | 0.36 | 0.42 |
| 8 | 1012890 | rs9650539 | 0.559 | 2.364 | 0.722 | 0.134 | -0.035 | 0.029 | 0.029 | 0.384 | 0.08 | 0.02 | 0.2 | 0.02 | 0.19 | 0.12 |
| 8 | 1014177 | rs17740775 | 0.96 | 2.387 | 0.847 | 0.07 | 0 | 0.027 | 0.027 | 0.03 | 0.2 | 0.14 | 0.11 | 0.02 | 0.16 | 0.25 |
| 8 | 1014668 | rs17668166 | 0.282 | 2.281 | 0.852 | 0.209 | -0.002 | -0.016 | -0.016 | 0 | 0.3 | 0.27 | 0.33 | 0.04 | 0.36 | 0.37 |
| 8 | 1016903 | rs2600487 | 1.342 | 2.177 | 0.918 | 0.354 | 0.032 | 0.033 | 0.033 | 0.167 | 0.46 | 0.14 | 0.37 | 0.47 | 0.36 | 0.43 |
| 8 | 1018332 | rs4735980 | 0.775 | 2.32 | 0.834 | 0.374 | 0.006 | 0.018 | 0.018 | 0.02 | 0.51 | 0.62 | 0.49 | 0.5 | 0.42 | 0.43 |
| 8 | 1020249 | rs7015529 | 0.846 | 1.754 | 0.768 | 0.417 | 0.021 | 0.025 | 0.025 | 0.046 | 0.51 | 0.6 | 0.43 | 0.45 | 0.36 | 0.42 |
| 8 | 1021250 | rs7818865 | -1.337 | 1.828 | 0.235 | 0.614 | 0.003 | 0.191 | 0.191 | 0 | 0.95 | 0.91 | 0.89 | 0.55 | 0.94 | 0.93 |
| 8 | 1024476 | rs9314422 | 0.481 | 1.885 | 0.012 | 0.368 | -0.042 | 0.046 | 0.046 | 0 | 0.15 | 0.02 | 0.09 | 0.03 | 0.19 | 0.14 |
| 8 | 1029500 | rs2123055 | 1.336 | 1.732 | 0.213 | 0.325 | 0.039 | 0.052 | 0.052 | 1.076 | 0.43 | 0.26 | 0.33 | 0.36 | 0.34 | 0.27 |
| 8 | 1032472 | rs727570 | -0.944 | 1.725 | 0.349 | 0.176 | 0.008 | 0.001 | 0.001 | 0.275 | 0.5 | 0.48 | 0.45 | 0.09 | 0.6 | 0.51 |
| 8 | 1033625 | rs2336695 | 0.124 | 1.64 | 0.372 | 0.206 | 0.007 | 0.008 | 0.008 | 0.716 | 0.62 | 0.44 | 0.54 | 0.55 | 0.57 | 0.58 |
| 8 | 1034497 | rs6982844 | 0.049 | 1.664 | -0.121 | -0.553 | 0.026 | -0.128 | -0.128 | 0.358 | 0.33 | 0.42 | 0.35 | 0.05 | 0.38 | 0.36 |
| 8 | 1035729 | rs6988649 | -0.147 | 1.203 | -0.683 | -0.079 | -0.002 | 0.008 | 0.008 | 0 | 0.16 | 0.16 | 0.1 | 0.02 | 0.19 | 0.13 |
| 8 | 1037264 | rs4735837 | -0.102 | 0.665 | -1.231 | 0.292 | 0.038 | 0.018 | 0.018 | 0 | 0.47 | 0.73 | 0.42 | 0.37 | 0.37 | 0.48 |
| 8 | 1039289 | rs17668341 | 0.196 | 0.183 | -1.267 | 0.225 | 0.004 | 0.003 | 0.003 | 1.44 | 0.15 | 0.13 | 0.06 | 0.18 | 0.08 | 0.07 |
| 8 | 1039740 | rs13268693 | -0.034 | 1.076 | -1.21 | 0.416 | 0.002 | -0.003 | -0.003 | 0 | 0.21 | 0.2 | 0.25 | 0.15 | 0.27 | 0.2 |
| 8 | 1061074 | rs1376786 | -0.266 | 0.579 | -1.596 | 0.812 | -0.007 | -0.05 | -0.05 | 0 | 0.14 | 0.05 | 0.05 | 0.22 | 0.11 | 0.16 |
| 8 | 1068910 | rs7842241 | 0.447 | 1.043 | -2.179 | 0.184 | -0.059 | 0.081 | 0.081 | 0 | 0.3 | 0.1 | 0.25 | 0.11 | 0.38 | 0.22 |
| 8 | 1069489 | rs12675380 | 1.142 | 0.577 | -2.75 | 0.186 | -0.057 | 0.029 | 0.029 | 0 | 0.25 | 0.05 | 0.35 | 0.19 | 0.31 | 0.24 |
| 8 | 1073519 | rs7012421 | 0.19 | -0.22 | -1.198 | 0.998 | -0.049 | 0.102 | 0.102 | 0 | 0.74 | 0.48 | 0.8 | 0.54 | 0.84 | 0.61 |
| 8 | 1074425 | rs11775878 | -0.493 | -0.096 | -1.009 | 0.798 | 0.005 | -0.003 | -0.003 | 0 | 0.42 | 0.39 | 0.34 | 0.5 | 0.33 | 0.37 |
| 8 | 1077688 | rs13259992 | -0.375 | 0.448 | -0.978 | 1.15 | 0.002 | 0.003 | 0.003 | 0 | 0.92 | 0.93 | 0.92 | 0.78 | 0.93 | 0.78 |
| 8 | 1082750 | rs2100321 | 0.116 | 0.808 | -1.314 | 0.703 | -0.053 | 0.002 | 0.002 | 0 | 0.11 | 0.01 | 0.05 | 0.09 | 0.13 | 0.14 |
| 8 | 1083364 | rs1451882 | 0.313 | 0.444 | -1.258 | 1.127 | -0.012 | -0.024 | -0.024 | 0 | 0.08 | 0.04 | 0.04 | 0.24 | 0.08 | 0.08 |
| 8 | 1086161 | rs717406 | -0.081 | 0.389 | -0.425 | 1.267 | 0.018 | 0.146 | 0.146 | 0.033 | 0.89 | 0.72 | 0.86 | 0.75 | 0.91 | 0.81 |
| 8 | 1087212 | rs11779020 | 0.093 | 0.705 | -1.284 | 0.608 | -0.11 | 0.082 | 0.082 | 0.195 | 0.29 | 0.08 | 0.4 | 0.15 | 0.42 | 0.35 |
| 8 | 1089766 | rs10104725 | -0.735 | 0.649 | -1.08 | 0.722 | -0.001 | 0.026 | 0.026 | 0 | 0.9 | 0.89 | 0.88 | 0.74 | 0.9 | 0.81 |
| 8 | 1091202 | rs7822337 | -0.669 | 0.593 | -0.806 | 0.345 | -0.028 | 0.087 | 0.087 | 0.56 | 0.38 | 0.2 | 0.47 | 0.18 | 0.57 | 0.42 |
| 8 | 1091262 | rs7839167 | 1.737 | -0.099 | -0.559 | 0.05 | -0.005 | -0.01 | -0.01 | 0 | 0.34 | 0.29 | 0.16 | 0.16 | 0.19 | 0.24 |
| 8 | 1094105 | rs10046782 | 0.17 | 0.264 | -0.806 | 0.046 | -0.101 | 0.087 | 0.087 | 0.817 | 0.3 | 0.1 | 0.41 | 0.15 | 0.44 | 0.34 |
| 8 | 1096519 | rs12716579 | -0.362 | 0.271 | -0.681 | 0.113 | -0.041 | 0.088 | 0.088 | 0.41 | 0.38 | 0.2 | 0.46 | 0.18 | 0.53 | 0.4 |
| 8 | 1101954 | rs7832117 | 1.259 | -0.152 | -0.295 | 0.481 | -0.01 | -0.003 | -0.003 | 0 | 0.37 | 0.32 | 0.18 | 0.33 | 0.24 | 0.31 |
| 8 | 1105431 | rs17065707 | 1.464 | 0.1 | 0.011 | 0.527 | 0.015 | 0.012 | 0.012 | 3.035 | 0.15 | 0.23 | 0.21 | 0.22 | 0.16 | 0.07 |
| 8 | 1105818 | rs7834689 | 1.473 | -0.006 | -0.022 | 0.58 | 0.029 | -0.005 | -0.005 | 0 | 0.35 | 0.22 | 0.2 | 0.38 | 0.2 | 0.34 |
| 8 | 1107201 | rs1451884 | -0.915 | 0.272 | -0.001 | 0.686 | 0.006 | 0.059 | 0.059 | 0 | 0.86 | 0.75 | 0.9 | 0.79 | 0.89 | 0.8 |
| 8 | 1108571 | rs13273049 | -0.226 | 0.568 | -0.238 | 0.563 | -0.021 | 0.024 | 0.024 | 0 | 0.34 | 0.26 | 0.5 | 0.21 | 0.49 | 0.39 |
| 8 | 1110914 | rs10093077 | 1.072 | 1.153 | -0.683 | 0.585 | -0.077 | -0.047 | -0.047 | 0.048 | 0.13 | 0.02 | 0.3 | 0.18 | 0.2 | 0.2 |
| 8 | 1111158 | rs7008181 | 1.325 | 0.278 | -0.345 | 0.876 | -0.002 | 0.061 | 0.061 | 0.592 | 0.05 | 0.07 | 0.03 | 0.21 | 0.04 | 0.08 |
| 8 | 1114849 | rs10096352 | -0.84 | 0.205 | -0.134 | 0.702 | 0.032 | 0.003 | 0.003 | 0.248 | 0.55 | 0.69 | 0.76 | 0.62 | 0.68 | 0.53 |
| 8 | 1115647 | rs4355799 | -1.148 | 0.099 | -0.245 | 0.702 | 0 | 0.021 | 0.021 | 0 | 0.87 | 0.81 | 0.91 | 0.77 | 0.88 | 0.82 |

| | | | | | | | | | | | | | | | | |
|---|---|---|---|---|---|---|---|---|---|---|---|---|---|---|---|---|
| 8 | 1116638 | rs7827800 | 1.779 | 0.078 | -0.477 | 0.431 | 0.05 | 0.08 | 0.08 | 0 | 0.32 | 0.11 | 0.15 | 0.16 | 0.2 | 0.29 |
| 8 | 1117739 | rs13260791 | 0.133 | 0.189 | 0.064 | 0.612 | -0.007 | 0.144 | 0.144 | 0 | 0.79 | 0.52 | 0.85 | 0.58 | 0.84 | 0.75 |
| 8 | 1124032 | rs13267567 | 0.31 | 0.409 | -0.632 | 0.464 | -0.095 | 0.062 | 0.062 | 0 | 0.3 | 0.09 | 0.47 | 0.16 | 0.48 | 0.38 |
| 8 | 1133247 | rs6558418 | 2.134 | -0.57 | -0.847 | 0.349 | 0.004 | 0.006 | 0.006 | 0.05 | 0.15 | 0.08 | 0.05 | 0.12 | 0.04 | 0.15 |
| 8 | 1133355 | rs12334702 | -0.033 | -0.59 | -0.748 | 0.19 | 0.029 | 0.002 | 0.002 | 1.649 | 0.5 | 0.63 | 0.73 | 0.56 | 0.65 | 0.53 |
| 8 | 1134511 | rs11998278 | 1.795 | -0.748 | -0.531 | 0.332 | -0.004 | 0.003 | 0.003 | 0.151 | 0.16 | 0.1 | 0.05 | 0.08 | 0.05 | 0.15 |
| 8 | 1138415 | rs7016616 | -0.908 | 0.267 | -0.271 | 0.953 | -0.011 | -0.052 | -0.052 | 0 | 0.91 | 1 | 0.98 | 0.85 | 0.93 | 0.88 |
| 8 | 1141930 | rs6989310 | -0.028 | 0.496 | -0.016 | 0.932 | 0.034 | 0.119 | 0.119 | 0 | 0.86 | 0.74 | 0.81 | 0.68 | 0.83 | 0.76 |
| 8 | 1143854 | rs11784898 | -0.056 | 0.231 | 0.128 | 0.756 | 0.009 | 0.034 | 0.034 | 0.594 | 0.45 | 0.31 | 0.63 | 0.39 | 0.61 | 0.4 |
| 8 | 1143930 | rs17815788 | 1.138 | -0.09 | -0.014 | 0.582 | 0.069 | 0.043 | 0.043 | 0.277 | 0.37 | 0.16 | 0.15 | 0.29 | 0.2 | 0.31 |
| 8 | 1144060 | rs17065769 | 0.858 | -0.086 | 0.004 | 0.584 | -0.022 | -0.013 | -0.013 | 0 | 0.41 | 0.44 | 0.18 | 0.29 | 0.21 | 0.35 |
| 8 | 1146100 | rs13273765 | -0.579 | 0.5 | -0.136 | 0.742 | 0.023 | 0.023 | 0.023 | 2.143 | 0.36 | 0.52 | 0.57 | 0.45 | 0.52 | 0.49 |
| 8 | 1146438 | rs12676200 | 0.413 | -0.072 | 0.031 | 0.59 | 0 | 0 | 0 | 1.874 | 0.59 | 0.52 | 0.4 | 0.51 | 0.43 | 0.44 |
| 8 | 1146999 | rs11136358 | 0.43 | -0.072 | 0.005 | 0.576 | -0.001 | 0.001 | 0.001 | 1.962 | 0.59 | 0.48 | 0.4 | 0.51 | 0.43 | 0.44 |
| 8 | 1148273 | rs7007815 | 1.292 | -0.252 | -0.107 | 0.481 | 0.076 | 0.073 | 0.073 | 2.129 | 0.36 | 0.16 | 0.15 | 0.18 | 0.18 | 0.21 |
| 8 | 1149028 | rs12547340 | 1.11 | -0.128 | -0.152 | 0.529 | -0.012 | -0.014 | -0.014 | 0 | 0.22 | 0.12 | 0.24 | 0.34 | 0.24 | 0.17 |
| 8 | 1149677 | rs7826946 | 0.456 | -0.074 | -0.116 | 0.417 | 0.024 | 0.011 | 0.011 | 1.017 | 0.63 | 0.42 | 0.43 | 0.52 | 0.49 | 0.43 |
| 8 | 1149907 | rs12543247 | 0.135 | -0.051 | -0.325 | 0.431 | 0.04 | 0.024 | 0.024 | 1.734 | 0.67 | 0.53 | 0.44 | 0.51 | 0.49 | 0.43 |
| 8 | 1150384 | rs12548097 | -0.096 | -0.057 | -0.826 | 0.358 | -0.016 | -0.021 | -0.021 | 0.71 | 0.35 | 0.28 | 0.43 | 0.5 | 0.4 | 0.33 |
| 8 | 1150582 | rs2044925 | 1.875 | -0.281 | -1.263 | 0.28 | 0.055 | 0.044 | 0.044 | 0 | 0.29 | 0.1 | 0.12 | 0.18 | 0.17 | 0.21 |
| 8 | 1150705 | rs17065805 | 0.576 | 0.322 | -1.003 | 0.411 | -0.007 | -0.038 | -0.038 | 0 | 0.1 | 0.13 | 0.05 | 0 | 0.07 | 0.06 |
| 8 | 1150848 | rs7006250 | -0.543 | 0.689 | -1.346 | 0.643 | -0.082 | -0.073 | -0.073 | 0 | 0.79 | 0.92 | 0.63 | 0.69 | 0.63 | 0.65 |
| 8 | 1151503 | rs1562920 | 0.299 | 0.182 | -0.805 | 0.291 | 0.001 | -0.017 | -0.017 | 0 | 0.23 | 0.39 | 0.24 | 0.21 | 0.21 | 0.15 |
| 8 | 1153719 | rs4735993 | -0.799 | 0.333 | -1.781 | 0.402 | 0.013 | -0.045 | -0.045 | 0 | 0.33 | 0.79 | 0.42 | 0.48 | 0.27 | 0.43 |
| 8 | 1153876 | rs4735994 | 0.296 | 1.236 | -1.692 | 0.851 | -0.001 | 0.026 | 0.026 | 0 | 0.13 | 0.18 | 0.3 | 0.37 | 0.19 | 0.25 |
| 8 | 1153941 | rs17669161 | 1.052 | 1.291 | -2.06 | 0.754 | -0.063 | -0.024 | -0.024 | 0 | 0.17 | 0.03 | 0.22 | 0.18 | 0.22 | 0.1 |
| 8 | 1154027 | rs2044924 | -0.058 | 1.077 | -2.122 | 0.656 | -0.051 | 0.032 | 0.032 | 0 | 0.59 | 0.89 | 0.55 | 0.7 | 0.43 | 0.6 |
| 8 | 1154309 | rs2044922 | -0.058 | 1.077 | -2.122 | 0.659 | -0.051 | 0.035 | 0.035 | 0 | 0.59 | 0.88 | 0.55 | 0.7 | 0.43 | 0.6 |
| 8 | 1156822 | rs13261391 | 0.32 | 1.136 | -1.898 | 0.866 | -0.018 | -0.02 | -0.02 | 0 | 0.13 | 0.11 | 0.25 | 0.34 | 0.21 | 0.26 |
| 8 | 1156972 | rs10094928 | 0.552 | 1.17 | -1.797 | 0.184 | 0 | 0.08 | 0.08 | 0 | 0.53 | 0.86 | 0.53 | 0.7 | 0.42 | 0.6 |
| 8 | 1161954 | rs10111597 | 0.32 | 1.134 | -1.133 | 0.251 | -0.007 | 0.01 | 0.01 | 1.074 | 0.7 | 0.77 | 0.73 | 0.88 | 0.64 | 0.72 |
| 8 | 1162454 | rs7829754 | 0.044 | 0.909 | -0.935 | 0.658 | -0.006 | 0.013 | 0.013 | 0 | 0.58 | 0.67 | 0.61 | 0.63 | 0.53 | 0.73 |
| 8 | 1163713 | rs4735997 | -0.733 | 0.885 | -0.135 | 1.274 | 0.015 | 0.111 | 0.111 | 0.066 | 0.08 | 0.16 | 0.23 | 0.22 | 0.11 | 0.11 |
| 8 | 1164482 | rs1463249 | -0.422 | 0.279 | 0.291 | 1.17 | -0.035 | 0.002 | 0.002 | 2.665 | 0.09 | 0.3 | 0.06 | 0.06 | 0.03 | 0.02 |
| 8 | 1165148 | rs7388163 | -0.054 | 0.346 | -0.574 | 1.001 | 0.002 | -0.034 | -0.034 | 0 | 0.28 | 0.16 | 0.33 | 0.45 | 0.28 | 0.38 |
| 8 | 1170382 | rs11136364 | -1.109 | 0.318 | -0.799 | 0.451 | 0.035 | 0.173 | 0.173 | 0 | 0.51 | 0.87 | 0.64 | 0.76 | 0.44 | 0.65 |
| 8 | 1174072 | rs2084803 | -0.426 | 0.247 | -0.35 | 0.918 | 0.014 | -0.032 | -0.032 | 5.546 | 0.28 | 0.26 | 0.34 | 0.53 | 0.24 | 0.34 |
| 8 | 1174950 | rs11776363 | -0.815 | 0.14 | 0.41 | 0.899 | 0.015 | -0.014 | -0.014 | 0 | 0.33 | 0.33 | 0.37 | 0.54 | 0.26 | 0.3 |
| 8 | 1178457 | rs4549802 | -1.547 | 0.411 | 0.312 | 0.504 | 0.007 | 0.157 | 0.157 | 0 | 0.5 | 0.82 | 0.65 | 0.77 | 0.48 | 0.65 |
| 8 | 1191927 | rs13277024 | 0.395 | 0.637 | 0.172 | 1.074 | 0.008 | -0.115 | -0.115 | 0 | 0.17 | 0.01 | 0.09 | 0.37 | 0.13 | 0.19 |
| 8 | 1192438 | rs4976875 | 0.485 | 2.514 | 0.754 | 0.649 | -0.024 | 0.003 | 0.003 | 0 | 0.87 | 0.9 | 0.8 | 0.85 | 0.7 | 0.77 |
| 8 | 1194935 | rs7821163 | -1.028 | 1.992 | 1.126 | 0.463 | 0.056 | -0.04 | -0.04 | 0.676 | 0.15 | 0.38 | 0.13 | 0.08 | 0.17 | 0.22 |
| 8 | 1214057 | rs6558434 | -0.114 | 2.023 | 0.822 | 0.267 | 0.058 | -0.023 | -0.023 | 0 | 0.14 | 0.39 | 0.14 | 0.07 | 0.18 | 0.15 |
| 8 | 1215445 | rs6558435 | -0.268 | 2.238 | 0.534 | 0.702 | -0.008 | -0.01 | -0.01 | 2.42 | 0.9 | 0.91 | 0.83 | 0.78 | 0.88 | 0.8 |
| 8 | 1216248 | rs7009764 | 1.047 | 2.607 | 0.543 | 0.719 | 0.039 | 0.036 | 0.036 | 0.333 | 0.69 | 0.47 | 0.6 | 0.59 | 0.59 | 0.49 |
| 8 | 1218258 | rs4272406 | -0.319 | 2.158 | 0.388 | 0.689 | 0.033 | 0.014 | 0.014 | 0 | 0.35 | 0.42 | 0.35 | 0.45 | 0.52 | 0.35 |
| 8 | 1220581 | rs4495453 | -0.212 | 1.771 | 0.018 | 0.478 | 0 | 0.009 | 0.009 | 0 | 0.82 | 0.77 | 0.68 | 0.77 | 0.83 | 0.83 |
| 8 | 1226530 | rs7000219 | 1.067 | 2.228 | 0.099 | 0.366 | 0.043 | 0.02 | 0.02 | 0 | 0.68 | 0.51 | 0.59 | 0.64 | 0.55 | 0.55 |
| 8 | 1227513 | rs11136377 | 1.476 | 1.851 | -0.608 | 0.055 | 0.025 | 0.009 | 0.009 | 1.14 | 0.44 | 0.24 | 0.23 | 0.44 | 0.36 | 0.28 |
| 8 | 1231248 | rs6995458 | -0.817 | 1.911 | -0.445 | -0.084 | -0.005 | 0.008 | 0.008 | 1.212 | 0.27 | 0.68 | 0.34 | 0.13 | 0.36 | 0.38 |
| 8 | 1236319 | rs7814902 | -1.253 | 1.79 | -0.11 | 0.045 | 0.007 | 0.002 | 0.002 | 1.12 | 0.4 | 0.47 | 0.49 | 0.4 | 0.56 | 0.58 |
| 8 | 1238564 | rs6998229 | -1.019 | 1.669 | -0.265 | 0.256 | 0.013 | 0.073 | 0.073 | 0 | 0.89 | 0.86 | 0.72 | 0.68 | 0.87 | 0.88 |
| 8 | 1240600 | rs760179 | 2.215 | 1.395 | -0.467 | 0.034 | 0.063 | 0.023 | 0.023 | 1.272 | 0.37 | 0.15 | 0.16 | 0.35 | 0.23 | 0.24 |
| 8 | 1240646 | rs17816317 | -0.004 | 1.567 | -0.518 | -0.032 | 0.006 | 0.01 | 0.01 | 0 | 0.2 | 0.27 | 0.2 | 0.26 | 0.23 | 0.25 |
| 8 | 1240819 | rs4875987 | -0.522 | 1.719 | -0.373 | 0.036 | 0.018 | -0.082 | -0.082 | 0.221 | 0.28 | 0.52 | 0.3 | 0.09 | 0.38 | 0.22 |
| 8 | 1241224 | rs7463743 | -1.362 | 1.871 | -0.9 | 0.233 | -0.05 | -0.063 | -0.063 | 0 | 0.9 | 0.99 | 0.73 | 0.76 | 0.87 | 0.89 |
| 8 | 1243574 | rs10283028 | 1.112 | 1.954 | -0.8 | 0.016 | -0.025 | 0.022 | 0.022 | 0.006 | 0.13 | 0.06 | 0.18 | 0.07 | 0.25 | 0.2 |
| 8 | 1244374 | rs10101376 | 0.857 | 1.609 | -0.512 | 0.094 | 0.027 | 0.022 | 0.022 | 0.714 | 0.67 | 0.44 | 0.45 | 0.45 | 0.64 | 0.52 |
| 8 | 1248400 | rs17747701 | 0.87 | 0.989 | -1.062 | 0.244 | -0.019 | -0.025 | -0.025 | 0 | 0.12 | 0.03 | 0.15 | 0.17 | 0.14 | 0.17 |
| 8 | 1248648 | rs12679463 | 1.466 | 0.934 | -1.277 | 0.168 | 0.013 | 0 | 0 | 0 | 0.39 | 0.07 | 0.17 | 0.36 | 0.26 | 0.28 |
| 8 | 1250263 | rs17754335 | 1.435 | 1.005 | -1.217 | 0.219 | 0.006 | 0.018 | 0.018 | 1.152 | 0.39 | 0.07 | 0.18 | 0.35 | 0.29 | 0.3 |
| 8 | 1251756 | rs7817445 | 0.45 | 1.226 | -0.94 | 0.134 | -0.015 | -0.022 | -0.022 | 0.272 | 0.21 | 0.33 | 0.26 | 0.13 | 0.23 | 0.34 |
| 8 | 1251835 | rs6558440 | -0.463 | 1.833 | -0.959 | 0.559 | 0.11 | 0.103 | 0.103 | 0 | 0.85 | 0.72 | 0.66 | 0.69 | 0.68 | 0.85 |
| 8 | 1252450 | rs2178855 | 0.642 | 1.264 | -0.938 | 0.346 | 0 | 0 | 0 | 0.181 | 0.6 | 0.63 | 0.45 | 0.54 | 0.5 | 0.64 |
| 8 | 1266814 | rs11775819 | 0.29 | 0.92 | -1.457 | 0.05 | 0.024 | 0.058 | 0.058 | 0 | 0.14 | 0.04 | 0.14 | 0.03 | 0.07 | 0.05 |
| 8 | 1266929 | rs10094861 | 0.206 | 0.302 | -1.786 | -0.405 | 0.015 | 0.052 | 0.052 | 0 | 0.3 | 0.17 | 0.27 | 0.14 | 0.3 | 0.37 |
| 8 | 1267212 | rs11782996 | 0.327 | 1.165 | -1.409 | 0.011 | 0.086 | 0.044 | 0.044 | 0 | 0.05 | 0.11 | 0.09 | 0.09 | 0.17 | 0.04 |
| 8 | 1270993 | rs6558446 | -1.722 | 0.682 | -2.438 | 0.046 | -0.002 | -0.067 | -0.067 | 0 | 0.79 | 0.95 | 0.9 | 0.65 | 0.84 | 0.84 |
| 8 | 1273426 | rs6994475 | 0.495 | 0.281 | -2.224 | 0.12 | -0.008 | 0.005 | 0.005 | 0 | 0.17 | 0.77 | 0.14 | 0.23 | 0.11 | 0.14 |

| | | | | | | | | | | | | | | | | |
|---|---|---|---|---|---|---|---|---|---|---|---|---|---|---|---|---|
| 8 | 1274056 | rs6558447 | -0.58 | 0.11 | -2.915 | -0.133 | -0.011 | -0.006 | -0.006 | 0 | 0.34 | 0.84 | 0.33 | 0.3 | 0.36 | 0.36 |
| 8 | 1282530 | rs11778753 | -0.098 | 0.467 | -3.147 | -0.115 | -0.049 | 0.031 | 0.031 | 0.017 | 0.45 | 0.9 | 0.46 | 0.34 | 0.5 | 0.39 |
| 8 | 1282843 | rs11778654 | -1.646 | 0.763 | -3.338 | -0.104 | -0.043 | -0.063 | -0.063 | 0 | 0.72 | 0.95 | 0.88 | 0.63 | 0.77 | 0.75 |
| 8 | 1283208 | rs13255561 | 1.705 | 0.688 | -2.893 | -0.391 | 0.008 | -0.004 | -0.004 | 0 | 0.13 | 0.18 | 0.2 | 0.1 | 0.18 | 0.17 |
| 8 | 1284332 | rs17816653 | 1.111 | 0.705 | -3.337 | -0.166 | -0.044 | -0.057 | -0.057 | 0 | 0.15 | 0.04 | 0.33 | 0.22 | 0.19 | 0.2 |
| 8 | 1286006 | rs12546588 | 0.127 | 0.297 | -2.938 | -0.078 | 0.009 | 0.031 | 0.031 | 0 | 0.33 | 0.83 | 0.27 | 0.24 | 0.32 | 0.22 |
| 8 | 1286920 | rs13256102 | -0.623 | 0.331 | -2.349 | 0.199 | -0.046 | -0.105 | -0.105 | 1.688 | 0.73 | 0.95 | 0.83 | 0.58 | 0.77 | 0.75 |
| 8 | 1292488 | rs5028021 | -0.154 | 0.005 | -1.485 | -0.807 | -0.021 | -0.12 | -0.12 | 0.103 | 0.76 | 0.45 | 0.84 | 0.99 | 0.84 | 0.85 |
| 8 | 1293061 | rs10093402 | 0.276 | 0.427 | -2.134 | -0.498 | 0.009 | 0.023 | 0.023 | 0.122 | 0.53 | 0.24 | 0.45 | 0.61 | 0.47 | 0.53 |
| 8 | 1293894 | rs12678087 | 1.12 | 0.551 | -2.298 | -0.881 | -0.037 | -0.083 | -0.083 | 0.198 | 0.12 | 0.02 | 0.07 | 0.33 | 0.12 | 0.19 |
| 8 | 1294073 | rs10110073 | 1.281 | 0.138 | -2.479 | -1.485 | -0.029 | 0.039 | 0.039 | 0.071 | 0.23 | 0.03 | 0.31 | 0.16 | 0.25 | 0.23 |
| 8 | 1297156 | rs1882801 | -0.434 | 0.805 | -1.962 | -1.416 | 0.01 | -0.059 | -0.059 | 0.475 | 0.71 | 0.27 | 0.71 | 0.92 | 0.67 | 0.71 |
| 8 | 1301097 | rs722524 | -1.241 | 0.779 | -1.407 | -0.928 | 0.035 | -0.033 | -0.033 | 0.609 | 0.94 | 0.85 | 0.95 | 1 | 0.92 | 0.98 |
| 8 | 1301110 | rs7823212 | 0.04 | 0.125 | -1.869 | -0.797 | 0.015 | -0.016 | -0.016 | 0.457 | 0.25 | 0.15 | 0.12 | 0.3 | 0.19 | 0.24 |
| 8 | 1304128 | rs4876218 | 0.086 | 0.338 | -1.369 | -1.524 | 0.001 | -0.036 | -0.036 | 0 | 0.43 | 0.55 | 0.51 | 0.25 | 0.45 | 0.37 |
| 8 | 1306071 | rs4454302 | -0.752 | 0.258 | -1.47 | -1.564 | -0.064 | 0.104 | 0.104 | 0.008 | 0.67 | 0.87 | 0.62 | 0.93 | 0.58 | 0.74 |
| 8 | 1308026 | rs12541553 | -0.362 | 0.226 | -0.825 | -1.5 | -0.003 | 0.023 | 0.023 | 0 | 0.48 | 0.63 | 0.55 | 0.91 | 0.52 | 0.65 |
| 8 | 1312469 | rs6989471 | -0.346 | 0.244 | -0.578 | -0.933 | -0.016 | -0.034 | -0.034 | 0 | 0.28 | 0.43 | 0.39 | 0.76 | 0.24 | 0.44 |
| 8 | 1316133 | rs2142138 | -0.86 | 0.314 | -1.066 | -1.181 | -0.04 | 0.185 | 0.185 | 0 | 0.6 | 0.87 | 0.67 | 0.91 | 0.59 | 0.77 |
| 8 | 1318693 | rs12114124 | -0.168 | 0.104 | -0.59 | -0.06 | -0.016 | 0.148 | 0.148 | 0 | 0.16 | 0.28 | 0.28 | 0.48 | 0.17 | 0.28 |
| 8 | 1319724 | rs10283337 | -0.345 | 0.283 | -0.466 | -0.081 | -0.007 | -0.004 | -0.004 | 0 | 0.4 | 0.3 | 0.34 | 0.43 | 0.39 | 0.49 |
| 8 | 1326916 | rs6558462 | -0.833 | 0.214 | -0.347 | 0.234 | 0.001 | -0.003 | -0.003 | 0 | 0.36 | 0.7 | 0.49 | 0.57 | 0.37 | 0.44 |
| 8 | 1329505 | rs10110777 | -0.086 | 0.059 | 0.026 | 0.92 | -0.036 | 0.037 | 0.037 | 0 | 0.12 | 0.17 | 0.19 | 0.23 | 0.06 | 0.18 |
| 8 | 1337849 | rs13276952 | -0.217 | 0.363 | 0.148 | 1.154 | -0.011 | -0.01 | -0.01 | 0 | 0.16 | 0.13 | 0.25 | 0.34 | 0.3 | 0.24 |
| 8 | 1338752 | rs17754782 | 0.4 | 0.201 | 0.298 | 1.156 | 0.004 | 0.002 | 0.002 | 0.354 | 0.09 | 0.06 | 0.18 | 0.24 | 0.04 | 0.18 |
| 8 | 1339130 | rs757888 | -0.448 | 0.153 | -0.342 | 0.421 | 0.007 | 0.019 | 0.019 | 0.373 | 0.66 | 0.87 | 0.77 | 0.64 | 0.74 | 0.76 |
| 8 | 1339570 | rs10866924 | -0.068 | 0.147 | -0.048 | 0.407 | 0.014 | 0.007 | 0.007 | 0.133 | 0.61 | 0.68 | 0.75 | 0.6 | 0.72 | 0.72 |
| 8 | 1339682 | rs10866925 | -0.064 | 0.14 | -0.035 | 0.403 | 0.012 | 0.006 | 0.006 | 0.138 | 0.61 | 0.67 | 0.75 | 0.6 | 0.72 | 0.72 |
| 8 | 1339829 | rs12677155 | 0.105 | 0.326 | 0.085 | 0.629 | -0.019 | -0.017 | -0.017 | 0 | 0.16 | 0.13 | 0.25 | 0.3 | 0.29 | 0.2 |
| 8 | 1340507 | rs7017774 | -0.69 | -0.125 | 0.087 | 0.44 | -0.001 | 0 | 0 | 0.027 | 0.46 | 0.5 | 0.45 | 0.65 | 0.5 | 0.42 |
| 8 | 1356333 | rs740237 | 0.19 | 0.273 | 0.056 | 0.687 | 0 | 0 | 0 | 0 | 0.26 | 0.28 | 0.22 | 0.35 | 0.26 | 0.21 |
| 8 | 1359156 | rs17680989 | 0.081 | 0.44 | -0.201 | 0.774 | 0.005 | 0.052 | 0.052 | 0 | 0.06 | 0.07 | 0.17 | 0.3 | 0.14 | 0.11 |
| 8 | 1362713 | rs10094948 | -0.409 | -0.039 | 0.185 | 0.595 | -0.014 | 0.02 | 0.02 | 0 | 0.35 | 0.49 | 0.41 | 0.61 | 0.31 | 0.4 |
| 8 | 1363967 | rs1019024 | 0.176 | -0.071 | -0.317 | 0.288 | -0.014 | -0.007 | -0.007 | 0 | 0.27 | 0.13 | 0.22 | 0.37 | 0.27 | 0.22 |
| 8 | 1366526 | rs6997056 | -0.589 | 0.004 | 0.423 | 0.749 | -0.038 | 0.219 | 0.219 | 0.098 | 0.19 | 0.42 | 0.25 | 0.36 | 0.14 | 0.24 |
| 8 | 1369636 | rs7816367 | -0.471 | 0.269 | 0.183 | 0.25 | 0.006 | -0.048 | -0.048 | 0 | 0.82 | 0.8 | 0.81 | 1 | 0.75 | 0.94 |
| 8 | 1370352 | rs4876039 | 0.208 | -0.075 | -0.679 | 0.058 | 0.035 | 0 | 0 | 0 | 0.21 | 0.07 | 0.24 | 0.18 | 0.17 | 0.27 |
| 8 | 1372223 | rs6983401 | 1.836 | -0.055 | -0.499 | -0.125 | 0.009 | 0.049 | 0.049 | 0 | 0.3 | 0.21 | 0.16 | 0.13 | 0.25 | 0.27 |
| 8 | 1373626 | rs1558027 | 0.209 | -0.127 | -0.433 | -0.022 | -0.016 | 0.047 | 0.047 | 0 | 0.59 | 0.32 | 0.6 | 0.65 | 0.5 | 0.75 |
| 8 | 1375610 | rs763869 | 0.25 | -0.184 | -0.163 | 0.028 | 0.006 | -0.016 | -0.016 | 0 | 0.6 | 0.5 | 0.59 | 0.64 | 0.49 | 0.75 |
| 8 | 1377915 | rs12674781 | 1.144 | -0.099 | -0.322 | 0.039 | -0.007 | -0.006 | -0.006 | 0 | 0.17 | 0.12 | 0.19 | 0.34 | 0.25 | 0.19 |
| 8 | 1379715 | rs7839262 | 1.592 | 0.045 | -0.389 | -0.445 | -0.004 | 0.091 | 0.091 | 0 | 0.29 | 0.11 | 0.16 | 0.12 | 0.25 | 0.27 |
| 8 | 1380394 | rs6998158 | 1.592 | 0.045 | -0.389 | -0.445 | -0.004 | 0.091 | 0.091 | 0 | 0.29 | 0.11 | 0.16 | 0.12 | 0.25 | 0.27 |
| 8 | 1385526 | rs2040988 | 0.06 | -0.032 | 0.081 | 0.004 | 0.008 | -0.003 | -0.003 | 0 | 0.18 | 0.15 | 0.22 | 0.19 | 0.15 | 0.24 |
| 8 | 1408448 | rs733597 | -0.669 | 0.391 | 0.366 | -0.297 | -0.001 | -0.022 | -0.022 | 0 | 0.51 | 0.5 | 0.57 | 0.83 | 0.47 | 0.61 |
| 8 | 1418574 | rs4876063 | 0.605 | 0.459 | 0.071 | -0.308 | -0.003 | 0.009 | 0.009 | 0 | 0.46 | 0.43 | 0.39 | 0.17 | 0.45 | 0.35 |
| 8 | 1421678 | rs2108122 | 0.893 | 0.625 | -0.694 | -0.215 | -0.042 | 0.14 | 0.14 | 0 | 0.4 | 0.13 | 0.33 | 0.22 | 0.4 | 0.29 |
| 8 | 1430681 | rs10106787 | 0.611 | 0.76 | 0.285 | 0.498 | 0.007 | 0.01 | 0.01 | 0 | 0.67 | 0.61 | 0.71 | 0.57 | 0.63 | 0.61 |
| 8 | 1439795 | rs12542312 | -1.639 | 0.814 | 0.076 | 0.179 | 0.004 | 0.015 | 0.015 | 0.921 | 0.2 | 0.81 | 0.16 | 0.11 | 0.13 | 0.2 |
| 8 | 1440839 | rs7845120 | 1.091 | 0.801 | 0.555 | 0.293 | -0.004 | -0.009 | -0.009 | 1.178 | 0.67 | 0.64 | 0.7 | 0.56 | 0.62 | 0.55 |
| 8 | 1444499 | rs4876067 | 0.35 | 0.913 | 0.543 | 0.352 | -0.022 | -0.049 | -0.049 | 0.453 | 0.77 | 0.85 | 0.78 | 0.56 | 0.71 | 0.74 |
| 8 | 1445111 | rs4875864 | 1.614 | 0.761 | 0.468 | 0.23 | -0.044 | -0.072 | -0.072 | 0 | 0.59 | 0.16 | 0.64 | 0.53 | 0.58 | 0.54 |
| 8 | 1445874 | rs4876069 | 1.524 | 0.869 | 0.834 | 0.337 | -0.018 | -0.051 | -0.051 | 0 | 0.65 | 0.25 | 0.67 | 0.53 | 0.64 | 0.72 |
| 8 | 1446009 | rs4876070 | 1.056 | 0.834 | 1.549 | 0.319 | 0.01 | 0.026 | 0.026 | 0 | 0.68 | 0.62 | 0.71 | 0.53 | 0.64 | 0.72 |
| 8 | 1449183 | rs2286216 | 0.575 | 0.714 | 1.723 | 0.377 | 0.007 | 0.047 | 0.047 | 0 | 0.75 | 0.61 | 0.8 | 0.56 | 0.73 | 0.77 |
| 8 | 1449268 | rs2286217 | -2.189 | 0.651 | 1.905 | 0.476 | 0.078 | 0.225 | 0.225 | 0.954 | 0.06 | 0.42 | 0.06 | 0.11 | 0.02 | 0.01 |
| 8 | 1453450 | rs4876074 | 0.387 | 0.739 | 1.756 | 0.442 | 0.02 | 0.145 | 0.145 | 0 | 0.78 | 0.46 | 0.78 | 0.45 | 0.74 | 0.78 |
| 8 | 1457884 | rs4875868 | 0.458 | 0.694 | 1.222 | -0.004 | -0.003 | -0.011 | -0.011 | 8.144 | 0.14 | 0.11 | 0.11 | 0.01 | 0.17 | 0.12 |
| 8 | 1460796 | rs4876080 | -0.031 | 1.135 | 1.611 | -0.042 | 0.055 | 0.115 | 0.115 | 0 | 0.76 | 0.57 | 0.72 | 0.46 | 0.68 | 0.81 |
| 8 | 1461822 | rs1076302 | 0.458 | 1.403 | 1.445 | -0.07 | -0.006 | -0.011 | -0.011 | 0.086 | 0.14 | 0.14 | 0.14 | 0.01 | 0.2 | 0.09 |
| 8 | 1463750 | rs4876082 | 1.161 | 0.927 | 1.63 | -0.12 | -0.006 | -0.019 | -0.019 | 0 | 0.58 | 0.3 | 0.56 | 0.44 | 0.46 | 0.71 |
| 8 | 1466044 | rs876664 | -0.186 | 0.993 | 1.722 | -0.063 | 0.077 | 0.093 | 0.093 | 0 | 0.74 | 0.58 | 0.69 | 0.45 | 0.64 | 0.8 |
| 8 | 1467648 | rs9314433 | -0.064 | 0.84 | 2.095 | -0.03 | 0.004 | -0.023 | -0.023 | 0.244 | 0.17 | 0.43 | 0.17 | 0.11 | 0.17 | 0.09 |
| 8 | 1468360 | rs17828263 | 0.036 | -0.193 | 1.645 | -0.597 | -0.052 | -0.037 | -0.037 | 0 | 0.2 | 0.32 | 0.11 | 0.12 | 0.09 | 0.2 |
| 8 | 1469213 | rs2020405 | -0.255 | 0.254 | 1.64 | -0.636 | 0.048 | 0.075 | 0.075 | 0 | 0.74 | 0.5 | 0.71 | 0.43 | 0.66 | 0.8 |
| 8 | 1470921 | rs13268384 | -0.566 | 0.295 | 1.319 | -0.582 | 0.033 | 0.048 | 0.048 | 0 | 0.77 | 0.74 | 0.73 | 0.44 | 0.65 | 0.81 |
| 8 | 1471389 | rs6558479 | -0.725 | 0.316 | 0.905 | -0.572 | -0.043 | -0.051 | -0.051 | 0 | 0.78 | 0.86 | 0.74 | 0.44 | 0.65 | 0.81 |
| 8 | 1480965 | rs17748677 | 0.724 | -0.587 | 0.357 | -0.299 | -0.05 | -0.059 | -0.059 | 0 | 0.1 | 0.02 | 0.15 | 0.25 | 0.16 | 0.23 |
| 8 | 1483050 | rs4876086 | 1.801 | -1.003 | 0.391 | -0.64 | 0.047 | 0.071 | 0.071 | 0 | 0.33 | 0.2 | 0.35 | 0.16 | 0.25 | 0.29 |

| | | | | | | | | | | | | | | | | |
|---|---|---|---|---|---|---|---|---|---|---|---|---|---|---|---|---|
| 8 | 1484510 | rs1990126 | 0.346 | -1.051 | 0.672 | -0.689 | 0.004 | -0.011 | -0.011 | 0 | 0.5 | 0.55 | 0.45 | 0.17 | 0.34 | 0.47 |
| 8 | 1484887 | rs7815979 | -0.072 | -1.018 | 0.962 | -0.359 | 0.002 | 0.003 | 0.003 | 0 | 0.62 | 0.65 | 0.61 | 0.45 | 0.5 | 0.71 |
| 8 | 1486112 | rs17064068 | 0.197 | -0.791 | 0.567 | -0.389 | -0.015 | -0.043 | -0.043 | 0 | 0.15 | 0.2 | 0.08 | 0.01 | 0.08 | 0.17 |
| 8 | 1489033 | rs11136406 | -0.134 | -1.316 | 0.709 | 0.054 | 0.022 | 0.012 | 0.012 | 0 | 0.35 | 0.54 | 0.39 | 0.6 | 0.43 | 0.31 |
| 8 | 1504155 | rs757890 | 0.117 | -1.813 | 0.303 | 0.013 | 0.018 | -0.014 | -0.014 | 7.31 | 0.58 | 0.79 | 0.58 | 0.4 | 0.42 | 0.48 |
| 8 | 1507069 | rs11136413 | 0.441 | -1.783 | 0.786 | -0.025 | 0.011 | 0.012 | 0.012 | 0 | 0.32 | 0.37 | 0.36 | 0.5 | 0.42 | 0.33 |
| 8 | 1512571 | rs2956914 | 0.086 | -1.924 | 0.942 | -0.164 | 0 | 0.005 | 0.005 | 0 | 0.59 | 0.45 | 0.59 | 0.45 | 0.46 | 0.56 |
| 8 | 1514009 | rs2301963 | 0.311 | -1.994 | 0.725 | -0.203 | 0.005 | -0.002 | -0.002 | 0 | 0.57 | 0.34 | 0.57 | 0.45 | 0.45 | 0.56 |
| 8 | 1514333 | rs11787306 | 0.861 | -1.964 | 0.873 | -0.198 | -0.076 | -0.068 | -0.068 | 0.49 | 0.29 | 0.16 | 0.33 | 0.52 | 0.42 | 0.3 |
| 8 | 1514749 | rs2956917 | -0.274 | -1.926 | 0.522 | -0.157 | 0.011 | 0.014 | 0.014 | 0 | 0.64 | 0.47 | 0.61 | 0.48 | 0.51 | 0.68 |
| 8 | 1517844 | rs929383 | -0.012 | -1.811 | 0.504 | -0.036 | -0.003 | 0.01 | 0.01 | 0 | 0.07 | 0.15 | 0.09 | 0.06 | 0.06 | 0.08 |
| 8 | 1518671 | rs2906590 | -0.254 | -1.774 | 0.179 | -0.069 | 0.002 | 0.003 | 0.003 | 0 | 0.64 | 0.41 | 0.6 | 0.48 | 0.51 | 0.65 |
| 8 | 1526557 | rs12546094 | -0.415 | -1.87 | -0.171 | -0.004 | -0.003 | 0.018 | 0.018 | 0.981 | 0.47 | 0.86 | 0.46 | 0.61 | 0.51 | 0.48 |
| 8 | 1534503 | rs10094463 | -0.545 | -1.803 | -0.029 | 0.03 | 0.047 | 0.012 | 0.012 | 0 | 0.53 | 0.8 | 0.47 | 0.58 | 0.56 | 0.48 |
| 8 | 1537126 | rs6558491 | 0.654 | -1.201 | 0.73 | 0.363 | 0.017 | 0.015 | 0.015 | 0 | 0.31 | 0.4 | 0.39 | 0.45 | 0.42 | 0.34 |
| 8 | 1548389 | rs2956943 | -0.455 | -1.988 | 0.714 | -0.03 | -0.006 | -0.046 | -0.046 | 0 | 0.55 | 0.61 | 0.75 | 0.92 | 0.73 | 0.7 |
| 8 | 1550644 | rs6999089 | 0.73 | -1.904 | 0.531 | 0.086 | 0.052 | 0.04 | 0.04 | 0 | 0.32 | 0.54 | 0.4 | 0.45 | 0.44 | 0.37 |
| 8 | 1551838 | rs17828747 | 0.123 | -1.859 | 0.46 | 0.158 | -0.003 | -0.019 | -0.019 | 4.57 | 0.25 | 0.2 | 0.37 | 0.48 | 0.3 | 0.35 |
| 8 | 1561756 | rs7819115 | -0.045 | -1.996 | 0.293 | 0.127 | 0.156 | 0.234 | 0.234 | 3.049 | 0.48 | 0.72 | 0.26 | 0.12 | 0.25 | 0.32 |
| 8 | 1566505 | rs2957086 | -1.04 | -1.863 | 0.403 | 0.119 | 0.024 | 0.017 | 0.017 | 0 | 0.69 | 0.56 | 0.6 | 0.55 | 0.54 | 0.64 |
| 8 | 1567920 | rs2957090 | 0.984 | -1.68 | 0.127 | 0.239 | -0.061 | -0.119 | -0.119 | 0 | 0.17 | 0.03 | 0.31 | 0.43 | 0.24 | 0.26 |
| 8 | 1570151 | rs2956964 | 1.113 | -1.736 | 0.127 | 0.238 | -0.068 | -0.101 | -0.101 | 0 | 0.16 | 0.03 | 0.26 | 0.38 | 0.24 | 0.25 |
| 8 | 1578073 | rs13280292 | 0.284 | -2.25 | 0.553 | -0.089 | -0.034 | -0.069 | -0.069 | 0.106 | 0.4 | 0.37 | 0.19 | 0.07 | 0.19 | 0.3 |
| 8 | 1581839 | rs6980953 | -0.105 | -2.242 | 0.451 | -0.047 | 0.152 | 0.162 | 0.162 | 0.323 | 0.47 | 0.76 | 0.29 | 0.18 | 0.24 | 0.36 |
| 8 | 1587877 | rs4875883 | 0.239 | -2.253 | 0.668 | -0.096 | 0.014 | 0.004 | 0.004 | 0.292 | 0.42 | 0.37 | 0.21 | 0.11 | 0.24 | 0.34 |
| 8 | 1595450 | rs4876115 | 0.366 | -2.239 | 0.297 | -0.082 | -0.057 | -0.057 | -0.057 | 1.606 | 0.39 | 0.45 | 0.26 | 0.14 | 0.19 | 0.3 |
| 8 | 1596082 | rs6558504 | -0.643 | -1.778 | 0.248 | -0.014 | -0.035 | -0.056 | -0.056 | 0.059 | 0.64 | 0.78 | 0.56 | 0.48 | 0.47 | 0.63 |
| 8 | 1599813 | rs4552929 | 0.997 | -2.028 | 0.088 | -0.194 | -0.029 | -0.046 | -0.046 | 0 | 0.3 | 0.35 | 0.16 | 0.13 | 0.16 | 0.33 |
| 8 | 1600382 | rs9644258 | 0.812 | -2.007 | 0.022 | -0.171 | -0.014 | -0.046 | -0.046 | 0 | 0.18 | 0.12 | 0.32 | 0.35 | 0.23 | 0.27 |
| 8 | 1600701 | rs9644315 | -0.664 | -1.901 | -0.323 | -0.108 | -0.032 | -0.057 | -0.057 | 0 | 0.63 | 0.77 | 0.55 | 0.49 | 0.47 | 0.63 |
| 8 | 1603043 | rs10448097 | -0.007 | -1.897 | -0.485 | -0.194 | -0.001 | 0.001 | 0.001 | 0 | 0.32 | 0.3 | 0.37 | 0.45 | 0.31 | 0.3 |
| 8 | 1613232 | rs7836020 | -0.888 | -1.329 | -1.528 | 0.106 | -0.074 | -0.057 | -0.057 | 0 | 0.71 | 0.82 | 0.61 | 0.59 | 0.52 | 0.67 |
| 8 | 1615694 | rs4471073 | -0.188 | -1.703 | -0.624 | 1.009 | -0.066 | 0.109 | 0.109 | 0 | 0.2 | 0.61 | 0.27 | 0.47 | 0.09 | 0.2 |
| 8 | 1618087 | rs11136419 | -0.035 | -1.63 | -0.693 | 0.976 | -0.077 | 0.146 | 0.146 | 0.128 | 0.19 | 0.52 | 0.27 | 0.42 | 0.09 | 0.2 |
| 8 | 1625104 | rs2235114 | -0.755 | -1.318 | -1.726 | 0.738 | 0.08 | -0.012 | -0.012 | 0.362 | 0.56 | 0.85 | 0.51 | 0.61 | 0.32 | 0.58 |
| 8 | 1625124 | rs4876121 | -0.264 | -0.665 | -1.132 | 0.413 | 0.017 | 0.017 | 0.017 | 0 | 0.31 | 0.23 | 0.15 | 0.15 | 0.16 | 0.35 |
| 8 | 1630469 | rs4875888 | 0.707 | 0.365 | -1.029 | 1.595 | -0.001 | 0.035 | 0.035 | 0 | 0.87 | 0.87 | 0.69 | 0.51 | 0.81 | 0.76 |
| 8 | 1632525 | rs4507794 | -0.428 | -0.085 | -1.506 | 1.508 | 0.014 | -0.04 | -0.04 | 1.495 | 0.13 | 0.03 | 0.22 | 0.21 | 0.09 | 0.16 |
| 8 | 1634089 | rs6986016 | -0.984 | -0.198 | -1.674 | 1.313 | 0.065 | -0.009 | -0.009 | 0 | 0.52 | 0.78 | 0.45 | 0.49 | 0.38 | 0.53 |
| 8 | 1640790 | rs6558511 | -0.895 | 0.182 | -0.863 | 1.234 | -0.001 | 0.001 | 0.001 | 0 | 0.55 | 0.52 | 0.69 | 0.69 | 0.56 | 0.67 |
| 8 | 1643051 | rs7835556 | -0.949 | -0.04 | -0.614 | 1.23 | 0.001 | -0.01 | -0.01 | 0.362 | 0.56 | 0.48 | 0.69 | 0.66 | 0.55 | 0.66 |
| 8 | 1644150 | rs7465223 | -0.403 | -0.236 | -0.605 | 1.362 | 0.021 | -0.034 | -0.034 | 0 | 0.27 | 0.16 | 0.32 | 0.35 | 0.21 | 0.34 |
| 8 | 1645536 | rs2293909 | -0.737 | -0.223 | -0.201 | 1.258 | 0.026 | 0.021 | 0.021 | 3.214 | 0.36 | 0.28 | 0.3 | 0.29 | 0.24 | 0.29 |
| 8 | 1648035 | rs7825359 | -1.083 | -0.272 | 0.218 | 1.227 | -0.011 | -0.005 | -0.005 | 0 | 0.39 | 0.53 | 0.35 | 0.36 | 0.27 | 0.35 |
| 8 | 1649132 | rs4876123 | -1.515 | -0.177 | 0.401 | 1.306 | 0.008 | 0.035 | 0.035 | 4.248 | 0.5 | 0.63 | 0.63 | 0.64 | 0.47 | 0.49 |
| 8 | 1649750 | rs17749622 | 0.031 | -0.554 | -0.148 | 1.184 | -0.032 | -0.044 | -0.044 | 0 | 0.14 | 0.01 | 0.21 | 0.14 | 0.11 | 0.14 |
| 8 | 1649960 | rs7005715 | -1.455 | -0.087 | 0.053 | 0.946 | -0.011 | 0.118 | 0.118 | 2.259 | 0.5 | 0.81 | 0.66 | 0.68 | 0.43 | 0.6 |
| 8 | 1651211 | rs11987205 | -0.846 | -0.118 | 0.797 | 1.128 | -0.002 | 0.076 | 0.076 | 0.115 | 0.09 | 0.28 | 0.14 | 0.14 | 0.08 | 0.09 |
| 8 | 1651275 | rs2977177 | -0.494 | -0.038 | 0.645 | 0.974 | -0.001 | 0.009 | 0.009 | 4.559 | 0.09 | 0.05 | 0.06 | 0.05 | 0.1 | 0.18 |
| 8 | 1652879 | rs2972175 | 0.149 | -0.454 | 0.884 | 1.355 | -0.001 | 0.001 | 0.001 | 1.059 | 0.21 | 0.15 | 0.25 | 0.39 | 0.16 | 0.27 |
| 8 | 1653690 | rs2957054 | -0.224 | -0.447 | 0.102 | 1.365 | -0.006 | 0.006 | 0.006 | 0.36 | 0.23 | 0.24 | 0.26 | 0.39 | 0.16 | 0.28 |
| 8 | 1654818 | rs2972174 | -0.557 | -0.168 | -0.548 | 1.069 | -0.003 | 0.003 | 0.003 | 0.13 | 0.34 | 0.59 | 0.41 | 0.59 | 0.25 | 0.49 |
| 8 | 1656704 | rs2972171 | -0.723 | -0.033 | -1.248 | 0.238 | 0.013 | 0.067 | 0.067 | 0.494 | 0.43 | 0.74 | 0.66 | 0.74 | 0.38 | 0.59 |
| 8 | 1658593 | rs2957055 | 0.593 | 0.301 | -1.297 | 0.389 | -0.043 | 0.099 | 0.099 | 0.405 | 0.44 | 0.18 | 0.31 | 0.27 | 0.55 | 0.37 |
| 8 | 1676789 | rs2957060 | 0.42 | 0.297 | -0.758 | 0.588 | 0 | 0.001 | 0.001 | 0 | 0.5 | 0.48 | 0.43 | 0.39 | 0.52 | 0.42 |
| 8 | 1678019 | rs12550582 | -0.066 | 0.202 | -1.635 | 0.112 | -0.031 | 0.125 | 0.125 | 0.929 | 0.3 | 0.12 | 0.22 | 0.11 | 0.27 | 0.15 |
| 8 | 1678986 | rs2977185 | 0.602 | -0.013 | -1.33 | 0.349 | 0.013 | 0.047 | 0.047 | 0.161 | 0.58 | 0.67 | 0.71 | 0.66 | 0.65 | 0.74 |
| 8 | 1679777 | rs2977187 | 0.092 | 0.252 | -1.32 | 0.59 | 0.007 | 0.038 | 0.038 | 0.095 | 0.94 | 0.87 | 0.94 | 0.76 | 0.92 | 0.89 |
| 8 | 1680901 | rs11990748 | 0.079 | 0.153 | -1.449 | 0.391 | 0.002 | 0.063 | 0.063 | 2.564 | 0.3 | 0.2 | 0.24 | 0.12 | 0.26 | 0.14 |
| 8 | 1681171 | rs2977189 | 0.45 | 0.388 | -1.21 | 0.593 | 0.004 | 0.025 | 0.025 | 1.675 | 0.92 | 0.89 | 0.95 | 0.76 | 0.91 | 0.87 |
| 8 | 1681733 | rs2972180 | 0.896 | 0.448 | -1.681 | 0.498 | -0.113 | 0.07 | 0.07 | 0.345 | 0.38 | 0.05 | 0.27 | 0.25 | 0.43 | 0.27 |
| 8 | 1686410 | rs2977195 | -0.969 | 0.256 | -0.155 | 1.608 | -0.022 | 0.202 | 0.202 | 0.013 | 0.14 | 0.42 | 0.35 | 0.3 | 0.1 | 0.26 |
| 8 | 1690346 | rs4876132 | 1.19 | 1.135 | -0.012 | 1.726 | -0.001 | -0.001 | -0.001 | 0 | 0.66 | 0.34 | 0.35 | 0.6 | 0.49 | 0.68 |
| 8 | 1692723 | rs7461883 | -0.424 | 2.204 | -0.159 | 1.912 | -0.029 | -0.032 | -0.032 | 0 | 0.05 | 0.01 | 0.14 | 0.13 | 0.14 | 0.06 |
| 8 | 1694236 | rs4242539 | 0.699 | 1.973 | 0.721 | 1.673 | 0.02 | 0.004 | 0.004 | 0 | 0.81 | 0.72 | 0.6 | 0.78 | 0.75 | 0.79 |
| 8 | 1697573 | rs6558520 | 0.916 | 1.901 | 1.061 | 1.641 | 0.069 | 0.021 | 0.021 | 0 | 0.8 | 0.51 | 0.58 | 0.78 | 0.76 | 0.78 |
| 8 | 1698911 | rs4623455 | -0.242 | 1.241 | 0.247 | 1.782 | 0.029 | -0.053 | -0.053 | 0 | 0.14 | 0.03 | 0.06 | 0.31 | 0.07 | 0.26 |
| 8 | 1700398 | rs7005592 | 0.566 | 1.713 | 0.469 | 1.265 | 0.011 | 0.009 | 0.009 | 0 | 0.86 | 0.84 | 0.62 | 0.78 | 0.81 | 0.79 |
| 8 | 1708044 | rs4639549 | 1.307 | 2.122 | 0.536 | 1.064 | 0.028 | 0.001 | 0.001 | 0 | 0.77 | 0.74 | 0.54 | 0.77 | 0.68 | 0.78 |

| | | | | | | | | | | | | | | | | |
|---|---|---|---|---|---|---|---|---|---|---|---|---|---|---|---|---|
| 8 | 1709918 | rs6558533 | 0.422 | 2.127 | 0.393 | 1.077 | 0.012 | 0.014 | 0.014 | 0 | 0.86 | 0.85 | 0.63 | 0.77 | 0.81 | 0.79 |
| 8 | 1717036 | rs11136424 | 1.419 | 1.931 | 0.381 | 0.942 | 0.036 | 0.003 | 0.003 | 0 | 0.76 | 0.68 | 0.54 | 0.76 | 0.7 | 0.79 |
| 8 | 1718389 | rs6993706 | 0.667 | 1.855 | 0.202 | 0.646 | -0.01 | -0.011 | -0.011 | 0.011 | 0.83 | 0.88 | 0.61 | 0.76 | 0.79 | 0.79 |
| 8 | 1721090 | rs4875958 | 1.558 | 1.829 | 0.247 | 0.636 | 0.012 | -0.003 | -0.003 | 0 | 0.74 | 0.7 | 0.54 | 0.76 | 0.68 | 0.77 |
| 8 | 1723749 | rs10100341 | 1.525 | 1.832 | 0.253 | 0.623 | 0.01 | -0.004 | -0.004 | 1.318 | 0.74 | 0.7 | 0.54 | 0.77 | 0.68 | 0.77 |
| 8 | 1728292 | rs12546965 | 0.832 | 1.887 | -0.174 | 0.734 | -0.037 | 0.04 | 0.04 | 0.41 | 0.07 | 0.01 | 0.07 | 0 | 0.11 | 0.01 |
| 8 | 1732756 | rs7827285 | 0.473 | 1.609 | -0.13 | 0.808 | -0.034 | -0.043 | -0.043 | 0 | 0.85 | 0.94 | 0.64 | 0.76 | 0.81 | 0.79 |
| 8 | 1748941 | rs6558541 | 1.198 | 0.995 | -0.073 | 0.768 | -0.004 | -0.009 | -0.009 | 0 | 0.72 | 0.83 | 0.59 | 0.7 | 0.72 | 0.6 |
| 8 | 1753497 | rs4327894 | 0.211 | 0.914 | -0.169 | 0.46 | -0.005 | 0 | 0 | 2.03 | 0.83 | 0.87 | 0.66 | 0.83 | 0.77 | 0.77 |
| 8 | 1756585 | rs6558545 | 1.41 | 0.966 | 0.111 | 0.361 | -0.002 | 0.001 | 0.001 | 0.237 | 0.61 | 0.61 | 0.54 | 0.71 | 0.59 | 0.55 |
| 8 | 1765371 | rs6980781 | -0.219 | 0.238 | 0.39 | 0.599 | -0.027 | 0.012 | 0.012 | 0 | 0.17 | 0.3 | 0.09 | 0.13 | 0.12 | 0.29 |
| 8 | 1768004 | rs6558550 | -0.497 | 0.449 | 0.401 | 0.222 | 0.019 | -0.004 | -0.004 | 0 | 0.2 | 0.28 | 0.32 | 0.17 | 0.3 | 0.28 |
| 8 | 1772641 | rs10090061 | -0.196 | -0.104 | -0.233 | -0.053 | -0.007 | 0.017 | 0.017 | 0 | 0.25 | 0.15 | 0.29 | 0.2 | 0.29 | 0.27 |
| 8 | 1777571 | rs17064297 | 0.051 | 0.768 | -0.261 | -0.387 | 0.041 | -0.087 | -0.087 | 0 | 0.2 | 0.31 | 0.18 | 0.03 | 0.25 | 0.03 |
| 8 | 1778697 | rs9802017 | 0.848 | 1.017 | -0.324 | 0.105 | -0.001 | -0.007 | -0.007 | 0 | 0.6 | 0.66 | 0.57 | 0.54 | 0.5 | 0.59 |
| 8 | 1782495 | rs6558551 | 0.458 | 0.93 | -0.056 | 0.067 | 0.002 | -0.003 | -0.003 | 0 | 0.67 | 0.55 | 0.56 | 0.38 | 0.67 | 0.49 |
| 8 | 1782876 | rs7387547 | 0.463 | 0.941 | -0.05 | 0.071 | 0.002 | -0.001 | -0.001 | 0 | 0.67 | 0.55 | 0.56 | 0.41 | 0.68 | 0.49 |
| 8 | 1784859 | rs6992502 | 0.79 | 0.452 | -0.234 | -0.243 | 0.024 | 0.103 | 0.103 | 0.269 | 0.47 | 0.27 | 0.24 | 0.24 | 0.39 | 0.22 |
| 8 | 1785264 | rs4875952 | -0.637 | 0.719 | -0.203 | -0.237 | 0 | 0.013 | 0.013 | 0.198 | 0.47 | 0.44 | 0.64 | 0.71 | 0.52 | 0.68 |
| 8 | 1789627 | rs7015432 | -0.859 | 0.835 | -0.466 | -0.161 | 0.022 | -0.041 | -0.041 | 0 | 0.33 | 0.55 | 0.27 | 0.16 | 0.35 | 0.3 |
| 8 | 1791433 | rs11136431 | -0.87 | 1.105 | -1.03 | 0.441 | 0.008 | -0.041 | -0.041 | 0.01 | 0.47 | 0.21 | 0.39 | 0.46 | 0.53 | 0.58 |
| 8 | 1794023 | rs6651438 | -0.725 | 1.129 | -1.034 | 0.442 | -0.009 | -0.034 | -0.034 | 0 | 0.46 | 0.18 | 0.36 | 0.4 | 0.52 | 0.52 |
| 8 | 1802388 | rs13278409 | -0.051 | 1.659 | -0.915 | 0.656 | -0.02 | -0.018 | -0.018 | 0 | 0.17 | 0.17 | 0.22 | 0.29 | 0.34 | 0.19 |
| 8 | 1802711 | rs13258335 | -0.186 | 0.912 | -1.019 | 0.521 | 0.05 | 0.048 | 0.048 | 0 | 0.28 | 0.12 | 0.15 | 0.13 | 0.19 | 0.33 |
| 8 | 1804388 | rs11136436 | -1.48 | 0.808 | -0.331 | 0.606 | 0.017 | 0.004 | 0.004 | 0 | 0.59 | 0.66 | 0.63 | 0.58 | 0.67 | 0.56 |
| 8 | 1806229 | rs4875950 | -0.609 | 0.894 | -0.14 | 0.264 | 0.001 | 0 | 0 | 1.514 | 0.35 | 0.31 | 0.29 | 0.24 | 0.27 | 0.42 |
| 8 | 1807756 | rs7007884 | -0.456 | 1.012 | -0.242 | 0.357 | 0.01 | 0.014 | 0.014 | 0 | 0.33 | 0.23 | 0.34 | 0.29 | 0.29 | 0.43 |
| 8 | 1811397 | rs11779733 | -1.005 | 1.226 | -0.127 | 0.542 | 0 | 0 | 0 | 0.392 | 0.44 | 0.6 | 0.42 | 0.64 | 0.5 | 0.44 |
| 8 | 1811873 | rs13277792 | -0.848 | 1.23 | -0.116 | 0.614 | -0.003 | 0.003 | 0.003 | 2.48 | 0.43 | 0.55 | 0.4 | 0.49 | 0.49 | 0.39 |
| 8 | 1812053 | rs2294030 | 0.517 | 1.056 | -0.172 | 0.463 | -0.001 | -0.023 | -0.023 | 0 | 0.17 | 0.13 | 0.19 | 0.02 | 0.17 | 0.2 |
| 8 | 1814142 | rs11136437 | -1.262 | 1.679 | -0.19 | 1.084 | -0.005 | -0.011 | -0.011 | 0.26 | 0.43 | 0.57 | 0.48 | 0.69 | 0.51 | 0.42 |
| 8 | 1817367 | rs7003969 | -0.587 | 1.994 | -0.11 | 1.476 | -0.028 | -0.029 | -0.029 | 0.427 | 0.15 | 0.1 | 0.21 | 0.52 | 0.22 | 0.17 |
| 8 | 1817961 | rs17829629 | -0.594 | 1.97 | -0.115 | 1.129 | 0.01 | 0.032 | 0.032 | 0 | 0.15 | 0.18 | 0.21 | 0.35 | 0.21 | 0.15 |
| 8 | 1818575 | rs7013410 | 0.286 | 2.155 | -0.42 | 1.205 | 0.02 | 0.078 | 0.078 | 0 | 0.1 | 0.14 | 0.18 | 0.31 | 0.15 | 0.13 |
| 8 | 1820656 | rs3735866 | -0.42 | 1.969 | -0.416 | 0.916 | 0.025 | 0.079 | 0.079 | 0.018 | 0.16 | 0.22 | 0.22 | 0.41 | 0.21 | 0.16 |
| 8 | 1821516 | rs12545104 | -0.85 | 1.53 | -1.033 | -0.265 | -0.005 | -0.023 | -0.023 | 0 | 0.39 | 0.43 | 0.54 | 0.88 | 0.49 | 0.59 |
| 8 | 1822423 | rs4875947 | -0.288 | 0.804 | -1.987 | 0.249 | 0.036 | -0.019 | -0.019 | 4.946 | 0.53 | 0.77 | 0.62 | 0.57 | 0.56 | 0.6 |
| 8 | 1822639 | rs3735869 | -0.382 | 0.694 | -2.002 | 0.184 | -0.082 | -0.118 | -0.118 | 0 | 0.26 | 0.08 | 0.27 | 0.38 | 0.32 | 0.16 |
| 8 | 1823481 | rs4875946 | -0.495 | 0.005 | -1.462 | -0.466 | 0 | -0.026 | -0.026 | 0.025 | 0.45 | 0.5 | 0.64 | 0.85 | 0.48 | 0.56 |
| 8 | 1823659 | rs7011997 | 0.807 | -0.515 | -1.316 | -0.832 | 0.016 | -0.128 | -0.128 | 0.056 | 0.27 | 0.4 | 0.21 | 0.04 | 0.32 | 0.1 |
| 8 | 1828807 | rs3779698 | 0.393 | -0.308 | -1.558 | 0.004 | -0.019 | -0.066 | -0.066 | 0 | 0.14 | 0.03 | 0.12 | 0.28 | 0.13 | 0.33 |
| 8 | 1830275 | rs2280885 | 0.447 | -0.312 | -1.565 | 0.042 | -0.031 | -0.087 | -0.087 | 0 | 0.14 | 0.01 | 0.12 | 0.29 | 0.13 | 0.34 |
| 8 | 1831436 | rs3758001 | -0.719 | -0.41 | -1.399 | -0.28 | 0.001 | -0.008 | -0.008 | 0.054 | 0.34 | 0.3 | 0.28 | 0.4 | 0.31 | 0.52 |
| 8 | 1832844 | rs7014895 | -0.879 | -0.444 | -1.384 | -0.218 | 0.002 | -0.009 | -0.009 | 0.142 | 0.35 | 0.34 | 0.28 | 0.48 | 0.32 | 0.53 |
| 8 | 1839973 | rs4292733 | -0.79 | -0.887 | -1.618 | -0.463 | -0.024 | 0.114 | 0.114 | 0 | 0.48 | 0.85 | 0.64 | 0.65 | 0.45 | 0.48 |
| 8 | 1840497 | rs6558560 | -0.754 | -0.951 | -1.105 | -0.46 | -0.036 | -0.023 | -0.023 | 0.017 | 0.14 | 0.51 | 0.1 | 0.12 | 0.1 | 0.09 |
| 8 | 1843091 | rs997378 | 0.323 | -1.096 | -1.099 | -0.289 | 0.063 | 0.085 | 0.085 | 0 | 0.26 | 0.57 | 0.53 | 0.43 | 0.35 | 0.22 |
| 8 | 1846010 | rs4474061 | 0.468 | -1.636 | -0.483 | 0.453 | 0.014 | 0.002 | 0.002 | 0 | 0.14 | 0.1 | 0.07 | 0.08 | 0.06 | 0.19 |
| 8 | 1846296 | rs2272712 | 0.75 | -1.552 | -0.682 | 0.473 | -0.046 | -0.078 | -0.078 | 8.483 | 0.11 | 0.01 | 0.12 | 0.33 | 0.14 | 0.29 |
| 8 | 1846368 | rs2272713 | -0.961 | -1.62 | -0.611 | 0.444 | -0.019 | 0.023 | 0.023 | 0 | 0.3 | 0.35 | 0.22 | 0.52 | 0.19 | 0.5 |
| 8 | 1847184 | rs3758012 | 0.978 | -1.427 | -0.657 | 0.528 | -0.034 | -0.069 | -0.069 | 0 | 0.1 | 0 | 0.11 | 0.32 | 0.12 | 0.28 |
| 8 | 1847564 | rs6990473 | -0.489 | -1.446 | -0.472 | 0.202 | -0.052 | -0.007 | -0.007 | 0.204 | 0.24 | 0.31 | 0.12 | 0.2 | 0.09 | 0.23 |
| 8 | 1848653 | rs7014554 | -0.437 | -0.822 | -0.346 | 0.592 | -0.018 | 0.105 | 0.105 | 0.031 | 0.19 | 0.39 | 0.32 | 0.38 | 0.19 | 0.34 |
| 8 | 1848860 | rs7013741 | -0.651 | -1.1 | -0.56 | 0.076 | -0.059 | -0.011 | -0.011 | 0 | 0.28 | 0.4 | 0.28 | 0.27 | 0.14 | 0.3 |
| 8 | 1848927 | rs7015000 | -0.915 | -0.594 | -0.839 | -0.109 | -0.012 | -0.008 | -0.008 | 0.616 | 0.35 | 0.39 | 0.32 | 0.32 | 0.25 | 0.35 |
| 8 | 1849275 | rs3758014 | -0.963 | 0.1 | -0.946 | -0.512 | -0.002 | 0.005 | 0.005 | 0 | 0.25 | 0.21 | 0.31 | 0.1 | 0.27 | 0.26 |
| 8 | 1849398 | rs3758015 | -0.26 | 0.095 | -0.607 | -0.854 | 0.006 | -0.051 | -0.051 | 0.02 | 0.32 | 0.53 | 0.37 | 0.17 | 0.39 | 0.24 |
| 8 | 1849530 | rs3758018 | -0.097 | 0.084 | -0.907 | -0.45 | 0.019 | 0.013 | 0.013 | 0 | 0.5 | 0.54 | 0.39 | 0.48 | 0.41 | 0.5 |
| 8 | 1849736 | rs13258210 | 0.144 | 0.549 | -1.189 | -1.023 | 0.002 | -0.038 | -0.038 | 0.394 | 0.09 | 0.14 | 0.05 | 0 | 0.06 | 0.04 |
| 8 | 1849882 | rs1006520 | -0.665 | 1.259 | -1.536 | -1.121 | -0.011 | 0.014 | 0.014 | 0 | 0.22 | 0.19 | 0.29 | 0.15 | 0.24 | 0.2 |
| 8 | 1850559 | rs28485777 | 0.197 | 1.041 | -1.354 | -1.301 | -0.001 | -0.022 | -0.022 | 0.534 | 0.12 | 0.09 | 0.09 | 0 | 0.16 | 0.06 |
| 8 | 1850622 | rs7836624 | 1.43 | 0.252 | -1.281 | -0.951 | -0.034 | 0.034 | 0.034 | 1.966 | 0.2 | 0.29 | 0.11 | 0.32 | 0.12 | 0.28 |
| 8 | 1850997 | rs12679580 | 1.152 | 1.116 | -1.534 | -0.937 | 0 | 0.025 | 0.025 | 0 | 0.12 | 0.11 | 0.21 | 0.4 | 0.18 | 0.22 |
| 8 | 1852425 | rs13280012 | -0.052 | 0.338 | -1.619 | -1.462 | 0.025 | 0.123 | 0.123 | 0.059 | 0.35 | 0.14 | 0.23 | 0.01 | 0.26 | 0.27 |
| 8 | 1857293 | rs2294035 | -0.517 | 0.711 | -0.597 | -0.416 | -0.005 | 0.001 | 0.001 | 0 | 0.57 | 0.68 | 0.53 | 0.58 | 0.58 | 0.57 |
| 8 | 1857367 | rs2294037 | -0.435 | 0.802 | -0.534 | -0.329 | 0.003 | -0.005 | -0.005 | 0 | 0.55 | 0.65 | 0.49 | 0.53 | 0.54 | 0.54 |
| 8 | 1857649 | rs2294041 | -0.715 | 0.611 | -0.399 | -0.589 | -0.004 | -0.024 | -0.024 | 0 | 0.27 | 0.37 | 0.31 | 0.16 | 0.29 | 0.18 |
| 8 | 1859248 | rs13261404 | -0.11 | 0.669 | -0.384 | -0.576 | 0 | 0 | 0 | 0 | 0.15 | 0.17 | 0.23 | 0.16 | 0.17 | 0.13 |
| 8 | 1860200 | rs12548089 | -0.789 | 0.67 | -0.343 | -0.577 | -0.003 | -0.028 | -0.028 | 0.715 | 0.26 | 0.38 | 0.3 | 0.16 | 0.29 | 0.18 |

| | | | | | | | | | | | | | | | | |
|---|---|---|---|---|---|---|---|---|---|---|---|---|---|---|---|---|
| 8 | 1860395 | rs6990129 | 1.154 | 0.494 | -0.568 | -0.525 | 0.002 | -0.003 | -0.003 | 1.177 | 0.28 | 0.28 | 0.18 | 0.38 | 0.24 | 0.31 |
| 8 | 1861971 | rs11995882 | -0.411 | 0.562 | -0.017 | -1.142 | -0.004 | -0.046 | -0.046 | 0 | 0.15 | 0.19 | 0.11 | 0 | 0.16 | 0.07 |
| 8 | 1865205 | rs4379489 | -0.01 | 0.611 | -0.234 | -1.013 | -0.007 | -0.026 | -0.026 | 0 | 0.22 | 0.3 | 0.28 | 0.12 | 0.25 | 0.15 |
| 8 | 1865887 | rs7004405 | 0.59 | 0.792 | -1.057 | -0.868 | -0.012 | 0.032 | 0.032 | 0 | 0.43 | 0.36 | 0.47 | 0.74 | 0.47 | 0.61 |
| 8 | 1866796 | rs13250852 | 0.169 | 1.119 | -1.187 | -0.353 | -0.006 | 0.029 | 0.029 | 0 | 0.72 | 0.81 | 0.76 | 0.85 | 0.74 | 0.79 |
| 8 | 1870734 | rs4609237 | 0.097 | 1.248 | -0.894 | -0.696 | 0.005 | -0.048 | -0.048 | 1.369 | 0.6 | 0.54 | 0.65 | 0.91 | 0.57 | 0.75 |
| 8 | 1870992 | rs17830000 | -0.141 | 0.702 | -0.601 | 0.233 | -0.02 | 0.11 | 0.11 | 0 | 0.14 | 0.31 | 0.09 | 0.37 | 0.11 | 0.2 |
| 8 | 1873540 | rs2272613 | 0.236 | 0.923 | -0.665 | 0.275 | 0.017 | -0.049 | -0.049 | 0 | 0.09 | 0.01 | 0.03 | 0.31 | 0.07 | 0.12 |
| 8 | 1874732 | rs17064390 | 0.322 | 0.77 | -0.935 | 0.516 | -0.004 | 0.006 | 0.006 | 0.136 | 0.13 | 0.15 | 0.16 | 0.15 | 0.07 | 0.1 |
| 8 | 1876631 | rs2272611 | 0.105 | 1.198 | -1.022 | 0.505 | 0.001 | -0.004 | -0.004 | 0 | 0.15 | 0.14 | 0.18 | 0.16 | 0.11 | 0.14 |
| 8 | 1879736 | rs13257924 | -0.974 | 1.688 | -1.118 | -0.225 | 0.017 | -0.022 | -0.022 | 0 | 0.94 | 0.9 | 0.92 | 1 | 0.9 | 0.99 |
| 8 | 1880530 | rs4242548 | -0.347 | 1.452 | -1.274 | -0.418 | 0.007 | 0.068 | 0.068 | 0.012 | 0.46 | 0.36 | 0.37 | 0.85 | 0.47 | 0.59 |
| 8 | 1886640 | rs3779707 | 0.264 | 1.085 | -0.839 | 0.326 | 0.002 | 0.004 | 0.004 | 0 | 0.13 | 0.13 | 0.06 | 0.35 | 0.09 | 0.12 |
| 8 | 1888447 | rs3824141 | 0.038 | 0.994 | -0.562 | 0.387 | -0.009 | 0.033 | 0.033 | 0 | 0.15 | 0.18 | 0.07 | 0.38 | 0.12 | 0.17 |
| 8 | 1890395 | rs9650470 | 0.206 | 1.235 | -0.524 | 0.475 | 0.002 | 0.009 | 0.009 | 0 | 0.15 | 0.17 | 0.21 | 0.19 | 0.13 | 0.16 |
| 8 | 1890533 | rs7386016 | -0.639 | 1.277 | -0.138 | 0.183 | 0.052 | -0.017 | -0.017 | 0 | 0.94 | 0.8 | 0.92 | 1 | 0.92 | 0.96 |
| 8 | 1897174 | rs17064407 | 0.974 | 1.238 | -0.077 | 0.044 | -0.005 | -0.024 | -0.024 | 0 | 0.26 | 0.39 | 0.19 | 0.25 | 0.26 | 0.27 |
| 8 | 1897249 | rs17830077 | 0.59 | 1.138 | -0.356 | 0.271 | 0.013 | -0.061 | -0.061 | 0 | 0.12 | 0.02 | 0.06 | 0.33 | 0.1 | 0.15 |
| 8 | 1897657 | rs2280823 | 0.6 | 1.388 | -0.21 | -0.135 | -0.001 | -0.003 | -0.003 | 0 | 0.14 | 0.14 | 0.19 | 0.23 | 0.13 | 0.21 |
| 8 | 1897727 | rs2280822 | 1.056 | 1.304 | -0.187 | -0.499 | 0.002 | -0.013 | -0.013 | 0.022 | 0.34 | 0.4 | 0.32 | 0.19 | 0.36 | 0.32 |
| 8 | 1899691 | rs17830107 | 0.906 | 1.587 | -0.106 | -0.515 | 0 | 0 | 0 | 0.751 | 0.35 | 0.31 | 0.23 | 0.59 | 0.39 | 0.42 |
| 8 | 1900370 | rs17750595 | -0.777 | 1.552 | -0.142 | -0.55 | -0.01 | -0.041 | -0.041 | 0 | 0.62 | 0.78 | 0.72 | 0.42 | 0.57 | 0.58 |
| 8 | 1900410 | rs3779713 | 1.107 | 1.541 | -0.134 | -0.568 | -0.009 | 0.043 | 0.043 | 0.847 | 0.33 | 0.16 | 0.42 | 0.2 | 0.33 | 0.34 |
| 8 | 1901930 | rs4876271 | 1.317 | 1.497 | -0.019 | -0.489 | 0.001 | 0.033 | 0.033 | 0.976 | 0.12 | 0.15 | 0.18 | 0.21 | 0.1 | 0.16 |
| 8 | 1902750 | rs7386942 | -0.433 | 1.24 | -0.166 | -0.665 | -0.003 | 0.015 | 0.015 | 0.216 | 0.61 | 0.67 | 0.49 | 0.79 | 0.58 | 0.62 |
| 8 | 1904157 | rs4876268 | 0.846 | 1.239 | -0.17 | -0.654 | -0.007 | -0.038 | -0.038 | 0 | 0.38 | 0.21 | 0.24 | 0.58 | 0.41 | 0.43 |
| 8 | 1905132 | rs3735876 | 0.916 | 1.227 | -0.176 | -0.659 | -0.009 | -0.039 | -0.039 | 0 | 0.37 | 0.21 | 0.23 | 0.57 | 0.4 | 0.43 |
| 8 | 1905823 | rs6991392 | 0.844 | 1.291 | 0.068 | -0.696 | -0.002 | -0.013 | -0.013 | 0 | 0.38 | 0.31 | 0.24 | 0.6 | 0.39 | 0.42 |
| 8 | 1907935 | rs7006457 | -0.811 | 1.197 | -0.127 | -1.113 | -0.004 | 0.004 | 0.004 | 0.137 | 0.65 | 0.66 | 0.51 | 0.8 | 0.58 | 0.68 |
| 8 | 1908052 | rs13281187 | -0.323 | 0.918 | -0.568 | -0.964 | -0.001 | 0.049 | 0.049 | 0 | 0.49 | 0.39 | 0.53 | 0.22 | 0.48 | 0.42 |
| 8 | 1911393 | rs17757154 | 1.16 | 1.007 | -1.058 | -1.068 | -0.038 | -0.034 | -0.034 | 0 | 0.46 | 0.12 | 0.31 | 0.58 | 0.52 | 0.5 |
| 8 | 1914752 | rs17830215 | 1.151 | 0.966 | -0.885 | -1.093 | -0.022 | -0.018 | -0.018 | 0 | 0.45 | 0.24 | 0.29 | 0.53 | 0.5 | 0.52 |
| 8 | 1916144 | rs4876266 | 0.239 | 1 | -0.99 | -1.508 | -0.012 | 0.058 | 0.058 | 0 | 0.8 | 0.9 | 0.76 | 0.92 | 0.78 | 0.8 |
| 8 | 1916347 | rs4876265 | -0.061 | 0.888 | -0.04 | -1.433 | -0.004 | 0.014 | 0.014 | 0 | 0.32 | 0.38 | 0.34 | 0.35 | 0.28 | 0.26 |
| 8 | 1918352 | rs6558568 | -0.7 | 0.59 | 0.648 | -2.303 | -0.016 | -0.118 | -0.118 | 0.914 | 0.34 | 0.52 | 0.39 | 0.07 | 0.26 | 0.24 |
| 8 | 1920461 | rs4242546 | 1.435 | 0.752 | 1.014 | -1.378 | 0.008 | 0.013 | 0.013 | 0 | 0.73 | 0.78 | 0.75 | 0.82 | 0.75 | 0.79 |
| 8 | 1921160 | rs4557743 | 0.59 | 0.769 | 1.226 | -1.345 | 0.018 | 0.038 | 0.038 | 0 | 0.86 | 0.73 | 0.8 | 0.83 | 0.84 | 0.88 |
| 8 | 1921428 | rs4633105 | 1.515 | 0.837 | 1.072 | -1.346 | -0.004 | -0.028 | -0.028 | 0 | 0.71 | 0.6 | 0.72 | 0.82 | 0.7 | 0.78 |
| 8 | 1926958 | rs13271843 | 1.209 | 0.86 | 0.689 | -0.909 | 0.024 | 0.006 | 0.006 | 0 | 0.81 | 0.69 | 0.76 | 0.83 | 0.76 | 0.85 |
| 8 | 1937286 | rs4410948 | 0.237 | 0.945 | 0.404 | -0.857 | 0.01 | -0.024 | -0.024 | 0 | 0.93 | 0.89 | 0.92 | 1 | 0.9 | 0.95 |
| 8 | 1938034 | rs4492421 | 1.709 | 0.814 | 0.217 | -1.173 | -0.021 | 0.089 | 0.089 | 2.348 | 0.6 | 0.17 | 0.57 | 0.83 | 0.59 | 0.72 |
| 8 | 1939557 | rs6983882 | -1.532 | 0.814 | -0.023 | -1.385 | -0.003 | 0.039 | 0.039 | 1.296 | 0.32 | 0.76 | 0.36 | 0.17 | 0.32 | 0.35 |
| 8 | 1947938 | rs3824142 | -0.696 | 0.216 | 0.846 | -1.358 | 0.03 | -0.025 | -0.025 | 0.056 | 0.11 | 0.43 | 0.24 | 0.04 | 0.11 | 0.16 |
| 8 | 1949126 | rs4322044 | 0.86 | 0.121 | 0.743 | -1.32 | 0.064 | -0.036 | -0.036 | 0 | 0.92 | 0.71 | 0.92 | 1 | 0.9 | 0.95 |
| 8 | 1952175 | rs7386994 | -0.5 | 0.165 | -0.153 | -1.376 | -0.039 | 0.059 | 0.059 | 0 | 0.1 | 0.01 | 0.09 | 0 | 0.12 | 0.06 |
| 8 | 1953042 | rs1043221 | -0.914 | 0.117 | -0.177 | -1.4 | -0.035 | 0.071 | 0.071 | 0 | 0.12 | 0.01 | 0.09 | 0 | 0.12 | 0.06 |
| 8 | 1953194 | rs3087819 | 0.712 | 0.215 | 0.057 | -1.383 | 0.008 | -0.036 | -0.036 | 0 | 0.92 | 0.84 | 0.9 | 1 | 0.9 | 0.99 |
| 8 | 1953223 | rs1043223 | 2.031 | -0.018 | -0.032 | -1.415 | -0.013 | 0.168 | 0.168 | 7.064 | 0.66 | 0.11 | 0.55 | 0.94 | 0.66 | 0.75 |
| 8 | 1953567 | rs1043225 | 1.005 | 0.194 | 0.015 | -1.303 | 0.003 | -0.051 | -0.051 | 0 | 0.8 | 0.77 | 0.81 | 1 | 0.77 | 0.93 |
| 8 | 1954464 | rs17064496 | -0.945 | -0.363 | 0.222 | -0.622 | -0.022 | -0.024 | -0.024 | 0 | 0.06 | 0.26 | 0.04 | 0 | 0.02 | 0.01 |
| 8 | 1955283 | rs12677235 | -0.925 | 0.231 | -0.359 | -0.332 | 0.016 | -0.011 | -0.011 | 2.017 | 0.11 | 0.38 | 0.23 | 0.05 | 0.11 | 0.17 |
| 8 | 1959336 | rs768753 | -0.342 | 0.162 | -1.94 | -0.09 | -0.026 | 0.002 | 0.002 | 0.821 | 0.34 | 0.03 | 0.24 | 0.27 | 0.31 | 0.3 |
| 8 | 1967924 | rs2280819 | 0.343 | -0.106 | -0.901 | 0.258 | -0.025 | 0.036 | 0.036 | 0 | 0.83 | 0.69 | 0.89 | 0.78 | 0.85 | 0.85 |
| 8 | 1968004 | rs6992819 | -0.061 | 0.682 | -0.9 | 0.25 | 0 | 0 | 0 | 0.297 | 0.17 | 0.16 | 0.23 | 0.16 | 0.2 | 0.17 |
| 8 | 1968128 | rs2280820 | -0.545 | 0.693 | -1.136 | -0.75 | 0.014 | 0.036 | 0.036 | 0.984 | 0.5 | 0.42 | 0.69 | 0.66 | 0.59 | 0.56 |
| 8 | 1969137 | rs9314445 | 0.289 | 1.079 | -0.58 | -0.851 | -0.031 | 0.031 | 0.031 | 0 | 0.14 | 0.07 | 0.19 | 0.01 | 0.2 | 0.08 |
| 8 | 1969726 | rs6558578 | 0.661 | 0.589 | -0.219 | -1.115 | 0.016 | -0.031 | -0.031 | 0.084 | 0.69 | 0.55 | 0.68 | 0.84 | 0.67 | 0.75 |
| 8 | 1975962 | rs11136456 | -0.857 | 0.241 | 0.242 | -0.961 | 0.002 | 0.02 | 0.02 | 0 | 0.15 | 0.26 | 0.13 | 0.18 | 0.15 | 0.2 |
| 8 | 1977401 | rs7464878 | 1.537 | 0.024 | 0.596 | -0.996 | -0.009 | 0.098 | 0.098 | 1.072 | 0.81 | 0.37 | 0.73 | 0.67 | 0.84 | 0.73 |
| 8 | 1978115 | rs4876252 | 1.301 | -0.084 | 0.563 | -1.03 | -0.023 | -0.026 | -0.026 | 0.082 | 0.77 | 0.61 | 0.84 | 0.85 | 0.82 | 0.77 |
| 8 | 1983582 | rs7814914 | 0.758 | -0.045 | -0.048 | -1.385 | 0.05 | 0.051 | 0.051 | 2.38 | 0.6 | 0.68 | 0.7 | 0.83 | 0.72 | 0.71 |
| 8 | 1987720 | rs7006192 | 1.002 | -0.208 | -0.025 | -1.418 | -0.034 | 0.043 | 0.043 | 0 | 0.73 | 0.47 | 0.71 | 0.67 | 0.8 | 0.69 |
| 8 | 1989980 | rs17683694 | 1.967 | 0.489 | -0.109 | -1.58 | -0.003 | -0.002 | -0.002 | 0 | 0.35 | 0.31 | 0.42 | 0.66 | 0.37 | 0.47 |
| 8 | 1991352 | rs4875929 | 0.432 | 0.448 | -0.073 | -1.565 | -0.021 | 0.054 | 0.054 | 0 | 0.82 | 0.51 | 0.86 | 0.79 | 0.87 | 0.83 |
| 8 | 1991386 | rs17750980 | 1.05 | 0.657 | -0.506 | -1.699 | -0.042 | 0.04 | 0.04 | 0.012 | 0.14 | 0.06 | 0.18 | 0.01 | 0.25 | 0.11 |
| 8 | 1991921 | rs4875928 | -0.231 | 0.049 | -0.415 | -1.452 | 0.061 | 0.092 | 0.092 | 0.027 | 0.32 | 0.15 | 0.25 | 0.12 | 0.25 | 0.24 |
| 8 | 1992937 | rs12678739 | -0.397 | 0.603 | 0.287 | -0.963 | 0.018 | 0.061 | 0.061 | 0.092 | 0.1 | 0.22 | 0.1 | 0.18 | 0.1 | 0.13 |
| 8 | 1997537 | rs17064574 | 0.492 | 0.298 | 0.164 | -1.036 | -0.013 | -0.03 | -0.03 | 0 | 0.06 | 0.13 | 0.01 | 0 | 0.02 | 0.02 |
| 8 | 1997993 | rs11136457 | -0.316 | 0.873 | 0.451 | -1.234 | 0.041 | -0.002 | -0.002 | 0 | 0.8 | 0.77 | 0.75 | 0.85 | 0.66 | 0.81 |

| | | | | | | | | | | | | | | | | |
|---|---|---|---|---|---|---|---|---|---|---|---|---|---|---|---|---|
| 8 | 2000587 | rs11136458 | 0.525 | 0.454 | 0.367 | -1.445 | 0.007 | 0.01 | 0.01 | 0 | 0.13 | 0.08 | 0.08 | 0 | 0.1 | 0.06 |
| 8 | 2000626 | rs11136459 | -0.503 | 0.375 | 0.551 | -1.537 | -0.029 | -0.1 | -0.1 | 0.054 | 0.2 | 0.37 | 0.14 | 0.03 | 0.14 | 0.08 |
| 8 | 2001244 | rs7835264 | -0.568 | 0.339 | 0.203 | -1.487 | -0.04 | -0.096 | -0.096 | 0.63 | 0.21 | 0.53 | 0.14 | 0.06 | 0.14 | 0.08 |
| 8 | 2001704 | rs1822036 | 1.753 | 0.332 | -0.115 | -1.639 | -0.026 | 0.003 | 0.003 | 1.305 | 0.39 | 0.22 | 0.46 | 0.69 | 0.42 | 0.58 |
| 8 | 2001755 | rs4876236 | -0.915 | 0.175 | -0.062 | -1.641 | 0.011 | 0.015 | 0.015 | 2.039 | 0.81 | 0.85 | 0.92 | 0.97 | 0.86 | 0.92 |
| 8 | 2001866 | rs7463722 | -0.913 | 0.18 | -0.059 | -1.638 | 0.009 | 0.01 | 0.01 | 2.039 | 0.81 | 0.84 | 0.91 | 0.97 | 0.86 | 0.92 |
| 8 | 2002029 | rs7464436 | 0.549 | 0.379 | -0.23 | -1.49 | 0.021 | 0.01 | 0.01 | 0.027 | 0.61 | 0.32 | 0.78 | 0.94 | 0.72 | 0.83 |
| 8 | 2003295 | rs2033909 | -0.004 | 0.792 | -0.795 | -1.193 | -0.042 | -0.028 | -0.028 | 0.932 | 0.23 | 0.09 | 0.33 | 0.25 | 0.32 | 0.28 |
| 8 | 2005346 | rs7833402 | 0.344 | 0.311 | -0.601 | -1.22 | 0.011 | -0.003 | -0.003 | 0.464 | 0.67 | 0.7 | 0.83 | 0.97 | 0.74 | 0.83 |
| 8 | 2010761 | rs7819286 | 0.652 | 0.877 | -0.454 | -0.989 | 0.019 | -0.038 | -0.038 | 0 | 0.62 | 0.55 | 0.54 | 0.82 | 0.54 | 0.72 |
| 8 | 2011010 | rs1078937 | -0.892 | 1.033 | -0.632 | -0.766 | -0.016 | -0.009 | -0.009 | 0 | 0.31 | 0.36 | 0.28 | 0.22 | 0.22 | 0.32 |
| 8 | 2011098 | rs893875 | -0.875 | 1.381 | -0.763 | -0.935 | -0.002 | 0.001 | 0.001 | 0 | 0.28 | 0.25 | 0.36 | 0.14 | 0.39 | 0.27 |
| 8 | 2011215 | rs12542139 | 0.243 | 1.459 | -1.172 | -0.564 | -0.041 | -0.015 | -0.015 | 0 | 0.15 | 0.08 | 0.29 | 0.14 | 0.27 | 0.17 |
| 8 | 2013234 | rs3779857 | 0.271 | 1.031 | -1.059 | -0.823 | -0.004 | 0.029 | 0.029 | 0.072 | 0.6 | 0.64 | 0.58 | 0.83 | 0.57 | 0.78 |
| 8 | 2014196 | rs13258621 | -0.158 | 1.057 | -1.972 | -0.508 | -0.06 | -0.033 | -0.033 | 0 | 0.25 | 0.06 | 0.19 | 0.25 | 0.29 | 0.21 |
| 8 | 2015700 | rs3779855 | 0.543 | 0.773 | -1.189 | 0.378 | 0.019 | 0.108 | 0.108 | 0 | 0.92 | 0.82 | 0.78 | 0.78 | 0.9 | 0.86 |
| 8 | 2016053 | rs3779853 | 0.819 | 0.624 | -2.192 | 0.763 | -0.093 | 0.055 | 0.055 | 7.183 | 0.48 | 0.04 | 0.41 | 0.33 | 0.43 | 0.42 |
| 8 | 2017014 | rs13263401 | 0.262 | 0.713 | -1.465 | 0.911 | 0.027 | 0.134 | 0.134 | 0 | 0.91 | 0.79 | 0.77 | 0.78 | 0.9 | 0.86 |
| 8 | 2020003 | rs11136462 | -0.438 | 0.654 | -1.839 | 0.747 | -0.017 | 0.024 | 0.024 | 4.88 | 0.43 | 0.82 | 0.32 | 0.38 | 0.46 | 0.4 |
| 8 | 2020751 | rs2294057 | -0.155 | 0.101 | -1.258 | 0.721 | -0.031 | 0.005 | 0.005 | 0.021 | 0.3 | 0.2 | 0.21 | 0.25 | 0.42 | 0.29 |
| 8 | 2021421 | rs2272720 | 1.369 | 0.089 | -1.651 | 0.852 | -0.107 | -0.028 | -0.028 | 1.941 | 0.44 | 0.03 | 0.38 | 0.36 | 0.4 | 0.34 |
| 8 | 2027154 | rs2235118 | -0.128 | 0.163 | -1.069 | 0.946 | 0.006 | 0.054 | 0.054 | 0 | 0.11 | 0.17 | 0.28 | 0.32 | 0.13 | 0.25 |
| 8 | 2028110 | rs6558595 | 0.703 | 0.481 | -0.424 | 0.7 | 0.01 | 0.062 | 0.062 | 0.271 | 0.75 | 0.59 | 0.64 | 0.61 | 0.74 | 0.64 |
| 8 | 2030094 | rs1440294 | -0.671 | 0.535 | -0.458 | 0.462 | -0.019 | 0.036 | 0.036 | 0.116 | 0.3 | 0.25 | 0.23 | 0.11 | 0.37 | 0.3 |
| 8 | 2031701 | rs7838972 | -0.454 | 0.735 | -0.215 | 0.944 | 0.001 | 0.015 | 0.015 | 0 | 0.09 | 0.13 | 0.09 | 0.14 | 0.11 | 0.03 |
| 8 | 2034776 | rs7815609 | -1.201 | 0.648 | -0.255 | 0.971 | -0.003 | -0.004 | -0.004 | 0 | 0.4 | 0.7 | 0.48 | 0.42 | 0.43 | 0.51 |
| 8 | 2035025 | rs7816051 | 0.454 | 0.546 | 0.094 | 1.396 | 0.024 | 0.025 | 0.025 | 0 | 0.86 | 0.8 | 0.84 | 0.8 | 0.77 | 0.82 |
| 8 | 2036476 | rs13269383 | -0.927 | 0.261 | 0.439 | 1.254 | -0.013 | 0.004 | 0.004 | 0 | 0.15 | 0.24 | 0.1 | 0.14 | 0.12 | 0.07 |
| 8 | 2037220 | rs1025798 | -1.594 | 0.125 | 0.53 | 1.238 | -0.033 | 0.008 | 0.008 | 0.097 | 0.22 | 0.62 | 0.17 | 0.18 | 0.16 | 0.08 |
| 8 | 2037281 | rs1025797 | -1.612 | 0.115 | 0.525 | 1.28 | -0.034 | 0.013 | 0.013 | 0.245 | 0.22 | 0.62 | 0.18 | 0.21 | 0.16 | 0.07 |
| 8 | 2039595 | rs1550454 | -1.08 | 0.258 | 0.507 | 1.156 | -0.009 | -0.018 | -0.018 | 0 | 0.13 | 0.31 | 0.12 | 0.06 | 0.12 | 0.04 |
| 8 | 2040298 | rs2235122 | 0.056 | 0.432 | -0.248 | 1.269 | 0.004 | 0.004 | 0.004 | 0 | 0.1 | 0.12 | 0.07 | 0.15 | 0.16 | 0.17 |
| 8 | 2040401 | rs2235123 | 1.221 | 0.375 | -1.17 | 0.967 | -0.024 | 0.112 | 0.112 | 0.6 | 0.39 | 0.11 | 0.21 | 0.25 | 0.37 | 0.27 |
| 8 | 2041466 | rs4559266 | -1.064 | 0.082 | -0.247 | 1.564 | 0.024 | 0.069 | 0.069 | 0.006 | 0.24 | 0.43 | 0.39 | 0.37 | 0.29 | 0.49 |
| 8 | 2042070 | rs7828132 | 0.755 | 0.16 | -0.272 | 1.225 | -0.011 | 0.093 | 0.093 | 1.573 | 0.77 | 0.51 | 0.85 | 0.68 | 0.79 | 0.8 |
| 8 | 2044363 | rs7834633 | -0.695 | 0.264 | -0.951 | 0.99 | -0.009 | 0.024 | 0.024 | 0 | 0.46 | 0.77 | 0.59 | 0.39 | 0.55 | 0.47 |
| 8 | 2044426 | rs7820357 | -0.695 | 0.192 | -0.538 | 0.987 | -0.008 | 0.019 | 0.019 | 0 | 0.45 | 0.7 | 0.58 | 0.38 | 0.54 | 0.47 |
| 8 | 2045725 | rs6558605 | -0.737 | 0.042 | -0.896 | 0.965 | -0.041 | 0.012 | 0.012 | 0 | 0.39 | 0.77 | 0.55 | 0.36 | 0.48 | 0.4 |
| 8 | 2046576 | rs2294065 | 0.008 | 0.32 | -0.036 | 0.897 | 0.021 | 0.088 | 0.088 | 0 | 0.13 | 0.31 | 0.21 | 0.26 | 0.17 | 0.11 |
| 8 | 2046700 | rs2294066 | 0.317 | 0.321 | 0.05 | 0.896 | 0.003 | 0.02 | 0.02 | 0 | 0.12 | 0.12 | 0.19 | 0.26 | 0.17 | 0.11 |
| 8 | 2047356 | rs10092320 | -1.488 | 0.24 | -0.265 | 1.317 | -0.01 | -0.017 | -0.017 | 0 | 0.47 | 0.79 | 0.74 | 0.48 | 0.55 | 0.48 |
| 8 | 2048480 | rs3779840 | -0.072 | 0.646 | -0.256 | 1.838 | -0.043 | -0.059 | -0.059 | 1.295 | 0.13 | 0.05 | 0.39 | 0.31 | 0.22 | 0.2 |
| 8 | 2048518 | rs3779839 | 0.547 | 0.253 | -0.173 | 1.255 | 0.004 | 0.008 | 0.008 | 1.444 | 0.72 | 0.66 | 0.51 | 0.58 | 0.7 | 0.67 |
| 8 | 2048650 | rs3779837 | 0.405 | -0.111 | -0.738 | 1.55 | 0.004 | -0.056 | -0.056 | 0 | 0.09 | 0 | 0.04 | 0.19 | 0.05 | 0.2 |
| 8 | 2048831 | rs968381 | -0.907 | -0.575 | -0.797 | 1.094 | 0.035 | 0.01 | 0.01 | 0 | 0.42 | 0.68 | 0.36 | 0.4 | 0.34 | 0.42 |
| 8 | 2050240 | rs2294069 | -1.252 | -0.353 | -0.72 | 1.143 | -0.005 | 0.018 | 0.018 | 0 | 0.48 | 0.62 | 0.7 | 0.54 | 0.54 | 0.54 |
| 8 | 2050519 | rs2294071 | 0.509 | -0.212 | -0.536 | 1.202 | -0.002 | -0.004 | -0.004 | 0 | 0.12 | 0.14 | 0.06 | 0.08 | 0.06 | 0.16 |
| 8 | 2050596 | rs2294072 | 0.107 | -0.075 | -0.551 | 1.292 | 0.008 | 0.154 | 0.154 | 0 | 0.16 | 0.39 | 0.23 | 0.28 | 0.19 | 0.13 |
| 8 | 2052656 | rs17684398 | 0.7 | 0.263 | -1.624 | 1.178 | -0.067 | -0.064 | -0.064 | 0 | 0.11 | 0.02 | 0.36 | 0.18 | 0.25 | 0.19 |
| 8 | 2054171 | rs3817705 | 1.415 | -0.199 | -2.074 | 0.791 | -0.078 | 0.121 | 0.121 | 0 | 0.41 | 0.1 | 0.31 | 0.23 | 0.41 | 0.29 |
| 8 | 2054569 | rs3817708 | 0.816 | 0.12 | -1.882 | 0.993 | -0.067 | -0.055 | -0.055 | 0 | 0.11 | 0.02 | 0.16 | 0.16 | 0.24 | 0.2 |
| 8 | 2055331 | rs6988125 | 0.1 | -0.205 | -1.262 | 1.037 | -0.017 | 0.015 | 0.015 | 0.001 | 0.78 | 0.65 | 0.8 | 0.75 | 0.86 | 0.69 |
| 8 | 2056569 | rs1978898 | 0.756 | 0.14 | -1.994 | 1.194 | -0.067 | -0.052 | -0.052 | 0 | 0.11 | 0.02 | 0.16 | 0.16 | 0.24 | 0.2 |
| 8 | 2056864 | rs1550458 | -0.12 | -0.13 | -2.043 | 1.071 | -0.05 | -0.091 | -0.091 | 0.877 | 0.28 | 0.06 | 0.37 | 0.4 | 0.33 | 0.44 |
| 8 | 2057298 | rs2280897 | 0.615 | -0.019 | -1.77 | 1.14 | -0.007 | -0.06 | -0.06 | 0.433 | 0.11 | 0.03 | 0.16 | 0.28 | 0.14 | 0.14 |
| 8 | 2058991 | rs7825233 | -0.913 | 0.184 | -1.649 | 0.393 | -0.063 | 0.078 | 0.078 | 0 | 0.59 | 0.83 | 0.78 | 0.74 | 0.54 | 0.72 |
| 8 | 2059994 | rs4876225 | -0.631 | 0.616 | -1.372 | -0.212 | -0.004 | -0.003 | -0.003 | 0 | 0.68 | 0.73 | 0.78 | 0.92 | 0.63 | 0.81 |
| 8 | 2060411 | rs3779834 | 0.147 | 1.138 | -1.153 | 0.528 | -0.001 | -0.001 | -0.001 | 0 | 0.15 | 0.18 | 0.18 | 0.23 | 0.18 | 0.17 |
| 8 | 2060471 | rs1866716 | -0.272 | 0.983 | -0.799 | 0.194 | 0.021 | 0.037 | 0.037 | 0.038 | 0.7 | 0.45 | 0.51 | 0.4 | 0.65 | 0.56 |
| 8 | 2060549 | rs1866717 | 1.173 | 0.863 | -1.012 | 0.242 | -0.003 | 0.032 | 0.032 | 0 | 0.42 | 0.26 | 0.26 | 0.28 | 0.43 | 0.36 |
| 8 | 2061006 | rs17684547 | 0.894 | 1.332 | -1.301 | -0.04 | -0.067 | 0.124 | 0.124 | 0 | 0.19 | 0.01 | 0.13 | 0 | 0.19 | 0.07 |
| 8 | 2061084 | rs17751791 | 0.918 | 1.328 | -1.303 | -0.042 | -0.068 | 0.122 | 0.122 | 0 | 0.19 | 0.01 | 0.13 | 0 | 0.19 | 0.07 |
| 8 | 2061155 | rs3779831 | 0.322 | 0.742 | -0.779 | 0.209 | 0.001 | 0.001 | 0.001 | 0 | 0.51 | 0.47 | 0.36 | 0.35 | 0.46 | 0.47 |
| 8 | 2063261 | rs7002629 | 0.507 | 0.824 | -0.824 | 0.312 | -0.007 | -0.023 | -0.023 | 0 | 0.16 | 0.07 | 0.17 | 0.18 | 0.18 | 0.16 |
| 8 | 2065092 | rs4876223 | -0.975 | 0.624 | -0.533 | 0.15 | 0.014 | 0.015 | 0.015 | 0.167 | 0.54 | 0.64 | 0.71 | 0.69 | 0.66 | 0.64 |
| 8 | 2066033 | rs10097327 | -0.804 | 0.593 | -0.434 | 0.203 | 0.033 | 0.038 | 0.038 | 0.126 | 0.53 | 0.72 | 0.72 | 0.71 | 0.65 | 0.61 |
| 8 | 2067475 | rs6990007 | -0.097 | 0.692 | -0.17 | 0.767 | -0.001 | -0.001 | -0.001 | 0 | 0.44 | 0.48 | 0.49 | 0.4 | 0.49 | 0.31 |
| 8 | 2067616 | rs10101799 | -0.22 | 0.642 | -0.243 | 0.848 | -0.003 | 0.023 | 0.023 | 0 | 0.1 | 0.12 | 0.19 | 0.2 | 0.08 | 0.25 |
| 8 | 2072342 | rs2280899 | -1.413 | 0.635 | -0.379 | 0.858 | 0.014 | 0.007 | 0.007 | 0 | 0.64 | 0.75 | 0.8 | 0.71 | 0.71 | 0.61 |

| | | | | | | | | | | | | | | | | |
|---|---|---|---|---|---|---|---|---|---|---|---|---|---|---|---|---|
| 8 | 2076757 | rs2280900 | -0.077 | 0.579 | 0.209 | 0.951 | 0.016 | 0.177 | 0.177 | 0 | 0.08 | 0.29 | 0.15 | 0.19 | 0.05 | 0.21 |
| 8 | 2077072 | rs2280902 | -0.386 | 0.788 | -0.032 | 0.771 | -0.016 | 0.014 | 0.014 | 0 | 0.65 | 0.7 | 0.59 | 0.69 | 0.58 | 0.71 |
| 8 | 2080289 | rs12680793 | -0.809 | 0.836 | -0.057 | 0.742 | -0.012 | 0.016 | 0.016 | 0 | 0.69 | 0.73 | 0.68 | 0.78 | 0.62 | 0.74 |
| 8 | 2086493 | rs3779817 | 1.964 | 0.698 | -0.129 | 0.837 | -0.004 | -0.027 | -0.027 | 0 | 0.3 | 0.06 | 0.16 | 0.27 | 0.28 | 0.34 |
| 8 | 2091139 | rs2280903 | 0.37 | 0.676 | 0.076 | 0.904 | 0.002 | 0.15 | 0.15 | 0.018 | 0.08 | 0.22 | 0.15 | 0.22 | 0.05 | 0.21 |
| 8 | 2091184 | rs2280904 | 1.946 | 0.755 | -0.288 | 0.93 | -0.006 | -0.075 | -0.075 | 6.523 | 0.3 | 0.06 | 0.17 | 0.36 | 0.28 | 0.35 |
| 8 | 2091227 | rs2280905 | 1.589 | 0.649 | -0.302 | 0.718 | -0.033 | -0.051 | -0.051 | 0.714 | 0.1 | 0.03 | 0.12 | 0.2 | 0.16 | 0.12 |
| 8 | 2093181 | rs3407 | 2.029 | 0.585 | -0.308 | 0.761 | -0.008 | 0.009 | 0.009 | 0 | 0.29 | 0.06 | 0.17 | 0.24 | 0.28 | 0.34 |
| 8 | 2094501 | rs6558610 | -0.664 | 0.53 | -0.159 | 0.75 | -0.006 | 0.015 | 0.015 | 0 | 0.34 | 0.53 | 0.39 | 0.42 | 0.31 | 0.36 |
| 8 | 2095011 | rs11774937 | 0.967 | 0.579 | -0.244 | 0.765 | -0.035 | -0.082 | -0.082 | 0 | 0.43 | 0.11 | 0.27 | 0.44 | 0.47 | 0.5 |
| 8 | 2095247 | rs9644319 | 1.94 | 0.58 | -0.242 | 0.709 | -0.006 | -0.008 | -0.008 | 0 | 0.3 | 0.07 | 0.17 | 0.26 | 0.29 | 0.35 |
| 8 | 2097007 | rs9314453 | -0.966 | 0.563 | -0.101 | 0.694 | -0.002 | 0.005 | 0.005 | 0 | 0.38 | 0.53 | 0.41 | 0.44 | 0.34 | 0.39 |
| 8 | 2097247 | rs17752155 | 1.088 | 0.575 | -0.132 | 0.651 | -0.037 | -0.051 | -0.051 | 0 | 0.13 | 0.03 | 0.11 | 0.17 | 0.19 | 0.15 |
| 8 | 2097827 | rs4529490 | -0.648 | 0.54 | -0.066 | 0.707 | -0.011 | 0.132 | 0.132 | 0.931 | 0.21 | 0.57 | 0.28 | 0.42 | 0.21 | 0.33 |
| 8 | 2097909 | rs4400411 | 0.003 | 0.553 | -0.031 | 0.702 | -0.024 | 0.022 | 0.022 | 0 | 0.25 | 0.48 | 0.27 | 0.25 | 0.15 | 0.24 |
| 8 | 2098770 | rs4876217 | -0.811 | 0.606 | 0.004 | 0.742 | -0.011 | 0.009 | 0.009 | 0 | 0.68 | 0.7 | 0.57 | 0.69 | 0.62 | 0.74 |
| 8 | 2101074 | rs11987927 | -0.12 | 0.52 | 0.294 | 0.947 | 0.007 | 0.226 | 0.226 | 1.088 | 0.08 | 0.28 | 0.15 | 0.23 | 0.05 | 0.2 |
| 8 | 2101655 | rs12676956 | 1.575 | 0.549 | 0.004 | 0.912 | -0.007 | 0.009 | 0.009 | 0 | 0.3 | 0.09 | 0.24 | 0.23 | 0.29 | 0.34 |
| 8 | 2102110 | rs7007697 | 0.574 | 0.597 | 0.095 | 0.875 | -0.012 | 0.007 | 0.007 | 0 | 0.4 | 0.49 | 0.42 | 0.5 | 0.34 | 0.55 |
| 8 | 2103379 | rs1866719 | -0.727 | 0.878 | 0.216 | 0.816 | 0.03 | 0.194 | 0.194 | 0.738 | 0.12 | 0.42 | 0.17 | 0.24 | 0.11 | 0.21 |
| 8 | 2103819 | rs17752323 | 1.234 | 1.037 | -0.337 | 0.789 | -0.028 | -0.05 | -0.05 | 0.061 | 0.38 | 0.11 | 0.22 | 0.43 | 0.38 | 0.48 |
| 8 | 2104726 | rs7828726 | -0.019 | 0.88 | 0.049 | 0.803 | 0.02 | 0.15 | 0.15 | 0.031 | 0.13 | 0.39 | 0.17 | 0.21 | 0.1 | 0.19 |
| 8 | 2104747 | rs7829787 | -0.019 | 0.88 | 0.069 | 0.803 | 0.019 | 0.148 | 0.148 | 0.031 | 0.13 | 0.39 | 0.17 | 0.21 | 0.1 | 0.19 |
| 8 | 2105181 | rs9644321 | 1.25 | 0.974 | -0.024 | 0.805 | -0.031 | -0.038 | -0.038 | 0 | 0.39 | 0.13 | 0.22 | 0.42 | 0.38 | 0.51 |
| 8 | 2105210 | rs10094743 | 0.01 | 0.97 | 0.033 | 0.814 | 0.02 | 0.15 | 0.15 | 0.042 | 0.13 | 0.39 | 0.17 | 0.21 | 0.11 | 0.19 |
| 8 | 2105621 | rs12543116 | 0.237 | 1.01 | -0.018 | 0.671 | -0.001 | 0.009 | 0.009 | 0 | 0.52 | 0.51 | 0.41 | 0.66 | 0.48 | 0.68 |
| 8 | 2106161 | rs6991191 | -1.415 | 0.844 | 0.132 | 0.386 | 0.013 | 0.018 | 0.018 | 0 | 0.67 | 0.8 | 0.82 | 0.76 | 0.69 | 0.6 |
| 8 | 2106764 | rs17752424 | 1.774 | 0.658 | -0.242 | -0.224 | -0.002 | 0.044 | 0.044 | 0 | 0.15 | 0.08 | 0.12 | 0 | 0.12 | 0.04 |
| 8 | 2106904 | rs5028635 | 0.248 | 0.668 | 0.111 | -0.323 | -0.005 | -0.106 | -0.106 | 0.573 | 0.26 | 0.35 | 0.16 | 0 | 0.2 | 0.08 |
| 8 | 2107653 | rs17752483 | 0.215 | 0.681 | 0.116 | -0.317 | -0.006 | -0.107 | -0.107 | 9.19 | 0.27 | 0.35 | 0.16 | 0 | 0.2 | 0.08 |
| 8 | 2107717 | rs13271597 | 1.442 | 0.684 | 0.009 | -0.139 | -0.052 | -0.022 | -0.022 | 0 | 0.36 | 0.11 | 0.21 | 0.4 | 0.39 | 0.51 |
| 8 | 2108188 | rs9314454 | 1.313 | 0.704 | 0.203 | -0.044 | 0.002 | 0.088 | 0.088 | 0 | 0.13 | 0.15 | 0.23 | 0.39 | 0.18 | 0.27 |
| 8 | 2109059 | rs9314455 | -1.059 | 0.665 | 0.366 | -0.107 | 0.008 | -0.02 | -0.02 | 0.244 | 0.41 | 0.74 | 0.44 | 0.46 | 0.38 | 0.36 |
| 8 | 2109376 | rs10217026 | 1.267 | 0.69 | 0.314 | 0.027 | 0.002 | 0.082 | 0.082 | 0 | 0.13 | 0.15 | 0.23 | 0.38 | 0.18 | 0.28 |
| 8 | 2110076 | rs7846356 | 0.569 | 0.682 | 0.368 | -0.056 | 0.016 | -0.058 | -0.058 | 0 | 0.1 | 0.37 | 0.06 | 0 | 0.06 | 0.01 |
| 8 | 2111830 | rs17064803 | 0.718 | 0.71 | 0.312 | -0.063 | -0.006 | -0.109 | -0.109 | 0.689 | 0.23 | 0.32 | 0.13 | 0 | 0.17 | 0.05 |
| 8 | 2112041 | rs1036688 | 1.197 | 0.766 | 0.309 | 0.048 | 0.002 | 0.09 | 0.09 | 0 | 0.13 | 0.18 | 0.23 | 0.41 | 0.18 | 0.29 |
| 8 | 2112130 | rs1036689 | 1.435 | 0.806 | 0.118 | -0.012 | -0.026 | -0.008 | -0.008 | 0.14 | 0.36 | 0.19 | 0.23 | 0.4 | 0.39 | 0.49 |
| 8 | 2112514 | rs17064809 | 0.621 | 0.74 | 0.243 | -0.215 | 0.014 | -0.074 | -0.074 | 0.518 | 0.12 | 0.32 | 0.04 | 0 | 0.09 | 0.04 |
| 8 | 2113776 | rs17685410 | 1.881 | 0.956 | -0.321 | 0.03 | -0.007 | 0.012 | 0.012 | 0 | 0.3 | 0.11 | 0.16 | 0.27 | 0.3 | 0.4 |
| 8 | 2117238 | rs10100885 | 0.972 | 0.874 | -0.124 | 0.272 | -0.003 | 0.016 | 0.016 | 0 | 0.13 | 0.11 | 0.26 | 0.39 | 0.19 | 0.23 |
| 8 | 2118784 | rs17064827 | 1.538 | 0.706 | 0.186 | 0.023 | 0.008 | -0.054 | -0.054 | 0.173 | 0.1 | 0.29 | 0.03 | 0 | 0.06 | 0.03 |
| 8 | 2118895 | rs2618851 | -1.457 | 0.707 | 0.158 | -0.078 | 0.001 | -0.042 | -0.042 | 0.164 | 0.79 | 0.65 | 0.62 | 0.87 | 0.77 | 0.88 |
| 8 | 2119582 | rs11784069 | 1.32 | 0.895 | -0.159 | -0.138 | -0.001 | -0.03 | -0.03 | 0.177 | 0.37 | 0.21 | 0.21 | 0.44 | 0.35 | 0.54 |
| 8 | 2120055 | rs2581570 | -1.764 | 0.859 | 0.008 | -0.142 | -0.001 | 0.009 | 0.009 | 0.115 | 0.8 | 0.84 | 0.65 | 0.87 | 0.77 | 0.88 |
| 8 | 2120235 | rs2618814 | 0.584 | 0.883 | 0.117 | -0.174 | 0 | 0.015 | 0.015 | 0.038 | 0.52 | 0.24 | 0.34 | 0.48 | 0.48 | 0.58 |
| 8 | 2120876 | rs17064834 | 1.139 | 0.838 | 0.234 | -0.217 | -0.008 | -0.079 | -0.079 | 0 | 0.13 | 0.31 | 0.07 | 0.01 | 0.09 | 0.06 |
| 8 | 2121238 | rs1371764 | 1.385 | 0.86 | 0.179 | -0.207 | -0.003 | -0.025 | -0.025 | 0 | 0.36 | 0.21 | 0.21 | 0.44 | 0.35 | 0.54 |
| 8 | 2121498 | rs899409 | -0.309 | 0.917 | 0.118 | -0.081 | 0 | 0.021 | 0.021 | 0.581 | 0.66 | 0.6 | 0.43 | 0.48 | 0.61 | 0.65 |
| 8 | 2121789 | rs899408 | 0.549 | 0.855 | 0.133 | -0.072 | 0.002 | 0.004 | 0.004 | 0 | 0.5 | 0.55 | 0.28 | 0.45 | 0.45 | 0.6 |
| 8 | 2121846 | rs10110395 | 0.215 | 0.868 | 0.144 | -0.039 | -0.014 | -0.044 | -0.044 | 2.499 | 0.29 | 0.18 | 0.37 | 0.42 | 0.33 | 0.28 |
| 8 | 2122689 | rs7462690 | -0.692 | 0.795 | 0.323 | 0.061 | 0 | -0.044 | -0.044 | 0.184 | 0.63 | 0.6 | 0.51 | 0.83 | 0.64 | 0.84 |
| 8 | 2123109 | rs731869 | 1.751 | 0.791 | 0.143 | 0.12 | 0.011 | -0.004 | -0.004 | 1.239 | 0.3 | 0.15 | 0.16 | 0.3 | 0.29 | 0.42 |
| 8 | 2123694 | rs2581578 | -0.737 | 0.835 | 0.166 | 0.054 | -0.001 | 0.003 | 0.003 | 0 | 0.64 | 0.68 | 0.52 | 0.83 | 0.64 | 0.84 |
| 8 | 2123928 | rs2581579 | -0.722 | 0.869 | -0.091 | 0.071 | 0 | -0.027 | -0.027 | 0 | 0.63 | 0.62 | 0.51 | 0.83 | 0.64 | 0.84 |
| 8 | 2124151 | rs2618806 | 0.314 | 1.005 | -0.202 | 0.382 | 0.012 | -0.013 | -0.013 | 0 | 0.19 | 0.32 | 0.1 | 0.15 | 0.16 | 0.18 |
| 8 | 2124338 | rs2119238 | 1.675 | 0.684 | -1.115 | 0.389 | -0.013 | -0.077 | -0.077 | 1.249 | 0.15 | 0.01 | 0.08 | 0.27 | 0.18 | 0.34 |
| 8 | 2128786 | rs11987758 | 0.796 | 1.119 | -0.223 | 0.559 | 0.013 | 0.025 | 0.025 | 0 | 0.09 | 0.13 | 0.09 | 0.13 | 0.13 | 0.09 |
| 8 | 2129547 | rs2581568 | 0.516 | 1.16 | -0.48 | 0.266 | -0.035 | 0.044 | 0.044 | 0 | 0.6 | 0.27 | 0.54 | 0.68 | 0.51 | 0.74 |
| 8 | 2131125 | rs11989864 | 0.259 | 1.133 | -0.216 | 0.24 | 0.025 | 0.024 | 0.024 | 0 | 0.13 | 0.28 | 0.13 | 0.14 | 0.17 | 0.13 |
| 8 | 2132561 | rs2581599 | 0.491 | 0.813 | -0.947 | 0.559 | -0.014 | -0.041 | -0.041 | 0.07 | 0.65 | 0.26 | 0.36 | 0.45 | 0.59 | 0.61 |
| 8 | 2133338 | rs4876213 | 1.186 | 0.699 | -0.285 | -0.227 | 0.03 | -0.053 | -0.053 | 0.027 | 0.4 | 0.15 | 0.37 | 0.5 | 0.36 | 0.55 |
| 8 | 2135762 | rs6558621 | -0.38 | 0.847 | 0.537 | -0.721 | 0.005 | -0.012 | -0.012 | 0.602 | 0.82 | 0.78 | 0.69 | 0.86 | 0.76 | 0.87 |
| 8 | 2143681 | rs1564289 | 0.81 | 0.833 | 0.52 | -0.607 | 0.015 | 0.02 | 0.02 | 0 | 0.39 | 0.24 | 0.42 | 0.67 | 0.4 | 0.58 |
| 8 | 2145379 | rs317220 | -0.145 | 1.151 | 1.694 | 0.264 | 0.007 | 0.012 | 0.012 | 0 | 0.87 | 0.81 | 0.59 | 0.78 | 0.81 | 0.88 |
| 8 | 2146093 | rs317221 | -0.026 | 1.176 | 1.933 | 0.129 | 0 | -0.034 | -0.034 | 0.798 | 0.13 | 0.16 | 0.07 | 0 | 0.1 | 0.06 |
| 8 | 2147744 | rs2244707 | -1.676 | 1.141 | 2.493 | 0.155 | 0.016 | 0.007 | 0.007 | 0 | 0.32 | 0.54 | 0.55 | 0.36 | 0.34 | 0.28 |
| 8 | 2149448 | rs717920 | -1.154 | 1.341 | 2.458 | -0.055 | 0 | -0.055 | -0.055 | 0 | 0.23 | 0.26 | 0.13 | 0.01 | 0.19 | 0.1 |
| 8 | 2150224 | rs2581592 | -0.769 | 1.32 | 2.482 | 0.107 | -0.013 | -0.066 | -0.066 | 0 | 0.18 | 0.42 | 0.15 | 0.06 | 0.1 | 0.06 |

| | | | | | | | | | | | | | | | | |
|---|---|---|---|---|---|---|---|---|---|---|---|---|---|---|---|---|
| 8 | 2151798 | rs11782440 | 0.426 | 1.435 | 2.409 | 0.246 | -0.051 | -0.059 | -0.059 | 0 | 0.11 | 0.03 | 0.26 | 0.22 | 0.18 | 0.08 |
| 8 | 2151954 | rs2618841 | 0.539 | 1.411 | 2.659 | 0.244 | 0.051 | 0.115 | 0.115 | 0.019 | 0.83 | 0.54 | 0.69 | 0.68 | 0.77 | 0.82 |
| 8 | 2163403 | rs10503174 | 1.326 | 1.337 | 2.144 | 0.147 | 0.046 | 0.145 | 0.145 | 1.239 | 0.93 | 0.48 | 0.92 | 0.87 | 0.95 | 0.9 |
| 8 | 2164005 | rs17064922 | -0.242 | 1.433 | 1.491 | 0.065 | 0.011 | 0.013 | 0.013 | 0 | 0.14 | 0.22 | 0.29 | 0.21 | 0.19 | 0.08 |
| 8 | 2164581 | rs1382608 | 1.72 | 1.444 | 1.136 | -0.1 | -0.049 | 0.03 | 0.03 | 0.138 | 0.56 | 0.12 | 0.51 | 0.61 | 0.55 | 0.73 |
| 8 | 2164712 | rs7838658 | -0.899 | 1.529 | 1.005 | -0.052 | 0.006 | -0.006 | -0.006 | 0.202 | 0.29 | 0.37 | 0.3 | 0.24 | 0.28 | 0.12 |
| 8 | 2167540 | rs10110313 | 1.416 | 1.118 | 0.088 | -0.032 | -0.052 | 0.04 | 0.04 | 0.012 | 0.53 | 0.07 | 0.46 | 0.61 | 0.52 | 0.68 |
| 8 | 2170955 | rs4876200 | 0.874 | 1.307 | 0.52 | -0.244 | -0.001 | -0.001 | -0.001 | 0 | 0.7 | 0.37 | 0.7 | 0.82 | 0.74 | 0.83 |
| 8 | 2171959 | rs11136479 | 0.878 | 1.385 | 0.935 | 0.42 | 0.031 | 0.059 | 0.059 | 0 | 0.77 | 0.55 | 0.68 | 0.68 | 0.7 | 0.8 |
| 8 | 2178596 | rs10107849 | 1.439 | 0.982 | 0.627 | 0.066 | 0.016 | -0.037 | -0.037 | 0 | 0.86 | 0.76 | 0.91 | 0.92 | 0.85 | 0.88 |
| 8 | 2181540 | rs35717865 | -1.17 | 0.889 | -0.067 | 0.11 | 0.001 | -0.031 | -0.031 | 0 | 0.09 | 0.2 | 0.06 | 0.01 | 0.08 | 0.03 |
| 8 | 2289839 | rs6558648 | 0.462 | 0.806 | -0.558 | 0.159 | 0.006 | 0.02 | 0.02 | 0 | 0.51 | 0.42 | 0.48 | 0.41 | 0.48 | 0.57 |
| 8 | 2290039 | rs7002201 | -0.49 | 1.248 | -0.181 | -0.135 | 0.024 | -0.04 | -0.04 | 0.355 | 0.92 | 0.86 | 0.92 | 1 | 0.88 | 0.99 |
| 8 | 2292024 | rs9644328 | -0.186 | 0.961 | 0.445 | 0.209 | -0.002 | -0.02 | -0.02 | 2.882 | 0.34 | 0.22 | 0.36 | 0.45 | 0.38 | 0.39 |
| 8 | 2292537 | rs7017375 | -1.198 | 0.84 | 0.917 | 0.038 | 0.07 | 0.016 | 0.016 | 0 | 0.05 | 0.31 | 0.05 | 0 | 0.03 | 0.02 |
| 8 | 2293408 | rs12675491 | -1.415 | 0.991 | 0.642 | 0.256 | 0.062 | 0.023 | 0.023 | 0 | 0.24 | 0.48 | 0.33 | 0.28 | 0.31 | 0.11 |
| 8 | 2295755 | rs897445 | 1.134 | 0.727 | 0.123 | -0.268 | 0.006 | 0.003 | 0.003 | 0 | 0.45 | 0.39 | 0.44 | 0.41 | 0.4 | 0.46 |
| 8 | 2304334 | rs4876153 | 0.281 | 0.843 | 0.051 | -0.275 | -0.067 | -0.135 | -0.135 | 0 | 0.31 | 0.05 | 0.32 | 0.43 | 0.34 | 0.38 |
| 8 | 2310313 | rs7812885 | -0.678 | 0.632 | 0.508 | -0.437 | -0.043 | -0.126 | -0.126 | 0 | 0.17 | 0.47 | 0.1 | 0 | 0.09 | 0.09 |
| 8 | 2311671 | rs6558654 | -0.717 | 0.628 | 0.54 | -0.439 | -0.042 | -0.124 | -0.124 | 0 | 0.17 | 0.46 | 0.1 | 0 | 0.09 | 0.09 |
| 8 | 2312429 | rs1455642 | -0.609 | 0.635 | 0.53 | -0.435 | -0.036 | -0.105 | -0.105 | 0 | 0.16 | 0.39 | 0.09 | 0 | 0.09 | 0.09 |
| 8 | 2314023 | rs7843995 | -0.307 | 0.441 | 0.15 | -0.443 | -0.005 | 0.003 | 0.003 | 0 | 0.89 | 0.8 | 0.89 | 0.86 | 0.91 | 0.95 |
| 8 | 2314556 | rs17072197 | -0.91 | 0.659 | 0.718 | -0.541 | 0.067 | -0.114 | -0.114 | 0 | 0.15 | 0.48 | 0.13 | 0 | 0.14 | 0.03 |
| 8 | 2319093 | rs12056750 | 1.517 | 0.186 | -0.413 | -0.435 | -0.008 | -0.063 | -0.063 | 0 | 0.35 | 0.1 | 0.39 | 0.4 | 0.36 | 0.41 |
| 8 | 2319271 | rs6991331 | 0.702 | 0.148 | -0.244 | -0.491 | 0.035 | 0.018 | 0.018 | 0 | 0.46 | 0.69 | 0.47 | 0.4 | 0.39 | 0.45 |
| 8 | 2319797 | rs11786540 | 1.36 | 0.086 | -0.378 | -0.422 | -0.02 | -0.058 | -0.058 | 2.824 | 0.37 | 0.1 | 0.39 | 0.4 | 0.38 | 0.44 |
| 8 | 2321337 | rs11136518 | -0.973 | -0.354 | 0.227 | -0.489 | -0.047 | -0.078 | -0.078 | 0.061 | 0.12 | 0.51 | 0.07 | 0 | 0.03 | 0.01 |
| 8 | 2346867 | rs4875971 | -0.408 | -0.517 | -0.549 | -0.195 | 0.025 | 0.034 | 0.034 | 0 | 0.5 | 0.34 | 0.34 | 0.41 | 0.41 | 0.54 |
| 8 | 2346899 | rs17692676 | -0.155 | -1.045 | -0.723 | -0.341 | -0.004 | 0.033 | 0.033 | 0 | 0.39 | 0.21 | 0.3 | 0.33 | 0.37 | 0.49 |
| 8 | 2350283 | rs6558659 | -0.787 | -1.173 | -0.244 | -0.111 | -0.022 | 0.062 | 0.062 | 0 | 0.87 | 0.63 | 0.87 | 0.8 | 0.92 | 0.92 |
| 8 | 2350511 | rs6998131 | -0.24 | -1.271 | -0.529 | -0.085 | 0.023 | -0.003 | -0.003 | 0 | 0.13 | 0.16 | 0.17 | 0.16 | 0.19 | 0.04 |
| 8 | 2352384 | rs11781947 | 0.636 | -1.37 | -0.643 | -0.111 | -0.006 | -0.01 | -0.01 | 0 | 0.71 | 0.25 | 0.67 | 0.64 | 0.72 | 0.87 |
| 8 | 2352452 | rs13261880 | -0.497 | -1.302 | -0.628 | -0.099 | -0.042 | 0.028 | 0.028 | 0 | 0.82 | 0.53 | 0.85 | 0.8 | 0.9 | 0.92 |
| 8 | 2354123 | rs1457449 | -0.506 | -1.596 | -0.938 | -0.011 | -0.003 | -0.006 | -0.006 | 0 | 0.47 | 0.47 | 0.34 | 0.38 | 0.39 | 0.53 |
| 8 | 2354194 | rs1993480 | -0.48 | -1.165 | -1.699 | 0.162 | 0.008 | 0.004 | 0.004 | 0.018 | 0.15 | 0.17 | 0.18 | 0.24 | 0.19 | 0.07 |
| 8 | 2355745 | rs13254407 | 1.081 | -0.806 | -1.848 | 0.258 | -0.135 | -0.148 | -0.148 | 0 | 0.36 | 0.08 | 0.51 | 0.42 | 0.51 | 0.39 |
| 8 | 2356921 | rs12549051 | 1.04 | -0.815 | -1.854 | 0.268 | -0.101 | -0.117 | -0.117 | 0 | 0.36 | 0.11 | 0.52 | 0.42 | 0.52 | 0.4 |
| 8 | 2361015 | rs7460386 | 0.332 | -0.951 | -1.971 | 0.066 | 0.145 | 0.145 | 0.145 | 0 | 0.5 | 0.16 | 0.71 | 0.71 | 0.72 | 0.58 |
| 8 | 2362084 | rs4332162 | 0.381 | -0.627 | -1.836 | -0.003 | 0.067 | 0.055 | 0.055 | 0 | 0.46 | 0.22 | 0.67 | 0.65 | 0.68 | 0.48 |
| 8 | 2362875 | rs11782214 | 0.655 | -0.418 | -2.134 | 0.301 | -0.129 | -0.158 | -0.158 | 2.826 | 0.34 | 0.06 | 0.33 | 0.43 | 0.48 | 0.37 |
| 8 | 2365648 | rs2100623 | -0.55 | -1.171 | -2.096 | -0.689 | 0.099 | 0.128 | 0.128 | 1.408 | 0.42 | 0.81 | 0.25 | 0.18 | 0.23 | 0.15 |
| 8 | 2366909 | rs7017120 | 0.025 | -1.415 | -1.534 | -0.766 | 0.004 | 0 | 0 | 0 | 0.17 | 0.16 | 0.12 | 0.18 | 0.11 | 0.08 |
| 8 | 2368638 | rs3849845 | -0.419 | -1.494 | -0.999 | -1.143 | 0.003 | 0.049 | 0.049 | 0 | 0.55 | 0.65 | 0.65 | 0.7 | 0.57 | 0.41 |
| 8 | 2370882 | rs4875825 | 0.541 | -1.506 | -1.248 | -1.159 | -0.068 | 0.09 | 0.09 | 0 | 0.38 | 0.19 | 0.56 | 0.71 | 0.5 | 0.38 |
| 8 | 2376531 | rs12545773 | 0.466 | -1.465 | -1.309 | -1.28 | -0.035 | 0.089 | 0.089 | 0 | 0.4 | 0.24 | 0.57 | 0.71 | 0.5 | 0.41 |
| 8 | 2378707 | rs4875828 | 0.764 | -1.486 | -0.32 | -1.279 | -0.002 | -0.03 | -0.03 | 0.15 | 0.07 | 0.19 | 0.03 | 0 | 0.04 | 0.03 |
| 8 | 2380877 | rs4875831 | 0.615 | -1.459 | -0.567 | -1.304 | -0.03 | 0.051 | 0.051 | 0 | 0.37 | 0.29 | 0.54 | 0.7 | 0.46 | 0.39 |
| 8 | 2381004 | rs4875832 | 1.81 | -1.558 | -0.834 | -1.175 | -0.114 | -0.101 | -0.101 | 0 | 0.24 | 0.05 | 0.5 | 0.52 | 0.39 | 0.33 |
| 8 | 2384721 | rs3894390 | -0.545 | -1.531 | -0.204 | -1.533 | -0.006 | 0.027 | 0.027 | 0 | 0.57 | 0.65 | 0.42 | 0.3 | 0.51 | 0.61 |
| 8 | 2386070 | rs12678432 | 0.909 | -1.601 | -0.671 | -1.62 | -0.107 | 0.034 | 0.034 | 0.039 | 0.32 | 0.02 | 0.29 | 0.3 | 0.36 | 0.49 |
| 8 | 2386695 | rs4875990 | -0.773 | -1.578 | -0.536 | -1.612 | 0.031 | 0.015 | 0.015 | 0 | 0.31 | 0.75 | 0.16 | 0 | 0.17 | 0.11 |
| 8 | 2386894 | rs9657389 | 0.53 | -1.472 | -0.656 | -1.513 | -0.02 | -0.091 | -0.091 | 0.11 | 0.16 | 0.33 | 0.07 | 0 | 0.09 | 0.06 |
| 8 | 2388930 | rs12674549 | -0.37 | -1.052 | -0.872 | -1.613 | -0.004 | 0.013 | 0.013 | 0 | 0.54 | 0.43 | 0.4 | 0.3 | 0.49 | 0.58 |
| 8 | 2391471 | rs10104906 | -0.763 | -0.523 | -1.515 | -1.367 | -0.014 | 0.036 | 0.036 | 0 | 0.65 | 0.76 | 0.46 | 0.28 | 0.57 | 0.56 |
| 8 | 2394776 | rs13248321 | 0.519 | -0.146 | -1.598 | -1.185 | -0.155 | 0.059 | 0.059 | 0 | 0.33 | 0.03 | 0.32 | 0.28 | 0.41 | 0.48 |
| 8 | 2396588 | rs6420206 | -0.433 | 0.578 | -0.604 | -0.416 | 0.002 | -0.007 | -0.007 | 0 | 0.68 | 0.67 | 0.47 | 0.28 | 0.6 | 0.6 |
| 8 | 2400236 | rs4451330 | -0.163 | 1.074 | -0.613 | 1.073 | -0.049 | -0.044 | -0.044 | 0.064 | 0.73 | 0.85 | 0.61 | 0.64 | 0.6 | 0.64 |
| 8 | 2403261 | rs10503177 | 0.216 | 1.178 | -0.735 | 1.195 | -0.01 | -0.012 | -0.012 | 0 | 0.1 | 0.04 | 0.14 | 0.26 | 0.13 | 0.13 |
| 8 | 2406818 | rs9314461 | 0.614 | 1.083 | -0.788 | 1.148 | -0.002 | 0.066 | 0.066 | 0 | 0.54 | 0.3 | 0.26 | 0.3 | 0.46 | 0.43 |
| 8 | 2407398 | rs10113359 | 0.087 | 1.185 | -0.316 | 1.281 | 0.05 | 0.172 | 0.172 | 0.092 | 0.11 | 0.28 | 0.3 | 0.34 | 0.14 | 0.16 |
| 8 | 2408327 | rs1457446 | 0.246 | 0.81 | -1.278 | 0.687 | -0.011 | 0.031 | 0.031 | 0 | 0.55 | 0.31 | 0.27 | 0.39 | 0.47 | 0.45 |
| 8 | 2410524 | rs4876000 | -0.244 | 0.263 | -1.735 | 0.558 | 0.007 | -0.042 | -0.042 | 0.031 | 0.25 | 0.12 | 0.4 | 0.31 | 0.18 | 0.39 |
| 8 | 2412280 | rs17070319 | 0.26 | -0.604 | -1.331 | -0.396 | -0.023 | -0.09 | -0.09 | 0.131 | 0.22 | 0.25 | 0.1 | 0 | 0.11 | 0.04 |
| 8 | 2415854 | rs17070399 | 0.872 | -0.301 | -0.953 | 0.102 | 0.008 | 0.046 | 0.046 | 0 | 0.05 | 0.12 | 0.05 | 0.2 | 0.08 | 0.04 |
| 8 | 2416120 | rs4455858 | -1.247 | -0.667 | -0.774 | -0.004 | -0.006 | -0.05 | -0.05 | 0 | 0.59 | 0.8 | 0.6 | 0.49 | 0.39 | 0.47 |
| 8 | 2417678 | rs6996532 | -0.641 | -0.668 | -0.241 | -0.027 | 0.021 | 0.147 | 0.147 | 0 | 0.11 | 0.41 | 0.1 | 0.22 | 0.1 | 0.07 |
| 8 | 2423004 | rs7016232 | 0.165 | -0.816 | -0.405 | -0.245 | -0.004 | -0.031 | -0.031 | 0.605 | 0.09 | 0.1 | 0.04 | 0 | 0.04 | 0.02 |
| 8 | 2433479 | rs7841009 | -1.223 | -1.247 | -0.659 | -0.58 | 0.056 | 0.066 | 0.066 | 5.046 | 0.42 | 0.66 | 0.24 | 0.24 | 0.28 | 0.17 |
| 8 | 2446418 | rs4876003 | -1.266 | -1.202 | -0.404 | -0.527 | 0.018 | 0.02 | 0.02 | 1.146 | 0.37 | 0.64 | 0.21 | 0.25 | 0.25 | 0.13 |

| | | | | | | | | | | | | | | | | |
|---|---|---|---|---|---|---|---|---|---|---|---|---|---|---|---|---|
| 8 | 2447630 | rs7827855 | -1.154 | -1.181 | -0.408 | -0.509 | 0.012 | 0.014 | 0.014 | 2.215 | 0.36 | 0.63 | 0.21 | 0.24 | 0.24 | 0.13 |
| 8 | 2449499 | rs1483809 | -0.442 | -1.038 | -0.147 | -0.461 | -0.002 | -0.005 | -0.005 | 0 | 0.14 | 0.12 | 0.08 | 0.22 | 0.14 | 0.07 |
| 8 | 2454978 | rs9785100 | 0.687 | -1.093 | -0.026 | -0.513 | -0.017 | -0.059 | -0.059 | 0 | 0.14 | 0.17 | 0.07 | 0 | 0.08 | 0.02 |
| 8 | 2466309 | rs4279621 | -0.561 | -1.084 | 0.504 | -0.459 | 0.012 | 0.023 | 0.023 | 0 | 0.14 | 0.3 | 0.11 | 0.22 | 0.14 | 0.06 |
| 8 | 2470160 | rs6990227 | 0.587 | -1.132 | 0.296 | -0.524 | -0.01 | -0.041 | -0.041 | 0 | 0.14 | 0.13 | 0.08 | 0 | 0.08 | 0.02 |
| 8 | 2475475 | rs12544337 | 1.127 | -0.294 | -0.646 | -0.167 | -0.118 | -0.054 | -0.054 | 0 | 0.32 | 0.03 | 0.31 | 0.24 | 0.36 | 0.43 |
| 8 | 2479259 | rs7830883 | -0.022 | 0.325 | 0.339 | 0.655 | 0.001 | 0.024 | 0.024 | 2.092 | 0.85 | 0.84 | 0.9 | 0.64 | 0.82 | 0.81 |
| 8 | 2479456 | rs7843177 | 1.205 | 0.356 | 0.512 | 0.623 | 0.017 | 0.064 | 0.064 | 0.136 | 0.7 | 0.48 | 0.8 | 0.41 | 0.66 | 0.75 |
| 8 | 2481966 | rs10282797 | -0.729 | 0.564 | 0.36 | 0.632 | 0.02 | 0.025 | 0.025 | 0 | 0.14 | 0.3 | 0.1 | 0.23 | 0.17 | 0.06 |
| 8 | 2482413 | rs4457356 | -0.672 | 0.585 | 0.371 | 0.6 | 0.023 | 0.019 | 0.019 | 0 | 0.14 | 0.3 | 0.1 | 0.22 | 0.17 | 0.06 |
| 8 | 2484674 | rs7465625 | 0.897 | 0.643 | 0.319 | 0.652 | 0.033 | 0.078 | 0.078 | 0.019 | 0.73 | 0.59 | 0.8 | 0.44 | 0.67 | 0.77 |
| 8 | 2487969 | rs9693133 | 0.83 | 0.59 | -0.071 | 0.678 | 0.007 | 0.02 | 0.02 | 0 | 0.74 | 0.7 | 0.82 | 0.44 | 0.67 | 0.77 |
| 8 | 2497827 | rs7814276 | 0.883 | 0.705 | -0.527 | 0.61 | -0.003 | -0.001 | -0.001 | 0 | 0.06 | 0.04 | 0.02 | 0 | 0.05 | 0.02 |
| 8 | 2500349 | rs13258322 | -0.541 | 0.656 | -0.403 | 0.72 | -0.001 | 0 | 0 | 3.538 | 0.33 | 0.36 | 0.48 | 0.38 | 0.36 | 0.4 |
| 8 | 2502867 | rs9644331 | -0.327 | 0.451 | -0.772 | 1.086 | -0.06 | -0.079 | -0.079 | 0.36 | 0.88 | 0.99 | 0.92 | 0.67 | 0.82 | 0.83 |
| 8 | 2505366 | rs13257520 | -0.678 | 0.246 | -0.388 | 1.047 | 0.004 | 0.04 | 0.04 | 0 | 0.08 | 0.15 | 0.07 | 0.21 | 0.12 | 0.04 |
| 8 | 2506899 | rs10097215 | -0.175 | 0.284 | -0.764 | 1.106 | -0.011 | -0.014 | -0.014 | 0 | 0.08 | 0.05 | 0.28 | 0.21 | 0.13 | 0.06 |
| 8 | 2509106 | rs10094826 | 1.089 | -0.073 | -0.826 | 1.154 | 0.011 | -0.023 | -0.023 | 0.088 | 0.64 | 0.74 | 0.59 | 0.36 | 0.65 | 0.59 |
| 8 | 2511536 | rs4876024 | -1.052 | -0.238 | -0.654 | 0.756 | -0.017 | 0.006 | 0.006 | 0.001 | 0.28 | 0.41 | 0.4 | 0.33 | 0.23 | 0.3 |
| 8 | 2517411 | rs4409422 | -0.457 | -0.505 | -1.352 | 0.336 | 0.048 | 0.046 | 0.046 | 0 | 0.28 | 0.15 | 0.18 | 0.18 | 0.17 | 0.23 |
| 8 | 2517882 | rs6982025 | 0.292 | -0.177 | -0.891 | 0.851 | -0.008 | 0.02 | 0.02 | 0 | 0.79 | 0.73 | 0.9 | 0.69 | 0.81 | 0.82 |
| 8 | 2519571 | rs4074184 | 0.01 | 0.496 | -1.486 | 0.533 | -0.033 | -0.005 | -0.005 | 0 | 0.16 | 0.09 | 0.37 | 0.2 | 0.24 | 0.18 |
| 8 | 2520024 | rs4075492 | -0.365 | 0.701 | -1.321 | 0.078 | 0.003 | -0.001 | -0.001 | 0 | 0.39 | 0.49 | 0.56 | 0.56 | 0.49 | 0.57 |
| 8 | 2522769 | rs10453151 | -0.94 | 0.426 | -1.964 | -0.179 | 0.002 | -0.039 | -0.039 | 1.18 | 0.81 | 0.9 | 0.82 | 0.58 | 0.79 | 0.86 |
| 8 | 2523398 | rs11777237 | 0.243 | 0.683 | -2.365 | -0.982 | -0.107 | 0.095 | 0.095 | 0 | 0.23 | 0.01 | 0.15 | 0.05 | 0.26 | 0.27 |
| 8 | 2527251 | rs10481364 | 0.508 | 0.326 | -1.662 | -0.605 | 0.002 | 0.006 | 0.006 | 0 | 0.2 | 0.2 | 0.15 | 0.14 | 0.19 | 0.17 |
| 8 | 2532368 | rs12675592 | -0.611 | 0.391 | -1.849 | -0.752 | -0.044 | -0.047 | -0.047 | 1.051 | 0.45 | 0.85 | 0.55 | 0.45 | 0.48 | 0.49 |
| 8 | 2535086 | rs13278616 | 0.357 | 0.405 | -1.855 | -1.161 | -0.125 | -0.053 | -0.053 | 0.071 | 0.23 | 0.02 | 0.2 | 0.18 | 0.32 | 0.34 |
| 8 | 2536024 | rs6997305 | -1.561 | 0.161 | -1.598 | -1.294 | 0.009 | -0.039 | -0.039 | 0 | 0.66 | 0.84 | 0.7 | 0.52 | 0.68 | 0.72 |
| 8 | 2537101 | rs4876030 | -0.2 | 0.139 | -0.66 | -1.291 | -0.014 | -0.049 | -0.049 | 0 | 0.3 | 0.48 | 0.2 | 0.16 | 0.25 | 0.32 |
| 8 | 2538086 | rs13273297 | -0.153 | 0.261 | -0.771 | -1.272 | -0.058 | 0.003 | 0.003 | 0 | 0.3 | 0.12 | 0.25 | 0.22 | 0.35 | 0.49 |
| 8 | 2540711 | rs11993221 | -0.07 | 0.334 | -0.694 | -1.242 | 0.01 | -0.01 | -0.01 | 0 | 0.28 | 0.33 | 0.19 | 0.2 | 0.28 | 0.41 |
| 8 | 2542045 | rs10111822 | 0.179 | -0.061 | -0.424 | -1.555 | -0.006 | 0.02 | 0.02 | 0.153 | 0.2 | 0.07 | 0.13 | 0.13 | 0.19 | 0.21 |
| 8 | 2546456 | rs10091941 | -0.744 | -0.027 | 0.092 | -1.454 | -0.015 | -0.02 | -0.02 | 0 | 0.24 | 0.48 | 0.15 | 0.2 | 0.21 | 0.14 |
| 8 | 2548916 | rs13439672 | -0.831 | -0.093 | -0.702 | -1.537 | 0.002 | 0.105 | 0.105 | 0 | 0.43 | 0.73 | 0.31 | 0.17 | 0.39 | 0.27 |
| 8 | 2549355 | rs17515 | -1.186 | 0.317 | -0.839 | -1.314 | 0.021 | 0.047 | 0.047 | 0.298 | 0.39 | 0.84 | 0.26 | 0.27 | 0.36 | 0.24 |
| 8 | 2549380 | rs4333626 | 0.181 | 1.023 | -0.499 | -0.898 | -0.02 | -0.006 | -0.006 | 0 | 0.12 | 0.02 | 0.1 | 0.1 | 0.16 | 0.08 |
| 8 | 2552402 | rs4350007 | 0.007 | 1.059 | 0.071 | -1.232 | 0.017 | -0.063 | -0.063 | 1.654 | 0.15 | 0.18 | 0.11 | 0 | 0.18 | 0.02 |
| 8 | 2552691 | rs10087989 | -1.16 | 0.959 | 0.396 | -1.355 | 0.013 | -0.055 | -0.055 | 0 | 0.34 | 0.49 | 0.23 | 0.19 | 0.38 | 0.17 |
| 8 | 2558468 | rs4367579 | 0.2 | 0.893 | 0.28 | -1.142 | 0.003 | 0.001 | 0.001 | 0 | 0.16 | 0.23 | 0.11 | 0.18 | 0.19 | 0.15 |
| 8 | 2561906 | rs4286971 | -0.221 | 0.865 | -0.232 | -1.553 | -0.006 | 0.043 | 0.043 | 0 | 0.18 | 0.07 | 0.13 | 0 | 0.19 | 0.09 |
| 8 | 2567831 | rs10109257 | -0.287 | 0.708 | 0.229 | -1.631 | -0.008 | -0.107 | -0.107 | 0.215 | 0.16 | 0.32 | 0.12 | 0 | 0.14 | 0.03 |
| 8 | 2568467 | rs17063062 | -0.225 | 0.666 | 0.233 | -1.533 | 0.009 | -0.036 | -0.036 | 0 | 0.22 | 0.5 | 0.16 | 0.16 | 0.24 | 0.15 |
| 8 | 2568679 | rs17063065 | -0.235 | 0.662 | -0.003 | -1.541 | 0.01 | -0.045 | -0.045 | 0.644 | 0.22 | 0.55 | 0.16 | 0.16 | 0.24 | 0.15 |
| 8 | 2568692 | rs10102441 | -0.441 | 0.988 | -0.874 | -1.394 | -0.103 | 0.173 | 0.173 | 0.466 | 0.27 | 0.04 | 0.22 | 0 | 0.33 | 0.05 |
| 8 | 2576054 | rs10106748 | -0.929 | 0.067 | -0.009 | -0.278 | 0.001 | 0.003 | 0.003 | 0 | 0.35 | 0.64 | 0.34 | 0.26 | 0.34 | 0.31 |
| 8 | 2581969 | rs10503181 | -0.565 | -0.073 | 0.569 | -0.229 | -0.001 | -0.01 | -0.01 | 0 | 0.27 | 0.3 | 0.27 | 0.19 | 0.3 | 0.23 |
| 8 | 2582764 | rs7005766 | -0.544 | -0.021 | 0.117 | -0.199 | 0 | -0.009 | -0.009 | 3.882 | 0.27 | 0.29 | 0.27 | 0.19 | 0.3 | 0.3 |
| 8 | 2589993 | rs7827228 | -0.501 | -0.256 | 0.437 | 0.462 | 0.002 | 0.042 | 0.042 | 0 | 0.32 | 0.54 | 0.24 | 0.5 | 0.34 | 0.34 |
| 8 | 2590103 | rs4457357 | -0.82 | -0.579 | 0.162 | 0.569 | -0.001 | 0 | 0 | 0 | 0.41 | 0.57 | 0.55 | 0.56 | 0.43 | 0.48 |
| 8 | 2593695 | rs11136540 | 0.258 | -0.845 | 0.038 | 0.932 | -0.09 | -0.092 | -0.092 | 0 | 0.24 | 0.07 | 0.17 | 0.34 | 0.33 | 0.39 |
| 8 | 2594419 | rs4875852 | -0.082 | -0.839 | 0.029 | 0.917 | 0.012 | 0.001 | 0.001 | 0.038 | 0.09 | 0.04 | 0.32 | 0.1 | 0.09 | 0.16 |
| 8 | 2595978 | rs11988178 | -1.423 | -0.739 | 0.127 | 0.899 | -0.048 | 0.107 | 0.107 | 0.004 | 0.16 | 0.49 | 0.07 | 0.29 | 0.08 | 0.15 |
| 8 | 2596106 | rs17755518 | 1.09 | -0.735 | 0.097 | 0.81 | -0.096 | -0.079 | -0.079 | 0.651 | 0.16 | 0.02 | 0.15 | 0.23 | 0.27 | 0.26 |
| 8 | 2597331 | rs6558685 | -0.969 | -0.725 | 0.118 | 0.878 | 0.007 | 0.02 | 0.02 | 0.083 | 0.33 | 0.53 | 0.23 | 0.63 | 0.36 | 0.42 |
| 8 | 2598677 | rs7836236 | 0.792 | -2.078 | 0.221 | 1.111 | -0.165 | -0.062 | -0.062 | 0 | 0.32 | 0.04 | 0.29 | 0.29 | 0.5 | 0.34 |
| 8 | 2604187 | rs6558687 | -2.271 | -2.079 | 1.701 | 1.496 | -0.052 | 0.163 | 0.163 | 0.013 | 0.16 | 0.5 | 0.08 | 0.38 | 0.08 | 0.16 |
| 8 | 2608183 | rs7824108 | -1.016 | -2.079 | 2.125 | 1.214 | -0.027 | -0.042 | -0.042 | 0.548 | 0.08 | 0.22 | 0.03 | 0 | 0.03 | 0.03 |
| 8 | 2610043 | rs17076772 | -1.782 | -2.055 | 2.024 | 1.249 | -0.071 | 0.141 | 0.141 | 0 | 0.14 | 0.48 | 0.06 | 0.28 | 0.06 | 0.05 |
| 8 | 2619210 | rs17833641 | -0.352 | -2.083 | 1.335 | 1.039 | 0.013 | 0.007 | 0.007 | 4.589 | 0.12 | 0.09 | 0.05 | 0.06 | 0.04 | 0.05 |
| 8 | 2622605 | rs4483185 | -1.687 | -2.206 | 1.796 | 1.124 | -0.035 | -0.008 | -0.008 | 0 | 0.23 | 0.3 | 0.43 | 0.17 | 0.15 | 0.17 |
| 8 | 2623635 | rs7017610 | -2.467 | -2.313 | 1.436 | 1.245 | 0.096 | -0.153 | -0.153 | 0.206 | 0.4 | 0.91 | 0.5 | 0.51 | 0.26 | 0.31 |
| 8 | 2626135 | rs969476 | -0.727 | -2.43 | 1.559 | 1.151 | -0.002 | 0 | 0 | 0 | 0.1 | 0.11 | 0.36 | 0.11 | 0.1 | 0.12 |
| 8 | 2626253 | rs2978340 | -2.231 | -2.408 | 1.5 | 1.176 | 0.107 | -0.058 | -0.058 | 0.358 | 0.38 | 0.8 | 0.49 | 0.44 | 0.23 | 0.25 |
| 8 | 2626600 | rs17833845 | -0.653 | -2.047 | 1.428 | 1.078 | -0.009 | 0.057 | 0.057 | 0 | 0.1 | 0.17 | 0.04 | 0.26 | 0.09 | 0.07 |
| 8 | 2635152 | rs9314465 | -1.627 | -2.094 | 0.778 | 0.94 | -0.009 | -0.001 | -0.001 | 0 | 0.29 | 0.66 | 0.13 | 0.33 | 0.14 | 0.11 |
| 8 | 2643793 | rs2124154 | -1.419 | -1.847 | 0.88 | 0.963 | 0.019 | -0.022 | -0.022 | 0 | 0.22 | 0.15 | 0.36 | 0.35 | 0.15 | 0.22 |
| 8 | 2646618 | rs974120 | -1.475 | -2.037 | 1.51 | 0.793 | -0.055 | -0.061 | -0.061 | 0 | 0.2 | 0.56 | 0.13 | 0.06 | 0.11 | 0.06 |
| 8 | 2650841 | rs12550109 | -0.543 | -1.896 | 1.212 | 0.929 | 0.013 | -0.052 | -0.052 | 0 | 0.11 | 0.04 | 0.33 | 0.29 | 0.1 | 0.18 |

| | | | | | | | | | | | | | | | | |
|---|---|---|---|---|---|---|---|---|---|---|---|---|---|---|---|---|
| 8 | 2653455 | rs1026377 | -0.577 | -1.918 | 1.22 | 0.943 | 0.017 | -0.054 | -0.054 | 0 | 0.11 | 0.04 | 0.33 | 0.29 | 0.1 | 0.18 |
| 8 | 2659537 | rs1378261 | -1.013 | -1.934 | 1.848 | 0.736 | -0.008 | -0.052 | -0.052 | 0.363 | 0.09 | 0.25 | 0.07 | 0 | 0.07 | 0.02 |
| 8 | 2662135 | rs1026378 | -1.049 | -2.032 | 1.655 | 0.753 | -0.045 | 0.007 | 0.007 | 0 | 0.16 | 0.47 | 0.09 | 0.13 | 0.09 | 0.16 |
| 8 | 2664858 | rs12542624 | -2.28 | -2.111 | 1.741 | 0.9 | 0.046 | -0.008 | -0.008 | 0 | 0.36 | 0.73 | 0.29 | 0.39 | 0.24 | 0.36 |
| 8 | 2667474 | rs10104883 | -0.798 | -2.102 | 1.766 | 0.813 | -0.048 | -0.022 | -0.022 | 0 | 0.14 | 0.4 | 0.08 | 0.06 | 0.07 | 0.11 |
| 8 | 2667530 | rs746917 | -1.783 | -2.008 | 1.747 | 0.885 | -0.039 | 0.062 | 0.062 | 0 | 0.22 | 0.38 | 0.42 | 0.33 | 0.17 | 0.24 |
| 8 | 2669518 | rs12676839 | -0.954 | -1.961 | 1.176 | 0.854 | 0.007 | -0.07 | -0.07 | 0 | 0.15 | 0.04 | 0.37 | 0.34 | 0.15 | 0.25 |
| 8 | 2669970 | rs17078492 | -0.323 | -1.929 | 1.28 | 0.727 | -0.004 | -0.022 | -0.022 | 2.355 | 0.07 | 0.1 | 0.04 | 0 | 0.05 | 0.02 |
| 8 | 2671510 | rs7010867 | -1.983 | -1.913 | 0.558 | 0.82 | 0.043 | -0.029 | -0.029 | 6.389 | 0.35 | 0.73 | 0.49 | 0.45 | 0.24 | 0.41 |
| 8 | 2671622 | rs17834306 | -0.947 | -1.644 | 0.222 | 0.933 | 0.012 | -0.071 | -0.071 | 0 | 0.14 | 0.04 | 0.36 | 0.39 | 0.14 | 0.3 |
| 8 | 2681051 | rs1470995 | -1.533 | -1.805 | 0.036 | 0.724 | 0.055 | -0.017 | -0.017 | 0 | 0.39 | 0.69 | 0.5 | 0.43 | 0.26 | 0.39 |
| 8 | 2682287 | rs10216332 | -0.604 | -1.366 | 0.249 | 0.592 | -0.037 | -0.024 | -0.024 | 0 | 0.12 | 0.22 | 0.05 | 0.05 | 0.05 | 0.08 |
| 8 | 2683929 | rs2978337 | -0.366 | -1.028 | -0.935 | 0.059 | -0.01 | 0.063 | 0.063 | 0 | 0.54 | 0.85 | 0.56 | 0.36 | 0.44 | 0.36 |
| 8 | 2685370 | rs17834523 | -0.503 | -1.019 | -0.824 | 0.666 | 0.015 | -0.034 | -0.034 | 0 | 0.15 | 0.07 | 0.16 | 0.22 | 0.12 | 0.18 |
| 8 | 2695795 | rs2100485 | -1.451 | -1.47 | 0.149 | 0.483 | -0.05 | 0.001 | 0.001 | 0 | 0.22 | 0.59 | 0.2 | 0.23 | 0.12 | 0.22 |
| 8 | 2696190 | rs2978319 | -1.653 | -1.702 | 0.076 | 0.371 | -0.005 | 0 | 0 | 0 | 0.25 | 0.72 | 0.22 | 0.25 | 0.13 | 0.2 |
| 8 | 2701836 | rs17761609 | -1.06 | -1.585 | 0.687 | 0.508 | -0.009 | 0.003 | 0.003 | 0 | 0.21 | 0.25 | 0.18 | 0.27 | 0.13 | 0.24 |
| 8 | 2704193 | rs11786908 | -0.024 | -1.057 | -0.272 | -0.38 | 0.005 | -0.006 | -0.006 | 0 | 0.42 | 0.44 | 0.57 | 0.35 | 0.49 | 0.35 |
| 8 | 2704274 | rs17078711 | 0.101 | -0.878 | -0.494 | -0.614 | 0.019 | -0.009 | -0.009 | 0 | 0.34 | 0.45 | 0.52 | 0.32 | 0.41 | 0.3 |
| 8 | 2706777 | rs10111815 | 0.305 | -0.872 | -0.604 | -0.612 | -0.01 | 0.024 | 0.024 | 0 | 0.09 | 0.02 | 0.05 | 0.03 | 0.07 | 0.05 |
| 8 | 2709107 | rs2124153 | -1.538 | -1.204 | -0.587 | -0.62 | 0.021 | 0.052 | 0.052 | 0 | 0.75 | 0.55 | 0.76 | 0.67 | 0.72 | 0.64 |
| 8 | 2709332 | rs13264309 | 0.422 | -1.373 | -0.726 | -0.518 | 0.043 | 0.005 | 0.005 | 0.256 | 0.33 | 0.07 | 0.2 | 0.23 | 0.23 | 0.29 |
| 8 | 2710392 | rs2924871 | -0.52 | -1.612 | -0.59 | -0.716 | -0.027 | 0.071 | 0.071 | 0 | 0.18 | 0.34 | 0.37 | 0.34 | 0.13 | 0.26 |
| 8 | 2712829 | rs341668 | 0.01 | -1.687 | -0.773 | -0.916 | -0.001 | -0.003 | -0.003 | 0 | 0.51 | 0.48 | 0.57 | 0.41 | 0.54 | 0.41 |
| 8 | 2712873 | rs17756764 | 1.432 | -1.62 | -0.799 | -0.899 | -0.016 | 0.014 | 0.014 | 0 | 0.24 | 0.19 | 0.2 | 0.1 | 0.35 | 0.1 |
| 8 | 2713726 | rs2978310 | -0.487 | -1.747 | -0.651 | -0.945 | 0.026 | 0.026 | 0.026 | 0 | 0.45 | 0.6 | 0.3 | 0.26 | 0.32 | 0.34 |
| 8 | 2715180 | rs2942800 | -0.358 | -1.684 | -0.719 | -0.825 | -0.004 | -0.003 | -0.003 | 0.792 | 0.58 | 0.51 | 0.62 | 0.44 | 0.61 | 0.42 |
| 8 | 2717835 | rs6992851 | 1.467 | -1.565 | -0.758 | -0.599 | -0.013 | 0.136 | 0.136 | 0 | 0.07 | 0.14 | 0.3 | 0.33 | 0.05 | 0.17 |
| 8 | 2723169 | rs9644266 | -2.33 | -1.567 | -1.16 | -0.741 | -0.08 | -0.11 | -0.11 | 0.06 | 0.79 | 0.98 | 0.81 | 0.68 | 0.74 | 0.63 |
| 8 | 2723686 | rs2978339 | -2.33 | -1.567 | -1.16 | -0.547 | -0.08 | -0.14 | -0.14 | 0.038 | 0.79 | 0.98 | 0.81 | 0.59 | 0.73 | 0.63 |
| 8 | 2724253 | rs1984864 | -0.315 | -1.677 | -0.783 | -0.693 | -0.009 | -0.063 | -0.063 | 0 | 0.14 | 0.45 | 0.11 | 0.03 | 0.09 | 0.04 |
| 8 | 2726469 | rs2978352 | -2.567 | -1.741 | -0.933 | -0.686 | -0.086 | -0.153 | -0.153 | 0 | 0.8 | 0.97 | 0.81 | 0.53 | 0.74 | 0.63 |
| 8 | 2727315 | rs17761793 | 1.07 | -1.835 | -0.893 | -0.674 | 0.034 | -0.055 | -0.055 | 0 | 0.15 | 0.05 | 0.13 | 0.23 | 0.11 | 0.18 |
| 8 | 2728207 | rs2978348 | -0.422 | -1.636 | -0.72 | -0.452 | -0.023 | 0.016 | 0.016 | 0.039 | 0.15 | 0.45 | 0.12 | 0.15 | 0.1 | 0.16 |
| 8 | 2731382 | rs2924907 | -2.795 | -1.302 | -1.249 | -0.307 | -0.074 | -0.108 | -0.108 | 0 | 0.85 | 0.99 | 0.88 | 0.63 | 0.78 | 0.72 |
| 8 | 2732886 | rs341696 | 1.278 | -1.366 | -1.093 | -0.342 | -0.021 | 0.023 | 0.023 | 0 | 0.09 | 0.01 | 0.05 | 0.02 | 0.09 | 0.08 |
| 8 | 2734202 | rs341672 | 0.244 | -1.691 | -0.988 | -0.445 | -0.075 | 0.103 | 0.103 | 0 | 0.21 | 0.48 | 0.18 | 0.36 | 0.1 | 0.27 |
| 8 | 2734771 | rs4875854 | 1.948 | -1.559 | -0.91 | -0.572 | 0.015 | -0.046 | -0.046 | 0 | 0.24 | 0.28 | 0.21 | 0.08 | 0.34 | 0.11 |
| 8 | 2734916 | rs2604146 | 0.6 | -1.586 | -0.95 | -0.582 | -0.009 | 0.005 | 0.005 | 0 | 0.45 | 0.24 | 0.55 | 0.54 | 0.46 | 0.55 |
| 8 | 2735767 | rs2721280 | 0.565 | -1.629 | -0.878 | -0.424 | -0.038 | 0.021 | 0.021 | 0 | 0.69 | 0.46 | 0.76 | 0.62 | 0.81 | 0.65 |
| 8 | 2738093 | rs11987190 | 1.577 | -1.525 | -0.892 | -0.309 | 0.016 | -0.022 | -0.022 | 1.622 | 0.07 | 0.03 | 0.29 | 0.12 | 0.06 | 0.07 |
| 8 | 2740099 | rs2721278 | 1.945 | -1.433 | -0.867 | -0.497 | 0.013 | 0.015 | 0.015 | 0 | 0.18 | 0.1 | 0.07 | 0 | 0.13 | 0.1 |
| 8 | 2740502 | rs641525 | 0.597 | -1.301 | -0.534 | -0.189 | -0.026 | 0.239 | 0.239 | 0 | 0.14 | 0.45 | 0.14 | 0.36 | 0.07 | 0.25 |
| 8 | 2742187 | rs7829261 | 1.879 | -0.744 | -0.918 | -0.553 | -0.012 | 0.009 | 0.009 | 0.209 | 0.22 | 0.19 | 0.38 | 0.09 | 0.33 | 0.11 |
| 8 | 2745751 | rs13265251 | 1.398 | -0.887 | -1.563 | -0.192 | -0.115 | -0.126 | -0.126 | 2.433 | 0.26 | 0.08 | 0.24 | 0.37 | 0.33 | 0.48 |
| 8 | 2745873 | rs341704 | 0.849 | -1.467 | -1.22 | -0.586 | 0 | 0.018 | 0.018 | 0.238 | 0.47 | 0.37 | 0.6 | 0.63 | 0.49 | 0.41 |
| 8 | 2747940 | rs341709 | -0.352 | -1.571 | -1.521 | -0.506 | -0.162 | -0.081 | -0.081 | 0 | 0.63 | 0.92 | 0.71 | 0.64 | 0.56 | 0.43 |
| 8 | 2748821 | rs17078780 | 1.762 | -1.086 | -1.237 | -0.538 | 0.031 | -0.038 | -0.038 | 0 | 0.07 | 0.14 | 0.09 | 0 | 0.15 | 0.02 |
| 8 | 2748965 | rs341712 | -1.615 | -1.6 | -1.221 | -0.472 | -0.023 | 0.054 | 0.054 | 0 | 0.89 | 0.99 | 0.95 | 0.99 | 0.9 | 0.92 |
| 8 | 2749070 | rs341714 | 1.174 | -1.906 | -0.996 | -0.65 | 0.008 | 0.002 | 0.002 | 0 | 0.38 | 0.43 | 0.62 | 0.63 | 0.48 | 0.37 |
| 8 | 2751839 | rs1869657 | -0.394 | -1.777 | -1.058 | -0.405 | 0.001 | 0.038 | 0.038 | 0 | 0.51 | 0.41 | 0.41 | 0.66 | 0.46 | 0.68 |
| 8 | 2753127 | rs2721266 | 0.285 | -1.748 | -1.027 | -0.245 | 0.006 | 0.021 | 0.021 | 0 | 0.42 | 0.3 | 0.27 | 0.33 | 0.36 | 0.5 |
| 8 | 2755070 | rs2627272 | 0.392 | -1.644 | -0.877 | -0.212 | -0.031 | 0.01 | 0.01 | 0 | 0.25 | 0.31 | 0.18 | 0.29 | 0.13 | 0.2 |
| 8 | 2757991 | rs12546362 | 1.335 | -1.617 | -1.154 | -0.201 | -0.105 | -0.142 | -0.142 | 0.168 | 0.26 | 0.07 | 0.24 | 0.43 | 0.33 | 0.48 |
| 8 | 2759300 | rs341750 | -0.803 | -1.824 | -0.878 | -0.315 | 0.033 | -0.019 | -0.019 | 0.084 | 0.38 | 0.66 | 0.3 | 0.53 | 0.22 | 0.32 |
| 8 | 2760061 | rs11997932 | 0.597 | -1.807 | -0.701 | -0.39 | -0.026 | 0.032 | 0.032 | 0.672 | 0.14 | 0.29 | 0.11 | 0.24 | 0.09 | 0.13 |
| 8 | 2760178 | rs10503185 | 2.785 | -1.557 | -0.861 | -0.67 | -0.073 | 0.102 | 0.102 | 0.021 | 0.23 | 0.05 | 0.39 | 0.04 | 0.35 | 0.12 |
| 8 | 2763334 | rs2670068 | 0.191 | -1.689 | -0.77 | -0.593 | 0.037 | -0.032 | -0.032 | 0 | 0.33 | 0.22 | 0.26 | 0.57 | 0.22 | 0.33 |
| 8 | 2763790 | rs12546807 | 0.396 | -1.481 | -0.71 | -0.498 | 0.031 | 0.005 | 0.005 | 0 | 0.55 | 0.65 | 0.67 | 0.42 | 0.68 | 0.6 |
| 8 | 2766689 | rs341764 | -2.089 | -1.258 | -1.001 | -0.674 | -0.097 | 0.135 | 0.135 | 0.081 | 0.73 | 0.93 | 0.58 | 0.96 | 0.59 | 0.85 |
| 8 | 2770357 | rs3907942 | 0.638 | -2.243 | -0.552 | -0.372 | -0.006 | -0.002 | -0.002 | 0 | 0.52 | 0.5 | 0.47 | 0.4 | 0.68 | 0.61 |
| 8 | 2770936 | rs636990 | 0.145 | -2.093 | -0.608 | -0.263 | 0.005 | -0.056 | -0.056 | 0 | 0.74 | 0.87 | 0.55 | 0.42 | 0.78 | 0.66 |
| 8 | 2771252 | rs341727 | 0.537 | -1.851 | -0.319 | -0.119 | 0.021 | 0.035 | 0.035 | 1.914 | 0.31 | 0.53 | 0.28 | 0.42 | 0.33 | 0.49 |
| 8 | 2773456 | rs6981972 | 0.633 | -1.908 | -0.347 | -0.124 | 0.032 | 0.006 | 0.006 | 0.326 | 0.54 | 0.6 | 0.47 | 0.45 | 0.69 | 0.58 |
| 8 | 2774057 | rs2604135 | -0.306 | -0.818 | 0.544 | 0.182 | -0.015 | 0.074 | 0.074 | 0 | 0.85 | 0.74 | 0.83 | 0.7 | 0.88 | 0.83 |
| 8 | 2774638 | rs921777 | 1.441 | -0.712 | 0.714 | -0.224 | -0.002 | -0.1 | -0.1 | 0 | 0.42 | 0.42 | 0.27 | 0.03 | 0.44 | 0.14 |
| 8 | 2776401 | rs341731 | 0.548 | 0.169 | 0.832 | 0.454 | 0.021 | -0.013 | -0.013 | 0 | 0.18 | 0.27 | 0.15 | 0.14 | 0.24 | 0.35 |
| 8 | 2777465 | rs12675661 | -0.121 | -0.116 | 0.858 | 0.575 | 0.035 | -0.015 | -0.015 | 0 | 0.27 | 0.4 | 0.34 | 0.25 | 0.32 | 0.48 |
| 8 | 2777552 | rs2604132 | 1.549 | -0.168 | 0.735 | 0.726 | 0.024 | 0.018 | 0.018 | 1.138 | 0.51 | 0.25 | 0.43 | 0.44 | 0.49 | 0.34 |

| | | | | | | | | | | | | | | | | |
|---|---|---|---|---|---|---|---|---|---|---|---|---|---|---|---|---|
| 8 | 2777814 | rs17078893 | -0.317 | -0.038 | 0.762 | 0.796 | 0.026 | -0.012 | -0.012 | 0 | 0.27 | 0.38 | 0.33 | 0.26 | 0.32 | 0.48 |
| 8 | 2781277 | rs2627283 | 0.379 | -0.77 | 0.888 | 1.118 | 0.031 | 0.057 | 0.057 | 1.423 | 0.88 | 0.52 | 0.88 | 0.87 | 0.92 | 0.86 |
| 8 | 2783183 | rs341742 | 0.176 | -0.531 | 0.08 | 1.195 | 0.003 | 0.004 | 0.004 | 0 | 0.87 | 0.88 | 0.84 | 0.69 | 0.9 | 0.83 |
| 8 | 2784418 | rs341744 | 0.691 | -0.509 | 0.176 | 1.129 | -0.015 | 0.094 | 0.094 | 0 | 0.81 | 0.63 | 0.81 | 0.68 | 0.86 | 0.78 |
| 8 | 2788108 | rs7840244 | 0.201 | -0.55 | -0.693 | 0.668 | 0.009 | 0.005 | 0.005 | 0 | 0.36 | 0.23 | 0.36 | 0.28 | 0.33 | 0.48 |
| 8 | 2789796 | rs12682123 | -0.128 | 0.315 | -0.458 | 0.27 | -0.001 | -0.012 | -0.012 | 0 | 0.59 | 0.57 | 0.56 | 0.69 | 0.41 | 0.67 |
| 8 | 2795835 | rs583087 | 0.704 | 0.437 | -0.109 | 0.216 | 0.012 | -0.016 | -0.016 | 0.094 | 0.92 | 0.79 | 0.94 | 0.93 | 0.92 | 0.92 |
| 8 | 2796810 | rs897275 | 0.159 | 0.273 | -1.263 | -0.032 | -0.039 | -0.024 | -0.024 | 0 | 0.39 | 0.08 | 0.37 | 0.61 | 0.39 | 0.48 |
| 8 | 2797776 | rs602123 | 0.094 | -1.019 | -1.267 | -0.706 | 0.016 | 0.014 | 0.014 | 5.45 | 0.81 | 0.87 | 0.93 | 0.93 | 0.9 | 0.9 |
| 8 | 2799369 | rs646181 | 0.67 | -1.097 | -1.606 | -0.979 | 0.01 | 0.005 | 0.005 | 3.524 | 0.71 | 0.77 | 0.73 | 0.91 | 0.82 | 0.71 |
| 8 | 2801788 | rs677637 | 1.099 | -1.155 | -0.509 | -1.228 | -0.006 | -0.041 | -0.041 | 0 | 0.62 | 0.44 | 0.67 | 0.89 | 0.74 | 0.68 |
| 8 | 2802745 | rs584541 | 0.036 | -1.135 | -0.408 | -1.151 | -0.005 | -0.033 | -0.033 | 0 | 0.8 | 0.77 | 0.91 | 0.93 | 0.84 | 0.9 |
| 8 | 2802860 | rs2721290 | 0.686 | -1.154 | 0.095 | -1.159 | -0.029 | -0.104 | -0.104 | 0 | 0.76 | 0.52 | 0.86 | 0.93 | 0.83 | 0.87 |
| 8 | 2808265 | rs637954 | -0.002 | -0.944 | 0.45 | -1.022 | -0.012 | -0.056 | -0.056 | 0 | 0.72 | 0.47 | 0.69 | 0.89 | 0.77 | 0.74 |
| 8 | 2808520 | rs688579 | -0.374 | -1.029 | 0.547 | -1.059 | 0.016 | 0.026 | 0.026 | 3.948 | 0.75 | 0.78 | 0.74 | 0.89 | 0.82 | 0.72 |
| 8 | 2811241 | rs624128 | -0.214 | -1.305 | 0.343 | -0.959 | 0.03 | 0.042 | 0.042 | 3.982 | 0.73 | 0.81 | 0.66 | 0.81 | 0.81 | 0.73 |
| 8 | 2811708 | rs1979509 | -0.052 | -1.104 | -0.149 | -0.548 | -0.004 | -0.03 | -0.03 | 0 | 0.22 | 0.18 | 0.2 | 0.55 | 0.25 | 0.36 |
| 8 | 2812271 | rs865303 | -0.983 | -1.018 | -0.18 | -0.539 | 0.015 | 0.021 | 0.021 | 3.739 | 0.75 | 0.79 | 0.65 | 0.78 | 0.79 | 0.71 |
| 8 | 2813376 | rs654126 | 0.155 | -1.679 | -0.185 | -0.414 | -0.008 | 0.018 | 0.018 | 0.009 | 0.21 | 0.24 | 0.12 | 0.3 | 0.17 | 0.31 |
| 8 | 2814698 | rs2670072 | 0.62 | -1.474 | -0.175 | -0.124 | 0.005 | 0.126 | 0.126 | 0.408 | 0.07 | 0.1 | 0.05 | 0.31 | 0.09 | 0.28 |
| 8 | 2817319 | rs1872232 | 1.013 | -1.459 | -0.039 | 0.019 | 0 | 0.003 | 0.003 | 0.27 | 0.12 | 0.12 | 0.11 | 0.16 | 0.16 | 0.09 |
| 8 | 2819809 | rs4875857 | -0.716 | -1.593 | -0.017 | -0.126 | -0.016 | 0.01 | 0.01 | 0.367 | 0.24 | 0.61 | 0.22 | 0.28 | 0.18 | 0.12 |
| 8 | 2826822 | rs648885 | -0.828 | -1.783 | 0.099 | -0.268 | -0.014 | 0.018 | 0.018 | 2.744 | 0.34 | 0.54 | 0.26 | 0.55 | 0.29 | 0.39 |
| 8 | 2827474 | rs12682210 | 1.034 | -0.999 | -0.474 | 0.392 | -0.031 | -0.034 | -0.034 | 0.044 | 0.08 | 0.02 | 0.07 | 0.12 | 0.11 | 0.09 |
| 8 | 2827657 | rs2670083 | 0.511 | -1.154 | -0.09 | 0.425 | -0.005 | -0.024 | -0.024 | 2.407 | 0.15 | 0.08 | 0.07 | 0.29 | 0.17 | 0.28 |
| 8 | 2832449 | rs666276 | 0.928 | -1.086 | -0.065 | 0.397 | 0.002 | -0.006 | -0.006 | 0.383 | 0.13 | 0.09 | 0.05 | 0.28 | 0.11 | 0.24 |
| 8 | 2833794 | rs623202 | -1.152 | -0.819 | -0.214 | -0.042 | 0 | -0.001 | -0.001 | 0 | 0.75 | 0.73 | 0.65 | 0.8 | 0.82 | 0.71 |
| 8 | 2837777 | rs671958 | -0.864 | -0.335 | 0.336 | 0.387 | -0.012 | 0.046 | 0.046 | 7.133 | 0.27 | 0.57 | 0.19 | 0.41 | 0.2 | 0.31 |
| 8 | 2847793 | rs675492 | -0.958 | -0.004 | 1.182 | -0.108 | 0.081 | 0.006 | 0.006 | 4.85 | 0.07 | 0.37 | 0.06 | 0 | 0.06 | 0 |
| 8 | 2854453 | rs649296 | -1.033 | -0.107 | 1.033 | -0.147 | 0.026 | -0.012 | -0.012 | 4.788 | 0.14 | 0.5 | 0.14 | 0.11 | 0.11 | 0.04 |
| 8 | 2855892 | rs664600 | -1.322 | -0.141 | 0.945 | -0.187 | 0.001 | -0.033 | -0.033 | 0.142 | 0.16 | 0.59 | 0.15 | 0.12 | 0.12 | 0.05 |
| 8 | 2865886 | rs11984719 | 0.716 | -0.201 | 0.559 | -0.101 | -0.002 | 0 | 0 | 0 | 0.07 | 0.09 | 0.06 | 0.08 | 0.04 | 0.03 |
| 8 | 2865938 | rs13277378 | 0.818 | 0.329 | -0.116 | -0.854 | -0.036 | 0.06 | 0.06 | 0 | 0.19 | 0.09 | 0.17 | 0 | 0.26 | 0.1 |
| 8 | 2868746 | rs11786445 | 0.173 | -0.006 | 0.157 | -0.55 | -0.003 | 0.001 | 0.001 | 0 | 0.1 | 0.14 | 0.07 | 0.13 | 0.06 | 0.04 |
| 8 | 2873996 | rs659370 | -1.276 | -0.69 | -0.29 | -0.88 | 0.016 | 0.017 | 0.017 | 6.28 | 0.74 | 0.79 | 0.67 | 0.8 | 0.84 | 0.73 |
| 8 | 2875501 | rs632014 | -1.183 | -0.556 | 0.026 | -0.761 | -0.042 | -0.042 | -0.042 | 2.502 | 0.19 | 0.55 | 0.13 | 0.13 | 0.08 | 0.05 |
| 8 | 2878632 | rs592319 | 0.729 | -0.274 | 0.03 | -0.254 | -0.004 | 0.033 | 0.033 | 2.126 | 0.14 | 0.22 | 0.18 | 0.26 | 0.13 | 0.16 |
| 8 | 2879070 | rs17079099 | -0.283 | -0.167 | -0.001 | -0.336 | -0.011 | 0.045 | 0.045 | 4.244 | 0.26 | 0.56 | 0.27 | 0.38 | 0.19 | 0.21 |
| 8 | 2884693 | rs681667 | -0.992 | -0.127 | -0.614 | -0.674 | -0.001 | 0.006 | 0.006 | 0 | 0.51 | 0.5 | 0.47 | 0.37 | 0.49 | 0.32 |
| 8 | 2885465 | rs603688 | 0.064 | -0.082 | -0.623 | -0.602 | -0.005 | 0.01 | 0.01 | 1.471 | 0.23 | 0.25 | 0.25 | 0.32 | 0.18 | 0.19 |
| 8 | 2887904 | rs13258764 | 0.506 | -0.23 | -0.828 | -0.833 | -0.055 | 0.119 | 0.119 | 0 | 0.24 | 0.07 | 0.19 | 0 | 0.3 | 0.1 |
| 8 | 2891804 | rs11784845 | 0.349 | -0.246 | -0.539 | -0.824 | -0.039 | 0.1 | 0.1 | 0 | 0.26 | 0.09 | 0.2 | 0.04 | 0.32 | 0.12 |
| 8 | 2900054 | rs7826843 | -0.217 | -0.476 | -0.768 | -0.944 | -0.005 | 0.014 | 0.014 | 0 | 0.34 | 0.23 | 0.28 | 0.12 | 0.39 | 0.2 |
| 8 | 2901470 | rs10089936 | -0.568 | -0.643 | -1.172 | -0.807 | 0.006 | -0.018 | -0.018 | 0 | 0.26 | 0.23 | 0.15 | 0.43 | 0.24 | 0.38 |
| 8 | 2902276 | rs10092843 | 0.687 | -0.619 | -1.091 | -0.858 | -0.015 | -0.009 | -0.009 | 0 | 0.11 | 0.2 | 0.1 | 0.08 | 0.09 | 0.08 |
| 8 | 2902570 | rs11136562 | 0.389 | -0.642 | -0.53 | -0.869 | -0.005 | 0.004 | 0.004 | 0.129 | 0.12 | 0.4 | 0.12 | 0.08 | 0.09 | 0.08 |
| 8 | 2903020 | rs7387065 | 0.224 | -0.632 | -0.656 | -0.918 | -0.005 | 0.013 | 0.013 | 0 | 0.32 | 0.28 | 0.29 | 0.09 | 0.39 | 0.18 |
| 8 | 2903458 | rs7388550 | -1.823 | -0.679 | -0.909 | -0.814 | -0.03 | -0.022 | -0.022 | 0.249 | 0.72 | 0.58 | 0.64 | 0.76 | 0.83 | 0.72 |
| 8 | 2905177 | rs7460441 | -2.008 | -0.859 | -0.937 | -0.97 | -0.029 | -0.038 | -0.038 | 0 | 0.75 | 0.6 | 0.64 | 0.8 | 0.83 | 0.73 |
| 8 | 2907031 | rs11990503 | 0.27 | -0.659 | -1.014 | -0.833 | -0.001 | -0.001 | -0.001 | 0 | 0.26 | 0.23 | 0.3 | 0.37 | 0.31 | 0.25 |
| 8 | 2907455 | rs7830499 | 0.353 | -0.654 | -1.107 | -0.831 | -0.069 | -0.09 | -0.09 | 0 | 0.25 | 0.07 | 0.28 | 0.37 | 0.31 | 0.25 |
| 8 | 2909351 | rs13249456 | -0.052 | -0.823 | -0.997 | -0.925 | 0.023 | 0.016 | 0.016 | 0.503 | 0.47 | 0.33 | 0.49 | 0.37 | 0.59 | 0.35 |
| 8 | 2909992 | rs6558702 | -1.623 | -0.986 | -0.917 | -0.998 | -0.029 | -0.05 | -0.05 | 0.907 | 0.71 | 0.58 | 0.62 | 0.79 | 0.79 | 0.71 |
| 8 | 2910289 | rs7004774 | -1.634 | -1.021 | -0.733 | -1.022 | -0.028 | -0.051 | -0.051 | 1.138 | 0.71 | 0.58 | 0.62 | 0.8 | 0.79 | 0.71 |
| 8 | 2911102 | rs10081578 | -0.028 | -0.917 | -0.728 | -0.913 | -0.008 | 0.03 | 0.03 | 0 | 0.24 | 0.27 | 0.13 | 0.41 | 0.19 | 0.35 |
| 8 | 2911314 | rs10081466 | -0.038 | -0.915 | -0.727 | -0.912 | -0.008 | 0.03 | 0.03 | 0 | 0.24 | 0.28 | 0.13 | 0.42 | 0.19 | 0.36 |
| 8 | 2913248 | rs7818200 | -1.588 | -0.89 | -0.789 | -0.954 | -0.016 | -0.029 | -0.029 | 0 | 0.71 | 0.62 | 0.62 | 0.76 | 0.78 | 0.7 |
| 8 | 2915398 | rs11988388 | -0.653 | -1.008 | -0.99 | -1.003 | -0.011 | 0.006 | 0.006 | 0 | 0.59 | 0.54 | 0.57 | 0.43 | 0.68 | 0.42 |
| 8 | 2915530 | rs13251747 | 1.723 | -0.806 | -1.307 | -1.009 | -0.071 | 0.127 | 0.127 | 0.113 | 0.22 | 0.04 | 0.18 | 0 | 0.29 | 0.1 |
| 8 | 2915862 | rs1993679 | 1.033 | -0.728 | -1.005 | -0.642 | -0.023 | -0.057 | -0.057 | 0 | 0.1 | 0 | 0.04 | 0.31 | 0.1 | 0.28 |
| 8 | 2919891 | rs10092374 | 0.265 | -0.713 | -0.65 | -0.695 | -0.024 | -0.007 | -0.007 | 0.111 | 0.14 | 0.33 | 0.25 | 0.12 | 0.1 | 0.08 |
| 8 | 2920183 | rs6995874 | 0.21 | -0.729 | -0.637 | -0.73 | -0.025 | -0.031 | -0.031 | 0.85 | 0.15 | 0.33 | 0.1 | 0.09 | 0.1 | 0.08 |
| 8 | 2923948 | rs11781078 | 2.095 | -0.161 | -1.055 | -0.961 | -0.042 | 0.037 | 0.037 | 0 | 0.1 | 0.04 | 0.1 | 0 | 0.19 | 0.11 |
| 8 | 2923969 | rs7013027 | -0.195 | -0.39 | -1.152 | -0.913 | 0.002 | 0.003 | 0.003 | 6.262 | 0.36 | 0.34 | 0.49 | 0.39 | 0.53 | 0.36 |
| 8 | 2924014 | rs6993396 | -0.548 | -0.619 | -1.717 | -0.595 | -0.027 | 0.057 | 0.057 | 0 | 0.45 | 0.85 | 0.47 | 0.34 | 0.42 | 0.37 |
| 8 | 2924238 | rs6992732 | -1.087 | -0.53 | -1.356 | -0.745 | 0.005 | -0.002 | -0.002 | 0.132 | 0.54 | 0.58 | 0.68 | 0.52 | 0.65 | 0.44 |
| 8 | 2924429 | rs7845867 | -2.48 | -0.477 | -1.142 | -0.744 | -0.016 | -0.028 | -0.028 | 0.024 | 0.77 | 0.72 | 0.82 | 0.83 | 0.84 | 0.71 |
| 8 | 2924890 | rs4335144 | 1.831 | 0.43 | -0.76 | -0.381 | 0.031 | -0.052 | -0.052 | 0 | 0.12 | 0.24 | 0.13 | 0 | 0.19 | 0.11 |
| 8 | 2926490 | rs2406583 | -0.127 | 0.217 | -0.894 | -0.331 | 0.042 | 0.015 | 0.015 | 3.762 | 0.32 | 0.48 | 0.44 | 0.34 | 0.49 | 0.31 |

| | | | | | | | | | | | | | | | | |
|---|---|---|---|---|---|---|---|---|---|---|---|---|---|---|---|---|
| 8 | 2930476 | rs1563863 | -0.939 | 0.044 | -0.905 | -0.355 | 0.002 | 0 | 0 | 0 | 0.48 | 0.54 | 0.53 | 0.46 | 0.61 | 0.42 |
| 8 | 2934938 | rs12681676 | -0.178 | 0.227 | -0.904 | -0.295 | -0.003 | -0.007 | -0.007 | 0 | 0.28 | 0.29 | 0.32 | 0.41 | 0.29 | 0.27 |
| 8 | 2935178 | rs11136568 | -1.139 | 0.168 | -0.792 | -0.339 | 0.018 | 0.005 | 0.005 | 0.049 | 0.5 | 0.58 | 0.53 | 0.46 | 0.58 | 0.41 |
| 8 | 2935525 | rs11986105 | 0.808 | 0.53 | -0.731 | -0.061 | -0.01 | 0.062 | 0.062 | 0.036 | 0.18 | 0.29 | 0.23 | 0.33 | 0.13 | 0.16 |
| 8 | 2936347 | rs10081479 | -0.323 | 0.313 | -0.779 | -0.14 | -0.003 | -0.004 | -0.004 | 0 | 0.31 | 0.29 | 0.35 | 0.46 | 0.33 | 0.28 |
| 8 | 2936461 | rs1020833 | -1.877 | 0.198 | -0.923 | 0.039 | -0.002 | -0.001 | -0.001 | 0 | 0.7 | 0.68 | 0.63 | 0.72 | 0.74 | 0.68 |
| 8 | 2941053 | rs1108095 | -0.618 | 0.315 | -1.183 | -0.026 | 0.002 | 0.003 | 0.003 | 0 | 0.39 | 0.45 | 0.39 | 0.43 | 0.41 | 0.29 |
| 8 | 2946196 | rs1870616 | -1.283 | 0.694 | -1.147 | 0.459 | 0.019 | 0.039 | 0.039 | 3.44 | 0.56 | 0.65 | 0.53 | 0.65 | 0.61 | 0.61 |
| 8 | 2950533 | rs17390162 | 0.798 | 0.888 | -2.116 | 0.179 | 0.02 | 0.007 | 0.007 | 0 | 0.2 | 0.1 | 0.13 | 0.18 | 0.16 | 0.31 |
| 8 | 2950798 | rs7818488 | -0.4 | 1.183 | -1.973 | 0.202 | -0.005 | 0.024 | 0.024 | 0.002 | 0.35 | 0.27 | 0.25 | 0.22 | 0.37 | 0.43 |
| 8 | 2951965 | rs7013604 | -0.185 | 1.864 | -1.685 | 0.593 | -0.008 | -0.012 | -0.012 | 0 | 0.19 | 0.14 | 0.28 | 0.39 | 0.25 | 0.19 |
| 8 | 2953622 | rs11776711 | 0.757 | 1.563 | -1.199 | -0.062 | -0.006 | -0.006 | -0.006 | 0 | 0.1 | 0.06 | 0.1 | 0 | 0.19 | 0.08 |
| 8 | 2955158 | rs7826548 | 0.015 | 1.025 | -1.317 | -0.035 | -0.008 | 0.041 | 0.041 | 0 | 0.53 | 0.35 | 0.59 | 0.73 | 0.53 | 0.68 |
| 8 | 2955341 | rs13253818 | 0.384 | 0.925 | -1.854 | 0.151 | -0.061 | -0.045 | -0.045 | 0.755 | 0.27 | 0.01 | 0.21 | 0.2 | 0.25 | 0.24 |
| 8 | 2956230 | rs11782360 | 0.456 | 1.572 | -1.067 | 0.107 | -0.026 | 0.02 | 0.02 | 0 | 0.09 | 0.03 | 0.11 | 0 | 0.18 | 0.06 |
| 8 | 2957974 | rs11787300 | 0.241 | 1.545 | -0.165 | 0.063 | 0.111 | -0.062 | -0.062 | 0 | 0.1 | 0.23 | 0.12 | 0 | 0.19 | 0.06 |
| 8 | 2959872 | rs4875731 | 1.382 | 1.268 | -1.192 | 0.118 | -0.053 | -0.064 | -0.064 | 0 | 0.46 | 0.12 | 0.54 | 0.46 | 0.48 | 0.57 |
| 8 | 2962919 | rs13277425 | 0.088 | 1.481 | -1.155 | 0.14 | -0.037 | -0.018 | -0.018 | 2.122 | 0.11 | 0.03 | 0.09 | 0.1 | 0.11 | 0.06 |
| 8 | 2964206 | rs4875742 | 0.23 | 1.525 | -0.707 | 0 | 0.027 | -0.051 | -0.051 | 0 | 0.12 | 0.22 | 0.13 | 0 | 0.19 | 0.07 |
| 8 | 2964705 | rs11993207 | -0.088 | 1.508 | -0.643 | 0.023 | -0.021 | 0.011 | 0.011 | 0 | 0.13 | 0.06 | 0.09 | 0.07 | 0.12 | 0.08 |
| 8 | 2965244 | rs3824271 | 0.142 | 1.391 | -0.459 | -0.068 | 0.048 | -0.063 | -0.063 | 0 | 0.86 | 0.74 | 0.85 | 0.95 | 0.8 | 0.91 |
| 8 | 2966328 | rs3802292 | 1.632 | 1.307 | -0.117 | 0.064 | 0 | -0.001 | -0.001 | 0 | 0.57 | 0.41 | 0.6 | 0.56 | 0.51 | 0.62 |
| 8 | 2966852 | rs750318 | -0.193 | 1.33 | 0.107 | 0.254 | 0.047 | -0.051 | -0.051 | 0.299 | 0.88 | 0.76 | 0.87 | 0.95 | 0.83 | 0.95 |
| 8 | 2967472 | rs4995273 | 0.09 | 1.229 | -0.084 | 0.414 | -0.007 | 0.004 | 0.004 | 0 | 0.16 | 0.23 | 0.14 | 0.14 | 0.15 | 0.11 |
| 8 | 2968777 | rs12542474 | 1.529 | 1.065 | -0.728 | 0.411 | -0.065 | -0.027 | -0.027 | 1.702 | 0.56 | 0.11 | 0.59 | 0.55 | 0.55 | 0.65 |
| 8 | 2970507 | rs10112871 | 0.485 | 1.073 | -0.108 | 0.299 | 0.027 | -0.002 | -0.002 | 0 | 0.92 | 0.66 | 0.94 | 0.95 | 0.95 | 0.97 |
| 8 | 2970539 | rs10503196 | -1.002 | 1.398 | -0.585 | 0.521 | 0.025 | -0.012 | -0.012 | 1.1 | 0.25 | 0.43 | 0.28 | 0.15 | 0.31 | 0.16 |
| 8 | 2971213 | rs10106035 | 0.579 | 0.496 | -0.837 | 0.463 | -0.001 | 0.024 | 0.024 | 2.613 | 0.73 | 0.42 | 0.59 | 0.43 | 0.76 | 0.7 |
| 8 | 2973853 | rs7002853 | -0.487 | 0.2 | -2.012 | -0.462 | -0.153 | 0.149 | 0.149 | 0 | 0.66 | 0.96 | 0.41 | 0.89 | 0.45 | 0.78 |
| 8 | 2975448 | rs12550459 | -0.155 | -0.132 | -1.905 | 0.5 | 0.018 | -0.03 | -0.03 | 0 | 0.1 | 0.03 | 0.03 | 0.35 | 0.04 | 0.12 |
| 8 | 2975578 | rs7841865 | 0.161 | -0.616 | -1.048 | 0.121 | 0.006 | -0.062 | -0.062 | 0 | 0.11 | 0.36 | 0.07 | 0 | 0.07 | 0.03 |
| 8 | 2975781 | rs2045637 | -1.324 | -0.736 | -1.272 | 0.053 | 0.039 | -0.061 | -0.061 | 0 | 0.23 | 0.8 | 0.15 | 0.36 | 0.12 | 0.15 |
| 8 | 2975905 | rs2045638 | -1.482 | -0.881 | -1.711 | 0.02 | -0.001 | -0.143 | -0.143 | 0 | 0.34 | 0.9 | 0.19 | 0.48 | 0.22 | 0.39 |
| 8 | 2977909 | rs2045640 | -1.335 | -0.907 | -1.745 | -0.018 | 0.011 | -0.12 | -0.12 | 0 | 0.36 | 0.88 | 0.19 | 0.42 | 0.23 | 0.31 |
| 8 | 2980976 | rs7837404 | -0.577 | -0.773 | -1.417 | -0.017 | 0.007 | 0.029 | 0.029 | 0.448 | 0.26 | 0.79 | 0.15 | 0.13 | 0.18 | 0.24 |
| 8 | 2982351 | rs7842818 | -0.77 | -0.592 | -1.438 | 0.063 | 0.009 | -0.062 | -0.062 | 1.145 | 0.37 | 0.85 | 0.19 | 0.49 | 0.26 | 0.37 |
| 8 | 2995761 | rs13273359 | 0.158 | -0.203 | -0.963 | -0.039 | -0.002 | 0.002 | 0.002 | 0.386 | 0.87 | 0.82 | 0.68 | 0.85 | 0.89 | 0.87 |
| 8 | 2996012 | rs10046663 | 0.4 | 0.053 | -1.357 | -0.484 | -0.139 | 0.212 | 0.212 | 0.831 | 0.34 | 0.07 | 0.34 | 0 | 0.53 | 0.19 |
| 8 | 3000462 | rs12545633 | 0.115 | -0.048 | -0.9 | 0.245 | -0.001 | 0.012 | 0.012 | 0.713 | 0.42 | 0.28 | 0.45 | 0.33 | 0.54 | 0.32 |
| 8 | 3002967 | rs2406358 | -1.318 | 0.432 | 0.076 | 1.19 | 0.112 | 0.447 | 0.447 | 0 | 0.07 | 0.34 | 0.14 | 0.37 | 0.08 | 0.12 |
| 8 | 3004567 | rs1545702 | 0.531 | -0.259 | 0.098 | 1.082 | -0.014 | 0.095 | 0.095 | 0.473 | 0.76 | 0.57 | 0.76 | 0.59 | 0.82 | 0.75 |
| 8 | 3013464 | rs13250365 | 0.228 | -0.235 | -0.376 | 0.923 | -0.02 | 0.043 | 0.043 | 0 | 0.08 | 0.01 | 0.06 | 0 | 0.1 | 0.03 |
| 8 | 3014491 | rs11989498 | -0.767 | -0.377 | 0.114 | 1.211 | 0.017 | 0.207 | 0.207 | 2.231 | 0.06 | 0.28 | 0.05 | 0.26 | 0.05 | 0.12 |
| 8 | 3018383 | rs2406305 | 0.366 | -0.293 | -0.426 | 1.658 | -0.06 | -0.081 | -0.081 | 1.733 | 0.83 | 0.99 | 0.73 | 0.78 | 0.8 | 0.87 |
| 8 | 3019592 | rs11998250 | -0.807 | -0.938 | 0.403 | 1.466 | 0.005 | 0.139 | 0.139 | 0 | 0.15 | 0.38 | 0.12 | 0.36 | 0.17 | 0.24 |
| 8 | 3021495 | rs1460404 | -1.18 | -0.941 | -0.148 | 1.088 | -0.019 | 0.004 | 0.004 | 1.346 | 0.26 | 0.66 | 0.17 | 0.3 | 0.19 | 0.25 |
| 8 | 3024326 | rs7822051 | -0.925 | -0.482 | -0.098 | 0.871 | -0.012 | -0.025 | -0.025 | 1.925 | 0.22 | 0.66 | 0.16 | 0.14 | 0.19 | 0.25 |
| 8 | 3025657 | rs17079527 | -0.364 | -0.43 | -0.453 | 1.063 | -0.003 | 0.002 | 0.002 | 0 | 0.22 | 0.24 | 0.39 | 0.61 | 0.15 | 0.42 |
| 8 | 3030335 | rs11136586 | -0.29 | -0.092 | -0.774 | 0.127 | 0.003 | 0.008 | 0.008 | 3.252 | 0.31 | 0.22 | 0.24 | 0.13 | 0.33 | 0.17 |
| 8 | 3030793 | rs4875090 | -0.56 | 0.216 | -1.573 | 0.547 | -0.076 | -0.047 | -0.047 | 0 | 0.65 | 0.91 | 0.49 | 0.66 | 0.61 | 0.75 |
| 8 | 3032815 | rs7827348 | -0.132 | -0.397 | -0.884 | 0.878 | -0.005 | 0.004 | 0.004 | 0 | 0.33 | 0.58 | 0.25 | 0.59 | 0.2 | 0.49 |
| 8 | 3032861 | rs2440721 | -0.104 | -0.251 | -0.641 | 0.784 | -0.032 | 0.027 | 0.027 | 0.157 | 0.12 | 0.3 | 0.08 | 0.16 | 0.07 | 0.2 |
| 8 | 3033518 | rs7828114 | -0.232 | -0.151 | -0.708 | 1.034 | 0.001 | 0.02 | 0.02 | 0.006 | 0.2 | 0.19 | 0.24 | 0.46 | 0.18 | 0.34 |
| 8 | 3035053 | rs7004125 | -0.653 | -0.272 | -0.769 | 1.074 | -0.021 | 0.024 | 0.024 | 0 | 0.31 | 0.42 | 0.42 | 0.44 | 0.2 | 0.39 |
| 8 | 3035124 | rs259856 | -1.128 | -0.072 | -1.915 | 0.444 | -0.127 | 0.188 | 0.188 | 0.123 | 0.55 | 0.95 | 0.5 | 0.84 | 0.45 | 0.7 |
| 8 | 3035669 | rs17079553 | -0.062 | 0.114 | -1.131 | 0.902 | 0 | 0.003 | 0.003 | 0 | 0.11 | 0.15 | 0.05 | 0.15 | 0.1 | 0.1 |
| 8 | 3035992 | rs17319073 | 0.41 | 0.291 | -1.126 | 0.692 | 0.002 | -0.017 | -0.017 | 0 | 0.05 | 0.06 | 0.06 | 0 | 0.06 | 0.05 |
| 8 | 3037530 | rs1460402 | 0.046 | 0.365 | -1.603 | 0.955 | -0.033 | -0.056 | -0.056 | 0.502 | 0.11 | 0.01 | 0.22 | 0.14 | 0.17 | 0.13 |
| 8 | 3042361 | rs10090467 | 1.142 | 0.63 | -2.313 | 0.465 | -0.072 | 0.016 | 0.016 | 0 | 0.25 | 0.11 | 0.29 | 0.19 | 0.33 | 0.15 |
| 8 | 3042665 | rs10099847 | 1.142 | 0.63 | -2.313 | 0.465 | -0.072 | 0.016 | 0.016 | 0 | 0.25 | 0.11 | 0.29 | 0.19 | 0.33 | 0.15 |
| 8 | 3043312 | rs2730048 | 0.317 | -0.347 | -1.645 | 0.325 | -0.001 | -0.002 | -0.002 | 0 | 0.46 | 0.53 | 0.37 | 0.52 | 0.42 | 0.6 |
| 8 | 3044961 | rs10216980 | 1.698 | -0.321 | -1.818 | 0.32 | -0.084 | 0.017 | 0.017 | 0 | 0.26 | 0.1 | 0.29 | 0.19 | 0.33 | 0.15 |
| 8 | 3045187 | rs2125952 | 0.348 | -0.402 | -1.452 | 0.331 | -0.001 | -0.005 | -0.005 | 0 | 0.46 | 0.54 | 0.37 | 0.52 | 0.42 | 0.6 |
| 8 | 3045230 | rs17319192 | 1.615 | -0.183 | -1.518 | 0.401 | -0.081 | 0.014 | 0.014 | 0 | 0.25 | 0.11 | 0.29 | 0.19 | 0.33 | 0.15 |
| 8 | 3049023 | rs922162 | 0.314 | -0.536 | -0.958 | 0.21 | 0 | 0.002 | 0.002 | 0 | 0.43 | 0.4 | 0.36 | 0.53 | 0.41 | 0.6 |
| 8 | 3049799 | rs10086940 | 0.009 | -1.204 | -0.787 | 0.438 | -0.002 | 0.007 | 0.007 | 0 | 0.68 | 0.64 | 0.68 | 0.59 | 0.73 | 0.55 |
| 8 | 3049850 | rs7016896 | -1.151 | -1.515 | -0.81 | 0.455 | 0.009 | 0.001 | 0.001 | 0 | 0.86 | 0.9 | 0.93 | 0.82 | 0.94 | 0.86 |
| 8 | 3050110 | rs17079568 | 0.705 | -1.591 | -0.861 | 0.458 | -0.039 | -0.008 | -0.008 | 2.776 | 0.17 | 0.19 | 0.08 | 0.17 | 0.06 | 0.18 |
| 8 | 3050524 | rs7816370 | 0.262 | -1.48 | -1.392 | 0.531 | 0.028 | 0.004 | 0.004 | 0 | 0.22 | 0.07 | 0.09 | 0.2 | 0.17 | 0.21 |

| | | | | | | | | | | | | | | | | |
|---|---|---|---|---|---|---|---|---|---|---|---|---|---|---|---|---|
| 8 | 3051115 | rs17079571 | 1.093 | -1.215 | -1.518 | 0.642 | -0.028 | -0.056 | -0.056 | 0 | 0.13 | 0.01 | 0.23 | 0.14 | 0.17 | 0.1 |
| 8 | 3051156 | rs13261550 | 1.143 | -1.297 | -1.656 | 0.62 | -0.026 | 0.027 | 0.027 | 0.045 | 0.42 | 0.22 | 0.4 | 0.36 | 0.55 | 0.26 |
| 8 | 3051367 | rs983779 | -0.374 | -1.229 | -1.375 | 0.663 | 0.035 | -0.005 | -0.005 | 0 | 0.36 | 0.19 | 0.35 | 0.34 | 0.34 | 0.29 |
| 8 | 3052978 | rs11775083 | 2.532 | -0.421 | -1.299 | 0.376 | 0.013 | -0.021 | -0.021 | 2.495 | 0.08 | 0.05 | 0.08 | 0 | 0.18 | 0.05 |
| 8 | 3053592 | rs1077153 | 1.899 | -0.699 | -1.17 | 0.635 | 0.007 | -0.012 | -0.012 | 0 | 0.09 | 0.03 | 0.04 | 0.08 | 0.05 | 0.1 |
| 8 | 3054780 | rs2098764 | -0.991 | -0.988 | -0.618 | 0.695 | -0.035 | -0.003 | -0.003 | 0 | 0.8 | 0.75 | 0.88 | 0.76 | 0.94 | 0.78 |
| 8 | 3054804 | rs7823098 | 0.124 | -1.002 | -0.749 | 0.85 | 0.006 | -0.008 | -0.008 | 0 | 0.63 | 0.41 | 0.67 | 0.52 | 0.77 | 0.46 |
| 8 | 3055301 | rs6558735 | 1.458 | -0.504 | -0.871 | 0.939 | 0.023 | 0.051 | 0.051 | 0 | 0.17 | 0.25 | 0.21 | 0.24 | 0.17 | 0.32 |
| 8 | 3058148 | rs12545450 | 2.066 | -0.445 | -1.166 | 0.783 | -0.032 | -0.002 | -0.002 | 0 | 0.18 | 0.11 | 0.25 | 0.16 | 0.25 | 0.11 |
| 8 | 3060078 | rs17319387 | 1.995 | -0.431 | -1.173 | 0.828 | -0.021 | -0.015 | -0.015 | 0 | 0.18 | 0.14 | 0.29 | 0.2 | 0.25 | 0.11 |
| 8 | 3061993 | rs1442398 | 0.596 | -0.726 | -0.955 | 0.769 | 0.017 | -0.01 | -0.01 | 0 | 0.37 | 0.44 | 0.43 | 0.29 | 0.43 | 0.45 |
| 8 | 3062695 | rs1867044 | -1.643 | -0.953 | -1.435 | 0.801 | 0.021 | 0.001 | 0.001 | 0 | 0.86 | 0.89 | 0.92 | 0.82 | 0.95 | 0.86 |
| 8 | 3064489 | rs10085969 | -0.137 | -0.963 | -1.552 | 0.68 | 0.052 | -0.003 | -0.003 | 0 | 0.64 | 0.74 | 0.66 | 0.64 | 0.74 | 0.66 |
| 8 | 3064568 | rs10086891 | -0.134 | -0.969 | -1.554 | 0.746 | 0.052 | -0.017 | -0.017 | 0.558 | 0.64 | 0.74 | 0.66 | 0.61 | 0.74 | 0.66 |
| 8 | 3065016 | rs4875110 | -0.245 | -0.97 | -1.553 | 0.668 | 0.053 | -0.003 | -0.003 | 0 | 0.64 | 0.74 | 0.66 | 0.64 | 0.75 | 0.66 |
| 8 | 3065125 | rs4875111 | -0.852 | -0.958 | -1.509 | 0.701 | 0.022 | -0.005 | -0.005 | 0 | 0.74 | 0.79 | 0.7 | 0.7 | 0.79 | 0.76 |
| 8 | 3066746 | rs6651442 | -0.083 | -0.44 | -1.253 | 0.842 | -0.021 | -0.024 | -0.024 | 0.563 | 0.44 | 0.66 | 0.39 | 0.51 | 0.48 | 0.5 |
| 8 | 3067724 | rs10101045 | -0.103 | -0.436 | -1.286 | 0.836 | -0.022 | -0.024 | -0.024 | 0 | 0.44 | 0.66 | 0.39 | 0.51 | 0.49 | 0.52 |
| 8 | 3069893 | rs10099364 | 0.62 | -0.181 | -1.142 | 0.937 | 0.003 | 0.032 | 0.032 | 0 | 0.26 | 0.53 | 0.23 | 0.35 | 0.31 | 0.2 |
| 8 | 3072322 | rs17079629 | 1.164 | 0.566 | -1.208 | 0.978 | -0.074 | -0.063 | -0.063 | 0 | 0.14 | 0.01 | 0.14 | 0.17 | 0.15 | 0.27 |
| 8 | 3072722 | rs11988224 | 0.642 | 0.188 | -0.481 | 0.998 | 0.005 | 0.032 | 0.032 | 0.288 | 0.26 | 0.42 | 0.35 | 0.32 | 0.27 | 0.18 |
| 8 | 3074711 | rs259866 | 0.078 | -0.07 | -0.465 | 0.756 | -0.002 | -0.003 | -0.003 | 0 | 0.13 | 0.17 | 0.23 | 0.09 | 0.17 | 0.09 |
| 8 | 3080184 | rs17079645 | -0.232 | -0.241 | -0.524 | 0.731 | -0.001 | 0.022 | 0.022 | 0.684 | 0.06 | 0.13 | 0.04 | 0.12 | 0.06 | 0.1 |
| 8 | 3081903 | rs17079652 | -1.339 | -0.12 | -0.527 | 0.908 | -0.008 | 0.032 | 0.032 | 0.93 | 0.24 | 0.29 | 0.27 | 0.32 | 0.25 | 0.23 |
| 8 | 3084216 | rs3943652 | 0.707 | -0.341 | -0.615 | 0.933 | -0.002 | 0.014 | 0.014 | 0.373 | 0.08 | 0.06 | 0.06 | 0.17 | 0.06 | 0.05 |
| 8 | 3086423 | rs13256014 | 0.488 | -0.546 | -0.82 | 0.418 | -0.001 | -0.004 | -0.004 | 0 | 0.27 | 0.28 | 0.21 | 0.13 | 0.26 | 0.44 |
| 8 | 3086808 | rs7845677 | 1.051 | -0.958 | -1.189 | 0.247 | -0.068 | 0.055 | 0.055 | 0.058 | 0.32 | 0.17 | 0.49 | 0.17 | 0.41 | 0.23 |
| 8 | 3087809 | rs1442403 | -0.7 | -1.343 | -1.101 | 0.322 | -0.023 | 0.11 | 0.11 | 0 | 0.84 | 0.65 | 0.86 | 0.6 | 0.87 | 0.86 |
| 8 | 3088422 | rs1477423 | -0.013 | -1.301 | -1.2 | 0.26 | -0.015 | 0.074 | 0.074 | 0 | 0.79 | 0.64 | 0.7 | 0.53 | 0.78 | 0.85 |
| 8 | 3088577 | rs12216867 | 1.864 | -1.253 | -1.266 | 0.105 | -0.046 | 0.047 | 0.047 | 0.022 | 0.27 | 0.17 | 0.34 | 0.11 | 0.32 | 0.22 |
| 8 | 3089598 | rs4546683 | 0.578 | -1.443 | -1.119 | 0.222 | -0.011 | -0.005 | -0.005 | 0 | 0.6 | 0.56 | 0.57 | 0.36 | 0.61 | 0.77 |
| 8 | 3089739 | rs1467980 | 0.354 | -1.614 | -1.207 | 0.045 | -0.017 | -0.008 | -0.008 | 0.381 | 0.19 | 0.24 | 0.16 | 0.13 | 0.13 | 0.29 |
| 8 | 3091084 | rs1562116 | 0.267 | -1.525 | -1.194 | 0.133 | -0.008 | 0.011 | 0.011 | 0 | 0.68 | 0.61 | 0.64 | 0.53 | 0.67 | 0.8 |
| 8 | 3091584 | rs1442407 | 0.221 | -1.45 | -1.127 | 0.123 | 0.003 | -0.007 | -0.007 | 0 | 0.15 | 0.09 | 0.23 | 0.23 | 0.16 | 0.09 |
| 8 | 3092375 | rs1442408 | 0.053 | -1.506 | -1.132 | 0.044 | -0.019 | 0.066 | 0.066 | 0 | 0.75 | 0.61 | 0.67 | 0.43 | 0.72 | 0.73 |
| 8 | 3092572 | rs1442410 | 0.642 | -1.487 | -1.143 | -0.019 | -0.032 | -0.018 | -0.018 | 0.011 | 0.66 | 0.53 | 0.6 | 0.25 | 0.67 | 0.7 |
| 8 | 3092889 | rs7002936 | 1.517 | -1.408 | -0.979 | -0.024 | -0.011 | -0.047 | -0.047 | 1.061 | 0.12 | 0.15 | 0.07 | 0 | 0.1 | 0.04 |
| 8 | 3093194 | rs7814326 | 0.576 | -1.314 | -0.888 | 0.084 | 0.002 | -0.045 | -0.045 | 0 | 0.28 | 0.35 | 0.21 | 0.11 | 0.25 | 0.43 |
| 8 | 3093985 | rs7819976 | 0.448 | -1.135 | -0.901 | 0.155 | -0.002 | 0.004 | 0.004 | 0.027 | 0.29 | 0.21 | 0.17 | 0.17 | 0.27 | 0.23 |
| 8 | 3101584 | rs1020347 | -0.233 | -1.128 | -0.538 | 0.367 | -0.038 | 0.116 | 0.116 | 0 | 0.89 | 0.64 | 0.87 | 0.77 | 0.93 | 0.89 |
| 8 | 3101916 | rs10104274 | 0.322 | -1.077 | -0.498 | 0.39 | -0.017 | 0.098 | 0.098 | 0 | 0.85 | 0.63 | 0.71 | 0.71 | 0.83 | 0.85 |
| 8 | 3105800 | rs17319624 | 0.34 | -0.838 | -1.046 | -0.247 | 0.004 | 0.033 | 0.033 | 0 | 0.17 | 0.1 | 0.24 | 0.08 | 0.21 | 0.06 |
| 8 | 3115165 | rs2081279 | 1.356 | -0.901 | -1.115 | -0.494 | -0.005 | 0.012 | 0.012 | 0 | 0.31 | 0.28 | 0.34 | 0.15 | 0.32 | 0.29 |
| 8 | 3117251 | rs17319673 | 0.287 | -0.867 | -1.234 | -0.433 | 0 | -0.003 | -0.003 | 0 | 0.21 | 0.2 | 0.15 | 0.06 | 0.19 | 0.06 |
| 8 | 3121966 | rs9650505 | 0.221 | -0.694 | -1.5 | -0.314 | -0.003 | 0.004 | 0.004 | 0 | 0.67 | 0.62 | 0.72 | 0.41 | 0.7 | 0.71 |
| 8 | 3128099 | rs6991834 | 0.63 | 0.073 | -1.018 | 0.224 | 0.03 | 0.057 | 0.057 | 0 | 0.23 | 0.34 | 0.31 | 0.28 | 0.24 | 0.33 |
| 8 | 3134810 | rs905502 | -0.499 | -0.085 | -1.319 | 0.126 | 0.016 | -0.008 | -0.008 | 0 | 0.58 | 0.62 | 0.68 | 0.48 | 0.63 | 0.65 |
| 8 | 3135398 | rs17391851 | 0.206 | 0.332 | -1.681 | -0.055 | -0.011 | -0.017 | -0.017 | 0 | 0.06 | 0.01 | 0.05 | 0.07 | 0.09 | 0.05 |
| 8 | 3135739 | rs10110184 | 0.276 | 0.513 | -1.357 | -0.11 | -0.019 | 0.025 | 0.025 | 0 | 0.27 | 0.21 | 0.29 | 0.12 | 0.3 | 0.27 |
| 8 | 3136580 | rs17391886 | -0.945 | 0.104 | -1.211 | 0.381 | -0.012 | 0.004 | 0.004 | 0.02 | 0.13 | 0.02 | 0.09 | 0.14 | 0.13 | 0.21 |
| 8 | 3136833 | rs7821628 | -1.517 | -0.164 | -0.23 | 0.274 | -0.017 | -0.036 | -0.036 | 0 | 0.18 | 0.38 | 0.13 | 0.17 | 0.14 | 0.2 |
| 8 | 3137646 | rs6558750 | 1.255 | -0.156 | 0.355 | 0.119 | -0.006 | -0.028 | -0.028 | 7.402 | 0.82 | 0.74 | 0.87 | 0.83 | 0.81 | 0.83 |
| 8 | 3139251 | rs13269823 | -0.528 | -0.127 | 0.357 | 0.085 | 0.001 | -0.03 | -0.03 | 0 | 0.07 | 0.14 | 0.04 | 0 | 0.07 | 0.03 |
| 8 | 3139579 | rs10105711 | 1.456 | -0.346 | 0.258 | 0.52 | -0.028 | 0.028 | 0.028 | 0 | 0.76 | 0.52 | 0.82 | 0.68 | 0.77 | 0.69 |
| 8 | 3140821 | rs9650506 | 1.563 | -0.224 | -0.053 | 1.068 | -0.006 | 0.005 | 0.005 | 0 | 0.63 | 0.39 | 0.54 | 0.42 | 0.65 | 0.45 |
| 8 | 3141447 | rs1542104 | 1.344 | -0.142 | 0.093 | 0.955 | -0.028 | 0.026 | 0.026 | 0 | 0.76 | 0.51 | 0.82 | 0.68 | 0.77 | 0.69 |
| 8 | 3153093 | rs6558756 | -0.34 | -0.115 | 0.103 | 1.447 | -0.007 | 0.181 | 0.181 | 0.091 | 0.92 | 0.87 | 0.81 | 0.6 | 0.93 | 0.85 |
| 8 | 3153107 | rs10108100 | 1.775 | 0.205 | 0.014 | 1.082 | 0 | -0.034 | -0.034 | 0 | 0.65 | 0.27 | 0.46 | 0.44 | 0.61 | 0.64 |
| 8 | 3154726 | rs6558759 | 0.563 | 0.326 | 0.468 | 1.062 | 0.003 | 0.062 | 0.062 | 0.108 | 0.86 | 0.85 | 0.74 | 0.61 | 0.8 | 0.79 |
| 8 | 3155871 | rs11777638 | 1.169 | 0.27 | 0.744 | 1 | -0.005 | 0.106 | 0.106 | 0.02 | 0.78 | 0.61 | 0.74 | 0.62 | 0.79 | 0.76 |
| 8 | 3157117 | rs4875568 | -0.943 | 0.354 | 0.564 | 1.075 | -0.001 | 0.162 | 0.162 | 0.213 | 0.92 | 0.86 | 0.84 | 0.64 | 0.93 | 0.84 |
| 8 | 3158741 | rs12678380 | -0.666 | 0.375 | 0.362 | 0.994 | -0.027 | -0.03 | -0.03 | 0.14 | 0.07 | 0.02 | 0.21 | 0.13 | 0.15 | 0.1 |
| 8 | 3159063 | rs1442399 | -0.193 | 0.378 | 0.618 | 0.896 | -0.002 | -0.01 | -0.01 | 0 | 0.12 | 0.13 | 0.09 | 0 | 0.14 | 0.08 |
| 8 | 3159095 | rs12676614 | -0.358 | 0.384 | 0.569 | 0.929 | -0.026 | -0.026 | -0.026 | 0 | 0.06 | 0.02 | 0.22 | 0.12 | 0.16 | 0.1 |
| 8 | 3159349 | rs718122 | 2.53 | -0.22 | 0.545 | 0.874 | 0.004 | -0.033 | -0.033 | 0 | 0.6 | 0.18 | 0.44 | 0.4 | 0.6 | 0.62 |
| 8 | 3160221 | rs4875574 | 0.976 | -0.208 | 0.927 | 0.933 | -0.052 | 0.172 | 0.172 | 1.334 | 0.81 | 0.48 | 0.77 | 0.52 | 0.89 | 0.8 |
| 8 | 3161362 | rs12545641 | 1.702 | 0.333 | -0.224 | -0.152 | 0.016 | 0.003 | 0.003 | 0.032 | 0.39 | 0.21 | 0.21 | 0.36 | 0.34 | 0.48 |
| 8 | 3161497 | rs17320105 | 1.351 | 0.472 | -0.301 | -0.354 | -0.046 | 0.092 | 0.092 | 0 | 0.23 | 0.09 | 0.24 | 0.02 | 0.27 | 0.14 |
| 8 | 3164019 | rs924730 | -0.277 | 0.2 | -0.338 | -0.432 | -0.019 | 0.099 | 0.099 | 0 | 0.41 | 0.27 | 0.36 | 0.1 | 0.42 | 0.28 |

| | | | | | | | | | | | | | | | | |
|---|---|---|---|---|---|---|---|---|---|---|---|---|---|---|---|---|
| 8 | 3164593 | rs1503277 | -0.758 | 0.049 | -0.41 | -0.541 | 0 | 0.073 | 0.073 | 0 | 0.45 | 0.4 | 0.44 | 0.1 | 0.53 | 0.3 |
| 8 | 3165207 | rs10103770 | -0.403 | 0.04 | -0.188 | -0.306 | -0.004 | 0.06 | 0.06 | 0 | 0.46 | 0.4 | 0.51 | 0.22 | 0.45 | 0.36 |
| 8 | 3165624 | rs2052599 | 1.582 | -0.03 | -0.202 | -0.103 | 0.003 | -0.002 | -0.002 | 1.972 | 0.4 | 0.21 | 0.23 | 0.38 | 0.37 | 0.42 |
| 8 | 3165814 | rs1393843 | -1.685 | 0.007 | -0.246 | 0.274 | 0.001 | 0.201 | 0.201 | 0.195 | 0.93 | 0.86 | 0.83 | 0.62 | 0.93 | 0.81 |
| 8 | 3167217 | rs10086567 | -1.685 | 0.007 | -0.252 | 0.262 | 0.001 | 0.161 | 0.161 | 0.195 | 0.93 | 0.86 | 0.84 | 0.65 | 0.93 | 0.81 |
| 8 | 3167898 | rs1007010 | -0.203 | -0.022 | -0.338 | 0.1 | -0.001 | 0.006 | 0.006 | 0 | 0.16 | 0.14 | 0.11 | 0 | 0.16 | 0.14 |
| 8 | 3168573 | rs4875592 | -1.596 | 0.064 | -0.375 | 0.304 | -0.001 | 0.156 | 0.156 | 0.156 | 0.93 | 0.86 | 0.84 | 0.65 | 0.93 | 0.81 |
| 8 | 3172225 | rs924735 | -1.438 | 0.039 | -0.396 | 0.333 | -0.003 | 0.178 | 0.178 | 0.123 | 0.92 | 0.85 | 0.84 | 0.64 | 0.93 | 0.81 |
| 8 | 3172404 | rs12156329 | 0.592 | 0.182 | -0.83 | -0.058 | -0.032 | 0.026 | 0.026 | 0 | 0.32 | 0.22 | 0.36 | 0.05 | 0.44 | 0.2 |
| 8 | 3174707 | rs894333 | 0.069 | 0.047 | -0.761 | 0.113 | -0.005 | 0.024 | 0.024 | 0.285 | 0.49 | 0.32 | 0.46 | 0.55 | 0.55 | 0.6 |
| 8 | 3175959 | rs2052600 | 1.446 | -0.113 | -0.854 | 0.176 | -0.031 | -0.048 | -0.048 | 0 | 0.34 | 0.09 | 0.18 | 0.36 | 0.32 | 0.39 |
| 8 | 3176299 | rs9644337 | 1.338 | -0.102 | -0.829 | 0.015 | -0.034 | -0.134 | -0.134 | 0.544 | 0.36 | 0.1 | 0.32 | 0.47 | 0.35 | 0.46 |
| 8 | 3177911 | rs4875610 | 0.688 | -0.028 | -0.44 | 0.045 | 0.009 | -0.008 | -0.008 | 0 | 0.43 | 0.39 | 0.37 | 0.47 | 0.4 | 0.53 |
| 8 | 3178349 | rs4875611 | 0.51 | -0.004 | -0.796 | 0.041 | 0.022 | 0.025 | 0.025 | 0 | 0.44 | 0.27 | 0.39 | 0.67 | 0.41 | 0.6 |
| 8 | 3179320 | rs11783178 | -0.681 | 0.103 | -0.209 | -0.267 | 0.028 | 0.011 | 0.011 | 2.785 | 0.1 | 0.34 | 0.09 | 0.07 | 0.07 | 0.01 |
| 8 | 3182062 | rs149756 | 1.734 | -0.001 | -0.861 | -0.802 | -0.046 | 0.114 | 0.114 | 1.405 | 0.24 | 0.05 | 0.23 | 0.01 | 0.24 | 0.19 |
| 8 | 3183640 | rs11775610 | 0.38 | 0.194 | -0.564 | -0.35 | 0.009 | -0.002 | -0.002 | 0 | 0.1 | 0.25 | 0.09 | 0.09 | 0.08 | 0.02 |
| 8 | 3185421 | rs2654478 | -1.305 | 0.164 | -0.617 | -0.368 | -0.005 | -0.024 | -0.024 | 0 | 0.66 | 0.85 | 0.84 | 0.67 | 0.68 | 0.55 |
| 8 | 3185484 | rs2551044 | 0.628 | 0.041 | -0.473 | -0.443 | -0.005 | -0.094 | -0.094 | 0.156 | 0.42 | 0.5 | 0.33 | 0.04 | 0.34 | 0.4 |
| 8 | 3186454 | rs17080008 | 1.707 | 0.425 | -0.313 | 0.186 | -0.015 | -0.04 | -0.04 | 0 | 0.07 | 0.01 | 0.08 | 0.2 | 0.06 | 0.08 |
| 8 | 3187806 | rs11776403 | -0.33 | 0.226 | -0.161 | 0.211 | 0.002 | 0.117 | 0.117 | 3.515 | 0.12 | 0.28 | 0.23 | 0.37 | 0.12 | 0.11 |
| 8 | 3189896 | rs2551043 | 0.535 | 0.05 | -0.412 | -0.08 | -0.007 | 0.033 | 0.033 | 0.686 | 0.44 | 0.2 | 0.31 | 0.34 | 0.51 | 0.48 |
| 8 | 3192042 | rs17080030 | 1.717 | 0.036 | -0.239 | -0.102 | -0.001 | 0.006 | 0.006 | 0 | 0.07 | 0.05 | 0.07 | 0.18 | 0.04 | 0.09 |
| 8 | 3192247 | rs7843496 | 0.017 | 0.172 | 0.088 | -0.396 | 0 | 0.016 | 0.016 | 0 | 0.45 | 0.63 | 0.37 | 0.16 | 0.38 | 0.37 |
| 8 | 3193698 | rs11777803 | 0.291 | 0.202 | 0.318 | -0.362 | 0.012 | 0.016 | 0.016 | 3.108 | 0.09 | 0.31 | 0.08 | 0.11 | 0.07 | 0.01 |
| 8 | 3194845 | rs963155 | -0.596 | 0.286 | 0.341 | -0.221 | -0.014 | 0.056 | 0.056 | 0 | 0.85 | 0.73 | 0.78 | 0.72 | 0.87 | 0.86 |
| 8 | 3196760 | rs13282938 | 1.786 | 0.329 | 0.148 | -0.648 | -0.051 | 0.097 | 0.097 | 0 | 0.22 | 0.05 | 0.21 | 0.07 | 0.21 | 0.31 |
| 8 | 3196871 | rs1444613 | 0.23 | 0.382 | 0.422 | -0.709 | -0.005 | -0.074 | -0.074 | 0 | 0.3 | 0.45 | 0.26 | 0.11 | 0.34 | 0.13 |
| 8 | 3199421 | rs7013730 | 0.334 | 0.493 | 0.212 | -0.664 | -0.014 | -0.069 | -0.069 | 0 | 0.3 | 0.58 | 0.21 | 0.11 | 0.27 | 0.14 |
| 8 | 3200119 | rs17320990 | 0.231 | 0.828 | -0.028 | -0.277 | -0.027 | 0.027 | 0.027 | 0 | 0.07 | 0.02 | 0.1 | 0.01 | 0.17 | 0.03 |
| 8 | 3201636 | rs13251425 | 0.959 | 0.617 | 0.262 | -0.172 | 0.024 | -0.042 | -0.042 | 0 | 0.09 | 0.27 | 0.07 | 0 | 0.09 | 0.04 |
| 8 | 3202945 | rs12548150 | 0.667 | 0.67 | -0.348 | -0.063 | 0 | -0.001 | -0.001 | 0.165 | 0.1 | 0.15 | 0.15 | 0.06 | 0.13 | 0.05 |
| 8 | 3206462 | rs1822243 | 0.218 | 0.475 | -0.185 | 0.017 | 0.002 | 0.011 | 0.011 | 0 | 0.59 | 0.53 | 0.65 | 0.5 | 0.55 | 0.52 |
| 8 | 3210699 | rs10102960 | 0.523 | 0.666 | -0.473 | 0.041 | -0.022 | 0.006 | 0.006 | 0 | 0.06 | 0.02 | 0.09 | 0.02 | 0.13 | 0.03 |
| 8 | 3212595 | rs1096505 | 0.05 | 0.323 | -0.097 | -0.125 | 0.002 | -0.004 | -0.004 | 0.017 | 0.36 | 0.33 | 0.26 | 0.48 | 0.29 | 0.39 |
| 8 | 3213288 | rs13280571 | 0.573 | 0.28 | 0.23 | -0.361 | 0.01 | -0.02 | -0.02 | 0.273 | 0.38 | 0.47 | 0.45 | 0.33 | 0.38 | 0.43 |
| 8 | 3214267 | rs1442401 | 0.517 | 0.249 | 0.148 | -0.405 | -0.009 | 0.013 | 0.013 | 0 | 0.38 | 0.27 | 0.47 | 0.31 | 0.41 | 0.48 |
| 8 | 3214344 | rs1442400 | 0.717 | 0.216 | 0.113 | -0.391 | 0.001 | 0.068 | 0.068 | 0 | 0.11 | 0.21 | 0.21 | 0.25 | 0.12 | 0.14 |
| 8 | 3214873 | rs11136609 | 0.543 | 0.091 | 0.471 | -0.434 | -0.002 | 0.018 | 0.018 | 0.787 | 0.52 | 0.44 | 0.46 | 0.23 | 0.45 | 0.42 |
| 8 | 3215497 | rs1348267 | -1.052 | 0.065 | 0.04 | -0.111 | 0.001 | -0.016 | -0.016 | 0 | 0.75 | 0.8 | 0.83 | 0.66 | 0.73 | 0.69 |
| 8 | 3217419 | rs1503283 | 0.203 | 0.417 | -0.124 | -0.244 | -0.002 | 0.001 | 0.001 | 0 | 0.44 | 0.44 | 0.61 | 0.37 | 0.47 | 0.48 |
| 8 | 3219559 | rs13268772 | -0.547 | 0.345 | 0.094 | -0.241 | -0.003 | 0.02 | 0.02 | 0 | 0.68 | 0.65 | 0.75 | 0.49 | 0.68 | 0.59 |
| 8 | 3221569 | rs13253492 | 2.26 | 0.49 | -0.432 | -0.2 | -0.035 | 0.08 | 0.08 | 0 | 0.23 | 0.08 | 0.24 | 0.05 | 0.23 | 0.33 |
| 8 | 3221795 | rs13272021 | 0.544 | 0.224 | -0.125 | 0.109 | 0.002 | -0.018 | -0.018 | 0 | 0.14 | 0.18 | 0.13 | 0.02 | 0.16 | 0.06 |
| 8 | 3223127 | rs10089434 | -0.469 | 0.216 | -0.145 | 0.132 | 0.019 | 0.06 | 0.06 | 0 | 0.19 | 0.32 | 0.37 | 0.33 | 0.25 | 0.16 |
| 8 | 3223829 | rs13266825 | -0.306 | 0.206 | -0.146 | 0.121 | 0.001 | 0.009 | 0.009 | 0 | 0.2 | 0.28 | 0.28 | 0.27 | 0.22 | 0.15 |
| 8 | 3224276 | rs17065961 | -0.35 | 0.205 | -0.146 | 0.121 | 0 | 0.009 | 0.009 | 0 | 0.2 | 0.28 | 0.28 | 0.26 | 0.22 | 0.15 |
| 8 | 3224430 | rs17065962 | 1.791 | 0.158 | -0.188 | 0.144 | 0.004 | -0.012 | -0.012 | 0 | 0.07 | 0.02 | 0.08 | 0.17 | 0.05 | 0.09 |
| 8 | 3224561 | rs4875703 | -0.706 | 0.246 | -0.136 | 0.137 | 0.005 | 0.032 | 0.032 | 0 | 0.23 | 0.33 | 0.38 | 0.33 | 0.25 | 0.16 |
| 8 | 3228908 | rs13278552 | 1.508 | 0.307 | -0.135 | 0.053 | 0 | -0.017 | -0.017 | 0 | 0.09 | 0.12 | 0.06 | 0 | 0.09 | 0.04 |
| 8 | 3228952 | rs7842576 | 0.52 | 0.191 | -0.191 | 0.002 | -0.002 | 0.002 | 0.002 | 0 | 0.57 | 0.58 | 0.55 | 0.39 | 0.51 | 0.61 |
| 8 | 3229812 | rs11985163 | 0.351 | 0.154 | -0.183 | 0.021 | -0.003 | -0.01 | -0.01 | 0 | 0.15 | 0.24 | 0.19 | 0.08 | 0.15 | 0.06 |
| 8 | 3230281 | rs13257496 | 0.561 | -0.168 | -0.251 | -0.087 | -0.008 | 0.015 | 0.015 | 0.087 | 0.42 | 0.37 | 0.45 | 0.15 | 0.45 | 0.47 |
| 8 | 3230582 | rs7015057 | 0.361 | -0.233 | -0.283 | -0.034 | -0.004 | 0.001 | 0.001 | 0.224 | 0.44 | 0.36 | 0.59 | 0.39 | 0.51 | 0.54 |
| 8 | 3231137 | rs10092265 | -0.976 | -0.131 | -0.468 | 0.069 | -0.013 | -0.057 | -0.057 | 0 | 0.77 | 0.91 | 0.83 | 0.66 | 0.74 | 0.77 |
| 8 | 3233443 | rs7011397 | 0.159 | -0.412 | -0.03 | -0.086 | -0.033 | -0.001 | -0.001 | 0 | 0.32 | 0.48 | 0.31 | 0.33 | 0.24 | 0.2 |
| 8 | 3244615 | rs895696 | 0.549 | -0.5 | -0.038 | -0.173 | -0.001 | -0.019 | -0.019 | 2.024 | 0.53 | 0.44 | 0.43 | 0.28 | 0.48 | 0.39 |
| 8 | 3248652 | rs1979830 | 0.619 | -0.518 | -0.132 | 0.012 | 0.005 | 0.052 | 0.052 | 0 | 0.49 | 0.59 | 0.44 | 0.26 | 0.43 | 0.52 |
| 8 | 3253768 | rs3802303 | 1.082 | -0.554 | 0.114 | -0.104 | -0.007 | 0.063 | 0.063 | 0 | 0.43 | 0.35 | 0.34 | 0.1 | 0.42 | 0.29 |
| 8 | 3255581 | rs270054 | 1.3 | -0.734 | 0.014 | 0.23 | -0.045 | -0.05 | -0.05 | 0 | 0.19 | 0.11 | 0.19 | 0.32 | 0.32 | 0.29 |
| 8 | 3257106 | rs13269697 | 1.711 | -0.771 | 0.199 | -0.074 | -0.001 | -0.032 | -0.032 | 0 | 0.11 | 0.13 | 0.07 | 0 | 0.11 | 0.05 |
| 8 | 3257401 | rs11785551 | 1.218 | -0.666 | 0.216 | -0.144 | 0.002 | -0.089 | -0.089 | 0.788 | 0.36 | 0.42 | 0.3 | 0.06 | 0.34 | 0.37 |
| 8 | 3257586 | rs11136621 | 0.836 | -0.428 | 0.755 | 0.77 | 0.001 | -0.021 | -0.021 | 0 | 0.59 | 0.55 | 0.48 | 0.17 | 0.54 | 0.5 |
| 8 | 3258715 | rs9969557 | 1.376 | -0.444 | 0.746 | 0.747 | 0.001 | -0.069 | -0.069 | 0.329 | 0.54 | 0.47 | 0.4 | 0.03 | 0.54 | 0.33 |
| 8 | 3260131 | rs1161539 | 0.272 | -0.113 | 0.696 | 1.096 | -0.015 | -0.028 | -0.028 | 0 | 0.23 | 0.22 | 0.34 | 0.66 | 0.33 | 0.37 |
| 8 | 3260534 | rs1178231 | -0.492 | -0.113 | 0.778 | 1.07 | 0.003 | 0.001 | 0.001 | 0.002 | 0.31 | 0.33 | 0.42 | 0.72 | 0.38 | 0.47 |
| 8 | 3261022 | rs1161536 | 0.929 | 0.137 | 0.717 | 0.913 | 0.005 | -0.014 | -0.014 | 0.014 | 0.78 | 0.72 | 0.8 | 0.81 | 0.8 | 0.83 |
| 8 | 3261483 | rs1161531 | -0.634 | 0.073 | 0.749 | 0.972 | -0.001 | -0.003 | -0.003 | 0 | 0.94 | 0.96 | 0.94 | 0.98 | 0.92 | 0.97 |
| 8 | 3262061 | rs1182739 | 0.685 | 0.058 | 0.709 | 0.977 | -0.019 | -0.004 | -0.004 | 0 | 0.81 | 0.76 | 0.82 | 0.83 | 0.87 | 0.86 |

| | | | | | | | | | | | | | | | | |
|---|---|---|---|---|---|---|---|---|---|---|---|---|---|---|---|---|
| 8 | 3263500 | rs10097933 | -0.066 | 0.056 | 0.617 | 0.982 | -0.027 | 0.019 | 0.019 | 1.213 | 0.12 | 0.26 | 0.12 | 0.16 | 0.05 | 0.11 |
| 8 | 3264509 | rs6990027 | -0.817 | 0.071 | 0.421 | 0.986 | 0.012 | -0.002 | -0.002 | 0.38 | 0.18 | 0.15 | 0.16 | 0.19 | 0.13 | 0.14 |
| 8 | 3265590 | rs10088378 | -0.182 | 0.12 | 0.436 | 0.985 | 0.024 | 0.054 | 0.054 | 0 | 0.77 | 0.7 | 0.65 | 0.34 | 0.67 | 0.64 |
| 8 | 3266909 | rs10094820 | -0.061 | 0.104 | -0.283 | 1.111 | 0 | 0.004 | 0.004 | 0 | 0.07 | 0.09 | 0.07 | 0.02 | 0.08 | 0.03 |
| 8 | 3267743 | rs17066034 | -0.289 | 0.113 | -0.214 | 1.172 | 0.007 | -0.009 | -0.009 | 0 | 0.13 | 0.15 | 0.18 | 0.26 | 0.06 | 0.12 |
| 8 | 3272206 | rs2654484 | 0.301 | 0.244 | -0.345 | 1.285 | -0.026 | 0.027 | 0.027 | 0 | 0.21 | 0.15 | 0.21 | 0.13 | 0.33 | 0.2 |
| 8 | 3274215 | rs6996823 | 1.01 | 0.157 | -0.11 | 1.079 | -0.002 | -0.046 | -0.046 | 0 | 0.11 | 0.16 | 0.08 | 0 | 0.11 | 0.05 |
| 8 | 3276592 | rs1553183 | 1.967 | 0.055 | 0.237 | 1.471 | 0.001 | -0.016 | -0.016 | 0 | 0.4 | 0.3 | 0.29 | 0.08 | 0.35 | 0.43 |
| 8 | 3278315 | rs270081 | 0.895 | 0.099 | 0.163 | 1.731 | 0.004 | 0.029 | 0.029 | 0 | 0.78 | 0.67 | 0.76 | 0.57 | 0.77 | 0.84 |
| 8 | 3281550 | rs270074 | 0.762 | 0.077 | 0.163 | 1.67 | -0.01 | 0.025 | 0.025 | 0 | 0.18 | 0.13 | 0.17 | 0.11 | 0.27 | 0.2 |
| 8 | 3281896 | rs270072 | 0.885 | 0.103 | 0.177 | 1.813 | 0.001 | 0.013 | 0.013 | 0 | 0.78 | 0.69 | 0.77 | 0.68 | 0.79 | 0.84 |
| 8 | 3282012 | rs17394143 | -0.09 | 0.09 | 0.128 | 1.771 | -0.029 | 0.031 | 0.031 | 0 | 0.1 | 0.02 | 0.09 | 0.02 | 0.12 | 0.04 |
| 8 | 3284183 | rs270066 | 1.419 | 0.063 | 0.215 | 1.864 | 0.005 | -0.006 | -0.006 | 0 | 0.6 | 0.48 | 0.59 | 0.43 | 0.61 | 0.68 |
| 8 | 3285570 | rs4461920 | 1.182 | 0.074 | 0.083 | 1.832 | -0.001 | 0.009 | 0.009 | 0.042 | 0.07 | 0.09 | 0.09 | 0.16 | 0.04 | 0.09 |
| 8 | 3286487 | rs270063 | 1.312 | 0.058 | 0.114 | 1.874 | 0.01 | 0.018 | 0.018 | 0 | 0.66 | 0.45 | 0.68 | 0.68 | 0.66 | 0.78 |
| 8 | 3286518 | rs1447355 | 2.436 | 0.033 | 0.094 | 1.492 | 0.002 | -0.004 | -0.004 | 0 | 0.39 | 0.25 | 0.28 | 0.06 | 0.34 | 0.4 |
| 8 | 3287064 | rs1393845 | -1.165 | 0.015 | 0.002 | 1.604 | -0.003 | 0.011 | 0.011 | 0 | 0.3 | 0.24 | 0.26 | 0.19 | 0.3 | 0.21 |
| 8 | 3289450 | rs1499693 | 1.079 | 0.04 | 0.003 | 1.727 | 0.012 | 0.055 | 0.055 | 0.299 | 0.73 | 0.57 | 0.75 | 0.52 | 0.73 | 0.75 |
| 8 | 3290339 | rs12544202 | -0.431 | -0.019 | -0.135 | 1.7 | 0.009 | -0.011 | -0.011 | 0 | 0.14 | 0.12 | 0.17 | 0.26 | 0.05 | 0.13 |
| 8 | 3292247 | rs7013161 | -1.486 | 0.04 | -0.021 | 1.525 | 0.002 | 0.015 | 0.015 | 0 | 0.29 | 0.4 | 0.32 | 0.57 | 0.26 | 0.26 |
| 8 | 3294730 | rs7820363 | 0.535 | 0.098 | -0.238 | 1.386 | 0.005 | 0.012 | 0.012 | 0 | 0.63 | 0.65 | 0.66 | 0.78 | 0.61 | 0.71 |
| 8 | 3295628 | rs11778675 | 0.527 | 0.102 | -0.232 | 1.388 | 0.005 | 0.011 | 0.011 | 0 | 0.63 | 0.65 | 0.66 | 0.78 | 0.61 | 0.71 |
| 8 | 3296720 | rs6982744 | 0.521 | 0.1 | -0.233 | 1.378 | 0.005 | 0.012 | 0.012 | 0 | 0.63 | 0.65 | 0.66 | 0.79 | 0.61 | 0.71 |
| 8 | 3303471 | rs7008713 | 0.093 | 0.179 | -0.514 | 1.388 | -0.019 | -0.009 | -0.009 | 0 | 0.73 | 0.87 | 0.75 | 0.72 | 0.68 | 0.66 |
| 8 | 3316099 | rs7007801 | -0.899 | 0.126 | -0.492 | 1.376 | -0.034 | 0.021 | 0.021 | 0 | 0.82 | 0.93 | 0.75 | 0.88 | 0.73 | 0.81 |
| 8 | 3316392 | rs12548247 | 0.052 | 0.034 | -0.512 | 1.415 | 0.014 | -0.014 | -0.014 | 0 | 0.11 | 0.04 | 0.09 | 0.17 | 0.05 | 0.13 |
| 8 | 3316475 | rs7464854 | -0.721 | -0.021 | -0.534 | 1.298 | -0.031 | 0.024 | 0.024 | 0 | 0.81 | 0.93 | 0.75 | 0.88 | 0.72 | 0.81 |
| 8 | 3316914 | rs6995462 | 0.429 | 0.019 | -0.128 | 1.574 | -0.003 | -0.001 | -0.001 | 0 | 0.66 | 0.55 | 0.61 | 0.63 | 0.64 | 0.69 |
| 8 | 3316986 | rs17394429 | 0.826 | 0.193 | -0.082 | 1.546 | 0.011 | 0.016 | 0.016 | 0 | 0.09 | 0.12 | 0.09 | 0.23 | 0.15 | 0.19 |
| 8 | 3317490 | rs1875070 | 1.521 | -0.006 | -0.025 | 1.01 | 0.003 | 0.01 | 0.01 | 0 | 0.44 | 0.33 | 0.32 | 0.12 | 0.39 | 0.37 |
| 8 | 3319748 | rs6558778 | -0.129 | -0.103 | 0.108 | 1.54 | -0.029 | 0 | 0 | 0.97 | 0.16 | 0.31 | 0.23 | 0.19 | 0.09 | 0.12 |
| 8 | 3322528 | rs1499686 | -1.418 | 0.181 | -0.122 | 1.797 | -0.006 | 0.042 | 0.042 | 0 | 0.21 | 0.29 | 0.27 | 0.38 | 0.19 | 0.21 |
| 8 | 3324801 | rs17394506 | 0.343 | 0.738 | -0.377 | 1.702 | 0.01 | 0.022 | 0.022 | 0 | 0.09 | 0.11 | 0.09 | 0.24 | 0.15 | 0.19 |
| 8 | 3328210 | rs4875719 | -0.615 | 0.042 | -0.516 | 1.609 | 0 | -0.015 | -0.015 | 0 | 0.1 | 0.03 | 0.14 | 0.18 | 0.08 | 0.09 |
| 8 | 3331675 | rs4875720 | -1.588 | -0.369 | -0.292 | 1.526 | -0.029 | 0.045 | 0.045 | 0.062 | 0.22 | 0.37 | 0.37 | 0.61 | 0.13 | 0.37 |
| 8 | 3332290 | rs11780580 | -2.012 | -0.325 | -0.192 | 1.41 | -0.026 | 0.017 | 0.017 | 0 | 0.3 | 0.45 | 0.44 | 0.64 | 0.21 | 0.41 |
| 8 | 3332750 | rs6558779 | 1.06 | -0.366 | -0.719 | 1.004 | 0.03 | 0.035 | 0.035 | 0 | 0.57 | 0.63 | 0.4 | 0.28 | 0.6 | 0.42 |
| 8 | 3332767 | rs6558780 | -0.513 | -0.401 | -0.698 | 1.179 | -0.002 | 0.029 | 0.029 | 0 | 0.78 | 0.89 | 0.76 | 0.89 | 0.72 | 0.8 |
| 8 | 3333127 | rs10106960 | 0.732 | -0.408 | -0.706 | 1.04 | 0.031 | 0.014 | 0.014 | 0 | 0.64 | 0.66 | 0.43 | 0.28 | 0.67 | 0.44 |
| 8 | 3336289 | rs7836865 | 0.274 | -0.125 | -0.578 | 1.557 | 0.011 | -0.027 | -0.027 | 0 | 0.72 | 0.73 | 0.59 | 0.52 | 0.7 | 0.65 |
| 8 | 3339898 | rs11783144 | -0.186 | 0.014 | -0.502 | 1.682 | 0 | -0.001 | -0.001 | 0 | 0.78 | 0.73 | 0.74 | 0.71 | 0.73 | 0.71 |
| 8 | 3342972 | rs7844574 | 0.285 | -0.049 | -0.299 | 1.809 | 0.006 | -0.009 | -0.009 | 0 | 0.72 | 0.67 | 0.54 | 0.36 | 0.71 | 0.61 |
| 8 | 3343547 | rs10087333 | -0.964 | 0.278 | -0.495 | 1.294 | -0.032 | -0.04 | -0.04 | 0 | 0.37 | 0.69 | 0.5 | 0.49 | 0.4 | 0.33 |
| 8 | 3344204 | rs11777685 | 0.606 | 0.538 | 0.061 | 1.124 | 0.012 | -0.044 | -0.044 | 0 | 0.14 | 0.23 | 0.14 | 0.04 | 0.18 | 0.04 |
| 8 | 3345029 | rs9314492 | -1.005 | 0.586 | 0.499 | 1.298 | 0.024 | 0.053 | 0.053 | 0.312 | 0.3 | 0.59 | 0.45 | 0.43 | 0.35 | 0.31 |
| 8 | 3346096 | rs17066250 | 0.762 | 0.742 | 0.454 | 1.332 | 0.04 | 0.119 | 0.119 | 0.485 | 0.08 | 0.19 | 0.13 | 0.23 | 0.11 | 0.16 |
| 8 | 3346459 | rs2055339 | -0.33 | 0.722 | 0.214 | 1.396 | 0.01 | 0.028 | 0.028 | 2.303 | 0.26 | 0.64 | 0.45 | 0.41 | 0.35 | 0.3 |
| 8 | 3347755 | rs17322708 | 0.444 | 0.763 | 0.272 | 1.326 | 0.037 | 0.029 | 0.029 | 1.4 | 0.19 | 0.31 | 0.23 | 0.22 | 0.28 | 0.19 |
| 8 | 3347822 | rs895704 | 0.953 | 0.49 | -0.133 | 1.121 | 0.036 | 0.071 | 0.071 | 0.256 | 0.5 | 0.33 | 0.44 | 0.35 | 0.38 | 0.42 |
| 8 | 3348108 | rs895701 | -1.367 | 0.145 | 0.133 | 1.765 | -0.008 | 0.047 | 0.047 | 0 | 0.14 | 0.21 | 0.22 | 0.32 | 0.08 | 0.17 |
| 8 | 3348684 | rs1532583 | -0.071 | 0.25 | -0.089 | 1.691 | -0.008 | -0.033 | -0.033 | 0.08 | 0.74 | 0.86 | 0.79 | 0.65 | 0.73 | 0.64 |
| 8 | 3349112 | rs6985693 | -0.714 | 0.338 | 0.003 | 1.697 | -0.001 | 0.013 | 0.013 | 0.153 | 0.1 | 0.17 | 0.17 | 0.16 | 0.06 | 0.12 |
| 8 | 3350962 | rs4875724 | -0.397 | 0.21 | -0.536 | 1.487 | 0.013 | 0.01 | 0.01 | 0 | 0.78 | 0.85 | 0.82 | 0.88 | 0.78 | 0.82 |
| 8 | 3351147 | rs17066296 | 0.39 | 0.38 | -0.647 | 1.495 | -0.008 | -0.017 | -0.017 | 0 | 0.08 | 0.04 | 0.05 | 0.23 | 0.07 | 0.16 |
| 8 | 3353059 | rs4493928 | 0.366 | 0.392 | -0.646 | 1.513 | -0.012 | -0.017 | -0.017 | 0 | 0.08 | 0.04 | 0.07 | 0.25 | 0.08 | 0.16 |
| 8 | 3357202 | rs4487801 | 0.903 | 0.074 | -0.419 | 1.442 | -0.017 | 0.025 | 0.025 | 0 | 0.62 | 0.54 | 0.57 | 0.56 | 0.65 | 0.66 |
| 8 | 3357255 | rs4471061 | 0.229 | 0.364 | -0.373 | 1.285 | -0.015 | 0.021 | 0.021 | 0.012 | 0.73 | 0.61 | 0.64 | 0.71 | 0.72 | 0.73 |
| 8 | 3360149 | rs4875727 | 0.181 | 0.618 | -0.788 | 1.322 | -0.029 | -0.024 | -0.024 | 0 | 0.73 | 0.85 | 0.77 | 0.69 | 0.66 | 0.71 |
| 8 | 3361471 | rs10106674 | -1.073 | 0.309 | -0.014 | 1.173 | -0.015 | 0.005 | 0.005 | 0 | 0.21 | 0.39 | 0.36 | 0.25 | 0.17 | 0.28 |
| 8 | 3363444 | rs6983016 | 0.258 | 0.496 | 0.004 | 1.318 | 0.005 | 0.002 | 0.002 | 0 | 0.4 | 0.52 | 0.45 | 0.44 | 0.47 | 0.43 |
| 8 | 3366303 | rs10090810 | -0.444 | 0.404 | -0.186 | 1.277 | 0.027 | 0.016 | 0.016 | 0 | 0.5 | 0.71 | 0.66 | 0.55 | 0.54 | 0.56 |
| 8 | 3366934 | rs12114535 | -0.879 | 0.722 | 0.221 | 1.387 | 0.006 | 0.074 | 0.074 | 0 | 0.15 | 0.25 | 0.28 | 0.28 | 0.16 | 0.28 |
| 8 | 3367210 | rs12114480 | 0.481 | 0.757 | 0.117 | 1.401 | 0.032 | 0.087 | 0.087 | 0 | 0.06 | 0.17 | 0.09 | 0.15 | 0.1 | 0.14 |
| 8 | 3367705 | rs4875728 | 0.355 | 0.608 | -0.336 | 1.061 | 0.003 | -0.013 | -0.013 | 1.787 | 0.08 | 0.03 | 0.18 | 0.13 | 0.06 | 0.14 |
| 8 | 3369812 | rs7839985 | -0.563 | 0.548 | -0.448 | 1.022 | 0.022 | -0.012 | -0.012 | 0 | 0.55 | 0.71 | 0.65 | 0.47 | 0.59 | 0.6 |
| 8 | 3373517 | rs7820873 | -0.651 | 0.4 | -0.167 | 1.214 | -0.001 | -0.002 | -0.002 | 0 | 0.15 | 0.15 | 0.26 | 0.18 | 0.14 | 0.21 |
| 8 | 3374958 | rs11778536 | -1.599 | 0.201 | 0.362 | 1.209 | -0.016 | 0.047 | 0.047 | 0 | 0.21 | 0.44 | 0.27 | 0.22 | 0.13 | 0.3 |
| 8 | 3378898 | rs10092000 | -1.611 | 0.228 | 0.28 | 1.244 | -0.016 | 0.053 | 0.053 | 0 | 0.21 | 0.41 | 0.27 | 0.23 | 0.14 | 0.3 |
| 8 | 3382982 | rs4413794 | -1.87 | 0.302 | -0.042 | 1.076 | -0.005 | 0.01 | 0.01 | 0 | 0.24 | 0.29 | 0.29 | 0.24 | 0.19 | 0.32 |

| | | | | | | | | | | | | | | | | |
|---|---|---|---|---|---|---|---|---|---|---|---|---|---|---|---|---|
| 8 | 3383253 | rs1026878 | -0.704 | 0.18 | -0.527 | 1.404 | -0.001 | -0.008 | -0.008 | 0 | 0.14 | 0.13 | 0.3 | 0.34 | 0.12 | 0.24 |
| 8 | 3385588 | rs13275302 | -0.244 | -0.129 | -1.082 | 0.631 | 0.014 | -0.014 | -0.014 | 0 | 0.19 | 0.06 | 0.17 | 0.26 | 0.15 | 0.2 |
| 8 | 3385757 | rs4875730 | -1.071 | -0.021 | -1.597 | 0.099 | -0.009 | 0.022 | 0.022 | 0 | 0.72 | 0.81 | 0.69 | 0.76 | 0.73 | 0.76 |
| 8 | 3390144 | rs6999415 | 0.842 | -0.08 | -1.524 | 0.016 | 0.001 | 0.009 | 0.009 | 0 | 0.13 | 0.07 | 0.08 | 0.02 | 0.13 | 0.04 |
| 8 | 3390607 | rs4503118 | 0.525 | -0.084 | -1.626 | 0.531 | -0.026 | 0.007 | 0.007 | 0 | 0.41 | 0.34 | 0.34 | 0.32 | 0.49 | 0.4 |
| 8 | 3391361 | rs10091895 | -0.932 | -0.488 | -1.085 | 0.799 | -0.003 | 0.001 | 0.001 | 1.484 | 0.31 | 0.63 | 0.4 | 0.43 | 0.21 | 0.38 |
| 8 | 3393402 | rs7844359 | -0.892 | -0.326 | -0.912 | 0.562 | -0.049 | -0.008 | -0.008 | 0 | 0.25 | 0.54 | 0.13 | 0.28 | 0.16 | 0.17 |
| 8 | 3393482 | rs13266649 | 0.516 | 0.134 | -1.795 | 0.528 | -0.115 | -0.058 | -0.058 | 0 | 0.45 | 0.05 | 0.42 | 0.4 | 0.53 | 0.37 |
| 8 | 3399578 | rs17066354 | -0.147 | -0.019 | -1.111 | 1.088 | 0.005 | -0.027 | -0.027 | 0 | 0.08 | 0.04 | 0.08 | 0.16 | 0.05 | 0.11 |
| 8 | 3404802 | rs6995564 | 0.328 | 0.095 | -0.858 | 0.902 | 0 | 0 | 0 | 0.017 | 0.46 | 0.51 | 0.39 | 0.52 | 0.45 | 0.44 |
| 8 | 3406285 | rs7459738 | 0.335 | 0.093 | -0.9 | 0.903 | 0.005 | 0.001 | 0.001 | 0 | 0.46 | 0.46 | 0.39 | 0.52 | 0.45 | 0.44 |
| 8 | 3407734 | rs9314498 | 0.588 | 0.14 | -1.413 | 0.87 | -0.013 | 0.01 | 0.01 | 0 | 0.42 | 0.3 | 0.37 | 0.38 | 0.45 | 0.39 |
| 8 | 3409744 | rs12056919 | 0.041 | 0.682 | -1.098 | 1.015 | -0.007 | 0.001 | 0.001 | 0 | 0.56 | 0.47 | 0.5 | 0.54 | 0.54 | 0.49 |
| 8 | 3414155 | rs7825970 | -0.084 | 1.002 | -1.661 | 0.873 | -0.047 | -0.047 | -0.047 | 0 | 0.63 | 0.83 | 0.53 | 0.51 | 0.56 | 0.56 |
| 8 | 3420815 | rs2170481 | 0.451 | 1.158 | -1.376 | 0.926 | -0.009 | -0.01 | -0.01 | 0 | 0.89 | 0.93 | 0.9 | 0.8 | 0.86 | 0.8 |
| 8 | 3421598 | rs13265032 | 0.682 | 1.009 | -2.055 | 0.328 | -0.027 | 0.04 | 0.04 | 1.053 | 0.33 | 0.16 | 0.28 | 0.27 | 0.37 | 0.25 |
| 8 | 3422332 | rs12675134 | 0.288 | 0.855 | -1.524 | 0.362 | 0.016 | 0.01 | 0.01 | 0 | 0.51 | 0.56 | 0.53 | 0.45 | 0.59 | 0.45 |
| 8 | 3423848 | rs13260637 | 0.974 | 0.673 | -0.791 | 0.354 | 0 | -0.011 | -0.011 | 0 | 0.41 | 0.46 | 0.35 | 0.31 | 0.42 | 0.26 |
| 8 | 3425381 | rs13279490 | -0.285 | -0.13 | -1.282 | 0.544 | 0.024 | -0.036 | -0.036 | 0 | 0.85 | 0.92 | 0.86 | 0.58 | 0.89 | 0.72 |
| 8 | 3426669 | rs10102248 | 0.359 | -0.226 | -1.354 | 0.438 | -0.027 | 0.012 | 0.012 | 0 | 0.21 | 0.13 | 0.24 | 0.16 | 0.28 | 0.25 |
| 8 | 3427306 | rs17395021 | 1.036 | -0.215 | -1.494 | 0.19 | -0.01 | 0.029 | 0.029 | 0 | 0.1 | 0.05 | 0.07 | 0 | 0.12 | 0.1 |
| 8 | 3427849 | rs11785740 | 2.392 | -0.319 | -1.89 | -0.072 | -0.143 | -0.065 | -0.065 | 0 | 0.28 | 0.02 | 0.25 | 0.3 | 0.36 | 0.26 |
| 8 | 3428652 | rs13273515 | 0.74 | -0.056 | -0.9 | 0.658 | -0.011 | -0.014 | -0.014 | 0 | 0.65 | 0.74 | 0.61 | 0.41 | 0.6 | 0.47 |
| 8 | 3432054 | rs7828513 | -0.613 | 0.086 | -0.597 | 0.694 | -0.014 | 0.018 | 0.018 | 0 | 0.31 | 0.24 | 0.33 | 0.17 | 0.39 | 0.33 |
| 8 | 3432105 | rs1564571 | 0.487 | 0.084 | -0.681 | 0.662 | -0.001 | 0 | 0 | 0 | 0.18 | 0.15 | 0.25 | 0.16 | 0.27 | 0.22 |
| 8 | 3432362 | rs1564573 | 2.184 | -0.107 | -0.694 | 0.642 | -0.086 | 0.061 | 0.061 | 1.131 | 0.4 | 0.09 | 0.3 | 0.3 | 0.45 | 0.31 |
| 8 | 3432860 | rs7816319 | 1.418 | -0.173 | 0.222 | 0.709 | -0.003 | 0.007 | 0.007 | 0.241 | 0.53 | 0.37 | 0.49 | 0.45 | 0.49 | 0.41 |
| 8 | 3433136 | rs17323204 | -0.577 | -0.088 | 0.101 | 0.606 | -0.016 | 0.022 | 0.022 | 0 | 0.31 | 0.24 | 0.33 | 0.14 | 0.39 | 0.34 |
| 8 | 3434657 | rs11780268 | 1.926 | -0.044 | -0.198 | 0.587 | -0.099 | 0.058 | 0.058 | 1.434 | 0.4 | 0.08 | 0.3 | 0.31 | 0.45 | 0.33 |
| 8 | 3437862 | rs13275823 | 1.926 | -0.085 | -0.237 | 0.587 | -0.129 | 0.047 | 0.047 | 1.434 | 0.4 | 0.05 | 0.3 | 0.31 | 0.45 | 0.33 |
| 8 | 3438733 | rs17395377 | -0.237 | -0.068 | -0.164 | 0.53 | 0.006 | 0.072 | 0.072 | 0 | 0.14 | 0.04 | 0.06 | 0 | 0.12 | 0.12 |
| 8 | 3441711 | rs17066457 | 0.201 | 0.073 | 0.134 | 0.561 | -0.001 | -0.028 | -0.028 | 0.005 | 0.12 | 0.13 | 0.09 | 0 | 0.13 | 0.11 |
| 8 | 3444008 | rs986183 | 1.938 | -0.017 | 0.496 | 0.736 | -0.033 | 0.059 | 0.059 | 0.742 | 0.42 | 0.2 | 0.31 | 0.3 | 0.45 | 0.33 |
| 8 | 3444748 | rs13258350 | 0.445 | -0.142 | 0.556 | 0.813 | -0.047 | 0.076 | 0.076 | 0 | 0.77 | 0.61 | 0.66 | 0.44 | 0.84 | 0.69 |
| 8 | 3444836 | rs13266061 | 2.132 | -0.083 | 0.306 | 0.817 | -0.067 | 0.054 | 0.054 | 1.137 | 0.41 | 0.15 | 0.3 | 0.31 | 0.45 | 0.33 |
| 8 | 3445665 | rs2469370 | -0.021 | 0.04 | 0.308 | 1.173 | 0.027 | -0.052 | -0.052 | 3.356 | 0.86 | 0.92 | 0.85 | 0.58 | 0.88 | 0.73 |
| 8 | 3448428 | rs2469368 | -0.036 | 0.106 | 0.312 | 1.148 | 0.013 | -0.048 | -0.048 | 6.363 | 0.85 | 0.91 | 0.84 | 0.6 | 0.87 | 0.72 |
| 8 | 3448547 | rs2469367 | 1.362 | 0.075 | 0.57 | 1.16 | 0.009 | 0.026 | 0.026 | 0.468 | 0.56 | 0.38 | 0.39 | 0.34 | 0.58 | 0.47 |
| 8 | 3448671 | rs2449179 | -0.05 | 0.097 | 0.39 | 1.171 | 0.014 | -0.053 | -0.053 | 0.382 | 0.85 | 0.91 | 0.84 | 0.6 | 0.87 | 0.72 |
| 8 | 3449198 | rs12541850 | -0.463 | 0.18 | 0.642 | 1.037 | 0.007 | 0.004 | 0.004 | 0.015 | 0.28 | 0.48 | 0.45 | 0.26 | 0.29 | 0.24 |
| 8 | 3450317 | rs9650507 | 0.242 | 0.366 | 0.274 | 1.123 | -0.005 | -0.003 | -0.003 | 0.019 | 0.19 | 0.16 | 0.27 | 0.19 | 0.26 | 0.21 |
| 8 | 3451057 | rs2469359 | 0.459 | 0.346 | 0.214 | 1.254 | -0.002 | 0.049 | 0.049 | 0.37 | 0.87 | 0.82 | 0.81 | 0.59 | 0.85 | 0.76 |
| 8 | 3451111 | rs2469358 | 1.213 | 0.428 | -0.075 | 0.962 | -0.005 | -0.044 | -0.044 | 0 | 0.71 | 0.77 | 0.71 | 0.58 | 0.68 | 0.63 |
| 8 | 3452058 | rs17066503 | 0.623 | 0.239 | -0.13 | 0.936 | 0.034 | -0.021 | -0.021 | 0 | 0.14 | 0.01 | 0.17 | 0.15 | 0.06 | 0.07 |
| 8 | 3452887 | rs2469354 | 0.391 | 0.212 | 0.653 | 1.101 | 0.008 | 0.301 | 0.301 | 0.19 | 0.87 | 0.58 | 0.79 | 0.49 | 0.87 | 0.71 |
| 8 | 3454412 | rs748304 | -1.531 | 0.242 | 0.578 | 0.943 | 0.014 | 0.084 | 0.084 | 2.989 | 0.46 | 0.81 | 0.64 | 0.67 | 0.41 | 0.55 |
| 8 | 3454736 | rs900217 | 0.385 | 0.216 | 0.619 | 1.456 | 0.001 | 0.22 | 0.22 | 0 | 0.07 | 0.25 | 0.11 | 0.23 | 0.03 | 0.19 |
| 8 | 3454880 | rs900213 | 2.249 | 0.222 | 0.232 | 1.262 | -0.067 | 0.01 | 0.01 | 3.253 | 0.39 | 0.05 | 0.25 | 0.32 | 0.41 | 0.32 |
| 8 | 3455664 | rs11998372 | -0.129 | 0.072 | 0.408 | 0.993 | 0.005 | -0.006 | -0.006 | 0 | 0.19 | 0.21 | 0.25 | 0.34 | 0.14 | 0.15 |
| 8 | 3455812 | rs931203 | 0.569 | 0.102 | 0.433 | 1.018 | 0.012 | 0.038 | 0.038 | 0 | 0.06 | 0.14 | 0.07 | 0.17 | 0.09 | 0.1 |
| 8 | 3455838 | rs931202 | -0.025 | 0.128 | 0.435 | 0.939 | 0.007 | -0.007 | -0.007 | 0 | 0.19 | 0.21 | 0.25 | 0.33 | 0.14 | 0.16 |
| 8 | 3457546 | rs2449174 | -0.178 | 0.17 | 0.317 | 0.671 | -0.03 | 0.069 | 0.069 | 0 | 0.85 | 0.97 | 0.91 | 1 | 0.83 | 0.88 |
| 8 | 3458866 | rs2449173 | 0.316 | -0.097 | 0.795 | 1.14 | -0.006 | 0.252 | 0.252 | 0 | 0.91 | 0.7 | 0.87 | 0.7 | 0.96 | 0.81 |
| 8 | 3459198 | rs2469350 | 1.366 | 0.089 | 0.344 | 1.078 | -0.002 | 0.094 | 0.094 | 2.119 | 0.53 | 0.17 | 0.34 | 0.29 | 0.57 | 0.45 |
| 8 | 3460063 | rs7815959 | -1.023 | 0.056 | 0.628 | 1.101 | -0.015 | 0.012 | 0.012 | 0.11 | 0.29 | 0.68 | 0.4 | 0.59 | 0.18 | 0.4 |
| 8 | 3460162 | rs17066560 | 0.366 | 0.173 | 0.57 | 1.087 | 0.016 | -0.006 | -0.006 | 0 | 0.19 | 0.24 | 0.27 | 0.12 | 0.25 | 0.17 |
| 8 | 3461624 | rs2469349 | 0.237 | 0.161 | 0.453 | 1.282 | 0.025 | -0.036 | -0.036 | 0 | 0.79 | 0.87 | 0.83 | 0.66 | 0.88 | 0.7 |
| 8 | 3461712 | rs9644274 | 0.214 | 0.234 | 0.575 | 1.264 | -0.001 | 0.032 | 0.032 | 0 | 0.07 | 0.12 | 0.21 | 0.25 | 0.09 | 0.14 |
| 8 | 3463535 | rs2449166 | 1.873 | 0.138 | 0.392 | 0.923 | -0.052 | 0.077 | 0.077 | 2.408 | 0.45 | 0.18 | 0.28 | 0.29 | 0.48 | 0.33 |
| 8 | 3463847 | rs2170479 | 0.797 | -0.01 | 0.616 | 1.137 | -0.008 | 0.015 | 0.015 | 0 | 0.72 | 0.64 | 0.61 | 0.42 | 0.79 | 0.58 |
| 8 | 3465842 | rs1471610 | 0.976 | 0.19 | 0.497 | 1.247 | -0.002 | 0.06 | 0.06 | 0.323 | 0.82 | 0.73 | 0.63 | 0.47 | 0.82 | 0.61 |
| 8 | 3474090 | rs2469319 | 0.108 | 0.378 | 0.531 | 1.02 | 0.014 | 0.03 | 0.03 | 0.543 | 0.8 | 0.77 | 0.56 | 0.57 | 0.72 | 0.74 |
| 8 | 3474590 | rs7840019 | -0.608 | -0.156 | 0.731 | 0.094 | -0.023 | -0.071 | -0.071 | 0.205 | 0.23 | 0.28 | 0.14 | 0.05 | 0.15 | 0.22 |
| 8 | 3476246 | rs10107351 | -0.26 | -0.066 | 0.722 | 0.126 | -0.002 | -0.027 | -0.027 | 0 | 0.19 | 0.18 | 0.12 | 0.05 | 0.14 | 0.15 |
| 8 | 3477129 | rs4577978 | 1.021 | -0.078 | 1.084 | 0.285 | 0.022 | 0.023 | 0.023 | 0.873 | 0.65 | 0.52 | 0.56 | 0.45 | 0.6 | 0.54 |
| 8 | 3478076 | rs2086586 | 0.94 | -0.083 | 1.191 | 0.279 | 0.006 | 0.005 | 0.005 | 0.702 | 0.64 | 0.42 | 0.54 | 0.44 | 0.6 | 0.52 |
| 8 | 3478336 | rs1600857 | -0.136 | 0.048 | 1.278 | 0.222 | 0.007 | 0.043 | 0.043 | 0.056 | 0.2 | 0.25 | 0.28 | 0.53 | 0.23 | 0.23 |
| 8 | 3478498 | rs3102099 | 0.456 | 0.044 | 1.262 | 0.204 | 0.002 | 0.005 | 0.005 | 1.938 | 0.75 | 0.75 | 0.62 | 0.45 | 0.68 | 0.63 |
| 8 | 3478958 | rs10091628 | 0.389 | -0.39 | 1.295 | 0.43 | 0.006 | 0.006 | 0.006 | 0 | 0.15 | 0.17 | 0.19 | 0.23 | 0.21 | 0.1 |

| | | | | | | | | | | | | | | | | |
|---|---|---|---|---|---|---|---|---|---|---|---|---|---|---|---|---|
| 8 | 3480906 | rs2469313 | 0.265 | -0.505 | 1.32 | 0.37 | -0.002 | -0.053 | -0.053 | 0 | 0.09 | 0.2 | 0.06 | 0.01 | 0.08 | 0.1 |
| 8 | 3481618 | rs922798 | 1.968 | -0.577 | 1.242 | 0.345 | -0.017 | 0.007 | 0.007 | 1.722 | 0.43 | 0.26 | 0.43 | 0.39 | 0.46 | 0.35 |
| 8 | 3482819 | rs2449223 | -0.249 | -0.549 | 1.067 | 0.334 | 0.003 | -0.027 | -0.027 | 0 | 0.85 | 0.84 | 0.81 | 0.97 | 0.81 | 0.73 |
| 8 | 3484731 | rs10108630 | 0.395 | -0.328 | 1.117 | -0.203 | 0.018 | 0.012 | 0.012 | 0.007 | 0.15 | 0.21 | 0.2 | 0.23 | 0.23 | 0.1 |
| 8 | 3485036 | rs2469312 | 0.306 | -0.396 | 0.956 | -0.317 | 0.007 | 0.043 | 0.043 | 0 | 0.73 | 0.83 | 0.79 | 0.92 | 0.77 | 0.68 |
| 8 | 3485700 | rs17396710 | 0.423 | -0.41 | 0.964 | -0.298 | -0.003 | -0.037 | -0.037 | 0 | 0.1 | 0.12 | 0.06 | 0 | 0.1 | 0.1 |
| 8 | 3486209 | rs2128219 | 1.156 | -0.43 | 1.204 | -0.267 | -0.018 | -0.062 | -0.062 | 0.475 | 0.62 | 0.48 | 0.72 | 0.76 | 0.68 | 0.57 |
| 8 | 3486233 | rs2469310 | 0.156 | 0.122 | 1.151 | -0.114 | -0.007 | -0.053 | -0.053 | 0 | 0.11 | 0.17 | 0.06 | 0.01 | 0.1 | 0.1 |
| 8 | 3486374 | rs13281285 | 0.268 | 0.214 | 0.918 | -0.059 | 0.006 | -0.008 | -0.008 | 0 | 0.14 | 0.12 | 0.21 | 0.06 | 0.22 | 0.08 |
| 8 | 3487401 | rs2469308 | -0.028 | -0.082 | 0.778 | -0.085 | 0.008 | 0 | 0 | 0 | 0.85 | 0.79 | 0.82 | 0.8 | 0.83 | 0.73 |
| 8 | 3487768 | rs11775463 | 0.659 | -0.149 | 0.649 | -0.103 | 0.02 | -0.055 | -0.055 | 0 | 0.75 | 0.53 | 0.75 | 0.8 | 0.73 | 0.63 |
| 8 | 3488338 | rs2449215 | -0.578 | -1.383 | -0.15 | -0.262 | 0.066 | 0.024 | 0.024 | 0.002 | 0.43 | 0.66 | 0.35 | 0.63 | 0.24 | 0.46 |
| 8 | 3488835 | rs2128216 | 0.598 | -1.208 | 0.028 | -0.547 | -0.002 | 0.019 | 0.019 | 4.845 | 0.47 | 0.34 | 0.44 | 0.37 | 0.45 | 0.4 |
| 8 | 3489030 | rs2406985 | -0.029 | -1.399 | 0.498 | -0.667 | 0.009 | -0.063 | -0.063 | 0 | 0.61 | 0.53 | 0.64 | 0.89 | 0.52 | 0.58 |
| 8 | 3489660 | rs17066698 | -0.351 | -1.391 | 0.524 | -0.581 | -0.019 | 0.015 | 0.015 | 0 | 0.09 | 0.12 | 0.02 | 0.17 | 0.03 | 0.05 |
| 8 | 3490806 | rs1482211 | 0.401 | -1.472 | 0.789 | -0.65 | -0.001 | -0.004 | -0.004 | 0 | 0.47 | 0.36 | 0.45 | 0.51 | 0.46 | 0.41 |
| 8 | 3490822 | rs10089138 | 0.003 | -1.671 | 1.059 | -0.787 | -0.019 | -0.077 | -0.077 | 0 | 0.18 | 0.29 | 0.1 | 0.01 | 0.12 | 0.15 |
| 8 | 3494062 | rs13279667 | 0.141 | -1.474 | 0.89 | -0.743 | -0.002 | -0.003 | -0.003 | 0.073 | 0.55 | 0.42 | 0.48 | 0.52 | 0.52 | 0.47 |
| 8 | 3494151 | rs11987743 | -0.65 | -1.51 | 0.8 | -0.731 | 0.005 | 0.019 | 0.019 | 4.457 | 0.66 | 0.63 | 0.56 | 0.52 | 0.62 | 0.56 |
| 8 | 3496633 | rs4523281 | -0.346 | -1.583 | 0.744 | -0.699 | -0.081 | -0.035 | -0.035 | 0.377 | 0.17 | 0.44 | 0.06 | 0.14 | 0.05 | 0.1 |
| 8 | 3498171 | rs11136645 | 0.022 | -1.327 | 0.533 | -0.6 | -0.003 | -0.003 | -0.003 | 0 | 0.56 | 0.58 | 0.5 | 0.52 | 0.52 | 0.47 |
| 8 | 3502239 | rs2449199 | -0.035 | -1.309 | 0.079 | -1.013 | -0.008 | 0.008 | 0.008 | 1.154 | 0.56 | 0.6 | 0.52 | 0.36 | 0.52 | 0.43 |
| 8 | 3502719 | rs13273027 | 0.778 | -0.773 | -0.667 | -0.898 | -0.066 | -0.032 | -0.032 | 0 | 0.33 | 0.14 | 0.41 | 0.34 | 0.4 | 0.35 |
| 8 | 3502796 | rs17066721 | 0.56 | -0.835 | -0.011 | -0.351 | -0.002 | -0.025 | -0.025 | 0 | 0.11 | 0.09 | 0.04 | 0 | 0.1 | 0.02 |
| 8 | 3503009 | rs7830816 | 0.385 | -0.659 | -0.117 | -0.208 | -0.001 | -0.043 | -0.043 | 0.045 | 0.11 | 0.15 | 0.06 | 0 | 0.1 | 0.1 |
| 8 | 3503182 | rs1482207 | 0.157 | -0.257 | -0.377 | 0.13 | 0 | 0.021 | 0.021 | 0.681 | 0.19 | 0.21 | 0.3 | 0.35 | 0.15 | 0.29 |
| 8 | 3504117 | rs17066733 | -0.168 | -0.527 | -0.352 | -0.542 | -0.021 | -0.056 | -0.056 | 0.039 | 0.14 | 0.16 | 0.05 | 0.01 | 0.1 | 0.04 |
| 8 | 3505575 | rs7003067 | 0.638 | -0.213 | -0.62 | -0.621 | -0.011 | 0.015 | 0.015 | 0 | 0.06 | 0.01 | 0.04 | 0 | 0.07 | 0.07 |
| 8 | 3505698 | rs6999124 | -0.625 | -0.493 | -0.684 | -0.603 | 0 | 0.016 | 0.016 | 0.067 | 0.38 | 0.5 | 0.58 | 0.54 | 0.35 | 0.58 |
| 8 | 3506404 | rs2469406 | 1.274 | -0.807 | -0.886 | -0.979 | -0.05 | 0.02 | 0.02 | 0 | 0.4 | 0.26 | 0.32 | 0.35 | 0.51 | 0.26 |
| 8 | 3506725 | rs17066769 | 1.51 | -0.918 | -1.108 | -0.729 | -0.037 | -0.057 | -0.057 | 0 | 0.13 | 0.04 | 0.34 | 0.33 | 0.21 | 0.34 |
| 8 | 3507129 | rs2449202 | 0.357 | -1.324 | -0.51 | -0.914 | -0.006 | -0.018 | -0.018 | 0 | 0.64 | 0.74 | 0.54 | 0.52 | 0.62 | 0.52 |
| 8 | 3508672 | rs2469400 | 1.201 | -0.872 | -0.701 | -1.059 | -0.002 | 0.03 | 0.03 | 0.091 | 0.48 | 0.41 | 0.33 | 0.35 | 0.53 | 0.29 |
| 8 | 3508977 | rs1586030 | 0.737 | -0.921 | -0.594 | -1.085 | -0.001 | -0.001 | -0.001 | 0 | 0.59 | 0.46 | 0.37 | 0.36 | 0.61 | 0.37 |
| 8 | 3510361 | rs12680010 | 1.206 | -0.831 | -0.918 | -1.479 | -0.002 | -0.004 | -0.004 | 0.011 | 0.27 | 0.3 | 0.19 | 0.36 | 0.26 | 0.23 |
| 8 | 3510462 | rs13439830 | 1.194 | -0.918 | -0.95 | -1.498 | -0.028 | 0.004 | 0.004 | 0 | 0.38 | 0.2 | 0.5 | 0.34 | 0.52 | 0.38 |
| 8 | 3510491 | rs11783239 | 2.317 | -0.818 | -0.889 | -1.298 | -0.017 | 0.06 | 0.06 | 0.042 | 0.1 | 0 | 0.03 | 0.01 | 0.06 | 0.08 |
| 8 | 3511506 | rs17066823 | 1.151 | -0.781 | -0.557 | -1.1 | -0.035 | 0.04 | 0.04 | 0 | 0.07 | 0.12 | 0.14 | 0.16 | 0 | 0.12 |
| 8 | 3511972 | rs2469395 | -0.749 | -0.719 | -0.261 | -1.118 | -0.023 | 0.065 | 0.065 | 0 | 0.86 | 0.65 | 0.84 | 0.78 | 0.93 | 0.87 |
| 8 | 3513337 | rs2469390 | -0.596 | -0.57 | -0.701 | -1.087 | -0.017 | 0.027 | 0.027 | 0 | 0.78 | 0.67 | 0.75 | 0.7 | 0.86 | 0.73 |
| 8 | 3514601 | rs1871886 | 0.838 | -0.3 | -1.888 | -1.073 | -0.064 | 0.036 | 0.036 | 0.262 | 0.44 | 0.19 | 0.32 | 0.36 | 0.53 | 0.28 |
| 8 | 3515413 | rs10103372 | 1.748 | -0.679 | -1.469 | -0.713 | -0.046 | -0.027 | -0.027 | 0.671 | 0.29 | 0.15 | 0.43 | 0.33 | 0.43 | 0.36 |
| 8 | 3515449 | rs2469385 | 0.719 | -0.944 | -0.513 | -0.086 | -0.021 | 0 | 0 | 0 | 0.69 | 0.48 | 0.68 | 0.7 | 0.78 | 0.72 |
| 8 | 3516792 | rs1905100 | 0.431 | -0.483 | -0.919 | 0.037 | 0.005 | 0.034 | 0.034 | 0 | 0.13 | 0.19 | 0.17 | 0.36 | 0.18 | 0.18 |
| 8 | 3516984 | rs922794 | 0.911 | -0.75 | -1.023 | -0.163 | 0.013 | 0.002 | 0.002 | 0 | 0.57 | 0.34 | 0.59 | 0.39 | 0.6 | 0.55 |
| 8 | 3517336 | rs17066851 | 0.548 | -0.831 | -0.654 | 0.164 | 0.007 | -0.002 | -0.002 | 2.803 | 0.08 | 0.06 | 0.04 | 0.11 | 0.06 | 0.03 |
| 8 | 3519630 | rs1482196 | 0.51 | -1.123 | -0.301 | 0.229 | -0.018 | 0.046 | 0.046 | 0.871 | 0.7 | 0.67 | 0.75 | 0.44 | 0.74 | 0.63 |
| 8 | 3519665 | rs1482195 | 1.026 | -1.463 | 0.034 | 0.276 | -0.016 | 0.004 | 0.004 | 1.614 | 0.61 | 0.51 | 0.66 | 0.39 | 0.7 | 0.62 |
| 8 | 3520763 | rs2623625 | 0.238 | -1.515 | 0.083 | 0.394 | 0.008 | -0.05 | -0.05 | 0 | 0.76 | 0.88 | 0.81 | 0.51 | 0.75 | 0.64 |
| 8 | 3520871 | rs2624087 | -0.164 | -1.258 | -0.055 | -0.187 | 0.008 | -0.073 | -0.073 | 0 | 0.31 | 0.38 | 0.25 | 0.12 | 0.31 | 0.3 |
| 8 | 3520923 | rs2624088 | -0.925 | -1.313 | -0.389 | -0.058 | -0.005 | -0.001 | -0.001 | 0.256 | 0.87 | 0.89 | 0.94 | 0.85 | 0.93 | 0.81 |
| 8 | 3521949 | rs13255284 | 2.45 | -1.392 | -0.425 | -0.008 | -0.067 | -0.026 | -0.026 | 0.2 | 0.32 | 0.1 | 0.46 | 0.27 | 0.4 | 0.31 |
| 8 | 3523100 | rs17325824 | 0.757 | -1.238 | -0.466 | -0.29 | 0.005 | 0.012 | 0.012 | 0 | 0.13 | 0.09 | 0.07 | 0 | 0.12 | 0.09 |
| 8 | 3527105 | rs1481704 | 0.292 | -1.334 | -0.139 | -0.303 | -0.012 | -0.019 | -0.019 | 0 | 0.75 | 0.75 | 0.88 | 0.85 | 0.81 | 0.72 |
| 8 | 3527234 | rs6998429 | 0.442 | -1.242 | -0.26 | -0.262 | -0.028 | 0.024 | 0.024 | 0 | 0.16 | 0.11 | 0.16 | 0.11 | 0.19 | 0.21 |
| 8 | 3529955 | rs2623612 | -0.844 | -1.284 | -0.256 | -0.224 | 0.012 | 0.005 | 0.005 | 1.598 | 0.87 | 0.94 | 0.95 | 0.85 | 0.91 | 0.78 |
| 8 | 3531913 | rs2624100 | -0.809 | -1.172 | -0.204 | -0.218 | -0.002 | -0.001 | -0.001 | 0 | 0.86 | 0.91 | 0.94 | 0.85 | 0.9 | 0.8 |
| 8 | 3532921 | rs4875252 | 2.224 | -1.016 | -0.367 | -0.011 | -0.135 | -0.103 | -0.103 | 1.043 | 0.3 | 0.03 | 0.45 | 0.31 | 0.4 | 0.27 |
| 8 | 3533558 | rs13261262 | 1.771 | -0.838 | -0.087 | 0.006 | -0.042 | -0.016 | -0.016 | 4.334 | 0.34 | 0.16 | 0.46 | 0.31 | 0.41 | 0.32 |
| 8 | 3534630 | rs12676229 | 1.145 | -0.725 | 0.253 | 0.138 | 0 | -0.004 | -0.004 | 1.903 | 0.46 | 0.45 | 0.57 | 0.44 | 0.45 | 0.34 |
| 8 | 3541190 | rs7844039 | 1.113 | -0.619 | 0.223 | 0.295 | -0.002 | 0.004 | 0.004 | 2.612 | 0.47 | 0.44 | 0.59 | 0.51 | 0.48 | 0.35 |
| 8 | 3546520 | rs17066905 | 0.798 | -0.623 | 0.43 | 0.105 | -0.029 | 0.116 | 0.116 | 0.485 | 0.12 | 0.33 | 0.12 | 0.24 | 0.07 | 0.07 |
| 8 | 3546812 | rs4307367 | 0.174 | -0.751 | 0.371 | -0.176 | -0.007 | -0.024 | -0.024 | 2.128 | 0.8 | 0.71 | 0.9 | 0.84 | 0.81 | 0.79 |
| 8 | 3547270 | rs2046200 | -0.301 | -0.783 | 0.481 | -0.171 | -0.034 | -0.034 | -0.034 | 0 | 0.18 | 0.41 | 0.1 | 0.16 | 0.13 | 0.11 |
| 8 | 3547954 | rs2101116 | -0.161 | -0.712 | 0.027 | -0.129 | -0.029 | 0.027 | 0.027 | 0.925 | 0.31 | 0.11 | 0.28 | 0.28 | 0.33 | 0.4 |
| 8 | 3547988 | rs17326146 | 0.457 | -0.723 | 0.083 | -0.133 | -0.009 | -0.009 | -0.009 | 0 | 0.14 | 0.21 | 0.09 | 0.15 | 0.12 | 0.11 |
| 8 | 3548027 | rs2086532 | -0.169 | -0.72 | 0.033 | -0.133 | -0.027 | 0.028 | 0.028 | 0.951 | 0.32 | 0.11 | 0.28 | 0.28 | 0.33 | 0.41 |
| 8 | 3548551 | rs2624074 | -0.846 | -0.578 | 0.058 | -0.043 | -0.002 | 0.024 | 0.024 | 0.721 | 0.46 | 0.34 | 0.36 | 0.42 | 0.44 | 0.51 |
| 8 | 3551408 | rs2624062 | 1.424 | -0.249 | -0.257 | -0.135 | -0.063 | -0.03 | -0.03 | 1.868 | 0.37 | 0.1 | 0.49 | 0.33 | 0.43 | 0.32 |

| | | | | | | | | | | | | | | | | |
|---|---|---|---|---|---|---|---|---|---|---|---|---|---|---|---|---|
| 8 | 3553750 | rs13264352 | 0.238 | -0.195 | 0.072 | -0.229 | -0.002 | 0.001 | 0.001 | 0.767 | 0.6 | 0.43 | 0.64 | 0.54 | 0.62 | 0.53 |
| 8 | 3555378 | rs2623589 | -0.597 | -0.076 | -0.098 | -0.11 | 0.036 | 0.001 | 0.001 | 0.387 | 0.7 | 0.86 | 0.83 | 0.76 | 0.78 | 0.65 |
| 8 | 3555522 | rs2407122 | 0.873 | -0.01 | -0.014 | -0.174 | 0 | 0 | 0 | 0 | 0.51 | 0.54 | 0.6 | 0.54 | 0.53 | 0.42 |
| 8 | 3555639 | rs2251850 | 0.332 | 0.163 | -0.021 | -0.092 | 0.045 | -0.048 | -0.048 | 0 | 0.09 | 0.25 | 0.07 | 0.01 | 0.09 | 0.12 |
| 8 | 3555657 | rs17066956 | 1.396 | 0.157 | -0.044 | -0.079 | -0.042 | -0.052 | -0.052 | 0 | 0.09 | 0.02 | 0.15 | 0.22 | 0.14 | 0.12 |
| 8 | 3556268 | rs2623587 | 0.026 | 0.173 | 0.075 | -0.108 | 0.04 | -0.057 | -0.057 | 0 | 0.1 | 0.26 | 0.07 | 0.01 | 0.11 | 0.12 |
| 8 | 3560723 | rs4875773 | 0.731 | -0.037 | 0.055 | -0.052 | 0.017 | 0.004 | 0.004 | 5.548 | 0.49 | 0.48 | 0.6 | 0.46 | 0.53 | 0.43 |
| 8 | 3560760 | rs4875774 | 2.302 | -0.018 | -0.139 | -0.12 | -0.095 | -0.104 | -0.104 | 0.114 | 0.29 | 0.03 | 0.43 | 0.32 | 0.34 | 0.28 |
| 8 | 3562041 | rs3739356 | 0.977 | 0.08 | 0.35 | 0.257 | -0.005 | 0.002 | 0.002 | 0 | 0.11 | 0.1 | 0.09 | 0.12 | 0.13 | 0.12 |
| 8 | 3562992 | rs7825205 | -0.525 | -0.377 | 0.237 | 0.386 | 0.003 | -0.017 | -0.017 | 1.481 | 0.95 | 0.86 | 0.97 | 0.99 | 0.93 | 0.87 |
| 8 | 3563509 | rs17067066 | -0.212 | -0.43 | 0.255 | 0.399 | 0.03 | -0.036 | -0.036 | 0 | 0.36 | 0.15 | 0.32 | 0.46 | 0.31 | 0.37 |
| 8 | 3564244 | rs17326685 | 1.134 | -0.389 | 0.321 | 0.432 | 0.008 | 0.002 | 0.002 | 5.911 | 0.49 | 0.58 | 0.62 | 0.46 | 0.56 | 0.44 |
| 8 | 3564568 | rs11986814 | 1.08 | -0.388 | 0.217 | 0.433 | 0.033 | 0.008 | 0.008 | 4.558 | 0.51 | 0.65 | 0.62 | 0.46 | 0.56 | 0.45 |
| 8 | 3565900 | rs12674750 | 0.477 | -0.327 | -0.076 | 0.395 | -0.025 | 0 | 0 | 0.079 | 0.27 | 0.22 | 0.16 | 0.24 | 0.15 | 0.25 |
| 8 | 3566615 | rs6981596 | 0.168 | 0.053 | 0.173 | 0.155 | 0.027 | -0.021 | -0.021 | 0.01 | 0.93 | 0.75 | 0.97 | 0.99 | 0.93 | 0.87 |
| 8 | 3568042 | rs4433172 | 0.084 | 0.073 | -0.055 | 0.218 | 0.009 | -0.01 | -0.01 | 0 | 0.37 | 0.25 | 0.34 | 0.47 | 0.31 | 0.38 |
| 8 | 3572855 | rs2720790 | 1.024 | 0.662 | 0.177 | 0.121 | -0.003 | -0.023 | -0.023 | 0 | 0.87 | 0.74 | 0.94 | 0.93 | 0.87 | 0.82 |
| 8 | 3574311 | rs2720797 | 0.544 | 0.561 | 0.128 | 0.292 | -0.011 | -0.001 | -0.001 | 0 | 0.92 | 0.81 | 0.95 | 0.94 | 0.93 | 0.92 |
| 8 | 3575929 | rs7843888 | -0.767 | 1.119 | 0.383 | 0.201 | 0.077 | -0.009 | -0.009 | 0 | 0.14 | 0.38 | 0.13 | 0.14 | 0.15 | 0.12 |
| 8 | 3576390 | rs11996752 | -0.36 | 0.93 | -0.15 | 0.057 | -0.001 | -0.015 | -0.015 | 0 | 0.3 | 0.4 | 0.21 | 0.24 | 0.31 | 0.14 |
| 8 | 3577522 | rs4875783 | 0.665 | 0.801 | -0.451 | 0.543 | 0.007 | 0.007 | 0.007 | 0 | 0.63 | 0.48 | 0.55 | 0.54 | 0.56 | 0.71 |
| 8 | 3578650 | rs11136653 | 0.199 | 0.375 | 0.1 | 0.941 | 0.019 | 0.088 | 0.088 | 0.005 | 0.89 | 0.73 | 0.78 | 0.73 | 0.85 | 0.86 |
| 8 | 3578731 | rs11136656 | 0.199 | 0.375 | 0.1 | 0.941 | 0.019 | 0.088 | 0.088 | 0.005 | 0.89 | 0.73 | 0.78 | 0.73 | 0.85 | 0.86 |
| 8 | 3580126 | rs17326768 | 1.37 | -0.106 | -0.413 | 0.673 | -0.015 | 0.155 | 0.155 | 0.113 | 0.46 | 0.1 | 0.37 | 0.29 | 0.56 | 0.37 |
| 8 | 3581019 | rs4990839 | 1.015 | 0.146 | 0.369 | 1.003 | 0.018 | 0.048 | 0.048 | 0 | 0.79 | 0.48 | 0.72 | 0.73 | 0.76 | 0.87 |
| 8 | 3582183 | rs1905030 | -0.323 | 0.105 | 0.169 | 1.092 | 0.047 | 0.093 | 0.093 | 0 | 0.93 | 0.78 | 0.84 | 0.83 | 0.88 | 0.93 |
| 8 | 3583503 | rs2623753 | -0.035 | -0.408 | -0.204 | 0.85 | 0.004 | -0.003 | -0.003 | 0 | 0.11 | 0.14 | 0.1 | 0.12 | 0.09 | 0.09 |
| 8 | 3584423 | rs2623749 | -0.085 | -0.034 | 0.037 | 1.107 | -0.037 | 0.128 | 0.128 | 0 | 0.79 | 0.61 | 0.7 | 0.67 | 0.88 | 0.67 |
| 8 | 3585282 | rs12682219 | -0.902 | 0.232 | -0.128 | 0.828 | 0.016 | 0.022 | 0.022 | 1.109 | 0.06 | 0.22 | 0.03 | 0.03 | 0.03 | 0.01 |
| 8 | 3585967 | rs13277081 | -0.421 | -0.19 | -0.496 | 0.998 | 0.009 | -0.011 | -0.011 | 0 | 0.28 | 0.16 | 0.36 | 0.33 | 0.18 | 0.38 |
| 8 | 3586010 | rs17063147 | -0.134 | -0.369 | -0.498 | 1.589 | -0.004 | 0.001 | 0.001 | 0 | 0.23 | 0.19 | 0.22 | 0.22 | 0.14 | 0.36 |
| 8 | 3586061 | rs2623746 | 1.016 | -0.435 | -1.307 | 1.304 | 0.035 | 0.119 | 0.119 | 0.976 | 0.48 | 0.12 | 0.39 | 0.33 | 0.66 | 0.37 |
| 8 | 3586074 | rs2623745 | 0.032 | -0.458 | -0.091 | 1.597 | -0.01 | 0.133 | 0.133 | 0.2 | 0.88 | 0.62 | 0.83 | 0.77 | 0.91 | 0.88 |
| 8 | 3586120 | rs2623744 | 0.718 | -0.401 | -0.168 | 1.612 | -0.032 | 0.139 | 0.139 | 0.837 | 0.83 | 0.48 | 0.81 | 0.68 | 0.9 | 0.86 |
| 8 | 3586818 | rs17067182 | 0.977 | -0.379 | -1.353 | 0.989 | -0.033 | -0.091 | -0.091 | 1.894 | 0.3 | 0.01 | 0.29 | 0.27 | 0.25 | 0.22 |
| 8 | 3587957 | rs2897573 | 0.898 | -0.132 | -0.857 | 0.812 | 0.039 | 0.026 | 0.026 | 2.42 | 0.62 | 0.18 | 0.72 | 0.64 | 0.71 | 0.63 |
| 8 | 3588100 | rs1031977 | -0.568 | -0.26 | -0.198 | 0.882 | -0.029 | 0.032 | 0.032 | 0.365 | 0.12 | 0.24 | 0.11 | 0.17 | 0.05 | 0.13 |
| 8 | 3589016 | rs10503215 | -0.775 | -0.178 | -0.069 | 0.875 | -0.022 | 0.016 | 0.016 | 0.041 | 0.12 | 0.24 | 0.12 | 0.15 | 0.06 | 0.14 |
| 8 | 3589501 | rs17067208 | -1.184 | -0.061 | 0.466 | 0.736 | -0.006 | -0.004 | -0.004 | 5.218 | 0.2 | 0.22 | 0.16 | 0.2 | 0.18 | 0.17 |
| 8 | 3589751 | rs6996403 | 0.66 | -0.142 | 0.282 | 0.562 | -0.001 | -0.033 | -0.033 | 0 | 0.1 | 0.11 | 0.04 | 0 | 0.08 | 0.09 |
| 8 | 3589776 | rs1026849 | 0.785 | 0.105 | 0.048 | 0.761 | -0.068 | -0.095 | -0.095 | 0.31 | 0.24 | 0.08 | 0.34 | 0.37 | 0.32 | 0.36 |
| 8 | 3589990 | rs1564552 | 1.731 | 0.003 | 0.256 | 0.687 | 0.007 | 0.038 | 0.038 | 3.05 | 0.41 | 0.23 | 0.38 | 0.33 | 0.38 | 0.32 |
| 8 | 3590610 | rs17067216 | -0.013 | -0.021 | 0.411 | 0.732 | -0.004 | 0.001 | 0.001 | 0 | 0.1 | 0.13 | 0.11 | 0.11 | 0.05 | 0.08 |
| 8 | 3592750 | rs17326950 | 0.833 | 0.071 | 0.33 | 0.755 | -0.071 | -0.103 | -0.103 | 0 | 0.25 | 0.07 | 0.33 | 0.37 | 0.32 | 0.37 |
| 8 | 3594287 | rs2981369 | 1.317 | -0.056 | 0.454 | 0.68 | 0.007 | 0.014 | 0.014 | 1.872 | 0.65 | 0.35 | 0.75 | 0.76 | 0.71 | 0.69 |
| 8 | 3595317 | rs2720823 | 0.329 | 0.131 | 0.78 | 0.661 | 0.005 | 0.004 | 0.004 | 0 | 0.6 | 0.55 | 0.55 | 0.42 | 0.53 | 0.46 |
| 8 | 3596746 | rs6981739 | 0.889 | 0.023 | 0.841 | 0.702 | -0.018 | -0.019 | -0.019 | 0.257 | 0.74 | 0.45 | 0.79 | 0.81 | 0.82 | 0.8 |
| 8 | 3598337 | rs2623713 | 0.32 | 0.047 | 0.787 | 0.623 | 0.01 | -0.027 | -0.027 | 1.687 | 0.13 | 0.19 | 0.08 | 0.09 | 0.13 | 0.11 |
| 8 | 3598374 | rs11136659 | 0.381 | 0.278 | 0.458 | 0.577 | -0.08 | -0.145 | -0.145 | 1.182 | 0.27 | 0.08 | 0.32 | 0.5 | 0.34 | 0.44 |
| 8 | 3600189 | rs9650510 | 1.883 | 0.331 | 0.585 | 0.54 | 0.017 | 0.05 | 0.05 | 1.06 | 0.41 | 0.27 | 0.36 | 0.26 | 0.36 | 0.31 |
| 8 | 3600743 | rs10096790 | -1.01 | 0.339 | 0.669 | 0.55 | 0.001 | -0.009 | -0.009 | 0 | 0.47 | 0.52 | 0.56 | 0.66 | 0.52 | 0.59 |
| 8 | 3603635 | rs12334508 | -0.375 | 0.31 | 0.436 | 0.542 | -0.031 | -0.017 | -0.017 | 0 | 0.37 | 0.21 | 0.49 | 0.53 | 0.4 | 0.47 |
| 8 | 3604955 | rs10503216 | 1.834 | 0.3 | 0.475 | 0.518 | 0.013 | 0.046 | 0.046 | 1.251 | 0.39 | 0.25 | 0.36 | 0.29 | 0.36 | 0.3 |
| 8 | 3605461 | rs12679480 | 0.274 | 0.182 | 0.275 | 0.669 | 0 | -0.002 | -0.002 | 3.672 | 0.15 | 0.13 | 0.2 | 0.14 | 0.16 | 0.1 |
| 8 | 3611169 | rs2623729 | 0.567 | 0.436 | 0.311 | 0.789 | -0.026 | -0.091 | -0.091 | 0 | 0.26 | 0.13 | 0.31 | 0.47 | 0.25 | 0.4 |
| 8 | 3612977 | rs2720865 | -0.659 | 0.327 | 0.754 | 0.631 | 0.004 | 0.01 | 0.01 | 0 | 0.41 | 0.48 | 0.4 | 0.47 | 0.39 | 0.53 |
| 8 | 3614229 | rs12543111 | 0.641 | 0.408 | 0.761 | 1.17 | -0.045 | 0.002 | 0.002 | 0 | 0.25 | 0.06 | 0.19 | 0.22 | 0.24 | 0.29 |
| 8 | 3614453 | rs1481720 | -0.149 | 0.459 | 0.875 | 1.118 | 0.012 | 0.045 | 0.045 | 0 | 0.31 | 0.42 | 0.36 | 0.47 | 0.29 | 0.41 |
| 8 | 3617569 | rs4363216 | 0.433 | 0.411 | 1.173 | 1.28 | 0.023 | 0.128 | 0.128 | 0 | 0.14 | 0.32 | 0.26 | 0.24 | 0.16 | 0.08 |
| 8 | 3617597 | rs13278570 | 1.514 | 0.368 | 1.268 | 1.137 | 0.009 | 0.039 | 0.039 | 0.606 | 0.44 | 0.29 | 0.35 | 0.3 | 0.42 | 0.32 |
| 8 | 3618117 | rs7826540 | 0.078 | 0.425 | 1.128 | 1.131 | 0.103 | 0.042 | 0.042 | 0.299 | 0.05 | 0.32 | 0.04 | 0 | 0.04 | 0.01 |
| 8 | 3618574 | rs2623734 | -1.251 | 0.437 | 0.871 | 1.157 | -0.002 | -0.003 | -0.003 | 0 | 0.93 | 0.89 | 0.96 | 0.91 | 0.92 | 0.91 |
| 8 | 3620123 | rs10093538 | -0.291 | 0.435 | 0.79 | 1.206 | -0.002 | -0.007 | -0.007 | 0 | 0.2 | 0.21 | 0.36 | 0.39 | 0.24 | 0.26 |
| 8 | 3620935 | rs12675982 | 0.755 | 0.452 | 0.761 | 1.081 | -0.037 | 0.056 | 0.056 | 0 | 0.24 | 0.1 | 0.2 | 0.14 | 0.23 | 0.27 |
| 8 | 3621844 | rs6558819 | 0.253 | 0.444 | 0.777 | 1.045 | 0.002 | -0.004 | -0.004 | 0 | 0.72 | 0.67 | 0.7 | 0.77 | 0.69 | 0.74 |
| 8 | 3623388 | rs11136663 | 1.673 | 0.435 | 0.479 | 1.086 | 0.01 | 0.032 | 0.032 | 0.191 | 0.38 | 0.23 | 0.32 | 0.29 | 0.38 | 0.3 |
| 8 | 3624046 | rs12678875 | 0.783 | 0.673 | 0.17 | 0.934 | 0 | 0.002 | 0.002 | 0 | 0.14 | 0.15 | 0.24 | 0.14 | 0.16 | 0.1 |
| 8 | 3624896 | rs10216475 | 0.422 | 0.601 | 0.124 | 0.685 | -0.011 | 0.024 | 0.024 | 0 | 0.24 | 0.14 | 0.2 | 0.14 | 0.24 | 0.26 |
| 8 | 3625284 | rs7816314 | 0.878 | 0.253 | 0.561 | 0.886 | -0.013 | 0 | 0 | 0 | 0.76 | 0.58 | 0.88 | 0.74 | 0.81 | 0.81 |

| | | | | | | | | | | | | | | | | |
|---|---|---|---|---|---|---|---|---|---|---|---|---|---|---|---|---|
| 8 | 3625780 | rs10095924 | -0.149 | 0.393 | 0.131 | 0.915 | -0.001 | -0.001 | -0.001 | 0 | 0.12 | 0.11 | 0.05 | 0.1 | 0.11 | 0.1 |
| 8 | 3630424 | rs2720770 | -0.496 | 0.358 | 0.241 | 0.997 | -0.002 | 0.023 | 0.023 | 0 | 0.13 | 0.14 | 0.06 | 0.3 | 0.11 | 0.11 |
| 8 | 3631634 | rs1994051 | 0.844 | 0.349 | 0.753 | 0.785 | 0.018 | -0.05 | -0.05 | 0 | 0.09 | 0.33 | 0.08 | 0 | 0.07 | 0.08 |
| 8 | 3631875 | rs1982696 | 0.281 | 0.39 | -0.522 | 0.706 | -0.079 | 0.03 | 0.03 | 0 | 0.37 | 0.06 | 0.38 | 0.25 | 0.41 | 0.37 |
| 8 | 3636066 | rs2623657 | -0.677 | 0.486 | -0.012 | 0.84 | 0.006 | 0.023 | 0.023 | 1.194 | 0.23 | 0.43 | 0.14 | 0.31 | 0.2 | 0.19 |
| 8 | 3636749 | rs1481713 | -0.789 | 0.443 | -0.065 | 1.31 | -0.001 | 0.123 | 0.123 | 1.162 | 0.24 | 0.43 | 0.26 | 0.48 | 0.21 | 0.31 |
| 8 | 3637074 | rs4875786 | 1.682 | 0.206 | -0.044 | 1.17 | -0.032 | 0.017 | 0.017 | 0.441 | 0.34 | 0.13 | 0.36 | 0.29 | 0.38 | 0.31 |
| 8 | 3637980 | rs2623662 | -1.077 | 0.348 | 0.097 | 1.524 | -0.001 | 0.124 | 0.124 | 1.266 | 0.24 | 0.44 | 0.27 | 0.48 | 0.21 | 0.32 |
| 8 | 3638047 | rs10503217 | 0.5 | 0.227 | -0.049 | 1.252 | 0.006 | 0.093 | 0.093 | 4.176 | 0.11 | 0.33 | 0.2 | 0.17 | 0.09 | 0.21 |
| 8 | 3641922 | rs13282787 | 0.431 | 0.425 | -0.966 | 0.899 | -0.017 | -0.024 | -0.024 | 0.022 | 0.17 | 0.02 | 0.21 | 0.16 | 0.2 | 0.11 |
| 8 | 3643487 | rs2720890 | -0.891 | 0.537 | -0.814 | 1.016 | 0.002 | -0.011 | -0.011 | 0 | 0.17 | 0.15 | 0.29 | 0.29 | 0.16 | 0.16 |
| 8 | 3643595 | rs2623665 | -0.837 | 1.153 | -0.791 | 0.523 | 0.004 | -0.006 | -0.006 | 0.109 | 0.1 | 0.1 | 0.1 | 0.06 | 0.16 | 0.17 |
| 8 | 3644651 | rs2720875 | -0.177 | 0.382 | -0.494 | 0.542 | 0.028 | 0.027 | 0.027 | 0 | 0.48 | 0.59 | 0.55 | 0.67 | 0.4 | 0.42 |
| 8 | 3644685 | rs2623668 | -0.916 | -0.139 | 0.053 | 1.042 | -0.044 | 0.045 | 0.045 | 0 | 0.18 | 0.27 | 0.18 | 0.39 | 0.08 | 0.21 |
| 8 | 3645747 | rs2623669 | -1.495 | -0.223 | 0.098 | 1.209 | -0.09 | 0.099 | 0.099 | 0.157 | 0.22 | 0.52 | 0.22 | 0.44 | 0.11 | 0.31 |
| 8 | 3645978 | rs2720851 | -0.473 | -0.075 | 0.064 | 0.956 | 0.022 | 0.018 | 0.018 | 0.077 | 0.42 | 0.65 | 0.43 | 0.67 | 0.33 | 0.55 |
| 8 | 3647890 | rs2623679 | 0.082 | -0.193 | 0.088 | 0.73 | -0.015 | 0.01 | 0.01 | 0 | 0.11 | 0.07 | 0.13 | 0.24 | 0.05 | 0.1 |
| 8 | 3648320 | rs2720811 | 0.114 | -0.179 | -0.142 | 0.701 | -0.002 | 0.005 | 0.005 | 0 | 0.33 | 0.3 | 0.36 | 0.52 | 0.28 | 0.43 |
| 8 | 3650502 | rs10093930 | 0.246 | -0.003 | -0.064 | 0.794 | 0.038 | -0.029 | -0.029 | 0 | 0.57 | 0.32 | 0.58 | 0.51 | 0.68 | 0.56 |
| 8 | 3650583 | rs2623683 | -0.243 | -0.001 | -0.039 | 0.761 | 0.011 | -0.01 | -0.01 | 0 | 0.42 | 0.59 | 0.41 | 0.48 | 0.32 | 0.44 |
| 8 | 3651181 | rs2623685 | -0.23 | 0.029 | -0.211 | 0.496 | 0.025 | 0.041 | 0.041 | 0.371 | 0.52 | 0.62 | 0.59 | 0.68 | 0.46 | 0.43 |
| 8 | 3655261 | rs12550117 | -0.082 | 0.102 | -0.247 | 0.509 | -0.023 | 0.063 | 0.063 | 0 | 0.29 | 0.22 | 0.21 | 0.11 | 0.34 | 0.3 |
| 8 | 3655372 | rs7017017 | -0.082 | 0.11 | -0.244 | 0.509 | -0.024 | 0.06 | 0.06 | 0 | 0.29 | 0.23 | 0.21 | 0.11 | 0.34 | 0.31 |
| 8 | 3660565 | rs4527894 | -1.3 | 0.215 | -0.289 | 0.572 | -0.004 | 0.011 | 0.011 | 2.055 | 0.73 | 0.88 | 0.75 | 0.78 | 0.74 | 0.63 |
| 8 | 3663940 | rs6998283 | 0.652 | 0.092 | -0.267 | 0.656 | 0.006 | -0.024 | -0.024 | 0 | 0.4 | 0.19 | 0.42 | 0.43 | 0.42 | 0.31 |
| 8 | 3663971 | rs7015612 | 0.658 | 0.099 | -0.268 | 0.626 | 0.01 | -0.013 | -0.013 | 0.003 | 0.41 | 0.3 | 0.4 | 0.47 | 0.42 | 0.34 |
| 8 | 3666456 | rs1871816 | 1.065 | 0.101 | -0.194 | 0.784 | -0.015 | 0.212 | 0.212 | 0.167 | 0.06 | 0.14 | 0.13 | 0.28 | 0.03 | 0.05 |
| 8 | 3668651 | rs2720887 | -0.288 | 0.078 | -0.213 | 0.805 | -0.037 | 0.1 | 0.1 | 0 | 0.11 | 0.15 | 0.15 | 0.35 | 0.06 | 0.1 |
| 8 | 3670381 | rs10102119 | -1.278 | 0.172 | -0.706 | 0.38 | -0.015 | -0.01 | -0.01 | 0 | 0.73 | 0.92 | 0.75 | 0.78 | 0.74 | 0.69 |
| 8 | 3672437 | rs1383945 | 1.091 | 0.59 | -0.628 | 0.018 | -0.005 | -0.024 | -0.024 | 1.142 | 0.35 | 0.21 | 0.36 | 0.39 | 0.35 | 0.3 |
| 8 | 3672470 | rs2929496 | -0.043 | 1.358 | -0.351 | -0.638 | -0.062 | 0.093 | 0.093 | 0.169 | 0.24 | 0.11 | 0.18 | 0.04 | 0.31 | 0.27 |
| 8 | 3674674 | rs17744884 | 0.679 | 1.64 | -0.19 | -0.523 | -0.02 | 0.019 | 0.019 | 0.035 | 0.18 | 0.16 | 0.16 | 0.03 | 0.25 | 0.17 |
| 8 | 3675565 | rs2720739 | -0.862 | 1.409 | 0.042 | -0.476 | 0.017 | -0.021 | -0.021 | 1.02 | 0.33 | 0.69 | 0.31 | 0.21 | 0.45 | 0.39 |
| 8 | 3677594 | rs17063168 | -0.871 | 1.379 | 0.296 | -0.449 | 0.079 | 0.126 | 0.126 | 0 | 0.08 | 0.37 | 0.09 | 0.2 | 0.11 | 0.14 |
| 8 | 3678006 | rs4607641 | 0.13 | 1.354 | 0.345 | -0.439 | 0.048 | -0.105 | -0.105 | 0 | 0.24 | 0.5 | 0.21 | 0.04 | 0.28 | 0.23 |
| 8 | 3678385 | rs9694077 | -1.233 | 1.314 | 0.33 | -0.397 | 0.056 | 0.089 | 0.089 | 0.494 | 0.13 | 0.58 | 0.11 | 0.22 | 0.13 | 0.17 |
| 8 | 3679446 | rs7007032 | -0.461 | 1.136 | -0.078 | -0.574 | 0.019 | 0.05 | 0.05 | 1.885 | 0.67 | 0.88 | 0.74 | 0.81 | 0.7 | 0.52 |
| 8 | 3679504 | rs7007070 | -1.049 | 1.343 | 0.051 | -0.472 | 0.002 | 0.016 | 0.016 | 1.026 | 0.73 | 0.91 | 0.77 | 0.81 | 0.76 | 0.6 |
| 8 | 3680182 | rs10110717 | 1.076 | 1.336 | 0.111 | -0.45 | 0.002 | 0 | 0 | 0 | 0.39 | 0.29 | 0.47 | 0.61 | 0.32 | 0.22 |
| 8 | 3681516 | rs17807654 | 0.692 | 1.413 | 0.058 | -0.43 | -0.038 | 0.056 | 0.056 | 0.005 | 0.2 | 0.11 | 0.17 | 0.04 | 0.26 | 0.18 |
| 8 | 3683879 | rs4437679 | -0.739 | 1.532 | 0.074 | -0.343 | 0.013 | 0.037 | 0.037 | 0.157 | 0.72 | 0.88 | 0.77 | 0.82 | 0.73 | 0.61 |
| 8 | 3686944 | rs4270991 | 1.091 | 1.561 | -0.345 | -0.25 | 0.023 | -0.017 | -0.017 | 0 | 0.39 | 0.21 | 0.48 | 0.57 | 0.32 | 0.26 |
| 8 | 3687399 | rs7824328 | -0.49 | 1.548 | -0.316 | -0.238 | 0.011 | 0.022 | 0.022 | 0.162 | 0.69 | 0.79 | 0.77 | 0.8 | 0.72 | 0.59 |
| 8 | 3687848 | rs11988752 | 1.237 | 1.407 | -0.564 | -0.111 | 0.015 | -0.015 | -0.015 | 0 | 0.37 | 0.21 | 0.45 | 0.55 | 0.29 | 0.25 |
| 8 | 3687917 | rs2975373 | 0.085 | 1.133 | 0.052 | 0.125 | 0.032 | 0.031 | 0.031 | 0 | 0.07 | 0.18 | 0.07 | 0.16 | 0.11 | 0.11 |
| 8 | 3694644 | rs9314506 | 1.087 | 1.142 | -0.223 | 0.01 | 0.005 | 0.012 | 0.012 | 0 | 0.46 | 0.2 | 0.53 | 0.6 | 0.44 | 0.34 |
| 8 | 3694817 | rs12542916 | -0.445 | 1.087 | -0.391 | -0.041 | 0.044 | -0.007 | -0.007 | 1.023 | 0.27 | 0.58 | 0.26 | 0.23 | 0.34 | 0.28 |
| 8 | 3695136 | rs11777359 | -0.011 | 1.108 | -0.646 | 0.117 | 0.016 | 0.038 | 0.038 | 0 | 0.09 | 0.17 | 0.08 | 0.18 | 0.11 | 0.12 |
| 8 | 3695148 | rs10216665 | -0.998 | 1.1 | -0.725 | -0.045 | 0.023 | 0.014 | 0.014 | 0.22 | 0.79 | 0.92 | 0.83 | 0.85 | 0.86 | 0.69 |
| 8 | 3698189 | rs7341563 | -1.45 | 1.792 | 0.126 | 0.218 | 0.015 | -0.017 | -0.017 | 0 | 0.34 | 0.65 | 0.28 | 0.22 | 0.43 | 0.35 |
| 8 | 3699398 | rs6991653 | -0.631 | 1.568 | 0.018 | 0.073 | 0.015 | -0.11 | -0.11 | 0 | 0.27 | 0.47 | 0.23 | 0.05 | 0.34 | 0.24 |
| 8 | 3703484 | rs10093889 | -0.949 | 1.755 | -0.54 | 0.855 | 0.01 | 0.025 | 0.025 | 0 | 0.09 | 0.17 | 0.05 | 0.18 | 0.1 | 0.11 |
| 8 | 3703884 | rs10086452 | -0.913 | 1.701 | -0.689 | 0.633 | -0.026 | 0.004 | 0.004 | 0 | 0.28 | 0.27 | 0.26 | 0.21 | 0.34 | 0.25 |
| 8 | 3706150 | rs2954219 | -1.344 | 1.134 | 0.001 | 1.338 | -0.003 | 0.069 | 0.069 | 0 | 0.15 | 0.22 | 0.17 | 0.42 | 0.14 | 0.26 |
| 8 | 3707645 | rs2954224 | -1.381 | 1.189 | -0.012 | 1.329 | 0 | 0.065 | 0.065 | 0 | 0.15 | 0.22 | 0.16 | 0.42 | 0.14 | 0.26 |
| 8 | 3708700 | rs2930362 | -1.951 | 0.78 | 0.463 | 0.77 | 0 | 0 | 0 | 0 | 0.34 | 0.68 | 0.39 | 0.49 | 0.33 | 0.48 |
| 8 | 3709049 | rs13253701 | -0.492 | 0.574 | 0.646 | 0.743 | -0.003 | 0.002 | 0.002 | 0 | 0.09 | 0.09 | 0.15 | 0.06 | 0.14 | 0.2 |
| 8 | 3709660 | rs2930357 | -2.258 | 0.387 | 1.047 | 0.829 | -0.018 | 0.053 | 0.053 | 0.008 | 0.28 | 0.56 | 0.35 | 0.52 | 0.27 | 0.47 |
| 8 | 3711034 | rs2930355 | -1.477 | 0.642 | 1.221 | 0.642 | -0.002 | 0.02 | 0.02 | 0 | 0.21 | 0.25 | 0.21 | 0.44 | 0.19 | 0.23 |
| 8 | 3717468 | rs2930353 | -2.566 | 0.425 | 1.569 | 0.547 | -0.011 | 0.008 | 0.008 | 0.168 | 0.39 | 0.51 | 0.4 | 0.49 | 0.35 | 0.48 |
| 8 | 3717935 | rs7012625 | -2.155 | 0.249 | 1.909 | 0.508 | 0.011 | -0.006 | -0.006 | 0 | 0.3 | 0.74 | 0.27 | 0.4 | 0.24 | 0.3 |
| 8 | 3717980 | rs17067779 | 0.114 | 0.356 | 1.606 | 0.569 | -0.007 | -0.011 | -0.011 | 0 | 0.07 | 0.03 | 0.13 | 0.06 | 0.08 | 0.14 |
| 8 | 3719161 | rs2975367 | -1.421 | 0.326 | 1.818 | 0.635 | 0 | 0 | 0 | 0 | 0.22 | 0.2 | 0.2 | 0.38 | 0.2 | 0.29 |
| 8 | 3726055 | rs13268401 | -0.413 | 0.51 | 1.312 | 0.657 | -0.004 | -0.011 | -0.011 | 0 | 0.08 | 0.04 | 0.13 | 0.07 | 0.09 | 0.19 |
| 8 | 3726907 | rs4146932 | -2.251 | 0.429 | 1.417 | 0.557 | 0.019 | -0.03 | -0.03 | 0 | 0.38 | 0.81 | 0.4 | 0.45 | 0.32 | 0.4 |
| 8 | 3728020 | rs2975359 | -1.356 | 0.426 | 2.124 | 0.566 | 0 | 0.04 | 0.04 | 0.273 | 0.26 | 0.35 | 0.24 | 0.41 | 0.23 | 0.24 |
| 8 | 3729496 | rs7830991 | -1.66 | 0.697 | 2.05 | 0.552 | 0 | 0.001 | 0.001 | 0.054 | 0.28 | 0.66 | 0.26 | 0.45 | 0.28 | 0.27 |
| 8 | 3731642 | rs13259787 | -0.396 | 0.647 | 1.8 | 0.54 | 0 | 0.002 | 0.002 | 0 | 0.09 | 0.07 | 0.14 | 0.06 | 0.09 | 0.19 |
| 8 | 3732601 | rs6996498 | -1.744 | 0.654 | 2.306 | 0.611 | 0 | 0.025 | 0.025 | 0.113 | 0.28 | 0.32 | 0.34 | 0.45 | 0.28 | 0.41 |

| | | | | | | | | | | | | | | | | |
|---|---|---|---|---|---|---|---|---|---|---|---|---|---|---|---|---|
| 8 | 3733003 | rs6998344 | -1.169 | 0.654 | 2.436 | 0.579 | 0.004 | -0.036 | -0.036 | 0 | 0.2 | 0.31 | 0.13 | 0.11 | 0.19 | 0.19 |
| 8 | 3734934 | rs2975348 | 1.597 | 1.359 | 2.697 | 0.901 | 0.034 | 0.056 | 0.056 | 0.04 | 0.75 | 0.55 | 0.81 | 0.66 | 0.72 | 0.76 |
| 8 | 3735032 | rs4388474 | 1.424 | 1.336 | 2.823 | 0.882 | 0.015 | 0.026 | 0.026 | 0.077 | 0.75 | 0.35 | 0.79 | 0.67 | 0.72 | 0.76 |
| 8 | 3736503 | rs13257055 | -1.956 | 1.118 | 2.995 | 0.816 | -0.014 | 0.007 | 0.007 | 0.042 | 0.16 | 0.27 | 0.12 | 0.23 | 0.12 | 0.24 |
| 8 | 3736913 | rs10089344 | -2.616 | 1.158 | 3.027 | 0.8 | 0.011 | -0.002 | -0.002 | 2.605 | 0.34 | 0.75 | 0.24 | 0.33 | 0.28 | 0.36 |
| 8 | 3737474 | rs11989166 | -1.147 | 1.155 | 3.018 | 0.808 | 0.003 | -0.05 | -0.05 | 0 | 0.2 | 0.48 | 0.14 | 0.09 | 0.18 | 0.13 |
| 8 | 3742245 | rs11786301 | 2.157 | 1.152 | 3.031 | 0.756 | -0.018 | -0.023 | -0.023 | 0 | 0.65 | 0.16 | 0.72 | 0.64 | 0.64 | 0.62 |
| 8 | 3743017 | rs2627359 | -1.256 | 1.077 | 2.838 | 0.771 | -0.009 | 0.083 | 0.083 | 0.161 | 0.21 | 0.48 | 0.15 | 0.3 | 0.18 | 0.16 |
| 8 | 3759146 | rs2930378 | -1.08 | 0.476 | 2.465 | 0.565 | -0.012 | 0.048 | 0.048 | 0.328 | 0.22 | 0.29 | 0.18 | 0.35 | 0.17 | 0.13 |
| 8 | 3759339 | rs2627425 | -2.213 | 0.375 | 2.444 | 0.495 | 0.044 | -0.031 | -0.031 | 2.271 | 0.35 | 0.86 | 0.29 | 0.37 | 0.25 | 0.31 |
| 8 | 3761373 | rs2948660 | -1.259 | 0.554 | 2.423 | 0.424 | -0.014 | 0.062 | 0.062 | 0.111 | 0.24 | 0.35 | 0.2 | 0.35 | 0.19 | 0.15 |
| 8 | 3761737 | rs10107710 | 2.527 | 0.611 | 2.351 | 0.38 | -0.043 | -0.089 | -0.089 | 0 | 0.67 | 0.1 | 0.68 | 0.59 | 0.65 | 0.66 |
| 8 | 3761816 | rs2975333 | -0.637 | 0.652 | 2.461 | 0.432 | -0.004 | 0.064 | 0.064 | 0.244 | 0.2 | 0.29 | 0.2 | 0.36 | 0.19 | 0.12 |
| 8 | 3763420 | rs11775224 | 2.063 | 0.663 | 1.988 | 0.451 | -0.037 | -0.066 | -0.066 | 0 | 0.63 | 0.1 | 0.68 | 0.6 | 0.65 | 0.66 |
| 8 | 3763463 | rs2688402 | 2.191 | 0.615 | 2.153 | 0.425 | 0.002 | -0.015 | -0.015 | 0 | 0.6 | 0.23 | 0.67 | 0.6 | 0.63 | 0.64 |
| 8 | 3763540 | rs17067892 | -0.304 | 0.603 | 2.157 | 0.382 | -0.001 | -0.012 | -0.012 | 0 | 0.07 | 0.1 | 0.05 | 0.03 | 0.08 | 0.15 |
| 8 | 3764615 | rs2688288 | -1.283 | 0.55 | 2.201 | 0.341 | -0.012 | -0.064 | -0.064 | 0.706 | 0.13 | 0.32 | 0.1 | 0.03 | 0.12 | 0.18 |
| 8 | 3766337 | rs2930372 | -0.518 | 0.552 | 2.248 | 0.373 | -0.018 | 0.156 | 0.156 | 0.847 | 0.19 | 0.41 | 0.18 | 0.35 | 0.14 | 0.13 |
| 8 | 3767210 | rs2688325 | 1.784 | 0.629 | 2.091 | 0.258 | 0 | 0.006 | 0.006 | 0.83 | 0.67 | 0.37 | 0.72 | 0.62 | 0.69 | 0.8 |
| 8 | 3768068 | rs2554507 | -0.495 | 0.629 | 2.078 | 0.253 | 0.013 | -0.028 | -0.028 | 2.057 | 0.08 | 0.25 | 0.06 | 0.03 | 0.07 | 0.16 |
| 8 | 3768466 | rs10216531 | 2.264 | 0.891 | 1.95 | 0.265 | -0.035 | -0.07 | -0.07 | 0 | 0.58 | 0.09 | 0.65 | 0.59 | 0.6 | 0.64 |
| 8 | 3769690 | rs2954625 | -1.081 | 0.92 | 2.23 | 0.253 | -0.016 | 0.055 | 0.055 | 0.348 | 0.26 | 0.46 | 0.22 | 0.35 | 0.21 | 0.15 |
| 8 | 3770471 | rs2554702 | -1.888 | 0.933 | 2.207 | 0.256 | 0.016 | -0.018 | -0.018 | 1.73 | 0.35 | 0.77 | 0.31 | 0.38 | 0.31 | 0.32 |
| 8 | 3771050 | rs2975331 | -0.407 | 0.927 | 2.362 | 0.325 | -0.009 | 0.063 | 0.063 | 0.057 | 0.2 | 0.33 | 0.18 | 0.35 | 0.17 | 0.13 |
| 8 | 3771236 | rs2912288 | -1.519 | 0.903 | 2.205 | 0.295 | -0.021 | 0.031 | 0.031 | 0 | 0.31 | 0.46 | 0.25 | 0.37 | 0.25 | 0.3 |
| 8 | 3772322 | rs17067967 | -0.486 | 1.017 | 1.562 | 0.301 | -0.003 | -0.037 | -0.037 | 0 | 0.12 | 0.16 | 0.08 | 0.03 | 0.12 | 0.19 |
| 8 | 3772648 | rs11779551 | 1.772 | 1.202 | 1.14 | 0.322 | -0.058 | -0.068 | -0.068 | 0 | 0.6 | 0.1 | 0.65 | 0.61 | 0.61 | 0.62 |
| 8 | 3773842 | rs2954621 | -0.621 | 1.084 | -0.269 | -0.063 | -0.042 | -0.046 | -0.046 | 0.959 | 0.69 | 0.83 | 0.63 | 0.55 | 0.63 | 0.78 |
| 8 | 3775951 | rs1464620 | 0.708 | 0.969 | -0.395 | -0.155 | -0.017 | -0.058 | -0.058 | 0 | 0.28 | 0.12 | 0.33 | 0.42 | 0.29 | 0.17 |
| 8 | 3778852 | rs2627395 | -1.221 | 0.613 | -0.351 | -0.24 | -0.011 | 0.04 | 0.04 | 0.167 | 0.59 | 0.67 | 0.58 | 0.78 | 0.54 | 0.5 |
| 8 | 3779090 | rs11136679 | 0.987 | 0.639 | -0.488 | 0.019 | -0.061 | -0.111 | -0.111 | 0 | 0.25 | 0.03 | 0.3 | 0.42 | 0.27 | 0.15 |
| 8 | 3779457 | rs1582766 | -0.192 | 0.734 | -0.292 | 0.123 | -0.016 | 0.101 | 0.101 | 0.031 | 0.24 | 0.44 | 0.21 | 0.33 | 0.18 | 0.17 |
| 8 | 3781893 | rs2954610 | -0.383 | 0.796 | -0.373 | -0.382 | -0.005 | 0.041 | 0.041 | 0.479 | 0.53 | 0.57 | 0.56 | 0.75 | 0.49 | 0.33 |
| 8 | 3783344 | rs2204223 | 0.345 | 0.473 | -1.102 | -0.357 | -0.008 | -0.035 | -0.035 | 0.792 | 0.63 | 0.52 | 0.66 | 0.76 | 0.68 | 0.74 |
| 8 | 3783471 | rs2627403 | 0.081 | 1.194 | -1.608 | 0.108 | -0.014 | -0.001 | -0.001 | 0 | 0.8 | 0.81 | 0.81 | 0.81 | 0.78 | 0.79 |
| 8 | 3784833 | rs17068023 | -0.394 | 0.368 | -1.721 | 0.673 | 0.016 | -0.034 | -0.034 | 0 | 0.25 | 0.15 | 0.2 | 0.37 | 0.21 | 0.23 |
| 8 | 3786543 | rs28407329 | 0.148 | 0.55 | -2.599 | 0.331 | -0.041 | -0.013 | -0.013 | 0 | 0.35 | 0.08 | 0.37 | 0.36 | 0.35 | 0.42 |
| 8 | 3786823 | rs2627336 | -0.407 | 0.771 | -2.07 | 0.235 | -0.021 | 0.019 | 0.019 | 0 | 0.66 | 0.87 | 0.66 | 0.71 | 0.63 | 0.71 |
| 8 | 3788431 | rs2948644 | -0.502 | 1.006 | -2.199 | 0.092 | -0.059 | -0.01 | -0.01 | 0 | 0.35 | 0.2 | 0.46 | 0.38 | 0.46 | 0.34 |
| 8 | 3788599 | rs2930336 | -0.37 | 0.96 | -2.038 | 0.035 | -0.057 | -0.013 | -0.013 | 0.123 | 0.35 | 0.18 | 0.45 | 0.38 | 0.45 | 0.34 |
| 8 | 3789272 | rs2554660 | -0.449 | 0.75 | -1.398 | 0.178 | -0.016 | -0.011 | -0.011 | 0 | 0.36 | 0.33 | 0.46 | 0.49 | 0.44 | 0.36 |
| 8 | 3791362 | rs2688266 | -0.592 | 0.265 | -1.218 | 0.35 | -0.003 | 0.018 | 0.018 | 0 | 0.82 | 0.77 | 0.82 | 0.76 | 0.84 | 0.73 |
| 8 | 3792481 | rs2975401 | 0.571 | 0.889 | -1.519 | 0.457 | -0.097 | -0.08 | -0.08 | 0 | 0.19 | 0.01 | 0.28 | 0.29 | 0.25 | 0.12 |
| 8 | 3792674 | rs11774238 | 1.613 | 0.439 | -0.943 | 0.107 | 0.005 | 0.006 | 0.006 | 0.484 | 0.36 | 0.32 | 0.24 | 0.22 | 0.29 | 0.28 |
| 8 | 3792688 | rs11784451 | 1.385 | 0.56 | -1.008 | 0.143 | -0.023 | -0.038 | -0.038 | 0 | 0.38 | 0.51 | 0.26 | 0.22 | 0.3 | 0.29 |
| 8 | 3793600 | rs2554653 | 0.128 | 0.665 | -0.801 | 0.228 | 0.001 | -0.006 | -0.006 | 0 | 0.71 | 0.73 | 0.68 | 0.69 | 0.72 | 0.61 |
| 8 | 3795203 | rs17068071 | 1.2 | 0.692 | -0.849 | 0.179 | -0.012 | -0.024 | -0.024 | 0.108 | 0.43 | 0.46 | 0.27 | 0.22 | 0.33 | 0.33 |
| 8 | 3795532 | rs2554622 | -0.334 | 0.742 | -0.948 | 0.286 | -0.007 | -0.006 | -0.006 | 0 | 0.33 | 0.28 | 0.42 | 0.48 | 0.41 | 0.31 |
| 8 | 3797372 | rs2975398 | 0.607 | 1.046 | -1.16 | 0.356 | -0.032 | 0.009 | 0.009 | 0 | 0.13 | 0.05 | 0.17 | 0.07 | 0.2 | 0.17 |
| 8 | 3797879 | rs2627525 | 0.867 | 1.107 | -1.069 | 0.521 | -0.011 | -0.007 | -0.007 | 0.361 | 0.16 | 0.13 | 0.23 | 0.27 | 0.19 | 0.12 |
| 8 | 3799327 | rs2258265 | 0.774 | 0.954 | -0.865 | 0.47 | -0.002 | -0.001 | -0.001 | 0 | 0.17 | 0.15 | 0.26 | 0.26 | 0.21 | 0.12 |
| 8 | 3799582 | rs1464617 | 0.526 | 0.709 | -0.915 | 0.53 | 0 | -0.008 | -0.008 | 0 | 0.56 | 0.6 | 0.46 | 0.46 | 0.49 | 0.56 |
| 8 | 3800820 | rs17068110 | -0.089 | 0.58 | -0.831 | 0.867 | 0.004 | 0.008 | 0.008 | 0 | 0.07 | 0.04 | 0.13 | 0.06 | 0.07 | 0.09 |
| 8 | 3801441 | rs1593487 | -1.284 | 0.646 | -0.845 | 0.888 | -0.009 | -0.003 | -0.003 | 0.918 | 0.91 | 0.97 | 0.89 | 0.95 | 0.9 | 0.88 |
| 8 | 3801831 | rs1523250 | 0.861 | 0.645 | -0.845 | 0.831 | -0.034 | -0.003 | -0.003 | 0 | 0.1 | 0.01 | 0.12 | 0.06 | 0.15 | 0.16 |
| 8 | 3801902 | rs1464618 | -0.842 | 0.101 | -0.459 | 0.792 | -0.027 | 0.049 | 0.049 | 0 | 0.33 | 0.46 | 0.32 | 0.42 | 0.27 | 0.48 |
| 8 | 3803623 | rs17068126 | 0.446 | -0.062 | -0.346 | 1.053 | -0.032 | 0.007 | 0.007 | 0 | 0.16 | 0.28 | 0.13 | 0.16 | 0.08 | 0.17 |
| 8 | 3806794 | rs2740868 | -0.671 | 0.006 | -0.653 | 0.997 | 0.001 | -0.013 | -0.013 | 0 | 0.86 | 0.92 | 0.78 | 0.76 | 0.85 | 0.78 |
| 8 | 3806885 | rs11779200 | -0.214 | -0.036 | -0.34 | 1.11 | -0.035 | 0.068 | 0.068 | 0.126 | 0.21 | 0.36 | 0.18 | 0.36 | 0.12 | 0.31 |
| 8 | 3807177 | rs2975395 | 0.393 | 0.689 | -0.822 | 0.548 | -0.024 | 0.021 | 0.021 | 0 | 0.12 | 0.03 | 0.14 | 0.06 | 0.16 | 0.15 |
| 8 | 3807528 | rs6984912 | -0.631 | 0.903 | 0.014 | 0.845 | 0.042 | 0.162 | 0.162 | 1.46 | 0.18 | 0.43 | 0.16 | 0.38 | 0.21 | 0.25 |
| 8 | 3807747 | rs10780168 | -0.201 | 0.859 | -0.322 | 0.715 | -0.01 | -0.045 | -0.045 | 0.465 | 0.18 | 0.11 | 0.19 | 0.38 | 0.22 | 0.3 |
| 8 | 3807826 | rs2740866 | -0.113 | 1.047 | -0.276 | 0.197 | 0.017 | -0.007 | -0.007 | 0.096 | 0.28 | 0.38 | 0.37 | 0.25 | 0.32 | 0.23 |
| 8 | 3808005 | rs756094 | -0.012 | 0.714 | 0.19 | -0.001 | -0.03 | -0.109 | -0.109 | 2.782 | 0.14 | 0.49 | 0.09 | 0 | 0.08 | 0.08 |
| 8 | 3808039 | rs756095 | 0.621 | 0.74 | -0.633 | -0.233 | 0.002 | -0.034 | -0.034 | 0 | 0.35 | 0.2 | 0.42 | 0.5 | 0.35 | 0.29 |
| 8 | 3808868 | rs2740862 | 0.11 | 0.568 | -0.5 | -0.553 | 0.002 | 0.025 | 0.025 | 0.553 | 0.55 | 0.62 | 0.45 | 0.28 | 0.46 | 0.42 |
| 8 | 3809397 | rs2554513 | 0.806 | 0.818 | -0.712 | -0.469 | -0.039 | -0.024 | -0.024 | 2.222 | 0.17 | 0.1 | 0.33 | 0.26 | 0.25 | 0.12 |
| 8 | 3809788 | rs2740860 | 0.734 | 0.295 | -0.673 | -0.801 | 0.006 | 0.006 | 0.006 | 0.002 | 0.18 | 0.15 | 0.12 | 0.14 | 0.14 | 0.16 |
| 8 | 3809910 | rs2980779 | 1.184 | 0.079 | -0.277 | -0.97 | 0.004 | 0.037 | 0.037 | 2.027 | 0.33 | 0.23 | 0.3 | 0.07 | 0.31 | 0.25 |

| | | | | | | | | | | | | | | | | |
|---|---|---|---|---|---|---|---|---|---|---|---|---|---|---|---|---|
| 8 | 3811284 | rs2554509 | 0.152 | 0.107 | -0.085 | -0.766 | 0.024 | 0.008 | 0.008 | 0 | 0.23 | 0.61 | 0.36 | 0.27 | 0.24 | 0.12 |
| 8 | 3811706 | rs2627508 | 1.035 | 0.169 | -0.378 | -0.761 | 0.012 | 0.003 | 0.003 | 0.187 | 0.15 | 0.09 | 0.1 | 0.13 | 0.12 | 0.12 |
| 8 | 3811812 | rs2930382 | 0.943 | 0.339 | -0.612 | -0.95 | -0.043 | 0.193 | 0.193 | 1.606 | 0.34 | 0.04 | 0.31 | 0.07 | 0.35 | 0.26 |
| 8 | 3811838 | rs2688334 | 0.945 | 0.078 | -0.022 | -0.677 | 0.067 | 0.051 | 0.051 | 0 | 0.15 | 0.32 | 0.31 | 0.25 | 0.18 | 0.1 |
| 8 | 3813310 | rs10503224 | 1.687 | -0.766 | -0.073 | -1.009 | -0.033 | -0.066 | -0.066 | 0.084 | 0.2 | 0.27 | 0.09 | 0.02 | 0.11 | 0.08 |
| 8 | 3813355 | rs2740854 | 0.711 | -0.994 | -0.048 | -1.032 | 0.007 | -0.002 | -0.002 | 0.138 | 0.16 | 0.13 | 0.1 | 0.19 | 0.11 | 0.14 |
| 8 | 3813731 | rs10503225 | 1.815 | -0.933 | 0.001 | -1.09 | -0.031 | -0.068 | -0.068 | 0 | 0.2 | 0.28 | 0.09 | 0.02 | 0.12 | 0.08 |
| 8 | 3814493 | rs2449821 | -1.253 | 0.094 | -0.072 | -1.127 | 0.017 | -0.026 | -0.026 | 0 | 0.82 | 0.82 | 0.72 | 0.96 | 0.71 | 0.81 |
| 8 | 3816121 | rs2740841 | 0.518 | -0.015 | -0.201 | -0.828 | 0.014 | 0.01 | 0.01 | 0.025 | 0.17 | 0.11 | 0.1 | 0.14 | 0.11 | 0.13 |
| 8 | 3819578 | rs2140340 | 1.014 | 0.007 | -0.345 | -0.831 | 0.011 | 0.036 | 0.036 | 0 | 0.14 | 0.03 | 0.07 | 0.07 | 0.1 | 0.11 |
| 8 | 3819733 | rs2740825 | 0.067 | 0.097 | -0.128 | -1.062 | -0.008 | 0.094 | 0.094 | 2.253 | 0.48 | 0.3 | 0.56 | 0.8 | 0.53 | 0.65 |
| 8 | 3822916 | rs12680491 | 1.077 | -0.001 | -0.407 | -0.765 | -0.002 | 0 | 0 | 1.073 | 0.3 | 0.23 | 0.23 | 0.55 | 0.28 | 0.48 |
| 8 | 3824773 | rs2627470 | -0.51 | -0.058 | -0.654 | -0.868 | -0.002 | -0.006 | -0.006 | 0 | 0.38 | 0.38 | 0.47 | 0.49 | 0.39 | 0.34 |
| 8 | 3824996 | rs7001755 | 0.189 | 0.016 | -0.556 | -0.916 | 0 | 0.004 | 0.004 | 1.36 | 0.43 | 0.57 | 0.53 | 0.71 | 0.45 | 0.53 |
| 8 | 3828077 | rs2627453 | 0.593 | 0.008 | -0.745 | -0.857 | 0.014 | 0.017 | 0.017 | 0.292 | 0.18 | 0.11 | 0.1 | 0.14 | 0.13 | 0.14 |
| 8 | 3828692 | rs2627451 | 0.169 | -0.367 | -1.07 | -0.688 | 0.023 | 0.011 | 0.011 | 0 | 0.27 | 0.22 | 0.13 | 0.2 | 0.18 | 0.24 |
| 8 | 3828722 | rs2740817 | 0.863 | -0.052 | -1.056 | -0.607 | 0.012 | 0.021 | 0.021 | 0.048 | 0.19 | 0.27 | 0.36 | 0.35 | 0.25 | 0.16 |
| 8 | 3829247 | rs2688353 | -1.331 | -0.217 | -1.305 | -0.785 | -0.064 | 0.149 | 0.149 | 2.38 | 0.68 | 0.89 | 0.68 | 0.94 | 0.64 | 0.75 |
| 8 | 3831350 | rs6558831 | 0.022 | -0.377 | -0.949 | -0.467 | -0.033 | 0.026 | 0.026 | 0.655 | 0.34 | 0.52 | 0.23 | 0.47 | 0.25 | 0.48 |
| 8 | 3832465 | rs1474319 | 1.671 | -0.286 | -0.995 | -0.476 | -0.009 | -0.009 | -0.009 | 1.974 | 0.16 | 0.16 | 0.32 | 0.24 | 0.21 | 0.11 |
| 8 | 3833181 | rs17068332 | 1.641 | -0.663 | -1.175 | 0.194 | -0.008 | -0.054 | -0.054 | 0 | 0.09 | 0.01 | 0.06 | 0.19 | 0.1 | 0.18 |
| 8 | 3833598 | rs11136689 | 0.583 | -0.967 | -0.888 | -0.223 | -0.026 | 0.047 | 0.047 | 0.095 | 0.25 | 0.29 | 0.19 | 0.4 | 0.22 | 0.34 |
| 8 | 3833708 | rs2554713 | -0.088 | -0.746 | -1.201 | -0.29 | 0 | -0.002 | -0.002 | 0.002 | 0.34 | 0.38 | 0.4 | 0.4 | 0.31 | 0.29 |
| 8 | 3835098 | rs11997340 | -0.387 | -0.406 | -0.933 | -0.004 | -0.026 | 0.018 | 0.018 | 0.72 | 0.34 | 0.55 | 0.23 | 0.47 | 0.25 | 0.43 |
| 8 | 3835166 | rs17068366 | 1.184 | 0.292 | -1.166 | -0.226 | -0.003 | 0 | 0 | 0 | 0.11 | 0.12 | 0.06 | 0.06 | 0.06 | 0.11 |
| 8 | 3838255 | rs2740810 | -1.465 | 0.526 | -1.422 | -0.346 | -0.051 | 0.042 | 0.042 | 0 | 0.88 | 0.98 | 0.77 | 0.97 | 0.77 | 0.81 |
| 8 | 3841168 | rs2688392 | -0.799 | 0.627 | -1.173 | -0.193 | 0 | -0.002 | -0.002 | 0 | 0.78 | 0.78 | 0.77 | 0.9 | 0.79 | 0.72 |
| 8 | 3842999 | rs2554728 | 0.513 | 0.062 | -1.546 | -0.187 | 0.005 | 0.013 | 0.013 | 0.763 | 0.29 | 0.24 | 0.21 | 0.18 | 0.26 | 0.19 |
| 8 | 3843797 | rs2554510 | 0.478 | 0.636 | -1.596 | 0.247 | -0.007 | -0.006 | -0.006 | 0 | 0.26 | 0.17 | 0.34 | 0.23 | 0.27 | 0.19 |
| 8 | 3846553 | rs2627383 | -0.118 | 0.556 | -1.611 | 0.325 | 0.011 | 0.056 | 0.056 | 0.015 | 0.51 | 0.28 | 0.56 | 0.33 | 0.55 | 0.36 |
| 8 | 3848025 | rs11995240 | 0.308 | 0.729 | -1.095 | 0.535 | 0.006 | 0.002 | 0.002 | 0 | 0.17 | 0.31 | 0.2 | 0.15 | 0.17 | 0.16 |
| 8 | 3848415 | rs2259576 | -0.36 | 0.786 | -1.447 | 0.503 | 0.004 | -0.005 | -0.005 | 0 | 0.39 | 0.36 | 0.42 | 0.31 | 0.42 | 0.34 |
| 8 | 3848750 | rs940155 | 1.055 | 0.498 | -1.585 | 0.597 | 0.041 | -0.044 | -0.044 | 0 | 0.32 | 0.19 | 0.36 | 0.48 | 0.26 | 0.39 |
| 8 | 3849209 | rs1523255 | 0.335 | 0.618 | -1.602 | 0.808 | -0.01 | 0.082 | 0.082 | 0 | 0.22 | 0.38 | 0.32 | 0.38 | 0.19 | 0.19 |
| 8 | 3849426 | rs940150 | -0.195 | 0.68 | -1.59 | 0.654 | 0.001 | 0 | 0 | 0 | 0.44 | 0.43 | 0.54 | 0.53 | 0.45 | 0.37 |
| 8 | 3854382 | rs1523254 | -0.409 | 0.618 | -1.812 | 0.524 | -0.015 | 0.055 | 0.055 | 0 | 0.57 | 0.79 | 0.66 | 0.68 | 0.52 | 0.64 |
| 8 | 3854987 | rs17068473 | 1.315 | 0.68 | -1.71 | 0.545 | 0.013 | -0.002 | -0.002 | 0 | 0.26 | 0.11 | 0.27 | 0.23 | 0.25 | 0.34 |
| 8 | 3855302 | rs1863475 | -0.092 | 0.848 | -1.333 | 0.362 | -0.009 | 0.019 | 0.019 | 0 | 0.66 | 0.66 | 0.68 | 0.77 | 0.64 | 0.73 |
| 8 | 3855838 | rs2059582 | -0.725 | 1.28 | -0.891 | 0.618 | 0.108 | 0.078 | 0.078 | 0 | 0.87 | 0.61 | 0.87 | 0.85 | 0.82 | 0.87 |
| 8 | 3857417 | rs7003719 | 0.008 | 1.175 | -1.319 | 0.075 | 0.013 | -0.091 | -0.091 | 0 | 0.28 | 0.45 | 0.28 | 0.09 | 0.29 | 0.21 |
| 8 | 3857596 | rs2688371 | 1.131 | 0.645 | -1.992 | 0.107 | -0.058 | -0.04 | -0.04 | 0.297 | 0.29 | 0.01 | 0.26 | 0.21 | 0.27 | 0.32 |
| 8 | 3858820 | rs2258845 | 0.194 | 0.709 | -1.817 | 0.139 | -0.003 | 0.012 | 0.012 | 1.291 | 0.44 | 0.34 | 0.41 | 0.26 | 0.52 | 0.41 |
| 8 | 3859012 | rs7013771 | 1.976 | 0.515 | -1.876 | 0.299 | -0.001 | -0.002 | -0.002 | 0 | 0.17 | 0.17 | 0.2 | 0.2 | 0.1 | 0.23 |
| 8 | 3859936 | rs2740973 | -0.664 | 0.583 | -1.732 | 0.508 | 0.004 | 0.006 | 0.006 | 0 | 0.69 | 0.62 | 0.64 | 0.68 | 0.69 | 0.74 |
| 8 | 3863111 | rs17068510 | 0.919 | 0.565 | -1.643 | 0.631 | 0.018 | 0.01 | 0.01 | 0.623 | 0.27 | 0.26 | 0.26 | 0.22 | 0.2 | 0.3 |
| 8 | 3863732 | rs2740968 | 0.598 | 0.48 | -1.691 | 0.738 | 0.003 | 0.002 | 0.002 | 2.051 | 0.36 | 0.32 | 0.34 | 0.58 | 0.42 | 0.41 |
| 8 | 3865488 | rs2740963 | 0.786 | 0.373 | -1.608 | 0.611 | 0.004 | 0.004 | 0.004 | 4.063 | 0.34 | 0.32 | 0.33 | 0.41 | 0.39 | 0.32 |
| 8 | 3865670 | rs2740962 | 0.183 | 0.369 | -1.625 | 0.581 | -0.001 | -0.002 | -0.002 | 0 | 0.49 | 0.46 | 0.5 | 0.6 | 0.47 | 0.56 |
| 8 | 3867513 | rs12674864 | 2.21 | 0.172 | -1.674 | 0.636 | 0.027 | -0.012 | -0.012 | 0 | 0.16 | 0.09 | 0.17 | 0.2 | 0.09 | 0.26 |
| 8 | 3867760 | rs2740955 | -0.384 | 0.295 | -1.621 | 0.56 | -0.007 | 0.005 | 0.005 | 0 | 0.61 | 0.66 | 0.57 | 0.66 | 0.58 | 0.66 |
| 8 | 3868458 | rs2740953 | 0.992 | 0.372 | -1.638 | 0.637 | 0.008 | -0.006 | -0.006 | 0 | 0.33 | 0.32 | 0.33 | 0.27 | 0.38 | 0.31 |
| 8 | 3869134 | rs17068540 | 1.709 | 0.54 | -1.639 | 0.661 | -0.013 | -0.013 | -0.013 | 0.261 | 0.12 | 0.06 | 0.07 | 0.14 | 0.12 | 0.09 |
| 8 | 3869858 | rs2740947 | -0.087 | 0.549 | -1.474 | 0.712 | 0.005 | -0.001 | -0.001 | 0 | 0.61 | 0.53 | 0.57 | 0.61 | 0.59 | 0.65 |
| 8 | 3871360 | rs2688276 | -0.222 | 0.595 | -1.516 | 0.777 | 0.001 | -0.002 | -0.002 | 0 | 0.61 | 0.57 | 0.57 | 0.63 | 0.59 | 0.66 |
| 8 | 3872513 | rs2740939 | 0.449 | 0.523 | -1.767 | 0.652 | 0.006 | -0.01 | -0.01 | 0 | 0.5 | 0.26 | 0.49 | 0.46 | 0.48 | 0.57 |
| 8 | 3872987 | rs17068550 | 2.586 | 0.141 | -1.881 | 0.625 | 0.024 | -0.013 | -0.013 | 0 | 0.14 | 0.09 | 0.18 | 0.2 | 0.08 | 0.27 |
| 8 | 3874384 | rs2688388 | -1.043 | 0.406 | -1.818 | 0.393 | -0.001 | 0.003 | 0.003 | 0.025 | 0.73 | 0.73 | 0.76 | 0.71 | 0.73 | 0.73 |
| 8 | 3875497 | rs11780696 | 2.223 | 0.384 | -2.012 | 0.215 | -0.002 | 0.022 | 0.022 | 0 | 0.12 | 0.09 | 0.07 | 0 | 0.11 | 0.08 |
| 8 | 3876265 | rs4875262 | 1.356 | 0.393 | -2.1 | 0.259 | 0.003 | -0.058 | -0.058 | 0 | 0.36 | 0.11 | 0.41 | 0.39 | 0.35 | 0.48 |
| 8 | 3877943 | rs2980796 | 0.863 | 0.53 | -2.036 | 0.334 | -0.109 | 0.073 | 0.073 | 1.085 | 0.39 | 0.1 | 0.34 | 0.19 | 0.44 | 0.35 |
| 8 | 3878109 | rs11994492 | 0.678 | 0.672 | -1.564 | 0.527 | 0.018 | -0.001 | -0.001 | 0 | 0.14 | 0.4 | 0.09 | 0.13 | 0.11 | 0.11 |
| 8 | 3879918 | rs2740929 | 0.702 | 0.731 | -1.859 | 0.524 | -0.005 | 0.009 | 0.009 | 0.236 | 0.54 | 0.22 | 0.49 | 0.39 | 0.47 | 0.64 |
| 8 | 3880839 | rs2688333 | -1.25 | 0.971 | -1.762 | 0.352 | 0.007 | 0.021 | 0.021 | 0 | 0.81 | 0.75 | 0.78 | 0.75 | 0.76 | 0.85 |
| 8 | 3881946 | rs2688397 | 0.223 | 0.464 | -1.657 | 0.178 | -0.002 | 0.002 | 0.002 | 3.349 | 0.4 | 0.44 | 0.34 | 0.45 | 0.4 | 0.39 |
| 8 | 3882868 | rs7812972 | 0.577 | 0.794 | -0.988 | -0.051 | 0.036 | -0.04 | -0.04 | 0 | 0.12 | 0.37 | 0.1 | 0.06 | 0.12 | 0.08 |
| 8 | 3883399 | rs2740924 | 0.317 | 0.767 | -1.477 | 0.23 | -0.018 | -0.009 | -0.009 | 0.783 | 0.67 | 0.73 | 0.57 | 0.64 | 0.59 | 0.71 |
| 8 | 3892017 | rs2554566 | 0.331 | 0.791 | -1.471 | -0.831 | 0 | 0.055 | 0.055 | 0 | 0.54 | 0.68 | 0.53 | 0.77 | 0.52 | 0.54 |
| 8 | 3895072 | rs2740914 | 0.167 | 0.596 | -1.721 | -1.34 | -0.039 | 0.037 | 0.037 | 0 | 0.75 | 0.93 | 0.7 | 0.8 | 0.74 | 0.77 |
| 8 | 3896625 | rs4379463 | -0.505 | 0.365 | -1.871 | -1.508 | 0.001 | -0.016 | -0.016 | 0 | 0.57 | 0.53 | 0.62 | 0.64 | 0.56 | 0.55 |

| | | | | | | | | | | | | | | | | |
|---|---|---|---|---|---|---|---|---|---|---|---|---|---|---|---|---|
| 8 | 3896853 | rs12677236 | 2.307 | 0.388 | -1.98 | -1.346 | 0.004 | -0.112 | -0.112 | 0 | 0.16 | 0.02 | 0.29 | 0.36 | 0.17 | 0.14 |
| 8 | 3897851 | rs2740911 | -0.587 | -0.734 | -1.944 | -1.309 | -0.059 | -0.027 | -0.027 | 0 | 0.84 | 0.99 | 0.75 | 0.83 | 0.83 | 0.84 |
| 8 | 3898221 | rs2980795 | -0.313 | -0.608 | -1.605 | -1.161 | -0.001 | -0.001 | -0.001 | 0 | 0.58 | 0.55 | 0.54 | 0.55 | 0.54 | 0.6 |
| 8 | 3899186 | rs2740910 | -0.86 | -0.505 | -1.664 | -1.23 | -0.067 | 0.068 | 0.068 | 0 | 0.77 | 0.87 | 0.82 | 0.98 | 0.65 | 0.75 |
| 8 | 3900305 | rs2897595 | -0.547 | -0.851 | -0.892 | -0.827 | 0.001 | 0.04 | 0.04 | 0 | 0.27 | 0.13 | 0.29 | 0.2 | 0.24 | 0.24 |
| 8 | 3900812 | rs7003625 | -0.653 | -0.897 | -0.386 | -0.844 | -0.005 | -0.02 | -0.02 | 0.203 | 0.28 | 0.32 | 0.37 | 0.21 | 0.24 | 0.21 |
| 8 | 3902552 | rs2740902 | -0.469 | -0.74 | -0.306 | -0.745 | -0.005 | 0.024 | 0.024 | 0 | 0.74 | 0.71 | 0.65 | 0.59 | 0.73 | 0.82 |
| 8 | 3902583 | rs4629888 | -0.49 | -0.676 | -0.185 | -0.875 | 0.002 | -0.02 | -0.02 | 0 | 0.27 | 0.33 | 0.3 | 0.21 | 0.24 | 0.19 |
| 8 | 3902849 | rs4421367 | -0.327 | -0.856 | -0.257 | -0.823 | -0.001 | -0.007 | -0.007 | 0 | 0.25 | 0.26 | 0.3 | 0.22 | 0.22 | 0.18 |
| 8 | 3903905 | rs17068727 | 0.733 | -1.097 | -0.244 | -0.781 | -0.031 | 0.045 | 0.045 | 0 | 0.21 | 0.23 | 0.29 | 0.48 | 0.13 | 0.19 |
| 8 | 3904006 | rs17068729 | 1.774 | -1.038 | -0.22 | -0.732 | 0.035 | -0.064 | -0.064 | 0.038 | 0.14 | 0.05 | 0.26 | 0.33 | 0.12 | 0.13 |
| 8 | 3904671 | rs3843921 | -0.425 | -1.058 | -0.051 | -0.842 | -0.021 | 0.013 | 0.013 | 0 | 0.34 | 0.33 | 0.46 | 0.5 | 0.24 | 0.27 |
| 8 | 3904874 | rs7004524 | 0.177 | -1.002 | -0.19 | -0.999 | -0.006 | -0.005 | -0.005 | 0 | 0.17 | 0.19 | 0.2 | 0.17 | 0.11 | 0.15 |
| 8 | 3904947 | rs2554555 | 0.32 | -0.777 | -0.592 | -0.885 | 0.005 | 0.01 | 0.01 | 0 | 0.55 | 0.46 | 0.44 | 0.33 | 0.6 | 0.61 |
| 8 | 3906162 | rs12548766 | 1.013 | -0.735 | -0.761 | -0.97 | -0.016 | 0.056 | 0.056 | 0 | 0.39 | 0.26 | 0.21 | 0.22 | 0.42 | 0.5 |
| 8 | 3906703 | rs2554549 | 0.497 | -0.799 | -0.667 | -0.68 | -0.006 | -0.012 | -0.012 | 0.376 | 0.56 | 0.5 | 0.44 | 0.33 | 0.61 | 0.63 |
| 8 | 3906969 | rs17068768 | 0.513 | -0.792 | -1.101 | -0.411 | 0.005 | -0.006 | -0.006 | 0 | 0.07 | 0.05 | 0.02 | 0.15 | 0.02 | 0.06 |
| 8 | 3918118 | rs2740878 | 0.35 | -0.543 | -1.1 | -0.638 | 0.063 | 0.072 | 0.072 | 0 | 0.51 | 0.29 | 0.44 | 0.33 | 0.64 | 0.56 |
| 8 | 3920312 | rs2554704 | -0.2 | -1.102 | -1.111 | -0.101 | 0.047 | -0.011 | -0.011 | 0 | 0.28 | 0.2 | 0.37 | 0.62 | 0.16 | 0.21 |
| 8 | 3920395 | rs2552166 | -0.081 | -0.863 | -1.083 | -0.033 | 0.039 | -0.015 | -0.015 | 0 | 0.27 | 0.19 | 0.4 | 0.61 | 0.2 | 0.19 |
| 8 | 3921821 | rs2740873 | -0.81 | -0.912 | -1.072 | -0.268 | -0.002 | 0 | 0 | 0.07 | 0.84 | 0.8 | 0.89 | 0.81 | 0.87 | 0.84 |
| 8 | 3921924 | rs6558843 | 1.033 | -0.747 | -0.901 | -0.541 | 0.002 | -0.063 | -0.063 | 0 | 0.38 | 0.41 | 0.4 | 0.03 | 0.52 | 0.43 |
| 8 | 3922303 | rs1365277 | 0.452 | -0.007 | -0.782 | -0.08 | -0.001 | -0.003 | -0.003 | 0 | 0.66 | 0.64 | 0.79 | 0.65 | 0.7 | 0.65 |
| 8 | 3923744 | rs1035633 | -0.519 | 0.174 | -0.858 | 0.476 | -0.001 | 0.001 | 0.001 | 0 | 0.35 | 0.33 | 0.47 | 0.49 | 0.33 | 0.32 |
| 8 | 3926858 | rs2113669 | -0.795 | 0.118 | -0.719 | 0.08 | -0.009 | -0.034 | -0.034 | 0 | 0.14 | 0.23 | 0.08 | 0.06 | 0.13 | 0.06 |
| 8 | 3928521 | rs2552122 | -0.269 | -0.273 | -1.435 | 0.137 | 0.012 | 0.002 | 0.002 | 0 | 0.59 | 0.41 | 0.65 | 0.67 | 0.69 | 0.73 |
| 8 | 3929795 | rs2552125 | 0.559 | -0.298 | -1.695 | 0.173 | 0.007 | 0 | 0 | 0.347 | 0.18 | 0.15 | 0.16 | 0.18 | 0.16 | 0.37 |
| 8 | 3935771 | rs2554682 | 0.288 | -1.285 | -1.849 | -0.694 | 0.006 | -0.01 | -0.01 | 0 | 0.28 | 0.25 | 0.13 | 0.06 | 0.18 | 0.16 |
| 8 | 3937191 | rs1541718 | 0.332 | -0.445 | -1.895 | -0.367 | 0.056 | 0.021 | 0.021 | 0 | 0.48 | 0.23 | 0.61 | 0.6 | 0.63 | 0.63 |
| 8 | 3937595 | rs1971078 | 0.444 | -0.451 | -1.892 | -0.366 | -0.026 | 0.08 | 0.08 | 0 | 0.44 | 0.2 | 0.41 | 0.31 | 0.54 | 0.61 |
| 8 | 3938349 | rs2161929 | -1.292 | -0.492 | -1.714 | -0.314 | -0.011 | 0.066 | 0.066 | 0 | 0.77 | 0.58 | 0.75 | 0.7 | 0.81 | 0.83 |
| 8 | 3938481 | rs2552145 | 0.589 | -0.535 | -1.858 | -0.432 | -0.004 | -0.011 | -0.011 | 0 | 0.41 | 0.25 | 0.58 | 0.59 | 0.57 | 0.62 |
| 8 | 3938612 | rs13267061 | 1.564 | -0.573 | -1.825 | -0.463 | 0.031 | 0.055 | 0.055 | 0 | 0.18 | 0.08 | 0.09 | 0.02 | 0.12 | 0.11 |
| 8 | 3942747 | rs2554646 | -0.165 | -0.5 | -1.697 | -0.39 | 0.01 | 0.037 | 0.037 | 0 | 0.53 | 0.31 | 0.43 | 0.41 | 0.6 | 0.66 |
| 8 | 3944168 | rs10503233 | 1.873 | 0.063 | -1.982 | -0.536 | -0.094 | 0.037 | 0.037 | 0 | 0.2 | 0.01 | 0.2 | 0.12 | 0.28 | 0.26 |
| 8 | 3944529 | rs17068867 | 1.369 | -0.006 | -1.834 | -0.46 | -0.007 | -0.026 | -0.026 | 0 | 0.2 | 0.11 | 0.28 | 0.31 | 0.2 | 0.15 |
| 8 | 3946186 | rs17404086 | 0.622 | -0.107 | -1.774 | -0.557 | -0.027 | 0.087 | 0.087 | 0.076 | 0.34 | 0.18 | 0.26 | 0.16 | 0.39 | 0.3 |
| 8 | 3946649 | rs17068887 | 1.387 | -0.451 | -1.828 | -0.709 | -0.007 | -0.116 | -0.116 | 2.232 | 0.2 | 0.42 | 0.12 | 0.03 | 0.17 | 0.08 |
| 8 | 3947046 | rs17068893 | 1.051 | -0.445 | -1.932 | -0.693 | -0.033 | -0.03 | -0.03 | 0.001 | 0.2 | 0.06 | 0.18 | 0.23 | 0.23 | 0.39 |
| 8 | 3947325 | rs11783062 | 0.547 | -0.401 | -1.9 | -0.67 | -0.06 | -0.032 | -0.032 | 0 | 0.38 | 0.21 | 0.46 | 0.45 | 0.48 | 0.33 |
| 8 | 3950508 | rs4875274 | 0.298 | -0.708 | -1.858 | -0.337 | 0.01 | 0.112 | 0.112 | 0 | 0.53 | 0.26 | 0.51 | 0.26 | 0.57 | 0.47 |
| 8 | 3950542 | rs10107611 | 0.748 | -0.651 | -1.759 | -0.449 | 0.031 | 0.061 | 0.061 | 0 | 0.24 | 0.43 | 0.32 | 0.32 | 0.25 | 0.1 |
| 8 | 3951689 | rs11786984 | 0.73 | -0.764 | -1.813 | -0.488 | -0.054 | 0.076 | 0.076 | 0.177 | 0.35 | 0.19 | 0.29 | 0.2 | 0.45 | 0.33 |
| 8 | 3952552 | rs10087392 | 0.183 | -0.784 | -1.635 | -0.36 | -0.014 | 0.099 | 0.099 | 0 | 0.09 | 0.2 | 0.17 | 0.27 | 0.04 | 0.11 |
| 8 | 3955025 | rs918917 | 1.261 | -0.888 | -1.429 | -0.673 | 0.014 | -0.129 | -0.129 | 0.238 | 0.17 | 0.4 | 0.13 | 0 | 0.17 | 0.08 |
| 8 | 3956233 | rs17404336 | 1.085 | -1.014 | -1.529 | -0.664 | 0.017 | 0.03 | 0.03 | 6.186 | 0.15 | 0.1 | 0.06 | 0.06 | 0.1 | 0.04 |
| 8 | 3956349 | rs17068970 | -1.466 | -0.978 | -1.291 | -0.516 | -0.001 | -0.005 | -0.005 | 0.456 | 0.79 | 0.86 | 0.72 | 0.66 | 0.8 | 0.85 |
| 8 | 3956964 | rs10503234 | 1.03 | -0.992 | -1.317 | -0.499 | 0.004 | -0.034 | -0.034 | 0.013 | 0.23 | 0.12 | 0.29 | 0.35 | 0.2 | 0.18 |
| 8 | 3957348 | rs2059398 | 0.821 | -0.814 | -1.373 | -0.513 | -0.008 | -0.009 | -0.009 | 0.276 | 0.31 | 0.32 | 0.4 | 0.43 | 0.4 | 0.36 |
| 8 | 3959597 | rs10107716 | -0.708 | -0.719 | -1.323 | -0.301 | 0.002 | -0.018 | -0.018 | 0 | 0.75 | 0.85 | 0.71 | 0.65 | 0.79 | 0.8 |
| 8 | 3964022 | rs2194604 | -0.17 | -0.891 | -0.518 | -0.422 | -0.021 | 0.004 | 0.004 | 0.729 | 0.64 | 0.54 | 0.67 | 0.62 | 0.74 | 0.81 |
| 8 | 3964905 | rs17069018 | 0.772 | -1.08 | -0.68 | -0.509 | -0.006 | 0 | 0 | 0 | 0.22 | 0.26 | 0.18 | 0.23 | 0.17 | 0.37 |
| 8 | 3965028 | rs7828949 | -0.192 | -0.937 | -0.507 | -0.694 | -0.007 | 0.04 | 0.04 | 0.245 | 0.47 | 0.31 | 0.46 | 0.38 | 0.46 | 0.28 |
| 8 | 3965436 | rs4372022 | 0.836 | -1.147 | -0.583 | -0.774 | 0.006 | -0.012 | -0.012 | 0 | 0.29 | 0.2 | 0.35 | 0.36 | 0.24 | 0.23 |
| 8 | 3965925 | rs4242493 | 0.11 | -0.964 | -0.292 | -0.739 | 0 | 0.001 | 0.001 | 0.039 | 0.45 | 0.39 | 0.5 | 0.38 | 0.56 | 0.48 |
| 8 | 3967931 | rs9314513 | -0.016 | -0.88 | -0.221 | -0.745 | -0.004 | -0.033 | -0.033 | 0 | 0.29 | 0.38 | 0.18 | 0.11 | 0.27 | 0.13 |
| 8 | 3969537 | rs7013590 | 0.447 | -0.877 | -0.664 | -0.651 | 0.01 | -0.033 | -0.033 | 0 | 0.38 | 0.22 | 0.42 | 0.51 | 0.34 | 0.43 |
| 8 | 3972304 | rs12549476 | 1.213 | -0.603 | -0.674 | -0.581 | -0.019 | 0.022 | 0.022 | 0 | 0.24 | 0.17 | 0.22 | 0.12 | 0.36 | 0.33 |
| 8 | 3975865 | rs12155988 | 1.411 | -0.657 | -0.795 | -0.412 | 0.009 | 0.008 | 0.008 | 4.296 | 0.11 | 0.1 | 0.04 | 0.06 | 0.07 | 0.03 |
| 8 | 3976837 | rs10094785 | 0.517 | -1.023 | -0.604 | -0.659 | -0.003 | -0.011 | -0.011 | 0 | 0.52 | 0.5 | 0.67 | 0.74 | 0.58 | 0.6 |
| 8 | 3978488 | rs1991139 | 1.391 | -0.813 | -0.487 | -0.589 | 0.034 | -0.07 | -0.07 | 0 | 0.12 | 0.19 | 0.11 | 0 | 0.17 | 0.08 |
| 8 | 3980013 | rs12549072 | 0.639 | -1.073 | -0.353 | -0.429 | 0.002 | 0.003 | 0.003 | 0.026 | 0.35 | 0.35 | 0.42 | 0.39 | 0.41 | 0.45 |
| 8 | 3980158 | rs10088637 | -0.454 | 0.26 | -0.753 | -0.541 | 0.015 | -0.025 | -0.025 | 0 | 0.76 | 0.75 | 0.72 | 0.93 | 0.6 | 0.73 |
| 8 | 3980317 | rs7816211 | 1.579 | -0.11 | -0.659 | -0.4 | 0.054 | -0.012 | -0.012 | 0 | 0.12 | 0.19 | 0.17 | 0.07 | 0.24 | 0.17 |
| 8 | 3980781 | rs3891533 | 0.568 | -1.132 | -0.464 | -0.098 | 0.017 | 0.012 | 0.012 | 0 | 0.23 | 0.09 | 0.17 | 0.2 | 0.2 | 0.3 |
| 8 | 3981876 | rs1647326 | 1.707 | -0.816 | -0.246 | -0.116 | 0.002 | -0.055 | -0.055 | 0 | 0.1 | 0.16 | 0.08 | 0 | 0.11 | 0.08 |
| 8 | 3984758 | rs1714789 | -1.059 | -0.899 | -0.257 | 0.047 | -0.007 | -0.005 | -0.005 | 0 | 0.63 | 0.38 | 0.55 | 0.61 | 0.55 | 0.58 |
| 8 | 3984856 | rs11785517 | 0.878 | -0.903 | -0.165 | 0.026 | 0.016 | 0.011 | 0.011 | 0.186 | 0.37 | 0.49 | 0.45 | 0.39 | 0.46 | 0.49 |
| 8 | 3987036 | rs17427521 | 1.146 | -0.989 | -0.352 | -0.045 | 0.027 | 0.036 | 0.036 | 0.027 | 0.25 | 0.16 | 0.11 | 0.11 | 0.16 | 0.16 |

| | | | | | | | | | | | | | | | | |
|---|---|---|---|---|---|---|---|---|---|---|---|---|---|---|---|---|
| 8 | 3987251 | rs4875288 | 1.406 | -0.946 | -0.188 | -0.005 | 0.003 | 0.002 | 0.002 | 0 | 0.33 | 0.31 | 0.43 | 0.39 | 0.45 | 0.41 |
| 8 | 3988435 | rs17069091 | 0.435 | -0.86 | -0.247 | 0.276 | 0 | -0.002 | -0.002 | 0 | 0.3 | 0.27 | 0.31 | 0.41 | 0.26 | 0.22 |
| 8 | 3989696 | rs12155841 | 0.358 | -0.869 | -0.273 | 0.271 | 0.002 | -0.012 | -0.012 | 0 | 0.31 | 0.24 | 0.31 | 0.41 | 0.29 | 0.22 |
| 8 | 3996660 | rs1613815 | 0.032 | -0.496 | -0.088 | 0.581 | -0.018 | -0.038 | -0.038 | 0 | 0.23 | 0.12 | 0.25 | 0.38 | 0.3 | 0.16 |
| 8 | 3997289 | rs13259316 | 1.666 | -0.365 | -0.121 | 0.768 | -0.132 | 0.075 | 0.075 | 0 | 0.29 | 0.04 | 0.34 | 0.13 | 0.41 | 0.35 |
| 8 | 3998888 | rs7817129 | 0.23 | -0.505 | 0.042 | 0.872 | 0.081 | 0.103 | 0.103 | 0.332 | 0.46 | 0.26 | 0.27 | 0.29 | 0.34 | 0.38 |
| 8 | 3999648 | rs7830462 | -0.088 | -0.286 | 0.545 | 1.031 | 0.005 | 0.252 | 0.252 | 0.196 | 0.09 | 0.3 | 0.17 | 0.29 | 0.03 | 0.17 |
| 8 | 4002031 | rs1613009 | 1.033 | -0.197 | 0.34 | 1.042 | -0.013 | -0.013 | -0.013 | 0 | 0.13 | 0.08 | 0.21 | 0.32 | 0.22 | 0.12 |
| 8 | 4002653 | rs10503236 | -0.24 | -0.412 | 0.463 | 0.998 | 0.018 | 0.282 | 0.282 | 0.217 | 0.09 | 0.35 | 0.17 | 0.28 | 0.03 | 0.17 |
| 8 | 4005375 | rs10112217 | 0.687 | -0.345 | 0.083 | 0.852 | 0.022 | 0.005 | 0.005 | 0.698 | 0.38 | 0.43 | 0.54 | 0.44 | 0.46 | 0.49 |
| 8 | 4005670 | rs1714820 | 0.896 | -0.222 | -0.22 | 0.932 | -0.011 | -0.01 | -0.01 | 0 | 0.14 | 0.08 | 0.17 | 0.29 | 0.2 | 0.13 |
| 8 | 4008566 | rs4875292 | 1.302 | -0.323 | -0.481 | 0.616 | -0.139 | 0.085 | 0.085 | 0 | 0.31 | 0.08 | 0.36 | 0.17 | 0.43 | 0.37 |
| 8 | 4008816 | rs3849825 | 1.3 | -0.389 | -0.571 | 0.615 | -0.182 | 0.072 | 0.072 | 0 | 0.31 | 0.03 | 0.37 | 0.17 | 0.44 | 0.38 |
| 8 | 4009376 | rs2552171 | -0.805 | -0.956 | -0.063 | 0.525 | -0.016 | -0.022 | -0.022 | 0 | 0.66 | 0.64 | 0.78 | 0.82 | 0.78 | 0.67 |
| 8 | 4011688 | rs17069139 | 0.382 | -0.684 | 0.336 | 0.895 | 0.01 | 0.205 | 0.205 | 0.036 | 0.07 | 0.21 | 0.16 | 0.26 | 0.03 | 0.13 |
| 8 | 4012166 | rs2407307 | 0.504 | -0.681 | 0.308 | 0.883 | 0 | 0.158 | 0.158 | 0.042 | 0.07 | 0.19 | 0.15 | 0.25 | 0.03 | 0.13 |
| 8 | 4012464 | rs2002195 | 0.636 | -0.87 | 0.169 | 0.462 | 0.037 | 0.046 | 0.046 | 0 | 0.34 | 0.21 | 0.22 | 0.16 | 0.23 | 0.33 |
| 8 | 4013345 | rs3843923 | 0.595 | -0.914 | 0.216 | 0.551 | -0.045 | -0.043 | -0.043 | 0 | 0.34 | 0.44 | 0.35 | 0.21 | 0.22 | 0.31 |
| 8 | 4014336 | rs3849827 | 1.376 | -0.273 | -0.768 | 0.437 | -0.176 | 0.086 | 0.086 | 0 | 0.31 | 0.02 | 0.37 | 0.16 | 0.44 | 0.37 |
| 8 | 4015930 | rs4875294 | 1.331 | -0.292 | -0.726 | 0.442 | -0.154 | 0.119 | 0.119 | 0 | 0.31 | 0.04 | 0.37 | 0.13 | 0.43 | 0.38 |
| 8 | 4016156 | rs17069167 | 0.452 | -0.498 | -0.421 | 0.617 | 0.009 | 0.003 | 0.003 | 0 | 0.39 | 0.54 | 0.37 | 0.36 | 0.26 | 0.41 |
| 8 | 4017692 | rs1647292 | 0.221 | -0.325 | -0.729 | 0.897 | -0.037 | -0.105 | -0.105 | 0.138 | 0.25 | 0.11 | 0.25 | 0.41 | 0.29 | 0.16 |
| 8 | 4019462 | rs17405197 | 0.707 | -0.467 | -0.718 | 0.97 | 0.005 | -0.022 | -0.022 | 0 | 0.13 | 0.05 | 0.06 | 0.13 | 0.11 | 0.04 |
| 8 | 4022182 | rs13280168 | -0.201 | -0.454 | -0.227 | 0.899 | 0.008 | -0.001 | -0.001 | 0 | 0.49 | 0.39 | 0.4 | 0.49 | 0.36 | 0.46 |
| 8 | 4026979 | rs3849831 | -0.363 | -0.448 | -0.361 | 0.789 | 0.031 | 0.014 | 0.014 | 0.023 | 0.52 | 0.55 | 0.43 | 0.6 | 0.36 | 0.52 |
| 8 | 4028497 | rs1568332 | 1.098 | -0.13 | -0.768 | 0.844 | -0.016 | -0.026 | -0.026 | 0 | 0.13 | 0.1 | 0.18 | 0.29 | 0.18 | 0.09 |
| 8 | 4029513 | rs1714757 | 0.393 | -0.369 | -0.811 | 0.755 | 0.041 | -0.015 | -0.015 | 0.303 | 0.44 | 0.15 | 0.55 | 0.4 | 0.63 | 0.47 |
| 8 | 4032622 | rs10113741 | 0.531 | -0.818 | -0.505 | 0.633 | -0.038 | 0.002 | 0.002 | 0 | 0.39 | 0.41 | 0.36 | 0.44 | 0.24 | 0.4 |
| 8 | 4038869 | rs3849834 | -0.31 | -0.567 | -0.334 | 0.514 | 0 | 0.007 | 0.007 | 0.341 | 0.74 | 0.7 | 0.74 | 0.59 | 0.71 | 0.85 |
| 8 | 4039852 | rs17069226 | 0.412 | -0.265 | -0.402 | 0.25 | -0.006 | -0.002 | -0.002 | 0 | 0.4 | 0.34 | 0.38 | 0.39 | 0.32 | 0.38 |
| 8 | 4040057 | rs10503238 | 1.344 | -0.206 | -0.676 | 0.154 | -0.178 | 0.161 | 0.161 | 0.463 | 0.32 | 0.02 | 0.34 | 0.09 | 0.43 | 0.39 |
| 8 | 4040129 | rs3849836 | 0.277 | 0.002 | -0.288 | 0.5 | 0 | 0.05 | 0.05 | 0 | 0.05 | 0.12 | 0.02 | 0.12 | 0.03 | 0.09 |
| 8 | 4041575 | rs4875303 | 0.583 | -0.364 | -0.507 | 0.396 | -0.001 | -0.004 | -0.004 | 0 | 0.28 | 0.2 | 0.22 | 0.17 | 0.27 | 0.25 |
| 8 | 4042746 | rs7018112 | 1.314 | -0.268 | -0.829 | 0.502 | -0.123 | 0.1 | 0.1 | 0 | 0.4 | 0.07 | 0.39 | 0.24 | 0.49 | 0.52 |
| 8 | 4043515 | rs17069257 | 1.011 | 0.199 | -0.305 | 0.959 | 0.013 | 0.063 | 0.063 | 0 | 0.11 | 0.2 | 0.18 | 0.29 | 0.14 | 0.09 |
| 8 | 4045835 | rs6996352 | 1.117 | 0.152 | -0.291 | 0.959 | 0.009 | 0.069 | 0.069 | 0 | 0.1 | 0.2 | 0.16 | 0.28 | 0.1 | 0.09 |
| 8 | 4047054 | rs4875304 | -0.153 | 0.024 | -0.277 | 0.627 | -0.001 | 0.003 | 0.003 | 1.053 | 0.83 | 0.82 | 0.9 | 0.77 | 0.81 | 0.89 |
| 8 | 4051243 | rs11985852 | 0.68 | -0.472 | -0.565 | 0.193 | -0.003 | -0.003 | -0.003 | 0 | 0.25 | 0.19 | 0.32 | 0.2 | 0.24 | 0.27 |
| 8 | 4051387 | rs779131 | 0.357 | -0.419 | -0.746 | 0.121 | -0.001 | 0.002 | 0.002 | 1.99 | 0.15 | 0.18 | 0.1 | 0.22 | 0.15 | 0.09 |
| 8 | 4052031 | rs779132 | 0.389 | -0.237 | -0.446 | -0.106 | -0.007 | -0.024 | -0.024 | 0.011 | 0.61 | 0.51 | 0.63 | 0.79 | 0.67 | 0.67 |
| 8 | 4052555 | rs2656298 | 0.224 | -0.276 | -0.386 | -0.153 | -0.006 | -0.003 | -0.003 | 0 | 0.63 | 0.46 | 0.63 | 0.66 | 0.71 | 0.61 |
| 8 | 4055073 | rs7841311 | 0.755 | -0.636 | -0.968 | -0.225 | -0.027 | -0.026 | -0.026 | 0 | 0.14 | 0.21 | 0.18 | 0.03 | 0.05 | 0.05 |
| 8 | 4059676 | rs710258 | -0.002 | -0.667 | -1.088 | -0.264 | -0.027 | -0.03 | -0.03 | 0.752 | 0.73 | 0.62 | 0.87 | 0.85 | 0.82 | 0.88 |
| 8 | 4060604 | rs4875308 | 0.944 | -0.465 | -1.395 | -0.285 | -0.033 | 0.168 | 0.168 | 0 | 0.47 | 0.17 | 0.51 | 0.21 | 0.5 | 0.54 |
| 8 | 4061058 | rs3914776 | 1.499 | -0.765 | -1.046 | 0.041 | 0.001 | 0.064 | 0.064 | 0 | 0.1 | 0.12 | 0.13 | 0.29 | 0.11 | 0.23 |
| 8 | 4061204 | rs1714708 | -0.288 | -0.786 | -1.107 | 0.001 | -0.05 | 0.024 | 0.024 | 0.581 | 0.75 | 0.47 | 0.81 | 0.7 | 0.84 | 0.69 |
| 8 | 4063488 | rs3860868 | 1.056 | -0.721 | -1.184 | -0.347 | 0.016 | -0.006 | -0.006 | 0.455 | 0.46 | 0.36 | 0.55 | 0.53 | 0.58 | 0.55 |
| 8 | 4065529 | rs1568330 | 0.66 | -0.129 | -1.807 | -0.607 | 0.01 | 0.114 | 0.114 | 0 | 0.52 | 0.19 | 0.42 | 0.3 | 0.56 | 0.56 |
| 8 | 4066831 | rs1647315 | -0.952 | -0.661 | -1.526 | -0.16 | -0.001 | -0.003 | -0.003 | 0 | 0.35 | 0.38 | 0.16 | 0.19 | 0.23 | 0.12 |
| 8 | 4067830 | rs6991758 | -0.15 | 0.016 | -1.211 | 0.095 | -0.021 | -0.056 | -0.056 | 0.568 | 0.17 | 0.24 | 0.07 | 0.01 | 0.07 | 0.07 |
| 8 | 4070415 | rs3889326 | -0.429 | 0.942 | -1.573 | 1.03 | -0.008 | -0.026 | -0.026 | 0 | 0.16 | 0.06 | 0.31 | 0.38 | 0.2 | 0.3 |
| 8 | 4071270 | rs7829915 | 0.06 | 0.833 | -1.557 | 0.173 | -0.005 | 0.122 | 0.122 | 1.768 | 0.52 | 0.18 | 0.59 | 0.72 | 0.52 | 0.51 |
| 8 | 4073225 | rs2407312 | 0.234 | 0.617 | -0.944 | 0.326 | 0.002 | 0.002 | 0.002 | 3.35 | 0.49 | 0.43 | 0.58 | 0.27 | 0.38 | 0.65 |
| 8 | 4073265 | rs1714717 | 0.095 | 1.08 | -1.341 | 0.814 | -0.105 | 0.029 | 0.029 | 0.123 | 0.29 | 0.06 | 0.23 | 0.24 | 0.43 | 0.16 |
| 8 | 4075796 | rs3849842 | 0.61 | 0.334 | -0.585 | 1.327 | 0.037 | 0.067 | 0.067 | 2.146 | 0.53 | 0.78 | 0.57 | 0.35 | 0.38 | 0.69 |
| 8 | 4080144 | rs939151 | -0.737 | 0.429 | -0.157 | 2.407 | 0 | 0.001 | 0.001 | 2.595 | 0.45 | 0.3 | 0.38 | 0.49 | 0.58 | 0.28 |
| 8 | 4081569 | rs1394481 | 0.965 | 0.385 | 0.314 | 2.391 | 0.001 | 0 | 0 | 1.432 | 0.5 | 0.44 | 0.53 | 0.4 | 0.38 | 0.67 |
| 8 | 4083087 | rs1700059 | -0.726 | 0.312 | 0.315 | 2.281 | -0.001 | -0.021 | -0.021 | 0 | 0.45 | 0.29 | 0.36 | 0.53 | 0.58 | 0.26 |
| 8 | 4084879 | rs1714821 | -0.915 | 0.444 | 0.294 | 2.314 | 0.052 | 0.07 | 0.07 | 2.101 | 0.48 | 0.36 | 0.42 | 0.3 | 0.63 | 0.26 |
| 8 | 4084970 | rs1700055 | -0.881 | 0.461 | 0.287 | 2.294 | 0.032 | 0.046 | 0.046 | 2.311 | 0.47 | 0.4 | 0.42 | 0.28 | 0.62 | 0.26 |
| 8 | 4087191 | rs7828683 | 0.952 | 0.479 | 0.329 | 2.402 | 0.004 | 0.005 | 0.005 | 1.981 | 0.5 | 0.46 | 0.54 | 0.35 | 0.38 | 0.69 |
| 8 | 4087405 | rs7829008 | 0.986 | 0.482 | 0.354 | 1.964 | 0.011 | 0.018 | 0.018 | 2.115 | 0.5 | 0.41 | 0.56 | 0.73 | 0.39 | 0.74 |
| 8 | 4089132 | rs11992223 | -0.715 | 0.303 | -0.082 | 1.732 | -0.014 | 0.017 | 0.017 | 0 | 0.35 | 0.28 | 0.36 | 0.28 | 0.57 | 0.25 |
| 8 | 4090790 | rs1714678 | 0.615 | 0.265 | -0.189 | 1.628 | 0.029 | 0.023 | 0.023 | 3.234 | 0.12 | 0.12 | 0.06 | 0 | 0.04 | 0.01 |
| 8 | 4091132 | rs7822226 | -0.566 | 0.797 | 0.005 | 1.857 | -0.056 | 0.046 | 0.046 | 0 | 0.29 | 0.17 | 0.28 | 0.18 | 0.53 | 0.19 |
| 8 | 4096011 | rs17334464 | -0.809 | 1.012 | 0.231 | 1.906 | -0.129 | 0.089 | 0.089 | 0.224 | 0.3 | 0.09 | 0.24 | 0.17 | 0.5 | 0.2 |
| 8 | 4096261 | rs1700113 | 1.594 | 0.585 | 0.999 | 1.748 | 0.054 | 0.132 | 0.132 | 0 | 0.53 | 0.2 | 0.55 | 0.27 | 0.38 | 0.56 |
| 8 | 4096546 | rs1620593 | 1.53 | 0.581 | 0.994 | 1.942 | 0.034 | 0.059 | 0.059 | 0 | 0.55 | 0.2 | 0.55 | 0.35 | 0.39 | 0.62 |
| 8 | 4097418 | rs1700112 | 0.768 | 0.551 | 1.715 | 1.916 | 0.039 | 0.004 | 0.004 | 0 | 0.66 | 0.49 | 0.67 | 0.39 | 0.45 | 0.67 |

| | | | | | | | | | | | | | | | | |
|---|---|---|---|---|---|---|---|---|---|---|---|---|---|---|---|---|
| 8 | 4097519 | rs1714689 | 0.779 | 0.595 | 1.725 | 1.929 | 0.03 | 0 | 0 | 0 | 0.65 | 0.49 | 0.67 | 0.4 | 0.46 | 0.65 |
| 8 | 4098594 | rs1700110 | 1.558 | 0.592 | 1.395 | 1.728 | 0.054 | 0.175 | 0.175 | 0 | 0.53 | 0.18 | 0.44 | 0.2 | 0.36 | 0.57 |
| 8 | 4101492 | rs1700106 | 1.162 | 0.752 | 1.078 | 1.948 | -0.008 | -0.007 | -0.007 | 0.022 | 0.6 | 0.3 | 0.45 | 0.57 | 0.46 | 0.71 |
| 8 | 4101558 | rs13248374 | -0.475 | 0.932 | 0.492 | 1.859 | -0.145 | 0.01 | 0.01 | 0.003 | 0.28 | 0.06 | 0.32 | 0.26 | 0.5 | 0.18 |
| 8 | 4103130 | rs4875310 | -0.9 | 0.887 | 0.704 | 1.693 | 0.012 | 0.009 | 0.009 | 0 | 0.32 | 0.36 | 0.4 | 0.6 | 0.53 | 0.42 |
| 8 | 4104656 | rs1700100 | 1.314 | 0.525 | 0.448 | 0.999 | -0.006 | -0.014 | -0.014 | 0.605 | 0.65 | 0.44 | 0.65 | 0.83 | 0.7 | 0.75 |
| 8 | 4105147 | rs1714746 | 1.547 | 0.536 | 0.565 | 1.479 | 0.012 | 0.009 | 0.009 | 0 | 0.6 | 0.32 | 0.59 | 0.46 | 0.65 | 0.6 |
| 8 | 4107869 | rs4404937 | -0.026 | 0.519 | 0.251 | 1.317 | -0.038 | 0.105 | 0.105 | 0.288 | 0.17 | 0.02 | 0.18 | 0 | 0.2 | 0.06 |
| 8 | 4108050 | rs6998882 | 0.143 | 0.579 | 0.624 | 1.406 | 0 | 0.019 | 0.019 | 0 | 0.09 | 0.11 | 0.09 | 0.18 | 0.06 | 0.18 |
| 8 | 4109163 | rs6984808 | 0.909 | -0.189 | -0.27 | 0.393 | -0.058 | 0.051 | 0.051 | 0 | 0.36 | 0.22 | 0.36 | 0.27 | 0.49 | 0.27 |
| 8 | 4110032 | rs13259289 | -0.706 | 0.77 | -0.169 | 0.508 | -0.069 | 0.107 | 0.107 | 0 | 0.24 | 0.07 | 0.29 | 0 | 0.41 | 0.08 |
| 8 | 4112258 | rs1700096 | 1.728 | 0.054 | 0.694 | 0.946 | -0.016 | 0.093 | 0.093 | 0.01 | 0.78 | 0.48 | 0.79 | 0.46 | 0.85 | 0.66 |
| 8 | 4113492 | rs17069764 | -0.584 | 0.315 | 0.38 | 1.22 | 0.014 | 0.146 | 0.146 | 0.21 | 0.13 | 0.27 | 0.16 | 0.52 | 0.11 | 0.33 |
| 8 | 4117538 | rs2656285 | 1.418 | -0.005 | -0.122 | 0.802 | 0.013 | -0.046 | -0.046 | 0 | 0.66 | 0.15 | 0.62 | 0.48 | 0.72 | 0.61 |
| 8 | 4119374 | rs1504761 | -0.003 | 0.215 | 0.228 | 1.107 | 0.006 | 0.255 | 0.255 | 0.015 | 0.05 | 0.15 | 0.07 | 0.4 | 0.05 | 0.17 |
| 8 | 4122003 | rs9693235 | 0.319 | 0.234 | 0.213 | 1.017 | 0.006 | 0.04 | 0.04 | 0 | 0.08 | 0.16 | 0.04 | 0.12 | 0.06 | 0.16 |
| 8 | 4122078 | rs779103 | 0.548 | 0.131 | 0.76 | 0.946 | 0.001 | 0.102 | 0.102 | 0.198 | 0.87 | 0.63 | 0.88 | 0.85 | 0.93 | 0.83 |
| 8 | 4122597 | rs4617179 | -0.136 | 0.418 | 0.325 | 0.878 | 0.026 | 0.111 | 0.111 | 0 | 0.22 | 0.4 | 0.4 | 0.42 | 0.3 | 0.24 |
| 8 | 4122628 | rs4288395 | -0.595 | 0.39 | 0.318 | 0.911 | 0.013 | 0.053 | 0.053 | 0 | 0.28 | 0.41 | 0.42 | 0.42 | 0.35 | 0.24 |
| 8 | 4125100 | rs17069790 | 0.013 | 0.605 | -0.111 | 1.035 | 0.011 | 0.07 | 0.07 | 0 | 0.08 | 0.17 | 0.1 | 0.15 | 0.06 | 0.18 |
| 8 | 4127845 | rs924864 | 0.135 | 0.353 | -0.772 | 0.13 | 0.021 | -0.052 | -0.052 | 0 | 0.27 | 0.13 | 0.18 | 0.54 | 0.18 | 0.42 |
| 8 | 4131490 | rs812941 | -0.548 | 1.403 | -0.162 | -0.416 | 0.028 | -0.049 | -0.049 | 0 | 0.76 | 0.68 | 0.78 | 0.91 | 0.69 | 0.81 |
| 8 | 4134874 | rs2407940 | 0.251 | 1.473 | -0.535 | -0.743 | -0.003 | -0.006 | -0.006 | 0 | 0.6 | 0.42 | 0.62 | 0.9 | 0.51 | 0.62 |
| 8 | 4135849 | rs17413156 | 0.191 | 1.6 | -0.847 | -0.647 | -0.03 | 0.029 | 0.029 | 0 | 0.09 | 0.03 | 0.09 | 0 | 0.13 | 0.07 |
| 8 | 4136051 | rs17413170 | 0.214 | 1.591 | -0.869 | -0.644 | -0.041 | 0.042 | 0.042 | 0 | 0.09 | 0.02 | 0.09 | 0 | 0.13 | 0.07 |
| 8 | 4137396 | rs17069831 | 1.156 | 1.109 | -0.939 | -0.435 | 0.006 | -0.034 | -0.034 | 0 | 0.28 | 0.21 | 0.31 | 0.44 | 0.27 | 0.23 |
| 8 | 4140685 | rs2680602 | 1.504 | 1.004 | -1.144 | -0.872 | 0.027 | 0.019 | 0.019 | 1.202 | 0.09 | 0 | 0.05 | 0.02 | 0.04 | 0.1 |
| 8 | 4143049 | rs10113352 | 1.19 | 1.047 | -0.825 | -0.858 | -0.002 | 0.005 | 0.005 | 0 | 0.27 | 0.31 | 0.33 | 0.37 | 0.28 | 0.22 |
| 8 | 4143740 | rs1504757 | 0.652 | 2.059 | -0.545 | -1.081 | -0.082 | 0.035 | 0.035 | 0.099 | 0.18 | 0.02 | 0.28 | 0.09 | 0.28 | 0.15 |
| 8 | 4144890 | rs17335036 | -0.229 | 2.205 | -0.468 | -1.013 | -0.046 | 0.045 | 0.045 | 0 | 0.1 | 0.02 | 0.1 | 0 | 0.15 | 0.07 |
| 8 | 4145622 | rs2680605 | 0.483 | 1.49 | -0.252 | -0.834 | 0.008 | 0.01 | 0.01 | 0.451 | 0.1 | 0.05 | 0.07 | 0.02 | 0.07 | 0.16 |
| 8 | 4145796 | rs11787185 | 0.954 | 1.483 | -0.137 | -0.898 | -0.001 | -0.02 | -0.02 | 0.097 | 0.57 | 0.42 | 0.53 | 0.88 | 0.51 | 0.62 |
| 8 | 4146236 | rs2680606 | -0.416 | 1.209 | -0.213 | -0.153 | -0.022 | -0.034 | -0.034 | 0 | 0.12 | 0.17 | 0.07 | 0.02 | 0.07 | 0.16 |
| 8 | 4157137 | rs7845852 | 0.978 | 2.122 | -0.172 | -0.141 | 0.028 | -0.064 | -0.064 | 4.086 | 0.75 | 0.61 | 0.69 | 0.85 | 0.71 | 0.82 |
| 8 | 4157782 | rs9314521 | -0.72 | 2.165 | -0.453 | -0.148 | 0.014 | -0.035 | -0.035 | 0.656 | 0.07 | 0.11 | 0.07 | 0 | 0.09 | 0.05 |
| 8 | 4158919 | rs10866965 | 1.344 | 2.661 | -0.647 | -0.319 | -0.02 | 0.02 | 0.02 | 0 | 0.73 | 0.82 | 0.78 | 0.82 | 0.62 | 0.77 |
| 8 | 4161539 | rs1910716 | 1.08 | 2.21 | -0.496 | -0.13 | -0.001 | -0.001 | -0.001 | 0 | 0.79 | 0.81 | 0.82 | 0.82 | 0.74 | 0.8 |
| 8 | 4162862 | rs4875319 | -0.037 | 2.007 | -0.584 | -0.035 | -0.001 | -0.001 | -0.001 | 0 | 0.13 | 0.1 | 0.1 | 0.16 | 0.15 | 0.08 |
| 8 | 4164975 | rs11784834 | 0.864 | 1.836 | -0.86 | 0.537 | -0.015 | -0.025 | -0.025 | 0 | 0.6 | 0.33 | 0.68 | 0.46 | 0.59 | 0.71 |
| 8 | 4165096 | rs12548881 | -0.063 | 1.411 | -0.864 | 0.92 | 0.001 | -0.003 | -0.003 | 0.154 | 0.12 | 0.12 | 0.06 | 0.25 | 0.1 | 0.07 |
| 8 | 4165460 | rs13258014 | -0.026 | 1.145 | -1.509 | 0.705 | -0.011 | 0.052 | 0.052 | 0.099 | 0.56 | 0.21 | 0.42 | 0.33 | 0.56 | 0.55 |
| 8 | 4166426 | rs10503249 | 0.614 | 0.587 | -1.821 | 0.265 | -0.02 | -0.005 | -0.005 | 0 | 0.62 | 0.11 | 0.73 | 0.62 | 0.66 | 0.65 |
| 8 | 4166522 | rs10503250 | 0.424 | 0.497 | -0.997 | -0.168 | 0.002 | -0.04 | -0.04 | 0 | 0.67 | 0.42 | 0.8 | 0.86 | 0.67 | 0.7 |
| 8 | 4167608 | rs4875322 | 0.023 | 0.454 | -1.092 | -0.239 | -0.048 | 0.062 | 0.062 | 2.976 | 0.77 | 0.92 | 0.87 | 0.86 | 0.75 | 0.75 |
| 8 | 4169706 | rs9785155 | 0.185 | 0.332 | -1.152 | -0.301 | -0.043 | 0.068 | 0.068 | 2.469 | 0.76 | 0.91 | 0.86 | 0.86 | 0.75 | 0.76 |
| 8 | 4171740 | rs7002661 | -0.509 | 0.348 | -0.933 | -0.379 | 0.069 | -0.026 | -0.026 | 0.411 | 0.16 | 0.48 | 0.12 | 0.13 | 0.22 | 0.19 |
| 8 | 4173675 | rs12679077 | 0.744 | 0.212 | -2.159 | -0.57 | 0 | 0.067 | 0.067 | 0 | 0.57 | 0.13 | 0.71 | 0.65 | 0.59 | 0.56 |
| 8 | 4174454 | rs4875329 | 0.499 | 0.02 | -2.339 | -0.996 | -0.071 | 0.114 | 0.114 | 0 | 0.44 | 0.14 | 0.58 | 0.73 | 0.47 | 0.53 |
| 8 | 4176941 | rs4875330 | 0.574 | 0.007 | -2.303 | -1.077 | -0.069 | 0.013 | 0.013 | 0 | 0.36 | 0.14 | 0.53 | 0.63 | 0.43 | 0.4 |
| 8 | 4178604 | rs10108628 | 0.058 | -0.421 | -1.315 | -1.361 | -0.005 | 0.024 | 0.024 | 0 | 0.2 | 0.22 | 0.12 | 0.34 | 0.2 | 0.29 |
| 8 | 4178983 | rs4875334 | 0.829 | -0.326 | -1.882 | -1.826 | -0.094 | 0 | 0 | 1.729 | 0.39 | 0.11 | 0.53 | 0.61 | 0.44 | 0.43 |
| 8 | 4179086 | rs4875335 | 0.096 | -0.454 | -1.684 | -1.992 | 0.057 | 0.15 | 0.15 | 0 | 0.56 | 0.21 | 0.66 | 0.99 | 0.67 | 0.82 |
| 8 | 4179859 | rs10100098 | 0.925 | -0.461 | -1.069 | -1.962 | 0.011 | -0.106 | -0.106 | 0.536 | 0.16 | 0.4 | 0.1 | 0 | 0.14 | 0.07 |
| 8 | 4180844 | rs10503253 | 0.584 | -0.437 | -1.249 | -1.806 | -0.005 | -0.044 | -0.044 | 0.179 | 0.19 | 0.08 | 0.1 | 0.35 | 0.22 | 0.4 |
| 8 | 4181864 | rs1566861 | -0.122 | -0.559 | -1.201 | -1.921 | 0.016 | 0.015 | 0.015 | 0 | 0.63 | 0.8 | 0.7 | 0.65 | 0.62 | 0.51 |
| 8 | 4182085 | rs17413763 | 1.261 | -0.062 | -1.473 | -1.825 | -0.077 | -0.121 | -0.121 | 0.224 | 0.23 | 0.02 | 0.32 | 0.6 | 0.27 | 0.29 |
| 8 | 4183209 | rs1394474 | 0.313 | -0.411 | -0.385 | -1.793 | 0.003 | -0.116 | -0.116 | 0.102 | 0.17 | 0.41 | 0.11 | 0 | 0.14 | 0.06 |
| 8 | 4183528 | rs12541979 | -0.927 | -0.345 | -1.007 | -1.524 | 0.018 | 0.041 | 0.041 | 0 | 0.82 | 0.91 | 0.81 | 0.99 | 0.84 | 0.92 |
| 8 | 4185116 | rs10503255 | 0.51 | -0.305 | -0.935 | -1.45 | 0.018 | 0.022 | 0.022 | 0 | 0.19 | 0.12 | 0.19 | 0 | 0.16 | 0.07 |
| 8 | 4187170 | rs2062879 | 0.934 | -0.397 | -1.403 | -1.537 | -0.125 | -0.016 | -0.016 | 0.185 | 0.4 | 0.09 | 0.54 | 0.58 | 0.45 | 0.4 |
| 8 | 4188239 | rs11776532 | 1.408 | -0.39 | -1.363 | -1.674 | -0.007 | 0.05 | 0.05 | 0 | 0.15 | 0.04 | 0.17 | 0.06 | 0.15 | 0.04 |
| 8 | 4188891 | rs1875897 | 0.305 | -0.533 | -0.83 | -1.701 | -0.004 | 0.002 | 0.002 | 0 | 0.24 | 0.19 | 0.14 | 0.27 | 0.3 | 0.45 |
| 8 | 4190222 | rs10091134 | -0.565 | -0.516 | -0.717 | -1.663 | 0.011 | 0.049 | 0.049 | 0.23 | 0.41 | 0.71 | 0.26 | 0.34 | 0.38 | 0.51 |
| 8 | 4190874 | rs10107472 | -0.56 | -0.704 | -0.87 | -1.473 | 0.018 | 0.04 | 0.04 | 2.607 | 0.33 | 0.87 | 0.23 | 0.31 | 0.27 | 0.47 |
| 8 | 4194916 | rs7010684 | -1.147 | -0.61 | -1.002 | -1.576 | -0.038 | 0.114 | 0.114 | 0.378 | 0.53 | 0.93 | 0.41 | 0.33 | 0.46 | 0.52 |
| 8 | 4195287 | rs10102768 | 0.491 | -0.697 | -0.887 | -1.567 | -0.017 | 0.012 | 0.012 | 0.044 | 0.17 | 0.07 | 0.1 | 0.16 | 0.21 | 0.22 |
| 8 | 4201268 | rs6558866 | -0.643 | -0.842 | -0.726 | -1.649 | 0.055 | 0.063 | 0.063 | 0 | 0.33 | 0.79 | 0.31 | 0.19 | 0.17 | 0.29 |
| 8 | 4201343 | rs6558867 | 1.197 | -0.919 | -0.536 | -1.757 | 0.008 | -0.005 | -0.005 | 0 | 0.16 | 0.14 | 0.19 | 0.01 | 0.11 | 0.03 |
| 8 | 4202182 | rs13253947 | -0.063 | -0.825 | -0.273 | -1.64 | -0.046 | -0.026 | -0.026 | 0.305 | 0.16 | 0.53 | 0.11 | 0.13 | 0.07 | 0.25 |

| | | | | | | | | | | | | | | | | |
|---|---|---|---|---|---|---|---|---|---|---|---|---|---|---|---|---|
| 8 | 4203603 | rs11985125 | 0.256 | -0.493 | -1.276 | -1.75 | -0.044 | 0.026 | 0.026 | 0 | 0.21 | 0.07 | 0.13 | 0.16 | 0.31 | 0.24 |
| 8 | 4204310 | rs17413882 | 0.72 | -0.493 | -1.162 | -1.771 | -0.007 | 0.006 | 0.006 | 0 | 0.16 | 0.12 | 0.09 | 0.16 | 0.25 | 0.2 |
| 8 | 4204987 | rs17069985 | 0.712 | -0.582 | -0.937 | -1.671 | -0.012 | 0.013 | 0.013 | 0 | 0.11 | 0.26 | 0.1 | 0.14 | 0.06 | 0.22 |
| 8 | 4205477 | rs7832549 | -0.848 | -0.608 | -2.113 | -1.74 | 0.017 | 0.116 | 0.116 | 0.377 | 0.46 | 0.87 | 0.4 | 0.33 | 0.42 | 0.49 |
| 8 | 4206548 | rs6558868 | -1.143 | -0.503 | -2.103 | -1.721 | 0.024 | 0.159 | 0.159 | 0.565 | 0.5 | 0.87 | 0.42 | 0.33 | 0.45 | 0.49 |
| 8 | 4206751 | rs7008571 | -0.683 | -0.037 | -2.442 | -1.395 | -0.031 | 0.085 | 0.085 | 1.5 | 0.75 | 0.99 | 0.75 | 0.94 | 0.81 | 0.86 |
| 8 | 4208391 | rs10105465 | 0.106 | 0.24 | -1.553 | -0.754 | -0.007 | 0.013 | 0.013 | 0.116 | 0.18 | 0.11 | 0.1 | 0.15 | 0.21 | 0.21 |
| 8 | 4212932 | rs4292712 | 0.523 | 0.293 | -2.081 | -0.128 | -0.096 | -0.055 | -0.055 | 0.059 | 0.44 | 0.08 | 0.45 | 0.54 | 0.43 | 0.45 |
| 8 | 4214179 | rs10503256 | 0.009 | 0.175 | -1.221 | -0.201 | 0.004 | 0.008 | 0.008 | 0.004 | 0.6 | 0.34 | 0.71 | 0.63 | 0.6 | 0.5 |
| 8 | 4216317 | rs2163308 | -0.781 | 0.073 | -1.143 | -0.306 | -0.011 | 0.033 | 0.033 | 0 | 0.76 | 0.9 | 0.86 | 0.83 | 0.7 | 0.7 |
| 8 | 4217304 | rs1036556 | 1.201 | -0.318 | -1.654 | -0.2 | -0.118 | -0.119 | -0.119 | 0 | 0.38 | 0.07 | 0.45 | 0.49 | 0.41 | 0.43 |
| 8 | 4217893 | rs10104910 | 0.293 | -0.341 | -0.327 | -0.173 | -0.015 | 0.004 | 0.004 | 1.264 | 0.12 | 0.37 | 0.1 | 0.11 | 0.05 | 0.18 |
| 8 | 4219026 | rs13270369 | -0.164 | -0.473 | 0.007 | -0.269 | -0.018 | -0.014 | -0.014 | 0 | 0.32 | 0.52 | 0.35 | 0.27 | 0.26 | 0.24 |
| 8 | 4219116 | rs7837578 | -0.219 | -0.496 | 0.006 | -0.27 | -0.023 | -0.016 | -0.016 | 0 | 0.32 | 0.52 | 0.36 | 0.27 | 0.26 | 0.24 |
| 8 | 4219551 | rs7814853 | -0.562 | -0.504 | 0.432 | -0.279 | -0.002 | -0.005 | -0.005 | 0 | 0.34 | 0.64 | 0.38 | 0.18 | 0.26 | 0.23 |
| 8 | 4220542 | rs11787410 | -1.164 | -0.504 | 0.332 | -0.281 | -0.034 | -0.012 | -0.012 | 0 | 0.58 | 0.76 | 0.51 | 0.44 | 0.56 | 0.53 |
| 8 | 4220622 | rs11778711 | 0.868 | -0.574 | 0.405 | -0.473 | 0.001 | 0 | 0 | 0 | 0.22 | 0.2 | 0.28 | 0.07 | 0.21 | 0.06 |
| 8 | 4224136 | rs10107898 | -1.318 | -0.618 | 0.415 | 0.332 | -0.016 | 0.03 | 0.03 | 0 | 0.61 | 0.73 | 0.61 | 0.7 | 0.63 | 0.6 |
| 8 | 4224893 | rs7839613 | 1.14 | -0.654 | 0.303 | 0.418 | -0.021 | -0.078 | -0.078 | 0.02 | 0.25 | 0.09 | 0.21 | 0.4 | 0.34 | 0.33 |
| 8 | 4225233 | rs1594353 | -0.558 | -0.489 | 0.589 | 0.375 | -0.001 | 0.002 | 0.002 | 0 | 0.39 | 0.61 | 0.35 | 0.53 | 0.43 | 0.53 |
| 8 | 4226590 | rs1583651 | -0.271 | -0.438 | 0.424 | 0.416 | 0.001 | 0.016 | 0.016 | 0 | 0.48 | 0.37 | 0.5 | 0.58 | 0.56 | 0.42 |
| 8 | 4227483 | rs1583130 | -1.299 | -0.347 | -0.088 | 0.428 | -0.077 | 0.037 | 0.037 | 0 | 0.61 | 0.94 | 0.63 | 0.71 | 0.64 | 0.59 |
| 8 | 4228083 | rs11993860 | -1.234 | -0.413 | -0.075 | 0.296 | -0.07 | -0.041 | -0.041 | 0 | 0.6 | 0.91 | 0.61 | 0.57 | 0.62 | 0.55 |
| 8 | 4228543 | rs11780622 | -0.379 | -0.624 | 0.767 | 0.084 | -0.014 | -0.063 | -0.063 | 0 | 0.3 | 0.42 | 0.34 | 0.15 | 0.26 | 0.14 |
| 8 | 4230614 | rs7843687 | 0.35 | -0.697 | 0.522 | 0.188 | -0.006 | -0.015 | -0.015 | 0 | 0.22 | 0.3 | 0.31 | 0.11 | 0.23 | 0.14 |
| 8 | 4231556 | rs13269315 | 0.156 | -0.691 | 0.199 | 0.096 | -0.002 | -0.009 | -0.009 | 0.608 | 0.23 | 0.22 | 0.28 | 0.07 | 0.21 | 0.05 |
| 8 | 4232568 | rs11786916 | 0.669 | -0.558 | -0.038 | 0.171 | 0.005 | 0.022 | 0.022 | 0 | 0.19 | 0.12 | 0.26 | 0.07 | 0.19 | 0.05 |
| 8 | 4232963 | rs11775921 | 0.671 | -0.623 | -0.039 | 0.17 | 0.006 | 0.015 | 0.015 | 0 | 0.2 | 0.15 | 0.26 | 0.08 | 0.2 | 0.05 |
| 8 | 4233891 | rs10098163 | -1.31 | -0.488 | -0.654 | -0.095 | -0.041 | 0.025 | 0.025 | 0.022 | 0.85 | 1 | 0.93 | 0.94 | 0.89 | 0.88 |
| 8 | 4234106 | rs11781210 | 0.247 | -0.7 | -0.051 | 0.05 | -0.019 | -0.014 | -0.014 | 3.4 | 0.25 | 0.32 | 0.29 | 0.18 | 0.21 | 0.11 |
| 8 | 4235701 | rs11785440 | 0.805 | -0.708 | -0.428 | -0.002 | 0.003 | 0.004 | 0.004 | 0 | 0.2 | 0.19 | 0.25 | 0.07 | 0.2 | 0.05 |
| 8 | 4235758 | rs1438208 | -2.033 | -0.373 | -1.037 | 0.115 | -0.027 | -0.024 | -0.024 | 2.221 | 0.79 | 0.92 | 0.74 | 0.72 | 0.78 | 0.74 |
| 8 | 4235897 | rs17414118 | 1.143 | -0.336 | -1.185 | 0.028 | -0.029 | 0.034 | 0.034 | 0.061 | 0.17 | 0.01 | 0.08 | 0.05 | 0.12 | 0.15 |
| 8 | 4236743 | rs1595468 | 0.863 | -0.435 | -0.837 | 0.134 | -0.052 | -0.065 | -0.065 | 0 | 0.34 | 0.15 | 0.31 | 0.47 | 0.46 | 0.37 |
| 8 | 4238139 | rs6558872 | 0.141 | -0.479 | -0.566 | 0.23 | 0.001 | 0.002 | 0.002 | 0 | 0.48 | 0.57 | 0.62 | 0.57 | 0.4 | 0.61 |
| 8 | 4238243 | rs6558873 | -1.718 | -0.489 | -1.037 | 0.43 | -0.029 | -0.052 | -0.052 | 0.351 | 0.77 | 0.93 | 0.69 | 0.67 | 0.76 | 0.7 |
| 8 | 4238712 | rs17070107 | 0.122 | -0.45 | -0.823 | 0.22 | 0.003 | -0.006 | -0.006 | 0 | 0.42 | 0.4 | 0.41 | 0.52 | 0.34 | 0.57 |
| 8 | 4240142 | rs4875101 | 0.163 | -0.61 | -1.54 | 0.117 | 0.042 | -0.015 | -0.015 | 0 | 0.46 | 0.72 | 0.61 | 0.51 | 0.4 | 0.41 |
| 8 | 4241132 | rs1430447 | 0.715 | -0.59 | -1.745 | 0.06 | -0.062 | -0.001 | -0.001 | 0 | 0.36 | 0.18 | 0.31 | 0.37 | 0.5 | 0.38 |
| 8 | 4241151 | rs13439862 | 0.501 | -0.484 | -1.473 | 0.101 | 0.01 | 0.02 | 0.02 | 0 | 0.19 | 0.12 | 0.09 | 0.08 | 0.09 | 0.2 |
| 8 | 4246903 | rs11774916 | 0.172 | -0.388 | -1.613 | 0.078 | 0.025 | 0.038 | 0.038 | 1.554 | 0.37 | 0.21 | 0.4 | 0.19 | 0.32 | 0.22 |
| 8 | 4247780 | rs1438214 | 0.525 | -0.295 | -1.647 | 0.283 | -0.009 | -0.003 | -0.003 | 0 | 0.27 | 0.2 | 0.22 | 0.28 | 0.39 | 0.31 |
| 8 | 4251284 | rs11988906 | -0.127 | -0.335 | -1.877 | 0.166 | -0.039 | 0.02 | 0.02 | 0.304 | 0.57 | 0.82 | 0.71 | 0.63 | 0.46 | 0.5 |
| 8 | 4257256 | rs1438210 | -0.688 | -0.34 | -1.594 | 0.245 | -0.009 | -0.02 | -0.02 | 0 | 0.77 | 0.84 | 0.8 | 0.65 | 0.69 | 0.58 |
| 8 | 4257291 | rs1864517 | -0.297 | -0.74 | -0.936 | 0.138 | 0 | 0.016 | 0.016 | 0.011 | 0.31 | 0.63 | 0.25 | 0.42 | 0.32 | 0.42 |
| 8 | 4258091 | rs1430448 | 0.021 | -0.608 | -1.769 | 0.02 | -0.014 | -0.049 | -0.049 | 0 | 0.6 | 0.83 | 0.73 | 0.53 | 0.54 | 0.56 |
| 8 | 4259048 | rs7006904 | -0.158 | -0.655 | -1.356 | -0.228 | 0.002 | -0.014 | -0.014 | 0 | 0.69 | 0.66 | 0.65 | 0.84 | 0.73 | 0.74 |
| 8 | 4261356 | rs13275996 | 0.697 | -0.803 | -1.397 | -0.244 | 0.016 | 0.029 | 0.029 | 0 | 0.08 | 0.02 | 0.02 | 0 | 0.03 | 0.01 |
| 8 | 4263579 | rs17336043 | 0.618 | -0.869 | -1.395 | -0.13 | 0.005 | -0.001 | -0.001 | 0.035 | 0.2 | 0.15 | 0.11 | 0.21 | 0.17 | 0.18 |
| 8 | 4264539 | rs7837956 | -0.548 | -0.954 | -1.199 | -0.527 | -0.014 | -0.058 | -0.058 | 0 | 0.77 | 0.66 | 0.68 | 0.92 | 0.81 | 0.84 |
| 8 | 4264676 | rs7838113 | -0.58 | -0.94 | -1.211 | -0.453 | -0.011 | -0.025 | -0.025 | 0.784 | 0.78 | 0.66 | 0.66 | 0.83 | 0.8 | 0.86 |
| 8 | 4265754 | rs7831167 | 1.446 | -0.727 | -1.564 | -0.302 | -0.004 | 0.039 | 0.039 | 0 | 0.11 | 0.16 | 0.07 | 0.25 | 0.08 | 0.18 |
| 8 | 4265784 | rs6558876 | 0.492 | -0.753 | -1.834 | -0.492 | -0.005 | 0.043 | 0.043 | 0.106 | 0.44 | 0.31 | 0.44 | 0.33 | 0.44 | 0.46 |
| 8 | 4266162 | rs13264886 | 0.495 | -0.028 | -2.461 | -0.334 | -0.004 | -0.015 | -0.015 | 0.675 | 0.21 | 0.04 | 0.12 | 0.14 | 0.18 | 0.15 |
| 8 | 4269777 | rs10092807 | 1.028 | 0.421 | -1.77 | -0.247 | -0.062 | -0.079 | -0.079 | 0.942 | 0.15 | 0.03 | 0.16 | 0.26 | 0.27 | 0.22 |
| 8 | 4270661 | rs13276921 | -0.367 | -0.23 | -0.936 | -0.834 | 0.01 | 0.004 | 0.004 | 0 | 0.1 | 0.08 | 0.04 | 0 | 0.04 | 0.01 |
| 8 | 4271213 | rs1217651 | 0.178 | 0.244 | -0.646 | -0.477 | 0.027 | -0.067 | -0.067 | 0 | 0.82 | 0.5 | 0.72 | 0.94 | 0.81 | 0.93 |
| 8 | 4272581 | rs13262986 | -0.196 | 0.38 | -0.908 | -0.201 | 0.01 | -0.012 | -0.012 | 0.067 | 0.22 | 0.13 | 0.13 | 0.3 | 0.19 | 0.22 |
| 8 | 4274957 | rs1658815 | 0.645 | 0.752 | -0.271 | -0.556 | 0.01 | -0.082 | -0.082 | 0.018 | 0.8 | 0.47 | 0.7 | 0.95 | 0.81 | 0.93 |
| 8 | 4275392 | rs1238056 | 0.83 | 0.901 | -0.931 | -0.352 | -0.005 | 0.019 | 0.019 | 0.042 | 0.87 | 0.93 | 0.79 | 0.94 | 0.85 | 0.85 |
| 8 | 4275981 | rs7816189 | -0.425 | 1.217 | -0.753 | -0.991 | 0.002 | -0.05 | -0.05 | 0.656 | 0.38 | 0.51 | 0.32 | 0.16 | 0.36 | 0.23 |
| 8 | 4276270 | rs11136742 | -1.192 | 2.634 | -0.285 | -0.987 | 0.085 | -0.055 | -0.055 | 0.018 | 0.1 | 0.25 | 0.14 | 0 | 0.22 | 0.07 |
| 8 | 4276303 | rs17414411 | -0.659 | 2.356 | -0.913 | -1.208 | -0.028 | 0.055 | 0.055 | 0 | 0.1 | 0.01 | 0.07 | 0.01 | 0.16 | 0.05 |
| 8 | 4276579 | rs11783587 | -0.128 | 2.937 | -1.708 | -1.431 | -0.062 | 0.023 | 0.023 | 0 | 0.22 | 0.01 | 0.16 | 0.13 | 0.22 | 0.13 |
| 8 | 4276719 | rs17070231 | -0.336 | 2.481 | -1.154 | -1.377 | -0.001 | -0.017 | -0.017 | 0 | 0.09 | 0.12 | 0.06 | 0.01 | 0.12 | 0.03 |
| 8 | 4277173 | rs1725132 | 0.756 | 2.386 | -0.944 | -0.351 | -0.008 | -0.008 | -0.008 | 0 | 0.86 | 0.83 | 0.78 | 0.72 | 0.75 | 0.77 |
| 8 | 4277861 | rs1217534 | -0.563 | 1.99 | 0.345 | -0.695 | 0.098 | 0 | 0 | 0 | 0.07 | 0.36 | 0.06 | 0 | 0.07 | 0.07 |
| 8 | 4278256 | rs1238884 | -0.542 | 1.964 | 0.505 | -0.71 | 0.095 | 0.012 | 0.012 | 0 | 0.06 | 0.33 | 0.05 | 0 | 0.06 | 0.07 |
| 8 | 4279582 | rs7015105 | -0.191 | 0.95 | 0.502 | -0.544 | -0.028 | -0.009 | -0.009 | 0 | 0.27 | 0.51 | 0.3 | 0.27 | 0.21 | 0.21 |

| | | | | | | | | | | | | | | | | |
|---|---|---|---|---|---|---|---|---|---|---|---|---|---|---|---|---|
| 8 | 4281327 | rs1154044 | 0.758 | 0.584 | -0.054 | -0.406 | 0 | 0.001 | 0.001 | 0 | 0.27 | 0.28 | 0.25 | 0.13 | 0.23 | 0.34 |
| 8 | 4281973 | rs1438197 | 1.471 | 0.359 | -0.223 | -0.422 | 0.026 | 0.001 | 0.001 | 0 | 0.22 | 0.09 | 0.2 | 0.13 | 0.17 | 0.28 |
| 8 | 4282088 | rs1154046 | 0.022 | 0.705 | 0.009 | -0.221 | 0.001 | 0.018 | 0.018 | 0 | 0.49 | 0.45 | 0.35 | 0.22 | 0.51 | 0.29 |
| 8 | 4282631 | rs1978844 | 0.409 | 0.615 | -0.455 | -0.413 | -0.049 | 0.116 | 0.116 | 0 | 0.38 | 0.19 | 0.25 | 0.1 | 0.42 | 0.19 |
| 8 | 4283673 | rs17414486 | 1.172 | 0.661 | -0.743 | -0.339 | -0.02 | 0.049 | 0.049 | 0 | 0.09 | 0 | 0.06 | 0.01 | 0.12 | 0.03 |
| 8 | 4285173 | rs7815734 | 1.465 | 0.352 | -0.747 | -0.397 | 0.012 | 0.005 | 0.005 | 0 | 0.2 | 0.15 | 0.2 | 0.12 | 0.15 | 0.29 |
| 8 | 4285608 | rs1438198 | 1.611 | 0.336 | -0.754 | -0.405 | 0.02 | 0.004 | 0.004 | 0 | 0.19 | 0.09 | 0.2 | 0.12 | 0.15 | 0.29 |
| 8 | 4285624 | rs1438199 | 1.473 | 0.32 | -0.662 | -0.048 | 0.005 | -0.006 | -0.006 | 0 | 0.21 | 0.2 | 0.21 | 0.23 | 0.15 | 0.3 |
| 8 | 4285663 | rs13251521 | 1.297 | 0.413 | -0.876 | -0.452 | -0.092 | 0.125 | 0.125 | 0 | 0.29 | 0.04 | 0.18 | 0.08 | 0.3 | 0.17 |
| 8 | 4287134 | rs7015706 | 0.023 | 0.423 | -0.283 | -0.133 | 0.018 | 0.018 | 0.018 | 0 | 0.41 | 0.33 | 0.35 | 0.25 | 0.35 | 0.42 |
| 8 | 4287640 | rs4478597 | 0.467 | 0.503 | -0.42 | -0.057 | 0 | 0.063 | 0.063 | 0 | 0.5 | 0.36 | 0.43 | 0.34 | 0.45 | 0.31 |
| 8 | 4288356 | rs17070283 | 1.393 | 0.499 | -0.788 | 0.051 | -0.001 | -0.021 | -0.021 | 0 | 0.1 | 0.01 | 0.09 | 0.13 | 0.09 | 0.1 |
| 8 | 4288538 | rs17414612 | 1.122 | 0.769 | -0.775 | -0.135 | -0.036 | 0.108 | 0.108 | 0 | 0.32 | 0.13 | 0.19 | 0.09 | 0.29 | 0.19 |
| 8 | 4288640 | rs1658821 | -0.749 | 1.071 | -0.535 | 0.443 | -0.008 | 0.049 | 0.049 | 0 | 0.93 | 0.93 | 0.8 | 0.58 | 0.82 | 0.74 |
| 8 | 4289266 | rs10088898 | 0.635 | 0.361 | -0.199 | 0.074 | -0.021 | -0.006 | -0.006 | 1.503 | 0.23 | 0.28 | 0.23 | 0.25 | 0.13 | 0.12 |
| 8 | 4290275 | rs7007659 | 1.717 | 0.483 | -0.627 | 0.029 | -0.001 | 0.011 | 0.011 | 0 | 0.08 | 0.1 | 0.09 | 0.09 | 0.08 | 0.16 |
| 8 | 4290305 | rs11781724 | 1.125 | 0.639 | -1.211 | -0.341 | -0.028 | 0.035 | 0.035 | 0.171 | 0.2 | 0.02 | 0.25 | 0.14 | 0.22 | 0.21 |
| 8 | 4290386 | rs2915065 | -0.781 | 1.041 | -0.817 | 0.104 | 0.018 | 0.071 | 0.071 | 0 | 0.91 | 0.88 | 0.78 | 0.57 | 0.79 | 0.69 |
| 8 | 4292032 | rs1217647 | -0.714 | 0.767 | -0.607 | -0.075 | 0.003 | 0.015 | 0.015 | 0 | 0.72 | 0.74 | 0.65 | 0.6 | 0.66 | 0.67 |
| 8 | 4293806 | rs6990173 | 1.432 | 0.731 | -0.597 | 0.004 | 0.007 | -0.013 | -0.013 | 0 | 0.2 | 0.11 | 0.24 | 0.28 | 0.19 | 0.16 |
| 8 | 4295123 | rs1154057 | -1.554 | 0.977 | -0.515 | 0.048 | 0.05 | 0.057 | 0.057 | 0 | 0.91 | 0.85 | 0.84 | 0.83 | 0.82 | 0.84 |
| 8 | 4295181 | rs6558883 | 1.895 | 0.772 | -0.805 | -0.401 | -0.026 | 0.085 | 0.085 | 0 | 0.27 | 0.1 | 0.16 | 0.08 | 0.25 | 0.18 |
| 8 | 4296478 | rs1154059 | -0.865 | 0.699 | -0.443 | -0.13 | 0.047 | 0.029 | 0.029 | 0 | 0.86 | 0.79 | 0.84 | 0.83 | 0.73 | 0.82 |
| 8 | 4297876 | rs732468 | -0.158 | 0.323 | -0.573 | -0.287 | 0.011 | 0.008 | 0.008 | 0 | 0.37 | 0.34 | 0.45 | 0.69 | 0.28 | 0.45 |
| 8 | 4298057 | rs732467 | 0.913 | 0.335 | -1.048 | -0.331 | -0.072 | -0.089 | -0.089 | 0.593 | 0.45 | 0.1 | 0.41 | 0.49 | 0.45 | 0.5 |
| 8 | 4299117 | rs1235700 | -0.445 | 0.508 | -0.51 | -0.255 | 0.106 | 0.055 | 0.055 | 0 | 0.84 | 0.59 | 0.82 | 0.84 | 0.72 | 0.81 |
| 8 | 4299883 | rs17070309 | 0.317 | 0.313 | -0.703 | -0.477 | -0.011 | -0.015 | -0.015 | 0.148 | 0.12 | 0.12 | 0.2 | 0.39 | 0.16 | 0.3 |
| 8 | 4300537 | rs1228295 | -0.728 | 0.14 | -0.739 | -0.601 | -0.013 | 0.011 | 0.011 | 0 | 0.36 | 0.64 | 0.44 | 0.49 | 0.3 | 0.45 |
| 8 | 4302217 | rs17336607 | 1.879 | 0.206 | -1.565 | -0.531 | 0 | 0.003 | 0.003 | 0.117 | 0.12 | 0.01 | 0.18 | 0.1 | 0.11 | 0.16 |
| 8 | 4303216 | rs17414883 | 1.135 | 0.334 | -1.804 | -0.733 | -0.109 | 0.202 | 0.202 | 0.2 | 0.36 | 0.05 | 0.21 | 0.09 | 0.36 | 0.22 |
| 8 | 4303398 | rs17070316 | 0.751 | 0.323 | -1.325 | -0.191 | -0.031 | -0.028 | -0.028 | 1.016 | 0.08 | 0.04 | 0.16 | 0.38 | 0.08 | 0.24 |
| 8 | 4303418 | rs17070317 | -0.081 | 0.242 | -1.114 | -0.227 | 0.007 | -0.035 | -0.035 | 0 | 0.22 | 0.22 | 0.35 | 0.47 | 0.19 | 0.42 |
| 8 | 4303634 | rs17070327 | 0.464 | 0.415 | -1.139 | -0.054 | -0.01 | 0.005 | 0.005 | 0 | 0.09 | 0.06 | 0.17 | 0.39 | 0.1 | 0.26 |
| 8 | 4308969 | rs1006583 | 0.101 | 0.144 | -0.849 | -1.176 | -0.023 | -0.054 | -0.054 | 0 | 0.27 | 0.39 | 0.2 | 0.09 | 0.18 | 0.2 |
| 8 | 4309702 | rs13278525 | 0.166 | 0.486 | -1.406 | -1.402 | -0.01 | 0.178 | 0.178 | 0 | 0.49 | 0.24 | 0.34 | 0.16 | 0.44 | 0.39 |
| 8 | 4312402 | rs7006288 | 0.365 | 0.638 | -1.575 | -1.303 | -0.068 | 0.2 | 0.2 | 0.113 | 0.37 | 0.09 | 0.3 | 0.08 | 0.39 | 0.22 |
| 8 | 4314917 | rs1217680 | 0.498 | 0.733 | -0.579 | -0.658 | 0.06 | 0.008 | 0.008 | 0 | 0.9 | 0.74 | 0.85 | 0.9 | 0.82 | 0.93 |
| 8 | 4315838 | rs1217677 | 0.849 | 0.704 | -0.666 | -0.643 | 0.039 | -0.002 | -0.002 | 0 | 0.87 | 0.73 | 0.84 | 0.85 | 0.8 | 0.9 |
| 8 | 4317346 | rs1217675 | -0.662 | 0.247 | -0.894 | -1.22 | -0.033 | -0.083 | -0.083 | 1.155 | 0.28 | 0.44 | 0.15 | 0.08 | 0.23 | 0.13 |
| 8 | 4317699 | rs4875347 | -0.682 | 0.356 | -1.579 | -1.146 | -0.007 | 0.073 | 0.073 | 0 | 0.23 | 0.07 | 0.12 | 0.09 | 0.24 | 0.16 |
| 8 | 4319850 | rs12545377 | 0.368 | 0.372 | -1.34 | -1.033 | 0.024 | 0.034 | 0.034 | 0.701 | 0.25 | 0.25 | 0.09 | 0.08 | 0.15 | 0.15 |
| 8 | 4320053 | rs2163306 | 1.203 | 0.648 | -1.771 | -1.058 | -0.062 | 0.017 | 0.017 | 0.553 | 0.21 | 0.01 | 0.21 | 0.13 | 0.23 | 0.26 |
| 8 | 4320301 | rs12676259 | -0.242 | 0.668 | -1.71 | -1.087 | 0.019 | 0.072 | 0.072 | 0 | 0.31 | 0.25 | 0.22 | 0.08 | 0.27 | 0.2 |
| 8 | 4320656 | rs11136745 | 0.406 | 0.665 | -1.518 | -0.938 | 0.024 | 0.036 | 0.036 | 0.648 | 0.26 | 0.25 | 0.17 | 0.08 | 0.15 | 0.16 |
| 8 | 4320749 | rs1217666 | -0.603 | 1.197 | -1.544 | -0.469 | 0.027 | 0.029 | 0.029 | 0 | 0.93 | 0.87 | 0.87 | 0.86 | 0.84 | 0.92 |
| 8 | 4321106 | rs13250119 | 2.082 | 1.278 | -1.777 | -0.34 | -0.068 | 0.017 | 0.017 | 0 | 0.14 | 0.01 | 0.14 | 0.09 | 0.18 | 0.1 |
| 8 | 4322434 | rs9644367 | 0.807 | 1.138 | -1.131 | 0.207 | -0.056 | 0.012 | 0.012 | 0.104 | 0.18 | 0.01 | 0.21 | 0.12 | 0.21 | 0.23 |
| 8 | 4322454 | rs9314531 | 1.041 | 0.993 | -0.764 | 0.695 | -0.009 | 0.016 | 0.016 | 0 | 0.4 | 0.15 | 0.36 | 0.35 | 0.39 | 0.34 |
| 8 | 4328106 | rs4875349 | 1.135 | 0.899 | -0.144 | 1.165 | -0.007 | -0.018 | -0.018 | 0 | 0.53 | 0.22 | 0.5 | 0.44 | 0.57 | 0.43 |
| 8 | 4335682 | rs4875350 | 0.087 | 1.106 | -0.203 | 0.941 | -0.023 | 0 | 0 | 0.954 | 0.19 | 0.02 | 0.29 | 0.14 | 0.21 | 0.24 |
| 8 | 4339207 | rs1725122 | 1.148 | 1.321 | 1.296 | 1.295 | 0.101 | 0.196 | 0.196 | 0 | 0.93 | 0.72 | 0.91 | 0.78 | 0.9 | 0.93 |
| 8 | 4341641 | rs1725111 | 0.968 | 1.399 | 1.246 | 0.961 | 0.021 | -0.002 | -0.002 | 0.931 | 0.84 | 0.77 | 0.85 | 0.82 | 0.8 | 0.81 |
| 8 | 4345284 | rs1626142 | 1.945 | 1.371 | 1.316 | 0.939 | 0.002 | 0.004 | 0.004 | 0 | 0.68 | 0.35 | 0.61 | 0.6 | 0.7 | 0.57 |
| 8 | 4345457 | rs1658825 | 1.264 | 1.329 | 1.615 | 0.619 | 0.068 | 0.057 | 0.057 | 0.159 | 0.88 | 0.53 | 0.89 | 0.86 | 0.89 | 0.92 |
| 8 | 4349983 | rs2915061 | -0.722 | 1.461 | 1.041 | 0.683 | 0 | -0.041 | -0.041 | 2.482 | 0.12 | 0.18 | 0.07 | 0.04 | 0.09 | 0.1 |
| 8 | 4350040 | rs2043804 | 0.826 | 1.494 | 0.675 | 0.874 | 0.092 | 0.091 | 0.091 | 0 | 0.9 | 0.54 | 0.89 | 0.87 | 0.89 | 0.93 |
| 8 | 4352157 | rs1725130 | -0.994 | 1.335 | 0.218 | 0.762 | 0.016 | -0.01 | -0.01 | 0 | 0.26 | 0.47 | 0.31 | 0.23 | 0.21 | 0.34 |
| 8 | 4352592 | rs12676659 | -0.857 | 1.64 | -0.16 | 1.008 | 0.006 | 0.007 | 0.007 | 0 | 0.13 | 0.16 | 0.12 | 0.16 | 0.1 | 0.29 |
| 8 | 4352751 | rs941006 | 0.143 | 1.453 | -0.952 | 1.02 | -0.01 | -0.056 | -0.056 | 0.177 | 0.29 | 0.16 | 0.26 | 0.36 | 0.3 | 0.25 |
| 8 | 4353992 | rs893457 | 0.289 | 1.442 | -0.608 | 0.92 | 0 | 0 | 0 | 0 | 0.83 | 0.79 | 0.76 | 0.8 | 0.78 | 0.78 |
| 8 | 4354653 | rs11997705 | -0.279 | 1.26 | -0.535 | 0.569 | -0.001 | -0.012 | -0.012 | 0 | 0.09 | 0.1 | 0.04 | 0.02 | 0.06 | 0.07 |
| 8 | 4356569 | rs1014333 | 0.378 | 1.41 | 0.045 | 0.86 | 0.023 | 0.02 | 0.02 | 0 | 0.82 | 0.69 | 0.71 | 0.8 | 0.77 | 0.79 |
| 8 | 4357525 | rs1217688 | 0.157 | 1.228 | -0.051 | 0.802 | 0.003 | 0.002 | 0.002 | 0 | 0.84 | 0.74 | 0.85 | 0.83 | 0.82 | 0.83 |
| 8 | 4361088 | rs1658826 | -0.475 | 0.818 | 0.275 | 0.769 | -0.006 | 0.023 | 0.023 | 0 | 0.15 | 0.31 | 0.15 | 0.18 | 0.13 | 0.17 |
| 8 | 4361441 | rs1013702 | -0.243 | 0.825 | -0.04 | 0.778 | -0.001 | 0 | 0 | 0 | 0.14 | 0.18 | 0.14 | 0.17 | 0.15 | 0.17 |
| 8 | 4365381 | rs1864516 | 0.519 | 0.532 | -0.952 | 0.595 | -0.029 | -0.044 | -0.044 | 0 | 0.16 | 0.01 | 0.24 | 0.14 | 0.14 | 0.2 |
| 8 | 4366666 | rs1985200 | 0.046 | 0.429 | -0.779 | 0.561 | 0 | -0.001 | -0.001 | 0 | 0.14 | 0.22 | 0.16 | 0.15 | 0.14 | 0.16 |
| 8 | 4369334 | rs4609213 | -0.454 | -0.101 | -1.272 | 0.036 | 0.017 | 0.084 | 0.084 | 0 | 0.48 | 0.25 | 0.45 | 0.33 | 0.49 | 0.44 |
| 8 | 4371051 | rs11994346 | 0.504 | -0.178 | -1.601 | -0.113 | 0 | -0.014 | -0.014 | 0 | 0.09 | 0.09 | 0.04 | 0.02 | 0.06 | 0.06 |

| | | | | | | | | | | | | | | | | |
|---|---|---|---|---|---|---|---|---|---|---|---|---|---|---|---|---|
| 8 | 4371169 | rs1112779 | -0.416 | -0.4 | -2.156 | -0.167 | -0.025 | -0.006 | -0.006 | 1.296 | 0.44 | 0.28 | 0.57 | 0.47 | 0.46 | 0.47 |
| 8 | 4372185 | rs10503264 | 0.031 | -0.675 | -2.345 | -0.2 | -0.039 | -0.002 | -0.002 | 0 | 0.35 | 0.07 | 0.43 | 0.33 | 0.33 | 0.33 |
| 8 | 4373184 | rs13275581 | 1.719 | -0.686 | -2.295 | 0.12 | -0.107 | -0.112 | -0.112 | 0 | 0.32 | 0.04 | 0.26 | 0.4 | 0.36 | 0.39 |
| 8 | 4377712 | rs1113226 | 0.214 | -0.86 | -2.092 | -0.113 | -0.053 | -0.044 | -0.044 | 0 | 0.35 | 0.04 | 0.39 | 0.33 | 0.34 | 0.34 |
| 8 | 4378215 | rs9650516 | 1.439 | -0.782 | -1.653 | -0.092 | -0.053 | -0.067 | -0.067 | 0 | 0.35 | 0.12 | 0.24 | 0.46 | 0.37 | 0.4 |
| 8 | 4379269 | rs10106920 | 0.69 | -0.555 | -1.432 | -0.091 | -0.001 | -0.002 | -0.002 | 2.559 | 0.15 | 0.2 | 0.17 | 0.11 | 0.14 | 0.17 |
| 8 | 4381436 | rs4875108 | -0.56 | -0.592 | -1.186 | -0.023 | -0.001 | -0.004 | -0.004 | 0 | 0.5 | 0.26 | 0.67 | 0.5 | 0.55 | 0.5 |
| 8 | 4383116 | rs11136747 | 0.798 | -0.499 | -1.189 | -0.064 | -0.024 | -0.054 | -0.054 | 0 | 0.13 | 0.28 | 0.14 | 0.03 | 0.06 | 0.05 |
| 8 | 4384861 | rs13273099 | 0.916 | -0.488 | -1.806 | 0.083 | -0.087 | -0.027 | -0.027 | 0 | 0.39 | 0.13 | 0.28 | 0.42 | 0.46 | 0.46 |
| 8 | 4388680 | rs4875357 | 0.04 | -0.526 | -1.712 | 0.054 | 0.011 | -0.043 | -0.043 | 0.025 | 0.43 | 0.14 | 0.5 | 0.43 | 0.4 | 0.37 |
| 8 | 4391037 | rs17070498 | 0.875 | -0.581 | -2 | 0.169 | 0.003 | -0.038 | -0.038 | 0 | 0.21 | 0.01 | 0.29 | 0.15 | 0.14 | 0.18 |
| 8 | 4391377 | rs12056740 | 1.009 | -0.532 | -1.913 | 0.113 | -0.065 | -0.014 | -0.014 | 0.018 | 0.4 | 0.16 | 0.28 | 0.43 | 0.48 | 0.48 |
| 8 | 4391719 | rs11994887 | 0.392 | -0.331 | -1.986 | -0.066 | -0.002 | -0.001 | -0.001 | 0 | 0.11 | 0.09 | 0.05 | 0.04 | 0.13 | 0.13 |
| 8 | 4395685 | rs2406706 | 1.188 | -0.5 | -1.976 | -0.117 | -0.067 | -0.026 | -0.026 | 0.014 | 0.38 | 0.13 | 0.23 | 0.41 | 0.44 | 0.48 |
| 8 | 4397019 | rs1464977 | 0.455 | -0.232 | -1.587 | -0.209 | -0.006 | 0.057 | 0.057 | 0 | 0.55 | 0.23 | 0.45 | 0.64 | 0.58 | 0.64 |
| 8 | 4397180 | rs1526341 | -0.213 | -0.092 | -0.953 | 0.021 | 0.004 | -0.004 | -0.004 | 0 | 0.36 | 0.43 | 0.41 | 0.35 | 0.39 | 0.33 |
| 8 | 4398511 | rs7010832 | -0.102 | 0.088 | -0.567 | 0.468 | -0.002 | -0.002 | -0.002 | 0 | 0.18 | 0.17 | 0.13 | 0.17 | 0.14 | 0.2 |
| 8 | 4398526 | rs2406707 | 0.284 | 0.525 | -0.577 | 0.765 | -0.016 | -0.019 | -0.019 | 0 | 0.05 | 0.01 | 0.09 | 0.1 | 0.08 | 0.14 |
| 8 | 4400460 | rs12547182 | 0.088 | 0.573 | -1.468 | 0.715 | -0.057 | -0.036 | -0.036 | 0.01 | 0.34 | 0.08 | 0.23 | 0.33 | 0.36 | 0.4 |
| 8 | 4401124 | rs11786902 | 0.033 | 0.611 | -1.423 | 0.612 | -0.03 | -0.078 | -0.078 | 0.065 | 0.35 | 0.15 | 0.24 | 0.43 | 0.36 | 0.4 |
| 8 | 4403586 | rs1983474 | -0.456 | 0.706 | -0.856 | 0.465 | 0 | 0.016 | 0.016 | 0 | 0.69 | 0.63 | 0.81 | 0.61 | 0.73 | 0.61 |
| 8 | 4405955 | rs7818189 | -0.332 | 0.781 | -0.913 | 0.459 | 0.001 | 0.013 | 0.013 | 0 | 0.67 | 0.63 | 0.76 | 0.59 | 0.71 | 0.63 |
| 8 | 4406368 | rs7831820 | -0.18 | 0.73 | -0.738 | 0.388 | -0.002 | 0.018 | 0.018 | 0 | 0.64 | 0.58 | 0.76 | 0.58 | 0.7 | 0.6 |
| 8 | 4410514 | rs7357469 | 0.63 | 0.714 | -1.227 | -0.047 | -0.005 | 0.022 | 0.022 | 0 | 0.18 | 0.06 | 0.25 | 0.13 | 0.18 | 0.13 |
| 8 | 4412051 | rs1526335 | -0.498 | 0.914 | -1.296 | -0.183 | 0.001 | 0.128 | 0.128 | 0 | 0.82 | 0.66 | 0.86 | 0.57 | 0.81 | 0.71 |
| 8 | 4412355 | rs7357519 | -0.753 | 0.95 | -1.1 | 0.053 | -0.004 | -0.007 | -0.007 | 0 | 0.63 | 0.64 | 0.71 | 0.51 | 0.58 | 0.58 |
| 8 | 4413282 | rs6992457 | -0.342 | 1.044 | -0.46 | -0.106 | 0.027 | 0.04 | 0.04 | 0 | 0.18 | 0.39 | 0.22 | 0.25 | 0.23 | 0.16 |
| 8 | 4414063 | rs9314532 | 0.561 | 1.023 | -0.788 | -0.173 | -0.002 | 0.026 | 0.026 | 0 | 0.17 | 0.06 | 0.25 | 0.13 | 0.18 | 0.14 |
| 8 | 4414316 | rs1997137 | 0.13 | 0.832 | -0.306 | -0.007 | -0.005 | 0.01 | 0.01 | 0 | 0.17 | 0.25 | 0.11 | 0.25 | 0.13 | 0.17 |
| 8 | 4415872 | rs17344322 | -0.23 | 0.826 | -0.243 | -0.288 | 0.001 | -0.037 | -0.037 | 0 | 0.07 | 0.17 | 0.06 | 0 | 0.09 | 0.02 |
| 8 | 4418410 | rs1949909 | 0.169 | 0.902 | -0.091 | -0.074 | 0.049 | 0.045 | 0.045 | 0.233 | 0.11 | 0.33 | 0.16 | 0.13 | 0.14 | 0.14 |
| 8 | 4433995 | rs2725060 | -0.239 | 0.525 | -0.43 | -0.243 | -0.002 | -0.006 | -0.006 | 0.227 | 0.18 | 0.2 | 0.22 | 0.15 | 0.15 | 0.17 |
| 8 | 4434168 | rs7843036 | -0.149 | 0.407 | -0.824 | -0.502 | 0.004 | 0.011 | 0.011 | 0 | 0.24 | 0.21 | 0.19 | 0.07 | 0.19 | 0.17 |
| 8 | 4435074 | rs17416403 | -0.333 | 0.408 | -0.888 | -0.525 | 0.003 | 0.035 | 0.035 | 0.501 | 0.28 | 0.19 | 0.19 | 0.07 | 0.23 | 0.2 |
| 8 | 4436656 | rs2466732 | -0.805 | 0.394 | -0.687 | -0.447 | 0.01 | 0.026 | 0.026 | 0 | 0.49 | 0.35 | 0.52 | 0.33 | 0.5 | 0.51 |
| 8 | 4437465 | rs10097669 | 0.257 | 0.138 | -0.849 | -0.556 | -0.059 | 0.023 | 0.023 | 0 | 0.28 | 0.01 | 0.25 | 0.17 | 0.32 | 0.29 |
| 8 | 4437566 | rs2725065 | -0.491 | 0.145 | -0.646 | -0.519 | 0.002 | 0.031 | 0.031 | 0 | 0.36 | 0.23 | 0.28 | 0.2 | 0.39 | 0.32 |
| 8 | 4438279 | rs17070658 | 0.344 | 0.061 | -0.335 | -0.091 | 0.001 | 0.043 | 0.043 | 0.033 | 0.15 | 0.22 | 0.17 | 0.27 | 0.12 | 0.17 |
| 8 | 4438543 | rs17070660 | 0.338 | 0.058 | -0.336 | -0.093 | 0.001 | 0.042 | 0.042 | 0.045 | 0.15 | 0.22 | 0.17 | 0.27 | 0.12 | 0.17 |
| 8 | 4438876 | rs2617069 | -1.027 | -0.029 | -0.752 | -0.181 | 0.012 | -0.01 | -0.01 | 0 | 0.27 | 0.79 | 0.22 | 0.31 | 0.19 | 0.2 |
| 8 | 4442963 | rs2725024 | -0.232 | -0.063 | -0.401 | -0.379 | -0.003 | -0.022 | -0.022 | 0.417 | 0.33 | 0.39 | 0.29 | 0.2 | 0.34 | 0.36 |
| 8 | 4444386 | rs4875363 | -0.691 | -0.088 | -0.694 | -0.444 | -0.059 | 0.075 | 0.075 | 0.813 | 0.44 | 0.1 | 0.45 | 0.27 | 0.48 | 0.52 |
| 8 | 4444535 | rs11784571 | -0.659 | 0.093 | -0.584 | -0.409 | -0.001 | -0.005 | -0.005 | 0 | 0.44 | 0.45 | 0.41 | 0.32 | 0.46 | 0.51 |
| 8 | 4444592 | rs4875364 | 0.397 | -0.057 | -0.784 | -0.278 | -0.003 | 0.011 | 0.011 | 0 | 0.15 | 0.09 | 0.19 | 0.11 | 0.14 | 0.15 |
| 8 | 4445174 | rs2617076 | -1.892 | -0.131 | -0.873 | -0.28 | -0.094 | -0.112 | -0.112 | 0.252 | 0.62 | 0.91 | 0.59 | 0.53 | 0.6 | 0.65 |
| 8 | 4445944 | rs2617078 | -0.367 | -0.815 | -0.268 | -0.184 | 0.001 | 0.002 | 0.002 | 0.231 | 0.37 | 0.34 | 0.34 | 0.27 | 0.38 | 0.45 |
| 8 | 4447199 | rs17070693 | 0.858 | -0.788 | -0.201 | -0.147 | -0.006 | 0.031 | 0.031 | 0.21 | 0.05 | 0.14 | 0.02 | 0.13 | 0.03 | 0 |
| 8 | 4447768 | rs2617084 | -0.296 | -0.95 | -0.194 | -0.246 | -0.007 | -0.007 | -0.007 | 3.107 | 0.35 | 0.43 | 0.36 | 0.29 | 0.38 | 0.43 |
| 8 | 4448387 | rs2724985 | -1.765 | -0.886 | -0.291 | -0.241 | -0.097 | -0.099 | -0.099 | 0 | 0.56 | 0.9 | 0.56 | 0.51 | 0.52 | 0.59 |
| 8 | 4448785 | rs7832831 | 0.102 | -0.684 | -0.097 | -0.101 | -0.007 | 0.025 | 0.025 | 0 | 0.12 | 0.21 | 0.11 | 0.19 | 0.09 | 0.09 |
| 8 | 4449297 | rs2617087 | -1.478 | -0.679 | -0.038 | -0.295 | 0.001 | 0.009 | 0.009 | 0 | 0.54 | 0.78 | 0.46 | 0.44 | 0.47 | 0.51 |
| 8 | 4449506 | rs17070710 | 0.143 | -0.891 | 0.181 | -0.235 | 0.04 | 0.027 | 0.027 | 3.361 | 0.05 | 0.26 | 0.02 | 0.02 | 0.02 | 0.01 |
| 8 | 4451048 | rs6998447 | -1.342 | -0.874 | -0.278 | -0.274 | 0.026 | -0.078 | -0.078 | 0 | 0.14 | 0.47 | 0.06 | 0.02 | 0.09 | 0.03 |
| 8 | 4453036 | rs13259583 | 1.119 | -0.649 | -0.453 | -0.15 | -0.042 | -0.046 | -0.046 | 0.846 | 0.44 | 0.13 | 0.52 | 0.53 | 0.56 | 0.48 |
| 8 | 4453244 | rs13249431 | -0.022 | -0.728 | -0.32 | -0.19 | 0.013 | 0.006 | 0.006 | 0.237 | 0.62 | 0.75 | 0.73 | 0.64 | 0.68 | 0.62 |
| 8 | 4453385 | rs7009652 | -0.541 | -0.723 | -0.288 | -0.195 | -0.012 | -0.034 | -0.034 | 0 | 0.17 | 0.59 | 0.2 | 0.11 | 0.13 | 0.14 |
| 8 | 4453655 | rs4875365 | 1.012 | -0.456 | -0.546 | -0.06 | -0.045 | -0.056 | -0.056 | 0 | 0.44 | 0.13 | 0.52 | 0.52 | 0.55 | 0.48 |
| 8 | 4454584 | rs2724952 | -0.042 | -0.394 | -0.067 | -0.045 | -0.002 | 0 | 0 | 0 | 0.93 | 0.95 | 0.97 | 0.97 | 0.94 | 0.95 |
| 8 | 4456167 | rs2617094 | 0.415 | -0.54 | -0.002 | -0.031 | -0.007 | -0.027 | -0.027 | 0 | 0.82 | 0.75 | 0.8 | 0.87 | 0.87 | 0.82 |
| 8 | 4458212 | rs1382249 | 0.145 | -0.596 | 0.063 | 0.094 | -0.007 | -0.03 | -0.03 | 0 | 0.83 | 0.61 | 0.78 | 0.86 | 0.87 | 0.83 |
| 8 | 4460277 | rs11781113 | 0.551 | -0.287 | -1.055 | 0.309 | 0.036 | -0.028 | -0.028 | 1.843 | 0.46 | 0.18 | 0.49 | 0.5 | 0.6 | 0.41 |
| 8 | 4461958 | rs6990838 | 0.262 | -0.276 | 0.111 | 0.304 | -0.035 | 0.013 | 0.013 | 0 | 0.88 | 0.75 | 0.9 | 0.89 | 0.95 | 0.88 |
| 8 | 4462599 | rs13269869 | -0.632 | 0.351 | -0.1 | 0.094 | 0.013 | -0.003 | -0.003 | 2.365 | 0.17 | 0.26 | 0.13 | 0.19 | 0.17 | 0.16 |
| 8 | 4463359 | rs17070773 | -0.16 | 0.46 | -0.565 | 0.285 | -0.018 | -0.038 | -0.038 | 0 | 0.08 | 0.01 | 0.17 | 0.14 | 0.07 | 0.12 |
| 8 | 4464167 | rs2725014 | -1.4 | -0.041 | -0.369 | -0.075 | 0.005 | -0.001 | -0.001 | 0.06 | 0.35 | 0.66 | 0.32 | 0.38 | 0.28 | 0.39 |
| 8 | 4464209 | rs2617102 | -0.331 | 0.09 | 0.257 | -0.147 | 0.012 | -0.023 | -0.023 | 0 | 0.14 | 0.17 | 0.09 | 0.09 | 0.15 | 0.16 |
| 8 | 4465474 | rs1382251 | -1.808 | -0.44 | -0.203 | -0.318 | 0.027 | 0.032 | 0.032 | 0 | 0.5 | 0.75 | 0.42 | 0.47 | 0.39 | 0.49 |
| 8 | 4467853 | rs2725046 | -1.71 | -0.45 | -0.122 | -0.352 | 0.014 | 0.012 | 0.012 | 0 | 0.48 | 0.7 | 0.41 | 0.48 | 0.38 | 0.5 |
| 8 | 4468173 | rs1478272 | -0.87 | -0.619 | 0.421 | -0.331 | -0.004 | 0.028 | 0.028 | 0 | 0.16 | 0.23 | 0.23 | 0.42 | 0.12 | 0.27 |

| | | | | | | | | | | | | | | | | |
|---|---|---|---|---|---|---|---|---|---|---|---|---|---|---|---|---|
| 8 | 4468542 | rs2725047 | 0.049 | -0.513 | 0.678 | -0.376 | -0.003 | -0.013 | -0.013 | 0 | 0.22 | 0.23 | 0.13 | 0.02 | 0.22 | 0.21 |
| 8 | 4469134 | rs7843182 | -0.416 | -0.423 | 0.745 | -0.208 | -0.007 | -0.018 | -0.018 | 0 | 0.14 | 0.29 | 0.09 | 0.11 | 0.12 | 0.14 |
| 8 | 4471486 | rs2725037 | -1.258 | -0.494 | 0.589 | -0.199 | 0.017 | 0.074 | 0.074 | 0.009 | 0.46 | 0.74 | 0.28 | 0.31 | 0.37 | 0.4 |
| 8 | 4471581 | rs17070808 | -1.004 | -0.323 | 0.928 | 0.148 | 0 | 0 | 0 | 3.323 | 0.13 | 0.13 | 0.21 | 0.26 | 0.11 | 0.23 |
| 8 | 4471851 | rs2617106 | -0.663 | -0.305 | 0.9 | 0.001 | -0.01 | -0.052 | -0.052 | 0 | 0.32 | 0.47 | 0.2 | 0.21 | 0.25 | 0.25 |
| 8 | 4475968 | rs2725012 | -0.314 | 0.092 | 0.136 | 0.056 | -0.013 | 0.03 | 0.03 | 0 | 0.34 | 0.25 | 0.23 | 0.02 | 0.38 | 0.26 |
| 8 | 4476870 | rs2616972 | -1.04 | 0.264 | -0.382 | 0.262 | 0 | 0.02 | 0.02 | 0 | 0.5 | 0.68 | 0.47 | 0.44 | 0.49 | 0.48 |
| 8 | 4476957 | rs17070836 | -0.369 | 0.39 | -0.46 | 0.904 | -0.008 | -0.037 | -0.037 | 0 | 0.06 | 0.01 | 0.15 | 0.15 | 0.05 | 0.14 |
| 8 | 4480667 | rs10100024 | -1.031 | 0.472 | 0.23 | 0.717 | 0.027 | 0.043 | 0.043 | 0 | 0.09 | 0.35 | 0.08 | 0.1 | 0.12 | 0.06 |
| 8 | 4480689 | rs2616982 | 0.137 | -0.35 | 0.027 | 0.814 | -0.029 | 0.011 | 0.011 | 0.027 | 0.75 | 0.69 | 0.86 | 0.61 | 0.84 | 0.87 |
| 8 | 4481209 | rs2725019 | -1.129 | 0.371 | 0.308 | 0.806 | 0.019 | 0.016 | 0.016 | 0 | 0.14 | 0.33 | 0.12 | 0.19 | 0.14 | 0.16 |
| 8 | 4481250 | rs2725020 | 0.131 | 0.599 | -0.193 | 0.411 | 0.007 | 0.062 | 0.062 | 0.163 | 0.5 | 0.33 | 0.66 | 0.63 | 0.48 | 0.51 |
| 8 | 4481386 | rs10113503 | -0.133 | 0.925 | -0.102 | 0.721 | 0.019 | 0.035 | 0.035 | 0 | 0.09 | 0.18 | 0.09 | 0.1 | 0.11 | 0.15 |
| 8 | 4481743 | rs17417225 | 0.013 | 0.669 | -0.58 | -0.456 | 0 | -0.043 | -0.043 | 0 | 0.29 | 0.24 | 0.19 | 0.01 | 0.26 | 0.17 |
| 8 | 4481970 | rs17070864 | -0.208 | 1.226 | -0.828 | 0.397 | -0.018 | -0.029 | -0.029 | 0.772 | 0.13 | 0.07 | 0.08 | 0.35 | 0.22 | 0.19 |
| 8 | 4482561 | rs1982628 | 0.283 | 0.923 | -0.883 | 0.445 | 0.013 | 0.02 | 0.02 | 0 | 0.67 | 0.51 | 0.76 | 0.62 | 0.61 | 0.68 |
| 8 | 4483059 | rs13259288 | 0.888 | 0.51 | -0.95 | -0.02 | 0.01 | -0.029 | -0.029 | 0.046 | 0.27 | 0.18 | 0.52 | 0.41 | 0.24 | 0.44 |
| 8 | 4483202 | rs2724958 | -0.489 | 0.433 | -0.889 | -0.006 | 0.039 | 0.104 | 0.104 | 0 | 0.76 | 0.52 | 0.78 | 0.51 | 0.63 | 0.75 |
| 8 | 4484362 | rs17070881 | 0.831 | 0.498 | -0.735 | 0.016 | 0.03 | 0.114 | 0.114 | 0 | 0.08 | 0.33 | 0.11 | 0.11 | 0.07 | 0.1 |
| 8 | 4484498 | rs10104818 | 1.601 | 0.282 | -1.131 | -0.359 | -0.001 | -0.042 | -0.042 | 0 | 0.29 | 0.23 | 0.17 | 0 | 0.23 | 0.16 |
| 8 | 4485185 | rs4875113 | 1.908 | 0.292 | -1.274 | -0.189 | -0.013 | -0.021 | -0.021 | 0 | 0.07 | 0.03 | 0.02 | 0.1 | 0.07 | 0.1 |
| 8 | 4486561 | rs13282923 | 0.668 | 0.141 | -1.345 | -0.227 | 0.018 | -0.031 | -0.031 | 1.448 | 0.32 | 0.17 | 0.53 | 0.44 | 0.29 | 0.44 |
| 8 | 4487895 | rs2616991 | 0.836 | 0.033 | -1.251 | 0.106 | -0.003 | 0.054 | 0.054 | 0.993 | 0.52 | 0.26 | 0.65 | 0.37 | 0.46 | 0.48 |
| 8 | 4490277 | rs1478281 | -0.766 | 0.174 | -1.013 | 0.091 | 0.003 | 0.007 | 0.007 | 0.076 | 0.73 | 0.56 | 0.79 | 0.72 | 0.71 | 0.74 |
| 8 | 4491502 | rs13261335 | 0.562 | 0.107 | -1.25 | 0.029 | -0.002 | -0.012 | -0.012 | 0 | 0.16 | 0.16 | 0.1 | 0.08 | 0.22 | 0.19 |
| 8 | 4492509 | rs4875371 | 1.088 | 0.19 | -1.213 | 0.234 | -0.005 | -0.016 | -0.016 | 0 | 0.1 | 0.07 | 0.08 | 0.13 | 0.13 | 0.12 |
| 8 | 4492520 | rs4875372 | 0.634 | 0.17 | -1.178 | 0.213 | -0.001 | -0.004 | -0.004 | 0 | 0.12 | 0.11 | 0.09 | 0.13 | 0.13 | 0.12 |
| 8 | 4493979 | rs2617004 | 1.227 | -0.009 | -1.236 | 0.132 | 0 | -0.003 | -0.003 | 0.09 | 0.32 | 0.28 | 0.19 | 0.2 | 0.27 | 0.22 |
| 8 | 4494048 | rs11991366 | 0.602 | 0.118 | -1.208 | 0.206 | 0.001 | -0.01 | -0.01 | 0 | 0.15 | 0.09 | 0.11 | 0.13 | 0.17 | 0.13 |
| 8 | 4494520 | rs2724974 | 0.007 | 0.315 | -0.791 | 0.235 | 0.006 | 0.015 | 0.015 | 1.291 | 0.76 | 0.6 | 0.82 | 0.72 | 0.73 | 0.76 |
| 8 | 4496463 | rs2617009 | -0.611 | 0.963 | -1.452 | 0.271 | -0.079 | -0.036 | -0.036 | 0 | 0.83 | 0.98 | 0.84 | 0.81 | 0.76 | 0.9 |
| 8 | 4500036 | rs2725007 | -0.193 | 0.853 | -1.258 | 0.149 | -0.039 | -0.002 | -0.002 | 0.007 | 0.8 | 0.86 | 0.85 | 0.77 | 0.66 | 0.82 |
| 8 | 4501278 | rs7009980 | -0.568 | 0.3 | -0.802 | 0.318 | -0.025 | 0.094 | 0.094 | 0 | 0.12 | 0.2 | 0.06 | 0.33 | 0.04 | 0.15 |
| 8 | 4501563 | rs7001148 | 0.318 | 0.287 | -0.786 | 0.327 | -0.004 | -0.001 | -0.001 | 0 | 0.07 | 0.08 | 0.02 | 0.13 | 0.03 | 0.12 |
| 8 | 4502531 | rs1038061 | 0.714 | 0.132 | -0.488 | 0.301 | 0.02 | 0.025 | 0.025 | 0.517 | 0.71 | 0.54 | 0.71 | 0.57 | 0.62 | 0.68 |
| 8 | 4503462 | rs2617044 | 0.702 | 0.112 | -0.471 | 0.294 | 0.019 | 0.022 | 0.022 | 0.566 | 0.71 | 0.54 | 0.71 | 0.6 | 0.62 | 0.69 |
| 8 | 4503828 | rs7841173 | -1.281 | -0.132 | -0.496 | 0.134 | -0.051 | 0.045 | 0.045 | 0 | 0.16 | 0.44 | 0.09 | 0.23 | 0.05 | 0.16 |
| 8 | 4504531 | rs11984453 | -0.119 | -0.447 | -1.085 | -0.288 | 0.032 | 0.07 | 0.07 | 0 | 0.17 | 0.04 | 0.04 | 0.03 | 0.08 | 0.19 |
| 8 | 4505914 | rs10094093 | 2.203 | -0.605 | -0.693 | -0.602 | -0.003 | 0.024 | 0.024 | 0 | 0.22 | 0.25 | 0.45 | 0.4 | 0.2 | 0.34 |
| 8 | 4506266 | rs2725000 | 0.156 | -0.476 | -0.881 | -0.749 | -0.001 | -0.002 | -0.002 | 0 | 0.63 | 0.59 | 0.73 | 0.56 | 0.59 | 0.56 |
| 8 | 4507076 | rs2724996 | 0.508 | -0.434 | -0.689 | -0.687 | -0.001 | -0.001 | -0.001 | 0 | 0.58 | 0.52 | 0.69 | 0.56 | 0.57 | 0.53 |
| 8 | 4508067 | rs10102117 | -0.025 | -0.394 | -0.686 | -0.667 | -0.016 | 0.019 | 0.019 | 0.16 | 0.66 | 0.77 | 0.75 | 0.73 | 0.6 | 0.62 |
| 8 | 4510670 | rs2725074 | 0.177 | -0.092 | -0.561 | -0.791 | 0.004 | -0.01 | -0.01 | 0 | 0.6 | 0.54 | 0.68 | 0.73 | 0.51 | 0.58 |
| 8 | 4511452 | rs12543670 | 1.605 | -0.33 | -0.54 | -0.809 | -0.024 | 0.058 | 0.058 | 0 | 0.27 | 0.41 | 0.48 | 0.43 | 0.22 | 0.37 |
| 8 | 4512135 | rs2617030 | 0.134 | -0.185 | -0.533 | -0.893 | 0.002 | -0.044 | -0.044 | 0.107 | 0.37 | 0.53 | 0.28 | 0.08 | 0.36 | 0.21 |
| 8 | 4516591 | rs12675498 | -0.276 | 0.018 | -0.816 | -0.517 | -0.023 | 0.046 | 0.046 | 0 | 0.09 | 0.22 | 0.05 | 0.22 | 0.03 | 0.12 |
| 8 | 4519946 | rs2724962 | 0.305 | 0.143 | -1.009 | -0.789 | -0.007 | 0.005 | 0.005 | 0 | 0.62 | 0.67 | 0.72 | 0.74 | 0.53 | 0.56 |
| 8 | 4521513 | rs17070996 | -0.598 | 0.501 | -0.848 | -0.541 | -0.007 | 0.018 | 0.018 | 0 | 0.13 | 0.24 | 0.08 | 0.23 | 0.08 | 0.14 |
| 8 | 4523667 | rs2617021 | -0.01 | 0.703 | -1.224 | -0.551 | -0.08 | -0.015 | -0.015 | 0.037 | 0.73 | 0.91 | 0.76 | 0.74 | 0.6 | 0.64 |
| 8 | 4527217 | rs2617020 | 0.994 | 0.746 | -1.315 | -0.731 | -0.001 | 0.001 | 0.001 | 0 | 0.51 | 0.48 | 0.62 | 0.48 | 0.47 | 0.55 |
| 8 | 4527398 | rs2617019 | 0.972 | 0.836 | -1.112 | -0.529 | 0.002 | 0.001 | 0.001 | 0.216 | 0.57 | 0.48 | 0.66 | 0.49 | 0.55 | 0.61 |
| 8 | 4529945 | rs1904119 | -0.4 | 1.098 | -1.243 | -0.396 | -0.073 | -0.041 | -0.041 | 0 | 0.78 | 0.94 | 0.79 | 0.76 | 0.67 | 0.76 |
| 8 | 4531835 | rs7843255 | 0.621 | 0.873 | -0.941 | -0.333 | 0.043 | 0.071 | 0.071 | 0 | 0.06 | 0.27 | 0.07 | 0.1 | 0.05 | 0.03 |
| 8 | 4535022 | rs17071021 | 1.141 | 0.836 | -1.425 | -0.368 | 0.007 | 0.01 | 0.01 | 0.362 | 0.13 | 0.05 | 0.07 | 0.16 | 0.09 | 0.1 |
| 8 | 4536253 | rs17071029 | -0.347 | 0.769 | -1.288 | -0.432 | 0.001 | 0.001 | 0.001 | 0 | 0.2 | 0.25 | 0.12 | 0.21 | 0.14 | 0.18 |
| 8 | 4536318 | rs10092221 | 0.56 | 0.91 | -1.401 | -0.471 | -0.014 | -0.005 | -0.005 | 0 | 0.68 | 0.67 | 0.72 | 0.65 | 0.6 | 0.71 |
| 8 | 4541063 | rs2248448 | 0.732 | 1.01 | -1.778 | -0.421 | -0.067 | -0.031 | -0.031 | 0.163 | 0.6 | 0.82 | 0.7 | 0.54 | 0.53 | 0.63 |
| 8 | 4546729 | rs1038064 | -0.12 | 1.428 | -1.813 | -0.375 | -0.115 | -0.039 | -0.039 | 0 | 0.76 | 0.94 | 0.76 | 0.75 | 0.61 | 0.73 |
| 8 | 4550458 | rs2617014 | 0.363 | 1.155 | -1.892 | -0.442 | -0.059 | -0.01 | -0.01 | 0.057 | 0.64 | 0.82 | 0.74 | 0.61 | 0.57 | 0.65 |
| 8 | 4555958 | rs12545434 | 1.985 | 0.651 | -2.473 | -0.535 | -0.025 | -0.151 | -0.151 | 0 | 0.24 | 0.01 | 0.43 | 0.41 | 0.21 | 0.33 |
| 8 | 4556148 | rs10110982 | 0.118 | 1.148 | -1.893 | -0.409 | -0.038 | -0.034 | -0.034 | 0 | 0.72 | 0.77 | 0.72 | 0.55 | 0.58 | 0.68 |
| 8 | 4557773 | rs10108771 | 0.411 | 1.181 | -1.61 | -0.449 | 0.009 | 0.01 | 0.01 | 0 | 0.69 | 0.62 | 0.72 | 0.55 | 0.62 | 0.63 |
| 8 | 4559317 | rs2897681 | -0.14 | 1.282 | -1.476 | -0.495 | -0.083 | -0.019 | -0.019 | 0 | 0.74 | 0.89 | 0.73 | 0.73 | 0.57 | 0.7 |
| 8 | 4560754 | rs1471419 | 0.01 | 1.171 | -1.343 | -0.548 | -0.008 | -0.013 | -0.013 | 0.207 | 0.8 | 0.61 | 0.88 | 0.78 | 0.85 | 0.87 |
| 8 | 4561933 | rs10503270 | 0.743 | 1.451 | -1.358 | -0.605 | -0.009 | -0.022 | -0.022 | 0 | 0.68 | 0.57 | 0.83 | 0.74 | 0.79 | 0.78 |
| 8 | 4563819 | rs12679634 | 0.676 | 1.463 | -1.332 | -0.523 | -0.002 | -0.003 | -0.003 | 0 | 0.07 | 0.07 | 0.05 | 0.1 | 0.06 | 0.03 |
| 8 | 4564077 | rs2617058 | 0.415 | 1.799 | -1.437 | -0.527 | -0.028 | -0.025 | -0.025 | 0 | 0.14 | 0.09 | 0.19 | 0.24 | 0.28 | 0.16 |
| 8 | 4564461 | rs748274 | 1.66 | 1.843 | -1.453 | -0.506 | -0.054 | -0.052 | -0.052 | 0.664 | 0.09 | 0.01 | 0.13 | 0.26 | 0.22 | 0.13 |
| 8 | 4566521 | rs974790 | 1.254 | 1.389 | -1.315 | -0.529 | 0 | 0 | 0 | 0 | 0.5 | 0.48 | 0.6 | 0.5 | 0.42 | 0.48 |

| | | | | | | | | | | | | | | | | |
|---|---|---|---|---|---|---|---|---|---|---|---|---|---|---|---|---|
| 8 | 4568613 | rs2617049 | -0.26 | 1.466 | -1.282 | -0.453 | -0.015 | -0.014 | -0.014 | 0.58 | 0.3 | 0.23 | 0.29 | 0.41 | 0.43 | 0.35 |
| 8 | 4571273 | rs10094500 | 0.444 | 1.465 | -1.374 | -0.55 | -0.051 | -0.008 | -0.008 | 0 | 0.63 | 0.79 | 0.7 | 0.61 | 0.52 | 0.61 |
| 8 | 4572113 | rs2725036 | -0.405 | 1.368 | -1.245 | -0.509 | 0.029 | 0.016 | 0.016 | 0.021 | 0.31 | 0.42 | 0.3 | 0.38 | 0.42 | 0.41 |
| 8 | 4577681 | rs1676972 | -0.672 | 1.252 | -1.464 | -0.528 | 0.043 | 0.041 | 0.041 | 2.096 | 0.38 | 0.84 | 0.27 | 0.27 | 0.3 | 0.35 |
| 8 | 4577746 | rs17071123 | 0.348 | 1.238 | -1.21 | -0.538 | 0 | 0.013 | 0.013 | 0 | 0.08 | 0.18 | 0.06 | 0.08 | 0.05 | 0.03 |
| 8 | 4579906 | rs1351107 | -0.01 | 1.223 | -1.066 | -0.558 | -0.02 | -0.047 | -0.047 | 3.21 | 0.29 | 0.56 | 0.19 | 0.11 | 0.24 | 0.28 |
| 8 | 4580814 | rs1673272 | -0.882 | 1.533 | -1.258 | -0.479 | -0.004 | 0.014 | 0.014 | 0.294 | 0.86 | 0.94 | 0.93 | 0.84 | 0.87 | 0.81 |
| 8 | 4582242 | rs293872 | -0.595 | 1.084 | -1.498 | -0.669 | 0.01 | 0.008 | 0.008 | 1.1 | 0.27 | 0.74 | 0.22 | 0.18 | 0.21 | 0.25 |
| 8 | 4585866 | rs1031610 | 1.232 | 1.243 | -1.759 | -0.841 | 0.04 | 0.209 | 0.209 | 1.537 | 0.53 | 0.19 | 0.67 | 0.8 | 0.59 | 0.56 |
| 8 | 4587592 | rs293881 | -0.265 | 1.445 | -1.386 | -0.126 | -0.03 | -0.035 | -0.035 | 2.934 | 0.25 | 0.13 | 0.34 | 0.55 | 0.37 | 0.23 |
| 8 | 4589168 | rs293884 | -1.092 | 1.102 | -0.871 | -0.219 | 0 | 0 | 0 | 0 | 0.4 | 0.56 | 0.44 | 0.53 | 0.44 | 0.34 |
| 8 | 4591177 | rs13257013 | 1.566 | 0.951 | -0.688 | -0.342 | 0.014 | 0.01 | 0.01 | 0 | 0.47 | 0.6 | 0.5 | 0.44 | 0.41 | 0.56 |
| 8 | 4593089 | rs10481373 | 1.866 | 0.908 | -0.97 | -0.29 | -0.033 | 0.034 | 0.034 | 0.072 | 0.43 | 0.11 | 0.48 | 0.38 | 0.42 | 0.47 |
| 8 | 4595814 | rs7826378 | -1.087 | 1.259 | -0.686 | -0.168 | -0.016 | -0.005 | -0.005 | 0 | 0.41 | 0.16 | 0.41 | 0.55 | 0.54 | 0.44 |
| 8 | 4596801 | rs1457186 | 0.18 | 0.443 | -0.545 | -0.375 | -0.022 | -0.108 | -0.108 | 0 | 0.72 | 0.46 | 0.82 | 0.95 | 0.76 | 0.76 |
| 8 | 4597645 | rs1676936 | 0.952 | 0.567 | -0.252 | -0.27 | 0.082 | -0.053 | -0.053 | 0 | 0.11 | 0.39 | 0.11 | 0 | 0.13 | 0.07 |
| 8 | 4599868 | rs2617048 | 0.027 | 0.623 | -0.536 | 0.373 | 0.031 | 0.166 | 0.166 | 0 | 0.07 | 0.16 | 0.15 | 0.31 | 0.1 | 0.09 |
| 8 | 4601284 | rs13266752 | 2.078 | 0.463 | -0.912 | 0.253 | -0.032 | 0.003 | 0.003 | 0.145 | 0.38 | 0.15 | 0.48 | 0.36 | 0.39 | 0.45 |
| 8 | 4604021 | rs1676941 | -0.317 | 0.535 | -0.769 | 0.092 | -0.009 | -0.071 | -0.071 | 0 | 0.29 | 0.65 | 0.19 | 0.05 | 0.23 | 0.23 |
| 8 | 4605176 | rs11136764 | 1.926 | 0.567 | -0.996 | 0.401 | -0.037 | -0.035 | -0.035 | 0 | 0.39 | 0.14 | 0.52 | 0.41 | 0.37 | 0.39 |
| 8 | 4605810 | rs10503274 | 2.169 | 0.578 | -0.969 | 0.401 | -0.038 | -0.016 | -0.016 | 0.178 | 0.37 | 0.14 | 0.48 | 0.37 | 0.36 | 0.38 |
| 8 | 4606409 | rs1676945 | 0.729 | 0.621 | -0.418 | 0.577 | 0.061 | 0.012 | 0.012 | 0 | 0.12 | 0.31 | 0.1 | 0.11 | 0.15 | 0.08 |
| 8 | 4606609 | rs1673257 | -0.901 | 0.621 | -0.516 | 0.341 | 0.035 | 0.071 | 0.071 | 0 | 0.21 | 0.46 | 0.22 | 0.28 | 0.26 | 0.16 |
| 8 | 4606955 | rs7831951 | 2.486 | 0.54 | -0.882 | 0.201 | -0.023 | -0.033 | -0.033 | 0 | 0.33 | 0.14 | 0.47 | 0.37 | 0.33 | 0.35 |
| 8 | 4609232 | rs12541886 | 1.235 | 0.366 | -0.329 | 0.368 | -0.004 | 0.011 | 0.011 | 0 | 0.47 | 0.44 | 0.66 | 0.63 | 0.48 | 0.53 |
| 8 | 4610345 | rs2407592 | 1.384 | 0.339 | -0.525 | 0.357 | -0.022 | 0.026 | 0.026 | 0 | 0.46 | 0.3 | 0.65 | 0.62 | 0.48 | 0.5 |
| 8 | 4611071 | rs17418933 | 2.501 | 0.396 | -0.693 | 0.401 | -0.024 | 0.023 | 0.023 | 0.108 | 0.34 | 0.11 | 0.46 | 0.28 | 0.33 | 0.37 |
| 8 | 4611924 | rs12677376 | 0.141 | 0.224 | -0.55 | 0.526 | -0.006 | 0.046 | 0.046 | 0 | 0.06 | 0.03 | 0.16 | 0.31 | 0.11 | 0.1 |
| 8 | 4613200 | rs4875115 | -0.109 | 0.219 | -0.54 | 0.522 | -0.001 | 0.06 | 0.06 | 0 | 0.07 | 0.05 | 0.17 | 0.34 | 0.11 | 0.1 |
| 8 | 4613391 | rs1350309 | 0.358 | 0.22 | -0.495 | 0.331 | -0.003 | 0.001 | 0.001 | 0 | 0.72 | 0.76 | 0.82 | 0.78 | 0.69 | 0.75 |
| 8 | 4613548 | rs1350308 | 0.624 | 0.209 | -0.442 | 0.299 | -0.002 | -0.002 | -0.002 | 0 | 0.61 | 0.61 | 0.77 | 0.68 | 0.65 | 0.67 |
| 8 | 4617810 | rs1379326 | -1.014 | 0.387 | -0.455 | 0.31 | 0.043 | 0.073 | 0.073 | 0 | 0.22 | 0.4 | 0.3 | 0.35 | 0.31 | 0.26 |
| 8 | 4618380 | rs1673243 | -0.405 | 0.313 | -0.628 | 0.174 | -0.004 | -0.007 | -0.007 | 0.225 | 0.8 | 0.84 | 0.87 | 0.78 | 0.78 | 0.81 |
| 8 | 4619105 | rs1676961 | -0.536 | 0.317 | -0.553 | -0.009 | 0.045 | -0.094 | -0.094 | 0 | 0.18 | 0.39 | 0.16 | 0.06 | 0.26 | 0.23 |
| 8 | 4619604 | rs17071221 | 1.39 | 0.211 | -0.804 | 0.002 | 0.015 | 0.073 | 0.073 | 0 | 0.15 | 0.02 | 0.08 | 0.05 | 0.09 | 0.1 |
| 8 | 4622661 | rs17080168 | -0.055 | 0.307 | -0.603 | 0.042 | -0.014 | -0.085 | -0.085 | 0 | 0.71 | 0.62 | 0.82 | 0.92 | 0.76 | 0.74 |
| 8 | 4622885 | rs11776753 | 0.305 | 0.357 | -0.88 | 0.187 | -0.041 | -0.077 | -0.077 | 0 | 0.19 | 0.03 | 0.13 | 0.29 | 0.25 | 0.19 |
| 8 | 4622964 | rs10866977 | 1.472 | 0.416 | -0.735 | 0.291 | -0.002 | -0.019 | -0.019 | 0 | 0.45 | 0.23 | 0.61 | 0.54 | 0.41 | 0.46 |
| 8 | 4623869 | rs6985100 | 2.24 | 0.408 | -0.847 | 0.309 | -0.028 | 0.071 | 0.071 | 0.217 | 0.37 | 0.11 | 0.47 | 0.28 | 0.35 | 0.4 |
| 8 | 4624983 | rs2164905 | 0.914 | 0.457 | -0.72 | 0.215 | 0.001 | 0.003 | 0.003 | 0 | 0.52 | 0.43 | 0.68 | 0.57 | 0.51 | 0.53 |
| 8 | 4627197 | rs17071260 | 1.227 | 0.516 | -0.75 | 0.245 | 0.003 | -0.007 | -0.007 | 0 | 0.43 | 0.34 | 0.6 | 0.54 | 0.39 | 0.43 |
| 8 | 4630930 | rs9644369 | 0.437 | 0.991 | -0.213 | 0.687 | 0.004 | 0.029 | 0.029 | 0 | 0.13 | 0.16 | 0.1 | 0.27 | 0.19 | 0.17 |
| 8 | 4631106 | rs10091902 | -0.895 | 0.956 | -0.092 | 0.887 | 0.005 | 0.048 | 0.048 | 0 | 0.13 | 0.17 | 0.19 | 0.37 | 0.15 | 0.14 |
| 8 | 4631249 | rs17347502 | 1.986 | 0.867 | -0.054 | 0.664 | -0.017 | 0.087 | 0.087 | 0 | 0.4 | 0.21 | 0.5 | 0.24 | 0.38 | 0.42 |
| 8 | 4633314 | rs7835351 | 0.158 | 0.803 | -0.014 | 0.583 | -0.002 | 0.01 | 0.01 | 0 | 0.93 | 0.68 | 0.95 | 0.96 | 0.98 | 0.95 |
| 8 | 4633387 | rs7835487 | -0.986 | 0.922 | -0.615 | 0.744 | 0.009 | 0.046 | 0.046 | 0 | 0.13 | 0.22 | 0.15 | 0.29 | 0.15 | 0.14 |
| 8 | 4634415 | rs6986423 | 1.037 | 0.879 | -0.696 | 0.675 | 0.012 | 0.016 | 0.016 | 0 | 0.51 | 0.44 | 0.64 | 0.59 | 0.53 | 0.53 |
| 8 | 4635553 | rs17432165 | 0.879 | 0.995 | -0.452 | 0.738 | 0.025 | 0.061 | 0.061 | 0 | 0.19 | 0.01 | 0.12 | 0.07 | 0.11 | 0.12 |
| 8 | 4636093 | rs17071304 | -0.733 | 1.048 | -0.402 | 0.788 | 0.012 | -0.02 | -0.02 | 0 | 0.32 | 0.21 | 0.27 | 0.38 | 0.27 | 0.28 |
| 8 | 4636715 | rs4369008 | -0.022 | 0.997 | -0.28 | 1.034 | 0.001 | 0.094 | 0.094 | 0 | 0.05 | 0.06 | 0.12 | 0.37 | 0.09 | 0.1 |
| 8 | 4637667 | rs4875117 | 0.095 | 1.025 | -0.198 | 1.012 | -0.005 | 0.007 | 0.007 | 0 | 0.75 | 0.49 | 0.82 | 0.76 | 0.75 | 0.76 |
| 8 | 4638706 | rs10503282 | 0.115 | 0.787 | -0.675 | 1.075 | 0.013 | 0.133 | 0.133 | 0 | 0.05 | 0.07 | 0.12 | 0.37 | 0.1 | 0.1 |
| 8 | 4640275 | rs17347703 | 0.843 | 0.652 | -0.912 | 0.777 | 0.022 | 0.058 | 0.058 | 0 | 0.18 | 0.02 | 0.13 | 0.07 | 0.12 | 0.1 |
| 8 | 4641574 | rs921954 | 1.472 | 0.736 | -0.871 | 0.772 | -0.002 | 0.089 | 0.089 | 0 | 0.47 | 0.27 | 0.5 | 0.31 | 0.43 | 0.49 |
| 8 | 4641941 | rs17071342 | 0.641 | 0.71 | -0.985 | 0.858 | 0.011 | 0.029 | 0.029 | 0 | 0.19 | 0.02 | 0.26 | 0.11 | 0.12 | 0.12 |
| 8 | 4644668 | rs721302 | 1.397 | 0.697 | -0.851 | 1.026 | -0.051 | 0.171 | 0.171 | 0.567 | 0.46 | 0.17 | 0.51 | 0.23 | 0.47 | 0.44 |
| 8 | 4644789 | rs11136774 | -0.353 | 0.812 | -0.591 | 1.127 | -0.011 | -0.008 | -0.008 | 0 | 0.24 | 0.14 | 0.14 | 0.22 | 0.29 | 0.24 |
| 8 | 4648804 | rs11136777 | 1.249 | 0.838 | -0.503 | 0.983 | -0.03 | 0.074 | 0.074 | 1.078 | 0.48 | 0.22 | 0.56 | 0.36 | 0.48 | 0.4 |
| 8 | 4649052 | rs6558918 | -0.091 | 0.979 | -0.033 | 0.826 | 0.039 | -0.036 | -0.036 | 0 | 0.07 | 0.27 | 0.05 | 0 | 0.06 | 0.05 |
| 8 | 4649142 | rs1457182 | 0.417 | 0.842 | 0.384 | 0.865 | 0.066 | 0.142 | 0.142 | 1.648 | 0.95 | 0.66 | 0.95 | 0.92 | 0.98 | 0.93 |
| 8 | 4649341 | rs2043811 | 0.113 | 0.841 | 0.051 | 0.749 | -0.001 | -0.023 | -0.023 | 0 | 0.25 | 0.34 | 0.19 | 0.06 | 0.19 | 0.19 |
| 8 | 4650764 | rs11786958 | 1.837 | 0.903 | -0.049 | 0.781 | -0.034 | 0.074 | 0.074 | 0.699 | 0.39 | 0.1 | 0.47 | 0.24 | 0.39 | 0.34 |
| 8 | 4652202 | rs6986167 | 0.697 | 0.721 | 0.367 | 0.824 | 0.048 | 0.237 | 0.237 | 0.621 | 0.94 | 0.67 | 0.95 | 0.81 | 0.98 | 0.93 |
| 8 | 4654246 | rs6992998 | 1.152 | 0.815 | 0.356 | 0.421 | 0.003 | 0.006 | 0.006 | 0.003 | 0.49 | 0.44 | 0.56 | 0.47 | 0.45 | 0.43 |
| 8 | 4654394 | rs7010443 | -1.202 | 0.86 | 0.428 | 0.601 | 0.036 | 0.222 | 0.222 | 0 | 0.09 | 0.32 | 0.17 | 0.26 | 0.09 | 0.11 |
| 8 | 4655995 | rs2028229 | 0.487 | 0.901 | -0.037 | 0.523 | -0.005 | -0.005 | -0.005 | 0.306 | 0.68 | 0.41 | 0.7 | 0.71 | 0.76 | 0.7 |
| 8 | 4656445 | rs1457179 | 1.429 | 0.87 | -0.346 | 0.72 | -0.022 | -0.04 | -0.04 | 0.021 | 0.43 | 0.14 | 0.51 | 0.46 | 0.43 | 0.42 |
| 8 | 4657295 | rs11136778 | -0.833 | 0.863 | -0.319 | 0.666 | -0.018 | 0.065 | 0.065 | 0 | 0.44 | 0.21 | 0.32 | 0.25 | 0.5 | 0.42 |
| 8 | 4660585 | rs17432531 | 0.037 | 0.872 | -0.33 | 0.758 | -0.001 | -0.009 | -0.009 | 0 | 0.19 | 0.12 | 0.12 | 0.23 | 0.22 | 0.14 |

| | | | | | | | | | | | | | | | | |
|---|---|---|---|---|---|---|---|---|---|---|---|---|---|---|---|---|
| 8 | 4662524 | rs12679612 | -1.126 | 0.824 | -0.03 | 0.661 | 0.04 | 0.239 | 0.239 | 0 | 0.07 | 0.3 | 0.16 | 0.26 | 0.07 | 0.11 |
| 8 | 4662898 | rs10106620 | -1.013 | 0.797 | 0.079 | 0.544 | -0.001 | 0.009 | 0.009 | 0.643 | 0.51 | 0.43 | 0.35 | 0.35 | 0.51 | 0.47 |
| 8 | 4663171 | rs7016863 | 1.765 | 0.781 | 0.054 | 0.428 | -0.005 | -0.019 | -0.019 | 0 | 0.39 | 0.12 | 0.48 | 0.44 | 0.38 | 0.4 |
| 8 | 4665649 | rs10086985 | -0.871 | 0.755 | 0.1 | 0.419 | -0.001 | 0.011 | 0.011 | 0.832 | 0.49 | 0.41 | 0.33 | 0.32 | 0.49 | 0.44 |
| 8 | 4672939 | rs2407596 | 2.028 | 0.724 | -0.101 | 0.422 | -0.013 | -0.043 | -0.043 | 0.115 | 0.35 | 0.09 | 0.49 | 0.39 | 0.36 | 0.34 |
| 8 | 4675393 | rs4875388 | 0.248 | 0.793 | -0.112 | 0.33 | 0.003 | -0.014 | -0.014 | 0 | 0.27 | 0.18 | 0.28 | 0.3 | 0.26 | 0.28 |
| 8 | 4679752 | rs1866282 | 0.936 | 0.81 | 0.274 | 0.521 | 0.013 | 0.126 | 0.126 | 0 | 0.87 | 0.68 | 0.82 | 0.7 | 0.91 | 0.78 |
| 8 | 4682422 | rs7833231 | 0.524 | 0.774 | 0.179 | 0.141 | -0.003 | -0.088 | -0.088 | 0.021 | 0.63 | 0.56 | 0.77 | 0.9 | 0.68 | 0.6 |
| 8 | 4682604 | rs1457191 | -0.033 | 0.763 | 0.109 | 0.174 | 0.001 | 0.026 | 0.026 | 0 | 0.29 | 0.26 | 0.2 | 0.07 | 0.3 | 0.28 |
| 8 | 4682675 | rs1457192 | 0.33 | 0.762 | 0.049 | 0.179 | -0.004 | 0.083 | 0.083 | 0 | 0.26 | 0.16 | 0.17 | 0.07 | 0.25 | 0.26 |
| 8 | 4682750 | rs11774281 | 0.229 | 0.791 | 0.054 | 0.371 | -0.002 | -0.061 | -0.061 | 1.871 | 0.27 | 0.16 | 0.33 | 0.38 | 0.26 | 0.29 |
| 8 | 4684203 | rs1992205 | -0.752 | 0.895 | 0.415 | -0.38 | 0.002 | 0.039 | 0.039 | 0.465 | 0.5 | 0.36 | 0.3 | 0.3 | 0.52 | 0.4 |
| 8 | 4688235 | rs1379324 | -0.921 | 0.9 | 0.395 | -0.405 | -0.001 | 0.02 | 0.02 | 1.021 | 0.52 | 0.63 | 0.33 | 0.27 | 0.54 | 0.39 |
| 8 | 4688296 | rs9657403 | 0.689 | 0.82 | 0.388 | -0.349 | 0.041 | -0.048 | -0.048 | 0 | 0.09 | 0.27 | 0.06 | 0 | 0.11 | 0.04 |
| 8 | 4692648 | rs10087766 | 1.341 | 0.633 | -0.439 | -0.248 | -0.058 | -0.022 | -0.022 | 0 | 0.38 | 0.1 | 0.65 | 0.68 | 0.44 | 0.49 |
| 8 | 4693023 | rs7012483 | 0.353 | 0.852 | 0.326 | -0.261 | 0.074 | -0.012 | -0.012 | 0 | 0.92 | 0.63 | 0.95 | 0.98 | 0.94 | 0.9 |
| 8 | 4697379 | rs6996405 | 0.735 | 0.924 | 0.053 | -0.227 | 0.011 | -0.015 | -0.015 | 1.325 | 0.05 | 0.09 | 0.03 | 0.02 | 0.06 | 0.1 |
| 8 | 4697447 | rs7000406 | 0.579 | 0.803 | -0.137 | -0.345 | 0.031 | 0.075 | 0.075 | 2.284 | 0.5 | 0.26 | 0.72 | 0.71 | 0.56 | 0.68 |
| 8 | 4699995 | rs4875396 | 0.142 | 0.853 | -0.073 | -0.543 | 0.001 | -0.056 | -0.056 | 0.35 | 0.64 | 0.62 | 0.76 | 0.95 | 0.61 | 0.72 |
| 8 | 4700647 | rs6988310 | -0.211 | 0.843 | 0.025 | -0.501 | -0.001 | 0.054 | 0.054 | 0 | 0.21 | 0.34 | 0.1 | 0.28 | 0.17 | 0.16 |
| 8 | 4701250 | rs9314546 | -1.166 | 0.897 | -0.034 | -0.598 | 0.016 | -0.053 | -0.053 | 2.114 | 0.9 | 0.66 | 0.94 | 1 | 0.93 | 0.95 |
| 8 | 4702233 | rs1561360 | 0.2 | 0.924 | -0.065 | -0.62 | 0 | -0.053 | -0.053 | 0.082 | 0.66 | 0.64 | 0.78 | 0.95 | 0.64 | 0.72 |
| 8 | 4705811 | rs4875398 | -0.878 | 0.876 | -0.039 | -0.489 | -0.002 | 0.007 | 0.007 | 0 | 0.45 | 0.35 | 0.27 | 0.33 | 0.46 | 0.4 |
| 8 | 4707429 | rs1580188 | 1.593 | 0.73 | -0.452 | -0.499 | -0.084 | -0.03 | -0.03 | 0.673 | 0.41 | 0.1 | 0.65 | 0.67 | 0.47 | 0.56 |
| 8 | 4707735 | rs7006552 | 0.484 | 0.815 | -0.09 | -0.999 | -0.004 | -0.07 | -0.07 | 0 | 0.63 | 0.61 | 0.76 | 0.95 | 0.65 | 0.73 |
| 8 | 4710970 | rs1585165 | -0.39 | 1.013 | -0.45 | -0.278 | -0.009 | 0.041 | 0.041 | 0 | 0.28 | 0.23 | 0.19 | 0.06 | 0.31 | 0.33 |
| 8 | 4723047 | rs11136785 | -0.296 | 0.934 | -0.412 | 0.394 | -0.024 | -0.003 | -0.003 | 4.776 | 0.4 | 0.32 | 0.45 | 0.53 | 0.5 | 0.35 |
| 8 | 4731577 | rs7840747 | 0.413 | 0.742 | 0.359 | 0.662 | -0.014 | -0.002 | -0.002 | 0 | 0.57 | 0.34 | 0.56 | 0.46 | 0.5 | 0.59 |
| 8 | 4734649 | rs11136786 | -0.103 | 0.819 | 0.027 | 0.608 | -0.054 | -0.019 | -0.019 | 0 | 0.36 | 0.19 | 0.45 | 0.57 | 0.46 | 0.32 |
| 8 | 4739608 | rs2043810 | -0.068 | 0.887 | -0.049 | 0.655 | -0.027 | -0.039 | -0.039 | 2.903 | 0.35 | 0.21 | 0.46 | 0.53 | 0.43 | 0.34 |
| 8 | 4740951 | rs12681969 | 1.225 | 0.297 | 0.275 | 0.57 | 0.051 | 0.01 | 0.01 | 1.734 | 0.92 | 0.56 | 0.93 | 0.98 | 0.96 | 0.95 |
| 8 | 4742724 | rs6990325 | -0.28 | 0.189 | -0.413 | 0.458 | -0.111 | -0.045 | -0.045 | 0 | 0.34 | 0.03 | 0.34 | 0.28 | 0.4 | 0.15 |
| 8 | 4742921 | rs6986644 | -0.65 | 0.439 | 0.194 | 0.186 | 0.002 | 0.011 | 0.011 | 0.512 | 0.52 | 0.44 | 0.48 | 0.58 | 0.56 | 0.39 |
| 8 | 4746885 | rs13266795 | 1.07 | 0.271 | 0.052 | 0.079 | -0.006 | -0.002 | -0.002 | 0.073 | 0.39 | 0.32 | 0.44 | 0.4 | 0.4 | 0.44 |
| 8 | 4746950 | rs13267077 | 0.729 | 0.106 | 0.165 | 0.116 | -0.004 | 0.006 | 0.006 | 0 | 0.45 | 0.39 | 0.48 | 0.4 | 0.46 | 0.56 |
| 8 | 4748952 | rs13251949 | 0.636 | 0.152 | 0.031 | 0.17 | 0 | 0.002 | 0.002 | 1.447 | 0.3 | 0.29 | 0.33 | 0.44 | 0.32 | 0.28 |
| 8 | 4751194 | rs17348582 | 0.839 | 0.158 | -0.646 | -0.256 | 0.005 | 0.044 | 0.044 | 0 | 0.12 | 0.04 | 0.05 | 0.03 | 0.11 | 0.09 |
| 8 | 4752877 | rs6558934 | -0.488 | 0.385 | -0.426 | -0.018 | 0.01 | -0.043 | -0.043 | 1.169 | 0.91 | 0.8 | 0.93 | 0.97 | 0.92 | 0.82 |
| 8 | 4753474 | rs731109 | -0.166 | 0.647 | -1.05 | 0.397 | -0.002 | 0 | 0 | 0.013 | 0.37 | 0.33 | 0.43 | 0.59 | 0.4 | 0.35 |
| 8 | 4755088 | rs7000541 | 0.126 | 0.921 | -1.179 | 0.759 | -0.023 | -0.032 | -0.032 | 0 | 0.33 | 0.27 | 0.42 | 0.5 | 0.39 | 0.47 |
| 8 | 4755645 | rs10081492 | -0.388 | 0.516 | -1.046 | 0.497 | 0 | 0.018 | 0.018 | 0 | 0.15 | 0.07 | 0.09 | 0.01 | 0.17 | 0.04 |
| 8 | 4755990 | rs9650522 | -0.901 | 0.377 | -0.607 | 1.569 | -0.008 | 0.093 | 0.093 | 0 | 0.14 | 0.22 | 0.19 | 0.48 | 0.09 | 0.25 |
| 8 | 4756609 | rs7816996 | -0.835 | 0.424 | -0.416 | 1.239 | -0.003 | 0.083 | 0.083 | 0.788 | 0.23 | 0.59 | 0.27 | 0.41 | 0.21 | 0.26 |
| 8 | 4761250 | rs12678783 | -0.114 | 0.266 | -1.252 | 0.883 | -0.006 | -0.099 | -0.099 | 0.041 | 0.22 | 0.06 | 0.42 | 0.53 | 0.18 | 0.29 |
| 8 | 4761765 | rs893465 | 0.41 | -0.371 | -0.44 | 0.272 | -0.021 | -0.041 | -0.041 | 0 | 0.84 | 0.59 | 0.88 | 0.88 | 0.83 | 0.81 |
| 8 | 4764417 | rs6982855 | 0.116 | -0.691 | -0.896 | -0.134 | 0.005 | -0.004 | -0.004 | 0.529 | 0.78 | 0.79 | 0.85 | 0.67 | 0.85 | 0.74 |
| 8 | 4764865 | rs7001869 | 0.493 | -0.501 | -0.852 | -0.374 | 0.012 | -0.044 | -0.044 | 0 | 0.27 | 0.15 | 0.45 | 0.52 | 0.26 | 0.28 |
| 8 | 4767236 | rs7829049 | -0.024 | -0.308 | -0.995 | -0.338 | 0.014 | -0.008 | -0.008 | 0 | 0.78 | 0.82 | 0.86 | 0.72 | 0.84 | 0.75 |
| 8 | 4772534 | rs4592061 | 0.99 | -0.107 | -1.141 | -0.759 | -0.101 | 0.147 | 0.147 | 0 | 0.37 | 0.12 | 0.31 | 0.14 | 0.46 | 0.38 |
| 8 | 4775412 | rs13248460 | -0.188 | -0.251 | -1.132 | -0.872 | 0.004 | 0.227 | 0.227 | 0 | 0.47 | 0.11 | 0.56 | 0.82 | 0.44 | 0.54 |
| 8 | 4778969 | rs4373538 | 0.612 | -0.464 | -0.182 | -0.858 | -0.02 | -0.083 | -0.083 | 0 | 0.85 | 0.36 | 0.9 | 0.95 | 0.91 | 0.93 |
| 8 | 4779452 | rs4607643 | 0.613 | -0.463 | -0.199 | -0.855 | -0.02 | -0.077 | -0.077 | 0 | 0.85 | 0.35 | 0.89 | 0.95 | 0.91 | 0.92 |
| 8 | 4779581 | rs11774735 | 0.111 | -0.114 | -0.978 | -0.652 | -0.001 | -0.055 | -0.055 | 0 | 0.26 | 0.04 | 0.13 | 0.32 | 0.21 | 0.31 |
| 8 | 4780877 | rs9657404 | -0.735 | 0.176 | -0.499 | -0.801 | -0.003 | -0.008 | -0.008 | 0 | 0.74 | 0.83 | 0.51 | 0.46 | 0.66 | 0.73 |
| 8 | 4782658 | rs9657371 | 0.97 | 0.164 | -0.569 | -1.134 | -0.036 | 0.164 | 0.164 | 0 | 0.38 | 0.15 | 0.24 | 0.04 | 0.39 | 0.26 |
| 8 | 4784894 | rs11985845 | 0.81 | 0.102 | -0.362 | -1.16 | -0.033 | 0.122 | 0.122 | 0 | 0.42 | 0.23 | 0.27 | 0.04 | 0.44 | 0.31 |
| 8 | 4787446 | rs4076510 | 0.14 | -0.019 | -0.104 | -1.061 | 0 | -0.002 | -0.002 | 0 | 0.32 | 0.32 | 0.53 | 0.61 | 0.37 | 0.39 |
| 8 | 4792894 | rs6990769 | 0.985 | 0.059 | -0.41 | -1.236 | -0.057 | 0.171 | 0.171 | 0 | 0.37 | 0.15 | 0.23 | 0.04 | 0.41 | 0.31 |
| 8 | 4795226 | rs10096103 | -0.135 | -0.26 | 0.527 | -0.951 | 0.011 | -0.009 | -0.009 | 0.02 | 0.3 | 0.24 | 0.25 | 0.36 | 0.21 | 0.29 |
| 8 | 4796145 | rs10089221 | -0.136 | -0.199 | 0.509 | -0.96 | 0.019 | -0.009 | -0.009 | 0 | 0.3 | 0.24 | 0.24 | 0.36 | 0.2 | 0.29 |
| 8 | 4802339 | rs4875129 | 1.188 | -0.175 | 0.692 | -1.109 | -0.052 | 0.132 | 0.132 | 0 | 0.36 | 0.14 | 0.23 | 0.04 | 0.41 | 0.29 |
| 8 | 4802438 | rs4875417 | 0.691 | -0.298 | 0.934 | -1.18 | -0.031 | -0.064 | -0.064 | 0 | 0.88 | 0.48 | 0.93 | 0.97 | 0.96 | 0.97 |
| 8 | 4803156 | rs12681508 | -0.215 | -0.332 | 0.869 | -0.515 | -0.007 | 0.139 | 0.139 | 0 | 0.49 | 0.27 | 0.69 | 0.78 | 0.53 | 0.55 |
| 8 | 4804768 | rs7012396 | -0.672 | -0.451 | 0.342 | -1.037 | 0.036 | -0.135 | -0.135 | 1.398 | 0.72 | 0.5 | 0.78 | 0.93 | 0.65 | 0.75 |
| 8 | 4807157 | rs6558946 | -0.506 | -0.63 | 0.211 | -0.688 | -0.008 | 0.084 | 0.084 | 0 | 0.91 | 0.82 | 0.87 | 0.78 | 0.96 | 0.91 |
| 8 | 4807818 | rs7003760 | 0.117 | -0.646 | 0.235 | -0.642 | 0 | 0.007 | 0.007 | 0 | 0.5 | 0.55 | 0.62 | 0.57 | 0.54 | 0.47 |
| 8 | 4809701 | rs10448109 | 0.414 | -0.341 | 0.198 | -0.568 | 0.017 | -0.027 | -0.027 | 0 | 0.05 | 0.14 | 0.04 | 0 | 0.05 | 0.05 |
| 8 | 4811010 | rs13274629 | -0.17 | -0.676 | 0.045 | -0.653 | -0.005 | -0.007 | -0.007 | 0 | 0.79 | 0.77 | 0.88 | 0.83 | 0.84 | 0.76 |
| 8 | 4811455 | rs6997388 | -0.085 | -0.542 | -0.014 | -1.061 | 0 | 0.003 | 0.003 | 0 | 0.52 | 0.44 | 0.33 | 0.36 | 0.51 | 0.53 |

| | | | | | | | | | | | | | | | | |
|---|---|---|---|---|---|---|---|---|---|---|---|---|---|---|---|---|
| 8 | 4812436 | rs7815832 | 0.609 | -0.585 | -0.056 | -1.126 | -0.001 | -0.008 | -0.008 | 0 | 0.69 | 0.35 | 0.79 | 0.81 | 0.75 | 0.73 |
| 8 | 4812710 | rs6558948 | -1.093 | -0.653 | -0.025 | -1.115 | 0.013 | 0.003 | 0.003 | 0.182 | 0.83 | 0.91 | 0.91 | 0.83 | 0.89 | 0.81 |
| 8 | 4812812 | rs10081536 | 0.369 | -0.688 | -0.202 | -1.03 | -0.004 | 0.017 | 0.017 | 0 | 0.16 | 0.26 | 0.18 | 0.22 | 0.12 | 0.14 |
| 8 | 4813459 | rs11782341 | 0.22 | -0.623 | -0.498 | -1.021 | -0.014 | -0.009 | -0.009 | 0.009 | 0.22 | 0.48 | 0.17 | 0.2 | 0.18 | 0.21 |
| 8 | 4814303 | rs4458898 | 0.784 | -0.622 | -0.542 | -1.159 | -0.005 | -0.031 | -0.031 | 0 | 0.35 | 0.32 | 0.23 | 0.03 | 0.4 | 0.23 |
| 8 | 4814896 | rs4295679 | 2.069 | -0.804 | -0.548 | -0.975 | -0.034 | -0.093 | -0.093 | 0.682 | 0.24 | 0.1 | 0.44 | 0.41 | 0.32 | 0.27 |
| 8 | 4815582 | rs7017888 | 0.479 | -0.916 | -0.46 | -1.092 | -0.026 | 0 | 0 | 0 | 0.61 | 0.5 | 0.75 | 0.62 | 0.79 | 0.66 |
| 8 | 4815661 | rs7000866 | 0.654 | -0.713 | -0.16 | -0.848 | -0.037 | 0.033 | 0.033 | 0 | 0.73 | 0.41 | 0.77 | 0.61 | 0.86 | 0.7 |
| 8 | 4817813 | rs12542818 | 1.501 | -0.647 | -0.47 | -0.957 | -0.125 | 0.194 | 0.194 | 0 | 0.29 | 0.02 | 0.18 | 0.03 | 0.35 | 0.23 |
| 8 | 4820177 | rs7011965 | -0.994 | -0.754 | 0.102 | -0.491 | -0.046 | 0.069 | 0.069 | 0 | 0.25 | 0.49 | 0.21 | 0.36 | 0.14 | 0.27 |
| 8 | 4822461 | rs12115171 | 0.664 | -0.493 | 0.121 | -0.322 | 0.002 | 0.062 | 0.062 | 0 | 0.06 | 0.16 | 0.03 | 0.14 | 0.03 | 0.03 |
| 8 | 4824108 | rs12674992 | -0.302 | -0.646 | 0.301 | -0.915 | -0.004 | -0.041 | -0.041 | 1.575 | 0.56 | 0.53 | 0.34 | 0.18 | 0.57 | 0.44 |
| 8 | 4824577 | rs17348854 | 0.757 | -0.492 | 0.395 | -0.932 | 0.037 | -0.026 | -0.026 | 0.178 | 0.06 | 0.17 | 0.05 | 0.02 | 0.09 | 0.03 |
| 8 | 4826467 | rs7823766 | 0.395 | -0.687 | 0.54 | -1.081 | -0.003 | 0.005 | 0.005 | 0 | 0.43 | 0.4 | 0.27 | 0.19 | 0.47 | 0.39 |
| 8 | 4826556 | rs6558951 | -1.702 | -0.478 | 0.267 | -0.771 | -0.001 | -0.012 | -0.012 | 0 | 0.75 | 0.77 | 0.46 | 0.54 | 0.69 | 0.72 |
| 8 | 4828118 | rs4593564 | 0.678 | -0.555 | 0.211 | -0.759 | -0.006 | 0.014 | 0.014 | 0 | 0.2 | 0.25 | 0.12 | 0.33 | 0.13 | 0.28 |
| 8 | 4828351 | rs7004026 | -0.219 | -0.547 | 0.176 | -0.837 | -0.001 | -0.008 | -0.008 | 0 | 0.54 | 0.55 | 0.34 | 0.23 | 0.57 | 0.44 |
| 8 | 4832237 | rs11778280 | 0.67 | -0.444 | -0.272 | -0.859 | -0.001 | 0 | 0 | 0.062 | 0.21 | 0.22 | 0.1 | 0.31 | 0.13 | 0.28 |
| 8 | 4834406 | rs9657374 | -0.91 | -0.327 | -0.506 | -0.91 | 0.048 | 0.053 | 0.053 | 0.753 | 0.46 | 0.75 | 0.25 | 0.38 | 0.3 | 0.47 |
| 8 | 4835344 | rs4517143 | -1.131 | -0.121 | -0.094 | -0.552 | 0 | 0.013 | 0.013 | 0 | 0.75 | 0.69 | 0.44 | 0.42 | 0.69 | 0.7 |
| 8 | 4838022 | rs4875427 | -1.325 | -0.047 | -0.317 | -0.553 | -0.013 | -0.056 | -0.056 | 0.399 | 0.77 | 0.89 | 0.46 | 0.44 | 0.69 | 0.73 |
| 8 | 4840032 | rs2977728 | -0.441 | -0.274 | 0.239 | -0.64 | 0.038 | -0.089 | -0.089 | 0 | 0.14 | 0.45 | 0.09 | 0.02 | 0.13 | 0.08 |
| 8 | 4852649 | rs3886807 | -0.024 | -0.385 | -0.056 | -0.624 | -0.015 | 0 | 0 | 0.273 | 0.36 | 0.5 | 0.2 | 0.38 | 0.22 | 0.36 |
| 8 | 4852925 | rs3886808 | 1.042 | -0.142 | 0.009 | -1.12 | 0.003 | -0.083 | -0.083 | 0.156 | 0.27 | 0.32 | 0.16 | 0 | 0.3 | 0.22 |
| 8 | 4854554 | rs4875428 | -1.307 | 0.306 | 0.045 | -0.499 | -0.01 | -0.027 | -0.027 | 0.113 | 0.78 | 0.84 | 0.49 | 0.52 | 0.66 | 0.75 |
| 8 | 4858104 | rs6558954 | 0.216 | -0.101 | 0.053 | -0.292 | 0.004 | -0.003 | -0.003 | 0.25 | 0.41 | 0.37 | 0.2 | 0.47 | 0.24 | 0.37 |
| 8 | 4863084 | rs10104215 | -1.117 | 0.14 | -0.081 | 0.124 | -0.008 | -0.007 | -0.007 | 1.899 | 0.81 | 0.81 | 0.46 | 0.59 | 0.67 | 0.78 |
| 8 | 4865315 | rs6982311 | -0.985 | 0.117 | -0.244 | 0.147 | -0.008 | -0.009 | -0.009 | 0 | 0.8 | 0.83 | 0.46 | 0.57 | 0.67 | 0.78 |
| 8 | 4868702 | rs12335274 | 0.615 | -0.046 | -0.059 | -0.3 | 0.002 | -0.042 | -0.042 | 0 | 0.14 | 0.31 | 0.1 | 0.1 | 0.14 | 0.19 |
| 8 | 4874850 | rs17434263 | 1.338 | -0.058 | -0.597 | -0.57 | -0.012 | 0.014 | 0.014 | 0.099 | 0.23 | 0.15 | 0.12 | 0 | 0.25 | 0.21 |
| 8 | 4875172 | rs10112481 | -1.507 | -0.187 | -0.8 | -0.413 | 0.021 | 0.058 | 0.058 | 0.118 | 0.81 | 0.77 | 0.46 | 0.59 | 0.65 | 0.76 |
| 8 | 4875997 | rs6992840 | 0.474 | -0.386 | -1.609 | -0.336 | 0.06 | -0.056 | -0.056 | 0 | 0.4 | 0.1 | 0.22 | 0.48 | 0.26 | 0.35 |
| 8 | 4880786 | rs6990880 | 0.147 | -0.397 | -1.232 | -0.044 | 0.005 | 0.011 | 0.011 | 0 | 0.61 | 0.5 | 0.34 | 0.53 | 0.5 | 0.56 |
| 8 | 4881304 | rs10095864 | 0.157 | -0.391 | -1.229 | -0.077 | 0.005 | 0.019 | 0.019 | 0 | 0.6 | 0.49 | 0.34 | 0.51 | 0.5 | 0.56 |
| 8 | 4883613 | rs11787515 | 0.623 | -0.223 | -2.436 | -0.09 | 0.018 | -0.117 | -0.117 | 0 | 0.28 | 0.06 | 0.19 | 0.54 | 0.19 | 0.29 |
| 8 | 4884403 | rs17072086 | 0.699 | -0.17 | -2.675 | 0.079 | -0.027 | -0.142 | -0.142 | 2.742 | 0.27 | 0.02 | 0.18 | 0.44 | 0.19 | 0.29 |
| 8 | 4884701 | rs6997954 | -0.235 | -0.345 | -2.654 | -0.017 | 0.041 | 0.045 | 0.045 | 0 | 0.46 | 0.75 | 0.31 | 0.68 | 0.31 | 0.53 |
| 8 | 4886316 | rs1454280 | -0.031 | -0.425 | -2.605 | -0.304 | -0.013 | 0.054 | 0.054 | 0 | 0.44 | 0.67 | 0.29 | 0.21 | 0.42 | 0.49 |
| 8 | 4887009 | rs11787076 | 0.844 | -0.142 | -3.122 | 0.002 | -0.101 | 0.161 | 0.161 | 0 | 0.27 | 0.03 | 0.18 | 0 | 0.32 | 0.26 |
| 8 | 4888974 | rs12677392 | -0.626 | -0.841 | -2.438 | 1.062 | 0.072 | -0.053 | -0.053 | 0.403 | 0.32 | 0.13 | 0.17 | 0.6 | 0.18 | 0.32 |
| 8 | 4889651 | rs17072101 | -0.116 | -0.158 | -2.506 | 0.998 | 0.018 | -0.148 | -0.148 | 2.5 | 0.25 | 0.03 | 0.15 | 0.47 | 0.17 | 0.27 |
| 8 | 4892834 | rs12544224 | -0.518 | 0.385 | -2.456 | 1.275 | -0.036 | -0.124 | -0.124 | 0.248 | 0.18 | 0.03 | 0.46 | 0.46 | 0.22 | 0.24 |
| 8 | 4893353 | rs10503290 | 0.471 | 0.765 | -3.526 | 0.007 | -0.01 | 0.006 | 0.006 | 0 | 0.14 | 0.08 | 0.09 | 0.1 | 0.19 | 0.19 |
| 8 | 4905437 | rs2977702 | -0.688 | -0.383 | -3.197 | -0.512 | -0.006 | 0 | 0 | 0.072 | 0.66 | 0.77 | 0.73 | 0.66 | 0.62 | 0.69 |
| 8 | 4910045 | rs2977709 | -0.001 | -0.452 | -3.51 | -0.987 | -0.144 | 0.11 | 0.11 | 2.492 | 0.28 | 0.03 | 0.22 | 0.08 | 0.39 | 0.3 |
| 8 | 4914546 | rs2924747 | 1.857 | -0.985 | -3.001 | -1.35 | -0.008 | -0.034 | -0.034 | 0 | 0.13 | 0.16 | 0.05 | 0 | 0.06 | 0.01 |
| 8 | 4915575 | rs13253703 | 1.01 | -0.824 | -2.522 | -1.41 | 0.007 | -0.008 | -0.008 | 0 | 0.17 | 0.18 | 0.13 | 0.08 | 0.25 | 0.2 |
| 8 | 4915797 | rs2977715 | -0.429 | -1.205 | -1.957 | -1.287 | 0.004 | 0.002 | 0.002 | 0 | 0.45 | 0.56 | 0.59 | 0.56 | 0.33 | 0.41 |
| 8 | 4918678 | rs17072176 | 1.571 | -0.998 | -2.034 | -1.113 | -0.023 | -0.114 | -0.114 | 0 | 0.15 | 0.02 | 0.44 | 0.53 | 0.15 | 0.24 |
| 8 | 4919521 | rs2168055 | -1.793 | -1.017 | -1.984 | -1.23 | -0.008 | 0.005 | 0.005 | 0.027 | 0.75 | 0.81 | 0.77 | 0.82 | 0.66 | 0.73 |
| 8 | 4922113 | rs2407739 | -1.006 | -1.107 | -1.354 | -1.375 | 0 | -0.01 | -0.01 | 0.472 | 0.6 | 0.49 | 0.71 | 0.75 | 0.61 | 0.62 |
| 8 | 4924197 | rs7843140 | -0.376 | -0.99 | -1.152 | -1.365 | 0 | 0.009 | 0.009 | 0 | 0.46 | 0.46 | 0.62 | 0.75 | 0.47 | 0.53 |
| 8 | 4924379 | rs17072200 | 0.653 | -0.973 | -1.272 | -1.065 | 0.023 | -0.055 | -0.055 | 2.465 | 0.22 | 0.14 | 0.46 | 0.52 | 0.19 | 0.26 |
| 8 | 4925260 | rs7817362 | -0.548 | -0.734 | -0.963 | -1.651 | -0.002 | 0.045 | 0.045 | 0 | 0.51 | 0.64 | 0.29 | 0.12 | 0.48 | 0.38 |
| 8 | 4927142 | rs1377888 | -0.67 | -0.542 | -0.913 | -1.305 | -0.013 | -0.001 | -0.001 | 0 | 0.63 | 0.8 | 0.65 | 0.59 | 0.59 | 0.55 |
| 8 | 4928491 | rs10086338 | 0.352 | -0.771 | -0.744 | -1.475 | -0.017 | -0.042 | -0.042 | 2.314 | 0.17 | 0.39 | 0.11 | 0.1 | 0.15 | 0.2 |
| 8 | 4930433 | rs4875138 | 0.865 | -0.861 | -1.2 | -1.696 | -0.043 | 0.106 | 0.106 | 0 | 0.27 | 0.11 | 0.14 | 0.07 | 0.28 | 0.22 |
| 8 | 4933779 | rs921703 | -0.634 | -0.927 | -0.968 | -1.484 | 0.044 | 0.039 | 0.039 | 0.021 | 0.46 | 0.73 | 0.59 | 0.59 | 0.41 | 0.44 |
| 8 | 4937165 | rs2123901 | 1.133 | -0.95 | -0.962 | -1.259 | -0.001 | 0.017 | 0.017 | 0 | 0.18 | 0.16 | 0.39 | 0.53 | 0.15 | 0.18 |
| 8 | 4937312 | rs17350657 | 1.192 | -0.858 | -1.156 | -1.342 | -0.001 | -0.012 | -0.012 | 7.285 | 0.1 | 0.14 | 0.07 | 0.03 | 0.1 | 0.11 |
| 8 | 4938002 | rs17435274 | 1.362 | -0.042 | -1.392 | -1.493 | -0.067 | 0.079 | 0.079 | 0.2 | 0.22 | 0.04 | 0.11 | 0.08 | 0.25 | 0.15 |
| 8 | 4938821 | rs17072323 | 1.059 | 0.296 | -0.729 | -1.18 | 0.019 | -0.02 | -0.02 | 0.09 | 0.11 | 0.27 | 0.08 | 0.04 | 0.11 | 0.07 |
| 8 | 4940320 | rs17072336 | 0.324 | 0.201 | -1.015 | -1.134 | 0.015 | -0.035 | -0.035 | 0 | 0.27 | 0.18 | 0.46 | 0.55 | 0.24 | 0.29 |
| 8 | 4942645 | rs17350817 | 1.812 | 0.35 | -1.011 | -1.078 | -0.009 | 0.02 | 0.02 | 0.134 | 0.09 | 0.03 | 0.07 | 0 | 0.11 | 0.07 |
| 8 | 4942981 | rs7818650 | -0.46 | 0.296 | -1.034 | -1.313 | -0.039 | 0.01 | 0.01 | 4.046 | 0.62 | 0.81 | 0.65 | 0.66 | 0.56 | 0.49 |
| 8 | 4943310 | rs12549047 | 0.173 | 0.406 | -0.84 | -1.371 | 0.002 | 0.003 | 0.003 | 2.277 | 0.5 | 0.52 | 0.53 | 0.63 | 0.43 | 0.43 |
| 8 | 4944714 | rs7827959 | -1.877 | 0.823 | -1.142 | -1.184 | -0.011 | -0.009 | -0.009 | 0 | 0.78 | 0.83 | 0.75 | 0.77 | 0.74 | 0.63 |
| 8 | 4948514 | rs2897754 | 0.143 | 0.436 | -0.888 | -1.302 | 0.004 | 0.034 | 0.034 | 0.68 | 0.42 | 0.65 | 0.25 | 0.1 | 0.4 | 0.27 |
| 8 | 4951542 | rs7014013 | -1.124 | 0.484 | -1.025 | -1.089 | -0.02 | -0.007 | -0.007 | 0 | 0.71 | 0.82 | 0.69 | 0.68 | 0.67 | 0.59 |

| | | | | | | | | | | | | | | | | |
|---|---|---|---|---|---|---|---|---|---|---|---|---|---|---|---|---|
| 8 | 4951610 | rs10100814 | 0.381 | 0.406 | -0.821 | -1.067 | -0.012 | 0.07 | 0.07 | 0 | 0.3 | 0.4 | 0.47 | 0.52 | 0.28 | 0.24 |
| 8 | 4955470 | rs2407749 | -0.916 | 0.492 | -1.363 | -0.908 | -0.091 | -0.083 | -0.083 | 0.015 | 0.7 | 0.94 | 0.69 | 0.64 | 0.66 | 0.58 |
| 8 | 4958358 | rs1377880 | 0.058 | 0.281 | -0.894 | -0.879 | 0.005 | 0.054 | 0.054 | 0.34 | 0.42 | 0.67 | 0.25 | 0.13 | 0.41 | 0.35 |
| 8 | 4958752 | rs7824955 | 1.068 | 0.234 | -1.073 | -0.609 | 0.013 | -0.102 | -0.102 | 0.099 | 0.18 | 0.05 | 0.35 | 0.48 | 0.15 | 0.13 |
| 8 | 4960050 | rs7838881 | 0.296 | 0.064 | -0.45 | -1.449 | -0.013 | -0.046 | -0.046 | 0 | 0.16 | 0.3 | 0.1 | 0.05 | 0.12 | 0.14 |
| 8 | 4960945 | rs7843651 | 0.124 | 0.126 | -0.359 | -1.548 | -0.009 | -0.067 | -0.067 | 0 | 0.36 | 0.55 | 0.19 | 0.12 | 0.3 | 0.21 |
| 8 | 4961439 | rs12546378 | -0.225 | 0.08 | -0.558 | -1.625 | 0.009 | 0.012 | 0.012 | 1.729 | 0.47 | 0.63 | 0.23 | 0.13 | 0.34 | 0.23 |
| 8 | 4968807 | rs17435487 | 0.551 | -0.003 | -0.653 | -1.589 | 0.019 | 0.055 | 0.055 | 0.498 | 0.32 | 0.24 | 0.13 | 0.1 | 0.24 | 0.19 |
| 8 | 4972000 | rs13274395 | 0.542 | -0.046 | -0.655 | -1.59 | 0.022 | 0.054 | 0.054 | 0.238 | 0.31 | 0.24 | 0.13 | 0.1 | 0.23 | 0.19 |
| 8 | 4975744 | rs11995687 | 1.614 | -0.018 | -0.307 | -1.489 | -0.024 | -0.056 | -0.056 | 0 | 0.1 | 0.19 | 0.04 | 0 | 0.04 | 0.01 |
| 8 | 4985361 | rs7011733 | -0.888 | 0.747 | -0.558 | -1.047 | -0.022 | -0.02 | -0.02 | 0.036 | 0.86 | 0.93 | 0.56 | 0.48 | 0.72 | 0.65 |
| 8 | 4987302 | rs17435605 | 0.64 | 0.543 | -0.698 | -1.635 | -0.006 | 0.013 | 0.013 | 0 | 0.19 | 0.13 | 0.13 | 0.03 | 0.19 | 0.17 |
| 8 | 4988269 | rs11778306 | 0.472 | 0.518 | -0.599 | -1.652 | -0.002 | -0.023 | -0.023 | 0.217 | 0.2 | 0.2 | 0.13 | 0.01 | 0.19 | 0.14 |
| 8 | 4989141 | rs11779265 | -0.229 | 0.457 | -0.389 | -1.684 | 0 | -0.018 | -0.018 | 0 | 0.51 | 0.53 | 0.26 | 0.12 | 0.42 | 0.28 |
| 8 | 4998837 | rs2168053 | 0.258 | 0.27 | -0.505 | -1.72 | 0.004 | 0.014 | 0.014 | 0 | 0.44 | 0.42 | 0.23 | 0.13 | 0.38 | 0.28 |
| 8 | 5000769 | rs4875441 | -0.249 | 0.231 | -0.495 | -1.754 | 0 | 0.036 | 0.036 | 0 | 0.56 | 0.7 | 0.28 | 0.13 | 0.42 | 0.28 |
| 8 | 5001614 | rs4875448 | -0.23 | 0.211 | -0.373 | -1.764 | 0 | -0.011 | -0.011 | 0.429 | 0.56 | 0.64 | 0.28 | 0.13 | 0.42 | 0.28 |
| 8 | 5003571 | rs6558971 | -0.643 | 0.616 | -0.447 | -1.29 | -0.03 | -0.052 | -0.052 | 0.286 | 0.82 | 0.93 | 0.58 | 0.48 | 0.74 | 0.58 |
| 8 | 5010329 | rs11775723 | -1.268 | 0.065 | -0.373 | -1.411 | 0.008 | 0.012 | 0.012 | 0 | 0.76 | 0.79 | 0.9 | 0.9 | 0.81 | 0.89 |
| 8 | 5015728 | rs7001682 | -1.178 | -0.015 | -0.5 | -1.451 | -0.003 | -0.009 | -0.009 | 0.031 | 0.75 | 0.73 | 0.89 | 0.9 | 0.82 | 0.89 |
| 8 | 5015861 | rs6997699 | -0.043 | 0.023 | -1.118 | -1.398 | 0.031 | -0.012 | -0.012 | 0 | 0.45 | 0.25 | 0.64 | 0.48 | 0.6 | 0.52 |
| 8 | 5024469 | rs6558973 | -0.138 | 0.108 | -1.003 | -1.259 | 0.006 | 0.008 | 0.008 | 0 | 0.39 | 0.4 | 0.58 | 0.5 | 0.47 | 0.4 |
| 8 | 5026520 | rs4130201 | 0.86 | -0.895 | -1.363 | -1.32 | -0.023 | -0.029 | -0.029 | 0 | 0.3 | 0.32 | 0.12 | 0.09 | 0.14 | 0.11 |
| 8 | 5029375 | rs4610773 | 0.791 | 0.249 | -1.305 | -0.766 | 0.005 | 0.024 | 0.024 | 1.73 | 0.26 | 0.29 | 0.27 | 0.44 | 0.26 | 0.47 |
| 8 | 5041417 | rs6989820 | -0.098 | 0.157 | -2.257 | -0.604 | 0 | 0.023 | 0.023 | 0 | 0.52 | 0.74 | 0.4 | 0.44 | 0.43 | 0.51 |
| 8 | 5044296 | rs7835499 | -0.843 | 1.149 | -1.152 | 0.34 | -0.065 | -0.058 | -0.058 | 0 | 0.9 | 0.97 | 0.62 | 0.58 | 0.72 | 0.84 |
| 8 | 5053822 | rs11990769 | -0.356 | 0.118 | -0.905 | -0.231 | -0.023 | -0.04 | -0.04 | 0 | 0.18 | 0.74 | 0.1 | 0.01 | 0.06 | 0.04 |
| 8 | 5053877 | rs11991556 | -0.275 | 0.139 | -0.903 | -0.22 | -0.034 | -0.054 | -0.054 | 0.047 | 0.18 | 0.7 | 0.1 | 0.01 | 0.07 | 0.04 |
| 8 | 5057818 | rs4305919 | 0.616 | 0.818 | -1.348 | 0.387 | -0.02 | -0.027 | -0.027 | 1.054 | 0.63 | 0.32 | 0.51 | 0.49 | 0.55 | 0.62 |
| 8 | 5059504 | rs4073014 | 0.201 | 1.379 | -1.702 | 1.367 | -0.098 | -0.129 | -0.129 | 0 | 0.16 | 0.02 | 0.12 | 0.5 | 0.26 | 0.22 |
| 8 | 5069057 | rs12549617 | 0.788 | 1.086 | -1.71 | 0.477 | -0.002 | 0.005 | 0.005 | 0 | 0.14 | 0.08 | 0.09 | 0 | 0.14 | 0.02 |
| 8 | 5071270 | rs12056538 | 0.169 | 1.209 | -1.889 | 0.932 | -0.078 | -0.1 | -0.1 | 0 | 0.19 | 0.06 | 0.14 | 0.43 | 0.29 | 0.25 |
| 8 | 5073294 | rs11136831 | 0.992 | 1.116 | -1.691 | 0.851 | 0 | -0.015 | -0.015 | 0.034 | 0.14 | 0.11 | 0.09 | 0 | 0.13 | 0.02 |
| 8 | 5076089 | rs3934968 | 0.348 | 0.696 | -1.093 | 0.354 | 0.01 | -0.068 | -0.068 | 0.022 | 0.17 | 0.33 | 0.21 | 0.07 | 0.18 | 0.2 |
| 8 | 5077899 | rs10435678 | 0.057 | 0.317 | -1.162 | 0.123 | -0.003 | 0.025 | 0.025 | 0 | 0.29 | 0.36 | 0.47 | 0.47 | 0.26 | 0.38 |
| 8 | 5078168 | rs7816277 | 1.305 | 0.23 | -1.104 | 0.316 | -0.012 | -0.061 | -0.061 | 0 | 0.14 | 0.04 | 0.3 | 0.39 | 0.13 | 0.3 |
| 8 | 5085053 | rs10089783 | 0.639 | 0.203 | -1.155 | 0.02 | 0.006 | 0.003 | 0.003 | 0 | 0.49 | 0.39 | 0.38 | 0.11 | 0.46 | 0.36 |
| 8 | 5088786 | rs4875149 | 0.107 | 0.161 | -1.097 | 0.16 | -0.018 | -0.016 | -0.016 | 0.157 | 0.34 | 0.27 | 0.23 | 0.46 | 0.42 | 0.29 |
| 8 | 5090707 | rs12679985 | -0.408 | 0.041 | -1.052 | -0.309 | 0.013 | 0.129 | 0.129 | 0 | 0.46 | 0.25 | 0.52 | 0.89 | 0.53 | 0.6 |
| 8 | 5092554 | rs4875475 | 0.493 | -0.182 | -0.57 | -0.153 | 0 | 0 | 0 | 0.014 | 0.55 | 0.57 | 0.68 | 0.47 | 0.55 | 0.52 |
| 8 | 5092970 | rs13275667 | -0.602 | -0.19 | -0.669 | -0.244 | -0.003 | -0.001 | -0.001 | 0.341 | 0.45 | 0.49 | 0.37 | 0.53 | 0.49 | 0.49 |
| 8 | 5101411 | rs10087168 | 0.068 | -0.413 | -0.75 | -0.423 | -0.001 | -0.022 | -0.022 | 0.132 | 0.29 | 0.57 | 0.34 | 0.08 | 0.25 | 0.26 |
| 8 | 5101520 | rs7832451 | 1.928 | -0.635 | -1.065 | -0.603 | 0.023 | 0.091 | 0.091 | 0 | 0.31 | 0.21 | 0.18 | 0.1 | 0.27 | 0.19 |
| 8 | 5101602 | rs4460395 | 0.386 | -0.392 | -0.858 | -0.631 | -0.009 | -0.043 | -0.043 | 0 | 0.63 | 0.28 | 0.42 | 0.56 | 0.63 | 0.6 |
| 8 | 5102909 | rs7838806 | 2.342 | -0.352 | -0.77 | -0.677 | -0.003 | -0.046 | -0.046 | 0 | 0.1 | 0.19 | 0.06 | 0 | 0.09 | 0.02 |
| 8 | 5104646 | rs10109186 | -0.82 | -0.362 | -0.808 | -0.622 | -0.005 | 0.014 | 0.014 | 0 | 0.44 | 0.67 | 0.69 | 0.58 | 0.43 | 0.6 |
| 8 | 5106642 | rs7016448 | 1.091 | -0.341 | -0.699 | -0.605 | 0.031 | -0.062 | -0.062 | 0.004 | 0.11 | 0.35 | 0.17 | 0 | 0.1 | 0.1 |
| 8 | 5106677 | rs7823489 | 1.46 | -0.346 | -0.851 | -0.518 | -0.003 | -0.005 | -0.005 | 0 | 0.1 | 0.13 | 0.06 | 0.07 | 0.08 | 0.15 |
| 8 | 5108569 | rs10111494 | 0.278 | -0.202 | -1.093 | -0.436 | 0.05 | 0.01 | 0.01 | 0 | 0.28 | 0.43 | 0.32 | 0.33 | 0.4 | 0.32 |
| 8 | 5112194 | rs10109571 | -1.859 | -0.5 | -1.602 | -0.545 | 0.052 | 0.142 | 0.142 | 0.842 | 0.68 | 0.89 | 0.87 | 0.96 | 0.77 | 0.84 |
| 8 | 5113692 | rs7831902 | 0.964 | -0.306 | -1.411 | -0.302 | -0.055 | -0.044 | -0.044 | 0.082 | 0.2 | 0.08 | 0.18 | 0.35 | 0.27 | 0.26 |
| 8 | 5119564 | rs6558990 | -0.384 | -0.624 | -0.694 | -0.003 | 0.012 | 0.023 | 0.023 | 0 | 0.51 | 0.57 | 0.48 | 0.4 | 0.56 | 0.51 |
| 8 | 5125606 | rs10866981 | -0.306 | -0.433 | -0.832 | -0.337 | 0 | -0.002 | -0.002 | 0 | 0.59 | 0.44 | 0.58 | 0.64 | 0.54 | 0.69 |
| 8 | 5125700 | rs12679472 | 0.084 | 0.171 | -1.256 | 0.093 | -0.048 | -0.1 | -0.1 | 0 | 0.18 | 0.07 | 0.37 | 0.46 | 0.26 | 0.33 |
| 8 | 5128247 | rs6651426 | -0.227 | -0.451 | -0.474 | -0.225 | -0.026 | -0.003 | -0.003 | 0 | 0.13 | 0.28 | 0.08 | 0.1 | 0.07 | 0.21 |
| 8 | 5128947 | rs7845021 | -0.264 | -0.032 | -0.342 | -0.416 | 0.015 | 0.015 | 0.015 | 0 | 0.49 | 0.54 | 0.62 | 0.65 | 0.6 | 0.57 |
| 8 | 5131892 | rs11783970 | 0.769 | -0.65 | -0.79 | -0.329 | 0.048 | -0.008 | -0.008 | 0 | 0.33 | 0.19 | 0.17 | 0.34 | 0.21 | 0.32 |
| 8 | 5133839 | rs12550676 | 1.092 | -0.311 | -0.959 | -0.505 | -0.057 | -0.127 | -0.127 | 0 | 0.25 | 0.06 | 0.36 | 0.46 | 0.33 | 0.27 |
| 8 | 5134838 | rs7001926 | -0.045 | -0.373 | -0.597 | -0.619 | -0.001 | 0.003 | 0.003 | 0 | 0.54 | 0.61 | 0.55 | 0.45 | 0.53 | 0.63 |
| 8 | 5137018 | rs11136848 | -0.009 | -0.355 | -0.584 | -0.662 | 0 | 0.005 | 0.005 | 0 | 0.54 | 0.61 | 0.54 | 0.45 | 0.53 | 0.62 |
| 8 | 5137057 | rs10112245 | 1.154 | -0.363 | -0.666 | -0.679 | 0.003 | 0.031 | 0.031 | 0 | 0.16 | 0.1 | 0.05 | 0.07 | 0.13 | 0.06 |
| 8 | 5137475 | rs918150 | -1.204 | -0.229 | -0.805 | -0.584 | -0.046 | -0.106 | -0.106 | 0 | 0.74 | 0.9 | 0.63 | 0.54 | 0.67 | 0.73 |
| 8 | 5138892 | rs17063324 | 1.198 | -0.23 | -0.794 | -0.748 | 0.053 | -0.011 | -0.011 | 0.477 | 0.31 | 0.17 | 0.16 | 0.36 | 0.19 | 0.29 |
| 8 | 5140102 | rs17351521 | 0.344 | 1.001 | -0.557 | -0.073 | -0.061 | -0.135 | -0.135 | 0.144 | 0.27 | 0.06 | 0.39 | 0.46 | 0.34 | 0.31 |
| 8 | 5142665 | rs7813806 | -0.656 | 1.156 | -0.223 | -0.203 | 0.077 | -0.105 | -0.105 | 0.058 | 0.25 | 0.47 | 0.39 | 0.08 | 0.35 | 0.37 |
| 8 | 5144783 | rs9314561 | 0.149 | 0.39 | -0.426 | 0.283 | 0.017 | 0.002 | 0.002 | 0 | 0.56 | 0.63 | 0.28 | 0.46 | 0.34 | 0.39 |
| 8 | 5146247 | rs7013564 | 0.48 | 0.46 | -0.07 | 0.114 | -0.016 | -0.036 | -0.036 | 0 | 0.73 | 0.38 | 0.65 | 0.84 | 0.77 | 0.68 |
| 8 | 5150113 | rs1420838 | 1.21 | 0.594 | 0.664 | 0.942 | -0.045 | 0.072 | 0.072 | 0.002 | 0.85 | 0.4 | 0.91 | 0.71 | 0.92 | 0.88 |
| 8 | 5154026 | rs17351653 | -0.73 | 0.556 | 0.641 | 0.901 | 0.002 | 0.036 | 0.036 | 0 | 0.07 | 0.01 | 0.01 | 0 | 0.05 | 0.05 |

| | | | | | | | | | | | | | | | | |
|---|---|---|---|---|---|---|---|---|---|---|---|---|---|---|---|---|
| 8 | 5154664 | rs2407956 | 1.282 | 0.549 | 1.047 | 0.922 | 0.026 | 0.305 | 0.305 | 0.924 | 0.93 | 0.53 | 0.94 | 0.71 | 0.97 | 0.94 |
| 8 | 5157922 | rs13273557 | 1.482 | 1.526 | 0.431 | 0.898 | -0.019 | -0.005 | -0.005 | 0 | 0.58 | 0.27 | 0.54 | 0.51 | 0.47 | 0.62 |
| 8 | 5158004 | rs17351786 | -0.32 | 1.586 | 0.43 | 0.888 | -0.088 | 0.034 | 0.034 | 0 | 0.26 | 0.14 | 0.37 | 0.19 | 0.43 | 0.26 |
| 8 | 5159593 | rs11136852 | 0.917 | 1.373 | 0.82 | 0.962 | 0.046 | 0.333 | 0.333 | 0.848 | 0.93 | 0.55 | 0.94 | 0.71 | 0.97 | 0.94 |
| 8 | 5160626 | rs7009701 | 1.749 | 1.438 | 0.724 | 0.978 | -0.022 | 0.089 | 0.089 | 0 | 0.86 | 0.47 | 0.91 | 0.71 | 0.9 | 0.88 |
| 8 | 5162491 | rs17436379 | -0.183 | 1.453 | 0.633 | 0.942 | -0.012 | 0.034 | 0.034 | 0 | 0.06 | 0.01 | 0.02 | 0 | 0.07 | 0.06 |
| 8 | 5165467 | rs6983919 | 1.785 | 1.417 | 0.711 | 0.974 | -0.039 | 0.065 | 0.065 | 0.485 | 0.84 | 0.41 | 0.91 | 0.71 | 0.9 | 0.88 |
| 8 | 5165806 | rs1946161 | 1.671 | 2.147 | 0.624 | 0.809 | -0.003 | -0.021 | -0.021 | 0 | 0.6 | 0.22 | 0.56 | 0.53 | 0.6 | 0.62 |
| 8 | 5165936 | rs10092857 | -0.335 | 2.177 | 0.623 | 0.819 | -0.019 | 0.017 | 0.017 | 0 | 0.23 | 0.15 | 0.34 | 0.19 | 0.31 | 0.26 |
| 8 | 5166336 | rs2111419 | 1.258 | 2.359 | 0.581 | 0.72 | -0.01 | -0.001 | -0.001 | 0.022 | 0.75 | 0.41 | 0.85 | 0.71 | 0.79 | 0.85 |
| 8 | 5169237 | rs11996707 | 0.448 | 2.409 | 1.037 | 0.907 | 0.016 | -0.03 | -0.03 | 0.662 | 0.83 | 0.51 | 0.88 | 0.93 | 0.85 | 0.92 |
| 8 | 5169388 | rs6997443 | 1.204 | 2.353 | 1.298 | 0.899 | 0.07 | 0.06 | 0.06 | 3.139 | 0.94 | 0.61 | 0.95 | 1 | 0.97 | 0.98 |
| 8 | 5172635 | rs11985327 | 1.808 | 2.619 | 0.993 | 1.573 | -0.015 | -0.039 | -0.039 | 0 | 0.69 | 0.24 | 0.51 | 0.59 | 0.63 | 0.64 |
| 8 | 5172992 | rs7007204 | -1.276 | 2.648 | 1.063 | 1.629 | -0.007 | 0.034 | 0.034 | 0 | 0.21 | 0.6 | 0.2 | 0.23 | 0.2 | 0.14 |
| 8 | 5173842 | rs11986426 | -0.721 | 2.639 | 1.066 | 1.467 | 0 | -0.034 | -0.034 | 2.782 | 0.17 | 0.49 | 0.12 | 0.07 | 0.14 | 0.08 |
| 8 | 5176066 | rs10866987 | -1.371 | 3.12 | 0.598 | 1.327 | -0.015 | 0 | 0 | 0 | 0.17 | 0.12 | 0.32 | 0.19 | 0.33 | 0.26 |
| 8 | 5177400 | rs6992382 | -1.371 | 2.977 | 0.72 | 1.398 | -0.004 | 0.004 | 0.004 | 0 | 0.24 | 0.2 | 0.35 | 0.18 | 0.3 | 0.28 |
| 8 | 5186047 | rs7840725 | -1.324 | 2.472 | 0.76 | 1.472 | -0.007 | 0.01 | 0.01 | 0 | 0.24 | 0.18 | 0.34 | 0.19 | 0.29 | 0.27 |
| 8 | 5187640 | rs4875509 | -1.044 | 2.659 | 0.737 | 1.427 | -0.004 | -0.003 | -0.003 | 0.092 | 0.14 | 0.11 | 0.38 | 0.19 | 0.21 | 0.28 |
| 8 | 5189722 | rs1592623 | -1.95 | 2.624 | 0.896 | 1.755 | 0.005 | 0.021 | 0.021 | 0 | 0.31 | 0.39 | 0.56 | 0.56 | 0.38 | 0.38 |
| 8 | 5189771 | rs4875157 | -1.225 | 2.624 | 0.892 | 1.743 | -0.007 | 0.009 | 0.009 | 0 | 0.25 | 0.2 | 0.46 | 0.19 | 0.32 | 0.31 |
| 8 | 5191869 | rs973660 | -1.334 | 2.499 | 0.863 | 1.779 | 0.005 | 0.185 | 0.185 | 0 | 0.06 | 0.19 | 0.11 | 0.37 | 0.06 | 0.07 |
| 8 | 5201210 | rs7845635 | -1.387 | 2.574 | 1.717 | 1.633 | 0.003 | -0.03 | -0.03 | 0 | 0.24 | 0.39 | 0.42 | 0.2 | 0.24 | 0.3 |
| 8 | 5201718 | rs7815392 | -1.393 | 2.605 | 1.717 | 1.633 | 0.008 | -0.03 | -0.03 | 0 | 0.24 | 0.4 | 0.43 | 0.2 | 0.24 | 0.3 |
| 8 | 5203682 | rs10104026 | -1.169 | 2.581 | 1.734 | 1.617 | 0 | -0.005 | -0.005 | 0 | 0.22 | 0.26 | 0.41 | 0.19 | 0.23 | 0.29 |
| 8 | 5204568 | rs7829940 | -1.632 | 2.634 | 2.087 | 1.593 | 0.015 | -0.046 | -0.046 | 0 | 0.27 | 0.43 | 0.47 | 0.2 | 0.31 | 0.31 |
| 8 | 5206751 | rs7843555 | -1.657 | 2.509 | 2.415 | 1.716 | -0.007 | 0.01 | 0.01 | 0 | 0.27 | 0.59 | 0.47 | 0.41 | 0.24 | 0.32 |
| 8 | 5207483 | rs11985153 | -0.736 | 2.509 | 2.383 | 1.717 | -0.002 | 0.002 | 0.002 | 0 | 0.12 | 0.12 | 0.34 | 0.18 | 0.12 | 0.27 |
| 8 | 5207530 | rs11988885 | -1.88 | 2.458 | 2.427 | 1.799 | 0 | -0.001 | -0.001 | 0 | 0.29 | 0.65 | 0.46 | 0.48 | 0.24 | 0.34 |
| 8 | 5209980 | rs6559005 | -1.578 | 2.443 | 2.334 | 1.775 | -0.002 | 0.03 | 0.03 | 0 | 0.25 | 0.44 | 0.45 | 0.4 | 0.25 | 0.33 |
| 8 | 5212268 | rs2052335 | -2.559 | 2.159 | 2.195 | 1.744 | 0.031 | 0.488 | 0.488 | 2.365 | 0.08 | 0.45 | 0.12 | 0.41 | 0.03 | 0.09 |
| 8 | 5214904 | rs10107399 | -1.479 | 2.601 | 1.633 | 2.108 | -0.014 | -0.003 | -0.003 | 0 | 0.28 | 0.24 | 0.47 | 0.31 | 0.4 | 0.34 |
| 8 | 5217751 | rs7814455 | 0.136 | 2.448 | 1.635 | 1.883 | 0.007 | -0.06 | -0.06 | 0 | 0.11 | 0.19 | 0.09 | 0 | 0.11 | 0.03 |
| 8 | 5218198 | rs7816093 | 0.365 | 2.449 | 1.594 | 1.884 | 0.003 | -0.041 | -0.041 | 0.02 | 0.1 | 0.1 | 0.08 | 0 | 0.11 | 0.03 |
| 8 | 5220524 | rs11781530 | -1.237 | 2.654 | 1.963 | 2.135 | 0.035 | 0.035 | 0.035 | 0 | 0.18 | 0.34 | 0.34 | 0.33 | 0.31 | 0.32 |
| 8 | 5222299 | rs4404939 | -2.221 | 1.928 | 2.093 | 1.823 | 0.019 | 0.183 | 0.183 | 0 | 0.06 | 0.35 | 0.11 | 0.16 | 0.05 | 0.08 |
| 8 | 5224855 | rs12678704 | -0.856 | 1.936 | 1.838 | 1.805 | 0.015 | 0.059 | 0.059 | 0 | 0.16 | 0.46 | 0.18 | 0.16 | 0.16 | 0.11 |
| 8 | 5226521 | rs7003556 | -1.363 | 1.971 | 1.671 | 1.63 | 0.046 | 0.015 | 0.015 | 0 | 0.08 | 0.2 | 0.09 | 0.05 | 0.08 | 0.05 |
| 8 | 5227409 | rs13272940 | 1.272 | 1.544 | 1.099 | 1.525 | 0.008 | 0.017 | 0.017 | 0.068 | 0.52 | 0.3 | 0.38 | 0.33 | 0.43 | 0.44 |
| 8 | 5228896 | rs11785007 | -1.125 | 1.462 | 1.279 | 1.66 | 0.021 | 0.003 | 0.003 | 0 | 0.38 | 0.5 | 0.52 | 0.51 | 0.54 | 0.46 |
| 8 | 5230140 | rs9314565 | 0.682 | 1.384 | 1.288 | 1.541 | 0.041 | 0.064 | 0.064 | 0.01 | 0.68 | 0.52 | 0.54 | 0.41 | 0.55 | 0.52 |
| 8 | 5237620 | rs12676390 | -1.973 | 1.133 | 0.571 | 1.491 | -0.017 | -0.007 | -0.007 | 0 | 0.54 | 0.87 | 0.65 | 0.66 | 0.58 | 0.56 |
| 8 | 5243099 | rs6994195 | 1.758 | 1.133 | 0.632 | 1.539 | -0.007 | 0.028 | 0.028 | 0.157 | 0.46 | 0.11 | 0.36 | 0.32 | 0.42 | 0.45 |
| 8 | 5245451 | rs6992051 | 0.827 | 1.029 | 1.005 | 1.549 | 0.002 | 0.003 | 0.003 | 0 | 0.6 | 0.44 | 0.46 | 0.46 | 0.47 | 0.53 |
| 8 | 5246298 | rs12676661 | 0.846 | 1.001 | 0.623 | 1.587 | -0.024 | -0.033 | -0.033 | 1.104 | 0.06 | 0 | 0.12 | 0.11 | 0.06 | 0.18 |
| 8 | 5246716 | rs4875530 | 1.623 | 1.06 | 0.85 | 1.541 | 0.012 | 0.006 | 0.006 | 0.002 | 0.48 | 0.18 | 0.38 | 0.39 | 0.44 | 0.46 |
| 8 | 5248533 | rs11136867 | 1.38 | 1.297 | 0.917 | 1.672 | -0.013 | -0.005 | -0.005 | 0 | 0.53 | 0.18 | 0.39 | 0.43 | 0.47 | 0.46 |
| 8 | 5249230 | rs7842355 | -0.638 | 1.248 | 1.069 | 1.677 | -0.013 | 0 | 0 | 0.35 | 0.06 | 0.15 | 0.07 | 0.07 | 0.02 | 0.07 |
| 8 | 5249754 | rs7015584 | 1.332 | 1.261 | 1.072 | 1.661 | -0.016 | -0.006 | -0.006 | 0.395 | 0.54 | 0.2 | 0.4 | 0.43 | 0.47 | 0.46 |
| 8 | 5252599 | rs7823997 | -0.847 | 1.157 | 0.701 | 1.234 | -0.019 | 0.004 | 0.004 | 0.13 | 0.06 | 0.2 | 0.08 | 0.07 | 0.02 | 0.07 |
| 8 | 5255155 | rs7817927 | 0.756 | 1.377 | 0.462 | 1.145 | 0.011 | -0.045 | -0.045 | 0 | 0.09 | 0.12 | 0.07 | 0 | 0.09 | 0.02 |
| 8 | 5255714 | rs6991139 | 0.36 | 1.346 | 0.504 | 1.129 | 0.008 | -0.056 | -0.056 | 0.795 | 0.1 | 0.18 | 0.08 | 0 | 0.09 | 0.02 |
| 8 | 5257588 | rs11136873 | 1.348 | 1.269 | 0.137 | 1.548 | -0.098 | -0.063 | -0.063 | 0.043 | 0.54 | 0.08 | 0.4 | 0.44 | 0.49 | 0.47 |
| 8 | 5258496 | rs7011009 | 0.371 | 1.263 | 0.461 | 1.426 | -0.009 | -0.041 | -0.041 | 2.936 | 0.84 | 0.66 | 0.87 | 0.93 | 0.9 | 0.9 |
| 8 | 5265851 | rs4146469 | -1.084 | 1.596 | 0.529 | 1.529 | 0 | 0.001 | 0.001 | 0 | 0.37 | 0.54 | 0.52 | 0.55 | 0.45 | 0.44 |
| 8 | 5269217 | rs1526364 | 1.6 | 1.439 | 0.093 | 1.464 | -0.039 | -0.023 | -0.023 | 0 | 0.54 | 0.08 | 0.42 | 0.4 | 0.53 | 0.48 |
| 8 | 5270693 | rs6980950 | 0.588 | 1.54 | -0.131 | 1.237 | 0.001 | 0.001 | 0.001 | 0 | 0.78 | 0.76 | 0.75 | 0.56 | 0.8 | 0.63 |
| 8 | 5270992 | rs6994611 | 1.73 | 1.438 | -0.086 | 0.954 | -0.039 | 0.015 | 0.015 | 0 | 0.91 | 0.68 | 0.89 | 0.91 | 0.98 | 0.84 |
| 8 | 5271716 | rs11988642 | 0.812 | 1.535 | -0.293 | 0.869 | 0.008 | -0.042 | -0.042 | 0.254 | 0.09 | 0.12 | 0.05 | 0 | 0.08 | 0.04 |
| 8 | 5272142 | rs7831446 | 0.813 | 1.254 | -0.912 | 1.082 | -0.024 | -0.014 | -0.014 | 0.233 | 0.93 | 0.8 | 0.92 | 0.96 | 0.98 | 0.91 |
| 8 | 5276592 | rs7841853 | -0.787 | 1.398 | -1.194 | 1.064 | -0.019 | -0.056 | -0.056 | 0.391 | 0.4 | 0.76 | 0.54 | 0.5 | 0.46 | 0.44 |
| 8 | 5279964 | rs4875544 | 0.308 | 1.234 | -1.268 | 0.949 | -0.006 | -0.015 | -0.015 | 4.47 | 0.94 | 0.9 | 0.96 | 0.99 | 0.97 | 0.9 |
| 8 | 5280169 | rs4875545 | 0.682 | 1.36 | -1.305 | 1.104 | -0.025 | -0.03 | -0.03 | 0 | 0.05 | 0 | 0.11 | 0.14 | 0.07 | 0.15 |
| 8 | 5283604 | rs1526355 | -0.652 | 1.384 | -1.321 | 0.964 | -0.007 | -0.015 | -0.015 | 0 | 0.38 | 0.64 | 0.53 | 0.43 | 0.43 | 0.37 |
| 8 | 5283728 | rs1358127 | 0.752 | 1.395 | -1.313 | 0.958 | 0.009 | -0.038 | -0.038 | 0 | 0.55 | 0.19 | 0.43 | 0.5 | 0.54 | 0.47 |
| 8 | 5288955 | rs4875552 | -0.932 | 1.27 | -1.333 | 0.941 | -0.021 | 0.001 | 0.001 | 0 | 0.45 | 0.71 | 0.6 | 0.44 | 0.51 | 0.51 |
| 8 | 5302917 | rs11776896 | 1.968 | 1.027 | -1.754 | 0.747 | -0.032 | -0.008 | -0.008 | 0 | 0.34 | 0.02 | 0.17 | 0.25 | 0.3 | 0.18 |
| 8 | 5304766 | rs4521783 | 1.21 | 1.04 | -1.6 | 0.818 | -0.046 | 0.046 | 0.046 | 0 | 0.46 | 0.09 | 0.26 | 0.35 | 0.44 | 0.31 |
| 8 | 5306942 | rs1880798 | 0.808 | 0.885 | -1.445 | 0.708 | -0.034 | 0.007 | 0.007 | 0.022 | 0.92 | 0.7 | 0.93 | 0.92 | 0.97 | 0.91 |

| | | | | | | | | | | | | | | | | |
|---|---|---|---|---|---|---|---|---|---|---|---|---|---|---|---|---|
| 8 | 5308224 | rs1464980 | -0.442 | 1.188 | -1.315 | 0.785 | -0.021 | -0.014 | -0.014 | 0.005 | 0.41 | 0.64 | 0.6 | 0.45 | 0.49 | 0.45 |
| 8 | 5317452 | rs4875174 | -0.399 | 1.16 | -0.948 | 0.761 | -0.003 | 0 | 0 | 0 | 0.4 | 0.5 | 0.56 | 0.41 | 0.49 | 0.46 |
| 8 | 5317617 | rs17073628 | -0.307 | 1.151 | -1.035 | 0.86 | 0 | 0.013 | 0.013 | 0 | 0.09 | 0.16 | 0.16 | 0.16 | 0.08 | 0.21 |
| 8 | 5323185 | rs11774646 | -0.51 | 1.159 | -1.086 | 0.841 | -0.003 | 0 | 0 | 0 | 0.4 | 0.5 | 0.55 | 0.41 | 0.49 | 0.45 |
| 8 | 5324981 | rs10503303 | 2.051 | 0.87 | -1.515 | 0.73 | -0.044 | -0.032 | -0.032 | 0 | 0.33 | 0.02 | 0.17 | 0.23 | 0.31 | 0.18 |
| 8 | 5334148 | rs10503306 | 0.179 | 0.997 | -1.271 | 0.612 | 0.006 | -0.055 | -0.055 | 0 | 0.22 | 0.08 | 0.28 | 0.39 | 0.21 | 0.33 |
| 8 | 5334655 | rs1526347 | 0.599 | 0.9 | -0.854 | 0.538 | -0.012 | -0.015 | -0.015 | 0 | 0.57 | 0.41 | 0.43 | 0.51 | 0.51 | 0.49 |
| 8 | 5335196 | rs12541107 | 1.28 | 0.954 | -1.164 | 0.539 | -0.02 | 0.051 | 0.051 | 0 | 0.47 | 0.1 | 0.26 | 0.35 | 0.45 | 0.34 |
| 8 | 5335374 | rs7016512 | 0.729 | 0.783 | -0.565 | 0.751 | 0.001 | 0.031 | 0.031 | 0 | 0.9 | 0.84 | 0.83 | 0.76 | 0.92 | 0.74 |
| 8 | 5339713 | rs1981578 | -0.214 | 0.26 | -0.77 | 0.344 | 0.003 | -0.002 | -0.002 | 0 | 0.47 | 0.62 | 0.41 | 0.48 | 0.4 | 0.53 |
| 8 | 5340264 | rs6997466 | 0.717 | 0.18 | -0.986 | 0.211 | 0 | 0.016 | 0.016 | 0 | 0.1 | 0.04 | 0.04 | 0.07 | 0.06 | 0.03 |
| 8 | 5344373 | rs7816614 | 1.327 | 0.005 | -1.228 | 0.201 | -0.021 | -0.03 | -0.03 | 0.99 | 0.25 | 0.02 | 0.16 | 0.2 | 0.25 | 0.3 |
| 8 | 5346663 | rs1526359 | -0.032 | -0.141 | -1.075 | 0.12 | -0.006 | -0.006 | -0.006 | 0 | 0.4 | 0.57 | 0.36 | 0.34 | 0.34 | 0.45 |
| 8 | 5348088 | rs10089077 | -0.683 | -0.098 | -1.024 | 0.148 | 0.006 | 0.007 | 0.007 | 0 | 0.5 | 0.59 | 0.4 | 0.44 | 0.4 | 0.48 |
| 8 | 5351668 | rs974995 | 0.635 | -0.058 | -0.735 | 0.068 | 0.003 | 0.007 | 0.007 | 0 | 0.31 | 0.31 | 0.28 | 0.19 | 0.3 | 0.37 |
| 8 | 5351965 | rs2408057 | -1.208 | -0.177 | -0.848 | 0.065 | -0.003 | -0.005 | -0.005 | 0 | 0.58 | 0.64 | 0.52 | 0.51 | 0.44 | 0.59 |
| 8 | 5357339 | rs7014409 | -1.078 | -0.466 | -0.904 | 0.035 | 0.008 | -0.003 | -0.003 | 0 | 0.52 | 0.63 | 0.44 | 0.5 | 0.39 | 0.58 |
| 8 | 5363039 | rs1918036 | -1.013 | -0.157 | -0.762 | 0.164 | 0.015 | -0.001 | -0.001 | 0.102 | 0.51 | 0.63 | 0.46 | 0.5 | 0.38 | 0.55 |
| 8 | 5368274 | rs17073828 | 0.709 | -0.058 | -0.91 | 0.258 | -0.04 | -0.038 | -0.038 | 0 | 0.1 | 0.01 | 0.12 | 0.13 | 0.1 | 0.2 |
| 8 | 5369076 | rs7824162 | 0.838 | -0.219 | -0.969 | 0.186 | 0.009 | 0.003 | 0.003 | 0 | 0.11 | 0.04 | 0.06 | 0.13 | 0.06 | 0.03 |
| 8 | 5369603 | rs1526343 | 1.243 | -0.295 | -1.106 | 0.13 | -0.029 | -0.039 | -0.039 | 1.273 | 0.23 | 0.04 | 0.21 | 0.19 | 0.24 | 0.3 |
| 8 | 5371613 | rs1011000 | -1.173 | -0.308 | -0.309 | 0.113 | 0.013 | 0.004 | 0.004 | 0.203 | 0.49 | 0.62 | 0.43 | 0.53 | 0.39 | 0.58 |
| 8 | 5372022 | rs1526371 | 1.052 | -0.006 | -0.288 | 0.199 | -0.003 | 0.004 | 0.004 | 0 | 0.25 | 0.16 | 0.24 | 0.2 | 0.29 | 0.33 |
| 8 | 5377735 | rs17073854 | 0.221 | -0.041 | -0.047 | 0.288 | -0.005 | 0.073 | 0.073 | 0 | 0.12 | 0.23 | 0.17 | 0.21 | 0.07 | 0.23 |
| 8 | 5380180 | rs7843051 | -0.957 | -0.131 | 0.024 | 0.233 | -0.028 | 0.038 | 0.038 | 0 | 0.24 | 0.41 | 0.24 | 0.34 | 0.13 | 0.26 |
| 8 | 5380363 | rs17073881 | 0.799 | -0.098 | -0.035 | 0.31 | -0.001 | 0.002 | 0.002 | 0 | 0.1 | 0.1 | 0.15 | 0.14 | 0.07 | 0.19 |
| 8 | 5380809 | rs1526352 | -0.311 | 0.023 | -0.093 | 0.253 | 0.005 | 0.021 | 0.021 | 0 | 0.15 | 0.2 | 0.2 | 0.22 | 0.13 | 0.24 |
| 8 | 5383666 | rs6995806 | -0.363 | -0.124 | -0.152 | 0.253 | -0.01 | -0.011 | -0.011 | 0 | 0.4 | 0.52 | 0.37 | 0.31 | 0.36 | 0.51 |
| 8 | 5383685 | rs7016608 | -0.32 | -0.132 | -0.142 | 0.224 | 0.007 | 0.013 | 0.013 | 0 | 0.41 | 0.41 | 0.27 | 0.3 | 0.36 | 0.37 |
| 8 | 5387418 | rs2732954 | -0.313 | -0.198 | 0.244 | 0.204 | 0.003 | -0.003 | -0.003 | 0.47 | 0.4 | 0.39 | 0.37 | 0.38 | 0.36 | 0.55 |
| 8 | 5387976 | rs2732995 | -0.47 | -0.013 | 0.471 | 0.314 | 0 | 0.002 | 0.002 | 0 | 0.4 | 0.44 | 0.4 | 0.39 | 0.4 | 0.56 |
| 8 | 5390967 | rs2527731 | -0.204 | -0.027 | 0.534 | 0.311 | -0.007 | 0.003 | 0.003 | 0 | 0.38 | 0.46 | 0.28 | 0.37 | 0.35 | 0.4 |
| 8 | 5392336 | rs2527737 | 1.067 | -0.015 | 0.515 | 0.307 | 0 | 0.004 | 0.004 | 0 | 0.25 | 0.29 | 0.22 | 0.25 | 0.29 | 0.37 |
| 8 | 5393748 | rs7833729 | 0.645 | -0.03 | 0.482 | 0.306 | -0.012 | -0.012 | -0.012 | 0 | 0.13 | 0.22 | 0.07 | 0.12 | 0.07 | 0.03 |
| 8 | 5393862 | rs2527743 | -1.518 | -0.033 | 0.385 | 0.303 | -0.01 | -0.004 | -0.004 | 0 | 0.54 | 0.73 | 0.48 | 0.51 | 0.46 | 0.59 |
| 8 | 5396199 | rs2527748 | 1.049 | -0.03 | 0.363 | 0.309 | 0.004 | -0.031 | -0.031 | 1.23 | 0.26 | 0.1 | 0.18 | 0.25 | 0.27 | 0.37 |
| 8 | 5397094 | rs1557982 | -0.406 | 0.03 | 0.436 | 0.296 | 0.007 | 0.001 | 0.001 | 0 | 0.41 | 0.64 | 0.28 | 0.38 | 0.34 | 0.4 |
| 8 | 5400926 | rs1319755 | -0.26 | 0.358 | 0.182 | 0.282 | -0.002 | -0.001 | -0.001 | 0 | 0.41 | 0.44 | 0.29 | 0.37 | 0.42 | 0.44 |
| 8 | 5405120 | rs2733016 | -0.235 | 0.254 | -0.203 | 0.352 | -0.012 | 0.017 | 0.017 | 0 | 0.42 | 0.25 | 0.41 | 0.34 | 0.46 | 0.56 |
| 8 | 5405145 | rs2733019 | -0.297 | 0.242 | -0.141 | 0.348 | -0.006 | 0.016 | 0.016 | 0 | 0.42 | 0.3 | 0.41 | 0.34 | 0.46 | 0.55 |
| 8 | 5407375 | rs17073983 | 1.115 | 0.577 | -0.561 | 0.873 | -0.061 | -0.076 | -0.076 | 0 | 0.11 | 0.01 | 0.25 | 0.31 | 0.16 | 0.26 |
| 8 | 5408321 | rs17073986 | 0.904 | -0.494 | -0.501 | 0.234 | -0.001 | 0 | 0 | 0.131 | 0.12 | 0.14 | 0.16 | 0.15 | 0.07 | 0.21 |
| 8 | 5408863 | rs2733027 | -1.075 | -0.516 | -0.764 | 0.235 | 0.013 | -0.012 | -0.012 | 0 | 0.53 | 0.71 | 0.44 | 0.49 | 0.46 | 0.63 |
| 8 | 5408994 | rs2733028 | 1.174 | -0.231 | -1.165 | -0.01 | -0.017 | 0.027 | 0.027 | 0 | 0.24 | 0.09 | 0.18 | 0.14 | 0.32 | 0.32 |
| 8 | 5409371 | rs767239 | 0.256 | -0.11 | -1.179 | 0.05 | -0.005 | -0.008 | -0.008 | 0 | 0.17 | 0.33 | 0.21 | 0.15 | 0.11 | 0.22 |
| 8 | 5409640 | rs960249 | -0.065 | -0.012 | -1.804 | 0.1 | -0.013 | 0.063 | 0.063 | 0 | 0.32 | 0.03 | 0.18 | 0.15 | 0.32 | 0.32 |
| 8 | 5410407 | rs11987584 | -0.022 | 0.698 | -1.169 | 0.532 | 0.001 | 0.02 | 0.02 | 0 | 0.36 | 0.23 | 0.24 | 0.29 | 0.38 | 0.35 |
| 8 | 5412336 | rs13271542 | -0.792 | 1.174 | -0.699 | 1.138 | 0.004 | 0.023 | 0.023 | 0 | 0.13 | 0.15 | 0.2 | 0.25 | 0.19 | 0.32 |
| 8 | 5413325 | rs4875580 | -1.27 | 1.306 | -0.159 | 0.421 | 0.017 | 0.003 | 0.003 | 0 | 0.3 | 0.42 | 0.44 | 0.25 | 0.37 | 0.39 |
| 8 | 5413358 | rs17354768 | -1.217 | 1.314 | -0.154 | 0.409 | 0.011 | 0.008 | 0.008 | 0 | 0.29 | 0.42 | 0.43 | 0.25 | 0.32 | 0.39 |
| 8 | 5415344 | rs17073999 | 0.283 | 1.2 | -0.199 | 0.819 | -0.025 | -0.05 | -0.05 | 2.09 | 0.1 | 0.03 | 0.1 | 0.23 | 0.14 | 0.27 |
| 8 | 5415799 | rs1981579 | -0.565 | 1.034 | -0.418 | 0.814 | -0.001 | -0.002 | -0.002 | 0.656 | 0.39 | 0.3 | 0.41 | 0.58 | 0.39 | 0.55 |
| 8 | 5416240 | rs2733055 | -1.902 | 1.103 | -0.902 | 0.503 | -0.004 | 0.026 | 0.026 | 0 | 0.73 | 0.9 | 0.81 | 0.78 | 0.77 | 0.76 |
| 8 | 5416295 | rs13271255 | 1.2 | 1.062 | -0.902 | 0.534 | -0.025 | -0.045 | -0.045 | 0 | 0.1 | 0.03 | 0.1 | 0.15 | 0.14 | 0.27 |
| 8 | 5417018 | rs4875582 | 1.21 | 1.01 | -0.381 | 0.301 | 0.007 | -0.047 | -0.047 | 0.009 | 0.12 | 0.15 | 0.21 | 0 | 0.15 | 0.05 |
| 8 | 5417839 | rs2733056 | 0.887 | 1.014 | -0.287 | 0.5 | -0.001 | -0.004 | -0.004 | 0.004 | 0.21 | 0.24 | 0.18 | 0.36 | 0.25 | 0.19 |
| 8 | 5421005 | rs2733059 | -1.804 | 0.625 | -0.267 | 0.5 | -0.043 | 0.003 | 0.003 | 0 | 0.74 | 0.98 | 0.83 | 0.78 | 0.78 | 0.76 |
| 8 | 5421206 | rs7835344 | 0.051 | 0.324 | 0.451 | 0.687 | 0 | 0 | 0 | 0.036 | 0.47 | 0.51 | 0.34 | 0.48 | 0.52 | 0.41 |
| 8 | 5421232 | rs17074025 | 1.222 | 0.117 | 0.272 | 0.543 | 0.013 | 0.015 | 0.015 | 2.068 | 0.19 | 0.07 | 0.08 | 0.12 | 0.15 | 0.12 |
| 8 | 5422146 | rs7012234 | 0.523 | 0.426 | 0.693 | 0.593 | -0.004 | -0.073 | -0.073 | 2.574 | 0.12 | 0.31 | 0.2 | 0 | 0.08 | 0.05 |
| 8 | 5423292 | rs994707 | 0.757 | 0.541 | 0.441 | 0.787 | -0.054 | -0.077 | -0.077 | 0 | 0.12 | 0.01 | 0.19 | 0.26 | 0.18 | 0.25 |
| 8 | 5423449 | rs7833139 | -0.51 | 0.484 | 0.444 | 0.697 | -0.001 | 0.038 | 0.038 | 0 | 0.44 | 0.78 | 0.45 | 0.38 | 0.43 | 0.38 |
| 8 | 5423742 | rs17074059 | 0.908 | 0.476 | 0.434 | 0.692 | -0.032 | -0.019 | -0.019 | 0 | 0.27 | 0.09 | 0.22 | 0.33 | 0.33 | 0.3 |
| 8 | 5423975 | rs1541423 | 0.098 | 0.475 | 0.595 | 0.672 | -0.022 | -0.033 | -0.033 | 0 | 0.3 | 0.52 | 0.28 | 0.16 | 0.22 | 0.2 |
| 8 | 5424408 | rs7008921 | 0.277 | 0.563 | 0.586 | 0.59 | 0.011 | -0.085 | -0.085 | 0.222 | 0.13 | 0.42 | 0.21 | 0 | 0.09 | 0.05 |
| 8 | 5425042 | rs2527760 | -0.547 | 0.708 | 0.402 | 0.825 | 0.001 | -0.004 | -0.004 | 0.724 | 0.43 | 0.57 | 0.44 | 0.66 | 0.5 | 0.58 |
| 8 | 5425625 | rs2527762 | 0.221 | 0.975 | -0.061 | 0.851 | -0.018 | -0.043 | -0.043 | 4.318 | 0.13 | 0.1 | 0.19 | 0.26 | 0.16 | 0.24 |
| 8 | 5427730 | rs2040943 | -0.419 | 0.816 | 0.268 | 0.391 | -0.002 | -0.023 | -0.023 | 0.959 | 0.59 | 0.5 | 0.56 | 0.75 | 0.61 | 0.66 |
| 8 | 5430301 | rs4875186 | -0.088 | 0.988 | 0.805 | 0.636 | 0.04 | 0.015 | 0.015 | 0 | 0.11 | 0.31 | 0.22 | 0.08 | 0.11 | 0.09 |

| | | | | | | | | | | | | | | | | |
|---|---|---|---|---|---|---|---|---|---|---|---|---|---|---|---|---|
| 8 | 5430697 | rs2725725 | -0.083 | 0.873 | 0.534 | 0.928 | 0.001 | 0.011 | 0.011 | 0 | 0.14 | 0.21 | 0.11 | 0.21 | 0.16 | 0.21 |
| 8 | 5431602 | rs10094006 | 0.448 | 0.549 | 0.317 | 1.016 | -0.02 | 0.016 | 0.016 | 0.929 | 0.2 | 0.26 | 0.22 | 0.24 | 0.13 | 0.14 |
| 8 | 5433699 | rs2733072 | 0.471 | 0.839 | 0.163 | 1.05 | 0 | 0 | 0 | 0 | 0.47 | 0.48 | 0.36 | 0.39 | 0.47 | 0.5 |
| 8 | 5436509 | rs1468385 | -0.596 | 0.837 | 0.212 | 1.161 | 0 | 0.001 | 0.001 | 0.185 | 0.51 | 0.53 | 0.69 | 0.57 | 0.5 | 0.48 |
| 8 | 5438130 | rs1468387 | -0.59 | 0.98 | -0.044 | 0.876 | 0.004 | 0.009 | 0.009 | 0.558 | 0.5 | 0.41 | 0.54 | 0.44 | 0.53 | 0.46 |
| 8 | 5443571 | rs727781 | 0.707 | 0.619 | -0.454 | 0.922 | 0.02 | 0.016 | 0.016 | 0.412 | 0.18 | 0.11 | 0.13 | 0.1 | 0.12 | 0.18 |
| 8 | 5444800 | rs7815374 | 0.562 | 0.794 | -0.793 | 0.813 | -0.071 | -0.075 | -0.075 | 0 | 0.25 | 0.06 | 0.34 | 0.35 | 0.33 | 0.26 |
| 8 | 5447686 | rs1859595 | 0.48 | 0.688 | -0.594 | 0.732 | -0.002 | -0.026 | -0.026 | 0.748 | 0.07 | 0.09 | 0.05 | 0.01 | 0.06 | 0.03 |
| 8 | 5448834 | rs7816199 | 0.574 | 0.769 | -0.874 | 0.822 | -0.063 | -0.076 | -0.076 | 0.995 | 0.25 | 0.05 | 0.33 | 0.34 | 0.32 | 0.23 |
| 8 | 5449630 | rs2107989 | -1.506 | 0.818 | -0.842 | 1.004 | -0.059 | 0.048 | 0.048 | 0.027 | 0.7 | 0.99 | 0.84 | 0.81 | 0.73 | 0.76 |
| 8 | 5452537 | rs2725664 | 0.083 | 0.841 | -0.279 | 1.141 | 0.007 | 0.003 | 0.003 | 0 | 0.15 | 0.24 | 0.12 | 0.14 | 0.2 | 0.24 |
| 8 | 5453453 | rs11136906 | 0.622 | 0.736 | -0.748 | 1.071 | -0.096 | -0.019 | -0.019 | 1.638 | 0.38 | 0.04 | 0.44 | 0.35 | 0.43 | 0.27 |
| 8 | 5454376 | rs2733052 | -1.314 | 0.737 | -0.47 | 1.137 | -0.035 | 0.069 | 0.069 | 0.052 | 0.68 | 0.95 | 0.82 | 0.8 | 0.71 | 0.74 |
| 8 | 5456893 | rs10092635 | -0.529 | 0.761 | 0.558 | 1.393 | 0.02 | 0.117 | 0.117 | 1.321 | 0.18 | 0.34 | 0.23 | 0.39 | 0.21 | 0.38 |
| 8 | 5457857 | rs2527720 | -0.398 | 0.882 | 0.317 | 1.201 | 0.001 | -0.002 | -0.002 | 0 | 0.56 | 0.66 | 0.65 | 0.59 | 0.56 | 0.43 |
| 8 | 5460166 | rs2527721 | -0.527 | 0.849 | -0.252 | 1.076 | 0.015 | 0.021 | 0.021 | 0 | 0.25 | 0.39 | 0.26 | 0.28 | 0.28 | 0.36 |
| 8 | 5460340 | rs17074186 | 0.281 | 0.82 | -1.348 | 0.815 | -0.111 | -0.04 | -0.04 | 4.078 | 0.38 | 0.04 | 0.44 | 0.34 | 0.42 | 0.26 |
| 8 | 5460669 | rs1015284 | 0.535 | 0.815 | -0.635 | 1.043 | 0.003 | 0 | 0 | 0 | 0.39 | 0.35 | 0.4 | 0.39 | 0.33 | 0.45 |
| 8 | 5460755 | rs1015283 | 0.194 | 1.197 | -0.417 | 0.954 | 0.021 | -0.015 | -0.015 | 0 | 0.78 | 0.69 | 0.83 | 0.84 | 0.71 | 0.7 |
| 8 | 5460838 | rs2408007 | -0.119 | 0.769 | -1.08 | 1.209 | -0.044 | -0.047 | -0.047 | 0 | 0.2 | 0.07 | 0.25 | 0.28 | 0.29 | 0.17 |
| 8 | 5461753 | rs4875598 | 0.263 | -0.083 | -0.854 | 1.028 | 0.003 | 0.001 | 0.001 | 0 | 0.66 | 0.6 | 0.77 | 0.65 | 0.63 | 0.58 |
| 8 | 5462398 | rs13272285 | 1.057 | -0.413 | -1.479 | 0.707 | -0.104 | 0.03 | 0.03 | 0 | 0.26 | 0.05 | 0.32 | 0.15 | 0.36 | 0.21 |
| 8 | 5462574 | rs10110186 | 0.185 | -0.671 | -0.931 | 1.023 | 0.033 | 0.019 | 0.019 | 0 | 0.33 | 0.48 | 0.55 | 0.35 | 0.41 | 0.43 |
| 8 | 5463238 | rs17074252 | 0.537 | -0.606 | -0.694 | 1.111 | 0.01 | 0.038 | 0.038 | 0.188 | 0.05 | 0.17 | 0.15 | 0.03 | 0.04 | 0.18 |
| 8 | 5466977 | rs6998217 | -0.267 | -1.148 | -1.154 | 0.82 | 0 | 0.001 | 0.001 | 0 | 0.58 | 0.46 | 0.66 | 0.44 | 0.56 | 0.63 |
| 8 | 5467156 | rs6981227 | 0.272 | -1.216 | -1.341 | 0.61 | -0.017 | -0.004 | -0.004 | 0 | 0.32 | 0.37 | 0.33 | 0.28 | 0.21 | 0.41 |
| 8 | 5468841 | rs11784618 | 1.526 | -1.054 | -1.972 | 0.363 | -0.103 | 0.078 | 0.078 | 0 | 0.25 | 0.04 | 0.32 | 0.12 | 0.35 | 0.21 |
| 8 | 5469190 | rs2527724 | -1.233 | -0.824 | -1.875 | 0.65 | -0.07 | -0.012 | -0.012 | 0 | 0.86 | 0.99 | 0.9 | 0.9 | 0.82 | 0.72 |
| 8 | 5470052 | rs17074275 | 0.805 | -1.009 | -1.481 | 0.467 | -0.018 | -0.014 | -0.014 | 0 | 0.12 | 0.15 | 0.17 | 0.04 | 0.07 | 0.21 |
| 8 | 5470116 | rs12155626 | 1.322 | -0.748 | -1.693 | 0.431 | 0.032 | 0.039 | 0.039 | 0 | 0.19 | 0.04 | 0.09 | 0.11 | 0.13 | 0.14 |
| 8 | 5473428 | rs2077535 | 0.703 | -0.517 | -1.419 | 0.59 | -0.048 | 0.021 | 0.021 | 0 | 0.41 | 0.2 | 0.49 | 0.39 | 0.46 | 0.23 |
| 8 | 5474720 | rs2725668 | 0.495 | -0.6 | -0.937 | 0.728 | -0.004 | 0.003 | 0.003 | 0 | 0.64 | 0.41 | 0.74 | 0.62 | 0.67 | 0.56 |
| 8 | 5476393 | rs2733033 | -0.44 | -0.744 | -0.628 | 0.771 | 0.017 | 0.006 | 0.006 | 0 | 0.9 | 0.83 | 0.92 | 0.92 | 0.86 | 0.91 |
| 8 | 5476479 | rs9644396 | 1.498 | -0.771 | -0.581 | 0.783 | -0.016 | 0.003 | 0.003 | 0 | 0.12 | 0.22 | 0.1 | 0.12 | 0.06 | 0.11 |
| 8 | 5477083 | rs2733032 | 1.062 | -0.722 | -1.166 | 0.795 | 0.022 | -0.005 | -0.005 | 0 | 0.56 | 0.25 | 0.54 | 0.45 | 0.59 | 0.41 |
| 8 | 5478843 | rs17074308 | 1.954 | -0.803 | -1.234 | 0.875 | 0.023 | -0.011 | -0.011 | 0 | 0.15 | 0.09 | 0.15 | 0.21 | 0.09 | 0.04 |
| 8 | 5479323 | rs2107772 | 0.188 | -0.72 | -1.321 | 0.72 | -0.006 | 0.005 | 0.005 | 0 | 0.47 | 0.4 | 0.66 | 0.56 | 0.52 | 0.39 |
| 8 | 5485006 | rs9314572 | 1.405 | -0.629 | -1.311 | 0.149 | 0.003 | 0.021 | 0.021 | 0 | 0.25 | 0.16 | 0.13 | 0.05 | 0.21 | 0.23 |
| 8 | 5487708 | rs2057860 | 1.207 | -0.592 | -1.077 | 0.461 | 0.001 | 0.065 | 0.065 | 0 | 0.31 | 0.16 | 0.16 | 0.08 | 0.27 | 0.3 |
| 8 | 5491128 | rs2733081 | -0.571 | -0.674 | -0.88 | 0.552 | 0.034 | 0.117 | 0.117 | 0.918 | 0.86 | 0.6 | 0.74 | 0.76 | 0.79 | 0.79 |
| 8 | 5491721 | rs2725685 | -0.188 | -0.531 | -1.544 | 0.469 | -0.022 | -0.025 | -0.025 | 0 | 0.39 | 0.25 | 0.44 | 0.47 | 0.43 | 0.44 |
| 8 | 5491824 | rs2725686 | 0.286 | -0.206 | -1.154 | 0.674 | 0.014 | 0.003 | 0.003 | 0.039 | 0.66 | 0.27 | 0.55 | 0.32 | 0.68 | 0.58 |
| 8 | 5492579 | rs2733083 | 0.697 | 0.462 | -1.458 | 0.479 | -0.113 | -0.018 | -0.018 | 1.118 | 0.2 | 0.02 | 0.29 | 0.15 | 0.31 | 0.15 |
| 8 | 5492707 | rs2527095 | 0.195 | 0.27 | -1.25 | 0.549 | -0.001 | 0 | 0 | 0 | 0.64 | 0.34 | 0.62 | 0.53 | 0.64 | 0.69 |
| 8 | 5493312 | rs2733084 | -0.166 | 0.629 | -1.262 | 0.519 | -0.033 | -0.033 | -0.033 | 0 | 0.31 | 0.24 | 0.51 | 0.43 | 0.47 | 0.31 |
| 8 | 5494638 | rs12680594 | 0.259 | 0.15 | -1.397 | 0.417 | 0.04 | 0.033 | 0.033 | 0 | 0.49 | 0.23 | 0.38 | 0.63 | 0.39 | 0.58 |
| 8 | 5495468 | rs2189889 | 0.187 | -0.457 | -1.393 | 0.285 | 0.039 | 0.05 | 0.05 | 0.089 | 0.49 | 0.3 | 0.6 | 0.63 | 0.59 | 0.31 |
| 8 | 5495670 | rs17452790 | 0.544 | -0.93 | -1.844 | -0.214 | 0.013 | 0.161 | 0.161 | 0 | 0.31 | 0.07 | 0.39 | 0.08 | 0.27 | 0.37 |
| 8 | 5495902 | rs2732949 | 0.147 | -0.251 | -1.556 | 0.365 | -0.051 | 0.057 | 0.057 | 0 | 0.37 | 0.16 | 0.39 | 0.74 | 0.52 | 0.5 |
| 8 | 5498411 | rs12541274 | 0.038 | -0.112 | -0.907 | 0.639 | 0.022 | 0.015 | 0.015 | 0 | 0.62 | 0.52 | 0.65 | 0.52 | 0.52 | 0.76 |
| 8 | 5498446 | rs11136911 | 0.036 | -0.109 | -0.905 | 0.641 | 0.022 | 0.014 | 0.014 | 0 | 0.62 | 0.52 | 0.65 | 0.52 | 0.52 | 0.76 |
| 8 | 5498487 | rs7838354 | -0.178 | -0.136 | -0.723 | 0.712 | -0.003 | -0.002 | -0.002 | 0.075 | 0.64 | 0.67 | 0.67 | 0.62 | 0.52 | 0.77 |
| 8 | 5499242 | rs11777608 | 0.481 | -0.256 | 0.022 | 0.545 | -0.012 | 0.061 | 0.061 | 0.873 | 0.89 | 0.54 | 0.92 | 0.9 | 0.95 | 0.98 |
| 8 | 5502080 | rs2732961 | 0.216 | -0.309 | -0.348 | 0.411 | 0.012 | -0.001 | -0.001 | 0.003 | 0.58 | 0.59 | 0.54 | 0.58 | 0.66 | 0.54 |
| 8 | 5503113 | rs2732962 | 0.232 | -0.845 | 0.347 | 0.241 | -0.01 | 0 | 0 | 0 | 0.64 | 0.46 | 0.81 | 0.56 | 0.73 | 0.68 |
| 8 | 5503800 | rs7824313 | -0.386 | -0.566 | 0.049 | 0.377 | -0.001 | 0.009 | 0.009 | 0 | 0.18 | 0.24 | 0.24 | 0.37 | 0.15 | 0.24 |
| 8 | 5504590 | rs10503314 | 0.59 | -0.49 | 0.001 | 0.391 | -0.012 | 0.013 | 0.013 | 0 | 0.07 | 0.03 | 0.28 | 0.03 | 0.07 | 0.16 |
| 8 | 5505067 | rs2527083 | -0.044 | -0.822 | 0.587 | 0.283 | 0 | -0.001 | -0.001 | 0.631 | 0.66 | 0.71 | 0.84 | 0.55 | 0.74 | 0.67 |
| 8 | 5505877 | rs2527081 | 1.773 | -0.576 | 0.711 | 0.031 | -0.004 | 0.002 | 0.002 | 0 | 0.38 | 0.32 | 0.3 | 0.17 | 0.47 | 0.18 |
| 8 | 5507168 | rs2057859 | 0.93 | -0.628 | 0.65 | -0.156 | -0.027 | -0.054 | -0.054 | 0 | 0.1 | 0.01 | 0.17 | 0.29 | 0.11 | 0.17 |
| 8 | 5508065 | rs2107770 | -0.967 | -0.696 | 1.059 | -0.396 | 0.017 | 0.012 | 0.012 | 0 | 0.46 | 0.33 | 0.45 | 0.4 | 0.37 | 0.57 |
| 8 | 5508110 | rs10106061 | 0.53 | -1.037 | 0.964 | -0.669 | 0.023 | 0.046 | 0.046 | 0 | 0.22 | 0.11 | 0.07 | 0 | 0.15 | 0.19 |
| 8 | 5509374 | rs7827127 | 0.886 | -0.939 | 0.997 | -0.673 | 0.006 | -0.006 | -0.006 | 0 | 0.19 | 0.13 | 0.07 | 0 | 0.13 | 0.16 |
| 8 | 5509449 | rs2527074 | 0.416 | -1.089 | 1.349 | -0.733 | -0.065 | -0.055 | -0.055 | 0 | 0.78 | 0.62 | 0.72 | 0.82 | 0.84 | 0.71 |
| 8 | 5509734 | rs7824017 | 0.096 | -1.098 | 1.479 | -0.717 | -0.034 | -0.06 | -0.06 | 0 | 0.25 | 0.38 | 0.14 | 0.16 | 0.15 | 0.27 |
| 8 | 5509961 | rs2732968 | 0.382 | -1.039 | 1.288 | -0.743 | -0.018 | -0.081 | -0.081 | 0 | 0.64 | 0.64 | 0.7 | 0.95 | 0.73 | 0.63 |
| 8 | 5511301 | rs10503315 | -0.692 | -0.301 | 1.162 | -0.551 | 0.028 | 0.061 | 0.061 | 0.058 | 0.23 | 0.36 | 0.37 | 0.47 | 0.25 | 0.4 |
| 8 | 5511450 | rs4875607 | -1.32 | -0.374 | 1.012 | -0.573 | -0.001 | -0.001 | -0.001 | 0 | 0.37 | 0.42 | 0.59 | 0.4 | 0.39 | 0.53 |
| 8 | 5514933 | rs6999868 | 0.9 | -0.451 | 0.65 | -0.709 | 0.005 | -0.011 | -0.011 | 0.027 | 0.2 | 0.18 | 0.08 | 0 | 0.14 | 0.16 |

| | | | | | | | | | | | | | | | | |
|---|---|---|---|---|---|---|---|---|---|---|---|---|---|---|---|---|
| 8 | 5515615 | rs17453068 | 0.121 | -0.294 | 0.716 | -0.692 | 0.01 | 0.01 | 0.01 | 0 | 0.16 | 0.09 | 0.21 | 0.03 | 0.15 | 0.19 |
| 8 | 5516171 | rs7005584 | -0.527 | -0.39 | 0.792 | -0.61 | -0.009 | -0.002 | -0.002 | 0.104 | 0.38 | 0.37 | 0.43 | 0.37 | 0.37 | 0.5 |
| 8 | 5518172 | rs6559035 | -1.73 | -0.367 | 0.316 | -0.586 | -0.001 | 0 | 0 | 0 | 0.58 | 0.56 | 0.64 | 0.49 | 0.52 | 0.75 |
| 8 | 5519029 | rs10503320 | 0.385 | -0.327 | -0.555 | -0.566 | 0.007 | 0.003 | 0.003 | 0 | 0.14 | 0.08 | 0.2 | 0.03 | 0.13 | 0.18 |
| 8 | 5521372 | rs6991017 | -0.094 | -0.147 | -0.873 | -0.48 | -0.006 | -0.022 | -0.022 | 0 | 0.22 | 0.17 | 0.32 | 0.34 | 0.18 | 0.32 |
| 8 | 5522149 | rs2725644 | 1.486 | 0.137 | -1.065 | -0.325 | -0.104 | -0.088 | -0.088 | 0.683 | 0.37 | 0.11 | 0.31 | 0.46 | 0.52 | 0.23 |
| 8 | 5526305 | rs7017768 | -0.019 | -0.112 | -0.349 | -0.683 | 0.002 | -0.03 | -0.03 | 0.41 | 0.27 | 0.22 | 0.11 | 0.02 | 0.23 | 0.2 |
| 8 | 5526951 | rs9987153 | 0.057 | -0.234 | -0.235 | -0.359 | -0.002 | 0.018 | 0.018 | 0.013 | 0.14 | 0.16 | 0.09 | 0.31 | 0.07 | 0.13 |
| 8 | 5527167 | rs17453333 | 0.846 | 0.008 | -0.421 | -1.132 | -0.001 | 0.005 | 0.005 | 0 | 0.15 | 0.13 | 0.06 | 0.02 | 0.13 | 0.16 |
| 8 | 5528773 | rs17074479 | 0.105 | -0.072 | -0.603 | -1.156 | 0.013 | 0.013 | 0.013 | 0 | 0.14 | 0.06 | 0.06 | 0 | 0.13 | 0.18 |
| 8 | 5528849 | rs6995190 | -0.4 | 0.121 | -0.198 | -0.982 | 0.001 | 0.022 | 0.022 | 0 | 0.11 | 0.23 | 0.14 | 0.17 | 0.08 | 0.21 |
| 8 | 5529622 | rs2725698 | 1.26 | 0.155 | -0.436 | -0.858 | -0.063 | 0.143 | 0.143 | 0 | 0.44 | 0.11 | 0.57 | 0.2 | 0.52 | 0.36 |
| 8 | 5532112 | rs10503322 | 0.071 | 0.256 | -0.102 | -0.657 | 0.035 | -0.005 | -0.005 | 0 | 0.06 | 0.16 | 0.03 | 0.04 | 0.07 | 0.11 |
| 8 | 5532124 | rs6559038 | -1.198 | 0.214 | -0.146 | -0.589 | -0.011 | -0.005 | -0.005 | 0 | 0.09 | 0.16 | 0.07 | 0.06 | 0.05 | 0.2 |
| 8 | 5532738 | rs10503324 | -1.195 | 0.373 | -0.165 | -0.63 | -0.005 | -0.022 | -0.022 | 0 | 0.09 | 0.16 | 0.06 | 0.03 | 0.06 | 0.18 |
| 8 | 5533388 | rs11136914 | -1.194 | 0.281 | 0.208 | -0.574 | -0.006 | 0.114 | 0.114 | 0.588 | 0.18 | 0.42 | 0.16 | 0.63 | 0.12 | 0.34 |
| 8 | 5533560 | rs17074516 | -2.053 | 0.215 | -0.055 | -0.616 | 0.001 | 0.001 | 0.001 | 0 | 0.15 | 0.19 | 0.16 | 0.14 | 0.11 | 0.29 |
| 8 | 5533605 | rs10503326 | -0.661 | 0.277 | -0.069 | -0.621 | -0.002 | -0.001 | -0.001 | 0.122 | 0.08 | 0.09 | 0.14 | 0.12 | 0.11 | 0.12 |
| 8 | 5533663 | rs2527165 | 1.786 | 0.18 | -0.305 | -0.872 | -0.11 | 0.114 | 0.114 | 0 | 0.4 | 0.04 | 0.55 | 0.18 | 0.48 | 0.23 |
| 8 | 5533702 | rs17453424 | -1.937 | 0.399 | -0.159 | -0.7 | -0.003 | 0.001 | 0.001 | 0 | 0.16 | 0.19 | 0.19 | 0.16 | 0.17 | 0.3 |
| 8 | 5533942 | rs17362595 | 1.363 | 0.736 | -0.846 | -1.035 | -0.035 | 0.033 | 0.033 | 0 | 0.1 | 0.01 | 0.1 | 0.01 | 0.13 | 0.08 |
| 8 | 5534086 | rs17362609 | -0.458 | 0.797 | -0.616 | -0.998 | 0.027 | 0.009 | 0.009 | 0 | 0.07 | 0.1 | 0.17 | 0.14 | 0.16 | 0.13 |
| 8 | 5534353 | rs7821325 | -1.455 | 0.713 | -0.539 | -1.035 | 0 | 0 | 0 | 0.169 | 0.14 | 0.19 | 0.18 | 0.14 | 0.17 | 0.29 |
| 8 | 5534617 | rs7011826 | -0.658 | 0.502 | -0.527 | -1.163 | 0.024 | 0.024 | 0.024 | 0.203 | 0.31 | 0.47 | 0.27 | 0.6 | 0.32 | 0.38 |
| 8 | 5534671 | rs2527163 | -0.23 | 0.267 | -0.628 | -1.136 | 0.028 | 0.051 | 0.051 | 2.365 | 0.73 | 0.84 | 0.78 | 0.84 | 0.79 | 0.75 |
| 8 | 5534849 | rs2527161 | -0.566 | 0.271 | -0.628 | -1.118 | 0.026 | 0.07 | 0.07 | 1.756 | 0.76 | 0.93 | 0.83 | 0.88 | 0.82 | 0.79 |
| 8 | 5535139 | rs10112126 | 0.484 | 0.374 | -0.253 | -1.031 | 0.064 | 0.292 | 0.292 | 0.05 | 0.12 | 0.37 | 0.12 | 0.56 | 0.12 | 0.16 |
| 8 | 5535501 | rs10113430 | -1.437 | 0.36 | -0.228 | -1.259 | 0.023 | 0.028 | 0.028 | 0 | 0.18 | 0.46 | 0.23 | 0.24 | 0.18 | 0.33 |
| 8 | 5536072 | rs2725672 | -1.041 | 0.145 | -0.459 | -1.328 | 0.024 | 0.008 | 0.008 | 1.639 | 0.8 | 0.85 | 0.94 | 1 | 0.88 | 0.92 |
| 8 | 5538424 | rs2527158 | 0.231 | 0.174 | -0.411 | -1.407 | -0.008 | -0.036 | -0.036 | 0 | 0.68 | 0.63 | 0.84 | 0.93 | 0.75 | 0.73 |
| 8 | 5539122 | rs2725673 | -0.213 | 0.171 | -0.479 | -1.414 | 0.012 | -0.002 | -0.002 | 0 | 0.72 | 0.76 | 0.9 | 0.97 | 0.78 | 0.81 |
| 8 | 5539878 | rs10503328 | 0.412 | 0.192 | -0.454 | -1.361 | -0.012 | -0.027 | -0.027 | 0.48 | 0.08 | 0.15 | 0.03 | 0.03 | 0.05 | 0.1 |
| 8 | 5540571 | rs7357494 | 0.021 | 0.267 | -0.433 | -1.324 | 0.074 | 0.21 | 0.21 | 1.487 | 0.15 | 0.37 | 0.15 | 0.56 | 0.18 | 0.17 |
| 8 | 5541143 | rs2732986 | 1.381 | 0.2 | -0.812 | -1.42 | -0.047 | 0.161 | 0.161 | 0 | 0.48 | 0.08 | 0.65 | 0.23 | 0.55 | 0.47 |
| 8 | 5542087 | rs2732987 | -0.671 | 0.245 | -0.538 | -1.356 | 0.015 | -0.006 | -0.006 | 0.669 | 0.78 | 0.81 | 0.92 | 1 | 0.85 | 0.89 |
| 8 | 5542779 | rs2107778 | 2.48 | 0.344 | -1.092 | -1.598 | -0.067 | -0.025 | -0.025 | 0 | 0.27 | 0.02 | 0.47 | 0.26 | 0.29 | 0.11 |
| 8 | 5544291 | rs17074583 | 0.039 | 0.353 | -0.854 | -1.509 | 0.039 | 0.143 | 0.143 | 0.282 | 0.16 | 0.32 | 0.17 | 0.55 | 0.18 | 0.17 |
| 8 | 5545090 | rs7831044 | 0.75 | 0.341 | -0.868 | -1.635 | -0.004 | 0.02 | 0.02 | 0.032 | 0.19 | 0.14 | 0.12 | 0.04 | 0.19 | 0.22 |
| 8 | 5545975 | rs12334664 | -1.029 | 0.317 | -0.788 | -1.663 | -0.005 | 0.015 | 0.015 | 0.45 | 0.42 | 0.63 | 0.36 | 0.68 | 0.44 | 0.58 |
| 8 | 5546739 | rs2258817 | 2.685 | 0.233 | -0.866 | -1.832 | -0.063 | 0 | 0 | 0 | 0.26 | 0.03 | 0.47 | 0.22 | 0.29 | 0.11 |
| 8 | 5546778 | rs6992802 | 1.147 | 0.312 | -0.777 | -1.831 | 0 | -0.002 | -0.002 | 0.041 | 0.08 | 0.08 | 0.05 | 0.06 | 0.07 | 0.18 |
| 8 | 5547185 | rs17074598 | -0.576 | 0.368 | -0.93 | -1.773 | 0.034 | -0.006 | -0.006 | 0 | 0.36 | 0.48 | 0.32 | 0.63 | 0.43 | 0.4 |
| 8 | 5547288 | rs17074600 | 0.011 | 0.267 | -0.868 | -1.671 | 0.037 | 0.116 | 0.116 | 0.174 | 0.16 | 0.32 | 0.17 | 0.58 | 0.18 | 0.18 |
| 8 | 5549438 | rs10090995 | -0.36 | -0.003 | -0.888 | -1.649 | 0.013 | 0.113 | 0.113 | 0 | 0.21 | 0.45 | 0.17 | 0.62 | 0.18 | 0.32 |
| 8 | 5549469 | rs4875190 | 1.369 | -0.02 | -1.164 | -1.741 | 0.038 | 0.062 | 0.062 | 0.361 | 0.17 | 0.04 | 0.05 | 0 | 0.11 | 0.08 |
| 8 | 5551530 | rs7006473 | -0.031 | 0.238 | -1.09 | -1.58 | 0.024 | -0.121 | -0.121 | 0.139 | 0.27 | 0.47 | 0.22 | 0.04 | 0.28 | 0.25 |
| 8 | 5553031 | rs6988955 | 0.051 | -0.095 | -0.694 | -0.919 | -0.011 | -0.026 | -0.026 | 0.738 | 0.08 | 0.15 | 0.04 | 0.05 | 0.07 | 0.15 |
| 8 | 5554620 | rs2732997 | 0.089 | 0.028 | -1.104 | -0.843 | -0.019 | -0.101 | -0.101 | 0 | 0.65 | 0.55 | 0.83 | 0.9 | 0.7 | 0.58 |
| 8 | 5555854 | rs2214688 | 0.654 | 0.337 | -2.195 | -0.479 | -0.029 | 0.051 | 0.051 | 0 | 0.45 | 0.17 | 0.67 | 0.33 | 0.52 | 0.35 |
| 8 | 5559875 | rs1707428 | 0.752 | 0.425 | -2.183 | -0.683 | -0.075 | 0.086 | 0.086 | 0 | 0.43 | 0.11 | 0.61 | 0.3 | 0.52 | 0.31 |
| 8 | 5563282 | rs1635657 | 0.598 | 0.358 | -2.124 | -0.595 | -0.018 | 0.085 | 0.085 | 0 | 0.45 | 0.17 | 0.63 | 0.31 | 0.54 | 0.31 |
| 8 | 5567808 | rs1615810 | 0.44 | 0.621 | -1.685 | -0.202 | 0.003 | 0.024 | 0.024 | 0 | 0.63 | 0.21 | 0.69 | 0.32 | 0.67 | 0.44 |
| 8 | 5569460 | rs1635649 | -0.533 | 1.045 | -1.494 | 0.681 | -0.006 | -0.029 | -0.029 | 0 | 0.25 | 0.19 | 0.24 | 0.5 | 0.29 | 0.31 |
| 8 | 5573346 | rs1567725 | 0.139 | 1.177 | -1.355 | 0.684 | 0.006 | 0.015 | 0.015 | 0 | 0.17 | 0.25 | 0.42 | 0.28 | 0.18 | 0.08 |
| 8 | 5573697 | rs263336 | -0.438 | 1.001 | -1.984 | 0.609 | -0.026 | -0.121 | -0.121 | 0.169 | 0.27 | 0.06 | 0.26 | 0.52 | 0.28 | 0.34 |
| 8 | 5575157 | rs263333 | 0.494 | 0.886 | -1.307 | 0.26 | 0 | 0.002 | 0.002 | 0 | 0.62 | 0.62 | 0.66 | 0.46 | 0.58 | 0.45 |
| 8 | 5576038 | rs10096897 | 0.141 | 0.746 | -1.041 | 0.168 | -0.001 | 0 | 0 | 0 | 0.34 | 0.25 | 0.55 | 0.67 | 0.35 | 0.42 |
| 8 | 5576744 | rs263332 | -0.138 | 0.338 | -0.257 | -0.144 | 0 | -0.088 | -0.088 | 0.438 | 0.22 | 0.43 | 0.16 | 0.07 | 0.2 | 0.15 |
| 8 | 5578558 | rs4644284 | 0.574 | 0.387 | -0.562 | 0.057 | 0.005 | 0.023 | 0.023 | 0 | 0.49 | 0.39 | 0.5 | 0.35 | 0.46 | 0.52 |
| 8 | 5578896 | rs2725728 | 0.13 | 0.375 | -0.322 | 0.008 | 0 | -0.001 | -0.001 | 0 | 0.59 | 0.55 | 0.55 | 0.35 | 0.53 | 0.53 |
| 8 | 5579875 | rs263328 | 0.803 | 0.34 | -0.427 | -0.324 | 0.006 | -0.056 | -0.056 | 0 | 0.1 | 0.21 | 0.05 | 0 | 0.07 | 0.02 |
| 8 | 5581059 | rs2725729 | -0.489 | 0.242 | -0.567 | -0.463 | -0.014 | -0.062 | -0.062 | 0.226 | 0.18 | 0.39 | 0.12 | 0.07 | 0.15 | 0.1 |
| 8 | 5581368 | rs13282034 | 0.745 | 0.577 | -0.971 | -0.131 | -0.078 | -0.085 | -0.085 | 0.023 | 0.2 | 0.02 | 0.25 | 0.31 | 0.28 | 0.28 |
| 8 | 5581690 | rs2733020 | 0.796 | 0.477 | -0.229 | -0.258 | 0.004 | -0.047 | -0.047 | 0.515 | 0.09 | 0.17 | 0.05 | 0 | 0.07 | 0.02 |
| 8 | 5581833 | rs4875613 | 0.119 | 0.565 | -0.576 | -0.005 | -0.002 | -0.021 | -0.021 | 0 | 0.14 | 0.2 | 0.11 | 0.04 | 0.15 | 0.09 |
| 8 | 5593810 | rs859812 | 0.339 | 0.625 | -0.936 | 0.109 | -0.004 | 0.038 | 0.038 | 0.221 | 0.54 | 0.36 | 0.54 | 0.29 | 0.5 | 0.45 |
| 8 | 5601191 | rs12543145 | 0.264 | 0.349 | -1.289 | 0.907 | -0.022 | -0.102 | -0.102 | 0 | 0.14 | 0.01 | 0.25 | 0.63 | 0.12 | 0.28 |
| 8 | 5607214 | rs1707418 | -0.474 | 0.203 | -0.939 | 0.577 | 0.001 | -0.005 | -0.005 | 0.059 | 0.25 | 0.22 | 0.32 | 0.59 | 0.19 | 0.27 |
| 8 | 5607797 | rs7820786 | 0.965 | 0.225 | -0.972 | 0.025 | -0.004 | -0.012 | -0.012 | 0 | 0.34 | 0.33 | 0.3 | 0.22 | 0.32 | 0.3 |

| | | | | | | | | | | | | | | | | |
|---|---|---|---|---|---|---|---|---|---|---|---|---|---|---|---|---|
| 8 | 5607876 | rs1707421 | -0.564 | 0.144 | -0.789 | 0.033 | -0.015 | 0.042 | 0.042 | 0.15 | 0.61 | 0.61 | 0.62 | 0.89 | 0.52 | 0.62 |
| 8 | 5608189 | rs1398476 | 0.089 | 0.1 | -0.856 | 0.631 | -0.01 | 0.053 | 0.053 | 0 | 0.22 | 0.27 | 0.29 | 0.48 | 0.17 | 0.18 |
| 8 | 5608667 | rs1635674 | -0.566 | 0.097 | -0.78 | 0.606 | -0.005 | 0.008 | 0.008 | 0.1 | 0.6 | 0.63 | 0.63 | 0.82 | 0.52 | 0.58 |
| 8 | 5608718 | rs1635671 | -0.331 | 0.131 | -0.984 | 0.607 | 0.003 | -0.007 | -0.007 | 2.688 | 0.27 | 0.24 | 0.32 | 0.67 | 0.19 | 0.32 |
| 8 | 5609719 | rs10112106 | -0.563 | 0.107 | -0.916 | 0.52 | -0.042 | 0.037 | 0.037 | 0 | 0.59 | 0.7 | 0.6 | 0.71 | 0.49 | 0.49 |
| 8 | 5615643 | rs870585 | 0.998 | -0.042 | -0.667 | 1.054 | -0.005 | -0.054 | -0.054 | 0 | 0.09 | 0.01 | 0.22 | 0.4 | 0.08 | 0.1 |
| 8 | 5617213 | rs10100001 | -0.551 | 0.119 | -0.608 | 0.671 | -0.001 | -0.001 | -0.001 | 0 | 0.68 | 0.66 | 0.68 | 0.71 | 0.65 | 0.52 |
| 8 | 5627509 | rs11992122 | 0.468 | -0.051 | -0.594 | 0.548 | 0.005 | 0.025 | 0.025 | 0 | 0.52 | 0.53 | 0.3 | 0.18 | 0.54 | 0.34 |
| 8 | 5628926 | rs6993631 | -0.501 | 0.033 | -0.687 | 0.685 | 0.002 | 0 | 0 | 0 | 0.67 | 0.62 | 0.69 | 0.71 | 0.64 | 0.51 |
| 8 | 5631607 | rs10092220 | -0.426 | 0.126 | -0.733 | 0.727 | -0.003 | -0.001 | -0.001 | 0 | 0.66 | 0.64 | 0.69 | 0.7 | 0.63 | 0.5 |
| 8 | 5637984 | rs11136920 | -0.749 | 0.051 | -0.777 | 0.6 | -0.003 | 0.025 | 0.025 | 0 | 0.58 | 0.66 | 0.76 | 0.82 | 0.55 | 0.72 |
| 8 | 5638005 | rs11783102 | 0.098 | 0.158 | -0.669 | 0.635 | -0.001 | 0 | 0 | 0 | 0.44 | 0.44 | 0.35 | 0.36 | 0.44 | 0.55 |
| 8 | 5638437 | rs7824014 | -0.834 | 0.298 | -0.806 | 0.62 | -0.004 | 0.014 | 0.014 | 0 | 0.59 | 0.66 | 0.76 | 0.79 | 0.55 | 0.71 |
| 8 | 5640236 | rs6559048 | -0.252 | 0.793 | -0.715 | 0.596 | 0.001 | -0.001 | -0.001 | 0.037 | 0.43 | 0.43 | 0.35 | 0.37 | 0.44 | 0.55 |
| 8 | 5641443 | rs7813894 | -0.579 | 1.167 | -0.815 | 0.374 | 0 | 0 | 0 | 0.882 | 0.32 | 0.39 | 0.23 | 0.24 | 0.35 | 0.44 |
| 8 | 5644695 | rs11136922 | -0.979 | 1.013 | -0.068 | 0.516 | -0.015 | 0.071 | 0.071 | 0.594 | 0.89 | 0.67 | 0.93 | 0.83 | 0.94 | 0.93 |
| 8 | 5644933 | rs1512817 | -0.63 | 0.641 | -0.081 | 0.477 | -0.009 | 0.014 | 0.014 | 0.305 | 0.36 | 0.29 | 0.38 | 0.28 | 0.4 | 0.51 |
| 8 | 5645521 | rs1512814 | 1.032 | 0.787 | -0.091 | 0.146 | -0.005 | -0.004 | -0.004 | 0 | 0.15 | 0.12 | 0.2 | 0.04 | 0.23 | 0.21 |
| 8 | 5646199 | rs4875623 | 0.002 | 0.69 | -0.101 | 0.122 | 0.017 | -0.021 | -0.021 | 5.535 | 0.72 | 0.52 | 0.71 | 0.79 | 0.69 | 0.72 |
| 8 | 5647593 | rs17364091 | 0.546 | 0.921 | -0.309 | 0.046 | -0.038 | 0.021 | 0.021 | 3.198 | 0.2 | 0.14 | 0.16 | 0.12 | 0.27 | 0.37 |
| 8 | 5648236 | rs1849397 | 0.041 | 0.846 | -0.52 | 0.288 | -0.024 | -0.038 | -0.038 | 0.795 | 0.25 | 0.13 | 0.38 | 0.46 | 0.33 | 0.33 |
| 8 | 5650784 | rs7012966 | -0.507 | 0.704 | -0.546 | 0.522 | -0.014 | -0.012 | -0.012 | 0.313 | 0.38 | 0.24 | 0.48 | 0.63 | 0.43 | 0.6 |
| 8 | 5651001 | rs1858459 | 0.24 | 0.632 | -0.513 | 0.423 | -0.015 | -0.026 | -0.026 | 0.418 | 0.25 | 0.2 | 0.39 | 0.47 | 0.33 | 0.33 |
| 8 | 5651425 | rs6980919 | 2.103 | 0.579 | -0.397 | 0.341 | 0.014 | 0.038 | 0.038 | 0.115 | 0.31 | 0.2 | 0.19 | 0.08 | 0.25 | 0.1 |
| 8 | 5651836 | rs12548568 | -0.121 | 0.634 | -0.479 | 0.31 | -0.007 | 0.006 | 0.006 | 0.735 | 0.32 | 0.26 | 0.35 | 0.28 | 0.36 | 0.5 |
| 8 | 5652215 | rs2048575 | -2.159 | 0.724 | -0.232 | 0.308 | -0.008 | -0.054 | -0.054 | 0 | 0.91 | 0.71 | 0.93 | 1 | 0.94 | 0.99 |
| 8 | 5652595 | rs2137452 | -1.026 | 0.796 | -0.426 | 0.47 | -0.004 | -0.001 | -0.001 | 0 | 0.47 | 0.48 | 0.56 | 0.59 | 0.61 | 0.7 |
| 8 | 5654482 | rs12156005 | 1.075 | 1.341 | -0.714 | 0.521 | -0.012 | -0.001 | -0.001 | 0 | 0.1 | 0.1 | 0.11 | 0.1 | 0.18 | 0.11 |
| 8 | 5656723 | rs4455854 | -0.608 | 1.317 | -0.657 | 0.533 | -0.045 | -0.043 | -0.043 | 0 | 0.36 | 0.24 | 0.46 | 0.56 | 0.5 | 0.6 |
| 8 | 5663038 | rs10866997 | 0.405 | 1.486 | -0.397 | 0.407 | 0.013 | -0.013 | -0.013 | 0 | 0.22 | 0.29 | 0.18 | 0.13 | 0.29 | 0.37 |
| 8 | 5666048 | rs2203207 | 1.121 | 1.149 | -0.495 | 0.212 | -0.001 | 0.003 | 0.003 | 0 | 0.43 | 0.35 | 0.36 | 0.26 | 0.42 | 0.28 |
| 8 | 5666474 | rs7812557 | -0.684 | 1.439 | -0.608 | 0.413 | 0 | -0.004 | -0.004 | 0 | 0.42 | 0.61 | 0.52 | 0.46 | 0.47 | 0.41 |
| 8 | 5666765 | rs4875628 | 1.757 | 1.136 | -0.58 | 0.454 | 0.004 | -0.005 | -0.005 | 0 | 0.09 | 0.09 | 0.14 | 0.07 | 0.15 | 0.1 |
| 8 | 5667902 | rs10095371 | 2.182 | 1.039 | -0.695 | 0.354 | 0.015 | 0.03 | 0.03 | 0.371 | 0.29 | 0.2 | 0.17 | 0.09 | 0.22 | 0.11 |
| 8 | 5668821 | rs9987113 | -0.649 | 0.692 | -0.522 | 0.588 | -0.006 | -0.002 | -0.002 | 0 | 0.34 | 0.33 | 0.35 | 0.63 | 0.42 | 0.58 |
| 8 | 5671701 | rs11776652 | 1.106 | 0.67 | -0.809 | 0.607 | -0.026 | -0.035 | -0.035 | 0.104 | 0.09 | 0 | 0.08 | 0.09 | 0.1 | 0.1 |
| 8 | 5674540 | rs1355665 | 0.073 | 0.665 | -0.658 | 0.532 | -0.036 | 0.071 | 0.071 | 0 | 0.72 | 0.45 | 0.68 | 0.57 | 0.8 | 0.72 |
| 8 | 5674575 | rs1355667 | 0.084 | 0.662 | -0.653 | 0.53 | -0.035 | 0.067 | 0.067 | 0 | 0.72 | 0.43 | 0.68 | 0.57 | 0.8 | 0.72 |
| 8 | 5676046 | rs10106213 | 0.362 | 0.832 | -0.655 | 0.558 | -0.041 | -0.055 | -0.055 | 0.062 | 0.26 | 0.18 | 0.31 | 0.47 | 0.35 | 0.48 |
| 8 | 5676058 | rs10092548 | 0.13 | 0.728 | -0.67 | 0.569 | -0.028 | 0.043 | 0.043 | 0 | 0.68 | 0.45 | 0.65 | 0.57 | 0.75 | 0.7 |
| 8 | 5677416 | rs12682166 | 1.653 | 0.796 | -0.794 | 0.669 | -0.016 | 0 | 0 | 0 | 0.06 | 0 | 0.14 | 0.35 | 0.07 | 0.11 |
| 8 | 5677494 | rs13275999 | 0.807 | 0.809 | -0.719 | 0.574 | -0.042 | 0.028 | 0.028 | 3.855 | 0.21 | 0.14 | 0.16 | 0.12 | 0.29 | 0.39 |
| 8 | 5678443 | rs7845160 | 0.542 | 0.267 | -0.375 | 1.008 | -0.027 | 0.044 | 0.044 | 0 | 0.68 | 0.43 | 0.65 | 0.58 | 0.75 | 0.72 |
| 8 | 5679269 | rs17074954 | -0.014 | 0.275 | -0.343 | 0.95 | 0.001 | -0.002 | -0.002 | 0.393 | 0.21 | 0.17 | 0.27 | 0.26 | 0.19 | 0.22 |
| 8 | 5689689 | rs10111249 | 0.759 | 0.052 | -0.349 | 0.79 | -0.007 | 0.003 | 0.003 | 0 | 0.66 | 0.37 | 0.65 | 0.6 | 0.75 | 0.66 |
| 8 | 5690484 | rs11136934 | -1.443 | 0.333 | -0.265 | 0.401 | -0.008 | -0.055 | -0.055 | 0 | 0.91 | 0.69 | 0.94 | 1 | 0.94 | 0.99 |
| 8 | 5691302 | rs4875202 | 0.562 | 0.216 | -0.636 | 0.449 | -0.02 | -0.022 | -0.022 | 0.195 | 0.62 | 0.46 | 0.53 | 0.24 | 0.7 | 0.59 |
| 8 | 5692720 | rs2408100 | 0.162 | 0.498 | -1.182 | 1.044 | -0.003 | 0.045 | 0.045 | 4.04 | 0.81 | 0.71 | 0.7 | 0.55 | 0.81 | 0.76 |
| 8 | 5693520 | rs4875633 | -1.659 | 0.554 | -1.03 | 0.898 | 0.016 | -0.034 | -0.034 | 0 | 0.93 | 0.76 | 0.95 | 1 | 0.94 | 0.99 |
| 8 | 5698503 | rs17364498 | 1.426 | 0.562 | -2.047 | 0.474 | -0.072 | 0.025 | 0.025 | 0.009 | 0.28 | 0.04 | 0.25 | 0.17 | 0.34 | 0.14 |
| 8 | 5700021 | rs2089961 | 0.045 | 0.353 | -1.16 | 1.024 | -0.038 | 0.111 | 0.111 | 1.941 | 0.85 | 0.55 | 0.82 | 0.67 | 0.91 | 0.89 |
| 8 | 5701562 | rs4875634 | -0.139 | 0.429 | -1.442 | 1.121 | -0.027 | 0.125 | 0.125 | 2.038 | 0.86 | 0.71 | 0.82 | 0.56 | 0.91 | 0.85 |
| 8 | 5703465 | rs2408147 | -0.012 | 0.384 | -1.512 | 1.123 | -0.036 | 0.178 | 0.178 | 2.709 | 0.85 | 0.64 | 0.82 | 0.55 | 0.91 | 0.85 |
| 8 | 5703760 | rs11136935 | -1.068 | 0.457 | -1.576 | 1.14 | -0.004 | 0.05 | 0.05 | 0.085 | 0.91 | 0.74 | 0.94 | 0.88 | 0.94 | 0.94 |
| 8 | 5704180 | rs12155543 | 1.034 | 0.875 | -2.079 | 0.972 | -0.059 | -0.017 | -0.017 | 0 | 0.11 | 0.01 | 0.11 | 0.07 | 0.17 | 0.09 |
| 8 | 5706993 | rs1512803 | -0.654 | 0.525 | -1.807 | 1.152 | -0.01 | 0.054 | 0.054 | 0.501 | 0.18 | 0.23 | 0.21 | 0.45 | 0.15 | 0.17 |
| 8 | 5713881 | rs6990815 | 0.148 | 0.528 | -1.749 | 1.257 | 0.02 | 0.03 | 0.03 | 0 | 0.07 | 0.03 | 0.13 | 0.46 | 0.04 | 0.13 |
| 8 | 5714823 | rs1581112 | 0.332 | 0.833 | -1.799 | 0.861 | 0.011 | 0.025 | 0.025 | 0 | 0.13 | 0.23 | 0.07 | 0.14 | 0.12 | 0.09 |
| 8 | 5719971 | rs12678020 | -0.293 | 0.955 | -1.878 | 0.599 | -0.003 | -0.008 | -0.008 | 0.387 | 0.32 | 0.29 | 0.38 | 0.57 | 0.37 | 0.28 |
| 8 | 5723448 | rs12678174 | -0.302 | 0.992 | -1.943 | 0.592 | -0.008 | -0.021 | -0.021 | 0 | 0.32 | 0.27 | 0.37 | 0.56 | 0.36 | 0.28 |
| 8 | 5739913 | rs7820953 | -0.426 | 0.764 | -2.026 | 0.519 | -0.006 | -0.011 | -0.011 | 0 | 0.34 | 0.32 | 0.42 | 0.6 | 0.4 | 0.3 |
| 8 | 5742291 | rs1567724 | 1.179 | 0.711 | -2.018 | 0.796 | -0.018 | -0.01 | -0.01 | 1.475 | 0.07 | 0.01 | 0.15 | 0.44 | 0.1 | 0.11 |
| 8 | 5742851 | rs1512820 | -1.228 | 0.528 | -2.372 | 0.181 | 0.001 | 0 | 0 | 0 | 0.58 | 0.43 | 0.58 | 0.64 | 0.68 | 0.52 |
| 8 | 5745690 | rs2840518 | -0.55 | 0.629 | -2.354 | 0.084 | 0.011 | 0.066 | 0.066 | 0 | 0.48 | 0.36 | 0.42 | 0.2 | 0.54 | 0.39 |
| 8 | 5745852 | rs2840399 | 1.083 | 0.68 | -2.651 | -0.063 | -0.052 | -0.014 | -0.014 | 0.421 | 0.2 | 0.06 | 0.24 | 0.16 | 0.25 | 0.18 |
| 8 | 5746669 | rs13277934 | 0.669 | 0.671 | -2.563 | -0.196 | -0.045 | 0.084 | 0.084 | 0.349 | 0.21 | 0.06 | 0.14 | 0.03 | 0.26 | 0.21 |
| 8 | 5749849 | rs2840460 | -0.918 | 0.604 | -2.227 | 0.4 | 0 | -0.005 | -0.005 | 0 | 0.37 | 0.35 | 0.4 | 0.57 | 0.4 | 0.31 |
| 8 | 5751093 | rs2246733 | -0.251 | 0.748 | -2.436 | 0.508 | -0.023 | -0.068 | -0.068 | 0.263 | 0.29 | 0.16 | 0.36 | 0.57 | 0.35 | 0.3 |
| 8 | 5752056 | rs2840492 | 0.717 | 0.769 | -2.44 | 0.358 | -0.063 | 0.129 | 0.129 | 4.424 | 0.2 | 0.03 | 0.2 | 0 | 0.25 | 0.14 |

| | | | | | | | | | | | | | | | | |
|---|---|---|---|---|---|---|---|---|---|---|---|---|---|---|---|---|
| 8 | 5755611 | rs11136938 | 0.707 | 0.722 | -2.141 | 0.782 | -0.001 | 0.109 | 0.109 | 0 | 0.08 | 0.12 | 0.16 | 0.59 | 0.1 | 0.16 |
| 8 | 5757479 | rs2703191 | 0.023 | 0.836 | -2.193 | 0.312 | -0.001 | -0.05 | -0.05 | 0 | 0.17 | 0.23 | 0.17 | 0.01 | 0.17 | 0.12 |
| 8 | 5763950 | rs2840443 | 1.479 | 1.024 | -2.151 | 0.368 | -0.028 | 0.034 | 0.034 | 0 | 0.09 | 0.03 | 0.13 | 0 | 0.13 | 0.1 |
| 8 | 5764839 | rs17075065 | 1.185 | 0.939 | -2.135 | 0.325 | -0.008 | -0.038 | -0.038 | 0 | 0.07 | 0.15 | 0.04 | 0 | 0.05 | 0.01 |
| 8 | 5764942 | rs2840445 | -0.729 | 0.93 | -2.152 | 0.652 | 0.002 | 0.037 | 0.037 | 0 | 0.24 | 0.33 | 0.32 | 0.52 | 0.27 | 0.29 |
| 8 | 5765381 | rs17075075 | 0.105 | 0.939 | -2.186 | 0.775 | 0.001 | -0.004 | -0.004 | 0 | 0.17 | 0.21 | 0.17 | 0.15 | 0.17 | 0.18 |
| 8 | 5765619 | rs2703249 | 0.185 | 1.039 | -2.234 | 0.878 | -0.029 | -0.079 | -0.079 | 0 | 0.17 | 0.07 | 0.28 | 0.52 | 0.23 | 0.28 |
| 8 | 5768423 | rs2703251 | 0.029 | 0.753 | -2.65 | 0.308 | -0.001 | 0 | 0 | 0 | 0.43 | 0.57 | 0.5 | 0.4 | 0.42 | 0.6 |
| 8 | 5769919 | rs2703253 | -0.53 | 0.738 | -2.827 | 0.276 | 0.006 | 0.1 | 0.1 | 0 | 0.52 | 0.71 | 0.66 | 0.8 | 0.53 | 0.71 |
| 8 | 5770873 | rs17075117 | 1.688 | 0.883 | -3.23 | 0.64 | -0.022 | -0.005 | -0.005 | 0 | 0.07 | 0.01 | 0.14 | 0.38 | 0.1 | 0.1 |
| 8 | 5772121 | rs13281936 | -0.042 | 1.075 | -3.721 | 0.306 | -0.142 | -0.185 | -0.185 | 0 | 0.33 | 0.02 | 0.35 | 0.48 | 0.45 | 0.26 |
| 8 | 5773051 | rs12679571 | 1.499 | 1.25 | -3.242 | 0.758 | -0.018 | 0.005 | 0.005 | 0 | 0.07 | 0.01 | 0.15 | 0.5 | 0.1 | 0.14 |
| 8 | 5773243 | rs2703257 | -0.151 | 1.049 | -3.56 | 0.392 | -0.045 | -0.037 | -0.037 | 0 | 0.35 | 0.1 | 0.5 | 0.68 | 0.36 | 0.4 |
| 8 | 5773530 | rs13275785 | 1.282 | 1.055 | -3.583 | 0.229 | 0.002 | -0.012 | -0.012 | 1.471 | 0.13 | 0.13 | 0.12 | 0.04 | 0.18 | 0.09 |
| 8 | 5773934 | rs2816512 | -0.717 | 0.997 | -3.597 | 0.248 | -0.018 | 0.058 | 0.058 | 0 | 0.46 | 0.23 | 0.62 | 0.7 | 0.53 | 0.48 |
| 8 | 5774771 | rs17456044 | 2.203 | 0.949 | -3.392 | 0.323 | -0.003 | -0.028 | -0.028 | 0 | 0.11 | 0 | 0.18 | 0.15 | 0.06 | 0.11 |
| 8 | 5776021 | rs17456092 | 2.159 | 0.956 | -3.383 | 0.315 | -0.004 | -0.021 | -0.021 | 0.721 | 0.11 | 0 | 0.18 | 0.15 | 0.06 | 0.11 |
| 8 | 5776116 | rs4512397 | 0.938 | 1.058 | -2.995 | 0.408 | -0.015 | 0.025 | 0.025 | 0 | 0.17 | 0.13 | 0.13 | 0.06 | 0.2 | 0.3 |
| 8 | 5776499 | rs11785532 | 1.398 | 0.864 | -2.921 | 1.071 | -0.001 | 0.017 | 0.017 | 0 | 0.07 | 0.05 | 0.15 | 0.36 | 0.1 | 0.09 |
| 8 | 5777585 | rs2816516 | -0.228 | 0.903 | -2.954 | 0.646 | -0.007 | -0.042 | -0.042 | 0 | 0.33 | 0.21 | 0.52 | 0.5 | 0.34 | 0.35 |
| 8 | 5778001 | rs2816517 | -0.054 | 0.926 | -3.072 | 0.556 | -0.032 | 0.108 | 0.108 | 6.821 | 0.36 | 0.16 | 0.46 | 0.18 | 0.37 | 0.29 |
| 8 | 5778519 | rs11787399 | 1.355 | 0.991 | -2.806 | 0.756 | -0.001 | 0.001 | 0.001 | 0 | 0.07 | 0.05 | 0.15 | 0.32 | 0.1 | 0.09 |
| 8 | 5778862 | rs2816518 | 0.422 | 1.014 | -2.836 | 0.351 | -0.008 | 0.041 | 0.041 | 0.459 | 0.26 | 0.16 | 0.35 | 0.18 | 0.25 | 0.29 |
| 8 | 5780487 | rs9792349 | -0.311 | 1.023 | -2.701 | 0.784 | -0.019 | -0.014 | -0.014 | 0 | 0.4 | 0.73 | 0.37 | 0.59 | 0.43 | 0.55 |
| 8 | 5780785 | rs2703263 | -0.08 | 1.038 | -2.229 | 0.587 | -0.007 | -0.032 | -0.032 | 0 | 0.37 | 0.21 | 0.47 | 0.59 | 0.43 | 0.37 |
| 8 | 5781984 | rs2816520 | 0.331 | 1.84 | -1.453 | 0.544 | 0.01 | 0.012 | 0.012 | 0 | 0.74 | 0.64 | 0.59 | 0.56 | 0.6 | 0.71 |
| 8 | 5782057 | rs12548359 | -0.509 | 2 | -1.395 | 0.341 | -0.009 | -0.022 | -0.022 | 0 | 0.19 | 0.16 | 0.28 | 0.44 | 0.24 | 0.22 |
| 8 | 5782377 | rs2816522 | 0.48 | 1.209 | -1.536 | -0.382 | -0.015 | 0 | 0 | 0 | 0.41 | 0.48 | 0.4 | 0.4 | 0.3 | 0.51 |
| 8 | 5783249 | rs4628275 | 0.443 | 1.004 | -1.012 | -0.43 | -0.001 | 0.054 | 0.054 | 0 | 0.15 | 0.3 | 0.28 | 0.22 | 0.15 | 0.31 |
| 8 | 5784490 | rs2816530 | 1.402 | 0.872 | -1.058 | -0.525 | 0.003 | 0.003 | 0.003 | 0 | 0.24 | 0.19 | 0.1 | 0.17 | 0.15 | 0.2 |
| 8 | 5785930 | rs10095454 | 0.762 | 1.298 | -1.243 | -0.703 | 0.005 | 0.032 | 0.032 | 0.004 | 0.39 | 0.32 | 0.26 | 0.14 | 0.35 | 0.29 |
| 8 | 5790102 | rs12548556 | -0.149 | 1.417 | -1.567 | -0.752 | -0.066 | -0.071 | -0.071 | 0 | 0.31 | 0.18 | 0.41 | 0.48 | 0.47 | 0.22 |
| 8 | 5792438 | rs12115050 | 1.063 | 1.249 | -1.47 | -0.832 | 0.009 | 0.025 | 0.025 | 0 | 0.32 | 0.28 | 0.21 | 0.15 | 0.24 | 0.28 |
| 8 | 5792884 | rs7826510 | 0.368 | 1.684 | -1.302 | -0.899 | 0.045 | 0.051 | 0.051 | 0 | 0.13 | 0.25 | 0.11 | 0.18 | 0.18 | 0.3 |
| 8 | 5793059 | rs12546419 | 0.116 | 1.208 | -1.256 | -0.916 | -0.02 | -0.03 | -0.03 | 0.08 | 0.25 | 0.21 | 0.53 | 0.51 | 0.39 | 0.22 |
| 8 | 5793209 | rs10503343 | -0.726 | 1.243 | -1.205 | -0.902 | 0.023 | -0.01 | -0.01 | 0 | 0.35 | 0.45 | 0.61 | 0.68 | 0.47 | 0.5 |
| 8 | 5793297 | rs10092006 | 0.46 | 1.403 | -1.141 | -0.85 | 0.013 | 0.071 | 0.071 | 0 | 0.45 | 0.33 | 0.47 | 0.18 | 0.42 | 0.32 |
| 8 | 5793832 | rs7831364 | -1.748 | 1.202 | -1.134 | -0.792 | -0.019 | -0.017 | -0.017 | 0.018 | 0.76 | 0.73 | 0.89 | 0.83 | 0.85 | 0.8 |
| 8 | 5794525 | rs17075202 | 1.897 | 1.168 | -1.143 | -0.817 | -0.027 | -0.017 | -0.017 | 0.142 | 0.05 | 0.02 | 0.07 | 0.18 | 0.1 | 0.14 |
| 8 | 5798050 | rs2840466 | 1.777 | 1.111 | -1.101 | -0.837 | -0.005 | -0.004 | -0.004 | 0 | 0.24 | 0.21 | 0.11 | 0.17 | 0.15 | 0.2 |
| 8 | 5798398 | rs2840467 | -1.835 | 1.167 | -1.041 | -0.812 | 0.003 | 0.027 | 0.027 | 0 | 0.94 | 0.94 | 0.93 | 0.83 | 0.89 | 0.87 |
| 8 | 5799576 | rs11136948 | 0.004 | 1.181 | -1.092 | -0.864 | -0.062 | -0.07 | -0.07 | 0.194 | 0.29 | 0.18 | 0.41 | 0.49 | 0.46 | 0.23 |
| 8 | 5801577 | rs2840474 | 1.961 | 1.065 | -1.22 | -0.919 | -0.013 | -0.006 | -0.006 | 0 | 0.22 | 0.21 | 0.1 | 0.17 | 0.13 | 0.12 |
| 8 | 5802228 | rs2703289 | 1.19 | 1.077 | -1.13 | -0.843 | 0.005 | -0.009 | -0.009 | 0 | 0.3 | 0.22 | 0.18 | 0.34 | 0.25 | 0.33 |
| 8 | 5807227 | rs2840485 | 0.952 | 1.021 | -1.061 | -0.895 | -0.003 | -0.016 | -0.016 | 0.189 | 0.58 | 0.5 | 0.31 | 0.32 | 0.39 | 0.61 |
| 8 | 5808134 | rs2840486 | 0.814 | 0.972 | -1.164 | -0.942 | 0.005 | -0.005 | -0.005 | 0.217 | 0.34 | 0.27 | 0.36 | 0.36 | 0.3 | 0.38 |
| 8 | 5809543 | rs2703301 | -0.372 | 0.949 | -1.097 | -0.952 | 0.002 | 0.004 | 0.004 | 0 | 0.58 | 0.42 | 0.52 | 0.68 | 0.63 | 0.46 |
| 8 | 5810703 | rs2703304 | -0.78 | 1.011 | -1.134 | -0.995 | 0.016 | 0.054 | 0.054 | 0.367 | 0.59 | 0.68 | 0.73 | 0.86 | 0.66 | 0.6 |
| 8 | 5810889 | rs2703305 | 0.208 | 1.009 | -1.094 | -0.977 | 0 | 0.026 | 0.026 | 0.017 | 0.48 | 0.39 | 0.45 | 0.66 | 0.51 | 0.44 |
| 8 | 5815774 | rs2816403 | -0.453 | 0.834 | -0.935 | -0.8 | 0.001 | 0.004 | 0.004 | 0 | 0.59 | 0.61 | 0.53 | 0.66 | 0.63 | 0.46 |
| 8 | 5816126 | rs2840495 | -1.225 | 0.836 | -0.945 | -0.849 | 0.003 | 0.001 | 0.001 | 0 | 0.68 | 0.68 | 0.79 | 0.86 | 0.77 | 0.61 |
| 8 | 5816490 | rs2816405 | -0.006 | 0.822 | -0.948 | -0.844 | 0.024 | 0.068 | 0.068 | 0 | 0.54 | 0.15 | 0.5 | 0.68 | 0.63 | 0.46 |
| 8 | 5817424 | rs2816410 | -0.842 | 0.9 | -0.978 | -0.8 | 0.001 | -0.009 | -0.009 | 0.03 | 0.64 | 0.63 | 0.6 | 0.85 | 0.72 | 0.6 |
| 8 | 5817790 | rs11998109 | -0.643 | 1.29 | -0.592 | -0.473 | 0.001 | -0.001 | -0.001 | 0 | 0.36 | 0.38 | 0.48 | 0.66 | 0.55 | 0.36 |
| 8 | 5818128 | rs2703312 | 0.594 | 1.349 | -0.572 | -0.432 | 0.056 | 0.139 | 0.139 | 0 | 0.07 | 0.21 | 0.09 | 0.17 | 0.1 | 0.13 |
| 8 | 5818306 | rs11780095 | -0.317 | 0.96 | -0.66 | -0.527 | 0.012 | 0.018 | 0.018 | 0 | 0.31 | 0.36 | 0.42 | 0.49 | 0.45 | 0.23 |
| 8 | 5820274 | rs2816416 | -0.62 | 0.982 | -1.013 | -0.496 | -0.008 | -0.019 | -0.019 | 0.217 | 0.36 | 0.66 | 0.6 | 0.5 | 0.44 | 0.22 |
| 8 | 5821708 | rs2222831 | 0.575 | 0.784 | -0.54 | -0.702 | -0.015 | -0.024 | -0.024 | 0 | 0.39 | 0.52 | 0.22 | 0.2 | 0.25 | 0.18 |
| 8 | 5822191 | rs10099750 | 1.641 | 0.948 | -0.552 | -0.782 | 0.013 | -0.028 | -0.028 | 0 | 0.07 | 0.09 | 0.11 | 0 | 0.1 | 0.11 |
| 8 | 5823186 | rs2816423 | 0.813 | 1.003 | -0.423 | -0.672 | -0.009 | 0.08 | 0.08 | 0.518 | 0.37 | 0.12 | 0.2 | 0.2 | 0.28 | 0.18 |
| 8 | 5823205 | rs2703318 | 0.021 | 1.03 | -0.413 | -0.649 | 0.026 | 0.032 | 0.032 | 0 | 0.47 | 0.6 | 0.3 | 0.3 | 0.33 | 0.31 |
| 8 | 5823890 | rs13257524 | 0.155 | 1.017 | -0.402 | -0.624 | 0.037 | 0.083 | 0.083 | 0 | 0.5 | 0.32 | 0.68 | 0.7 | 0.63 | 0.68 |
| 8 | 5824905 | rs13259125 | 0.565 | 1.014 | -0.653 | -0.493 | -0.077 | -0.129 | -0.129 | 0 | 0.17 | 0.04 | 0.45 | 0.48 | 0.32 | 0.17 |
| 8 | 5825194 | rs2700797 | -0.128 | 0.861 | -0.519 | -0.544 | 0.022 | 0.096 | 0.096 | 0.835 | 0.54 | 0.22 | 0.66 | 0.67 | 0.6 | 0.34 |
| 8 | 5825487 | rs13269249 | 1.021 | 0.903 | -0.552 | -0.525 | -0.006 | -0.028 | -0.028 | 0 | 0.06 | 0 | 0.04 | 0.11 | 0.06 | 0.14 |
| 8 | 5826979 | rs9314586 | 1.029 | 0.912 | -0.364 | -0.419 | -0.008 | 0.033 | 0.033 | 0.085 | 0.32 | 0.19 | 0.23 | 0.23 | 0.32 | 0.51 |
| 8 | 5827045 | rs9314587 | 1.066 | 0.908 | -0.391 | -0.421 | -0.015 | 0.038 | 0.038 | 0.245 | 0.32 | 0.15 | 0.23 | 0.23 | 0.32 | 0.52 |
| 8 | 5827383 | rs2840507 | 0.06 | 0.937 | -0.009 | -0.379 | 0.005 | 0.012 | 0.012 | 0.533 | 0.44 | 0.57 | 0.32 | 0.35 | 0.4 | 0.66 |
| 8 | 5828855 | rs2816434 | 0.799 | 1.055 | -0.256 | -0.666 | 0.002 | 0.083 | 0.083 | 1.036 | 0.38 | 0.12 | 0.21 | 0.2 | 0.29 | 0.21 |

| | | | | | | | | | | | | | | | | |
|---|---|---|---|---|---|---|---|---|---|---|---|---|---|---|---|---|
| 8 | 5829376 | rs2816435 | -0.794 | 1.025 | 0.4 | -0.582 | -0.004 | -0.068 | -0.068 | 0 | 0.64 | 0.5 | 0.72 | 0.83 | 0.67 | 0.5 |
| 8 | 5830891 | rs2700791 | 0.399 | 1.031 | 0.105 | -0.618 | -0.006 | -0.051 | -0.051 | 0 | 0.39 | 0.5 | 0.27 | 0.17 | 0.34 | 0.5 |
| 8 | 5832448 | rs2493169 | -0.372 | 1.248 | 0.004 | -0.309 | 0.033 | 0.077 | 0.077 | 0 | 0.8 | 0.72 | 0.51 | 0.5 | 0.64 | 0.85 |
| 8 | 5835671 | rs2703335 | -1.036 | 1.084 | -0.163 | -0.557 | 0 | -0.013 | -0.013 | 0 | 0.67 | 0.65 | 0.76 | 0.83 | 0.67 | 0.51 |
| 8 | 5843106 | rs2816443 | 0.833 | 0.834 | -0.888 | -0.481 | 0.026 | 0.053 | 0.053 | 0.1 | 0.34 | 0.1 | 0.18 | 0.2 | 0.22 | 0.19 |
| 8 | 5846580 | rs17075533 | 0.57 | 0.918 | -0.772 | -0.337 | -0.003 | 0.006 | 0.006 | 0 | 0.18 | 0.15 | 0.45 | 0.51 | 0.3 | 0.15 |
| 8 | 5848670 | rs2816456 | 0.234 | 0.923 | -0.753 | -0.429 | -0.005 | 0.053 | 0.053 | 0.06 | 0.43 | 0.7 | 0.33 | 0.27 | 0.42 | 0.62 |
| 8 | 5850153 | rs4413793 | -0.336 | 0.901 | -0.681 | -0.271 | -0.001 | -0.003 | -0.003 | 0 | 0.25 | 0.22 | 0.52 | 0.64 | 0.37 | 0.29 |
| 8 | 5850799 | rs13271355 | 0.247 | 0.701 | -0.423 | -0.273 | -0.004 | 0.034 | 0.034 | 0 | 0.08 | 0.17 | 0.07 | 0.13 | 0.07 | 0.14 |
| 8 | 5852560 | rs2141643 | 0.348 | 0.882 | -0.485 | -0.395 | 0.005 | -0.057 | -0.057 | 0.053 | 0.36 | 0.53 | 0.26 | 0.16 | 0.35 | 0.47 |
| 8 | 5852840 | rs2141641 | 0.315 | 0.882 | -0.478 | -0.375 | 0.005 | -0.055 | -0.055 | 0.019 | 0.36 | 0.54 | 0.26 | 0.18 | 0.35 | 0.47 |
| 8 | 5853743 | rs2552240 | 0.597 | 0.675 | -0.754 | -0.099 | -0.023 | -0.024 | -0.024 | 0 | 0.61 | 0.3 | 0.44 | 0.49 | 0.53 | 0.68 |
| 8 | 5857354 | rs890027 | -0.53 | 0.048 | -0.165 | -0.387 | -0.017 | -0.027 | -0.027 | 0 | 0.68 | 0.4 | 0.81 | 0.81 | 0.8 | 0.84 |
| 8 | 5859894 | rs2141639 | 0.61 | 0.175 | -0.43 | -0.279 | -0.03 | -0.061 | -0.061 | 0 | 0.51 | 0.08 | 0.59 | 0.59 | 0.62 | 0.42 |
| 8 | 5862887 | rs17366343 | 0.858 | 0.149 | -0.256 | -0.241 | 0.019 | 0.053 | 0.053 | 0 | 0.29 | 0.07 | 0.15 | 0.19 | 0.2 | 0.16 |
| 8 | 5864122 | rs2700736 | -0.48 | 0.231 | -0.353 | -0.268 | -0.018 | 0.017 | 0.017 | 0 | 0.67 | 0.39 | 0.72 | 0.61 | 0.78 | 0.78 |
| 8 | 5864854 | rs10108017 | -0.127 | 0.304 | -0.016 | -0.308 | 0.015 | -0.046 | -0.046 | 0 | 0.14 | 0.19 | 0.13 | 0.02 | 0.16 | 0.36 |
| 8 | 5864927 | rs1834201 | -0.389 | 0.278 | 0.042 | -0.322 | 0.043 | -0.086 | -0.086 | 0.195 | 0.15 | 0.32 | 0.13 | 0.02 | 0.16 | 0.36 |
| 8 | 5865883 | rs2700737 | -0.409 | 0.219 | -0.216 | -0.29 | 0.024 | -0.024 | -0.024 | 0 | 0.66 | 0.28 | 0.71 | 0.61 | 0.79 | 0.78 |
| 8 | 5866081 | rs2703212 | -0.379 | 0.22 | -0.228 | -0.29 | 0.027 | -0.025 | -0.025 | 3.55 | 0.66 | 0.27 | 0.71 | 0.61 | 0.79 | 0.78 |
| 8 | 5866221 | rs17075693 | 0.53 | 0.317 | -0.402 | -0.177 | -0.105 | -0.154 | -0.154 | 0 | 0.21 | 0.02 | 0.42 | 0.47 | 0.33 | 0.21 |
| 8 | 5871134 | rs28759338 | -0.267 | 0.157 | 0.044 | -0.054 | -0.008 | 0.056 | 0.056 | 0 | 0.4 | 0.23 | 0.26 | 0.22 | 0.35 | 0.52 |
| 8 | 5874500 | rs6987785 | -0.269 | 0.155 | 0.282 | -0.053 | -0.002 | 0.012 | 0.012 | 0 | 0.42 | 0.32 | 0.3 | 0.22 | 0.38 | 0.49 |
| 8 | 5874901 | rs2840426 | 1.147 | 0.38 | 0.287 | -0.146 | 0.011 | 0.183 | 0.183 | 0.003 | 0.39 | 0.13 | 0.2 | 0.12 | 0.39 | 0.24 |
| 8 | 5875308 | rs12164148 | -0.12 | 0.394 | 0.256 | -0.143 | -0.002 | 0.012 | 0.012 | 0 | 0.42 | 0.31 | 0.3 | 0.23 | 0.38 | 0.49 |
| 8 | 5875495 | rs1424724 | 0.842 | 0.466 | 0.203 | 0.06 | -0.076 | -0.106 | -0.106 | 0 | 0.14 | 0.02 | 0.36 | 0.46 | 0.21 | 0.21 |
| 8 | 5881382 | rs12545850 | 0.087 | 0.523 | 0.328 | -0.042 | 0.012 | -0.048 | -0.048 | 0 | 0.58 | 0.22 | 0.63 | 0.6 | 0.66 | 0.67 |
| 8 | 5883129 | rs17075807 | 0.712 | 0.586 | 0.275 | -0.038 | -0.036 | -0.058 | -0.058 | 0.331 | 0.15 | 0.08 | 0.36 | 0.47 | 0.21 | 0.21 |
| 8 | 5888705 | rs4875678 | -0.528 | 0.411 | 0.754 | -0.034 | 0.041 | 0.017 | 0.017 | 0.02 | 0.5 | 0.24 | 0.59 | 0.59 | 0.52 | 0.46 |
| 8 | 5897635 | rs2052647 | -0.848 | 0.502 | 0.449 | -0.001 | 0.02 | 0.005 | 0.005 | 0 | 0.52 | 0.7 | 0.61 | 0.6 | 0.52 | 0.48 |
| 8 | 5899514 | rs7016104 | -0.82 | 0.466 | 0.433 | -0.006 | 0.02 | 0.002 | 0.002 | 0 | 0.52 | 0.69 | 0.61 | 0.59 | 0.51 | 0.47 |
| 8 | 5901623 | rs7821175 | 0.136 | 0.564 | 0.362 | -0.044 | 0 | -0.001 | -0.001 | 0 | 0.35 | 0.3 | 0.24 | 0.12 | 0.31 | 0.27 |
| 8 | 5909125 | rs17367195 | 1.5 | 0.522 | -0.109 | -0.087 | -0.059 | 0.177 | 0.177 | 0 | 0.36 | 0.05 | 0.21 | 0.1 | 0.4 | 0.23 |
| 8 | 5911367 | rs7010123 | 0.766 | 0.597 | -0.089 | -0.001 | -0.084 | -0.116 | -0.116 | 0 | 0.15 | 0.02 | 0.36 | 0.48 | 0.22 | 0.22 |
| 8 | 5911777 | rs13253909 | 1.487 | 0.511 | -0.149 | -0.091 | -0.079 | 0.186 | 0.186 | 0 | 0.37 | 0.06 | 0.22 | 0.1 | 0.41 | 0.23 |
| 8 | 5912100 | rs2408151 | -0.381 | 0.614 | 0.328 | 0.011 | 0.002 | 0.023 | 0.023 | 0 | 0.42 | 0.32 | 0.29 | 0.21 | 0.36 | 0.5 |
| 8 | 5912591 | rs11783989 | 0.718 | 0.666 | 0.114 | 0.029 | -0.012 | -0.054 | -0.054 | 0 | 0.52 | 0.09 | 0.57 | 0.59 | 0.62 | 0.44 |
| 8 | 5913047 | rs10503351 | 1.044 | 0.657 | 0.037 | 0.017 | -0.072 | -0.092 | -0.092 | 0.824 | 0.13 | 0.02 | 0.35 | 0.48 | 0.21 | 0.21 |
| 8 | 5913205 | rs1343923 | 1.359 | 0.582 | 0.158 | 0.073 | -0.053 | 0.199 | 0.199 | 0 | 0.38 | 0.05 | 0.2 | 0.1 | 0.4 | 0.23 |
| 8 | 5913360 | rs13269415 | -1.129 | 0.884 | 0.923 | 0.584 | 0.062 | 0.171 | 0.171 | 0 | 0.15 | 0.48 | 0.25 | 0.27 | 0.17 | 0.38 |
| 8 | 5920480 | rs4875695 | -0.685 | 0.915 | 0.676 | 0.332 | 0.059 | -0.037 | -0.037 | 0 | 0.12 | 0.28 | 0.14 | 0.07 | 0.14 | 0.35 |
| 8 | 5923150 | rs1071978 | -0.267 | 0.704 | 0.708 | 0.377 | -0.036 | 0.021 | 0.021 | 0 | 0.81 | 0.74 | 0.85 | 0.77 | 0.88 | 0.85 |
| 8 | 5924556 | rs4875219 | -0.446 | 0.773 | 0.644 | 0.487 | 0.059 | -0.014 | -0.014 | 0.547 | 0.12 | 0.29 | 0.23 | 0.09 | 0.15 | 0.35 |
| 8 | 5925196 | rs1364553 | -0.274 | 0.663 | 0.353 | 0.624 | -0.016 | 0.012 | 0.012 | 0 | 0.81 | 0.76 | 0.84 | 0.77 | 0.86 | 0.85 |
| 8 | 5926632 | rs12547659 | 0.246 | 0.899 | 0.558 | 0.808 | 0.021 | 0.085 | 0.085 | 0.03 | 0.22 | 0.4 | 0.34 | 0.46 | 0.23 | 0.22 |
| 8 | 5926852 | rs13273456 | 1.294 | 0.713 | 0.388 | 0.547 | 0.021 | 0.093 | 0.093 | 0.022 | 0.52 | 0.24 | 0.33 | 0.27 | 0.49 | 0.38 |
| 8 | 5926872 | rs17075926 | -0.741 | 0.739 | 0.721 | 0.743 | 0.008 | -0.01 | -0.01 | 0 | 0.11 | 0.15 | 0.12 | 0.07 | 0.13 | 0.32 |
| 8 | 5929378 | rs17075934 | -1.441 | 0.611 | 0.829 | 0.607 | -0.022 | 0.007 | 0.007 | 0.154 | 0.3 | 0.39 | 0.36 | 0.31 | 0.25 | 0.34 |
| 8 | 5929952 | rs17075940 | -0.752 | 0.749 | 0.473 | 0.601 | -0.001 | 0.001 | 0.001 | 0 | 0.18 | 0.15 | 0.19 | 0.13 | 0.19 | 0.17 |
| 8 | 5933204 | rs17075950 | -0.311 | 0.849 | 0.542 | 0.588 | 0.007 | -0.023 | -0.023 | 0 | 0.09 | 0.15 | 0.19 | 0.02 | 0.1 | 0.14 |
| 8 | 5935576 | rs17075959 | 0.013 | 0.853 | 0.551 | 0.584 | 0 | -0.006 | -0.006 | 0 | 0.08 | 0.09 | 0.18 | 0.02 | 0.1 | 0.15 |
| 8 | 5936536 | rs1157252 | 0.083 | 0.854 | 0.54 | 0.584 | -0.003 | 0.001 | 0.001 | 0 | 0.08 | 0.08 | 0.18 | 0.02 | 0.1 | 0.15 |
| 8 | 5939560 | rs890021 | -0.331 | 0.795 | 0.98 | 0.645 | -0.025 | 0.029 | 0.029 | 2.807 | 0.15 | 0.3 | 0.22 | 0.24 | 0.09 | 0.38 |
| 8 | 5939849 | rs890026 | 0.714 | 0.791 | 0.918 | 0.653 | -0.033 | 0.022 | 0.022 | 1.226 | 0.1 | 0.21 | 0.17 | 0.17 | 0.05 | 0.16 |
| 8 | 5941490 | rs2222834 | -1.833 | 0.832 | 0.87 | 0.664 | -0.005 | 0.006 | 0.006 | 0.544 | 0.33 | 0.4 | 0.39 | 0.37 | 0.28 | 0.55 |
| 8 | 5942863 | rs9987312 | -0.583 | 0.829 | 0.867 | 0.677 | -0.014 | -0.005 | -0.005 | 0 | 0.18 | 0.3 | 0.25 | 0.18 | 0.15 | 0.3 |
| 8 | 5942924 | rs17076003 | 0.92 | 0.856 | 0.659 | 0.674 | -0.014 | -0.043 | -0.043 | 0 | 0.09 | 0 | 0.1 | 0.13 | 0.08 | 0.03 |
| 8 | 5942981 | rs13257249 | 0.194 | 0.872 | 0.691 | 0.662 | 0.002 | -0.022 | -0.022 | 0 | 0.08 | 0.09 | 0.08 | 0.01 | 0.09 | 0.14 |
| 8 | 5949442 | rs754252 | -0.755 | 1.093 | 0.236 | 0.801 | -0.034 | -0.061 | -0.061 | 0 | 0.71 | 0.87 | 0.78 | 0.53 | 0.69 | 0.78 |
| 8 | 5950645 | rs17367507 | 1.922 | 1.038 | 0.203 | 0.709 | -0.004 | 0.061 | 0.061 | 0 | 0.35 | 0.15 | 0.21 | 0.17 | 0.4 | 0.22 |
| 8 | 5950845 | rs989538 | 0.809 | 0.894 | -0.203 | 0.809 | -0.053 | -0.14 | -0.14 | 0.108 | 0.28 | 0.01 | 0.27 | 0.45 | 0.23 | 0.21 |
| 8 | 5951565 | rs7842740 | -1.786 | 0.921 | 0.865 | 0.957 | -0.014 | 0.022 | 0.022 | 0.619 | 0.33 | 0.5 | 0.56 | 0.37 | 0.28 | 0.53 |
| 8 | 5952944 | rs10503354 | -0.787 | 1.203 | 0.6 | 0.89 | 0.001 | 0.017 | 0.017 | 0 | 0.15 | 0.21 | 0.23 | 0.21 | 0.13 | 0.25 |
| 8 | 5955405 | rs4875699 | -1.166 | 1.506 | 0.267 | 0.506 | 0.038 | -0.013 | -0.013 | 0 | 0.08 | 0.18 | 0.25 | 0.04 | 0.14 | 0.25 |
| 8 | 5956277 | rs1424720 | -1.212 | 1.667 | 0.163 | 0.568 | -0.018 | 0.002 | 0.002 | 0 | 0.08 | 0.06 | 0.31 | 0.06 | 0.17 | 0.31 |
| 8 | 5957456 | rs17076034 | -0.55 | 1.345 | 0.455 | 1.283 | -0.007 | 0.155 | 0.155 | 0 | 0.08 | 0.21 | 0.06 | 0.24 | 0.05 | 0.09 |
| 8 | 5958595 | rs12677147 | 0.246 | 1.074 | 0.459 | 1.72 | -0.002 | 0.038 | 0.038 | 1.309 | 0.1 | 0.13 | 0.11 | 0.4 | 0.07 | 0.17 |
| 8 | 5959744 | rs10107799 | -0.565 | 0.985 | -0.155 | 0.842 | 0.009 | 0.007 | 0.007 | 0 | 0.49 | 0.45 | 0.67 | 0.42 | 0.61 | 0.62 |
| 8 | 5960845 | rs12545754 | -0.076 | 0.712 | -0.43 | 0.59 | -0.02 | 0.196 | 0.196 | 0.255 | 0.46 | 0.16 | 0.63 | 0.18 | 0.54 | 0.51 |

| | | | | | | | | | | | | | | | | |
|---|---|---|---|---|---|---|---|---|---|---|---|---|---|---|---|---|
| 8 | 5964052 | rs7011696 | 0.604 | 0.464 | -0.44 | 0.575 | -0.061 | 0.149 | 0.149 | 2.353 | 0.36 | 0.05 | 0.38 | 0.14 | 0.36 | 0.21 |
| 8 | 5967474 | rs6980961 | -0.662 | 0.381 | 0.202 | 0.655 | 0.002 | 0.008 | 0.008 | 0.669 | 0.52 | 0.5 | 0.54 | 0.39 | 0.56 | 0.42 |
| 8 | 5968366 | rs11985779 | -0.348 | 0.84 | 0.295 | 0.844 | -0.007 | 0.014 | 0.014 | 0 | 0.44 | 0.3 | 0.49 | 0.38 | 0.54 | 0.41 |
| 8 | 5973842 | rs11136971 | -1.107 | 0.774 | 0.273 | 0.842 | 0.012 | 0.033 | 0.033 | 0 | 0.54 | 0.38 | 0.59 | 0.79 | 0.62 | 0.61 |
| 8 | 5981454 | rs750835 | -0.499 | 0.859 | 0.21 | 0.854 | 0 | -0.001 | -0.001 | 0 | 0.45 | 0.53 | 0.51 | 0.4 | 0.53 | 0.43 |
| 8 | 5981709 | rs17076074 | -0.235 | 0.938 | -0.029 | 0.828 | 0.055 | -0.029 | -0.029 | 0.12 | 0.06 | 0.16 | 0.08 | 0 | 0.12 | 0.1 |
| 8 | 5981856 | rs17076075 | 0.644 | 0.892 | -0.221 | 0.904 | -0.01 | -0.05 | -0.05 | 0 | 0.09 | 0.01 | 0.1 | 0.39 | 0.08 | 0.18 |
| 8 | 5984917 | rs10088851 | -0.034 | 1.061 | 0.007 | 0.408 | -0.021 | 0.028 | 0.028 | 0 | 0.37 | 0.28 | 0.47 | 0.15 | 0.5 | 0.3 |
| 8 | 5990270 | rs2570704 | -0.543 | 1.04 | -0.366 | 0.327 | 0 | -0.014 | -0.014 | 0 | 0.44 | 0.46 | 0.4 | 0.35 | 0.55 | 0.39 |
| 8 | 5990329 | rs1971079 | 0.386 | 0.939 | -1.183 | 0.127 | -0.119 | 0.111 | 0.111 | 0 | 0.29 | 0.02 | 0.26 | 0.13 | 0.36 | 0.19 |
| 8 | 5994410 | rs7012076 | -0.211 | 1.019 | -0.595 | 0.456 | 0.01 | 0.048 | 0.048 | 0 | 0.47 | 0.3 | 0.51 | 0.74 | 0.48 | 0.4 |
| 8 | 5994598 | rs2005213 | -0.632 | 0.74 | -0.13 | 0.28 | -0.001 | -0.009 | -0.009 | 0 | 0.51 | 0.52 | 0.68 | 0.87 | 0.47 | 0.63 |
| 8 | 5994671 | rs2005212 | -0.285 | 0.771 | 0.059 | 0.691 | 0.003 | 0.015 | 0.015 | 0 | 0.45 | 0.4 | 0.64 | 0.66 | 0.4 | 0.55 |
| 8 | 5995309 | rs2700709 | -0.709 | 0.628 | -0.036 | 0.945 | -0.006 | 0.174 | 0.174 | 0.045 | 0.18 | 0.39 | 0.35 | 0.61 | 0.17 | 0.2 |
| 8 | 5995912 | rs6983976 | 0.067 | 0.736 | -0.067 | 0.793 | -0.011 | 0.026 | 0.026 | 0 | 0.58 | 0.36 | 0.58 | 0.69 | 0.57 | 0.49 |
| 8 | 5998754 | rs2007793 | 0.446 | 0.959 | -0.259 | 0.568 | -0.015 | -0.034 | -0.034 | 0 | 0.07 | 0.01 | 0.03 | 0.17 | 0.07 | 0.06 |
| 8 | 5999311 | rs2552220 | 1.24 | 0.899 | -0.261 | 0.498 | -0.001 | 0.091 | 0.091 | 0 | 0.41 | 0.21 | 0.24 | 0.09 | 0.41 | 0.26 |
| 8 | 5999421 | rs2700714 | 0.61 | 1.032 | -0.179 | 0.531 | -0.002 | 0.073 | 0.073 | 0 | 0.49 | 0.35 | 0.29 | 0.26 | 0.49 | 0.33 |
| 8 | 6001071 | rs2700716 | -1.017 | 0.941 | -0.017 | 0.793 | 0.013 | 0.172 | 0.172 | 1.294 | 0.22 | 0.53 | 0.39 | 0.52 | 0.2 | 0.27 |
| 8 | 6001772 | rs2552224 | -0.062 | 0.388 | 0.049 | 0.84 | -0.001 | 0.225 | 0.225 | 0.686 | 0.15 | 0.45 | 0.33 | 0.34 | 0.11 | 0.17 |
| 8 | 6002470 | rs918924 | 0.308 | 0.422 | 0.036 | 0.856 | -0.004 | 0.044 | 0.044 | 0.949 | 0.13 | 0.2 | 0.33 | 0.33 | 0.11 | 0.2 |
| 8 | 6005877 | rs2700724 | -0.036 | 0.45 | 0.028 | 0.802 | -0.002 | 0.01 | 0.01 | 0 | 0.63 | 0.6 | 0.62 | 0.6 | 0.63 | 0.52 |
| 8 | 6006443 | rs11136975 | 0.379 | 0.288 | 0.026 | 1.086 | -0.003 | 0.045 | 0.045 | 0.686 | 0.12 | 0.18 | 0.31 | 0.34 | 0.11 | 0.16 |
| 8 | 6009600 | rs2700821 | -0.146 | 0.445 | 0.042 | 0.966 | 0 | -0.019 | -0.019 | 2.157 | 0.68 | 0.76 | 0.64 | 0.6 | 0.68 | 0.6 |
| 8 | 6015416 | rs2552266 | -0.059 | 0.212 | -0.448 | 1.011 | -0.019 | 0.14 | 0.14 | 0 | 0.23 | 0.48 | 0.36 | 0.5 | 0.19 | 0.27 |
| 8 | 6016142 | rs10503358 | 1.078 | 0.275 | -0.49 | 0.874 | -0.01 | 0.044 | 0.044 | 0 | 0.44 | 0.36 | 0.27 | 0.15 | 0.47 | 0.32 |
| 8 | 6016650 | rs4875228 | 0.434 | 0.378 | -1.183 | 0.853 | -0.086 | -0.126 | -0.126 | 0.92 | 0.23 | 0.01 | 0.26 | 0.32 | 0.27 | 0.3 |
| 8 | 6016801 | rs2570689 | -0.276 | 0.33 | -0.543 | 1.013 | -0.019 | 0.08 | 0.08 | 0 | 0.22 | 0.45 | 0.3 | 0.41 | 0.18 | 0.26 |
| 8 | 6018779 | rs2552279 | 0.088 | 0.611 | -1 | 1.063 | 0.002 | -0.016 | -0.016 | 3.226 | 0.71 | 0.77 | 0.61 | 0.63 | 0.7 | 0.62 |
| 8 | 6020683 | rs10091677 | -0.397 | 0.382 | -0.899 | 1.042 | 0.001 | -0.005 | -0.005 | 0.942 | 0.27 | 0.29 | 0.38 | 0.43 | 0.24 | 0.26 |
| 8 | 6021203 | rs2570683 | -0.301 | -0.42 | -0.6 | 0.998 | -0.039 | 0.04 | 0.04 | 0.363 | 0.32 | 0.47 | 0.35 | 0.49 | 0.21 | 0.39 |
| 8 | 6023022 | rs11779054 | -0.286 | -0.06 | -0.369 | 0.568 | 0.01 | -0.003 | -0.003 | 0 | 0.07 | 0.1 | 0.06 | 0.04 | 0.09 | 0.06 |
| 8 | 6024175 | rs890032 | 0.2 | 0.485 | -0.841 | -0.223 | -0.018 | 0.064 | 0.064 | 0 | 0.79 | 0.89 | 0.86 | 0.99 | 0.76 | 0.88 |
| 8 | 6025119 | rs10867007 | 0.345 | 0.388 | -0.651 | -0.154 | -0.005 | 0.026 | 0.026 | 0 | 0.78 | 0.82 | 0.85 | 0.95 | 0.76 | 0.82 |
| 8 | 6026858 | rs17076166 | 0.501 | 0.176 | -0.148 | 0.08 | 0.001 | 0.038 | 0.038 | 0 | 0.25 | 0.3 | 0.42 | 0.61 | 0.27 | 0.44 |
| 8 | 6030568 | rs2570668 | -0.4 | -0.214 | 0.324 | 0.203 | -0.018 | 0.023 | 0.023 | 0.022 | 0.32 | 0.53 | 0.26 | 0.35 | 0.21 | 0.3 |
| 8 | 6031012 | rs918925 | -0.14 | 0.326 | 0.346 | 0.078 | 0.029 | 0.004 | 0.004 | 0 | 0.07 | 0.2 | 0.07 | 0.04 | 0.09 | 0.06 |
| 8 | 6031212 | rs918926 | 0.967 | 0.355 | 0.033 | -0.024 | 0.001 | -0.027 | -0.027 | 0 | 0.86 | 0.78 | 0.91 | 0.96 | 0.85 | 0.9 |
| 8 | 6032731 | rs4543562 | -0.137 | 0.51 | -0.059 | 0.134 | 0.012 | 0.002 | 0.002 | 0 | 0.06 | 0.12 | 0.05 | 0.04 | 0.08 | 0.06 |
| 8 | 6033545 | rs4599833 | 0.29 | 0.001 | -0.023 | 0.232 | 0.009 | -0.002 | -0.002 | 0 | 0.55 | 0.38 | 0.65 | 0.52 | 0.7 | 0.58 |
| 8 | 6033899 | rs13260564 | -0.141 | 0.249 | -0.239 | -0.081 | 0.002 | 0.001 | 0.001 | 0 | 0.62 | 0.43 | 0.52 | 0.51 | 0.59 | 0.52 |
| 8 | 6037337 | rs12676543 | 0.348 | 0.254 | -0.733 | -0.168 | -0.039 | -0.075 | -0.075 | 1.713 | 0.35 | 0.07 | 0.41 | 0.62 | 0.33 | 0.35 |
| 8 | 6037491 | rs2570660 | -0.029 | 0.298 | -0.922 | -0.555 | 0.006 | -0.001 | -0.001 | 0.502 | 0.6 | 0.17 | 0.61 | 0.64 | 0.66 | 0.52 |
| 8 | 6039117 | rs10108062 | 0.133 | 0.283 | -0.439 | -0.552 | 0 | 0 | 0 | 0.094 | 0.37 | 0.37 | 0.36 | 0.33 | 0.34 | 0.41 |
| 8 | 6044874 | rs17368375 | 1.055 | 0.395 | -0.855 | -1.035 | -0.071 | 0.132 | 0.132 | 0.055 | 0.21 | 0.03 | 0.17 | 0.03 | 0.29 | 0.17 |
| 8 | 6055638 | rs6559108 | 0.594 | 0.678 | -0.31 | -1.03 | 0.007 | -0.036 | -0.036 | 0 | 0.26 | 0.33 | 0.26 | 0.07 | 0.35 | 0.23 |
| 8 | 6059955 | rs4311675 | 0.883 | 0.383 | -1.149 | -0.892 | -0.05 | -0.002 | -0.002 | 0 | 0.36 | 0.06 | 0.41 | 0.68 | 0.32 | 0.36 |
| 8 | 6060429 | rs11775757 | 0.086 | 0.621 | -0.662 | -0.648 | -0.005 | -0.016 | -0.016 | 0 | 0.31 | 0.35 | 0.32 | 0.24 | 0.3 | 0.28 |
| 8 | 6061170 | rs11136985 | 0.09 | 1.39 | -1.042 | -0.146 | -0.015 | -0.012 | -0.012 | 0 | 0.1 | 0.05 | 0.2 | 0.18 | 0.18 | 0.15 |
| 8 | 6062800 | rs17368494 | 0.209 | 0.825 | -1.285 | -0.351 | -0.072 | 0.076 | 0.076 | 0 | 0.23 | 0.09 | 0.24 | 0.09 | 0.35 | 0.21 |
| 8 | 6063569 | rs7007868 | 0.166 | 0.93 | -1.501 | -0.204 | -0.021 | -0.006 | -0.006 | 0 | 0.24 | 0.19 | 0.31 | 0.28 | 0.33 | 0.23 |
| 8 | 6065210 | rs7015351 | 0.686 | 0.804 | -1.451 | -0.192 | -0.052 | 0.09 | 0.09 | 0 | 0.19 | 0.03 | 0.15 | 0.05 | 0.25 | 0.12 |
| 8 | 6069941 | rs7012285 | 0.476 | -0.075 | -0.504 | 0.451 | -0.009 | 0.031 | 0.031 | 0 | 0.24 | 0.26 | 0.27 | 0.54 | 0.18 | 0.35 |
| 8 | 6071321 | rs17368536 | 0.83 | -0.577 | -1.138 | -0.14 | 0.059 | 0.086 | 0.086 | 0 | 0.24 | 0.06 | 0.12 | 0.05 | 0.12 | 0.15 |
| 8 | 6071352 | rs4506242 | 0.343 | -0.835 | -0.344 | -0.1 | 0.01 | 0.009 | 0.009 | 0 | 0.46 | 0.41 | 0.57 | 0.37 | 0.66 | 0.41 |
| 8 | 6072076 | rs12546953 | 0.535 | -0.779 | -0.041 | 0.101 | -0.019 | 0.051 | 0.051 | 0.217 | 0.28 | 0.41 | 0.3 | 0.54 | 0.23 | 0.38 |
| 8 | 6073852 | rs4480139 | -0.132 | -0.542 | 0.194 | 0.013 | 0.004 | -0.008 | -0.008 | 0.137 | 0.65 | 0.33 | 0.65 | 0.47 | 0.72 | 0.55 |
| 8 | 6075093 | rs11783067 | 0.964 | -0.636 | -0.051 | -0.099 | 0.021 | 0.121 | 0.121 | 0 | 0.27 | 0.05 | 0.15 | 0.05 | 0.17 | 0.17 |
| 8 | 6076066 | rs7820429 | 0.679 | -0.324 | -0.023 | -0.024 | -0.018 | -0.019 | -0.019 | 0 | 0.4 | 0.19 | 0.51 | 0.42 | 0.56 | 0.37 |
| 8 | 6080043 | rs4604465 | 1.176 | -0.514 | 0.273 | -0.146 | 0.018 | 0.016 | 0.016 | 0 | 0.26 | 0.19 | 0.13 | 0.05 | 0.13 | 0.15 |
| 8 | 6080742 | rs7005733 | 0.548 | -0.472 | 0.024 | -0.028 | -0.016 | -0.015 | -0.015 | 0.277 | 0.4 | 0.18 | 0.51 | 0.41 | 0.57 | 0.37 |
| 8 | 6082233 | rs6993405 | 0.467 | -0.623 | 0.278 | 0.331 | -0.006 | 0.014 | 0.014 | 0.066 | 0.28 | 0.3 | 0.28 | 0.51 | 0.21 | 0.38 |
| 8 | 6084048 | rs7815835 | 0.267 | -0.269 | 0.239 | 0.368 | -0.018 | 0.079 | 0.079 | 0.211 | 0.28 | 0.41 | 0.3 | 0.48 | 0.23 | 0.43 |
| 8 | 6084755 | rs7008774 | 0.686 | -0.221 | -0.785 | -0.439 | -0.073 | 0.154 | 0.154 | 0.043 | 0.4 | 0.11 | 0.28 | 0.18 | 0.45 | 0.35 |
| 8 | 6086549 | rs11136987 | -0.56 | -0.228 | -0.629 | -0.767 | 0.003 | 0.087 | 0.087 | 0.287 | 0.49 | 0.67 | 0.66 | 0.85 | 0.57 | 0.64 |
| 8 | 6086905 | rs17368655 | 1.883 | -0.271 | -0.661 | -0.857 | -0.045 | 0.1 | 0.1 | 0 | 0.27 | 0.07 | 0.18 | 0.07 | 0.31 | 0.14 |
| 8 | 6088783 | rs17076292 | 1.637 | -0.388 | -0.968 | -0.272 | -0.04 | -0.098 | -0.098 | 0 | 0.16 | 0.01 | 0.13 | 0.3 | 0.15 | 0.14 |
| 8 | 6088885 | rs7826963 | -0.247 | -0.411 | -0.842 | -0.351 | 0.011 | 0.002 | 0.002 | 0 | 0.43 | 0.59 | 0.43 | 0.38 | 0.31 | 0.41 |
| 8 | 6091254 | rs6992649 | 0.447 | -0.209 | -0.587 | -0.13 | 0.005 | 0.051 | 0.051 | 0 | 0.25 | 0.52 | 0.3 | 0.32 | 0.23 | 0.28 |

| | | | | | | | | | | | | | | | | |
|---|---|---|---|---|---|---|---|---|---|---|---|---|---|---|---|---|
| 8 | 6094406 | rs7822472 | -0.864 | -0.153 | -0.652 | -0.285 | 0.006 | 0.052 | 0.052 | 0.408 | 0.49 | 0.62 | 0.68 | 0.89 | 0.57 | 0.67 |
| 8 | 6095927 | rs13269139 | -0.121 | 0.15 | -0.738 | 0.047 | -0.033 | -0.103 | -0.103 | 0 | 0.24 | 0.11 | 0.36 | 0.53 | 0.34 | 0.36 |
| 8 | 6100733 | rs4313178 | -0.534 | 0.135 | -0.25 | 0.074 | -0.001 | -0.001 | -0.001 | 0 | 0.29 | 0.32 | 0.39 | 0.52 | 0.37 | 0.39 |
| 8 | 6103090 | rs5025213 | -0.628 | 0.214 | 0.035 | -0.498 | -0.013 | 0.061 | 0.061 | 0.14 | 0.63 | 0.29 | 0.74 | 0.93 | 0.61 | 0.69 |
| 8 | 6103913 | rs4457353 | -0.847 | -0.039 | 0.258 | -0.4 | 0.008 | -0.085 | -0.085 | 0 | 0.63 | 0.51 | 0.73 | 0.92 | 0.61 | 0.75 |
| 8 | 6104157 | rs6559122 | -0.909 | -0.036 | 0.238 | -0.398 | 0.01 | -0.086 | -0.086 | 0 | 0.64 | 0.6 | 0.73 | 0.93 | 0.61 | 0.76 |
| 8 | 6106751 | rs4840555 | -1.361 | -0.006 | 0.09 | -0.471 | -0.009 | 0.007 | 0.007 | 0.021 | 0.68 | 0.78 | 0.76 | 0.95 | 0.65 | 0.78 |
| 8 | 6108423 | rs6984741 | -0.523 | 0.024 | 0.3 | -0.433 | 0.017 | -0.039 | -0.039 | 0 | 0.08 | 0.27 | 0.05 | 0.01 | 0.07 | 0.11 |
| 8 | 6112580 | rs4454296 | -0.791 | 0.021 | 0.088 | -0.424 | -0.003 | -0.015 | -0.015 | 0 | 0.63 | 0.62 | 0.72 | 0.92 | 0.57 | 0.68 |
| 8 | 6114142 | rs6983190 | -1.322 | 0.04 | 0.075 | -0.421 | -0.006 | 0.005 | 0.005 | 0.262 | 0.68 | 0.77 | 0.76 | 0.93 | 0.65 | 0.77 |
| 8 | 6114273 | rs4537324 | 0.818 | -0.194 | -0.112 | -0.205 | -0.001 | 0.025 | 0.025 | 0.389 | 0.2 | 0.48 | 0.18 | 0.26 | 0.17 | 0.22 |
| 8 | 6114621 | rs4240683 | -1.045 | -0.121 | -0.433 | -0.181 | 0.018 | -0.081 | -0.081 | 0 | 0.63 | 0.56 | 0.73 | 0.91 | 0.61 | 0.71 |
| 8 | 6116071 | rs6994867 | 1.204 | -0.356 | -1.393 | -0.015 | 0.045 | -0.026 | -0.026 | 0 | 0.18 | 0.05 | 0.1 | 0.14 | 0.09 | 0.11 |
| 8 | 6117252 | rs13257360 | -0.695 | -0.351 | -1.384 | -0.028 | 0.006 | 0.064 | 0.064 | 0.461 | 0.44 | 0.2 | 0.56 | 0.66 | 0.45 | 0.49 |
| 8 | 6118677 | rs12682045 | 0.547 | -0.024 | -1.202 | 0.092 | -0.017 | 0.033 | 0.033 | 0.352 | 0.11 | 0.08 | 0.07 | 0.06 | 0.15 | 0.17 |
| 8 | 6118694 | rs13274647 | -0.414 | 0.132 | -1.127 | 0.175 | -0.006 | -0.077 | -0.077 | 0 | 0.26 | 0.15 | 0.44 | 0.5 | 0.33 | 0.34 |
| 8 | 6119926 | rs17381669 | 0.949 | 0.311 | -1.216 | 0.221 | -0.142 | 0.125 | 0.125 | 0.181 | 0.3 | 0.03 | 0.22 | 0.07 | 0.36 | 0.19 |
| 8 | 6120950 | rs4570185 | -0.81 | 0.305 | -1.172 | 0.238 | 0.05 | -0.027 | -0.027 | 0.685 | 0.58 | 0.21 | 0.69 | 0.59 | 0.71 | 0.6 |
| 8 | 6123137 | rs17466210 | 0.857 | 0.288 | -1.174 | 0.37 | -0.141 | 0.123 | 0.123 | 0.275 | 0.31 | 0.02 | 0.23 | 0.07 | 0.36 | 0.21 |
| 8 | 6123393 | rs17063413 | -1.076 | 0.226 | 0.192 | 1.394 | -0.009 | -0.089 | -0.089 | 0 | 0.25 | 0.14 | 0.44 | 0.52 | 0.31 | 0.39 |
| 8 | 6128828 | rs4841788 | -1.125 | 0.272 | 0.542 | 1.673 | 0.001 | 0.001 | 0.001 | 0 | 0.21 | 0.19 | 0.39 | 0.45 | 0.18 | 0.27 |
| 8 | 6136512 | rs7827867 | -1.085 | 0.737 | -0.716 | 1.547 | -0.003 | 0.017 | 0.017 | 0 | 0.23 | 0.21 | 0.4 | 0.46 | 0.22 | 0.35 |
| 8 | 6138130 | rs11777993 | -1.09 | 0.655 | -0.77 | 1.555 | -0.008 | 0.028 | 0.028 | 0 | 0.23 | 0.23 | 0.39 | 0.46 | 0.22 | 0.35 |
| 8 | 6142331 | rs11136998 | -0.685 | 1.406 | -1.941 | 1.313 | -0.104 | -0.139 | -0.139 | 2.075 | 0.29 | 0.02 | 0.29 | 0.42 | 0.34 | 0.35 |
| 8 | 6150756 | rs4406414 | -0.859 | 1.248 | -1.193 | 1.713 | 0.008 | 0.088 | 0.088 | 0 | 0.09 | 0.15 | 0.16 | 0.34 | 0.11 | 0.15 |
| 8 | 6155658 | rs12679001 | 0.127 | 1.285 | -2.102 | 1.21 | -0.035 | -0.033 | -0.033 | 0 | 0.24 | 0.08 | 0.18 | 0.28 | 0.25 | 0.19 |
| 8 | 6159718 | rs12675721 | 0.066 | 0.932 | -1.971 | 0.359 | -0.006 | 0.043 | 0.043 | 0 | 0.59 | 0.26 | 0.46 | 0.31 | 0.6 | 0.42 |
| 8 | 6164560 | rs7840430 | -0.127 | 0.941 | -0.612 | 0.166 | 0.033 | -0.056 | -0.056 | 0.83 | 0.9 | 0.71 | 0.83 | 0.99 | 0.92 | 0.93 |
| 8 | 6167610 | rs10105162 | -0.264 | 0.805 | -0.399 | 0.664 | -0.004 | 0.043 | 0.043 | 0 | 0.16 | 0.27 | 0.19 | 0.28 | 0.17 | 0.31 |
| 8 | 6168912 | rs4587352 | -0.595 | 0.691 | -0.096 | 0.589 | 0 | 0.022 | 0.022 | 0 | 0.08 | 0.27 | 0.04 | 0.05 | 0.04 | 0.11 |
| 8 | 6169072 | rs13261309 | 0.848 | 0.457 | -0.948 | 0.122 | 0.011 | 0.077 | 0.077 | 0 | 0.44 | 0.11 | 0.3 | 0.32 | 0.39 | 0.26 |
| 8 | 6169577 | rs2166008 | -0.463 | 0.503 | -0.477 | 0.123 | -0.017 | -0.021 | -0.021 | 0 | 0.45 | 0.72 | 0.51 | 0.45 | 0.48 | 0.51 |
| 8 | 6169780 | rs2120482 | 0.766 | 0.479 | 0.104 | 0.268 | -0.001 | 0.062 | 0.062 | 0 | 0.13 | 0.41 | 0.14 | 0.11 | 0.07 | 0.13 |
| 8 | 6170683 | rs7845004 | -1.042 | 0.677 | 0.118 | 0.198 | 0.001 | 0.02 | 0.02 | 0 | 0.36 | 0.47 | 0.54 | 0.41 | 0.36 | 0.54 |
| 8 | 6171324 | rs6999296 | -0.195 | 0.635 | 0.239 | 0.243 | -0.022 | 0.029 | 0.029 | 0.014 | 0.19 | 0.48 | 0.12 | 0.14 | 0.12 | 0.21 |
| 8 | 6173075 | rs10086903 | 1.261 | 0.529 | -0.134 | 0.082 | 0.015 | 0.044 | 0.044 | 0 | 0.4 | 0.13 | 0.29 | 0.33 | 0.34 | 0.28 |
| 8 | 6173651 | rs17076428 | 1.106 | 0.567 | -0.132 | 0.076 | -0.001 | -0.005 | -0.005 | 2.254 | 0.12 | 0.07 | 0.34 | 0.26 | 0.12 | 0.23 |
| 8 | 6173868 | rs17597848 | 0.027 | 0.564 | -0.105 | 0.008 | -0.006 | -0.008 | -0.008 | 0 | 0.22 | 0.25 | 0.12 | 0.09 | 0.16 | 0.21 |
| 8 | 6173931 | rs7843394 | -0.338 | 0.609 | -0.032 | 0.004 | -0.015 | -0.017 | -0.017 | 0.758 | 0.25 | 0.49 | 0.17 | 0.14 | 0.2 | 0.22 |
| 8 | 6175430 | rs7005128 | 1.075 | 0.535 | 0.114 | 0.114 | -0.003 | -0.004 | -0.004 | 0 | 0.19 | 0.2 | 0.11 | 0.08 | 0.15 | 0.08 |
| 8 | 6175643 | rs7005444 | -0.481 | 0.651 | -0.075 | 0.017 | -0.002 | -0.033 | -0.033 | 0 | 0.35 | 0.5 | 0.25 | 0.16 | 0.32 | 0.27 |
| 8 | 6176137 | rs17076436 | -0.655 | 0.786 | -0.13 | 0.139 | 0.008 | 0.002 | 0.002 | 0 | 0.49 | 0.58 | 0.39 | 0.44 | 0.55 | 0.5 |
| 8 | 6176935 | rs9650534 | 1.499 | 0.701 | -0.436 | 0.271 | -0.002 | -0.002 | -0.002 | 0 | 0.18 | 0.19 | 0.24 | 0.33 | 0.28 | 0.25 |
| 8 | 6178302 | rs13265841 | 1.29 | 0.538 | -0.413 | 0.185 | 0.019 | 0.026 | 0.026 | 0 | 0.36 | 0.24 | 0.26 | 0.29 | 0.32 | 0.25 |
| 8 | 6179115 | rs10110812 | 1.01 | 0.651 | -0.612 | 0.275 | -0.003 | 0.038 | 0.038 | 0.525 | 0.13 | 0.18 | 0.35 | 0.25 | 0.13 | 0.2 |
| 8 | 6180270 | rs2120485 | 0.545 | 0.757 | -0.965 | 0.265 | -0.012 | 0.01 | 0.01 | 1.725 | 0.12 | 0.06 | 0.09 | 0.08 | 0.12 | 0.21 |
| 8 | 6182107 | rs12679827 | -0.015 | 0.757 | -0.95 | 0.272 | -0.025 | -0.009 | -0.009 | 3.48 | 0.3 | 0.25 | 0.25 | 0.37 | 0.39 | 0.44 |
| 8 | 6184017 | rs17598589 | 0.173 | 0.517 | -0.967 | 0.518 | -0.01 | 0.011 | 0.011 | 1.665 | 0.12 | 0.07 | 0.09 | 0.08 | 0.12 | 0.21 |
| 8 | 6184254 | rs2053615 | 0.744 | 0.496 | -0.932 | 0.424 | 0.006 | -0.003 | -0.003 | 0 | 0.54 | 0.41 | 0.43 | 0.59 | 0.6 | 0.51 |
| 8 | 6188792 | rs13251950 | 0.081 | 0.802 | -1.181 | 0.653 | -0.013 | 0.025 | 0.025 | 2.238 | 0.11 | 0.05 | 0.09 | 0.03 | 0.11 | 0.11 |
| 8 | 6192831 | rs12550228 | 0.33 | 0.435 | -1.17 | 1.245 | 0.025 | -0.099 | -0.099 | 1.008 | 0.19 | 0.01 | 0.13 | 0.34 | 0.11 | 0.18 |
| 8 | 6194816 | rs9693244 | 0.547 | 0.738 | -0.893 | 1.04 | 0.001 | -0.007 | -0.007 | 0.1 | 0.67 | 0.71 | 0.59 | 0.51 | 0.68 | 0.62 |
| 8 | 6200413 | rs9657422 | -0.311 | 1.034 | -0.49 | 1.017 | 0.008 | 0.057 | 0.057 | 0 | 0.13 | 0.18 | 0.17 | 0.3 | 0.15 | 0.21 |
| 8 | 6204395 | rs2920648 | 0.17 | 1.024 | -0.755 | 0.157 | -0.001 | -0.003 | -0.003 | 0 | 0.41 | 0.36 | 0.27 | 0.16 | 0.45 | 0.17 |
| 8 | 6208138 | rs2916708 | -0.743 | 0.548 | -1.105 | 0.143 | 0.018 | -0.009 | -0.009 | 2.885 | 0.74 | 0.81 | 0.82 | 0.69 | 0.8 | 0.78 |
| 8 | 6209425 | rs4840830 | 0.488 | 0.939 | -1.337 | -0.72 | -0.007 | 0.036 | 0.036 | 0.211 | 0.29 | 0.17 | 0.28 | 0.16 | 0.3 | 0.27 |
| 8 | 6210377 | rs17076494 | 0.595 | 0.597 | -1.027 | -0.95 | -0.001 | -0.039 | -0.039 | 0.405 | 0.2 | 0.21 | 0.23 | 0.02 | 0.27 | 0.15 |
| 8 | 6210460 | rs2916705 | 0.623 | 0.275 | -1.451 | -1.142 | 0.033 | 0.039 | 0.039 | 0 | 0.3 | 0.12 | 0.16 | 0.2 | 0.25 | 0.26 |
| 8 | 6211879 | rs17599378 | -0.853 | 0.831 | -1.286 | -0.593 | -0.005 | -0.001 | -0.001 | 0 | 0.15 | 0.1 | 0.21 | 0.19 | 0.16 | 0.22 |
| 8 | 6214573 | rs4840843 | 0.16 | 0.25 | -1.386 | -0.935 | -0.005 | 0.114 | 0.114 | 0.053 | 0.53 | 0.27 | 0.35 | 0.26 | 0.5 | 0.5 |
| 8 | 6214753 | rs2916701 | -0.763 | -0.508 | -0.953 | -0.624 | -0.023 | -0.04 | -0.04 | 0 | 0.24 | 0.41 | 0.22 | 0.17 | 0.19 | 0.24 |
| 8 | 6214856 | rs11775766 | -0.722 | -0.334 | -1.499 | -0.066 | -0.048 | 0.014 | 0.014 | 0 | 0.2 | 0.05 | 0.17 | 0.12 | 0.21 | 0.15 |
| 8 | 6215165 | rs17076502 | -0.775 | -1.358 | -0.769 | 0.089 | -0.008 | -0.001 | -0.001 | 0 | 0.23 | 0.23 | 0.19 | 0.28 | 0.08 | 0.19 |
| 8 | 6216811 | rs2916698 | 0.6 | -1.831 | -0.189 | -0.399 | -0.099 | -0.098 | -0.098 | 0 | 0.74 | 0.62 | 0.94 | 0.94 | 0.95 | 0.88 |
| 8 | 6220077 | rs1470348 | 1.016 | -2.294 | 0.663 | 0.541 | 0.008 | 0.326 | 0.326 | 0 | 0.93 | 0.55 | 0.81 | 0.69 | 0.98 | 0.83 |
| 8 | 6223125 | rs7001462 | 1.159 | -2.282 | 0.824 | 0.588 | -0.016 | -0.003 | -0.003 | 0 | 0.71 | 0.25 | 0.76 | 0.66 | 0.93 | 0.75 |
| 8 | 6229808 | rs2916674 | -1.465 | -2.184 | 1.195 | 0.533 | -0.081 | -0.017 | -0.017 | 0 | 0.29 | 0.59 | 0.19 | 0.27 | 0.1 | 0.23 |
| 8 | 6232009 | rs17534653 | 1.51 | -1.771 | 0.26 | 0.224 | 0.009 | -0.04 | -0.04 | 0 | 0.4 | 0.17 | 0.25 | 0.45 | 0.4 | 0.49 |
| 8 | 6232277 | rs2920617 | -0.104 | -2.342 | 0.666 | 0.39 | -0.092 | 0.084 | 0.084 | 1.755 | 0.16 | 0.5 | 0.15 | 0.25 | 0.04 | 0.16 |

| | | | | | | | | | | | | | | | | |
|---|---|---|---|---|---|---|---|---|---|---|---|---|---|---|---|---|
| 8 | 6236764 | rs2034140 | 0.88 | -2.368 | 0.808 | 0.39 | -0.01 | 0.246 | 0.246 | 0.217 | 0.9 | 0.47 | 0.85 | 0.71 | 0.94 | 0.83 |
| 8 | 6237397 | rs6997285 | -0.943 | -2.417 | 0.74 | 0.397 | -0.043 | -0.075 | -0.075 | 0 | 0.26 | 0.28 | 0.07 | 0.05 | 0.09 | 0.11 |
| 8 | 6248958 | rs11137015 | 1.183 | -2.386 | 0.436 | 0.431 | 0.022 | 0.004 | 0.004 | 0.728 | 0.69 | 0.24 | 0.79 | 0.7 | 0.91 | 0.75 |
| 8 | 6251986 | rs11997958 | 0.44 | -2.378 | 0.375 | 0.374 | -0.006 | -0.013 | -0.013 | 0 | 0.13 | 0.11 | 0.03 | 0.02 | 0.03 | 0.05 |
| 8 | 6255426 | rs7010345 | 0.249 | -1.816 | -0.079 | 0.406 | -0.01 | -0.019 | -0.019 | 1.155 | 0.1 | 0.03 | 0.23 | 0.22 | 0.11 | 0.14 |
| 8 | 6260430 | rs10104726 | 1.15 | -1.813 | -0.126 | 0.398 | 0.088 | -0.036 | -0.036 | 0 | 0.59 | 0.21 | 0.55 | 0.46 | 0.8 | 0.62 |
| 8 | 6263083 | rs1057187 | 1.336 | -1.757 | -0.155 | 0.409 | 0.103 | -0.009 | -0.009 | 0 | 0.56 | 0.21 | 0.5 | 0.44 | 0.78 | 0.62 |
| 8 | 6264879 | rs2440452 | 0.996 | -1.571 | 0.097 | -0.013 | 0.056 | 0.018 | 0.018 | 1.746 | 0.64 | 0.25 | 0.78 | 0.68 | 0.86 | 0.77 |
| 8 | 6266774 | rs1550697 | -1.183 | -1.333 | 0.205 | -0.154 | 0.033 | 0.024 | 0.024 | 3.153 | 0.41 | 0.67 | 0.26 | 0.32 | 0.22 | 0.25 |
| 8 | 6270096 | rs6995735 | -1.144 | -1.359 | 0.676 | -0.1 | -0.038 | -0.048 | -0.048 | 0 | 0.39 | 0.48 | 0.17 | 0.22 | 0.21 | 0.26 |
| 8 | 6272399 | rs2305022 | 1.058 | -1.365 | 0.579 | -0.103 | 0.005 | 0.004 | 0.004 | 2.56 | 0.6 | 0.38 | 0.83 | 0.78 | 0.79 | 0.76 |
| 8 | 6279379 | rs1530408 | 0.957 | -1.423 | 0.475 | -0.18 | 0.079 | 0.083 | 0.083 | 4.748 | 0.6 | 0.2 | 0.82 | 0.76 | 0.8 | 0.76 |
| 8 | 6283479 | rs17536777 | -0.1 | -1.43 | 0.327 | -0.299 | -0.009 | 0.14 | 0.14 | 2.04 | 0.24 | 0.03 | 0.1 | 0.03 | 0.18 | 0.07 |
| 8 | 6285539 | rs2515564 | 0.804 | -1.489 | 1.039 | -0.283 | -0.043 | -0.056 | -0.056 | 0 | 0.65 | 0.45 | 0.84 | 0.82 | 0.82 | 0.81 |
| 8 | 6286733 | rs2440427 | 0.879 | -1.482 | 1.087 | -0.282 | -0.008 | -0.008 | -0.008 | 0 | 0.64 | 0.36 | 0.83 | 0.77 | 0.82 | 0.81 |
| 8 | 6288113 | rs2442524 | 0.837 | -1.472 | 1.132 | -0.278 | -0.02 | -0.019 | -0.019 | 0.707 | 0.64 | 0.38 | 0.83 | 0.77 | 0.81 | 0.8 |
| 8 | 6290206 | rs2440423 | 0.83 | -1.481 | 1.133 | -0.277 | -0.038 | -0.053 | -0.053 | 0.035 | 0.64 | 0.45 | 0.84 | 0.81 | 0.8 | 0.81 |
| 8 | 6291524 | rs2442517 | 1.643 | -0.367 | 0.802 | -0.583 | 0.019 | 0.12 | 0.12 | 0.277 | 0.46 | 0.19 | 0.58 | 0.74 | 0.46 | 0.66 |
| 8 | 6292473 | rs6559162 | -0.809 | -0.38 | 0.912 | -0.544 | -0.026 | -0.033 | -0.033 | 0.007 | 0.34 | 0.41 | 0.15 | 0.19 | 0.19 | 0.19 |
| 8 | 6296550 | rs2442513 | 1.722 | -0.388 | 0.828 | -0.596 | -0.01 | -0.012 | -0.012 | 0 | 0.94 | 0.74 | 0.98 | 0.99 | 0.99 | 0.97 |
| 8 | 6302154 | rs2083914 | 0.428 | -0.356 | 0.604 | -0.592 | 0.003 | 0.128 | 0.128 | 1.879 | 0.22 | 0.03 | 0.09 | 0.03 | 0.16 | 0.06 |
| 8 | 6302671 | rs2920676 | -0.175 | -0.384 | 0.702 | -0.586 | -0.064 | 0.033 | 0.033 | 0 | 0.14 | 0.48 | 0.06 | 0.17 | 0.03 | 0.16 |
| 8 | 6303025 | rs2584 | 0.875 | -0.212 | 0.601 | -0.531 | -0.108 | 0.064 | 0.064 | 0 | 0.14 | 0.02 | 0.25 | 0.03 | 0.34 | 0.1 |
| 8 | 6303409 | rs2916750 | -1.02 | -0.954 | 1.137 | -0.101 | -0.031 | -0.041 | -0.041 | 0 | 0.36 | 0.57 | 0.15 | 0.2 | 0.2 | 0.22 |
| 8 | 6304694 | rs1129703 | 0.985 | -0.952 | 0.921 | -0.1 | 0.045 | 0.051 | 0.051 | 4.059 | 0.62 | 0.25 | 0.83 | 0.8 | 0.8 | 0.78 |
| 8 | 6308056 | rs2440413 | 0.935 | -0.951 | 0.919 | -0.086 | 0.043 | 0.043 | 0.043 | 3.62 | 0.62 | 0.25 | 0.83 | 0.77 | 0.81 | 0.78 |
| 8 | 6310738 | rs2034143 | 0.977 | -0.949 | 0.928 | -0.084 | 0.036 | 0.037 | 0.037 | 3.96 | 0.62 | 0.27 | 0.84 | 0.77 | 0.81 | 0.78 |
| 8 | 6322703 | rs2916734 | -1.275 | -1.29 | 0.944 | 0.436 | -0.014 | 0 | 0 | 1.637 | 0.39 | 0.6 | 0.26 | 0.46 | 0.2 | 0.33 |
| 8 | 6325975 | rs2440399 | 1.191 | -1.284 | 0.717 | 0.444 | 0.096 | -0.021 | -0.021 | 0 | 0.58 | 0.21 | 0.73 | 0.5 | 0.79 | 0.66 |
| 8 | 6329410 | rs2515585 | 0.192 | -1.306 | 1.106 | 0.405 | -0.047 | -0.036 | -0.036 | 0 | 0.73 | 0.62 | 0.88 | 0.78 | 0.85 | 0.81 |
| 8 | 6331758 | rs2979656 | 1.598 | -1.345 | 0.891 | 0.258 | 0.092 | 0.002 | 0.002 | 1.961 | 0.54 | 0.21 | 0.69 | 0.5 | 0.7 | 0.64 |
| 8 | 6334101 | rs3020285 | 0.337 | -0.986 | 1.003 | 0.465 | -0.016 | 0.034 | 0.034 | 0 | 0.11 | 0.19 | 0.13 | 0.27 | 0.04 | 0.11 |
| 8 | 6334619 | rs11137033 | -0.216 | -0.964 | 0.957 | 0.19 | 0.018 | 0.015 | 0.015 | 1.849 | 0.25 | 0.15 | 0.11 | 0.22 | 0.15 | 0.19 |
| 8 | 6343519 | rs2440445 | -0.99 | -0.905 | 0.847 | 0.185 | 0.003 | -0.088 | -0.088 | 0 | 0.15 | 0.42 | 0.1 | 0.02 | 0.15 | 0.06 |
| 8 | 6347219 | rs13282577 | -0.329 | -1.191 | 0.557 | 0.311 | 0.007 | 0.007 | 0.007 | 1.625 | 0.28 | 0.23 | 0.13 | 0.2 | 0.17 | 0.23 |
| 8 | 6351252 | rs17077154 | 0.306 | -0.869 | 0.078 | 0.619 | -0.019 | 0.035 | 0.035 | 0 | 0.1 | 0.21 | 0.13 | 0.21 | 0.05 | 0.11 |
| 8 | 6352358 | rs1031309 | 0.686 | -0.843 | -0.25 | 0.51 | 0.055 | -0.005 | -0.005 | 0.088 | 0.61 | 0.23 | 0.71 | 0.56 | 0.78 | 0.66 |
| 8 | 6354159 | rs2922806 | 0.663 | -0.259 | 0.074 | 0.753 | -0.011 | -0.004 | -0.004 | 0 | 0.69 | 0.52 | 0.74 | 0.73 | 0.77 | 0.72 |
| 8 | 6354270 | rs2442477 | -1.01 | 0.501 | 0.048 | 0.487 | 0.002 | 0.004 | 0.004 | 0 | 0.48 | 0.42 | 0.67 | 0.69 | 0.55 | 0.59 |
| 8 | 6355321 | rs2916718 | 0.108 | 0.765 | 0.082 | 0.702 | 0.003 | -0.012 | -0.012 | 0 | 0.27 | 0.23 | 0.32 | 0.47 | 0.21 | 0.37 |
| 8 | 6355460 | rs2122279 | -0.314 | 0.936 | -0.113 | 0.551 | -0.037 | -0.039 | -0.039 | 0 | 0.83 | 0.91 | 0.69 | 0.76 | 0.72 | 0.86 |
| 8 | 6356645 | rs2916717 | -0.535 | 0.751 | 0.72 | 0.362 | -0.019 | 0.031 | 0.031 | 0 | 0.35 | 0.47 | 0.37 | 0.54 | 0.27 | 0.48 |
| 8 | 6356937 | rs2442475 | -0.485 | 0.844 | 0.511 | 0.322 | -0.042 | -0.041 | -0.041 | 6.174 | 0.84 | 0.91 | 0.7 | 0.77 | 0.74 | 0.88 |
| 8 | 6359480 | rs2167071 | 1.384 | 0.617 | 0.965 | -0.302 | 0.041 | 0.137 | 0.137 | 0 | 0.43 | 0.2 | 0.27 | 0.21 | 0.39 | 0.28 |
| 8 | 6364922 | rs2515409 | -0.707 | 0.667 | 1.471 | -0.237 | 0.037 | -0.039 | -0.039 | 0 | 0.15 | 0.4 | 0.09 | 0.08 | 0.14 | 0.2 |
| 8 | 6365620 | rs2515413 | -1.866 | 0.229 | 1.108 | -0.512 | -0.017 | -0.026 | -0.026 | 0 | 0.25 | 0.64 | 0.15 | 0.23 | 0.16 | 0.29 |
| 8 | 6367302 | rs2442635 | 0.56 | 0.231 | 1.248 | -0.547 | -0.001 | -0.018 | -0.018 | 1.8 | 0.52 | 0.5 | 0.34 | 0.22 | 0.47 | 0.4 |
| 8 | 6368222 | rs2515418 | 1.316 | 0.194 | 1.49 | -0.6 | 0.001 | 0.008 | 0.008 | 0 | 0.42 | 0.37 | 0.3 | 0.07 | 0.44 | 0.29 |
| 8 | 6368776 | rs2442631 | 0.618 | 0.232 | 1.482 | -0.595 | 0.007 | 0.014 | 0.014 | 0 | 0.51 | 0.45 | 0.33 | 0.22 | 0.45 | 0.4 |
| 8 | 6369896 | rs3780088 | 1.465 | 0.058 | 1.498 | -0.542 | -0.007 | -0.017 | -0.017 | 0 | 0.08 | 0.14 | 0.03 | 0.06 | 0.05 | 0.1 |
| 8 | 6371291 | rs2916747 | -1.27 | 0.035 | 1.632 | -0.566 | -0.043 | 0.013 | 0.013 | 0 | 0.13 | 0.35 | 0.08 | 0.16 | 0.04 | 0.15 |
| 8 | 6373705 | rs4276711 | -0.489 | 0.106 | 1.602 | -0.555 | -0.013 | -0.038 | -0.038 | 0 | 0.16 | 0.23 | 0.06 | 0.07 | 0.09 | 0.16 |
| 8 | 6374065 | rs10102851 | -0.409 | 0.113 | 1.395 | -0.53 | -0.023 | 0.054 | 0.054 | 0.278 | 0.09 | 0.31 | 0.06 | 0.17 | 0.03 | 0.15 |
| 8 | 6374773 | rs2515432 | -0.093 | 0.443 | 1.272 | -0.618 | -0.02 | 0.052 | 0.052 | 0 | 0.6 | 0.72 | 0.38 | 0.28 | 0.51 | 0.49 |
| 8 | 6376877 | rs13281117 | 0.879 | 0.377 | 0.817 | -0.481 | 0.043 | 0.091 | 0.091 | 0 | 0.45 | 0.28 | 0.27 | 0.23 | 0.39 | 0.33 |
| 8 | 6377433 | rs1961222 | 1.759 | 0.385 | 0.71 | -0.488 | -0.027 | 0.206 | 0.206 | 0 | 0.35 | 0.01 | 0.21 | 0.05 | 0.34 | 0.2 |
| 8 | 6378763 | rs3020221 | 0.92 | 0.507 | 0.86 | -0.394 | 0.036 | 0.073 | 0.073 | 0 | 0.45 | 0.31 | 0.28 | 0.23 | 0.38 | 0.33 |
| 8 | 6383599 | rs2515462 | 0.407 | 0.787 | 0.876 | -0.397 | 0.046 | -0.098 | -0.098 | 2.362 | 0.75 | 0.57 | 0.68 | 0.93 | 0.68 | 0.77 |
| 8 | 6386204 | rs6990020 | 0.831 | 0.466 | 0.617 | -0.376 | 0.007 | -0.024 | -0.024 | 2.867 | 0.64 | 0.42 | 0.6 | 0.79 | 0.62 | 0.66 |
| 8 | 6386250 | rs3824310 | 1.718 | 0.706 | -0.16 | 0.146 | 0.009 | 0.086 | 0.086 | 0 | 0.34 | 0.07 | 0.31 | 0.27 | 0.32 | 0.22 |
| 8 | 6386286 | rs2515466 | -0.621 | 0.765 | -0.003 | 0.126 | -0.032 | 0.015 | 0.015 | 0 | 0.84 | 0.9 | 0.7 | 0.89 | 0.76 | 0.88 |
| 8 | 6386620 | rs2442608 | 0.364 | 0.737 | 0.227 | 0.171 | 0.006 | 0.04 | 0.04 | 0 | 0.52 | 0.57 | 0.42 | 0.35 | 0.46 | 0.37 |
| 8 | 6386824 | rs1868553 | -0.44 | 0.712 | 0.325 | 0.148 | 0.005 | -0.09 | -0.09 | 0.736 | 0.18 | 0.51 | 0.12 | 0.04 | 0.14 | 0.16 |
| 8 | 6388306 | rs734704 | -0.569 | 0.706 | 0.224 | 0.178 | -0.002 | 0.002 | 0.002 | 0 | 0.67 | 0.64 | 0.71 | 0.46 | 0.7 | 0.48 |
| 8 | 6388640 | rs2515477 | -0.106 | 0.712 | 0.434 | 0.065 | 0.022 | -0.079 | -0.079 | 0 | 0.16 | 0.38 | 0.1 | 0.04 | 0.13 | 0.16 |
| 8 | 6389216 | rs12674822 | 0.569 | 0.731 | 0.389 | 0.527 | 0.016 | -0.001 | -0.001 | 0.195 | 0.51 | 0.26 | 0.62 | 0.51 | 0.57 | 0.33 |
| 8 | 6390027 | rs2515481 | -0.58 | 0.635 | 0.498 | 0.617 | 0.033 | -0.062 | -0.062 | 1.663 | 0.31 | 0.43 | 0.39 | 0.14 | 0.37 | 0.28 |
| 8 | 6390079 | rs2515482 | -0.476 | 0.633 | 0.516 | 0.637 | 0 | 0.001 | 0.001 | 0 | 0.67 | 0.66 | 0.71 | 0.55 | 0.7 | 0.49 |
| 8 | 6390530 | rs17077419 | 1.511 | 0.654 | 0.421 | 0.658 | 0.022 | -0.013 | -0.013 | 0 | 0.36 | 0.21 | 0.31 | 0.42 | 0.33 | 0.21 |

| | | | | | | | | | | | | | | | | |
|---|---|---|---|---|---|---|---|---|---|---|---|---|---|---|---|---|
| 8 | 6391302 | rs2959812 | -0.484 | 0.754 | 0.228 | 0.722 | 0.001 | -0.008 | -0.008 | 0 | 0.68 | 0.69 | 0.71 | 0.57 | 0.7 | 0.51 |
| 8 | 6393780 | rs2959809 | -0.701 | 0.775 | 0.117 | 0.47 | 0.009 | -0.019 | -0.019 | 0 | 0.3 | 0.3 | 0.38 | 0.14 | 0.36 | 0.27 |
| 8 | 6394821 | rs2442598 | -0.141 | 0.666 | -0.085 | 0.992 | -0.002 | -0.019 | -0.019 | 0.268 | 0.7 | 0.74 | 0.8 | 0.55 | 0.69 | 0.62 |
| 8 | 6394876 | rs12541780 | -0.104 | 0.332 | 0.007 | 0.963 | -0.003 | 0.001 | 0.001 | 0 | 0.15 | 0.16 | 0.25 | 0.16 | 0.11 | 0.12 |
| 8 | 6395909 | rs11989215 | 0.734 | 0.01 | -0.071 | 1.281 | -0.004 | 0.002 | 0.002 | 0 | 0.65 | 0.42 | 0.57 | 0.62 | 0.68 | 0.62 |
| 8 | 6396182 | rs11137037 | -0.395 | -0.084 | -0.425 | 1.04 | -0.017 | -0.022 | -0.022 | 0 | 0.37 | 0.55 | 0.37 | 0.29 | 0.3 | 0.39 |
| 8 | 6396341 | rs17623313 | 1.45 | -0.013 | -0.783 | 0.753 | -0.054 | 0.069 | 0.069 | 0 | 0.42 | 0.17 | 0.38 | 0.22 | 0.48 | 0.31 |
| 8 | 6396696 | rs2922876 | 0.638 | -0.143 | -0.379 | 0.936 | -0.009 | -0.022 | -0.022 | 0 | 0.15 | 0.23 | 0.24 | 0.07 | 0.13 | 0.13 |
| 8 | 6396763 | rs13269021 | 0.721 | 0.166 | -1.054 | 1.142 | -0.011 | -0.026 | -0.026 | 0 | 0.27 | 0.19 | 0.25 | 0.42 | 0.27 | 0.31 |
| 8 | 6396870 | rs1375668 | 1.013 | -0.382 | -0.021 | 1.062 | 0.01 | 0.049 | 0.049 | 0 | 0.69 | 0.24 | 0.62 | 0.91 | 0.66 | 0.76 |
| 8 | 6396986 | rs2897911 | 1.133 | -0.314 | -0.017 | 1.213 | 0.032 | 0.015 | 0.015 | 0 | 0.62 | 0.25 | 0.59 | 0.64 | 0.68 | 0.58 |
| 8 | 6398327 | rs2922873 | 0.481 | 0.39 | -0.399 | 1.523 | -0.008 | 0.011 | 0.011 | 0 | 0.82 | 0.73 | 0.86 | 0.8 | 0.84 | 0.78 |
| 8 | 6402366 | rs4478599 | 0.223 | 0.596 | -0.685 | 1.057 | 0.003 | 0 | 0 | 0 | 0.51 | 0.53 | 0.5 | 0.46 | 0.4 | 0.3 |
| 8 | 6402367 | rs13250248 | -0.271 | 1.406 | -0.518 | 0.757 | 0.007 | -0.003 | -0.003 | 4.314 | 0.19 | 0.21 | 0.11 | 0.16 | 0.21 | 0.13 |
| 8 | 6403006 | rs2515488 | -0.054 | 0.605 | -0.416 | 0.625 | -0.003 | -0.003 | -0.003 | 0 | 0.44 | 0.58 | 0.45 | 0.51 | 0.49 | 0.56 |
| 8 | 6404947 | rs17553089 | -1.138 | 0.344 | -0.151 | 0.452 | -0.009 | -0.02 | -0.02 | 2.638 | 0.14 | 0.02 | 0.14 | 0.14 | 0.11 | 0.25 |
| 8 | 6405608 | rs2442592 | 1.741 | 0.004 | 0.387 | 0.308 | -0.004 | -0.001 | -0.001 | 0 | 0.72 | 0.31 | 0.76 | 0.73 | 0.8 | 0.61 |
| 8 | 6406572 | rs2515493 | 1.42 | 0.132 | 1.087 | 0.193 | 0.007 | 0.003 | 0.003 | 0 | 0.85 | 0.61 | 0.85 | 0.84 | 0.87 | 0.71 |
| 8 | 6413948 | rs12543837 | -1.619 | 0.144 | 0.88 | 0.083 | 0.005 | 0.005 | 0.005 | 0 | 0.12 | 0.07 | 0.14 | 0.11 | 0.12 | 0.18 |
| 8 | 6420534 | rs3739391 | 2.211 | 0.059 | 1.105 | 0.065 | 0.002 | 0.001 | 0.001 | 0 | 0.79 | 0.24 | 0.79 | 0.83 | 0.85 | 0.73 |
| 8 | 6420574 | rs3739392 | -1.542 | 0.049 | 1.148 | 0.072 | -0.004 | -0.002 | -0.002 | 14.752 | 0.1 | 0.06 | 0.13 | 0.11 | 0.12 | 0.17 |
| 8 | 6421927 | rs2442509 | 1.307 | 0.078 | 1.414 | 0.075 | 0.009 | -0.024 | -0.024 | 0 | 0.87 | 0.36 | 0.87 | 0.93 | 0.9 | 0.82 |
| 8 | 6421998 | rs2515508 | 1.307 | 0.078 | 1.414 | 0.074 | 0.009 | -0.027 | -0.027 | 2.339 | 0.87 | 0.36 | 0.87 | 0.94 | 0.9 | 0.82 |
| 8 | 6423978 | rs7817104 | -1.129 | -0.02 | 1.094 | -0.035 | -0.004 | -0.001 | -0.001 | 0 | 0.08 | 0.07 | 0.06 | 0.11 | 0.05 | 0.09 |
| 8 | 6427868 | rs2959776 | -1.681 | -0.067 | 1.682 | -0.075 | -0.051 | 0.009 | 0.009 | 0 | 0.13 | 0.35 | 0.1 | 0.11 | 0.07 | 0.1 |
| 8 | 6431666 | rs7833581 | -1.412 | 0.201 | 1.408 | -0.079 | 0.006 | -0.046 | -0.046 | 1.513 | 0.09 | 0.21 | 0.1 | 0.01 | 0.09 | 0.18 |
| 8 | 6433935 | rs2959777 | -1.424 | 0.814 | 0.781 | -0.583 | 0.005 | -0.046 | -0.046 | 0.47 | 0.31 | 0.58 | 0.31 | 0.12 | 0.35 | 0.36 |
| 8 | 6435402 | rs2442589 | -0.6 | 1.505 | 0.062 | -0.875 | -0.046 | 0.06 | 0.06 | 2.226 | 0.22 | 0.11 | 0.18 | 0.08 | 0.29 | 0.2 |
| 8 | 6435875 | rs2922861 | 0.54 | 0.864 | 0.225 | -1.648 | -0.001 | -0.01 | -0.01 | 0 | 0.48 | 0.54 | 0.69 | 0.85 | 0.55 | 0.48 |
| 8 | 6438242 | rs2515521 | 1.104 | 0.873 | 0.517 | -1.716 | -0.004 | 0.03 | 0.03 | 0.303 | 0.36 | 0.31 | 0.53 | 0.83 | 0.44 | 0.33 |
| 8 | 6444567 | rs17077621 | 0.383 | 0.401 | 0.628 | -1.505 | 0.045 | 0.091 | 0.091 | 0 | 0.34 | 0.19 | 0.12 | 0.07 | 0.2 | 0.33 |
| 8 | 6445786 | rs2442581 | 0.496 | 0.695 | 0.608 | -0.319 | -0.037 | -0.063 | -0.063 | 0.348 | 0.2 | 0.13 | 0.25 | 0.43 | 0.29 | 0.23 |
| 8 | 6445827 | rs3020246 | 0.529 | 0.622 | 1.099 | -0.258 | -0.001 | -0.002 | -0.002 | 0 | 0.43 | 0.42 | 0.61 | 0.5 | 0.49 | 0.42 |
| 8 | 6449368 | rs9657427 | -0.483 | 0.732 | 0.828 | -0.193 | 0.023 | -0.038 | -0.038 | 0.432 | 0.08 | 0.18 | 0.1 | 0.01 | 0.11 | 0.14 |
| 8 | 6453371 | rs11777882 | 1.348 | 0.599 | 0.172 | -0.078 | -0.072 | -0.068 | -0.068 | 0 | 0.3 | 0.02 | 0.41 | 0.3 | 0.32 | 0.22 |
| 8 | 6454104 | rs3020249 | 0.968 | 0.509 | 0.832 | -0.143 | 0.001 | -0.012 | -0.012 | 0.423 | 0.34 | 0.29 | 0.49 | 0.49 | 0.35 | 0.32 |
| 8 | 6454442 | rs2442573 | 0.624 | 0.682 | 0.386 | -0.217 | 0.004 | -0.019 | -0.019 | 2.896 | 0.2 | 0.21 | 0.17 | 0.08 | 0.25 | 0.19 |
| 8 | 6454559 | rs2442572 | -0.569 | 0.623 | 0.509 | -0.267 | 0.003 | -0.002 | -0.002 | 0 | 0.56 | 0.38 | 0.43 | 0.5 | 0.58 | 0.59 |
| 8 | 6455279 | rs4841336 | 0.422 | 0.499 | 0.381 | -0.157 | -0.002 | 0.09 | 0.09 | 0 | 0.36 | 0.17 | 0.19 | 0.08 | 0.31 | 0.35 |
| 8 | 6455556 | rs2922852 | 0.207 | 0.436 | 0.376 | -0.148 | 0.002 | -0.011 | -0.011 | 0 | 0.37 | 0.26 | 0.27 | 0.42 | 0.34 | 0.41 |
| 8 | 6455701 | rs2922851 | 0.298 | 0.295 | -0.015 | 0.057 | 0 | -0.001 | -0.001 | 0.899 | 0.32 | 0.62 | 0.47 | 0.46 | 0.28 | 0.33 |
| 8 | 6456370 | rs2442570 | -0.287 | 0.14 | 0.453 | 0.91 | 0.014 | 0.43 | 0.43 | 0 | 0.89 | 0.57 | 0.88 | 0.47 | 0.94 | 0.84 |
| 8 | 6456430 | rs2442569 | -0.679 | 0.362 | -0.297 | 0.177 | -0.026 | 0.053 | 0.053 | 1.072 | 0.35 | 0.79 | 0.26 | 0.22 | 0.35 | 0.28 |
| 8 | 6458438 | rs2442562 | 0.069 | 0.204 | -1.158 | 0.479 | 0.039 | -0.011 | -0.011 | 0 | 0.29 | 0.43 | 0.3 | 0.27 | 0.38 | 0.29 |
| 8 | 6459642 | rs17631450 | 0.742 | 0.029 | -2.728 | -0.335 | -0.026 | 0.011 | 0.011 | 0 | 0.13 | 0.02 | 0.08 | 0.07 | 0.12 | 0.28 |
| 8 | 6460877 | rs11785507 | 0.063 | 0.207 | -2.627 | -0.161 | -0.053 | 0.06 | 0.06 | 0.304 | 0.59 | 0.18 | 0.55 | 0.26 | 0.52 | 0.62 |
| 8 | 6461802 | rs17077658 | 0.415 | 0.137 | -2.291 | 0.499 | -0.047 | 0.019 | 0.019 | 0 | 0.32 | 0.02 | 0.42 | 0.2 | 0.31 | 0.33 |
| 8 | 6464034 | rs2454518 | 0.554 | 0.012 | -0.954 | 1.102 | -0.002 | -0.001 | -0.001 | 0 | 0.66 | 0.35 | 0.47 | 0.26 | 0.77 | 0.69 |
| 8 | 6469600 | rs1077701 | 0.676 | 0.814 | -0.897 | 1.234 | 0.001 | 0.055 | 0.055 | 0 | 0.49 | 0.38 | 0.51 | 0.22 | 0.55 | 0.5 |
| 8 | 6471182 | rs3020258 | -0.082 | 0.712 | -1.312 | 0.962 | -0.082 | 0.074 | 0.074 | 0 | 0.18 | 0.01 | 0.1 | 0.03 | 0.21 | 0.14 |
| 8 | 6471526 | rs13276344 | 1.114 | 0.601 | -0.746 | 0.969 | 0.001 | 0.048 | 0.048 | 0 | 0.51 | 0.42 | 0.53 | 0.23 | 0.58 | 0.53 |
| 8 | 6472105 | rs2959801 | -0.673 | 0.529 | -0.214 | 0.637 | -0.012 | 0.063 | 0.063 | 0 | 0.31 | 0.18 | 0.17 | 0.05 | 0.3 | 0.21 |
| 8 | 6472795 | rs3020259 | -1.376 | 0.136 | 0.841 | 0.971 | 0.001 | 0.002 | 0.002 | 0 | 0.44 | 0.55 | 0.36 | 0.36 | 0.36 | 0.41 |
| 8 | 6473288 | rs2959800 | -1.202 | 0.202 | 1.205 | 1.322 | 0.004 | 0.086 | 0.086 | 0.372 | 0.12 | 0.18 | 0.3 | 0.64 | 0.13 | 0.25 |
| 8 | 6473715 | rs2959799 | -1.411 | 0.203 | 1.244 | 1.358 | -0.003 | -0.003 | -0.003 | 0.569 | 0.45 | 0.54 | 0.38 | 0.4 | 0.37 | 0.42 |
| 8 | 6473746 | rs2959798 | -1.461 | 0.211 | 1.243 | 1.361 | 0.003 | 0.003 | 0.003 | 0 | 0.46 | 0.57 | 0.38 | 0.4 | 0.37 | 0.42 |
| 8 | 6473758 | rs2454517 | -0.495 | 0.145 | 1.325 | 1.337 | -0.022 | 0.128 | 0.128 | 0.614 | 0.32 | 0.06 | 0.15 | 0.09 | 0.29 | 0.21 |
| 8 | 6474136 | rs2936501 | -1.403 | 0.169 | 1.326 | 1.35 | -0.002 | -0.002 | -0.002 | 0.462 | 0.44 | 0.46 | 0.37 | 0.4 | 0.37 | 0.42 |
| 8 | 6475904 | rs2013938 | -1.745 | 0.433 | 1.279 | 1.006 | 0.007 | 0.042 | 0.042 | 0 | 0.49 | 0.65 | 0.5 | 0.74 | 0.42 | 0.46 |
| 8 | 6477605 | rs1971314 | -1.437 | 0.475 | 1.384 | 0.998 | 0.005 | 0.027 | 0.027 | 0 | 0.46 | 0.67 | 0.48 | 0.71 | 0.41 | 0.45 |
| 8 | 6478986 | rs2912010 | -1.338 | 0.496 | 1.619 | 0.988 | -0.002 | 0.007 | 0.007 | 0 | 0.44 | 0.34 | 0.35 | 0.68 | 0.41 | 0.49 |
| 8 | 6479042 | rs1057090 | -1.467 | 0.456 | 1.696 | 0.995 | 0 | 0 | 0 | 0 | 0.46 | 0.45 | 0.36 | 0.67 | 0.41 | 0.48 |
| 8 | 6479178 | rs2912016 | -1.243 | 0.42 | 1.659 | 1.111 | -0.001 | 0 | 0 | 0 | 0.43 | 0.43 | 0.29 | 0.44 | 0.36 | 0.4 |
| 8 | 6479432 | rs2515559 | -0.434 | 0.443 | 1.147 | 1.083 | -0.02 | 0.156 | 0.156 | 0.465 | 0.31 | 0.05 | 0.14 | 0.07 | 0.28 | 0.23 |
| 8 | 6481697 | rs17077721 | -0.838 | 0.333 | 1.174 | 1.235 | -0.027 | -0.03 | -0.03 | 0.019 | 0.08 | 0.01 | 0.12 | 0.35 | 0.08 | 0.15 |
| 8 | 6484968 | rs3020261 | -1.581 | 0.528 | 1.538 | 1.101 | 0.004 | 0.02 | 0.02 | 0 | 0.48 | 0.65 | 0.46 | 0.68 | 0.41 | 0.49 |
| 8 | 6487449 | rs3020263 | -0.577 | 0.554 | 1.748 | 1.025 | 0 | -0.011 | -0.011 | 2.124 | 0.31 | 0.28 | 0.15 | 0.09 | 0.28 | 0.21 |
| 8 | 6488577 | rs3020264 | -0.493 | 0.539 | 1.666 | 1.025 | 0 | 0.064 | 0.064 | 0 | 0.31 | 0.17 | 0.14 | 0.09 | 0.28 | 0.21 |
| 8 | 6488921 | rs3020265 | -1.379 | 0.54 | 1.539 | 1.159 | 0.002 | 0.008 | 0.008 | 0 | 0.44 | 0.6 | 0.43 | 0.71 | 0.38 | 0.48 |

| | | | | | | | | | | | | | | | | |
|---|---|---|---|---|---|---|---|---|---|---|---|---|---|---|---|---|
| 8 | 6491035 | rs2911974 | -1.614 | 0.326 | 1.205 | 0.69 | -0.003 | 0.036 | 0.036 | 0 | 0.53 | 0.63 | 0.3 | 0.33 | 0.44 | 0.44 |
| 8 | 6493910 | rs2042670 | -0.673 | 0.413 | 0.76 | 0.357 | -0.012 | 0.093 | 0.093 | 0 | 0.35 | 0.16 | 0.16 | 0.09 | 0.32 | 0.26 |
| 8 | 6494383 | rs2042668 | -1.626 | 0.346 | 1.031 | 0.65 | -0.005 | 0.006 | 0.006 | 0 | 0.53 | 0.67 | 0.52 | 0.59 | 0.47 | 0.58 |
| 8 | 6494655 | rs2911991 | -1.455 | 0.244 | 0.96 | 0.637 | 0.007 | 0.005 | 0.005 | 0 | 0.51 | 0.55 | 0.47 | 0.59 | 0.42 | 0.57 |
| 8 | 6500544 | rs1057091 | -1.246 | 0.155 | 0.447 | -0.117 | 0.027 | 0.069 | 0.069 | 0.001 | 0.36 | 0.17 | 0.18 | 0.21 | 0.3 | 0.36 |
| 8 | 6502152 | rs2433150 | -0.707 | 0.121 | 0.481 | -0.174 | 0.008 | 0.005 | 0.005 | 0 | 0.26 | 0.23 | 0.11 | 0.09 | 0.18 | 0.2 |
| 8 | 6514185 | rs6997076 | -0.388 | 0.071 | 0.229 | -0.014 | 0.02 | 0.022 | 0.022 | 0 | 0.14 | 0.06 | 0.04 | 0.07 | 0.07 | 0.07 |
| 8 | 6518191 | rs10867020 | 1.191 | 0.003 | 0.333 | -0.348 | 0.008 | 0.092 | 0.092 | 0 | 0.56 | 0.3 | 0.47 | 0.18 | 0.61 | 0.5 |
| 8 | 6520628 | rs13281444 | 0.668 | -0.108 | 1.039 | -0.306 | -0.007 | -0.027 | -0.027 | 0 | 0.64 | 0.53 | 0.65 | 0.32 | 0.7 | 0.6 |
| 8 | 6521432 | rs4841493 | -1.277 | -0.283 | 1.203 | -0.356 | -0.053 | -0.02 | -0.02 | 0.885 | 0.27 | 0.45 | 0.23 | 0.22 | 0.16 | 0.2 |
| 8 | 6527846 | rs11775232 | 1.501 | -0.294 | 1.13 | -0.398 | 0.023 | 0.125 | 0.125 | 0 | 0.53 | 0.33 | 0.46 | 0.17 | 0.61 | 0.45 |
| 8 | 6529048 | rs17077835 | 0.159 | -0.295 | 0.954 | -0.364 | 0.004 | -0.026 | -0.026 | 4.671 | 0.07 | 0.14 | 0.1 | 0.01 | 0.08 | 0.06 |
| 8 | 6531541 | rs11137053 | -1.554 | -0.287 | 0.979 | -0.42 | -0.015 | -0.005 | -0.005 | 0 | 0.3 | 0.63 | 0.42 | 0.26 | 0.19 | 0.26 |
| 8 | 6533492 | rs6559174 | -0.169 | -0.282 | 0.709 | -0.407 | -0.043 | -0.034 | -0.034 | 0 | 0.17 | 0.33 | 0.06 | 0.09 | 0.06 | 0.05 |
| 8 | 6533966 | rs12542460 | 1.876 | -0.186 | -0.108 | -0.435 | -0.142 | 0.274 | 0.274 | 0 | 0.45 | 0.03 | 0.32 | 0.14 | 0.55 | 0.37 |
| 8 | 6534000 | rs6559175 | 0.036 | 0.238 | 0.585 | 0.137 | 0.013 | 0.032 | 0.032 | 0 | 0.79 | 0.72 | 0.75 | 0.4 | 0.76 | 0.65 |
| 8 | 6535046 | rs7825811 | -1.324 | -0.039 | 0.331 | -0.127 | -0.025 | -0.03 | -0.03 | 0.342 | 0.12 | 0.2 | 0.2 | 0.05 | 0.05 | 0.17 |
| 8 | 6539289 | rs6559176 | -0.407 | 0.418 | -0.596 | -0.89 | -0.002 | 0.016 | 0.016 | 0 | 0.47 | 0.6 | 0.58 | 0.83 | 0.45 | 0.5 |
| 8 | 6539987 | rs1014199 | -0.216 | 0.409 | -1.173 | -0.558 | -0.018 | -0.014 | -0.014 | 0 | 0.05 | 0.03 | 0.21 | 0.06 | 0.06 | 0.08 |
| 8 | 6554249 | rs11137055 | 0.585 | 0.086 | -1.539 | -1.014 | -0.101 | 0.227 | 0.227 | 0 | 0.42 | 0.07 | 0.37 | 0.13 | 0.5 | 0.46 |
| 8 | 6554615 | rs11783611 | 0.585 | 0.1 | -1.539 | -1.014 | -0.104 | 0.227 | 0.227 | 0 | 0.42 | 0.07 | 0.38 | 0.14 | 0.5 | 0.46 |
| 8 | 6562092 | rs10108451 | -0.244 | -0.736 | -0.847 | -0.319 | 0.003 | -0.005 | -0.005 | 0 | 0.65 | 0.54 | 0.69 | 0.66 | 0.63 | 0.58 |
| 8 | 6564609 | rs1834450 | -0.458 | -0.759 | -0.723 | -0.287 | -0.004 | 0.046 | 0.046 | 3.678 | 0.09 | 0.36 | 0.06 | 0.06 | 0.05 | 0.02 |
| 8 | 6572165 | rs10503368 | 1.063 | -1.355 | -1.614 | 0.092 | -0.106 | -0.043 | -0.043 | 2.877 | 0.34 | 0.07 | 0.3 | 0.33 | 0.44 | 0.24 |
| 8 | 6580426 | rs10110103 | -0.417 | -1.58 | -1.73 | -0.219 | -0.047 | 0.059 | 0.059 | 0 | 0.63 | 0.89 | 0.72 | 0.72 | 0.61 | 0.57 |
| 8 | 6582485 | rs2980686 | -1.148 | -1.164 | -1.251 | 0.084 | -0.049 | 0.02 | 0.02 | 0 | 0.74 | 0.94 | 0.83 | 0.79 | 0.7 | 0.59 |
| 8 | 6585864 | rs2980684 | -1.082 | -1.177 | -1.23 | 0.116 | -0.086 | 0.015 | 0.015 | 0 | 0.73 | 0.97 | 0.82 | 0.79 | 0.68 | 0.58 |
| 8 | 6594940 | rs12676235 | 1.054 | -1.363 | -0.772 | 0.128 | -0.014 | -0.01 | -0.01 | 0 | 0.4 | 0.32 | 0.48 | 0.42 | 0.48 | 0.32 |
| 8 | 6599005 | rs2936512 | -1.213 | -1.209 | -0.802 | 0.034 | -0.117 | -0.031 | -0.031 | 0 | 0.74 | 0.98 | 0.77 | 0.74 | 0.69 | 0.58 |
| 8 | 6604403 | rs7012528 | 0.836 | -1.488 | -0.548 | -0.035 | 0.006 | -0.006 | -0.006 | 0 | 0.42 | 0.46 | 0.58 | 0.67 | 0.5 | 0.33 |
| 8 | 6605662 | rs2912078 | 1.116 | -1.333 | -0.494 | -0.018 | 0.017 | 0.03 | 0.03 | 0 | 0.37 | 0.47 | 0.44 | 0.48 | 0.44 | 0.27 |
| 8 | 6610163 | rs2928579 | -1.102 | -1.641 | -0.436 | 0.35 | -0.044 | -0.035 | -0.035 | 0 | 0.74 | 0.99 | 0.8 | 0.73 | 0.77 | 0.6 |
| 8 | 6612219 | rs3824307 | 0.347 | -1.599 | -0.358 | 0.466 | 0.004 | -0.017 | -0.017 | 0.553 | 0.18 | 0.05 | 0.2 | 0.17 | 0.14 | 0.19 |
| 8 | 6613787 | rs1559746 | 0.551 | -1.655 | -0.145 | 0.453 | 0.004 | -0.001 | -0.001 | 2.456 | 0.48 | 0.51 | 0.54 | 0.48 | 0.59 | 0.32 |
| 8 | 6615754 | rs1046449 | -0.081 | -1.011 | -0.387 | 0.577 | 0.013 | -0.003 | -0.003 | 3.578 | 0.18 | 0.08 | 0.21 | 0.18 | 0.15 | 0.21 |
| 8 | 6622685 | rs13281818 | -0.072 | -0.795 | -0.632 | 0.657 | 0.005 | -0.003 | -0.003 | 0 | 0.18 | 0.15 | 0.21 | 0.19 | 0.17 | 0.18 |
| 8 | 6623285 | rs2928596 | -1.083 | -0.535 | -0.975 | 0.794 | -0.115 | -0.039 | -0.039 | 0 | 0.69 | 0.91 | 0.67 | 0.72 | 0.52 | 0.77 |
| 8 | 6628841 | rs3990015 | -0.108 | -0.642 | -0.246 | 0.8 | 0.02 | -0.019 | -0.019 | 0.041 | 0.15 | 0.12 | 0.11 | 0.23 | 0.12 | 0.2 |
| 8 | 6628927 | rs17078076 | -0.458 | -0.706 | 0.287 | 0.778 | -0.039 | 0.025 | 0.025 | 0 | 0.17 | 0.31 | 0.2 | 0.24 | 0.12 | 0.2 |
| 8 | 6631138 | rs2936524 | -0.678 | -0.17 | 0.374 | 0.761 | 0.011 | 0.007 | 0.007 | 0 | 0.31 | 0.58 | 0.4 | 0.34 | 0.34 | 0.28 |
| 8 | 6631197 | rs2936523 | -0.3 | -0.124 | 0.369 | 0.786 | 0.009 | 0.004 | 0.004 | 0 | 0.25 | 0.34 | 0.24 | 0.26 | 0.31 | 0.19 |
| 8 | 6633702 | rs17637986 | 0.253 | -0.53 | -0.191 | 0.614 | 0.027 | 0.026 | 0.026 | 0 | 0.17 | 0.02 | 0.05 | 0.08 | 0.09 | 0.24 |
| 8 | 6634008 | rs17078102 | -0.647 | -0.77 | 0.375 | 0.699 | 0.01 | 0.088 | 0.088 | 2.904 | 0.08 | 0.32 | 0.13 | 0.09 | 0.01 | 0.07 |
| 8 | 6634129 | rs13277331 | 0.444 | -0.512 | -0.058 | 0.716 | -0.005 | -0.001 | -0.001 | 0.014 | 0.14 | 0.12 | 0.19 | 0.13 | 0.21 | 0.13 |
| 8 | 6634373 | rs12549951 | 0.542 | -0.49 | -0.216 | 0.735 | -0.054 | -0.017 | -0.017 | 0 | 0.14 | 0.04 | 0.19 | 0.13 | 0.21 | 0.13 |
| 8 | 6634774 | rs2912089 | -1.135 | -1.142 | 0.505 | 0.829 | -0.028 | -0.003 | -0.003 | 0.142 | 0.31 | 0.57 | 0.23 | 0.28 | 0.15 | 0.34 |
| 8 | 6635669 | rs1077489 | -0.569 | -0.003 | 0.382 | 0.988 | 0.003 | 0.022 | 0.022 | 0 | 0.07 | 0.14 | 0.05 | 0.11 | 0.05 | 0.03 |
| 8 | 6637188 | rs17078124 | -1.451 | -0.007 | 0.837 | 1.247 | 0.035 | 0.084 | 0.084 | 1.004 | 0.11 | 0.31 | 0.11 | 0.21 | 0.12 | 0.11 |
| 8 | 6637851 | rs2912045 | -1.622 | -0.283 | 1.362 | 1.207 | 0.049 | 0.159 | 0.159 | 2.402 | 0.06 | 0.43 | 0.03 | 0.08 | 0 | 0.14 |
| 8 | 6638081 | rs2978931 | -1.361 | -0.639 | 0.976 | 1.202 | -0.026 | 0.069 | 0.069 | 0 | 0.16 | 0.37 | 0.15 | 0.28 | 0.11 | 0.23 |
| 8 | 6638899 | rs2912024 | -1.755 | -0.663 | 1.041 | 1.181 | -0.039 | 0.064 | 0.064 | 0 | 0.19 | 0.55 | 0.15 | 0.28 | 0.11 | 0.24 |
| 8 | 6639240 | rs2936519 | -1.567 | -0.612 | 1.067 | 1.182 | -0.029 | 0.089 | 0.089 | 0 | 0.18 | 0.46 | 0.15 | 0.28 | 0.11 | 0.24 |
| 8 | 6640209 | rs2912023 | -1.913 | -1.339 | 0.598 | 0.675 | -0.028 | -0.005 | -0.005 | 2.433 | 0.35 | 0.58 | 0.2 | 0.35 | 0.19 | 0.43 |
| 8 | 6640355 | rs1974947 | 0.054 | -1.32 | 0.538 | 0.742 | 0.03 | 0.03 | 0.03 | 0 | 0.17 | 0.1 | 0.05 | 0.07 | 0.09 | 0.19 |
| 8 | 6640527 | rs1974949 | -1.41 | -1.241 | 0.389 | 0.773 | -0.033 | -0.001 | -0.001 | 0 | 0.33 | 0.59 | 0.21 | 0.35 | 0.2 | 0.33 |
| 8 | 6645394 | rs1541749 | -1.494 | -1.043 | 0.125 | 0.738 | -0.001 | -0.001 | -0.001 | 1.078 | 0.37 | 0.63 | 0.22 | 0.27 | 0.24 | 0.43 |
| 8 | 6645836 | rs2912052 | -1.587 | -1.096 | 0.075 | 0.885 | 0.003 | 0 | 0 | 0 | 0.39 | 0.64 | 0.25 | 0.38 | 0.25 | 0.44 |
| 8 | 6648542 | rs9693942 | 0.205 | -0.436 | -0.472 | 0.812 | -0.002 | 0.001 | 0.001 | 0 | 0.54 | 0.5 | 0.5 | 0.54 | 0.46 | 0.4 |
| 8 | 6655391 | rs966561 | -0.325 | -0.513 | -0.488 | 0.772 | -0.001 | 0 | 0 | 0 | 0.45 | 0.66 | 0.53 | 0.46 | 0.56 | 0.55 |
| 8 | 6656981 | rs10503365 | 0.432 | -1.178 | -0.731 | 0.673 | 0.021 | 0.01 | 0.01 | 0 | 0.16 | 0.11 | 0.05 | 0.11 | 0.05 | 0.16 |
| 8 | 6659754 | rs1469272 | -1.043 | -0.356 | -1.068 | 0.759 | -0.064 | -0.006 | -0.006 | 1.357 | 0.64 | 0.84 | 0.63 | 0.67 | 0.57 | 0.76 |
| 8 | 6661587 | rs12381471 | -0.374 | -0.598 | -0.599 | 0.778 | -0.041 | 0.024 | 0.024 | 0 | 0.17 | 0.26 | 0.18 | 0.25 | 0.04 | 0.25 |
| 8 | 6664969 | rs2553723 | 0.338 | -0.3 | -0.86 | 0.596 | 0.011 | 0.011 | 0.011 | 0 | 0.36 | 0.48 | 0.47 | 0.41 | 0.45 | 0.34 |
| 8 | 6665227 | rs17078194 | -0.311 | -0.529 | -0.566 | 0.812 | -0.022 | 0.028 | 0.028 | 0 | 0.06 | 0.14 | 0.15 | 0.14 | 0.05 | 0.08 |
| 8 | 6665742 | rs17078199 | -0.922 | -0.617 | -0.623 | 0.769 | -0.028 | 0.069 | 0.069 | 0.785 | 0.1 | 0.38 | 0.17 | 0.14 | 0.04 | 0.1 |
| 8 | 6666488 | rs17078203 | 0.238 | -0.35 | -1.557 | 0.752 | 0.018 | -0.038 | -0.038 | 0 | 0.13 | 0.01 | 0.04 | 0.14 | 0.05 | 0.17 |
| 8 | 6669949 | rs2553728 | -0.574 | 0.189 | -1.642 | 0.369 | 0.029 | 0.062 | 0.062 | 0 | 0.5 | 0.68 | 0.44 | 0.62 | 0.38 | 0.63 |
| 8 | 6674458 | rs9773025 | -0.812 | 0.886 | -1.576 | 0.172 | -0.032 | 0.106 | 0.106 | 0 | 0.58 | 0.73 | 0.64 | 0.94 | 0.47 | 0.75 |
| 8 | 6675458 | rs12541503 | -0.325 | 2.212 | -1.169 | 1.629 | -0.042 | -0.045 | -0.045 | 0.011 | 0.16 | 0.11 | 0.2 | 0.28 | 0.22 | 0.26 |

| | | | | | | | | | | | | | | | | |
|---|---|---|---|---|---|---|---|---|---|---|---|---|---|---|---|---|
| 8 | 6677842 | rs12546593 | -0.418 | 1.852 | -1.038 | 1.877 | -0.008 | 0.107 | 0.107 | 0.522 | 0.07 | 0.16 | 0.3 | 0.34 | 0.08 | 0.07 |
| 8 | 6679031 | rs10104503 | -0.763 | 2.075 | -1.403 | 1.373 | -0.079 | -0.073 | -0.073 | 0.794 | 0.19 | 0.07 | 0.24 | 0.3 | 0.28 | 0.26 |
| 8 | 6684137 | rs2741083 | -0.818 | 1.77 | -0.899 | 1.072 | 0.011 | 0.005 | 0.005 | 0.378 | 0.36 | 0.49 | 0.59 | 0.6 | 0.41 | 0.52 |
| 8 | 6684819 | rs2980957 | 1.014 | 1.756 | -0.606 | 1.154 | 0.003 | 0.003 | 0.003 | 0 | 0.78 | 0.7 | 0.82 | 0.71 | 0.75 | 0.73 |
| 8 | 6686791 | rs2741087 | -1.353 | 1.464 | -0.804 | 1.274 | 0.004 | -0.006 | -0.006 | 0 | 0.21 | 0.16 | 0.17 | 0.23 | 0.23 | 0.27 |
| 8 | 6690276 | rs2741098 | 0.708 | 1.731 | -1.483 | 1.423 | -0.076 | -0.096 | -0.096 | 0 | 0.68 | 0.91 | 0.51 | 0.55 | 0.54 | 0.71 |
| 8 | 6697450 | rs2977798 | 0.782 | 1.67 | -0.66 | 1.257 | -0.013 | -0.002 | -0.002 | 6.111 | 0.63 | 0.74 | 0.56 | 0.62 | 0.51 | 0.69 |
| 8 | 6707885 | rs2702934 | -0.103 | 1.66 | -0.454 | 2.074 | -0.016 | -0.039 | -0.039 | 0 | 0.91 | 0.99 | 0.78 | 0.58 | 0.87 | 0.88 |
| 8 | 6709095 | rs11989153 | -0.283 | 0.832 | 0.432 | 1.127 | -0.002 | 0.061 | 0.061 | 0 | 0.07 | 0.31 | 0.04 | 0.09 | 0.01 | 0.18 |
| 8 | 6709805 | rs2977795 | 0.976 | 0.218 | 0.356 | 0.616 | 0.006 | 0.156 | 0.156 | 0 | 0.48 | 0.2 | 0.48 | 0.15 | 0.51 | 0.42 |
| 8 | 6710502 | rs2951863 | 0.76 | 0.173 | 0.825 | 0.718 | 0.007 | 0.037 | 0.037 | 0 | 0.52 | 0.27 | 0.48 | 0.32 | 0.51 | 0.5 |
| 8 | 6713695 | rs2980948 | 0.864 | 0.238 | 0.689 | 0.226 | 0.009 | 0.12 | 0.12 | 0 | 0.5 | 0.25 | 0.47 | 0.16 | 0.5 | 0.42 |
| 8 | 6714699 | rs11777699 | 0.602 | 0.229 | 0.631 | 0.178 | -0.002 | 0.058 | 0.058 | 0.446 | 0.54 | 0.31 | 0.52 | 0.24 | 0.52 | 0.59 |
| 8 | 6714902 | rs11774240 | 0.85 | -0.059 | 0.76 | -0.128 | 0.003 | 0.057 | 0.057 | 0.351 | 0.49 | 0.4 | 0.52 | 0.16 | 0.48 | 0.38 |
| 8 | 6715057 | rs4841777 | 0.332 | -0.078 | 0.636 | -0.39 | 0 | -0.001 | -0.001 | 0 | 0.57 | 0.63 | 0.57 | 0.31 | 0.51 | 0.61 |
| 8 | 6715751 | rs2980945 | 0.477 | -0.126 | 0.47 | -0.865 | -0.001 | -0.019 | -0.019 | 0 | 0.44 | 0.43 | 0.47 | 0.14 | 0.43 | 0.32 |
| 8 | 6716312 | rs2977790 | 0.258 | -0.555 | 0.657 | -0.85 | -0.001 | 0 | 0 | 0 | 0.47 | 0.5 | 0.5 | 0.25 | 0.49 | 0.56 |
| 8 | 6716958 | rs2741101 | -1.082 | -0.365 | 1.141 | -1.067 | -0.019 | -0.043 | -0.043 | 0 | 0.15 | 0.46 | 0.08 | 0.09 | 0.07 | 0.12 |
| 8 | 6717079 | rs7018192 | -0.558 | -0.338 | 0.896 | -1.057 | -0.015 | -0.027 | -0.027 | 0 | 0.13 | 0.31 | 0.06 | 0.09 | 0.06 | 0.12 |
| 8 | 6717107 | rs6989229 | -0.708 | -0.335 | 0.563 | -1.053 | -0.015 | -0.028 | -0.028 | 8.058 | 0.14 | 0.3 | 0.06 | 0.09 | 0.07 | 0.12 |
| 8 | 6719487 | rs2980942 | -0.233 | -0.44 | -0.075 | -1.395 | -0.001 | -0.013 | -0.013 | 2.134 | 0.51 | 0.46 | 0.34 | 0.23 | 0.48 | 0.4 |
| 8 | 6721326 | rs2738050 | -0.057 | -0.45 | -0.445 | -1.594 | -0.001 | -0.025 | -0.025 | 0 | 0.39 | 0.4 | 0.38 | 0.17 | 0.4 | 0.32 |
| 8 | 6721402 | rs2702842 | -1.293 | -0.363 | -0.914 | -1.574 | -0.102 | 0.006 | 0.006 | 1.128 | 0.62 | 0.94 | 0.51 | 0.26 | 0.55 | 0.63 |
| 8 | 6722082 | rs2702839 | -0.593 | -0.331 | -0.734 | -1.801 | -0.005 | 0.135 | 0.135 | 0.27 | 0.41 | 0.7 | 0.24 | 0.11 | 0.37 | 0.31 |
| 8 | 6722382 | rs7834740 | -0.506 | -0.508 | -0.516 | -1.778 | 0 | -0.024 | -0.024 | 0 | 0.37 | 0.58 | 0.22 | 0.11 | 0.33 | 0.29 |
| 8 | 6722662 | rs2980938 | -0.779 | -0.563 | -0.286 | -1.691 | 0 | -0.033 | -0.033 | 0 | 0.44 | 0.46 | 0.42 | 0.17 | 0.44 | 0.39 |
| 8 | 6723033 | rs2980937 | 0.325 | -0.434 | -0.42 | -1.311 | -0.007 | -0.015 | -0.015 | 0.474 | 0.16 | 0.08 | 0.26 | 0.16 | 0.17 | 0.22 |
| 8 | 6724306 | rs2978880 | -0.25 | -0.49 | -0.258 | -1.194 | 0.003 | 0.012 | 0.012 | 0 | 0.19 | 0.23 | 0.27 | 0.28 | 0.18 | 0.24 |
| 8 | 6724576 | rs2741108 | -0.259 | -0.756 | -0.569 | -0.897 | 0.017 | 0.009 | 0.009 | 0 | 0.48 | 0.3 | 0.47 | 0.45 | 0.51 | 0.52 |
| 8 | 6724615 | rs2741109 | -1.372 | -0.736 | -0.527 | -0.909 | 0.007 | -0.028 | -0.028 | 0 | 0.66 | 0.54 | 0.74 | 0.75 | 0.68 | 0.76 |
| 8 | 6725056 | rs2702833 | -0.909 | -0.774 | -0.224 | -0.905 | -0.001 | -0.001 | -0.001 | 0 | 0.53 | 0.53 | 0.48 | 0.42 | 0.52 | 0.54 |
| 8 | 6727082 | rs2741121 | -0.237 | -0.895 | -0.507 | -0.965 | 0.006 | -0.007 | -0.007 | 0.42 | 0.39 | 0.32 | 0.44 | 0.43 | 0.44 | 0.42 |
| 8 | 6728116 | rs1800971 | -0.243 | -0.875 | -0.499 | -0.806 | -0.025 | -0.009 | -0.009 | 0.012 | 0.07 | 0.25 | 0.03 | 0.04 | 0.02 | 0.06 |
| 8 | 6728198 | rs1047031 | 1.237 | -0.759 | -1.057 | -0.827 | -0.037 | -0.076 | -0.076 | 0 | 0.11 | 0.01 | 0.26 | 0.41 | 0.17 | 0.21 |
| 8 | 6731487 | rs2741127 | 1.928 | -0.614 | -0.971 | -0.865 | -0.048 | -0.057 | -0.057 | 0 | 0.3 | 0.14 | 0.24 | 0.44 | 0.31 | 0.28 |
| 8 | 6731993 | rs2980924 | -1.273 | -0.158 | -0.682 | -0.749 | -0.005 | -0.004 | -0.004 | 0 | 0.88 | 0.91 | 0.74 | 0.9 | 0.81 | 0.86 |
| 8 | 6735399 | rs11362 | -0.339 | -0.464 | -0.414 | -0.838 | 0.017 | -0.008 | -0.008 | 0.599 | 0.42 | 0.3 | 0.44 | 0.43 | 0.44 | 0.42 |
| 8 | 6735423 | rs1800972 | -1.374 | -0.456 | -0.444 | -0.848 | -0.035 | -0.016 | -0.016 | 0 | 0.88 | 0.95 | 0.74 | 0.9 | 0.8 | 0.85 |
| 8 | 6735431 | rs1799946 | 0.71 | -0.692 | -0.196 | -0.758 | 0.008 | 0.006 | 0.006 | 0 | 0.46 | 0.59 | 0.3 | 0.47 | 0.36 | 0.43 |
| 8 | 6737121 | rs5743404 | -0.801 | -0.565 | -0.334 | -0.675 | 0.018 | 0.016 | 0.016 | 0 | 0.51 | 0.42 | 0.71 | 0.52 | 0.65 | 0.57 |
| 8 | 6740074 | rs2977826 | -0.09 | -0.293 | -0.411 | -0.809 | 0.024 | -0.018 | -0.018 | 0 | 0.75 | 0.69 | 0.63 | 0.86 | 0.68 | 0.81 |
| 8 | 6740938 | rs2738173 | -0.164 | -0.582 | -0.555 | -0.986 | -0.011 | -0.067 | -0.067 | 0 | 0.24 | 0.25 | 0.08 | 0.01 | 0.18 | 0.21 |
| 8 | 6742110 | rs2951850 | -0.031 | 0.058 | -0.703 | -0.459 | -0.014 | 0.007 | 0.007 | 0 | 0.76 | 0.81 | 0.63 | 0.85 | 0.68 | 0.81 |
| 8 | 6742558 | rs2738169 | -0.131 | -0.254 | -0.195 | 0.235 | 0.002 | 0.091 | 0.091 | 0 | 0.14 | 0.21 | 0.28 | 0.41 | 0.17 | 0.21 |
| 8 | 6743029 | rs2738166 | -1.175 | -0.328 | -0.278 | 0.052 | -0.001 | 0.044 | 0.044 | 0 | 0.33 | 0.45 | 0.46 | 0.49 | 0.36 | 0.39 |
| 8 | 6743108 | rs7826831 | -0.718 | -0.509 | -0.733 | -0.24 | 0.016 | 0.047 | 0.047 | 0 | 0.26 | 0.17 | 0.15 | 0.07 | 0.25 | 0.16 |
| 8 | 6743568 | rs2738165 | 1.587 | -0.832 | -1.087 | -0.175 | 0.05 | -0.067 | -0.067 | 0 | 0.52 | 0.1 | 0.44 | 0.51 | 0.61 | 0.42 |
| 8 | 6745485 | rs17581175 | 0.901 | -0.897 | -1.043 | -0.121 | -0.015 | 0.035 | 0.035 | 0 | 0.07 | 0.01 | 0.07 | 0 | 0.08 | 0.11 |
| 8 | 6746641 | rs2741718 | 1.877 | -0.613 | -1.099 | -0.113 | 0.022 | -0.068 | -0.068 | 0 | 0.49 | 0.08 | 0.43 | 0.5 | 0.59 | 0.42 |
| 8 | 6749770 | rs7842755 | -1.08 | -0.671 | -0.195 | -0.093 | -0.043 | 0.129 | 0.129 | 0 | 0.25 | 0.51 | 0.36 | 0.44 | 0.18 | 0.28 |
| 8 | 6750251 | rs2738157 | -0.173 | -0.512 | -0.779 | -0.295 | -0.046 | 0.104 | 0.104 | 0 | 0.18 | 0.02 | 0.13 | 0.02 | 0.2 | 0.14 |
| 8 | 6754548 | rs751009 | 0.822 | -0.497 | 0.002 | 0.054 | -0.013 | 0.139 | 0.139 | 0.07 | 0.77 | 0.59 | 0.6 | 0.51 | 0.78 | 0.64 |
| 8 | 6755820 | rs2741056 | -1.769 | -0.443 | -0.83 | 0.056 | 0.032 | -0.028 | -0.028 | 0 | 0.51 | 0.9 | 0.57 | 0.51 | 0.42 | 0.6 |
| 8 | 6760142 | rs2702933 | 1.601 | -0.289 | -0.773 | 0.098 | -0.004 | -0.044 | -0.044 | 0.54 | 0.48 | 0.12 | 0.43 | 0.51 | 0.55 | 0.42 |
| 8 | 6761571 | rs20527 | -1.08 | -0.337 | -0.004 | -0.338 | -0.015 | -0.135 | -0.135 | 0 | 0.31 | 0.56 | 0.18 | 0.02 | 0.28 | 0.24 |
| 8 | 6763241 | rs2741073 | 1.185 | -0.147 | -0.394 | -0.242 | -0.002 | -0.002 | -0.002 | 0.924 | 0.39 | 0.59 | 0.33 | 0.33 | 0.34 | 0.3 |
| 8 | 6763986 | rs2741710 | -1.003 | 0.014 | -0.712 | 0.004 | -0.026 | 0.038 | 0.038 | 0 | 0.76 | 0.84 | 0.66 | 0.84 | 0.68 | 0.87 |
| 8 | 6765136 | rs2702889 | -1.645 | -0.128 | -0.488 | 0.132 | 0.03 | -0.001 | -0.001 | 0.35 | 0.45 | 0.71 | 0.37 | 0.52 | 0.36 | 0.59 |
| 8 | 6767192 | rs2702898 | -1.148 | 0.083 | 0.006 | 0.431 | -0.003 | 0.025 | 0.025 | 0.138 | 0.32 | 0.35 | 0.28 | 0.47 | 0.29 | 0.41 |
| 8 | 6770015 | rs2741706 | 0.186 | 0.116 | -0.207 | 0.71 | -0.001 | -0.002 | -0.002 | 0.818 | 0.12 | 0.09 | 0.09 | 0.11 | 0.12 | 0.2 |
| 8 | 6772913 | rs7461956 | 1.171 | 0.032 | -0.084 | 0.478 | 0 | 0.003 | 0.003 | 0 | 0.42 | 0.54 | 0.41 | 0.76 | 0.39 | 0.43 |
| 8 | 6774414 | rs2738133 | -1.243 | -0.099 | -0.164 | 0.997 | -0.021 | 0.012 | 0.012 | 0.087 | 0.3 | 0.45 | 0.21 | 0.31 | 0.26 | 0.38 |
| 8 | 6777051 | rs2741694 | -0.062 | -0.055 | -0.666 | 1.049 | 0.003 | 0.002 | 0.002 | 0.056 | 0.85 | 0.88 | 0.72 | 0.62 | 0.88 | 0.75 |
| 8 | 6780107 | rs2741692 | 0.403 | 0.051 | -1.244 | 0.858 | 0.006 | 0.012 | 0.012 | 0 | 0.24 | 0.1 | 0.32 | 0.25 | 0.27 | 0.2 |
| 8 | 6780244 | rs13275170 | 0.99 | 0.312 | -1.337 | 0.63 | -0.02 | 0.072 | 0.072 | 0 | 0.45 | 0.25 | 0.35 | 0.32 | 0.49 | 0.53 |
| 8 | 6782271 | rs712276 | -0.593 | 0.838 | -1.164 | 0.147 | -0.024 | 0.01 | 0.01 | 0 | 0.14 | 0.05 | 0.08 | 0.06 | 0.18 | 0.14 |
| 8 | 6784971 | rs2738118 | 0.109 | 0.434 | -1.017 | 0.292 | 0 | -0.003 | -0.003 | 1.152 | 0.84 | 0.84 | 0.78 | 0.85 | 0.81 | 0.81 |
| 8 | 6785352 | rs2741688 | -0.238 | 0.385 | -1.136 | -0.456 | 0.004 | 0.007 | 0.007 | 0.22 | 0.3 | 0.22 | 0.28 | 0.09 | 0.29 | 0.1 |
| 8 | 6786417 | rs2738116 | 0.13 | 0.407 | -0.971 | -0.273 | -0.005 | -0.02 | -0.02 | 0.799 | 0.22 | 0.26 | 0.26 | 0.11 | 0.23 | 0.09 |

| | | | | | | | | | | | | | | | | |
|---|---|---|---|---|---|---|---|---|---|---|---|---|---|---|---|---|
| 8 | 6787075 | rs2702938 | -0.011 | 0.767 | -1.867 | -0.07 | -0.044 | -0.084 | -0.084 | 0.496 | 0.65 | 0.87 | 0.55 | 0.54 | 0.59 | 0.45 |
| 8 | 6788102 | rs2003880 | 0.868 | 1.061 | -1.796 | -0.1 | -0.038 | 0.038 | 0.038 | 0 | 0.17 | 0.02 | 0.22 | 0.1 | 0.2 | 0.09 |
| 8 | 6789368 | rs4526381 | 0.516 | 1.309 | -1.345 | 0.311 | 0.009 | 0.01 | 0.01 | 0 | 0.16 | 0.21 | 0.09 | 0.22 | 0.15 | 0.25 |
| 8 | 6790242 | rs2702860 | 1.123 | 1.294 | -2.099 | 0.081 | -0.06 | 0.134 | 0.134 | 0 | 0.29 | 0.03 | 0.28 | 0.1 | 0.29 | 0.18 |
| 8 | 6790710 | rs13251447 | 1.157 | 1.16 | -1.779 | 0.064 | -0.001 | -0.006 | -0.006 | 0.025 | 0.22 | 0.19 | 0.18 | 0.31 | 0.21 | 0.22 |
| 8 | 6793540 | rs736227 | -0.204 | 0.893 | -1.969 | 0.08 | 0.005 | -0.022 | -0.022 | 0 | 0.69 | 0.76 | 0.58 | 0.62 | 0.67 | 0.66 |
| 8 | 6793581 | rs2738100 | -0.059 | 1.279 | -1.425 | 0.479 | -0.001 | -0.001 | -0.001 | 0 | 0.39 | 0.62 | 0.29 | 0.46 | 0.36 | 0.48 |
| 8 | 6797532 | rs2741675 | 0.383 | 1.464 | -1.573 | 0.577 | -0.036 | -0.03 | -0.03 | 0 | 0.87 | 0.96 | 0.78 | 0.86 | 0.8 | 0.82 |
| 8 | 6801712 | rs2981398 | -0.1 | 1.276 | -2.029 | 0.098 | -0.024 | 0.063 | 0.063 | 0 | 0.34 | 0.12 | 0.44 | 0.22 | 0.4 | 0.33 |
| 8 | 6802073 | rs2741669 | 0.356 | 1.186 | -1.442 | -0.19 | -0.002 | -0.014 | -0.014 | 0 | 0.41 | 0.53 | 0.42 | 0.6 | 0.39 | 0.41 |
| 8 | 6811559 | rs2741658 | -0.622 | 1.464 | -0.574 | -0.403 | 0.002 | -0.028 | -0.028 | 0 | 0.36 | 0.55 | 0.38 | 0.29 | 0.38 | 0.36 |
| 8 | 6812198 | rs2741657 | -1.108 | 1.187 | -0.906 | -0.568 | 0 | 0 | 0 | 0 | 0.58 | 0.63 | 0.51 | 0.33 | 0.61 | 0.49 |
| 8 | 6813501 | rs2702852 | -0.522 | 1.154 | -0.816 | -0.74 | 0.001 | -0.023 | -0.023 | 0 | 0.44 | 0.55 | 0.3 | 0.22 | 0.45 | 0.44 |
| 8 | 6820464 | rs12680018 | 0.537 | 1.172 | -0.671 | -0.084 | -0.002 | -0.003 | -0.003 | 0.018 | 0.06 | 0.1 | 0.21 | 0.15 | 0.12 | 0.18 |
| 8 | 6821419 | rs2977793 | -1.004 | 0.773 | -0.631 | -0.192 | -0.003 | -0.003 | -0.003 | 0 | 0.54 | 0.5 | 0.65 | 0.49 | 0.63 | 0.59 |
| 8 | 6822785 | rs2738048 | 0.052 | 1.041 | -0.424 | -0.113 | -0.004 | -0.005 | -0.005 | 0 | 0.28 | 0.18 | 0.33 | 0.34 | 0.32 | 0.3 |
| 8 | 6823542 | rs2738046 | -0.669 | 1.066 | -0.003 | -0.132 | 0.006 | 0.007 | 0.007 | 0 | 0.36 | 0.55 | 0.55 | 0.51 | 0.45 | 0.49 |
| 8 | 6823891 | rs2702910 | 0.093 | 0.922 | -0.005 | -0.146 | -0.005 | 0.006 | 0.006 | 0 | 0.28 | 0.16 | 0.32 | 0.32 | 0.31 | 0.29 |
| 8 | 6825295 | rs6996047 | 0.352 | 0.322 | -0.087 | -0.311 | 0.002 | -0.022 | -0.022 | 0 | 0.17 | 0.18 | 0.11 | 0 | 0.15 | 0.08 |
| 8 | 6880925 | rs7826487 | -0.199 | 0.79 | 0.131 | 0.657 | 0.048 | 0.013 | 0.013 | 0 | 0.1 | 0.34 | 0.11 | 0.09 | 0.13 | 0.07 |
| 8 | 6883291 | rs9774483 | 0.243 | 0.869 | -0.53 | 0.833 | -0.003 | 0.001 | 0.001 | 0 | 0.93 | 0.9 | 0.82 | 0.83 | 0.95 | 0.83 |
| 8 | 6897900 | rs4546682 | 0.304 | 0.591 | -0.732 | 1.474 | -0.001 | 0.06 | 0.06 | 0 | 0.93 | 0.87 | 0.82 | 0.71 | 0.96 | 0.86 |
| 8 | 6898731 | rs4358823 | 0.016 | 0.343 | -0.865 | 1.386 | -0.002 | 0.018 | 0.018 | 0 | 0.93 | 0.89 | 0.82 | 0.79 | 0.96 | 0.87 |
| 8 | 8266397 | rs2976921 | 0.104 | 0.961 | -0.153 | -0.431 | -0.011 | 0.017 | 0.017 | 0.206 | 0.3 | 0.36 | 0.29 | 0.58 | 0.28 | 0.49 |
| 8 | 8275062 | rs13256028 | -1.157 | 1.455 | -0.85 | -0.265 | -0.003 | 0.005 | 0.005 | 0 | 0.34 | 0.68 | 0.34 | 0.27 | 0.34 | 0.2 |
| 8 | 8276626 | rs2976941 | 0.096 | 1.17 | -1.128 | 0.237 | -0.035 | -0.081 | -0.081 | 0 | 0.32 | 0.16 | 0.43 | 0.49 | 0.39 | 0.49 |
| 8 | 8283706 | rs13252804 | -0.291 | 1.311 | -0.448 | 0.423 | -0.003 | 0.042 | 0.042 | 0.588 | 0.15 | 0.17 | 0.26 | 0.39 | 0.11 | 0.15 |
| 8 | 8285302 | rs10099571 | 0.32 | 1.628 | -0.153 | 0.305 | 0.083 | -0.029 | -0.029 | 2.349 | 0.07 | 0.26 | 0.07 | 0 | 0.13 | 0.05 |
| 8 | 8291357 | rs2979164 | -0.319 | 1.471 | -0.388 | 0.274 | -0.031 | -0.04 | -0.04 | 0.92 | 0.37 | 0.18 | 0.43 | 0.51 | 0.43 | 0.5 |
| 8 | 8296071 | rs2976846 | -0.359 | 1.457 | -0.315 | 0.247 | -0.027 | -0.015 | -0.015 | 0 | 0.37 | 0.21 | 0.47 | 0.55 | 0.43 | 0.51 |
| 8 | 8304502 | rs2921077 | -1.346 | 1.064 | -0.263 | 0.8 | -0.022 | 0.054 | 0.054 | 5.718 | 0.4 | 0.25 | 0.62 | 0.83 | 0.46 | 0.56 |
| 8 | 8307128 | rs2976862 | -0.738 | 1.007 | -0.364 | 0.852 | 0.001 | -0.028 | -0.028 | 0 | 0.14 | 0.06 | 0.24 | 0.32 | 0.1 | 0.13 |
| 8 | 8313010 | rs2976871 | -0.393 | 1.101 | -0.219 | 0.743 | -0.023 | -0.032 | -0.032 | 1.391 | 0.26 | 0.19 | 0.54 | 0.55 | 0.34 | 0.44 |
| 8 | 8318850 | rs2976876 | -1.344 | 0.938 | 0.116 | 0.319 | -0.001 | -0.034 | -0.034 | 0 | 0.39 | 0.39 | 0.61 | 0.91 | 0.41 | 0.68 |
| 8 | 8319925 | rs2921054 | -1.028 | 1.055 | 0.029 | 0.509 | 0 | 0.004 | 0.004 | 1.615 | 0.17 | 0.16 | 0.13 | 0.34 | 0.22 | 0.17 |
| 8 | 8321836 | rs2980771 | -0.884 | 1.07 | -0.008 | 0.561 | -0.003 | -0.008 | -0.008 | 0 | 0.16 | 0.12 | 0.13 | 0.34 | 0.22 | 0.17 |
| 8 | 8323042 | rs2980772 | -0.41 | 1.162 | 0.09 | 0.692 | 0 | -0.003 | -0.003 | 2.215 | 0.25 | 0.24 | 0.5 | 0.58 | 0.22 | 0.53 |
| 8 | 8325753 | rs2921044 | -0.231 | 1.048 | 0.206 | 0.766 | -0.003 | 0.018 | 0.018 | 0 | 0.22 | 0.27 | 0.48 | 0.56 | 0.19 | 0.51 |
| 8 | 8335210 | rs2976890 | -1.745 | 1.204 | 0.58 | 0.884 | 0.033 | 0.115 | 0.115 | 0.893 | 0.17 | 0.38 | 0.21 | 0.38 | 0.22 | 0.15 |
| 8 | 8341678 | rs2980742 | -1.784 | 1.216 | 0.531 | 0.686 | 0.042 | 0.106 | 0.106 | 0.481 | 0.17 | 0.35 | 0.23 | 0.35 | 0.23 | 0.14 |
| 8 | 8342506 | rs2921027 | -1.385 | 1.323 | -0.116 | 0.722 | 0.004 | 0.015 | 0.015 | 0.319 | 0.14 | 0.13 | 0.21 | 0.34 | 0.23 | 0.14 |
| 8 | 8344163 | rs2979203 | -0.578 | 1.285 | -0.238 | 0.795 | -0.002 | 0.013 | 0.013 | 0.016 | 0.11 | 0.1 | 0.2 | 0.35 | 0.18 | 0.09 |
| 8 | 8349087 | rs2921024 | 0.059 | 0.64 | -0.484 | 0.162 | -0.006 | -0.005 | -0.005 | 0 | 0.91 | 0.95 | 0.8 | 0.76 | 0.83 | 0.93 |
| 8 | 8349757 | rs6997361 | 0.461 | 0.239 | -0.046 | -0.189 | -0.005 | -0.025 | -0.025 | 1.288 | 0.67 | 0.55 | 0.48 | 0.26 | 0.74 | 0.38 |
| 8 | 8350370 | rs2921023 | 0 | -0.014 | 0.196 | 0.078 | 0.001 | 0.095 | 0.095 | 0 | 0.23 | 0.49 | 0.32 | 0.6 | 0.21 | 0.45 |
| 8 | 8351068 | rs2980748 | 0.236 | 0.041 | 0.246 | -0.061 | 0.004 | 0.015 | 0.015 | 0 | 0.87 | 0.86 | 0.72 | 0.72 | 0.81 | 0.83 |
| 8 | 8351677 | rs2921016 | -0.8 | -0.227 | 0.186 | -0.429 | 0 | 0 | 0 | 0 | 0.37 | 0.62 | 0.41 | 0.63 | 0.3 | 0.52 |
| 8 | 8351697 | rs2980751 | 0.378 | -0.194 | 0.49 | -0.151 | -0.001 | 0.014 | 0.014 | 0 | 0.22 | 0.26 | 0.23 | 0.61 | 0.21 | 0.45 |
| 8 | 8353587 | rs2921004 | 0.167 | -0.192 | 0.315 | 0.155 | -0.006 | 0.121 | 0.121 | 0 | 0.23 | 0.45 | 0.26 | 0.53 | 0.21 | 0.46 |
| 8 | 8354436 | rs4840344 | 0.985 | -0.214 | -0.348 | -0.762 | -0.004 | -0.015 | -0.015 | 0.018 | 0.4 | 0.37 | 0.25 | 0.02 | 0.38 | 0.14 |
| 8 | 8357183 | rs2921102 | -0.433 | -0.02 | -0.603 | -0.469 | -0.036 | -0.058 | -0.058 | 0 | 0.89 | 0.98 | 0.66 | 0.73 | 0.83 | 0.85 |
| 8 | 8359387 | rs2921092 | -0.292 | -0.183 | -0.524 | -0.644 | -0.003 | -0.002 | -0.002 | 0 | 0.35 | 0.6 | 0.31 | 0.63 | 0.3 | 0.53 |
| 8 | 8361139 | rs2921074 | -0.488 | -0.027 | -0.472 | -0.211 | -0.045 | -0.057 | -0.057 | 0 | 0.9 | 0.98 | 0.75 | 0.73 | 0.81 | 0.83 |
| 8 | 8361254 | rs2980754 | -0.144 | -0.103 | -0.446 | -0.314 | 0.002 | -0.004 | -0.004 | 0 | 0.66 | 0.75 | 0.61 | 0.62 | 0.62 | 0.63 |
| 8 | 8365421 | rs7000205 | -0.493 | -0.652 | 0.028 | -1.061 | -0.015 | -0.091 | -0.091 | 0 | 0.17 | 0.37 | 0.12 | 0.02 | 0.1 | 0.1 |
| 8 | 8366246 | rs2921098 | 0.684 | -0.485 | -0.032 | -0.863 | -0.02 | 0.007 | 0.007 | 0 | 0.67 | 0.79 | 0.59 | 0.65 | 0.6 | 0.72 |
| 8 | 8374342 | rs4375003 | -0.425 | -0.249 | 0.16 | -0.652 | 0.044 | 0.021 | 0.021 | 0.153 | 0.24 | 0.45 | 0.4 | 0.33 | 0.33 | 0.21 |
| 8 | 8374439 | rs7007897 | 0.857 | -0.441 | 0.257 | -0.729 | 0.007 | -0.012 | -0.012 | 0 | 0.24 | 0.2 | 0.28 | 0.58 | 0.16 | 0.49 |
| 8 | 8384479 | rs11777086 | -0.328 | -0.184 | -0.656 | -0.678 | 0.02 | 0.047 | 0.047 | 1.739 | 0.31 | 0.19 | 0.2 | 0.09 | 0.25 | 0.22 |
| 8 | 8390067 | rs6994806 | 0.221 | 0.089 | -0.491 | 0.009 | -0.004 | -0.002 | -0.002 | 0.551 | 0.66 | 0.69 | 0.59 | 0.61 | 0.53 | 0.61 |
| 8 | 8390411 | rs4840980 | 0.253 | -0.023 | 0.088 | -0.033 | 0.005 | 0.002 | 0.002 | 0 | 0.65 | 0.52 | 0.6 | 0.61 | 0.53 | 0.61 |
| 8 | 8394297 | rs13249144 | -0.573 | -0.317 | -0.406 | -0.271 | -0.136 | -0.017 | -0.017 | 0 | 0.42 | 0.05 | 0.44 | 0.39 | 0.55 | 0.4 |
| 8 | 8394877 | rs4840986 | 0.93 | -0.518 | -0.222 | 0.298 | 0.009 | 0.061 | 0.061 | 0 | 0.49 | 0.64 | 0.31 | 0.23 | 0.41 | 0.44 |
| 8 | 8396676 | rs17153465 | -0.511 | -0.534 | 0.065 | 0.28 | 0.003 | 0.083 | 0.083 | 0 | 0.1 | 0.21 | 0.29 | 0.32 | 0.09 | 0.15 |
| 8 | 8397785 | rs11782884 | 1.545 | -0.631 | 0.19 | 0.183 | -0.009 | 0.06 | 0.06 | 0 | 0.41 | 0.16 | 0.23 | 0.23 | 0.36 | 0.41 |
| 8 | 8400723 | rs10903346 | 0.282 | -0.725 | -0.182 | 0.137 | -0.124 | -0.065 | -0.065 | 0 | 0.31 | 0.02 | 0.36 | 0.33 | 0.45 | 0.34 |
| 8 | 8402347 | rs7837584 | 0.718 | -0.673 | 0.085 | 0.176 | -0.017 | 0.07 | 0.07 | 0 | 0.74 | 0.45 | 0.62 | 0.55 | 0.82 | 0.74 |
| 8 | 8403142 | rs4240619 | -0.24 | -0.634 | 0.132 | 0.139 | -0.013 | -0.02 | -0.02 | 0 | 0.82 | 0.69 | 0.9 | 0.87 | 0.89 | 0.87 |
| 8 | 8407886 | rs2001445 | 0.349 | -0.572 | 0.299 | 0.337 | -0.002 | 0.006 | 0.006 | 0 | 0.53 | 0.59 | 0.53 | 0.55 | 0.44 | 0.54 |

| | | | | | | | | | | | | | | | | |
|---|---|---|---|---|---|---|---|---|---|---|---|---|---|---|---|---|
| 8 | 8413380 | rs2294143 | 0.65 | -0.458 | 0.343 | 0.34 | 0.002 | 0.004 | 0.004 | 0 | 0.49 | 0.45 | 0.5 | 0.55 | 0.41 | 0.52 |
| 8 | 8415810 | rs6986647 | 0.424 | -0.438 | 0.273 | 0.464 | -0.003 | 0.081 | 0.081 | 0 | 0.07 | 0.09 | 0.24 | 0.32 | 0.04 | 0.13 |
| 8 | 8419612 | rs10108910 | 0.371 | -0.455 | 0.224 | 0.407 | -0.005 | 0.016 | 0.016 | 0 | 0.53 | 0.65 | 0.53 | 0.55 | 0.45 | 0.53 |
| 8 | 8426753 | rs1986346 | 0.523 | -0.702 | -0.66 | -0.153 | -0.015 | -0.007 | -0.007 | 1.556 | 0.3 | 0.34 | 0.22 | 0.22 | 0.22 | 0.34 |
| 8 | 8429600 | rs12544932 | 0.451 | -0.134 | -0.878 | 0.544 | -0.006 | 0.086 | 0.086 | 0 | 0.06 | 0.11 | 0.2 | 0.28 | 0.04 | 0.1 |
| 8 | 8429687 | rs4840991 | 0.099 | -0.007 | -1.02 | 0.413 | -0.036 | -0.044 | -0.044 | 0 | 0.64 | 0.77 | 0.61 | 0.53 | 0.56 | 0.6 |
| 8 | 8432208 | rs7813863 | -0.045 | -0.297 | -0.525 | 0.027 | 0.026 | 0.056 | 0.056 | 0.068 | 0.7 | 0.81 | 0.79 | 0.88 | 0.74 | 0.74 |
| 8 | 8433876 | rs4595128 | -0.957 | 0.343 | 0.116 | 0.793 | 0.002 | 0.085 | 0.085 | 0 | 0.1 | 0.25 | 0.15 | 0.26 | 0.08 | 0.14 |
| 8 | 8437123 | rs10099072 | -0.107 | 0.713 | 0.054 | 0.108 | 0.025 | -0.066 | -0.066 | 0 | 0.23 | 0.46 | 0.16 | 0.13 | 0.25 | 0.15 |
| 8 | 8439068 | rs7843504 | -0.831 | 0.442 | -0.372 | 0.207 | -0.019 | 0.075 | 0.075 | 0.688 | 0.53 | 0.75 | 0.65 | 0.7 | 0.51 | 0.59 |
| 8 | 8444891 | rs4268132 | 0.536 | 0.302 | -0.511 | 0.808 | 0.005 | 0.017 | 0.017 | 0.273 | 0.07 | 0.04 | 0.24 | 0.35 | 0.06 | 0.11 |
| 8 | 8448429 | rs13280653 | 1.865 | 0.305 | -1.115 | 0.113 | -0.037 | 0.14 | 0.14 | 0 | 0.23 | 0.04 | 0.17 | 0 | 0.23 | 0.07 |
| 8 | 8450798 | rs7826228 | 0.044 | 0.224 | -0.135 | 0.418 | 0.007 | 0.078 | 0.078 | 0 | 0.3 | 0.52 | 0.39 | 0.52 | 0.31 | 0.28 |
| 8 | 8460285 | rs17154314 | 0.01 | 0.204 | -0.292 | 0.596 | -0.003 | 0.069 | 0.069 | 0 | 0.08 | 0.12 | 0.24 | 0.41 | 0.06 | 0.18 |
| 8 | 8461280 | rs17154329 | 0.344 | 0.212 | -0.328 | 0.603 | 0.002 | 0.033 | 0.033 | 0.131 | 0.07 | 0.07 | 0.24 | 0.36 | 0.06 | 0.17 |
| 8 | 8462928 | rs2980758 | 1.022 | 0.255 | -0.177 | 0.405 | 0.012 | 0.005 | 0.005 | 0.289 | 0.24 | 0.16 | 0.28 | 0.15 | 0.18 | 0.32 |
| 8 | 8472504 | rs9329273 | 1.02 | 0.431 | 0.106 | 0.223 | -0.008 | -0.033 | -0.033 | 0 | 0.21 | 0.58 | 0.11 | 0.15 | 0.2 | 0.09 |
| 8 | 8472561 | rs9329274 | -0.313 | 0.376 | -0.182 | 0.119 | -0.006 | 0.007 | 0.007 | 0 | 0.66 | 0.68 | 0.45 | 0.5 | 0.72 | 0.57 |
| 8 | 8472758 | rs10216570 | 0.764 | 0.469 | -0.09 | 0.178 | -0.029 | 0.043 | 0.043 | 1.126 | 0.43 | 0.22 | 0.33 | 0.35 | 0.49 | 0.43 |
| 8 | 8482211 | rs10109515 | -1.717 | 0.424 | -0.321 | 0.078 | 0.011 | 0.007 | 0.007 | 0.31 | 0.78 | 0.86 | 0.72 | 0.84 | 0.82 | 0.71 |
| 8 | 8483293 | rs12675573 | 0.514 | 0.506 | 0.123 | 0.181 | 0.008 | 0.01 | 0.01 | 0 | 0.27 | 0.34 | 0.34 | 0.32 | 0.28 | 0.25 |
| 8 | 8484187 | rs13256641 | 0.962 | 0.514 | 0.062 | 0.164 | -0.031 | -0.044 | -0.044 | 0.007 | 0.23 | 0.1 | 0.17 | 0.37 | 0.31 | 0.28 |
| 8 | 8485387 | rs10903347 | 0.545 | 0.519 | 0.184 | 0.296 | 0.015 | 0.023 | 0.023 | 0 | 0.26 | 0.46 | 0.36 | 0.3 | 0.27 | 0.18 |
| 8 | 8487486 | rs17682402 | 1.561 | 0.487 | -0.327 | 0.034 | -0.03 | 0.209 | 0.209 | 1.344 | 0.29 | 0.01 | 0.19 | 0.01 | 0.26 | 0.17 |
| 8 | 8491382 | rs4841003 | -0.287 | 0.72 | -0.062 | 0.19 | -0.001 | 0.004 | 0.004 | 4.373 | 0.55 | 0.39 | 0.57 | 0.69 | 0.54 | 0.6 |
| 8 | 8493783 | rs12549726 | 1.114 | 0.576 | -0.194 | 0.268 | -0.003 | 0.072 | 0.072 | 0 | 0.07 | 0.07 | 0.24 | 0.31 | 0.06 | 0.13 |
| 8 | 8496077 | rs11250215 | 1.267 | 0.641 | -0.174 | 0.086 | -0.002 | 0 | 0 | 0.731 | 0.19 | 0.18 | 0.1 | 0.11 | 0.22 | 0.06 |
| 8 | 8500990 | rs2140246 | -1.539 | 0.563 | -0.296 | 0.092 | 0.012 | 0.007 | 0.007 | 0.342 | 0.78 | 0.85 | 0.73 | 0.85 | 0.83 | 0.69 |
| 8 | 8501230 | rs4841005 | 0.002 | 0.625 | -0.086 | 0.198 | 0.002 | 0.003 | 0.003 | 1.21 | 0.57 | 0.51 | 0.56 | 0.46 | 0.56 | 0.43 |
| 8 | 8501322 | rs7013418 | 0.625 | 0.615 | -0.065 | 0.228 | 0.027 | 0.042 | 0.042 | 1.784 | 0.21 | 0.28 | 0.17 | 0.36 | 0.27 | 0.26 |
| 8 | 8503233 | rs6985949 | -1.372 | 0.283 | 0.204 | 0.498 | 0.011 | 0.007 | 0.007 | 0.285 | 0.78 | 0.86 | 0.73 | 0.85 | 0.83 | 0.71 |
| 8 | 8503437 | rs6986287 | -0.568 | 0.322 | 0.23 | 0.457 | -0.003 | 0.001 | 0.001 | 0.564 | 0.4 | 0.58 | 0.41 | 0.44 | 0.37 | 0.37 |
| 8 | 8506404 | rs12550829 | -0.357 | 0.142 | 0.111 | 0.409 | -0.018 | 0.013 | 0.013 | 1.757 | 0.35 | 0.49 | 0.36 | 0.42 | 0.31 | 0.29 |
| 8 | 8514059 | rs17627505 | 0.931 | 0.048 | -0.4 | 0.089 | 0.016 | 0.011 | 0.011 | 0.015 | 0.14 | 0.11 | 0.06 | 0.02 | 0.09 | 0.17 |
| 8 | 8515205 | rs2177254 | -0.971 | -0.024 | -0.564 | 0.142 | 0.019 | 0.014 | 0.014 | 0.017 | 0.76 | 0.86 | 0.68 | 0.85 | 0.8 | 0.68 |
| 8 | 8517919 | rs1522838 | -0.119 | -0.012 | -0.341 | -0.126 | -0.001 | -0.084 | -0.084 | 0 | 0.32 | 0.45 | 0.18 | 0.13 | 0.32 | 0.24 |
| 8 | 8520051 | rs7813831 | -0.613 | -0.331 | -0.795 | 0.346 | 0.022 | 0.009 | 0.009 | 0.178 | 0.77 | 0.87 | 0.69 | 0.86 | 0.82 | 0.71 |
| 8 | 8523158 | rs9693402 | -0.497 | -0.139 | -0.344 | 0.703 | -0.026 | 0.009 | 0.009 | 0 | 0.2 | 0.25 | 0.23 | 0.26 | 0.13 | 0.24 |
| 8 | 8534192 | rs1112331 | -0.688 | 0.264 | -0.695 | 0.528 | 0 | -0.001 | -0.001 | 0.644 | 0.46 | 0.48 | 0.44 | 0.47 | 0.52 | 0.49 |
| 8 | 8539241 | rs17154584 | -0.639 | 0.883 | -0.696 | 0.35 | 0 | -0.017 | -0.017 | 0 | 0.33 | 0.31 | 0.36 | 0.43 | 0.31 | 0.32 |
| 8 | 8540189 | rs17149208 | -0.053 | 0.888 | -1.078 | 0.472 | -0.077 | -0.124 | -0.124 | 0.891 | 0.16 | 0.01 | 0.28 | 0.35 | 0.22 | 0.21 |
| 8 | 8542718 | rs13267387 | 0.863 | 0.88 | -1.291 | 0.616 | -0.021 | -0.049 | -0.049 | 0.772 | 0.09 | 0.01 | 0.1 | 0.23 | 0.11 | 0.14 |
| 8 | 8543324 | rs7011229 | 0.431 | 0.55 | -0.905 | -0.551 | -0.004 | 0.076 | 0.076 | 0.106 | 0.53 | 0.68 | 0.35 | 0.12 | 0.5 | 0.4 |
| 8 | 8547642 | rs17684514 | 2.857 | 0.586 | -0.696 | -0.584 | 0.004 | 0.027 | 0.027 | 0 | 0.22 | 0.12 | 0.14 | 0 | 0.21 | 0.07 |
| 8 | 8550255 | rs11249878 | 0.439 | 0.652 | -0.628 | -0.314 | 0.001 | -0.011 | -0.011 | 0.195 | 0.27 | 0.25 | 0.34 | 0.49 | 0.26 | 0.41 |
| 8 | 8557220 | rs10104950 | 0.424 | 0.603 | -0.357 | -0.476 | 0 | 0.004 | 0.004 | 0.489 | 0.66 | 0.65 | 0.59 | 0.53 | 0.64 | 0.67 |
| 8 | 8559832 | rs9644774 | 1.802 | 0.627 | -0.566 | -0.507 | -0.023 | 0.062 | 0.062 | 0 | 0.36 | 0.19 | 0.24 | 0.03 | 0.4 | 0.24 |
| 8 | 8561452 | rs1060106 | 0.744 | 0.587 | -0.651 | -0.172 | 0.017 | -0.055 | -0.055 | 0 | 0.25 | 0.14 | 0.3 | 0.49 | 0.2 | 0.4 |
| 8 | 8568875 | rs885000 | 0.781 | 0.452 | -0.652 | -0.173 | 0.013 | -0.056 | -0.056 | 1.363 | 0.25 | 0.14 | 0.3 | 0.48 | 0.21 | 0.39 |
| 8 | 8580453 | rs1522839 | 0.747 | 0.534 | -0.57 | -0.098 | 0.006 | -0.032 | -0.032 | 4.406 | 0.24 | 0.15 | 0.31 | 0.45 | 0.2 | 0.38 |
| 8 | 8583109 | rs777710 | -0.124 | 0.581 | -0.645 | -0.19 | -0.019 | -0.035 | -0.035 | 0.202 | 0.84 | 0.92 | 0.73 | 0.61 | 0.77 | 0.8 |
| 8 | 8583872 | rs777709 | 0.478 | 0.626 | -0.363 | -0.204 | -0.014 | 0.061 | 0.061 | 3.605 | 0.59 | 0.73 | 0.39 | 0.13 | 0.55 | 0.41 |
| 8 | 8584153 | rs11780672 | 1.583 | 0.693 | -0.618 | -0.355 | -0.066 | 0.056 | 0.056 | 0 | 0.12 | 0.02 | 0.1 | 0.03 | 0.17 | 0.19 |
| 8 | 8586016 | rs12678414 | 0.956 | 0.571 | -0.644 | -0.394 | -0.001 | -0.064 | -0.064 | 0.292 | 0.26 | 0.11 | 0.33 | 0.46 | 0.23 | 0.38 |
| 8 | 8587589 | rs17628982 | 2.818 | 0.622 | -0.263 | -0.738 | -0.001 | -0.049 | -0.049 | 1.284 | 0.25 | 0.21 | 0.15 | 0 | 0.22 | 0.14 |
| 8 | 8589783 | rs11781985 | 0.087 | 0.64 | -0.111 | -0.735 | 0.008 | -0.038 | -0.038 | 0 | 0.18 | 0.23 | 0.13 | 0.03 | 0.21 | 0.17 |
| 8 | 8593638 | rs570364 | -0.395 | 0.664 | -0.172 | -0.697 | -0.001 | -0.01 | -0.01 | 0.569 | 0.31 | 0.32 | 0.23 | 0.13 | 0.3 | 0.26 |
| 8 | 8596181 | rs559534 | 1.019 | 0.572 | -0.11 | -0.659 | -0.005 | -0.04 | -0.04 | 0 | 0.43 | 0.56 | 0.27 | 0.1 | 0.34 | 0.23 |
| 8 | 8596325 | rs17154677 | 0.755 | 0.819 | -0.155 | -0.15 | -0.045 | -0.017 | -0.017 | 0.205 | 0.07 | 0.01 | 0.24 | 0.35 | 0.16 | 0.16 |
| 8 | 8596562 | rs473118 | 0.412 | 0.423 | 0.036 | -0.69 | -0.033 | -0.036 | -0.036 | 0.619 | 0.18 | 0.33 | 0.12 | 0.1 | 0.13 | 0.11 |
| 8 | 8601350 | rs4841020 | 2.778 | 0.337 | -0.098 | -0.843 | 0 | -0.033 | -0.033 | 0 | 0.19 | 0.15 | 0.13 | 0 | 0.19 | 0.12 |
| 8 | 8603182 | rs4841026 | 0.486 | 0.315 | -0.11 | -0.858 | 0.001 | 0.019 | 0.019 | 0.465 | 0.5 | 0.59 | 0.35 | 0.12 | 0.49 | 0.38 |
| 8 | 8605632 | rs4841029 | 0.952 | 0.174 | -0.261 | -1.055 | -0.015 | 0.011 | 0.011 | 0 | 0.4 | 0.3 | 0.28 | 0.04 | 0.44 | 0.31 |
| 8 | 8606482 | rs484074 | -0.167 | 0.351 | -0.377 | -0.515 | -0.001 | 0.005 | 0.005 | 0.027 | 0.2 | 0.19 | 0.34 | 0.35 | 0.25 | 0.19 |
| 8 | 8606967 | rs568927 | -1.134 | 0.556 | -0.089 | -0.243 | 0.001 | -0.02 | -0.02 | 4.229 | 0.75 | 0.82 | 0.7 | 0.56 | 0.77 | 0.69 |
| 8 | 8607764 | rs11774744 | 0.124 | 0.505 | 0.276 | -0.32 | 0.002 | -0.033 | -0.033 | 0 | 0.22 | 0.23 | 0.16 | 0.03 | 0.26 | 0.18 |
| 8 | 8613387 | rs635594 | -0.819 | 0.627 | -0.036 | -0.045 | -0.009 | -0.028 | -0.028 | 0 | 0.68 | 0.8 | 0.62 | 0.53 | 0.65 | 0.63 |
| 8 | 8618430 | rs517394 | -0.436 | 0.663 | 0.361 | -0.175 | -0.003 | 0.112 | 0.112 | 0 | 0.13 | 0.2 | 0.3 | 0.34 | 0.13 | 0.15 |
| 8 | 8620141 | rs17154709 | 0.722 | 0.45 | 0.458 | -1.49 | 0.024 | -0.028 | -0.028 | 0 | 0.1 | 0.18 | 0.08 | 0.03 | 0.18 | 0.14 |

| | | | | | | | | | | | | | | | | |
|---|---|---|---|---|---|---|---|---|---|---|---|---|---|---|---|---|
| 8 | 8623637 | rs686189 | 0.169 | 0.303 | 0.227 | -1.586 | 0.008 | 0.19 | 0.19 | 0.001 | 0.47 | 0.81 | 0.3 | 0.12 | 0.45 | 0.35 |
| 8 | 8625041 | rs659256 | -0.876 | 0.489 | 0.482 | -1.092 | -0.025 | -0.012 | -0.012 | 0 | 0.13 | 0.4 | 0.08 | 0.08 | 0.09 | 0.03 |
| 8 | 8629936 | rs677039 | -0.675 | 0.523 | 0.374 | -1.115 | -0.004 | -0.016 | -0.016 | 0 | 0.23 | 0.23 | 0.14 | 0.08 | 0.17 | 0.08 |
| 8 | 8632957 | rs565203 | -1.124 | 0.738 | 0.205 | -0.971 | 0.03 | -0.089 | -0.089 | 0 | 0.3 | 0.52 | 0.22 | 0.12 | 0.33 | 0.2 |
| 8 | 8637957 | rs565169 | -0.789 | 0.504 | 0.245 | -0.928 | -0.021 | -0.09 | -0.09 | 0.433 | 0.23 | 0.48 | 0.16 | 0.08 | 0.18 | 0.08 |
| 8 | 8641125 | rs2427 | 1.559 | 0.809 | 0.194 | -0.812 | -0.037 | 0.023 | 0.023 | 0 | 0.07 | 0.01 | 0.06 | 0.03 | 0.15 | 0.11 |
| 8 | 8641145 | rs2428 | 0.128 | 0.527 | 0.494 | -1.299 | 0.002 | 0.04 | 0.04 | 0.601 | 0.45 | 0.36 | 0.63 | 0.88 | 0.46 | 0.67 |
| 8 | 8643387 | rs10903311 | -0.486 | 0.399 | 0.75 | -1.373 | -0.006 | -0.065 | -0.065 | 0 | 0.53 | 0.5 | 0.7 | 0.91 | 0.6 | 0.77 |
| 8 | 8644322 | rs12677550 | -1.023 | 0.408 | 0.91 | -1.437 | -0.012 | 0.065 | 0.065 | 0 | 0.63 | 0.74 | 0.74 | 0.89 | 0.59 | 0.74 |
| 8 | 8646301 | rs10093872 | 1.61 | 0.507 | 0.803 | -1.217 | -0.024 | 0.019 | 0.019 | 0 | 0.07 | 0.03 | 0.06 | 0.03 | 0.15 | 0.11 |
| 8 | 8649023 | rs10092600 | -0.508 | 0.42 | 1.099 | -1.282 | 0.03 | -0.118 | -0.118 | 0.016 | 0.26 | 0.42 | 0.18 | 0.03 | 0.29 | 0.19 |
| 8 | 8650939 | rs964771 | -0.593 | 0.411 | 1.355 | -1.199 | 0.033 | -0.109 | -0.109 | 0.113 | 0.26 | 0.52 | 0.18 | 0.05 | 0.29 | 0.19 |
| 8 | 8651879 | rs11774860 | 0.295 | 0.396 | 1.503 | -1.23 | 0.019 | -0.043 | -0.043 | 0 | 0.16 | 0.25 | 0.11 | 0.04 | 0.2 | 0.11 |
| 8 | 8652504 | rs11985640 | 0.079 | 0.396 | 1.531 | -1.238 | 0.021 | -0.051 | -0.051 | 0 | 0.17 | 0.32 | 0.12 | 0.05 | 0.2 | 0.11 |
| 8 | 8653478 | rs10093805 | 0.307 | 0.367 | 1.461 | -1.221 | 0.018 | -0.059 | -0.059 | 0 | 0.1 | 0.22 | 0.06 | 0 | 0.09 | 0.07 |
| 8 | 8655023 | rs3748144 | -1.819 | 0.253 | 1.364 | -1.307 | 0.017 | 0.038 | 0.038 | 0 | 0.7 | 0.81 | 0.81 | 0.92 | 0.74 | 0.85 |
| 8 | 8656647 | rs7831224 | -0.265 | 0.047 | 1.267 | -1.236 | -0.006 | 0.019 | 0.019 | 0.757 | 0.21 | 0.18 | 0.12 | 0.11 | 0.22 | 0.25 |
| 8 | 8659038 | rs3827811 | -1.303 | 0.031 | 1.231 | -1.261 | 0.007 | -0.054 | -0.054 | 0 | 0.32 | 0.44 | 0.19 | 0.11 | 0.31 | 0.39 |
| 8 | 8659559 | rs17154769 | -1.435 | -0.014 | 1.315 | -1.223 | -0.025 | -0.071 | -0.071 | 1.077 | 0.2 | 0.58 | 0.1 | 0.06 | 0.12 | 0.28 |
| 8 | 8664622 | rs4841042 | 0.448 | -0.021 | 1.236 | -1.24 | -0.003 | 0.006 | 0.006 | 5.521 | 0.38 | 0.33 | 0.61 | 0.8 | 0.42 | 0.52 |
| 8 | 8664940 | rs4841044 | -0.89 | 0.051 | 0.996 | -1.175 | -0.011 | 0.133 | 0.133 | 1.566 | 0.57 | 0.81 | 0.7 | 0.86 | 0.54 | 0.74 |
| 8 | 8669622 | rs10503390 | -0.497 | 0.139 | 1.164 | -1.129 | 0.037 | -0.093 | -0.093 | 0 | 0.13 | 0.49 | 0.08 | 0 | 0.1 | 0.14 |
| 8 | 8673601 | rs13259216 | 0.335 | 0.111 | 1.022 | -1.05 | -0.021 | 0.095 | 0.095 | 3.203 | 0.41 | 0.2 | 0.61 | 0.8 | 0.44 | 0.54 |
| 8 | 8676490 | rs950721 | -0.98 | 0.45 | 0.815 | -0.606 | 0 | -0.005 | -0.005 | 0.213 | 0.61 | 0.61 | 0.57 | 0.69 | 0.59 | 0.76 |
| 8 | 8681756 | rs1876836 | -1.706 | 0.323 | 0.451 | -1.038 | 0.013 | 0.029 | 0.029 | 0 | 0.71 | 0.82 | 0.81 | 0.91 | 0.74 | 0.85 |
| 8 | 8686845 | rs3789850 | -0.787 | 0.466 | 0.087 | -0.942 | 0.004 | -0.016 | -0.016 | 0 | 0.32 | 0.4 | 0.24 | 0.15 | 0.32 | 0.42 |
| 8 | 8687325 | rs7013471 | 0.371 | -0.279 | 0.018 | -0.611 | 0.014 | 0.11 | 0.11 | 0.652 | 0.52 | 0.34 | 0.38 | 0.15 | 0.53 | 0.45 |
| 8 | 8693386 | rs9650614 | 0.624 | -0.099 | -0.281 | -0.571 | 0 | -0.002 | -0.002 | 0.992 | 0.16 | 0.12 | 0.1 | 0 | 0.14 | 0.16 |
| 8 | 8695574 | rs6601265 | -1.448 | -0.098 | -0.277 | -0.503 | -0.038 | 0.121 | 0.121 | 0 | 0.64 | 0.83 | 0.75 | 0.87 | 0.59 | 0.84 |
| 8 | 8699161 | rs4841055 | -1.439 | -0.089 | -0.316 | -0.533 | -0.026 | 0.107 | 0.107 | 0 | 0.64 | 0.83 | 0.75 | 0.88 | 0.59 | 0.8 |
| 8 | 8702827 | rs907180 | -0.225 | -0.136 | -0.036 | -0.447 | 0 | 0.005 | 0.005 | 0.129 | 0.39 | 0.57 | 0.58 | 0.74 | 0.42 | 0.5 |
| 8 | 8702875 | rs907181 | -0.247 | -0.138 | -0.037 | -0.447 | 0 | 0.006 | 0.006 | 0.178 | 0.4 | 0.57 | 0.58 | 0.74 | 0.42 | 0.51 |
| 8 | 8705180 | rs11249895 | 0.047 | -0.18 | -0.237 | -0.382 | -0.004 | 0.008 | 0.008 | 0.577 | 0.19 | 0.11 | 0.16 | 0.13 | 0.16 | 0.19 |
| 8 | 8707970 | rs387706 | -1.008 | -0.291 | 0.001 | -0.525 | -0.014 | -0.044 | -0.044 | 0 | 0.31 | 0.59 | 0.23 | 0.14 | 0.23 | 0.38 |
| 8 | 8712895 | rs4348501 | 1.142 | -0.052 | -0.581 | -0.253 | -0.099 | 0.104 | 0.104 | 0 | 0.39 | 0.08 | 0.36 | 0.22 | 0.48 | 0.38 |
| 8 | 8713038 | rs4523255 | -0.806 | -0.008 | 0.235 | 1.33 | -0.012 | 0 | 0 | 1.341 | 0.37 | 0.17 | 0.43 | 0.67 | 0.36 | 0.51 |
| 8 | 8716166 | rs332029 | -1.352 | -0.2 | 0.751 | 1.308 | -0.004 | 0.076 | 0.076 | 0.907 | 0.1 | 0.34 | 0.17 | 0.15 | 0.06 | 0.17 |
| 8 | 8716971 | rs5023278 | 0.501 | -0.053 | 0.205 | 1.878 | -0.002 | -0.039 | -0.039 | 0 | 0.19 | 0.07 | 0.15 | 0.32 | 0.22 | 0.26 |
| 8 | 8718850 | rs1039917 | 0.126 | -0.304 | 0.343 | 1.564 | -0.063 | -0.119 | -0.119 | 0.898 | 0.28 | 0.07 | 0.36 | 0.49 | 0.35 | 0.38 |
| 8 | 8722378 | rs10108954 | -1.538 | -0.2 | 0.836 | 1.399 | 0.053 | 0.015 | 0.015 | 1.518 | 0.07 | 0.42 | 0.11 | 0 | 0.03 | 0.07 |
| 8 | 8724365 | rs7841466 | -1.753 | -0.201 | 0.401 | 1.539 | -0.007 | 0.179 | 0.179 | 3.654 | 0.1 | 0.55 | 0.21 | 0.2 | 0.05 | 0.15 |
| 8 | 8729193 | rs13274028 | 0.026 | 0.175 | -0.464 | 1.42 | -0.071 | -0.102 | -0.102 | 0 | 0.27 | 0.06 | 0.41 | 0.4 | 0.35 | 0.37 |
| 8 | 8729900 | rs10098667 | -1.576 | -0.204 | 0.607 | 1.152 | -0.024 | 0.029 | 0.029 | 0.62 | 0.13 | 0.29 | 0.17 | 0.18 | 0.05 | 0.27 |
| 8 | 8736386 | rs12679021 | 0.972 | -0.036 | 0.744 | 0.785 | -0.018 | 0.007 | 0.007 | 0.528 | 0.87 | 0.51 | 0.86 | 0.91 | 0.93 | 0.85 |
| 8 | 8737605 | rs410487 | 1.195 | 0.187 | 0.64 | 1.22 | -0.011 | 0.094 | 0.094 | 0 | 0.87 | 0.65 | 0.79 | 0.75 | 0.9 | 0.93 |
| 8 | 8738868 | rs13249564 | 0.201 | 0.285 | 0.444 | 0.485 | -0.012 | -0.029 | -0.029 | 0 | 0.22 | 0.43 | 0.21 | 0.16 | 0.18 | 0.14 |
| 8 | 8744240 | rs7818276 | 0.038 | 1.101 | -0.569 | 0.867 | -0.088 | -0.138 | -0.138 | 0.882 | 0.2 | 0.06 | 0.36 | 0.49 | 0.28 | 0.27 |
| 8 | 8747245 | rs4841067 | 0.89 | 0.742 | -0.21 | 0.447 | 0.001 | -0.125 | -0.125 | 0 | 0.19 | 0.35 | 0.13 | 0 | 0.16 | 0.15 |
| 8 | 8747537 | rs399123 | -0.52 | 0.363 | -0.966 | -0.24 | -0.045 | 0.166 | 0.166 | 3.67 | 0.41 | 0.18 | 0.63 | 0.88 | 0.48 | 0.52 |
| 8 | 8752684 | rs3762032 | 1.312 | 0.599 | -0.397 | -0.096 | -0.004 | -0.01 | -0.01 | 0.148 | 0.2 | 0.34 | 0.18 | 0.19 | 0.17 | 0.15 |
| 8 | 8758058 | rs1077951 | 0.221 | 0.737 | -0.425 | 0.324 | -0.034 | -0.06 | -0.06 | 0.672 | 0.21 | 0.15 | 0.38 | 0.44 | 0.28 | 0.28 |
| 8 | 8759261 | rs332013 | 0.432 | 0.686 | -0.182 | 0.032 | 0.025 | 0.004 | 0.004 | 0.82 | 0.71 | 0.5 | 0.5 | 0.32 | 0.65 | 0.67 |
| 8 | 8760138 | rs13282211 | 0.261 | 0.859 | -0.663 | 0.277 | -0.092 | -0.157 | -0.157 | 0 | 0.23 | 0.05 | 0.38 | 0.47 | 0.28 | 0.32 |
| 8 | 8761295 | rs7004322 | -0.06 | 0.574 | -0.873 | -1.229 | 0.005 | -0.018 | -0.018 | 0 | 0.32 | 0.35 | 0.32 | 0.24 | 0.37 | 0.37 |
| 8 | 8762040 | rs10093912 | 1.148 | 0.676 | -0.849 | -1.89 | -0.021 | 0.056 | 0.056 | 0.33 | 0.2 | 0.1 | 0.12 | 0.03 | 0.22 | 0.27 |
| 8 | 8764824 | rs10097305 | -0.195 | 0.668 | -0.714 | -1.845 | 0.027 | -0.045 | -0.045 | 0 | 0.09 | 0.15 | 0.11 | 0 | 0.13 | 0.07 |
| 8 | 8768160 | rs433724 | -0.845 | 0.532 | -0.533 | -1.846 | -0.013 | -0.084 | -0.084 | 0.359 | 0.57 | 0.51 | 0.74 | 0.9 | 0.63 | 0.76 |
| 8 | 8768319 | rs433960 | -0.797 | 0.554 | -0.859 | -1.835 | 0.004 | -0.015 | -0.015 | 4.687 | 0.56 | 0.41 | 0.74 | 0.9 | 0.63 | 0.75 |
| 8 | 8770454 | rs400404 | 0.493 | 0.358 | -0.985 | -1.765 | -0.005 | 0.056 | 0.056 | 4.812 | 0.28 | 0.17 | 0.52 | 0.86 | 0.28 | 0.42 |
| 8 | 8771402 | rs373204 | -0.587 | 0.207 | -0.851 | -1.732 | 0.009 | 0.038 | 0.038 | 0 | 0.56 | 0.39 | 0.71 | 0.91 | 0.59 | 0.74 |
| 8 | 8779328 | rs11775257 | 0.407 | 0.415 | -0.988 | -1.466 | -0.039 | 0.109 | 0.109 | 0.427 | 0.25 | 0.1 | 0.16 | 0.03 | 0.27 | 0.26 |
| 8 | 8779418 | rs10104541 | 0.366 | 0.387 | -0.906 | -1.467 | -0.015 | 0.053 | 0.053 | 0.462 | 0.25 | 0.16 | 0.16 | 0.03 | 0.27 | 0.26 |
| 8 | 8783465 | rs332022 | -1.242 | 0.025 | -0.279 | -1.564 | -0.002 | -0.164 | -0.164 | 0.127 | 0.77 | 0.6 | 0.86 | 1 | 0.79 | 0.92 |
| 8 | 8786490 | rs2941658 | -0.415 | -0.137 | -0.422 | -1.589 | -0.008 | -0.054 | -0.054 | 0.404 | 0.66 | 0.48 | 0.79 | 0.89 | 0.7 | 0.82 |
| 8 | 8793654 | rs17700611 | 0.94 | 0.026 | -0.329 | -1.596 | -0.051 | 0.132 | 0.132 | 0.212 | 0.4 | 0.18 | 0.27 | 0 | 0.46 | 0.18 |
| 8 | 8794251 | rs6601272 | -0.229 | 0.049 | -0.015 | -1.675 | -0.011 | -0.016 | -0.016 | 0 | 0.15 | 0.41 | 0.08 | 0.1 | 0.1 | 0.1 |
| 8 | 8794454 | rs891570 | -0.689 | -0.232 | -0.197 | -1.63 | -0.019 | 0.072 | 0.072 | 4.063 | 0.63 | 0.73 | 0.38 | 0.18 | 0.58 | 0.47 |
| 8 | 8802067 | rs7007974 | -0.166 | -0.391 | 0.074 | -1.463 | -0.016 | -0.02 | -0.02 | 0.546 | 0.15 | 0.47 | 0.09 | 0.1 | 0.11 | 0.1 |
| 8 | 8802082 | rs6987224 | -1.122 | -0.446 | -0.009 | -1.445 | -0.032 | -0.017 | -0.017 | 0 | 0.23 | 0.6 | 0.12 | 0.18 | 0.12 | 0.26 |

| | | | | | | | | | | | | | | | | |
|---|---|---|---|---|---|---|---|---|---|---|---|---|---|---|---|---|
| 8 | 8805969 | rs17154995 | -0.342 | -0.127 | 0.063 | -1.613 | 0.006 | -0.006 | -0.006 | 0.007 | 0.62 | 0.62 | 0.41 | 0.05 | 0.66 | 0.33 |
| 8 | 8807259 | rs6601276 | -0.002 | -0.125 | 0.15 | -1.568 | -0.012 | -0.047 | -0.047 | 0 | 0.15 | 0.41 | 0.08 | 0.05 | 0.1 | 0.1 |
| 8 | 8808695 | rs11785268 | -0.142 | -0.036 | 0.407 | -1.488 | -0.01 | -0.072 | -0.072 | 1.517 | 0.59 | 0.54 | 0.4 | 0.05 | 0.65 | 0.29 |
| 8 | 8809696 | rs11776020 | -0.526 | 0.055 | 0.218 | -1.198 | 0.001 | 0.002 | 0.002 | 1.209 | 0.65 | 0.63 | 0.43 | 0.18 | 0.67 | 0.33 |
| 8 | 8814890 | rs11778913 | -0.629 | -0.077 | 0.139 | -1.2 | 0.038 | 0.051 | 0.051 | 0.325 | 0.61 | 0.74 | 0.81 | 0.77 | 0.75 | 0.63 |
| 8 | 8821318 | rs6981573 | 0.268 | -0.025 | 0.205 | -0.277 | 0.006 | 0.004 | 0.004 | 0.148 | 0.39 | 0.38 | 0.55 | 0.69 | 0.35 | 0.56 |
| 8 | 8821637 | rs13259143 | 0.316 | -0.029 | 0.208 | -0.258 | 0.006 | 0.014 | 0.014 | 0.221 | 0.38 | 0.34 | 0.54 | 0.74 | 0.36 | 0.56 |
| 8 | 8824858 | rs12542733 | 0.082 | -0.047 | 0.185 | -0.385 | -0.002 | 0.12 | 0.12 | 2.015 | 0.43 | 0.3 | 0.63 | 0.93 | 0.46 | 0.6 |
| 8 | 8831662 | rs11778747 | 1.212 | 0.147 | 0.6 | -0.345 | 0.072 | -0.127 | -0.127 | 0 | 0.18 | 0.36 | 0.14 | 0 | 0.25 | 0.06 |
| 8 | 8832086 | rs12677456 | 0.65 | -0.409 | 0.409 | 0.21 | 0.005 | -0.008 | -0.008 | 0.06 | 0.11 | 0.11 | 0.12 | 0.23 | 0.08 | 0.11 |
| 8 | 8834240 | rs10103953 | -0.167 | -0.46 | 0.487 | 0.445 | -0.017 | 0.008 | 0.008 | 0.139 | 0.6 | 0.55 | 0.73 | 0.52 | 0.7 | 0.59 |
| 8 | 8837354 | rs7015461 | -0.881 | -0.521 | 0.275 | 0.321 | 0.021 | 0.043 | 0.043 | 0 | 0.68 | 0.75 | 0.82 | 0.95 | 0.77 | 0.8 |
| 8 | 8840042 | rs9644674 | 0.719 | -0.44 | 0.32 | 0.449 | -0.002 | 0.02 | 0.02 | 0 | 0.11 | 0.13 | 0.12 | 0.23 | 0.08 | 0.11 |
| 8 | 8840583 | rs10107215 | -0.066 | -0.535 | 0.448 | 0.438 | -0.007 | 0 | 0 | 0 | 0.6 | 0.6 | 0.74 | 0.6 | 0.71 | 0.6 |
| 8 | 8841162 | rs4840384 | -1.323 | -0.501 | 0.386 | 0.478 | 0.02 | 0.004 | 0.004 | 0 | 0.73 | 0.82 | 0.87 | 0.77 | 0.8 | 0.85 |
| 8 | 8849443 | rs4472520 | -0.687 | -0.293 | 0.548 | 0.418 | -0.019 | 0.061 | 0.061 | 0 | 0.17 | 0.25 | 0.15 | 0.26 | 0.11 | 0.22 |
| 8 | 8859747 | rs2956242 | 0.304 | -0.358 | 0.627 | 0.436 | -0.036 | 0.087 | 0.087 | 0 | 0.84 | 0.67 | 0.85 | 0.74 | 0.91 | 0.78 |
| 8 | 8861184 | rs2979261 | -0.259 | -0.327 | 0.487 | 0.444 | -0.005 | 0.016 | 0.016 | 0 | 0.87 | 0.81 | 0.87 | 0.76 | 0.91 | 0.88 |
| 8 | 8867471 | rs17701980 | 1.168 | -0.065 | 0.472 | -0.118 | -0.033 | 0.038 | 0.038 | 0.474 | 0.22 | 0.09 | 0.16 | 0 | 0.3 | 0.14 |
| 8 | 8869199 | rs9650616 | 0.57 | -0.366 | 0.973 | -0.421 | 0.003 | -0.114 | -0.114 | 0.684 | 0.44 | 0.44 | 0.28 | 0 | 0.47 | 0.26 |
| 8 | 8873293 | rs7017868 | 0.44 | -0.387 | 0.857 | -0.433 | -0.018 | 0.137 | 0.137 | 0.194 | 0.46 | 0.68 | 0.3 | 0 | 0.47 | 0.26 |
| 8 | 8875128 | rs11781203 | 1.196 | -0.448 | 0.835 | -0.47 | 0.007 | -0.063 | -0.063 | 0 | 0.23 | 0.22 | 0.16 | 0 | 0.31 | 0.13 |
| 8 | 8875238 | rs13250919 | 2.324 | -0.504 | 0.99 | -0.428 | 0 | -0.032 | -0.032 | 0 | 0.2 | 0.19 | 0.11 | 0 | 0.17 | 0.12 |
| 8 | 8881820 | rs1016512 | 2.352 | -0.501 | 0.948 | -0.426 | 0.004 | 0.017 | 0.017 | 0 | 0.19 | 0.1 | 0.11 | 0 | 0.16 | 0.12 |
| 8 | 8888948 | rs2979247 | 0.948 | -0.769 | 0.651 | -0.014 | -0.08 | -0.095 | -0.095 | 0 | 0.35 | 0.11 | 0.51 | 0.49 | 0.41 | 0.42 |
| 8 | 8890041 | rs1045527 | 0.606 | -0.725 | 0.849 | -0.244 | -0.017 | 0.084 | 0.084 | 0.063 | 0.46 | 0.63 | 0.3 | 0 | 0.48 | 0.25 |
| 8 | 8894700 | rs2979258 | -0.878 | -0.757 | 0.798 | -0.171 | -0.012 | -0.018 | -0.018 | 0 | 0.87 | 0.79 | 0.91 | 0.94 | 0.93 | 0.83 |
| 8 | 8895470 | rs7016139 | 0.513 | -0.746 | 0.642 | -0.117 | -0.079 | -0.007 | -0.007 | 0 | 0.4 | 0.11 | 0.57 | 0.7 | 0.45 | 0.54 |
| 8 | 8899621 | rs7828944 | 1.042 | -0.641 | 0.889 | -0.155 | 0.032 | -0.154 | -0.154 | 0.275 | 0.26 | 0.42 | 0.18 | 0 | 0.29 | 0.14 |
| 8 | 8899815 | rs4841087 | 0.885 | -0.56 | 0.451 | 0.424 | -0.001 | 0.104 | 0.104 | 2.125 | 0.52 | 0.21 | 0.51 | 0.3 | 0.54 | 0.46 |
| 8 | 8901140 | rs2979269 | 0.815 | -0.534 | 0.477 | 0.365 | -0.006 | 0.095 | 0.095 | 1.503 | 0.52 | 0.22 | 0.5 | 0.32 | 0.53 | 0.47 |
| 8 | 8902424 | rs1562211 | 0.389 | -0.456 | 0.178 | 0.151 | -0.069 | -0.001 | -0.001 | 0.632 | 0.4 | 0.12 | 0.58 | 0.7 | 0.46 | 0.53 |
| 8 | 8907557 | rs13273769 | 2.276 | -0.291 | 0.393 | -0.266 | 0.005 | -0.003 | -0.003 | 0 | 0.2 | 0.16 | 0.11 | 0 | 0.17 | 0.1 |
| 8 | 8908596 | rs17702602 | 0.449 | -0.207 | 0.258 | -0.242 | -0.005 | 0.103 | 0.103 | 0.758 | 0.47 | 0.66 | 0.29 | 0.03 | 0.45 | 0.37 |
| 8 | 8911308 | rs3989373 | 0.439 | -0.29 | 0.249 | -0.256 | -0.01 | 0.093 | 0.093 | 0.537 | 0.47 | 0.68 | 0.29 | 0.03 | 0.45 | 0.37 |
| 8 | 8912705 | rs2100021 | 0.51 | -0.403 | 0.413 | 0.423 | -0.012 | 0.015 | 0.015 | 0 | 0.32 | 0.39 | 0.36 | 0.44 | 0.27 | 0.39 |
| 8 | 8915740 | rs4841092 | 1.971 | -0.231 | 0.339 | -0.275 | -0.001 | -0.038 | -0.038 | 0.015 | 0.2 | 0.19 | 0.11 | 0 | 0.17 | 0.12 |
| 8 | 8923585 | rs1900102 | -0.284 | -0.166 | 0.371 | -0.069 | 0.017 | -0.011 | -0.011 | 0 | 0.64 | 0.54 | 0.7 | 0.73 | 0.6 | 0.8 |
| 8 | 8928744 | rs920974 | 0.155 | -0.379 | 0.064 | -0.167 | 0.036 | 0.009 | 0.009 | 0.776 | 0.33 | 0.16 | 0.52 | 0.73 | 0.27 | 0.47 |
| 8 | 8929596 | rs11249900 | -1.077 | -0.182 | 0.614 | 0.332 | 0.002 | 0.257 | 0.257 | 0.158 | 0.05 | 0.24 | 0.07 | 0.25 | 0.02 | 0.14 |
| 8 | 8930740 | rs10087078 | 1.261 | -0.363 | 0.597 | -0.5 | -0.027 | -0.116 | -0.116 | 0.284 | 0.23 | 0.42 | 0.14 | 0.03 | 0.17 | 0.19 |
| 8 | 8934720 | rs6997088 | 0.178 | 0.016 | 0.346 | -1.121 | 0.003 | -0.038 | -0.038 | 1.116 | 0.25 | 0.32 | 0.22 | 0.05 | 0.28 | 0.17 |
| 8 | 8936101 | rs17155176 | 1.679 | 0.217 | -0.192 | -1.419 | -0.025 | 0.02 | 0.02 | 2.41 | 0.18 | 0.12 | 0.12 | 0 | 0.23 | 0.11 |
| 8 | 8936749 | rs6990464 | -1.516 | -0.277 | -0.801 | -0.643 | -0.038 | -0.108 | -0.108 | 1.475 | 0.85 | 0.98 | 0.74 | 0.55 | 0.81 | 0.69 |
| 8 | 8936962 | rs10090227 | -0.429 | -0.113 | -0.413 | -1.379 | 0.03 | 0.171 | 0.171 | 0 | 0.5 | 0.76 | 0.38 | 0.05 | 0.53 | 0.34 |
| 8 | 8939373 | rs2979245 | -1.01 | -0.205 | -0.531 | -1.061 | 0.006 | -0.03 | -0.03 | 0.875 | 0.78 | 0.86 | 0.57 | 0.27 | 0.79 | 0.58 |
| 8 | 8941044 | rs17703328 | 0.663 | 0.283 | -0.266 | -1.711 | -0.014 | 0.013 | 0.013 | 0 | 0.26 | 0.19 | 0.25 | 0.03 | 0.35 | 0.19 |
| 8 | 8945102 | rs7009435 | 0.111 | 0.165 | -0.376 | -1.761 | -0.013 | 0.213 | 0.213 | 0 | 0.52 | 0.75 | 0.41 | 0.06 | 0.53 | 0.4 |
| 8 | 8946163 | rs4841097 | -0.95 | 0.177 | -0.283 | -1.677 | 0.012 | 0.024 | 0.024 | 2.24 | 0.76 | 0.82 | 0.56 | 0.12 | 0.78 | 0.55 |
| 8 | 8946391 | rs7825188 | 1.792 | 0.569 | -0.267 | -1.706 | -0.021 | 0.042 | 0.042 | 2.927 | 0.18 | 0.09 | 0.1 | 0 | 0.19 | 0.1 |
| 8 | 8948469 | rs6992247 | -0.002 | 0.452 | 0.041 | -1.849 | -0.012 | -0.082 | -0.082 | 1.793 | 0.63 | 0.62 | 0.52 | 0.03 | 0.72 | 0.36 |
| 8 | 8948751 | rs11784750 | 0.971 | 0.862 | 0.741 | -1.3 | 0.001 | -0.052 | -0.052 | 0 | 0.2 | 0.21 | 0.12 | 0 | 0.19 | 0.11 |
| 8 | 8952019 | rs4841098 | -0.102 | 0.786 | 0.637 | -1.31 | 0.023 | 0.065 | 0.065 | 0 | 0.39 | 0.68 | 0.26 | 0.12 | 0.32 | 0.31 |
| 8 | 8959386 | rs13264785 | 0.298 | 0.901 | 0.787 | -1.243 | -0.001 | -0.029 | -0.029 | 0 | 0.34 | 0.62 | 0.23 | 0.11 | 0.31 | 0.17 |
| 8 | 8961017 | rs1809006 | 0.893 | 0.869 | 0.985 | -1.263 | 0 | -0.02 | -0.02 | 0.018 | 0.3 | 0.34 | 0.2 | 0.08 | 0.31 | 0.15 |
| 8 | 8961149 | rs895251 | 0.283 | 0.906 | 0.749 | -1.236 | -0.002 | -0.025 | -0.025 | 0 | 0.34 | 0.61 | 0.23 | 0.13 | 0.31 | 0.17 |
| 8 | 8964626 | rs17155236 | 1.6 | 1.277 | 0.467 | -1.163 | -0.002 | -0.015 | -0.015 | 0 | 0.16 | 0.14 | 0.09 | 0 | 0.15 | 0.08 |
| 8 | 8964985 | rs2898208 | 0.562 | 1.066 | 0.593 | -0.864 | 0.006 | -0.008 | -0.008 | 1.667 | 0.13 | 0.21 | 0.1 | 0.08 | 0.14 | 0.1 |
| 8 | 8966606 | rs17155242 | 0.037 | 1.274 | 0.174 | -0.868 | -0.003 | -0.061 | -0.061 | 0 | 0.35 | 0.53 | 0.23 | 0.1 | 0.33 | 0.17 |
| 8 | 8966972 | rs2979248 | -0.425 | 1.141 | 0.251 | -0.452 | -0.004 | 0.034 | 0.034 | 0 | 0.44 | 0.62 | 0.36 | 0.24 | 0.48 | 0.42 |
| 8 | 8967105 | rs17155244 | -0.271 | 1.185 | 0.364 | -0.436 | 0.014 | -0.034 | -0.034 | 0.025 | 0.19 | 0.46 | 0.14 | 0.11 | 0.16 | 0.23 |
| 8 | 8971100 | rs7822664 | -0.161 | 1.197 | 0.267 | -0.407 | 0.011 | -0.023 | -0.023 | 0 | 0.18 | 0.46 | 0.14 | 0.11 | 0.16 | 0.23 |
| 8 | 8972515 | rs7011104 | 1.137 | 1.382 | 0.006 | -0.265 | -0.002 | 0.001 | 0.001 | 3.25 | 0.19 | 0.19 | 0.18 | 0.19 | 0.22 | 0.11 |
| 8 | 8973846 | rs13439673 | -0.213 | 1.211 | -0.54 | -0.162 | -0.012 | 0.037 | 0.037 | 3.188 | 0.37 | 0.66 | 0.27 | 0.11 | 0.38 | 0.34 |
| 8 | 8973858 | rs12155586 | -0.685 | 1.183 | -0.865 | -0.151 | 0.031 | -0.036 | -0.036 | 0 | 0.63 | 0.83 | 0.5 | 0.4 | 0.67 | 0.53 |
| 8 | 8979776 | rs12334631 | -0.004 | 1.203 | -0.632 | -1.01 | 0.007 | -0.039 | -0.039 | 0.015 | 0.34 | 0.56 | 0.26 | 0.1 | 0.36 | 0.23 |
| 8 | 8982396 | rs753012 | -0.894 | 1.093 | -0.91 | -1.013 | 0.033 | 0.03 | 0.03 | 1.041 | 0.64 | 0.9 | 0.5 | 0.24 | 0.7 | 0.56 |
| 8 | 8984302 | rs10111820 | -0.008 | 1.085 | -0.686 | -1.041 | 0.001 | -0.017 | -0.017 | 0 | 0.35 | 0.64 | 0.26 | 0.1 | 0.36 | 0.23 |
| 8 | 8986613 | rs7836377 | -0.288 | 1.053 | -0.793 | -0.937 | -0.018 | 0.041 | 0.041 | 1.892 | 0.39 | 0.71 | 0.3 | 0.17 | 0.4 | 0.38 |

| | | | | | | | | | | | | | | | | |
|---|---|---|---|---|---|---|---|---|---|---|---|---|---|---|---|---|
| 8 | 8990577 | rs189798 | -0.491 | 0.71 | -0.98 | -0.637 | 0.044 | -0.009 | -0.009 | 0 | 0.63 | 0.76 | 0.55 | 0.33 | 0.72 | 0.57 |
| 8 | 8994512 | rs9949 | 0.417 | 0.654 | -0.881 | -1.184 | 0.042 | 0.019 | 0.019 | 0.133 | 0.36 | 0.19 | 0.38 | 0.73 | 0.28 | 0.42 |
| 8 | 8995166 | rs330908 | 0.13 | 0.626 | -0.806 | -1.216 | 0.001 | 0.001 | 0.001 | 0 | 0.4 | 0.37 | 0.4 | 0.77 | 0.29 | 0.62 |
| 8 | 8996273 | rs330911 | 0.449 | 0.566 | -1.394 | -1.246 | 0.042 | 0.037 | 0.037 | 0.857 | 0.35 | 0.14 | 0.38 | 0.76 | 0.29 | 0.53 |
| 8 | 8997415 | rs330919 | 0.254 | 0.708 | -1.31 | -1.155 | 0.055 | 0.053 | 0.053 | 0.029 | 0.38 | 0.14 | 0.38 | 0.77 | 0.3 | 0.54 |
| 8 | 8997614 | rs330920 | 0.207 | 0.706 | -1.308 | -1.162 | 0.057 | 0.052 | 0.052 | 0 | 0.38 | 0.14 | 0.38 | 0.76 | 0.3 | 0.54 |
| 8 | 8997623 | rs330921 | 0.442 | 0.714 | -1.326 | -1.148 | 0.043 | -0.005 | -0.005 | 0 | 0.34 | 0.08 | 0.37 | 0.71 | 0.28 | 0.38 |
| 8 | 8998022 | rs17653841 | 0.936 | 1.119 | -1.239 | -1.974 | -0.075 | 0.105 | 0.105 | 1.808 | 0.18 | 0.03 | 0.12 | 0 | 0.22 | 0.06 |
| 8 | 9009906 | rs19334 | 0.09 | 0.652 | -1.138 | -1.655 | 0.038 | 0.06 | 0.06 | 2.791 | 0.39 | 0.28 | 0.54 | 0.84 | 0.31 | 0.48 |
| 8 | 9017979 | rs330938 | 0.066 | 0.87 | -1.087 | -1.611 | 0.037 | 0.079 | 0.079 | 3.123 | 0.41 | 0.23 | 0.49 | 0.8 | 0.38 | 0.5 |
| 8 | 9021026 | rs330943 | 0.056 | 0.766 | -1.394 | -1.622 | 0.048 | 0.11 | 0.11 | 2.788 | 0.4 | 0.16 | 0.54 | 0.83 | 0.33 | 0.48 |
| 8 | 9026639 | rs378974 | 0.163 | 0.625 | -1.604 | -1.673 | 0.056 | 0.091 | 0.091 | 2.998 | 0.39 | 0.09 | 0.45 | 0.81 | 0.32 | 0.5 |
| 8 | 9028261 | rs1530483 | 1.56 | 0.346 | -1.927 | -2.438 | -0.038 | 0.098 | 0.098 | 0.325 | 0.19 | 0 | 0.09 | 0.04 | 0.18 | 0.21 |
| 8 | 9029413 | rs435581 | -0.55 | 0.256 | -0.83 | -1.916 | 0.006 | 0.032 | 0.032 | 3.012 | 0.48 | 0.57 | 0.68 | 0.92 | 0.48 | 0.61 |
| 8 | 9034148 | rs367543 | -0.766 | 0.263 | -0.863 | -1.844 | -0.002 | 0.049 | 0.049 | 1.857 | 0.52 | 0.58 | 0.72 | 0.91 | 0.54 | 0.59 |
| 8 | 9038596 | rs1994020 | -0.534 | 0.307 | -1.595 | -1.528 | -0.005 | 0.048 | 0.048 | 0.021 | 0.4 | 0.28 | 0.6 | 0.92 | 0.41 | 0.5 |
| 8 | 9042454 | rs11249905 | -0.58 | 0.19 | -2.061 | -1.309 | -0.006 | 0.198 | 0.198 | 1.432 | 0.39 | 0.11 | 0.52 | 0.91 | 0.33 | 0.52 |
| 8 | 9044856 | rs4841106 | -0.647 | 0.216 | -1.704 | -0.887 | -0.001 | 0.024 | 0.024 | 1.622 | 0.29 | 0.2 | 0.48 | 0.84 | 0.32 | 0.62 |
| 8 | 9046312 | rs1968853 | -0.922 | 0.123 | -1.697 | -0.767 | 0.004 | 0.016 | 0.016 | 5.711 | 0.47 | 0.4 | 0.53 | 0.63 | 0.44 | 0.57 |
| 8 | 9048104 | rs2929301 | 0.727 | -0.128 | -1.133 | -0.752 | -0.008 | -0.011 | -0.011 | 0 | 0.66 | 0.57 | 0.56 | 0.28 | 0.67 | 0.39 |
| 8 | 9049381 | rs2929478 | 0.315 | -0.171 | -0.913 | -0.284 | -0.012 | 0.041 | 0.041 | 6.137 | 0.72 | 0.55 | 0.63 | 0.36 | 0.71 | 0.51 |
| 8 | 9056151 | rs2242281 | 0.972 | -0.151 | -0.89 | -0.418 | -0.026 | -0.032 | -0.032 | 0 | 0.66 | 0.5 | 0.58 | 0.24 | 0.74 | 0.38 |
| 8 | 9057175 | rs17654950 | -0.267 | 0.02 | -0.791 | -0.419 | -0.001 | 0.001 | 0.001 | 4.4 | 0.26 | 0.21 | 0.38 | 0.77 | 0.25 | 0.51 |
| 8 | 9062490 | rs2929470 | -1.428 | -0.104 | -0.984 | -1.128 | -0.001 | 0.021 | 0.021 | 0.172 | 0.62 | 0.75 | 0.75 | 1 | 0.56 | 0.79 |
| 8 | 9064580 | rs2929464 | 0.013 | -0.095 | -0.886 | -0.989 | -0.027 | -0.036 | -0.036 | 0.547 | 0.93 | 0.82 | 0.97 | 1 | 0.98 | 0.89 |
| 8 | 9066272 | rs2929462 | -0.103 | 0.067 | -0.898 | -0.915 | -0.001 | -0.018 | -0.018 | 0 | 0.33 | 0.44 | 0.35 | 0.25 | 0.26 | 0.26 |
| 8 | 9069476 | rs2929292 | 0.186 | 0.107 | -1.252 | 0.079 | -0.012 | 0.003 | 0.003 | 0.011 | 0.91 | 0.93 | 0.74 | 0.71 | 0.79 | 0.83 |
| 8 | 9070973 | rs2929290 | -0.21 | -0.023 | -1.149 | 0.115 | 0.005 | -0.002 | -0.002 | 0 | 0.34 | 0.54 | 0.44 | 0.31 | 0.28 | 0.27 |
| 8 | 9071416 | rs17149544 | 0.422 | -0.166 | -1.131 | 0.411 | 0.001 | -0.001 | -0.001 | 0 | 0.17 | 0.15 | 0.07 | 0.35 | 0.08 | 0.26 |
| 8 | 9075279 | rs17149548 | 0.25 | -0.18 | -1.182 | 0.331 | 0.015 | -0.023 | -0.023 | 0.533 | 0.17 | 0.1 | 0.11 | 0.36 | 0.12 | 0.27 |
| 8 | 9075705 | rs2929286 | 0.176 | -0.037 | -1.116 | 0.462 | -0.023 | -0.021 | -0.021 | 0 | 0.91 | 0.94 | 0.77 | 0.75 | 0.79 | 0.84 |
| 8 | 9079297 | rs10087768 | 0.271 | -0.227 | -1.041 | 0.386 | -0.002 | 0.015 | 0.015 | 2.838 | 0.18 | 0.16 | 0.1 | 0.47 | 0.1 | 0.29 |
| 8 | 9085217 | rs2929305 | 0.656 | -0.124 | -1.123 | -0.419 | -0.009 | 0.042 | 0.042 | 2.079 | 0.55 | 0.61 | 0.33 | 0.03 | 0.52 | 0.23 |
| 8 | 9087278 | rs330048 | 0.731 | -0.045 | -1.102 | -0.379 | -0.013 | 0.051 | 0.051 | 2.479 | 0.56 | 0.63 | 0.33 | 0.03 | 0.53 | 0.27 |
| 8 | 9088245 | rs330052 | 0.205 | 0.578 | -0.67 | -0.319 | -0.001 | -0.064 | -0.064 | 0 | 0.24 | 0.28 | 0.14 | 0.03 | 0.22 | 0.05 |
| 8 | 9094288 | rs4840399 | -0.792 | 0.344 | -0.276 | 0.341 | -0.001 | 0.082 | 0.082 | 0 | 0.15 | 0.34 | 0.28 | 0.26 | 0.15 | 0.14 |
| 8 | 9095896 | rs17149592 | 0.146 | 0.859 | -0.861 | 0.764 | -0.057 | -0.075 | -0.075 | 0 | 0.12 | 0.04 | 0.19 | 0.21 | 0.16 | 0.16 |
| 8 | 9097767 | rs17707436 | 1.597 | 0.708 | -0.607 | -0.081 | -0.013 | 0.038 | 0.038 | 2.405 | 0.32 | 0.19 | 0.22 | 0 | 0.39 | 0.13 |
| 8 | 9097793 | rs419477 | 0.775 | 0.948 | -0.186 | 1.068 | 0.067 | 0.178 | 0.178 | 0 | 0.92 | 0.7 | 0.77 | 0.74 | 0.87 | 0.86 |
| 8 | 9100576 | rs17656431 | -0.445 | 1.163 | -1.064 | 0.678 | -0.065 | -0.04 | -0.04 | 0 | 0.15 | 0.01 | 0.12 | 0.15 | 0.18 | 0.13 |
| 8 | 9101842 | rs1551790 | 0.167 | 0.622 | -0.436 | 0.139 | -0.037 | -0.08 | -0.08 | 9.141 | 0.82 | 0.6 | 0.93 | 0.94 | 0.9 | 0.84 |
| 8 | 9103000 | rs330037 | 1.113 | 0.401 | -0.778 | -0.036 | -0.024 | 0.078 | 0.078 | 2.886 | 0.61 | 0.83 | 0.47 | 0.23 | 0.61 | 0.47 |
| 8 | 9104301 | rs172957 | -0.582 | 0.284 | -0.505 | -0.196 | -0.016 | -0.024 | -0.024 | 0 | 0.87 | 0.84 | 0.96 | 0.94 | 0.92 | 0.89 |
| 8 | 9104831 | rs1011158 | -1.744 | 0.317 | -0.539 | -0.154 | -0.025 | 0.219 | 0.219 | 0.022 | 0.48 | 0.17 | 0.58 | 0.9 | 0.46 | 0.64 |
| 8 | 9105172 | rs330042 | -0.856 | 0.406 | -0.469 | -0.128 | 0 | -0.017 | -0.017 | 0.024 | 0.29 | 0.37 | 0.16 | 0.21 | 0.25 | 0.26 |
| 8 | 9106638 | rs17149626 | 0.443 | 0.381 | -1.167 | 0.611 | -0.045 | -0.029 | -0.029 | 0 | 0.08 | 0.01 | 0.28 | 0.3 | 0.13 | 0.09 |
| 8 | 9109216 | rs17149632 | -0.12 | 0.467 | -1.261 | -0.482 | -0.05 | -0.074 | -0.074 | 0 | 0.2 | 0.04 | 0.5 | 0.68 | 0.26 | 0.41 |
| 8 | 9110822 | rs6984551 | -1.118 | 0.715 | -0.833 | -0.805 | -0.011 | 0.019 | 0.019 | 0 | 0.33 | 0.25 | 0.2 | 0.22 | 0.32 | 0.3 |
| 8 | 9112572 | rs4841112 | 2.555 | 0.762 | -1.133 | -1.129 | -0.052 | 0.124 | 0.124 | 2.069 | 0.29 | 0.09 | 0.2 | 0 | 0.4 | 0.11 |
| 8 | 9121398 | rs329997 | 0.809 | 0.319 | -0.716 | -0.837 | 0.001 | -0.006 | -0.006 | 0 | 0.71 | 0.3 | 0.75 | 0.83 | 0.74 | 0.78 |
| 8 | 9132309 | rs330023 | 0.664 | 0.231 | -0.819 | -0.781 | 0.005 | -0.006 | -0.006 | 1.069 | 0.72 | 0.69 | 0.83 | 0.79 | 0.75 | 0.76 |
| 8 | 9133798 | rs1017760 | 0.338 | 0.356 | -1.079 | -0.695 | 0.018 | 0.007 | 0.007 | 0 | 0.14 | 0.18 | 0.13 | 0.15 | 0.17 | 0.12 |
| 8 | 9134180 | rs11249912 | -0.355 | 0.279 | -2.216 | -0.929 | 0.005 | 0.009 | 0.009 | 1.133 | 0.31 | 0.08 | 0.54 | 0.77 | 0.3 | 0.56 |
| 8 | 9134637 | rs11783632 | 0.231 | 0.627 | -1.873 | -0.473 | -0.063 | -0.046 | -0.046 | 0.646 | 0.14 | 0.04 | 0.13 | 0.16 | 0.18 | 0.14 |
| 8 | 9135565 | rs17657675 | 0.877 | 0.339 | -1.481 | -0.676 | 0.005 | 0.02 | 0.02 | 0.31 | 0.11 | 0.03 | 0.04 | 0.06 | 0.07 | 0.12 |
| 8 | 9139629 | rs329985 | -1.216 | 0.612 | -1.235 | -0.929 | 0.001 | -0.071 | -0.071 | 0.007 | 0.6 | 0.44 | 0.74 | 0.97 | 0.55 | 0.82 |
| 8 | 9145082 | rs7839174 | -0.816 | 0.389 | -1.424 | -0.041 | 0.002 | 0.032 | 0.032 | 2.053 | 0.47 | 0.28 | 0.58 | 0.6 | 0.53 | 0.59 |
| 8 | 9151226 | rs719409 | -0.047 | 0.839 | -1.834 | 0.513 | -0.028 | -0.055 | -0.055 | 0 | 0.25 | 0.13 | 0.39 | 0.43 | 0.36 | 0.31 |
| 8 | 9151237 | rs330085 | -0.002 | 0.749 | -1.875 | 0.407 | -0.001 | 0.026 | 0.026 | 0.437 | 0.1 | 0.1 | 0.17 | 0.37 | 0.08 | 0.21 |
| 8 | 9152782 | rs747749 | 2.166 | 0.584 | -1.983 | 0.151 | -0.022 | 0.096 | 0.096 | 1.744 | 0.3 | 0.13 | 0.2 | 0 | 0.37 | 0.1 |
| 8 | 9153127 | rs11774353 | -0.695 | 0.831 | -1.334 | 1.291 | -0.027 | -0.059 | -0.059 | 2.106 | 0.25 | 0.12 | 0.42 | 0.42 | 0.36 | 0.3 |
| 8 | 9154394 | rs17149709 | -1.258 | 0.464 | -1.169 | 1.953 | -0.022 | -0.065 | -0.065 | 0 | 0.13 | 0.08 | 0.22 | 0.46 | 0.16 | 0.2 |
| 8 | 9154537 | rs10109698 | -1.569 | 0.218 | -0.631 | 1.754 | -0.015 | 0.082 | 0.082 | 0.049 | 0.12 | 0.24 | 0.15 | 0.24 | 0.08 | 0.12 |
| 8 | 9154694 | rs7820048 | -1.352 | -0.024 | 0.028 | 1.04 | -0.024 | -0.028 | -0.028 | 0 | 0.12 | 0.48 | 0.16 | 0.05 | 0.07 | 0.13 |
| 8 | 9155547 | rs4606064 | -0.919 | 0.299 | -0.685 | 1.467 | -0.006 | 0.009 | 0.009 | 0 | 0.09 | 0.09 | 0.3 | 0.45 | 0.13 | 0.27 |
| 8 | 9156468 | rs69526 | -0.032 | 0.303 | -0.76 | 0.919 | -0.007 | 0.015 | 0.015 | 0 | 0.63 | 0.7 | 0.65 | 0.77 | 0.61 | 0.64 |
| 8 | 9157884 | rs330075 | -0.243 | 0.259 | -0.89 | 0.774 | -0.004 | 0.001 | 0.001 | 0 | 0.68 | 0.69 | 0.67 | 0.76 | 0.65 | 0.68 |
| 8 | 9158475 | rs10086260 | -0.535 | -0.07 | -0.714 | 0.872 | -0.018 | 0.093 | 0.093 | 0 | 0.2 | 0.4 | 0.37 | 0.51 | 0.17 | 0.37 |
| 8 | 9159895 | rs330071 | 2.083 | 0.013 | -1.4 | 0.654 | -0.073 | 0.089 | 0.089 | 0.602 | 0.29 | 0.09 | 0.22 | 0.06 | 0.37 | 0.15 |

| | | | | | | | | | | | | | | | | |
|---|---|---|---|---|---|---|---|---|---|---|---|---|---|---|---|---|
| 8 | 9161873 | rs330069 | 0.123 | 0.742 | -0.747 | 1.112 | -0.037 | 0.013 | 0.013 | 0 | 0.65 | 0.88 | 0.66 | 0.76 | 0.61 | 0.74 |
| 8 | 9165380 | rs405001 | 0.435 | 0.76 | -0.554 | 0.38 | -0.003 | 0.008 | 0.008 | 0 | 0.55 | 0.58 | 0.52 | 0.29 | 0.51 | 0.43 |
| 8 | 9167175 | rs3102070 | -0.172 | 1.287 | -0.136 | 0.27 | 0.04 | 0.018 | 0.018 | 0 | 0.05 | 0.13 | 0.07 | 0.04 | 0.06 | 0.02 |
| 8 | 9171735 | rs713286 | -0.984 | 1.033 | 0.395 | -0.514 | 0.004 | -0.041 | -0.041 | 0.352 | 0.11 | 0.38 | 0.23 | 0.01 | 0.08 | 0.2 |
| 8 | 9172718 | rs330095 | -2.143 | 1.367 | 0.803 | -0.635 | 0.028 | -0.139 | -0.139 | 0 | 0.23 | 0.53 | 0.32 | 0.02 | 0.24 | 0.3 |
| 8 | 9175706 | rs7006947 | -0.115 | 1.465 | 0.43 | -0.301 | -0.006 | -0.019 | -0.019 | 0 | 0.23 | 0.2 | 0.29 | 0.66 | 0.27 | 0.35 |
| 8 | 9177732 | rs983309 | 0.971 | 1.182 | 0.672 | -0.48 | -0.003 | -0.057 | -0.057 | 0 | 0.88 | 0.7 | 0.77 | 0.99 | 0.9 | 0.9 |
| 8 | 9178135 | rs7002551 | 1.701 | 1.391 | 0.582 | -0.578 | 0.004 | -0.068 | -0.068 | 0 | 0.76 | 0.7 | 0.71 | 0.99 | 0.77 | 0.81 |
| 8 | 9178921 | rs12543276 | 1.701 | 1.322 | 0.743 | -0.576 | -0.007 | -0.12 | -0.12 | 0.017 | 0.74 | 0.44 | 0.69 | 0.98 | 0.78 | 0.7 |
| 8 | 9184099 | rs12546815 | 0.082 | 1.835 | 0.189 | -0.634 | -0.071 | -0.075 | -0.075 | 0 | 0.11 | 0.01 | 0.23 | 0.27 | 0.23 | 0.17 |
| 8 | 9184529 | rs11778929 | 2.098 | 1.818 | 0.389 | -0.688 | 0.03 | 0.011 | 0.011 | 0 | 0.66 | 0.49 | 0.48 | 0.71 | 0.57 | 0.53 |
| 8 | 9184628 | rs10106829 | 0.819 | 1.904 | 0.467 | -0.707 | 0.004 | -0.048 | -0.048 | 0 | 0.81 | 0.78 | 0.68 | 0.99 | 0.8 | 0.83 |
| 8 | 9187142 | rs1461730 | -1.091 | 2.101 | 0.506 | -0.55 | 0.028 | 0.082 | 0.082 | 0 | 0.15 | 0.29 | 0.2 | 0.28 | 0.23 | 0.29 |
| 8 | 9189971 | rs17149797 | -0.91 | 2.113 | 0.424 | -0.557 | -0.016 | -0.023 | -0.023 | 0 | 0.14 | 0.08 | 0.18 | 0.28 | 0.22 | 0.27 |
| 8 | 9192250 | rs6993236 | 0.32 | 2.108 | 0.348 | -0.593 | -0.04 | 0.042 | 0.042 | 0 | 0.08 | 0 | 0.06 | 0 | 0.12 | 0.08 |
| 8 | 9192423 | rs11781511 | 0.339 | 2.138 | 0.486 | -0.664 | 0.01 | -0.029 | -0.029 | 0 | 0.92 | 0.84 | 0.74 | 0.99 | 0.91 | 0.89 |
| 8 | 9192927 | rs17149807 | -0.181 | 2.067 | 0.346 | -0.651 | -0.065 | -0.079 | -0.079 | 0 | 0.11 | 0.01 | 0.18 | 0.3 | 0.21 | 0.17 |
| 8 | 9193501 | rs6999153 | 2.243 | 2.013 | 0.475 | -0.641 | 0.033 | 0.022 | 0.022 | 0 | 0.66 | 0.48 | 0.48 | 0.69 | 0.57 | 0.51 |
| 8 | 9197585 | rs717595 | 0.337 | 2.524 | 0.373 | -0.441 | -0.043 | -0.031 | -0.031 | 0.467 | 0.08 | 0.02 | 0.15 | 0.26 | 0.19 | 0.15 |
| 8 | 9197661 | rs958374 | 2.501 | 2.484 | 0.492 | -0.526 | 0.034 | 0.007 | 0.007 | 0 | 0.65 | 0.44 | 0.44 | 0.68 | 0.51 | 0.52 |
| 8 | 9200472 | rs6990912 | 1.761 | 2.405 | 0.485 | -0.541 | 0.036 | 0.059 | 0.059 | 0 | 0.75 | 0.5 | 0.53 | 0.7 | 0.68 | 0.71 |
| 8 | 9201041 | rs2199401 | 1.465 | 2.429 | 0.499 | -0.582 | 0.056 | -0.073 | -0.073 | 0.05 | 0.85 | 0.55 | 0.86 | 0.95 | 0.82 | 0.79 |
| 8 | 9203101 | rs930991 | -0.075 | 1.885 | 0.432 | -0.949 | 0 | -0.002 | -0.002 | 0 | 0.3 | 0.26 | 0.28 | 0.66 | 0.26 | 0.37 |
| 8 | 9204425 | rs9329185 | 1.357 | 2.283 | 0.6 | -0.507 | 0.037 | 0.086 | 0.086 | 0 | 0.83 | 0.69 | 0.79 | 0.71 | 0.74 | 0.72 |
| 8 | 9205584 | rs7018002 | -1.175 | 2.16 | 0.598 | -0.567 | 0.013 | 0.011 | 0.011 | 0 | 0.06 | 0.25 | 0.06 | 0.05 | 0.06 | 0.13 |
| 8 | 9206425 | rs10102703 | 1.676 | 2.223 | 0.604 | -0.586 | 0.055 | 0.04 | 0.04 | 0 | 0.78 | 0.55 | 0.77 | 0.75 | 0.67 | 0.75 |
| 8 | 9206678 | rs2169385 | 1.423 | 1.961 | 0.658 | -0.721 | 0.007 | -0.088 | -0.088 | 1.578 | 0.85 | 0.54 | 0.71 | 0.99 | 0.89 | 0.88 |
| 8 | 9215012 | rs10101673 | 0.812 | 1.899 | 0.414 | -0.611 | 0.009 | -0.015 | -0.015 | 0.019 | 0.95 | 0.8 | 0.98 | 1 | 0.98 | 0.99 |
| 8 | 9216371 | rs6989535 | 2.057 | 1.901 | 0.447 | -0.628 | -0.004 | -0.025 | -0.025 | 0 | 0.9 | 0.75 | 0.94 | 0.95 | 0.93 | 0.86 |
| 8 | 9216584 | rs9329191 | -1.208 | 2.132 | 0.163 | -0.359 | -0.014 | -0.008 | -0.008 | 0.528 | 0.13 | 0.09 | 0.2 | 0.34 | 0.26 | 0.31 |
| 8 | 9217374 | rs4538888 | -0.187 | 2.128 | 0.276 | -0.324 | 0.026 | 0.089 | 0.089 | 0 | 0.09 | 0.16 | 0.16 | 0.28 | 0.2 | 0.18 |
| 8 | 9221219 | rs17661379 | -0.506 | 1.815 | 0.03 | -0.597 | -0.001 | -0.014 | -0.014 | 0 | 0.09 | 0.09 | 0.07 | 0 | 0.1 | 0.1 |
| 8 | 9223144 | rs1993596 | 1.257 | 1.548 | 0.227 | -0.851 | -0.001 | 0.001 | 0.001 | 0.277 | 0.55 | 0.4 | 0.44 | 0.38 | 0.56 | 0.43 |
| 8 | 9223532 | rs13262846 | -0.675 | 1.567 | 0.142 | -0.871 | -0.043 | 0.071 | 0.071 | 0 | 0.14 | 0.04 | 0.29 | 0.01 | 0.17 | 0.17 |
| 8 | 9225872 | rs10091799 | -1.533 | 1.417 | 0.085 | -0.963 | 0 | -0.012 | -0.012 | 0 | 0.24 | 0.22 | 0.37 | 0.06 | 0.24 | 0.37 |
| 8 | 9226991 | rs13271803 | 0.175 | 1.616 | -0.199 | -0.793 | -0.036 | -0.011 | -0.011 | 1.05 | 0.07 | 0.02 | 0.14 | 0.29 | 0.17 | 0.13 |
| 8 | 9231148 | rs2409571 | -1.752 | 1.407 | -0.164 | -0.94 | -0.001 | 0.021 | 0.021 | 0.266 | 0.52 | 0.63 | 0.61 | 0.69 | 0.49 | 0.75 |
| 8 | 9234648 | rs17718351 | -0.51 | 1.579 | -0.758 | -0.718 | -0.002 | 0 | 0 | 0.078 | 0.05 | 0.05 | 0.04 | 0.03 | 0.05 | 0.16 |
| 8 | 9237403 | rs12235038 | 2.078 | 1.611 | -0.853 | -0.795 | -0.019 | 0.023 | 0.023 | 0.106 | 0.57 | 0.16 | 0.49 | 0.67 | 0.58 | 0.43 |
| 8 | 9239458 | rs6601302 | 1.118 | 1.473 | -0.753 | -0.729 | 0.011 | 0.029 | 0.029 | 0 | 0.86 | 0.83 | 0.79 | 0.74 | 0.78 | 0.69 |
| 8 | 9242557 | rs4240625 | 1.884 | 1.296 | -1.127 | -0.635 | -0.039 | 0.009 | 0.009 | 0.021 | 0.56 | 0.15 | 0.46 | 0.65 | 0.54 | 0.44 |
| 8 | 9243428 | rs7841407 | 0.764 | 0.847 | -0.696 | -0.594 | -0.006 | 0.019 | 0.019 | 0 | 0.94 | 0.85 | 0.94 | 0.97 | 0.95 | 0.84 |
| 8 | 9246837 | rs17662123 | -0.734 | 0.919 | -1.235 | -0.759 | -0.07 | -0.012 | -0.012 | 0 | 0.33 | 0.19 | 0.55 | 0.72 | 0.5 | 0.46 |
| 8 | 9250346 | rs922053 | 1.41 | 0.224 | -0.925 | -0.659 | 0.053 | -0.003 | -0.003 | 2.864 | 0.72 | 0.21 | 0.76 | 0.75 | 0.86 | 0.74 |
| 8 | 9253175 | rs7828601 | -1.181 | 0.417 | -0.514 | -0.544 | -0.034 | -0.087 | -0.087 | 0 | 0.18 | 0.6 | 0.13 | 0.04 | 0.11 | 0.06 |
| 8 | 9255250 | rs10435712 | 1.482 | 0.443 | -0.727 | -0.52 | -0.004 | 0 | 0 | 0.415 | 0.76 | 0.29 | 0.8 | 0.75 | 0.87 | 0.81 |
| 8 | 9263290 | rs7836066 | -0.894 | 1.053 | -0.664 | -0.664 | 0.001 | 0.005 | 0.005 | 0.211 | 0.12 | 0.13 | 0.09 | 0.18 | 0.14 | 0.26 |
| 8 | 9263460 | rs7818396 | -0.609 | 1.043 | -0.676 | -0.664 | 0 | -0.001 | -0.001 | 0 | 0.1 | 0.11 | 0.08 | 0.09 | 0.12 | 0.17 |
| 8 | 9264711 | rs6601306 | 0.963 | 0.848 | -0.193 | -0.938 | -0.005 | -0.069 | -0.069 | 0 | 0.88 | 0.47 | 0.9 | 0.96 | 0.93 | 0.97 |
| 8 | 9268764 | rs17149982 | -0.981 | 0.969 | -0.528 | -0.802 | 0.001 | 0.005 | 0.005 | 0.108 | 0.12 | 0.15 | 0.09 | 0.16 | 0.14 | 0.23 |
| 8 | 9269676 | rs17149987 | -1.04 | 0.964 | -0.522 | -0.826 | 0.001 | -0.004 | -0.004 | 0 | 0.12 | 0.16 | 0.09 | 0.1 | 0.14 | 0.25 |
| 8 | 9271136 | rs17149991 | -0.961 | 0.97 | -0.545 | -0.81 | -0.001 | 0 | 0 | 0 | 0.12 | 0.12 | 0.09 | 0.12 | 0.14 | 0.24 |
| 8 | 9272507 | rs13261949 | -1.048 | 1.792 | -0.961 | -1.394 | -0.066 | 0.015 | 0.015 | 0.027 | 0.26 | 0.09 | 0.55 | 0.79 | 0.38 | 0.36 |
| 8 | 9273247 | rs11776114 | -0.562 | 1.857 | -1.019 | -1.392 | -0.106 | -0.035 | -0.035 | 0 | 0.21 | 0.03 | 0.52 | 0.77 | 0.33 | 0.31 |
| 8 | 9276163 | rs17730481 | -1.652 | 2.201 | -1.052 | -1.514 | -0.04 | 0.029 | 0.029 | 0 | 0.22 | 0.09 | 0.19 | 0.12 | 0.3 | 0.27 |
| 8 | 9276286 | rs6989109 | 1.837 | 1.654 | -0.972 | -2.325 | -0.004 | 0.028 | 0.028 | 0.174 | 0.53 | 0.33 | 0.34 | 0.05 | 0.5 | 0.48 |
| 8 | 9276702 | rs17730541 | -0.557 | 1.69 | -1.063 | -2.249 | -0.021 | 0.031 | 0.031 | 0 | 0.1 | 0.03 | 0.11 | 0.01 | 0.16 | 0.07 |
| 8 | 9279553 | rs13281098 | 2.832 | 1.546 | -1.082 | -2.347 | 0.035 | 0.202 | 0.202 | 1.336 | 0.41 | 0.17 | 0.23 | 0.04 | 0.33 | 0.4 |
| 8 | 9280516 | rs17663071 | -0.737 | 1.597 | -1.078 | -2.159 | -0.097 | -0.02 | -0.02 | 0 | 0.19 | 0.04 | 0.52 | 0.79 | 0.32 | 0.31 |
| 8 | 9287209 | rs17150024 | -0.296 | 1.136 | -0.71 | -2.114 | -0.025 | -0.141 | -0.141 | 0.026 | 0.19 | 0.39 | 0.12 | 0 | 0.15 | 0.05 |
| 8 | 9288937 | rs17663167 | -0.689 | 1.156 | -0.761 | -2.031 | -0.001 | -0.005 | -0.005 | 6.051 | 0.16 | 0.1 | 0.09 | 0.16 | 0.16 | 0.29 |
| 8 | 9294503 | rs11997395 | -1.421 | 1.175 | -0.743 | -2.066 | -0.012 | -0.081 | -0.081 | 1.09 | 0.32 | 0.5 | 0.2 | 0.09 | 0.3 | 0.27 |
| 8 | 9294638 | rs1458939 | -0.173 | 1.206 | -1.026 | -2.18 | -0.027 | 0.044 | 0.044 | 0 | 0.11 | 0.02 | 0.11 | 0.05 | 0.16 | 0.08 |
| 8 | 9294964 | rs7817527 | 2.028 | 1.28 | -0.933 | -2.116 | -0.022 | 0.212 | 0.212 | 0.552 | 0.53 | 0.23 | 0.36 | 0.1 | 0.52 | 0.48 |
| 8 | 9310266 | rs17675053 | -0.378 | 1.139 | -1.195 | -1.97 | -0.063 | -0.048 | -0.048 | 0.076 | 0.17 | 0.04 | 0.45 | 0.74 | 0.22 | 0.25 |
| 8 | 9310511 | rs17150066 | 0.437 | 1.188 | -1.196 | -1.952 | -0.049 | -0.052 | -0.052 | 0.377 | 0.11 | 0.03 | 0.42 | 0.74 | 0.18 | 0.21 |
| 8 | 9313017 | rs6995907 | -1.655 | 0.765 | -1.016 | -2.186 | -0.017 | -0.057 | -0.057 | 0 | 0.35 | 0.54 | 0.21 | 0.13 | 0.28 | 0.3 |
| 8 | 9314784 | rs10503382 | -0.065 | 0.664 | -0.996 | -2.268 | -0.024 | -0.121 | -0.121 | 0.302 | 0.18 | 0.37 | 0.11 | 0 | 0.15 | 0.05 |
| 8 | 9315518 | rs4841154 | 1.913 | 0.676 | -1.116 | -2.257 | 0.051 | 0.169 | 0.169 | 1.2 | 0.59 | 0.12 | 0.73 | 0.82 | 0.7 | 0.62 |

| | | | | | | | | | | | | | | | | |
|---|---|---|---|---|---|---|---|---|---|---|---|---|---|---|---|---|
| 8 | 9315807 | rs17150086 | -1.547 | 0.737 | -1.097 | -2.236 | -0.02 | -0.065 | -0.065 | 0 | 0.28 | 0.52 | 0.22 | 0.12 | 0.23 | 0.29 |
| 8 | 9321780 | rs10100706 | -2.849 | 0.835 | -1.453 | -2.346 | -0.008 | 0.217 | 0.217 | 0.679 | 0.65 | 0.92 | 0.8 | 0.97 | 0.7 | 0.7 |
| 8 | 9323515 | rs10105543 | -1.402 | 0.756 | -1.297 | -2.383 | -0.008 | -0.039 | -0.039 | 0 | 0.31 | 0.39 | 0.26 | 0.19 | 0.3 | 0.35 |
| 8 | 9328926 | rs7007562 | -1.464 | 0.8 | -1.338 | -2.39 | -0.009 | -0.044 | -0.044 | 0 | 0.31 | 0.42 | 0.28 | 0.21 | 0.31 | 0.36 |
| 8 | 9331557 | rs11783183 | -1.102 | 0.809 | -1.367 | -2.296 | 0.008 | -0.013 | -0.013 | 0.749 | 0.34 | 0.37 | 0.53 | 0.68 | 0.38 | 0.39 |
| 8 | 9335968 | rs6999852 | -2.598 | 0.859 | -1.477 | -2.336 | 0.005 | 0.176 | 0.176 | 0 | 0.62 | 0.84 | 0.65 | 0.94 | 0.67 | 0.7 |
| 8 | 9336065 | rs17150106 | -0.267 | 0.846 | -1.38 | -2.377 | 0.028 | 0.012 | 0.012 | 0 | 0.23 | 0.36 | 0.34 | 0.68 | 0.26 | 0.26 |
| 8 | 9336714 | rs7819054 | -1.292 | 1.049 | -1.32 | -2.077 | 0.001 | -0.024 | -0.024 | 0.007 | 0.29 | 0.41 | 0.26 | 0.26 | 0.3 | 0.32 |
| 8 | 9340932 | rs6994287 | -1.258 | 1.082 | -1.048 | -1.628 | 0.008 | -0.026 | -0.026 | 0 | 0.38 | 0.44 | 0.56 | 0.72 | 0.41 | 0.45 |
| 8 | 9343978 | rs11249925 | -0.89 | 0.59 | -1.057 | -1.448 | 0.005 | -0.018 | -0.018 | 0.013 | 0.33 | 0.42 | 0.39 | 0.69 | 0.34 | 0.38 |
| 8 | 9351983 | rs6990812 | -0.497 | 0.623 | -0.939 | -1.3 | 0.007 | 0.005 | 0.005 | 0 | 0.11 | 0.14 | 0.12 | 0.14 | 0.13 | 0.23 |
| 8 | 9357874 | rs4841161 | -0.554 | 0.665 | -0.968 | -1.184 | 0.007 | 0.017 | 0.017 | 0.007 | 0.12 | 0.18 | 0.12 | 0.17 | 0.13 | 0.24 |
| 8 | 9365967 | rs7829843 | -1.893 | 0.543 | -1.24 | -1.459 | 0.003 | 0.132 | 0.132 | 0.465 | 0.49 | 0.73 | 0.65 | 0.8 | 0.5 | 0.53 |
| 8 | 9366203 | rs7831172 | -0.786 | 0.539 | -1.083 | -1.443 | -0.005 | -0.005 | -0.005 | 0 | 0.33 | 0.41 | 0.37 | 0.63 | 0.31 | 0.38 |
| 8 | 9367866 | rs4841165 | 1.012 | 1.434 | -0.26 | -0.452 | -0.008 | -0.013 | -0.013 | 0.156 | 0.94 | 0.93 | 0.98 | 0.99 | 0.98 | 0.91 |
| 8 | 9370559 | rs10100760 | -2.247 | 1.559 | -0.058 | -0.481 | 0.003 | -0.097 | -0.097 | 0 | 0.42 | 0.48 | 0.53 | 0.92 | 0.54 | 0.56 |
| 8 | 9374506 | rs10503381 | -1.434 | 1.194 | 0.662 | 0.283 | 0.027 | -0.024 | -0.024 | 0 | 0.13 | 0.29 | 0.11 | 0.1 | 0.16 | 0.09 |
| 8 | 9375944 | rs17733293 | -1.905 | 1.214 | 0.611 | 0.345 | 0.03 | -0.009 | -0.009 | 0 | 0.15 | 0.31 | 0.28 | 0.18 | 0.19 | 0.15 |
| 8 | 9376657 | rs6601324 | 2.115 | 1.206 | 0.585 | 0.445 | 0.013 | -0.025 | -0.025 | 0 | 0.81 | 0.62 | 0.7 | 0.82 | 0.8 | 0.76 |
| 8 | 9380132 | rs7830939 | 2.369 | 1.186 | 0.258 | 0.548 | -0.002 | -0.002 | -0.002 | 0.511 | 0.68 | 0.4 | 0.6 | 0.66 | 0.71 | 0.52 |
| 8 | 9384159 | rs11249929 | 2.181 | 1.278 | -0.362 | 0.557 | -0.001 | -0.001 | -0.001 | 1.622 | 0.93 | 0.93 | 0.82 | 0.93 | 0.96 | 0.84 |
| 8 | 9386210 | rs7013746 | 2.204 | 1.369 | -0.389 | 0.579 | 0.001 | 0.001 | 0.001 | 0 | 0.93 | 0.94 | 0.82 | 0.92 | 0.96 | 0.84 |
| 8 | 9388022 | rs2100700 | 1.487 | 1.227 | -0.325 | 1.051 | -0.001 | 0.005 | 0.005 | 0 | 0.93 | 0.9 | 0.9 | 0.86 | 0.92 | 0.81 |
| 8 | 9389929 | rs7016636 | 1.271 | 1.201 | -0.29 | 1.051 | 0 | 0.02 | 0.02 | 0 | 0.93 | 0.88 | 0.9 | 0.86 | 0.94 | 0.8 |
| 8 | 9394053 | rs6987057 | 2.628 | 1.247 | 0.034 | 1.567 | 0.004 | 0.119 | 0.119 | 0.414 | 0.83 | 0.69 | 0.65 | 0.6 | 0.84 | 0.73 |
| 8 | 9394331 | rs1471203 | -2.147 | 1.262 | 0.019 | 1.561 | 0.011 | 0.079 | 0.079 | 0.649 | 0.1 | 0.18 | 0.29 | 0.25 | 0.1 | 0.14 |
| 8 | 9398118 | rs17150211 | -1.326 | 1.551 | -0.145 | 2.329 | -0.028 | -0.027 | -0.027 | 0 | 0.08 | 0.03 | 0.11 | 0.24 | 0.16 | 0.14 |
| 8 | 9404663 | rs7013369 | -1.124 | 1.436 | -0.128 | 2.228 | 0 | 0 | 0 | 0 | 0.07 | 0.09 | 0.08 | 0.11 | 0.05 | 0.2 |
| 8 | 9411808 | rs9644677 | 2.82 | 1.437 | 0.303 | 2.312 | 0.044 | 0.139 | 0.139 | 0.111 | 0.79 | 0.57 | 0.79 | 0.58 | 0.75 | 0.66 |
| 8 | 9412153 | rs9329203 | 2.453 | 1.446 | 0.347 | 2.325 | 0.037 | 0.142 | 0.142 | 0.06 | 0.81 | 0.63 | 0.8 | 0.57 | 0.77 | 0.67 |
| 8 | 9412857 | rs6990097 | -2.742 | 1.424 | 0.351 | 2.325 | 0.037 | 0.116 | 0.116 | 0.026 | 0.18 | 0.32 | 0.19 | 0.43 | 0.25 | 0.34 |
| 8 | 9421789 | rs6986386 | -2.29 | 1.321 | 0.311 | 2.398 | 0.021 | 0.085 | 0.085 | 0 | 0.16 | 0.26 | 0.18 | 0.4 | 0.22 | 0.31 |
| 8 | 9422926 | rs13265363 | -2.748 | 1.291 | 0.298 | 2.466 | 0.002 | 0.056 | 0.056 | 1.552 | 0.11 | 0.22 | 0.29 | 0.21 | 0.12 | 0.13 |
| 8 | 9427048 | rs6601333 | -2.358 | 1.304 | 0.251 | 2.483 | 0.018 | 0.087 | 0.087 | 0.009 | 0.16 | 0.26 | 0.18 | 0.41 | 0.22 | 0.32 |
| 8 | 9428688 | rs11984585 | -2.339 | 1.343 | 0.252 | 2.426 | 0.022 | 0.086 | 0.086 | 0 | 0.16 | 0.26 | 0.18 | 0.4 | 0.23 | 0.32 |
| 8 | 9437352 | rs7834823 | -2.999 | 1.334 | 0.272 | 2.388 | 0.029 | 0.125 | 0.125 | 0.037 | 0.19 | 0.38 | 0.2 | 0.39 | 0.23 | 0.33 |
| 8 | 9438928 | rs6601338 | -3.602 | 1.328 | 0.277 | 2.385 | 0.027 | 0.047 | 0.047 | 1.947 | 0.27 | 0.45 | 0.47 | 0.62 | 0.33 | 0.45 |
| 8 | 9442027 | rs12549064 | -1.65 | 1.351 | 0.291 | 2.461 | 0.03 | 0.12 | 0.12 | 0 | 0.1 | 0.18 | 0.11 | 0.32 | 0.16 | 0.12 |
| 8 | 9445429 | rs12679892 | -3.043 | 1.318 | 0.192 | 2.247 | 0.002 | 0.012 | 0.012 | 1.499 | 0.22 | 0.23 | 0.19 | 0.4 | 0.28 | 0.37 |
| 8 | 9449132 | rs7002917 | -1.079 | 1.281 | 0.406 | 2.417 | 0.002 | 0.012 | 0.012 | 0 | 0.06 | 0.12 | 0.08 | 0.09 | 0.06 | 0.2 |
| 8 | 9450886 | rs10090277 | -1.932 | 1.279 | 0.442 | 2.419 | 0.014 | 0.037 | 0.037 | 0 | 0.08 | 0.26 | 0.09 | 0.09 | 0.07 | 0.21 |
| 8 | 9452088 | rs4610751 | -3.084 | 1.296 | 0.431 | 2.442 | 0.023 | 0.13 | 0.13 | 5.33 | 0.18 | 0.27 | 0.37 | 0.52 | 0.26 | 0.22 |
| 8 | 9467171 | rs5002815 | -2.023 | 1.226 | 0.426 | 2.377 | -0.004 | 0.007 | 0.007 | 0 | 0.09 | 0.21 | 0.11 | 0.1 | 0.07 | 0.21 |
| 8 | 9471265 | rs4629869 | -2.116 | 1.261 | 0.406 | 2.377 | 0.001 | 0.011 | 0.011 | 0 | 0.1 | 0.21 | 0.11 | 0.1 | 0.07 | 0.21 |
| 8 | 9472445 | rs11249930 | 1.576 | 1.276 | 0.465 | 2.438 | 0.039 | 0.151 | 0.151 | 0.094 | 0.89 | 0.75 | 0.89 | 0.7 | 0.84 | 0.89 |
| 8 | 9508879 | rs4128324 | -2.808 | 1.202 | 0.274 | 2.325 | -0.005 | 0.035 | 0.035 | 1.146 | 0.11 | 0.15 | 0.28 | 0.23 | 0.1 | 0.13 |
| 8 | 9526345 | rs13250838 | -2.728 | 1.206 | 0.277 | 2.335 | -0.002 | 0.036 | 0.036 | 1.148 | 0.11 | 0.15 | 0.27 | 0.22 | 0.1 | 0.13 |
| 8 | 9527707 | rs7015700 | -3.194 | 1.188 | 0.278 | 2.323 | -0.003 | 0.042 | 0.042 | 1.505 | 0.18 | 0.28 | 0.35 | 0.33 | 0.17 | 0.32 |
| 8 | 9533399 | rs4427176 | 1.43 | 1.158 | 0.244 | 2.466 | 0.024 | 0.054 | 0.054 | 0.096 | 0.91 | 0.84 | 0.9 | 0.8 | 0.84 | 0.9 |
| 8 | 9540693 | rs13273033 | 2.063 | 1.162 | 0.253 | 2.506 | 0.018 | 0.055 | 0.055 | 0.062 | 0.83 | 0.72 | 0.82 | 0.67 | 0.78 | 0.7 |
| 8 | 9547083 | rs11773928 | -2.934 | 1.119 | 0.352 | 2.474 | -0.024 | 0.056 | 0.056 | 0.167 | 0.09 | 0.13 | 0.24 | 0.22 | 0.05 | 0.12 |
| 8 | 9555092 | rs4841196 | 2.858 | 1.204 | 0.344 | 2.541 | 0.006 | 0.046 | 0.046 | 1.579 | 0.74 | 0.61 | 0.57 | 0.38 | 0.72 | 0.57 |
| 8 | 9577096 | rs6990116 | 2.02 | 1.2 | 0.418 | 2.115 | -0.007 | 0.014 | 0.014 | 0 | 0.9 | 0.71 | 0.93 | 0.89 | 0.94 | 0.81 |
| 8 | 9578982 | rs13276086 | 3.03 | 1.239 | 0.388 | 2.146 | 0.02 | 0.076 | 0.076 | 2.172 | 0.72 | 0.55 | 0.54 | 0.46 | 0.67 | 0.57 |
| 8 | 9593924 | rs7846023 | -1.424 | 1.242 | 0.349 | 2.173 | -0.004 | -0.002 | -0.002 | 0 | 0.08 | 0.16 | 0.05 | 0.08 | 0.06 | 0.19 |
| 8 | 9600917 | rs9644708 | -0.267 | 1.17 | -0.245 | 1.618 | -0.059 | -0.157 | -0.157 | 0 | 0.23 | 0.01 | 0.23 | 0.42 | 0.22 | 0.31 |
| 8 | 9601699 | rs12545912 | 2.337 | 0.896 | 0.475 | 2.225 | 0.017 | 0.043 | 0.043 | 1.002 | 0.8 | 0.65 | 0.84 | 0.72 | 0.77 | 0.69 |
| 8 | 9607863 | rs11249943 | -1.499 | 0.949 | 0.448 | 2.163 | 0.015 | 0.038 | 0.038 | 1.559 | 0.11 | 0.16 | 0.11 | 0.21 | 0.18 | 0.11 |
| 8 | 9611001 | rs7014007 | -2.606 | 0.933 | 0.496 | 2.115 | 0.026 | 0.083 | 0.083 | 2.372 | 0.18 | 0.25 | 0.4 | 0.41 | 0.28 | 0.24 |
| 8 | 9611470 | rs10102170 | -3.4 | 0.936 | 0.502 | 2.122 | 0.023 | 0.098 | 0.098 | 2.605 | 0.27 | 0.47 | 0.47 | 0.5 | 0.33 | 0.43 |
| 8 | 9616553 | rs10101132 | -2.571 | 0.965 | 0.471 | 2.121 | 0.014 | 0.05 | 0.05 | 2.852 | 0.18 | 0.21 | 0.4 | 0.41 | 0.28 | 0.24 |
| 8 | 9622559 | rs6986755 | -1.816 | 0.839 | 0.465 | 2.048 | -0.002 | 0.024 | 0.024 | 0 | 0.08 | 0.23 | 0.05 | 0.12 | 0.06 | 0.19 |
| 8 | 9634874 | rs7010154 | -1.206 | 0.845 | 0.425 | 2.049 | -0.002 | 0.011 | 0.011 | 0 | 0.07 | 0.13 | 0.04 | 0.12 | 0.06 | 0.2 |
| 8 | 9635770 | rs17734024 | -1.568 | 0.856 | 0.404 | 2.163 | -0.011 | 0.007 | 0.007 | 0 | 0.05 | 0.02 | 0.27 | 0.2 | 0.1 | 0.07 |
| 8 | 9638308 | rs4841215 | -1.36 | 0.81 | 0.399 | 1.848 | -0.002 | 0.013 | 0.013 | 0 | 0.07 | 0.16 | 0.05 | 0.12 | 0.06 | 0.17 |
| 8 | 9640400 | rs11785739 | -2.397 | 0.789 | 0.409 | 1.837 | -0.018 | 0.003 | 0.003 | 0 | 0.11 | 0.23 | 0.07 | 0.12 | 0.06 | 0.17 |
| 8 | 9646560 | rs7814507 | -2.403 | 0.863 | 0.353 | 2.034 | 0.002 | 0.019 | 0.019 | 1.721 | 0.19 | 0.21 | 0.43 | 0.38 | 0.3 | 0.22 |
| 8 | 9647603 | rs2898223 | -1.386 | 0.839 | 0.419 | 2.019 | -0.001 | 0.045 | 0.045 | 0 | 0.07 | 0.23 | 0.05 | 0.15 | 0.05 | 0.12 |
| 8 | 9647796 | rs7838571 | -3.377 | 0.824 | 0.377 | 2.019 | 0.018 | 0.055 | 0.055 | 2.042 | 0.29 | 0.5 | 0.51 | 0.52 | 0.35 | 0.35 |

| | | | | | | | | | | | | | | | | |
|---|---|---|---|---|---|---|---|---|---|---|---|---|---|---|---|---|
| 8 | 9649769 | rs7830613 | -2.342 | 0.967 | 0.251 | 2.063 | -0.006 | -0.003 | -0.003 | 0 | 0.18 | 0.18 | 0.42 | 0.37 | 0.29 | 0.2 |
| 8 | 9652735 | rs7837054 | 1.208 | 0.982 | 0.257 | 2.106 | 0.009 | 0.005 | 0.005 | 1.793 | 0.88 | 0.82 | 0.91 | 0.85 | 0.83 | 0.89 |
| 8 | 9653502 | rs13271796 | -2.622 | 0.987 | 0.234 | 2.108 | 0.002 | 0.018 | 0.018 | 1.773 | 0.19 | 0.22 | 0.43 | 0.38 | 0.3 | 0.22 |
| 8 | 9672946 | rs10107086 | -0.95 | 0.944 | 0.233 | 1.865 | 0.006 | 0.028 | 0.028 | 0 | 0.07 | 0.14 | 0.04 | 0.15 | 0.06 | 0.12 |
| 8 | 9673870 | rs4840440 | -1.35 | 0.984 | 0.193 | 1.925 | -0.014 | 0.014 | 0.014 | 0 | 0.06 | 0.03 | 0.31 | 0.22 | 0.11 | 0.08 |
| 8 | 9680223 | rs17734290 | -2.491 | 0.873 | 0.152 | 1.863 | -0.005 | -0.009 | -0.009 | 0 | 0.21 | 0.23 | 0.41 | 0.37 | 0.29 | 0.21 |
| 8 | 9681257 | rs2102658 | 1.243 | 0.838 | 0.189 | 1.843 | 0.003 | 0.029 | 0.029 | 0 | 0.92 | 0.81 | 0.95 | 0.85 | 0.94 | 0.88 |
| 8 | 9684014 | rs13279794 | -2.949 | 1.041 | 0.127 | 1.838 | 0.016 | 0.032 | 0.032 | 0.243 | 0.28 | 0.38 | 0.47 | 0.52 | 0.39 | 0.33 |
| 8 | 9685362 | rs10110146 | -2.592 | 0.976 | 0.145 | 1.204 | 0.009 | 0.006 | 0.006 | 0 | 0.23 | 0.33 | 0.36 | 0.29 | 0.29 | 0.24 |
| 8 | 9688094 | rs12547274 | 1.953 | 0.398 | 0.606 | 1.017 | -0.054 | -0.035 | -0.035 | 0 | 0.82 | 0.37 | 0.9 | 0.86 | 0.93 | 0.74 |
| 8 | 9692541 | rs998019 | -0.934 | 0.486 | 0.643 | 1.058 | 0.006 | 0.014 | 0.014 | 0.117 | 0.08 | 0.16 | 0.07 | 0.14 | 0.12 | 0.06 |
| 8 | 9692825 | rs4841239 | 2.544 | 0.453 | 0.613 | 1.051 | -0.004 | 0 | 0 | 0.019 | 0.76 | 0.24 | 0.85 | 0.71 | 0.81 | 0.7 |
| 8 | 9708433 | rs17734541 | 2.321 | 0.15 | 0.092 | 0.335 | -0.025 | 0.06 | 0.06 | 4.17 | 0.41 | 0.22 | 0.28 | 0.01 | 0.46 | 0.15 |
| 8 | 9711456 | rs7821773 | 1.827 | 0.226 | 1.022 | 1.262 | -0.04 | -0.019 | -0.019 | 1.255 | 0.85 | 0.38 | 0.92 | 0.83 | 0.93 | 0.73 |
| 8 | 9718278 | rs7015101 | -2.402 | 0.285 | 1.071 | 1.312 | 0.008 | 0.03 | 0.03 | 0.378 | 0.18 | 0.57 | 0.12 | 0.24 | 0.19 | 0.18 |
| 8 | 9729939 | rs1022258 | 1.357 | 0.371 | 0.868 | 1.313 | 0.009 | 0.013 | 0.013 | 0.251 | 0.9 | 0.77 | 0.93 | 0.87 | 0.88 | 0.94 |
| 8 | 9731188 | rs7003172 | -1.163 | 0.22 | 0.837 | 1.308 | 0.002 | 0.004 | 0.004 | 0.083 | 0.08 | 0.12 | 0.07 | 0.13 | 0.12 | 0.06 |
| 8 | 9731470 | rs13268133 | 2.738 | 0.252 | 0.869 | 1.639 | 0.005 | -0.03 | -0.03 | 0.013 | 0.7 | 0.22 | 0.7 | 0.53 | 0.7 | 0.6 |
| 8 | 9740636 | rs6601389 | 2.281 | 0.35 | 0.473 | 1.722 | -0.002 | -0.006 | -0.006 | 1.306 | 0.81 | 0.76 | 0.76 | 0.65 | 0.77 | 0.75 |
| 8 | 9744650 | rs6989793 | 2.092 | 0.339 | 0.431 | 1.738 | -0.002 | -0.014 | -0.014 | 0.168 | 0.82 | 0.78 | 0.77 | 0.65 | 0.78 | 0.76 |
| 8 | 9758230 | rs491364 | -1.954 | 0.562 | 0.424 | 1.938 | -0.016 | -0.014 | -0.014 | 0 | 0.13 | 0.12 | 0.22 | 0.36 | 0.22 | 0.14 |
| 8 | 9760420 | rs592420 | -2.336 | 0.555 | 0.657 | 1.915 | 0.007 | 0.049 | 0.049 | 0.417 | 0.14 | 0.2 | 0.22 | 0.37 | 0.23 | 0.18 |
| 8 | 9774864 | rs556819 | 2.155 | 0.546 | 0.686 | 1.798 | 0.013 | 0.027 | 0.027 | 0 | 0.92 | 0.88 | 0.85 | 0.82 | 0.89 | 0.89 |
| 8 | 9775516 | rs473034 | 2.624 | 0.532 | 0.768 | 1.791 | 0.015 | 0.034 | 0.034 | 0 | 0.9 | 0.83 | 0.84 | 0.82 | 0.89 | 0.86 |
| 8 | 9781492 | rs12548180 | -0.572 | 0.501 | 0.647 | 1.899 | -0.015 | 0 | 0 | 0 | 0.05 | 0 | 0.07 | 0.21 | 0.12 | 0.03 |
| 8 | 9781855 | rs17150788 | -1.148 | 0.526 | 0.665 | 1.862 | -0.026 | -0.024 | -0.024 | 0 | 0.06 | 0.01 | 0.14 | 0.18 | 0.11 | 0.1 |
| 8 | 9783789 | rs17150793 | -1.315 | 0.449 | 1.042 | 1.873 | -0.001 | -0.009 | -0.009 | 0.161 | 0.08 | 0.15 | 0.04 | 0.08 | 0.06 | 0.16 |
| 8 | 9785503 | rs1399054 | 2.803 | 0.456 | 1.151 | 1.869 | 0.025 | 0.04 | 0.04 | 0 | 0.82 | 0.59 | 0.79 | 0.74 | 0.82 | 0.7 |
| 8 | 9786494 | rs586123 | 2.059 | 0.427 | 0.663 | 1.824 | -0.012 | -0.014 | -0.014 | 0 | 0.92 | 0.93 | 0.85 | 0.82 | 0.88 | 0.89 |
| 8 | 9787398 | rs10503397 | -1.105 | 0.454 | 0.717 | 1.84 | 0 | -0.006 | -0.006 | 0 | 0.08 | 0.13 | 0.04 | 0.08 | 0.05 | 0.16 |
| 8 | 9787808 | rs661310 | -2.607 | 0.399 | 0.967 | 1.854 | -0.009 | 0.063 | 0.063 | 0.183 | 0.18 | 0.69 | 0.15 | 0.3 | 0.18 | 0.22 |
| 8 | 9791143 | rs4841254 | 2.69 | 0.377 | 0.856 | 1.768 | -0.001 | -0.016 | -0.016 | 0 | 0.72 | 0.25 | 0.69 | 0.52 | 0.69 | 0.65 |
| 8 | 9794093 | rs509820 | 1.792 | 0.498 | 0.607 | 1.809 | 0 | 0.018 | 0.018 | 0 | 0.88 | 0.87 | 0.88 | 0.72 | 0.83 | 0.81 |
| 8 | 9796189 | rs615171 | 1.679 | 0.509 | 0.624 | 1.781 | 0.004 | 0.046 | 0.046 | 0 | 0.88 | 0.85 | 0.81 | 0.73 | 0.84 | 0.82 |
| 8 | 9800153 | rs17150826 | -0.512 | 0.533 | 0.296 | 1.791 | -0.011 | 0.018 | 0.018 | 0.284 | 0.05 | 0.01 | 0.18 | 0.22 | 0.11 | 0.03 |
| 8 | 9812462 | rs471480 | 1.256 | -0.022 | 0.245 | 1.169 | 0.01 | -0.001 | -0.001 | 0 | 0.6 | 0.32 | 0.44 | 0.33 | 0.69 | 0.35 |
| 8 | 9813317 | rs671370 | 1.269 | -0.191 | 0.23 | 1.089 | 0.007 | -0.002 | -0.002 | 0.67 | 0.6 | 0.34 | 0.52 | 0.34 | 0.69 | 0.38 |
| 8 | 9818065 | rs591346 | 2.032 | -0.139 | 0.27 | 0.867 | 0.011 | 0.011 | 0.011 | 0.024 | 0.5 | 0.43 | 0.39 | 0.04 | 0.53 | 0.32 |
| 8 | 9819890 | rs609792 | 0.449 | -0.307 | 0.195 | 1.091 | 0.005 | -0.08 | -0.08 | 1.023 | 0.75 | 0.87 | 0.67 | 0.43 | 0.79 | 0.64 |
| 8 | 9821024 | rs11774592 | -0.195 | -0.248 | 0.254 | 1.19 | -0.008 | -0.008 | -0.008 | 0 | 0.07 | 0 | 0.08 | 0.31 | 0.08 | 0.08 |
| 8 | 9825154 | rs1521197 | -0.94 | -0.318 | 0.25 | 1.034 | 0.007 | 0.017 | 0.017 | 0 | 0.11 | 0.03 | 0.07 | 0.07 | 0.08 | 0.18 |
| 8 | 9826340 | rs7820910 | -1.969 | -0.313 | 0.591 | 1.018 | 0.067 | 0.041 | 0.041 | 0.324 | 0.06 | 0.38 | 0.04 | 0 | 0.03 | 0.03 |
| 8 | 9831664 | rs587059 | 1.79 | -0.089 | -0.22 | 0.666 | -0.032 | -0.031 | -0.031 | 0 | 0.61 | 0.82 | 0.58 | 0.49 | 0.55 | 0.57 |
| 8 | 9831907 | rs13273152 | -0.998 | -0.045 | -0.203 | 0.589 | -0.024 | 0.019 | 0.019 | 0 | 0.24 | 0.11 | 0.22 | 0.2 | 0.26 | 0.28 |
| 8 | 9834684 | rs12680244 | -0.325 | -0.235 | -0.249 | 0.756 | 0.014 | -0.035 | -0.035 | 0.032 | 0.19 | 0.15 | 0.35 | 0.45 | 0.14 | 0.47 |
| 8 | 9834862 | rs7813389 | -1.754 | -0.296 | -0.133 | 0.764 | 0.002 | -0.003 | -0.003 | 0 | 0.44 | 0.6 | 0.62 | 0.63 | 0.39 | 0.71 |
| 8 | 9838936 | rs17737355 | -0.593 | -0.236 | -0.247 | 0.733 | -0.008 | 0.008 | 0.008 | 0 | 0.21 | 0.13 | 0.25 | 0.18 | 0.21 | 0.27 |
| 8 | 9839731 | rs537231 | 0.511 | -0.254 | -0.198 | 0.665 | -0.002 | 0.004 | 0.004 | 0 | 0.79 | 0.84 | 0.68 | 0.81 | 0.79 | 0.73 |
| 8 | 9844384 | rs576405 | 1.993 | -0.344 | -0.166 | 0.632 | -0.001 | 0 | 0 | 0 | 0.6 | 0.48 | 0.54 | 0.46 | 0.58 | 0.6 |
| 8 | 9847606 | rs7815168 | 1.769 | 0.166 | -0.369 | 0.349 | -0.001 | 0.009 | 0.009 | 0 | 0.52 | 0.56 | 0.36 | 0.35 | 0.55 | 0.19 |
| 8 | 9847775 | rs6601403 | 0.473 | 0.212 | -0.327 | 0.325 | -0.001 | -0.016 | -0.016 | 0 | 0.73 | 0.72 | 0.67 | 0.53 | 0.76 | 0.47 |
| 8 | 9856072 | rs6601405 | 2.425 | 0.222 | -0.307 | 0.071 | 0 | -0.039 | -0.039 | 0 | 0.41 | 0.55 | 0.27 | 0 | 0.41 | 0.09 |
| 8 | 9862430 | rs4517127 | -1.554 | -0.094 | 0.081 | 0.808 | -0.001 | 0.136 | 0.136 | 0 | 0.2 | 0.42 | 0.14 | 0.32 | 0.19 | 0.18 |
| 8 | 9864228 | rs6601407 | 0.768 | -0.114 | -0.01 | 0.926 | -0.002 | -0.004 | -0.004 | 0 | 0.77 | 0.68 | 0.68 | 0.52 | 0.77 | 0.54 |
| 8 | 9864345 | rs7000335 | -1.506 | -0.143 | -0.16 | 0.921 | 0.006 | -0.01 | -0.01 | 0 | 0.14 | 0.09 | 0.23 | 0.2 | 0.1 | 0.22 |
| 8 | 9871971 | rs1962448 | -0.336 | -0.064 | -0.222 | 1.061 | -0.041 | -0.096 | -0.096 | 0.329 | 0.86 | 0.97 | 0.73 | 0.56 | 0.84 | 0.73 |
| 8 | 9875204 | rs1396186 | -1.189 | -0.089 | -0.312 | 0.992 | -0.039 | -0.113 | -0.113 | 0.181 | 0.17 | 0.01 | 0.14 | 0.32 | 0.19 | 0.21 |
| 8 | 9875444 | rs560835 | 1.642 | -0.196 | -0.032 | 0.937 | -0.001 | 0 | 0 | 0.48 | 0.69 | 0.39 | 0.49 | 0.36 | 0.74 | 0.54 |
| 8 | 9883177 | rs6987670 | -2.321 | -0.149 | 0.192 | 1.041 | -0.005 | 0.011 | 0.011 | 0 | 0.29 | 0.59 | 0.36 | 0.51 | 0.26 | 0.33 |
| 8 | 9888973 | rs657086 | 1.101 | -0.111 | 0.123 | 1.059 | 0 | -0.028 | -0.028 | 0 | 0.76 | 0.81 | 0.51 | 0.35 | 0.77 | 0.59 |
| 8 | 9892529 | rs670044 | 1.232 | 0.007 | 0.251 | 1.006 | 0.002 | -0.011 | -0.011 | 2.117 | 0.69 | 0.53 | 0.59 | 0.25 | 0.67 | 0.58 |
| 8 | 9895230 | rs1521191 | 2.012 | -0.121 | 0.362 | 1.12 | 0.003 | -0.003 | -0.003 | 0.015 | 0.71 | 0.27 | 0.79 | 0.71 | 0.74 | 0.56 |
| 8 | 9895811 | rs12216712 | -1.155 | -0.209 | 0.359 | 1.336 | -0.001 | 0.012 | 0.012 | 0.335 | 0.1 | 0.09 | 0.23 | 0.18 | 0.07 | 0.12 |
| 8 | 9897041 | rs7823928 | 1.991 | -0.288 | 0.373 | 1.284 | 0.004 | 0 | 0 | 0.235 | 0.71 | 0.27 | 0.81 | 0.73 | 0.74 | 0.56 |
| 8 | 9899255 | rs1521194 | -1.815 | -0.298 | 0.476 | 1.15 | -0.02 | -0.077 | -0.077 | 0 | 0.19 | 0.64 | 0.11 | 0.04 | 0.14 | 0.26 |
| 8 | 9902198 | rs6987910 | -0.569 | -0.382 | 0.262 | 1.133 | -0.011 | -0.042 | -0.042 | 0.813 | 0.1 | 0.04 | 0.09 | 0.26 | 0.12 | 0.15 |
| 8 | 9904597 | rs11775290 | -1.352 | -0.449 | 0.463 | 1.005 | -0.004 | -0.029 | -0.029 | 0.604 | 0.17 | 0.25 | 0.09 | 0.04 | 0.13 | 0.27 |
| 8 | 9906271 | rs1980262 | -0.79 | -0.343 | 0.454 | 1.185 | -0.003 | 0.008 | 0.008 | 0.394 | 0.08 | 0.08 | 0.2 | 0.16 | 0.05 | 0.1 |
| 8 | 9911617 | rs10098474 | 2.268 | -0.264 | 0.77 | 1.129 | -0.014 | 0.051 | 0.051 | 0 | 0.75 | 0.39 | 0.67 | 0.58 | 0.81 | 0.7 |

| | | | | | | | | | | | | | | | | |
|---|---|---|---|---|---|---|---|---|---|---|---|---|---|---|---|---|
| 8 | 9919062 | rs556049 | -2.047 | -0.503 | 0.736 | 0.984 | -0.004 | 0.003 | 0.003 | 0 | 0.2 | 0.25 | 0.28 | 0.18 | 0.17 | 0.23 |
| 8 | 9919269 | rs557835 | -2.047 | -0.503 | 0.736 | 0.984 | -0.004 | 0.003 | 0.003 | 0 | 0.2 | 0.25 | 0.29 | 0.18 | 0.17 | 0.23 |
| 8 | 9923902 | rs728877 | -2.079 | -0.516 | 0.752 | 0.985 | -0.005 | 0.009 | 0.009 | 0 | 0.2 | 0.29 | 0.29 | 0.19 | 0.16 | 0.24 |
| 8 | 9931941 | rs6983566 | 2.664 | -0.566 | 0.66 | 1.071 | 0.024 | -0.027 | -0.027 | 0 | 0.64 | 0.3 | 0.64 | 0.52 | 0.71 | 0.46 |
| 8 | 9932797 | rs17151026 | -0.751 | -0.563 | 0.679 | 1.076 | -0.001 | -0.031 | -0.031 | 0.052 | 0.1 | 0.02 | 0.07 | 0.27 | 0.1 | 0.18 |
| 8 | 9933528 | rs844668 | -2.481 | -0.47 | 1.103 | 1.057 | -0.016 | -0.009 | -0.009 | 0 | 0.26 | 0.63 | 0.28 | 0.2 | 0.18 | 0.35 |
| 8 | 9937217 | rs17739179 | -0.63 | -0.359 | 0.965 | 0.894 | -0.001 | -0.027 | -0.027 | 0 | 0.09 | 0.15 | 0.05 | 0.02 | 0.08 | 0.11 |
| 8 | 9942924 | rs814424 | -1.578 | -0.367 | 1.078 | 1.011 | 0.001 | -0.006 | -0.006 | 2.583 | 0.18 | 0.12 | 0.27 | 0.42 | 0.18 | 0.26 |
| 8 | 9956739 | rs10098342 | -2.31 | -0.39 | 1.24 | 0.819 | -0.024 | -0.094 | -0.094 | 0 | 0.17 | 0.61 | 0.09 | 0.04 | 0.11 | 0.23 |
| 8 | 9961451 | rs4294187 | -2.135 | -0.461 | 1.296 | 0.854 | -0.01 | 0.051 | 0.051 | 0 | 0.24 | 0.31 | 0.29 | 0.43 | 0.22 | 0.43 |
| 8 | 9963646 | rs4841281 | 0.789 | -0.47 | 1.335 | 0.795 | -0.007 | -0.033 | -0.033 | 0 | 0.91 | 0.82 | 0.95 | 0.98 | 0.94 | 0.87 |
| 8 | 9969692 | rs11774411 | -0.404 | -0.508 | 1.031 | 0.601 | -0.003 | 0.113 | 0.113 | 0 | 0.2 | 0.3 | 0.36 | 0.55 | 0.21 | 0.29 |
| 8 | 9971054 | rs1160369 | 2.067 | 0.03 | 1.187 | 0.573 | -0.001 | 0.036 | 0.036 | 0 | 0.65 | 0.53 | 0.63 | 0.55 | 0.64 | 0.44 |
| 8 | 9975212 | rs17151117 | -0.241 | 0.002 | 0.997 | 0.723 | -0.024 | -0.063 | -0.063 | 0 | 0.1 | 0 | 0.05 | 0.29 | 0.11 | 0.23 |
| 8 | 9977918 | rs6983332 | -1.992 | 0.052 | 1.237 | 0.405 | -0.001 | -0.126 | -0.126 | 0 | 0.4 | 0.44 | 0.62 | 0.96 | 0.41 | 0.59 |
| 8 | 9979144 | rs12674871 | -0.567 | 0.058 | 1.172 | 0.534 | -0.005 | 0.063 | 0.063 | 0.019 | 0.21 | 0.28 | 0.35 | 0.56 | 0.2 | 0.29 |
| 8 | 9983369 | rs17151147 | -1.016 | 0.112 | 1.012 | 0.572 | -0.011 | -0.067 | -0.067 | 0.172 | 0.18 | 0.05 | 0.26 | 0.4 | 0.21 | 0.31 |
| 8 | 9990098 | rs7005350 | 0.643 | 0.228 | 1.025 | 0.517 | 0.014 | -0.039 | -0.039 | 0 | 0.9 | 0.76 | 0.94 | 0.98 | 0.91 | 0.86 |
| 8 | 9990358 | rs2046400 | 0.614 | 0.23 | 0.957 | 0.518 | 0.005 | -0.033 | -0.033 | 0.045 | 0.91 | 0.81 | 0.94 | 0.98 | 0.91 | 0.86 |
| 8 | 9994504 | rs9644681 | 1.551 | 0.18 | 0.637 | 0.609 | -0.003 | 0.015 | 0.015 | 0 | 0.68 | 0.43 | 0.66 | 0.58 | 0.7 | 0.55 |
| 8 | 9995515 | rs12547884 | -0.581 | 0.2 | 0.193 | 0.989 | -0.054 | -0.122 | -0.122 | 0.215 | 0.18 | 0.04 | 0.39 | 0.55 | 0.23 | 0.29 |
| 8 | 10000966 | rs1484645 | -0.478 | 0.15 | 0.209 | 0.983 | -0.04 | -0.112 | -0.112 | 0 | 0.18 | 0.04 | 0.36 | 0.55 | 0.21 | 0.29 |
| 8 | 10007018 | rs12677675 | -0.42 | -0.169 | 0.503 | 1.122 | -0.007 | 0.144 | 0.144 | 0 | 0.11 | 0.2 | 0.2 | 0.47 | 0.1 | 0.19 |
| 8 | 10007227 | rs4841287 | 1.602 | -0.351 | 0.746 | 1.324 | -0.015 | 0.009 | 0.009 | 0.157 | 0.7 | 0.38 | 0.73 | 0.63 | 0.76 | 0.55 |
| 8 | 10016634 | rs7842835 | -1.895 | -0.436 | 0.76 | 1.075 | 0 | 0 | 0 | 0.3 | 0.3 | 0.7 | 0.32 | 0.52 | 0.24 | 0.43 |
| 8 | 10016969 | rs12678938 | -2.487 | -0.609 | 0.046 | 0.445 | -0.067 | 0.158 | 0.158 | 0.209 | 0.61 | 0.84 | 0.72 | 1 | 0.54 | 0.88 |
| 8 | 10018850 | rs1385256 | -1.257 | -0.589 | 0.748 | 0.889 | -0.023 | 0.089 | 0.089 | 2.392 | 0.21 | 0.38 | 0.27 | 0.53 | 0.17 | 0.35 |
| 8 | 10024748 | rs4255142 | -0.514 | -0.541 | 0.413 | 0.948 | 0.002 | -0.05 | -0.05 | 0 | 0.14 | 0.05 | 0.32 | 0.48 | 0.16 | 0.26 |
| 8 | 10030255 | rs7823314 | -0.353 | -0.504 | 0.868 | 0.861 | -0.003 | -0.013 | -0.013 | 0 | 0.06 | 0.14 | 0.04 | 0.03 | 0.05 | 0.08 |
| 8 | 10030902 | rs11787370 | -0.831 | -0.801 | 0.499 | 0.487 | -0.006 | -0.052 | -0.052 | 0 | 0.31 | 0.11 | 0.3 | 0.52 | 0.3 | 0.39 |
| 8 | 10032894 | rs11993663 | -1.294 | -0.827 | 0.74 | 0.385 | 0.004 | -0.01 | -0.01 | 0.012 | 0.35 | 0.36 | 0.32 | 0.48 | 0.31 | 0.48 |
| 8 | 10033264 | rs1905630 | -1.739 | -0.844 | 0.638 | 0.414 | 0.022 | -0.012 | -0.012 | 0.048 | 0.46 | 0.78 | 0.4 | 0.48 | 0.41 | 0.58 |
| 8 | 10035151 | rs11783143 | -0.131 | -0.742 | 0.296 | 0.034 | 0.003 | -0.009 | -0.009 | 0 | 0.77 | 0.8 | 0.63 | 0.51 | 0.78 | 0.7 |
| 8 | 10044801 | rs7820236 | -1.569 | -0.498 | 0.863 | 0.572 | 0.008 | -0.002 | -0.002 | 0 | 0.43 | 0.79 | 0.41 | 0.51 | 0.4 | 0.6 |
| 8 | 10046638 | rs11249973 | -1.267 | -0.56 | 1.206 | 0.562 | -0.002 | 0.001 | 0.001 | 0.439 | 0.4 | 0.45 | 0.39 | 0.51 | 0.39 | 0.6 |
| 8 | 10050618 | rs17748464 | -1.177 | -0.532 | 1.047 | 0.533 | -0.006 | -0.04 | -0.04 | 0 | 0.37 | 0.29 | 0.34 | 0.46 | 0.37 | 0.55 |
| 8 | 10056664 | rs6982806 | -1.447 | -0.485 | 0.921 | 0.554 | -0.001 | 0 | 0 | 0 | 0.42 | 0.63 | 0.4 | 0.48 | 0.39 | 0.59 |
| 8 | 10060638 | rs10097315 | -1.825 | -0.57 | 0.515 | 0.588 | 0.012 | -0.024 | -0.024 | 0 | 0.45 | 0.81 | 0.41 | 0.48 | 0.39 | 0.61 |
| 8 | 10065961 | rs7832976 | -0.414 | -0.406 | 0.587 | 1.068 | 0 | 0.004 | 0.004 | 0.008 | 0.18 | 0.15 | 0.17 | 0.25 | 0.14 | 0.26 |
| 8 | 10066119 | rs7836718 | -1.407 | -0.4 | 0.331 | 1.054 | -0.007 | -0.003 | -0.003 | 2.943 | 0.29 | 0.67 | 0.24 | 0.28 | 0.21 | 0.43 |
| 8 | 10067157 | rs6991853 | -1.186 | -0.233 | -0.055 | 1.127 | 0 | -0.007 | -0.007 | 0 | 0.24 | 0.68 | 0.22 | 0.21 | 0.19 | 0.35 |
| 8 | 10068397 | rs7829161 | -0.437 | 0.051 | -0.62 | 1.39 | -0.007 | 0 | 0 | 0 | 0.08 | 0.03 | 0.09 | 0.25 | 0.13 | 0.15 |
| 8 | 10069485 | rs10903321 | 1.628 | -0.2 | -1.698 | -0.016 | -0.168 | 0.234 | 0.234 | 0.156 | 0.31 | 0.02 | 0.27 | 0 | 0.42 | 0.11 |
| 8 | 10071620 | rs11775334 | 0.444 | -0.931 | -1.278 | 0.64 | 0.01 | -0.015 | -0.015 | 0.688 | 0.63 | 0.35 | 0.59 | 0.44 | 0.69 | 0.56 |
| 8 | 10072350 | rs7840347 | 2.011 | -0.996 | -1.392 | 0.456 | -0.129 | 0.202 | 0.202 | 0.004 | 0.35 | 0.12 | 0.28 | 0 | 0.46 | 0.11 |
| 8 | 10072855 | rs7012244 | -0.038 | -1.07 | -1.345 | 0.622 | 0.017 | -0.019 | -0.019 | 0 | 0.27 | 0.21 | 0.28 | 0.41 | 0.18 | 0.41 |
| 8 | 10073962 | rs4240634 | 1.239 | -1.286 | -1.561 | 0.896 | 0.117 | 0.176 | 0.176 | 0.528 | 0.52 | 0.16 | 0.58 | 0.18 | 0.69 | 0.38 |
| 8 | 10074778 | rs4474025 | 0.367 | -1.248 | -0.881 | 1.068 | -0.001 | 0.017 | 0.017 | 0 | 0.15 | 0.16 | 0.08 | 0.29 | 0.09 | 0.19 |
| 8 | 10077565 | rs4075952 | -0.193 | -1.208 | -0.958 | 1.167 | -0.045 | -0.072 | -0.072 | 0 | 0.19 | 0.08 | 0.35 | 0.28 | 0.26 | 0.26 |
| 8 | 10077806 | rs10110380 | -1.445 | -1.253 | -0.601 | 1.19 | 0.001 | 0 | 0 | 0 | 0.38 | 0.43 | 0.52 | 0.62 | 0.36 | 0.47 |
| 8 | 10078102 | rs10110731 | 0.309 | -1.245 | -0.517 | 1.278 | -0.036 | 0.012 | 0.012 | 0 | 0.69 | 0.44 | 0.78 | 0.63 | 0.79 | 0.57 |
| 8 | 10078853 | rs12543841 | 0.351 | -0.967 | -1.019 | 1.331 | -0.042 | -0.018 | -0.018 | 0 | 0.16 | 0.06 | 0.38 | 0.17 | 0.23 | 0.25 |
| 8 | 10079278 | rs12546137 | 0.526 | -0.952 | -1.042 | 1.341 | -0.065 | -0.049 | -0.049 | 0.933 | 0.16 | 0.02 | 0.38 | 0.17 | 0.21 | 0.24 |
| 8 | 10082204 | rs4529465 | 0.001 | -1.111 | -0.964 | 1.237 | 0.027 | -0.024 | -0.024 | 1.883 | 0.25 | 0.17 | 0.18 | 0.34 | 0.18 | 0.38 |
| 8 | 10084886 | rs13256234 | -0.124 | -1.072 | -0.964 | 1.272 | -0.012 | -0.041 | -0.041 | 0 | 0.2 | 0.1 | 0.43 | 0.33 | 0.25 | 0.3 |
| 8 | 10085359 | rs10503404 | 0.397 | -1.132 | -0.944 | 1.392 | 0.031 | -0.03 | -0.03 | 0.736 | 0.21 | 0.13 | 0.16 | 0.29 | 0.16 | 0.29 |
| 8 | 10086946 | rs4240635 | -1.342 | -1.058 | -0.65 | 1.086 | 0.086 | 0.045 | 0.045 | 0 | 0.48 | 0.78 | 0.3 | 0.66 | 0.31 | 0.61 |
| 8 | 10090680 | rs2975732 | 1.391 | -0.531 | -0.067 | 0.898 | -0.007 | 0.017 | 0.017 | 0 | 0.43 | 0.31 | 0.37 | 0.28 | 0.47 | 0.25 |
| 8 | 10091194 | rs2952248 | 1.291 | -0.61 | 0.053 | 0.873 | -0.005 | -0.033 | -0.033 | 0 | 0.07 | 0.1 | 0.05 | 0 | 0.04 | 0.03 |
| 8 | 10093228 | rs17151405 | -0.637 | -0.834 | 0.078 | 0.985 | -0.001 | 0.007 | 0.007 | 0 | 0.19 | 0.18 | 0.2 | 0.44 | 0.22 | 0.3 |
| 8 | 10094377 | rs2975726 | -0.653 | -1.463 | 0.188 | 0.878 | -0.011 | 0.009 | 0.009 | 0 | 0.24 | 0.21 | 0.17 | 0.31 | 0.14 | 0.29 |
| 8 | 10095049 | rs12680355 | -0.291 | -1.244 | 0.335 | 0.921 | 0.001 | 0.008 | 0.008 | 0 | 0.17 | 0.18 | 0.17 | 0.41 | 0.15 | 0.28 |
| 8 | 10095328 | rs2952238 | 0.61 | -1.159 | 0.23 | 0.873 | 0 | 0.109 | 0.109 | 0 | 0.1 | 0.18 | 0.06 | 0.21 | 0.05 | 0.1 |
| 8 | 10095707 | rs7842910 | -0.924 | -1.135 | 0.328 | 0.664 | 0.02 | 0.006 | 0.006 | 0 | 0.4 | 0.26 | 0.48 | 0.41 | 0.36 | 0.52 |
| 8 | 10097271 | rs2975722 | -0.043 | -1.055 | 0.735 | 1.076 | -0.008 | 0.104 | 0.104 | 0 | 0.86 | 0.68 | 0.84 | 0.64 | 0.87 | 0.76 |
| 8 | 10097511 | rs2975720 | -0.467 | -0.875 | 0.837 | 0.937 | -0.006 | 0.141 | 0.141 | 0 | 0.13 | 0.32 | 0.08 | 0.23 | 0.07 | 0.11 |
| 8 | 10104092 | rs2952223 | 0.714 | -0.984 | 0.34 | 0.722 | -0.044 | 0.048 | 0.048 | 0 | 0.68 | 0.52 | 0.73 | 0.51 | 0.77 | 0.55 |
| 8 | 10105427 | rs4295669 | 0.328 | -0.878 | 0.06 | 0.443 | 0 | 0.007 | 0.007 | 0.93 | 0.23 | 0.25 | 0.37 | 0.33 | 0.24 | 0.3 |
| 8 | 10106503 | rs13258235 | 0.798 | -0.863 | -0.094 | 0.437 | -0.062 | -0.08 | -0.08 | 0 | 0.21 | 0.02 | 0.36 | 0.31 | 0.23 | 0.29 |

| | | | | | | | | | | | | | | | | |
|---|---|---|---|---|---|---|---|---|---|---|---|---|---|---|---|---|
| 8 | 10107022 | rs12549696 | 0.574 | -0.84 | -0.031 | 0.441 | -0.001 | -0.002 | -0.002 | 0.003 | 0.22 | 0.2 | 0.36 | 0.32 | 0.23 | 0.29 |
| 8 | 10108972 | rs10503406 | 2.842 | -0.738 | -0.264 | 0.09 | -0.075 | 0.129 | 0.129 | 0 | 0.29 | 0.09 | 0.24 | 0 | 0.39 | 0.09 |
| 8 | 10110457 | rs2975701 | 0.756 | -0.767 | 0.11 | 0.585 | -0.006 | 0.055 | 0.055 | 0 | 0.09 | 0.12 | 0.06 | 0.22 | 0.05 | 0.11 |
| 8 | 10111571 | rs4841297 | 1.226 | -0.916 | 0.159 | 0.612 | 0.031 | 0.012 | 0.012 | 0 | 0.58 | 0.28 | 0.69 | 0.42 | 0.68 | 0.46 |
| 8 | 10112034 | rs10099182 | 0.615 | -0.942 | 0.249 | 0.644 | 0.024 | 0.082 | 0.082 | 0 | 0.1 | 0.16 | 0.16 | 0.32 | 0.17 | 0.2 |
| 8 | 10112706 | rs2975696 | 1.021 | -0.912 | 0.173 | 0.632 | 0.001 | -0.002 | -0.002 | 0 | 0.65 | 0.33 | 0.68 | 0.54 | 0.7 | 0.53 |
| 8 | 10113604 | rs11781781 | 1.139 | -1.246 | -0.339 | 0.333 | 0.013 | 0.011 | 0.011 | 0 | 0.58 | 0.37 | 0.65 | 0.32 | 0.71 | 0.43 |
| 8 | 10115960 | rs4258008 | 0.215 | -1.015 | -0.986 | 0.652 | -0.054 | -0.04 | -0.04 | 0 | 0.22 | 0.02 | 0.34 | 0.28 | 0.23 | 0.28 |
| 8 | 10117293 | rs10098383 | -0.692 | -1.107 | -0.286 | 0.713 | -0.008 | 0.014 | 0.014 | 0 | 0.19 | 0.18 | 0.16 | 0.32 | 0.15 | 0.36 |
| 8 | 10118004 | rs2952194 | -1.247 | -0.98 | -0.512 | -0.2 | -0.002 | -0.038 | -0.038 | 0.271 | 0.2 | 0.18 | 0.09 | 0 | 0.15 | 0.08 |
| 8 | 10118346 | rs2975684 | -0.756 | -0.507 | -0.44 | 0.307 | 0 | -0.001 | -0.001 | 0 | 0.1 | 0.14 | 0.09 | 0.06 | 0.08 | 0.06 |
| 8 | 10119523 | rs4305909 | 0.285 | -0.818 | -0.427 | -0.089 | -0.023 | -0.032 | -0.032 | 0 | 0.72 | 0.51 | 0.56 | 0.84 | 0.81 | 0.77 |
| 8 | 10122315 | rs4841306 | -0.269 | 0.182 | -0.855 | -0.879 | 0.013 | -0.038 | -0.038 | 0 | 0.73 | 0.6 | 0.54 | 0.85 | 0.75 | 0.79 |
| 8 | 10122423 | rs6997416 | 0.027 | 0.096 | -0.753 | -0.91 | 0.004 | 0.012 | 0.012 | 0 | 0.56 | 0.48 | 0.32 | 0.85 | 0.44 | 0.7 |
| 8 | 10123027 | rs12679687 | 1.166 | -0.36 | -1.478 | -0.27 | 0.046 | -0.122 | -0.122 | 0.181 | 0.25 | 0.02 | 0.1 | 0.4 | 0.14 | 0.4 |
| 8 | 10123974 | rs13250969 | 0.693 | -0.475 | -1.422 | -0.151 | -0.051 | -0.065 | -0.065 | 2.807 | 0.46 | 0.13 | 0.32 | 0.46 | 0.46 | 0.51 |
| 8 | 10125195 | rs2952184 | -1.244 | -0.616 | -1.012 | -0.336 | 0.025 | -0.056 | -0.056 | 0.965 | 0.84 | 0.73 | 0.88 | 0.97 | 0.84 | 0.91 |
| 8 | 10125535 | rs6994059 | 1.433 | -0.682 | -0.819 | -0.159 | -0.002 | -0.006 | -0.006 | 0 | 0.22 | 0.18 | 0.19 | 0.34 | 0.27 | 0.27 |
| 8 | 10126348 | rs4260895 | -0.482 | -0.963 | -0.978 | -0.34 | 0.006 | -0.016 | -0.016 | 0 | 0.64 | 0.72 | 0.83 | 0.96 | 0.74 | 0.73 |
| 8 | 10128539 | rs2975668 | 1.481 | -0.87 | -0.898 | -0.159 | 0 | -0.003 | -0.003 | 0.082 | 0.11 | 0.08 | 0.36 | 0.12 | 0.1 | 0.13 |
| 8 | 10129757 | rs4240637 | -0.295 | -0.785 | -0.858 | -0.222 | 0.004 | -0.029 | -0.029 | 0.557 | 0.74 | 0.64 | 0.61 | 0.85 | 0.77 | 0.82 |
| 8 | 10129773 | rs7001821 | 0.816 | -0.738 | -0.657 | 0.104 | 0.012 | 0.008 | 0.008 | 0 | 0.46 | 0.42 | 0.49 | 0.44 | 0.63 | 0.38 |
| 8 | 10131459 | rs11249979 | 0.185 | -0.701 | -0.797 | 0.493 | 0.009 | 0.011 | 0.011 | 0.285 | 0.53 | 0.63 | 0.48 | 0.7 | 0.64 | 0.58 |
| 8 | 10131799 | rs12681109 | 0.462 | -0.685 | -0.779 | 0.597 | -0.007 | 0.059 | 0.059 | 0 | 0.08 | 0.12 | 0.06 | 0.34 | 0.05 | 0.22 |
| 8 | 10132114 | rs2952166 | 0.643 | -0.58 | -0.853 | 0.386 | -0.003 | 0.002 | 0.002 | 0 | 0.14 | 0.08 | 0.35 | 0.12 | 0.11 | 0.16 |
| 8 | 10133477 | rs2975654 | -0.783 | -0.797 | -0.763 | 0.424 | -0.001 | -0.001 | -0.001 | 0.015 | 0.67 | 0.72 | 0.83 | 0.83 | 0.75 | 0.75 |
| 8 | 10135235 | rs2004678 | 1.044 | -0.62 | -0.825 | 0.413 | 0.004 | -0.001 | -0.001 | 0 | 0.12 | 0.07 | 0.27 | 0.12 | 0.08 | 0.12 |
| 8 | 10140875 | rs10102621 | 1.046 | -0.556 | -0.434 | 0.201 | 0.021 | -0.01 | -0.01 | 0 | 0.29 | 0.4 | 0.29 | 0.31 | 0.36 | 0.34 |
| 8 | 10145860 | rs11784779 | 0.336 | -0.565 | -0.54 | 0.073 | -0.02 | -0.014 | -0.014 | 0 | 0.38 | 0.25 | 0.53 | 0.45 | 0.43 | 0.41 |
| 8 | 10148197 | rs4840471 | -1.271 | -1.099 | -0.378 | -0.651 | 0.006 | -0.027 | -0.027 | 1.008 | 0.75 | 0.72 | 0.92 | 1 | 0.88 | 0.81 |
| 8 | 10149567 | rs10903323 | 0.049 | -0.864 | -0.588 | -0.311 | 0.016 | -0.034 | -0.034 | 0 | 0.2 | 0.11 | 0.33 | 0.6 | 0.11 | 0.38 |
| 8 | 10153082 | rs17151637 | 0.423 | -0.661 | -0.339 | -0.801 | -0.002 | -0.07 | -0.07 | 0.12 | 0.34 | 0.34 | 0.16 | 0 | 0.28 | 0.25 |
| 8 | 10153600 | rs11249982 | -0.124 | -0.636 | -0.434 | -0.543 | -0.012 | 0.014 | 0.014 | 0 | 0.22 | 0.24 | 0.35 | 0.65 | 0.12 | 0.4 |
| 8 | 10160977 | rs11249984 | -1.319 | -0.735 | -0.441 | -0.849 | 0.006 | -0.028 | -0.028 | 16.387 | 0.74 | 0.77 | 0.93 | 1 | 0.88 | 0.79 |
| 8 | 10162381 | rs13257313 | 2.511 | -0.31 | -0.832 | -0.907 | -0.098 | 0.109 | 0.109 | 0 | 0.18 | 0.04 | 0.2 | 0 | 0.29 | 0.09 |
| 8 | 10165439 | rs11782868 | -0.126 | -0.315 | -0.719 | -0.702 | 0.001 | -0.017 | -0.017 | 0 | 0.25 | 0.15 | 0.4 | 0.65 | 0.21 | 0.44 |
| 8 | 10168930 | rs17151700 | 0.008 | -0.428 | -0.741 | -0.645 | 0.02 | -0.043 | -0.043 | 0.041 | 0.23 | 0.08 | 0.35 | 0.64 | 0.15 | 0.39 |
| 8 | 10170545 | rs6601431 | -1.099 | -0.475 | -0.377 | -0.802 | -0.007 | -0.009 | -0.009 | 0 | 0.37 | 0.52 | 0.47 | 0.65 | 0.3 | 0.54 |
| 8 | 10172841 | rs7014570 | 1.326 | -0.362 | -0.28 | -0.794 | 0.047 | -0.054 | -0.054 | 0 | 0.1 | 0.27 | 0.1 | 0 | 0.14 | 0.07 |
| 8 | 10175492 | rs17151728 | 1.153 | -0.328 | -0.337 | -0.812 | 0.063 | -0.064 | -0.064 | 0.264 | 0.11 | 0.29 | 0.11 | 0 | 0.17 | 0.11 |
| 8 | 10175545 | rs17151731 | -0.221 | -0.416 | -0.663 | -0.725 | 0.021 | -0.025 | -0.025 | 0 | 0.24 | 0.14 | 0.36 | 0.66 | 0.16 | 0.45 |
| 8 | 10176467 | rs10089244 | 1.338 | -0.337 | -0.886 | -0.773 | -0.005 | -0.013 | -0.013 | 0 | 0.19 | 0.14 | 0.26 | 0.36 | 0.26 | 0.18 |
| 8 | 10179776 | rs17151743 | 1.3 | -0.252 | -0.993 | -0.309 | -0.009 | -0.066 | -0.066 | 0.474 | 0.12 | 0.01 | 0.27 | 0.18 | 0.07 | 0.12 |
| 8 | 10183347 | rs7006402 | -0.587 | -0.328 | -0.268 | -0.327 | -0.042 | 0.057 | 0.057 | 0 | 0.24 | 0.44 | 0.37 | 0.61 | 0.13 | 0.35 |
| 8 | 10183540 | rs12680918 | 1.077 | -0.124 | -0.603 | -0.156 | -0.013 | 0.052 | 0.052 | 0 | 0.06 | 0 | 0.06 | 0.46 | 0.05 | 0.22 |
| 8 | 10183576 | rs17707988 | 1.163 | -0.105 | -0.458 | -0.917 | 0.019 | -0.045 | -0.045 | 0.796 | 0.1 | 0.13 | 0.09 | 0.01 | 0.14 | 0.11 |
| 8 | 10186135 | rs4841317 | -0.474 | -0.412 | -0.718 | -1.087 | 0.005 | -0.01 | -0.01 | 0 | 0.44 | 0.32 | 0.61 | 0.99 | 0.42 | 0.59 |
| 8 | 10186789 | rs7816713 | -0.412 | 0.106 | -1.454 | -1.024 | -0.034 | 0.13 | 0.13 | 0 | 0.49 | 0.14 | 0.5 | 0.34 | 0.49 | 0.36 |
| 8 | 10187200 | rs4841318 | -0.714 | 0.3 | -0.92 | -1.158 | 0.038 | -0.123 | -0.123 | 0.141 | 0.72 | 0.66 | 0.76 | 1 | 0.62 | 0.81 |
| 8 | 10188617 | rs13249929 | 0.615 | 0.576 | -1.169 | -0.24 | -0.019 | -0.041 | -0.041 | 0.444 | 0.23 | 0.16 | 0.26 | 0.41 | 0.32 | 0.24 |
| 8 | 10197718 | rs7460226 | 0.152 | 0.472 | -0.984 | -0.622 | 0 | -0.007 | -0.007 | 0 | 0.32 | 0.32 | 0.38 | 0.15 | 0.39 | 0.31 |
| 8 | 10199548 | rs17708276 | 0.652 | 0.596 | -1.089 | -0.27 | -0.047 | 0.016 | 0.016 | 0 | 0.08 | 0 | 0.07 | 0.03 | 0.13 | 0.07 |
| 8 | 10201532 | rs4841324 | -0.576 | 0.535 | -0.471 | 0.176 | 0.006 | 0.147 | 0.147 | 0 | 0.9 | 0.74 | 0.92 | 0.67 | 0.91 | 0.94 |
| 8 | 10207464 | rs4260896 | 0.05 | 0.168 | -0.929 | -0.995 | 0.032 | 0.075 | 0.075 | 0 | 0.51 | 0.35 | 0.56 | 0.14 | 0.65 | 0.4 |
| 8 | 10225159 | rs7015455 | -0.537 | 0.219 | -0.625 | -0.824 | 0.004 | 0.012 | 0.012 | 0 | 0.49 | 0.58 | 0.43 | 0.28 | 0.56 | 0.39 |
| 8 | 10226355 | rs3088186 | 0.96 | 0.542 | -0.599 | -0.813 | -0.017 | -0.001 | -0.001 | 0 | 0.23 | 0.14 | 0.26 | 0.21 | 0.28 | 0.19 |
| 8 | 10234303 | rs13273009 | 0.126 | 0.059 | -0.748 | -1.16 | 0.014 | -0.001 | -0.001 | 5.096 | 0.73 | 0.81 | 0.56 | 0.23 | 0.81 | 0.59 |
| 8 | 10236559 | rs17151854 | -0.091 | -0.026 | 0.075 | -1.182 | 0.052 | -0.064 | -0.064 | 0 | 0.14 | 0.36 | 0.1 | 0.03 | 0.16 | 0.13 |
| 8 | 10236815 | rs11249994 | -0.487 | -0.143 | -0.026 | -1.288 | 0 | -0.001 | -0.001 | 0.109 | 0.43 | 0.45 | 0.42 | 0.28 | 0.5 | 0.39 |
| 8 | 10238922 | rs11249995 | 1.043 | -0.35 | -0.338 | -1.122 | -0.06 | -0.035 | -0.035 | 0.055 | 0.27 | 0.07 | 0.29 | 0.25 | 0.34 | 0.26 |
| 8 | 10243101 | rs6601450 | -0.153 | -0.486 | -0.054 | -0.529 | -0.006 | -0.015 | -0.015 | 0 | 0.38 | 0.37 | 0.42 | 0.52 | 0.42 | 0.35 |
| 8 | 10244946 | rs11783514 | 0.523 | -0.71 | 0.063 | -0.607 | -0.02 | 0.001 | 0.001 | 0 | 0.74 | 0.68 | 0.61 | 0.25 | 0.85 | 0.59 |
| 8 | 10245095 | rs13249940 | -1.64 | -0.621 | -0.014 | -0.423 | -0.016 | 0.043 | 0.043 | 0 | 0.88 | 0.92 | 0.74 | 0.59 | 0.97 | 0.75 |
| 8 | 10245463 | rs11249996 | 1.303 | -0.453 | -0.136 | -0.859 | -0.025 | -0.015 | -0.015 | 0 | 0.26 | 0.12 | 0.27 | 0.26 | 0.32 | 0.24 |
| 8 | 10245788 | rs11774221 | 0.904 | 0.044 | -0.308 | -1.012 | -0.036 | 0.03 | 0.03 | 0 | 0.13 | 0.06 | 0.14 | 0.03 | 0.28 | 0.17 |
| 8 | 10247816 | rs4841328 | -0.208 | -0.59 | 0.28 | -0.663 | -0.037 | 0.039 | 0.039 | 1.016 | 0.8 | 0.69 | 0.62 | 0.28 | 0.89 | 0.61 |
| 8 | 10248035 | rs17151885 | 0.671 | -0.618 | 0.449 | -0.833 | -0.019 | -0.04 | -0.04 | 0 | 0.08 | 0.25 | 0.03 | 0 | 0.03 | 0.02 |
| 8 | 10249480 | rs17765901 | 1.033 | -0.619 | 0.119 | -0.967 | 0.002 | 0.065 | 0.065 | 0 | 0.45 | 0.28 | 0.33 | 0.03 | 0.53 | 0.37 |
| 8 | 10250159 | rs12544727 | 0.696 | -0.696 | 0.149 | -0.89 | -0.05 | -0.006 | -0.006 | 0.038 | 0.7 | 0.37 | 0.6 | 0.25 | 0.85 | 0.6 |

| | | | | | | | | | | | | | | | | |
|---|---|---|---|---|---|---|---|---|---|---|---|---|---|---|---|---|
| 8 | 10250758 | rs17151892 | 0.135 | -0.608 | -0.01 | -0.789 | -0.005 | 0 | 0 | 0.891 | 0.2 | 0.24 | 0.37 | 0.75 | 0.12 | 0.39 |
| 8 | 10251217 | rs2001338 | 1.178 | -0.533 | -0.042 | -0.798 | -0.025 | 0.002 | 0.002 | 0 | 0.25 | 0.12 | 0.27 | 0.22 | 0.33 | 0.22 |
| 8 | 10253537 | rs17151903 | 0.058 | -0.579 | -0.19 | -0.752 | -0.004 | 0 | 0 | 0 | 0.21 | 0.24 | 0.37 | 0.75 | 0.13 | 0.39 |
| 8 | 10255402 | rs11777976 | 1.016 | -0.526 | -0.503 | -0.883 | -0.008 | 0.149 | 0.149 | 0 | 0.44 | 0.22 | 0.32 | 0.03 | 0.52 | 0.39 |
| 8 | 10256054 | rs11989640 | -1.148 | -0.369 | -0.493 | -0.649 | -0.031 | 0.006 | 0.006 | 0 | 0.75 | 0.86 | 0.74 | 0.79 | 0.67 | 0.79 |
| 8 | 10260553 | rs1962073 | 0.846 | -0.602 | -0.657 | -0.332 | 0.036 | 0.093 | 0.093 | 0.012 | 0.48 | 0.13 | 0.4 | 0.34 | 0.59 | 0.49 |
| 8 | 10260837 | rs12547997 | 1.067 | -0.43 | -0.643 | -0.208 | -0.042 | -0.031 | -0.031 | 0 | 0.26 | 0.11 | 0.29 | 0.27 | 0.33 | 0.21 |
| 8 | 10261068 | rs7459532 | -0.54 | -0.556 | -0.071 | -0.303 | -0.004 | -0.016 | -0.016 | 0 | 0.45 | 0.53 | 0.61 | 0.71 | 0.41 | 0.5 |
| 8 | 10265240 | rs12549419 | 0.417 | -0.57 | -0.179 | -0.071 | -0.029 | 0.049 | 0.049 | 0 | 0.16 | 0.2 | 0.3 | 0.44 | 0.05 | 0.28 |
| 8 | 10268736 | rs1986971 | 0.668 | -0.564 | -0.734 | -0.654 | 0.013 | 0.008 | 0.008 | 0 | 0.36 | 0.2 | 0.21 | 0.33 | 0.31 | 0.42 |
| 8 | 10268983 | rs1986973 | 0.294 | -0.55 | -0.247 | -0.251 | -0.025 | 0.056 | 0.056 | 0 | 0.16 | 0.23 | 0.31 | 0.45 | 0.06 | 0.29 |
| 8 | 10270958 | rs11250004 | 0.356 | -0.763 | -0.343 | -0.351 | 0.055 | 0.077 | 0.077 | 0 | 0.55 | 0.26 | 0.44 | 0.29 | 0.66 | 0.53 |
| 8 | 10271318 | rs4311657 | 0.527 | -0.548 | -0.195 | -0.022 | -0.044 | 0.082 | 0.082 | 0 | 0.14 | 0.2 | 0.29 | 0.46 | 0.04 | 0.25 |
| 8 | 10277494 | rs17767186 | 1.191 | -0.369 | -0.016 | -0.119 | -0.003 | 0 | 0 | 1.223 | 0.22 | 0.16 | 0.24 | 0.2 | 0.27 | 0.17 |
| 8 | 10278678 | rs11250005 | 0.225 | -0.565 | -0.068 | -0.081 | 0.034 | 0.04 | 0.04 | 0 | 0.58 | 0.3 | 0.45 | 0.29 | 0.68 | 0.53 |
| 8 | 10279503 | rs10112584 | 0.064 | -0.706 | 0.015 | 0.257 | -0.044 | 0.061 | 0.061 | 0 | 0.17 | 0.23 | 0.3 | 0.5 | 0.05 | 0.28 |
| 8 | 10280818 | rs6985267 | -0.848 | -0.551 | -0.226 | -0.053 | 0.004 | 0.001 | 0.001 | 0 | 0.28 | 0.28 | 0.42 | 0.76 | 0.22 | 0.44 |
| 8 | 10284051 | rs7009513 | -0.684 | -0.818 | 0.133 | 0.208 | -0.037 | 0.051 | 0.051 | 0 | 0.85 | 0.8 | 0.74 | 0.55 | 0.97 | 0.74 |
| 8 | 10285810 | rs3750314 | 0.337 | -0.779 | 0.224 | 0.324 | -0.057 | 0.127 | 0.127 | 0 | 0.13 | 0.26 | 0.27 | 0.42 | 0.03 | 0.23 |
| 8 | 10286050 | rs3750315 | 0.027 | -0.816 | 0.418 | 0.338 | -0.084 | 0.255 | 0.255 | 0.71 | 0.14 | 0.5 | 0.27 | 0.4 | 0.03 | 0.24 |
| 8 | 10286273 | rs14879 | 0.162 | -0.796 | 0.108 | 0.028 | 0.053 | 0.006 | 0.006 | 0.82 | 0.62 | 0.19 | 0.49 | 0.36 | 0.73 | 0.57 |
| 8 | 10286511 | rs10429305 | 1.077 | -0.741 | -0.385 | -0.744 | 0.007 | 0.035 | 0.035 | 0.13 | 0.31 | 0.24 | 0.18 | 0.08 | 0.26 | 0.34 |
| 8 | 10286615 | rs6996446 | -0.416 | -0.114 | -0.136 | -0.241 | 0.007 | -0.002 | -0.002 | 2.198 | 0.69 | 0.69 | 0.66 | 0.72 | 0.63 | 0.76 |
| 8 | 10287438 | rs10095339 | -0.061 | 0.327 | -0.154 | 0.209 | 0.03 | 0.029 | 0.029 | 0 | 0.75 | 0.7 | 0.63 | 0.61 | 0.63 | 0.66 |
| 8 | 10288461 | rs10100203 | 0.785 | 0.185 | 0.023 | 0.175 | 0.003 | 0.018 | 0.018 | 0.011 | 0.05 | 0.2 | 0.05 | 0.05 | 0.02 | 0.05 |
| 8 | 10289663 | rs10096610 | -0.645 | 0.537 | -0.111 | 0.317 | 0.007 | 0.002 | 0.002 | 0 | 0.92 | 0.91 | 0.9 | 0.87 | 0.83 | 0.86 |
| 8 | 10290314 | rs10107098 | -0.204 | 0.423 | -0.327 | 0.257 | 0.003 | 0.002 | 0.002 | 0.03 | 0.62 | 0.62 | 0.77 | 0.63 | 0.56 | 0.61 |
| 8 | 10290566 | rs7000959 | 1.195 | 0.462 | -0.457 | 0.178 | 0 | 0 | 0 | 0 | 0.28 | 0.22 | 0.14 | 0.25 | 0.27 | 0.25 |
| 8 | 10290650 | rs10903327 | 0.303 | 0.143 | -0.827 | 0.04 | -0.027 | 0.022 | 0.022 | 0 | 0.34 | 0.25 | 0.41 | 0.28 | 0.46 | 0.24 |
| 8 | 10294330 | rs4841339 | 1.318 | 0.241 | -1.055 | -0.114 | -0.055 | 0.077 | 0.077 | 0 | 0.18 | 0.07 | 0.18 | 0 | 0.26 | 0.05 |
| 8 | 10295378 | rs11996714 | 1.057 | -0.589 | -0.782 | 0.222 | 0.007 | -0.007 | -0.007 | 0.297 | 0.26 | 0.2 | 0.33 | 0.33 | 0.1 | 0.33 |
| 8 | 10300132 | rs7837520 | 1.28 | 0.14 | -0.372 | 0.263 | 0.026 | 0.047 | 0.047 | 0 | 0.15 | 0.24 | 0.24 | 0.29 | 0.21 | 0.19 |
| 8 | 10301469 | rs17151969 | 1.78 | -0.408 | -0.459 | -0.139 | -0.006 | -0.046 | -0.046 | 2.741 | 0.23 | 0.22 | 0.06 | 0 | 0.11 | 0.18 |
| 8 | 10302213 | rs10095788 | 0.575 | -0.151 | -0.558 | -0.222 | 0.01 | 0.005 | 0.005 | 0 | 0.32 | 0.27 | 0.41 | 0.69 | 0.26 | 0.43 |
| 8 | 10303265 | rs10110333 | 1.115 | 0 | -0.378 | 0.088 | 0.003 | 0.085 | 0.085 | 0.006 | 0.05 | 0.2 | 0.05 | 0.11 | 0.02 | 0.06 |
| 8 | 10305317 | rs10086673 | -0.706 | -0.068 | -0.716 | 0.034 | 0.053 | 0.023 | 0.023 | 0 | 0.51 | 0.66 | 0.6 | 0.44 | 0.67 | 0.47 |
| 8 | 10308074 | rs6993523 | 1.025 | -0.218 | -0.863 | 0.128 | 0.099 | 0.036 | 0.036 | 0.01 | 0.44 | 0.25 | 0.34 | 0.32 | 0.19 | 0.49 |
| 8 | 10309262 | rs6999196 | 1.067 | -0.215 | -0.891 | 0.13 | 0.105 | 0.037 | 0.037 | 0 | 0.44 | 0.24 | 0.34 | 0.32 | 0.19 | 0.48 |
| 8 | 10312694 | rs7011215 | 0.464 | 0.053 | -0.739 | 0.132 | 0.058 | 0.05 | 0.05 | 0 | 0.52 | 0.28 | 0.44 | 0.62 | 0.37 | 0.56 |
| 8 | 10314986 | rs17711111 | 0.425 | 0.465 | -0.511 | 0.112 | 0 | -0.029 | -0.029 | 0 | 0.26 | 0.35 | 0.24 | 0.13 | 0.26 | 0.12 |
| 8 | 10315460 | rs10102613 | 0.56 | 0.465 | -0.524 | -0.036 | 0.006 | 0.073 | 0.073 | 0 | 0.06 | 0.3 | 0.07 | 0.1 | 0.03 | 0.06 |
| 8 | 10316170 | rs1986358 | 1.596 | 0.713 | -0.854 | 0.01 | 0.006 | 0.013 | 0.013 | 0 | 0.15 | 0.18 | 0.23 | 0.27 | 0.2 | 0.17 |
| 8 | 10316236 | rs1986357 | 0.021 | 0.686 | -1.036 | 0.064 | -0.009 | 0.018 | 0.018 | 0 | 0.58 | 0.33 | 0.52 | 0.63 | 0.54 | 0.7 |
| 8 | 10328677 | rs7007193 | 0.747 | 0.56 | -0.897 | 0.061 | 0.001 | -0.005 | -0.005 | 0 | 0.29 | 0.21 | 0.14 | 0.29 | 0.28 | 0.25 |
| 8 | 10331754 | rs9329223 | 0.241 | 0.749 | -0.534 | -0.438 | 0.038 | -0.079 | -0.079 | 0 | 0.66 | 0.51 | 0.65 | 0.9 | 0.56 | 0.74 |
| 8 | 10334694 | rs4841356 | 0.786 | 1.365 | -0.641 | -0.558 | -0.054 | 0.117 | 0.117 | 4.302 | 0.33 | 0.12 | 0.21 | 0.01 | 0.34 | 0.13 |
| 8 | 10334723 | rs12546910 | -0.027 | 1.569 | 0.009 | -0.027 | 0.009 | 0.031 | 0.031 | 0 | 0.45 | 0.37 | 0.5 | 0.69 | 0.4 | 0.62 |
| 8 | 10334781 | rs11782819 | 0.69 | 2.144 | 0.61 | 0.787 | 0.023 | 0.011 | 0.011 | 0 | 0.69 | 0.59 | 0.44 | 0.58 | 0.53 | 0.64 |
| 8 | 10337449 | rs4342552 | -0.037 | 1.522 | 0.844 | 0.589 | 0.058 | -0.051 | -0.051 | 0 | 0.88 | 0.77 | 0.87 | 0.98 | 0.77 | 0.86 |
| 8 | 10338924 | rs4380891 | 1.004 | 1.589 | 0.871 | 0.963 | -0.001 | 0.008 | 0.008 | 0 | 0.12 | 0.14 | 0.26 | 0.3 | 0.2 | 0.15 |
| 8 | 10339254 | rs4840487 | 0.861 | 1.518 | 1.237 | 0.895 | 0.027 | 0.008 | 0.008 | 0.177 | 0.73 | 0.34 | 0.59 | 0.67 | 0.57 | 0.68 |
| 8 | 10340111 | rs11250018 | 0.093 | 1.013 | 1.077 | 1.15 | -0.001 | -0.002 | -0.002 | 0.028 | 0.6 | 0.59 | 0.6 | 0.63 | 0.6 | 0.47 |
| 8 | 10341903 | rs4531008 | -1.106 | 1.208 | 1.137 | 0.767 | 0.067 | 0.006 | 0.006 | 0.037 | 0.29 | 0.46 | 0.44 | 0.33 | 0.49 | 0.35 |
| 8 | 10343863 | rs6601469 | 0.055 | 1.216 | 0.962 | 1.068 | -0.005 | 0 | 0 | 0 | 0.57 | 0.5 | 0.58 | 0.6 | 0.59 | 0.47 |
| 8 | 10345753 | rs13282174 | 0.446 | 1.348 | 0.722 | 0.997 | 0.007 | 0.024 | 0.024 | 0 | 0.14 | 0.2 | 0.26 | 0.31 | 0.2 | 0.17 |
| 8 | 10352230 | rs4268093 | 0.959 | 1.242 | 0.692 | 0.814 | 0.001 | 0.03 | 0.03 | 0 | 0.43 | 0.29 | 0.33 | 0.32 | 0.39 | 0.31 |
| 8 | 10355378 | rs10086950 | 0.01 | 1.493 | 0.773 | 0.939 | 0.001 | 0.008 | 0.008 | 0 | 0.17 | 0.16 | 0.3 | 0.34 | 0.29 | 0.2 |
| 8 | 10356449 | rs4392860 | 0.736 | 1.43 | 0.739 | 0.972 | 0.002 | 0.023 | 0.023 | 0 | 0.13 | 0.15 | 0.26 | 0.32 | 0.2 | 0.17 |
| 8 | 10359394 | rs4588813 | -0.025 | 1.262 | 0.769 | 0.704 | -0.008 | 0 | 0 | 0.097 | 0.38 | 0.46 | 0.37 | 0.38 | 0.32 | 0.52 |
| 8 | 10372068 | rs4473996 | 0.265 | 1.25 | 0.7 | 0.607 | 0.015 | -0.022 | -0.022 | 0 | 0.35 | 0.17 | 0.35 | 0.37 | 0.32 | 0.45 |
| 8 | 10377861 | rs6601479 | -0.265 | 1.039 | 0.942 | 0.543 | 0.034 | -0.034 | -0.034 | 0 | 0.93 | 0.75 | 0.95 | 1 | 0.91 | 0.97 |
| 8 | 10378405 | rs7834351 | -0.669 | 1.063 | 0.829 | 0.562 | -0.001 | 0.005 | 0.005 | 0 | 0.52 | 0.53 | 0.64 | 0.69 | 0.53 | 0.68 |
| 8 | 10378485 | rs7846328 | 0.796 | 0.825 | 0.244 | 0.686 | 0.003 | 0.017 | 0.017 | 0 | 0.44 | 0.35 | 0.32 | 0.3 | 0.39 | 0.28 |
| 8 | 10380962 | rs7002304 | 0.962 | 0.654 | 0.012 | 0.687 | 0.053 | -0.036 | -0.036 | 0 | 0.26 | 0.15 | 0.24 | 0.35 | 0.13 | 0.35 |
| 8 | 10381128 | rs4531010 | -0.568 | 0.999 | 0.173 | 0.503 | 0.002 | 0.011 | 0.011 | 0 | 0.56 | 0.63 | 0.67 | 0.69 | 0.61 | 0.72 |
| 8 | 10389301 | rs4400345 | -0.76 | 1.076 | -0.066 | 0.226 | 0.004 | 0.041 | 0.041 | 0 | 0.62 | 0.77 | 0.67 | 0.75 | 0.66 | 0.68 |
| 8 | 10390452 | rs4406360 | 0.466 | 1.327 | 0.248 | 0.414 | 0.016 | -0.024 | -0.024 | 0 | 0.37 | 0.22 | 0.33 | 0.43 | 0.33 | 0.43 |
| 8 | 10390972 | rs4840489 | 0.456 | 1.616 | 0.776 | 0.486 | 0 | 0.018 | 0.018 | 0 | 0.13 | 0.14 | 0.23 | 0.32 | 0.17 | 0.19 |
| 8 | 10391917 | rs10086002 | -0.571 | 1.631 | 1.127 | 0.465 | 0.027 | 0.083 | 0.083 | 0.055 | 0.21 | 0.4 | 0.29 | 0.32 | 0.23 | 0.2 |

| | | | | | | | | | | | | | | | | |
|---|---|---|---|---|---|---|---|---|---|---|---|---|---|---|---|---|
| 8 | 10395495 | rs7820225 | -0.616 | 1.557 | 0.877 | 0.479 | 0.045 | 0.108 | 0.108 | 0.207 | 0.2 | 0.54 | 0.29 | 0.32 | 0.26 | 0.23 |
| 8 | 10403088 | rs4404876 | 0.937 | 1.576 | 0.516 | 0.449 | 0.005 | 0.004 | 0.004 | 0.318 | 0.72 | 0.32 | 0.65 | 0.67 | 0.68 | 0.77 |
| 8 | 10404019 | rs4566993 | 1.168 | 1.899 | 0.16 | 0.228 | -0.044 | 0.023 | 0.023 | 0 | 0.07 | 0.01 | 0.1 | 0.02 | 0.16 | 0.08 |
| 8 | 10406533 | rs6981847 | 0.041 | 1.829 | 0.703 | 0.23 | 0.08 | -0.093 | -0.093 | 0.182 | 0.86 | 0.53 | 0.89 | 1 | 0.85 | 0.96 |
| 8 | 10407928 | rs7011457 | 0.254 | 1.38 | -0.073 | 0.339 | -0.008 | 0.017 | 0.017 | 0 | 0.48 | 0.29 | 0.56 | 0.57 | 0.51 | 0.55 |
| 8 | 10411975 | rs6601489 | 1.098 | 1.461 | -0.202 | 0.362 | 0.006 | 0.038 | 0.038 | 0.213 | 0.34 | 0.13 | 0.32 | 0.24 | 0.33 | 0.32 |
| 8 | 10412116 | rs7004553 | 1.089 | 1.463 | -0.201 | 0.356 | 0.006 | 0.036 | 0.036 | 0.218 | 0.34 | 0.13 | 0.32 | 0.24 | 0.33 | 0.32 |
| 8 | 10418039 | rs4841380 | -0.091 | 1.382 | 0.212 | 0.332 | 0.005 | 0.059 | 0.059 | 0 | 0.14 | 0.21 | 0.24 | 0.33 | 0.16 | 0.18 |
| 8 | 10421967 | rs11785067 | -0.324 | 1.359 | 0.203 | -0.143 | 0.018 | -0.065 | -0.065 | 3.485 | 0.2 | 0.25 | 0.16 | 0.03 | 0.25 | 0.15 |
| 8 | 10422108 | rs11250031 | -0.683 | 1.112 | -0.008 | -0.216 | -0.007 | -0.011 | -0.011 | 0 | 0.66 | 0.48 | 0.57 | 0.69 | 0.65 | 0.7 |
| 8 | 10422207 | rs11787406 | -0.392 | 1.104 | -0.353 | 0.103 | -0.03 | 0.034 | 0.034 | 0.002 | 0.13 | 0.07 | 0.1 | 0 | 0.17 | 0.07 |
| 8 | 10436353 | rs11993892 | 0.101 | 0.252 | -0.811 | 0.412 | 0.032 | -0.07 | -0.07 | 0 | 0.31 | 0.1 | 0.25 | 0.6 | 0.26 | 0.44 |
| 8 | 10437868 | rs13250449 | -0.515 | 0.391 | -0.206 | 0.771 | 0 | 0.002 | 0.002 | 0 | 0.18 | 0.16 | 0.33 | 0.35 | 0.15 | 0.18 |
| 8 | 10438796 | rs12546950 | -0.577 | 0.47 | 0.004 | 0.776 | -0.007 | 0.045 | 0.045 | 0 | 0.19 | 0.23 | 0.36 | 0.35 | 0.18 | 0.18 |
| 8 | 10442013 | rs4458841 | 0.975 | 0.158 | 0.087 | 0.643 | 0.007 | -0.005 | -0.005 | 0 | 0.57 | 0.34 | 0.64 | 0.46 | 0.66 | 0.58 |
| 8 | 10444137 | rs10110911 | 0.79 | 0.159 | 0.261 | 0.794 | 0.023 | 0.006 | 0.006 | 0.372 | 0.22 | 0.11 | 0.14 | 0.18 | 0.14 | 0.39 |
| 8 | 10444179 | rs4841386 | 0.801 | 0.289 | 0.248 | 0.715 | 0.022 | 0.011 | 0.011 | 1.525 | 0.54 | 0.35 | 0.6 | 0.41 | 0.65 | 0.46 |
| 8 | 10448142 | rs12681639 | 0.265 | 0.6 | -0.307 | 0.827 | -0.029 | -0.041 | -0.041 | 0 | 0.13 | 0.05 | 0.21 | 0.33 | 0.2 | 0.17 |
| 8 | 10451011 | rs10089537 | 1.206 | 0.359 | 0.026 | 0.778 | 0.001 | 0.002 | 0.002 | 0.12 | 0.48 | 0.44 | 0.46 | 0.39 | 0.58 | 0.44 |
| 8 | 10451585 | rs4841389 | -0.155 | 0.382 | -0.028 | 0.838 | -0.028 | -0.066 | -0.066 | 0.061 | 0.19 | 0.08 | 0.34 | 0.41 | 0.21 | 0.13 |
| 8 | 10452070 | rs7009985 | 1.355 | 0.398 | -0.014 | 0.786 | 0.001 | 0.004 | 0.004 | 0.039 | 0.48 | 0.43 | 0.44 | 0.38 | 0.58 | 0.43 |
| 8 | 10453419 | rs4841390 | 0.959 | 0.677 | -0.623 | 0.917 | 0 | -0.013 | -0.013 | 0 | 0.07 | 0.11 | 0.03 | 0 | 0.05 | 0 |
| 8 | 10456111 | rs9644729 | -0.56 | 0.692 | -0.69 | 1.003 | -0.005 | -0.011 | -0.011 | 0 | 0.2 | 0.21 | 0.37 | 0.4 | 0.23 | 0.15 |
| 8 | 10456823 | rs4841394 | 0.873 | 0.484 | -0.853 | 0.991 | -0.012 | 0.034 | 0.034 | 0 | 0.07 | 0.02 | 0.03 | 0 | 0.06 | 0 |
| 8 | 10459305 | rs11784353 | 0.496 | 0.519 | -0.805 | 1.279 | -0.014 | -0.061 | -0.061 | 0 | 0.1 | 0.01 | 0.15 | 0.41 | 0.09 | 0.08 |
| 8 | 10460373 | rs12679862 | 0.22 | 0.535 | -0.612 | 1.147 | 0.011 | 0.054 | 0.054 | 0 | 0.13 | 0.17 | 0.33 | 0.38 | 0.2 | 0.13 |
| 8 | 10462806 | rs4841398 | 0.043 | 0.381 | -0.294 | 1.112 | -0.014 | 0.091 | 0.091 | 0 | 0.85 | 0.77 | 0.85 | 0.56 | 0.89 | 0.87 |
| 8 | 10463944 | rs7816990 | 0.637 | -0.049 | -0.345 | 0.754 | -0.026 | 0.008 | 0.008 | 0 | 0.63 | 0.53 | 0.67 | 0.41 | 0.76 | 0.47 |
| 8 | 10464161 | rs12542071 | 0.754 | 0.126 | -0.575 | 0.86 | 0.03 | -0.002 | -0.002 | 0 | 0.21 | 0.04 | 0.18 | 0.18 | 0.14 | 0.38 |
| 8 | 10464206 | rs12542075 | 0.741 | 0.128 | -0.574 | 0.861 | 0.03 | -0.002 | -0.002 | 2.188 | 0.21 | 0.04 | 0.17 | 0.18 | 0.14 | 0.38 |
| 8 | 10464307 | rs12542104 | -0.839 | 0.181 | -0.577 | 0.527 | 0.004 | -0.002 | -0.002 | 0.598 | 0.42 | 0.34 | 0.53 | 0.55 | 0.4 | 0.57 |
| 8 | 10468172 | rs4840502 | 1.148 | -0.085 | -0.45 | 0.886 | -0.002 | 0.005 | 0.005 | 0 | 0.71 | 0.65 | 0.76 | 0.6 | 0.73 | 0.84 |
| 8 | 10470915 | rs11773913 | 0.825 | -0.722 | -0.3 | 0.834 | -0.029 | 0.017 | 0.017 | 2.115 | 0.65 | 0.38 | 0.68 | 0.43 | 0.77 | 0.46 |
| 8 | 10471850 | rs7018397 | 1.154 | -0.628 | -0.308 | 0.696 | -0.013 | -0.002 | -0.002 | 0.011 | 0.6 | 0.44 | 0.65 | 0.4 | 0.71 | 0.41 |
| 8 | 10472468 | rs11775769 | 0.713 | -0.489 | -0.247 | 0.827 | -0.032 | 0.042 | 0.042 | 2.549 | 0.69 | 0.43 | 0.69 | 0.42 | 0.77 | 0.5 |
| 8 | 10472638 | rs4382429 | -1.369 | -0.114 | -0.783 | 0.89 | 0.006 | -0.047 | -0.047 | 0 | 0.19 | 0.1 | 0.16 | 0.48 | 0.16 | 0.09 |
| 8 | 10474042 | rs4388421 | -0.793 | 0.257 | -0.601 | 0.739 | -0.001 | 0.067 | 0.067 | 0 | 0.14 | 0.16 | 0.24 | 0.46 | 0.14 | 0.11 |
| 8 | 10482372 | rs9329228 | 0.213 | 0.199 | -0.659 | -0.672 | 0.016 | 0.03 | 0.03 | 0 | 0.56 | 0.65 | 0.56 | 0.22 | 0.6 | 0.47 |
| 8 | 10494675 | rs6981218 | 0.044 | -0.105 | -0.293 | -1.594 | -0.028 | -0.125 | -0.125 | 0.914 | 0.72 | 0.53 | 0.75 | 0.95 | 0.77 | 0.8 |
| 8 | 10497483 | rs4240662 | -0.14 | -0.336 | -0.44 | -1.578 | 0.017 | 0.023 | 0.023 | 4.949 | 0.77 | 0.83 | 0.76 | 0.89 | 0.83 | 0.89 |
| 8 | 10503525 | rs7386213 | 0.206 | 0.041 | -0.814 | -2.123 | 0.007 | 0.138 | 0.138 | 0 | 0.4 | 0.23 | 0.52 | 0.92 | 0.4 | 0.61 |
| 8 | 10506113 | rs7463228 | -0.689 | -0.275 | -0.629 | -2.261 | -0.011 | -0.067 | -0.067 | 0.221 | 0.14 | 0.27 | 0.07 | 0 | 0.07 | 0.15 |
| 8 | 10506207 | rs4841404 | -0.593 | 0.488 | -1.318 | -2.327 | -0.07 | 0.249 | 0.249 | 3.267 | 0.46 | 0.93 | 0.4 | 0.11 | 0.4 | 0.4 |
| 8 | 10519285 | rs7814795 | -0.339 | 0.567 | -0.26 | -2.07 | -0.003 | -0.002 | -0.002 | 0 | 0.65 | 0.66 | 0.49 | 0.1 | 0.56 | 0.53 |
| 8 | 10520255 | rs7815184 | -1.074 | 0.315 | 0.981 | -1.456 | 0.065 | -0.075 | -0.075 | 2.798 | 0.14 | 0.51 | 0.1 | 0.02 | 0.16 | 0.31 |
| 8 | 10520913 | rs13269921 | -1.605 | 0.234 | 0.627 | -1.425 | -0.007 | 0.007 | 0.007 | 0.249 | 0.08 | 0.32 | 0.1 | 0.07 | 0.06 | 0.07 |
| 8 | 10527144 | rs4373491 | 1.587 | 0.286 | 0.344 | -1.418 | -0.001 | -0.004 | -0.004 | 0 | 0.79 | 0.24 | 0.81 | 0.9 | 0.8 | 0.62 |
| 8 | 10530037 | rs11250057 | 0.507 | 0.63 | 0.421 | -1.204 | 0.082 | -0.013 | -0.013 | 0 | 0.87 | 0.48 | 0.91 | 0.94 | 0.87 | 0.65 |
| 8 | 10530750 | rs11778725 | 0.562 | 0.633 | 0.321 | -1.205 | 0.067 | -0.021 | -0.021 | 0 | 0.87 | 0.46 | 0.91 | 0.94 | 0.87 | 0.65 |
| 8 | 10535377 | rs7011192 | -0.489 | 0.526 | -0.53 | -1.101 | -0.006 | -0.005 | -0.005 | 0 | 0.06 | 0.01 | 0.18 | 0.08 | 0.1 | 0.05 |
| 8 | 10535826 | rs6601501 | 0.964 | 0.249 | -0.098 | -1.546 | 0.001 | -0.065 | -0.065 | 0 | 0.75 | 0.38 | 0.87 | 0.94 | 0.78 | 0.64 |
| 8 | 10540767 | rs6601503 | 0.399 | -0.457 | -0.961 | -2.067 | 0.051 | 0.182 | 0.182 | 0.073 | 0.61 | 0.17 | 0.8 | 0.91 | 0.75 | 0.67 |
| 8 | 10540931 | rs7827623 | 0.366 | -0.591 | -0.113 | -1.912 | -0.016 | -0.056 | -0.056 | 0 | 0.26 | 0.3 | 0.12 | 0.08 | 0.18 | 0.23 |
| 8 | 10541651 | rs13259547 | -0.078 | -0.461 | -0.379 | -1.683 | 0.003 | 0.003 | 0.003 | 0 | 0.09 | 0.06 | 0.05 | 0 | 0.07 | 0.11 |
| 8 | 10547758 | rs7823200 | -0.828 | -0.404 | -1.205 | -0.957 | -0.142 | -0.094 | -0.094 | 0.181 | 0.6 | 0.94 | 0.66 | 0.45 | 0.47 | 0.5 |
| 8 | 10549767 | rs11994746 | -1.019 | -0.289 | -0.016 | -0.734 | 0.013 | 0.11 | 0.11 | 2.743 | 0.05 | 0.66 | 0.1 | 0.1 | 0.01 | 0.05 |
| 8 | 10555734 | rs11996203 | -1.329 | -0.343 | -0.075 | -0.78 | -0.022 | 0.047 | 0.047 | 4.874 | 0.07 | 0.74 | 0.1 | 0.1 | 0.01 | 0.07 |
| 8 | 10556109 | rs11996321 | -0.231 | -0.393 | -0.65 | -1.268 | 0.013 | 0.068 | 0.068 | 0 | 0.32 | 0.79 | 0.25 | 0.1 | 0.24 | 0.29 |
| 8 | 10558748 | rs10087914 | -0.323 | -0.73 | -0.359 | -0.886 | 0.021 | 0.026 | 0.026 | 2.511 | 0.3 | 0.12 | 0.4 | 0.23 | 0.26 | 0.21 |
| 8 | 10560297 | rs11774792 | -0.302 | -0.904 | 0.387 | -0.542 | -0.006 | -0.003 | -0.003 | 0 | 0.16 | 0.49 | 0.15 | 0.1 | 0.11 | 0.09 |
| 8 | 10563114 | rs4840508 | -1.207 | -1.089 | -0.487 | -1.353 | 0.003 | 0.008 | 0.008 | 0 | 0.31 | 0.67 | 0.3 | 0.14 | 0.23 | 0.13 |
| 8 | 10564067 | rs4840510 | -0.564 | -1.099 | -0.213 | -1.085 | -0.012 | -0.01 | -0.01 | 0 | 0.18 | 0.7 | 0.25 | 0.17 | 0.12 | 0.1 |
| 8 | 10565219 | rs6601509 | -0.879 | -1.079 | -0.244 | -1.128 | -0.02 | 0.004 | 0.004 | 0 | 0.12 | 0.68 | 0.22 | 0.14 | 0.09 | 0.1 |
| 8 | 10567630 | rs4841428 | -0.925 | -1.103 | -0.152 | -1.145 | -0.015 | -0.019 | -0.019 | 0.083 | 0.17 | 0.56 | 0.23 | 0.14 | 0.12 | 0.1 |
| 8 | 10576464 | rs7834573 | -0.143 | -1.292 | -1.37 | -1.12 | 0.021 | 0.003 | 0.003 | 0.02 | 0.35 | 0.19 | 0.27 | 0.3 | 0.28 | 0.39 |
| 8 | 10579072 | rs12544976 | -0.382 | -0.931 | -1.552 | -1.225 | -0.022 | 0.035 | 0.035 | 0.263 | 0.57 | 0.29 | 0.41 | 0.3 | 0.51 | 0.55 |
| 8 | 10579392 | rs4503064 | -1.232 | -0.678 | -1.726 | -1.065 | -0.05 | -0.075 | -0.075 | 0.192 | 0.68 | 0.88 | 0.63 | 0.45 | 0.62 | 0.62 |
| 8 | 10583506 | rs4841432 | -0.412 | -0.851 | -1.738 | -1.415 | 0.007 | 0.234 | 0.234 | 0 | 0.46 | 0.84 | 0.38 | 0.14 | 0.38 | 0.28 |
| 8 | 10584212 | rs10100209 | -1.142 | -1.179 | -1.659 | -1.25 | -0.02 | -0.065 | -0.065 | 0 | 0.65 | 0.86 | 0.63 | 0.43 | 0.61 | 0.59 |

| | | | | | | | | | | | | | | | | |
|---|---|---|---|---|---|---|---|---|---|---|---|---|---|---|---|---|
| 8 | 10593065 | rs4841435 | 1.022 | -0.817 | -0.452 | -1.43 | 0.032 | 0.012 | 0.012 | 0.975 | 0.07 | 0.22 | 0.03 | 0.04 | 0.03 | 0.02 |
| 8 | 10595878 | rs2409674 | -1.024 | -1.259 | -0.822 | -1.663 | 0.001 | 0.007 | 0.007 | 1.043 | 0.58 | 0.68 | 0.49 | 0.35 | 0.54 | 0.45 |
| 8 | 10601281 | rs6601514 | -1.06 | -1.283 | -0.591 | -1.691 | 0.001 | 0.003 | 0.003 | 0.768 | 0.59 | 0.65 | 0.5 | 0.35 | 0.54 | 0.47 |
| 8 | 10605234 | rs6601518 | -0.008 | -1.425 | -0.813 | -1.845 | -0.062 | 0.007 | 0.007 | 0 | 0.39 | 0.2 | 0.4 | 0.32 | 0.43 | 0.33 |
| 8 | 10611882 | rs7014285 | -0.963 | -1.516 | -0.79 | -1.979 | -0.002 | 0.001 | 0.001 | 0 | 0.34 | 0.38 | 0.33 | 0.32 | 0.28 | 0.3 |
| 8 | 10620026 | rs17152302 | -0.37 | -1.377 | -0.866 | -1.878 | 0.011 | -0.022 | -0.022 | 0 | 0.3 | 0.23 | 0.31 | 0.33 | 0.26 | 0.25 |
| 8 | 10621061 | rs2898250 | 0.047 | -1.452 | -0.69 | -1.99 | -0.016 | -0.118 | -0.118 | 0 | 0.19 | 0.43 | 0.11 | 0.03 | 0.14 | 0.12 |
| 8 | 10621981 | rs733396 | -0.616 | -1.434 | -0.883 | -2.038 | -0.024 | 0.052 | 0.052 | 2.216 | 0.47 | 0.25 | 0.43 | 0.33 | 0.46 | 0.36 |
| 8 | 10622812 | rs12750 | 2.018 | -1.302 | -0.839 | -2.01 | -0.048 | 0.082 | 0.082 | 0 | 0.18 | 0.05 | 0.13 | 0 | 0.21 | 0.16 |
| 8 | 10626333 | rs17774023 | 0.895 | -1.271 | -0.612 | -2.011 | 0.014 | -0.093 | -0.093 | 1.712 | 0.27 | 0.34 | 0.18 | 0.03 | 0.28 | 0.2 |
| 8 | 10627844 | rs7826180 | -0.261 | -1.361 | -0.631 | -2.02 | 0.008 | -0.015 | -0.015 | 0 | 0.3 | 0.22 | 0.3 | 0.32 | 0.26 | 0.2 |
| 8 | 10627860 | rs7826189 | -0.766 | -1.357 | -0.508 | -2.007 | -0.002 | 0.025 | 0.025 | 1.35 | 0.49 | 0.4 | 0.43 | 0.32 | 0.46 | 0.41 |
| 8 | 10628947 | rs6601521 | -1.197 | -1.233 | -0.677 | -1.992 | -0.001 | -0.001 | -0.001 | 0 | 0.6 | 0.58 | 0.5 | 0.35 | 0.56 | 0.41 |
| 8 | 10629061 | rs6601522 | -2.234 | -1.129 | -0.943 | -1.889 | -0.027 | -0.041 | -0.041 | 0 | 0.71 | 0.83 | 0.55 | 0.35 | 0.61 | 0.53 |
| 8 | 10632684 | rs6993841 | -2.106 | -1.102 | -0.939 | -1.841 | 0 | 0.001 | 0.001 | 2.059 | 0.69 | 0.68 | 0.55 | 0.35 | 0.6 | 0.48 |
| 8 | 10635222 | rs6601524 | -0.223 | -0.991 | -0.831 | -1.674 | -0.001 | -0.039 | -0.039 | 0.705 | 0.28 | 0.15 | 0.28 | 0.31 | 0.25 | 0.19 |
| 8 | 10643164 | rs9657541 | -0.012 | -0.862 | -0.846 | -1.789 | 0.003 | 0.061 | 0.061 | 2.771 | 0.26 | 0.16 | 0.14 | 0.02 | 0.21 | 0.16 |
| 8 | 10645738 | rs6984094 | -0.144 | -0.992 | -0.635 | -1.854 | -0.043 | -0.13 | -0.13 | 0 | 0.22 | 0.48 | 0.14 | 0.03 | 0.14 | 0.12 |
| 8 | 10647823 | rs11250076 | -1.952 | -0.872 | -0.839 | -1.873 | -0.03 | 0.113 | 0.113 | 1.63 | 0.66 | 0.82 | 0.42 | 0.05 | 0.57 | 0.48 |
| 8 | 10656437 | rs11250077 | -2.242 | -0.811 | -0.844 | -1.758 | -0.026 | -0.04 | -0.04 | 0.03 | 0.69 | 0.82 | 0.56 | 0.35 | 0.6 | 0.53 |
| 8 | 10659517 | rs11998382 | 2.084 | -0.721 | -0.79 | -1.761 | -0.002 | -0.03 | -0.03 | 2.014 | 0.17 | 0.15 | 0.13 | 0 | 0.2 | 0.16 |
| 8 | 10661734 | rs7838341 | -0.321 | -0.791 | -0.836 | -1.756 | 0.004 | -0.032 | -0.032 | 0 | 0.3 | 0.19 | 0.29 | 0.32 | 0.26 | 0.25 |
| 8 | 10666813 | rs11779094 | 0.796 | -0.823 | -0.344 | -1.736 | 0.026 | -0.048 | -0.048 | 0 | 0.11 | 0.4 | 0.06 | 0.03 | 0.08 | 0.04 |
| 8 | 10666993 | rs7002282 | -2.256 | -0.765 | -0.734 | -1.574 | -0.038 | -0.064 | -0.064 | 0.063 | 0.7 | 0.86 | 0.56 | 0.37 | 0.6 | 0.53 |
| 8 | 10670108 | rs6989930 | 1.173 | -0.723 | -0.655 | -1.648 | -0.022 | -0.052 | -0.052 | 0 | 0.11 | 0.18 | 0.08 | 0 | 0.06 | 0.07 |
| 8 | 10674467 | rs11250080 | -0.843 | -0.659 | -0.725 | -1.57 | -0.003 | -0.003 | -0.003 | 0 | 0.39 | 0.45 | 0.35 | 0.34 | 0.34 | 0.26 |
| 8 | 10676497 | rs17776622 | 1.67 | -0.352 | -1.056 | -1.822 | -0.038 | 0.074 | 0.074 | 0 | 0.18 | 0.06 | 0.13 | 0 | 0.2 | 0.16 |
| 8 | 10685851 | rs17152584 | 1.352 | -0.265 | -1.108 | -1.879 | -0.027 | 0.023 | 0.023 | 0 | 0.08 | 0.03 | 0.05 | 0.02 | 0.13 | 0.08 |
| 8 | 10689212 | rs12375394 | 2.02 | -0.377 | -1.231 | -2.012 | -0.055 | 0.09 | 0.09 | 0 | 0.17 | 0.04 | 0.13 | 0 | 0.2 | 0.16 |
| 8 | 10690319 | rs2292370 | -1.213 | -0.94 | -1.202 | -2.1 | 0.009 | 0.088 | 0.088 | 0.12 | 0.41 | 0.73 | 0.28 | 0.22 | 0.35 | 0.32 |
| 8 | 10690509 | rs6993753 | 1.093 | -0.973 | -0.985 | -2.075 | -0.014 | -0.034 | -0.034 | 0 | 0.12 | 0.13 | 0.07 | 0 | 0.06 | 0.12 |
| 8 | 10701959 | rs7010709 | -0.237 | -0.903 | -1.13 | -2.099 | -0.015 | 0.005 | 0.005 | 2.17 | 0.2 | 0.15 | 0.14 | 0.2 | 0.22 | 0.16 |
| 8 | 10702534 | rs6990589 | -0.201 | -0.895 | -1.022 | -2.095 | -0.006 | 0.001 | 0.001 | 2.905 | 0.19 | 0.18 | 0.14 | 0.19 | 0.22 | 0.15 |
| 8 | 10703909 | rs1055329 | -1.406 | -0.968 | -1.016 | -2.168 | 0 | 0.017 | 0.017 | 0.884 | 0.39 | 0.61 | 0.27 | 0.19 | 0.35 | 0.25 |
| 8 | 10705360 | rs6989160 | -2.392 | -0.973 | -1.22 | -2.199 | -0.021 | 0.066 | 0.066 | 0.719 | 0.68 | 0.84 | 0.46 | 0.19 | 0.61 | 0.53 |
| 8 | 10706801 | rs1469557 | -0.259 | -0.79 | -1.043 | -1.951 | -0.013 | 0.013 | 0.013 | 2.833 | 0.19 | 0.16 | 0.14 | 0.16 | 0.21 | 0.15 |
| 8 | 10707974 | rs10089153 | -1.287 | -1.014 | -1.189 | -1.999 | 0.04 | 0.054 | 0.054 | 3.448 | 0.43 | 0.66 | 0.28 | 0.24 | 0.28 | 0.26 |
| 8 | 10718061 | rs17777835 | -0.077 | -0.789 | -1.122 | -1.928 | -0.003 | -0.07 | -0.07 | 0 | 0.64 | 0.47 | 0.71 | 0.89 | 0.7 | 0.73 |
| 8 | 10722058 | rs2409676 | -1.668 | -0.932 | -1.065 | -2.083 | 0 | 0.051 | 0.051 | 0 | 0.58 | 0.68 | 0.44 | 0.25 | 0.56 | 0.56 |
| 8 | 10726464 | rs7840344 | -2.413 | -0.844 | -1.448 | -2.115 | -0.027 | 0.04 | 0.04 | 0 | 0.68 | 0.85 | 0.49 | 0.23 | 0.61 | 0.57 |
| 8 | 10727072 | rs17152632 | 1.746 | -0.826 | -1.299 | -2.087 | 0.015 | 0.016 | 0.016 | 0.455 | 0.1 | 0.06 | 0.04 | 0 | 0.05 | 0.01 |
| 8 | 10729177 | rs10095927 | -0.799 | -0.734 | -1.165 | -2.028 | 0 | -0.044 | -0.044 | 0 | 0.45 | 0.45 | 0.37 | 0.14 | 0.46 | 0.47 |
| 8 | 10729623 | rs2409667 | -0.188 | -0.478 | -1.051 | -1.8 | -0.006 | 0.031 | 0.031 | 0.205 | 0.2 | 0.39 | 0.18 | 0.23 | 0.17 | 0.21 |
| 8 | 10730558 | rs17721866 | 1.876 | -0.041 | -1.659 | -1.715 | -0.053 | 0.085 | 0.085 | 0 | 0.14 | 0.02 | 0.08 | 0 | 0.17 | 0.2 |
| 8 | 10732050 | rs2163379 | -1.737 | 0.019 | -1.615 | -0.942 | -0.043 | -0.086 | -0.086 | 0 | 0.72 | 0.87 | 0.53 | 0.54 | 0.61 | 0.62 |
| 8 | 10732905 | rs1864585 | -0.417 | -0.068 | -0.899 | -2.481 | 0 | -0.069 | -0.069 | 0.237 | 0.79 | 0.73 | 0.86 | 1 | 0.8 | 0.72 |
| 8 | 10742059 | rs7843666 | -1.508 | -0.075 | -1.099 | -3.283 | -0.006 | -0.015 | -0.015 | 1.789 | 0.54 | 0.66 | 0.34 | 0 | 0.48 | 0.42 |
| 8 | 10745586 | rs2898249 | 2.47 | 0.318 | -1.534 | -3.332 | -0.056 | -0.059 | -0.059 | 0.129 | 0.25 | 0.02 | 0.23 | 0.75 | 0.29 | 0.35 |
| 8 | 10750204 | rs11784061 | -0.246 | -0.268 | -0.595 | -3.687 | -0.047 | -0.132 | -0.132 | 5.635 | 0.21 | 0.44 | 0.12 | 0.01 | 0.12 | 0.04 |
| 8 | 10751709 | rs1529598 | -0.98 | -0.362 | -0.551 | -3.709 | -0.014 | -0.132 | -0.132 | 0 | 0.37 | 0.51 | 0.22 | 0.02 | 0.3 | 0.25 |
| 8 | 10761585 | rs6985109 | -2.213 | -0.306 | -0.554 | -4.047 | 0.004 | 0.073 | 0.073 | 1.024 | 0.56 | 0.73 | 0.38 | 0.02 | 0.53 | 0.44 |
| 8 | 10767748 | rs4240671 | -2.157 | -0.265 | -0.431 | -3.986 | -0.003 | 0.023 | 0.023 | 2.083 | 0.53 | 0.65 | 0.34 | 0.02 | 0.49 | 0.43 |
| 8 | 10778546 | rs7823296 | -2.404 | -0.3 | -0.525 | -3.983 | -0.003 | 0.123 | 0.123 | 2.235 | 0.57 | 0.77 | 0.36 | 0.03 | 0.53 | 0.48 |
| 8 | 10781688 | rs4841458 | -0.786 | -0.142 | -0.1 | -3.946 | -0.012 | -0.139 | -0.139 | 0.142 | 0.33 | 0.51 | 0.2 | 0.02 | 0.28 | 0.3 |
| 8 | 10783223 | rs7821438 | -0.854 | -0.544 | -0.504 | -3.901 | -0.009 | 0.024 | 0.024 | 0 | 0.2 | 0.11 | 0.12 | 0 | 0.21 | 0.15 |
| 8 | 10788875 | rs11783247 | -2.08 | -0.418 | -0.8 | -3.78 | -0.007 | 0.134 | 0.134 | 3.194 | 0.56 | 0.75 | 0.36 | 0.03 | 0.53 | 0.46 |
| 8 | 10796093 | rs6984496 | -2.178 | -0.434 | -0.8 | -3.776 | -0.005 | 0.146 | 0.146 | 3.019 | 0.57 | 0.8 | 0.37 | 0.03 | 0.54 | 0.46 |
| 8 | 10797831 | rs11990550 | -1.946 | -0.501 | -0.49 | -3.473 | -0.001 | -0.063 | -0.063 | 0 | 0.39 | 0.58 | 0.24 | 0.02 | 0.32 | 0.21 |
| 8 | 10802102 | rs6988281 | -1.503 | -0.87 | -0.4 | -3.207 | -0.023 | -0.074 | -0.074 | 0 | 0.31 | 0.65 | 0.15 | 0.02 | 0.16 | 0.14 |
| 8 | 10803617 | rs4841462 | -0.552 | -0.384 | -0.916 | -2.87 | 0.008 | 0.047 | 0.047 | 2.865 | 0.14 | 0.05 | 0.07 | 0.03 | 0.1 | 0.09 |
| 8 | 10804843 | rs7000276 | -0.464 | -0.265 | -0.619 | -3.145 | -0.002 | 0.013 | 0.013 | 0 | 0.38 | 0.29 | 0.35 | 0.09 | 0.39 | 0.3 |
| 8 | 10807105 | rs4841463 | -0.151 | -0.276 | -0.857 | -3.116 | 0.006 | 0.033 | 0.033 | 1.623 | 0.16 | 0.09 | 0.05 | 0 | 0.1 | 0.11 |
| 8 | 10815515 | rs10095390 | -0.475 | -0.137 | -0.466 | -2.991 | 0.005 | -0.11 | -0.11 | 0.283 | 0.19 | 0.38 | 0.09 | 0.03 | 0.14 | 0.13 |
| 8 | 10816174 | rs4645525 | -0.475 | -0.137 | -0.466 | -2.991 | 0.005 | -0.11 | -0.11 | 0 | 0.19 | 0.38 | 0.08 | 0.03 | 0.14 | 0.13 |
| 8 | 10819854 | rs4841465 | -1.286 | -0.219 | -1.486 | -2.966 | -0.047 | 0.215 | 0.215 | 0.983 | 0.57 | 0.89 | 0.4 | 0.14 | 0.52 | 0.38 |
| 8 | 10830743 | rs6601541 | -0.423 | -0.057 | -1.169 | -2.74 | 0 | 0.057 | 0.057 | 0 | 0.38 | 0.77 | 0.38 | 0.17 | 0.36 | 0.31 |
| 8 | 10834431 | rs4078186 | -1.021 | -0.017 | -1.664 | -2.748 | -0.026 | 0.097 | 0.097 | 0 | 0.66 | 0.86 | 0.51 | 0.17 | 0.6 | 0.52 |
| 8 | 10836436 | rs6996943 | -1.224 | 0.201 | -1.622 | -2.734 | -0.075 | 0.267 | 0.267 | 0.087 | 0.56 | 0.89 | 0.42 | 0.09 | 0.51 | 0.39 |
| 8 | 10836472 | rs7010246 | -0.621 | 0.279 | -1.59 | -2.978 | -0.011 | 0.122 | 0.122 | 0.971 | 0.36 | 0.19 | 0.21 | 0.01 | 0.34 | 0.24 |

| | | | | | | | | | | | | | | | | |
|---|---|---|---|---|---|---|---|---|---|---|---|---|---|---|---|---|
| 8 | 10852164 | rs4841471 | -1.306 | 0.344 | -1.629 | -2.603 | -0.118 | 0.271 | 0.271 | 1.466 | 0.43 | 0.94 | 0.29 | 0.07 | 0.37 | 0.38 |
| 8 | 10861385 | rs7350066 | -0.841 | 0.52 | -0.577 | -2.727 | 0.014 | -0.079 | -0.079 | 0.795 | 0.25 | 0.36 | 0.14 | 0.01 | 0.23 | 0.31 |
| 8 | 10867349 | rs7016361 | -1.382 | 0.369 | -0.775 | -2.721 | -0.062 | 0.143 | 0.143 | 1.142 | 0.34 | 0.14 | 0.22 | 0.02 | 0.36 | 0.37 |
| 8 | 10879611 | rs4074356 | -1.192 | 0.38 | 0.36 | -2.437 | -0.003 | -0.008 | -0.008 | 0.328 | 0.16 | 0.16 | 0.15 | 0.01 | 0.19 | 0.11 |
| 8 | 10880517 | rs7822400 | -1.042 | 0.387 | 0.191 | -2.433 | -0.031 | 0.058 | 0.058 | 0.056 | 0.16 | 0.06 | 0.14 | 0.01 | 0.19 | 0.12 |
| 8 | 10881408 | rs10111872 | -0.36 | 0.504 | 0.444 | -2.554 | 0.012 | -0.028 | -0.028 | 0.372 | 0.07 | 0.1 | 0.04 | 0 | 0.09 | 0.13 |
| 8 | 10888096 | rs6601548 | -1.58 | 0.162 | 0.702 | -2.709 | 0.006 | 0.028 | 0.028 | 1.641 | 0.16 | 0.83 | 0.09 | 0.02 | 0.1 | 0.14 |
| 8 | 10889623 | rs10110123 | -2.201 | 0.108 | 0.477 | -2.775 | -0.074 | 0.209 | 0.209 | 0 | 0.32 | 0.91 | 0.23 | 0.03 | 0.29 | 0.26 |
| 8 | 10896697 | rs10104571 | -0.578 | 0.058 | 0.803 | -2.745 | -0.009 | -0.05 | -0.05 | 0.984 | 0.11 | 0.17 | 0.09 | 0 | 0.1 | 0.05 |
| 8 | 10899288 | rs10100449 | -2.226 | 0.082 | 0.663 | -2.788 | -0.006 | 0.053 | 0.053 | 0.067 | 0.32 | 0.8 | 0.22 | 0.03 | 0.28 | 0.25 |
| 8 | 10907051 | rs7462014 | -0.989 | -0.191 | -0.15 | -2.574 | -0.016 | -0.041 | -0.041 | 0.044 | 0.92 | 0.99 | 0.9 | 0.83 | 0.93 | 0.78 |
| 8 | 10907821 | rs6601552 | 0.97 | -0.454 | -0.112 | -2.834 | 0.001 | 0.073 | 0.073 | 0 | 0.72 | 0.89 | 0.76 | 0.86 | 0.74 | 0.61 |
| 8 | 10914233 | rs10096332 | -1.736 | -0.652 | -0.157 | -2.913 | 0.048 | 0.113 | 0.113 | 0.239 | 0.39 | 0.81 | 0.31 | 0.2 | 0.27 | 0.39 |
| 8 | 10924369 | rs11780798 | 0.232 | -0.501 | -0.681 | -3.307 | -0.049 | 0.136 | 0.136 | 0.062 | 0.21 | 0.03 | 0.12 | 0 | 0.22 | 0.14 |
| 8 | 10937019 | rs6999969 | -1.186 | -0.494 | 0.443 | -3.289 | -0.02 | -0.151 | -0.151 | 3.365 | 0.26 | 0.47 | 0.18 | 0.05 | 0.2 | 0.11 |
| 8 | 10937853 | rs11986748 | -1.099 | -0.527 | 0.633 | -3.305 | -0.004 | -0.097 | -0.097 | 0 | 0.23 | 0.33 | 0.17 | 0.05 | 0.2 | 0.11 |
| 8 | 10942742 | rs13248264 | -0.247 | -0.658 | -0.106 | -3.602 | -0.078 | 0.096 | 0.096 | 0.082 | 0.19 | 0.07 | 0.14 | 0.01 | 0.27 | 0.29 |
| 8 | 10943276 | rs2409691 | -2.076 | -0.75 | 0.306 | -3.66 | -0.02 | 0.224 | 0.224 | 0 | 0.48 | 0.79 | 0.33 | 0.06 | 0.47 | 0.41 |
| 8 | 10954187 | rs7840068 | -1.059 | -0.825 | 0.907 | -3.569 | -0.02 | -0.107 | -0.107 | 0 | 0.22 | 0.53 | 0.15 | 0.04 | 0.16 | 0.11 |
| 8 | 10958539 | rs2898261 | -1.808 | -1.194 | 0.224 | -3.953 | 0.002 | 0.182 | 0.182 | 0 | 0.42 | 0.8 | 0.28 | 0.1 | 0.41 | 0.41 |
| 8 | 10959666 | rs7000695 | -0.156 | -1.078 | 0.815 | -3.647 | -0.015 | -0.022 | -0.022 | 0 | 0.05 | 0.08 | 0.02 | 0.04 | 0.02 | 0.04 |
| 8 | 10963902 | rs11991271 | -1.518 | -1.169 | 1.522 | -3.686 | 0.022 | 0.019 | 0.019 | 0 | 0.06 | 0.37 | 0.03 | 0 | 0 | 0.02 |
| 8 | 10965442 | rs7832722 | -1.346 | -1.26 | 1.367 | -3.87 | -0.03 | 0.029 | 0.029 | 0 | 0.3 | 0.19 | 0.21 | 0.04 | 0.4 | 0.27 |
| 8 | 10969075 | rs7460507 | -2.289 | -1.305 | 1.072 | -3.866 | -0.059 | 0.229 | 0.229 | 0 | 0.44 | 0.84 | 0.29 | 0.07 | 0.45 | 0.45 |
| 8 | 10983534 | rs3021494 | -2.196 | -1.251 | 0.884 | -3.847 | -0.095 | 0.264 | 0.264 | 0.361 | 0.6 | 0.95 | 0.37 | 0.08 | 0.54 | 0.47 |
| 8 | 10984294 | rs3736015 | 0.079 | -1.238 | 1.077 | -3.795 | -0.015 | -0.019 | -0.019 | 2.374 | 0.05 | 0.07 | 0.01 | 0.04 | 0.02 | 0.04 |
| 8 | 10987651 | rs10106914 | -1.14 | -1.168 | 1.101 | -3.875 | 0.004 | -0.089 | -0.089 | 0 | 0.4 | 0.55 | 0.29 | 0.04 | 0.43 | 0.32 |
| 8 | 10987967 | rs10088853 | -0.545 | -1.285 | 1.277 | -3.875 | 0.005 | -0.094 | -0.094 | 0.093 | 0.33 | 0.34 | 0.23 | 0 | 0.37 | 0.15 |
| 8 | 10990164 | rs4291231 | -0.619 | -1.285 | 1.232 | -3.875 | 0.012 | -0.157 | -0.157 | 0 | 0.33 | 0.46 | 0.23 | 0 | 0.37 | 0.15 |
| 8 | 10995060 | rs3021496 | -1.08 | -1.303 | 1.309 | -3.82 | 0.02 | -0.072 | -0.072 | 0 | 0.12 | 0.33 | 0.05 | 0 | 0.09 | 0.11 |
| 8 | 10996693 | rs9329237 | -0.846 | -1.318 | 1.374 | -3.84 | 0.033 | -0.021 | -0.021 | 0 | 0.09 | 0.43 | 0.06 | 0.02 | 0.04 | 0.14 |
| 8 | 10998445 | rs7845493 | -1.432 | -1.364 | 1.384 | -3.809 | -0.01 | -0.057 | -0.057 | 0.005 | 0.16 | 0.23 | 0.07 | 0.04 | 0.12 | 0.14 |
| 8 | 11001301 | rs7834295 | -2.604 | -1.299 | 1.038 | -3.841 | 0.028 | 0.044 | 0.044 | 0.057 | 0.29 | 0.79 | 0.15 | 0.05 | 0.17 | 0.28 |
| 8 | 11002093 | rs6986002 | -2.365 | -1.451 | 0.493 | -3.709 | -0.021 | 0.062 | 0.062 | 0.023 | 0.61 | 0.77 | 0.37 | 0.19 | 0.48 | 0.6 |
| 8 | 11009317 | rs17725480 | 0.115 | -1.557 | 0.69 | -3.736 | -0.01 | 0.021 | 0.021 | 1.823 | 0.27 | 0.17 | 0.16 | 0.01 | 0.28 | 0.15 |
| 8 | 11009797 | rs2409713 | -1.823 | -1.344 | 0.524 | -3.531 | -0.013 | 0.097 | 0.097 | 0.796 | 0.54 | 0.73 | 0.3 | 0.08 | 0.44 | 0.46 |
| 8 | 11012977 | rs2409718 | -2.045 | -1.315 | -0.139 | -3.403 | -0.108 | 0.259 | 0.259 | 0.691 | 0.6 | 0.95 | 0.38 | 0.08 | 0.52 | 0.46 |
| 8 | 11024663 | rs4841504 | -1.386 | -1.351 | 0.063 | -3.421 | 0 | 0.051 | 0.051 | 0 | 0.52 | 0.66 | 0.34 | 0.04 | 0.5 | 0.43 |
| 8 | 11031472 | rs7004825 | -2.033 | -1.127 | -0.537 | -3.214 | -0.129 | 0.186 | 0.186 | 0 | 0.63 | 0.95 | 0.39 | 0.11 | 0.54 | 0.47 |
| 8 | 11037034 | rs11250119 | -1.293 | -1.175 | -0.265 | -3.283 | -0.005 | 0.302 | 0.302 | 1.618 | 0.5 | 0.82 | 0.31 | 0.06 | 0.45 | 0.33 |
| 8 | 11039816 | rs2409722 | -2.028 | -1.136 | -0.376 | -3.117 | -0.107 | 0.223 | 0.223 | 0.194 | 0.6 | 0.94 | 0.37 | 0.12 | 0.51 | 0.46 |
| 8 | 11045161 | rs7819412 | -1.952 | -1.113 | -0.349 | -3.207 | -0.099 | 0.267 | 0.267 | 0.396 | 0.59 | 0.94 | 0.37 | 0.07 | 0.51 | 0.45 |
| 8 | 11060638 | rs2409730 | -0.351 | -1.313 | 0.233 | -3.262 | 0 | -0.06 | -0.06 | 0 | 0.33 | 0.29 | 0.23 | 0.03 | 0.35 | 0.2 |
| 8 | 11064269 | rs6996752 | -1.745 | -1.401 | 0.187 | -3.233 | -0.067 | -0.096 | -0.096 | 0 | 0.19 | 0.55 | 0.11 | 0.07 | 0.07 | 0.09 |
| 8 | 11068950 | rs2409732 | 0.702 | -1.264 | -0.634 | -3.168 | 0.001 | 0.014 | 0.014 | 2.244 | 0.07 | 0.07 | 0.06 | 0.01 | 0.05 | 0.02 |
| 8 | 11071057 | rs6987059 | -0.546 | -1.299 | -0.493 | -3.065 | -0.023 | 0.209 | 0.209 | 0.373 | 0.41 | 0.16 | 0.29 | 0.05 | 0.42 | 0.21 |
| 8 | 11072549 | rs2409744 | -0.747 | -0.904 | -0.645 | -2.933 | 0.001 | -0.055 | -0.055 | 0 | 0.13 | 0.2 | 0.06 | 0 | 0.11 | 0.09 |
| 8 | 11073578 | rs6601569 | -1.668 | -0.825 | -0.861 | -2.586 | -0.135 | 0.14 | 0.14 | 0.016 | 0.64 | 0.97 | 0.44 | 0.14 | 0.54 | 0.55 |
| 8 | 11076635 | rs2409745 | -1.509 | -0.78 | -0.582 | -2.558 | -0.078 | 0.227 | 0.227 | 0.223 | 0.6 | 0.9 | 0.39 | 0.08 | 0.52 | 0.46 |
| 8 | 11087694 | rs2409750 | -1.122 | -0.984 | -0.373 | -2.297 | 0.008 | -0.028 | -0.028 | 0.317 | 0.29 | 0.71 | 0.15 | 0.01 | 0.17 | 0.3 |
| 8 | 11088131 | rs10103062 | -0.766 | -0.523 | 0.003 | -2.298 | -0.015 | -0.118 | -0.118 | 0 | 0.28 | 0.6 | 0.19 | 0.01 | 0.25 | 0.24 |
| 8 | 11094751 | rs6601573 | -1.439 | -0.179 | -0.564 | -0.827 | -0.028 | -0.059 | -0.059 | 1.95 | 0.65 | 0.84 | 0.69 | 0.48 | 0.58 | 0.54 |
| 8 | 11096054 | rs11774569 | -0.64 | -0.634 | -0.03 | -1.285 | 0.004 | -0.062 | -0.062 | 0 | 0.3 | 0.3 | 0.22 | 0.02 | 0.35 | 0.24 |
| 8 | 11097804 | rs6601575 | -1.695 | -0.622 | -0.258 | -0.886 | -0.049 | 0.15 | 0.15 | 0.166 | 0.6 | 0.85 | 0.63 | 0.15 | 0.53 | 0.45 |
| 8 | 11099612 | rs2245257 | -0.923 | -0.825 | 0.188 | -0.902 | 0.003 | -0.005 | -0.005 | 0.992 | 0.39 | 0.61 | 0.51 | 0.5 | 0.22 | 0.26 |
| 8 | 11101723 | rs2572432 | -0.547 | -0.775 | 0.089 | -1.132 | -0.13 | 0.156 | 0.156 | 0.436 | 0.24 | 0.03 | 0.17 | 0 | 0.35 | 0.17 |
| 8 | 11103895 | rs958648 | -1.677 | -0.643 | 0.821 | -0.515 | 0 | 0 | 0 | 0.25 | 0.59 | 0.43 | 0.62 | 0.54 | 0.57 | 0.39 |
| 8 | 11110503 | rs2082775 | -0.031 | -0.013 | 0.276 | -1.108 | -0.022 | 0.012 | 0.012 | 0 | 0.12 | 0.07 | 0.11 | 0 | 0.2 | 0.1 |
| 8 | 11111892 | rs4504596 | -0.225 | -0.219 | 0.505 | -0.69 | -0.004 | 0.043 | 0.043 | 0 | 0.06 | 0.1 | 0.02 | 0.23 | 0.04 | 0.07 |
| 8 | 11113828 | rs2736374 | -1.286 | -0.33 | 0.418 | -0.998 | -0.058 | 0.061 | 0.061 | 0.153 | 0.29 | 0.18 | 0.23 | 0.01 | 0.4 | 0.26 |
| 8 | 11114566 | rs2196295 | -1.178 | -0.638 | 0.914 | -1.126 | 0.005 | 0.005 | 0.005 | 0.406 | 0.43 | 0.59 | 0.46 | 0.35 | 0.35 | 0.4 |
| 8 | 11116661 | rs17728678 | -0.353 | -0.588 | 0.743 | -1.133 | -0.014 | -0.004 | -0.004 | 1.495 | 0.17 | 0.14 | 0.14 | 0 | 0.24 | 0.11 |
| 8 | 11118983 | rs10503418 | -2.328 | -0.501 | 0.659 | -0.982 | -0.02 | -0.007 | -0.007 | 0 | 0.64 | 0.82 | 0.62 | 0.39 | 0.58 | 0.53 |
| 8 | 11119037 | rs2251301 | 0.222 | -0.38 | 0.49 | -0.935 | -0.04 | 0.052 | 0.052 | 0 | 0.11 | 0.03 | 0.09 | 0 | 0.17 | 0.14 |
| 8 | 11121260 | rs6990463 | -0.415 | -0.395 | 0.846 | -0.942 | 0.022 | -0.055 | -0.055 | 0.318 | 0.13 | 0.25 | 0.12 | 0.01 | 0.18 | 0.15 |
| 8 | 11125262 | rs2736379 | -0.982 | -0.485 | 0.323 | -0.748 | -0.004 | -0.037 | -0.037 | 3.55 | 0.2 | 0.14 | 0.33 | 0.38 | 0.22 | 0.28 |
| 8 | 11135974 | rs7004362 | -1.953 | -0.739 | 0.376 | -1.489 | -0.011 | 0.05 | 0.05 | 3.383 | 0.64 | 0.74 | 0.62 | 0.16 | 0.59 | 0.47 |
| 8 | 11143272 | rs3808518 | 2.09 | -0.714 | 0.819 | -1.516 | -0.037 | 0.024 | 0.024 | 2.245 | 0.35 | 0.14 | 0.35 | 0.7 | 0.4 | 0.45 |
| 8 | 11147123 | rs2572426 | 0.834 | -0.898 | 0.818 | -1.665 | -0.088 | 0.11 | 0.11 | 1.903 | 0.67 | 0.87 | 0.52 | 0.87 | 0.55 | 0.67 |

| | | | | | | | | | | | | | | | | |
|---|---|---|---|---|---|---|---|---|---|---|---|---|---|---|---|---|
| 8 | 11152957 | rs2245937 | 0.909 | -0.945 | 0.849 | -1.687 | -0.109 | 0.125 | 0.125 | 1.874 | 0.67 | 0.89 | 0.53 | 0.87 | 0.55 | 0.67 |
| 8 | 11156041 | rs3808515 | 0.818 | -0.958 | 0.861 | -1.698 | -0.113 | 0.125 | 0.125 | 2.093 | 0.67 | 0.89 | 0.53 | 0.87 | 0.54 | 0.67 |
| 8 | 11156163 | rs3824211 | -0.08 | -1.063 | 1.199 | -1.739 | -0.04 | -0.037 | -0.037 | 0.002 | 0.28 | 0.38 | 0.15 | 0.17 | 0.15 | 0.21 |
| 8 | 11157371 | rs2736387 | 1.085 | -1.113 | 1.301 | -1.768 | -0.017 | 0.026 | 0.026 | 0.934 | 0.63 | 0.68 | 0.51 | 0.86 | 0.53 | 0.59 |
| 8 | 11157460 | rs3808513 | 2.509 | -1.12 | 1.18 | -1.757 | -0.032 | 0.058 | 0.058 | 2.713 | 0.34 | 0.14 | 0.35 | 0.77 | 0.39 | 0.46 |
| 8 | 11160752 | rs10503420 | 0.551 | -1.165 | 1.144 | -1.78 | 0.028 | 0.147 | 0.147 | 2.119 | 0.23 | 0.02 | 0.11 | 0.02 | 0.13 | 0.04 |
| 8 | 11162496 | rs2164272 | 2.479 | -1.07 | 1.21 | -1.732 | -0.029 | 0.022 | 0.022 | 2.631 | 0.34 | 0.15 | 0.35 | 0.71 | 0.39 | 0.46 |
| 8 | 11165009 | rs11782742 | 1.065 | -1.036 | 1.383 | -1.713 | 0.017 | -0.047 | -0.047 | 0.36 | 0.63 | 0.56 | 0.5 | 0.84 | 0.53 | 0.67 |
| 8 | 11169983 | rs2409755 | 0.876 | -1.048 | 1.183 | -1.717 | -0.016 | 0.02 | 0.02 | 0.896 | 0.65 | 0.71 | 0.51 | 0.83 | 0.54 | 0.67 |
| 8 | 11173444 | rs978149 | 0.831 | -1.029 | 1.065 | -1.56 | 0.025 | 0.284 | 0.284 | 0.017 | 0.51 | 0.17 | 0.67 | 0.9 | 0.62 | 0.64 |
| 8 | 11176147 | rs17153128 | -0.493 | -1.037 | 1.222 | -1.358 | 0.001 | -0.003 | -0.003 | 0 | 0.08 | 0.09 | 0.23 | 0.16 | 0.05 | 0.06 |
| 8 | 11182455 | rs7016671 | 1.534 | -0.938 | 1.211 | -1.381 | -0.071 | 0.062 | 0.062 | 0.55 | 0.37 | 0.15 | 0.38 | 0.74 | 0.46 | 0.5 |
| 8 | 11183505 | rs3779891 | 1.534 | -0.855 | 1.211 | -1.405 | -0.066 | 0.059 | 0.059 | 0.55 | 0.37 | 0.14 | 0.37 | 0.73 | 0.46 | 0.5 |
| 8 | 11191537 | rs11774673 | 1.491 | -0.825 | 1.227 | -1.378 | -0.065 | 0.062 | 0.062 | 0.813 | 0.38 | 0.15 | 0.37 | 0.74 | 0.46 | 0.5 |
| 8 | 11196970 | rs1435277 | 0.545 | -0.836 | 1.317 | -1.444 | -0.001 | -0.013 | -0.013 | 0 | 0.49 | 0.57 | 0.64 | 0.98 | 0.52 | 0.63 |
| 8 | 11202154 | rs11250129 | 1.291 | -0.718 | 1.023 | -1.372 | -0.012 | 0.025 | 0.025 | 0.285 | 0.39 | 0.31 | 0.38 | 0.76 | 0.46 | 0.5 |
| 8 | 11213092 | rs17741537 | 1.37 | -0.744 | 0.493 | -1.386 | -0.07 | 0.058 | 0.058 | 0.782 | 0.38 | 0.15 | 0.37 | 0.72 | 0.47 | 0.51 |
| 8 | 11213589 | rs2250028 | 0.297 | -0.856 | 0.901 | -1.512 | -0.003 | 0.033 | 0.033 | 0 | 0.53 | 0.64 | 0.64 | 0.98 | 0.53 | 0.66 |
| 8 | 11214455 | rs11250130 | 0.587 | -0.789 | 0.433 | -1.476 | -0.022 | 0.231 | 0.231 | 0 | 0.49 | 0.18 | 0.4 | 0.84 | 0.52 | 0.64 |
| 8 | 11214972 | rs11781637 | 0.773 | -0.816 | 0.424 | -1.473 | -0.032 | 0.212 | 0.212 | 0 | 0.47 | 0.16 | 0.4 | 0.83 | 0.52 | 0.63 |
| 8 | 11219386 | rs2293860 | 1.4 | -0.73 | 0.33 | -1.59 | -0.069 | 0.079 | 0.079 | 0.405 | 0.38 | 0.15 | 0.36 | 0.75 | 0.47 | 0.5 |
| 8 | 11223022 | rs2736280 | 0.971 | -0.806 | 0.45 | -1.999 | -0.001 | -0.02 | -0.02 | 3.223 | 0.43 | 0.59 | 0.62 | 0.86 | 0.49 | 0.57 |
| 8 | 11229638 | rs2572450 | -1.056 | -0.512 | -1.035 | -2.209 | -0.074 | 0.356 | 0.356 | 0 | 0.53 | 0.92 | 0.38 | 0.02 | 0.47 | 0.37 |
| 8 | 11236975 | rs2572386 | -0.846 | -0.284 | -0.839 | -2.099 | 0.005 | 0.162 | 0.162 | 0 | 0.5 | 0.71 | 0.36 | 0.02 | 0.47 | 0.27 |
| 8 | 11248036 | rs2043507 | -0.062 | -0.386 | -0.668 | -1.276 | -0.002 | -0.009 | -0.009 | 0 | 0.07 | 0.07 | 0.04 | 0.01 | 0.04 | 0.01 |
| 8 | 11251440 | rs2736310 | -1.154 | -0.514 | -1.026 | -1.657 | 0.033 | 0.038 | 0.038 | 1.533 | 0.27 | 0.8 | 0.19 | 0.02 | 0.18 | 0.16 |
| 8 | 11253879 | rs10105588 | -0.914 | 0.047 | -0.84 | -1.806 | -0.001 | -0.074 | -0.074 | 0.916 | 0.23 | 0.31 | 0.16 | 0.03 | 0.21 | 0.12 |
| 8 | 11257495 | rs10100187 | -1.093 | 0.617 | -1.965 | -2.338 | -0.079 | 0.142 | 0.142 | 0 | 0.69 | 0.91 | 0.42 | 0.09 | 0.6 | 0.59 |
| 8 | 11258378 | rs4841526 | 0.843 | 0.589 | -2.008 | -2.381 | -0.093 | 0.169 | 0.169 | 0 | 0.31 | 0.07 | 0.58 | 0.93 | 0.41 | 0.41 |
| 8 | 11269005 | rs9644737 | 1.487 | 0.772 | -2.184 | -1.625 | -0.077 | -0.116 | -0.116 | 0.997 | 0.22 | 0.02 | 0.39 | 0.66 | 0.3 | 0.33 |
| 8 | 11272079 | rs11775511 | 0.04 | 0.44 | -1.405 | -2.437 | 0.025 | -0.043 | -0.043 | 0.021 | 0.08 | 0.2 | 0.06 | 0 | 0.07 | 0.07 |
| 8 | 11272164 | rs7836059 | -0.161 | 0.952 | -1.479 | -2.54 | -0.019 | 0.049 | 0.049 | 0 | 0.6 | 0.71 | 0.31 | 0.06 | 0.48 | 0.48 |
| 8 | 11274122 | rs7018324 | -0.428 | 0.501 | -1.781 | -3.02 | 0.122 | 0.276 | 0.276 | 0.347 | 0.5 | 0.79 | 0.25 | 0.06 | 0.31 | 0.35 |
| 8 | 11276542 | rs2002030 | -0.01 | 0.197 | -1.792 | -2.828 | -0.025 | -0.065 | -0.065 | 1.474 | 0.13 | 0.24 | 0.05 | 0.02 | 0.07 | 0.18 |
| 8 | 11278146 | rs11250135 | -1.008 | 0.675 | -2.323 | -2.62 | -0.09 | 0.181 | 0.181 | 0 | 0.68 | 0.91 | 0.41 | 0.07 | 0.59 | 0.57 |
| 8 | 11281273 | rs2409764 | -0.477 | 0.545 | -1.394 | -2.818 | 0 | -0.102 | -0.102 | 0.002 | 0.46 | 0.47 | 0.29 | 0.03 | 0.45 | 0.29 |
| 8 | 11282893 | rs11780980 | 0.621 | 0.836 | -1.785 | -2.399 | -0.061 | 0.074 | 0.074 | 0 | 0.12 | 0.02 | 0.09 | 0 | 0.2 | 0.07 |
| 8 | 11285186 | rs10503422 | -0.075 | 0.108 | -1.564 | -2.921 | 0.026 | 0.055 | 0.055 | 0.707 | 0.36 | 0.21 | 0.2 | 0.03 | 0.26 | 0.24 |
| 8 | 11285384 | rs10102600 | -1.226 | 0.448 | -2.066 | -1.994 | -0.038 | -0.104 | -0.104 | 0 | 0.85 | 0.98 | 0.86 | 0.69 | 0.84 | 0.75 |
| 8 | 11286069 | rs7831885 | 0.606 | 0.115 | -1.449 | -2.385 | 0.001 | -0.054 | -0.054 | 0 | 0.31 | 0.25 | 0.16 | 0 | 0.21 | 0.15 |
| 8 | 11286146 | rs7844834 | 0.076 | 0.23 | -0.819 | -2.687 | 0.004 | -0.115 | -0.115 | 0 | 0.3 | 0.54 | 0.2 | 0 | 0.3 | 0.2 |
| 8 | 11287371 | rs10099909 | 0.792 | 0.403 | -0.362 | -2.571 | 0.013 | -0.043 | -0.043 | 1.175 | 0.08 | 0.18 | 0.06 | 0.01 | 0.07 | 0.12 |
| 8 | 11289146 | rs4841530 | -1.297 | 0.36 | -0.173 | -2.5 | -0.003 | 0.064 | 0.064 | 0 | 0.85 | 0.75 | 0.84 | 0.71 | 0.86 | 0.71 |
| 8 | 11289685 | rs12115021 | 1.206 | 0.218 | -0.743 | -2.775 | -0.003 | -0.005 | -0.005 | 0 | 0.17 | 0.12 | 0.1 | 0 | 0.19 | 0.06 |
| 8 | 11289697 | rs12549796 | 1.367 | 0.109 | -0.996 | -2.868 | -0.061 | -0.07 | -0.07 | 0 | 0.23 | 0.08 | 0.52 | 0.7 | 0.36 | 0.39 |
| 8 | 11289769 | rs10448145 | 1.493 | 0.126 | -0.985 | -2.857 | -0.048 | -0.058 | -0.058 | 0.493 | 0.23 | 0.1 | 0.52 | 0.7 | 0.36 | 0.38 |
| 8 | 11290966 | rs6984212 | -1.795 | -0.397 | -0.936 | -2.648 | -0.038 | -0.096 | -0.096 | 0.132 | 0.86 | 0.97 | 0.86 | 0.72 | 0.86 | 0.78 |
| 8 | 11295925 | rs3021517 | 1.368 | -0.262 | -0.848 | -2.646 | -0.039 | 0.065 | 0.065 | 0 | 0.16 | 0.03 | 0.11 | 0 | 0.21 | 0.06 |
| 8 | 11296600 | rs7001675 | 0.151 | -0.062 | -0.514 | -2.816 | 0.006 | 0.004 | 0.004 | 0.622 | 0.33 | 0.41 | 0.56 | 0.72 | 0.41 | 0.58 |
| 8 | 11302445 | rs11779875 | 1.573 | -0.007 | -0.587 | -2.808 | -0.037 | -0.075 | -0.075 | 0 | 0.21 | 0.07 | 0.49 | 0.69 | 0.33 | 0.36 |
| 8 | 11303011 | rs2003423 | -0.346 | -0.102 | -0.432 | -2.867 | -0.001 | -0.002 | -0.002 | 2.154 | 0.47 | 0.42 | 0.65 | 0.7 | 0.59 | 0.55 |
| 8 | 11306805 | rs2409775 | -0.18 | -0.286 | -0.534 | -2.683 | 0.01 | -0.005 | -0.005 | 0 | 0.38 | 0.65 | 0.57 | 0.68 | 0.4 | 0.49 |
| 8 | 11309893 | rs13439411 | 0.452 | -0.29 | -0.443 | -2.509 | -0.007 | 0.005 | 0.005 | 0 | 0.67 | 0.57 | 0.76 | 0.72 | 0.73 | 0.66 |
| 8 | 11310290 | rs10903338 | -0.721 | -0.217 | -1.061 | -2.263 | 0.013 | -0.064 | -0.064 | 0.241 | 0.81 | 0.88 | 0.85 | 0.72 | 0.85 | 0.75 |
| 8 | 11320152 | rs4840563 | 0.794 | -0.285 | -1.271 | -2.754 | -0.003 | 0.113 | 0.113 | 0.267 | 0.32 | 0.11 | 0.18 | 0.03 | 0.29 | 0.19 |
| 8 | 11321292 | rs2898280 | -0.111 | -0.491 | -0.656 | -2.575 | -0.001 | -0.006 | -0.006 | 0 | 0.45 | 0.48 | 0.6 | 0.69 | 0.46 | 0.44 |
| 8 | 11323025 | rs9644687 | 0.442 | -0.337 | -0.861 | -2.3 | 0.002 | -0.028 | -0.028 | 0.402 | 0.24 | 0.2 | 0.47 | 0.68 | 0.25 | 0.36 |
| 8 | 11329544 | rs899364 | 0.235 | -0.4 | -0.337 | -2.949 | 0.012 | -0.141 | -0.141 | 0 | 0.33 | 0.47 | 0.25 | 0.03 | 0.33 | 0.23 |
| 8 | 11330364 | rs6989817 | -0.041 | -0.375 | -0.326 | -2.96 | -0.006 | 0.036 | 0.036 | 0 | 0.52 | 0.69 | 0.67 | 0.97 | 0.53 | 0.57 |
| 8 | 11332247 | rs2736328 | -1.095 | -0.127 | -0.359 | -2.562 | -0.001 | -0.01 | -0.01 | 3.202 | 0.87 | 0.81 | 0.89 | 0.98 | 0.88 | 0.69 |
| 8 | 11333521 | rs17799348 | -0.258 | -0.283 | -0.24 | -2.518 | 0 | -0.032 | -0.032 | 1.592 | 0.52 | 0.57 | 0.59 | 0.87 | 0.57 | 0.49 |
| 8 | 11339372 | rs7830501 | -0.19 | -0.501 | 0.305 | -2.236 | 0.036 | 0.022 | 0.022 | 0.474 | 0.06 | 0.33 | 0.02 | 0 | 0.02 | 0.01 |
| 8 | 11343973 | rs2736340 | 0.654 | -0.245 | -0.721 | -2.053 | -0.01 | -0.011 | -0.011 | 0 | 0.19 | 0.12 | 0.49 | 0.72 | 0.24 | 0.35 |
| 8 | 11349186 | rs13277113 | 0.737 | -0.256 | -0.726 | -2.033 | -0.01 | -0.013 | -0.013 | 0.23 | 0.19 | 0.12 | 0.49 | 0.71 | 0.24 | 0.35 |
| 8 | 11350899 | rs1382567 | 1.168 | -1.02 | -1.069 | -2.837 | 0.001 | 0.118 | 0.118 | 0.013 | 0.43 | 0.21 | 0.34 | 0.01 | 0.54 | 0.4 |
| 8 | 11352541 | rs2618476 | 1.067 | -0.913 | -1.295 | -2.895 | -0.01 | -0.011 | -0.011 | 0.497 | 0.2 | 0.12 | 0.5 | 0.72 | 0.25 | 0.36 |
| 8 | 11354933 | rs4606022 | 1.13 | -0.179 | -1.578 | -2.954 | -0.047 | 0.179 | 0.179 | 0 | 0.37 | 0.11 | 0.26 | 0 | 0.39 | 0.19 |
| 8 | 11355821 | rs2618479 | -0.721 | -0.712 | -1.189 | -2.752 | 0.016 | -0.023 | -0.023 | 0.022 | 0.82 | 0.9 | 0.91 | 0.75 | 0.84 | 0.83 |
| 8 | 11359557 | rs2409781 | -0.13 | -0.898 | -0.549 | -2.893 | -0.05 | -0.009 | -0.009 | 0.003 | 0.29 | 0.51 | 0.54 | 0.69 | 0.23 | 0.42 |

| | | | | | | | | | | | | | | | | |
|---|---|---|---|---|---|---|---|---|---|---|---|---|---|---|---|---|
| 8 | 11359638 | rs1600249 | 1.06 | -0.603 | -1.109 | -2.737 | -0.021 | -0.069 | -0.069 | 0 | 0.17 | 0.05 | 0.5 | 0.69 | 0.21 | 0.41 |
| 8 | 11369989 | rs6993775 | -0.093 | -0.878 | -0.535 | -3.13 | -0.012 | -0.004 | -0.004 | 0.054 | 0.74 | 0.66 | 0.9 | 0.73 | 0.82 | 0.8 |
| 8 | 11370878 | rs2618455 | -0.42 | -0.903 | -0.595 | -3.18 | 0.019 | -0.002 | -0.002 | 0 | 0.78 | 0.87 | 0.92 | 0.74 | 0.83 | 0.8 |
| 8 | 11372141 | rs11250141 | 0.974 | -0.808 | -1.187 | -3.778 | 0 | 0.169 | 0.169 | 2.974 | 0.49 | 0.21 | 0.76 | 0.71 | 0.58 | 0.53 |
| 8 | 11373264 | rs17153385 | 1.56 | -0.736 | -1.102 | -3.739 | -0.004 | -0.025 | -0.025 | 0 | 0.14 | 0.07 | 0.05 | 0.26 | 0.15 | 0.18 |
| 8 | 11373467 | rs2736360 | 1.376 | -0.77 | -1.114 | -3.754 | -0.001 | 0.01 | 0.01 | 0 | 0.15 | 0.16 | 0.06 | 0.26 | 0.15 | 0.18 |
| 8 | 11374779 | rs4841548 | 1.514 | -0.784 | -1.026 | -3.952 | -0.022 | 0.094 | 0.094 | 0.814 | 0.32 | 0.17 | 0.26 | 0 | 0.36 | 0.11 |
| 8 | 11376266 | rs2618451 | 0.127 | -0.773 | -0.887 | -3.94 | 0.01 | 0.015 | 0.015 | 0.538 | 0.39 | 0.32 | 0.57 | 0.97 | 0.38 | 0.64 |
| 8 | 11376790 | rs2618450 | -0.699 | -0.475 | -0.577 | -4.015 | 0.028 | -0.082 | -0.082 | 0 | 0.79 | 0.71 | 0.86 | 0.97 | 0.77 | 0.75 |
| 8 | 11384556 | rs2618443 | -0.722 | -0.879 | -0.712 | -4.01 | 0.022 | 0.033 | 0.033 | 0 | 0.52 | 0.67 | 0.62 | 0.97 | 0.44 | 0.63 |
| 8 | 11384713 | rs2252534 | -1.513 | -0.527 | -1.266 | -4.212 | -0.081 | 0.079 | 0.079 | 0.452 | 0.84 | 0.99 | 0.87 | 0.97 | 0.77 | 0.86 |
| 8 | 11387861 | rs7814834 | -0.037 | -0.751 | -1.118 | -4.267 | -0.001 | 0.008 | 0.008 | 0 | 0.46 | 0.43 | 0.67 | 0.73 | 0.51 | 0.56 |
| 8 | 11388449 | rs2264306 | 1.301 | -0.681 | -1.238 | -4.289 | 0.001 | -0.067 | -0.067 | 0 | 0.32 | 0.3 | 0.24 | 0 | 0.33 | 0.23 |
| 8 | 11388764 | rs2249259 | 0.306 | -0.792 | -1.231 | -4.276 | 0 | 0.018 | 0.018 | 0 | 0.48 | 0.43 | 0.31 | 0.27 | 0.47 | 0.42 |
| 8 | 11391650 | rs2248932 | 0.042 | -1.79 | -0.59 | -3.92 | -0.01 | -0.021 | -0.021 | 0 | 0.64 | 0.57 | 0.44 | 0.27 | 0.67 | 0.48 |
| 8 | 11394233 | rs17153419 | 1.084 | -1.786 | -1.097 | -3.957 | -0.034 | -0.037 | -0.037 | 0.295 | 0.23 | 0.05 | 0.52 | 0.72 | 0.27 | 0.43 |
| 8 | 11396818 | rs7014565 | 0.857 | -1.452 | -0.848 | -3.742 | -0.002 | -0.026 | -0.026 | 0.154 | 0.15 | 0.08 | 0.06 | 0.25 | 0.14 | 0.19 |
| 8 | 11398865 | rs2618434 | 0.312 | -1.23 | -0.57 | -4.292 | -0.001 | 0 | 0 | 0 | 0.44 | 0.34 | 0.35 | 0 | 0.52 | 0.28 |
| 8 | 11404514 | rs7820492 | -0.094 | -1.014 | -0.393 | -4.09 | -0.004 | 0.098 | 0.098 | 0 | 0.46 | 0.26 | 0.35 | 0.03 | 0.52 | 0.36 |
| 8 | 11405141 | rs11779613 | -0.357 | -0.981 | -0.155 | -4.061 | 0.006 | -0.001 | -0.001 | 0 | 0.51 | 0.39 | 0.37 | 0.06 | 0.56 | 0.43 |
| 8 | 11406593 | rs3816668 | -0.659 | -0.961 | 0.054 | -4.01 | 0 | -0.039 | -0.039 | 0 | 0.55 | 0.54 | 0.39 | 0.08 | 0.57 | 0.46 |
| 8 | 11408401 | rs922484 | 0.079 | -0.934 | -0.33 | -3.805 | 0.007 | 0.048 | 0.048 | 0.659 | 0.14 | 0.05 | 0.09 | 0.03 | 0.13 | 0.18 |
| 8 | 11410987 | rs936550 | -0.585 | -0.905 | -0.235 | -3.82 | -0.006 | -0.037 | -0.037 | 0.644 | 0.19 | 0.19 | 0.09 | 0.03 | 0.17 | 0.22 |
| 8 | 11411823 | rs17744726 | 0.811 | -0.41 | -0.567 | -3.575 | -0.046 | 0.083 | 0.083 | 0.433 | 0.27 | 0.12 | 0.24 | 0 | 0.36 | 0.17 |
| 8 | 11423439 | rs755320 | 0.522 | -0.898 | 0.358 | -3.031 | -0.024 | -0.069 | -0.069 | 0 | 0.81 | 0.39 | 0.87 | 0.96 | 0.87 | 0.85 |
| 8 | 11425077 | rs4841562 | 0.064 | -1.061 | -0.166 | -3.428 | 0.008 | 0.102 | 0.102 | 0.546 | 0.35 | 0.23 | 0.53 | 0.94 | 0.35 | 0.58 |
| 8 | 11429147 | rs732947 | -0.393 | -0.891 | 0.181 | -3.284 | -0.003 | -0.028 | -0.028 | 0 | 0.87 | 0.86 | 0.92 | 0.97 | 0.88 | 0.87 |
| 8 | 11433909 | rs2898290 | 0.213 | -1.069 | 0.586 | -3.252 | 0.005 | -0.05 | -0.05 | 0.814 | 0.54 | 0.58 | 0.4 | 0.01 | 0.55 | 0.29 |
| 8 | 11444516 | rs10097870 | -0.461 | -0.929 | 0.175 | -3.111 | 0 | -0.094 | -0.094 | 0 | 0.46 | 0.55 | 0.62 | 0.99 | 0.44 | 0.7 |
| 8 | 11448529 | rs1478892 | 0.609 | -0.765 | -0.817 | -2.849 | -0.025 | 0.066 | 0.066 | 0 | 0.27 | 0.17 | 0.52 | 0.93 | 0.34 | 0.48 |
| 8 | 11448738 | rs11783137 | 0.619 | -0.737 | -1.132 | -2.8 | 0.016 | 0.017 | 0.017 | 4.429 | 0.08 | 0.02 | 0.06 | 0.03 | 0.06 | 0.08 |
| 8 | 11450587 | rs10903340 | -0.252 | -0.774 | -0.858 | -2.901 | 0.001 | -0.017 | -0.017 | 3.578 | 0.38 | 0.38 | 0.61 | 0.97 | 0.4 | 0.57 |
| 8 | 11455106 | rs11777273 | 0.619 | -0.428 | -2.031 | -3.009 | -0.1 | 0.168 | 0.168 | 1.356 | 0.35 | 0.09 | 0.26 | 0.01 | 0.41 | 0.29 |
| 8 | 11463015 | rs17807624 | 0.02 | 0.148 | -1.456 | -2.019 | -0.029 | 0.102 | 0.102 | 0 | 0.29 | 0.11 | 0.54 | 0.92 | 0.34 | 0.48 |
| 8 | 11469622 | rs10106995 | -0.766 | 0.139 | -0.922 | -1.676 | -0.004 | -0.03 | -0.03 | 0.111 | 0.1 | 0.15 | 0.06 | 0.01 | 0.07 | 0.02 |
| 8 | 11470175 | rs2448074 | 0.02 | 0.147 | -0.926 | -2.557 | -0.008 | 0.062 | 0.062 | 1.395 | 0.46 | 0.69 | 0.35 | 0.06 | 0.48 | 0.26 |
| 8 | 11470460 | rs2250562 | 0.05 | 0.559 | -0.372 | -2.956 | -0.001 | -0.116 | -0.116 | 0.279 | 0.3 | 0.5 | 0.2 | 0.05 | 0.28 | 0.22 |
| 8 | 11470573 | rs2250556 | 0.3 | 0.488 | 0.009 | -3.127 | 0.04 | -0.082 | -0.082 | 0 | 0.22 | 0.42 | 0.19 | 0.05 | 0.27 | 0.19 |
| 8 | 11474651 | rs2250056 | -0.908 | -0.165 | -0.486 | -2.68 | 0 | 0.002 | 0.002 | 0.164 | 0.84 | 0.9 | 0.9 | 0.85 | 0.82 | 0.84 |
| 8 | 11476145 | rs936549 | 0.712 | -0.66 | 0.046 | -1.922 | 0.005 | 0 | 0 | 1.509 | 0.67 | 0.34 | 0.82 | 0.7 | 0.76 | 0.7 |
| 8 | 11476595 | rs2249831 | -0.873 | -0.645 | 0.143 | -1.917 | 0 | -0.001 | -0.001 | 0 | 0.82 | 0.83 | 0.9 | 0.84 | 0.8 | 0.79 |
| 8 | 11477956 | rs1017804 | 0.593 | -0.673 | 0.258 | -1.883 | -0.031 | 0.002 | 0.002 | 0 | 0.67 | 0.44 | 0.82 | 0.71 | 0.76 | 0.7 |
| 8 | 11479356 | rs10111627 | -0.237 | -0.567 | 0.307 | -1.884 | 0.027 | 0.231 | 0.231 | 0 | 0.07 | 0.37 | 0.06 | 0.13 | 0.01 | 0.1 |
| 8 | 11480457 | rs2243407 | 0.674 | -0.546 | -0.68 | -1.467 | 0.028 | 0.001 | 0.001 | 0 | 0.57 | 0.33 | 0.72 | 0.63 | 0.68 | 0.7 |
| 8 | 11481253 | rs2246521 | -1.2 | -0.144 | -1.192 | -1.218 | 0 | 0.011 | 0.011 | 0.271 | 0.88 | 0.9 | 0.92 | 0.94 | 0.84 | 0.89 |
| 8 | 11483870 | rs12681079 | 0.72 | -0.128 | -1.474 | -1.134 | -0.009 | -0.033 | -0.033 | 0.009 | 0.25 | 0.09 | 0.49 | 0.71 | 0.29 | 0.41 |
| 8 | 11492153 | rs2256612 | -0.522 | 0.085 | -1.689 | -1.694 | -0.035 | 0.182 | 0.182 | 0 | 0.56 | 0.84 | 0.68 | 0.88 | 0.51 | 0.68 |
| 8 | 11496032 | rs11786272 | 0.847 | -0.121 | -1.714 | -1.949 | -0.025 | 0.076 | 0.076 | 0 | 0.2 | 0.06 | 0.13 | 0 | 0.18 | 0.03 |
| 8 | 11496193 | rs2256087 | 0.235 | -0.759 | -1.738 | -2.276 | 0.003 | -0.001 | -0.001 | 0 | 0.69 | 0.36 | 0.7 | 0.71 | 0.78 | 0.72 |
| 8 | 11496686 | rs11250154 | -0.092 | -0.825 | -1.476 | -2.207 | 0.004 | 0.004 | 0.004 | 0 | 0.64 | 0.73 | 0.66 | 0.71 | 0.69 | 0.74 |
| 8 | 11497458 | rs2253061 | -0.895 | -0.818 | -1.454 | -2.166 | -0.01 | 0.034 | 0.034 | 1.431 | 0.84 | 0.79 | 0.79 | 0.71 | 0.88 | 0.77 |
| 8 | 11499277 | rs2252902 | -0.2 | -0.87 | -1.203 | -2.192 | 0.004 | 0.004 | 0.004 | 0 | 0.64 | 0.75 | 0.66 | 0.71 | 0.69 | 0.74 |
| 8 | 11499967 | rs10903341 | 1.437 | -0.482 | -1.682 | -2.253 | 0.037 | -0.099 | -0.099 | 0.051 | 0.23 | 0.06 | 0.31 | 0.68 | 0.18 | 0.52 |
| 8 | 11501265 | rs1809356 | 0.248 | -0.258 | -1.132 | -2.007 | 0.046 | 0.253 | 0.253 | 0 | 0.08 | 0.39 | 0.19 | 0.19 | 0.05 | 0.08 |
| 8 | 11501447 | rs2100965 | 0.375 | -0.145 | -2.411 | -1.198 | -0.124 | -0.081 | -0.081 | 0 | 0.39 | 0.1 | 0.52 | 0.61 | 0.51 | 0.41 |
| 8 | 11502515 | rs12114452 | 0.83 | -0.135 | -1.739 | -0.775 | 0.002 | 0.001 | 0.001 | 2.697 | 0.14 | 0.15 | 0.06 | 0.08 | 0.11 | 0.18 |
| 8 | 11503333 | rs11782574 | 0.349 | 0.265 | -2.205 | -0.834 | -0.074 | -0.13 | -0.13 | 1.048 | 0.31 | 0.04 | 0.37 | 0.58 | 0.33 | 0.26 |
| 8 | 11503979 | rs1564265 | 0.642 | 0.42 | -1.792 | -0.665 | -0.041 | 0.141 | 0.141 | 0.291 | 0.4 | 0.15 | 0.29 | 0.14 | 0.46 | 0.49 |
| 8 | 11504919 | rs13267835 | 0.842 | 0.547 | -1.717 | -0.379 | -0.04 | -0.049 | -0.049 | 0.056 | 0.1 | 0.01 | 0.26 | 0.58 | 0.15 | 0.23 |
| 8 | 11508688 | rs7838703 | 0.782 | 0.508 | -1.718 | -0.705 | -0.04 | 0.102 | 0.102 | 0 | 0.37 | 0.17 | 0.27 | 0.14 | 0.44 | 0.45 |
| 8 | 11514071 | rs2248334 | -0.252 | 0.455 | -0.668 | -0.031 | 0.004 | 0.135 | 0.135 | 0.128 | 0.13 | 0.24 | 0.3 | 0.59 | 0.15 | 0.22 |
| 8 | 11515193 | rs17153607 | -0.017 | 0.216 | -0.786 | -0.106 | 0.004 | 0.003 | 0.003 | 0 | 0.3 | 0.27 | 0.26 | 0.25 | 0.24 | 0.24 |
| 8 | 11526986 | rs12675779 | 0.041 | 0.767 | -0.051 | 0.052 | 0.066 | 0.204 | 0.204 | 0 | 0.13 | 0.28 | 0.29 | 0.54 | 0.25 | 0.22 |
| 8 | 11527910 | rs4240677 | -0.648 | 0.395 | -0.665 | -1.406 | -0.035 | 0.099 | 0.099 | 1.044 | 0.58 | 0.76 | 0.64 | 0.95 | 0.54 | 0.8 |
| 8 | 11533384 | rs4510815 | -1.293 | 0.69 | -0.634 | -1.333 | 0.016 | -0.098 | -0.098 | 0.273 | 0.75 | 0.61 | 0.75 | 0.97 | 0.69 | 0.83 |
| 8 | 11536638 | rs4840575 | -0.241 | 0.004 | -1.162 | -0.815 | -0.026 | -0.053 | -0.053 | 0 | 0.19 | 0.07 | 0.3 | 0.73 | 0.26 | 0.29 |
| 8 | 11541697 | rs11784691 | 0.25 | -0.437 | -1.119 | -1.064 | 0.002 | 0.002 | 0.002 | 0.058 | 0.15 | 0.11 | 0.06 | 0.1 | 0.12 | 0.14 |
| 8 | 11542996 | rs2409805 | 1.024 | -1.019 | -1.449 | -1.304 | 0.081 | 0.139 | 0.139 | 4.577 | 0.57 | 0.13 | 0.61 | 0.17 | 0.66 | 0.46 |
| 8 | 11546264 | rs4841583 | -1.037 | -0.779 | -0.552 | -0.973 | -0.011 | 0.095 | 0.095 | 0 | 0.08 | 0.29 | 0.06 | 0.13 | 0.02 | 0.18 |

| | | | | | | | | | | | | | | | | |
|---|---|---|---|---|---|---|---|---|---|---|---|---|---|---|---|---|
| 8 | 11547150 | rs2409809 | 0.478 | -0.791 | -1.278 | -1.567 | 0.075 | 0.154 | 0.154 | 0 | 0.51 | 0.18 | 0.6 | 0.78 | 0.66 | 0.56 |
| 8 | 11551470 | rs34838488 | 0.438 | -0.56 | -0.65 | -1.117 | -0.003 | -0.002 | -0.002 | 1.314 | 0.17 | 0.16 | 0.28 | 0.74 | 0.24 | 0.29 |
| 8 | 11570036 | rs6990313 | 0.607 | -0.842 | -0.308 | -1.231 | -0.013 | 0.013 | 0.013 | 0.158 | 0.11 | 0.28 | 0.07 | 0.09 | 0.09 | 0.14 |
| 8 | 11575518 | rs6601604 | 0.563 | -1.277 | -0.483 | -1.803 | 0.015 | 0.009 | 0.009 | 0 | 0.5 | 0.53 | 0.52 | 0.14 | 0.7 | 0.3 |
| 8 | 11579802 | rs10112596 | -1.116 | -1.668 | -1.03 | -1.926 | 0.005 | 0.007 | 0.007 | 0 | 0.8 | 0.88 | 0.78 | 0.85 | 0.85 | 0.7 |
| 8 | 11579954 | rs12550668 | 1.825 | -1.615 | -1.269 | -2.018 | -0.097 | 0.219 | 0.219 | 0.444 | 0.35 | 0.03 | 0.42 | 0.04 | 0.61 | 0.17 |
| 8 | 11582929 | rs2898292 | -0.244 | -2.078 | -0.493 | -1.777 | -0.063 | 0.009 | 0.009 | 0 | 0.2 | 0.31 | 0.22 | 0.84 | 0.1 | 0.4 |
| 8 | 11584425 | rs4840579 | -0.081 | -1.46 | -0.646 | -1.991 | 0.03 | -0.051 | -0.051 | 0 | 0.68 | 0.45 | 0.63 | 0.86 | 0.64 | 0.65 |
| 8 | 11587234 | rs11250159 | 0.358 | -1.446 | -0.761 | -1.767 | -0.048 | 0.053 | 0.053 | 0 | 0.16 | 0.22 | 0.19 | 0.72 | 0.09 | 0.37 |
| 8 | 11588481 | rs17153694 | 0.67 | -1.444 | -0.77 | -1.76 | 0.019 | -0.022 | -0.022 | 0.071 | 0.15 | 0.09 | 0.18 | 0.74 | 0.08 | 0.37 |
| 8 | 11590008 | rs17153698 | -0.395 | -1.534 | -0.841 | -1.853 | -0.059 | 0.026 | 0.026 | 0 | 0.25 | 0.34 | 0.23 | 0.72 | 0.17 | 0.5 |
| 8 | 11591136 | rs6983129 | 0.131 | -1.083 | -0.907 | -2.049 | -0.009 | 0.014 | 0.014 | 0 | 0.61 | 0.67 | 0.48 | 0.11 | 0.55 | 0.3 |
| 8 | 11595969 | rs2898295 | 0.576 | -0.787 | -1.265 | -1.761 | 0.003 | 0.036 | 0.036 | 1.289 | 0.48 | 0.4 | 0.42 | 0.12 | 0.5 | 0.17 |
| 8 | 11596437 | rs11250163 | 0.941 | -0.395 | -2.218 | -1.723 | -0.154 | 0.259 | 0.259 | 2.936 | 0.39 | 0.04 | 0.38 | 0.03 | 0.48 | 0.15 |
| 8 | 11601075 | rs13275657 | 0.79 | 0.111 | -1.633 | -0.375 | -0.052 | -0.062 | -0.062 | 0 | 0.1 | 0.02 | 0.29 | 0.58 | 0.19 | 0.27 |
| 8 | 11604858 | rs2029969 | -0.88 | -0.392 | -1.482 | -1.05 | 0.011 | 0.008 | 0.008 | 0.14 | 0.35 | 0.21 | 0.4 | 0.71 | 0.36 | 0.45 |
| 8 | 11605130 | rs2173117 | -0.204 | -0.416 | -1.402 | -1.096 | 0.002 | -0.007 | -0.007 | 2.308 | 0.27 | 0.22 | 0.37 | 0.71 | 0.32 | 0.43 |
| 8 | 11607048 | rs3779664 | 0.919 | -0.364 | -1.747 | -1.202 | -0.017 | -0.068 | -0.068 | 0.676 | 0.17 | 0.05 | 0.23 | 0.26 | 0.14 | 0.3 |
| 8 | 11608905 | rs2740434 | -1.327 | -0.086 | -1.354 | -1.662 | -0.004 | -0.003 | -0.003 | 2.409 | 0.71 | 0.7 | 0.72 | 0.92 | 0.65 | 0.83 |
| 8 | 11609169 | rs2645399 | -1.091 | -0.437 | -0.643 | -1.717 | 0.029 | -0.093 | -0.093 | 0.629 | 0.67 | 0.48 | 0.69 | 0.89 | 0.63 | 0.81 |
| 8 | 11610674 | rs11784693 | 1.27 | -0.528 | -0.653 | -1.822 | -0.019 | 0.02 | 0.02 | 0.487 | 0.29 | 0.23 | 0.24 | 0.02 | 0.32 | 0.11 |
| 8 | 11611043 | rs804283 | -1.538 | -0.545 | -0.885 | -1.841 | -0.083 | 0.138 | 0.138 | 1.039 | 0.73 | 0.93 | 0.77 | 0.99 | 0.68 | 0.9 |
| 8 | 11611353 | rs17153747 | 1.842 | -0.721 | -0.977 | -2.146 | -0.004 | 0.018 | 0.018 | 0.173 | 0.12 | 0.06 | 0.07 | 0 | 0.11 | 0.04 |
| 8 | 11611743 | rs804282 | -0.557 | -1.248 | -0.79 | -2.276 | 0.002 | 0.023 | 0.023 | 0 | 0.48 | 0.56 | 0.5 | 0.28 | 0.52 | 0.52 |
| 8 | 11612315 | rs13264774 | 1.075 | -1.246 | -0.822 | -2.101 | -0.027 | 0.055 | 0.055 | 0 | 0.17 | 0.25 | 0.22 | 0.62 | 0.15 | 0.38 |
| 8 | 11612381 | rs13273672 | 0.446 | -0.929 | -0.883 | -2.219 | -0.002 | 0.058 | 0.058 | 0 | 0.23 | 0.37 | 0.34 | 0.63 | 0.3 | 0.41 |
| 8 | 11612698 | rs804280 | -0.721 | -1.141 | -0.855 | -2.373 | -0.003 | -0.06 | -0.06 | 0.011 | 0.57 | 0.6 | 0.69 | 0.99 | 0.56 | 0.87 |
| 8 | 11616338 | rs867858 | 0.507 | -0.971 | -1.105 | -1.817 | 0.008 | -0.024 | -0.024 | 0 | 0.3 | 0.24 | 0.3 | 0.58 | 0.32 | 0.39 |
| 8 | 11616836 | rs804290 | 1.831 | -0.88 | -1.476 | -2.198 | -0.078 | 0.141 | 0.141 | 1.865 | 0.23 | 0.02 | 0.15 | 0 | 0.25 | 0.21 |
| 8 | 11617142 | rs11785481 | 2.072 | -0.721 | -1.2 | -1.892 | -0.056 | 0.047 | 0.047 | 0 | 0.11 | 0.01 | 0.12 | 0.01 | 0.16 | 0.03 |
| 8 | 11617240 | rs12458 | -0.295 | -0.91 | -0.809 | -1.837 | -0.011 | 0.005 | 0.005 | 0 | 0.39 | 0.39 | 0.32 | 0.55 | 0.32 | 0.39 |
| 8 | 11618107 | rs9329248 | 0.818 | -0.821 | -0.503 | -1.573 | -0.076 | 0.162 | 0.162 | 0 | 0.22 | 0.5 | 0.2 | 0.56 | 0.16 | 0.37 |
| 8 | 11620315 | rs2686206 | 0.025 | -0.518 | -1.099 | -1.434 | -0.004 | 0.005 | 0.005 | 0 | 0.45 | 0.28 | 0.43 | 0.62 | 0.5 | 0.65 |
| 8 | 11622865 | rs17153785 | 1.144 | -0.398 | -0.917 | -0.748 | 0.001 | -0.046 | -0.046 | 0 | 0.12 | 0.06 | 0.2 | 0.22 | 0.09 | 0.1 |
| 8 | 11623212 | rs10101195 | 0.098 | -0.58 | -0.509 | -0.844 | -0.037 | -0.058 | -0.058 | 0.002 | 0.32 | 0.53 | 0.27 | 0.17 | 0.27 | 0.15 |
| 8 | 11628162 | rs2010628 | 0.891 | -0.335 | -1.018 | -1.183 | -0.004 | 0.077 | 0.077 | 0 | 0.23 | 0.08 | 0.14 | 0.01 | 0.24 | 0.15 |
| 8 | 11629241 | rs804267 | 0.123 | -0.351 | -0.437 | -1.275 | -0.003 | -0.071 | -0.071 | 0 | 0.57 | 0.41 | 0.75 | 0.97 | 0.65 | 0.79 |
| 8 | 11635946 | rs4840583 | 0.053 | -0.128 | -0.857 | -0.755 | 0.018 | 0.004 | 0.004 | 0.152 | 0.49 | 0.22 | 0.48 | 0.48 | 0.54 | 0.47 |
| 8 | 11636017 | rs4840584 | 1.476 | -0.169 | -1.267 | -0.911 | -0.006 | 0.067 | 0.067 | 0 | 0.13 | 0.02 | 0.04 | 0 | 0.08 | 0.03 |
| 8 | 11636862 | rs804256 | 0.798 | 0.092 | -1.505 | -0.993 | -0.096 | -0.055 | -0.055 | 0 | 0.23 | 0.06 | 0.37 | 0.26 | 0.33 | 0.38 |
| 8 | 11636884 | rs8191604 | 0.966 | 0.148 | -0.882 | -0.964 | 0.005 | 0.016 | 0.016 | 0.012 | 0.28 | 0.25 | 0.18 | 0.01 | 0.27 | 0.15 |
| 8 | 11639991 | rs8180912 | 1.277 | 0.259 | -0.995 | -0.596 | -0.004 | -0.023 | -0.023 | 0 | 0.14 | 0.08 | 0.28 | 0.47 | 0.19 | 0.23 |
| 8 | 11641871 | rs6982453 | 0.217 | 0.256 | -0.818 | -0.98 | 0.003 | -0.018 | -0.018 | 0.216 | 0.42 | 0.48 | 0.49 | 0.76 | 0.48 | 0.45 |
| 8 | 11643803 | rs1534862 | 0.825 | 0.229 | -0.919 | -1.146 | -0.008 | -0.004 | -0.004 | 1.341 | 0.3 | 0.35 | 0.19 | 0.25 | 0.26 | 0.16 |
| 8 | 11643915 | rs804292 | -1.451 | 0.273 | -0.793 | -1.326 | 0.007 | -0.086 | -0.086 | 0 | 0.76 | 0.76 | 0.84 | 1 | 0.75 | 0.81 |
| 8 | 11646934 | rs904015 | 0.832 | 0.493 | -0.712 | -0.846 | 0.009 | 0.035 | 0.035 | 0.11 | 0.41 | 0.3 | 0.28 | 0.25 | 0.38 | 0.19 |
| 8 | 11648789 | rs2740444 | 1.237 | 0.533 | -0.766 | -0.932 | -0.007 | -0.029 | -0.029 | 0 | 0.19 | 0.1 | 0.29 | 0.23 | 0.18 | 0.31 |
| 8 | 11650475 | rs7001819 | 0.442 | 0.324 | -0.442 | -1.185 | 0.005 | 0.015 | 0.015 | 0.015 | 0.44 | 0.34 | 0.32 | 0.32 | 0.42 | 0.23 |
| 8 | 11651553 | rs2645444 | -0.573 | 0.447 | -0.46 | -1.363 | 0.002 | -0.073 | -0.073 | 0 | 0.79 | 0.77 | 0.89 | 1 | 0.81 | 0.84 |
| 8 | 11662885 | rs10098874 | 0.678 | -0.009 | -0.106 | -1.035 | -0.001 | -0.155 | -0.155 | 0.006 | 0.22 | 0.49 | 0.13 | 0.01 | 0.18 | 0.11 |
| 8 | 11665518 | rs1736067 | -0.385 | -0.264 | -0.455 | -0.816 | -0.011 | 0.004 | 0.004 | 0 | 0.7 | 0.67 | 0.77 | 0.73 | 0.73 | 0.84 |
| 8 | 11667179 | rs8417 | 1.094 | 0.186 | -0.69 | -0.023 | -0.002 | -0.046 | -0.046 | 0 | 0.15 | 0.09 | 0.24 | 0.45 | 0.19 | 0.26 |
| 8 | 11670932 | rs6991234 | 0.083 | 0.013 | -0.428 | -0.312 | -0.008 | -0.104 | -0.104 | 0.009 | 0.32 | 0.39 | 0.17 | 0.02 | 0.27 | 0.16 |
| 8 | 11671041 | rs1736058 | 0.586 | 0.353 | -0.694 | 0.758 | 0.007 | -0.055 | -0.055 | 0 | 0.16 | 0.09 | 0.23 | 0.45 | 0.18 | 0.27 |
| 8 | 11682566 | rs2645427 | -0.706 | 0.261 | -0.637 | 0.324 | 0.008 | -0.017 | -0.017 | 0 | 0.83 | 0.87 | 0.71 | 0.23 | 0.79 | 0.7 |
| 8 | 11688148 | rs4841600 | -0.061 | 0.295 | -0.337 | -0.712 | -0.006 | 0.013 | 0.013 | 0.332 | 0.54 | 0.38 | 0.66 | 0.99 | 0.53 | 0.75 |
| 8 | 11694604 | rs6995787 | 0.996 | 0.209 | -0.329 | -0.215 | 0.007 | -0.033 | -0.033 | 0.738 | 0.17 | 0.11 | 0.23 | 0.5 | 0.19 | 0.23 |
| 8 | 11696352 | rs3258 | 0.844 | 0.077 | -0.079 | -0.386 | -0.001 | 0.01 | 0.01 | 0.541 | 0.36 | 0.38 | 0.42 | 0.51 | 0.33 | 0.46 |
| 8 | 11696492 | rs8944 | 1.026 | 0.14 | -0.212 | -0.345 | 0.004 | -0.082 | -0.082 | 0 | 0.17 | 0.07 | 0.23 | 0.49 | 0.19 | 0.24 |
| 8 | 11698747 | rs1296028 | 0.556 | 0.201 | -0.237 | -0.628 | -0.042 | 0.048 | 0.048 | 0 | 0.22 | 0.07 | 0.19 | 0.05 | 0.27 | 0.16 |
| 8 | 11699730 | rs4841601 | 1.059 | 0.055 | 0.285 | -0.81 | 0 | -0.003 | -0.003 | 0.148 | 0.37 | 0.32 | 0.42 | 0.51 | 0.34 | 0.47 |
| 8 | 11700101 | rs6731 | 1.134 | 0.061 | 0.286 | -0.826 | -0.001 | 0.006 | 0.006 | 1.055 | 0.36 | 0.35 | 0.42 | 0.5 | 0.34 | 0.47 |
| 8 | 11701198 | rs12898 | -1.039 | -0.002 | 0.325 | -0.863 | 0.008 | -0.024 | -0.024 | 0.702 | 0.59 | 0.31 | 0.56 | 0.48 | 0.65 | 0.47 |
| 8 | 11701933 | rs4839 | 0.614 | 0.134 | 0.248 | -0.757 | -0.034 | 0.069 | 0.069 | 0 | 0.21 | 0.07 | 0.18 | 0 | 0.26 | 0.1 |
| 8 | 11711059 | rs2272767 | -0.019 | 0.182 | -0.046 | -0.606 | -0.009 | -0.049 | -0.049 | 0 | 0.32 | 0.17 | 0.37 | 0.47 | 0.37 | 0.34 |
| 8 | 11713852 | rs17154027 | -0.312 | 0.13 | 0.019 | -0.711 | -0.002 | -0.035 | -0.035 | 0.336 | 0.35 | 0.26 | 0.39 | 0.51 | 0.38 | 0.38 |
| 8 | 11718528 | rs1293288 | -0.789 | 0.094 | 0.197 | -0.804 | 0.001 | -0.001 | -0.001 | 0 | 0.54 | 0.53 | 0.51 | 0.52 | 0.57 | 0.55 |
| 8 | 11722046 | rs2645415 | -0.432 | 0.245 | 0.081 | -0.773 | -0.001 | 0.054 | 0.054 | 0 | 0.29 | 0.43 | 0.36 | 0.48 | 0.31 | 0.37 |
| 8 | 11726817 | rs1296023 | -0.04 | 0.424 | -0.289 | -0.809 | 0.009 | 0.047 | 0.047 | 0.623 | 0.24 | 0.31 | 0.33 | 0.5 | 0.3 | 0.39 |

| | | | | | | | | | | | | | | | | |
|---|---|---|---|---|---|---|---|---|---|---|---|---|---|---|---|---|
| 8 | 11734701 | rs1619200 | 0.334 | 0.452 | -0.851 | -0.817 | -0.036 | -0.087 | -0.087 | 0.289 | 0.22 | 0.11 | 0.3 | 0.49 | 0.28 | 0.35 |
| 8 | 11735939 | rs880632 | -0.113 | 0.359 | -1.046 | -1.247 | 0.011 | 0.065 | 0.065 | 0.109 | 0.33 | 0.22 | 0.18 | 0.03 | 0.3 | 0.21 |
| 8 | 11745820 | rs1630828 | -0.788 | 0.43 | -0.979 | -0.893 | -0.016 | -0.03 | -0.03 | 0 | 0.61 | 0.77 | 0.53 | 0.54 | 0.56 | 0.55 |
| 8 | 11746122 | rs11783694 | -0.654 | 0.67 | -1.136 | -0.539 | -0.008 | -0.011 | -0.011 | 0.341 | 0.27 | 0.2 | 0.34 | 0.69 | 0.3 | 0.48 |
| 8 | 11746358 | rs2645408 | -0.418 | 0.489 | -0.539 | -0.89 | -0.005 | 0.062 | 0.062 | 0.023 | 0.31 | 0.5 | 0.34 | 0.5 | 0.32 | 0.4 |
| 8 | 11748645 | rs1692793 | -0.251 | 0.435 | -0.934 | -0.104 | 0.012 | 0.001 | 0.001 | 0 | 0.49 | 0.44 | 0.5 | 0.5 | 0.59 | 0.39 |
| 8 | 12633235 | rs6530980 | -0.006 | -0.456 | -1.178 | 0.192 | -0.057 | 0.034 | 0.034 | 0 | 0.86 | 0.71 | 0.82 | 0.8 | 0.96 | 0.64 |
| 8 | 12661835 | rs12548697 | -0.155 | -0.495 | -0.608 | 0.223 | 0.031 | 0.037 | 0.037 | 0 | 0.46 | 0.31 | 0.62 | 0.6 | 0.57 | 0.42 |
| 8 | 12673698 | rs6980603 | -0.684 | -0.803 | 0.019 | 0.013 | 0.016 | -0.017 | -0.017 | 0 | 0.2 | 0.3 | 0.1 | 0.16 | 0.22 | 0.23 |
| 8 | 12679589 | rs4240185 | 0.185 | -1.01 | 0.153 | -0.019 | 0.016 | 0.007 | 0.007 | 0 | 0.45 | 0.37 | 0.64 | 0.54 | 0.57 | 0.39 |
| 8 | 12680107 | rs17830696 | 0.025 | -0.937 | 0.286 | 0.013 | 0 | 0 | 0 | 0 | 0.15 | 0.15 | 0.09 | 0.17 | 0.18 | 0.19 |
| 8 | 12681715 | rs10503429 | 0.114 | -0.929 | 0.253 | 0.015 | -0.01 | -0.004 | -0.004 | 0 | 0.15 | 0.1 | 0.09 | 0.18 | 0.19 | 0.2 |
| 8 | 12685783 | rs6994104 | 1.397 | -0.733 | -0.069 | 0.24 | 0.001 | 0.003 | 0.003 | 0 | 0.08 | 0.09 | 0.04 | 0 | 0.07 | 0.07 |
| 8 | 12687498 | rs4831834 | 0.304 | -0.779 | -0.036 | 0.275 | -0.017 | -0.01 | -0.01 | 0 | 0.37 | 0.28 | 0.6 | 0.52 | 0.49 | 0.31 |
| 8 | 12688558 | rs11203841 | -0.447 | -0.85 | 0.048 | 0.256 | 0.002 | -0.007 | -0.007 | 0.109 | 0.25 | 0.28 | 0.13 | 0.19 | 0.27 | 0.27 |
| 8 | 12690464 | rs4609193 | 0.614 | -1.175 | 0.222 | 0.255 | -0.001 | -0.006 | -0.006 | 0 | 0.71 | 0.67 | 0.86 | 0.78 | 0.72 | 0.71 |
| 8 | 12692084 | rs2068281 | 0.951 | -1.18 | 0.293 | 0.293 | -0.002 | 0.015 | 0.015 | 0 | 0.14 | 0.17 | 0.26 | 0.27 | 0.11 | 0.23 |
| 8 | 12692882 | rs7014965 | 0.48 | -1.177 | 0.276 | 0.206 | -0.008 | -0.054 | -0.054 | 0 | 0.11 | 0.19 | 0.05 | 0.02 | 0.09 | 0.13 |
| 8 | 12695190 | rs6981317 | -0.868 | -1.448 | 0.618 | 0.1 | 0.015 | -0.006 | -0.006 | 0 | 0.55 | 0.58 | 0.5 | 0.51 | 0.68 | 0.42 |
| 8 | 12696125 | rs4361764 | 0.046 | -1.329 | 0.729 | 0.129 | -0.004 | 0.012 | 0.012 | 0 | 0.57 | 0.38 | 0.57 | 0.65 | 0.48 | 0.73 |
| 8 | 12697557 | rs4602886 | -0.412 | -0.978 | 1.047 | -0.01 | -0.036 | -0.083 | -0.083 | 7.543 | 0.16 | 0.28 | 0.07 | 0 | 0.07 | 0.11 |
| 8 | 12698174 | rs11774014 | 0.607 | -0.87 | 0.984 | 0.01 | 0 | -0.015 | -0.015 | 0 | 0.12 | 0.08 | 0.06 | 0.16 | 0.11 | 0.12 |
| 8 | 12698470 | rs6985529 | -0.123 | -0.737 | 1.081 | 0.096 | -0.001 | -0.005 | -0.005 | 0 | 0.93 | 0.85 | 0.98 | 0.95 | 0.94 | 0.92 |
| 8 | 12700069 | rs17120657 | 1.003 | -0.769 | 1.061 | 0.101 | 0.001 | -0.002 | -0.002 | 0 | 0.16 | 0.16 | 0.26 | 0.25 | 0.11 | 0.23 |
| 8 | 12700974 | rs10102302 | 0.738 | -0.777 | 1.11 | 0.067 | -0.025 | -0.029 | -0.029 | 1.423 | 0.75 | 0.59 | 0.86 | 0.83 | 0.82 | 0.77 |
| 8 | 12701140 | rs13282410 | 1.465 | -0.773 | 0.972 | 0.069 | -0.028 | -0.037 | -0.037 | 0 | 0.68 | 0.44 | 0.84 | 0.78 | 0.76 | 0.7 |
| 8 | 12701924 | rs12548516 | 0.177 | -0.425 | 0.501 | 0.23 | -0.003 | 0.009 | 0.009 | 0 | 0.84 | 0.78 | 0.91 | 0.79 | 0.84 | 0.8 |
| 8 | 12703263 | rs4612342 | 0.641 | -0.434 | 0.051 | 0.076 | -0.001 | -0.018 | -0.018 | 0.664 | 0.09 | 0.11 | 0.04 | 0 | 0.07 | 0.07 |
| 8 | 12703642 | rs10113673 | -0.174 | -0.001 | 0.119 | 0.504 | -0.003 | -0.025 | -0.025 | 0 | 0.26 | 0.24 | 0.34 | 0.44 | 0.26 | 0.38 |
| 8 | 12706130 | rs6984565 | 0.546 | -0.094 | 0.425 | 0.179 | 0 | 0.006 | 0.006 | 0.277 | 0.94 | 0.78 | 0.98 | 0.96 | 0.97 | 0.97 |
| 8 | 12706446 | rs4527872 | -0.704 | -0.382 | 0.266 | -0.094 | -0.002 | -0.015 | -0.015 | 0 | 0.65 | 0.67 | 0.83 | 0.86 | 0.73 | 0.55 |
| 8 | 12707900 | rs12677284 | 0.046 | -0.379 | 0.085 | -0.091 | 0.007 | 0 | 0 | 0 | 0.54 | 0.43 | 0.77 | 0.81 | 0.7 | 0.49 |
| 8 | 12710800 | rs13273810 | 0.355 | 0.039 | 0.063 | 0.352 | -0.002 | 0.006 | 0.006 | 0 | 0.79 | 0.73 | 0.82 | 0.7 | 0.81 | 0.76 |
| 8 | 12710960 | rs4574847 | 0.089 | 0.639 | -0.091 | 0.689 | -0.002 | -0.002 | -0.002 | 0.624 | 0.46 | 0.46 | 0.66 | 0.55 | 0.55 | 0.36 |
| 8 | 12711863 | rs4831837 | -0.158 | 0.738 | -0.79 | 0.693 | 0.012 | 0.012 | 0.012 | 3.632 | 0.49 | 0.4 | 0.66 | 0.6 | 0.58 | 0.39 |
| 8 | 12714790 | rs17770189 | -1.233 | 1.754 | -1.024 | 0.886 | -0.044 | 0.004 | 0.004 | 0 | 0.12 | 0.04 | 0.27 | 0.14 | 0.28 | 0.16 |
| 8 | 12715139 | rs17120947 | -1.303 | 0.783 | -0.592 | 1.532 | -0.011 | 0.053 | 0.053 | 0 | 0.11 | 0.2 | 0.3 | 0.3 | 0.07 | 0.1 |
| 8 | 12715744 | rs17120965 | -1.129 | 0.629 | -1.169 | 1.74 | -0.006 | -0.1 | -0.1 | 0 | 0.16 | 0.04 | 0.17 | 0.47 | 0.14 | 0.12 |
| 8 | 12717076 | rs10091521 | -0.88 | 0.186 | -0.132 | 1.817 | 0.015 | 0.367 | 0.367 | 0.423 | 0.06 | 0.33 | 0.23 | 0.3 | 0.02 | 0.08 |
| 8 | 12718090 | rs6997599 | 0.637 | 0.395 | -1.609 | 0.578 | -0.156 | 0.138 | 0.138 | 0 | 0.32 | 0.05 | 0.27 | 0.11 | 0.45 | 0.41 |
| 8 | 12718500 | rs17832679 | 0.269 | -0.156 | -1.354 | 0.488 | 0.021 | -0.006 | -0.006 | 0 | 0.37 | 0.16 | 0.43 | 0.32 | 0.35 | 0.19 |
| 8 | 12723282 | rs17121134 | -0.147 | -0.327 | -0.904 | 0.131 | -0.004 | -0.038 | -0.038 | 0.02 | 0.14 | 0.16 | 0.06 | 0 | 0.1 | 0.07 |
| 8 | 12725720 | rs2956768 | -0.077 | -0.285 | -0.865 | 0.002 | 0 | -0.014 | -0.014 | 0 | 0.09 | 0.09 | 0.06 | 0 | 0.1 | 0.07 |
| 8 | 12726547 | rs4501576 | 0.124 | -0.35 | -0.79 | 0.801 | -0.005 | 0.118 | 0.118 | 0 | 0.09 | 0.12 | 0.21 | 0.5 | 0.06 | 0.12 |
| 8 | 12728419 | rs7823534 | 1.203 | -0.681 | -0.486 | -0.038 | 0.095 | 0.144 | 0.144 | 0.161 | 0.54 | 0.22 | 0.41 | 0.13 | 0.71 | 0.58 |
| 8 | 12728705 | rs7828391 | 1.206 | -0.692 | -0.491 | -0.044 | 0.095 | 0.145 | 0.145 | 0.203 | 0.54 | 0.21 | 0.41 | 0.13 | 0.71 | 0.57 |
| 8 | 12732754 | rs6991079 | 1.234 | 0.083 | -0.057 | -0.222 | 0.017 | 0.086 | 0.086 | 0 | 0.54 | 0.32 | 0.36 | 0.1 | 0.63 | 0.55 |
| 8 | 12733077 | rs2977173 | 0.142 | 0.117 | 0.379 | 0.226 | 0 | 0.097 | 0.097 | 0 | 0.92 | 0.68 | 0.86 | 0.81 | 0.95 | 0.93 |
| 8 | 12733321 | rs2977171 | -0.33 | 0.108 | -0.086 | 0.098 | -0.08 | 0.074 | 0.074 | 0 | 0.13 | 0.01 | 0.13 | 0.01 | 0.21 | 0.13 |
| 8 | 12754267 | rs17121552 | -0.122 | -0.137 | 0.31 | 0.878 | -0.006 | 0.052 | 0.052 | 0 | 0.12 | 0.13 | 0.27 | 0.49 | 0.05 | 0.1 |
| 8 | 12769518 | rs2947381 | 0.956 | -0.403 | 0.579 | 0.418 | -0.08 | 0.101 | 0.101 | 0 | 0.78 | 0.44 | 0.59 | 0.34 | 0.91 | 0.87 |
| 8 | 12770006 | rs2017656 | 0.576 | -0.105 | 0.645 | 0.497 | -0.04 | 0.109 | 0.109 | 0 | 0.81 | 0.53 | 0.6 | 0.34 | 0.89 | 0.87 |
| 8 | 12772901 | rs2977124 | -1.075 | 0.01 | 0.51 | 0.269 | 0.017 | -0.01 | -0.01 | 3.92 | 0.3 | 0.43 | 0.23 | 0.23 | 0.3 | 0.32 |
| 8 | 12773413 | rs2956755 | -1.924 | 0.7 | -0.336 | -0.18 | -0.059 | 0.29 | 0.29 | 0.612 | 0.53 | 0.92 | 0.7 | 0.9 | 0.48 | 0.51 |
| 8 | 12773958 | rs1915480 | -1.15 | 1.225 | 0.479 | -0.056 | 0.015 | 0.023 | 0.023 | 0 | 0.28 | 0.48 | 0.53 | 0.67 | 0.36 | 0.23 |
| 8 | 12780892 | rs2977139 | -0.865 | 1.453 | 0.705 | 0.595 | 0.002 | 0.01 | 0.01 | 0 | 0.23 | 0.29 | 0.44 | 0.48 | 0.34 | 0.16 |
| 8 | 12786594 | rs2947384 | -0.829 | 1.451 | 0.721 | 0.662 | 0.004 | 0.016 | 0.016 | 0 | 0.23 | 0.3 | 0.44 | 0.53 | 0.35 | 0.16 |
| 8 | 12792352 | rs10749624 | 1.472 | 1.367 | 1.124 | 0.358 | 0.009 | -0.026 | -0.026 | 0 | 0.6 | 0.54 | 0.45 | 0.28 | 0.55 | 0.57 |
| 8 | 12804101 | rs2977097 | -0.619 | 1.235 | 0.948 | 0.243 | 0.002 | -0.026 | -0.026 | 0 | 0.27 | 0.22 | 0.44 | 0.54 | 0.32 | 0.19 |
| 8 | 12807305 | rs2946511 | -1.029 | 1.112 | 1.218 | 0.16 | -0.012 | 0.003 | 0.003 | 0 | 0.3 | 0.42 | 0.53 | 0.7 | 0.32 | 0.21 |
| 8 | 12808110 | rs7824233 | -0.416 | 1.152 | 1.699 | 0.157 | 0.033 | 0.004 | 0.004 | 1.216 | 0.09 | 0.39 | 0.05 | 0.04 | 0.04 | 0.05 |
| 8 | 12811152 | rs2946505 | 1.4 | 1.325 | 1.441 | 0.169 | 0.031 | 0.14 | 0.14 | 0 | 0.8 | 0.56 | 0.74 | 0.66 | 0.84 | 0.66 |
| 8 | 12817506 | rs2977078 | -1.144 | 1.421 | 1.355 | -0.416 | -0.004 | -0.001 | -0.001 | 0 | 0.32 | 0.37 | 0.58 | 0.7 | 0.34 | 0.29 |
| 8 | 12824893 | rs2203834 | 0.568 | 2.386 | 1.702 | -0.024 | 0.01 | 0.026 | 0.026 | 0 | 0.92 | 0.91 | 0.9 | 0.8 | 0.87 | 0.73 |
| 8 | 12826813 | rs2954170 | -0.513 | 2.664 | 1.822 | 0.024 | 0.02 | 0.046 | 0.046 | 0.113 | 0.14 | 0.22 | 0.46 | 0.64 | 0.29 | 0.15 |
| 8 | 12828161 | rs2977102 | -1.534 | 2.799 | 2.039 | 0.063 | 0.037 | 0.077 | 0.077 | 0.477 | 0.19 | 0.35 | 0.49 | 0.63 | 0.35 | 0.17 |
| 8 | 12829038 | rs2977098 | 0.336 | 2.858 | 1.963 | 0.064 | -0.003 | 0.034 | 0.034 | 0.012 | 0.1 | 0.12 | 0.44 | 0.62 | 0.23 | 0.14 |
| 8 | 12832522 | rs2460349 | -1.638 | 2.829 | 2.005 | 0.043 | 0.028 | 0.066 | 0.066 | 0.122 | 0.19 | 0.32 | 0.48 | 0.63 | 0.32 | 0.17 |
| 8 | 12832636 | rs2460901 | -1.584 | 2.825 | 2.022 | 0.041 | 0.025 | 0.059 | 0.059 | 0.142 | 0.18 | 0.3 | 0.48 | 0.63 | 0.32 | 0.17 |

| | | | | | | | | | | | | | | | | |
|---|---|---|---|---|---|---|---|---|---|---|---|---|---|---|---|---|
| 8 | 12834320 | rs11203996 | 2.157 | 2.727 | 1.737 | -0.336 | -0.022 | 0.036 | 0.036 | 0.222 | 0.6 | 0.34 | 0.68 | 0.76 | 0.58 | 0.52 |
| 8 | 12837011 | rs11786846 | -0.86 | 2.817 | 1.914 | -0.014 | 0.015 | 0.052 | 0.052 | 0.06 | 0.15 | 0.23 | 0.46 | 0.62 | 0.29 | 0.15 |
| 8 | 12846907 | rs6531017 | 1.893 | 2.596 | 2.244 | -0.188 | 0.049 | 0.024 | 0.024 | 0 | 0.79 | 0.48 | 0.79 | 0.8 | 0.76 | 0.72 |
| 8 | 12847210 | rs7821427 | 2.217 | 2.644 | 2.337 | -0.237 | 0.002 | -0.002 | -0.002 | 0 | 0.73 | 0.38 | 0.75 | 0.79 | 0.72 | 0.69 |
| 8 | 12847873 | rs2460341 | 0.001 | 2.494 | 2.397 | -0.218 | 0.002 | 0.013 | 0.013 | 0 | 0.89 | 0.88 | 0.85 | 0.81 | 0.87 | 0.76 |
| 8 | 12850442 | rs2954179 | 0.759 | 2.482 | 2.56 | -0.206 | 0.019 | 0.043 | 0.043 | 0 | 0.86 | 0.74 | 0.83 | 0.8 | 0.85 | 0.76 |
| 8 | 12850918 | rs2460911 | 1.75 | 2.479 | 2.746 | -0.223 | 0.028 | 0.028 | 0.028 | 0 | 0.79 | 0.56 | 0.79 | 0.79 | 0.8 | 0.72 |
| 8 | 12851669 | rs2954184 | -1.43 | 2.478 | 2.721 | -0.221 | 0.009 | -0.067 | -0.067 | 0.287 | 0.11 | 0.26 | 0.07 | 0.01 | 0.08 | 0.07 |
| 8 | 12851817 | rs10888160 | 0.09 | 2.655 | 2.529 | -0.129 | -0.013 | 0.005 | 0.005 | 0 | 0.11 | 0.11 | 0.42 | 0.62 | 0.28 | 0.16 |
| 8 | 12853625 | rs12375378 | 0.846 | 2.256 | 2.58 | -0.234 | -0.015 | 0.009 | 0.009 | 0 | 0.06 | 0 | 0.07 | 0.02 | 0.06 | 0.1 |
| 8 | 12854628 | rs7841451 | 2.581 | 2.281 | 2.724 | -0.249 | 0.002 | -0.002 | -0.002 | 0 | 0.68 | 0.37 | 0.69 | 0.76 | 0.7 | 0.59 |
| 8 | 12855473 | rs2460893 | 0.993 | 2.285 | 2.729 | -0.213 | 0.04 | 0.058 | 0.058 | 0 | 0.85 | 0.69 | 0.83 | 0.79 | 0.84 | 0.75 |
| 8 | 12856457 | rs7846534 | -0.897 | 2.215 | 2.475 | -0.278 | 0 | -0.017 | -0.017 | 0 | 0.12 | 0.12 | 0.12 | 0.04 | 0.11 | 0.14 |
| 8 | 12857756 | rs10503430 | -0.87 | 2.215 | 2.415 | -0.278 | 0 | -0.009 | -0.009 | 0 | 0.12 | 0.11 | 0.12 | 0.04 | 0.11 | 0.14 |
| 8 | 12857798 | rs10113105 | -1.166 | 2.217 | 2.449 | -0.277 | 0.003 | -0.054 | -0.054 | 0.008 | 0.13 | 0.21 | 0.12 | 0.04 | 0.11 | 0.14 |
| 8 | 12858916 | rs17784878 | 0.124 | 2.218 | 2.137 | -0.206 | 0 | -0.022 | -0.022 | 0 | 0.06 | 0.08 | 0.04 | 0.01 | 0.05 | 0.04 |
| 8 | 12860528 | rs2466270 | 1.31 | 2.267 | 2.268 | -0.136 | 0.035 | 0.018 | 0.018 | 0 | 0.8 | 0.67 | 0.81 | 0.8 | 0.77 | 0.75 |
| 8 | 12866004 | rs2466273 | -1.54 | 2.102 | 2.139 | -0.203 | 0.061 | -0.039 | -0.039 | 0 | 0.09 | 0.37 | 0.06 | 0 | 0.08 | 0.03 |
| 8 | 12868122 | rs13278029 | -1.417 | 2.186 | 1.969 | -0.115 | 0.011 | 0.017 | 0.017 | 0.236 | 0.21 | 0.29 | 0.29 | 0.22 | 0.2 | 0.34 |
| 8 | 12870928 | rs10091369 | -1.912 | 2.056 | 1.801 | -0.253 | 0.002 | -0.003 | -0.003 | 0 | 0.23 | 0.3 | 0.21 | 0.2 | 0.22 | 0.29 |
| 8 | 12873414 | rs2466247 | -0.405 | 1.937 | 1.406 | -0.064 | 0.003 | -0.027 | -0.027 | 0 | 0.06 | 0.11 | 0.03 | 0.01 | 0.05 | 0.05 |
| 8 | 12874915 | rs11779957 | -0.047 | 2.188 | 1.286 | -0.055 | 0.002 | 0.024 | 0.024 | 0.011 | 0.12 | 0.13 | 0.44 | 0.64 | 0.29 | 0.14 |
| 8 | 12876558 | rs10109178 | 1.828 | 1.295 | 1.638 | -0.381 | -0.001 | -0.001 | -0.001 | 0 | 0.71 | 0.32 | 0.71 | 0.78 | 0.77 | 0.59 |
| 8 | 12878843 | rs505480 | -0.427 | 1.401 | 1.495 | -0.328 | 0.001 | -0.019 | -0.019 | 0.103 | 0.06 | 0.08 | 0.03 | 0.01 | 0.05 | 0.05 |
| 8 | 12879062 | rs503550 | 1 | 1.374 | 1.617 | -0.271 | 0.008 | 0.025 | 0.025 | 0.372 | 0.78 | 0.59 | 0.76 | 0.79 | 0.82 | 0.65 |
| 8 | 12879334 | rs608909 | 1.029 | 1.373 | 1.616 | -0.272 | 0.007 | 0.024 | 0.024 | 0.394 | 0.78 | 0.58 | 0.76 | 0.79 | 0.82 | 0.65 |
| 8 | 12879539 | rs608052 | 1.029 | 1.373 | 1.625 | -0.272 | 0.008 | 0.025 | 0.025 | 0.394 | 0.78 | 0.56 | 0.76 | 0.79 | 0.82 | 0.65 |
| 8 | 12879777 | rs475181 | 1.073 | 1.38 | 1.58 | -0.268 | 0.009 | 0.027 | 0.027 | 0.554 | 0.78 | 0.56 | 0.76 | 0.79 | 0.82 | 0.65 |
| 8 | 12880105 | rs472347 | 0.982 | 1.401 | 1.454 | -0.258 | 0.01 | 0.026 | 0.026 | 0.326 | 0.79 | 0.59 | 0.76 | 0.79 | 0.82 | 0.65 |
| 8 | 12880521 | rs579024 | 0.999 | 1.403 | 1.456 | -0.256 | 0.008 | 0.025 | 0.025 | 0.372 | 0.78 | 0.59 | 0.76 | 0.79 | 0.82 | 0.65 |
| 8 | 12884804 | rs579253 | -0.658 | 1.134 | 0.76 | -0.574 | 0 | -0.029 | -0.029 | 0 | 0.13 | 0.14 | 0.06 | 0.01 | 0.09 | 0.1 |
| 8 | 12890377 | rs709829 | -0.9 | 0.778 | 0.962 | -0.487 | -0.027 | -0.038 | -0.038 | 0 | 0.17 | 0.27 | 0.15 | 0.06 | 0.08 | 0.28 |
| 8 | 12890974 | rs7816868 | 1.204 | 0.67 | 1.15 | -0.526 | -0.052 | -0.07 | -0.07 | 0 | 0.78 | 0.6 | 0.82 | 0.91 | 0.9 | 0.69 |
| 8 | 12895780 | rs7840927 | 1.345 | 0.629 | 1.179 | -0.575 | -0.034 | -0.054 | -0.054 | 0 | 0.72 | 0.42 | 0.77 | 0.85 | 0.85 | 0.64 |
| 8 | 12898184 | rs529983 | -1.427 | 0.805 | 1.002 | -0.456 | 0.02 | 0.018 | 0.018 | 0 | 0.05 | 0.16 | 0.05 | 0.08 | 0.04 | 0.09 |
| 8 | 12898806 | rs713536 | 1.652 | 1.124 | 0.102 | -0.291 | -0.03 | 0.044 | 0.044 | 0 | 0.49 | 0.05 | 0.32 | 0.3 | 0.47 | 0.4 |
| 8 | 12899416 | rs601101 | 0.405 | 0.86 | 0.663 | -0.316 | 0.003 | 0.003 | 0.003 | 0.209 | 0.88 | 0.88 | 0.9 | 0.94 | 0.92 | 0.88 |
| 8 | 12900843 | rs1474140 | -0.755 | 0.781 | 0.423 | -0.425 | -0.075 | -0.147 | -0.147 | 0 | 0.27 | 0.05 | 0.47 | 0.53 | 0.33 | 0.28 |
| 8 | 12901247 | rs544302 | -0.04 | 0.773 | 0.53 | -0.53 | 0 | -0.009 | -0.009 | 0 | 0.89 | 0.91 | 0.93 | 0.96 | 0.92 | 0.89 |
| 8 | 12908074 | rs542548 | -1.734 | 0.592 | 1.855 | -0.597 | -0.058 | -0.087 | -0.087 | 0 | 0.14 | 0.52 | 0.08 | 0.03 | 0.04 | 0.09 |
| 8 | 12908899 | rs643378 | -1.768 | 0.633 | 1.519 | -0.52 | -0.009 | -0.069 | -0.069 | 0 | 0.4 | 0.85 | 0.6 | 0.61 | 0.4 | 0.44 |
| 8 | 12913259 | rs2460362 | -2.408 | 0.627 | 1.716 | -0.667 | -0.037 | 0.003 | 0.003 | 1.687 | 0.08 | 0.68 | 0.08 | 0.08 | 0.01 | 0.12 |
| 8 | 12916186 | rs2460365 | 0.295 | 0.605 | 1.701 | -0.701 | -0.017 | -0.025 | -0.025 | 0.03 | 0.87 | 0.74 | 0.92 | 0.9 | 0.92 | 0.82 |
| 8 | 12917220 | rs606181 | 0.244 | 0.612 | 1.698 | -0.697 | -0.016 | -0.023 | -0.023 | 0 | 0.87 | 0.74 | 0.92 | 0.9 | 0.93 | 0.82 |
| 8 | 12918912 | rs634228 | 0.22 | 0.613 | 1.69 | -0.697 | -0.015 | -0.023 | -0.023 | 0 | 0.88 | 0.75 | 0.92 | 0.9 | 0.93 | 0.82 |
| 8 | 12919518 | rs2119910 | 1.952 | 0.652 | 1.231 | -0.75 | -0.046 | 0.019 | 0.019 | 0.062 | 0.51 | 0.07 | 0.37 | 0.29 | 0.53 | 0.45 |
| 8 | 12926048 | rs630498 | 0.138 | 0.411 | 1.991 | -0.969 | 0.006 | -0.043 | -0.043 | 0 | 0.88 | 0.66 | 0.92 | 0.92 | 0.91 | 0.87 |
| 8 | 12927294 | rs2584561 | -1.04 | 0.533 | 2.001 | -0.996 | 0.03 | 0.075 | 0.075 | 0 | 0.3 | 0.52 | 0.52 | 0.59 | 0.36 | 0.34 |
| 8 | 12929162 | rs543265 | 0.049 | 0.528 | 2.032 | -0.956 | 0.004 | -0.045 | -0.045 | 0.859 | 0.88 | 0.76 | 0.92 | 0.93 | 0.91 | 0.87 |
| 8 | 12933568 | rs492974 | -0.477 | 0.538 | 1.76 | -0.952 | 0 | -0.027 | -0.027 | 0.242 | 0.9 | 0.86 | 0.94 | 0.96 | 0.91 | 0.98 |
| 8 | 12933581 | rs633979 | -0.734 | 0.523 | 1.833 | -0.982 | -0.023 | -0.042 | -0.042 | 0 | 0.27 | 0.21 | 0.52 | 0.6 | 0.37 | 0.32 |
| 8 | 12941709 | rs1044011 | -0.004 | 0.367 | 1.441 | -0.758 | -0.06 | -0.103 | -0.103 | 0 | 0.2 | 0.07 | 0.47 | 0.59 | 0.33 | 0.29 |
| 8 | 12942510 | rs486596 | -0.167 | 0.308 | 1.656 | -0.851 | 0.002 | -0.038 | -0.038 | 0 | 0.89 | 0.77 | 0.93 | 0.95 | 0.91 | 0.98 |
| 8 | 12945508 | rs2584560 | -0.986 | 0.252 | 1.492 | -0.867 | -0.029 | -0.07 | -0.07 | 0 | 0.13 | 0.32 | 0.06 | 0.03 | 0.05 | 0.15 |
| 8 | 12948275 | rs2466274 | -0.456 | 0.336 | 1.485 | -0.852 | -0.002 | 0.001 | 0.001 | 2.362 | 0.89 | 0.9 | 0.93 | 0.95 | 0.89 | 0.97 |
| 8 | 12950315 | rs568182 | -0.704 | 0.359 | 1.254 | -0.907 | -0.022 | -0.059 | -0.059 | 0 | 0.12 | 0.25 | 0.06 | 0.03 | 0.06 | 0.12 |
| 8 | 12952230 | rs658948 | 0.977 | 0.372 | 1.313 | -0.896 | 0.022 | 0.019 | 0.019 | 0 | 0.66 | 0.46 | 0.42 | 0.3 | 0.57 | 0.62 |
| 8 | 12952293 | rs658856 | -0.726 | 0.515 | 1.05 | -0.89 | -0.022 | -0.06 | -0.06 | 0 | 0.12 | 0.25 | 0.06 | 0.03 | 0.06 | 0.11 |
| 8 | 12957475 | rs532841 | 1.119 | 0.728 | 0.441 | -0.691 | -0.005 | 0 | 0 | 0.496 | 0.57 | 0.61 | 0.38 | 0.34 | 0.5 | 0.59 |
| 8 | 12966739 | rs1372707 | -0.281 | 0.937 | -0.904 | -0.59 | -0.033 | -0.08 | -0.08 | 0 | 0.23 | 0.11 | 0.49 | 0.57 | 0.33 | 0.31 |
| 8 | 12969055 | rs13274488 | 1.346 | 0.903 | -0.905 | -0.688 | -0.023 | 0.025 | 0.025 | 0 | 0.52 | 0.23 | 0.34 | 0.31 | 0.49 | 0.51 |
| 8 | 12969754 | rs2203838 | 0.758 | 0.451 | -0.021 | -1.192 | -0.007 | -0.052 | -0.052 | 0 | 0.74 | 0.41 | 0.82 | 0.9 | 0.81 | 0.84 |
| 8 | 12973575 | rs2410025 | 0.673 | 0.557 | -0.884 | -1.08 | 0.007 | 0.01 | 0.01 | 2.024 | 0.74 | 0.28 | 0.82 | 0.9 | 0.82 | 0.84 |
| 8 | 12979400 | rs12541254 | -0.726 | 0.917 | -0.948 | -1.117 | 0.011 | -0.088 | -0.088 | 0 | 0.33 | 0.54 | 0.2 | 0.14 | 0.35 | 0.36 |
| 8 | 12979786 | rs10108207 | -0.632 | 0.391 | -1.78 | -1.308 | 0.009 | 0.077 | 0.077 | 0 | 0.26 | 0.12 | 0.11 | 0.08 | 0.2 | 0.12 |
| 8 | 12980647 | rs9325866 | -0.478 | 0.123 | -1.82 | -0.526 | 0.08 | 0.045 | 0.045 | 0.081 | 0.45 | 0.73 | 0.45 | 0.32 | 0.3 | 0.47 |
| 8 | 12986826 | rs4831399 | -0.418 | 0.194 | -1.424 | -0.035 | 0.02 | -0.016 | -0.016 | 0 | 0.36 | 0.26 | 0.51 | 0.43 | 0.31 | 0.38 |
| 8 | 12988714 | rs7824944 | -0.699 | 0.639 | -1.494 | -0.344 | -0.002 | -0.002 | -0.002 | 0 | 0.39 | 0.27 | 0.6 | 0.4 | 0.4 | 0.37 |
| 8 | 12993628 | rs12334589 | -0.927 | 0.93 | -1.341 | -0.702 | 0.001 | 0.008 | 0.008 | 0.22 | 0.51 | 0.45 | 0.68 | 0.44 | 0.54 | 0.52 |

| | | | | | | | | | | | | | | | | |
|---|---|---|---|---|---|---|---|---|---|---|---|---|---|---|---|---|
| 8 | 12995743 | rs7015013 | -0.226 | 0.705 | -0.696 | -0.851 | -0.012 | -0.001 | -0.001 | 0 | 0.39 | 0.44 | 0.55 | 0.39 | 0.32 | 0.43 |
| 8 | 12996384 | rs7015031 | -0.432 | 0.723 | -0.729 | -0.825 | 0.006 | 0.003 | 0.003 | 0.301 | 0.46 | 0.41 | 0.62 | 0.44 | 0.46 | 0.51 |
| 8 | 13000610 | rs17126052 | -0.834 | 0.077 | -0.441 | -0.165 | 0 | 0.002 | 0.002 | 0 | 0.19 | 0.17 | 0.3 | 0.33 | 0.2 | 0.29 |
| 8 | 13000629 | rs12677920 | 1.498 | 0.031 | -0.957 | -0.792 | -0.067 | -0.07 | -0.07 | 0.34 | 0.39 | 0.12 | 0.29 | 0.56 | 0.46 | 0.35 |
| 8 | 13002472 | rs10109292 | 0.23 | -0.173 | -0.526 | -1.077 | -0.001 | -0.008 | -0.008 | 0 | 0.14 | 0.15 | 0.11 | 0.03 | 0.17 | 0.16 |
| 8 | 13002824 | rs6981656 | 0.817 | -0.356 | -0.212 | -1.116 | -0.013 | -0.026 | -0.026 | 0 | 0.22 | 0.25 | 0.26 | 0.09 | 0.17 | 0.21 |
| 8 | 13005559 | rs10093276 | -0.582 | -0.095 | -0.417 | -0.95 | -0.003 | -0.006 | -0.006 | 0 | 0.18 | 0.18 | 0.12 | 0.03 | 0.18 | 0.16 |
| 8 | 13007470 | rs1020248 | 0.715 | -0.208 | -0.235 | -0.772 | 0.037 | 0.002 | 0.002 | 0 | 0.57 | 0.3 | 0.41 | 0.59 | 0.63 | 0.52 |
| 8 | 13019554 | rs1348280 | 0.329 | 0.091 | 0.196 | -1.108 | -0.025 | 0.13 | 0.13 | 0 | 0.61 | 0.8 | 0.64 | 0.85 | 0.54 | 0.65 |
| 8 | 13024625 | rs7827434 | 0.443 | -0.111 | 0.764 | -1.249 | -0.003 | -0.062 | -0.062 | 0 | 0.58 | 0.52 | 0.61 | 0.87 | 0.54 | 0.65 |
| 8 | 13025409 | rs17126375 | -0.653 | -0.006 | 0.786 | -1.378 | 0.06 | -0.08 | -0.08 | 0 | 0.16 | 0.41 | 0.13 | 0.02 | 0.15 | 0.16 |
| 8 | 13025800 | rs10099359 | 1.365 | -0.058 | 0.892 | -1.353 | 0.016 | 0.08 | 0.08 | 0 | 0.46 | 0.31 | 0.49 | 0.85 | 0.39 | 0.49 |
| 8 | 13027912 | rs7828964 | -0.949 | 0.052 | 1.063 | -0.707 | 0.007 | 0.252 | 0.252 | 0.013 | 0.14 | 0.44 | 0.31 | 0.31 | 0.11 | 0.14 |
| 8 | 13034369 | rs1431204 | 1.036 | -0.216 | 0.619 | -1.23 | 0.006 | 0.113 | 0.113 | 0 | 0.49 | 0.32 | 0.52 | 0.85 | 0.44 | 0.5 |
| 8 | 13036405 | rs9644024 | 0.174 | -0.247 | 0.777 | -1.286 | 0.002 | -0.084 | -0.084 | 0 | 0.61 | 0.5 | 0.61 | 0.87 | 0.55 | 0.71 |
| 8 | 13043878 | rs7004673 | 0 | -0.263 | 0.731 | -1.414 | 0.01 | -0.081 | -0.081 | 0 | 0.64 | 0.54 | 0.64 | 0.87 | 0.57 | 0.72 |
| 8 | 13044729 | rs10099039 | 1.415 | -0.268 | 0.744 | -1.403 | 0.006 | 0.04 | 0.04 | 0 | 0.49 | 0.4 | 0.52 | 0.85 | 0.42 | 0.49 |
| 8 | 13050887 | rs6988111 | 0.741 | -0.233 | 0.641 | -1.437 | 0.003 | -0.045 | -0.045 | 0.558 | 0.56 | 0.53 | 0.57 | 0.86 | 0.5 | 0.61 |
| 8 | 13061713 | rs919590 | 0.352 | -0.228 | 0.751 | -1.465 | -0.009 | 0.042 | 0.042 | 0 | 0.62 | 0.29 | 0.61 | 0.87 | 0.54 | 0.7 |
| 8 | 13062363 | rs6995936 | -0.922 | -0.303 | 0.661 | -1.057 | -0.018 | 0.104 | 0.104 | 0 | 0.16 | 0.28 | 0.3 | 0.34 | 0.12 | 0.25 |
| 8 | 13065646 | rs6997092 | 1.402 | -0.301 | 0.275 | -1.098 | 0 | 0.182 | 0.182 | 0 | 0.48 | 0.21 | 0.51 | 0.85 | 0.43 | 0.49 |
| 8 | 13066409 | rs1188306 | -1.616 | -0.294 | 0.243 | -1.085 | -0.006 | 0.02 | 0.02 | 0 | 0.2 | 0.71 | 0.34 | 0.35 | 0.13 | 0.31 |
| 8 | 13068051 | rs1188309 | 0.069 | -0.097 | 0.318 | -1.468 | -0.017 | 0.065 | 0.065 | 0 | 0.63 | 0.75 | 0.75 | 0.94 | 0.54 | 0.71 |
| 8 | 13072177 | rs3739300 | 0.471 | -0.138 | -0.011 | -1.432 | -0.041 | 0.176 | 0.176 | 0.021 | 0.55 | 0.79 | 0.55 | 0.85 | 0.47 | 0.56 |
| 8 | 13082745 | rs1209107 | 1.047 | -0.097 | -0.105 | -1.284 | 0.006 | 0.033 | 0.033 | 0.564 | 0.44 | 0.64 | 0.46 | 0.83 | 0.38 | 0.41 |
| 8 | 13086762 | rs1653032 | 0.368 | 0.022 | -0.312 | -1.407 | -0.005 | 0.038 | 0.038 | 0 | 0.62 | 0.68 | 0.61 | 0.91 | 0.57 | 0.6 |
| 8 | 13089647 | rs1729161 | 0.98 | 0.028 | -0.156 | -1.342 | 0 | 0.014 | 0.014 | 0.014 | 0.48 | 0.41 | 0.52 | 0.86 | 0.49 | 0.41 |
| 8 | 13096264 | rs1188267 | 0.932 | 0.034 | -0.102 | -1.264 | -0.001 | -0.026 | -0.026 | 0 | 0.49 | 0.44 | 0.53 | 0.84 | 0.52 | 0.38 |
| 8 | 13097749 | rs17188894 | 1.517 | -0.136 | -0.538 | -0.584 | -0.087 | -0.124 | -0.124 | 0 | 0.39 | 0.03 | 0.22 | 0.48 | 0.38 | 0.32 |
| 8 | 13098348 | rs17127548 | 1.027 | -0.072 | -0.231 | -1.033 | 0.002 | 0.002 | 0.002 | 0 | 0.1 | 0.12 | 0.07 | 0.05 | 0.05 | 0.16 |
| 8 | 13098707 | rs17188985 | 1.297 | -0.043 | -0.284 | -1.03 | 0.007 | 0.039 | 0.039 | 0 | 0.09 | 0.02 | 0.07 | 0.02 | 0.05 | 0.16 |
| 8 | 13100245 | rs17127579 | -0.089 | -0.092 | -0.023 | -1.228 | 0.038 | -0.042 | -0.042 | 0 | 0.14 | 0.35 | 0.1 | 0.04 | 0.14 | 0.2 |
| 8 | 13106308 | rs7834940 | -0.608 | -0.134 | -0.061 | -1.256 | -0.001 | -0.049 | -0.049 | 0 | 0.23 | 0.3 | 0.17 | 0.07 | 0.2 | 0.35 |
| 8 | 13108897 | rs2410043 | -0.563 | -0.125 | -0.143 | -1.297 | -0.001 | -0.054 | -0.054 | 0 | 0.22 | 0.37 | 0.17 | 0.1 | 0.19 | 0.35 |
| 8 | 13109645 | rs17793145 | 2.61 | -0.323 | -0.981 | -1.591 | -0.055 | 0.024 | 0.024 | 2.792 | 0.18 | 0.02 | 0.1 | 0.11 | 0.22 | 0.11 |
| 8 | 13112976 | rs1528628 | 0.901 | -0.424 | 0.093 | -1.325 | 0.002 | -0.048 | -0.048 | 0 | 0.56 | 0.49 | 0.53 | 0.88 | 0.53 | 0.49 |
| 8 | 13114705 | rs17127848 | 1.01 | -0.399 | 0.069 | -1.286 | -0.003 | -0.009 | -0.009 | 2.549 | 0.1 | 0.12 | 0.08 | 0.05 | 0.06 | 0.2 |
| 8 | 13115090 | rs17127858 | 0.984 | -0.4 | 0.172 | -1.286 | -0.004 | -0.023 | -0.023 | 0.298 | 0.1 | 0.18 | 0.09 | 0.05 | 0.06 | 0.2 |
| 8 | 13117733 | rs10503444 | 0.234 | -0.399 | 0.071 | -1.294 | 0.001 | 0.002 | 0.002 | 0 | 0.14 | 0.13 | 0.11 | 0.06 | 0.1 | 0.27 |
| 8 | 13118537 | rs6991346 | -0.485 | -0.423 | 0.102 | -1.336 | -0.012 | 0.016 | 0.016 | 0 | 0.73 | 0.78 | 0.66 | 0.93 | 0.65 | 0.81 |
| 8 | 13120332 | rs1203096 | 0.928 | -0.52 | 0.128 | -1.276 | 0.003 | -0.035 | -0.035 | 0 | 0.57 | 0.53 | 0.55 | 0.88 | 0.5 | 0.59 |
| 8 | 13124314 | rs1209992 | -0.126 | -0.331 | 0.072 | -1.229 | -0.02 | 0.025 | 0.025 | 0 | 0.7 | 0.75 | 0.61 | 0.89 | 0.59 | 0.72 |
| 8 | 13128100 | rs884058 | 1.731 | -0.253 | 0.2 | -1.216 | 0.013 | 0.07 | 0.07 | 1.034 | 0.44 | 0.3 | 0.45 | 0.81 | 0.4 | 0.35 |
| 8 | 13131656 | rs1188284 | 0.378 | -0.075 | -0.191 | -1.188 | -0.026 | 0.077 | 0.077 | 0 | 0.65 | 0.79 | 0.64 | 0.93 | 0.59 | 0.76 |
| 8 | 13146630 | rs13262128 | 0.236 | 0.086 | -0.172 | -1.141 | -0.016 | 0.017 | 0.017 | 0 | 0.27 | 0.21 | 0.35 | 0.04 | 0.34 | 0.2 |
| 8 | 13147465 | rs13250216 | -1.263 | -0.05 | 0.08 | -1.225 | -0.001 | -0.026 | -0.026 | 0 | 0.47 | 0.47 | 0.52 | 0.11 | 0.53 | 0.48 |
| 8 | 13147657 | rs17128435 | 1.177 | -0.06 | 0.092 | -1.237 | -0.001 | -0.009 | -0.009 | 0.653 | 0.1 | 0.08 | 0.07 | 0.02 | 0.05 | 0.13 |
| 8 | 13148221 | rs3842964 | -0.549 | -0.02 | 0.06 | -1.222 | -0.032 | 0.065 | 0.065 | 0 | 0.34 | 0.21 | 0.39 | 0.05 | 0.41 | 0.25 |
| 8 | 13149874 | rs7836264 | 2.275 | -0.213 | -0.085 | -0.848 | -0.051 | -0.068 | -0.068 | 0 | 0.39 | 0.03 | 0.22 | 0.44 | 0.38 | 0.31 |
| 8 | 13150955 | rs10888181 | -1.431 | -0.171 | 0.1 | -0.869 | -0.001 | -0.025 | -0.025 | 0 | 0.5 | 0.52 | 0.54 | 0.11 | 0.55 | 0.53 |
| 8 | 13153938 | rs13279835 | 0.395 | -0.112 | -0.002 | -0.926 | -0.015 | 0.015 | 0.015 | 0 | 0.27 | 0.21 | 0.35 | 0.06 | 0.34 | 0.2 |
| 8 | 13159120 | rs1653024 | 0.015 | -0.154 | 0.46 | -0.837 | 0.011 | -0.034 | -0.034 | 0 | 0.84 | 0.57 | 0.85 | 0.92 | 0.86 | 0.72 |
| 8 | 13163596 | rs2583106 | 2.137 | -0.235 | 0.298 | -0.824 | 0.007 | -0.012 | -0.012 | 0 | 0.39 | 0.22 | 0.24 | 0.47 | 0.38 | 0.32 |
| 8 | 13165729 | rs10091996 | -0.643 | -0.244 | 0.358 | -0.844 | -0.007 | -0.004 | -0.004 | 0.055 | 0.37 | 0.32 | 0.4 | 0.08 | 0.42 | 0.25 |
| 8 | 13167115 | rs1188133 | 2.119 | -0.286 | 0.306 | -0.799 | 0.008 | -0.007 | -0.007 | 0 | 0.39 | 0.23 | 0.23 | 0.47 | 0.38 | 0.32 |
| 8 | 13171496 | rs10102530 | 1.938 | -0.486 | 0.382 | -0.831 | -0.01 | -0.011 | -0.011 | 0.091 | 0.4 | 0.33 | 0.29 | 0.48 | 0.44 | 0.44 |
| 8 | 13173084 | rs1188130 | -0.532 | -0.603 | 0.357 | -0.987 | -0.039 | -0.086 | -0.086 | 0 | 0.25 | 0.3 | 0.14 | 0.03 | 0.14 | 0.2 |
| 8 | 13174184 | rs13282920 | -0.857 | -0.528 | 0.123 | -0.959 | -0.008 | 0.041 | 0.041 | 0 | 0.42 | 0.31 | 0.4 | 0.11 | 0.43 | 0.32 |
| 8 | 13176407 | rs1671400 | 0.473 | -0.561 | 0.05 | -0.848 | 0.026 | 0.019 | 0.019 | 0.002 | 0.74 | 0.81 | 0.87 | 0.97 | 0.87 | 0.8 |
| 8 | 13179510 | rs3911339 | -1.261 | -0.54 | 0.044 | -0.65 | 0.074 | -0.063 | -0.063 | 0 | 0.31 | 0.12 | 0.32 | 0.4 | 0.18 | 0.31 |
| 8 | 13181518 | rs1671389 | -0.322 | -0.105 | 0.609 | -0.431 | 0.065 | 0.024 | 0.024 | 0 | 0.93 | 0.7 | 0.92 | 0.96 | 0.91 | 0.86 |
| 8 | 13183399 | rs12543681 | -0.719 | -0.183 | 0.49 | -0.445 | 0.016 | -0.027 | -0.027 | 0 | 0.18 | 0.13 | 0.28 | 0.39 | 0.1 | 0.26 |
| 8 | 13185325 | rs1671382 | 0.885 | -0.092 | 0.6 | -0.342 | 0.017 | 0.243 | 0.243 | 0 | 0.87 | 0.66 | 0.71 | 0.58 | 0.86 | 0.74 |
| 8 | 13186949 | rs1406345 | -1.297 | -0.06 | 0.567 | -0.604 | -0.004 | 0 | 0 | 0 | 0.45 | 0.56 | 0.62 | 0.53 | 0.48 | 0.43 |
| 8 | 13187829 | rs1454949 | -0.308 | -0.017 | 0.603 | -0.576 | 0.062 | 0.021 | 0.021 | 0 | 0.93 | 0.68 | 0.92 | 0.97 | 0.91 | 0.87 |
| 8 | 13188361 | rs4492382 | 2.325 | 0.045 | 0.599 | -0.594 | -0.025 | -0.052 | -0.052 | 3.275 | 0.37 | 0.09 | 0.24 | 0.42 | 0.38 | 0.3 |
| 8 | 13188804 | rs13266051 | 0.991 | 0.251 | 0.64 | -0.712 | 0.001 | -0.004 | -0.004 | 2.569 | 0.18 | 0.17 | 0.3 | 0.08 | 0.28 | 0.12 |
| 8 | 13190938 | rs1671365 | 2.176 | 0.198 | 0.659 | -0.642 | 0.002 | 0 | 0 | 0.058 | 0.39 | 0.33 | 0.26 | 0.41 | 0.38 | 0.3 |
| 8 | 13194328 | rs1729109 | -0.213 | 0.218 | 0.705 | -0.613 | 0.038 | 0.004 | 0.004 | 0 | 0.93 | 0.71 | 0.92 | 0.96 | 0.91 | 0.86 |

| | | | | | | | | | | | | | | | | |
|---|---|---|---|---|---|---|---|---|---|---|---|---|---|---|---|---|
| 8 | 13194983 | rs1528624 | -1.839 | 0.177 | 0.613 | -0.641 | 0.003 | -0.004 | -0.004 | 0 | 0.54 | 0.34 | 0.66 | 0.5 | 0.54 | 0.53 |
| 8 | 13195741 | rs1454937 | 0.022 | 0.104 | 0.656 | -0.709 | -0.034 | -0.093 | -0.093 | 0 | 0.84 | 0.6 | 0.92 | 0.98 | 0.91 | 0.81 |
| 8 | 13198040 | rs17091779 | 2.06 | 0.135 | 0.45 | -0.692 | -0.044 | -0.075 | -0.075 | 0 | 0.4 | 0.1 | 0.27 | 0.48 | 0.42 | 0.35 |
| 8 | 13199151 | rs2278945 | -1.288 | 0.324 | 0.477 | -0.611 | -0.01 | -0.009 | -0.009 | 0 | 0.43 | 0.34 | 0.64 | 0.5 | 0.53 | 0.52 |
| 8 | 13200021 | rs767763 | -0.429 | 0.354 | 0.582 | -0.625 | 0.055 | -0.075 | -0.075 | 0 | 0.28 | 0.4 | 0.37 | 0.12 | 0.38 | 0.23 |
| 8 | 13202119 | rs7013038 | -0.681 | 0.368 | 0.406 | -0.572 | 0.004 | -0.023 | -0.023 | 0 | 0.2 | 0.15 | 0.3 | 0.42 | 0.18 | 0.35 |
| 8 | 13202138 | rs1671339 | 0.591 | 0.436 | 0.451 | -0.714 | 0.036 | -0.03 | -0.03 | 0 | 0.2 | 0.29 | 0.32 | 0.12 | 0.29 | 0.17 |
| 8 | 13203803 | rs7018361 | -1.294 | 0.41 | -0.141 | -0.693 | 0.004 | 0.002 | 0.002 | 0.178 | 0.46 | 0.32 | 0.65 | 0.51 | 0.55 | 0.55 |
| 8 | 13204116 | rs12681573 | -0.477 | 0.631 | -0.322 | -0.555 | -0.02 | -0.029 | -0.029 | 0 | 0.31 | 0.26 | 0.54 | 0.46 | 0.42 | 0.31 |
| 8 | 13205853 | rs17091974 | 1.734 | 0.52 | -0.972 | -0.441 | 0 | -0.012 | -0.012 | 0 | 0.38 | 0.14 | 0.25 | 0.43 | 0.38 | 0.3 |
| 8 | 13209661 | rs17802857 | 0.285 | 0.878 | -0.652 | -0.415 | -0.002 | 0.003 | 0.003 | 0.61 | 0.11 | 0.07 | 0.06 | 0.03 | 0.08 | 0.07 |
| 8 | 13209681 | rs1729133 | 0.231 | 0.847 | -0.58 | -0.491 | 0 | -0.012 | -0.012 | 0 | 0.9 | 0.88 | 0.94 | 1 | 0.9 | 0.87 |
| 8 | 13211769 | rs1729159 | 0.294 | 0.84 | -0.493 | -0.495 | 0 | -0.038 | -0.038 | 0 | 0.9 | 0.83 | 0.93 | 1 | 0.9 | 0.87 |
| 8 | 13212584 | rs4831408 | 0.086 | 0.569 | -0.643 | -0.083 | 0.015 | -0.016 | -0.016 | 1.095 | 0.1 | 0.05 | 0.21 | 0.47 | 0.05 | 0.14 |
| 8 | 13213144 | rs1671431 | -0.402 | 0.693 | -0.483 | -1.119 | 0.001 | -0.022 | -0.022 | 0 | 0.89 | 0.86 | 0.94 | 0.98 | 0.92 | 0.93 |
| 8 | 13216803 | rs1454959 | 2.093 | 0.721 | -0.671 | -1.495 | 0.012 | 0.018 | 0.018 | 0.441 | 0.41 | 0.6 | 0.47 | 0.79 | 0.37 | 0.35 |
| 8 | 13216874 | rs10503450 | -1.448 | 0.96 | -0.812 | -1.655 | -0.024 | 0.062 | 0.062 | 0 | 0.22 | 0.14 | 0.13 | 0.03 | 0.21 | 0.16 |
| 8 | 13220951 | rs41468845 | -1.199 | 0.954 | -0.815 | -1.659 | -0.008 | 0.051 | 0.051 | 0.345 | 0.19 | 0.11 | 0.1 | 0.02 | 0.15 | 0.13 |
| 8 | 13224672 | rs897265 | 0.473 | 1.06 | -0.818 | -1.674 | -0.058 | 0.08 | 0.08 | 0.109 | 0.68 | 0.83 | 0.61 | 0.85 | 0.6 | 0.63 |
| 8 | 13224806 | rs897266 | 1.976 | 1.012 | -0.828 | -1.713 | 0.041 | 0.105 | 0.105 | 2.03 | 0.48 | 0.66 | 0.5 | 0.81 | 0.42 | 0.36 |
| 8 | 13224865 | rs4102144 | -0.872 | 1.07 | -0.785 | -1.684 | 0 | -0.026 | -0.026 | 0.647 | 0.18 | 0.16 | 0.09 | 0.01 | 0.15 | 0.17 |
| 8 | 13226588 | rs1671413 | 0.241 | 1.027 | -0.73 | -1.64 | -0.036 | 0.067 | 0.067 | 0.014 | 0.71 | 0.84 | 0.64 | 0.87 | 0.67 | 0.65 |
| 8 | 13228164 | rs11991613 | 0.487 | 1.015 | -0.677 | -1.612 | -0.01 | 0.006 | 0.006 | 0 | 0.12 | 0.09 | 0.08 | 0.01 | 0.14 | 0.08 |
| 8 | 13230921 | rs921773 | 0.25 | 1.265 | -0.76 | -1.672 | -0.052 | 0.032 | 0.032 | 0.446 | 0.21 | 0.07 | 0.31 | 0.12 | 0.29 | 0.25 |
| 8 | 13233321 | rs1729181 | 1.824 | 1.194 | -0.601 | -1.724 | 0.018 | 0.05 | 0.05 | 0.356 | 0.51 | 0.58 | 0.5 | 0.83 | 0.45 | 0.46 |
| 8 | 13233591 | rs1729175 | 0.967 | 1.181 | -0.7 | -1.733 | -0.024 | 0.062 | 0.062 | 0.327 | 0.66 | 0.78 | 0.6 | 0.86 | 0.61 | 0.56 |
| 8 | 13234234 | rs921774 | 0.377 | 1.18 | -0.709 | -1.742 | -0.043 | 0.028 | 0.028 | 0.431 | 0.2 | 0.08 | 0.3 | 0.12 | 0.27 | 0.25 |
| 8 | 13234600 | rs1729163 | 1.008 | 1.176 | -0.677 | -1.74 | -0.022 | 0.055 | 0.055 | 0.364 | 0.66 | 0.78 | 0.6 | 0.86 | 0.61 | 0.55 |
| 8 | 13238878 | rs7837580 | -1.709 | 0.997 | 0.04 | -1.802 | -0.004 | 0.018 | 0.018 | 0 | 0.44 | 0.6 | 0.43 | 0.19 | 0.46 | 0.44 |
| 8 | 13241026 | rs6994536 | -0.213 | 0.999 | -0.385 | -1.712 | -0.007 | 0.013 | 0.013 | 0 | 0.16 | 0.14 | 0.09 | 0.03 | 0.15 | 0.1 |
| 8 | 13241664 | rs289510 | 0.683 | 1.002 | -0.523 | -1.702 | -0.002 | -0.023 | -0.023 | 0 | 0.86 | 0.8 | 0.89 | 0.98 | 0.87 | 0.8 |
| 8 | 13242061 | rs167042 | -1.005 | 1.003 | -0.53 | -1.704 | -0.001 | 0.014 | 0.014 | 0.007 | 0.93 | 0.94 | 0.93 | 0.98 | 0.92 | 0.88 |
| 8 | 13242215 | rs289528 | 0.14 | 1.014 | -0.519 | -1.682 | 0 | -0.011 | -0.011 | 0.769 | 0.87 | 0.86 | 0.9 | 0.98 | 0.88 | 0.81 |
| 8 | 13242413 | rs289527 | 1.731 | 1.031 | -0.52 | -1.685 | -0.003 | 0.041 | 0.041 | 0.824 | 0.67 | 0.72 | 0.79 | 0.91 | 0.69 | 0.63 |
| 8 | 13245917 | rs2444947 | 1.369 | 0.955 | -0.504 | -1.589 | -0.02 | 0.059 | 0.059 | 0.445 | 0.58 | 0.67 | 0.55 | 0.81 | 0.52 | 0.43 |
| 8 | 13246338 | rs289553 | 0.825 | 0.927 | -0.597 | -1.481 | 0.006 | 0.01 | 0.01 | 0.103 | 0.82 | 0.85 | 0.87 | 0.98 | 0.84 | 0.74 |
| 8 | 13251235 | rs17805176 | -0.088 | 1.179 | -0.798 | -1.429 | -0.058 | 0.077 | 0.077 | 0.021 | 0.16 | 0.03 | 0.09 | 0.03 | 0.18 | 0.1 |
| 8 | 13251758 | rs1021094 | -1.345 | 1.202 | -0.757 | -1.52 | -0.028 | 0.096 | 0.096 | 0 | 0.39 | 0.17 | 0.43 | 0.19 | 0.51 | 0.38 |
| 8 | 13253898 | rs7842439 | 0.407 | 1.269 | -0.777 | -1.53 | -0.046 | 0.025 | 0.025 | 0.145 | 0.18 | 0.07 | 0.31 | 0.11 | 0.29 | 0.19 |
| 8 | 13256290 | rs7008120 | 0.028 | 1.195 | -0.746 | -1.507 | -0.058 | 0.076 | 0.076 | 0.011 | 0.16 | 0.03 | 0.09 | 0.03 | 0.18 | 0.1 |
| 8 | 13257778 | rs1454957 | 0.034 | 1.119 | -0.524 | -1.543 | -0.06 | 0.023 | 0.023 | 0 | 0.22 | 0.08 | 0.33 | 0.16 | 0.33 | 0.28 |
| 8 | 13259865 | rs289584 | 2.135 | 0.578 | -0.264 | -1.551 | 0.042 | 0.073 | 0.073 | 0.821 | 0.4 | 0.23 | 0.43 | 0.79 | 0.34 | 0.35 |
| 8 | 13259928 | rs6996724 | -0.792 | 0.78 | -0.539 | -1.424 | -0.077 | 0.11 | 0.11 | 0 | 0.38 | 0.1 | 0.42 | 0.19 | 0.51 | 0.38 |
| 8 | 13260536 | rs289585 | 0.678 | 0.633 | 0.188 | -0.995 | -0.001 | -0.016 | -0.016 | 0 | 0.88 | 0.84 | 0.9 | 0.98 | 0.86 | 0.8 |
| 8 | 13261721 | rs289587 | 0.599 | 0.721 | 0.156 | -0.829 | -0.001 | -0.066 | -0.066 | 0.574 | 0.88 | 0.55 | 0.92 | 0.98 | 0.93 | 0.86 |
| 8 | 13264691 | rs289603 | 0.021 | 0.901 | -0.541 | -0.524 | -0.037 | 0.065 | 0.065 | 0 | 0.86 | 0.99 | 0.93 | 0.97 | 0.9 | 0.93 |
| 8 | 13267353 | rs289609 | 0.881 | 1.141 | -0.71 | -0.587 | -0.046 | 0.059 | 0.059 | 0 | 0.75 | 0.95 | 0.73 | 0.88 | 0.74 | 0.82 |
| 8 | 13267974 | rs11203485 | -1.738 | 0.932 | -0.008 | -0.065 | -0.002 | 0.017 | 0.017 | 0 | 0.34 | 0.3 | 0.35 | 0.13 | 0.35 | 0.31 |
| 8 | 13269694 | rs17817461 | 0.078 | 0.754 | 0.068 | 0.441 | -0.004 | -0.002 | -0.002 | 0.013 | 0.1 | 0.06 | 0.22 | 0.1 | 0.14 | 0.11 |
| 8 | 13271982 | rs7826429 | -0.694 | 0.651 | 0.216 | 0.947 | -0.005 | 0.178 | 0.178 | 0 | 0.09 | 0.18 | 0.25 | 0.42 | 0.08 | 0.17 |
| 8 | 13273425 | rs7822301 | 1.208 | 0.752 | 0.369 | 0.472 | 0.001 | 0.025 | 0.025 | 0 | 0.56 | 0.37 | 0.43 | 0.38 | 0.56 | 0.58 |
| 8 | 13273477 | rs7834383 | 0.301 | 0.924 | 0.385 | 0.271 | -0.008 | -0.004 | -0.004 | 0 | 0.23 | 0.2 | 0.25 | 0.04 | 0.29 | 0.28 |
| 8 | 13282821 | rs370730 | -0.682 | 0.614 | 0.854 | 0.543 | 0.002 | 0.002 | 0.002 | 0 | 0.48 | 0.59 | 0.59 | 0.52 | 0.51 | 0.49 |
| 8 | 13285702 | rs7003283 | 0.54 | 0.649 | 1.042 | 0.597 | 0.022 | 0.036 | 0.036 | 0 | 0.52 | 0.65 | 0.44 | 0.39 | 0.48 | 0.47 |
| 8 | 13286160 | rs10503451 | -0.6 | 0.679 | 1.338 | 0.587 | 0.004 | 0.016 | 0.016 | 0 | 0.13 | 0.17 | 0.2 | 0.14 | 0.14 | 0.19 |
| 8 | 13290539 | rs17116045 | 1.335 | 0.725 | 1.207 | 0.616 | -0.012 | -0.015 | -0.015 | 0 | 0.05 | 0.02 | 0.13 | 0.09 | 0.07 | 0.09 |
| 8 | 13294635 | rs289620 | 0.108 | 0.628 | 1.313 | 0.631 | -0.019 | 0.008 | 0.008 | 0 | 0.58 | 0.72 | 0.46 | 0.43 | 0.52 | 0.53 |
| 8 | 13294808 | rs289621 | -0.28 | 0.652 | 1.525 | 0.738 | 0.004 | -0.042 | -0.042 | 1.685 | 0.08 | 0.17 | 0.05 | 0 | 0.08 | 0.05 |
| 8 | 13297265 | rs192627 | -1.31 | 0.683 | 1.398 | 0.778 | 0.033 | 0.059 | 0.059 | 0 | 0.15 | 0.4 | 0.22 | 0.14 | 0.15 | 0.2 |
| 8 | 13304484 | rs289634 | 0.148 | 0.565 | 0.97 | 0.826 | -0.019 | 0.005 | 0.005 | 0 | 0.57 | 0.73 | 0.47 | 0.41 | 0.52 | 0.55 |
| 8 | 13309547 | rs430888 | 0.547 | 0.682 | 1.059 | 0.763 | 0.013 | 0.027 | 0.027 | 0 | 0.52 | 0.66 | 0.43 | 0.39 | 0.48 | 0.54 |
| 8 | 13314628 | rs289543 | -0.802 | 0.528 | 1.28 | 0.875 | 0.036 | 0.131 | 0.131 | 0 | 0.11 | 0.5 | 0.19 | 0.14 | 0.09 | 0.12 |
| 8 | 13315533 | rs289542 | 0.192 | 0.399 | 1.178 | 0.895 | -0.016 | 0.016 | 0.016 | 0 | 0.56 | 0.72 | 0.47 | 0.38 | 0.52 | 0.52 |
| 8 | 13330640 | rs4376497 | 0.564 | 0.213 | 1.23 | 1.029 | -0.002 | -0.001 | -0.001 | 0 | 0.49 | 0.5 | 0.42 | 0.44 | 0.47 | 0.48 |
| 8 | 13333763 | rs2410058 | -0.93 | 0.295 | 1.015 | 0.894 | 0.007 | 0.011 | 0.011 | 0 | 0.54 | 0.59 | 0.7 | 0.8 | 0.57 | 0.68 |
| 8 | 13333857 | rs4831439 | 1.095 | 0.302 | 0.476 | 0.929 | 0.031 | 0.133 | 0.133 | 0 | 0.41 | 0.1 | 0.26 | 0.2 | 0.39 | 0.31 |
| 8 | 13348868 | rs17094050 | -0.337 | 0.303 | 0.876 | 1.441 | -0.005 | 0.078 | 0.078 | 0 | 0.08 | 0.14 | 0.15 | 0.22 | 0.06 | 0.25 |
| 8 | 13349605 | rs12549364 | 1.064 | 0.179 | 1.192 | 1.675 | -0.003 | 0.235 | 0.235 | 0.054 | 0.92 | 0.75 | 0.73 | 0.6 | 0.93 | 0.93 |
| 8 | 13349951 | rs12542629 | 1.447 | 0.172 | 1.332 | 1.671 | -0.001 | 0.279 | 0.279 | 0.025 | 0.91 | 0.69 | 0.73 | 0.6 | 0.93 | 0.85 |

| | | | | | | | | | | | | | | | | |
|---|---|---|---|---|---|---|---|---|---|---|---|---|---|---|---|---|
| 8 | 13356802 | rs3816747 | 0.632 | 0.175 | 1.335 | 1.685 | -0.002 | 0.165 | 0.165 | 0.066 | 0.93 | 0.84 | 0.73 | 0.61 | 0.94 | 0.92 |
| 8 | 13356819 | rs11203495 | 1.294 | 0.178 | 1.293 | 1.686 | -0.004 | 0.264 | 0.264 | 0.01 | 0.91 | 0.68 | 0.72 | 0.61 | 0.94 | 0.85 |
| 8 | 13357587 | rs1383942 | 1.167 | 0.183 | 1.328 | 1.7 | 0.034 | 0.433 | 0.433 | 0.003 | 0.91 | 0.57 | 0.72 | 0.61 | 0.94 | 0.93 |
| 8 | 13369061 | rs17222010 | 0.062 | 0.371 | 0.53 | 1.272 | 0.005 | 0.036 | 0.036 | 0.808 | 0.21 | 0.13 | 0.1 | 0.12 | 0.16 | 0.12 |
| 8 | 13382080 | rs12234951 | 0.791 | 0.408 | -0.382 | 1.682 | -0.051 | 0.007 | 0.007 | 0.059 | 0.36 | 0.03 | 0.24 | 0.26 | 0.38 | 0.26 |
| 8 | 13390938 | rs979728 | 0.466 | 0.422 | -0.105 | 1.717 | 0.018 | -0.015 | -0.015 | 0 | 0.41 | 0.22 | 0.47 | 0.59 | 0.41 | 0.36 |
| 8 | 13392754 | rs1481687 | -1.49 | 0.395 | 0.399 | 2.22 | 0.022 | 0.475 | 0.475 | 0.021 | 0.09 | 0.49 | 0.19 | 0.44 | 0.03 | 0.13 |
| 8 | 13395085 | rs2101109 | 0.398 | 0.793 | -0.504 | 1.457 | 0 | -0.01 | -0.01 | 0 | 0.61 | 0.64 | 0.6 | 0.46 | 0.65 | 0.48 |
| 8 | 13395662 | rs10089628 | -0.204 | 0.599 | -1.39 | 1.12 | 0.044 | 0.006 | 0.006 | 0 | 0.51 | 0.17 | 0.42 | 0.39 | 0.63 | 0.51 |
| 8 | 13396623 | rs1160220 | -0.193 | 0.559 | -1.393 | 0.818 | 0.003 | -0.004 | -0.004 | 0 | 0.46 | 0.44 | 0.61 | 0.48 | 0.47 | 0.41 |
| 8 | 13397098 | rs17094556 | 0.472 | 0.331 | -1.517 | 0.887 | 0.007 | -0.006 | -0.006 | 0 | 0.19 | 0.16 | 0.14 | 0.22 | 0.15 | 0.24 |
| 8 | 13397806 | rs966213 | -0.119 | 0.551 | -1.558 | 0.824 | -0.006 | 0 | 0 | 0.698 | 0.27 | 0.25 | 0.23 | 0.24 | 0.32 | 0.31 |
| 8 | 13401816 | rs1160554 | 0.714 | 0.478 | -1.672 | 0.834 | -0.026 | -0.01 | -0.01 | 0 | 0.14 | 0.08 | 0.23 | 0.14 | 0.24 | 0.07 |
| 8 | 13401972 | rs1160555 | -1.291 | 0.299 | -1.54 | 0.594 | 0 | 0 | 0 | 0 | 0.5 | 0.49 | 0.63 | 0.48 | 0.51 | 0.49 |
| 8 | 13402197 | rs10503453 | 1.213 | 0.426 | -2.013 | 0.256 | -0.063 | -0.045 | -0.045 | 0 | 0.25 | 0.09 | 0.22 | 0.36 | 0.34 | 0.32 |
| 8 | 13402465 | rs7822459 | 2.348 | 0.323 | -1.55 | 0.164 | 0.003 | -0.031 | -0.031 | 0.383 | 0.06 | 0.19 | 0.04 | 0 | 0.04 | 0.08 |
| 8 | 13404010 | rs12549031 | 2.303 | 0.339 | -1.685 | 0.237 | -0.005 | -0.037 | -0.037 | 0.745 | 0.09 | 0 | 0.16 | 0.14 | 0.08 | 0.05 |
| 8 | 13404835 | rs12541022 | -0.822 | 0.189 | -1.002 | 0.102 | -0.001 | -0.002 | -0.002 | 0.415 | 0.52 | 0.6 | 0.64 | 0.49 | 0.51 | 0.48 |
| 8 | 13405241 | rs10503454 | 0.762 | 0.018 | -1.107 | -0.141 | 0.003 | -0.003 | -0.003 | 0 | 0.22 | 0.24 | 0.27 | 0.14 | 0.32 | 0.08 |
| 8 | 13406688 | rs7842394 | -0.109 | -0.063 | -1.126 | -0.089 | -0.037 | 0.02 | 0.02 | 0 | 0.3 | 0.39 | 0.37 | 0.37 | 0.19 | 0.4 |
| 8 | 13407297 | rs1905014 | -0.045 | 0.25 | -1.245 | -0.277 | 0 | 0 | 0 | 0.743 | 0.56 | 0.45 | 0.5 | 0.41 | 0.4 | 0.65 |
| 8 | 13414154 | rs1481597 | 0.162 | 0.203 | -0.816 | -0.181 | 0.022 | 0.007 | 0.007 | 0.126 | 0.33 | 0.19 | 0.29 | 0.28 | 0.24 | 0.29 |
| 8 | 13414280 | rs1481596 | -2.502 | 0.312 | -0.579 | -0.169 | -0.009 | -0.057 | -0.057 | 2.215 | 0.78 | 0.72 | 0.89 | 0.97 | 0.85 | 0.83 |
| 8 | 13416666 | rs6981209 | 1.673 | 0.504 | -0.575 | 0.077 | -0.015 | -0.019 | -0.019 | 1.133 | 0.2 | 0.17 | 0.19 | 0.36 | 0.29 | 0.27 |
| 8 | 13417869 | rs6985737 | 1.885 | 0.449 | -0.461 | 0.163 | 0.048 | 0.152 | 0.152 | 0.094 | 0.07 | 0.19 | 0.31 | 0.28 | 0.16 | 0.09 |
| 8 | 13418788 | rs7821950 | -0.102 | -0.019 | -0.522 | -0.374 | -0.003 | -0.007 | -0.007 | 0.287 | 0.42 | 0.58 | 0.59 | 0.64 | 0.61 | 0.47 |
| 8 | 13419965 | rs7006225 | -0.787 | -0.037 | -0.508 | -0.363 | 0.001 | 0.001 | 0.001 | 0 | 0.56 | 0.62 | 0.65 | 0.69 | 0.67 | 0.67 |
| 8 | 13423179 | rs6989489 | 0.34 | 0.055 | -0.38 | -0.317 | -0.02 | -0.002 | -0.002 | 0 | 0.34 | 0.33 | 0.53 | 0.67 | 0.52 | 0.45 |
| 8 | 13424091 | rs17225114 | 0.618 | 0.048 | -0.486 | -0.352 | 0.005 | 0.001 | 0.001 | 0 | 0.3 | 0.37 | 0.51 | 0.64 | 0.5 | 0.42 |
| 8 | 13424583 | rs13273355 | -1.966 | -0.04 | -0.668 | -0.418 | -0.009 | -0.065 | -0.065 | 3.021 | 0.74 | 0.69 | 0.86 | 0.97 | 0.82 | 0.77 |
| 8 | 13426508 | rs10090349 | 1.91 | -0.076 | -0.873 | -0.413 | 0.004 | 0.001 | 0.001 | 0 | 0.26 | 0.23 | 0.14 | 0.03 | 0.19 | 0.23 |
| 8 | 13426832 | rs4145815 | 1.185 | -0.282 | -0.939 | -0.056 | 0.007 | -0.009 | -0.009 | 0 | 0.18 | 0.09 | 0.23 | 0.24 | 0.15 | 0.1 |
| 8 | 13430578 | rs11783751 | 0.526 | -0.063 | -1.052 | -0.274 | 0.024 | 0.095 | 0.095 | 0 | 0.24 | 0.05 | 0.12 | 0.06 | 0.17 | 0.23 |
| 8 | 13434686 | rs10086825 | -0.572 | -0.065 | -0.53 | 0.21 | -0.032 | 0.029 | 0.029 | 0 | 0.76 | 0.57 | 0.74 | 0.69 | 0.81 | 0.83 |
| 8 | 13434921 | rs11785331 | 1.33 | -0.224 | -0.497 | 0.073 | 0 | 0 | 0 | 0 | 0.15 | 0.13 | 0.08 | 0.06 | 0.07 | 0.21 |
| 8 | 13435046 | rs2410092 | -0.393 | -0.105 | -0.491 | -0.312 | 0.001 | 0.001 | 0.001 | 0 | 0.57 | 0.45 | 0.64 | 0.66 | 0.67 | 0.64 |
| 8 | 13438708 | rs7817551 | 2.052 | -0.075 | -0.518 | -0.323 | -0.011 | -0.051 | -0.051 | 0.63 | 0.16 | 0.26 | 0.1 | 0.03 | 0.1 | 0.12 |
| 8 | 13444412 | rs900065 | 0.317 | 0.189 | -0.771 | -0.048 | -0.001 | -0.001 | -0.001 | 1.564 | 0.35 | 0.75 | 0.45 | 0.32 | 0.35 | 0.3 |
| 8 | 13448112 | rs6530649 | 1.163 | 0.256 | -1.229 | 0.089 | -0.079 | -0.062 | -0.062 | 0.128 | 0.24 | 0.07 | 0.21 | 0.35 | 0.35 | 0.33 |
| 8 | 13452449 | rs12677258 | 1.072 | 0.218 | -1.141 | 0.125 | -0.007 | 0.008 | 0.008 | 0 | 0.27 | 0.36 | 0.37 | 0.31 | 0.21 | 0.25 |
| 8 | 13454418 | rs1481590 | 0.169 | 0.304 | -1.376 | 0.025 | 0.01 | -0.013 | -0.013 | 0.276 | 0.5 | 0.19 | 0.44 | 0.53 | 0.48 | 0.62 |
| 8 | 13456131 | rs1021087 | -1.067 | 0.189 | -1.422 | -0.057 | -0.028 | 0.006 | 0.006 | 0 | 0.84 | 0.89 | 0.7 | 0.87 | 0.73 | 0.9 |
| 8 | 13458794 | rs17227506 | 0.502 | 0 | -1.677 | 0.006 | -0.035 | -0.069 | -0.069 | 0 | 0.4 | 0.13 | 0.47 | 0.5 | 0.43 | 0.56 |
| 8 | 13460704 | rs17094707 | 1.329 | -0.025 | -1.37 | 0.084 | -0.017 | -0.011 | -0.011 | 0.125 | 0.22 | 0.31 | 0.14 | 0.17 | 0.13 | 0.17 |
| 8 | 13462286 | rs9325661 | -1.91 | 0.01 | -1.351 | 0.112 | -0.001 | 0.097 | 0.097 | 0 | 0.88 | 0.6 | 0.9 | 0.82 | 0.91 | 0.89 |
| 8 | 13462398 | rs9650348 | 0.036 | 0.068 | -1.641 | -0.045 | 0.011 | 0.059 | 0.059 | 0 | 0.5 | 0.28 | 0.35 | 0.37 | 0.54 | 0.59 |
| 8 | 13463024 | rs7843188 | 1.59 | 0.086 | -1.485 | -0.53 | 0 | 0.068 | 0.068 | 0 | 0.14 | 0.21 | 0.13 | 0.46 | 0.14 | 0.22 |
| 8 | 13468201 | rs10091421 | 1.129 | 0.162 | -2.06 | -0.646 | -0.002 | -0.003 | -0.003 | 0 | 0.27 | 0.27 | 0.26 | 0.19 | 0.2 | 0.19 |
| 8 | 13475278 | rs12155866 | 0.577 | 0.224 | -2.02 | -0.607 | 0.022 | 0.035 | 0.035 | 0 | 0.42 | 0.31 | 0.38 | 0.25 | 0.3 | 0.41 |
| 8 | 13475785 | rs7013716 | -0.625 | 0.482 | -2.198 | -0.501 | 0.042 | 0.105 | 0.105 | 0.223 | 0.78 | 0.48 | 0.79 | 0.6 | 0.75 | 0.73 |
| 8 | 13484121 | rs11776158 | 1 | 0.705 | -2.336 | -0.446 | -0.084 | 0.03 | 0.03 | 0 | 0.2 | 0.03 | 0.17 | 0.1 | 0.29 | 0.22 |
| 8 | 13485034 | rs7816852 | -0.966 | 0.66 | -2.307 | -0.408 | -0.012 | -0.015 | -0.015 | 0.576 | 0.8 | 0.92 | 0.68 | 0.78 | 0.78 | 0.71 |
| 8 | 13486906 | rs7006666 | 0.357 | 0.308 | -1.976 | -0.494 | -0.005 | -0.002 | -0.002 | 0 | 0.41 | 0.55 | 0.27 | 0.42 | 0.36 | 0.39 |
| 8 | 13487294 | rs10503460 | 0.994 | 0.262 | -1.738 | 0.182 | -0.001 | -0.036 | -0.036 | 0 | 0.13 | 0.01 | 0.16 | 0.14 | 0.09 | 0.06 |
| 8 | 13487425 | rs6992740 | 0.441 | 0.736 | -1.303 | 0.339 | 0.014 | 0.059 | 0.059 | 0 | 0.08 | 0.22 | 0.07 | 0.11 | 0.04 | 0.03 |
| 8 | 13489891 | rs11203501 | 0.324 | 1.121 | -1.611 | 0.345 | -0.003 | -0.004 | -0.004 | 0.001 | 0.71 | 0.73 | 0.76 | 0.59 | 0.61 | 0.69 |
| 8 | 13490323 | rs4629867 | 0.625 | 1.089 | -2.052 | 1.102 | -0.019 | -0.018 | -0.018 | 0.045 | 0.17 | 0.02 | 0.21 | 0.14 | 0.15 | 0.08 |
| 8 | 13490463 | rs1383879 | -0.07 | 1.518 | -1.118 | 1.399 | -0.008 | -0.044 | -0.044 | 0 | 0.12 | 0.03 | 0.12 | 0.4 | 0.17 | 0.14 |
| 8 | 13490995 | rs10503462 | 0.178 | 1.565 | -0.62 | 0.598 | 0.016 | -0.023 | -0.023 | 0 | 0.74 | 0.65 | 0.78 | 0.85 | 0.71 | 0.8 |
| 8 | 13492542 | rs7002898 | -0.793 | 1.618 | -0.987 | 1.068 | -0.024 | -0.022 | -0.022 | 0 | 0.05 | 0.02 | 0.22 | 0.25 | 0.15 | 0.03 |
| 8 | 13493056 | rs10089460 | -0.378 | 1.589 | -0.695 | 0.705 | 0.015 | 0 | 0 | 0 | 0.39 | 0.49 | 0.49 | 0.42 | 0.51 | 0.3 |
| 8 | 13495289 | rs17116929 | 0.063 | 1.42 | -0.248 | 0.09 | -0.001 | -0.012 | -0.012 | 0.601 | 0.13 | 0.15 | 0.05 | 0.03 | 0.12 | 0.37 |
| 8 | 13496786 | rs6530659 | 0.412 | 0.951 | -0.218 | 0.384 | -0.009 | 0.043 | 0.043 | 0 | 0.71 | 0.5 | 0.65 | 0.6 | 0.77 | 0.49 |
| 8 | 13509864 | rs17116959 | 0.349 | 0.952 | -0.131 | 0.32 | 0 | -0.02 | -0.02 | 0.909 | 0.13 | 0.17 | 0.05 | 0.04 | 0.12 | 0.38 |
| 8 | 13510293 | rs7008403 | 0.687 | 1.069 | -0.478 | -0.039 | -0.003 | 0.014 | 0.014 | 0.335 | 0.62 | 0.44 | 0.57 | 0.55 | 0.65 | 0.38 |
| 8 | 13513544 | rs6986816 | 0.62 | 1.014 | -0.004 | -0.646 | -0.001 | 0.003 | 0.003 | 0 | 0.41 | 0.62 | 0.21 | 0.27 | 0.36 | 0.25 |
| 8 | 13515457 | rs1481608 | 1.036 | 1.087 | 0.038 | -1.123 | -0.001 | 0 | 0 | 0 | 0.24 | 0.24 | 0.09 | 0.21 | 0.14 | 0.15 |
| 8 | 13516023 | rs7843942 | 0.027 | 1.085 | 0.077 | -1.122 | 0.002 | -0.015 | -0.015 | 0 | 0.41 | 0.54 | 0.21 | 0.26 | 0.36 | 0.25 |
| 8 | 13516584 | rs17236421 | 1.65 | 1.118 | -0.092 | -1.049 | -0.024 | 0.001 | 0.001 | 0 | 0.09 | 0 | 0.08 | 0.04 | 0.1 | 0.1 |

| | | | | | | | | | | | | | | | | |
|---|---|---|---|---|---|---|---|---|---|---|---|---|---|---|---|---|
| 8 | 13520160 | rs1481604 | 0.013 | 1.068 | 0.008 | -1.061 | 0 | -0.001 | -0.001 | 0 | 0.41 | 0.6 | 0.21 | 0.26 | 0.36 | 0.26 |
| 8 | 13521103 | rs2062045 | -0.465 | 0.94 | 0.091 | -0.655 | 0 | -0.002 | -0.002 | 0 | 0.47 | 0.53 | 0.27 | 0.4 | 0.4 | 0.32 |
| 8 | 13524802 | rs12677921 | -0.731 | 0.956 | 0.06 | -0.711 | -0.011 | 0.001 | 0.001 | 0.089 | 0.71 | 0.47 | 0.87 | 0.78 | 0.79 | 0.85 |
| 8 | 13526692 | rs13253300 | 0.783 | 0.882 | -0.084 | -0.876 | -0.001 | -0.023 | -0.023 | 0 | 0.1 | 0.1 | 0.05 | 0.02 | 0.1 | 0.2 |
| 8 | 13527166 | rs10503464 | 1.901 | 0.9 | 0.01 | -0.799 | -0.004 | -0.034 | -0.034 | 0.237 | 0.06 | 0 | 0.07 | 0.14 | 0.04 | 0.06 |
| 8 | 13527276 | rs1947905 | 1.899 | 0.963 | -0.041 | -0.904 | -0.001 | -0.015 | -0.015 | 5.818 | 0.11 | 0.1 | 0.04 | 0.02 | 0.11 | 0.27 |
| 8 | 13527569 | rs11782494 | -0.601 | 0.733 | -0.002 | -0.996 | -0.019 | 0.023 | 0.023 | 0 | 0.7 | 0.43 | 0.83 | 0.64 | 0.82 | 0.8 |
| 8 | 13527801 | rs13265604 | 0.267 | 0.641 | -0.101 | -1.085 | 0.018 | 0.123 | 0.123 | 0 | 0.81 | 0.44 | 0.84 | 0.63 | 0.82 | 0.85 |
| 8 | 13529635 | rs17237212 | -0.108 | 0.63 | -0.25 | -1.177 | -0.033 | 0.139 | 0.139 | 0 | 0.38 | 0.16 | 0.23 | 0.1 | 0.38 | 0.48 |
| 8 | 13530357 | rs3849006 | 1.402 | 0.636 | -0.235 | -1.153 | -0.001 | -0.002 | -0.002 | 0 | 0.31 | 0.33 | 0.58 | 0.54 | 0.38 | 0.32 |
| 8 | 13531381 | rs17237541 | -0.678 | 0.723 | -0.243 | -1.134 | -0.006 | 0.071 | 0.071 | 0 | 0.45 | 0.32 | 0.31 | 0.25 | 0.42 | 0.53 |
| 8 | 13535612 | rs17237821 | 0.274 | 0.751 | -0.409 | -1.174 | -0.058 | 0.027 | 0.027 | 0 | 0.25 | 0.1 | 0.2 | 0.19 | 0.26 | 0.17 |
| 8 | 13536002 | rs11785905 | 1.531 | 0.814 | -0.391 | -1.184 | -0.051 | 0.043 | 0.043 | 0 | 0.15 | 0.07 | 0.12 | 0.06 | 0.22 | 0.11 |
| 8 | 13537443 | rs7845073 | -0.277 | 0.702 | -0.414 | -1.197 | -0.062 | 0.181 | 0.181 | 0 | 0.37 | 0.14 | 0.23 | 0.1 | 0.39 | 0.48 |
| 8 | 13539917 | rs4831454 | 1.383 | 0.72 | -0.334 | -1.164 | 0 | -0.035 | -0.035 | 0 | 0.14 | 0.19 | 0.07 | 0.02 | 0.13 | 0.27 |
| 8 | 13540001 | rs4831455 | -0.362 | 0.699 | -0.269 | -1.177 | -0.003 | -0.024 | -0.024 | 0 | 0.3 | 0.34 | 0.22 | 0.19 | 0.26 | 0.44 |
| 8 | 13541779 | rs7821547 | 0.553 | 0.711 | -0.485 | -1.191 | 0 | -0.008 | -0.008 | 1.429 | 0.22 | 0.22 | 0.14 | 0.05 | 0.21 | 0.37 |
| 8 | 13542084 | rs12542520 | 1.086 | 0.747 | -0.506 | -1.183 | -0.046 | 0.075 | 0.075 | 0 | 0.21 | 0.05 | 0.16 | 0.07 | 0.25 | 0.2 |
| 8 | 13542432 | rs12542609 | 1.988 | 0.617 | -0.439 | -1.096 | -0.017 | 0.015 | 0.015 | 0 | 0.09 | 0 | 0.07 | 0.03 | 0.09 | 0.1 |
| 8 | 13543195 | rs13282358 | -0.739 | 0.487 | -0.328 | -1.007 | -0.005 | 0.001 | 0.001 | 0.649 | 0.71 | 0.45 | 0.85 | 0.78 | 0.79 | 0.85 |
| 8 | 13543520 | rs13248289 | 1.35 | 0.493 | -0.417 | -0.951 | 0.001 | -0.046 | -0.046 | 3.051 | 0.3 | 0.21 | 0.56 | 0.55 | 0.37 | 0.31 |
| 8 | 13543579 | rs13256918 | -0.251 | 0.479 | 0.07 | -0.772 | -0.007 | 0.005 | 0.005 | 0.539 | 0.72 | 0.54 | 0.86 | 0.77 | 0.79 | 0.85 |
| 8 | 13544506 | rs1947919 | -0.091 | 0.428 | 0.344 | -0.839 | -0.03 | -0.053 | -0.053 | 0.236 | 0.7 | 0.51 | 0.8 | 0.81 | 0.83 | 0.85 |
| 8 | 13544610 | rs12676987 | 1.662 | 0.586 | -0.296 | -0.963 | -0.024 | -0.131 | -0.131 | 1.435 | 0.35 | 0.07 | 0.52 | 0.56 | 0.36 | 0.42 |
| 8 | 13544718 | rs1947917 | -0.828 | 0.409 | 0.239 | -1.015 | -0.01 | -0.066 | -0.066 | 0 | 0.77 | 0.68 | 0.88 | 0.93 | 0.85 | 0.9 |
| 8 | 13545280 | rs17239169 | 1.503 | 0.664 | -0.236 | -0.98 | -0.014 | 0.035 | 0.035 | 0 | 0.12 | 0.05 | 0.08 | 0.03 | 0.15 | 0.07 |
| 8 | 13556813 | rs11784733 | -0.071 | 0.728 | -0.066 | -0.904 | 0.003 | 0.009 | 0.009 | 0 | 0.08 | 0.14 | 0.06 | 0.14 | 0.07 | 0.06 |
| 8 | 13557493 | rs2410094 | 0.81 | 0.741 | -0.173 | -1.042 | -0.011 | 0.142 | 0.142 | 0 | 0.49 | 0.2 | 0.67 | 0.75 | 0.55 | 0.66 |
| 8 | 13560688 | rs4474022 | -0.241 | 0.699 | -0.033 | -1.071 | 0.003 | 0.011 | 0.011 | 0 | 0.41 | 0.36 | 0.25 | 0.11 | 0.37 | 0.29 |
| 8 | 13562564 | rs12544492 | 1.198 | 1.322 | 0.378 | -0.47 | -0.006 | 0.005 | 0.005 | 0 | 0.07 | 0.06 | 0.08 | 0.02 | 0.12 | 0.11 |
| 8 | 13565045 | rs10103102 | 1.024 | 1.309 | 0.384 | -0.453 | -0.009 | 0.013 | 0.013 | 0 | 0.09 | 0.06 | 0.1 | 0.02 | 0.13 | 0.12 |
| 8 | 13568143 | rs13282636 | 1.509 | 1.293 | 0.383 | -0.414 | 0.002 | -0.01 | -0.01 | 0 | 0.53 | 0.23 | 0.63 | 0.59 | 0.46 | 0.51 |
| 8 | 13572288 | rs10094983 | -1.388 | 1.163 | 0.89 | -0.051 | 0.053 | 0.074 | 0.074 | 0.655 | 0.19 | 0.58 | 0.19 | 0.22 | 0.24 | 0.24 |
| 8 | 13574437 | rs11775652 | -0.656 | 1.136 | 0.793 | -0.124 | -0.004 | -0.005 | -0.005 | 0 | 0.19 | 0.3 | 0.13 | 0.15 | 0.17 | 0.17 |
| 8 | 13574646 | rs12681268 | -0.264 | 1.535 | 0.863 | 0.099 | 0.007 | 0.028 | 0.028 | 0 | 0.13 | 0.15 | 0.12 | 0.19 | 0.19 | 0.23 |
| 8 | 13582367 | rs1481897 | -1.615 | 1.463 | 0.443 | 0.054 | 0.006 | 0.007 | 0.007 | 0.268 | 0.33 | 0.66 | 0.29 | 0.37 | 0.41 | 0.41 |
| 8 | 13593642 | rs6985907 | -0.846 | 1.468 | 0.038 | 0.012 | 0.008 | 0.002 | 0.002 | 0 | 0.19 | 0.28 | 0.16 | 0.18 | 0.21 | 0.18 |
| 8 | 13594258 | rs10092409 | -0.629 | 1.3 | 0.027 | 0.013 | 0.001 | 0.01 | 0.01 | 0 | 0.08 | 0.13 | 0.05 | 0.16 | 0.06 | 0.06 |
| 8 | 13597430 | rs4483156 | 1.308 | 1.07 | 0.066 | -0.506 | 0.017 | 0.006 | 0.006 | 0 | 0.47 | 0.55 | 0.49 | 0.61 | 0.38 | 0.46 |
| 8 | 13600416 | rs1973227 | 0.931 | 1.218 | -0.06 | -0.671 | -0.003 | 0.029 | 0.029 | 0 | 0.16 | 0.09 | 0.08 | 0.04 | 0.17 | 0.1 |
| 8 | 13601481 | rs5012676 | 1.461 | 1.123 | 0.063 | -0.539 | 0.013 | 0.003 | 0.003 | 0 | 0.46 | 0.51 | 0.48 | 0.61 | 0.38 | 0.47 |
| 8 | 13604114 | rs1384043 | -0.077 | 1.326 | 0.174 | -0.481 | 0.009 | 0.036 | 0.036 | 0 | 0.8 | 0.72 | 0.67 | 0.69 | 0.72 | 0.78 |
| 8 | 13608309 | rs13256272 | 1.754 | 0.596 | -0.182 | -0.524 | -0.028 | 0.007 | 0.007 | 0.22 | 0.14 | 0.16 | 0.05 | 0.16 | 0.04 | 0.08 |
| 8 | 13609377 | rs11780002 | 1.909 | 0.948 | -0.621 | -0.687 | -0.08 | -0.104 | -0.104 | 0.981 | 0.3 | 0.05 | 0.41 | 0.46 | 0.34 | 0.38 |
| 8 | 13609518 | rs11783610 | 1.632 | 0.783 | -0.535 | -0.618 | -0.031 | 0.004 | 0.004 | 0 | 0.1 | 0.03 | 0.22 | 0.07 | 0.14 | 0.13 |
| 8 | 13610731 | rs12678711 | 1.937 | 0.709 | -0.597 | -0.609 | 0 | -0.062 | -0.062 | 0 | 0.28 | 0.09 | 0.38 | 0.46 | 0.27 | 0.34 |
| 8 | 13613335 | rs6984781 | -0.166 | 0.307 | 0.08 | -0.53 | -0.016 | -0.036 | -0.036 | 0 | 0.3 | 0.54 | 0.17 | 0.17 | 0.21 | 0.25 |
| 8 | 13623820 | rs13258997 | 1.369 | 0.188 | -0.643 | -0.302 | 0.054 | -0.013 | -0.013 | 0 | 0.16 | 0.04 | 0.05 | 0.15 | 0.06 | 0.1 |
| 8 | 13625816 | rs17117434 | 1.238 | 0.187 | -0.665 | -0.305 | 0.065 | -0.007 | -0.007 | 0 | 0.18 | 0.04 | 0.05 | 0.15 | 0.06 | 0.1 |
| 8 | 13628490 | rs17254321 | 1.449 | 0.574 | -0.665 | -0.366 | -0.029 | 0.012 | 0.012 | 0 | 0.1 | 0.03 | 0.22 | 0.08 | 0.14 | 0.13 |
| 8 | 13631473 | rs7814407 | 0.245 | 0.449 | -0.253 | -0.39 | 0.006 | -0.001 | -0.001 | 0 | 0.54 | 0.5 | 0.73 | 0.54 | 0.63 | 0.66 |
| 8 | 13638207 | rs1384048 | 0.077 | 0.424 | -1.01 | -0.341 | 0.062 | -0.042 | -0.042 | 0.113 | 0.57 | 0.25 | 0.76 | 0.56 | 0.69 | 0.74 |
| 8 | 13653625 | rs11774781 | 1.4 | 0.757 | -1.438 | -0.248 | -0.109 | -0.079 | -0.079 | 0 | 0.31 | 0.04 | 0.42 | 0.43 | 0.39 | 0.38 |
| 8 | 13653983 | rs6530673 | 0.887 | 0.698 | -1.427 | -0.298 | -0.079 | -0.038 | -0.038 | 0.637 | 0.37 | 0.1 | 0.46 | 0.47 | 0.43 | 0.44 |
| 8 | 13656961 | rs12164168 | -0.299 | 0.877 | -1.102 | -0.108 | -0.005 | 0.023 | 0.023 | 0 | 0.92 | 0.92 | 0.92 | 0.76 | 0.87 | 0.95 |
| 8 | 13657315 | rs11203526 | 1.408 | 0.87 | -1.251 | -0.327 | -0.029 | 0.01 | 0.01 | 0 | 0.11 | 0.02 | 0.21 | 0.07 | 0.13 | 0.13 |
| 8 | 13659502 | rs7014031 | 0.847 | 0.661 | -0.89 | -0.503 | 0.009 | 0.012 | 0.012 | 0.019 | 0.26 | 0.22 | 0.11 | 0.16 | 0.17 | 0.23 |
| 8 | 13660787 | rs7015855 | 0.477 | 0.705 | -0.249 | -0.417 | 0.005 | -0.002 | -0.002 | 0 | 0.52 | 0.38 | 0.7 | 0.56 | 0.58 | 0.61 |
| 8 | 13665492 | rs4831476 | 0.323 | 0.436 | -0.14 | -0.345 | 0.011 | -0.008 | -0.008 | 1.807 | 0.56 | 0.61 | 0.59 | 0.54 | 0.62 | 0.62 |
| 8 | 13667268 | rs2054608 | 0.923 | 0.402 | 0.005 | -0.377 | 0.008 | -0.003 | -0.003 | 0 | 0.47 | 0.15 | 0.52 | 0.52 | 0.58 | 0.54 |
| 8 | 13669693 | rs13275974 | 0.506 | 0.293 | 0.148 | -0.405 | 0.008 | 0.008 | 0.008 | 0.184 | 0.31 | 0.29 | 0.14 | 0.18 | 0.16 | 0.26 |
| 8 | 13670677 | rs10086085 | 0.715 | 0.596 | 0.455 | -0.365 | 0.001 | 0.001 | 0.001 | 0.127 | 0.49 | 0.38 | 0.52 | 0.52 | 0.56 | 0.56 |
| 8 | 13673447 | rs2410116 | -0.378 | 0.658 | 0.385 | -0.393 | -0.001 | 0.001 | 0.001 | 0 | 0.8 | 0.82 | 0.84 | 0.72 | 0.72 | 0.85 |
| 8 | 13673943 | rs17117636 | 0.164 | 0.634 | 0.364 | -0.373 | -0.03 | -0.027 | -0.027 | 0 | 0.08 | 0.02 | 0.07 | 0.23 | 0.14 | 0.04 |
| 8 | 13674437 | rs13252414 | -0.205 | 1.248 | 0.475 | -0.189 | 0.002 | 0.004 | 0.004 | 0 | 0.43 | 0.45 | 0.44 | 0.35 | 0.38 | 0.46 |
| 8 | 13674554 | rs9325677 | -0.305 | 0.75 | 0.626 | -0.471 | -0.017 | -0.027 | -0.027 | 0 | 0.44 | 0.52 | 0.39 | 0.25 | 0.31 | 0.38 |
| 8 | 13676425 | rs7827109 | 0.896 | 0.768 | 0.496 | -0.467 | 0.007 | 0.003 | 0.003 | 0.22 | 0.47 | 0.2 | 0.51 | 0.52 | 0.56 | 0.57 |
| 8 | 13677952 | rs2128127 | 1.598 | 0.824 | 0.389 | -0.424 | -0.105 | -0.098 | -0.098 | 0 | 0.34 | 0.05 | 0.44 | 0.47 | 0.41 | 0.46 |
| 8 | 13680840 | rs6994400 | 0.496 | 0.931 | 0.739 | -0.316 | 0.128 | 0.298 | 0.298 | 0.884 | 0.93 | 0.59 | 0.91 | 0.78 | 0.89 | 0.95 |

| | | | | | | | | | | | | | | | | |
|---|---|---|---|---|---|---|---|---|---|---|---|---|---|---|---|---|
| 8 | 13684419 | rs12718360 | 0.542 | 0.781 | 0.36 | -0.505 | 0.016 | 0.029 | 0.029 | 0.07 | 0.72 | 0.6 | 0.76 | 0.72 | 0.66 | 0.76 |
| 8 | 13685474 | rs7016635 | 2.09 | 0.787 | 0.158 | -0.546 | -0.055 | -0.071 | -0.071 | 0.054 | 0.29 | 0.11 | 0.42 | 0.46 | 0.35 | 0.44 |
| 8 | 13686240 | rs17117713 | 0.363 | 0.79 | 0.126 | -0.495 | -0.028 | -0.022 | -0.022 | 0 | 0.08 | 0.02 | 0.07 | 0.25 | 0.13 | 0.05 |
| 8 | 13686259 | rs10282775 | 1.306 | 0.715 | 0.222 | -0.565 | 0.002 | -0.026 | -0.026 | 0 | 0.14 | 0.2 | 0.2 | 0.09 | 0.15 | 0.11 |
| 8 | 13686816 | rs10503477 | 1.57 | 0.514 | -0.056 | -0.632 | 0.078 | 0.01 | 0.01 | 0.613 | 0.19 | 0.04 | 0.05 | 0.14 | 0.06 | 0.1 |
| 8 | 13687215 | rs10216855 | 0.612 | 0.926 | 0.208 | -0.449 | 0.017 | 0.037 | 0.037 | 0.049 | 0.72 | 0.58 | 0.76 | 0.71 | 0.66 | 0.76 |
| 8 | 13688369 | rs7013031 | -0.243 | 0.907 | 0.19 | -0.484 | -0.002 | -0.054 | -0.054 | 0 | 0.81 | 0.73 | 0.84 | 0.96 | 0.8 | 0.82 |
| 8 | 13691166 | rs10107281 | 1.539 | 0.947 | -0.615 | -0.397 | -0.009 | -0.022 | -0.022 | 0 | 0.06 | 0.1 | 0.03 | 0.02 | 0.07 | 0.05 |
| 8 | 13694986 | rs12675887 | 1.941 | 1.027 | -0.756 | -0.361 | -0.099 | -0.113 | -0.113 | 1.837 | 0.29 | 0.04 | 0.4 | 0.45 | 0.34 | 0.43 |
| 8 | 13696026 | rs1157338 | 1.443 | 0.82 | -0.262 | -0.242 | -0.028 | -0.021 | -0.021 | 0 | 0.41 | 0.25 | 0.61 | 0.53 | 0.48 | 0.51 |
| 8 | 13697822 | rs17117781 | 1.412 | 0.771 | -0.477 | -0.238 | -0.064 | -0.052 | -0.052 | 0 | 0.41 | 0.15 | 0.61 | 0.53 | 0.49 | 0.51 |
| 8 | 13699894 | rs7009172 | 0.272 | 0.746 | -0.526 | -0.108 | -0.046 | -0.056 | -0.056 | 0.2 | 0.29 | 0.48 | 0.15 | 0.17 | 0.16 | 0.22 |
| 8 | 13703387 | rs13265387 | -0.412 | 1.512 | -0.823 | 0.44 | -0.033 | -0.049 | -0.049 | 0 | 0.09 | 0.01 | 0.09 | 0.24 | 0.13 | 0.05 |
| 8 | 13711241 | rs1480859 | -0.537 | 1.488 | -0.779 | 0.355 | -0.005 | 0.009 | 0.009 | 0 | 0.88 | 0.89 | 0.88 | 0.98 | 0.87 | 0.87 |
| 8 | 13711549 | rs1480858 | -0.811 | 1.425 | -0.754 | 0.602 | -0.015 | -0.017 | -0.017 | 0 | 0.1 | 0.05 | 0.11 | 0.25 | 0.17 | 0.09 |
| 8 | 13715911 | rs1600646 | -0.704 | 1.172 | -0.7 | 0.128 | -0.007 | 0.012 | 0.012 | 0.029 | 0.45 | 0.4 | 0.3 | 0.37 | 0.49 | 0.42 |
| 8 | 13716524 | rs1383492 | 0.051 | 1.23 | -0.069 | -0.149 | 0.019 | -0.08 | -0.08 | 0 | 0.74 | 0.52 | 0.84 | 0.86 | 0.68 | 0.81 |
| 8 | 13719796 | rs10087180 | 0.335 | 1.001 | -0.203 | 0.189 | 0 | 0 | 0 | 1.283 | 0.64 | 0.59 | 0.66 | 0.66 | 0.59 | 0.59 |
| 8 | 13721994 | rs4831486 | 0.42 | 0.295 | -0.462 | 0.032 | -0.006 | -0.001 | -0.001 | 0.021 | 0.64 | 0.39 | 0.75 | 0.63 | 0.73 | 0.7 |
| 8 | 13726384 | rs4831495 | -0.106 | 0.276 | -0.464 | 0.068 | 0.009 | 0.001 | 0.001 | 0 | 0.54 | 0.42 | 0.55 | 0.47 | 0.56 | 0.5 |
| 8 | 13726729 | rs13249149 | 1.71 | 0.337 | -0.718 | -0.064 | -0.111 | 0.015 | 0.015 | 0 | 0.21 | 0.02 | 0.24 | 0.11 | 0.3 | 0.21 |
| 8 | 13727528 | rs13259601 | -0.203 | 0.219 | -0.491 | -0.005 | -0.007 | 0.013 | 0.013 | 0 | 0.38 | 0.44 | 0.33 | 0.53 | 0.3 | 0.34 |
| 8 | 13729850 | rs6530685 | -0.194 | 0.235 | -0.499 | -0.012 | -0.007 | 0.013 | 0.013 | 0 | 0.38 | 0.44 | 0.33 | 0.52 | 0.3 | 0.35 |
| 8 | 13731228 | rs10888072 | -0.459 | -0.049 | -0.253 | -0.017 | -0.001 | -0.001 | -0.001 | 0.34 | 0.59 | 0.6 | 0.58 | 0.61 | 0.6 | 0.56 |
| 8 | 13733048 | rs13265025 | 0.979 | -0.106 | -0.128 | 0.126 | -0.001 | 0.028 | 0.028 | 0 | 0.21 | 0.08 | 0.19 | 0.14 | 0.17 | 0.17 |
| 8 | 13734903 | rs13250045 | -0.234 | -0.126 | 0.087 | 0.168 | -0.019 | -0.012 | -0.012 | 0 | 0.33 | 0.49 | 0.31 | 0.31 | 0.25 | 0.26 |
| 8 | 13736874 | rs6988928 | 1.794 | -0.316 | -0.225 | -0.034 | -0.138 | 0.066 | 0.066 | 0 | 0.27 | 0.02 | 0.26 | 0.11 | 0.36 | 0.23 |
| 8 | 13736961 | rs13269960 | -0.42 | -0.366 | -0.079 | 0.011 | -0.001 | 0 | 0 | 0 | 0.37 | 0.37 | 0.32 | 0.47 | 0.28 | 0.32 |
| 8 | 13737259 | rs7016993 | -0.582 | -0.366 | -0.134 | 0.023 | -0.003 | 0.001 | 0.001 | 0 | 0.66 | 0.63 | 0.61 | 0.62 | 0.66 | 0.57 |
| 8 | 13737515 | rs7013408 | -0.673 | -0.363 | 0.171 | 0.19 | 0.001 | -0.002 | -0.002 | 0.502 | 0.38 | 0.59 | 0.33 | 0.47 | 0.29 | 0.32 |
| 8 | 13738781 | rs10888073 | -0.416 | -0.325 | 0.108 | 0.017 | -0.007 | -0.001 | -0.001 | 0 | 0.33 | 0.59 | 0.32 | 0.35 | 0.25 | 0.28 |
| 8 | 13739012 | rs969365 | 0.035 | -0.359 | 0.032 | -0.045 | 0.003 | 0.007 | 0.007 | 0 | 0.7 | 0.75 | 0.75 | 0.84 | 0.76 | 0.71 |
| 8 | 13739069 | rs13271017 | -0.513 | -0.271 | -0.059 | 0.206 | -0.003 | -0.001 | -0.001 | 0.419 | 0.33 | 0.38 | 0.31 | 0.35 | 0.28 | 0.3 |
| 8 | 13741178 | rs11203543 | 0.337 | -0.317 | -0.237 | 0.518 | -0.012 | 0.096 | 0.096 | 0 | 0.13 | 0.52 | 0.14 | 0.21 | 0.08 | 0.12 |
| 8 | 13742298 | rs12547223 | 0.617 | -0.055 | -1.294 | 0.48 | -0.013 | -0.075 | -0.075 | 0 | 0.22 | 0.08 | 0.31 | 0.36 | 0.24 | 0.3 |
| 8 | 13743940 | rs6530688 | -0.752 | -0.409 | -1.326 | -0.015 | -0.058 | 0.069 | 0.069 | 0 | 0.66 | 0.79 | 0.65 | 0.9 | 0.53 | 0.76 |
| 8 | 13748391 | rs10089158 | 0.107 | -1.098 | -1.064 | 0.105 | -0.002 | -0.001 | -0.001 | 1.439 | 0.18 | 0.17 | 0.21 | 0.22 | 0.15 | 0.2 |
| 8 | 13748948 | rs7836269 | 0.797 | -1.043 | -0.286 | 0.738 | 0.014 | 0.117 | 0.117 | 0 | 0.07 | 0.18 | 0.1 | 0.17 | 0.04 | 0.04 |
| 8 | 13749772 | rs1383498 | 0.463 | -1.435 | -0.107 | 0.542 | -0.036 | 0.061 | 0.061 | 0.575 | 0.87 | 0.64 | 0.86 | 0.83 | 0.96 | 0.94 |
| 8 | 13750535 | rs10448132 | 0.224 | -1.728 | -0.371 | 0.326 | -0.012 | 0.011 | 0.011 | 0 | 0.14 | 0.16 | 0.12 | 0.17 | 0.07 | 0.09 |
| 8 | 13750744 | rs10156329 | 0.155 | -1.551 | -0.256 | 0.972 | -0.021 | 0.022 | 0.022 | 0 | 0.13 | 0.26 | 0.07 | 0.18 | 0.05 | 0.19 |
| 8 | 13752511 | rs7845848 | -0.714 | -1.523 | -0.267 | 0.918 | -0.029 | 0.098 | 0.098 | 0.307 | 0.18 | 0.34 | 0.16 | 0.35 | 0.09 | 0.23 |
| 8 | 13758130 | rs10096982 | 0.309 | -1.638 | -0.583 | 0.742 | 0.005 | 0.001 | 0.001 | 0 | 0.46 | 0.37 | 0.46 | 0.46 | 0.49 | 0.48 |
| 8 | 13760419 | rs10888075 | 1.076 | -1.609 | -0.456 | 0.802 | -0.025 | 0.08 | 0.08 | 0.176 | 0.77 | 0.54 | 0.8 | 0.65 | 0.85 | 0.76 |
| 8 | 13762061 | rs10102306 | 0.777 | -1.472 | -0.578 | 0.802 | 0.008 | -0.029 | -0.029 | 0 | 0.05 | 0.14 | 0.04 | 0 | 0.06 | 0.01 |
| 8 | 13764930 | rs1904875 | -0.556 | -1.512 | -0.413 | 0.869 | -0.001 | 0.08 | 0.08 | 0.444 | 0.18 | 0.3 | 0.16 | 0.36 | 0.15 | 0.23 |
| 8 | 13766875 | rs4831250 | -0.151 | -1.631 | -0.412 | 0.764 | 0.021 | 0.005 | 0.005 | 0 | 0.49 | 0.55 | 0.49 | 0.54 | 0.45 | 0.51 |
| 8 | 13777214 | rs10888076 | -0.917 | -1.616 | -0.032 | 0.782 | 0.012 | -0.009 | -0.009 | 0 | 0.59 | 0.49 | 0.61 | 0.71 | 0.54 | 0.55 |
| 8 | 13786824 | rs1520754 | -1.256 | -1.619 | -0.007 | 0.691 | -0.022 | 0.036 | 0.036 | 0 | 0.64 | 0.7 | 0.64 | 0.9 | 0.59 | 0.74 |
| 8 | 13787726 | rs17118033 | 1.169 | -1.703 | -0.316 | 0.974 | -0.021 | -0.04 | -0.04 | 0 | 0.08 | 0.01 | 0.06 | 0.16 | 0.12 | 0.13 |
| 8 | 13789061 | rs1356945 | 2.338 | -1.473 | -0.112 | 0.822 | -0.013 | -0.035 | -0.035 | 0 | 0.06 | 0.01 | 0.03 | 0.18 | 0.05 | 0.19 |
| 8 | 13790074 | rs1402331 | -1.174 | -1.436 | 0.385 | 0.625 | 0.006 | -0.018 | -0.018 | 0.126 | 0.66 | 0.59 | 0.65 | 0.89 | 0.6 | 0.75 |
| 8 | 13790151 | rs1829561 | 0.234 | -1.365 | 0.553 | 0.726 | 0.057 | 0.134 | 0.134 | 0 | 0.09 | 0.31 | 0.14 | 0.16 | 0.1 | 0.04 |
| 8 | 13791180 | rs6991638 | 2.178 | -1.342 | -0.044 | 0.545 | -0.157 | 0.013 | 0.013 | 1.118 | 0.23 | 0.01 | 0.28 | 0.1 | 0.37 | 0.22 |
| 8 | 13792292 | rs2622478 | 0.85 | -1.3 | 0.645 | 0.758 | -0.033 | 0.034 | 0.034 | 0.041 | 0.83 | 0.55 | 0.82 | 0.83 | 0.9 | 0.93 |
| 8 | 13798005 | rs2682665 | -0.414 | -1.276 | 0.731 | 0.612 | -0.021 | -0.024 | -0.024 | 0 | 0.93 | 0.7 | 0.94 | 1 | 1 | 0.97 |
| 8 | 13799783 | rs9643963 | 0.095 | -1.446 | 0.537 | 0.537 | -0.035 | -0.046 | -0.046 | 0 | 0.84 | 0.34 | 0.78 | 0.92 | 0.92 | 0.86 |
| 8 | 13800461 | rs4562326 | 1.355 | -1.511 | 0.23 | 0.712 | 0 | -0.045 | -0.045 | 0 | 0.09 | 0.02 | 0.04 | 0.19 | 0.06 | 0.17 |
| 8 | 13802354 | rs11203550 | -0.323 | -1.504 | 0.826 | 0.606 | -0.03 | -0.049 | -0.049 | 0.039 | 0.86 | 0.5 | 0.79 | 0.92 | 0.92 | 0.86 |
| 8 | 13803253 | rs12677281 | -0.034 | -1.514 | 0.755 | 0.601 | -0.039 | -0.055 | -0.055 | 0 | 0.85 | 0.43 | 0.79 | 0.92 | 0.92 | 0.86 |
| 8 | 13803949 | rs17202944 | 2.256 | -1.212 | 0.253 | 0.556 | -0.161 | 0.046 | 0.046 | 0 | 0.27 | 0.02 | 0.29 | 0.1 | 0.37 | 0.23 |
| 8 | 13804128 | rs17118128 | -0.438 | -1.284 | 0.562 | 0.798 | 0.037 | 0.117 | 0.117 | 0 | 0.15 | 0.39 | 0.17 | 0.27 | 0.16 | 0.1 |
| 8 | 13807670 | rs10092922 | 0.673 | -1.257 | 0.491 | 0.78 | 0.007 | 0.001 | 0.001 | 0 | 0.08 | 0.13 | 0.06 | 0.09 | 0.12 | 0.05 |
| 8 | 13810061 | rs2139722 | -0.525 | -1.418 | 0.606 | 0.714 | -0.018 | -0.045 | -0.045 | 0 | 0.86 | 0.67 | 0.8 | 0.92 | 0.9 | 0.86 |
| 8 | 13810119 | rs2176903 | 0.684 | -1.376 | 0.362 | 0.831 | 0.044 | 0.002 | 0.002 | 0 | 0.37 | 0.16 | 0.29 | 0.38 | 0.3 | 0.38 |
| 8 | 13815335 | rs1385620 | 0.116 | -1.365 | 0.337 | 0.827 | -0.011 | 0.003 | 0.003 | 0 | 0.43 | 0.42 | 0.4 | 0.55 | 0.34 | 0.39 |
| 8 | 13816796 | rs12681911 | 0.767 | -1.122 | 0.429 | 0.962 | 0.009 | 0.119 | 0.119 | 0 | 0.06 | 0.19 | 0.11 | 0.17 | 0.04 | 0.03 |
| 8 | 13823380 | rs12678513 | 0.746 | -1.123 | 0.378 | 0.945 | 0.016 | 0.166 | 0.166 | 0 | 0.06 | 0.22 | 0.11 | 0.18 | 0.04 | 0.03 |
| 8 | 13823917 | rs991885 | 0.514 | -1.044 | 0.19 | 0.721 | 0.001 | -0.004 | -0.004 | 0 | 0.79 | 0.72 | 0.77 | 0.82 | 0.78 | 0.8 |

| | | | | | | | | | | | | | | | | |
|---|---|---|---|---|---|---|---|---|---|---|---|---|---|---|---|---|
| 8 | 13825467 | rs1485532 | 0.513 | -1.057 | 0.19 | 0.69 | 0.001 | -0.004 | -0.004 | 0 | 0.79 | 0.72 | 0.77 | 0.82 | 0.78 | 0.8 |
| 8 | 13827505 | rs2410132 | 1.192 | -1.078 | 0.072 | 0.707 | -0.002 | 0.012 | 0.012 | 2.09 | 0.11 | 0.1 | 0.06 | 0.18 | 0.1 | 0.18 |
| 8 | 13829314 | rs13258175 | 0.577 | -1.49 | -0.175 | 0.429 | 0.006 | 0.012 | 0.012 | 0.077 | 0.49 | 0.47 | 0.56 | 0.23 | 0.56 | 0.37 |
| 8 | 13832082 | rs10503478 | 0.358 | -1.496 | -0.368 | 0.561 | -0.028 | -0.031 | -0.031 | 0 | 0.39 | 0.16 | 0.42 | 0.51 | 0.42 | 0.53 |
| 8 | 13832286 | rs1485534 | 0.624 | -1.62 | -0.231 | 0.504 | -0.074 | 0.024 | 0.024 | 0.772 | 0.77 | 0.52 | 0.77 | 0.78 | 0.87 | 0.91 |
| 8 | 13834371 | rs4831257 | 1.462 | -1.27 | -0.539 | 0.389 | -0.075 | 0.007 | 0.007 | 0 | 0.36 | 0.17 | 0.33 | 0.27 | 0.46 | 0.38 |
| 8 | 13843667 | rs4831518 | 0.096 | -1.216 | -0.119 | 0.521 | -0.013 | 0.033 | 0.033 | 1.069 | 0.17 | 0.32 | 0.14 | 0.21 | 0.16 | 0.19 |
| 8 | 13847960 | rs11780201 | 0.874 | -1.127 | -0.638 | 0.653 | -0.007 | 0.031 | 0.031 | 0 | 0.07 | 0.12 | 0.07 | 0.16 | 0.06 | 0.03 |
| 8 | 13851562 | rs10503482 | 0.742 | -1.319 | -0.759 | 0.322 | 0.005 | -0.004 | -0.004 | 0.2 | 0.14 | 0.09 | 0.05 | 0.17 | 0.06 | 0.11 |
| 8 | 13864063 | rs10503473 | 1.865 | -0.813 | -0.66 | 0.289 | -0.054 | 0.071 | 0.071 | 0 | 0.29 | 0.13 | 0.29 | 0.11 | 0.37 | 0.22 |
| 8 | 13868456 | rs17118297 | 0.333 | -0.858 | -0.572 | 0.442 | -0.014 | -0.011 | -0.011 | 0 | 0.1 | 0.18 | 0.09 | 0.06 | 0.06 | 0.03 |
| 8 | 13868530 | rs17118305 | 0.333 | -0.86 | -0.617 | 0.441 | -0.011 | -0.009 | -0.009 | 0.048 | 0.1 | 0.16 | 0.09 | 0.06 | 0.07 | 0.03 |
| 8 | 13870210 | rs7828991 | -0.155 | -0.972 | -0.597 | 0.477 | 0.011 | 0.026 | 0.026 | 0 | 0.46 | 0.3 | 0.53 | 0.68 | 0.44 | 0.56 |
| 8 | 13871034 | rs12674595 | 1.197 | -0.874 | -0.618 | 0.702 | 0.012 | -0.028 | -0.028 | 0.267 | 0.29 | 0.22 | 0.29 | 0.5 | 0.23 | 0.4 |
| 8 | 13874153 | rs10503468 | 0.348 | -0.304 | -0.813 | 0.626 | 0 | -0.014 | -0.014 | 0 | 0.12 | 0.11 | 0.09 | 0.18 | 0.12 | 0.19 |
| 8 | 13877465 | rs12678107 | 0.038 | -0.426 | -0.92 | 0.586 | 0.014 | -0.014 | -0.014 | 0 | 0.47 | 0.25 | 0.49 | 0.56 | 0.46 | 0.53 |
| 8 | 13885513 | rs1843360 | 0.389 | -0.265 | -1.255 | 0.603 | 0.001 | 0.015 | 0.015 | 0.13 | 0.71 | 0.61 | 0.75 | 0.65 | 0.69 | 0.65 |
| 8 | 13891058 | rs4398915 | 0.325 | -0.161 | -2.431 | 0.353 | -0.031 | 0.049 | 0.049 | 0 | 0.43 | 0.11 | 0.37 | 0.27 | 0.53 | 0.42 |
| 8 | 13891431 | rs12542625 | 0.987 | -0.088 | -2.691 | 0.31 | -0.143 | 0.113 | 0.113 | 0 | 0.33 | 0.04 | 0.31 | 0.1 | 0.4 | 0.25 |
| 8 | 13891562 | rs2046436 | 0.401 | -0.496 | -2.246 | 0.662 | 0.045 | -0.025 | -0.025 | 0.017 | 0.33 | 0.15 | 0.34 | 0.43 | 0.26 | 0.37 |
| 8 | 13894700 | rs901117 | 0.059 | 0.322 | -2.303 | 0.624 | 0.02 | 0.077 | 0.077 | 0 | 0.47 | 0.09 | 0.39 | 0.27 | 0.57 | 0.44 |
| 8 | 13895191 | rs11203567 | 0.31 | 0.136 | -1.577 | 0.516 | -0.029 | 0.011 | 0.011 | 0 | 0.11 | 0 | 0.06 | 0.05 | 0.1 | 0.12 |
| 8 | 13897999 | rs1872465 | 0.005 | 0.246 | -0.869 | 0.867 | -0.007 | 0 | 0 | 0.005 | 0.45 | 0.52 | 0.48 | 0.52 | 0.34 | 0.49 |
| 8 | 13898663 | rs13271261 | -0.064 | 0.229 | -0.483 | 0.721 | -0.005 | 0.001 | 0.001 | 0 | 0.73 | 0.6 | 0.71 | 0.74 | 0.74 | 0.71 |
| 8 | 13899644 | rs12544126 | 0.062 | 0.801 | -0.416 | 0.554 | -0.04 | 0.021 | 0.021 | 0 | 0.31 | 0.18 | 0.31 | 0.21 | 0.42 | 0.25 |
| 8 | 13902880 | rs11784230 | 1.015 | 0.435 | -0.829 | 0.496 | -0.039 | -0.118 | -0.118 | 0 | 0.3 | 0.05 | 0.26 | 0.47 | 0.33 | 0.4 |
| 8 | 13903756 | rs13261147 | -0.162 | 0.472 | -0.784 | 0.095 | 0.082 | 0.058 | 0.058 | 0.472 | 0.6 | 0.14 | 0.56 | 0.73 | 0.75 | 0.69 |
| 8 | 13905487 | rs11784664 | 0.698 | 0.229 | -0.417 | 0.306 | -0.018 | -0.106 | -0.106 | 0 | 0.3 | 0.1 | 0.26 | 0.5 | 0.33 | 0.4 |
| 8 | 13905760 | rs1948039 | 0.57 | 0.221 | -0.419 | 0.325 | -0.008 | -0.036 | -0.036 | 0 | 0.31 | 0.12 | 0.27 | 0.64 | 0.33 | 0.46 |
| 8 | 13918143 | rs10100940 | 0.507 | 0.477 | -0.599 | 0.345 | -0.03 | -0.077 | -0.077 | 0.735 | 0.38 | 0.15 | 0.31 | 0.49 | 0.42 | 0.51 |
| 8 | 13921358 | rs6990047 | 0.426 | 0.468 | -0.385 | 0.34 | -0.007 | -0.032 | -0.032 | 0 | 0.39 | 0.22 | 0.31 | 0.49 | 0.42 | 0.5 |
| 8 | 13923923 | rs6981128 | 0.535 | 0.621 | -0.359 | 0.375 | -0.003 | -0.034 | -0.034 | 0.5 | 0.4 | 0.2 | 0.31 | 0.49 | 0.42 | 0.5 |
| 8 | 13924510 | rs11779074 | 0.554 | 0.458 | -0.72 | 0.413 | -0.065 | -0.1 | -0.1 | 0.058 | 0.4 | 0.09 | 0.31 | 0.49 | 0.46 | 0.51 |
| 8 | 13937395 | rs10106631 | 0.323 | -0.325 | -0.172 | -0.289 | -0.052 | -0.1 | -0.1 | 0.203 | 0.72 | 0.55 | 0.77 | 0.92 | 0.86 | 0.79 |
| 8 | 13947515 | rs17118615 | -1.288 | 0.996 | -0.002 | 0.243 | 0.03 | -0.038 | -0.038 | 0 | 0.09 | 0.2 | 0.1 | 0.03 | 0.13 | 0.06 |
| 8 | 13950615 | rs17118622 | -1.052 | 0.102 | -0.28 | -0.009 | -0.013 | -0.019 | -0.019 | 0 | 0.31 | 0.3 | 0.37 | 0.79 | 0.24 | 0.48 |
| 8 | 13950829 | rs11775245 | 0.574 | 0.507 | -0.731 | -1.183 | -0.018 | 0.034 | 0.034 | 0.851 | 0.23 | 0.18 | 0.27 | 0.02 | 0.26 | 0.14 |
| 8 | 13955943 | rs12678173 | -0.391 | 0.695 | -0.494 | -1.131 | 0 | 0.004 | 0.004 | 0 | 0.3 | 0.22 | 0.37 | 0.8 | 0.26 | 0.49 |
| 8 | 13956109 | rs1478039 | 0.128 | 0.687 | -0.513 | -0.727 | 0.015 | -0.049 | -0.049 | 0.125 | 0.24 | 0.06 | 0.27 | 0.71 | 0.17 | 0.43 |
| 8 | 13958463 | rs7007117 | -0.509 | 0.619 | -0.108 | -0.734 | 0.069 | 0.065 | 0.065 | 0.862 | 0.07 | 0.25 | 0.09 | 0.06 | 0.09 | 0.05 |
| 8 | 13961133 | rs10503484 | 1.399 | 0.226 | 0.265 | -1.06 | -0.057 | 0.071 | 0.071 | 0.04 | 0.37 | 0.16 | 0.3 | 0.19 | 0.45 | 0.33 |
| 8 | 13963152 | rs13266182 | 1.345 | 0.171 | 0.167 | -1.106 | -0.164 | 0.046 | 0.046 | 0.307 | 0.37 | 0.02 | 0.33 | 0.2 | 0.47 | 0.34 |
| 8 | 13966546 | rs10503485 | -0.567 | 0.359 | 0.617 | -1.15 | 0.053 | 0.05 | 0.05 | 0.059 | 0.08 | 0.24 | 0.14 | 0.09 | 0.11 | 0.06 |
| 8 | 13967618 | rs9643972 | 1.332 | 0.287 | 0.641 | -1.181 | -0.075 | 0.079 | 0.079 | 0.219 | 0.37 | 0.1 | 0.3 | 0.19 | 0.47 | 0.34 |
| 8 | 13967792 | rs11993263 | 0.79 | -0.113 | 0.515 | -1.217 | 0 | 0.009 | 0.009 | 0 | 0.15 | 0.13 | 0.25 | 0.68 | 0.12 | 0.39 |
| 8 | 13968169 | rs10503486 | 0.915 | -0.327 | 1.172 | -1.299 | -0.045 | -0.054 | -0.054 | 0 | 0.78 | 0.6 | 0.86 | 0.92 | 0.85 | 0.9 |
| 8 | 13969304 | rs7010173 | 0.509 | -0.38 | 1.169 | -1.309 | 0.005 | 0.036 | 0.036 | 0 | 0.53 | 0.36 | 0.38 | 0.25 | 0.59 | 0.42 |
| 8 | 13971646 | rs13259591 | 1.758 | -0.366 | 0.676 | -1.462 | -0.128 | 0.168 | 0.168 | 0 | 0.36 | 0.02 | 0.2 | 0.07 | 0.42 | 0.3 |
| 8 | 13973465 | rs10109252 | 0.776 | -0.319 | 0.856 | -1.348 | -0.014 | 0.07 | 0.07 | 0.123 | 0.25 | 0.17 | 0.31 | 0.05 | 0.26 | 0.16 |
| 8 | 13979289 | rs17118702 | 0.695 | -0.264 | 0.847 | -0.98 | -0.001 | 0.015 | 0.015 | 0 | 0.16 | 0.14 | 0.26 | 0.67 | 0.15 | 0.39 |
| 8 | 13980629 | rs17118706 | 0.696 | -0.249 | 1.133 | -0.984 | 0.001 | 0.04 | 0.04 | 0 | 0.16 | 0.18 | 0.27 | 0.69 | 0.15 | 0.4 |
| 8 | 13981100 | rs12675733 | 0.656 | -0.245 | 1.024 | -0.979 | -0.001 | -0.015 | -0.015 | 1.247 | 0.16 | 0.12 | 0.26 | 0.69 | 0.15 | 0.41 |
| 8 | 14002020 | rs13253386 | 1.44 | -0.306 | 1.002 | -1.058 | -0.039 | 0.052 | 0.052 | 0 | 0.4 | 0.2 | 0.35 | 0.22 | 0.48 | 0.36 |
| 8 | 14002309 | rs2126809 | 0.507 | -0.106 | 0.791 | -1.027 | -0.001 | -0.003 | -0.003 | 0 | 0.17 | 0.14 | 0.27 | 0.67 | 0.16 | 0.4 |
| 8 | 14012279 | rs3888260 | -0.405 | -0.151 | 0.806 | -0.978 | -0.001 | 0.001 | 0.001 | 0 | 0.43 | 0.62 | 0.6 | 0.75 | 0.42 | 0.57 |
| 8 | 14012782 | rs7817792 | -0.467 | 0.009 | 0.211 | -0.94 | 0.024 | 0.01 | 0.01 | 0 | 0.17 | 0.07 | 0.07 | 0.08 | 0.13 | 0.1 |
| 8 | 14015338 | rs1382147 | -0.35 | -0.002 | 0.228 | -0.93 | 0.022 | 0.008 | 0.008 | 0 | 0.16 | 0.1 | 0.07 | 0.08 | 0.13 | 0.1 |
| 8 | 14017547 | rs7841810 | -0.187 | 0.062 | 0.495 | -0.854 | 0.015 | 0.047 | 0.047 | 0.052 | 0.41 | 0.37 | 0.36 | 0.14 | 0.37 | 0.25 |
| 8 | 14019665 | rs12547822 | -0.161 | 0.101 | 0.34 | -0.83 | 0.021 | 0.078 | 0.078 | 0.046 | 0.4 | 0.3 | 0.36 | 0.14 | 0.37 | 0.25 |
| 8 | 14021456 | rs6992644 | 0.208 | 0.217 | 0.079 | -0.504 | -0.002 | 0.003 | 0.003 | 0 | 0.17 | 0.15 | 0.28 | 0.66 | 0.17 | 0.4 |
| 8 | 14025722 | rs7839595 | -0.809 | 0.363 | 0.067 | -0.773 | 0.004 | -0.045 | -0.045 | 0.6 | 0.61 | 0.58 | 0.7 | 0.82 | 0.58 | 0.68 |
| 8 | 14035832 | rs12680272 | -0.345 | 0.637 | -0.324 | -0.776 | -0.019 | -0.002 | -0.002 | 1.937 | 0.32 | 0.2 | 0.34 | 0.71 | 0.34 | 0.49 |
| 8 | 14036072 | rs4831279 | -0.614 | 0.252 | 0.203 | -1.138 | -0.021 | 0.021 | 0.021 | 0.068 | 0.18 | 0.48 | 0.28 | 0.14 | 0.15 | 0.19 |
| 8 | 14037030 | rs10888085 | -0.016 | 0.374 | -0.216 | -0.862 | -0.018 | -0.019 | -0.019 | 0 | 0.29 | 0.2 | 0.34 | 0.63 | 0.32 | 0.48 |
| 8 | 14037301 | rs11203582 | -0.368 | 0.234 | -0.238 | -0.901 | -0.035 | -0.007 | -0.007 | 0 | 0.36 | 0.2 | 0.38 | 0.63 | 0.39 | 0.51 |
| 8 | 14043262 | rs1904078 | -0.356 | 0.123 | 0.511 | -0.657 | -0.01 | 0.055 | 0.055 | 0 | 0.12 | 0.25 | 0.25 | 0.17 | 0.11 | 0.1 |
| 8 | 14044475 | rs7828061 | 0.312 | -0.127 | -0.506 | -0.513 | -0.008 | 0.038 | 0.038 | 0.134 | 0.52 | 0.24 | 0.45 | 0.63 | 0.48 | 0.72 |
| 8 | 14044795 | rs13251919 | -0.191 | 0.003 | -0.412 | -0.075 | -0.012 | 0.082 | 0.082 | 0 | 0.13 | 0.28 | 0.26 | 0.23 | 0.12 | 0.11 |
| 8 | 14045251 | rs7002819 | -0.469 | -0.014 | -1.199 | 0.114 | -0.084 | 0.003 | 0.003 | 0 | 0.58 | 0.88 | 0.66 | 0.71 | 0.51 | 0.77 |

| | | | | | | | | | | | | | | | | |
|---|---|---|---|---|---|---|---|---|---|---|---|---|---|---|---|---|
| 8 | 14053276 | rs1478014 | 0.45 | -0.08 | -0.408 | 0.114 | -0.003 | -0.002 | -0.002 | 0 | 0.42 | 0.28 | 0.5 | 0.56 | 0.41 | 0.66 |
| 8 | 14064049 | rs11996588 | -1.637 | 0.013 | -0.698 | -0.182 | -0.053 | 0.032 | 0.032 | 0 | 0.74 | 0.9 | 0.77 | 0.84 | 0.66 | 0.87 |
| 8 | 14074348 | rs7016966 | 0.029 | -0.298 | -0.638 | 0.139 | 0.037 | 0.037 | 0.037 | 0 | 0.47 | 0.76 | 0.44 | 0.48 | 0.39 | 0.56 |
| 8 | 14078154 | rs13278543 | 2.075 | -0.235 | -0.753 | 0.03 | -0.131 | 0.003 | 0.003 | 0 | 0.24 | 0.02 | 0.22 | 0.16 | 0.34 | 0.13 |
| 8 | 14082230 | rs981905 | 1.211 | -0.527 | -0.542 | -0.122 | -0.066 | 0.073 | 0.073 | 0.763 | 0.32 | 0.1 | 0.26 | 0.18 | 0.37 | 0.17 |
| 8 | 14088127 | rs4637829 | 1.088 | -0.295 | -0.272 | -0.103 | -0.023 | 0.032 | 0.032 | 0 | 0.32 | 0.26 | 0.27 | 0.24 | 0.37 | 0.2 |
| 8 | 14091774 | rs17118836 | 2.236 | -0.338 | -0.163 | -0.053 | -0.046 | -0.061 | -0.061 | 0.853 | 0.25 | 0.03 | 0.26 | 0.34 | 0.27 | 0.42 |
| 8 | 14092342 | rs17292180 | -0.168 | -0.474 | -0.056 | 0.02 | 0.004 | 0.043 | 0.043 | 0.022 | 0.19 | 0.2 | 0.35 | 0.39 | 0.2 | 0.27 |
| 8 | 14092667 | rs13266960 | -1.234 | -0.48 | -0.083 | -0.207 | -0.018 | -0.022 | -0.022 | 0.41 | 0.92 | 0.88 | 0.98 | 1 | 0.98 | 0.98 |
| 8 | 14093929 | rs1382142 | -1.631 | -0.354 | 0.225 | -0.012 | 0.026 | -0.012 | -0.012 | 0.098 | 0.44 | 0.71 | 0.49 | 0.46 | 0.36 | 0.39 |
| 8 | 14098105 | rs17118860 | 0.098 | -0.472 | 0.388 | 0.081 | -0.007 | 0.038 | 0.038 | 0.167 | 0.26 | 0.42 | 0.19 | 0.29 | 0.21 | 0.27 |
| 8 | 14098306 | rs17118863 | 2.032 | -0.371 | 0.128 | 0.137 | 0.005 | -0.029 | -0.029 | 0 | 0.29 | 0.13 | 0.26 | 0.38 | 0.28 | 0.42 |
| 8 | 14099307 | rs11988897 | 0.469 | -0.596 | 0.013 | -0.006 | -0.001 | -0.001 | -0.001 | 0 | 0.13 | 0.16 | 0.27 | 0.1 | 0.16 | 0.13 |
| 8 | 14105056 | rs4521757 | 1.832 | -0.187 | 0.025 | 0.14 | 0.006 | -0.022 | -0.022 | 0 | 0.3 | 0.13 | 0.27 | 0.38 | 0.28 | 0.44 |
| 8 | 14107396 | rs17293289 | 1.858 | 0.114 | -0.647 | 0.016 | -0.134 | 0.022 | 0.022 | 0 | 0.24 | 0.02 | 0.23 | 0.14 | 0.35 | 0.14 |
| 8 | 14108434 | rs9325698 | 0.56 | -0.003 | -0.262 | 0.138 | -0.028 | -0.009 | -0.009 | 0.019 | 0.92 | 0.75 | 0.96 | 0.91 | 0.97 | 0.94 |
| 8 | 14113816 | rs9886428 | 0.63 | 0.107 | -0.826 | 0.159 | -0.074 | 0.097 | 0.097 | 0 | 0.41 | 0.14 | 0.33 | 0.24 | 0.51 | 0.22 |
| 8 | 14119944 | rs7014940 | 0.123 | -0.032 | -0.576 | 0.193 | -0.003 | -0.002 | -0.002 | 0 | 0.15 | 0.18 | 0.28 | 0.1 | 0.15 | 0.11 |
| 8 | 14122672 | rs12542368 | 0.819 | -0.107 | -0.461 | 0.333 | -0.002 | 0.004 | 0.004 | 0 | 0.72 | 0.28 | 0.62 | 0.57 | 0.8 | 0.7 |
| 8 | 14122745 | rs12549721 | 0.882 | -0.056 | -0.839 | 0.124 | 0.009 | 0.044 | 0.044 | 0 | 0.17 | 0.08 | 0.09 | 0.05 | 0.16 | 0.09 |
| 8 | 14124809 | rs17294565 | 1.992 | 0.112 | -0.748 | 0.153 | -0.136 | -0.011 | -0.011 | 0.471 | 0.24 | 0.02 | 0.25 | 0.18 | 0.37 | 0.13 |
| 8 | 14126134 | rs17228952 | 1.646 | 0.09 | -0.741 | 0.14 | -0.089 | 0.042 | 0.042 | 0 | 0.27 | 0.08 | 0.27 | 0.2 | 0.4 | 0.16 |
| 8 | 14131509 | rs6651379 | -1.067 | -0.095 | -0.093 | 0.2 | -0.016 | -0.006 | -0.006 | 0 | 0.63 | 0.59 | 0.7 | 0.66 | 0.71 | 0.5 |
| 8 | 14132428 | rs12545677 | 1.709 | -0.115 | -0.156 | 0.202 | -0.001 | -0.045 | -0.045 | 0 | 0.3 | 0.07 | 0.26 | 0.34 | 0.28 | 0.49 |
| 8 | 14135277 | rs6530746 | 1.574 | -0.089 | 0.012 | 0.195 | 0.008 | -0.01 | -0.01 | 0 | 0.31 | 0.16 | 0.27 | 0.34 | 0.29 | 0.48 |
| 8 | 14139307 | rs6530747 | -1.163 | -0.11 | 0.277 | 0.256 | 0 | 0 | 0 | 0 | 0.64 | 0.68 | 0.71 | 0.66 | 0.71 | 0.5 |
| 8 | 14141283 | rs6999172 | -1.31 | -0.19 | 0.353 | 0.242 | 0.023 | 0.007 | 0.007 | 0 | 0.65 | 0.82 | 0.72 | 0.68 | 0.71 | 0.5 |
| 8 | 14141326 | rs6999312 | -1.32 | -0.18 | 0.357 | 0.247 | 0.022 | 0.006 | 0.006 | 0 | 0.65 | 0.82 | 0.72 | 0.68 | 0.71 | 0.5 |
| 8 | 14142455 | rs10503490 | 0.283 | -0.237 | 0.36 | 0.216 | 0.001 | 0.023 | 0.023 | 0 | 0.14 | 0.18 | 0.14 | 0.3 | 0.15 | 0.23 |
| 8 | 14143471 | rs11775244 | 1.733 | -0.279 | 0.281 | 0.115 | 0.002 | 0.01 | 0.01 | 0 | 0.08 | 0.06 | 0.04 | 0.02 | 0.06 | 0.03 |
| 8 | 14144867 | rs890542 | 0.216 | -0.296 | 0.357 | 0.07 | -0.013 | 0.001 | 0.001 | 0 | 0.16 | 0.29 | 0.27 | 0.16 | 0.13 | 0.1 |
| 8 | 14145876 | rs12548539 | 0.792 | -0.287 | 0.304 | 0.077 | -0.023 | 0.026 | 0.026 | 0.021 | 0.72 | 0.5 | 0.79 | 0.68 | 0.78 | 0.72 |
| 8 | 14151079 | rs12674914 | 0.022 | -0.05 | -0.344 | 0.281 | -0.001 | 0.007 | 0.007 | 0 | 0.17 | 0.18 | 0.17 | 0.36 | 0.18 | 0.3 |
| 8 | 14152719 | rs7838948 | -0.261 | -0.041 | -0.391 | 0.301 | -0.003 | 0.011 | 0.011 | 0 | 0.46 | 0.59 | 0.55 | 0.35 | 0.5 | 0.25 |
| 8 | 14153650 | rs10097743 | 0.302 | 0.428 | -0.248 | 0.344 | 0 | 0.007 | 0.007 | 0 | 0.74 | 0.67 | 0.8 | 0.68 | 0.77 | 0.67 |
| 8 | 14157277 | rs12678379 | -0.453 | 0.327 | -0.405 | 0.383 | 0.041 | 0.002 | 0.002 | 0.099 | 0.48 | 0.26 | 0.46 | 0.57 | 0.59 | 0.44 |
| 8 | 14159065 | rs17119018 | -0.328 | 0.327 | -0.404 | 0.368 | -0.001 | -0.005 | -0.005 | 1.886 | 0.24 | 0.24 | 0.22 | 0.38 | 0.25 | 0.31 |
| 8 | 14161038 | rs17231019 | 1.647 | 0.475 | -0.629 | 0.086 | -0.12 | 0.118 | 0.118 | 0.001 | 0.23 | 0.02 | 0.18 | 0.05 | 0.33 | 0.09 |
| 8 | 14164118 | rs1030423 | -0.519 | 0.147 | -0.194 | 0.261 | 0.012 | 0.001 | 0.001 | 0.118 | 0.5 | 0.48 | 0.45 | 0.49 | 0.6 | 0.43 |
| 8 | 14168180 | rs1820528 | 0.727 | 0.053 | -0.213 | 0.197 | 0.008 | -0.024 | -0.024 | 0 | 0.3 | 0.32 | 0.31 | 0.14 | 0.43 | 0.14 |
| 8 | 14168197 | rs1820529 | -0.296 | -0.017 | -0.145 | 0.279 | 0.003 | 0.02 | 0.02 | 0 | 0.95 | 0.84 | 0.77 | 0.89 | 0.95 | 0.96 |
| 8 | 14170563 | rs17297046 | 1.779 | -0.255 | -0.397 | 0.218 | -0.017 | -0.015 | -0.015 | 0 | 0.34 | 0.11 | 0.25 | 0.32 | 0.31 | 0.51 |
| 8 | 14171703 | rs13277849 | 0.296 | -0.304 | -0.134 | 0.233 | -0.008 | 0.01 | 0.01 | 0 | 0.9 | 0.71 | 0.76 | 0.9 | 0.94 | 0.94 |
| 8 | 14172535 | rs12546389 | 0.675 | -0.082 | -0.099 | 0.388 | -0.02 | 0.052 | 0.052 | 0 | 0.68 | 0.56 | 0.57 | 0.49 | 0.73 | 0.69 |
| 8 | 14173974 | rs6990042 | -0.408 | 0.366 | -0.156 | 0.26 | 0.013 | -0.008 | -0.008 | 2.133 | 0.66 | 0.64 | 0.48 | 0.7 | 0.52 | 0.8 |
| 8 | 14191967 | rs7003304 | -0.023 | 0.281 | 0.345 | 0.684 | -0.005 | -0.014 | -0.014 | 0 | 0.16 | 0.26 | 0.12 | 0.12 | 0.16 | 0.07 |
| 8 | 14194119 | rs1593626 | 1.653 | 0.438 | -0.185 | 0.444 | -0.024 | 0.077 | 0.077 | 0 | 0.5 | 0.09 | 0.36 | 0.63 | 0.44 | 0.74 |
| 8 | 14194356 | rs1426993 | 0.2 | 0.402 | -0.025 | 0.427 | 0.012 | 0.001 | 0.001 | 0 | 0.79 | 0.65 | 0.64 | 0.79 | 0.76 | 0.88 |
| 8 | 14194620 | rs7000497 | 0.214 | 0.381 | 0.411 | 0.403 | 0.01 | -0.027 | -0.027 | 0 | 0.79 | 0.47 | 0.75 | 0.82 | 0.78 | 0.89 |
| 8 | 14198260 | rs2898389 | 2.012 | 0.258 | 0.122 | 0.795 | -0.006 | 0.01 | 0.01 | 0 | 0.45 | 0.27 | 0.42 | 0.39 | 0.42 | 0.58 |
| 8 | 14199014 | rs9643974 | -0.787 | 0.229 | 0.728 | 1.028 | -0.011 | 0.068 | 0.068 | 0 | 0.13 | 0.18 | 0.12 | 0.37 | 0.07 | 0.24 |
| 8 | 14201728 | rs11997612 | 0.321 | 0.242 | 0.77 | 1.009 | 0.001 | 0.044 | 0.044 | 0 | 0.06 | 0.17 | 0.07 | 0.07 | 0.02 | 0.04 |
| 8 | 14203326 | rs7000337 | -1.365 | 0.153 | 0.774 | 0.945 | -0.003 | 0.01 | 0.01 | 0 | 0.36 | 0.56 | 0.35 | 0.42 | 0.37 | 0.32 |
| 8 | 14215025 | rs989870 | 1.274 | 0.039 | 0.808 | 0.713 | 0 | 0.021 | 0.021 | 0 | 0.66 | 0.28 | 0.59 | 0.73 | 0.64 | 0.81 |
| 8 | 14220732 | rs6997359 | -0.866 | 0.076 | 0.914 | 0.655 | -0.001 | -0.058 | -0.058 | 0 | 0.29 | 0.56 | 0.28 | 0.15 | 0.31 | 0.12 |
| 8 | 14242443 | rs12155793 | 0.432 | 0.021 | 0.307 | 0.593 | -0.095 | 0.061 | 0.061 | 0 | 0.18 | 0.01 | 0.19 | 0.06 | 0.3 | 0.08 |
| 8 | 14243131 | rs7463769 | -1.815 | -0.096 | 0.306 | 0.757 | 0.001 | 0.008 | 0.008 | 0 | 0.42 | 0.73 | 0.51 | 0.58 | 0.41 | 0.39 |
| 8 | 14252372 | rs6988872 | -0.98 | 0.028 | 0.499 | 0.444 | 0.005 | -0.054 | -0.054 | 0 | 0.3 | 0.56 | 0.25 | 0.14 | 0.32 | 0.16 |
| 8 | 14253354 | rs2054356 | -1.459 | -0.012 | 0.499 | 0.465 | -0.002 | 0.018 | 0.018 | 0 | 0.36 | 0.71 | 0.37 | 0.25 | 0.37 | 0.23 |
| 8 | 14256574 | rs1037934 | -1.067 | -0.02 | 0.604 | 0.456 | -0.047 | 0.015 | 0.015 | 0 | 0.12 | 0.55 | 0.11 | 0.08 | 0.03 | 0.05 |
| 8 | 14263619 | rs6530767 | -0.762 | 0.09 | 0.735 | 0.509 | 0.012 | 0.08 | 0.08 | 0.068 | 0.09 | 0.45 | 0.11 | 0.08 | 0.03 | 0.05 |
| 8 | 14265602 | rs7827220 | -0.783 | 0.086 | 0.733 | 0.508 | 0.01 | 0.078 | 0.078 | 0.071 | 0.09 | 0.45 | 0.11 | 0.08 | 0.03 | 0.05 |
| 8 | 14272269 | rs12543569 | -0.15 | 0.166 | 0.432 | 0.368 | -0.032 | 0.06 | 0.06 | 0 | 0.22 | 0.16 | 0.21 | 0.06 | 0.31 | 0.12 |
| 8 | 14275001 | rs11987419 | 0.454 | 0.138 | 0.475 | 0.409 | -0.006 | 0.008 | 0.008 | 0 | 0.74 | 0.73 | 0.67 | 0.83 | 0.65 | 0.82 |
| 8 | 14275444 | rs11776823 | 0.303 | 0.14 | 0.366 | 0.413 | -0.012 | 0.013 | 0.013 | 0 | 0.76 | 0.79 | 0.68 | 0.83 | 0.66 | 0.83 |
| 8 | 14275561 | rs11781105 | 1.372 | 0.092 | 0.581 | 0.459 | 0.003 | -0.013 | -0.013 | 0 | 0.65 | 0.38 | 0.58 | 0.75 | 0.62 | 0.77 |
| 8 | 14276337 | rs11782022 | 1.725 | 0.024 | 0.076 | 0.633 | 0.004 | 0.017 | 0.017 | 0 | 0.57 | 0.21 | 0.5 | 0.41 | 0.58 | 0.58 |
| 8 | 14282382 | rs10503497 | -0.012 | 0.061 | 0.133 | 0.267 | -0.053 | 0.101 | 0.101 | 0.323 | 0.23 | 0.06 | 0.2 | 0.06 | 0.3 | 0.11 |
| 8 | 14282776 | rs7837188 | -0.95 | -0.054 | 0.271 | 0.16 | 0.004 | -0.112 | -0.112 | 0 | 0.3 | 0.56 | 0.26 | 0.06 | 0.32 | 0.14 |

| | | | | | | | | | | | | | | | | |
|---|---|---|---|---|---|---|---|---|---|---|---|---|---|---|---|---|
| 8 | 14286167 | rs17242772 | 0.232 | 0.082 | -0.423 | 0.218 | -0.099 | 0.092 | 0.092 | 0.595 | 0.21 | 0.01 | 0.19 | 0.06 | 0.3 | 0.11 |
| 8 | 14286552 | rs728845 | 1.464 | -0.002 | -0.4 | 0.401 | 0.001 | -0.007 | -0.007 | 0 | 0.58 | 0.25 | 0.44 | 0.48 | 0.59 | 0.59 |
| 8 | 14292205 | rs13265496 | -0.147 | 0.464 | -0.541 | -0.226 | -0.053 | 0.088 | 0.088 | 0.696 | 0.22 | 0.07 | 0.21 | 0.06 | 0.3 | 0.11 |
| 8 | 14297969 | rs4644256 | -0.074 | 1.044 | -0.574 | -0.136 | -0.087 | 0.029 | 0.029 | 0.992 | 0.18 | 0.03 | 0.35 | 0.13 | 0.3 | 0.12 |
| 8 | 14308744 | rs10089641 | -0.912 | 0.781 | 0.339 | -0.264 | 0.033 | -0.065 | -0.065 | 0.47 | 0.23 | 0.42 | 0.36 | 0.13 | 0.3 | 0.14 |
| 8 | 14308995 | rs7830759 | -1.543 | 1.053 | 0.309 | -0.435 | -0.021 | 0.028 | 0.028 | 0 | 0.32 | 0.27 | 0.44 | 0.16 | 0.44 | 0.21 |
| 8 | 14312131 | rs12543902 | 0.274 | 0.981 | 0.279 | -0.358 | -0.088 | 0.003 | 0.003 | 0.419 | 0.17 | 0.03 | 0.35 | 0.15 | 0.3 | 0.12 |
| 8 | 14313293 | rs6530778 | -1.278 | 1.002 | 0.707 | -0.248 | 0.029 | -0.012 | -0.012 | 0 | 0.3 | 0.6 | 0.46 | 0.25 | 0.41 | 0.17 |
| 8 | 14318124 | rs1903597 | -2.139 | 0.991 | 0.704 | -0.297 | -0.009 | 0.01 | 0.01 | 0 | 0.4 | 0.74 | 0.55 | 0.31 | 0.46 | 0.36 |
| 8 | 14324437 | rs10110727 | -0.118 | 0.875 | 0.951 | -0.408 | -0.042 | 0.046 | 0.046 | 0.038 | 0.21 | 0.08 | 0.35 | 0.12 | 0.29 | 0.13 |
| 8 | 14325742 | rs7816029 | -0.298 | 0.883 | 0.895 | -0.405 | 0.013 | -0.025 | -0.025 | 0 | 0.22 | 0.25 | 0.36 | 0.12 | 0.3 | 0.13 |
| 8 | 14326637 | rs11203612 | -0.946 | 0.879 | 0.831 | -0.407 | 0.022 | -0.028 | -0.028 | 0 | 0.27 | 0.59 | 0.41 | 0.2 | 0.32 | 0.17 |
| 8 | 14328522 | rs4831597 | -1.48 | 0.739 | 0.799 | -0.343 | 0.002 | -0.006 | -0.006 | 0 | 0.32 | 0.71 | 0.45 | 0.25 | 0.35 | 0.25 |
| 8 | 14337080 | rs7834728 | 0.379 | 0.509 | 0.814 | -0.193 | 0.002 | -0.001 | -0.001 | 0 | 0.93 | 0.66 | 0.97 | 1 | 0.99 | 0.98 |
| 8 | 14339483 | rs3848991 | 0.171 | 0.234 | 0.182 | -0.866 | 0 | 0.029 | 0.029 | 0 | 0.43 | 0.29 | 0.28 | 0.68 | 0.41 | 0.59 |
| 8 | 14339987 | rs5023503 | 0.559 | 0.234 | -0.029 | -1.003 | -0.076 | 0.075 | 0.075 | 4.652 | 0.24 | 0.04 | 0.39 | 0.12 | 0.32 | 0.13 |
| 8 | 14343159 | rs17119369 | -1.226 | 0.354 | 0.341 | -0.602 | -0.011 | -0.027 | -0.027 | 0.831 | 0.71 | 0.42 | 0.7 | 0.8 | 0.76 | 0.75 |
| 8 | 14346985 | rs17303140 | 0.061 | 0.26 | 0.122 | -0.567 | -0.029 | 0.053 | 0.053 | 0.203 | 0.2 | 0.08 | 0.33 | 0.12 | 0.26 | 0.13 |
| 8 | 14347293 | rs7836622 | 0.186 | 0.309 | 0.161 | -0.556 | -0.027 | 0.04 | 0.04 | 0 | 0.19 | 0.09 | 0.33 | 0.12 | 0.26 | 0.13 |
| 8 | 14348505 | rs1542503 | 2.322 | 0.38 | 0.313 | -0.56 | -0.001 | -0.01 | -0.01 | 0.206 | 0.2 | 0.23 | 0.25 | 0.13 | 0.2 | 0.23 |
| 8 | 14349013 | rs13259013 | 2.257 | 0.378 | 0.331 | -0.561 | -0.001 | -0.018 | -0.018 | 1.02 | 0.21 | 0.26 | 0.25 | 0.13 | 0.2 | 0.23 |
| 8 | 14352641 | rs1947709 | 0.181 | 0.366 | 0.396 | -0.544 | -0.003 | 0.007 | 0.007 | 0 | 0.41 | 0.62 | 0.62 | 0.32 | 0.48 | 0.37 |
| 8 | 14355629 | rs13252317 | 0.201 | 0.373 | 0.178 | -0.531 | -0.001 | 0 | 0 | 0 | 0.41 | 0.48 | 0.6 | 0.32 | 0.48 | 0.38 |
| 8 | 14358350 | rs7828975 | 0.015 | 0.434 | 0.015 | -0.498 | -0.002 | 0.001 | 0.001 | 0.292 | 0.2 | 0.25 | 0.35 | 0.12 | 0.26 | 0.14 |
| 8 | 14362238 | rs12682277 | -0.568 | 0.166 | 0.155 | -0.407 | -0.026 | 0.005 | 0.005 | 0 | 0.08 | 0.18 | 0.04 | 0.07 | 0.03 | 0.05 |
| 8 | 14362930 | rs13270111 | 2.084 | 0.234 | 0.355 | -0.599 | 0.002 | -0.095 | -0.095 | 0 | 0.2 | 0.32 | 0.25 | 0.05 | 0.19 | 0.21 |
| 8 | 14369890 | rs2054357 | 0.018 | 0.247 | 0.258 | -0.449 | -0.009 | 0.001 | 0.001 | 0 | 0.07 | 0.11 | 0.04 | 0.07 | 0.03 | 0.05 |
| 8 | 14371006 | rs12056456 | 0.559 | 0.344 | 0.182 | -0.251 | -0.001 | -0.004 | -0.004 | 0.252 | 0.75 | 0.7 | 0.63 | 0.72 | 0.72 | 0.82 |
| 8 | 14373990 | rs17119539 | -0.01 | 0.317 | 0.211 | -0.23 | -0.021 | 0.006 | 0.006 | 0 | 0.07 | 0.17 | 0.04 | 0.07 | 0.03 | 0.05 |
| 8 | 14374513 | rs7839802 | 0.39 | 0.319 | 0.411 | -0.251 | 0.006 | 0.026 | 0.026 | 0 | 0.75 | 0.47 | 0.61 | 0.72 | 0.72 | 0.79 |
| 8 | 14376095 | rs17119601 | 0.447 | 0.293 | 0.401 | -0.236 | 0.002 | 0.021 | 0.021 | 0 | 0.74 | 0.43 | 0.61 | 0.72 | 0.72 | 0.79 |
| 8 | 14377392 | rs17119609 | 0.451 | 0.351 | 0.056 | 0.106 | -0.003 | -0.012 | -0.012 | 0 | 0.22 | 0.21 | 0.18 | 0.45 | 0.23 | 0.31 |
| 8 | 14380663 | rs7812643 | -0.983 | 0.498 | -0.208 | 0.255 | -0.077 | -0.023 | -0.023 | 0.149 | 0.85 | 0.98 | 0.8 | 0.87 | 0.75 | 0.9 |
| 8 | 14384443 | rs1461865 | 0.005 | 0.369 | -0.074 | 0.284 | 0 | 0.011 | 0.011 | 0 | 0.55 | 0.38 | 0.38 | 0.66 | 0.53 | 0.6 |
| 8 | 14385320 | rs1381410 | 0.018 | 0.456 | -0.062 | 0.753 | 0 | 0.054 | 0.054 | 0 | 0.06 | 0.24 | 0.04 | 0.07 | 0.03 | 0.03 |
| 8 | 14387833 | rs10113046 | 0.091 | 0.986 | -1.027 | 1.132 | -0.021 | -0.055 | -0.055 | 0 | 0.78 | 0.83 | 0.65 | 0.55 | 0.71 | 0.85 |
| 8 | 14388093 | rs11987549 | 0.153 | 0.549 | -0.649 | 0.783 | -0.005 | 0.02 | 0.02 | 0 | 0.21 | 0.29 | 0.2 | 0.29 | 0.19 | 0.21 |
| 8 | 14389640 | rs1461854 | 0.012 | 0.409 | -0.726 | 0.031 | 0.03 | 0.058 | 0.058 | 1.173 | 0.42 | 0.22 | 0.22 | 0.2 | 0.31 | 0.35 |
| 8 | 14389709 | rs1903592 | 0.136 | 0.697 | -0.927 | 0.224 | -0.046 | -0.051 | -0.051 | 0 | 0.24 | 0.18 | 0.4 | 0.54 | 0.42 | 0.22 |
| 8 | 14390350 | rs10111879 | -0.511 | 0.7 | -1.1 | 0.084 | 0.018 | 0.037 | 0.037 | 0.437 | 0.55 | 0.31 | 0.46 | 0.32 | 0.61 | 0.47 |
| 8 | 14391475 | rs13280583 | 0.397 | 0.669 | -0.927 | 0.295 | 0.007 | -0.008 | -0.008 | 0 | 0.45 | 0.27 | 0.62 | 0.54 | 0.58 | 0.44 |
| 8 | 14391844 | rs13254243 | 1.064 | 0.664 | -0.96 | 0.257 | -0.075 | 0.13 | 0.13 | 0 | 0.37 | 0.17 | 0.46 | 0.16 | 0.47 | 0.32 |
| 8 | 14392130 | rs1461849 | -0.384 | 0.702 | -0.458 | 0.605 | 0.028 | 0.176 | 0.176 | 0.37 | 0.15 | 0.37 | 0.23 | 0.42 | 0.14 | 0.24 |
| 8 | 14392244 | rs17119718 | 0.76 | 0.615 | -0.426 | 0.593 | 0.027 | 0.092 | 0.092 | 0.202 | 0.06 | 0.24 | 0.06 | 0.1 | 0.03 | 0.12 |
| 8 | 14392321 | rs17119719 | -0.606 | 0.636 | -0.499 | 0.508 | 0.053 | 0.066 | 0.066 | 0 | 0.3 | 0.47 | 0.46 | 0.57 | 0.45 | 0.34 |
| 8 | 14397466 | rs17119762 | 1.233 | 0.484 | -0.188 | 0.646 | -0.014 | -0.003 | -0.003 | 0.083 | 0.29 | 0.61 | 0.34 | 0.22 | 0.24 | 0.29 |
| 8 | 14397689 | rs17119771 | 0.11 | 0.347 | -0.484 | 0.447 | 0.002 | 0.006 | 0.006 | 0 | 0.38 | 0.29 | 0.22 | 0.3 | 0.31 | 0.48 |
| 8 | 14397963 | rs13271560 | 1.478 | 0.319 | -0.44 | 0.43 | -0.005 | 0.003 | 0.003 | 0 | 0.3 | 0.27 | 0.39 | 0.43 | 0.24 | 0.32 |
| 8 | 14400367 | rs2410192 | -0.43 | 0.28 | -0.399 | 0.288 | -0.002 | 0.013 | 0.013 | 0 | 0.68 | 0.63 | 0.8 | 0.59 | 0.71 | 0.61 |
| 8 | 14400664 | rs2410193 | -0.124 | 0.298 | -0.492 | 0.345 | 0.003 | 0.015 | 0.015 | 0 | 0.38 | 0.25 | 0.22 | 0.29 | 0.31 | 0.48 |
| 8 | 14400878 | rs4831602 | -0.775 | 0.315 | -0.603 | 0.509 | 0.016 | 0.184 | 0.184 | 0 | 0.93 | 0.78 | 0.91 | 0.78 | 0.93 | 0.9 |
| 8 | 14411773 | rs825908 | -1.627 | 0.248 | -0.657 | 0.345 | -0.004 | 0.003 | 0.003 | 0.109 | 0.73 | 0.65 | 0.69 | 0.73 | 0.76 | 0.7 |
| 8 | 14417189 | rs1510427 | 0.923 | 0.783 | -1.175 | 0.067 | -0.056 | 0.005 | 0.005 | 0 | 0.25 | 0.05 | 0.15 | 0.2 | 0.31 | 0.29 |
| 8 | 14418331 | rs1510428 | 2.624 | 0.781 | -1.14 | -0.063 | 0.016 | 0.066 | 0.066 | 0 | 0.21 | 0.12 | 0.22 | 0.06 | 0.16 | 0.21 |
| 8 | 14423424 | rs13265072 | 1.238 | 0.825 | -1.414 | -0.079 | -0.113 | 0.053 | 0.053 | 0 | 0.23 | 0.02 | 0.26 | 0.1 | 0.34 | 0.11 |
| 8 | 14428679 | rs1510432 | 1.72 | -0.029 | -0.835 | -0.021 | -0.001 | -0.002 | -0.002 | 0.038 | 0.28 | 0.28 | 0.27 | 0.22 | 0.25 | 0.31 |
| 8 | 14429287 | rs10503502 | -0.002 | -0.235 | -0.87 | 0.083 | -0.037 | -0.05 | -0.05 | 0 | 0.36 | 0.12 | 0.42 | 0.43 | 0.39 | 0.26 |
| 8 | 14430153 | rs1912470 | 0.58 | -0.077 | -0.687 | 0.227 | -0.006 | -0.015 | -0.015 | 0.723 | 0.12 | 0.09 | 0.18 | 0.35 | 0.13 | 0.14 |
| 8 | 14433115 | rs17249839 | 2.841 | -0.131 | -0.729 | 0.02 | -0.024 | -0.028 | -0.028 | 0 | 0.18 | 0.07 | 0.22 | 0.16 | 0.19 | 0.21 |
| 8 | 14433716 | rs13274454 | 2.061 | -0.092 | -0.54 | 0.148 | 0.002 | -0.004 | -0.004 | 0 | 0.26 | 0.27 | 0.27 | 0.18 | 0.23 | 0.24 |
| 8 | 14437388 | rs17119850 | 1.857 | -0.108 | -1.326 | 0.169 | 0.016 | -0.022 | -0.022 | 0 | 0.06 | 0.01 | 0.06 | 0.1 | 0.03 | 0.15 |
| 8 | 14437498 | rs17309528 | 0.999 | 0.08 | -1.228 | 0.053 | -0.061 | -0.012 | -0.012 | 0 | 0.24 | 0.07 | 0.14 | 0.2 | 0.3 | 0.29 |
| 8 | 14447094 | rs11203633 | -0.496 | 0.176 | -0.655 | 0.201 | 0 | -0.006 | -0.006 | 0.105 | 0.57 | 0.52 | 0.48 | 0.34 | 0.61 | 0.46 |
| 8 | 14450366 | rs1510424 | 0.848 | 0.332 | -1.335 | 0.379 | -0.017 | -0.015 | -0.015 | 0.137 | 0.09 | 0.05 | 0.18 | 0.39 | 0.12 | 0.13 |
| 8 | 14451068 | rs17250457 | 0.832 | 0.14 | -1.26 | -0.248 | 0.005 | 0.028 | 0.028 | 0.004 | 0.23 | 0.17 | 0.27 | 0.09 | 0.23 | 0.09 |
| 8 | 14451137 | rs17119866 | -0.639 | 0.222 | -1.107 | -0.116 | -0.005 | 0.02 | 0.02 | 0 | 0.57 | 0.39 | 0.47 | 0.34 | 0.61 | 0.48 |
| 8 | 14452324 | rs10091580 | 0.173 | 0.261 | -0.954 | -0.301 | 0.004 | 0.019 | 0.019 | 0 | 0.25 | 0.19 | 0.23 | 0.09 | 0.25 | 0.11 |
| 8 | 14453097 | rs10453158 | -1.887 | 0.39 | -0.574 | -0.232 | 0.008 | -0.019 | -0.019 | 0 | 0.95 | 0.9 | 0.97 | 0.97 | 0.97 | 0.99 |
| 8 | 14453288 | rs9325717 | 0.066 | 0.312 | -0.514 | -0.207 | 0.045 | 0.035 | 0.035 | 0 | 0.3 | 0.5 | 0.29 | 0.34 | 0.41 | 0.36 |

| | | | | | | | | | | | | | | | | |
|---|---|---|---|---|---|---|---|---|---|---|---|---|---|---|---|---|
| 8 | 14453644 | rs17250568 | 0.851 | -0.14 | -1.076 | -0.879 | -0.004 | 0.193 | 0.193 | 0 | 0.52 | 0.21 | 0.49 | 0.18 | 0.49 | 0.45 |
| 8 | 14461355 | rs2175278 | -0.017 | -0.054 | -0.771 | -1.137 | -0.003 | 0.015 | 0.015 | 3.102 | 0.54 | 0.61 | 0.62 | 0.78 | 0.5 | 0.72 |
| 8 | 14461831 | rs2898393 | 0.788 | 0.118 | -0.259 | -0.696 | 0.049 | 0.049 | 0.049 | 0.163 | 0.11 | 0.28 | 0.13 | 0.13 | 0.14 | 0.19 |
| 8 | 14463388 | rs1355303 | 1.47 | -0.039 | -0.592 | -0.868 | -0.046 | 0.056 | 0.056 | 0.058 | 0.18 | 0.02 | 0.21 | 0.07 | 0.21 | 0.09 |
| 8 | 14463704 | rs1397279 | 0.639 | -0.272 | -0.686 | -1.043 | -0.013 | -0.012 | -0.012 | 3.622 | 0.27 | 0.21 | 0.37 | 0.66 | 0.33 | 0.33 |
| 8 | 14464147 | rs12675018 | 2.125 | -0.54 | -0.949 | -1.321 | -0.004 | 0.004 | 0.004 | 0.255 | 0.27 | 0.21 | 0.24 | 0.08 | 0.27 | 0.24 |
| 8 | 14464669 | rs17119892 | 1.873 | -0.59 | -0.948 | -1.256 | 0.001 | -0.001 | -0.001 | 0 | 0.07 | 0.06 | 0.05 | 0.08 | 0.05 | 0.15 |
| 8 | 14465018 | rs4240170 | 0.479 | -0.611 | -0.781 | -1.318 | 0.005 | 0.01 | 0.01 | 0 | 0.56 | 0.62 | 0.64 | 0.74 | 0.6 | 0.59 |
| 8 | 14465315 | rs12542406 | 1.954 | -0.52 | -0.884 | -1.137 | 0.001 | -0.001 | -0.001 | 0 | 0.08 | 0.01 | 0.22 | 0.56 | 0.06 | 0.14 |
| 8 | 14468227 | rs10088391 | 1.568 | -0.602 | -0.963 | -1.313 | -0.013 | 0.05 | 0.05 | 0.603 | 0.18 | 0.09 | 0.21 | 0.07 | 0.2 | 0.11 |
| 8 | 14471820 | rs1355304 | 0.954 | -0.657 | -0.92 | -1.408 | -0.004 | 0.051 | 0.051 | 0 | 0.47 | 0.36 | 0.47 | 0.15 | 0.48 | 0.35 |
| 8 | 14473500 | rs4831619 | 1.903 | -0.652 | -1.043 | -1.413 | -0.073 | 0.02 | 0.02 | 0.035 | 0.18 | 0.04 | 0.14 | 0.11 | 0.27 | 0.19 |
| 8 | 14474514 | rs10109020 | 2.05 | -0.648 | -1.038 | -1.392 | -0.046 | 0.066 | 0.066 | 0.879 | 0.16 | 0.02 | 0.2 | 0.04 | 0.19 | 0.08 |
| 8 | 14474682 | rs10091736 | 0.678 | -0.68 | -1.019 | -1.355 | -0.002 | -0.058 | -0.058 | 0.139 | 0.26 | 0.15 | 0.43 | 0.59 | 0.25 | 0.25 |
| 8 | 14476252 | rs6981363 | -0.538 | -0.735 | -0.757 | -1.241 | -0.019 | -0.033 | -0.033 | 0 | 0.81 | 0.7 | 0.89 | 0.87 | 0.86 | 0.83 |
| 8 | 14479404 | rs13258861 | 2.417 | -0.681 | -0.934 | -1.275 | -0.026 | 0.127 | 0.127 | 0.104 | 0.25 | 0.01 | 0.2 | 0.05 | 0.2 | 0.23 |
| 8 | 14480628 | rs1397282 | -0.484 | -0.349 | -0.443 | -0.938 | 0.007 | 0.073 | 0.073 | 0.484 | 0.5 | 0.4 | 0.63 | 0.79 | 0.57 | 0.56 |
| 8 | 14481802 | rs17119925 | 1.461 | -0.084 | -0.492 | -0.605 | 0.005 | 0.006 | 0.006 | 0.751 | 0.08 | 0.03 | 0.23 | 0.56 | 0.06 | 0.14 |
| 8 | 14489024 | rs6530791 | -0.76 | -0.231 | -0.455 | -0.906 | 0.012 | 0.024 | 0.024 | 0.461 | 0.55 | 0.65 | 0.69 | 0.81 | 0.65 | 0.49 |
| 8 | 14500655 | rs964337 | 0.393 | 0.075 | -0.334 | -0.28 | 0.054 | 0.192 | 0.192 | 1.824 | 0.11 | 0.25 | 0.24 | 0.61 | 0.14 | 0.16 |
| 8 | 14505407 | rs12542237 | 0.394 | 0.085 | -0.352 | -0.282 | 0.052 | 0.18 | 0.18 | 2.369 | 0.11 | 0.24 | 0.24 | 0.6 | 0.15 | 0.15 |
| 8 | 14508593 | rs12546155 | 0.116 | -0.6 | -0.654 | -0.914 | 0.005 | 0.062 | 0.062 | 0 | 0.62 | 0.25 | 0.6 | 0.82 | 0.63 | 0.7 |
| 8 | 14508908 | rs10503504 | 1.061 | -0.545 | -0.525 | -0.982 | 0.017 | 0.108 | 0.108 | 0.502 | 0.32 | 0.11 | 0.25 | 0.11 | 0.25 | 0.35 |
| 8 | 14513442 | rs11778242 | -0.547 | -0.492 | -0.842 | -0.571 | -0.026 | 0.046 | 0.046 | 0 | 0.43 | 0.19 | 0.27 | 0.31 | 0.48 | 0.46 |
| 8 | 14513469 | rs6982610 | 0.262 | -0.565 | -0.766 | -0.534 | -0.003 | -0.013 | -0.013 | 0 | 0.62 | 0.48 | 0.73 | 0.69 | 0.61 | 0.58 |
| 8 | 14513780 | rs9987304 | -0.647 | -0.798 | -1.255 | -0.125 | -0.001 | -0.001 | -0.001 | 0.152 | 0.21 | 0.22 | 0.15 | 0.18 | 0.16 | 0.25 |
| 8 | 14518540 | rs12216821 | 0.735 | -0.798 | -1.743 | -0.14 | 0.01 | 0.015 | 0.015 | 0.066 | 0.31 | 0.72 | 0.25 | 0.27 | 0.23 | 0.26 |
| 8 | 14519517 | rs12541555 | 0.162 | -0.706 | -2.167 | -0.048 | 0.007 | -0.042 | -0.042 | 0 | 0.23 | 0.17 | 0.42 | 0.49 | 0.24 | 0.28 |
| 8 | 14522350 | rs17119984 | 0.311 | -0.708 | -2.121 | -0.054 | -0.016 | -0.004 | -0.004 | 0 | 0.06 | 0.11 | 0.02 | 0.05 | 0.02 | 0.1 |
| 8 | 14529851 | rs10097849 | -0.028 | -0.685 | -2.169 | -0.054 | 0.028 | 0.104 | 0.104 | 1.739 | 0.49 | 0.79 | 0.65 | 0.71 | 0.45 | 0.43 |
| 8 | 14543582 | rs920616 | 0.316 | -0.527 | -2.132 | 0.057 | -0.027 | 0.088 | 0.088 | 1.114 | 0.43 | 0.16 | 0.33 | 0.25 | 0.51 | 0.43 |
| 8 | 14544079 | rs6987164 | -1.241 | -0.336 | -1.973 | 0.352 | -0.019 | -0.092 | -0.092 | 0 | 0.83 | 0.94 | 0.62 | 0.55 | 0.77 | 0.79 |
| 8 | 14547284 | rs6989198 | 1.044 | -0.413 | -1.984 | 0.405 | 0.018 | -0.027 | -0.027 | 0 | 0.31 | 0.82 | 0.39 | 0.37 | 0.24 | 0.26 |
| 8 | 14550156 | rs11989868 | 1.131 | -0.41 | -1.851 | 0.424 | 0.011 | -0.008 | -0.008 | 0 | 0.31 | 0.73 | 0.38 | 0.37 | 0.24 | 0.25 |
| 8 | 14552065 | rs13282394 | -0.31 | -0.331 | -2.025 | 0.213 | -0.004 | 0.123 | 0.123 | 1.071 | 0.5 | 0.11 | 0.33 | 0.28 | 0.53 | 0.53 |
| 8 | 14554261 | rs10503513 | 0.605 | -0.056 | -1.764 | 0.229 | -0.02 | 0.006 | 0.006 | 0 | 0.1 | 0.02 | 0.09 | 0.05 | 0.12 | 0.12 |
| 8 | 14566827 | rs957607 | -0.484 | -0.19 | -0.859 | 0.252 | -0.021 | -0.004 | -0.004 | 0 | 0.07 | 0.19 | 0.02 | 0.05 | 0.02 | 0.1 |
| 8 | 14573221 | rs1372420 | 1.496 | 0.281 | -1.478 | 0.117 | -0.085 | -0.023 | -0.023 | 0.021 | 0.16 | 0.01 | 0.19 | 0.13 | 0.21 | 0.21 |
| 8 | 14573575 | rs2036829 | 1.279 | -0.225 | -1.312 | 0.306 | 0.002 | -0.003 | -0.003 | 0 | 0.31 | 0.68 | 0.4 | 0.42 | 0.24 | 0.27 |
| 8 | 14578221 | rs2250166 | 0.13 | -0.324 | -1.303 | 0.209 | -0.009 | 0.012 | 0.012 | 0.036 | 0.71 | 0.69 | 0.77 | 0.66 | 0.72 | 0.69 |
| 8 | 14580231 | rs1010895 | -0.083 | -0.318 | -1.293 | 0.187 | -0.015 | -0.009 | -0.009 | 0 | 0.07 | 0.23 | 0.02 | 0.02 | 0.02 | 0.08 |
| 8 | 14582913 | rs2165591 | 1.034 | -0.285 | -1.769 | 0.305 | -0.048 | -0.098 | -0.098 | 0.788 | 0.19 | 0.02 | 0.18 | 0.3 | 0.24 | 0.22 |
| 8 | 14585560 | rs2466921 | -0.989 | -0.331 | -1.126 | 0.258 | -0.001 | 0.008 | 0.008 | 0 | 0.8 | 0.76 | 0.8 | 0.69 | 0.75 | 0.75 |
| 8 | 14587716 | rs13263700 | 2.05 | -0.335 | -1.31 | 0.307 | 0.006 | 0.017 | 0.017 | 0 | 0.25 | 0.11 | 0.29 | 0.22 | 0.19 | 0.23 |
| 8 | 14589550 | rs2254300 | 0.182 | -0.225 | -1.577 | 0.134 | -0.087 | 0.066 | 0.066 | 0.528 | 0.42 | 0.1 | 0.38 | 0.27 | 0.49 | 0.43 |
| 8 | 14596241 | rs10087547 | 0.188 | -0.207 | -1.174 | 0.151 | -0.019 | -0.014 | -0.014 | 0.081 | 0.06 | 0.18 | 0.03 | 0.02 | 0.02 | 0.08 |
| 8 | 14599592 | rs2055857 | 1.344 | -0.268 | -1.154 | 0.266 | 0.002 | -0.004 | -0.004 | 0.318 | 0.3 | 0.65 | 0.39 | 0.41 | 0.24 | 0.26 |
| 8 | 14600390 | rs2244476 | -0.745 | -0.15 | -0.974 | 0.036 | 0.015 | 0.086 | 0.086 | 0.502 | 0.51 | 0.3 | 0.43 | 0.31 | 0.52 | 0.5 |
| 8 | 14604752 | rs2247583 | 0.383 | -0.025 | -1.149 | -0.084 | -0.105 | 0.074 | 0.074 | 2.698 | 0.33 | 0.08 | 0.3 | 0.2 | 0.38 | 0.32 |
| 8 | 14610232 | rs1965816 | 0.015 | 0.104 | -0.581 | -0.169 | -0.001 | -0.003 | -0.003 | 0 | 0.62 | 0.61 | 0.65 | 0.77 | 0.62 | 0.65 |
| 8 | 14612067 | rs2248499 | -0.676 | 0.09 | -0.499 | -0.134 | -0.023 | 0.034 | 0.034 | 2.943 | 0.68 | 0.8 | 0.68 | 0.79 | 0.64 | 0.71 |
| 8 | 14612100 | rs2248500 | -0.533 | 0.085 | -0.4 | -0.135 | -0.006 | 0.011 | 0.011 | 3.173 | 0.67 | 0.72 | 0.67 | 0.79 | 0.64 | 0.71 |
| 8 | 14614134 | rs894259 | 0.317 | 0.028 | -0.391 | -0.079 | -0.004 | 0.01 | 0.01 | 1.077 | 0.57 | 0.61 | 0.64 | 0.69 | 0.57 | 0.54 |
| 8 | 14619817 | rs1441944 | 1.023 | -0.126 | -0.684 | -0.013 | 0.007 | -0.031 | -0.031 | 0.006 | 0.46 | 0.17 | 0.46 | 0.5 | 0.45 | 0.5 |
| 8 | 14620320 | rs2165594 | -0.883 | -0.083 | -0.503 | 0.076 | 0.023 | 0.062 | 0.062 | 0.392 | 0.14 | 0.54 | 0.19 | 0.19 | 0.13 | 0.05 |
| 8 | 14625850 | rs2255685 | -0.699 | -0.204 | -0.399 | 0.013 | -0.002 | -0.002 | -0.002 | 0 | 0.13 | 0.15 | 0.09 | 0.06 | 0.1 | 0.1 |
| 8 | 14626452 | rs10105241 | -0.262 | -0.259 | -0.125 | 0.066 | 0.001 | 0.144 | 0.144 | 0 | 0.08 | 0.29 | 0.09 | 0.22 | 0.06 | 0.09 |
| 8 | 14628659 | rs6980965 | -0.918 | -0.279 | 0.438 | 0.044 | 0.007 | 0.169 | 0.169 | 0 | 0.11 | 0.51 | 0.12 | 0.22 | 0.09 | 0.12 |
| 8 | 14628891 | rs1031052 | -1.483 | -0.287 | 0.277 | 0.036 | -0.031 | 0.079 | 0.079 | 0.001 | 0.18 | 0.44 | 0.18 | 0.28 | 0.15 | 0.17 |
| 8 | 14632825 | rs2256560 | -0.364 | -0.182 | -0.186 | -0.287 | 0.005 | 0.035 | 0.035 | 0 | 0.2 | 0.14 | 0.17 | 0.06 | 0.2 | 0.11 |
| 8 | 14634691 | rs2252460 | 0.182 | -0.27 | -0.507 | -0.391 | -0.024 | -0.036 | -0.036 | 0 | 0.62 | 0.17 | 0.63 | 0.57 | 0.65 | 0.63 |
| 8 | 14637987 | rs7846701 | -0.359 | -0.445 | -0.363 | -0.448 | 0.011 | -0.037 | -0.037 | 0.696 | 0.15 | 0.32 | 0.15 | 0.07 | 0.18 | 0.11 |
| 8 | 14639668 | rs2253107 | -0.206 | -0.453 | -0.609 | -0.455 | -0.002 | 0.009 | 0.009 | 0 | 0.14 | 0.12 | 0.13 | 0.07 | 0.16 | 0.09 |
| 8 | 14640051 | rs2253225 | 0.863 | -0.316 | -0.796 | -0.499 | -0.013 | 0.012 | 0.012 | 3.354 | 0.52 | 0.15 | 0.58 | 0.56 | 0.55 | 0.54 |
| 8 | 14642903 | rs2253458 | -0.7 | -0.246 | -0.957 | -0.482 | -0.066 | 0.045 | 0.045 | 4.002 | 0.68 | 0.94 | 0.73 | 0.78 | 0.68 | 0.68 |
| 8 | 14646549 | rs1899371 | 1.727 | -0.323 | -0.864 | -0.391 | -0.136 | -0.176 | -0.176 | 0 | 0.39 | 0.02 | 0.45 | 0.48 | 0.44 | 0.47 |
| 8 | 14652812 | rs1441933 | -0.586 | -0.245 | -1.288 | -0.391 | -0.112 | 0.062 | 0.062 | 1.895 | 0.66 | 0.93 | 0.69 | 0.79 | 0.62 | 0.7 |
| 8 | 14654116 | rs2246433 | 1.519 | -0.027 | -1.109 | -0.221 | -0.108 | -0.174 | -0.174 | 0 | 0.38 | 0.04 | 0.45 | 0.47 | 0.41 | 0.45 |
| 8 | 14660575 | rs2247142 | 1.122 | -0.04 | -0.938 | -0.214 | -0.025 | -0.035 | -0.035 | 0.257 | 0.42 | 0.25 | 0.49 | 0.49 | 0.46 | 0.51 |

| | | | | | | | | | | | | | | | | |
|---|---|---|---|---|---|---|---|---|---|---|---|---|---|---|---|---|
| 8 | 14667283 | rs6995984 | 0.297 | -0.132 | -0.902 | -0.154 | 0.006 | 0.007 | 0.007 | 0 | 0.14 | 0.1 | 0.11 | 0.07 | 0.13 | 0.11 |
| 8 | 14668859 | rs7002469 | 0.515 | 0.112 | -0.551 | -0.054 | -0.001 | 0.019 | 0.019 | 1.932 | 0.51 | 0.37 | 0.58 | 0.56 | 0.55 | 0.59 |
| 8 | 14669834 | rs2250991 | -0.969 | 0.424 | -0.718 | 0.486 | -0.066 | -0.019 | -0.019 | 0 | 0.79 | 0.95 | 0.79 | 0.79 | 0.75 | 0.77 |
| 8 | 14670425 | rs17120307 | -1.192 | 0.048 | 0.315 | 0.946 | 0.029 | 0.197 | 0.197 | 0 | 0.1 | 0.52 | 0.1 | 0.22 | 0.07 | 0.12 |
| 8 | 14672184 | rs2251303 | 0.033 | 0.311 | -0.837 | 1.311 | -0.059 | -0.114 | -0.114 | 0 | 0.19 | 0.04 | 0.28 | 0.44 | 0.26 | 0.24 |
| 8 | 14672930 | rs17120311 | 0.36 | -0.571 | -0.425 | 1.63 | -0.024 | 0.033 | 0.033 | 0.059 | 0.15 | 0.14 | 0.17 | 0.32 | 0.06 | 0.21 |
| 8 | 14674205 | rs17278826 | 0.433 | -0.421 | -0.714 | 1.58 | -0.018 | -0.056 | -0.056 | 3.746 | 0.22 | 0.04 | 0.2 | 0.25 | 0.23 | 0.21 |
| 8 | 14676812 | rs4831651 | -0.836 | -0.557 | 0.324 | 1.861 | -0.044 | 0.134 | 0.134 | 0 | 0.22 | 0.42 | 0.36 | 0.57 | 0.15 | 0.31 |
| 8 | 14678519 | rs7823866 | -0.697 | -0.466 | 0.354 | 1.839 | 0.055 | 0.233 | 0.233 | 0 | 0.05 | 0.27 | 0.17 | 0.26 | 0.08 | 0.1 |
| 8 | 14681936 | rs7826396 | 0.364 | -0.546 | 0.052 | 1.39 | -0.021 | 0.033 | 0.033 | 0 | 0.15 | 0.14 | 0.2 | 0.32 | 0.07 | 0.21 |
| 8 | 14688728 | rs7824647 | -1.592 | -0.473 | 0.831 | 0.918 | 0.074 | 0.052 | 0.052 | 0.386 | 0.07 | 0.45 | 0.13 | 0.03 | 0.06 | 0.09 |
| 8 | 14694373 | rs4240174 | -0.49 | -0.429 | 0.694 | 1.02 | 0.011 | 0.042 | 0.042 | 0 | 0.33 | 0.5 | 0.44 | 0.54 | 0.38 | 0.32 |
| 8 | 14695745 | rs4596659 | -0.53 | -0.52 | 0.696 | 1.025 | -0.004 | 0.017 | 0.017 | 0 | 0.35 | 0.56 | 0.34 | 0.46 | 0.35 | 0.32 |
| 8 | 14696388 | rs11780862 | 0.35 | -0.516 | 0.384 | 1.024 | -0.087 | -0.153 | -0.153 | 0 | 0.27 | 0.02 | 0.29 | 0.44 | 0.32 | 0.24 |
| 8 | 14696964 | rs4262322 | -0.186 | -0.407 | 0.52 | 0.758 | -0.012 | 0.024 | 0.024 | 0 | 0.19 | 0.28 | 0.31 | 0.26 | 0.15 | 0.24 |
| 8 | 14697130 | rs10107666 | -1.471 | -0.289 | -0.131 | 0.31 | -0.036 | 0.099 | 0.099 | 0 | 0.57 | 0.74 | 0.66 | 0.79 | 0.52 | 0.57 |
| 8 | 14697181 | rs10104703 | -1.712 | -0.263 | -0.509 | 0.324 | -0.092 | 0.107 | 0.107 | 0.008 | 0.59 | 0.87 | 0.67 | 0.79 | 0.52 | 0.57 |
| 8 | 14698830 | rs4831658 | -0.322 | 0.002 | 0.035 | 0.425 | 0.017 | -0.024 | -0.024 | 0 | 0.06 | 0.09 | 0.15 | 0 | 0.1 | 0.06 |
| 8 | 14711886 | rs6530810 | -1.077 | -0.8 | 0.162 | 0.361 | -0.019 | -0.008 | -0.008 | 0 | 0.29 | 0.65 | 0.34 | 0.22 | 0.15 | 0.27 |
| 8 | 14746634 | rs1480696 | -1.141 | -0.642 | -0.014 | 0.372 | 0.012 | 0.007 | 0.007 | 0 | 0.3 | 0.7 | 0.36 | 0.24 | 0.18 | 0.28 |
| 8 | 14755579 | rs9643991 | 0.497 | -0.373 | -0.439 | 0.431 | -0.078 | -0.141 | -0.141 | 0.396 | 0.26 | 0.01 | 0.29 | 0.63 | 0.3 | 0.27 |
| 8 | 14756351 | rs17120493 | -0.587 | -0.069 | 0.596 | 0.368 | 0.013 | -0.038 | -0.038 | 0 | 0.07 | 0.18 | 0.05 | 0 | 0.07 | 0.04 |
| 8 | 14762991 | rs4831661 | -0.492 | -0.441 | 1.229 | 0.41 | -0.059 | 0.018 | 0.018 | 0 | 0.21 | 0.49 | 0.21 | 0.22 | 0.1 | 0.2 |
| 8 | 14768548 | rs17601874 | -0.082 | -0.351 | 1.077 | 0.442 | -0.014 | -0.002 | -0.002 | 0 | 0.17 | 0.27 | 0.21 | 0.18 | 0.12 | 0.25 |
| 8 | 14773143 | rs1564449 | -0.866 | -0.364 | 1.083 | 0.41 | 0.026 | -0.006 | -0.006 | 0 | 0.08 | 0.39 | 0.05 | 0 | 0.03 | 0.01 |
| 8 | 14777031 | rs10099062 | 0.4 | -0.339 | 0.93 | 0.508 | -0.019 | 0.004 | 0.004 | 0 | 0.14 | 0.17 | 0.1 | 0.17 | 0.07 | 0.18 |
| 8 | 14779330 | rs922657 | -0.785 | -0.24 | 0.685 | 0.509 | -0.016 | 0.004 | 0.004 | 0 | 0.2 | 0.64 | 0.3 | 0.23 | 0.14 | 0.27 |
| 8 | 14785484 | rs17120612 | -0.54 | -0.149 | 0.484 | 1.062 | 0.003 | 0.021 | 0.021 | 0 | 0.36 | 0.51 | 0.46 | 0.54 | 0.41 | 0.39 |
| 8 | 14790181 | rs11992231 | -0.194 | -0.515 | -0.491 | 0.909 | 0.015 | -0.099 | -0.099 | 1.166 | 0.73 | 0.94 | 0.77 | 0.65 | 0.82 | 0.78 |
| 8 | 14791491 | rs11203669 | -0.508 | -0.088 | -0.569 | 0.87 | -0.004 | -0.015 | -0.015 | 0 | 0.42 | 0.7 | 0.5 | 0.61 | 0.5 | 0.46 |
| 8 | 14794917 | rs12679928 | -0.559 | -0.324 | -0.334 | 0.749 | 0 | 0 | 0 | 0.095 | 0.42 | 0.51 | 0.46 | 0.43 | 0.41 | 0.48 |
| 8 | 14795799 | rs1480695 | -0.387 | -0.33 | -0.194 | 0.655 | -0.034 | -0.062 | -0.062 | 0 | 0.18 | 0.45 | 0.19 | 0.11 | 0.11 | 0.2 |
| 8 | 14796949 | rs7813529 | -0.12 | -0.332 | -0.381 | 0.589 | -0.02 | -0.119 | -0.119 | 0.141 | 0.71 | 0.95 | 0.77 | 0.6 | 0.76 | 0.82 |
| 8 | 14797292 | rs11781498 | -0.375 | -0.399 | -0.22 | 0.287 | -0.034 | -0.062 | -0.062 | 0.478 | 0.18 | 0.45 | 0.19 | 0.11 | 0.11 | 0.2 |
| 8 | 14807910 | rs4831665 | 0.271 | -0.311 | -0.135 | 0.312 | 0.026 | 0.031 | 0.031 | 0.616 | 0.67 | 0.79 | 0.77 | 0.81 | 0.81 | 0.8 |
| 8 | 14808002 | rs17470444 | 0.732 | -0.159 | -0.172 | 0.334 | -0.01 | -0.003 | -0.003 | 0 | 0.24 | 0.2 | 0.27 | 0.29 | 0.31 | 0.27 |
| 8 | 14809219 | rs962727 | 1.175 | -0.301 | -0.279 | 0.366 | -0.049 | -0.061 | -0.061 | 0 | 0.22 | 0.03 | 0.24 | 0.28 | 0.27 | 0.24 |
| 8 | 14809796 | rs1383411 | 0.36 | -0.377 | -0.011 | 0.329 | -0.008 | -0.01 | -0.01 | 0 | 0.63 | 0.64 | 0.74 | 0.73 | 0.75 | 0.8 |
| 8 | 14811169 | rs17608413 | 0.095 | -0.23 | 0.312 | 0.399 | -0.015 | -0.012 | -0.012 | 0.049 | 0.43 | 0.22 | 0.44 | 0.47 | 0.45 | 0.38 |
| 8 | 14811233 | rs13439657 | -1.21 | -0.159 | 0.37 | 0.443 | -0.018 | -0.038 | -0.038 | 0 | 0.62 | 0.74 | 0.66 | 0.57 | 0.57 | 0.6 |
| 8 | 14814308 | rs10101010 | 0.551 | -0.197 | 0.853 | 0.231 | -0.026 | -0.007 | -0.007 | 0.875 | 0.09 | 0.16 | 0.04 | 0.09 | 0.03 | 0.06 |
| 8 | 14822104 | rs10081602 | 0.335 | -0.418 | 0.664 | 0.229 | -0.024 | -0.008 | -0.008 | 0 | 0.1 | 0.18 | 0.05 | 0.09 | 0.03 | 0.06 |
| 8 | 14822757 | rs7827121 | -0.196 | -0.435 | 1.072 | 0.21 | -0.021 | 0.005 | 0.005 | 0 | 0.12 | 0.46 | 0.08 | 0.09 | 0.04 | 0.08 |
| 8 | 14824575 | rs7008305 | 0.128 | -0.479 | 0.728 | 0.168 | -0.022 | -0.018 | -0.018 | 0 | 0.86 | 0.79 | 0.93 | 0.9 | 0.95 | 0.82 |
| 8 | 14830796 | rs13267348 | 1.566 | -0.221 | 0.208 | 0.206 | -0.02 | -0.023 | -0.023 | 0 | 0.3 | 0.25 | 0.38 | 0.37 | 0.41 | 0.39 |
| 8 | 14831991 | rs5028286 | -0.58 | -0.391 | 0.03 | 0.161 | -0.022 | -0.003 | -0.003 | 0 | 0.16 | 0.35 | 0.2 | 0.14 | 0.1 | 0.21 |
| 8 | 14837778 | rs7012038 | -0.183 | -0.378 | -0.447 | 0.233 | 0.003 | 0.096 | 0.096 | 0 | 0.51 | 0.2 | 0.51 | 0.69 | 0.5 | 0.52 |
| 8 | 14839006 | rs7016798 | 0.997 | -0.291 | -0.552 | 0.278 | -0.058 | 0.051 | 0.051 | 0 | 0.36 | 0.17 | 0.38 | 0.22 | 0.44 | 0.39 |
| 8 | 14839946 | rs10109779 | 1.339 | -0.099 | -0.449 | 0.446 | -0.06 | -0.066 | -0.066 | 0.454 | 0.3 | 0.19 | 0.38 | 0.38 | 0.43 | 0.39 |
| 8 | 14841762 | rs12681103 | 0.966 | -0.078 | -0.647 | 0.541 | -0.066 | -0.125 | -0.125 | 0.641 | 0.19 | 0.02 | 0.26 | 0.44 | 0.26 | 0.25 |
| 8 | 14848938 | rs1480694 | 0.187 | -0.186 | -0.054 | 0.466 | -0.012 | -0.004 | -0.004 | 0 | 0.11 | 0.17 | 0.05 | 0.09 | 0.03 | 0.1 |
| 8 | 14850978 | rs9657247 | 0.158 | -0.188 | -0.039 | 0.465 | -0.013 | -0.005 | -0.005 | 0 | 0.11 | 0.18 | 0.05 | 0.09 | 0.03 | 0.1 |
| 8 | 14853394 | rs13251088 | 1.193 | 0.056 | -0.062 | 0.339 | 0.002 | -0.005 | -0.005 | 0 | 0.31 | 0.35 | 0.39 | 0.23 | 0.37 | 0.42 |
| 8 | 14855673 | rs17120703 | -0.644 | 0.062 | -0.168 | 0.277 | 0.001 | 0.001 | 0.001 | 1.928 | 0.4 | 0.57 | 0.32 | 0.25 | 0.3 | 0.36 |
| 8 | 14858666 | rs7840310 | -0.623 | 0.004 | -0.16 | 0.712 | -0.036 | -0.061 | -0.061 | 0.098 | 0.7 | 0.88 | 0.69 | 0.62 | 0.64 | 0.7 |
| 8 | 14860953 | rs11203672 | 0.495 | -0.297 | 0.202 | 0.572 | -0.026 | -0.004 | -0.004 | 1.322 | 0.1 | 0.19 | 0.06 | 0.08 | 0.03 | 0.08 |
| 8 | 14861409 | rs6530824 | -0.711 | -0.013 | 0.175 | 0.603 | 0.017 | -0.038 | -0.038 | 0.653 | 0.7 | 0.81 | 0.72 | 0.61 | 0.71 | 0.7 |
| 8 | 14862366 | rs10888094 | 0.495 | -0.039 | 0.148 | 0.359 | -0.001 | 0.009 | 0.009 | 0.114 | 0.47 | 0.53 | 0.46 | 0.32 | 0.5 | 0.55 |
| 8 | 14863434 | rs1125265 | 0.678 | -0.057 | 0.196 | 0.363 | -0.003 | 0.013 | 0.013 | 0.318 | 0.45 | 0.4 | 0.45 | 0.31 | 0.49 | 0.51 |
| 8 | 14868055 | rs11786914 | 0.091 | -0.098 | 0.15 | 0.359 | 0 | 0.014 | 0.014 | 0.104 | 0.12 | 0.33 | 0.07 | 0.11 | 0.05 | 0.12 |
| 8 | 14868162 | rs2127774 | -0.78 | -0.106 | 0.083 | 0.374 | -0.024 | 0.041 | 0.041 | 2.053 | 0.28 | 0.41 | 0.27 | 0.38 | 0.18 | 0.25 |
| 8 | 14868516 | rs13251017 | 1.306 | -0.068 | -0.286 | 0.364 | -0.061 | 0.067 | 0.067 | 0.49 | 0.36 | 0.17 | 0.39 | 0.22 | 0.45 | 0.41 |
| 8 | 14868636 | rs10111834 | -0.547 | -0.003 | -0.387 | 0.381 | 0.003 | -0.013 | -0.013 | 0 | 0.65 | 0.68 | 0.67 | 0.61 | 0.63 | 0.7 |
| 8 | 14868808 | rs13251961 | 1.037 | 0.01 | -0.544 | 0.417 | -0.016 | 0.03 | 0.03 | 0.274 | 0.38 | 0.3 | 0.4 | 0.22 | 0.45 | 0.41 |
| 8 | 14870508 | rs10088098 | -0.409 | 0.17 | -1.272 | 0.567 | -0.025 | 0.017 | 0.017 | 0.541 | 0.21 | 0.08 | 0.25 | 0.15 | 0.21 | 0.16 |
| 8 | 14870622 | rs10283038 | 0.733 | -0.137 | -1.049 | 0.788 | -0.001 | -0.01 | -0.01 | 0 | 0.25 | 0.28 | 0.27 | 0.38 | 0.3 | 0.24 |
| 8 | 14873709 | rs7015648 | -0.628 | -0.941 | -1.073 | 0.538 | -0.004 | -0.002 | -0.002 | 0.423 | 0.29 | 0.64 | 0.25 | 0.35 | 0.12 | 0.25 |
| 8 | 14874924 | rs2170239 | 0.647 | -0.084 | -1.663 | 0.31 | -0.077 | 0.053 | 0.053 | 1.811 | 0.31 | 0.07 | 0.38 | 0.18 | 0.36 | 0.44 |
| 8 | 14875723 | rs10216426 | -0.129 | -0.762 | -0.439 | 0.563 | -0.027 | 0.004 | 0.004 | 0.009 | 0.14 | 0.29 | 0.11 | 0.17 | 0.06 | 0.15 |

| | | | | | | | | | | | | | | | | |
|---|---|---|---|---|---|---|---|---|---|---|---|---|---|---|---|---|
| 8 | 14877288 | rs7815606 | -0.696 | -0.482 | -0.21 | 0.516 | -0.032 | 0.043 | 0.043 | 0 | 0.19 | 0.29 | 0.17 | 0.2 | 0.11 | 0.1 |
| 8 | 14879360 | rs7830454 | 0.252 | -1.04 | -0.579 | -0.346 | 0.043 | 0.054 | 0.054 | 0 | 0.51 | 0.47 | 0.73 | 0.82 | 0.72 | 0.55 |
| 8 | 14879525 | rs7834376 | -0.621 | -1.392 | 0.045 | -0.4 | 0.062 | -0.003 | -0.003 | 0 | 0.11 | 0.21 | 0.1 | 0.08 | 0.16 | 0.06 |
| 8 | 14879659 | rs7830867 | -1.098 | -2.959 | -0.259 | -0.852 | 0.012 | 0.016 | 0.016 | 0.465 | 0.52 | 0.49 | 0.35 | 0.32 | 0.38 | 0.48 |
| 8 | 14880448 | rs936631 | -0.709 | -3.229 | 0.205 | -1.047 | 0.021 | 0.037 | 0.037 | 0 | 0.36 | 0.27 | 0.28 | 0.25 | 0.31 | 0.48 |
| 8 | 14881568 | rs12678080 | -0.28 | -2.784 | 0.584 | -0.17 | 0.004 | 0.077 | 0.077 | 0 | 0.88 | 0.76 | 0.88 | 0.74 | 0.9 | 0.89 |
| 8 | 14888292 | rs746836 | -0.674 | -3.241 | 0.65 | -0.443 | -0.001 | -0.002 | -0.002 | 1.418 | 0.33 | 0.29 | 0.25 | 0.25 | 0.28 | 0.46 |
| 8 | 14892661 | rs9650384 | 1.347 | -3.168 | 0.762 | -0.375 | 0.065 | -0.008 | -0.008 | 0 | 0.49 | 0.3 | 0.59 | 0.48 | 0.63 | 0.44 |
| 8 | 14897211 | rs7007999 | -1.526 | -3.259 | 0.932 | -0.452 | 0.031 | -0.002 | -0.002 | 1.372 | 0.52 | 0.59 | 0.37 | 0.5 | 0.36 | 0.55 |
| 8 | 14897267 | rs6983207 | -1.243 | -3.245 | 0.937 | -0.444 | 0.017 | -0.002 | -0.002 | 0 | 0.47 | 0.55 | 0.35 | 0.5 | 0.34 | 0.55 |
| 8 | 14899788 | rs7004485 | -1.115 | -3.204 | 0.905 | -0.387 | 0.045 | -0.009 | -0.009 | 0.402 | 0.47 | 0.35 | 0.33 | 0.49 | 0.35 | 0.54 |
| 8 | 14900324 | rs6995750 | 1.201 | -3.209 | 1.144 | -0.206 | 0.046 | -0.005 | -0.005 | 0.032 | 0.52 | 0.33 | 0.62 | 0.49 | 0.6 | 0.42 |
| 8 | 14902400 | rs7002722 | -0.8 | -3.05 | 1.15 | -0.428 | 0.026 | 0.043 | 0.043 | 0 | 0.36 | 0.3 | 0.28 | 0.19 | 0.29 | 0.36 |
| 8 | 14902794 | rs10888101 | 1.004 | -2.646 | 1.219 | -0.524 | -0.017 | -0.026 | -0.026 | 0 | 0.92 | 0.71 | 0.97 | 1 | 0.97 | 0.98 |
| 8 | 14909339 | rs7844246 | -0.77 | -2.402 | 1.067 | -0.455 | 0.033 | 0.008 | 0.008 | 0 | 0.07 | 0.31 | 0.04 | 0 | 0.04 | 0.01 |
| 8 | 14909909 | rs1869578 | 1.351 | -2.749 | 0.852 | -0.453 | 0.04 | -0.001 | -0.001 | 0 | 0.5 | 0.33 | 0.61 | 0.53 | 0.59 | 0.44 |
| 8 | 14912769 | rs1454598 | 0.5 | -2.569 | 1.268 | -0.121 | -0.034 | -0.078 | -0.078 | 0.095 | 0.71 | 0.57 | 0.77 | 0.89 | 0.8 | 0.77 |
| 8 | 14916925 | rs17655231 | -0.244 | -2.575 | 1.426 | -0.002 | -0.008 | 0.031 | 0.031 | 1.046 | 0.11 | 0.12 | 0.07 | 0.29 | 0.08 | 0.14 |
| 8 | 14917224 | rs4831675 | 0.948 | -2.596 | 1.611 | -0.039 | -0.034 | -0.042 | -0.042 | 0 | 0.65 | 0.55 | 0.73 | 0.8 | 0.77 | 0.64 |
| 8 | 14919478 | rs9643995 | 0.641 | -2.517 | 1.471 | -0.044 | -0.014 | 0.035 | 0.035 | 0.099 | 0.07 | 0.19 | 0.04 | 0.11 | 0.03 | 0.13 |
| 8 | 14919557 | rs10111470 | 1.666 | -2.507 | 1.366 | -0.049 | 0.065 | -0.01 | -0.01 | 0 | 0.53 | 0.28 | 0.63 | 0.51 | 0.64 | 0.48 |
| 8 | 14921494 | rs17608964 | -0.068 | -2.022 | 1.317 | 0.138 | 0.002 | 0.013 | 0.013 | 2.739 | 0.1 | 0.06 | 0.06 | 0.29 | 0.08 | 0.13 |
| 8 | 14922477 | rs13281953 | 0.614 | -2.107 | 1.316 | 0.026 | -0.019 | -0.075 | -0.075 | 0.67 | 0.71 | 0.45 | 0.76 | 0.91 | 0.79 | 0.79 |
| 8 | 14928878 | rs2898407 | 0.26 | -2.077 | 0.819 | 0.303 | -0.005 | 0.021 | 0.021 | 4.422 | 0.89 | 0.86 | 0.92 | 0.72 | 0.92 | 0.86 |
| 8 | 14932048 | rs17609053 | -0.805 | -1.984 | 0.83 | 0.233 | -0.005 | -0.023 | -0.023 | 1.327 | 0.17 | 0.14 | 0.14 | 0.29 | 0.22 | 0.18 |
| 8 | 14932322 | rs17609074 | -0.703 | -1.981 | 0.825 | 0.234 | -0.004 | -0.017 | -0.017 | 0 | 0.17 | 0.14 | 0.14 | 0.29 | 0.22 | 0.19 |
| 8 | 14933399 | rs7011953 | -0.469 | -2.032 | 1.499 | 0.544 | -0.009 | -0.012 | -0.012 | 0 | 0.21 | 0.21 | 0.17 | 0.09 | 0.11 | 0.16 |
| 8 | 14934535 | rs17120855 | -0.509 | -2.061 | 1.581 | 0.532 | -0.025 | -0.031 | -0.031 | 0.848 | 0.21 | 0.27 | 0.18 | 0.1 | 0.12 | 0.16 |
| 8 | 14935227 | rs1454592 | 0.432 | -2.016 | 1.477 | 0.746 | 0.009 | 0.036 | 0.036 | 0 | 0.06 | 0.03 | 0.03 | 0.28 | 0.05 | 0.12 |
| 8 | 14938404 | rs2410229 | 1.575 | -2.173 | 1.422 | 0.752 | 0.013 | -0.011 | -0.011 | 0 | 0.63 | 0.33 | 0.72 | 0.51 | 0.76 | 0.56 |
| 8 | 14940146 | rs1378033 | 0.648 | -2.088 | 0.977 | 0.763 | 0.026 | 0.014 | 0.014 | 0 | 0.74 | 0.82 | 0.78 | 0.8 | 0.82 | 0.67 |
| 8 | 14944692 | rs1378032 | 0.552 | -1.983 | 0.879 | 0.87 | 0.008 | 0.035 | 0.035 | 0 | 0.05 | 0.03 | 0.02 | 0.28 | 0.05 | 0.11 |
| 8 | 14944908 | rs10503531 | 0.053 | -1.941 | 0.91 | 0.744 | -0.001 | 0 | 0 | 0.009 | 0.92 | 0.91 | 0.94 | 0.89 | 0.91 | 0.82 |
| 8 | 14946168 | rs4831677 | 0.093 | -1.979 | 0.95 | 0.744 | -0.001 | 0.011 | 0.011 | 0.007 | 0.92 | 0.89 | 0.94 | 0.89 | 0.91 | 0.82 |
| 8 | 14948644 | rs17120898 | 0.057 | -2.038 | 0.756 | 0.708 | 0.027 | 0.01 | 0.01 | 0 | 0.18 | 0.01 | 0.13 | 0.12 | 0.11 | 0.16 |
| 8 | 14950895 | rs4831679 | 1.17 | -2.075 | 1.155 | 0.771 | -0.032 | 0.122 | 0.122 | 0 | 0.82 | 0.58 | 0.89 | 0.61 | 0.85 | 0.7 |
| 8 | 14951436 | rs7821053 | 1.058 | -1.934 | 1.214 | 0.65 | -0.01 | 0.003 | 0.003 | 0 | 0.64 | 0.55 | 0.76 | 0.49 | 0.75 | 0.54 |
| 8 | 14951943 | rs6530833 | -1.072 | -1.664 | 1.098 | 0.692 | -0.062 | 0.045 | 0.045 | 0 | 0.31 | 0.52 | 0.22 | 0.4 | 0.22 | 0.31 |
| 8 | 14962494 | rs1026349 | -0.705 | -1.561 | -0.067 | 0.476 | 0.054 | -0.061 | -0.061 | 0 | 0.3 | 0.14 | 0.19 | 0.48 | 0.17 | 0.35 |
| 8 | 14962552 | rs6651385 | 0.914 | -1.57 | 0.079 | 0.42 | 0.011 | -0.006 | -0.006 | 0 | 0.62 | 0.37 | 0.75 | 0.5 | 0.77 | 0.61 |
| 8 | 14965403 | rs2168123 | -0.001 | -0.709 | -0.496 | 0.014 | -0.047 | 0.059 | 0.059 | 0 | 0.16 | 0.03 | 0.11 | 0.06 | 0.18 | 0.14 |
| 8 | 14969057 | rs13265356 | 1.351 | -1.369 | -1.157 | -0.17 | -0.051 | -0.025 | -0.025 | 0 | 0.45 | 0.1 | 0.63 | 0.58 | 0.6 | 0.51 |
| 8 | 14972402 | rs17574959 | 0.198 | -1.26 | -0.767 | -0.068 | -0.027 | 0.026 | 0.026 | 0 | 0.17 | 0.09 | 0.12 | 0.11 | 0.18 | 0.14 |
| 8 | 14975143 | rs1454583 | 0.274 | -1.275 | -0.773 | -0.094 | -0.035 | 0.047 | 0.047 | 0 | 0.17 | 0.07 | 0.12 | 0.07 | 0.18 | 0.13 |
| 8 | 14979482 | rs7844593 | 0.179 | -1.917 | -0.071 | -0.088 | -0.046 | -0.02 | -0.02 | 0 | 0.74 | 0.48 | 0.86 | 0.83 | 0.84 | 0.78 |
| 8 | 14981439 | rs1378030 | -1.411 | -2.051 | 0.49 | 0.207 | 0.007 | 0.012 | 0.012 | 0 | 0.44 | 0.68 | 0.35 | 0.38 | 0.35 | 0.36 |
| 8 | 14982273 | rs17575042 | 0.493 | -2.033 | 0.408 | 0.091 | -0.073 | 0.083 | 0.083 | 0 | 0.17 | 0.01 | 0.12 | 0.06 | 0.18 | 0.13 |
| 8 | 14982457 | rs1454580 | -1.191 | -2.038 | 0.586 | 0.115 | 0.001 | 0.002 | 0.002 | 0 | 0.41 | 0.6 | 0.35 | 0.38 | 0.33 | 0.35 |
| 8 | 14985667 | rs10503533 | 1.111 | -2.153 | 0.414 | -0.131 | 0.029 | -0.005 | -0.005 | 0 | 0.6 | 0.32 | 0.76 | 0.64 | 0.77 | 0.64 |
| 8 | 14987658 | rs12056855 | -0.125 | -2.135 | 0.431 | -0.125 | 0.035 | -0.01 | -0.01 | 0.519 | 0.22 | 0.1 | 0.11 | 0.18 | 0.14 | 0.21 |
| 8 | 14989966 | rs4831685 | 1.068 | -2.1 | 0.241 | -0.181 | -0.002 | -0.004 | -0.004 | 0 | 0.58 | 0.44 | 0.74 | 0.66 | 0.67 | 0.65 |
| 8 | 14990069 | rs13249540 | 1.753 | -1.911 | -0.251 | -0.117 | -0.017 | -0.016 | -0.016 | 0 | 0.42 | 0.27 | 0.56 | 0.55 | 0.58 | 0.51 |
| 8 | 14991599 | rs17471747 | 0.532 | -1.365 | -0.346 | -0.213 | -0.074 | 0.093 | 0.093 | 0 | 0.16 | 0.01 | 0.11 | 0.01 | 0.18 | 0.13 |
| 8 | 14996912 | rs268428 | -0.629 | -1.651 | 0.024 | -0.085 | -0.028 | 0.09 | 0.09 | 0.878 | 0.2 | 0.53 | 0.13 | 0.31 | 0.13 | 0.16 |
| 8 | 14997275 | rs268425 | 0.556 | -1.941 | -0.148 | -0.353 | 0.011 | 0.079 | 0.079 | 0 | 0.63 | 0.76 | 0.77 | 0.9 | 0.72 | 0.67 |
| 8 | 14997519 | rs268422 | 0.038 | -2.111 | 0.035 | -0.168 | 0.008 | 0.008 | 0.008 | 0 | 0.79 | 0.79 | 0.87 | 0.94 | 0.9 | 0.8 |
| 8 | 14997611 | rs1869577 | 1.189 | -2.138 | 0.004 | -0.143 | 0.048 | -0.006 | -0.006 | 0 | 0.59 | 0.29 | 0.75 | 0.62 | 0.76 | 0.65 |
| 8 | 14999256 | rs7832745 | -1.005 | -2.105 | 0.242 | -0.096 | -0.033 | 0.003 | 0.003 | 0 | 0.25 | 0.41 | 0.13 | 0.2 | 0.16 | 0.2 |
| 8 | 14999559 | rs7833187 | 0.644 | -1.842 | -0.196 | -0.069 | -0.01 | -0.015 | -0.015 | 0 | 0.77 | 0.72 | 0.86 | 0.93 | 0.88 | 0.8 |
| 8 | 15000112 | rs17575278 | 0.298 | -1.898 | -0.274 | -0.383 | 0.024 | 0.006 | 0.006 | 0 | 0.14 | 0.01 | 0.1 | 0.07 | 0.07 | 0.13 |
| 8 | 15001979 | rs478019 | -0.724 | -2.069 | -0.49 | -0.488 | 0.066 | 0.07 | 0.07 | 0 | 0.24 | 0.12 | 0.13 | 0.07 | 0.13 | 0.21 |
| 8 | 15003364 | rs554891 | -0.773 | -0.413 | -0.334 | -0.323 | -0.017 | 0.012 | 0.012 | 0.937 | 0.42 | 0.34 | 0.28 | 0.37 | 0.49 | 0.38 |
| 8 | 15003951 | rs198529 | 0.455 | -0.13 | 0.416 | -0.459 | 0.035 | -0.105 | -0.105 | 0.068 | 0.84 | 0.63 | 0.85 | 1 | 0.86 | 0.9 |
| 8 | 15005820 | rs17471917 | -0.236 | -0.118 | -0.029 | -0.472 | -0.034 | 0.045 | 0.045 | 0 | 0.08 | 0 | 0.06 | 0.01 | 0.08 | 0.07 |
| 8 | 15006154 | rs13250784 | 0.766 | -0.171 | -1.048 | -0.604 | -0.122 | 0.02 | 0.02 | 0 | 0.33 | 0.03 | 0.27 | 0.19 | 0.43 | 0.36 |
| 8 | 15006325 | rs1457219 | 0.773 | -0.02 | -0.7 | -0.393 | -0.002 | 0.003 | 0.003 | 0 | 0.06 | 0.06 | 0.08 | 0.04 | 0.08 | 0.06 |
| 8 | 15008979 | rs268379 | -0.496 | -0.144 | -0.468 | 0.086 | -0.002 | 0.014 | 0.014 | 0 | 0.88 | 0.84 | 0.84 | 0.82 | 0.89 | 0.89 |
| 8 | 15009878 | rs268377 | 0.839 | -0.07 | -0.028 | 0.048 | -0.003 | 0.083 | 0.083 | 0.135 | 0.8 | 0.46 | 0.71 | 0.71 | 0.8 | 0.81 |
| 8 | 15011716 | rs12548503 | 1.042 | -0.065 | -0.766 | 0.208 | 0.005 | -0.096 | -0.096 | 0 | 0.28 | 0.12 | 0.32 | 0.49 | 0.22 | 0.32 |

| | | | | | | | | | | | | | | | | |
|---|---|---|---|---|---|---|---|---|---|---|---|---|---|---|---|---|
| 8 | 15015581 | rs12678727 | 0.366 | -0.08 | -0.703 | 0.15 | 0.002 | -0.021 | -0.021 | 0 | 0.33 | 0.28 | 0.42 | 0.6 | 0.3 | 0.39 |
| 8 | 15016172 | rs10503539 | 0.683 | -0.038 | -0.809 | -0.04 | 0 | -0.002 | -0.002 | 0 | 0.06 | 0.08 | 0.08 | 0.04 | 0.08 | 0.06 |
| 8 | 15024850 | rs2410233 | 0.699 | -0.31 | -1.129 | 0.257 | 0.014 | -0.123 | -0.123 | 1.158 | 0.31 | 0.11 | 0.36 | 0.61 | 0.24 | 0.32 |
| 8 | 15031027 | rs7840960 | 1.019 | -0.23 | -0.954 | -0.093 | -0.039 | 0.063 | 0.063 | 0 | 0.34 | 0.18 | 0.3 | 0.15 | 0.44 | 0.33 |
| 8 | 15032119 | rs268386 | 0.48 | -0.331 | -0.243 | -0.205 | -0.004 | 0.011 | 0.011 | 0 | 0.42 | 0.36 | 0.35 | 0.15 | 0.49 | 0.37 |
| 8 | 15035322 | rs750895 | 0.876 | -0.428 | -0.656 | 0.053 | -0.078 | -0.005 | -0.005 | 0 | 0.37 | 0.13 | 0.38 | 0.29 | 0.5 | 0.37 |
| 8 | 15036752 | rs268353 | 0.651 | -0.425 | -0.527 | 0.019 | -0.001 | -0.013 | -0.013 | 0.141 | 0.07 | 0.09 | 0.05 | 0 | 0.05 | 0.05 |
| 8 | 15037002 | rs1457223 | 0.805 | -0.502 | -0.496 | 0.003 | -0.066 | -0.051 | -0.051 | 0.04 | 0.38 | 0.14 | 0.37 | 0.36 | 0.5 | 0.39 |
| 8 | 15037929 | rs17609949 | 1.449 | -0.77 | -0.632 | -0.002 | -0.022 | -0.114 | -0.114 | 0 | 0.28 | 0.07 | 0.33 | 0.46 | 0.24 | 0.33 |
| 8 | 15038435 | rs268359 | 0.293 | -0.347 | -0.218 | 0.186 | 0.006 | 0.003 | 0.003 | 0 | 0.46 | 0.32 | 0.43 | 0.37 | 0.55 | 0.43 |
| 8 | 15039864 | rs7820870 | 0.164 | -0.554 | 0.066 | 0.163 | 0.036 | 0.015 | 0.015 | 0 | 0.45 | 0.65 | 0.47 | 0.62 | 0.35 | 0.49 |
| 8 | 15040697 | rs268365 | 0.111 | -0.574 | 0.116 | 0.033 | 0.013 | -0.001 | -0.001 | 0 | 0.44 | 0.58 | 0.43 | 0.48 | 0.32 | 0.44 |
| 8 | 15045575 | rs268349 | -0.628 | -0.628 | -0.253 | -0.111 | -0.028 | 0.095 | 0.095 | 0 | 0.55 | 0.82 | 0.59 | 0.69 | 0.42 | 0.53 |
| 8 | 15045825 | rs17575734 | 0.151 | -0.6 | -0.227 | -0.088 | 0.004 | 0.005 | 0.005 | 0 | 0.09 | 0.08 | 0.07 | 0.05 | 0.08 | 0.07 |
| 8 | 15047014 | rs9325744 | -0.244 | -0.54 | 0.102 | -0.088 | 0.004 | -0.019 | -0.019 | 0 | 0.13 | 0.18 | 0.13 | 0.05 | 0.12 | 0.1 |
| 8 | 15047485 | rs552993 | -0.221 | -0.538 | 0.098 | -0.087 | 0.004 | -0.018 | -0.018 | 0 | 0.13 | 0.18 | 0.13 | 0.05 | 0.11 | 0.1 |
| 8 | 15050793 | rs573856 | 0.662 | -0.589 | -0.115 | -0.183 | -0.06 | 0.1 | 0.1 | 0 | 0.4 | 0.14 | 0.34 | 0.2 | 0.53 | 0.39 |
| 8 | 15051158 | rs7840505 | 1.304 | -0.565 | -0.229 | -0.192 | -0.153 | 0.022 | 0.022 | 0 | 0.34 | 0.03 | 0.28 | 0.2 | 0.46 | 0.34 |
| 8 | 15051872 | rs17472351 | -0.696 | -0.762 | -0.027 | -0.22 | -0.009 | -0.011 | -0.011 | 0.049 | 0.13 | 0.18 | 0.11 | 0.07 | 0.11 | 0.08 |
| 8 | 15052361 | rs2410239 | -0.067 | -0.841 | 0.708 | -0.029 | -0.004 | 0 | 0 | 0 | 0.47 | 0.55 | 0.46 | 0.51 | 0.37 | 0.47 |
| 8 | 15057078 | rs17656261 | 1.175 | -0.789 | 0.877 | -0.016 | 0.009 | -0.024 | -0.024 | 0.077 | 0.32 | 0.2 | 0.33 | 0.42 | 0.25 | 0.38 |
| 8 | 15058103 | rs959358 | 0.321 | -0.663 | 1.163 | 0.033 | -0.006 | 0.053 | 0.053 | 0 | 0.44 | 0.32 | 0.37 | 0.22 | 0.55 | 0.41 |
| 8 | 15060890 | rs6990575 | 0.513 | -0.663 | 1.079 | 0.145 | -0.004 | 0.021 | 0.021 | 0 | 0.87 | 0.84 | 0.83 | 0.76 | 0.89 | 0.88 |
| 8 | 15061346 | rs17610165 | 0.341 | -0.699 | 1.077 | 0.124 | 0.017 | -0.001 | -0.001 | 0 | 0.48 | 0.39 | 0.47 | 0.55 | 0.4 | 0.49 |
| 8 | 15067980 | rs9325747 | 1.26 | -0.58 | 1.413 | 0.378 | 0.003 | 0.066 | 0.066 | 0 | 0.82 | 0.67 | 0.77 | 0.74 | 0.85 | 0.85 |
| 8 | 15075310 | rs268335 | 0.752 | -0.395 | 1.105 | 0.489 | 0.042 | 0.142 | 0.142 | 1.093 | 0.9 | 0.68 | 0.84 | 0.79 | 0.93 | 0.91 |
| 8 | 15086109 | rs17121158 | -0.598 | -0.361 | 1.136 | -0.004 | 0.055 | -0.048 | -0.048 | 1.071 | 0.09 | 0.25 | 0.1 | 0 | 0.09 | 0.06 |
| 8 | 15091086 | rs7010086 | -0.323 | -0.39 | 1.364 | -0.037 | 0.054 | -0.042 | -0.042 | 0 | 0.08 | 0.24 | 0.07 | 0 | 0.07 | 0.05 |
| 8 | 15094597 | rs7820168 | -0.215 | -0.683 | 0.525 | -0.157 | 0.006 | 0.009 | 0.009 | 0.022 | 0.5 | 0.55 | 0.43 | 0.3 | 0.61 | 0.37 |
| 8 | 15109424 | rs4618695 | -0.539 | -1.234 | -0.164 | -0.164 | 0.038 | 0.084 | 0.084 | 0 | 0.48 | 0.75 | 0.47 | 0.34 | 0.36 | 0.54 |
| 8 | 15112267 | rs17576070 | 0.818 | -1.355 | -0.214 | -0.188 | 0.009 | -0.025 | -0.025 | 0.01 | 0.3 | 0.09 | 0.31 | 0.34 | 0.24 | 0.42 |
| 8 | 15113977 | rs11987274 | 0.086 | -1.415 | 0.381 | -0.188 | 0.017 | -0.072 | -0.072 | 0 | 0.13 | 0.34 | 0.09 | 0.01 | 0.09 | 0.11 |
| 8 | 15117476 | rs2410246 | -0.757 | -1.825 | 0.362 | -0.409 | 0.019 | 0.047 | 0.047 | 0 | 0.44 | 0.73 | 0.45 | 0.36 | 0.35 | 0.55 |
| 8 | 15118628 | rs17121207 | 0.333 | -1.819 | 0.605 | -0.42 | 0.006 | -0.055 | -0.055 | 0 | 0.13 | 0.24 | 0.09 | 0.01 | 0.09 | 0.11 |
| 8 | 15121774 | rs6530848 | -0.705 | -1.8 | 0.312 | -0.384 | 0.016 | 0.044 | 0.044 | 0 | 0.44 | 0.72 | 0.45 | 0.36 | 0.35 | 0.55 |
| 8 | 15131170 | rs1457221 | -0.818 | -2.037 | -0.391 | -0.362 | -0.04 | -0.06 | -0.06 | 0 | 0.58 | 0.8 | 0.48 | 0.49 | 0.39 | 0.64 |
| 8 | 15148242 | rs11776929 | 0.329 | -1.937 | 0.113 | -0.189 | -0.017 | -0.003 | -0.003 | 0 | 0.64 | 0.51 | 0.66 | 0.69 | 0.75 | 0.7 |
| 8 | 15151390 | rs11203689 | 1.24 | -1.819 | 0.015 | -0.369 | -0.032 | -0.073 | -0.073 | 0 | 0.22 | 0.04 | 0.28 | 0.35 | 0.23 | 0.34 |
| 8 | 15152395 | rs7822797 | -0.607 | -1.842 | 0.65 | -0.472 | -0.006 | -0.12 | -0.12 | 0 | 0.19 | 0.56 | 0.15 | 0.02 | 0.12 | 0.14 |
| 8 | 15152460 | rs7840581 | 0.081 | -1.846 | 0.851 | -0.472 | 0.019 | -0.095 | -0.095 | 0 | 0.15 | 0.4 | 0.11 | 0.01 | 0.1 | 0.11 |
| 8 | 15152926 | rs6991671 | 0.6 | -1.83 | 0.688 | -0.468 | 0.002 | -0.045 | -0.045 | 0 | 0.13 | 0.19 | 0.09 | 0 | 0.09 | 0.11 |
| 8 | 15157843 | rs12548410 | 0.692 | -1.879 | 0.198 | -0.417 | 0.001 | -0.03 | -0.03 | 1.257 | 0.27 | 0.12 | 0.3 | 0.36 | 0.25 | 0.45 |
| 8 | 15169007 | rs12114950 | -0.608 | -1.771 | 0.342 | -0.67 | -0.027 | -0.154 | -0.154 | 0 | 0.24 | 0.46 | 0.15 | 0.02 | 0.14 | 0.14 |
| 8 | 15171989 | rs7003979 | -1.237 | -1.871 | 0.058 | -0.576 | 0.008 | 0.035 | 0.035 | 0.313 | 0.45 | 0.7 | 0.45 | 0.37 | 0.39 | 0.56 |
| 8 | 15173109 | rs6991005 | 0.33 | -1.698 | -0.333 | -0.763 | -0.001 | -0.038 | -0.038 | 0 | 0.15 | 0.2 | 0.09 | 0.01 | 0.1 | 0.11 |
| 8 | 15178106 | rs12678297 | 1.62 | -1.893 | -0.73 | -0.715 | -0.177 | -0.069 | -0.069 | 0 | 0.34 | 0.03 | 0.32 | 0.32 | 0.5 | 0.25 |
| 8 | 15182925 | rs10093109 | -0.822 | -1.467 | 0.062 | -0.718 | -0.027 | -0.152 | -0.152 | 0 | 0.24 | 0.5 | 0.14 | 0.02 | 0.14 | 0.13 |
| 8 | 15184667 | rs11992115 | 0.219 | -1.438 | -0.035 | -0.703 | -0.003 | -0.079 | -0.079 | 0 | 0.15 | 0.32 | 0.09 | 0.01 | 0.1 | 0.11 |
| 8 | 15186378 | rs11990343 | -0.237 | -1.383 | 0.016 | -0.651 | -0.006 | -0.064 | -0.064 | 0 | 0.23 | 0.32 | 0.13 | 0.02 | 0.13 | 0.13 |
| 8 | 15190987 | rs4265187 | -0.301 | -1.385 | -0.098 | -0.655 | -0.008 | -0.073 | -0.073 | 0.315 | 0.23 | 0.32 | 0.13 | 0.02 | 0.14 | 0.13 |
| 8 | 15191015 | rs4383971 | -0.391 | -1.32 | 0.013 | -0.622 | -0.019 | -0.128 | -0.128 | 0 | 0.24 | 0.45 | 0.14 | 0.02 | 0.14 | 0.13 |
| 8 | 15195565 | rs11985409 | -0.508 | -1.33 | -0.191 | -0.627 | -0.027 | -0.158 | -0.158 | 0 | 0.25 | 0.53 | 0.15 | 0.02 | 0.14 | 0.13 |
| 8 | 15197417 | rs12676170 | 0.536 | -0.862 | -1.095 | 0.064 | -0.059 | -0.115 | -0.115 | 0.015 | 0.18 | 0.02 | 0.29 | 0.43 | 0.23 | 0.35 |
| 8 | 15202493 | rs11779599 | -0.98 | -1.133 | -0.742 | -0.025 | 0.003 | -0.003 | -0.003 | 4.844 | 0.85 | 0.86 | 0.86 | 0.89 | 0.88 | 0.83 |
| 8 | 15214144 | rs4831699 | 0.516 | -1.252 | -0.904 | -0.149 | 0.068 | 0.058 | 0.058 | 0.362 | 0.56 | 0.27 | 0.62 | 0.77 | 0.73 | 0.71 |
| 8 | 15226444 | rs12544670 | -0.439 | -0.882 | -0.999 | -0.14 | -0.012 | 0.014 | 0.014 | 0.162 | 0.4 | 0.72 | 0.5 | 0.65 | 0.38 | 0.52 |
| 8 | 15229295 | rs4240179 | 0.406 | -1.095 | -0.709 | -0.213 | 0.109 | -0.018 | -0.018 | 0 | 0.49 | 0.13 | 0.51 | 0.5 | 0.65 | 0.48 |
| 8 | 15236462 | rs6530854 | 0.159 | -1.118 | -0.207 | -0.213 | 0.013 | 0.001 | 0.001 | 0 | 0.56 | 0.4 | 0.53 | 0.54 | 0.67 | 0.59 |
| 8 | 15236867 | rs11781165 | 3.009 | -0.745 | -0.326 | -0.052 | -0.039 | -0.016 | -0.016 | 0 | 0.07 | 0 | 0.04 | 0.05 | 0.11 | 0.04 |
| 8 | 15237056 | rs1994950 | 0.593 | -0.987 | 0.238 | -0.078 | 0 | 0.06 | 0.06 | 0 | 0.18 | 0.28 | 0.32 | 0.42 | 0.2 | 0.24 |
| 8 | 15238678 | rs1565919 | 0.081 | -1.053 | 0.344 | -0.209 | -0.003 | 0.062 | 0.062 | 0.024 | 0.77 | 0.6 | 0.65 | 0.58 | 0.75 | 0.76 |
| 8 | 15240589 | rs6530859 | 0.636 | -0.895 | 0.366 | -0.131 | 0.001 | 0.045 | 0.045 | 0 | 0.17 | 0.25 | 0.32 | 0.43 | 0.2 | 0.24 |
| 8 | 15242455 | rs7001851 | 0.306 | -1.167 | 0.544 | -0.227 | -0.036 | -0.04 | -0.04 | 2.55 | 0.3 | 0.39 | 0.18 | 0.12 | 0.17 | 0.24 |
| 8 | 15242806 | rs1994952 | -0.948 | -1.081 | 0.528 | -0.144 | 0.002 | 0.06 | 0.06 | 0 | 0.82 | 0.72 | 0.68 | 0.57 | 0.8 | 0.77 |
| 8 | 15246658 | rs13263367 | 2.047 | -0.666 | -0.038 | -0.309 | -0.14 | -0.113 | -0.113 | 0.64 | 0.3 | 0.03 | 0.33 | 0.35 | 0.4 | 0.32 |
| 8 | 15246970 | rs12546480 | 1.559 | -0.705 | 0.257 | -0.28 | 0.003 | 0.042 | 0.042 | 0 | 0.14 | 0.17 | 0.29 | 0.43 | 0.18 | 0.21 |
| 8 | 15249056 | rs4831713 | -0.995 | -0.829 | 0.464 | -0.367 | -0.048 | -0.045 | -0.045 | 0 | 0.74 | 0.6 | 0.84 | 0.89 | 0.86 | 0.84 |
| 8 | 15249492 | rs11783448 | -0.922 | -0.824 | 0.273 | -0.364 | -0.067 | -0.063 | -0.063 | 0 | 0.73 | 0.55 | 0.83 | 0.89 | 0.86 | 0.84 |
| 8 | 15250768 | rs1495097 | 0.603 | -0.859 | 0.212 | -0.397 | 0.022 | -0.013 | -0.013 | 0 | 0.19 | 0.24 | 0.13 | 0.1 | 0.2 | 0.15 |

| | | | | | | | | | | | | | | | | |
|---|---|---|---|---|---|---|---|---|---|---|---|---|---|---|---|---|
| 8 | 15251170 | rs2101809 | 0.278 | -0.834 | 0.048 | -0.372 | 0.026 | 0.09 | 0.09 | 0.025 | 0.53 | 0.26 | 0.7 | 0.79 | 0.65 | 0.69 |
| 8 | 15257883 | rs2201476 | 1.96 | -0.763 | -0.397 | -0.26 | -0.088 | -0.067 | -0.067 | 0.133 | 0.29 | 0.06 | 0.33 | 0.35 | 0.4 | 0.31 |
| 8 | 15258706 | rs1495075 | -1.046 | -0.88 | -0.228 | -0.376 | -0.046 | -0.092 | -0.092 | 0 | 0.74 | 0.64 | 0.82 | 0.97 | 0.85 | 0.84 |
| 8 | 15259345 | rs12542950 | -0.869 | -0.678 | -0.144 | -0.251 | -0.002 | 0.001 | 0.001 | 0.116 | 0.44 | 0.59 | 0.48 | 0.61 | 0.43 | 0.52 |
| 8 | 15260357 | rs10503543 | 0.038 | -0.663 | -0.069 | -0.168 | -0.003 | 0.001 | 0.001 | 0 | 0.26 | 0.25 | 0.16 | 0.18 | 0.24 | 0.25 |
| 8 | 15262565 | rs2410269 | 0.017 | -0.675 | 0.017 | -0.167 | -0.006 | 0.007 | 0.007 | 0 | 0.25 | 0.21 | 0.15 | 0.1 | 0.24 | 0.24 |
| 8 | 15264125 | rs7835718 | 0.087 | -0.853 | 0 | -0.139 | -0.002 | -0.003 | -0.003 | 0 | 0.25 | 0.25 | 0.22 | 0.24 | 0.22 | 0.24 |
| 8 | 15265778 | rs2132135 | -0.779 | -0.986 | 0.038 | -0.287 | 0.008 | -0.014 | -0.014 | 0 | 0.7 | 0.74 | 0.77 | 0.96 | 0.79 | 0.79 |
| 8 | 15265818 | rs17657086 | 1.018 | -0.959 | -0.089 | -0.243 | 0.028 | 0.184 | 0.184 | 0 | 0.16 | 0.42 | 0.3 | 0.42 | 0.18 | 0.24 |
| 8 | 15265901 | rs17657098 | 1.333 | -0.948 | -0.064 | -0.215 | 0.026 | 0.205 | 0.205 | 0 | 0.14 | 0.4 | 0.28 | 0.42 | 0.15 | 0.22 |
| 8 | 15266218 | rs2132134 | 0.236 | -0.968 | -0.55 | -0.292 | -0.013 | 0.009 | 0.009 | 0 | 0.24 | 0.15 | 0.21 | 0.14 | 0.22 | 0.24 |
| 8 | 15268160 | rs4831719 | 1.701 | -0.982 | -0.607 | -0.659 | -0.144 | -0.05 | -0.05 | 0 | 0.3 | 0.03 | 0.27 | 0.3 | 0.42 | 0.31 |
| 8 | 15270545 | rs13264819 | 1.676 | -0.965 | -0.606 | -0.649 | -0.143 | -0.049 | -0.049 | 0 | 0.3 | 0.03 | 0.27 | 0.3 | 0.42 | 0.31 |
| 8 | 15271666 | rs11990887 | -0.823 | -0.428 | 0.284 | -0.094 | -0.006 | 0.016 | 0.016 | 0.107 | 0.4 | 0.66 | 0.5 | 0.66 | 0.37 | 0.48 |
| 8 | 15273777 | rs2029795 | -0.131 | -0.593 | 0.235 | -0.086 | 0.001 | 0.012 | 0.012 | 0 | 0.18 | 0.27 | 0.09 | 0.19 | 0.15 | 0.16 |
| 8 | 15276707 | rs6985514 | -0.691 | -1.443 | 0.287 | -0.846 | 0.007 | 0.005 | 0.005 | 0.016 | 0.4 | 0.57 | 0.25 | 0.15 | 0.3 | 0.43 |
| 8 | 15278285 | rs6981134 | -0.16 | -1.439 | 0.424 | -0.995 | 0.004 | -0.007 | -0.007 | 0.029 | 0.25 | 0.16 | 0.17 | 0 | 0.22 | 0.23 |
| 8 | 15280085 | rs7827410 | -0.096 | -1.613 | 0.594 | -1.005 | -0.002 | -0.086 | -0.086 | 0 | 0.19 | 0.27 | 0.17 | 0 | 0.19 | 0.23 |
| 8 | 15281140 | rs1495088 | -1.352 | -1.926 | 0.323 | -1.011 | -0.006 | -0.011 | -0.011 | 0 | 0.32 | 0.6 | 0.22 | 0.16 | 0.24 | 0.4 |
| 8 | 15282409 | rs10090209 | -0.698 | -1.815 | 0.422 | -0.876 | -0.025 | -0.007 | -0.007 | 0 | 0.16 | 0.33 | 0.07 | 0.13 | 0.1 | 0.17 |
| 8 | 15284720 | rs1110854 | 1.253 | -2.394 | -0.322 | -0.754 | 0.139 | 0.089 | 0.089 | 0.264 | 0.52 | 0.16 | 0.76 | 0.7 | 0.73 | 0.53 |
| 8 | 15289066 | rs7837098 | 0.042 | -2.752 | -0.029 | -0.689 | 0.029 | 0.029 | 0.029 | 0 | 0.23 | 0.16 | 0.12 | 0.02 | 0.13 | 0.17 |
| 8 | 15290119 | rs10282994 | 0.04 | -2.616 | -0.024 | -0.59 | 0.024 | 0.03 | 0.03 | 0 | 0.24 | 0.16 | 0.14 | 0.02 | 0.16 | 0.18 |
| 8 | 15297137 | rs12155687 | 1.435 | -3.003 | -0.668 | -0.725 | 0.162 | 0.089 | 0.089 | 0 | 0.48 | 0.14 | 0.74 | 0.71 | 0.71 | 0.49 |
| 8 | 15299298 | rs17657296 | 0.088 | -2.284 | -0.169 | -0.311 | 0 | 0.002 | 0.002 | 0 | 0.09 | 0.1 | 0.04 | 0.12 | 0.07 | 0.1 |
| 8 | 15302403 | rs17611350 | 0.057 | -2.282 | -0.143 | -0.316 | -0.001 | 0.005 | 0.005 | 0 | 0.09 | 0.11 | 0.03 | 0.12 | 0.07 | 0.1 |
| 8 | 15312275 | rs10098376 | -0.476 | -2.464 | 0.239 | -0.596 | -0.053 | -0.112 | -0.112 | 0 | 0.16 | 0.35 | 0.07 | 0.01 | 0.06 | 0.21 |
| 8 | 15313497 | rs7818555 | 0.131 | -2.267 | 0.156 | -0.254 | -0.025 | 0.095 | 0.095 | 0 | 0.89 | 0.73 | 0.95 | 0.78 | 0.93 | 0.89 |
| 8 | 15314352 | rs990544 | 0.833 | -2.74 | -0.37 | -0.592 | 0.057 | 0.029 | 0.029 | 0.077 | 0.6 | 0.3 | 0.79 | 0.75 | 0.76 | 0.58 |
| 8 | 15319937 | rs10093209 | 0.184 | -2.733 | -0.834 | -0.055 | -0.002 | 0.004 | 0.004 | 0 | 0.26 | 0.26 | 0.3 | 0.4 | 0.25 | 0.38 |
| 8 | 15322770 | rs1390051 | -0.662 | -2.513 | -1.287 | -0.095 | 0.036 | -0.008 | -0.008 | 0 | 0.49 | 0.76 | 0.61 | 0.56 | 0.45 | 0.46 |
| 8 | 15324518 | rs12674702 | -0.628 | -2.394 | -1.283 | -0.181 | 0.017 | 0.026 | 0.026 | 0 | 0.47 | 0.76 | 0.62 | 0.6 | 0.47 | 0.57 |
| 8 | 15324559 | rs17576922 | 0.913 | -2.319 | -1.133 | -0.204 | 0.001 | 0.012 | 0.012 | 0.014 | 0.19 | 0.12 | 0.14 | 0.15 | 0.22 | 0.19 |
| 8 | 15327144 | rs1587099 | -1.259 | -2.441 | -1.328 | -0.308 | -0.031 | 0.083 | 0.083 | 0 | 0.58 | 0.77 | 0.67 | 0.73 | 0.54 | 0.66 |
| 8 | 15328797 | rs4563886 | 2.345 | -2.292 | -0.886 | 0.131 | -0.06 | -0.084 | -0.084 | 0 | 0.12 | 0.01 | 0.38 | 0.39 | 0.18 | 0.1 |
| 8 | 15331514 | rs1845609 | -0.783 | -2.179 | -0.758 | 0.138 | 0.017 | 0.01 | 0.01 | 0 | 0.45 | 0.68 | 0.55 | 0.6 | 0.41 | 0.55 |
| 8 | 15332551 | rs10095315 | 0.779 | -2.126 | -0.586 | 0.198 | -0.018 | -0.004 | -0.004 | 0.385 | 0.18 | 0.26 | 0.24 | 0.19 | 0.16 | 0.21 |
| 8 | 15337889 | rs1353319 | 2.829 | -2.047 | -0.741 | 0.253 | -0.038 | -0.038 | -0.038 | 0.056 | 0.09 | 0.01 | 0.22 | 0.34 | 0.16 | 0.07 |
| 8 | 15339941 | rs10087154 | -0.778 | -2.123 | -0.713 | 0.188 | 0.023 | 0.012 | 0.012 | 0 | 0.44 | 0.69 | 0.55 | 0.6 | 0.41 | 0.55 |
| 8 | 15341953 | rs17576978 | 0.75 | -1.774 | -0.74 | 0.259 | -0.003 | -0.003 | -0.003 | 1.164 | 0.17 | 0.11 | 0.26 | 0.2 | 0.21 | 0.18 |
| 8 | 15344980 | rs17576999 | 0.414 | -1.773 | -0.775 | 0.393 | -0.006 | -0.02 | -0.02 | 0.027 | 0.19 | 0.09 | 0.29 | 0.22 | 0.22 | 0.2 |
| 8 | 15345301 | rs10110222 | 1.035 | -1.75 | -0.857 | -0.048 | -0.007 | -0.035 | -0.035 | 0 | 0.07 | 0.15 | 0.04 | 0 | 0.05 | 0.1 |
| 8 | 15345669 | rs1976289 | -1.189 | -2.04 | -0.985 | 0.009 | -0.015 | 0.033 | 0.033 | 0 | 0.59 | 0.73 | 0.62 | 0.67 | 0.52 | 0.63 |
| 8 | 15347819 | rs870712 | 0.957 | -1.627 | -1.35 | 0.373 | -0.015 | -0.007 | -0.007 | 0 | 0.14 | 0.01 | 0.11 | 0.15 | 0.16 | 0.15 |
| 8 | 15349153 | rs1495084 | -0.333 | -1.352 | -0.835 | 0.557 | 0.061 | -0.013 | -0.013 | 0 | 0.22 | 0.09 | 0.07 | 0.18 | 0.1 | 0.25 |
| 8 | 15354208 | rs7828050 | 1.618 | -0.656 | -0.723 | 0.495 | -0.05 | 0.023 | 0.023 | 0.408 | 0.14 | 0.08 | 0.23 | 0.09 | 0.29 | 0.15 |
| 8 | 15356234 | rs7820988 | 1.356 | -0.748 | -0.656 | 0.515 | -0.039 | 0.003 | 0.003 | 0.498 | 0.16 | 0.13 | 0.24 | 0.16 | 0.31 | 0.15 |
| 8 | 15357992 | rs11203708 | 0.591 | -1.409 | -0.096 | 1.014 | -0.016 | 0.041 | 0.041 | 0 | 0.82 | 0.73 | 0.85 | 0.65 | 0.85 | 0.76 |
| 8 | 15358801 | rs17611665 | -0.358 | -1.475 | -0.441 | 1.062 | 0.049 | 0.001 | 0.001 | 0 | 0.23 | 0.09 | 0.1 | 0.14 | 0.15 | 0.25 |
| 8 | 15359174 | rs10095606 | -0.004 | -0.679 | 0.036 | 1.358 | 0.008 | 0.123 | 0.123 | 0.201 | 0.93 | 0.85 | 0.9 | 0.67 | 0.93 | 0.92 |
| 8 | 15359838 | rs12550443 | 0.394 | -0.659 | -0.371 | 0.84 | -0.013 | -0.057 | -0.057 | 0 | 0.31 | 0.08 | 0.41 | 0.31 | 0.28 | 0.26 |
| 8 | 15360737 | rs2132129 | 0.671 | -0.384 | 1.027 | 0.794 | -0.024 | 0.051 | 0.051 | 0 | 0.82 | 0.59 | 0.89 | 0.72 | 0.86 | 0.78 |
| 8 | 15365443 | rs6989485 | 0.581 | 0.121 | 0.801 | 0.252 | 0.033 | -0.058 | -0.058 | 0.011 | 0.89 | 0.65 | 0.85 | 0.99 | 0.91 | 0.77 |
| 8 | 15375002 | rs17657638 | -1.036 | -0.385 | -0.187 | 0.27 | 0.016 | 0.016 | 0.016 | 0.728 | 0.16 | 0.12 | 0.08 | 0.12 | 0.12 | 0.05 |
| 8 | 15376049 | rs2132131 | 0.681 | -0.447 | 0.258 | -0.022 | -0.002 | -0.02 | -0.02 | 0 | 0.78 | 0.72 | 0.81 | 0.87 | 0.77 | 0.69 |
| 8 | 15378002 | rs1495087 | 1.315 | -0.74 | 0.406 | 0.152 | 0.037 | 0.029 | 0.029 | 2.038 | 0.63 | 0.18 | 0.57 | 0.65 | 0.66 | 0.56 |
| 8 | 15386048 | rs4430095 | 1.172 | -0.868 | 0.753 | 0.164 | 0 | 0.049 | 0.049 | 0 | 0.91 | 0.78 | 0.95 | 0.81 | 0.95 | 0.97 |
| 8 | 15388566 | rs7813286 | -1.156 | -0.985 | 1.077 | -0.092 | -0.013 | -0.036 | -0.036 | 0 | 0.25 | 0.27 | 0.16 | 0.11 | 0.21 | 0.39 |
| 8 | 15395144 | rs11784308 | 0.476 | -1.604 | 0.162 | 0.163 | 0.018 | 0.006 | 0.006 | 0 | 0.29 | 0.2 | 0.19 | 0.25 | 0.2 | 0.26 |
| 8 | 15400574 | rs1429087 | 1.105 | -0.653 | -0.453 | 0.121 | -0.05 | 0.014 | 0.014 | 0 | 0.21 | 0.13 | 0.38 | 0.11 | 0.35 | 0.15 |
| 8 | 15402936 | rs1346593 | 1.118 | -1.15 | -0.494 | 0.529 | 0.034 | -0.023 | -0.023 | 0 | 0.29 | 0.16 | 0.24 | 0.37 | 0.2 | 0.25 |
| 8 | 15405959 | rs7821056 | -0.291 | -1.289 | -0.023 | 0.374 | -0.001 | -0.001 | -0.001 | 0 | 0.64 | 0.66 | 0.4 | 0.63 | 0.43 | 0.44 |
| 8 | 15407216 | rs919401 | -0.665 | -1.201 | 0.168 | 0.35 | -0.006 | -0.006 | -0.006 | 0 | 0.84 | 0.82 | 0.87 | 0.87 | 0.87 | 0.59 |
| 8 | 15409462 | rs10100797 | 1.222 | -1.298 | 0.323 | 0.413 | -0.033 | -0.012 | -0.012 | 0 | 0.23 | 0.2 | 0.47 | 0.26 | 0.44 | 0.16 |
| 8 | 15410250 | rs7010988 | -0.841 | -1.265 | 0.373 | 0.363 | -0.007 | -0.009 | -0.009 | 0 | 0.85 | 0.81 | 0.87 | 0.88 | 0.87 | 0.6 |
| 8 | 15419777 | rs12549184 | -0.592 | -1.258 | 0.324 | 0.323 | -0.017 | -0.026 | -0.026 | 0.597 | 0.83 | 0.68 | 0.74 | 0.88 | 0.86 | 0.61 |
| 8 | 15420894 | rs12677906 | 1.968 | -1.447 | 0.307 | 0.311 | 0.021 | -0.001 | -0.001 | 0 | 0.26 | 0.17 | 0.18 | 0.24 | 0.19 | 0.24 |
| 8 | 15422664 | rs7009698 | -0.622 | -1.392 | 0.29 | 0.304 | -0.014 | -0.026 | -0.026 | 0 | 0.83 | 0.68 | 0.74 | 0.88 | 0.86 | 0.62 |
| 8 | 15424898 | rs1560216 | -0.938 | -1.384 | 0.275 | 0.308 | -0.001 | -0.002 | -0.002 | 0 | 0.85 | 0.86 | 0.76 | 0.88 | 0.86 | 0.61 |

| | | | | | | | | | | | | | | | | |
|---|---|---|---|---|---|---|---|---|---|---|---|---|---|---|---|---|
| 8 | 15425610 | rs2410277 | 0.241 | -1.567 | 0.126 | 0.404 | 0.042 | -0.05 | -0.05 | 0 | 0.3 | 0.05 | 0.17 | 0.35 | 0.23 | 0.19 |
| 8 | 15426417 | rs35349964 | 0.43 | -1.814 | -0.101 | 0.52 | 0.011 | 0.047 | 0.047 | 0 | 0.24 | 0.05 | 0.15 | 0.11 | 0.21 | 0.11 |
| 8 | 15427155 | rs1429088 | 0.438 | -1.812 | -0.1 | 0.503 | 0.011 | 0.069 | 0.069 | 0.006 | 0.24 | 0.04 | 0.14 | 0.07 | 0.21 | 0.11 |
| 8 | 15431475 | rs2898420 | -0.525 | -1.635 | -0.11 | 0.638 | -0.002 | 0 | 0 | 0 | 0.65 | 0.67 | 0.44 | 0.68 | 0.52 | 0.48 |
| 8 | 15442613 | rs11203713 | -0.07 | -1.727 | -0.501 | 0.369 | 0.011 | 0.008 | 0.008 | 0.969 | 0.58 | 0.64 | 0.35 | 0.66 | 0.35 | 0.44 |
| 8 | 15447243 | rs994641 | -0.93 | -1.675 | -0.433 | 0.419 | -0.02 | -0.017 | -0.017 | 0 | 0.85 | 0.66 | 0.74 | 0.88 | 0.87 | 0.63 |
| 8 | 15447478 | rs2042993 | 1.396 | -1.713 | -0.452 | 0.455 | 0.049 | -0.05 | -0.05 | 0 | 0.33 | 0.17 | 0.2 | 0.48 | 0.22 | 0.34 |
| 8 | 15450409 | rs1579450 | 0.583 | -1.543 | -0.632 | 0.402 | -0.081 | 0.046 | 0.046 | 0 | 0.19 | 0.01 | 0.17 | 0.05 | 0.27 | 0.09 |
| 8 | 15452518 | rs13253051 | 1.7 | -1.328 | -0.303 | 0.531 | 0.015 | 0.001 | 0.001 | 0 | 0.2 | 0.21 | 0.32 | 0.22 | 0.34 | 0.13 |
| 8 | 15461996 | rs7009035 | 0.171 | -1.447 | -0.171 | 0.556 | 0.001 | -0.001 | -0.001 | 0.358 | 0.54 | 0.54 | 0.36 | 0.45 | 0.41 | 0.38 |
| 8 | 15465489 | rs6530890 | -1.016 | -1.532 | -0.194 | 0.584 | -0.013 | -0.005 | -0.005 | 0 | 0.86 | 0.8 | 0.78 | 0.88 | 0.89 | 0.65 |
| 8 | 15469237 | rs1366667 | 1.139 | -1.5 | -0.21 | 0.68 | -0.043 | -0.033 | -0.033 | 0 | 0.25 | 0.15 | 0.28 | 0.39 | 0.37 | 0.21 |
| 8 | 15470543 | rs13256392 | -0.398 | -1.594 | -0.055 | 0.698 | -0.004 | -0.003 | -0.003 | 0 | 0.83 | 0.82 | 0.74 | 0.88 | 0.87 | 0.64 |
| 8 | 15472430 | rs1991531 | -0.092 | -1.629 | -0.061 | 0.846 | -0.004 | 0.002 | 0.002 | 0 | 0.81 | 0.79 | 0.68 | 0.71 | 0.86 | 0.63 |
| 8 | 15475667 | rs2410280 | -0.676 | -1.504 | -0.065 | 0.81 | -0.002 | 0 | 0 | 0 | 0.84 | 0.78 | 0.74 | 0.82 | 0.85 | 0.64 |
| 8 | 15476868 | rs4831757 | -0.218 | -1.705 | -0.099 | 0.794 | -0.002 | -0.001 | -0.001 | 0 | 0.82 | 0.78 | 0.74 | 0.82 | 0.86 | 0.61 |
| 8 | 15477945 | rs11203714 | 1.818 | -1.845 | -0.207 | 0.765 | 0.026 | 0.005 | 0.005 | 0.006 | 0.28 | 0.17 | 0.19 | 0.23 | 0.19 | 0.26 |
| 8 | 15478125 | rs10107831 | 0.599 | -1.833 | -0.008 | 0.724 | -0.002 | -0.003 | -0.003 | 0 | 0.6 | 0.49 | 0.49 | 0.59 | 0.58 | 0.51 |
| 8 | 15480039 | rs1429079 | 0.6 | -1.863 | 0.014 | 0.724 | -0.003 | -0.002 | -0.002 | 0 | 0.3 | 0.26 | 0.3 | 0.37 | 0.38 | 0.23 |
| 8 | 15482653 | rs1835141 | 0.2 | -1.794 | -0.015 | 0.526 | -0.014 | 0.028 | 0.028 | 0 | 0.24 | 0.13 | 0.22 | 0.07 | 0.3 | 0.13 |
| 8 | 15488238 | rs1366671 | 0.566 | -1.87 | 0.081 | 0.559 | -0.001 | -0.016 | -0.016 | 0.768 | 0.58 | 0.49 | 0.54 | 0.72 | 0.56 | 0.49 |
| 8 | 15488656 | rs1864236 | 0.6 | -1.866 | 0.041 | 0.598 | -0.027 | -0.022 | -0.022 | 0 | 0.3 | 0.24 | 0.34 | 0.41 | 0.38 | 0.23 |
| 8 | 15488774 | rs4623427 | 1.866 | -1.682 | -0.116 | 0.578 | -0.031 | -0.011 | -0.011 | 0.006 | 0.19 | 0.12 | 0.3 | 0.22 | 0.33 | 0.14 |
| 8 | 15491440 | rs1429091 | -0.108 | -1.731 | 0.114 | 0.506 | -0.034 | -0.006 | -0.006 | 0 | 0.78 | 0.58 | 0.72 | 0.78 | 0.85 | 0.6 |
| 8 | 15491989 | rs12541025 | 1.148 | -1.693 | 0.008 | 0.548 | -0.057 | -0.034 | -0.034 | 3.518 | 0.25 | 0.15 | 0.33 | 0.35 | 0.37 | 0.2 |
| 8 | 15493417 | rs4258002 | 0.335 | -1.651 | 0.07 | 0.58 | -0.014 | 0.008 | 0.008 | 0 | 0.18 | 0.28 | 0.28 | 0.25 | 0.15 | 0.4 |
| 8 | 15494159 | rs4645564 | 2.206 | -1.585 | -0.241 | 0.57 | 0.035 | -0.005 | -0.005 | 0 | 0.27 | 0.16 | 0.18 | 0.25 | 0.18 | 0.25 |
| 8 | 15497331 | rs10081561 | 0.811 | -1.65 | -0.248 | 0.429 | 0 | 0.003 | 0.003 | 0 | 0.57 | 0.43 | 0.53 | 0.61 | 0.58 | 0.46 |
| 8 | 15500967 | rs12541402 | 0.423 | -1.389 | -0.681 | 0.333 | -0.088 | 0.064 | 0.064 | 0 | 0.23 | 0.01 | 0.2 | 0.06 | 0.33 | 0.11 |
| 8 | 15508200 | rs1035972 | 0.719 | -1.137 | -0.491 | 0.597 | -0.049 | -0.052 | -0.052 | 0 | 0.28 | 0.16 | 0.34 | 0.43 | 0.38 | 0.22 |
| 8 | 15509105 | rs11777115 | 0.269 | -1.156 | -0.48 | 0.546 | -0.043 | 0.065 | 0.065 | 0 | 0.24 | 0.04 | 0.18 | 0.07 | 0.29 | 0.13 |
| 8 | 15511767 | rs7011443 | 0.173 | -1.235 | -0.161 | 0.584 | 0 | 0.002 | 0.002 | 0 | 0.55 | 0.53 | 0.4 | 0.37 | 0.47 | 0.4 |
| 8 | 15523366 | rs7838101 | 0.606 | -1.213 | -0.142 | 0.605 | 0 | -0.006 | -0.006 | 0 | 0.58 | 0.58 | 0.55 | 0.71 | 0.56 | 0.48 |
| 8 | 15525501 | rs9650390 | 2.503 | -1.277 | -0.122 | 0.615 | 0.033 | -0.007 | -0.007 | 0 | 0.24 | 0.13 | 0.17 | 0.23 | 0.17 | 0.24 |
| 8 | 15526849 | rs354519 | 0.138 | -0.902 | 0.01 | 0.627 | 0.019 | -0.018 | -0.018 | 0 | 0.73 | 0.82 | 0.69 | 0.66 | 0.79 | 0.52 |
| 8 | 15527376 | rs10093681 | -0.345 | -0.992 | 0.17 | 0.664 | -0.052 | 0.09 | 0.09 | 1.977 | 0.13 | 0.38 | 0.07 | 0.24 | 0.05 | 0.08 |
| 8 | 15527984 | rs354517 | -0.475 | -0.9 | 0.089 | 0.647 | -0.007 | 0.022 | 0.022 | 3.931 | 0.33 | 0.1 | 0.21 | 0.24 | 0.35 | 0.21 |
| 8 | 15529890 | rs354515 | -0.807 | -0.853 | 0.026 | 0.534 | -0.013 | -0.002 | -0.002 | 0.953 | 0.37 | 0.46 | 0.25 | 0.35 | 0.35 | 0.21 |
| 8 | 15535226 | rs354508 | -0.552 | -0.785 | -0.04 | 0.574 | -0.001 | 0 | 0 | 0 | 0.81 | 0.78 | 0.72 | 0.78 | 0.84 | 0.58 |
| 8 | 15545152 | rs7841137 | 1.632 | -0.956 | 0.02 | 0.467 | 0.001 | 0.003 | 0.003 | 0.069 | 0.43 | 0.45 | 0.61 | 0.44 | 0.5 | 0.37 |
| 8 | 15557421 | rs1477207 | -0.118 | -1.122 | -0.222 | 0.434 | 0.071 | 0.039 | 0.039 | 0 | 0.45 | 0.26 | 0.62 | 0.31 | 0.65 | 0.31 |
| 8 | 15558404 | rs12682254 | 0.257 | -1.122 | -0.301 | 0.403 | 0.024 | 0.021 | 0.021 | 0 | 0.39 | 0.19 | 0.59 | 0.3 | 0.62 | 0.25 |
| 8 | 15561260 | rs1421260 | -1.244 | -1.16 | -0.204 | 0.447 | 0.004 | -0.002 | -0.002 | 0.381 | 0.85 | 0.84 | 0.87 | 0.81 | 0.88 | 0.6 |
| 8 | 15561631 | rs12680995 | -1.005 | -1.169 | -0.073 | 0.442 | -0.016 | 0.003 | 0.003 | 0 | 0.84 | 0.72 | 0.87 | 0.8 | 0.87 | 0.6 |
| 8 | 15562284 | rs4273855 | 2.742 | -1.189 | -0.135 | 0.438 | 0.016 | -0.013 | -0.013 | 0.823 | 0.23 | 0.12 | 0.18 | 0.22 | 0.19 | 0.22 |
| 8 | 15565126 | rs10104063 | 0.141 | -1.157 | -0.045 | 0.512 | -0.007 | 0.001 | 0.001 | 0 | 0.72 | 0.68 | 0.82 | 0.58 | 0.82 | 0.51 |
| 8 | 15566164 | rs12679528 | -0.123 | -1.122 | -0.011 | 0.481 | -0.009 | -0.005 | -0.005 | 0.05 | 0.46 | 0.49 | 0.62 | 0.36 | 0.61 | 0.27 |
| 8 | 15571187 | rs10109890 | 0.059 | -0.859 | -0.039 | 0.437 | -0.011 | 0.049 | 0.049 | 0 | 0.65 | 0.8 | 0.73 | 0.76 | 0.63 | 0.52 |
| 8 | 15581392 | rs1421246 | 1.514 | -0.928 | -0.175 | 0.456 | -0.01 | -0.011 | -0.011 | 1.002 | 0.45 | 0.62 | 0.62 | 0.44 | 0.5 | 0.37 |
| 8 | 15583586 | rs352763 | 1.116 | -0.962 | -0.364 | 0.439 | 0.021 | 0.005 | 0.005 | 0.662 | 0.51 | 0.78 | 0.65 | 0.48 | 0.53 | 0.4 |
| 8 | 15585452 | rs2543139 | -0.238 | -0.983 | -0.507 | 0.467 | 0.05 | -0.064 | -0.064 | 2.129 | 0.77 | 0.91 | 0.84 | 0.6 | 0.85 | 0.54 |
| 8 | 15587568 | rs2721216 | 1.735 | -0.825 | -0.363 | 0.499 | 0.016 | 0.003 | 0.003 | 0 | 0.21 | 0.25 | 0.44 | 0.26 | 0.35 | 0.15 |
| 8 | 15590011 | rs352750 | -0.329 | -0.916 | -0.572 | 0.395 | 0.027 | -0.027 | -0.027 | 5.921 | 0.43 | 0.25 | 0.61 | 0.57 | 0.61 | 0.31 |
| 8 | 15595515 | rs11998428 | -0.989 | 0.411 | -0.841 | 0.144 | -0.112 | 0.021 | 0.021 | 3.022 | 0.38 | 0.08 | 0.3 | 0.34 | 0.52 | 0.25 |
| 8 | 15596087 | rs918245 | -1.004 | 0.147 | -0.689 | 0.196 | 0.003 | -0.002 | -0.002 | 0 | 0.81 | 0.83 | 0.87 | 0.74 | 0.87 | 0.6 |
| 8 | 15598202 | rs13274016 | -0.55 | 0.245 | -0.859 | 0.129 | -0.053 | 0.038 | 0.038 | 1.9 | 0.3 | 0.1 | 0.23 | 0.26 | 0.38 | 0.21 |
| 8 | 15598482 | rs7826764 | 0.905 | -0.018 | -0.396 | 0.128 | -0.002 | 0.071 | 0.071 | 0 | 0.06 | 0.16 | 0.05 | 0.15 | 0.04 | 0.15 |
| 8 | 15598698 | rs7827167 | 0.165 | -0.094 | -0.339 | 0.127 | -0.029 | -0.011 | -0.011 | 0 | 0.35 | 0.26 | 0.54 | 0.65 | 0.42 | 0.37 |
| 8 | 15599492 | rs17121874 | 3.183 | -0.202 | -0.32 | 0.075 | 0.028 | -0.016 | -0.016 | 0.13 | 0.23 | 0.15 | 0.17 | 0.22 | 0.16 | 0.23 |
| 8 | 15599587 | rs168474 | -0.276 | -0.158 | -0.145 | 0.092 | -0.011 | 0.002 | 0.002 | 0 | 0.39 | 0.36 | 0.56 | 0.66 | 0.44 | 0.39 |
| 8 | 15604455 | rs2168247 | 1.234 | -0.049 | 0.003 | 0.178 | 0.007 | 0.051 | 0.051 | 0.073 | 0.49 | 0.62 | 0.65 | 0.67 | 0.51 | 0.46 |
| 8 | 15606182 | rs7820648 | 1.666 | -0.028 | 0.085 | 0.158 | -0.005 | 0.012 | 0.012 | 0 | 0.44 | 0.6 | 0.64 | 0.62 | 0.49 | 0.44 |
| 8 | 15606598 | rs11986266 | 1.969 | 0.122 | 0.14 | 0.313 | 0.038 | 0.02 | 0.02 | 0.036 | 0.19 | 0.24 | 0.45 | 0.26 | 0.32 | 0.14 |
| 8 | 15608896 | rs751687 | 0.93 | -0.061 | -0.068 | 0.225 | 0.005 | 0.072 | 0.072 | 0 | 0.48 | 0.18 | 0.34 | 0.3 | 0.49 | 0.36 |
| 8 | 15612603 | rs352794 | 1.339 | 0.287 | 0.252 | 0.304 | 0.013 | 0.104 | 0.104 | 0.068 | 0.23 | 0.52 | 0.4 | 0.42 | 0.21 | 0.22 |
| 8 | 15615689 | rs352797 | -0.034 | 0.191 | 0.073 | 0.183 | 0.024 | 0.004 | 0.004 | 0.088 | 0.74 | 0.83 | 0.82 | 0.73 | 0.81 | 0.6 |
| 8 | 15616806 | rs352801 | -1.171 | 0.328 | -0.008 | 0.06 | -0.026 | 0.018 | 0.018 | 3.866 | 0.35 | 0.24 | 0.29 | 0.28 | 0.49 | 0.22 |
| 8 | 15617084 | rs352804 | -1.122 | 0.524 | -0.07 | 0.065 | 0.002 | 0.028 | 0.028 | 4.313 | 0.34 | 0.21 | 0.22 | 0.28 | 0.36 | 0.2 |
| 8 | 15617917 | rs1421240 | -1.007 | 0.275 | -0.014 | 0.18 | -0.025 | -0.013 | -0.013 | 0 | 0.34 | 0.31 | 0.3 | 0.48 | 0.49 | 0.32 |

| | | | | | | | | | | | | | | | | |
|---|---|---|---|---|---|---|---|---|---|---|---|---|---|---|---|---|
| 8 | 15618380 | rs352805 | -1.095 | 0.174 | -0.089 | 0.108 | -0.004 | -0.026 | -0.026 | 0.346 | 0.83 | 0.83 | 0.9 | 0.98 | 0.89 | 0.69 |
| 8 | 15618407 | rs352806 | 1.807 | 0.293 | 0.063 | 0.345 | 0.003 | 0.005 | 0.005 | 0 | 0.22 | 0.23 | 0.48 | 0.29 | 0.25 | 0.14 |
| 8 | 15622519 | rs352810 | -0.521 | 0.213 | -0.078 | 0.295 | -0.012 | 0.008 | 0.008 | 0 | 0.77 | 0.85 | 0.79 | 0.76 | 0.74 | 0.61 |
| 8 | 15622858 | rs987926 | 2.065 | 0.235 | -0.095 | 0.264 | -0.018 | -0.023 | -0.023 | 0 | 0.2 | 0.12 | 0.39 | 0.27 | 0.23 | 0.14 |
| 8 | 15624843 | rs2721230 | 1.035 | 0.196 | 0.006 | 0.24 | 0.012 | 0.086 | 0.086 | 0 | 0.28 | 0.47 | 0.45 | 0.44 | 0.26 | 0.25 |
| 8 | 15625017 | rs2543112 | -0.873 | 0.123 | -0.037 | 0.17 | 0.023 | 0.001 | 0.001 | 0 | 0.51 | 0.65 | 0.61 | 0.53 | 0.58 | 0.38 |
| 8 | 15628324 | rs7842027 | 0.632 | 0.188 | 0.009 | 0.251 | 0.016 | 0.065 | 0.065 | 0 | 0.5 | 0.36 | 0.35 | 0.3 | 0.49 | 0.38 |
| 8 | 15632028 | rs1584963 | 0.653 | -0.539 | 0.247 | 0.472 | 0.008 | 0.065 | 0.065 | 0 | 0.3 | 0.49 | 0.53 | 0.43 | 0.28 | 0.26 |
| 8 | 15642213 | rs2604376 | 0.662 | -0.69 | 0.149 | 0.316 | 0.023 | -0.006 | -0.006 | 0 | 0.44 | 0.62 | 0.31 | 0.47 | 0.31 | 0.43 |
| 8 | 15643179 | rs12678466 | 2.618 | -0.767 | 0.332 | 0.531 | -0.022 | -0.006 | -0.006 | 0.354 | 0.24 | 0.28 | 0.18 | 0.18 | 0.17 | 0.24 |
| 8 | 15643201 | rs2543130 | 0.91 | -0.613 | 0.218 | 0.595 | 0.002 | 0.091 | 0.091 | 0 | 0.06 | 0.16 | 0.05 | 0.17 | 0.04 | 0.1 |
| 8 | 15643760 | rs352765 | -0.564 | -0.605 | -0.276 | 0.462 | 0.003 | -0.013 | -0.013 | 0 | 0.44 | 0.65 | 0.54 | 0.52 | 0.4 | 0.32 |
| 8 | 15644559 | rs352766 | 0.719 | -0.327 | -0.553 | 0.334 | -0.002 | -0.01 | -0.01 | 0 | 0.15 | 0.18 | 0.09 | 0.08 | 0.13 | 0.09 |
| 8 | 15646216 | rs352771 | 0.261 | 0.018 | -0.909 | 0.486 | -0.057 | 0.027 | 0.027 | 0.189 | 0.68 | 0.83 | 0.72 | 0.72 | 0.56 | 0.56 |
| 8 | 15646620 | rs187089 | 1.136 | 0.046 | -0.784 | 0.716 | -0.007 | -0.012 | -0.012 | 0.112 | 0.23 | 0.16 | 0.41 | 0.28 | 0.25 | 0.13 |
| 8 | 15648936 | rs373806 | 0.972 | -0.264 | -0.472 | 0.689 | 0.002 | 0.021 | 0.021 | 0 | 0.24 | 0.32 | 0.39 | 0.3 | 0.24 | 0.13 |
| 8 | 15649416 | rs13253786 | 0.871 | 0.144 | -0.882 | 0.645 | -0.071 | -0.013 | -0.013 | 0 | 0.14 | 0.06 | 0.18 | 0.2 | 0.34 | 0.14 |
| 8 | 15649922 | rs17612884 | 0.433 | -1.435 | -1.05 | 0.437 | 0.006 | 0.006 | 0.006 | 0.109 | 0.15 | 0.02 | 0.08 | 0.13 | 0.1 | 0.18 |
| 8 | 15650858 | rs2736062 | -0.398 | -1.945 | -0.369 | 0.549 | 0.058 | 0.011 | 0.011 | 0.123 | 0.55 | 0.34 | 0.65 | 0.62 | 0.69 | 0.37 |
| 8 | 15655081 | rs429004 | 1.019 | -2.522 | -0.429 | 0.392 | -0.036 | -0.03 | -0.03 | 0.319 | 0.29 | 0.5 | 0.22 | 0.21 | 0.2 | 0.23 |
| 8 | 15655686 | rs2205236 | -0.344 | -1.967 | -0.377 | 0.11 | -0.028 | -0.085 | -0.085 | 0.38 | 0.85 | 0.55 | 0.85 | 0.98 | 0.89 | 0.7 |
| 8 | 15657119 | rs2410284 | 0.046 | -1.774 | -1.69 | 0.525 | 0.019 | -0.022 | -0.022 | 0 | 0.3 | 0.12 | 0.36 | 0.32 | 0.23 | 0.44 |
| 8 | 15660239 | rs240664 | 1.338 | -1.753 | -1.38 | 0.081 | 0.026 | 0.061 | 0.061 | 0 | 0.19 | 0 | 0.1 | 0.08 | 0.16 | 0.13 |
| 8 | 15666727 | rs240654 | -0.522 | -2.17 | -1.281 | 0.3 | 0.025 | 0.009 | 0.009 | 0 | 0.8 | 0.9 | 0.79 | 0.78 | 0.88 | 0.85 |
| 8 | 15666899 | rs240653 | 0.241 | -2.572 | -0.597 | -0.007 | -0.018 | -0.038 | -0.038 | 0 | 0.21 | 0.33 | 0.16 | 0.07 | 0.16 | 0.28 |
| 8 | 15667827 | rs240652 | 0.868 | -2.814 | -0.11 | 0.207 | 0.003 | 0.005 | 0.005 | 0 | 0.32 | 0.31 | 0.43 | 0.48 | 0.56 | 0.35 |
| 8 | 15668481 | rs240650 | -0.235 | -2.748 | -0.058 | 0.105 | 0.004 | 0 | 0 | 0 | 0.7 | 0.71 | 0.67 | 0.67 | 0.84 | 0.73 |
| 8 | 15668830 | rs240648 | 0.761 | -2.788 | -0.181 | 0.008 | 0.059 | 0.085 | 0.085 | 0 | 0.49 | 0.27 | 0.51 | 0.67 | 0.68 | 0.46 |
| 8 | 15669311 | rs240647 | -0.514 | -2.808 | -0.109 | 0.028 | 0.03 | 0.012 | 0.012 | 0 | 0.75 | 0.82 | 0.71 | 0.76 | 0.88 | 0.77 |
| 8 | 15669454 | rs10110697 | -0.307 | -2.362 | 0.26 | 0.157 | -0.024 | -0.082 | -0.082 | 0 | 0.28 | 0.53 | 0.2 | 0.07 | 0.2 | 0.32 |
| 8 | 15672486 | rs7812326 | 1.353 | -2.204 | 0.037 | 0.455 | -0.017 | -0.005 | -0.005 | 0.382 | 0.21 | 0.3 | 0.1 | 0.2 | 0.12 | 0.11 |
| 8 | 15672534 | rs7812350 | 0.043 | -2.159 | 0.254 | 0.992 | -0.048 | 0.032 | 0.032 | 0 | 0.36 | 0.5 | 0.19 | 0.44 | 0.23 | 0.26 |
| 8 | 15673075 | rs7828611 | -0.674 | -2.372 | 0.266 | 0.896 | 0.04 | 0.056 | 0.056 | 0 | 0.58 | 0.75 | 0.48 | 0.7 | 0.36 | 0.53 |
| 8 | 15673430 | rs17613164 | 0.441 | -2.377 | 0.273 | 0.907 | 0.01 | -0.044 | -0.044 | 8.828 | 0.13 | 0.04 | 0.08 | 0.24 | 0.11 | 0.15 |
| 8 | 15673980 | rs2736050 | 0.919 | -2.401 | 0.236 | 0.958 | -0.007 | 0.08 | 0.08 | 0 | 0.36 | 0.19 | 0.41 | 0.23 | 0.6 | 0.43 |
| 8 | 15674616 | rs1455069 | 0.125 | -2.393 | 0.285 | 0.974 | -0.019 | 0.073 | 0.073 | 0 | 0.8 | 0.74 | 0.82 | 0.69 | 0.85 | 0.82 |
| 8 | 15675537 | rs1455067 | 0.28 | -2.388 | 0.272 | 0.959 | -0.025 | -0.002 | -0.002 | 0 | 0.44 | 0.55 | 0.41 | 0.47 | 0.25 | 0.4 |
| 8 | 15676266 | rs2736046 | -1.596 | -2.324 | 0.311 | 0.936 | -0.005 | 0.005 | 0.005 | 0 | 0.93 | 0.87 | 0.9 | 0.94 | 0.96 | 0.96 |
| 8 | 15677287 | rs2604308 | -0.581 | -2.386 | 0.274 | 0.98 | -0.023 | 0.004 | 0.004 | 0 | 0.73 | 0.7 | 0.62 | 0.71 | 0.85 | 0.73 |
| 8 | 15679463 | rs1569334 | 1.874 | -2.281 | 0.001 | 0.916 | -0.16 | 0.028 | 0.028 | 0 | 0.23 | 0.04 | 0.31 | 0.19 | 0.46 | 0.21 |
| 8 | 15679475 | rs7003942 | -0.229 | -2.158 | 0.526 | 1.32 | -0.015 | 0.064 | 0.064 | 0 | 0.14 | 0.21 | 0.09 | 0.25 | 0.11 | 0.15 |
| 8 | 15682305 | rs2720625 | 0.201 | -2.245 | 0.526 | 1.301 | 0.041 | 0.001 | 0.001 | 0 | 0.51 | 0.4 | 0.5 | 0.49 | 0.71 | 0.57 |
| 8 | 15682639 | rs6981193 | 0.808 | -2.172 | 0.333 | 1.318 | -0.004 | 0 | 0 | 0.125 | 0.22 | 0.21 | 0.29 | 0.22 | 0.13 | 0.28 |
| 8 | 15686349 | rs1869676 | 0.032 | -2.044 | 0.101 | 1.347 | 0.008 | -0.018 | -0.018 | 0 | 0.13 | 0.06 | 0.08 | 0.25 | 0.11 | 0.15 |
| 8 | 15689365 | rs2205235 | 0.221 | -2.089 | 0.116 | 1.267 | 0.093 | -0.01 | -0.01 | 0 | 0.49 | 0.25 | 0.49 | 0.48 | 0.7 | 0.57 |
| 8 | 15692532 | rs2720630 | 0.827 | -1.2 | 0.045 | 1.475 | -0.031 | -0.011 | -0.011 | 0.81 | 0.4 | 0.23 | 0.38 | 0.4 | 0.49 | 0.49 |
| 8 | 15695725 | rs17613567 | 0.788 | -2.896 | -0.108 | 1.471 | 0.022 | -0.007 | -0.007 | 0 | 0.21 | 0.14 | 0.29 | 0.24 | 0.14 | 0.28 |
| 8 | 15698099 | rs17675663 | 0.372 | -3.113 | -0.25 | 1.523 | 0.072 | 0.062 | 0.062 | 0.131 | 0.37 | 0.16 | 0.3 | 0.31 | 0.25 | 0.29 |
| 8 | 15698639 | rs1997573 | 1.262 | -3.137 | -0.139 | 1.576 | -0.005 | 0.033 | 0.033 | 0 | 0.38 | 0.22 | 0.41 | 0.31 | 0.58 | 0.45 |
| 8 | 15699845 | rs2205233 | -0.244 | -3.042 | 0.038 | 1.654 | 0.024 | -0.006 | -0.006 | 0.054 | 0.72 | 0.84 | 0.61 | 0.71 | 0.82 | 0.7 |
| 8 | 15700683 | rs2720632 | 0.003 | -3.08 | -0.578 | 1.293 | 0.091 | 0.071 | 0.071 | 0.225 | 0.55 | 0.81 | 0.6 | 0.63 | 0.7 | 0.65 |
| 8 | 15701508 | rs17122092 | 0.87 | -3.056 | -0.582 | 1.287 | 0.02 | 0.024 | 0.024 | 0 | 0.2 | 0.13 | 0.22 | 0.16 | 0.13 | 0.18 |
| 8 | 15702143 | rs17613658 | 1.621 | -2.969 | -0.577 | 1.323 | 0.048 | 0.019 | 0.019 | 0 | 0.19 | 0.02 | 0.09 | 0.16 | 0.13 | 0.14 |
| 8 | 15702272 | rs2736012 | 1 | -3.043 | -0.504 | 1.324 | 0.063 | 0.057 | 0.057 | 0 | 0.56 | 0.26 | 0.49 | 0.39 | 0.71 | 0.55 |
| 8 | 15702433 | rs2604316 | -2.102 | -2.925 | -0.66 | 1.387 | 0 | 0 | 0 | 0 | 0.94 | 0.96 | 0.9 | 0.94 | 0.96 | 0.97 |
| 8 | 15702633 | rs2736009 | -0.038 | -2.941 | -0.679 | 1.404 | 0.068 | 0.004 | 0.004 | 0 | 0.57 | 0.76 | 0.53 | 0.56 | 0.7 | 0.57 |
| 8 | 15703097 | rs1017654 | -0.843 | -1.461 | -0.118 | 1.659 | -0.001 | 0 | 0 | 0.011 | 0.38 | 0.59 | 0.4 | 0.55 | 0.25 | 0.42 |
| 8 | 15703250 | rs1017660 | -0.719 | -1.429 | -0.097 | 1.559 | -0.004 | 0.001 | 0.001 | 0 | 0.37 | 0.59 | 0.32 | 0.47 | 0.24 | 0.34 |
| 8 | 15704894 | rs1017665 | 0.154 | -0.881 | -0.802 | 1.167 | -0.01 | 0.146 | 0.146 | 0 | 0.07 | 0.21 | 0.25 | 0.19 | 0.03 | 0.11 |
| 8 | 15706243 | rs2736003 | 0.227 | -0.548 | -1.564 | 0.808 | 0.009 | -0.01 | -0.01 | 0.113 | 0.65 | 0.71 | 0.76 | 0.55 | 0.75 | 0.66 |
| 8 | 15709114 | rs2736070 | -0.368 | 0.269 | -2.287 | 0.205 | -0.088 | 0.121 | 0.121 | 0.015 | 0.35 | 0.06 | 0.41 | 0.18 | 0.52 | 0.28 |
| 8 | 15713330 | rs2736043 | -0.382 | 0.192 | -1.585 | 0.744 | 0 | 0.001 | 0.001 | 0 | 0.45 | 0.55 | 0.47 | 0.47 | 0.49 | 0.53 |
| 8 | 15714725 | rs2736017 | -0.132 | -0.231 | -2.352 | 0.326 | 0.007 | -0.05 | -0.05 | 0 | 0.71 | 0.58 | 0.81 | 0.82 | 0.77 | 0.78 |
| 8 | 15715499 | rs1349973 | 0.079 | -0.454 | -2.207 | 0.416 | -0.023 | -0.041 | -0.041 | 0.148 | 0.56 | 0.21 | 0.5 | 0.49 | 0.51 | 0.4 |
| 8 | 15716196 | rs2736068 | 1.275 | -0.426 | -2.269 | 0.518 | 0.016 | -0.031 | -0.031 | 0 | 0.34 | 0.04 | 0.32 | 0.29 | 0.31 | 0.25 |
| 8 | 15716896 | rs2720644 | 0.46 | -0.576 | -2.177 | 0.607 | -0.072 | -0.036 | -0.036 | 0 | 0.24 | 0.12 | 0.23 | 0.3 | 0.39 | 0.22 |
| 8 | 15720845 | rs11203722 | 0.42 | -0.84 | -1.624 | 0.054 | -0.007 | -0.01 | -0.01 | 0.058 | 0.07 | 0.12 | 0.04 | 0.03 | 0.05 | 0.11 |
| 8 | 15721809 | rs12680012 | 1.65 | -1.062 | -2.051 | 0.067 | -0.01 | -0.16 | -0.16 | 0 | 0.36 | 0.03 | 0.34 | 0.52 | 0.32 | 0.29 |
| 8 | 15723927 | rs11785189 | -0.274 | -0.938 | -1.764 | -0.258 | -0.003 | -0.002 | -0.002 | 0 | 0.09 | 0.11 | 0.1 | 0.05 | 0.06 | 0.27 |

| | | | | | | | | | | | | | | | | |
|---|---|---|---|---|---|---|---|---|---|---|---|---|---|---|---|---|
| 8 | 15725282 | rs2604329 | -0.273 | -0.954 | -1.21 | -0.289 | -0.051 | -0.067 | -0.067 | 0.476 | 0.84 | 0.62 | 0.94 | 0.94 | 0.92 | 0.78 |
| 8 | 15732889 | rs2410283 | 0.393 | -0.574 | -1.393 | -0.304 | -0.04 | 0.037 | 0.037 | 0 | 0.19 | 0.13 | 0.22 | 0.04 | 0.37 | 0.18 |
| 8 | 15738015 | rs1563297 | 0.595 | -0.388 | -1.32 | -0.281 | 0.006 | 0.016 | 0.016 | 0 | 0.54 | 0.44 | 0.59 | 0.63 | 0.61 | 0.61 |
| 8 | 15742366 | rs9325770 | -0.404 | -0.523 | -0.608 | -0.382 | -0.023 | -0.069 | -0.069 | 0 | 0.22 | 0.34 | 0.13 | 0.06 | 0.18 | 0.21 |
| 8 | 15743366 | rs1968723 | 0.119 | -0.5 | -0.437 | -0.388 | -0.022 | -0.072 | -0.072 | 0 | 0.79 | 0.67 | 0.89 | 0.94 | 0.83 | 0.8 |
| 8 | 15743626 | rs2720611 | 1.019 | -0.638 | -0.524 | -0.42 | 0.003 | 0.008 | 0.008 | 0 | 0.49 | 0.49 | 0.51 | 0.6 | 0.58 | 0.45 |
| 8 | 15747067 | rs2596576 | -0.809 | -0.403 | -0.447 | -0.452 | 0 | 0.007 | 0.007 | 0 | 0.45 | 0.61 | 0.55 | 0.38 | 0.52 | 0.49 |
| 8 | 15748729 | rs2604333 | 0.886 | -0.317 | -0.228 | -0.486 | -0.021 | 0.018 | 0.018 | 0 | 0.12 | 0.12 | 0.13 | 0.04 | 0.26 | 0.15 |
| 8 | 15748902 | rs2596577 | -0.377 | -0.407 | 0.234 | -0.53 | -0.006 | 0.005 | 0.005 | 0 | 0.38 | 0.33 | 0.44 | 0.35 | 0.5 | 0.34 |
| 8 | 15751090 | rs2720606 | -0.23 | -0.392 | 0.419 | -0.561 | 0.107 | -0.088 | -0.088 | 0 | 0.18 | 0.5 | 0.19 | 0.05 | 0.3 | 0.18 |
| 8 | 15752089 | rs17613901 | 0.897 | -0.241 | 0.079 | -0.505 | -0.018 | -0.019 | -0.019 | 0.011 | 0.23 | 0.09 | 0.27 | 0.28 | 0.21 | 0.18 |
| 8 | 15752275 | rs9650357 | 0.192 | -0.996 | 0.13 | -0.582 | -0.012 | -0.054 | -0.054 | 0.622 | 0.2 | 0.27 | 0.11 | 0.06 | 0.18 | 0.21 |
| 8 | 15754033 | rs17675975 | 2.107 | -1.039 | -0.074 | -0.514 | -0.016 | -0.129 | -0.129 | 0 | 0.33 | 0.01 | 0.31 | 0.58 | 0.27 | 0.27 |
| 8 | 15755789 | rs11984634 | 0.498 | -1.047 | 0.262 | -0.545 | -0.041 | -0.024 | -0.024 | 0 | 0.85 | 0.43 | 0.84 | 0.92 | 0.93 | 0.78 |
| 8 | 15757977 | rs10503553 | -0.794 | -0.953 | 0.234 | -0.365 | 0.005 | 0.004 | 0.004 | 0 | 0.51 | 0.5 | 0.53 | 0.4 | 0.62 | 0.48 |
| 8 | 15763818 | rs10503555 | 0.501 | -0.639 | 0.089 | -0.381 | -0.088 | -0.023 | -0.023 | 0 | 0.29 | 0.1 | 0.42 | 0.33 | 0.44 | 0.27 |
| 8 | 15765112 | rs10503559 | -0.218 | -0.709 | 0.13 | -0.406 | -0.01 | -0.006 | -0.006 | 0 | 0.07 | 0.13 | 0.11 | 0.03 | 0.03 | 0.17 |
| 8 | 15765369 | rs6982582 | -0.266 | -0.689 | 0.423 | -0.401 | 0.007 | -0.003 | -0.003 | 0 | 0.37 | 0.4 | 0.55 | 0.34 | 0.47 | 0.45 |
| 8 | 15767521 | rs4831778 | -0.741 | -0.704 | 0.232 | -0.539 | 0.018 | 0.028 | 0.028 | 0 | 0.46 | 0.64 | 0.63 | 0.36 | 0.58 | 0.51 |
| 8 | 15767978 | rs10503560 | 1.277 | -1.26 | -0.528 | -0.916 | 0.025 | 0.051 | 0.051 | 4.86 | 0.13 | 0 | 0.03 | 0.03 | 0.07 | 0.2 |
| 8 | 15770661 | rs12676637 | 1.343 | -1.24 | -0.461 | -0.9 | 0.002 | 0.005 | 0.005 | 0 | 0.08 | 0.08 | 0.05 | 0.04 | 0.07 | 0.09 |
| 8 | 15771612 | rs6987931 | -2.018 | -1.226 | -0.595 | -0.772 | -0.034 | -0.109 | -0.109 | 0 | 0.68 | 0.95 | 0.71 | 0.45 | 0.69 | 0.78 |
| 8 | 15772118 | rs6991301 | -0.865 | -1.357 | -0.24 | -0.894 | 0.033 | 0.055 | 0.055 | 1.188 | 0.35 | 0.79 | 0.2 | 0.09 | 0.25 | 0.34 |
| 8 | 15772489 | rs10108727 | 0.106 | -1.379 | -0.067 | -0.802 | 0.016 | 0.016 | 0.016 | 0.346 | 0.18 | 0.12 | 0.14 | 0.08 | 0.08 | 0.37 |
| 8 | 15773120 | rs7827166 | 0.471 | -1.414 | 0.172 | -0.45 | -0.084 | 0 | 0 | 2.83 | 0.17 | 0.5 | 0.06 | 0.11 | 0.05 | 0.2 |
| 8 | 15773592 | rs17579480 | 0.189 | -1.296 | 0.174 | -0.531 | -0.002 | -0.031 | -0.031 | 0 | 0.19 | 0.26 | 0.13 | 0.06 | 0.19 | 0.16 |
| 8 | 15773679 | rs13280081 | 0.829 | -1.188 | 0.068 | -0.455 | -0.051 | -0.008 | -0.008 | 0.379 | 0.28 | 0.13 | 0.42 | 0.29 | 0.43 | 0.28 |
| 8 | 15774587 | rs11203730 | 0.817 | -1.188 | 0.037 | -0.265 | -0.079 | -0.018 | -0.018 | 0 | 0.28 | 0.1 | 0.42 | 0.3 | 0.43 | 0.28 |
| 8 | 15776837 | rs7000884 | -1.004 | -1.289 | 0.063 | -0.381 | -0.001 | -0.001 | -0.001 | 0.524 | 0.53 | 0.47 | 0.64 | 0.38 | 0.63 | 0.59 |
| 8 | 15784174 | rs4831355 | 0.561 | -1.303 | 0.267 | -0.274 | -0.022 | -0.003 | -0.003 | 0 | 0.72 | 0.37 | 0.66 | 0.75 | 0.8 | 0.66 |
| 8 | 15786340 | rs4291272 | 1.043 | -1.139 | 0.003 | -0.501 | 0.014 | 0.003 | 0.003 | 0 | 0.51 | 0.38 | 0.4 | 0.52 | 0.57 | 0.32 |
| 8 | 15786889 | rs7832401 | 0.611 | -1.096 | 0.148 | -0.575 | 0 | -0.005 | -0.005 | 0 | 0.26 | 0.27 | 0.28 | 0.23 | 0.23 | 0.33 |
| 8 | 15788332 | rs7012411 | 0.682 | -1.181 | 0.066 | -0.64 | 0 | 0.073 | 0.073 | 3.734 | 0.53 | 0.35 | 0.54 | 0.76 | 0.51 | 0.55 |
| 8 | 15788551 | rs4831356 | -0.206 | -1.2 | 0.028 | -0.558 | -0.001 | 0 | 0 | 0.44 | 0.25 | 0.3 | 0.36 | 0.27 | 0.21 | 0.37 |
| 8 | 15789001 | rs2410368 | -1.044 | -1.009 | 0.105 | -0.686 | 0.006 | 0.092 | 0.092 | 1.115 | 0.46 | 0.21 | 0.51 | 0.28 | 0.52 | 0.61 |
| 8 | 15789762 | rs2410365 | -0.564 | -0.776 | 0.01 | -0.64 | 0.003 | -0.036 | -0.036 | 0.132 | 0.24 | 0.3 | 0.25 | 0.05 | 0.29 | 0.26 |
| 8 | 15791633 | rs2410363 | -1.477 | -0.889 | -0.365 | -0.559 | 0.006 | 0.023 | 0.023 | 0.097 | 0.55 | 0.6 | 0.55 | 0.28 | 0.55 | 0.61 |
| 8 | 15792451 | rs12548645 | -1.448 | -0.902 | -0.367 | -0.566 | 0.007 | 0.024 | 0.024 | 0 | 0.55 | 0.6 | 0.55 | 0.28 | 0.55 | 0.61 |
| 8 | 15795560 | rs35422228 | 1.18 | 0.792 | -0.341 | -0.589 | -0.037 | 0.097 | 0.097 | 0 | 0.6 | 0.18 | 0.43 | 0.78 | 0.56 | 0.52 |
| 8 | 15798908 | rs10503561 | -1.703 | 1.394 | 0.675 | -0.054 | 0.006 | -0.011 | -0.011 | 0 | 0.4 | 0.59 | 0.6 | 0.31 | 0.44 | 0.52 |
| 8 | 15800500 | rs2410342 | -1.659 | 1.365 | 1.04 | -0.021 | 0.002 | -0.02 | -0.02 | 0.021 | 0.41 | 0.56 | 0.65 | 0.3 | 0.48 | 0.54 |
| 8 | 15801843 | rs6998493 | 0.906 | 1.529 | 1.12 | -0.194 | 0.008 | -0.103 | -0.103 | 0.109 | 0.75 | 0.36 | 0.8 | 1 | 0.73 | 0.87 |
| 8 | 15801934 | rs6985396 | 0.183 | 1.773 | 0.832 | 0.198 | -0.001 | -0.002 | -0.002 | 0 | 0.15 | 0.15 | 0.39 | 0.23 | 0.19 | 0.27 |
| 8 | 15805361 | rs13281682 | -0.436 | 1.897 | 0.837 | 0.127 | 0.052 | -0.04 | -0.04 | 0 | 0.08 | 0.18 | 0.13 | 0 | 0.21 | 0.07 |
| 8 | 15807735 | rs2410341 | -0.487 | 1.947 | 1.102 | 0.396 | 0.046 | 0.01 | 0.01 | 0 | 0.86 | 0.72 | 0.84 | 0.86 | 0.82 | 0.86 |
| 8 | 15808002 | rs10112114 | 0.655 | 1.765 | 0.974 | 0.372 | 0.004 | -0.016 | -0.016 | 0.834 | 0.72 | 0.63 | 0.69 | 0.76 | 0.78 | 0.71 |
| 8 | 15808593 | rs2410339 | 1.069 | 1.74 | 0.979 | 0.42 | 0 | 0.018 | 0.018 | 0 | 0.68 | 0.57 | 0.6 | 0.62 | 0.75 | 0.54 |
| 8 | 15809365 | rs13271046 | 1.887 | 1.226 | 1.032 | 0.443 | 0.008 | -0.003 | -0.003 | 0.327 | 0.6 | 0.43 | 0.47 | 0.62 | 0.54 | 0.45 |
| 8 | 15809821 | rs7829952 | -0.107 | 1.21 | 1.062 | 0.304 | -0.064 | 0.007 | 0.007 | 0 | 0.85 | 0.65 | 0.83 | 0.85 | 0.96 | 0.85 |
| 8 | 15809938 | rs9643930 | 1.864 | 1.241 | 0.921 | 0.333 | -0.001 | 0 | 0 | 0 | 0.6 | 0.37 | 0.46 | 0.67 | 0.54 | 0.45 |
| 8 | 15810092 | rs1074844 | -0.906 | 1.308 | 0.057 | 0.352 | -0.045 | -0.013 | -0.013 | 0.724 | 0.25 | 0.19 | 0.49 | 0.33 | 0.44 | 0.49 |
| 8 | 15813380 | rs7016422 | 0.426 | 0.902 | 0.058 | 0.563 | -0.029 | 0.007 | 0.007 | 0 | 0.82 | 0.74 | 0.74 | 0.76 | 0.93 | 0.67 |
| 8 | 15820384 | rs2410326 | 1.671 | 1.333 | -0.97 | 0.516 | -0.044 | -0.015 | -0.015 | 0 | 0.41 | 0.05 | 0.51 | 0.68 | 0.43 | 0.38 |
| 8 | 15822369 | rs9325777 | 1.441 | 1.996 | 1.012 | 1.013 | 0.012 | -0.005 | -0.005 | 0 | 0.47 | 0.41 | 0.32 | 0.54 | 0.45 | 0.3 |
| 8 | 15823912 | rs10503566 | 0.375 | 2.026 | 1.43 | 0.958 | -0.005 | -0.006 | -0.006 | 0.023 | 0.72 | 0.27 | 0.56 | 0.69 | 0.71 | 0.58 |
| 8 | 15825165 | rs12056902 | 0.015 | 2.013 | 1.485 | 0.954 | -0.063 | -0.062 | -0.062 | 0.483 | 0.23 | 0.07 | 0.39 | 0.31 | 0.29 | 0.42 |
| 8 | 15826667 | rs13256281 | 1.485 | 1.977 | 1.76 | 1.086 | 0.001 | 0 | 0 | 0.627 | 0.51 | 0.45 | 0.46 | 0.54 | 0.55 | 0.33 |
| 8 | 15830136 | rs17676346 | -1.076 | 1.914 | 2.086 | 1.119 | 0.002 | -0.006 | -0.006 | 0 | 0.31 | 0.28 | 0.39 | 0.43 | 0.26 | 0.51 |
| 8 | 15830643 | rs7845736 | 0.161 | 1.98 | 2.137 | 1.15 | 0.01 | 0.018 | 0.018 | 0 | 0.75 | 0.36 | 0.65 | 0.69 | 0.74 | 0.59 |
| 8 | 15832617 | rs13252630 | -0.408 | 1.967 | 2.01 | 1.14 | -0.008 | 0.01 | 0.01 | 0.081 | 0.19 | 0.1 | 0.21 | 0.15 | 0.22 | 0.21 |
| 8 | 15833227 | rs1080226 | -0.868 | 1.944 | 2.175 | 1.153 | 0.031 | -0.052 | -0.052 | 0.052 | 0.83 | 0.76 | 0.85 | 0.96 | 0.8 | 0.88 |
| 8 | 15837977 | rs2898432 | -0.966 | 2.023 | 2.253 | 1.304 | 0.017 | 0.046 | 0.046 | 1.036 | 0.25 | 0.32 | 0.44 | 0.47 | 0.35 | 0.53 |
| 8 | 15838886 | rs4831789 | 1.64 | 1.902 | 2.142 | 0.968 | 0.05 | -0.005 | -0.005 | 0 | 0.49 | 0.33 | 0.37 | 0.52 | 0.41 | 0.26 |
| 8 | 15839117 | rs2410319 | -1.05 | 1.809 | 1.833 | 1.114 | -0.001 | 0.001 | 0.001 | 0.836 | 0.24 | 0.25 | 0.42 | 0.43 | 0.33 | 0.51 |
| 8 | 15849729 | rs10089515 | 1.562 | 1.615 | 2.076 | 0.818 | 0.058 | -0.003 | -0.003 | 0 | 0.49 | 0.24 | 0.36 | 0.45 | 0.42 | 0.33 |
| 8 | 15853724 | rs6983454 | -1.014 | 1.596 | 2.567 | 0.778 | 0.014 | 0.016 | 0.016 | 0 | 0.34 | 0.41 | 0.45 | 0.48 | 0.44 | 0.51 |
| 8 | 15854454 | rs13263124 | -0.072 | 1.596 | 2.378 | 0.62 | 0.002 | 0.033 | 0.033 | 0.349 | 0.16 | 0.06 | 0.18 | 0.08 | 0.14 | 0.2 |
| 8 | 15855142 | rs4455832 | 1.738 | 1.656 | 2.334 | 0.661 | 0.049 | -0.006 | -0.006 | 15.536 | 0.49 | 0.23 | 0.34 | 0.45 | 0.42 | 0.28 |
| 8 | 15856708 | rs10503567 | -0.672 | 1.695 | 2.367 | 0.608 | 0.021 | -0.007 | -0.007 | 0.669 | 0.31 | 0.37 | 0.26 | 0.22 | 0.37 | 0.43 |

| | | | | | | | | | | | | | | | | |
|---|---|---|---|---|---|---|---|---|---|---|---|---|---|---|---|---|
| 8 | 15857397 | rs10481437 | 0.143 | 1.701 | 2.544 | 0.832 | 0.085 | 0.036 | 0.036 | 0 | 0.19 | 0.36 | 0.21 | 0.22 | 0.32 | 0.33 |
| 8 | 15872029 | rs4623424 | 0.019 | 1.728 | 2.412 | 0.841 | 0 | 0 | 0 | 1.083 | 0.85 | 0.8 | 0.8 | 0.81 | 0.84 | 0.7 |
| 8 | 15872655 | rs17483163 | -0.749 | 1.841 | 2.883 | 0.814 | -0.008 | 0.007 | 0.007 | 0 | 0.3 | 0.23 | 0.34 | 0.17 | 0.37 | 0.38 |
| 8 | 15872769 | rs2898430 | -1.34 | 1.903 | 3.147 | 0.98 | 0.084 | 0.213 | 0.213 | 0 | 0.1 | 0.32 | 0.27 | 0.31 | 0.19 | 0.17 |
| 8 | 15873133 | rs2410305 | 0.941 | 1.869 | 2.84 | 0.926 | -0.031 | 0.008 | 0.008 | 0 | 0.15 | 0.07 | 0.15 | 0.09 | 0.22 | 0.17 |
| 8 | 15874259 | rs10089993 | 0.482 | 1.676 | 2.978 | 0.94 | 0.042 | 0.081 | 0.081 | 0 | 0.75 | 0.57 | 0.61 | 0.47 | 0.69 | 0.63 |
| 8 | 15874680 | rs17580332 | 0.99 | 1.798 | 2.504 | 0.859 | -0.022 | 0.025 | 0.025 | 0 | 0.15 | 0.07 | 0.14 | 0.03 | 0.19 | 0.16 |
| 8 | 15874858 | rs7819568 | -0.013 | 1.505 | 2.483 | 0.875 | -0.011 | 0.03 | 0.03 | 0.438 | 0.13 | 0.06 | 0.17 | 0.03 | 0.15 | 0.2 |
| 8 | 15877689 | rs9942835 | -1.542 | 1.276 | 2.722 | 0.891 | -0.029 | -0.014 | -0.014 | 0.066 | 0.32 | 0.53 | 0.27 | 0.29 | 0.26 | 0.42 |
| 8 | 15885126 | rs8181023 | -0.005 | 1.434 | 1.726 | 0.699 | -0.042 | 0.052 | 0.052 | 0 | 0.26 | 0.07 | 0.18 | 0.1 | 0.26 | 0.26 |
| 8 | 15898666 | rs1484767 | -0.367 | 1.184 | 1.838 | 0.717 | 0.002 | 0.033 | 0.033 | 0 | 0.15 | 0.06 | 0.18 | 0.08 | 0.14 | 0.2 |
| 8 | 15899688 | rs765262 | -0.134 | 1.25 | 1.968 | 0.69 | -0.018 | 0.037 | 0.037 | 0 | 0.28 | 0.17 | 0.2 | 0.12 | 0.29 | 0.29 |
| 8 | 15901155 | rs2596583 | -1.888 | 1.278 | 2.232 | 0.827 | 0.122 | 0.197 | 0.197 | 0.235 | 0.05 | 0.26 | 0.24 | 0.17 | 0.12 | 0.15 |
| 8 | 15901219 | rs11203752 | -0.309 | 1.278 | 2.071 | 0.783 | -0.017 | 0.036 | 0.036 | 0 | 0.31 | 0.15 | 0.24 | 0.16 | 0.33 | 0.33 |
| 8 | 15905116 | rs9987282 | -0.432 | 1.011 | 1.737 | 0.879 | -0.01 | 0.01 | 0.01 | 1.818 | 0.12 | 0.21 | 0.16 | 0.16 | 0.09 | 0.26 |
| 8 | 15916389 | rs2628289 | -1.623 | 0.927 | 1.551 | 0.779 | 0.06 | 0.045 | 0.045 | 1.947 | 0.49 | 0.76 | 0.66 | 0.6 | 0.61 | 0.69 |
| 8 | 15924506 | rs2410373 | -0.861 | 1.031 | 1.661 | 0.731 | -0.046 | 0.112 | 0.112 | 0.013 | 0.4 | 0.16 | 0.39 | 0.19 | 0.43 | 0.48 |
| 8 | 15926382 | rs1484744 | -0.427 | 1.094 | 1.627 | 0.16 | 0 | 0.02 | 0.02 | 0.158 | 0.14 | 0.07 | 0.18 | 0.07 | 0.14 | 0.2 |
| 8 | 15928246 | rs2170939 | 0.61 | 1.016 | 2.132 | 0.209 | 0.004 | -0.026 | -0.026 | 0 | 0.65 | 0.59 | 0.73 | 0.82 | 0.64 | 0.71 |
| 8 | 15928415 | rs4577960 | 0.894 | 1.11 | 2.088 | 0.288 | -0.006 | -0.006 | -0.006 | 0 | 0.61 | 0.35 | 0.52 | 0.54 | 0.51 | 0.46 |
| 8 | 15928918 | rs13263158 | 1.322 | 0.122 | 1.666 | 0.276 | -0.093 | 0.05 | 0.05 | 0.005 | 0.15 | 0.01 | 0.17 | 0.05 | 0.23 | 0.19 |
| 8 | 15929038 | rs10106273 | 0.361 | 0.179 | 1.942 | 0.287 | 0.001 | 0.029 | 0.029 | 0 | 0.92 | 0.81 | 0.94 | 0.87 | 0.91 | 0.98 |
| 8 | 15931037 | rs11786905 | 0.917 | 0.087 | 1.645 | 0.194 | -0.002 | -0.013 | -0.013 | 0 | 0.6 | 0.33 | 0.59 | 0.56 | 0.63 | 0.51 |
| 8 | 15932333 | rs11784460 | 1.504 | 0.011 | 0.629 | 0.044 | 0.059 | 0.014 | 0.014 | 0.624 | 0.66 | 0.26 | 0.7 | 0.71 | 0.8 | 0.69 |
| 8 | 15933011 | rs13259717 | 1.774 | 0.045 | 0.45 | 0.035 | 0.079 | 0.008 | 0.008 | 1.711 | 0.6 | 0.19 | 0.55 | 0.63 | 0.73 | 0.6 |
| 8 | 15934868 | rs4831803 | 1.965 | 0.039 | 0.231 | -0.19 | -0.06 | 0.004 | 0.004 | 0 | 0.41 | 0.18 | 0.38 | 0.6 | 0.48 | 0.4 |
| 8 | 15935790 | rs10099900 | 0.795 | 0.048 | 0.755 | -0.2 | 0.002 | -0.006 | -0.006 | 1.356 | 0.59 | 0.46 | 0.56 | 0.69 | 0.58 | 0.56 |
| 8 | 15939182 | rs10094036 | 0.735 | 0.014 | 0.779 | -0.203 | -0.016 | -0.022 | -0.022 | 0 | 0.82 | 0.38 | 0.89 | 0.88 | 0.85 | 0.85 |
| 8 | 15939341 | rs10107034 | 1.634 | 0.003 | 0.772 | -0.21 | 0.007 | 0.008 | 0.008 | 5.445 | 0.75 | 0.28 | 0.74 | 0.8 | 0.8 | 0.78 |
| 8 | 15939879 | rs7012344 | 0.869 | 0.021 | 0.806 | -0.21 | -0.014 | 0.087 | 0.087 | 0 | 0.59 | 0.3 | 0.71 | 0.84 | 0.57 | 0.65 |
| 8 | 15942311 | rs4831368 | 2.184 | -0.55 | 0.552 | -0.252 | -0.107 | -0.122 | -0.122 | 0 | 0.38 | 0.06 | 0.35 | 0.5 | 0.42 | 0.25 |
| 8 | 15943072 | rs7846392 | 0.413 | -1.062 | 1.077 | -0.578 | 0.033 | 0.072 | 0.072 | 0.856 | 0.91 | 0.66 | 0.94 | 0.88 | 0.92 | 0.99 |
| 8 | 15944307 | rs1872342 | -0.004 | -1.024 | 0.51 | -0.588 | -0.001 | -0.075 | -0.075 | 0 | 0.72 | 0.53 | 0.77 | 0.86 | 0.71 | 0.78 |
| 8 | 15949771 | rs13280294 | 2.31 | -1.636 | 1.188 | 0.152 | 0.073 | 0.017 | 0.017 | 5.305 | 0.6 | 0.24 | 0.56 | 0.64 | 0.76 | 0.6 |
| 8 | 15950878 | rs11988785 | -0.431 | -1.656 | 1.233 | 0.128 | -0.005 | -0.028 | -0.028 | 0 | 0.12 | 0.11 | 0.04 | 0 | 0.07 | 0.12 |
| 8 | 15950899 | rs4409402 | 0.692 | -1.666 | 1.37 | 0.126 | 0.009 | 0.01 | 0.01 | 0 | 0.83 | 0.8 | 0.82 | 0.93 | 0.9 | 0.82 |
| 8 | 15951562 | rs10096072 | -1.867 | -1.657 | 1.023 | 0.139 | -0.043 | 0.025 | 0.025 | 0 | 0.27 | 0.57 | 0.38 | 0.36 | 0.16 | 0.27 |
| 8 | 15952199 | rs1038463 | 2.201 | -1.724 | 0.813 | 0.196 | 0.016 | 0.005 | 0.005 | 5.052 | 0.61 | 0.32 | 0.57 | 0.64 | 0.77 | 0.6 |
| 8 | 15957442 | rs2898437 | -0.238 | -1.6 | 0.124 | 0.174 | 0.022 | 0.017 | 0.017 | 0 | 0.89 | 0.94 | 0.97 | 1 | 0.97 | 0.92 |
| 8 | 15957615 | rs2410379 | 1.906 | -1.55 | -0.189 | 0.217 | 0.037 | 0.014 | 0.014 | 4.998 | 0.63 | 0.29 | 0.68 | 0.71 | 0.79 | 0.68 |
| 8 | 15966542 | rs9650391 | 1.769 | -1.197 | -0.032 | 0.47 | 0.018 | 0.002 | 0.002 | 0.018 | 0.71 | 0.28 | 0.7 | 0.72 | 0.84 | 0.75 |
| 8 | 15966855 | rs6986419 | 1.674 | -1.005 | 0.061 | 0.571 | 0.016 | -0.001 | -0.001 | 0.911 | 0.71 | 0.27 | 0.68 | 0.7 | 0.83 | 0.74 |
| 8 | 15968147 | rs7015682 | -1.212 | -0.752 | -0.393 | 0.496 | -0.01 | 0.032 | 0.032 | 0 | 0.08 | 0.2 | 0.11 | 0.13 | 0.05 | 0.15 |
| 8 | 15968313 | rs6982066 | -1.212 | -0.752 | -0.393 | 0.496 | -0.01 | 0.032 | 0.032 | 0 | 0.08 | 0.2 | 0.11 | 0.13 | 0.05 | 0.15 |
| 8 | 15981879 | rs10503575 | 1.102 | -0.967 | -0.925 | 0.532 | 0.045 | -0.006 | -0.006 | 0 | 0.74 | 0.19 | 0.7 | 0.75 | 0.86 | 0.75 |
| 8 | 15984662 | rs12681382 | -0.633 | -0.603 | -0.9 | 0.541 | 0.039 | -0.032 | -0.032 | 0.774 | 0.11 | 0.01 | 0.11 | 0.13 | 0.04 | 0.1 |
| 8 | 15997974 | rs4333601 | -0.643 | -0.898 | -0.457 | 0.511 | -0.023 | 0.033 | 0.033 | 0 | 0.33 | 0.55 | 0.46 | 0.56 | 0.22 | 0.37 |
| 8 | 15998145 | rs12718376 | 0.645 | -1.085 | -0.063 | 0.441 | 0.028 | -0.03 | -0.03 | 0 | 0.65 | 0.27 | 0.53 | 0.44 | 0.78 | 0.63 |
| 8 | 15999039 | rs351562 | -0.467 | -0.995 | 0.71 | 0.377 | -0.009 | 0.034 | 0.034 | 0 | 0.3 | 0.37 | 0.46 | 0.55 | 0.22 | 0.36 |
| 8 | 16000235 | rs17484273 | 0.663 | -0.72 | -0.185 | -0.6 | -0.021 | -0.06 | -0.06 | 0 | 0.35 | 0.18 | 0.26 | 0.42 | 0.34 | 0.46 |
| 8 | 16003484 | rs10503573 | -1.127 | -0.842 | 0.592 | 0.099 | -0.014 | 0.122 | 0.122 | 0 | 0.07 | 0.16 | 0.19 | 0.35 | 0.05 | 0.22 |
| 8 | 16007395 | rs351554 | -0.716 | -1.151 | 0.873 | -0.176 | -0.048 | 0.057 | 0.057 | 1.748 | 0.26 | 0.62 | 0.43 | 0.56 | 0.12 | 0.35 |
| 8 | 16008635 | rs17620811 | 0.749 | -1.228 | 0.874 | -0.108 | 0.027 | -0.002 | -0.002 | 0 | 0.14 | 0.04 | 0.22 | 0.11 | 0.07 | 0.13 |
| 8 | 16019957 | rs1484751 | 0.737 | -1.653 | 0.37 | -0.502 | -0.119 | -0.086 | -0.086 | 1.299 | 0.42 | 0.07 | 0.32 | 0.42 | 0.44 | 0.47 |
| 8 | 16021468 | rs351572 | -1.24 | -1.404 | 0.973 | -0.382 | 0.002 | -0.056 | -0.056 | 0.508 | 0.7 | 0.72 | 0.75 | 0.97 | 0.57 | 0.82 |
| 8 | 16029684 | rs17677449 | -0.959 | -1.086 | 1.844 | 0.629 | -0.002 | 0.091 | 0.091 | 0 | 0.05 | 0.05 | 0.17 | 0.34 | 0.04 | 0.19 |
| 8 | 16039922 | rs386310 | -1.323 | -1.128 | 2.331 | 0.626 | -0.049 | 0.08 | 0.08 | 0 | 0.24 | 0.36 | 0.41 | 0.54 | 0.13 | 0.36 |
| 8 | 16040632 | rs383836 | -0.694 | -1.117 | 2.325 | 0.637 | -0.055 | 0.007 | 0.007 | 0.407 | 0.2 | 0.34 | 0.24 | 0.22 | 0.09 | 0.16 |
| 8 | 16042713 | rs3789015 | -0.829 | -1.106 | 2.587 | 0.623 | -0.004 | 0.184 | 0.184 | 0 | 0.06 | 0.09 | 0.18 | 0.43 | 0.05 | 0.23 |
| 8 | 16045949 | rs381111 | -0.685 | -1.154 | 2.702 | 0.573 | -0.073 | -0.037 | -0.037 | 0 | 0.2 | 0.44 | 0.24 | 0.11 | 0.08 | 0.13 |
| 8 | 16050240 | rs433235 | -0.699 | -1.095 | 2.711 | 0.569 | -0.067 | -0.038 | -0.038 | 0 | 0.2 | 0.44 | 0.25 | 0.11 | 0.08 | 0.13 |
| 8 | 16054928 | rs6530946 | 1.25 | -1.069 | 2.446 | 0.576 | -0.038 | 0.044 | 0.044 | 0.999 | 0.72 | 0.4 | 0.55 | 0.46 | 0.85 | 0.63 |
| 8 | 16059926 | rs7815674 | 1.217 | -0.963 | 2.24 | 0.549 | -0.036 | 0.039 | 0.039 | 1.146 | 0.72 | 0.4 | 0.57 | 0.44 | 0.86 | 0.61 |
| 8 | 16066481 | rs17677666 | -1.335 | -0.993 | 2.048 | 0.642 | -0.015 | 0.176 | 0.176 | 0 | 0.07 | 0.16 | 0.18 | 0.43 | 0.05 | 0.25 |
| 8 | 16067292 | rs12176618 | 1.662 | -0.954 | 1.817 | 0.572 | -0.017 | 0.008 | 0.008 | 0.962 | 0.67 | 0.36 | 0.54 | 0.44 | 0.84 | 0.59 |
| 8 | 16069524 | rs899683 | -0.72 | -0.664 | 1.302 | 0.564 | -0.041 | 0.053 | 0.053 | 0.018 | 0.2 | 0.32 | 0.27 | 0.37 | 0.1 | 0.31 |
| 8 | 16074001 | rs12546189 | 1.329 | -0.662 | 1.185 | -0.216 | -0.011 | -0.027 | -0.027 | 0 | 0.68 | 0.36 | 0.79 | 0.84 | 0.79 | 0.74 |
| 8 | 16074294 | rs12545741 | 0.916 | -0.743 | 1.973 | -0.262 | -0.008 | -0.077 | -0.077 | 0.042 | 0.73 | 0.67 | 0.82 | 0.97 | 0.79 | 0.79 |
| 8 | 16074573 | rs13280216 | 1.141 | -0.9 | 1.951 | -0.352 | -0.036 | -0.077 | -0.077 | 0.095 | 0.73 | 0.32 | 0.84 | 0.97 | 0.88 | 0.82 |

| | | | | | | | | | | | | | | | | |
|---|---|---|---|---|---|---|---|---|---|---|---|---|---|---|---|---|
| 8 | 16078712 | rs11780850 | 1.511 | -0.695 | 2.042 | -0.262 | -0.02 | -0.056 | -0.056 | 0 | 0.68 | 0.51 | 0.79 | 0.81 | 0.76 | 0.73 |
| 8 | 16081313 | rs4352853 | 1.607 | -0.675 | 2.153 | -0.375 | -0.016 | -0.046 | -0.046 | 0 | 0.65 | 0.42 | 0.78 | 0.8 | 0.73 | 0.73 |
| 8 | 16081596 | rs1600543 | 1.172 | -0.632 | 2.028 | -0.427 | 0.003 | -0.033 | -0.033 | 0 | 0.71 | 0.73 | 0.81 | 0.97 | 0.75 | 0.79 |
| 8 | 16087631 | rs10101109 | 1.214 | -0.39 | 1.932 | -0.433 | 0.001 | -0.117 | -0.117 | 5.915 | 0.71 | 0.53 | 0.76 | 0.92 | 0.74 | 0.79 |
| 8 | 16087729 | rs10100384 | 1.656 | -0.001 | 1.842 | -0.453 | 0.002 | -0.13 | -0.13 | 0.756 | 0.63 | 0.48 | 0.7 | 0.91 | 0.66 | 0.76 |
| 8 | 16090854 | rs9643931 | 2.332 | -0.494 | 2.17 | -0.199 | -0.045 | 0.054 | 0.054 | 4.748 | 0.9 | 0.33 | 0.94 | 0.86 | 0.98 | 0.93 |
| 8 | 16092656 | rs1480138 | 2.503 | -0.383 | 2.626 | -0.22 | 0.019 | 0.117 | 0.117 | 1.61 | 0.91 | 0.43 | 0.94 | 0.87 | 0.96 | 0.93 |
| 8 | 16097847 | rs13281928 | 2.41 | -0.395 | 2.6 | -0.188 | -0.022 | 0.063 | 0.063 | 4.824 | 0.9 | 0.33 | 0.93 | 0.86 | 0.96 | 0.93 |
| 8 | 16121167 | rs2163120 | 2.421 | -0.477 | 3.152 | -0.228 | -0.015 | 0.078 | 0.078 | 1.65 | 0.9 | 0.41 | 0.93 | 0.85 | 0.96 | 0.93 |
| 8 | 16124298 | rs958697 | 2.582 | -0.527 | 3.048 | 0.4 | -0.041 | 0.106 | 0.106 | 4.723 | 0.88 | 0.29 | 0.88 | 0.73 | 0.95 | 0.83 |
| 8 | 16125987 | rs1480147 | -1.276 | -0.538 | 2.619 | 0.445 | 0.003 | 0.116 | 0.116 | 0 | 0.3 | 0.12 | 0.31 | 0.09 | 0.27 | 0.21 |
| 8 | 16135246 | rs10503576 | -1.053 | -0.716 | 2.279 | 0.941 | -0.046 | 0.069 | 0.069 | 0 | 0.19 | 0.01 | 0.13 | 0.06 | 0.18 | 0.17 |
| 8 | 16162446 | rs11998457 | 1.975 | 0.425 | 2.315 | 0.95 | -0.008 | 0.009 | 0.009 | 0.974 | 0.62 | 0.13 | 0.59 | 0.62 | 0.67 | 0.57 |
| 8 | 16166495 | rs5008211 | -1.94 | 0.535 | 2.294 | 0.608 | 0.007 | -0.045 | -0.045 | 0 | 0.33 | 0.45 | 0.39 | 0.22 | 0.34 | 0.3 |
| 8 | 16168390 | rs1366960 | 1.808 | 0.232 | 1.675 | 0.587 | -0.006 | 0.046 | 0.046 | 0.064 | 0.62 | 0.13 | 0.59 | 0.67 | 0.64 | 0.55 |
| 8 | 16169243 | rs1541837 | -1.562 | 0.227 | 2.031 | 0.531 | -0.006 | 0.006 | 0.006 | 0.552 | 0.31 | 0.24 | 0.28 | 0.08 | 0.33 | 0.2 |
| 8 | 16172469 | rs1560307 | 2.295 | -0.109 | 2.174 | 0.627 | -0.038 | 0.119 | 0.119 | 4.591 | 0.87 | 0.34 | 0.85 | 0.74 | 0.94 | 0.73 |
| 8 | 16173694 | rs12545451 | -1.613 | -0.108 | 2.282 | 0.636 | -0.011 | 0.059 | 0.059 | 0.855 | 0.08 | 0.24 | 0.11 | 0.15 | 0.04 | 0.2 |
| 8 | 16175171 | rs1366957 | -0.974 | -0.047 | 2.113 | 0.636 | -0.004 | 0.014 | 0.014 | 0.778 | 0.06 | 0.07 | 0.1 | 0.15 | 0.04 | 0.19 |
| 8 | 16175495 | rs17582024 | -1.121 | -0.061 | 2.04 | 0.63 | 0.006 | 0.008 | 0.008 | 0.852 | 0.27 | 0.17 | 0.18 | 0.03 | 0.24 | 0.17 |
| 8 | 16175982 | rs1366956 | 0.414 | -0.326 | 1.446 | 0.845 | -0.127 | -0.118 | -0.118 | 0 | 0.22 | 0.02 | 0.26 | 0.37 | 0.3 | 0.2 |
| 8 | 16179167 | rs12675862 | 2.393 | -1.265 | 2.248 | 0.572 | 0.052 | 0.135 | 0.135 | 1.829 | 0.54 | 0.09 | 0.63 | 0.71 | 0.64 | 0.55 |
| 8 | 16188310 | rs1030517 | -0.899 | -0.811 | 2.916 | 0.585 | 0.017 | 0.029 | 0.029 | 0 | 0.21 | 0.12 | 0.2 | 0.07 | 0.18 | 0.16 |
| 8 | 16197551 | rs1429652 | 2.817 | -0.768 | 2.872 | 0.679 | -0.037 | 0.057 | 0.057 | 4.857 | 0.87 | 0.27 | 0.85 | 0.74 | 0.94 | 0.74 |
| 8 | 16211810 | rs2115696 | 2.022 | -0.751 | 2.561 | 0.649 | 0.019 | 0.056 | 0.056 | 0.974 | 0.64 | 0.1 | 0.67 | 0.71 | 0.7 | 0.58 |
| 8 | 16224716 | rs34484299 | -1.224 | -1.779 | 3.098 | 0.926 | -0.061 | -0.055 | -0.055 | 0 | 0.2 | 0.54 | 0.15 | 0.14 | 0.13 | 0.2 |
| 8 | 16236680 | rs1429667 | 1.812 | -1.819 | 2.674 | 0.974 | 0.083 | -0.029 | -0.029 | 0 | 0.72 | 0.1 | 0.67 | 0.66 | 0.81 | 0.64 |
| 8 | 16243135 | rs7003671 | 1.487 | -1.65 | 3.29 | 0.874 | -0.063 | -0.024 | -0.024 | 0 | 0.77 | 0.48 | 0.72 | 0.8 | 0.84 | 0.7 |
| 8 | 16245654 | rs891320 | 1.855 | -1.66 | 2.743 | 0.857 | 0.074 | -0.004 | -0.004 | 0 | 0.72 | 0.11 | 0.68 | 0.68 | 0.82 | 0.64 |
| 8 | 16259612 | rs10099016 | 0.912 | -1.594 | 2.925 | 0.848 | -0.004 | -0.033 | -0.033 | 0 | 0.94 | 0.84 | 0.97 | 1 | 0.96 | 0.98 |
| 8 | 16260166 | rs1429660 | 2.29 | -1.625 | 3.121 | 0.846 | -0.022 | 0.016 | 0.016 | 2.809 | 0.88 | 0.53 | 0.93 | 0.88 | 0.94 | 0.86 |
| 8 | 16262509 | rs12542835 | -1.193 | -1.615 | 2.528 | 0.871 | -0.036 | 0.024 | 0.024 | 0 | 0.2 | 0.23 | 0.26 | 0.32 | 0.14 | 0.3 |
| 8 | 16264714 | rs12544354 | -0.931 | -1.613 | 2.557 | 0.91 | -0.037 | 0.036 | 0.036 | 0 | 0.18 | 0.23 | 0.26 | 0.33 | 0.14 | 0.3 |
| 8 | 16282259 | rs17621828 | -1.449 | -1.419 | 1.426 | 0.919 | 0.077 | -0.014 | -0.014 | 0 | 0.28 | 0.9 | 0.32 | 0.34 | 0.18 | 0.38 |
| 8 | 16331191 | rs11781732 | 1.24 | -1.36 | 1.313 | 0.916 | 0.013 | 0.001 | 0.001 | 0.02 | 0.67 | 0.27 | 0.67 | 0.62 | 0.79 | 0.54 |
| 8 | 16336416 | rs10110883 | 0.892 | -1.284 | 1.017 | 0.947 | -0.002 | 0.001 | 0.001 | 0 | 0.7 | 0.35 | 0.73 | 0.71 | 0.81 | 0.61 |
| 8 | 16337635 | rs9325791 | 0.911 | -1.223 | 0.728 | 0.831 | -0.002 | 0 | 0 | 0 | 0.69 | 0.29 | 0.74 | 0.65 | 0.81 | 0.55 |
| 8 | 16350300 | rs2141026 | 1.224 | -0.976 | 1.32 | 0.825 | -0.021 | 0.022 | 0.022 | 1.355 | 0.88 | 0.45 | 0.92 | 0.92 | 0.94 | 0.87 |
| 8 | 16356260 | rs10503578 | -1.413 | -1.213 | 1.368 | 1.561 | -0.015 | 0.087 | 0.087 | 0.265 | 0.06 | 0.09 | 0.05 | 0.25 | 0.01 | 0.12 |
| 8 | 16364918 | rs17123050 | -1.945 | -1.211 | 1.901 | 0.904 | -0.038 | -0.076 | -0.076 | 2.443 | 0.12 | 0.73 | 0.09 | 0.01 | 0.05 | 0.09 |
| 8 | 16366723 | rs17123069 | -1.868 | -1.192 | 1.883 | 0.912 | -0.027 | -0.077 | -0.077 | 0.914 | 0.12 | 0.67 | 0.09 | 0.01 | 0.06 | 0.11 |
| 8 | 16370057 | rs17123099 | -0.26 | -1.102 | 1.562 | 0.96 | 0 | -0.01 | -0.01 | 0 | 0.05 | 0.06 | 0.04 | 0.01 | 0.05 | 0.05 |
| 8 | 16399163 | rs11987842 | -2.267 | -1.342 | 2.436 | 0.728 | -0.014 | -0.085 | -0.085 | 0 | 0.13 | 0.42 | 0.08 | 0.01 | 0.08 | 0.13 |
| 8 | 16401836 | rs3942029 | -2.242 | -1.32 | 2.897 | 1.857 | 0.016 | 0.31 | 0.31 | 0 | 0.09 | 0.28 | 0.23 | 0.37 | 0.06 | 0.23 |
| 8 | 16402524 | rs7823625 | 0.472 | -1.289 | 1.811 | 0.534 | -0.016 | 0.05 | 0.05 | 1.603 | 0.37 | 0.23 | 0.29 | 0.09 | 0.41 | 0.28 |
| 8 | 16406371 | rs12681937 | -2.275 | -1.273 | 2.006 | 0.635 | -0.036 | 0.203 | 0.203 | 0.484 | 0.16 | 0.58 | 0.26 | 0.37 | 0.09 | 0.27 |
| 8 | 16409107 | rs11780907 | -0.694 | -1.249 | 1.831 | 0.6 | -0.012 | 0.206 | 0.206 | 0.02 | 0.51 | 0.79 | 0.38 | 0.1 | 0.49 | 0.41 |
| 8 | 16410046 | rs1474449 | -0.956 | -1.353 | 1.696 | 0.626 | -0.049 | -0.055 | -0.055 | 1.712 | 0.53 | 0.88 | 0.55 | 0.48 | 0.49 | 0.57 |
| 8 | 16416409 | rs4461937 | -0.421 | -1.361 | 1.788 | 0.314 | -0.001 | -0.033 | -0.033 | 0 | 0.6 | 0.65 | 0.72 | 0.95 | 0.6 | 0.69 |
| 8 | 16416711 | rs4301485 | 1.006 | -1.484 | 1.053 | 0.193 | -0.097 | -0.111 | -0.111 | 0.5 | 0.42 | 0.05 | 0.41 | 0.54 | 0.45 | 0.5 |
| 8 | 16417482 | rs7017704 | -1.236 | -1.088 | 1.438 | 0.527 | -0.002 | 0.133 | 0.133 | 0 | 0.08 | 0.13 | 0.22 | 0.41 | 0.07 | 0.17 |
| 8 | 16417588 | rs7007310 | -1.008 | -1.073 | 1.422 | 0.534 | -0.001 | 0.114 | 0.114 | 0 | 0.08 | 0.1 | 0.22 | 0.41 | 0.07 | 0.17 |
| 8 | 16427536 | rs2037727 | -0.811 | -1.998 | 0.636 | -0.385 | -0.026 | 0.031 | 0.031 | 0.894 | 0.56 | 0.81 | 0.63 | 0.34 | 0.53 | 0.5 |
| 8 | 16429770 | rs17123254 | -0.592 | -2.109 | 0.776 | -0.502 | -0.01 | -0.036 | -0.036 | 0 | 0.1 | 0.15 | 0.04 | 0 | 0.05 | 0.06 |
| 8 | 16444725 | rs10503583 | 0.877 | -2.128 | 0.962 | -0.503 | -0.016 | 0.023 | 0.023 | 0.44 | 0.42 | 0.24 | 0.37 | 0.67 | 0.47 | 0.47 |
| 8 | 16445870 | rs4921630 | -0.515 | -2.063 | 0.976 | -0.322 | 0.001 | 0.026 | 0.026 | 0 | 0.08 | 0.05 | 0.31 | 0.29 | 0.09 | 0.15 |
| 8 | 16447075 | rs822226 | 2.116 | -2.228 | 1.251 | -0.456 | -0.054 | 0.066 | 0.066 | 0 | 0.78 | 0.58 | 0.64 | 0.71 | 0.86 | 0.77 |
| 8 | 16460684 | rs2687581 | -0.889 | -2.291 | 0.683 | -0.577 | -0.027 | 0.063 | 0.063 | 1.006 | 0.53 | 0.82 | 0.65 | 0.34 | 0.54 | 0.5 |
| 8 | 16467525 | rs1654781 | -0.528 | -2.299 | 0.8 | -0.694 | 0.002 | 0.122 | 0.122 | 0.36 | 0.51 | 0.67 | 0.37 | 0.06 | 0.5 | 0.37 |
| 8 | 16474041 | rs822304 | -0.421 | -2.313 | 0.628 | -0.409 | -0.042 | 0.001 | 0.001 | 1.184 | 0.48 | 0.82 | 0.64 | 0.41 | 0.51 | 0.47 |
| 8 | 16476498 | rs822301 | 0.173 | -2.377 | 0.831 | -0.431 | -0.004 | -0.035 | -0.035 | 1.598 | 0.41 | 0.55 | 0.33 | 0.06 | 0.44 | 0.31 |
| 8 | 16478760 | rs822298 | -1.099 | -2.34 | 0.447 | -0.347 | -0.073 | -0.048 | -0.048 | 0 | 0.57 | 0.88 | 0.66 | 0.4 | 0.54 | 0.51 |
| 8 | 16487401 | rs822272 | -0.47 | -1.994 | 1.168 | 0.006 | -0.008 | 0.094 | 0.094 | 0 | 0.07 | 0.08 | 0.34 | 0.29 | 0.07 | 0.18 |
| 8 | 16489491 | rs822277 | -1.316 | -1.621 | 1.046 | -0.236 | -0.008 | 0.022 | 0.022 | 2.291 | 0.55 | 0.84 | 0.7 | 0.36 | 0.58 | 0.53 |
| 8 | 16491849 | rs822278 | -2.551 | -1.23 | 1.68 | -0.205 | -0.016 | 0.124 | 0.124 | 0.011 | 0.22 | 0.48 | 0.42 | 0.37 | 0.18 | 0.34 |
| 8 | 16500728 | rs10046696 | -2.119 | -1.253 | 1.627 | -0.211 | -0.017 | 0.04 | 0.04 | 0 | 0.18 | 0.24 | 0.39 | 0.36 | 0.12 | 0.24 |
| 8 | 16502722 | rs1719421 | -0.747 | -1.023 | 1.632 | -0.056 | -0.004 | 0.097 | 0.097 | 0 | 0.08 | 0.08 | 0.35 | 0.35 | 0.09 | 0.18 |
| 8 | 16503082 | rs10110604 | -0.346 | -0.983 | 1.791 | -0.145 | -0.001 | -0.062 | -0.062 | 0 | 0.38 | 0.61 | 0.32 | 0.06 | 0.46 | 0.33 |
| 8 | 16510222 | rs822284 | -0.803 | -0.905 | 1.935 | 0.066 | 0.034 | 0.374 | 0.374 | 0 | 0.06 | 0.26 | 0.3 | 0.31 | 0.06 | 0.21 |

| | | | | | | | | | | | | | | | | |
|---|---|---|---|---|---|---|---|---|---|---|---|---|---|---|---|---|
| 8 | 16512119 | rs822286 | -1.019 | -0.937 | 1.483 | 0.004 | -0.021 | 0.049 | 0.049 | 2.466 | 0.48 | 0.8 | 0.63 | 0.36 | 0.52 | 0.51 |
| 8 | 16513105 | rs7814890 | -1.373 | -1.069 | 1.638 | -0.303 | -0.018 | -0.088 | -0.088 | 0 | 0.16 | 0.29 | 0.04 | 0.02 | 0.09 | 0.17 |
| 8 | 16517137 | rs822293 | -0.138 | -0.933 | 1.52 | -0.176 | -0.005 | 0.168 | 0.168 | 0 | 0.05 | 0.07 | 0.3 | 0.35 | 0.07 | 0.2 |
| 8 | 16517562 | rs822295 | -0.92 | -0.932 | 1.419 | -0.206 | -0.012 | 0 | 0 | 0.032 | 0.46 | 0.66 | 0.63 | 0.42 | 0.52 | 0.5 |
| 8 | 16517981 | rs6997598 | -0.961 | -1.018 | 1.084 | -0.132 | -0.004 | -0.053 | -0.053 | 0.083 | 0.13 | 0.15 | 0.04 | 0.02 | 0.09 | 0.16 |
| 8 | 16518066 | rs17487009 | -0.323 | -0.993 | 1.03 | -0.06 | 0 | 0.007 | 0.007 | 0.066 | 0.1 | 0.05 | 0.03 | 0.02 | 0.07 | 0.15 |
| 8 | 16520858 | rs1654801 | 0.209 | -0.936 | 1.077 | -0.02 | -0.002 | -0.035 | -0.035 | 0 | 0.9 | 0.87 | 0.96 | 0.98 | 0.93 | 0.87 |
| 8 | 16525028 | rs4521804 | 0.534 | -0.967 | 0.068 | 0.089 | -0.039 | -0.021 | -0.021 | 0 | 0.53 | 0.16 | 0.36 | 0.58 | 0.51 | 0.49 |
| 8 | 16526958 | rs17679694 | -0.829 | -0.952 | 0.775 | 1.054 | -0.005 | 0.138 | 0.138 | 0 | 0.05 | 0.05 | 0.3 | 0.36 | 0.04 | 0.22 |
| 8 | 16538391 | rs17123445 | -0.948 | -0.967 | 1.248 | 0.158 | 0.013 | 0.011 | 0.011 | 0.125 | 0.05 | 0.3 | 0.02 | 0 | 0 | 0.01 |
| 8 | 16543188 | rs1719442 | 0.449 | -0.536 | 0.456 | -0.479 | -0.141 | 0.17 | 0.17 | 0 | 0.28 | 0.06 | 0.29 | 0.01 | 0.44 | 0.17 |
| 8 | 16546204 | rs1654808 | 0.501 | -0.375 | 0.731 | -0.526 | -0.068 | 0.092 | 0.092 | 0 | 0.27 | 0.11 | 0.26 | 0.01 | 0.38 | 0.17 |
| 8 | 16547647 | rs1719437 | 1.314 | -1.097 | 1.617 | -0.648 | -0.057 | -0.078 | -0.078 | 0 | 0.88 | 0.44 | 0.94 | 1 | 0.98 | 0.93 |
| 8 | 16552997 | rs822259 | -0.264 | -1.036 | 1.682 | -0.639 | -0.054 | 0.082 | 0.082 | 0 | 0.35 | 0.23 | 0.31 | 0.02 | 0.47 | 0.22 |
| 8 | 16553165 | rs822258 | 0.428 | -1.035 | 1.653 | -0.624 | -0.097 | 0.12 | 0.12 | 0 | 0.29 | 0.11 | 0.29 | 0.01 | 0.45 | 0.18 |
| 8 | 16553779 | rs13264607 | 1.311 | -1.19 | 1.437 | -0.003 | -0.015 | -0.012 | -0.012 | 0 | 0.46 | 0.05 | 0.35 | 0.4 | 0.42 | 0.46 |
| 8 | 16556033 | rs7012141 | -1.993 | -1.096 | 1.988 | -0.504 | -0.043 | -0.03 | -0.03 | 0 | 0.13 | 0.33 | 0.05 | 0.09 | 0.05 | 0.16 |
| 8 | 16558730 | rs10097146 | 0.614 | -1.068 | 1.688 | -0.512 | -0.017 | -0.019 | -0.019 | 0 | 0.29 | 0.21 | 0.3 | 0.01 | 0.45 | 0.13 |
| 8 | 16568328 | rs12546337 | -1.319 | -1.06 | 1.504 | -0.104 | 0.006 | 0.011 | 0.011 | 0 | 0.11 | 0.09 | 0.06 | 0.02 | 0.06 | 0.19 |
| 8 | 16573252 | rs13258727 | 1.864 | -1.199 | 1.686 | -0.191 | -0.063 | -0.076 | -0.076 | 0.113 | 0.82 | 0.5 | 0.93 | 0.91 | 0.91 | 0.85 |
| 8 | 16573891 | rs11992032 | -0.752 | -1.073 | 1.702 | -0.174 | -0.016 | -0.028 | -0.028 | 0 | 0.07 | 0.31 | 0.04 | 0 | 0.02 | 0.05 |
| 8 | 16576557 | rs822241 | -0.103 | -1.034 | 1.452 | -0.153 | -0.009 | -0.031 | -0.031 | 0 | 0.93 | 0.87 | 0.96 | 1 | 0.96 | 1 |
| 8 | 16586768 | rs822231 | 1.551 | -0.952 | 1.064 | -0.161 | -0.037 | -0.143 | -0.143 | 0 | 0.82 | 0.5 | 0.91 | 0.98 | 0.92 | 0.78 |
| 8 | 16592601 | rs13251770 | 0.469 | -0.832 | 0.445 | 0.341 | 0.027 | 0.009 | 0.009 | 0 | 0.47 | 0.21 | 0.35 | 0.5 | 0.4 | 0.57 |
| 8 | 16596098 | rs11985010 | 0.563 | -0.851 | 0.349 | 0.331 | 0.023 | 0.002 | 0.002 | 0 | 0.46 | 0.17 | 0.35 | 0.5 | 0.4 | 0.56 |
| 8 | 16607394 | rs1697648 | -0.051 | -0.748 | 0.682 | 0.317 | -0.024 | -0.015 | -0.015 | 0 | 0.42 | 0.32 | 0.58 | 0.39 | 0.51 | 0.44 |
| 8 | 16608872 | rs7827906 | 0.418 | -0.806 | 0.452 | 0.335 | 0.013 | -0.005 | -0.005 | 0.123 | 0.52 | 0.23 | 0.37 | 0.53 | 0.46 | 0.45 |
| 8 | 16610997 | rs6996349 | 0.158 | -0.767 | 0.144 | 0.306 | -0.001 | -0.001 | -0.001 | 0 | 0.55 | 0.49 | 0.41 | 0.62 | 0.48 | 0.55 |
| 8 | 16622351 | rs13249535 | 1.106 | -0.634 | -0.902 | 0.38 | -0.039 | -0.096 | -0.096 | 0 | 0.4 | 0.05 | 0.32 | 0.48 | 0.37 | 0.42 |
| 8 | 16625172 | rs6586877 | -0.391 | -0.194 | -0.385 | 0.176 | -0.027 | -0.022 | -0.022 | 0.12 | 0.67 | 0.72 | 0.61 | 0.55 | 0.56 | 0.71 |
| 8 | 16632702 | rs13262310 | 0.086 | 0.187 | -1.853 | 0.558 | 0.024 | 0.005 | 0.005 | 0 | 0.51 | 0.15 | 0.51 | 0.54 | 0.47 | 0.72 |
| 8 | 16634751 | rs971797 | 0.089 | 0.04 | -1.823 | 0.576 | -0.007 | 0.049 | 0.049 | 0.053 | 0.47 | 0.12 | 0.52 | 0.57 | 0.56 | 0.48 |
| 8 | 16643823 | rs631290 | -0.919 | -1.127 | -0.144 | 0.766 | -0.094 | -0.01 | -0.01 | 1.895 | 0.23 | 0.4 | 0.11 | 0.18 | 0.08 | 0.16 |
| 8 | 16644426 | rs521882 | 0.258 | -0.535 | -0.844 | 0.527 | -0.032 | 0.038 | 0.038 | 0.106 | 0.61 | 0.77 | 0.76 | 0.77 | 0.6 | 0.73 |
| 8 | 16645869 | rs11203801 | 0.545 | -0.85 | 0.025 | 0.242 | -0.041 | 0.021 | 0.021 | 0 | 0.88 | 0.71 | 0.94 | 0.85 | 0.97 | 0.92 |
| 8 | 16648391 | rs17487711 | 0.905 | -0.071 | 0.252 | -0.104 | 0.013 | 0.185 | 0.185 | 0 | 0.52 | 0.22 | 0.67 | 0.81 | 0.53 | 0.69 |
| 8 | 16652106 | rs540481 | -0.665 | -0.116 | 0.794 | -0.133 | -0.051 | -0.076 | -0.076 | 3.79 | 0.12 | 0.37 | 0.02 | 0 | 0.01 | 0.04 |
| 8 | 16655774 | rs1604375 | 1.458 | 0.225 | 0.67 | -0.437 | -0.023 | -0.051 | -0.051 | 0 | 0.79 | 0.5 | 0.85 | 0.85 | 0.86 | 0.88 |
| 8 | 16657321 | rs652078 | 0.414 | 0.255 | 0.669 | -0.453 | 0.004 | 0.001 | 0.001 | 0.133 | 0.57 | 0.58 | 0.63 | 0.82 | 0.45 | 0.71 |
| 8 | 16659905 | rs1521357 | 0.188 | 0.28 | 0.658 | -0.477 | 0.002 | -0.046 | -0.046 | 0.255 | 0.34 | 0.33 | 0.25 | 0.04 | 0.44 | 0.23 |
| 8 | 16668990 | rs6992801 | -0.467 | 0.211 | 0.464 | -0.357 | 0.007 | 0 | 0 | 0.086 | 0.43 | 0.34 | 0.33 | 0.05 | 0.57 | 0.28 |
| 8 | 16669657 | rs1846949 | -0.615 | 0.205 | 0.397 | -0.35 | -0.006 | -0.071 | -0.071 | 0 | 0.44 | 0.45 | 0.36 | 0.1 | 0.57 | 0.28 |
| 8 | 16672922 | rs633944 | 0.182 | 0.12 | 0.66 | -0.356 | -0.034 | -0.044 | -0.044 | 0 | 0.08 | 0.14 | 0.01 | 0 | 0.01 | 0.03 |
| 8 | 16674096 | rs823422 | -1.451 | 0.468 | 0.356 | -0.43 | -0.041 | 0.183 | 0.183 | 0.132 | 0.55 | 0.96 | 0.43 | 0.2 | 0.54 | 0.33 |
| 8 | 16674526 | rs823424 | -1.805 | 0.652 | 0.512 | -0.425 | -0.004 | -0.043 | -0.043 | 6.479 | 0.23 | 0.43 | 0.21 | 0.16 | 0.21 | 0.14 |
| 8 | 16676099 | rs6992579 | 0.224 | 0.984 | 0.553 | -0.496 | 0.004 | -0.044 | -0.044 | 0 | 0.32 | 0.3 | 0.22 | 0.03 | 0.43 | 0.23 |
| 8 | 16679361 | rs17585055 | -0.417 | 1.112 | 0.127 | -0.597 | -0.022 | 0.038 | 0.038 | 4.987 | 0.08 | 0.03 | 0.08 | 0.01 | 0.12 | 0.04 |
| 8 | 16681701 | rs1717322 | 0.248 | 0.756 | -0.17 | -0.817 | -0.007 | -0.001 | -0.001 | 0 | 0.77 | 0.8 | 0.74 | 0.75 | 0.75 | 0.82 |
| 8 | 16690648 | rs475738 | 0.525 | 0.387 | -0.378 | -1.174 | 0.001 | -0.054 | -0.054 | 0 | 0.31 | 0.31 | 0.2 | 0.05 | 0.32 | 0.16 |
| 8 | 16691422 | rs611863 | 0.525 | 0.514 | -0.099 | -0.52 | 0.025 | -0.01 | -0.01 | 0 | 0.74 | 0.61 | 0.7 | 0.76 | 0.68 | 0.81 |
| 8 | 16698609 | rs635070 | 0.348 | 0.256 | -0.432 | -0.65 | -0.006 | 0.038 | 0.038 | 0.165 | 0.54 | 0.65 | 0.55 | 0.67 | 0.48 | 0.75 |
| 8 | 16700937 | rs666028 | 1.386 | 0.039 | -0.093 | -0.41 | -0.002 | -0.012 | -0.012 | 0.218 | 0.31 | 0.24 | 0.4 | 0.42 | 0.29 | 0.59 |
| 8 | 16702047 | rs6982320 | 0.524 | 0.099 | -0.234 | -0.699 | 0.028 | -0.001 | -0.001 | 1.973 | 0.32 | 0.22 | 0.15 | 0.01 | 0.22 | 0.07 |
| 8 | 16702455 | rs644062 | -0.672 | 0.169 | -0.243 | -0.649 | -0.02 | -0.02 | -0.02 | 0 | 0.86 | 0.89 | 0.7 | 0.67 | 0.69 | 0.83 |
| 8 | 16703588 | rs572151 | 1.263 | 0.057 | 0.225 | -0.719 | 0.003 | 0.005 | 0.005 | 0.468 | 0.38 | 0.38 | 0.53 | 0.46 | 0.53 | 0.66 |
| 8 | 16703648 | rs2037436 | 0.581 | 0.053 | 0.105 | -0.741 | 0.046 | 0.047 | 0.047 | 3.394 | 0.32 | 0.17 | 0.14 | 0.01 | 0.22 | 0.07 |
| 8 | 16704939 | rs1697632 | -0.132 | 0.017 | 0.201 | -0.663 | -0.013 | 0.111 | 0.111 | 0 | 0.88 | 0.75 | 0.79 | 0.47 | 0.92 | 0.78 |
| 8 | 16709730 | rs630757 | -0.12 | -0.004 | 0.067 | -0.718 | 0.04 | 0.033 | 0.033 | 0.622 | 0.61 | 0.73 | 0.68 | 0.71 | 0.72 | 0.82 |
| 8 | 16713686 | rs12234982 | 2.214 | 0.065 | -0.075 | -0.779 | -0.049 | -0.076 | -0.076 | 0 | 0.28 | 0.06 | 0.39 | 0.39 | 0.28 | 0.56 |
| 8 | 16713991 | rs477516 | 1.934 | 0.042 | 0.124 | -0.802 | 0.006 | 0.015 | 0.015 | 0 | 0.3 | 0.35 | 0.4 | 0.39 | 0.28 | 0.56 |
| 8 | 16714197 | rs10503585 | 0.924 | -0.342 | -0.17 | -0.975 | 0.052 | 0.096 | 0.096 | 0 | 0.46 | 0.22 | 0.58 | 0.7 | 0.6 | 0.78 |
| 8 | 16714318 | rs666336 | -0.746 | -0.322 | -0.042 | -0.88 | -0.018 | 0.093 | 0.093 | 0 | 0.88 | 0.77 | 0.79 | 0.67 | 0.92 | 0.86 |
| 8 | 16716348 | rs1521365 | 0.692 | -0.096 | -0.366 | -0.827 | -0.004 | 0.022 | 0.022 | 0 | 0.11 | 0.13 | 0.06 | 0.27 | 0.06 | 0.13 |
| 8 | 16718206 | rs11986428 | -0.045 | -0.156 | -0.558 | -0.919 | -0.005 | -0.001 | -0.001 | 0.069 | 0.33 | 0.36 | 0.17 | 0.28 | 0.23 | 0.17 |
| 8 | 16718818 | rs588564 | -1.16 | 0.184 | -0.644 | -0.748 | -0.005 | 0.007 | 0.007 | 0.859 | 0.61 | 0.72 | 0.42 | 0.33 | 0.64 | 0.32 |
| 8 | 16725205 | rs883355 | 0.194 | 0.167 | -0.797 | -0.814 | -0.033 | 0.086 | 0.086 | 0 | 0.3 | 0.15 | 0.24 | 0.05 | 0.41 | 0.13 |
| 8 | 16725613 | rs622595 | 0.096 | 0.109 | -1.077 | -0.786 | -0.031 | 0 | 0 | 0 | 0.3 | 0.51 | 0.17 | 0.27 | 0.22 | 0.18 |
| 8 | 16725835 | rs17585543 | 0.411 | 0.315 | -1.247 | -1.071 | 0.033 | 0.055 | 0.055 | 0.092 | 0.2 | 0.08 | 0.1 | 0 | 0.16 | 0.04 |
| 8 | 16726107 | rs11778346 | -0.232 | 1.045 | -0.677 | -0.081 | -0.071 | 0.038 | 0.038 | 0 | 0.11 | 0.02 | 0.14 | 0.04 | 0.25 | 0.1 |

| | | | | | | | | | | | | | | | | |
|---|---|---|---|---|---|---|---|---|---|---|---|---|---|---|---|---|
| 8 | 16735361 | rs17624390 | 0.775 | -0.067 | -0.004 | 0.419 | -0.03 | 0.086 | 0.086 | 0 | 0.84 | 0.71 | 0.78 | 0.66 | 0.89 | 0.86 |
| 8 | 16743181 | rs6990377 | -0.964 | 0.518 | -0.629 | -0.391 | -0.013 | 0.041 | 0.041 | 0.119 | 0.15 | 0.1 | 0.07 | 0.03 | 0.18 | 0.08 |
| 8 | 16748627 | rs10503586 | -0.767 | 0.101 | -0.295 | -0.5 | -0.001 | -0.022 | -0.022 | 0.012 | 0.12 | 0.13 | 0.06 | 0.01 | 0.08 | 0.03 |
| 8 | 16754703 | rs6987784 | 1.687 | -0.512 | 0.93 | 0.137 | -0.004 | 0.009 | 0.009 | 0.522 | 0.76 | 0.26 | 0.68 | 0.61 | 0.81 | 0.82 |
| 8 | 16755485 | rs1565741 | 1.597 | -0.515 | 1.493 | 0.14 | -0.032 | 0.073 | 0.073 | 0.568 | 0.77 | 0.39 | 0.7 | 0.62 | 0.83 | 0.84 |
| 8 | 16758938 | rs823580 | 0.785 | -0.484 | 1.568 | 0.045 | -0.018 | -0.081 | -0.081 | 1.371 | 0.85 | 0.73 | 0.91 | 0.99 | 0.89 | 0.96 |
| 8 | 16760196 | rs13270470 | 0.699 | -0.278 | 0.3 | -0.746 | -0.098 | 0.246 | 0.246 | 0.013 | 0.36 | 0.02 | 0.21 | 0.05 | 0.34 | 0.38 |
| 8 | 16764673 | rs923854 | -1.397 | -0.347 | 1.876 | 0.318 | -0.014 | 0.201 | 0.201 | 0.17 | 0.1 | 0.18 | 0.21 | 0.38 | 0.09 | 0.14 |
| 8 | 16774970 | rs823616 | 1.258 | -0.449 | 1.521 | 0.323 | -0.029 | 0.05 | 0.05 | 0 | 0.78 | 0.35 | 0.73 | 0.61 | 0.89 | 0.76 |
| 8 | 16790472 | rs10104915 | -1.437 | 0.072 | 1.513 | -0.227 | 0.048 | 0.033 | 0.033 | 0.937 | 0.06 | 0.4 | 0.04 | 0 | 0.01 | 0.07 |
| 8 | 16796681 | rs4406442 | 0.635 | 0.779 | 1 | -0.549 | 0.053 | -0.084 | -0.084 | 0 | 0.77 | 0.55 | 0.81 | 0.92 | 0.71 | 0.83 |
| 8 | 16799989 | rs1523655 | 1.186 | 0.661 | 1.059 | -0.951 | -0.006 | -0.03 | -0.03 | 0 | 0.79 | 0.32 | 0.83 | 0.92 | 0.81 | 0.9 |
| 8 | 16817064 | rs823602 | -0.129 | 0.686 | 0.319 | -2.187 | 0.006 | 0.033 | 0.033 | 0.041 | 0.49 | 0.44 | 0.62 | 0.87 | 0.42 | 0.68 |
| 8 | 16817416 | rs823601 | -0.419 | 0.969 | -0.024 | -0.925 | -0.001 | 0.007 | 0.007 | 0.168 | 0.24 | 0.27 | 0.35 | 0.44 | 0.21 | 0.35 |
| 8 | 16817909 | rs12674530 | 0.547 | 0.616 | -0.388 | -1.213 | -0.037 | 0.003 | 0.003 | 0 | 0.13 | 0.01 | 0.08 | 0.07 | 0.14 | 0.12 |
| 8 | 16821484 | rs823595 | -0.747 | 0.533 | -0.319 | -1.21 | -0.006 | 0.026 | 0.026 | 0 | 0.53 | 0.42 | 0.67 | 0.88 | 0.51 | 0.72 |
| 8 | 16822979 | rs1721073 | -0.775 | -0.099 | -0.213 | -0.93 | 0.004 | -0.015 | -0.015 | 0 | 0.29 | 0.24 | 0.39 | 0.44 | 0.29 | 0.38 |
| 8 | 16823648 | rs10503587 | 0.776 | -0.082 | -0.316 | -0.598 | 0.007 | -0.019 | -0.019 | 0 | 0.09 | 0.03 | 0.15 | 0.4 | 0.06 | 0.27 |
| 8 | 16826297 | rs7813540 | -0.17 | -0.086 | -0.268 | -0.592 | 0.003 | -0.048 | -0.048 | 0 | 0.14 | 0.06 | 0.2 | 0.4 | 0.15 | 0.3 |
| 8 | 16830161 | rs1689856 | 0.046 | -0.033 | -0.164 | -0.661 | 0 | -0.002 | -0.002 | 0.021 | 0.15 | 0.15 | 0.2 | 0.04 | 0.15 | 0.08 |
| 8 | 16830934 | rs1721078 | 0.713 | 0.089 | 0.752 | -0.777 | 0.073 | 0.097 | 0.097 | 0.007 | 0.94 | 0.71 | 0.95 | 0.93 | 0.96 | 0.99 |
| 8 | 16831026 | rs1908145 | 1.688 | 0.151 | 0.795 | -1.204 | -0.003 | 0.005 | 0.005 | 0 | 0.35 | 0.3 | 0.25 | 0.04 | 0.37 | 0.18 |
| 8 | 16831284 | rs1689871 | -0.096 | 0.295 | 0.921 | -0.13 | -0.025 | -0.051 | -0.051 | 0 | 0.13 | 0.07 | 0.29 | 0.34 | 0.18 | 0.24 |
| 8 | 16838292 | rs2055092 | 1.141 | 0.224 | 1.105 | -0.135 | 0.03 | 0.057 | 0.057 | 0.871 | 0.72 | 0.61 | 0.66 | 0.53 | 0.63 | 0.58 |
| 8 | 16840672 | rs1689888 | 1.11 | 0.217 | 1.174 | -0.041 | 0.035 | -0.004 | -0.004 | 0 | 0.87 | 0.75 | 0.86 | 0.92 | 0.8 | 0.91 |
| 8 | 16841292 | rs950549 | 0.486 | 0.177 | 0.716 | 0.013 | -0.002 | 0.053 | 0.053 | 0.978 | 0.06 | 0.06 | 0.24 | 0.36 | 0.06 | 0.15 |
| 8 | 16842573 | rs1721105 | -0.036 | 0.208 | 0.88 | -0.063 | 0.067 | -0.019 | -0.019 | 0 | 0.93 | 0.79 | 0.93 | 0.99 | 0.87 | 0.94 |
| 8 | 16843277 | rs1615196 | -0.903 | 0.106 | 0.856 | -0.061 | 0.031 | 0.198 | 0.198 | 1.679 | 0.11 | 0.35 | 0.29 | 0.37 | 0.1 | 0.16 |
| 8 | 16844315 | rs2517037 | -0.619 | 0.095 | 0.554 | -0.227 | -0.013 | -0.034 | -0.034 | 0 | 0.21 | 0.11 | 0.29 | 0.4 | 0.28 | 0.33 |
| 8 | 16845975 | rs1721101 | 0.946 | 0.263 | 0.794 | -0.101 | -0.007 | 0.111 | 0.111 | 0.021 | 0.76 | 0.58 | 0.74 | 0.55 | 0.73 | 0.62 |
| 8 | 16854803 | rs10106536 | 0.165 | 0.258 | 0.184 | -0.548 | -0.009 | 0.055 | 0.055 | 0 | 0.19 | 0.24 | 0.28 | 0.44 | 0.12 | 0.21 |
| 8 | 16862202 | rs2188014 | -0.51 | 0.247 | -1.541 | -0.952 | -0.027 | 0.113 | 0.113 | 0.07 | 0.78 | 0.94 | 0.86 | 1 | 0.76 | 0.84 |
| 8 | 16864155 | rs7822029 | 1.218 | 0.074 | -1.648 | -0.836 | -0.021 | -0.048 | -0.048 | 0 | 0.42 | 0.12 | 0.43 | 0.5 | 0.42 | 0.49 |
| 8 | 16867469 | rs6985266 | 0.65 | -0.102 | -1.646 | -0.833 | -0.013 | -0.001 | -0.001 | 0 | 0.5 | 0.14 | 0.49 | 0.49 | 0.53 | 0.57 |
| 8 | 16870536 | rs709844 | -0.218 | -0.23 | -1.339 | -0.738 | -0.013 | 0 | 0 | 0.531 | 0.68 | 0.72 | 0.78 | 0.99 | 0.63 | 0.76 |
| 8 | 16871448 | rs12056820 | -0.701 | -0.508 | -0.892 | 0.559 | 0.016 | -0.058 | -0.058 | 0.017 | 0.29 | 0.79 | 0.38 | 0.48 | 0.23 | 0.26 |
| 8 | 16877472 | rs850429 | -0.846 | -0.395 | -0.21 | 0.316 | -0.041 | 0.019 | 0.019 | 0 | 0.22 | 0.46 | 0.35 | 0.71 | 0.19 | 0.21 |
| 8 | 16881104 | rs850408 | 0.265 | -1.143 | -0.457 | -0.649 | -0.044 | -0.1 | -0.1 | 0.354 | 0.64 | 0.53 | 0.84 | 0.89 | 0.8 | 0.68 |
| 8 | 16881740 | rs2517010 | 0.012 | -0.861 | -1.042 | -0.559 | -0.001 | -0.002 | -0.002 | 0 | 0.48 | 0.48 | 0.4 | 0.41 | 0.48 | 0.46 |
| 8 | 16889380 | rs7812526 | -0.523 | -0.515 | -0.616 | 0.279 | 0.053 | 0.119 | 0.119 | 0.006 | 0.21 | 0.49 | 0.26 | 0.6 | 0.23 | 0.4 |
| 8 | 16892136 | rs11203825 | 0.111 | -0.493 | -0.501 | 0.173 | 0.014 | -0.031 | -0.031 | 0 | 0.09 | 0.1 | 0.06 | 0.01 | 0.13 | 0.27 |
| 8 | 16899180 | rs11203826 | -1.078 | -0.466 | -0.439 | 0.252 | 0.05 | 0.107 | 0.107 | 0 | 0.19 | 0.38 | 0.25 | 0.59 | 0.23 | 0.4 |
| 8 | 16905525 | rs6988267 | -0.949 | -0.14 | -0.587 | 0.355 | 0.109 | 0.367 | 0.367 | 0.012 | 0.06 | 0.48 | 0.09 | 0.22 | 0.05 | 0.09 |
| 8 | 16912655 | rs17488918 | -1.345 | -0.455 | -1.055 | 0.038 | -0.038 | -0.02 | -0.02 | 0 | 0.41 | 0.88 | 0.37 | 0.63 | 0.35 | 0.49 |
| 8 | 16915648 | rs3850756 | -0.368 | -0.329 | -0.98 | -0.01 | -0.004 | 0.008 | 0.008 | 0 | 0.25 | 0.29 | 0.27 | 0.62 | 0.2 | 0.2 |
| 8 | 16915980 | rs17624782 | 0.631 | -0.349 | -1.018 | 0.006 | -0.002 | -0.082 | -0.082 | 0 | 0.15 | 0.01 | 0.11 | 0.25 | 0.1 | 0.12 |
| 8 | 16916363 | rs3915070 | -1.978 | -0.622 | -1.493 | -0.291 | -0.094 | 0.17 | 0.17 | 0 | 0.55 | 0.89 | 0.61 | 0.81 | 0.51 | 0.61 |
| 8 | 16919086 | rs17624806 | 1.097 | -0.665 | -1.557 | -0.333 | -0.029 | -0.05 | -0.05 | 2.878 | 0.1 | 0.01 | 0.21 | 0.18 | 0.13 | 0.09 |
| 8 | 16922013 | rs2285261 | -1.289 | -0.671 | -1.514 | -0.312 | 0.004 | 0.002 | 0.002 | 0 | 0.44 | 0.58 | 0.44 | 0.44 | 0.41 | 0.54 |
| 8 | 16925693 | rs4145471 | 0.709 | -0.692 | -1.437 | -0.262 | 0.021 | 0.214 | 0.214 | 0.117 | 0.1 | 0.27 | 0.16 | 0.37 | 0.1 | 0.07 |
| 8 | 16929635 | rs10090288 | 0.096 | -0.743 | -1.376 | -0.29 | -0.062 | 0.049 | 0.049 | 0 | 0.2 | 0.46 | 0.15 | 0.25 | 0.1 | 0.12 |
| 8 | 16930341 | rs6998957 | 0.362 | -0.661 | -1.331 | -0.25 | 0.009 | 0.178 | 0.178 | 0.227 | 0.11 | 0.27 | 0.16 | 0.38 | 0.1 | 0.07 |
| 8 | 16931954 | rs11787157 | -0.639 | -0.614 | -1.356 | -0.276 | -0.013 | 0.004 | 0.004 | 0 | 0.24 | 0.09 | 0.29 | 0.23 | 0.26 | 0.38 |
| 8 | 16947295 | rs6587015 | -0.538 | -0.636 | -1.359 | -0.296 | -0.001 | 0.046 | 0.046 | 0 | 0.23 | 0.29 | 0.38 | 0.57 | 0.25 | 0.16 |
| 8 | 16949718 | rs7839562 | -1.306 | -0.773 | -1.558 | -0.568 | 0.009 | 0.011 | 0.011 | 0 | 0.4 | 0.49 | 0.45 | 0.56 | 0.45 | 0.5 |
| 8 | 16957500 | rs17124087 | 0.16 | -0.68 | -1.482 | -0.599 | 0 | -0.049 | -0.049 | 0 | 0.16 | 0.16 | 0.11 | 0.01 | 0.16 | 0.3 |
| 8 | 16960541 | rs10503589 | -0.38 | -0.674 | -1.467 | -0.61 | 0.004 | -0.061 | -0.061 | 0 | 0.19 | 0.21 | 0.13 | 0.01 | 0.21 | 0.33 |
| 8 | 16962762 | rs3862113 | -0.199 | -0.729 | -1.539 | -0.616 | 0.002 | -0.004 | -0.004 | 0 | 0.3 | 0.3 | 0.32 | 0.42 | 0.25 | 0.23 |
| 8 | 16963329 | rs11989486 | 0.345 | -0.711 | -1.535 | -0.617 | -0.001 | -0.016 | -0.016 | 0.821 | 0.16 | 0.11 | 0.11 | 0.01 | 0.16 | 0.32 |
| 8 | 16965977 | rs3988342 | -0.774 | -0.755 | -1.175 | -0.326 | 0.01 | -0.011 | -0.011 | 0 | 0.35 | 0.68 | 0.36 | 0.42 | 0.3 | 0.25 |
| 8 | 16967584 | rs7835868 | -0.892 | -0.737 | -1.338 | -0.336 | 0.011 | 0.038 | 0.038 | 0 | 0.38 | 0.29 | 0.46 | 0.81 | 0.35 | 0.3 |
| 8 | 16973070 | rs12680013 | 0.23 | -0.401 | -1.265 | 0.276 | -0.01 | -0.034 | -0.034 | 1.445 | 0.07 | 0.01 | 0.08 | 0.26 | 0.06 | 0.07 |
| 8 | 16975580 | rs2904647 | -0.922 | -0.484 | -0.773 | 0.263 | 0.004 | 0.038 | 0.038 | 0 | 0.21 | 0.24 | 0.31 | 0.41 | 0.25 | 0.17 |
| 8 | 16975802 | rs2517042 | -1.554 | -0.064 | -0.836 | 0.727 | -0.001 | -0.001 | -0.001 | 0 | 0.58 | 0.54 | 0.66 | 0.66 | 0.66 | 0.65 |
| 8 | 16978400 | rs13253571 | 0.298 | 0.018 | -0.87 | 0.632 | -0.023 | -0.027 | -0.027 | 0 | 0.13 | 0.05 | 0.27 | 0.19 | 0.19 | 0.11 |
| 8 | 16982443 | rs2904649 | 0.154 | 0.02 | -0.803 | 0.629 | -0.003 | -0.003 | -0.003 | 0 | 0.14 | 0.11 | 0.28 | 0.19 | 0.19 | 0.11 |
| 8 | 16983693 | rs2213901 | -0.725 | -0.117 | -0.534 | 0.388 | -0.001 | 0.001 | 0.001 | 0 | 0.44 | 0.42 | 0.44 | 0.5 | 0.44 | 0.54 |
| 8 | 16985962 | rs1158577 | -0.389 | 0.058 | -0.568 | -0.37 | -0.011 | 0.028 | 0.028 | 0.08 | 0.37 | 0.3 | 0.37 | 0.26 | 0.42 | 0.48 |
| 8 | 16988955 | rs10093037 | 0.061 | -0.426 | -1.824 | -0.501 | 0.091 | 0.129 | 0.129 | 2.327 | 0.42 | 0.22 | 0.18 | 0.15 | 0.22 | 0.41 |

| | | | | | | | | | | | | | | | | |
|---|---|---|---|---|---|---|---|---|---|---|---|---|---|---|---|---|
| 8 | 16989032 | rs11203833 | 0.361 | 0.712 | -2.634 | -0.235 | -0.102 | -0.103 | -0.103 | 0.64 | 0.52 | 0.07 | 0.45 | 0.59 | 0.51 | 0.52 |
| 8 | 16989568 | rs2952134 | -0.105 | 0.983 | -1.786 | -0.783 | 0.005 | 0.001 | 0.001 | 0 | 0.27 | 0.29 | 0.17 | 0 | 0.23 | 0.27 |
| 8 | 16991254 | rs10106054 | -0.43 | 1.483 | -1.07 | 0.302 | 0.063 | 0.114 | 0.114 | 0 | 0.27 | 0.4 | 0.43 | 0.55 | 0.35 | 0.41 |
| 8 | 16998449 | rs6984752 | -0.466 | 0.931 | -0.86 | -0.057 | -0.001 | 0.006 | 0.006 | 0.18 | 0.48 | 0.35 | 0.61 | 0.63 | 0.49 | 0.7 |
| 8 | 17000463 | rs2705215 | 0.077 | 1.066 | -0.561 | -0.6 | 0.008 | -0.159 | -0.159 | 2.495 | 0.69 | 0.41 | 0.67 | 1 | 0.69 | 0.68 |
| 8 | 17012250 | rs2904658 | -0.179 | 0.697 | -0.434 | 0.048 | 0.006 | -0.003 | -0.003 | 0 | 0.78 | 0.62 | 0.69 | 0.8 | 0.75 | 0.9 |
| 8 | 17018849 | rs2705204 | 0.682 | 0.561 | -0.981 | 0.161 | -0.08 | 0.135 | 0.135 | 1.419 | 0.45 | 0.13 | 0.31 | 0.23 | 0.5 | 0.31 |
| 8 | 17025169 | rs17586654 | -0.685 | 0.302 | -0.444 | 0.404 | 0 | -0.004 | -0.004 | 1.017 | 0.53 | 0.49 | 0.58 | 0.36 | 0.49 | 0.42 |
| 8 | 17025769 | rs13267854 | 0.185 | 0.382 | -0.628 | 0.053 | -0.009 | 0.062 | 0.062 | 0.793 | 0.29 | 0.18 | 0.23 | 0.09 | 0.26 | 0.31 |
| 8 | 17026852 | rs2959634 | -1.139 | -0.829 | -0.7 | -0.027 | 0.053 | 0.118 | 0.118 | 0 | 0.52 | 0.66 | 0.58 | 0.86 | 0.41 | 0.68 |
| 8 | 17028091 | rs730243 | 1.066 | -0.858 | -0.98 | 0.337 | -0.04 | -0.062 | -0.062 | 0 | 0.16 | 0.04 | 0.32 | 0.23 | 0.18 | 0.08 |
| 8 | 17029975 | rs7827043 | -0.683 | -1.121 | -0.579 | 0.237 | 0.066 | 0.046 | 0.046 | 0.587 | 0.39 | 0.74 | 0.52 | 0.74 | 0.25 | 0.57 |
| 8 | 17032687 | rs17586696 | 0.857 | -0.983 | -0.604 | 0.316 | -0.011 | -0.011 | -0.011 | 0 | 0.06 | 0 | 0.04 | 0.05 | 0.08 | 0.05 |
| 8 | 17034509 | rs2959631 | 0.469 | -1.07 | -0.236 | 0.608 | 0.001 | 0.001 | 0.001 | 0.009 | 0.17 | 0.2 | 0.13 | 0.15 | 0.07 | 0.19 |
| 8 | 17035927 | rs2904665 | -0.815 | -1.202 | 0.141 | 0.689 | -0.031 | 0.012 | 0.012 | 0.526 | 0.28 | 0.59 | 0.23 | 0.62 | 0.15 | 0.56 |
| 8 | 17036261 | rs2517082 | 0.38 | -1.004 | 0.277 | 0.555 | -0.01 | -0.007 | -0.007 | 0 | 0.73 | 0.68 | 0.79 | 0.81 | 0.75 | 0.75 |
| 8 | 17041823 | rs2904668 | 0.422 | -0.925 | 0.109 | 0.53 | 0.06 | 0.039 | 0.039 | 0.047 | 0.18 | 0.07 | 0.07 | 0.1 | 0.08 | 0.28 |
| 8 | 17046726 | rs2023627 | 0.276 | -0.851 | 0.25 | 0.744 | -0.001 | 0.225 | 0.225 | 0 | 0.84 | 0.67 | 0.88 | 0.56 | 0.84 | 0.69 |
| 8 | 17047134 | rs2023628 | 1.008 | -1.285 | -0.159 | 0.412 | -0.007 | -0.001 | -0.001 | 0 | 0.59 | 0.47 | 0.75 | 0.42 | 0.69 | 0.37 |
| 8 | 17052601 | rs10094988 | -0.732 | -1.186 | -1.204 | -0.112 | 0.032 | 0.07 | 0.07 | 1.36 | 0.44 | 0.75 | 0.52 | 0.7 | 0.42 | 0.43 |
| 8 | 17052840 | rs17124184 | 0.105 | -0.742 | -1.663 | -0.264 | -0.001 | 0.008 | 0.008 | 0 | 0.13 | 0.09 | 0.08 | 0.03 | 0.16 | 0.1 |
| 8 | 17053322 | rs2904669 | 0.949 | -0.406 | -1.787 | -0.116 | -0.132 | 0.135 | 0.135 | 2.134 | 0.37 | 0.03 | 0.4 | 0.14 | 0.5 | 0.29 |
| 8 | 17056529 | rs2904670 | 0.48 | -0.567 | -1.212 | 0.622 | 0.064 | -0.037 | -0.037 | 0 | 0.65 | 0.21 | 0.76 | 0.44 | 0.77 | 0.44 |
| 8 | 17056651 | rs2904671 | 0.337 | -0.204 | -0.83 | 0.14 | 0.023 | 0.039 | 0.039 | 0.195 | 0.5 | 0.57 | 0.72 | 0.79 | 0.62 | 0.58 |
| 8 | 17063335 | rs2904674 | 0.404 | -0.681 | -0.863 | 0.71 | 0.106 | 0.198 | 0.198 | 0.037 | 0.55 | 0.09 | 0.49 | 0.23 | 0.66 | 0.46 |
| 8 | 17064853 | rs6997380 | -0.126 | -0.929 | -0.755 | 0.685 | -0.024 | 0.016 | 0.016 | 0 | 0.33 | 0.34 | 0.39 | 0.38 | 0.25 | 0.32 |
| 8 | 17069284 | rs17124241 | 0.516 | -0.886 | -0.269 | 0.205 | -0.046 | 0.011 | 0.011 | 0.172 | 0.1 | 0.3 | 0.03 | 0.06 | 0.03 | 0.06 |
| 8 | 17073919 | rs9644663 | -0.198 | -1.125 | -0.474 | 0.216 | -0.044 | -0.012 | -0.012 | 0 | 0.26 | 0.3 | 0.1 | 0.17 | 0.12 | 0.24 |
| 8 | 17079358 | rs3750249 | -0.475 | -0.996 | -0.508 | 0.198 | -0.045 | -0.028 | -0.028 | 0.054 | 0.35 | 0.44 | 0.45 | 0.71 | 0.26 | 0.48 |
| 8 | 17081010 | rs2285268 | -0.642 | -0.993 | -0.505 | 0.209 | -0.045 | -0.027 | -0.027 | 0 | 0.35 | 0.44 | 0.44 | 0.7 | 0.26 | 0.48 |
| 8 | 17083856 | rs7460815 | 1.372 | 0.326 | -0.866 | -0.059 | -0.051 | 0.291 | 0.291 | 0.051 | 0.48 | 0.09 | 0.37 | 0.17 | 0.52 | 0.4 |
| 8 | 17084866 | rs7845103 | -0.497 | -0.135 | -0.746 | -0.004 | -0.046 | -0.046 | -0.046 | 0 | 0.35 | 0.45 | 0.49 | 0.75 | 0.27 | 0.48 |
| 8 | 17085769 | rs13267258 | -0.968 | -0.023 | -0.738 | 0.033 | 0.091 | 0.171 | 0.171 | 0.146 | 0.45 | 0.75 | 0.56 | 0.8 | 0.35 | 0.53 |
| 8 | 17089564 | rs2904696 | 0.372 | 0.01 | -0.533 | 0.041 | 0.002 | 0.002 | 0.002 | 0.063 | 0.63 | 0.37 | 0.49 | 0.26 | 0.68 | 0.52 |
| 8 | 17090798 | rs2904699 | -0.777 | -0.198 | -0.577 | -0.026 | -0.002 | 0.001 | 0.001 | 0 | 0.8 | 0.79 | 0.91 | 0.89 | 0.89 | 0.79 |
| 8 | 17110788 | rs7818793 | 0.167 | -0.295 | -0.349 | 0.036 | -0.036 | 0.152 | 0.152 | 0 | 0.19 | 0.38 | 0.4 | 0.59 | 0.18 | 0.28 |
| 8 | 17111633 | rs11991741 | 0.735 | 0.015 | -0.62 | -0.139 | -0.01 | -0.006 | -0.006 | 0 | 0.11 | 0.23 | 0.04 | 0.06 | 0.06 | 0.08 |
| 8 | 17130582 | rs10094736 | -1.49 | 0.254 | -1.182 | -0.407 | -0.038 | 0.192 | 0.192 | 0.042 | 0.56 | 0.92 | 0.56 | 0.76 | 0.48 | 0.65 |
| 8 | 17133045 | rs10087602 | 0.168 | 0.325 | -0.817 | -0.229 | -0.044 | 0.118 | 0.118 | 0 | 0.21 | 0.39 | 0.37 | 0.58 | 0.17 | 0.27 |
| 8 | 17137687 | rs3213601 | 1.698 | 0.578 | -1.034 | -0.369 | -0.167 | 0.147 | 0.147 | 0.014 | 0.42 | 0.03 | 0.38 | 0.24 | 0.47 | 0.34 |
| 8 | 17138567 | rs2188015 | 0.327 | 0.503 | -0.877 | -0.403 | -0.021 | -0.009 | -0.009 | 0 | 0.26 | 0.34 | 0.11 | 0.17 | 0.15 | 0.29 |
| 8 | 17140378 | rs10110023 | -0.896 | 0.524 | -0.793 | -0.394 | 0.022 | 0.002 | 0.002 | 0.066 | 0.32 | 0.72 | 0.42 | 0.65 | 0.24 | 0.38 |
| 8 | 17143246 | rs7815340 | -0.879 | 0.608 | -0.567 | -0.419 | 0.014 | 0.032 | 0.032 | 0.048 | 0.44 | 0.6 | 0.51 | 0.69 | 0.42 | 0.51 |
| 8 | 17155432 | rs1053903 | 1.452 | 1.295 | -0.953 | -0.727 | -0.011 | 0.004 | 0.004 | 0 | 0.11 | 0.14 | 0.12 | 0.01 | 0.13 | 0.09 |
| 8 | 17156393 | rs28434234 | 1.116 | 1.084 | -0.856 | -0.702 | 0.009 | 0.016 | 0.016 | 0 | 0.07 | 0.18 | 0.02 | 0.07 | 0.05 | 0.09 |
| 8 | 17156904 | rs6981281 | 0.165 | 0.896 | -0.98 | -0.684 | -0.051 | 0.085 | 0.085 | 0 | 0.23 | 0.4 | 0.39 | 0.6 | 0.2 | 0.31 |
| 8 | 17157259 | rs6992513 | -2.024 | 0.963 | -1.088 | -0.826 | 0.014 | 0.092 | 0.092 | 0 | 0.65 | 0.9 | 0.72 | 0.77 | 0.69 | 0.74 |
| 8 | 17157677 | rs3764796 | 0.998 | 0.791 | -0.924 | -0.648 | 0.04 | 0.006 | 0.006 | 0 | 0.19 | 0.01 | 0.1 | 0.11 | 0.12 | 0.23 |
| 8 | 17164074 | rs11779713 | 0.028 | 1.267 | -0.251 | -0.354 | 0.005 | 0.086 | 0.086 | 0 | 0.22 | 0.36 | 0.35 | 0.3 | 0.23 | 0.14 |
| 8 | 17165768 | rs7000979 | -0.438 | 1.212 | -0.081 | -0.315 | 0.081 | 0.106 | 0.106 | 0 | 0.2 | 0.54 | 0.4 | 0.29 | 0.32 | 0.13 |
| 8 | 17166765 | rs1548353 | -0.872 | 0.844 | -0.355 | -0.689 | 0.007 | 0.005 | 0.005 | 0 | 0.46 | 0.67 | 0.62 | 0.33 | 0.58 | 0.23 |
| 8 | 17171599 | rs2285275 | 0.542 | 0.742 | -0.172 | -0.618 | 0.004 | 0.253 | 0.253 | 0 | 0.13 | 0.37 | 0.31 | 0.3 | 0.15 | 0.06 |
| 8 | 17171737 | rs4921750 | 1.444 | 0.883 | -0.327 | -0.766 | -0.011 | 0.009 | 0.009 | 0 | 0.14 | 0.17 | 0.16 | 0.01 | 0.17 | 0.07 |
| 8 | 17174343 | rs12680752 | 0.445 | 0.702 | 0.718 | -0.416 | -0.01 | 0.099 | 0.099 | 0 | 0.12 | 0.23 | 0.3 | 0.26 | 0.14 | 0.06 |
| 8 | 17174534 | rs1858709 | -0.345 | 0.729 | 0.789 | -0.619 | -0.001 | 0.01 | 0.01 | 0 | 0.46 | 0.59 | 0.56 | 0.3 | 0.47 | 0.24 |
| 8 | 17174912 | rs6587031 | -0.551 | 0.434 | 0.691 | -0.494 | 0.001 | 0.12 | 0.12 | 0 | 0.21 | 0.48 | 0.34 | 0.3 | 0.21 | 0.12 |
| 8 | 17174994 | rs2269688 | 0.831 | 0.25 | 0.127 | -0.49 | 0.061 | -0.104 | -0.104 | 0 | 0.21 | 0.05 | 0.12 | 0.47 | 0.11 | 0.51 |
| 8 | 17176644 | rs10090731 | 1.902 | 0.503 | 0.184 | -0.856 | -0.001 | -0.049 | -0.049 | 1.506 | 0.24 | 0.25 | 0.21 | 0 | 0.27 | 0.09 |
| 8 | 17177934 | rs756604 | -1.036 | 0.904 | 0.603 | -0.487 | 0.021 | 0.029 | 0.029 | 0.035 | 0.42 | 0.65 | 0.42 | 0.76 | 0.32 | 0.63 |
| 8 | 17178199 | rs756606 | 2.351 | 0.872 | 0.501 | -0.51 | -0.036 | -0.015 | -0.015 | 0 | 0.21 | 0.01 | 0.27 | 0.24 | 0.2 | 0.2 |
| 8 | 17178237 | rs756607 | -1.499 | 1.324 | 2.071 | 0.861 | -0.056 | -0.048 | -0.048 | 0 | 0.34 | 0.5 | 0.38 | 0.72 | 0.25 | 0.57 |
| 8 | 17179288 | rs4593587 | -2.302 | 1.39 | 1.613 | 0.844 | -0.01 | 0.102 | 0.102 | 0 | 0.48 | 0.7 | 0.5 | 0.75 | 0.5 | 0.69 |
| 8 | 17179695 | rs2051663 | -1.806 | 1.13 | 1.769 | 0.928 | 0.011 | 0.016 | 0.016 | 0 | 0.4 | 0.6 | 0.41 | 0.72 | 0.32 | 0.62 |
| 8 | 17181772 | rs7003503 | -1.317 | 1.172 | 1.741 | 1.153 | 0 | 0.124 | 0.124 | 0.425 | 0.21 | 0.56 | 0.33 | 0.38 | 0.21 | 0.12 |
| 8 | 17182541 | rs2239865 | -1.688 | 1.124 | 1.716 | 1.193 | 0.012 | -0.013 | -0.013 | 0 | 0.37 | 0.64 | 0.42 | 0.4 | 0.32 | 0.31 |
| 8 | 17182965 | rs2239868 | -0.08 | 1.208 | 1.751 | 1.216 | 0.027 | -0.031 | -0.031 | 0 | 0.21 | 0.12 | 0.1 | 0.39 | 0.12 | 0.51 |
| 8 | 17183714 | rs2239869 | -0.904 | 1.205 | 1.986 | 1.243 | 0.021 | 0.013 | 0.013 | 0.769 | 0.14 | 0.4 | 0.1 | 0.08 | 0.1 | 0.09 |
| 8 | 17187235 | rs10453161 | -1.968 | 1.382 | 1.558 | 1.18 | -0.004 | 0.096 | 0.096 | 0 | 0.47 | 0.65 | 0.5 | 0.76 | 0.52 | 0.69 |
| 8 | 17193011 | rs2269692 | -0.566 | 2.101 | 2.194 | 2.231 | 0.004 | 0.188 | 0.188 | 0 | 0.08 | 0.21 | 0.24 | 0.28 | 0.11 | 0.03 |

| | | | | | | | | | | | | | | | | |
|---|---|---|---|---|---|---|---|---|---|---|---|---|---|---|---|---|
| 8 | 17193274 | rs2269693 | -1.122 | 2.177 | 1.9 | 2.162 | -0.033 | -0.027 | -0.027 | 0 | 0.09 | 0.04 | 0.09 | 0.36 | 0.2 | 0.35 |
| 8 | 17197967 | rs17124423 | -2.015 | 1.536 | 2.623 | 2.281 | 0.007 | 0.257 | 0.257 | 0.507 | 0.15 | 0.56 | 0.12 | 0.44 | 0.1 | 0.34 |
| 8 | 17202871 | rs739337 | -1.317 | 1.341 | 2.349 | 2.376 | -0.015 | 0.166 | 0.166 | 0 | 0.07 | 0.21 | 0.24 | 0.3 | 0.07 | 0.03 |
| 8 | 17218629 | rs1024355 | -2.53 | 1.431 | 2.688 | 2.328 | 0.233 | 0.128 | 0.128 | 0.103 | 0.07 | 0.49 | 0.13 | 0.07 | 0.15 | 0.02 |
| 8 | 17222410 | rs7822451 | -0.365 | 1.442 | 2.4 | 2.445 | 0.017 | 0.269 | 0.269 | 0 | 0.06 | 0.19 | 0.16 | 0.3 | 0.09 | 0.03 |
| 8 | 17222590 | rs11203846 | -2.394 | 1.431 | 2.407 | 2.427 | 0.069 | 0.224 | 0.224 | 0.731 | 0.14 | 0.64 | 0.29 | 0.37 | 0.24 | 0.05 |
| 8 | 17226856 | rs4921763 | -1.51 | 1.703 | 1.402 | 2.07 | 0.074 | 0.216 | 0.216 | 0 | 0.06 | 0.15 | 0.1 | 0.4 | 0.15 | 0.32 |
| 8 | 17236268 | rs10087746 | -1.184 | 0.864 | 1.19 | 1.081 | -0.011 | -0.042 | -0.042 | 0 | 0.13 | 0.22 | 0.06 | 0.02 | 0.09 | 0.1 |
| 8 | 17237608 | rs10089145 | -2.161 | 1.1 | 1.849 | 2.024 | 0.143 | 0.433 | 0.433 | 0 | 0.08 | 0.42 | 0.17 | 0.33 | 0.16 | 0.07 |
| 8 | 17240957 | rs7823989 | 0.784 | 1.015 | 2.115 | 1.647 | -0.001 | 0 | 0 | 0.03 | 0.63 | 0.62 | 0.67 | 0.36 | 0.52 | 0.44 |
| 8 | 17241040 | rs7828510 | -1.304 | 0.96 | 2.331 | 1.604 | 0.039 | 0.083 | 0.083 | 0 | 0.05 | 0.24 | 0.05 | 0.05 | 0.03 | 0.02 |
| 8 | 17244051 | rs4921768 | 1.299 | 0.966 | 2.192 | 1.655 | 0.014 | 0.043 | 0.043 | 0.002 | 0.54 | 0.35 | 0.59 | 0.32 | 0.44 | 0.42 |
| 8 | 17244248 | rs2898461 | 0.674 | 0.822 | 2.081 | 1.811 | -0.013 | -0.009 | -0.009 | 0.529 | 0.65 | 0.69 | 0.71 | 0.49 | 0.55 | 0.63 |
| 8 | 17247311 | rs4921546 | -0.76 | 0.931 | 1.299 | 1.139 | -0.029 | 0.023 | 0.023 | 0 | 0.32 | 0.23 | 0.2 | 0.21 | 0.37 | 0.34 |
| 8 | 17253619 | rs6992452 | -1.014 | 0.888 | 1.08 | 0.94 | 0.012 | -0.027 | -0.027 | 1.043 | 0.37 | 0.42 | 0.22 | 0.24 | 0.38 | 0.44 |
| 8 | 17257720 | rs1002810 | 0.518 | 0.757 | 0.539 | 0.411 | -0.013 | 0.047 | 0.047 | 0 | 0.67 | 0.81 | 0.84 | 0.99 | 0.66 | 0.69 |
| 8 | 17258798 | rs2898463 | -0.899 | 0.596 | 0.831 | 0.939 | 0.043 | 0.206 | 0.206 | 0.107 | 0.11 | 0.29 | 0.15 | 0.32 | 0.13 | 0.06 |
| 8 | 17258906 | rs2157640 | 1.675 | 0.46 | -0.247 | 0.865 | -0.102 | -0.054 | -0.054 | 0 | 0.46 | 0.04 | 0.61 | 0.42 | 0.49 | 0.49 |
| 8 | 17266143 | rs12548066 | 0.176 | 0.427 | 0.037 | 0.385 | -0.008 | 0.006 | 0.006 | 0 | 0.7 | 0.77 | 0.86 | 0.99 | 0.69 | 0.69 |
| 8 | 17267569 | rs13254505 | -0.233 | 0.5 | -0.421 | 0.387 | -0.048 | 0.139 | 0.139 | 0 | 0.28 | 0.08 | 0.12 | 0.02 | 0.3 | 0.31 |
| 8 | 17268961 | rs9802064 | 1.052 | 0.461 | -0.346 | 0.721 | 0.042 | 0.176 | 0.176 | 0 | 0.54 | 0.14 | 0.74 | 0.8 | 0.63 | 0.52 |
| 8 | 17269633 | rs1003299 | 1.372 | 0.441 | -0.491 | 1.069 | -0.064 | -0.018 | -0.018 | 0.354 | 0.5 | 0.07 | 0.62 | 0.51 | 0.51 | 0.49 |
| 8 | 17271136 | rs2073695 | -0.457 | 0.187 | 0.324 | 0.9 | -0.008 | 0.024 | 0.024 | 0 | 0.08 | 0.16 | 0.07 | 0.11 | 0.08 | 0.15 |
| 8 | 17271478 | rs7817826 | -0.97 | 0.202 | 0.011 | 0.769 | -0.021 | -0.006 | -0.006 | 0.111 | 0.28 | 0.18 | 0.19 | 0.29 | 0.3 | 0.33 |
| 8 | 17274709 | rs6981038 | -0.129 | -0.378 | 0.267 | 0.741 | -0.018 | 0.011 | 0.011 | 0 | 0.12 | 0.22 | 0.05 | 0.21 | 0.05 | 0.16 |
| 8 | 17282411 | rs13271465 | 1.509 | -0.86 | 0.492 | 0.731 | -0.046 | 0.054 | 0.054 | 0.605 | 0.69 | 0.41 | 0.78 | 0.51 | 0.83 | 0.76 |
| 8 | 17283970 | rs10503595 | 0.395 | -0.704 | -0.033 | 0.469 | -0.05 | 0.064 | 0.064 | 0 | 0.16 | 0.04 | 0.07 | 0 | 0.22 | 0.18 |
| 8 | 17284564 | rs10086318 | 2.071 | -0.726 | 0.292 | 0.456 | 0.027 | -0.005 | -0.005 | 0 | 0.51 | 0.29 | 0.7 | 0.5 | 0.6 | 0.58 |
| 8 | 17287780 | rs7462219 | 0.827 | -0.733 | 0.519 | 0.409 | -0.003 | -0.014 | -0.014 | 0 | 0.68 | 0.6 | 0.87 | 0.78 | 0.72 | 0.66 |
| 8 | 17290124 | rs4921772 | 2.071 | -0.717 | 0.413 | 0.406 | 0.025 | 0.017 | 0.017 | 2.18 | 0.49 | 0.25 | 0.7 | 0.45 | 0.6 | 0.56 |
| 8 | 17290628 | rs7387693 | 1.145 | -0.717 | 0.486 | 0.378 | 0.001 | 0.049 | 0.049 | 3.678 | 0.61 | 0.71 | 0.85 | 0.78 | 0.68 | 0.65 |
| 8 | 17292283 | rs17587994 | 2.13 | -0.723 | 0.227 | 0.517 | -0.005 | 0.054 | 0.054 | 1.496 | 0.45 | 0.19 | 0.68 | 0.34 | 0.59 | 0.53 |
| 8 | 17292466 | rs11203855 | -0.884 | -0.623 | 0.42 | 0.497 | -0.029 | 0.101 | 0.101 | 0 | 0.19 | 0.4 | 0.18 | 0.32 | 0.12 | 0.12 |
| 8 | 17298699 | rs4921773 | -0.029 | -0.309 | -0.869 | 0.356 | 0.012 | -0.123 | -0.123 | 0.07 | 0.15 | 0.01 | 0.15 | 0.52 | 0.1 | 0.18 |
| 8 | 17298825 | rs4921774 | 1.213 | -0.122 | -0.732 | 0.062 | 0.028 | 0.006 | 0.006 | 3.119 | 0.49 | 0.23 | 0.7 | 0.47 | 0.62 | 0.54 |
| 8 | 17299625 | rs6986498 | -0.894 | -0.601 | -0.527 | 0.372 | -0.009 | 0.017 | 0.017 | 0 | 0.24 | 0.26 | 0.19 | 0.54 | 0.14 | 0.18 |
| 8 | 17299950 | rs2027922 | 0.811 | -0.495 | -0.696 | 0.178 | 0.051 | 0.001 | 0.001 | 0 | 0.59 | 0.77 | 0.75 | 0.43 | 0.73 | 0.68 |
| 8 | 17301152 | rs12546964 | 0.888 | -0.065 | -0.102 | 0.292 | -0.001 | 0.007 | 0.007 | 0.065 | 0.64 | 0.39 | 0.61 | 0.54 | 0.64 | 0.64 |
| 8 | 17302931 | rs11203858 | 0.908 | -0.08 | 0.173 | 0.297 | 0 | -0.001 | -0.001 | 0 | 0.67 | 0.36 | 0.61 | 0.54 | 0.68 | 0.65 |
| 8 | 17303403 | rs1557591 | -1.632 | -0.156 | 0.212 | -0.113 | 0.005 | 0.006 | 0.006 | 0.019 | 0.32 | 0.61 | 0.17 | 0.3 | 0.27 | 0.29 |
| 8 | 17303717 | rs17632013 | 0.708 | -0.192 | 0.167 | -0.104 | 0.014 | 0.007 | 0.007 | 0 | 0.11 | 0.08 | 0.05 | 0.09 | 0.08 | 0.12 |
| 8 | 17309151 | rs973986 | -0.966 | -0.137 | 0.567 | -0.109 | -0.016 | 0.02 | 0.02 | 0.02 | 0.15 | 0.27 | 0.08 | 0.22 | 0.13 | 0.18 |
| 8 | 17312407 | rs1012640 | -0.941 | -0.228 | 0.585 | -0.105 | -0.022 | 0.01 | 0.01 | 0.136 | 0.15 | 0.23 | 0.07 | 0.21 | 0.13 | 0.2 |
| 8 | 17314613 | rs4554505 | 2.002 | -0.278 | 0.225 | -0.024 | -0.122 | -0.104 | -0.104 | 0.741 | 0.48 | 0.04 | 0.57 | 0.47 | 0.55 | 0.49 |
| 8 | 17319881 | rs2517102 | -0.274 | -0.308 | 0.562 | 0.04 | -0.01 | 0.055 | 0.055 | 0 | 0.12 | 0.3 | 0.07 | 0.2 | 0.11 | 0.15 |
| 8 | 17320152 | rs12543516 | 1.395 | -0.428 | 0.588 | -0.011 | 0.025 | 0.009 | 0.009 | 1.704 | 0.72 | 0.24 | 0.86 | 0.71 | 0.82 | 0.67 |
| 8 | 17322180 | rs2705044 | 0.49 | -0.333 | 0.629 | -0.033 | -0.019 | -0.059 | -0.059 | 0.42 | 0.82 | 0.49 | 0.91 | 0.91 | 0.87 | 0.84 |
| 8 | 17322899 | rs17632120 | 1.009 | -0.324 | 0.608 | -0.01 | 0.012 | 0.004 | 0.004 | 0 | 0.09 | 0.07 | 0.03 | 0.09 | 0.05 | 0.12 |
| 8 | 17325343 | rs2517105 | -1.913 | -0.309 | 0.605 | -0.011 | 0.008 | -0.026 | -0.026 | 0 | 0.24 | 0.35 | 0.14 | 0.23 | 0.26 | 0.25 |
| 8 | 17327208 | rs17632167 | 0.936 | -0.303 | 0.408 | -0.06 | -0.045 | -0.055 | -0.055 | 0.439 | 0.16 | 0.06 | 0.26 | 0.22 | 0.19 | 0.15 |
| 8 | 17328731 | rs2517111 | -1.074 | -0.252 | 0.461 | -0.063 | -0.029 | 0.006 | 0.006 | 0 | 0.18 | 0.35 | 0.1 | 0.22 | 0.17 | 0.22 |
| 8 | 17330523 | rs2588240 | -0.837 | -0.357 | 0.541 | -0.112 | -0.001 | -0.002 | -0.002 | 0 | 0.34 | 0.27 | 0.33 | 0.42 | 0.33 | 0.32 |
| 8 | 17331692 | rs2517120 | -1.968 | -0.353 | 0.307 | -0.116 | 0.013 | 0.009 | 0.009 | 0 | 0.49 | 0.57 | 0.42 | 0.53 | 0.45 | 0.48 |
| 8 | 17332465 | rs10448129 | 0.875 | -0.192 | 0.411 | -0.093 | -0.048 | -0.06 | -0.06 | 0 | 0.17 | 0.06 | 0.26 | 0.22 | 0.2 | 0.15 |
| 8 | 17333462 | rs2517124 | 0.75 | -0.431 | 0.597 | -0.126 | 0.04 | -0.07 | -0.07 | 0.02 | 0.88 | 0.61 | 0.93 | 1 | 0.9 | 0.91 |
| 8 | 17334887 | rs11985222 | -0.52 | -0.363 | 0.052 | 0.065 | -0.031 | -0.007 | -0.007 | 0 | 0.11 | 0.19 | 0.05 | 0.1 | 0.07 | 0.1 |
| 8 | 17341027 | rs2720524 | -0.4 | -0.176 | -0.261 | 0.132 | 0.012 | 0.013 | 0.013 | 0 | 0.08 | 0.15 | 0.04 | 0.11 | 0.06 | 0.09 |
| 8 | 17344517 | rs1007691 | 1.42 | -0.283 | -1.239 | 0.055 | -0.105 | -0.118 | -0.118 | 3.594 | 0.39 | 0.06 | 0.52 | 0.47 | 0.46 | 0.44 |
| 8 | 17357290 | rs2517217 | -0.053 | -0.709 | -0.697 | -0.402 | -0.044 | 0.002 | 0.002 | 0 | 0.79 | 0.7 | 0.92 | 0.74 | 0.88 | 0.71 |
| 8 | 17357438 | rs4921552 | 0.485 | -0.599 | -1.139 | -0.374 | -0.1 | 0.016 | 0.016 | 0 | 0.25 | 0.04 | 0.3 | 0.11 | 0.31 | 0.15 |
| 8 | 17359171 | rs11203867 | 0.743 | -0.443 | -0.956 | -0.444 | 0.011 | 0.013 | 0.013 | 0 | 0.64 | 0.39 | 0.8 | 0.75 | 0.72 | 0.62 |
| 8 | 17360318 | rs2720540 | -0.734 | 0.444 | -0.745 | -0.12 | 0.012 | -0.075 | -0.075 | 0 | 0.18 | 0.24 | 0.13 | 0.04 | 0.19 | 0.14 |
| 8 | 17363239 | rs17632417 | 0.559 | 0.628 | -1.741 | 0.284 | -0.022 | -0.023 | -0.023 | 0.613 | 0.36 | 0.22 | 0.48 | 0.6 | 0.38 | 0.46 |
| 8 | 17369376 | rs11203868 | 0.334 | -0.115 | -1.239 | 0.577 | 0.046 | 0.01 | 0.01 | 0.04 | 0.54 | 0.24 | 0.4 | 0.48 | 0.63 | 0.49 |
| 8 | 17371287 | rs17124738 | -0.763 | -0.545 | -0.265 | 1.082 | 0 | 0.196 | 0.196 | 0 | 0.1 | 0.41 | 0.15 | 0.24 | 0.02 | 0.33 |
| 8 | 17372560 | rs17124742 | -0.644 | -0.95 | -0.007 | 0.829 | 0 | -0.001 | -0.001 | 0 | 0.19 | 0.18 | 0.15 | 0.19 | 0.08 | 0.25 |
| 8 | 17374960 | rs2188020 | 0.169 | -0.955 | 0.352 | 0.876 | -0.006 | 0.047 | 0.047 | 0 | 0.76 | 0.57 | 0.59 | 0.68 | 0.72 | 0.75 |
| 8 | 17375959 | rs2517237 | -0.408 | -1.006 | 0.488 | 1.05 | 0.006 | 0.016 | 0.016 | 0.013 | 0.84 | 0.71 | 0.64 | 0.84 | 0.8 | 0.88 |
| 8 | 17377242 | rs2188021 | 1.371 | -1.038 | 0.51 | 1.067 | 0.019 | -0.005 | -0.005 | 3.991 | 0.58 | 0.62 | 0.45 | 0.48 | 0.65 | 0.49 |

| | | | | | | | | | | | | | | | | |
|---|---|---|---|---|---|---|---|---|---|---|---|---|---|---|---|---|
| 8 | 17380076 | rs10110252 | 1.159 | -1.056 | 0.183 | 1.119 | 0.013 | 0.019 | 0.019 | 1.53 | 0.58 | 0.31 | 0.39 | 0.35 | 0.59 | 0.35 |
| 8 | 17383503 | rs17124760 | -0.261 | -1.044 | 0.044 | 1.158 | -0.021 | 0.091 | 0.091 | 0 | 0.15 | 0.3 | 0.22 | 0.33 | 0.11 | 0.25 |
| 8 | 17383689 | rs2720493 | 1.064 | -0.99 | 0.008 | 1.126 | -0.01 | -0.014 | -0.014 | 0.56 | 0.59 | 0.45 | 0.4 | 0.35 | 0.6 | 0.35 |
| 8 | 17384262 | rs2588209 | 0.199 | -0.581 | -0.219 | 1.121 | -0.006 | -0.013 | -0.013 | 0.435 | 0.76 | 0.81 | 0.6 | 0.71 | 0.7 | 0.7 |
| 8 | 17386831 | rs2705084 | 0.652 | -0.313 | -0.412 | 0.796 | 0.005 | 0.003 | 0.003 | 0 | 0.47 | 0.38 | 0.36 | 0.5 | 0.56 | 0.43 |
| 8 | 17390063 | rs2517244 | 0.622 | -1.116 | -0.906 | 0.631 | 0.08 | 0.073 | 0.073 | 0 | 0.53 | 0.26 | 0.63 | 0.27 | 0.63 | 0.32 |
| 8 | 17390525 | rs2588228 | 0.593 | -1.358 | 0.141 | 1.351 | -0.006 | 0 | 0 | 0 | 0.6 | 0.41 | 0.68 | 0.51 | 0.7 | 0.44 |
| 8 | 17390689 | rs2517245 | 0.211 | -1.882 | 0.067 | 1.419 | -0.086 | -0.041 | -0.041 | 0 | 0.79 | 0.55 | 0.87 | 0.83 | 0.92 | 0.75 |
| 8 | 17391277 | rs2720498 | 1.209 | -1.892 | 0.091 | 1.586 | -0.02 | -0.005 | -0.005 | 0 | 0.61 | 0.44 | 0.72 | 0.6 | 0.71 | 0.52 |
| 8 | 17391913 | rs2588232 | 1.173 | -1.611 | 0.695 | 1.404 | -0.009 | -0.005 | -0.005 | 0 | 0.59 | 0.51 | 0.68 | 0.6 | 0.65 | 0.5 |
| 8 | 17392357 | rs2720499 | 0.869 | -1.71 | 1.378 | 1.414 | 0.012 | 0.124 | 0.124 | 1.15 | 0.86 | 0.62 | 0.93 | 0.73 | 0.9 | 0.82 |
| 8 | 17392603 | rs2588233 | 2.339 | -2.029 | 0.658 | 0.981 | 0.034 | 0.097 | 0.097 | 0 | 0.46 | 0.18 | 0.57 | 0.23 | 0.56 | 0.29 |
| 8 | 17396224 | rs11993109 | -1.096 | -2.379 | 0.407 | 0.504 | 0.02 | -0.033 | -0.033 | 1.914 | 0.14 | 0.04 | 0.1 | 0.2 | 0.09 | 0.2 |
| 8 | 17397822 | rs2705017 | 0.735 | -2.335 | 0.804 | 0.22 | 0 | -0.007 | -0.007 | 0 | 0.84 | 0.77 | 0.83 | 0.91 | 0.77 | 0.88 |
| 8 | 17399149 | rs2517250 | 0.474 | -2.329 | 0.668 | 0.129 | -0.006 | 0.001 | 0.001 | 0 | 0.86 | 0.88 | 0.89 | 1 | 0.79 | 0.94 |
| 8 | 17399762 | rs2237817 | 2.362 | -2.542 | 0.753 | 0.145 | 0.07 | 0.05 | 0.05 | 0 | 0.52 | 0.27 | 0.64 | 0.57 | 0.65 | 0.43 |
| 8 | 17401878 | rs2517252 | 0.978 | -2.305 | 0.962 | 0.118 | -0.031 | -0.11 | -0.11 | 0 | 0.72 | 0.57 | 0.8 | 0.89 | 0.77 | 0.71 |
| 8 | 17405336 | rs2015840 | -0.361 | -2.183 | 0.585 | 0.173 | -0.002 | 0.001 | 0.001 | 0 | 0.13 | 0.11 | 0.15 | 0.1 | 0.2 | 0.15 |
| 8 | 17406183 | rs737039 | 1.616 | -2.011 | 0.433 | 0.348 | 0.014 | 0.012 | 0.012 | 0 | 0.58 | 0.35 | 0.66 | 0.59 | 0.65 | 0.45 |
| 8 | 17412333 | rs2301539 | -1.197 | -1.356 | 0.019 | 1.065 | -0.008 | 0.031 | 0.031 | 0 | 0.13 | 0.2 | 0.14 | 0.32 | 0.1 | 0.25 |
| 8 | 17414450 | rs2720588 | -0.616 | -0.986 | -0.35 | 0.952 | 0.008 | -0.012 | -0.012 | 0 | 0.33 | 0.2 | 0.25 | 0.37 | 0.31 | 0.3 |
| 8 | 17416478 | rs2248010 | -0.589 | -0.982 | -0.562 | 0.926 | 0.012 | -0.012 | -0.012 | 1.276 | 0.33 | 0.16 | 0.24 | 0.32 | 0.3 | 0.29 |
| 8 | 17417329 | rs2588252 | -0.865 | -0.377 | -0.662 | 0.577 | -0.092 | -0.011 | -0.011 | 0 | 0.7 | 0.87 | 0.45 | 0.75 | 0.5 | 0.71 |
| 8 | 17418671 | rs3780121 | -1.544 | -0.286 | 0.665 | 0.521 | 0.045 | 0.035 | 0.035 | 0 | 0.18 | 0.42 | 0.21 | 0.27 | 0.21 | 0.18 |
| 8 | 17422717 | rs7838611 | -1.198 | -0.659 | 0.639 | 0.033 | -0.062 | -0.066 | -0.066 | 0 | 0.14 | 0.38 | 0.13 | 0.05 | 0.02 | 0.03 |
| 8 | 17423065 | rs7842456 | -1.198 | -0.659 | 0.639 | 0.033 | -0.062 | -0.066 | -0.066 | 0 | 0.14 | 0.38 | 0.13 | 0.05 | 0.02 | 0.03 |
| 8 | 17423513 | rs6586610 | -1.36 | -0.365 | 0.46 | 0.103 | 0 | -0.029 | -0.029 | 0 | 0.17 | 0.34 | 0.19 | 0.09 | 0.11 | 0.1 |
| 8 | 17433755 | rs2237819 | -0.565 | -0.902 | 0.276 | -0.226 | -0.044 | -0.061 | -0.061 | 0 | 0.28 | 0.37 | 0.11 | 0.25 | 0.18 | 0.37 |
| 8 | 17434339 | rs2517262 | 0.754 | -0.766 | 0.314 | -0.038 | -0.024 | 0.003 | 0.003 | 0.067 | 0.77 | 0.46 | 0.71 | 0.8 | 0.79 | 0.6 |
| 8 | 17438439 | rs9792282 | 0.01 | -1.059 | 0 | -0.253 | 0.019 | 0.033 | 0.033 | 0.037 | 0.17 | 0.08 | 0.06 | 0.13 | 0.12 | 0.15 |
| 8 | 17438490 | rs9792165 | -1.328 | -1.191 | 0.274 | -0.38 | 0.025 | 0.032 | 0.032 | 0.098 | 0.38 | 0.33 | 0.23 | 0.22 | 0.26 | 0.25 |
| 8 | 17439187 | rs2517273 | 0.179 | -1.178 | -0.323 | -0.718 | -0.003 | -0.012 | -0.012 | 0 | 0.64 | 0.51 | 0.77 | 0.76 | 0.7 | 0.52 |
| 8 | 17439560 | rs2517275 | 1.789 | -0.935 | -0.747 | -0.643 | -0.057 | -0.062 | -0.062 | 0 | 0.34 | 0.15 | 0.53 | 0.54 | 0.45 | 0.31 |
| 8 | 17442136 | rs2720565 | 0.768 | -1.39 | -0.358 | -0.58 | 0.052 | 0.025 | 0.025 | 0 | 0.64 | 0.29 | 0.73 | 0.71 | 0.78 | 0.54 |
| 8 | 17444642 | rs3850741 | -0.992 | -1.531 | 0.114 | -0.864 | -0.001 | -0.03 | -0.03 | 0.003 | 0.47 | 0.43 | 0.22 | 0.18 | 0.33 | 0.49 |
| 8 | 17446444 | rs2588156 | 1.056 | -1.472 | 0.258 | -0.676 | 0.038 | 0.123 | 0.123 | 3.203 | 0.57 | 0.25 | 0.66 | 0.87 | 0.7 | 0.61 |
| 8 | 17448800 | rs2720582 | -0.772 | -1.139 | 0.69 | 0.361 | 0.006 | 0.049 | 0.049 | 0 | 0.29 | 0.46 | 0.34 | 0.41 | 0.27 | 0.3 |
| 8 | 17449657 | rs2588153 | 0.818 | -0.824 | 0.196 | 0.025 | 0.122 | 0.186 | 0.186 | 2.253 | 0.94 | 0.47 | 0.88 | 0.85 | 0.95 | 0.81 |
| 8 | 17449710 | rs2588152 | 0.627 | -1.015 | -0.047 | -0.134 | -0.022 | -0.022 | -0.022 | 0.695 | 0.82 | 0.44 | 0.77 | 0.85 | 0.91 | 0.77 |
| 8 | 17450013 | rs2588150 | 0.969 | -0.92 | -0.93 | -0.064 | 0.051 | 0.079 | 0.079 | 0 | 0.52 | 0.21 | 0.63 | 0.6 | 0.71 | 0.5 |
| 8 | 17454300 | rs2246488 | 0.9 | -0.432 | -1.015 | 0.121 | -0.044 | -0.013 | -0.013 | 0 | 0.54 | 0.15 | 0.56 | 0.52 | 0.59 | 0.4 |
| 8 | 17455169 | rs2246376 | -1.255 | -0.671 | 0.175 | 0.232 | -0.001 | 0.07 | 0.07 | 3.703 | 0.08 | 0.51 | 0.13 | 0.11 | 0.01 | 0.2 |
| 8 | 17455860 | rs2588139 | -1.597 | -0.362 | -0.078 | 0.135 | 0.002 | 0.021 | 0.021 | 1.579 | 0.3 | 0.53 | 0.32 | 0.39 | 0.29 | 0.49 |
| 8 | 17456739 | rs2588137 | 1.218 | -0.418 | 0.952 | -0.046 | 0.036 | 0.021 | 0.021 | 0.092 | 0.87 | 0.74 | 0.86 | 0.87 | 0.85 | 0.71 |
| 8 | 17457420 | rs2517168 | 2.304 | -1.248 | -0.136 | -0.537 | -0.11 | -0.039 | -0.039 | 0.838 | 0.39 | 0.08 | 0.55 | 0.57 | 0.55 | 0.34 |
| 8 | 17458727 | rs7007653 | -0.125 | -0.945 | 1.147 | -0.3 | 0.01 | -0.091 | -0.091 | 0 | 0.18 | 0.35 | 0.09 | 0.02 | 0.15 | 0.11 |
| 8 | 17460245 | rs2517177 | 2.021 | -1.168 | 0.864 | -0.342 | 0.002 | 0.018 | 0.018 | 0 | 0.56 | 0.21 | 0.62 | 0.57 | 0.64 | 0.48 |
| 8 | 17461421 | rs13256095 | 0.153 | -0.804 | 0.912 | -0.144 | -0.012 | -0.01 | -0.01 | 0 | 0.06 | 0.04 | 0.09 | 0.06 | 0.12 | 0.12 |
| 8 | 17465862 | rs2517199 | 1.746 | -0.964 | 1.314 | -0.216 | -0.003 | 0 | 0 | 0 | 0.61 | 0.37 | 0.64 | 0.57 | 0.67 | 0.48 |
| 8 | 17466212 | rs2427715 | 0.669 | -1.003 | 1.654 | -0.23 | -0.015 | -0.072 | -0.072 | 0.179 | 0.73 | 0.47 | 0.81 | 0.83 | 0.81 | 0.82 |
| 8 | 17467347 | rs3862079 | -1.375 | -0.895 | 1.499 | -0.213 | -0.017 | -0.019 | -0.019 | 0 | 0.13 | 0.08 | 0.15 | 0.21 | 0.15 | 0.26 |
| 8 | 17468605 | rs17632963 | -1.989 | -0.898 | 1.706 | -0.206 | -0.003 | -0.005 | -0.005 | 0.02 | 0.17 | 0.15 | 0.24 | 0.27 | 0.16 | 0.37 |
| 8 | 17469167 | rs13249125 | -1.184 | -0.85 | 1.579 | -0.197 | -0.016 | -0.019 | -0.019 | 0 | 0.12 | 0.08 | 0.15 | 0.21 | 0.15 | 0.25 |
| 8 | 17469788 | rs2517211 | 0.336 | -0.962 | 1.735 | -0.333 | 0.034 | -0.094 | -0.094 | 0.412 | 0.84 | 0.66 | 0.9 | 0.98 | 0.85 | 0.88 |
| 8 | 17470286 | rs2588115 | 1.27 | -1.021 | 1.446 | -0.26 | -0.002 | 0.004 | 0.004 | 0.001 | 0.77 | 0.78 | 0.82 | 0.72 | 0.83 | 0.6 |
| 8 | 17470693 | rs3862082 | 2.4 | -0.833 | 0.979 | -0.156 | -0.039 | 0.029 | 0.029 | 0 | 0.54 | 0.14 | 0.55 | 0.58 | 0.61 | 0.44 |
| 8 | 17471838 | rs17124938 | -1.01 | -0.391 | 1.35 | 0.003 | -0.001 | 0.002 | 0.002 | 0 | 0.11 | 0.16 | 0.1 | 0.09 | 0.08 | 0.27 |
| 8 | 17474391 | rs2237825 | 2.344 | -0.334 | 1.468 | 0.192 | -0.005 | -0.002 | -0.002 | 0.216 | 0.59 | 0.25 | 0.5 | 0.57 | 0.66 | 0.46 |
| 8 | 17474915 | rs2237826 | 2.411 | -0.363 | 1.513 | 0.213 | -0.003 | -0.003 | -0.003 | 0 | 0.59 | 0.28 | 0.5 | 0.56 | 0.65 | 0.44 |
| 8 | 17475155 | rs2237830 | 1.61 | -0.366 | 2.114 | 0.217 | 0.011 | -0.003 | -0.003 | 0 | 0.7 | 0.6 | 0.6 | 0.67 | 0.71 | 0.64 |
| 8 | 17475999 | rs2283105 | 2.024 | -0.378 | 2.077 | 0.217 | 0.002 | 0 | 0 | 0 | 0.65 | 0.37 | 0.52 | 0.62 | 0.66 | 0.56 |
| 8 | 17476704 | rs2237831 | 0.689 | -0.346 | 2.232 | 0.074 | 0 | 0.006 | 0.006 | 0 | 0.82 | 0.73 | 0.68 | 0.76 | 0.83 | 0.69 |
| 8 | 17478086 | rs2588108 | -2.33 | -0.291 | 2.3 | 0.107 | 0.001 | -0.004 | -0.004 | 0 | 0.33 | 0.4 | 0.4 | 0.3 | 0.29 | 0.44 |
| 8 | 17479066 | rs2237833 | -2.653 | -0.282 | 2.258 | 0.075 | 0.004 | -0.003 | -0.003 | 0.012 | 0.37 | 0.6 | 0.48 | 0.37 | 0.35 | 0.46 |
| 8 | 17480971 | rs2588104 | 0.695 | -0.259 | 2.327 | 0.126 | 0.072 | 0.057 | 0.057 | 0.2 | 0.95 | 0.62 | 0.97 | 1 | 0.99 | 0.99 |
| 8 | 17483028 | rs13259157 | 1.902 | -0.265 | 2.231 | 0.143 | 0.004 | 0.039 | 0.039 | 0.07 | 0.83 | 0.36 | 0.62 | 0.74 | 0.85 | 0.69 |
| 8 | 17483118 | rs11993165 | -0.964 | -0.245 | 2.205 | 0.108 | 0.02 | 0.026 | 0.026 | 0 | 0.11 | 0.41 | 0.1 | 0.06 | 0.06 | 0.14 |
| 8 | 17484137 | rs2237836 | 2.336 | -0.553 | 1.059 | 0.397 | -0.089 | -0.014 | -0.014 | 0 | 0.37 | 0.08 | 0.33 | 0.35 | 0.51 | 0.29 |
| 8 | 17487184 | rs2237839 | 0.941 | -0.515 | 2.174 | 0.693 | -0.012 | -0.053 | -0.053 | 0 | 0.8 | 0.64 | 0.84 | 0.93 | 0.86 | 0.84 |

| | | | | | | | | | | | | | | | | |
|---|---|---|---|---|---|---|---|---|---|---|---|---|---|---|---|---|
| 8 | 17488610 | rs2237842 | -1.441 | -0.573 | 1.854 | 0.794 | -0.004 | 0.043 | 0.043 | 0 | 0.13 | 0.22 | 0.24 | 0.26 | 0.12 | 0.26 |
| 8 | 17488845 | rs2237843 | 2.082 | -0.505 | 0.831 | 0.743 | 0.06 | -0.013 | -0.013 | 0 | 0.65 | 0.16 | 0.58 | 0.63 | 0.8 | 0.5 |
| 8 | 17491144 | rs2237846 | 2.052 | -0.502 | 0.856 | 0.69 | 0.046 | -0.002 | -0.002 | 0 | 0.66 | 0.19 | 0.59 | 0.63 | 0.8 | 0.57 |
| 8 | 17491730 | rs7006613 | -1.867 | -0.552 | 0.765 | 0.732 | 0.01 | -0.002 | -0.002 | 0 | 0.3 | 0.75 | 0.39 | 0.35 | 0.14 | 0.39 |
| 8 | 17497874 | rs4921797 | 0.58 | -0.12 | 1.122 | 0.902 | -0.018 | 0.065 | 0.065 | 0 | 0.84 | 0.63 | 0.72 | 0.74 | 0.91 | 0.68 |
| 8 | 17500223 | rs4705 | 1.678 | 0.08 | 0.034 | 0.869 | -0.023 | -0.047 | -0.047 | 0 | 0.6 | 0.2 | 0.41 | 0.46 | 0.58 | 0.45 |
| 8 | 17503125 | rs3203733 | 1.665 | -0.075 | 0.35 | 0.731 | -0.001 | 0.058 | 0.058 | 0 | 0.59 | 0.22 | 0.51 | 0.7 | 0.62 | 0.52 |
| 8 | 17507964 | rs4921799 | 1.15 | 0.323 | 1.673 | 0.905 | 0.04 | -0.04 | -0.04 | 0 | 0.79 | 0.61 | 0.79 | 0.82 | 0.78 | 0.78 |
| 8 | 17509722 | rs2410493 | 0.329 | 0.486 | 1.951 | 0.783 | 0.073 | -0.042 | -0.042 | 0 | 0.91 | 0.82 | 0.95 | 0.99 | 0.86 | 0.94 |
| 8 | 17513414 | rs12224 | 0.192 | 0.301 | 1.591 | 0.843 | 0 | -0.012 | -0.012 | 0 | 0.09 | 0.13 | 0.05 | 0.02 | 0.07 | 0.09 |
| 8 | 17518957 | rs10094030 | -0.868 | 0.171 | 1.942 | 1.218 | 0.016 | 0.272 | 0.272 | 0 | 0.06 | 0.24 | 0.15 | 0.25 | 0.05 | 0.18 |
| 8 | 17520999 | rs6586619 | 1.025 | 0.255 | 2.132 | 1.165 | 0.049 | -0.003 | -0.003 | 0 | 0.85 | 0.73 | 0.79 | 0.83 | 0.79 | 0.73 |
| 8 | 17523646 | rs4388484 | 1.009 | 0.27 | 2.163 | 0.866 | 0.089 | -0.071 | -0.071 | 0 | 0.85 | 0.61 | 0.85 | 0.97 | 0.8 | 0.86 |
| 8 | 17528777 | rs3862089 | 1.081 | 0.184 | 2.071 | 0.986 | 0.06 | -0.006 | -0.006 | 0 | 0.84 | 0.67 | 0.76 | 0.86 | 0.82 | 0.69 |
| 8 | 17535483 | rs7010721 | -0.44 | 0.254 | 1.547 | 0.932 | 0 | -0.001 | -0.001 | 0 | 0.13 | 0.18 | 0.13 | 0.15 | 0.11 | 0.21 |
| 8 | 17535659 | rs12114081 | -0.404 | 0.177 | 1.532 | 0.939 | -0.023 | 0.004 | 0.004 | 0 | 0.07 | 0.13 | 0.12 | 0.1 | 0.01 | 0.19 |
| 8 | 17536347 | rs7462063 | 1.666 | 0.31 | 1.607 | 1.113 | 0.012 | 0.023 | 0.023 | 0.17 | 0.71 | 0.39 | 0.59 | 0.56 | 0.69 | 0.49 |
| 8 | 17540324 | rs7464791 | -0.037 | 0.367 | 0.857 | 0.457 | 0.029 | 0.022 | 0.022 | 0 | 0.71 | 0.53 | 0.62 | 0.71 | 0.65 | 0.68 |
| 8 | 17541999 | rs2979792 | 1.553 | 0.019 | 0.059 | -0.049 | -0.084 | -0.067 | -0.067 | 0 | 0.37 | 0.07 | 0.28 | 0.37 | 0.41 | 0.19 |
| 8 | 17542215 | rs6421404 | -0.578 | 0.099 | 0.433 | 0.106 | 0.025 | -0.108 | -0.108 | 0 | 0.71 | 0.54 | 0.73 | 0.96 | 0.66 | 0.84 |
| 8 | 17542548 | rs7000487 | 0.584 | 0.088 | 0.2 | 0.178 | -0.012 | -0.005 | -0.005 | 0 | 0.55 | 0.34 | 0.48 | 0.55 | 0.51 | 0.47 |
| 8 | 17549493 | rs182363 | 0.305 | 0.306 | 1.127 | 0.503 | 0.086 | -0.047 | -0.047 | 0 | 0.91 | 0.73 | 0.94 | 0.99 | 0.87 | 0.94 |
| 8 | 17550623 | rs4921804 | 2.036 | -0.072 | 0.569 | 0.292 | -0.06 | 0.149 | 0.149 | 3.721 | 0.35 | 0.1 | 0.3 | 0.04 | 0.42 | 0.14 |
| 8 | 17550770 | rs186022 | -0.374 | -0.199 | 0.72 | 0.745 | 0.073 | -0.043 | -0.043 | 0 | 0.38 | 0.15 | 0.42 | 0.64 | 0.31 | 0.52 |
| 8 | 17550787 | rs4921805 | 1.925 | -0.172 | 0.847 | 0.592 | -0.014 | 0.036 | 0.036 | 2.615 | 0.37 | 0.28 | 0.32 | 0.04 | 0.42 | 0.14 |
| 8 | 17550854 | rs209973 | -0.43 | -0.335 | 0.673 | 0.91 | 0.053 | -0.062 | -0.062 | 0 | 0.35 | 0.14 | 0.32 | 0.53 | 0.3 | 0.34 |
| 8 | 17555422 | rs2979790 | 0.521 | 0.4 | 1.87 | 1.724 | 0.025 | 0.07 | 0.07 | 0.097 | 0.85 | 0.74 | 0.91 | 0.72 | 0.85 | 0.86 |
| 8 | 17555736 | rs454100 | 0.063 | 0.136 | 1.96 | 1.879 | -0.002 | 0.002 | 0.002 | 0 | 0.25 | 0.24 | 0.34 | 0.46 | 0.2 | 0.38 |
| 8 | 17555952 | rs415088 | 0.428 | 0.146 | 1.87 | 1.996 | 0.032 | -0.032 | -0.032 | 0.049 | 0.22 | 0.09 | 0.24 | 0.31 | 0.17 | 0.19 |
| 8 | 17556514 | rs17506026 | 2.431 | 0.326 | 1.81 | 1.826 | -0.073 | 0.2 | 0.2 | 2.77 | 0.38 | 0.09 | 0.31 | 0.06 | 0.46 | 0.14 |
| 8 | 17558519 | rs374386 | 1.072 | -0.257 | 2.315 | 2.274 | -0.009 | 0.012 | 0.012 | 0 | 0.64 | 0.58 | 0.67 | 0.4 | 0.68 | 0.66 |
| 8 | 17560234 | rs371997 | 1.386 | -0.261 | 2.336 | 2.265 | -0.003 | 0.002 | 0.002 | 0 | 0.6 | 0.59 | 0.64 | 0.4 | 0.63 | 0.64 |
| 8 | 17563062 | rs10503603 | -0.841 | -0.261 | 2.088 | 2.292 | -0.001 | -0.003 | -0.003 | 0.331 | 0.06 | 0.04 | 0.12 | 0.17 | 0.09 | 0.21 |
| 8 | 17563762 | rs4504675 | 1.726 | -0.26 | 2.266 | 2.3 | 0.021 | 0.074 | 0.074 | 0.914 | 0.54 | 0.25 | 0.52 | 0.25 | 0.58 | 0.45 |
| 8 | 17564973 | rs4921808 | -1.476 | -0.256 | 2.286 | 2.385 | 0.002 | 0 | 0 | 0.032 | 0.4 | 0.65 | 0.31 | 0.39 | 0.37 | 0.26 |
| 8 | 17565118 | rs7386698 | 0.906 | -0.2 | 1.961 | 2.355 | -0.003 | 0.019 | 0.019 | 0 | 0.94 | 0.92 | 0.9 | 0.8 | 0.93 | 0.87 |
| 8 | 17565556 | rs380129 | -1.432 | -0.135 | 1.739 | 2.306 | -0.004 | 0.007 | 0.007 | 0.02 | 0.38 | 0.45 | 0.37 | 0.55 | 0.34 | 0.41 |
| 8 | 17569104 | rs12678396 | -1.309 | 0.008 | 1.103 | 2.366 | 0.001 | 0.061 | 0.061 | 0 | 0.07 | 0.19 | 0.12 | 0.16 | 0.09 | 0.18 |
| 8 | 17569838 | rs376600 | -0.06 | 0.075 | 0.98 | 1.934 | -0.008 | -0.011 | -0.011 | 0 | 0.22 | 0.22 | 0.18 | 0.11 | 0.15 | 0.08 |
| 8 | 17573107 | rs2937891 | -1.835 | 0.897 | 1.109 | 2.559 | 0.019 | 0.055 | 0.055 | 0.019 | 0.15 | 0.26 | 0.09 | 0.28 | 0.15 | 0.15 |
| 8 | 17578652 | rs2979774 | 1.784 | 0.906 | 0.715 | 2.255 | 0 | 0.1 | 0.1 | 0.168 | 0.77 | 0.58 | 0.79 | 0.49 | 0.81 | 0.71 |
| 8 | 17580791 | rs2937886 | -1.55 | 0.388 | 0.112 | 2.386 | 0.004 | 0.016 | 0.016 | 0 | 0.12 | 0.12 | 0.14 | 0.49 | 0.04 | 0.22 |
| 8 | 17581700 | rs2904724 | -1.779 | 1.274 | 0.586 | 1.915 | 0.001 | 0.005 | 0.005 | 0.011 | 0.13 | 0.14 | 0.11 | 0.24 | 0.18 | 0.11 |
| 8 | 17582137 | rs2979775 | 1.039 | 1.006 | 0.32 | 1.034 | 0.013 | 0.022 | 0.022 | 1.346 | 0.59 | 0.35 | 0.54 | 0.19 | 0.68 | 0.29 |
| 8 | 17583151 | rs6586637 | 0.64 | 1.541 | 0.409 | 1.729 | -0.009 | -0.006 | -0.006 | 0 | 0.81 | 0.81 | 0.72 | 0.48 | 0.74 | 0.57 |
| 8 | 17585129 | rs7842088 | 1.161 | 1.622 | 0.295 | 1.589 | -0.038 | -0.062 | -0.062 | 0.436 | 0.77 | 0.84 | 0.7 | 0.47 | 0.68 | 0.49 |
| 8 | 17586321 | rs2979776 | 1.697 | 1.759 | 0.945 | 1.303 | 0.011 | -0.003 | -0.003 | 0 | 0.61 | 0.41 | 0.51 | 0.35 | 0.54 | 0.36 |
| 8 | 17589960 | rs6586639 | 1.057 | 1.733 | 1.165 | 1.062 | 0.002 | 0 | 0 | 1.287 | 0.7 | 0.61 | 0.66 | 0.31 | 0.66 | 0.42 |
| 8 | 17599136 | rs2937885 | -0.758 | 1.413 | 1.079 | 0.904 | 0.003 | -0.029 | -0.029 | 0 | 0.19 | 0.21 | 0.19 | 0.07 | 0.2 | 0.14 |
| 8 | 17599574 | rs2979784 | -1.093 | 1.248 | 1.133 | 0.928 | -0.001 | -0.019 | -0.019 | 0.384 | 0.22 | 0.25 | 0.19 | 0.11 | 0.2 | 0.18 |
| 8 | 17600912 | rs2904748 | 1.36 | 1.019 | 0.965 | 1.131 | -0.002 | 0.004 | 0.004 | 0.08 | 0.55 | 0.35 | 0.51 | 0.27 | 0.52 | 0.34 |
| 8 | 17604586 | rs17690761 | -2.155 | 0.316 | 1.37 | 1.614 | -0.006 | 0.08 | 0.08 | 0 | 0.08 | 0.14 | 0.04 | 0.19 | 0.05 | 0.12 |
| 8 | 17607488 | rs3862101 | 2.403 | 0.2 | 1.67 | 1.775 | -0.024 | 0.058 | 0.058 | 2.122 | 0.81 | 0.35 | 0.83 | 0.63 | 0.86 | 0.64 |
| 8 | 17610093 | rs9325821 | 1.988 | 0.387 | 1.555 | 1.666 | 0.007 | 0.11 | 0.11 | 0 | 0.93 | 0.84 | 0.96 | 0.8 | 0.95 | 0.94 |
| 8 | 17611886 | rs2979788 | 2.122 | 0.134 | 1.744 | 1.827 | -0.021 | 0.089 | 0.089 | 0 | 0.81 | 0.37 | 0.82 | 0.59 | 0.85 | 0.85 |
| 8 | 17612639 | rs209568 | -0.885 | 0.28 | 0.876 | 2.124 | -0.003 | -0.088 | -0.088 | 0.007 | 0.24 | 0.07 | 0.2 | 0.45 | 0.22 | 0.57 |
| 8 | 17612846 | rs3739408 | -0.698 | 0.268 | 0.89 | 1.988 | 0.001 | -0.014 | -0.014 | 0 | 0.15 | 0.06 | 0.23 | 0.15 | 0.15 | 0.11 |
| 8 | 17612875 | rs3739407 | 1.971 | 0.148 | 1.672 | 1.945 | -0.017 | 0.076 | 0.076 | 0 | 0.76 | 0.56 | 0.7 | 0.68 | 0.8 | 0.86 |
| 8 | 17613606 | rs7387668 | 1.32 | 0.174 | 1.832 | 1.978 | -0.002 | 0.057 | 0.057 | 0 | 0.8 | 0.73 | 0.71 | 0.66 | 0.82 | 0.87 |
| 8 | 17616191 | rs17690879 | -0.823 | 0.112 | 1.843 | 1.699 | 0.001 | -0.003 | -0.003 | 1.519 | 0.16 | 0.1 | 0.24 | 0.15 | 0.15 | 0.11 |
| 8 | 17620890 | rs10097099 | 1.686 | 0.029 | 1.774 | 1.594 | -0.032 | -0.009 | -0.009 | 0 | 0.77 | 0.57 | 0.74 | 0.86 | 0.83 | 0.83 |
| 8 | 17620942 | rs451322 | 1.951 | -0.011 | 1.722 | 1.521 | 0 | -0.015 | -0.015 | 0 | 0.54 | 0.44 | 0.5 | 0.22 | 0.58 | 0.27 |
| 8 | 17630019 | rs6998733 | 2.191 | 0.073 | 1.727 | 1.459 | 0.018 | 0.08 | 0.08 | 0.137 | 0.5 | 0.33 | 0.47 | 0.16 | 0.51 | 0.18 |
| 8 | 17631418 | rs17125338 | -1.635 | 0.109 | 1.353 | 1.465 | -0.025 | 0.007 | 0.007 | 0 | 0.22 | 0.35 | 0.26 | 0.23 | 0.18 | 0.21 |
| 8 | 17631429 | rs1910110 | -0.296 | 0.18 | 1.196 | 1.63 | -0.008 | -0.054 | -0.054 | 0 | 0.19 | 0.09 | 0.18 | 0.45 | 0.2 | 0.53 |
| 8 | 17632737 | rs447516 | -1.352 | 0.183 | 1.188 | 1.637 | 0 | -0.038 | -0.038 | 1.768 | 0.22 | 0.15 | 0.32 | 0.39 | 0.26 | 0.23 |
| 8 | 17632943 | rs385586 | -0.65 | -0.055 | 1.184 | 1.764 | -0.001 | 0.028 | 0.028 | 0.018 | 0.22 | 0.24 | 0.19 | 0.45 | 0.22 | 0.58 |
| 8 | 17636728 | rs430238 | -0.385 | -0.379 | 0.757 | 1.912 | 0.002 | -0.117 | -0.117 | 0 | 0.19 | 0.03 | 0.18 | 0.47 | 0.15 | 0.46 |
| 8 | 17637674 | rs6986061 | 2.232 | -0.667 | 0.979 | 1.66 | 0.078 | 0.17 | 0.17 | 1.319 | 0.49 | 0.14 | 0.47 | 0.2 | 0.59 | 0.31 |

| | | | | | | | | | | | | | | | | |
|---|---|---|---|---|---|---|---|---|---|---|---|---|---|---|---|---|
| 8 | 17638034 | rs7460414 | -1.331 | -0.82 | 2.156 | 1.739 | -0.041 | 0.068 | 0.068 | 0 | 0.28 | 0.61 | 0.31 | 0.52 | 0.17 | 0.48 |
| 8 | 17638826 | rs414703 | -1.607 | -0.847 | 2.2 | 1.999 | -0.002 | 0 | 0 | 0 | 0.3 | 0.71 | 0.32 | 0.48 | 0.2 | 0.51 |
| 8 | 17643685 | rs10099690 | 0.959 | -0.709 | 2.899 | 2.034 | -0.057 | 0.047 | 0.047 | 0 | 0.73 | 0.49 | 0.78 | 0.57 | 0.84 | 0.55 |
| 8 | 17645028 | rs2073700 | -0.632 | -0.676 | 2.932 | 2.036 | 0.007 | -0.01 | -0.01 | 0 | 0.21 | 0.18 | 0.19 | 0.43 | 0.13 | 0.44 |
| 8 | 17648127 | rs2036425 | 0.668 | -0.681 | 2.95 | 2.074 | 0.01 | -0.016 | -0.016 | 0 | 0.77 | 0.82 | 0.8 | 0.56 | 0.85 | 0.55 |
| 8 | 17655406 | rs10098719 | 0.349 | -0.658 | 2.852 | 2.069 | 0.017 | -0.059 | -0.059 | 1.297 | 0.8 | 0.89 | 0.81 | 0.55 | 0.85 | 0.55 |
| 8 | 17666107 | rs4406440 | -2.716 | -0.639 | 2.929 | 1.802 | 0.002 | 0.055 | 0.055 | 0 | 0.05 | 0.18 | 0.12 | 0.06 | 0.04 | 0.06 |
| 8 | 17670845 | rs395737 | 1.334 | -0.665 | 2.806 | 1.84 | -0.021 | 0.06 | 0.06 | 0 | 0.71 | 0.46 | 0.62 | 0.45 | 0.78 | 0.43 |
| 8 | 17672670 | rs403930 | 0.113 | -0.7 | 2.196 | 1.859 | 0.012 | -0.055 | -0.055 | 0.097 | 0.81 | 0.9 | 0.78 | 0.51 | 0.85 | 0.57 |
| 8 | 17673299 | rs7816561 | 1.395 | -0.698 | 2.312 | 1.788 | -0.011 | 0.026 | 0.026 | 0.23 | 0.69 | 0.41 | 0.61 | 0.45 | 0.74 | 0.45 |
| 8 | 17674988 | rs429412 | 0.2 | -0.746 | 1.872 | 1.882 | 0.003 | -0.02 | -0.02 | 0 | 0.8 | 0.84 | 0.66 | 0.56 | 0.83 | 0.54 |
| 8 | 17681402 | rs414677 | 0.842 | -0.933 | 2.229 | 1.763 | -0.013 | 0.027 | 0.027 | 0 | 0.73 | 0.66 | 0.63 | 0.55 | 0.79 | 0.47 |
| 8 | 17691690 | rs3900397 | 1.163 | -1.168 | 2.68 | 1.727 | -0.017 | 0.037 | 0.037 | 0 | 0.69 | 0.39 | 0.48 | 0.44 | 0.75 | 0.46 |
| 8 | 17692414 | rs376247 | 0 | -1.165 | 2.719 | 1.758 | 0.005 | -0.058 | -0.058 | 0 | 0.8 | 0.9 | 0.66 | 0.55 | 0.83 | 0.55 |
| 8 | 17693964 | rs434532 | 0.607 | -1.165 | 2.723 | 1.758 | -0.021 | 0.06 | 0.06 | 0 | 0.77 | 0.64 | 0.63 | 0.55 | 0.83 | 0.55 |
| 8 | 17697376 | rs10503644 | 2.784 | -1.568 | 2.528 | 1.731 | 0.037 | 0.202 | 0.202 | 0.895 | 0.46 | 0.1 | 0.32 | 0.15 | 0.57 | 0.3 |
| 8 | 17698128 | rs379407 | -0.177 | -1.568 | 2.654 | 1.751 | 0.002 | -0.009 | -0.009 | 0.82 | 0.8 | 0.82 | 0.65 | 0.55 | 0.83 | 0.55 |
| 8 | 17698251 | rs687105 | -1.545 | -1.589 | 2.749 | 1.755 | 0.007 | 0.067 | 0.067 | 0.087 | 0.13 | 0.46 | 0.18 | 0.12 | 0.09 | 0.13 |
| 8 | 17701629 | rs10100795 | 1.714 | -1.594 | 2.503 | 1.693 | 0.061 | -0.038 | -0.038 | 0 | 0.59 | 0.14 | 0.44 | 0.43 | 0.7 | 0.36 |
| 8 | 17702161 | rs11778291 | 2.8 | -1.437 | 2.473 | 1.625 | 0.021 | 0.189 | 0.189 | 1.021 | 0.46 | 0.11 | 0.32 | 0.15 | 0.56 | 0.29 |
| 8 | 17719243 | rs7004717 | -0.064 | -1.285 | 2.7 | 1.618 | 0.006 | -0.028 | -0.028 | 0 | 0.8 | 0.84 | 0.56 | 0.5 | 0.83 | 0.56 |
| 8 | 17721720 | rs2517330 | -1.01 | -1.289 | 3.269 | 1.537 | -0.011 | -0.014 | -0.014 | 0 | 0.1 | 0.26 | 0.08 | 0.07 | 0.07 | 0.1 |
| 8 | 17724142 | rs2517287 | -1.186 | -1.252 | 3.297 | 1.588 | -0.013 | 0.041 | 0.041 | 0 | 0.11 | 0.29 | 0.17 | 0.13 | 0.07 | 0.11 |
| 8 | 17725423 | rs2653417 | 0.212 | -1.283 | 3.34 | 1.625 | 0.006 | -0.036 | -0.036 | 0.038 | 0.76 | 0.81 | 0.5 | 0.5 | 0.77 | 0.55 |
| 8 | 17725863 | rs2653416 | 0.246 | -1.282 | 3.325 | 1.625 | 0.006 | -0.035 | -0.035 | 0 | 0.75 | 0.81 | 0.5 | 0.5 | 0.77 | 0.55 |
| 8 | 17726805 | rs34259886 | -0.928 | -1.279 | 3.456 | 1.623 | -0.005 | -0.03 | -0.03 | 0 | 0.06 | 0.16 | 0.03 | 0 | 0.04 | 0.06 |
| 8 | 17728689 | rs2276473 | 1.223 | -1.312 | 3.533 | 1.615 | -0.027 | 0.036 | 0.036 | 0.004 | 0.66 | 0.46 | 0.45 | 0.5 | 0.73 | 0.49 |
| 8 | 17729478 | rs2653409 | 0.408 | -1.315 | 3.525 | 1.613 | -0.01 | 0.034 | 0.034 | 0 | 0.72 | 0.66 | 0.49 | 0.49 | 0.77 | 0.55 |
| 8 | 17730338 | rs7815785 | 1.874 | -1.319 | 3.518 | 1.609 | 0.068 | -0.022 | -0.022 | 0.044 | 0.56 | 0.18 | 0.4 | 0.43 | 0.66 | 0.38 |
| 8 | 17731324 | rs2653408 | 1.059 | -1.314 | 3.532 | 1.617 | -0.014 | 0.011 | 0.011 | 0 | 0.67 | 0.59 | 0.58 | 0.54 | 0.73 | 0.49 |
| 8 | 17731961 | rs2653406 | -1.105 | -1.261 | 3.436 | 1.598 | -0.013 | 0.036 | 0.036 | 0 | 0.1 | 0.26 | 0.17 | 0.12 | 0.07 | 0.11 |
| 8 | 17733479 | rs2517296 | -1.883 | -1.257 | 3.464 | 1.634 | -0.026 | -0.017 | -0.017 | 0 | 0.13 | 0.28 | 0.16 | 0.06 | 0.07 | 0.11 |
| 8 | 17737589 | rs2427731 | -1.552 | -1.265 | 3.48 | 1.651 | 0.005 | 0.069 | 0.069 | 0 | 0.11 | 0.4 | 0.18 | 0.13 | 0.08 | 0.11 |
| 8 | 17737700 | rs2467073 | -2.252 | -1.365 | 3.477 | 1.65 | -0.03 | 0.002 | 0.002 | 0 | 0.16 | 0.42 | 0.2 | 0.13 | 0.11 | 0.17 |
| 8 | 17738125 | rs2517319 | -2.306 | -1.37 | 3.529 | 1.647 | -0.032 | 0.003 | 0.003 | 0.007 | 0.17 | 0.44 | 0.2 | 0.13 | 0.11 | 0.16 |
| 8 | 17739538 | rs3739406 | 1.846 | -1.41 | 3.475 | 1.613 | 0.05 | -0.016 | -0.016 | 1.531 | 0.56 | 0.23 | 0.4 | 0.43 | 0.65 | 0.39 |
| 8 | 17742072 | rs403732 | 0.308 | -1.403 | 3.33 | 1.589 | 0.003 | -0.014 | -0.014 | 0 | 0.73 | 0.76 | 0.47 | 0.45 | 0.73 | 0.49 |
| 8 | 17743020 | rs484373 | 0.166 | -1.4 | 3.133 | 1.6 | 0.007 | -0.051 | -0.051 | 0.032 | 0.76 | 0.83 | 0.5 | 0.5 | 0.77 | 0.53 |
| 8 | 17744768 | rs693812 | 0.678 | -1.394 | 3.089 | 1.607 | -0.023 | 0.056 | 0.056 | 0 | 0.7 | 0.58 | 0.45 | 0.43 | 0.73 | 0.5 |
| 8 | 17746785 | rs402124 | 0.374 | -1.395 | 3.072 | 1.583 | -0.001 | 0.005 | 0.005 | 0 | 0.72 | 0.72 | 0.49 | 0.49 | 0.77 | 0.56 |
| 8 | 17747717 | rs401153 | 1.151 | -1.345 | 2.828 | 1.572 | -0.023 | 0.03 | 0.03 | 0 | 0.65 | 0.56 | 0.44 | 0.44 | 0.69 | 0.46 |
| 8 | 17747909 | rs3913558 | 1.789 | -1.363 | 2.522 | 1.5 | 0.07 | -0.018 | -0.018 | 6.862 | 0.55 | 0.17 | 0.4 | 0.43 | 0.66 | 0.41 |
| 8 | 17754461 | rs2653426 | -1.281 | -1.24 | 2.527 | 1.344 | -0.007 | -0.043 | -0.043 | 0 | 0.1 | 0.38 | 0.03 | 0.01 | 0.04 | 0.05 |
| 8 | 17755571 | rs452075 | -1.224 | -1.24 | 2.553 | 1.559 | -0.01 | 0.005 | 0.005 | 0.009 | 0.1 | 0.34 | 0.07 | 0.06 | 0.04 | 0.05 |
| 8 | 17758989 | rs432070 | -1.086 | -1.241 | 2.552 | 1.441 | -0.016 | -0.037 | -0.037 | 0 | 0.09 | 0.25 | 0.03 | 0.01 | 0.04 | 0.05 |
| 8 | 17759924 | rs537700 | -2.044 | -1.293 | 2.533 | 1.38 | -0.027 | -0.08 | -0.08 | 0 | 0.14 | 0.29 | 0.05 | 0.01 | 0.08 | 0.1 |
| 8 | 17760348 | rs407833 | 0.514 | -1.32 | 2.541 | 1.606 | 0.002 | -0.008 | -0.008 | 0 | 0.71 | 0.71 | 0.46 | 0.51 | 0.73 | 0.51 |
| 8 | 17766207 | rs6992489 | -2.543 | -1.297 | 2.767 | 1.563 | 0.025 | 0.042 | 0.042 | 2.211 | 0.07 | 0.34 | 0.06 | 0.05 | 0.04 | 0.05 |
| 8 | 17769193 | rs439968 | -0.594 | -1.316 | 2.68 | 1.541 | -0.016 | -0.036 | -0.036 | 0 | 0.08 | 0.15 | 0.02 | 0.01 | 0.04 | 0.05 |
| 8 | 17771154 | rs366298 | -2.427 | -1.225 | 2.824 | 1.532 | -0.008 | -0.049 | -0.049 | 0 | 0.14 | 0.45 | 0.05 | 0.07 | 0.07 | 0.11 |
| 8 | 17773482 | rs382667 | -0.604 | -1.205 | 2.499 | 1.541 | -0.015 | -0.034 | -0.034 | 0.906 | 0.08 | 0.14 | 0.02 | 0.01 | 0.04 | 0.05 |
| 8 | 17776084 | rs6586671 | 1.132 | -1.015 | 2.514 | 1.648 | -0.008 | 0.015 | 0.015 | 0 | 0.64 | 0.57 | 0.46 | 0.53 | 0.69 | 0.48 |
| 8 | 17781257 | rs12718385 | 0.859 | -1.137 | 2.478 | 1.634 | -0.003 | 0.009 | 0.009 | 0 | 0.68 | 0.65 | 0.48 | 0.54 | 0.73 | 0.52 |
| 8 | 17782692 | rs106294 | -0.671 | -1.113 | 2.459 | 1.724 | -0.014 | 0.003 | 0.003 | 0 | 0.08 | 0.13 | 0.14 | 0.06 | 0.04 | 0.05 |
| 8 | 17788511 | rs386075 | -1.718 | -0.876 | 2.503 | 1.713 | -0.021 | -0.012 | -0.012 | 0 | 0.13 | 0.22 | 0.17 | 0.06 | 0.07 | 0.1 |
| 8 | 17793084 | rs916550 | -1.517 | -0.967 | 2.451 | 1.765 | -0.024 | 0.152 | 0.152 | 0.124 | 0.15 | 0.35 | 0.14 | 0.41 | 0.14 | 0.11 |
| 8 | 17793093 | rs445422 | -0.435 | -0.938 | 2.303 | 1.714 | -0.014 | 0.004 | 0.004 | 0 | 0.08 | 0.13 | 0.14 | 0.06 | 0.04 | 0.05 |
| 8 | 17795496 | rs404186 | 0.046 | -0.994 | 2.129 | 1.749 | 0.002 | -0.017 | -0.017 | 0 | 0.76 | 0.79 | 0.63 | 0.62 | 0.76 | 0.57 |
| 8 | 17799241 | rs10108545 | 2.274 | -1.018 | 2.2 | 1.726 | 0.038 | 0.197 | 0.197 | 0.552 | 0.48 | 0.15 | 0.32 | 0.16 | 0.55 | 0.36 |
| 8 | 17800769 | rs2299587 | 2.238 | -1.013 | 2.201 | 1.721 | 0.051 | 0.201 | 0.201 | 0.441 | 0.48 | 0.16 | 0.33 | 0.16 | 0.58 | 0.37 |
| 8 | 17800896 | rs208770 | 1.677 | -0.904 | 2.355 | 1.741 | 0.008 | 0.019 | 0.019 | 0.031 | 0.57 | 0.34 | 0.46 | 0.22 | 0.58 | 0.41 |
| 8 | 17801725 | rs2299588 | 1.079 | -1.017 | 2.355 | 1.793 | -0.005 | 0.009 | 0.009 | 0 | 0.64 | 0.58 | 0.46 | 0.56 | 0.69 | 0.48 |
| 8 | 17807728 | rs17126008 | -1.54 | -1.003 | 2.457 | 1.804 | -0.029 | 0.111 | 0.111 | 0 | 0.14 | 0.31 | 0.13 | 0.38 | 0.11 | 0.11 |
| 8 | 17808730 | rs2299596 | -2.072 | -1.004 | 2.464 | 1.789 | -0.027 | 0.115 | 0.115 | 0.382 | 0.19 | 0.43 | 0.16 | 0.39 | 0.18 | 0.16 |
| 8 | 17810141 | rs7837060 | 2.273 | -1.008 | 2.402 | 1.751 | 0.039 | 0.15 | 0.15 | 0.434 | 0.48 | 0.22 | 0.32 | 0.18 | 0.55 | 0.31 |
| 8 | 17811536 | rs208756 | -2.52 | -1.024 | 2.396 | 1.76 | -0.035 | 0.061 | 0.061 | 0.63 | 0.27 | 0.56 | 0.31 | 0.43 | 0.21 | 0.21 |
| 8 | 17814915 | rs208753 | 0.034 | -1.008 | 2.416 | 1.907 | 0.001 | -0.023 | -0.023 | 0 | 0.76 | 0.81 | 0.63 | 0.63 | 0.76 | 0.57 |
| 8 | 17819572 | rs3780103 | 2.174 | -1.012 | 2.345 | 1.842 | 0.052 | 0.163 | 0.163 | 0.177 | 0.5 | 0.21 | 0.32 | 0.17 | 0.56 | 0.31 |
| 8 | 17823299 | rs12056521 | -1.85 | -1.026 | 2.414 | 1.864 | -0.017 | 0.071 | 0.071 | 0.139 | 0.17 | 0.3 | 0.16 | 0.37 | 0.17 | 0.16 |

| | | | | | | | | | | | | | | | | |
|---|---|---|---|---|---|---|---|---|---|---|---|---|---|---|---|---|
| 8 | 17827597 | rs2285310 | -1.714 | -1.046 | 2.375 | 1.864 | -0.034 | 0.15 | 0.15 | 0.047 | 0.15 | 0.38 | 0.14 | 0.37 | 0.11 | 0.11 |
| 8 | 17830459 | rs10112514 | 2.187 | -1.134 | 2.238 | 1.773 | 0.054 | 0.157 | 0.157 | 0.218 | 0.5 | 0.22 | 0.33 | 0.18 | 0.58 | 0.32 |
| 8 | 17844353 | rs17126060 | -1.856 | -1.117 | 2.226 | 2.005 | -0.031 | 0.046 | 0.046 | 0.064 | 0.17 | 0.33 | 0.15 | 0.32 | 0.14 | 0.16 |
| 8 | 17845447 | rs439511 | -0.883 | -1.113 | 2.252 | 2.02 | -0.008 | 0.058 | 0.058 | 0 | 0.09 | 0.21 | 0.15 | 0.11 | 0.04 | 0.05 |
| 8 | 17849147 | rs370429 | -0.672 | -1.112 | 2.246 | 2.02 | -0.007 | 0.052 | 0.052 | 0 | 0.09 | 0.18 | 0.14 | 0.11 | 0.04 | 0.05 |
| 8 | 17851248 | rs429633 | 1.399 | -1.13 | 2.185 | 1.976 | 0.007 | 0.015 | 0.015 | 0 | 0.6 | 0.58 | 0.48 | 0.23 | 0.63 | 0.35 |
| 8 | 17860863 | rs12056629 | -2.025 | -1.111 | 2.172 | 1.963 | -0.033 | 0.047 | 0.047 | 0.116 | 0.18 | 0.42 | 0.17 | 0.31 | 0.17 | 0.22 |
| 8 | 17864832 | rs437707 | -2.108 | -1.127 | 2.102 | 1.949 | -0.02 | -0.002 | -0.002 | 0 | 0.13 | 0.28 | 0.19 | 0.11 | 0.09 | 0.12 |
| 8 | 17874330 | rs11777465 | -0.2 | -1.099 | 2.058 | 2 | 0.017 | -0.044 | -0.044 | 0.024 | 0.08 | 0.02 | 0.1 | 0.25 | 0.09 | 0.05 |
| 8 | 17874480 | rs10503607 | -1.877 | -1.102 | 2.144 | 1.975 | -0.006 | -0.023 | -0.023 | 0 | 0.09 | 0.29 | 0.06 | 0.05 | 0.07 | 0.11 |
| 8 | 17874604 | rs2237848 | 1.051 | -1.13 | 2.163 | 1.962 | -0.007 | 0.018 | 0.018 | 0 | 0.63 | 0.57 | 0.47 | 0.49 | 0.69 | 0.43 |
| 8 | 17879392 | rs208021 | -1.592 | -1.143 | 2.161 | 1.949 | -0.024 | 0.04 | 0.04 | 0 | 0.17 | 0.26 | 0.23 | 0.31 | 0.12 | 0.09 |
| 8 | 17880243 | rs2283113 | 1.422 | -1.122 | 2.189 | 1.938 | -0.012 | 0.018 | 0.018 | 0 | 0.6 | 0.48 | 0.42 | 0.49 | 0.65 | 0.37 |
| 8 | 17880616 | rs3886398 | 2.44 | -1.479 | 1.991 | 1.773 | 0.021 | 0.118 | 0.118 | 0.327 | 0.46 | 0.22 | 0.31 | 0.17 | 0.55 | 0.32 |
| 8 | 17882483 | rs2299603 | -0.3 | -1.27 | 2.012 | 1.876 | -0.004 | -0.047 | -0.047 | 0.051 | 0.09 | 0.02 | 0.1 | 0.32 | 0.1 | 0.06 |
| 8 | 17887205 | rs444181 | -0.063 | -1.263 | 2.107 | 1.866 | -0.013 | 0.012 | 0.012 | 0 | 0.08 | 0.17 | 0.15 | 0.06 | 0.04 | 0.04 |
| 8 | 17888770 | rs492069 | -0.054 | -1.263 | 2.059 | 1.88 | -0.001 | -0.026 | -0.026 | 0 | 0.77 | 0.84 | 0.63 | 0.61 | 0.76 | 0.53 |
| 8 | 17889343 | rs7832527 | 1.416 | -1.288 | 2.114 | 1.864 | -0.008 | 0.015 | 0.015 | 0 | 0.61 | 0.47 | 0.44 | 0.5 | 0.67 | 0.37 |
| 8 | 17891869 | rs17692377 | -0.327 | -1.93 | 1.289 | 1.188 | -0.012 | 0.008 | 0.008 | 0 | 0.23 | 0.29 | 0.15 | 0.32 | 0.19 | 0.07 |
| 8 | 17899172 | rs408466 | -0.903 | -1.993 | 1.096 | 1.145 | 0.019 | -0.019 | -0.019 | 1.564 | 0.3 | 0.23 | 0.32 | 0.39 | 0.25 | 0.12 |
| 8 | 17903046 | rs441606 | -0.352 | -2.024 | 1.13 | 1.192 | 0.001 | -0.005 | -0.005 | 0.069 | 0.75 | 0.75 | 0.6 | 0.59 | 0.75 | 0.48 |
| 8 | 17907843 | rs4087059 | -0.39 | -1.918 | 1.047 | 1.296 | -0.003 | -0.111 | -0.111 | 0.115 | 0.8 | 0.97 | 0.85 | 0.66 | 0.81 | 0.92 |
| 8 | 17908116 | rs441517 | 1.072 | -2.009 | 1.127 | 1.195 | 0.016 | -0.018 | -0.018 | 0.085 | 0.65 | 0.7 | 0.65 | 0.52 | 0.7 | 0.81 |
| 8 | 17909492 | rs7844114 | 1.353 | -1.796 | 1.327 | 1.1 | 0.001 | -0.025 | -0.025 | 0.191 | 0.44 | 0.5 | 0.3 | 0.16 | 0.49 | 0.35 |
| 8 | 17910777 | rs10112108 | 1.68 | -1.79 | 1.285 | 1.079 | 0.017 | -0.053 | -0.053 | 0 | 0.4 | 0.47 | 0.27 | 0.16 | 0.46 | 0.3 |
| 8 | 17911822 | rs543449 | 0.928 | -1.991 | 1.118 | 1.131 | -0.002 | -0.008 | -0.008 | 0 | 0.49 | 0.45 | 0.43 | 0.23 | 0.53 | 0.37 |
| 8 | 17913970 | rs7508 | -0.386 | -2.112 | 0.827 | 1.391 | -0.03 | -0.142 | -0.142 | 0.256 | 0.72 | 0.95 | 0.65 | 0.55 | 0.69 | 0.64 |
| 8 | 17914799 | rs405308 | 0.193 | -2.01 | 1.156 | 1.344 | -0.008 | 0.001 | 0.001 | 0 | 0.09 | 0.2 | 0.15 | 0.07 | 0.04 | 0.05 |
| 8 | 17914883 | rs574774 | 0.193 | -2.01 | 1.156 | 1.344 | -0.008 | 0.001 | 0.001 | 0 | 0.09 | 0.2 | 0.15 | 0.07 | 0.04 | 0.05 |
| 8 | 17916224 | rs420610 | 1.658 | -2.201 | 0.998 | 1.459 | -0.021 | 0.034 | 0.034 | 0.814 | 0.78 | 0.26 | 0.71 | 0.52 | 0.94 | 0.73 |
| 8 | 17916553 | rs7465164 | 2.069 | -2.196 | 1.234 | 1.487 | 0.03 | 0.215 | 0.215 | 0.128 | 0.92 | 0.55 | 0.97 | 0.86 | 0.98 | 0.97 |
| 8 | 17916752 | rs10107973 | 0.883 | -2.163 | 1.164 | 1.497 | -0.019 | 0.149 | 0.149 | 0 | 0.05 | 0.18 | 0.12 | 0.28 | 0.01 | 0.18 |
| 8 | 17918527 | rs1234367 | -0.623 | -2.173 | 1.166 | 1.484 | -0.03 | 0.081 | 0.081 | 0 | 0.14 | 0.3 | 0.26 | 0.34 | 0.04 | 0.24 |
| 8 | 17918934 | rs10103355 | 1.721 | -2.184 | 1.262 | 1.504 | 0.038 | 0.215 | 0.215 | 0.015 | 0.93 | 0.59 | 0.97 | 0.86 | 0.98 | 0.97 |
| 8 | 17918986 | rs17515291 | 0.335 | -2.172 | 0.915 | 1.48 | -0.001 | -0.017 | -0.017 | 0.522 | 0.27 | 0.09 | 0.34 | 0.3 | 0.27 | 0.38 |
| 8 | 17919470 | rs17692671 | 0.802 | -2.082 | 0.797 | 1.492 | -0.019 | 0.159 | 0.159 | 0 | 0.05 | 0.18 | 0.12 | 0.29 | 0.01 | 0.18 |
| 8 | 17921921 | rs2073574 | 0.914 | -2.056 | 0.571 | 1.39 | 0.025 | -0.016 | -0.016 | 1.315 | 0.57 | 0.68 | 0.49 | 0.49 | 0.64 | 0.46 |
| 8 | 17925151 | rs12155668 | -0.7 | -1.667 | 0.576 | 1.276 | -0.005 | 0.074 | 0.074 | 0 | 0.52 | 0.23 | 0.57 | 0.37 | 0.48 | 0.53 |
| 8 | 17927327 | rs1049874 | -0.752 | -1.669 | 0.616 | 1.292 | -0.001 | 0.063 | 0.063 | 0 | 0.52 | 0.25 | 0.57 | 0.37 | 0.48 | 0.53 |
| 8 | 17930772 | rs3753115 | -0.689 | -1.668 | 0.616 | 1.246 | 0.002 | 0.076 | 0.076 | 0 | 0.51 | 0.25 | 0.57 | 0.35 | 0.48 | 0.52 |
| 8 | 17934029 | rs7462991 | -0.837 | -1.218 | 0.871 | 0.611 | -0.001 | -0.002 | -0.002 | 0 | 0.13 | 0.14 | 0.16 | 0.06 | 0.07 | 0.12 |
| 8 | 17937364 | rs7825389 | 0.561 | -1.373 | 0.679 | 0.633 | 0 | 0 | 0 | 0 | 0.4 | 0.41 | 0.4 | 0.43 | 0.52 | 0.48 |
| 8 | 17937530 | rs11986226 | 1.048 | -1.351 | 0.681 | 0.639 | -0.028 | 0.019 | 0.019 | 0 | 0.35 | 0.24 | 0.28 | 0.25 | 0.49 | 0.31 |
| 8 | 17938574 | rs2283120 | 0.297 | -1.309 | 0.744 | 0.612 | -0.001 | 0 | 0 | 0 | 0.43 | 0.39 | 0.42 | 0.46 | 0.54 | 0.54 |
| 8 | 17943036 | rs7387252 | -1.281 | -0.955 | 0.725 | 0.395 | -0.019 | -0.008 | -0.008 | 0 | 0.26 | 0.65 | 0.3 | 0.29 | 0.13 | 0.32 |
| 8 | 17943183 | rs7388634 | -0.329 | -0.496 | -0.441 | 0.721 | -0.006 | 0.022 | 0.022 | 0 | 0.09 | 0.2 | 0.11 | 0.18 | 0.06 | 0.19 |
| 8 | 17944663 | rs2010799 | -0.015 | -0.565 | -1.148 | 0.566 | 0.036 | 0.001 | 0.001 | 0 | 0.55 | 0.72 | 0.57 | 0.49 | 0.57 | 0.6 |
| 8 | 17948298 | rs17126209 | -0.506 | -0.601 | -1.381 | 0.149 | -0.012 | 0.051 | 0.051 | 0 | 0.14 | 0.01 | 0.07 | 0.06 | 0.11 | 0.19 |
| 8 | 17951737 | rs7829634 | -0.922 | -0.79 | -1.364 | 0.494 | 0.004 | -0.009 | -0.009 | 0 | 0.74 | 0.78 | 0.78 | 0.49 | 0.8 | 0.79 |
| 8 | 17951763 | rs11986482 | 0.147 | -1.092 | -0.556 | -0.157 | 0.005 | -0.079 | -0.079 | 0.103 | 0.12 | 0.4 | 0.13 | 0 | 0.08 | 0.07 |
| 8 | 17953439 | rs11774166 | 0.5 | -0.965 | -0.936 | -0.433 | 0.035 | 0.027 | 0.027 | 0 | 0.49 | 0.1 | 0.56 | 0.37 | 0.61 | 0.61 |
| 8 | 17953808 | rs9325825 | -1.951 | -1.032 | -0.803 | -0.381 | 0.016 | -0.058 | -0.058 | 0.677 | 0.77 | 0.87 | 0.79 | 0.48 | 0.8 | 0.82 |
| 8 | 17954298 | rs916551 | 1.53 | -1.072 | -0.875 | -0.088 | -0.176 | 0.046 | 0.046 | 0 | 0.36 | 0.03 | 0.41 | 0.2 | 0.5 | 0.43 |
| 8 | 17955256 | rs2739684 | 0.644 | -1.04 | -0.488 | -0.195 | -0.002 | -0.008 | -0.008 | 1.064 | 0.09 | 0.15 | 0.07 | 0.06 | 0.07 | 0.06 |
| 8 | 17957107 | rs11203939 | 0.173 | -1.091 | 0.411 | -0.218 | 0.015 | 0.008 | 0.008 | 0 | 0.55 | 0.41 | 0.6 | 0.38 | 0.65 | 0.65 |
| 8 | 17958628 | rs7825369 | 0.357 | -0.76 | 0.545 | -0.193 | -0.005 | -0.05 | -0.05 | 0.731 | 0.1 | 0.15 | 0.1 | 0 | 0.07 | 0.08 |
| 8 | 17964559 | rs7844251 | 0.684 | -0.744 | 0.613 | -0.185 | -0.001 | -0.033 | -0.033 | 0 | 0.08 | 0.1 | 0.1 | 0 | 0.07 | 0.07 |
| 8 | 17972040 | rs2517343 | -0.388 | -0.706 | 0.926 | -0.153 | 0.067 | 0.048 | 0.048 | 0 | 0.07 | 0.36 | 0.09 | 0.06 | 0.07 | 0.02 |
| 8 | 17975812 | rs2006919 | -0.535 | -0.707 | 0.477 | -0.26 | -0.002 | -0.009 | -0.009 | 0 | 0.06 | 0.11 | 0.04 | 0.03 | 0.04 | 0.03 |
| 8 | 17977230 | rs1858720 | -1.169 | -0.753 | 0.419 | -0.322 | -0.01 | -0.061 | -0.061 | 0 | 0.2 | 0.3 | 0.14 | 0.04 | 0.13 | 0.16 |
| 8 | 17977650 | rs17126232 | -0.185 | -0.708 | -0.001 | -0.284 | 0.005 | -0.006 | -0.006 | 0 | 0.14 | 0.12 | 0.12 | 0 | 0.1 | 0.1 |
| 8 | 17981877 | rs7388381 | -0.876 | -0.706 | -0.317 | -0.147 | 0.027 | -0.027 | -0.027 | 0 | 0.64 | 0.78 | 0.72 | 0.4 | 0.7 | 0.62 |
| 8 | 17982155 | rs2739679 | -1.615 | -0.563 | 0.25 | -0.338 | 0.004 | -0.019 | -0.019 | 0.017 | 0.29 | 0.71 | 0.22 | 0.05 | 0.21 | 0.21 |
| 8 | 17983723 | rs2027924 | -1.35 | -0.593 | 0.351 | -0.293 | -0.023 | -0.118 | -0.118 | 0.586 | 0.26 | 0.56 | 0.2 | 0.05 | 0.19 | 0.2 |
| 8 | 17984690 | rs2739685 | 0.697 | -0.914 | 0.024 | 0.322 | 0 | -0.001 | -0.001 | 2.438 | 0.74 | 0.76 | 0.87 | 0.68 | 0.83 | 0.7 |
| 8 | 17987012 | rs2898455 | 1.27 | -1.137 | 0.27 | 1.08 | 0.001 | -0.001 | -0.001 | 0.05 | 0.72 | 0.27 | 0.69 | 0.49 | 0.79 | 0.76 |
| 8 | 17987186 | rs7830223 | 2.106 | -0.516 | -0.57 | 0.719 | -0.083 | -0.041 | -0.041 | 0 | 0.25 | 0.03 | 0.24 | 0.23 | 0.28 | 0.17 |
| 8 | 17990273 | rs2739696 | -1.101 | -0.343 | 0.544 | 1.286 | -0.021 | 0.016 | 0.016 | 0.27 | 0.2 | 0.69 | 0.27 | 0.28 | 0.14 | 0.18 |
| 8 | 17992833 | rs714131 | 0.709 | -0.311 | 0.965 | 1.328 | -0.012 | -0.04 | -0.04 | 0 | 0.9 | 0.97 | 0.94 | 0.77 | 0.91 | 0.94 |

| | | | | | | | | | | | | | | | | |
|---|---|---|---|---|---|---|---|---|---|---|---|---|---|---|---|---|
| 8 | 17995760 | rs2517355 | -1.567 | -0.387 | 1.385 | 1.036 | -0.006 | 0.013 | 0.013 | 0 | 0.23 | 0.32 | 0.27 | 0.34 | 0.2 | 0.25 |
| 8 | 17996495 | rs992617 | -2.152 | 0.205 | 0.94 | 0.878 | -0.039 | -0.029 | -0.029 | 0 | 0.4 | 0.93 | 0.37 | 0.4 | 0.36 | 0.45 |
| 8 | 17998436 | rs4921865 | -0.831 | 0.163 | 1.236 | 0.783 | -0.048 | -0.115 | -0.115 | 0 | 0.19 | 0.39 | 0.08 | 0.04 | 0.1 | 0.2 |
| 8 | 17998577 | rs2739726 | -0.17 | 0.241 | 0.828 | 0.846 | -0.005 | 0.006 | 0.006 | 0 | 0.08 | 0.03 | 0.05 | 0.06 | 0.06 | 0.05 |
| 8 | 17999052 | rs12544705 | 1.051 | 0.274 | 0.791 | 0.968 | -0.035 | -0.034 | -0.034 | 0 | 0.06 | 0.01 | 0.18 | 0.15 | 0.13 | 0.05 |
| 8 | 18004224 | rs2739734 | -2.073 | 0.183 | 0.019 | 0.717 | -0.092 | -0.015 | -0.015 | 0 | 0.51 | 0.93 | 0.61 | 0.6 | 0.52 | 0.72 |
| 8 | 18005973 | rs11203940 | 1.32 | 0.282 | 0.137 | 0.734 | -0.035 | -0.034 | -0.034 | 1.144 | 0.06 | 0.01 | 0.18 | 0.15 | 0.13 | 0.05 |
| 8 | 18006561 | rs2739737 | -1.776 | 0.084 | 0.678 | 0.684 | -0.033 | 0.006 | 0.006 | 0.037 | 0.2 | 0.56 | 0.12 | 0.2 | 0.14 | 0.19 |
| 8 | 18009367 | rs987341 | 0.514 | 0.116 | 0.162 | 0.712 | 0.009 | -0.012 | -0.012 | 0.54 | 0.07 | 0.04 | 0.02 | 0.11 | 0.05 | 0.12 |
| 8 | 18020696 | rs6586704 | -2.489 | 0.076 | 0.041 | 0.723 | -0.036 | -0.01 | -0.01 | 0.256 | 0.58 | 0.89 | 0.65 | 0.64 | 0.59 | 0.78 |
| 8 | 18028739 | rs17593996 | -1.582 | 0.127 | 0.223 | 0.627 | -0.028 | -0.03 | -0.03 | 0 | 0.43 | 0.2 | 0.56 | 0.51 | 0.49 | 0.65 |
| 8 | 18029277 | rs2157651 | -1.266 | 0.21 | 0.179 | 0.575 | -0.014 | 0.016 | 0.016 | 0 | 0.28 | 0.17 | 0.29 | 0.24 | 0.29 | 0.3 |
| 8 | 18029977 | rs6992951 | -2.131 | 0.072 | -0.038 | 0.518 | -0.002 | 0.048 | 0.048 | 0 | 0.51 | 0.93 | 0.47 | 0.39 | 0.41 | 0.66 |
| 8 | 18032110 | rs7003927 | -0.76 | -0.159 | 0.891 | 0.539 | -0.028 | 0.021 | 0.021 | 0 | 0.24 | 0.4 | 0.31 | 0.3 | 0.2 | 0.41 |
| 8 | 18032419 | rs2040475 | -1.764 | -0.214 | 0.947 | 0.491 | 0.041 | 0.003 | 0.003 | 0.762 | 0.36 | 0.86 | 0.39 | 0.34 | 0.3 | 0.49 |
| 8 | 18032510 | rs2040476 | -0.447 | -0.182 | 1.265 | 0.507 | 0.01 | -0.026 | -0.026 | 0 | 0.09 | 0.19 | 0.07 | 0.03 | 0.09 | 0.07 |
| 8 | 18033462 | rs10094013 | -1.669 | -0.202 | 1.145 | 0.501 | -0.002 | -0.001 | -0.001 | 0 | 0.41 | 0.42 | 0.43 | 0.34 | 0.33 | 0.62 |
| 8 | 18036404 | rs11996662 | -0.746 | -0.154 | 1.103 | 0.581 | -0.03 | 0.016 | 0.016 | 0 | 0.25 | 0.41 | 0.32 | 0.3 | 0.21 | 0.43 |
| 8 | 18039231 | rs2188023 | -1.283 | -0.185 | 0.775 | 0.57 | -0.033 | -0.003 | -0.003 | 0 | 0.31 | 0.53 | 0.33 | 0.31 | 0.24 | 0.46 |
| 8 | 18042767 | rs6990966 | -0.46 | -0.038 | 0.934 | 0.69 | -0.002 | 0.07 | 0.07 | 0 | 0.2 | 0.52 | 0.3 | 0.28 | 0.2 | 0.35 |
| 8 | 18044101 | rs7818834 | -1.833 | 0.119 | 0.646 | 0.653 | -0.012 | 0.017 | 0.017 | 0 | 0.39 | 0.27 | 0.32 | 0.34 | 0.39 | 0.42 |
| 8 | 18044222 | rs7831983 | -2.509 | 0.181 | 0.563 | 0.711 | -0.008 | 0.003 | 0.003 | 0 | 0.58 | 0.77 | 0.62 | 0.61 | 0.57 | 0.77 |
| 8 | 18045349 | rs3901866 | -1.309 | -0.3 | 0.062 | 0.592 | 0.061 | 0.006 | 0.006 | 0.007 | 0.35 | 0.81 | 0.34 | 0.3 | 0.23 | 0.38 |
| 8 | 18051545 | rs7823976 | 1.79 | -0.458 | 0.272 | 0.588 | -0.004 | 0.009 | 0.009 | 0 | 0.49 | 0.34 | 0.42 | 0.41 | 0.47 | 0.27 |
| 8 | 18059563 | rs6586711 | 0.994 | -0.151 | 1.023 | 0.576 | 0.023 | 0.065 | 0.065 | 0 | 0.7 | 0.51 | 0.64 | 0.53 | 0.7 | 0.57 |
| 8 | 18061043 | rs6998240 | 1.697 | -0.288 | 0.737 | 0.355 | -0.046 | -0.049 | -0.049 | 0 | 0.52 | 0.12 | 0.46 | 0.48 | 0.52 | 0.41 |
| 8 | 18062769 | rs7003666 | -0.091 | -0.416 | 1.071 | 0.421 | -0.02 | -0.023 | -0.023 | 0 | 0.09 | 0.37 | 0.03 | 0.04 | 0.02 | 0.02 |
| 8 | 18064030 | rs7843046 | 0.026 | -0.415 | 0.936 | 0.422 | -0.019 | -0.022 | -0.022 | 0 | 0.09 | 0.33 | 0.03 | 0.04 | 0.02 | 0.02 |
| 8 | 18066126 | rs10888150 | 1.486 | -0.348 | 0.22 | 0.473 | 0.002 | -0.025 | -0.025 | 0 | 0.55 | 0.18 | 0.5 | 0.49 | 0.57 | 0.42 |
| 8 | 18067395 | rs7845127 | -0.334 | -0.005 | -0.108 | 0.517 | -0.068 | -0.077 | -0.077 | 0 | 0.8 | 0.94 | 0.66 | 0.6 | 0.68 | 0.63 |
| 8 | 18068835 | rs2410545 | 0.237 | 0.022 | -0.164 | 0.485 | -0.071 | -0.057 | -0.057 | 0 | 0.73 | 0.91 | 0.64 | 0.61 | 0.65 | 0.6 |
| 8 | 18070927 | rs11203943 | 1.025 | -0.644 | -0.235 | 0.807 | 0.012 | -0.05 | -0.05 | 0 | 0.11 | 0.06 | 0.18 | 0.37 | 0.11 | 0.25 |
| 8 | 18071095 | rs4921581 | 1.252 | -0.841 | -0.26 | 0.693 | 0.061 | 0.007 | 0.007 | 0 | 0.58 | 0.24 | 0.51 | 0.38 | 0.68 | 0.44 |
| 8 | 18071907 | rs13253389 | 1.498 | -0.742 | -0.264 | 0.676 | 0.087 | 0.037 | 0.037 | 0 | 0.53 | 0.25 | 0.51 | 0.35 | 0.66 | 0.4 |
| 8 | 18073182 | rs9325827 | -0.014 | -0.215 | -0.363 | 0.897 | 0.002 | -0.084 | -0.084 | 0.017 | 0.18 | 0.07 | 0.21 | 0.39 | 0.17 | 0.26 |
| 8 | 18074322 | rs17126350 | -0.774 | -0.045 | 0.637 | 0.593 | 0.024 | 0.026 | 0.026 | 0.164 | 0.06 | 0.23 | 0.11 | 0.04 | 0.06 | 0.06 |
| 8 | 18076731 | rs28359520 | 0.005 | -0.267 | -0.042 | 0.254 | 0.022 | 0.049 | 0.049 | 0.011 | 0.09 | 0 | 0.01 | 0.01 | 0.02 | 0.04 |
| 8 | 18076751 | rs28359521 | 0.005 | -0.267 | -0.042 | 0.254 | 0.022 | 0.049 | 0.049 | 0.011 | 0.09 | 0 | 0.01 | 0.01 | 0.02 | 0.04 |
| 8 | 18077045 | rs28359526 | 0.005 | -0.221 | -0.075 | 0.223 | 0.022 | 0.049 | 0.049 | 0.011 | 0.09 | 0 | 0.01 | 0.01 | 0.02 | 0.04 |
| 8 | 18077310 | rs7003890 | 1.553 | -0.266 | 0.629 | 0.176 | -0.003 | 0.009 | 0.009 | 0 | 0.53 | 0.35 | 0.49 | 0.42 | 0.51 | 0.53 |
| 8 | 18077320 | rs28359529 | 0.443 | -0.444 | 0.636 | 0.193 | 0.022 | 0.048 | 0.048 | 0 | 0.09 | 0 | 0.01 | 0.01 | 0.02 | 0.04 |
| 8 | 18077576 | rs28359531 | 0.443 | -0.444 | 0.636 | 0.193 | 0.022 | 0.048 | 0.048 | 0 | 0.09 | 0 | 0.01 | 0.01 | 0.02 | 0.04 |
| 8 | 18077987 | rs8190837 | 0.132 | -0.458 | 0.871 | 0.05 | -0.002 | 0.008 | 0.008 | 0 | 0.08 | 0.05 | 0.05 | 0.01 | 0.1 | 0.1 |
| 8 | 18078171 | rs28359532 | 0.524 | -0.461 | 0.863 | 0.051 | 0.021 | 0.048 | 0.048 | 0.017 | 0.09 | 0 | 0.01 | 0.01 | 0.02 | 0.04 |
| 8 | 18078357 | rs8190841 | 0.522 | -0.46 | 0.864 | 0.051 | 0.021 | 0.048 | 0.048 | 0 | 0.09 | 0 | 0.01 | 0.01 | 0.02 | 0.04 |
| 8 | 18078445 | rs8190842 | 0.386 | -0.461 | 0.869 | 0.051 | 0.024 | 0.029 | 0.029 | 0 | 0.09 | 0.03 | 0.02 | 0.01 | 0.02 | 0.04 |
| 8 | 18078628 | rs8190845 | -1.648 | -0.508 | 1.245 | 0.228 | -0.032 | -0.08 | -0.08 | 0 | 0.21 | 0.43 | 0.1 | 0.07 | 0.15 | 0.14 |
| 8 | 18078689 | rs8190846 | 0.543 | -0.498 | 1.088 | 0.234 | 0.021 | 0.048 | 0.048 | 0 | 0.09 | 0 | 0.01 | 0.01 | 0.02 | 0.04 |
| 8 | 18078713 | rs8190847 | 0.543 | -0.498 | 1.088 | 0.234 | 0.021 | 0.048 | 0.048 | 0 | 0.09 | 0 | 0.01 | 0.01 | 0.02 | 0.04 |
| 8 | 18078731 | rs8190848 | 0.543 | -0.498 | 1.088 | 0.234 | 0.021 | 0.048 | 0.048 | 0 | 0.09 | 0 | 0.01 | 0.01 | 0.02 | 0.04 |
| 8 | 18078851 | rs8190851 | 0.543 | -0.498 | 1.088 | 0.234 | 0.021 | 0.048 | 0.048 | 0 | 0.09 | 0 | 0.01 | 0.01 | 0.02 | 0.04 |
| 8 | 18078872 | rs8190853 | 0.543 | -0.498 | 1.088 | 0.234 | 0.021 | 0.048 | 0.048 | 0 | 0.09 | 0 | 0.01 | 0.01 | 0.02 | 0.04 |
| 8 | 18079124 | rs8190856 | 0.543 | -0.498 | 1.103 | 0.234 | 0.021 | 0.048 | 0.048 | 0 | 0.09 | 0 | 0.01 | 0.01 | 0.02 | 0.04 |
| 8 | 18079517 | rs4986989 | 0.509 | -0.498 | 1.103 | 0.234 | 0.022 | 0.048 | 0.048 | 0 | 0.09 | 0 | 0.01 | 0.01 | 0.02 | 0.04 |
| 8 | 18080015 | rs4986990 | 0.51 | -0.5 | 1.117 | 0.243 | 0.022 | 0.043 | 0.043 | 0 | 0.09 | 0 | 0.01 | 0.01 | 0.02 | 0.04 |
| 8 | 18080196 | rs4986783 | 0.535 | -0.511 | 1.12 | 0.23 | 0.021 | 0.048 | 0.048 | 0.018 | 0.09 | 0 | 0.01 | 0.01 | 0.02 | 0.04 |
| 8 | 18080651 | rs15561 | 1.759 | -0.544 | 1.117 | 0.218 | -0.031 | 0.015 | 0.015 | 0 | 0.62 | 0.48 | 0.62 | 0.45 | 0.72 | 0.59 |
| 8 | 18080747 | rs4986993 | 1.796 | -0.55 | 1.114 | 0.215 | -0.031 | 0.014 | 0.014 | 0 | 0.61 | 0.48 | 0.62 | 0.45 | 0.72 | 0.59 |
| 8 | 18080933 | rs8190863 | 0.623 | -0.503 | 0.794 | 0.203 | 0.021 | 0.048 | 0.048 | 0 | 0.09 | 0 | 0.01 | 0.01 | 0.02 | 0.04 |
| 8 | 18081272 | rs8190870 | -1.296 | -0.505 | 0.879 | 0.199 | -0.005 | -0.05 | -0.05 | 0 | 0.19 | 0.25 | 0.08 | 0.02 | 0.15 | 0.14 |
| 8 | 18081829 | rs9650592 | -0.401 | -0.515 | 0.627 | 0.249 | 0.003 | -0.147 | -0.147 | 0.003 | 0.83 | 0.99 | 0.72 | 0.48 | 0.86 | 0.72 |
| 8 | 18082354 | rs13270034 | -0.749 | -0.583 | 0.83 | 0.214 | 0.017 | 0.04 | 0.04 | 0 | 0.16 | 0.08 | 0.06 | 0.02 | 0.13 | 0.12 |
| 8 | 18083413 | rs13248278 | 0.153 | -0.533 | 1.004 | 0.327 | -0.001 | -0.011 | -0.011 | 1.111 | 0.08 | 0.08 | 0.05 | 0.01 | 0.1 | 0.08 |
| 8 | 18086096 | rs13277177 | 0.144 | -0.531 | 0.913 | 0.328 | -0.015 | 0.038 | 0.038 | 0 | 0.08 | 0.03 | 0.05 | 0.01 | 0.1 | 0.1 |
| 8 | 18086217 | rs13277481 | -0.881 | -0.57 | 0.907 | 0.302 | 0.015 | 0.067 | 0.067 | 0 | 0.17 | 0.05 | 0.06 | 0.02 | 0.13 | 0.14 |
| 8 | 18091656 | rs1389110 | 1.324 | -0.463 | 1.161 | 0.37 | 0.01 | 0.009 | 0.009 | 0 | 0.62 | 0.43 | 0.49 | 0.42 | 0.52 | 0.52 |
| 8 | 18092958 | rs11203945 | 0.928 | -0.516 | 1.108 | 0.349 | -0.011 | -0.047 | -0.047 | 0 | 0.65 | 0.79 | 0.58 | 0.52 | 0.64 | 0.61 |
| 8 | 18093176 | rs11203948 | 1.012 | -0.52 | 1.226 | 0.363 | -0.002 | -0.011 | -0.011 | 0.356 | 0.63 | 0.69 | 0.55 | 0.42 | 0.63 | 0.6 |
| 8 | 18095343 | rs4921884 | -1.119 | -0.672 | 1.304 | 0.238 | -0.011 | -0.047 | -0.047 | 0 | 0.08 | 0.28 | 0.04 | 0 | 0.05 | 0.08 |

| | | | | | | | | | | | | | | | | |
|---|---|---|---|---|---|---|---|---|---|---|---|---|---|---|---|---|
| 8 | 18096128 | rs6586722 | 1.123 | -0.639 | 1.288 | 0.267 | -0.012 | -0.034 | -0.034 | 0 | 0.58 | 0.75 | 0.54 | 0.47 | 0.55 | 0.5 |
| 8 | 18097080 | rs5026870 | -1.577 | -0.666 | 1.316 | 0.269 | -0.018 | -0.008 | -0.008 | 0.824 | 0.1 | 0.38 | 0.06 | 0.04 | 0.05 | 0.08 |
| 8 | 18099148 | rs7009832 | -1.826 | -0.662 | 1.319 | 0.274 | 0.077 | 0.097 | 0.097 | 1.502 | 0.05 | 0.37 | 0.05 | 0.04 | 0.02 | 0 |
| 8 | 18099509 | rs3959118 | 2.196 | -0.786 | 1.704 | 0.293 | -0.036 | 0.018 | 0.018 | 0 | 0.45 | 0.22 | 0.43 | 0.37 | 0.46 | 0.42 |
| 8 | 18099877 | rs6586725 | 1.244 | -0.734 | 1.614 | 0.28 | -0.004 | -0.001 | -0.001 | 0 | 0.57 | 0.65 | 0.51 | 0.4 | 0.53 | 0.5 |
| 8 | 18100220 | rs6586726 | 0.769 | -0.752 | 1.639 | 0.266 | -0.009 | -0.01 | -0.01 | 0.016 | 0.65 | 0.69 | 0.52 | 0.42 | 0.56 | 0.53 |
| 8 | 18101807 | rs4921593 | -0.391 | -0.73 | 1.607 | 0.27 | -0.009 | -0.038 | -0.038 | 0 | 0.07 | 0.18 | 0.03 | 0 | 0.05 | 0.07 |
| 8 | 18105556 | rs7834428 | 1.585 | -0.707 | 1.571 | 0.296 | -0.002 | 0.001 | 0.001 | 0 | 0.53 | 0.4 | 0.49 | 0.43 | 0.51 | 0.49 |
| 8 | 18109256 | rs2172427 | 0.819 | -0.704 | 1.616 | 0.304 | -0.001 | 0 | 0 | 3.108 | 0.65 | 0.61 | 0.53 | 0.47 | 0.55 | 0.52 |
| 8 | 18111750 | rs6985224 | 0.101 | -0.714 | 1.403 | 0.11 | -0.02 | -0.039 | -0.039 | 0 | 0.1 | 0.14 | 0.02 | 0 | 0.03 | 0.03 |
| 8 | 18114563 | rs17126406 | -0.855 | -0.712 | 1.419 | 0.115 | -0.035 | -0.034 | -0.034 | 0 | 0.14 | 0.31 | 0.06 | 0.06 | 0.05 | 0.03 |
| 8 | 18121307 | rs6586730 | 0.485 | -0.697 | 1.239 | 0.423 | -0.091 | -0.127 | -0.127 | 1.709 | 0.68 | 0.92 | 0.56 | 0.48 | 0.57 | 0.54 |
| 8 | 18123076 | rs2172425 | 0.872 | -0.697 | 1.343 | 0.483 | -0.012 | -0.018 | -0.018 | 0 | 0.65 | 0.73 | 0.53 | 0.44 | 0.57 | 0.55 |
| 8 | 18124099 | rs7817820 | -0.575 | -0.774 | 1.277 | 0.42 | -0.044 | -0.025 | -0.025 | 0 | 0.12 | 0.24 | 0.04 | 0.06 | 0.03 | 0.03 |
| 8 | 18125900 | rs12542228 | 2.241 | -0.835 | 1.3 | 0.384 | -0.032 | 0.013 | 0.013 | 0 | 0.45 | 0.23 | 0.44 | 0.38 | 0.46 | 0.42 |
| 8 | 18134632 | rs2131422 | -0.46 | -0.732 | 0.92 | 0.297 | -0.043 | -0.023 | -0.023 | 0 | 0.12 | 0.24 | 0.04 | 0.06 | 0.03 | 0.04 |
| 8 | 18139163 | rs13270961 | 0.076 | -0.661 | 0.687 | 0.456 | -0.007 | -0.01 | -0.01 | 0 | 0.07 | 0.04 | 0.06 | 0.13 | 0.11 | 0.09 |
| 8 | 18139427 | rs1493040 | -0.716 | -0.673 | 0.755 | 0.492 | -0.016 | -0.036 | -0.036 | 0.044 | 0.32 | 0.2 | 0.44 | 0.43 | 0.37 | 0.45 |
| 8 | 18140633 | rs1353035 | 1.066 | -0.7 | 0.895 | 0.454 | 0 | 0 | 0 | 0 | 0.81 | 0.8 | 0.91 | 0.87 | 0.83 | 0.86 |
| 8 | 18141181 | rs11787299 | -0.7 | -0.731 | 0.893 | 0.463 | -0.02 | -0.043 | -0.043 | 0 | 0.32 | 0.2 | 0.43 | 0.43 | 0.37 | 0.45 |
| 8 | 18142353 | rs7357578 | -2.004 | -0.74 | 0.922 | 0.456 | -0.002 | 0.007 | 0.007 | 0 | 0.5 | 0.4 | 0.52 | 0.56 | 0.53 | 0.59 |
| 8 | 18144874 | rs7834791 | -0.841 | -0.746 | 0.992 | 0.464 | 0.008 | 0.012 | 0.012 | 0 | 0.16 | 0.12 | 0.07 | 0.13 | 0.14 | 0.14 |
| 8 | 18146650 | rs11776179 | -0.741 | -0.764 | 0.966 | 0.448 | -0.009 | -0.024 | -0.024 | 0 | 0.32 | 0.26 | 0.43 | 0.43 | 0.37 | 0.42 |
| 8 | 18148015 | rs13282737 | -2.302 | -0.751 | 1.042 | 0.425 | -0.002 | 0.005 | 0.005 | 0 | 0.52 | 0.6 | 0.54 | 0.62 | 0.54 | 0.56 |
| 8 | 18159185 | rs6986036 | -1.215 | -0.729 | 0.86 | 0.157 | -0.003 | -0.007 | -0.007 | 0 | 0.35 | 0.34 | 0.47 | 0.45 | 0.39 | 0.41 |
| 8 | 18164113 | rs4440669 | -0.405 | -0.544 | 0.694 | 0.131 | -0.01 | -0.043 | -0.043 | 0 | 0.28 | 0.21 | 0.4 | 0.46 | 0.31 | 0.35 |
| 8 | 18169443 | rs7839590 | -0.658 | -0.633 | 0.673 | 0.117 | -0.03 | -0.035 | -0.035 | 0 | 0.07 | 0.13 | 0.04 | 0 | 0.03 | 0.07 |
| 8 | 18172011 | rs17642444 | -0.204 | -0.524 | 0.583 | 0.152 | -0.012 | -0.054 | -0.054 | 0 | 0.26 | 0.18 | 0.35 | 0.45 | 0.29 | 0.3 |
| 8 | 18173967 | rs7828904 | -1.649 | -0.465 | 0.603 | 0.145 | -0.005 | 0 | 0 | 0 | 0.44 | 0.54 | 0.43 | 0.44 | 0.38 | 0.43 |
| 8 | 18175677 | rs7834906 | -0.312 | -0.207 | 0.133 | 0.357 | 0.004 | 0.019 | 0.019 | 0 | 0.24 | 0.26 | 0.36 | 0.44 | 0.29 | 0.31 |
| 8 | 18181069 | rs12541267 | -1.004 | 0.057 | -0.155 | 0.48 | -0.007 | -0.044 | -0.044 | 0 | 0.43 | 0.17 | 0.51 | 0.46 | 0.47 | 0.56 |
| 8 | 18192279 | rs969189 | -0.866 | -0.172 | -0.226 | 0.412 | -0.045 | -0.015 | -0.015 | 0 | 0.43 | 0.16 | 0.52 | 0.55 | 0.5 | 0.53 |
| 8 | 18193096 | rs7834039 | -1.138 | -0.189 | 0.014 | 0.446 | -0.002 | 0.013 | 0.013 | 0 | 0.49 | 0.62 | 0.58 | 0.56 | 0.54 | 0.59 |
| 8 | 18193852 | rs1353037 | 1.103 | -1.013 | 0.361 | 0.649 | -0.065 | 0.045 | 0.045 | 0.168 | 0.71 | 0.54 | 0.8 | 0.58 | 0.79 | 0.65 |
| 8 | 18194387 | rs6586734 | 0.644 | -0.591 | 0.496 | 0.736 | -0.01 | 0.02 | 0.02 | 0 | 0.72 | 0.58 | 0.81 | 0.31 | 0.76 | 0.63 |
| 8 | 18196020 | rs7827099 | -0.987 | -0.682 | 0.708 | 0.572 | -0.008 | 0.008 | 0.008 | 0 | 0.32 | 0.33 | 0.47 | 0.63 | 0.28 | 0.32 |
| 8 | 18200916 | rs7812408 | -1.181 | -0.726 | 0.837 | 0.558 | -0.012 | 0.01 | 0.01 | 0 | 0.34 | 0.37 | 0.48 | 0.64 | 0.3 | 0.33 |
| 8 | 18201818 | rs1602782 | 1.309 | -0.823 | 1.152 | 0.521 | 0.023 | 0 | 0 | 0 | 0.61 | 0.31 | 0.48 | 0.36 | 0.67 | 0.67 |
| 8 | 18205455 | rs11203956 | 1.975 | -0.965 | 0.199 | 0.134 | -0.156 | 0.131 | 0.131 | 0 | 0.32 | 0.03 | 0.3 | 0.09 | 0.42 | 0.3 |
| 8 | 18205573 | rs10095674 | 0.382 | -0.849 | 1.151 | 0.888 | -0.005 | 0.012 | 0.012 | 0 | 0.15 | 0.15 | 0.36 | 0.31 | 0.14 | 0.17 |
| 8 | 18208003 | rs6586740 | 0.183 | -0.842 | 1.394 | 0.872 | -0.002 | 0.015 | 0.015 | 0 | 0.86 | 0.88 | 0.92 | 0.68 | 0.9 | 0.89 |
| 8 | 18209487 | rs10093448 | -0.282 | -0.718 | 1.427 | 0.803 | -0.049 | 0.103 | 0.103 | 0 | 0.18 | 0.39 | 0.39 | 0.31 | 0.15 | 0.17 |
| 8 | 18209776 | rs11775054 | 0.041 | -0.674 | 0.879 | 0.826 | -0.023 | 0.039 | 0.039 | 0 | 0.17 | 0.23 | 0.38 | 0.31 | 0.15 | 0.18 |
| 8 | 18209795 | rs11785281 | 0.041 | -0.674 | 0.879 | 0.826 | -0.023 | 0.039 | 0.039 | 0 | 0.17 | 0.23 | 0.37 | 0.31 | 0.14 | 0.18 |
| 8 | 18209976 | rs10094010 | 0.239 | -0.633 | 0.473 | 0.849 | 0.004 | -0.005 | -0.005 | 0 | 0.16 | 0.12 | 0.36 | 0.31 | 0.15 | 0.17 |
| 8 | 18212381 | rs2410552 | 0.382 | -0.613 | 0.287 | 0.896 | 0.017 | -0.03 | -0.03 | 0.217 | 0.14 | 0.07 | 0.34 | 0.3 | 0.14 | 0.15 |
| 8 | 18213874 | rs6983849 | -1.184 | -0.658 | 0.905 | 0.813 | -0.028 | 0.056 | 0.056 | 0 | 0.2 | 0.42 | 0.15 | 0.33 | 0.14 | 0.15 |
| 8 | 18214566 | rs11993911 | -0.643 | -0.66 | 0.626 | 0.352 | -0.017 | -0.025 | -0.025 | 0 | 0.1 | 0.2 | 0.04 | 0.03 | 0.05 | 0.07 |
| 8 | 18215738 | rs13256693 | 1.322 | -0.72 | 0.214 | -0.178 | -0.074 | 0.17 | 0.17 | 0 | 0.4 | 0.15 | 0.34 | 0.1 | 0.48 | 0.31 |
| 8 | 18217658 | rs1495754 | 0.81 | -0.751 | 0.56 | 0.12 | 0.008 | 0.01 | 0.01 | 0 | 0.55 | 0.39 | 0.7 | 0.41 | 0.62 | 0.46 |
| 8 | 18217814 | rs971770 | 1.12 | -0.433 | 1.112 | -0.25 | 0.075 | -0.016 | -0.016 | 1.261 | 0.92 | 0.64 | 0.93 | 0.99 | 0.92 | 0.97 |
| 8 | 18233582 | rs7006687 | 0.609 | -0.651 | 0.609 | 0.074 | 0.008 | 0.006 | 0.006 | 0 | 0.52 | 0.57 | 0.62 | 0.52 | 0.55 | 0.45 |
| 8 | 18241486 | rs13277723 | -0.336 | -0.276 | 0.329 | -0.345 | 0.02 | 0.09 | 0.09 | 0.035 | 0.52 | 0.32 | 0.47 | 0.74 | 0.49 | 0.58 |
| 8 | 18241771 | rs34546534 | 1.413 | -0.318 | 0.582 | -0.504 | -0.026 | 0.067 | 0.067 | 0 | 0.31 | 0.23 | 0.33 | 0.04 | 0.38 | 0.28 |
| 8 | 18243113 | rs7016785 | 0.523 | -0.054 | 0.585 | -0.441 | 0.005 | 0.051 | 0.051 | 0 | 0.37 | 0.23 | 0.2 | 0.26 | 0.32 | 0.37 |
| 8 | 18243468 | rs7013253 | 0.44 | -0.053 | 0.611 | -0.439 | 0.005 | 0.045 | 0.045 | 0 | 0.38 | 0.25 | 0.21 | 0.26 | 0.32 | 0.4 |
| 8 | 18243603 | rs10109286 | 0.22 | -0.13 | 1.255 | -0.263 | 0.044 | 0.169 | 0.169 | 0 | 0.16 | 0.29 | 0.27 | 0.48 | 0.17 | 0.18 |
| 8 | 18246664 | rs1565684 | 0.423 | 0.085 | 1.395 | -0.642 | 0.005 | 0.108 | 0.108 | 0 | 0.51 | 0.33 | 0.53 | 0.26 | 0.54 | 0.45 |
| 8 | 18246696 | rs4646241 | 0.887 | 0.285 | 1.381 | -0.572 | -0.009 | 0.024 | 0.024 | 0 | 0.3 | 0.24 | 0.18 | 0.26 | 0.29 | 0.35 |
| 8 | 18247449 | rs4646242 | 0.887 | 0.275 | 1.381 | -0.572 | -0.007 | 0.024 | 0.024 | 0 | 0.3 | 0.24 | 0.18 | 0.26 | 0.28 | 0.35 |
| 8 | 18247913 | rs4646267 | 0.728 | 0.228 | 1.35 | -0.504 | -0.023 | -0.025 | -0.025 | 0 | 0.14 | 0.08 | 0.26 | 0.47 | 0.16 | 0.15 |
| 8 | 18248208 | rs4345600 | 1.003 | 0.163 | 1.249 | -0.553 | 0.025 | -0.039 | -0.039 | 0 | 0.13 | 0.03 | 0.18 | 0.22 | 0.13 | 0.14 |
| 8 | 18248268 | rs4271002 | 0.959 | 0.161 | 1.292 | -0.554 | 0.014 | -0.013 | -0.013 | 0.293 | 0.14 | 0.08 | 0.19 | 0.21 | 0.13 | 0.14 |
| 8 | 18248661 | rs4646246 | 0.33 | 0.203 | 1.391 | -0.539 | 0.049 | 0.158 | 0.158 | 0 | 0.16 | 0.32 | 0.27 | 0.47 | 0.17 | 0.18 |
| 8 | 18249652 | rs11780272 | 1.161 | 0.262 | 1.248 | -0.708 | -0.022 | 0.125 | 0.125 | 0 | 0.38 | 0.23 | 0.35 | 0.04 | 0.43 | 0.34 |
| 8 | 18250592 | rs13277605 | 1.181 | 0.262 | 1.248 | -0.708 | -0.023 | 0.127 | 0.127 | 0 | 0.38 | 0.22 | 0.35 | 0.04 | 0.43 | 0.31 |
| 8 | 18250922 | rs9987109 | 0.912 | 0.264 | 1.282 | -0.708 | -0.006 | 0.043 | 0.043 | 0 | 0.4 | 0.29 | 0.37 | 0.04 | 0.45 | 0.36 |
| 8 | 18251926 | rs2087852 | 0.885 | 0.205 | 1.173 | -0.691 | -0.008 | 0.036 | 0.036 | 0 | 0.3 | 0.2 | 0.17 | 0.26 | 0.27 | 0.36 |
| 8 | 18252755 | rs1390358 | 1.177 | 0.212 | 1.195 | -0.692 | -0.008 | 0.107 | 0.107 | 0 | 0.38 | 0.24 | 0.35 | 0.04 | 0.42 | 0.33 |

| | | | | | | | | | | | | | | | | |
|---|---|---|---|---|---|---|---|---|---|---|---|---|---|---|---|---|
| 8 | 18254575 | rs11996129 | 0.742 | 0.547 | 1.183 | -0.445 | -0.005 | 0.035 | 0.035 | 0 | 0.3 | 0.2 | 0.16 | 0.25 | 0.27 | 0.35 |
| 8 | 18255410 | rs1115784 | -0.591 | 0.611 | 1.421 | -0.385 | 0.002 | -0.008 | -0.008 | 0 | 0.29 | 0.32 | 0.45 | 0.69 | 0.29 | 0.3 |
| 8 | 18255709 | rs1961456 | -0.614 | 0.61 | 1.465 | -0.385 | 0.004 | -0.012 | -0.012 | 0.141 | 0.29 | 0.35 | 0.45 | 0.69 | 0.29 | 0.3 |
| 8 | 18255876 | rs1112005 | 0.87 | 0.527 | 1.381 | -0.454 | -0.006 | 0.028 | 0.028 | 0 | 0.28 | 0.18 | 0.16 | 0.25 | 0.27 | 0.35 |
| 8 | 18257280 | rs7832071 | 1.049 | 0.657 | 1.851 | -0.432 | 0.001 | -0.04 | -0.04 | 0 | 0.41 | 0.41 | 0.36 | 0.04 | 0.43 | 0.32 |
| 8 | 18257795 | rs1041983 | 0.215 | 0.521 | 2.005 | -0.342 | -0.016 | 0.001 | 0.001 | 0 | 0.36 | 0.47 | 0.29 | 0.44 | 0.31 | 0.43 |
| 8 | 18257854 | rs1801280 | 1.216 | 0.532 | 1.934 | -0.357 | -0.009 | 0.082 | 0.082 | 0 | 0.41 | 0.29 | 0.36 | 0.04 | 0.45 | 0.35 |
| 8 | 18257994 | rs1799929 | 1.319 | 0.539 | 1.961 | -0.359 | -0.022 | 0.129 | 0.129 | 0 | 0.39 | 0.24 | 0.34 | 0.04 | 0.44 | 0.32 |
| 8 | 18258103 | rs1799930 | 0.642 | 0.585 | 1.869 | -0.261 | 0.001 | 0.029 | 0.029 | 0 | 0.31 | 0.24 | 0.17 | 0.26 | 0.28 | 0.36 |
| 8 | 18258908 | rs4646247 | 0.642 | 0.583 | 1.857 | -0.258 | 0.001 | 0.027 | 0.027 | 0 | 0.31 | 0.24 | 0.17 | 0.26 | 0.28 | 0.36 |
| 8 | 18259305 | rs721398 | 0.653 | 0.587 | 1.836 | -0.264 | 0.001 | 0.027 | 0.027 | 0 | 0.31 | 0.23 | 0.17 | 0.26 | 0.28 | 0.36 |
| 8 | 18259366 | rs721399 | 0.694 | 0.593 | 1.868 | -0.285 | 0.012 | -0.019 | -0.019 | 0 | 0.72 | 0.54 | 0.53 | 0.3 | 0.72 | 0.7 |
| 8 | 18260013 | rs11784251 | 0.671 | 0.594 | 1.862 | -0.284 | 0.013 | -0.019 | -0.019 | 0 | 0.72 | 0.54 | 0.53 | 0.3 | 0.72 | 0.7 |
| 8 | 18260355 | rs4646248 | 0.426 | 0.593 | 1.892 | -0.285 | 0.002 | -0.003 | -0.003 | 0 | 0.73 | 0.75 | 0.54 | 0.3 | 0.72 | 0.7 |
| 8 | 18260431 | rs4646249 | 0.426 | 0.593 | 1.892 | -0.285 | 0.002 | -0.003 | -0.003 | 0 | 0.73 | 0.75 | 0.54 | 0.3 | 0.72 | 0.7 |
| 8 | 18260598 | rs4646250 | 0.414 | 0.593 | 1.892 | -0.285 | 0.002 | -0.003 | -0.003 | 0 | 0.73 | 0.75 | 0.54 | 0.3 | 0.72 | 0.7 |
| 8 | 18261493 | rs1390360 | 0.693 | 0.595 | 1.892 | -0.285 | 0.011 | -0.019 | -0.019 | 0.04 | 0.72 | 0.55 | 0.53 | 0.3 | 0.72 | 0.71 |
| 8 | 18261497 | rs1908586 | 1.348 | 0.596 | 1.892 | -0.239 | 0 | 0.101 | 0.101 | 0 | 0.4 | 0.28 | 0.35 | 0.09 | 0.43 | 0.35 |
| 8 | 18261679 | rs1908587 | 1.274 | 0.598 | 1.884 | -0.239 | 0.003 | 0.084 | 0.084 | 0 | 0.41 | 0.3 | 0.36 | 0.09 | 0.43 | 0.35 |
| 8 | 18262723 | rs1495745 | 0.414 | 0.6 | 1.88 | -0.235 | 0.001 | -0.002 | -0.002 | 0 | 0.73 | 0.76 | 0.54 | 0.3 | 0.72 | 0.71 |
| 8 | 18262845 | rs4646259 | 0.655 | 0.599 | 1.881 | -0.238 | 0 | 0.038 | 0.038 | 0 | 0.31 | 0.24 | 0.17 | 0.21 | 0.29 | 0.36 |
| 8 | 18262861 | rs1495747 | 0.683 | 0.644 | 1.885 | -0.236 | 0.013 | -0.019 | -0.019 | 0 | 0.72 | 0.54 | 0.53 | 0.3 | 0.72 | 0.71 |
| 8 | 18263484 | rs2410558 | 0.678 | 0.628 | 1.884 | -0.237 | 0.011 | -0.019 | -0.019 | 0.194 | 0.72 | 0.55 | 0.53 | 0.3 | 0.72 | 0.71 |
| 8 | 18263603 | rs4540438 | 1.044 | 0.666 | 1.745 | -0.219 | -0.004 | -0.08 | -0.08 | 0.774 | 0.43 | 0.52 | 0.37 | 0.09 | 0.43 | 0.35 |
| 8 | 18263663 | rs12674710 | 0.372 | 0.677 | 1.64 | -0.132 | -0.008 | 0 | 0 | 0 | 0.15 | 0.11 | 0.26 | 0.48 | 0.17 | 0.17 |
| 8 | 18267338 | rs10103029 | 0.207 | 0.546 | 1.572 | -0.143 | 0.028 | 0.148 | 0.148 | 0 | 0.16 | 0.29 | 0.26 | 0.48 | 0.15 | 0.17 |
| 8 | 18267878 | rs10088333 | 0.443 | 0.551 | 1.446 | -0.14 | 0.012 | 0.077 | 0.077 | 0 | 0.15 | 0.18 | 0.26 | 0.48 | 0.15 | 0.17 |
| 8 | 18271912 | rs12543818 | -0.164 | 0.539 | 1.057 | -0.19 | 0.013 | 0.021 | 0.021 | 0 | 0.42 | 0.3 | 0.36 | 0.44 | 0.4 | 0.47 |
| 8 | 18271948 | rs12545528 | -0.173 | 0.538 | 1.072 | -0.19 | 0.012 | 0.019 | 0.019 | 0 | 0.42 | 0.33 | 0.36 | 0.44 | 0.4 | 0.47 |
| 8 | 18272107 | rs4921911 | 0.209 | 0.53 | 1.051 | -0.201 | 0.008 | 0.006 | 0.006 | 0 | 0.36 | 0.3 | 0.29 | 0.4 | 0.31 | 0.43 |
| 8 | 18272377 | rs4921913 | 0.354 | 0.572 | 1.127 | -0.194 | 0.029 | 0.1 | 0.1 | 0 | 0.76 | 0.58 | 0.65 | 0.49 | 0.76 | 0.78 |
| 8 | 18272438 | rs4921914 | 0.354 | 0.572 | 1.127 | -0.194 | 0.029 | 0.1 | 0.1 | 0 | 0.76 | 0.58 | 0.65 | 0.49 | 0.76 | 0.78 |
| 8 | 18272881 | rs1495741 | 0.348 | 0.578 | 1.131 | -0.209 | 0.028 | 0.097 | 0.097 | 0 | 0.76 | 0.58 | 0.65 | 0.48 | 0.76 | 0.78 |
| 8 | 18272901 | rs1495742 | 0.888 | 0.523 | 0.837 | -0.227 | 0.004 | 0.066 | 0.066 | 0 | 0.44 | 0.32 | 0.47 | 0.27 | 0.47 | 0.42 |
| 8 | 18273996 | rs6998197 | 0.377 | 0.589 | 0.84 | -0.208 | -0.002 | 0.078 | 0.078 | 0 | 0.51 | 0.34 | 0.53 | 0.32 | 0.55 | 0.47 |
| 8 | 18275189 | rs7816847 | 0.432 | 0.613 | 0.737 | -0.05 | 0.009 | 0.057 | 0.057 | 0 | 0.15 | 0.19 | 0.27 | 0.47 | 0.15 | 0.18 |
| 8 | 18275845 | rs2410564 | 1.932 | 0.59 | 0.096 | -0.084 | -0.004 | 0.015 | 0.015 | 0 | 0.07 | 0.02 | 0.07 | 0.04 | 0.09 | 0.04 |
| 8 | 18281867 | rs17126633 | 1.044 | 0.69 | 0.033 | -0.475 | -0.015 | 0.091 | 0.091 | 0 | 0.4 | 0.26 | 0.37 | 0.09 | 0.46 | 0.33 |
| 8 | 18283339 | rs6989506 | 0.054 | 0.693 | -0.161 | -0.299 | 0.005 | 0.031 | 0.031 | 0 | 0.34 | 0.3 | 0.2 | 0.21 | 0.3 | 0.39 |
| 8 | 18286997 | rs6997340 | 0.473 | 0.516 | -0.072 | 0.657 | 0.002 | 0.056 | 0.056 | 0 | 0.69 | 0.49 | 0.61 | 0.49 | 0.71 | 0.58 |
| 8 | 18307161 | rs7817770 | -0.388 | 0.166 | -0.318 | 0.303 | 0.008 | 0.013 | 0.013 | 0 | 0.51 | 0.58 | 0.45 | 0.7 | 0.37 | 0.51 |
| 8 | 18309074 | rs13262005 | -0.625 | 0.155 | 0.08 | -0.212 | 0.024 | -0.068 | -0.068 | 0 | 0.64 | 0.58 | 0.6 | 0.86 | 0.53 | 0.76 |
| 8 | 18310071 | rs9325829 | -1.072 | 0.369 | -0.106 | 0.284 | -0.003 | 0 | 0 | 0 | 0.7 | 0.75 | 0.6 | 0.77 | 0.57 | 0.77 |
| 8 | 18310705 | rs12544627 | -0.179 | -0.986 | -0.482 | 0.268 | -0.013 | 0.011 | 0.011 | 0 | 0.38 | 0.55 | 0.33 | 0.44 | 0.29 | 0.28 |
| 8 | 18318883 | rs7017310 | -0.862 | -1.001 | -0.559 | 0.272 | -0.018 | 0.043 | 0.043 | 0 | 0.63 | 0.79 | 0.55 | 0.82 | 0.39 | 0.75 |
| 8 | 18320873 | rs7817217 | -0.674 | -0.845 | 0.138 | 0.172 | -0.012 | 0.027 | 0.027 | 0 | 0.14 | 0.2 | 0.15 | 0.27 | 0.11 | 0.17 |
| 8 | 18332127 | rs901940 | 0.771 | -1.221 | 0.381 | 0.119 | -0.083 | 0.03 | 0.03 | 0.964 | 0.85 | 0.39 | 0.93 | 0.8 | 0.98 | 0.64 |
| 8 | 18332971 | rs1995002 | 0.142 | -1.047 | 0.154 | 0.11 | 0.008 | 0.014 | 0.014 | 4.206 | 0.08 | 0.14 | 0.13 | 0.19 | 0.13 | 0.16 |
| 8 | 18333109 | rs1995003 | -0.741 | -1.03 | -0.066 | 0.109 | -0.013 | -0.009 | -0.009 | 0 | 0.16 | 0.14 | 0.25 | 0.29 | 0.24 | 0.27 |
| 8 | 18333171 | rs17595454 | 0.577 | -0.973 | -0.089 | 0.145 | -0.043 | -0.017 | -0.017 | 0 | 0.08 | 0.01 | 0.13 | 0.1 | 0.11 | 0.1 |
| 8 | 18335423 | rs4921928 | -0.768 | -1.406 | -0.383 | -0.199 | 0.023 | 0.045 | 0.045 | 0 | 0.25 | 0.1 | 0.21 | 0.2 | 0.21 | 0.2 |
| 8 | 18339293 | rs2035686 | 0.072 | -1.536 | 0.492 | -0.247 | -0.06 | -0.047 | -0.047 | 0 | 0.88 | 0.55 | 0.96 | 0.94 | 0.98 | 0.72 |
| 8 | 18342941 | rs13250636 | 1.795 | -1.087 | -0.077 | -0.197 | 0.01 | -0.012 | -0.012 | 0 | 0.48 | 0.18 | 0.46 | 0.46 | 0.59 | 0.32 |
| 8 | 18342961 | rs13259283 | 1.783 | -1.081 | -0.074 | -0.221 | 0.011 | -0.007 | -0.007 | 1.499 | 0.48 | 0.18 | 0.46 | 0.47 | 0.59 | 0.32 |
| 8 | 18348902 | rs17126664 | 0.353 | -0.32 | 0.405 | -0.129 | -0.006 | 0.002 | 0.002 | 0 | 0.06 | 0.03 | 0.06 | 0.07 | 0.08 | 0.11 |
| 8 | 18350362 | rs12546055 | -1.131 | -0.342 | 1.219 | -0.159 | 0.015 | -0.012 | -0.012 | 0 | 0.26 | 0.44 | 0.38 | 0.3 | 0.25 | 0.51 |
| 8 | 18358093 | rs17595636 | -1.035 | 0.009 | 1.424 | -0.388 | 0.044 | -0.022 | -0.022 | 0 | 0.11 | 0.24 | 0.09 | 0.07 | 0.15 | 0.19 |
| 8 | 18361385 | rs12546956 | 0.515 | 0.113 | 1.483 | -0.529 | -0.033 | 0.016 | 0.016 | 0 | 0.07 | 0.02 | 0.06 | 0.02 | 0.1 | 0.1 |
| 8 | 18362239 | rs11203964 | 1.213 | -0.238 | 1.734 | -0.963 | -0.007 | -0.04 | -0.04 | 0.011 | 0.69 | 0.37 | 0.83 | 0.89 | 0.78 | 0.59 |
| 8 | 18363313 | rs10098661 | 0.334 | 0.061 | 1.627 | -0.783 | 0.011 | -0.029 | -0.029 | 0 | 0.08 | 0.12 | 0.06 | 0 | 0.12 | 0.09 |
| 8 | 18363607 | rs17126683 | -0.158 | 0.04 | 1.685 | -0.596 | 0.029 | 0.02 | 0.02 | 0 | 0.09 | 0.17 | 0.07 | 0.07 | 0.12 | 0.11 |
| 8 | 18364774 | rs901938 | 0.722 | 0.042 | 1.713 | -0.594 | 0.036 | 0.035 | 0.035 | 0 | 0.88 | 0.67 | 0.9 | 0.87 | 0.86 | 0.87 |
| 8 | 18367759 | rs7836260 | 0.589 | 0.08 | 1.699 | -0.588 | 0.038 | 0.037 | 0.037 | 0 | 0.88 | 0.66 | 0.9 | 0.87 | 0.86 | 0.89 |
| 8 | 18372093 | rs11787522 | 1.439 | 0.073 | 1.002 | -0.957 | -0.032 | 0.094 | 0.094 | 0.365 | 0.6 | 0.21 | 0.7 | 0.79 | 0.61 | 0.5 |
| 8 | 18374592 | rs7461797 | 1.729 | 0.189 | 0.99 | -0.915 | -0.011 | 0.148 | 0.148 | 0 | 0.52 | 0.21 | 0.66 | 0.78 | 0.54 | 0.47 |
| 8 | 18379513 | rs766143 | 1.162 | 0.393 | 1.724 | -0.4 | 0.028 | -0.033 | -0.033 | 0.107 | 0.72 | 0.42 | 0.78 | 0.86 | 0.68 | 0.76 |
| 8 | 18380024 | rs6995993 | -0.427 | 0.408 | 1.412 | -0.397 | 0.015 | -0.024 | -0.024 | 0 | 0.93 | 0.85 | 0.93 | 0.96 | 0.9 | 0.97 |
| 8 | 18381235 | rs10090117 | -0.738 | 0.284 | 1.363 | -0.326 | 0.005 | -0.005 | -0.005 | 0.377 | 0.2 | 0.23 | 0.13 | 0.1 | 0.2 | 0.25 |
| 8 | 18385056 | rs9741 | -1.346 | 0.118 | 0.906 | -0.398 | -0.001 | -0.026 | -0.026 | 0.121 | 0.29 | 0.58 | 0.17 | 0.16 | 0.24 | 0.5 |

| | | | | | | | | | | | | | | | | |
|---|---|---|---|---|---|---|---|---|---|---|---|---|---|---|---|---|
| 8 | 18386237 | rs3739398 | -1.417 | 0.167 | 1.02 | -0.35 | 0.01 | -0.008 | -0.008 | 0 | 0.21 | 0.6 | 0.13 | 0.16 | 0.18 | 0.46 |
| 8 | 18387374 | rs13275817 | 1.474 | 0.224 | 1.048 | -0.427 | 0.002 | -0.003 | -0.003 | 0 | 0.73 | 0.28 | 0.81 | 0.8 | 0.73 | 0.5 |
| 8 | 18388041 | rs3739396 | 1.171 | 0.257 | 1.212 | -0.461 | -0.005 | -0.07 | -0.07 | 0.068 | 0.81 | 0.42 | 0.89 | 0.9 | 0.86 | 0.65 |
| 8 | 18388254 | rs8695 | 0.943 | 0.279 | 1.143 | -0.442 | -0.022 | 0.012 | 0.012 | 0 | 0.06 | 0.01 | 0.06 | 0.03 | 0.11 | 0.12 |
| 8 | 18389112 | rs17126760 | 0.77 | 0.201 | 1.567 | -0.432 | -0.026 | -0.033 | -0.033 | 0 | 0.07 | 0.21 | 0.02 | 0 | 0.01 | 0.2 |
| 8 | 18390691 | rs901933 | 1.513 | 0.23 | 1.337 | -0.48 | -0.007 | 0.017 | 0.017 | 0 | 0.65 | 0.3 | 0.77 | 0.81 | 0.67 | 0.47 |
| 8 | 18394992 | rs13279660 | -0.531 | 0.455 | 1.437 | -0.306 | 0.044 | -0.008 | -0.008 | 0 | 0.93 | 0.82 | 0.92 | 0.94 | 0.91 | 0.96 |
| 8 | 18396405 | rs7844662 | 1.169 | -0.183 | 0.873 | 0.131 | 0.003 | 0.016 | 0.016 | 0.069 | 0.48 | 0.3 | 0.7 | 0.46 | 0.62 | 0.38 |
| 8 | 18399242 | rs3780102 | 0.173 | 0.012 | 0.609 | 0.122 | 0.017 | -0.002 | -0.002 | 0.733 | 0.84 | 0.82 | 0.84 | 0.87 | 0.78 | 0.76 |
| 8 | 18399723 | rs3780099 | 0.813 | -0.251 | 0.5 | 0.18 | 0.038 | -0.063 | -0.063 | 0 | 0.25 | 0.12 | 0.11 | 0.38 | 0.14 | 0.19 |
| 8 | 18400819 | rs7013675 | 0.378 | -0.2 | 0.598 | 0.155 | 0.072 | -0.002 | -0.002 | 0 | 0.83 | 0.66 | 0.83 | 0.86 | 0.76 | 0.75 |
| 8 | 18404638 | rs10888156 | 0.23 | -0.143 | 0.525 | 0.342 | 0 | 0.02 | 0.02 | 0.474 | 0.64 | 0.41 | 0.74 | 0.53 | 0.68 | 0.7 |
| 8 | 18404837 | rs10888157 | 0.107 | -0.144 | 0.524 | 0.364 | 0.001 | 0.012 | 0.012 | 0.024 | 0.66 | 0.41 | 0.79 | 0.61 | 0.69 | 0.76 |
| 8 | 18407858 | rs876983 | 1.581 | -0.196 | -0.148 | 0.605 | -0.056 | 0.069 | 0.069 | 0.335 | 0.43 | 0.09 | 0.47 | 0.24 | 0.54 | 0.3 |
| 8 | 18409460 | rs17694569 | 0.641 | 0.384 | -0.151 | 0.905 | 0.027 | -0.064 | -0.064 | 0 | 0.23 | 0.06 | 0.1 | 0.31 | 0.14 | 0.17 |
| 8 | 18412006 | rs12545132 | 0.151 | 0.557 | -0.231 | 0.895 | -0.017 | -0.011 | -0.011 | 0 | 0.1 | 0.04 | 0.08 | 0.11 | 0.12 | 0.21 |
| 8 | 18412742 | rs17517680 | -0.432 | 0.563 | -0.222 | 0.886 | -0.018 | -0.021 | -0.021 | 0 | 0.12 | 0.06 | 0.13 | 0.19 | 0.13 | 0.28 |
| 8 | 18419589 | rs12546918 | 0.178 | 0.588 | -0.177 | 0.782 | -0.007 | -0.003 | -0.003 | 0 | 0.09 | 0.06 | 0.08 | 0.08 | 0.1 | 0.21 |
| 8 | 18420041 | rs1979597 | 0.916 | 0.497 | -0.083 | 0.778 | 0.004 | -0.002 | -0.002 | 0.597 | 0.58 | 0.49 | 0.57 | 0.43 | 0.62 | 0.51 |
| 8 | 18422211 | rs12681068 | 0.389 | 0.472 | -0.131 | 0.771 | 0.021 | -0.055 | -0.055 | 0 | 0.24 | 0.12 | 0.21 | 0.36 | 0.14 | 0.17 |
| 8 | 18423590 | rs2634436 | 0.096 | 0.448 | -0.118 | 0.846 | 0.001 | -0.013 | -0.013 | 0 | 0.3 | 0.27 | 0.29 | 0.4 | 0.24 | 0.21 |
| 8 | 18429663 | rs2717756 | 0.397 | 0.916 | -0.461 | 0.779 | 0.012 | 0.013 | 0.013 | 0 | 0.86 | 0.81 | 0.86 | 0.83 | 0.87 | 0.78 |
| 8 | 18429782 | rs6984438 | 0.401 | 1.251 | -1.185 | 0.872 | -0.03 | -0.019 | -0.019 | 0 | 0.19 | 0.14 | 0.37 | 0.26 | 0.37 | 0.26 |
| 8 | 18429869 | rs923391 | -0.216 | -0.259 | -1.249 | 0.235 | 0.045 | 0.032 | 0.032 | 0.399 | 0.64 | 0.33 | 0.8 | 0.78 | 0.83 | 0.63 |
| 8 | 18430473 | rs10503617 | 0.305 | 0.997 | -1.248 | 1.226 | -0.044 | -0.118 | -0.118 | 0 | 0.23 | 0.02 | 0.26 | 0.36 | 0.22 | 0.11 |
| 8 | 18430556 | rs11780899 | -0.565 | 1.287 | -0.659 | 1.233 | -0.007 | -0.017 | -0.017 | 0 | 0.33 | 0.23 | 0.37 | 0.47 | 0.35 | 0.31 |
| 8 | 18431478 | rs1546001 | -0.497 | 1.484 | -0.083 | 1.263 | -0.025 | -0.059 | -0.059 | 0 | 0.31 | 0.19 | 0.35 | 0.46 | 0.33 | 0.32 |
| 8 | 18432308 | rs2634431 | -0.209 | 1.658 | 0.084 | 1.148 | 0.024 | 0.01 | 0.01 | 0 | 0.9 | 0.73 | 0.93 | 0.88 | 0.89 | 0.96 |
| 8 | 18433183 | rs1546003 | 0.945 | 1.632 | 0.08 | 1.129 | -0.005 | -0.001 | -0.001 | 0.008 | 0.6 | 0.69 | 0.62 | 0.43 | 0.63 | 0.63 |
| 8 | 18435816 | rs12675535 | 0.995 | 1.606 | 0.019 | 1.097 | 0.008 | -0.001 | -0.001 | 0.114 | 0.58 | 0.52 | 0.58 | 0.44 | 0.56 | 0.64 |
| 8 | 18437729 | rs17643851 | 0.995 | 1.449 | -0.121 | 1.099 | 0.003 | -0.02 | -0.02 | 0 | 0.07 | 0.01 | 0.02 | 0.11 | 0.04 | 0.03 |
| 8 | 18438866 | rs2634444 | 1.191 | 1.815 | 0.353 | 1.144 | -0.003 | 0.008 | 0.008 | 0 | 0.56 | 0.38 | 0.56 | 0.38 | 0.51 | 0.62 |
| 8 | 18442351 | rs2634447 | 0.788 | 1.567 | 0.635 | 1.036 | 0.032 | 0.027 | 0.027 | 0 | 0.87 | 0.6 | 0.9 | 0.81 | 0.86 | 0.94 |
| 8 | 18443677 | rs13256522 | 0.313 | 1.807 | 0.249 | 0.932 | -0.031 | 0.024 | 0.024 | 0 | 0.1 | 0.02 | 0.07 | 0.03 | 0.13 | 0.17 |
| 8 | 18444450 | rs2634449 | 0.794 | 1.869 | 0.486 | 0.923 | 0.12 | 0.055 | 0.055 | 0.115 | 0.94 | 0.66 | 0.88 | 0.93 | 0.89 | 0.98 |
| 8 | 18444535 | rs10110600 | 0.724 | 2.215 | 0.506 | 1.196 | 0.006 | -0.01 | -0.01 | 0 | 0.61 | 0.55 | 0.59 | 0.69 | 0.48 | 0.66 |
| 8 | 18445966 | rs17518171 | 0.964 | 2.17 | 0.403 | 1.401 | -0.099 | 0.018 | 0.018 | 0 | 0.22 | 0.02 | 0.18 | 0.13 | 0.27 | 0.1 |
| 8 | 18446080 | rs2717743 | -0.538 | 2.097 | 0.445 | 1.282 | -0.06 | -0.058 | -0.058 | 0 | 0.36 | 0.15 | 0.4 | 0.46 | 0.44 | 0.36 |
| 8 | 18447453 | rs7839376 | 0.782 | 2.094 | 0.433 | 1.302 | -0.064 | -0.031 | -0.031 | 0 | 0.23 | 0.06 | 0.21 | 0.21 | 0.28 | 0.16 |
| 8 | 18452156 | rs2465884 | 0.53 | 2.201 | 0.471 | 1.237 | 0.005 | 0.007 | 0.007 | 0 | 0.64 | 0.54 | 0.56 | 0.39 | 0.6 | 0.75 |
| 8 | 18452272 | rs7007390 | 0.467 | 2.194 | 0.42 | 1.25 | 0.016 | -0.004 | -0.004 | 0 | 0.64 | 0.55 | 0.64 | 0.61 | 0.56 | 0.78 |
| 8 | 18452628 | rs2717751 | 0.5 | 2.219 | 0.24 | 1.13 | 0.025 | 0.171 | 0.171 | 0 | 0.85 | 0.58 | 0.74 | 0.6 | 0.83 | 0.91 |
| 8 | 18456179 | rs10091332 | 1.267 | 2.109 | -0.142 | 1.065 | 0.022 | 0.022 | 0.022 | 0 | 0.52 | 0.37 | 0.45 | 0.35 | 0.41 | 0.61 |
| 8 | 18457679 | rs11203969 | 1.024 | 1.85 | -0.184 | 1.11 | -0.059 | -0.042 | -0.042 | 0 | 0.2 | 0.03 | 0.18 | 0.21 | 0.23 | 0.14 |
| 8 | 18458608 | rs11203970 | 0.779 | 1.499 | 0.028 | 0.802 | 0.007 | 0.015 | 0.015 | 0 | 0.53 | 0.37 | 0.48 | 0.36 | 0.47 | 0.61 |
| 8 | 18460837 | rs13251595 | 0.977 | 0.208 | -0.286 | -0.356 | 0.052 | 0.057 | 0.057 | 0 | 0.31 | 0.17 | 0.17 | 0.01 | 0.2 | 0.32 |
| 8 | 18463969 | rs2717763 | -0.458 | 0.114 | -0.067 | -0.367 | 0.021 | 0.017 | 0.017 | 0.21 | 0.58 | 0.61 | 0.74 | 0.83 | 0.75 | 0.66 |
| 8 | 18464798 | rs10101988 | -0.901 | 0.172 | -0.032 | -0.376 | 0.001 | -0.038 | -0.038 | 0.08 | 0.65 | 0.64 | 0.77 | 0.93 | 0.78 | 0.67 |
| 8 | 18466123 | rs9325833 | -0.203 | 0.259 | -0.068 | 0.109 | 0.057 | 0.058 | 0.058 | 0 | 0.53 | 0.35 | 0.44 | 0.35 | 0.34 | 0.52 |
| 8 | 18467291 | rs17126921 | 0.986 | 0.044 | -0.014 | -0.125 | 0.037 | 0.039 | 0.039 | 0 | 0.36 | 0.24 | 0.22 | 0.07 | 0.22 | 0.33 |
| 8 | 18467318 | rs17126924 | 0.681 | 0.019 | -0.065 | -0.098 | 0.004 | 0.019 | 0.019 | 0 | 0.1 | 0.01 | 0.07 | 0.04 | 0.06 | 0.14 |
| 8 | 18468245 | rs7013632 | 0.466 | 0.252 | 0.189 | 0.037 | 0.062 | 0.095 | 0.095 | 0 | 0.45 | 0.27 | 0.29 | 0.1 | 0.28 | 0.47 |
| 8 | 18470613 | rs17596751 | 1.153 | 0.851 | 0.423 | 0.262 | -0.006 | 0.001 | 0.001 | 0 | 0.19 | 0.13 | 0.15 | 0.12 | 0.24 | 0.09 |
| 8 | 18471234 | rs10108055 | 0.348 | 0.856 | 0.5 | 0.309 | 0.086 | 0.126 | 0.126 | 0 | 0.49 | 0.31 | 0.32 | 0.2 | 0.31 | 0.51 |
| 8 | 18474596 | rs13266017 | -0.031 | 1.05 | 0.578 | 0.32 | 0.016 | -0.007 | -0.007 | 0 | 0.64 | 0.49 | 0.52 | 0.29 | 0.59 | 0.54 |
| 8 | 18479776 | rs13263545 | 0.438 | 0.648 | 0.612 | 0.091 | 0.017 | 0.016 | 0.016 | 0 | 0.49 | 0.46 | 0.36 | 0.18 | 0.33 | 0.46 |
| 8 | 18482596 | rs2129703 | -0.059 | 0.707 | 0.832 | 0.505 | -0.003 | 0.089 | 0.089 | 0 | 0.06 | 0.21 | 0.07 | 0.17 | 0.05 | 0.07 |
| 8 | 18484094 | rs6991100 | -1.413 | 0.931 | 0.879 | 0.989 | 0.001 | 0.014 | 0.014 | 0 | 0.83 | 0.84 | 0.74 | 0.65 | 0.77 | 0.8 |
| 8 | 18492827 | rs1487742 | 1.116 | 0.713 | 0.856 | 0.739 | -0.006 | 0.028 | 0.028 | 0 | 0.08 | 0.01 | 0.07 | 0.03 | 0.06 | 0.11 |
| 8 | 18493652 | rs1487743 | 0.613 | 0.722 | 1.082 | 0.724 | 0.034 | 0.047 | 0.047 | 0 | 0.47 | 0.39 | 0.35 | 0.17 | 0.33 | 0.45 |
| 8 | 18497616 | rs722311 | -0.588 | 0.755 | 1.149 | 0.772 | 0.062 | 0.138 | 0.138 | 0 | 0.77 | 0.58 | 0.63 | 0.5 | 0.69 | 0.71 |
| 8 | 18497842 | rs2018849 | 0.424 | 0.654 | 0.861 | 0.928 | 0.095 | 0.113 | 0.113 | 0 | 0.47 | 0.26 | 0.31 | 0.2 | 0.31 | 0.47 |
| 8 | 18500939 | rs4921943 | 0.841 | 0.834 | 0.834 | 0.895 | -0.022 | -0.014 | -0.014 | 0 | 0.23 | 0.18 | 0.28 | 0.29 | 0.33 | 0.26 |
| 8 | 18505600 | rs9325837 | -0.485 | -0.113 | 0.929 | 0.631 | 0.019 | 0.019 | 0.019 | 0 | 0.5 | 0.42 | 0.35 | 0.25 | 0.31 | 0.49 |
| 8 | 18507535 | rs10112384 | 0.199 | -0.118 | 0.736 | 0.627 | 0.095 | 0.121 | 0.121 | 0 | 0.42 | 0.2 | 0.26 | 0.12 | 0.27 | 0.45 |
| 8 | 18509985 | rs13258758 | 0.851 | 0.192 | 0.874 | 0.782 | 0.06 | 0.099 | 0.099 | 0 | 0.29 | 0.11 | 0.17 | 0.02 | 0.21 | 0.34 |
| 8 | 18510321 | rs4504676 | -0.409 | 0.576 | 1.126 | 1.092 | 0.006 | 0.007 | 0.007 | 0 | 0.47 | 0.56 | 0.42 | 0.34 | 0.38 | 0.54 |
| 8 | 18511097 | rs13268366 | 0.464 | 0.491 | 1.121 | 0.898 | 0.06 | 0.093 | 0.093 | 0 | 0.37 | 0.18 | 0.22 | 0.03 | 0.26 | 0.45 |
| 8 | 18512245 | rs4921944 | 0.461 | 0.528 | 1.109 | 0.935 | 0.053 | 0.073 | 0.073 | 0 | 0.37 | 0.21 | 0.26 | 0.11 | 0.27 | 0.44 |

| | | | | | | | | | | | | | | | | |
|---|---|---|---|---|---|---|---|---|---|---|---|---|---|---|---|---|
| 8 | 18514100 | rs10106032 | -0.039 | 0.704 | 0.781 | 1.095 | -0.001 | -0.006 | -0.006 | 0 | 0.47 | 0.54 | 0.4 | 0.26 | 0.34 | 0.52 |
| 8 | 18514675 | rs2410591 | -0.031 | 0.794 | 0.82 | 1.157 | -0.001 | -0.006 | -0.006 | 0 | 0.47 | 0.53 | 0.39 | 0.26 | 0.35 | 0.52 |
| 8 | 18515902 | rs11990818 | -0.242 | 0.844 | 0.235 | 1.402 | 0.022 | 0.026 | 0.026 | 0 | 0.49 | 0.56 | 0.39 | 0.29 | 0.36 | 0.51 |
| 8 | 18517026 | rs7824406 | -1.517 | 0.444 | 0.176 | 2.666 | 0.018 | -0.033 | -0.033 | 0 | 0.13 | 0.07 | 0.15 | 0.47 | 0.08 | 0.24 |
| 8 | 18519032 | rs1121922 | -2.198 | 0.03 | 0.994 | 2.538 | -0.066 | 0.055 | 0.055 | 0 | 0.23 | 0.51 | 0.33 | 0.68 | 0.12 | 0.4 |
| 8 | 18520079 | rs7015570 | -2.294 | 0.092 | 1.017 | 2.508 | -0.041 | 0.027 | 0.027 | 0 | 0.26 | 0.57 | 0.35 | 0.68 | 0.14 | 0.43 |
| 8 | 18524337 | rs17695622 | -0.104 | 0.151 | 0.772 | 2.413 | -0.013 | -0.003 | -0.003 | 0.021 | 0.05 | 0.09 | 0.1 | 0.05 | 0.02 | 0.05 |
| 8 | 18527302 | rs2046668 | -0.096 | 0.162 | 0.721 | 2.432 | 0 | 0.001 | 0.001 | 0 | 0.05 | 0.05 | 0.09 | 0.05 | 0.02 | 0.05 |
| 8 | 18527459 | rs7844180 | -1.21 | 0.104 | 0.966 | 2.335 | -0.006 | -0.022 | -0.022 | 0 | 0.11 | 0.16 | 0.06 | 0.02 | 0.06 | 0.15 |
| 8 | 18528135 | rs17695695 | -0.54 | 0.113 | 0.951 | 2.841 | 0.01 | 0.063 | 0.063 | 0 | 0.06 | 0.03 | 0.13 | 0.4 | 0.02 | 0.2 |
| 8 | 18530834 | rs11995995 | -1.667 | 0.705 | 0.973 | 2.293 | 0.018 | 0 | 0 | 0.42 | 0.39 | 0.46 | 0.3 | 0.37 | 0.47 | 0.57 |
| 8 | 18531247 | rs11992396 | -0.995 | 0.525 | 1.031 | 2.118 | 0.006 | -0.014 | -0.014 | 1.159 | 0.23 | 0.27 | 0.19 | 0.17 | 0.29 | 0.3 |
| 8 | 18534285 | rs10087403 | -0.722 | 0.657 | 0.982 | 2.448 | -0.029 | -0.038 | -0.038 | 0 | 0.16 | 0.13 | 0.25 | 0.3 | 0.25 | 0.33 |
| 8 | 18534417 | rs10090559 | -1.03 | 0.121 | 1.742 | 1.978 | 0.01 | -0.057 | -0.057 | 0 | 0.1 | 0.3 | 0.09 | 0 | 0.06 | 0.04 |
| 8 | 18534520 | rs2046670 | 1.527 | -0.026 | 2.106 | 1.925 | 0.047 | 0.043 | 0.043 | 1.445 | 0.92 | 0.63 | 0.97 | 0.94 | 0.95 | 0.98 |
| 8 | 18534708 | rs7834242 | -1.961 | -0.149 | 2.123 | 2.253 | -0.03 | 0.101 | 0.101 | 1.002 | 0.14 | 0.55 | 0.11 | 0.23 | 0.08 | 0.14 |
| 8 | 18535065 | rs721562 | 1.267 | -0.134 | 2.312 | 2.58 | 0.007 | 0.065 | 0.065 | 0.802 | 0.74 | 0.41 | 0.64 | 0.49 | 0.71 | 0.53 |
| 8 | 18535156 | rs2220352 | -1.603 | -0.145 | 2.04 | 2.546 | -0.002 | -0.002 | -0.002 | 0 | 0.29 | 0.32 | 0.41 | 0.38 | 0.33 | 0.43 |
| 8 | 18535768 | rs4921951 | 1.83 | -0.624 | 2.53 | 2.776 | -0.034 | 0.141 | 0.141 | 0 | 0.81 | 0.63 | 0.71 | 0.62 | 0.9 | 0.72 |
| 8 | 18537560 | rs11787358 | 2.608 | -0.498 | 2.683 | 2.901 | 0.014 | 0.325 | 0.325 | 3.105 | 0.92 | 0.4 | 0.78 | 0.75 | 0.97 | 0.96 |
| 8 | 18539039 | rs4921607 | 2.537 | -0.51 | 2.704 | 2.884 | 0.011 | 0.312 | 0.312 | 3.323 | 0.93 | 0.37 | 0.77 | 0.75 | 0.97 | 0.96 |
| 8 | 18539768 | rs4921608 | 2.674 | -0.665 | 2.121 | 2.772 | 0.006 | -0.008 | -0.008 | 2.043 | 0.78 | 0.22 | 0.73 | 0.63 | 0.9 | 0.69 |
| 8 | 18546783 | rs12546585 | -1.64 | -0.456 | 1.728 | 2.823 | 0.007 | -0.02 | -0.02 | 0 | 0.11 | 0.07 | 0.25 | 0.45 | 0.07 | 0.24 |
| 8 | 18548407 | rs1982801 | 0.615 | -0.545 | 1.681 | 3.009 | 0.003 | 0.025 | 0.025 | 0 | 0.92 | 0.94 | 0.76 | 0.68 | 0.95 | 0.9 |
| 8 | 18549142 | rs13259970 | 1.712 | -0.583 | 1.806 | 3.032 | 0.003 | 0.002 | 0.002 | 0 | 0.85 | 0.83 | 0.68 | 0.44 | 0.92 | 0.74 |
| 8 | 18550278 | rs6996073 | 1.13 | -0.538 | 2.207 | 2.857 | -0.007 | 0.223 | 0.223 | 0 | 0.9 | 0.75 | 0.69 | 0.67 | 0.95 | 0.9 |
| 8 | 18554798 | rs972433 | 2.401 | -0.665 | 2.541 | 2.826 | -0.048 | 0.299 | 0.299 | 1.699 | 0.86 | 0.45 | 0.67 | 0.65 | 0.94 | 0.9 |
| 8 | 18555053 | rs972431 | 0.094 | -0.655 | 2.431 | 2.87 | 0 | -0.001 | -0.001 | 0 | 0.94 | 0.94 | 0.97 | 0.89 | 0.98 | 0.93 |
| 8 | 18556281 | rs10089319 | 2.369 | -0.696 | 2.615 | 2.804 | -0.051 | 0.251 | 0.251 | 0 | 0.86 | 0.58 | 0.69 | 0.65 | 0.95 | 0.88 |
| 8 | 18557985 | rs11998342 | -2.808 | -0.68 | 2.656 | 2.72 | 0.05 | 0.057 | 0.057 | 2.362 | 0.07 | 0.45 | 0.04 | 0.03 | 0.02 | 0 |
| 8 | 18559060 | rs9650595 | 1.653 | -0.682 | 2.553 | 2.841 | -0.025 | 0.317 | 0.317 | 0 | 0.9 | 0.6 | 0.69 | 0.65 | 0.96 | 0.89 |
| 8 | 18560687 | rs755683 | 2.869 | -0.868 | 2.121 | 2.689 | 0.106 | -0.075 | -0.075 | 0.026 | 0.66 | 0.11 | 0.63 | 0.45 | 0.85 | 0.64 |
| 8 | 18565844 | rs17127078 | -1.889 | -0.694 | 2.618 | 2.826 | -0.02 | 0.182 | 0.182 | 0.983 | 0.11 | 0.45 | 0.05 | 0.23 | 0.05 | 0.15 |
| 8 | 18567975 | rs6987756 | -0.921 | -0.667 | 2.211 | 2.855 | 0.013 | -0.014 | -0.014 | 0 | 0.09 | 0.06 | 0.02 | 0.23 | 0.04 | 0.2 |
| 8 | 18568985 | rs2171443 | 0.41 | -0.5 | 2.162 | 2.778 | 0.001 | -0.004 | -0.004 | 0 | 0.93 | 0.94 | 0.88 | 0.87 | 0.95 | 0.93 |
| 8 | 18574874 | rs17696125 | -1.095 | -0.502 | 2.28 | 2.734 | 0.013 | 0.151 | 0.151 | 0 | 0.07 | 0.2 | 0.16 | 0.17 | 0.04 | 0.1 |
| 8 | 18576156 | rs13254772 | 2.472 | -0.597 | 2.193 | 2.87 | -0.015 | 0.013 | 0.013 | 0.326 | 0.74 | 0.32 | 0.69 | 0.48 | 0.89 | 0.7 |
| 8 | 18579638 | rs7815682 | 1.085 | -0.446 | 1.92 | 2.609 | -0.01 | 0.001 | 0.001 | 0 | 0.91 | 0.88 | 0.87 | 0.91 | 0.95 | 0.91 |
| 8 | 18585728 | rs1487747 | 1.455 | -0.501 | 1.789 | 2.732 | 0.008 | -0.024 | -0.024 | 0 | 0.88 | 0.96 | 0.86 | 0.84 | 0.92 | 0.87 |
| 8 | 18586732 | rs10090564 | -0.85 | -0.765 | 1.777 | 2.912 | 0.021 | -0.026 | -0.026 | 0 | 0.08 | 0.03 | 0.02 | 0.25 | 0.03 | 0.16 |
| 8 | 18587676 | rs10094532 | -2.646 | -0.881 | 2.249 | 2.958 | -0.053 | 0.112 | 0.112 | 0.864 | 0.18 | 0.64 | 0.21 | 0.44 | 0.07 | 0.24 |
| 8 | 18588911 | rs17127110 | -1.217 | -0.708 | 2.183 | 3.044 | -0.001 | 0.051 | 0.051 | 0 | 0.1 | 0.11 | 0.15 | 0.42 | 0.05 | 0.23 |
| 8 | 18589437 | rs13265655 | 0.782 | -0.71 | 2.149 | 3.046 | 0.01 | -0.013 | -0.013 | 0 | 0.92 | 0.96 | 0.87 | 0.84 | 0.96 | 0.92 |
| 8 | 18590892 | rs2410586 | 2.359 | -0.724 | 2.675 | 3.003 | -0.054 | 0.085 | 0.085 | 0.202 | 0.8 | 0.66 | 0.7 | 0.43 | 0.91 | 0.7 |
| 8 | 18592842 | rs13439396 | -1.692 | -0.758 | 2.714 | 2.615 | 0.008 | -0.05 | -0.05 | 1.458 | 0.1 | 0.42 | 0.06 | 0.03 | 0.06 | 0.04 |
| 8 | 18593329 | rs6996474 | 0.977 | -0.508 | 2.331 | 2.943 | 0.01 | -0.009 | -0.009 | 0 | 0.92 | 0.97 | 0.87 | 0.92 | 0.95 | 0.93 |
| 8 | 18595103 | rs923315 | 1.88 | -0.542 | 2.599 | 3.054 | -0.043 | 0.085 | 0.085 | 0 | 0.84 | 0.73 | 0.84 | 0.64 | 0.92 | 0.76 |
| 8 | 18596143 | rs4244449 | 1.45 | -0.5 | 2.304 | 3.036 | -0.019 | 0.012 | 0.012 | 0 | 0.89 | 0.82 | 0.86 | 0.91 | 0.94 | 0.93 |
| 8 | 18596762 | rs1473482 | 2.029 | -0.432 | 2.297 | 3.13 | -0.012 | 0.054 | 0.054 | 0 | 0.86 | 0.81 | 0.72 | 0.75 | 0.92 | 0.87 |
| 8 | 18598597 | rs10096702 | -1.04 | -0.303 | 2.276 | 2.785 | -0.007 | 0.083 | 0.083 | 0 | 0.1 | 0.12 | 0.16 | 0.43 | 0.05 | 0.22 |
| 8 | 18601063 | rs12548270 | -1.469 | -0.216 | 1.946 | 2.49 | -0.012 | 0.065 | 0.065 | 0 | 0.21 | 0.32 | 0.23 | 0.44 | 0.16 | 0.24 |
| 8 | 18602354 | rs2410584 | 1.082 | -0.073 | 1.451 | 2.337 | 0.001 | 0.001 | 0.001 | 0 | 0.93 | 0.96 | 0.88 | 0.87 | 0.95 | 0.91 |
| 8 | 18612464 | rs7833606 | 1.621 | -0.063 | 1.929 | 2.305 | 0 | 0.101 | 0.101 | 0 | 0.91 | 0.73 | 0.88 | 0.87 | 0.95 | 0.92 |
| 8 | 18612612 | rs10110943 | -1.768 | -0.133 | 2.451 | 2.86 | 0.015 | 0.228 | 0.228 | 0.258 | 0.1 | 0.39 | 0.14 | 0.28 | 0.08 | 0.1 |
| 8 | 18618427 | rs7012496 | -2.093 | -0.313 | 2.574 | 3.098 | -0.032 | 0.259 | 0.259 | 0.292 | 0.17 | 0.56 | 0.34 | 0.47 | 0.1 | 0.14 |
| 8 | 18618764 | rs6994455 | -2.142 | -0.361 | 2.736 | 3.15 | -0.015 | 0.132 | 0.132 | 0 | 0.15 | 0.48 | 0.19 | 0.27 | 0.09 | 0.11 |
| 8 | 18621332 | rs1487745 | -2.407 | -0.366 | 2.668 | 3.094 | -0.037 | 0.207 | 0.207 | 0.104 | 0.17 | 0.6 | 0.34 | 0.48 | 0.1 | 0.15 |
| 8 | 18629355 | rs335231 | 1.258 | 0.433 | 2.293 | 2.881 | 0.021 | 0.076 | 0.076 | 0.168 | 0.78 | 0.68 | 0.71 | 0.59 | 0.77 | 0.49 |
| 8 | 18632093 | rs6586764 | -1.233 | 0.597 | 2.31 | 2.907 | 0.044 | 0.027 | 0.027 | 0 | 0.14 | 0.24 | 0.23 | 0.18 | 0.21 | 0.35 |
| 8 | 18632736 | rs7845297 | -2.5 | 0.612 | 2.381 | 3.054 | 0.025 | 0.016 | 0.016 | 0.059 | 0.24 | 0.31 | 0.49 | 0.66 | 0.3 | 0.51 |
| 8 | 18636126 | rs335232 | -2.401 | 0.613 | 2.351 | 3.031 | 0.039 | -0.01 | -0.01 | 0.169 | 0.25 | 0.43 | 0.3 | 0.19 | 0.31 | 0.45 |
| 8 | 18642757 | rs10100475 | -1.174 | 0.552 | 1.678 | 2.857 | 0.006 | 0.212 | 0.212 | 0 | 0.06 | 0.18 | 0.13 | 0.34 | 0.05 | 0.08 |
| 8 | 18643985 | rs335243 | -0.648 | 0.691 | 1.298 | 2.621 | 0.004 | 0.008 | 0.008 | 0 | 0.06 | 0.09 | 0.03 | 0 | 0.06 | 0.03 |
| 8 | 18644651 | rs17383284 | -0.567 | 0.795 | 1.23 | 2.694 | -0.062 | -0.005 | -0.005 | 0 | 0.09 | 0.01 | 0.11 | 0.08 | 0.18 | 0.34 |
| 8 | 18645679 | rs335246 | 0.462 | 0.458 | 1.339 | 2.75 | 0.019 | -0.023 | -0.023 | 0 | 0.92 | 0.95 | 0.95 | 0.78 | 0.98 | 0.8 |
| 8 | 18652623 | rs2469946 | 0.773 | 0.443 | 1.335 | 2.666 | 0.021 | -0.026 | -0.026 | 0 | 0.91 | 0.95 | 0.93 | 0.75 | 0.97 | 0.82 |
| 8 | 18652812 | rs13250313 | -1.234 | 0.736 | 1.733 | 2.679 | 0.027 | 0.219 | 0.219 | 0.008 | 0.11 | 0.3 | 0.17 | 0.36 | 0.1 | 0.15 |
| 8 | 18653742 | rs10503633 | 2.416 | 0.663 | 1.849 | 2.597 | -0.042 | 0.044 | 0.044 | 0 | 0.72 | 0.46 | 0.72 | 0.37 | 0.82 | 0.63 |
| 8 | 18654293 | rs10503634 | 2.364 | 0.67 | 1.879 | 2.601 | -0.048 | 0.056 | 0.056 | 0 | 0.72 | 0.49 | 0.72 | 0.37 | 0.82 | 0.62 |

| | | | | | | | | | | | | | | | | |
|---|---|---|---|---|---|---|---|---|---|---|---|---|---|---|---|---|
| 8 | 18654478 | rs12549858 | -2.007 | 1.462 | 1.65 | 2.456 | 0.082 | 0.083 | 0.083 | 0 | 0.19 | 0.38 | 0.41 | 0.32 | 0.3 | 0.41 |
| 8 | 18655330 | rs13254153 | -1.605 | 1.197 | 1.637 | 2.453 | 0.055 | 0.233 | 0.233 | 0.18 | 0.13 | 0.35 | 0.18 | 0.37 | 0.12 | 0.16 |
| 8 | 18656601 | rs2248631 | 0.513 | 0.908 | 1.191 | 2.501 | 0.002 | 0.012 | 0.012 | 0 | 0.92 | 0.94 | 0.97 | 0.71 | 0.97 | 0.82 |
| 8 | 18656805 | rs2306825 | -1.553 | 0.954 | 1.511 | 2.32 | -0.011 | 0.029 | 0.029 | 0 | 0.17 | 0.24 | 0.17 | 0.3 | 0.16 | 0.17 |
| 8 | 18659019 | rs1386690 | 0.582 | 0.928 | 1.279 | 2.251 | -0.012 | 0.019 | 0.019 | 0 | 0.88 | 0.84 | 0.89 | 0.75 | 0.91 | 0.81 |
| 8 | 18661844 | rs13255143 | -0.875 | 1.004 | 1.4 | 2.392 | 0.011 | 0.058 | 0.058 | 0.368 | 0.1 | 0.16 | 0.15 | 0.3 | 0.14 | 0.16 |
| 8 | 18664022 | rs1386689 | 0.44 | 0.965 | 1.354 | 2.334 | -0.009 | 0.017 | 0.017 | 0 | 0.88 | 0.86 | 0.88 | 0.76 | 0.91 | 0.82 |
| 8 | 18665140 | rs7837572 | -1.795 | 0.932 | 1.435 | 2.315 | 0 | 0.069 | 0.069 | 0 | 0.19 | 0.4 | 0.2 | 0.31 | 0.19 | 0.17 |
| 8 | 18665523 | rs1487753 | -1.596 | 1.089 | 0.825 | 2.095 | 0.007 | 0.018 | 0.018 | 0 | 0.17 | 0.23 | 0.32 | 0.33 | 0.22 | 0.41 |
| 8 | 18665597 | rs11778064 | 2.347 | 1.025 | 0.394 | 1.944 | 0.002 | 0.03 | 0.03 | 0.136 | 0.57 | 0.22 | 0.42 | 0.36 | 0.6 | 0.35 |
| 8 | 18665836 | rs11203999 | -0.868 | 1.186 | -0.116 | 1.952 | 0 | 0.017 | 0.017 | 0 | 0.15 | 0.27 | 0.15 | 0.13 | 0.13 | 0.08 |
| 8 | 18668608 | rs2172290 | 1.246 | 1.329 | -1.789 | 1.398 | 0.018 | 0.026 | 0.026 | 0.425 | 0.63 | 0.2 | 0.57 | 0.3 | 0.66 | 0.57 |
| 8 | 18670645 | rs17467958 | -0.6 | 1.59 | -1.518 | 1.954 | -0.035 | -0.016 | -0.016 | 0.262 | 0.06 | 0.01 | 0.19 | 0.24 | 0.13 | 0.3 |
| 8 | 18670908 | rs2244320 | 1.056 | 0.831 | -1.421 | 1.51 | 0.024 | -0.061 | -0.061 | 0.15 | 0.87 | 0.96 | 0.85 | 0.65 | 0.9 | 0.78 |
| 8 | 18671742 | rs2638659 | -0.498 | 1.388 | -1.941 | -0.394 | -0.043 | 0.083 | 0.083 | 0 | 0.28 | 0.16 | 0.18 | 0 | 0.35 | 0.15 |
| 8 | 18671783 | rs2632841 | 0.483 | 0.835 | -2.784 | -0.912 | 0.004 | 0.193 | 0.193 | 0.112 | 0.46 | 0.12 | 0.6 | 0.8 | 0.6 | 0.53 |
| 8 | 18674816 | rs2632838 | 0.493 | 0.251 | -1.678 | -0.98 | -0.002 | 0.008 | 0.008 | 0 | 0.35 | 0.3 | 0.29 | 0.73 | 0.35 | 0.35 |
| 8 | 18675150 | rs2632837 | -0.249 | -0.997 | -0.98 | -1.446 | -0.04 | -0.043 | -0.043 | 0 | 0.83 | 0.7 | 0.86 | 0.88 | 0.89 | 0.79 |
| 8 | 18676521 | rs871762 | 0.288 | -1.167 | -0.149 | -1.552 | -0.059 | -0.069 | -0.069 | 0 | 0.8 | 0.61 | 0.85 | 0.9 | 0.89 | 0.79 |
| 8 | 18679795 | rs4921964 | -0.03 | -0.963 | -0.183 | -1.82 | -0.001 | 0.177 | 0.177 | 0 | 0.45 | 0.22 | 0.56 | 0.09 | 0.48 | 0.3 |
| 8 | 18682532 | rs11204000 | -0.967 | -1.29 | 0.299 | -1.985 | -0.02 | -0.034 | -0.034 | 0 | 0.31 | 0.48 | 0.43 | 0.27 | 0.28 | 0.38 |
| 8 | 18691956 | rs747555 | -0.132 | -1.23 | 0.369 | -1.969 | 0 | 0.03 | 0.03 | 0 | 0.37 | 0.3 | 0.29 | 0.17 | 0.43 | 0.37 |
| 8 | 18693453 | rs11782622 | 0.263 | -1.227 | 0.461 | -1.978 | 0.012 | 0.076 | 0.076 | 0 | 0.34 | 0.2 | 0.22 | 0.07 | 0.34 | 0.17 |
| 8 | 18694209 | rs6586769 | 0.224 | -1.228 | 0.492 | -1.968 | 0.008 | 0.026 | 0.026 | 0 | 0.34 | 0.27 | 0.23 | 0.08 | 0.34 | 0.2 |
| 8 | 18695756 | rs6982585 | -0.999 | -1.289 | 0.494 | -1.837 | 0 | -0.005 | -0.005 | 0 | 0.5 | 0.49 | 0.4 | 0.24 | 0.51 | 0.54 |
| 8 | 18695831 | rs6586773 | -2.206 | -1.383 | 0.67 | -1.582 | -0.02 | -0.017 | -0.017 | 0 | 0.69 | 0.76 | 0.73 | 0.41 | 0.65 | 0.68 |
| 8 | 18698031 | rs11785914 | -1.599 | -1.274 | 0.512 | -1.619 | -0.011 | 0.015 | 0.015 | 0 | 0.63 | 0.68 | 0.64 | 0.2 | 0.55 | 0.47 |
| 8 | 18698418 | rs4921966 | -0.017 | -1.379 | 0.348 | -1.681 | -0.01 | 0.197 | 0.197 | 0 | 0.41 | 0.15 | 0.47 | 0.07 | 0.44 | 0.25 |
| 8 | 18699659 | rs11204002 | -1.682 | -1.315 | 0.459 | -1.59 | -0.024 | 0.029 | 0.029 | 0 | 0.64 | 0.72 | 0.65 | 0.2 | 0.56 | 0.48 |
| 8 | 18699820 | rs11779137 | 0.288 | -1.062 | 0.424 | -1.405 | 0.006 | 0.018 | 0.018 | 0 | 0.16 | 0.14 | 0.3 | 0.05 | 0.14 | 0.11 |
| 8 | 18712513 | rs748208 | -0.259 | -1.003 | 0.462 | -1.328 | -0.014 | 0.131 | 0.131 | 0 | 0.68 | 0.88 | 0.79 | 0.93 | 0.66 | 0.84 |
| 8 | 18713148 | rs1492289 | 0.69 | -1.094 | 0.985 | -1.354 | 0.014 | 0.051 | 0.051 | 1.056 | 0.55 | 0.38 | 0.61 | 0.83 | 0.53 | 0.53 |
| 8 | 18714766 | rs17127341 | 0.474 | -1.201 | 0.857 | -1.304 | 0.002 | 0.003 | 0.003 | 0 | 0.13 | 0.14 | 0.27 | 0.05 | 0.09 | 0.1 |
| 8 | 18717366 | rs1116017 | -0.446 | -1.23 | 1.174 | -1.234 | 0.002 | 0.112 | 0.112 | 0 | 0.12 | 0.24 | 0.13 | 0.3 | 0.12 | 0.24 |
| 8 | 18720624 | rs2131181 | -0.738 | -1.248 | 1.125 | -1.42 | -0.004 | -0.006 | -0.006 | 0 | 0.74 | 0.75 | 0.78 | 0.98 | 0.65 | 0.85 |
| 8 | 18721215 | rs6998100 | 0.22 | -1.242 | 1.025 | -1.345 | -0.002 | -0.008 | -0.008 | 0 | 0.1 | 0.08 | 0.1 | 0.17 | 0.11 | 0.19 |
| 8 | 18722312 | rs768073 | 0.092 | -1.322 | 1.162 | -1.384 | -0.012 | -0.049 | -0.049 | 0.587 | 0.65 | 0.54 | 0.65 | 0.85 | 0.61 | 0.7 |
| 8 | 18725072 | rs1009482 | 0.158 | -1.332 | 0.832 | -1.35 | 0.007 | 0.021 | 0.021 | 0 | 0.14 | 0.12 | 0.27 | 0.03 | 0.1 | 0.11 |
| 8 | 18728941 | rs6993670 | -1.976 | -1.429 | 0.936 | -1.313 | -0.047 | 0.018 | 0.018 | 0.964 | 0.61 | 0.79 | 0.72 | 0.35 | 0.6 | 0.61 |
| 8 | 18729459 | rs13273954 | -1.202 | -1.42 | 0.7 | -1.309 | -0.037 | 0.056 | 0.056 | 0.038 | 0.22 | 0.37 | 0.24 | 0.31 | 0.17 | 0.34 |
| 8 | 18729818 | rs7003060 | -1.304 | -1.327 | 0.159 | -1.415 | -0.047 | 0.111 | 0.111 | 0.001 | 0.78 | 0.94 | 0.8 | 0.98 | 0.67 | 0.85 |
| 8 | 18730581 | rs17127379 | -0.26 | -1.195 | 0.496 | -1.022 | 0.011 | 0.02 | 0.02 | 0 | 0.14 | 0.12 | 0.27 | 0.05 | 0.1 | 0.11 |
| 8 | 18731495 | rs2029611 | -0.135 | -1.161 | 0.513 | -1.007 | 0.008 | 0.018 | 0.018 | 0 | 0.14 | 0.12 | 0.27 | 0.05 | 0.1 | 0.11 |
| 8 | 18741038 | rs7010288 | -1.89 | -1.172 | 0.588 | -0.993 | -0.031 | -0.008 | -0.008 | 0 | 0.34 | 0.5 | 0.5 | 0.32 | 0.26 | 0.43 |
| 8 | 18742049 | rs7843239 | -0.911 | -1.179 | 0.663 | -1.061 | 0.001 | 0.001 | 0.001 | 0 | 0.18 | 0.17 | 0.19 | 0.16 | 0.16 | 0.29 |
| 8 | 18742713 | rs12156368 | -1.368 | -0.951 | 0.321 | -0.911 | -0.022 | -0.011 | -0.011 | 0 | 0.31 | 0.55 | 0.5 | 0.29 | 0.26 | 0.43 |
| 8 | 18749120 | rs17127410 | -0.847 | -1.023 | -0.316 | -0.798 | 0.012 | 0.009 | 0.009 | 0.158 | 0.21 | 0.2 | 0.33 | 0.05 | 0.12 | 0.22 |
| 8 | 18763357 | rs2555619 | -1.326 | -0.478 | -0.398 | -0.585 | -0.024 | 0.002 | 0.002 | 0.095 | 0.6 | 0.68 | 0.65 | 0.4 | 0.59 | 0.6 |
| 8 | 18776114 | rs6999221 | -0.689 | -0.307 | -0.95 | -0.638 | 0.006 | 0.007 | 0.007 | 0.082 | 0.37 | 0.29 | 0.47 | 0.15 | 0.37 | 0.35 |
| 8 | 18777006 | rs1157546 | -0.451 | 0.037 | -1.066 | 0.129 | -0.001 | 0.001 | 0.001 | 0 | 0.11 | 0.09 | 0.22 | 0.06 | 0.06 | 0.09 |
| 8 | 18779024 | rs334748 | -0.475 | 0.866 | -1.697 | 0.666 | -0.108 | -0.079 | -0.079 | 0.037 | 0.76 | 0.96 | 0.75 | 0.74 | 0.71 | 0.75 |
| 8 | 18779065 | rs12156073 | -1.091 | 0.588 | -1.358 | 1.09 | 0.003 | -0.003 | -0.003 | 0 | 0.12 | 0.08 | 0.14 | 0.15 | 0.09 | 0.15 |
| 8 | 18779126 | rs12155975 | -1.193 | -0.367 | -1.518 | 1.209 | 0.037 | -0.006 | -0.006 | 0.889 | 0.26 | 0.15 | 0.26 | 0.31 | 0.14 | 0.25 |
| 8 | 18779787 | rs17127442 | -0.755 | -0.191 | -0.994 | 1.304 | 0.012 | -0.028 | -0.028 | 0.615 | 0.24 | 0.19 | 0.22 | 0.41 | 0.23 | 0.22 |
| 8 | 18785476 | rs6586786 | -1.731 | -0.235 | 0.228 | 1.252 | 0.037 | 0.035 | 0.035 | 0 | 0.46 | 0.7 | 0.53 | 0.66 | 0.38 | 0.48 |
| 8 | 18786682 | rs11996563 | 0.145 | -0.291 | 0.212 | 1.486 | -0.03 | 0.09 | 0.09 | 0 | 0.11 | 0.21 | 0.07 | 0.32 | 0.05 | 0.1 |
| 8 | 18787784 | rs17384209 | 0.372 | -0.313 | 0.087 | 1.491 | 0.005 | 0.009 | 0.009 | 0 | 0.14 | 0.11 | 0.14 | 0.06 | 0.15 | 0.12 |
| 8 | 18787795 | rs11786514 | -0.917 | -0.286 | 0.205 | 1.514 | -0.038 | 0.054 | 0.054 | 0 | 0.28 | 0.46 | 0.29 | 0.38 | 0.24 | 0.31 |
| 8 | 18789374 | rs932649 | -1.006 | -0.311 | 0.085 | 1.469 | -0.04 | 0.024 | 0.024 | 0.095 | 0.26 | 0.4 | 0.2 | 0.36 | 0.17 | 0.19 |
| 8 | 18790341 | rs7839913 | -1.171 | 0.038 | -0.573 | 1.277 | 0.045 | 0.04 | 0.04 | 0.047 | 0.46 | 0.77 | 0.52 | 0.65 | 0.38 | 0.48 |
| 8 | 18791682 | rs17468779 | -1.023 | 0.654 | -0.385 | 1.483 | -0.003 | 0.013 | 0.013 | 0 | 0.14 | 0.09 | 0.12 | 0.1 | 0.13 | 0.07 |
| 8 | 18793884 | rs13254740 | -0.935 | 0.54 | -0.359 | 1.509 | 0.001 | 0.012 | 0.012 | 0 | 0.14 | 0.08 | 0.12 | 0.1 | 0.11 | 0.07 |
| 8 | 18801513 | rs11784244 | -0.359 | 0.598 | -0.31 | 1.146 | 0.006 | 0.006 | 0.006 | 0.187 | 0.51 | 0.6 | 0.54 | 0.67 | 0.63 | 0.6 |
| 8 | 18804631 | rs6982196 | -0.418 | 1.143 | 0.036 | 1.139 | 0.025 | 0.026 | 0.026 | 0.238 | 0.52 | 0.64 | 0.54 | 0.65 | 0.63 | 0.58 |
| 8 | 18807198 | rs7001786 | 0.052 | 1.592 | 0.442 | 1.497 | 0.014 | 0.028 | 0.028 | 0.013 | 0.29 | 0.37 | 0.28 | 0.4 | 0.37 | 0.36 |
| 8 | 18812020 | rs1025229 | 1.031 | 1.506 | 0.841 | 1.568 | 0.042 | 0.04 | 0.04 | 0.237 | 0.48 | 0.3 | 0.46 | 0.38 | 0.37 | 0.41 |
| 8 | 18817819 | rs17469073 | -0.598 | 1.482 | 0.934 | 1.662 | 0.01 | 0 | 0 | 0 | 0.4 | 0.62 | 0.46 | 0.58 | 0.47 | 0.53 |
| 8 | 18819473 | rs11780765 | 0.582 | 2.007 | 0.949 | 2.073 | -0.016 | -0.043 | -0.043 | 0 | 0.18 | 0.06 | 0.16 | 0.21 | 0.21 | 0.19 |
| 8 | 18820391 | rs1019704 | 1.857 | 1.916 | 1.188 | 2.046 | 0.048 | 0.055 | 0.055 | 0.281 | 0.48 | 0.21 | 0.45 | 0.36 | 0.37 | 0.41 |

| | | | | | | | | | | | | | | | | |
|---|---|---|---|---|---|---|---|---|---|---|---|---|---|---|---|---|
| 8 | 18821054 | rs11775276 | -0.533 | 1.906 | 1.303 | 2.183 | 0.019 | 0.021 | 0.021 | 0.024 | 0.29 | 0.42 | 0.29 | 0.37 | 0.37 | 0.36 |
| 8 | 18821398 | rs11780329 | 0.607 | 1.865 | 1.264 | 2.169 | -0.023 | -0.043 | -0.043 | 0 | 0.17 | 0.06 | 0.15 | 0.2 | 0.22 | 0.18 |
| 8 | 18828343 | rs4921622 | 1.184 | 1.867 | 1.537 | 2.25 | -0.005 | -0.002 | -0.002 | 0.044 | 0.58 | 0.41 | 0.59 | 0.55 | 0.43 | 0.58 |
| 8 | 18829714 | rs6586789 | 0.583 | 1.867 | 1.519 | 2.387 | 0.013 | 0.049 | 0.049 | 0 | 0.78 | 0.6 | 0.74 | 0.68 | 0.77 | 0.78 |
| 8 | 18829923 | rs11783114 | 0.348 | 1.878 | 1.508 | 2.326 | -0.042 | -0.011 | -0.011 | 0 | 0.19 | 0.04 | 0.16 | 0.15 | 0.25 | 0.18 |
| 8 | 18830828 | rs720266 | 0.653 | 1.878 | 1.497 | 2.229 | 0.012 | 0 | 0 | 0 | 0.78 | 0.6 | 0.74 | 0.77 | 0.77 | 0.76 |
| 8 | 18831269 | rs12548880 | 0.43 | 1.363 | 1.499 | 2.048 | 0.067 | 0.041 | 0.041 | 0.122 | 0.11 | 0.3 | 0.13 | 0.17 | 0.16 | 0.18 |
| 8 | 18836545 | rs4921978 | -0.447 | 1.278 | 1.34 | 2.056 | 0.019 | 0.013 | 0.013 | 0 | 0.88 | 0.75 | 0.84 | 0.85 | 0.84 | 0.9 |
| 8 | 18839091 | rs1426919 | -0.872 | 1.267 | 1.115 | 2.175 | -0.119 | -0.064 | -0.064 | 0 | 0.36 | 0.13 | 0.36 | 0.38 | 0.46 | 0.46 |
| 8 | 18842968 | rs6981220 | -0.256 | 1.342 | 1.089 | 2.184 | -0.055 | -0.037 | -0.037 | 4.102 | 0.27 | 0.09 | 0.25 | 0.27 | 0.33 | 0.3 |
| 8 | 18844382 | rs10216388 | 0.911 | 1.381 | 1.514 | 2.17 | 0.008 | 0.002 | 0.002 | 0.011 | 0.62 | 0.58 | 0.68 | 0.62 | 0.54 | 0.54 |
| 8 | 18846401 | rs17384769 | 0.845 | 1.404 | 1.396 | 2.203 | -0.04 | 0.006 | 0.006 | 0 | 0.08 | 0.03 | 0.05 | 0.06 | 0.13 | 0.14 |
| 8 | 18846534 | rs7017835 | -0.251 | 1.384 | 1.413 | 2.182 | -0.02 | -0.031 | -0.031 | 2.975 | 0.26 | 0.2 | 0.25 | 0.31 | 0.31 | 0.31 |
| 8 | 18847646 | rs2059657 | -0.105 | 1.373 | 1.492 | 2.197 | 0.008 | 0.03 | 0.03 | 0 | 0.86 | 0.73 | 0.85 | 0.8 | 0.84 | 0.9 |
| 8 | 18848091 | rs2059660 | -0.285 | 1.375 | 1.338 | 2.169 | -0.029 | -0.023 | -0.023 | 0 | 0.25 | 0.15 | 0.24 | 0.26 | 0.31 | 0.29 |
| 8 | 18852474 | rs890518 | -0.181 | 1.379 | 1.293 | 2.182 | -0.059 | -0.071 | -0.071 | 0.774 | 0.24 | 0.09 | 0.23 | 0.31 | 0.3 | 0.28 |
| 8 | 18852851 | rs10429350 | 1.623 | 1.694 | 1.659 | 2.09 | 0.033 | 0.015 | 0.015 | 0.777 | 0.5 | 0.4 | 0.53 | 0.45 | 0.37 | 0.46 |
| 8 | 18853687 | rs7824049 | -0.472 | 1.766 | 1.788 | 2.102 | 0.001 | 0.005 | 0.005 | 0.056 | 0.28 | 0.27 | 0.29 | 0.41 | 0.34 | 0.29 |
| 8 | 18853940 | rs11988880 | -0.446 | 1.14 | 1.45 | 1.639 | 0.021 | 0.028 | 0.028 | 0 | 0.16 | 0.36 | 0.14 | 0.23 | 0.17 | 0.2 |
| 8 | 18854678 | rs11204014 | -1.221 | 1.092 | 1.23 | 1.526 | -0.009 | -0.008 | -0.008 | 0 | 0.43 | 0.67 | 0.4 | 0.5 | 0.48 | 0.49 |
| 8 | 18856363 | rs1365600 | 0.451 | 1.111 | 1.3 | 1.157 | 0.035 | -0.058 | -0.058 | 0 | 0.78 | 0.61 | 0.83 | 0.89 | 0.72 | 0.81 |
| 8 | 18857090 | rs13256382 | 0.096 | 1.364 | 0.993 | 1.529 | -0.004 | -0.006 | -0.006 | 1.799 | 0.22 | 0.17 | 0.24 | 0.36 | 0.29 | 0.3 |
| 8 | 18858239 | rs6994627 | -0.137 | 1.262 | 1.174 | 1.344 | 0.008 | 0.012 | 0.012 | 0.572 | 0.24 | 0.22 | 0.25 | 0.4 | 0.33 | 0.31 |
| 8 | 18866368 | rs7010496 | -0.413 | 1.443 | 1.178 | 1.164 | 0.067 | 0 | 0 | 0 | 0.05 | 0.2 | 0.07 | 0.05 | 0.11 | 0.03 |
| 8 | 18870349 | rs7002540 | -0.82 | 1.37 | 1.133 | 1.116 | 0.077 | -0.038 | -0.038 | 0 | 0.16 | 0.4 | 0.14 | 0.1 | 0.24 | 0.17 |
| 8 | 18873422 | rs4921982 | 1.386 | 0.606 | -0.053 | 0.943 | 0.025 | -0.092 | -0.092 | 3.358 | 0.48 | 0.05 | 0.5 | 0.43 | 0.36 | 0.42 |
| 8 | 18892291 | rs2048090 | -1.032 | 0.776 | 0.641 | 0.761 | 0.008 | -0.009 | -0.009 | 0 | 0.27 | 0.42 | 0.29 | 0.31 | 0.3 | 0.3 |
| 8 | 18896703 | rs6984468 | 0.808 | 0.799 | 0.607 | 0.845 | -0.003 | -0.016 | -0.016 | 0.012 | 0.62 | 0.27 | 0.48 | 0.47 | 0.43 | 0.54 |
| 8 | 18898040 | rs10104732 | -0.683 | 0.835 | 0.874 | 0.849 | -0.047 | -0.062 | -0.062 | 0 | 0.8 | 0.9 | 0.64 | 0.57 | 0.67 | 0.71 |
| 8 | 18898198 | rs170794 | 0.445 | 0.824 | 0.853 | 0.844 | -0.08 | -0.103 | -0.103 | 0 | 0.2 | 0.06 | 0.36 | 0.43 | 0.33 | 0.3 |
| 8 | 18899170 | rs764511 | -1.029 | 0.878 | 1.027 | 0.794 | 0.102 | 0.033 | 0.033 | 0.03 | 0.09 | 0.47 | 0.09 | 0.05 | 0.11 | 0.05 |
| 8 | 18901644 | rs1529348 | -0.653 | 0.946 | 0.682 | 0.86 | -0.051 | -0.072 | -0.072 | 0 | 0.8 | 0.9 | 0.64 | 0.57 | 0.67 | 0.71 |
| 8 | 18903018 | rs7841005 | -0.745 | 0.776 | 0.43 | 0.772 | 0 | -0.002 | -0.002 | 1.047 | 0.26 | 0.26 | 0.25 | 0.27 | 0.24 | 0.31 |
| 8 | 18904368 | rs919149 | 1.341 | 0.416 | -0.227 | 0.662 | 0.089 | 0.083 | 0.083 | 2.705 | 0.46 | 0.18 | 0.3 | 0.25 | 0.33 | 0.37 |
| 8 | 18914420 | rs6985805 | -0.383 | 0.319 | -0.161 | 0.215 | 0.012 | -0.003 | -0.003 | 0.194 | 0.17 | 0.09 | 0.19 | 0.22 | 0.12 | 0.18 |
| 8 | 18919720 | rs1027803 | 1.041 | 0.084 | 0.07 | 0.044 | 0.008 | 0.014 | 0.014 | 0.115 | 0.71 | 0.32 | 0.81 | 0.87 | 0.78 | 0.78 |
| 8 | 18919857 | rs1027804 | 0.41 | 0.463 | -0.072 | 0.217 | -0.008 | -0.008 | -0.008 | 0 | 0.2 | 0.2 | 0.38 | 0.61 | 0.31 | 0.23 |
| 8 | 18922377 | rs2410599 | 0.88 | -0.557 | -0.335 | -0.946 | -0.013 | -0.105 | -0.105 | 0 | 0.35 | 0.48 | 0.23 | 0.09 | 0.31 | 0.4 |
| 8 | 18924056 | rs4921984 | 0.518 | -0.169 | -0.554 | -0.957 | -0.016 | -0.01 | -0.01 | 0 | 0.24 | 0.25 | 0.49 | 0.61 | 0.44 | 0.27 |
| 8 | 18926800 | rs36110451 | 0.407 | -0.175 | -0.322 | -1.153 | 0.03 | 0.175 | 0.175 | 0 | 0.5 | 0.2 | 0.32 | 0.18 | 0.44 | 0.47 |
| 8 | 18926827 | rs5026830 | 0.362 | -0.158 | -0.254 | -1.145 | 0.027 | 0.151 | 0.151 | 0 | 0.51 | 0.22 | 0.32 | 0.18 | 0.44 | 0.47 |
| 8 | 18928440 | rs1472834 | 0.808 | -0.194 | -0.295 | -1.066 | 0.008 | 0.048 | 0.048 | 0 | 0.33 | 0.2 | 0.19 | 0.08 | 0.29 | 0.36 |
| 8 | 18930380 | rs7000610 | -0.106 | -0.102 | 0.163 | -0.258 | -0.041 | -0.034 | -0.034 | 0 | 0.76 | 0.57 | 0.85 | 0.84 | 0.86 | 0.81 |
| 8 | 18936250 | rs17127791 | -0.372 | 0.859 | -0.55 | 0.065 | 0.002 | -0.004 | -0.004 | 0 | 0.36 | 0.37 | 0.26 | 0.29 | 0.37 | 0.37 |
| 8 | 18936350 | rs1506896 | 0.18 | 0.866 | -0.514 | -0.387 | 0 | -0.006 | -0.006 | 0.274 | 0.68 | 0.38 | 0.5 | 0.26 | 0.66 | 0.65 |
| 8 | 18937033 | rs17469836 | 0.658 | 0.902 | -1.423 | -0.645 | -0.176 | 0.088 | 0.088 | 0 | 0.37 | 0.05 | 0.39 | 0.19 | 0.5 | 0.45 |
| 8 | 18938601 | rs10101681 | 0.19 | 0.739 | -0.527 | -0.721 | 0.001 | 0 | 0 | 0 | 0.47 | 0.44 | 0.42 | 0.19 | 0.61 | 0.61 |
| 8 | 18939267 | rs6651477 | 0.191 | 0.48 | -0.848 | -1.04 | 0.004 | 0.017 | 0.017 | 0.329 | 0.76 | 0.84 | 0.88 | 0.92 | 0.8 | 0.8 |
| 8 | 18940973 | rs17385318 | 0.057 | 0.312 | -0.791 | -0.956 | -0.002 | -0.007 | -0.007 | 0 | 0.11 | 0.11 | 0.07 | 0.06 | 0.12 | 0.12 |
| 8 | 18941341 | rs6651480 | -0.541 | 0.377 | -1.051 | -1.141 | 0 | 0.156 | 0.156 | 0 | 0.54 | 0.81 | 0.6 | 0.85 | 0.42 | 0.47 |
| 8 | 18941488 | rs9657493 | 1.056 | 0.478 | -1.142 | -1.038 | -0.164 | 0.09 | 0.09 | 0 | 0.34 | 0.05 | 0.32 | 0.12 | 0.46 | 0.41 |
| 8 | 18942656 | rs7464855 | 0.86 | 0.256 | -0.334 | -0.899 | 0.011 | -0.024 | -0.024 | 0 | 0.4 | 0.46 | 0.37 | 0.2 | 0.49 | 0.44 |
| 8 | 18943802 | rs17469898 | 0.121 | 0.077 | -0.334 | -0.691 | 0 | -0.003 | -0.003 | 0.29 | 0.34 | 0.21 | 0.51 | 0.68 | 0.32 | 0.32 |
| 8 | 18945519 | rs7839794 | -0.015 | 0.249 | -0.318 | -0.699 | -0.003 | -0.021 | -0.021 | 2.045 | 0.59 | 0.54 | 0.46 | 0.16 | 0.63 | 0.5 |
| 8 | 18946962 | rs12545227 | -0.148 | -0.476 | -0.156 | -0.222 | -0.032 | 0.002 | 0.002 | 0.45 | 0.69 | 0.44 | 0.67 | 0.6 | 0.78 | 0.65 |
| 8 | 18947138 | rs11204020 | -0.503 | -0.262 | -0.56 | -0.278 | -0.021 | -0.01 | -0.01 | 0 | 0.73 | 0.59 | 0.67 | 0.7 | 0.77 | 0.7 |
| 8 | 18948957 | rs11784798 | 1.595 | -0.159 | -1.458 | -0.419 | -0.05 | 0.027 | 0.027 | 0 | 0.26 | 0.04 | 0.19 | 0.1 | 0.31 | 0.32 |
| 8 | 18951822 | rs10108465 | 0.138 | 0.148 | -1.015 | -0.283 | 0.065 | 0.066 | 0.066 | 0 | 0.27 | 0.38 | 0.41 | 0.49 | 0.38 | 0.22 |
| 8 | 18954802 | rs4921994 | -0.49 | -0.701 | -1.413 | -0.158 | -0.016 | -0.003 | -0.003 | 0 | 0.81 | 0.77 | 0.89 | 0.82 | 0.89 | 0.87 |
| 8 | 18955169 | rs7014714 | 0.061 | -0.081 | -2.103 | 0.043 | 0.104 | 0.042 | 0.042 | 0 | 0.54 | 0.18 | 0.56 | 0.57 | 0.68 | 0.58 |
| 8 | 18959366 | rs2048091 | -0.626 | 0.529 | -2.097 | 0.011 | -0.019 | 0.017 | 0.017 | 0 | 0.3 | 0.13 | 0.27 | 0.22 | 0.33 | 0.26 |
| 8 | 18960448 | rs13266438 | -0.219 | 0.403 | -2.218 | -0.509 | -0.001 | 0.069 | 0.069 | 0.243 | 0.36 | 0.25 | 0.26 | 0.18 | 0.37 | 0.16 |
| 8 | 18962087 | rs10102187 | -0.799 | 0.221 | -2.994 | -0.032 | -0.122 | -0.129 | -0.129 | 0 | 0.63 | 0.9 | 0.53 | 0.59 | 0.54 | 0.4 |
| 8 | 18962544 | rs10088192 | -0.329 | -0.027 | -2.317 | -0.353 | 0.019 | 0.088 | 0.088 | 0.968 | 0.5 | 0.31 | 0.33 | 0.37 | 0.45 | 0.21 |
| 8 | 18962643 | rs10102778 | -0.427 | -0.053 | -2.079 | -0.364 | 0.009 | 0.064 | 0.064 | 0.783 | 0.51 | 0.35 | 0.34 | 0.37 | 0.45 | 0.22 |
| 8 | 18967822 | rs11779160 | 1.087 | -0.159 | -2.564 | -0.519 | -0.025 | 0.001 | 0.001 | 0 | 0.21 | 0.01 | 0.14 | 0.19 | 0.2 | 0.08 |
| 8 | 18969786 | rs7817975 | 0.239 | -0.158 | -2.312 | -0.502 | 0.017 | -0.023 | -0.023 | 0 | 0.14 | 0.03 | 0.1 | 0.18 | 0.09 | 0.05 |
| 8 | 18973729 | rs10111009 | -0.766 | -0.07 | -1.251 | -0.588 | 0.008 | -0.032 | -0.032 | 0 | 0.22 | 0.54 | 0.19 | 0.18 | 0.18 | 0.12 |
| 8 | 18973808 | rs17127839 | 1.067 | 0.274 | -1.969 | -0.881 | -0.026 | -0.059 | -0.059 | 0 | 0.3 | 0.16 | 0.34 | 0.47 | 0.36 | 0.5 |

| | | | | | | | | | | | | | | | | |
|---|---|---|---|---|---|---|---|---|---|---|---|---|---|---|---|---|
| 8 | 18975171 | rs1027802 | 0.41 | 0.313 | -1.479 | -1.116 | 0.001 | 0.001 | 0.001 | 2.645 | 0.12 | 0.13 | 0.09 | 0.1 | 0.09 | 0.08 |
| 8 | 18976000 | rs13262483 | 0.554 | 0.469 | -1.564 | -1.131 | -0.031 | -0.012 | -0.012 | 3.498 | 0.24 | 0.02 | 0.25 | 0.19 | 0.22 | 0.11 |
| 8 | 18979716 | rs925037 | 1.181 | 0.744 | -1.189 | -1.328 | -0.015 | -0.011 | -0.011 | 1.193 | 0.31 | 0.27 | 0.48 | 0.58 | 0.42 | 0.38 |
| 8 | 18981713 | rs13268591 | 0.191 | 0.571 | -1.178 | -1.584 | 0.014 | 0.045 | 0.045 | 1.984 | 0.32 | 0.17 | 0.3 | 0.22 | 0.3 | 0.17 |
| 8 | 18985382 | rs333041 | -1.514 | 1.086 | -0.719 | -1.709 | -0.001 | 0.001 | 0.001 | 0 | 0.68 | 0.64 | 0.54 | 0.4 | 0.65 | 0.61 |
| 8 | 18986961 | rs6985201 | 0.148 | 1.35 | -1.142 | -1.771 | 0.013 | -0.02 | -0.02 | 0 | 0.13 | 0.17 | 0.12 | 0.03 | 0.24 | 0.14 |
| 8 | 18989735 | rs692512 | -0.055 | 0.562 | -0.806 | -1.879 | 0.016 | 0.058 | 0.058 | 1.069 | 0.34 | 0.22 | 0.21 | 0.2 | 0.27 | 0.43 |
| 8 | 18991089 | rs13264451 | 1.419 | 0.663 | -1.087 | -1.954 | -0.01 | -0.022 | -0.022 | 0 | 0.21 | 0.01 | 0.22 | 0.18 | 0.17 | 0.07 |
| 8 | 18991838 | rs17470093 | 0.509 | 0.95 | -0.673 | -1.822 | 0.013 | -0.027 | -0.027 | 0.236 | 0.11 | 0.17 | 0.08 | 0.04 | 0.17 | 0.11 |
| 8 | 18996630 | rs17385577 | 0.407 | 0.191 | -0.494 | -1.856 | 0.006 | 0.024 | 0.024 | 0 | 0.46 | 0.44 | 0.57 | 0.65 | 0.61 | 0.5 |
| 8 | 18997571 | rs333029 | 0.365 | 0.229 | -0.217 | -1.206 | -0.003 | -0.001 | -0.001 | 0 | 0.09 | 0.07 | 0.1 | 0.1 | 0.1 | 0.1 |
| 8 | 19003304 | rs679287 | -0.72 | 0.14 | -0.307 | -1.32 | -0.007 | -0.022 | -0.022 | 0.271 | 0.41 | 0.53 | 0.37 | 0.3 | 0.36 | 0.22 |
| 8 | 19007442 | rs11785844 | -0.311 | 0.052 | -0.26 | -1.118 | 0.001 | -0.025 | -0.025 | 1.863 | 0.13 | 0.17 | 0.08 | 0.02 | 0.15 | 0.08 |
| 8 | 19007783 | rs11775805 | -1.166 | 0.016 | -0.046 | -1.32 | 0.005 | -0.091 | -0.091 | 0 | 0.21 | 0.35 | 0.12 | 0.02 | 0.22 | 0.1 |
| 8 | 19008265 | rs170773 | -1.365 | -0.047 | -0.337 | -1.299 | -0.001 | -0.009 | -0.009 | 0 | 0.48 | 0.52 | 0.4 | 0.36 | 0.44 | 0.24 |
| 8 | 19012671 | rs2410603 | 0.926 | -0.313 | -0.749 | -1.234 | 0.003 | -0.043 | -0.043 | 0 | 0.16 | 0.01 | 0.16 | 0.22 | 0.11 | 0.03 |
| 8 | 19018044 | rs10503648 | -0.189 | -0.106 | -0.567 | -1.513 | -0.019 | 0.058 | 0.058 | 2.85 | 0.16 | 0.06 | 0.09 | 0.02 | 0.19 | 0.12 |
| 8 | 19018434 | rs691589 | 0.291 | -0.177 | 0.073 | -1.403 | 0.008 | -0.043 | -0.043 | 0 | 0.72 | 0.72 | 0.79 | 0.85 | 0.7 | 0.78 |
| 8 | 19018974 | rs590640 | 0.288 | -0.176 | -0.004 | -1.403 | -0.002 | -0.003 | -0.003 | 1.117 | 0.72 | 0.76 | 0.79 | 0.85 | 0.69 | 0.78 |
| 8 | 19019943 | rs2672275 | 0.346 | -0.201 | -0.059 | -1.302 | -0.001 | -0.006 | -0.006 | 0.82 | 0.7 | 0.7 | 0.79 | 0.81 | 0.69 | 0.78 |
| 8 | 19020143 | rs11204022 | 1.028 | -0.176 | -0.259 | -1.086 | 0.002 | 0.092 | 0.092 | 0.146 | 0.15 | 0.01 | 0.07 | 0 | 0.1 | 0.02 |
| 8 | 19028649 | rs691605 | 0.528 | 0.017 | -0.05 | -0.564 | -0.017 | -0.029 | -0.029 | 0.053 | 0.69 | 0.51 | 0.79 | 0.71 | 0.73 | 0.77 |
| 8 | 19030337 | rs12544783 | -0.113 | 0.426 | -0.383 | -0.13 | -0.001 | -0.002 | -0.002 | 0 | 0.66 | 0.33 | 0.52 | 0.5 | 0.65 | 0.38 |
| 8 | 19036043 | rs333056 | 0.578 | -0.118 | -0.075 | -0.222 | 0.008 | 0.003 | 0.003 | 2.589 | 0.62 | 0.34 | 0.53 | 0.63 | 0.7 | 0.65 |
| 8 | 19037928 | rs6998937 | -0.149 | -0.234 | 0.472 | -0.304 | -0.003 | 0.021 | 0.021 | 3.269 | 0.19 | 0.26 | 0.19 | 0.28 | 0.18 | 0.19 |
| 8 | 19039163 | rs333052 | 0.745 | -0.255 | 0.731 | -0.736 | -0.015 | -0.071 | -0.071 | 0 | 0.78 | 0.5 | 0.8 | 0.93 | 0.82 | 0.81 |
| 8 | 19039642 | rs184565 | 0.702 | -0.21 | 0.99 | -0.644 | 0 | -0.004 | -0.004 | 0 | 0.68 | 0.65 | 0.72 | 0.74 | 0.67 | 0.75 |
| 8 | 19041987 | rs1395524 | -0.441 | -0.158 | 0.34 | -0.651 | -0.014 | -0.014 | -0.014 | 0 | 0.22 | 0.28 | 0.36 | 0.16 | 0.15 | 0.28 |
| 8 | 19044189 | rs6982478 | -1.58 | -0.189 | 0.388 | -0.671 | 0 | 0 | 0 | 0 | 0.48 | 0.52 | 0.54 | 0.36 | 0.43 | 0.41 |
| 8 | 19044253 | rs17127960 | 0.657 | -0.071 | 0.215 | -0.394 | -0.004 | 0.016 | 0.016 | 0 | 0.06 | 0.16 | 0.11 | 0.05 | 0.05 | 0.13 |
| 8 | 19044862 | rs17127963 | -0.089 | 0.133 | 0.108 | -0.446 | -0.014 | -0.03 | -0.03 | 1.78 | 0.16 | 0.22 | 0.26 | 0.09 | 0.1 | 0.16 |
| 8 | 19047187 | rs968001 | 0.119 | 0.577 | 0.344 | 0.211 | 0.019 | 0.069 | 0.069 | 2.23 | 0.12 | 0.2 | 0.17 | 0.28 | 0.18 | 0.06 |
| 8 | 19047343 | rs4922001 | 0.451 | 0.356 | 0.836 | -0.328 | -0.006 | -0.058 | -0.058 | 1.691 | 0.86 | 0.69 | 0.9 | 0.97 | 0.86 | 0.86 |
| 8 | 19047771 | rs333042 | -0.816 | 0.356 | 0.274 | -1.187 | 0 | -0.019 | -0.019 | 0 | 0.5 | 0.49 | 0.66 | 0.89 | 0.49 | 0.61 |
| 8 | 19049811 | rs333047 | -0.046 | 0.446 | 0.166 | -0.876 | 0.002 | 0.004 | 0.004 | 0.143 | 0.34 | 0.27 | 0.54 | 0.68 | 0.3 | 0.49 |
| 8 | 19052474 | rs12678474 | -0.835 | 0.978 | -0.009 | -0.199 | 0.003 | 0.012 | 0.012 | 0.035 | 0.25 | 0.33 | 0.45 | 0.61 | 0.3 | 0.39 |
| 8 | 19054974 | rs7825941 | -0.696 | 1.215 | 0.18 | -0.189 | 0.02 | -0.004 | -0.004 | 0.6 | 0.3 | 0.41 | 0.36 | 0.65 | 0.41 | 0.34 |
| 8 | 19056642 | rs1567926 | -0.579 | 1.001 | -0.195 | -0.19 | -0.001 | 0.003 | 0.003 | 3.619 | 0.84 | 0.85 | 0.91 | 0.8 | 0.83 | 0.91 |
| 8 | 19058430 | rs11204024 | 0.491 | 0.987 | -0.331 | -0.44 | -0.001 | 0 | 0 | 0 | 0.6 | 0.34 | 0.54 | 0.39 | 0.49 | 0.52 |
| 8 | 19067320 | rs11986238 | 0.833 | 1.123 | -0.458 | 0.106 | -0.023 | 0.045 | 0.045 | 0 | 0.07 | 0.02 | 0.19 | 0.41 | 0.11 | 0.12 |
| 8 | 19067578 | rs17127976 | 1.037 | 0.735 | -0.489 | -0.669 | 0.002 | -0.016 | -0.016 | 0 | 0.34 | 0.32 | 0.21 | 0.08 | 0.26 | 0.2 |
| 8 | 19070373 | rs1193327 | 0.058 | 0.408 | -0.486 | -0.63 | 0.001 | 0.002 | 0.002 | 0 | 0.55 | 0.51 | 0.47 | 0.51 | 0.48 | 0.49 |
| 8 | 19071911 | rs1193324 | 0.144 | 0.22 | -0.406 | -0.44 | 0.014 | -0.015 | -0.015 | 0 | 0.81 | 0.89 | 0.69 | 0.75 | 0.84 | 0.72 |
| 8 | 19072536 | rs12544391 | 0.031 | 0.18 | -0.477 | -0.426 | 0.022 | -0.03 | -0.03 | 0.188 | 0.15 | 0.07 | 0.27 | 0.27 | 0.09 | 0.32 |
| 8 | 19072672 | rs4922003 | -0.147 | 0.3 | -0.173 | -0.351 | 0.002 | 0.075 | 0.075 | 0 | 0.1 | 0.11 | 0.21 | 0.46 | 0.12 | 0.22 |
| 8 | 19074836 | rs2683300 | 0.399 | 0.382 | -0.045 | -0.555 | 0.003 | 0.003 | 0.003 | 0.95 | 0.74 | 0.73 | 0.49 | 0.3 | 0.74 | 0.6 |
| 8 | 19076743 | rs11204025 | 1.164 | 0.324 | -0.431 | -0.825 | -0.04 | 0 | 0 | 0 | 0.24 | 0.15 | 0.24 | 0.22 | 0.37 | 0.15 |
| 8 | 19076995 | rs2947255 | -0.272 | 0.019 | -0.062 | -0.968 | -0.002 | 0.001 | 0.001 | 1.347 | 0.46 | 0.47 | 0.58 | 0.46 | 0.56 | 0.48 |
| 8 | 19077526 | rs1915411 | 0.637 | 0.084 | -0.302 | -0.959 | 0 | 0.028 | 0.028 | 0 | 0.15 | 0.07 | 0.09 | 0 | 0.17 | 0.16 |
| 8 | 19079284 | rs6993460 | 1.158 | 0.091 | -0.122 | -0.656 | -0.039 | 0.009 | 0.009 | 0.613 | 0.24 | 0.15 | 0.24 | 0.21 | 0.36 | 0.15 |
| 8 | 19079796 | rs2090789 | -0.364 | -0.067 | 0.138 | -0.624 | 0.012 | 0.005 | 0.005 | 0.097 | 0.62 | 0.54 | 0.47 | 0.56 | 0.54 | 0.62 |
| 8 | 19080151 | rs12056334 | -0.162 | -0.056 | 0.236 | 0.194 | 0.001 | 0.179 | 0.179 | 0 | 0.12 | 0.15 | 0.23 | 0.44 | 0.11 | 0.11 |
| 8 | 19081781 | rs977262 | 0.389 | 0.113 | -0.423 | -0.63 | -0.001 | -0.067 | -0.067 | 0 | 0.71 | 0.71 | 0.86 | 0.99 | 0.71 | 0.72 |
| 8 | 19085172 | rs991359 | 0.321 | 0.072 | -0.539 | 0.111 | -0.002 | 0.016 | 0.016 | 0 | 0.63 | 0.52 | 0.64 | 0.51 | 0.64 | 0.47 |
| 8 | 19090103 | rs7817195 | -0.011 | 0.014 | -0.95 | 0.276 | -0.021 | 0.015 | 0.015 | 0 | 0.14 | 0.2 | 0.23 | 0.19 | 0.09 | 0.1 |
| 8 | 19093006 | rs1915418 | 0.14 | 0.77 | -1.481 | 0.349 | -0.035 | -0.03 | -0.03 | 0 | 0.58 | 0.75 | 0.55 | 0.52 | 0.56 | 0.36 |
| 8 | 19106346 | rs1877856 | 0.73 | 0.642 | -0.945 | 0.006 | -0.002 | -0.026 | -0.026 | 0.021 | 0.86 | 0.85 | 0.93 | 0.97 | 0.89 | 0.79 |
| 8 | 19107092 | rs7010676 | -0.765 | 0.403 | -1.066 | -0.314 | 0.001 | 0.001 | 0.001 | 1.985 | 0.3 | 0.24 | 0.17 | 0.13 | 0.28 | 0.17 |
| 8 | 19108021 | rs1915413 | -0.236 | 0.306 | -0.737 | 0.034 | -0.002 | 0 | 0 | 0 | 0.48 | 0.48 | 0.35 | 0.46 | 0.37 | 0.38 |
| 8 | 19109719 | rs10092053 | -0.103 | 0.269 | -0.827 | 0.104 | 0.001 | 0.018 | 0.018 | 0 | 0.15 | 0.16 | 0.32 | 0.33 | 0.16 | 0.28 |
| 8 | 19111578 | rs1399395 | 0.526 | -0.213 | -0.536 | 0.025 | 0.006 | -0.005 | -0.005 | 0.022 | 0.34 | 0.27 | 0.3 | 0.32 | 0.3 | 0.19 |
| 8 | 19112868 | rs2672230 | -0.026 | -0.169 | -0.533 | 0.078 | 0 | 0 | 0 | 0.012 | 0.48 | 0.5 | 0.35 | 0.43 | 0.38 | 0.39 |
| 8 | 19113393 | rs11786215 | 0.286 | 0.562 | -0.846 | -0.137 | -0.033 | 0.051 | 0.051 | 0 | 0.1 | 0.05 | 0.08 | 0 | 0.15 | 0.1 |
| 8 | 19120154 | rs11780720 | 0.256 | 0.46 | -0.903 | -0.128 | -0.046 | 0.041 | 0.041 | 0 | 0.45 | 0.1 | 0.35 | 0.3 | 0.45 | 0.28 |
| 8 | 19121318 | rs967509 | 0.262 | 0.133 | -0.223 | -0.215 | -0.063 | -0.06 | -0.06 | 0 | 0.22 | 0.38 | 0.08 | 0.13 | 0.11 | 0.29 |
| 8 | 19121924 | rs9693033 | -1.163 | 0.202 | 0.127 | -0.2 | -0.005 | 0.034 | 0.034 | 0 | 0.26 | 0.42 | 0.4 | 0.34 | 0.26 | 0.35 |
| 8 | 19122763 | rs2244965 | -1.271 | 0.311 | 0.204 | -0.137 | 0.001 | -0.001 | -0.001 | 0.097 | 0.75 | 0.73 | 0.72 | 0.78 | 0.64 | 0.76 |
| 8 | 19128303 | rs9644625 | 0.952 | 0.301 | 0.238 | 0.026 | -0.001 | 0.061 | 0.061 | 0 | 0.06 | 0.05 | 0.28 | 0.32 | 0.08 | 0.19 |
| 8 | 19128886 | rs2683279 | -0.509 | 0.203 | 0.24 | -0.089 | 0.023 | -0.047 | -0.047 | 0 | 0.66 | 0.5 | 0.68 | 0.75 | 0.6 | 0.7 |

| | | | | | | | | | | | | | | | | |
|---|---|---|---|---|---|---|---|---|---|---|---|---|---|---|---|---|
| 8 | 19137542 | rs2410608 | -0.356 | 0.206 | 0.424 | 0.05 | -0.006 | -0.009 | -0.009 | 0.4 | 0.16 | 0.13 | 0.37 | 0.32 | 0.23 | 0.26 |
| 8 | 19144414 | rs7831738 | 0.829 | 0.207 | 0.746 | 0.148 | 0.007 | 0.125 | 0.125 | 0 | 0.06 | 0.08 | 0.28 | 0.32 | 0.08 | 0.18 |
| 8 | 19145032 | rs1193293 | -1.234 | 0.227 | 0.758 | 0.024 | -0.004 | 0.001 | 0.001 | 0 | 0.75 | 0.75 | 0.72 | 0.76 | 0.64 | 0.76 |
| 8 | 19145800 | rs1193288 | -1.549 | 0.265 | 0.777 | 0.008 | -0.014 | 0.005 | 0.005 | 0 | 0.78 | 0.81 | 0.75 | 0.78 | 0.69 | 0.77 |
| 8 | 19150829 | rs2683321 | -0.112 | 0.04 | 0.766 | -0.022 | 0.028 | 0.018 | 0.018 | 0 | 0.08 | 0.25 | 0.06 | 0.08 | 0.08 | 0.1 |
| 8 | 19150880 | rs10503656 | 0.925 | -0.08 | 0.732 | 0.028 | 0.052 | -0.012 | -0.012 | 2.844 | 0.46 | 0.18 | 0.29 | 0.4 | 0.34 | 0.34 |
| 8 | 19151139 | rs9325854 | 0.657 | -0.087 | 0.799 | 0.023 | -0.025 | -0.049 | -0.049 | 0 | 0.09 | 0.21 | 0.03 | 0.01 | 0.03 | 0.08 |
| 8 | 19152067 | rs871668 | 0.262 | -0.089 | 0.999 | 0.128 | 0 | -0.008 | -0.008 | 0 | 0.55 | 0.38 | 0.6 | 0.74 | 0.45 | 0.61 |
| 8 | 19152556 | rs7826320 | 0.587 | -0.165 | 0.698 | 0.208 | 0.056 | 0 | 0 | 0 | 0.49 | 0.19 | 0.32 | 0.46 | 0.37 | 0.43 |
| 8 | 19153728 | rs1193329 | -1.174 | -0.218 | 0.991 | 0.254 | -0.051 | 0.001 | 0.001 | 0 | 0.77 | 0.89 | 0.76 | 0.76 | 0.68 | 0.76 |
| 8 | 19156100 | rs1915416 | 0.165 | -1.029 | 0.827 | 0.216 | 0.011 | -0.004 | -0.004 | 0 | 0.16 | 0.09 | 0.04 | 0.16 | 0.08 | 0.26 |
| 8 | 19156367 | rs10109833 | -0.769 | -0.471 | 0.827 | 0.087 | -0.008 | -0.055 | -0.055 | 0 | 0.12 | 0.34 | 0.06 | 0.03 | 0.08 | 0.07 |
| 8 | 19169851 | rs10091966 | -0.743 | -0.437 | 0.523 | 0.098 | -0.012 | -0.056 | -0.056 | 0.261 | 0.13 | 0.3 | 0.06 | 0.03 | 0.08 | 0.07 |
| 8 | 19174166 | rs7005298 | -0.297 | 0.287 | -0.077 | 0.59 | 0.007 | 0.019 | 0.019 | 0 | 0.8 | 0.72 | 0.73 | 0.65 | 0.74 | 0.75 |
| 8 | 19177772 | rs6997589 | -0.572 | 0.106 | -0.39 | 0.739 | -0.003 | -0.006 | -0.006 | 0 | 0.82 | 0.81 | 0.75 | 0.68 | 0.76 | 0.75 |
| 8 | 19189125 | rs3824328 | -0.908 | -0.689 | -0.385 | 0.094 | 0.019 | 0.019 | 0.019 | 0 | 0.34 | 0.68 | 0.18 | 0.21 | 0.24 | 0.25 |
| 8 | 19189757 | rs7845057 | -0.459 | -0.841 | 0.147 | -0.015 | -0.038 | -0.048 | -0.048 | 0 | 0.23 | 0.33 | 0.12 | 0.11 | 0.16 | 0.12 |
| 8 | 19190491 | rs1574288 | -0.886 | -0.84 | 0.161 | -0.014 | -0.031 | -0.052 | -0.052 | 0.538 | 0.29 | 0.39 | 0.15 | 0.14 | 0.23 | 0.13 |
| 8 | 19197531 | rs10448137 | -0.438 | -0.912 | -0.643 | 0.036 | -0.012 | -0.046 | -0.046 | 0 | 0.72 | 0.83 | 0.54 | 0.35 | 0.68 | 0.57 |
| 8 | 19200063 | rs5018145 | -1.91 | -0.881 | -0.684 | 0.455 | -0.036 | -0.066 | -0.066 | 0.203 | 0.89 | 0.99 | 0.78 | 0.82 | 0.87 | 0.78 |
| 8 | 19200609 | rs1316901 | -0.669 | -1.254 | -0.124 | -0.52 | -0.028 | -0.059 | -0.059 | 0.13 | 0.16 | 0.2 | 0.05 | 0 | 0.09 | 0.09 |
| 8 | 19204945 | rs13271942 | -1.167 | -1.279 | -0.03 | -0.56 | -0.05 | -0.076 | -0.076 | 0 | 0.29 | 0.38 | 0.17 | 0.07 | 0.18 | 0.23 |
| 8 | 19206063 | rs17128246 | -0.905 | -1.284 | -0.064 | -0.583 | -0.028 | -0.086 | -0.086 | 0.209 | 0.25 | 0.25 | 0.1 | 0.01 | 0.17 | 0.14 |
| 8 | 19207820 | rs6999618 | -0.414 | -1.175 | -0.187 | -0.401 | -0.011 | -0.029 | -0.029 | 0 | 0.68 | 0.81 | 0.53 | 0.34 | 0.64 | 0.53 |
| 8 | 19211220 | rs13279141 | -1.312 | -1.08 | 0.336 | -0.059 | -0.029 | -0.013 | -0.013 | 0.535 | 0.27 | 0.32 | 0.18 | 0.2 | 0.17 | 0.23 |
| 8 | 19213714 | rs17128272 | 0.085 | -0.811 | 0.234 | 0.128 | -0.003 | 0.001 | 0.001 | 0 | 0.13 | 0.14 | 0.08 | 0.13 | 0.12 | 0.07 |
| 8 | 19215011 | rs11995233 | -1.293 | -0.712 | -0.037 | 0.112 | -0.025 | -0.022 | -0.022 | 0 | 0.3 | 0.52 | 0.22 | 0.21 | 0.25 | 0.29 |
| 8 | 19218766 | rs4921637 | -0.639 | -0.478 | 0.299 | 0.382 | -0.041 | 0.004 | 0.004 | 0 | 0.2 | 0.47 | 0.17 | 0.18 | 0.14 | 0.17 |
| 8 | 19221997 | rs3824319 | 0.409 | -0.112 | -1.334 | -0.099 | -0.075 | 0.157 | 0.157 | 0 | 0.42 | 0.1 | 0.3 | 0.14 | 0.48 | 0.36 |
| 8 | 19228197 | rs10105819 | -0.095 | -0.197 | -0.181 | -0.31 | 0.008 | -0.048 | -0.048 | 2.393 | 0.09 | 0.33 | 0.06 | 0 | 0.08 | 0 |
| 8 | 19235381 | rs10094541 | -0.547 | -0.547 | -0.308 | -0.974 | -0.012 | -0.067 | -0.067 | 0 | 0.2 | 0.27 | 0.12 | 0 | 0.17 | 0.2 |
| 8 | 19237690 | rs13265589 | 1.667 | -0.657 | -0.328 | -0.324 | -0.001 | 0.017 | 0.017 | 1.718 | 0.11 | 0.11 | 0.27 | 0.31 | 0.1 | 0.15 |
| 8 | 19241970 | rs1320326 | -0.433 | -0.899 | -0.282 | -1.104 | -0.015 | -0.09 | -0.09 | 0 | 0.17 | 0.32 | 0.1 | 0 | 0.16 | 0.18 |
| 8 | 19248361 | rs2410611 | -0.037 | -1.191 | -0.386 | -1.304 | 0.016 | 0.012 | 0.012 | 6.049 | 0.27 | 0.17 | 0.42 | 0.8 | 0.2 | 0.31 |
| 8 | 19248579 | rs17406794 | 0.606 | -1.184 | -0.393 | -1.375 | -0.009 | -0.056 | -0.056 | 0 | 0.13 | 0.22 | 0.09 | 0 | 0.14 | 0.17 |
| 8 | 19249622 | rs1915422 | 2.349 | -1.062 | -0.816 | -1.866 | -0.043 | 0.068 | 0.068 | 0.069 | 0.31 | 0.15 | 0.24 | 0.15 | 0.32 | 0.25 |
| 8 | 19252486 | rs7357583 | 0.64 | -1.014 | -0.639 | -1.786 | 0.003 | 0.056 | 0.056 | 0.609 | 0.54 | 0.36 | 0.37 | 0.12 | 0.53 | 0.48 |
| 8 | 19252602 | rs2280443 | 1.348 | -0.966 | -0.774 | -1.758 | -0.003 | 0.152 | 0.152 | 1.66 | 0.47 | 0.23 | 0.35 | 0.12 | 0.5 | 0.44 |
| 8 | 19255000 | rs7839555 | 0.488 | -0.775 | -0.734 | -1.514 | 0.004 | 0.004 | 0.004 | 0.902 | 0.23 | 0.15 | 0.42 | 0.81 | 0.23 | 0.33 |
| 8 | 19260599 | rs4308750 | -0.279 | -0.552 | -0.166 | -1.506 | 0.005 | -0.114 | -0.114 | 0 | 0.8 | 0.53 | 0.89 | 0.97 | 0.82 | 0.82 |
| 8 | 19261730 | rs7001231 | 1.829 | -0.512 | -0.316 | -1.448 | -0.007 | -0.01 | -0.01 | 0 | 0.06 | 0.1 | 0.02 | 0.03 | 0.03 | 0.04 |
| 8 | 19263003 | rs3802330 | 1.08 | -0.821 | -0.519 | -1.803 | 0.028 | 0.046 | 0.046 | 0 | 0.57 | 0.35 | 0.77 | 0.96 | 0.65 | 0.66 |
| 8 | 19264789 | rs12545119 | -0.483 | -0.233 | -0.749 | -1.597 | 0.005 | -0.019 | -0.019 | 0.344 | 0.3 | 0.27 | 0.13 | 0.03 | 0.23 | 0.19 |
| 8 | 19266727 | rs3802329 | 0.178 | 0.664 | -1.249 | -1.421 | -0.047 | 0.001 | 0.001 | 0 | 0.25 | 0.09 | 0.58 | 0.82 | 0.34 | 0.45 |
| 8 | 19274742 | rs6980861 | 1.892 | 0.74 | -1.371 | -2.46 | -0.005 | -0.002 | -0.002 | 0.03 | 0.31 | 0.33 | 0.18 | 0.04 | 0.29 | 0.21 |
| 8 | 19274848 | rs17407149 | 0.256 | 0.682 | -0.963 | -1.917 | -0.005 | -0.005 | -0.005 | 0 | 0.33 | 0.22 | 0.52 | 0.69 | 0.4 | 0.45 |
| 8 | 19277056 | rs11204038 | 2.132 | 0.669 | -1.088 | -2.051 | -0.007 | 0.018 | 0.018 | 0.006 | 0.08 | 0.05 | 0.03 | 0 | 0.06 | 0.06 |
| 8 | 19277109 | rs17479085 | 3.966 | 0.815 | -1.28 | -1.776 | -0.036 | -0.018 | -0.018 | 0 | 0.08 | 0.01 | 0.22 | 0.11 | 0.11 | 0.05 |
| 8 | 19279199 | rs10481426 | 1.059 | 0.508 | -0.862 | -1.75 | 0 | -0.022 | -0.022 | 0 | 0.44 | 0.54 | 0.26 | 0.17 | 0.37 | 0.42 |
| 8 | 19280152 | rs12678270 | 1.364 | 0.694 | -1.018 | -1.241 | -0.013 | 0.069 | 0.069 | 0 | 0.07 | 0.04 | 0.18 | 0.49 | 0.13 | 0.29 |
| 8 | 19280442 | rs11204039 | -0.403 | 0.626 | -0.965 | -1.511 | -0.003 | 0.01 | 0.01 | 0 | 0.41 | 0.31 | 0.66 | 0.85 | 0.49 | 0.52 |
| 8 | 19285726 | rs13258090 | -0.38 | 0.62 | -0.994 | -1.513 | -0.008 | 0.041 | 0.041 | 0 | 0.41 | 0.28 | 0.66 | 0.84 | 0.49 | 0.52 |
| 8 | 19293565 | rs4431609 | 0.163 | 0.78 | -0.827 | -1.435 | -0.006 | -0.004 | -0.004 | 0 | 0.33 | 0.22 | 0.52 | 0.68 | 0.4 | 0.45 |
| 8 | 19295831 | rs11985575 | 3.013 | 0.736 | -0.946 | -1.687 | -0.026 | 0.09 | 0.09 | 0 | 0.21 | 0.06 | 0.13 | 0 | 0.18 | 0.15 |
| 8 | 19297051 | rs3802328 | -0.058 | 0.653 | -0.8 | -1.655 | -0.005 | -0.026 | -0.026 | 0.03 | 0.61 | 0.54 | 0.8 | 0.85 | 0.67 | 0.66 |
| 8 | 19303954 | rs4307360 | 0.244 | 0.689 | -0.874 | -1.955 | 0.002 | -0.028 | -0.028 | 0 | 0.57 | 0.5 | 0.74 | 0.86 | 0.56 | 0.6 |
| 8 | 19307051 | rs7835461 | -0.328 | 0.081 | -0.885 | -1.681 | -0.009 | 0.024 | 0.024 | 0 | 0.38 | 0.28 | 0.66 | 0.87 | 0.49 | 0.52 |
| 8 | 19313404 | rs4486225 | 0.162 | 0.187 | -0.842 | -1.373 | -0.001 | -0.014 | -0.014 | 0 | 0.52 | 0.48 | 0.68 | 0.86 | 0.51 | 0.57 |
| 8 | 19315845 | rs17128441 | 0.44 | 0.031 | -0.853 | -0.861 | 0.008 | 0.009 | 0.009 | 0 | 0.13 | 0.07 | 0.15 | 0.48 | 0.09 | 0.36 |
| 8 | 19321634 | rs4072131 | 0.091 | 0.178 | -0.536 | -1.805 | -0.002 | -0.014 | -0.014 | 0 | 0.66 | 0.69 | 0.8 | 0.87 | 0.7 | 0.69 |
| 8 | 19323985 | rs7002357 | -0.744 | 0.146 | -0.468 | -1.8 | -0.012 | -0.031 | -0.031 | 0 | 0.73 | 0.7 | 0.84 | 0.87 | 0.79 | 0.79 |
| 8 | 19326083 | rs6994666 | 0.259 | 0.262 | -0.43 | -1.662 | -0.001 | 0 | 0 | 1.984 | 0.1 | 0.13 | 0.13 | 0.11 | 0.13 | 0.13 |
| 8 | 19327959 | rs6984449 | 0.366 | 0.2 | -0.242 | -1.959 | 0.01 | 0.076 | 0.076 | 0.396 | 0.53 | 0.38 | 0.68 | 0.86 | 0.58 | 0.62 |
| 8 | 19330215 | rs13261732 | 0.464 | 0.318 | -0.367 | -1.956 | -0.018 | 0.049 | 0.049 | 0.096 | 0.34 | 0.2 | 0.58 | 0.86 | 0.44 | 0.49 |
| 8 | 19341239 | rs4922040 | 0.195 | 0.762 | 0.157 | -1.604 | 0 | -0.083 | -0.083 | 0 | 0.38 | 0.45 | 0.24 | 0.02 | 0.35 | 0.29 |
| 8 | 19348020 | rs6998334 | -0.619 | 0.714 | 0.228 | -1.509 | -0.004 | -0.013 | -0.013 | 0 | 0.76 | 0.74 | 0.87 | 0.86 | 0.79 | 0.7 |
| 8 | 19353933 | rs7818871 | 1.13 | 0.748 | 0.204 | -1.547 | 0.001 | -0.023 | -0.023 | 0 | 0.17 | 0.24 | 0.13 | 0.02 | 0.2 | 0.14 |
| 8 | 19355978 | rs11204050 | -0.261 | 0.703 | 0.166 | -1.608 | 0.011 | 0.078 | 0.078 | 1.362 | 0.57 | 0.36 | 0.75 | 0.86 | 0.66 | 0.58 |
| 8 | 19357642 | rs4317590 | 0.711 | 0.731 | 0.27 | -1.558 | -0.003 | 0.006 | 0.006 | 1.612 | 0.38 | 0.32 | 0.64 | 0.84 | 0.46 | 0.44 |

| | | | | | | | | | | | | | | | | |
|---|---|---|---|---|---|---|---|---|---|---|---|---|---|---|---|---|
| 8 | 19358961 | rs10888165 | 0.177 | 0.696 | 0.139 | -1.561 | 0 | -0.023 | -0.023 | 0.034 | 0.58 | 0.49 | 0.74 | 0.84 | 0.59 | 0.57 |
| 8 | 19359184 | rs3923920 | 0.547 | 0.684 | 0.002 | -1.456 | -0.001 | 0.079 | 0.079 | 3.425 | 0.4 | 0.24 | 0.63 | 0.84 | 0.45 | 0.45 |
| 8 | 19362768 | rs7017776 | -0.488 | 0.561 | 0.419 | -1.206 | -0.011 | -0.044 | -0.044 | 0 | 0.75 | 0.63 | 0.86 | 0.86 | 0.8 | 0.7 |
| 8 | 19363425 | rs6984644 | 0.694 | 0.53 | 0.14 | -1.338 | 0.009 | 0.126 | 0.126 | 1.723 | 0.54 | 0.25 | 0.72 | 0.84 | 0.59 | 0.53 |
| 8 | 19363963 | rs4922049 | -0.426 | 0.546 | 0.343 | -1.308 | -0.009 | -0.023 | -0.023 | 0 | 0.75 | 0.72 | 0.87 | 0.86 | 0.8 | 0.7 |
| 8 | 19364515 | rs13266857 | 0.207 | 0.508 | 0.341 | -1.27 | -0.002 | 0.041 | 0.041 | 1.791 | 0.59 | 0.37 | 0.74 | 0.84 | 0.6 | 0.53 |
| 8 | 19371767 | rs7816777 | 0.375 | 0.544 | 0.223 | -1.261 | 0.005 | 0.008 | 0.008 | 0.949 | 0.58 | 0.58 | 0.73 | 0.84 | 0.63 | 0.59 |
| 8 | 19377911 | rs2975469 | -0.144 | 0.387 | 0.352 | -0.896 | -0.002 | 0.007 | 0.007 | 0 | 0.39 | 0.75 | 0.37 | 0.28 | 0.37 | 0.4 |
| 8 | 19379421 | rs10107533 | 0.995 | 0.542 | -0.082 | -1.019 | -0.001 | -0.005 | -0.005 | 0 | 0.17 | 0.19 | 0.16 | 0.02 | 0.19 | 0.11 |
| 8 | 19379676 | rs4922053 | -1.93 | 0.608 | -0.326 | -0.976 | -0.003 | 0 | 0 | 0 | 0.82 | 0.86 | 0.82 | 0.86 | 0.82 | 0.89 |
| 8 | 19381519 | rs1014482 | 1.163 | 0.641 | -0.194 | -0.917 | -0.019 | 0.046 | 0.046 | 0 | 0.15 | 0.09 | 0.15 | 0.02 | 0.19 | 0.11 |
| 8 | 19383697 | rs6586842 | -0.618 | 0.543 | -0.195 | -0.797 | -0.018 | 0.061 | 0.061 | 0.408 | 0.79 | 0.93 | 0.88 | 0.99 | 0.78 | 0.83 |
| 8 | 19389865 | rs7002643 | -0.425 | 0.587 | 0.26 | -0.244 | -0.002 | 0.045 | 0.045 | 0 | 0.08 | 0.18 | 0.06 | 0.13 | 0.07 | 0.16 |
| 8 | 19398267 | rs13282951 | -0.28 | 0.54 | 0.332 | -0.311 | 0.006 | 0.069 | 0.069 | 0 | 0.07 | 0.2 | 0.06 | 0.12 | 0.07 | 0.16 |
| 8 | 19400018 | rs4921659 | 0.698 | 0.494 | 0.523 | -0.627 | -0.003 | -0.036 | -0.036 | 0 | 0.16 | 0.21 | 0.17 | 0.02 | 0.17 | 0.12 |
| 8 | 19401179 | rs10282938 | -0.71 | 0.296 | 0.816 | -0.706 | -0.016 | -0.026 | -0.026 | 0 | 0.27 | 0.6 | 0.23 | 0.14 | 0.24 | 0.35 |
| 8 | 19411717 | rs10464993 | 0.662 | -0.093 | 0.271 | -0.312 | -0.006 | -0.098 | -0.098 | 0 | 0.16 | 0.02 | 0.16 | 0.38 | 0.14 | 0.35 |
| 8 | 19418028 | rs6996732 | -0.441 | 0.041 | 0.878 | -0.478 | -0.031 | -0.005 | -0.005 | 0 | 0.23 | 0.53 | 0.21 | 0.17 | 0.2 | 0.34 |
| 8 | 19419494 | rs10106622 | -0.256 | 0.123 | 0.837 | -0.358 | 0.023 | 0.141 | 0.141 | 0 | 0.07 | 0.31 | 0.06 | 0.13 | 0.06 | 0.16 |
| 8 | 19429049 | rs6980558 | -0.667 | 0.092 | 0.795 | -0.363 | -0.007 | 0.107 | 0.107 | 0 | 0.08 | 0.29 | 0.06 | 0.13 | 0.06 | 0.16 |
| 8 | 19432198 | rs17128665 | 1.783 | 0.321 | 0.284 | -0.63 | -0.028 | -0.002 | -0.002 | 2.643 | 0.16 | 0.09 | 0.17 | 0.13 | 0.18 | 0.11 |
| 8 | 19434698 | rs12334858 | -0.463 | 0.155 | 0.66 | -0.828 | -0.029 | -0.087 | -0.087 | 0 | 0.25 | 0.62 | 0.22 | 0.04 | 0.21 | 0.24 |
| 8 | 19437153 | rs4922065 | -0.852 | 0.245 | 0.465 | -1.035 | 0.003 | -0.078 | -0.078 | 0.144 | 0.8 | 0.76 | 0.88 | 0.99 | 0.8 | 0.87 |
| 8 | 19439548 | rs7818368 | 0.878 | 0.295 | 0.344 | -1.105 | 0.001 | 0.02 | 0.02 | 0 | 0.18 | 0.16 | 0.17 | 0.04 | 0.18 | 0.13 |
| 8 | 19440069 | rs4298501 | 0.731 | 0.291 | 0.105 | -1.147 | 0.022 | 0.18 | 0.18 | 0 | 0.53 | 0.23 | 0.64 | 0.95 | 0.55 | 0.61 |
| 8 | 19443165 | rs13255998 | 0.039 | 0.423 | 0.308 | -0.826 | 0.011 | 0.02 | 0.02 | 0 | 0.54 | 0.19 | 0.57 | 0.63 | 0.57 | 0.52 |
| 8 | 19445170 | rs10100557 | -1.034 | 0.589 | 1.156 | -0.202 | 0.034 | 0.189 | 0.189 | 0 | 0.08 | 0.36 | 0.11 | 0.16 | 0.07 | 0.25 |
| 8 | 19446147 | rs1013438 | -0.381 | 0.701 | 0.564 | -0.348 | 0.011 | -0.101 | -0.101 | 0 | 0.81 | 0.71 | 0.87 | 0.99 | 0.81 | 0.79 |
| 8 | 19463823 | rs4472533 | -1.278 | 0.772 | 1.149 | 0.416 | 0.013 | 0.103 | 0.103 | 0.205 | 0.09 | 0.42 | 0.13 | 0.12 | 0.07 | 0.15 |
| 8 | 19465619 | rs12542189 | 1.041 | 0.96 | 1.043 | 0.403 | -0.003 | 0.015 | 0.015 | 0.064 | 0.71 | 0.33 | 0.64 | 0.63 | 0.7 | 0.71 |
| 8 | 19472837 | rs6988825 | -1.2 | 1.05 | 1.205 | 0.268 | 0.041 | 0.198 | 0.198 | 0.171 | 0.09 | 0.47 | 0.13 | 0.17 | 0.08 | 0.12 |
| 8 | 19474715 | rs4922069 | 0.006 | 1.266 | 0.535 | -0.569 | -0.029 | 0.072 | 0.072 | 0.138 | 0.18 | 0.04 | 0.13 | 0.05 | 0.2 | 0.13 |
| 8 | 19483259 | rs4922070 | 0.67 | 1.274 | 0.906 | -0.563 | 0.006 | -0.071 | -0.071 | 0 | 0.73 | 0.52 | 0.69 | 0.9 | 0.7 | 0.75 |
| 8 | 19484954 | rs2957624 | -0.438 | 1.484 | 0.785 | -0.534 | 0.001 | 0.005 | 0.005 | 0 | 0.5 | 0.55 | 0.59 | 0.36 | 0.52 | 0.35 |
| 8 | 19489046 | rs2975416 | -0.459 | 1.25 | 0.258 | -0.599 | 0.008 | 0.03 | 0.03 | 0 | 0.51 | 0.66 | 0.66 | 0.41 | 0.57 | 0.38 |
| 8 | 19489763 | rs2975418 | -0.424 | 1.194 | 0.248 | -0.618 | 0.009 | 0.034 | 0.034 | 0 | 0.51 | 0.65 | 0.65 | 0.39 | 0.57 | 0.38 |
| 8 | 19489948 | rs2975419 | -0.242 | 1.197 | -0.864 | -0.54 | -0.036 | 0.101 | 0.101 | 0 | 0.46 | 0.2 | 0.51 | 0.31 | 0.54 | 0.35 |
| 8 | 19491705 | rs4921662 | -0.475 | 1.165 | -0.353 | -0.227 | 0.021 | 0.01 | 0.01 | 0 | 0.28 | 0.34 | 0.5 | 0.36 | 0.4 | 0.13 |
| 8 | 19492840 | rs10503658 | -0.335 | -0.324 | -0.746 | -1.285 | 0.028 | -0.005 | -0.005 | 0.184 | 0.31 | 0.25 | 0.14 | 0.01 | 0.16 | 0.18 |
| 8 | 19493529 | rs2979874 | -0.007 | 0.698 | -0.373 | -0.668 | 0.012 | 0.012 | 0.012 | 0 | 0.69 | 0.65 | 0.41 | 0.35 | 0.64 | 0.53 |
| 8 | 19494153 | rs17480734 | -0.899 | 0.507 | -0.559 | -0.831 | 0.001 | -0.015 | -0.015 | 0 | 0.1 | 0.14 | 0.05 | 0.03 | 0.12 | 0.19 |
| 8 | 19495610 | rs13252345 | -1.152 | 0.439 | -0.493 | -1.272 | 0.002 | -0.084 | -0.084 | 0 | 0.27 | 0.4 | 0.14 | 0.03 | 0.26 | 0.33 |
| 8 | 19502855 | rs10283283 | -1.099 | 0.511 | -0.589 | -1.263 | -0.01 | 0.008 | 0.008 | 0.554 | 0.13 | 0.11 | 0.07 | 0.02 | 0.16 | 0.2 |
| 8 | 19506410 | rs4922073 | 0.583 | 0.529 | -0.34 | -1.13 | 0 | 0.004 | 0.004 | 0 | 0.6 | 0.44 | 0.49 | 0.41 | 0.57 | 0.41 |
| 8 | 19507270 | rs7001715 | -0.125 | 0.56 | -0.266 | -1.143 | -0.001 | -0.016 | -0.016 | 0 | 0.83 | 0.83 | 0.9 | 0.97 | 0.81 | 0.82 |
| 8 | 19508835 | rs4535751 | -0.868 | 0.553 | -0.31 | -1.135 | 0.002 | -0.003 | -0.003 | 0 | 0.87 | 0.93 | 0.93 | 0.99 | 0.87 | 0.92 |
| 8 | 19510286 | rs17128793 | 0.867 | 0.48 | -0.257 | -1.049 | 0.003 | 0.069 | 0.069 | 0 | 0.21 | 0.28 | 0.4 | 0.57 | 0.23 | 0.41 |
| 8 | 19513321 | rs12543931 | 0.802 | 0.478 | -0.411 | -0.961 | -0.001 | 0.02 | 0.02 | 0.293 | 0.51 | 0.32 | 0.42 | 0.38 | 0.48 | 0.29 |
| 8 | 19514868 | rs7828491 | -0.11 | 0.856 | -0.306 | -0.388 | 0.002 | 0.038 | 0.038 | 0 | 0.28 | 0.44 | 0.46 | 0.62 | 0.29 | 0.47 |
| 8 | 19518908 | rs4921664 | -0.191 | -0.073 | -0.674 | -0.488 | -0.02 | -0.004 | -0.004 | 0.572 | 0.59 | 0.61 | 0.83 | 0.67 | 0.7 | 0.7 |
| 8 | 19529575 | rs2172350 | 0.134 | 0.227 | -0.757 | -0.435 | 0.004 | 0.008 | 0.008 | 0 | 0.57 | 0.33 | 0.44 | 0.42 | 0.55 | 0.52 |
| 8 | 19530403 | rs17128820 | -0.091 | 0.137 | 0.05 | -0.705 | 0.027 | -0.041 | -0.041 | 0 | 0.09 | 0.3 | 0.05 | 0.01 | 0.08 | 0.1 |
| 8 | 19530963 | rs923768 | 0.589 | 0.825 | -0.229 | -0.713 | -0.002 | 0.014 | 0.014 | 0 | 0.55 | 0.32 | 0.4 | 0.41 | 0.52 | 0.56 |
| 8 | 19530983 | rs958416 | 0.268 | 0.838 | 0.199 | -0.733 | 0.009 | -0.01 | -0.01 | 0.157 | 0.59 | 0.47 | 0.43 | 0.42 | 0.53 | 0.55 |
| 8 | 19534545 | rs11780937 | 0.644 | 1.303 | 0.308 | -1.269 | -0.001 | 0.057 | 0.057 | 0 | 0.48 | 0.37 | 0.27 | 0.03 | 0.43 | 0.39 |
| 8 | 19535134 | rs9987322 | 0.674 | 0.946 | 0.19 | -1.569 | -0.041 | -0.086 | -0.086 | 0 | 0.13 | 0.28 | 0.05 | 0 | 0.06 | 0.07 |
| 8 | 19536790 | rs4922081 | -0.349 | 1.346 | 0.229 | -0.504 | 0 | 0.006 | 0.006 | 0 | 0.38 | 0.31 | 0.56 | 0.65 | 0.39 | 0.42 |
| 8 | 19537084 | rs4922084 | 0.418 | 1.32 | 0.214 | -0.262 | 0.009 | 0.004 | 0.004 | 0 | 0.29 | 0.25 | 0.35 | 0.23 | 0.23 | 0.21 |
| 8 | 19538319 | rs11993323 | 1.126 | 1.355 | 0.124 | -0.24 | -0.023 | -0.021 | -0.021 | 0.039 | 0.11 | 0.09 | 0.22 | 0.42 | 0.16 | 0.22 |
| 8 | 19540122 | rs11985450 | 0.145 | 1.359 | 0.179 | -0.293 | 0.045 | 0.136 | 0.136 | 0 | 0.14 | 0.3 | 0.24 | 0.41 | 0.2 | 0.27 |
| 8 | 19540262 | rs11204064 | 0.689 | 1.043 | -0.008 | -0.749 | -0.001 | 0.001 | 0.001 | 0 | 0.5 | 0.49 | 0.43 | 0.44 | 0.48 | 0.58 |
| 8 | 19540437 | rs4921665 | 0.132 | 1.048 | 0.032 | -0.747 | -0.023 | -0.019 | -0.019 | 0 | 0.34 | 0.41 | 0.37 | 0.26 | 0.28 | 0.27 |
| 8 | 19540966 | rs4548199 | 0.98 | 1.127 | -0.107 | -0.737 | -0.008 | -0.002 | -0.002 | 0 | 0.1 | 0.09 | 0.07 | 0 | 0.1 | 0.05 |
| 8 | 19543862 | rs7827153 | 0.149 | 0.996 | 0.051 | -0.376 | 0.013 | 0.001 | 0.001 | 0 | 0.59 | 0.31 | 0.48 | 0.6 | 0.59 | 0.64 |
| 8 | 19544135 | rs17090706 | -0.171 | 1.04 | -0.01 | -0.392 | -0.005 | -0.005 | -0.005 | 0.225 | 0.23 | 0.24 | 0.27 | 0.42 | 0.31 | 0.31 |
| 8 | 19544426 | rs17481249 | 0.869 | 0.709 | 0.067 | -0.391 | 0.018 | 0.005 | 0.005 | 0 | 0.27 | 0.17 | 0.36 | 0.24 | 0.23 | 0.21 |
| 8 | 19547541 | rs4922087 | 0.134 | 0.681 | -0.004 | -0.397 | -0.012 | -0.013 | -0.013 | 0 | 0.35 | 0.56 | 0.41 | 0.25 | 0.28 | 0.26 |
| 8 | 19547919 | rs4921667 | 0.351 | 0.747 | -0.017 | -0.351 | -0.027 | -0.022 | -0.022 | 0.605 | 0.71 | 0.73 | 0.6 | 0.43 | 0.57 | 0.59 |
| 8 | 19550298 | rs17090735 | 1.851 | 0.008 | -0.599 | -0.972 | -0.01 | -0.02 | -0.02 | 2.478 | 0.18 | 0.04 | 0.12 | 0.18 | 0.17 | 0.17 |

| | | | | | | | | | | | | | | | | |
|---|---|---|---|---|---|---|---|---|---|---|---|---|---|---|---|---|
| 8 | 19552218 | rs1552783 | 1.143 | 0.135 | -0.458 | -0.568 | 0.034 | 0.047 | 0.047 | 0 | 0.36 | 0.17 | 0.2 | 0.22 | 0.28 | 0.31 |
| 8 | 19553497 | rs7017723 | 0.004 | 0.363 | -0.301 | -0.451 | -0.007 | 0.01 | 0.01 | 0 | 0.54 | 0.37 | 0.44 | 0.62 | 0.51 | 0.61 |
| 8 | 19555762 | rs10090551 | -0.025 | 0.345 | -0.022 | -0.325 | 0.021 | 0.024 | 0.024 | 0 | 0.73 | 0.65 | 0.59 | 0.46 | 0.59 | 0.63 |
| 8 | 19561997 | rs1907800 | 0.814 | 0.197 | 0.055 | -0.544 | 0.034 | 0.057 | 0.057 | 0 | 0.42 | 0.27 | 0.22 | 0.19 | 0.32 | 0.35 |
| 8 | 19563900 | rs923765 | 1.181 | 0.201 | 0.173 | -0.559 | 0.009 | 0.008 | 0.008 | 0 | 0.27 | 0.19 | 0.37 | 0.22 | 0.24 | 0.22 |
| 8 | 19564742 | rs7836255 | -0.752 | 0.52 | 0.266 | -0.626 | 0.016 | -0.002 | -0.002 | 0 | 0.84 | 0.81 | 0.8 | 0.86 | 0.75 | 0.83 |
| 8 | 19569489 | rs7815576 | -0.329 | 0.458 | 0.391 | -0.63 | 0.092 | -0.014 | -0.014 | 0 | 0.82 | 0.55 | 0.78 | 0.86 | 0.75 | 0.83 |
| 8 | 19575066 | rs7002401 | 0.193 | 0.365 | -0.343 | -0.683 | -0.002 | -0.011 | -0.011 | 0.666 | 0.42 | 0.54 | 0.58 | 0.66 | 0.43 | 0.48 |
| 8 | 19576471 | rs10503660 | 0.753 | 0.304 | -0.413 | -0.783 | 0.008 | 0.017 | 0.017 | 0 | 0.29 | 0.2 | 0.17 | 0.19 | 0.24 | 0.26 |
| 8 | 19576883 | rs1021436 | 0.115 | 0.805 | -0.31 | -0.371 | 0.017 | -0.009 | -0.009 | 0.14 | 0.8 | 0.76 | 0.78 | 0.86 | 0.72 | 0.78 |
| 8 | 19577136 | rs10503661 | 0.482 | 0.624 | -0.539 | 0.16 | -0.009 | 0.004 | 0.004 | 0 | 0.14 | 0.12 | 0.23 | 0.51 | 0.19 | 0.28 |
| 8 | 19580291 | rs1388943 | -0.19 | 0.674 | -0.572 | 0.055 | 0.005 | 0.004 | 0.004 | 0.84 | 0.45 | 0.62 | 0.65 | 0.63 | 0.5 | 0.48 |
| 8 | 19580824 | rs7823121 | 1.161 | 1.197 | -0.239 | 0.034 | 0.032 | -0.053 | -0.053 | 0 | 0.11 | 0.23 | 0.1 | 0 | 0.14 | 0.07 |
| 8 | 19582144 | rs6999473 | -0.913 | 0.87 | -0.591 | -0.342 | 0 | 0.019 | 0.019 | 0 | 0.73 | 0.81 | 0.79 | 0.82 | 0.69 | 0.73 |
| 8 | 19582804 | rs17091113 | 0.972 | 0.839 | -0.528 | -0.313 | 0.01 | 0.012 | 0.012 | 0.765 | 0.33 | 0.37 | 0.54 | 0.63 | 0.37 | 0.41 |
| 8 | 19585176 | rs10100391 | 1.42 | 0.794 | -0.704 | -0.465 | 0.013 | 0.005 | 0.005 | 0.017 | 0.2 | 0.14 | 0.07 | 0.01 | 0.15 | 0.16 |
| 8 | 19586205 | rs10104473 | 0.364 | 0.733 | -0.651 | -0.473 | 0.103 | 0.111 | 0.111 | 0.459 | 0.47 | 0.21 | 0.21 | 0.21 | 0.34 | 0.4 |
| 8 | 19589706 | rs1107613 | 3.008 | 1.005 | -0.872 | -0.022 | -0.025 | -0.015 | -0.015 | 0 | 0.09 | 0.03 | 0.22 | 0.35 | 0.14 | 0.21 |
| 8 | 19590012 | rs7825914 | -0.977 | 1.163 | -0.628 | 0.003 | 0 | 0 | 0 | 0 | 0.83 | 0.8 | 0.77 | 0.82 | 0.71 | 0.82 |
| 8 | 19590098 | rs11204065 | 2.199 | 1.187 | -0.8 | 0.027 | 0.005 | -0.011 | -0.011 | 0.044 | 0.2 | 0.1 | 0.2 | 0.23 | 0.17 | 0.19 |
| 8 | 19590800 | rs1546100 | 1.759 | 0.943 | -0.782 | 0.007 | 0.005 | 0.032 | 0.032 | 0.013 | 0.21 | 0.25 | 0.35 | 0.39 | 0.22 | 0.27 |
| 8 | 19593584 | rs17091237 | -1.108 | 0.861 | -0.963 | -0.031 | 0.026 | -0.037 | -0.037 | 0 | 0.66 | 0.87 | 0.71 | 0.62 | 0.65 | 0.65 |
| 8 | 19600329 | rs6983139 | 0.582 | 1.101 | -0.317 | 0.042 | 0.004 | 0.002 | 0.002 | 0 | 0.49 | 0.51 | 0.5 | 0.58 | 0.41 | 0.51 |
| 8 | 19605966 | rs12544497 | 0.783 | 0.619 | -0.115 | 0.341 | -0.02 | -0.005 | -0.005 | 0 | 0.22 | 0.36 | 0.22 | 0.22 | 0.17 | 0.19 |
| 8 | 19607876 | rs2172349 | 0.141 | 0.509 | -0.292 | 0.01 | 0.002 | -0.017 | -0.017 | 1.709 | 0.24 | 0.34 | 0.21 | 0.15 | 0.19 | 0.15 |
| 8 | 19609029 | rs11204068 | -0.028 | 0.858 | -0.171 | 0.353 | 0.013 | 0.029 | 0.029 | 0.022 | 0.19 | 0.33 | 0.22 | 0.23 | 0.19 | 0.18 |
| 8 | 19614339 | rs10105057 | -1.423 | 1.186 | -0.683 | 0.643 | 0.013 | 0.02 | 0.02 | 0 | 0.86 | 0.83 | 0.76 | 0.67 | 0.72 | 0.6 |
| 8 | 19619751 | rs4262337 | 0.382 | 0.542 | -0.643 | 0.238 | -0.003 | 0.004 | 0.004 | 0.343 | 0.2 | 0.3 | 0.21 | 0.24 | 0.17 | 0.13 |
| 8 | 19623055 | rs17091325 | 1.408 | -0.053 | -1.179 | -0.069 | 0.042 | -0.005 | -0.005 | 0 | 0.16 | 0.03 | 0.14 | 0.14 | 0.08 | 0.15 |
| 8 | 19623129 | rs11782067 | -0.443 | 0.616 | -0.861 | 0.248 | 0.02 | 0.023 | 0.023 | 0 | 0.75 | 0.56 | 0.71 | 0.67 | 0.67 | 0.61 |
| 8 | 19623911 | rs6996964 | 1.468 | 0.658 | -0.76 | 0.246 | -0.014 | -0.068 | -0.068 | 0.011 | 0.11 | 0.35 | 0.08 | 0.01 | 0.07 | 0.04 |
| 8 | 19625464 | rs13282247 | -0.501 | 0.421 | -0.886 | 0.158 | 0 | 0.004 | 0.004 | 0.092 | 0.3 | 0.47 | 0.33 | 0.33 | 0.3 | 0.25 |
| 8 | 19625912 | rs7836144 | -0.302 | 0.607 | -0.791 | 0.436 | 0.032 | 0.045 | 0.045 | 0 | 0.76 | 0.55 | 0.68 | 0.64 | 0.64 | 0.61 |
| 8 | 19626120 | rs7837390 | -0.241 | 0.524 | -0.822 | 0.209 | 0.026 | 0.011 | 0.011 | 0 | 0.74 | 0.54 | 0.69 | 0.67 | 0.62 | 0.61 |
| 8 | 19626356 | rs13254708 | 0.127 | -0.274 | -0.727 | -0.163 | -0.01 | -0.041 | -0.041 | 0 | 0.18 | 0.36 | 0.16 | 0.06 | 0.17 | 0.19 |
| 8 | 19626439 | rs1994786 | 0.088 | -0.402 | -0.91 | -0.271 | -0.048 | -0.034 | -0.034 | 0 | 0.3 | 0.52 | 0.22 | 0.2 | 0.22 | 0.2 |
| 8 | 19628529 | rs6981472 | 0.571 | -0.573 | -1.29 | -0.539 | 0.024 | 0.001 | 0.001 | 0 | 0.29 | 0.19 | 0.35 | 0.27 | 0.22 | 0.31 |
| 8 | 19628809 | rs7014781 | 0.23 | -0.404 | -1.242 | -0.563 | 0.02 | -0.022 | -0.022 | 0.124 | 0.36 | 0.24 | 0.48 | 0.42 | 0.32 | 0.39 |
| 8 | 19632090 | rs7010610 | -0.654 | 0.159 | -1.038 | -0.236 | 0.002 | 0.014 | 0.014 | 0 | 0.74 | 0.73 | 0.76 | 0.44 | 0.71 | 0.56 |
| 8 | 19634864 | rs7820520 | 0.147 | -0.585 | -1.501 | -0.591 | 0.092 | 0.089 | 0.089 | 0 | 0.43 | 0.19 | 0.35 | 0.25 | 0.3 | 0.31 |
| 8 | 19635098 | rs10788700 | 0.211 | -0.654 | -1.489 | -0.625 | 0.101 | 0.098 | 0.098 | 0 | 0.43 | 0.14 | 0.35 | 0.25 | 0.3 | 0.31 |
| 8 | 19635117 | rs10749625 | -0.152 | -0.446 | -1.453 | -0.562 | 0.07 | 0.038 | 0.038 | 0 | 0.51 | 0.25 | 0.49 | 0.42 | 0.39 | 0.39 |
| 8 | 19639397 | rs7815086 | 0.232 | -0.905 | -1.681 | -0.851 | 0.027 | 0.02 | 0.02 | 0 | 0.29 | 0.19 | 0.28 | 0.24 | 0.23 | 0.27 |
| 8 | 19642668 | rs12375354 | 0.571 | -0.911 | -1.689 | -0.887 | 0.036 | 0.004 | 0.004 | 0.01 | 0.23 | 0.07 | 0.21 | 0.18 | 0.16 | 0.12 |
| 8 | 19642778 | rs7002897 | 1.321 | -0.112 | -0.97 | -0.941 | 0.131 | -0.032 | -0.032 | 0 | 0.07 | 0.21 | 0.08 | 0 | 0.13 | 0.06 |
| 8 | 19644764 | rs13261437 | 1.598 | -1.472 | -1.611 | -0.878 | 0.003 | 0.02 | 0.02 | 0 | 0.08 | 0.06 | 0.05 | 0 | 0.08 | 0.05 |
| 8 | 19646515 | rs10112900 | 0.796 | -1.714 | -1.341 | -0.743 | 0.001 | -0.001 | -0.001 | 0.134 | 0.18 | 0.16 | 0.2 | 0.18 | 0.12 | 0.12 |
| 8 | 19650333 | rs11204072 | 0.401 | -2.038 | -1.697 | -1.179 | 0.039 | 0.079 | 0.079 | 0 | 0.26 | 0.14 | 0.15 | 0.07 | 0.17 | 0.18 |
| 8 | 19651581 | rs17482069 | 1.677 | -1.984 | -1.807 | -1.078 | -0.025 | -0.034 | -0.034 | 0 | 0.07 | 0.01 | 0.13 | 0.15 | 0.09 | 0.09 |
| 8 | 19652410 | rs7006468 | -0.502 | -2.653 | -1.626 | -0.746 | 0.016 | -0.005 | -0.005 | 0 | 0.39 | 0.34 | 0.41 | 0.54 | 0.27 | 0.35 |
| 8 | 19652710 | rs6986179 | 0.968 | -2.735 | -1.641 | -0.791 | 0.044 | 0.055 | 0.055 | 0 | 0.23 | 0.15 | 0.13 | 0.07 | 0.13 | 0.22 |
| 8 | 19654041 | rs6996994 | -0.074 | -2.748 | -1.602 | -0.745 | 0.003 | -0.003 | -0.003 | 0 | 0.35 | 0.34 | 0.39 | 0.54 | 0.24 | 0.35 |
| 8 | 19656648 | rs7008703 | -1.113 | -2.762 | -1.622 | -0.78 | -0.005 | -0.004 | -0.004 | 0 | 0.62 | 0.5 | 0.54 | 0.61 | 0.4 | 0.57 |
| 8 | 19656859 | rs17091499 | 0.176 | -2.611 | -1.37 | -0.76 | 0.018 | -0.019 | -0.019 | 0 | 0.29 | 0.22 | 0.27 | 0.38 | 0.17 | 0.25 |
| 8 | 19662458 | rs6993714 | 0.849 | -2.858 | -1.565 | -1.168 | 0.025 | 0.03 | 0.03 | 0 | 0.21 | 0.14 | 0.09 | 0 | 0.13 | 0.07 |
| 8 | 19662498 | rs7012358 | -0.093 | -2.56 | -1.405 | -0.909 | 0.055 | 0.028 | 0.028 | 0 | 0.45 | 0.34 | 0.34 | 0.29 | 0.27 | 0.31 |
| 8 | 19669333 | rs6982084 | 0.772 | -2.561 | -1.453 | -0.875 | -0.003 | 0 | 0 | 0.023 | 0.43 | 0.53 | 0.51 | 0.55 | 0.63 | 0.54 |
| 8 | 19669503 | rs10283069 | -1.083 | -2.533 | -1.445 | -0.837 | 0.002 | 0.015 | 0.015 | 0 | 0.92 | 0.87 | 0.86 | 0.85 | 0.9 | 0.9 |
| 8 | 19671104 | rs6993525 | -0.082 | -2.607 | -1.488 | -0.912 | 0.099 | 0.037 | 0.037 | 0.167 | 0.43 | 0.25 | 0.34 | 0.29 | 0.27 | 0.29 |
| 8 | 19673750 | rs10090638 | 0.557 | -2.208 | -1.321 | -0.772 | -0.004 | 0 | 0 | 0 | 0.44 | 0.52 | 0.52 | 0.55 | 0.64 | 0.55 |
| 8 | 19681647 | rs2410612 | 0.59 | -1.236 | -1.399 | -0.462 | 0.026 | 0.013 | 0.013 | 0 | 0.56 | 0.64 | 0.56 | 0.66 | 0.65 | 0.67 |
| 8 | 19682545 | rs3816246 | 0.143 | -1.001 | -0.871 | 0.07 | -0.001 | -0.001 | -0.001 | 0 | 0.69 | 0.66 | 0.67 | 0.66 | 0.78 | 0.72 |
| 8 | 19686177 | rs11777843 | -0.38 | -0.106 | -0.816 | 0.086 | -0.001 | -0.005 | -0.005 | 0 | 0.9 | 0.93 | 0.87 | 0.86 | 0.9 | 0.89 |
| 8 | 19709575 | rs4599828 | -0.062 | -0.414 | -0.678 | -0.307 | 0.007 | 0.053 | 0.053 | 0 | 0.45 | 0.64 | 0.61 | 0.72 | 0.54 | 0.49 |
| 8 | 19709775 | rs6586870 | 0.517 | -0.222 | -0.66 | 0.045 | 0.098 | 0.06 | 0.06 | 0 | 0.24 | 0.5 | 0.35 | 0.41 | 0.38 | 0.36 |
| 8 | 19710388 | rs4922108 | 0.746 | -0.252 | -0.851 | -0.019 | 0.013 | -0.021 | -0.021 | 0.659 | 0.22 | 0.21 | 0.25 | 0.31 | 0.17 | 0.14 |
| 8 | 19710468 | rs4244456 | -0.082 | -0.278 | -0.996 | -0.174 | 0.017 | 0.115 | 0.115 | 0 | 0.47 | 0.76 | 0.62 | 0.73 | 0.54 | 0.51 |
| 8 | 19717091 | rs6586874 | 0.051 | 0.3 | -0.376 | 0.067 | 0.025 | -0.01 | -0.01 | 0 | 0.84 | 0.79 | 0.85 | 0.87 | 0.78 | 0.86 |
| 8 | 19720833 | rs10096561 | 0.059 | 0.354 | -0.733 | 0.069 | -0.058 | -0.003 | -0.003 | 0.472 | 0.85 | 0.96 | 0.86 | 0.87 | 0.78 | 0.86 |

| | | | | | | | | | | | | | | | | |
|---|---|---|---|---|---|---|---|---|---|---|---|---|---|---|---|---|
| 8 | 19723503 | rs2044061 | 0.2 | 1.291 | -0.339 | 0.333 | 0.016 | 0.032 | 0.032 | 0 | 0.2 | 0.22 | 0.36 | 0.49 | 0.32 | 0.47 |
| 8 | 19726130 | rs767145 | 0.096 | 0.669 | -0.872 | -0.198 | 0.002 | 0 | 0 | 1.429 | 0.47 | 0.47 | 0.54 | 0.5 | 0.58 | 0.59 |
| 8 | 19726742 | rs6988864 | 0.436 | 0.288 | -1.044 | -0.79 | 0.033 | 0.148 | 0.148 | 0 | 0.4 | 0.16 | 0.28 | 0.07 | 0.34 | 0.25 |
| 8 | 19727047 | rs1441778 | 0.409 | 0.726 | -0.033 | -0.414 | 0.016 | -0.004 | -0.004 | 0 | 0.83 | 0.69 | 0.89 | 0.85 | 0.84 | 0.89 |
| 8 | 19731375 | rs7821631 | -1.515 | 1.275 | -0.519 | 0.201 | -0.034 | -0.043 | -0.043 | 0 | 0.88 | 0.96 | 0.76 | 0.64 | 0.8 | 0.68 |
| 8 | 19733568 | rs1561747 | 0.538 | 0.871 | -0.317 | -0.31 | 0.005 | 0.013 | 0.013 | 0 | 0.43 | 0.37 | 0.33 | 0.15 | 0.42 | 0.23 |
| 8 | 19737157 | rs2197088 | 0.83 | 0.833 | 0.144 | -0.334 | -0.002 | -0.023 | -0.023 | 0 | 0.4 | 0.47 | 0.32 | 0.15 | 0.42 | 0.22 |
| 8 | 19738260 | rs6586876 | -0.419 | 0.953 | -0.344 | -0.402 | -0.005 | 0.059 | 0.059 | 0 | 0.58 | 0.72 | 0.43 | 0.27 | 0.61 | 0.47 |
| 8 | 19746876 | rs17482310 | -0.016 | 1.683 | 0.123 | -0.399 | -0.039 | -0.004 | -0.004 | 0.449 | 0.1 | 0.03 | 0.09 | 0.09 | 0.17 | 0.19 |
| 8 | 19748856 | rs7822518 | 1.397 | 1.497 | 0.14 | -0.389 | -0.016 | -0.022 | -0.022 | 0 | 0.52 | 0.2 | 0.53 | 0.43 | 0.51 | 0.35 |
| 8 | 19749390 | rs3898938 | 0.646 | 1.58 | 0.059 | -0.527 | -0.02 | 0.001 | 0.001 | 0 | 0.57 | 0.61 | 0.5 | 0.53 | 0.46 | 0.47 |
| 8 | 19751939 | rs1441773 | -0.518 | 1.528 | 0.117 | -0.457 | -0.001 | -0.018 | -0.018 | 0.042 | 0.72 | 0.73 | 0.67 | 0.54 | 0.72 | 0.6 |
| 8 | 19753602 | rs1441772 | 0.363 | 1.61 | 1.017 | -0.881 | -0.001 | -0.022 | -0.022 | 1.116 | 0.13 | 0.14 | 0.14 | 0 | 0.23 | 0.11 |
| 8 | 19756813 | rs10102717 | -1.62 | 1.536 | 0.887 | -0.877 | 0.002 | -0.018 | -0.018 | 0 | 0.3 | 0.66 | 0.27 | 0.1 | 0.42 | 0.29 |
| 8 | 19758457 | rs6586878 | -0.707 | 1.915 | 1.19 | -0.856 | 0.071 | -0.113 | -0.113 | 0 | 0.18 | 0.62 | 0.2 | 0 | 0.3 | 0.18 |
| 8 | 19776981 | rs4466415 | -0.465 | 1.625 | 0.999 | -0.764 | -0.01 | -0.006 | -0.006 | 0 | 0.13 | 0.08 | 0.09 | 0.19 | 0.15 | 0.25 |
| 8 | 19784574 | rs3900537 | -1.438 | 1.332 | 1.731 | -0.878 | 0.022 | -0.001 | -0.001 | 1.176 | 0.08 | 0.49 | 0.04 | 0 | 0.01 | 0.03 |
| 8 | 19786891 | rs7816032 | -2.094 | 1.463 | 1.856 | -0.963 | 0.001 | 0.008 | 0.008 | 0.312 | 0.26 | 0.76 | 0.17 | 0.18 | 0.22 | 0.3 |
| 8 | 19787528 | rs7009128 | -1.482 | 1.484 | 1.741 | -0.842 | 0.012 | -0.002 | -0.002 | 0.266 | 0.07 | 0.37 | 0.02 | 0 | 0.01 | 0.03 |
| 8 | 19795789 | rs1470186 | -0.586 | 1.49 | 1.749 | -0.839 | 0.007 | -0.005 | -0.005 | 0 | 0.06 | 0.23 | 0.02 | 0 | 0.01 | 0.03 |
| 8 | 19796671 | rs1800590 | -1.458 | 1.521 | 1.641 | -0.839 | 0.028 | 0.007 | 0.007 | 2.655 | 0.07 | 0.46 | 0.03 | 0 | 0.01 | 0.03 |
| 8 | 19799641 | rs1534649 | -2.082 | 1.618 | 0.904 | -0.932 | -0.086 | 0.122 | 0.122 | 1.29 | 0.35 | 0.9 | 0.31 | 0.18 | 0.45 | 0.38 |
| 8 | 19803802 | rs7000460 | -1.407 | 1.363 | 1.421 | -0.482 | -0.005 | -0.017 | -0.017 | 0 | 0.07 | 0.28 | 0.02 | 0 | 0.01 | 0.03 |
| 8 | 19809435 | rs1121923 | 0.318 | 1.434 | 0.598 | -0.533 | -0.011 | -0.036 | -0.036 | 0 | 0.07 | 0.14 | 0.03 | 0 | 0.04 | 0 |
| 8 | 19810787 | rs343 | 0.024 | 1.624 | 0.151 | -0.263 | -0.008 | -0.024 | -0.024 | 0 | 0.09 | 0.04 | 0.12 | 0.16 | 0.09 | 0.12 |
| 8 | 19811417 | rs253 | -2.328 | 2.075 | 0.275 | -0.21 | -0.038 | 0.001 | 0.001 | 0 | 0.34 | 0.77 | 0.42 | 0.35 | 0.48 | 0.43 |
| 8 | 19811967 | rs256 | -0.35 | 2.049 | -0.03 | 0.107 | -0.017 | -0.038 | -0.038 | 0 | 0.11 | 0.04 | 0.15 | 0.25 | 0.15 | 0.18 |
| 8 | 19813676 | rs270 | -0.06 | 2.11 | 0.287 | 0.022 | 0.026 | 0.015 | 0.015 | 0.081 | 0.06 | 0.07 | 0.16 | 0.11 | 0.18 | 0.14 |
| 8 | 19818436 | rs316 | -0.534 | 1.558 | 0.758 | -0.216 | -0.004 | -0.009 | -0.009 | 0 | 0.14 | 0.24 | 0.15 | 0.11 | 0.12 | 0.12 |
| 8 | 19818976 | rs319 | -0.235 | 2.156 | 0.647 | -0.304 | 0.006 | -0.002 | -0.002 | 0.033 | 0.1 | 0.07 | 0.18 | 0.1 | 0.24 | 0.17 |
| 8 | 19819536 | rs327 | -1.213 | 1.877 | 0.868 | -0.686 | -0.003 | -0.024 | -0.024 | 0 | 0.31 | 0.42 | 0.25 | 0.24 | 0.29 | 0.21 |
| 8 | 19822830 | rs4922115 | 0.219 | 1.773 | 0.965 | -0.736 | 0.007 | 0.026 | 0.026 | 0 | 0.17 | 0.11 | 0.17 | 0.11 | 0.14 | 0.12 |
| 8 | 19823617 | rs11570892 | -0.272 | 1.746 | 1.214 | -0.75 | -0.008 | -0.02 | -0.02 | 0 | 0.19 | 0.23 | 0.18 | 0.11 | 0.15 | 0.15 |
| 8 | 19824492 | rs13702 | -1.709 | 1.581 | 1.611 | -0.918 | -0.012 | -0.038 | -0.038 | 0 | 0.37 | 0.54 | 0.28 | 0.23 | 0.32 | 0.22 |
| 8 | 19824563 | rs1059611 | -0.345 | 1.52 | 1.659 | -0.913 | 0.005 | -0.003 | -0.003 | 2.586 | 0.11 | 0.18 | 0.07 | 0.12 | 0.14 | 0.09 |
| 8 | 19824667 | rs15285 | -1.634 | 1.46 | 1.625 | -0.913 | -0.013 | -0.038 | -0.038 | 0.002 | 0.37 | 0.53 | 0.28 | 0.23 | 0.32 | 0.22 |
| 8 | 19826373 | rs2197089 | -2.468 | 1.608 | 1.796 | -0.97 | 0.024 | 0.106 | 0.106 | 0 | 0.51 | 0.81 | 0.49 | 0.32 | 0.58 | 0.42 |
| 8 | 19827848 | rs10105606 | -1.831 | 1.379 | 1.866 | -0.971 | 0.022 | 0.067 | 0.067 | 0.032 | 0.41 | 0.71 | 0.3 | 0.23 | 0.34 | 0.25 |
| 8 | 19831977 | rs17091872 | -0.305 | 1.464 | 1.343 | -0.913 | -0.001 | -0.007 | -0.007 | 0 | 0.22 | 0.21 | 0.2 | 0.12 | 0.18 | 0.15 |
| 8 | 19832646 | rs17482753 | 0.206 | 1.524 | 0.867 | -0.785 | -0.013 | -0.004 | -0.004 | 0 | 0.09 | 0.05 | 0.06 | 0.12 | 0.13 | 0.08 |
| 8 | 19841801 | rs7843426 | 0.215 | 1.502 | 0.966 | -0.843 | -0.022 | -0.043 | -0.043 | 0.642 | 0.08 | 0.15 | 0.04 | 0 | 0.04 | 0.01 |
| 8 | 19845376 | rs7841189 | -0.66 | 1.625 | 0.949 | -0.798 | 0.005 | -0.017 | -0.017 | 0 | 0.14 | 0.23 | 0.09 | 0.12 | 0.16 | 0.09 |
| 8 | 19849757 | rs17091905 | -0.876 | 1.614 | 0.878 | -0.803 | 0.005 | -0.031 | -0.031 | 0 | 0.16 | 0.31 | 0.1 | 0.12 | 0.17 | 0.09 |
| 8 | 19854773 | rs2410622 | -1.676 | 2.329 | -0.261 | -0.277 | -0.04 | 0.012 | 0.012 | 0 | 0.93 | 0.99 | 0.7 | 0.5 | 0.86 | 0.66 |
| 8 | 19856539 | rs4523270 | -0.93 | 2.09 | -0.082 | -0.577 | -0.009 | 0.035 | 0.035 | 0 | 0.3 | 0.22 | 0.27 | 0.24 | 0.31 | 0.25 |
| 8 | 19858330 | rs10108800 | 0.052 | 2.257 | -0.326 | -0.483 | 0.016 | 0.002 | 0.002 | 0.121 | 0.1 | 0.1 | 0.19 | 0.1 | 0.23 | 0.17 |
| 8 | 19867220 | rs9644637 | -0.643 | 1.504 | -0.467 | -0.49 | -0.025 | 0.029 | 0.029 | 0 | 0.26 | 0.13 | 0.23 | 0.23 | 0.28 | 0.21 |
| 8 | 19870693 | rs7005359 | -0.373 | 0.944 | 0.267 | -0.193 | -0.006 | 0.024 | 0.024 | 0 | 0.25 | 0.45 | 0.26 | 0.31 | 0.23 | 0.19 |
| 8 | 19878154 | rs1534650 | -0.459 | 1.124 | 0.144 | -0.847 | -0.022 | -0.039 | -0.039 | 0 | 0.74 | 0.87 | 0.57 | 0.36 | 0.73 | 0.51 |
| 8 | 19890612 | rs10103634 | -1.091 | 0.935 | 0.473 | -1.283 | 0.03 | -0.008 | -0.008 | 0 | 0.26 | 0.55 | 0.25 | 0.26 | 0.34 | 0.27 |
| 8 | 19899046 | rs4244457 | -0.71 | 0.957 | -0.516 | -1.332 | -0.069 | -0.053 | -0.053 | 0 | 0.72 | 0.94 | 0.56 | 0.34 | 0.68 | 0.49 |
| 8 | 19907912 | rs11991231 | 0.135 | 0.912 | -0.38 | -1.113 | -0.01 | -0.006 | -0.006 | 0.096 | 0.1 | 0.06 | 0.07 | 0.13 | 0.13 | 0.09 |
| 8 | 19919486 | rs10086580 | 0.414 | 0.651 | -0.265 | -1.581 | 0.002 | 0.153 | 0.153 | 0.018 | 0.49 | 0.2 | 0.42 | 0.22 | 0.48 | 0.37 |
| 8 | 19923758 | rs13254929 | 0.737 | 0.187 | -0.261 | -1.814 | 0.019 | 0.059 | 0.059 | 0.573 | 0.4 | 0.26 | 0.27 | 0.09 | 0.3 | 0.19 |
| 8 | 19923902 | rs12541270 | 0.862 | 0.224 | -0.631 | -1.796 | 0.02 | 0.127 | 0.127 | 0.507 | 0.38 | 0.16 | 0.27 | 0.09 | 0.3 | 0.19 |
| 8 | 19936355 | rs12544393 | 0.958 | 0.275 | -0.898 | -1.906 | 0.018 | 0.066 | 0.066 | 0 | 0.32 | 0.15 | 0.13 | 0 | 0.25 | 0.13 |
| 8 | 19938044 | rs12545731 | 0.401 | -0.123 | -0.72 | -2.113 | -0.001 | -0.07 | -0.07 | 0 | 0.41 | 0.39 | 0.18 | 0.03 | 0.31 | 0.26 |
| 8 | 19941448 | rs6989064 | 0.002 | -0.063 | -0.964 | -1.86 | -0.011 | 0.061 | 0.061 | 0.028 | 0.61 | 0.74 | 0.52 | 0.19 | 0.58 | 0.36 |
| 8 | 19943027 | rs13265868 | 0.863 | 0.14 | -1.196 | -2.055 | -0.008 | 0.016 | 0.016 | 0 | 0.4 | 0.29 | 0.41 | 0.09 | 0.46 | 0.29 |
| 8 | 19943308 | rs13276972 | -0.233 | 0.628 | -1.507 | -1.266 | -0.09 | 0.106 | 0.106 | 0.024 | 0.72 | 0.95 | 0.58 | 0.13 | 0.64 | 0.48 |
| 8 | 19946066 | rs4401880 | 1.012 | -0.091 | -0.78 | -0.407 | 0.001 | 0 | 0 | 0 | 0.78 | 0.22 | 0.85 | 0.81 | 0.88 | 0.82 |
| 8 | 19955813 | rs10888177 | -0.294 | 0.344 | -0.62 | -0.631 | -0.002 | 0.067 | 0.067 | 0 | 0.52 | 0.61 | 0.67 | 0.79 | 0.52 | 0.79 |
| 8 | 19957089 | rs7008150 | -0.296 | 0.593 | -0.416 | 0.08 | 0.005 | -0.002 | -0.002 | 0 | 0.64 | 0.44 | 0.5 | 0.36 | 0.56 | 0.38 |
| 8 | 19958166 | rs4559256 | -0.761 | 0.547 | -0.215 | 0.154 | 0.046 | 0.029 | 0.029 | 0 | 0.7 | 0.53 | 0.54 | 0.38 | 0.61 | 0.46 |
| 8 | 19961274 | rs17092008 | -0.567 | 0.183 | -0.279 | 0.043 | -0.007 | -0.034 | -0.034 | 0 | 0.08 | 0.17 | 0.03 | 0.01 | 0.06 | 0.07 |
| 8 | 19966205 | rs17092018 | -0.243 | 0.311 | -0.28 | 0.117 | -0.012 | -0.048 | -0.048 | 0 | 0.14 | 0.38 | 0.07 | 0.03 | 0.11 | 0.15 |
| 8 | 19968929 | rs16842 | -1.029 | 0.223 | -0.781 | 0.282 | -0.012 | 0.018 | 0.018 | 0.679 | 0.31 | 0.48 | 0.26 | 0.38 | 0.26 | 0.28 |
| 8 | 19970337 | rs1566122 | 1.013 | 0.399 | -0.852 | 0.6 | 0.01 | 0.009 | 0.009 | 0 | 0.28 | 0.33 | 0.31 | 0.35 | 0.36 | 0.52 |
| 8 | 19970840 | rs1018078 | -1.065 | 0.453 | -0.941 | -0.397 | 0.014 | 0.013 | 0.013 | 0 | 0.71 | 0.63 | 0.56 | 0.37 | 0.64 | 0.47 |

| | | | | | | | | | | | | | | | | |
|---|---|---|---|---|---|---|---|---|---|---|---|---|---|---|---|---|
| 8 | 19970889 | rs1018077 | 0.285 | 0.397 | -1.019 | -0.468 | -0.004 | -0.026 | -0.026 | 0 | 0.17 | 0.08 | 0.18 | 0.36 | 0.15 | 0.13 |
| 8 | 19971767 | rs7838269 | 0.515 | 0.579 | -0.498 | 0.142 | -0.023 | 0.046 | 0.046 | 0 | 0.57 | 0.23 | 0.48 | 0.37 | 0.53 | 0.32 |
| 8 | 19972368 | rs1039122 | -1.168 | 0.265 | 0.097 | 0.113 | -0.012 | 0.004 | 0.004 | 0 | 0.14 | 0.4 | 0.08 | 0.08 | 0.11 | 0.17 |
| 8 | 19977043 | rs17411601 | -0.282 | 0.271 | -0.755 | -0.165 | -0.005 | -0.02 | -0.02 | 0 | 0.41 | 0.4 | 0.38 | 0.21 | 0.36 | 0.34 |
| 8 | 19983859 | rs7001825 | 0.397 | 0.02 | -1.243 | 0.587 | 0.008 | -0.024 | -0.024 | 0.748 | 0.8 | 0.83 | 0.69 | 0.63 | 0.86 | 0.74 |
| 8 | 19985382 | rs10503670 | 0.578 | 0.545 | -1.549 | -0.062 | -0.021 | 0.015 | 0.015 | 0 | 0.44 | 0.26 | 0.38 | 0.61 | 0.5 | 0.45 |
| 8 | 19987703 | rs10503672 | 0.122 | 0.61 | -2.202 | -0.495 | -0.095 | 0.103 | 0.103 | 0 | 0.22 | 0.02 | 0.25 | 0.05 | 0.28 | 0.16 |
| 8 | 19987889 | rs10503673 | -0.251 | 0.502 | -2.207 | 0.037 | -0.004 | -0.039 | -0.039 | 0.014 | 0.82 | 0.9 | 0.7 | 0.68 | 0.81 | 0.78 |
| 8 | 19987914 | rs17489871 | 0.498 | 1.08 | -2.95 | -0.697 | -0.066 | 0.094 | 0.094 | 0.194 | 0.21 | 0.04 | 0.15 | 0.08 | 0.25 | 0.05 |
| 8 | 19988211 | rs11204094 | 0.189 | 0.419 | -2.348 | 0.035 | 0.019 | 0.023 | 0.023 | 0 | 0.49 | 0.65 | 0.46 | 0.42 | 0.44 | 0.49 |
| 8 | 19989194 | rs11783715 | -0.03 | -0.092 | -2.262 | 1.029 | -0.032 | -0.147 | -0.147 | 0 | 0.74 | 0.95 | 0.71 | 0.56 | 0.78 | 0.57 |
| 8 | 19992955 | rs11997907 | 0.347 | 0.139 | -1.22 | 1.321 | -0.005 | 0.064 | 0.064 | 0.583 | 0.13 | 0.23 | 0.15 | 0.29 | 0.12 | 0.12 |
| 8 | 19995409 | rs2047360 | -0.725 | 0.238 | -1.491 | 1.461 | -0.072 | -0.109 | -0.109 | 0.978 | 0.82 | 0.97 | 0.74 | 0.7 | 0.79 | 0.71 |
| 8 | 19995717 | rs4921691 | -0.148 | 0.461 | -1.542 | 1.262 | -0.105 | -0.159 | -0.159 | 0 | 0.75 | 0.96 | 0.68 | 0.52 | 0.7 | 0.61 |
| 8 | 19997540 | rs4922131 | 0.798 | 0.283 | -0.711 | 0.924 | 0.004 | 0.001 | 0.001 | 0.327 | 0.51 | 0.55 | 0.49 | 0.46 | 0.45 | 0.4 |
| 8 | 20001454 | rs11204097 | 1 | 0.367 | -1.098 | 0.633 | -0.007 | 0.001 | 0.001 | 0.523 | 0.2 | 0.14 | 0.17 | 0.02 | 0.26 | 0.2 |
| 8 | 20001995 | rs7003858 | 0.211 | -0.001 | -0.78 | 1.049 | 0.003 | 0.014 | 0.014 | 0 | 0.7 | 0.7 | 0.64 | 0.56 | 0.7 | 0.61 |
| 8 | 20002426 | rs1497020 | 0.056 | 0.133 | -2.085 | 1.115 | -0.092 | -0.15 | -0.15 | 0 | 0.72 | 0.96 | 0.65 | 0.56 | 0.7 | 0.6 |
| 8 | 20004641 | rs6586896 | 1.293 | 0.479 | -1.64 | 0.914 | -0.018 | 0.016 | 0.016 | 0.161 | 0.06 | 0.02 | 0.06 | 0.05 | 0.09 | 0.02 |
| 8 | 20008012 | rs2270650 | 0.869 | 0.256 | -1.704 | 0.959 | -0.023 | 0.093 | 0.093 | 0 | 0.35 | 0.12 | 0.37 | 0.18 | 0.38 | 0.31 |
| 8 | 20010158 | rs4922132 | -0.645 | 0.171 | -1.341 | 0.918 | -0.013 | -0.027 | -0.027 | 0 | 0.8 | 0.94 | 0.73 | 0.74 | 0.83 | 0.73 |
| 8 | 20021657 | rs7013199 | -0.373 | 0.122 | -0.627 | 1.038 | -0.013 | 0.062 | 0.062 | 0 | 0.26 | 0.39 | 0.16 | 0.4 | 0.21 | 0.3 |
| 8 | 20028474 | rs7841346 | -0.066 | 0.196 | -0.773 | 0.915 | 0 | -0.004 | -0.004 | 0.384 | 0.21 | 0.24 | 0.16 | 0.14 | 0.21 | 0.23 |
| 8 | 20032103 | rs3779672 | 0.637 | 0.136 | -1.687 | 0.643 | -0.009 | 0.022 | 0.022 | 0 | 0.16 | 0.09 | 0.14 | 0.11 | 0.18 | 0.1 |
| 8 | 20032587 | rs952859 | -0.758 | 0.488 | -2.269 | 0.242 | -0.019 | -0.021 | -0.021 | 2.722 | 0.77 | 0.9 | 0.56 | 0.71 | 0.74 | 0.67 |
| 8 | 20036713 | rs1390938 | -0.745 | 0.504 | -2.337 | 0.25 | -0.028 | -0.03 | -0.03 | 0 | 0.76 | 0.93 | 0.56 | 0.71 | 0.75 | 0.66 |
| 8 | 20037017 | rs2270638 | 0.664 | 0.739 | -1.84 | -0.404 | -0.042 | 0.076 | 0.076 | 0 | 0.33 | 0.1 | 0.24 | 0.17 | 0.37 | 0.28 |
| 8 | 20040712 | rs1390939 | 0.301 | 0.468 | -1.533 | -0.098 | -0.019 | 0.177 | 0.177 | 0 | 0.47 | 0.11 | 0.31 | 0.19 | 0.55 | 0.39 |
| 8 | 20049524 | rs11783752 | 0.112 | 0.783 | -0.699 | -0.013 | -0.002 | 0.002 | 0.002 | 0 | 0.67 | 0.71 | 0.79 | 0.73 | 0.69 | 0.65 |
| 8 | 20058757 | rs6586899 | -1.352 | 1.035 | -0.196 | -0.252 | 0.035 | 0.028 | 0.028 | 0 | 0.13 | 0.33 | 0.28 | 0.15 | 0.13 | 0.21 |
| 8 | 20060856 | rs2410633 | -0.493 | 1.005 | -0.407 | -0.347 | -0.029 | 0.056 | 0.056 | 0 | 0.73 | 0.88 | 0.87 | 0.88 | 0.72 | 0.71 |
| 8 | 20066618 | rs11778205 | -0.582 | 0.861 | -0.198 | -0.722 | 0.002 | -0.02 | -0.02 | 0 | 0.34 | 0.6 | 0.45 | 0.22 | 0.33 | 0.29 |
| 8 | 20071503 | rs11204099 | 0.941 | 0.597 | -0.393 | -0.675 | 0.016 | 0.017 | 0.017 | 0 | 0.39 | 0.28 | 0.43 | 0.65 | 0.38 | 0.42 |
| 8 | 20078643 | rs1042426 | -0.111 | 0.643 | -0.548 | -0.595 | -0.002 | -0.002 | -0.002 | 0.544 | 0.06 | 0.06 | 0.17 | 0.08 | 0.09 | 0.14 |
| 8 | 20080322 | rs13252207 | 0.769 | 0.624 | -0.526 | -0.697 | -0.001 | -0.019 | -0.019 | 2.664 | 0.21 | 0.22 | 0.17 | 0.08 | 0.2 | 0.06 |
| 8 | 20081890 | rs1390943 | -0.633 | 0.633 | -1.339 | -0.683 | 0.005 | 0.021 | 0.021 | 2.279 | 0.37 | 0.69 | 0.45 | 0.23 | 0.32 | 0.29 |
| 8 | 20088118 | rs10095373 | -0.482 | 0.789 | -1.227 | -0.581 | 0 | 0.008 | 0.008 | 1.49 | 0.34 | 0.69 | 0.41 | 0.23 | 0.31 | 0.26 |
| 8 | 20091363 | rs952149 | 0.401 | 0.687 | -1.218 | -0.113 | 0.029 | 0.028 | 0.028 | 0 | 0.4 | 0.2 | 0.43 | 0.66 | 0.38 | 0.44 |
| 8 | 20091519 | rs952148 | -0.535 | 1.105 | -1.048 | 0.257 | -0.014 | -0.009 | -0.009 | 0.058 | 0.07 | 0.06 | 0.21 | 0.11 | 0.1 | 0.18 |
| 8 | 20097325 | rs17092176 | 0.12 | 0.56 | -1.091 | 0.47 | 0.031 | -0.016 | -0.016 | 0 | 0.41 | 0.26 | 0.43 | 0.51 | 0.37 | 0.37 |
| 8 | 20101079 | rs12682557 | -0.522 | 1.245 | -1.18 | 0.693 | -0.045 | -0.066 | -0.066 | 0 | 0.29 | 0.17 | 0.21 | 0.5 | 0.35 | 0.33 |
| 8 | 20102878 | rs1982959 | -0.755 | 0.812 | -1.213 | 0.064 | 0.007 | 0.02 | 0.02 | 0 | 0.78 | 0.81 | 0.77 | 0.93 | 0.8 | 0.86 |
| 8 | 20104985 | rs1075 | 0.298 | 1.252 | -0.905 | 0.477 | -0.088 | -0.041 | -0.041 | 0.371 | 0.19 | 0.01 | 0.12 | 0.18 | 0.21 | 0.17 |
| 8 | 20106874 | rs904002 | -0.769 | 1.034 | -0.773 | 0.258 | 0.011 | 0.076 | 0.076 | 0 | 0.77 | 0.91 | 0.78 | 0.93 | 0.8 | 0.86 |
| 8 | 20109187 | rs2935312 | -0.493 | 1.118 | 0.123 | 0.413 | 0.03 | 0.037 | 0.037 | 0 | 0.29 | 0.38 | 0.22 | 0.46 | 0.32 | 0.35 |
| 8 | 20109809 | rs903999 | 0.794 | 0.853 | -0.119 | 0.303 | -0.006 | -0.024 | -0.024 | 0 | 0.41 | 0.27 | 0.53 | 0.47 | 0.45 | 0.49 |
| 8 | 20116621 | rs4921697 | 0.932 | 0.752 | -0.554 | 0.279 | -0.065 | -0.06 | -0.06 | 0.468 | 0.35 | 0.08 | 0.51 | 0.37 | 0.42 | 0.44 |
| 8 | 20120139 | rs13259742 | -0.795 | 0.903 | 0.145 | 0.468 | 0.032 | 0.095 | 0.095 | 0.431 | 0.21 | 0.44 | 0.2 | 0.44 | 0.25 | 0.28 |
| 8 | 20127595 | rs1497017 | -0.379 | 0.632 | 0.082 | -0.106 | -0.001 | -0.004 | -0.004 | 0.412 | 0.56 | 0.62 | 0.55 | 0.51 | 0.45 | 0.45 |
| 8 | 20128078 | rs4436141 | -0.135 | 0.408 | 0.156 | -0.159 | 0.006 | -0.021 | -0.021 | 0.058 | 0.31 | 0.22 | 0.35 | 0.52 | 0.26 | 0.36 |
| 8 | 20130379 | rs4922145 | 0.493 | 0.47 | 0.029 | -0.354 | -0.006 | -0.03 | -0.03 | 0 | 0.08 | 0.09 | 0.04 | 0 | 0.05 | 0.02 |
| 8 | 20131697 | rs6984493 | -0.203 | 0.56 | 0.153 | -0.392 | 0 | -0.045 | -0.045 | 0.26 | 0.89 | 0.76 | 0.94 | 0.98 | 0.9 | 0.93 |
| 8 | 20131799 | rs1532445 | 0.505 | 0.53 | 0.647 | -0.361 | 0.005 | -0.017 | -0.017 | 0 | 0.59 | 0.34 | 0.58 | 0.48 | 0.62 | 0.62 |
| 8 | 20132255 | rs13280649 | 0.309 | 0.54 | 0.533 | -0.357 | 0 | -0.033 | -0.033 | 0 | 0.29 | 0.19 | 0.34 | 0.5 | 0.27 | 0.32 |
| 8 | 20132391 | rs11995187 | 0.673 | 0.607 | 0.172 | -0.312 | -0.031 | -0.086 | -0.086 | 0 | 0.29 | 0.4 | 0.21 | 0.05 | 0.2 | 0.21 |
| 8 | 20134283 | rs11204108 | 0 | 0.639 | 0.032 | -0.299 | -0.004 | 0.014 | 0.014 | 0 | 0.59 | 0.51 | 0.6 | 0.49 | 0.63 | 0.62 |
| 8 | 20134377 | rs11786102 | 0.645 | 1.21 | -0.488 | 0.266 | -0.068 | -0.074 | -0.074 | 0 | 0.31 | 0.14 | 0.38 | 0.44 | 0.43 | 0.42 |
| 8 | 20138047 | rs10107110 | 0.175 | 0.904 | 0.438 | 0.144 | 0.019 | -0.006 | -0.006 | 0 | 0.86 | 0.53 | 0.85 | 0.91 | 0.87 | 0.83 |
| 8 | 20138477 | rs17092214 | 0.586 | 1.452 | 0.02 | 0.36 | -0.01 | -0.006 | -0.006 | 0.145 | 0.63 | 0.35 | 0.46 | 0.6 | 0.53 | 0.65 |
| 8 | 20139567 | rs12675554 | 1.078 | 1.253 | -0.239 | -0.031 | 0 | -0.001 | -0.001 | 0 | 0.4 | 0.39 | 0.32 | 0.41 | 0.35 | 0.43 |
| 8 | 20139680 | rs2898500 | 0.78 | 1.422 | -0.538 | 0.164 | -0.009 | -0.008 | -0.008 | 2.372 | 0.21 | 0.15 | 0.34 | 0.26 | 0.29 | 0.16 |
| 8 | 20142119 | rs12677666 | 1.311 | 1.343 | -0.248 | 0.054 | 0.004 | -0.007 | -0.007 | 0 | 0.38 | 0.24 | 0.3 | 0.41 | 0.32 | 0.43 |
| 8 | 20143440 | rs13270944 | 0.23 | 1.545 | -0.571 | 0.078 | -0.024 | -0.027 | -0.027 | 0 | 0.26 | 0.17 | 0.48 | 0.35 | 0.37 | 0.22 |
| 8 | 20144440 | rs10098648 | -0.523 | 1.196 | -0.959 | -0.237 | -0.01 | -0.001 | -0.001 | 0.602 | 0.82 | 0.86 | 0.9 | 1 | 0.78 | 0.91 |
| 8 | 20146182 | rs12114422 | 1.441 | 0.8 | -1.098 | -0.086 | 0.016 | -0.017 | -0.017 | 0 | 0.35 | 0.21 | 0.26 | 0.4 | 0.25 | 0.41 |
| 8 | 20146260 | rs7012277 | 0.66 | 0.483 | -1.187 | 0.008 | 0.053 | 0.009 | 0.009 | 0.001 | 0.15 | 0.05 | 0.08 | 0.1 | 0.07 | 0.11 |
| 8 | 20146804 | rs6998516 | 0.258 | 1.373 | -0.645 | 0.166 | 0.028 | -0.011 | -0.011 | 0 | 0.72 | 0.67 | 0.69 | 0.8 | 0.61 | 0.73 |
| 8 | 20147170 | rs11204109 | 0.851 | 0.961 | -1.137 | 0.155 | 0.02 | -0.014 | -0.014 | 0 | 0.39 | 0.17 | 0.28 | 0.41 | 0.28 | 0.41 |
| 8 | 20150536 | rs7835387 | -0.386 | 0.662 | -0.751 | -0.141 | 0.031 | 0.003 | 0.003 | 0 | 0.49 | 0.62 | 0.65 | 0.5 | 0.63 | 0.67 |

| | | | | | | | | | | | | | | | | |
|---|---|---|---|---|---|---|---|---|---|---|---|---|---|---|---|---|
| 8 | 20151095 | rs868223 | -0.22 | 0.58 | -0.623 | -0.059 | 0.012 | 0.022 | 0.022 | 0 | 0.47 | 0.56 | 0.56 | 0.39 | 0.63 | 0.65 |
| 8 | 20151152 | rs11782092 | 1.299 | 0.775 | -0.918 | -0.339 | 0.013 | 0.024 | 0.024 | 0 | 0.21 | 0.12 | 0.1 | 0.09 | 0.18 | 0.11 |
| 8 | 20152143 | rs923057 | 0.384 | 0.848 | -0.641 | -0.179 | -0.022 | 0.038 | 0.038 | 0 | 0.37 | 0.26 | 0.46 | 0.21 | 0.52 | 0.43 |
| 8 | 20153776 | rs4922155 | -0.203 | 0.786 | -1.189 | -0.135 | 0.048 | 0.012 | 0.012 | 0 | 0.5 | 0.63 | 0.66 | 0.56 | 0.65 | 0.68 |
| 8 | 20153950 | rs4922156 | -0.403 | 1.174 | -1.208 | -0.468 | 0 | -0.008 | -0.008 | 0.802 | 0.86 | 0.88 | 0.94 | 1 | 0.85 | 0.94 |
| 8 | 20167753 | rs931292 | 0.148 | 1.206 | -1.419 | -0.385 | -0.029 | 0.039 | 0.039 | 0 | 0.8 | 0.88 | 0.81 | 0.88 | 0.75 | 0.85 |
| 8 | 20184824 | rs7821344 | 0.79 | 0.693 | -1.337 | -0.545 | 0.003 | 0.029 | 0.029 | 0 | 0.21 | 0.13 | 0.1 | 0.09 | 0.2 | 0.11 |
| 8 | 20185105 | rs17092240 | 0.728 | 0.021 | -1.307 | -0.546 | 0.055 | -0.039 | -0.039 | 0.281 | 0.36 | 0.21 | 0.26 | 0.52 | 0.2 | 0.3 |
| 8 | 20185310 | rs4921702 | -0.666 | 0.017 | -0.656 | -0.551 | -0.001 | -0.046 | -0.046 | 0 | 0.86 | 0.79 | 0.89 | 1 | 0.85 | 0.86 |
| 8 | 20186028 | rs13265570 | 1.104 | 0.046 | -0.749 | -0.537 | 0.005 | 0.014 | 0.014 | 0 | 0.19 | 0.13 | 0.09 | 0.08 | 0.18 | 0.1 |
| 8 | 20189277 | rs7827808 | -0.347 | 0.25 | -0.251 | -0.107 | -0.002 | -0.053 | -0.053 | 0 | 0.86 | 0.77 | 0.9 | 0.99 | 0.86 | 0.85 |
| 8 | 20190224 | rs10503676 | -0.178 | -0.344 | -0.236 | 0.348 | -0.032 | 0.041 | 0.041 | 0 | 0.23 | 0.32 | 0.3 | 0.59 | 0.12 | 0.26 |
| 8 | 20191012 | rs7838163 | 0.459 | 0.483 | -0.131 | 0.373 | 0.035 | 0.006 | 0.006 | 0 | 0.07 | 0.11 | 0.07 | 0.1 | 0.1 | 0.06 |
| 8 | 20191087 | rs7007963 | -0.418 | 0.148 | -0.024 | 0.664 | 0.002 | 0 | 0 | 0 | 0.43 | 0.39 | 0.43 | 0.46 | 0.35 | 0.46 |
| 8 | 20192835 | rs13276304 | -0.111 | 0.419 | -0.374 | 0.673 | -0.04 | -0.081 | -0.081 | 0.186 | 0.36 | 0.08 | 0.44 | 0.46 | 0.36 | 0.42 |
| 8 | 20196093 | rs1021201 | -0.718 | 0.354 | 0.271 | 0.591 | 0.003 | 0.058 | 0.058 | 0 | 0.46 | 0.63 | 0.5 | 0.73 | 0.43 | 0.52 |
| 8 | 20204044 | rs10503677 | -0.706 | 0.301 | 0.334 | 0.625 | 0.011 | 0.041 | 0.041 | 0 | 0.45 | 0.62 | 0.49 | 0.74 | 0.42 | 0.53 |
| 8 | 20211529 | rs900955 | -0.199 | 0.328 | 0.97 | 0.636 | -0.022 | -0.003 | -0.003 | 0.055 | 0.35 | 0.2 | 0.39 | 0.63 | 0.4 | 0.56 |
| 8 | 20212403 | rs1872346 | -0.745 | 0.303 | 1.097 | 0.619 | -0.001 | 0.03 | 0.03 | 0 | 0.4 | 0.63 | 0.47 | 0.79 | 0.41 | 0.57 |
| 8 | 20215762 | rs2062348 | -0.344 | 0.433 | 1.668 | 0.682 | -0.038 | -0.009 | -0.009 | 0 | 0.35 | 0.16 | 0.39 | 0.63 | 0.4 | 0.56 |
| 8 | 20221905 | rs12549080 | -0.204 | 0.401 | 1.614 | 0.651 | -0.037 | -0.002 | -0.002 | 0.074 | 0.35 | 0.15 | 0.39 | 0.65 | 0.4 | 0.62 |
| 8 | 20224079 | rs6985844 | -1.216 | 0.307 | 1.421 | 0.44 | 0.015 | 0.158 | 0.158 | 0 | 0.5 | 0.71 | 0.53 | 0.84 | 0.47 | 0.68 |
| 8 | 20233146 | rs1038465 | 0.272 | 0.24 | 0.849 | 0.551 | -0.044 | -0.129 | -0.129 | 0.187 | 0.3 | 0.07 | 0.34 | 0.52 | 0.33 | 0.45 |
| 8 | 20237266 | rs1385309 | -1.443 | 0.459 | 0.785 | 0.396 | -0.038 | 0.2 | 0.2 | 0 | 0.6 | 0.93 | 0.69 | 0.88 | 0.58 | 0.74 |
| 8 | 20251128 | rs4922184 | -1.153 | 0.071 | 1.542 | 0.914 | 0.013 | 0.075 | 0.075 | 0 | 0.47 | 0.62 | 0.51 | 0.8 | 0.45 | 0.64 |
| 8 | 20260729 | rs4921709 | -0.142 | 0.337 | 0.785 | 1.188 | 0.023 | 0.156 | 0.156 | 0 | 0.06 | 0.18 | 0.12 | 0.29 | 0.07 | 0.07 |
| 8 | 20274088 | rs12547668 | -0.46 | 0.289 | -0.118 | 0.879 | -0.008 | -0.073 | -0.073 | 0 | 0.34 | 0.15 | 0.37 | 0.5 | 0.37 | 0.37 |
| 8 | 20280942 | rs13248317 | -0.367 | 0.307 | -0.13 | 0.868 | -0.005 | -0.076 | -0.076 | 3.16 | 0.35 | 0.15 | 0.37 | 0.5 | 0.37 | 0.46 |
| 8 | 20283164 | rs2410644 | -0.956 | 0.36 | 0.292 | 0.365 | -0.002 | 0.047 | 0.047 | 0.009 | 0.42 | 0.58 | 0.42 | 0.79 | 0.45 | 0.63 |
| 8 | 20283918 | rs6995343 | -0.923 | 0.233 | 1.176 | 0.508 | 0.003 | 0.021 | 0.021 | 0 | 0.43 | 0.39 | 0.39 | 0.82 | 0.42 | 0.64 |
| 8 | 20284121 | rs12677076 | -0.514 | 0.252 | 1.366 | 0.718 | 0.007 | 0.021 | 0.021 | 0 | 0.36 | 0.25 | 0.33 | 0.75 | 0.36 | 0.51 |
| 8 | 20285163 | rs4430111 | -1.357 | 0.257 | 1.546 | 0.953 | 0.029 | 0.168 | 0.168 | 0 | 0.09 | 0.32 | 0.11 | 0.22 | 0.06 | 0.15 |
| 8 | 20288391 | rs4922193 | -1.323 | 0.154 | 1.543 | 0.633 | 0.017 | 0.093 | 0.093 | 0 | 0.47 | 0.64 | 0.5 | 0.8 | 0.43 | 0.71 |
| 8 | 20290327 | rs7817914 | -0.934 | 0.196 | 1.66 | 0.725 | 0 | -0.003 | -0.003 | 0 | 0.42 | 0.39 | 0.49 | 0.69 | 0.42 | 0.65 |
| 8 | 20290539 | rs6982875 | -0.646 | 0.22 | 1.092 | 0.747 | 0.001 | -0.024 | -0.024 | 0 | 0.37 | 0.15 | 0.34 | 0.61 | 0.37 | 0.52 |
| 8 | 20300074 | rs13282008 | -0.493 | -0.19 | 1.189 | 0.734 | 0.019 | -0.035 | -0.035 | 0 | 0.32 | 0.17 | 0.3 | 0.61 | 0.27 | 0.47 |
| 8 | 20309866 | rs13266471 | -1.076 | -0.197 | 1.418 | 0.647 | 0.01 | 0.023 | 0.023 | 0 | 0.42 | 0.35 | 0.41 | 0.67 | 0.38 | 0.55 |
| 8 | 20315601 | rs12550821 | -0.021 | -0.226 | 0.935 | 0.794 | -0.022 | -0.141 | -0.141 | 1.139 | 0.28 | 0.08 | 0.23 | 0.46 | 0.27 | 0.4 |
| 8 | 20320695 | rs13279091 | -1.166 | -0.339 | 1.14 | 0.832 | 0.002 | -0.001 | -0.001 | 0 | 0.11 | 0.11 | 0.13 | 0.15 | 0.06 | 0.1 |
| 8 | 20322808 | rs17092514 | -2.085 | -0.297 | 1.639 | 0.877 | -0.02 | 0.044 | 0.044 | 0 | 0.16 | 0.38 | 0.26 | 0.22 | 0.12 | 0.23 |
| 8 | 20328614 | rs10101153 | -1.858 | -0.333 | 1.604 | 0.855 | -0.046 | -0.001 | -0.001 | 0 | 0.13 | 0.29 | 0.21 | 0.14 | 0.07 | 0.18 |
| 8 | 20331946 | rs17092536 | -0.538 | -0.342 | 1.329 | 0.72 | -0.007 | -0.033 | -0.033 | 0.031 | 0.36 | 0.14 | 0.33 | 0.6 | 0.36 | 0.48 |
| 8 | 20332653 | rs4921719 | -0.191 | -0.272 | 1.32 | 0.853 | -0.03 | -0.084 | -0.084 | 0 | 0.31 | 0.08 | 0.31 | 0.57 | 0.33 | 0.4 |
| 8 | 20335626 | rs2127649 | -1.275 | -0.266 | 1.388 | 0.756 | 0.01 | 0.055 | 0.055 | 0 | 0.46 | 0.37 | 0.5 | 0.67 | 0.45 | 0.63 |
| 8 | 20339332 | rs10103232 | -1.798 | -0.133 | 2.049 | 0.768 | -0.014 | 0.086 | 0.086 | 0 | 0.08 | 0.33 | 0.16 | 0.14 | 0.02 | 0.16 |
| 8 | 20347457 | rs1480347 | -2.413 | -0.155 | 2.05 | 0.756 | 0.008 | -0.003 | -0.003 | 0 | 0.19 | 0.75 | 0.28 | 0.2 | 0.12 | 0.23 |
| 8 | 20351911 | rs11204117 | -2.325 | -0.139 | 2.078 | 0.763 | -0.014 | 0.004 | 0.004 | 0.021 | 0.19 | 0.67 | 0.28 | 0.2 | 0.12 | 0.23 |
| 8 | 20353398 | rs4921720 | -2.121 | -0.106 | 2.014 | 0.78 | -0.027 | 0.026 | 0.026 | 0 | 0.18 | 0.6 | 0.27 | 0.2 | 0.12 | 0.23 |
| 8 | 20356843 | rs11990984 | -1.845 | -0.225 | 1.512 | 0.539 | -0.066 | -0.039 | -0.039 | 0 | 0.15 | 0.5 | 0.22 | 0.06 | 0.07 | 0.19 |
| 8 | 20366515 | rs10283119 | -1.184 | 0.015 | 0.497 | 0.37 | 0.009 | 0.167 | 0.167 | 0.561 | 0.51 | 0.83 | 0.57 | 0.76 | 0.55 | 0.71 |
| 8 | 20369496 | rs6993760 | -0.804 | 0.073 | 1.501 | 0.568 | 0.001 | -0.001 | -0.001 | 0 | 0.39 | 0.56 | 0.49 | 0.59 | 0.44 | 0.5 |
| 8 | 20369873 | rs1564427 | 0.251 | 0.077 | 1.446 | 0.656 | -0.019 | -0.05 | -0.05 | 0.282 | 0.26 | 0.17 | 0.28 | 0.53 | 0.33 | 0.39 |
| 8 | 20370176 | rs1564425 | -0.917 | 0.16 | 1.237 | 0.696 | -0.023 | -0.001 | -0.001 | 0 | 0.39 | 0.77 | 0.49 | 0.59 | 0.44 | 0.5 |
| 8 | 20391160 | rs9325871 | -0.163 | -0.156 | 1.564 | 0.899 | -0.028 | -0.035 | -0.035 | 0 | 0.06 | 0.01 | 0.05 | 0.08 | 0.06 | 0.11 |
| 8 | 20393206 | rs7000921 | -1.674 | -0.343 | 1.868 | 0.98 | 0.009 | 0.031 | 0.031 | 0 | 0.21 | 0.56 | 0.27 | 0.26 | 0.18 | 0.32 |
| 8 | 20400341 | rs12543862 | -1.168 | -0.313 | 1.735 | 0.815 | 0.002 | 0.01 | 0.01 | 0 | 0.17 | 0.39 | 0.21 | 0.2 | 0.12 | 0.29 |
| 8 | 20412168 | rs4921723 | -1.429 | -0.515 | 1.806 | 0.982 | 0.015 | 0.072 | 0.072 | 0.001 | 0.16 | 0.49 | 0.23 | 0.22 | 0.12 | 0.19 |
| 8 | 20413142 | rs4454287 | -1.989 | -1.189 | 1.291 | 0.86 | 0.02 | 0.001 | 0.001 | 0 | 0.32 | 0.71 | 0.35 | 0.38 | 0.19 | 0.31 |
| 8 | 20419397 | rs7824498 | -1.674 | -1.035 | 1.003 | 0.932 | 0.001 | -0.002 | -0.002 | 0 | 0.31 | 0.69 | 0.39 | 0.59 | 0.26 | 0.39 |
| 8 | 20425772 | rs1351381 | 0.335 | -0.896 | 1.194 | 0.778 | 0.024 | -0.02 | -0.02 | 0 | 0.05 | 0.17 | 0.06 | 0 | 0.05 | 0.01 |
| 8 | 20437022 | rs6586919 | -2.032 | -1.361 | 0.841 | 0.716 | 0.052 | 0.066 | 0.066 | 0 | 0.43 | 0.73 | 0.49 | 0.76 | 0.29 | 0.43 |
| 8 | 20437702 | rs7816876 | -1.22 | -1.059 | 1.325 | 0.749 | 0.02 | -0.071 | -0.071 | 0 | 0.12 | 0.52 | 0.12 | 0 | 0.09 | 0.03 |
| 8 | 20441965 | rs1979433 | -0.957 | -0.914 | 0.977 | 1.054 | -0.005 | -0.04 | -0.04 | 0.13 | 0.21 | 0.15 | 0.35 | 0.58 | 0.22 | 0.33 |
| 8 | 20442258 | rs970293 | -0.97 | -0.916 | 0.951 | 1.077 | -0.007 | -0.05 | -0.05 | 0 | 0.22 | 0.15 | 0.35 | 0.57 | 0.22 | 0.33 |
| 8 | 20457201 | rs7827575 | -0.799 | -1.073 | 1.537 | 0.468 | 0.039 | -0.034 | -0.034 | 0 | 0.09 | 0.32 | 0.11 | 0.01 | 0.08 | 0.03 |
| 8 | 20459712 | rs17420770 | 0.617 | -1.098 | 1.336 | 0.555 | -0.016 | -0.008 | -0.008 | 0 | 0.06 | 0.01 | 0.16 | 0.04 | 0.06 | 0.14 |
| 8 | 20464805 | rs12547174 | -0.505 | -1.19 | 1.512 | 0.627 | -0.006 | -0.061 | -0.061 | 0 | 0.17 | 0.03 | 0.31 | 0.74 | 0.16 | 0.38 |
| 8 | 20465952 | rs4592052 | -1.442 | -1.187 | 1.577 | 0.635 | -0.003 | 0.009 | 0.009 | 0 | 0.26 | 0.42 | 0.42 | 0.74 | 0.25 | 0.4 |
| 8 | 20468247 | rs11204121 | -1.141 | -1.15 | 1.486 | 0.658 | 0 | 0.014 | 0.014 | 0 | 0.23 | 0.34 | 0.37 | 0.75 | 0.21 | 0.39 |

| | | | | | | | | | | | | | | | | |
|---|---|---|---|---|---|---|---|---|---|---|---|---|---|---|---|---|
| 8 | 20468502 | rs4329288 | 1.554 | -1.261 | 0.636 | -0.29 | 0.052 | 0.168 | 0.168 | 0 | 0.5 | 0.27 | 0.42 | 0.13 | 0.59 | 0.44 |
| 8 | 20474703 | rs4922223 | -0.87 | -0.629 | 0.755 | 0.368 | 0.003 | 0.05 | 0.05 | 0 | 0.22 | 0.38 | 0.36 | 0.68 | 0.21 | 0.39 |
| 8 | 20482878 | rs1450299 | 0.384 | -0.44 | 0.488 | 0.121 | 0.033 | -0.001 | -0.001 | 0 | 0.06 | 0.25 | 0.04 | 0.01 | 0.05 | 0.03 |
| 8 | 20483935 | rs7841532 | -0.06 | -0.789 | 0.045 | 0.17 | -0.001 | 0.009 | 0.009 | 0 | 0.17 | 0.21 | 0.18 | 0.18 | 0.18 | 0.14 |
| 8 | 20484131 | rs1031317 | 0.05 | -0.878 | -0.772 | -0.182 | -0.025 | -0.061 | -0.061 | 0.01 | 0.8 | 0.95 | 0.69 | 0.25 | 0.78 | 0.74 |
| 8 | 20484381 | rs1562786 | -0.091 | -0.269 | -1.032 | -1.007 | -0.037 | 0.03 | 0.03 | 0 | 0.71 | 0.82 | 0.53 | 0.19 | 0.59 | 0.65 |
| 8 | 20486184 | rs1868660 | 0.107 | -0.213 | -0.896 | -1.539 | 0.012 | -0.133 | -0.133 | 1.039 | 0.2 | 0.54 | 0.22 | 0.01 | 0.21 | 0.3 |
| 8 | 20486240 | rs1868661 | 1.251 | -0.268 | -0.938 | -1.597 | -0.014 | 0.04 | 0.04 | 0.043 | 0.12 | 0.04 | 0.07 | 0 | 0.15 | 0.21 |
| 8 | 20487616 | rs4922225 | 2.54 | -0.381 | -1.494 | -1.913 | -0.099 | 0.011 | 0.011 | 1.193 | 0.22 | 0.01 | 0.14 | 0.11 | 0.23 | 0.19 |
| 8 | 20487794 | rs4922226 | -0.355 | -0.361 | -1.231 | -2.111 | 0.014 | -0.136 | -0.136 | 0 | 0.8 | 0.46 | 0.77 | 0.99 | 0.79 | 0.68 |
| 8 | 20489396 | rs10106155 | 0.289 | -0.402 | -1.097 | -2.051 | -0.017 | 0.091 | 0.091 | 0.293 | 0.45 | 0.72 | 0.38 | 0.13 | 0.44 | 0.52 |
| 8 | 20492735 | rs2198011 | -0.332 | -0.487 | -1.139 | -2.19 | 0.028 | 0.148 | 0.148 | 1.213 | 0.54 | 0.77 | 0.48 | 0.16 | 0.54 | 0.64 |
| 8 | 20494172 | rs969825 | -0.245 | -0.528 | -1.316 | -1.43 | 0.014 | 0.016 | 0.016 | 0.586 | 0.58 | 0.74 | 0.56 | 0.63 | 0.59 | 0.68 |
| 8 | 20494871 | rs2898513 | -0.44 | -0.447 | -1.225 | -1.487 | 0.005 | 0.038 | 0.038 | 0 | 0.32 | 0.58 | 0.4 | 0.51 | 0.35 | 0.45 |
| 8 | 20494897 | rs2410657 | 0.032 | -0.482 | -0.608 | -1.55 | 0 | 0.028 | 0.028 | 0 | 0.27 | 0.33 | 0.35 | 0.51 | 0.31 | 0.41 |
| 8 | 20496648 | rs1947260 | -0.05 | -0.474 | -0.779 | -1.372 | 0.02 | 0.038 | 0.038 | 1.583 | 0.57 | 0.73 | 0.66 | 0.68 | 0.6 | 0.67 |
| 8 | 20498604 | rs17420979 | 0.712 | -0.218 | -0.683 | -1.322 | -0.015 | 0.024 | 0.024 | 0 | 0.08 | 0.02 | 0.09 | 0.03 | 0.09 | 0.12 |
| 8 | 20499038 | rs12543515 | 2.101 | -0.145 | -0.746 | -1.222 | -0.01 | 0.02 | 0.02 | 0.091 | 0.07 | 0.02 | 0.14 | 0.49 | 0.07 | 0.08 |
| 8 | 20499758 | rs7018113 | -0.24 | -0.195 | -0.422 | -1.348 | 0.014 | -0.118 | -0.118 | 0 | 0.27 | 0.57 | 0.32 | 0.04 | 0.3 | 0.43 |
| 8 | 20500940 | rs6988396 | -0.999 | -0.356 | -0.748 | -1.167 | 0.011 | -0.069 | -0.069 | 0 | 0.75 | 0.93 | 0.7 | 0.64 | 0.77 | 0.78 |
| 8 | 20511029 | rs7013866 | 1.855 | 0.087 | -0.714 | -1.433 | -0.009 | 0.025 | 0.025 | 0 | 0.24 | 0.16 | 0.16 | 0.08 | 0.23 | 0.18 |
| 8 | 20519467 | rs11204123 | -0.222 | 0.16 | -0.125 | -1.476 | -0.011 | -0.149 | -0.149 | 0 | 0.24 | 0.45 | 0.15 | 0.01 | 0.22 | 0.28 |
| 8 | 20521191 | rs17421063 | 0.23 | -0.072 | -0.442 | -1.498 | 0.002 | -0.003 | -0.003 | 0.102 | 0.58 | 0.4 | 0.57 | 0.84 | 0.6 | 0.62 |
| 8 | 20526310 | rs6586925 | -0.536 | 0.07 | -0.519 | -1.08 | -0.007 | -0.029 | -0.029 | 0 | 0.82 | 0.88 | 0.66 | 0.47 | 0.76 | 0.75 |
| 8 | 20529012 | rs11781541 | 0.153 | 0.103 | -0.561 | -1.167 | 0.001 | 0.006 | 0.006 | 0.751 | 0.27 | 0.26 | 0.29 | 0.51 | 0.34 | 0.33 |
| 8 | 20532503 | rs11990986 | -0.139 | 0.09 | -0.244 | -1.153 | -0.015 | -0.059 | -0.059 | 1.569 | 0.66 | 0.5 | 0.61 | 0.85 | 0.66 | 0.67 |
| 8 | 20544296 | rs2597415 | 0.76 | 0.204 | -0.252 | -1.066 | 0.025 | 0.041 | 0.041 | 0 | 0.71 | 0.51 | 0.55 | 0.42 | 0.65 | 0.58 |
| 8 | 20545032 | rs12335299 | 1.798 | 0.212 | -0.202 | -1.243 | 0.002 | -0.019 | -0.019 | 1.048 | 0.06 | 0.12 | 0.04 | 0.01 | 0.05 | 0.08 |
| 8 | 20548817 | rs7839655 | 0.851 | -0.029 | -0.242 | -1.342 | 0.02 | 0.032 | 0.032 | 0 | 0.26 | 0.22 | 0.19 | 0.06 | 0.21 | 0.15 |
| 8 | 20549450 | rs1868662 | -0.42 | -0.062 | 0.102 | -1.183 | 0.022 | 0.008 | 0.008 | 0.05 | 0.88 | 0.74 | 0.78 | 0.9 | 0.87 | 0.77 |
| 8 | 20550536 | rs2597384 | 0.816 | -0.059 | 0.02 | -1.149 | 0.006 | 0.045 | 0.045 | 0 | 0.5 | 0.35 | 0.4 | 0.34 | 0.47 | 0.48 |
| 8 | 20551783 | rs2597385 | -1.587 | 0.054 | 0.182 | -1.117 | -0.002 | 0.002 | 0.002 | 0 | 0.94 | 0.87 | 0.83 | 0.96 | 0.94 | 0.87 |
| 8 | 20553125 | rs2597388 | 0.187 | 0.057 | 0.083 | -1.149 | 0.018 | -0.029 | -0.029 | 0 | 0.82 | 0.54 | 0.73 | 0.89 | 0.83 | 0.7 |
| 8 | 20559099 | rs13273516 | 1.326 | -0.037 | -0.426 | -1.143 | 0.016 | 0.027 | 0.027 | 0.024 | 0.18 | 0.14 | 0.14 | 0.06 | 0.14 | 0.13 |
| 8 | 20562642 | rs896044 | 0.567 | 0.069 | -0.1 | -1.093 | 0.007 | 0.005 | 0.005 | 0 | 0.66 | 0.5 | 0.53 | 0.4 | 0.59 | 0.6 |
| 8 | 20563929 | rs17093001 | 0.944 | 0.194 | -0.134 | -1.269 | 0.019 | -0.097 | -0.097 | 0 | 0.14 | 0.34 | 0.09 | 0 | 0.15 | 0.21 |
| 8 | 20564192 | rs13260745 | -0.619 | -0.046 | -0.407 | -1.447 | -0.002 | 0.002 | 0.002 | 0 | 0.51 | 0.59 | 0.42 | 0.49 | 0.56 | 0.49 |
| 8 | 20564798 | rs2198012 | 0.298 | -0.044 | -0.488 | -1.394 | -0.011 | -0.017 | -0.017 | 0 | 0.67 | 0.79 | 0.56 | 0.41 | 0.61 | 0.62 |
| 8 | 20566971 | rs2442976 | -0.868 | -0.008 | -0.462 | -1.347 | 0.002 | -0.001 | -0.001 | 0 | 0.89 | 0.87 | 0.79 | 0.9 | 0.86 | 0.79 |
| 8 | 20569247 | rs2616231 | 1.292 | 0.005 | -0.363 | -1.487 | -0.008 | 0.014 | 0.014 | 0.329 | 0.41 | 0.37 | 0.36 | 0.33 | 0.41 | 0.43 |
| 8 | 20571619 | rs1868664 | 0.355 | -0.104 | -0.452 | -1.545 | -0.017 | 0.008 | 0.008 | 0 | 0.58 | 0.68 | 0.49 | 0.42 | 0.45 | 0.45 |
| 8 | 20571970 | rs2597406 | 2.324 | -0.097 | -0.507 | -1.516 | -0.057 | -0.044 | -0.044 | 0 | 0.27 | 0.08 | 0.28 | 0.34 | 0.26 | 0.25 |
| 8 | 20575099 | rs2616241 | 1.442 | -0.161 | -0.23 | -1.551 | -0.001 | -0.01 | -0.01 | 0 | 0.34 | 0.46 | 0.32 | 0.34 | 0.27 | 0.26 |
| 8 | 20575213 | rs2616243 | 1.606 | -0.236 | -0.418 | -1.656 | -0.052 | 0.053 | 0.053 | 0.203 | 0.42 | 0.12 | 0.38 | 0.35 | 0.38 | 0.36 |
| 8 | 20575467 | rs6984463 | 0.273 | -0.2 | -0.295 | -1.587 | -0.028 | -0.06 | -0.06 | 0.309 | 0.29 | 0.34 | 0.19 | 0.07 | 0.18 | 0.2 |
| 8 | 20575493 | rs6984590 | 1.698 | -0.186 | -0.282 | -1.58 | -0.017 | -0.073 | -0.073 | 0.793 | 0.17 | 0.23 | 0.11 | 0.01 | 0.12 | 0.13 |
| 8 | 20576344 | rs17603934 | 0.958 | 0.062 | -0.766 | -1.321 | -0.005 | -0.002 | -0.002 | 0 | 0.11 | 0.11 | 0.08 | 0.07 | 0.06 | 0.08 |
| 8 | 20576761 | rs13258625 | 0.172 | 0.125 | -0.825 | -1.32 | -0.033 | -0.066 | -0.066 | 0 | 0.31 | 0.17 | 0.3 | 0.49 | 0.42 | 0.33 |
| 8 | 20578934 | rs11786346 | 1.497 | 0.136 | -0.76 | -1.291 | 0.004 | -0.047 | -0.047 | 1.337 | 0.13 | 0.16 | 0.09 | 0 | 0.16 | 0.19 |
| 8 | 20579843 | rs1530783 | 1.281 | 0.092 | -0.695 | -1.304 | 0 | 0.003 | 0.003 | 0.162 | 0.47 | 0.57 | 0.42 | 0.39 | 0.39 | 0.37 |
| 8 | 20580756 | rs1530780 | 0.661 | 0.153 | -1.034 | -1.084 | -0.014 | 0.004 | 0.004 | 1.853 | 0.55 | 0.69 | 0.48 | 0.45 | 0.46 | 0.48 |
| 8 | 20582438 | rs1530779 | -0.459 | 0.406 | -1.211 | -0.721 | -0.016 | -0.057 | -0.057 | 0 | 0.76 | 0.86 | 0.63 | 0.52 | 0.67 | 0.75 |
| 8 | 20589713 | rs2616178 | 1.229 | 0.306 | -1.498 | -0.791 | -0.085 | -0.115 | -0.115 | 0 | 0.32 | 0.03 | 0.32 | 0.41 | 0.29 | 0.29 |
| 8 | 20590319 | rs2442983 | 0.556 | 0.069 | -0.71 | -1.102 | -0.001 | -0.001 | -0.001 | 0.15 | 0.54 | 0.43 | 0.46 | 0.38 | 0.47 | 0.49 |
| 8 | 20590386 | rs2442982 | 0.856 | 0.071 | -0.389 | -1.315 | -0.001 | -0.067 | -0.067 | 0 | 0.24 | 0.33 | 0.19 | 0.05 | 0.24 | 0.27 |
| 8 | 20590474 | rs1450292 | -0.607 | 0.159 | -0.315 | -1.457 | -0.005 | 0.037 | 0.037 | 0 | 0.79 | 0.72 | 0.67 | 0.44 | 0.78 | 0.76 |
| 8 | 20590669 | rs2410659 | 1.208 | -0.062 | -0.642 | -1.482 | 0.002 | -0.025 | -0.025 | 0 | 0.33 | 0.1 | 0.3 | 0.4 | 0.24 | 0.32 |
| 8 | 20592077 | rs11204129 | -0.079 | 0.171 | -0.447 | -1.541 | -0.008 | 0.032 | 0.032 | 0 | 0.54 | 0.69 | 0.43 | 0.45 | 0.43 | 0.55 |
| 8 | 20592509 | rs11985951 | 0.675 | -0.084 | 0.097 | -1.328 | 0.007 | -0.042 | -0.042 | 0.06 | 0.15 | 0.21 | 0.13 | 0.05 | 0.18 | 0.24 |
| 8 | 20592764 | rs1376010 | -1.255 | -0.116 | 0.004 | -0.979 | 0.025 | 0.051 | 0.051 | 0 | 0.93 | 0.73 | 0.83 | 0.95 | 0.95 | 0.88 |
| 8 | 20594439 | rs2616175 | 0.019 | -0.068 | -0.074 | -1.197 | 0.003 | 0.025 | 0.025 | 0 | 0.73 | 0.64 | 0.57 | 0.46 | 0.68 | 0.74 |
| 8 | 20594766 | rs2597419 | 0.721 | -0.273 | -0.262 | -1.288 | -0.007 | -0.015 | -0.015 | 0 | 0.63 | 0.59 | 0.51 | 0.36 | 0.62 | 0.65 |
| 8 | 20595218 | rs6984510 | 0.331 | -0.357 | -0.374 | -1.35 | -0.005 | 0 | 0 | 0 | 0.11 | 0.1 | 0.07 | 0.1 | 0.06 | 0.09 |
| 8 | 20607908 | rs13281313 | -0.84 | -0.362 | -0.494 | -1.705 | -0.003 | -0.019 | -0.019 | 0 | 0.26 | 0.71 | 0.17 | 0.07 | 0.22 | 0.25 |
| 8 | 20613337 | rs2616217 | 0.386 | -0.348 | -0.729 | -2.254 | -0.001 | -0.021 | -0.021 | 0.071 | 0.37 | 0.36 | 0.27 | 0.07 | 0.4 | 0.28 |
| 8 | 20615434 | rs13269596 | 0.55 | -0.344 | -0.592 | -2.053 | -0.006 | -0.006 | -0.006 | 3.244 | 0.1 | 0.08 | 0.05 | 0.01 | 0.15 | 0.11 |
| 8 | 20615678 | rs10110077 | 0.931 | -1.022 | -1.264 | -1.961 | -0.005 | -0.029 | -0.029 | 0 | 0.29 | 0.05 | 0.21 | 0.28 | 0.2 | 0.27 |
| 8 | 20619641 | rs10090500 | 0.073 | -1.141 | -0.789 | -2.053 | -0.039 | 0 | 0 | 0 | 0.33 | 0.43 | 0.24 | 0.31 | 0.15 | 0.32 |
| 8 | 20625091 | rs10503692 | 0.096 | -1.13 | -1.004 | -2.058 | 0.069 | -0.008 | -0.008 | 0 | 0.32 | 0.21 | 0.23 | 0.31 | 0.16 | 0.34 |

| | | | | | | | | | | | | | | | | |
|---|---|---|---|---|---|---|---|---|---|---|---|---|---|---|---|---|
| 8 | 20625232 | rs17576339 | 1.862 | -1.065 | -1.312 | -2.16 | -0.054 | -0.104 | -0.104 | 0.134 | 0.15 | 0.02 | 0.25 | 0.59 | 0.19 | 0.18 |
| 8 | 20625484 | rs2597418 | -1.378 | -0.948 | -0.829 | -2.242 | -0.001 | 0 | 0 | 0 | 0.76 | 0.79 | 0.54 | 0.36 | 0.74 | 0.68 |
| 8 | 20626321 | rs2122469 | 1.549 | -1.108 | -1.111 | -2.043 | 0.029 | -0.003 | -0.003 | 0 | 0.11 | 0.04 | 0.06 | 0.06 | 0.06 | 0.12 |
| 8 | 20629602 | rs17576395 | 0.661 | -1.006 | -1.523 | -2.052 | 0.015 | -0.009 | -0.009 | 0.012 | 0.26 | 0.08 | 0.2 | 0.27 | 0.18 | 0.25 |
| 8 | 20629958 | rs17604209 | 1.245 | -1.014 | -1.52 | -2.05 | 0.051 | -0.03 | -0.03 | 0 | 0.21 | 0.07 | 0.17 | 0.27 | 0.11 | 0.21 |
| 8 | 20630963 | rs11204131 | -0.378 | -1.02 | -1.478 | -2.047 | 0.057 | 0.001 | 0.001 | 0 | 0.38 | 0.22 | 0.26 | 0.34 | 0.24 | 0.37 |
| 8 | 20632022 | rs2616143 | 2.15 | -0.824 | -1.847 | -2.129 | -0.104 | 0.152 | 0.152 | 0 | 0.24 | 0.04 | 0.18 | 0.03 | 0.33 | 0.19 |
| 8 | 20634750 | rs13328379 | 0.808 | -0.881 | -1.677 | -1.948 | 0.003 | -0.016 | -0.016 | 0 | 0.24 | 0.08 | 0.19 | 0.25 | 0.16 | 0.24 |
| 8 | 20640478 | rs10089927 | 0.165 | -1.261 | -1.497 | -1.813 | 0.082 | -0.05 | -0.05 | 0 | 0.31 | 0.11 | 0.23 | 0.35 | 0.17 | 0.32 |
| 8 | 20647375 | rs7007919 | -1.513 | -0.811 | -1.182 | -1.833 | -0.059 | 0.057 | 0.057 | 0 | 0.71 | 0.83 | 0.78 | 0.96 | 0.58 | 0.78 |
| 8 | 20647438 | rs4352868 | -0.579 | -0.399 | -0.651 | -1.343 | 0.01 | 0.037 | 0.037 | 0 | 0.41 | 0.67 | 0.58 | 0.69 | 0.41 | 0.49 |
| 8 | 20654010 | rs2410666 | -0.096 | 0.035 | -0.951 | -1.049 | 0.002 | 0.002 | 0.002 | 0 | 0.3 | 0.29 | 0.51 | 0.64 | 0.38 | 0.41 |
| 8 | 20656633 | rs7013899 | -0.415 | -0.036 | -1.33 | -1.156 | -0.003 | 0.025 | 0.025 | 0 | 0.42 | 0.68 | 0.66 | 0.69 | 0.44 | 0.48 |
| 8 | 20658230 | rs7000415 | 0.166 | 0.156 | -1.917 | -1.492 | 0.007 | 0.043 | 0.043 | 0 | 0.37 | 0.19 | 0.52 | 0.67 | 0.34 | 0.44 |
| 8 | 20659561 | rs11204137 | -0.17 | 0.155 | -1.938 | -1.513 | -0.033 | 0.065 | 0.065 | 0 | 0.4 | 0.19 | 0.56 | 0.68 | 0.45 | 0.5 |
| 8 | 20665979 | rs2583693 | -0.926 | -0.094 | -1.467 | -1.362 | 0.055 | 0.046 | 0.046 | 0 | 0.37 | 0.78 | 0.27 | 0.3 | 0.28 | 0.36 |
| 8 | 20669205 | rs2616191 | -0.783 | 0.321 | -1.404 | -1.31 | 0.015 | 0.066 | 0.066 | 0 | 0.39 | 0.78 | 0.3 | 0.28 | 0.32 | 0.34 |
| 8 | 20673550 | rs17494322 | 0.557 | 0.309 | -1.403 | -0.816 | -0.018 | -0.015 | -0.015 | 0.572 | 0.08 | 0 | 0.06 | 0.06 | 0.07 | 0.09 |
| 8 | 20680983 | rs1530288 | -0.299 | 0.275 | -0.885 | -0.893 | 0.008 | 0.001 | 0.001 | 1.346 | 0.26 | 0.13 | 0.2 | 0.27 | 0.19 | 0.24 |
| 8 | 20683523 | rs6985254 | -0.45 | 0.316 | -0.005 | -1.348 | -0.015 | -0.077 | -0.077 | 0.002 | 0.23 | 0.32 | 0.13 | 0.09 | 0.17 | 0.22 |
| 8 | 20683558 | rs2616155 | -1.183 | 0.344 | -0.133 | -1.669 | 0.013 | 0.13 | 0.13 | 0.614 | 0.37 | 0.77 | 0.22 | 0.1 | 0.34 | 0.31 |
| 8 | 20684477 | rs2616160 | -1.175 | 0.329 | 0.023 | -1.678 | 0.017 | 0.065 | 0.065 | 0.475 | 0.35 | 0.73 | 0.18 | 0.1 | 0.29 | 0.3 |
| 8 | 20685467 | rs10103263 | 0.327 | 0.357 | 0.35 | -1.666 | 0.007 | -0.052 | -0.052 | 0.067 | 0.12 | 0.2 | 0.08 | 0.01 | 0.17 | 0.1 |
| 8 | 20688643 | rs2410669 | -1.247 | 0.359 | 0.244 | -1.696 | 0.014 | 0.1 | 0.1 | 0.832 | 0.38 | 0.74 | 0.22 | 0.1 | 0.34 | 0.32 |
| 8 | 20695239 | rs2583680 | -1.163 | 0.728 | 0.397 | -1.745 | 0 | -0.005 | -0.005 | 0 | 0.38 | 0.64 | 0.26 | 0.1 | 0.39 | 0.38 |
| 8 | 20697695 | rs7823871 | 0.143 | 0.686 | 0.525 | -1.724 | -0.02 | -0.058 | -0.058 | 0 | 0.2 | 0.3 | 0.1 | 0.09 | 0.12 | 0.2 |
| 8 | 20700431 | rs2583684 | -1.166 | 0.767 | 0.23 | -1.762 | 0.003 | 0.168 | 0.168 | 0.236 | 0.41 | 0.78 | 0.25 | 0.11 | 0.37 | 0.31 |
| 8 | 20714686 | rs2583691 | -1.305 | 0.732 | 0.265 | -1.735 | -0.009 | 0.158 | 0.158 | 0.567 | 0.42 | 0.76 | 0.3 | 0.1 | 0.44 | 0.42 |
| 8 | 20721412 | rs2410670 | 0.144 | 0.721 | 0.477 | -1.701 | 0.015 | 0.051 | 0.051 | 0.847 | 0.2 | 0.07 | 0.09 | 0.09 | 0.13 | 0.2 |
| 8 | 20724468 | rs2616186 | -0.734 | 1.105 | 0.863 | -1.434 | 0.013 | -0.112 | -0.112 | 0 | 0.19 | 0.6 | 0.15 | 0.01 | 0.23 | 0.15 |
| 8 | 20731418 | rs4961396 | -1.577 | 0.951 | 0.389 | -1.776 | -0.028 | 0.242 | 0.242 | 0 | 0.47 | 0.81 | 0.5 | 0.11 | 0.49 | 0.53 |
| 8 | 20732231 | rs6586933 | -1.162 | 0.969 | 0.603 | -1.777 | -0.001 | -0.06 | -0.06 | 0 | 0.34 | 0.38 | 0.41 | 0.01 | 0.42 | 0.37 |
| 8 | 20733771 | rs6996901 | -1.021 | 1.169 | 0.717 | -1.651 | -0.001 | -0.051 | -0.051 | 0 | 0.28 | 0.68 | 0.23 | 0.01 | 0.35 | 0.25 |
| 8 | 20742407 | rs11779820 | 0.736 | 1.095 | 0.743 | -1.548 | -0.001 | -0.03 | -0.03 | 1.448 | 0.06 | 0.12 | 0.2 | 0 | 0.06 | 0.11 |
| 8 | 20742927 | rs1866959 | -1.592 | 1.572 | 0.62 | -1.685 | -0.053 | 0.217 | 0.217 | 0 | 0.45 | 0.81 | 0.51 | 0.1 | 0.49 | 0.52 |
| 8 | 20743401 | rs6984780 | -1.688 | 1.155 | 0.604 | -1.704 | -0.028 | 0.217 | 0.217 | 0 | 0.45 | 0.81 | 0.48 | 0.1 | 0.49 | 0.51 |
| 8 | 20747139 | rs7004724 | -1.358 | 1.118 | 0.626 | -1.664 | -0.009 | 0.117 | 0.117 | 0 | 0.39 | 0.76 | 0.28 | 0.1 | 0.39 | 0.39 |
| 8 | 20748890 | rs17093306 | -0.112 | 1.041 | 0.712 | -1.63 | -0.002 | -0.051 | -0.051 | 0 | 0.19 | 0.22 | 0.09 | 0.01 | 0.18 | 0.12 |
| 8 | 20753081 | rs1441782 | 0.438 | 1.027 | 0.939 | -0.733 | -0.006 | 0.065 | 0.065 | 0.225 | 0.14 | 0.2 | 0.08 | 0.32 | 0.09 | 0.17 |
| 8 | 20755975 | rs17093318 | -0.474 | 1.042 | 0.765 | -0.59 | -0.011 | 0.043 | 0.043 | 0.861 | 0.2 | 0.29 | 0.28 | 0.35 | 0.15 | 0.28 |
| 8 | 20759417 | rs2165562 | 0.316 | 0.938 | 0.607 | -0.514 | 0.007 | -0.031 | -0.031 | 0 | 0.2 | 0.15 | 0.19 | 0.49 | 0.14 | 0.26 |
| 8 | 20759704 | rs11775933 | -0.963 | 0.971 | 0.554 | -0.584 | -0.01 | 0.034 | 0.034 | 1.46 | 0.33 | 0.45 | 0.45 | 0.5 | 0.26 | 0.4 |
| 8 | 20762445 | rs7840513 | -0.861 | 0.875 | 0.75 | -1.059 | -0.01 | -0.018 | -0.018 | 0 | 0.28 | 0.62 | 0.4 | 0.24 | 0.2 | 0.37 |
| 8 | 20764624 | rs6586940 | -1.901 | 0.854 | 0.628 | -1.15 | -0.025 | 0.223 | 0.223 | 0 | 0.49 | 0.87 | 0.6 | 0.24 | 0.55 | 0.62 |
| 8 | 20765290 | rs10102368 | -0.754 | 0.84 | 0.683 | -1.141 | -0.016 | -0.026 | -0.026 | 0 | 0.28 | 0.58 | 0.4 | 0.23 | 0.2 | 0.37 |
| 8 | 20767229 | rs4961412 | -0.589 | 0.894 | 0.539 | -1.173 | 0.017 | -0.099 | -0.099 | 0.034 | 0.19 | 0.35 | 0.31 | 0.01 | 0.26 | 0.26 |
| 8 | 20770917 | rs4961409 | -2.189 | 0.83 | 0.512 | -1.169 | -0.029 | 0.177 | 0.177 | 0 | 0.53 | 0.86 | 0.6 | 0.24 | 0.56 | 0.62 |
| 8 | 20770926 | rs17093344 | -0.369 | 0.811 | 0.589 | -1.083 | 0.002 | 0.004 | 0.004 | 0.903 | 0.25 | 0.23 | 0.38 | 0.23 | 0.2 | 0.38 |
| 8 | 20771752 | rs17093356 | -0.123 | 0.806 | 0.6 | -1.069 | 0.053 | -0.057 | -0.057 | 0.188 | 0.1 | 0.29 | 0.1 | 0 | 0.15 | 0.1 |
| 8 | 20779697 | rs985559 | -0.161 | 0.733 | 0.479 | -0.894 | -0.001 | -0.022 | -0.022 | 0.319 | 0.28 | 0.35 | 0.28 | 0.2 | 0.25 | 0.35 |
| 8 | 20781471 | rs7826784 | -0.754 | 0.742 | 0.397 | -0.851 | 0 | -0.006 | -0.006 | 0.047 | 0.33 | 0.34 | 0.47 | 0.22 | 0.34 | 0.48 |
| 8 | 20783432 | rs7836543 | -0.687 | 0.775 | 0.44 | -1.093 | -0.002 | -0.098 | -0.098 | 0.287 | 0.22 | 0.34 | 0.28 | 0.01 | 0.24 | 0.26 |
| 8 | 20785504 | rs17605349 | -0.768 | 1.232 | -0.072 | -0.502 | 0.001 | 0 | 0 | 0 | 0.32 | 0.33 | 0.33 | 0.38 | 0.32 | 0.42 |
| 8 | 20787009 | rs9325877 | -0.377 | 0.793 | -0.246 | -0.151 | -0.004 | 0.006 | 0.006 | 0 | 0.41 | 0.41 | 0.38 | 0.22 | 0.47 | 0.49 |
| 8 | 20787381 | rs7834527 | -0.515 | 0.647 | -1.126 | -0.058 | -0.003 | 0.031 | 0.031 | 0 | 0.46 | 0.64 | 0.45 | 0.38 | 0.51 | 0.55 |
| 8 | 20795744 | rs11780068 | -0.543 | 0.389 | -1.099 | 0.364 | 0.001 | 0.003 | 0.003 | 0 | 0.11 | 0.07 | 0.29 | 0.15 | 0.1 | 0.2 |
| 8 | 20803341 | rs502740 | 0.245 | -0.034 | -1.113 | 0.517 | 0.007 | -0.004 | -0.004 | 0 | 0.58 | 0.61 | 0.56 | 0.49 | 0.65 | 0.43 |
| 8 | 20809616 | rs483654 | 0.283 | -0.774 | -1.992 | -0.138 | 0.106 | 0.064 | 0.064 | 0 | 0.51 | 0.15 | 0.5 | 0.4 | 0.68 | 0.63 |
| 8 | 20810194 | rs510395 | 0.133 | -1.386 | -1.294 | -0.31 | -0.025 | -0.024 | -0.024 | 0 | 0.64 | 0.52 | 0.57 | 0.24 | 0.73 | 0.49 |
| 8 | 20812112 | rs569950 | 0.857 | -1.231 | -1.487 | -0.374 | 0.02 | 0.004 | 0.004 | 0.073 | 0.59 | 0.31 | 0.51 | 0.41 | 0.63 | 0.55 |
| 8 | 20825287 | rs1441780 | 0.261 | -1.258 | -1.437 | -0.37 | 0.033 | -0.008 | -0.008 | 0 | 0.69 | 0.25 | 0.61 | 0.35 | 0.81 | 0.66 |
| 8 | 20825873 | rs568433 | 1.015 | -1.353 | -1.408 | -0.505 | 0.044 | 0.041 | 0.041 | 0.782 | 0.57 | 0.24 | 0.5 | 0.35 | 0.63 | 0.54 |
| 8 | 20832794 | rs2280488 | -0.499 | -0.954 | -1.291 | -0.385 | -0.014 | -0.003 | -0.003 | 0 | 0.12 | 0.06 | 0.1 | 0.15 | 0.19 | 0.22 |
| 8 | 20836701 | rs573980 | 0.027 | -1.045 | -1.118 | -0.31 | -0.024 | 0.105 | 0.105 | 0 | 0.77 | 0.52 | 0.55 | 0.42 | 0.79 | 0.75 |
| 8 | 20843176 | rs536554 | 0.666 | -1.024 | -1.732 | -0.689 | 0.03 | 0.137 | 0.137 | 0 | 0.54 | 0.1 | 0.4 | 0.26 | 0.6 | 0.48 |
| 8 | 20856234 | rs1402649 | -0.878 | -0.913 | -1.229 | -1.24 | 0.007 | 0 | 0 | 0.494 | 0.5 | 0.51 | 0.24 | 0.16 | 0.36 | 0.44 |
| 8 | 20857659 | rs1402648 | -0.171 | -0.028 | -0.375 | -0.896 | 0.044 | -0.075 | -0.075 | 0 | 0.13 | 0.25 | 0.14 | 0 | 0.19 | 0.08 |
| 8 | 20858217 | rs1474240 | -0.571 | -0.012 | -0.686 | -0.856 | -0.01 | -0.136 | -0.136 | 0 | 0.27 | 0.43 | 0.19 | 0.01 | 0.25 | 0.11 |
| 8 | 20875901 | rs7815422 | 0.153 | 0.435 | -0.721 | -0.028 | 0 | 0 | 0 | 0 | 0.66 | 0.67 | 0.67 | 0.64 | 0.62 | 0.63 |

| | | | | | | | | | | | | | | | | |
|---|---|---|---|---|---|---|---|---|---|---|---|---|---|---|---|---|
| 8 | 20879350 | rs7462165 | -0.267 | 0.268 | -0.408 | 0.087 | -0.002 | 0.105 | 0.105 | 1.261 | 0.83 | 0.37 | 0.66 | 0.62 | 0.84 | 0.8 |
| 8 | 20880472 | rs1521455 | 0.443 | 0.431 | -0.64 | 0.266 | -0.022 | 0.033 | 0.033 | 0 | 0.31 | 0.16 | 0.29 | 0.2 | 0.38 | 0.35 |
| 8 | 20881134 | rs9644648 | 0.861 | 0.397 | -0.777 | -0.11 | -0.013 | -0.04 | -0.04 | 0 | 0.27 | 0.13 | 0.29 | 0.37 | 0.29 | 0.32 |
| 8 | 20881539 | rs7821595 | -0.048 | 0.39 | -0.625 | -0.14 | 0.003 | -0.016 | -0.016 | 0 | 0.41 | 0.57 | 0.61 | 0.74 | 0.44 | 0.43 |
| 8 | 20882453 | rs6586963 | -1.177 | 0.556 | -0.439 | -0.023 | 0.001 | -0.007 | -0.007 | 0 | 0.78 | 0.83 | 0.92 | 0.95 | 0.86 | 0.87 |
| 8 | 20883105 | rs2204051 | 0.976 | 0.576 | -0.295 | 0.167 | 0.046 | 0.242 | 0.242 | 0.961 | 0.11 | 0.38 | 0.27 | 0.37 | 0.1 | 0.19 |
| 8 | 20886454 | rs9692729 | 0.603 | 0.719 | -0.959 | -0.17 | 0.001 | 0.068 | 0.068 | 0.449 | 0.37 | 0.15 | 0.28 | 0.22 | 0.4 | 0.35 |
| 8 | 20893039 | rs11998635 | 0.869 | 0.893 | -0.902 | -0.117 | -0.012 | -0.032 | -0.032 | 0.042 | 0.26 | 0.17 | 0.37 | 0.39 | 0.3 | 0.33 |
| 8 | 20894128 | rs11204153 | 0.023 | 1.152 | -0.551 | -0.284 | -0.001 | 0 | 0 | 0.698 | 0.61 | 0.4 | 0.58 | 0.58 | 0.67 | 0.64 |
| 8 | 20894811 | rs13265344 | -0.066 | 0.909 | -0.616 | -0.491 | -0.005 | 0.021 | 0.021 | 0 | 0.24 | 0.42 | 0.42 | 0.35 | 0.2 | 0.28 |
| 8 | 20895883 | rs7814769 | 0.674 | 0.804 | -1.134 | -0.988 | 0.044 | 0.11 | 0.11 | 1.034 | 0.41 | 0.24 | 0.21 | 0.15 | 0.36 | 0.34 |
| 8 | 20900090 | rs7014506 | 0.344 | 0.684 | -1.352 | -0.917 | 0.002 | 0.065 | 0.065 | 0 | 0.31 | 0.16 | 0.2 | 0.06 | 0.26 | 0.14 |
| 8 | 20907847 | rs1546710 | 0.516 | 0.404 | -1.878 | -1.003 | 0.039 | 0.088 | 0.088 | 0 | 0.33 | 0.05 | 0.11 | 0.15 | 0.21 | 0.34 |
| 8 | 20911783 | rs2177011 | 0.863 | 0.844 | -1.844 | -1.367 | -0.062 | 0.105 | 0.105 | 0 | 0.24 | 0.11 | 0.16 | 0.06 | 0.28 | 0.14 |
| 8 | 20912550 | rs12677687 | 0.77 | 0.501 | -1.495 | -0.962 | -0.013 | 0.008 | 0.008 | 0 | 0.35 | 0.35 | 0.39 | 0.5 | 0.22 | 0.53 |
| 8 | 20914161 | rs1004711 | 0.057 | 0.773 | -1.524 | -1.141 | 0.004 | -0.035 | -0.035 | 0.074 | 0.52 | 0.51 | 0.61 | 0.93 | 0.42 | 0.8 |
| 8 | 20914898 | rs12114973 | -1.179 | 1.086 | -1.983 | -1.055 | -0.007 | 0.009 | 0.009 | 0 | 0.75 | 0.83 | 0.81 | 0.94 | 0.73 | 0.86 |
| 8 | 20915316 | rs1473634 | 0.459 | 1.3 | -1.842 | -0.521 | -0.052 | 0.072 | 0.072 | 0 | 0.19 | 0.08 | 0.17 | 0 | 0.3 | 0.05 |
| 8 | 20915962 | rs1004933 | 0.762 | 0.604 | -1.329 | 0.225 | -0.012 | -0.028 | -0.028 | 0 | 0.11 | 0.04 | 0.18 | 0.44 | 0.16 | 0.27 |
| 8 | 20917469 | rs4961399 | 0.18 | 0.31 | -1.281 | -0.704 | -0.004 | 0.014 | 0.014 | 0 | 0.63 | 0.69 | 0.5 | 0.22 | 0.63 | 0.56 |
| 8 | 20927542 | rs1015380 | -0.201 | -0.056 | -1.394 | -0.941 | 0.033 | 0.021 | 0.021 | 0.582 | 0.25 | 0.18 | 0.17 | 0 | 0.13 | 0.08 |
| 8 | 20928675 | rs4961390 | -0.79 | 0.125 | -1.717 | -0.369 | -0.057 | -0.116 | -0.116 | 0 | 0.86 | 0.99 | 0.75 | 0.51 | 0.82 | 0.75 |
| 8 | 20930420 | rs4961400 | -0.106 | -0.076 | -1.782 | -0.542 | 0.024 | 0.017 | 0.017 | 1.488 | 0.21 | 0.13 | 0.11 | 0 | 0.12 | 0.05 |
| 8 | 20934631 | rs12678610 | -1.106 | -0.079 | -1.394 | -0.501 | 0.007 | -0.009 | -0.009 | 0 | 0.38 | 0.35 | 0.37 | 0.5 | 0.3 | 0.34 |
| 8 | 20938375 | rs1464426 | -0.61 | 0.156 | -1.625 | -0.29 | -0.015 | -0.074 | -0.074 | 0 | 0.23 | 0.13 | 0.32 | 0.52 | 0.24 | 0.33 |
| 8 | 20939367 | rs1949640 | -0.421 | 0.188 | -1.771 | -0.324 | -0.038 | -0.117 | -0.117 | 0 | 0.22 | 0.08 | 0.31 | 0.5 | 0.24 | 0.3 |
| 8 | 20944386 | rs6986709 | -0.171 | -0.153 | -1.774 | -0.682 | 0.034 | 0.1 | 0.1 | 1.914 | 0.21 | 0.06 | 0.1 | 0 | 0.12 | 0.03 |
| 8 | 20947906 | rs1471295 | 0.408 | -0.092 | -1.61 | -0.691 | 0.025 | 0.02 | 0.02 | 4.108 | 0.16 | 0.1 | 0.08 | 0 | 0.09 | 0.02 |
| 8 | 20949171 | rs11785073 | -0.169 | -0.097 | -1.62 | -0.689 | 0.033 | 0.094 | 0.094 | 0.135 | 0.2 | 0.06 | 0.1 | 0.01 | 0.12 | 0.05 |
| 8 | 20952060 | rs7813936 | 0.738 | 0.256 | -1.61 | -1.107 | 0.003 | -0.005 | -0.005 | 0 | 0.52 | 0.54 | 0.66 | 0.89 | 0.62 | 0.59 |
| 8 | 20960490 | rs898648 | -1.069 | 0.218 | -1.643 | -0.979 | 0.021 | -0.001 | -0.001 | 0 | 0.47 | 0.41 | 0.46 | 0.51 | 0.36 | 0.37 |
| 8 | 20962541 | rs7012438 | -0.284 | 0.434 | -1.418 | -0.821 | -0.014 | 0.016 | 0.016 | 0 | 0.73 | 0.8 | 0.78 | 0.86 | 0.7 | 0.64 |
| 8 | 20990398 | rs12674821 | -1.068 | -0.166 | -1.295 | -0.128 | 0.01 | -0.015 | -0.015 | 0 | 0.35 | 0.3 | 0.41 | 0.49 | 0.27 | 0.37 |
| 8 | 21000894 | rs12549890 | -0.655 | -0.095 | -1.221 | 0.072 | -0.01 | 0.095 | 0.095 | 0 | 0.07 | 0.12 | 0.23 | 0.32 | 0.04 | 0.16 |
| 8 | 21001379 | rs11990863 | 0.643 | 0.039 | -1.164 | -0.1 | -0.002 | 0.027 | 0.027 | 0 | 0.13 | 0.07 | 0.27 | 0.49 | 0.16 | 0.27 |
| 8 | 21005710 | rs12676503 | 0.908 | 0.064 | -1.346 | -0.126 | -0.043 | -0.084 | -0.084 | 0 | 0.12 | 0.01 | 0.23 | 0.48 | 0.15 | 0.27 |
| 8 | 21006870 | rs2126108 | -1.022 | -0.113 | -0.855 | -0.996 | -0.003 | -0.028 | -0.028 | 0 | 0.1 | 0.12 | 0.11 | 0 | 0.06 | 0.08 |
| 8 | 21016194 | rs10283289 | -0.67 | -0.198 | -0.859 | -1.055 | -0.043 | -0.12 | -0.12 | 0 | 0.24 | 0.35 | 0.17 | 0 | 0.13 | 0.07 |
| 8 | 21019822 | rs898647 | 0.572 | 0.731 | -1.051 | -0.613 | 0.013 | -0.018 | -0.018 | 0 | 0.62 | 0.49 | 0.68 | 0.72 | 0.6 | 0.53 |
| 8 | 21025253 | rs9942827 | -0.315 | 0.831 | -0.708 | -0.268 | 0.044 | 0.086 | 0.086 | 1.707 | 0.19 | 0.44 | 0.24 | 0.3 | 0.24 | 0.4 |
| 8 | 21026005 | rs2613713 | -0.575 | 0.606 | -1.562 | -0.504 | 0.015 | 0.03 | 0.03 | 0.309 | 0.4 | 0.77 | 0.36 | 0.31 | 0.34 | 0.51 |
| 8 | 21028044 | rs13257229 | 0.084 | 0.726 | -2.044 | -0.263 | -0.058 | -0.106 | -0.106 | 0.023 | 0.26 | 0.03 | 0.46 | 0.67 | 0.32 | 0.34 |
| 8 | 21032148 | rs11991767 | -0.036 | -0.449 | -1.57 | -0.875 | 0.03 | 0.051 | 0.051 | 0.248 | 0.47 | 0.38 | 0.25 | 0.25 | 0.32 | 0.42 |
| 8 | 21034986 | rs13274146 | 0.405 | -1.484 | -1.141 | -1.494 | -0.047 | -0.111 | -0.111 | 0 | 0.26 | 0.33 | 0.08 | 0.01 | 0.12 | 0.07 |
| 8 | 21035642 | rs2126107 | 0.677 | -1.372 | -0.95 | -1.822 | -0.002 | -0.046 | -0.046 | 0.358 | 0.1 | 0.17 | 0.1 | 0 | 0.09 | 0.07 |
| 8 | 21037511 | rs17606149 | 0.679 | -1.302 | -1.211 | -1.859 | -0.078 | 0.034 | 0.034 | 0 | 0.2 | 0.02 | 0.16 | 0.16 | 0.22 | 0.33 |
| 8 | 21038301 | rs2631874 | -0.579 | -1.403 | -0.456 | -1.969 | 0.001 | -0.021 | -0.021 | 0 | 0.36 | 0.43 | 0.32 | 0.27 | 0.35 | 0.46 |
| 8 | 21038785 | rs2613671 | -1.199 | -1.449 | -0.361 | -1.998 | -0.018 | 0.03 | 0.03 | 0 | 0.62 | 0.7 | 0.38 | 0.27 | 0.47 | 0.53 |
| 8 | 21039424 | rs17093664 | -0.151 | -2.816 | -0.373 | -1.379 | 0.005 | -0.001 | -0.001 | 0 | 0.25 | 0.28 | 0.2 | 0.24 | 0.14 | 0.33 |
| 8 | 21039961 | rs10503699 | -0.182 | -2.799 | -0.522 | -1.678 | 0.018 | 0.034 | 0.034 | 0 | 0.19 | 0.1 | 0.11 | 0 | 0.14 | 0.11 |
| 8 | 21040212 | rs10503700 | -0.378 | -2.806 | -0.538 | -1.688 | 0.019 | 0.043 | 0.043 | 0 | 0.21 | 0.1 | 0.13 | 0.01 | 0.17 | 0.14 |
| 8 | 21040466 | rs17578394 | 1.176 | -2.697 | -0.599 | -1.54 | 0.015 | 0.026 | 0.026 | 0 | 0.14 | 0.01 | 0.1 | 0.14 | 0.08 | 0.28 |
| 8 | 21041621 | rs2631857 | -1.501 | -3.07 | -0.583 | -1.554 | 0.08 | 0.161 | 0.161 | 0 | 0.54 | 0.76 | 0.31 | 0.23 | 0.33 | 0.45 |
| 8 | 21041896 | rs1381003 | 0.332 | -3.115 | -0.378 | -1.305 | -0.04 | -0.029 | -0.029 | 0.035 | 0.28 | 0.37 | 0.21 | 0.22 | 0.2 | 0.36 |
| 8 | 21043949 | rs2631856 | 0.658 | -2.666 | 0.295 | -0.942 | 0.022 | -0.06 | -0.06 | 0 | 0.1 | 0.32 | 0.12 | 0 | 0.1 | 0.06 |
| 8 | 21046469 | rs2631855 | -1.639 | -3.024 | 0.039 | -0.838 | 0.051 | 0.006 | 0.006 | 0 | 0.36 | 0.75 | 0.28 | 0.28 | 0.21 | 0.4 |
| 8 | 21047759 | rs17606303 | -0.119 | -3.041 | 0.006 | -0.853 | 0.004 | -0.002 | -0.002 | 0 | 0.13 | 0.09 | 0.07 | 0.16 | 0.07 | 0.29 |
| 8 | 21047865 | rs2613674 | -1.652 | -3.027 | -0.065 | -0.847 | 0.081 | -0.015 | -0.015 | 0.229 | 0.36 | 0.88 | 0.29 | 0.27 | 0.21 | 0.4 |
| 8 | 21051235 | rs2613675 | -2.126 | -3.141 | -0.144 | -0.916 | 0.125 | 0.043 | 0.043 | 0 | 0.46 | 0.9 | 0.31 | 0.28 | 0.26 | 0.45 |
| 8 | 21053295 | rs2631933 | -1.814 | -2.918 | 0.041 | -0.76 | 0.091 | -0.013 | -0.013 | 0 | 0.35 | 0.87 | 0.28 | 0.27 | 0.19 | 0.39 |
| 8 | 21058813 | rs7836199 | 0.477 | -2.776 | 0.102 | -0.659 | -0.025 | 0.043 | 0.043 | 0 | 0.11 | 0.16 | 0.1 | 0.16 | 0.04 | 0.06 |
| 8 | 21060367 | rs7464772 | 0.524 | -2.648 | -0.169 | -0.647 | 0.001 | 0 | 0 | 0 | 0.1 | 0.06 | 0.03 | 0.11 | 0.04 | 0.23 |
| 8 | 21072402 | rs2613696 | -1.616 | -2.639 | -0.542 | -0.818 | 0.055 | 0.024 | 0.024 | 0 | 0.37 | 0.74 | 0.28 | 0.25 | 0.23 | 0.42 |
| 8 | 21075277 | rs2610934 | -0.567 | -2.656 | -0.433 | -0.773 | -0.006 | -0.001 | -0.001 | 2.688 | 0.23 | 0.69 | 0.28 | 0.18 | 0.14 | 0.13 |
| 8 | 21077378 | rs2613702 | -0.442 | -2.639 | -0.33 | -0.762 | -0.019 | -0.024 | -0.024 | 0 | 0.22 | 0.61 | 0.18 | 0.1 | 0.14 | 0.12 |
| 8 | 21077655 | rs2610969 | -0.899 | -2.613 | -0.459 | -0.769 | -0.013 | -0.014 | -0.014 | 0 | 0.25 | 0.65 | 0.18 | 0.1 | 0.15 | 0.16 |
| 8 | 21082177 | rs2631914 | -0.889 | -2.623 | -0.425 | -0.773 | -0.015 | -0.024 | -0.024 | 0 | 0.24 | 0.6 | 0.2 | 0.1 | 0.16 | 0.16 |
| 8 | 21083752 | rs2613706 | -0.964 | -2.635 | -0.512 | -0.768 | -0.003 | -0.005 | -0.005 | 0 | 0.25 | 0.67 | 0.2 | 0.11 | 0.16 | 0.17 |
| 8 | 21085331 | rs1563916 | -2.25 | -2.952 | -0.978 | -0.801 | -0.007 | -0.02 | -0.02 | 0 | 0.55 | 0.96 | 0.32 | 0.35 | 0.35 | 0.49 |

| | | | | | | | | | | | | | | | | |
|---|---|---|---|---|---|---|---|---|---|---|---|---|---|---|---|---|
| 8 | 21087783 | rs989329 | -1.79 | -2.882 | -1.013 | -0.915 | 0.052 | 0.08 | 0.08 | 0 | 0.39 | 0.86 | 0.27 | 0.19 | 0.26 | 0.43 |
| 8 | 21093911 | rs2610975 | -1.965 | -2.83 | -0.624 | -0.864 | 0.044 | 0.012 | 0.012 | 0.136 | 0.48 | 0.73 | 0.4 | 0.43 | 0.36 | 0.49 |
| 8 | 21105345 | rs2446614 | -1.783 | -2.696 | -0.903 | -0.974 | 0.127 | 0.162 | 0.162 | 0 | 0.41 | 0.81 | 0.24 | 0.12 | 0.22 | 0.45 |
| 8 | 21106064 | rs17093744 | 0.149 | -2.659 | -0.829 | -0.959 | -0.002 | -0.02 | -0.02 | 0.022 | 0.12 | 0.11 | 0.04 | 0 | 0.07 | 0.08 |
| 8 | 21107763 | rs17606653 | -0.662 | -2.662 | -0.907 | -0.99 | -0.005 | -0.016 | -0.016 | 0 | 0.07 | 0.04 | 0.15 | 0.14 | 0.07 | 0.05 |
| 8 | 21109328 | rs2631865 | -1.85 | -2.562 | -0.756 | -0.943 | 0.129 | 0.17 | 0.17 | 0 | 0.41 | 0.83 | 0.24 | 0.12 | 0.23 | 0.42 |
| 8 | 21112084 | rs2631864 | -0.593 | -2.289 | -0.516 | -0.845 | -0.021 | -0.031 | -0.031 | 0.872 | 0.21 | 0.64 | 0.17 | 0.06 | 0.11 | 0.1 |
| 8 | 21119617 | rs10101710 | -0.893 | -2.227 | -0.712 | -1 | 0.026 | 0.016 | 0.016 | 0 | 0.25 | 0.83 | 0.25 | 0.14 | 0.14 | 0.14 |
| 8 | 21123464 | rs10089199 | -1.537 | -2.175 | -0.732 | -1.348 | 0.081 | 0.229 | 0.229 | 0 | 0.36 | 0.9 | 0.21 | 0.03 | 0.18 | 0.24 |
| 8 | 21124021 | rs2631896 | -1.76 | -2.171 | -0.734 | -1.337 | 0.075 | 0.193 | 0.193 | 0 | 0.38 | 0.91 | 0.22 | 0.06 | 0.22 | 0.26 |
| 8 | 21124700 | rs2446620 | -1.456 | -2.148 | -0.698 | -1.334 | 0.081 | 0.16 | 0.16 | 0 | 0.35 | 0.84 | 0.2 | 0.03 | 0.18 | 0.2 |
| 8 | 21124811 | rs3934909 | -0.663 | -2.046 | -0.589 | -1.271 | -0.011 | -0.04 | -0.04 | 0 | 0.23 | 0.72 | 0.16 | 0.03 | 0.11 | 0.11 |
| 8 | 21130702 | rs2631879 | -0.502 | -1.977 | -0.567 | -1.261 | -0.016 | -0.026 | -0.026 | 0 | 0.22 | 0.69 | 0.19 | 0.07 | 0.11 | 0.11 |
| 8 | 21137028 | rs1459320 | -1.208 | -1.742 | -1.132 | -1.124 | 0.084 | 0.164 | 0.164 | 0.058 | 0.41 | 0.89 | 0.28 | 0.13 | 0.25 | 0.42 |
| 8 | 21138357 | rs6998973 | -1.28 | -1.368 | 0.142 | -0.828 | 0.001 | 0.015 | 0.015 | 0 | 0.08 | 0.25 | 0.05 | 0.05 | 0.05 | 0.23 |
| 8 | 21138621 | rs10110485 | -1.341 | -1.829 | 0.32 | -1.096 | -0.029 | -0.041 | -0.041 | 0 | 0.21 | 0.66 | 0.1 | 0.02 | 0.08 | 0.11 |
| 8 | 21138707 | rs10110590 | -1.092 | -1.813 | 0.679 | -0.936 | -0.071 | -0.049 | -0.049 | 0 | 0.16 | 0.35 | 0.09 | 0.04 | 0.06 | 0.14 |
| 8 | 21138916 | rs10109981 | -1.459 | -1.037 | -0.23 | -1.348 | 0.012 | 0.026 | 0.026 | 0 | 0.38 | 0.63 | 0.33 | 0.16 | 0.28 | 0.47 |
| 8 | 21139032 | rs2610946 | 0.702 | -0.838 | -0.228 | -1.396 | -0.033 | -0.063 | -0.063 | 0.043 | 0.74 | 0.42 | 0.86 | 0.88 | 0.83 | 0.68 |
| 8 | 21151009 | rs2610941 | -0.508 | 0.218 | -0.8 | -1.236 | -0.006 | 0.062 | 0.062 | 1.992 | 0.39 | 0.76 | 0.53 | 0.74 | 0.4 | 0.35 |
| 8 | 21153190 | rs12676298 | -0.507 | 0.152 | -0.531 | -1.519 | -0.001 | 0.003 | 0.003 | 0 | 0.42 | 0.41 | 0.6 | 0.78 | 0.49 | 0.42 |
| 8 | 21157621 | rs10086856 | -0.958 | 0.109 | -0.38 | -1.631 | -0.027 | -0.061 | -0.061 | 0 | 0.13 | 0.69 | 0.16 | 0.01 | 0.11 | 0.07 |
| 8 | 21162980 | rs10086198 | -0.934 | 0.105 | -0.224 | -1.613 | 0.037 | 0.024 | 0.024 | 5.337 | 0.05 | 0.6 | 0.06 | 0 | 0.01 | 0 |
| 8 | 21164751 | rs2631889 | 0.009 | -0.095 | -0.694 | -1.926 | 0.002 | 0.001 | 0.001 | 0.04 | 0.15 | 0.12 | 0.07 | 0.11 | 0.11 | 0.2 |
| 8 | 21173660 | rs898249 | -1.688 | 0.019 | -1.083 | -2.049 | -0.099 | 0.195 | 0.195 | 0.085 | 0.64 | 0.93 | 0.74 | 0.93 | 0.63 | 0.67 |
| 8 | 21176295 | rs2610983 | -0.523 | 0.127 | -0.915 | -1.914 | -0.007 | 0.085 | 0.085 | 0.037 | 0.43 | 0.3 | 0.61 | 0.82 | 0.51 | 0.47 |
| 8 | 21176834 | rs2959369 | -0.007 | 0.216 | -1.13 | -1.871 | -0.032 | 0.057 | 0.057 | 0.051 | 0.36 | 0.12 | 0.5 | 0.79 | 0.4 | 0.38 |
| 8 | 21180872 | rs2926427 | 0.6 | 0.253 | -1.165 | -1.802 | -0.083 | -0.063 | -0.063 | 0.253 | 0.25 | 0.02 | 0.41 | 0.74 | 0.33 | 0.31 |
| 8 | 21183761 | rs881914 | -0.343 | -0.141 | -0.059 | -1.359 | -0.017 | -0.009 | -0.009 | 0.017 | 0.15 | 0.46 | 0.11 | 0.09 | 0.11 | 0.18 |
| 8 | 21183993 | rs898252 | -0.372 | 0.189 | 0.115 | -1.057 | 0.008 | -0.037 | -0.037 | 0.192 | 0.09 | 0.24 | 0.12 | 0.01 | 0.11 | 0.08 |
| 8 | 21190268 | rs2066940 | 0.932 | 0.623 | -0.28 | -1 | 0.022 | 0.104 | 0.104 | 0.024 | 0.43 | 0.2 | 0.19 | 0.18 | 0.36 | 0.35 |
| 8 | 21191742 | rs2631887 | -0.238 | 0.633 | 0.21 | -1.016 | 0.094 | 0.033 | 0.033 | 0.695 | 0.07 | 0.43 | 0.07 | 0 | 0.04 | 0.05 |
| 8 | 21199381 | rs2125649 | -0.558 | 0.742 | -0.055 | -0.8 | 0.001 | 0.033 | 0.033 | 0 | 0.11 | 0.27 | 0.12 | 0.17 | 0.09 | 0.24 |
| 8 | 21199398 | rs999012 | 0.36 | 0.762 | -0.808 | -0.847 | 0.009 | 0.01 | 0.01 | 0 | 0.11 | 0.08 | 0.09 | 0 | 0.09 | 0.12 |
| 8 | 21199670 | rs1466349 | -0.704 | 0.885 | -0.858 | -0.755 | -0.152 | -0.113 | -0.113 | 0.001 | 0.72 | 0.98 | 0.45 | 0.34 | 0.59 | 0.77 |
| 8 | 21199998 | rs7009171 | -0.704 | 0.885 | -0.858 | -0.755 | -0.152 | -0.113 | -0.113 | 0 | 0.72 | 0.98 | 0.45 | 0.33 | 0.59 | 0.77 |
| 8 | 21202578 | rs7007767 | -0.59 | 0.591 | 0.01 | -0.336 | -0.018 | 0.022 | 0.022 | 0.518 | 0.12 | 0.55 | 0.08 | 0.11 | 0.07 | 0.19 |
| 8 | 21203442 | rs12114901 | -0.257 | 0.906 | -0.774 | -0.36 | 0.014 | 0.003 | 0.003 | 0 | 0.06 | 0.12 | 0.1 | 0.06 | 0.08 | 0.11 |
| 8 | 21209281 | rs2045563 | 0.009 | 0.978 | -1.372 | -0.442 | -0.079 | 0.083 | 0.083 | 0.992 | 0.56 | 0.89 | 0.33 | 0.25 | 0.44 | 0.64 |
| 8 | 21209945 | rs6996307 | 0.284 | 1.321 | -0.72 | -0.153 | 0.011 | 0.04 | 0.04 | 0.898 | 0.45 | 0.34 | 0.32 | 0.23 | 0.43 | 0.4 |
| 8 | 21215152 | rs13282264 | -1.169 | 1.469 | -0.335 | 1.214 | -0.002 | 0.04 | 0.04 | 0.241 | 0.09 | 0.16 | 0.18 | 0.22 | 0.08 | 0.05 |
| 8 | 21217118 | rs9693351 | 0.641 | 1.149 | -0.605 | 0.55 | 0.026 | 0.067 | 0.067 | 1.19 | 0.39 | 0.25 | 0.21 | 0.16 | 0.34 | 0.31 |
| 8 | 21217238 | rs17606997 | -0.29 | 0.797 | -0.397 | 1.322 | -0.02 | 0.033 | 0.033 | 0.035 | 0.13 | 0.17 | 0.35 | 0.26 | 0.1 | 0.2 |
| 8 | 21217677 | rs17579232 | -0.399 | 0.812 | -0.184 | 1.315 | -0.027 | 0.052 | 0.052 | 0.176 | 0.14 | 0.24 | 0.36 | 0.26 | 0.1 | 0.2 |
| 8 | 21217838 | rs11780424 | 0.157 | 1.721 | -0.726 | 1.445 | -0.037 | -0.039 | -0.039 | 0.407 | 0.67 | 0.92 | 0.71 | 0.65 | 0.66 | 0.67 |
| 8 | 21219361 | rs9694615 | -0.722 | 1.684 | -0.756 | 1.573 | -0.079 | -0.147 | -0.147 | 0 | 0.75 | 0.98 | 0.69 | 0.58 | 0.75 | 0.62 |
| 8 | 21221776 | rs4237068 | 0.041 | 1.599 | -0.402 | 1.405 | 0.001 | -0.01 | -0.01 | 0 | 0.69 | 0.73 | 0.47 | 0.38 | 0.73 | 0.55 |
| 8 | 21222143 | rs6995784 | -0.227 | 1.858 | -0.045 | 1.375 | 0.193 | 0.054 | 0.054 | 0 | 0.09 | 0.54 | 0.1 | 0.06 | 0.15 | 0.09 |
| 8 | 21222933 | rs10108672 | 0.582 | 1.688 | -0.116 | 1.421 | 0.006 | 0.003 | 0.003 | 0.452 | 0.6 | 0.66 | 0.42 | 0.37 | 0.66 | 0.44 |
| 8 | 21224174 | rs4739199 | 1.221 | 1.713 | 0.077 | 1.298 | -0.001 | -0.002 | -0.002 | 0 | 0.48 | 0.55 | 0.3 | 0.23 | 0.53 | 0.39 |
| 8 | 21226852 | rs4237069 | -0.823 | 1.93 | -0.07 | 1.885 | -0.077 | -0.079 | -0.079 | 0 | 0.79 | 0.99 | 0.8 | 0.72 | 0.79 | 0.76 |
| 8 | 21227151 | rs4606049 | -1.457 | 2.727 | 0.569 | 2.03 | 0.084 | 0.107 | 0.107 | 0 | 0.14 | 0.42 | 0.15 | 0.21 | 0.21 | 0.12 |
| 8 | 21228286 | rs7009887 | -0.432 | 2.274 | 0.057 | 1.966 | -0.058 | -0.079 | -0.079 | 0 | 0.77 | 0.99 | 0.79 | 0.72 | 0.77 | 0.75 |
| 8 | 21233872 | rs2410685 | 1.844 | 2.514 | 0.258 | 1.982 | 0.063 | 0.025 | 0.025 | 0.623 | 0.47 | 0.13 | 0.39 | 0.4 | 0.38 | 0.37 |
| 8 | 21238653 | rs1035746 | -1.059 | 2.653 | 0.565 | 1.986 | 0.129 | 0.029 | 0.029 | 0 | 0.12 | 0.56 | 0.12 | 0.09 | 0.18 | 0.22 |
| 8 | 21238695 | rs1964328 | 0.606 | 2.641 | 0.464 | 2.043 | 0.008 | 0.005 | 0.005 | 0 | 0.64 | 0.38 | 0.67 | 0.57 | 0.59 | 0.5 |
| 8 | 21238755 | rs2959389 | -0.488 | 2.485 | 0.393 | 2.102 | -0.046 | -0.092 | -0.092 | 0 | 0.77 | 0.96 | 0.79 | 0.66 | 0.77 | 0.73 |
| 8 | 21239232 | rs1549467 | -0.324 | 2.43 | 0.505 | 2.077 | -0.002 | -0.002 | -0.002 | 0 | 0.11 | 0.08 | 0.13 | 0.16 | 0.15 | 0.06 |
| 8 | 21240387 | rs13271607 | -1.24 | 2.427 | 0.513 | 2.064 | 0 | -0.001 | -0.001 | 0.74 | 0.18 | 0.16 | 0.19 | 0.16 | 0.24 | 0.1 |
| 8 | 21243200 | rs17497685 | 0.58 | 2.459 | 0.552 | 2.065 | 0 | -0.012 | -0.012 | 0 | 0.07 | 0.07 | 0.05 | 0 | 0.09 | 0.04 |
| 8 | 21257195 | rs2912236 | -0.846 | 2.642 | 0.561 | 2.084 | -0.004 | -0.003 | -0.003 | 0 | 0.34 | 0.67 | 0.33 | 0.42 | 0.37 | 0.45 |
| 8 | 21258876 | rs1365216 | -0.916 | 2.52 | 0.471 | 2.04 | -0.008 | 0.004 | 0.004 | 0 | 0.17 | 0.15 | 0.18 | 0.15 | 0.24 | 0.14 |
| 8 | 21259722 | rs7005839 | -0.888 | 2.65 | 0.851 | 2.017 | 0.141 | 0.062 | 0.062 | 0 | 0.1 | 0.47 | 0.11 | 0.07 | 0.14 | 0.18 |
| 8 | 21260356 | rs6992081 | -1.039 | 2.645 | 0.724 | 2.013 | 0.079 | 0.018 | 0.018 | 0 | 0.11 | 0.61 | 0.12 | 0.07 | 0.14 | 0.19 |
| 8 | 21260780 | rs1346147 | 0.233 | 2.682 | 0.376 | 2.079 | -0.061 | -0.102 | -0.102 | 2.624 | 0.23 | 0.03 | 0.21 | 0.35 | 0.23 | 0.26 |
| 8 | 21263978 | rs9325882 | -1.137 | 2.646 | 0.609 | 2.082 | -0.014 | 0.031 | 0.031 | 0 | 0.42 | 0.28 | 0.59 | 0.71 | 0.46 | 0.43 |
| 8 | 21266578 | rs11995191 | -1.362 | 2.616 | 0.598 | 1.963 | 0.006 | -0.008 | -0.008 | 0 | 0.46 | 0.68 | 0.43 | 0.56 | 0.52 | 0.51 |
| 8 | 21268290 | rs890315 | -0.702 | 2.573 | 0.784 | 1.968 | 0.007 | 0.002 | 0.002 | 0.678 | 0.36 | 0.52 | 0.31 | 0.39 | 0.37 | 0.44 |
| 8 | 21270062 | rs12679362 | -0.319 | 2.818 | 0.837 | 2.169 | -0.007 | -0.005 | -0.005 | 0 | 0.09 | 0.06 | 0.1 | 0.14 | 0.14 | 0.06 |

| | | | | | | | | | | | | | | | | |
|---|---|---|---|---|---|---|---|---|---|---|---|---|---|---|---|---|
| 8 | 21273654 | rs13265523 | -1.674 | 2.88 | 1.355 | 2.327 | 0.007 | -0.009 | -0.009 | 0 | 0.36 | 0.37 | 0.63 | 0.76 | 0.4 | 0.58 |
| 8 | 21273776 | rs13263769 | -1.3 | 2.873 | 1.503 | 2.54 | -0.008 | -0.01 | -0.01 | 0 | 0.29 | 0.25 | 0.3 | 0.46 | 0.34 | 0.44 |
| 8 | 21275297 | rs13249365 | -1.731 | 2.881 | 1.478 | 2.479 | 0.014 | -0.016 | -0.016 | 0 | 0.35 | 0.42 | 0.63 | 0.77 | 0.4 | 0.57 |
| 8 | 21277933 | rs9325883 | -0.955 | 3.041 | 1.544 | 2.372 | 0.23 | 0.13 | 0.13 | 0 | 0.08 | 0.51 | 0.12 | 0.08 | 0.15 | 0.04 |
| 8 | 21278526 | rs10098621 | -0.26 | 3.146 | 1.731 | 2.383 | 0.103 | 0.14 | 0.14 | 0 | 0.06 | 0.38 | 0.08 | 0.08 | 0.07 | 0.03 |
| 8 | 21283876 | rs13275716 | -1.367 | 3.038 | 1.819 | 2.381 | 0.009 | 0.013 | 0.013 | 0 | 0.3 | 0.37 | 0.32 | 0.46 | 0.34 | 0.45 |
| 8 | 21284771 | rs12155764 | -1.27 | 3.029 | 1.423 | 2.385 | -0.016 | -0.019 | -0.019 | 0 | 0.29 | 0.23 | 0.3 | 0.46 | 0.34 | 0.45 |
| 8 | 21298535 | rs734364 | -0.449 | 3.039 | 1.23 | 2.42 | -0.005 | -0.011 | -0.011 | 0 | 0.21 | 0.14 | 0.19 | 0.26 | 0.21 | 0.4 |
| 8 | 21298827 | rs1549471 | -2.246 | 2.853 | 2.052 | 2.745 | 0.042 | 0.165 | 0.165 | 0.284 | 0.17 | 0.35 | 0.45 | 0.44 | 0.21 | 0.2 |
| 8 | 21299092 | rs13280180 | -1.129 | 2.852 | 2.083 | 2.818 | 0.015 | 0.006 | 0.006 | 0 | 0.11 | 0.15 | 0.18 | 0.18 | 0.15 | 0.11 |
| 8 | 21299827 | rs17607826 | -1.193 | 2.84 | 2.072 | 2.822 | 0.033 | 0.262 | 0.262 | 0.113 | 0.06 | 0.23 | 0.27 | 0.26 | 0.06 | 0.09 |
| 8 | 21301000 | rs1549470 | -1.206 | 3.039 | 1.96 | 2.841 | 0.223 | 0.208 | 0.208 | 0 | 0.07 | 0.45 | 0.11 | 0.15 | 0.14 | 0.05 |
| 8 | 21301625 | rs1426182 | -1.119 | 3.009 | 1.804 | 2.825 | 0.005 | 0.002 | 0.002 | 0 | 0.11 | 0.1 | 0.18 | 0.18 | 0.15 | 0.11 |
| 8 | 21308489 | rs7011968 | -1.815 | 2.641 | 1.816 | 2.317 | 0.163 | 0.128 | 0.128 | 0 | 0.15 | 0.5 | 0.19 | 0.27 | 0.26 | 0.11 |
| 8 | 21311364 | rs13275264 | -0.407 | 2.631 | 1.495 | 2.316 | -0.012 | -0.008 | -0.008 | 0 | 0.08 | 0.03 | 0.08 | 0.12 | 0.12 | 0.06 |
| 8 | 21311470 | rs6982234 | -2.252 | 2.619 | 1.512 | 2.318 | 0.041 | 0.035 | 0.035 | 0.025 | 0.2 | 0.65 | 0.37 | 0.33 | 0.28 | 0.27 |
| 8 | 21324811 | rs9644577 | -1.271 | 2.421 | 1.092 | 2.322 | 0.163 | 0.233 | 0.233 | 0 | 0.07 | 0.31 | 0.27 | 0.23 | 0.14 | 0.06 |
| 8 | 21326878 | rs6586986 | -1.19 | 2.409 | 1.096 | 2.068 | 0.163 | 0.151 | 0.151 | 0 | 0.06 | 0.31 | 0.09 | 0.14 | 0.14 | 0.05 |
| 8 | 21333478 | rs11204173 | -0.225 | 1.903 | -0.016 | 2.018 | -0.035 | -0.08 | -0.08 | 0 | 0.19 | 0.04 | 0.29 | 0.29 | 0.2 | 0.32 |
| 8 | 21336738 | rs7002607 | -0.027 | 1.466 | -0.568 | 1.506 | -0.009 | 0.015 | 0.015 | 0 | 0.18 | 0.08 | 0.1 | 0.13 | 0.18 | 0.19 |
| 8 | 21341744 | rs1426194 | 0.114 | 0.867 | 0.135 | 1.96 | -0.019 | 0.252 | 0.252 | 0.021 | 0.09 | 0.23 | 0.08 | 0.46 | 0.08 | 0.09 |
| 8 | 21341837 | rs1426193 | -1.024 | 0.18 | 0.374 | 1.428 | -0.062 | 0.052 | 0.052 | 0.617 | 0.23 | 0.64 | 0.12 | 0.39 | 0.13 | 0.11 |
| 8 | 21341924 | rs7003897 | 1.15 | 0.444 | 0.008 | 1.359 | -0.027 | -0.059 | -0.059 | 0 | 0.17 | 0.04 | 0.28 | 0.19 | 0.18 | 0.24 |
| 8 | 21348352 | rs1426192 | -0.565 | 0.367 | 0.588 | 1.023 | -0.005 | -0.004 | -0.004 | 0 | 0.32 | 0.64 | 0.42 | 0.69 | 0.27 | 0.33 |
| 8 | 21348442 | rs1426190 | -0.708 | 0.791 | 0.347 | 0.768 | -0.014 | 0.022 | 0.022 | 0.02 | 0.38 | 0.68 | 0.5 | 0.69 | 0.43 | 0.39 |
| 8 | 21349334 | rs13277366 | -1.838 | 0.628 | 0.462 | 0.526 | 0 | 0.031 | 0.031 | 0 | 0.64 | 0.73 | 0.6 | 0.8 | 0.64 | 0.62 |
| 8 | 21352198 | rs919072 | -1.832 | 0.571 | 0.436 | 0.568 | -0.001 | 0.074 | 0.074 | 0 | 0.63 | 0.83 | 0.6 | 0.76 | 0.64 | 0.59 |
| 8 | 21354617 | rs7820789 | -0.617 | 0.571 | 0.855 | 0.818 | -0.008 | 0.002 | 0.002 | 0 | 0.42 | 0.39 | 0.46 | 0.55 | 0.45 | 0.39 |
| 8 | 21364298 | rs11781875 | 1.924 | 0.648 | 0.551 | 0.811 | -0.027 | 0.031 | 0.031 | 0 | 0.14 | 0.01 | 0.23 | 0.08 | 0.14 | 0.18 |
| 8 | 21368779 | rs4739207 | 0.944 | 0.613 | 0.78 | 0.707 | -0.009 | 0.004 | 0.004 | 0 | 0.17 | 0.14 | 0.11 | 0.16 | 0.18 | 0.2 |
| 8 | 21370386 | rs11786869 | 1.335 | 0.675 | 0.932 | 0.975 | -0.005 | -0.06 | -0.06 | 0 | 0.18 | 0.08 | 0.28 | 0.23 | 0.15 | 0.25 |
| 8 | 21370888 | rs7816543 | -0.28 | 0.607 | 1.161 | 0.837 | -0.003 | -0.006 | -0.006 | 1.148 | 0.23 | 0.18 | 0.19 | 0.39 | 0.29 | 0.16 |
| 8 | 21373577 | rs12549377 | 0.441 | 0.698 | 1.04 | 0.902 | 0.001 | 0.005 | 0.005 | 0 | 0.15 | 0.12 | 0.16 | 0.31 | 0.24 | 0.14 |
| 8 | 21377986 | rs1466478 | -2.074 | 0.63 | 0.831 | 0.922 | 0.005 | 0.04 | 0.04 | 0 | 0.63 | 0.83 | 0.6 | 0.73 | 0.64 | 0.62 |
| 8 | 21381296 | rs1481003 | -0.029 | 1 | 0.867 | 1.059 | -0.001 | 0.05 | 0.05 | 0 | 0.3 | 0.44 | 0.39 | 0.52 | 0.28 | 0.37 |
| 8 | 21382927 | rs11775524 | 1.102 | 1.191 | 0.635 | 0.944 | -0.004 | -0.016 | -0.016 | 0 | 0.18 | 0.09 | 0.28 | 0.19 | 0.17 | 0.22 |
| 8 | 21383205 | rs13272802 | 0.046 | 0.902 | 0.836 | 0.917 | -0.002 | -0.005 | -0.005 | 0 | 0.17 | 0.14 | 0.1 | 0.2 | 0.16 | 0.22 |
| 8 | 21384428 | rs17498886 | 1.019 | 0.836 | 0.52 | 0.889 | -0.025 | 0.023 | 0.023 | 0.064 | 0.05 | 0.02 | 0.07 | 0 | 0.13 | 0.03 |
| 8 | 21395310 | rs1010999 | -0.61 | 0.379 | 0.406 | 0.851 | -0.012 | 0.039 | 0.039 | 0 | 0.74 | 0.73 | 0.71 | 0.58 | 0.75 | 0.62 |
| 8 | 21400565 | rs6992005 | 0.229 | 0.444 | 0.286 | 0.608 | 0.055 | -0.069 | -0.069 | 0 | 0.11 | 0.28 | 0.1 | 0 | 0.13 | 0.05 |
| 8 | 21401976 | rs1481002 | 0.229 | 0.338 | -0.132 | 0.767 | -0.001 | -0.001 | -0.001 | 0 | 0.22 | 0.31 | 0.13 | 0.21 | 0.25 | 0.2 |
| 8 | 21402658 | rs1564480 | 0.42 | 0.419 | -0.108 | 0.532 | 0.001 | 0.002 | 0.002 | 0.852 | 0.52 | 0.52 | 0.58 | 0.4 | 0.51 | 0.45 |
| 8 | 21404381 | rs13255302 | -0.495 | 0.467 | -0.066 | 0.111 | 0.023 | -0.102 | -0.102 | 0.01 | 0.32 | 0.59 | 0.22 | 0.09 | 0.37 | 0.22 |
| 8 | 21404961 | rs6994223 | 0.023 | -0.312 | 0.005 | 1.278 | 0.005 | -0.022 | -0.022 | 0.815 | 0.69 | 0.78 | 0.65 | 0.53 | 0.71 | 0.56 |
| 8 | 21407392 | rs3902428 | -0.824 | -0.785 | 0.495 | 1.592 | 0.016 | -0.005 | -0.005 | 0 | 0.46 | 0.64 | 0.38 | 0.47 | 0.38 | 0.56 |
| 8 | 21408218 | rs17094172 | -0.492 | -0.921 | 1.017 | 1.545 | 0.001 | 0.003 | 0.003 | 0 | 0.08 | 0.24 | 0.05 | 0.06 | 0.06 | 0.14 |
| 8 | 21409341 | rs17580809 | -0.264 | -1.006 | 0.721 | 1.691 | -0.011 | 0.012 | 0.012 | 0.994 | 0.35 | 0.17 | 0.31 | 0.34 | 0.32 | 0.4 |
| 8 | 21416030 | rs899958 | -0.052 | -1.074 | 0.655 | 1.657 | -0.062 | -0.023 | -0.023 | 0 | 0.33 | 0.08 | 0.31 | 0.33 | 0.32 | 0.4 |
| 8 | 21416971 | rs13256459 | -1.243 | -1.274 | 0.666 | 1.998 | 0.034 | 0.049 | 0.049 | 0.458 | 0.48 | 0.69 | 0.39 | 0.67 | 0.39 | 0.56 |
| 8 | 21424405 | rs11777093 | 1.766 | -1.196 | -0.061 | 1.952 | -0.192 | -0.022 | -0.022 | 0 | 0.37 | 0.03 | 0.53 | 0.27 | 0.48 | 0.26 |
| 8 | 21428969 | rs4739257 | 0.674 | -1.184 | 0.384 | 1.975 | 0.01 | 0.015 | 0.015 | 0.015 | 0.55 | 0.59 | 0.64 | 0.6 | 0.62 | 0.47 |
| 8 | 21431520 | rs6993164 | 0.029 | -0.83 | 0.515 | 2.017 | 0.003 | 0.02 | 0.02 | 0.12 | 0.65 | 0.79 | 0.68 | 0.67 | 0.68 | 0.6 |
| 8 | 21438783 | rs13260877 | -0.03 | -0.501 | 0.5 | 1.926 | -0.029 | -0.073 | -0.073 | 3.664 | 0.31 | 0.16 | 0.3 | 0.53 | 0.32 | 0.38 |
| 8 | 21447276 | rs1481008 | 0.208 | -0.592 | 0.706 | 1.785 | -0.039 | -0.054 | -0.054 | 0.832 | 0.81 | 0.45 | 0.9 | 0.89 | 0.87 | 0.81 |
| 8 | 21448663 | rs1383576 | 1.079 | -0.536 | 0.647 | 1.956 | 0 | -0.019 | -0.019 | 0.337 | 0.49 | 0.18 | 0.58 | 0.42 | 0.55 | 0.43 |
| 8 | 21451894 | rs4739261 | 1.078 | -0.534 | 0.648 | 1.956 | 0 | -0.019 | -0.019 | 0 | 0.49 | 0.18 | 0.58 | 0.42 | 0.55 | 0.43 |
| 8 | 21462456 | rs7844095 | -0.193 | -0.381 | 0.663 | 1.918 | -0.006 | -0.014 | -0.014 | 0 | 0.32 | 0.24 | 0.31 | 0.46 | 0.33 | 0.37 |
| 8 | 21466249 | rs1383570 | -0.214 | -0.419 | 0.853 | 1.838 | -0.024 | -0.068 | -0.068 | 0 | 0.83 | 0.71 | 0.94 | 0.94 | 0.88 | 0.77 |
| 8 | 21469061 | rs6986039 | 0.921 | -0.296 | 0.631 | 1.991 | 0 | 0.005 | 0.005 | 1.057 | 0.52 | 0.45 | 0.63 | 0.37 | 0.54 | 0.43 |
| 8 | 21470887 | rs2010282 | -0.521 | -0.514 | 0.531 | 1.831 | -0.017 | -0.041 | -0.041 | 0 | 0.85 | 0.75 | 0.94 | 0.92 | 0.88 | 0.81 |
| 8 | 21472776 | rs12680722 | 0.073 | -0.485 | 0.413 | 1.933 | -0.016 | -0.059 | -0.059 | 0.028 | 0.29 | 0.16 | 0.3 | 0.53 | 0.31 | 0.36 |
| 8 | 21473209 | rs6990471 | 0.649 | -0.438 | 0.179 | 2.104 | 0 | -0.001 | -0.001 | 0 | 0.55 | 0.41 | 0.63 | 0.47 | 0.57 | 0.45 |
| 8 | 21474049 | rs10503709 | 0.645 | -0.378 | 0.137 | 2.059 | -0.001 | 0 | 0 | 0 | 0.55 | 0.51 | 0.62 | 0.38 | 0.56 | 0.45 |
| 8 | 21477345 | rs1871743 | 0.66 | -0.38 | 0.006 | 2.041 | 0.002 | 0 | 0 | 0 | 0.56 | 0.59 | 0.64 | 0.46 | 0.57 | 0.47 |
| 8 | 21479255 | rs4739274 | 0.709 | -0.343 | -0.621 | 1.905 | 0 | 0 | 0 | 0.365 | 0.56 | 0.4 | 0.64 | 0.51 | 0.57 | 0.45 |
| 8 | 21481330 | rs1979450 | -0.087 | -0.089 | -1.591 | 1.776 | -0.039 | -0.024 | -0.024 | 0 | 0.28 | 0.12 | 0.3 | 0.35 | 0.3 | 0.34 |
| 8 | 21483253 | rs13270674 | -0.243 | -0.086 | -1.564 | 1.604 | -0.054 | -0.091 | -0.091 | 1.356 | 0.31 | 0.13 | 0.32 | 0.5 | 0.33 | 0.38 |
| 8 | 21485754 | rs7001002 | -0.77 | -0.32 | -0.671 | 1.354 | -0.034 | 0.01 | 0.01 | 1.882 | 0.12 | 0.7 | 0.12 | 0.13 | 0.08 | 0.18 |
| 8 | 21487597 | rs10788703 | 0.754 | -0.414 | -1.813 | 1.316 | -0.034 | -0.039 | -0.039 | 0 | 0.47 | 0.03 | 0.59 | 0.29 | 0.59 | 0.34 |

| | | | | | | | | | | | | | | | | |
|---|---|---|---|---|---|---|---|---|---|---|---|---|---|---|---|---|
| 8 | 21488273 | rs1106570 | 0.956 | -0.224 | -0.502 | 1.108 | -0.021 | 0.002 | 0.002 | 1.105 | 0.87 | 0.3 | 0.82 | 0.84 | 0.91 | 0.7 |
| 8 | 21500728 | rs10503711 | -0.102 | 0.158 | -0.597 | 1.059 | 0.009 | 0.004 | 0.004 | 0 | 0.5 | 0.38 | 0.39 | 0.43 | 0.45 | 0.44 |
| 8 | 21512427 | rs1383573 | -0.004 | 0.053 | 0.473 | 1.192 | -0.001 | -0.001 | -0.001 | 0 | 0.49 | 0.47 | 0.49 | 0.57 | 0.55 | 0.57 |
| 8 | 21525358 | rs12678835 | -0.123 | -0.058 | 0.705 | 1.087 | 0 | 0 | 0 | 0 | 0.52 | 0.51 | 0.36 | 0.4 | 0.41 | 0.43 |
| 8 | 21541084 | rs4443656 | -0.216 | -0.109 | 0.267 | 1.307 | -0.005 | -0.024 | -0.024 | 0 | 0.56 | 0.84 | 0.39 | 0.46 | 0.43 | 0.56 |
| 8 | 21547340 | rs10090737 | 0.147 | -0.445 | -0.126 | 0.967 | 0.009 | 0.051 | 0.051 | 0 | 0.45 | 0.23 | 0.31 | 0.37 | 0.39 | 0.35 |
| 8 | 21550800 | rs1128397 | -0.308 | -0.37 | 0.101 | 0.81 | 0.011 | 0.017 | 0.017 | 0.092 | 0.54 | 0.34 | 0.32 | 0.43 | 0.4 | 0.36 |
| 8 | 21557285 | rs11990425 | 0.189 | -0.499 | -0.017 | 0.526 | 0.036 | 0.002 | 0.002 | 0.885 | 0.42 | 0.3 | 0.3 | 0.46 | 0.34 | 0.35 |
| 8 | 21557443 | rs10109522 | -0.46 | -0.902 | 0.587 | 0.487 | -0.053 | 0.006 | 0.006 | 0.02 | 0.12 | 0.24 | 0.06 | 0.07 | 0.03 | 0.19 |
| 8 | 21563222 | rs7813735 | -0.504 | -0.122 | -0.738 | 0.404 | -0.113 | -0.136 | -0.136 | 0.194 | 0.62 | 0.95 | 0.43 | 0.53 | 0.44 | 0.59 |
| 8 | 21565402 | rs7823726 | -0.576 | -0.295 | -0.189 | -0.041 | -0.004 | 0.002 | 0.002 | 0.084 | 0.07 | 0.24 | 0.06 | 0.02 | 0.04 | 0.14 |
| 8 | 21565623 | rs11997721 | -1.604 | 0.726 | -0.288 | -0.001 | 0.193 | 0.083 | 0.083 | 4.632 | 0.11 | 0.47 | 0.11 | 0.11 | 0.21 | 0.07 |
| 8 | 21569104 | rs4237073 | 0.457 | 0.707 | -2.489 | -0.299 | -0.124 | 0.05 | 0.05 | 0.682 | 0.4 | 0.09 | 0.64 | 0.68 | 0.53 | 0.44 |
| 8 | 21573959 | rs4464960 | 0.242 | 0.733 | -2.382 | -0.484 | 0.034 | 0.106 | 0.106 | 0.886 | 0.42 | 0.18 | 0.71 | 0.71 | 0.65 | 0.47 |
| 8 | 21577185 | rs4739285 | 0.407 | 0.975 | -2.895 | -0.405 | -0.124 | 0.019 | 0.019 | 0 | 0.41 | 0.08 | 0.63 | 0.6 | 0.49 | 0.41 |
| 8 | 21578387 | rs4739286 | -0.599 | 0.364 | -2.78 | -0.692 | 0.05 | 0.183 | 0.183 | 0.063 | 0.6 | 0.89 | 0.8 | 0.82 | 0.72 | 0.64 |
| 8 | 21587997 | rs7014143 | -0.283 | 0.812 | -2.899 | 0.036 | -0.04 | -0.078 | -0.078 | 0 | 0.63 | 0.96 | 0.43 | 0.4 | 0.59 | 0.61 |
| 8 | 21598077 | rs4506202 | -0.291 | 1.057 | -2.747 | 0.099 | -0.131 | -0.021 | -0.021 | 0 | 0.58 | 0.94 | 0.37 | 0.39 | 0.48 | 0.46 |
| 8 | 21602192 | rs6587004 | -0.727 | 1.21 | -0.739 | 0.228 | 0.12 | 0.017 | 0.017 | 4.816 | 0.08 | 0.51 | 0.07 | 0.03 | 0.1 | 0.12 |
| 8 | 21615817 | rs11786696 | -1.201 | 0.986 | -1.467 | 0.54 | 0 | 0.003 | 0.003 | 0.254 | 0.18 | 0.23 | 0.14 | 0.24 | 0.12 | 0.24 |
| 8 | 21616063 | rs6587006 | -0.592 | 0.957 | -2.508 | 0.823 | -0.085 | -0.093 | -0.093 | 0 | 0.63 | 0.97 | 0.63 | 0.61 | 0.56 | 0.48 |
| 8 | 21618317 | rs11775754 | -0.472 | 0.706 | -1.253 | 1.312 | -0.001 | 0 | 0 | 0 | 0.31 | 0.34 | 0.28 | 0.37 | 0.26 | 0.4 |
| 8 | 21618967 | rs4739218 | 0.494 | 0.825 | -1.901 | 1.027 | -0.009 | -0.022 | -0.022 | 0 | 0.56 | 0.11 | 0.53 | 0.36 | 0.67 | 0.51 |
| 8 | 21662445 | rs17581368 | -0.894 | 0.994 | -0.769 | 1.333 | 0.005 | 0.026 | 0.026 | 0 | 0.32 | 0.43 | 0.45 | 0.55 | 0.33 | 0.44 |
| 8 | 21668241 | rs10095228 | -1.281 | 0.528 | -1.208 | 1.19 | -0.036 | 0.007 | 0.007 | 0.053 | 0.41 | 0.88 | 0.56 | 0.62 | 0.38 | 0.46 |
| 8 | 21668901 | rs12548155 | -0.716 | 0.517 | -0.738 | 0.916 | -0.006 | -0.018 | -0.018 | 2.477 | 0.16 | 0.16 | 0.11 | 0.03 | 0.09 | 0.05 |
| 8 | 21670092 | rs899420 | -0.938 | 0.567 | -0.541 | 0.845 | -0.003 | -0.022 | -0.022 | 0 | 0.13 | 0.18 | 0.12 | 0.03 | 0.11 | 0.05 |
| 8 | 21673695 | rs7013204 | -2.332 | 0.17 | 0.182 | 0.554 | 0.002 | -0.023 | -0.023 | 0 | 0.34 | 0.61 | 0.26 | 0.05 | 0.27 | 0.12 |
| 8 | 21673762 | rs17060577 | -0.843 | 0.16 | 0.425 | 0.594 | -0.006 | 0.028 | 0.028 | 0.195 | 0.15 | 0.12 | 0.15 | 0.01 | 0.17 | 0.08 |
| 8 | 21674737 | rs17615281 | -1.024 | 0.147 | 0.403 | 1.396 | -0.011 | 0.078 | 0.078 | 0 | 0.06 | 0.04 | 0.27 | 0.46 | 0.1 | 0.1 |
| 8 | 21678103 | rs748065 | -1.771 | 0.386 | 1.288 | 1.456 | 0.045 | 0.084 | 0.084 | 0 | 0.25 | 0.43 | 0.45 | 0.59 | 0.31 | 0.34 |
| 8 | 21679414 | rs4871993 | -0.862 | 0.314 | 1.426 | 1.379 | -0.004 | -0.013 | -0.013 | 0 | 0.13 | 0.18 | 0.07 | 0.05 | 0.1 | 0.04 |
| 8 | 21680506 | rs6992129 | -1.671 | 0.299 | 1.507 | 1.351 | -0.036 | -0.118 | -0.118 | 0 | 0.26 | 0.49 | 0.14 | 0.06 | 0.18 | 0.12 |
| 8 | 21682044 | rs922504 | -2.833 | 0.272 | 1.314 | 1.37 | -0.046 | 0.016 | 0.016 | 0 | 0.43 | 0.87 | 0.54 | 0.65 | 0.41 | 0.45 |
| 8 | 21692229 | rs4872498 | -1.896 | -0.017 | 1.493 | 1.317 | -0.054 | -0.1 | -0.1 | 0 | 0.2 | 0.51 | 0.11 | 0.06 | 0.11 | 0.1 |
| 8 | 21696657 | rs9644053 | -2.636 | 0.024 | 1.112 | 1.817 | 0.021 | -0.035 | -0.035 | 0 | 0.39 | 0.9 | 0.35 | 0.63 | 0.29 | 0.34 |
| 8 | 21699960 | rs7003813 | -1.989 | -0.03 | 1.825 | 2.421 | -0.008 | 0.012 | 0.012 | 0 | 0.29 | 0.45 | 0.23 | 0.25 | 0.26 | 0.29 |
| 8 | 21701375 | rs4872508 | 1.495 | 0.012 | 1.857 | 2.464 | -0.005 | 0.002 | 0.002 | 0.436 | 0.79 | 0.3 | 0.8 | 0.67 | 0.88 | 0.94 |
| 8 | 21705910 | rs17581453 | -1.302 | 0.132 | 1.641 | 2.497 | -0.021 | -0.027 | -0.027 | 0 | 0.07 | 0.03 | 0.08 | 0.16 | 0.11 | 0.09 |
| 8 | 21706074 | rs4872002 | 0.499 | -0.026 | 2.149 | 2.256 | -0.016 | -0.08 | -0.08 | 0 | 0.86 | 0.42 | 0.89 | 0.99 | 0.92 | 0.98 |
| 8 | 21711431 | rs17428041 | -1.695 | 0.973 | 1.778 | 2.476 | 0.001 | 0.007 | 0.007 | 0 | 0.12 | 0.12 | 0.23 | 0.31 | 0.25 | 0.1 |
| 8 | 21712293 | rs6990439 | -1.17 | 0.859 | 2.011 | 2.484 | -0.011 | 0 | 0 | 0 | 0.12 | 0.16 | 0.07 | 0.11 | 0.06 | 0.18 |
| 8 | 21712401 | rs12545534 | -1.325 | 0.878 | 1.976 | 2.505 | -0.067 | -0.072 | -0.072 | 0 | 0.11 | 0.03 | 0.22 | 0.31 | 0.25 | 0.09 |
| 8 | 21712590 | rs13249393 | -2.59 | 0.868 | 2.039 | 2.458 | 0.004 | 0.012 | 0.012 | 1.919 | 0.23 | 0.27 | 0.3 | 0.41 | 0.32 | 0.26 |
| 8 | 21713119 | rs17499933 | 3.023 | 0.808 | 2.048 | 2.529 | -0.013 | 0.008 | 0.008 | 0 | 0.6 | 0.21 | 0.6 | 0.4 | 0.57 | 0.56 |
| 8 | 21716861 | rs12550574 | 1.475 | 0.885 | 2.117 | 2.489 | -0.007 | 0.035 | 0.035 | 0.486 | 0.79 | 0.31 | 0.81 | 0.51 | 0.84 | 0.67 |
| 8 | 21717529 | rs12545347 | 2.478 | 0.901 | 2.149 | 2.456 | -0.004 | -0.01 | -0.01 | 0.589 | 0.7 | 0.32 | 0.65 | 0.52 | 0.62 | 0.6 |
| 8 | 21717790 | rs1000443 | 1.995 | 0.911 | 2.186 | 2.442 | 0 | 0 | 0 | 0 | 0.73 | 0.35 | 0.66 | 0.67 | 0.65 | 0.83 |
| 8 | 21717841 | rs11780601 | -1.22 | 1.003 | 1.734 | 2.295 | -0.027 | 0.025 | 0.025 | 0.117 | 0.08 | 0.03 | 0.16 | 0 | 0.22 | 0.06 |
| 8 | 21728788 | rs1564307 | -0.234 | 0.898 | 1.939 | 2.293 | -0.007 | -0.058 | -0.058 | 0.104 | 0.89 | 0.77 | 0.93 | 1 | 0.91 | 0.94 |
| 8 | 21733768 | rs4872033 | -2.514 | 0.956 | 1.409 | 2.517 | -0.031 | -0.045 | -0.045 | 0.03 | 0.17 | 0.13 | 0.26 | 0.51 | 0.33 | 0.35 |
| 8 | 21734560 | rs4871840 | -2.543 | 0.955 | 1.373 | 2.627 | -0.03 | -0.035 | -0.035 | 0 | 0.14 | 0.08 | 0.18 | 0.26 | 0.29 | 0.31 |
| 8 | 21737730 | rs11778338 | -2.609 | 0.917 | 1.415 | 2.575 | -0.017 | -0.034 | -0.034 | 0 | 0.18 | 0.13 | 0.25 | 0.52 | 0.3 | 0.36 |
| 8 | 21750834 | rs11783175 | -1.305 | 0.872 | 1.574 | 1.97 | 0.02 | -0.02 | -0.02 | 0.001 | 0.09 | 0.15 | 0.14 | 0.04 | 0.21 | 0.07 |
| 8 | 21751006 | rs11774438 | -1.332 | 0.872 | 1.626 | 1.97 | 0.038 | -0.029 | -0.029 | 0.018 | 0.09 | 0.21 | 0.14 | 0.04 | 0.21 | 0.06 |
| 8 | 21752742 | rs899427 | 0.529 | 0.615 | 1.545 | 1.657 | -0.034 | -0.086 | -0.086 | 0 | 0.81 | 0.43 | 0.87 | 0.95 | 0.87 | 0.89 |
| 8 | 21753869 | rs899425 | 1.484 | 0.734 | 1.429 | 1.629 | -0.002 | 0.012 | 0.012 | 0 | 0.72 | 0.26 | 0.72 | 0.92 | 0.64 | 0.83 |
| 8 | 21757661 | rs17500192 | 0.643 | 0.132 | 0.694 | 1.234 | 0.026 | 0.019 | 0.019 | 0 | 0.13 | 0.03 | 0.17 | 0.06 | 0.05 | 0.11 |
| 8 | 21761443 | rs11135701 | 1.567 | 0.526 | 1.303 | 1.384 | -0.001 | -0.003 | -0.003 | 0.053 | 0.64 | 0.39 | 0.69 | 0.8 | 0.57 | 0.64 |
| 8 | 21761603 | rs4872056 | 0.462 | 0.494 | 1.218 | 1.316 | -0.04 | -0.155 | -0.155 | 0 | 0.75 | 0.61 | 0.83 | 0.99 | 0.8 | 0.89 |
| 8 | 21764007 | rs7461577 | 1.577 | 0.597 | 0.646 | 1.43 | -0.002 | 0.049 | 0.049 | 0 | 0.62 | 0.22 | 0.67 | 0.76 | 0.55 | 0.62 |
| 8 | 21766101 | rs17581764 | -0.937 | 0.991 | 0.693 | 1.847 | -0.013 | -0.022 | -0.022 | 1.732 | 0.1 | 0.08 | 0.07 | 0.37 | 0.14 | 0.27 |
| 8 | 21770796 | rs3758156 | -2.123 | 1.371 | 0.455 | 1.351 | 0.032 | 0.011 | 0.011 | 0.255 | 0.51 | 0.75 | 0.43 | 0.54 | 0.59 | 0.45 |
| 8 | 21773384 | rs11783414 | -2.105 | 1.492 | 0.457 | 1.338 | -0.012 | 0.002 | 0.002 | 0.175 | 0.51 | 0.74 | 0.35 | 0.49 | 0.55 | 0.44 |
| 8 | 21781629 | rs10866823 | -2.307 | 1.205 | 0.561 | 1.398 | 0.013 | -0.007 | -0.007 | 0.576 | 0.54 | 0.7 | 0.37 | 0.49 | 0.59 | 0.45 |
| 8 | 21799737 | rs13278159 | -1.5 | 1.182 | 0.669 | 1.304 | -0.028 | -0.028 | -0.028 | 0.002 | 0.31 | 0.24 | 0.21 | 0.42 | 0.45 | 0.18 |
| 8 | 21803735 | rs7846485 | -0.417 | 1.027 | 0.779 | 1.297 | -0.048 | -0.068 | -0.068 | 0.055 | 0.22 | 0.5 | 0.16 | 0.09 | 0.15 | 0.25 |
| 8 | 21833965 | rs2306641 | -1.476 | 1.105 | 0.734 | 1.344 | -0.03 | -0.026 | -0.026 | 1.28 | 0.3 | 0.22 | 0.21 | 0.4 | 0.46 | 0.18 |
| 8 | 21845619 | rs3943207 | -0.342 | 1.033 | 0.757 | 1.321 | -0.048 | -0.076 | -0.076 | 0 | 0.21 | 0.51 | 0.16 | 0.08 | 0.14 | 0.25 |

| | | | | | | | | | | | | | | | | |
|---|---|---|---|---|---|---|---|---|---|---|---|---|---|---|---|---|
| 8 | 21850656 | rs10503714 | -0.43 | 1.021 | 0.871 | 1.316 | -0.048 | -0.069 | -0.069 | 0 | 0.22 | 0.5 | 0.14 | 0.09 | 0.14 | 0.2 |
| 8 | 21862551 | rs1058486 | -2.584 | 1.036 | 0.787 | 1.36 | 0.011 | -0.01 | -0.01 | 0 | 0.54 | 0.76 | 0.44 | 0.53 | 0.6 | 0.46 |
| 8 | 21866933 | rs17060733 | -0.457 | 0.922 | 0.885 | 1.563 | -0.046 | -0.01 | -0.01 | 0 | 0.22 | 0.52 | 0.16 | 0.19 | 0.15 | 0.25 |
| 8 | 21872328 | rs10097055 | -2.34 | 1.012 | 0.622 | 1.238 | 0.009 | 0.001 | 0.001 | 0.007 | 0.53 | 0.75 | 0.43 | 0.53 | 0.59 | 0.44 |
| 8 | 21878607 | rs11135789 | 1.317 | 0.5 | 1.083 | 1.072 | 0 | -0.026 | -0.026 | 0.494 | 0.65 | 0.64 | 0.77 | 0.85 | 0.61 | 0.83 |
| 8 | 21884544 | rs11776272 | -1.512 | 0.438 | 0.769 | 1.059 | -0.021 | 0.107 | 0.107 | 0 | 0.35 | 0.14 | 0.21 | 0.15 | 0.39 | 0.17 |
| 8 | 21888570 | rs12550716 | -0.217 | 0.512 | 0.617 | 1.281 | -0.012 | -0.08 | -0.08 | 0 | 0.21 | 0.04 | 0.17 | 0.35 | 0.19 | 0.08 |
| 8 | 21888853 | rs7818819 | 1.809 | 0.376 | 0.854 | 1.168 | -0.002 | 0.004 | 0.004 | 0 | 0.7 | 0.34 | 0.77 | 0.82 | 0.66 | 0.65 |
| 8 | 21890589 | rs1947986 | -1.515 | 0.619 | 0.809 | 1.051 | 0.016 | -0.008 | -0.008 | 0.727 | 0.39 | 0.31 | 0.28 | 0.45 | 0.31 | 0.39 |
| 8 | 21892263 | rs748468 | -0.616 | 1.04 | 0.378 | 0.889 | -0.003 | 0.013 | 0.013 | 0 | 0.26 | 0.65 | 0.28 | 0.43 | 0.23 | 0.34 |
| 8 | 21895146 | rs11992609 | 1.338 | 0.558 | -0.121 | 0.653 | 0.013 | 0.01 | 0.01 | 0.635 | 0.52 | 0.26 | 0.63 | 0.41 | 0.51 | 0.55 |
| 8 | 21896097 | rs7813392 | -0.987 | 1.588 | -0.096 | 0.929 | 0.002 | 0.005 | 0.005 | 0 | 0.32 | 0.35 | 0.34 | 0.58 | 0.42 | 0.39 |
| 8 | 21903316 | rs3176292 | -1.031 | 1.52 | 0.269 | 0.435 | 0.008 | 0.012 | 0.012 | 0 | 0.2 | 0.22 | 0.35 | 0.19 | 0.24 | 0.29 |
| 8 | 21904212 | rs1078363 | 0.992 | 1.818 | 0.535 | 0.424 | 0.033 | 0.021 | 0.021 | 0 | 0.78 | 0.71 | 0.63 | 0.8 | 0.7 | 0.8 |
| 8 | 21904891 | rs3176300 | -0.785 | 2.098 | 0.564 | -0.258 | -0.01 | 0.007 | 0.007 | 0 | 0.1 | 0.09 | 0.16 | 0.11 | 0.14 | 0.15 |
| 8 | 21911231 | rs12541126 | -1.643 | 1.895 | 1.703 | -0.576 | 0.073 | -0.058 | -0.058 | 0.201 | 0.13 | 0.47 | 0.2 | 0.02 | 0.15 | 0.15 |
| 8 | 21911601 | rs12549516 | -0.328 | 1.924 | 1.336 | -0.575 | -0.043 | 0.031 | 0.031 | 0 | 0.08 | 0.01 | 0.16 | 0.01 | 0.13 | 0.13 |
| 8 | 21920052 | rs12543764 | -1.997 | 1.819 | 1.393 | -0.759 | 0.019 | -0.077 | -0.077 | 0 | 0.18 | 0.38 | 0.21 | 0.05 | 0.19 | 0.18 |
| 8 | 21930280 | rs10091692 | 1.862 | 1.664 | 1.855 | -0.882 | 0.008 | -0.104 | -0.104 | 0 | 0.79 | 0.57 | 0.79 | 0.98 | 0.82 | 0.82 |
| 8 | 21940121 | rs1872224 | 1.319 | 1.562 | 1.722 | -0.8 | 0.019 | -0.028 | -0.028 | 1.919 | 0.92 | 0.7 | 0.96 | 0.99 | 0.95 | 0.98 |
| 8 | 21949070 | rs4872276 | -0.794 | 1.777 | 1.052 | -0.801 | -0.001 | 0.002 | 0.002 | 0.011 | 0.29 | 0.59 | 0.51 | 0.62 | 0.31 | 0.39 |
| 8 | 21951598 | rs2309308 | -0.907 | 2.085 | 1.263 | -0.861 | 0.033 | 0.021 | 0.021 | 0 | 0.29 | 0.3 | 0.64 | 0.6 | 0.44 | 0.53 |
| 8 | 21959877 | rs7812435 | 0.95 | 2.29 | 1.221 | -1.079 | 0.027 | -0.072 | -0.072 | 2.488 | 0.82 | 0.71 | 0.78 | 0.99 | 0.7 | 0.8 |
| 8 | 21961834 | rs7490 | -1.011 | 2.13 | 1.066 | -1.071 | 0.023 | 0.01 | 0.01 | 0 | 0.32 | 0.31 | 0.65 | 0.6 | 0.45 | 0.53 |
| 8 | 21964802 | rs1552286 | 0.242 | 2.214 | 1.12 | -1.04 | 0.003 | -0.02 | -0.02 | 0 | 0.91 | 0.88 | 0.93 | 0.99 | 0.84 | 0.97 |
| 8 | 21970230 | rs7831711 | 0.603 | 2.058 | 0.991 | -1.009 | 0.022 | 0.012 | 0.012 | 0 | 0.69 | 0.71 | 0.36 | 0.4 | 0.56 | 0.48 |
| 8 | 21971141 | rs6557826 | 0.707 | 2.069 | 0.992 | -1.005 | 0.021 | 0.009 | 0.009 | 0 | 0.68 | 0.7 | 0.35 | 0.4 | 0.55 | 0.47 |
| 8 | 21978072 | rs11135859 | 0.142 | 1.795 | 0.246 | -1.045 | 0.008 | 0.011 | 0.011 | 0 | 0.72 | 0.69 | 0.48 | 0.4 | 0.6 | 0.64 |
| 8 | 21984488 | rs12681529 | 0.913 | 1.094 | -0.817 | -0.943 | 0.071 | 0.058 | 0.058 | 0.208 | 0.47 | 0.16 | 0.21 | 0.37 | 0.3 | 0.43 |
| 8 | 21994951 | rs12678092 | -0.592 | 1.346 | 0.125 | -0.825 | 0 | -0.008 | -0.008 | 0 | 0.18 | 0.3 | 0.12 | 0.09 | 0.18 | 0.1 |
| 8 | 21995097 | rs12676497 | -0.592 | 1.346 | 0.119 | -0.825 | 0 | -0.007 | -0.007 | 0 | 0.18 | 0.3 | 0.12 | 0.09 | 0.18 | 0.1 |
| 8 | 22001072 | rs4872320 | -1.274 | 1.493 | -0.057 | -0.896 | -0.021 | 0.062 | 0.062 | 0 | 0.36 | 0.75 | 0.47 | 0.13 | 0.4 | 0.32 |
| 8 | 22007293 | rs12679786 | -0.944 | 0.942 | 0.447 | -0.409 | -0.001 | -0.084 | -0.084 | 0.852 | 0.22 | 0.48 | 0.13 | 0.09 | 0.18 | 0.12 |
| 8 | 22008480 | rs7838257 | -0.88 | 1.057 | 0.36 | -0.376 | 0.004 | -0.062 | -0.062 | 0.28 | 0.2 | 0.38 | 0.11 | 0.09 | 0.18 | 0.12 |
| 8 | 22019893 | rs8192325 | -0.499 | 0.702 | -0.361 | -0.154 | -0.005 | -0.007 | -0.007 | 0 | 0.29 | 0.34 | 0.22 | 0.18 | 0.27 | 0.15 |
| 8 | 22020294 | rs2070684 | 1.109 | 0.397 | 0.133 | 0.276 | -0.019 | 0.106 | 0.106 | 0.733 | 0.86 | 0.35 | 0.7 | 0.71 | 0.89 | 0.85 |
| 8 | 22021037 | rs4715 | -0.049 | 0.373 | -0.98 | 0.605 | -0.032 | -0.047 | -0.047 | 0 | 0.24 | 0.03 | 0.25 | 0.27 | 0.26 | 0.24 |
| 8 | 22021207 | rs2070685 | 0.601 | 0.996 | -1.246 | 1.031 | -0.037 | -0.032 | -0.032 | 0.605 | 0.07 | 0.01 | 0.12 | 0.29 | 0.1 | 0.26 |
| 8 | 22021517 | rs1124 | -0.047 | 1.141 | -1.18 | 0.835 | -0.029 | -0.032 | -0.032 | 0.593 | 0.28 | 0.08 | 0.27 | 0.3 | 0.32 | 0.26 |
| 8 | 22021883 | rs7592 | -0.376 | 1.058 | -0.281 | 0.789 | -0.005 | -0.003 | -0.003 | 0 | 0.31 | 0.44 | 0.3 | 0.3 | 0.33 | 0.26 |
| 8 | 22025644 | rs7812993 | 0.292 | 0.646 | -0.458 | 0.578 | 0.002 | 0.017 | 0.017 | 0 | 0.73 | 0.71 | 0.79 | 0.6 | 0.79 | 0.57 |
| 8 | 22033435 | rs11777932 | -0.033 | 0.684 | -0.888 | 0.412 | -0.015 | -0.009 | -0.009 | 0 | 0.05 | 0.02 | 0.05 | 0.11 | 0.09 | 0.09 |
| 8 | 22038952 | rs7838961 | -0.605 | 0.142 | -1.182 | 0.129 | -0.022 | 0.033 | 0.033 | 0 | 0.57 | 0.69 | 0.63 | 0.69 | 0.52 | 0.53 |
| 8 | 22042151 | rs7819541 | -0.132 | -0.112 | -0.755 | 0.339 | -0.044 | -0.034 | -0.034 | 1.414 | 0.16 | 0.58 | 0.17 | 0.07 | 0.09 | 0.06 |
| 8 | 22042197 | rs4507761 | 1.178 | -0.007 | -1.237 | 0.385 | -0.023 | -0.022 | -0.022 | 1.484 | 0.1 | 0 | 0.05 | 0.09 | 0.08 | 0.03 |
| 8 | 22049780 | rs3924231 | -0.941 | 0.047 | -0.461 | 0.46 | -0.008 | -0.034 | -0.034 | 0 | 0.23 | 0.56 | 0.23 | 0.16 | 0.19 | 0.15 |
| 8 | 22049892 | rs3924229 | -0.788 | -0.077 | -0.476 | 0.085 | -0.017 | -0.049 | -0.049 | 0 | 0.19 | 0.28 | 0.08 | 0.1 | 0.11 | 0.12 |
| 8 | 22053123 | rs4075478 | -0.14 | 0.725 | -1.49 | 0.509 | -0.001 | -0.006 | -0.006 | 0.497 | 0.37 | 0.67 | 0.56 | 0.54 | 0.4 | 0.4 |
| 8 | 22055348 | rs3857979 | 0.499 | 0.64 | -1.582 | 0.276 | -0.008 | 0.094 | 0.094 | 0.559 | 0.48 | 0.27 | 0.38 | 0.28 | 0.48 | 0.45 |
| 8 | 22057449 | rs7827382 | -1.202 | 0.876 | -1.337 | 0.665 | -0.03 | -0.016 | -0.016 | 0 | 0.25 | 0.91 | 0.28 | 0.27 | 0.22 | 0.22 |
| 8 | 22063438 | rs6983732 | -1.263 | 0.746 | -0.274 | 0.747 | 0.006 | 0.029 | 0.029 | 0 | 0.13 | 0.32 | 0.11 | 0.16 | 0.11 | 0.1 |
| 8 | 22067769 | rs4242430 | -0.427 | 0.62 | -1.55 | 0.676 | -0.067 | -0.189 | -0.189 | 3.347 | 0.68 | 0.97 | 0.58 | 0.49 | 0.67 | 0.58 |
| 8 | 22067774 | rs4076873 | -0.425 | 0.849 | -0.309 | -0.091 | -0.004 | -0.056 | -0.056 | 0 | 0.27 | 0.52 | 0.15 | 0.11 | 0.26 | 0.13 |
| 8 | 22069018 | rs11778448 | -0.405 | 0.999 | -0.581 | -0.341 | 0.006 | -0.134 | -0.134 | 1.399 | 0.25 | 0.37 | 0.14 | 0.01 | 0.26 | 0.09 |
| 8 | 22084653 | rs4871975 | 0.392 | 0.83 | -1.687 | 0.107 | -0.07 | -0.06 | -0.06 | 1.228 | 0.36 | 0.04 | 0.39 | 0.6 | 0.38 | 0.47 |
| 8 | 22085260 | rs10100426 | -0.18 | 0.772 | -0.708 | -0.281 | -0.003 | 0.013 | 0.013 | 0 | 0.54 | 0.61 | 0.6 | 0.73 | 0.5 | 0.56 |
| 8 | 22086491 | rs4871976 | 1.038 | 0.728 | -1 | -0.505 | 0.001 | 0.009 | 0.009 | 0.552 | 0.76 | 0.21 | 0.68 | 0.81 | 0.76 | 0.89 |
| 8 | 22086830 | rs4872428 | -0.006 | 0.859 | -1.589 | -0.435 | -0.096 | 0.016 | 0.016 | 0 | 0.35 | 0.11 | 0.41 | 0.29 | 0.41 | 0.47 |
| 8 | 22087184 | rs6557917 | 0.425 | 1.083 | -1.68 | -0.768 | -0.039 | 0.03 | 0.03 | 0 | 0.61 | 0.84 | 0.62 | 0.66 | 0.55 | 0.48 |
| 8 | 22088432 | rs12541335 | -0.788 | 1.139 | -1.94 | -1.068 | -0.035 | 0.246 | 0.246 | 0.635 | 0.43 | 0.16 | 0.34 | 0.04 | 0.43 | 0.33 |
| 8 | 22090263 | rs3888282 | 1.648 | 0.9 | -1.422 | -0.574 | 0.029 | -0.015 | -0.015 | 2.707 | 0.39 | 0.18 | 0.37 | 0.6 | 0.35 | 0.38 |
| 8 | 22100833 | rs12546622 | 0.547 | 1.757 | -1.199 | -0.23 | 0.011 | -0.011 | -0.011 | 0.846 | 0.73 | 0.36 | 0.75 | 0.82 | 0.71 | 0.89 |
| 8 | 22104691 | rs34392777 | 1.7 | 1.618 | -1.361 | -0.062 | 0.095 | -0.103 | -0.103 | 0 | 0.47 | 0.03 | 0.4 | 0.51 | 0.39 | 0.42 |
| 8 | 22108347 | rs2645 | 1.088 | 1.848 | -1.38 | -0.287 | -0.012 | 0.06 | 0.06 | 0.091 | 0.57 | 0.31 | 0.66 | 0.8 | 0.52 | 0.68 |
| 8 | 22108611 | rs9173 | 1.628 | 1.629 | -1.399 | -0.262 | 0.083 | -0.032 | -0.032 | 0 | 0.48 | 0.03 | 0.4 | 0.58 | 0.4 | 0.45 |
| 8 | 22113430 | rs12541504 | 0.439 | 1.686 | -1.298 | -0.219 | 0.008 | -0.013 | -0.013 | 0 | 0.72 | 0.37 | 0.75 | 0.82 | 0.72 | 0.88 |
| 8 | 22113565 | rs13249656 | -0.262 | 1.607 | -1.207 | -0.239 | -0.017 | -0.009 | -0.009 | 0.002 | 0.26 | 0.15 | 0.23 | 0.31 | 0.28 | 0.44 |
| 8 | 22121448 | rs4872461 | 0.772 | 1.707 | -0.811 | 0.175 | -0.018 | -0.016 | -0.016 | 0 | 0.08 | 0.04 | 0.14 | 0.24 | 0.1 | 0.23 |
| 8 | 22132033 | rs4872462 | 0.205 | 1.643 | -0.873 | -0.041 | -0.094 | 0.128 | 0.128 | 1.919 | 0.61 | 0.9 | 0.71 | 0.86 | 0.57 | 0.7 |

| | | | | | | | | | | | | | | | | |
|---|---|---|---|---|---|---|---|---|---|---|---|---|---|---|---|---|
| 8 | 22132837 | rs4871990 | 0.279 | 1.657 | -0.824 | -0.034 | -0.074 | 0.131 | 0.131 | 1.571 | 0.61 | 0.87 | 0.7 | 0.85 | 0.57 | 0.69 |
| 8 | 22133744 | rs13259097 | 0.554 | 1.687 | -0.548 | -0.003 | -0.017 | 0.077 | 0.077 | 1.221 | 0.55 | 0.29 | 0.64 | 0.81 | 0.55 | 0.68 |
| 8 | 22134826 | rs6557952 | 1.208 | 1.614 | -0.625 | 0.038 | 0.041 | 0.003 | 0.003 | 0 | 0.45 | 0.13 | 0.51 | 0.61 | 0.45 | 0.44 |
| 8 | 22147333 | rs4872463 | 0.61 | 1.62 | -0.502 | -0.086 | -0.016 | 0.121 | 0.121 | 0.388 | 0.55 | 0.18 | 0.55 | 0.81 | 0.5 | 0.69 |
| 8 | 22154133 | rs4565458 | 0.279 | 1.64 | -0.315 | -0.076 | -0.01 | 0.045 | 0.045 | 0.062 | 0.58 | 0.63 | 0.58 | 0.8 | 0.5 | 0.69 |
| 8 | 22167613 | rs6557986 | -0.375 | 1.646 | -0.283 | -0.045 | 0.069 | 0.094 | 0.094 | 2.914 | 0.06 | 0.54 | 0.06 | 0.04 | 0.02 | 0.01 |
| 8 | 22169087 | rs4266653 | 0.104 | 1.698 | -0.375 | -0.053 | -0.018 | 0.049 | 0.049 | 0.002 | 0.6 | 0.66 | 0.6 | 0.85 | 0.52 | 0.69 |
| 8 | 22172702 | rs4397403 | 1.151 | 1.843 | -1.196 | 0.217 | 0.053 | -0.124 | -0.124 | 0 | 0.42 | 0.03 | 0.38 | 0.53 | 0.38 | 0.41 |
| 8 | 22188330 | rs2404332 | 0.22 | 1.947 | -1.373 | -0.41 | 0 | 0.047 | 0.047 | 0 | 0.62 | 0.23 | 0.65 | 0.81 | 0.65 | 0.71 |
| 8 | 22192692 | rs4872471 | 0.04 | 1.707 | -1.408 | -0.403 | 0.016 | 0.029 | 0.029 | 0 | 0.64 | 0.29 | 0.76 | 0.82 | 0.71 | 0.73 |
| 8 | 22200148 | rs11135995 | 0.245 | 0.974 | -1.884 | -0.127 | -0.031 | 0.154 | 0.154 | 0.337 | 0.54 | 0.09 | 0.55 | 0.81 | 0.5 | 0.69 |
| 8 | 22227522 | rs6998654 | 0.027 | 1.349 | -1.553 | 0.494 | -0.045 | -0.052 | -0.052 | 1.969 | 0.9 | 0.99 | 0.79 | 0.83 | 0.89 | 0.97 |
| 8 | 22238030 | rs11136017 | -0.023 | 0.942 | -1.784 | -0.31 | -0.041 | 0.096 | 0.096 | 0.26 | 0.58 | 0.8 | 0.56 | 0.8 | 0.5 | 0.61 |
| 8 | 22239868 | rs12545568 | -0.065 | 0.844 | -0.891 | 0.189 | 0.063 | 0.101 | 0.101 | 0 | 0.15 | 0.45 | 0.18 | 0.28 | 0.21 | 0.25 |
| 8 | 22244897 | rs11783730 | 1 | 0.323 | -1.243 | -0.009 | 0.018 | -0.118 | -0.118 | 0.562 | 0.33 | 0.08 | 0.36 | 0.51 | 0.27 | 0.37 |
| 8 | 22249943 | rs1568569 | -0.709 | 0.382 | -1.208 | -0.057 | -0.048 | 0.079 | 0.079 | 0.722 | 0.59 | 0.82 | 0.56 | 0.78 | 0.51 | 0.59 |
| 8 | 22256310 | rs7833302 | 0.357 | 0.456 | -0.655 | 0.114 | 0.014 | 0.022 | 0.022 | 0 | 0.1 | 0.13 | 0.1 | 0.17 | 0.15 | 0.14 |
| 8 | 22256359 | rs4872486 | 0.79 | 0.289 | -0.655 | 0.03 | 0.036 | -0.063 | -0.063 | 0 | 0.36 | 0.15 | 0.4 | 0.6 | 0.29 | 0.41 |
| 8 | 22262418 | rs2293144 | -0.794 | 0.272 | -0.556 | -0.022 | -0.05 | 0.089 | 0.089 | 0.366 | 0.58 | 0.83 | 0.56 | 0.78 | 0.51 | 0.61 |
| 8 | 22262630 | rs10113026 | -0.768 | 0.216 | 0.181 | -0.04 | 0.004 | -0.003 | -0.003 | 0 | 0.21 | 0.58 | 0.15 | 0.18 | 0.21 | 0.18 |
| 8 | 22263172 | rs17060826 | -0.941 | 0.216 | 0.018 | -0.04 | -0.001 | -0.003 | -0.003 | 0 | 0.22 | 0.69 | 0.16 | 0.18 | 0.21 | 0.18 |
| 8 | 22264334 | rs7828089 | -0.58 | 0.265 | -0.004 | -0.009 | -0.033 | 0.068 | 0.068 | 0.493 | 0.56 | 0.76 | 0.53 | 0.7 | 0.51 | 0.58 |
| 8 | 22266292 | rs13259873 | 0.009 | 0.088 | 0.648 | 0.036 | 0.003 | 0.003 | 0.003 | 0 | 0.45 | 0.55 | 0.44 | 0.61 | 0.34 | 0.51 |
| 8 | 22267199 | rs13256933 | 0.819 | 0.103 | 0.309 | 0.049 | -0.003 | -0.131 | -0.131 | 0 | 0.34 | 0.07 | 0.36 | 0.52 | 0.29 | 0.4 |
| 8 | 22267385 | rs3802199 | 0.819 | 0.103 | 0.309 | 0.049 | -0.003 | -0.131 | -0.131 | 2.814 | 0.34 | 0.07 | 0.36 | 0.52 | 0.29 | 0.4 |
| 8 | 22270652 | rs11989798 | 0.711 | 0.296 | 0.742 | 0.119 | 0.005 | 0.01 | 0.01 | 0 | 0.07 | 0.15 | 0.06 | 0.09 | 0.05 | 0.14 |
| 8 | 22271995 | rs2280521 | -0.537 | -0.28 | 0.495 | -0.258 | 0.082 | 0.097 | 0.097 | 2.094 | 0.48 | 0.83 | 0.37 | 0.69 | 0.31 | 0.5 |
| 8 | 22272226 | rs2280522 | -1.744 | -0.31 | 0.791 | -0.154 | 0.016 | 0.079 | 0.079 | 2.933 | 0.06 | 0.62 | 0.06 | 0.11 | 0 | 0.01 |
| 8 | 22273687 | rs6558052 | -1.201 | -0.344 | 0.825 | -0.166 | -0.007 | 0.011 | 0.011 | 1.965 | 0.12 | 0.72 | 0.12 | 0.19 | 0.05 | 0.15 |
| 8 | 22277040 | rs10101909 | 0.059 | -0.377 | 0.756 | -0.193 | 0.038 | 0.011 | 0.011 | 1.916 | 0.41 | 0.15 | 0.3 | 0.58 | 0.3 | 0.49 |
| 8 | 22278781 | rs1051638 | -0.819 | -0.255 | 0.987 | -0.059 | 0.013 | 0.094 | 0.094 | 0.469 | 0.56 | 0.82 | 0.5 | 0.72 | 0.39 | 0.56 |
| 8 | 22279398 | rs1051647 | 1.582 | -0.272 | 0.82 | -0.096 | -0.001 | 0 | 0 | 0.015 | 0.05 | 0.04 | 0.05 | 0.08 | 0.05 | 0.13 |
| 8 | 22279863 | rs1051708 | -0.3 | -0.272 | 0.968 | -0.109 | 0.056 | 0.065 | 0.065 | 0.555 | 0.5 | 0.2 | 0.42 | 0.59 | 0.38 | 0.55 |
| 8 | 22282732 | rs3943520 | -0.88 | -0.235 | 0.874 | -0.113 | 0.005 | 0.066 | 0.066 | 0.992 | 0.56 | 0.81 | 0.49 | 0.68 | 0.39 | 0.56 |
| 8 | 22282748 | rs10113731 | -0.284 | -0.203 | 0.749 | -0.098 | 0.057 | 0.071 | 0.071 | 0.409 | 0.5 | 0.16 | 0.43 | 0.59 | 0.38 | 0.55 |
| 8 | 22288995 | rs7833266 | 0.016 | -0.189 | 0.814 | -0.1 | 0.066 | -0.008 | -0.008 | 1.117 | 0.46 | 0.24 | 0.36 | 0.51 | 0.33 | 0.41 |
| 8 | 22293306 | rs4872499 | 0.876 | -0.501 | 0.426 | 0.13 | -0.031 | -0.167 | -0.167 | 0 | 0.27 | 0.02 | 0.29 | 0.47 | 0.24 | 0.3 |
| 8 | 22295228 | rs12546391 | -1.195 | -0.16 | 1.235 | 0.256 | 0.033 | 0.095 | 0.095 | 0.539 | 0.11 | 0.51 | 0.12 | 0.17 | 0.05 | 0.15 |
| 8 | 22297849 | rs17060857 | 1.444 | -0.11 | 1.099 | 0.272 | -0.002 | -0.002 | -0.002 | 0 | 0.05 | 0.04 | 0.06 | 0.08 | 0.05 | 0.13 |
| 8 | 22303439 | rs10096012 | 0.401 | -0.171 | 1.171 | 0.382 | 0 | 0.004 | 0.004 | 0 | 0.07 | 0.06 | 0.06 | 0.09 | 0.05 | 0.17 |
| 8 | 22305501 | rs7821470 | -1.528 | -0.215 | 1.175 | 0.367 | -0.009 | 0.066 | 0.066 | 0.022 | 0.14 | 0.5 | 0.11 | 0.18 | 0.06 | 0.19 |
| 8 | 22320806 | rs10108011 | -0.773 | 0.136 | 0.588 | 0.013 | -0.047 | 0.066 | 0.066 | 0.065 | 0.62 | 0.82 | 0.56 | 0.78 | 0.57 | 0.61 |
| 8 | 22334610 | rs7827093 | -0.34 | 0.211 | 0.757 | 0.066 | -0.004 | 0.025 | 0.025 | 0.481 | 0.56 | 0.65 | 0.49 | 0.6 | 0.47 | 0.55 |
| 8 | 22353569 | rs2461493 | -0.157 | 0.182 | 1.012 | 0.033 | 0.003 | 0.015 | 0.015 | 0.184 | 0.53 | 0.37 | 0.47 | 0.6 | 0.46 | 0.54 |
| 8 | 22357598 | rs2469749 | -0.634 | 0.742 | 0.557 | -0.377 | -0.011 | -0.006 | -0.006 | 0 | 0.68 | 0.74 | 0.65 | 0.62 | 0.65 | 0.69 |
| 8 | 22374200 | rs896379 | -0.466 | 0.578 | 0.703 | 0.263 | -0.028 | -0.004 | -0.004 | 0 | 0.59 | 0.8 | 0.52 | 0.64 | 0.45 | 0.53 |
| 8 | 22374551 | rs2100291 | -0.01 | 0.556 | 0.967 | 0.255 | -0.001 | -0.001 | -0.001 | 0 | 0.53 | 0.43 | 0.47 | 0.54 | 0.45 | 0.51 |
| 8 | 22374794 | rs2449329 | -0.135 | 0.555 | 1.054 | 0.255 | 0 | -0.001 | -0.001 | 0 | 0.54 | 0.54 | 0.47 | 0.55 | 0.45 | 0.51 |
| 8 | 22378118 | rs1600877 | 0.041 | 0.558 | 0.955 | 0.296 | 0.006 | 0.003 | 0.003 | 0.508 | 0.53 | 0.35 | 0.46 | 0.54 | 0.45 | 0.51 |
| 8 | 22386479 | rs2461483 | -0.429 | 0.62 | 0.616 | 0.455 | -0.033 | -0.026 | -0.026 | 0.248 | 0.58 | 0.81 | 0.52 | 0.62 | 0.46 | 0.53 |
| 8 | 22394920 | rs1116085 | 0.011 | 0.302 | 0.737 | 0.547 | -0.002 | -0.005 | -0.005 | 0.371 | 0.53 | 0.45 | 0.46 | 0.58 | 0.45 | 0.52 |
| 8 | 22395309 | rs1116084 | 0.077 | 0.298 | 0.703 | 0.535 | 0.006 | -0.001 | -0.001 | 0.015 | 0.52 | 0.36 | 0.46 | 0.52 | 0.45 | 0.52 |
| 8 | 22398462 | rs7431 | -0.448 | 0.302 | 0.33 | 0.54 | -0.03 | -0.012 | -0.012 | 0 | 0.59 | 0.81 | 0.52 | 0.61 | 0.46 | 0.53 |
| 8 | 22403451 | rs2469776 | -0.182 | 0.453 | 0.335 | 0.374 | 0 | 0.009 | 0.009 | 0.254 | 0.54 | 0.53 | 0.47 | 0.61 | 0.46 | 0.52 |
| 8 | 22409882 | rs754270 | 0.254 | 0.53 | -0.26 | 0.306 | 0.002 | -0.001 | -0.001 | 0 | 0.46 | 0.54 | 0.5 | 0.52 | 0.42 | 0.43 |
| 8 | 22410157 | rs2005080 | -0.076 | 0.799 | -0.873 | 0.601 | 0.03 | 0.008 | 0.008 | 1.323 | 0.51 | 0.77 | 0.52 | 0.54 | 0.37 | 0.54 |
| 8 | 22414741 | rs2449347 | 0.402 | 0.771 | -0.407 | 0.326 | 0.002 | 0.006 | 0.006 | 0 | 0.47 | 0.38 | 0.38 | 0.4 | 0.46 | 0.5 |
| 8 | 22415992 | rs2449346 | 0.464 | 0.918 | -0.879 | 0.428 | -0.009 | 0.006 | 0.006 | 0 | 0.48 | 0.28 | 0.38 | 0.44 | 0.49 | 0.47 |
| 8 | 22422150 | rs3758038 | -0.339 | 0.794 | -0.778 | 0.06 | -0.017 | 0.019 | 0.019 | 0.45 | 0.38 | 0.3 | 0.37 | 0.22 | 0.52 | 0.46 |
| 8 | 22424489 | rs11779261 | -0.516 | 0.321 | -0.57 | 0.011 | -0.001 | -0.002 | -0.002 | 0.361 | 0.52 | 0.46 | 0.65 | 0.36 | 0.62 | 0.61 |
| 8 | 22428658 | rs2449331 | -0.255 | -0.609 | -0.289 | -0.244 | -0.019 | -0.051 | -0.051 | 0 | 0.91 | 0.73 | 0.95 | 1 | 0.96 | 0.97 |
| 8 | 22428708 | rs1047030 | -0.143 | -0.148 | -0.663 | -0.249 | -0.023 | 0.042 | 0.042 | 0.139 | 0.14 | 0.06 | 0.14 | 0.03 | 0.21 | 0.21 |
| 8 | 22430766 | rs10088686 | -0.291 | -0.092 | -0.316 | -0.244 | 0.037 | -0.043 | -0.043 | 0 | 0.16 | 0.21 | 0.2 | 0.03 | 0.29 | 0.24 |
| 8 | 22433508 | rs2013211 | -0.821 | -1.426 | -0.766 | 0.21 | 0.005 | -0.002 | -0.002 | 0 | 0.55 | 0.62 | 0.67 | 0.52 | 0.62 | 0.64 |
| 8 | 22439442 | rs4872512 | -0.319 | -1.541 | -0.695 | 0.167 | 0.005 | -0.01 | -0.01 | 0 | 0.31 | 0.18 | 0.37 | 0.32 | 0.28 | 0.34 |
| 8 | 22442661 | rs2294047 | -0.397 | -1.453 | -0.574 | 0.419 | -0.002 | 0 | 0 | 1.379 | 0.55 | 0.59 | 0.65 | 0.47 | 0.61 | 0.62 |
| 8 | 22452357 | rs11782130 | 0.731 | -1.46 | -0.195 | 0.377 | -0.003 | 0.005 | 0.005 | 0.051 | 0.28 | 0.3 | 0.41 | 0.26 | 0.34 | 0.41 |
| 8 | 22453192 | rs7012299 | 1.11 | -1.687 | -0.14 | 0.268 | -0.009 | -0.016 | -0.016 | 0 | 0.07 | 0.06 | 0.02 | 0 | 0.02 | 0 |
| 8 | 22453223 | rs11785755 | 1.183 | -1.639 | -0.08 | 0.324 | -0.049 | 0.035 | 0.035 | 0.137 | 0.25 | 0.07 | 0.39 | 0.2 | 0.34 | 0.4 |

| | | | | | | | | | | | | | | | | |
|---|---|---|---|---|---|---|---|---|---|---|---|---|---|---|---|---|
| 8 | 22454826 | rs3735894 | 0.835 | -1.625 | -0.04 | 0.301 | -0.007 | 0.008 | 0.008 | 0.236 | 0.28 | 0.23 | 0.4 | 0.26 | 0.34 | 0.42 |
| 8 | 22463623 | rs7843828 | 0.842 | -1.66 | -0.013 | 0.273 | -0.003 | 0.01 | 0.01 | 0.111 | 0.28 | 0.26 | 0.33 | 0.21 | 0.34 | 0.4 |
| 8 | 22466488 | rs11136092 | 1.192 | -1.667 | -0.054 | 0.27 | -0.046 | 0.022 | 0.022 | 0.247 | 0.25 | 0.07 | 0.31 | 0.21 | 0.34 | 0.4 |
| 8 | 22471147 | rs720745 | -0.187 | -1.827 | 0.396 | 0.26 | -0.012 | -0.018 | -0.018 | 0 | 0.39 | 0.61 | 0.37 | 0.28 | 0.37 | 0.48 |
| 8 | 22471824 | rs3736147 | 1.214 | -1.812 | 0.277 | 0.208 | -0.029 | -0.006 | -0.006 | 0.057 | 0.26 | 0.11 | 0.3 | 0.26 | 0.34 | 0.4 |
| 8 | 22473224 | rs6558165 | -0.236 | -1.813 | 0.471 | 0.092 | -0.069 | -0.067 | -0.067 | 0.327 | 0.13 | 0.38 | 0.05 | 0.02 | 0.03 | 0.09 |
| 8 | 22477585 | rs1869 | 1.201 | -1.984 | 0.142 | 0.027 | -0.102 | -0.115 | -0.115 | 0 | 0.29 | 0.02 | 0.41 | 0.63 | 0.3 | 0.32 |
| 8 | 22477778 | rs1059592 | 0.058 | -2.01 | 0.267 | 0.051 | -0.009 | -0.018 | -0.018 | 0 | 0.38 | 0.51 | 0.36 | 0.25 | 0.37 | 0.49 |
| 8 | 22477807 | rs1545837 | 1.071 | -2.002 | 0.229 | 0.027 | -0.04 | 0.052 | 0.052 | 0.831 | 0.28 | 0.11 | 0.32 | 0.18 | 0.37 | 0.46 |
| 8 | 22478018 | rs13439542 | 0.071 | -1.947 | 0.297 | 0.055 | -0.064 | -0.062 | -0.062 | 1.038 | 0.13 | 0.36 | 0.05 | 0.02 | 0.03 | 0.09 |
| 8 | 22483836 | rs878051 | -0.055 | -1.943 | 0.096 | 0.058 | 0.002 | 0.002 | 0.002 | 0 | 0.42 | 0.48 | 0.38 | 0.26 | 0.41 | 0.54 |
| 8 | 22485375 | rs7830206 | 0.094 | -1.898 | 0.144 | 0.055 | 0.018 | 0.063 | 0.063 | 0 | 0.4 | 0.33 | 0.35 | 0.21 | 0.41 | 0.54 |
| 8 | 22487087 | rs3758040 | -0.669 | -1.856 | 0.147 | 0.346 | 0.013 | -0.019 | -0.019 | 0 | 0.73 | 0.5 | 0.79 | 0.78 | 0.73 | 0.87 |
| 8 | 22491518 | rs2003908 | -0.452 | -1.817 | 0.316 | 0.392 | 0.015 | -0.033 | -0.033 | 0 | 0.7 | 0.56 | 0.78 | 0.8 | 0.69 | 0.78 |
| 8 | 22492708 | rs4872525 | 0.684 | -1.781 | -0.02 | 0.397 | -0.005 | -0.019 | -0.019 | 0.603 | 0.34 | 0.27 | 0.45 | 0.56 | 0.33 | 0.32 |
| 8 | 22493124 | rs11776549 | 1.382 | -1.715 | -0.011 | -0.075 | -0.028 | 0.029 | 0.029 | 0.24 | 0.24 | 0.14 | 0.3 | 0.18 | 0.33 | 0.39 |
| 8 | 22496731 | rs12334545 | 0.849 | -1.63 | 0.03 | 0.104 | -0.004 | 0.001 | 0.001 | 0 | 0.09 | 0.05 | 0.03 | 0.04 | 0.03 | 0.08 |
| 8 | 22506408 | rs7388243 | 0.677 | -1.643 | -0.083 | -0.088 | -0.009 | -0.023 | -0.023 | 0 | 0.33 | 0.24 | 0.37 | 0.62 | 0.32 | 0.34 |
| 8 | 22509478 | rs10216447 | -0.183 | -1.284 | 0.177 | 0.033 | -0.063 | -0.016 | -0.016 | 0 | 0.13 | 0.37 | 0.05 | 0.06 | 0.03 | 0.06 |
| 8 | 22522502 | rs6558176 | 0.027 | -1.166 | 0.171 | 0.115 | -0.003 | -0.009 | -0.009 | 0.243 | 0.42 | 0.48 | 0.27 | 0.2 | 0.31 | 0.21 |
| 8 | 22525980 | rs2280104 | -0.368 | -1.307 | 0.715 | 0.432 | -0.051 | 0.034 | 0.034 | 1.15 | 0.73 | 0.93 | 0.68 | 0.8 | 0.63 | 0.55 |
| 8 | 22531463 | rs11985023 | 0.66 | -1.244 | 0.8 | 0.483 | 0.02 | 0.021 | 0.021 | 0 | 0.55 | 0.63 | 0.61 | 0.66 | 0.57 | 0.43 |
| 8 | 22536084 | rs17088531 | 0.101 | -0.875 | -0.603 | 0.772 | -0.041 | 0.015 | 0.015 | 0 | 0.15 | 0.23 | 0.05 | 0.18 | 0.04 | 0.09 |
| 8 | 22536502 | rs4872006 | 0.072 | -0.204 | -0.898 | 0.725 | -0.009 | -0.004 | -0.004 | 0.49 | 0.58 | 0.43 | 0.63 | 0.63 | 0.62 | 0.52 |
| 8 | 22538384 | rs10104039 | 0.112 | 0.036 | -1.101 | 0.425 | -0.011 | -0.009 | -0.009 | 0 | 0.56 | 0.49 | 0.63 | 0.66 | 0.61 | 0.51 |
| 8 | 22539641 | rs12114462 | 0.147 | 0.071 | -1.119 | 0.421 | -0.007 | -0.006 | -0.006 | 0 | 0.55 | 0.5 | 0.63 | 0.67 | 0.61 | 0.51 |
| 8 | 22546561 | rs1877670 | 0.158 | 0.704 | -1.812 | 0.396 | -0.057 | -0.025 | -0.025 | 1.673 | 0.57 | 0.17 | 0.68 | 0.39 | 0.54 | 0.51 |
| 8 | 22553621 | rs1008949 | -0.122 | 0.989 | -1.257 | 0.194 | -0.012 | -0.035 | -0.035 | 0 | 0.62 | 0.79 | 0.78 | 0.5 | 0.61 | 0.56 |
| 8 | 22559693 | rs4872533 | 0.192 | 0.872 | -0.508 | -0.238 | 0.002 | 0.005 | 0.005 | 0 | 0.52 | 0.57 | 0.57 | 0.22 | 0.47 | 0.44 |
| 8 | 22562092 | rs4295666 | 0.2 | 0.83 | -0.685 | -1.114 | 0.03 | 0.053 | 0.053 | 0 | 0.36 | 0.18 | 0.23 | 0 | 0.26 | 0.16 |
| 8 | 22562352 | rs7823327 | -0.407 | 0.911 | -0.547 | -1.011 | 0.009 | 0.159 | 0.159 | 0 | 0.5 | 0.22 | 0.41 | 0.18 | 0.5 | 0.45 |
| 8 | 22563095 | rs2090052 | 0.669 | 0.881 | -0.381 | -0.947 | 0.009 | 0.009 | 0.009 | 0 | 0.16 | 0.21 | 0.2 | 0.18 | 0.23 | 0.28 |
| 8 | 22564999 | rs10103576 | -0.612 | 1.222 | -0.104 | -1.077 | 0 | -0.007 | -0.007 | 1.685 | 0.56 | 0.45 | 0.49 | 0.19 | 0.62 | 0.51 |
| 8 | 22567354 | rs11775096 | 0.315 | 1.236 | -0.398 | -1.296 | -0.017 | 0.183 | 0.183 | 0 | 0.36 | 0.1 | 0.27 | 0.01 | 0.34 | 0.18 |
| 8 | 22569211 | rs7388527 | -0.375 | 1.696 | 0.652 | -0.529 | 0.066 | -0.025 | -0.025 | 1.104 | 0.21 | 0.34 | 0.28 | 0.15 | 0.35 | 0.34 |
| 8 | 22570935 | rs1047406 | -0.944 | 0.972 | 0.913 | -0.303 | 0.004 | 0.001 | 0.001 | 0 | 0.26 | 0.24 | 0.46 | 0.3 | 0.32 | 0.17 |
| 8 | 22571018 | rs1047398 | -0.945 | 0.97 | 0.912 | -0.304 | 0.004 | 0.001 | 0.001 | 0 | 0.26 | 0.24 | 0.46 | 0.3 | 0.32 | 0.17 |
| 8 | 22571466 | rs2280107 | -1.31 | 0.908 | 1.189 | -0.292 | 0.057 | 0.208 | 0.208 | 0.1 | 0.1 | 0.32 | 0.26 | 0.29 | 0.13 | 0.1 |
| 8 | 22572116 | rs10091402 | -1.195 | 0.906 | 1.182 | -0.293 | 0.052 | 0.199 | 0.199 | 0.068 | 0.1 | 0.31 | 0.26 | 0.29 | 0.13 | 0.1 |
| 8 | 22575108 | rs907891 | -1.706 | 0.715 | 1.265 | -0.429 | 0.008 | -0.001 | -0.001 | 0.091 | 0.32 | 0.57 | 0.47 | 0.31 | 0.34 | 0.2 |
| 8 | 22578003 | rs17088571 | 0.018 | 0.744 | 1.266 | -0.648 | -0.001 | -0.019 | -0.019 | 0 | 0.06 | 0.01 | 0.18 | 0.23 | 0.06 | 0.08 |
| 8 | 22578041 | rs6558180 | -1.624 | 0.738 | 1.294 | -0.664 | 0.007 | -0.025 | -0.025 | 0.716 | 0.3 | 0.55 | 0.45 | 0.25 | 0.32 | 0.18 |
| 8 | 22578660 | rs17088580 | 0.053 | 0.792 | 0.823 | -0.615 | -0.001 | -0.018 | -0.018 | 0.102 | 0.06 | 0.01 | 0.18 | 0.23 | 0.06 | 0.08 |
| 8 | 22578872 | rs10503717 | -0.952 | 0.806 | 0.958 | -0.609 | -0.002 | 0.002 | 0.002 | 0 | 0.24 | 0.16 | 0.39 | 0.23 | 0.25 | 0.16 |
| 8 | 22587587 | rs7812900 | -0.744 | 0.934 | 0.933 | -0.699 | 0.015 | -0.069 | -0.069 | 0.014 | 0.25 | 0.58 | 0.37 | 0.08 | 0.26 | 0.1 |
| 8 | 22590779 | rs7001279 | -0.464 | 0.968 | 0.153 | -0.59 | -0.003 | 0.013 | 0.013 | 0 | 0.22 | 0.27 | 0.33 | 0.08 | 0.22 | 0.09 |
| 8 | 22595828 | rs13255912 | -0.733 | 0.328 | -0.221 | -0.472 | 0.027 | 0.025 | 0.025 | 0 | 0.18 | 0.07 | 0.13 | 0.13 | 0.13 | 0.17 |
| 8 | 22596518 | rs7841894 | 0.71 | 0.448 | -0.143 | -0.518 | -0.004 | -0.01 | -0.01 | 0 | 0.78 | 0.82 | 0.81 | 0.86 | 0.78 | 0.82 |
| 8 | 22597858 | rs17088602 | -1.356 | 1.073 | 0.107 | -0.162 | 0.043 | 0.005 | 0.005 | 0 | 0.07 | 0.25 | 0.05 | 0.03 | 0.07 | 0.08 |
| 8 | 22599421 | rs7845221 | -0.616 | 1.862 | -0.412 | -0.505 | -0.037 | 0.031 | 0.031 | 0 | 0.19 | 0.11 | 0.16 | 0.1 | 0.25 | 0.16 |
| 8 | 22600953 | rs2048651 | 0.223 | 1.495 | -0.284 | -0.308 | -0.001 | -0.004 | -0.004 | 1.007 | 0.68 | 0.47 | 0.69 | 0.73 | 0.66 | 0.65 |
| 8 | 22601787 | rs11781095 | 1.318 | 0.572 | -0.987 | -0.067 | 0.015 | -0.086 | -0.086 | 0.257 | 0.23 | 0.1 | 0.18 | 0.46 | 0.18 | 0.23 |
| 8 | 22606292 | rs1877674 | 0.198 | 0.146 | -0.93 | -0.584 | 0.024 | 0.013 | 0.013 | 0 | 0.48 | 0.38 | 0.47 | 0.38 | 0.57 | 0.47 |
| 8 | 22606613 | rs4872545 | -0.946 | 0.073 | -0.372 | -0.536 | 0.019 | -0.016 | -0.016 | 0.767 | 0.93 | 0.73 | 0.96 | 1 | 0.95 | 0.95 |
| 8 | 22608029 | rs11781835 | 0.068 | 0.001 | -1.081 | -0.425 | 0.048 | 0.067 | 0.067 | 0 | 0.53 | 0.23 | 0.52 | 0.31 | 0.6 | 0.48 |
| 8 | 22610888 | rs17088616 | 0.5 | 0.386 | -0.878 | 0.104 | 0.006 | 0.037 | 0.037 | 0 | 0.09 | 0.17 | 0.27 | 0.14 | 0.11 | 0.21 |
| 8 | 22613172 | rs12682145 | 1.445 | 0.359 | -1.297 | 0.071 | 0.001 | -0.041 | -0.041 | 2.408 | 0.22 | 0.14 | 0.14 | 0.52 | 0.2 | 0.24 |
| 8 | 22617312 | rs7846693 | 0.089 | 0.694 | -1.579 | -0.604 | -0.011 | 0.039 | 0.039 | 0.318 | 0.18 | 0.11 | 0.24 | 0.06 | 0.25 | 0.29 |
| 8 | 22617365 | rs17676811 | -0.337 | 0.78 | -2.1 | -0.769 | -0.062 | 0.093 | 0.093 | 0 | 0.32 | 0.09 | 0.35 | 0.13 | 0.42 | 0.34 |
| 8 | 22617903 | rs10503719 | 1.107 | 0.87 | -1.993 | -1.071 | -0.012 | 0.016 | 0.016 | 0 | 0.17 | 0.1 | 0.15 | 0.01 | 0.18 | 0.11 |
| 8 | 22627471 | rs12546464 | 0.811 | 0.88 | -1.832 | -0.554 | 0.004 | 0.008 | 0.008 | 0 | 0.16 | 0.18 | 0.2 | 0.26 | 0.2 | 0.2 |
| 8 | 22628469 | rs4872010 | 0.191 | 0.659 | -2.074 | -0.926 | -0.001 | 0.116 | 0.116 | 0 | 0.49 | 0.28 | 0.48 | 0.14 | 0.52 | 0.45 |
| 8 | 22629084 | rs11135676 | 0.052 | 0.803 | -1.871 | -0.617 | -0.008 | 0.042 | 0.042 | 0 | 0.59 | 0.28 | 0.6 | 0.2 | 0.57 | 0.56 |
| 8 | 22635335 | rs12676845 | 1.087 | 0.818 | -1.818 | -0.289 | -0.011 | -0.05 | -0.05 | 0.024 | 0.21 | 0.13 | 0.18 | 0.52 | 0.24 | 0.2 |
| 8 | 22635576 | rs4872011 | 0.439 | 0.781 | -1.769 | -0.555 | -0.029 | 0.162 | 0.162 | 0 | 0.54 | 0.23 | 0.57 | 0.2 | 0.49 | 0.5 |
| 8 | 22637220 | rs2055822 | 0.66 | 0.886 | -1.475 | -0.089 | 0.01 | -0.002 | -0.002 | 0 | 0.55 | 0.2 | 0.47 | 0.57 | 0.56 | 0.39 |
| 8 | 22641627 | rs11986963 | 0.163 | 0.945 | -1.091 | -0.029 | -0.003 | -0.021 | -0.021 | 0.104 | 0.08 | 0.15 | 0.12 | 0.02 | 0.09 | 0.06 |
| 8 | 22643323 | rs11991347 | 0.437 | 1.07 | -1.472 | 0.271 | -0.026 | -0.041 | -0.041 | 0.319 | 0.56 | 0.14 | 0.47 | 0.51 | 0.53 | 0.35 |
| 8 | 22647850 | rs17088625 | 0.34 | 1.003 | -1.251 | 0.272 | -0.005 | 0.018 | 0.018 | 0 | 0.42 | 0.29 | 0.42 | 0.37 | 0.38 | 0.38 |

| | | | | | | | | | | | | | | | | |
|---|---|---|---|---|---|---|---|---|---|---|---|---|---|---|---|---|
| 8 | 22648756 | rs10102337 | -0.409 | 0.899 | -0.608 | 0.621 | 0.002 | -0.01 | -0.01 | 0 | 0.61 | 0.58 | 0.64 | 0.63 | 0.57 | 0.72 |
| 8 | 22650358 | rs28414783 | 0.56 | 1.085 | -0.203 | 0.613 | 0 | 0.075 | 0.075 | 1.481 | 0.19 | 0.4 | 0.29 | 0.26 | 0.2 | 0.36 |
| 8 | 22653377 | rs4871836 | 0.619 | 1.165 | -0.242 | 0.584 | 0 | 0.02 | 0.02 | 0 | 0.42 | 0.26 | 0.35 | 0.36 | 0.37 | 0.37 |
| 8 | 22655188 | rs6557594 | 0.604 | 1.106 | -0.194 | 0.667 | 0.002 | 0.081 | 0.081 | 2.688 | 0.18 | 0.35 | 0.29 | 0.27 | 0.2 | 0.36 |
| 8 | 22661286 | rs10216846 | 0.33 | 1.048 | -0.328 | 0.513 | -0.002 | 0.008 | 0.008 | 0 | 0.43 | 0.37 | 0.36 | 0.37 | 0.39 | 0.39 |
| 8 | 22662664 | rs4872028 | 0.272 | 1.024 | -0.403 | 0.528 | -0.011 | 0.053 | 0.053 | 0 | 0.25 | 0.37 | 0.23 | 0.37 | 0.18 | 0.33 |
| 8 | 22664798 | rs11985147 | -0.474 | 1.297 | -0.761 | 0.483 | -0.003 | 0.023 | 0.023 | 0.261 | 0.43 | 0.69 | 0.5 | 0.62 | 0.43 | 0.7 |
| 8 | 22667639 | rs12681784 | 1.234 | 1.376 | -0.773 | 0.656 | -0.019 | -0.029 | -0.029 | 0.265 | 0.15 | 0.09 | 0.27 | 0.27 | 0.23 | 0.3 |
| 8 | 22669393 | rs1533308 | -0.221 | 1.372 | -0.761 | 0.297 | 0.001 | -0.006 | -0.006 | 0 | 0.37 | 0.38 | 0.49 | 0.67 | 0.39 | 0.67 |
| 8 | 22671505 | rs1996148 | 0.096 | 1.018 | -1.182 | 0.411 | 0.001 | -0.005 | -0.005 | 0 | 0.36 | 0.31 | 0.39 | 0.46 | 0.32 | 0.3 |
| 8 | 22673716 | rs10503720 | -1.744 | 0.702 | -1.14 | -0.432 | 0 | -0.089 | -0.089 | 2.148 | 0.81 | 0.74 | 0.88 | 1 | 0.84 | 0.95 |
| 8 | 22676135 | rs12677017 | 0.5 | 0.556 | -1.105 | -0.306 | -0.001 | -0.002 | -0.002 | 0 | 0.43 | 0.66 | 0.46 | 0.55 | 0.44 | 0.69 |
| 8 | 22685153 | rs12386970 | 0.644 | 0.61 | -1.013 | -0.796 | -0.001 | -0.01 | -0.01 | 0 | 0.39 | 0.46 | 0.51 | 0.7 | 0.4 | 0.7 |
| 8 | 22687572 | rs2872210 | 0.194 | 0.77 | -0.547 | -0.943 | 0.001 | -0.062 | -0.062 | 0 | 0.61 | 0.56 | 0.71 | 0.88 | 0.63 | 0.79 |
| 8 | 22697090 | rs7815300 | 0.68 | 0.508 | -0.424 | -0.901 | -0.008 | -0.062 | -0.062 | 0 | 0.2 | 0.25 | 0.11 | 0 | 0.13 | 0.05 |
| 8 | 22698090 | rs2063688 | 0.67 | 0.423 | 0.124 | -1.052 | 0.002 | -0.091 | -0.091 | 0.012 | 0.19 | 0.31 | 0.11 | 0.01 | 0.17 | 0.09 |
| 8 | 22698106 | rs4273853 | -0.132 | 0.421 | 0.005 | -1.036 | -0.003 | 0.003 | 0.003 | 0 | 0.37 | 0.35 | 0.38 | 0.34 | 0.4 | 0.21 |
| 8 | 22698209 | rs11986200 | 0.809 | 0.347 | 0.056 | -1.023 | -0.005 | 0.03 | 0.03 | 0 | 0.44 | 0.26 | 0.51 | 0.65 | 0.45 | 0.72 |
| 8 | 22699219 | rs1040053 | -0.308 | 0.525 | 0.108 | -1.07 | -0.007 | 0.082 | 0.082 | 0 | 0.8 | 0.61 | 0.68 | 0.73 | 0.81 | 0.87 |
| 8 | 22703333 | rs11135681 | 0.741 | 0.567 | -0.518 | -1.153 | 0.005 | 0.039 | 0.039 | 0 | 0.55 | 0.25 | 0.55 | 0.69 | 0.6 | 0.76 |
| 8 | 22703557 | rs12676524 | 1.76 | 0.384 | -0.902 | -1.32 | -0.022 | -0.07 | -0.07 | 0.065 | 0.27 | 0.14 | 0.35 | 0.59 | 0.33 | 0.63 |
| 8 | 22707074 | rs13268364 | 1.256 | 0.333 | -0.925 | -1.432 | -0.06 | 0.095 | 0.095 | 0 | 0.26 | 0.02 | 0.19 | 0.1 | 0.25 | 0.11 |
| 8 | 22707565 | rs1553777 | 0.811 | 0.176 | -0.41 | -1.429 | -0.01 | -0.01 | -0.01 | 0.731 | 0.87 | 0.51 | 0.93 | 0.92 | 0.93 | 0.89 |
| 8 | 22708838 | rs7816775 | 0.819 | 0.262 | -0.286 | -1.399 | 0.015 | -0.011 | -0.011 | 0 | 0.86 | 0.45 | 0.91 | 0.92 | 0.91 | 0.9 |
| 8 | 22709663 | rs1876875 | 0.869 | 0.205 | -0.382 | -1.39 | -0.003 | 0.001 | 0.001 | 0 | 0.44 | 0.39 | 0.44 | 0.6 | 0.48 | 0.74 |
| 8 | 22712077 | rs4284046 | 1.498 | 0.115 | -0.099 | -1.354 | 0.002 | -0.06 | -0.06 | 0.573 | 0.17 | 0.21 | 0.09 | 0.01 | 0.15 | 0.09 |
| 8 | 22713012 | rs7007235 | 0.782 | 0.048 | -0.072 | -1.192 | -0.005 | 0.004 | 0.004 | 0 | 0.7 | 0.34 | 0.64 | 0.7 | 0.75 | 0.85 |
| 8 | 22718237 | rs7005929 | 0.506 | -0.305 | 0.073 | -1.114 | -0.027 | 0.037 | 0.037 | 0 | 0.73 | 0.49 | 0.7 | 0.67 | 0.81 | 0.85 |
| 8 | 22720023 | rs1010795 | 0.412 | -0.175 | 0.086 | -1.372 | 0.047 | -0.013 | -0.013 | 0 | 0.91 | 0.69 | 0.95 | 0.97 | 0.94 | 0.89 |
| 8 | 22720683 | rs7823247 | 0.491 | -0.279 | -0.458 | -1.171 | 0.008 | -0.011 | -0.011 | 0.021 | 0.16 | 0.14 | 0.24 | 0.26 | 0.12 | 0.06 |
| 8 | 22730082 | rs2457436 | 0.41 | -0.085 | -0.8 | -1.274 | -0.054 | -0.028 | -0.028 | 0 | 0.29 | 0.06 | 0.32 | 0.25 | 0.3 | 0.15 |
| 8 | 22733383 | rs1104872 | -0.505 | -0.222 | -0.234 | -1.1 | 0.024 | 0.016 | 0.016 | 0.062 | 0.94 | 0.81 | 0.98 | 0.98 | 0.99 | 0.96 |
| 8 | 22736969 | rs11785590 | 1.111 | -0.14 | -1.2 | -1.213 | 0.004 | 0.078 | 0.078 | 0 | 0.22 | 0.06 | 0.17 | 0.08 | 0.19 | 0.09 |
| 8 | 22737971 | rs2466177 | -0.299 | 0.416 | -0.653 | -0.926 | -0.006 | 0.007 | 0.007 | 0 | 0.34 | 0.34 | 0.37 | 0.25 | 0.35 | 0.18 |
| 8 | 22738040 | rs2466176 | -0.328 | 0.434 | -0.677 | -0.917 | -0.002 | 0.002 | 0.002 | 0 | 0.34 | 0.39 | 0.37 | 0.25 | 0.35 | 0.17 |
| 8 | 22740325 | rs11784354 | 0.404 | 0.879 | -1.042 | -0.881 | -0.05 | 0.079 | 0.079 | 0 | 0.2 | 0.09 | 0.1 | 0.01 | 0.2 | 0.15 |
| 8 | 22744108 | rs2466241 | 0.965 | 0.585 | -1.122 | -0.528 | -0.011 | -0.064 | -0.064 | 0.027 | 0.34 | 0.2 | 0.44 | 0.55 | 0.4 | 0.64 |
| 8 | 22744564 | rs2246578 | 0.714 | 0.731 | -0.672 | -0.686 | -0.018 | 0.127 | 0.127 | 0 | 0.52 | 0.25 | 0.61 | 0.79 | 0.57 | 0.7 |
| 8 | 22749961 | rs2457434 | -0.645 | 1.065 | -0.13 | -0.715 | -0.002 | -0.12 | -0.12 | 0.102 | 0.81 | 0.44 | 0.9 | 0.99 | 0.87 | 0.95 |
| 8 | 22750571 | rs1858192 | -1.301 | 1.086 | 0.154 | -0.696 | 0.018 | -0.117 | -0.117 | 0 | 0.85 | 0.58 | 0.93 | 0.99 | 0.88 | 0.96 |
| 8 | 22751594 | rs13270026 | 0.012 | 1.187 | -0.827 | -0.526 | -0.036 | 0.008 | 0.008 | 0 | 0.23 | 0.12 | 0.27 | 0.17 | 0.26 | 0.2 |
| 8 | 22756126 | rs11135684 | 0.392 | 0.715 | -0.083 | -0.537 | 0.01 | -0.117 | -0.117 | 0.342 | 0.17 | 0.46 | 0.1 | 0.01 | 0.13 | 0.05 |
| 8 | 22761463 | rs17088737 | 0.435 | 0.917 | -0.434 | 0.281 | -0.012 | -0.031 | -0.031 | 0.506 | 0.17 | 0.07 | 0.23 | 0.2 | 0.17 | 0.13 |
| 8 | 22763248 | rs6557600 | -0.072 | 1.134 | -0.341 | 0.309 | 0.002 | 0.003 | 0.003 | 0.2 | 0.2 | 0.29 | 0.17 | 0.22 | 0.22 | 0.1 |
| 8 | 22766549 | rs2466208 | 0.894 | 0.825 | -0.222 | 0.129 | 0 | 0.015 | 0.015 | 0.009 | 0.5 | 0.25 | 0.53 | 0.52 | 0.53 | 0.68 |
| 8 | 22770385 | rs17088769 | 0.102 | 0.843 | -0.185 | 0.149 | -0.011 | -0.017 | -0.017 | 0.258 | 0.21 | 0.1 | 0.14 | 0.25 | 0.24 | 0.12 |
| 8 | 22770526 | rs13258613 | -1.274 | 0.952 | 0.289 | 0.164 | -0.001 | 0.012 | 0.012 | 0 | 0.49 | 0.65 | 0.46 | 0.48 | 0.47 | 0.31 |
| 8 | 22771004 | rs2457422 | -0.775 | 0.982 | 0.34 | 0.146 | -0.012 | -0.015 | -0.015 | 0 | 0.39 | 0.31 | 0.38 | 0.47 | 0.42 | 0.23 |
| 8 | 22772384 | rs17757261 | 0.374 | 1.135 | -0.15 | 0.131 | -0.021 | -0.017 | -0.017 | 0.012 | 0.22 | 0.13 | 0.24 | 0.21 | 0.21 | 0.21 |
| 8 | 22774103 | rs7829314 | -1.176 | 1.107 | -0.091 | 0.171 | 0.01 | -0.01 | -0.01 | 0 | 0.54 | 0.66 | 0.46 | 0.45 | 0.52 | 0.38 |
| 8 | 22774110 | rs7817198 | 0.188 | 1.015 | 0.022 | 0.12 | -0.004 | -0.035 | -0.035 | 0 | 0.09 | 0.22 | 0.07 | 0.02 | 0.07 | 0.08 |
| 8 | 22774668 | rs2466213 | -0.421 | 1.221 | -0.706 | 0.474 | -0.006 | 0.002 | 0.002 | 0.172 | 0.47 | 0.65 | 0.52 | 0.41 | 0.44 | 0.43 |
| 8 | 22775716 | rs2466216 | 0.9 | 1.534 | -1.137 | 0.273 | -0.066 | -0.01 | -0.01 | 0 | 0.13 | 0.01 | 0.11 | 0.08 | 0.2 | 0.05 |
| 8 | 22776796 | rs10216634 | 1.173 | 1.304 | -1.023 | 0.289 | 0 | 0 | 0 | 0.065 | 0.18 | 0.2 | 0.16 | 0.2 | 0.19 | 0.2 |
| 8 | 22778646 | rs2466219 | -0.521 | 1.236 | -1.298 | 0.109 | 0.026 | 0.054 | 0.054 | 0 | 0.54 | 0.36 | 0.52 | 0.76 | 0.64 | 0.54 |
| 8 | 22782633 | rs11135686 | 1.506 | 1.408 | -1.96 | 0.673 | -0.046 | -0.092 | -0.092 | 0 | 0.14 | 0.02 | 0.13 | 0.37 | 0.14 | 0.15 |
| 8 | 22782783 | rs17088800 | 0.546 | 1.304 | -2.14 | 0.439 | -0.03 | -0.054 | -0.054 | 0 | 0.31 | 0.05 | 0.18 | 0.35 | 0.28 | 0.3 |
| 8 | 22784735 | rs11135687 | 2.019 | 0.878 | -2.105 | 0.177 | -0.007 | -0.002 | -0.002 | 0 | 0.17 | 0.16 | 0.23 | 0.15 | 0.13 | 0.27 |
| 8 | 22785386 | rs2466225 | -0.671 | 1.531 | -2.497 | 0.657 | -0.029 | -0.006 | -0.006 | 0 | 0.39 | 0.23 | 0.37 | 0.58 | 0.46 | 0.32 |
| 8 | 22785401 | rs17685420 | 1.319 | 1.533 | -2.508 | 0.645 | 0.002 | -0.006 | -0.006 | 0 | 0.21 | 0.04 | 0.14 | 0.2 | 0.18 | 0.26 |
| 8 | 22786150 | rs2457426 | -0.69 | 1.492 | -2.491 | 0.687 | -0.031 | -0.006 | -0.006 | 0 | 0.39 | 0.24 | 0.36 | 0.58 | 0.45 | 0.32 |
| 8 | 22788370 | rs7846053 | 1.076 | 1.461 | -2.375 | 0.648 | 0.001 | 0.002 | 0.002 | 0.038 | 0.39 | 0.58 | 0.49 | 0.22 | 0.36 | 0.42 |
| 8 | 22793451 | rs12550367 | 1.206 | 1.618 | -2.583 | 0.799 | 0.008 | 0.022 | 0.022 | 0.129 | 0.4 | 0.65 | 0.5 | 0.29 | 0.36 | 0.43 |
| 8 | 22793954 | rs9644027 | 1.071 | 1.566 | -2.578 | 0.804 | 0.007 | 0.022 | 0.022 | 0.088 | 0.4 | 0.65 | 0.48 | 0.29 | 0.36 | 0.42 |
| 8 | 22794462 | rs7012193 | 1.305 | 1.657 | -2.653 | 0.467 | -0.005 | -0.003 | -0.003 | 0 | 0.11 | 0.1 | 0.07 | 0.13 | 0.13 | 0.07 |
| 8 | 22796318 | rs6557604 | -1.085 | 1.622 | -2.837 | 0.478 | 0.007 | 0.008 | 0.008 | 0 | 0.77 | 0.91 | 0.84 | 0.77 | 0.8 | 0.74 |
| 8 | 22797597 | rs1006333 | -0.157 | 1.974 | -2.233 | 1.065 | -0.023 | -0.079 | -0.079 | 0.751 | 0.25 | 0.09 | 0.22 | 0.41 | 0.31 | 0.23 |
| 8 | 22797720 | rs6986719 | 0.515 | 1.608 | -2.373 | 0.884 | -0.019 | -0.011 | -0.011 | 0 | 0.11 | 0.06 | 0.07 | 0.13 | 0.13 | 0.08 |
| 8 | 22801913 | rs2466206 | -0.603 | 1.285 | -2.111 | 1.141 | 0.007 | -0.009 | -0.009 | 0 | 0.1 | 0.09 | 0.1 | 0.09 | 0.12 | 0.13 |

| | | | | | | | | | | | | | | | | |
|---|---|---|---|---|---|---|---|---|---|---|---|---|---|---|---|---|
| 8 | 22802380 | rs1123828 | -0.635 | 1.553 | -2.246 | 1.284 | -0.066 | -0.088 | -0.088 | 0.039 | 0.83 | 0.97 | 0.84 | 0.64 | 0.78 | 0.86 |
| 8 | 22806848 | rs7010858 | 0.488 | 1.446 | -2.084 | 0.717 | -0.03 | -0.007 | -0.007 | 0 | 0.63 | 0.69 | 0.65 | 0.4 | 0.56 | 0.66 |
| 8 | 22816684 | rs2466195 | 0.938 | 1.08 | -2.046 | 0.471 | -0.008 | 0.004 | 0.004 | 0.07 | 0.81 | 0.85 | 0.81 | 0.81 | 0.79 | 0.78 |
| 8 | 22818588 | rs2466238 | 0.691 | 0.848 | -1.957 | 0.295 | 0.003 | 0.013 | 0.013 | 0.276 | 0.92 | 0.96 | 0.91 | 0.95 | 0.91 | 0.84 |
| 8 | 22819138 | rs17686010 | 0.322 | 0.617 | -2.565 | 0.074 | -0.018 | -0.008 | -0.008 | 0 | 0.21 | 0.02 | 0.1 | 0.17 | 0.17 | 0.16 |
| 8 | 22830059 | rs6989664 | -0.025 | 0.471 | -2.039 | -0.276 | 0.007 | 0.011 | 0.011 | 0.079 | 0.36 | 0.62 | 0.41 | 0.67 | 0.36 | 0.45 |
| 8 | 22831150 | rs11784679 | 0.845 | 0.683 | -2.7 | -0.818 | -0.07 | 0.098 | 0.098 | 0 | 0.23 | 0.03 | 0.13 | 0.09 | 0.23 | 0.09 |
| 8 | 22832115 | rs17088859 | 1.207 | 0.439 | -2.26 | -0.7 | -0.01 | -0.029 | -0.029 | 0.295 | 0.22 | 0.16 | 0.28 | 0.53 | 0.29 | 0.26 |
| 8 | 22837121 | rs2430802 | 0.981 | 0.186 | -2.329 | -0.876 | -0.044 | 0.036 | 0.036 | 0 | 0.29 | 0.14 | 0.35 | 0.18 | 0.31 | 0.28 |
| 8 | 22837613 | rs2466235 | -0.024 | -0.102 | -1.461 | -0.741 | -0.014 | -0.029 | -0.029 | 0.105 | 0.88 | 0.79 | 0.81 | 0.94 | 0.88 | 0.82 |
| 8 | 22844211 | rs2466212 | 0.513 | -0.158 | -1.598 | -0.278 | 0.018 | -0.028 | -0.028 | 0 | 0.82 | 0.88 | 0.77 | 0.72 | 0.82 | 0.62 |
| 8 | 22844981 | rs2251588 | 0.983 | -0.17 | -0.707 | -0.627 | -0.01 | -0.056 | -0.056 | 0 | 0.25 | 0.34 | 0.31 | 0.14 | 0.19 | 0.29 |
| 8 | 22846335 | rs13251475 | 1.6 | 0.158 | -1.042 | -0.715 | -0.063 | -0.128 | -0.128 | 3.643 | 0.23 | 0.05 | 0.28 | 0.42 | 0.3 | 0.23 |
| 8 | 22849422 | rs7826882 | 0.397 | -0.259 | -0.662 | -0.71 | 0.055 | 0.047 | 0.047 | 0 | 0.53 | 0.31 | 0.45 | 0.56 | 0.62 | 0.33 |
| 8 | 22851074 | rs2466189 | 0.127 | 0.038 | -0.103 | -0.564 | 0.001 | -0.003 | -0.003 | 0.279 | 0.87 | 0.87 | 0.79 | 0.95 | 0.87 | 0.8 |
| 8 | 22859307 | rs2430807 | 0.696 | 0.372 | -0.227 | -0.624 | 0.031 | 0.112 | 0.112 | 0.265 | 0.55 | 0.24 | 0.72 | 0.68 | 0.65 | 0.58 |
| 8 | 22861242 | rs2430809 | -0.859 | 0.295 | 0.072 | -0.595 | 0.004 | 0.023 | 0.023 | 0 | 0.87 | 0.74 | 0.86 | 0.79 | 0.85 | 0.65 |
| 8 | 22863247 | rs2430813 | 0.707 | 0.387 | 0.101 | -0.649 | -0.022 | -0.045 | -0.045 | 0 | 0.73 | 0.49 | 0.83 | 0.79 | 0.78 | 0.64 |
| 8 | 22864622 | rs2466178 | -0.09 | 0.564 | -0.355 | -0.566 | 0.006 | -0.028 | -0.028 | 0.094 | 0.88 | 0.86 | 0.89 | 0.95 | 0.85 | 0.86 |
| 8 | 22865866 | rs2430816 | -0.175 | 0.573 | -0.469 | -0.562 | 0.002 | -0.017 | -0.017 | 0 | 0.88 | 0.89 | 0.89 | 0.95 | 0.85 | 0.86 |
| 8 | 22873533 | rs876435 | -0.354 | 0.279 | -0.681 | -0.332 | -0.002 | 0.03 | 0.03 | 0 | 0.66 | 0.62 | 0.5 | 0.53 | 0.59 | 0.59 |
| 8 | 22873601 | rs888342 | 1.205 | 0.496 | -1.778 | -0.26 | -0.044 | -0.015 | -0.015 | 0 | 0.3 | 0.13 | 0.34 | 0.28 | 0.33 | 0.3 |
| 8 | 22875985 | rs2241260 | 0.935 | 0.606 | -1.692 | -0.014 | -0.004 | -0.057 | -0.057 | 0 | 0.26 | 0.14 | 0.39 | 0.46 | 0.32 | 0.28 |
| 8 | 22876739 | rs2241261 | 0.184 | 0.759 | -2.207 | -0.038 | 0.033 | 0.051 | 0.051 | 0 | 0.51 | 0.36 | 0.45 | 0.37 | 0.52 | 0.36 |
| 8 | 22892274 | rs11785599 | 0.017 | 1.172 | -2.106 | 0.749 | -0.09 | -0.081 | -0.081 | 0 | 0.43 | 0.12 | 0.42 | 0.44 | 0.5 | 0.35 |
| 8 | 22898404 | rs7834266 | -0.633 | 1.507 | -1.117 | 1.264 | 0.023 | 0.045 | 0.045 | 0 | 0.29 | 0.38 | 0.26 | 0.49 | 0.38 | 0.28 |
| 8 | 22900949 | rs1001793 | -0.436 | 0.475 | -2.053 | 1.313 | 0.05 | 0.061 | 0.061 | 0 | 0.38 | 0.78 | 0.22 | 0.28 | 0.3 | 0.24 |
| 8 | 22901202 | rs1001792 | -0.444 | 0.573 | -2.065 | 1.281 | 0.048 | 0.073 | 0.073 | 0 | 0.39 | 0.81 | 0.23 | 0.28 | 0.32 | 0.25 |
| 8 | 22901568 | rs4259415 | -0.959 | 0.44 | -2.798 | 1.143 | -0.075 | 0.091 | 0.091 | 0.267 | 0.59 | 0.92 | 0.58 | 0.77 | 0.51 | 0.46 |
| 8 | 22911073 | rs12677679 | 0.508 | -0.049 | -2.413 | 0.935 | 0.002 | 0.015 | 0.015 | 0 | 0.23 | 0.18 | 0.15 | 0.22 | 0.26 | 0.15 |
| 8 | 22919322 | rs4424253 | 0.126 | 0.027 | -1.895 | 1.148 | -0.001 | 0.01 | 0.01 | 2.128 | 0.21 | 0.16 | 0.33 | 0.49 | 0.18 | 0.27 |
| 8 | 22920149 | rs9644062 | 0.08 | 0.69 | -2.144 | 0.935 | 0.039 | 0.041 | 0.041 | 0.143 | 0.1 | 0.17 | 0.09 | 0.16 | 0.15 | 0.22 |
| 8 | 22924031 | rs4872047 | 0.348 | 0.604 | -2.354 | 0.961 | -0.01 | -0.093 | -0.093 | 0.588 | 0.2 | 0.06 | 0.33 | 0.51 | 0.18 | 0.28 |
| 8 | 22925154 | rs11135693 | -1.562 | 0.377 | -1.769 | 0.428 | -0.019 | 0.079 | 0.079 | 0 | 0.66 | 0.81 | 0.65 | 0.92 | 0.66 | 0.74 |
| 8 | 22929350 | rs11778166 | 1.252 | -0.073 | -1.706 | 0.329 | -0.034 | 0.077 | 0.077 | 0 | 0.39 | 0.23 | 0.27 | 0.05 | 0.42 | 0.3 |
| 8 | 22932122 | rs11784599 | -1.021 | -0.009 | -1.804 | 0.526 | -0.024 | 0.012 | 0.012 | 0 | 0.81 | 0.99 | 0.89 | 0.82 | 0.79 | 0.92 |
| 8 | 22944695 | rs7830593 | -0.349 | -0.087 | -1.252 | 1.008 | 0 | 0.015 | 0.015 | 0.222 | 0.12 | 0.13 | 0.1 | 0.27 | 0.16 | 0.26 |
| 8 | 22950389 | rs12678837 | 0.348 | -0.269 | -1.45 | 1.443 | -0.005 | -0.064 | -0.064 | 0.382 | 0.11 | 0.02 | 0.23 | 0.35 | 0.1 | 0.14 |
| 8 | 22954194 | rs11781924 | 1.223 | -0.23 | -1.425 | 0.764 | -0.001 | -0.024 | -0.024 | 0 | 0.07 | 0.01 | 0.21 | 0.08 | 0.06 | 0.06 |
| 8 | 22954205 | rs4518666 | -1.23 | -0.045 | -0.724 | 0.896 | -0.006 | -0.008 | -0.008 | 0 | 0.38 | 0.55 | 0.38 | 0.68 | 0.32 | 0.58 |
| 8 | 22957606 | rs7009522 | -0.199 | 0.725 | -0.638 | 1.162 | -0.018 | -0.001 | -0.001 | 0 | 0.8 | 0.96 | 0.87 | 0.8 | 0.77 | 0.89 |
| 8 | 22962553 | rs12546238 | -0.579 | 0.917 | -0.532 | 1.803 | -0.011 | -0.001 | -0.001 | 0 | 0.07 | 0.04 | 0.19 | 0.32 | 0.11 | 0.15 |
| 8 | 22963602 | rs12681965 | -0.082 | 0.965 | -0.534 | 1.912 | -0.027 | -0.003 | -0.003 | 1.118 | 0.08 | 0.05 | 0.15 | 0.29 | 0.16 | 0.17 |
| 8 | 22966200 | rs11779793 | -0.609 | 0.972 | -0.138 | 1.877 | 0.007 | 0.089 | 0.089 | 0.36 | 0.08 | 0.15 | 0.16 | 0.33 | 0.08 | 0.16 |
| 8 | 22966769 | rs12550612 | 0.164 | 0.884 | -0.024 | 1.75 | -0.004 | -0.003 | -0.003 | 0 | 0.86 | 0.86 | 0.89 | 0.8 | 0.8 | 0.86 |
| 8 | 22967438 | rs9314261 | -1.058 | 0.652 | 0.551 | 1.517 | -0.036 | 0.07 | 0.07 | 2.174 | 0.23 | 0.62 | 0.45 | 0.46 | 0.16 | 0.28 |
| 8 | 22968219 | rs11785284 | -0.423 | 1.207 | 0.26 | 1.628 | 0.008 | 0.077 | 0.077 | 0.37 | 0.08 | 0.14 | 0.15 | 0.32 | 0.08 | 0.15 |
| 8 | 22969091 | rs10111172 | 0.226 | 1.16 | 0.16 | 1.527 | -0.022 | -0.01 | -0.01 | 0 | 0.87 | 0.92 | 0.9 | 0.82 | 0.8 | 0.86 |
| 8 | 22977170 | rs10866820 | -0.991 | 0.581 | 0.937 | 1.633 | 0.108 | 0.297 | 0.297 | 1.132 | 0.11 | 0.36 | 0.19 | 0.32 | 0.17 | 0.17 |
| 8 | 22985399 | rs4242390 | -0.717 | 0.636 | 0.351 | 1.46 | 0.009 | 0.103 | 0.103 | 0.667 | 0.1 | 0.23 | 0.17 | 0.29 | 0.09 | 0.06 |
| 8 | 22987837 | rs7843320 | -0.755 | 0.696 | -0.095 | 1.101 | -0.034 | -0.059 | -0.059 | 0 | 0.13 | 0.05 | 0.19 | 0.4 | 0.19 | 0.28 |
| 8 | 22989954 | rs4872058 | -0.802 | 0.63 | 0.1 | 1.035 | -0.002 | 0.014 | 0.014 | 0.942 | 0.16 | 0.12 | 0.19 | 0.4 | 0.18 | 0.29 |
| 8 | 22996779 | rs4278155 | -0.909 | -0.09 | 0.078 | 0.904 | -0.018 | 0.033 | 0.033 | 0.127 | 0.24 | 0.3 | 0.41 | 0.46 | 0.18 | 0.21 |
| 8 | 23000183 | rs4242391 | 0.597 | 0.901 | -0.019 | -0.064 | -0.003 | 0.033 | 0.033 | 0 | 0.63 | 0.32 | 0.7 | 0.88 | 0.59 | 0.79 |
| 8 | 23001988 | rs1133782 | 0.509 | 1.082 | -0.116 | -0.05 | 0.011 | -0.094 | -0.094 | 0 | 0.66 | 0.55 | 0.71 | 0.91 | 0.6 | 0.79 |
| 8 | 23006176 | rs4871850 | 0.266 | 1.113 | -0.401 | 0.714 | 0.001 | 0.033 | 0.033 | 0 | 0.7 | 0.46 | 0.55 | 0.41 | 0.72 | 0.58 |
| 8 | 23007178 | rs13257094 | -0.029 | 1.309 | -1.266 | 0.409 | -0.052 | 0.076 | 0.076 | 0 | 0.27 | 0.02 | 0.26 | 0.1 | 0.25 | 0.19 |
| 8 | 23007793 | rs7464769 | -1.231 | 1.61 | 0.077 | 1.685 | 0.083 | 0.34 | 0.34 | 0 | 0.08 | 0.23 | 0.32 | 0.57 | 0.12 | 0.14 |
| 8 | 23010298 | rs7462795 | -0.937 | 1.663 | -0.262 | 1.649 | 0.054 | 0.267 | 0.267 | 0 | 0.07 | 0.17 | 0.32 | 0.53 | 0.12 | 0.12 |
| 8 | 23041082 | rs4871854 | -0.304 | 1.41 | -0.824 | 0.233 | -0.009 | 0.007 | 0.007 | 0 | 0.29 | 0.23 | 0.36 | 0.73 | 0.35 | 0.4 |
| 8 | 23049292 | rs2230229 | -0.966 | 1.814 | -0.865 | -0.321 | 0.005 | -0.015 | -0.015 | 0 | 0.9 | 0.88 | 0.9 | 0.98 | 0.86 | 0.95 |
| 8 | 23051485 | rs11775256 | 0.768 | 1.534 | -1.372 | -1.211 | -0.006 | 0.065 | 0.065 | 0 | 0.24 | 0.11 | 0.19 | 0.05 | 0.24 | 0.14 |
| 8 | 23054779 | rs3824218 | -0.545 | 1.687 | -1.484 | -1.41 | -0.009 | 0.052 | 0.052 | 0 | 0.69 | 0.8 | 0.74 | 0.9 | 0.66 | 0.81 |
| 8 | 23055729 | rs3808530 | 0.396 | 1.551 | -2.028 | -1.447 | -0.035 | 0.049 | 0.049 | 0 | 0.25 | 0.09 | 0.25 | 0.14 | 0.23 | 0.38 |
| 8 | 23057181 | rs2235126 | 1.038 | 1.338 | -1.761 | -1.222 | -0.009 | -0.028 | -0.028 | 0.18 | 0.23 | 0.17 | 0.32 | 0.69 | 0.27 | 0.33 |
| 8 | 23058188 | rs4872077 | 0.48 | 1.125 | -1.3 | -2.006 | 0.005 | -0.066 | -0.066 | 0.036 | 0.28 | 0.33 | 0.36 | 0.06 | 0.31 | 0.21 |
| 8 | 23058220 | rs20576 | 0.896 | 0.981 | -2.012 | -2.162 | -0.073 | 0.095 | 0.095 | 0 | 0.16 | 0.01 | 0.14 | 0.01 | 0.22 | 0.07 |
| 8 | 23061792 | rs6982233 | 1.716 | 1.026 | -1.872 | -1.901 | -0.001 | -0.004 | -0.004 | 1.62 | 0.2 | 0.22 | 0.28 | 0.67 | 0.24 | 0.31 |
| 8 | 23061878 | rs6995408 | -0.217 | 0.781 | -1.842 | -2.264 | -0.005 | 0.073 | 0.073 | 0.354 | 0.52 | 0.35 | 0.62 | 0.97 | 0.47 | 0.69 |

| | | | | | | | | | | | | | | | | |
|---|---|---|---|---|---|---|---|---|---|---|---|---|---|---|---|---|
| 8 | 23072256 | rs11785328 | 1.141 | 0.594 | -1.863 | -2.184 | -0.016 | 0.157 | 0.157 | 1.827 | 0.27 | 0.05 | 0.21 | 0.01 | 0.3 | 0.2 |
| 8 | 23075040 | rs4532600 | 1.027 | 0.858 | -1.379 | -1.601 | 0.003 | 0.01 | 0.01 | 0 | 0.21 | 0.27 | 0.27 | 0.67 | 0.26 | 0.32 |
| 8 | 23083546 | rs13255394 | 0.36 | 0.592 | -1.856 | -1.862 | -0.048 | 0.066 | 0.066 | 0.114 | 0.59 | 0.14 | 0.56 | 0.37 | 0.54 | 0.71 |
| 8 | 23085868 | rs4872091 | 0.14 | 0.541 | -1.171 | -1.977 | 0.005 | -0.073 | -0.073 | 0 | 0.72 | 0.33 | 0.81 | 0.98 | 0.74 | 0.92 |
| 8 | 23088245 | rs4242394 | 0.158 | 0.542 | -0.978 | -1.977 | 0.009 | -0.096 | -0.096 | 0 | 0.72 | 0.35 | 0.81 | 0.98 | 0.74 | 0.91 |
| 8 | 23094395 | rs11997323 | 0.329 | 1.167 | -1.037 | -1.864 | -0.048 | 0.061 | 0.061 | 0 | 0.18 | 0.07 | 0.14 | 0.01 | 0.24 | 0.07 |
| 8 | 23094417 | rs4872092 | -1.295 | 0.678 | -0.647 | -2.172 | -0.027 | -0.016 | -0.016 | 0 | 0.86 | 0.7 | 0.89 | 0.87 | 0.91 | 0.97 |
| 8 | 23100963 | rs4242397 | -0.179 | 0.595 | -1.196 | -2.264 | -0.029 | -0.051 | -0.051 | 0 | 0.78 | 0.39 | 0.84 | 0.88 | 0.86 | 0.93 |
| 8 | 23101620 | rs7463256 | 1.48 | 0.3 | -2.026 | -1.188 | -0.007 | 0.054 | 0.054 | 0.134 | 0.4 | 0.11 | 0.47 | 0.37 | 0.41 | 0.48 |
| 8 | 23104195 | rs2294123 | 1.495 | 0.298 | -2.027 | -1.189 | -0.008 | 0.053 | 0.053 | 0.128 | 0.4 | 0.11 | 0.47 | 0.37 | 0.41 | 0.48 |
| 8 | 23104612 | rs6981400 | 0.596 | 0.406 | -1.767 | -0.948 | -0.003 | 0.024 | 0.024 | 0 | 0.19 | 0.21 | 0.25 | 0.49 | 0.18 | 0.34 |
| 8 | 23107578 | rs12544038 | 0.498 | 0.594 | -2.181 | -0.924 | 0.025 | 0.097 | 0.097 | 0 | 0.54 | 0.13 | 0.61 | 0.38 | 0.63 | 0.53 |
| 8 | 23112662 | rs3736025 | -0.295 | 0.766 | -1.93 | -0.434 | -0.019 | -0.004 | -0.004 | 0.594 | 0.35 | 0.29 | 0.4 | 0.51 | 0.45 | 0.45 |
| 8 | 23113908 | rs4439140 | 0.68 | 0.73 | -2.479 | 0.463 | -0.08 | -0.024 | -0.024 | 0 | 0.33 | 0.07 | 0.37 | 0.33 | 0.41 | 0.37 |
| 8 | 23115646 | rs3736022 | -0.617 | 0.637 | -1.917 | 1.433 | -0.02 | 0.02 | 0.02 | 0 | 0.06 | 0.01 | 0.14 | 0.33 | 0.1 | 0.07 |
| 8 | 23116392 | rs4581063 | -1.336 | 0.656 | -2.057 | 1.421 | -0.026 | -0.046 | -0.046 | 0 | 0.08 | 0.01 | 0.14 | 0.18 | 0.12 | 0.07 |
| 8 | 23116431 | rs4872098 | 0.89 | 0.732 | -1.429 | 1.013 | 0.01 | 0.004 | 0.004 | 0 | 0.8 | 0.67 | 0.8 | 0.8 | 0.77 | 0.82 |
| 8 | 23117643 | rs2294125 | 0.855 | 0.314 | -2.405 | 0.39 | -0.127 | 0.329 | 0.329 | 0.362 | 0.42 | 0.05 | 0.4 | 0.06 | 0.46 | 0.4 |
| 8 | 23118155 | rs9828 | 0.922 | 0.33 | -2.403 | 0.392 | -0.13 | 0.296 | 0.296 | 0.225 | 0.4 | 0.05 | 0.39 | 0.06 | 0.44 | 0.37 |
| 8 | 23118503 | rs12194 | 0.922 | 0.33 | -2.403 | 0.392 | -0.13 | 0.296 | 0.296 | 0.225 | 0.4 | 0.05 | 0.39 | 0.06 | 0.44 | 0.37 |
| 8 | 23119205 | rs7361 | -0.243 | 0.846 | -1.626 | 1.077 | -0.008 | 0.024 | 0.024 | 0.962 | 0.13 | 0.24 | 0.09 | 0.15 | 0.09 | 0.32 |
| 8 | 23120358 | rs11778746 | 0.303 | 0.658 | -1.771 | 0.097 | -0.1 | 0.129 | 0.129 | 0 | 0.25 | 0.02 | 0.19 | 0.04 | 0.31 | 0.1 |
| 8 | 23131396 | rs10095543 | 0.429 | -0.04 | -0.443 | 0.947 | -0.067 | 0.143 | 0.143 | 1.442 | 0.83 | 0.56 | 0.8 | 0.41 | 0.95 | 0.68 |
| 8 | 23134374 | rs7836184 | 0.15 | 0.535 | -0.539 | 0.316 | 0.014 | -0.021 | -0.021 | 0 | 0.17 | 0.21 | 0.12 | 0.07 | 0.22 | 0.15 |
| 8 | 23135398 | rs11786959 | -0.343 | 0.351 | -0.376 | 0.684 | -0.02 | 0.021 | 0.021 | 0.027 | 0.94 | 0.9 | 0.98 | 0.89 | 0.98 | 0.94 |
| 8 | 23139436 | rs10091786 | -1.342 | 0.439 | -0.93 | -0.071 | -0.014 | 0.098 | 0.098 | 0.115 | 0.68 | 0.86 | 0.67 | 0.96 | 0.66 | 0.66 |
| 8 | 23140413 | rs7826635 | 0.871 | 0.338 | -0.8 | -0.055 | -0.036 | 0.115 | 0.115 | 0 | 0.28 | 0.09 | 0.25 | 0.06 | 0.31 | 0.12 |
| 8 | 23142869 | rs9918794 | -1.573 | 0.333 | -0.884 | -0.156 | -0.007 | 0.057 | 0.057 | 0 | 0.73 | 0.85 | 0.69 | 0.96 | 0.7 | 0.69 |
| 8 | 23148930 | rs2272762 | -0.476 | 0.27 | -0.19 | 0.111 | -0.001 | -0.016 | -0.016 | 0 | 0.58 | 0.43 | 0.46 | 0.8 | 0.54 | 0.56 |
| 8 | 23148940 | rs2272761 | -0.406 | 0.245 | -0.203 | 0.098 | -0.001 | -0.012 | -0.012 | 0 | 0.58 | 0.43 | 0.46 | 0.8 | 0.54 | 0.55 |
| 8 | 23151236 | rs3924018 | 1.305 | 0.189 | -0.257 | 0.335 | -0.026 | 0.032 | 0.032 | 0 | 0.27 | 0.08 | 0.27 | 0.19 | 0.3 | 0.12 |
| 8 | 23153022 | rs1002792 | -0.682 | 0.175 | -0.028 | 0.326 | 0 | -0.056 | -0.056 | 0 | 0.59 | 0.52 | 0.46 | 0.8 | 0.54 | 0.55 |
| 8 | 23153260 | rs1002791 | -0.137 | 0.172 | -0.116 | 0.325 | -0.002 | 0.003 | 0.003 | 0 | 0.32 | 0.35 | 0.19 | 0.61 | 0.24 | 0.44 |
| 8 | 23154486 | rs7834641 | -0.5 | 0.193 | -0.072 | 0.362 | -0.001 | -0.014 | -0.014 | 0 | 0.58 | 0.43 | 0.46 | 0.8 | 0.54 | 0.56 |
| 8 | 23155130 | rs2280935 | 0.253 | 0.237 | -0.062 | 1.1 | -0.005 | 0.053 | 0.053 | 0.04 | 0.11 | 0.13 | 0.05 | 0.35 | 0.05 | 0.28 |
| 8 | 23156837 | rs2280937 | 0.036 | 0.277 | 0.019 | 1.145 | -0.009 | 0.059 | 0.059 | 0.081 | 0.12 | 0.14 | 0.06 | 0.33 | 0.05 | 0.29 |
| 8 | 23156880 | rs2280939 | 0.051 | 0.226 | 0.137 | 0.62 | -0.001 | 0.006 | 0.006 | 0 | 0.4 | 0.33 | 0.49 | 0.21 | 0.47 | 0.23 |
| 8 | 23158775 | rs3808528 | 0.33 | 0.515 | 0.124 | 0.94 | -0.021 | -0.031 | -0.031 | 0 | 0.23 | 0.19 | 0.28 | 0.45 | 0.31 | 0.23 |
| 8 | 23158812 | rs3808527 | -0.449 | 0.514 | 0.366 | 0.253 | 0.011 | -0.029 | -0.029 | 0.003 | 0.36 | 0.51 | 0.32 | 0.72 | 0.36 | 0.52 |
| 8 | 23160155 | rs17760913 | 1.239 | 0.62 | -0.134 | 0.006 | -0.061 | 0.055 | 0.055 | 0 | 0.2 | 0.03 | 0.18 | 0.09 | 0.23 | 0.09 |
| 8 | 23160187 | rs11135722 | -0.72 | 0.211 | -0.11 | -0.209 | 0.016 | 0.153 | 0.153 | 0 | 0.51 | 0.72 | 0.5 | 0.81 | 0.47 | 0.55 |
| 8 | 23160433 | rs17760943 | 1.158 | 0.672 | 0.099 | 0.292 | -0.073 | 0.047 | 0.047 | 0.012 | 0.2 | 0.03 | 0.19 | 0.09 | 0.26 | 0.09 |
| 8 | 23161204 | rs11135723 | 0.003 | 0.214 | 0.482 | 0.368 | -0.025 | 0.048 | 0.048 | 0 | 0.29 | 0.41 | 0.27 | 0.58 | 0.21 | 0.46 |
| 8 | 23161933 | rs3808526 | -0.493 | 0.235 | 0.551 | 0.275 | 0.004 | 0.013 | 0.013 | 0 | 0.48 | 0.53 | 0.49 | 0.83 | 0.43 | 0.55 |
| 8 | 23162176 | rs3808522 | -0.547 | 0.246 | 0.593 | 0.282 | 0.009 | 0.047 | 0.047 | 0 | 0.49 | 0.57 | 0.5 | 0.83 | 0.43 | 0.56 |
| 8 | 23164367 | rs3808521 | -0.069 | 0.33 | 1.184 | 0.621 | 0.012 | 0.006 | 0.006 | 0.748 | 0.75 | 0.63 | 0.77 | 0.74 | 0.74 | 0.88 |
| 8 | 23167353 | rs1063582 | -0.274 | 0.334 | 1.686 | 0.724 | 0.01 | 0.028 | 0.028 | 0 | 0.78 | 0.67 | 0.82 | 0.74 | 0.8 | 0.9 |
| 8 | 23167701 | rs11135724 | 0.11 | 0.373 | 1.741 | 0.69 | 0.029 | -0.06 | -0.06 | 0 | 0.76 | 0.47 | 0.82 | 0.9 | 0.73 | 0.88 |
| 8 | 23168650 | rs13261574 | -0.526 | 0.272 | 1.344 | 0.702 | 0.009 | -0.03 | -0.03 | 0.171 | 0.8 | 0.77 | 0.87 | 0.9 | 0.8 | 0.91 |
| 8 | 23173314 | rs4288380 | -0.301 | 0.287 | 1.185 | 0.749 | 0.011 | 0.025 | 0.025 | 0 | 0.79 | 0.67 | 0.83 | 0.75 | 0.8 | 0.9 |
| 8 | 23174883 | rs13282766 | -0.091 | 0.037 | 0.959 | 1.31 | -0.014 | 0.067 | 0.067 | 0 | 0.73 | 0.54 | 0.68 | 0.54 | 0.79 | 0.66 |
| 8 | 23175252 | rs13248926 | 0.673 | -0.132 | 0.839 | 1.266 | 0.023 | -0.071 | -0.071 | 0.305 | 0.64 | 0.22 | 0.5 | 0.48 | 0.7 | 0.61 |
| 8 | 23175732 | rs10111946 | -1.494 | -0.059 | 0.816 | 1.36 | 0.007 | 0.123 | 0.123 | 0.136 | 0.09 | 0.35 | 0.21 | 0.22 | 0.08 | 0.06 |
| 8 | 23180616 | rs9792292 | -0.24 | -0.206 | 0.922 | 1.369 | -0.016 | 0.022 | 0.022 | 0 | 0.75 | 0.63 | 0.7 | 0.72 | 0.8 | 0.67 |
| 8 | 23181742 | rs4323477 | 0.774 | -0.29 | 1.042 | 1.401 | 0.028 | -0.066 | -0.066 | 0 | 0.65 | 0.24 | 0.51 | 0.5 | 0.74 | 0.63 |
| 8 | 23182268 | rs11775841 | -1.638 | -0.296 | 1.07 | 1.35 | 0.007 | 0.16 | 0.16 | 0.027 | 0.1 | 0.37 | 0.19 | 0.22 | 0.05 | 0.05 |
| 8 | 23183869 | rs12676713 | 0.607 | -0.214 | 0.484 | 1.289 | 0.022 | -0.054 | -0.054 | 3.185 | 0.65 | 0.26 | 0.51 | 0.52 | 0.73 | 0.62 |
| 8 | 23185202 | rs9792307 | 0.624 | -0.212 | 0.463 | 1.334 | 0.024 | 0.003 | 0.003 | 0.011 | 0.65 | 0.24 | 0.53 | 0.66 | 0.73 | 0.62 |
| 8 | 23185580 | rs17088178 | 0.621 | -0.209 | 0.465 | 1.367 | 0.024 | -0.065 | -0.065 | 0.014 | 0.65 | 0.24 | 0.51 | 0.5 | 0.73 | 0.62 |
| 8 | 23186007 | rs1051146 | 0.119 | -0.101 | 0.582 | 1.452 | -0.008 | 0.028 | 0.028 | 0 | 0.69 | 0.45 | 0.53 | 0.51 | 0.75 | 0.62 |
| 8 | 23190941 | rs1010156 | 0.094 | -0.095 | -0.252 | 1.238 | 0.007 | -0.025 | -0.025 | 0.266 | 0.44 | 0.32 | 0.39 | 0.47 | 0.55 | 0.34 |
| 8 | 23194591 | rs883524 | 0.369 | -0.62 | -0.486 | -0.067 | 0.05 | 0.076 | 0.076 | 0 | 0.27 | 0.14 | 0.19 | 0.07 | 0.18 | 0.37 |
| 8 | 23197978 | rs7828979 | 0.881 | -0.404 | -0.408 | -0.613 | 0.008 | -0.001 | -0.001 | 0 | 0.25 | 0.2 | 0.11 | 0.06 | 0.19 | 0.21 |
| 8 | 23198330 | rs4301460 | -0.116 | -0.21 | -0.736 | -0.331 | 0.006 | -0.01 | -0.01 | 0.355 | 0.34 | 0.68 | 0.43 | 0.41 | 0.3 | 0.23 |
| 8 | 23198732 | rs11781891 | 1.807 | -0.194 | -0.566 | -0.371 | -0.001 | 0.001 | 0.001 | 0.861 | 0.15 | 0.17 | 0.25 | 0.06 | 0.16 | 0.08 |
| 8 | 23201103 | rs7013955 | 0.513 | -0.174 | -0.068 | -0.158 | 0.011 | 0.017 | 0.017 | 0 | 0.07 | 0.16 | 0.05 | 0.11 | 0.08 | 0.14 |
| 8 | 23201313 | rs9785120 | 1.768 | -0.188 | -0.373 | -0.355 | 0.012 | 0.022 | 0.022 | 0.988 | 0.25 | 0.19 | 0.12 | 0.06 | 0.21 | 0.24 |
| 8 | 23202245 | rs11774789 | -0.665 | 0.177 | -0.781 | -0.565 | -0.032 | -0.041 | -0.041 | 0 | 0.64 | 0.79 | 0.63 | 0.47 | 0.57 | 0.56 |
| 8 | 23202401 | rs7832355 | -0.708 | 0.279 | -0.627 | -0.469 | 0.004 | 0.018 | 0.018 | 0 | 0.39 | 0.65 | 0.5 | 0.28 | 0.35 | 0.3 |

| | | | | | | | | | | | | | | | | |
|---|---|---|---|---|---|---|---|---|---|---|---|---|---|---|---|---|
| 8 | 23204355 | rs10866824 | -0.732 | 0.281 | -0.582 | -0.467 | 0.003 | 0.014 | 0.014 | 0 | 0.39 | 0.61 | 0.5 | 0.28 | 0.36 | 0.3 |
| 8 | 23206698 | rs7829982 | -0.91 | 0.211 | -0.673 | -0.518 | -0.007 | 0.009 | 0.009 | 0 | 0.25 | 0.64 | 0.28 | 0.33 | 0.19 | 0.23 |
| 8 | 23207583 | rs7018424 | 1.27 | 0.285 | -0.823 | -0.668 | -0.025 | 0.184 | 0.184 | 0 | 0.4 | 0.15 | 0.36 | 0.12 | 0.4 | 0.33 |
| 8 | 23208792 | rs7010657 | -0.789 | 0.197 | -0.754 | -0.622 | -0.007 | 0.009 | 0.009 | 0.004 | 0.26 | 0.64 | 0.28 | 0.34 | 0.19 | 0.23 |
| 8 | 23212224 | rs954705 | -0.284 | 0.329 | -0.863 | -0.541 | 0.035 | 0.033 | 0.033 | 1.82 | 0.51 | 0.78 | 0.41 | 0.41 | 0.42 | 0.52 |
| 8 | 23218101 | rs7010362 | 1.319 | 0.483 | -0.54 | -0.557 | 0.009 | -0.024 | -0.024 | 0 | 0.06 | 0.05 | 0.08 | 0 | 0.05 | 0.07 |
| 8 | 23218136 | rs6997194 | 0.764 | 0.362 | -0.45 | -0.614 | -0.015 | 0.104 | 0.104 | 0 | 0.46 | 0.3 | 0.52 | 0.22 | 0.46 | 0.41 |
| 8 | 23220509 | rs11782783 | 0.866 | 1.809 | 0.024 | -0.035 | 0.026 | 0.125 | 0.125 | 0.111 | 0.12 | 0.17 | 0.26 | 0.43 | 0.19 | 0.2 |
| 8 | 23223506 | rs6986654 | 0.883 | 1.922 | 0.106 | -0.106 | -0.031 | 0.035 | 0.035 | 0.056 | 0.2 | 0.09 | 0.33 | 0.06 | 0.26 | 0.15 |
| 8 | 23224014 | rs7818494 | -0.732 | 1.822 | 0.285 | -0.144 | -0.045 | 0.029 | 0.029 | 0 | 0.19 | 0.47 | 0.11 | 0.21 | 0.13 | 0.15 |
| 8 | 23224153 | rs7004710 | 0.862 | 1.992 | 0.22 | -0.087 | -0.029 | 0.033 | 0.033 | 0.048 | 0.2 | 0.09 | 0.32 | 0.06 | 0.26 | 0.15 |
| 8 | 23225745 | rs2294133 | 0.19 | 1.82 | 0.511 | -0.006 | 0.017 | 0.099 | 0.099 | 0 | 0.08 | 0.24 | 0.06 | 0.22 | 0.08 | 0.15 |
| 8 | 23226014 | rs6999447 | -0.552 | 2.058 | 0.171 | -0.139 | 0.009 | 0.01 | 0.01 | 0 | 0.24 | 0.27 | 0.44 | 0.55 | 0.34 | 0.29 |
| 8 | 23226464 | rs10087981 | -0.77 | 2.054 | 0.256 | -0.098 | -0.01 | -0.004 | -0.004 | 0 | 0.33 | 0.59 | 0.34 | 0.28 | 0.31 | 0.23 |
| 8 | 23227177 | rs11135728 | -0.319 | 1.343 | 0.423 | 0.033 | -0.02 | 0.049 | 0.049 | 0 | 0.16 | 0.39 | 0.1 | 0.21 | 0.13 | 0.17 |
| 8 | 23230137 | rs6983061 | -0.398 | 1.448 | 0.268 | 0.204 | 0.049 | 0.141 | 0.141 | 0 | 0.2 | 0.38 | 0.3 | 0.46 | 0.28 | 0.21 |
| 8 | 23237547 | rs4493910 | -1.463 | 1.227 | 0.004 | -0.173 | -0.012 | 0.097 | 0.097 | 0.611 | 0.43 | 0.69 | 0.57 | 0.76 | 0.51 | 0.52 |
| 8 | 23237983 | rs7845265 | -1.763 | 1.081 | -0.099 | -0.31 | 0.055 | 0.118 | 0.118 | 0.325 | 0.57 | 0.72 | 0.8 | 0.82 | 0.68 | 0.59 |
| 8 | 23239445 | rs6557665 | -1.393 | 1.026 | 0.203 | -0.372 | 0.068 | 0.118 | 0.118 | 0.756 | 0.51 | 0.71 | 0.76 | 0.84 | 0.63 | 0.58 |
| 8 | 23240061 | rs8180974 | -0.892 | 1.166 | 0.343 | -0.172 | 0.019 | 0.001 | 0.001 | 0.29 | 0.33 | 0.41 | 0.44 | 0.64 | 0.44 | 0.5 |
| 8 | 23240389 | rs17768910 | 1.782 | 0.912 | 0.151 | -0.185 | -0.039 | 0.024 | 0.024 | 0.385 | 0.14 | 0.02 | 0.23 | 0.05 | 0.16 | 0.08 |
| 8 | 23242026 | rs7017737 | -1.318 | 0.974 | 0.233 | -0.123 | -0.021 | 0.095 | 0.095 | 0 | 0.47 | 0.74 | 0.69 | 0.63 | 0.54 | 0.34 |
| 8 | 23245783 | rs10096495 | 0.341 | 1.072 | 0.637 | -0.1 | -0.018 | -0.04 | -0.04 | 0 | 0.09 | 0.15 | 0.07 | 0 | 0.07 | 0.01 |
| 8 | 23249947 | rs7824768 | 0.37 | 1.402 | 0.675 | 0.635 | 0.059 | 0.211 | 0.211 | 0 | 0.1 | 0.23 | 0.25 | 0.27 | 0.15 | 0.12 |
| 8 | 23257034 | rs4871874 | 0.349 | 0.953 | 0.167 | 0.49 | -0.002 | 0 | 0 | 0 | 0.11 | 0.12 | 0.06 | 0.1 | 0.1 | 0.19 |
| 8 | 23259377 | rs17696480 | 0.024 | 0.965 | 0.051 | 0.459 | -0.025 | 0.029 | 0.029 | 0.075 | 0.25 | 0.12 | 0.29 | 0.14 | 0.27 | 0.26 |
| 8 | 23275784 | rs7823585 | -1.091 | 0.81 | 0.333 | 0.439 | 0 | -0.002 | -0.002 | 0 | 0.35 | 0.67 | 0.29 | 0.44 | 0.31 | 0.41 |
| 8 | 23280443 | rs4872131 | -1.091 | 0.794 | 0.331 | 0.437 | 0 | -0.002 | -0.002 | 0 | 0.35 | 0.67 | 0.29 | 0.44 | 0.31 | 0.41 |
| 8 | 23282951 | rs11779953 | 1.644 | 0.802 | 0.224 | 0.375 | -0.043 | 0.032 | 0.032 | 0.121 | 0.15 | 0.01 | 0.23 | 0.06 | 0.18 | 0.08 |
| 8 | 23283300 | rs17089234 | 0.576 | 0.787 | 0.292 | 0.509 | 0.021 | 0.136 | 0.136 | 0 | 0.06 | 0.15 | 0.21 | 0.13 | 0.04 | 0.09 |
| 8 | 23288895 | rs7842651 | -1.018 | 0.79 | 0.308 | 0.296 | 0 | -0.001 | -0.001 | 2.424 | 0.35 | 0.67 | 0.29 | 0.44 | 0.31 | 0.41 |
| 8 | 23289536 | rs7843529 | -1.464 | 0.779 | 0.285 | 0.383 | -0.009 | 0.078 | 0.078 | 0.619 | 0.41 | 0.83 | 0.5 | 0.71 | 0.37 | 0.5 |
| 8 | 23292136 | rs12678121 | -0.993 | 0.771 | 0.734 | 0.384 | 0.007 | 0.023 | 0.023 | 0 | 0.33 | 0.53 | 0.27 | 0.44 | 0.31 | 0.4 |
| 8 | 23292985 | rs3736652 | 0.943 | 0.856 | 0.92 | 0.427 | 0.011 | 0.002 | 0.002 | 0 | 0.1 | 0.17 | 0.07 | 0.12 | 0.1 | 0.19 |
| 8 | 23294761 | rs2272641 | 0.471 | 0.789 | 0.891 | 0.411 | 0.017 | 0.266 | 0.266 | 0 | 0.06 | 0.16 | 0.21 | 0.27 | 0.06 | 0.09 |
| 8 | 23301340 | rs11787224 | -0.388 | 0.752 | 0.946 | 0.355 | 0.002 | 0.007 | 0.007 | 0 | 0.25 | 0.29 | 0.23 | 0.44 | 0.26 | 0.4 |
| 8 | 23307253 | rs2293936 | 1.654 | 0.752 | 0.915 | 0.331 | -0.014 | 0.034 | 0.034 | 0.041 | 0.15 | 0.04 | 0.23 | 0.06 | 0.18 | 0.07 |
| 8 | 23307580 | rs11777391 | -0.964 | 0.725 | 1.011 | 0.268 | 0.008 | 0.023 | 0.023 | 0 | 0.31 | 0.42 | 0.26 | 0.44 | 0.31 | 0.41 |
| 8 | 23310133 | rs10111621 | 1.091 | 0.676 | 0.87 | 0.275 | -0.01 | -0.033 | -0.033 | 0 | 0.06 | 0.13 | 0.04 | 0 | 0.04 | 0.01 |
| 8 | 23310478 | rs6557673 | -1.047 | 0.99 | 0.872 | 0.418 | -0.002 | 0.016 | 0.016 | 0 | 0.3 | 0.39 | 0.22 | 0.44 | 0.27 | 0.4 |
| 8 | 23310904 | rs10088178 | 0.49 | 1.148 | 0.938 | 0.805 | 0.001 | 0.037 | 0.037 | 0 | 0.21 | 0.2 | 0.44 | 0.33 | 0.24 | 0.18 |
| 8 | 23326033 | rs13248759 | -0.936 | 1.3 | 0.713 | 0.725 | 0.006 | 0.011 | 0.011 | 0 | 0.25 | 0.31 | 0.23 | 0.41 | 0.27 | 0.4 |
| 8 | 23337410 | rs2235116 | -0.411 | 1.176 | 0.523 | 0.813 | 0.014 | 0.074 | 0.074 | 0 | 0.17 | 0.34 | 0.15 | 0.32 | 0.16 | 0.2 |
| 8 | 23340546 | rs6557675 | -0.928 | 0.937 | 0.255 | 0.679 | -0.005 | -0.001 | -0.001 | 0 | 0.33 | 0.57 | 0.53 | 0.29 | 0.32 | 0.36 |
| 8 | 23344403 | rs10216695 | 0.688 | 0.837 | -0.217 | 0.51 | 0.001 | 0.025 | 0.025 | 0 | 0.16 | 0.07 | 0.11 | 0.06 | 0.16 | 0.04 |
| 8 | 23345526 | rs732563 | -1.092 | 0.876 | -0.34 | 0.323 | 0 | 0.004 | 0.004 | 0 | 0.51 | 0.44 | 0.67 | 0.62 | 0.53 | 0.64 |
| 8 | 23347851 | rs17770799 | 1.158 | 0.938 | -0.74 | 0.301 | -0.009 | 0.042 | 0.042 | 0 | 0.16 | 0.04 | 0.11 | 0.06 | 0.16 | 0.04 |
| 8 | 23349637 | rs17089296 | 1.035 | 0.913 | -0.805 | 0.554 | 0.004 | -0.051 | -0.051 | 0.576 | 0.15 | 0.05 | 0.11 | 0.31 | 0.14 | 0.24 |
| 8 | 23351559 | rs2017650 | -0.41 | 0.992 | -0.728 | 0.334 | 0 | -0.032 | -0.032 | 0 | 0.31 | 0.2 | 0.25 | 0.62 | 0.3 | 0.44 |
| 8 | 23351759 | rs1010848 | -2.1 | 0.908 | -0.401 | 0.113 | -0.007 | -0.106 | -0.106 | 0 | 0.67 | 0.52 | 0.8 | 0.93 | 0.69 | 0.83 |
| 8 | 23355459 | rs7813840 | -1.051 | 1.065 | -0.319 | 0.347 | 0 | 0 | 0 | 0 | 0.35 | 0.37 | 0.57 | 0.55 | 0.37 | 0.54 |
| 8 | 23355620 | rs7832926 | 0.022 | 1.127 | -0.373 | 0.35 | -0.023 | 0.051 | 0.051 | 0 | 0.28 | 0.17 | 0.22 | 0.18 | 0.34 | 0.24 |
| 8 | 23356800 | rs4554474 | -1.546 | 1.063 | -0.299 | 0.374 | 0.006 | 0.011 | 0.011 | 0 | 0.51 | 0.48 | 0.68 | 0.62 | 0.53 | 0.61 |
| 8 | 23359831 | rs13250356 | -0.689 | 0.995 | -0.088 | 0.615 | -0.001 | 0.048 | 0.048 | 0 | 0.22 | 0.31 | 0.45 | 0.43 | 0.19 | 0.33 |
| 8 | 23360383 | rs12546432 | 0.413 | 0.982 | -0.054 | 0.702 | -0.007 | 0.068 | 0.068 | 0 | 0.05 | 0.1 | 0.3 | 0.12 | 0.03 | 0.13 |
| 8 | 23364738 | rs6988938 | -1.698 | 1.013 | -0.207 | 0.678 | -0.001 | 0 | 0 | 0 | 0.44 | 0.47 | 0.61 | 0.6 | 0.46 | 0.59 |
| 8 | 23365884 | rs17089312 | 1.731 | 1.027 | -0.275 | 0.668 | -0.003 | 0.006 | 0.006 | 0 | 0.08 | 0.04 | 0.05 | 0.06 | 0.1 | 0.06 |
| 8 | 23367757 | rs11135740 | -0.663 | 1.032 | -0.265 | 0.686 | -0.001 | 0.043 | 0.043 | 0 | 0.22 | 0.3 | 0.45 | 0.42 | 0.2 | 0.33 |
| 8 | 23367883 | rs7824718 | 0.695 | 1.06 | -0.624 | 0.616 | -0.015 | 0.017 | 0.017 | 0 | 0.13 | 0.09 | 0.11 | 0.11 | 0.16 | 0.2 |
| 8 | 23370177 | rs11775345 | -1.123 | 1.105 | -0.563 | 0.6 | -0.024 | -0.028 | -0.028 | 0 | 0.36 | 0.22 | 0.28 | 0.49 | 0.39 | 0.46 |
| 8 | 23374454 | rs7009973 | -1.136 | 0.829 | -0.228 | 0.783 | -0.004 | -0.01 | -0.01 | 0.429 | 0.29 | 0.31 | 0.24 | 0.48 | 0.31 | 0.43 |
| 8 | 23377161 | rs11784090 | 0.256 | 0.604 | -0.437 | 0.818 | -0.013 | -0.056 | -0.056 | 0.144 | 0.13 | 0.05 | 0.12 | 0.31 | 0.12 | 0.16 |
| 8 | 23382594 | rs972575 | -1.847 | 0.572 | -0.186 | 0.333 | -0.005 | -0.045 | -0.045 | 0.227 | 0.71 | 0.71 | 0.85 | 0.93 | 0.75 | 0.85 |
| 8 | 23382985 | rs6557678 | -1.799 | 0.536 | -0.294 | 1.112 | -0.007 | 0.02 | 0.02 | 0 | 0.41 | 0.43 | 0.61 | 0.67 | 0.45 | 0.58 |
| 8 | 23383204 | rs7008867 | -0.461 | 0.702 | 0.173 | 1.456 | 0.005 | 0.002 | 0.002 | 0 | 0.16 | 0.24 | 0.16 | 0.22 | 0.24 | 0.21 |
| 8 | 23383743 | rs6557679 | -1.155 | 0.712 | 0.063 | 1.467 | 0.007 | 0.011 | 0.011 | 0.077 | 0.21 | 0.37 | 0.29 | 0.29 | 0.28 | 0.3 |
| 8 | 23394908 | rs1008856 | 0.344 | 0.137 | 0.559 | 1.373 | 0.005 | 0.001 | 0.001 | 0.01 | 0.88 | 0.64 | 0.93 | 0.9 | 0.92 | 0.97 |
| 8 | 23396101 | rs17698981 | 0.094 | 0.16 | -0.018 | 1.761 | -0.049 | -0.063 | -0.063 | 0.003 | 0.14 | 0.01 | 0.29 | 0.17 | 0.19 | 0.17 |
| 8 | 23396537 | rs10503725 | -0.555 | -0.014 | 0.147 | 1.845 | -0.02 | 0.003 | 0.003 | 0 | 0.05 | 0.12 | 0.15 | 0.08 | 0.01 | 0.06 |

| | | | | | | | | | | | | | | | |
|---|---|---|---|---|---|---|---|---|---|---|---|---|---|---|---|
| 8 | 23396732 | rs4593549 | -2.389 | -0.299 | 0.278 | 2.145 | -0.059 | 0.2 | 0.2 | 0.438 | 0.18 | 0.58 | 0.2 | 0.43 | 0.06 | 0.28 |
| 8 | 23397035 | rs9314268 | -1.951 | -0.274 | 0.272 | 2.022 | -0.015 | -0.005 | -0.005 | 0.938 | 0.32 | 0.61 | 0.49 | 0.6 | 0.25 | 0.46 |
| 8 | 23397393 | rs4872144 | -1.976 | -0.294 | 0.292 | 2.014 | -0.013 | -0.004 | -0.004 | 0 | 0.32 | 0.62 | 0.49 | 0.6 | 0.25 | 0.46 |
| 8 | 23397750 | rs4872145 | -1.128 | -0.272 | 0.443 | 2.209 | -0.017 | 0.109 | 0.109 | 0 | 0.1 | 0.16 | 0.16 | 0.32 | 0.03 | 0.12 |
| 8 | 23402070 | rs7015818 | -1.024 | -0.154 | 0.398 | 2.054 | -0.006 | 0.07 | 0.07 | 0 | 0.22 | 0.44 | 0.33 | 0.28 | 0.22 | 0.34 |
| 8 | 23411260 | rs2978473 | 0.263 | -0.028 | 0.031 | 1.651 | -0.002 | 0.001 | 0.001 | 0 | 0.44 | 0.6 | 0.4 | 0.53 | 0.41 | 0.36 |
| 8 | 23411798 | rs7816824 | -0.335 | -0.144 | 0.209 | 1.885 | -0.018 | 0.208 | 0.208 | 0 | 0.06 | 0.17 | 0.16 | 0.32 | 0.01 | 0.06 |
| 8 | 23414369 | rs10098490 | -0.898 | -0.133 | 0.057 | 1.535 | 0.001 | 0.036 | 0.036 | 0 | 0.61 | 0.71 | 0.67 | 0.73 | 0.62 | 0.68 |
| 8 | 23414743 | rs13254494 | -0.339 | -0.081 | 0.097 | 1.555 | 0.001 | 0.017 | 0.017 | 0 | 0.57 | 0.64 | 0.66 | 0.68 | 0.6 | 0.5 |
| 8 | 23414822 | rs9644032 | -1.08 | -0.071 | 0.045 | 1.372 | 0 | 0.023 | 0.023 | 0 | 0.64 | 0.74 | 0.68 | 0.76 | 0.64 | 0.69 |
| 8 | 23415240 | rs2137304 | 0.897 | 0.012 | -0.025 | 1.356 | 0 | 0.003 | 0.003 | 0 | 0.1 | 0.08 | 0.19 | 0.09 | 0.13 | 0.11 |
| 8 | 23415524 | rs11784870 | 0.505 | 0.01 | 0.015 | 1.325 | -0.01 | -0.012 | -0.012 | 1.954 | 0.16 | 0.08 | 0.27 | 0.16 | 0.19 | 0.19 |
| 8 | 23416034 | rs2978487 | 1.26 | -0.076 | 0.195 | 1.45 | -0.003 | 0.012 | 0.012 | 0 | 0.34 | 0.33 | 0.28 | 0.26 | 0.36 | 0.23 |
| 8 | 23418444 | rs2942202 | 0.087 | -0.015 | 0.19 | 1.382 | -0.002 | 0.022 | 0.022 | 0 | 0.51 | 0.63 | 0.48 | 0.38 | 0.51 | 0.42 |
| 8 | 23419176 | rs2942201 | 0.153 | 0.144 | 0.191 | 1.181 | 0.001 | -0.006 | -0.006 | 0 | 0.43 | 0.51 | 0.34 | 0.37 | 0.42 | 0.44 |
| 8 | 23419869 | rs7830129 | 0.864 | 0.227 | 0.247 | 1.106 | -0.002 | -0.033 | -0.033 | 0.664 | 0.34 | 0.45 | 0.28 | 0.22 | 0.32 | 0.23 |
| 8 | 23421696 | rs2928686 | 1.537 | 0.265 | 0.021 | 1.136 | -0.001 | -0.001 | -0.001 | 0 | 0.28 | 0.29 | 0.26 | 0.2 | 0.29 | 0.16 |
| 8 | 23423373 | rs2978477 | -0.252 | 0.069 | -0.459 | 1.369 | 0.007 | -0.012 | -0.012 | 0 | 0.94 | 0.96 | 0.84 | 0.86 | 0.97 | 0.93 |
| 8 | 23423669 | rs2942194 | 2.041 | 0.093 | -0.574 | 1.21 | -0.095 | -0.034 | -0.034 | 0.426 | 0.24 | 0.02 | 0.24 | 0.22 | 0.29 | 0.16 |
| 8 | 23423758 | rs10992 | 1.267 | -0.091 | -0.664 | 1.136 | 0.045 | 0.048 | 0.048 | 0.821 | 0.58 | 0.1 | 0.53 | 0.37 | 0.66 | 0.5 |
| 8 | 23427919 | rs2942213 | 0.458 | 0.244 | -0.319 | 0.952 | -0.037 | -0.017 | -0.017 | 0.004 | 0.42 | 0.24 | 0.34 | 0.4 | 0.46 | 0.42 |
| 8 | 23429329 | rs1047384 | -0.356 | 0.024 | -0.01 | 0.823 | -0.015 | 0.132 | 0.132 | 0.117 | 0.07 | 0.17 | 0.16 | 0.21 | 0.04 | 0.07 |
| 8 | 23431045 | rs1810193 | -0.778 | 0.033 | -0.345 | 0.637 | -0.039 | -0.043 | -0.043 | 0.051 | 0.56 | 0.86 | 0.58 | 0.45 | 0.54 | 0.61 |
| 8 | 23431407 | rs3174040 | -1.472 | 0.008 | 0.117 | 0.608 | -0.009 | 0.047 | 0.047 | 0 | 0.2 | 0.52 | 0.26 | 0.24 | 0.19 | 0.26 |
| 8 | 23432213 | rs10098411 | 0.737 | 0.106 | 0.175 | 0.305 | 0.024 | -0.004 | -0.004 | 0 | 0.05 | 0.17 | 0.02 | 0.04 | 0.06 | 0.12 |
| 8 | 23432332 | rs8534 | 0.587 | 0.136 | 0.079 | 0.296 | 0 | -0.008 | -0.008 | 0 | 0.36 | 0.34 | 0.32 | 0.21 | 0.34 | 0.35 |
| 8 | 23436752 | rs2314686 | 0.655 | -0.031 | 0.153 | 0.326 | 0.01 | 0.068 | 0.068 | 0.437 | 0.08 | 0.32 | 0.07 | 0.09 | 0.05 | 0.05 |
| 8 | 23438975 | rs2928679 | -0.382 | 0.003 | -0.19 | 0.195 | -0.029 | 0.117 | 0.117 | 0 | 0.6 | 0.75 | 0.68 | 0.85 | 0.58 | 0.52 |
| 8 | 23439308 | rs2942219 | 2.177 | 0.019 | -0.488 | 0.365 | -0.075 | -0.092 | -0.092 | 0 | 0.25 | 0.02 | 0.22 | 0.28 | 0.28 | 0.19 |
| 8 | 23440655 | rs7818607 | -1.441 | 0.029 | -0.296 | 0.43 | -0.078 | 0.113 | 0.113 | 0 | 0.73 | 0.97 | 0.76 | 0.89 | 0.73 | 0.7 |
| 8 | 23441319 | rs7009429 | 0.744 | 0.029 | 0.111 | 0.588 | -0.01 | 0.005 | 0.005 | 0 | 0.45 | 0.31 | 0.44 | 0.57 | 0.49 | 0.39 |
| 8 | 23447471 | rs17700954 | 0.266 | 0.003 | -0.059 | 0.578 | -0.001 | -0.009 | -0.009 | 0 | 0.08 | 0.09 | 0.06 | 0 | 0.09 | 0.06 |
| 8 | 23449483 | rs17089399 | 0.213 | -0.089 | 0.113 | 0.721 | -0.01 | 0.091 | 0.091 | 0 | 0.06 | 0.1 | 0.17 | 0.18 | 0.04 | 0.07 |
| 8 | 23451893 | rs6988205 | -0.643 | -0.083 | 0.213 | 0.663 | 0 | 0 | 0 | 1.103 | 0.29 | 0.33 | 0.27 | 0.33 | 0.3 | 0.34 |
| 8 | 23452968 | rs11780097 | -0.531 | -0.081 | 0.212 | 0.663 | -0.001 | -0.002 | -0.002 | 0.036 | 0.28 | 0.26 | 0.27 | 0.33 | 0.31 | 0.35 |
| 8 | 23455686 | rs6980743 | -0.595 | -0.159 | 0.406 | 0.692 | -0.044 | 0.195 | 0.195 | 0 | 0.15 | 0.42 | 0.23 | 0.28 | 0.09 | 0.13 |
| 8 | 23458802 | rs6557696 | -0.43 | -0.067 | -0.026 | 0.562 | -0.083 | 0.176 | 0.176 | 0 | 0.62 | 0.91 | 0.68 | 0.85 | 0.58 | 0.52 |
| 8 | 23461175 | rs12549100 | 1.535 | -0.021 | 0.386 | 0.565 | 0.015 | 0.002 | 0.002 | 0 | 0.3 | 0.45 | 0.27 | 0.28 | 0.34 | 0.23 |
| 8 | 23463823 | rs10503731 | 0.42 | 0.003 | 0.265 | 0.637 | -0.006 | 0.083 | 0.083 | 0 | 0.06 | 0.11 | 0.17 | 0.18 | 0.04 | 0.07 |
| 8 | 23464770 | rs725173 | 0.687 | -0.047 | 0.261 | 0.625 | 0.008 | -0.023 | -0.023 | 0 | 0.17 | 0.14 | 0.19 | 0.29 | 0.15 | 0.17 |
| 8 | 23464789 | rs13272418 | 1.225 | -0.157 | 0.45 | 0.68 | 0.004 | -0.001 | -0.001 | 0.634 | 0.32 | 0.38 | 0.28 | 0.28 | 0.34 | 0.23 |
| 8 | 23465745 | rs992531 | 0.72 | -0.064 | 0.42 | 0.73 | -0.019 | 0.017 | 0.017 | 0.082 | 0.08 | 0.18 | 0.05 | 0.1 | 0.05 | 0.06 |
| 8 | 23465916 | rs992530 | -1.379 | -0.032 | 0.287 | 0.648 | -0.071 | 0.113 | 0.113 | 0 | 0.73 | 0.97 | 0.76 | 0.89 | 0.73 | 0.7 |
| 8 | 23467398 | rs7841597 | 0.05 | -0.022 | 0.309 | 0.675 | 0 | 0.001 | 0.001 | 0 | 0.29 | 0.29 | 0.27 | 0.32 | 0.3 | 0.35 |
| 8 | 23469088 | rs11135759 | -0.418 | -0.039 | 1.178 | 1.139 | 0.007 | 0.02 | 0.02 | 0 | 0.44 | 0.68 | 0.49 | 0.56 | 0.43 | 0.35 |
| 8 | 23470246 | rs12549446 | -1.192 | 0.052 | 0.738 | 1.182 | -0.03 | 0.117 | 0.117 | 0 | 0.14 | 0.28 | 0.22 | 0.28 | 0.09 | 0.13 |
| 8 | 23470562 | rs12550045 | -1.192 | 0.052 | 0.738 | 1.182 | -0.03 | 0.117 | 0.117 | 0 | 0.14 | 0.28 | 0.22 | 0.28 | 0.09 | 0.13 |
| 8 | 23477195 | rs10098347 | -0.254 | 0.196 | 0.63 | 0.484 | -0.016 | 0.041 | 0.041 | 0.06 | 0.09 | 0.25 | 0.05 | 0.11 | 0.05 | 0.06 |
| 8 | 23477981 | rs4872167 | -0.786 | 0.196 | 0.61 | 0.435 | -0.004 | -0.003 | -0.003 | 0.235 | 0.71 | 0.72 | 0.73 | 0.72 | 0.66 | 0.77 |
| 8 | 23478587 | rs907586 | -0.074 | 0.404 | 0.077 | 0.344 | 0 | -0.002 | -0.002 | 0 | 0.6 | 0.36 | 0.66 | 0.58 | 0.58 | 0.71 |
| 8 | 23478809 | rs11986046 | -0.209 | 0.418 | -0.184 | 0.4 | -0.001 | 0.012 | 0.012 | 0 | 0.62 | 0.48 | 0.68 | 0.61 | 0.61 | 0.71 |
| 8 | 23481910 | rs1810195 | 0.648 | 0.313 | -0.415 | -0.063 | 0.004 | -0.002 | -0.002 | 0 | 0.92 | 0.93 | 0.94 | 1 | 0.93 | 0.95 |
| 8 | 23482464 | rs7003364 | 0.152 | 0.533 | -0.24 | -0.369 | 0.004 | 0.017 | 0.017 | 0 | 0.51 | 0.53 | 0.63 | 0.7 | 0.58 | 0.58 |
| 8 | 23485112 | rs907588 | -0.288 | 0.227 | 0.512 | -0.762 | 0.019 | -0.061 | -0.061 | 0 | 0.1 | 0.31 | 0.1 | 0.01 | 0.11 | 0.03 |
| 8 | 23490172 | rs4871896 | 0.081 | 0.324 | 0.14 | -0.83 | -0.005 | -0.003 | -0.003 | 0 | 0.62 | 0.53 | 0.64 | 0.7 | 0.65 | 0.58 |
| 8 | 23509858 | rs17296662 | 0.833 | 0.24 | 0.28 | -0.876 | -0.006 | -0.025 | -0.025 | 1.313 | 0.25 | 0.17 | 0.3 | 0.3 | 0.28 | 0.34 |
| 8 | 23514289 | rs4872171 | 0.904 | 0.164 | 0.106 | -0.943 | 0.026 | 0.069 | 0.069 | 0.036 | 0.51 | 0.18 | 0.54 | 0.69 | 0.54 | 0.56 |
| 8 | 23514823 | rs2137303 | 1.195 | 0.157 | -0.096 | -0.973 | -0.09 | -0.158 | -0.158 | 0 | 0.24 | 0.02 | 0.3 | 0.48 | 0.3 | 0.28 |
| 8 | 23520429 | rs907584 | -0.826 | 0.165 | 0.519 | -0.819 | -0.017 | -0.053 | -0.053 | 0 | 0.88 | 0.83 | 0.95 | 1 | 0.93 | 0.93 |
| 8 | 23526463 | rs1512268 | 0.779 | 0.259 | 0.46 | -0.812 | 0.012 | 0.027 | 0.027 | 0 | 0.51 | 0.31 | 0.55 | 0.7 | 0.53 | 0.55 |
| 8 | 23528230 | rs10503734 | 0.7 | 0.275 | 0.375 | -0.829 | -0.002 | 0.003 | 0.003 | 0 | 0.08 | 0.1 | 0.07 | 0.01 | 0.11 | 0.03 |
| 8 | 23534018 | rs10503733 | 0.598 | 0.287 | 0.316 | -0.825 | -0.001 | -0.006 | -0.006 | 0 | 0.28 | 0.29 | 0.32 | 0.28 | 0.29 | 0.34 |
| 8 | 23534802 | rs4872175 | 0.712 | 0.233 | 0.324 | -0.825 | 0.017 | 0.062 | 0.062 | 0 | 0.52 | 0.24 | 0.54 | 0.72 | 0.53 | 0.64 |
| 8 | 23536098 | rs7005368 | 0.533 | 0.219 | 0.34 | -0.464 | -0.015 | -0.063 | -0.063 | 0.636 | 0.11 | 0.21 | 0.06 | 0.01 | 0.07 | 0.07 |
| 8 | 23538533 | rs1567669 | 0.476 | 0.216 | -0.113 | -0.466 | -0.011 | -0.009 | -0.009 | 0 | 0.61 | 0.48 | 0.62 | 0.71 | 0.63 | 0.58 |
| 8 | 23540417 | rs11781886 | -0.356 | 0.349 | -0.811 | -0.377 | -0.001 | 0.004 | 0.004 | 0 | 0.72 | 0.67 | 0.69 | 0.71 | 0.7 | 0.66 |
| 8 | 23547214 | rs11774063 | 0.843 | 0.752 | -1.287 | -0.534 | -0.049 | -0.013 | -0.013 | 0 | 0.23 | 0.02 | 0.18 | 0.21 | 0.19 | 0.28 |
| 8 | 23547510 | rs1866347 | -0.915 | 0.973 | -0.207 | -0.377 | 0.019 | -0.112 | -0.112 | 0 | 0.21 | 0.51 | 0.26 | 0.02 | 0.23 | 0.28 |
| 8 | 23549767 | rs11135767 | -1.114 | 0.777 | -0.996 | 0.132 | 0.004 | -0.014 | -0.014 | 0 | 0.2 | 0.1 | 0.2 | 0.15 | 0.19 | 0.24 |

| | | | | | | | | | | | | | | | | |
|---|---|---|---|---|---|---|---|---|---|---|---|---|---|---|---|---|
| 8 | 23551671 | rs7844736 | -0.142 | 0.222 | -1.388 | -0.772 | 0.009 | 0.029 | 0.029 | 0 | 0.27 | 0.17 | 0.24 | 0.13 | 0.19 | 0.33 |
| 8 | 23551904 | rs4872179 | 0.245 | 0.108 | -1.365 | -0.641 | 0 | 0 | 0 | 0 | 0.42 | 0.64 | 0.56 | 0.58 | 0.55 | 0.42 |
| 8 | 23552773 | rs10106897 | -0.255 | 0.839 | -1.139 | -1.048 | 0.003 | 0.014 | 0.014 | 0 | 0.53 | 0.6 | 0.52 | 0.71 | 0.42 | 0.62 |
| 8 | 23553455 | rs13267142 | 0.695 | 1.148 | -0.954 | -1.163 | 0.075 | -0.067 | -0.067 | 0.151 | 0.14 | 0.24 | 0.17 | 0 | 0.28 | 0.06 |
| 8 | 23554565 | rs920300 | 0.482 | 0.98 | -1.059 | -1.325 | 0.011 | 0.051 | 0.051 | 0 | 0.33 | 0.19 | 0.29 | 0.14 | 0.26 | 0.37 |
| 8 | 23556615 | rs12543268 | 0.419 | 0.484 | -1.011 | -1.766 | 0.006 | 0.001 | 0.001 | 0 | 0.64 | 0.37 | 0.61 | 0.3 | 0.63 | 0.62 |
| 8 | 23559370 | rs749145 | 0.148 | 0.017 | -0.8 | -1.851 | 0.018 | 0.039 | 0.039 | 0 | 0.43 | 0.39 | 0.45 | 0.82 | 0.38 | 0.44 |
| 8 | 23562262 | rs13265089 | 0.591 | 0.076 | -0.765 | -1.783 | -0.001 | -0.002 | -0.002 | 0 | 0.48 | 0.43 | 0.52 | 0.29 | 0.48 | 0.54 |
| 8 | 23565878 | rs7834389 | -0.438 | -0.134 | -0.628 | -1.679 | -0.019 | 0.094 | 0.094 | 0 | 0.78 | 0.62 | 0.75 | 0.39 | 0.84 | 0.79 |
| 8 | 23568464 | rs7844903 | 0.093 | -0.119 | -0.877 | -1.768 | 0.038 | 0.076 | 0.076 | 0 | 0.5 | 0.24 | 0.43 | 0.31 | 0.58 | 0.42 |
| 8 | 23570489 | rs17782264 | 0.946 | 0.002 | -0.863 | -1.557 | -0.002 | -0.014 | -0.014 | 0 | 0.27 | 0.26 | 0.31 | 0.11 | 0.25 | 0.37 |
| 8 | 23572332 | rs17782270 | 1.358 | 0.217 | -1.193 | -1.413 | -0.026 | -0.066 | -0.066 | 0.776 | 0.2 | 0.01 | 0.2 | 0.19 | 0.21 | 0.21 |
| 8 | 23580010 | rs9785115 | -0.335 | 0.406 | -0.649 | -1.885 | -0.009 | -0.031 | -0.031 | 0 | 0.75 | 0.79 | 0.82 | 0.99 | 0.65 | 0.86 |
| 8 | 23584226 | rs13273073 | 0.604 | 0.435 | -0.223 | -1.786 | 0 | -0.013 | -0.013 | 1.317 | 0.28 | 0.31 | 0.19 | 0.07 | 0.25 | 0.37 |
| 8 | 23590795 | rs6557711 | -0.842 | 0.288 | 0.049 | -1.474 | 0.015 | 0.041 | 0.041 | 0 | 0.61 | 0.71 | 0.58 | 0.21 | 0.66 | 0.65 |
| 8 | 23595940 | rs7004766 | 0.613 | 0.389 | 0.072 | -1.405 | -0.015 | -0.082 | -0.082 | 0 | 0.29 | 0.52 | 0.21 | 0.11 | 0.26 | 0.38 |
| 8 | 23598040 | rs6991937 | 0.69 | 0.622 | -0.048 | -1.343 | 0.03 | -0.058 | -0.058 | 0.607 | 0.18 | 0.24 | 0.16 | 0.03 | 0.3 | 0.08 |
| 8 | 23599043 | rs1438455 | -1.589 | 0.44 | -0.112 | -1.047 | 0.014 | -0.002 | -0.002 | 0 | 0.7 | 0.74 | 0.75 | 0.75 | 0.76 | 0.73 |
| 8 | 23602802 | rs364675 | -0.283 | 1.123 | 0.211 | -0.528 | 0.018 | 0.022 | 0.022 | 0.062 | 0.29 | 0.33 | 0.34 | 0.58 | 0.41 | 0.22 |
| 8 | 23607733 | rs10866835 | 0.395 | 1.013 | 0.189 | -0.53 | 0 | 0.133 | 0.133 | 0 | 0.15 | 0.26 | 0.19 | 0.53 | 0.12 | 0.16 |
| 8 | 23610292 | rs218870 | -1.163 | 1.223 | -0.193 | -0.705 | -0.033 | 0.025 | 0.025 | 0 | 0.86 | 0.93 | 0.87 | 1 | 0.73 | 0.95 |
| 8 | 23610799 | rs1561105 | 0.416 | 1.124 | -0.149 | -0.507 | -0.001 | -0.008 | -0.008 | 0 | 0.26 | 0.29 | 0.22 | 0.11 | 0.26 | 0.3 |
| 8 | 23612956 | rs17089603 | -0.019 | 1.086 | -0.187 | -0.092 | 0.002 | -0.009 | -0.009 | 0 | 0.2 | 0.16 | 0.28 | 0.51 | 0.16 | 0.22 |
| 8 | 23614369 | rs310282 | 0.079 | 1.186 | -0.186 | -0.395 | -0.012 | 0.024 | 0.024 | 0.142 | 0.62 | 0.34 | 0.63 | 0.88 | 0.48 | 0.75 |
| 8 | 23615779 | rs10097518 | -0.682 | 1.266 | 0.09 | -0.407 | 0 | -0.001 | -0.001 | 0 | 0.42 | 0.55 | 0.5 | 0.62 | 0.41 | 0.4 |
| 8 | 23616084 | rs1212399 | 1.154 | 1.276 | -0.966 | -0.355 | 0.07 | -0.002 | -0.002 | 0 | 0.43 | 0.19 | 0.35 | 0.41 | 0.32 | 0.53 |
| 8 | 23617532 | rs6985626 | 0.376 | 1.132 | -0.817 | 0.402 | 0.001 | -0.002 | -0.002 | 0 | 0.21 | 0.24 | 0.22 | 0.12 | 0.27 | 0.18 |
| 8 | 23617540 | rs6985631 | -0.122 | 1.064 | -0.857 | 0.415 | 0.003 | -0.007 | -0.007 | 0.088 | 0.2 | 0.16 | 0.27 | 0.47 | 0.15 | 0.22 |
| 8 | 23617879 | rs6985798 | -0.154 | 1.216 | -0.859 | 0.178 | -0.002 | 0.001 | 0.001 | 0 | 0.21 | 0.24 | 0.22 | 0.1 | 0.26 | 0.2 |
| 8 | 23624725 | rs7832345 | 0.28 | 1.002 | -0.927 | 0.513 | -0.001 | -0.002 | -0.002 | 0 | 0.71 | 0.68 | 0.72 | 0.43 | 0.71 | 0.71 |
| 8 | 23625062 | rs192841 | -0.07 | -0.247 | -0.545 | 0.123 | -0.022 | -0.029 | -0.029 | 0 | 0.17 | 0.21 | 0.29 | 0.11 | 0.1 | 0.21 |
| 8 | 23626461 | rs188699 | -0.493 | -0.323 | -0.749 | 0.203 | 0.002 | -0.005 | -0.005 | 0 | 0.4 | 0.33 | 0.59 | 0.55 | 0.44 | 0.66 |
| 8 | 23629252 | rs1972844 | 0.797 | -0.644 | -0.957 | 0.329 | 0.025 | -0.073 | -0.073 | 0 | 0.13 | 0.03 | 0.08 | 0.29 | 0.12 | 0.24 |
| 8 | 23633291 | rs13272072 | 0.098 | -0.66 | -1.083 | 0.193 | 0.003 | 0.003 | 0.003 | 0 | 0.13 | 0.06 | 0.19 | 0.07 | 0.1 | 0.1 |
| 8 | 23638600 | rs450336 | 0.198 | -0.614 | -0.494 | 0.17 | 0.042 | -0.004 | -0.004 | 0 | 0.22 | 0.38 | 0.4 | 0.24 | 0.28 | 0.34 |
| 8 | 23647494 | rs310302 | -0.725 | -1.247 | -0.312 | -0.21 | 0.001 | -0.011 | -0.011 | 0 | 0.39 | 0.51 | 0.46 | 0.25 | 0.39 | 0.47 |
| 8 | 23649551 | rs167456 | 0.407 | -1.203 | -0.244 | -0.214 | 0.041 | -0.017 | -0.017 | 0.157 | 0.22 | 0.38 | 0.38 | 0.23 | 0.29 | 0.36 |
| 8 | 23652909 | rs6988827 | 0.774 | -1.259 | -0.308 | -0.183 | 0.014 | -0.001 | -0.001 | 14.491 | 0.12 | 0.05 | 0.17 | 0.06 | 0.06 | 0.09 |
| 8 | 23655784 | rs218872 | -0.751 | -1.258 | -0.178 | -0.206 | 0.005 | -0.025 | -0.025 | 0 | 0.39 | 0.51 | 0.46 | 0.25 | 0.4 | 0.48 |
| 8 | 23659511 | rs310291 | -0.916 | -1.508 | -0.405 | -0.102 | 0.007 | 0.007 | 0.007 | 0 | 0.44 | 0.58 | 0.48 | 0.56 | 0.39 | 0.67 |
| 8 | 23663101 | rs310292 | -0.706 | -1.539 | -0.367 | -0.011 | -0.001 | 0.001 | 0.001 | 0 | 0.39 | 0.55 | 0.49 | 0.3 | 0.4 | 0.48 |
| 8 | 23664441 | rs310296 | 0.46 | -1.55 | -0.437 | -0.02 | 0.028 | -0.016 | -0.016 | 0 | 0.23 | 0.35 | 0.38 | 0.23 | 0.28 | 0.36 |
| 8 | 23671498 | rs310299 | -1.077 | -1.552 | -0.51 | -0.016 | -0.002 | -0.006 | -0.006 | 0 | 0.47 | 0.49 | 0.51 | 0.6 | 0.43 | 0.71 |
| 8 | 23678000 | rs12678365 | 1.35 | -1.504 | -0.581 | 0.069 | 0.032 | -0.008 | -0.008 | 0 | 0.07 | 0.01 | 0.03 | 0.3 | 0.04 | 0.23 |
| 8 | 23687444 | rs17089666 | -0.565 | -1.556 | -0.53 | -0.233 | 0.02 | 0.028 | 0.028 | 0 | 0.29 | 0.19 | 0.27 | 0.13 | 0.18 | 0.21 |
| 8 | 23689103 | rs13267264 | -0.526 | -1.543 | -0.521 | -0.225 | 0.018 | 0.033 | 0.033 | 0 | 0.28 | 0.19 | 0.25 | 0.09 | 0.18 | 0.2 |
| 8 | 23689182 | rs10503732 | 0.388 | -1.562 | -0.536 | -0.239 | 0.009 | 0.046 | 0.046 | 0 | 0.17 | 0.08 | 0.08 | 0.03 | 0.11 | 0.11 |
| 8 | 23689919 | rs310318 | -2.272 | -1.56 | -0.577 | -0.337 | -0.077 | 0.001 | 0.001 | 0 | 0.67 | 0.9 | 0.75 | 0.62 | 0.6 | 0.81 |
| 8 | 23690631 | rs310319 | -1.706 | -1.521 | -0.615 | -0.212 | -0.007 | 0.011 | 0.011 | 0 | 0.59 | 0.59 | 0.68 | 0.66 | 0.5 | 0.8 |
| 8 | 23692067 | rs4872196 | -0.357 | -1.774 | -0.826 | -0.229 | 0.029 | 0.032 | 0.032 | 0 | 0.26 | 0.16 | 0.2 | 0.07 | 0.11 | 0.18 |
| 8 | 23695903 | rs409783 | -0.665 | -1.648 | -0.471 | 0.131 | 0.005 | 0.005 | 0.005 | 0 | 0.35 | 0.46 | 0.54 | 0.32 | 0.34 | 0.5 |
| 8 | 23696543 | rs566429 | 0.316 | -1.677 | -0.241 | 0.069 | -0.001 | 0 | 0 | 0 | 0.23 | 0.24 | 0.36 | 0.17 | 0.27 | 0.38 |
| 8 | 23698746 | rs182564 | -1.67 | -1.816 | -0.248 | 0.236 | -0.001 | 0 | 0 | 0.778 | 0.64 | 0.62 | 0.67 | 0.54 | 0.52 | 0.78 |
| 8 | 23701630 | rs3824220 | -0.23 | -1.585 | -0.26 | -0.238 | -0.009 | -0.006 | -0.006 | 0 | 0.18 | 0.19 | 0.24 | 0.09 | 0.16 | 0.15 |
| 8 | 23702072 | rs1062998 | -0.22 | -1.582 | -0.258 | -0.237 | -0.009 | -0.006 | -0.006 | 0 | 0.18 | 0.19 | 0.24 | 0.09 | 0.16 | 0.15 |
| 8 | 23702220 | rs2936 | -0.22 | -1.582 | -0.258 | -0.237 | -0.009 | -0.006 | -0.006 | 0 | 0.18 | 0.19 | 0.24 | 0.09 | 0.16 | 0.15 |
| 8 | 23702603 | rs11135775 | 1.733 | -1.581 | -0.235 | -0.24 | -0.001 | -0.004 | -0.004 | 0 | 0.07 | 0.04 | 0.05 | 0 | 0.1 | 0.01 |
| 8 | 23702685 | rs7010237 | -0.138 | -1.657 | -0.255 | -0.088 | 0.005 | 0.026 | 0.026 | 0 | 0.37 | 0.32 | 0.38 | 0.2 | 0.31 | 0.43 |
| 8 | 23703716 | rs9969426 | 0.061 | -1.054 | -0.081 | 0.251 | -0.013 | 0.021 | 0.021 | 0 | 0.11 | 0.14 | 0.19 | 0.09 | 0.06 | 0.14 |
| 8 | 23714122 | rs1369836 | -1.171 | -1.299 | -0.167 | 0.067 | 0.005 | 0.004 | 0.004 | 0.211 | 0.34 | 0.26 | 0.4 | 0.12 | 0.26 | 0.33 |
| 8 | 23714609 | rs3758087 | -0.895 | -1.486 | -0.2 | -0.245 | 0.004 | 0.022 | 0.022 | 0 | 0.65 | 0.69 | 0.72 | 0.77 | 0.71 | 0.64 |
| 8 | 23724908 | rs11989588 | 0.726 | -1.29 | 0.257 | 0.216 | 0.029 | -0.034 | -0.034 | 3.101 | 0.36 | 0.51 | 0.37 | 0.32 | 0.53 | 0.12 |
| 8 | 23728271 | rs6999484 | 1.686 | -1.285 | 0.352 | 0.178 | 0.012 | -0.004 | -0.004 | 0 | 0.27 | 0.32 | 0.28 | 0.29 | 0.43 | 0.11 |
| 8 | 23732888 | rs1436149 | 0.045 | -1.451 | 0.228 | 0.169 | -0.002 | -0.019 | -0.019 | 0 | 0.45 | 0.54 | 0.51 | 0.32 | 0.61 | 0.31 |
| 8 | 23742960 | rs819198 | -1.082 | -1.398 | 0.209 | 0.174 | 0.022 | -0.037 | -0.037 | 0 | 0.65 | 0.54 | 0.65 | 0.71 | 0.44 | 0.88 |
| 8 | 23743178 | rs12546342 | 0.745 | -1.388 | 0.014 | 0.157 | 0.005 | -0.004 | -0.004 | 0 | 0.11 | 0.06 | 0.17 | 0.08 | 0.07 | 0.13 |
| 8 | 23767867 | rs1705691 | 0.699 | -1.396 | 0.004 | 0.2 | 0.02 | 0.036 | 0.036 | 0 | 0.41 | 0.32 | 0.29 | 0.27 | 0.28 | 0.42 |
| 8 | 23772530 | rs9650405 | -0.436 | -0.965 | 0.233 | 0.507 | 0.011 | 0.019 | 0.019 | 0.085 | 0.48 | 0.59 | 0.54 | 0.65 | 0.64 | 0.51 |
| 8 | 23776392 | rs2089830 | -0.192 | -0.884 | 0.024 | 0.507 | 0.016 | -0.02 | -0.02 | 1.008 | 0.12 | 0.06 | 0.2 | 0.37 | 0.1 | 0.39 |

| | | | | | | | | | | | | | | | | |
|---|---|---|---|---|---|---|---|---|---|---|---|---|---|---|---|---|
| 8 | 23776938 | rs1511387 | -0.181 | -0.881 | 0.308 | 0.658 | 0.024 | 0.072 | 0.072 | 0.068 | 0.48 | 0.65 | 0.58 | 0.72 | 0.63 | 0.5 |
| 8 | 23786333 | rs1631430 | -0.219 | -0.692 | -0.077 | 0.027 | -0.001 | -0.003 | -0.003 | 0 | 0.56 | 0.46 | 0.43 | 0.62 | 0.37 | 0.69 |
| 8 | 23787759 | rs17787051 | 0.576 | -0.28 | -0.109 | -0.013 | -0.001 | -0.015 | -0.015 | 0.278 | 0.08 | 0.05 | 0.05 | 0 | 0.11 | 0.01 |
| 8 | 23796319 | rs7826970 | -0.573 | -0.808 | 0.412 | -0.107 | -0.069 | -0.078 | -0.078 | 0.961 | 0.13 | 0.47 | 0.05 | 0 | 0.02 | 0.07 |
| 8 | 23797763 | rs1731263 | -0.377 | -0.267 | -0.571 | 0.161 | -0.053 | -0.012 | -0.012 | 0.307 | 0.36 | 0.17 | 0.44 | 0.32 | 0.48 | 0.4 |
| 8 | 23858018 | rs10104259 | -0.543 | -0.728 | 0.861 | 0.026 | -0.062 | -0.099 | -0.099 | 0.255 | 0.14 | 0.38 | 0.04 | 0 | 0.02 | 0.08 |
| 8 | 23858386 | rs1705704 | -1.094 | -0.571 | 0.523 | 0.452 | -0.001 | -0.003 | -0.003 | 1.737 | 0.8 | 0.82 | 0.9 | 0.54 | 0.82 | 0.68 |
| 8 | 23878352 | rs1320881 | -0.798 | -0.41 | 0.649 | 0.356 | -0.004 | -0.015 | -0.015 | 0.161 | 0.78 | 0.83 | 0.91 | 0.54 | 0.81 | 0.66 |
| 8 | 23879104 | rs4407889 | -0.107 | -0.115 | 0.573 | 0.257 | -0.006 | 0.014 | 0.014 | 0 | 0.19 | 0.1 | 0.19 | 0.13 | 0.22 | 0.28 |
| 8 | 23892625 | rs10503742 | 0.459 | 0.213 | 1.376 | 0.661 | -0.026 | -0.037 | -0.037 | 0 | 0.08 | 0.22 | 0.02 | 0 | 0.01 | 0.04 |
| 8 | 23923170 | rs11989470 | -0.166 | 0.185 | 1.44 | 0.976 | -0.006 | 0.08 | 0.08 | 0.038 | 0.08 | 0.3 | 0.04 | 0.1 | 0.02 | 0.09 |
| 8 | 23927713 | rs4872211 | -0.096 | 0.205 | 1.145 | 1.261 | -0.003 | -0.001 | -0.001 | 0.923 | 0.1 | 0.1 | 0.29 | 0.26 | 0.14 | 0.2 |
| 8 | 23932243 | rs7845836 | -0.849 | 0.488 | 1.409 | 1.331 | -0.027 | 0.079 | 0.079 | 0.576 | 0.1 | 0.47 | 0.04 | 0.11 | 0.02 | 0.09 |
| 8 | 23936884 | rs7016438 | -1.049 | 0.473 | 1.29 | 1.306 | -0.031 | 0.062 | 0.062 | 0 | 0.12 | 0.43 | 0.05 | 0.11 | 0.04 | 0.1 |
| 8 | 23938133 | rs900655 | -1.195 | 0.839 | 0.606 | 1.731 | -0.015 | -0.024 | -0.024 | 0 | 0.85 | 0.93 | 0.83 | 0.68 | 0.85 | 0.8 |
| 8 | 23962408 | rs17727499 | -0.233 | 0.288 | 1.332 | 0.761 | 0.007 | -0.002 | -0.002 | 0 | 0.13 | 0.1 | 0.07 | 0 | 0.12 | 0.02 |
| 8 | 23991643 | rs17051661 | 0.672 | 0.246 | 1.526 | 0.92 | -0.015 | -0.017 | -0.017 | 4.401 | 0.05 | 0.19 | 0.01 | 0 | 0.01 | 0.02 |
| 8 | 23999082 | rs10503746 | -0.369 | 0.2 | 1.979 | 0.897 | 0.019 | 0.014 | 0.014 | 2.334 | 0.07 | 0.4 | 0.03 | 0 | 0.01 | 0.02 |
| 8 | 23999321 | rs10108392 | -0.26 | 0.214 | 1.924 | 0.904 | 0.01 | 0.006 | 0.006 | 1.764 | 0.07 | 0.35 | 0.03 | 0 | 0.01 | 0.02 |
| 8 | 24022239 | rs10101419 | -0.93 | 0.172 | 1.943 | 1.036 | -0.027 | 0.068 | 0.068 | 0.154 | 0.1 | 0.47 | 0.04 | 0.09 | 0.01 | 0.09 |
| 8 | 24049157 | rs17796497 | -0.304 | 0.097 | 1.263 | 0.954 | -0.008 | 0.002 | 0.002 | 0 | 0.12 | 0.03 | 0.14 | 0.08 | 0.14 | 0.14 |
| 8 | 24049625 | rs3106318 | 0.861 | -0.017 | 1.833 | 0.736 | -0.05 | -0.001 | -0.001 | 0 | 0.88 | 0.56 | 0.97 | 0.94 | 0.98 | 0.89 |
| 8 | 24053148 | rs17051701 | -0.657 | 0.009 | 0.966 | 0.772 | 0.01 | 0.005 | 0.005 | 0.428 | 0.09 | 0.06 | 0.15 | 0.07 | 0.01 | 0.11 |
| 8 | 24058590 | rs13254021 | 0.032 | 0.655 | 0.316 | 0.729 | -0.086 | -0.005 | -0.005 | 0.468 | 0.2 | 0.03 | 0.29 | 0.22 | 0.28 | 0.23 |
| 8 | 24071487 | rs17051717 | -0.702 | 0.116 | 0.997 | 0.876 | -0.04 | -0.009 | -0.009 | 0.727 | 0.1 | 0.18 | 0.02 | 0.06 | 0.01 | 0.11 |
| 8 | 24079693 | rs12156213 | -0.684 | 0.501 | 0.476 | 1.073 | -0.07 | -0.021 | -0.021 | 0.945 | 0.16 | 0.03 | 0.38 | 0.18 | 0.31 | 0.2 |
| 8 | 24088374 | rs11991223 | -1.449 | 0.368 | 1.233 | 1.282 | -0.012 | -0.035 | -0.035 | 0 | 0.22 | 0.35 | 0.17 | 0.14 | 0.21 | 0.24 |
| 8 | 24090176 | rs4311659 | -1.457 | 0.366 | 1.243 | 1.297 | -0.014 | -0.042 | -0.042 | 0.444 | 0.23 | 0.43 | 0.18 | 0.14 | 0.21 | 0.23 |
| 8 | 24094711 | rs10097956 | -1.697 | 0.369 | 1.121 | 1.204 | -0.01 | -0.048 | -0.048 | 0 | 0.29 | 0.43 | 0.19 | 0.17 | 0.27 | 0.28 |
| 8 | 24101897 | rs6989000 | -2.341 | 0.998 | 1.669 | 2.335 | 0.028 | 0.139 | 0.139 | 2.43 | 0.22 | 0.45 | 0.34 | 0.4 | 0.26 | 0.42 |
| 8 | 24105762 | rs17051743 | -2.345 | 1.255 | 1.545 | 2.37 | 0.044 | 0.123 | 0.123 | 1.232 | 0.21 | 0.35 | 0.34 | 0.4 | 0.27 | 0.41 |
| 8 | 24107465 | rs870033 | -1.557 | 1.29 | 1.491 | 2.439 | 0.003 | 0.053 | 0.053 | 0.72 | 0.15 | 0.22 | 0.23 | 0.3 | 0.12 | 0.33 |
| 8 | 24108631 | rs12547212 | -1.502 | 1.289 | 1.492 | 2.511 | 0.002 | 0.006 | 0.006 | 0 | 0.13 | 0.14 | 0.23 | 0.18 | 0.12 | 0.28 |
| 8 | 24109055 | rs10096579 | -2.631 | 1.379 | 1.567 | 2.459 | 0.066 | 0.158 | 0.158 | 0.719 | 0.08 | 0.28 | 0.13 | 0.22 | 0.15 | 0.12 |
| 8 | 24110439 | rs11135787 | -2.211 | 1.276 | 1.31 | 2.502 | 0.014 | 0.047 | 0.047 | 1.707 | 0.2 | 0.3 | 0.36 | 0.35 | 0.28 | 0.24 |
| 8 | 24111330 | rs17088190 | -1.507 | 1.252 | 0.913 | 2.56 | -0.024 | -0.055 | -0.055 | 3.836 | 0.16 | 0.07 | 0.33 | 0.34 | 0.26 | 0.23 |
| 8 | 24116722 | rs1013210 | -1.491 | 1.232 | 0.929 | 2.565 | -0.008 | -0.018 | -0.018 | 0 | 0.16 | 0.11 | 0.33 | 0.34 | 0.25 | 0.23 |
| 8 | 24133535 | rs17051819 | -2.107 | 1.52 | 1.571 | 2.591 | 0.099 | 0.205 | 0.205 | 0.231 | 0.09 | 0.33 | 0.14 | 0.24 | 0.18 | 0.14 |
| 8 | 24138128 | rs6980834 | -1.779 | 1.417 | 1.555 | 3.116 | 0.068 | 0.302 | 0.302 | 0 | 0.08 | 0.28 | 0.11 | 0.36 | 0.16 | 0.14 |
| 8 | 24144598 | rs436760 | 2.026 | 1.255 | 2.085 | 3.247 | 0.021 | 0.077 | 0.077 | 1.68 | 0.76 | 0.44 | 0.76 | 0.45 | 0.7 | 0.62 |
| 8 | 24151305 | rs7004152 | -2.014 | 1.212 | 1.686 | 3.073 | 0.1 | 0.381 | 0.381 | 0.046 | 0.08 | 0.33 | 0.11 | 0.36 | 0.16 | 0.15 |
| 8 | 24171051 | rs6996616 | -1.554 | 1.212 | 1.635 | 3.065 | 0.04 | 0.21 | 0.21 | 0 | 0.07 | 0.15 | 0.1 | 0.36 | 0.16 | 0.14 |
| 8 | 24174324 | rs11991134 | -1.803 | 1.208 | 1.646 | 3.047 | 0.061 | 0.268 | 0.268 | 0 | 0.07 | 0.21 | 0.1 | 0.36 | 0.16 | 0.15 |
| 8 | 24189188 | rs1369078 | -1.434 | 1.281 | 0.879 | 3.047 | 0.027 | 0.175 | 0.175 | 0 | 0.07 | 0.12 | 0.1 | 0.37 | 0.16 | 0.15 |
| 8 | 24208847 | rs7009516 | -0.093 | 0.846 | 0.042 | 1.38 | -0.003 | 0.027 | 0.027 | 0 | 0.52 | 0.66 | 0.43 | 0.67 | 0.5 | 0.48 |
| 8 | 24211716 | rs2289040 | -0.127 | 0.408 | -0.018 | 1.285 | -0.022 | 0.056 | 0.056 | 0 | 0.9 | 0.81 | 0.92 | 0.87 | 0.96 | 0.9 |
| 8 | 24212040 | rs1058914 | 0.128 | -0.351 | -0.72 | 0.99 | 0.033 | -0.081 | -0.081 | 0.02 | 0.21 | 0.1 | 0.14 | 0.48 | 0.16 | 0.22 |
| 8 | 24216331 | rs4242406 | -0.238 | 0.145 | -0.376 | 0.877 | -0.008 | 0.041 | 0.041 | 0 | 0.89 | 0.78 | 0.9 | 0.8 | 0.92 | 0.83 |
| 8 | 24218697 | rs13257546 | 0.651 | 0.15 | -0.534 | 0.75 | -0.083 | 0.093 | 0.093 | 0 | 0.37 | 0.09 | 0.49 | 0.2 | 0.46 | 0.42 |
| 8 | 24221015 | rs4559243 | -0.497 | -0.018 | -0.161 | 0.67 | -0.015 | 0.03 | 0.03 | 0 | 0.58 | 0.65 | 0.5 | 0.74 | 0.53 | 0.48 |
| 8 | 24222025 | rs13279576 | 0.324 | 0.114 | -0.212 | 0.675 | -0.055 | 0.025 | 0.025 | 0 | 0.42 | 0.16 | 0.51 | 0.32 | 0.47 | 0.45 |
| 8 | 24223031 | rs12542012 | 0.706 | 0.197 | -0.324 | 0.611 | -0.11 | 0.08 | 0.08 | 0 | 0.36 | 0.07 | 0.46 | 0.2 | 0.45 | 0.41 |
| 8 | 24240704 | rs9650409 | 0.535 | 0.198 | -0.265 | 0.654 | -0.033 | 0.068 | 0.068 | 0 | 0.38 | 0.2 | 0.47 | 0.21 | 0.44 | 0.4 |
| 8 | 24249668 | rs2291574 | 1.12 | 0.398 | -0.779 | 0.199 | -0.068 | 0.219 | 0.219 | 0 | 0.35 | 0.06 | 0.3 | 0.05 | 0.39 | 0.29 |
| 8 | 24256579 | rs2291578 | 1.321 | 0.139 | -0.93 | 0.565 | -0.003 | -0.061 | -0.061 | 0 | 0.13 | 0.02 | 0.12 | 0.39 | 0.13 | 0.18 |
| 8 | 24261526 | rs3765124 | 1.056 | 0.139 | -0.949 | 0.372 | -0.066 | 0.073 | 0.073 | 0 | 0.37 | 0.1 | 0.46 | 0.2 | 0.43 | 0.4 |
| 8 | 24263345 | rs1056222 | -0.816 | 0.135 | -0.852 | 0.374 | -0.007 | 0.047 | 0.047 | 0 | 0.6 | 0.69 | 0.53 | 0.79 | 0.57 | 0.54 |
| 8 | 24273719 | rs964331 | -0.826 | 0.135 | -0.449 | 0.786 | 0 | 0.024 | 0.024 | 0 | 0.86 | 0.83 | 0.87 | 0.61 | 0.87 | 0.8 |
| 8 | 24275679 | rs17737459 | -0.051 | 0.527 | -0.928 | 0.36 | -0.055 | -0.008 | -0.008 | 0.954 | 0.19 | 0.07 | 0.22 | 0.17 | 0.25 | 0.21 |
| 8 | 24275828 | rs11135802 | 0.975 | 0.021 | -0.755 | -0.315 | 0.008 | 0.012 | 0.012 | 0.13 | 0.56 | 0.33 | 0.59 | 0.38 | 0.6 | 0.5 |
| 8 | 24283237 | rs7012896 | 0.65 | 0.296 | -0.956 | -0.211 | -0.004 | -0.004 | -0.004 | 0 | 0.62 | 0.43 | 0.69 | 0.61 | 0.67 | 0.62 |
| 8 | 24285553 | rs6557746 | -0.704 | 0.268 | -0.411 | -0.865 | -0.001 | -0.007 | -0.007 | 0 | 0.78 | 0.82 | 0.79 | 0.3 | 0.74 | 0.67 |
| 8 | 24287743 | rs9314283 | -0.425 | 0.251 | -0.512 | -1.001 | 0 | 0.004 | 0.004 | 0 | 0.78 | 0.77 | 0.84 | 0.53 | 0.8 | 0.71 |
| 8 | 24292266 | rs10112054 | -1.099 | 0.288 | -0.468 | -1.072 | 0.001 | -0.013 | -0.013 | 0.198 | 0.39 | 0.53 | 0.32 | 0.28 | 0.35 | 0.23 |
| 8 | 24293499 | rs7846188 | -0.126 | 0.288 | -0.487 | -1.056 | 0.002 | 0.009 | 0.009 | 0 | 0.69 | 0.75 | 0.75 | 0.3 | 0.69 | 0.65 |
| 8 | 24297351 | rs1492421 | -0.876 | 0.056 | -0.324 | -1.065 | 0 | -0.013 | -0.013 | 0.597 | 0.41 | 0.46 | 0.32 | 0.28 | 0.36 | 0.28 |
| 8 | 24305307 | rs7845024 | -0.459 | -0.06 | -0.058 | -2.102 | -0.002 | 0.005 | 0.005 | 0 | 0.66 | 0.67 | 0.73 | 0.3 | 0.65 | 0.61 |
| 8 | 24327201 | rs7816257 | 1.315 | 0.081 | -0.079 | -2.428 | 0.006 | 0.015 | 0.015 | 0 | 0.35 | 0.33 | 0.44 | 0.19 | 0.31 | 0.4 |
| 8 | 24331204 | rs17738359 | 1.83 | 0.099 | -0.255 | -2.43 | -0.061 | 0.028 | 0.028 | 1.384 | 0.12 | 0.01 | 0.12 | 0.06 | 0.2 | 0.1 |

| | | | | | | | | | | | | | | | | |
|---|---|---|---|---|---|---|---|---|---|---|---|---|---|---|---|---|
| 8 | 24332030 | rs13275040 | 1.615 | 0.091 | -0.122 | -2.388 | 0.014 | 0.177 | 0.177 | 0 | 0.32 | 0.07 | 0.27 | 0.05 | 0.27 | 0.28 |
| 8 | 24339410 | rs7006845 | -1.613 | 0.074 | -0.052 | -2.383 | -0.001 | 0 | 0 | 0 | 0.75 | 0.72 | 0.77 | 0.3 | 0.69 | 0.63 |
| 8 | 24339679 | rs13255694 | 1.522 | 0.072 | -0.055 | -2.359 | 0.004 | 0.064 | 0.064 | 0 | 0.33 | 0.1 | 0.43 | 0.19 | 0.3 | 0.39 |
| 8 | 24345959 | rs990463 | 1.349 | 0.075 | -0.008 | -2.357 | 0.014 | 0.039 | 0.039 | 0 | 0.34 | 0.24 | 0.43 | 0.19 | 0.3 | 0.41 |
| 8 | 24348121 | rs1492400 | -0.845 | 0.08 | 0.013 | -2.387 | -0.001 | -0.039 | -0.039 | 0.597 | 0.4 | 0.56 | 0.33 | 0.11 | 0.39 | 0.22 |
| 8 | 24348241 | rs2242354 | 1.373 | 0.079 | -0.022 | -2.363 | 0.015 | 0.06 | 0.06 | 0 | 0.34 | 0.15 | 0.43 | 0.19 | 0.31 | 0.41 |
| 8 | 24350715 | rs12547971 | 1.359 | -0.001 | 0.031 | -2.36 | -0.013 | 0.016 | 0.016 | 0 | 0.07 | 0.04 | 0.12 | 0 | 0.09 | 0.05 |
| 8 | 24356818 | rs13259668 | 1.278 | -0.042 | 0.225 | -2.359 | 0.006 | 0.011 | 0.011 | 0 | 0.35 | 0.32 | 0.45 | 0.19 | 0.31 | 0.41 |
| 8 | 24362810 | rs1492418 | -1.512 | 0.083 | 0.113 | -2.257 | -0.012 | -0.001 | -0.001 | 0.222 | 0.75 | 0.81 | 0.77 | 0.29 | 0.69 | 0.63 |
| 8 | 24366894 | rs3173956 | -1.384 | -0.015 | 0.405 | -2.01 | -0.007 | 0.009 | 0.009 | 0 | 0.69 | 0.74 | 0.73 | 0.3 | 0.64 | 0.61 |
| 8 | 24367198 | rs902847 | -0.904 | 0.15 | 0.54 | -1.97 | 0.014 | -0.127 | -0.127 | 0 | 0.33 | 0.47 | 0.29 | 0.11 | 0.34 | 0.21 |
| 8 | 24368940 | rs17052096 | -0.292 | 0.156 | 0.56 | -1.962 | 0.002 | -0.123 | -0.123 | 0 | 0.26 | 0.48 | 0.18 | 0.11 | 0.25 | 0.17 |
| 8 | 24370834 | rs10503759 | 0.354 | 0.127 | 0.67 | -1.959 | -0.025 | -0.069 | -0.069 | 0 | 0.13 | 0.27 | 0.05 | 0.05 | 0.07 | 0.07 |
| 8 | 24370878 | rs7016551 | -0.803 | 0.141 | 0.406 | -1.978 | -0.018 | -0.143 | -0.143 | 0 | 0.22 | 0.55 | 0.19 | 0.05 | 0.16 | 0.11 |
| 8 | 24371091 | rs17812926 | 1.172 | 0.198 | 0.21 | -1.939 | -0.013 | 0.015 | 0.015 | 0 | 0.07 | 0.04 | 0.12 | 0 | 0.09 | 0.05 |
| 8 | 24377862 | rs4472524 | 0.132 | 0.208 | 0.23 | -1.98 | 0.005 | 0.061 | 0.061 | 0 | 0.49 | 0.32 | 0.55 | 0.26 | 0.49 | 0.5 |
| 8 | 24378741 | rs11781622 | 0.095 | 0.04 | 0.312 | -1.79 | 0.001 | -0.123 | -0.123 | 0 | 0.81 | 0.53 | 0.81 | 0.95 | 0.84 | 0.89 |
| 8 | 24381375 | rs12546913 | 0.501 | 0.073 | -0.199 | -1.433 | 0.033 | 0.066 | 0.066 | 0 | 0.37 | 0.21 | 0.44 | 0.2 | 0.31 | 0.41 |
| 8 | 24398854 | rs7831504 | 0.91 | 0.681 | 0.364 | -1.497 | -0.01 | 0.027 | 0.027 | 1.58 | 0.68 | 0.29 | 0.68 | 0.88 | 0.65 | 0.83 |
| 8 | 24400773 | rs12543494 | -0.146 | 0.783 | 1.03 | -1.527 | 0.058 | -0.057 | -0.057 | 0.297 | 0.1 | 0.33 | 0.14 | 0 | 0.1 | 0.05 |
| 8 | 24401779 | rs11991530 | 0.636 | 0.431 | 1.108 | -1.243 | 0.031 | -0.009 | -0.009 | 0.019 | 0.12 | 0.19 | 0.15 | 0.1 | 0.2 | 0.1 |
| 8 | 24410033 | rs12678642 | 0.707 | -0.817 | 1.401 | -1.152 | -0.007 | -0.017 | -0.017 | 0 | 0.58 | 0.52 | 0.49 | 0.81 | 0.68 | 0.57 |
| 8 | 24410561 | rs1492415 | 0.153 | -0.748 | 0.579 | -1.24 | 0.019 | 0.154 | 0.154 | 0 | 0.29 | 0.06 | 0.26 | 0.05 | 0.23 | 0.29 |
| 8 | 24412159 | rs969456 | 0.519 | -0.388 | 1.579 | -0.755 | 0.024 | -0.05 | -0.05 | 0 | 0.9 | 0.69 | 0.93 | 0.99 | 0.91 | 0.99 |
| 8 | 24416733 | rs13250765 | -0.663 | -0.593 | 1.75 | -0.898 | -0.025 | -0.04 | -0.04 | 0 | 0.33 | 0.5 | 0.45 | 0.2 | 0.28 | 0.45 |
| 8 | 24426949 | rs4871919 | -0.186 | -0.665 | 1.831 | -0.874 | 0.018 | 0.025 | 0.025 | 0 | 0.29 | 0.15 | 0.42 | 0.22 | 0.27 | 0.45 |
| 8 | 24427039 | rs11784810 | 1.349 | -0.681 | 1.67 | -0.89 | 0.052 | 0.119 | 0.119 | 0 | 0.57 | 0.19 | 0.49 | 0.77 | 0.64 | 0.54 |
| 8 | 24430258 | rs9314293 | -0.447 | -0.683 | 1.917 | -0.888 | 0.006 | 0.006 | 0.006 | 0 | 0.31 | 0.29 | 0.43 | 0.22 | 0.27 | 0.45 |
| 8 | 24430924 | rs9792163 | 1.204 | -0.664 | 1.635 | -0.877 | 0.03 | 0.066 | 0.066 | 0 | 0.57 | 0.29 | 0.5 | 0.77 | 0.63 | 0.53 |
| 8 | 24432087 | rs6991250 | -1.041 | -0.65 | 1.267 | -0.724 | -0.016 | -0.028 | -0.028 | 0 | 0.37 | 0.53 | 0.46 | 0.21 | 0.3 | 0.43 |
| 8 | 24433531 | rs11773926 | -0.134 | -0.633 | 1.212 | -0.596 | 0.008 | 0.007 | 0.007 | 0 | 0.26 | 0.25 | 0.41 | 0.21 | 0.22 | 0.42 |
| 8 | 24450423 | rs7828068 | -0.622 | -0.529 | 0.744 | -0.512 | -0.027 | -0.025 | -0.025 | 0.294 | 0.31 | 0.55 | 0.44 | 0.21 | 0.22 | 0.41 |
| 8 | 24456913 | rs11782327 | 0.355 | -0.104 | 0.48 | -0.175 | -0.016 | -0.032 | -0.032 | 0 | 0.69 | 0.46 | 0.55 | 0.78 | 0.71 | 0.68 |
| 8 | 24457899 | rs17052218 | -1.455 | -0.159 | 1.111 | 0.379 | -0.033 | 0.019 | 0.019 | 0.158 | 0.16 | 0.58 | 0.13 | 0.18 | 0.08 | 0.13 |
| 8 | 24473215 | rs17052249 | -0.99 | -0.042 | 2.318 | 0.417 | 0.002 | 0.086 | 0.086 | 0.25 | 0.12 | 0.46 | 0.11 | 0.16 | 0.07 | 0.12 |
| 8 | 24476252 | rs17052263 | -0.719 | -0.045 | 2.642 | 0.417 | -0.012 | 0.038 | 0.038 | 0.078 | 0.1 | 0.18 | 0.09 | 0.16 | 0.07 | 0.12 |
| 8 | 24484612 | rs6981398 | -1.479 | -0.071 | 2.658 | 0.404 | -0.021 | 0.051 | 0.051 | 1.463 | 0.13 | 0.58 | 0.12 | 0.16 | 0.07 | 0.12 |
| 8 | 24486153 | rs7001105 | -1.094 | -0.06 | 2.624 | 0.41 | -0.007 | 0.068 | 0.068 | 0.286 | 0.11 | 0.34 | 0.1 | 0.16 | 0.07 | 0.12 |
| 8 | 24510378 | rs7464587 | 1.088 | 0.121 | 1.332 | 0.162 | -0.018 | 0.154 | 0.154 | 0 | 0.45 | 0.2 | 0.37 | 0.13 | 0.49 | 0.29 |
| 8 | 24523301 | rs6557756 | 0.25 | 0.371 | 0.76 | 0.18 | -0.037 | -0.002 | -0.002 | 0 | 0.7 | 0.9 | 0.57 | 0.29 | 0.68 | 0.48 |
| 8 | 24538247 | rs4612351 | 0.31 | 0.483 | 1.022 | 1.365 | 0.001 | 0.095 | 0.095 | 0 | 0.92 | 0.94 | 0.7 | 0.59 | 0.87 | 0.87 |
| 8 | 24553382 | rs17744454 | 0.118 | 0.415 | 0.808 | 0.744 | -0.037 | 0.069 | 0.069 | 0 | 0.16 | 0.06 | 0.08 | 0 | 0.15 | 0.11 |
| 8 | 24562232 | rs4377969 | -0.033 | 0.726 | 1.082 | 0.889 | 0.013 | 0.101 | 0.101 | 0 | 0.87 | 0.86 | 0.66 | 0.55 | 0.83 | 0.85 |
| 8 | 24564992 | rs4872257 | 0.133 | 0.718 | 1.126 | 0.912 | 0.017 | 0.123 | 0.123 | 0 | 0.86 | 0.84 | 0.66 | 0.55 | 0.83 | 0.85 |
| 8 | 24566766 | rs4461905 | -0.747 | 0.477 | 1.449 | 0.693 | -0.002 | 0.021 | 0.021 | 0.223 | 0.31 | 0.27 | 0.16 | 0.22 | 0.28 | 0.34 |
| 8 | 24580132 | rs11990473 | -0.843 | 0.469 | 1.619 | 0.67 | 0.001 | -0.042 | -0.042 | 0 | 0.32 | 0.45 | 0.18 | 0.22 | 0.28 | 0.34 |
| 8 | 24580756 | rs11783712 | -0.745 | 0.453 | 1.533 | 0.662 | 0 | -0.001 | -0.001 | 8.503 | 0.31 | 0.28 | 0.17 | 0.22 | 0.28 | 0.34 |
| 8 | 24583700 | rs17052453 | 0.623 | 0.464 | 1.454 | 0.71 | 0.001 | -0.002 | -0.002 | 0 | 0.16 | 0.15 | 0.09 | 0.22 | 0.13 | 0.26 |
| 8 | 24587657 | rs11993163 | 1.491 | 0.446 | 1.484 | 0.72 | -0.003 | 0.048 | 0.048 | 0 | 0.52 | 0.39 | 0.39 | 0.19 | 0.53 | 0.35 |
| 8 | 24588610 | rs10095932 | 1.006 | 0.388 | 1.434 | 0.711 | 0.02 | 0.001 | 0.001 | 0.005 | 0.67 | 0.55 | 0.49 | 0.4 | 0.66 | 0.61 |
| 8 | 24589864 | rs6995471 | 0.347 | 0.405 | 1.379 | 0.759 | -0.014 | -0.004 | -0.004 | 0 | 0.16 | 0.12 | 0.17 | 0.16 | 0.17 | 0.24 |
| 8 | 24591705 | rs11992743 | -0.136 | 0.407 | 1.391 | 0.767 | -0.013 | -0.034 | -0.034 | 0 | 0.86 | 0.94 | 0.67 | 0.57 | 0.83 | 0.85 |
| 8 | 24591784 | rs11997946 | 0.914 | 0.386 | 1.405 | 0.74 | 0.008 | -0.001 | -0.001 | 0 | 0.67 | 0.59 | 0.49 | 0.4 | 0.66 | 0.61 |
| 8 | 24596387 | rs10103344 | 0.803 | 0.42 | 1.182 | 0.757 | -0.016 | -0.003 | -0.003 | 0.485 | 0.68 | 0.7 | 0.5 | 0.4 | 0.66 | 0.61 |
| 8 | 24604793 | rs1824190 | 1.346 | 0.38 | 1.18 | 0.641 | -0.001 | -0.016 | -0.016 | 0 | 0.52 | 0.47 | 0.4 | 0.2 | 0.53 | 0.34 |
| 8 | 24606874 | rs2607611 | 1.431 | 0.373 | 1.32 | 0.633 | 0.002 | 0.074 | 0.074 | 0 | 0.51 | 0.32 | 0.39 | 0.2 | 0.54 | 0.37 |
| 8 | 24612421 | rs2712773 | 1.004 | 0.385 | 1.289 | 0.806 | -0.003 | -0.008 | -0.008 | 5.783 | 0.62 | 0.7 | 0.56 | 0.41 | 0.62 | 0.59 |
| 8 | 24624746 | rs2607620 | 0.264 | 0.544 | 0.758 | 1.01 | -0.063 | -0.127 | -0.127 | 0.022 | 0.8 | 0.97 | 0.65 | 0.61 | 0.75 | 0.82 |
| 8 | 24628464 | rs2720723 | -0.079 | 0.236 | 0.828 | 1.21 | -0.013 | -0.069 | -0.069 | 0 | 0.83 | 0.94 | 0.7 | 0.67 | 0.82 | 0.84 |
| 8 | 24632351 | rs10503748 | 0.26 | 0.069 | 0.845 | 1.085 | 0.009 | -0.006 | -0.006 | 0 | 0.18 | 0.14 | 0.11 | 0.22 | 0.15 | 0.25 |
| 8 | 24634243 | rs17826587 | 0.242 | 0.06 | 1.135 | 1.081 | -0.002 | -0.001 | -0.001 | 0.022 | 0.18 | 0.2 | 0.11 | 0.22 | 0.15 | 0.25 |
| 8 | 24634770 | rs984923 | -0.962 | -0.066 | 0.725 | 0.774 | 0.011 | 0.011 | 0.011 | 0 | 0.41 | 0.87 | 0.33 | 0.39 | 0.33 | 0.51 |
| 8 | 24642112 | rs2245382 | 0.241 | -0.025 | 0.965 | 0.736 | 0.005 | -0.105 | -0.105 | 0 | 0.2 | 0.32 | 0.1 | 0.01 | 0.18 | 0.11 |
| 8 | 24644724 | rs2607604 | -0.176 | 0.182 | 0.72 | 0.794 | -0.027 | -0.093 | -0.093 | 0.4 | 0.82 | 0.97 | 0.69 | 0.66 | 0.79 | 0.85 |
| 8 | 24644749 | rs2607605 | 0.177 | 0.245 | 0.988 | 0.79 | 0.019 | -0.128 | -0.128 | 0.514 | 0.2 | 0.38 | 0.1 | 0.01 | 0.18 | 0.11 |
| 8 | 24649816 | rs6557770 | -0.116 | 0.232 | 0.325 | 1.01 | 0.005 | 0.017 | 0.017 | 0.013 | 0.23 | 0.58 | 0.22 | 0.26 | 0.2 | 0.29 |
| 8 | 24649968 | rs7835162 | 0.4 | 0.267 | 0.134 | 1.224 | 0.01 | 0.02 | 0.02 | 0 | 0.17 | 0.13 | 0.11 | 0.14 | 0.15 | 0.24 |
| 8 | 24650399 | rs17052625 | -0.046 | 0.381 | 0.112 | 1.201 | -0.006 | 0.08 | 0.08 | 0 | 0.2 | 0.34 | 0.23 | 0.42 | 0.18 | 0.43 |
| 8 | 24656085 | rs17754434 | 1.349 | 0.395 | -1.028 | 1.039 | -0.129 | 0.027 | 0.027 | 0 | 0.39 | 0.04 | 0.36 | 0.26 | 0.45 | 0.34 |

| | | | | | | | | | | | | | | | | |
|---|---|---|---|---|---|---|---|---|---|---|---|---|---|---|---|---|
| 8 | 24659652 | rs7838471 | 0.189 | 0.408 | -0.459 | 0.458 | 0 | -0.002 | -0.002 | 0.862 | 0.17 | 0.12 | 0.09 | 0.24 | 0.15 | 0.26 |
| 8 | 24663706 | rs7010000 | -0.059 | 0.496 | -0.712 | -0.002 | 0.004 | -0.173 | -0.173 | 0 | 0.72 | 0.53 | 0.85 | 0.99 | 0.75 | 0.87 |
| 8 | 24666099 | rs12545900 | 0.338 | 0.606 | -1.783 | 0.523 | -0.005 | 0.06 | 0.06 | 0.064 | 0.52 | 0.15 | 0.64 | 0.6 | 0.59 | 0.48 |
| 8 | 24677132 | rs7010085 | 0.133 | 0.57 | -1.386 | 0.349 | -0.009 | 0.026 | 0.026 | 1.06 | 0.53 | 0.3 | 0.47 | 0.35 | 0.53 | 0.45 |
| 8 | 24677490 | rs1016446 | 0.098 | 0.171 | -2.03 | -0.185 | 0.006 | 0.082 | 0.082 | 0.356 | 0.68 | 0.91 | 0.86 | 0.84 | 0.73 | 0.71 |
| 8 | 24681600 | rs11135827 | 0.505 | -0.525 | -2.331 | -0.593 | 0.055 | 0.089 | 0.089 | 0.03 | 0.63 | 0.1 | 0.79 | 0.76 | 0.73 | 0.74 |
| 8 | 24688776 | rs196800 | 0.2 | -0.619 | -0.919 | -0.683 | 0.011 | 0.003 | 0.003 | 0 | 0.25 | 0.21 | 0.12 | 0.25 | 0.18 | 0.22 |
| 8 | 24709967 | rs11775313 | 0.561 | -0.61 | -0.151 | -0.72 | 0.053 | 0.068 | 0.068 | 0.043 | 0.65 | 0.12 | 0.8 | 0.75 | 0.74 | 0.78 |
| 8 | 24716594 | rs196814 | -0.358 | -0.63 | -0.139 | -0.713 | 0.003 | 0.004 | 0.004 | 0 | 0.3 | 0.72 | 0.15 | 0.27 | 0.18 | 0.21 |
| 8 | 24722527 | rs17052718 | -0.73 | -0.698 | 0.274 | -0.759 | 0.01 | -0.017 | -0.017 | 0 | 0.1 | 0.53 | 0.08 | 0.01 | 0.07 | 0.06 |
| 8 | 24723726 | rs376886 | 0.337 | -0.724 | 0.252 | -0.776 | 0.024 | 0.007 | 0.007 | 0 | 0.23 | 0.15 | 0.07 | 0.25 | 0.13 | 0.2 |
| 8 | 24725705 | rs7003271 | -0.753 | -0.659 | 0.42 | -0.955 | 0.014 | -0.013 | -0.013 | 0 | 0.1 | 0.52 | 0.08 | 0.01 | 0.07 | 0.06 |
| 8 | 24726623 | rs2320562 | 1.732 | -0.592 | -0.355 | -1.349 | -0.109 | 0.16 | 0.16 | 1.453 | 0.33 | 0.05 | 0.42 | 0.08 | 0.43 | 0.29 |
| 8 | 24729914 | rs7844382 | -0.531 | -0.69 | -0.106 | -1.262 | -0.01 | -0.033 | -0.033 | 0.232 | 0.1 | 0.61 | 0.09 | 0.01 | 0.07 | 0.05 |
| 8 | 24730318 | rs196896 | -0.576 | -0.582 | -0.169 | -1.222 | 0.072 | -0.017 | -0.017 | 0 | 0.95 | 0.83 | 0.96 | 1 | 0.92 | 0.99 |
| 8 | 24736607 | rs196886 | 0.983 | -0.709 | -0.329 | -0.489 | 0.017 | -0.024 | -0.024 | 4.85 | 0.21 | 0.15 | 0.08 | 0.44 | 0.11 | 0.17 |
| 8 | 24739169 | rs926104 | 0.415 | -0.691 | 0.064 | -0.502 | -0.002 | 0.001 | 0.001 | 0 | 0.66 | 0.43 | 0.81 | 0.54 | 0.73 | 0.72 |
| 8 | 24745196 | rs11786728 | 1.487 | -0.621 | -0.263 | -0.523 | -0.158 | 0.118 | 0.118 | 1.392 | 0.35 | 0.05 | 0.47 | 0.17 | 0.5 | 0.31 |
| 8 | 24745451 | rs2975186 | 0.753 | -0.699 | -0.455 | -0.794 | 0.036 | -0.028 | -0.028 | 0.276 | 0.55 | 0.13 | 0.73 | 0.54 | 0.64 | 0.71 |
| 8 | 24754180 | rs196875 | -0.721 | -0.431 | -0.18 | -0.859 | 0.091 | -0.008 | -0.008 | 0 | 0.95 | 0.81 | 0.95 | 1 | 0.93 | 0.99 |
| 8 | 24759930 | rs9650410 | 1.425 | -0.442 | -0.751 | -0.871 | -0.127 | 0.112 | 0.112 | 1.482 | 0.31 | 0.05 | 0.42 | 0.13 | 0.44 | 0.31 |
| 8 | 24769595 | rs196868 | 0.482 | -0.788 | -0.21 | -0.678 | 0.041 | -0.045 | -0.045 | 0.306 | 0.24 | 0.18 | 0.08 | 0.45 | 0.12 | 0.18 |
| 8 | 24769852 | rs1457266 | 0.569 | -0.591 | -0.485 | -0.659 | 0.057 | -0.018 | -0.018 | 0.009 | 0.55 | 0.1 | 0.74 | 0.57 | 0.65 | 0.69 |
| 8 | 24778956 | rs13251967 | 0.648 | -0.725 | -0.482 | -0.698 | 0.075 | -0.009 | -0.009 | 0 | 0.53 | 0.11 | 0.73 | 0.55 | 0.65 | 0.67 |
| 8 | 24783920 | rs11135833 | 0.66 | -0.769 | -0.223 | -0.815 | -0.01 | -0.041 | -0.041 | 0.044 | 0.12 | 0.1 | 0.06 | 0 | 0.09 | 0.04 |
| 8 | 24786803 | rs10097219 | 0.672 | -0.99 | -0.278 | -0.733 | 0.027 | -0.05 | -0.05 | 0 | 0.21 | 0.11 | 0.08 | 0.42 | 0.13 | 0.17 |
| 8 | 24790573 | rs118727 | -0.296 | -0.56 | 0.005 | -0.881 | 0.006 | -0.019 | -0.019 | 0.137 | 0.92 | 0.89 | 0.94 | 0.99 | 0.88 | 0.9 |
| 8 | 24791552 | rs196839 | 0.349 | -0.589 | 0.352 | -0.899 | 0.053 | -0.034 | -0.034 | 0 | 0.9 | 0.61 | 0.91 | 0.99 | 0.88 | 0.91 |
| 8 | 24795851 | rs169061 | -0.64 | -0.3 | -1.116 | -1.326 | -0.035 | 0.161 | 0.161 | 0.028 | 0.63 | 0.86 | 0.5 | 0.93 | 0.42 | 0.6 |
| 8 | 24799742 | rs6557786 | -0.095 | -0.242 | -0.985 | -1.331 | 0.009 | -0.023 | -0.023 | 0 | 0.9 | 0.83 | 0.92 | 0.98 | 0.85 | 0.9 |
| 8 | 24800190 | rs10100189 | 1.279 | -0.676 | -1.103 | -1.26 | 0.027 | -0.048 | -0.048 | 0.078 | 0.2 | 0.11 | 0.08 | 0.43 | 0.13 | 0.16 |
| 8 | 24802235 | rs11994760 | 0.023 | -0.839 | -0.955 | -1.379 | -0.029 | 0.002 | 0.002 | 0 | 0.62 | 0.42 | 0.83 | 0.58 | 0.79 | 0.79 |
| 8 | 24809067 | rs2979704 | -0.758 | -0.84 | -0.863 | -1.369 | -0.073 | 0.094 | 0.094 | 0 | 0.75 | 0.53 | 0.9 | 0.58 | 0.87 | 0.84 |
| 8 | 24816437 | rs3757985 | 1.867 | -0.637 | -1.007 | -1.417 | -0.148 | 0.146 | 0.146 | 1.081 | 0.27 | 0.05 | 0.43 | 0.05 | 0.43 | 0.28 |
| 8 | 24817145 | rs17830286 | 0.096 | -0.818 | -0.639 | -1.255 | 0.015 | -0.034 | -0.034 | 0 | 0.1 | 0.19 | 0.08 | 0.02 | 0.15 | 0.1 |
| 8 | 24818901 | rs17830392 | 2.535 | -0.53 | -1.044 | -1.467 | -0.004 | 0.03 | 0.03 | 8.662 | 0.08 | 0.02 | 0.05 | 0.03 | 0.06 | 0.04 |
| 8 | 24819116 | rs2979682 | 0.798 | -0.602 | -0.653 | -1.122 | 0.067 | 0.019 | 0.019 | 0 | 0.52 | 0.23 | 0.74 | 0.56 | 0.64 | 0.69 |
| 8 | 24838662 | rs11779795 | -0.507 | -0.519 | -0.93 | -1.24 | -0.027 | -0.133 | -0.133 | 0.183 | 0.23 | 0.57 | 0.15 | 0.02 | 0.2 | 0.13 |
| 8 | 24844891 | rs11135836 | 0.469 | -0.336 | -2.328 | -1.134 | 0.021 | 0.041 | 0.041 | 0 | 0.19 | 0.14 | 0.09 | 0 | 0.15 | 0.11 |
| 8 | 24853843 | rs13264292 | 0.849 | -0.011 | -2.71 | -0.778 | 0.011 | -0.021 | -0.021 | 4.557 | 0.46 | 0.08 | 0.71 | 0.57 | 0.58 | 0.64 |
| 8 | 24855584 | rs4872278 | 1.328 | -0.083 | -2.957 | -0.51 | -0.136 | -0.092 | -0.092 | 0 | 0.34 | 0.07 | 0.51 | 0.36 | 0.43 | 0.39 |
| 8 | 24856504 | rs17763685 | 0.436 | 0.275 | -2.041 | 0.402 | -0.01 | 0.014 | 0.014 | 0 | 0.06 | 0.05 | 0.07 | 0.02 | 0.11 | 0.08 |
| 8 | 24857168 | rs17052966 | 0.919 | 0.217 | -1.97 | 0.555 | -0.03 | -0.051 | -0.051 | 0 | 0.13 | 0 | 0.16 | 0.19 | 0.13 | 0.22 |
| 8 | 24858551 | rs2976408 | -0.63 | 0.125 | -1.627 | 0.569 | 0.024 | -0.054 | -0.054 | 0.19 | 0.83 | 0.93 | 0.9 | 0.67 | 0.88 | 0.87 |
| 8 | 24861815 | rs2976411 | 0.336 | 0.572 | -1.069 | 0.372 | -0.004 | -0.003 | -0.003 | 0 | 0.54 | 0.55 | 0.7 | 0.39 | 0.57 | 0.53 |
| 8 | 24862709 | rs2950347 | -0.143 | 0.514 | -1.416 | 0.558 | -0.01 | -0.016 | -0.016 | 0 | 0.8 | 0.93 | 0.88 | 0.78 | 0.81 | 0.77 |
| 8 | 24865502 | rs35842725 | 0.643 | 1.025 | -1.719 | 0.334 | -0.03 | 0.025 | 0.025 | 0.711 | 0.25 | 0.11 | 0.37 | 0.16 | 0.32 | 0.2 |
| 8 | 24869436 | rs2976456 | -0.618 | 0.603 | -1.662 | 0.213 | 0.007 | -0.003 | -0.003 | 0.026 | 0.75 | 0.75 | 0.85 | 0.77 | 0.86 | 0.65 |
| 8 | 24873978 | rs2950324 | -0.04 | 0.535 | -1.156 | 0.369 | 0 | 0.006 | 0.006 | 0.349 | 0.48 | 0.51 | 0.45 | 0.62 | 0.53 | 0.48 |
| 8 | 24889912 | rs1527033 | -0.683 | 0.636 | -0.563 | 0.671 | 0.003 | -0.018 | -0.018 | 0.095 | 0.65 | 0.77 | 0.76 | 0.5 | 0.61 | 0.68 |
| 8 | 24896797 | rs4872285 | -1.187 | 0.605 | -0.66 | 0.925 | -0.001 | -0.003 | -0.003 | 0 | 0.74 | 0.79 | 0.82 | 0.76 | 0.74 | 0.81 |
| 8 | 24900556 | rs7002363 | 0.005 | 0.741 | -0.227 | 0.594 | -0.001 | 0 | 0 | 0 | 0.32 | 0.32 | 0.32 | 0.3 | 0.3 | 0.5 |
| 8 | 24901750 | rs11135839 | -0.096 | 0.618 | -0.118 | 0.591 | 0 | 0.001 | 0.001 | 0 | 0.32 | 0.36 | 0.31 | 0.33 | 0.29 | 0.45 |
| 8 | 24903745 | rs11773978 | -0.128 | 0.634 | -0.096 | 0.435 | 0 | -0.003 | -0.003 | 0 | 0.32 | 0.4 | 0.32 | 0.26 | 0.29 | 0.38 |
| 8 | 24915917 | rs13265334 | 1.475 | 0.742 | -0.27 | 0.373 | -0.063 | 0.021 | 0.021 | 5.063 | 0.28 | 0.07 | 0.45 | 0.2 | 0.33 | 0.17 |
| 8 | 24920430 | rs13272587 | 0.958 | 0.772 | 0.021 | 0.388 | 0.007 | -0.019 | -0.019 | 1.339 | 0.3 | 0.39 | 0.46 | 0.2 | 0.33 | 0.17 |
| 8 | 24921780 | rs1358269 | 0.183 | 0.943 | 0.02 | 0.567 | -0.011 | -0.003 | -0.003 | 0 | 0.27 | 0.22 | 0.29 | 0.26 | 0.28 | 0.37 |
| 8 | 24924178 | rs11135840 | -0.603 | 0.743 | -0.056 | 0.44 | 0.014 | -0.024 | -0.024 | 0 | 0.6 | 0.76 | 0.77 | 0.48 | 0.6 | 0.56 |
| 8 | 24924326 | rs17053065 | -0.061 | 0.721 | 0.13 | 0.411 | -0.001 | 0 | 0 | 0 | 0.3 | 0.37 | 0.3 | 0.27 | 0.28 | 0.38 |
| 8 | 24925058 | rs7017203 | -0.667 | 0.725 | -0.008 | 0.466 | 0.017 | -0.025 | -0.025 | 0 | 0.6 | 0.75 | 0.77 | 0.48 | 0.6 | 0.55 |
| 8 | 24934368 | rs4260904 | -0.318 | 1.198 | 0.943 | 0.485 | -0.003 | 0.008 | 0.008 | 0 | 0.24 | 0.34 | 0.25 | 0.27 | 0.17 | 0.34 |
| 8 | 24935155 | rs11994900 | -0.173 | 1.16 | 0.945 | 0.469 | 0 | -0.004 | -0.004 | 2.099 | 0.23 | 0.25 | 0.24 | 0.27 | 0.17 | 0.33 |
| 8 | 24936863 | rs6996803 | -1.133 | 1.221 | 1.144 | 0.457 | 0.011 | -0.009 | -0.009 | 0 | 0.32 | 0.54 | 0.33 | 0.27 | 0.3 | 0.35 |
| 8 | 24943384 | rs17053092 | -0.178 | 1.283 | 1.051 | 0.46 | 0.058 | -0.046 | -0.046 | 0 | 0.08 | 0.21 | 0.08 | 0 | 0.13 | 0.02 |
| 8 | 24944411 | rs6984035 | -0.386 | 1.291 | 1.064 | 0.463 | 0.064 | -0.048 | -0.048 | 0.545 | 0.09 | 0.26 | 0.08 | 0 | 0.13 | 0.02 |
| 8 | 24952142 | rs17053106 | -0.329 | 1.31 | 1.024 | 0.474 | 0.073 | -0.045 | -0.045 | 0.209 | 0.09 | 0.28 | 0.08 | 0 | 0.13 | 0.02 |
| 8 | 24954959 | rs10503768 | -0.667 | 1.21 | 0.774 | 0.543 | 0.016 | 0.001 | 0.001 | 0 | 0.29 | 0.4 | 0.31 | 0.28 | 0.3 | 0.35 |
| 8 | 24957951 | rs7014893 | -0.834 | 1.229 | 0.687 | 0.537 | 0.013 | -0.004 | -0.004 | 1.431 | 0.31 | 0.57 | 0.33 | 0.27 | 0.3 | 0.35 |
| 8 | 24959167 | rs7833099 | -0.655 | 1.211 | 1.488 | 0.21 | 0.09 | -0.038 | -0.038 | 0 | 0.09 | 0.36 | 0.1 | 0 | 0.12 | 0.02 |

| | | | | | | | | | | | | | | | | |
|---|---|---|---|---|---|---|---|---|---|---|---|---|---|---|---|---|
| 8 | 24967220 | rs1358266 | -1.328 | 1.049 | 1.453 | 0 | 0.009 | -0.09 | -0.09 | 0.1 | 0.15 | 0.29 | 0.13 | 0 | 0.18 | 0.09 |
| 8 | 24967468 | rs2874355 | -2.242 | 1.484 | 1.501 | -0.139 | 0.019 | -0.075 | -0.075 | 1.264 | 0.23 | 0.66 | 0.3 | 0.05 | 0.29 | 0.17 |
| 8 | 24970519 | rs7843688 | -1.568 | 1.304 | 1.6 | -0.049 | -0.005 | -0.042 | -0.042 | 0.306 | 0.12 | 0.4 | 0.18 | 0.05 | 0.09 | 0.11 |
| 8 | 24971005 | rs879958 | -0.929 | 1.301 | 0.793 | 0.14 | -0.034 | 0.05 | 0.05 | 1.022 | 0.42 | 0.76 | 0.44 | 0.27 | 0.52 | 0.36 |
| 8 | 24991003 | rs13250561 | 1.875 | 1.119 | 0.232 | 0.376 | -0.035 | -0.084 | -0.084 | 1.412 | 0.35 | 0.03 | 0.43 | 0.35 | 0.32 | 0.26 |
| 8 | 24991235 | rs2950289 | -0.693 | 1.318 | 0.789 | -0.126 | -0.022 | 0.027 | 0.027 | 0.524 | 0.4 | 0.71 | 0.43 | 0.28 | 0.52 | 0.36 |
| 8 | 24992463 | rs2950291 | 0.084 | 1.345 | 0.846 | -0.154 | -0.029 | 0.011 | 0.011 | 0 | 0.3 | 0.26 | 0.25 | 0.22 | 0.43 | 0.25 |
| 8 | 24993705 | rs2979831 | -0.117 | 1.291 | 0.967 | -0.14 | 0.001 | -0.004 | -0.004 | 0.055 | 0.32 | 0.28 | 0.26 | 0.23 | 0.4 | 0.29 |
| 8 | 24999429 | rs4872291 | -0.697 | 1.307 | 1.1 | -0.098 | 0.064 | -0.005 | -0.005 | 0 | 0.07 | 0.23 | 0.16 | 0.06 | 0.12 | 0.07 |
| 8 | 25006118 | rs1358268 | -0.931 | 1.226 | 1.176 | -0.107 | 0.001 | 0.028 | 0.028 | 0 | 0.5 | 0.53 | 0.35 | 0.35 | 0.5 | 0.56 |
| 8 | 25006687 | rs1554766 | -1.655 | 1.238 | 1.024 | -0.096 | -0.006 | -0.043 | -0.043 | 0.484 | 0.59 | 0.92 | 0.51 | 0.41 | 0.62 | 0.63 |
| 8 | 25009089 | rs2320783 | -0.325 | 1.176 | 1.101 | -0.128 | -0.009 | -0.022 | -0.022 | 0 | 0.24 | 0.26 | 0.1 | 0.12 | 0.15 | 0.34 |
| 8 | 25009616 | rs9694084 | -0.349 | 1.178 | 1.102 | -0.127 | -0.009 | -0.021 | -0.021 | 0.379 | 0.24 | 0.26 | 0.1 | 0.12 | 0.15 | 0.34 |
| 8 | 25013751 | rs2979813 | 0.54 | 1.194 | 1.092 | -0.058 | 0.019 | 0.004 | 0.004 | 0 | 0.27 | 0.34 | 0.25 | 0.24 | 0.36 | 0.21 |
| 8 | 25021361 | rs10283360 | -0.892 | 0.773 | 1.302 | -0.139 | -0.002 | -0.041 | -0.041 | 0.023 | 0.28 | 0.38 | 0.13 | 0.12 | 0.22 | 0.37 |
| 8 | 25025177 | rs2941606 | 1.043 | 0.827 | 1.449 | -0.034 | 0.004 | -0.053 | -0.053 | 0 | 0.72 | 0.39 | 0.73 | 0.83 | 0.77 | 0.6 |
| 8 | 25028671 | rs11985184 | -1.199 | 0.909 | 1.427 | 0.05 | 0.045 | -0.006 | -0.006 | 0 | 0.09 | 0.38 | 0.16 | 0.04 | 0.09 | 0.07 |
| 8 | 25037492 | rs2979823 | -0.8 | 1.027 | 0.618 | 0.048 | -0.019 | 0.057 | 0.057 | 0.199 | 0.46 | 0.82 | 0.59 | 0.32 | 0.55 | 0.37 |
| 8 | 25038234 | rs11774796 | 0.372 | 0.956 | 1.408 | 0.147 | 0.004 | -0.011 | -0.011 | 0.097 | 0.06 | 0.07 | 0.06 | 0 | 0.1 | 0.01 |
| 8 | 25052151 | rs11986821 | 0.929 | -0.01 | 1.695 | 0.595 | -0.022 | -0.002 | -0.002 | 0.133 | 0.72 | 0.53 | 0.83 | 0.64 | 0.81 | 0.59 |
| 8 | 25052372 | rs6996838 | 0.35 | -0.03 | 1.311 | 0.57 | -0.008 | 0.036 | 0.036 | 0 | 0.77 | 0.66 | 0.75 | 0.59 | 0.85 | 0.63 |
| 8 | 25054065 | rs17053187 | -1.123 | 0.083 | 1.297 | 0.602 | 0.005 | -0.063 | -0.063 | 0 | 0.11 | 0.36 | 0.07 | 0 | 0.08 | 0.11 |
| 8 | 25061807 | rs1510758 | 0.894 | 0.171 | 0.305 | 0.76 | 0.007 | 0 | 0 | 0 | 0.68 | 0.28 | 0.68 | 0.59 | 0.77 | 0.52 |
| 8 | 25079502 | rs11135856 | -1.446 | -0.181 | 1.414 | 1.194 | -0.023 | 0.09 | 0.09 | 0 | 0.16 | 0.5 | 0.31 | 0.17 | 0.13 | 0.11 |
| 8 | 25115730 | rs12549196 | -1.864 | -0.586 | 2.175 | 1.128 | -0.034 | 0.051 | 0.051 | 0 | 0.17 | 0.55 | 0.31 | 0.14 | 0.11 | 0.18 |
| 8 | 25116072 | rs11135858 | -0.96 | -0.561 | 2.228 | 1.139 | 0.008 | 0.084 | 0.084 | 0 | 0.11 | 0.23 | 0.3 | 0.14 | 0.11 | 0.12 |
| 8 | 25123492 | rs17053273 | -0.944 | -0.561 | 2.232 | 1.147 | 0.008 | 0.074 | 0.074 | 0 | 0.11 | 0.23 | 0.3 | 0.13 | 0.11 | 0.12 |
| 8 | 25129948 | rs1122450 | 2.126 | -1.096 | 2.35 | 1.18 | -0.044 | 0.027 | 0.027 | 0.104 | 0.76 | 0.36 | 0.65 | 0.77 | 0.85 | 0.69 |
| 8 | 25138403 | rs2666160 | 1.566 | -1.022 | 2.79 | 1.161 | -0.021 | 0.071 | 0.071 | 0 | 0.84 | 0.44 | 0.69 | 0.85 | 0.88 | 0.82 |
| 8 | 25141468 | rs2709620 | 0.739 | -1.028 | 2.95 | 2.055 | -0.029 | 0.084 | 0.084 | 0 | 0.89 | 0.77 | 0.94 | 0.73 | 0.96 | 0.85 |
| 8 | 25141983 | rs6982431 | -1.22 | -1.026 | 3.158 | 2.055 | -0.001 | 0.07 | 0.07 | 0 | 0.12 | 0.33 | 0.3 | 0.14 | 0.12 | 0.18 |
| 8 | 25142069 | rs2666163 | 1.988 | -1.001 | 2.696 | 1.93 | -0.005 | 0.008 | 0.008 | 0 | 0.74 | 0.27 | 0.62 | 0.59 | 0.84 | 0.67 |
| 8 | 25159885 | rs907157 | 1.094 | -1.044 | 0.734 | 0.797 | -0.135 | 0.131 | 0.131 | 0 | 0.39 | 0.05 | 0.36 | 0.24 | 0.5 | 0.33 |
| 8 | 25162736 | rs17053371 | -1.533 | -1.25 | 1.928 | 0.703 | -0.05 | 0.001 | 0.001 | 0 | 0.14 | 0.45 | 0.05 | 0.09 | 0.03 | 0.18 |
| 8 | 25163078 | rs17053372 | -0.703 | -1.247 | 1.682 | 0.706 | -0.003 | 0 | 0 | 6.228 | 0.09 | 0.1 | 0.03 | 0.09 | 0.03 | 0.13 |
| 8 | 25163802 | rs2956654 | 1.732 | -1.173 | 1.787 | 0.738 | -0.022 | 0.015 | 0.015 | 0 | 0.75 | 0.38 | 0.73 | 0.78 | 0.83 | 0.7 |
| 8 | 25172949 | rs2465006 | 1.251 | -1.152 | 1.429 | 0.437 | -0.01 | 0.009 | 0.009 | 0.018 | 0.78 | 0.75 | 0.84 | 0.77 | 0.86 | 0.67 |
| 8 | 25186301 | rs2960173 | 1.018 | -1.106 | 1.254 | 0.378 | -0.001 | -0.001 | -0.001 | 0 | 0.8 | 0.8 | 0.7 | 0.84 | 0.86 | 0.77 |
| 8 | 25190752 | rs2468896 | 1.835 | -1.235 | 1.428 | 0.168 | 0 | 0 | 0 | 0.256 | 0.67 | 0.34 | 0.76 | 0.77 | 0.8 | 0.68 |
| 8 | 25193338 | rs17197236 | 0.011 | -1.036 | 1.348 | 0.474 | 0.023 | 0.064 | 0.064 | 0 | 0.12 | 0.02 | 0.03 | 0 | 0.05 | 0.12 |
| 8 | 25198091 | rs13252932 | -0.636 | -0.979 | 1.576 | 0.52 | 0.023 | 0.025 | 0.025 | 0 | 0.17 | 0.11 | 0.04 | 0.07 | 0.07 | 0.21 |
| 8 | 25209144 | rs7012335 | -1.346 | -0.938 | 1.861 | 0.514 | -0.032 | 0.03 | 0.03 | 0 | 0.24 | 0.47 | 0.18 | 0.23 | 0.15 | 0.31 |
| 8 | 25213299 | rs2465004 | 1.043 | -1.132 | 1.301 | -0.005 | -0.019 | -0.018 | -0.018 | 0.296 | 0.66 | 0.4 | 0.85 | 0.77 | 0.81 | 0.63 |
| 8 | 25217938 | rs10086399 | 0.892 | -0.735 | 0.451 | 0.137 | 0.004 | 0.004 | 0.004 | 0 | 0.23 | 0.23 | 0.32 | 0.34 | 0.37 | 0.21 |
| 8 | 25218092 | rs10096386 | 0.7 | -0.763 | 0.484 | 0.107 | 0.016 | 0.015 | 0.015 | 0 | 0.24 | 0.29 | 0.32 | 0.34 | 0.37 | 0.22 |
| 8 | 25221000 | rs6557837 | -0.481 | -0.878 | 0.041 | -0.149 | -0.039 | -0.1 | -0.1 | 0.034 | 0.32 | 0.38 | 0.16 | 0.04 | 0.14 | 0.26 |
| 8 | 25225804 | rs2257081 | 0.765 | -0.318 | 0.397 | 0.393 | -0.003 | -0.01 | -0.01 | 0.178 | 0.38 | 0.29 | 0.46 | 0.41 | 0.39 | 0.41 |
| 8 | 25226595 | rs10103395 | -0.45 | -0.169 | -0.489 | 0.313 | 0 | 0 | 0 | 0.087 | 0.54 | 0.45 | 0.48 | 0.4 | 0.53 | 0.48 |
| 8 | 25230168 | rs2271108 | -0.303 | -0.13 | -0.852 | 0.335 | -0.003 | 0.015 | 0.015 | 0 | 0.52 | 0.38 | 0.46 | 0.4 | 0.51 | 0.47 |
| 8 | 25231756 | rs6993060 | -0.378 | -0.139 | -0.871 | 0.331 | -0.004 | 0.011 | 0.011 | 0 | 0.53 | 0.39 | 0.47 | 0.4 | 0.51 | 0.47 |
| 8 | 25232630 | rs7813096 | -0.334 | -0.086 | -0.842 | 0.338 | -0.003 | 0.015 | 0.015 | 0 | 0.52 | 0.39 | 0.47 | 0.39 | 0.51 | 0.47 |
| 8 | 25238462 | rs17053529 | 0.204 | 0.138 | -0.991 | 0.581 | -0.01 | -0.009 | -0.009 | 0 | 0.06 | 0.03 | 0.06 | 0.22 | 0.09 | 0.11 |
| 8 | 25240100 | rs7837477 | -0.137 | 0.05 | -0.959 | 0.451 | -0.001 | -0.004 | -0.004 | 0.138 | 0.21 | 0.19 | 0.17 | 0.2 | 0.18 | 0.28 |
| 8 | 25240437 | rs11779690 | -0.181 | 0.517 | -1.314 | 0.6 | -0.006 | -0.008 | -0.008 | 0 | 0.38 | 0.24 | 0.35 | 0.38 | 0.41 | 0.3 |
| 8 | 25242601 | rs925030 | 0.091 | 0.562 | -1.005 | 0.599 | -0.006 | -0.006 | -0.006 | 0 | 0.58 | 0.45 | 0.67 | 0.41 | 0.61 | 0.68 |
| 8 | 25242876 | rs925031 | -1.082 | 0.285 | -0.733 | 1.071 | -0.005 | 0.056 | 0.056 | 0 | 0.07 | 0.16 | 0.07 | 0.22 | 0.06 | 0.06 |
| 8 | 25245398 | rs931874 | 0.166 | 0.136 | -1.763 | -0.047 | 0.014 | 0.166 | 0.166 | 0 | 0.38 | 0.16 | 0.4 | 0.08 | 0.36 | 0.47 |
| 8 | 25245432 | rs931875 | 0.479 | 0.668 | -2.136 | 0.564 | -0.058 | -0.087 | -0.087 | 0 | 0.16 | 0.04 | 0.22 | 0.55 | 0.27 | 0.12 |
| 8 | 25247587 | rs1000798 | -0.217 | 0.434 | -1.914 | 0.57 | 0.027 | 0.025 | 0.025 | 0 | 0.54 | 0.21 | 0.62 | 0.63 | 0.63 | 0.58 |
| 8 | 25250957 | rs11774502 | 0.24 | 0.135 | -0.713 | 0.418 | 0.005 | 0.006 | 0.006 | 0 | 0.41 | 0.32 | 0.36 | 0.35 | 0.36 | 0.36 |
| 8 | 25256561 | rs17053593 | 0.841 | -0.041 | -0.751 | 0.154 | -0.001 | -0.017 | -0.017 | 0 | 0.12 | 0.12 | 0.06 | 0.05 | 0.09 | 0.08 |
| 8 | 25257341 | rs3763519 | 0.72 | -0.092 | -0.8 | 0.111 | 0.001 | -0.005 | -0.005 | 0 | 0.05 | 0.1 | 0.1 | 0.05 | 0.05 | 0.11 |
| 8 | 25264962 | rs2248185 | 0.447 | -0.058 | -0.611 | 0.273 | 0.035 | -0.007 | -0.007 | 0 | 0.54 | 0.25 | 0.62 | 0.59 | 0.63 | 0.58 |
| 8 | 25267622 | rs2709618 | 0.438 | -0.057 | -0.611 | 0.015 | 0.031 | 0.029 | 0.029 | 0 | 0.54 | 0.25 | 0.62 | 0.64 | 0.63 | 0.58 |
| 8 | 25269993 | rs2659582 | 0.487 | -0.089 | -0.197 | -0.028 | 0.02 | 0.019 | 0.019 | 0 | 0.55 | 0.32 | 0.63 | 0.64 | 0.64 | 0.6 |
| 8 | 25270508 | rs7008627 | -0.436 | -0.089 | -0.202 | -0.028 | 0.005 | 0.005 | 0.005 | 0 | 0.43 | 0.38 | 0.36 | 0.36 | 0.36 | 0.39 |
| 8 | 25275274 | rs2659590 | 0.458 | -0.091 | -0.213 | -0.029 | 0.027 | 0.027 | 0.027 | 0 | 0.54 | 0.26 | 0.63 | 0.64 | 0.64 | 0.56 |
| 8 | 25275527 | rs17201112 | 0.963 | -0.132 | -0.173 | -0.053 | 0.004 | 0.032 | 0.032 | 0.105 | 0.12 | 0.06 | 0.05 | 0.05 | 0.09 | 0.08 |
| 8 | 25287556 | rs965130 | 0.51 | -0.14 | -0.196 | -0.086 | 0.03 | 0.028 | 0.028 | 0 | 0.54 | 0.26 | 0.63 | 0.64 | 0.64 | 0.56 |

| | | | | | | | | | | | | | | | | |
|---|---|---|---|---|---|---|---|---|---|---|---|---|---|---|---|---|
| 8 | 25294779 | rs6557842 | 0.549 | -0.185 | -0.204 | -0.089 | 0.03 | 0.028 | 0.028 | 0 | 0.54 | 0.26 | 0.63 | 0.64 | 0.64 | 0.56 |
| 8 | 25296466 | rs3750198 | 0.667 | -0.191 | -0.194 | -0.12 | 0.001 | -0.008 | -0.008 | 0 | 0.13 | 0.12 | 0.06 | 0.05 | 0.09 | 0.08 |
| 8 | 25304906 | rs11787383 | -0.568 | -0.21 | 0.342 | -0.113 | -0.003 | -0.003 | -0.003 | 0 | 0.44 | 0.53 | 0.36 | 0.36 | 0.36 | 0.42 |
| 8 | 25306521 | rs11781066 | -0.239 | -0.213 | 0.34 | -0.115 | -0.004 | -0.003 | -0.003 | 0.769 | 0.39 | 0.43 | 0.27 | 0.31 | 0.31 | 0.31 |
| 8 | 25307798 | rs1567121 | 0.471 | -0.198 | 0.266 | -0.106 | 0.027 | 0.027 | 0.027 | 0 | 0.54 | 0.27 | 0.63 | 0.64 | 0.64 | 0.56 |
| 8 | 25313456 | rs13277090 | 0.465 | -0.24 | 0.237 | -0.111 | 0.026 | 0.026 | 0.026 | 0 | 0.55 | 0.26 | 0.64 | 0.64 | 0.64 | 0.57 |
| 8 | 25314860 | rs4872313 | 0.484 | -0.243 | 0.227 | -0.136 | 0.011 | 0.024 | 0.024 | 0 | 0.56 | 0.31 | 0.64 | 0.66 | 0.64 | 0.59 |
| 8 | 25316661 | rs3812422 | -0.493 | -0.284 | 0.218 | -0.143 | -0.002 | -0.004 | -0.004 | 0 | 0.43 | 0.54 | 0.36 | 0.34 | 0.35 | 0.39 |
| 8 | 25316789 | rs3812421 | -0.503 | -0.286 | 0.24 | -0.144 | -0.002 | -0.003 | -0.003 | 0 | 0.43 | 0.54 | 0.36 | 0.34 | 0.35 | 0.39 |
| 8 | 25318253 | rs17053743 | 0.087 | -0.269 | 0.241 | -0.137 | 0 | 0.009 | 0.009 | 0 | 0.36 | 0.22 | 0.26 | 0.3 | 0.31 | 0.24 |
| 8 | 25321076 | rs11775903 | 1.352 | -0.191 | 0.12 | -0.212 | -0.034 | 0.219 | 0.219 | 0 | 0.4 | 0.06 | 0.39 | 0.09 | 0.39 | 0.45 |
| 8 | 25321281 | rs9644041 | -0.62 | 0.015 | 0.266 | -0.297 | -0.002 | -0.003 | -0.003 | 0 | 0.43 | 0.53 | 0.36 | 0.36 | 0.35 | 0.41 |
| 8 | 25323241 | rs17053762 | 1.064 | -0.011 | 0.177 | -0.281 | -0.007 | -0.014 | -0.014 | 0 | 0.23 | 0.16 | 0.2 | 0.23 | 0.22 | 0.16 |
| 8 | 25325730 | rs11996213 | -0.158 | -0.017 | 0.187 | -0.275 | 0.003 | 0.008 | 0.008 | 0.004 | 0.37 | 0.3 | 0.27 | 0.3 | 0.31 | 0.24 |
| 8 | 25364960 | rs6990278 | 0.97 | -0.025 | 0.232 | -0.207 | -0.005 | -0.015 | -0.015 | 0 | 0.24 | 0.17 | 0.19 | 0.24 | 0.21 | 0.16 |
| 8 | 25371087 | rs4871944 | 0.28 | -0.033 | 0.187 | -0.219 | 0.014 | 0.015 | 0.015 | 0.549 | 0.61 | 0.29 | 0.73 | 0.7 | 0.71 | 0.73 |
| 8 | 25390946 | rs17053894 | 0.886 | -0.014 | 0.134 | -0.208 | -0.001 | -0.007 | -0.007 | 0 | 0.12 | 0.13 | 0.06 | 0.06 | 0.07 | 0.07 |
| 8 | 25394154 | rs9644098 | -0.175 | -0.002 | -0.043 | -0.185 | -0.005 | -0.006 | -0.006 | 0.538 | 0.37 | 0.56 | 0.27 | 0.3 | 0.29 | 0.25 |
| 8 | 25395597 | rs2979742 | -1.434 | -0.45 | -0.516 | -1.172 | -0.096 | 0.218 | 0.218 | 0.021 | 0.68 | 0.98 | 0.66 | 0.97 | 0.64 | 0.66 |
| 8 | 25398162 | rs11135872 | 1.022 | -0.707 | 0.165 | -0.377 | -0.002 | 0.006 | 0.006 | 0.009 | 0.23 | 0.21 | 0.19 | 0.21 | 0.21 | 0.15 |
| 8 | 25407292 | rs7832226 | -0.057 | -0.693 | 0.351 | -0.389 | -0.008 | -0.049 | -0.049 | 0 | 0.36 | 0.46 | 0.25 | 0.29 | 0.29 | 0.28 |
| 8 | 25421567 | rs4427183 | 0.177 | -0.339 | -0.142 | -0.032 | -0.002 | -0.05 | -0.05 | 0 | 0.27 | 0.51 | 0.21 | 0.22 | 0.22 | 0.2 |
| 8 | 25422536 | rs6993305 | 0.177 | -0.339 | -0.142 | -0.032 | -0.002 | -0.05 | -0.05 | 0.021 | 0.27 | 0.51 | 0.21 | 0.22 | 0.22 | 0.2 |
| 8 | 25434568 | rs10107581 | 0.296 | -0.284 | -0.366 | 0 | -0.001 | -0.038 | -0.038 | 0 | 0.26 | 0.41 | 0.2 | 0.22 | 0.22 | 0.2 |
| 8 | 25444716 | rs2979737 | -0.586 | -0.289 | -0.975 | 0.47 | 0.008 | -0.05 | -0.05 | 0 | 0.79 | 0.89 | 0.85 | 0.77 | 0.78 | 0.84 |
| 8 | 25445375 | rs4610756 | 0.53 | -0.34 | -0.938 | 0.381 | 0.022 | 0 | 0 | 0.207 | 0.64 | 0.33 | 0.76 | 0.73 | 0.71 | 0.71 |
| 8 | 25455330 | rs2979740 | -0.234 | 0.537 | -1.267 | 0.882 | 0.007 | -0.045 | -0.045 | 0.867 | 0.79 | 0.86 | 0.85 | 0.75 | 0.78 | 0.84 |
| 8 | 25461315 | rs7817569 | -0.164 | 0.808 | -0.66 | 0.679 | 0.009 | 0.006 | 0.006 | 0 | 0.73 | 0.61 | 0.58 | 0.69 | 0.7 | 0.67 |
| 8 | 25464092 | rs4425764 | -0.044 | 0.202 | -0.255 | 0.225 | -0.007 | -0.035 | -0.035 | 0 | 0.28 | 0.41 | 0.24 | 0.2 | 0.25 | 0.2 |
| 8 | 25464895 | rs11135873 | -0.37 | -0.532 | -0.94 | 0.522 | 0.042 | -0.064 | -0.064 | 0 | 0.26 | 0.09 | 0.24 | 0.37 | 0.18 | 0.23 |
| 8 | 25474711 | rs4596664 | -0.971 | -0.793 | 0.047 | -0.033 | -0.044 | 0.006 | 0.006 | 0 | 0.26 | 0.5 | 0.15 | 0.31 | 0.17 | 0.27 |
| 8 | 25477344 | rs11135875 | -0.956 | -0.882 | 0.237 | -0.077 | -0.055 | 0.013 | 0.013 | 0 | 0.26 | 0.42 | 0.14 | 0.31 | 0.16 | 0.27 |
| 8 | 25477382 | rs11135876 | -1.05 | -0.867 | 0.133 | -0.07 | -0.035 | 0.003 | 0.003 | 0 | 0.26 | 0.56 | 0.15 | 0.31 | 0.17 | 0.27 |
| 8 | 25478871 | rs11782477 | 1.313 | -0.619 | -0.757 | -0.472 | -0.015 | 0.067 | 0.067 | 0 | 0.47 | 0.21 | 0.61 | 0.59 | 0.51 | 0.52 |
| 8 | 25479295 | rs17054017 | -0.322 | -0.092 | -0.335 | 0.233 | 0.01 | 0.006 | 0.006 | 0 | 0.14 | 0.11 | 0.04 | 0.13 | 0.11 | 0.21 |
| 8 | 25480061 | rs7845031 | -0.733 | 0.309 | 0.124 | 0.757 | -0.013 | 0.015 | 0.015 | 0 | 0.09 | 0.14 | 0.09 | 0.17 | 0.05 | 0.06 |
| 8 | 25482766 | rs6990076 | -0.991 | 0.55 | 0.075 | 0.609 | 0.019 | 0.043 | 0.043 | 0 | 0.23 | 0.54 | 0.22 | 0.35 | 0.22 | 0.22 |
| 8 | 25484685 | rs17054045 | -0.542 | 0.699 | -0.13 | 0.562 | 0.002 | 0.01 | 0.01 | 0 | 0.3 | 0.61 | 0.3 | 0.63 | 0.29 | 0.32 |
| 8 | 25485084 | rs7812504 | 0.148 | 0.598 | 0.477 | 0.07 | -0.008 | -0.063 | -0.063 | 11.099 | 0.13 | 0.22 | 0.06 | 0 | 0.09 | 0.09 |
| 8 | 25487993 | rs4424264 | -0.256 | 0.931 | -0.276 | -0.661 | -0.036 | 0.035 | 0.035 | 0 | 0.38 | 0.27 | 0.42 | 0.76 | 0.43 | 0.48 |
| 8 | 25488542 | rs10081452 | 0.807 | 1.188 | -0.39 | 0.113 | 0.001 | -0.083 | -0.083 | 0.018 | 0.22 | 0.08 | 0.19 | 0.42 | 0.2 | 0.26 |
| 8 | 25488629 | rs2467732 | -1.226 | 0.809 | -0.571 | -0.351 | -0.006 | -0.017 | -0.017 | 0 | 0.9 | 0.84 | 0.97 | 0.98 | 0.93 | 0.9 |
| 8 | 25489096 | rs2433109 | -1.038 | 0.838 | -0.575 | -0.355 | -0.003 | -0.016 | -0.016 | 0 | 0.89 | 0.83 | 0.96 | 0.98 | 0.91 | 0.9 |
| 8 | 25490633 | rs1425741 | 1.681 | 0.692 | -1.236 | -0.773 | -0.001 | 0.052 | 0.052 | 1.502 | 0.21 | 0.03 | 0.12 | 0.1 | 0.19 | 0.04 |
| 8 | 25491742 | rs1425740 | 0.816 | 0.64 | -1.241 | -0.868 | -0.028 | 0.021 | 0.021 | 0.183 | 0.32 | 0.09 | 0.45 | 0.21 | 0.38 | 0.3 |
| 8 | 25494162 | rs2433094 | 0.295 | 0.401 | -1.06 | -0.945 | 0.01 | 0 | 0 | 0 | 0.46 | 0.21 | 0.67 | 0.56 | 0.59 | 0.49 |
| 8 | 25495087 | rs4872336 | 1.491 | 0.632 | -0.947 | -0.752 | -0.002 | -0.087 | -0.087 | 0 | 0.22 | 0.08 | 0.19 | 0.42 | 0.2 | 0.27 |
| 8 | 25498256 | rs11135879 | 0.465 | 0.598 | -0.735 | -0.629 | -0.003 | 0 | 0 | 0 | 0.41 | 0.49 | 0.29 | 0.42 | 0.34 | 0.37 |
| 8 | 25510851 | rs1593406 | -1.002 | 0.609 | -1.134 | -0.885 | 0.02 | 0.053 | 0.053 | 0.229 | 0.73 | 0.85 | 0.69 | 0.88 | 0.77 | 0.91 |
| 8 | 25519212 | rs7828851 | 0.798 | 0.671 | -0.82 | -0.969 | -0.01 | -0.105 | -0.105 | 0 | 0.22 | 0.27 | 0.11 | 0 | 0.16 | 0.15 |
| 8 | 25522134 | rs17801625 | 1.654 | 1.134 | -1.436 | -0.515 | -0.039 | -0.038 | -0.038 | 0.691 | 0.11 | 0.02 | 0.14 | 0.12 | 0.18 | 0.24 |
| 8 | 25524005 | rs2467715 | -1.444 | 0.787 | -1.601 | -0.71 | -0.026 | 0.022 | 0.022 | 0.02 | 0.81 | 0.97 | 0.76 | 0.88 | 0.8 | 0.91 |
| 8 | 25524300 | rs2467713 | -0.655 | 0.995 | -1.816 | -0.768 | -0.076 | -0.036 | -0.036 | 0.907 | 0.72 | 0.96 | 0.6 | 0.76 | 0.62 | 0.73 |
| 8 | 25524964 | rs2467712 | 0.613 | 0.726 | -1.636 | -0.852 | -0.012 | -0.012 | -0.012 | 0.487 | 0.32 | 0.3 | 0.41 | 0.56 | 0.41 | 0.48 |
| 8 | 25525396 | rs7835922 | -0.454 | 0.432 | -1.407 | -0.752 | 0.024 | 0.094 | 0.094 | 0.635 | 0.49 | 0.68 | 0.35 | 0.33 | 0.39 | 0.43 |
| 8 | 25526317 | rs4872342 | 0.632 | 0.661 | -1.552 | -0.527 | -0.052 | -0.057 | -0.057 | 1.893 | 0.21 | 0.05 | 0.22 | 0.32 | 0.23 | 0.22 |
| 8 | 25529047 | rs2433084 | 0.761 | 0.789 | -0.996 | 0.033 | 0 | 0.021 | 0.021 | 0 | 0.2 | 0.24 | 0.2 | 0.43 | 0.19 | 0.29 |
| 8 | 25529106 | rs11783334 | 0.245 | 0.929 | -1.045 | -0.219 | 0.014 | 0.067 | 0.067 | 0.791 | 0.47 | 0.34 | 0.36 | 0.32 | 0.4 | 0.37 |
| 8 | 25529797 | rs9644100 | -0.168 | 1.178 | -0.645 | -0.206 | 0.027 | -0.01 | -0.01 | 0 | 0.7 | 0.58 | 0.59 | 0.78 | 0.6 | 0.69 |
| 8 | 25530161 | rs2433086 | 0.304 | 1.159 | -1.192 | -0.272 | -0.016 | 0.004 | 0.004 | 0 | 0.66 | 0.75 | 0.66 | 0.66 | 0.58 | 0.76 |
| 8 | 25532110 | rs17802005 | 0.309 | 0.676 | -0.988 | -0.378 | 0.003 | 0.032 | 0.032 | 0 | 0.11 | 0.03 | 0.07 | 0.02 | 0.06 | 0.02 |
| 8 | 25532328 | rs17054141 | -0.574 | 0.988 | -0.289 | -0.309 | 0.001 | -0.003 | -0.003 | 1.516 | 0.21 | 0.28 | 0.2 | 0.15 | 0.19 | 0.14 |
| 8 | 25534668 | rs993047 | -1.329 | 0.728 | -0.93 | -0.428 | 0.039 | 0.118 | 0.118 | 0.685 | 0.49 | 0.85 | 0.41 | 0.32 | 0.41 | 0.52 |
| 8 | 25536691 | rs17054147 | -0.165 | 0.002 | -0.556 | -0.48 | 0.024 | 0.049 | 0.049 | 0 | 0.22 | 0.11 | 0.09 | 0.07 | 0.17 | 0.14 |
| 8 | 25536852 | rs1025157 | -0.857 | 0.24 | 0.224 | -0.287 | -0.009 | -0.018 | -0.018 | 0 | 0.39 | 0.56 | 0.33 | 0.22 | 0.31 | 0.25 |
| 8 | 25537518 | rs881366 | -1.83 | 0.984 | 1.08 | 0.614 | 0.008 | 0.003 | 0.003 | 1.28 | 0.25 | 0.81 | 0.31 | 0.27 | 0.2 | 0.3 |
| 8 | 25539830 | rs1365016 | -1.755 | 1.101 | 1.59 | 0.969 | 0.049 | 0.156 | 0.156 | 0.578 | 0.09 | 0.36 | 0.18 | 0.28 | 0.08 | 0.16 |
| 8 | 25544268 | rs11782600 | 1.42 | 1.121 | 1.422 | 0.436 | 0.003 | -0.005 | -0.005 | 0 | 0.67 | 0.42 | 0.67 | 0.71 | 0.7 | 0.51 |
| 8 | 25545756 | rs2433093 | -0.29 | 1.107 | 1.119 | 0.432 | -0.003 | -0.001 | -0.001 | 0 | 0.13 | 0.12 | 0.12 | 0.1 | 0.17 | 0.18 |

| | | | | | | | | | | | | | | | | |
|---|---|---|---|---|---|---|---|---|---|---|---|---|---|---|---|---|
| 8 | 25546083 | rs2467703 | 0.129 | 1.082 | 1.067 | 0.424 | 0.001 | 0.002 | 0.002 | 0 | 0.85 | 0.87 | 0.81 | 0.89 | 0.88 | 0.8 |
| 8 | 25547166 | rs4872343 | 1.599 | 1.046 | 0.681 | 0.392 | -0.005 | 0.011 | 0.011 | 0.654 | 0.65 | 0.16 | 0.65 | 0.71 | 0.7 | 0.51 |
| 8 | 25549528 | rs2433101 | 0.515 | 1.095 | 0.503 | 0.578 | -0.014 | -0.002 | -0.002 | 0 | 0.81 | 0.73 | 0.8 | 0.8 | 0.87 | 0.7 |
| 8 | 25557962 | rs11997651 | -1.039 | 1.805 | -0.636 | 0.641 | -0.045 | 0.001 | 0.001 | 0.078 | 0.19 | 0.09 | 0.16 | 0.13 | 0.27 | 0.27 |
| 8 | 25572870 | rs12056519 | -1.112 | 1.3 | 0.151 | 0.983 | 0.03 | 0.104 | 0.104 | 0 | 0.09 | 0.19 | 0.11 | 0.21 | 0.08 | 0.14 |
| 8 | 25595265 | rs1425726 | -0.428 | 0.826 | 0.03 | 0.1 | 0.018 | 0.021 | 0.021 | 0 | 0.46 | 0.43 | 0.72 | 0.69 | 0.64 | 0.55 |
| 8 | 25598401 | rs1346090 | -0.277 | 1.911 | -0.133 | 0.915 | 0.001 | 0 | 0 | 1.083 | 0.27 | 0.28 | 0.35 | 0.26 | 0.34 | 0.24 |
| 8 | 25599783 | rs17054249 | 0.404 | 1.907 | -0.14 | 0.916 | -0.028 | -0.043 | -0.043 | 0 | 0.19 | 0.15 | 0.37 | 0.44 | 0.3 | 0.31 |
| 8 | 25601529 | rs2467680 | -1.213 | 0.686 | -0.735 | 0.036 | 0.03 | 0.048 | 0.048 | 0 | 0.81 | 0.9 | 0.89 | 0.95 | 0.84 | 0.87 |
| 8 | 25603811 | rs10095777 | 1.316 | 0.498 | -0.655 | -0.021 | -0.003 | -0.002 | -0.002 | 0 | 0.33 | 0.22 | 0.16 | 0.25 | 0.2 | 0.32 |
| 8 | 25606163 | rs10108173 | -0.021 | 0.295 | -0.978 | -0.393 | 0.015 | -0.011 | -0.011 | 1.628 | 0.52 | 0.64 | 0.49 | 0.51 | 0.51 | 0.57 |
| 8 | 25608710 | rs13253626 | 0.72 | 0.173 | -1.094 | -0.313 | 0.008 | -0.002 | -0.002 | 0 | 0.26 | 0.28 | 0.3 | 0.23 | 0.35 | 0.25 |
| 8 | 25609836 | rs1365008 | -1.101 | -0.446 | -1.347 | -0.564 | 0 | 0.071 | 0.071 | 0 | 0.82 | 0.99 | 0.92 | 0.96 | 0.86 | 0.89 |
| 8 | 25615466 | rs7845807 | 1.579 | -0.389 | -1.324 | -0.526 | -0.042 | -0.002 | -0.002 | 0 | 0.25 | 0.37 | 0.14 | 0.25 | 0.15 | 0.31 |
| 8 | 25616023 | rs13279614 | -0.438 | -0.351 | -1.525 | -0.525 | 0.052 | 0.111 | 0.111 | 0 | 0.71 | 0.9 | 0.86 | 0.94 | 0.79 | 0.86 |
| 8 | 25625754 | rs12542834 | 0.19 | -0.132 | -1.416 | -0.392 | -0.039 | -0.053 | -0.053 | 0 | 0.32 | 0.22 | 0.45 | 0.5 | 0.42 | 0.4 |
| 8 | 25633877 | rs2278145 | 1.546 | 0.075 | -1.503 | 0.102 | -0.014 | -0.002 | -0.002 | 0.659 | 0.17 | 0.15 | 0.27 | 0.18 | 0.27 | 0.27 |
| 8 | 25634077 | rs2278144 | 0.295 | 0.13 | -1.456 | 0.259 | -0.01 | -0.04 | -0.04 | 0 | 0.15 | 0.09 | 0.18 | 0.34 | 0.16 | 0.13 |
| 8 | 25639329 | rs4872356 | 1.118 | 0.307 | -1.085 | 0.373 | -0.032 | -0.029 | -0.029 | 0.378 | 0.13 | 0.05 | 0.26 | 0.19 | 0.22 | 0.14 |
| 8 | 25643577 | rs4382486 | -0.07 | 0.02 | -0.784 | 0.274 | -0.003 | 0.003 | 0.003 | 0.043 | 0.63 | 0.63 | 0.62 | 0.76 | 0.57 | 0.6 |
| 8 | 25648792 | rs1978544 | -1.383 | 0.463 | -0.587 | 0.518 | -0.008 | 0.042 | 0.042 | 0 | 0.91 | 0.99 | 0.96 | 0.99 | 0.92 | 0.89 |
| 8 | 25649192 | rs13257008 | 1.357 | 0.265 | -0.483 | 0.546 | 0.028 | 0.063 | 0.063 | 1.027 | 0.41 | 0.19 | 0.18 | 0.18 | 0.31 | 0.39 |
| 8 | 25651493 | rs6557859 | -0.057 | 0.223 | -0.449 | 0.614 | 0.039 | -0.014 | -0.014 | 0 | 0.72 | 0.53 | 0.6 | 0.77 | 0.63 | 0.61 |
| 8 | 25657921 | rs1425717 | 0.57 | 0.198 | -0.374 | 0.317 | 0.087 | 0.034 | 0.034 | 0 | 0.19 | 0.49 | 0.36 | 0.22 | 0.29 | 0.28 |
| 8 | 25660392 | rs1365004 | -0.267 | 0.029 | -0.317 | 0.426 | 0.046 | -0.016 | -0.016 | 0.015 | 0.72 | 0.44 | 0.59 | 0.76 | 0.63 | 0.61 |
| 8 | 25665302 | rs17054357 | 0.97 | 0.118 | -0.545 | -0.151 | -0.037 | -0.046 | -0.046 | 1.894 | 0.14 | 0.03 | 0.24 | 0.26 | 0.2 | 0.12 |
| 8 | 25669550 | rs17054371 | 0.533 | -1.507 | -0.113 | -1.223 | -0.024 | -0.056 | -0.056 | 0.047 | 0.14 | 0.2 | 0.09 | 0.02 | 0.09 | 0.04 |
| 8 | 25673973 | rs11135892 | 0.533 | -1.534 | -0.082 | -1.222 | -0.024 | -0.056 | -0.056 | 0 | 0.14 | 0.2 | 0.09 | 0.02 | 0.09 | 0.04 |
| 8 | 25676188 | rs6990437 | -0.192 | -1.718 | -0.116 | -1.327 | -0.046 | 0.097 | 0.097 | 1.909 | 0.61 | 0.9 | 0.47 | 0.23 | 0.41 | 0.44 |
| 8 | 25681515 | rs1365017 | 0.815 | -1.756 | 0.096 | -1.361 | 0.012 | 0.024 | 0.024 | 1.763 | 0.43 | 0.32 | 0.3 | 0.21 | 0.31 | 0.39 |
| 8 | 25691031 | rs11135894 | 0.825 | -1.757 | 0.095 | -1.362 | 0.016 | 0.035 | 0.035 | 1.859 | 0.43 | 0.3 | 0.3 | 0.21 | 0.31 | 0.4 |
| 8 | 25697059 | rs6983496 | 0.825 | -1.758 | 0.094 | -1.362 | 0.016 | 0.035 | 0.035 | 1.836 | 0.43 | 0.3 | 0.3 | 0.21 | 0.31 | 0.4 |
| 8 | 25701661 | rs3829011 | 0.822 | -1.751 | 0.082 | -1.359 | 0.011 | 0.021 | 0.021 | 1.771 | 0.43 | 0.32 | 0.3 | 0.21 | 0.31 | 0.39 |
| 8 | 25708267 | rs10866845 | -0.202 | -1.739 | 0.013 | -1.36 | -0.012 | 0.034 | 0.034 | 2.77 | 0.62 | 0.7 | 0.41 | 0.23 | 0.45 | 0.47 |
| 8 | 25719803 | rs10448076 | 0.65 | -1.686 | -0.326 | -1.354 | 0.009 | 0.005 | 0.005 | 0 | 0.15 | 0.11 | 0.09 | 0.02 | 0.13 | 0.07 |
| 8 | 25726838 | rs4871957 | -0.016 | -1.687 | -0.377 | -1.35 | -0.019 | -0.036 | -0.036 | 0 | 0.18 | 0.22 | 0.21 | 0.05 | 0.14 | 0.08 |
| 8 | 25727286 | rs4263777 | 0.883 | -1.632 | -0.952 | -1.571 | 0.012 | 0.028 | 0.028 | 1.132 | 0.42 | 0.29 | 0.19 | 0.17 | 0.32 | 0.38 |
| 8 | 25728494 | rs17233617 | 1.255 | -1.473 | -1.61 | -1.5 | 0.003 | 0.058 | 0.058 | 0 | 0.12 | 0.01 | 0.07 | 0.02 | 0.12 | 0.05 |
| 8 | 25729752 | rs4568629 | -0.564 | -1.237 | -1.555 | -1.339 | -0.118 | -0.01 | -0.01 | 0.066 | 0.69 | 0.91 | 0.53 | 0.36 | 0.49 | 0.56 |
| 8 | 25733066 | rs7013854 | -0.177 | -1.413 | -1.229 | -1.562 | -0.053 | -0.066 | -0.066 | 2.383 | 0.26 | 0.6 | 0.31 | 0.11 | 0.17 | 0.17 |
| 8 | 25740411 | rs11781208 | -0.003 | -0.618 | -1.926 | -1.087 | -0.106 | 0.081 | 0.081 | 2.171 | 0.66 | 0.9 | 0.37 | 0.21 | 0.48 | 0.54 |
| 8 | 25743428 | rs10503779 | 0.034 | -1.234 | -1.922 | -1.232 | 0.006 | 0.123 | 0.123 | 0.185 | 0.63 | 0.89 | 0.41 | 0.2 | 0.34 | 0.53 |
| 8 | 25748553 | rs13278074 | 0.055 | -1.631 | -2.224 | -1.782 | 0.044 | 0.133 | 0.133 | 0.157 | 0.47 | 0.19 | 0.61 | 0.73 | 0.61 | 0.59 |
| 8 | 25748695 | rs11783974 | 0.567 | -1.506 | -2.214 | -1.825 | -0.054 | 0.006 | 0.006 | 0 | 0.27 | 0.18 | 0.55 | 0.72 | 0.51 | 0.42 |
| 8 | 25768391 | rs7819105 | 0.045 | -2.269 | -0.412 | -1.865 | -0.001 | -0.002 | -0.002 | 0 | 0.79 | 0.74 | 0.85 | 0.88 | 0.83 | 0.83 |
| 8 | 25769324 | rs4570167 | 0.962 | -2.542 | -0.937 | -2.073 | 0.096 | 0.156 | 0.156 | 0 | 0.48 | 0.18 | 0.64 | 0.75 | 0.66 | 0.46 |
| 8 | 25777363 | rs11786343 | 0.906 | -2.547 | -0.943 | -2.052 | 0.102 | 0.162 | 0.162 | 0 | 0.49 | 0.18 | 0.64 | 0.74 | 0.67 | 0.47 |
| 8 | 25778954 | rs17054598 | 0.751 | -2.632 | -1.256 | -2.15 | -0.006 | -0.01 | -0.01 | 0 | 0.09 | 0.01 | 0.05 | 0.11 | 0.06 | 0.16 |
| 8 | 25780152 | rs6985478 | 0.332 | -2.47 | -0.901 | -2.106 | -0.031 | -0.056 | -0.056 | 0 | 0.6 | 0.5 | 0.72 | 0.86 | 0.73 | 0.62 |
| 8 | 25787121 | rs17054638 | 0.428 | -2.392 | -0.777 | -1.951 | -0.035 | -0.105 | -0.105 | 0 | 0.24 | 0.32 | 0.12 | 0 | 0.13 | 0.21 |
| 8 | 25787232 | rs17054639 | 0.071 | -1.922 | -1.161 | -1.711 | -0.001 | -0.004 | -0.004 | 0.326 | 0.1 | 0.26 | 0.07 | 0.11 | 0.07 | 0.13 |
| 8 | 25789667 | rs9792201 | 0.22 | -1.403 | -2.08 | -1.566 | 0.08 | 0.159 | 0.159 | 0 | 0.54 | 0.13 | 0.61 | 0.74 | 0.66 | 0.47 |
| 8 | 25796376 | rs4872380 | -0.126 | -0.94 | -1.514 | -1.315 | 0.025 | 0.012 | 0.012 | 0 | 0.82 | 0.92 | 0.93 | 0.85 | 0.87 | 0.86 |
| 8 | 25805224 | rs17237257 | 0.07 | 0.024 | -1.574 | -1.905 | -0.028 | 0.077 | 0.077 | 0.059 | 0.39 | 0.26 | 0.42 | 0.15 | 0.47 | 0.36 |
| 8 | 25806471 | rs17237292 | 1.496 | 0.12 | -1.456 | -1.98 | -0.001 | 0.001 | 0.001 | 10.518 | 0.06 | 0.04 | 0.05 | 0 | 0.07 | 0.11 |
| 8 | 25808664 | rs7830333 | 0.963 | 0.457 | -0.616 | -2.45 | 0.016 | -0.083 | -0.083 | 0 | 0.13 | 0.3 | 0.07 | 0 | 0.12 | 0.02 |
| 8 | 25808945 | rs1485750 | -0.079 | 0.623 | -0.768 | -2.536 | -0.017 | 0.02 | 0.02 | 0 | 0.28 | 0.2 | 0.36 | 0.08 | 0.37 | 0.39 |
| 8 | 25812543 | rs9314316 | -0.704 | 0.493 | -0.598 | -2.937 | 0.001 | -0.027 | -0.027 | 0 | 0.5 | 0.54 | 0.67 | 0.88 | 0.54 | 0.62 |
| 8 | 25814150 | rs11779334 | 0.88 | 0.309 | -0.325 | -2.918 | 0.026 | -0.04 | -0.04 | 3.082 | 0.2 | 0.27 | 0.29 | 0.1 | 0.28 | 0.26 |
| 8 | 25814533 | rs11775931 | 0.247 | 0.315 | -0.309 | -2.95 | 0.015 | -0.075 | -0.075 | 0.75 | 0.27 | 0.37 | 0.32 | 0.1 | 0.3 | 0.37 |
| 8 | 25816376 | rs4872381 | -0.821 | 0.385 | -0.389 | -2.879 | 0 | 0.036 | 0.036 | 0 | 0.5 | 0.62 | 0.63 | 0.88 | 0.48 | 0.61 |
| 8 | 25817084 | rs1542654 | -0.041 | 0.416 | -0.538 | -2.742 | -0.002 | 0.007 | 0.007 | 0 | 0.33 | 0.25 | 0.34 | 0.12 | 0.34 | 0.29 |
| 8 | 25820971 | rs12543589 | 0.68 | 0.466 | -0.653 | -2.396 | -0.017 | 0.031 | 0.031 | 0 | 0.36 | 0.32 | 0.49 | 0.8 | 0.41 | 0.47 |
| 8 | 25821871 | rs11135904 | -0.2 | 0.582 | -0.498 | -2.568 | 0.006 | -0.007 | -0.007 | 0 | 0.5 | 0.58 | 0.61 | 0.89 | 0.53 | 0.66 |
| 8 | 25822022 | rs2171078 | 0.185 | 0.644 | -0.461 | -2.543 | 0 | -0.046 | -0.046 | 0.846 | 0.46 | 0.53 | 0.51 | 0.85 | 0.44 | 0.58 |
| 8 | 25825649 | rs11135905 | 0.852 | 0.694 | -0.483 | -2.471 | -0.007 | 0.013 | 0.013 | 0 | 0.36 | 0.36 | 0.47 | 0.8 | 0.4 | 0.46 |
| 8 | 25825680 | rs12541364 | 0.286 | 0.577 | -0.537 | -2.114 | 0.002 | -0.023 | -0.023 | 0 | 0.5 | 0.57 | 0.6 | 0.88 | 0.52 | 0.67 |
| 8 | 25829691 | rs6991575 | -0.098 | 0.327 | -0.589 | -1.901 | 0.004 | -0.048 | -0.048 | 0.01 | 0.3 | 0.25 | 0.2 | 0 | 0.2 | 0.07 |
| 8 | 25834968 | rs11984569 | 0.201 | 1.287 | -0.537 | -0.633 | 0.007 | 0.013 | 0.013 | 0 | 0.74 | 0.66 | 0.64 | 0.4 | 0.68 | 0.7 |

| | | | | | | | | | | | | | | | | |
|---|---|---|---|---|---|---|---|---|---|---|---|---|---|---|---|---|
| 8 | 25835942 | rs2171079 | -0.448 | 1.783 | -0.034 | -0.762 | 0.099 | -0.004 | -0.004 | 0 | 0.09 | 0.31 | 0.11 | 0.08 | 0.11 | 0.1 |
| 8 | 25836281 | rs7000178 | 0.948 | 1.189 | -1.021 | -1.326 | 0.027 | 0.055 | 0.055 | 0.506 | 0.42 | 0.26 | 0.31 | 0.07 | 0.33 | 0.46 |
| 8 | 25838780 | rs17817279 | -0.441 | 2.076 | -1.255 | -0.674 | -0.057 | 0.025 | 0.025 | 0 | 0.17 | 0.08 | 0.18 | 0.12 | 0.29 | 0.26 |
| 8 | 25839043 | rs11990648 | -0.248 | 1.186 | -1.306 | -0.06 | 0.008 | -0.009 | -0.009 | 0.489 | 0.09 | 0.06 | 0.25 | 0.3 | 0.09 | 0.15 |
| 8 | 25844226 | rs7001120 | 0.592 | 1.112 | -0.845 | -0.07 | 0.001 | -0.001 | -0.001 | 2.542 | 0.54 | 0.53 | 0.53 | 0.44 | 0.39 | 0.55 |
| 8 | 25844999 | rs17238923 | 0.104 | 1.764 | -1.061 | -0.128 | -0.069 | 0.037 | 0.037 | 0 | 0.16 | 0.03 | 0.18 | 0.06 | 0.27 | 0.28 |
| 8 | 25845675 | rs10866846 | -0.738 | 1.593 | -0.921 | -0.143 | -0.035 | -0.048 | -0.048 | 0.212 | 0.25 | 0.13 | 0.44 | 0.37 | 0.36 | 0.43 |
| 8 | 25846604 | rs6987894 | -0.283 | 1.58 | -0.653 | -0.091 | 0.01 | 0.025 | 0.025 | 0.456 | 0.71 | 0.62 | 0.71 | 0.5 | 0.66 | 0.83 |
| 8 | 25846741 | rs7006324 | -0.264 | 1.588 | -0.663 | -0.087 | 0.016 | 0.042 | 0.042 | 0.56 | 0.7 | 0.54 | 0.71 | 0.5 | 0.66 | 0.83 |
| 8 | 25858692 | rs4269551 | -1.085 | 1.487 | -0.998 | -0.21 | 0.059 | 0.032 | 0.032 | 0.43 | 0.49 | 0.3 | 0.62 | 0.37 | 0.66 | 0.48 |
| 8 | 25870252 | rs1485748 | -0.691 | 1.906 | -1.209 | -0.938 | -0.026 | 0.039 | 0.039 | 0 | 0.26 | 0.21 | 0.28 | 0.09 | 0.36 | 0.34 |
| 8 | 25870277 | rs17818083 | 1.906 | 0.954 | -2.098 | -1.624 | -0.014 | 0.066 | 0.066 | 3.679 | 0.23 | 0.01 | 0.11 | 0.07 | 0.17 | 0.16 |
| 8 | 25871350 | rs7003404 | -0.202 | 1.354 | -1.626 | -1.584 | 0.022 | -0.056 | -0.056 | 2.084 | 0.15 | 0.23 | 0.18 | 0.04 | 0.2 | 0.22 |
| 8 | 25871721 | rs11998023 | 0.305 | 1.513 | -1.685 | -1.536 | -0.015 | 0.017 | 0.017 | 0 | 0.11 | 0.09 | 0.16 | 0.04 | 0.17 | 0.18 |
| 8 | 25875310 | rs1485749 | -0.022 | 1.022 | -1.536 | -1.624 | -0.006 | -0.027 | -0.027 | 3.959 | 0.33 | 0.59 | 0.48 | 0.79 | 0.34 | 0.27 |
| 8 | 25875505 | rs17818275 | 1.073 | 1.272 | -1.367 | -1.339 | -0.015 | 0.021 | 0.021 | 3.774 | 0.11 | 0.05 | 0.11 | 0.04 | 0.17 | 0.1 |
| 8 | 25879726 | rs10503781 | 0.358 | 0.789 | -0.728 | -0.707 | 0.009 | 0.173 | 0.173 | 0 | 0.08 | 0.19 | 0.1 | 0.44 | 0.08 | 0.17 |
| 8 | 25888552 | rs17818622 | 0.54 | 0.853 | -0.825 | -1.003 | -0.004 | 0.006 | 0.006 | 0 | 0.08 | 0.06 | 0.12 | 0.04 | 0.08 | 0.15 |
| 8 | 25889446 | rs7821832 | -0.04 | 0.628 | -0.419 | -1.152 | 0.01 | 0.049 | 0.049 | 0 | 0.25 | 0.47 | 0.36 | 0.32 | 0.27 | 0.19 |
| 8 | 25891653 | rs6997996 | -0.057 | 0.623 | -0.355 | -1.145 | 0.002 | 0.023 | 0.023 | 0 | 0.26 | 0.39 | 0.36 | 0.33 | 0.27 | 0.19 |
| 8 | 25892142 | rs11135910 | 1.19 | 0.502 | -0.507 | -1.156 | -0.008 | 0.017 | 0.017 | 0 | 0.13 | 0.09 | 0.15 | 0.04 | 0.13 | 0.16 |
| 8 | 25894943 | rs17818981 | 1.474 | 0.457 | -0.567 | -1.194 | -0.009 | -0.004 | -0.004 | 0 | 0.23 | 0.16 | 0.2 | 0.23 | 0.29 | 0.34 |
| 8 | 25896410 | rs17054793 | 0.679 | 0.512 | 0.28 | -0.17 | 0.01 | 0.172 | 0.172 | 0 | 0.07 | 0.19 | 0.07 | 0.4 | 0.07 | 0.16 |
| 8 | 25901579 | rs1466562 | -0.377 | 0.511 | 0.417 | -0.277 | -0.002 | 0.02 | 0.02 | 0 | 0.26 | 0.38 | 0.35 | 0.32 | 0.27 | 0.19 |
| 8 | 25904764 | rs1008975 | 1.319 | 0.484 | 0.206 | -0.362 | 0.001 | 0.032 | 0.032 | 0 | 0.59 | 0.31 | 0.52 | 0.27 | 0.61 | 0.57 |
| 8 | 25910975 | rs4872390 | -1.292 | 0.838 | 0.411 | -0.36 | 0 | 0.002 | 0.002 | 0 | 0.39 | 0.59 | 0.5 | 0.34 | 0.4 | 0.35 |
| 8 | 25913258 | rs11778076 | -0.547 | 0.821 | 0.177 | -0.322 | 0 | 0.004 | 0.004 | 0 | 0.25 | 0.31 | 0.34 | 0.32 | 0.26 | 0.18 |
| 8 | 25916964 | rs7824253 | 0.721 | 0.834 | 0.041 | -0.37 | -0.002 | 0.019 | 0.019 | 0 | 0.14 | 0.1 | 0.14 | 0.03 | 0.13 | 0.18 |
| 8 | 25917711 | rs4872393 | 0 | 0.845 | 0.135 | -0.296 | 0 | 0.013 | 0.013 | 0 | 0.21 | 0.26 | 0.33 | 0.33 | 0.24 | 0.16 |
| 8 | 25918507 | rs980723 | 0.815 | 1.097 | 0.133 | -0.069 | 0.022 | -0.015 | -0.015 | 1.94 | 0.68 | 0.48 | 0.54 | 0.26 | 0.64 | 0.6 |
| 8 | 25920295 | rs6557880 | 1.754 | 1.007 | 0.101 | -0.079 | 0.006 | 0.128 | 0.128 | 0.15 | 0.49 | 0.22 | 0.39 | 0.23 | 0.5 | 0.4 |
| 8 | 25921494 | rs1601468 | 1.048 | 1.321 | -0.462 | 0.026 | -0.04 | -0.02 | -0.02 | 0 | 0.07 | 0.01 | 0.26 | 0.33 | 0.14 | 0.14 |
| 8 | 25922911 | rs7836754 | -0.298 | 1.138 | -0.253 | 0.105 | 0.004 | 0.018 | 0.018 | 0 | 0.78 | 0.71 | 0.66 | 0.67 | 0.75 | 0.77 |
| 8 | 25924029 | rs6993325 | -1.467 | 0.856 | -0.406 | 0.396 | 0.003 | 0.021 | 0.021 | 0 | 0.26 | 0.35 | 0.26 | 0.44 | 0.25 | 0.36 |
| 8 | 25929365 | rs6985329 | -0.675 | 0.52 | -0.675 | -0.713 | -0.003 | 0.019 | 0.019 | 0 | 0.17 | 0.12 | 0.15 | 0.03 | 0.15 | 0.19 |
| 8 | 25935389 | rs10104004 | -0.885 | 0.338 | -0.241 | -0.663 | -0.021 | -0.085 | -0.085 | 0.187 | 0.22 | 0.43 | 0.2 | 0.07 | 0.16 | 0.2 |
| 8 | 25938238 | rs13269936 | -0.447 | 0.431 | -0.582 | -0.998 | 0.001 | 0.009 | 0.009 | 0 | 0.27 | 0.25 | 0.14 | 0.12 | 0.22 | 0.21 |
| 8 | 25939846 | rs12548228 | 0.246 | 0.898 | -0.587 | -1.052 | -0.002 | -0.002 | -0.002 | 0 | 0.27 | 0.21 | 0.4 | 0.72 | 0.24 | 0.39 |
| 8 | 25940605 | rs11780628 | 0.941 | 0.888 | -0.297 | -0.786 | -0.006 | 0.001 | 0.001 | 0 | 0.06 | 0.06 | 0.02 | 0 | 0.07 | 0.06 |
| 8 | 25941193 | rs7818429 | 0.334 | 0.583 | -0.235 | -0.902 | 0 | -0.036 | -0.036 | 0 | 0.15 | 0.41 | 0.15 | 0.08 | 0.15 | 0.15 |
| 8 | 25942560 | rs1872498 | -0.666 | 0.897 | -1.107 | -0.791 | -0.014 | 0.003 | 0.003 | 0.652 | 0.86 | 0.92 | 0.87 | 0.99 | 0.81 | 0.88 |
| 8 | 25950732 | rs4872403 | 0.2 | 0.54 | -0.915 | 0.377 | -0.006 | 0.158 | 0.158 | 0 | 0.07 | 0.12 | 0.21 | 0.48 | 0.04 | 0.18 |
| 8 | 25952339 | rs11787090 | 0.064 | 0.292 | -1.084 | -0.353 | 0.004 | 0.031 | 0.031 | 0 | 0.63 | 0.73 | 0.63 | 0.74 | 0.6 | 0.65 |
| 8 | 25954604 | rs7460527 | 0.188 | 0.18 | -1.084 | -0.487 | 0.023 | 0.04 | 0.04 | 0 | 0.52 | 0.31 | 0.49 | 0.34 | 0.56 | 0.45 |
| 8 | 25965198 | rs10113210 | -0.252 | 0.135 | -1.181 | -0.244 | 0.002 | 0.054 | 0.054 | 0 | 0.44 | 0.23 | 0.48 | 0.66 | 0.42 | 0.48 |
| 8 | 25967898 | rs4463438 | -0.451 | 0.008 | -0.735 | -0.252 | -0.001 | 0.001 | 0.001 | 0 | 0.52 | 0.5 | 0.58 | 0.6 | 0.54 | 0.55 |
| 8 | 25974894 | rs2952051 | 0.897 | 0.085 | -1.124 | -0.204 | -0.022 | -0.079 | -0.079 | 0.785 | 0.16 | 0.06 | 0.25 | 0.44 | 0.15 | 0.27 |
| 8 | 25976442 | rs6999088 | -0.426 | -0.101 | -0.886 | -0.533 | -0.001 | 0.008 | 0.008 | 0 | 0.38 | 0.33 | 0.39 | 0.66 | 0.34 | 0.47 |
| 8 | 25978851 | rs4872412 | -2.13 | 0.286 | -1.073 | -0.91 | -0.023 | 0.033 | 0.033 | 0 | 0.82 | 0.91 | 0.85 | 0.95 | 0.77 | 0.86 |
| 8 | 25980730 | rs6987909 | 0.168 | -0.026 | -0.491 | -0.592 | 0.001 | 0.01 | 0.01 | 0 | 0.46 | 0.44 | 0.61 | 0.68 | 0.48 | 0.5 |
| 8 | 25982665 | rs6557882 | -1.394 | -0.008 | -0.389 | -0.624 | -0.015 | 0.027 | 0.027 | 0 | 0.7 | 0.81 | 0.77 | 0.94 | 0.66 | 0.7 |
| 8 | 25992682 | rs6557885 | 0.819 | 0.144 | -0.686 | -0.496 | 0.003 | -0.007 | -0.007 | 0.056 | 0.22 | 0.16 | 0.14 | 0.25 | 0.18 | 0.2 |
| 8 | 25994238 | rs7011753 | 1.293 | 0.118 | -0.721 | -0.479 | 0.016 | -0.026 | -0.026 | 0 | 0.17 | 0.1 | 0.1 | 0.24 | 0.11 | 0.16 |
| 8 | 25995356 | rs7828041 | 1.546 | 0.32 | -0.674 | -0.613 | -0.002 | -0.021 | -0.021 | 0 | 0.2 | 0.3 | 0.19 | 0.15 | 0.21 | 0.22 |
| 8 | 26000422 | rs2951987 | 1.576 | 0.298 | -0.767 | -0.651 | -0.006 | -0.031 | -0.031 | 0 | 0.08 | 0.11 | 0.11 | 0.01 | 0.1 | 0.01 |
| 8 | 26002358 | rs2976285 | 0.695 | 0.335 | -0.972 | -0.536 | -0.009 | -0.03 | -0.03 | 0 | 0.28 | 0.4 | 0.29 | 0.2 | 0.31 | 0.24 |
| 8 | 26007894 | rs2951996 | -0.666 | 0.529 | -1.25 | -0.334 | -0.003 | 0.008 | 0.008 | 0 | 0.66 | 0.37 | 0.67 | 0.82 | 0.65 | 0.7 |
| 8 | 26021965 | rs4872419 | 0.787 | 0.54 | -1.539 | 0.292 | -0.032 | -0.017 | -0.017 | 0 | 0.08 | 0.01 | 0.23 | 0.44 | 0.08 | 0.23 |
| 8 | 26023906 | rs16860 | -0.445 | 0.305 | -1.703 | -0.117 | 0.025 | -0.051 | -0.051 | 0 | 0.44 | 0.16 | 0.45 | 0.53 | 0.35 | 0.45 |
| 8 | 26026958 | rs2976282 | -0.259 | 0.345 | -1.297 | -1.253 | -0.003 | -0.069 | -0.069 | 1.445 | 0.67 | 0.67 | 0.77 | 0.96 | 0.71 | 0.78 |
| 8 | 26027462 | rs2976281 | 0.44 | 0.524 | -1.558 | -1.53 | 0.033 | -0.018 | -0.018 | 2.22 | 0.59 | 0.28 | 0.49 | 0.47 | 0.64 | 0.57 |
| 8 | 26027842 | rs2952020 | -0.092 | 0.641 | -0.909 | -2.487 | -0.016 | -0.011 | -0.011 | 0 | 0.7 | 0.51 | 0.55 | 0.24 | 0.78 | 0.59 |
| 8 | 26029798 | rs9650417 | 1.664 | 1.502 | -0.738 | -2.254 | -0.014 | -0.022 | -0.022 | 0 | 0.15 | 0.09 | 0.21 | 0.23 | 0.24 | 0.26 |
| 8 | 26034528 | rs10098922 | -0.42 | 1.359 | -0.607 | -2.368 | -0.003 | 0.015 | 0.015 | 0 | 0.47 | 0.53 | 0.38 | 0.24 | 0.5 | 0.44 |
| 8 | 26036343 | rs2951977 | 0.55 | 1.494 | -1.119 | -2.278 | -0.031 | -0.001 | -0.001 | 0 | 0.28 | 0.17 | 0.27 | 0.24 | 0.38 | 0.33 |
| 8 | 26038663 | rs11782690 | 1.292 | 1.63 | -0.97 | -2.085 | -0.025 | 0.042 | 0.042 | 0.402 | 0.1 | 0.03 | 0.03 | 0.01 | 0.11 | 0.07 |
| 8 | 26039903 | rs4347000 | -0.654 | 1.679 | -0.739 | -2.047 | -0.002 | -0.001 | -0.001 | 0 | 0.84 | 0.84 | 0.58 | 0.24 | 0.8 | 0.65 |
| 8 | 26043064 | rs13280608 | 0.962 | 1.751 | -0.918 | -1.536 | -0.017 | -0.012 | -0.012 | 0 | 0.23 | 0.13 | 0.25 | 0.23 | 0.3 | 0.27 |
| 8 | 26044276 | rs17055028 | 0.685 | 1.87 | -0.948 | -1.455 | 0.014 | -0.028 | -0.028 | 0 | 0.07 | 0.07 | 0.05 | 0 | 0.09 | 0.1 |

| | | | | | | | | | | | | | | | | |
|---|---|---|---|---|---|---|---|---|---|---|---|---|---|---|---|---|
| 8 | 26050530 | rs12056386 | 0.766 | 1.536 | -1.235 | -0.994 | -0.035 | -0.005 | -0.005 | 0 | 0.08 | 0.01 | 0.29 | 0.48 | 0.11 | 0.19 |
| 8 | 26051071 | rs2976295 | -0.94 | 1.289 | -0.72 | -2.057 | -0.004 | 0.022 | 0.022 | 0 | 0.43 | 0.36 | 0.41 | 0.25 | 0.54 | 0.37 |
| 8 | 26053562 | rs13250131 | -0.197 | 1.298 | -0.75 | -2.087 | -0.01 | 0.016 | 0.016 | 0 | 0.32 | 0.16 | 0.37 | 0.25 | 0.43 | 0.3 |
| 8 | 26053962 | rs11779143 | 0.35 | 1.1 | -0.748 | -2.027 | 0 | -0.026 | -0.026 | 0 | 0.11 | 0.14 | 0.04 | 0.01 | 0.11 | 0.07 |
| 8 | 26065901 | rs4355773 | 0.685 | 1.367 | -0.53 | -1.787 | -0.002 | 0.007 | 0.007 | 0 | 0.47 | 0.5 | 0.53 | 0.75 | 0.39 | 0.62 |
| 8 | 26069942 | rs7845865 | 0.173 | 1.018 | -0.235 | -1.828 | -0.022 | -0.089 | -0.089 | 0 | 0.81 | 0.72 | 0.91 | 0.99 | 0.87 | 0.92 |
| 8 | 26072964 | rs11775982 | -0.427 | 1.053 | -0.484 | -1.81 | -0.016 | -0.071 | -0.071 | 0 | 0.18 | 0.23 | 0.09 | 0.01 | 0.13 | 0.08 |
| 8 | 26076419 | rs7815658 | -0.628 | 1.527 | -0.88 | -1.607 | 0.013 | -0.068 | -0.068 | 1.119 | 0.34 | 0.37 | 0.32 | 0.09 | 0.46 | 0.34 |
| 8 | 26078020 | rs4531067 | -0.258 | 1.328 | -0.76 | -1.437 | 0.013 | -0.074 | -0.074 | 0.062 | 0.29 | 0.34 | 0.24 | 0.09 | 0.36 | 0.23 |
| 8 | 26081505 | rs4275231 | -0.21 | 0.335 | -1.055 | -1.281 | 0.01 | 0.002 | 0.002 | 2.57 | 0.29 | 0.21 | 0.4 | 0.9 | 0.16 | 0.39 |
| 8 | 26088624 | rs6996845 | 0.386 | 0.782 | -1.338 | -0.689 | 0.004 | 0.002 | 0.002 | 0 | 0.49 | 0.72 | 0.43 | 0.39 | 0.49 | 0.29 |
| 8 | 26089068 | rs6986899 | 0.374 | 0.758 | -1.34 | -0.701 | 0.005 | 0.003 | 0.003 | 0 | 0.49 | 0.74 | 0.43 | 0.39 | 0.49 | 0.29 |
| 8 | 26094767 | rs17834024 | 0.749 | 0.802 | -1.631 | -0.86 | -0.011 | 0.046 | 0.046 | 0 | 0.08 | 0.01 | 0.02 | 0 | 0.06 | 0.15 |
| 8 | 26108104 | rs1825183 | 0.523 | 0.659 | -1.863 | -1.086 | 0.021 | 0.069 | 0.069 | 0.001 | 0.12 | 0.01 | 0.03 | 0 | 0.05 | 0.01 |
| 8 | 26109041 | rs11135928 | -0.08 | 1.246 | -1.744 | -1.128 | -0.01 | -0.023 | -0.023 | 0 | 0.22 | 0.17 | 0.39 | 0.57 | 0.3 | 0.5 |
| 8 | 26116732 | rs17245857 | -0.973 | 1.037 | -1.469 | -1.156 | -0.008 | -0.01 | -0.01 | 0 | 0.41 | 0.32 | 0.55 | 0.59 | 0.47 | 0.69 |
| 8 | 26118439 | rs10102733 | -1.058 | 1.31 | -0.895 | -1.211 | 0.009 | 0.007 | 0.007 | 0.064 | 0.42 | 0.44 | 0.57 | 0.59 | 0.52 | 0.69 |
| 8 | 26122035 | rs4602893 | -0.478 | 1.446 | -0.89 | -1.23 | 0 | 0 | 0 | 0 | 0.19 | 0.13 | 0.16 | 0.04 | 0.18 | 0.19 |
| 8 | 26124172 | rs7009683 | -0.776 | 1.359 | -1.054 | -1.109 | 0.013 | 0.047 | 0.047 | 0 | 0.88 | 0.98 | 0.95 | 0.96 | 0.93 | 0.99 |
| 8 | 26124195 | rs17055073 | 0.274 | 1.546 | -0.811 | -0.916 | -0.002 | 0.011 | 0.011 | 0 | 0.11 | 0.07 | 0.09 | 0.03 | 0.1 | 0.15 |
| 8 | 26126137 | rs11135930 | -0.31 | 1.52 | -0.698 | -0.661 | 0.026 | 0.068 | 0.068 | 1.29 | 0.24 | 0.31 | 0.42 | 0.55 | 0.34 | 0.51 |
| 8 | 26141765 | rs12681736 | 0.432 | 1.918 | -0.832 | -0.373 | -0.013 | 0.03 | 0.03 | 0 | 0.7 | 0.79 | 0.72 | 0.84 | 0.58 | 0.66 |
| 8 | 26142995 | rs10093390 | 0.928 | 0.927 | -1.146 | -1.241 | 0.002 | -0.015 | -0.015 | 0.047 | 0.41 | 0.49 | 0.49 | 0.41 | 0.39 | 0.4 |
| 8 | 26144146 | rs984324 | -1.546 | 0.953 | -1.088 | -1.404 | -0.002 | -0.022 | -0.022 | 0 | 0.9 | 0.89 | 0.96 | 1 | 0.94 | 0.99 |
| 8 | 26148748 | rs7840855 | -0.132 | 1.474 | -1.308 | -1.296 | -0.091 | 0.044 | 0.044 | 0 | 0.28 | 0.05 | 0.29 | 0.19 | 0.45 | 0.41 |
| 8 | 26159509 | rs9650418 | -0.725 | 0.775 | -1.462 | -1.417 | -0.006 | 0.003 | 0.003 | 0 | 0.42 | 0.42 | 0.43 | 0.43 | 0.51 | 0.49 |
| 8 | 26160015 | rs7005099 | 1.085 | 0.676 | -1.442 | -1.382 | 0.004 | 0.003 | 0.003 | 0 | 0.06 | 0.08 | 0.1 | 0.25 | 0.03 | 0.1 |
| 8 | 26160739 | rs17055108 | -1.245 | 0.63 | -1.439 | -1.489 | -0.001 | -0.002 | -0.002 | 0 | 0.53 | 0.56 | 0.51 | 0.45 | 0.58 | 0.64 |
| 8 | 26162364 | rs11135933 | -1.252 | 0.632 | -1.439 | -1.488 | -0.001 | -0.002 | -0.002 | 0 | 0.53 | 0.56 | 0.51 | 0.45 | 0.58 | 0.64 |
| 8 | 26170246 | rs11784209 | -1.252 | 0.632 | -1.439 | -1.462 | -0.001 | -0.002 | -0.002 | 0.37 | 0.53 | 0.56 | 0.51 | 0.46 | 0.58 | 0.64 |
| 8 | 26178438 | rs17174717 | 1.859 | 0.745 | -1.627 | -1.521 | -0.004 | 0.011 | 0.011 | 0 | 0.07 | 0.04 | 0.03 | 0 | 0.04 | 0.14 |
| 8 | 26179066 | rs11135934 | 1.495 | 0.885 | -1.801 | -1.626 | -0.089 | -0.028 | -0.028 | 0 | 0.15 | 0.02 | 0.19 | 0.15 | 0.31 | 0.29 |
| 8 | 26187537 | rs7838409 | -0.449 | 0.755 | -1.949 | -2.034 | -0.001 | -0.033 | -0.033 | 0 | 0.27 | 0.24 | 0.2 | 0.02 | 0.24 | 0.29 |
| 8 | 26190212 | rs13271042 | -1.547 | 0.333 | -1.355 | -1.773 | 0.046 | 0.122 | 0.122 | 1.971 | 0.61 | 0.81 | 0.62 | 0.88 | 0.71 | 0.76 |
| 8 | 26206077 | rs1594829 | 2.04 | 0.207 | -1.318 | -1.902 | 0.005 | -0.03 | -0.03 | 0 | 0.3 | 0.26 | 0.19 | 0 | 0.24 | 0.13 |
| 8 | 26212413 | rs2046222 | -2.828 | 0.296 | -1.355 | -1.765 | 0.02 | 0.042 | 0.042 | 0.067 | 0.92 | 0.99 | 0.97 | 1 | 0.95 | 0.99 |
| 8 | 26221945 | rs17055163 | 0.43 | 0.87 | -0.812 | -1.391 | 0.005 | 0.024 | 0.024 | 0.023 | 0.15 | 0.31 | 0.16 | 0.28 | 0.14 | 0.18 |
| 8 | 26223458 | rs3808566 | 0.626 | 0.853 | -0.843 | -1.353 | 0.027 | 0.235 | 0.235 | 0 | 0.14 | 0.31 | 0.12 | 0.43 | 0.14 | 0.15 |
| 8 | 26224958 | rs7823021 | -1.858 | 0.826 | -0.675 | -1.418 | 0.036 | 0.076 | 0.076 | 1.32 | 0.6 | 0.74 | 0.6 | 0.88 | 0.7 | 0.75 |
| 8 | 26236978 | rs1064807 | -1.972 | 0.895 | -0.932 | -1.321 | 0.027 | 0.063 | 0.063 | 1.032 | 0.59 | 0.7 | 0.59 | 0.88 | 0.69 | 0.72 |
| 8 | 26238949 | rs3814484 | 1.53 | 0.813 | -0.906 | -1.273 | 0.004 | 0.004 | 0.004 | 0 | 0.06 | 0.08 | 0.1 | 0.25 | 0.03 | 0.1 |
| 8 | 26239383 | rs3758098 | 0.785 | 0.838 | -0.798 | -1.298 | 0.002 | 0.146 | 0.146 | 0 | 0.14 | 0.2 | 0.11 | 0.43 | 0.13 | 0.11 |
| 8 | 26245658 | rs2874670 | -0.011 | 0.996 | -0.825 | -1.297 | -0.016 | -0.013 | -0.013 | 0.384 | 0.3 | 0.33 | 0.35 | 0.45 | 0.45 | 0.43 |
| 8 | 26250906 | rs3779916 | -1.383 | 1.005 | -0.699 | -1.2 | 0.014 | 0.03 | 0.03 | 0.277 | 0.48 | 0.59 | 0.51 | 0.86 | 0.62 | 0.59 |
| 8 | 26251422 | rs3808578 | 0.898 | 0.946 | -0.67 | -1.155 | 0.006 | 0.17 | 0.17 | 0 | 0.13 | 0.21 | 0.11 | 0.41 | 0.13 | 0.11 |
| 8 | 26262744 | rs3808577 | -1.281 | 0.999 | -0.84 | -1.163 | 0.019 | 0.035 | 0.035 | 0.307 | 0.49 | 0.59 | 0.52 | 0.86 | 0.63 | 0.6 |
| 8 | 26263260 | rs7813520 | -1.613 | 0.988 | -0.806 | -0.918 | 0.029 | 0.066 | 0.066 | 1.302 | 0.6 | 0.71 | 0.59 | 0.88 | 0.69 | 0.73 |
| 8 | 26267362 | rs17055196 | 0.727 | 1.285 | -0.368 | -0.111 | 0.02 | -0.022 | -0.022 | 0 | 0.09 | 0.2 | 0.06 | 0.03 | 0.1 | 0.06 |
| 8 | 26269032 | rs17310286 | -0.576 | 0.813 | -0.478 | 0.256 | -0.003 | 0.138 | 0.138 | 0 | 0.19 | 0.25 | 0.16 | 0.41 | 0.17 | 0.13 |
| 8 | 26269191 | rs1042992 | -0.547 | 0.831 | -0.469 | 0.265 | -0.002 | 0.141 | 0.141 | 0 | 0.18 | 0.25 | 0.16 | 0.41 | 0.17 | 0.13 |
| 8 | 26279865 | rs17055205 | -0.491 | 1.007 | -0.862 | -0.059 | 0.008 | -0.005 | -0.005 | 0 | 0.36 | 0.42 | 0.38 | 0.61 | 0.49 | 0.44 |
| 8 | 26280472 | rs4871977 | -1.976 | 0.801 | -1.229 | -0.391 | -0.01 | 0.028 | 0.028 | 0 | 0.95 | 1 | 0.97 | 1 | 0.95 | 0.99 |
| 8 | 26280723 | rs10282829 | -0.093 | 0.618 | -1.503 | -0.55 | 0.002 | 0.013 | 0.013 | 0 | 0.79 | 0.89 | 0.69 | 0.83 | 0.82 | 0.66 |
| 8 | 26280815 | rs9694042 | -0.196 | 0.338 | -1.334 | -0.874 | 0.021 | 0.064 | 0.064 | 0 | 0.52 | 0.67 | 0.53 | 0.83 | 0.61 | 0.54 |
| 8 | 26285835 | rs1469756 | 1.54 | 0.963 | -1.475 | -0.2 | -0.057 | -0.01 | -0.01 | 0 | 0.14 | 0.07 | 0.2 | 0.17 | 0.29 | 0.15 |
| 8 | 26287852 | rs11987934 | -0.05 | 0.601 | -0.996 | -0.275 | 0.007 | 0.005 | 0.005 | 0 | 0.76 | 0.63 | 0.71 | 0.73 | 0.77 | 0.62 |
| 8 | 26288369 | rs4872433 | -0.104 | 0.596 | -0.837 | -0.277 | 0.004 | 0.003 | 0.003 | 0 | 0.77 | 0.73 | 0.72 | 0.73 | 0.77 | 0.62 |
| 8 | 26288441 | rs4872434 | 0.436 | 0.165 | -1.42 | -0.265 | -0.013 | 0.074 | 0.074 | 0 | 0.19 | 0.33 | 0.21 | 0.36 | 0.15 | 0.15 |
| 8 | 26288526 | rs17401856 | 0.926 | -0.131 | -1.886 | -0.988 | 0.014 | 0.127 | 0.127 | 0.735 | 0.3 | 0.09 | 0.15 | 0 | 0.22 | 0.28 |
| 8 | 26290268 | rs17310684 | 1.637 | 0.577 | -1.539 | -0.529 | -0.052 | -0.006 | -0.006 | 0.019 | 0.14 | 0.07 | 0.21 | 0.16 | 0.28 | 0.15 |
| 8 | 26290773 | rs6557919 | -0.554 | -0.108 | -1.561 | -1.026 | 0.039 | 0.078 | 0.078 | 0 | 0.54 | 0.68 | 0.76 | 0.9 | 0.69 | 0.59 |
| 8 | 26295234 | rs11786499 | -0.872 | 0.593 | -1.286 | 0.183 | -0.008 | -0.006 | -0.006 | 0.028 | 0.81 | 0.81 | 0.75 | 0.74 | 0.72 | 0.77 |
| 8 | 26298209 | rs11781866 | -0.065 | 0.861 | -0.848 | 0.283 | 0.019 | 0.001 | 0.001 | 0 | 0.23 | 0.31 | 0.26 | 0.17 | 0.33 | 0.18 |
| 8 | 26303033 | rs4407886 | 0.61 | 0.507 | -0.984 | 0.21 | 0.01 | -0.007 | -0.007 | 0 | 0.63 | 0.52 | 0.65 | 0.76 | 0.59 | 0.71 |
| 8 | 26303417 | rs7823542 | -0.612 | 0.124 | -1.353 | 0.238 | -0.016 | -0.004 | -0.004 | 0 | 0.87 | 0.85 | 0.93 | 0.92 | 0.93 | 0.9 |
| 8 | 26305735 | rs13276734 | -0.283 | 0.223 | -2.08 | 0.19 | 0.017 | -0.007 | -0.007 | 0.001 | 0.29 | 0.1 | 0.16 | 0.25 | 0.24 | 0.13 |
| 8 | 26313736 | rs11135938 | 0.314 | 0.669 | -1.706 | 0.073 | 0.017 | 0.099 | 0.099 | 0 | 0.15 | 0.21 | 0.26 | 0.42 | 0.19 | 0.23 |
| 8 | 26313832 | rs17310789 | 0.171 | 1.287 | -2.341 | -0.677 | -0.07 | 0.111 | 0.111 | 0.031 | 0.18 | 0.02 | 0.14 | 0 | 0.24 | 0.13 |
| 8 | 26315727 | rs2034039 | 0.838 | 1.172 | -2.048 | -0.161 | -0.067 | -0.046 | -0.046 | 0.074 | 0.18 | 0.03 | 0.17 | 0.27 | 0.29 | 0.13 |

| | | | | | | | | | | | | | | | | |
|---|---|---|---|---|---|---|---|---|---|---|---|---|---|---|---|---|
| 8 | 26317214 | rs2570638 | 0.326 | 0.208 | -1.528 | -0.196 | -0.024 | 0.041 | 0.041 | 0.539 | 0.41 | 0.29 | 0.38 | 0.39 | 0.47 | 0.36 |
| 8 | 26322970 | rs167871 | -0.41 | -0.125 | -1.025 | -0.678 | -0.003 | -0.058 | -0.058 | 0.24 | 0.31 | 0.4 | 0.19 | 0.07 | 0.29 | 0.21 |
| 8 | 26326383 | rs4872439 | 0.506 | -0.891 | -0.661 | -0.4 | 0.002 | -0.018 | -0.018 | 0 | 0.67 | 0.63 | 0.61 | 0.73 | 0.66 | 0.7 |
| 8 | 26326695 | rs17402279 | 1.294 | -1.037 | -1.046 | -0.667 | -0.009 | 0.035 | 0.035 | 0.051 | 0.1 | 0.04 | 0.08 | 0.02 | 0.11 | 0.24 |
| 8 | 26326894 | rs4872440 | 0.667 | -1.329 | -1.092 | -0.045 | -0.018 | 0.086 | 0.086 | 0 | 0.12 | 0.2 | 0.23 | 0.28 | 0.11 | 0.23 |
| 8 | 26328656 | rs328076 | 1.068 | -1.29 | -1.185 | -0.395 | -0.016 | 0.011 | 0.011 | 0 | 0.13 | 0.09 | 0.12 | 0 | 0.2 | 0.07 |
| 8 | 26329205 | rs328072 | 1.068 | -1.29 | -1.185 | -0.395 | -0.016 | 0.011 | 0.011 | 0 | 0.13 | 0.09 | 0.12 | 0 | 0.2 | 0.07 |
| 8 | 26332723 | rs4872441 | 0.622 | -1.502 | -1.156 | -0.145 | -0.018 | 0.122 | 0.122 | 0 | 0.13 | 0.21 | 0.37 | 0.41 | 0.11 | 0.23 |
| 8 | 26337515 | rs189025 | 0.365 | -1.639 | -1.256 | -0.407 | 0.02 | 0.023 | 0.023 | 0.436 | 0.67 | 0.3 | 0.78 | 0.81 | 0.78 | 0.62 |
| 8 | 26339256 | rs6984046 | 1.322 | -1.676 | -1.105 | -0.429 | -0.041 | -0.021 | -0.021 | 1.571 | 0.1 | 0.32 | 0.03 | 0.05 | 0.03 | 0.02 |
| 8 | 26339759 | rs328094 | 0.067 | -1.645 | -1.09 | -0.338 | -0.009 | -0.051 | -0.051 | 0 | 0.24 | 0.18 | 0.48 | 0.41 | 0.31 | 0.3 |
| 8 | 26340103 | rs328095 | -0.262 | -1.5 | -1.104 | -0.285 | -0.022 | -0.059 | -0.059 | 0 | 0.77 | 0.57 | 0.81 | 0.86 | 0.81 | 0.64 |
| 8 | 26340833 | rs328096 | -1.217 | -1.506 | -0.976 | -0.245 | -0.014 | 0.022 | 0.022 | 3.203 | 0.87 | 0.66 | 0.9 | 0.87 | 0.92 | 0.84 |
| 8 | 26343095 | rs328097 | -0.829 | -1.351 | -0.901 | -0.444 | -0.008 | 0.005 | 0.005 | 0 | 0.42 | 0.55 | 0.55 | 0.48 | 0.39 | 0.43 |
| 8 | 26359157 | rs719587 | -0.721 | -1.514 | -0.901 | -0.743 | -0.002 | -0.005 | -0.005 | 0 | 0.41 | 0.46 | 0.4 | 0.26 | 0.41 | 0.49 |
| 8 | 26361499 | rs11780047 | -0.345 | -1.24 | -0.93 | -0.76 | -0.014 | 0.008 | 0.008 | 0 | 0.31 | 0.22 | 0.37 | 0.22 | 0.41 | 0.48 |
| 8 | 26365716 | rs2233701 | 0.84 | -1.258 | -0.915 | -0.719 | -0.009 | 0.038 | 0.038 | 0.133 | 0.09 | 0.12 | 0.18 | 0.21 | 0.1 | 0.22 |
| 8 | 26369707 | rs9886448 | 0.451 | -1.283 | -0.888 | -0.731 | -0.006 | 0.063 | 0.063 | 0.287 | 0.1 | 0.2 | 0.19 | 0.21 | 0.12 | 0.22 |
| 8 | 26370514 | rs4872449 | 2.165 | -1.724 | -1.421 | -1.09 | -0.147 | -0.118 | -0.118 | 0 | 0.31 | 0.03 | 0.48 | 0.67 | 0.42 | 0.17 |
| 8 | 26374298 | rs10866851 | 0.77 | -1.599 | -1.458 | -1.179 | 0.07 | 0.155 | 0.155 | 0 | 0.51 | 0.28 | 0.77 | 0.89 | 0.69 | 0.58 |
| 8 | 26374888 | rs1442887 | -0.056 | -1.871 | -1.391 | -1.35 | -0.021 | -0.094 | -0.094 | 0.134 | 0.7 | 0.48 | 0.84 | 0.94 | 0.77 | 0.75 |
| 8 | 26376855 | rs4872450 | 0.354 | -1.878 | -1.362 | -1.253 | 0.001 | 0.004 | 0.004 | 0 | 0.21 | 0.24 | 0.28 | 0.22 | 0.26 | 0.41 |
| 8 | 26380059 | rs12680365 | 0.339 | -1.877 | -1.366 | -1.253 | -0.001 | -0.003 | -0.003 | 0 | 0.21 | 0.22 | 0.28 | 0.23 | 0.26 | 0.41 |
| 8 | 26380458 | rs1542114 | 0.526 | -2.017 | -1.278 | -1.33 | -0.005 | -0.014 | -0.014 | 0 | 0.63 | 0.4 | 0.86 | 0.89 | 0.83 | 0.65 |
| 8 | 26387784 | rs3866466 | -1.532 | -1.976 | -1.139 | -1.329 | 0.024 | -0.061 | -0.061 | 0.732 | 0.89 | 0.63 | 0.94 | 0.99 | 0.93 | 0.83 |
| 8 | 26387909 | rs17403886 | 1.623 | -1.914 | -1.483 | -1.351 | -0.002 | -0.002 | -0.002 | 0 | 0.1 | 0.07 | 0.07 | 0 | 0.12 | 0.06 |
| 8 | 26391147 | rs17055424 | 1.707 | -1.964 | -1.486 | -1.342 | -0.018 | -0.008 | -0.008 | 1.599 | 0.1 | 0.11 | 0.02 | 0.04 | 0.02 | 0.02 |
| 8 | 26393114 | rs17319277 | 0.564 | -2.026 | -1.561 | -1.379 | 0.026 | 0.047 | 0.047 | 1.741 | 0.18 | 0.04 | 0.06 | 0.05 | 0.09 | 0.21 |
| 8 | 26398643 | rs1899494 | -0.387 | -2.113 | -0.861 | -1.4 | -0.02 | -0.001 | -0.001 | 0.003 | 0.31 | 0.57 | 0.34 | 0.28 | 0.28 | 0.39 |
| 8 | 26405024 | rs13279683 | 2.616 | -2.126 | -0.932 | -1.44 | -0.149 | -0.115 | -0.115 | 0 | 0.3 | 0.04 | 0.47 | 0.66 | 0.43 | 0.17 |
| 8 | 26405253 | rs9314325 | -0.838 | -2.144 | -0.911 | -1.457 | 0.01 | 0.017 | 0.017 | 0 | 0.42 | 0.66 | 0.42 | 0.28 | 0.41 | 0.45 |
| 8 | 26405508 | rs7831864 | -1.66 | -2.053 | -0.847 | -1.406 | 0.006 | -0.056 | -0.056 | 0.013 | 0.9 | 0.76 | 0.95 | 0.99 | 0.93 | 0.83 |
| 8 | 26413989 | rs4872453 | 0.262 | -1.977 | -0.911 | -1.34 | 0.012 | 0.035 | 0.035 | 0 | 0.22 | 0.45 | 0.32 | 0.23 | 0.26 | 0.37 |
| 8 | 26414551 | rs4872454 | 0.508 | -1.947 | -0.892 | -1.353 | -0.027 | -0.048 | -0.048 | 0.677 | 0.62 | 0.57 | 0.87 | 0.89 | 0.82 | 0.6 |
| 8 | 26423632 | rs17404749 | 1.79 | -1.752 | -0.652 | -1.101 | -0.033 | 0.052 | 0.052 | 0.568 | 0.1 | 0.01 | 0.09 | 0.01 | 0.14 | 0.16 |
| 8 | 26424236 | rs9772104 | 1.433 | -1.725 | -0.536 | -0.995 | -0.002 | -0.009 | -0.009 | 0.448 | 0.1 | 0.09 | 0.09 | 0 | 0.13 | 0.06 |
| 8 | 26426579 | rs431217 | -0.378 | -1.831 | -0.532 | -1.016 | -0.023 | 0.005 | 0.005 | 3.025 | 0.27 | 0.51 | 0.32 | 0.23 | 0.27 | 0.33 |
| 8 | 26431466 | rs401004 | 0.327 | -1.693 | -0.38 | -0.957 | -0.022 | 0.062 | 0.062 | 8.167 | 0.16 | 0.3 | 0.22 | 0.22 | 0.14 | 0.26 |
| 8 | 26433886 | rs6983262 | 1.102 | -1.744 | -0.42 | -1.083 | -0.001 | -0.019 | -0.019 | 4.911 | 0.13 | 0.14 | 0.1 | 0 | 0.13 | 0.06 |
| 8 | 26434014 | rs2585456 | 0.264 | -1.743 | -0.572 | -1.089 | -0.004 | -0.006 | -0.006 | 0.586 | 0.25 | 0.29 | 0.24 | 0.2 | 0.26 | 0.4 |
| 8 | 26440229 | rs388047 | -0.686 | -1.761 | -0.889 | -1.089 | 0.006 | 0.008 | 0.008 | 0 | 0.39 | 0.65 | 0.33 | 0.25 | 0.33 | 0.55 |
| 8 | 26441275 | rs415524 | -1.293 | -1.68 | -0.925 | -0.971 | -0.02 | 0.013 | 0.013 | 0 | 0.6 | 0.75 | 0.43 | 0.29 | 0.48 | 0.63 |
| 8 | 26444669 | rs327234 | -1.273 | -1.682 | -0.91 | -0.958 | -0.012 | 0.008 | 0.008 | 0 | 0.6 | 0.71 | 0.43 | 0.3 | 0.48 | 0.63 |
| 8 | 26446923 | rs12678418 | 1.779 | -1.53 | -1.395 | -0.894 | -0.127 | -0.063 | -0.063 | 0 | 0.3 | 0.07 | 0.45 | 0.68 | 0.46 | 0.22 |
| 8 | 26449411 | rs17055486 | 1.592 | -1.514 | -1.065 | -0.793 | 0.024 | 0.016 | 0.016 | 0.03 | 0.08 | 0.07 | 0.02 | 0.04 | 0.02 | 0.02 |
| 8 | 26453830 | rs12550663 | 0.607 | -1.4 | -1.046 | -0.746 | 0.01 | -0.001 | -0.001 | 0 | 0.1 | 0.05 | 0.14 | 0.07 | 0.08 | 0.2 |
| 8 | 26461110 | rs7835006 | -0.288 | -1.617 | -0.877 | -0.985 | 0.029 | 0.054 | 0.054 | 0.154 | 0.33 | 0.2 | 0.16 | 0.09 | 0.23 | 0.24 |
| 8 | 26462161 | rs327229 | -0.765 | -1.843 | -0.54 | -1.12 | -0.037 | -0.072 | -0.072 | 0 | 0.85 | 0.78 | 0.9 | 0.97 | 0.93 | 0.78 |
| 8 | 26463344 | rs13249543 | 1.927 | -0.88 | -1.319 | -0.464 | -0.055 | -0.085 | -0.085 | 0 | 0.18 | 0.08 | 0.32 | 0.42 | 0.3 | 0.19 |
| 8 | 26466860 | rs11775967 | 0.088 | -1.583 | -0.58 | -0.656 | 0.01 | 0.011 | 0.011 | 0 | 0.29 | 0.27 | 0.14 | 0.09 | 0.2 | 0.19 |
| 8 | 26468630 | rs327214 | 0.471 | -1.135 | -0.688 | -0.238 | 0.02 | 0.019 | 0.019 | 0 | 0.16 | 0.27 | 0.16 | 0.18 | 0.21 | 0.24 |
| 8 | 26472037 | rs17321828 | 0.149 | -1.346 | -1.147 | -0.217 | 0.008 | 0.021 | 0.021 | 0.013 | 0.25 | 0.17 | 0.12 | 0.14 | 0.19 | 0.18 |
| 8 | 26472655 | rs4733013 | -0.176 | -1.602 | -1.022 | -0.123 | 0.016 | -0.016 | -0.016 | 0 | 0.75 | 0.82 | 0.61 | 0.72 | 0.8 | 0.67 |
| 8 | 26473403 | rs2584187 | -1.287 | -1.667 | -0.889 | -0.111 | -0.011 | 0.004 | 0.004 | 0 | 0.85 | 0.84 | 0.87 | 0.81 | 0.9 | 0.78 |
| 8 | 26473624 | rs2585459 | 0.192 | -1.659 | -0.756 | -0.14 | 0.026 | 0.007 | 0.007 | 0 | 0.6 | 0.71 | 0.73 | 0.68 | 0.69 | 0.59 |
| 8 | 26476721 | rs403185 | 0.644 | -1.592 | -0.622 | -0.234 | 0.04 | 0.03 | 0.03 | 0 | 0.48 | 0.32 | 0.66 | 0.66 | 0.62 | 0.45 |
| 8 | 26481035 | rs327221 | -0.53 | -1.657 | -0.187 | -0.421 | -0.017 | -0.038 | -0.038 | 0.531 | 0.22 | 0.36 | 0.32 | 0.1 | 0.16 | 0.28 |
| 8 | 26483190 | rs327225 | -0.446 | -1.96 | -0.382 | -0.789 | -0.001 | -0.024 | -0.024 | 0 | 0.8 | 0.81 | 0.83 | 0.93 | 0.83 | 0.8 |
| 8 | 26486255 | rs11995227 | -0.227 | -2.693 | -0.439 | -1.054 | -0.077 | -0.038 | -0.038 | 0 | 0.29 | 0.44 | 0.12 | 0.13 | 0.14 | 0.31 |
| 8 | 26486742 | rs7832576 | 0.129 | -2.626 | -0.16 | -1.157 | -0.063 | -0.102 | -0.102 | 1.446 | 0.17 | 0.4 | 0.07 | 0.01 | 0.07 | 0.14 |
| 8 | 26488246 | rs17322528 | 0.537 | -2.449 | -0.325 | -1.086 | 0.001 | 0.013 | 0.013 | 0 | 0.19 | 0.14 | 0.1 | 0 | 0.16 | 0.17 |
| 8 | 26491463 | rs327216 | 1.615 | -2.313 | -0.375 | -0.801 | -0.022 | -0.008 | -0.008 | 0 | 0.09 | 0.02 | 0.27 | 0.08 | 0.1 | 0.11 |
| 8 | 26495740 | rs1972921 | 0.535 | -2.577 | -0.476 | -0.898 | -0.043 | -0.046 | -0.046 | 0 | 0.67 | 0.47 | 0.59 | 0.81 | 0.81 | 0.71 |
| 8 | 26496387 | rs1442337 | 1.304 | -2.637 | -0.682 | -0.794 | 0.007 | 0 | 0 | 0 | 0.15 | 0.11 | 0.06 | 0.11 | 0.07 | 0.16 |
| 8 | 26498013 | rs13277175 | 1.548 | -2.537 | -0.791 | -0.962 | -0.006 | 0.006 | 0.006 | 1.047 | 0.37 | 0.25 | 0.43 | 0.7 | 0.59 | 0.38 |
| 8 | 26499089 | rs4733048 | -1.014 | -2.687 | -0.769 | -1.045 | 0.005 | 0.005 | 0.005 | 1.781 | 0.84 | 0.92 | 0.63 | 0.85 | 0.86 | 0.73 |
| 8 | 26499118 | rs10088957 | 0.007 | -2.696 | -0.508 | -1.273 | 0.031 | 0.057 | 0.057 | 0 | 0.31 | 0.15 | 0.15 | 0.11 | 0.22 | 0.28 |
| 8 | 26500090 | rs17055593 | 1.785 | -2.697 | -0.547 | -1.229 | 0.01 | -0.004 | -0.004 | 1.083 | 0.14 | 0.11 | 0.08 | 0.11 | 0.09 | 0.13 |
| 8 | 26505992 | rs9644118 | 1.229 | -2.566 | -0.635 | -0.807 | 0 | -0.001 | -0.001 | 0 | 0.36 | 0.36 | 0.36 | 0.36 | 0.56 | 0.35 |

| | | | | | | | | | | | | | | | | |
|---|---|---|---|---|---|---|---|---|---|---|---|---|---|---|---|---|
| 8 | 26508767 | rs6992095 | 0.21 | -2.466 | -0.475 | -0.776 | -0.038 | -0.107 | -0.107 | 0.207 | 0.24 | 0.48 | 0.12 | 0.04 | 0.14 | 0.2 |
| 8 | 26510792 | rs708621 | 0.363 | -2.393 | -0.846 | -0.649 | 0.069 | 0.002 | 0.002 | 1.287 | 0.55 | 0.78 | 0.46 | 0.4 | 0.67 | 0.54 |
| 8 | 26514293 | rs1058332 | 1.87 | -2.282 | -0.687 | -0.68 | -0.002 | -0.022 | -0.022 | 0 | 0.14 | 0.11 | 0.07 | 0 | 0.08 | 0.11 |
| 8 | 26514779 | rs920633 | -1.366 | -2.135 | -0.889 | -0.554 | 0.003 | -0.035 | -0.035 | 0.315 | 0.82 | 0.92 | 0.56 | 0.52 | 0.83 | 0.78 |
| 8 | 26515458 | rs17666 | -0.533 | -2.066 | -0.654 | -0.695 | -0.003 | -0.013 | -0.013 | 0 | 0.46 | 0.5 | 0.24 | 0.12 | 0.3 | 0.43 |
| 8 | 26518751 | rs7845740 | 0.088 | -2.218 | -0.667 | -0.819 | 0.012 | -0.002 | -0.002 | 0 | 0.59 | 0.72 | 0.45 | 0.5 | 0.64 | 0.59 |
| 8 | 26519881 | rs13249662 | 1.601 | -2.012 | -0.71 | -0.835 | 0 | 0 | 0 | 0 | 0.3 | 0.28 | 0.31 | 0.45 | 0.49 | 0.35 |
| 8 | 26524262 | rs13258612 | 0.967 | -2.114 | -0.859 | -0.973 | -0.006 | 0.11 | 0.11 | 0 | 0.18 | 0.03 | 0.09 | 0.01 | 0.13 | 0.18 |
| 8 | 26524421 | rs17055673 | 0.77 | -2.248 | -0.719 | -1.023 | -0.061 | -0.125 | -0.125 | 0 | 0.27 | 0.4 | 0.14 | 0 | 0.14 | 0.24 |
| 8 | 26530079 | rs13251131 | -1.013 | -2.206 | -0.755 | -0.921 | -0.004 | 0.016 | 0.016 | 0 | 0.76 | 0.77 | 0.55 | 0.51 | 0.79 | 0.76 |
| 8 | 26536443 | rs6984375 | 0.564 | -2.194 | -0.946 | -0.92 | 0.057 | 0.021 | 0.021 | 0.266 | 0.48 | 0.35 | 0.4 | 0.52 | 0.65 | 0.52 |
| 8 | 26537428 | rs11990945 | 2.057 | -1.705 | -0.953 | -0.704 | -0.009 | -0.005 | -0.005 | 0 | 0.27 | 0.22 | 0.26 | 0.29 | 0.4 | 0.23 |
| 8 | 26541693 | rs11783368 | -0.874 | -1.833 | -0.893 | -0.606 | -0.008 | 0.03 | 0.03 | 0 | 0.76 | 0.74 | 0.55 | 0.52 | 0.78 | 0.76 |
| 8 | 26548643 | rs11785701 | 0.744 | -1.525 | -1.17 | -0.62 | 0.01 | 0.105 | 0.105 | 0 | 0.17 | 0.03 | 0.08 | 0.01 | 0.11 | 0.17 |
| 8 | 26556867 | rs9693984 | 1.591 | -1.655 | -1.047 | -0.686 | 0.015 | 0.001 | 0.001 | 0 | 0.16 | 0.11 | 0.07 | 0 | 0.1 | 0.13 |
| 8 | 26558479 | rs7818301 | 2.08 | -0.626 | -1.084 | -0.676 | 0.038 | 0.104 | 0.104 | 0.075 | 0.14 | 0.27 | 0.12 | 0.26 | 0.16 | 0.09 |
| 8 | 26561030 | rs7820310 | 0.17 | -0.618 | -1.144 | -0.478 | 0.005 | 0.008 | 0.008 | 0 | 0.46 | 0.4 | 0.28 | 0.3 | 0.37 | 0.36 |
| 8 | 26565770 | rs7001655 | 1.111 | 0.165 | -1.782 | -0.353 | -0.06 | -0.021 | -0.021 | 0 | 0.19 | 0.13 | 0.21 | 0.21 | 0.38 | 0.24 |
| 8 | 26567555 | rs17055767 | 1.347 | -0.735 | -1.481 | -0.592 | 0.013 | 0.048 | 0.048 | 0 | 0.15 | 0.06 | 0.1 | 0 | 0.13 | 0.13 |
| 8 | 26569237 | rs1122214 | 0.346 | -1.097 | -1.484 | -0.7 | 0.029 | 0.042 | 0.042 | 0.209 | 0.27 | 0.16 | 0.16 | 0.06 | 0.18 | 0.31 |
| 8 | 26571605 | rs2028554 | 1.662 | -1.13 | -1.876 | -0.566 | -0.113 | -0.122 | -0.122 | 0.017 | 0.36 | 0.1 | 0.32 | 0.41 | 0.49 | 0.29 |
| 8 | 26573417 | rs6557943 | 1.812 | -1.087 | -1.791 | -0.565 | -0.004 | -0.004 | -0.004 | 0.629 | 0.11 | 0.1 | 0.06 | 0.06 | 0.05 | 0.17 |
| 8 | 26574853 | rs4073055 | 0.191 | -1.166 | -1.369 | -0.565 | -0.014 | -0.029 | -0.029 | 3.573 | 0.26 | 0.59 | 0.15 | 0.16 | 0.18 | 0.34 |
| 8 | 26576125 | rs11779314 | 0.543 | -0.992 | -1.189 | -0.455 | 0.095 | 0.089 | 0.089 | 0.054 | 0.09 | 0.46 | 0.08 | 0.1 | 0.09 | 0.11 |
| 8 | 26578017 | rs1036813 | -0.886 | -0.824 | -1.628 | -0.227 | -0.002 | 0.002 | 0.002 | 0 | 0.77 | 0.75 | 0.57 | 0.57 | 0.8 | 0.72 |
| 8 | 26578762 | rs1036814 | -0.198 | -0.75 | -1.264 | -0.712 | 0.024 | 0.052 | 0.052 | 0 | 0.41 | 0.66 | 0.25 | 0.16 | 0.31 | 0.43 |
| 8 | 26582313 | rs4732626 | 0.232 | -0.851 | -1.04 | -0.258 | 0.013 | 0.014 | 0.014 | 0 | 0.63 | 0.32 | 0.51 | 0.81 | 0.73 | 0.62 |
| 8 | 26583103 | rs17088261 | 0.545 | -0.621 | -0.791 | -0.283 | -0.003 | -0.029 | -0.029 | 0 | 0.1 | 0.14 | 0.08 | 0.01 | 0.08 | 0.11 |
| 8 | 26583589 | rs7819786 | -0.838 | -1.256 | -0.665 | 0.03 | -0.006 | 0.005 | 0.005 | 0 | 0.86 | 0.84 | 0.68 | 0.75 | 0.91 | 0.71 |
| 8 | 26585149 | rs6998714 | -0.339 | -1.09 | -1.12 | 0.002 | 0.002 | 0.002 | 0.002 | 0.517 | 0.56 | 0.7 | 0.35 | 0.39 | 0.45 | 0.37 |
| 8 | 26585268 | rs10503796 | 0.833 | -0.768 | -0.842 | 0.087 | -0.002 | -0.024 | -0.024 | 0 | 0.1 | 0.14 | 0.07 | 0.01 | 0.09 | 0.1 |
| 8 | 26588975 | rs6992526 | 0.598 | -0.926 | -0.458 | 0.597 | 0.02 | 0.184 | 0.184 | 0.018 | 0.1 | 0.25 | 0.09 | 0.3 | 0.1 | 0.1 |
| 8 | 26590630 | rs13249152 | -0.452 | -0.999 | -0.152 | 0.485 | 0.037 | 0.19 | 0.19 | 0.237 | 0.17 | 0.47 | 0.17 | 0.36 | 0.19 | 0.15 |
| 8 | 26591302 | rs1965328 | 0.448 | -1.824 | -0.556 | -0.097 | -0.005 | 0.002 | 0.002 | 0.033 | 0.63 | 0.43 | 0.57 | 0.57 | 0.69 | 0.6 |
| 8 | 26592410 | rs1016733 | -0.527 | -2.304 | -1.087 | -0.666 | 0.036 | 0.011 | 0.011 | 0.063 | 0.51 | 0.64 | 0.5 | 0.44 | 0.39 | 0.49 |
| 8 | 26593427 | rs17055842 | 0.973 | -2.166 | -1.009 | -0.789 | -0.001 | -0.065 | -0.065 | 0.005 | 0.24 | 0.28 | 0.16 | 0.01 | 0.2 | 0.18 |
| 8 | 26593933 | rs1390516 | 2.336 | -2.136 | -0.956 | -0.727 | -0.001 | -0.006 | -0.006 | 0 | 0.07 | 0.07 | 0.06 | 0.01 | 0.09 | 0.06 |
| 8 | 26594912 | rs1020232 | -0.299 | -2.12 | -0.925 | -0.647 | -0.028 | 0.035 | 0.035 | 0 | 0.71 | 0.65 | 0.64 | 0.57 | 0.78 | 0.65 |
| 8 | 26600068 | rs10105278 | 0.682 | -1.963 | -1.065 | -0.779 | -0.001 | -0.015 | -0.015 | 0 | 0.42 | 0.35 | 0.47 | 0.5 | 0.55 | 0.43 |
| 8 | 26601394 | rs1472346 | -0.07 | -1.96 | -0.97 | -0.699 | -0.019 | 0.025 | 0.025 | 0 | 0.68 | 0.67 | 0.63 | 0.52 | 0.75 | 0.59 |
| 8 | 26605110 | rs4732639 | -0.276 | -1.931 | -0.961 | -0.923 | -0.021 | -0.058 | -0.058 | 0 | 0.62 | 0.6 | 0.59 | 0.87 | 0.71 | 0.56 |
| 8 | 26605660 | rs9314328 | 2.891 | -1.736 | -1.418 | -0.841 | -0.008 | 0.027 | 0.027 | 0 | 0.13 | 0.07 | 0.07 | 0 | 0.11 | 0.12 |
| 8 | 26610427 | rs1442341 | -1.287 | -1.844 | -1.323 | -0.664 | 0.03 | 0.028 | 0.028 | 0 | 0.81 | 0.92 | 0.74 | 0.93 | 0.88 | 0.73 |
| 8 | 26610651 | rs4732853 | 1.497 | -1.027 | -0.797 | -0.611 | 0.01 | -0.052 | -0.052 | 2.005 | 0.15 | 0.03 | 0.11 | 0.22 | 0.12 | 0.1 |
| 8 | 26611846 | rs17055923 | -0.018 | -0.851 | -0.721 | -0.84 | 0.03 | -0.023 | -0.023 | 0 | 0.31 | 0.27 | 0.25 | 0.39 | 0.23 | 0.28 |
| 8 | 26616784 | rs2036109 | 0.043 | -0.478 | -0.211 | -0.984 | 0.023 | -0.005 | -0.005 | 0 | 0.46 | 0.42 | 0.4 | 0.52 | 0.35 | 0.46 |
| 8 | 26617129 | rs12674917 | 1.343 | -0.45 | -0.007 | -0.972 | 0.037 | 0.255 | 0.255 | 8.37 | 0.08 | 0.25 | 0.05 | 0.33 | 0.06 | 0.08 |
| 8 | 26618366 | rs4236678 | -0.533 | -0.469 | -0.065 | -1.006 | 0.008 | 0.123 | 0.123 | 0.298 | 0.55 | 0.78 | 0.48 | 0.89 | 0.42 | 0.54 |
| 8 | 26623384 | rs3802241 | -0.524 | -0.469 | -0.065 | -1.012 | 0.009 | 0.122 | 0.122 | 0.293 | 0.55 | 0.78 | 0.48 | 0.89 | 0.42 | 0.54 |
| 8 | 26623401 | rs17333700 | 1.799 | -0.528 | -0.073 | -1.056 | 0.01 | 0.049 | 0.049 | 0.128 | 0.25 | 0.11 | 0.16 | 0.02 | 0.2 | 0.21 |
| 8 | 26623868 | rs17055954 | 2.055 | -0.535 | -0.05 | -1.044 | 0.002 | -0.004 | -0.004 | 0.777 | 0.07 | 0.07 | 0.06 | 0 | 0.07 | 0.03 |
| 8 | 26626557 | rs4236679 | 0.918 | -0.526 | 0.012 | -0.993 | -0.015 | 0.124 | 0.124 | 0 | 0.11 | 0.22 | 0.15 | 0.44 | 0.06 | 0.22 |
| 8 | 26627249 | rs3739216 | 0.918 | -0.568 | 0.012 | -0.993 | -0.017 | 0.124 | 0.124 | 0 | 0.11 | 0.22 | 0.15 | 0.44 | 0.06 | 0.22 |
| 8 | 26628028 | rs1048101 | -0.571 | -0.304 | -0.072 | -1.021 | 0.001 | 0.116 | 0.116 | 0.532 | 0.56 | 0.78 | 0.49 | 0.89 | 0.43 | 0.54 |
| 8 | 26628318 | rs13261597 | 1.557 | -0.307 | 0.017 | -1.014 | 0.01 | 0.161 | 0.161 | 9.06 | 0.07 | 0.17 | 0.04 | 0.32 | 0.06 | 0.08 |
| 8 | 26628765 | rs13277287 | -0.541 | -0.3 | -0.025 | -1.085 | 0.003 | 0.093 | 0.093 | 0.384 | 0.56 | 0.73 | 0.48 | 0.85 | 0.43 | 0.54 |
| 8 | 26629336 | rs7842829 | -0.541 | -0.3 | -0.025 | -1.084 | 0.003 | 0.101 | 0.101 | 0.384 | 0.56 | 0.73 | 0.49 | 0.89 | 0.43 | 0.54 |
| 8 | 26633797 | rs12543356 | -0.249 | -0.292 | 0.01 | -1.097 | -0.014 | 0.009 | 0.009 | 0.544 | 0.22 | 0.42 | 0.24 | 0.79 | 0.18 | 0.33 |
| 8 | 26634589 | rs6557946 | -0.456 | -0.237 | -0.353 | -1.089 | -0.003 | 0.124 | 0.124 | 0.415 | 0.56 | 0.77 | 0.49 | 0.89 | 0.43 | 0.54 |
| 8 | 26635744 | rs10086077 | 2.152 | -0.219 | -0.422 | -1.066 | 0.014 | 0.009 | 0.009 | 0 | 0.21 | 0.1 | 0.11 | 0.01 | 0.14 | 0.17 |
| 8 | 26638910 | rs11135955 | 0.439 | -0.015 | -0.364 | -0.815 | -0.004 | 0.06 | 0.06 | 0 | 0.14 | 0.2 | 0.19 | 0.46 | 0.12 | 0.25 |
| 8 | 26641025 | rs2055195 | -0.249 | 0.005 | -0.417 | -1.018 | -0.015 | 0.168 | 0.168 | 0.171 | 0.56 | 0.78 | 0.5 | 0.89 | 0.44 | 0.55 |
| 8 | 26645612 | rs4732874 | -0.349 | 0.051 | -0.696 | -0.92 | -0.016 | -0.039 | -0.039 | 0 | 0.29 | 0.48 | 0.29 | 0.82 | 0.25 | 0.37 |
| 8 | 26652010 | rs11135957 | -0.349 | 0.051 | -0.696 | -0.92 | -0.016 | -0.039 | -0.039 | 0 | 0.29 | 0.48 | 0.29 | 0.82 | 0.25 | 0.37 |
| 8 | 26662246 | rs7835853 | 1.509 | 0.39 | -0.965 | -0.859 | -0.027 | -0.091 | -0.091 | 0 | 0.29 | 0.34 | 0.21 | 0.07 | 0.2 | 0.2 |
| 8 | 26662926 | rs12547707 | -0.011 | 0.699 | -1.629 | -0.46 | -0.008 | 0.089 | 0.089 | 0 | 0.16 | 0.24 | 0.2 | 0.5 | 0.12 | 0.26 |
| 8 | 26667045 | rs13261054 | 0.963 | -0.006 | -2.073 | -0.409 | 0.04 | 0.008 | 0.008 | 0.005 | 0.42 | 0.7 | 0.32 | 0.43 | 0.33 | 0.29 |
| 8 | 26667066 | rs7817265 | -0.549 | 0.559 | -1.392 | 0.402 | 0 | 0.02 | 0.02 | 0 | 0.15 | 0.18 | 0.19 | 0.47 | 0.12 | 0.25 |
| 8 | 26670795 | rs4732897 | -0.663 | 1.172 | -1.347 | 0.182 | -0.05 | 0.04 | 0.04 | 0 | 0.85 | 0.94 | 0.86 | 0.97 | 0.66 | 0.88 |

| | | | | | | | | | | | | | | | | |
|---|---|---|---|---|---|---|---|---|---|---|---|---|---|---|---|---|
| 8 | 26672929 | rs7017961 | -0.581 | 1.167 | -1.748 | -0.147 | 0.071 | -0.052 | -0.052 | 0.005 | 0.83 | 0.74 | 0.83 | 0.97 | 0.65 | 0.88 |
| 8 | 26673902 | rs13257637 | -0.526 | 0.394 | -3.046 | -0.112 | -0.003 | 0.035 | 0.035 | 0 | 0.36 | 0.15 | 0.37 | 0.28 | 0.35 | 0.48 |
| 8 | 26674936 | rs11779546 | -0.06 | 0.67 | -3.112 | 0.319 | -0.03 | 0.04 | 0.04 | 0 | 0.21 | 0.01 | 0.17 | 0.07 | 0.19 | 0.3 |
| 8 | 26678206 | rs13282836 | -0.034 | 0.653 | -2.95 | 0.354 | -0.052 | -0.046 | -0.046 | 0 | 0.25 | 0.07 | 0.31 | 0.29 | 0.28 | 0.46 |
| 8 | 26679578 | rs4732902 | 0.195 | 0.235 | -2.747 | 0.348 | 0.05 | 0.012 | 0.012 | 0 | 0.33 | 0.15 | 0.32 | 0.24 | 0.2 | 0.29 |
| 8 | 26681450 | rs11781115 | -0.484 | 0.942 | -2.705 | 0.331 | -0.056 | -0.039 | -0.039 | 0 | 0.31 | 0.08 | 0.38 | 0.34 | 0.34 | 0.51 |
| 8 | 26682677 | rs4416829 | -0.722 | 1.242 | -3 | -0.35 | -0.104 | 0.109 | 0.109 | 0 | 0.27 | 0.03 | 0.28 | 0.07 | 0.33 | 0.39 |
| 8 | 26683792 | rs7820633 | -0.13 | 0.429 | -2.533 | -0.309 | -0.004 | -0.003 | -0.003 | 3.086 | 0.55 | 0.46 | 0.54 | 0.7 | 0.63 | 0.6 |
| 8 | 26688826 | rs17334323 | 0.172 | 0.775 | -2.741 | -0.27 | -0.045 | -0.038 | -0.038 | 0 | 0.39 | 0.24 | 0.33 | 0.51 | 0.43 | 0.43 |
| 8 | 26690695 | rs526302 | 0.827 | 1.405 | -2.966 | 0.009 | 0.008 | 0 | 0 | 0 | 0.16 | 0.16 | 0.14 | 0.13 | 0.25 | 0.09 |
| 8 | 26692367 | rs472151 | -0.286 | 0.415 | -3.09 | -0.693 | -0.007 | 0.091 | 0.091 | 3.024 | 0.43 | 0.74 | 0.36 | 0.19 | 0.56 | 0.25 |
| 8 | 26695255 | rs2322333 | -0.412 | -0.185 | -2.969 | -1.015 | -0.001 | -0.001 | -0.001 | 4.273 | 0.63 | 0.61 | 0.55 | 0.53 | 0.59 | 0.63 |
| 8 | 26695617 | rs10503800 | 0.576 | -0.48 | -2.258 | -1.168 | -0.02 | -0.01 | -0.01 | 0 | 0.4 | 0.44 | 0.35 | 0.47 | 0.29 | 0.44 |
| 8 | 26697057 | rs574647 | -0.644 | -0.447 | -2.333 | -1.113 | 0.011 | -0.03 | -0.03 | 0.421 | 0.71 | 0.74 | 0.53 | 0.58 | 0.72 | 0.58 |
| 8 | 26697365 | rs577366 | 2.216 | -0.362 | -2.376 | -1.097 | 0.006 | -0.001 | -0.001 | 0.659 | 0.15 | 0.15 | 0.13 | 0.14 | 0.25 | 0.07 |
| 8 | 26698047 | rs1079078 | 2.073 | -0.459 | -2.427 | -1.139 | -0.003 | -0.041 | -0.041 | 0 | 0.24 | 0.1 | 0.19 | 0.38 | 0.19 | 0.35 |
| 8 | 26699067 | rs556793 | 0.398 | -0.53 | -2.382 | -0.945 | 0.033 | 0.008 | 0.008 | 0 | 0.48 | 0.74 | 0.41 | 0.48 | 0.53 | 0.46 |
| 8 | 26699937 | rs3102087 | 0.48 | -0.552 | -2.358 | -0.956 | 0.012 | 0.005 | 0.005 | 0 | 0.47 | 0.39 | 0.4 | 0.48 | 0.53 | 0.47 |
| 8 | 26701444 | rs2036107 | -0.232 | -0.53 | -2.369 | -0.962 | 0.074 | -0.042 | -0.042 | 0.097 | 0.56 | 0.77 | 0.56 | 0.56 | 0.64 | 0.59 |
| 8 | 26701818 | rs13274679 | 0.938 | -0.513 | -2.355 | -0.955 | 0.006 | -0.029 | -0.029 | 0 | 0.06 | 0.09 | 0.05 | 0 | 0.07 | 0.04 |
| 8 | 26703221 | rs498194 | -0.859 | -0.554 | -2.351 | -0.945 | 0.021 | -0.075 | -0.075 | 0 | 0.67 | 0.79 | 0.6 | 0.57 | 0.68 | 0.6 |
| 8 | 26704875 | rs10093667 | 2.338 | -0.461 | -2.449 | -0.909 | -0.01 | 0.022 | 0.022 | 0.6 | 0.12 | 0.07 | 0.09 | 0.06 | 0.16 | 0.11 |
| 8 | 26705581 | rs558455 | -0.004 | -0.142 | -2.179 | -0.773 | 0.067 | 0.002 | 0.002 | 0 | 0.5 | 0.67 | 0.44 | 0.52 | 0.57 | 0.49 |
| 8 | 26708986 | rs12541572 | 0.689 | 0.116 | -2.274 | -0.674 | -0.004 | 0.011 | 0.011 | 0 | 0.08 | 0.06 | 0.16 | 0.09 | 0.13 | 0.12 |
| 8 | 26711990 | rs544104 | 0.868 | 0.177 | -2.322 | -0.632 | -0.019 | 0.023 | 0.023 | 0.102 | 0.27 | 0.17 | 0.19 | 0.13 | 0.3 | 0.09 |
| 8 | 26712028 | rs544215 | 0.192 | 0.139 | -2.267 | -0.586 | 0.024 | 0.025 | 0.025 | 0.333 | 0.5 | 0.27 | 0.37 | 0.48 | 0.49 | 0.43 |
| 8 | 26712981 | rs11782159 | -0.072 | 0.089 | -1.838 | -0.342 | -0.001 | 0.023 | 0.023 | 0 | 0.4 | 0.62 | 0.42 | 0.43 | 0.39 | 0.5 |
| 8 | 26714874 | rs13278849 | 0.509 | -0.348 | -1.432 | -0.101 | -0.001 | -0.001 | -0.001 | 0 | 0.74 | 0.34 | 0.61 | 0.77 | 0.75 | 0.78 |
| 8 | 26716622 | rs489790 | 0.408 | -0.363 | -1.224 | -0.116 | 0 | -0.005 | -0.005 | 2.643 | 0.13 | 0.14 | 0.22 | 0.13 | 0.11 | 0.11 |
| 8 | 26717817 | rs17426222 | 1.324 | -0.053 | -1.98 | -0.038 | -0.065 | -0.049 | -0.049 | 0 | 0.23 | 0.05 | 0.15 | 0.23 | 0.3 | 0.17 |
| 8 | 26720449 | rs10503801 | 2.5 | -0.133 | -1.842 | -0.142 | 0.004 | 0 | 0 | 0 | 0.06 | 0.06 | 0.05 | 0 | 0.05 | 0.01 |
| 8 | 26720490 | rs580644 | -0.719 | -0.134 | -0.722 | -0.122 | 0.063 | 0.009 | 0.009 | 0 | 0.91 | 0.65 | 0.86 | 0.91 | 0.89 | 0.89 |
| 8 | 26720515 | rs17056112 | 2.055 | -0.574 | -1.049 | 0.243 | 0.022 | -0.042 | -0.042 | 0.495 | 0.07 | 0.01 | 0.07 | 0.18 | 0.03 | 0.06 |
| 8 | 26721281 | rs573514 | 0.537 | -0.493 | -1.053 | 0.193 | -0.035 | -0.047 | -0.047 | 0.044 | 0.37 | 0.22 | 0.39 | 0.36 | 0.41 | 0.29 |
| 8 | 26723049 | rs1383914 | 0.181 | -0.421 | -1.068 | 0.242 | 0 | 0 | 0 | 0 | 0.52 | 0.46 | 0.47 | 0.55 | 0.48 | 0.61 |
| 8 | 26724326 | rs3808585 | 0.686 | -0.444 | -1.228 | 0.007 | 0.008 | -0.012 | -0.012 | 0 | 0.35 | 0.17 | 0.33 | 0.4 | 0.29 | 0.54 |
| 8 | 26727370 | rs922733 | 1.829 | -0.233 | -1.245 | -0.124 | -0.036 | 0.032 | 0.032 | 0 | 0.17 | 0.07 | 0.19 | 0.09 | 0.2 | 0.32 |
| 8 | 26728237 | rs472585 | -0.868 | -0.411 | -0.501 | 0.171 | 0.019 | -0.011 | -0.011 | 0.179 | 0.91 | 0.77 | 0.86 | 0.91 | 0.89 | 0.89 |
| 8 | 26731706 | rs7830143 | -0.738 | -0.91 | -0.592 | 0.283 | -0.003 | -0.001 | -0.001 | 0 | 0.4 | 0.59 | 0.39 | 0.41 | 0.33 | 0.37 |
| 8 | 26743306 | rs10503802 | -0.098 | -1.031 | -0.517 | 0.654 | 0 | 0.021 | 0.021 | 0 | 0.12 | 0.12 | 0.29 | 0.35 | 0.13 | 0.26 |
| 8 | 26743789 | rs478360 | 1.799 | -1.053 | -0.922 | 0.345 | -0.05 | 0.053 | 0.053 | 0.058 | 0.32 | 0.18 | 0.32 | 0.13 | 0.45 | 0.24 |
| 8 | 26743933 | rs1842646 | -0.692 | -1.083 | -0.907 | 0.334 | 0.014 | 0.007 | 0.007 | 1.255 | 0.44 | 0.62 | 0.55 | 0.63 | 0.37 | 0.5 |
| 8 | 26749304 | rs2322315 | 1.399 | -1.304 | -0.651 | 0.162 | -0.01 | 0 | 0 | 0 | 0.2 | 0.2 | 0.16 | 0.27 | 0.09 | 0.22 |
| 8 | 26752716 | rs1481649 | 2.369 | -1.455 | -0.847 | 0.089 | 0.03 | 0.032 | 0.032 | 0 | 0.12 | 0.01 | 0.07 | 0.09 | 0.04 | 0.16 |
| 8 | 26755162 | rs839201 | -2.552 | -1.436 | -0.808 | 0.021 | 0.002 | -0.033 | -0.033 | 0 | 0.89 | 0.88 | 0.94 | 0.98 | 0.88 | 0.8 |
| 8 | 26764947 | rs1600790 | -0.774 | -1.634 | -0.907 | 0.037 | 0.01 | 0.004 | 0.004 | 1.255 | 0.44 | 0.6 | 0.56 | 0.63 | 0.37 | 0.5 |
| 8 | 26765621 | rs6982177 | 2.382 | -1.652 | -0.848 | -0.118 | -0.021 | 0.027 | 0.027 | 0 | 0.1 | 0.05 | 0.09 | 0 | 0.15 | 0.03 |
| 8 | 26765831 | rs6982580 | -1.314 | -1.643 | -0.868 | -0.099 | 0.011 | 0.045 | 0.045 | 1.196 | 0.57 | 0.68 | 0.61 | 0.85 | 0.43 | 0.55 |
| 8 | 26767065 | rs12549158 | 0.245 | -1.316 | -0.724 | 0.101 | -0.027 | -0.023 | -0.023 | 0 | 0.31 | 0.43 | 0.25 | 0.27 | 0.24 | 0.25 |
| 8 | 26773511 | rs1481650 | 0.197 | -1.316 | -0.729 | 0.107 | -0.031 | -0.023 | -0.023 | 0 | 0.31 | 0.44 | 0.25 | 0.28 | 0.25 | 0.25 |
| 8 | 26776134 | rs7816179 | 1.093 | -1.118 | -0.805 | 0.128 | 0.004 | 0.022 | 0.022 | 0 | 0.07 | 0.07 | 0.22 | 0.27 | 0.07 | 0.16 |
| 8 | 26776330 | rs13258346 | 0.06 | -1.168 | -0.703 | 0.053 | 0.002 | 0.032 | 0.032 | 0 | 0.21 | 0.33 | 0.35 | 0.27 | 0.24 | 0.22 |
| 8 | 26781104 | rs1481653 | 2.18 | -0.977 | -0.929 | -0.107 | -0.103 | 0.077 | 0.077 | 0.274 | 0.29 | 0.07 | 0.27 | 0.13 | 0.41 | 0.22 |
| 8 | 26785371 | rs10102024 | 1.046 | -1.323 | -0.868 | 0.018 | 0.002 | 0 | 0 | 0 | 0.23 | 0.22 | 0.18 | 0.26 | 0.12 | 0.25 |
| 8 | 26786291 | rs6557957 | -0.549 | -1.274 | -0.898 | -0.051 | 0.02 | -0.015 | -0.015 | 0.863 | 0.4 | 0.32 | 0.49 | 0.53 | 0.34 | 0.44 |
| 8 | 26800625 | rs9886536 | 0.088 | -1.265 | -0.849 | 0.002 | 0.01 | -0.014 | -0.014 | 0.099 | 0.3 | 0.28 | 0.4 | 0.52 | 0.18 | 0.41 |
| 8 | 26812027 | rs11135963 | 1.8 | -0.999 | -0.708 | -0.487 | -0.05 | 0.054 | 0.054 | 0.157 | 0.32 | 0.18 | 0.32 | 0.14 | 0.45 | 0.24 |
| 8 | 26812434 | rs2046187 | -0.644 | -1.004 | -0.878 | -0.38 | 0.012 | 0.019 | 0.019 | 0.41 | 0.52 | 0.47 | 0.53 | 0.7 | 0.39 | 0.48 |
| 8 | 26814879 | rs7825001 | 0.244 | -1 | -0.888 | -0.492 | -0.005 | -0.016 | -0.016 | 0 | 0.55 | 0.54 | 0.51 | 0.23 | 0.67 | 0.39 |
| 8 | 26819163 | rs12547764 | 1.144 | -0.635 | -0.657 | -0.149 | -0.018 | 0.012 | 0.012 | 0 | 0.23 | 0.3 | 0.23 | 0.31 | 0.16 | 0.29 |
| 8 | 26819362 | rs12549537 | 1.343 | -0.642 | -0.608 | -0.13 | -0.003 | 0 | 0 | 0 | 0.22 | 0.23 | 0.19 | 0.24 | 0.15 | 0.24 |
| 8 | 26821013 | rs6557962 | 0.039 | -0.565 | -0.62 | -0.167 | 0 | 0 | 0 | 0.118 | 0.36 | 0.35 | 0.34 | 0.32 | 0.34 | 0.34 |
| 8 | 26831305 | rs1481646 | 0.007 | -0.568 | -0.649 | -0.305 | -0.002 | -0.001 | -0.001 | 0 | 0.37 | 0.38 | 0.34 | 0.35 | 0.34 | 0.34 |
| 8 | 26833198 | rs12547343 | 0.752 | -0.615 | -0.607 | -0.258 | -0.021 | 0.015 | 0.015 | 0.069 | 0.27 | 0.34 | 0.25 | 0.35 | 0.19 | 0.31 |
| 8 | 26834246 | rs6420159 | 0.29 | -0.526 | -0.682 | -0.288 | -0.033 | 0.023 | 0.023 | 0.405 | 0.32 | 0.44 | 0.34 | 0.44 | 0.24 | 0.36 |
| 8 | 26834297 | rs900073 | -1.728 | -0.281 | -0.892 | -0.017 | -0.004 | -0.002 | -0.002 | 0.688 | 0.89 | 0.95 | 0.95 | 0.99 | 0.89 | 0.79 |
| 8 | 26838170 | rs17430706 | 1.416 | -0.203 | -0.932 | 0.066 | -0.026 | 0.034 | 0.034 | 0 | 0.1 | 0.04 | 0.09 | 0 | 0.15 | 0.03 |
| 8 | 26838759 | rs10095546 | -0.27 | -0.169 | -0.948 | 0.109 | 0.002 | -0.013 | -0.013 | 0 | 0.7 | 0.75 | 0.69 | 0.6 | 0.75 | 0.58 |
| 8 | 26843131 | rs1871807 | 0.76 | -0.051 | -0.833 | 0.065 | 0.009 | -0.009 | -0.009 | 0 | 0.12 | 0.16 | 0.13 | 0.08 | 0.16 | 0.07 |

| | | | | | | | | | | | | | | | | |
|---|---|---|---|---|---|---|---|---|---|---|---|---|---|---|---|---|
| 8 | 26843152 | rs1871808 | 0.76 | -0.051 | -0.833 | 0.065 | 0.009 | -0.009 | -0.009 | 0.006 | 0.12 | 0.16 | 0.13 | 0.08 | 0.16 | 0.07 |
| 8 | 26844116 | rs17346184 | 0.797 | 0.186 | -1.313 | -0.243 | -0.006 | 0.054 | 0.054 | 0 | 0.22 | 0.11 | 0.18 | 0.07 | 0.21 | 0.21 |
| 8 | 26844269 | rs10087840 | -0.271 | 0.11 | -1.281 | -0.025 | -0.013 | -0.025 | -0.025 | 0 | 0.73 | 0.8 | 0.62 | 0.54 | 0.66 | 0.56 |
| 8 | 26844293 | rs12334593 | 1.058 | -0.169 | -1.219 | -0.241 | -0.009 | -0.004 | -0.004 | 0 | 0.39 | 0.5 | 0.33 | 0.4 | 0.36 | 0.32 |
| 8 | 26847397 | rs12681793 | 1.329 | -0.286 | -1.14 | 0.109 | -0.03 | 0.101 | 0.101 | 0 | 0.08 | 0.17 | 0.07 | 0.36 | 0.04 | 0.1 |
| 8 | 26849975 | rs10503804 | 1.106 | -0.324 | -1.219 | -0.062 | 0.002 | 0.054 | 0.054 | 0 | 0.23 | 0.11 | 0.19 | 0.07 | 0.21 | 0.19 |
| 8 | 26857295 | rs9314331 | -0.055 | -0.142 | -1.272 | -0.154 | -0.004 | 0.008 | 0.008 | 2.102 | 0.69 | 0.63 | 0.79 | 0.57 | 0.74 | 0.65 |
| 8 | 26857693 | rs7842266 | 0.454 | -0.141 | -1.21 | -0.109 | -0.024 | 0.008 | 0.008 | 0 | 0.31 | 0.18 | 0.46 | 0.22 | 0.38 | 0.33 |
| 8 | 26858439 | rs6980493 | 1.391 | -0.127 | -1.203 | -0.116 | -0.006 | -0.006 | -0.006 | 0.396 | 0.38 | 0.46 | 0.33 | 0.35 | 0.36 | 0.32 |
| 8 | 26859730 | rs10096027 | -0.662 | -0.09 | -1.315 | -0.073 | -0.002 | -0.006 | -0.006 | 0 | 0.78 | 0.81 | 0.86 | 0.93 | 0.78 | 0.75 |
| 8 | 26867904 | rs2874755 | 0.944 | -0.184 | -1.082 | 0.134 | -0.052 | 0.091 | 0.091 | 0 | 0.18 | 0.4 | 0.12 | 0.37 | 0.09 | 0.11 |
| 8 | 26873319 | rs12545634 | 0.849 | -0.39 | -1.505 | -0.09 | -0.013 | 0.036 | 0.036 | 0.008 | 0.31 | 0.41 | 0.47 | 0.4 | 0.29 | 0.38 |
| 8 | 26873380 | rs13279967 | 2.158 | -0.392 | -1.554 | -0.055 | -0.044 | -0.049 | -0.049 | 0 | 0.08 | 0.01 | 0.27 | 0.16 | 0.16 | 0.13 |
| 8 | 26877573 | rs10107927 | 1.785 | -0.423 | -1.506 | -0.071 | -0.044 | 0.013 | 0.013 | 0 | 0.23 | 0.35 | 0.2 | 0.24 | 0.13 | 0.23 |
| 8 | 26877998 | rs12543619 | -0.609 | -0.399 | -1.567 | -0.127 | -0.032 | 0.046 | 0.046 | 0.15 | 0.73 | 0.52 | 0.82 | 0.57 | 0.79 | 0.71 |
| 8 | 26883003 | rs10503806 | 2.69 | 0.096 | -2.078 | -0.26 | -0.038 | 0.033 | 0.033 | 0 | 0.06 | 0.01 | 0.08 | 0 | 0.14 | 0.03 |
| 8 | 26892214 | rs2874779 | 1.382 | 0.104 | -2.151 | -0.334 | -0.093 | 0.054 | 0.054 | 0 | 0.19 | 0.04 | 0.16 | 0.1 | 0.29 | 0.14 |
| 8 | 26893734 | rs9314338 | 0.5 | -0.156 | -1.731 | -0.124 | -0.065 | 0.187 | 0.187 | 0 | 0.42 | 0.1 | 0.35 | 0.17 | 0.49 | 0.34 |
| 8 | 26894544 | rs7461897 | -1.464 | 0.642 | -0.911 | 0.525 | 0.005 | -0.004 | -0.004 | 0 | 0.91 | 0.92 | 0.95 | 0.92 | 0.88 | 0.81 |
| 8 | 26894971 | rs13251780 | 1.124 | 0.49 | -1.025 | 0.626 | -0.011 | -0.01 | -0.01 | 1.227 | 0.08 | 0.07 | 0.27 | 0.16 | 0.16 | 0.13 |
| 8 | 26899943 | rs10086635 | -0.073 | 0.144 | -0.913 | 0.287 | 0.005 | 0.008 | 0.008 | 0.714 | 0.48 | 0.39 | 0.64 | 0.48 | 0.45 | 0.57 |
| 8 | 26901426 | rs7459991 | 0.685 | -0.227 | -0.815 | 0.021 | -0.013 | 0.002 | 0.002 | 0 | 0.21 | 0.28 | 0.17 | 0.23 | 0.14 | 0.23 |
| 8 | 26907844 | rs17362506 | 0.828 | -0.099 | -1.26 | -0.15 | -0.025 | 0.02 | 0.02 | 0 | 0.21 | 0.05 | 0.16 | 0.13 | 0.18 | 0.17 |
| 8 | 26907937 | rs7823902 | -1.087 | -0.257 | -1.352 | -0.344 | -0.012 | -0.047 | -0.047 | 0 | 0.78 | 0.86 | 0.61 | 0.53 | 0.72 | 0.56 |
| 8 | 26908599 | rs7461503 | 0.032 | -0.459 | -1.262 | -0.714 | 0.007 | 0.032 | 0.032 | 0.001 | 0.33 | 0.26 | 0.23 | 0.13 | 0.26 | 0.21 |
| 8 | 26910150 | rs7834004 | 1.877 | -0.453 | -1.206 | -0.703 | -0.005 | -0.029 | -0.029 | 0 | 0.1 | 0.12 | 0.04 | 0 | 0.05 | 0.01 |
| 8 | 26912993 | rs17056735 | 1.877 | -0.453 | -1.206 | -0.703 | -0.005 | -0.029 | -0.029 | 0 | 0.1 | 0.12 | 0.04 | 0 | 0.05 | 0.01 |
| 8 | 26913183 | rs11778045 | 1.265 | -0.376 | -1.312 | -0.642 | -0.025 | 0.017 | 0.017 | 0 | 0.21 | 0.05 | 0.16 | 0.13 | 0.18 | 0.17 |
| 8 | 26921594 | rs1878447 | 0.666 | -0.469 | -1.17 | -0.58 | -0.114 | 0.116 | 0.116 | 0.03 | 0.4 | 0.08 | 0.3 | 0.21 | 0.47 | 0.28 |
| 8 | 26923058 | rs4623440 | -0.469 | -0.424 | -0.823 | -0.468 | 0.009 | 0.075 | 0.075 | 0 | 0.72 | 0.56 | 0.55 | 0.44 | 0.68 | 0.55 |
| 8 | 26933124 | rs10503809 | -0.354 | -0.471 | -1.201 | -0.583 | 0.033 | 0.131 | 0.131 | 0 | 0.49 | 0.75 | 0.58 | 0.77 | 0.46 | 0.68 |
| 8 | 26941432 | rs4732708 | -0.406 | -0.589 | -0.985 | -0.604 | 0.004 | 0.004 | 0.004 | 0 | 0.68 | 0.66 | 0.81 | 0.79 | 0.79 | 0.74 |
| 8 | 26942659 | rs11776692 | 1.872 | -0.447 | -1.307 | -0.817 | -0.086 | 0.096 | 0.096 | 0.128 | 0.2 | 0.03 | 0.17 | 0.08 | 0.29 | 0.14 |
| 8 | 26943632 | rs935408 | 0.283 | -0.552 | -0.904 | -0.493 | 0.001 | 0.024 | 0.024 | 0 | 0.29 | 0.58 | 0.47 | 0.56 | 0.32 | 0.45 |
| 8 | 26944966 | rs2028909 | 0.9 | -0.538 | -0.964 | -0.493 | -0.117 | 0.098 | 0.098 | 0.225 | 0.4 | 0.07 | 0.35 | 0.24 | 0.47 | 0.28 |
| 8 | 26948163 | rs2121400 | 1.891 | -0.572 | -1.001 | -0.52 | -0.008 | 0.002 | 0.002 | 0.005 | 0.15 | 0.16 | 0.1 | 0.21 | 0.1 | 0.18 |
| 8 | 26950586 | rs12675131 | 0.964 | -0.649 | -0.629 | -0.505 | 0.001 | 0.017 | 0.017 | 0 | 0.23 | 0.24 | 0.39 | 0.47 | 0.28 | 0.43 |
| 8 | 26951968 | rs17056778 | 0.483 | -0.745 | -0.081 | -0.294 | 0.006 | -0.003 | -0.003 | 0.107 | 0.2 | 0.21 | 0.12 | 0.21 | 0.14 | 0.23 |
| 8 | 26953656 | rs3906912 | 0.102 | -0.806 | -0.038 | -0.257 | 0.006 | 0.005 | 0.005 | 0 | 0.69 | 0.68 | 0.82 | 0.79 | 0.79 | 0.73 |
| 8 | 26953801 | rs3906913 | -1.447 | -0.119 | 0.027 | 0.063 | -0.005 | -0.028 | -0.028 | 0 | 0.92 | 0.91 | 0.96 | 1 | 0.95 | 0.99 |
| 8 | 26962037 | rs17056786 | -0.807 | 0.2 | -0.012 | 0.453 | 0.01 | 0.007 | 0.007 | 2.974 | 0.1 | 0.31 | 0.09 | 0.09 | 0.08 | 0.06 |
| 8 | 26964212 | rs10110980 | -0.332 | 0.848 | -0.623 | 0.513 | 0.004 | 0.007 | 0.007 | 0 | 0.29 | 0.34 | 0.47 | 0.3 | 0.32 | 0.31 |
| 8 | 26967488 | rs10102339 | 0.378 | 0.668 | -0.735 | 0.359 | 0.002 | -0.014 | -0.014 | 0 | 0.58 | 0.32 | 0.59 | 0.42 | 0.64 | 0.62 |
| 8 | 26968532 | rs12541830 | 0.706 | 0.198 | -1.208 | 0.048 | 0.006 | -0.035 | -0.035 | 0 | 0.23 | 0.16 | 0.34 | 0.32 | 0.23 | 0.26 |
| 8 | 26969601 | rs4621816 | 0.094 | 0.17 | -1.096 | -0.118 | 0.007 | -0.041 | -0.041 | 0.291 | 0.34 | 0.27 | 0.33 | 0.58 | 0.27 | 0.32 |
| 8 | 26970565 | rs17436930 | 1.624 | 0.144 | -1.15 | -0.213 | -0.11 | 0.135 | 0.135 | 0 | 0.31 | 0.02 | 0.22 | 0.1 | 0.39 | 0.34 |
| 8 | 26977478 | rs4510876 | -1.371 | 0.323 | -0.374 | -0.089 | 0.005 | -0.062 | -0.062 | 0 | 0.65 | 0.63 | 0.74 | 0.9 | 0.59 | 0.65 |
| 8 | 26979015 | rs13282907 | -0.136 | 0.369 | -0.365 | 0.112 | -0.001 | 0.012 | 0.012 | 0 | 0.31 | 0.35 | 0.41 | 0.32 | 0.32 | 0.33 |
| 8 | 26986250 | rs6557981 | 1.384 | 0.226 | -0.579 | 0.186 | 0.036 | -0.036 | -0.036 | 0 | 0.17 | 0.08 | 0.13 | 0.34 | 0.1 | 0.19 |
| 8 | 26991054 | rs9644048 | -0.261 | 0.335 | -0.514 | 0.097 | -0.011 | -0.006 | -0.006 | 0.014 | 0.42 | 0.44 | 0.48 | 0.66 | 0.34 | 0.45 |
| 8 | 26996374 | rs2322509 | -0.616 | 0.3 | -0.488 | 0.121 | 0.004 | 0.006 | 0.006 | 0 | 0.47 | 0.44 | 0.54 | 0.66 | 0.42 | 0.51 |
| 8 | 27001175 | rs11775507 | -0.798 | 0.36 | -0.604 | 0.163 | 0 | 0 | 0 | 0 | 0.51 | 0.54 | 0.56 | 0.66 | 0.45 | 0.52 |
| 8 | 27001846 | rs12544600 | 1.179 | 0.408 | -0.99 | 0.492 | 0.032 | -0.034 | -0.034 | 0 | 0.17 | 0.08 | 0.14 | 0.34 | 0.1 | 0.19 |
| 8 | 27002114 | rs17437785 | 1.445 | 0.527 | -1.006 | 0.445 | -0.01 | -0.012 | -0.012 | 5.614 | 0.12 | 0.09 | 0.18 | 0.22 | 0.15 | 0.12 |
| 8 | 27002663 | rs17056852 | 1.21 | 0.377 | -0.992 | 0.466 | 0.024 | -0.032 | -0.032 | 0 | 0.16 | 0.08 | 0.13 | 0.34 | 0.11 | 0.19 |
| 8 | 27005955 | rs2166536 | 1.192 | 0.761 | -1.011 | 0.479 | 0.009 | -0.014 | -0.014 | 0 | 0.16 | 0.11 | 0.11 | 0.19 | 0.16 | 0.1 |
| 8 | 27011201 | rs2322516 | 1.168 | 0.686 | -0.932 | 0.24 | 0.025 | -0.026 | -0.026 | 0.885 | 0.16 | 0.09 | 0.13 | 0.34 | 0.1 | 0.19 |
| 8 | 27016165 | rs12544814 | 1.188 | 0.683 | -0.933 | 0.238 | 0.024 | -0.026 | -0.026 | 0.053 | 0.16 | 0.09 | 0.13 | 0.34 | 0.1 | 0.19 |
| 8 | 27017642 | rs13274189 | 1.197 | 0.817 | -1.036 | 0.266 | -0.055 | -0.055 | -0.055 | 0 | 0.11 | 0.02 | 0.26 | 0.14 | 0.17 | 0.17 |
| 8 | 27030123 | rs10107664 | -0.882 | 0.522 | -0.587 | -0.169 | 0.047 | -0.031 | -0.031 | 0 | 0.94 | 0.81 | 0.93 | 1 | 0.91 | 0.93 |
| 8 | 27036631 | rs10110140 | -0.023 | 0.318 | -0.387 | -0.083 | -0.037 | 0.008 | 0.008 | 0 | 0.4 | 0.25 | 0.55 | 0.55 | 0.48 | 0.39 |
| 8 | 27040155 | rs10097427 | -0.02 | 0.385 | -0.735 | -0.049 | -0.026 | 0.007 | 0.007 | 0 | 0.39 | 0.28 | 0.56 | 0.55 | 0.48 | 0.39 |
| 8 | 27050300 | rs7341551 | -0.02 | 0.385 | -0.708 | -0.049 | -0.023 | 0.007 | 0.007 | 0 | 0.39 | 0.3 | 0.56 | 0.55 | 0.48 | 0.39 |
| 8 | 27057879 | rs12548107 | 0.843 | 0.129 | -0.585 | 0.113 | -0.028 | 0.017 | 0.017 | 0 | 0.16 | 0.21 | 0.13 | 0.33 | 0.08 | 0.18 |
| 8 | 27066369 | rs10099964 | -0.249 | 1.013 | -0.888 | 0.228 | -0.031 | 0.006 | 0.006 | 0 | 0.38 | 0.27 | 0.55 | 0.56 | 0.49 | 0.35 |
| 8 | 27073268 | rs1446684 | -0.289 | 1.154 | -0.329 | 0.644 | 0.031 | 0.065 | 0.065 | 0 | 0.24 | 0.38 | 0.32 | 0.37 | 0.27 | 0.24 |
| 8 | 27075395 | rs1077739 | 1.064 | 0.588 | -1.344 | -0.07 | -0.011 | 0.09 | 0.09 | 0.26 | 0.25 | 0.09 | 0.15 | 0.11 | 0.22 | 0.1 |
| 8 | 27076127 | rs10503811 | 1.466 | 0.512 | -1.541 | -0.167 | -0.025 | 0.033 | 0.033 | 0.337 | 0.06 | 0 | 0.06 | 0 | 0.09 | 0.05 |
| 8 | 27079609 | rs13282363 | -0.213 | 0.283 | -1.225 | -0.053 | 0.026 | 0.155 | 0.155 | 1.809 | 0.19 | 0.45 | 0.35 | 0.4 | 0.22 | 0.24 |

| | | | | | | | | | | | | | | | | |
|---|---|---|---|---|---|---|---|---|---|---|---|---|---|---|---|---|
| 8 | 27080408 | rs2084227 | -1.602 | -0.13 | -1.147 | -0.251 | -0.014 | -0.003 | -0.003 | 0.754 | 0.93 | 0.87 | 0.96 | 0.96 | 0.95 | 0.88 |
| 8 | 27081237 | rs2322411 | 1.505 | -0.053 | -1.173 | -0.352 | 0.007 | 0.04 | 0.04 | 0.559 | 0.26 | 0.22 | 0.16 | 0.11 | 0.22 | 0.11 |
| 8 | 27084275 | rs9314343 | 1.438 | -0.038 | -1.132 | -0.338 | 0.014 | 0.055 | 0.055 | 0.291 | 0.27 | 0.2 | 0.15 | 0.12 | 0.21 | 0.11 |
| 8 | 27088607 | rs10099974 | 1.434 | -0.033 | -1.132 | -0.336 | 0.015 | 0.058 | 0.058 | 0.222 | 0.27 | 0.2 | 0.15 | 0.12 | 0.21 | 0.11 |
| 8 | 27093495 | rs4733037 | 0.387 | -0.012 | -0.937 | -0.219 | 0.011 | -0.03 | -0.03 | 0 | 0.37 | 0.27 | 0.4 | 0.49 | 0.34 | 0.3 |
| 8 | 27093924 | rs4733038 | -0.866 | -0.392 | -0.813 | -0.39 | -0.01 | 0.021 | 0.021 | 0.135 | 0.92 | 0.7 | 0.97 | 0.96 | 0.97 | 0.96 |
| 8 | 27096634 | rs1542185 | -0.382 | -0.347 | -0.886 | -0.434 | 0.009 | -0.001 | -0.001 | 0.583 | 0.51 | 0.57 | 0.48 | 0.53 | 0.45 | 0.39 |
| 8 | 27098436 | rs1562330 | 1.402 | -0.6 | -0.741 | 0.03 | -0.046 | 0.062 | 0.062 | 0.939 | 0.37 | 0.17 | 0.29 | 0.28 | 0.38 | 0.3 |
| 8 | 27099879 | rs3739212 | 0.804 | -0.543 | -0.458 | 0.033 | 0.01 | 0.027 | 0.027 | 1.206 | 0.48 | 0.3 | 0.42 | 0.42 | 0.51 | 0.35 |
| 8 | 27100014 | rs3739213 | 1.343 | -0.537 | -0.471 | 0.055 | -0.008 | -0.011 | -0.011 | 0 | 0.1 | 0.05 | 0.25 | 0.42 | 0.13 | 0.2 |
| 8 | 27101206 | rs7830662 | -1.155 | -0.535 | -0.477 | 0.03 | -0.007 | -0.046 | -0.046 | 0.191 | 0.72 | 0.63 | 0.75 | 0.88 | 0.75 | 0.64 |
| 8 | 27103621 | rs7844646 | -1.155 | -0.535 | -0.477 | 0.03 | -0.007 | -0.046 | -0.046 | 2.79 | 0.72 | 0.63 | 0.75 | 0.88 | 0.75 | 0.64 |
| 8 | 27105384 | rs7831609 | 0.219 | -0.439 | -0.765 | 0.024 | 0.039 | 0.012 | 0.012 | 0.316 | 0.54 | 0.29 | 0.47 | 0.42 | 0.6 | 0.38 |
| 8 | 27111154 | rs7819130 | 0.56 | -0.464 | 0.276 | 0.344 | -0.026 | -0.053 | -0.053 | 0 | 0.85 | 0.48 | 0.9 | 0.96 | 0.89 | 0.91 |
| 8 | 27115405 | rs11779435 | 1.215 | -0.106 | -0.22 | 0.434 | -0.024 | 0.03 | 0.03 | 0.22 | 0.06 | 0 | 0.06 | 0 | 0.09 | 0.04 |
| 8 | 27116494 | rs7005224 | -0.659 | -0.348 | 0.2 | 0.824 | -0.03 | 0.119 | 0.119 | 0 | 0.23 | 0.42 | 0.33 | 0.45 | 0.18 | 0.22 |
| 8 | 27120575 | rs1867887 | -0.028 | -0.067 | 0.795 | 0.136 | -0.003 | -0.026 | -0.026 | 0.009 | 0.88 | 0.61 | 0.89 | 0.95 | 0.88 | 0.96 |
| 8 | 27125867 | rs7003044 | 0.398 | -0.077 | 0.781 | 0.397 | 0.005 | 0.101 | 0.101 | 0 | 0.12 | 0.18 | 0.27 | 0.63 | 0.13 | 0.21 |
| 8 | 27125887 | rs7832778 | 1.14 | -0.144 | 1.103 | 0.335 | -0.013 | 0.019 | 0.019 | 0.01 | 0.75 | 0.42 | 0.62 | 0.32 | 0.75 | 0.75 |
| 8 | 27126551 | rs7003027 | 0.898 | 0.061 | 0.941 | 0.707 | 0.039 | 0.266 | 0.266 | 0 | 0.08 | 0.17 | 0.24 | 0.44 | 0.12 | 0.11 |
| 8 | 27134100 | rs957008 | 1.171 | -0.367 | 0.615 | -0.174 | 0.019 | 0.064 | 0.064 | 0 | 0.61 | 0.29 | 0.49 | 0.21 | 0.65 | 0.54 |
| 8 | 27141452 | rs2322601 | 1.176 | -0.369 | 0.614 | -0.175 | 0.019 | 0.065 | 0.065 | 0 | 0.61 | 0.29 | 0.49 | 0.2 | 0.65 | 0.54 |
| 8 | 27146346 | rs6557990 | 0.177 | -0.378 | 0.746 | -0.117 | 0.002 | -0.003 | -0.003 | 0 | 0.73 | 0.79 | 0.63 | 0.33 | 0.75 | 0.58 |
| 8 | 27147716 | rs12386801 | 1.001 | -0.347 | 0.902 | -0.184 | 0 | -0.003 | -0.003 | 0 | 0.63 | 0.42 | 0.5 | 0.21 | 0.65 | 0.54 |
| 8 | 27156502 | rs2292979 | 1.595 | -0.211 | 0.772 | -0.09 | -0.014 | -0.021 | -0.021 | 1.021 | 0.07 | 0.01 | 0.09 | 0.09 | 0.08 | 0.03 |
| 8 | 27157307 | rs2322602 | 1.335 | -0.241 | 0.423 | -0.848 | -0.073 | 0.101 | 0.101 | 0 | 0.35 | 0.13 | 0.37 | 0.17 | 0.44 | 0.44 |
| 8 | 27159459 | rs7813625 | 0.341 | -0.294 | 0.753 | -0.849 | 0.003 | 0.039 | 0.039 | 0 | 0.47 | 0.68 | 0.51 | 0.29 | 0.54 | 0.48 |
| 8 | 27159836 | rs11996459 | 1.011 | -0.297 | 0.982 | -0.854 | 0.016 | -0.039 | -0.039 | 2.822 | 0.38 | 0.49 | 0.4 | 0.21 | 0.45 | 0.44 |
| 8 | 27163123 | rs2322603 | 0.343 | -0.299 | 0.942 | -0.821 | 0.003 | 0.037 | 0.037 | 0 | 0.47 | 0.68 | 0.51 | 0.3 | 0.54 | 0.48 |
| 8 | 27167942 | rs17446091 | 1.219 | -0.375 | 0.946 | -0.988 | 0.022 | 0.07 | 0.07 | 0.061 | 0.27 | 0.13 | 0.12 | 0.06 | 0.2 | 0.11 |
| 8 | 27168030 | rs13253058 | 1.364 | -0.352 | 0.901 | -0.995 | -0.111 | 0.116 | 0.116 | 0 | 0.35 | 0.09 | 0.37 | 0.17 | 0.44 | 0.44 |
| 8 | 27168820 | rs7006183 | 1.19 | -0.342 | 0.965 | -0.966 | -0.001 | -0.001 | -0.001 | 0 | 0.08 | 0.11 | 0.1 | 0.09 | 0.09 | 0.04 |
| 8 | 27174900 | rs6986075 | 0.645 | -0.375 | 0.909 | -1.007 | -0.009 | 0.051 | 0.051 | 0 | 0.43 | 0.34 | 0.47 | 0.26 | 0.52 | 0.47 |
| 8 | 27175308 | rs6557991 | 1.189 | -0.239 | 0.88 | -0.857 | -0.039 | 0.053 | 0.053 | 0 | 0.36 | 0.2 | 0.37 | 0.21 | 0.45 | 0.44 |
| 8 | 27181065 | rs1316370 | -0.084 | -0.232 | 0.86 | -0.765 | 0.002 | -0.003 | -0.003 | 0 | 0.73 | 0.81 | 0.64 | 0.32 | 0.75 | 0.58 |
| 8 | 27182276 | rs10095624 | -0.084 | -0.148 | 0.75 | -0.707 | 0.002 | -0.003 | -0.003 | 0.114 | 0.73 | 0.81 | 0.64 | 0.32 | 0.75 | 0.58 |
| 8 | 27183794 | rs7005866 | 0.076 | 0.318 | 1.26 | -0.527 | -0.026 | -0.086 | -0.086 | 1.672 | 0.35 | 0.48 | 0.24 | 0.12 | 0.28 | 0.13 |
| 8 | 27184577 | rs4291226 | 0.152 | 0.357 | 0.867 | -0.495 | 0.024 | 0.082 | 0.082 | 0.342 | 0.33 | 0.17 | 0.21 | 0.11 | 0.29 | 0.13 |
| 8 | 27186923 | rs2322606 | -0.14 | 0.21 | 1.012 | 0.034 | 0.003 | 0.056 | 0.056 | 0 | 0.84 | 0.79 | 0.7 | 0.4 | 0.81 | 0.74 |
| 8 | 27188518 | rs10097861 | 0.506 | 0.145 | 0.998 | -0.04 | 0.001 | 0.048 | 0.048 | 0 | 0.8 | 0.74 | 0.69 | 0.4 | 0.79 | 0.74 |
| 8 | 27193923 | rs1879188 | 0.611 | 0.304 | 1.096 | 0.108 | 0.014 | 0.122 | 0.122 | 0 | 0.83 | 0.65 | 0.67 | 0.38 | 0.82 | 0.74 |
| 8 | 27198884 | rs1879189 | 0.376 | 0.349 | 0.707 | 0.132 | 0.003 | 0.045 | 0.045 | 0 | 0.84 | 0.82 | 0.68 | 0.38 | 0.82 | 0.74 |
| 8 | 27203039 | rs3735759 | -1.564 | 0.349 | 0.547 | 0.296 | -0.035 | 0.045 | 0.045 | 0 | 0.94 | 0.99 | 0.7 | 0.53 | 0.85 | 0.88 |
| 8 | 27205777 | rs2322608 | 0.478 | 0.335 | 0.603 | 0.198 | 0.002 | 0.044 | 0.044 | 0 | 0.84 | 0.87 | 0.68 | 0.42 | 0.83 | 0.74 |
| 8 | 27211333 | rs7834529 | 0.301 | 0.369 | 0.574 | 0.268 | -0.002 | 0.012 | 0.012 | 0 | 0.86 | 0.9 | 0.68 | 0.42 | 0.82 | 0.74 |
| 8 | 27212276 | rs10109834 | 0.018 | 1.324 | 0.497 | -0.226 | 0.017 | -0.036 | -0.036 | 0 | 0.36 | 0.43 | 0.4 | 0.17 | 0.48 | 0.21 |
| 8 | 27217763 | rs10097651 | 0.047 | 1.32 | 0.487 | -0.286 | -0.045 | 0.07 | 0.07 | 0 | 0.35 | 0.21 | 0.39 | 0.09 | 0.47 | 0.21 |
| 8 | 27220310 | rs17057043 | 0.591 | 1.43 | 0.575 | -0.295 | 0.014 | 0.02 | 0.02 | 0 | 0.49 | 0.47 | 0.28 | 0.26 | 0.35 | 0.51 |
| 8 | 27226790 | rs755951 | 0.264 | 1.404 | 0.517 | -0.302 | -0.007 | -0.023 | -0.023 | 0 | 0.55 | 0.53 | 0.3 | 0.26 | 0.39 | 0.53 |
| 8 | 27230819 | rs12679874 | 0.322 | 1.453 | 0.214 | -0.232 | 0.05 | 0.065 | 0.065 | 0 | 0.53 | 0.71 | 0.29 | 0.3 | 0.34 | 0.51 |
| 8 | 27237708 | rs4732720 | -0.077 | 1.56 | 0.159 | -0.265 | -0.036 | 0.108 | 0.108 | 0 | 0.41 | 0.2 | 0.41 | 0.24 | 0.51 | 0.37 |
| 8 | 27238826 | rs11785606 | 0.012 | 1.562 | 0.13 | -0.263 | -0.049 | 0.111 | 0.111 | 0 | 0.4 | 0.17 | 0.41 | 0.24 | 0.52 | 0.36 |
| 8 | 27243074 | rs17057065 | 1.145 | 2.183 | -0.062 | -0.263 | -0.044 | -0.041 | -0.041 | 0 | 0.07 | 0.01 | 0.06 | 0.18 | 0.16 | 0.12 |
| 8 | 27244300 | rs11776858 | -0.076 | 1.682 | 0.286 | -0.093 | -0.002 | 0.05 | 0.05 | 0 | 0.41 | 0.27 | 0.39 | 0.28 | 0.43 | 0.35 |
| 8 | 27244919 | rs17447286 | -0.002 | 1.682 | 0.28 | -0.104 | -0.016 | 0.092 | 0.092 | 0 | 0.4 | 0.18 | 0.38 | 0.27 | 0.43 | 0.35 |
| 8 | 27245790 | rs11783759 | -0.09 | 1.681 | 0.279 | -0.104 | -0.021 | 0.106 | 0.106 | 0 | 0.39 | 0.17 | 0.38 | 0.24 | 0.43 | 0.35 |
| 8 | 27248435 | rs11987089 | -1.552 | 1.716 | 0.324 | -0.079 | 0.031 | 0.187 | 0.187 | 0 | 0.94 | 0.92 | 0.69 | 0.54 | 0.85 | 0.87 |
| 8 | 27251325 | rs13260543 | 0.659 | 1.685 | 0.404 | -0.127 | 0.001 | 0.001 | 0.001 | 0 | 0.49 | 0.5 | 0.3 | 0.27 | 0.4 | 0.52 |
| 8 | 27255128 | rs1045510 | 0.022 | 1.61 | 0.336 | -0.156 | -0.011 | 0.084 | 0.084 | 0 | 0.4 | 0.21 | 0.38 | 0.27 | 0.43 | 0.35 |
| 8 | 27257787 | rs2322718 | -0.338 | 1.605 | 0.294 | -0.125 | 0.001 | 0.058 | 0.058 | 0 | 0.44 | 0.29 | 0.39 | 0.27 | 0.45 | 0.35 |
| 8 | 27260484 | rs7005183 | 0.577 | 1.402 | 0.206 | -0.293 | 0.002 | 0.004 | 0.004 | 0 | 0.49 | 0.48 | 0.29 | 0.27 | 0.4 | 0.52 |
| 8 | 27262195 | rs1367088 | 0.081 | 1.357 | -0.109 | -0.353 | 0 | -0.002 | -0.002 | 0 | 0.59 | 0.54 | 0.54 | 0.39 | 0.6 | 0.56 |
| 8 | 27263433 | rs7000615 | -0.086 | 1.81 | 0.016 | -0.229 | 0.033 | 0.131 | 0.131 | 0 | 0.88 | 0.82 | 0.68 | 0.53 | 0.81 | 0.85 |
| 8 | 27263900 | rs11135991 | 0.954 | 1.748 | 0.284 | -0.441 | 0.009 | 0.019 | 0.019 | 0 | 0.49 | 0.46 | 0.28 | 0.27 | 0.41 | 0.51 |
| 8 | 27264952 | rs11986969 | 0.512 | 1.784 | 0.186 | -0.521 | 0.016 | 0.047 | 0.047 | 0 | 0.29 | 0.19 | 0.15 | 0.17 | 0.22 | 0.28 |
| 8 | 27265476 | rs725787 | 0.357 | 1.837 | 0.118 | -0.438 | -0.028 | 0 | 0 | 0 | 0.11 | 0.05 | 0.24 | 0.09 | 0.2 | 0.06 |
| 8 | 27268750 | rs3757908 | -1.526 | 1.808 | 0.126 | -0.383 | -0.039 | 0.021 | 0.021 | 0 | 0.94 | 0.99 | 0.7 | 0.54 | 0.84 | 0.85 |
| 8 | 27269884 | rs11775958 | -0.728 | 1.806 | 0.212 | -0.386 | 0.053 | 0.187 | 0.187 | 0.142 | 0.92 | 0.84 | 0.69 | 0.54 | 0.84 | 0.85 |
| 8 | 27274931 | rs2322714 | 0.954 | 1.712 | 0.268 | -0.48 | 0.014 | 0.035 | 0.035 | 0 | 0.49 | 0.42 | 0.27 | 0.27 | 0.41 | 0.51 |

| | | | | | | | | | | | | | | | | |
|---|---|---|---|---|---|---|---|---|---|---|---|---|---|---|---|---|
| 8 | 27275509 | rs891391 | -1.346 | 1.786 | 0.147 | -0.442 | 0.066 | 0.222 | 0.222 | 0 | 0.93 | 0.87 | 0.69 | 0.55 | 0.84 | 0.85 |
| 8 | 27275794 | rs750539 | -0.157 | 1.769 | 0.146 | -0.463 | -0.006 | 0.042 | 0.042 | 0 | 0.37 | 0.25 | 0.39 | 0.25 | 0.41 | 0.34 |
| 8 | 27279705 | rs2241652 | -0.821 | 1.804 | 0.158 | -0.424 | 0.05 | 0.189 | 0.189 | 0.142 | 0.92 | 0.84 | 0.69 | 0.55 | 0.84 | 0.85 |
| 8 | 27280860 | rs3779634 | 0.899 | 1.826 | 0.137 | -0.436 | -0.001 | -0.006 | -0.006 | 0 | 0.51 | 0.53 | 0.29 | 0.29 | 0.4 | 0.52 |
| 8 | 27281896 | rs4733058 | 1.183 | 1.86 | 0.124 | -0.452 | -0.035 | -0.018 | -0.018 | 0 | 0.08 | 0.03 | 0.24 | 0.09 | 0.18 | 0.06 |
| 8 | 27282063 | rs13268201 | 0.656 | 1.613 | 0.261 | -0.375 | -0.001 | -0.006 | -0.006 | 1.215 | 0.51 | 0.53 | 0.29 | 0.29 | 0.41 | 0.52 |
| 8 | 27284394 | rs7846568 | 0.656 | 1.613 | 0.261 | -0.375 | -0.001 | -0.006 | -0.006 | 0 | 0.51 | 0.53 | 0.29 | 0.29 | 0.41 | 0.52 |
| 8 | 27284692 | rs3779635 | -0.111 | 1.495 | -0.068 | -0.473 | -0.002 | 0.048 | 0.048 | 0 | 0.41 | 0.28 | 0.39 | 0.25 | 0.43 | 0.34 |
| 8 | 27288033 | rs7827965 | 0.417 | 1.49 | 0.269 | -0.275 | -0.001 | -0.022 | -0.022 | 1.769 | 0.55 | 0.5 | 0.44 | 0.27 | 0.45 | 0.55 |
| 8 | 27291557 | rs919493 | -1.116 | 1.63 | 0.422 | -0.083 | 0.089 | 0.335 | 0.335 | 0 | 0.93 | 0.79 | 0.81 | 0.56 | 0.85 | 0.84 |
| 8 | 27291749 | rs12056620 | 1.025 | 1.41 | 0.299 | -0.411 | 0.019 | 0.075 | 0.075 | 0 | 0.45 | 0.31 | 0.39 | 0.2 | 0.39 | 0.43 |
| 8 | 27293865 | rs7005936 | 0.966 | 1.415 | 0.115 | -0.407 | -0.003 | -0.023 | -0.023 | 0.58 | 0.46 | 0.46 | 0.38 | 0.2 | 0.39 | 0.4 |
| 8 | 27299277 | rs9773817 | 0.567 | 1.546 | -0.009 | -0.487 | 0.006 | -0.024 | -0.024 | 0.147 | 0.13 | 0.17 | 0.1 | 0.05 | 0.13 | 0.07 |
| 8 | 27299553 | rs3736525 | 0.701 | 1.513 | -0.006 | -0.501 | 0 | 0.017 | 0.017 | 0 | 0.55 | 0.62 | 0.44 | 0.26 | 0.45 | 0.54 |
| 8 | 27302433 | rs9657295 | 0.753 | 1.511 | -0.204 | -0.478 | -0.002 | -0.018 | -0.018 | 0.082 | 0.53 | 0.49 | 0.43 | 0.26 | 0.45 | 0.54 |
| 8 | 27303015 | rs11135993 | 0.753 | 1.511 | -0.204 | -0.502 | -0.002 | -0.019 | -0.019 | 0 | 0.53 | 0.49 | 0.43 | 0.26 | 0.45 | 0.54 |
| 8 | 27307072 | rs1019832 | 0.929 | 1.498 | -0.251 | -0.452 | -0.041 | -0.005 | -0.005 | 0 | 0.24 | 0.05 | 0.28 | 0.24 | 0.27 | 0.24 |
| 8 | 27307095 | rs6557999 | 0.039 | 1.501 | -0.215 | -0.472 | 0 | -0.004 | -0.004 | 0 | 0.2 | 0.2 | 0.13 | 0.11 | 0.19 | 0.17 |
| 8 | 27308538 | rs751018 | 1.199 | 1.499 | -0.136 | -0.433 | -0.006 | -0.033 | -0.033 | 0 | 0.46 | 0.48 | 0.39 | 0.2 | 0.39 | 0.43 |
| 8 | 27308585 | rs751019 | -0.662 | 1.568 | -0.601 | -0.391 | -0.012 | 0.056 | 0.056 | 0 | 0.45 | 0.24 | 0.41 | 0.36 | 0.46 | 0.41 |
| 8 | 27308849 | rs2251430 | -1.578 | 1.567 | -0.586 | -0.388 | 0.037 | 0.192 | 0.192 | 0 | 0.93 | 0.89 | 0.82 | 0.55 | 0.85 | 0.84 |
| 8 | 27309339 | rs1106359 | 0.996 | 1.561 | -0.587 | -0.397 | 0.023 | 0.06 | 0.06 | 0 | 0.49 | 0.64 | 0.41 | 0.2 | 0.39 | 0.43 |
| 8 | 27309379 | rs891393 | -1.482 | 1.544 | -0.571 | -0.372 | 0.044 | 0.214 | 0.214 | 0 | 0.93 | 0.88 | 0.82 | 0.56 | 0.85 | 0.84 |
| 8 | 27311703 | rs1879184 | -1.796 | 1.544 | -0.571 | -0.372 | 0.04 | 0.204 | 0.204 | 0 | 0.94 | 0.9 | 0.82 | 0.56 | 0.85 | 0.84 |
| 8 | 27312040 | rs10093964 | 0.953 | 1.563 | -0.588 | -0.398 | 0.021 | 0.063 | 0.063 | 0 | 0.49 | 0.64 | 0.41 | 0.2 | 0.4 | 0.43 |
| 8 | 27315876 | rs1879182 | -1.577 | 1.564 | -0.587 | -0.396 | 0.037 | 0.194 | 0.194 | 0 | 0.93 | 0.89 | 0.82 | 0.56 | 0.85 | 0.84 |
| 8 | 27315900 | rs1879181 | -1.492 | 1.564 | -0.588 | -0.396 | 0.044 | 0.214 | 0.214 | 0 | 0.93 | 0.88 | 0.82 | 0.56 | 0.85 | 0.84 |
| 8 | 27316117 | rs2271920 | 0.962 | 1.543 | -0.601 | -0.408 | 0.021 | 0.062 | 0.062 | 0 | 0.49 | 0.64 | 0.41 | 0.19 | 0.4 | 0.43 |
| 8 | 27316980 | rs748282 | -1.521 | 1.758 | -0.619 | -0.237 | 0.035 | 0.191 | 0.191 | 0 | 0.93 | 0.9 | 0.82 | 0.56 | 0.85 | 0.84 |
| 8 | 27317337 | rs2280375 | -1.711 | 1.764 | -0.616 | -0.022 | 0.04 | 0.198 | 0.198 | 0 | 0.94 | 0.9 | 0.82 | 0.6 | 0.85 | 0.84 |
| 8 | 27318391 | rs2292974 | -0.522 | 1.276 | -0.831 | -0.49 | 0.005 | 0.084 | 0.084 | 0 | 0.48 | 0.21 | 0.42 | 0.36 | 0.47 | 0.42 |
| 8 | 27319292 | rs735421 | 0.779 | 1.351 | -0.853 | -0.42 | -0.022 | 0.031 | 0.031 | 0 | 0.29 | 0.1 | 0.31 | 0.26 | 0.33 | 0.24 |
| 8 | 27319805 | rs9314347 | 1.246 | 1.52 | -0.714 | -0.285 | -0.002 | 0.002 | 0.002 | 0 | 0.08 | 0.06 | 0.04 | 0.06 | 0.08 | 0.12 |
| 8 | 27323860 | rs2163177 | -1.253 | 1.414 | -0.303 | 0.086 | 0.116 | 0.39 | 0.39 | 0 | 0.93 | 0.76 | 0.8 | 0.57 | 0.85 | 0.85 |
| 8 | 27324822 | rs891398 | -0.474 | 1.053 | -1.033 | -0.31 | -0.049 | 0.038 | 0.038 | 0 | 0.43 | 0.83 | 0.56 | 0.65 | 0.51 | 0.41 |
| 8 | 27325473 | rs747111 | 0.852 | 1.014 | -1.104 | -0.35 | -0.078 | 0.048 | 0.048 | 0 | 0.29 | 0.05 | 0.31 | 0.2 | 0.34 | 0.26 |
| 8 | 27326127 | rs2741343 | -0.234 | 1.25 | -0.906 | -0.227 | -0.038 | 0.043 | 0.043 | 0 | 0.43 | 0.81 | 0.55 | 0.64 | 0.51 | 0.41 |
| 8 | 27328021 | rs2565065 | -0.823 | 1.14 | -1.058 | -0.001 | -0.03 | 0.073 | 0.073 | 0.816 | 0.66 | 0.98 | 0.71 | 0.79 | 0.72 | 0.63 |
| 8 | 27328511 | rs2472553 | 0.337 | 1.422 | 0.239 | 0.804 | 0.091 | 0.352 | 0.352 | 0 | 0.07 | 0.19 | 0.19 | 0.43 | 0.14 | 0.15 |
| 8 | 27330096 | rs2741342 | -0.395 | 1.573 | 0.308 | 0.762 | 0.173 | 0.334 | 0.334 | 0.538 | 0.11 | 0.35 | 0.25 | 0.41 | 0.23 | 0.19 |
| 8 | 27330937 | rs7819756 | -0.185 | -0.12 | -0.191 | -0.172 | -0.005 | 0.003 | 0.003 | 0 | 0.39 | 0.37 | 0.44 | 0.61 | 0.48 | 0.44 |
| 8 | 27331119 | rs2565067 | -0.713 | -0.612 | -0.364 | -0.204 | 0.029 | 0.018 | 0.018 | 0 | 0.74 | 0.84 | 0.76 | 0.77 | 0.85 | 0.76 |
| 8 | 27334969 | rs2741339 | -0.504 | -0.59 | -0.387 | -0.286 | 0.032 | 0.061 | 0.061 | 2.879 | 0.35 | 0.77 | 0.27 | 0.24 | 0.29 | 0.23 |
| 8 | 27336978 | rs2565059 | 0.833 | -0.665 | 0.045 | -0.353 | 0.013 | -0.006 | -0.006 | 0 | 0.14 | 0.21 | 0.18 | 0.09 | 0.22 | 0.09 |
| 8 | 27337380 | rs2565055 | -0.045 | -0.807 | 0.389 | -0.413 | 0.003 | -0.011 | -0.011 | 1.228 | 0.23 | 0.31 | 0.22 | 0.23 | 0.27 | 0.22 |
| 8 | 27337507 | rs7823854 | -0.116 | -0.897 | 0.772 | -0.441 | -0.016 | -0.019 | -0.019 | 0.872 | 0.07 | 0.23 | 0.01 | 0 | 0 | 0.01 |
| 8 | 27337791 | rs2741335 | -1.172 | -0.816 | 0.56 | -0.467 | 0.02 | 0.047 | 0.047 | 3.087 | 0.34 | 0.76 | 0.26 | 0.24 | 0.29 | 0.23 |
| 8 | 27344360 | rs12334538 | -2.016 | -1.184 | 0.824 | -0.55 | -0.064 | -0.075 | -0.075 | 4.635 | 0.13 | 0.66 | 0.05 | 0 | 0.02 | 0.02 |
| 8 | 27345305 | rs7017417 | 1.501 | -1.423 | 0.908 | -0.636 | -0.004 | -0.038 | -0.038 | 0 | 0.13 | 0.17 | 0.09 | 0 | 0.09 | 0.07 |
| 8 | 27347194 | rs7003694 | 1.738 | -1.435 | 0.92 | -0.624 | -0.008 | -0.04 | -0.04 | 0 | 0.12 | 0.17 | 0.08 | 0 | 0.09 | 0.07 |
| 8 | 27361241 | rs17057255 | 0.535 | -1.485 | 1.454 | -0.637 | -0.022 | -0.022 | -0.022 | 0.059 | 0.06 | 0.1 | 0.02 | 0 | 0 | 0.01 |
| 8 | 27362142 | rs7846038 | 1.558 | -1.501 | 1.447 | -0.656 | -0.014 | -0.056 | -0.056 | 0 | 0.13 | 0.21 | 0.08 | 0 | 0.09 | 0.07 |
| 8 | 27363136 | rs4149239 | 0.085 | -1.568 | 1.444 | -0.657 | -0.033 | -0.067 | -0.067 | 0 | 0.23 | 0.54 | 0.14 | 0.16 | 0.15 | 0.22 |
| 8 | 27364787 | rs2741334 | -2.096 | -1.837 | 0.958 | -0.506 | 0.01 | -0.048 | -0.048 | 0.564 | 0.87 | 0.99 | 0.81 | 0.82 | 0.92 | 0.89 |
| 8 | 27365430 | rs7839733 | -0.571 | -1.579 | 1.292 | -0.5 | 0.05 | 0.065 | 0.065 | 0.954 | 0.3 | 0.77 | 0.17 | 0.16 | 0.15 | 0.22 |
| 8 | 27366152 | rs17057284 | 1.96 | -1.539 | 1.28 | -0.482 | 0.002 | -0.009 | -0.009 | 1.246 | 0.11 | 0.09 | 0.08 | 0 | 0.09 | 0.07 |
| 8 | 27371546 | rs7000388 | -0.463 | -1.557 | 1.275 | -0.488 | -0.001 | -0.011 | -0.011 | 0 | 0.29 | 0.66 | 0.16 | 0.16 | 0.15 | 0.22 |
| 8 | 27372189 | rs7018249 | -0.463 | -1.557 | 1.275 | -0.488 | -0.001 | -0.011 | -0.011 | 0 | 0.29 | 0.66 | 0.16 | 0.16 | 0.15 | 0.22 |
| 8 | 27373923 | rs4149243 | -0.387 | -1.564 | 1.263 | -0.451 | -0.006 | -0.018 | -0.018 | 0.469 | 0.29 | 0.66 | 0.16 | 0.16 | 0.15 | 0.21 |
| 8 | 27377162 | rs4149244 | 1.102 | -1.537 | 1.201 | -0.411 | -0.004 | -0.002 | -0.002 | 0 | 0.12 | 0.11 | 0.04 | 0.15 | 0.06 | 0.14 |
| 8 | 27381996 | rs721619 | 1.37 | -1.145 | 1.05 | -0.119 | -0.057 | 0.036 | 0.036 | 0.571 | 0.29 | 0.09 | 0.36 | 0.22 | 0.32 | 0.32 |
| 8 | 27383901 | rs4149246 | 0.719 | -1.61 | 2.207 | 0.42 | -0.005 | 0.001 | 0.001 | 0 | 0.11 | 0.11 | 0.06 | 0.18 | 0.06 | 0.14 |
| 8 | 27385415 | rs4149247 | -1.004 | -1.618 | 2.174 | 0.412 | 0.03 | 0.033 | 0.033 | 1.192 | 0.29 | 0.76 | 0.19 | 0.19 | 0.15 | 0.21 |
| 8 | 27385450 | rs4149248 | -0.572 | -1.618 | 2.336 | 0.412 | -0.052 | -0.063 | -0.063 | 0.002 | 0.26 | 0.44 | 0.16 | 0.19 | 0.15 | 0.21 |
| 8 | 27388199 | rs10503812 | -0.638 | -1.575 | 2.237 | 0.408 | -0.049 | -0.06 | -0.06 | 0.004 | 0.27 | 0.54 | 0.17 | 0.19 | 0.15 | 0.21 |
| 8 | 27389457 | rs7843805 | -1.048 | -1.575 | 2.089 | 0.38 | 0.033 | 0.037 | 0.037 | 1.513 | 0.3 | 0.76 | 0.19 | 0.19 | 0.15 | 0.21 |
| 8 | 27389542 | rs6558002 | -1.018 | -1.554 | 2.096 | 0.396 | 0.034 | 0.036 | 0.036 | 1.585 | 0.3 | 0.76 | 0.19 | 0.2 | 0.15 | 0.21 |
| 8 | 27390192 | rs6558003 | -1.205 | -1.536 | 2.114 | 0.386 | 0.053 | 0.057 | 0.057 | 5.055 | 0.32 | 0.78 | 0.19 | 0.19 | 0.15 | 0.22 |
| 8 | 27391620 | rs747276 | -1.205 | -1.535 | 2.097 | 0.386 | 0.045 | 0.048 | 0.048 | 5.055 | 0.32 | 0.76 | 0.19 | 0.2 | 0.15 | 0.22 |

| | | | | | | | | | | | | | | | | |
|---|---|---|---|---|---|---|---|---|---|---|---|---|---|---|---|---|
| 8 | 27393066 | rs7828349 | -0.653 | -1.532 | 1.996 | 0.388 | -0.054 | -0.062 | -0.062 | 0 | 0.27 | 0.46 | 0.16 | 0.19 | 0.15 | 0.22 |
| 8 | 27393220 | rs6558004 | -1.211 | -1.537 | 1.966 | 0.385 | 0.036 | 0.039 | 0.039 | 5.055 | 0.32 | 0.75 | 0.19 | 0.19 | 0.15 | 0.22 |
| 8 | 27394366 | rs13439459 | 1.371 | -1.543 | 1.897 | 0.38 | 0.005 | -0.003 | -0.003 | 0 | 0.12 | 0.08 | 0.08 | 0 | 0.08 | 0.06 |
| 8 | 27394957 | rs10282967 | -0.633 | -1.524 | 1.83 | 0.393 | -0.061 | -0.071 | -0.071 | 0 | 0.27 | 0.48 | 0.17 | 0.2 | 0.15 | 0.22 |
| 8 | 27395249 | rs10283378 | -0.648 | -1.523 | 1.831 | 0.393 | -0.061 | -0.071 | -0.071 | 0.582 | 0.27 | 0.48 | 0.17 | 0.19 | 0.15 | 0.22 |
| 8 | 27396111 | rs4149252 | -0.648 | -1.523 | 1.831 | 0.393 | -0.061 | -0.071 | -0.071 | 1.03 | 0.27 | 0.48 | 0.17 | 0.19 | 0.15 | 0.22 |
| 8 | 27396523 | rs729609 | -1.157 | -1.53 | 1.844 | 0.39 | 0.049 | 0.053 | 0.053 | 5.055 | 0.32 | 0.77 | 0.19 | 0.19 | 0.15 | 0.22 |
| 8 | 27397157 | rs4149255 | -1.098 | -1.506 | 1.39 | 0.392 | 0.048 | 0.052 | 0.052 | 4.751 | 0.31 | 0.77 | 0.19 | 0.19 | 0.15 | 0.22 |
| 8 | 27398432 | rs2071575 | -1.425 | -1.392 | 1.138 | 0.579 | 0.03 | -0.017 | -0.017 | 0 | 0.37 | 0.87 | 0.27 | 0.4 | 0.26 | 0.34 |
| 8 | 27399704 | rs4149257 | -1.445 | -1.375 | 1.245 | 0.686 | 0.03 | -0.027 | -0.027 | 0 | 0.35 | 0.87 | 0.27 | 0.4 | 0.24 | 0.33 |
| 8 | 27400604 | rs13269963 | -0.136 | -1.307 | 1.541 | 0.688 | 0.002 | 0.01 | 0.01 | 0 | 0.19 | 0.28 | 0.17 | 0.22 | 0.18 | 0.19 |
| 8 | 27401142 | rs2291635 | 0.759 | -1.248 | 1.33 | 0.716 | 0.029 | 0.09 | 0.09 | 0 | 0.07 | 0.19 | 0.08 | 0.2 | 0.09 | 0.11 |
| 8 | 27401964 | rs1126452 | -1.086 | -1.297 | 1.405 | 0.69 | 0.013 | -0.017 | -0.017 | 0 | 0.32 | 0.77 | 0.26 | 0.41 | 0.24 | 0.32 |
| 8 | 27402132 | rs1042064 | -0.992 | -1.299 | 1.49 | 0.689 | 0.005 | -0.008 | -0.008 | 0 | 0.31 | 0.72 | 0.26 | 0.41 | 0.24 | 0.32 |
| 8 | 27402494 | rs4149259 | -0.484 | -1.298 | 1.555 | 0.644 | -0.048 | -0.047 | -0.047 | 0 | 0.24 | 0.52 | 0.17 | 0.2 | 0.15 | 0.21 |
| 8 | 27402664 | rs4149260 | -0.261 | -1.296 | 1.663 | 0.647 | -0.025 | -0.023 | -0.023 | 0.84 | 0.23 | 0.32 | 0.16 | 0.2 | 0.15 | 0.21 |
| 8 | 27402777 | rs7341557 | 0.685 | -1.199 | 1.515 | 0.625 | 0.062 | 0.147 | 0.147 | 0 | 0.07 | 0.25 | 0.09 | 0.2 | 0.09 | 0.11 |
| 8 | 27403730 | rs6990074 | 0.23 | -1.69 | 0.387 | 0.011 | -0.012 | 0.006 | 0.006 | 2.534 | 0.07 | 0.16 | 0.04 | 0.08 | 0.05 | 0.01 |
| 8 | 27403777 | rs10096092 | 0.683 | -1.304 | 0.074 | 0.077 | -0.008 | -0.041 | -0.041 | 0.033 | 0.13 | 0.17 | 0.09 | 0 | 0.09 | 0.06 |
| 8 | 27404137 | rs6990733 | -0.695 | -1.423 | -0.489 | 0.13 | -0.035 | -0.032 | -0.032 | 0 | 0.24 | 0.58 | 0.17 | 0.2 | 0.14 | 0.21 |
| 8 | 27404354 | rs2565050 | 0.214 | -1.099 | -1.338 | 0.616 | 0.027 | -0.017 | -0.017 | 0 | 0.77 | 0.89 | 0.88 | 0.68 | 0.86 | 0.74 |
| 8 | 27405207 | rs34319290 | -0.426 | -1.347 | -1.48 | 0.28 | 0.042 | 0.002 | 0.002 | 0 | 0.15 | 0.01 | 0.05 | 0.1 | 0.06 | 0.05 |
| 8 | 27405809 | rs2640726 | -0.125 | -1.547 | -1.18 | 0.008 | -0.002 | -0.002 | -0.002 | 0 | 0.55 | 0.47 | 0.67 | 0.35 | 0.59 | 0.55 |
| 8 | 27417024 | rs2741353 | -0.545 | -2.311 | -0.608 | 0.38 | -0.004 | 0.029 | 0.029 | 0 | 0.81 | 0.81 | 0.83 | 0.53 | 0.81 | 0.84 |
| 8 | 27417422 | rs1873933 | 1.294 | -2.581 | -0.595 | -0.364 | -0.049 | 0.039 | 0.039 | 0 | 0.3 | 0.22 | 0.57 | 0.16 | 0.46 | 0.38 |
| 8 | 27418325 | rs2640723 | -0.266 | -2.396 | -0.378 | 0.139 | -0.005 | 0.01 | 0.01 | 0 | 0.78 | 0.77 | 0.81 | 0.4 | 0.79 | 0.73 |
| 8 | 27418747 | rs11780592 | 1.547 | -2.168 | -0.609 | -0.045 | 0.005 | -0.002 | -0.002 | 0.446 | 0.17 | 0.13 | 0.11 | 0 | 0.17 | 0.1 |
| 8 | 27420782 | rs2741348 | 0.401 | -1.642 | -0.443 | 0.01 | 0.041 | 0.11 | 0.11 | 0 | 0.49 | 0.3 | 0.61 | 0.25 | 0.53 | 0.44 |
| 8 | 27422250 | rs7824574 | -0.503 | -1.214 | 0.019 | 0.581 | 0.009 | 0.045 | 0.045 | 0 | 0.1 | 0.21 | 0.09 | 0.19 | 0.09 | 0.18 |
| 8 | 27423338 | rs4732725 | 0.44 | -1.337 | 0.036 | 0.761 | -0.008 | 0.031 | 0.031 | 0.11 | 0.08 | 0.13 | 0.04 | 0.15 | 0.08 | 0.14 |
| 8 | 27425117 | rs9632863 | -0.176 | -1.996 | 0.248 | 0.92 | -0.017 | 0.003 | 0.003 | 0 | 0.42 | 0.57 | 0.32 | 0.45 | 0.31 | 0.44 |
| 8 | 27429228 | rs1316801 | -1.056 | -1.97 | 0.119 | 0.814 | 0 | 0.014 | 0.014 | 0 | 0.57 | 0.81 | 0.39 | 0.62 | 0.43 | 0.59 |
| 8 | 27437388 | rs17057419 | -0.179 | -2.311 | 0.479 | 0.768 | -0.024 | 0.032 | 0.032 | 0 | 0.26 | 0.36 | 0.21 | 0.44 | 0.15 | 0.36 |
| 8 | 27440879 | rs10216623 | 0.832 | -2.566 | 0.517 | 0.858 | 0.006 | -0.016 | -0.016 | 0.614 | 0.14 | 0.11 | 0.14 | 0.23 | 0.1 | 0.14 |
| 8 | 27441010 | rs7828131 | -1.143 | -2.597 | 0.644 | 0.941 | 0.061 | 0.118 | 0.118 | 0 | 0.51 | 0.71 | 0.36 | 0.86 | 0.39 | 0.61 |
| 8 | 27441800 | rs12114305 | 0.325 | -2.509 | 0.28 | 0.794 | -0.027 | 0.168 | 0.168 | 0 | 0.06 | 0.19 | 0.05 | 0.23 | 0.01 | 0.13 |
| 8 | 27442064 | rs7844965 | 0.598 | -2.47 | 0.244 | 0.688 | 0.005 | 0.009 | 0.009 | 0 | 0.24 | 0.18 | 0.14 | 0.15 | 0.23 | 0.22 |
| 8 | 27447024 | rs12680584 | 0.855 | -1.564 | 0.254 | 0.914 | 0 | 0.043 | 0.043 | 0 | 0.28 | 0.12 | 0.23 | 0.2 | 0.26 | 0.24 |
| 8 | 27448729 | rs7012010 | 0.189 | -1.403 | 0.375 | 0.781 | 0.012 | -0.008 | -0.008 | 0.071 | 0.36 | 0.3 | 0.27 | 0.39 | 0.28 | 0.38 |
| 8 | 27450073 | rs17057438 | -0.097 | -1.366 | 0.245 | 0.773 | -0.019 | 0.001 | 0.001 | 0 | 0.42 | 0.54 | 0.32 | 0.4 | 0.31 | 0.45 |
| 8 | 27455442 | rs3087554 | -0.644 | 0.213 | -0.33 | 0.636 | -0.002 | 0.004 | 0.004 | 0 | 0.18 | 0.2 | 0.29 | 0.31 | 0.17 | 0.28 |
| 8 | 27464519 | rs11136000 | 0.058 | 0.513 | -0.656 | -0.07 | -0.001 | -0.012 | -0.012 | 0 | 0.61 | 0.43 | 0.64 | 0.78 | 0.61 | 0.72 |
| 8 | 27466315 | rs1532278 | -0.138 | 0.718 | -1.652 | 0.04 | -0.02 | 0.065 | 0.065 | 0.104 | 0.66 | 0.84 | 0.67 | 0.78 | 0.61 | 0.72 |
| 8 | 27474202 | rs9314349 | -0.535 | 0.978 | -1.95 | -0.065 | -0.052 | 0.063 | 0.063 | 0 | 0.3 | 0.12 | 0.31 | 0.14 | 0.38 | 0.28 |
| 8 | 27476815 | rs538181 | 0.27 | 0.862 | -2.385 | 0.439 | -0.09 | -0.057 | -0.057 | 0 | 0.56 | 0.89 | 0.53 | 0.47 | 0.55 | 0.48 |
| 8 | 27479227 | rs492638 | -0.925 | 0.949 | -2.384 | 0.574 | -0.098 | -0.16 | -0.16 | 0 | 0.74 | 0.97 | 0.63 | 0.55 | 0.69 | 0.67 |
| 8 | 27483267 | rs1493521 | 0.445 | 0.805 | -1.171 | 0.578 | 0.022 | 0.243 | 0.243 | 0 | 0.08 | 0.2 | 0.14 | 0.38 | 0.06 | 0.2 |
| 8 | 27489421 | rs520769 | 0.121 | 0.84 | -1.098 | 0.456 | 0.011 | 0.144 | 0.144 | 0 | 0.86 | 0.45 | 0.82 | 0.62 | 0.91 | 0.79 |
| 8 | 27494612 | rs894019 | -0.078 | 0.801 | -1.828 | 0.251 | 0.064 | -0.038 | -0.038 | 0 | 0.46 | 0.23 | 0.41 | 0.46 | 0.37 | 0.47 |
| 8 | 27497790 | rs500048 | 0.625 | 0.901 | -2.001 | 0.495 | -0.025 | 0.048 | 0.048 | 0 | 0.34 | 0.03 | 0.33 | 0.19 | 0.33 | 0.33 |
| 8 | 27500427 | rs12542107 | 1.124 | 0.99 | -2.106 | 0.503 | -0.141 | 0.144 | 0.144 | 0 | 0.38 | 0.02 | 0.33 | 0.16 | 0.45 | 0.28 |
| 8 | 27503608 | rs4732736 | 0.038 | 0.678 | -2.101 | 0.917 | -0.015 | -0.055 | -0.055 | 0 | 0.13 | 0.05 | 0.14 | 0.29 | 0.12 | 0.18 |
| 8 | 27505756 | rs2582369 | 0.705 | 0.638 | -2.17 | 0.829 | 0.034 | 0.056 | 0.056 | 0.051 | 0.31 | 0.02 | 0.26 | 0.17 | 0.25 | 0.28 |
| 8 | 27508543 | rs2582370 | -0.212 | 0.759 | -2.028 | 0.829 | 0.048 | -0.065 | -0.065 | 0.014 | 0.44 | 0.12 | 0.4 | 0.48 | 0.37 | 0.49 |
| 8 | 27520266 | rs516125 | -0.302 | 0.853 | -2.035 | 0.869 | -0.011 | -0.053 | -0.053 | 0 | 0.46 | 0.11 | 0.48 | 0.49 | 0.45 | 0.55 |
| 8 | 27528446 | rs17057523 | 0.272 | 0.724 | -1.66 | 1.147 | -0.009 | 0.019 | 0.019 | 0 | 0.06 | 0.06 | 0.11 | 0.23 | 0.08 | 0.15 |
| 8 | 27529639 | rs1036708 | 0.405 | 0.729 | -1.635 | 1.14 | -0.006 | 0.021 | 0.021 | 0 | 0.06 | 0.06 | 0.11 | 0.21 | 0.08 | 0.12 |
| 8 | 27530475 | rs1036710 | 0.298 | 0.731 | -1.569 | 1.196 | -0.004 | 0.035 | 0.035 | 0 | 0.06 | 0.06 | 0.12 | 0.25 | 0.08 | 0.12 |
| 8 | 27532654 | rs2640734 | 0.501 | 0.404 | -1.836 | 0.898 | 0.064 | 0.096 | 0.096 | 0.023 | 0.32 | 0.05 | 0.28 | 0.13 | 0.26 | 0.31 |
| 8 | 27534727 | rs735277 | 0.91 | 0.462 | -2.477 | 0.543 | -0.166 | -0.005 | -0.005 | 3.939 | 0.24 | 0.03 | 0.3 | 0.22 | 0.43 | 0.24 |
| 8 | 27539370 | rs559251 | -0.025 | 0.224 | -1.112 | 0.557 | -0.006 | -0.012 | -0.012 | 0 | 0.59 | 0.43 | 0.6 | 0.35 | 0.69 | 0.55 |
| 8 | 27542849 | rs512941 | -0.571 | 0.137 | -1.058 | 0.582 | -0.013 | 0.008 | 0.008 | 0 | 0.67 | 0.57 | 0.72 | 0.59 | 0.76 | 0.68 |
| 8 | 27546148 | rs11997515 | 0.583 | 0.298 | -1.077 | 0.793 | -0.004 | 0.029 | 0.029 | 0 | 0.06 | 0.06 | 0.09 | 0.21 | 0.07 | 0.14 |
| 8 | 27549421 | rs3103908 | -0.118 | -0.278 | -0.61 | 0.248 | 0.013 | 0.03 | 0.03 | 0 | 0.64 | 0.49 | 0.57 | 0.54 | 0.48 | 0.58 |
| 8 | 27558358 | rs7823813 | 0.019 | -0.312 | -0.559 | 0.207 | -0.013 | -0.048 | -0.048 | 0 | 0.36 | 0.17 | 0.43 | 0.44 | 0.54 | 0.4 |
| 8 | 27560630 | rs13253543 | 0.496 | 0.069 | -0.691 | 0.079 | -0.086 | -0.077 | -0.077 | 0 | 0.28 | 0.11 | 0.39 | 0.44 | 0.45 | 0.34 |
| 8 | 27560651 | rs17384485 | 0.12 | -0.214 | -0.64 | -0.094 | 0.002 | 0 | 0 | 0.205 | 0.38 | 0.35 | 0.31 | 0.43 | 0.31 | 0.34 |
| 8 | 27562211 | rs566398 | 0.952 | -0.259 | -1.012 | -0.358 | 0.048 | 0.098 | 0.098 | 0.33 | 0.3 | 0.05 | 0.27 | 0.11 | 0.25 | 0.32 |
| 8 | 27564087 | rs17469423 | 1.28 | 0.133 | -0.823 | -0.49 | -0.035 | 0.043 | 0.043 | 0 | 0.08 | 0.01 | 0.03 | 0 | 0.09 | 0.06 |

| | | | | | | | | | | | | | | | | |
|---|---|---|---|---|---|---|---|---|---|---|---|---|---|---|---|---|
| 8 | 27564846 | rs500458 | 0.913 | -0.233 | -0.794 | -0.483 | 0.002 | -0.034 | -0.034 | 0 | 0.25 | 0.24 | 0.25 | 0.01 | 0.21 | 0.17 |
| 8 | 27567885 | rs11780158 | 0.275 | 0.917 | -1.599 | 0.503 | -0.149 | -0.098 | -0.098 | 0.263 | 0.23 | 0.04 | 0.36 | 0.32 | 0.41 | 0.28 |
| 8 | 27577392 | rs2014617 | 0.203 | -0.459 | -0.957 | -0.249 | -0.024 | -0.041 | -0.041 | 0.058 | 0.66 | 0.44 | 0.84 | 0.85 | 0.77 | 0.68 |
| 8 | 27580039 | rs10216990 | -0.096 | -0.337 | -0.949 | -0.394 | -0.002 | -0.019 | -0.019 | 0 | 0.41 | 0.57 | 0.56 | 0.75 | 0.51 | 0.56 |
| 8 | 27581042 | rs11984514 | 0.544 | -0.282 | -0.644 | -0.209 | 0.008 | -0.001 | -0.001 | 0 | 0.08 | 0.18 | 0.04 | 0.1 | 0.09 | 0.11 |
| 8 | 27590027 | rs2010776 | 0.299 | -0.79 | -0.72 | -0.718 | 0.021 | 0.156 | 0.156 | 0.158 | 0.45 | 0.25 | 0.37 | 0.12 | 0.44 | 0.39 |
| 8 | 27591164 | rs7109 | 0.687 | -0.894 | -0.314 | -0.567 | 0.058 | 0.006 | 0.006 | 0 | 0.32 | 0.42 | 0.52 | 0.63 | 0.42 | 0.45 |
| 8 | 27591506 | rs2010525 | 0.687 | -0.894 | -0.314 | -0.567 | 0.058 | 0.006 | 0.006 | 0 | 0.32 | 0.42 | 0.52 | 0.63 | 0.42 | 0.45 |
| 8 | 27591849 | rs9797 | -0.069 | -0.936 | -0.205 | -0.605 | 0.02 | 0.1 | 0.1 | 0 | 0.49 | 0.36 | 0.39 | 0.12 | 0.47 | 0.39 |
| 8 | 27593150 | rs1048887 | 0.734 | -0.927 | -0.143 | -0.6 | 0.059 | 0.007 | 0.007 | 0 | 0.32 | 0.42 | 0.52 | 0.63 | 0.42 | 0.45 |
| 8 | 27595880 | rs11784867 | 0.734 | -0.927 | -0.143 | -0.6 | 0.059 | 0.007 | 0.007 | 0 | 0.32 | 0.42 | 0.52 | 0.63 | 0.42 | 0.45 |
| 8 | 27599514 | rs17477221 | 1.309 | -1.192 | -0.38 | -0.92 | 0.01 | 0.205 | 0.205 | 1.589 | 0.3 | 0.01 | 0.28 | 0.01 | 0.29 | 0.26 |
| 8 | 27600450 | rs6987111 | -1.4 | -1.278 | -0.292 | -0.84 | 0.002 | -0.003 | -0.003 | 1.031 | 0.82 | 0.83 | 0.91 | 0.77 | 0.89 | 0.84 |
| 8 | 27603200 | rs17477326 | 0.745 | -1.188 | -0.352 | -0.848 | -0.004 | 0.163 | 0.163 | 0.593 | 0.37 | 0.1 | 0.33 | 0.12 | 0.39 | 0.32 |
| 8 | 27607934 | rs10216403 | -0.655 | -1.135 | -0.219 | -0.745 | -0.037 | 0.008 | 0.008 | 0 | 0.77 | 0.63 | 0.88 | 0.74 | 0.86 | 0.84 |
| 8 | 27614138 | rs2280884 | 0.829 | -1.043 | -0.264 | -0.636 | 0.055 | 0.015 | 0.015 | 0 | 0.32 | 0.43 | 0.51 | 0.62 | 0.42 | 0.45 |
| 8 | 27623209 | rs10102309 | -0.003 | -1.135 | -0.158 | -0.781 | 0.017 | 0.075 | 0.075 | 0.176 | 0.51 | 0.62 | 0.62 | 0.95 | 0.53 | 0.65 |
| 8 | 27625201 | rs13262948 | 0.941 | -1.366 | -0.346 | -0.98 | 0.002 | 0.135 | 0.135 | 0.009 | 0.33 | 0.1 | 0.31 | 0.02 | 0.35 | 0.21 |
| 8 | 27628886 | rs10093956 | 0.221 | -1.351 | -0.316 | -0.98 | 0.015 | 0.048 | 0.048 | 0.106 | 0.5 | 0.58 | 0.61 | 0.95 | 0.53 | 0.65 |
| 8 | 27630152 | rs7842666 | 0.238 | -1.352 | -0.316 | -0.942 | 0.015 | 0.065 | 0.065 | 0.098 | 0.5 | 0.58 | 0.61 | 0.86 | 0.53 | 0.61 |
| 8 | 27634064 | rs4732748 | 1.43 | -1.372 | -0.368 | -0.913 | 0.02 | -0.01 | -0.01 | 0 | 0.17 | 0.1 | 0.09 | 0.24 | 0.11 | 0.16 |
| 8 | 27640383 | rs12677603 | 1.433 | -1.371 | -0.373 | -0.912 | 0.02 | -0.01 | -0.01 | 0 | 0.17 | 0.1 | 0.09 | 0.25 | 0.11 | 0.16 |
| 8 | 27644513 | rs4732750 | 0.898 | -1.35 | -0.399 | -0.964 | 0.007 | 0.105 | 0.105 | 0 | 0.34 | 0.14 | 0.31 | 0.02 | 0.35 | 0.2 |
| 8 | 27645704 | rs6558014 | -0.361 | -1.304 | -0.322 | -0.84 | -0.002 | -0.042 | -0.042 | 0 | 0.85 | 0.82 | 0.94 | 1 | 0.89 | 0.86 |
| 8 | 27658491 | rs12675119 | 0.841 | -1.305 | -0.386 | -0.744 | -0.022 | 0.015 | 0.015 | 0 | 0.2 | 0.28 | 0.11 | 0.36 | 0.11 | 0.2 |
| 8 | 27664686 | rs4732756 | 0.971 | -1.305 | -0.385 | -0.742 | -0.01 | 0.003 | 0.003 | 0 | 0.19 | 0.22 | 0.1 | 0.33 | 0.11 | 0.2 |
| 8 | 27667793 | rs1052873 | 1.851 | -1.659 | -0.743 | -1.049 | 0.004 | 0.127 | 0.127 | 0.086 | 0.22 | 0.01 | 0.23 | 0.01 | 0.22 | 0.13 |
| 8 | 27668726 | rs17479454 | 0.758 | -1.493 | -0.335 | -0.995 | 0.003 | 0.108 | 0.108 | 0.009 | 0.33 | 0.11 | 0.31 | 0.02 | 0.35 | 0.2 |
| 8 | 27676378 | rs6991862 | 1.08 | -1.533 | -0.435 | -0.991 | -0.022 | 0.015 | 0.015 | 0 | 0.2 | 0.28 | 0.11 | 0.36 | 0.11 | 0.2 |
| 8 | 27680576 | rs727813 | -0.298 | -1.519 | -0.398 | -0.995 | 0.016 | 0.021 | 0.021 | 0 | 0.53 | 0.43 | 0.42 | 0.38 | 0.46 | 0.4 |
| 8 | 27681725 | rs10087558 | 0.927 | -1.6 | -0.297 | -0.775 | -0.02 | 0.013 | 0.013 | 0 | 0.2 | 0.28 | 0.11 | 0.36 | 0.11 | 0.2 |
| 8 | 27689305 | rs12682017 | 0.927 | -1.6 | -0.3 | -0.775 | -0.022 | 0.015 | 0.015 | 0 | 0.2 | 0.28 | 0.11 | 0.36 | 0.11 | 0.2 |
| 8 | 27702453 | rs11136015 | 0.459 | -1.564 | -0.165 | -0.833 | -0.005 | -0.013 | -0.013 | 0.066 | 0.43 | 0.53 | 0.67 | 0.65 | 0.54 | 0.5 |
| 8 | 27711673 | rs4732776 | -0.266 | -1.476 | -0.045 | -1.011 | 0.017 | 0.015 | 0.015 | 2.726 | 0.64 | 0.71 | 0.77 | 0.98 | 0.67 | 0.69 |
| 8 | 27715116 | rs1870587 | -0.716 | -1.392 | 0.057 | -0.947 | -0.004 | -0.043 | -0.043 | 0 | 0.89 | 0.83 | 0.95 | 1 | 0.91 | 0.86 |
| 8 | 27719703 | rs10866866 | -0.389 | -1.454 | 0.719 | -0.908 | -0.001 | -0.081 | -0.081 | 0.427 | 0.87 | 0.66 | 0.93 | 1 | 0.91 | 0.86 |
| 8 | 27720824 | rs898460 | -0.141 | -1.592 | 0.565 | -1.02 | -0.012 | -0.106 | -0.106 | 0 | 0.63 | 0.55 | 0.75 | 0.98 | 0.66 | 0.69 |
| 8 | 27728151 | rs4383927 | -0.446 | -1.47 | 0.489 | -0.863 | -0.012 | -0.045 | -0.045 | 0 | 0.91 | 0.78 | 0.96 | 1 | 0.94 | 0.9 |
| 8 | 27731779 | rs6558023 | 0.485 | -1.349 | 0.378 | -0.527 | 0.007 | 0.015 | 0.015 | 0 | 0.52 | 0.5 | 0.65 | 0.72 | 0.54 | 0.51 |
| 8 | 27734464 | rs7827647 | 0.904 | -0.815 | 0.181 | -0.625 | -0.003 | 0.095 | 0.095 | 2.884 | 0.29 | 0.1 | 0.22 | 0.04 | 0.3 | 0.21 |
| 8 | 27740066 | rs6999359 | 1.091 | -0.986 | 0.32 | -0.26 | -0.022 | -0.008 | -0.008 | 0 | 0.42 | 0.25 | 0.52 | 0.6 | 0.44 | 0.48 |
| 8 | 27740237 | rs11778759 | -1.019 | -0.989 | 0.235 | -0.265 | 0 | 0.019 | 0.019 | 0.126 | 0.88 | 0.87 | 0.85 | 0.68 | 0.84 | 0.78 |
| 8 | 27740417 | rs11774576 | 0.508 | -0.996 | 0.289 | -0.264 | -0.001 | 0.003 | 0.003 | 0.297 | 0.49 | 0.53 | 0.56 | 0.65 | 0.47 | 0.48 |
| 8 | 27740693 | rs10103507 | 0.508 | -0.996 | 0.289 | -0.264 | -0.001 | 0.003 | 0.003 | 0.297 | 0.49 | 0.53 | 0.56 | 0.65 | 0.47 | 0.48 |
| 8 | 27741555 | rs4545047 | 0.227 | -0.839 | 0.122 | -0.15 | 0 | 0.005 | 0.005 | 0 | 0.8 | 0.75 | 0.8 | 0.68 | 0.78 | 0.68 |
| 8 | 27742959 | rs13281095 | 0.209 | -0.769 | 0.153 | -0.29 | -0.014 | -0.132 | -0.132 | 1.926 | 0.37 | 0.51 | 0.27 | 0.03 | 0.34 | 0.2 |
| 8 | 27743401 | rs2726928 | -0.488 | -0.763 | 0.129 | -0.226 | -0.001 | -0.009 | -0.009 | 0 | 0.48 | 0.51 | 0.38 | 0.28 | 0.49 | 0.41 |
| 8 | 27745205 | rs1879678 | 0.929 | -0.737 | -0.077 | -0.179 | -0.042 | -0.066 | -0.066 | 0.057 | 0.11 | 0.01 | 0.11 | 0.25 | 0.16 | 0.21 |
| 8 | 27745776 | rs2685379 | -0.485 | -0.604 | 0.099 | -0.142 | -0.001 | 0.054 | 0.054 | 0.02 | 0.49 | 0.64 | 0.38 | 0.28 | 0.5 | 0.41 |
| 8 | 27749344 | rs2726959 | 0.478 | -1.125 | 0.051 | 0.244 | -0.012 | 0.049 | 0.049 | 0 | 0.79 | 0.56 | 0.79 | 0.77 | 0.83 | 0.85 |
| 8 | 27749394 | rs2726960 | 0.579 | -1.217 | -0.362 | 0.299 | 0.076 | -0.007 | -0.007 | 0 | 0.54 | 0.24 | 0.68 | 0.53 | 0.68 | 0.64 |
| 8 | 27750732 | rs939711 | 1.114 | -1.363 | -0.39 | 0.093 | 0.027 | 0.06 | 0.06 | 0 | 0.27 | 0.07 | 0.18 | 0.14 | 0.27 | 0.27 |
| 8 | 27753688 | rs7817722 | -0.239 | -1.228 | 0.479 | 0.372 | -0.001 | 0.035 | 0.035 | 0 | 0.13 | 0.22 | 0.15 | 0.19 | 0.12 | 0.05 |
| 8 | 27754827 | rs2726972 | 0.807 | -1.169 | -0.16 | 0.584 | -0.037 | 0.093 | 0.093 | 0.853 | 0.85 | 0.4 | 0.87 | 0.81 | 0.93 | 0.87 |
| 8 | 27757789 | rs17058183 | 0.114 | -0.677 | -0.693 | 0.604 | -0.01 | 0.008 | 0.008 | 7.029 | 0.06 | 0.12 | 0.03 | 0.07 | 0.02 | 0.03 |
| 8 | 27758376 | rs7002829 | 0.114 | -0.677 | -0.705 | 0.657 | -0.01 | 0.029 | 0.029 | 7.029 | 0.06 | 0.12 | 0.05 | 0.11 | 0.02 | 0.03 |
| 8 | 27758922 | rs17058190 | -0.161 | -0.977 | -1.21 | 0.264 | 0.043 | 0.027 | 0.027 | 0.885 | 0.32 | 0.16 | 0.16 | 0.21 | 0.22 | 0.23 |
| 8 | 27759126 | rs2685372 | 0.031 | -0.11 | -1.146 | 0.558 | 0.052 | 0 | 0 | 0 | 0.47 | 0.22 | 0.66 | 0.58 | 0.64 | 0.62 |
| 8 | 27760475 | rs6999364 | -0.677 | 0.724 | -0.291 | 0.796 | 0.005 | 0.009 | 0.009 | 0 | 0.4 | 0.48 | 0.44 | 0.55 | 0.36 | 0.45 |
| 8 | 27763129 | rs4515501 | -0.023 | 0.687 | 0.378 | 1.024 | 0 | -0.007 | -0.007 | 0 | 0.72 | 0.67 | 0.62 | 0.83 | 0.75 | 0.75 |
| 8 | 27764576 | rs7826640 | -0.267 | 0.715 | 0.797 | 1.019 | 0.015 | 0.12 | 0.12 | 0.042 | 0.12 | 0.23 | 0.07 | 0.37 | 0.09 | 0.19 |
| 8 | 27767066 | rs17058204 | -0.114 | 0.725 | 0.825 | 1.089 | 0.011 | 0.088 | 0.088 | 0 | 0.11 | 0.16 | 0.07 | 0.38 | 0.09 | 0.19 |
| 8 | 27767171 | rs2726991 | 0.786 | 0.681 | 0.684 | 0.84 | -0.003 | -0.008 | -0.008 | 0 | 0.6 | 0.39 | 0.55 | 0.46 | 0.65 | 0.56 |
| 8 | 27774299 | rs756187 | 0.835 | 0.66 | 0.576 | 0.819 | -0.003 | -0.013 | -0.013 | 0 | 0.59 | 0.35 | 0.55 | 0.46 | 0.65 | 0.56 |
| 8 | 27775117 | rs2685325 | 0.213 | 0.79 | 0.882 | 0.246 | -0.005 | -0.094 | -0.094 | 0 | 0.73 | 0.48 | 0.8 | 0.92 | 0.77 | 0.78 |
| 8 | 27778842 | rs2685320 | -0.679 | 1.511 | 0.935 | 0.811 | 0.044 | 0.05 | 0.05 | 0 | 0.23 | 0.4 | 0.28 | 0.37 | 0.35 | 0.25 |
| 8 | 27790436 | rs2727014 | -0.346 | 1.472 | 1.206 | 0.842 | 0.022 | 0.027 | 0.027 | 0 | 0.21 | 0.28 | 0.28 | 0.36 | 0.34 | 0.24 |
| 8 | 27791766 | rs10104217 | 0.89 | 1.392 | 1.401 | 0.804 | 0.019 | 0.026 | 0.026 | 0 | 0.72 | 0.34 | 0.67 | 0.57 | 0.6 | 0.71 |
| 8 | 27808994 | rs10866867 | 0.278 | 1.691 | 1.514 | 0.768 | 0.122 | 0.022 | 0.022 | 0.007 | 0.85 | 0.57 | 0.84 | 0.82 | 0.73 | 0.9 |

| | | | | | | | | | | | | | | | | |
|---|---|---|---|---|---|---|---|---|---|---|---|---|---|---|---|---|
| 8 | 27810577 | rs2685413 | -0.167 | 1.78 | 1.301 | 0.75 | 0.065 | 0.034 | 0.034 | 0 | 0.13 | 0.25 | 0.15 | 0.24 | 0.28 | 0.14 |
| 8 | 27812107 | rs884829 | -0.298 | 1.777 | 1.248 | 1.023 | 0.016 | 0.073 | 0.073 | 0 | 0.15 | 0.22 | 0.23 | 0.44 | 0.28 | 0.3 |
| 8 | 27819749 | rs2726942 | -0.017 | 1.11 | 0.508 | -0.161 | 0.001 | 0.007 | 0.007 | 0 | 0.49 | 0.45 | 0.62 | 0.56 | 0.55 | 0.65 |
| 8 | 27823832 | rs11780263 | 0.539 | 0.89 | 0.352 | -0.896 | 0.019 | -0.022 | -0.022 | 0.014 | 0.28 | 0.17 | 0.21 | 0.43 | 0.18 | 0.25 |
| 8 | 27824470 | rs2726939 | -0.47 | 0.976 | -0.113 | -0.827 | -0.037 | 0.032 | 0.032 | 0 | 0.68 | 0.62 | 0.77 | 0.57 | 0.8 | 0.75 |
| 8 | 27825560 | rs17393091 | 0.631 | 1.117 | -0.245 | -0.85 | -0.01 | -0.008 | -0.008 | 0 | 0.2 | 0.18 | 0.15 | 0.01 | 0.26 | 0.09 |
| 8 | 27827391 | rs9650426 | 0.74 | 1.034 | -0.455 | -0.516 | 0.03 | 0.116 | 0.116 | 0 | 0.34 | 0.15 | 0.34 | 0.04 | 0.29 | 0.3 |
| 8 | 27828280 | rs2726930 | -0.285 | 1.021 | -0.643 | -0.467 | 0 | 0.004 | 0.004 | 0 | 0.51 | 0.47 | 0.62 | 0.57 | 0.56 | 0.65 |
| 8 | 27829251 | rs4732617 | 0.395 | 0.603 | -1.485 | -1.28 | 0.005 | 0.203 | 0.203 | 0 | 0.33 | 0.06 | 0.34 | 0.04 | 0.29 | 0.3 |
| 8 | 27836417 | rs1001866 | 0.204 | 0.995 | -0.915 | -1.259 | 0.013 | -0.037 | -0.037 | 0 | 0.67 | 0.63 | 0.52 | 0.14 | 0.56 | 0.49 |
| 8 | 27839246 | rs11786962 | 0.732 | 0.164 | -1.987 | -2.166 | -0.001 | 0.192 | 0.192 | 0.061 | 0.35 | 0.03 | 0.33 | 0.12 | 0.31 | 0.36 |
| 8 | 27840616 | rs11780951 | 0.107 | -0.018 | -1.442 | -2.411 | -0.004 | -0.052 | -0.052 | 0.02 | 0.52 | 0.5 | 0.41 | 0.1 | 0.39 | 0.41 |
| 8 | 27844622 | rs9650427 | -0.033 | 0.121 | -0.717 | -2.208 | 0.007 | 0.063 | 0.063 | 0 | 0.54 | 0.64 | 0.43 | 0.1 | 0.46 | 0.43 |
| 8 | 27848226 | rs2685391 | -0.213 | 0.954 | 0.062 | -0.786 | 0.067 | 0.072 | 0.072 | 0 | 0.2 | 0.3 | 0.25 | 0.35 | 0.32 | 0.28 |
| 8 | 27850161 | rs2685390 | -0.566 | 0.883 | 0.114 | -0.773 | 0.054 | 0.056 | 0.056 | 0 | 0.23 | 0.33 | 0.26 | 0.36 | 0.34 | 0.29 |
| 8 | 27853359 | rs2726961 | -0.353 | 1.082 | 0.225 | -0.646 | 0.079 | 0.128 | 0.128 | 0.016 | 0.18 | 0.53 | 0.21 | 0.36 | 0.26 | 0.24 |
| 8 | 27858737 | rs4732797 | 0.217 | 0.745 | -0.217 | -1.515 | -0.001 | -0.026 | -0.026 | 0 | 0.59 | 0.63 | 0.41 | 0.11 | 0.48 | 0.46 |
| 8 | 27871594 | rs10090626 | 0.395 | 0.451 | -0.339 | -1.721 | -0.001 | -0.067 | -0.067 | 3.193 | 0.48 | 0.54 | 0.37 | 0.11 | 0.44 | 0.38 |
| 8 | 27875050 | rs7840091 | 0.287 | 0.396 | -0.578 | -0.848 | -0.005 | -0.028 | -0.028 | 1.036 | 0.26 | 0.27 | 0.39 | 0.51 | 0.28 | 0.34 |
| 8 | 27875620 | rs4732802 | -0.064 | 0.147 | -0.192 | -0.949 | -0.016 | 0.026 | 0.026 | 0.214 | 0.72 | 0.57 | 0.77 | 0.66 | 0.73 | 0.73 |
| 8 | 27877269 | rs4732619 | 0.099 | 0.218 | -0.158 | -0.924 | 0.01 | -0.033 | -0.033 | 0 | 0.74 | 0.16 | 0.77 | 0.68 | 0.75 | 0.77 |
| 8 | 27878672 | rs2084918 | -0.069 | 0.303 | 0.112 | -0.893 | 0.059 | 0.016 | 0.016 | 0 | 0.07 | 0.39 | 0.05 | 0 | 0.05 | 0.01 |
| 8 | 27878996 | rs2100365 | 0.72 | 0.281 | 0.009 | -0.868 | -0.05 | -0.126 | -0.126 | 0 | 0.23 | 0.07 | 0.36 | 0.53 | 0.24 | 0.34 |
| 8 | 27879595 | rs997549 | -0.017 | 0.291 | 0.321 | -1.202 | 0.002 | -0.052 | -0.052 | 2.279 | 0.58 | 0.53 | 0.47 | 0.15 | 0.57 | 0.45 |
| 8 | 27880155 | rs10046724 | 0.226 | 0.28 | -0.14 | -1.176 | -0.003 | 0.052 | 0.052 | 0 | 0.56 | 0.35 | 0.46 | 0.15 | 0.55 | 0.44 |
| 8 | 27880826 | rs921597 | 0.137 | 0.307 | -0.505 | -1.163 | -0.009 | 0.146 | 0.146 | 0 | 0.55 | 0.23 | 0.45 | 0.15 | 0.56 | 0.44 |
| 8 | 27883101 | rs7815173 | 0.339 | 0.458 | -0.783 | -1.2 | -0.03 | 0.252 | 0.252 | 0 | 0.46 | 0.15 | 0.34 | 0.12 | 0.51 | 0.35 |
| 8 | 27885140 | rs10097488 | 0.253 | 0.379 | -0.217 | -1.216 | -0.003 | -0.061 | -0.061 | 0.061 | 0.46 | 0.53 | 0.3 | 0.12 | 0.43 | 0.33 |
| 8 | 27885843 | rs11993800 | -0.381 | 0.447 | -0.265 | -0.722 | -0.004 | 0.094 | 0.094 | 0 | 0.86 | 0.76 | 0.8 | 0.63 | 0.87 | 0.74 |
| 8 | 27886903 | rs9314357 | 0.339 | 0.467 | -0.511 | -0.823 | 0 | -0.04 | -0.04 | 0.015 | 0.44 | 0.43 | 0.29 | 0.12 | 0.43 | 0.32 |
| 8 | 27887575 | rs4732803 | -0.241 | 0.168 | -1.008 | -0.092 | 0.029 | -0.064 | -0.064 | 0.204 | 0.2 | 0.05 | 0.25 | 0.36 | 0.19 | 0.31 |
| 8 | 27894741 | rs4732807 | 0.32 | 0.818 | -0.295 | 0.047 | 0.022 | 0.116 | 0.116 | 0 | 0.25 | 0.49 | 0.42 | 0.47 | 0.29 | 0.31 |
| 8 | 27898580 | rs17058502 | 0.546 | 0.889 | -0.338 | -0.218 | 0.002 | -0.02 | -0.02 | 0 | 0.13 | 0.17 | 0.11 | 0.05 | 0.12 | 0.08 |
| 8 | 27900827 | rs4054868 | -0.965 | 0.509 | -0.386 | 0.146 | 0.007 | 0.028 | 0.028 | 0 | 0.45 | 0.8 | 0.61 | 0.61 | 0.45 | 0.56 |
| 8 | 27903069 | rs13279787 | 0.412 | 0.605 | -0.172 | 0.214 | 0.019 | 0.078 | 0.078 | 0 | 0.26 | 0.43 | 0.41 | 0.44 | 0.3 | 0.31 |
| 8 | 27905599 | rs1000658 | 0.101 | -0.04 | -0.856 | 0.355 | -0.002 | 0 | 0 | 0 | 0.16 | 0.2 | 0.2 | 0.17 | 0.14 | 0.24 |
| 8 | 27905861 | rs1984880 | -0.247 | 0.443 | -0.76 | -0.1 | -0.009 | -0.004 | -0.004 | 0.483 | 0.43 | 0.52 | 0.59 | 0.61 | 0.37 | 0.55 |
| 8 | 27918368 | rs12549429 | -0.355 | 0.594 | -1.312 | 0.417 | -0.057 | -0.099 | -0.099 | 2.677 | 0.17 | 0 | 0.14 | 0.33 | 0.17 | 0.19 |
| 8 | 27922574 | rs2045029 | 0.155 | 1.063 | -0.556 | 0.272 | 0.018 | 0.004 | 0.004 | 0 | 0.79 | 0.78 | 0.7 | 0.69 | 0.61 | 0.89 |
| 8 | 27923388 | rs4732812 | 0.934 | -0.084 | -0.549 | -0.237 | 0.044 | 0.063 | 0.063 | 0.059 | 0.35 | 0.25 | 0.32 | 0.15 | 0.26 | 0.24 |
| 8 | 27923551 | rs4732813 | 0.59 | -0.4 | -0.444 | -0.371 | 0.008 | -0.012 | -0.012 | 0.219 | 0.22 | 0.17 | 0.25 | 0.39 | 0.15 | 0.34 |
| 8 | 27924924 | rs17408912 | 0.838 | -0.064 | 0.054 | 0.024 | -0.011 | 0.011 | 0.011 | 0.098 | 0.15 | 0.06 | 0.09 | 0.11 | 0.16 | 0.26 |
| 8 | 27925204 | rs4732620 | -0.409 | -0.154 | 0.177 | -0.116 | 0.022 | 0.01 | 0.01 | 0 | 0.64 | 0.61 | 0.61 | 0.61 | 0.52 | 0.58 |
| 8 | 27926117 | rs10107497 | -1.259 | -0.009 | 1.116 | 0.82 | 0.011 | 0.075 | 0.075 | 2.197 | 0.28 | 0.56 | 0.23 | 0.6 | 0.27 | 0.4 |
| 8 | 27929927 | rs13271784 | 0.174 | 0.365 | 0.233 | 0.047 | -0.027 | 0.018 | 0.018 | 0.374 | 0.15 | 0.07 | 0.1 | 0.1 | 0.19 | 0.24 |
| 8 | 27930510 | rs9644050 | 0.018 | -0.231 | 0.268 | 0.051 | -0.033 | 0.013 | 0.013 | 0 | 0.09 | 0.22 | 0.11 | 0.14 | 0.04 | 0.12 |
| 8 | 27931751 | rs17058573 | -0.168 | 0.461 | -0.119 | 0.106 | 0.001 | -0.009 | -0.009 | 0 | 0.16 | 0.24 | 0.1 | 0.11 | 0.16 | 0.24 |
| 8 | 27933097 | rs13274329 | -0.461 | 0.424 | -0.089 | 0.087 | 0.005 | -0.016 | -0.016 | 0 | 0.19 | 0.28 | 0.14 | 0.12 | 0.21 | 0.25 |
| 8 | 27934199 | rs1901744 | 0.193 | 0.16 | 0.171 | -0.106 | -0.005 | -0.017 | -0.017 | 0.041 | 0.7 | 0.62 | 0.64 | 0.16 | 0.74 | 0.61 |
| 8 | 27936248 | rs4732815 | -0.738 | -0.123 | 1.164 | 0.761 | -0.043 | 0.235 | 0.235 | 0 | 0.16 | 0.46 | 0.32 | 0.63 | 0.11 | 0.21 |
| 8 | 27950670 | rs2305451 | -0.311 | 0.057 | 0.751 | 0.193 | -0.04 | -0.02 | -0.02 | 0 | 0.28 | 0.1 | 0.32 | 0.25 | 0.34 | 0.33 |
| 8 | 27951556 | rs2123472 | -1.181 | -0.04 | 1.58 | 0.894 | 0.004 | 0.01 | 0.01 | 0 | 0.42 | 0.68 | 0.46 | 0.58 | 0.44 | 0.53 |
| 8 | 27954465 | rs1563055 | -0.308 | -0.054 | 2.152 | 0.892 | 0.008 | 0.001 | 0.001 | 0 | 0.29 | 0.31 | 0.33 | 0.25 | 0.34 | 0.33 |
| 8 | 27957529 | rs2290370 | 0.725 | -0.055 | 2.261 | 0.905 | -0.01 | 0.064 | 0.064 | 0 | 0.1 | 0.16 | 0.12 | 0.31 | 0.1 | 0.19 |
| 8 | 27957556 | rs2290369 | -1.233 | -0.044 | 2.242 | 0.837 | 0.006 | 0.018 | 0.018 | 0 | 0.42 | 0.71 | 0.46 | 0.59 | 0.44 | 0.53 |
| 8 | 27967856 | rs3213997 | -1.394 | -0.041 | 1.735 | 0.811 | 0.003 | 0.006 | 0.006 | 0 | 0.45 | 0.61 | 0.66 | 0.89 | 0.47 | 0.57 |
| 8 | 28001680 | rs10104739 | -1.347 | -0.498 | 1.157 | 0.55 | -0.019 | 0.023 | 0.023 | 0 | 0.31 | 0.48 | 0.41 | 0.65 | 0.29 | 0.32 |
| 8 | 28003817 | rs17058631 | 0.077 | -0.501 | 1.168 | 0.562 | 0.01 | -0.012 | -0.012 | 0.81 | 0.15 | 0.12 | 0.12 | 0.31 | 0.13 | 0.2 |
| 8 | 28004531 | rs4377920 | -1.561 | -0.33 | 1.118 | -0.131 | 0.002 | -0.026 | -0.026 | 0.615 | 0.49 | 0.58 | 0.67 | 0.91 | 0.51 | 0.59 |
| 8 | 28022926 | rs2015443 | -0.052 | 0.046 | 0.825 | 0.09 | 0.003 | -0.016 | -0.016 | 0 | 0.28 | 0.21 | 0.36 | 0.56 | 0.31 | 0.47 |
| 8 | 28027006 | rs2061587 | -0.677 | -0.062 | 0.925 | 0.242 | -0.002 | 0.013 | 0.013 | 0 | 0.42 | 0.4 | 0.65 | 0.9 | 0.48 | 0.59 |
| 8 | 28031201 | rs6558041 | -0.682 | -0.051 | 0.737 | 0.248 | 0 | -0.037 | -0.037 | 0.184 | 0.42 | 0.45 | 0.65 | 0.9 | 0.48 | 0.59 |
| 8 | 28055236 | rs13268953 | -0.44 | 0.317 | 0.025 | 0.177 | 0.002 | 0.002 | 0.002 | 0.262 | 0.51 | 0.46 | 0.56 | 0.63 | 0.61 | 0.59 |
| 8 | 28058868 | rs13257057 | 0.464 | 0.359 | -0.077 | -0.218 | 0.014 | 0.014 | 0.014 | 1.166 | 0.56 | 0.31 | 0.58 | 0.61 | 0.62 | 0.51 |
| 8 | 28059541 | rs4054817 | 0.73 | 0.314 | 0.613 | -0.375 | 0.009 | -0.077 | -0.077 | 0 | 0.83 | 0.59 | 0.78 | 0.94 | 0.85 | 0.82 |
| 8 | 28059645 | rs4319045 | 0.044 | 0.295 | 0.814 | -0.378 | 0.042 | -0.045 | -0.045 | 0 | 0.88 | 0.59 | 0.92 | 0.95 | 0.9 | 0.89 |
| 8 | 28060032 | rs17411703 | -0.228 | 0.805 | 0.589 | -0.641 | 0 | -0.009 | -0.009 | 0 | 0.15 | 0.17 | 0.31 | 0.06 | 0.22 | 0.18 |
| 8 | 28060690 | rs7821358 | 1.38 | 0.706 | 0.641 | -0.691 | -0.032 | 0.008 | 0.008 | 1.715 | 0.45 | 0.14 | 0.43 | 0.57 | 0.46 | 0.42 |
| 8 | 28060783 | rs7840565 | 0.623 | 0.672 | 0.751 | -0.695 | 0.016 | 0.031 | 0.031 | 1.145 | 0.54 | 0.23 | 0.48 | 0.62 | 0.56 | 0.52 |

| | | | | | | | | | | | | | | | | |
|---|---|---|---|---|---|---|---|---|---|---|---|---|---|---|---|---|
| 8 | 28064115 | rs17411752 | 0.903 | 0.674 | 0.686 | -0.65 | -0.001 | 0.002 | 0.002 | 0 | 0.05 | 0.05 | 0.15 | 0.02 | 0.05 | 0.08 |
| 8 | 28064417 | rs12675992 | 0.679 | 0.591 | 0.764 | -0.659 | 0.014 | -0.017 | -0.017 | 0 | 0.28 | 0.21 | 0.18 | 0.32 | 0.22 | 0.31 |
| 8 | 28064546 | rs13271596 | 1.096 | 0.623 | 0.997 | -0.685 | 0.007 | -0.064 | -0.064 | 0 | 0.73 | 0.37 | 0.61 | 0.89 | 0.68 | 0.71 |
| 8 | 28069862 | rs4732839 | -0.481 | 0.732 | 0.73 | -0.64 | -0.022 | 0.031 | 0.031 | 0 | 0.19 | 0.12 | 0.21 | 0.09 | 0.28 | 0.21 |
| 8 | 28074892 | rs11784982 | 0.079 | 0.549 | 1.016 | -0.484 | 0.002 | 0.005 | 0.005 | 0.498 | 0.35 | 0.41 | 0.25 | 0.37 | 0.35 | 0.4 |
| 8 | 28076271 | rs9314362 | 0.069 | 0.549 | 1.016 | -0.484 | 0.002 | 0.005 | 0.005 | 0.552 | 0.35 | 0.41 | 0.25 | 0.37 | 0.35 | 0.41 |
| 8 | 28076422 | rs6985192 | 0.555 | 0.787 | 1.263 | -0.601 | 0.015 | -0.064 | -0.064 | 0 | 0.78 | 0.62 | 0.65 | 0.9 | 0.73 | 0.78 |
| 8 | 28095939 | rs11994255 | 0.406 | 0.723 | 0.959 | -0.551 | -0.026 | -0.089 | -0.089 | 2.641 | 0.27 | 0.08 | 0.2 | 0.39 | 0.29 | 0.35 |
| 8 | 28096383 | rs7010836 | -0.218 | 0.754 | 1.272 | -0.63 | 0.012 | -0.027 | -0.027 | 0.377 | 0.22 | 0.21 | 0.24 | 0.11 | 0.31 | 0.25 |
| 8 | 28100234 | rs13277389 | 1.301 | 0.727 | 1.126 | -0.557 | 0.03 | 0.005 | 0.005 | 2.697 | 0.44 | 0.19 | 0.39 | 0.52 | 0.37 | 0.37 |
| 8 | 28103444 | rs9314364 | -0.415 | 0.753 | 1.266 | -0.632 | 0.055 | -0.093 | -0.093 | 0 | 0.23 | 0.44 | 0.26 | 0.11 | 0.3 | 0.25 |
| 8 | 28104473 | rs4732844 | 1.378 | 0.716 | 1.137 | -0.618 | 0.02 | -0.046 | -0.046 | 0 | 0.44 | 0.13 | 0.38 | 0.48 | 0.37 | 0.37 |
| 8 | 28106784 | rs2614114 | -0.64 | 0.916 | 1.187 | -0.624 | 0.054 | -0.092 | -0.092 | 0 | 0.24 | 0.53 | 0.27 | 0.11 | 0.31 | 0.25 |
| 8 | 28118130 | rs2100814 | -0.254 | 0.47 | 0.961 | -0.456 | -0.042 | -0.06 | -0.06 | 0 | 0.35 | 0.19 | 0.27 | 0.48 | 0.41 | 0.43 |
| 8 | 28119260 | rs2722887 | 0.13 | 0.487 | 1.201 | -0.675 | 0.032 | -0.051 | -0.051 | 1.046 | 0.16 | 0.32 | 0.22 | 0.08 | 0.19 | 0.17 |
| 8 | 28121178 | rs1531383 | 0.081 | 0.498 | 1.035 | -0.616 | 0.014 | -0.017 | -0.017 | 0 | 0.16 | 0.27 | 0.22 | 0.1 | 0.19 | 0.19 |
| 8 | 28129261 | rs7830271 | -0.151 | 0.511 | 0.673 | -0.587 | -0.032 | -0.008 | -0.008 | 0 | 0.34 | 0.2 | 0.26 | 0.41 | 0.41 | 0.41 |
| 8 | 28131298 | rs2722924 | -0.534 | 0.22 | 0.745 | -0.869 | -0.02 | -0.083 | -0.083 | 0.412 | 0.85 | 0.63 | 0.81 | 0.97 | 0.91 | 0.96 |
| 8 | 28135494 | rs1841552 | 0.536 | 0.287 | 0.664 | -0.854 | -0.006 | 0.003 | 0.003 | 0 | 0.15 | 0.11 | 0.21 | 0.11 | 0.19 | 0.2 |
| 8 | 28136109 | rs2614046 | -0.587 | 0.172 | 0.722 | -0.894 | -0.02 | -0.085 | -0.085 | 0 | 0.85 | 0.62 | 0.81 | 0.97 | 0.91 | 0.96 |
| 8 | 28139238 | rs1461223 | 1.783 | 0.124 | 0.684 | -0.933 | 0.016 | -0.068 | -0.068 | 0 | 0.35 | 0.15 | 0.34 | 0.46 | 0.31 | 0.36 |
| 8 | 28143259 | rs7009996 | 0.24 | 0.155 | 0.73 | -0.936 | -0.019 | -0.005 | -0.005 | 0 | 0.32 | 0.21 | 0.23 | 0.39 | 0.35 | 0.39 |
| 8 | 28144790 | rs2722939 | -0.708 | 0.1 | 0.761 | -0.823 | -0.018 | -0.083 | -0.083 | 0 | 0.86 | 0.63 | 0.81 | 0.97 | 0.91 | 0.96 |
| 8 | 28146297 | rs17414182 | 1.187 | 0.181 | -0.144 | -0.894 | -0.003 | -0.017 | -0.017 | 0 | 0.45 | 0.2 | 0.48 | 0.47 | 0.57 | 0.45 |
| 8 | 28146753 | rs1381114 | -0.213 | 0.229 | -0.426 | -0.501 | 0.003 | -0.011 | -0.011 | 0.082 | 0.16 | 0.13 | 0.11 | 0.36 | 0.12 | 0.14 |
| 8 | 28146771 | rs1381115 | 0.262 | 0.282 | -0.291 | -0.334 | -0.005 | 0.047 | 0.047 | 0 | 0.1 | 0.13 | 0.08 | 0.38 | 0.07 | 0.14 |
| 8 | 28147240 | rs2722937 | -0.318 | 0.553 | -0.221 | -0.632 | 0.004 | -0.012 | -0.012 | 0.074 | 0.34 | 0.35 | 0.27 | 0.14 | 0.45 | 0.43 |
| 8 | 28150727 | rs2614053 | -0.934 | 0.322 | 0.145 | 0.048 | -0.002 | -0.001 | -0.001 | 1.041 | 0.41 | 0.58 | 0.54 | 0.46 | 0.36 | 0.38 |
| 8 | 28151129 | rs2614054 | -0.762 | 0.39 | 0.217 | 0.579 | -0.006 | -0.026 | -0.026 | 0 | 0.22 | 0.19 | 0.42 | 0.51 | 0.25 | 0.28 |
| 8 | 28151145 | rs13263030 | 0.747 | 0.608 | 0.206 | 0.23 | -0.056 | 0.103 | 0.103 | 0.232 | 0.44 | 0.17 | 0.32 | 0.24 | 0.46 | 0.31 |
| 8 | 28155890 | rs2614056 | -1.083 | 0.742 | 1.182 | 0.677 | 0.019 | 0.013 | 0.013 | 0 | 0.11 | 0.2 | 0.12 | 0.16 | 0.17 | 0.17 |
| 8 | 28156896 | rs2614059 | -1.692 | 0.743 | 1.287 | 0.673 | 0.085 | 0.065 | 0.065 | 0.153 | 0.14 | 0.43 | 0.14 | 0.17 | 0.17 | 0.17 |
| 8 | 28156978 | rs2614060 | -0.679 | 0.736 | 1.1 | 0.694 | 0.018 | 0.015 | 0.015 | 0 | 0.1 | 0.14 | 0.12 | 0.17 | 0.17 | 0.17 |
| 8 | 28157226 | rs2614062 | -1.266 | 0.707 | 0.765 | 0.711 | 0.073 | 0.062 | 0.062 | 0.146 | 0.13 | 0.41 | 0.13 | 0.17 | 0.17 | 0.17 |
| 8 | 28158138 | rs2614064 | -1.252 | 0.714 | 0.72 | 0.715 | 0.079 | 0.068 | 0.068 | 0.116 | 0.13 | 0.44 | 0.14 | 0.17 | 0.17 | 0.17 |
| 8 | 28163935 | rs2722892 | 1.543 | -0.385 | 0.623 | 0.584 | -0.024 | 0.036 | 0.036 | 2.681 | 0.91 | 0.46 | 0.95 | 0.93 | 0.99 | 0.92 |
| 8 | 28163980 | rs2614067 | 1.484 | -0.388 | 1.072 | 0.578 | 0.008 | 0.072 | 0.072 | 2.106 | 0.92 | 0.53 | 0.95 | 0.94 | 0.99 | 0.92 |
| 8 | 28165985 | rs2614071 | 1.081 | -0.701 | 0.5 | 0.825 | -0.043 | 0.032 | 0.032 | 1.051 | 0.8 | 0.5 | 0.8 | 0.77 | 0.89 | 0.8 |
| 8 | 28166098 | rs7826140 | 1.124 | -0.624 | -0.52 | 0.787 | 0.043 | -0.015 | -0.015 | 0 | 0.53 | 0.28 | 0.45 | 0.5 | 0.46 | 0.49 |
| 8 | 28166870 | rs2614075 | 0.084 | -0.612 | -0.008 | 0.763 | -0.024 | 0.006 | 0.006 | 1.254 | 0.83 | 0.64 | 0.83 | 0.83 | 0.89 | 0.88 |
| 8 | 28167580 | rs4732636 | 1.812 | -1.704 | -1.306 | 0.375 | -0.011 | -0.109 | -0.109 | 0.031 | 0.3 | 0.06 | 0.34 | 0.43 | 0.32 | 0.31 |
| 8 | 28167726 | rs10503827 | 0.85 | -0.929 | -0.848 | 0.411 | -0.127 | 0.034 | 0.034 | 0.846 | 0.18 | 0.04 | 0.23 | 0.11 | 0.39 | 0.19 |
| 8 | 28168659 | rs2614079 | 0.283 | -0.988 | -0.306 | 0.641 | -0.03 | -0.011 | -0.011 | 0 | 0.81 | 0.45 | 0.81 | 0.83 | 0.89 | 0.87 |
| 8 | 28168886 | rs2722910 | 0.824 | -0.987 | -0.334 | 0.603 | -0.016 | 0.005 | 0.005 | 0.476 | 0.73 | 0.41 | 0.7 | 0.68 | 0.85 | 0.76 |
| 8 | 28173751 | rs3808366 | 0.174 | -0.768 | -0.63 | 0.85 | 0.001 | -0.029 | -0.029 | 0 | 0.07 | 0.02 | 0.09 | 0.16 | 0.05 | 0.09 |
| 8 | 28174354 | rs17058952 | 0.759 | -1.315 | -0.114 | 0.552 | -0.012 | 0.001 | 0.001 | 0 | 0.47 | 0.51 | 0.47 | 0.52 | 0.37 | 0.52 |
| 8 | 28178513 | rs7825003 | 1.078 | -1.463 | -0.228 | 0.578 | 0.048 | -0.025 | -0.025 | 0 | 0.67 | 0.24 | 0.74 | 0.61 | 0.81 | 0.71 |
| 8 | 28179013 | rs7825480 | 0.031 | -1.447 | -0.07 | 0.682 | 0.008 | -0.016 | -0.016 | 0 | 0.17 | 0.04 | 0.15 | 0.17 | 0.15 | 0.17 |
| 8 | 28183219 | rs2614091 | 0.505 | -1.296 | -0.182 | 0.645 | -0.16 | 0.084 | 0.084 | 0.78 | 0.32 | 0.05 | 0.28 | 0.14 | 0.44 | 0.29 |
| 8 | 28184548 | rs2722901 | 0.57 | -1.256 | -0.184 | 0.643 | -0.163 | 0.078 | 0.078 | 0.791 | 0.31 | 0.04 | 0.28 | 0.14 | 0.43 | 0.28 |
| 8 | 28189387 | rs2722931 | 0.495 | -1.243 | -0.158 | 0.65 | -0.148 | 0.088 | 0.088 | 0.629 | 0.32 | 0.05 | 0.28 | 0.14 | 0.43 | 0.29 |
| 8 | 28191306 | rs351776 | -0.09 | -1.3 | 0.438 | 0.633 | -0.02 | 0.086 | 0.086 | 0.743 | 0.39 | 0.16 | 0.33 | 0.18 | 0.52 | 0.33 |
| 8 | 28191706 | rs189509 | -0.489 | -1.288 | 1.028 | 0.699 | -0.038 | 0.024 | 0.024 | 2.554 | 0.86 | 0.74 | 0.87 | 0.82 | 0.93 | 0.86 |
| 8 | 28193846 | rs4732849 | 1.521 | -1.54 | 0.44 | 0.705 | 0.001 | -0.047 | -0.047 | 0 | 0.33 | 0.17 | 0.46 | 0.42 | 0.35 | 0.34 |
| 8 | 28195931 | rs17058979 | 0.669 | -1.357 | 0.628 | 0.764 | 0.022 | -0.003 | -0.003 | 0.648 | 0.06 | 0.1 | 0.05 | 0.04 | 0.09 | 0.04 |
| 8 | 28198159 | rs351783 | 0.804 | -1.845 | 1.164 | 1.025 | -0.064 | 0.046 | 0.046 | 2.431 | 0.73 | 0.43 | 0.79 | 0.62 | 0.87 | 0.65 |
| 8 | 28198716 | rs4732850 | -0.003 | -1.443 | 0.749 | 1.07 | 0.008 | -0.034 | -0.034 | 0.933 | 0.13 | 0.05 | 0.16 | 0.2 | 0.11 | 0.21 |
| 8 | 28201643 | rs4732638 | -0.132 | -1.437 | 0.888 | 1.075 | 0.009 | -0.022 | -0.022 | 0.541 | 0.13 | 0.09 | 0.16 | 0.21 | 0.11 | 0.21 |
| 8 | 28204264 | rs351779 | 0.35 | -1.435 | 0.739 | 1.069 | -0.037 | 0.095 | 0.095 | 0.559 | 0.37 | 0.14 | 0.3 | 0.19 | 0.51 | 0.29 |
| 8 | 28209837 | rs2272750 | 0.479 | -1.031 | 1.263 | 1.619 | 0.012 | -0.024 | -0.024 | 0 | 0.11 | 0.06 | 0.06 | 0.22 | 0.07 | 0.15 |
| 8 | 28214096 | rs2272749 | -0.71 | -1.111 | 1.972 | 1.64 | -0.046 | 0.105 | 0.105 | 0 | 0.16 | 0.58 | 0.09 | 0.28 | 0.08 | 0.17 |
| 8 | 28226017 | rs17059034 | 0.69 | -1.209 | 1.824 | 1.519 | 0.006 | -0.01 | -0.01 | 0.182 | 0.34 | 0.29 | 0.49 | 0.53 | 0.3 | 0.44 |
| 8 | 28240092 | rs12056907 | -0.445 | -1.172 | 1.749 | 1.511 | -0.028 | 0.171 | 0.171 | 0 | 0.14 | 0.48 | 0.09 | 0.28 | 0.08 | 0.16 |
| 8 | 28244724 | rs757 | -0.362 | -1.111 | 1.038 | 1.48 | 0.086 | 0.209 | 0.209 | 0 | 0.49 | 0.89 | 0.56 | 0.8 | 0.37 | 0.6 |
| 8 | 28272574 | rs7832040 | -0.988 | -0.967 | 1.852 | 1.737 | -0.029 | 0.084 | 0.084 | 0.114 | 0.15 | 0.58 | 0.09 | 0.23 | 0.08 | 0.17 |
| 8 | 28286071 | rs11895 | -0.686 | -0.696 | 1.484 | 1.374 | -0.018 | 0.029 | 0.029 | 0.019 | 0.18 | 0.39 | 0.13 | 0.2 | 0.12 | 0.29 |
| 8 | 28288715 | rs11784510 | -0.958 | -0.723 | 0.704 | 1.355 | -0.022 | 0.033 | 0.033 | 0.468 | 0.22 | 0.65 | 0.19 | 0.4 | 0.13 | 0.39 |
| 8 | 28296171 | rs924184 | 0.044 | 0.491 | -0.636 | 0.522 | -0.014 | 0.025 | 0.025 | 0 | 0.56 | 0.27 | 0.58 | 0.62 | 0.56 | 0.71 |
| 8 | 28300460 | rs17059088 | -0.419 | 0.586 | -0.434 | 0.444 | 0 | -0.003 | -0.003 | 1.51 | 0.21 | 0.18 | 0.13 | 0.26 | 0.21 | 0.3 |

| | | | | | | | | | | | | | | | | |
|---|---|---|---|---|---|---|---|---|---|---|---|---|---|---|---|---|
| 8 | 28302765 | rs1060411 | -0.199 | 0.538 | 0.116 | 0.385 | -0.004 | 0.002 | 0.002 | 0 | 0.36 | 0.54 | 0.46 | 0.6 | 0.34 | 0.39 |
| 8 | 28304767 | rs7016831 | -0.796 | 0.735 | 0.461 | 0.215 | 0.02 | -0.027 | -0.027 | 0 | 0.07 | 0.3 | 0.03 | 0 | 0.04 | 0.02 |
| 8 | 28307911 | rs11993007 | -0.302 | 0.684 | 0.218 | 0.404 | 0.001 | -0.004 | -0.004 | 0 | 0.4 | 0.66 | 0.43 | 0.56 | 0.36 | 0.38 |
| 8 | 28308594 | rs17059112 | -0.432 | 0.744 | 0.129 | 0.436 | -0.001 | 0.002 | 0.002 | 0 | 0.39 | 0.59 | 0.42 | 0.55 | 0.36 | 0.38 |
| 8 | 28319438 | rs35107433 | -0.399 | 0.698 | -0.488 | 0.464 | 0.007 | -0.02 | -0.02 | 0 | 0.39 | 0.32 | 0.39 | 0.5 | 0.31 | 0.38 |
| 8 | 28325411 | rs6992329 | 0.095 | 0.636 | -0.864 | 0.358 | 0.015 | -0.001 | -0.001 | 0 | 0.37 | 0.11 | 0.43 | 0.62 | 0.31 | 0.38 |
| 8 | 28333294 | rs10503829 | -0.052 | 0.745 | 0.296 | 0.234 | -0.001 | -0.002 | -0.002 | 0 | 0.46 | 0.47 | 0.49 | 0.65 | 0.38 | 0.43 |
| 8 | 28336041 | rs11136047 | 0.753 | 1.1 | 0.524 | -0.153 | -0.003 | -0.022 | -0.022 | 0.887 | 0.28 | 0.26 | 0.25 | 0.09 | 0.25 | 0.24 |
| 8 | 28337560 | rs7011772 | 0.093 | 1.301 | 0.877 | 0.549 | -0.026 | 0.009 | 0.009 | 0 | 0.75 | 0.84 | 0.74 | 0.74 | 0.64 | 0.67 |
| 8 | 28347283 | rs1908916 | 1.84 | 0.852 | 1.836 | 0.994 | 0.058 | 0.08 | 0.08 | 0 | 0.93 | 0.73 | 0.91 | 0.9 | 0.89 | 0.94 |
| 8 | 28350753 | rs1946583 | 1.547 | 0.889 | 2.119 | 1.144 | 0.047 | 0.058 | 0.058 | 0 | 0.93 | 0.84 | 0.92 | 0.88 | 0.87 | 0.93 |
| 8 | 28359611 | rs6997072 | 0.919 | 0.566 | 2.546 | 1.482 | -0.003 | -0.009 | -0.009 | 0.007 | 0.62 | 0.34 | 0.6 | 0.53 | 0.59 | 0.62 |
| 8 | 28362459 | rs7842884 | 0.692 | 0.53 | 2.864 | 1.536 | 0.003 | -0.005 | -0.005 | 0 | 0.66 | 0.69 | 0.63 | 0.69 | 0.6 | 0.63 |
| 8 | 28362792 | rs7833751 | 0.949 | 0.539 | 2.828 | 1.537 | -0.002 | -0.004 | -0.004 | 0 | 0.62 | 0.34 | 0.6 | 0.57 | 0.61 | 0.62 |
| 8 | 28369097 | rs6984655 | 0.697 | 0.496 | 2.847 | 1.533 | 0.009 | -0.009 | -0.009 | 0.034 | 0.64 | 0.54 | 0.61 | 0.65 | 0.59 | 0.63 |
| 8 | 28384712 | rs2241802 | 0.898 | 0.541 | 2.619 | 1.563 | 0.003 | 0.005 | 0.005 | 0 | 0.63 | 0.39 | 0.6 | 0.57 | 0.59 | 0.62 |
| 8 | 28394701 | rs352203 | 0.78 | 0.546 | 2.649 | 1.417 | -0.006 | 0.003 | 0.003 | 0 | 0.65 | 0.7 | 0.62 | 0.65 | 0.59 | 0.63 |
| 8 | 28416551 | rs6980605 | 0.887 | 0.556 | 2.471 | 1.4 | 0.003 | 0.011 | 0.011 | 0 | 0.63 | 0.39 | 0.6 | 0.53 | 0.59 | 0.62 |
| 8 | 28439437 | rs12681261 | -0.277 | 0.298 | 1.657 | 1.364 | -0.003 | -0.008 | -0.008 | 0 | 0.28 | 0.15 | 0.2 | 0.28 | 0.26 | 0.31 |
| 8 | 28441331 | rs352199 | 0.392 | 0.281 | 1.731 | 1.376 | -0.001 | -0.01 | -0.01 | 0.557 | 0.69 | 0.74 | 0.75 | 0.63 | 0.67 | 0.62 |
| 8 | 28446734 | rs352423 | -0.637 | 0.117 | 0.971 | 1.174 | -0.014 | 0.034 | 0.034 | 0.017 | 0.32 | 0.54 | 0.23 | 0.45 | 0.27 | 0.32 |
| 8 | 28452799 | rs352416 | -1.945 | 0.109 | 1.382 | 0.747 | 0.011 | -0.03 | -0.03 | 2.067 | 0.09 | 0.57 | 0.04 | 0 | 0.02 | 0.01 |
| 8 | 28454347 | rs6983363 | -1.805 | 0.144 | 1.724 | 0.761 | 0.03 | 0.004 | 0.004 | 0.082 | 0.06 | 0.33 | 0.04 | 0 | 0.03 | 0.03 |
| 8 | 28454447 | rs6996714 | -1.754 | -0.708 | 1.338 | 1.394 | -0.011 | 0.006 | 0.006 | 0.008 | 0.23 | 0.76 | 0.13 | 0.35 | 0.09 | 0.19 |
| 8 | 28461258 | rs6992319 | -1.678 | -0.783 | 1.364 | 1.423 | -0.013 | 0.005 | 0.005 | 0.359 | 0.22 | 0.77 | 0.12 | 0.36 | 0.08 | 0.14 |
| 8 | 28461524 | rs6558068 | -1.678 | -0.783 | 1.33 | 1.423 | -0.007 | 0.001 | 0.001 | 0.359 | 0.22 | 0.8 | 0.13 | 0.36 | 0.08 | 0.14 |
| 8 | 28464593 | rs7008984 | -1.589 | -0.454 | 1.794 | 1.839 | 0.023 | 0.174 | 0.174 | 0 | 0.06 | 0.24 | 0.16 | 0.23 | 0.04 | 0.08 |
| 8 | 28472973 | rs9644143 | -0.526 | 0.272 | 1.44 | 1.078 | 0.014 | -0.002 | -0.002 | 0.218 | 0.26 | 0.39 | 0.29 | 0.23 | 0.3 | 0.23 |
| 8 | 28478892 | rs352428 | -0.531 | -0.193 | 0.217 | 0.878 | -0.033 | -0.019 | -0.019 | 0 | 0.84 | 0.99 | 0.91 | 0.87 | 0.85 | 0.8 |
| 8 | 28479611 | rs1441706 | -0.808 | -0.352 | 1.036 | 1.061 | 0 | -0.002 | -0.002 | 0.239 | 0.34 | 0.58 | 0.32 | 0.27 | 0.3 | 0.26 |
| 8 | 28487460 | rs6985395 | -0.468 | -0.388 | 1.161 | 1.029 | 0.009 | -0.004 | -0.004 | 0.021 | 0.3 | 0.58 | 0.31 | 0.24 | 0.31 | 0.25 |
| 8 | 28489068 | rs6994291 | -0.962 | -0.392 | -0.189 | 1.134 | -0.064 | -0.074 | -0.074 | 0 | 0.47 | 0.91 | 0.62 | 0.46 | 0.43 | 0.42 |
| 8 | 28491418 | rs12235009 | 0.784 | -0.208 | -0.034 | 1.002 | -0.084 | -0.05 | -0.05 | 0 | 0.21 | 0.05 | 0.27 | 0.23 | 0.29 | 0.24 |
| 8 | 28516077 | rs352427 | 0.395 | -0.736 | 0.131 | 0.523 | 0.017 | 0.005 | 0.005 | 5.364 | 0.86 | 0.9 | 0.96 | 1 | 0.95 | 0.98 |
| 8 | 28516514 | rs240748 | -0.331 | -0.17 | -0.058 | 0.763 | -0.019 | -0.023 | -0.023 | 0.44 | 0.35 | 0.24 | 0.4 | 0.41 | 0.38 | 0.34 |
| 8 | 28519849 | rs7813692 | -1.41 | -0.493 | 0.716 | 0.282 | -0.021 | -0.066 | -0.066 | 0 | 0.12 | 0.5 | 0.05 | 0.03 | 0.06 | 0.14 |
| 8 | 28521410 | rs164664 | -1.276 | -1.186 | 0.553 | -0.068 | -0.011 | -0.03 | -0.03 | 2.251 | 0.21 | 0.68 | 0.25 | 0.09 | 0.11 | 0.21 |
| 8 | 28522426 | rs171302 | -0.758 | -0.986 | 0.484 | 0.053 | 0.028 | -0.008 | -0.008 | 0.987 | 0.09 | 0.32 | 0.21 | 0.06 | 0.06 | 0.07 |
| 8 | 28523675 | rs164661 | -0.291 | -0.947 | 0.004 | 0.073 | 0.008 | -0.015 | -0.015 | 0.814 | 0.08 | 0.17 | 0.2 | 0.06 | 0.06 | 0.06 |
| 8 | 28528915 | rs4732865 | 0.533 | -1.07 | -1.818 | 0.131 | -0.062 | -0.029 | -0.029 | 0 | 0.48 | 0.07 | 0.4 | 0.36 | 0.58 | 0.39 |
| 8 | 28535241 | rs240920 | 0.363 | -0.216 | -1.467 | 0.437 | -0.017 | -0.065 | -0.065 | 0 | 0.28 | 0.1 | 0.38 | 0.41 | 0.31 | 0.26 |
| 8 | 28541640 | rs2856802 | 0.045 | 0.009 | -1.226 | 0.521 | 0.006 | -0.021 | -0.021 | 0 | 0.31 | 0.15 | 0.28 | 0.35 | 0.29 | 0.25 |
| 8 | 28557627 | rs240940 | -0.307 | -0.181 | 0.045 | 0.448 | -0.021 | -0.018 | -0.018 | 0 | 0.36 | 0.5 | 0.3 | 0.3 | 0.29 | 0.25 |
| 8 | 28557762 | rs240941 | -0.377 | -0.215 | 0.332 | 0.431 | -0.008 | -0.007 | -0.007 | 0 | 0.37 | 0.57 | 0.3 | 0.3 | 0.3 | 0.25 |
| 8 | 28566677 | rs2299563 | 0.589 | -0.435 | 0.164 | -0.128 | -0.004 | 0.026 | 0.026 | 0 | 0.18 | 0.11 | 0.21 | 0.11 | 0.15 | 0.23 |
| 8 | 28569393 | rs2299561 | -0.119 | -0.02 | 0.017 | -0.177 | -0.018 | 0.036 | 0.036 | 0 | 0.37 | 0.24 | 0.43 | 0.25 | 0.39 | 0.34 |
| 8 | 28585204 | rs2237814 | 0.828 | 0.041 | 0.39 | -0.136 | -0.009 | 0.001 | 0.001 | 0 | 0.36 | 0.39 | 0.29 | 0.35 | 0.29 | 0.25 |
| 8 | 28585452 | rs2237813 | -2.234 | -0.08 | 0.309 | -0.158 | -0.053 | -0.07 | -0.07 | 0.103 | 0.78 | 0.95 | 0.75 | 0.62 | 0.69 | 0.62 |
| 8 | 28586036 | rs2237812 | 1.252 | -0.528 | 0.38 | -0.202 | 0.012 | -0.033 | -0.033 | 0 | 0.31 | 0.11 | 0.2 | 0.35 | 0.28 | 0.25 |
| 8 | 28593399 | rs7015836 | -1.774 | -0.365 | 0.496 | -0.149 | -0.006 | -0.005 | -0.005 | 0 | 0.74 | 0.77 | 0.62 | 0.59 | 0.63 | 0.52 |
| 8 | 28597765 | rs743676 | 0.637 | -0.375 | 0.332 | -0.238 | -0.016 | 0.002 | 0.002 | 0 | 0.35 | 0.56 | 0.3 | 0.35 | 0.29 | 0.26 |
| 8 | 28601554 | rs4732873 | 1.019 | -0.285 | 0.384 | -0.191 | -0.003 | 0.002 | 0.002 | 0 | 0.32 | 0.36 | 0.28 | 0.35 | 0.29 | 0.26 |
| 8 | 28609136 | rs10091907 | -0.393 | -0.178 | -0.104 | -0.292 | 0.006 | 0.016 | 0.016 | 0 | 0.37 | 0.32 | 0.38 | 0.22 | 0.33 | 0.25 |
| 8 | 28612724 | rs241149 | 0.245 | 0.395 | -0.151 | 0.051 | -0.001 | 0 | 0 | 0 | 0.29 | 0.27 | 0.13 | 0.32 | 0.24 | 0.24 |
| 8 | 28613447 | rs10095449 | -0.389 | 0.3 | -0.7 | 0.036 | -0.003 | 0.013 | 0.013 | 0 | 0.27 | 0.24 | 0.23 | 0.12 | 0.3 | 0.14 |
| 8 | 28614068 | rs6995174 | -1.131 | 0.279 | -1.402 | 0.13 | 0 | 0.007 | 0.007 | 0 | 0.51 | 0.73 | 0.42 | 0.45 | 0.54 | 0.33 |
| 8 | 28615433 | rs2213306 | 0.493 | 0.666 | -1.49 | 0.656 | -0.065 | -0.103 | -0.103 | 0 | 0.22 | 0.02 | 0.14 | 0.35 | 0.24 | 0.24 |
| 8 | 28621169 | rs11786308 | -0.786 | 0.654 | -0.429 | 0.306 | -0.001 | -0.029 | -0.029 | 0.016 | 0.27 | 0.4 | 0.16 | 0.14 | 0.27 | 0.15 |
| 8 | 28622089 | rs241179 | -1.386 | 0.59 | 0.272 | 0.948 | 0.04 | 0.143 | 0.143 | 0 | 0.19 | 0.47 | 0.21 | 0.32 | 0.24 | 0.23 |
| 8 | 28623666 | rs17443293 | -0.532 | 0.888 | 0.137 | 0.337 | -0.033 | 0.032 | 0.032 | 0 | 0.11 | 0.04 | 0.1 | 0 | 0.17 | 0.05 |
| 8 | 28626418 | rs241178 | -0.591 | 0.215 | 0.379 | 0.479 | 0.004 | -0.009 | -0.009 | 0 | 0.28 | 0.23 | 0.19 | 0.52 | 0.25 | 0.34 |
| 8 | 28626470 | rs241177 | 1.983 | -0.469 | 0.402 | 0.38 | -0.027 | 0.021 | 0.021 | 0 | 0.44 | 0.18 | 0.54 | 0.33 | 0.51 | 0.48 |
| 8 | 28629160 | rs241174 | -1.295 | -0.274 | 0.761 | 0.235 | -0.003 | -0.007 | -0.007 | 0 | 0.39 | 0.35 | 0.3 | 0.53 | 0.43 | 0.34 |
| 8 | 28635787 | rs7828890 | -0.076 | -0.383 | 0.871 | 0.069 | 0.017 | -0.065 | -0.065 | 0 | 0.11 | 0.37 | 0.06 | 0 | 0.11 | 0.04 |
| 8 | 28640370 | rs7814699 | -0.259 | -0.328 | 0.801 | 0.288 | 0.003 | 0.139 | 0.139 | 0 | 0.14 | 0.27 | 0.17 | 0.47 | 0.16 | 0.15 |
| 8 | 28643060 | rs241189 | 0.917 | -0.397 | 1.092 | 0.176 | -0.017 | 0.065 | 0.065 | 0 | 0.69 | 0.55 | 0.74 | 0.44 | 0.71 | 0.7 |
| 8 | 28643722 | rs171313 | 1.815 | -0.698 | 0.859 | 0.152 | 0.079 | 0.116 | 0.116 | 0.615 | 0.53 | 0.24 | 0.55 | 0.18 | 0.65 | 0.39 |
| 8 | 28647832 | rs241181 | 0.403 | -0.628 | 1.02 | 0.128 | 0.046 | -0.029 | -0.029 | 0 | 0.72 | 0.88 | 0.78 | 0.65 | 0.82 | 0.58 |
| 8 | 28651801 | rs17522554 | 0.39 | -0.371 | 0.871 | 0.011 | -0.037 | 0.036 | 0.036 | 0 | 0.11 | 0.03 | 0.09 | 0 | 0.17 | 0.05 |

| | | | | | | | | | | | | | | | | |
|---|---|---|---|---|---|---|---|---|---|---|---|---|---|---|---|---|
| 8 | 28657761 | rs241167 | 1.551 | -0.428 | 0.978 | 0.037 | 0.029 | 0.037 | 0.037 | 0 | 0.55 | 0.39 | 0.57 | 0.18 | 0.66 | 0.4 |
| 8 | 28657998 | rs241166 | 0.576 | -0.427 | 0.552 | 0.077 | 0.024 | -0.008 | -0.008 | 0 | 0.69 | 0.78 | 0.74 | 0.64 | 0.82 | 0.54 |
| 8 | 28682594 | rs10102198 | 2.512 | -0.431 | 0.424 | 0.054 | -0.062 | 0.104 | 0.104 | 0 | 0.4 | 0.17 | 0.45 | 0.18 | 0.47 | 0.33 |
| 8 | 28683829 | rs17059489 | 0.372 | -0.625 | 0.26 | 0.171 | 0.049 | -0.054 | -0.054 | 0 | 0.14 | 0.04 | 0.17 | 0.27 | 0.06 | 0.25 |
| 8 | 28694549 | rs907133 | -0.299 | -0.501 | 0.407 | -0.167 | -0.007 | -0.023 | -0.023 | 0 | 0.82 | 0.81 | 0.91 | 0.91 | 0.88 | 0.81 |
| 8 | 28696395 | rs4732885 | 1.656 | -0.565 | 0.471 | -0.204 | 0.089 | 0.136 | 0.136 | 0.682 | 0.53 | 0.27 | 0.64 | 0.18 | 0.65 | 0.41 |
| 8 | 28698588 | rs10096475 | -0.4 | -0.817 | 0.442 | -0.098 | -0.002 | -0.012 | -0.012 | 0 | 0.83 | 0.84 | 0.91 | 0.91 | 0.88 | 0.81 |
| 8 | 28707570 | rs4732887 | 2.024 | -0.732 | 0.144 | -0.115 | 0.012 | 0.103 | 0.103 | 0 | 0.45 | 0.2 | 0.57 | 0.26 | 0.55 | 0.41 |
| 8 | 28719284 | rs10503833 | -0.282 | -0.467 | 0.557 | -0.135 | 0.008 | -0.05 | -0.05 | 0 | 0.12 | 0.2 | 0.09 | 0 | 0.14 | 0.05 |
| 8 | 28741584 | rs12547794 | -0.725 | -0.653 | 0.417 | 0.089 | 0.02 | -0.014 | -0.014 | 0 | 0.26 | 0.08 | 0.17 | 0.21 | 0.2 | 0.29 |
| 8 | 28746324 | rs11782284 | 2.302 | -0.914 | 0.436 | -0.088 | -0.045 | 0.067 | 0.067 | 0 | 0.38 | 0.19 | 0.55 | 0.2 | 0.51 | 0.36 |
| 8 | 28751381 | rs4732890 | 1.475 | -1.11 | 0.859 | -0.088 | -0.001 | -0.002 | -0.002 | 0 | 0.52 | 0.49 | 0.72 | 0.67 | 0.64 | 0.53 |
| 8 | 28756738 | rs13280242 | 1.285 | -1.116 | 0.914 | -0.095 | 0.018 | 0.027 | 0.027 | 0 | 0.54 | 0.63 | 0.72 | 0.74 | 0.65 | 0.55 |
| 8 | 28791159 | rs2221894 | 1.346 | -1.16 | 1.024 | -0.092 | 0.006 | 0.008 | 0.008 | 0 | 0.54 | 0.58 | 0.71 | 0.74 | 0.65 | 0.55 |
| 8 | 28802932 | rs6558085 | 1.214 | -1.166 | 1.031 | -0.1 | 0.042 | 0.071 | 0.071 | 0 | 0.55 | 0.77 | 0.73 | 0.74 | 0.65 | 0.55 |
| 8 | 28805908 | rs17524918 | 0.325 | -1.228 | 0.982 | -0.158 | -0.031 | 0.032 | 0.032 | 0 | 0.1 | 0.05 | 0.1 | 0 | 0.16 | 0.05 |
| 8 | 28816852 | rs4559202 | 1.393 | -1.24 | 1.279 | -0.153 | 0.013 | 0.018 | 0.018 | 0 | 0.54 | 0.63 | 0.71 | 0.74 | 0.64 | 0.54 |
| 8 | 28843721 | rs12674777 | 1.484 | -1.247 | 1.308 | -0.158 | 0.001 | -0.001 | -0.001 | 1.543 | 0.53 | 0.49 | 0.71 | 0.74 | 0.64 | 0.54 |
| 8 | 28851177 | rs7823358 | 0.108 | -1.261 | 1.373 | -0.113 | -0.009 | 0.187 | 0.187 | 0 | 0.12 | 0.23 | 0.16 | 0.47 | 0.13 | 0.17 |
| 8 | 28855526 | rs11136055 | 1.734 | -1.279 | 1.352 | -0.143 | 0.042 | 0.052 | 0.052 | 0 | 0.5 | 0.34 | 0.71 | 0.67 | 0.64 | 0.53 |
| 8 | 28892090 | rs12546597 | 0.578 | -1.391 | 1.166 | -0.147 | 0.054 | -0.053 | -0.053 | 0 | 0.15 | 0.03 | 0.09 | 0.2 | 0.07 | 0.24 |
| 8 | 28902759 | rs17526069 | 0.329 | -1.354 | 1.204 | -0.176 | 0 | -0.013 | -0.013 | 0 | 0.2 | 0.15 | 0.08 | 0.05 | 0.13 | 0.16 |
| 8 | 28909523 | rs10088428 | -0.82 | -1.363 | 1.172 | -0.2 | -0.001 | -0.098 | -0.098 | 0.026 | 0.3 | 0.39 | 0.19 | 0.05 | 0.28 | 0.21 |
| 8 | 28910397 | rs7004667 | -0.578 | -1.494 | 1.088 | -0.089 | -0.007 | 0.099 | 0.099 | 0 | 0.88 | 0.87 | 0.85 | 0.53 | 0.88 | 0.83 |
| 8 | 28916392 | rs2279446 | -0.086 | -1.604 | 0.969 | -0.096 | 0.015 | -0.015 | -0.015 | 1.091 | 0.18 | 0.12 | 0.1 | 0.29 | 0.08 | 0.27 |
| 8 | 28923898 | rs936920 | 0.684 | -1.579 | 1.071 | -0.096 | 0.027 | -0.031 | -0.031 | 0 | 0.69 | 0.27 | 0.77 | 0.42 | 0.81 | 0.66 |
| 8 | 28924858 | rs2693 | -1.666 | -1.542 | 1.137 | -0.132 | 0.005 | 0.005 | 0.005 | 0 | 0.47 | 0.57 | 0.29 | 0.36 | 0.35 | 0.47 |
| 8 | 28935001 | rs6999630 | 2.197 | -1.392 | 0.722 | -0.297 | -0.002 | 0.078 | 0.078 | 0 | 0.42 | 0.24 | 0.34 | 0.21 | 0.55 | 0.35 |
| 8 | 28935457 | rs7000444 | 1.329 | -1.418 | 0.822 | -0.283 | -0.013 | -0.014 | -0.014 | 0 | 0.62 | 0.44 | 0.44 | 0.27 | 0.69 | 0.53 |
| 8 | 28939010 | rs7462138 | 1.142 | -1.135 | 0.899 | -0.203 | -0.004 | -0.004 | -0.004 | 0 | 0.65 | 0.63 | 0.47 | 0.27 | 0.69 | 0.57 |
| 8 | 28941305 | rs1471939 | 0.608 | -1.191 | 0.858 | -0.229 | -0.017 | -0.001 | -0.001 | 0 | 0.71 | 0.69 | 0.53 | 0.26 | 0.79 | 0.6 |
| 8 | 28943734 | rs7816628 | 1.461 | -1.221 | 0.845 | -0.252 | 0.119 | 0.114 | 0.114 | 0 | 0.53 | 0.27 | 0.48 | 0.21 | 0.72 | 0.46 |
| 8 | 28954892 | rs7844537 | -1.038 | -1.194 | 0.873 | -0.093 | -0.027 | 0.215 | 0.215 | 0 | 0.9 | 0.77 | 0.87 | 0.57 | 0.94 | 0.84 |
| 8 | 28958517 | rs10091515 | 0.097 | -1.081 | 0.93 | -0.297 | -0.044 | -0.101 | -0.101 | 0 | 0.21 | 0.45 | 0.11 | 0.06 | 0.14 | 0.17 |
| 8 | 28960219 | rs9987365 | 2.218 | -0.848 | 0.578 | -0.405 | -0.045 | 0.077 | 0.077 | 0 | 0.4 | 0.19 | 0.32 | 0.2 | 0.49 | 0.38 |
| 8 | 28971558 | rs11778430 | -0.273 | -0.96 | 0.699 | -0.379 | -0.003 | -0.003 | -0.003 | 0 | 0.21 | 0.16 | 0.37 | 0.24 | 0.21 | 0.23 |
| 8 | 28975073 | rs936922 | 0.3 | -0.478 | 0.615 | -0.452 | 0.004 | 0.064 | 0.064 | 0 | 0.2 | 0.07 | 0.09 | 0.02 | 0.16 | 0.17 |
| 8 | 28977112 | rs7836759 | 1.113 | -0.503 | 0.769 | -0.469 | -0.001 | 0 | 0 | 0 | 0.59 | 0.61 | 0.68 | 0.45 | 0.64 | 0.6 |
| 8 | 28984042 | rs3793435 | 1.826 | -0.36 | 0.498 | -0.902 | 0.023 | 0.163 | 0.163 | 0 | 0.45 | 0.23 | 0.37 | 0.15 | 0.58 | 0.4 |
| 8 | 28984572 | rs3763566 | -0.818 | -0.419 | 0.633 | -0.816 | -0.016 | 0.076 | 0.076 | 0 | 0.85 | 0.77 | 0.84 | 0.4 | 0.87 | 0.76 |
| 8 | 28986357 | rs11786952 | 2.224 | -0.341 | 0.539 | -0.817 | -0.037 | 0.096 | 0.096 | 0 | 0.38 | 0.23 | 0.3 | 0.14 | 0.45 | 0.37 |
| 8 | 28992540 | rs6558096 | 0.05 | -0.143 | 0.751 | -0.603 | -0.018 | 0.024 | 0.024 | 0 | 0.19 | 0.23 | 0.37 | 0.24 | 0.15 | 0.2 |
| 8 | 29004822 | rs4732909 | 0.055 | -0.127 | 0.689 | -0.595 | -0.023 | 0.036 | 0.036 | 0 | 0.19 | 0.27 | 0.37 | 0.23 | 0.15 | 0.2 |
| 8 | 29006677 | rs6981956 | -0.62 | -0.087 | 0.619 | -0.478 | -0.009 | 0.052 | 0.052 | 0 | 0.85 | 0.8 | 0.84 | 0.41 | 0.87 | 0.76 |
| 8 | 29006708 | rs7002710 | 2.134 | 0.048 | 0.302 | -0.752 | -0.039 | 0.1 | 0.1 | 0 | 0.37 | 0.19 | 0.3 | 0.14 | 0.43 | 0.37 |
| 8 | 29017087 | rs6994211 | 1.086 | -0.17 | 0.535 | -0.586 | -0.002 | 0.032 | 0.032 | 0 | 0.65 | 0.5 | 0.71 | 0.41 | 0.67 | 0.62 |
| 8 | 29021362 | rs11778025 | 1.161 | -0.106 | 0.508 | -0.631 | 0.007 | 0.013 | 0.013 | 0 | 0.62 | 0.47 | 0.68 | 0.41 | 0.6 | 0.61 |
| 8 | 29025431 | rs876058 | 1.03 | -0.126 | 0.587 | -0.63 | 0.02 | -0.017 | -0.017 | 0 | 0.66 | 0.28 | 0.65 | 0.33 | 0.73 | 0.63 |
| 8 | 29027141 | rs4732912 | -0.698 | -0.154 | 0.798 | -0.662 | -0.018 | 0.111 | 0.111 | 0 | 0.85 | 0.75 | 0.84 | 0.43 | 0.89 | 0.81 |
| 8 | 29033258 | rs4461945 | 0.534 | -0.116 | 0.871 | -0.89 | -0.002 | 0.007 | 0.007 | 0 | 0.15 | 0.08 | 0.08 | 0.02 | 0.15 | 0.17 |
| 8 | 29041440 | rs10102538 | -0.606 | -0.124 | 0.88 | -0.888 | 0.019 | -0.065 | -0.065 | 0 | 0.22 | 0.25 | 0.16 | 0.02 | 0.3 | 0.2 |
| 8 | 29042453 | rs7007878 | -1.542 | -0.264 | 0.768 | -0.826 | -0.009 | 0.122 | 0.122 | 0 | 0.48 | 0.73 | 0.55 | 0.21 | 0.46 | 0.4 |
| 8 | 29048477 | rs1551479 | -0.387 | -0.121 | 1.096 | -0.149 | 0.001 | -0.049 | -0.049 | 0 | 0.17 | 0.28 | 0.18 | 0.09 | 0.16 | 0.18 |
| 8 | 29054302 | rs10448081 | 1.242 | -0.137 | 0.698 | -0.125 | 0.025 | -0.009 | -0.009 | 0.035 | 0.66 | 0.26 | 0.65 | 0.31 | 0.73 | 0.63 |
| 8 | 29060581 | rs6989058 | 1.091 | -0.11 | 0.605 | -0.134 | -0.004 | -0.001 | -0.001 | 0.21 | 0.65 | 0.39 | 0.65 | 0.31 | 0.71 | 0.6 |
| 8 | 29079558 | rs12682543 | 1.253 | -0.069 | 0.035 | -0.16 | 0.015 | 0.001 | 0.001 | 0 | 0.63 | 0.24 | 0.62 | 0.31 | 0.65 | 0.57 |
| 8 | 29082965 | rs10100409 | -0.624 | -0.213 | 0.24 | -0.279 | 0.01 | -0.029 | -0.029 | 0 | 0.12 | 0.14 | 0.1 | 0.01 | 0.17 | 0.1 |
| 8 | 29091635 | rs2925210 | -1.217 | -0.215 | 0.341 | -0.23 | 0.015 | -0.058 | -0.058 | 0 | 0.19 | 0.35 | 0.21 | 0.09 | 0.21 | 0.19 |
| 8 | 29100294 | rs1378703 | 0.527 | -0.292 | -0.216 | -0.039 | 0.009 | 0.003 | 0.003 | 0.02 | 0.07 | 0.04 | 0.11 | 0.07 | 0.05 | 0.1 |
| 8 | 29107388 | rs2954794 | -0.24 | -0.591 | -0.515 | -0.006 | -0.004 | -0.032 | -0.032 | 0 | 0.91 | 1 | 0.96 | 0.91 | 0.93 | 0.92 |
| 8 | 29109665 | rs17059855 | 0.392 | -0.478 | -0.135 | 0.01 | -0.003 | -0.001 | -0.001 | 0 | 0.08 | 0.09 | 0.11 | 0.08 | 0.05 | 0.11 |
| 8 | 29110154 | rs4732922 | -1.016 | -0.853 | 0.204 | -0.004 | -0.049 | 0.081 | 0.081 | 0.018 | 0.25 | 0.51 | 0.28 | 0.68 | 0.17 | 0.29 |
| 8 | 29124351 | rs7009885 | -0.055 | -0.571 | 0.202 | -0.292 | -0.001 | -0.006 | -0.006 | 0 | 0.11 | 0.09 | 0.15 | 0.07 | 0.1 | 0.1 |
| 8 | 29140616 | rs677485 | 0.68 | -0.694 | -0.299 | -0.652 | -0.049 | 0.077 | 0.077 | 0 | 0.14 | 0.01 | 0.1 | 0.01 | 0.16 | 0.1 |
| 8 | 29144515 | rs597695 | -1.701 | -0.899 | -0.078 | -0.643 | 0.011 | 0.03 | 0.03 | 0 | 0.48 | 0.63 | 0.49 | 0.67 | 0.44 | 0.44 |
| 8 | 29146988 | rs599531 | -1.18 | -0.873 | -0.033 | -0.603 | -0.001 | 0 | 0 | 0 | 0.38 | 0.59 | 0.36 | 0.61 | 0.33 | 0.33 |
| 8 | 29150284 | rs649959 | 0.617 | -0.84 | -0.078 | -0.638 | -0.003 | -0.001 | -0.001 | 0.2 | 0.16 | 0.12 | 0.11 | 0.01 | 0.17 | 0.09 |
| 8 | 29151173 | rs4732924 | -0.543 | -1.214 | 0.58 | 0.081 | -0.014 | 0.127 | 0.127 | 0 | 0.2 | 0.38 | 0.23 | 0.59 | 0.16 | 0.23 |
| 8 | 29153777 | rs643472 | 1.191 | -1.668 | 0.175 | -0.17 | -0.004 | -0.015 | -0.015 | 1.538 | 0.72 | 0.7 | 0.58 | 0.85 | 0.75 | 0.77 |

| | | | | | | | | | | | | | | | | |
|---|---|---|---|---|---|---|---|---|---|---|---|---|---|---|---|---|
| 8 | 29159473 | rs471214 | -1.376 | -1.324 | -0.123 | 0.049 | 0.011 | -0.007 | -0.007 | 0 | 0.37 | 0.65 | 0.36 | 0.62 | 0.31 | 0.3 |
| 8 | 29176919 | rs592382 | -1.675 | -1.235 | -0.34 | -0.233 | 0.047 | 0.065 | 0.065 | 0 | 0.46 | 0.73 | 0.59 | 0.71 | 0.39 | 0.46 |
| 8 | 29188937 | rs7812684 | -0.575 | -0.873 | -0.309 | -0.307 | -0.002 | -0.026 | -0.026 | 0 | 0.12 | 0.13 | 0.03 | 0 | 0.07 | 0.03 |
| 8 | 29198256 | rs2341674 | -0.892 | -1.063 | -0.29 | -0.389 | 0.009 | 0.01 | 0.01 | 0 | 0.22 | 0.2 | 0.21 | 0.04 | 0.17 | 0.15 |
| 8 | 29199062 | rs3824133 | -0.69 | -1.611 | -0.678 | 0.277 | -0.027 | -0.112 | -0.112 | 0.919 | 0.21 | 0.06 | 0.38 | 0.61 | 0.22 | 0.31 |
| 8 | 29205803 | rs7831728 | -0.594 | -1.848 | 0.324 | 0.07 | -0.025 | -0.061 | -0.061 | 0.031 | 0.26 | 0.37 | 0.16 | 0.04 | 0.15 | 0.17 |
| 8 | 29206126 | rs1455915 | -0.698 | -1.87 | 0.455 | 0.057 | -0.051 | -0.102 | -0.102 | 0.198 | 0.27 | 0.43 | 0.17 | 0.04 | 0.15 | 0.17 |
| 8 | 29207872 | rs2637819 | -1.039 | -1.834 | 0.461 | 0.088 | -0.003 | 0.057 | 0.057 | 0 | 0.24 | 0.32 | 0.41 | 0.61 | 0.24 | 0.3 |
| 8 | 29223864 | rs10108738 | -0.76 | -1.781 | 0.436 | 0.06 | -0.041 | -0.088 | -0.088 | 0.015 | 0.27 | 0.39 | 0.16 | 0.04 | 0.16 | 0.17 |
| 8 | 29228241 | rs10097270 | -0.995 | -1.754 | 0.395 | 0.05 | -0.011 | -0.07 | -0.07 | 0.045 | 0.32 | 0.39 | 0.2 | 0.04 | 0.24 | 0.21 |
| 8 | 29230135 | rs4732934 | -1.985 | -1.806 | 0.416 | 0.254 | 0.003 | 0.003 | 0.003 | 0 | 0.49 | 0.51 | 0.54 | 0.71 | 0.4 | 0.46 |
| 8 | 29231270 | rs734131 | -0.843 | -1.783 | -0.347 | 0.338 | 0.006 | -0.03 | -0.03 | 0 | 0.16 | 0.11 | 0.33 | 0.66 | 0.16 | 0.25 |
| 8 | 29231378 | rs746959 | -0.194 | -1.724 | -0.528 | 0.475 | 0.015 | -0.004 | -0.004 | 0.042 | 0.08 | 0.04 | 0.21 | 0.09 | 0.05 | 0.11 |
| 8 | 29235497 | rs2637822 | 1.399 | -1.779 | -0.568 | 0.452 | -0.006 | 0.001 | 0.001 | 0.002 | 0.55 | 0.51 | 0.61 | 0.43 | 0.6 | 0.58 |
| 8 | 29244053 | rs611302 | 0.622 | -1.668 | -0.478 | 0.137 | -0.003 | 0.04 | 0.04 | 0 | 0.71 | 0.57 | 0.75 | 0.39 | 0.72 | 0.66 |
| 8 | 29246441 | rs650026 | 0.694 | -1.259 | -0.456 | 0.348 | -0.011 | -0.001 | -0.001 | 0 | 0.59 | 0.52 | 0.64 | 0.39 | 0.66 | 0.65 |
| 8 | 29250546 | rs601962 | 0.601 | -0.843 | -1.395 | 0.592 | 0.1 | 0.066 | 0.066 | 0 | 0.53 | 0.25 | 0.47 | 0.31 | 0.69 | 0.52 |
| 8 | 29253667 | rs4732936 | -1.331 | 0.286 | -1.266 | 1.129 | -0.021 | -0.05 | -0.05 | 0 | 0.2 | 0.11 | 0.21 | 0.66 | 0.24 | 0.25 |
| 8 | 29253809 | rs4732937 | -0.962 | 0.059 | -1.218 | 1.243 | 0.002 | -0.042 | -0.042 | 0 | 0.16 | 0.1 | 0.15 | 0.68 | 0.15 | 0.21 |
| 8 | 29255584 | rs10503841 | -0.786 | 0.204 | -1.175 | 1.341 | -0.014 | 0.008 | 0.008 | 0.16 | 0.07 | 0.02 | 0.08 | 0.5 | 0.09 | 0.15 |
| 8 | 29266126 | rs653125 | -0.076 | -0.186 | -1.003 | -0.253 | 0.038 | 0.062 | 0.062 | 0 | 0.44 | 0.27 | 0.6 | 0.12 | 0.66 | 0.43 |
| 8 | 29272247 | rs2670683 | 0.036 | 0.532 | -0.833 | -0.028 | -0.025 | 0.03 | 0.03 | 3.048 | 0.37 | 0.3 | 0.5 | 0.21 | 0.48 | 0.39 |
| 8 | 29276435 | rs680913 | 0.4 | -0.392 | -0.313 | 0.055 | -0.038 | -0.043 | -0.043 | 0.168 | 0.77 | 0.65 | 0.89 | 0.9 | 0.85 | 0.79 |
| 8 | 29278868 | rs608321 | -0.901 | 0.5 | -0.695 | 0.467 | -0.017 | -0.012 | -0.012 | 0 | 0.07 | 0.04 | 0.08 | 0.16 | 0.11 | 0.11 |
| 8 | 29284161 | rs624461 | 0.123 | -0.718 | -0.855 | 0.391 | 0.024 | 0.003 | 0.003 | 2.91 | 0.82 | 0.86 | 0.95 | 0.83 | 0.94 | 0.84 |
| 8 | 29287001 | rs10110940 | -0.124 | -1.096 | -0.398 | -0.017 | -0.014 | -0.039 | -0.039 | 0 | 0.11 | 0.21 | 0.17 | 0.01 | 0.07 | 0.06 |
| 8 | 29291973 | rs6995249 | 1.457 | -0.432 | -1.179 | 0.046 | -0.075 | 0.022 | 0.022 | 3.603 | 0.51 | 0.05 | 0.56 | 0.59 | 0.62 | 0.49 |
| 8 | 29295712 | rs13278369 | 0.938 | -0.55 | 0.112 | 0.075 | 0.065 | 0.039 | 0.039 | 3.322 | 0.63 | 0.26 | 0.72 | 0.7 | 0.81 | 0.63 |
| 8 | 29300685 | rs11136066 | 0.32 | -0.313 | 0.078 | 0.043 | -0.034 | -0.006 | -0.006 | 0 | 0.09 | 0.04 | 0.12 | 0.11 | 0.18 | 0.14 |
| 8 | 29303526 | rs4732948 | -0.53 | -1.485 | 0.603 | 0.173 | 0.012 | 0.001 | 0.001 | 3.866 | 0.82 | 0.83 | 0.95 | 0.83 | 0.94 | 0.86 |
| 8 | 29304358 | rs13279037 | 0.465 | -1.548 | 0.789 | 0.596 | -0.04 | 0.015 | 0.015 | 6.439 | 0.73 | 0.4 | 0.88 | 0.65 | 0.88 | 0.73 |
| 8 | 29305308 | rs11136067 | 1.416 | -1.568 | 0.788 | 0.585 | 0.117 | -0.024 | -0.024 | 0 | 0.6 | 0.1 | 0.68 | 0.64 | 0.8 | 0.62 |
| 8 | 29305953 | rs2341803 | 0.427 | -1.425 | 0.995 | 0.67 | -0.061 | 0.031 | 0.031 | 6.099 | 0.74 | 0.47 | 0.88 | 0.65 | 0.88 | 0.73 |
| 8 | 29319875 | rs7834608 | -0.199 | -1.08 | 1.438 | 0.519 | 0.065 | 0.181 | 0.181 | 0.584 | 0.09 | 0.4 | 0.07 | 0.18 | 0.06 | 0.15 |
| 8 | 29321482 | rs6991662 | -0.618 | -1.024 | 0.73 | 0.459 | -0.017 | 0.013 | 0.013 | 0.309 | 0.18 | 0.4 | 0.22 | 0.18 | 0.13 | 0.22 |
| 8 | 29323625 | rs4732954 | 0.446 | -1.263 | 0.051 | 0.915 | -0.119 | 0.101 | 0.101 | 2.217 | 0.75 | 0.49 | 0.92 | 0.6 | 0.92 | 0.82 |
| 8 | 29327431 | rs11786009 | -0.213 | -0.951 | -0.692 | -0.017 | 0.014 | 0.031 | 0.031 | 0 | 0.27 | 0.18 | 0.12 | 0.07 | 0.25 | 0.28 |
| 8 | 29327924 | rs7823751 | -0.017 | -1.456 | -0.63 | -0.509 | -0.006 | -0.03 | -0.03 | 0.014 | 0.1 | 0.14 | 0.04 | 0 | 0.08 | 0.04 |
| 8 | 29331251 | rs2341797 | 0.713 | -1.638 | -0.845 | -0.735 | 0.052 | 0.093 | 0.093 | 0 | 0.48 | 0.23 | 0.75 | 0.69 | 0.6 | 0.7 |
| 8 | 29336008 | rs11987193 | 0.486 | -1.721 | -0.328 | -0.971 | -0.015 | -0.045 | -0.045 | 0 | 0.66 | 0.57 | 0.82 | 0.77 | 0.73 | 0.86 |
| 8 | 29339824 | rs4732956 | 0.223 | -1.668 | -0.48 | -0.862 | -0.014 | 0.034 | 0.034 | 0.268 | 0.23 | 0.07 | 0.13 | 0.2 | 0.2 | 0.08 |
| 8 | 29340948 | rs11777087 | 0.186 | -1.665 | -0.275 | -0.861 | 0.001 | 0.02 | 0.02 | 0.198 | 0.23 | 0.16 | 0.13 | 0.2 | 0.2 | 0.08 |
| 8 | 29344426 | rs11136072 | -0.569 | -1.754 | -0.243 | -0.83 | 0.01 | 0.004 | 0.004 | 0 | 0.81 | 0.86 | 0.93 | 0.89 | 0.85 | 0.72 |
| 8 | 29344462 | rs12548021 | 1.498 | -0.603 | -0.64 | -0.405 | -0.029 | -0.002 | -0.002 | 0 | 0.27 | 0.16 | 0.45 | 0.21 | 0.35 | 0.25 |
| 8 | 29345678 | rs7828717 | 0.434 | -0.47 | -0.174 | -0.296 | 0.002 | -0.021 | -0.021 | 0.018 | 0.22 | 0.35 | 0.15 | 0.21 | 0.19 | 0.09 |
| 8 | 29353675 | rs17552942 | 0.248 | -0.342 | -0.418 | 0.334 | -0.003 | 0.001 | 0.001 | 0 | 0.18 | 0.2 | 0.15 | 0.17 | 0.16 | 0.29 |
| 8 | 29353919 | rs4236681 | -0.465 | 0.67 | -0.823 | 0.822 | -0.033 | -0.043 | -0.043 | 0 | 0.86 | 0.91 | 0.73 | 0.63 | 0.78 | 0.65 |
| 8 | 29355430 | rs10106093 | 0.282 | 0.363 | -0.504 | 0.309 | -0.018 | 0.043 | 0.043 | 0 | 0.19 | 0.41 | 0.13 | 0.21 | 0.13 | 0.12 |
| 8 | 29355491 | rs10095540 | 1.352 | -0.004 | -0.956 | 0.106 | -0.035 | 0.091 | 0.091 | 0.009 | 0.42 | 0.22 | 0.45 | 0.25 | 0.47 | 0.21 |
| 8 | 29355530 | rs17060256 | 0.711 | -0.155 | -0.544 | 0.144 | -0.005 | 0.03 | 0.03 | 0 | 0.17 | 0.25 | 0.12 | 0.21 | 0.13 | 0.12 |
| 8 | 29359592 | rs4732961 | -0.748 | -0.33 | -0.801 | -0.023 | 0.008 | 0.004 | 0.004 | 0 | 0.91 | 0.9 | 0.98 | 1 | 0.98 | 0.95 |
| 8 | 29360305 | rs10091038 | -0.658 | 0.306 | -1.038 | -0.198 | -0.001 | -0.002 | -0.002 | 0 | 0.44 | 0.56 | 0.55 | 0.77 | 0.59 | 0.7 |
| 8 | 29361255 | rs10503848 | 0.2 | 0.724 | -1.427 | -0.09 | -0.004 | -0.01 | -0.01 | 0 | 0.3 | 0.25 | 0.33 | 0.57 | 0.36 | 0.58 |
| 8 | 29367054 | rs13277135 | 1.784 | 1.051 | -1.869 | -0.207 | -0.017 | 0.141 | 0.141 | 0.526 | 0.29 | 0.05 | 0.31 | 0.04 | 0.25 | 0.11 |
| 8 | 29370590 | rs7838475 | 0.276 | 1.225 | -1.341 | 0.351 | 0.015 | 0.003 | 0.003 | 0 | 0.11 | 0.15 | 0.21 | 0.21 | 0.23 | 0.1 |
| 8 | 29383034 | rs7012981 | 0.913 | 0.362 | -1.992 | 0.225 | -0.124 | -0.075 | -0.075 | 0 | 0.44 | 0.05 | 0.6 | 0.42 | 0.52 | 0.5 |
| 8 | 29387385 | rs3824106 | -0.256 | -0.002 | -2.263 | 0.156 | -0.071 | -0.12 | -0.12 | 1.075 | 0.27 | 0.08 | 0.19 | 0.44 | 0.29 | 0.19 |
| 8 | 29389615 | rs13250654 | -0.258 | -0.011 | -1.956 | 0.153 | -0.071 | -0.048 | -0.048 | 0 | 0.27 | 0.08 | 0.18 | 0.29 | 0.29 | 0.16 |
| 8 | 29392146 | rs17060323 | 0.306 | -0.291 | -1.265 | 0.06 | -0.012 | 0.088 | 0.088 | 0 | 0.11 | 0.21 | 0.12 | 0.27 | 0.09 | 0.22 |
| 8 | 29393711 | rs13255288 | -0.048 | -0.233 | -1.009 | 0.038 | -0.069 | -0.04 | -0.04 | 1.66 | 0.27 | 0.09 | 0.18 | 0.3 | 0.29 | 0.16 |
| 8 | 29394261 | rs4732686 | -0.083 | 0.112 | -0.932 | 0.034 | -0.022 | -0.017 | -0.017 | 0 | 0.3 | 0.23 | 0.39 | 0.39 | 0.41 | 0.48 |
| 8 | 29394468 | rs10503847 | 1.066 | 0.106 | -1.208 | 0.004 | -0.095 | -0.098 | -0.098 | 0 | 0.18 | 0.03 | 0.33 | 0.37 | 0.31 | 0.4 |
| 8 | 29394781 | rs7822830 | 1.085 | 0.241 | -1.024 | -0.516 | -0.005 | -0.017 | -0.017 | 0 | 0.12 | 0.14 | 0.06 | 0.02 | 0.1 | 0.08 |
| 8 | 29395511 | rs4295704 | 0.445 | 0.291 | -1.17 | -0.368 | -0.005 | 0.033 | 0.033 | 0.027 | 0.52 | 0.32 | 0.46 | 0.34 | 0.48 | 0.29 |
| 8 | 29396185 | rs11778860 | -0.414 | 0.75 | -1.214 | 0.312 | -0.054 | -0.031 | -0.031 | 0.154 | 0.22 | 0.14 | 0.24 | 0.3 | 0.33 | 0.24 |
| 8 | 29404084 | rs4732689 | 0.126 | 0.569 | -1.265 | -0.17 | -0.029 | 0.069 | 0.069 | 0 | 0.8 | 0.94 | 0.79 | 1 | 0.73 | 0.82 |
| 8 | 29404109 | rs4732690 | 0.574 | 0.526 | -1.737 | 0.114 | -0.071 | -0.056 | -0.056 | 0.43 | 0.25 | 0.1 | 0.43 | 0.37 | 0.34 | 0.46 |
| 8 | 29406768 | rs10503845 | 1.184 | -0.119 | -1.087 | 0.09 | 0.012 | 0.003 | 0.003 | 0 | 0.35 | 0.29 | 0.24 | 0.31 | 0.27 | 0.25 |
| 8 | 29409159 | rs13273498 | 1.26 | -0.051 | -0.781 | 0.38 | -0.001 | -0.014 | -0.014 | 0 | 0.11 | 0.08 | 0.07 | 0.32 | 0.12 | 0.08 |

| | | | | | | | | | | | | | | | | |
|---|---|---|---|---|---|---|---|---|---|---|---|---|---|---|---|---|
| 8 | 29409939 | rs12056634 | 0.724 | -0.061 | -0.882 | 0.275 | -0.073 | -0.033 | -0.033 | 0 | 0.25 | 0.1 | 0.43 | 0.32 | 0.34 | 0.45 |
| 8 | 29410072 | rs10099976 | -0.841 | -0.888 | -0.374 | 0.323 | -0.06 | 0.057 | 0.057 | 0 | 0.28 | 0.46 | 0.13 | 0.61 | 0.19 | 0.27 |
| 8 | 29412004 | rs4133117 | 0.999 | -0.293 | -0.742 | 0.6 | 0 | -0.02 | -0.02 | 3.225 | 0.12 | 0.07 | 0.08 | 0.36 | 0.12 | 0.07 |
| 8 | 29413436 | rs17060351 | 0.644 | 0.345 | -0.665 | 0.105 | -0.019 | -0.012 | -0.012 | 0 | 0.25 | 0.16 | 0.38 | 0.32 | 0.26 | 0.45 |
| 8 | 29416526 | rs6986919 | -0.027 | -0.277 | -0.139 | -0.24 | 0.013 | -0.057 | -0.057 | 0 | 0.79 | 0.65 | 0.78 | 0.94 | 0.76 | 0.86 |
| 8 | 29416684 | rs10503844 | 0.972 | -0.645 | -0.373 | -0.271 | 0.025 | 0.082 | 0.082 | 0 | 0.53 | 0.24 | 0.62 | 0.33 | 0.57 | 0.59 |
| 8 | 29419564 | rs10090180 | 0.251 | -0.519 | -0.499 | 0.019 | -0.007 | 0.027 | 0.027 | 0 | 0.17 | 0.22 | 0.13 | 0.61 | 0.13 | 0.22 |
| 8 | 29422893 | rs13250512 | 0.265 | -0.514 | -0.604 | 0.022 | 0 | 0.005 | 0.005 | 0.006 | 0.17 | 0.17 | 0.13 | 0.61 | 0.13 | 0.22 |
| 8 | 29427765 | rs17555250 | 0.805 | 0.191 | -1.047 | 0.007 | -0.14 | -0.037 | -0.037 | 0 | 0.26 | 0.02 | 0.42 | 0.31 | 0.38 | 0.49 |
| 8 | 29441006 | rs4259448 | 1.033 | -0.108 | -0.459 | 0.454 | 0 | 0.047 | 0.047 | 0.347 | 0.13 | 0.13 | 0.13 | 0.38 | 0.12 | 0.07 |
| 8 | 29441961 | rs9314379 | 0.303 | -0.492 | -0.311 | 0.623 | -0.019 | 0.052 | 0.052 | 0.093 | 0.12 | 0.16 | 0.05 | 0.24 | 0.08 | 0.23 |
| 8 | 29443591 | rs4732972 | 1.448 | -0.668 | -0.41 | -0.213 | 0.023 | 0.03 | 0.03 | 0.149 | 0.31 | 0.24 | 0.22 | 0.03 | 0.22 | 0.1 |
| 8 | 29449996 | rs17470763 | 0.763 | -0.378 | -0.521 | -0.238 | -0.033 | 0.043 | 0.043 | 0 | 0.09 | 0.02 | 0.13 | 0 | 0.14 | 0.11 |
| 8 | 29451004 | rs11996657 | 0.075 | -0.585 | -0.157 | -0.16 | -0.031 | 0.035 | 0.035 | 0 | 0.73 | 0.45 | 0.81 | 0.44 | 0.8 | 0.72 |
| 8 | 29451488 | rs12546462 | 0.13 | -0.434 | -0.48 | -0.113 | -0.024 | -0.019 | -0.019 | 0 | 0.41 | 0.11 | 0.59 | 0.42 | 0.57 | 0.62 |
| 8 | 29453239 | rs4732976 | 0.819 | -0.371 | -0.49 | -0.082 | -0.144 | -0.088 | -0.088 | 6.214 | 0.31 | 0.04 | 0.45 | 0.42 | 0.43 | 0.51 |
| 8 | 29454051 | rs6558124 | 0.421 | -0.505 | 0.163 | -0.01 | -0.041 | 0.043 | 0.043 | 0.25 | 0.13 | 0.26 | 0.05 | 0.18 | 0.08 | 0.21 |
| 8 | 29464437 | rs7821268 | -0.216 | -0.458 | 0.322 | 0.019 | -0.012 | -0.008 | -0.008 | 0.321 | 0.24 | 0.35 | 0.21 | 0.18 | 0.22 | 0.32 |
| 8 | 29469606 | rs6558129 | -0.202 | -0.575 | 0.172 | -0.071 | 0.002 | 0.022 | 0.022 | 0 | 0.56 | 0.65 | 0.42 | 0.2 | 0.46 | 0.43 |
| 8 | 29470596 | rs6994244 | -0.209 | -0.573 | 0.169 | -0.069 | 0.003 | 0.026 | 0.026 | 0 | 0.56 | 0.65 | 0.41 | 0.2 | 0.46 | 0.43 |
| 8 | 29473431 | rs4732980 | 0.673 | -0.311 | -0.259 | 0.223 | 0.023 | -0.01 | -0.01 | 0 | 0.62 | 0.27 | 0.68 | 0.44 | 0.66 | 0.59 |
| 8 | 29485927 | rs4732986 | 0.861 | -0.187 | -0.914 | -0.035 | -0.117 | -0.157 | -0.157 | 0.101 | 0.3 | 0.04 | 0.46 | 0.49 | 0.4 | 0.49 |
| 8 | 29489388 | rs12682644 | 0.084 | -0.336 | -0.414 | -0.194 | 0.017 | 0.012 | 0.012 | 0 | 0.21 | 0.35 | 0.21 | 0.21 | 0.24 | 0.27 |
| 8 | 29491696 | rs12675660 | -0.236 | -0.397 | -0.655 | -0.211 | 0.005 | -0.006 | -0.006 | 0 | 0.25 | 0.54 | 0.23 | 0.21 | 0.26 | 0.3 |
| 8 | 29494941 | rs4732987 | 1.092 | -0.33 | -0.925 | -0.208 | -0.115 | -0.157 | -0.157 | 0 | 0.3 | 0.04 | 0.46 | 0.48 | 0.39 | 0.48 |
| 8 | 29502909 | rs4073220 | -1.283 | -0.955 | -0.878 | -0.744 | -0.017 | 0.173 | 0.173 | 0.233 | 0.71 | 0.96 | 0.82 | 0.94 | 0.77 | 0.9 |
| 8 | 29505165 | rs7465364 | -0.311 | -0.682 | -0.38 | -0.656 | 0.038 | 0.059 | 0.059 | 3.658 | 0.54 | 0.61 | 0.69 | 0.7 | 0.65 | 0.78 |
| 8 | 29527339 | rs12546747 | -0.091 | -0.684 | -0.482 | -0.69 | 0.024 | 0.035 | 0.035 | 5.619 | 0.51 | 0.53 | 0.69 | 0.71 | 0.65 | 0.76 |
| 8 | 29536211 | rs4732990 | 0.14 | -0.926 | -0.761 | -1.298 | -0.029 | -0.106 | -0.106 | 0 | 0.8 | 0.62 | 0.83 | 1 | 0.87 | 0.87 |
| 8 | 29536945 | rs7460831 | -0.5 | -0.912 | -0.797 | -1.255 | -0.017 | -0.08 | -0.08 | 1.224 | 0.84 | 0.72 | 0.85 | 1 | 0.9 | 0.9 |
| 8 | 29537217 | rs11776937 | 0.081 | -1.18 | -1.095 | -1.529 | 0.04 | 0.026 | 0.026 | 0.133 | 0.6 | 0.28 | 0.76 | 1 | 0.76 | 0.82 |
| 8 | 29540751 | rs7839112 | -0.333 | -0.391 | -0.944 | -0.624 | 0.002 | 0.002 | 0.002 | 0 | 0.24 | 0.25 | 0.36 | 0.33 | 0.27 | 0.4 |
| 8 | 29541902 | rs7002486 | -0.863 | -0.474 | -0.902 | -0.493 | -0.012 | -0.01 | -0.01 | 0 | 0.63 | 0.66 | 0.5 | 0.57 | 0.47 | 0.57 |
| 8 | 29542521 | rs7016719 | 0.088 | -0.61 | -1.149 | -0.356 | -0.01 | -0.027 | -0.027 | 0 | 0.19 | 0.09 | 0.24 | 0.36 | 0.2 | 0.31 |
| 8 | 29545470 | rs10112805 | 0.584 | -1.803 | -0.958 | -0.673 | 0.04 | 0.022 | 0.022 | 0.016 | 0.3 | 0.19 | 0.18 | 0.19 | 0.18 | 0.22 |
| 8 | 29551058 | rs7459435 | 0.73 | -1.734 | -0.713 | -0.579 | 0.061 | 0.041 | 0.041 | 0.4 | 0.31 | 0.2 | 0.13 | 0.2 | 0.15 | 0.21 |
| 8 | 29552383 | rs7003746 | -0.818 | -1.799 | -0.396 | -0.52 | 0.014 | 0.01 | 0.01 | 3.163 | 0.59 | 0.71 | 0.43 | 0.35 | 0.37 | 0.58 |
| 8 | 29552712 | rs7009155 | 0.107 | -1.717 | -0.128 | -0.488 | -0.031 | -0.032 | -0.032 | 0 | 0.25 | 0.36 | 0.26 | 0.14 | 0.19 | 0.3 |
| 8 | 29558453 | rs10283295 | 0.124 | -1.684 | -0.1 | -0.471 | -0.012 | -0.014 | -0.014 | 0.014 | 0.26 | 0.27 | 0.27 | 0.14 | 0.21 | 0.32 |
| 8 | 29569127 | rs12216758 | 0.313 | -1.91 | -0.164 | -0.206 | 0.025 | 0.023 | 0.023 | 0 | 0.32 | 0.28 | 0.16 | 0.2 | 0.18 | 0.26 |
| 8 | 29587962 | rs10435648 | -0.051 | -1.551 | -0.293 | -0.07 | 0.008 | 0.003 | 0.003 | 0.342 | 0.16 | 0.29 | 0.15 | 0.2 | 0.13 | 0.14 |
| 8 | 29588279 | rs6993885 | -0.554 | -1.6 | -0.949 | -0.267 | 0.042 | 0.063 | 0.063 | 0.106 | 0.68 | 0.82 | 0.73 | 0.82 | 0.78 | 0.74 |
| 8 | 29598009 | rs12676320 | -1.802 | -1.499 | 0.155 | 0.454 | -0.033 | 0 | 0 | 0 | 0.39 | 0.47 | 0.39 | 0.39 | 0.26 | 0.42 |
| 8 | 29598297 | rs12678225 | -1.187 | -1.509 | -0.447 | 0.211 | 0.008 | -0.001 | -0.001 | 0 | 0.22 | 0.18 | 0.25 | 0.19 | 0.13 | 0.3 |
| 8 | 29598591 | rs12682519 | -1.843 | -1.459 | -0.315 | 0.259 | 0.013 | 0.005 | 0.005 | 0 | 0.35 | 0.3 | 0.38 | 0.33 | 0.26 | 0.41 |
| 8 | 29600080 | rs7006016 | -0.236 | -1.15 | -0.02 | 0.553 | 0.001 | 0.003 | 0.003 | 0 | 0.12 | 0.12 | 0.12 | 0.14 | 0.12 | 0.09 |
| 8 | 29603201 | rs6983144 | -0.929 | -1.051 | -0.124 | 0.609 | -0.013 | 0.016 | 0.016 | 0 | 0.14 | 0.18 | 0.18 | 0.19 | 0.09 | 0.25 |
| 8 | 29608755 | rs7836482 | -1.041 | -0.929 | 0.117 | 0.635 | -0.017 | 0.088 | 0.088 | 0 | 0.26 | 0.55 | 0.22 | 0.6 | 0.22 | 0.35 |
| 8 | 29613316 | rs7388052 | 0.081 | -0.734 | -0.225 | 0.179 | 0 | -0.031 | -0.031 | 0 | 0.82 | 0.81 | 0.74 | 0.98 | 0.84 | 0.81 |
| 8 | 29616242 | rs6983033 | -0.391 | -1.115 | -0.665 | 0.323 | 0.019 | -0.086 | -0.086 | 0 | 0.19 | 0.08 | 0.17 | 0.63 | 0.14 | 0.24 |
| 8 | 29619600 | rs6558150 | 1.09 | -1.16 | -0.17 | 0.241 | -0.002 | 0 | 0 | 0.654 | 0.61 | 0.43 | 0.55 | 0.35 | 0.69 | 0.57 |
| 8 | 29630942 | rs6982080 | -1.076 | -1.475 | -0.921 | -0.351 | 0.049 | 0.147 | 0.147 | 0.462 | 0.43 | 0.79 | 0.4 | 0.2 | 0.31 | 0.31 |
| 8 | 29631661 | rs3923822 | -0.207 | -1.135 | -1.018 | -0.081 | -0.001 | -0.001 | -0.001 | 0 | 0.21 | 0.23 | 0.16 | 0.23 | 0.2 | 0.15 |
| 8 | 29633694 | rs6980701 | -0.809 | -0.895 | -0.924 | -0.317 | 0.022 | 0.046 | 0.046 | 0 | 0.4 | 0.69 | 0.35 | 0.22 | 0.27 | 0.39 |
| 8 | 29635910 | rs7813195 | -0.614 | -0.446 | -0.704 | 0.17 | -0.011 | -0.001 | -0.001 | 0 | 0.38 | 0.59 | 0.34 | 0.4 | 0.26 | 0.34 |
| 8 | 29639419 | rs7827693 | -0.197 | 0.482 | -0.614 | 0.004 | 0.001 | -0.027 | -0.027 | 0 | 0.45 | 0.39 | 0.38 | 0.11 | 0.45 | 0.39 |
| 8 | 29648515 | rs9694300 | 0.603 | 0.688 | -0.17 | 1.5 | 0.046 | 0.196 | 0.196 | 0.007 | 0.94 | 0.77 | 0.79 | 0.7 | 0.94 | 0.91 |
| 8 | 29671207 | rs10090311 | -0.875 | 0.447 | -1.706 | 0.913 | -0.142 | 0.009 | 0.009 | 0.042 | 0.56 | 0.96 | 0.41 | 0.67 | 0.5 | 0.61 |
| 8 | 29676487 | rs4130873 | -0.459 | 0.157 | -0.816 | 0.807 | 0.021 | 0.013 | 0.013 | 0.972 | 0.42 | 0.72 | 0.33 | 0.62 | 0.36 | 0.51 |
| 8 | 29679740 | rs7387323 | 0.118 | 0.2 | -0.789 | 0.705 | -0.004 | -0.028 | -0.028 | 0 | 0.85 | 0.77 | 0.91 | 0.95 | 0.86 | 0.92 |
| 8 | 29682635 | rs7459662 | -0.464 | -0.532 | 0.035 | 1.779 | 0 | 0 | 0 | 0 | 0.36 | 0.61 | 0.3 | 0.32 | 0.35 | 0.32 |
| 8 | 29687858 | rs6989695 | -0.761 | -0.642 | 0.123 | 2.002 | -0.002 | 0 | 0 | 0.996 | 0.37 | 0.56 | 0.3 | 0.4 | 0.35 | 0.37 |
| 8 | 29691214 | rs11989810 | -0.557 | -0.724 | 0.524 | 1.915 | -0.015 | 0.032 | 0.032 | 0.116 | 0.24 | 0.37 | 0.21 | 0.38 | 0.2 | 0.28 |
| 8 | 29692897 | rs11774732 | -1.219 | -0.747 | 0.544 | 1.902 | -0.002 | 0.011 | 0.011 | 0.41 | 0.35 | 0.45 | 0.3 | 0.4 | 0.34 | 0.37 |
| 8 | 29694470 | rs13278786 | -1.697 | -0.791 | 0.506 | 2.083 | 0.02 | -0.035 | -0.035 | 0 | 0.42 | 0.76 | 0.34 | 0.52 | 0.37 | 0.48 |
| 8 | 29697805 | rs13270786 | 0.217 | -0.709 | 0.445 | 1.987 | 0 | -0.002 | -0.002 | 0 | 0.11 | 0.08 | 0.09 | 0.02 | 0.15 | 0.08 |
| 8 | 29702442 | rs6992654 | -0.361 | -0.771 | 0.466 | 2.053 | -0.006 | 0.004 | 0.004 | 0.289 | 0.23 | 0.35 | 0.21 | 0.28 | 0.2 | 0.22 |
| 8 | 29707868 | rs13275481 | -0.624 | -0.805 | 0.244 | 1.859 | -0.001 | 0 | 0 | 0 | 0.14 | 0.12 | 0.1 | 0.12 | 0.16 | 0.14 |
| 8 | 29718019 | rs1545405 | -0.669 | -0.75 | 0.22 | 1.758 | -0.013 | 0.086 | 0.086 | 1.43 | 0.24 | 0.58 | 0.21 | 0.47 | 0.2 | 0.26 |

| | | | | | | | | | | | | | | | | |
|---|---|---|---|---|---|---|---|---|---|---|---|---|---|---|---|---|
| 8 | 29731466 | rs7818468 | -1.32 | -0.804 | -0.2 | 1.616 | 0.01 | -0.004 | -0.004 | 0 | 0.41 | 0.77 | 0.35 | 0.61 | 0.38 | 0.44 |
| 8 | 29734572 | rs7814169 | 1.025 | -1.125 | 0.272 | 1.681 | -0.004 | -0.002 | -0.002 | 0.876 | 0.69 | 0.63 | 0.76 | 0.74 | 0.71 | 0.67 |
| 8 | 29740955 | rs10808343 | 0.625 | -1.155 | 0.611 | 1.837 | 0.004 | 0.184 | 0.184 | 0.001 | 0.81 | 0.52 | 0.81 | 0.51 | 0.8 | 0.73 |
| 8 | 29741748 | rs12547428 | 1.774 | -1.47 | -0.293 | 1.62 | 0.059 | 0.045 | 0.045 | 0.427 | 0.51 | 0.06 | 0.53 | 0.33 | 0.62 | 0.5 |
| 8 | 29742361 | rs6468351 | -1.044 | -0.789 | 0.556 | 1.992 | -0.004 | 0.139 | 0.139 | 0 | 0.2 | 0.34 | 0.18 | 0.49 | 0.2 | 0.27 |
| 8 | 29743305 | rs7387246 | 0.504 | -0.83 | 0.608 | 1.905 | -0.006 | 0.021 | 0.021 | 0 | 0.89 | 0.86 | 0.92 | 0.88 | 0.88 | 0.91 |
| 8 | 29743361 | rs7013957 | -1.25 | -0.804 | 1.097 | 2.2 | -0.011 | 0.056 | 0.056 | 0.24 | 0.27 | 0.59 | 0.25 | 0.57 | 0.23 | 0.29 |
| 8 | 29744518 | rs6981562 | -1.754 | -0.977 | 1.509 | 2.189 | -0.022 | 0.021 | 0.021 | 0.011 | 0.3 | 0.65 | 0.18 | 0.63 | 0.28 | 0.41 |
| 8 | 29745193 | rs2122068 | -2.03 | -1.148 | 1.425 | 2.538 | -0.021 | 0.029 | 0.029 | 0 | 0.24 | 0.62 | 0.19 | 0.37 | 0.24 | 0.26 |
| 8 | 29754597 | rs13267681 | 2.559 | -1.28 | 1.52 | 2.596 | 0.02 | -0.022 | -0.022 | 0 | 0.65 | 0.25 | 0.78 | 0.56 | 0.73 | 0.57 |
| 8 | 29760101 | rs7386992 | 1.347 | -1.372 | 1.562 | 2.54 | 0 | 0.004 | 0.004 | 0 | 0.85 | 0.87 | 0.92 | 0.74 | 0.92 | 0.8 |
| 8 | 29780940 | rs7005253 | -1.014 | -1.305 | 2.527 | 2.566 | -0.013 | -0.028 | -0.028 | 0 | 0.16 | 0.31 | 0.07 | 0.1 | 0.14 | 0.2 |
| 8 | 29786701 | rs6468011 | 1.568 | -1.364 | 2.731 | 2.488 | -0.026 | 0.043 | 0.043 | 0 | 0.8 | 0.36 | 0.85 | 0.62 | 0.82 | 0.77 |
| 8 | 29792105 | rs10503858 | -0.785 | -1.329 | 2.469 | 2.489 | -0.003 | -0.044 | -0.044 | 0 | 0.07 | 0.01 | 0.05 | 0.23 | 0.05 | 0.06 |
| 8 | 29799481 | rs7006983 | -0.368 | -1.347 | 2.445 | 2.599 | -0.001 | -0.033 | -0.033 | 0.012 | 0.07 | 0.02 | 0.02 | 0.2 | 0.04 | 0.16 |
| 8 | 29801968 | rs2762077 | 2.381 | -1.492 | 2.546 | 2.544 | 0.069 | -0.008 | -0.008 | 0 | 0.64 | 0.17 | 0.78 | 0.37 | 0.76 | 0.55 |
| 8 | 29802727 | rs10101117 | -2.242 | -1.421 | 2.278 | 2.1 | -0.052 | 0.034 | 0.034 | 0 | 0.28 | 0.58 | 0.18 | 0.44 | 0.2 | 0.28 |
| 8 | 29811542 | rs6997575 | -1.153 | -1.336 | 1.976 | 1.923 | -0.016 | -0.015 | -0.015 | 0.753 | 0.18 | 0.29 | 0.05 | 0.14 | 0.13 | 0.21 |
| 8 | 29812451 | rs16876237 | -1.397 | -1.307 | 1.565 | 1.967 | -0.048 | -0.007 | -0.007 | 0 | 0.19 | 0.42 | 0.12 | 0.2 | 0.12 | 0.22 |
| 8 | 29814414 | rs755926 | -0.89 | -1.395 | 1.251 | 1.954 | -0.009 | -0.005 | -0.005 | 0 | 0.17 | 0.22 | 0.05 | 0.14 | 0.11 | 0.21 |
| 8 | 29818527 | rs7838594 | -0.287 | -1.329 | 0.656 | 2.127 | -0.002 | -0.039 | -0.039 | 0.196 | 0.07 | 0.02 | 0.02 | 0.16 | 0.05 | 0.15 |
| 8 | 29830431 | rs6468038 | -2.214 | -1.352 | 0.667 | 2.018 | -0.004 | 0.009 | 0.009 | 0 | 0.22 | 0.25 | 0.16 | 0.43 | 0.15 | 0.22 |
| 8 | 29830855 | rs7845816 | -1.44 | -1.34 | 0.703 | 1.885 | -0.023 | 0.074 | 0.074 | 0 | 0.11 | 0.23 | 0.11 | 0.31 | 0.07 | 0.06 |
| 8 | 29831265 | rs10096447 | -2.419 | -1.42 | 0.737 | 1.856 | -0.046 | 0.028 | 0.028 | 0.039 | 0.37 | 0.46 | 0.16 | 0.51 | 0.26 | 0.43 |
| 8 | 29834020 | rs7821925 | -1.933 | -1.397 | 0.704 | 1.831 | -0.043 | 0.001 | 0.001 | 0.01 | 0.3 | 0.41 | 0.11 | 0.28 | 0.19 | 0.37 |
| 8 | 29836592 | rs10108668 | -1.173 | -1.068 | 1.104 | 2.05 | -0.056 | 0.036 | 0.036 | 0 | 0.18 | 0.34 | 0.06 | 0.23 | 0.11 | 0.23 |
| 8 | 29838800 | rs16876298 | -1.318 | -0.263 | 0.489 | 1.688 | -0.005 | 0.012 | 0.012 | 0 | 0.1 | 0.14 | 0.06 | 0.23 | 0.08 | 0.06 |
| 8 | 29840089 | rs10503860 | -1.622 | -0.226 | 0.434 | 1.622 | -0.025 | 0.068 | 0.068 | 0 | 0.13 | 0.28 | 0.13 | 0.29 | 0.08 | 0.07 |
| 8 | 29846153 | rs2341175 | 1.148 | 0.65 | -0.272 | 0.522 | 0.003 | 0.13 | 0.13 | 0.008 | 0.62 | 0.26 | 0.74 | 0.97 | 0.62 | 0.76 |
| 8 | 29848666 | rs309843 | -0.609 | 0.799 | -0.532 | 0.374 | -0.054 | 0.062 | 0.062 | 0.031 | 0.16 | 0.02 | 0.12 | 0.03 | 0.21 | 0.1 |
| 8 | 29850729 | rs6987138 | 1.665 | 1.138 | 0.132 | 1.507 | -0.007 | -0.013 | -0.013 | 0 | 0.57 | 0.36 | 0.67 | 0.5 | 0.55 | 0.65 |
| 8 | 29860405 | rs2595042 | -1.868 | 1.219 | 0.829 | 1.594 | -0.049 | 0.023 | 0.023 | 0 | 0.44 | 0.82 | 0.61 | 0.64 | 0.48 | 0.45 |
| 8 | 29873603 | rs309853 | -1.376 | 0.685 | 1.307 | 1.925 | -0.003 | 0.032 | 0.032 | 0 | 0.28 | 0.52 | 0.27 | 0.32 | 0.27 | 0.15 |
| 8 | 29875964 | rs4107736 | 2.169 | 0.453 | 0.732 | 2.198 | -0.057 | 0.184 | 0.184 | 0.033 | 0.47 | 0.05 | 0.53 | 0.2 | 0.53 | 0.38 |
| 8 | 29876605 | rs9297181 | -2.041 | 0.285 | 1.813 | 2.38 | -0.003 | 0.063 | 0.063 | 0.421 | 0.19 | 0.33 | 0.15 | 0.29 | 0.19 | 0.16 |
| 8 | 29877729 | rs991124 | -1.518 | 0.346 | 1.848 | 2.416 | 0.048 | 0.327 | 0.327 | 0.647 | 0.07 | 0.31 | 0.09 | 0.28 | 0.06 | 0.04 |
| 8 | 29883905 | rs10086650 | -2.073 | 0.359 | 1.815 | 2.414 | 0 | 0.064 | 0.064 | 0.52 | 0.19 | 0.34 | 0.14 | 0.3 | 0.19 | 0.16 |
| 8 | 29885706 | rs17489292 | -0.892 | 0.495 | 1.125 | 1.744 | -0.015 | 0.056 | 0.056 | 0 | 0.12 | 0.03 | 0.05 | 0 | 0.14 | 0.12 |
| 8 | 29894450 | rs7011477 | 1.557 | 0.339 | 1.886 | 1.843 | 0.008 | -0.034 | -0.034 | 0.072 | 0.93 | 0.82 | 0.97 | 0.98 | 0.95 | 0.91 |
| 8 | 29905051 | rs16876366 | -1.295 | 0.172 | 2.147 | 1.954 | -0.016 | -0.007 | -0.007 | 0 | 0.08 | 0.1 | 0.05 | 0.09 | 0.05 | 0.04 |
| 8 | 29910706 | rs2175408 | -0.888 | 0.172 | 2.726 | 2.964 | 0.001 | 0.036 | 0.036 | 0 | 0.17 | 0.15 | 0.14 | 0.36 | 0.16 | 0.32 |
| 8 | 29918642 | rs9656754 | -2.169 | 0.222 | 3.561 | 3.472 | 0.004 | 0.025 | 0.025 | 0.013 | 0.1 | 0.11 | 0.09 | 0.2 | 0.12 | 0.13 |
| 8 | 29918902 | rs9656755 | -1.933 | 0.24 | 3.56 | 3.473 | -0.003 | -0.001 | -0.001 | 0.044 | 0.09 | 0.05 | 0.09 | 0.2 | 0.12 | 0.13 |
| 8 | 29920825 | rs9130 | 1.482 | 0.098 | 3.699 | 3.425 | -0.042 | 0.043 | 0.043 | 0.667 | 0.89 | 0.74 | 0.88 | 0.85 | 0.94 | 0.95 |
| 8 | 29932284 | rs11777119 | 2.726 | 0.089 | 3.71 | 3.466 | -0.036 | 0.127 | 0.127 | 0 | 0.79 | 0.48 | 0.79 | 0.64 | 0.82 | 0.81 |
| 8 | 29937404 | rs10090988 | 2.629 | 0.11 | 3.714 | 3.413 | -0.03 | 0.086 | 0.086 | 0.006 | 0.8 | 0.48 | 0.79 | 0.74 | 0.83 | 0.85 |
| 8 | 29955587 | rs7845133 | 2.809 | 0.049 | 3.895 | 3.583 | -0.034 | 0.065 | 0.065 | 0.473 | 0.81 | 0.39 | 0.81 | 0.7 | 0.85 | 0.86 |
| 8 | 29956313 | rs4733354 | 1.301 | 0.046 | 4.043 | 3.552 | -0.026 | -0.051 | -0.051 | 0 | 0.91 | 0.75 | 0.94 | 1 | 0.96 | 0.99 |
| 8 | 29957664 | rs16876445 | -1.761 | 0.047 | 4.038 | 3.563 | 0 | 0.017 | 0.017 | 0 | 0.07 | 0.05 | 0.07 | 0.23 | 0.09 | 0.12 |
| 8 | 29957816 | rs12386822 | 2.48 | 0.192 | 4.1 | 3.719 | 0.011 | 0.016 | 0.016 | 0.147 | 0.64 | 0.26 | 0.7 | 0.41 | 0.68 | 0.56 |
| 8 | 29961951 | rs11557703 | -2.111 | 0.071 | 3.922 | 3.466 | 0.003 | 0.04 | 0.04 | 0 | 0.07 | 0.08 | 0.07 | 0.23 | 0.09 | 0.12 |
| 8 | 29965403 | rs2657 | 1.519 | 0.03 | 3.947 | 3.237 | -0.025 | -0.051 | -0.051 | 0 | 0.91 | 0.72 | 0.94 | 1 | 0.96 | 0.99 |
| 8 | 29965519 | rs2656 | 1.622 | 0.002 | 3.927 | 3.223 | -0.038 | -0.062 | -0.062 | 0 | 0.9 | 0.72 | 0.94 | 1 | 0.96 | 0.99 |
| 8 | 29969029 | rs11984896 | 3.067 | 0.109 | 3.842 | 3.298 | -0.034 | 0.061 | 0.061 | 0.473 | 0.81 | 0.39 | 0.81 | 0.71 | 0.85 | 0.86 |
| 8 | 29971290 | rs16876489 | -2.742 | 0.089 | 3.839 | 3.284 | 0.006 | 0.069 | 0.069 | 0.458 | 0.09 | 0.19 | 0.07 | 0.23 | 0.09 | 0.12 |
| 8 | 29973421 | rs12682122 | 1.806 | 0.048 | 3.611 | 3.27 | -0.019 | 0.06 | 0.06 | 0.653 | 0.9 | 0.58 | 0.88 | 0.94 | 0.95 | 0.98 |
| 8 | 29982518 | rs12681963 | 1.868 | 0.089 | 3.125 | 3.112 | -0.021 | 0.026 | 0.026 | 0 | 0.84 | 0.7 | 0.88 | 0.76 | 0.87 | 0.87 |
| 8 | 29996824 | rs7838088 | 2.706 | 0.284 | 3.204 | 3.442 | 0.006 | 0.241 | 0.241 | 0.38 | 0.88 | 0.56 | 0.84 | 0.71 | 0.91 | 0.93 |
| 8 | 30000346 | rs12334730 | 2.948 | 0.263 | 3.204 | 3.411 | 0.028 | 0.305 | 0.305 | 1.792 | 0.92 | 0.46 | 0.86 | 0.72 | 0.94 | 0.93 |
| 8 | 30004493 | rs2341177 | 2.517 | 0.264 | 3.245 | 3.427 | 0.007 | 0.12 | 0.12 | 0.175 | 0.89 | 0.64 | 0.89 | 0.77 | 0.92 | 0.94 |
| 8 | 30012514 | rs4733400 | 2.798 | 0.266 | 2.948 | 3.416 | -0.015 | 0.157 | 0.157 | 0.768 | 0.88 | 0.42 | 0.83 | 0.72 | 0.91 | 0.93 |
| 8 | 30041916 | rs12548744 | -3.226 | 0.316 | 2.5 | 3.895 | -0.02 | 0.25 | 0.25 | 0 | 0.13 | 0.29 | 0.24 | 0.48 | 0.1 | 0.19 |
| 8 | 30042658 | rs2242457 | -0.697 | 0.474 | 2.305 | 4.166 | -0.043 | -0.071 | -0.071 | 0 | 0.14 | 0.03 | 0.11 | 0.27 | 0.19 | 0.26 |
| 8 | 30062814 | rs7817838 | -2.592 | 0.477 | 2.316 | 4.161 | -0.009 | 0.062 | 0.062 | 0 | 0.1 | 0.16 | 0.1 | 0.28 | 0.09 | 0.14 |
| 8 | 30068583 | rs2958751 | -1.058 | 0.491 | 2.316 | 4.179 | 0.008 | 0.066 | 0.066 | 1.062 | 0.16 | 0.19 | 0.21 | 0.42 | 0.19 | 0.29 |
| 8 | 30073503 | rs2860330 | -0.663 | 0.586 | 1.859 | 4.229 | -0.045 | -0.064 | -0.064 | 0 | 0.14 | 0.03 | 0.12 | 0.23 | 0.19 | 0.26 |
| 8 | 30080900 | rs6987718 | -2.757 | 0.037 | 2.246 | 3.906 | -0.01 | 0.12 | 0.12 | 0 | 0.07 | 0.13 | 0.11 | 0.32 | 0.06 | 0.14 |
| 8 | 30099822 | rs12677295 | -2.577 | 0.05 | 2.144 | 3.885 | -0.007 | 0.1 | 0.1 | 0 | 0.09 | 0.14 | 0.15 | 0.32 | 0.08 | 0.08 |
| 8 | 30112318 | rs16876741 | -2.173 | 0.121 | 2.119 | 3.869 | -0.006 | 0.181 | 0.181 | 0.014 | 0.05 | 0.15 | 0.11 | 0.29 | 0.05 | 0.08 |

| | | | | | | | | | | | | | | | | |
|---|---|---|---|---|---|---|---|---|---|---|---|---|---|---|---|---|
| 8 | 30126708 | rs7822547 | -2.009 | 0.178 | 1.896 | 3.194 | -0.005 | 0.126 | 0.126 | 0 | 0.07 | 0.22 | 0.08 | 0.23 | 0.05 | 0.12 |
| 8 | 30131119 | rs7008022 | -1.88 | 0.137 | 2.052 | 2.61 | 0.05 | 0.29 | 0.29 | 0 | 0.06 | 0.32 | 0.11 | 0.25 | 0.03 | 0.11 |
| 8 | 30146429 | rs12542640 | -1.776 | 0.026 | 1.878 | 2.364 | -0.016 | 0.129 | 0.129 | 0.778 | 0.12 | 0.57 | 0.2 | 0.24 | 0.08 | 0.2 |
| 8 | 30162683 | rs6468273 | 0.308 | -0.362 | 0.149 | 2.145 | 0.026 | -0.063 | -0.063 | 0 | 0.73 | 0.85 | 0.64 | 0.54 | 0.75 | 0.61 |
| 8 | 30166202 | rs12679495 | -0.675 | -0.582 | -0.496 | 2.106 | -0.027 | -0.105 | -0.105 | 0 | 0.23 | 0.02 | 0.33 | 0.34 | 0.23 | 0.22 |
| 8 | 30168589 | rs9297225 | -0.369 | 0.283 | -1.527 | 1.286 | -0.114 | -0.039 | -0.039 | 0 | 0.77 | 0.98 | 0.74 | 0.79 | 0.69 | 0.75 |
| 8 | 30169122 | rs4733476 | -1.028 | -0.215 | -1.255 | 1.124 | 0.03 | 0 | 0 | 1.429 | 0.44 | 0.79 | 0.55 | 0.53 | 0.38 | 0.43 |
| 8 | 30169551 | rs4584112 | -0.494 | -0.206 | -0.507 | 0.543 | -0.018 | -0.062 | -0.062 | 0 | 0.14 | 0.24 | 0.09 | 0.01 | 0.09 | 0.06 |
| 8 | 30171254 | rs10454342 | -0.481 | -0.475 | -0.61 | 0.624 | -0.004 | 0.002 | 0.002 | 0 | 0.23 | 0.28 | 0.29 | 0.23 | 0.21 | 0.15 |
| 8 | 30171796 | rs4733181 | -0.505 | -0.184 | -0.642 | 0.684 | 0.001 | 0.015 | 0.015 | 0 | 0.75 | 0.69 | 0.74 | 0.61 | 0.72 | 0.72 |
| 8 | 30174964 | rs11996929 | 1.745 | -0.06 | -1.139 | -0.01 | -0.027 | 0.026 | 0.026 | 0 | 0.26 | 0.13 | 0.24 | 0.18 | 0.29 | 0.33 |
| 8 | 30175381 | rs12674731 | 0.113 | 0.006 | -0.996 | -0.227 | 0.019 | -0.029 | -0.029 | 0 | 0.3 | 0.19 | 0.33 | 0.39 | 0.25 | 0.23 |
| 8 | 30177562 | rs4434584 | 0.557 | 0.054 | -1.159 | -0.375 | -0.012 | -0.051 | -0.051 | 5.882 | 0.22 | 0.07 | 0.28 | 0.32 | 0.21 | 0.09 |
| 8 | 30177969 | rs7829473 | -0.823 | 0.035 | -1.018 | -0.441 | -0.003 | 0.017 | 0.017 | 0 | 0.68 | 0.61 | 0.72 | 0.54 | 0.68 | 0.6 |
| 8 | 30179213 | rs7015785 | -0.09 | 0.023 | -1.076 | -0.537 | 0.002 | -0.005 | -0.005 | 0.103 | 0.58 | 0.58 | 0.65 | 0.5 | 0.58 | 0.48 |
| 8 | 30182338 | rs4260861 | 0.453 | 0.175 | -1.015 | -0.527 | -0.001 | -0.003 | -0.003 | 0 | 0.45 | 0.52 | 0.44 | 0.22 | 0.46 | 0.51 |
| 8 | 30185572 | rs12541729 | -0.125 | 0.127 | -1.026 | -0.39 | -0.012 | -0.023 | -0.023 | 0 | 0.27 | 0.15 | 0.37 | 0.32 | 0.27 | 0.14 |
| 8 | 30188487 | rs7002660 | -1.861 | 0.472 | -0.531 | -0.378 | -0.005 | 0.012 | 0.012 | 0 | 0.74 | 0.79 | 0.73 | 0.85 | 0.7 | 0.66 |
| 8 | 30202711 | rs6993943 | 0.015 | 0.712 | 0.041 | 0.182 | 0.001 | 0.031 | 0.031 | 0 | 0.11 | 0.13 | 0.18 | 0.31 | 0.11 | 0.07 |
| 8 | 30202885 | rs4398866 | -1.567 | 0.626 | -0.127 | -0.273 | -0.001 | 0 | 0 | 0.157 | 0.41 | 0.57 | 0.41 | 0.37 | 0.42 | 0.3 |
| 8 | 30202954 | rs12678882 | 0.72 | 0.569 | -0.077 | -0.283 | -0.035 | 0.082 | 0.082 | 1.647 | 0.22 | 0.06 | 0.13 | 0.05 | 0.25 | 0.16 |
| 8 | 30203555 | rs13262664 | -1.26 | 0.586 | -0.079 | -0.269 | -0.001 | 0.001 | 0.001 | 0.297 | 0.36 | 0.56 | 0.34 | 0.37 | 0.36 | 0.25 |
| 8 | 30217449 | rs4509276 | -0.467 | 0.615 | 0.194 | 0.146 | 0.02 | 0.11 | 0.11 | 0 | 0.14 | 0.33 | 0.21 | 0.32 | 0.13 | 0.08 |
| 8 | 30218245 | rs9642710 | -0.599 | 0.612 | 0.196 | 0.145 | 0.026 | 0.15 | 0.15 | 0 | 0.15 | 0.43 | 0.21 | 0.32 | 0.13 | 0.08 |
| 8 | 30221351 | rs7000773 | -1.549 | 0.486 | 0.136 | 0.091 | 0 | 0.056 | 0.056 | 0 | 0.22 | 0.44 | 0.29 | 0.3 | 0.19 | 0.13 |
| 8 | 30230319 | rs7009669 | -0.165 | 0.875 | -0.476 | 0.365 | 0.004 | 0.024 | 0.024 | 0 | 0.13 | 0.16 | 0.18 | 0.31 | 0.16 | 0.07 |
| 8 | 30237882 | rs10087801 | 0.223 | 1.053 | -0.761 | -0.049 | -0.016 | 0.041 | 0.041 | 0 | 0.22 | 0.11 | 0.19 | 0.08 | 0.27 | 0.14 |
| 8 | 30240549 | rs12677454 | -0.89 | 0.898 | -0.59 | 0.094 | 0.013 | 0.002 | 0.002 | 0 | 0.39 | 0.46 | 0.42 | 0.41 | 0.47 | 0.33 |
| 8 | 30248320 | rs4259380 | -1.282 | 0.456 | -0.785 | -0.176 | -0.013 | -0.034 | -0.034 | 0 | 0.57 | 0.52 | 0.67 | 0.8 | 0.65 | 0.47 |
| 8 | 30259921 | rs2915621 | 0.428 | 1.178 | -1.715 | -0.318 | -0.039 | 0.066 | 0.066 | 0.435 | 0.22 | 0.1 | 0.2 | 0.04 | 0.3 | 0.1 |
| 8 | 30261558 | rs10435524 | -0.126 | 1.148 | -1.965 | 0.11 | -0.084 | -0.079 | -0.079 | 1.225 | 0.3 | 0.06 | 0.39 | 0.69 | 0.34 | 0.3 |
| 8 | 30267599 | rs3116010 | -1.162 | 0.455 | -1.444 | -0.15 | 0.006 | 0.042 | 0.042 | 0 | 0.64 | 0.85 | 0.69 | 0.76 | 0.68 | 0.45 |
| 8 | 30275342 | rs952774 | -0.585 | 0.788 | -1.31 | -0.403 | -0.022 | 0.127 | 0.127 | 0 | 0.47 | 0.77 | 0.66 | 0.77 | 0.5 | 0.46 |
| 8 | 30286174 | rs7011660 | 0.449 | 1.004 | -0.789 | -0.259 | 0.011 | -0.03 | -0.03 | 0.001 | 0.17 | 0.2 | 0.13 | 0.05 | 0.22 | 0.06 |
| 8 | 30288258 | rs11988241 | 0.826 | 0.913 | -1.393 | -0.06 | -0.02 | -0.012 | -0.012 | 0 | 0.28 | 0.1 | 0.23 | 0.31 | 0.28 | 0.15 |
| 8 | 30288272 | rs2979470 | -1.056 | 0.921 | -1.157 | -0.027 | -0.021 | 0.004 | 0.004 | 0 | 0.45 | 0.82 | 0.58 | 0.52 | 0.51 | 0.39 |
| 8 | 30300365 | rs2915603 | -0.335 | 0.895 | -0.983 | 0.065 | 0 | 0.006 | 0.006 | 0 | 0.28 | 0.72 | 0.46 | 0.48 | 0.3 | 0.33 |
| 8 | 30310983 | rs12679764 | 1.616 | 0.959 | -1.172 | 0.259 | 0 | -0.003 | -0.003 | 0.702 | 0.06 | 0.04 | 0.03 | 0.06 | 0.04 | 0.11 |
| 8 | 30319299 | rs7010539 | -0.253 | 0.888 | -0.5 | 0.298 | 0.045 | 0.143 | 0.143 | 0.048 | 0.8 | 0.53 | 0.6 | 0.59 | 0.75 | 0.76 |
| 8 | 30331968 | rs3736645 | 0.468 | 0.832 | -0.478 | 0.135 | -0.023 | -0.041 | -0.041 | 0 | 0.17 | 0.14 | 0.38 | 0.42 | 0.25 | 0.22 |
| 8 | 30332809 | rs7813272 | 0.078 | 0.912 | -0.447 | 0.176 | 0.019 | -0.004 | -0.004 | 0.81 | 0.24 | 0.35 | 0.17 | 0.19 | 0.27 | 0.17 |
| 8 | 30346544 | rs12548554 | 2.034 | 0.745 | -0.467 | 0.064 | 0.038 | 0.056 | 0.056 | 0 | 0.28 | 0.07 | 0.18 | 0.17 | 0.21 | 0.43 |
| 8 | 30352258 | rs7812836 | -0.333 | 0.755 | -0.485 | 0.08 | 0.001 | 0.001 | 0.001 | 0.338 | 0.51 | 0.46 | 0.4 | 0.43 | 0.54 | 0.32 |
| 8 | 30358824 | rs7459974 | 0.762 | 0.675 | -0.812 | -0.098 | 0.046 | 0.124 | 0.124 | 0 | 0.41 | 0.16 | 0.27 | 0.17 | 0.32 | 0.56 |
| 8 | 30364388 | rs16877096 | 1.307 | 1.085 | -0.247 | 0.205 | -0.017 | -0.021 | -0.021 | 0.233 | 0.31 | 0.33 | 0.2 | 0.18 | 0.2 | 0.43 |
| 8 | 30369438 | rs10503868 | 1.19 | 1.169 | -0.32 | 0.27 | -0.024 | 0.005 | 0.005 | 1.938 | 0.29 | 0.1 | 0.24 | 0.24 | 0.28 | 0.14 |
| 8 | 30371917 | rs13251659 | 1.403 | 1.188 | -0.977 | 0.243 | -0.077 | -0.046 | -0.046 | 0 | 0.28 | 0.02 | 0.23 | 0.25 | 0.28 | 0.14 |
| 8 | 30383013 | rs2979531 | -0.143 | 1.244 | -0.639 | 0.363 | -0.001 | -0.001 | -0.001 | 0.204 | 0.48 | 0.5 | 0.59 | 0.56 | 0.45 | 0.68 |
| 8 | 30384436 | rs4733498 | 0.36 | 1.279 | -0.641 | 0.508 | -0.075 | -0.123 | -0.123 | 0.117 | 0.16 | 0.03 | 0.38 | 0.37 | 0.25 | 0.22 |
| 8 | 30392824 | rs7018462 | 0.264 | 1.288 | -0.606 | 0.512 | -0.028 | -0.058 | -0.058 | 0 | 0.17 | 0.11 | 0.38 | 0.37 | 0.25 | 0.22 |
| 8 | 30397935 | rs11776064 | 0.124 | 1.284 | -0.586 | 0.511 | -0.013 | -0.028 | -0.028 | 1.652 | 0.18 | 0.15 | 0.38 | 0.38 | 0.25 | 0.24 |
| 8 | 30410795 | rs1984838 | -0.343 | 1.269 | -0.496 | 0.504 | 0.007 | 0.045 | 0.045 | 0 | 0.2 | 0.31 | 0.4 | 0.38 | 0.25 | 0.25 |
| 8 | 30419334 | rs17554116 | 1.369 | 1.18 | -0.661 | -0.027 | 0.033 | 0.101 | 0.101 | 0 | 0.29 | 0.07 | 0.2 | 0.12 | 0.23 | 0.41 |
| 8 | 30421455 | rs3779641 | 1.247 | 1.308 | -0.563 | -0.047 | 0 | 0 | 0 | 0 | 0.31 | 0.36 | 0.27 | 0.29 | 0.3 | 0.15 |
| 8 | 30422888 | rs7833229 | 1.774 | 0.878 | -0.557 | 0.13 | -0.032 | -0.034 | -0.034 | 3.104 | 0.11 | 0.02 | 0.07 | 0.16 | 0.15 | 0.15 |
| 8 | 30425905 | rs3757916 | -0.794 | 0.905 | -0.63 | 0.127 | -0.021 | -0.06 | -0.06 | 0.041 | 0.84 | 0.91 | 0.64 | 0.59 | 0.8 | 0.78 |
| 8 | 30427531 | rs17554408 | 1.3 | 0.898 | -0.623 | 0.125 | 0.036 | 0.092 | 0.092 | 0 | 0.3 | 0.1 | 0.19 | 0.11 | 0.23 | 0.41 |
| 8 | 30434903 | rs6990642 | -0.839 | 0.919 | -0.564 | 0.237 | -0.015 | -0.053 | -0.053 | 0 | 0.85 | 0.91 | 0.65 | 0.59 | 0.81 | 0.79 |
| 8 | 30437008 | rs11783765 | -0.26 | 0.918 | -0.509 | 0.235 | 0.007 | 0.088 | 0.088 | 0 | 0.82 | 0.7 | 0.62 | 0.59 | 0.8 | 0.78 |
| 8 | 30450114 | rs2979502 | 0.408 | 0.825 | -0.722 | 0.296 | 0 | -0.009 | -0.009 | 0 | 0.4 | 0.55 | 0.35 | 0.22 | 0.4 | 0.19 |
| 8 | 30455200 | rs7833529 | 0.453 | 0.839 | -0.852 | 0.278 | -0.001 | -0.001 | -0.001 | 0 | 0.45 | 0.47 | 0.39 | 0.47 | 0.52 | 0.35 |
| 8 | 30462149 | rs10503873 | 1.226 | 0.829 | -0.823 | 0.275 | 0.009 | 0.136 | 0.136 | 0 | 0.32 | 0.06 | 0.21 | 0.13 | 0.27 | 0.42 |
| 8 | 30471443 | rs2979534 | -0.614 | 0.889 | -0.526 | 0.645 | -0.045 | -0.109 | -0.109 | 0 | 0.85 | 0.96 | 0.64 | 0.5 | 0.81 | 0.78 |
| 8 | 30498859 | rs2978263 | -0.567 | 0.913 | -0.496 | 0.386 | -0.011 | -0.046 | -0.046 | 0 | 0.84 | 0.9 | 0.64 | 0.59 | 0.81 | 0.78 |
| 8 | 30500715 | rs2978257 | 0.278 | 0.925 | -0.476 | 0.34 | -0.002 | -0.011 | -0.011 | 0 | 0.51 | 0.49 | 0.33 | 0.29 | 0.46 | 0.63 |
| 8 | 30503371 | rs16877238 | 0.601 | 0.955 | -0.634 | 0.37 | 0.009 | -0.002 | -0.002 | 0 | 0.33 | 0.42 | 0.32 | 0.3 | 0.36 | 0.16 |
| 8 | 30509469 | rs9649911 | 0.209 | 0.618 | -1.137 | -0.391 | 0.016 | 0.077 | 0.077 | 0.001 | 0.44 | 0.25 | 0.29 | 0.28 | 0.42 | 0.59 |
| 8 | 30516496 | rs6981869 | -0.101 | 0.807 | -1.007 | 0.133 | -0.002 | -0.022 | -0.022 | 0 | 0.94 | 0.93 | 0.98 | 0.99 | 0.97 | 0.97 |
| 8 | 30535660 | rs3594 | 0.07 | 0.792 | -1.651 | -0.028 | -0.028 | 0.07 | 0.07 | 0 | 0.36 | 0.14 | 0.23 | 0.26 | 0.41 | 0.4 |

| | | | | | | | | | | | | | | | | |
|---|---|---|---|---|---|---|---|---|---|---|---|---|---|---|---|---|
| 8 | 30546636 | rs2551715 | -0.333 | 0.56 | -1.407 | 0.129 | -0.019 | -0.024 | -0.024 | 0.062 | 0.69 | 0.75 | 0.5 | 0.58 | 0.63 | 0.59 |
| 8 | 30554006 | rs8190996 | 0.487 | 0.571 | -1.306 | 0.087 | -0.022 | -0.032 | -0.032 | 0 | 0.37 | 0.29 | 0.49 | 0.49 | 0.45 | 0.47 |
| 8 | 30560887 | rs3779647 | 0.612 | 0.236 | -1.121 | -0.111 | -0.012 | -0.017 | -0.017 | 0.973 | 0.47 | 0.13 | 0.53 | 0.43 | 0.57 | 0.48 |
| 8 | 30565945 | rs2978663 | 0.151 | 0.072 | -1.059 | -0.207 | 0.042 | -0.029 | -0.029 | 0.132 | 0.55 | 0.24 | 0.65 | 0.48 | 0.64 | 0.72 |
| 8 | 30569827 | rs2978662 | 1.34 | -0.103 | 0.342 | -0.808 | 0.014 | 0.011 | 0.011 | 0 | 0.72 | 0.26 | 0.74 | 0.82 | 0.82 | 0.63 |
| 8 | 30575799 | rs2911670 | 1.418 | -0.129 | 1.286 | -0.818 | -0.009 | 0 | 0 | 0 | 0.91 | 0.7 | 0.86 | 0.94 | 0.94 | 0.79 |
| 8 | 30575838 | rs17557435 | 0.082 | -0.159 | 1.36 | -0.835 | 0.002 | 0.011 | 0.011 | 0 | 0.13 | 0.05 | 0.09 | 0.08 | 0.11 | 0.11 |
| 8 | 30578282 | rs16877292 | -1.944 | -0.235 | 1.882 | -0.848 | 0.047 | 0.083 | 0.083 | 1.001 | 0.05 | 0.34 | 0.03 | 0.04 | 0.01 | 0.04 |
| 8 | 30580577 | rs2551698 | 2.58 | -0.262 | 1.843 | -0.862 | -0.032 | -0.016 | -0.016 | 0.524 | 0.84 | 0.27 | 0.82 | 0.9 | 0.93 | 0.74 |
| 8 | 30581501 | rs10088455 | -2.085 | -0.056 | 1.857 | -0.751 | 0.055 | 0.093 | 0.093 | 1.059 | 0.06 | 0.4 | 0.03 | 0.04 | 0.01 | 0.04 |
| 8 | 30583581 | rs8190898 | -2.151 | -0.049 | 1.841 | -0.748 | 0.042 | 0.078 | 0.078 | 1.718 | 0.06 | 0.37 | 0.03 | 0.04 | 0.01 | 0.04 |
| 8 | 30585738 | rs1002149 | -0.91 | -0.047 | 1.537 | -0.651 | -0.007 | 0.005 | 0.005 | 0 | 0.2 | 0.28 | 0.2 | 0.15 | 0.16 | 0.3 |
| 8 | 30593869 | rs11986256 | -1.083 | -0.039 | 1.548 | -0.647 | -0.022 | 0.001 | 0.001 | 0 | 0.22 | 0.44 | 0.25 | 0.16 | 0.17 | 0.33 |
| 8 | 30600304 | rs1469175 | -1.083 | -0.039 | 1.523 | -0.696 | -0.022 | -0.005 | -0.005 | 0.182 | 0.22 | 0.46 | 0.25 | 0.16 | 0.17 | 0.33 |
| 8 | 30609061 | rs2911689 | 1.002 | -0.003 | 0.944 | -0.71 | -0.024 | -0.016 | -0.016 | 1.048 | 0.77 | 0.45 | 0.75 | 0.85 | 0.83 | 0.66 |
| 8 | 30627027 | rs10105118 | -0.815 | 0.02 | 0.933 | -0.829 | -0.033 | -0.016 | -0.016 | 0 | 0.19 | 0.54 | 0.14 | 0.14 | 0.11 | 0.24 |
| 8 | 30630150 | rs6468455 | -1.761 | 0.2 | 0.665 | -0.932 | 0.016 | 0.016 | 0.016 | 0.474 | 0.29 | 0.88 | 0.44 | 0.24 | 0.21 | 0.37 |
| 8 | 30632191 | rs12541220 | -1.538 | 0.191 | 0.714 | -0.953 | 0.009 | 0.011 | 0.011 | 0.988 | 0.27 | 0.86 | 0.43 | 0.22 | 0.21 | 0.37 |
| 8 | 30635443 | rs7017734 | -0.854 | 0.152 | 1.3 | -0.952 | -0.048 | -0.025 | -0.025 | 0 | 0.21 | 0.45 | 0.15 | 0.14 | 0.12 | 0.23 |
| 8 | 30636140 | rs10216603 | -0.513 | 0.125 | 1.458 | -0.968 | -0.027 | -0.018 | -0.018 | 0 | 0.17 | 0.28 | 0.11 | 0.1 | 0.1 | 0.11 |
| 8 | 30647243 | rs10089206 | 0.268 | 0.661 | 1.349 | -1.223 | 0.013 | 0.003 | 0.003 | 0 | 0.13 | 0.1 | 0.06 | 0.11 | 0.08 | 0.11 |
| 8 | 30651335 | rs10087700 | 0.19 | 0.65 | 1.467 | -1.213 | 0.003 | 0.001 | 0.001 | 0 | 0.13 | 0.15 | 0.08 | 0.1 | 0.09 | 0.11 |
| 8 | 30655363 | rs1116003 | -1.781 | 0.725 | 1.812 | -1.185 | 0.077 | 0.04 | 0.04 | 0 | 0.12 | 0.49 | 0.33 | 0.11 | 0.12 | 0.26 |
| 8 | 30665278 | rs4733515 | -1.781 | 0.73 | 1.737 | -1.18 | 0.077 | 0.051 | 0.051 | 0 | 0.12 | 0.49 | 0.32 | 0.11 | 0.12 | 0.27 |
| 8 | 30674390 | rs4733202 | -1.88 | 0.729 | 1.627 | -1.218 | 0.056 | 0.024 | 0.024 | 0.047 | 0.12 | 0.52 | 0.33 | 0.11 | 0.12 | 0.26 |
| 8 | 30687895 | rs16877392 | 0.032 | 0.538 | 1.241 | -1.436 | -0.011 | -0.023 | -0.023 | 0 | 0.09 | 0.21 | 0.07 | 0.06 | 0.06 | 0.19 |
| 8 | 30694153 | rs6980502 | 0.052 | 0.475 | 1.108 | -1.57 | -0.009 | -0.024 | -0.024 | 0 | 0.1 | 0.3 | 0.07 | 0.06 | 0.06 | 0.19 |
| 8 | 30702602 | rs323345 | -1.619 | 0.472 | 1.135 | -1.621 | 0.009 | 0.016 | 0.016 | 2.152 | 0.17 | 0.85 | 0.16 | 0.07 | 0.13 | 0.21 |
| 8 | 30703431 | rs323346 | -1.831 | 0.512 | 1.156 | -1.628 | 0.002 | 0.008 | 0.008 | 0.011 | 0.2 | 0.8 | 0.37 | 0.11 | 0.17 | 0.33 |
| 8 | 30706224 | rs323347 | -1.953 | 0.514 | 1.113 | -1.628 | 0.007 | 0.034 | 0.034 | 0.534 | 0.21 | 0.85 | 0.38 | 0.11 | 0.17 | 0.33 |
| 8 | 30713343 | rs323352 | -1.922 | 0.51 | 1.202 | -1.629 | 0.006 | 0.033 | 0.033 | 0.467 | 0.21 | 0.85 | 0.37 | 0.11 | 0.17 | 0.33 |
| 8 | 30719034 | rs323362 | 1.645 | 0.28 | 0.555 | -1.923 | -0.062 | 0.177 | 0.177 | 0 | 0.38 | 0.03 | 0.23 | 0.11 | 0.37 | 0.35 |
| 8 | 30725122 | rs323368 | -1.587 | 0.255 | 0.668 | -1.873 | 0.004 | 0.015 | 0.015 | 0.453 | 0.21 | 0.82 | 0.38 | 0.11 | 0.16 | 0.33 |
| 8 | 30726911 | rs2467542 | -0.955 | 0.4 | 0.631 | -1.756 | -0.136 | -0.076 | -0.076 | 0 | 0.73 | 0.98 | 0.68 | 0.32 | 0.62 | 0.77 |
| 8 | 30731171 | rs323375 | -1.818 | 0.13 | 1.753 | -0.944 | -0.019 | 0.067 | 0.067 | 0.416 | 0.36 | 0.95 | 0.47 | 0.21 | 0.26 | 0.42 |
| 8 | 30746492 | rs10109208 | -2.057 | 0.246 | 1.974 | -0.875 | 0.082 | -0.009 | -0.009 | 0.968 | 0.1 | 0.52 | 0.31 | 0.06 | 0.1 | 0.14 |
| 8 | 30784071 | rs7822191 | -1.689 | 0.248 | 1.892 | -0.862 | 0.119 | 0.033 | 0.033 | 4.195 | 0.07 | 0.48 | 0.1 | 0.01 | 0.06 | 0.02 |
| 8 | 30787906 | rs4733206 | -0.716 | 0.187 | 1.894 | -0.904 | -0.009 | -0.014 | -0.014 | 0 | 0.26 | 0.32 | 0.36 | 0.2 | 0.19 | 0.4 |
| 8 | 30794336 | rs6986926 | -1.571 | 0.234 | 1.376 | -0.902 | 0.011 | 0.046 | 0.046 | 0.185 | 0.32 | 0.78 | 0.46 | 0.21 | 0.25 | 0.43 |
| 8 | 30797840 | rs10100241 | -1.355 | 0.214 | 1.15 | -0.925 | -0.003 | -0.006 | -0.006 | 0.043 | 0.3 | 0.66 | 0.4 | 0.19 | 0.19 | 0.4 |
| 8 | 30802695 | rs2880773 | -0.453 | 0.25 | 0.175 | -0.909 | -0.022 | -0.026 | -0.026 | 0 | 0.22 | 0.35 | 0.15 | 0.16 | 0.16 | 0.29 |
| 8 | 30804490 | rs1978402 | -1.555 | 0.258 | -0.853 | -0.897 | 0.014 | 0.074 | 0.074 | 0.138 | 0.35 | 0.87 | 0.46 | 0.23 | 0.26 | 0.43 |
| 8 | 30809614 | rs11990181 | -1.33 | 0.256 | -0.823 | -0.893 | 0.026 | 0.084 | 0.084 | 0.432 | 0.31 | 0.86 | 0.25 | 0.17 | 0.23 | 0.32 |
| 8 | 30816027 | rs4733210 | -1.685 | 0.537 | -0.18 | -0.509 | -0.003 | -0.012 | -0.012 | 0.802 | 0.15 | 0.78 | 0.32 | 0.07 | 0.11 | 0.14 |
| 8 | 30817510 | rs16877581 | -1.405 | 0.289 | 0.208 | -0.325 | -0.021 | 0.033 | 0.033 | 0.558 | 0.06 | 0.22 | 0.05 | 0.07 | 0.02 | 0.04 |
| 8 | 30820305 | rs17647942 | 0.521 | 0.736 | -0.171 | -0.139 | -0.077 | -0.057 | -0.057 | 0 | 0.18 | 0.01 | 0.2 | 0.24 | 0.25 | 0.07 |
| 8 | 30821036 | rs4733212 | -1.39 | 0.895 | 0.815 | 0.268 | 0.066 | 0.143 | 0.143 | 0.024 | 0.09 | 0.32 | 0.23 | 0.2 | 0.09 | 0.15 |
| 8 | 30822094 | rs6985028 | -1.008 | 0.884 | 0.486 | -0.353 | 0.023 | 0.15 | 0.15 | 0 | 0.51 | 0.73 | 0.7 | 0.89 | 0.57 | 0.53 |
| 8 | 30829623 | rs6981541 | -0.536 | 0.7 | 0.575 | -0.174 | 0.028 | 0.074 | 0.074 | 0 | 0.07 | 0.17 | 0.22 | 0.2 | 0.09 | 0.15 |
| 8 | 30838546 | rs11783562 | 0.136 | 1.053 | 0.438 | -0.415 | -0.152 | -0.014 | -0.014 | 0 | 0.37 | 0.04 | 0.44 | 0.68 | 0.52 | 0.33 |
| 8 | 30853102 | rs10503879 | -0.686 | 1.031 | 0.901 | -0.414 | 0.048 | 0.118 | 0.118 | 0 | 0.08 | 0.26 | 0.22 | 0.2 | 0.09 | 0.15 |
| 8 | 30854033 | rs2725371 | -1.494 | 0.679 | 0.392 | -0.918 | 0.026 | 0.192 | 0.192 | 0 | 0.64 | 0.9 | 0.79 | 0.95 | 0.71 | 0.61 |
| 8 | 30856464 | rs1362910 | -0.334 | 0.682 | 0.451 | -0.914 | 0.027 | 0.164 | 0.164 | 0.202 | 0.52 | 0.1 | 0.53 | 0.76 | 0.61 | 0.46 |
| 8 | 30858054 | rs1125841 | -0.827 | 0.738 | 0.673 | -0.802 | 0.03 | 0.07 | 0.07 | 0 | 0.52 | 0.7 | 0.72 | 0.95 | 0.62 | 0.53 |
| 8 | 30859050 | rs10954771 | -0.691 | 0.753 | 0.214 | -0.794 | 0.028 | 0.036 | 0.036 | 0 | 0.5 | 0.4 | 0.71 | 0.95 | 0.61 | 0.53 |
| 8 | 30864339 | rs11775287 | 0.359 | 0.741 | -0.366 | -0.704 | -0.124 | 0.007 | 0.007 | 1.488 | 0.37 | 0.08 | 0.43 | 0.68 | 0.5 | 0.33 |
| 8 | 30864944 | rs1362911 | -1.567 | 0.755 | 0.238 | -0.49 | -0.004 | 0.102 | 0.102 | 1.607 | 0.1 | 0.66 | 0.13 | 0.19 | 0.07 | 0.07 |
| 8 | 30871656 | rs7827210 | 0.433 | 0.472 | -0.661 | -0.394 | 0.009 | 0.226 | 0.226 | 0.217 | 0.48 | 0.08 | 0.37 | 0.18 | 0.39 | 0.48 |
| 8 | 30896151 | rs13269094 | -0.114 | 0.452 | 0.074 | -0.41 | 0.011 | -0.033 | -0.033 | 0 | 0.15 | 0.27 | 0.08 | 0.1 | 0.14 | 0.15 |
| 8 | 30900890 | rs4733220 | 0.238 | 0.166 | 0.265 | -0.318 | 0.011 | 0.028 | 0.028 | 0.191 | 0.53 | 0.25 | 0.6 | 0.58 | 0.59 | 0.45 |
| 8 | 30921935 | rs2230009 | -0.295 | 0.103 | 0.266 | -0.441 | 0.034 | -0.019 | -0.019 | 0 | 0.06 | 0.15 | 0.02 | 0.02 | 0.06 | 0.08 |
| 8 | 30928146 | rs2725385 | -0.232 | -0.078 | 0.572 | -0.391 | -0.017 | -0.061 | -0.061 | 0 | 0.62 | 0.5 | 0.79 | 0.88 | 0.71 | 0.67 |
| 8 | 30936356 | rs2553268 | -0.244 | -0.062 | 0.711 | -0.393 | -0.016 | -0.061 | -0.061 | 2.662 | 0.62 | 0.5 | 0.79 | 0.87 | 0.7 | 0.66 |
| 8 | 30946403 | rs3087409 | -0.401 | 0.248 | 0.664 | -0.281 | 0.037 | -0.017 | -0.017 | 0.283 | 0.06 | 0.15 | 0.02 | 0.02 | 0.06 | 0.08 |
| 8 | 30947203 | rs11574245 | 1.068 | 0.338 | 0.602 | -0.859 | -0.005 | -0.038 | -0.038 | 1.842 | 0.1 | 0.12 | 0.06 | 0.01 | 0.09 | 0.07 |
| 8 | 30957765 | rs2737327 | 1.063 | 0.184 | 0.635 | -1.053 | -0.004 | -0.053 | -0.053 | 0 | 0.77 | 0.27 | 0.88 | 0.97 | 0.82 | 0.87 |
| 8 | 30962693 | rs9649886 | -0.549 | 0.63 | 0.764 | -0.713 | 0.008 | 0.02 | 0.02 | 0 | 0.39 | 0.3 | 0.28 | 0.27 | 0.36 | 0.41 |
| 8 | 30967369 | rs4733224 | -0.394 | 0.604 | 0.772 | -0.75 | 0.013 | 0.029 | 0.029 | 0 | 0.37 | 0.23 | 0.28 | 0.27 | 0.35 | 0.41 |
| 8 | 30967673 | rs17651080 | 1.537 | 0.826 | 0.582 | -0.858 | -0.017 | 0.057 | 0.057 | 0.568 | 0.22 | 0.07 | 0.12 | 0.1 | 0.2 | 0.26 |

| | | | | | | | | | | | | | | | | |
|---|---|---|---|---|---|---|---|---|---|---|---|---|---|---|---|---|
| 8 | 30973957 | rs1800392 | 0.694 | 0.823 | 0.973 | -0.852 | 0.001 | -0.001 | -0.001 | 3.129 | 0.39 | 0.42 | 0.58 | 0.62 | 0.45 | 0.33 |
| 8 | 30974348 | rs11574306 | 1.526 | 0.82 | 0.813 | -0.855 | -0.006 | 0.041 | 0.041 | 0.665 | 0.22 | 0.12 | 0.12 | 0.1 | 0.2 | 0.26 |
| 8 | 30977392 | rs4733225 | 1.225 | 0.766 | 0.694 | -0.911 | 0.006 | -0.058 | -0.058 | 0.111 | 0.21 | 0.03 | 0.2 | 0.24 | 0.2 | 0.24 |
| 8 | 30977696 | rs4987034 | 1.482 | 0.836 | 0.656 | -0.851 | -0.016 | 0.06 | 0.06 | 0.494 | 0.23 | 0.09 | 0.13 | 0.1 | 0.2 | 0.26 |
| 8 | 30981201 | rs2725353 | 1.027 | 0.781 | 0.866 | -0.793 | -0.054 | -0.021 | -0.021 | 0 | 0.36 | 0.14 | 0.56 | 0.62 | 0.45 | 0.34 |
| 8 | 30990110 | rs2252480 | 0.976 | 0.861 | 0.9 | -0.877 | 0 | -0.001 | -0.001 | 1.237 | 0.36 | 0.38 | 0.54 | 0.64 | 0.38 | 0.3 |
| 8 | 31008747 | rs10954778 | 0.08 | 0.773 | 0.944 | -0.935 | 0.014 | 0.015 | 0.015 | 0 | 0.34 | 0.22 | 0.25 | 0.23 | 0.3 | 0.33 |
| 8 | 31018685 | rs6994361 | -1.366 | 0.798 | 0.939 | -0.932 | 0.001 | 0.001 | 0.001 | 0.009 | 0.66 | 0.62 | 0.45 | 0.36 | 0.62 | 0.7 |
| 8 | 31020159 | rs4398867 | 1.429 | 0.843 | 0.603 | -0.914 | -0.004 | -0.038 | -0.038 | 0 | 0.11 | 0.1 | 0.06 | 0.01 | 0.09 | 0.09 |
| 8 | 31024638 | rs1801196 | 0.363 | 1.09 | 0.715 | -0.665 | 0.003 | 0.003 | 0.003 | 0 | 0.34 | 0.26 | 0.25 | 0.23 | 0.29 | 0.33 |
| 8 | 31024654 | rs1346044 | 1.032 | 1.126 | 0.434 | -0.617 | -0.029 | 0.063 | 0.063 | 0.134 | 0.25 | 0.15 | 0.17 | 0.1 | 0.27 | 0.28 |
| 8 | 31025343 | rs11574398 | -0.417 | 0.764 | 0.647 | -0.478 | -0.014 | -0.028 | -0.028 | 0 | 0.4 | 0.46 | 0.29 | 0.26 | 0.35 | 0.42 |
| 8 | 31042409 | rs2553244 | -0.404 | 0.962 | -0.099 | -0.452 | 0.018 | 0.057 | 0.057 | 0 | 0.49 | 0.35 | 0.37 | 0.29 | 0.47 | 0.42 |
| 8 | 31052018 | rs2737374 | 0.067 | 0.867 | 0.061 | -0.557 | 0.057 | -0.042 | -0.042 | 0 | 0.91 | 0.63 | 0.92 | 0.98 | 0.89 | 0.91 |
| 8 | 31054787 | rs7827455 | 0.474 | 0.983 | -0.308 | -0.559 | 0.008 | 0.043 | 0.043 | 0 | 0.49 | 0.32 | 0.59 | 0.72 | 0.5 | 0.56 |
| 8 | 31057097 | rs2553239 | 0.225 | 0.907 | -1.248 | -0.42 | 0.02 | -0.02 | -0.02 | 0 | 0.39 | 0.18 | 0.3 | 0.36 | 0.37 | 0.35 |
| 8 | 31062127 | rs17653630 | 0.658 | 1.448 | -1.436 | -0.613 | -0.09 | 0.007 | 0.007 | 1.776 | 0.23 | 0.01 | 0.18 | 0.12 | 0.27 | 0.23 |
| 8 | 31066640 | rs6981774 | 0.301 | 1.531 | -0.957 | -1.382 | 0.005 | -0.013 | -0.013 | 0 | 0.07 | 0.12 | 0.05 | 0.02 | 0.07 | 0.07 |
| 8 | 31070598 | rs7837872 | 0.549 | 1.405 | -1.196 | -1.747 | -0.047 | 0.052 | 0.052 | 0 | 0.17 | 0.03 | 0.12 | 0.06 | 0.21 | 0.13 |
| 8 | 31072165 | rs7843551 | -0.165 | 1.052 | -0.827 | -1.591 | 0.008 | 0.102 | 0.102 | 0 | 0.52 | 0.67 | 0.33 | 0.15 | 0.53 | 0.38 |
| 8 | 31076640 | rs10954782 | 0.166 | 1.011 | -1.67 | -1.433 | -0.008 | 0.068 | 0.068 | 0 | 0.43 | 0.32 | 0.25 | 0.14 | 0.43 | 0.37 |
| 8 | 31079161 | rs11998012 | -0.331 | 0.832 | -1.921 | -0.881 | -0.03 | -0.096 | -0.096 | 2.186 | 0.27 | 0.03 | 0.37 | 0.3 | 0.25 | 0.27 |
| 8 | 31087429 | rs16877840 | 0.1 | 0.903 | -0.771 | -1.624 | 0.045 | -0.081 | -0.081 | 0.112 | 0.12 | 0.41 | 0.1 | 0 | 0.13 | 0.04 |
| 8 | 31097266 | rs10447952 | -0.066 | 0.746 | -1.061 | -1.702 | 0.019 | -0.119 | -0.119 | 3.456 | 0.16 | 0.55 | 0.1 | 0 | 0.13 | 0.04 |
| 8 | 31107532 | rs1559681 | -0.213 | 0.694 | -1.854 | -2.325 | -0.017 | 0.154 | 0.154 | 0 | 0.65 | 0.93 | 0.82 | 0.89 | 0.66 | 0.79 |
| 8 | 31120019 | rs1425293 | -0.924 | 0.742 | -2.146 | -2.383 | -0.004 | 0.022 | 0.022 | 0 | 0.88 | 0.97 | 0.95 | 0.94 | 0.92 | 0.94 |
| 8 | 31120862 | rs6468029 | -0.705 | 0.668 | -1.778 | -2.301 | -0.001 | -0.056 | -0.056 | 1.244 | 0.39 | 0.63 | 0.23 | 0.06 | 0.4 | 0.2 |
| 8 | 31121769 | rs7812391 | 0.03 | 0.664 | -2.056 | -1.585 | -0.047 | 0.115 | 0.115 | 0 | 0.22 | 0.02 | 0.13 | 0.06 | 0.27 | 0.16 |
| 8 | 31123341 | rs11993516 | 0.099 | 0.177 | -1.72 | -1.373 | 0.004 | -0.014 | -0.014 | 0 | 0.2 | 0.13 | 0.14 | 0.01 | 0.16 | 0.13 |
| 8 | 31132850 | rs6997014 | 0.165 | 0.536 | -1.936 | -1.022 | -0.008 | 0.108 | 0.108 | 0 | 0.77 | 0.95 | 0.86 | 0.92 | 0.74 | 0.83 |
| 8 | 31133658 | rs4609156 | -0.6 | 1.031 | -1.424 | -0.439 | 0.016 | 0.007 | 0.007 | 0.647 | 0.34 | 0.4 | 0.64 | 0.44 | 0.55 | 0.48 |
| 8 | 31135931 | rs10110117 | 0.259 | 0.266 | -1.143 | -0.728 | -0.001 | -0.011 | -0.011 | 0 | 0.63 | 0.56 | 0.62 | 0.76 | 0.62 | 0.66 |
| 8 | 31140870 | rs4733071 | 0.378 | -0.505 | -0.688 | -0.94 | 0.011 | -0.044 | -0.044 | 0.16 | 0.67 | 0.54 | 0.61 | 0.79 | 0.65 | 0.7 |
| 8 | 31144815 | rs10282922 | 1.259 | -1.46 | -1.296 | -1.375 | 0.045 | 0.066 | 0.066 | 0.434 | 0.14 | 0.04 | 0.03 | 0 | 0.05 | 0.03 |
| 8 | 31146458 | rs7005168 | 0.516 | -1.337 | -1.412 | -1.363 | -0.001 | 0.002 | 0.002 | 0 | 0.56 | 0.53 | 0.56 | 0.7 | 0.52 | 0.62 |
| 8 | 31148512 | rs4733242 | 1.127 | -1.679 | -1.166 | -1.569 | -0.058 | 0.04 | 0.04 | 0 | 0.38 | 0.19 | 0.49 | 0.72 | 0.47 | 0.59 |
| 8 | 31159012 | rs4400341 | 0.928 | -1.188 | -1.162 | -1.356 | -0.06 | 0.06 | 0.06 | 0 | 0.42 | 0.17 | 0.41 | 0.67 | 0.51 | 0.52 |
| 8 | 31159672 | rs2881123 | 0.628 | -1.119 | -0.687 | -1.301 | -0.001 | 0.009 | 0.009 | 0 | 0.48 | 0.44 | 0.45 | 0.66 | 0.52 | 0.53 |
| 8 | 31160864 | rs17654713 | 1.41 | -1.112 | -1.256 | -1.411 | 0.047 | 0.069 | 0.069 | 0.577 | 0.14 | 0.04 | 0.03 | 0 | 0.05 | 0.05 |
| 8 | 31166101 | rs4336565 | -0.927 | -0.921 | -0.951 | -1.299 | -0.031 | 0.002 | 0.002 | 0 | 0.79 | 0.87 | 0.66 | 0.79 | 0.72 | 0.76 |
| 8 | 31166328 | rs11780547 | 1.056 | -1.099 | -0.589 | -1.268 | -0.018 | -0.025 | -0.025 | 0.275 | 0.09 | 0.13 | 0.02 | 0.01 | 0.04 | 0.06 |
| 8 | 31167530 | rs10100049 | -0.211 | -1.24 | -0.742 | -0.766 | -0.011 | -0.018 | -0.018 | 0 | 0.68 | 0.83 | 0.61 | 0.62 | 0.62 | 0.63 |
| 8 | 31173834 | rs16877873 | 0.122 | -1.222 | -0.768 | -0.772 | -0.029 | -0.031 | -0.031 | 0 | 0.31 | 0.12 | 0.39 | 0.36 | 0.38 | 0.35 |
| 8 | 31174830 | rs2575029 | -0.397 | -0.966 | -0.773 | -0.4 | 0.005 | -0.02 | -0.02 | 0.001 | 0.87 | 0.83 | 0.84 | 0.92 | 0.84 | 0.84 |
| 8 | 31175862 | rs1347564 | 0.785 | -0.934 | -1.362 | -0.466 | 0.07 | 0.115 | 0.115 | 1.439 | 0.19 | 0.03 | 0.03 | 0 | 0.05 | 0.06 |
| 8 | 31177347 | rs16877899 | 0.418 | -0.977 | -1.199 | -0.445 | -0.029 | 0.037 | 0.037 | 0 | 0.5 | 0.14 | 0.42 | 0.56 | 0.49 | 0.53 |
| 8 | 31208088 | rs6997892 | -0.41 | -1.076 | -0.244 | -0.579 | -0.029 | -0.063 | -0.063 | 0.364 | 0.27 | 0.39 | 0.18 | 0.07 | 0.15 | 0.15 |
| 8 | 31209774 | rs7017656 | 0.396 | -0.385 | -0.189 | -0.808 | -0.001 | 0 | 0 | 0 | 0.14 | 0.15 | 0.2 | 0.09 | 0.19 | 0.11 |
| 8 | 31218482 | rs1836851 | -1.218 | -0.633 | 0.463 | -0.479 | 0.005 | 0.005 | 0.005 | 3.107 | 0.42 | 0.62 | 0.3 | 0.23 | 0.31 | 0.27 |
| 8 | 31230777 | rs1583528 | -1.107 | -0.606 | 0.298 | -0.482 | -0.006 | -0.016 | -0.016 | 10.038 | 0.41 | 0.51 | 0.3 | 0.22 | 0.31 | 0.27 |
| 8 | 31237833 | rs6998531 | -0.635 | -0.63 | 0.248 | -0.47 | 0.001 | -0.001 | -0.001 | 0 | 0.22 | 0.34 | 0.23 | 0.22 | 0.21 | 0.18 |
| 8 | 31248913 | rs2681595 | -0.206 | -0.658 | 0.172 | -0.484 | 0.012 | 0.018 | 0.018 | 0.525 | 0.71 | 0.77 | 0.86 | 0.97 | 0.79 | 0.88 |
| 8 | 31249003 | rs10503885 | 1.294 | -0.634 | 0.161 | -0.506 | -0.015 | 0.019 | 0.019 | 0 | 0.11 | 0.1 | 0.1 | 0.02 | 0.15 | 0.05 |
| 8 | 31249870 | rs745957 | 0.141 | -0.68 | 0.319 | -0.389 | -0.052 | -0.013 | -0.013 | 3.515 | 0.72 | 0.62 | 0.82 | 0.77 | 0.89 | 0.78 |
| 8 | 31253149 | rs2681591 | -0.111 | -0.463 | -0.015 | -0.684 | 0.005 | -0.031 | -0.031 | 0.577 | 0.69 | 0.67 | 0.85 | 0.98 | 0.79 | 0.87 |
| 8 | 31254697 | rs2575069 | 0.577 | -0.21 | -0.371 | -0.448 | -0.002 | -0.004 | -0.004 | 0 | 0.37 | 0.6 | 0.4 | 0.38 | 0.39 | 0.41 |
| 8 | 31258539 | rs10954784 | 0.339 | -0.149 | -0.218 | -0.24 | -0.008 | -0.031 | -0.031 | 0.116 | 0.33 | 0.28 | 0.44 | 0.54 | 0.34 | 0.44 |
| 8 | 31261439 | rs2681590 | -0.32 | -0.363 | 0.067 | -0.718 | -0.009 | -0.04 | -0.04 | 0 | 0.81 | 0.8 | 0.92 | 0.97 | 0.87 | 0.89 |
| 8 | 31265972 | rs10282849 | -0.317 | 0.248 | -0.208 | -0.206 | -0.014 | -0.038 | -0.038 | 0 | 0.13 | 0.19 | 0.22 | 0.01 | 0.08 | 0.16 |
| 8 | 31272592 | rs2681600 | 0.992 | 0.832 | -0.257 | -0.598 | 0.005 | -0.072 | -0.072 | 0 | 0.69 | 0.63 | 0.63 | 0.89 | 0.68 | 0.64 |
| 8 | 31272835 | rs11781075 | 0.933 | 0.872 | -1.363 | -0.387 | -0.038 | -0.07 | -0.07 | 0.013 | 0.28 | 0.04 | 0.3 | 0.33 | 0.28 | 0.26 |
| 8 | 31278876 | rs13276104 | 1.005 | 1.305 | -0.76 | -0.016 | -0.024 | -0.015 | -0.015 | 0 | 0.33 | 0.2 | 0.23 | 0.3 | 0.41 | 0.24 |
| 8 | 31280717 | rs6983355 | 1.139 | 1.108 | -0.743 | 0.038 | -0.004 | -0.012 | -0.012 | 0 | 0.32 | 0.19 | 0.2 | 0.28 | 0.34 | 0.23 |
| 8 | 31283253 | rs12543574 | 0.339 | 1.012 | -0.643 | -0.11 | -0.023 | -0.032 | -0.032 | 0 | 0.13 | 0.21 | 0.23 | 0.06 | 0.08 | 0.19 |
| 8 | 31283473 | rs6981897 | 0.965 | 1.125 | -0.919 | -0.04 | -0.004 | -0.016 | -0.016 | 0 | 0.33 | 0.19 | 0.2 | 0.3 | 0.34 | 0.24 |
| 8 | 31297286 | rs2681599 | -0.508 | 1.38 | -0.774 | -0.796 | -0.005 | -0.004 | -0.004 | 0 | 0.18 | 0.15 | 0.3 | 0.04 | 0.2 | 0.24 |
| 8 | 31297579 | rs10107214 | 1.237 | 1.217 | -0.737 | -0.773 | 0 | -0.007 | -0.007 | 0 | 0.33 | 0.3 | 0.21 | 0.48 | 0.34 | 0.26 |
| 8 | 31321243 | rs2681619 | -0.31 | 1.241 | -0.54 | -0.928 | 0.004 | -0.003 | -0.003 | 0 | 0.62 | 0.72 | 0.62 | 0.57 | 0.65 | 0.53 |
| 8 | 31330263 | rs7813593 | 0.406 | 1.137 | 0.061 | -0.911 | 0 | 0.017 | 0.017 | 0.137 | 0.1 | 0.26 | 0.13 | 0.09 | 0.08 | 0.07 |

| | | | | | | | | | | | | | | | | |
|---|---|---|---|---|---|---|---|---|---|---|---|---|---|---|---|---|
| 8 | 31333436 | rs7828595 | -0.854 | 1.382 | 0.012 | -1.045 | 0.027 | -0.08 | -0.08 | 0 | 0.25 | 0.39 | 0.37 | 0.1 | 0.27 | 0.33 |
| 8 | 31350562 | rs10954791 | 1.663 | 1.199 | -0.065 | -1.037 | 0.001 | -0.057 | -0.057 | 0 | 0.34 | 0.19 | 0.21 | 0.48 | 0.34 | 0.28 |
| 8 | 31352499 | rs10101482 | 0.424 | 1.125 | 0.05 | -1.148 | -0.007 | 0.01 | 0.01 | 0.002 | 0.11 | 0.26 | 0.13 | 0.09 | 0.08 | 0.08 |
| 8 | 31353463 | rs2681615 | -1.814 | 1.1 | 0.055 | -1.349 | -0.01 | 0.03 | 0.03 | 0.172 | 0.43 | 0.63 | 0.54 | 0.18 | 0.48 | 0.45 |
| 8 | 31354360 | rs2681611 | -2.285 | 1.027 | -0.168 | -1.436 | -0.02 | 0.003 | 0.003 | 0 | 0.94 | 1 | 0.97 | 0.97 | 0.96 | 0.96 |
| 8 | 31354452 | rs10094967 | 0.604 | 1.02 | -0.11 | -1.407 | -0.007 | 0.011 | 0.011 | 0.012 | 0.11 | 0.26 | 0.13 | 0.09 | 0.08 | 0.08 |
| 8 | 31354758 | rs2247267 | -0.411 | 1.085 | -0.298 | -1.439 | -0.064 | 0.044 | 0.044 | 0.147 | 0.17 | 0.07 | 0.32 | 0.1 | 0.26 | 0.31 |
| 8 | 31355172 | rs2880958 | 0.205 | 1.137 | -0.298 | -1.438 | -0.026 | 0.023 | 0.023 | 0 | 0.13 | 0.07 | 0.12 | 0.09 | 0.16 | 0.17 |
| 8 | 31369158 | rs4326344 | -0.413 | 1.108 | -0.239 | -1.452 | 0.014 | -0.066 | -0.066 | 0.492 | 0.86 | 0.78 | 0.75 | 1 | 0.84 | 0.83 |
| 8 | 31370455 | rs13253857 | 1.865 | 1.105 | -0.355 | -1.448 | -0.032 | 0.115 | 0.115 | 0.532 | 0.56 | 0.26 | 0.45 | 0.81 | 0.52 | 0.55 |
| 8 | 31384518 | rs12676976 | -0.567 | 1.23 | -0.209 | -1.279 | 0.02 | 0.005 | 0.005 | 0 | 0.16 | 0.23 | 0.23 | 0.18 | 0.21 | 0.19 |
| 8 | 31398121 | rs4489285 | 1.342 | 1.15 | -0.174 | -1.34 | 0.029 | -0.018 | -0.018 | 2.498 | 0.68 | 0.61 | 0.5 | 0.8 | 0.57 | 0.59 |
| 8 | 31398144 | rs4478530 | 0.582 | 1.178 | -0.229 | -1.341 | -0.075 | 0.058 | 0.058 | 0 | 0.77 | 0.91 | 0.65 | 0.89 | 0.7 | 0.64 |
| 8 | 31400219 | rs10954797 | 1.861 | 0.921 | -0.182 | -1.353 | 0.018 | -0.027 | -0.027 | 0 | 0.41 | 0.29 | 0.23 | 0.48 | 0.37 | 0.34 |
| 8 | 31401339 | rs9792249 | 1.484 | 0.978 | -0.151 | -1.365 | -0.002 | 0.011 | 0.011 | 0 | 0.59 | 0.39 | 0.44 | 0.79 | 0.51 | 0.57 |
| 8 | 31401802 | rs11777352 | 2.428 | 0.923 | -0.148 | -1.398 | 0.001 | -0.057 | -0.057 | 0 | 0.34 | 0.19 | 0.21 | 0.48 | 0.34 | 0.29 |
| 8 | 31403217 | rs10105518 | 1.412 | 1.596 | 0.132 | -1.222 | 0.051 | -0.016 | -0.016 | 0.454 | 0.75 | 0.61 | 0.57 | 0.84 | 0.67 | 0.64 |
| 8 | 31403563 | rs10095141 | 1.252 | 1.481 | 0.008 | -1.24 | 0.01 | -0.005 | -0.005 | 0.123 | 0.77 | 0.73 | 0.62 | 0.84 | 0.75 | 0.67 |
| 8 | 31414246 | rs7003512 | 1.317 | 1.526 | -0.254 | -1.09 | 0.029 | -0.041 | -0.041 | 0 | 0.63 | 0.51 | 0.48 | 0.79 | 0.59 | 0.6 |
| 8 | 31434385 | rs1557800 | -0.033 | 1.652 | -0.583 | -0.996 | -0.016 | 0.01 | 0.01 | 0 | 0.1 | 0.08 | 0.23 | 0.05 | 0.16 | 0.26 |
| 8 | 31447143 | rs11991032 | -0.435 | 1.369 | -0.725 | -1.039 | -0.003 | 0.002 | 0.002 | 0 | 0.2 | 0.33 | 0.11 | 0.16 | 0.2 | 0.18 |
| 8 | 31448469 | rs13258147 | 1.527 | 1.356 | -1.473 | -1.477 | 0.046 | 0.15 | 0.15 | 0.031 | 0.46 | 0.15 | 0.38 | 0.81 | 0.41 | 0.46 |
| 8 | 31449769 | rs10489349 | 0.494 | 2.033 | -1.309 | -0.816 | -0.047 | 0.043 | 0.043 | 0 | 0.82 | 0.88 | 0.7 | 0.93 | 0.77 | 0.67 |
| 8 | 31453914 | rs7817546 | 0.816 | 1.904 | -1.212 | -0.79 | -0.027 | 0.023 | 0.023 | 0 | 0.8 | 0.85 | 0.61 | 0.9 | 0.69 | 0.82 |
| 8 | 31469479 | rs4513929 | 1.067 | 1.696 | -0.876 | -0.892 | 0 | -0.001 | -0.001 | 0.04 | 0.75 | 0.26 | 0.52 | 0.86 | 0.68 | 0.62 |
| 8 | 31474141 | rs35753505 | 1.501 | 1.04 | -1.822 | -0.871 | -0.006 | -0.096 | -0.096 | 0.415 | 0.4 | 0.05 | 0.28 | 0.57 | 0.35 | 0.35 |
| 8 | 31486788 | rs10109424 | -0.724 | 1.082 | -1.251 | -0.814 | -0.003 | -0.001 | -0.001 | 0 | 0.41 | 0.61 | 0.59 | 0.41 | 0.51 | 0.57 |
| 8 | 31487450 | rs4452759 | -0.518 | 1.294 | -0.939 | -0.778 | -0.037 | -0.016 | -0.016 | 0 | 0.34 | 0.24 | 0.54 | 0.41 | 0.46 | 0.4 |
| 8 | 31490474 | rs4733263 | -0.84 | 0.902 | -0.521 | -0.949 | 0.011 | 0.007 | 0.007 | 0 | 0.48 | 0.42 | 0.65 | 0.41 | 0.62 | 0.5 |
| 8 | 31495374 | rs4281084 | 0.385 | 1.183 | -0.685 | -0.893 | -0.002 | -0.008 | -0.008 | 0 | 0.2 | 0.18 | 0.24 | 0.31 | 0.25 | 0.25 |
| 8 | 31495581 | rs6994992 | 0.794 | 0.969 | -0.445 | -1.018 | 0.025 | 0.019 | 0.019 | 0 | 0.5 | 0.39 | 0.34 | 0.59 | 0.38 | 0.49 |
| 8 | 31496006 | rs28401439 | -0.281 | 0.449 | 0.231 | -1.117 | -0.024 | -0.049 | -0.049 | 0 | 0.09 | 0.28 | 0.06 | 0 | 0.04 | 0.15 |
| 8 | 31500207 | rs1081064 | -0.309 | 0.578 | 0.069 | -0.834 | -0.001 | 0.038 | 0.038 | 0 | 0.22 | 0.39 | 0.26 | 0.3 | 0.25 | 0.22 |
| 8 | 31500222 | rs1081063 | 1.559 | 0.591 | -1.187 | -0.985 | -0.096 | -0.163 | -0.163 | 0.05 | 0.26 | 0.02 | 0.23 | 0.59 | 0.27 | 0.26 |
| 8 | 31504639 | rs4733267 | 1.582 | 0.591 | -1.187 | -0.984 | -0.097 | -0.163 | -0.163 | 1.949 | 0.26 | 0.02 | 0.23 | 0.59 | 0.27 | 0.26 |
| 8 | 31506771 | rs6987996 | 0.432 | 0.452 | -1.219 | -1.334 | 0.049 | 0.077 | 0.077 | 0.02 | 0.52 | 0.28 | 0.52 | 0.68 | 0.52 | 0.52 |
| 8 | 31523972 | rs7463426 | 1.628 | 0.408 | -1.418 | -1.275 | -0.047 | -0.119 | -0.119 | 1.384 | 0.28 | 0.08 | 0.22 | 0.6 | 0.26 | 0.29 |
| 8 | 31524553 | rs7009043 | -0.267 | 0.328 | -1.205 | -1.468 | -0.006 | 0.045 | 0.045 | 0 | 0.39 | 0.2 | 0.45 | 0.28 | 0.47 | 0.31 |
| 8 | 31539942 | rs12677942 | 0.745 | 0.393 | -1.22 | -1.379 | -0.016 | -0.039 | -0.039 | 0.001 | 0.17 | 0.05 | 0.24 | 0.27 | 0.24 | 0.2 |
| 8 | 31555139 | rs4552856 | 1.734 | 0.484 | -1.238 | -1.744 | 0.005 | -0.02 | -0.02 | 0.03 | 0.1 | 0.14 | 0.23 | 0.01 | 0.14 | 0.09 |
| 8 | 31555743 | rs10954810 | 0.933 | 0.231 | -1.426 | -1.859 | 0.005 | 0.015 | 0.015 | 0 | 0.21 | 0.18 | 0.21 | 0.01 | 0.23 | 0.1 |
| 8 | 31573695 | rs4566990 | -0.828 | 0.088 | -1.651 | -1.946 | -0.027 | 0.042 | 0.042 | 0 | 0.56 | 0.24 | 0.52 | 0.33 | 0.59 | 0.47 |
| 8 | 31575884 | rs7006674 | 0.767 | 0.069 | -1.661 | -1.972 | 0.009 | 0.048 | 0.048 | 0 | 0.23 | 0.18 | 0.22 | 0.01 | 0.24 | 0.12 |
| 8 | 31585000 | rs13260061 | -0.061 | 0.151 | -1.857 | -1.87 | -0.039 | -0.023 | -0.023 | 1.35 | 0.32 | 0.06 | 0.3 | 0.31 | 0.35 | 0.34 |
| 8 | 31597593 | rs4733094 | 2.192 | 0.157 | -1.835 | -1.826 | -0.074 | -0.11 | -0.11 | 0 | 0.25 | 0.02 | 0.21 | 0.65 | 0.26 | 0.28 |
| 8 | 31609048 | rs16878243 | 1.008 | 0.294 | -1.692 | -1.688 | -0.022 | -0.03 | -0.03 | 0.883 | 0.16 | 0.02 | 0.12 | 0.21 | 0.18 | 0.16 |
| 8 | 31633447 | rs10503887 | 1.539 | 0.244 | -1.758 | -1.915 | -0.033 | 0.077 | 0.077 | 0 | 0.16 | 0.02 | 0.18 | 0.01 | 0.2 | 0.1 |
| 8 | 31638990 | rs13279876 | 0.717 | 0.321 | -1.327 | -1.845 | -0.002 | 0.002 | 0.002 | 0 | 0.16 | 0.08 | 0.17 | 0.12 | 0.16 | 0.18 |
| 8 | 31650609 | rs2221349 | 0.367 | 0.338 | -0.782 | -1.969 | -0.009 | 0.016 | 0.016 | 0 | 0.22 | 0.17 | 0.26 | 0.08 | 0.24 | 0.24 |
| 8 | 31655026 | rs1503499 | 0.353 | 0.337 | -0.783 | -1.97 | -0.009 | 0.017 | 0.017 | 0 | 0.23 | 0.17 | 0.26 | 0.08 | 0.24 | 0.23 |
| 8 | 31663923 | rs11775517 | 0.947 | -0.239 | -0.57 | -2.061 | 0.006 | 0.002 | 0.002 | 0 | 0.22 | 0.2 | 0.23 | 0.01 | 0.23 | 0.11 |
| 8 | 31667034 | rs17603821 | 1.115 | -0.26 | -0.453 | -2.074 | -0.002 | -0.003 | -0.003 | 0 | 0.15 | 0.08 | 0.12 | 0.19 | 0.18 | 0.16 |
| 8 | 31668404 | rs17603876 | 1.866 | -0.233 | -0.637 | -2.146 | -0.036 | 0.038 | 0.038 | 0 | 0.07 | 0 | 0.08 | 0 | 0.12 | 0.02 |
| 8 | 31675630 | rs6987310 | 1.189 | -0.258 | -0.537 | -2.083 | -0.019 | 0.118 | 0.118 | 0 | 0.21 | 0.05 | 0.21 | 0.01 | 0.23 | 0.11 |
| 8 | 31680070 | rs1354334 | 1.023 | -0.277 | -0.055 | -2.079 | -0.004 | 0.003 | 0.003 | 0 | 0.42 | 0.44 | 0.47 | 0.68 | 0.41 | 0.53 |
| 8 | 31700703 | rs975199 | -0.538 | -0.433 | 0.177 | -2.235 | -0.035 | -0.141 | -0.141 | 0 | 0.73 | 0.55 | 0.76 | 0.99 | 0.76 | 0.88 |
| 8 | 31703542 | rs1383895 | 0.734 | -0.295 | 0.006 | -2.34 | -0.004 | 0.127 | 0.127 | 0 | 0.47 | 0.24 | 0.39 | 0.8 | 0.47 | 0.61 |
| 8 | 31706695 | rs776404 | -0.153 | -0.292 | -0.24 | -2.335 | 0.005 | 0.018 | 0.018 | 0 | 0.39 | 0.34 | 0.36 | 0.19 | 0.43 | 0.29 |
| 8 | 31716962 | rs776401 | -0.787 | -0.059 | 0.089 | -2.076 | 0.011 | 0.044 | 0.044 | 0 | 0.66 | 0.76 | 0.74 | 0.93 | 0.67 | 0.78 |
| 8 | 31718367 | rs16878368 | 1.87 | 0.009 | -0.14 | -2.034 | 0.002 | -0.025 | -0.025 | 0.422 | 0.1 | 0.13 | 0.23 | 0.01 | 0.11 | 0.1 |
| 8 | 31721207 | rs1383890 | -0.188 | -0.015 | -0.05 | -2.029 | -0.01 | -0.056 | -0.056 | 0 | 0.58 | 0.54 | 0.6 | 0.89 | 0.6 | 0.7 |
| 8 | 31743113 | rs2062055 | 0.901 | 0.142 | -0.764 | -1.865 | -0.037 | 0.019 | 0.019 | 0 | 0.34 | 0.16 | 0.36 | 0.74 | 0.37 | 0.49 |
| 8 | 31758989 | rs1383888 | 1.741 | 0.115 | -0.803 | -1.801 | -0.105 | -0.088 | -0.088 | 0.011 | 0.27 | 0.02 | 0.22 | 0.69 | 0.3 | 0.28 |
| 8 | 31764280 | rs13266217 | 0.796 | 0.222 | -0.75 | -1.907 | -0.013 | -0.008 | -0.008 | 0.678 | 0.18 | 0.08 | 0.25 | 0.22 | 0.25 | 0.23 |
| 8 | 31765337 | rs776398 | 0.178 | 0.264 | -0.685 | -1.762 | 0 | -0.008 | -0.008 | 0.547 | 0.46 | 0.5 | 0.48 | 0.71 | 0.43 | 0.55 |
| 8 | 31768217 | rs796549 | -0.167 | -0.069 | -0.167 | -1.501 | -0.001 | -0.108 | -0.108 | 0 | 0.17 | 0.44 | 0.26 | 0.01 | 0.12 | 0.12 |
| 8 | 31779956 | rs1481624 | 0.089 | -0.135 | -0.805 | -1.104 | 0.015 | 0.07 | 0.07 | 0 | 0.7 | 0.82 | 0.79 | 0.92 | 0.71 | 0.78 |
| 8 | 31789101 | rs435904 | -0.449 | -0.253 | -0.458 | -0.433 | -0.006 | 0.038 | 0.038 | 0.064 | 0.84 | 0.82 | 0.85 | 0.4 | 0.88 | 0.85 |
| 8 | 31791685 | rs4373486 | 0.489 | -0.781 | 0.38 | -0.795 | -0.052 | 0.068 | 0.068 | 1.88 | 0.74 | 0.52 | 0.83 | 0.39 | 0.85 | 0.71 |

| | | | | | | | | | | | | | | | | |
|---|---|---|---|---|---|---|---|---|---|---|---|---|---|---|---|---|
| 8 | 31801518 | rs16878500 | 0.352 | -0.233 | -0.044 | -1.019 | 0.022 | -0.006 | -0.006 | 0.031 | 0.05 | 0.08 | 0.15 | 0.05 | 0.09 | 0.05 |
| 8 | 31803534 | rs7002063 | 1.302 | -0.721 | -0.234 | -1.059 | 0.035 | 0.056 | 0.056 | 2.785 | 0.62 | 0.13 | 0.65 | 0.23 | 0.76 | 0.62 |
| 8 | 31815427 | rs13251778 | 0.56 | -0.771 | -0.593 | -1.252 | -0.025 | -0.034 | -0.034 | 0 | 0.11 | 0 | 0.05 | 0.09 | 0.13 | 0.14 |
| 8 | 31816746 | rs10808317 | -0.173 | -0.623 | 0.114 | -1.416 | -0.007 | 0.014 | 0.014 | 0.73 | 0.73 | 0.74 | 0.79 | 0.4 | 0.78 | 0.81 |
| 8 | 31824555 | rs1016165 | 2.262 | -0.989 | -0.55 | -1.768 | -0.148 | 0.134 | 0.134 | 0.276 | 0.33 | 0.03 | 0.41 | 0.14 | 0.42 | 0.4 |
| 8 | 31826219 | rs327416 | -1.158 | -0.96 | -0.022 | -1.334 | -0.054 | -0.126 | -0.126 | 0 | 0.81 | 0.99 | 0.81 | 0.39 | 0.8 | 0.83 |
| 8 | 31830933 | rs1462891 | 0.07 | -0.897 | 0.536 | -1.298 | -0.024 | 0.033 | 0.033 | 1.616 | 0.71 | 0.52 | 0.78 | 0.39 | 0.78 | 0.81 |
| 8 | 31830996 | rs1462892 | 0.812 | -0.896 | 0.032 | -1.304 | 0.064 | -0.015 | -0.015 | 0 | 0.63 | 0.21 | 0.74 | 0.38 | 0.77 | 0.72 |
| 8 | 31835829 | rs13276200 | 0.853 | -0.681 | 0.146 | -1.203 | 0.02 | 0.003 | 0.003 | 0 | 0.05 | 0.09 | 0.15 | 0.08 | 0.08 | 0.05 |
| 8 | 31846380 | rs2010243 | 0.368 | -0.882 | -0.22 | -1.196 | 0.021 | -0.012 | -0.012 | 0 | 0.68 | 0.3 | 0.75 | 0.38 | 0.78 | 0.81 |
| 8 | 31887758 | rs13266401 | 1.047 | -1.102 | -0.26 | -1.323 | -0.035 | -0.043 | -0.043 | 0 | 0.87 | 0.27 | 0.94 | 0.99 | 0.97 | 0.96 |
| 8 | 31888494 | rs7827456 | 1.096 | -1.16 | -0.649 | -1.361 | 0.017 | 0.082 | 0.082 | 2.136 | 0.55 | 0.11 | 0.57 | 0.29 | 0.68 | 0.57 |
| 8 | 31919990 | rs13271344 | 0.976 | -1.144 | -0.422 | -1.178 | 0.036 | 0.063 | 0.063 | 1.417 | 0.57 | 0.19 | 0.65 | 0.33 | 0.68 | 0.66 |
| 8 | 31926881 | rs1462904 | 0.702 | -1.146 | 0.011 | -1.179 | 0.002 | 0.003 | 0.003 | 0.677 | 0.6 | 0.42 | 0.67 | 0.34 | 0.68 | 0.66 |
| 8 | 31927638 | rs1462900 | 0.416 | -1.035 | -0.38 | -1.145 | -0.03 | -0.052 | -0.052 | 0 | 0.26 | 0.11 | 0.27 | 0.64 | 0.26 | 0.19 |
| 8 | 31935302 | rs1381871 | 0.644 | -1.056 | -0.446 | -1.115 | -0.066 | -0.088 | -0.088 | 0 | 0.24 | 0.03 | 0.27 | 0.64 | 0.26 | 0.19 |
| 8 | 31936552 | rs16878764 | -0.181 | -1.175 | -0.712 | -1.335 | -0.064 | -0.032 | -0.032 | 0.369 | 0.35 | 0.09 | 0.38 | 0.65 | 0.38 | 0.25 |
| 8 | 31941211 | rs16878772 | -0.251 | -0.902 | -0.234 | -1.154 | -0.017 | 0.035 | 0.035 | 1.17 | 0.07 | 0.3 | 0.03 | 0.09 | 0.01 | 0.09 |
| 8 | 31942557 | rs1381874 | -0.395 | -0.888 | -0.339 | -1.166 | -0.027 | -0.007 | -0.007 | 0 | 0.11 | 0.41 | 0.05 | 0.09 | 0.04 | 0.13 |
| 8 | 31943664 | rs7841220 | 0.392 | -0.526 | -1.303 | -1.19 | 0.029 | 0.117 | 0.117 | 1.513 | 0.55 | 0.1 | 0.65 | 0.25 | 0.68 | 0.58 |
| 8 | 31947234 | rs10503899 | 1.026 | -0.038 | -1.22 | -1.106 | -0.151 | 0.127 | 0.127 | 0 | 0.34 | 0.03 | 0.41 | 0.15 | 0.44 | 0.4 |
| 8 | 31947880 | rs7838546 | -1.669 | 0.012 | 0.322 | -0.425 | 0.061 | 0.086 | 0.086 | 0 | 0.1 | 0.4 | 0.27 | 0.09 | 0.07 | 0.08 |
| 8 | 31948864 | rs11776959 | 0.561 | -0.345 | -0.75 | -1.166 | 0.036 | 0.202 | 0.202 | 0.012 | 0.48 | 0.15 | 0.69 | 0.23 | 0.62 | 0.43 |
| 8 | 31952710 | rs10954827 | 0.545 | -0.348 | -0.752 | -1.162 | 0.037 | 0.23 | 0.23 | 0.007 | 0.48 | 0.15 | 0.69 | 0.16 | 0.62 | 0.44 |
| 8 | 31952745 | rs7007662 | -0.405 | -0.372 | 0.06 | -0.419 | 0.002 | -0.001 | -0.001 | 0.109 | 0.47 | 0.58 | 0.18 | 0.52 | 0.33 | 0.45 |
| 8 | 31955149 | rs13252152 | -0.424 | -0.65 | 0.015 | -0.774 | 0.018 | -0.001 | -0.001 | 0 | 0.48 | 0.28 | 0.24 | 0.49 | 0.45 | 0.39 |
| 8 | 31959943 | rs1542517 | 1.077 | -0.563 | 0.666 | -0.719 | 0.026 | 0.325 | 0.325 | 0 | 0.9 | 0.62 | 0.86 | 0.66 | 0.95 | 0.79 |
| 8 | 31960376 | rs9297186 | 0.953 | -0.525 | 0.681 | -0.695 | 0.059 | 0.284 | 0.284 | 0.042 | 0.9 | 0.61 | 0.84 | 0.72 | 0.92 | 0.79 |
| 8 | 31964189 | rs7815226 | -0.26 | -0.884 | 0.362 | -1.013 | -0.037 | -0.091 | -0.091 | 0.019 | 0.42 | 0.11 | 0.24 | 0.51 | 0.44 | 0.34 |
| 8 | 31965188 | rs6993043 | 1.003 | -1.033 | 0.487 | -1.1 | 0.059 | 0.215 | 0.215 | 0 | 0.5 | 0.18 | 0.69 | 0.22 | 0.63 | 0.44 |
| 8 | 31966653 | rs12056821 | 0.843 | -1.029 | 0.581 | -1.098 | -0.001 | -0.1 | -0.1 | 0.456 | 0.28 | 0.12 | 0.11 | 0.5 | 0.26 | 0.28 |
| 8 | 31967245 | rs1481756 | 0.936 | -1.052 | 0.621 | -1.11 | 0.036 | 0.093 | 0.093 | 0 | 0.51 | 0.32 | 0.71 | 0.22 | 0.63 | 0.44 |
| 8 | 31970568 | rs6995821 | -0.842 | -0.973 | 0.461 | -1.078 | -0.025 | -0.118 | -0.118 | 0.065 | 0.72 | 0.91 | 0.89 | 0.5 | 0.71 | 0.72 |
| 8 | 31980642 | rs1481736 | 0.465 | -0.929 | 0.404 | -1.074 | 0.032 | -0.043 | -0.043 | 0.378 | 0.52 | 0.79 | 0.75 | 0.5 | 0.58 | 0.62 |
| 8 | 31986192 | rs981817 | 1.709 | -0.803 | 0.402 | -1.098 | -0.038 | 0.062 | 0.062 | 0 | 0.36 | 0.22 | 0.57 | 0.14 | 0.48 | 0.39 |
| 8 | 31987432 | rs6989889 | -0.109 | -0.874 | 0.294 | -1.071 | 0.034 | -0.029 | -0.029 | 0.417 | 0.63 | 0.74 | 0.87 | 0.46 | 0.71 | 0.58 |
| 8 | 31991227 | rs1481762 | 0.413 | -1.032 | 0.18 | -1.166 | 0.019 | 0.001 | 0.001 | 0 | 0.54 | 0.7 | 0.77 | 0.45 | 0.56 | 0.56 |
| 8 | 31993810 | rs1481763 | 1.599 | -0.771 | 0.258 | -1.076 | -0.025 | 0.104 | 0.104 | 0.003 | 0.39 | 0.21 | 0.61 | 0.14 | 0.54 | 0.43 |
| 8 | 31995958 | rs7835014 | -0.463 | -0.823 | 0.417 | -0.978 | -0.005 | -0.048 | -0.048 | 0 | 0.18 | 0.2 | 0.14 | 0 | 0.14 | 0.07 |
| 8 | 32000665 | rs6468085 | -0.194 | -0.909 | 0.378 | -0.829 | 0.014 | -0.01 | -0.01 | 0 | 0.2 | 0.16 | 0.19 | 0.23 | 0.13 | 0.2 |
| 8 | 32008253 | rs1481761 | 1.666 | -0.481 | 0.427 | -0.795 | -0.059 | 0.162 | 0.162 | 0 | 0.41 | 0.17 | 0.61 | 0.15 | 0.54 | 0.42 |
| 8 | 32008543 | rs34442697 | 1.219 | -0.35 | 0.704 | -0.562 | -0.003 | 0.084 | 0.084 | 0.556 | 0.09 | 0.15 | 0.12 | 0.35 | 0.06 | 0.16 |
| 8 | 32010506 | rs2068226 | 1.303 | -0.33 | 0.883 | -0.607 | 0 | 0.024 | 0.024 | 0 | 0.47 | 0.36 | 0.66 | 0.18 | 0.58 | 0.48 |
| 8 | 32019288 | rs4733303 | 1.41 | -0.294 | 0.9 | -0.652 | -0.002 | 0.021 | 0.021 | 0 | 0.46 | 0.36 | 0.65 | 0.24 | 0.58 | 0.49 |
| 8 | 32019682 | rs16878885 | 0.137 | -0.287 | 0.679 | -0.662 | 0.022 | 0.044 | 0.044 | 0.033 | 0.66 | 0.85 | 0.8 | 0.25 | 0.72 | 0.63 |
| 8 | 32020891 | rs16878890 | -0.326 | -0.245 | 0.853 | -0.662 | 0.025 | -0.048 | -0.048 | 0.036 | 0.09 | 0.26 | 0.07 | 0 | 0.07 | 0.07 |
| 8 | 32021186 | rs1481742 | -0.523 | -0.119 | 0.336 | -0.403 | 0.021 | -0.081 | -0.081 | 0 | 0.75 | 0.87 | 0.92 | 0.54 | 0.8 | 0.81 |
| 8 | 32066467 | rs7001060 | 0.047 | 0.173 | 0.214 | -0.226 | -0.001 | 0.015 | 0.015 | 0.806 | 0.12 | 0.12 | 0.14 | 0.29 | 0.09 | 0.17 |
| 8 | 32068180 | rs7836913 | -1.162 | 0.187 | 0.007 | -0.239 | 0.017 | 0.045 | 0.045 | 0 | 0.23 | 0.49 | 0.25 | 0.28 | 0.2 | 0.27 |
| 8 | 32078855 | rs7841677 | -0.058 | 0.347 | -0.985 | -0.591 | -0.045 | 0.07 | 0.07 | 0 | 0.62 | 0.81 | 0.66 | 0.27 | 0.61 | 0.64 |
| 8 | 32082436 | rs4733308 | 0.661 | -0.157 | -1.239 | -0.918 | -0.03 | -0.043 | -0.043 | 0 | 0.45 | 0.17 | 0.71 | 0.52 | 0.59 | 0.65 |
| 8 | 32085171 | rs10503901 | -0.665 | -0.178 | -1.001 | -0.96 | 0.004 | -0.03 | -0.03 | 0 | 0.23 | 0.73 | 0.14 | 0 | 0.1 | 0.13 |
| 8 | 32100541 | rs1481734 | 1.201 | -0.204 | -0.712 | -0.925 | -0.005 | 0.01 | 0.01 | 0 | 0.37 | 0.25 | 0.5 | 0.2 | 0.47 | 0.43 |
| 8 | 32105334 | rs989465 | -0.055 | -0.263 | -1.365 | -0.827 | 0.024 | -0.029 | -0.029 | 0 | 0.58 | 0.78 | 0.81 | 0.51 | 0.65 | 0.72 |
| 8 | 32112409 | rs2881544 | -0.226 | -0.234 | -1.323 | -0.822 | 0.023 | -0.027 | -0.027 | 0 | 0.59 | 0.79 | 0.82 | 0.5 | 0.66 | 0.69 |
| 8 | 32116927 | rs1481728 | -0.975 | 0.063 | -1.243 | -0.451 | -0.014 | 0.031 | 0.031 | 0 | 0.18 | 0.29 | 0.23 | 0.26 | 0.13 | 0.16 |
| 8 | 32117412 | rs6992010 | -0.988 | 0.059 | -1.245 | -0.453 | -0.014 | 0.03 | 0.03 | 0 | 0.18 | 0.29 | 0.23 | 0.26 | 0.13 | 0.16 |
| 8 | 32118296 | rs16879060 | -0.677 | 0.221 | -1.178 | -0.351 | -0.005 | 0.064 | 0.064 | 0 | 0.15 | 0.26 | 0.21 | 0.26 | 0.12 | 0.15 |
| 8 | 32118416 | rs4733313 | 0.158 | 0.275 | -1.129 | -0.434 | 0.019 | 0.018 | 0.018 | 0 | 0.5 | 0.3 | 0.7 | 0.47 | 0.59 | 0.6 |
| 8 | 32120188 | rs900102 | -0.056 | 0.745 | -1.196 | -1.022 | 0 | -0.019 | -0.019 | 0 | 0.64 | 0.42 | 0.54 | 0.82 | 0.66 | 0.55 |
| 8 | 32121424 | rs16879067 | 0.67 | 0.535 | -1.976 | -0.737 | 0.005 | -0.087 | -0.087 | 0 | 0.18 | 0.05 | 0.21 | 0.53 | 0.19 | 0.24 |
| 8 | 32122424 | rs17624592 | 0.004 | 1.039 | -2.051 | -0.84 | -0.02 | 0.02 | 0.02 | 0 | 0.05 | 0.03 | 0.05 | 0.01 | 0.09 | 0.07 |
| 8 | 32126374 | rs17624670 | 1.316 | 0.951 | -2.319 | -1.212 | -0.002 | -0.016 | -0.016 | 0 | 0.23 | 0.1 | 0.18 | 0.26 | 0.22 | 0.16 |
| 8 | 32131798 | rs1481758 | 0.963 | 1.206 | -2.27 | -0.903 | -0.01 | -0.098 | -0.098 | 0 | 0.2 | 0.05 | 0.23 | 0.52 | 0.21 | 0.25 |
| 8 | 32133284 | rs16879088 | -0.262 | 1.16 | -2.15 | -1.062 | 0.012 | -0.022 | -0.022 | 1.165 | 0.06 | 0.13 | 0.04 | 0.01 | 0.09 | 0.08 |
| 8 | 32134420 | rs1487146 | 0.474 | 1.034 | -2.39 | -0.835 | -0.012 | -0.096 | -0.096 | 2.917 | 0.24 | 0.08 | 0.26 | 0.56 | 0.23 | 0.28 |
| 8 | 32134461 | rs1487145 | -1.226 | 1.138 | -2.052 | -1.038 | -0.061 | 0.024 | 0.024 | 0 | 0.75 | 0.93 | 0.58 | 0.77 | 0.67 | 0.62 |
| 8 | 32137313 | rs16879099 | -0.246 | 1.085 | -1.056 | -0.944 | 0.09 | -0.005 | -0.005 | 0 | 0.05 | 0.23 | 0.04 | 0.02 | 0.08 | 0.06 |
| 8 | 32141515 | rs7824399 | -1.13 | 1.201 | -1.751 | -0.99 | -0.103 | 0.044 | 0.044 | 0 | 0.75 | 0.95 | 0.68 | 0.83 | 0.67 | 0.62 |

| | | | | | | | | | | | | | | | |
|---|---|---|---|---|---|---|---|---|---|---|---|---|---|---|---|
| 8 | 32143462 | rs901561 | -1.063 | 1.189 | -1.757 | -0.807 | -0.102 | 0.019 | 0.019 | 0 | 0.74 | 0.95 | 0.68 | 0.8 | 0.66 | 0.61 |
| 8 | 32147740 | rs7836407 | 1.305 | 0.128 | -1.652 | -0.969 | -0.035 | 0.003 | 0.003 | 0.674 | 0.24 | 0.35 | 0.17 | 0.22 | 0.16 | 0.12 |
| 8 | 32153164 | rs7818821 | 0.212 | 0.051 | -1.789 | -1.111 | -0.016 | -0.025 | -0.025 | 0.715 | 0.37 | 0.43 | 0.28 | 0.19 | 0.33 | 0.21 |
| 8 | 32156724 | rs11778887 | 1.025 | 0.217 | -1.976 | -1.013 | -0.025 | -0.07 | -0.07 | 0.29 | 0.23 | 0.09 | 0.35 | 0.6 | 0.24 | 0.29 |
| 8 | 32164476 | rs6468095 | -0.998 | 0.763 | -1.572 | -0.596 | 0.004 | -0.001 | -0.001 | 0 | 0.76 | 0.79 | 0.82 | 0.79 | 0.8 | 0.87 |
| 8 | 32167730 | rs7005124 | 0.417 | 1.081 | -1.831 | -0.65 | -0.071 | 0.078 | 0.078 | 0 | 0.3 | 0.19 | 0.33 | 0.18 | 0.39 | 0.49 |
| 8 | 32170767 | rs4317533 | 0.12 | 0.838 | -1.527 | -0.457 | 0.03 | 0.024 | 0.024 | 0 | 0.46 | 0.66 | 0.39 | 0.59 | 0.41 | 0.37 |
| 8 | 32172010 | rs10503907 | 0.381 | 1.055 | -1.05 | -1.333 | -0.004 | 0.002 | 0.002 | 0 | 0.25 | 0.29 | 0.16 | 0.15 | 0.35 | 0.42 |
| 8 | 32177531 | rs1386440 | 0.257 | 1.193 | -0.456 | -1.519 | 0.054 | 0.016 | 0.016 | 0.013 | 0.81 | 0.7 | 0.63 | 0.71 | 0.68 | 0.65 |
| 8 | 32184932 | rs1948098 | 0.15 | 1.313 | -0.338 | -1.75 | 0.022 | -0.008 | -0.008 | 0 | 0.21 | 0.28 | 0.23 | 0.2 | 0.34 | 0.38 |
| 8 | 32190881 | rs7822917 | 1.388 | 0.733 | -0.788 | -2.065 | -0.082 | -0.013 | -0.013 | 0 | 0.2 | 0.01 | 0.2 | 0.14 | 0.24 | 0.24 |
| 8 | 32192397 | rs16879153 | 1.227 | -0.226 | -0.475 | -2.445 | -0.021 | -0.09 | -0.09 | 0 | 0.26 | 0.31 | 0.13 | 0 | 0.18 | 0.11 |
| 8 | 32195259 | rs1487155 | -0.522 | -0.234 | -0.394 | -2.452 | -0.005 | -0.046 | -0.046 | 0 | 0.66 | 0.6 | 0.56 | 0.88 | 0.67 | 0.76 |
| 8 | 32195932 | rs17705398 | 1.996 | 0.086 | -0.395 | -2.266 | -0.003 | -0.006 | -0.006 | 0 | 0.12 | 0.15 | 0.08 | 0.02 | 0.17 | 0.15 |
| 8 | 32197182 | rs1685117 | 1.387 | -0.521 | -0.242 | -2.058 | -0.023 | 0.019 | 0.019 | 1.074 | 0.23 | 0.22 | 0.14 | 0.55 | 0.09 | 0.14 |
| 8 | 32197913 | rs1685101 | 1.596 | -0.533 | -0.215 | -2.06 | -0.002 | -0.003 | -0.003 | 0 | 0.21 | 0.16 | 0.14 | 0.56 | 0.09 | 0.14 |
| 8 | 32201084 | rs970998 | -0.012 | -0.757 | -0.394 | -2.228 | -0.006 | -0.007 | -0.007 | 0.113 | 0.59 | 0.56 | 0.49 | 0.33 | 0.65 | 0.69 |
| 8 | 32201135 | rs1565031 | -0.841 | -0.353 | -0.493 | -2.115 | 0.033 | 0.007 | 0.007 | 0 | 0.91 | 0.81 | 0.69 | 0.88 | 0.85 | 0.88 |
| 8 | 32214317 | rs10106329 | 1.289 | -1.153 | -0.501 | -2.27 | -0.014 | -0.054 | -0.054 | 0 | 0.22 | 0.2 | 0.11 | 0 | 0.11 | 0.08 |
| 8 | 32215602 | rs1623372 | 0.939 | -0.989 | -0.216 | -2.454 | 0.062 | -0.02 | -0.02 | 0 | 0.21 | 0.34 | 0.22 | 0.19 | 0.34 | 0.37 |
| 8 | 32217225 | rs1487152 | -0.264 | -1.203 | -0.368 | -2.469 | 0 | 0.031 | 0.031 | 0.029 | 0.52 | 0.62 | 0.42 | 0.72 | 0.51 | 0.54 |
| 8 | 32220852 | rs1626771 | 0.115 | -1.174 | -0.198 | -2.351 | 0.045 | 0.011 | 0.011 | 1.521 | 0.53 | 0.57 | 0.31 | 0.53 | 0.28 | 0.24 |
| 8 | 32227926 | rs6992907 | -0.343 | -1.073 | -0.333 | -2.399 | 0.003 | -0.02 | -0.02 | 0 | 0.57 | 0.51 | 0.64 | 0.28 | 0.58 | 0.49 |
| 8 | 32229351 | rs7011088 | 0.676 | -0.7 | -0.314 | -2.127 | 0.004 | 0 | 0 | 0 | 0.27 | 0.28 | 0.47 | 0.28 | 0.38 | 0.39 |
| 8 | 32232827 | rs2347503 | -1.079 | -0.8 | -0.263 | -1.772 | -0.009 | 0.004 | 0.004 | 1.253 | 0.78 | 0.79 | 0.86 | 0.47 | 0.91 | 0.86 |
| 8 | 32234782 | rs2347505 | -0.331 | -0.547 | 0.463 | -1.799 | -0.05 | 0.034 | 0.034 | 0 | 0.71 | 0.62 | 0.77 | 0.46 | 0.8 | 0.81 |
| 8 | 32234929 | rs2347506 | -0.75 | -0.487 | 0.227 | -1.785 | 0.029 | 0.032 | 0.032 | 0 | 0.6 | 0.8 | 0.66 | 0.36 | 0.68 | 0.57 |
| 8 | 32235025 | rs2347507 | 0.374 | 0.022 | 0.275 | -1.601 | -0.047 | -0.048 | -0.048 | 0.237 | 0.08 | 0.01 | 0.29 | 0.19 | 0.16 | 0.11 |
| 8 | 32235881 | rs4733319 | 0.675 | 0.122 | 0.412 | -1.577 | -0.014 | -0.014 | -0.014 | 0 | 0.07 | 0.04 | 0.29 | 0.19 | 0.15 | 0.11 |
| 8 | 32238916 | rs4352806 | -1.29 | 0.02 | 0.453 | -1.565 | -0.015 | 0.103 | 0.103 | 0 | 0.91 | 0.88 | 0.87 | 0.44 | 0.94 | 0.87 |
| 8 | 32240840 | rs16879304 | 1.159 | 0.021 | 0.63 | -1.731 | -0.009 | -0.043 | -0.043 | 1.104 | 0.08 | 0.12 | 0.07 | 0 | 0.08 | 0.03 |
| 8 | 32242498 | rs13276181 | 0.383 | 0.339 | 0.556 | -1.724 | -0.002 | -0.006 | -0.006 | 0 | 0.39 | 0.35 | 0.23 | 0 | 0.39 | 0.23 |
| 8 | 32245538 | rs939077 | 0.623 | 0.342 | 0.585 | -1.711 | -0.017 | 0.072 | 0.072 | 0 | 0.41 | 0.23 | 0.51 | 0.28 | 0.43 | 0.42 |
| 8 | 32247220 | rs4733325 | 0.54 | 0.364 | 0.619 | -1.7 | -0.004 | 0.035 | 0.035 | 0 | 0.42 | 0.36 | 0.51 | 0.28 | 0.43 | 0.42 |
| 8 | 32247739 | rs6986716 | -0.063 | 0.375 | 0.491 | -1.672 | -0.009 | 0.029 | 0.029 | 0 | 0.8 | 0.7 | 0.74 | 0.29 | 0.82 | 0.65 |
| 8 | 32250029 | rs6986789 | 0.535 | 0.413 | 0.037 | -1.628 | -0.009 | 0.054 | 0.054 | 0 | 0.42 | 0.3 | 0.51 | 0.28 | 0.43 | 0.42 |
| 8 | 32254397 | rs4733327 | 0.535 | 0.413 | 0.037 | -1.628 | -0.009 | 0.054 | 0.054 | 0 | 0.42 | 0.3 | 0.51 | 0.28 | 0.43 | 0.42 |
| 8 | 32254835 | rs10100501 | -1.177 | 0.047 | -0.214 | -1.293 | -0.015 | 0.101 | 0.101 | 0 | 0.91 | 0.88 | 0.87 | 0.45 | 0.94 | 0.88 |
| 8 | 32257621 | rs16879327 | 0.766 | 0.379 | -0.501 | -1.539 | 0.009 | 0.014 | 0.014 | 0 | 0.09 | 0.13 | 0.13 | 0.16 | 0.12 | 0.21 |
| 8 | 32257673 | rs4276645 | 0.459 | 0.271 | -0.965 | -1.682 | 0.017 | 0.108 | 0.108 | 0 | 0.34 | 0.16 | 0.22 | 0.09 | 0.28 | 0.31 |
| 8 | 32282019 | rs3847131 | -0.286 | 0.696 | -0.32 | -1.931 | 0.002 | -0.088 | -0.088 | 0.014 | 0.66 | 0.61 | 0.78 | 0.97 | 0.64 | 0.75 |
| 8 | 32282597 | rs9918832 | 0.399 | 0.65 | -0.141 | -1.923 | 0.003 | 0.005 | 0.005 | 0 | 0.2 | 0.23 | 0.29 | 0.71 | 0.22 | 0.31 |
| 8 | 32283471 | rs9297191 | -0.235 | 0.547 | -0.186 | -2.078 | 0 | -0.003 | -0.003 | 0 | 0.27 | 0.26 | 0.35 | 0.71 | 0.3 | 0.33 |
| 8 | 32283517 | rs9297192 | 1.167 | 0.607 | -0.133 | -1.808 | 0.02 | 0.007 | 0.007 | 0 | 0.11 | 0.19 | 0.19 | 0.13 | 0.16 | 0.18 |
| 8 | 32285177 | rs10503915 | 1.453 | 0.374 | -0.372 | -1.963 | 0.003 | 0.065 | 0.065 | 0 | 0.28 | 0.07 | 0.18 | 0.16 | 0.23 | 0.31 |
| 8 | 32286041 | rs16879366 | 0.842 | 0.54 | -0.03 | -1.848 | -0.001 | 0 | 0 | 0 | 0.08 | 0.1 | 0.26 | 0.09 | 0.12 | 0.1 |
| 8 | 32290601 | rs11782671 | -0.646 | 0.512 | 0.185 | -1.854 | 0.026 | -0.027 | -0.027 | 0.4 | 0.28 | 0.47 | 0.51 | 0.24 | 0.37 | 0.3 |
| 8 | 32292231 | rs17642273 | 1.595 | 0.853 | -0.345 | -2.056 | -0.064 | 0.062 | 0.062 | 0 | 0.11 | 0.02 | 0.07 | 0.02 | 0.16 | 0.18 |
| 8 | 32294151 | rs2347501 | 0.082 | 0.573 | 0.021 | -1.973 | 0.022 | -0.033 | -0.033 | 0 | 0.21 | 0.41 | 0.26 | 0.15 | 0.24 | 0.2 |
| 8 | 32298100 | rs4129581 | 1.249 | 0.21 | -0.16 | -2.113 | 0.011 | 0.013 | 0.013 | 0 | 0.2 | 0.14 | 0.18 | 0.01 | 0.19 | 0.08 |
| 8 | 32298476 | rs6981660 | 0.501 | 0.262 | 0.309 | -2.178 | 0.019 | -0.018 | -0.018 | 0 | 0.26 | 0.33 | 0.23 | 0.17 | 0.33 | 0.36 |
| 8 | 32299316 | rs4584111 | -0.415 | 0.218 | 0.526 | -2.2 | -0.015 | -0.108 | -0.108 | 1.883 | 0.71 | 0.57 | 0.56 | 0.9 | 0.68 | 0.83 |
| 8 | 32305720 | rs11993611 | -0.951 | 0.116 | 0.428 | -2.189 | -0.001 | -0.011 | -0.011 | 1.721 | 0.62 | 0.66 | 0.57 | 0.79 | 0.65 | 0.59 |
| 8 | 32306984 | rs6996957 | -0.806 | 0.099 | 0.433 | -2.134 | -0.011 | -0.03 | -0.03 | 0 | 0.6 | 0.57 | 0.69 | 0.28 | 0.67 | 0.55 |
| 8 | 32317917 | rs17716295 | 0.831 | 0.12 | 0.433 | -2.169 | -0.02 | 0.014 | 0.014 | 0 | 0.24 | 0.2 | 0.21 | 0.17 | 0.33 | 0.36 |
| 8 | 32319538 | rs6985581 | -1.194 | 0.054 | 0.482 | -2.211 | 0.002 | -0.024 | -0.024 | 0.639 | 0.5 | 0.57 | 0.64 | 0.88 | 0.57 | 0.61 |
| 8 | 32325220 | rs10096770 | 0.612 | 0.118 | 0.479 | -2.157 | 0.021 | -0.02 | -0.02 | 0 | 0.26 | 0.33 | 0.22 | 0.17 | 0.33 | 0.36 |
| 8 | 32332633 | rs10092307 | -0.568 | 0.051 | 0.511 | -2.157 | -0.005 | -0.03 | -0.03 | 0 | 0.55 | 0.62 | 0.54 | 0.79 | 0.63 | 0.58 |
| 8 | 32340967 | rs12216802 | 1.749 | 0.035 | 0.402 | -2.142 | 0 | 0.093 | 0.093 | 0 | 0.29 | 0.07 | 0.17 | 0.11 | 0.24 | 0.33 |
| 8 | 32345693 | rs7844698 | -0.619 | 0.067 | 0.438 | -2.115 | 0.004 | 0.009 | 0.009 | 0 | 0.56 | 0.68 | 0.55 | 0.78 | 0.62 | 0.58 |
| 8 | 32348088 | rs13254149 | 1.593 | 0.062 | 0.422 | -2.172 | 0.009 | 0.016 | 0.016 | 0 | 0.19 | 0.12 | 0.17 | 0.01 | 0.19 | 0.07 |
| 8 | 32349236 | rs4733342 | 0.556 | 0.051 | 0.465 | -2.137 | 0.001 | 0.01 | 0.01 | 0 | 0.14 | 0.07 | 0.28 | 0.11 | 0.15 | 0.1 |
| 8 | 32351333 | rs17645417 | 0.166 | 0.065 | 0.483 | -1.974 | -0.01 | 0.061 | 0.061 | 0 | 0.43 | 0.33 | 0.4 | 0.21 | 0.52 | 0.43 |
| 8 | 32363288 | rs7013361 | -1.154 | 0.063 | 0.411 | -2.12 | -0.006 | -0.032 | -0.032 | 0 | 0.63 | 0.71 | 0.81 | 0.89 | 0.75 | 0.66 |
| 8 | 32364970 | rs11777461 | 1.492 | 0.093 | 0.247 | -2.061 | 0 | 0 | 0 | 0 | 0.08 | 0.07 | 0.19 | 0.11 | 0.12 | 0.09 |
| 8 | 32368723 | rs13253155 | 1.569 | 0.039 | 0.178 | -2.12 | 0.01 | 0.033 | 0.033 | 0 | 0.19 | 0.09 | 0.17 | 0.01 | 0.19 | 0.08 |
| 8 | 32370520 | rs4422736 | 0.456 | -0.005 | 0.346 | -2.103 | -0.03 | 0.14 | 0.14 | 0 | 0.12 | 0.32 | 0.12 | 0.61 | 0.05 | 0.13 |
| 8 | 32375617 | rs7844425 | 0.904 | 0.061 | 0.261 | -2.149 | -0.019 | 0.013 | 0.013 | 0 | 0.23 | 0.19 | 0.21 | 0.16 | 0.32 | 0.36 |
| 8 | 32376010 | rs4733347 | -1.809 | 0.077 | 0.183 | -2.153 | -0.005 | -0.028 | -0.028 | 0.16 | 0.69 | 0.75 | 0.82 | 0.88 | 0.77 | 0.68 |

| | | | | | | | | | | | | | | | | |
|---|---|---|---|---|---|---|---|---|---|---|---|---|---|---|---|---|
| 8 | 32377064 | rs13282123 | 0.397 | 0.151 | 0.076 | -2.018 | -0.045 | 0.005 | 0.005 | 0 | 0.14 | 0.02 | 0.31 | 0.11 | 0.2 | 0.11 |
| 8 | 32382668 | rs6651144 | -1.184 | 0.126 | 0.232 | -1.984 | 0.014 | 0.056 | 0.056 | 0 | 0.51 | 0.7 | 0.66 | 0.82 | 0.56 | 0.6 |
| 8 | 32383339 | rs12548924 | 0.377 | 0.306 | 0.187 | -1.977 | -0.01 | 0.013 | 0.013 | 0 | 0.14 | 0.07 | 0.31 | 0.11 | 0.21 | 0.1 |
| 8 | 32383863 | rs7000397 | 0.13 | 0.279 | 0.265 | -1.987 | -0.011 | 0.019 | 0.019 | 0 | 0.31 | 0.28 | 0.24 | 0.16 | 0.33 | 0.39 |
| 8 | 32397189 | rs12546350 | 0.399 | 0.269 | 0.108 | -1.953 | 0.005 | 0.012 | 0.012 | 0 | 0.15 | 0.13 | 0.28 | 0.11 | 0.14 | 0.09 |
| 8 | 32399662 | rs4489283 | -0.652 | 0.234 | 0.155 | -2.027 | -0.001 | -0.006 | -0.006 | 0.02 | 0.54 | 0.56 | 0.53 | 0.72 | 0.61 | 0.59 |
| 8 | 32399742 | rs10503919 | 1.546 | 0.173 | 0.002 | -1.96 | 0.009 | 0.068 | 0.068 | 0 | 0.18 | 0.06 | 0.16 | 0.03 | 0.19 | 0.08 |
| 8 | 32400628 | rs7000590 | -1.523 | 0.356 | 0.362 | -1.938 | -0.002 | -0.024 | -0.024 | 0 | 0.69 | 0.67 | 0.8 | 0.83 | 0.75 | 0.67 |
| 8 | 32401561 | rs6468119 | -0.615 | 0.21 | 0.08 | -2.011 | 0 | -0.008 | -0.008 | 0 | 0.42 | 0.46 | 0.62 | 0.8 | 0.6 | 0.58 |
| 8 | 32406994 | rs4733130 | 0.114 | 0.266 | 0.172 | -1.972 | -0.043 | 0.043 | 0.043 | 0 | 0.31 | 0.22 | 0.47 | 0.18 | 0.47 | 0.42 |
| 8 | 32412359 | rs2439312 | -1.644 | 0.698 | -0.362 | -1.969 | -0.001 | -0.002 | -0.002 | 0 | 0.7 | 0.77 | 0.85 | 0.78 | 0.81 | 0.64 |
| 8 | 32414166 | rs17719687 | 2.08 | 1.005 | -0.66 | -2.019 | -0.046 | 0.048 | 0.048 | 0 | 0.09 | 0.02 | 0.08 | 0.01 | 0.17 | 0.17 |
| 8 | 32416274 | rs7825175 | 1.286 | 0.629 | -0.491 | -1.944 | -0.001 | 0 | 0 | 0 | 0.14 | 0.15 | 0.14 | 0.12 | 0.19 | 0.2 |
| 8 | 32417600 | rs13260545 | 1.238 | 0.588 | -0.708 | -1.845 | -0.026 | 0.093 | 0.093 | 0 | 0.34 | 0.05 | 0.19 | 0.19 | 0.27 | 0.39 |
| 8 | 32419119 | rs11776203 | 0.978 | 0.901 | -0.622 | -1.793 | -0.095 | 0.049 | 0.049 | 0 | 0.17 | 0.02 | 0.33 | 0.06 | 0.28 | 0.22 |
| 8 | 32428968 | rs4733362 | -0.202 | 0.737 | -0.349 | -1.744 | -0.002 | -0.047 | -0.047 | 0 | 0.64 | 0.43 | 0.63 | 0.85 | 0.61 | 0.73 |
| 8 | 32436875 | rs17721043 | 0.69 | 0.881 | -0.731 | -1.52 | -0.067 | 0.062 | 0.062 | 0 | 0.17 | 0.06 | 0.33 | 0.06 | 0.27 | 0.21 |
| 8 | 32437994 | rs2466067 | 0.073 | 0.45 | -1.011 | -1.967 | 0.017 | 0.047 | 0.047 | 0 | 0.6 | 0.25 | 0.64 | 0.27 | 0.69 | 0.55 |
| 8 | 32438416 | rs2466066 | 1.452 | 0.429 | -1.206 | -2.173 | -0.043 | 0.034 | 0.034 | 0 | 0.25 | 0.04 | 0.15 | 0.16 | 0.22 | 0.26 |
| 8 | 32443025 | rs17721282 | 0.623 | 0.67 | -0.389 | -1.739 | 0 | -0.002 | -0.002 | 0 | 0.24 | 0.24 | 0.23 | 0.18 | 0.26 | 0.37 |
| 8 | 32443090 | rs2466062 | -0.255 | 0.877 | -0.586 | -1.517 | -0.01 | -0.07 | -0.07 | 1.523 | 0.64 | 0.5 | 0.73 | 0.82 | 0.71 | 0.61 |
| 8 | 32446882 | rs2439292 | 0.322 | 0.442 | -1.077 | -1.322 | 0.019 | 0.109 | 0.109 | 0 | 0.43 | 0.19 | 0.32 | 0.21 | 0.41 | 0.34 |
| 8 | 32451898 | rs4602844 | 0.51 | 0.296 | -0.093 | -0.962 | 0.056 | 0.04 | 0.04 | 0 | 0.91 | 0.54 | 0.84 | 0.93 | 0.91 | 0.96 |
| 8 | 32453358 | rs3924999 | 0.336 | 0.364 | -1.882 | -1.064 | -0.128 | 0.028 | 0.028 | 0 | 0.35 | 0.07 | 0.37 | 0.76 | 0.38 | 0.45 |
| 8 | 32455027 | rs17665441 | 0.38 | -0.362 | -2.162 | -1.63 | -0.073 | 0.129 | 0.129 | 0 | 0.38 | 0.11 | 0.4 | 0.17 | 0.48 | 0.47 |
| 8 | 32466409 | rs2439279 | -0.955 | -0.391 | -2.117 | -1.526 | -0.086 | 0.105 | 0.105 | 0 | 0.6 | 0.84 | 0.67 | 0.73 | 0.53 | 0.58 |
| 8 | 32472617 | rs2466059 | 1.288 | -0.459 | -2.177 | -1.255 | -0.01 | -0.015 | -0.015 | 0 | 0.08 | 0.03 | 0.13 | 0.13 | 0.1 | 0.09 |
| 8 | 32474622 | rs4535704 | -0.17 | -1.035 | -1.958 | -1.338 | -0.021 | 0.055 | 0.055 | 0 | 0.32 | 0.22 | 0.31 | 0.13 | 0.37 | 0.36 |
| 8 | 32476816 | rs2919370 | -1.497 | -0.735 | -1.439 | -1.146 | -0.044 | 0.055 | 0.055 | 0 | 0.64 | 0.71 | 0.68 | 0.73 | 0.56 | 0.64 |
| 8 | 32487053 | rs12547858 | 0.798 | -0.422 | -0.997 | -0.949 | -0.049 | -0.026 | -0.026 | 0 | 0.1 | 0.01 | 0.19 | 0.11 | 0.19 | 0.16 |
| 8 | 32490310 | rs2439273 | 0.123 | -0.675 | -0.157 | -0.936 | 0.008 | -0.024 | -0.024 | 0 | 0.16 | 0.25 | 0.18 | 0.12 | 0.17 | 0.1 |
| 8 | 32499740 | rs10095694 | -0.851 | -0.771 | -0.658 | -0.963 | -0.058 | 0.048 | 0.048 | 0.241 | 0.6 | 0.83 | 0.62 | 0.62 | 0.55 | 0.5 |
| 8 | 32501987 | rs4531002 | 0.561 | -0.643 | -0.451 | -0.835 | -0.026 | 0.022 | 0.022 | 0.607 | 0.21 | 0.17 | 0.33 | 0.11 | 0.37 | 0.24 |
| 8 | 32504401 | rs7825588 | 1.108 | -1.033 | -0.402 | -1.071 | 0 | -0.029 | -0.029 | 0 | 0.17 | 0.15 | 0.06 | 0 | 0.12 | 0.06 |
| 8 | 32507149 | rs2466058 | 0.576 | -0.912 | -0.192 | -0.59 | -0.013 | -0.005 | -0.005 | 1.012 | 0.1 | 0.2 | 0.13 | 0.11 | 0.08 | 0.14 |
| 8 | 32511006 | rs13259346 | 0.971 | -1.067 | -0.481 | -0.822 | 0.058 | 0.205 | 0.205 | 0 | 0.51 | 0.16 | 0.38 | 0.25 | 0.55 | 0.43 |
| 8 | 32512309 | rs2439327 | 0.721 | -1.143 | -0.304 | -0.776 | -0.011 | -0.003 | -0.003 | 0 | 0.09 | 0.15 | 0.13 | 0.11 | 0.08 | 0.14 |
| 8 | 32512823 | rs2466051 | -1.137 | -1.254 | -0.16 | 0.436 | -0.017 | 0.048 | 0.048 | 0.239 | 0.87 | 0.87 | 0.81 | 0.66 | 0.93 | 0.8 |
| 8 | 32514916 | rs2466049 | 0.813 | -0.995 | -0.494 | 0.298 | -0.008 | 0.001 | 0.001 | 0 | 0.09 | 0.15 | 0.13 | 0.12 | 0.08 | 0.09 |
| 8 | 32520472 | rs2439322 | 0.42 | -0.78 | -0.517 | 0.364 | 0.008 | -0.006 | -0.006 | 0 | 0.6 | 0.36 | 0.63 | 0.63 | 0.59 | 0.62 |
| 8 | 32524451 | rs2919392 | 0.433 | -0.816 | -0.55 | 0.313 | 0.018 | 0.058 | 0.058 | 0 | 0.59 | 0.31 | 0.5 | 0.28 | 0.59 | 0.52 |
| 8 | 32525041 | rs2919391 | -0.569 | -1.129 | 0.105 | 0.353 | -0.048 | 0.192 | 0.192 | 0.575 | 0.17 | 0.38 | 0.31 | 0.49 | 0.11 | 0.21 |
| 8 | 32527281 | rs723811 | 0.172 | -1.251 | 0.11 | -0.185 | -0.07 | -0.173 | -0.173 | 0 | 0.74 | 0.54 | 0.88 | 1 | 0.87 | 0.85 |
| 8 | 32535920 | rs12546380 | 0.513 | -0.56 | -0.456 | 0.151 | -0.042 | -0.06 | -0.06 | 0.425 | 0.16 | 0.08 | 0.22 | 0.37 | 0.29 | 0.26 |
| 8 | 32537908 | rs10099273 | 0.983 | -0.714 | -0.112 | -0.478 | 0.002 | -0.028 | -0.028 | 0 | 0.08 | 0.09 | 0.07 | 0 | 0.1 | 0.08 |
| 8 | 32539327 | rs6468122 | -0.688 | -1.399 | -1.07 | -1.247 | 0.103 | 0.269 | 0.269 | 0.05 | 0.52 | 0.83 | 0.33 | 0.12 | 0.32 | 0.31 |
| 8 | 32541216 | rs10503926 | -0.805 | -1.715 | -0.391 | -1.181 | 0.01 | 0.012 | 0.012 | 3.042 | 0.59 | 0.5 | 0.49 | 0.5 | 0.52 | 0.48 |
| 8 | 32545001 | rs12541309 | 1.496 | -1.48 | -0.38 | -1.182 | 0.002 | 0.045 | 0.045 | 0 | 0.14 | 0.14 | 0.19 | 0.38 | 0.14 | 0.24 |
| 8 | 32549845 | rs7835765 | -1.784 | -1.492 | -0.481 | -1.375 | -0.109 | -0.136 | -0.136 | 2.944 | 0.72 | 0.94 | 0.59 | 0.5 | 0.63 | 0.57 |
| 8 | 32559943 | rs2919384 | 0.029 | -1.712 | -0.374 | -1.462 | -0.014 | 0.001 | 0.001 | 0 | 0.63 | 0.46 | 0.68 | 0.65 | 0.78 | 0.76 |
| 8 | 32563363 | rs10092449 | 0.921 | -1.743 | -0.395 | -1.489 | -0.001 | -0.042 | -0.042 | 0 | 0.26 | 0.25 | 0.09 | 0 | 0.11 | 0.06 |
| 8 | 32563924 | rs2919381 | 1.122 | -1.695 | -0.508 | -1.461 | -0.036 | -0.071 | -0.071 | 3.904 | 0.44 | 0.12 | 0.44 | 0.55 | 0.5 | 0.6 |
| 8 | 32569669 | rs2919378 | -0.752 | -1.084 | -0.156 | -1.033 | 0.069 | 0.326 | 0.326 | 0.511 | 0.89 | 0.65 | 0.77 | 0.66 | 0.89 | 0.82 |
| 8 | 32570493 | rs16879801 | 1.201 | -1.058 | -0.433 | -1.086 | 0.006 | -0.036 | -0.036 | 0.068 | 0.08 | 0.12 | 0.07 | 0 | 0.1 | 0.07 |
| 8 | 32572873 | rs16879809 | 1.455 | -1.089 | -0.715 | -1.019 | -0.043 | -0.104 | -0.104 | 0 | 0.14 | 0.02 | 0.15 | 0.39 | 0.13 | 0.26 |
| 8 | 32572983 | rs2976525 | -0.983 | -1.299 | -0.344 | -1.04 | -0.027 | 0.132 | 0.132 | 0 | 0.81 | 0.49 | 0.77 | 0.65 | 0.9 | 0.88 |
| 8 | 32573583 | rs16879814 | -0.786 | -0.777 | -0.181 | -1.141 | -0.003 | 0.015 | 0.015 | 0 | 0.27 | 0.24 | 0.33 | 0.11 | 0.38 | 0.26 |
| 8 | 32579109 | rs17669617 | -0.439 | -0.607 | -0.309 | -0.932 | -0.004 | 0.037 | 0.037 | 0 | 0.19 | 0.09 | 0.23 | 0.11 | 0.22 | 0.19 |
| 8 | 32584857 | rs16879886 | 0.393 | -0.305 | 0.451 | -0.318 | 0.002 | 0.08 | 0.08 | 0 | 0.1 | 0.13 | 0.22 | 0.38 | 0.13 | 0.16 |
| 8 | 32585434 | rs3735776 | -0.61 | -0.952 | 0.373 | -0.614 | -0.031 | 0.053 | 0.053 | 0 | 0.2 | 0.24 | 0.25 | 0.38 | 0.15 | 0.24 |
| 8 | 32585992 | rs11785668 | 0.565 | -0.938 | 0.26 | -0.605 | -0.026 | -0.031 | -0.031 | 0.049 | 0.1 | 0.08 | 0.03 | 0 | 0.03 | 0.07 |
| 8 | 32588901 | rs2919389 | 0.006 | -1.077 | -0.392 | -1.111 | -0.013 | -0.043 | -0.043 | 3.038 | 0.35 | 0.21 | 0.44 | 0.42 | 0.42 | 0.45 |
| 8 | 32589118 | rs3757930 | 0.023 | -0.869 | -0.217 | -1.276 | 0 | 0 | 0 | 0 | 0.47 | 0.44 | 0.38 | 0.39 | 0.34 | 0.31 |
| 8 | 32592129 | rs7844475 | 0.967 | -0.606 | -0.125 | -0.888 | -0.027 | -0.031 | -0.031 | 0 | 0.1 | 0.08 | 0.03 | 0 | 0.03 | 0.06 |
| 8 | 32596998 | rs10103750 | -0.517 | -0.615 | -0.207 | -1.021 | -0.041 | 0.05 | 0.05 | 0 | 0.23 | 0.57 | 0.28 | 0.39 | 0.16 | 0.23 |
| 8 | 32605388 | rs4733372 | 0.951 | 0.237 | -0.114 | -0.637 | -0.002 | 0.028 | 0.028 | 0 | 0.09 | 0.08 | 0.22 | 0.39 | 0.12 | 0.17 |
| 8 | 32611221 | rs16879922 | 1.122 | 0.27 | -0.341 | -0.883 | 0.052 | -0.107 | -0.107 | 1.629 | 0.25 | 0.05 | 0.24 | 0.41 | 0.16 | 0.33 |
| 8 | 32612705 | rs16879927 | 0.77 | 0.381 | 0.091 | -0.942 | 0.018 | -0.033 | -0.033 | 3.595 | 0.1 | 0.3 | 0.13 | 0.02 | 0.11 | 0.09 |
| 8 | 32615098 | rs6982890 | 0.968 | 0.569 | 0.259 | -0.668 | -0.041 | 0.008 | 0.008 | 0 | 0.17 | 0.12 | 0.16 | 0.18 | 0.22 | 0.23 |

| | | | | | | | | | | | | | | | | |
|---|---|---|---|---|---|---|---|---|---|---|---|---|---|---|---|---|
| 8 | 32624387 | rs6992642 | 0.47 | 0.454 | 0.624 | -0.209 | -0.009 | 0.099 | 0.099 | 0 | 0.48 | 0.35 | 0.3 | 0.17 | 0.42 | 0.3 |
| 8 | 32633362 | rs17671468 | -0.286 | 0.566 | 1.078 | 0.695 | -0.002 | -0.024 | -0.024 | 0 | 0.09 | 0.1 | 0.11 | 0.01 | 0.11 | 0.09 |
| 8 | 32670264 | rs4556053 | 0.687 | 0.569 | 1.064 | 0.832 | 0.014 | 0.281 | 0.281 | 0 | 0.88 | 0.61 | 0.77 | 0.67 | 0.89 | 0.84 |
| 8 | 32679661 | rs10217030 | -0.492 | 0.775 | 0.306 | 0.848 | -0.001 | 0.021 | 0.021 | 0 | 0.1 | 0.12 | 0.22 | 0.32 | 0.12 | 0.16 |
| 8 | 32695333 | rs6990963 | 0.324 | 0.779 | 0.336 | 0.85 | 0.017 | 0.181 | 0.181 | 0 | 0.89 | 0.75 | 0.78 | 0.68 | 0.88 | 0.84 |
| 8 | 32696905 | rs17673099 | 0.001 | 0.85 | -0.317 | 0.702 | -0.027 | 0.029 | 0.029 | 0 | 0.08 | 0.01 | 0.12 | 0.01 | 0.11 | 0.09 |
| 8 | 32697912 | rs6983065 | 0.395 | 0.822 | -0.242 | 0.764 | -0.002 | 0.032 | 0.032 | 0 | 0.86 | 0.84 | 0.77 | 0.68 | 0.87 | 0.83 |
| 8 | 32697958 | rs17673138 | -0.354 | 0.712 | -0.435 | 0.358 | 0.007 | 0.007 | 0.007 | 0 | 0.12 | 0.08 | 0.11 | 0 | 0.08 | 0.09 |
| 8 | 32700343 | rs10503931 | -0.979 | 0.836 | -0.508 | 0.515 | 0.006 | -0.018 | -0.018 | 1.104 | 0.19 | 0.14 | 0.3 | 0.27 | 0.15 | 0.23 |
| 8 | 32703798 | rs17746038 | -0.743 | 0.82 | -0.505 | 0.545 | 0.004 | 0.048 | 0.048 | 0 | 0.17 | 0.08 | 0.19 | 0.06 | 0.15 | 0.16 |
| 8 | 32703920 | rs12546782 | -0.932 | 0.82 | -0.501 | 0.557 | 0.007 | -0.018 | -0.018 | 0.895 | 0.18 | 0.14 | 0.3 | 0.27 | 0.15 | 0.23 |
| 8 | 32704845 | rs4330646 | -1 | 0.819 | -0.501 | 0.556 | 0.008 | -0.018 | -0.018 | 2.878 | 0.19 | 0.14 | 0.3 | 0.27 | 0.15 | 0.23 |
| 8 | 32705069 | rs10109272 | -0.003 | 1.659 | -0.688 | -0.151 | -0.028 | 0.015 | 0.015 | 0 | 0.18 | 0.13 | 0.21 | 0.01 | 0.32 | 0.14 |
| 8 | 32716483 | rs9656744 | 1.225 | 1.464 | 0.351 | -0.206 | 0.018 | -0.029 | -0.029 | 2.735 | 0.62 | 0.5 | 0.49 | 0.75 | 0.54 | 0.62 |
| 8 | 32748779 | rs4557662 | 1.11 | 1.609 | 0.648 | -0.384 | 0.027 | -0.057 | -0.057 | 0 | 0.66 | 0.52 | 0.55 | 0.81 | 0.59 | 0.67 |
| 8 | 32750127 | rs10954876 | 0.174 | 1.668 | 0.669 | -0.256 | 0.038 | -0.074 | -0.074 | 0 | 0.15 | 0.25 | 0.16 | 0 | 0.27 | 0.08 |
| 8 | 32761948 | rs16880153 | 0.956 | 1.296 | 0.04 | -0.304 | -0.011 | -0.026 | -0.026 | 0 | 0.12 | 0.01 | 0.06 | 0.12 | 0.11 | 0.15 |
| 8 | 32803573 | rs13255644 | 0.636 | 1.699 | 0.308 | -0.53 | 0.016 | -0.044 | -0.044 | 0.369 | 0.14 | 0.15 | 0.15 | 0 | 0.27 | 0.08 |
| 8 | 32842853 | rs4236722 | -0.162 | 0.838 | 0.244 | -0.885 | -0.008 | -0.035 | -0.035 | 0 | 0.42 | 0.48 | 0.44 | 0.82 | 0.38 | 0.59 |
| 8 | 32844577 | rs6996680 | -0.033 | 0.717 | 0.268 | -0.926 | 0.018 | 0.058 | 0.058 | 0 | 0.38 | 0.23 | 0.39 | 0.81 | 0.34 | 0.61 |
| 8 | 32863768 | rs17749288 | 0.766 | 0.681 | 0.353 | -0.215 | -0.005 | -0.011 | -0.011 | 0 | 0.12 | 0.11 | 0.18 | 0.24 | 0.13 | 0.21 |
| 8 | 32918302 | rs4493872 | -0.951 | 0.752 | 0.744 | -1.007 | 0.002 | -0.096 | -0.096 | 0.008 | 0.59 | 0.58 | 0.71 | 0.99 | 0.5 | 0.82 |
| 8 | 32947353 | rs7014840 | -0.012 | 0.389 | 0.339 | -0.945 | 0.006 | 0.034 | 0.034 | 0 | 0.45 | 0.31 | 0.33 | 0.72 | 0.41 | 0.64 |
| 8 | 32959945 | rs7460258 | -0.226 | 0.62 | 0.246 | -0.923 | 0.019 | 0.04 | 0.04 | 0 | 0.5 | 0.36 | 0.35 | 0.71 | 0.42 | 0.59 |
| 8 | 32962851 | rs6468141 | -0.351 | 0.51 | 0.187 | -0.932 | 0.027 | 0.066 | 0.066 | 0 | 0.52 | 0.75 | 0.37 | 0.69 | 0.4 | 0.59 |
| 8 | 32969507 | rs11785345 | 0.91 | 0.396 | -0.161 | -1.068 | -0.047 | -0.086 | -0.086 | 0.398 | 0.14 | 0.01 | 0.15 | 0.24 | 0.14 | 0.17 |
| 8 | 32979314 | rs7833961 | -0.289 | 0.19 | 0.313 | -0.969 | 0.017 | 0.038 | 0.038 | 0 | 0.5 | 0.36 | 0.42 | 0.68 | 0.45 | 0.6 |
| 8 | 33005939 | rs16880419 | 0.528 | -0.202 | 0.526 | -0.749 | 0.01 | -0.087 | -0.087 | 0 | 0.22 | 0.06 | 0.18 | 0.57 | 0.2 | 0.38 |
| 8 | 33042526 | rs17685354 | -0.048 | -0.107 | -0.239 | -1.09 | 0.002 | -0.001 | -0.001 | 1.174 | 0.36 | 0.32 | 0.26 | 0.1 | 0.3 | 0.25 |
| 8 | 33059828 | rs7815128 | -0.451 | -0.656 | 0.07 | -0.457 | -0.005 | -0.014 | -0.014 | 0 | 0.46 | 0.51 | 0.52 | 0.73 | 0.4 | 0.64 |
| 8 | 33067307 | rs4131222 | -0.289 | -1.082 | 0.189 | -0.071 | -0.014 | -0.001 | -0.001 | 0 | 0.4 | 0.47 | 0.42 | 0.61 | 0.26 | 0.45 |
| 8 | 33068303 | rs10808333 | 1.202 | -1.226 | -0.17 | -0.268 | -0.063 | 0.105 | 0.105 | 0.004 | 0.4 | 0.06 | 0.38 | 0.17 | 0.55 | 0.28 |
| 8 | 33070458 | rs4413730 | -0.25 | -0.875 | 0.442 | 0.219 | -0.008 | -0.03 | -0.03 | 0.576 | 0.06 | 0 | 0.08 | 0.14 | 0.06 | 0.11 |
| 8 | 33096139 | rs12676973 | -0.32 | -1.453 | 1.194 | 0.174 | 0.001 | 0 | 0 | 0 | 0.35 | 0.4 | 0.42 | 0.7 | 0.25 | 0.52 |
| 8 | 33096300 | rs10954888 | -0.142 | -1.47 | 1.137 | 0.165 | 0.032 | -0.002 | -0.002 | 0 | 0.33 | 0.21 | 0.42 | 0.7 | 0.25 | 0.52 |
| 8 | 33115544 | rs7843085 | -0.792 | -1.496 | 1.227 | 0.123 | -0.019 | -0.034 | -0.034 | 0 | 0.42 | 0.49 | 0.5 | 0.83 | 0.31 | 0.63 |
| 8 | 33116943 | rs13253577 | -0.625 | -1.469 | 1.226 | 0.139 | -0.006 | 0.031 | 0.031 | 0 | 0.07 | 0.16 | 0.08 | 0.13 | 0.06 | 0.11 |
| 8 | 33116988 | rs4587290 | -0.908 | -1.514 | 1.4 | 0.116 | 0.031 | 0.043 | 0.043 | 0 | 0.43 | 0.68 | 0.51 | 0.83 | 0.31 | 0.63 |
| 8 | 33133775 | rs7833088 | -0.242 | -1.517 | 1.343 | 0.132 | 0.036 | 0.014 | 0.014 | 0 | 0.36 | 0.27 | 0.43 | 0.7 | 0.26 | 0.52 |
| 8 | 33137029 | rs7844346 | -0.327 | -1.519 | 1.475 | 0.131 | -0.009 | -0.004 | -0.004 | 0 | 0.37 | 0.44 | 0.44 | 0.7 | 0.26 | 0.52 |
| 8 | 33143930 | rs4427132 | -1.038 | -1.49 | 1.387 | 0.156 | 0.032 | 0.046 | 0.046 | 0 | 0.48 | 0.6 | 0.35 | 0.82 | 0.35 | 0.67 |
| 8 | 33155972 | rs11988745 | -1.032 | -1.415 | 1.172 | 0.228 | 0.047 | 0.072 | 0.072 | 0 | 0.48 | 0.67 | 0.35 | 0.83 | 0.35 | 0.67 |
| 8 | 33167056 | rs13250346 | -0.936 | -1.522 | 1.192 | 0.257 | 0.048 | 0.064 | 0.064 | 0 | 0.46 | 0.67 | 0.33 | 0.83 | 0.33 | 0.65 |
| 8 | 33168975 | rs3923524 | -0.94 | -1.541 | 1.19 | 0.254 | 0.047 | 0.064 | 0.064 | 0 | 0.46 | 0.67 | 0.33 | 0.83 | 0.33 | 0.65 |
| 8 | 33178687 | rs16880728 | -0.358 | -1.567 | 1.343 | 0.283 | 0.003 | 0.074 | 0.074 | 0 | 0.06 | 0.24 | 0.06 | 0.13 | 0.04 | 0.09 |
| 8 | 33178822 | rs10954905 | -0.479 | -1.583 | 1.219 | 0.255 | -0.004 | -0.002 | -0.002 | 0 | 0.41 | 0.43 | 0.27 | 0.7 | 0.29 | 0.56 |
| 8 | 33181829 | rs11986601 | -0.898 | -1.086 | 1.211 | 0.942 | -0.006 | -0.002 | -0.002 | 0.056 | 0.41 | 0.61 | 0.32 | 0.7 | 0.33 | 0.62 |
| 8 | 33187102 | rs12546032 | -0.566 | -1.004 | 1.321 | 1.411 | -0.001 | -0.001 | -0.001 | 0 | 0.31 | 0.32 | 0.2 | 0.42 | 0.2 | 0.51 |
| 8 | 33192353 | rs12541516 | -0.667 | -0.993 | 1.386 | 1.549 | -0.035 | 0.041 | 0.041 | 0.008 | 0.32 | 0.45 | 0.2 | 0.59 | 0.21 | 0.58 |
| 8 | 33195455 | rs954009 | -0.418 | -0.915 | 1.223 | 1.367 | 0.014 | -0.026 | -0.026 | 0 | 0.29 | 0.22 | 0.19 | 0.43 | 0.21 | 0.52 |
| 8 | 33202506 | rs11785888 | -0.356 | -1.166 | 1.055 | 1.372 | 0.042 | -0.088 | -0.088 | 0.164 | 0.28 | 0.12 | 0.18 | 0.46 | 0.21 | 0.53 |
| 8 | 33203726 | rs1996363 | -0.267 | -0.95 | 1.101 | 1.189 | -0.007 | 0.003 | 0.003 | 0 | 0.08 | 0.06 | 0.11 | 0 | 0.14 | 0.09 |
| 8 | 33206572 | rs1474034 | -0.459 | -0.967 | 1.291 | 1.573 | 0.041 | -0.088 | -0.088 | 0 | 0.28 | 0.12 | 0.18 | 0.46 | 0.21 | 0.53 |
| 8 | 33228162 | rs1474035 | -0.323 | -0.868 | 1.269 | 1.601 | 0.04 | -0.11 | -0.11 | 0.197 | 0.26 | 0.1 | 0.17 | 0.47 | 0.2 | 0.49 |
| 8 | 33235671 | rs7837460 | -1.116 | -1.001 | 1.859 | 1.604 | -0.029 | 0.023 | 0.023 | 0.333 | 0.33 | 0.58 | 0.22 | 0.58 | 0.21 | 0.55 |
| 8 | 33246538 | rs16880849 | -0.48 | -0.944 | 1.635 | 1.652 | 0.036 | -0.054 | -0.054 | 0 | 0.28 | 0.18 | 0.17 | 0.47 | 0.19 | 0.48 |
| 8 | 33246685 | rs16880852 | -0.48 | -0.944 | 1.635 | 1.651 | 0.036 | -0.053 | -0.053 | 0.176 | 0.28 | 0.18 | 0.17 | 0.47 | 0.19 | 0.48 |
| 8 | 33260047 | rs11987164 | -0.761 | -0.707 | 1.205 | 1.646 | -0.009 | 0.011 | 0.011 | 0.022 | 0.29 | 0.35 | 0.22 | 0.48 | 0.23 | 0.45 |
| 8 | 33264711 | rs12549316 | -0.991 | -0.537 | 1.182 | 1.699 | -0.019 | -0.005 | -0.005 | 0 | 0.33 | 0.44 | 0.27 | 0.32 | 0.29 | 0.46 |
| 8 | 33274426 | rs9656765 | -0.808 | -0.383 | 0.407 | 1.744 | 0.01 | -0.003 | -0.003 | 0 | 0.31 | 0.23 | 0.25 | 0.33 | 0.29 | 0.46 |
| 8 | 33275793 | rs2676434 | 0.565 | -0.273 | 0.173 | 1.924 | 0.001 | -0.046 | -0.046 | 0.062 | 0.7 | 0.76 | 0.73 | 0.49 | 0.7 | 0.53 |
| 8 | 33286370 | rs1533470 | 0.773 | -0.314 | 0.367 | 1.888 | 0.004 | -0.014 | -0.014 | 0 | 0.67 | 0.74 | 0.52 | 0.56 | 0.67 | 0.47 |
| 8 | 33287802 | rs12155581 | 1.12 | -0.468 | 0.542 | 1.861 | 0.002 | -0.007 | -0.007 | 0 | 0.61 | 0.64 | 0.48 | 0.46 | 0.62 | 0.38 |
| 8 | 33292469 | rs6982159 | 1.566 | -0.853 | 0.596 | 1.923 | -0.003 | -0.003 | -0.003 | 0 | 0.52 | 0.48 | 0.43 | 0.31 | 0.57 | 0.27 |
| 8 | 33294524 | rs2348613 | 0.569 | -0.707 | 0.614 | 1.755 | -0.006 | -0.053 | -0.053 | 0 | 0.67 | 0.89 | 0.53 | 0.58 | 0.67 | 0.47 |
| 8 | 33297428 | rs2732255 | -0.476 | -0.713 | 0.569 | 1.685 | -0.01 | 0.024 | 0.024 | 0 | 0.94 | 1 | 0.97 | 1 | 0.94 | 0.99 |
| 8 | 33303493 | rs6995454 | 0.466 | -0.598 | 0.686 | 1.761 | -0.004 | -0.047 | -0.047 | 0.535 | 0.66 | 0.88 | 0.52 | 0.57 | 0.67 | 0.47 |
| 8 | 33304557 | rs2732286 | -1.013 | -0.561 | 0.712 | 1.717 | -0.011 | 0.01 | 0.01 | 0 | 0.07 | 0.14 | 0.04 | 0.07 | 0.05 | 0.05 |
| 8 | 33325055 | rs7018447 | 1.202 | -0.564 | 0.72 | 1.755 | -0.01 | 0.006 | 0.006 | 0.676 | 0.57 | 0.48 | 0.45 | 0.49 | 0.6 | 0.41 |

| | | | | | | | | | | | | | | | | |
|---|---|---|---|---|---|---|---|---|---|---|---|---|---|---|---|---|
| 8 | 33342018 | rs2037213 | 1.314 | -0.623 | 0.797 | 1.723 | -0.005 | 0.001 | 0.001 | 0.486 | 0.56 | 0.43 | 0.46 | 0.49 | 0.6 | 0.41 |
| 8 | 33356074 | rs6468171 | 0.959 | -0.506 | 0.803 | 1.725 | 0.016 | -0.023 | -0.023 | 0 | 0.6 | 0.68 | 0.49 | 0.49 | 0.63 | 0.42 |
| 8 | 33358270 | rs8685 | 1.369 | -0.511 | 0.809 | 1.722 | -0.007 | 0.002 | 0.002 | 0.274 | 0.55 | 0.45 | 0.46 | 0.49 | 0.6 | 0.41 |
| 8 | 33358730 | rs7848 | 0.468 | -0.504 | 0.81 | 1.701 | 0.006 | -0.01 | -0.01 | 0 | 0.64 | 0.7 | 0.5 | 0.56 | 0.66 | 0.46 |
| 8 | 33369944 | rs2304748 | 0.957 | -0.481 | 0.799 | 1.713 | -0.01 | 0 | 0 | 0 | 0.59 | 0.45 | 0.47 | 0.56 | 0.63 | 0.45 |
| 8 | 33371296 | rs12682118 | 0.624 | -0.421 | 0.529 | 1.765 | 0.006 | -0.006 | -0.006 | 3.984 | 0.63 | 0.64 | 0.5 | 0.56 | 0.66 | 0.46 |
| 8 | 33384027 | rs12386778 | 0.774 | -0.277 | 0.556 | 1.768 | 0.008 | -0.008 | -0.008 | 0 | 0.58 | 0.59 | 0.5 | 0.49 | 0.64 | 0.42 |
| 8 | 33391380 | rs7842778 | 0.549 | -0.066 | 0.494 | 1.676 | 0.021 | -0.017 | -0.017 | 0.028 | 0.58 | 0.8 | 0.5 | 0.54 | 0.61 | 0.46 |
| 8 | 33397032 | rs2043967 | 1.291 | -0.007 | 0.857 | 1.618 | 0.012 | 0.015 | 0.015 | 0.035 | 0.48 | 0.34 | 0.42 | 0.36 | 0.56 | 0.39 |
| 8 | 33412036 | rs13439478 | -2.222 | 0.082 | 0.787 | 1.609 | 0.002 | 0.123 | 0.123 | 0.01 | 0.08 | 0.43 | 0.07 | 0.13 | 0.03 | 0.08 |
| 8 | 33414494 | rs1440530 | 1.167 | 0.145 | 0.715 | 1.602 | 0.007 | 0.006 | 0.006 | 0.093 | 0.51 | 0.39 | 0.45 | 0.38 | 0.57 | 0.46 |
| 8 | 33416189 | rs3735950 | 0.969 | 0.156 | 0.209 | 1.611 | 0.022 | 0.022 | 0.022 | 0.303 | 0.53 | 0.62 | 0.45 | 0.38 | 0.56 | 0.42 |
| 8 | 33416222 | rs3735951 | 0.549 | 0.183 | 0.055 | 1.642 | 0.003 | -0.052 | -0.052 | 0 | 0.58 | 0.85 | 0.5 | 0.5 | 0.59 | 0.5 |
| 8 | 33417258 | rs9297209 | 0.593 | 0.203 | 0.056 | 1.652 | 0.007 | -0.048 | -0.048 | 0.083 | 0.57 | 0.84 | 0.5 | 0.5 | 0.58 | 0.49 |
| 8 | 33421253 | rs6468175 | 0.739 | 0.43 | -0.001 | 1.671 | -0.008 | -0.044 | -0.044 | 0 | 0.56 | 0.83 | 0.48 | 0.51 | 0.56 | 0.49 |
| 8 | 33421534 | rs754437 | 0.995 | 0.437 | 0.101 | 1.563 | 0.017 | 0.028 | 0.028 | 0.121 | 0.51 | 0.61 | 0.43 | 0.39 | 0.54 | 0.42 |
| 8 | 33443076 | rs1530344 | 0.924 | 0.558 | -0.085 | 1.601 | 0.018 | 0.019 | 0.019 | 0.17 | 0.52 | 0.66 | 0.43 | 0.42 | 0.54 | 0.42 |
| 8 | 33457369 | rs1044115 | -0.864 | 1.226 | 0.074 | 1.837 | -0.048 | -0.089 | -0.089 | 0.055 | 0.31 | 0.06 | 0.5 | 0.58 | 0.35 | 0.51 |
| 8 | 33466398 | rs7838624 | -1.06 | 1.496 | 0.503 | 1.232 | -0.042 | 0.017 | 0.017 | 0 | 0.33 | 0.12 | 0.51 | 0.77 | 0.39 | 0.62 |
| 8 | 33466724 | rs11786051 | -1.043 | 1.506 | 0.508 | 1.237 | -0.043 | 0.016 | 0.016 | 0 | 0.33 | 0.12 | 0.51 | 0.77 | 0.39 | 0.61 |
| 8 | 33467010 | rs10092750 | -1.518 | 1.354 | 1.294 | 1.166 | 0.007 | -0.027 | -0.027 | 0.7 | 0.39 | 0.48 | 0.54 | 0.77 | 0.41 | 0.68 |
| 8 | 33470899 | rs7814270 | -1.188 | 1.359 | 1.284 | 1.168 | -0.019 | 0.023 | 0.023 | 0 | 0.36 | 0.23 | 0.53 | 0.77 | 0.4 | 0.67 |
| 8 | 33484845 | rs7387564 | -1.52 | 1.327 | 1.261 | 1.181 | 0.002 | -0.026 | -0.026 | 0 | 0.39 | 0.47 | 0.55 | 0.77 | 0.4 | 0.67 |
| 8 | 33495185 | rs11782575 | -0.921 | 1.34 | 1.116 | 1.188 | -0.047 | 0.012 | 0.012 | 0 | 0.33 | 0.07 | 0.52 | 0.77 | 0.39 | 0.61 |
| 8 | 33506910 | rs13272880 | -1.417 | 1.314 | 1.339 | 1.165 | -0.001 | -0.006 | -0.006 | 0 | 0.41 | 0.57 | 0.55 | 0.77 | 0.39 | 0.67 |
| 8 | 33524179 | rs10107668 | -1.46 | 1.375 | 1.225 | 1.185 | 0.003 | -0.026 | -0.026 | 0.008 | 0.39 | 0.46 | 0.54 | 0.77 | 0.39 | 0.67 |
| 8 | 33530099 | rs10954922 | -1.018 | 1.446 | 0.985 | 1.238 | -0.002 | 0.003 | 0.003 | 0 | 0.35 | 0.28 | 0.52 | 0.77 | 0.38 | 0.62 |
| 8 | 33548107 | rs1443080 | -1.142 | 1.928 | 0.483 | 1.057 | -0.03 | 0.021 | 0.021 | 0 | 0.3 | 0.23 | 0.5 | 0.82 | 0.39 | 0.59 |
| 8 | 33571496 | rs4733166 | -0.636 | 1.944 | 0.424 | 1.518 | -0.039 | -0.075 | -0.075 | 0 | 0.23 | 0.11 | 0.44 | 0.64 | 0.33 | 0.53 |
| 8 | 33574235 | rs12056344 | 0.715 | 1.969 | 0.317 | 1.918 | -0.024 | -0.03 | -0.03 | 0 | 0.08 | 0.01 | 0.33 | 0.42 | 0.11 | 0.2 |
| 8 | 33578917 | rs4733168 | -1.349 | 1.664 | 0.664 | 1.432 | -0.004 | 0.008 | 0.008 | 0 | 0.33 | 0.29 | 0.5 | 0.81 | 0.38 | 0.62 |
| 8 | 33587898 | rs10454341 | -0.127 | 1.623 | 0.353 | 1.416 | -0.126 | -0.145 | -0.145 | 0.328 | 0.27 | 0.04 | 0.46 | 0.64 | 0.37 | 0.53 |
| 8 | 33600806 | rs16881180 | -0.023 | 1.739 | 0.479 | 1.181 | -0.011 | 0.006 | 0.006 | 0 | 0.07 | 0.04 | 0.12 | 0.06 | 0.11 | 0.07 |
| 8 | 33638554 | rs6997661 | -1.221 | 1.563 | 0.818 | 0.951 | 0.001 | -0.006 | -0.006 | 0.078 | 0.4 | 0.4 | 0.57 | 0.82 | 0.49 | 0.62 |
| 8 | 33656359 | rs4733456 | -0.795 | 1.595 | 0.801 | 0.927 | -0.015 | 0.02 | 0.02 | 0 | 0.35 | 0.27 | 0.54 | 0.82 | 0.45 | 0.62 |
| 8 | 33662884 | rs7833512 | -0.92 | 1.589 | 1.116 | 1.14 | 0.022 | 0.028 | 0.028 | 0 | 0.35 | 0.42 | 0.4 | 0.59 | 0.42 | 0.57 |
| 8 | 33667153 | rs7836762 | -1.089 | 1.532 | 1.129 | 0.817 | 0.016 | -0.04 | -0.04 | 0 | 0.4 | 0.5 | 0.44 | 0.82 | 0.48 | 0.65 |
| 8 | 33713451 | rs4612298 | -0.516 | 1.566 | 1.204 | 0.992 | 0.019 | 0.044 | 0.044 | 0.075 | 0.32 | 0.36 | 0.36 | 0.58 | 0.4 | 0.56 |
| 8 | 33718321 | rs7465686 | 0.269 | 1.573 | 1.194 | 0.985 | -0.04 | -0.067 | -0.067 | 0.384 | 0.25 | 0.14 | 0.33 | 0.58 | 0.38 | 0.51 |
| 8 | 33744877 | rs7388666 | -0.091 | 1.595 | 1.107 | 0.949 | -0.109 | -0.119 | -0.119 | 1.057 | 0.28 | 0.07 | 0.34 | 0.52 | 0.42 | 0.52 |
| 8 | 33765126 | rs11776207 | -0.813 | 1.599 | 1.283 | 0.913 | 0.028 | -0.023 | -0.023 | 0.498 | 0.35 | 0.38 | 0.41 | 0.79 | 0.46 | 0.66 |
| 8 | 33779934 | rs7824990 | 0.001 | 1.656 | 1.363 | 0.97 | -0.004 | -0.006 | -0.006 | 2.564 | 0.3 | 0.26 | 0.35 | 0.52 | 0.39 | 0.54 |
| 8 | 33782471 | rs7386384 | -1.32 | 1.767 | 1.527 | 0.828 | 0.018 | 0.035 | 0.035 | 0 | 0.47 | 0.6 | 0.56 | 0.81 | 0.59 | 0.74 |
| 8 | 33786830 | rs10105616 | -0.983 | 1.796 | 1.544 | 0.84 | -0.002 | -0.027 | -0.027 | 1.137 | 0.43 | 0.47 | 0.52 | 0.81 | 0.55 | 0.73 |
| 8 | 33806910 | rs11780927 | -1.368 | 1.403 | 1.767 | 0.844 | 0.018 | 0.054 | 0.054 | 0 | 0.12 | 0.15 | 0.17 | 0.3 | 0.16 | 0.18 |
| 8 | 33829906 | rs10087815 | -0.44 | 1.022 | 1.31 | 0.57 | -0.004 | 0.003 | 0.003 | 0 | 0.08 | 0.06 | 0.11 | 0.03 | 0.11 | 0.08 |
| 8 | 33853868 | rs7465297 | 2.173 | 0.865 | 1.521 | 0.466 | 0.007 | 0.017 | 0.017 | 0.723 | 0.78 | 0.31 | 0.76 | 0.71 | 0.79 | 0.8 |
| 8 | 33864647 | rs4493901 | 1.158 | 0.837 | 1.467 | 0.455 | 0.05 | 0.211 | 0.211 | 0.901 | 0.87 | 0.47 | 0.88 | 0.73 | 0.91 | 0.89 |
| 8 | 33896448 | rs12682471 | -2.162 | 0.786 | 1.445 | 0.437 | 0.032 | 0.065 | 0.065 | 0.402 | 0.21 | 0.61 | 0.23 | 0.29 | 0.2 | 0.18 |
| 8 | 33903459 | rs10954947 | -0.813 | 0.794 | 1.358 | 0.407 | 0.051 | -0.026 | -0.026 | 0.016 | 0.09 | 0.51 | 0.06 | 0 | 0.04 | 0.01 |
| 8 | 33927648 | rs4357281 | -0.374 | 0.826 | -0.242 | 0.453 | -0.01 | 0.009 | 0.009 | 0 | 0.08 | 0.05 | 0.11 | 0.02 | 0.11 | 0.07 |
| 8 | 33953787 | rs10101028 | -1.539 | 0.695 | -0.468 | 0.441 | 0 | -0.013 | -0.013 | 0 | 0.3 | 0.82 | 0.26 | 0.29 | 0.25 | 0.21 |
| 8 | 33970202 | rs2643282 | -1.232 | 1.031 | -0.248 | 0.72 | -0.003 | -0.006 | -0.006 | 0 | 0.23 | 0.75 | 0.24 | 0.29 | 0.2 | 0.2 |
| 8 | 33982039 | rs7843751 | -0.713 | 0.816 | -0.457 | -0.026 | 0.025 | 0.093 | 0.093 | 0 | 0.53 | 0.7 | 0.53 | 0.82 | 0.59 | 0.78 |
| 8 | 33988414 | rs7014743 | -0.155 | 0.815 | -0.173 | -0.183 | -0.001 | -0.019 | -0.019 | 1.334 | 0.08 | 0.08 | 0.04 | 0 | 0.06 | 0.07 |
| 8 | 33994378 | rs16875745 | -0.764 | 1.106 | 0.499 | 0.1 | 0.033 | 0.018 | 0.018 | 0 | 0.19 | 0.31 | 0.24 | 0.22 | 0.27 | 0.15 |
| 8 | 33996986 | rs924008 | 0.023 | 1.23 | 0.628 | 0.155 | -0.018 | -0.056 | -0.056 | 0.962 | 0.14 | 0.01 | 0.07 | 0.2 | 0.15 | 0.09 |
| 8 | 34001156 | rs1495216 | -0.81 | 1.092 | 0.8 | -0.133 | -0.003 | -0.029 | -0.029 | 0 | 0.71 | 0.82 | 0.7 | 0.38 | 0.72 | 0.43 |
| 8 | 34004175 | rs12541339 | -0.115 | 1.091 | 0.905 | -0.243 | 0.134 | 0.019 | 0.019 | 0 | 0.05 | 0.26 | 0.17 | 0.03 | 0.12 | 0.05 |
| 8 | 34005746 | rs7839708 | 0.329 | 1.064 | 0.935 | -0.256 | 0.002 | 0.099 | 0.099 | 0 | 0.56 | 0.76 | 0.63 | 0.18 | 0.57 | 0.33 |
| 8 | 34011204 | rs17283209 | 1.861 | 0.784 | 0.884 | -0.484 | -0.033 | 0.142 | 0.142 | 0 | 0.37 | 0.07 | 0.39 | 0.14 | 0.37 | 0.2 |
| 8 | 34011345 | rs1495219 | -0.83 | 0.913 | 0.918 | -0.315 | -0.002 | -0.016 | -0.016 | 0 | 0.7 | 0.82 | 0.7 | 0.35 | 0.72 | 0.42 |
| 8 | 34013305 | rs7818844 | 0.608 | 0.91 | 0.888 | -0.294 | 0 | -0.02 | -0.02 | 0 | 0.51 | 0.5 | 0.46 | 0.15 | 0.45 | 0.28 |
| 8 | 34022494 | rs16881601 | 0.443 | 0.937 | 0.65 | -0.133 | -0.013 | -0.026 | -0.026 | 0.027 | 0.06 | 0.01 | 0.07 | 0.17 | 0.1 | 0.06 |
| 8 | 34041162 | rs10094759 | 0.055 | 0.904 | 1.039 | -0.198 | -0.003 | -0.047 | -0.047 | 4.06 | 0.09 | 0.16 | 0.04 | 0 | 0.06 | 0.09 |
| 8 | 34044529 | rs2730003 | 0.917 | 0.79 | 1.08 | 0.692 | 0.071 | 0.185 | 0.185 | 0.946 | 0.89 | 0.65 | 0.78 | 0.69 | 0.86 | 0.65 |
| 8 | 34061676 | rs2729958 | 0.352 | 0.792 | 0.225 | 0.595 | -0.012 | -0.056 | -0.056 | 0 | 0.33 | 0.11 | 0.26 | 0.39 | 0.32 | 0.31 |
| 8 | 34068459 | rs12679346 | 0.297 | 0.964 | 0.451 | 0.495 | -0.002 | -0.001 | -0.001 | 3.76 | 0.07 | 0.04 | 0.06 | 0.17 | 0.08 | 0.06 |
| 8 | 34083175 | rs7835367 | -0.477 | 1.159 | 0.564 | 0.714 | 0.003 | 0.047 | 0.047 | 0 | 0.06 | 0.06 | 0.08 | 0.23 | 0.07 | 0.39 |

| | | | | | | | | | | | | | | | | |
|---|---|---|---|---|---|---|---|---|---|---|---|---|---|---|---|---|
| 8 | 34083654 | rs17264354 | 0.986 | 0.481 | 0.047 | -0.186 | -0.041 | 0.2 | 0.2 | 0 | 0.42 | 0.11 | 0.39 | 0.12 | 0.44 | 0.19 |
| 8 | 34087160 | rs2729976 | 0.401 | 0.505 | 0.474 | -0.209 | 0 | -0.001 | -0.001 | 0.539 | 0.32 | 0.31 | 0.43 | 0.69 | 0.33 | 0.66 |
| 8 | 34089670 | rs2643261 | 1.307 | 0.52 | 0.891 | -0.154 | 0.093 | -0.022 | -0.022 | 0 | 0.91 | 0.59 | 0.9 | 0.98 | 0.89 | 0.96 |
| 8 | 34102353 | rs2643270 | -1.006 | 0.673 | 0.516 | -0.242 | 0 | -0.018 | -0.018 | 0 | 0.63 | 0.68 | 0.63 | 0.84 | 0.6 | 0.81 |
| 8 | 34108334 | rs2132167 | 0.598 | 0.501 | 0.7 | -0.258 | 0.002 | -0.015 | -0.015 | 2.718 | 0.75 | 0.46 | 0.85 | 0.79 | 0.77 | 0.79 |
| 8 | 34111195 | rs12681698 | -0.135 | 0.475 | 0.831 | -0.249 | -0.006 | -0.003 | -0.003 | 3.275 | 0.2 | 0.26 | 0.11 | 0.18 | 0.14 | 0.17 |
| 8 | 34116194 | rs6998084 | 0.782 | 0.439 | 0.733 | -0.054 | -0.026 | 0.077 | 0.077 | 0 | 0.46 | 0.18 | 0.48 | 0.33 | 0.49 | 0.49 |
| 8 | 34118035 | rs12677511 | -0.204 | 0.56 | 1.122 | 0.085 | -0.003 | -0.001 | -0.001 | 0 | 0.2 | 0.25 | 0.11 | 0.18 | 0.15 | 0.18 |
| 8 | 34119062 | rs1495235 | 0.6 | 0.672 | 1.147 | 0.083 | 0.181 | 0.086 | 0.086 | 0 | 0.95 | 0.69 | 0.96 | 0.97 | 0.91 | 0.96 |
| 8 | 34120699 | rs2978836 | 1.241 | 0.705 | 0.794 | 0.053 | -0.03 | 0.052 | 0.052 | 0.069 | 0.09 | 0.01 | 0.04 | 0 | 0.1 | 0.06 |
| 8 | 34131943 | rs10091208 | -2.19 | 0.884 | 1.395 | 0.636 | 0.064 | 0.112 | 0.112 | 0.072 | 0.13 | 0.52 | 0.18 | 0.25 | 0.18 | 0.3 |
| 8 | 34132075 | rs10091344 | -1.985 | 0.8 | 1.283 | 0.629 | 0.006 | -0.018 | -0.018 | 0 | 0.35 | 0.76 | 0.37 | 0.44 | 0.32 | 0.47 |
| 8 | 34167955 | rs2953923 | -2.156 | 0.705 | 1.415 | 0.551 | 0.056 | 0.138 | 0.138 | 0 | 0.12 | 0.53 | 0.18 | 0.25 | 0.16 | 0.3 |
| 8 | 34176431 | rs2609622 | -1.023 | 0.818 | 1.384 | 0.576 | 0.118 | 0.206 | 0.206 | 0.623 | 0.08 | 0.27 | 0.16 | 0.25 | 0.15 | 0.3 |
| 8 | 34197587 | rs10503960 | -0.247 | 0.741 | 1.135 | 0.55 | -0.001 | 0 | 0 | 0 | 0.2 | 0.24 | 0.18 | 0.16 | 0.14 | 0.17 |
| 8 | 34199010 | rs1458903 | -2.281 | 0.831 | 1.336 | 0.579 | 0.078 | 0.113 | 0.113 | 0.049 | 0.13 | 0.5 | 0.2 | 0.25 | 0.19 | 0.31 |
| 8 | 34202968 | rs2034976 | -1.624 | 0.799 | 1.155 | 0.546 | 0.008 | -0.01 | -0.01 | 0 | 0.35 | 0.77 | 0.39 | 0.41 | 0.33 | 0.48 |
| 8 | 34227159 | rs2590468 | 0.669 | 0.843 | 1.394 | 0.579 | 0.141 | 0.046 | 0.046 | 0.071 | 0.93 | 0.73 | 0.87 | 0.94 | 0.85 | 0.85 |
| 8 | 34228681 | rs978439 | 0.669 | 0.843 | 1.397 | 0.579 | 0.139 | 0.044 | 0.044 | 0.071 | 0.93 | 0.74 | 0.87 | 0.94 | 0.85 | 0.85 |
| 8 | 34249110 | rs2028774 | 0.887 | 0.811 | 1.071 | 0.408 | 0.003 | -0.037 | -0.037 | 0 | 0.71 | 0.46 | 0.66 | 0.79 | 0.69 | 0.64 |
| 8 | 34255851 | rs1445190 | 1.259 | 0.737 | 0.718 | 0.273 | -0.073 | -0.113 | -0.113 | 0 | 0.34 | 0.05 | 0.29 | 0.45 | 0.35 | 0.36 |
| 8 | 34316909 | rs4294176 | -1.229 | 0.865 | 1.409 | 0.689 | 0.125 | 0.005 | 0.005 | 0.092 | 0.09 | 0.29 | 0.18 | 0.06 | 0.19 | 0.19 |
| 8 | 34317465 | rs4739565 | -0.231 | 0.849 | 1.448 | 0.684 | 0.008 | 0.006 | 0.006 | 0 | 0.21 | 0.24 | 0.18 | 0.16 | 0.14 | 0.18 |
| 8 | 34320094 | rs7840167 | -1.23 | 0.872 | 1.406 | 0.68 | 0.125 | 0.005 | 0.005 | 0.092 | 0.09 | 0.29 | 0.18 | 0.06 | 0.19 | 0.19 |
| 8 | 34324240 | rs1551653 | -1.761 | 0.849 | 1.292 | 0.674 | 0.004 | -0.001 | -0.001 | 0.023 | 0.37 | 0.75 | 0.43 | 0.43 | 0.36 | 0.52 |
| 8 | 34325200 | rs16882257 | -2.039 | 0.864 | 1.207 | 0.682 | 0.009 | -0.006 | -0.006 | 0 | 0.4 | 0.88 | 0.44 | 0.44 | 0.38 | 0.53 |
| 8 | 34331257 | rs10954965 | -0.961 | 0.818 | 1.402 | 0.653 | -0.014 | -0.035 | -0.035 | 0.178 | 0.28 | 0.57 | 0.27 | 0.17 | 0.23 | 0.35 |
| 8 | 34331822 | rs10088476 | -1.485 | 0.898 | 1.473 | 0.637 | 0.077 | -0.034 | -0.034 | 0 | 0.09 | 0.3 | 0.09 | 0.01 | 0.1 | 0.17 |
| 8 | 34353811 | rs11994160 | -1.576 | 0.463 | 1.468 | 0.666 | 0.056 | -0.041 | -0.041 | 0 | 0.1 | 0.42 | 0.08 | 0.01 | 0.08 | 0.17 |
| 8 | 34371604 | rs6468265 | -1.962 | 0.368 | 1.403 | 0.801 | 0.028 | -0.008 | -0.008 | 0 | 0.35 | 0.78 | 0.31 | 0.4 | 0.26 | 0.5 |
| 8 | 34372293 | rs7000467 | -0.602 | 0.453 | 1.553 | 0.811 | 0.062 | -0.012 | -0.012 | 0 | 0.07 | 0.24 | 0.06 | 0.01 | 0.07 | 0.18 |
| 8 | 34384758 | rs12541775 | -0.295 | 0.431 | 1.492 | 0.838 | -0.012 | -0.005 | -0.005 | 0.03 | 0.2 | 0.29 | 0.18 | 0.17 | 0.15 | 0.18 |
| 8 | 34415528 | rs4291266 | -0.256 | 0.435 | 1.499 | 0.839 | -0.002 | -0.001 | -0.001 | 0 | 0.19 | 0.26 | 0.17 | 0.17 | 0.15 | 0.18 |
| 8 | 34436475 | rs6995321 | -1.486 | 0.505 | 1.538 | 0.978 | -0.007 | 0.011 | 0.011 | 0.512 | 0.3 | 0.62 | 0.29 | 0.41 | 0.24 | 0.5 |
| 8 | 34443390 | rs16882479 | -1.169 | 0.59 | 1.616 | 1.022 | 0.045 | 0.155 | 0.155 | 0 | 0.08 | 0.25 | 0.09 | 0.24 | 0.08 | 0.32 |
| 8 | 34459723 | rs10481034 | -1.146 | 0.591 | 1.621 | 1.015 | 0.046 | 0.153 | 0.153 | 0 | 0.08 | 0.25 | 0.09 | 0.23 | 0.08 | 0.32 |
| 8 | 34460767 | rs10217066 | -0.53 | 0.603 | 1.916 | 0.892 | 0.029 | -0.028 | -0.028 | 0 | 0.06 | 0.18 | 0.06 | 0.01 | 0.07 | 0.16 |
| 8 | 34470646 | rs7003000 | -1.047 | 0.604 | 1.943 | 0.893 | 0.061 | -0.016 | -0.016 | 0 | 0.08 | 0.29 | 0.06 | 0.01 | 0.07 | 0.16 |
| 8 | 34478701 | rs4739367 | -0.646 | 0.489 | 1.818 | 0.882 | -0.01 | 0.025 | 0.025 | 0.015 | 0.23 | 0.32 | 0.23 | 0.41 | 0.17 | 0.35 |
| 8 | 34497650 | rs7819762 | -1.357 | 0.509 | 1.822 | 0.906 | -0.015 | 0.03 | 0.03 | 0.033 | 0.3 | 0.56 | 0.29 | 0.41 | 0.25 | 0.5 |
| 8 | 34501546 | rs4342589 | -2.156 | 0.632 | 1.471 | 0.804 | 0.004 | -0.013 | -0.013 | 0 | 0.44 | 0.87 | 0.46 | 0.46 | 0.41 | 0.57 |
| 8 | 34503776 | rs7844647 | -1.802 | 0.608 | 1.679 | 0.945 | 0.025 | -0.019 | -0.019 | 0 | 0.34 | 0.79 | 0.32 | 0.41 | 0.26 | 0.51 |
| 8 | 34515169 | rs7817181 | -1.071 | 0.56 | 1.754 | 0.866 | 0.024 | -0.035 | -0.035 | 0 | 0.07 | 0.21 | 0.06 | 0.01 | 0.07 | 0.16 |
| 8 | 34524003 | rs6987004 | -0.318 | 0.327 | 1.558 | 1.238 | -0.016 | 0.042 | 0.042 | 0 | 0.21 | 0.29 | 0.23 | 0.45 | 0.13 | 0.28 |
| 8 | 34524021 | rs7005465 | -0.289 | 0.326 | 1.557 | 1.08 | -0.016 | 0.024 | 0.024 | 0 | 0.2 | 0.29 | 0.23 | 0.35 | 0.13 | 0.26 |
| 8 | 34524479 | rs4455823 | -1.667 | 0.544 | 1.304 | 0.943 | 0 | 0 | 0 | 0.003 | 0.4 | 0.69 | 0.45 | 0.42 | 0.36 | 0.5 |
| 8 | 34524513 | rs16882722 | -0.993 | 0.118 | 2.021 | 0.55 | 0.031 | -0.043 | -0.043 | 0 | 0.11 | 0.36 | 0.13 | 0.04 | 0.12 | 0.04 |
| 8 | 34532101 | rs7005808 | -1.235 | -0.34 | 1.869 | 0.679 | 0.032 | 0.049 | 0.049 | 0.565 | 0.3 | 0.82 | 0.23 | 0.07 | 0.16 | 0.17 |
| 8 | 34537363 | rs10109248 | -0.491 | -0.251 | 1.446 | 0.697 | -0.016 | -0.057 | -0.057 | 0 | 0.23 | 0.38 | 0.19 | 0.07 | 0.15 | 0.17 |
| 8 | 34550799 | rs7819985 | -1.382 | -0.287 | 0.55 | 0.786 | -0.122 | -0.09 | -0.09 | 0 | 0.57 | 0.97 | 0.63 | 0.65 | 0.44 | 0.51 |
| 8 | 34568015 | rs7826148 | -1.002 | -0.469 | 2.159 | 0.802 | -0.054 | -0.088 | -0.088 | 1.39 | 0.25 | 0.49 | 0.2 | 0.09 | 0.14 | 0.2 |
| 8 | 34591853 | rs10094381 | -1.021 | -0.475 | 2.21 | 0.793 | -0.025 | -0.047 | -0.047 | 1.195 | 0.26 | 0.64 | 0.21 | 0.09 | 0.14 | 0.19 |
| 8 | 34607039 | rs13263701 | -1.238 | -0.435 | 2.172 | 0.814 | -0.044 | -0.076 | -0.076 | 0 | 0.28 | 0.6 | 0.22 | 0.09 | 0.15 | 0.2 |
| 8 | 34631702 | rs7003480 | -1.472 | -0.449 | 2.102 | 0.807 | -0.009 | -0.021 | -0.021 | 0 | 0.29 | 0.74 | 0.23 | 0.09 | 0.15 | 0.2 |
| 8 | 34636144 | rs2685607 | -1.472 | -0.449 | 2.102 | 0.808 | -0.009 | -0.023 | -0.023 | 0 | 0.29 | 0.74 | 0.23 | 0.09 | 0.15 | 0.2 |
| 8 | 34641284 | rs13254921 | -1.097 | -0.329 | 1.611 | 0.868 | -0.042 | -0.08 | -0.08 | 0 | 0.27 | 0.49 | 0.21 | 0.09 | 0.15 | 0.2 |
| 8 | 34661961 | rs2687548 | -0.999 | -0.319 | 1.612 | 0.449 | -0.027 | -0.053 | -0.053 | 0.478 | 0.26 | 0.41 | 0.21 | 0.09 | 0.15 | 0.2 |
| 8 | 34700488 | rs10096577 | -1.973 | -0.116 | 1.999 | 0.748 | -0.023 | -0.054 | -0.054 | 0 | 0.11 | 0.5 | 0.05 | 0 | 0.03 | 0.05 |
| 8 | 34721328 | rs2685608 | 0.285 | -0.1 | 2.473 | 0.637 | 0.008 | 0.003 | 0.003 | 0 | 0.9 | 0.8 | 0.83 | 0.91 | 0.91 | 0.91 |
| 8 | 34723346 | rs10089466 | -2.304 | -0.12 | 2.425 | 0.596 | -0.042 | -0.074 | -0.074 | 0 | 0.12 | 0.59 | 0.06 | 0 | 0.03 | 0.05 |
| 8 | 34762526 | rs7813448 | -1.945 | 0.023 | 2.319 | 0.733 | 0.01 | 0 | 0 | 0.713 | 0.07 | 0.46 | 0.04 | 0 | 0.02 | 0.06 |
| 8 | 34762640 | rs7813623 | -2.033 | 0.017 | 2.281 | 0.73 | 0.014 | 0.004 | 0.004 | 0.765 | 0.08 | 0.55 | 0.05 | 0 | 0.02 | 0.06 |
| 8 | 34770257 | rs6995738 | -1.504 | 0.134 | 1.328 | 0.789 | -0.014 | -0.02 | -0.02 | 0 | 0.05 | 0.26 | 0.03 | 0 | 0.02 | 0.05 |
| 8 | 34808188 | rs11996585 | -0.656 | -0.12 | 1.379 | 1.013 | -0.043 | -0.099 | -0.099 | 0 | 0.17 | 0.36 | 0.09 | 0 | 0.1 | 0.14 |
| 8 | 34819174 | rs4324915 | -1.707 | -0.456 | 1.972 | 0.842 | 0.002 | -0.004 | -0.004 | 0 | 0.27 | 0.77 | 0.23 | 0.42 | 0.2 | 0.36 |
| 8 | 34838513 | rs10095892 | -0.591 | -0.496 | 2.059 | 0.801 | -0.05 | -0.108 | -0.108 | 0 | 0.18 | 0.39 | 0.1 | 0 | 0.1 | 0.14 |
| 8 | 34843712 | rs6996639 | -1.224 | -0.338 | 0.945 | -0.618 | 0.001 | -0.001 | -0.001 | 1.014 | 0.55 | 0.6 | 0.37 | 0.43 | 0.5 | 0.46 |
| 8 | 34854498 | rs4739383 | -0.055 | -0.3 | 0.752 | -0.573 | -0.02 | -0.019 | -0.019 | 0.185 | 0.37 | 0.25 | 0.27 | 0.42 | 0.39 | 0.33 |
| 8 | 34866368 | rs10503970 | 0.799 | -0.319 | 0.324 | -0.6 | -0.104 | 0.199 | 0.199 | 0 | 0.28 | 0.02 | 0.15 | 0 | 0.3 | 0.13 |

| | | | | | | | | | | | | | | | | |
|---|---|---|---|---|---|---|---|---|---|---|---|---|---|---|---|---|
| 8 | 34882693 | rs4520153 | -0.921 | -0.256 | 0.841 | -0.454 | -0.011 | 0.009 | 0.009 | 0 | 0.54 | 0.66 | 0.36 | 0.42 | 0.5 | 0.46 |
| 8 | 34887877 | rs13273022 | 1.01 | 0.324 | -0.086 | -0.371 | -0.031 | -0.037 | -0.037 | 0 | 0.23 | 0.06 | 0.24 | 0.27 | 0.26 | 0.36 |
| 8 | 34889071 | rs4269538 | -0.267 | 0.069 | -0.112 | -1.207 | 0.016 | -0.101 | -0.101 | 0 | 0.75 | 0.68 | 0.82 | 1 | 0.69 | 0.87 |
| 8 | 34892197 | rs4613990 | -0.112 | 0.537 | -0.343 | -1.176 | 0.004 | 0.021 | 0.021 | 0 | 0.39 | 0.34 | 0.58 | 0.28 | 0.45 | 0.38 |
| 8 | 34897803 | rs7004239 | 0.125 | 0.468 | 0.121 | -1.015 | -0.003 | -0.006 | -0.006 | 0.225 | 0.38 | 0.42 | 0.5 | 0.74 | 0.35 | 0.48 |
| 8 | 34899169 | rs10503972 | 0.052 | 0.507 | 0.096 | -1.014 | 0.005 | -0.027 | -0.027 | 0 | 0.05 | 0.11 | 0.07 | 0.01 | 0.08 | 0.04 |
| 8 | 34900118 | rs10087728 | -0.083 | 0.513 | -0.082 | -0.973 | -0.006 | -0.026 | -0.026 | 0 | 0.41 | 0.49 | 0.57 | 0.74 | 0.43 | 0.52 |
| 8 | 34906448 | rs12678629 | 1.454 | 0.618 | -1.005 | -1.343 | -0.01 | 0.012 | 0.012 | 0.812 | 0.32 | 0.09 | 0.26 | 0.26 | 0.28 | 0.37 |
| 8 | 34933127 | rs4314649 | -1.085 | 0.611 | -0.953 | -1.353 | 0.013 | 0.044 | 0.044 | 0.689 | 0.63 | 0.85 | 0.72 | 0.74 | 0.7 | 0.63 |
| 8 | 34935896 | rs1376515 | -1.085 | 0.611 | -0.953 | -1.353 | 0.013 | 0.044 | 0.044 | 0.689 | 0.63 | 0.85 | 0.72 | 0.74 | 0.7 | 0.63 |
| 8 | 34941640 | rs2923661 | -1.085 | 0.611 | -0.953 | -1.353 | 0.013 | 0.044 | 0.044 | 0.689 | 0.63 | 0.85 | 0.72 | 0.74 | 0.7 | 0.63 |
| 8 | 34953826 | rs6468303 | 0.278 | 1.023 | -0.634 | -1.193 | 0.047 | -0.129 | -0.129 | 0 | 0.22 | 0.36 | 0.15 | 0 | 0.26 | 0.11 |
| 8 | 34955107 | rs2981319 | -1.174 | 0.948 | -1.098 | -1.172 | -0.001 | 0.012 | 0.012 | 0.534 | 0.64 | 0.89 | 0.64 | 0.71 | 0.67 | 0.58 |
| 8 | 34957138 | rs7013529 | -0.214 | 1.056 | -0.756 | -1.179 | -0.023 | 0.055 | 0.055 | 0 | 0.26 | 0.51 | 0.34 | 0.64 | 0.2 | 0.43 |
| 8 | 34960258 | rs2860367 | -1.293 | 1.029 | -0.356 | -0.735 | 0.005 | 0.001 | 0.001 | 1.367 | 0.66 | 0.78 | 0.6 | 0.69 | 0.64 | 0.6 |
| 8 | 34962205 | rs1376508 | -1.754 | 0.997 | -0.387 | -0.741 | -0.071 | -0.09 | -0.09 | 0.831 | 0.71 | 0.98 | 0.64 | 0.69 | 0.68 | 0.62 |
| 8 | 34967179 | rs921449 | 0.039 | 1.637 | -0.402 | -0.773 | 0.004 | 0.008 | 0.008 | 1.278 | 0.47 | 0.54 | 0.46 | 0.69 | 0.39 | 0.51 |
| 8 | 34976128 | rs6468304 | -0.242 | 1.709 | -0.562 | -0.717 | 0.018 | 0.041 | 0.041 | 2.202 | 0.51 | 0.64 | 0.5 | 0.69 | 0.43 | 0.52 |
| 8 | 34978091 | rs930705 | -1.843 | 1.716 | -0.561 | -0.723 | -0.003 | -0.017 | -0.017 | 1.505 | 0.69 | 0.87 | 0.63 | 0.69 | 0.68 | 0.61 |
| 8 | 34978269 | rs2981342 | -0.182 | 1.724 | -0.568 | -0.713 | 0.015 | -0.059 | -0.059 | 0.017 | 0.19 | 0.23 | 0.13 | 0 | 0.25 | 0.09 |
| 8 | 34982768 | rs1993093 | -1.693 | 1.657 | -0.797 | -0.467 | -0.009 | -0.028 | -0.028 | 0 | 0.7 | 0.89 | 0.63 | 0.69 | 0.68 | 0.61 |
| 8 | 35004235 | rs1451354 | -1.608 | 1.711 | -0.688 | -0.383 | 0.002 | -0.004 | -0.004 | 0.027 | 0.67 | 0.86 | 0.61 | 0.7 | 0.65 | 0.59 |
| 8 | 35008702 | rs1451362 | -1.363 | 1.406 | -0.415 | -0.188 | 0.028 | 0.008 | 0.008 | 0.104 | 0.62 | 0.84 | 0.61 | 0.69 | 0.67 | 0.53 |
| 8 | 35009268 | rs7009205 | -0.327 | 1.325 | 0.047 | -0.108 | -0.001 | 0.001 | 0.001 | 0.298 | 0.45 | 0.53 | 0.48 | 0.69 | 0.45 | 0.45 |
| 8 | 35019465 | rs16883597 | -0.108 | 1.469 | 0.455 | -0.033 | 0.032 | -0.075 | -0.075 | 0 | 0.13 | 0.27 | 0.1 | 0 | 0.17 | 0.09 |
| 8 | 35021424 | rs7003452 | -1.594 | 1.364 | 0.162 | 0.045 | -0.042 | -0.002 | -0.002 | 0.983 | 0.62 | 0.93 | 0.59 | 0.69 | 0.63 | 0.55 |
| 8 | 35041361 | rs7825763 | 0.487 | 1.501 | 0.331 | 0.003 | 0.033 | -0.056 | -0.056 | 0.962 | 0.1 | 0.19 | 0.06 | 0 | 0.14 | 0.08 |
| 8 | 35067119 | rs6997817 | -0.765 | 1.637 | 0.683 | 0.139 | -0.002 | -0.011 | -0.011 | 0.842 | 0.48 | 0.48 | 0.49 | 0.73 | 0.46 | 0.45 |
| 8 | 35075632 | rs1947300 | 0.383 | 0.901 | 0.667 | 0.181 | 0.029 | -0.052 | -0.052 | 0.153 | 0.1 | 0.19 | 0.06 | 0 | 0.14 | 0.08 |
| 8 | 35102627 | rs2056761 | 1.407 | 0.882 | 0.516 | 0.24 | -0.006 | 0.066 | 0.066 | 5.012 | 0.43 | 0.21 | 0.44 | 0.27 | 0.41 | 0.45 |
| 8 | 35111603 | rs2056769 | 1.475 | 0.881 | 0.365 | 0.247 | -0.028 | 0.072 | 0.072 | 5.318 | 0.42 | 0.13 | 0.43 | 0.28 | 0.41 | 0.45 |
| 8 | 35133122 | rs2037840 | 0.233 | 0.771 | 0.963 | 0.858 | -0.002 | 0.186 | 0.186 | 0 | 0.08 | 0.18 | 0.11 | 0.35 | 0.07 | 0.15 |
| 8 | 35137310 | rs2950916 | 1.534 | 0.803 | 0.384 | 0.98 | -0.097 | -0.104 | -0.104 | 0.978 | 0.43 | 0.05 | 0.43 | 0.44 | 0.41 | 0.46 |
| 8 | 35169020 | rs1528714 | -0.157 | 0.893 | 0.644 | 0.819 | 0.067 | -0.083 | -0.083 | 0.237 | 0.87 | 0.7 | 0.9 | 1 | 0.81 | 0.91 |
| 8 | 35182128 | rs1528707 | 1.095 | 0.732 | 0.548 | 0.963 | 0 | 0 | 0 | 0.037 | 0.09 | 0.09 | 0.19 | 0.23 | 0.07 | 0.14 |
| 8 | 35184361 | rs12680114 | -0.576 | 0.909 | 0.123 | 0.507 | -0.002 | -0.006 | -0.006 | 0 | 0.43 | 0.44 | 0.48 | 0.72 | 0.4 | 0.46 |
| 8 | 35218676 | rs12056409 | -0.567 | 1.045 | 0.229 | 0.092 | -0.009 | -0.024 | -0.024 | 0 | 0.43 | 0.49 | 0.48 | 0.72 | 0.4 | 0.46 |
| 8 | 35244430 | rs1950045 | -0.543 | 1.121 | 0.259 | 0.04 | 0.027 | -0.046 | -0.046 | 0 | 0.91 | 0.84 | 0.95 | 1 | 0.86 | 0.92 |
| 8 | 35247297 | rs4521753 | -0.094 | 1.257 | 0.204 | 0.345 | -0.021 | 0.043 | 0.043 | 0.692 | 0.25 | 0.4 | 0.35 | 0.65 | 0.19 | 0.35 |
| 8 | 35273031 | rs4739406 | -0.97 | 1.306 | 0.028 | -0.193 | 0.006 | 0.006 | 0.006 | 3.234 | 0.46 | 0.63 | 0.52 | 0.57 | 0.46 | 0.47 |
| 8 | 35280730 | rs1919344 | -1.042 | 1.325 | -0.214 | -0.183 | 0.018 | 0.015 | 0.015 | 3.327 | 0.47 | 0.68 | 0.53 | 0.57 | 0.46 | 0.47 |
| 8 | 35305866 | rs12545139 | -0.07 | 1.027 | -1.179 | 0.163 | -0.016 | 0.096 | 0.096 | 0.337 | 0.23 | 0.39 | 0.34 | 0.5 | 0.19 | 0.37 |
| 8 | 35312380 | rs4739408 | -0.438 | 0.968 | -2.131 | 0.258 | -0.116 | -0.072 | -0.072 | 0.053 | 0.54 | 0.95 | 0.57 | 0.54 | 0.56 | 0.54 |
| 8 | 35316657 | rs9297240 | -0.956 | 1.051 | -0.566 | -0.069 | 0.007 | -0.015 | -0.015 | 0 | 0.33 | 0.61 | 0.3 | 0.22 | 0.36 | 0.3 |
| 8 | 35316788 | rs10113333 | -1.252 | 0.188 | 0.291 | -0.337 | -0.016 | -0.099 | -0.099 | 0 | 0.24 | 0.57 | 0.15 | 0.05 | 0.22 | 0.13 |
| 8 | 35319481 | rs4577954 | 1.015 | -0.065 | 0.793 | 0.443 | -0.018 | 0.056 | 0.056 | 0 | 0.86 | 0.73 | 0.73 | 0.78 | 0.91 | 0.8 |
| 8 | 35328477 | rs2579880 | -1.266 | -0.133 | 1.545 | 0.072 | 0.014 | -0.068 | -0.068 | 0 | 0.21 | 0.41 | 0.18 | 0.11 | 0.23 | 0.21 |
| 8 | 35331411 | rs2589332 | -1.351 | -0.133 | 1.685 | 0.084 | 0.021 | -0.094 | -0.094 | 0 | 0.21 | 0.49 | 0.18 | 0.09 | 0.22 | 0.2 |
| 8 | 35341484 | rs10503980 | -1.246 | -0.251 | 1.591 | -0.013 | -0.03 | 0.055 | 0.055 | 0 | 0.17 | 0.31 | 0.28 | 0.24 | 0.1 | 0.2 |
| 8 | 35375058 | rs12544701 | -0.959 | -0.259 | 1.519 | -0.04 | -0.013 | 0.02 | 0.02 | 0 | 0.16 | 0.19 | 0.28 | 0.24 | 0.09 | 0.19 |
| 8 | 35378030 | rs2579891 | -1.201 | -0.385 | 1.6 | -0.187 | 0 | -0.01 | -0.01 | 2.387 | 0.14 | 0.17 | 0.1 | 0.08 | 0.12 | 0.14 |
| 8 | 35380121 | rs2579892 | -2.541 | -0.492 | 1.323 | -0.243 | 0.022 | 0.05 | 0.05 | 0 | 0.41 | 0.9 | 0.46 | 0.29 | 0.32 | 0.4 |
| 8 | 35397152 | rs2589344 | -1.885 | -0.478 | 1.517 | -0.28 | 0.006 | -0.075 | -0.075 | 0 | 0.22 | 0.35 | 0.17 | 0.06 | 0.21 | 0.2 |
| 8 | 35412398 | rs6980691 | -1.617 | -0.169 | 1.769 | 0.011 | -0.042 | 0.045 | 0.045 | 0 | 0.19 | 0.54 | 0.29 | 0.24 | 0.1 | 0.19 |
| 8 | 35423882 | rs13260765 | -0.776 | -0.041 | 1.699 | -0.095 | 0.02 | -0.051 | -0.051 | 3.052 | 0.09 | 0.22 | 0.07 | 0 | 0.1 | 0.06 |
| 8 | 35424457 | rs2579897 | -1.919 | -0.032 | 1.324 | 0.019 | 0.006 | -0.007 | -0.007 | 0 | 0.31 | 0.56 | 0.45 | 0.27 | 0.31 | 0.36 |
| 8 | 35432266 | rs2053343 | -2.275 | -0.319 | 0.704 | 0.313 | 0.012 | 0.004 | 0.004 | 0 | 0.39 | 0.79 | 0.41 | 0.35 | 0.33 | 0.39 |
| 8 | 35438155 | rs2589763 | -0.736 | -0.253 | 0.757 | 0.3 | -0.001 | -0.012 | -0.012 | 0 | 0.09 | 0.17 | 0.05 | 0.04 | 0.09 | 0.07 |
| 8 | 35441893 | rs1837347 | -1.216 | -0.286 | 0.924 | 0.314 | -0.035 | -0.038 | -0.038 | 0 | 0.13 | 0.57 | 0.07 | 0.05 | 0.07 | 0.05 |
| 8 | 35467287 | rs7826624 | -2.102 | -0.291 | 0.844 | 0.367 | 0.01 | -0.01 | -0.01 | 0 | 0.3 | 0.7 | 0.36 | 0.32 | 0.24 | 0.31 |
| 8 | 35479110 | rs2978575 | -2.278 | -0.282 | 0.76 | 0.31 | 0.017 | 0.024 | 0.024 | 0.047 | 0.23 | 0.82 | 0.14 | 0.09 | 0.16 | 0.12 |
| 8 | 35479468 | rs17353731 | -0.9 | -0.28 | 0.748 | 0.332 | -0.017 | -0.035 | -0.035 | 0 | 0.17 | 0.07 | 0.28 | 0.26 | 0.17 | 0.27 |
| 8 | 35479911 | rs2978576 | -1.097 | -0.266 | 0.667 | 0.304 | -0.004 | -0.023 | -0.023 | 0 | 0.11 | 0.21 | 0.06 | 0.04 | 0.09 | 0.08 |
| 8 | 35491784 | rs4739420 | 2.44 | -0.3 | 0.497 | 0.312 | 0.001 | -0.009 | -0.009 | 0.54 | 0.6 | 0.1 | 0.58 | 0.66 | 0.67 | 0.63 |
| 8 | 35512394 | rs7845368 | -1.06 | -0.58 | 1.232 | 0.146 | -0.004 | -0.013 | -0.013 | 0.006 | 0.11 | 0.48 | 0.07 | 0.04 | 0.07 | 0.05 |
| 8 | 35522772 | rs1946745 | 0.786 | -0.588 | 0.179 | 0.168 | -0.003 | 0.006 | 0.006 | 0 | 0.91 | 0.88 | 0.94 | 0.95 | 0.89 | 0.92 |
| 8 | 35524881 | rs1898259 | 1.968 | -0.603 | 0.183 | 0.16 | 0.025 | 0.039 | 0.039 | 0.282 | 0.77 | 0.16 | 0.87 | 0.92 | 0.84 | 0.87 |
| 8 | 35528279 | rs1821746 | 1.049 | 0.201 | 0.236 | 0.585 | -0.003 | 0.006 | 0.006 | 0 | 0.91 | 0.88 | 0.94 | 0.95 | 0.89 | 0.92 |
| 8 | 35534138 | rs17270119 | -1.112 | -0.008 | 0.197 | 1.016 | -0.002 | 0.013 | 0.013 | 0 | 0.06 | 0.05 | 0.17 | 0.14 | 0.05 | 0.08 |

| | | | | | | | | | | | | | | | | |
|---|---|---|---|---|---|---|---|---|---|---|---|---|---|---|---|---|
| 8 | 35549181 | rs11985782 | -1.498 | -0.525 | 1.207 | 1.031 | -0.027 | 0.091 | 0.091 | 1.671 | 0.1 | 0.45 | 0.14 | 0.1 | 0.02 | 0.15 |
| 8 | 35558646 | rs6998842 | -1.63 | -0.626 | 2.163 | 0.978 | -0.016 | -0.032 | -0.032 | 0 | 0.07 | 0.27 | 0.03 | 0 | 0.01 | 0.05 |
| 8 | 35561981 | rs16884271 | -1.751 | -1.649 | 2.422 | 1.67 | 0.046 | -0.042 | -0.042 | 0 | 0.3 | 0.76 | 0.25 | 0.38 | 0.1 | 0.31 |
| 8 | 35572305 | rs12675805 | -2.499 | -1.661 | 2.94 | 2.063 | -0.062 | 0.093 | 0.093 | 3 | 0.13 | 0.65 | 0.16 | 0.19 | 0.02 | 0.22 |
| 8 | 35576360 | rs7842132 | -0.606 | -1.654 | 2.885 | 1.982 | -0.02 | -0.033 | -0.033 | 0 | 0.06 | 0.19 | 0.02 | 0 | 0.01 | 0.05 |
| 8 | 35579664 | rs11988094 | -1.143 | -1.656 | 2.899 | 1.981 | -0.021 | -0.035 | -0.035 | 0.029 | 0.07 | 0.25 | 0.03 | 0 | 0.01 | 0.05 |
| 8 | 35583692 | rs6983275 | -2.236 | -1.662 | 2.978 | 1.995 | -0.041 | 0.081 | 0.081 | 2.073 | 0.11 | 0.5 | 0.15 | 0.1 | 0.02 | 0.15 |
| 8 | 35587621 | rs16884338 | -0.618 | -1.89 | 2.729 | 1.946 | 0.033 | -0.004 | -0.004 | 0.27 | 0.18 | 0.03 | 0.05 | 0.14 | 0.09 | 0.09 |
| 8 | 35591808 | rs6998018 | -1.451 | -1.725 | 2.831 | 1.947 | 0.002 | -0.022 | -0.022 | 0 | 0.08 | 0.37 | 0.04 | 0 | 0.01 | 0.05 |
| 8 | 35600202 | rs11993299 | -1.235 | -1.725 | 2.777 | 1.95 | -0.012 | -0.028 | -0.028 | 0 | 0.07 | 0.31 | 0.04 | 0 | 0.01 | 0.06 |
| 8 | 35606168 | rs1436615 | -1.107 | -1.723 | 2.97 | 1.952 | -0.001 | -0.018 | -0.018 | 0.674 | 0.07 | 0.3 | 0.03 | 0 | 0.01 | 0.05 |
| 8 | 35613060 | rs6468336 | -1.944 | -1.732 | 3.189 | 1.947 | -0.015 | -0.035 | -0.035 | 0 | 0.09 | 0.42 | 0.04 | 0 | 0.01 | 0.05 |
| 8 | 35624434 | rs16884404 | -1.226 | -1.739 | 3.202 | 2.017 | -0.003 | -0.02 | -0.02 | 0 | 0.07 | 0.32 | 0.04 | 0 | 0.01 | 0.05 |
| 8 | 35626517 | rs16884409 | -0.792 | -1.737 | 3.082 | 2.018 | -0.022 | -0.034 | -0.034 | 0.016 | 0.06 | 0.18 | 0.02 | 0 | 0.01 | 0.05 |
| 8 | 35641942 | rs7836205 | -1.948 | -2.326 | 3.077 | 1.947 | 0.087 | -0.014 | -0.014 | 0 | 0.35 | 0.76 | 0.15 | 0.3 | 0.12 | 0.26 |
| 8 | 35648941 | rs16884441 | -0.482 | -2.178 | 3.328 | 1.987 | -0.014 | -0.029 | -0.029 | 0 | 0.06 | 0.2 | 0.02 | 0 | 0.01 | 0.05 |
| 8 | 35648980 | rs16884444 | -1.611 | -2.18 | 3.421 | 1.986 | -0.006 | -0.027 | -0.027 | 0 | 0.09 | 0.41 | 0.04 | 0 | 0.01 | 0.05 |
| 8 | 35649884 | rs6987079 | -1.167 | -2.179 | 3.392 | 1.987 | -0.025 | -0.041 | -0.041 | 0.271 | 0.08 | 0.24 | 0.02 | 0 | 0.01 | 0.05 |
| 8 | 35680145 | rs7824105 | -1.59 | -2.179 | 3.177 | 1.978 | -0.007 | -0.029 | -0.029 | 0 | 0.09 | 0.43 | 0.04 | 0 | 0.01 | 0.05 |
| 8 | 35721159 | rs9642758 | -0.276 | -1.494 | 2.168 | 2.67 | -0.049 | -0.054 | -0.054 | 0 | 0.09 | 0.01 | 0.12 | 0.22 | 0.14 | 0.09 |
| 8 | 35754100 | rs17293069 | 0.488 | -0.94 | 0.759 | 1.882 | -0.076 | -0.013 | -0.013 | 0.414 | 0.14 | 0.02 | 0.23 | 0.13 | 0.22 | 0.23 |
| 8 | 35769316 | rs6468343 | -1.744 | -1.201 | 1.113 | 1.851 | -0.046 | 0.052 | 0.052 | 2.556 | 0.09 | 0.72 | 0.09 | 0.19 | 0.01 | 0.17 |
| 8 | 35801618 | rs7014119 | -1.327 | -1.597 | 0.783 | 1.619 | 0.134 | 0.04 | 0.04 | 0 | 0.36 | 0.81 | 0.19 | 0.26 | 0.14 | 0.29 |
| 8 | 35825293 | rs2226323 | -0.062 | -2.237 | 0.497 | 1.258 | -0.05 | -0.059 | -0.059 | 0 | 0.74 | 0.85 | 0.55 | 0.38 | 0.67 | 0.6 |
| 8 | 35831046 | rs3015820 | -0.302 | -2.337 | 0.457 | 1.263 | 0.055 | 0.08 | 0.08 | 0.508 | 0.26 | 0.02 | 0.09 | 0.1 | 0.13 | 0.12 |
| 8 | 35855973 | rs7820891 | -1.244 | -2.306 | 0.662 | 1.314 | -0.046 | 0.043 | 0.043 | 1.083 | 0.09 | 0.7 | 0.09 | 0.18 | 0.01 | 0.19 |
| 8 | 35883816 | rs2979574 | -0.007 | -2.285 | 0.353 | 1.386 | -0.071 | -0.089 | -0.089 | 0 | 0.74 | 0.9 | 0.56 | 0.39 | 0.68 | 0.6 |
| 8 | 35897989 | rs11992651 | 1.507 | -2.091 | 0.464 | 1.256 | -0.015 | 0.118 | 0.118 | 0 | 0.46 | 0.78 | 0.46 | 0.29 | 0.56 | 0.49 |
| 8 | 35920869 | rs12114308 | 1.711 | -2.344 | 0.562 | 1.019 | -0.009 | 0.03 | 0.03 | 0 | 0.41 | 0.31 | 0.39 | 0.13 | 0.56 | 0.3 |
| 8 | 35922813 | rs4739429 | 1.711 | -2.388 | 0.684 | 0.989 | -0.003 | 0 | 0 | 0 | 0.41 | 0.34 | 0.39 | 0.12 | 0.56 | 0.26 |
| 8 | 35928167 | rs3015796 | 1.084 | -2.391 | 0.466 | 1.112 | -0.007 | 0.128 | 0.128 | 0 | 0.5 | 0.85 | 0.47 | 0.29 | 0.57 | 0.5 |
| 8 | 35943847 | rs2951810 | 1.055 | -2.362 | 0.379 | 1.087 | -0.033 | 0.12 | 0.12 | 0 | 0.5 | 0.88 | 0.46 | 0.29 | 0.56 | 0.49 |
| 8 | 35971841 | rs11998466 | 1.212 | -2.297 | 0.45 | 1.188 | -0.043 | 0.147 | 0.147 | 0.355 | 0.46 | 0.85 | 0.43 | 0.23 | 0.56 | 0.48 |
| 8 | 35999591 | rs6415407 | 0.683 | -1.896 | 0.82 | 1.359 | -0.036 | -0.006 | -0.006 | 0 | 0.69 | 0.84 | 0.5 | 0.3 | 0.64 | 0.59 |
| 8 | 36019172 | rs7832435 | 1.017 | -1.22 | 0.76 | 1.825 | -0.04 | -0.069 | -0.069 | 0 | 0.65 | 0.84 | 0.52 | 0.51 | 0.63 | 0.6 |
| 8 | 36025847 | rs2843894 | 1.547 | -1.372 | 1.287 | 1.874 | -0.002 | -0.007 | -0.007 | 0 | 0.56 | 0.43 | 0.47 | 0.27 | 0.62 | 0.39 |
| 8 | 36049909 | rs2843886 | -1.464 | -0.742 | 1.784 | 1.978 | -0.024 | 0.266 | 0.266 | 2.528 | 0.09 | 0.54 | 0.07 | 0.25 | 0 | 0.2 |
| 8 | 36104136 | rs602382 | 0.851 | -0.683 | 1.799 | 2.312 | -0.039 | 0.072 | 0.072 | 0.094 | 0.73 | 0.5 | 0.62 | 0.5 | 0.81 | 0.58 |
| 8 | 36114239 | rs1413875 | 0.53 | -0.612 | 1.165 | 2.03 | 0.006 | 0.004 | 0.004 | 0 | 0.77 | 0.8 | 0.66 | 0.75 | 0.81 | 0.74 |
| 8 | 36169691 | rs604756 | 0.673 | -0.355 | 1.007 | 2.451 | -0.002 | 0.007 | 0.007 | 0 | 0.87 | 0.87 | 0.87 | 0.59 | 0.88 | 0.71 |
| 8 | 36217127 | rs4739439 | -0.454 | -0.327 | 1.165 | 2.238 | -0.002 | -0.022 | -0.022 | 0.041 | 0.14 | 0.07 | 0.31 | 0.24 | 0.16 | 0.17 |
| 8 | 36272346 | rs2843871 | 0.748 | -0.288 | 1.208 | 2.286 | 0 | 0.001 | 0.001 | 0 | 0.86 | 0.86 | 0.87 | 0.67 | 0.89 | 0.73 |
| 8 | 36406388 | rs1018739 | 1.463 | 0.33 | -0.73 | 0.728 | -0.029 | 0.191 | 0.191 | 0.186 | 0.34 | 0.01 | 0.28 | 0.04 | 0.31 | 0.24 |
| 8 | 36410648 | rs10503990 | -1.42 | 0.391 | -0.114 | 1.181 | -0.046 | 0.066 | 0.066 | 1.093 | 0.19 | 0.44 | 0.12 | 0.25 | 0.1 | 0.23 |
| 8 | 36418643 | rs4739319 | -1.869 | 0.772 | -0.09 | 1.506 | 0.004 | 0.002 | 0.002 | 0.107 | 0.37 | 0.64 | 0.47 | 0.66 | 0.31 | 0.56 |
| 8 | 36419196 | rs7009451 | -1.538 | 1.263 | 0.037 | 1.79 | -0.101 | -0.058 | -0.058 | 0 | 0.78 | 0.96 | 0.78 | 0.7 | 0.64 | 0.81 |
| 8 | 36425166 | rs6468370 | -1.453 | 1.155 | 0.512 | 1.641 | -0.043 | 0.076 | 0.076 | 1.39 | 0.19 | 0.47 | 0.12 | 0.25 | 0.1 | 0.3 |
| 8 | 36459009 | rs7813659 | -1.585 | 1.24 | 0.506 | 1.511 | -0.017 | 0.001 | 0.001 | 1.579 | 0.21 | 0.65 | 0.15 | 0.19 | 0.12 | 0.3 |
| 8 | 36459454 | rs7814403 | -1.549 | 2.047 | 0.325 | 1.643 | 0.023 | 0.009 | 0.009 | 0 | 0.36 | 0.42 | 0.41 | 0.43 | 0.46 | 0.39 |
| 8 | 36486830 | rs16885250 | -1.186 | 1.624 | 0.452 | 1.066 | 0.021 | -0.06 | -0.06 | 0.015 | 0.31 | 0.47 | 0.24 | 0.17 | 0.31 | 0.29 |
| 8 | 36492258 | rs2211743 | -1.17 | 1.634 | 0.468 | 1.064 | 0.024 | -0.052 | -0.052 | 2.655 | 0.31 | 0.4 | 0.23 | 0.17 | 0.33 | 0.3 |
| 8 | 36494238 | rs16885261 | -1.363 | 1.617 | 0.425 | 1.091 | 0.015 | -0.06 | -0.06 | 0 | 0.32 | 0.49 | 0.24 | 0.17 | 0.31 | 0.31 |
| 8 | 36547313 | rs6982811 | -1.292 | 1.727 | 0.271 | 0.879 | 0.013 | -0.097 | -0.097 | 0.502 | 0.3 | 0.6 | 0.19 | 0.05 | 0.3 | 0.19 |
| 8 | 36552965 | rs13438817 | -0.886 | 1.796 | 0.197 | 0.913 | 0.027 | -0.123 | -0.123 | 0.029 | 0.29 | 0.5 | 0.17 | 0.05 | 0.3 | 0.18 |
| 8 | 36565705 | rs3890448 | -0.269 | 2.107 | 0.003 | 1.995 | -0.019 | -0.049 | -0.049 | 0.047 | 0.12 | 0.04 | 0.26 | 0.49 | 0.17 | 0.21 |
| 8 | 36603585 | rs4739325 | 0.116 | 2.066 | -0.373 | 1.917 | -0.015 | -0.021 | -0.021 | 0 | 0.1 | 0.04 | 0.27 | 0.51 | 0.16 | 0.21 |
| 8 | 36628405 | rs17305805 | 0.143 | 1.264 | 0.424 | 2.019 | -0.067 | -0.067 | -0.067 | 0 | 0.11 | 0.01 | 0.2 | 0.2 | 0.19 | 0.12 |
| 8 | 36629798 | rs1981323 | -2.079 | 0.138 | 1.249 | 1.575 | -0.035 | -0.029 | -0.029 | 0 | 0.08 | 0.66 | 0.07 | 0.01 | 0.01 | 0.11 |
| 8 | 36634382 | rs10094607 | -1.643 | 0.27 | 1.238 | 1.704 | -0.004 | 0.137 | 0.137 | 4.555 | 0.07 | 0.57 | 0.1 | 0.11 | 0.01 | 0.12 |
| 8 | 36646217 | rs6989810 | -0.661 | 0.521 | 0.411 | 2.084 | -0.002 | 0.201 | 0.201 | 0.247 | 0.18 | 0.5 | 0.41 | 0.43 | 0.17 | 0.31 |
| 8 | 36659001 | rs7016533 | -0.409 | 0.487 | 0.681 | 2.486 | 0.009 | 0.02 | 0.02 | 0.018 | 0.17 | 0.42 | 0.39 | 0.17 | 0.17 | 0.25 |
| 8 | 36660957 | rs2211730 | -0.446 | 0.455 | 1.018 | 2.45 | 0.01 | 0.016 | 0.016 | 0.012 | 0.17 | 0.45 | 0.39 | 0.17 | 0.17 | 0.25 |
| 8 | 36661686 | rs12543523 | -0.024 | 0.448 | 0.975 | 2.847 | -0.002 | -0.004 | -0.004 | 0.801 | 0.14 | 0.14 | 0.38 | 0.24 | 0.17 | 0.24 |
| 8 | 36662680 | rs16885430 | -0.651 | 0.449 | 1.035 | 2.722 | 0.009 | 0.014 | 0.014 | 0.093 | 0.17 | 0.45 | 0.39 | 0.17 | 0.17 | 0.25 |
| 8 | 36686343 | rs16885468 | 0.082 | 0.666 | 0.297 | 2.787 | -0.053 | -0.09 | -0.09 | 0.029 | 0.14 | 0.02 | 0.12 | 0.36 | 0.2 | 0.2 |
| 8 | 36690323 | rs13439887 | -0.514 | 0.319 | 0.345 | 2.405 | 0.001 | -0.009 | -0.009 | 0 | 0.22 | 0.23 | 0.42 | 0.43 | 0.22 | 0.32 |
| 8 | 36707921 | rs10107777 | -1.196 | 0.797 | 0.304 | 2.141 | 0.012 | 0.016 | 0.016 | 0 | 0.3 | 0.38 | 0.48 | 0.39 | 0.4 | 0.36 |
| 8 | 36727529 | rs10100186 | -1.152 | 0.794 | 0.268 | 2.093 | -0.011 | -0.012 | -0.012 | 0.02 | 0.29 | 0.27 | 0.49 | 0.38 | 0.4 | 0.36 |
| 8 | 36729594 | rs2211727 | -0.761 | 0.799 | 0.267 | 2.096 | -0.033 | -0.036 | -0.036 | 0.117 | 0.28 | 0.19 | 0.48 | 0.38 | 0.4 | 0.36 |

| | | | | | | | | | | | | | | | | |
|---|---|---|---|---|---|---|---|---|---|---|---|---|---|---|---|---|
| 8 | 36741559 | rs10099214 | -0.845 | 0.826 | 0.287 | 2.102 | -0.01 | -0.011 | -0.011 | 0 | 0.28 | 0.28 | 0.47 | 0.38 | 0.4 | 0.36 |
| 8 | 36745664 | rs35727025 | -0.864 | 0.802 | 0.311 | 2.069 | 0.004 | 0.004 | 0.004 | 0.293 | 0.28 | 0.32 | 0.48 | 0.38 | 0.4 | 0.36 |
| 8 | 36754130 | rs7460480 | -0.864 | 0.802 | 0.311 | 2.069 | 0.004 | 0.004 | 0.004 | 0.293 | 0.28 | 0.32 | 0.48 | 0.38 | 0.4 | 0.36 |
| 8 | 36786379 | rs17400167 | 0.354 | 0.917 | 0.123 | 2.098 | -0.041 | -0.016 | -0.016 | 0 | 0.17 | 0.05 | 0.4 | 0.18 | 0.25 | 0.26 |
| 8 | 36789591 | rs7386222 | -0.186 | -0.151 | 0.01 | 1.443 | 0.023 | 0.01 | 0.01 | 0 | 0.65 | 0.58 | 0.64 | 0.57 | 0.53 | 0.65 |
| 8 | 36825080 | rs10090367 | 1.505 | -0.21 | -0.051 | 1.42 | 0.046 | 0.04 | 0.04 | 0.54 | 0.37 | 0.25 | 0.16 | 0.18 | 0.13 | 0.32 |
| 8 | 36827102 | rs2970663 | 0.031 | -0.51 | -0.221 | 1.237 | 0.006 | -0.013 | -0.013 | 0.038 | 0.58 | 0.5 | 0.63 | 0.69 | 0.52 | 0.65 |
| 8 | 36834333 | rs2970646 | -0.037 | 0.403 | 0.198 | 1.538 | -0.073 | -0.033 | -0.033 | 0.654 | 0.26 | 0.13 | 0.47 | 0.3 | 0.42 | 0.32 |
| 8 | 36839221 | rs2970653 | 0.188 | 0.368 | 0.22 | 1.52 | 0.027 | 0.01 | 0.01 | 0 | 0.64 | 0.5 | 0.63 | 0.59 | 0.54 | 0.64 |
| 8 | 36848357 | rs16885613 | 1.26 | 0.018 | 0.153 | 1.36 | 0.003 | 0.005 | 0.005 | 0.007 | 0.4 | 0.35 | 0.2 | 0.31 | 0.15 | 0.33 |
| 8 | 36861905 | rs7464912 | -0.042 | 0.037 | 0.633 | 1.522 | 0.038 | 0.032 | 0.032 | 0 | 0.7 | 0.6 | 0.55 | 0.58 | 0.61 | 0.66 |
| 8 | 36865149 | rs11997488 | -0.087 | -0.021 | 0.639 | 1.522 | 0.024 | 0.021 | 0.021 | 0 | 0.7 | 0.65 | 0.56 | 0.57 | 0.61 | 0.66 |
| 8 | 36869192 | rs3015758 | 1.755 | -0.069 | 0.083 | 1.447 | 0.002 | 0.009 | 0.009 | 0 | 0.38 | 0.34 | 0.23 | 0.28 | 0.32 | 0.34 |
| 8 | 36872094 | rs1119539 | -1.169 | -0.505 | 0.617 | 2.13 | 0.016 | -0.025 | -0.025 | 0.092 | 0.31 | 0.17 | 0.33 | 0.3 | 0.29 | 0.33 |
| 8 | 36880802 | rs2970636 | -1.821 | 0.474 | 1.494 | 2.631 | 0.004 | 0.011 | 0.011 | 0.204 | 0.34 | 0.41 | 0.56 | 0.56 | 0.43 | 0.47 |
| 8 | 36884712 | rs2970627 | 1.601 | 0.462 | 1.7 | 2.657 | 0.009 | 0.023 | 0.023 | 0.254 | 0.66 | 0.54 | 0.44 | 0.43 | 0.57 | 0.53 |
| 8 | 36899857 | rs3015775 | 1.502 | 0.406 | 1.94 | 2.598 | 0.003 | 0.015 | 0.015 | 0.363 | 0.65 | 0.44 | 0.43 | 0.44 | 0.57 | 0.53 |
| 8 | 36908104 | rs2040115 | 1.504 | 0.43 | 1.733 | 2.624 | -0.008 | -0.019 | -0.019 | 0.117 | 0.67 | 0.67 | 0.45 | 0.44 | 0.57 | 0.53 |
| 8 | 36911307 | rs3015729 | 1.107 | 0.456 | 1.762 | 2.753 | -0.01 | -0.021 | -0.021 | 0.046 | 0.71 | 0.78 | 0.62 | 0.67 | 0.67 | 0.63 |
| 8 | 36939390 | rs2970708 | 3.023 | -0.224 | 2.759 | 2.929 | 0.01 | 0.262 | 0.262 | 0.816 | 0.88 | 0.6 | 0.77 | 0.67 | 0.89 | 0.84 |
| 8 | 36939955 | rs16885876 | -2.186 | -0.195 | 2.752 | 2.917 | 0.053 | 0.2 | 0.2 | 0 | 0.06 | 0.21 | 0.18 | 0.23 | 0.1 | 0.1 |
| 8 | 36953646 | rs11776875 | -2.543 | -0.248 | 2.791 | 2.733 | -0.008 | -0.072 | -0.072 | 0 | 0.12 | 0.38 | 0.09 | 0.02 | 0.07 | 0.03 |
| 8 | 36961089 | rs11996325 | -1.517 | -0.242 | 2.549 | 2.736 | -0.002 | -0.019 | -0.019 | 0 | 0.09 | 0.12 | 0.05 | 0.02 | 0.07 | 0.04 |
| 8 | 36965264 | rs3015666 | 2.512 | -0.235 | 2.363 | 2.806 | -0.026 | 0.083 | 0.083 | 0.481 | 0.8 | 0.46 | 0.71 | 0.65 | 0.83 | 0.8 |
| 8 | 36981730 | rs502322 | 2.615 | -0.166 | 2.391 | 2.91 | -0.016 | 0.102 | 0.102 | 0.742 | 0.78 | 0.42 | 0.68 | 0.45 | 0.79 | 0.78 |
| 8 | 36987916 | rs1089432 | 1.975 | -0.143 | 2.515 | 2.821 | 0.01 | 0.418 | 0.418 | 0.667 | 0.89 | 0.57 | 0.8 | 0.45 | 0.91 | 0.84 |
| 8 | 36990733 | rs2013020 | -2.745 | -0.186 | 2.199 | 2.5 | -0.003 | 0.237 | 0.237 | 0.792 | 0.16 | 0.57 | 0.22 | 0.51 | 0.15 | 0.21 |
| 8 | 36991553 | rs6468392 | -1.961 | 0.028 | 1.787 | 1.354 | 0.068 | 0.002 | 0.002 | 0.027 | 0.07 | 0.36 | 0.05 | 0.02 | 0.06 | 0.06 |
| 8 | 36994316 | rs7814066 | -1.66 | -0.189 | 1.16 | 1.689 | -0.009 | 0.119 | 0.119 | 0 | 0.24 | 0.43 | 0.2 | 0.52 | 0.23 | 0.23 |
| 8 | 36995874 | rs543946 | 0.818 | -0.102 | 0.88 | 1.577 | 0.001 | 0.002 | 0.002 | 0 | 0.93 | 0.94 | 0.94 | 0.97 | 0.94 | 0.95 |
| 8 | 37003868 | rs489583 | 1.637 | -0.129 | 1.335 | 1.71 | 0.001 | -0.005 | -0.005 | 0 | 0.75 | 0.25 | 0.76 | 0.47 | 0.77 | 0.76 |
| 8 | 37004569 | rs604302 | 1.218 | -0.265 | 1.446 | 1.833 | -0.004 | 0.049 | 0.049 | 0 | 0.81 | 0.71 | 0.83 | 0.61 | 0.81 | 0.76 |
| 8 | 37006731 | rs16886043 | -0.495 | -0.204 | 1.499 | 1.359 | 0.001 | 0.011 | 0.011 | 0 | 0.12 | 0.12 | 0.14 | 0.24 | 0.19 | 0.14 |
| 8 | 37008299 | rs492548 | 0.083 | -0.195 | 1.286 | 1.303 | 0.011 | 0.019 | 0.019 | 0 | 0.91 | 0.97 | 0.95 | 0.97 | 0.96 | 0.91 |
| 8 | 37011661 | rs508635 | 1.604 | -0.236 | 1.37 | 1.343 | 0.001 | -0.003 | -0.003 | 2.739 | 0.71 | 0.27 | 0.77 | 0.58 | 0.75 | 0.72 |
| 8 | 37012275 | rs481779 | 1.305 | -0.123 | 1.336 | 1.387 | -0.001 | 0.011 | 0.011 | 0 | 0.76 | 0.67 | 0.8 | 0.72 | 0.77 | 0.73 |
| 8 | 37013573 | rs508295 | 0.516 | 0.203 | 0.745 | 1.212 | 0.021 | -0.031 | -0.031 | 0 | 0.38 | 0.19 | 0.34 | 0.47 | 0.33 | 0.55 |
| 8 | 37027184 | rs6998075 | -1.718 | 0.572 | 1.322 | 1.855 | 0.021 | 0.175 | 0.175 | 1.02 | 0.17 | 0.61 | 0.18 | 0.39 | 0.2 | 0.19 |
| 8 | 37030919 | rs7005136 | -1.336 | 0.826 | 0.302 | 1.157 | 0.029 | 0.081 | 0.081 | 0.186 | 0.14 | 0.29 | 0.16 | 0.23 | 0.18 | 0.16 |
| 8 | 37044187 | rs576019 | -1.021 | 0.485 | 0.296 | 1.37 | -0.005 | 0.065 | 0.065 | 0 | 0.09 | 0.18 | 0.11 | 0.22 | 0.07 | 0.18 |
| 8 | 37045176 | rs7841889 | -1.746 | 0.301 | 1.46 | 1.257 | 0 | 0.167 | 0.167 | 0 | 0.13 | 0.43 | 0.13 | 0.22 | 0.09 | 0.2 |
| 8 | 37048234 | rs9642769 | -0.386 | 0.224 | 0.981 | 0.626 | -0.004 | 0.035 | 0.035 | 0 | 0.57 | 0.63 | 0.6 | 0.78 | 0.57 | 0.72 |
| 8 | 37048751 | rs7826398 | 0.396 | 0.209 | 0.859 | 0.61 | -0.001 | -0.001 | -0.001 | 0 | 0.45 | 0.4 | 0.49 | 0.56 | 0.49 | 0.54 |
| 8 | 37050143 | rs2843837 | 0.681 | 0.291 | 0.076 | 0.242 | -0.029 | -0.015 | -0.015 | 0 | 0.16 | 0.05 | 0.25 | 0.18 | 0.23 | 0.18 |
| 8 | 37053824 | rs493477 | 0.145 | -0.172 | 0.324 | 0.321 | -0.008 | 0.019 | 0.019 | 0 | 0.85 | 0.78 | 0.87 | 0.78 | 0.89 | 0.75 |
| 8 | 37054785 | rs519676 | 0.212 | -0.203 | 0.491 | 0.305 | -0.024 | 0.051 | 0.051 | 0 | 0.84 | 0.69 | 0.86 | 0.78 | 0.89 | 0.75 |
| 8 | 37055719 | rs510561 | 0.044 | -0.203 | 0.487 | 0.305 | -0.008 | 0.018 | 0.018 | 0 | 0.85 | 0.78 | 0.87 | 0.78 | 0.89 | 0.75 |
| 8 | 37057541 | rs531990 | 0.228 | -0.207 | 0.409 | 0.304 | -0.026 | 0.053 | 0.053 | 0 | 0.84 | 0.68 | 0.86 | 0.78 | 0.89 | 0.75 |
| 8 | 37065784 | rs1850650 | 1.124 | -0.417 | -0.744 | 0.025 | -0.102 | -0.098 | -0.098 | 0.627 | 0.38 | 0.05 | 0.44 | 0.52 | 0.42 | 0.49 |
| 8 | 37073040 | rs472223 | 0.266 | -0.265 | -0.698 | 0.056 | -0.092 | 0.039 | 0.039 | 0.785 | 0.33 | 0.06 | 0.32 | 0.23 | 0.4 | 0.2 |
| 8 | 37074938 | rs532022 | 0.727 | -0.308 | 0.292 | 0.391 | -0.049 | 0.068 | 0.068 | 0.022 | 0.81 | 0.38 | 0.84 | 0.78 | 0.89 | 0.74 |
| 8 | 37076504 | rs511544 | 0.743 | -0.304 | 0.294 | 0.376 | -0.05 | 0.058 | 0.058 | 0.047 | 0.81 | 0.38 | 0.84 | 0.79 | 0.89 | 0.74 |
| 8 | 37096059 | rs1015003 | 0.013 | 0.356 | -0.4 | -0.13 | 0.001 | -0.003 | -0.003 | 0 | 0.63 | 0.66 | 0.52 | 0.73 | 0.54 | 0.7 |
| 8 | 37098052 | rs408109 | 0.384 | 0.844 | -0.619 | -0.36 | -0.067 | -0.026 | -0.026 | 0 | 0.27 | 0.06 | 0.37 | 0.27 | 0.3 | 0.19 |
| 8 | 37100928 | rs6985083 | 0.69 | -0.092 | -1.533 | -1.038 | 0.056 | 0.089 | 0.089 | 0.656 | 0.22 | 0.1 | 0.08 | 0.02 | 0.13 | 0.1 |
| 8 | 37109147 | rs405501 | -0.957 | 0.311 | -1.318 | -0.81 | -0.005 | -0.024 | -0.024 | 0.145 | 0.74 | 0.8 | 0.7 | 0.44 | 0.68 | 0.58 |
| 8 | 37118078 | rs6468400 | -0.283 | 0.224 | -0.96 | -0.837 | 0.011 | 0.017 | 0.017 | 0.052 | 0.5 | 0.6 | 0.42 | 0.64 | 0.44 | 0.53 |
| 8 | 37120864 | rs6468401 | -0.962 | 0.32 | -0.896 | -0.423 | -0.005 | 0.022 | 0.022 | 0 | 0.54 | 0.66 | 0.52 | 0.72 | 0.56 | 0.7 |
| 8 | 37121241 | rs3015657 | 0.154 | 0.28 | -1.257 | -0.448 | -0.005 | 0.071 | 0.071 | 0.053 | 0.26 | 0.13 | 0.28 | 0.12 | 0.26 | 0.27 |
| 8 | 37127023 | rs402003 | 0.765 | 0.299 | -1.248 | -0.194 | -0.008 | -0.009 | -0.009 | 1.12 | 0.15 | 0.1 | 0.23 | 0.18 | 0.18 | 0.21 |
| 8 | 37127143 | rs433689 | -0.366 | 0.291 | -1.093 | -0.233 | -0.009 | 0.012 | 0.012 | 0 | 0.39 | 0.32 | 0.51 | 0.7 | 0.49 | 0.63 |
| 8 | 37128540 | rs10504003 | 0.413 | 0.085 | -1.177 | -0.273 | -0.016 | -0.05 | -0.05 | 1.302 | 0.31 | 0.19 | 0.34 | 0.52 | 0.41 | 0.44 |
| 8 | 37129297 | rs407732 | 0.927 | 0.419 | -1.017 | -0.048 | -0.004 | -0.009 | -0.009 | 0 | 0.14 | 0.1 | 0.22 | 0.17 | 0.15 | 0.2 |
| 8 | 37144505 | rs398625 | 1.058 | 0.422 | -1.027 | -0.08 | -0.01 | 0.026 | 0.026 | 0 | 0.14 | 0.06 | 0.21 | 0.1 | 0.15 | 0.19 |
| 8 | 37146598 | rs441155 | -0.4 | 0.237 | -0.652 | 0.062 | 0.001 | -0.019 | -0.019 | 2.749 | 0.58 | 0.51 | 0.63 | 0.72 | 0.65 | 0.72 |
| 8 | 37146819 | rs454228 | -0.355 | 0.278 | -0.759 | -0.021 | 0 | 0 | 0 | 0 | 0.58 | 0.45 | 0.62 | 0.63 | 0.65 | 0.7 |
| 8 | 37151564 | rs7002677 | 0.458 | 0.22 | -0.569 | -0.311 | 0.008 | 0.015 | 0.015 | 0 | 0.42 | 0.36 | 0.26 | 0.26 | 0.31 | 0.25 |
| 8 | 37158002 | rs384875 | 0.373 | 0.451 | -0.686 | -0.501 | -0.001 | 0.012 | 0.012 | 0 | 0.26 | 0.27 | 0.3 | 0.08 | 0.28 | 0.33 |
| 8 | 37159154 | rs10955016 | 0.43 | 0.251 | -0.416 | -0.421 | 0.013 | 0.019 | 0.019 | 0 | 0.49 | 0.58 | 0.41 | 0.43 | 0.43 | 0.35 |

| | | | | | | | | | | | | | | | | |
|---|---|---|---|---|---|---|---|---|---|---|---|---|---|---|---|---|
| 8 | 37159346 | rs364583 | 0.629 | 0.311 | -0.329 | -0.353 | -0.009 | 0.051 | 0.051 | 0 | 0.25 | 0.16 | 0.29 | 0.08 | 0.28 | 0.33 |
| 8 | 37161773 | rs11783184 | 0.531 | 0.088 | -0.568 | -0.378 | -0.002 | -0.002 | -0.002 | 0 | 0.41 | 0.53 | 0.36 | 0.41 | 0.38 | 0.31 |
| 8 | 37161801 | rs11785575 | 0.642 | 0.125 | -0.428 | -0.377 | 0.002 | 0.005 | 0.005 | 0 | 0.07 | 0.08 | 0.14 | 0.15 | 0.12 | 0.1 |
| 8 | 37162062 | rs3896235 | 1.176 | 0.128 | -0.617 | -0.467 | -0.012 | -0.019 | -0.019 | 0 | 0.34 | 0.46 | 0.23 | 0.26 | 0.26 | 0.21 |
| 8 | 37167934 | rs422549 | 0.596 | 0.618 | -1.273 | -0.35 | -0.013 | 0.061 | 0.061 | 0 | 0.21 | 0.07 | 0.24 | 0.07 | 0.22 | 0.31 |
| 8 | 37169244 | rs390287 | -0.165 | 0.443 | -1.393 | -0.13 | -0.046 | -0.086 | -0.086 | 2.414 | 0.7 | 0.84 | 0.63 | 0.48 | 0.64 | 0.64 |
| 8 | 37173317 | rs641384 | -0.425 | 0.209 | -0.601 | -0.256 | 0 | -0.017 | -0.017 | 0.006 | 0.31 | 0.35 | 0.4 | 0.22 | 0.35 | 0.36 |
| 8 | 37178897 | rs391583 | 0.715 | 0.146 | -0.627 | -0.33 | -0.014 | -0.033 | -0.033 | 0.583 | 0.65 | 0.55 | 0.6 | 0.75 | 0.7 | 0.66 |
| 8 | 37181510 | rs368641 | 0.565 | 0.574 | -0.121 | 0.131 | 0.034 | 0.059 | 0.059 | 0.554 | 0.87 | 0.57 | 0.82 | 0.81 | 0.87 | 0.88 |
| 8 | 37183686 | rs446276 | -0.871 | 0.997 | -0.362 | 0.332 | 0.031 | 0.052 | 0.052 | 0 | 0.23 | 0.31 | 0.41 | 0.5 | 0.33 | 0.3 |
| 8 | 37188456 | rs392552 | -0.867 | 0.48 | -0.573 | 0.431 | -0.047 | -0.096 | -0.096 | 0 | 0.24 | 0.09 | 0.37 | 0.48 | 0.3 | 0.33 |
| 8 | 37194555 | rs462931 | 0.068 | 0.335 | 0.029 | 0.06 | -0.013 | -0.082 | -0.082 | 0 | 0.86 | 0.57 | 0.91 | 0.97 | 0.89 | 0.96 |
| 8 | 37200241 | rs461858 | -0.441 | 0.355 | -0.867 | 0.169 | -0.087 | -0.137 | -0.137 | 0.193 | 0.2 | 0.03 | 0.3 | 0.48 | 0.27 | 0.33 |
| 8 | 37200741 | rs7818467 | 1.116 | 0.396 | -0.644 | 0.197 | -0.008 | -0.017 | -0.017 | 0 | 0.63 | 0.28 | 0.56 | 0.49 | 0.61 | 0.61 |
| 8 | 37209300 | rs11785595 | 0.982 | 0.171 | -0.662 | 0.273 | 0.002 | -0.029 | -0.029 | 0 | 0.62 | 0.24 | 0.57 | 0.5 | 0.65 | 0.61 |
| 8 | 37217380 | rs406494 | 0.386 | 0.203 | -0.11 | 0.674 | -0.007 | -0.011 | -0.011 | 0 | 0.71 | 0.74 | 0.64 | 0.54 | 0.65 | 0.57 |
| 8 | 37219077 | rs10096200 | 1.052 | -0.328 | -0.33 | 0.144 | -0.06 | 0.163 | 0.163 | 1.64 | 0.41 | 0.09 | 0.25 | 0.18 | 0.43 | 0.34 |
| 8 | 37220942 | rs239451 | -0.253 | -0.157 | 0.21 | 0.495 | -0.025 | -0.071 | -0.071 | 4.573 | 0.24 | 0.12 | 0.38 | 0.46 | 0.27 | 0.3 |
| 8 | 37237491 | rs413003 | 0.096 | -0.687 | 0.53 | 0.433 | 0.009 | -0.057 | -0.057 | 0.348 | 0.23 | 0.13 | 0.26 | 0.52 | 0.18 | 0.22 |
| 8 | 37246921 | rs16886686 | -0.508 | -0.779 | 1.182 | 0.33 | 0 | -0.007 | -0.007 | 0 | 0.08 | 0.32 | 0.05 | 0.03 | 0.06 | 0.02 |
| 8 | 37259045 | rs2843807 | 0.053 | -0.583 | 1.378 | 0.452 | -0.018 | 0.051 | 0.051 | 0 | 0.74 | 0.64 | 0.73 | 0.48 | 0.82 | 0.75 |
| 8 | 37286161 | rs2541454 | 0.67 | -0.442 | 1.382 | 0.175 | 0.003 | -0.001 | -0.001 | 0.221 | 0.65 | 0.32 | 0.68 | 0.39 | 0.76 | 0.72 |
| 8 | 37305509 | rs1121262 | 0.917 | -0.681 | 0.818 | 0.02 | -0.078 | 0.14 | 0.14 | 0 | 0.35 | 0.09 | 0.24 | 0.11 | 0.42 | 0.31 |
| 8 | 37317145 | rs6980823 | -0.801 | -0.949 | 1.737 | 1.017 | 0.023 | -0.037 | -0.037 | 0 | 0.27 | 0.2 | 0.26 | 0.58 | 0.19 | 0.23 |
| 8 | 37331826 | rs16886781 | -0.758 | -0.949 | 1.221 | 1.036 | 0.016 | -0.038 | -0.038 | 0 | 0.28 | 0.16 | 0.27 | 0.58 | 0.23 | 0.23 |
| 8 | 37336212 | rs16886796 | -0.587 | -0.595 | 1.234 | -0.007 | -0.008 | -0.018 | -0.018 | 0.167 | 0.05 | 0.25 | 0.03 | 0.01 | 0.04 | 0.03 |
| 8 | 37336363 | rs2407275 | 0.932 | -0.59 | 0.346 | 0.076 | 0.037 | 0.051 | 0.051 | 0 | 0.67 | 0.22 | 0.8 | 0.85 | 0.75 | 0.77 |
| 8 | 37337950 | rs11777137 | 0.582 | -0.539 | 0.309 | 0.042 | -0.017 | 0.015 | 0.015 | 0 | 0.09 | 0.05 | 0.07 | 0 | 0.16 | 0.03 |
| 8 | 37339153 | rs11986415 | 1.164 | -0.939 | 0.94 | 0.689 | 0.016 | 0.021 | 0.021 | 0 | 0.68 | 0.27 | 0.81 | 0.85 | 0.75 | 0.77 |
| 8 | 37360288 | rs7387040 | 1.209 | -0.919 | 0.585 | 0.699 | 0.036 | 0.05 | 0.05 | 0 | 0.67 | 0.23 | 0.8 | 0.85 | 0.75 | 0.78 |
| 8 | 37372363 | rs16886842 | -0.759 | 0.049 | -0.977 | 1.27 | 0 | 0.029 | 0.029 | 0.189 | 0.2 | 0.07 | 0.15 | 0.12 | 0.19 | 0.16 |
| 8 | 37381976 | rs2211704 | -1.098 | 0.235 | -0.282 | 2.045 | 0 | 0.309 | 0.309 | 0 | 0.07 | 0.25 | 0.21 | 0.5 | 0.06 | 0.15 |
| 8 | 37394958 | rs16886859 | -0.869 | -0.263 | -1.185 | 1.271 | 0.013 | 0.011 | 0.011 | 0 | 0.13 | 0.01 | 0.07 | 0.05 | 0.09 | 0.05 |
| 8 | 37404478 | rs1467846 | -1.587 | -1.466 | 0.085 | 1.695 | -0.093 | 0.003 | 0.003 | 0 | 0.24 | 0.5 | 0.12 | 0.17 | 0.11 | 0.07 |
| 8 | 37405129 | rs12548714 | 1.315 | -1.889 | -0.064 | 1.108 | 0.01 | 0.009 | 0.009 | 0.025 | 0.68 | 0.32 | 0.75 | 0.83 | 0.81 | 0.91 |
| 8 | 37409288 | rs7003946 | -1.376 | -1.861 | -0.07 | 1.557 | -0.088 | 0.116 | 0.116 | 0 | 0.26 | 0.48 | 0.29 | 0.49 | 0.12 | 0.17 |
| 8 | 37444536 | rs7386256 | -0.538 | -1.418 | -1.11 | 0.655 | -0.027 | 0.092 | 0.092 | 0 | 0.73 | 0.93 | 0.84 | 0.96 | 0.75 | 0.75 |
| 8 | 37444583 | rs4739523 | -0.771 | -1.74 | -0.949 | 0.508 | -0.068 | 0.125 | 0.125 | 1.69 | 0.17 | 0.68 | 0.27 | 0.44 | 0.07 | 0.13 |
| 8 | 37445273 | rs1014998 | -0.042 | -1.716 | -0.898 | 0.544 | -0.064 | 0.324 | 0.324 | 0.579 | 0.12 | 0.53 | 0.24 | 0.44 | 0.03 | 0.09 |
| 8 | 37445639 | rs4739525 | -0.942 | -1.828 | -1.032 | 0.396 | -0.071 | 0.108 | 0.108 | 1.5 | 0.19 | 0.69 | 0.28 | 0.52 | 0.05 | 0.12 |
| 8 | 37450001 | rs7821974 | -0.619 | -1.387 | -1.256 | 0.32 | -0.088 | 0.208 | 0.208 | 0.065 | 0.18 | 0.54 | 0.28 | 0.52 | 0.06 | 0.13 |
| 8 | 37456304 | rs2298321 | 0.597 | -0.292 | -1.835 | 0.43 | -0.025 | -0.004 | -0.004 | 0 | 0.14 | 0 | 0.07 | 0.07 | 0.11 | 0.22 |
| 8 | 37459524 | rs4739526 | -0.092 | -0.19 | -1.576 | 0.324 | -0.028 | -0.022 | -0.022 | 0 | 0.39 | 0.27 | 0.46 | 0.42 | 0.48 | 0.45 |
| 8 | 37469049 | rs6468418 | -0.218 | -0.104 | -0.986 | 0.257 | -0.002 | 0.005 | 0.005 | 0 | 0.47 | 0.47 | 0.5 | 0.61 | 0.51 | 0.49 |
| 8 | 37469921 | rs10100061 | -0.717 | -0.277 | -0.42 | 0.36 | 0.01 | 0.154 | 0.154 | 0.137 | 0.12 | 0.38 | 0.19 | 0.25 | 0.12 | 0.21 |
| 8 | 37470110 | rs10481048 | 0.067 | -0.439 | -0.657 | -0.517 | -0.017 | -0.002 | -0.002 | 0.141 | 0.17 | 0.21 | 0.08 | 0.13 | 0.05 | 0.15 |
| 8 | 37470129 | rs4739348 | -0.302 | 0.314 | -1.02 | -0.789 | 0.002 | 0.014 | 0.014 | 0 | 0.27 | 0.25 | 0.2 | 0.16 | 0.2 | 0.27 |
| 8 | 37472911 | rs713189 | -0.403 | 0.376 | -1.363 | -1.149 | 0.001 | -0.001 | -0.001 | 0 | 0.84 | 0.92 | 0.92 | 1 | 0.83 | 0.92 |
| 8 | 37488996 | rs7386601 | 0.027 | 0.466 | -1.505 | -1.289 | -0.008 | 0.028 | 0.028 | 0 | 0.79 | 0.92 | 0.8 | 1 | 0.77 | 0.89 |
| 8 | 37489299 | rs2507710 | -0.053 | 0.502 | -1.447 | -1.271 | -0.007 | 0.009 | 0.009 | 0 | 0.79 | 0.89 | 0.8 | 1 | 0.77 | 0.89 |
| 8 | 37492872 | rs2407531 | 0.136 | 0.438 | -1.301 | -1.303 | 0 | -0.039 | -0.039 | 0 | 0.78 | 0.8 | 0.8 | 1 | 0.77 | 0.89 |
| 8 | 37493505 | rs7832424 | 0.873 | 0.576 | -1.475 | -1.814 | -0.007 | 0.008 | 0.008 | 0 | 0.59 | 0.67 | 0.67 | 0.96 | 0.54 | 0.75 |
| 8 | 37495521 | rs6992720 | 0.248 | 0.602 | -1.234 | -1.184 | -0.021 | 0.033 | 0.033 | 0.015 | 0.13 | 0.05 | 0.1 | 0.03 | 0.18 | 0.05 |
| 8 | 37532411 | rs10088732 | 0.203 | 0.133 | -0.996 | -1.02 | -0.016 | -0.029 | -0.029 | 0 | 0.09 | 0.18 | 0.03 | 0.01 | 0.05 | 0.09 |
| 8 | 37538022 | rs16887011 | -0.407 | 0.237 | -1.146 | -0.864 | -0.005 | -0.017 | -0.017 | 0 | 0.1 | 0.05 | 0.06 | 0.18 | 0.07 | 0.13 |
| 8 | 37556987 | rs36107033 | -0.074 | 0.054 | -1.121 | -0.989 | 0.01 | 0.041 | 0.041 | 0 | 0.08 | 0.01 | 0.05 | 0.01 | 0.06 | 0.13 |
| 8 | 37571392 | rs7814863 | 0.186 | 0.302 | -1.212 | -1.05 | -0.001 | 0.003 | 0.003 | 0 | 0.25 | 0.21 | 0.2 | 0.07 | 0.27 | 0.14 |
| 8 | 37589772 | rs7831942 | 0.621 | -0.223 | -1.772 | -0.023 | 0.037 | -0.083 | -0.083 | 13.054 | 0.22 | 0.07 | 0.18 | 0.32 | 0.14 | 0.16 |
| 8 | 37590446 | rs6468437 | -0.44 | 0.732 | -1.486 | -0.047 | -0.054 | 0.01 | 0.01 | 0 | 0.85 | 0.98 | 0.89 | 0.9 | 0.8 | 0.83 |
| 8 | 37635649 | rs7463174 | -0.181 | 0.887 | -0.976 | 0.682 | -0.007 | -0.015 | -0.015 | 1.667 | 0.13 | 0.08 | 0.24 | 0.31 | 0.14 | 0.3 |
| 8 | 37645466 | rs6985987 | -0.178 | 0.327 | 0.427 | 0.367 | 0.034 | 0.064 | 0.064 | 4.062 | 0.06 | 0.43 | 0.04 | 0.03 | 0 | 0.11 |
| 8 | 37646650 | rs10955032 | 0.105 | 0.451 | -0.153 | 0.333 | 0.007 | -0.011 | -0.011 | 0 | 0.57 | 0.62 | 0.6 | 0.49 | 0.61 | 0.59 |
| 8 | 37651477 | rs7008274 | 0.227 | 0.258 | -0.183 | 0.31 | -0.029 | 0.032 | 0.032 | 0 | 0.3 | 0.45 | 0.3 | 0.46 | 0.19 | 0.42 |
| 8 | 37656183 | rs7827048 | 0.154 | 1.198 | -0.263 | 0.209 | -0.069 | -0.073 | -0.073 | 0.788 | 0.66 | 0.83 | 0.56 | 0.54 | 0.54 | 0.55 |
| 8 | 37657526 | rs915650 | 0.513 | 0.838 | -0.202 | -0.087 | 0.027 | 0.02 | 0.02 | 0 | 0.5 | 0.36 | 0.36 | 0.49 | 0.38 | 0.33 |
| 8 | 37660948 | rs4976893 | 0.272 | 1.203 | 0.193 | 0.302 | -0.001 | 0 | 0 | 0.788 | 0.61 | 0.66 | 0.46 | 0.61 | 0.48 | 0.52 |
| 8 | 37667018 | rs4976891 | 0.41 | 0.88 | 0.558 | 0.101 | 0.026 | 0.052 | 0.052 | 0.747 | 0.49 | 0.64 | 0.25 | 0.22 | 0.31 | 0.24 |
| 8 | 37667471 | rs7013416 | -1.139 | 0.658 | 0.285 | 0.636 | 0.036 | 0.414 | 0.414 | 3.018 | 0.08 | 0.52 | 0.09 | 0.31 | 0.05 | 0.14 |
| 8 | 37673542 | rs7823249 | 0.684 | 0.889 | -0.628 | -0.737 | -0.011 | -0.065 | -0.065 | 0.004 | 0.43 | 0.43 | 0.18 | 0.14 | 0.26 | 0.18 |

| | | | | | | | | | | | | | | | | |
|---|---|---|---|---|---|---|---|---|---|---|---|---|---|---|---|---|
| 8 | 37691711 | rs11785789 | -0.605 | 1.481 | -0.551 | 0.581 | 0.018 | 0.042 | 0.042 | 1.401 | 0.26 | 0.56 | 0.24 | 0.54 | 0.34 | 0.42 |
| 8 | 37696874 | rs6998793 | -0.695 | 1.671 | -0.805 | 0.156 | -0.023 | -0.01 | -0.01 | 0 | 0.27 | 0.28 | 0.5 | 0.66 | 0.4 | 0.46 |
| 8 | 37706267 | rs12541064 | -0.436 | 1.664 | -0.633 | 0.068 | -0.016 | -0.005 | -0.005 | 0 | 0.27 | 0.25 | 0.52 | 0.65 | 0.42 | 0.45 |
| 8 | 37711398 | rs10808346 | -0.434 | 1.16 | -0.594 | -0.355 | -0.008 | 0.008 | 0.008 | 0 | 0.37 | 0.65 | 0.54 | 0.7 | 0.43 | 0.45 |
| 8 | 37712037 | rs1977912 | -0.293 | 1.411 | -0.711 | -0.498 | 0.003 | -0.006 | -0.006 | 0.783 | 0.22 | 0.28 | 0.23 | 0.22 | 0.28 | 0.23 |
| 8 | 37727281 | rs6468443 | -0.803 | 0.695 | -0.513 | -0.581 | 0.045 | 0.082 | 0.082 | 0 | 0.62 | 0.8 | 0.78 | 0.98 | 0.74 | 0.7 |
| 8 | 37728750 | rs2070493 | -0.801 | 1.462 | 0.151 | 0.788 | -0.009 | -0.049 | -0.049 | 0 | 0.08 | 0 | 0.05 | 0.17 | 0.06 | 0.14 |
| 8 | 37730017 | rs16887092 | 0.298 | 1.25 | 1.326 | 0.356 | 0.035 | 0.025 | 0.025 | 1.243 | 0.06 | 0.37 | 0.04 | 0.01 | 0.01 | 0.01 |
| 8 | 37730456 | rs7341564 | -1 | 2.366 | 1.729 | 0.351 | 0.038 | -0.092 | -0.092 | 0 | 0.27 | 0.5 | 0.27 | 0.12 | 0.38 | 0.15 |
| 8 | 37738616 | rs10955035 | -1.463 | 2.402 | 1.967 | 0.904 | -0.007 | 0.004 | 0.004 | 0 | 0.36 | 0.28 | 0.32 | 0.15 | 0.48 | 0.33 |
| 8 | 37750289 | rs7812866 | -1.811 | 2.055 | 2.331 | 1.65 | 0.016 | -0.021 | -0.021 | 0 | 0.39 | 0.53 | 0.38 | 0.26 | 0.48 | 0.38 |
| 8 | 37784204 | rs28490371 | -1.807 | 1.22 | 3.273 | 2.218 | 0.019 | 0.001 | 0.001 | 0 | 0.23 | 0.28 | 0.24 | 0.23 | 0.26 | 0.37 |
| 8 | 37786457 | rs28380463 | -2.1 | 1.446 | 3.398 | 2.325 | 0.046 | 0.004 | 0.004 | 0.847 | 0.23 | 0.57 | 0.27 | 0.23 | 0.26 | 0.37 |
| 8 | 37788799 | rs28427316 | -2.114 | 1.447 | 3.398 | 2.326 | 0.045 | 0.003 | 0.003 | 1.092 | 0.23 | 0.57 | 0.27 | 0.23 | 0.26 | 0.36 |
| 8 | 37791109 | rs28655870 | -1.953 | 1.385 | 3.387 | 2.33 | 0.058 | 0.011 | 0.011 | 0.473 | 0.22 | 0.55 | 0.27 | 0.23 | 0.26 | 0.36 |
| 8 | 37809609 | rs7004842 | -2.573 | 1.061 | 3.371 | 2.782 | 0.01 | 0.016 | 0.016 | 0 | 0.1 | 0.25 | 0.12 | 0.12 | 0.08 | 0.22 |
| 8 | 37821486 | rs4998 | -1.915 | 1.173 | 2.532 | 3.093 | -0.005 | -0.013 | -0.013 | 5.773 | 0.08 | 0.09 | 0.12 | 0.13 | 0.08 | 0.15 |
| 8 | 37823798 | rs4994 | -1.975 | 1.172 | 2.477 | 3.093 | -0.004 | -0.013 | -0.013 | 0.009 | 0.09 | 0.09 | 0.12 | 0.13 | 0.08 | 0.16 |
| 8 | 37834312 | rs2298423 | -2.394 | 1.511 | 2.628 | 3.244 | 0.049 | 0.064 | 0.064 | 0 | 0.06 | 0.23 | 0.13 | 0.12 | 0.08 | 0.14 |
| 8 | 37883438 | rs28813686 | 1.449 | 1.753 | 2.528 | 3.4 | 0.088 | 0.171 | 0.171 | 0 | 0.81 | 0.57 | 0.78 | 0.61 | 0.76 | 0.66 |
| 8 | 37884310 | rs28797500 | 1.449 | 1.753 | 2.528 | 3.4 | 0.088 | 0.171 | 0.171 | 0 | 0.81 | 0.57 | 0.78 | 0.61 | 0.76 | 0.65 |
| 8 | 37887422 | rs3813521 | 0.894 | 1.801 | 2.274 | 3.462 | -0.008 | -0.007 | -0.007 | 0.014 | 0.84 | 0.88 | 0.8 | 0.65 | 0.76 | 0.71 |
| 8 | 37897855 | rs28368989 | 1.338 | 1.804 | 2.051 | 3.441 | 0.074 | 0.159 | 0.159 | 0 | 0.81 | 0.59 | 0.79 | 0.6 | 0.76 | 0.64 |
| 8 | 37899702 | rs28377666 | 1.225 | 1.804 | 2.069 | 3.441 | 0.06 | 0.126 | 0.126 | 0 | 0.81 | 0.67 | 0.79 | 0.6 | 0.76 | 0.64 |
| 8 | 37902453 | rs6605631 | 1.297 | 1.835 | 2.063 | 3.428 | 0.074 | 0.163 | 0.163 | 0 | 0.81 | 0.59 | 0.78 | 0.6 | 0.76 | 0.64 |
| 8 | 37977732 | rs2517388 | -1.046 | 1.089 | 0.467 | 2.325 | 0.009 | 0.064 | 0.064 | 0 | 0.18 | 0.31 | 0.29 | 0.65 | 0.21 | 0.31 |
| 8 | 38013848 | rs6474491 | -0.248 | 0.266 | -1.015 | 1.353 | 0.031 | -0.017 | -0.017 | 0 | 0.31 | 0.23 | 0.21 | 0.35 | 0.23 | 0.38 |
| 8 | 38020325 | rs2843727 | -0.953 | 1.138 | -0.062 | 1.857 | 0.077 | 0.173 | 0.173 | 0 | 0.07 | 0.23 | 0.17 | 0.28 | 0.11 | 0.21 |
| 8 | 38025175 | rs2843744 | -1.003 | 1.791 | 0.079 | 2.123 | 0.06 | 0.064 | 0.064 | 0 | 0.27 | 0.47 | 0.37 | 0.6 | 0.37 | 0.3 |
| 8 | 38027519 | rs16887236 | -0.372 | 1.79 | 0.078 | 2.123 | 0.085 | 0.185 | 0.185 | 0 | 0.07 | 0.23 | 0.16 | 0.28 | 0.11 | 0.21 |
| 8 | 38028195 | rs7823144 | -0.56 | 1.778 | 0.206 | 2.119 | 0.116 | 0.243 | 0.243 | 0 | 0.07 | 0.31 | 0.18 | 0.28 | 0.11 | 0.2 |
| 8 | 38031345 | rs16887244 | -0.216 | 1.71 | 0.186 | 2.07 | -0.018 | -0.03 | -0.03 | 0 | 0.19 | 0.13 | 0.19 | 0.33 | 0.26 | 0.09 |
| 8 | 38033004 | rs17435276 | -0.182 | 1.867 | 0.884 | 1.864 | -0.028 | -0.035 | -0.035 | 0.112 | 0.2 | 0.12 | 0.2 | 0.32 | 0.29 | 0.1 |
| 8 | 38036165 | rs11783967 | -0.214 | 1.801 | 1.127 | 1.866 | 0.005 | 0.012 | 0.012 | 0 | 0.2 | 0.25 | 0.27 | 0.32 | 0.26 | 0.09 |
| 8 | 38063235 | rs7836805 | -0.362 | 1.801 | 1.125 | 1.867 | 0.004 | 0.011 | 0.011 | 0 | 0.2 | 0.24 | 0.26 | 0.32 | 0.26 | 0.09 |
| 8 | 38095662 | rs2306899 | 0.136 | 1.824 | 0.997 | 1.832 | -0.081 | -0.099 | -0.099 | 2.564 | 0.16 | 0.02 | 0.24 | 0.32 | 0.25 | 0.09 |
| 8 | 38101003 | rs16887273 | 0.044 | 1.823 | 1.022 | 1.831 | -0.018 | -0.022 | -0.022 | 0 | 0.16 | 0.12 | 0.25 | 0.32 | 0.25 | 0.09 |
| 8 | 38110215 | rs2898674 | 0.028 | 1.811 | 1.017 | 1.826 | -0.018 | -0.022 | -0.022 | 0.347 | 0.16 | 0.12 | 0.25 | 0.32 | 0.25 | 0.09 |
| 8 | 38120276 | rs6981405 | -0.22 | 1.842 | 1.047 | 1.818 | -0.002 | 0.001 | 0.001 | 0 | 0.18 | 0.19 | 0.25 | 0.32 | 0.24 | 0.09 |
| 8 | 38130025 | rs1906672 | -0.123 | 1.815 | 1.006 | 1.851 | -0.01 | -0.013 | -0.013 | 0 | 0.17 | 0.16 | 0.25 | 0.34 | 0.24 | 0.09 |
| 8 | 38133612 | rs1488934 | -0.182 | 1.82 | 1.045 | 1.873 | -0.008 | -0.01 | -0.01 | 2.39 | 0.17 | 0.18 | 0.25 | 0.31 | 0.24 | 0.09 |
| 8 | 38133793 | rs1488935 | -0.192 | 1.821 | 1.044 | 1.854 | -0.005 | -0.006 | -0.006 | 0.272 | 0.18 | 0.18 | 0.25 | 0.32 | 0.24 | 0.09 |
| 8 | 38135412 | rs7845911 | -0.007 | 1.909 | 1.018 | 1.828 | -0.001 | 0.006 | 0.006 | 0 | 0.16 | 0.18 | 0.24 | 0.32 | 0.23 | 0.08 |
| 8 | 38193518 | rs10101168 | -0.146 | 1.739 | 1.692 | 1.871 | -0.005 | -0.006 | -0.006 | 0 | 0.18 | 0.17 | 0.25 | 0.32 | 0.24 | 0.09 |
| 8 | 38212413 | rs11774214 | -0.273 | 1.719 | 1.64 | 1.832 | -0.008 | -0.012 | -0.012 | 0 | 0.18 | 0.18 | 0.26 | 0.32 | 0.26 | 0.09 |
| 8 | 38231874 | rs2932005 | -1.38 | 1.516 | 1.895 | 1.822 | 0.018 | 0.015 | 0.015 | 0 | 0.3 | 0.57 | 0.38 | 0.61 | 0.39 | 0.31 |
| 8 | 38232738 | rs10107763 | -0.27 | 1.53 | 1.492 | 1.843 | -0.007 | -0.01 | -0.01 | 0 | 0.18 | 0.18 | 0.26 | 0.32 | 0.26 | 0.09 |
| 8 | 38236890 | rs10095753 | -0.111 | 1.533 | 1.493 | 1.844 | -0.006 | -0.006 | -0.006 | 0.835 | 0.18 | 0.18 | 0.25 | 0.32 | 0.24 | 0.09 |
| 8 | 38259481 | rs11986274 | -0.251 | 1.533 | 1.493 | 2.046 | -0.007 | -0.009 | -0.009 | 0 | 0.18 | 0.18 | 0.28 | 0.35 | 0.25 | 0.09 |
| 8 | 38260815 | rs13248631 | 1.091 | 1.846 | 1.338 | 1.566 | -0.023 | 0 | 0 | 0 | 0.62 | 0.33 | 0.51 | 0.39 | 0.49 | 0.68 |
| 8 | 38269514 | rs13317 | -0.117 | 1.703 | 0.744 | 1.643 | -0.007 | -0.01 | -0.01 | 0 | 0.18 | 0.18 | 0.28 | 0.35 | 0.25 | 0.09 |
| 8 | 38281658 | rs3925 | 0.158 | 1.744 | 0.676 | 1.67 | -0.035 | -0.048 | -0.048 | 1.228 | 0.17 | 0.09 | 0.25 | 0.32 | 0.26 | 0.07 |
| 8 | 38283208 | rs3779921 | 0.22 | 1.644 | 0.662 | 1.729 | -0.026 | -0.038 | -0.038 | 0 | 0.17 | 0.09 | 0.25 | 0.32 | 0.24 | 0.07 |
| 8 | 38286225 | rs2288696 | 0.839 | 1.045 | -0.548 | 0.456 | -0.054 | 0.051 | 0.051 | 0 | 0.24 | 0.01 | 0.26 | 0.1 | 0.19 | 0.2 |
| 8 | 38299419 | rs6983315 | -0.86 | 1.3 | 0.316 | 0.708 | 0.029 | 0.014 | 0.014 | 0 | 0.34 | 0.43 | 0.47 | 0.6 | 0.48 | 0.3 |
| 8 | 38302965 | rs6474354 | -0.24 | 0.981 | 0.198 | 0.609 | 0.017 | 0 | 0 | 0 | 0.17 | 0.28 | 0.21 | 0.16 | 0.22 | 0.21 |
| 8 | 38317398 | rs7012413 | 0.12 | 0.063 | -0.182 | 0.557 | 0.002 | 0 | 0 | 0 | 0.67 | 0.33 | 0.65 | 0.66 | 0.71 | 0.63 |
| 8 | 38319349 | rs4733946 | -0.151 | -0.148 | -0.008 | 0.588 | 0 | -0.001 | -0.001 | 0.355 | 0.1 | 0.08 | 0.23 | 0.09 | 0.08 | 0.12 |
| 8 | 38322346 | rs6996321 | -1.001 | -0.113 | 0.09 | 0.367 | 0 | 0 | 0 | 0 | 0.62 | 0.61 | 0.62 | 0.6 | 0.59 | 0.86 |
| 8 | 38327287 | rs7825208 | -0.009 | 0.134 | 0.26 | 0.257 | 0.023 | 0.022 | 0.022 | 0 | 0.13 | 0.25 | 0.13 | 0.16 | 0.12 | 0.25 |
| 8 | 38332318 | rs881301 | 0.006 | -0.093 | 0.046 | 0.113 | -0.012 | 0.003 | 0.003 | 0.342 | 0.41 | 0.25 | 0.26 | 0.39 | 0.4 | 0.51 |
| 8 | 38350315 | rs7830964 | -0.609 | -0.039 | -0.236 | 0.252 | -0.004 | 0.008 | 0.008 | 0.751 | 0.4 | 0.63 | 0.5 | 0.28 | 0.4 | 0.53 |
| 8 | 38359381 | rs13262495 | 0.209 | 0.203 | 0.069 | 0.179 | 0.032 | 0.034 | 0.034 | 0.006 | 0.07 | 0.13 | 0.06 | 0.12 | 0.08 | 0.23 |
| 8 | 38359468 | rs7845393 | -0.045 | 0.097 | -0.178 | 0.09 | -0.026 | 0.003 | 0.003 | 0 | 0.73 | 0.42 | 0.86 | 0.75 | 0.81 | 0.66 |
| 8 | 38361379 | rs328294 | 0.701 | 0.25 | -0.182 | 0.232 | 0.006 | -0.013 | -0.013 | 0 | 0.61 | 0.34 | 0.77 | 0.58 | 0.69 | 0.38 |
| 8 | 38362521 | rs7825829 | -0.56 | 0.379 | -0.334 | 0.623 | 0.055 | 0.102 | 0.102 | 1.486 | 0.09 | 0.28 | 0.1 | 0.17 | 0.09 | 0.25 |
| 8 | 38363472 | rs1541897 | 0.369 | 0.095 | -1.417 | 0.445 | -0.071 | -0.104 | -0.104 | 0 | 0.43 | 0.08 | 0.61 | 0.55 | 0.5 | 0.25 |
| 8 | 38369493 | rs6474384 | -0.74 | -0.695 | -0.177 | 0.219 | -0.008 | -0.002 | -0.002 | 0 | 0.18 | 0.24 | 0.12 | 0.14 | 0.07 | 0.29 |
| 8 | 38369831 | rs4733951 | 1.176 | -1.301 | -0.753 | 0.01 | 0.095 | 0.042 | 0.042 | 0.084 | 0.62 | 0.14 | 0.78 | 0.74 | 0.82 | 0.58 |

| | | | | | | | | | | | | | | | | |
|---|---|---|---|---|---|---|---|---|---|---|---|---|---|---|---|---|
| 8 | 38374393 | rs2461328 | -0.517 | -1.416 | 0.243 | 0.105 | -0.055 | -0.004 | -0.004 | 0 | 0.17 | 0.37 | 0.07 | 0.1 | 0.07 | 0.07 |
| 8 | 38378345 | rs2411271 | 0.07 | -1.427 | 0.481 | 0.07 | -0.025 | -0.046 | -0.046 | 0.649 | 0.13 | 0.21 | 0.03 | 0.02 | 0.07 | 0.05 |
| 8 | 38379631 | rs2461333 | 0.139 | -1.467 | 0.742 | 0.106 | -0.041 | 0.059 | 0.059 | 0 | 0.88 | 0.61 | 0.93 | 0.87 | 0.96 | 0.9 |
| 8 | 38381884 | rs12682402 | 1.075 | -1.48 | 0.662 | 0.053 | -0.041 | -0.024 | -0.024 | 3.442 | 0.76 | 0.48 | 0.84 | 0.83 | 0.88 | 0.63 |
| 8 | 38384366 | rs2445019 | 1.12 | -1.431 | 0.598 | 0.075 | -0.023 | -0.029 | -0.029 | 0 | 0.69 | 0.45 | 0.83 | 0.82 | 0.83 | 0.59 |
| 8 | 38399485 | rs7813370 | -0.686 | -1.357 | 0.628 | 0.081 | -0.068 | 0.052 | 0.052 | 0 | 0.19 | 0.48 | 0.11 | 0.21 | 0.1 | 0.17 |
| 8 | 38407075 | rs4733918 | 1.177 | -1.269 | -0.067 | 0.124 | 0.052 | 0.008 | 0.008 | 0.165 | 0.63 | 0.23 | 0.77 | 0.71 | 0.81 | 0.6 |
| 8 | 38410557 | rs17581781 | 0.158 | -0.177 | -0.846 | -0.187 | -0.048 | 0 | 0 | 0 | 0.12 | 0.02 | 0.19 | 0.09 | 0.17 | 0.16 |
| 8 | 38419633 | rs11985941 | -0.453 | -0.641 | -0.58 | -0.27 | 0.008 | 0.017 | 0.017 | 0 | 0.16 | 0.17 | 0.08 | 0.05 | 0.09 | 0.21 |
| 8 | 38423868 | rs10958731 | -0.282 | -0.927 | -0.567 | -0.347 | -0.006 | -0.012 | -0.012 | 0 | 0.16 | 0.19 | 0.05 | 0.05 | 0.11 | 0.14 |
| 8 | 38430234 | rs7817828 | 0.593 | -0.683 | -0.953 | -0.867 | -0.013 | 0.024 | 0.024 | 0 | 0.4 | 0.37 | 0.46 | 0.31 | 0.49 | 0.4 |
| 8 | 38432037 | rs16887465 | 1.346 | -0.5 | -1.291 | -1.165 | -0.005 | 0.024 | 0.024 | 0 | 0.16 | 0.07 | 0.13 | 0.1 | 0.15 | 0.11 |
| 8 | 38433937 | rs921802 | -0.347 | -1.024 | -0.759 | -0.638 | 0.001 | 0.001 | 0.001 | 0.361 | 0.49 | 0.6 | 0.54 | 0.47 | 0.5 | 0.44 |
| 8 | 38437040 | rs4733883 | 1.99 | -0.99 | -1.245 | -0.711 | -0.084 | -0.157 | -0.157 | 0.152 | 0.23 | 0.02 | 0.4 | 0.44 | 0.32 | 0.19 |
| 8 | 38438589 | rs4733924 | -0.075 | -0.76 | -0.46 | -0.575 | -0.049 | 0.016 | 0.016 | 0 | 0.78 | 0.67 | 0.9 | 0.79 | 0.86 | 0.8 |
| 8 | 38440791 | rs11776404 | -0.534 | -0.688 | -0.726 | -0.382 | -0.015 | 0.024 | 0.024 | 0 | 0.13 | 0.17 | 0.08 | 0.21 | 0.09 | 0.19 |
| 8 | 38444229 | rs7004885 | 0.506 | -0.537 | -0.646 | -0.494 | -0.003 | 0 | 0 | 0 | 0.33 | 0.31 | 0.34 | 0.21 | 0.39 | 0.29 |
| 8 | 38445425 | rs1665529 | -0.554 | -0.292 | -0.948 | -0.142 | -0.004 | -0.002 | -0.002 | 0 | 0.93 | 0.94 | 0.95 | 0.95 | 0.94 | 0.88 |
| 8 | 38445999 | rs1466050 | 0.101 | -0.376 | -0.999 | -0.165 | 0.008 | -0.008 | -0.008 | 0 | 0.72 | 0.8 | 0.8 | 0.69 | 0.76 | 0.71 |
| 8 | 38446937 | rs1550879 | -0.074 | 0.089 | -0.906 | -0.22 | -0.004 | -0.002 | -0.002 | 0 | 0.93 | 0.94 | 0.95 | 0.95 | 0.94 | 0.88 |
| 8 | 38450131 | rs6474427 | -0.19 | 0.154 | -1.103 | -0.395 | 0.004 | -0.009 | -0.009 | 0.675 | 0.37 | 0.45 | 0.34 | 0.25 | 0.41 | 0.25 |
| 8 | 38457123 | rs327931 | 0.89 | 0.186 | -1.851 | -0.363 | 0.025 | 0.058 | 0.058 | 2.715 | 0.27 | 0.07 | 0.31 | 0.13 | 0.23 | 0.15 |
| 8 | 38460284 | rs327937 | 0.445 | 0.737 | -1.403 | -0.101 | 0.001 | -0.001 | -0.001 | 0 | 0.73 | 0.25 | 0.81 | 0.72 | 0.81 | 0.61 |
| 8 | 38465928 | rs4733887 | -0.078 | 0.89 | -1.706 | 0.164 | -0.102 | -0.128 | -0.128 | 0 | 0.22 | 0.06 | 0.32 | 0.54 | 0.36 | 0.2 |
| 8 | 38466443 | rs2445001 | 0.134 | 0.774 | -0.884 | -0.267 | -0.053 | -0.12 | -0.12 | 0 | 0.75 | 0.5 | 0.87 | 0.96 | 0.89 | 0.57 |
| 8 | 38468605 | rs2593095 | 1.321 | 1.092 | -0.835 | -0.182 | -0.005 | -0.001 | -0.001 | 0.089 | 0.52 | 0.53 | 0.5 | 0.48 | 0.4 | 0.41 |
| 8 | 38469303 | rs7832232 | -0.483 | 1.25 | -0.774 | -0.09 | 0.028 | 0.018 | 0.018 | 0 | 0.29 | 0.32 | 0.4 | 0.48 | 0.5 | 0.25 |
| 8 | 38470624 | rs16887551 | 0.382 | 1.446 | -0.848 | -0.018 | -0.06 | -0.078 | -0.078 | 0.025 | 0.2 | 0.08 | 0.28 | 0.47 | 0.3 | 0.18 |
| 8 | 38471714 | rs7845862 | -0.756 | 1.455 | -0.68 | -0.65 | 0.03 | -0.095 | -0.095 | 0 | 0.28 | 0.39 | 0.22 | 0.05 | 0.3 | 0.39 |
| 8 | 38474484 | rs1000236 | 1.487 | 0.897 | -0.634 | -0.332 | -0.009 | -0.049 | -0.049 | 0.744 | 0.53 | 0.14 | 0.51 | 0.4 | 0.45 | 0.47 |
| 8 | 38476208 | rs7844617 | 0.235 | 0.944 | -0.535 | -0.304 | 0.056 | 0.088 | 0.088 | 0.074 | 0.22 | 0.28 | 0.34 | 0.56 | 0.36 | 0.29 |
| 8 | 38480529 | rs99733 | -0.267 | 0.979 | -0.38 | -0.312 | 0.012 | -0.066 | -0.066 | 0.025 | 0.88 | 0.78 | 0.92 | 0.99 | 0.86 | 0.9 |
| 8 | 38484612 | rs12335077 | 0.329 | 1.113 | -0.454 | -0.171 | 0.049 | 0.094 | 0.094 | 0.531 | 0.2 | 0.27 | 0.31 | 0.57 | 0.31 | 0.28 |
| 8 | 38484726 | rs6992470 | -0.371 | 0.695 | -0.927 | -0.424 | -0.028 | -0.08 | -0.08 | 0.314 | 0.15 | 0.34 | 0.09 | 0.02 | 0.1 | 0.15 |
| 8 | 38486970 | rs6474458 | -0.627 | 1.227 | -0.849 | 0.059 | 0.056 | 0.021 | 0.021 | 0.283 | 0.33 | 0.47 | 0.41 | 0.63 | 0.45 | 0.39 |
| 8 | 38488601 | rs7818839 | -0.701 | 0.987 | -1.461 | -0.644 | -0.004 | 0.122 | 0.122 | 2.304 | 0.32 | 0.85 | 0.28 | 0.06 | 0.31 | 0.45 |
| 8 | 38497650 | rs13253867 | 0.064 | 0.698 | -1.552 | -1.295 | -0.095 | 0.082 | 0.082 | 0 | 0.33 | 0.09 | 0.51 | 0.16 | 0.44 | 0.32 |
| 8 | 38498637 | rs6474464 | -0.365 | 0.537 | -1.256 | -1.203 | -0.015 | -0.009 | -0.009 | 0 | 0.8 | 0.37 | 0.87 | 0.81 | 0.86 | 0.69 |
| 8 | 38508357 | rs895621 | 0.671 | 0.65 | -1.383 | -1.255 | -0.105 | 0.076 | 0.076 | 0 | 0.32 | 0.09 | 0.51 | 0.16 | 0.44 | 0.32 |
| 8 | 38514745 | rs1447057 | 0.263 | 0.511 | -1.106 | -1.046 | 0.023 | 0.068 | 0.068 | 0.193 | 0.47 | 0.26 | 0.37 | 0.66 | 0.42 | 0.37 |
| 8 | 38518131 | rs11782681 | 0.968 | 0.196 | -1.135 | -0.631 | 0.003 | -0.01 | -0.01 | 0.015 | 0.14 | 0.01 | 0.1 | 0.1 | 0.14 | 0.09 |
| 8 | 38532918 | rs10755988 | -0.471 | 0.42 | -1.013 | -1.164 | 0.013 | 0.034 | 0.034 | 0.925 | 0.61 | 0.72 | 0.58 | 0.82 | 0.66 | 0.5 |
| 8 | 38535185 | rs2411279 | -0.166 | 0.885 | -0.94 | -1.59 | 0.01 | 0.059 | 0.059 | 0.468 | 0.47 | 0.27 | 0.32 | 0.29 | 0.45 | 0.37 |
| 8 | 38536625 | rs7011775 | -0.687 | 0.949 | -0.689 | -1.563 | -0.003 | -0.036 | -0.036 | 0 | 0.55 | 0.52 | 0.51 | 0.85 | 0.57 | 0.48 |
| 8 | 38539742 | rs11777840 | 0.293 | 1.107 | -0.615 | -1.678 | 0.006 | 0.013 | 0.013 | 1.633 | 0.3 | 0.2 | 0.21 | 0.23 | 0.29 | 0.19 |
| 8 | 38539754 | rs6651370 | -1.83 | 1.125 | -0.621 | -2.069 | 0.004 | -0.099 | -0.099 | 0 | 0.79 | 0.74 | 0.86 | 1 | 0.78 | 0.79 |
| 8 | 38540944 | rs10958805 | 1.286 | 1.03 | -0.715 | -1.935 | 0.008 | -0.022 | -0.022 | 0 | 0.19 | 0.11 | 0.12 | 0.32 | 0.17 | 0.15 |
| 8 | 38546707 | rs1010588 | 1.676 | 1.139 | -0.796 | -1.996 | -0.009 | 0.061 | 0.061 | 0.794 | 0.17 | 0.05 | 0.1 | 0.05 | 0.16 | 0.1 |
| 8 | 38547033 | rs16887675 | 1.263 | 1.187 | -0.841 | -2.106 | -0.001 | 0.002 | 0.002 | 0 | 0.13 | 0.11 | 0.09 | 0.07 | 0.12 | 0.17 |
| 8 | 38548619 | rs4733942 | 1.267 | 1.186 | -0.935 | -2.217 | 0.009 | 0.093 | 0.093 | 0 | 0.25 | 0.1 | 0.34 | 0.07 | 0.22 | 0.29 |
| 8 | 38549335 | rs1010589 | -0.04 | 1.22 | -0.829 | -2.221 | -0.005 | -0.084 | -0.084 | 0.506 | 0.25 | 0.37 | 0.2 | 0.07 | 0.23 | 0.23 |
| 8 | 38552532 | rs7837897 | 1.371 | 1.212 | -0.839 | -2.21 | -0.001 | 0.002 | 0.002 | 0 | 0.13 | 0.11 | 0.09 | 0.07 | 0.13 | 0.17 |
| 8 | 38552757 | rs7816405 | -0.739 | 1.283 | -0.858 | -2.233 | 0.021 | 0.014 | 0.014 | 0 | 0.34 | 0.49 | 0.4 | 0.61 | 0.39 | 0.33 |
| 8 | 38554723 | rs13279807 | 0.311 | 1.301 | -1.045 | -2.14 | -0.009 | -0.01 | -0.01 | 4.869 | 0.19 | 0.22 | 0.29 | 0.48 | 0.25 | 0.14 |
| 8 | 38555872 | rs7001342 | 1.237 | 1.215 | -1.089 | -2.193 | 0.01 | 0.094 | 0.094 | 0 | 0.25 | 0.1 | 0.34 | 0.07 | 0.22 | 0.29 |
| 8 | 38558577 | rs10282916 | 1.426 | 1.307 | -0.896 | -1.957 | 0 | -0.015 | -0.015 | 0 | 0.11 | 0.1 | 0.1 | 0 | 0.11 | 0.05 |
| 8 | 38559013 | rs4364602 | -0.29 | 1.298 | -0.819 | -1.967 | 0.004 | -0.071 | -0.071 | 0.033 | 0.41 | 0.53 | 0.41 | 0.93 | 0.44 | 0.45 |
| 8 | 38559362 | rs11997272 | 1.654 | 1.383 | -1.073 | -1.749 | -0.004 | 0.072 | 0.072 | 0 | 0.08 | 0.12 | 0.19 | 0.47 | 0.14 | 0.1 |
| 8 | 38561633 | rs11785498 | 1.786 | 0.974 | -1.073 | -1.854 | 0.008 | 0.054 | 0.054 | 1.493 | 0.16 | 0.05 | 0.09 | 0.05 | 0.14 | 0.09 |
| 8 | 38572422 | rs6983695 | -0.317 | 1.063 | -0.802 | -1.657 | 0.009 | 0.017 | 0.017 | 0.269 | 0.5 | 0.44 | 0.48 | 0.66 | 0.51 | 0.41 |
| 8 | 38574366 | rs10755989 | -1.198 | 1.032 | -0.624 | -2.42 | -0.001 | -0.097 | -0.097 | 2.072 | 0.78 | 0.73 | 0.86 | 1 | 0.79 | 0.79 |
| 8 | 38576759 | rs6474495 | -0.362 | 1.361 | -0.683 | -2.199 | -0.014 | 0.019 | 0.019 | 0 | 0.28 | 0.26 | 0.4 | 0.89 | 0.38 | 0.46 |
| 8 | 38582053 | rs16887717 | 0.433 | 1.331 | -0.779 | -2.311 | -0.001 | -0.013 | -0.013 | 0 | 0.14 | 0.1 | 0.08 | 0 | 0.1 | 0.15 |
| 8 | 38582476 | rs17506431 | 0.676 | 1.469 | -0.955 | -2.102 | -0.016 | -0.02 | -0.02 | 0.386 | 0.14 | 0.13 | 0.26 | 0.72 | 0.2 | 0.33 |
| 8 | 38588313 | rs6985355 | -0.532 | 0.964 | -1.004 | -2.444 | -0.025 | 0.073 | 0.073 | 0 | 0.42 | 0.34 | 0.49 | 0.95 | 0.49 | 0.59 |
| 8 | 38592135 | rs7821911 | 0.001 | 0.892 | -1.11 | -2.08 | 0.003 | 0.082 | 0.082 | 0 | 0.3 | 0.14 | 0.17 | 0.06 | 0.22 | 0.26 |
| 8 | 38596395 | rs4733947 | -1.187 | 0.885 | -0.912 | -2.14 | 0.002 | 0.036 | 0.036 | 0 | 0.6 | 0.73 | 0.6 | 0.95 | 0.6 | 0.75 |
| 8 | 38597727 | rs13278546 | 0.455 | 0.893 | -1.079 | -1.402 | 0.012 | -0.016 | -0.016 | 0 | 0.17 | 0.15 | 0.25 | 0.74 | 0.17 | 0.33 |
| 8 | 38599218 | rs7844381 | 0.179 | 1.493 | -0.779 | -0.791 | 0.003 | 0 | 0 | 0 | 0.12 | 0.16 | 0.14 | 0.15 | 0.19 | 0.13 |

| | | | | | | | | | | | | | | | | |
|---|---|---|---|---|---|---|---|---|---|---|---|---|---|---|---|---|
| 8 | 38599893 | rs7814409 | 0 | 0.799 | -0.777 | -0.955 | 0 | -0.008 | -0.008 | 0.295 | 0.18 | 0.17 | 0.12 | 0.05 | 0.14 | 0.18 |
| 8 | 38602323 | rs7845067 | 0.037 | 1.01 | -1.003 | -1.104 | -0.002 | 0.009 | 0.009 | 0 | 0.58 | 0.65 | 0.57 | 0.88 | 0.61 | 0.7 |
| 8 | 38610096 | rs7816768 | -0.149 | 0.517 | -1.277 | -0.795 | 0.016 | 0.019 | 0.019 | 0 | 0.49 | 0.6 | 0.3 | 0.31 | 0.37 | 0.44 |
| 8 | 38628758 | rs10958810 | 0.256 | 0.723 | -1.195 | -1.182 | -0.003 | -0.106 | -0.106 | 0.587 | 0.27 | 0.39 | 0.19 | 0 | 0.24 | 0.15 |
| 8 | 38656652 | rs13267903 | 1.08 | 0.108 | -1.551 | -1.272 | 0.036 | -0.033 | -0.033 | 0.328 | 0.25 | 0.02 | 0.12 | 0.23 | 0.17 | 0.29 |
| 8 | 38657695 | rs6997835 | -0.077 | 0.3 | -1.474 | -0.831 | 0 | 0.001 | 0.001 | 0 | 0.34 | 0.39 | 0.25 | 0.27 | 0.34 | 0.26 |
| 8 | 38658394 | rs4733950 | 1.235 | 0.907 | -1.888 | -0.536 | -0.054 | -0.09 | -0.09 | 4.344 | 0.14 | 0.01 | 0.28 | 0.26 | 0.15 | 0.12 |
| 8 | 38684853 | rs2037204 | -0.128 | 0.869 | -1.506 | -0.781 | -0.001 | -0.001 | -0.001 | 0 | 0.36 | 0.4 | 0.28 | 0.27 | 0.39 | 0.27 |
| 8 | 38695996 | rs2292741 | 0.762 | 0.903 | -1.536 | -0.73 | -0.031 | -0.037 | -0.037 | 1.809 | 0.08 | 0.04 | 0.1 | 0.27 | 0.15 | 0.12 |
| 8 | 38701329 | rs7009521 | -0.584 | 0.819 | -1.45 | -0.769 | 0.017 | 0.017 | 0.017 | 0 | 0.61 | 0.56 | 0.41 | 0.5 | 0.56 | 0.57 |
| 8 | 38706369 | rs1802267 | 1.251 | 0.194 | -1.55 | -0.687 | 0.032 | -0.036 | -0.036 | 0.312 | 0.25 | 0.03 | 0.11 | 0.23 | 0.17 | 0.29 |
| 8 | 38709553 | rs1135990 | 1.242 | 0.229 | -1.811 | -0.664 | 0.032 | -0.036 | -0.036 | 0.18 | 0.25 | 0.03 | 0.11 | 0.23 | 0.17 | 0.29 |
| 8 | 38717029 | rs9298642 | 1.365 | 0.398 | -1.768 | -0.665 | -0.009 | 0.064 | 0.064 | 0 | 0.22 | 0.09 | 0.14 | 0 | 0.21 | 0.15 |
| 8 | 38727464 | rs7003661 | -0.444 | 0.441 | -1.72 | -0.509 | 0 | 0.006 | 0.006 | 0 | 0.58 | 0.56 | 0.36 | 0.5 | 0.48 | 0.6 |
| 8 | 38727972 | rs16887849 | 0.541 | 0.363 | -1.662 | 0.327 | -0.015 | -0.027 | -0.027 | 1.106 | 0.07 | 0.02 | 0.07 | 0.27 | 0.1 | 0.11 |
| 8 | 38748612 | rs13248839 | 0.717 | 0.239 | -1.691 | 0.086 | 0.021 | -0.049 | -0.049 | 0.421 | 0.22 | 0.03 | 0.12 | 0.22 | 0.15 | 0.32 |
| 8 | 38754530 | rs3847141 | 0.085 | 0.359 | -1.234 | 0.03 | -0.01 | 0.012 | 0.012 | 0 | 0.31 | 0.35 | 0.22 | 0.46 | 0.26 | 0.43 |
| 8 | 38762056 | rs11776027 | 0.35 | 0.365 | -1.891 | -0.257 | -0.013 | -0.064 | -0.064 | 0 | 0.3 | 0.04 | 0.2 | 0.38 | 0.26 | 0.34 |
| 8 | 38763716 | rs2056170 | 0.103 | 0.22 | -1.665 | -0.208 | 0.038 | 0.009 | 0.009 | 0.729 | 0.48 | 0.34 | 0.64 | 0.4 | 0.59 | 0.47 |
| 8 | 38764942 | rs17515835 | 1.285 | 0.542 | -1.753 | -0.886 | -0.001 | 0.088 | 0.088 | 0.489 | 0.22 | 0.05 | 0.15 | 0.06 | 0.2 | 0.15 |
| 8 | 38766074 | rs10110562 | 0.166 | 1.328 | -1.027 | -0.35 | 0.034 | -0.07 | -0.07 | 0.853 | 0.16 | 0.27 | 0.19 | 0.01 | 0.23 | 0.12 |
| 8 | 38771581 | rs7386365 | -0.228 | 1.051 | -1.602 | -0.188 | 0.002 | -0.012 | -0.012 | 0 | 0.8 | 0.83 | 0.86 | 0.8 | 0.81 | 0.8 |
| 8 | 38777146 | rs13254784 | -0.615 | 0.926 | -1.621 | -0.135 | -0.007 | 0.032 | 0.032 | 0 | 0.35 | 0.79 | 0.35 | 0.22 | 0.36 | 0.31 |
| 8 | 38779647 | rs9643901 | 0.575 | 0.883 | -2.008 | 0.356 | -0.026 | -0.108 | -0.108 | 0 | 0.79 | 0.96 | 0.63 | 0.6 | 0.81 | 0.76 |
| 8 | 38780606 | rs6474512 | 0.217 | 0.756 | -1.729 | -0.384 | -0.001 | 0.062 | 0.062 | 0 | 0.47 | 0.45 | 0.39 | 0.23 | 0.55 | 0.46 |
| 8 | 38781040 | rs11782824 | -0.168 | 0.723 | -1.661 | -0.305 | 0 | 0 | 0 | 0.05 | 0.53 | 0.58 | 0.42 | 0.42 | 0.56 | 0.55 |
| 8 | 38781696 | rs11785248 | 1.337 | 0.394 | -2.107 | -0.895 | 0.015 | -0.05 | -0.05 | 0.249 | 0.18 | 0.03 | 0.08 | 0.21 | 0.16 | 0.22 |
| 8 | 38782627 | rs7835508 | 0.101 | 0.998 | -1.427 | -0.309 | -0.02 | 0.001 | 0.001 | 0 | 0.29 | 0.32 | 0.3 | 0.03 | 0.4 | 0.24 |
| 8 | 38790786 | rs9792206 | 0.37 | 1.009 | -1.249 | -0.244 | 0.001 | 0.002 | 0.002 | 0 | 0.78 | 0.73 | 0.58 | 0.28 | 0.76 | 0.61 |
| 8 | 38793682 | rs4633027 | 0.538 | 1.095 | -1.402 | -0.299 | -0.026 | 0.032 | 0.032 | 0 | 0.24 | 0.23 | 0.15 | 0.01 | 0.29 | 0.15 |
| 8 | 38794457 | rs4076625 | 0.777 | 0.849 | -1.339 | 0.048 | -0.007 | -0.019 | -0.019 | 0 | 0.47 | 0.38 | 0.28 | 0.24 | 0.41 | 0.35 |
| 8 | 38804069 | rs6995018 | -0.438 | 1.111 | -1.666 | 0.072 | -0.008 | 0.017 | 0.017 | 0 | 0.4 | 0.43 | 0.36 | 0.18 | 0.44 | 0.38 |
| 8 | 38804226 | rs6474513 | 0.509 | 1.161 | -1.838 | 0.973 | -0.025 | -0.059 | -0.059 | 0.01 | 0.84 | 0.92 | 0.62 | 0.45 | 0.8 | 0.71 |
| 8 | 38808657 | rs6984245 | 0.285 | 0.509 | -2.214 | -0.018 | 0.016 | 0.105 | 0.105 | 0 | 0.77 | 0.98 | 0.94 | 0.93 | 0.85 | 0.85 |
| 8 | 38808752 | rs6998933 | 0.239 | 0.961 | -2.034 | 0.246 | -0.011 | 0.002 | 0.002 | 0 | 0.38 | 0.35 | 0.47 | 0.39 | 0.4 | 0.4 |
| 8 | 38813046 | rs7007530 | 0.145 | 0.565 | -2.39 | 0.084 | 0.008 | 0.043 | 0.043 | 0 | 0.76 | 0.97 | 0.92 | 0.89 | 0.85 | 0.82 |
| 8 | 38817412 | rs10958812 | 0.414 | 1.359 | -2.053 | -0.275 | -0.008 | 0.034 | 0.034 | 0 | 0.69 | 0.8 | 0.73 | 0.85 | 0.67 | 0.72 |
| 8 | 38820437 | rs7015219 | 0.933 | 1.898 | -2.098 | 0.094 | -0.007 | 0.024 | 0.024 | 0 | 0.58 | 0.65 | 0.44 | 0.31 | 0.46 | 0.6 |
| 8 | 38821327 | rs10808999 | -0.915 | 1.681 | -2.073 | -0.293 | -0.022 | 0.043 | 0.043 | 0 | 0.81 | 0.98 | 0.93 | 0.92 | 0.86 | 0.88 |
| 8 | 38823999 | rs4404880 | -0.023 | 1.745 | -1.856 | -0.132 | -0.016 | 0.039 | 0.039 | 0 | 0.69 | 0.81 | 0.7 | 0.81 | 0.66 | 0.71 |
| 8 | 38826365 | rs6985572 | -0.038 | 1.716 | -1.841 | -0.116 | -0.018 | 0.03 | 0.03 | 0 | 0.69 | 0.82 | 0.69 | 0.79 | 0.64 | 0.71 |
| 8 | 38831059 | rs7295 | 0.054 | 1.717 | -1.774 | -0.097 | -0.006 | 0.016 | 0.016 | 0 | 0.68 | 0.71 | 0.68 | 0.79 | 0.65 | 0.72 |
| 8 | 38835269 | rs6474514 | 0.382 | 1.444 | -1.113 | -0.523 | -0.011 | 0.013 | 0.013 | 0 | 0.39 | 0.35 | 0.55 | 0.79 | 0.45 | 0.33 |
| 8 | 38835666 | rs4733961 | -1.366 | 1.241 | -0.596 | -0.601 | -0.02 | -0.05 | -0.05 | 0.036 | 0.74 | 0.68 | 0.86 | 0.95 | 0.8 | 0.68 |
| 8 | 38840300 | rs4733962 | 0.445 | 1.294 | -0.714 | -0.582 | -0.01 | 0.01 | 0.01 | 0 | 0.39 | 0.36 | 0.53 | 0.75 | 0.45 | 0.33 |
| 8 | 38840591 | rs7814264 | 0.616 | 1.282 | -0.718 | -0.53 | -0.02 | 0.018 | 0.018 | 0 | 0.37 | 0.31 | 0.51 | 0.75 | 0.43 | 0.32 |
| 8 | 38845716 | rs7823125 | 0.529 | 1.256 | -0.706 | -0.537 | -0.019 | 0.022 | 0.022 | 0 | 0.38 | 0.31 | 0.52 | 0.75 | 0.44 | 0.32 |
| 8 | 38848850 | rs4733895 | -1.362 | 1.178 | -0.525 | -0.569 | 0.023 | 0.028 | 0.028 | 0 | 0.74 | 0.84 | 0.87 | 0.95 | 0.8 | 0.66 |
| 8 | 38853983 | rs7840270 | 0.581 | 1.161 | -0.293 | -0.553 | -0.083 | 0.041 | 0.041 | 0 | 0.37 | 0.18 | 0.51 | 0.75 | 0.44 | 0.32 |
| 8 | 38855847 | rs10093975 | 0.315 | 1.113 | -0.355 | -0.576 | -0.023 | 0.027 | 0.027 | 0 | 0.39 | 0.3 | 0.52 | 0.75 | 0.45 | 0.33 |
| 8 | 38858484 | rs10089487 | -0.018 | 1.036 | -0.331 | -0.559 | -0.005 | -0.035 | -0.035 | 2.192 | 0.36 | 0.55 | 0.33 | 0.21 | 0.33 | 0.34 |
| 8 | 38872186 | rs7833611 | -1.466 | 1.056 | -0.509 | -0.612 | 0.016 | 0.019 | 0.019 | 0 | 0.75 | 0.84 | 0.86 | 0.95 | 0.79 | 0.69 |
| 8 | 38873945 | rs10958813 | 0.331 | 1.147 | -0.57 | -0.539 | -0.008 | 0.054 | 0.054 | 0 | 0.31 | 0.11 | 0.26 | 0.18 | 0.28 | 0.33 |
| 8 | 38897470 | rs13282163 | 1.609 | 0.946 | -0.706 | -0.611 | -0.027 | 0.008 | 0.008 | 0 | 0.12 | 0 | 0.06 | 0.09 | 0.09 | 0.16 |
| 8 | 38901877 | rs7015505 | 0.114 | 1.142 | -0.305 | -0.367 | -0.021 | 0.022 | 0.022 | 0 | 0.39 | 0.29 | 0.54 | 0.73 | 0.47 | 0.35 |
| 8 | 38912806 | rs6990299 | -1.343 | 1.015 | -0.721 | -0.506 | 0.018 | 0.108 | 0.108 | 0 | 0.76 | 0.98 | 0.87 | 0.95 | 0.78 | 0.67 |
| 8 | 38940922 | rs6474529 | -1.248 | 0.993 | -0.599 | -0.505 | 0.016 | 0.025 | 0.025 | 0 | 0.74 | 0.84 | 0.86 | 0.94 | 0.78 | 0.67 |
| 8 | 38940934 | rs4130393 | 1.561 | 1.231 | -0.577 | -0.421 | 0.019 | 0.057 | 0.057 | 0 | 0.2 | 0.09 | 0.12 | 0.06 | 0.15 | 0.07 |
| 8 | 38966013 | rs11985898 | 0.221 | 1.291 | -0.531 | -0.188 | -0.002 | 0.003 | 0.003 | 0 | 0.58 | 0.39 | 0.42 | 0.27 | 0.51 | 0.64 |
| 8 | 38968798 | rs12542595 | -0.133 | 1.646 | 0.104 | 0.569 | -0.001 | 0.008 | 0.008 | 0.026 | 0.75 | 0.28 | 0.63 | 0.47 | 0.76 | 0.9 |
| 8 | 38971606 | rs12549232 | 0.758 | 1.61 | 0.275 | 0.351 | 0.042 | 0.059 | 0.059 | 0 | 0.37 | 0.2 | 0.28 | 0.18 | 0.25 | 0.3 |
| 8 | 38975368 | rs11775433 | 0.134 | 1.689 | 0.162 | 0.355 | -0.034 | -0.029 | -0.029 | 0.014 | 0.12 | 0.07 | 0.24 | 0.2 | 0.28 | 0.24 |
| 8 | 38989813 | rs4733980 | 0.762 | 1.672 | 0.11 | 0.361 | 0.036 | 0.08 | 0.08 | 0 | 0.38 | 0.15 | 0.32 | 0.19 | 0.29 | 0.31 |
| 8 | 38993605 | rs6994035 | 0.206 | 1.404 | 0.46 | 0.391 | -0.001 | -0.001 | -0.001 | 0.218 | 0.47 | 0.5 | 0.43 | 0.6 | 0.43 | 0.43 |
| 8 | 39021272 | rs4585712 | 0.966 | 2.232 | 1.728 | 0.796 | 0.101 | 0.064 | 0.064 | 0.058 | 0.85 | 0.71 | 0.75 | 0.79 | 0.72 | 0.76 |
| 8 | 39042042 | rs4273822 | 1.36 | 2.314 | 1.844 | 0.741 | -0.037 | -0.001 | -0.001 | 0 | 0.67 | 0.24 | 0.56 | 0.66 | 0.53 | 0.72 |
| 8 | 39052223 | rs11781014 | -0.815 | 2.27 | 1.862 | 0.796 | 0.018 | 0.009 | 0.009 | 0 | 0.13 | 0.12 | 0.24 | 0.24 | 0.28 | 0.23 |
| 8 | 39089941 | rs10958542 | 1.355 | 2.255 | 1.829 | 0.819 | -0.047 | -0.028 | -0.028 | 0.007 | 0.67 | 0.16 | 0.58 | 0.62 | 0.58 | 0.72 |
| 8 | 39091526 | rs4515515 | -0.443 | 2.269 | 1.836 | 0.813 | 0.017 | 0.023 | 0.023 | 0 | 0.11 | 0.12 | 0.21 | 0.29 | 0.24 | 0.24 |

| | | | | | | | | | | | | | | | | |
|---|---|---|---|---|---|---|---|---|---|---|---|---|---|---|---|---|
| 8 | 39123820 | rs4422746 | 0.184 | 2.236 | 1.545 | 0.727 | -0.013 | 0.038 | 0.038 | 0 | 0.17 | 0.09 | 0.15 | 0.07 | 0.2 | 0.08 |
| 8 | 39128387 | rs4733994 | -0.5 | 2.292 | 1.713 | 0.887 | 0.021 | 0.029 | 0.029 | 0 | 0.11 | 0.13 | 0.21 | 0.29 | 0.24 | 0.23 |
| 8 | 39135121 | rs6988280 | -1.257 | 2.335 | 1.502 | 0.891 | 0.101 | 0.116 | 0.116 | 0.469 | 0.15 | 0.61 | 0.26 | 0.29 | 0.24 | 0.23 |
| 8 | 39136219 | rs6994939 | -0.924 | 2.198 | 1.194 | 0.764 | 0.078 | 0.039 | 0.039 | 0 | 0.28 | 0.54 | 0.38 | 0.34 | 0.41 | 0.26 |
| 8 | 39142264 | rs7008808 | 0.887 | 1.164 | 0.738 | 0.315 | -0.001 | -0.061 | -0.061 | 0 | 0.42 | 0.49 | 0.29 | 0.13 | 0.37 | 0.33 |
| 8 | 39144098 | rs10100075 | 0.288 | 1.245 | 0.758 | 0.369 | -0.001 | 0.001 | 0.001 | 0 | 0.17 | 0.17 | 0.18 | 0.1 | 0.21 | 0.07 |
| 8 | 39147294 | rs4324894 | 1.501 | 1.107 | 0.73 | 0.295 | -0.097 | 0.178 | 0.178 | 0 | 0.37 | 0.03 | 0.25 | 0.09 | 0.37 | 0.32 |
| 8 | 39148173 | rs13439218 | -0.087 | 1.266 | 0.726 | 0.538 | -0.008 | -0.009 | -0.009 | 0 | 0.2 | 0.19 | 0.31 | 0.31 | 0.31 | 0.23 |
| 8 | 39148247 | rs12708189 | -0.181 | 1.201 | 0.733 | 0.551 | 0.021 | -0.022 | -0.022 | 0 | 0.63 | 0.72 | 0.62 | 0.44 | 0.71 | 0.58 |
| 8 | 39163716 | rs12681970 | -0.347 | 1.193 | 0.761 | 0.579 | 0.022 | -0.021 | -0.021 | 0.155 | 0.64 | 0.72 | 0.62 | 0.44 | 0.72 | 0.58 |
| 8 | 39170959 | rs10087894 | -0.548 | 1.058 | 1.022 | 0.795 | 0.006 | 0.008 | 0.008 | 0 | 0.22 | 0.22 | 0.33 | 0.32 | 0.33 | 0.25 |
| 8 | 39182188 | rs4484654 | 0.198 | 0.725 | 0.736 | 0.755 | 0.013 | 0.009 | 0.009 | 0 | 0.51 | 0.56 | 0.69 | 0.41 | 0.59 | 0.55 |
| 8 | 39187256 | rs7009846 | -0.002 | 0.653 | 0.862 | 0.719 | 0.022 | 0.012 | 0.012 | 0 | 0.54 | 0.66 | 0.69 | 0.41 | 0.59 | 0.55 |
| 8 | 39188834 | rs3935233 | 0.072 | 0.65 | 0.939 | 0.728 | 0.004 | 0.002 | 0.002 | 0 | 0.53 | 0.53 | 0.69 | 0.41 | 0.59 | 0.55 |
| 8 | 39191670 | rs4595090 | -0.005 | 0.66 | 0.642 | 0.744 | 0.021 | 0.01 | 0.01 | 0 | 0.54 | 0.66 | 0.69 | 0.41 | 0.59 | 0.55 |
| 8 | 39192481 | rs13252535 | 0.989 | 0.63 | 0.14 | 0.371 | -0.109 | 0.153 | 0.153 | 0.917 | 0.41 | 0.07 | 0.4 | 0.15 | 0.43 | 0.36 |
| 8 | 39194984 | rs6985917 | 0.358 | 0.539 | 0.613 | 0.641 | 0.004 | 0.061 | 0.061 | 0 | 0.5 | 0.64 | 0.65 | 0.36 | 0.5 | 0.52 |
| 8 | 39195489 | rs7000147 | -1.373 | 0.709 | 1.511 | 0.697 | 0.032 | 0.063 | 0.063 | 0.076 | 0.1 | 0.37 | 0.22 | 0.11 | 0.06 | 0.16 |
| 8 | 39199543 | rs6474129 | -1.479 | 0.703 | 1.546 | 0.695 | 0.03 | 0.068 | 0.068 | 0.092 | 0.1 | 0.4 | 0.22 | 0.12 | 0.06 | 0.17 |
| 8 | 39206995 | rs4379414 | -1.281 | 0.697 | 2.035 | 0.692 | 0.006 | 0.032 | 0.032 | 0 | 0.09 | 0.25 | 0.22 | 0.12 | 0.06 | 0.17 |
| 8 | 39219286 | rs7461947 | 0.192 | 0.634 | 1.777 | 0.683 | -0.003 | 0.048 | 0.048 | 0 | 0.52 | 0.62 | 0.65 | 0.33 | 0.49 | 0.52 |
| 8 | 39224178 | rs13270080 | 0.878 | 0.7 | 1.706 | 0.519 | -0.052 | 0.188 | 0.188 | 0.104 | 0.44 | 0.12 | 0.42 | 0.15 | 0.45 | 0.39 |
| 8 | 39224968 | rs6474131 | 0.394 | 0.689 | 1.678 | 0.561 | 0 | 0.011 | 0.011 | 0 | 0.51 | 0.5 | 0.64 | 0.26 | 0.49 | 0.52 |
| 8 | 39390219 | rs7829181 | 0.423 | 0.379 | 1.239 | 0.626 | -0.023 | 0.011 | 0.011 | 0.211 | 0.11 | 0.18 | 0.08 | 0.12 | 0.06 | 0.13 |
| 8 | 39397015 | rs7463768 | -0.049 | 0.266 | 1.114 | 0.651 | -0.015 | 0.039 | 0.039 | 0 | 0.77 | 0.85 | 0.84 | 0.9 | 0.76 | 0.68 |
| 8 | 39399057 | rs10108531 | 0.341 | 0.326 | 1.13 | 0.69 | -0.019 | 0.005 | 0.005 | 0.049 | 0.12 | 0.19 | 0.1 | 0.12 | 0.08 | 0.14 |
| 8 | 39405337 | rs6474156 | -0.608 | 0.338 | 1.136 | 1.302 | 0.004 | 0.001 | 0.001 | 0 | 0.85 | 0.92 | 0.93 | 0.89 | 0.89 | 0.74 |
| 8 | 39408586 | rs10095849 | 0.266 | 0.244 | 1.675 | 1.227 | -0.013 | 0.059 | 0.059 | 0 | 0.2 | 0.35 | 0.14 | 0.34 | 0.18 | 0.12 |
| 8 | 39410399 | rs12056485 | 1.002 | 0.248 | 1.494 | 1.208 | 0.013 | -0.009 | -0.009 | 0.586 | 0.1 | 0.06 | 0.07 | 0.12 | 0.06 | 0.12 |
| 8 | 39429071 | rs7830114 | 0.399 | 0.343 | 1.301 | 1.128 | 0.004 | 0.006 | 0.006 | 0 | 0.15 | 0.08 | 0.07 | 0.1 | 0.11 | 0.26 |
| 8 | 39434544 | rs4733916 | 0.87 | 0.287 | 1.473 | 1.203 | 0.085 | 0.331 | 0.331 | 0 | 0.93 | 0.64 | 0.8 | 0.71 | 0.91 | 0.91 |
| 8 | 39438921 | rs11985201 | 2.061 | 0.305 | 1.182 | 1.071 | -0.064 | 0.16 | 0.16 | 0.507 | 0.4 | 0.13 | 0.41 | 0.15 | 0.42 | 0.35 |
| 8 | 39439518 | rs10108019 | 0.304 | 0.336 | 1.174 | 1.139 | -0.006 | -0.004 | -0.004 | 0 | 0.69 | 0.66 | 0.63 | 0.25 | 0.68 | 0.68 |
| 8 | 39449187 | rs13270163 | 1.96 | 0.393 | 0.948 | 1.12 | -0.051 | 0.163 | 0.163 | 1.605 | 0.42 | 0.12 | 0.41 | 0.14 | 0.42 | 0.33 |
| 8 | 39454677 | rs13248954 | -0.191 | 0.277 | 1.096 | 1.139 | 0 | 0 | 0 | 0 | 0.27 | 0.26 | 0.19 | 0.46 | 0.24 | 0.23 |
| 8 | 39463823 | rs13268577 | 1.953 | 0.311 | 0.934 | 1.137 | -0.058 | 0.17 | 0.17 | 1.618 | 0.42 | 0.12 | 0.41 | 0.14 | 0.44 | 0.34 |
| 8 | 39480366 | rs11988623 | -0.374 | 0.157 | 0.974 | 1.362 | 0 | 0 | 0 | 0 | 0.27 | 0.26 | 0.19 | 0.46 | 0.24 | 0.23 |
| 8 | 39495325 | rs3824144 | 0.457 | 0.536 | 0.879 | 0.679 | -0.001 | 0.003 | 0.003 | 0.407 | 0.69 | 0.74 | 0.6 | 0.25 | 0.64 | 0.67 |
| 8 | 39502217 | rs2060802 | 0.628 | 0.572 | 1.291 | 0.629 | -0.008 | -0.009 | -0.009 | 1.77 | 0.68 | 0.45 | 0.59 | 0.25 | 0.67 | 0.67 |
| 8 | 39503079 | rs6474161 | 0.598 | 0.531 | 1.21 | 0.631 | 0 | -0.002 | -0.002 | 0.307 | 0.68 | 0.66 | 0.6 | 0.25 | 0.64 | 0.67 |
| 8 | 39521419 | rs6988389 | 0.203 | 0.361 | 1.265 | 0.943 | 0.017 | 0.108 | 0.108 | 0 | 0.85 | 0.75 | 0.74 | 0.36 | 0.82 | 0.82 |
| 8 | 39522613 | rs1993120 | 0.203 | 0.361 | 1.265 | 0.943 | 0.017 | 0.108 | 0.108 | 0 | 0.85 | 0.75 | 0.74 | 0.36 | 0.82 | 0.82 |
| 8 | 39535829 | rs7011098 | 0.313 | 0.338 | 0.947 | 0.455 | 0.004 | 0.001 | 0.001 | 0 | 0.14 | 0.09 | 0.07 | 0.1 | 0.1 | 0.28 |
| 8 | 39539916 | rs6474166 | -0.512 | 0.432 | 0.949 | 0.379 | -0.004 | 0.014 | 0.014 | 0 | 0.22 | 0.15 | 0.15 | 0.1 | 0.21 | 0.33 |
| 8 | 39543458 | rs10105183 | -0.498 | 0.438 | 0.767 | 0.382 | -0.004 | 0.014 | 0.014 | 0 | 0.22 | 0.15 | 0.15 | 0.1 | 0.21 | 0.33 |
| 8 | 39553628 | rs11991456 | -0.563 | 0.386 | 1.013 | 0.483 | -0.005 | 0.002 | 0.002 | 0 | 0.27 | 0.36 | 0.35 | 0.75 | 0.26 | 0.3 |
| 8 | 39555708 | rs7464542 | -0.595 | 0.431 | 0.092 | 0.5 | 0.004 | 0.001 | 0.001 | 0 | 0.86 | 0.9 | 0.93 | 0.9 | 0.91 | 0.72 |
| 8 | 39557645 | rs1451736 | 1.378 | 0.373 | 0.137 | 0.459 | 0.034 | 0.136 | 0.136 | 1.514 | 0.56 | 0.22 | 0.56 | 0.14 | 0.65 | 0.42 |
| 8 | 39564151 | rs7460786 | -0.291 | 0.318 | -0.083 | 0.641 | 0.004 | 0.001 | 0.001 | 0 | 0.86 | 0.9 | 0.93 | 0.9 | 0.91 | 0.72 |
| 8 | 39572749 | rs7386927 | -0.219 | 0.11 | -0.266 | 0.534 | 0.01 | 0.006 | 0.006 | 0 | 0.84 | 0.9 | 0.93 | 0.9 | 0.91 | 0.73 |
| 8 | 39575853 | rs7460182 | -0.205 | 0.095 | -0.22 | 0.561 | 0.005 | 0.003 | 0.003 | 0 | 0.84 | 0.9 | 0.93 | 0.9 | 0.91 | 0.73 |
| 8 | 39577963 | rs6474169 | 1.025 | -0.146 | -0.29 | 0.804 | 0.004 | 0 | 0 | 0 | 0.55 | 0.62 | 0.56 | 0.5 | 0.62 | 0.37 |
| 8 | 39587864 | rs4440676 | -0.358 | -0.155 | -0.5 | 0.833 | -0.005 | 0.051 | 0.051 | 0.571 | 0.24 | 0.36 | 0.32 | 0.32 | 0.2 | 0.36 |
| 8 | 39587888 | rs10958563 | -0.991 | -0.101 | -0.504 | 0.832 | 0.003 | 0.015 | 0.015 | 1.119 | 0.32 | 0.37 | 0.4 | 0.42 | 0.31 | 0.42 |
| 8 | 39588338 | rs12545052 | -0.11 | 0.006 | -0.741 | 0.93 | 0.001 | 0.018 | 0.018 | 0 | 0.11 | 0.1 | 0.23 | 0.26 | 0.13 | 0.19 |
| 8 | 39588519 | rs201993 | 1.438 | -0.12 | -0.935 | 0.599 | -0.077 | 0.183 | 0.183 | 0.324 | 0.45 | 0.1 | 0.48 | 0.22 | 0.54 | 0.34 |
| 8 | 39591821 | rs201986 | 1.422 | -0.201 | -0.957 | 0.558 | -0.08 | 0.182 | 0.182 | 0.473 | 0.44 | 0.1 | 0.48 | 0.22 | 0.54 | 0.34 |
| 8 | 39594979 | rs201982 | -0.162 | -0.097 | 0.196 | 0.597 | 0.005 | 0.21 | 0.21 | 0.057 | 0.09 | 0.33 | 0.06 | 0.27 | 0.07 | 0.03 |
| 8 | 39596120 | rs12682366 | 0.132 | -0.089 | 0.4 | 0.714 | 0.015 | 0.105 | 0.105 | 1.734 | 0.19 | 0.33 | 0.32 | 0.4 | 0.19 | 0.33 |
| 8 | 39599660 | rs2122987 | -0.291 | -0.231 | 0.398 | 0.451 | 0.009 | 0.031 | 0.031 | 4.009 | 0.26 | 0.32 | 0.4 | 0.5 | 0.31 | 0.4 |
| 8 | 39608631 | rs673382 | -1.058 | 0.203 | 0.175 | -0.523 | 0 | 0.001 | 0.001 | 0.229 | 0.55 | 0.6 | 0.55 | 0.86 | 0.56 | 0.65 |
| 8 | 39611992 | rs12541801 | 0.051 | 0.725 | 0.007 | -0.521 | -0.008 | 0.006 | 0.006 | 0 | 0.58 | 0.38 | 0.53 | 0.61 | 0.54 | 0.66 |
| 8 | 39620015 | rs12546336 | -1.46 | 0.567 | -0.284 | -0.656 | -0.069 | 0.038 | 0.038 | 0 | 0.79 | 0.89 | 0.64 | 0.88 | 0.66 | 0.73 |
| 8 | 39628986 | rs202006 | 0.393 | 0.504 | 0.13 | -0.66 | -0.007 | 0.011 | 0.011 | 0 | 0.19 | 0.27 | 0.13 | 0.27 | 0.16 | 0.07 |
| 8 | 39633967 | rs2290301 | 0.076 | 0.464 | 0.32 | -0.584 | 0.003 | -0.003 | -0.003 | 0 | 0.58 | 0.39 | 0.5 | 0.61 | 0.5 | 0.67 |
| 8 | 39648002 | rs1451745 | 0.793 | 0.49 | 0.302 | -0.759 | 0.035 | 0.046 | 0.046 | 0 | 0.48 | 0.24 | 0.42 | 0.61 | 0.4 | 0.59 |
| 8 | 39671926 | rs10113392 | -0.378 | 0.987 | 0.49 | -0.142 | 0 | -0.033 | -0.033 | 0 | 0.11 | 0.12 | 0.09 | 0 | 0.09 | 0.08 |
| 8 | 39687113 | rs6415698 | 0.12 | 1.722 | 0.806 | 0.063 | 0.167 | 0.061 | 0.061 | 0.778 | 0.92 | 0.68 | 0.72 | 0.88 | 0.79 | 0.84 |
| 8 | 39694610 | rs13255763 | 0.118 | 1.55 | 0.739 | -0.012 | 0.008 | 0.005 | 0.005 | 0.667 | 0.3 | 0.28 | 0.18 | 0.24 | 0.26 | 0.07 |

| | | | | | | | | | | | | | | | | |
|---|---|---|---|---|---|---|---|---|---|---|---|---|---|---|---|---|
| 8 | 39695535 | rs7844819 | 0.364 | 1.561 | 0.826 | 0.007 | 0.13 | 0.046 | 0.046 | 0.271 | 0.07 | 0.21 | 0.27 | 0.11 | 0.2 | 0.16 |
| 8 | 39697212 | rs6984521 | -1.458 | 1.444 | 0.846 | -0.079 | 0.022 | 0.037 | 0.037 | 0 | 0.49 | 0.77 | 0.52 | 0.38 | 0.55 | 0.34 |
| 8 | 39703449 | rs7836577 | -0.723 | 1.433 | 0.769 | -0.069 | -0.001 | -0.003 | -0.003 | 0 | 0.41 | 0.44 | 0.25 | 0.27 | 0.35 | 0.19 |
| 8 | 39710324 | rs10096637 | -0.062 | 1.441 | 0.894 | -0.119 | 0.19 | 0.075 | 0.075 | 1.126 | 0.08 | 0.38 | 0.28 | 0.11 | 0.2 | 0.16 |
| 8 | 39720982 | rs6998587 | -1.43 | 1.409 | 0.259 | -0.136 | 0.037 | 0.063 | 0.063 | 0 | 0.5 | 0.83 | 0.54 | 0.37 | 0.56 | 0.34 |
| 8 | 39721031 | rs6998739 | -1.348 | 1.419 | 0.376 | -0.164 | 0.086 | -0.036 | -0.036 | 0 | 0.2 | 0.55 | 0.34 | 0.14 | 0.29 | 0.27 |
| 8 | 39722425 | rs7004755 | 0.161 | 1.322 | 0.367 | -0.15 | 0.008 | 0.006 | 0.006 | 0.635 | 0.3 | 0.28 | 0.19 | 0.22 | 0.27 | 0.07 |
| 8 | 39727430 | rs6988758 | -0.708 | 1.594 | 0.194 | 0.012 | 0.016 | -0.053 | -0.053 | 0 | 0.1 | 0.23 | 0.1 | 0 | 0.1 | 0.08 |
| 8 | 39746386 | rs7839240 | -1.646 | 1.533 | -0.812 | 0.058 | 0.032 | 0.007 | 0.007 | 1.184 | 0.54 | 0.84 | 0.55 | 0.41 | 0.59 | 0.35 |
| 8 | 39751406 | rs7812568 | 0.134 | 1.392 | -0.476 | 0.296 | -0.007 | -0.005 | -0.005 | 0 | 0.29 | 0.36 | 0.17 | 0.22 | 0.24 | 0.1 |
| 8 | 39769375 | rs3808606 | -1.088 | 1.429 | -0.793 | 0.705 | -0.01 | -0.021 | -0.021 | 0 | 0.48 | 0.8 | 0.51 | 0.49 | 0.51 | 0.3 |
| 8 | 39769939 | rs10089078 | -0.543 | 0.613 | -1.268 | 0.224 | 0.005 | 0.033 | 0.033 | 0.074 | 0.38 | 0.25 | 0.22 | 0.23 | 0.34 | 0.17 |
| 8 | 39777529 | rs7820268 | -0.467 | 0.284 | -1.531 | 0.237 | 0.018 | 0.05 | 0.05 | 0.288 | 0.37 | 0.2 | 0.21 | 0.22 | 0.32 | 0.17 |
| 8 | 39779989 | rs10108662 | -0.373 | 0.311 | -1.464 | 0.177 | 0.005 | 0.042 | 0.042 | 0 | 0.35 | 0.2 | 0.21 | 0.22 | 0.33 | 0.17 |
| 8 | 39792914 | rs16888361 | 0.495 | -0.342 | -1.827 | 0.6 | 0.093 | -0.017 | -0.017 | 0.636 | 0.32 | 0.07 | 0.21 | 0.28 | 0.19 | 0.31 |
| 8 | 39793368 | rs13278237 | 0.451 | 0.53 | -2.363 | 0.933 | -0.137 | -0.092 | -0.092 | 1.537 | 0.2 | 0.03 | 0.5 | 0.31 | 0.45 | 0.34 |
| 8 | 39796413 | rs4737159 | -1.087 | 0.34 | -1.576 | 0.716 | 0.01 | 0.04 | 0.04 | 0 | 0.85 | 0.58 | 0.85 | 0.83 | 0.85 | 0.86 |
| 8 | 39797458 | rs11773909 | 0.189 | 0.277 | -1.405 | 0.859 | 0.016 | 0.014 | 0.014 | 0.006 | 0.64 | 0.45 | 0.46 | 0.55 | 0.39 | 0.52 |
| 8 | 39798956 | rs13248917 | 0.699 | 0.575 | -1.703 | 0.81 | -0.138 | -0.076 | -0.076 | 0.005 | 0.2 | 0.03 | 0.38 | 0.29 | 0.46 | 0.34 |
| 8 | 39805797 | rs3927941 | 0.403 | 0.419 | -1.485 | 0.791 | -0.051 | -0.017 | -0.017 | 0 | 0.24 | 0.15 | 0.4 | 0.31 | 0.46 | 0.37 |
| 8 | 39816468 | rs6990252 | 0.023 | 0.63 | -0.852 | 0.503 | -0.002 | 0.007 | 0.007 | 0 | 0.27 | 0.2 | 0.12 | 0.24 | 0.19 | 0.17 |
| 8 | 39824406 | rs28631334 | -0.148 | 0.794 | -0.21 | 0.515 | 0.033 | 0.023 | 0.023 | 0 | 0.15 | 0.47 | 0.15 | 0.14 | 0.15 | 0.14 |
| 8 | 39831647 | rs2543061 | -0.052 | 1.006 | -0.974 | 0.882 | -0.038 | -0.054 | -0.054 | 2.327 | 0.82 | 0.92 | 0.84 | 0.63 | 0.8 | 0.84 |
| 8 | 39834819 | rs10504014 | 0.242 | 0.05 | -1.115 | 0.575 | 0.007 | 0.029 | 0.029 | 0 | 0.6 | 0.69 | 0.57 | 0.3 | 0.65 | 0.64 |
| 8 | 39835582 | rs2729467 | 0.072 | 0.021 | -0.896 | 0.594 | 0.006 | 0.022 | 0.022 | 0 | 0.81 | 0.73 | 0.83 | 0.63 | 0.79 | 0.84 |
| 8 | 39836128 | rs7002522 | 0.561 | -0.589 | -0.569 | 0.069 | 0.044 | 0.07 | 0.07 | 0 | 0.5 | 0.34 | 0.52 | 0.3 | 0.61 | 0.6 |
| 8 | 39843295 | rs12541238 | 1.239 | -0.336 | -0.907 | -0.169 | -0.019 | 0.045 | 0.045 | 0 | 0.1 | 0.01 | 0.05 | 0 | 0.13 | 0.02 |
| 8 | 39845265 | rs16888478 | 0.758 | -0.27 | -0.537 | 0.347 | 0.046 | -0.061 | -0.061 | 0.016 | 0.22 | 0.09 | 0.27 | 0.67 | 0.17 | 0.24 |
| 8 | 39848935 | rs2955877 | -0.701 | -0.234 | -0.391 | 0.303 | -0.035 | -0.01 | -0.01 | 0 | 0.39 | 0.5 | 0.4 | 0.67 | 0.25 | 0.31 |
| 8 | 39850058 | rs2543073 | 0.311 | 0.095 | -0.35 | 0.371 | -0.002 | 0.012 | 0.012 | 0 | 0.62 | 0.54 | 0.6 | 0.6 | 0.64 | 0.66 |
| 8 | 39855954 | rs888444 | 0.601 | 0.153 | -0.291 | 0.314 | -0.009 | 0.031 | 0.031 | 0.759 | 0.71 | 0.76 | 0.77 | 0.93 | 0.7 | 0.79 |
| 8 | 39858026 | rs10104711 | -0.551 | 0.146 | -0.206 | 0.311 | 0.003 | -0.003 | -0.003 | 0 | 0.85 | 0.82 | 0.85 | 0.93 | 0.76 | 0.86 |
| 8 | 39858740 | rs7837357 | 1.098 | 0.027 | -0.208 | 0.264 | -0.04 | 0.069 | 0.069 | 0 | 0.2 | 0.26 | 0.24 | 0.34 | 0.12 | 0.18 |
| 8 | 39862881 | rs10109853 | 1.044 | -0.008 | -0.513 | 0.089 | -0.007 | 0.053 | 0.053 | 0 | 0.47 | 0.33 | 0.46 | 0.29 | 0.51 | 0.55 |
| 8 | 39864479 | rs2909333 | 0.417 | 0.608 | -0.396 | 0.546 | 0.007 | -0.018 | -0.018 | 0 | 0.61 | 0.49 | 0.54 | 0.29 | 0.58 | 0.61 |
| 8 | 39867827 | rs10958579 | -0.296 | 1.117 | -0.722 | 0.811 | -0.018 | -0.019 | -0.019 | 0.464 | 0.15 | 0.11 | 0.19 | 0.3 | 0.28 | 0.17 |
| 8 | 39869083 | rs6415701 | -0.666 | 0.533 | -0.568 | 0.818 | 0.003 | -0.005 | -0.005 | 0 | 0.18 | 0.18 | 0.12 | 0.37 | 0.18 | 0.15 |
| 8 | 39869633 | rs2981156 | -0.259 | 0.38 | -0.137 | 0.422 | -0.012 | 0.016 | 0.016 | 0 | 0.63 | 0.54 | 0.61 | 0.61 | 0.73 | 0.74 |
| 8 | 39869841 | rs2981157 | 0.04 | 0.53 | 0.127 | 0.349 | -0.002 | -0.006 | -0.006 | 0.018 | 0.6 | 0.38 | 0.41 | 0.6 | 0.63 | 0.72 |
| 8 | 39872129 | rs4617138 | -0.239 | 0.306 | 0.527 | 0.138 | 0.037 | -0.058 | -0.058 | 0 | 0.87 | 0.64 | 0.75 | 0.98 | 0.84 | 0.93 |
| 8 | 39872362 | rs7008343 | -0.18 | 0.231 | 0.426 | 0.337 | 0.001 | 0.002 | 0.002 | 0 | 0.37 | 0.38 | 0.56 | 0.38 | 0.36 | 0.28 |
| 8 | 39873304 | rs2955903 | 1.359 | 0.088 | 0.344 | 0.438 | -0.033 | 0.048 | 0.048 | 0.001 | 0.4 | 0.22 | 0.3 | 0.35 | 0.45 | 0.57 |
| 8 | 39880766 | rs2955914 | 1.359 | 0.091 | 0.329 | 0.204 | -0.031 | 0.1 | 0.1 | 0.001 | 0.4 | 0.22 | 0.28 | 0.22 | 0.45 | 0.57 |
| 8 | 39880881 | rs2981165 | 0.034 | 0.102 | 0.293 | 0.225 | 0.004 | 0.009 | 0.009 | 0 | 0.62 | 0.55 | 0.43 | 0.62 | 0.64 | 0.72 |
| 8 | 39884165 | rs16888648 | 0.66 | 0.087 | 0.31 | 0.22 | -0.025 | -0.033 | -0.033 | 0 | 0.1 | 0.19 | 0.06 | 0.01 | 0.05 | 0.05 |
| 8 | 39884574 | rs13270833 | -0.153 | 0.1 | 0.27 | 0.262 | 0.012 | -0.026 | -0.026 | 0.244 | 0.21 | 0.17 | 0.13 | 0.4 | 0.19 | 0.15 |
| 8 | 39887665 | rs12544923 | 0.747 | 0.248 | 0.242 | 0.273 | -0.022 | -0.029 | -0.029 | 1.707 | 0.27 | 0.17 | 0.5 | 0.37 | 0.31 | 0.24 |
| 8 | 39892501 | rs16888662 | 0.007 | 0.458 | 0.441 | 0.15 | -0.009 | -0.019 | -0.019 | 0 | 0.13 | 0.18 | 0.06 | 0.05 | 0.12 | 0.08 |
| 8 | 39900189 | rs6997214 | 0.016 | 0.675 | -0.582 | -0.067 | 0.001 | 0.01 | 0.01 | 0 | 0.43 | 0.57 | 0.45 | 0.68 | 0.38 | 0.33 |
| 8 | 39901373 | rs6983223 | 0.412 | 0.374 | -0.741 | -0.086 | 0.016 | 0.019 | 0.019 | 4.005 | 0.15 | 0.08 | 0.11 | 0.05 | 0.12 | 0.11 |
| 8 | 39901761 | rs2955864 | 0.05 | 0.596 | -0.545 | 0.152 | -0.011 | 0.045 | 0.045 | 0 | 0.55 | 0.35 | 0.37 | 0.31 | 0.52 | 0.64 |
| 8 | 39909152 | rs2929106 | 0.086 | 0.804 | -0.789 | 0.105 | -0.004 | 0.004 | 0.004 | 0 | 0.48 | 0.71 | 0.53 | 0.57 | 0.5 | 0.6 |
| 8 | 39910724 | rs13268381 | -0.356 | 0.764 | -0.559 | -0.003 | 0.002 | 0.011 | 0.011 | 0 | 0.18 | 0.17 | 0.15 | 0.07 | 0.21 | 0.11 |
| 8 | 39911650 | rs2929104 | 0.858 | 0.821 | -0.486 | 0.188 | 0.006 | -0.008 | -0.008 | 0.519 | 0.43 | 0.29 | 0.41 | 0.45 | 0.39 | 0.54 |
| 8 | 39913064 | rs2170306 | 0.263 | 0.685 | 0.105 | 0.535 | -0.003 | -0.002 | -0.002 | 0 | 0.59 | 0.53 | 0.52 | 0.54 | 0.52 | 0.65 |
| 8 | 39928050 | rs7821033 | -1.256 | 0.74 | 0.375 | 0.693 | 0.043 | 0.098 | 0.098 | 0 | 0.08 | 0.33 | 0.11 | 0.13 | 0.1 | 0.03 |
| 8 | 39938945 | rs13275833 | -0.022 | 0.679 | 0.086 | 0.549 | -0.09 | -0.09 | -0.09 | 4.453 | 0.33 | 0.07 | 0.39 | 0.43 | 0.4 | 0.31 |
| 8 | 39942372 | rs1904906 | -0.597 | 0.705 | 0.44 | 0.402 | 0.012 | -0.025 | -0.025 | 0 | 0.06 | 0.2 | 0.05 | 0 | 0.05 | 0.01 |
| 8 | 39955252 | rs7828920 | 0.779 | 0.664 | 0.149 | 0.562 | -0.002 | 0.005 | 0.005 | 0 | 0.53 | 0.29 | 0.46 | 0.46 | 0.46 | 0.66 |
| 8 | 39956020 | rs16888751 | -0.094 | 0.547 | 0.38 | 0.643 | -0.012 | 0.001 | 0.001 | 0 | 0.07 | 0.16 | 0.07 | 0.05 | 0.05 | 0.03 |
| 8 | 39959393 | rs16888778 | -0.166 | 0.52 | 0.491 | 0.586 | -0.01 | -0.006 | -0.006 | 1.653 | 0.07 | 0.13 | 0.06 | 0.03 | 0.05 | 0.03 |
| 8 | 39967058 | rs1471559 | 0.107 | 0.67 | 0.343 | 0.682 | -0.028 | -0.027 | -0.027 | 1.803 | 0.65 | 0.76 | 0.6 | 0.59 | 0.58 | 0.73 |
| 8 | 39969006 | rs12682158 | 0.639 | 0.626 | 0.186 | 0.685 | 0.005 | 0.023 | 0.023 | 0 | 0.55 | 0.26 | 0.48 | 0.42 | 0.43 | 0.63 |
| 8 | 39969392 | rs4370521 | -0.591 | 1.201 | 0.325 | 1.149 | -0.055 | -0.095 | -0.095 | 0.23 | 0.27 | 0.09 | 0.34 | 0.44 | 0.35 | 0.25 |
| 8 | 39976970 | rs10113242 | 1.017 | 1.242 | 0.129 | 0.882 | -0.003 | -0.002 | -0.002 | 2.613 | 0.66 | 0.28 | 0.59 | 0.44 | 0.57 | 0.73 |
| 8 | 39984467 | rs4737047 | 0.232 | 1.448 | -0.13 | 0.902 | -0.009 | -0.004 | -0.004 | 0 | 0.73 | 0.75 | 0.77 | 0.74 | 0.65 | 0.73 |
| 8 | 39993166 | rs7825010 | -0.7 | 0.974 | 0.225 | 0.88 | 0.036 | 0.056 | 0.056 | 0.071 | 0.14 | 0.49 | 0.16 | 0.15 | 0.17 | 0.13 |
| 8 | 40006096 | rs4398898 | 1.019 | 0.861 | -0.033 | 0.951 | 0 | 0.04 | 0.04 | 0 | 0.94 | 0.77 | 0.83 | 0.93 | 0.95 | 0.97 |
| 8 | 40010486 | rs10504018 | -0.138 | 1.348 | -0.706 | 0.738 | -0.005 | 0.003 | 0.003 | 0.236 | 0.94 | 0.93 | 0.94 | 1 | 0.9 | 0.94 |

| | | | | | | | | | | | | | | | |
|---|---|---|---|---|---|---|---|---|---|---|---|---|---|---|---|
| 8 | 40010820 | rs2242277 | -1.046 | 1.166 | -0.501 | 0.88 | 0.006 | -0.013 | -0.013 | 0 | 0.2 | 0.8 | 0.33 | 0.22 | 0.21 | 0.15 |
| 8 | 40012063 | rs10199 | 1.153 | 1.008 | -1.251 | 0.931 | 0.025 | 0.014 | 0.014 | 0 | 0.49 | 0.09 | 0.42 | 0.47 | 0.4 | 0.56 |
| 8 | 40013811 | rs10958604 | 1.101 | 0.905 | -0.306 | 0.98 | 0.005 | -0.023 | -0.023 | 0.486 | 0.8 | 0.15 | 0.67 | 0.78 | 0.78 | 0.85 |
| 8 | 40020831 | rs7822661 | -0.951 | 0.781 | 0.334 | 1.122 | -0.015 | 0.079 | 0.079 | 0 | 0.15 | 0.3 | 0.25 | 0.24 | 0.14 | 0.11 |
| 8 | 40023334 | rs6994498 | -1.173 | 1.129 | 0.454 | 0.782 | 0.012 | -0.033 | -0.033 | 2.752 | 0.2 | 0.32 | 0.25 | 0.13 | 0.24 | 0.16 |
| 8 | 40024606 | rs6474228 | -1.17 | 1.123 | 0.456 | 0.776 | 0.011 | -0.037 | -0.037 | 0 | 0.2 | 0.31 | 0.24 | 0.12 | 0.24 | 0.16 |
| 8 | 40028306 | rs2980821 | 1.121 | 0.812 | -0.199 | 0.953 | 0.02 | -0.033 | -0.033 | 0.556 | 0.51 | 0.16 | 0.45 | 0.53 | 0.44 | 0.49 |
| 8 | 40033476 | rs10097923 | -0.624 | 0.644 | 0.294 | 0.972 | -0.017 | 0.156 | 0.156 | 2.254 | 0.09 | 0.5 | 0.09 | 0.14 | 0.01 | 0.07 |
| 8 | 40039431 | rs13270718 | -0.723 | 0.663 | -0.522 | 0.989 | -0.035 | 0.022 | 0.022 | 0.059 | 0.38 | 0.15 | 0.45 | 0.33 | 0.55 | 0.44 |
| 8 | 40046361 | rs2925034 | 0.868 | 1.059 | -0.93 | 1.084 | -0.046 | -0.02 | -0.02 | 0 | 0.52 | 0.15 | 0.47 | 0.48 | 0.48 | 0.5 |
| 8 | 40053605 | rs10958605 | 0.954 | 0.584 | -1.224 | 0.921 | 0.019 | 0.013 | 0.013 | 0 | 0.47 | 0.13 | 0.42 | 0.42 | 0.43 | 0.48 |
| 8 | 40055056 | rs2929079 | 0.285 | 0.168 | -0.481 | 0.607 | -0.008 | -0.005 | -0.005 | 0.086 | 0.75 | 0.7 | 0.86 | 0.79 | 0.84 | 0.74 |
| 8 | 40059650 | rs10091442 | 0.022 | -0.567 | -1.241 | 0.256 | 0.077 | 0.082 | 0.082 | 1.784 | 0.5 | 0.21 | 0.29 | 0.36 | 0.4 | 0.38 |
| 8 | 40064509 | rs16888961 | -0.146 | -0.576 | -0.914 | 0.189 | -0.005 | -0.004 | -0.004 | 0 | 0.61 | 0.35 | 0.36 | 0.47 | 0.48 | 0.45 |
| 8 | 40065457 | rs10086086 | 0.632 | -1.514 | -1.383 | -0.312 | 0.04 | 0.057 | 0.057 | 1.296 | 0.25 | 0.12 | 0.11 | 0.03 | 0.14 | 0.22 |
| 8 | 40065987 | rs6987930 | -0.147 | -1.157 | -1.153 | -0.693 | 0.014 | 0.035 | 0.035 | 2.822 | 0.38 | 0.25 | 0.34 | 0.03 | 0.31 | 0.4 |
| 8 | 40066888 | rs2980806 | -0.975 | -1.165 | -0.645 | -0.681 | -0.028 | -0.025 | -0.025 | 0 | 0.75 | 0.79 | 0.58 | 0.45 | 0.61 | 0.63 |
| 8 | 40069791 | rs3116102 | 0.788 | -1.217 | 0.015 | -0.689 | 0.019 | -0.003 | -0.003 | 0 | 0.36 | 0.3 | 0.23 | 0.42 | 0.3 | 0.23 |
| 8 | 40071825 | rs10504019 | 0.928 | -1.196 | 0.068 | -0.729 | 0.036 | 0.048 | 0.048 | 1.344 | 0.25 | 0.14 | 0.11 | 0.03 | 0.14 | 0.22 |
| 8 | 40084203 | rs7822183 | 1.799 | -1.32 | 0.017 | -0.731 | 0.001 | -0.001 | -0.001 | 0 | 0.26 | 0.24 | 0.19 | 0.31 | 0.25 | 0.16 |
| 8 | 40085270 | rs726921 | 1.416 | -1.395 | -0.065 | -0.619 | -0.004 | -0.009 | -0.009 | 0 | 0.13 | 0.11 | 0.24 | 0 | 0.17 | 0.19 |
| 8 | 40096615 | rs1584989 | 0.035 | -1.218 | -0.017 | -0.596 | -0.014 | 0.01 | 0.01 | 2.783 | 0.61 | 0.33 | 0.49 | 0.34 | 0.52 | 0.55 |
| 8 | 40106630 | rs3116105 | 0.544 | -1.278 | 0.036 | -0.587 | -0.067 | -0.004 | -0.004 | 0 | 0.14 | 0.39 | 0.08 | 0.12 | 0.05 | 0.08 |
| 8 | 40107855 | rs2925014 | -0.963 | -1.022 | -0.236 | -0.488 | -0.026 | -0.02 | -0.02 | 0 | 0.75 | 0.79 | 0.57 | 0.45 | 0.57 | 0.63 |
| 8 | 40124799 | rs2980830 | 0.715 | -1.337 | -0.109 | -0.425 | -0.046 | -0.014 | -0.014 | 0 | 0.16 | 0.23 | 0.06 | 0.11 | 0.05 | 0.08 |
| 8 | 40131362 | rs2925002 | -0.462 | -1.502 | -0.58 | -0.194 | 0.01 | -0.014 | -0.014 | 0.366 | 0.63 | 0.86 | 0.4 | 0.66 | 0.41 | 0.51 |
| 8 | 40138358 | rs2925010 | -0.021 | -1.175 | -0.12 | 0.314 | -0.093 | 0.114 | 0.114 | 0 | 0.19 | 0.43 | 0.14 | 0.32 | 0.07 | 0.16 |
| 8 | 40146549 | rs970675 | 1.358 | -0.689 | -0.907 | 0.289 | 0.021 | -0.052 | -0.052 | 0 | 0.23 | 0.06 | 0.15 | 0.31 | 0.22 | 0.15 |
| 8 | 40152297 | rs13276767 | 0.587 | -0.327 | -0.599 | 0.57 | 0.003 | 0.023 | 0.023 | 0 | 0.08 | 0.04 | 0.03 | 0.34 | 0.09 | 0.12 |
| 8 | 40154240 | rs10504021 | -0.118 | -0.219 | -0.492 | 0.677 | 0.016 | 0.177 | 0.177 | 0.631 | 0.06 | 0.19 | 0.07 | 0.26 | 0.05 | 0.09 |
| 8 | 40154794 | rs1378414 | 0.842 | -0.014 | -0.679 | -0.189 | 0.006 | -0.037 | -0.037 | 0 | 0.14 | 0.2 | 0.24 | 0 | 0.16 | 0.18 |
| 8 | 40164860 | rs2168282 | 0.632 | 0.742 | -1.068 | -0.121 | 0.043 | -0.003 | -0.003 | 0 | 0.67 | 0.47 | 0.58 | 0.35 | 0.55 | 0.55 |
| 8 | 40167696 | rs1455387 | 0.533 | 0.79 | -1.175 | -0.205 | -0.041 | -0.01 | -0.01 | 0 | 0.07 | 0.02 | 0.08 | 0.07 | 0.2 | 0.08 |
| 8 | 40171900 | rs1598355 | -0.018 | 0.054 | -0.891 | 0.073 | 0.013 | 0.073 | 0.073 | 0.647 | 0.73 | 0.54 | 0.67 | 0.44 | 0.74 | 0.65 |
| 8 | 40172507 | rs2341978 | 0.331 | 0.186 | -0.794 | -0.252 | -0.018 | -0.01 | -0.01 | 0 | 0.6 | 0.53 | 0.64 | 0.34 | 0.7 | 0.55 |
| 8 | 40177668 | rs16889144 | 0.603 | 0.749 | -0.991 | -0.193 | 0.002 | 0.004 | 0.004 | 0 | 0.19 | 0.22 | 0.31 | 0.24 | 0.24 | 0.17 |
| 8 | 40179383 | rs4146443 | 0.907 | 0.403 | -1.646 | -0.354 | -0.022 | -0.114 | -0.114 | 0 | 0.27 | 0.05 | 0.19 | 0.57 | 0.3 | 0.23 |
| 8 | 40179891 | rs210665 | 0.603 | 0.742 | -1.024 | -0.068 | 0.011 | 0.001 | 0.001 | 0 | 0.45 | 0.39 | 0.3 | 0.57 | 0.44 | 0.4 |
| 8 | 40180752 | rs909293 | -0.376 | 0.917 | -1.283 | 0.025 | -0.002 | -0.001 | -0.001 | 1.611 | 0.39 | 0.36 | 0.61 | 0.42 | 0.44 | 0.52 |
| 8 | 40181183 | rs210662 | 0.182 | 0.428 | -1.452 | 0.02 | 0.001 | 0 | 0 | 2.012 | 0.68 | 0.72 | 0.67 | 0.82 | 0.77 | 0.62 |
| 8 | 40190876 | rs3736646 | -0.459 | 1.041 | -1.803 | 0.816 | -0.042 | -0.063 | -0.063 | 0 | 0.11 | 0.02 | 0.11 | 0.28 | 0.17 | 0.14 |
| 8 | 40193414 | rs210676 | -0.24 | 0.613 | -2.726 | 0.219 | -0.032 | 0.039 | 0.039 | 2.581 | 0.56 | 0.9 | 0.45 | 0.7 | 0.62 | 0.59 |
| 8 | 40194456 | rs10108825 | 0.176 | 0.313 | -2.996 | 0.153 | -0.053 | -0.073 | -0.073 | 0 | 0.33 | 0.1 | 0.36 | 0.39 | 0.28 | 0.39 |
| 8 | 40200810 | rs7013912 | 0.97 | 0.286 | -2.758 | -0.251 | 0.003 | -0.026 | -0.026 | 0 | 0.12 | 0.16 | 0.09 | 0.04 | 0.1 | 0.14 |
| 8 | 40204544 | rs996015 | -0.334 | 0.017 | -1.799 | -0.403 | 0 | 0.004 | 0.004 | 0 | 0.5 | 0.54 | 0.48 | 0.43 | 0.41 | 0.55 |
| 8 | 40208140 | rs11776037 | 1.219 | -0.023 | -2.208 | -0.415 | -0.015 | -0.007 | -0.007 | 0.077 | 0.1 | 0.01 | 0.08 | 0.07 | 0.08 | 0.14 |
| 8 | 40221041 | rs16889230 | 1.428 | -0.097 | -2.381 | -0.569 | -0.086 | -0.136 | -0.136 | 2.825 | 0.26 | 0.02 | 0.33 | 0.38 | 0.25 | 0.26 |
| 8 | 40221754 | rs210705 | -0.63 | -0.128 | -1.486 | -0.489 | 0 | 0.002 | 0.002 | 0 | 0.68 | 0.76 | 0.6 | 0.55 | 0.54 | 0.8 |
| 8 | 40229762 | rs10100953 | -0.434 | -0.357 | -1.264 | -0.725 | -0.003 | -0.009 | -0.009 | 0.913 | 0.3 | 0.31 | 0.19 | 0.12 | 0.19 | 0.33 |
| 8 | 40230928 | rs210840 | -0.412 | -0.131 | -1.127 | -0.657 | 0.005 | 0.012 | 0.012 | 0.109 | 0.35 | 0.64 | 0.24 | 0.15 | 0.26 | 0.41 |
| 8 | 40243954 | rs760349 | 0.125 | 0.558 | -1.301 | -0.284 | -0.029 | -0.023 | -0.023 | 0 | 0.25 | 0.16 | 0.43 | 0.7 | 0.37 | 0.45 |
| 8 | 40245200 | rs125676 | -1.105 | -0.376 | -0.863 | -0.961 | -0.038 | -0.118 | -0.118 | 0.047 | 0.76 | 0.72 | 0.86 | 1 | 0.83 | 0.95 |
| 8 | 40246274 | rs13265404 | 0.519 | -0.354 | -0.794 | -1.232 | 0 | -0.005 | -0.005 | 0 | 0.2 | 0.19 | 0.16 | 0.26 | 0.2 | 0.27 |
| 8 | 40251102 | rs210860 | -1.401 | -0.403 | -0.645 | -1.266 | -0.02 | -0.083 | -0.083 | 0 | 0.77 | 0.76 | 0.87 | 1 | 0.83 | 0.95 |
| 8 | 40252983 | rs16889302 | 1.242 | -0.379 | -0.807 | -1.197 | 0.014 | 0.011 | 0.011 | 0 | 0.14 | 0.08 | 0.08 | 0.13 | 0.1 | 0.24 |
| 8 | 40253726 | rs1367101 | 1.638 | 0.061 | -0.851 | -0.573 | -0.064 | -0.069 | -0.069 | 0 | 0.12 | 0.01 | 0.19 | 0.28 | 0.23 | 0.07 |
| 8 | 40254456 | rs210866 | 0.423 | -0.535 | -0.438 | -1.131 | 0.001 | 0.001 | 0.001 | 0 | 0.56 | 0.48 | 0.7 | 0.74 | 0.63 | 0.68 |
| 8 | 40261358 | rs169788 | -0.252 | -0.325 | -0.239 | -0.828 | -0.011 | 0.039 | 0.039 | 0 | 0.23 | 0.4 | 0.18 | 0.27 | 0.2 | 0.27 |
| 8 | 40264512 | rs1981999 | -0.21 | -0.311 | -0.128 | -0.831 | 0.057 | 0.152 | 0.152 | 1.024 | 0.93 | 0.71 | 0.91 | 0.86 | 0.9 | 0.97 |
| 8 | 40265646 | rs16889358 | 0.812 | -0.35 | -0.221 | -0.84 | 0.006 | 0.006 | 0.006 | 0 | 0.16 | 0.13 | 0.09 | 0.13 | 0.11 | 0.24 |
| 8 | 40267939 | rs2565116 | 1.09 | -0.364 | -0.499 | -1.084 | -0.012 | 0.056 | 0.056 | 0 | 0.29 | 0.2 | 0.26 | 0.04 | 0.25 | 0.2 |
| 8 | 40270318 | rs10111027 | 0.481 | -0.247 | -0.37 | -0.925 | 0.004 | 0.028 | 0.028 | 0.388 | 0.07 | 0.09 | 0.07 | 0.14 | 0.1 | 0.03 |
| 8 | 40276391 | rs891404 | 0.434 | -0.342 | -0.351 | -0.941 | -0.002 | 0.017 | 0.017 | 0 | 0.48 | 0.39 | 0.57 | 0.7 | 0.55 | 0.52 |
| 8 | 40278197 | rs2730136 | 0.204 | -0.007 | -0.565 | -0.722 | -0.002 | 0 | 0 | 0 | 0.24 | 0.26 | 0.27 | 0.17 | 0.28 | 0.12 |
| 8 | 40283141 | rs2730142 | -1.032 | -0.268 | -0.642 | -0.693 | -0.031 | -0.007 | -0.007 | 0 | 0.64 | 0.75 | 0.54 | 0.37 | 0.56 | 0.5 |
| 8 | 40285421 | rs7824285 | 0.854 | -0.666 | -0.112 | -1.089 | 0.009 | -0.022 | -0.022 | 0 | 0.23 | 0.15 | 0.13 | 0 | 0.18 | 0.05 |
| 8 | 40288067 | rs11783883 | 0.412 | -0.687 | 0.165 | -1.154 | 0.03 | 0.092 | 0.092 | 0 | 0.09 | 0.22 | 0.14 | 0.27 | 0.13 | 0.18 |
| 8 | 40288625 | rs2565125 | 1.089 | -0.761 | 0.077 | -1.295 | -0.002 | -0.005 | -0.005 | 0 | 0.15 | 0.15 | 0.19 | 0.03 | 0.15 | 0.08 |
| 8 | 40303200 | rs2730152 | -0.451 | -0.733 | 0.002 | -1.23 | 0.027 | 0.031 | 0.031 | 0 | 0.26 | 0.55 | 0.35 | 0.32 | 0.28 | 0.24 |

| | | | | | | | | | | | | | | | | |
|---|---|---|---|---|---|---|---|---|---|---|---|---|---|---|---|---|
| 8 | 40312565 | rs9298596 | 0.068 | -0.693 | 0.042 | -1.24 | -0.005 | -0.006 | -0.006 | 0.299 | 0.32 | 0.34 | 0.27 | 0.28 | 0.31 | 0.23 |
| 8 | 40313583 | rs2730194 | -0.746 | -0.647 | -0.195 | -1.149 | 0.012 | 0.05 | 0.05 | 0.008 | 0.48 | 0.7 | 0.47 | 0.33 | 0.46 | 0.3 |
| 8 | 40314725 | rs2730198 | -0.498 | -0.792 | -0.261 | -1.212 | 0.004 | -0.003 | -0.003 | 0 | 0.34 | 0.55 | 0.39 | 0.34 | 0.37 | 0.26 |
| 8 | 40314789 | rs4549753 | 1.479 | -0.938 | -0.4 | -1.367 | -0.06 | 0.042 | 0.042 | 1.241 | 0.43 | 0.12 | 0.48 | 0.66 | 0.48 | 0.59 |
| 8 | 40315004 | rs7823502 | 0.568 | -0.823 | -0.155 | -1.274 | -0.023 | -0.09 | -0.09 | 0 | 0.17 | 0.23 | 0.1 | 0 | 0.11 | 0.05 |
| 8 | 40316691 | rs13277698 | 0.533 | -0.703 | -0.207 | -1.177 | -0.006 | -0.027 | -0.027 | 0 | 0.07 | 0.11 | 0.04 | 0 | 0.06 | 0.02 |
| 8 | 40319197 | rs4736865 | 1.315 | -0.668 | -0.122 | -0.927 | -0.004 | -0.024 | -0.024 | 0 | 0.05 | 0.03 | 0.08 | 0.14 | 0.05 | 0.15 |
| 8 | 40319744 | rs4737131 | -0.124 | -0.684 | 0.058 | -0.942 | 0.001 | 0.024 | 0.024 | 0 | 0.11 | 0.17 | 0.14 | 0.28 | 0.14 | 0.18 |
| 8 | 40319953 | rs2565068 | 0.695 | -0.843 | 0.044 | -1.107 | -0.01 | -0.014 | -0.014 | 0 | 0.6 | 0.45 | 0.68 | 0.71 | 0.63 | 0.66 |
| 8 | 40330071 | rs7819777 | -0.233 | -0.825 | -0.185 | -1.184 | -0.025 | -0.107 | -0.107 | 0 | 0.73 | 0.68 | 0.84 | 0.99 | 0.78 | 0.84 |
| 8 | 40333321 | rs13278529 | 0.985 | -0.71 | -0.339 | -1.161 | 0.003 | -0.02 | -0.02 | 0.01 | 0.15 | 0.2 | 0.2 | 0.03 | 0.15 | 0.06 |
| 8 | 40334268 | rs1429969 | -0.943 | -0.76 | -0.584 | -1.06 | -0.006 | 0.022 | 0.022 | 0 | 0.93 | 0.87 | 0.92 | 0.81 | 0.95 | 0.85 |
| 8 | 40335724 | rs10102300 | 0.092 | -0.675 | -0.582 | -1.046 | 0.009 | -0.003 | -0.003 | 0 | 0.21 | 0.3 | 0.25 | 0.17 | 0.24 | 0.09 |
| 8 | 40336076 | rs11782411 | 0.108 | -0.843 | -0.5 | -1.139 | -0.024 | -0.04 | -0.04 | 2.927 | 0.69 | 0.46 | 0.76 | 0.81 | 0.76 | 0.79 |
| 8 | 40343490 | rs13269280 | 0.142 | -0.832 | -0.483 | -1.121 | -0.022 | -0.039 | -0.039 | 0.306 | 0.68 | 0.44 | 0.73 | 0.81 | 0.75 | 0.59 |
| 8 | 40351859 | rs4736869 | -1.818 | -0.612 | -0.451 | -1.008 | -0.005 | -0.019 | -0.019 | 0 | 0.95 | 0.92 | 0.97 | 0.99 | 0.96 | 0.89 |
| 8 | 40356700 | rs2195588 | 1.691 | -0.169 | -0.728 | -0.701 | -0.039 | -0.027 | -0.027 | 0.284 | 0.08 | 0.01 | 0.22 | 0.41 | 0.13 | 0.23 |
| 8 | 40359895 | rs1367104 | 0.026 | -0.533 | -0.178 | -1.251 | -0.027 | -0.152 | -0.152 | 0 | 0.73 | 0.49 | 0.82 | 1 | 0.77 | 0.83 |
| 8 | 40362168 | rs2342618 | 0.269 | -0.54 | -0.147 | -1.25 | -0.021 | -0.134 | -0.134 | 0.986 | 0.22 | 0.35 | 0.15 | 0 | 0.19 | 0.06 |
| 8 | 40362878 | rs10091962 | -0.687 | -0.558 | -0.207 | -1.303 | 0.009 | 0.057 | 0.057 | 0 | 0.37 | 0.71 | 0.35 | 0.04 | 0.34 | 0.12 |
| 8 | 40373509 | rs2565085 | 1.299 | -0.466 | 0.215 | -0.633 | -0.003 | -0.001 | -0.001 | 0.11 | 0.66 | 0.45 | 0.64 | 0.62 | 0.68 | 0.66 |
| 8 | 40382843 | rs7018220 | 0.492 | -0.378 | -0.35 | -0.683 | 0.011 | -0.045 | -0.045 | 0 | 0.15 | 0.3 | 0.2 | 0.04 | 0.15 | 0.06 |
| 8 | 40383512 | rs2115809 | 1.465 | -0.358 | -0.412 | -0.625 | 0.004 | 0.002 | 0.002 | 0.016 | 0.62 | 0.29 | 0.62 | 0.61 | 0.67 | 0.64 |
| 8 | 40385028 | rs11786438 | -0.483 | -0.115 | -0.686 | -0.169 | -0.014 | -0.081 | -0.081 | 1.949 | 0.14 | 0.03 | 0.12 | 0.34 | 0.13 | 0.18 |
| 8 | 40387791 | rs13274527 | 0.363 | 0.033 | -0.344 | -0.733 | -0.044 | 0.068 | 0.068 | 0 | 0.13 | 0.03 | 0.17 | 0.01 | 0.13 | 0.05 |
| 8 | 40388318 | rs753133 | -0.505 | -0.01 | -0.221 | -0.748 | 0.005 | -0.04 | -0.04 | 0.155 | 0.17 | 0.27 | 0.2 | 0.04 | 0.15 | 0.09 |
| 8 | 40399069 | rs721176 | -0.574 | -0.2 | -0.093 | -0.468 | 0.084 | 0.062 | 0.062 | 3.099 | 0.05 | 0.45 | 0.04 | 0 | 0.01 | 0.03 |
| 8 | 40401035 | rs1429948 | 0.46 | -0.071 | -1.493 | -0.294 | -0.042 | -0.064 | -0.064 | 0.005 | 0.11 | 0 | 0.09 | 0.19 | 0.14 | 0.07 |
| 8 | 40406406 | rs11996359 | -0.215 | -0.613 | -1.104 | -0.89 | 0.016 | 0.012 | 0.012 | 0.295 | 0.2 | 0.09 | 0.08 | 0 | 0.13 | 0.06 |
| 8 | 40411787 | rs2599675 | 0.657 | -0.435 | -1.647 | -0.66 | 0.06 | 0.083 | 0.083 | 2.241 | 0.67 | 0.2 | 0.82 | 0.84 | 0.81 | 0.77 |
| 8 | 40417433 | rs1429949 | 0.649 | 0.299 | -1.229 | -0.246 | 0.014 | 0.04 | 0.04 | 0.727 | 0.65 | 0.27 | 0.79 | 0.77 | 0.76 | 0.78 |
| 8 | 40420851 | rs10093578 | -0.601 | 0.53 | -0.794 | 0.027 | 0.003 | -0.028 | -0.028 | 0 | 0.07 | 0.1 | 0.05 | 0.01 | 0.06 | 0.04 |
| 8 | 40423517 | rs2722413 | -0.337 | 1.161 | -1.493 | 0.449 | -0.097 | -0.077 | -0.077 | 0.361 | 0.17 | 0.02 | 0.19 | 0.25 | 0.3 | 0.23 |
| 8 | 40431991 | rs10088250 | 0.163 | 0.252 | -0.991 | 0.631 | -0.013 | 0.003 | 0.003 | 0 | 0.14 | 0.15 | 0.22 | 0.15 | 0.06 | 0.13 |
| 8 | 40436774 | rs10098129 | 0.175 | 0.251 | -0.987 | 0.659 | -0.011 | 0.003 | 0.003 | 0 | 0.14 | 0.15 | 0.22 | 0.16 | 0.06 | 0.13 |
| 8 | 40437853 | rs2589934 | -0.96 | -0.196 | -1.506 | 0.51 | -0.034 | 0.009 | 0.009 | 0.202 | 0.65 | 0.8 | 0.66 | 0.65 | 0.6 | 0.62 |
| 8 | 40438757 | rs17851751 | 1.483 | -0.342 | -1.64 | -0.053 | -0.072 | 0.108 | 0.108 | 0 | 0.23 | 0.07 | 0.21 | 0 | 0.31 | 0.16 |
| 8 | 40439463 | rs1451687 | -0.115 | -0.741 | -0.821 | 0.018 | 0.001 | -0.006 | -0.006 | 0 | 0.56 | 0.33 | 0.61 | 0.42 | 0.57 | 0.57 |
| 8 | 40440413 | rs7016856 | 0.714 | -0.81 | -0.37 | 0.066 | -0.03 | 0.066 | 0.066 | 0 | 0.13 | 0.32 | 0.06 | 0.17 | 0.06 | 0.08 |
| 8 | 40442565 | rs10958623 | 1.71 | -0.68 | -0.483 | 0.082 | -0.071 | 0.11 | 0.11 | 0 | 0.23 | 0.07 | 0.22 | 0 | 0.31 | 0.16 |
| 8 | 40442938 | rs1562906 | 0.603 | -0.7 | -0.267 | 0.365 | -0.007 | 0.077 | 0.077 | 0 | 0.07 | 0.1 | 0.12 | 0.42 | 0.05 | 0.19 |
| 8 | 40446494 | rs10504028 | 1.211 | -0.564 | -0.109 | 0.29 | -0.001 | -0.001 | -0.001 | 0.02 | 0.12 | 0.08 | 0.21 | 0.12 | 0.06 | 0.13 |
| 8 | 40451400 | rs4737151 | -0.391 | -0.462 | 0.187 | 0.367 | -0.014 | 0.178 | 0.178 | 0 | 0.11 | 0.22 | 0.32 | 0.57 | 0.08 | 0.26 |
| 8 | 40452558 | rs16889640 | 0.966 | -0.65 | 0.472 | 0.28 | 0.005 | -0.006 | -0.006 | 0 | 0.13 | 0.08 | 0.05 | 0.17 | 0.07 | 0.08 |
| 8 | 40456385 | rs6995580 | 1.251 | -0.754 | 0.112 | 0.1 | 0.006 | 0.023 | 0.023 | 0 | 0.26 | 0.18 | 0.16 | 0.03 | 0.2 | 0.19 |
| 8 | 40456995 | rs4736874 | 0.025 | -0.75 | 0.123 | 0.103 | -0.006 | 0.004 | 0.004 | 0 | 0.37 | 0.42 | 0.49 | 0.59 | 0.28 | 0.46 |
| 8 | 40462615 | rs1111280 | 0.638 | -0.725 | 0.157 | 0.089 | 0.053 | 0.159 | 0.159 | 1.001 | 0.47 | 0.15 | 0.44 | 0.25 | 0.65 | 0.46 |
| 8 | 40465121 | rs7843803 | 0.969 | -0.694 | 0.182 | 0.05 | -0.041 | 0.059 | 0.059 | 0.343 | 0.14 | 0.31 | 0.06 | 0.17 | 0.07 | 0.08 |
| 8 | 40466720 | rs2254577 | -1.222 | -0.577 | 0.048 | 0.012 | -0.001 | -0.024 | -0.024 | 2.911 | 0.73 | 0.72 | 0.82 | 0.97 | 0.79 | 0.81 |
| 8 | 40467685 | rs1157732 | -1.187 | -0.565 | 0.077 | 0.011 | 0 | -0.024 | -0.024 | 0 | 0.73 | 0.72 | 0.82 | 0.97 | 0.8 | 0.81 |
| 8 | 40478096 | rs2599693 | 0.856 | -0.314 | -0.209 | 0.121 | -0.067 | -0.05 | -0.05 | 0.184 | 0.15 | 0.03 | 0.21 | 0.25 | 0.29 | 0.26 |
| 8 | 40479828 | rs10429444 | 0.089 | -0.511 | -0.104 | 0.172 | -0.003 | 0.003 | 0.003 | 0 | 0.08 | 0.1 | 0.21 | 0.1 | 0.05 | 0.15 |
| 8 | 40481476 | rs1823642 | -0.27 | -0.578 | -0.041 | 0.161 | 0.025 | 0.03 | 0.03 | 0.779 | 0.59 | 0.38 | 0.75 | 0.8 | 0.73 | 0.74 |
| 8 | 40482871 | rs2722429 | -0.444 | -0.562 | 0.04 | 0.189 | -0.049 | 0.149 | 0.149 | 0.008 | 0.82 | 0.55 | 0.73 | 0.73 | 0.9 | 0.8 |
| 8 | 40484239 | rs2722425 | -0.437 | -0.565 | 0.091 | 0.115 | -0.045 | 0.118 | 0.118 | 0 | 0.82 | 0.59 | 0.72 | 0.74 | 0.9 | 0.79 |
| 8 | 40488149 | rs2722423 | 1.258 | -0.103 | -0.106 | 0.268 | -0.027 | -0.025 | -0.025 | 0 | 0.06 | 0.01 | 0.08 | 0.07 | 0.12 | 0.08 |
| 8 | 40489691 | rs2599644 | -0.151 | -0.445 | 0.02 | 0.181 | 0.043 | 0.021 | 0.021 | 0.514 | 0.55 | 0.34 | 0.55 | 0.71 | 0.7 | 0.6 |
| 8 | 40490793 | rs10504033 | 1.197 | -0.439 | -0.211 | 0.11 | -0.039 | 0.158 | 0.158 | 0 | 0.32 | 0.11 | 0.21 | 0 | 0.33 | 0.19 |
| 8 | 40494700 | rs10092513 | 0.437 | -0.535 | 0.331 | 0.789 | 0.007 | 0.034 | 0.034 | 0 | 0.31 | 0.22 | 0.39 | 0.12 | 0.25 | 0.33 |
| 8 | 40499624 | rs12114099 | 0.461 | -0.547 | 0.339 | 0.822 | -0.035 | 0.04 | 0.04 | 0 | 0.14 | 0.25 | 0.06 | 0.17 | 0.07 | 0.09 |
| 8 | 40499961 | rs2599656 | 0.25 | -0.53 | -0.074 | 0.811 | 0.017 | 0.011 | 0.011 | 0.056 | 0.54 | 0.43 | 0.54 | 0.72 | 0.68 | 0.58 |
| 8 | 40501168 | rs2599658 | 0.375 | -0.456 | -0.43 | 0.897 | 0.07 | 0.132 | 0.132 | 0.005 | 0.5 | 0.25 | 0.43 | 0.27 | 0.65 | 0.46 |
| 8 | 40515088 | rs1823643 | 0.159 | -0.052 | -0.12 | 0.439 | -0.039 | 0.092 | 0.092 | 0.022 | 0.85 | 0.63 | 0.92 | 0.8 | 0.93 | 0.91 |
| 8 | 40517786 | rs2589893 | -0.141 | 0.326 | -0.998 | -0.018 | 0.004 | 0.072 | 0.072 | 0 | 0.42 | 0.21 | 0.28 | 0.18 | 0.43 | 0.24 |
| 8 | 40519227 | rs6474264 | -0.386 | -0.594 | -0.743 | -0.017 | -0.005 | 0 | 0 | 0 | 0.24 | 0.22 | 0.13 | 0.19 | 0.12 | 0.17 |
| 8 | 40519461 | rs2599664 | -0.43 | -0.963 | -0.357 | -0.374 | -0.013 | -0.031 | -0.031 | 0 | 0.83 | 0.71 | 0.87 | 0.92 | 0.87 | 0.84 |
| 8 | 40525615 | rs2122943 | 1.496 | -0.245 | -0.397 | -0.164 | -0.044 | 0.059 | 0.059 | 0 | 0.35 | 0.15 | 0.35 | 0.24 | 0.43 | 0.26 |
| 8 | 40527778 | rs2599672 | 0.44 | -0.256 | 0.157 | -0.056 | 0.006 | 0.002 | 0.002 | 0 | 0.59 | 0.49 | 0.47 | 0.42 | 0.55 | 0.41 |
| 8 | 40530253 | rs4374978 | -2.305 | -0.129 | 0.158 | -0.006 | 0.008 | -0.002 | -0.002 | 0 | 0.9 | 0.9 | 0.89 | 0.97 | 0.81 | 0.89 |

| | | | | | | | | | | | | | | | | |
|---|---|---|---|---|---|---|---|---|---|---|---|---|---|---|---|---|
| 8 | 40532142 | rs10504034 | 0.296 | -0.394 | 0.453 | -0.136 | -0.038 | 0.013 | 0.013 | 0 | 0.27 | 0.42 | 0.2 | 0.26 | 0.17 | 0.27 |
| 8 | 40532665 | rs4541895 | -0.459 | 0.013 | 0.373 | 0.033 | -0.029 | 0.042 | 0.042 | 0 | 0.78 | 0.86 | 0.85 | 0.97 | 0.74 | 0.85 |
| 8 | 40535838 | rs7007113 | 0.742 | -0.02 | 0.454 | 0.054 | -0.029 | -0.056 | -0.056 | 2.821 | 0.65 | 0.79 | 0.55 | 0.5 | 0.61 | 0.53 |
| 8 | 40539356 | rs4301441 | -0.075 | 0.071 | 0.965 | 0.359 | -0.015 | 0.124 | 0.124 | 0 | 0.14 | 0.22 | 0.31 | 0.47 | 0.13 | 0.32 |
| 8 | 40541533 | rs10958629 | -0.152 | 0.156 | 0.92 | 0.351 | 0.008 | -0.003 | -0.003 | 0 | 0.78 | 0.72 | 0.79 | 0.83 | 0.68 | 0.71 |
| 8 | 40542986 | rs4576415 | 0.137 | -0.182 | 1.034 | 0.325 | 0.025 | -0.031 | -0.031 | 0 | 0.71 | 0.56 | 0.72 | 0.83 | 0.62 | 0.67 |
| 8 | 40555742 | rs4496956 | -0.319 | 0.197 | 0.862 | 0.385 | 0.05 | -0.031 | -0.031 | 0 | 0.31 | 0.45 | 0.31 | 0.19 | 0.48 | 0.36 |
| 8 | 40562776 | rs6474267 | 0.001 | 0.036 | 1.123 | 0.947 | -0.007 | 0.269 | 0.269 | 0 | 0.09 | 0.29 | 0.28 | 0.37 | 0.07 | 0.22 |
| 8 | 40569004 | rs10089331 | 0.613 | -0.386 | 0.777 | 0.661 | 0.014 | 0.01 | 0.01 | 0 | 0.47 | 0.4 | 0.62 | 0.62 | 0.44 | 0.54 |
| 8 | 40572816 | rs9643878 | -0.888 | -0.618 | 0.653 | 0.648 | -0.004 | 0.05 | 0.05 | 0 | 0.56 | 0.77 | 0.56 | 0.66 | 0.55 | 0.6 |
| 8 | 40573190 | rs6988966 | 0.469 | -0.815 | 1.119 | 0.928 | 0.014 | 0.303 | 0.303 | 0 | 0.08 | 0.28 | 0.26 | 0.37 | 0.07 | 0.2 |
| 8 | 40582822 | rs4535727 | -0.334 | -0.897 | 1.075 | 0.808 | -0.038 | 0.041 | 0.041 | 0 | 0.32 | 0.43 | 0.35 | 0.57 | 0.19 | 0.36 |
| 8 | 40594029 | rs11785191 | 1.364 | -0.724 | 0.71 | 0.832 | 0 | 0.082 | 0.082 | 1.336 | 0.39 | 0.13 | 0.33 | 0.23 | 0.36 | 0.3 |
| 8 | 40596858 | rs6474274 | -0.552 | -0.833 | 0.881 | 0.937 | 0.012 | 0.012 | 0.012 | 0 | 0.78 | 0.85 | 0.91 | 0.81 | 0.87 | 0.83 |
| 8 | 40606053 | rs10092049 | 0.145 | -0.796 | 0.936 | 0.829 | -0.036 | 0.03 | 0.03 | 0 | 0.7 | 0.59 | 0.65 | 0.62 | 0.81 | 0.63 |
| 8 | 40620539 | rs11781404 | 0.683 | -0.498 | 0.446 | 0.525 | -0.012 | 0.018 | 0.018 | 0 | 0.2 | 0.17 | 0.2 | 0.06 | 0.26 | 0.21 |
| 8 | 40636042 | rs6987241 | 0.805 | -0.534 | 0.455 | 0.577 | -0.012 | 0.015 | 0.015 | 0 | 0.2 | 0.21 | 0.2 | 0.07 | 0.26 | 0.21 |
| 8 | 40637448 | rs4487749 | -0.527 | -0.958 | 0.051 | 0.12 | -0.018 | 0.247 | 0.247 | 0 | 0.49 | 0.22 | 0.49 | 0.08 | 0.48 | 0.44 |
| 8 | 40640413 | rs7820759 | -0.885 | -0.604 | 0.532 | 0.787 | 0.02 | 0.023 | 0.023 | 0 | 0.79 | 0.71 | 0.6 | 0.7 | 0.75 | 0.73 |
| 8 | 40666189 | rs11777437 | -0.34 | -0.689 | 0.862 | 0.754 | 0.01 | 0.084 | 0.084 | 0 | 0.74 | 0.57 | 0.57 | 0.55 | 0.7 | 0.59 |
| 8 | 40671114 | rs7835606 | 0.52 | -0.913 | 0.645 | 0.65 | -0.028 | 0.046 | 0.046 | 0 | 0.54 | 0.24 | 0.46 | 0.33 | 0.5 | 0.42 |
| 8 | 40673248 | rs17561583 | 0.701 | -0.922 | 0.539 | 0.587 | 0.001 | 0.022 | 0.022 | 0 | 0.13 | 0.02 | 0.09 | 0.08 | 0.11 | 0.15 |
| 8 | 40673946 | rs10481311 | 0.442 | -0.849 | 0.563 | 0.596 | -0.027 | 0.045 | 0.045 | 0 | 0.54 | 0.25 | 0.46 | 0.33 | 0.5 | 0.42 |
| 8 | 40676775 | rs16889988 | -0.109 | -0.811 | 0.269 | 0.407 | 0.001 | -0.007 | -0.007 | 0 | 0.24 | 0.21 | 0.11 | 0.32 | 0.25 | 0.3 |
| 8 | 40683338 | rs872054 | 1.657 | -0.827 | -0.302 | -0.001 | -0.041 | 0.129 | 0.129 | 0.034 | 0.33 | 0.13 | 0.3 | 0 | 0.35 | 0.12 |
| 8 | 40686754 | rs13270397 | 0.124 | -0.471 | 0.09 | 0.751 | -0.001 | 0.004 | 0.004 | 0 | 0.14 | 0.12 | 0.11 | 0.09 | 0.14 | 0.15 |
| 8 | 40687433 | rs10113804 | -0.32 | -0.318 | 0.192 | 0.611 | -0.016 | 0.018 | 0.018 | 0.832 | 0.23 | 0.32 | 0.19 | 0.3 | 0.18 | 0.3 |
| 8 | 40693883 | rs16890016 | 0.194 | 0.14 | -0.059 | 0.893 | -0.024 | 0.164 | 0.164 | 0 | 0.05 | 0.12 | 0.02 | 0.37 | 0.03 | 0.16 |
| 8 | 40696404 | rs870478 | 0.264 | 0.658 | -0.348 | -0.459 | 0.003 | 0.057 | 0.057 | 0 | 0.51 | 0.42 | 0.62 | 0.76 | 0.53 | 0.66 |
| 8 | 40698232 | rs1028014 | -0.251 | 0.649 | -1.007 | -0.174 | 0.004 | 0.01 | 0.01 | 0 | 0.82 | 0.79 | 0.92 | 0.76 | 0.82 | 0.79 |
| 8 | 40699841 | rs17562411 | 0.28 | 0.732 | -1.467 | -0.338 | -0.026 | 0.121 | 0.121 | 0 | 0.25 | 0.07 | 0.25 | 0 | 0.23 | 0.12 |
| 8 | 40701766 | rs6992115 | -0.393 | 0.518 | -0.812 | 0.574 | -0.001 | 0 | 0 | 0 | 0.35 | 0.38 | 0.25 | 0.58 | 0.32 | 0.36 |
| 8 | 40703367 | rs7837405 | -1.114 | 0.973 | -0.357 | 0.602 | 0.021 | 0.041 | 0.041 | 0.002 | 0.12 | 0.22 | 0.21 | 0.24 | 0.12 | 0.19 |
| 8 | 40704851 | rs868586 | -0.061 | 0.994 | -1.097 | -0.104 | -0.004 | 0.126 | 0.126 | 1.375 | 0.5 | 0.73 | 0.55 | 0.2 | 0.54 | 0.54 |
| 8 | 40717349 | rs1467108 | -0.499 | 0.95 | -1.755 | 0.318 | 0.009 | -0.008 | -0.008 | 0 | 0.7 | 0.76 | 0.81 | 0.6 | 0.81 | 0.77 |
| 8 | 40717778 | rs998609 | -0.295 | 0.999 | -1.504 | -0.337 | 0.008 | 0.004 | 0.004 | 0.216 | 0.52 | 0.63 | 0.63 | 0.57 | 0.64 | 0.59 |
| 8 | 40723038 | rs16890054 | 0.855 | 0.824 | -0.77 | -0.55 | 0.008 | -0.019 | -0.019 | 0.868 | 0.16 | 0.25 | 0.24 | 0.1 | 0.23 | 0.09 |
| 8 | 40727444 | rs10504036 | 1.095 | 0.924 | -0.774 | -0.491 | -0.101 | -0.117 | -0.117 | 0 | 0.2 | 0.02 | 0.27 | 0.4 | 0.31 | 0.24 |
| 8 | 40739249 | rs1533178 | 1.206 | 0.764 | -0.596 | -0.649 | -0.008 | -0.004 | -0.004 | 0 | 0.08 | 0.13 | 0.14 | 0.08 | 0.08 | 0.05 |
| 8 | 40740473 | rs9298606 | 0.684 | 0.778 | -0.289 | -0.633 | -0.006 | 0.001 | 0.001 | 0 | 0.11 | 0.17 | 0.07 | 0.07 | 0.08 | 0.17 |
| 8 | 40756407 | rs11985600 | 0.664 | 0.769 | -0.198 | -0.601 | 0.002 | 0.016 | 0.016 | 0 | 0.06 | 0.16 | 0.05 | 0.04 | 0.04 | 0.12 |
| 8 | 40766191 | rs7015608 | -0.024 | 0.593 | -0.862 | -0.879 | -0.034 | -0.022 | -0.022 | 0 | 0.36 | 0.31 | 0.49 | 0.46 | 0.49 | 0.35 |
| 8 | 40773508 | rs7817426 | 0.376 | 0.462 | -0.639 | -0.804 | 0.005 | -0.033 | -0.033 | 0 | 0.15 | 0.29 | 0.22 | 0.04 | 0.18 | 0.12 |
| 8 | 40781558 | rs16890097 | -0.337 | 0.133 | -0.619 | -0.742 | 0.047 | -0.002 | -0.002 | 0 | 0.51 | 0.41 | 0.61 | 0.52 | 0.66 | 0.55 |
| 8 | 40781583 | rs16890099 | 0.872 | 0.343 | -0.566 | -0.904 | -0.001 | -0.014 | -0.014 | 0 | 0.17 | 0.12 | 0.18 | 0.03 | 0.14 | 0.15 |
| 8 | 40783739 | rs11786553 | 0.286 | 0.427 | -1.083 | -0.942 | -0.032 | 0.034 | 0.034 | 0 | 0.15 | 0.01 | 0.32 | 0.06 | 0.13 | 0.17 |
| 8 | 40786645 | rs2055949 | -0.824 | 0.676 | -0.891 | -1.611 | -0.001 | 0.001 | 0.001 | 0 | 0.42 | 0.63 | 0.54 | 0.1 | 0.41 | 0.43 |
| 8 | 40787440 | rs10109724 | -0.453 | 0.594 | -0.625 | -1.694 | 0 | -0.004 | -0.004 | 0 | 0.33 | 0.32 | 0.47 | 0.09 | 0.31 | 0.4 |
| 8 | 40789256 | rs11783346 | -1.125 | 0.708 | -1.089 | -1.764 | -0.028 | 0.218 | 0.218 | 0 | 0.48 | 0.82 | 0.59 | 0.1 | 0.49 | 0.48 |
| 8 | 40793533 | rs10112489 | -0.975 | 0.892 | -1.691 | -1.002 | -0.01 | 0.06 | 0.06 | 0 | 0.54 | 0.8 | 0.67 | 0.62 | 0.54 | 0.6 |
| 8 | 40801951 | rs963520 | -1.733 | 0.892 | -2.322 | -1.044 | -0.041 | -0.028 | -0.028 | 0 | 0.74 | 0.98 | 0.77 | 0.74 | 0.74 | 0.81 |
| 8 | 40805151 | rs7827529 | 0.746 | 1.131 | -1.576 | -0.969 | 0.001 | -0.024 | -0.024 | 0 | 0.22 | 0.3 | 0.14 | 0.11 | 0.23 | 0.22 |
| 8 | 40805213 | rs7826645 | 0.442 | 1.103 | -1.502 | -0.984 | 0.041 | -0.033 | -0.033 | 0.245 | 0.07 | 0.13 | 0.07 | 0 | 0.12 | 0.09 |
| 8 | 40806515 | rs6988809 | 0.561 | 1.042 | -1.541 | -0.953 | 0.004 | -0.019 | -0.019 | 2.849 | 0.25 | 0.3 | 0.17 | 0.11 | 0.27 | 0.26 |
| 8 | 40808054 | rs4736895 | 0.666 | 0.813 | -1.352 | -0.198 | 0 | 0.013 | 0.013 | 0 | 0.19 | 0.18 | 0.14 | 0.36 | 0.2 | 0.22 |
| 8 | 40809189 | rs10091158 | 1.398 | 0.932 | -1.284 | -0.046 | -0.001 | -0.036 | -0.036 | 0.033 | 0.16 | 0.04 | 0.32 | 0.16 | 0.12 | 0.22 |
| 8 | 40811968 | rs6995294 | -0.11 | 0.837 | -0.783 | 0.123 | -0.02 | 0.044 | 0.044 | 0.286 | 0.05 | 0.1 | 0.03 | 0.18 | 0.02 | 0.11 |
| 8 | 40821132 | rs10089590 | 0.678 | 0.808 | -0.895 | 0.362 | 0.003 | -0.009 | -0.009 | 0 | 0.1 | 0.06 | 0.16 | 0.31 | 0.1 | 0.14 |
| 8 | 40821738 | rs7836545 | -0.037 | 0.688 | -0.277 | -0.305 | -0.013 | -0.033 | -0.033 | 0 | 0.32 | 0.52 | 0.24 | 0.23 | 0.28 | 0.36 |
| 8 | 40831222 | rs2353240 | -2.305 | 0.92 | -0.505 | -0.492 | -0.057 | 0.095 | 0.095 | 0 | 0.84 | 0.99 | 0.87 | 1 | 0.78 | 0.95 |
| 8 | 40831294 | rs10094981 | 0.443 | 0.877 | -0.391 | -0.373 | -0.021 | -0.025 | -0.025 | 0 | 0.29 | 0.51 | 0.22 | 0.23 | 0.26 | 0.32 |
| 8 | 40834880 | rs11987006 | 2.077 | 0.778 | -0.276 | -0.382 | 0.005 | 0.014 | 0.014 | 0 | 0.17 | 0.16 | 0.1 | 0.04 | 0.16 | 0.15 |
| 8 | 40836593 | rs3847180 | 0.263 | 0.863 | -0.184 | -0.299 | -0.002 | -0.006 | -0.006 | 0.449 | 0.39 | 0.34 | 0.49 | 0.53 | 0.41 | 0.42 |
| 8 | 40837275 | rs3842923 | -1.002 | 0.98 | 0.077 | -0.384 | -0.001 | 0 | 0 | 0 | 0.7 | 0.73 | 0.73 | 0.76 | 0.69 | 0.78 |
| 8 | 40844800 | rs10958646 | -1.313 | 1.012 | -0.358 | -0.361 | -0.008 | -0.002 | -0.002 | 0 | 0.72 | 0.86 | 0.75 | 0.76 | 0.72 | 0.78 |
| 8 | 40845908 | rs3847182 | -0.057 | 1.078 | -0.488 | -0.383 | -0.007 | 0.021 | 0.021 | 0 | 0.43 | 0.32 | 0.52 | 0.7 | 0.45 | 0.54 |
| 8 | 40855553 | rs10958648 | 0.057 | 1.156 | -0.719 | -0.481 | -0.026 | 0.055 | 0.055 | 0 | 0.58 | 0.33 | 0.58 | 0.73 | 0.54 | 0.67 |
| 8 | 40858257 | rs1379137 | -1.327 | 1.189 | -0.965 | -0.497 | -0.009 | 0.017 | 0.017 | 0 | 0.73 | 0.89 | 0.78 | 0.78 | 0.73 | 0.78 |
| 8 | 40866575 | rs10504037 | 0.564 | 1.23 | -0.924 | -0.412 | -0.019 | 0.016 | 0.016 | 0.333 | 0.11 | 0.08 | 0.14 | 0.05 | 0.16 | 0.11 |

| | | | | | | | | | | | | | | | | |
|---|---|---|---|---|---|---|---|---|---|---|---|---|---|---|---|---|
| 8 | 40880501 | rs1456767 | 0.808 | 1.092 | -1.307 | -0.35 | -0.017 | -0.136 | -0.136 | 0 | 0.29 | 0.06 | 0.41 | 0.52 | 0.28 | 0.43 |
| 8 | 40881755 | rs11995320 | 1.583 | 1.021 | -1.273 | -0.504 | 0.027 | 0.003 | 0.003 | 0 | 0.23 | 0.18 | 0.12 | 0.21 | 0.17 | 0.23 |
| 8 | 40883577 | rs7826932 | 0.48 | 1.128 | -1.168 | -0.36 | -0.014 | -0.112 | -0.112 | 0 | 0.3 | 0.06 | 0.31 | 0.6 | 0.29 | 0.43 |
| 8 | 40885578 | rs1379134 | -2.189 | 1.367 | -1.144 | -0.602 | -0.043 | 0.093 | 0.093 | 0 | 0.84 | 0.99 | 0.87 | 1 | 0.82 | 0.96 |
| 8 | 40889727 | rs6474296 | -0.002 | 1.329 | -0.497 | -0.306 | -0.009 | 0.081 | 0.081 | 0 | 0.44 | 0.73 | 0.35 | 0.17 | 0.44 | 0.36 |
| 8 | 40895774 | rs7011811 | -1.436 | 1.243 | -0.455 | -0.204 | -0.001 | 0.007 | 0.007 | 0 | 0.72 | 0.79 | 0.65 | 0.77 | 0.72 | 0.78 |
| 8 | 40896048 | rs4736790 | 0.203 | 1.489 | -0.171 | -0.169 | 0.01 | -0.013 | -0.013 | 0 | 0.11 | 0.12 | 0.14 | 0.05 | 0.16 | 0.12 |
| 8 | 40914727 | rs7813807 | -0.586 | 1.491 | -0.147 | -0.143 | -0.005 | 0.004 | 0.004 | 0 | 0.39 | 0.34 | 0.47 | 0.68 | 0.45 | 0.54 |
| 8 | 40931683 | rs10097162 | -1.365 | 1.415 | 0.091 | -0.233 | 0 | 0.026 | 0.026 | 0 | 0.69 | 0.79 | 0.66 | 0.8 | 0.72 | 0.76 |
| 8 | 40933986 | rs13257857 | 1.557 | 1.5 | -0.349 | -0.239 | 0.006 | 0.054 | 0.054 | 0 | 0.26 | 0.19 | 0.16 | 0.06 | 0.24 | 0.16 |
| 8 | 40934691 | rs11775247 | 0.35 | 2.027 | -0.107 | 0.967 | -0.017 | -0.039 | -0.039 | 0 | 0.08 | 0.01 | 0.12 | 0.3 | 0.1 | 0.14 |
| 8 | 40937651 | rs10504039 | -0.578 | 2.089 | -0.057 | 0.87 | 0.031 | -0.03 | -0.03 | 0 | 0.12 | 0.21 | 0.14 | 0.05 | 0.17 | 0.1 |
| 8 | 40949654 | rs16890282 | 0.252 | 1.821 | -0.767 | 1.054 | -0.063 | -0.111 | -0.111 | 0.512 | 0.19 | 0.01 | 0.19 | 0.35 | 0.2 | 0.27 |
| 8 | 40951638 | rs4736917 | -0.347 | 1.829 | -1.212 | 0.223 | -0.044 | -0.082 | -0.082 | 0 | 0.28 | 0.06 | 0.31 | 0.65 | 0.3 | 0.42 |
| 8 | 40954561 | rs4554449 | -0.382 | 1.899 | -1.164 | 0.197 | -0.048 | -0.058 | -0.058 | 0 | 0.29 | 0.09 | 0.33 | 0.65 | 0.32 | 0.44 |
| 8 | 40957887 | rs17652249 | -0.307 | 1.669 | -1.102 | 0.228 | -0.035 | -0.055 | -0.055 | 0 | 0.28 | 0.09 | 0.29 | 0.65 | 0.3 | 0.43 |
| 8 | 40965681 | rs6474300 | -1.359 | 1.515 | -0.278 | 0.027 | 0.029 | -0.048 | -0.048 | 0.438 | 0.14 | 0.28 | 0.19 | 0.05 | 0.19 | 0.12 |
| 8 | 40967164 | rs12550040 | -0.995 | 1.702 | -1.154 | 0.104 | -0.011 | 0.012 | 0.012 | 0.119 | 0.13 | 0.08 | 0.18 | 0.05 | 0.19 | 0.12 |
| 8 | 40978059 | rs7007458 | -0.866 | 1.774 | -1.084 | 0.26 | 0.001 | -0.001 | -0.001 | 4.516 | 0.14 | 0.14 | 0.14 | 0.1 | 0.19 | 0.16 |
| 8 | 40978302 | rs13276016 | -0.16 | 2.294 | -1.738 | 0.924 | -0.059 | -0.093 | -0.093 | 0 | 0.16 | 0.05 | 0.29 | 0.46 | 0.24 | 0.37 |
| 8 | 40978925 | rs6474309 | -0.616 | 1.456 | -1.984 | 0.207 | -0.012 | -0.033 | -0.033 | 0 | 0.3 | 0.15 | 0.38 | 0.37 | 0.31 | 0.35 |
| 8 | 40980390 | rs737505 | 0.041 | 0.746 | -1.845 | -0.878 | 0 | -0.034 | -0.034 | 0 | 0.28 | 0.2 | 0.2 | 0.04 | 0.31 | 0.16 |
| 8 | 40981047 | rs13269996 | 0.445 | 0.84 | -1.587 | -0.305 | -0.003 | 0.009 | 0.009 | 0 | 0.08 | 0.08 | 0.09 | 0.18 | 0.07 | 0.14 |
| 8 | 40984060 | rs13266712 | -0.202 | 0.278 | -1.299 | -1.314 | -0.004 | -0.084 | -0.084 | 0 | 0.54 | 0.5 | 0.7 | 0.96 | 0.58 | 0.71 |
| 8 | 40988523 | rs7827040 | -0.169 | 0.304 | -1.351 | -0.987 | -0.029 | 0.077 | 0.077 | 0.007 | 0.59 | 0.71 | 0.38 | 0.23 | 0.51 | 0.39 |
| 8 | 40995139 | rs4736795 | 0.238 | 0.418 | -1.409 | -0.212 | 0.008 | -0.08 | -0.08 | 0 | 0.25 | 0.15 | 0.23 | 0.48 | 0.22 | 0.39 |
| 8 | 40995372 | rs4736796 | -0.787 | 0.046 | -0.862 | -0.646 | -0.001 | 0.013 | 0.013 | 0.139 | 0.56 | 0.64 | 0.59 | 0.68 | 0.56 | 0.67 |
| 8 | 40996160 | rs11996523 | 0.852 | 0.221 | -1.061 | -0.717 | 0 | -0.001 | -0.001 | 0.437 | 0.16 | 0.2 | 0.23 | 0.17 | 0.18 | 0.2 |
| 8 | 41001296 | rs10958660 | -1.222 | 0.323 | -0.829 | -0.779 | -0.002 | -0.012 | -0.012 | 0 | 0.59 | 0.52 | 0.63 | 0.68 | 0.63 | 0.69 |
| 8 | 41003003 | rs12156199 | 0.941 | 0.516 | -0.765 | -0.69 | -0.003 | -0.028 | -0.028 | 0.172 | 0.14 | 0.16 | 0.1 | 0.03 | 0.14 | 0.07 |
| 8 | 41003359 | rs7812672 | 1.084 | 0.52 | -0.852 | -0.682 | 0 | -0.004 | -0.004 | 0 | 0.13 | 0.1 | 0.09 | 0.03 | 0.14 | 0.06 |
| 8 | 41004159 | rs7818335 | -0.098 | 0.528 | -0.938 | -0.541 | 0.001 | -0.025 | -0.025 | 0 | 0.34 | 0.39 | 0.38 | 0.19 | 0.39 | 0.28 |
| 8 | 41004362 | rs4736927 | -0.369 | 0.304 | -0.95 | -0.558 | 0 | 0.002 | 0.002 | 0.257 | 0.56 | 0.6 | 0.6 | 0.67 | 0.62 | 0.67 |
| 8 | 41006048 | rs11986992 | 0.047 | -0.234 | -1.055 | -1.089 | -0.007 | -0.041 | -0.041 | 0.438 | 0.44 | 0.45 | 0.27 | 0.15 | 0.4 | 0.21 |
| 8 | 41007120 | rs10958662 | 0.606 | -0.312 | -1.073 | -0.536 | -0.001 | 0.039 | 0.039 | 0 | 0.19 | 0.29 | 0.19 | 0.31 | 0.21 | 0.19 |
| 8 | 41007462 | rs12541730 | -0.319 | -0.723 | -1.294 | -0.814 | -0.011 | 0.005 | 0.005 | 0 | 0.34 | 0.39 | 0.3 | 0.42 | 0.31 | 0.39 |
| 8 | 41008081 | rs6474312 | -1.102 | -0.952 | -1.822 | -1.134 | -0.139 | -0.102 | -0.102 | 0.213 | 0.62 | 0.9 | 0.49 | 0.46 | 0.55 | 0.53 |
| 8 | 41010900 | rs4736932 | -1.032 | -0.482 | -1.67 | -0.551 | -0.089 | -0.033 | -0.033 | 0.052 | 0.7 | 0.94 | 0.66 | 0.75 | 0.63 | 0.77 |
| 8 | 41011531 | rs4392880 | -0.043 | -0.873 | -1.055 | -0.486 | 0 | 0.002 | 0.002 | 0.239 | 0.43 | 0.57 | 0.41 | 0.33 | 0.39 | 0.4 |
| 8 | 41017797 | rs10101921 | -0.081 | -1.944 | 0.051 | -1.148 | -0.024 | -0.138 | -0.138 | 0 | 0.22 | 0.52 | 0.16 | 0 | 0.2 | 0.12 |
| 8 | 41025313 | rs10102223 | -0.223 | -1.999 | 0.493 | -1.175 | -0.027 | -0.143 | -0.143 | 1.089 | 0.23 | 0.5 | 0.17 | 0 | 0.2 | 0.12 |
| 8 | 41028520 | rs6983729 | 0.394 | -2.28 | 0.276 | -1.182 | -0.009 | 0.018 | 0.018 | 0 | 0.19 | 0.21 | 0.16 | 0.33 | 0.14 | 0.31 |
| 8 | 41028701 | rs4540396 | 0.412 | -2.278 | 0.798 | -1.385 | -0.005 | -0.082 | -0.082 | 0.078 | 0.19 | 0.22 | 0.15 | 0 | 0.2 | 0.12 |
| 8 | 41033553 | rs10112327 | -0.31 | -2.253 | 0.721 | -1.384 | -0.023 | -0.121 | -0.121 | 0 | 0.23 | 0.57 | 0.18 | 0 | 0.2 | 0.12 |
| 8 | 41035938 | rs11997968 | 0.159 | -2.263 | 0.812 | -1.389 | -0.012 | -0.141 | -0.141 | 0 | 0.21 | 0.41 | 0.16 | 0 | 0.2 | 0.12 |
| 8 | 41037709 | rs7820359 | -0.778 | -2.361 | 0.248 | -1.417 | 0.011 | 0.002 | 0.002 | 1.145 | 0.29 | 0.73 | 0.21 | 0.1 | 0.22 | 0.24 |
| 8 | 41064077 | rs4736943 | -0.198 | -2.168 | 0.387 | -1.198 | -0.027 | 0.024 | 0.024 | 0 | 0.25 | 0.31 | 0.16 | 0.39 | 0.17 | 0.41 |
| 8 | 41080046 | rs4736949 | -0.727 | -1.975 | 0.257 | -1.18 | 0.014 | 0.008 | 0.008 | 0 | 0.41 | 0.64 | 0.43 | 0.31 | 0.35 | 0.43 |
| 8 | 41102437 | rs10108063 | -0.254 | -1.901 | 0.364 | -1.123 | -0.041 | 0.042 | 0.042 | 0 | 0.26 | 0.44 | 0.28 | 0.31 | 0.18 | 0.32 |
| 8 | 41106582 | rs4412355 | 0.055 | -1.999 | 0.346 | -1.284 | 0.003 | 0.036 | 0.036 | 0 | 0.31 | 0.19 | 0.2 | 0.03 | 0.34 | 0.18 |
| 8 | 41108648 | rs7823929 | -1.023 | -2.064 | 0.407 | -1.226 | 0.045 | 0.034 | 0.034 | 0 | 0.53 | 0.75 | 0.47 | 0.4 | 0.44 | 0.55 |
| 8 | 41118021 | rs4236914 | -0.106 | -1.969 | 0.807 | -1.068 | -0.002 | 0.239 | 0.239 | 0 | 0.09 | 0.27 | 0.21 | 0.28 | 0.05 | 0.24 |
| 8 | 41121280 | rs1127379 | -1.021 | -1.984 | 0.429 | -1.282 | 0.007 | 0.005 | 0.005 | 0 | 0.54 | 0.58 | 0.48 | 0.45 | 0.45 | 0.58 |
| 8 | 41128410 | rs11781990 | -0.019 | -1.972 | 0.024 | -1.381 | 0.013 | 0.181 | 0.181 | 0 | 0.38 | 0.13 | 0.2 | 0.05 | 0.37 | 0.23 |
| 8 | 41131170 | rs13254926 | -0.213 | -1.928 | 0.073 | -1.299 | 0.011 | 0.033 | 0.033 | 0 | 0.4 | 0.34 | 0.23 | 0.05 | 0.37 | 0.23 |
| 8 | 41135769 | rs7843510 | -0.808 | -1.559 | -0.182 | -0.937 | 0.017 | -0.01 | -0.01 | 0.121 | 0.55 | 0.77 | 0.45 | 0.43 | 0.44 | 0.58 |
| 8 | 41143382 | rs11783547 | -0.209 | -1.422 | -0.318 | -1.312 | 0.027 | 0.081 | 0.081 | 0 | 0.43 | 0.29 | 0.23 | 0.05 | 0.38 | 0.26 |
| 8 | 41144413 | rs6651363 | 0.153 | -1.179 | -0.594 | -1.04 | 0.022 | 0.107 | 0.107 | 0 | 0.37 | 0.13 | 0.24 | 0.17 | 0.36 | 0.21 |
| 8 | 41147578 | rs10098898 | -1.196 | -0.649 | 0.053 | -0.137 | -0.045 | 0.134 | 0.134 | 0 | 0.16 | 0.51 | 0.32 | 0.29 | 0.11 | 0.29 |
| 8 | 41150652 | rs730648 | -0.154 | -0.192 | -0.514 | -0.08 | -0.043 | -0.018 | -0.018 | 0 | 0.6 | 0.81 | 0.6 | 0.41 | 0.61 | 0.58 |
| 8 | 41155341 | rs10958672 | 0.562 | 0.113 | -0.332 | 0.142 | 0 | -0.01 | -0.01 | 0.954 | 0.51 | 0.42 | 0.32 | 0.12 | 0.53 | 0.32 |
| 8 | 41158004 | rs747417 | -0.12 | 0.7 | -0.919 | 0.517 | -0.007 | -0.006 | -0.006 | 0 | 0.64 | 0.69 | 0.61 | 0.43 | 0.62 | 0.6 |
| 8 | 41159859 | rs7832767 | -1.256 | 0.526 | -0.894 | 1.115 | -0.004 | 0.031 | 0.031 | 0 | 0.09 | 0.12 | 0.21 | 0.25 | 0.07 | 0.22 |
| 8 | 41165794 | rs968427 | -0.535 | 1.269 | -1.445 | 1.143 | -0.053 | -0.066 | -0.066 | 6.218 | 0.3 | 0.16 | 0.49 | 0.42 | 0.45 | 0.47 |
| 8 | 41174462 | rs2121605 | -0.441 | 1.38 | -1.564 | 1.125 | -0.105 | -0.131 | -0.131 | 0 | 0.25 | 0.05 | 0.39 | 0.43 | 0.4 | 0.48 |
| 8 | 41189599 | rs16890461 | -0.745 | 0.923 | -0.416 | 1.059 | -0.005 | 0.119 | 0.119 | 0 | 0.14 | 0.42 | 0.12 | 0.26 | 0.08 | 0.19 |
| 8 | 41190199 | rs16890462 | -0.775 | 0.91 | -0.409 | 1.053 | -0.007 | 0.111 | 0.111 | 0 | 0.14 | 0.39 | 0.11 | 0.25 | 0.08 | 0.19 |
| 8 | 41190971 | rs6993927 | 0.642 | 0.755 | -0.883 | 0.381 | -0.005 | -0.025 | -0.025 | 0 | 0.66 | 0.48 | 0.68 | 0.74 | 0.69 | 0.71 |
| 8 | 41203570 | rs11779994 | 0.468 | 0.301 | -1.716 | 0.221 | -0.075 | 0.048 | 0.048 | 0.019 | 0.37 | 0.17 | 0.38 | 0.25 | 0.43 | 0.24 |

| | | | | | | | | | | | | | | | |
|---|---|---|---|---|---|---|---|---|---|---|---|---|---|---|---|
| 8 | 41204825 | rs7841046 | -0.674 | -0.22 | -1.153 | -0.262 | -0.017 | -0.113 | -0.113 | 0 | 0.82 | 0.44 | 0.86 | 1 | 0.85 | 0.96 |
| 8 | 41204995 | rs12549180 | -0.077 | -0.362 | -1.469 | 0.101 | 0.002 | 0.001 | 0.001 | 0 | 0.76 | 0.31 | 0.8 | 0.79 | 0.83 | 0.79 |
| 8 | 41215593 | rs4546646 | 0.884 | -0.342 | -1.539 | 0.151 | -0.056 | 0.048 | 0.048 | 0.009 | 0.38 | 0.21 | 0.39 | 0.25 | 0.43 | 0.26 |
| 8 | 41218751 | rs16890476 | 0.942 | -0.213 | -1.667 | 0.563 | -0.064 | -0.095 | -0.095 | 0 | 0.15 | 0.01 | 0.28 | 0.32 | 0.21 | 0.32 |
| 8 | 41220162 | rs717218 | 0.577 | -0.301 | -1.656 | 0.376 | 0.016 | -0.025 | -0.025 | 0 | 0.54 | 0.22 | 0.49 | 0.46 | 0.57 | 0.42 |
| 8 | 41223011 | rs10958679 | 0.285 | -0.226 | -0.655 | 0.262 | -0.014 | 0.001 | 0.001 | 0 | 0.73 | 0.52 | 0.74 | 0.74 | 0.74 | 0.73 |
| 8 | 41228458 | rs10958680 | 0.233 | -0.167 | -0.901 | 0.151 | 0.005 | 0.002 | 0.002 | 0.243 | 0.74 | 0.23 | 0.57 | 0.77 | 0.77 | 0.85 |
| 8 | 41231323 | rs4736805 | 0.323 | 0.094 | -0.557 | 0.222 | 0.021 | 0.064 | 0.064 | 0.166 | 0.92 | 0.77 | 0.89 | 0.81 | 0.9 | 0.88 |
| 8 | 41231950 | rs2883471 | 0.855 | 0.205 | -1.489 | -0.338 | -0.057 | 0.011 | 0.011 | 0 | 0.3 | 0.11 | 0.26 | 0.2 | 0.34 | 0.21 |
| 8 | 41232360 | rs2069122 | 0.709 | 0.225 | -1.264 | -0.501 | -0.004 | -0.006 | -0.006 | 0.09 | 0.08 | 0.06 | 0.11 | 0 | 0.1 | 0.09 |
| 8 | 41252090 | rs6474334 | 0.319 | -0.364 | -1.152 | -0.955 | -0.012 | -0.103 | -0.103 | 0.743 | 0.64 | 0.4 | 0.53 | 0.97 | 0.68 | 0.85 |
| 8 | 41256225 | rs7830124 | 0.962 | -0.568 | -0.865 | -1.101 | 0.006 | -0.064 | -0.064 | 0.308 | 0.11 | 0.31 | 0.1 | 0 | 0.1 | 0.03 |
| 8 | 41260259 | rs973441 | 1.008 | -0.662 | -1.228 | -0.833 | 0.01 | 0.024 | 0.024 | 0 | 0.57 | 0.36 | 0.44 | 0.76 | 0.6 | 0.68 |
| 8 | 41261544 | rs10097592 | 1.155 | -0.609 | -1.568 | -0.703 | 0.009 | 0.018 | 0.018 | 0.197 | 0.29 | 0.31 | 0.3 | 0.44 | 0.32 | 0.25 |
| 8 | 41273233 | rs1447214 | 0.219 | -0.693 | -1.204 | -0.86 | -0.058 | -0.13 | -0.13 | 0 | 0.77 | 0.43 | 0.66 | 0.97 | 0.83 | 0.93 |
| 8 | 41292579 | rs11774152 | -0.03 | -0.581 | -1.153 | -0.842 | -0.009 | -0.137 | -0.137 | 0 | 0.71 | 0.56 | 0.57 | 0.97 | 0.71 | 0.87 |
| 8 | 41298716 | rs1425778 | 0.108 | -0.624 | -0.823 | -0.724 | -0.003 | -0.02 | -0.02 | 0.531 | 0.94 | 0.81 | 0.78 | 0.97 | 0.95 | 0.96 |
| 8 | 41302561 | rs13260111 | 1.255 | -0.661 | -0.942 | -0.739 | -0.002 | -0.009 | -0.009 | 0 | 0.06 | 0.06 | 0.04 | 0 | 0.02 | 0.02 |
| 8 | 41303896 | rs7018403 | -0.128 | -0.7 | -0.981 | -0.878 | -0.025 | -0.171 | -0.171 | 1.457 | 0.29 | 0.5 | 0.26 | 0 | 0.25 | 0.11 |
| 8 | 41308421 | rs4130040 | 1.407 | -0.609 | -1.383 | -0.746 | 0.015 | -0.009 | -0.009 | 0 | 0.26 | 0.31 | 0.2 | 0.14 | 0.29 | 0.18 |
| 8 | 41308927 | rs6474342 | 0.792 | -0.608 | -1.269 | -0.438 | -0.004 | -0.008 | -0.008 | 0.21 | 0.61 | 0.41 | 0.44 | 0.68 | 0.66 | 0.79 |
| 8 | 41310010 | rs1030370 | 0.219 | -0.653 | -1.209 | -0.444 | -0.036 | 0.016 | 0.016 | 0 | 0.74 | 0.57 | 0.57 | 0.68 | 0.8 | 0.87 |
| 8 | 41312419 | rs6474346 | 0.197 | -0.653 | -1.205 | -0.444 | -0.028 | 0.012 | 0.012 | 0 | 0.74 | 0.6 | 0.57 | 0.68 | 0.8 | 0.87 |
| 8 | 41314993 | rs1425781 | -0.954 | -0.653 | -1.22 | -0.431 | 0.003 | 0.088 | 0.088 | 0 | 0.94 | 0.9 | 0.9 | 0.71 | 0.96 | 0.92 |
| 8 | 41316426 | rs7018401 | -0.936 | -0.662 | -1.228 | -0.482 | 0.014 | 0.027 | 0.027 | 0 | 0.81 | 0.71 | 0.77 | 0.71 | 0.81 | 0.85 |
| 8 | 41321076 | rs2042667 | 0.96 | -0.693 | -1.252 | -0.51 | 0.001 | 0.001 | 0.001 | 0.133 | 0.61 | 0.39 | 0.44 | 0.67 | 0.65 | 0.79 |
| 8 | 41322336 | rs16890610 | 1.447 | -0.864 | -1.434 | -0.656 | -0.009 | -0.055 | -0.055 | 0.166 | 0.1 | 0.01 | 0.09 | 0.21 | 0.08 | 0.16 |
| 8 | 41324400 | rs11986663 | 0.762 | -0.798 | -1.097 | -0.817 | -0.036 | -0.071 | -0.071 | 0 | 0.14 | 0.28 | 0.28 | 0.03 | 0.12 | 0.04 |
| 8 | 41328522 | rs6998193 | 1.962 | -0.706 | -1.126 | -0.765 | 0.007 | -0.051 | -0.051 | 0.804 | 0.13 | 0.18 | 0.13 | 0 | 0.15 | 0.08 |
| 8 | 41347396 | rs3824277 | -0.944 | -0.688 | -1.156 | -0.571 | 0.004 | 0.055 | 0.055 | 0.007 | 0.95 | 0.89 | 0.89 | 0.77 | 0.96 | 0.92 |
| 8 | 41348534 | rs3780023 | 0.104 | -0.741 | -1.057 | -0.641 | -0.038 | -0.071 | -0.071 | 0 | 0.19 | 0.3 | 0.32 | 0.03 | 0.14 | 0.04 |
| 8 | 41352075 | rs12679812 | 1.938 | -0.931 | -1.363 | -0.749 | -0.062 | -0.107 | -0.107 | 0 | 0.2 | 0.01 | 0.14 | 0.32 | 0.26 | 0.41 |
| 8 | 41355096 | rs17656952 | 1.56 | -1.058 | -0.458 | -1.038 | -0.003 | -0.017 | -0.017 | 0 | 0.13 | 0.12 | 0.13 | 0 | 0.15 | 0.07 |
| 8 | 41363149 | rs4736814 | -0.365 | -1.079 | -0.497 | -1.043 | -0.04 | -0.074 | -0.074 | 0 | 0.2 | 0.32 | 0.32 | 0.03 | 0.14 | 0.04 |
| 8 | 41367608 | rs1043869 | -0.288 | -1.09 | -0.431 | -1.048 | -0.019 | -0.043 | -0.043 | 0 | 0.19 | 0.24 | 0.32 | 0.03 | 0.14 | 0.04 |
| 8 | 41367719 | rs1043873 | -0.003 | -0.963 | -0.626 | -0.93 | 0.016 | 0.074 | 0.074 | 0 | 0.92 | 0.8 | 0.88 | 0.77 | 0.94 | 0.91 |
| 8 | 41367882 | rs7442 | -0.511 | -0.966 | -0.611 | -0.94 | -0.02 | -0.117 | -0.117 | 0.625 | 0.32 | 0.42 | 0.46 | 0.03 | 0.29 | 0.11 |
| 8 | 41368500 | rs7014054 | 0.482 | -0.241 | -0.113 | -0.156 | 0.016 | 0.074 | 0.074 | 0 | 0.92 | 0.8 | 0.88 | 0.77 | 0.94 | 0.91 |
| 8 | 41376476 | rs6990366 | -0.017 | -0.422 | -0.273 | -0.472 | -0.035 | -0.084 | -0.084 | 4.446 | 0.16 | 0.33 | 0.11 | 0.02 | 0.09 | 0.03 |
| 8 | 41385848 | rs4257991 | 0.47 | 0.087 | -0.561 | -0.287 | 0.001 | -0.023 | -0.023 | 0 | 0.17 | 0.19 | 0.15 | 0.05 | 0.2 | 0.09 |
| 8 | 41387921 | rs7819949 | -0.678 | -0.059 | -0.806 | 0.015 | 0.003 | 0.005 | 0.005 | 0 | 0.39 | 0.55 | 0.36 | 0.37 | 0.35 | 0.33 |
| 8 | 41402037 | rs13265966 | -0.318 | 0.577 | -0.37 | 0.44 | 0.047 | 0.046 | 0.046 | 0.016 | 0.1 | 0.32 | 0.14 | 0.13 | 0.13 | 0.16 |
| 8 | 41406773 | rs6994462 | 0.493 | 0.508 | -1.09 | 0.909 | -0.038 | -0.082 | -0.082 | 0.236 | 0.15 | 0.01 | 0.12 | 0.23 | 0.17 | 0.17 |
| 8 | 41416952 | rs7002916 | -1.332 | -0.279 | -0.826 | 0.1 | 0.004 | 0 | 0 | 0 | 0.45 | 0.67 | 0.39 | 0.4 | 0.47 | 0.41 |
| 8 | 41424636 | rs11986407 | -0.075 | -0.223 | -0.756 | 0.091 | 0.025 | 0.013 | 0.013 | 0 | 0.13 | 0.33 | 0.16 | 0.13 | 0.16 | 0.16 |
| 8 | 41424806 | rs11994079 | -0.867 | -0.335 | -0.539 | 0.011 | 0.014 | -0.002 | -0.002 | 0 | 0.2 | 0.41 | 0.18 | 0.17 | 0.21 | 0.2 |
| 8 | 41432893 | rs11785763 | 2.188 | -0.077 | -1.063 | -0.084 | -0.055 | 0.005 | 0.005 | 0 | 0.13 | 0.01 | 0.09 | 0.1 | 0.17 | 0.08 |
| 8 | 41435795 | rs999188 | -0.933 | -0.385 | -1.122 | -0.199 | 0.001 | 0 | 0 | 2.053 | 0.5 | 0.5 | 0.36 | 0.47 | 0.44 | 0.39 |
| 8 | 41447470 | rs7837006 | -0.874 | -0.403 | -0.967 | -0.208 | 0.002 | 0 | 0 | 0.24 | 0.5 | 0.59 | 0.37 | 0.47 | 0.45 | 0.39 |
| 8 | 41457094 | rs2977844 | 2.731 | -0.874 | -1.466 | -0.179 | -0.105 | -0.057 | -0.057 | 1.846 | 0.19 | 0.02 | 0.35 | 0.18 | 0.26 | 0.32 |
| 8 | 41460187 | rs11996226 | 1.133 | -1.104 | -1.287 | -0.297 | -0.012 | -0.015 | -0.015 | 0 | 0.25 | 0.24 | 0.2 | 0.29 | 0.26 | 0.2 |
| 8 | 41462955 | rs2354576 | -1.212 | -0.801 | -1.347 | -0.127 | -0.007 | -0.007 | -0.007 | 0 | 0.59 | 0.7 | 0.44 | 0.47 | 0.54 | 0.43 |
| 8 | 41467827 | rs919035 | 1.517 | -0.741 | -1.467 | -0.301 | 0.004 | -0.006 | -0.006 | 0 | 0.08 | 0.07 | 0.07 | 0 | 0.09 | 0.04 |
| 8 | 41476991 | rs11786297 | 1.517 | -0.741 | -1.467 | -0.301 | 0.004 | -0.006 | -0.006 | 1.066 | 0.08 | 0.07 | 0.07 | 0 | 0.09 | 0.04 |
| 8 | 41478361 | rs890220 | 0.055 | -0.519 | -1.414 | -0.137 | 0.011 | 0.005 | 0.005 | 0 | 0.18 | 0.12 | 0.14 | 0.14 | 0.14 | 0.16 |
| 8 | 41496007 | rs2920126 | 1.556 | -0.052 | -1.004 | -0.12 | -0.005 | 0.001 | 0.001 | 2.008 | 0.23 | 0.16 | 0.38 | 0.17 | 0.28 | 0.34 |
| 8 | 41501562 | rs13254454 | 1.253 | -0.23 | -0.951 | -0.231 | 0.004 | -0.003 | -0.003 | 0 | 0.08 | 0.08 | 0.07 | 0 | 0.1 | 0.04 |
| 8 | 41503293 | rs2241895 | -0.17 | -0.409 | -1.333 | 0.317 | 0.008 | -0.103 | -0.103 | 0 | 0.21 | 0.06 | 0.25 | 0.34 | 0.22 | 0.18 |
| 8 | 41503727 | rs755162 | 1.547 | 0.039 | -0.932 | -0.102 | -0.003 | 0.004 | 0.004 | 1.497 | 0.23 | 0.16 | 0.37 | 0.16 | 0.27 | 0.33 |
| 8 | 41506380 | rs12549294 | 0.039 | 0.026 | -0.711 | -0.076 | 0.005 | -0.005 | -0.005 | 0.01 | 0.6 | 0.71 | 0.62 | 0.38 | 0.58 | 0.6 |
| 8 | 41507237 | rs7825337 | 0.476 | 0.017 | -0.454 | -0.098 | 0.001 | 0 | 0 | 0 | 0.56 | 0.6 | 0.59 | 0.38 | 0.55 | 0.57 |
| 8 | 41509259 | rs12549902 | 0.076 | 0.094 | -0.652 | -0.054 | 0.008 | -0.009 | -0.009 | 0 | 0.59 | 0.73 | 0.63 | 0.39 | 0.57 | 0.56 |
| 8 | 41516790 | rs492083 | 1.265 | 0.143 | -0.77 | 0.055 | -0.025 | -0.081 | -0.081 | 0 | 0.22 | 0.08 | 0.2 | 0.46 | 0.23 | 0.28 |
| 8 | 41516915 | rs6981587 | 0.098 | -0.023 | -0.372 | -0.274 | 0.001 | 0.003 | 0.003 | 3.25 | 0.37 | 0.62 | 0.22 | 0.2 | 0.26 | 0.27 |
| 8 | 41519248 | rs516946 | 0.089 | 0.06 | -0.791 | 0.025 | 0.016 | 0.023 | 0.023 | 0.611 | 0.77 | 0.79 | 0.79 | 0.86 | 0.77 | 0.81 |
| 8 | 41522991 | rs508419 | 0.33 | 0.052 | -0.717 | 0.021 | 0.007 | 0.009 | 0.009 | 0.221 | 0.75 | 0.73 | 0.78 | 0.85 | 0.76 | 0.81 |
| 8 | 41523745 | rs6989203 | -0.379 | 0.057 | -0.785 | 0.023 | 0.009 | 0.012 | 0.012 | 0.156 | 0.24 | 0.22 | 0.21 | 0.14 | 0.24 | 0.19 |
| 8 | 41524287 | rs475831 | -1.249 | 0.007 | -0.809 | -0.096 | -0.001 | 0 | 0 | 0 | 0.75 | 0.66 | 0.82 | 0.76 | 0.79 | 0.79 |
| 8 | 41525914 | rs750625 | -0.344 | 0.064 | -1.206 | -0.139 | 0.017 | 0.023 | 0.023 | 0.16 | 0.24 | 0.19 | 0.21 | 0.14 | 0.24 | 0.19 |

| | | | | | | | | | | | | | | | | |
|---|---|---|---|---|---|---|---|---|---|---|---|---|---|---|---|---|
| 8 | 41531478 | rs12543292 | 0.841 | 0.574 | -0.904 | 0.071 | -0.051 | -0.015 | -0.015 | 0 | 0.23 | 0.02 | 0.16 | 0.17 | 0.21 | 0.19 |
| 8 | 41538286 | rs2355254 | -0.723 | 0.605 | -0.491 | 0.067 | -0.003 | -0.003 | -0.003 | 0 | 0.72 | 0.63 | 0.81 | 0.79 | 0.78 | 0.76 |
| 8 | 41540884 | rs4737004 | 1.13 | 1.315 | -1.08 | -0.042 | -0.076 | 0.002 | 0.002 | 0.448 | 0.23 | 0.02 | 0.37 | 0.13 | 0.32 | 0.31 |
| 8 | 41541049 | rs7844297 | -0.033 | 1.406 | -0.298 | -0.118 | 0.001 | -0.006 | -0.006 | 2.75 | 0.61 | 0.56 | 0.61 | 0.29 | 0.59 | 0.6 |
| 8 | 41547386 | rs7831590 | -0.779 | 1.412 | 0.147 | -0.28 | 0.087 | 0.082 | 0.082 | 4.214 | 0.08 | 0.54 | 0.06 | 0.08 | 0.06 | 0.04 |
| 8 | 41547660 | rs2278621 | 1.06 | 1.334 | -0.641 | -0.448 | -0.042 | 0.062 | 0.062 | 0 | 0.29 | 0.04 | 0.19 | 0.13 | 0.23 | 0.24 |
| 8 | 41555473 | rs2241896 | -0.035 | 1.633 | 0.136 | -0.085 | -0.004 | 0.001 | 0.001 | 0 | 0.38 | 0.26 | 0.41 | 0.65 | 0.37 | 0.39 |
| 8 | 41555954 | rs490157 | -0.423 | 1.982 | 0.923 | -0.102 | -0.003 | -0.088 | -0.088 | 1.183 | 0.72 | 0.58 | 0.67 | 0.89 | 0.69 | 0.67 |
| 8 | 41558898 | rs2304881 | -0.466 | 2.065 | 0.731 | -0.138 | -0.003 | 0.046 | 0.046 | 0.014 | 0.35 | 0.75 | 0.38 | 0.15 | 0.37 | 0.37 |
| 8 | 41559609 | rs2304880 | 0.639 | 2.074 | 0.256 | -0.111 | -0.065 | 0.122 | 0.122 | 0 | 0.25 | 0.02 | 0.3 | 0.08 | 0.28 | 0.32 |
| 8 | 41563685 | rs1137177 | 0.518 | 2.002 | 0.135 | -0.145 | -0.062 | 0.122 | 0.122 | 0 | 0.25 | 0.02 | 0.3 | 0.08 | 0.28 | 0.32 |
| 8 | 41573488 | rs473235 | 1.365 | 2.242 | 0.17 | 0.363 | -0.018 | -0.058 | -0.058 | 0 | 0.2 | 0.1 | 0.28 | 0.46 | 0.23 | 0.2 |
| 8 | 41576746 | rs12549656 | 1.357 | 1.992 | 0.476 | -0.172 | 0.033 | 0.021 | 0.021 | 0 | 0.34 | 0.22 | 0.21 | 0.25 | 0.28 | 0.27 |
| 8 | 41580607 | rs7826127 | 0.701 | 2.117 | 0.691 | -0.191 | -0.01 | -0.022 | -0.022 | 0 | 0.4 | 0.55 | 0.29 | 0.26 | 0.35 | 0.3 |
| 8 | 41585384 | rs2304872 | 0.67 | 2.16 | 0.565 | -0.392 | -0.034 | 0.096 | 0.096 | 0 | 0.23 | 0.08 | 0.31 | 0.08 | 0.29 | 0.3 |
| 8 | 41585571 | rs1872877 | 0.034 | 2.222 | 0.685 | -0.265 | 0.003 | -0.006 | -0.006 | 0 | 0.63 | 0.64 | 0.6 | 0.34 | 0.64 | 0.6 |
| 8 | 41586356 | rs11989909 | 0.247 | 2.225 | 0.851 | -0.266 | -0.004 | 0.004 | 0.004 | 0 | 0.61 | 0.43 | 0.54 | 0.31 | 0.58 | 0.59 |
| 8 | 41586822 | rs7016707 | -1.554 | 2.302 | 0.578 | -0.223 | 0.162 | 0.033 | 0.033 | 0.141 | 0.08 | 0.43 | 0.09 | 0.03 | 0.11 | 0.04 |
| 8 | 41591643 | rs2304869 | -0.238 | 1.994 | -0.201 | -0.013 | -0.032 | -0.033 | -0.033 | 0.662 | 0.14 | 0.03 | 0.1 | 0.15 | 0.12 | 0.16 |
| 8 | 41593802 | rs7820101 | 0.758 | 2.066 | -0.244 | -0.294 | -0.046 | 0.09 | 0.09 | 0 | 0.22 | 0.02 | 0.29 | 0.08 | 0.26 | 0.29 |
| 8 | 41595613 | rs516606 | 0.727 | 1.868 | 0.247 | -0.238 | 0 | -0.019 | -0.019 | 0.16 | 0.6 | 0.45 | 0.56 | 0.75 | 0.61 | 0.55 |
| 8 | 41599417 | rs497561 | -0.164 | 1.735 | -0.38 | -0.15 | 0.003 | -0.001 | -0.001 | 0.222 | 0.87 | 0.91 | 0.9 | 0.86 | 0.9 | 0.86 |
| 8 | 41600125 | rs7836809 | 0.697 | 1.842 | -0.542 | -0.172 | -0.049 | 0.065 | 0.065 | 0.004 | 0.22 | 0.04 | 0.3 | 0.11 | 0.28 | 0.31 |
| 8 | 41601685 | rs521184 | 0.659 | 1.935 | -0.19 | -0.085 | 0.005 | 0.092 | 0.092 | 0 | 0.59 | 0.23 | 0.55 | 0.75 | 0.61 | 0.56 |
| 8 | 41602668 | rs11784487 | 1.297 | 1.922 | -0.402 | -0.118 | 0.025 | 0.056 | 0.056 | 0 | 0.36 | 0.1 | 0.21 | 0.24 | 0.32 | 0.32 |
| 8 | 41605979 | rs510889 | 0.774 | 2.082 | -0.122 | 0.31 | -0.056 | -0.075 | -0.075 | 0.461 | 0.23 | 0.11 | 0.34 | 0.52 | 0.3 | 0.25 |
| 8 | 41607715 | rs488758 | 0.599 | 1.848 | 0.564 | -0.1 | 0.015 | 0.012 | 0.012 | 0 | 0.26 | 0.29 | 0.52 | 0.64 | 0.29 | 0.28 |
| 8 | 41611745 | rs515791 | 0.666 | 1.777 | 0.975 | -0.237 | -0.006 | -0.011 | -0.011 | 0 | 0.85 | 0.68 | 0.88 | 0.88 | 0.88 | 0.86 |
| 8 | 41612255 | rs6981979 | 0.458 | 1.856 | 0.497 | -0.233 | -0.005 | 0.01 | 0.01 | 0 | 0.59 | 0.39 | 0.36 | 0.23 | 0.58 | 0.58 |
| 8 | 41614503 | rs7831492 | 1.15 | 1.837 | 0.248 | -0.443 | 0.002 | 0.004 | 0.004 | 0 | 0.41 | 0.37 | 0.28 | 0.14 | 0.4 | 0.32 |
| 8 | 41621903 | rs11997827 | -0.831 | 1.4 | 0.187 | -0.372 | 0.005 | 0.013 | 0.013 | 0 | 0.32 | 0.25 | 0.18 | 0.21 | 0.3 | 0.4 |
| 8 | 41623602 | rs11780780 | -0.762 | 1.42 | 0.092 | -0.062 | 0 | 0.001 | 0.001 | 0.013 | 0.12 | 0.13 | 0.1 | 0.12 | 0.13 | 0.14 |
| 8 | 41630405 | rs4737009 | -0.032 | 1.56 | 0.33 | 0.246 | -0.003 | 0.037 | 0.037 | 0.403 | 0.29 | 0.62 | 0.27 | 0.51 | 0.25 | 0.16 |
| 8 | 41636071 | rs13255458 | -0.243 | 1.963 | 0.371 | -0.683 | -0.011 | 0.056 | 0.056 | 0 | 0.28 | 0.19 | 0.16 | 0.11 | 0.3 | 0.27 |
| 8 | 41639266 | rs2032736 | 0.7 | 1.608 | 0.291 | -0.716 | 0.001 | -0.056 | -0.056 | 0 | 0.25 | 0.07 | 0.19 | 0.7 | 0.23 | 0.23 |
| 8 | 41640125 | rs9650332 | 0.246 | 2.12 | 0.429 | -0.682 | -0.092 | 0.039 | 0.039 | 0.28 | 0.23 | 0.06 | 0.23 | 0.15 | 0.35 | 0.4 |
| 8 | 41640657 | rs12156072 | 0.036 | 2.038 | 0.4 | -0.714 | -0.094 | 0.059 | 0.059 | 0 | 0.27 | 0.07 | 0.24 | 0.15 | 0.36 | 0.41 |
| 8 | 41641075 | rs10958698 | -0.059 | 2.052 | 0.537 | -0.719 | -0.074 | 0.044 | 0.044 | 0 | 0.28 | 0.12 | 0.25 | 0.18 | 0.38 | 0.41 |
| 8 | 41641624 | rs13273224 | -0.447 | 2.089 | 0.784 | -0.666 | -0.01 | 0.009 | 0.009 | 0 | 0.33 | 0.33 | 0.3 | 0.19 | 0.4 | 0.42 |
| 8 | 41648358 | rs10958699 | 1.053 | 2.047 | 0.898 | -0.941 | 0.005 | 0.016 | 0.016 | 0.016 | 0.4 | 0.31 | 0.49 | 0.13 | 0.37 | 0.36 |
| 8 | 41660854 | rs6999814 | -1.017 | 2.072 | 0.86 | -0.848 | -0.008 | -0.053 | -0.053 | 0 | 0.77 | 0.9 | 0.82 | 0.31 | 0.77 | 0.79 |
| 8 | 41661716 | rs13272350 | -0.449 | 2.162 | 1.099 | -0.951 | 0.012 | -0.04 | -0.04 | 0 | 0.36 | 0.5 | 0.33 | 0.17 | 0.4 | 0.42 |
| 8 | 41666355 | rs12680139 | 0.99 | 1.936 | 0.378 | -0.995 | 0.011 | 0.023 | 0.023 | 0.037 | 0.4 | 0.26 | 0.48 | 0.12 | 0.36 | 0.37 |
| 8 | 41670952 | rs7006871 | -0.919 | 1.969 | -0.135 | -0.898 | 0 | -0.057 | -0.057 | 0.122 | 0.76 | 0.92 | 0.82 | 0.32 | 0.77 | 0.78 |
| 8 | 41674525 | rs17662607 | 0.897 | 1.786 | -0.026 | -0.874 | -0.021 | 0.124 | 0.124 | 0.081 | 0.34 | 0.02 | 0.43 | 0.11 | 0.28 | 0.3 |
| 8 | 41684488 | rs1549064 | 0.679 | 1.884 | 0.198 | -0.774 | 0.009 | 0.108 | 0.108 | 0.039 | 0.38 | 0.16 | 0.47 | 0.14 | 0.35 | 0.39 |
| 8 | 41688828 | rs2102360 | -1.063 | 2.119 | 1.194 | 0.154 | 0.081 | -0.031 | -0.031 | 0 | 0.07 | 0.18 | 0.16 | 0.02 | 0.14 | 0.09 |
| 8 | 41697078 | rs6989331 | -1.293 | 1.611 | 0.585 | 0.124 | -0.013 | -0.06 | -0.06 | 0 | 0.34 | 0.57 | 0.33 | 0.84 | 0.38 | 0.41 |
| 8 | 41704910 | rs7833494 | -1.303 | 0.95 | -0.141 | 0.172 | 0.004 | 0.047 | 0.047 | 1.931 | 0.33 | 0.84 | 0.31 | 0.19 | 0.36 | 0.5 |
| 8 | 41706484 | rs12156316 | -1.006 | 0.886 | -0.196 | 0.215 | 0.006 | 0.024 | 0.024 | 1.327 | 0.29 | 0.2 | 0.25 | 0.18 | 0.32 | 0.51 |
| 8 | 41714362 | rs4737018 | -0.839 | 0.767 | 0.057 | 0.297 | -0.003 | -0.025 | -0.025 | 0 | 0.26 | 0.41 | 0.27 | 0.17 | 0.3 | 0.42 |
| 8 | 41727195 | rs10101217 | -1.481 | 0.586 | 0.597 | 0.93 | 0.019 | 0.162 | 0.162 | 0.099 | 0.07 | 0.3 | 0.06 | 0.18 | 0.04 | 0.15 |
| 8 | 41744094 | rs7000214 | -0.717 | 1.083 | 0.468 | 0.085 | 0.003 | -0.029 | -0.029 | 0 | 0.21 | 0.24 | 0.16 | 0.06 | 0.28 | 0.13 |
| 8 | 41746303 | rs10958702 | -0.362 | 0.419 | -0.017 | 0.618 | 0.017 | -0.047 | -0.047 | 0 | 0.19 | 0.12 | 0.15 | 0.41 | 0.19 | 0.35 |
| 8 | 41751299 | rs4737021 | -1.831 | -0.171 | -0.712 | 0.815 | -0.124 | -0.001 | -0.001 | 0 | 0.55 | 0.98 | 0.53 | 0.74 | 0.53 | 0.73 |
| 8 | 41761807 | rs13276409 | -0.473 | -0.043 | 0.612 | 0.618 | 0.019 | -0.13 | -0.13 | 0 | 0.25 | 0.47 | 0.18 | 0.07 | 0.28 | 0.13 |
| 8 | 41791283 | rs3739365 | -0.248 | 0.051 | 0.839 | 0.859 | -0.001 | 0.002 | 0.002 | 0.01 | 0.11 | 0.1 | 0.18 | 0.23 | 0.07 | 0.26 |
| 8 | 41794934 | rs13748 | -0.335 | -0.156 | 1.673 | 0.594 | 0.05 | -0.138 | -0.138 | 0 | 0.2 | 0.48 | 0.15 | 0 | 0.27 | 0.03 |
| 8 | 41797694 | rs6474373 | -0.048 | -0.193 | 2.021 | 0.576 | 0.043 | -0.099 | -0.099 | 1.716 | 0.18 | 0.38 | 0.13 | 0 | 0.27 | 0.03 |
| 8 | 41797902 | rs1318481 | 0.018 | -0.589 | 1.09 | 0.008 | -0.016 | 0 | 0 | 0 | 0.64 | 0.67 | 0.39 | 0.38 | 0.58 | 0.41 |
| 8 | 41798202 | rs2929896 | 1.049 | -0.59 | 1.088 | 0.008 | 0.098 | 0.038 | 0.038 | 0.891 | 0.44 | 0.18 | 0.24 | 0.38 | 0.31 | 0.37 |
| 8 | 41798417 | rs3824275 | 0.496 | -0.569 | 1.041 | 0.019 | -0.003 | 0.006 | 0.006 | 0.009 | 0.11 | 0.1 | 0.19 | 0.23 | 0.07 | 0.26 |
| 8 | 41805249 | rs10504048 | 0.731 | -0.548 | 1.259 | 0.013 | 0.014 | -0.053 | -0.053 | 0 | 0.17 | 0.25 | 0.13 | 0 | 0.27 | 0.03 |
| 8 | 41807375 | rs13260358 | -0.941 | -0.458 | 0.867 | 0.089 | -0.062 | -0.067 | -0.067 | 0 | 0.75 | 0.85 | 0.58 | 0.61 | 0.65 | 0.66 |
| 8 | 41823538 | rs1871991 | 0.056 | -0.48 | 0.588 | 0.051 | -0.026 | 0 | 0 | 0 | 0.64 | 0.71 | 0.39 | 0.38 | 0.58 | 0.41 |
| 8 | 41846687 | rs7008906 | 1.005 | -0.549 | 0.594 | -0.011 | 0.082 | 0.038 | 0.038 | 1.003 | 0.44 | 0.22 | 0.25 | 0.37 | 0.31 | 0.37 |
| 8 | 41854101 | rs10958705 | -0.775 | -0.219 | 0.438 | 0.322 | -0.08 | -0.089 | -0.089 | 1.344 | 0.75 | 0.89 | 0.58 | 0.61 | 0.65 | 0.67 |
| 8 | 41868921 | rs10112548 | 0.266 | -0.202 | 0.451 | 0.33 | -0.028 | 0 | 0 | 0.75 | 0.64 | 0.7 | 0.39 | 0.38 | 0.58 | 0.4 |
| 8 | 41877317 | rs6987070 | -1.043 | -0.128 | -0.006 | 0.356 | -0.15 | -0.146 | -0.146 | 0 | 0.78 | 0.97 | 0.59 | 0.68 | 0.65 | 0.75 |

| | | | | | | | | | | | | | | | | |
|---|---|---|---|---|---|---|---|---|---|---|---|---|---|---|---|---|
| 8 | 41906095 | rs3824276 | -0.131 | -0.091 | 0.735 | 0.244 | -0.003 | 0.005 | 0.005 | 0.01 | 0.11 | 0.1 | 0.19 | 0.23 | 0.07 | 0.26 |
| 8 | 41908148 | rs12546107 | -0.283 | -0.09 | 0.724 | 0.244 | -0.006 | 0.009 | 0.009 | 0 | 0.11 | 0.13 | 0.19 | 0.23 | 0.07 | 0.26 |
| 8 | 41951881 | rs10103250 | -1.154 | 0.276 | 0.663 | -0.117 | 0.006 | -0.025 | -0.025 | 0.511 | 0.19 | 0.69 | 0.14 | 0.1 | 0.2 | 0.23 |
| 8 | 41969930 | rs7842131 | -1.978 | 0.277 | 0.469 | -0.127 | 0.001 | 0.02 | 0.02 | 0 | 0.34 | 0.72 | 0.19 | 0.25 | 0.31 | 0.38 |
| 8 | 41974805 | rs4266635 | 1.792 | 0.256 | 0.098 | -0.137 | -0.004 | 0.067 | 0.067 | 0 | 0.66 | 0.17 | 0.79 | 0.75 | 0.69 | 0.62 |
| 8 | 41983076 | rs12544318 | -1.156 | 0.015 | 0.03 | -0.12 | 0.034 | 0.005 | 0.005 | 0.759 | 0.17 | 0.02 | 0.05 | 0.15 | 0.09 | 0.16 |
| 8 | 41994579 | rs6980818 | -1.505 | -0.098 | 1.142 | -0.18 | 0.058 | 0.053 | 0.053 | 4.173 | 0.06 | 0.5 | 0.04 | 0 | 0 | 0 |
| 8 | 41997694 | rs10958707 | -2.292 | 0.073 | 1.255 | -0.086 | -0.039 | -0.025 | -0.025 | 0 | 0.24 | 0.63 | 0.12 | 0.21 | 0.13 | 0.21 |
| 8 | 42001906 | rs13270574 | -0.88 | 0.559 | 0.354 | -0.08 | 0.008 | -0.007 | -0.007 | 0.343 | 0.54 | 0.67 | 0.43 | 0.5 | 0.41 | 0.38 |
| 8 | 42002818 | rs4641052 | -0.709 | 0.535 | 0.568 | -0.088 | 0.012 | 0.001 | 0.001 | 0 | 0.52 | 0.61 | 0.42 | 0.46 | 0.4 | 0.32 |
| 8 | 42003124 | rs11998613 | 1.023 | 0.501 | 0.767 | -0.081 | -0.011 | -0.041 | -0.041 | 0.016 | 0.31 | 0.11 | 0.31 | 0.31 | 0.29 | 0.17 |
| 8 | 42010466 | rs6982888 | -0.955 | 0.477 | 1.004 | -0.103 | 0.008 | -0.01 | -0.01 | 0 | 0.54 | 0.66 | 0.43 | 0.5 | 0.41 | 0.37 |
| 8 | 42013295 | rs11778382 | -0.659 | 0.509 | 0.961 | -0.087 | 0.01 | -0.019 | -0.019 | 0 | 0.17 | 0.02 | 0.08 | 0.18 | 0.11 | 0.14 |
| 8 | 42014996 | rs13277203 | 1.206 | 0.514 | 0.966 | -0.085 | -0.06 | -0.08 | -0.08 | 0.004 | 0.3 | 0.02 | 0.31 | 0.28 | 0.28 | 0.17 |
| 8 | 42026987 | rs1050262 | -0.956 | 0.515 | 1.008 | -0.131 | 0.009 | -0.009 | -0.009 | 0 | 0.54 | 0.66 | 0.42 | 0.5 | 0.41 | 0.38 |
| 8 | 42028077 | rs10958709 | -0.244 | 0.512 | 0.943 | -0.131 | -0.032 | -0.092 | -0.092 | 0 | 0.47 | 0.04 | 0.38 | 0.46 | 0.4 | 0.32 |
| 8 | 42032026 | rs4581040 | 0.757 | 0.541 | 0.998 | -0.015 | -0.01 | -0.01 | -0.01 | 0 | 0.33 | 0.43 | 0.33 | 0.28 | 0.28 | 0.18 |
| 8 | 42032379 | rs8178895 | -0.787 | 0.481 | 0.7 | 0.013 | 0.015 | -0.033 | -0.033 | 0.025 | 0.17 | 0.02 | 0.08 | 0.19 | 0.12 | 0.15 |
| 8 | 42036577 | rs1136159 | -0.737 | 0.564 | 0.687 | -0.051 | 0.01 | -0.038 | -0.038 | 0 | 0.17 | 0.02 | 0.07 | 0.19 | 0.11 | 0.14 |
| 8 | 42044954 | rs1058720 | 0.606 | 0.846 | 0.776 | -0.245 | 0.007 | -0.009 | -0.009 | 0.327 | 0.46 | 0.35 | 0.57 | 0.5 | 0.59 | 0.62 |
| 8 | 42045655 | rs2070713 | 1.088 | 0.778 | 0.338 | -0.206 | -0.047 | -0.084 | -0.084 | 0.062 | 0.34 | 0.1 | 0.46 | 0.46 | 0.41 | 0.42 |
| 8 | 42051908 | rs8178721 | -0.418 | 0.746 | 0.067 | -0.179 | -0.005 | 0.052 | 0.052 | 0.001 | 0.11 | 0.02 | 0.05 | 0.02 | 0.08 | 0.07 |
| 8 | 42057370 | rs7837156 | -0.001 | 0.861 | 0.556 | -0.231 | 0.024 | 0.03 | 0.03 | 0 | 0.54 | 0.78 | 0.71 | 0.69 | 0.63 | 0.73 |
| 8 | 42065209 | rs2020919 | -2.207 | 0.973 | 1.512 | -0.254 | 0.029 | -0.053 | -0.053 | 0 | 0.11 | 0.6 | 0.08 | 0.01 | 0.06 | 0.08 |
| 8 | 42072438 | rs2020918 | -0.303 | 0.925 | 0.789 | -0.243 | 0 | 0.028 | 0.028 | 0 | 0.61 | 0.79 | 0.75 | 0.72 | 0.66 | 0.8 |
| 8 | 42086472 | rs7820274 | -0.722 | 0.969 | 0.609 | -0.141 | -0.021 | -0.076 | -0.076 | 0 | 0.66 | 0.95 | 0.77 | 0.67 | 0.7 | 0.8 |
| 8 | 42087576 | rs10094577 | -2.439 | 0.92 | 1.798 | -0.268 | 0.053 | -0.028 | -0.028 | 0 | 0.11 | 0.54 | 0.07 | 0.05 | 0.06 | 0.25 |
| 8 | 42100552 | rs12115174 | -0.533 | 1.323 | 1.915 | -0.437 | 0.05 | -0.057 | -0.057 | 1.091 | 0.14 | 0.31 | 0.11 | 0.05 | 0.19 | 0.25 |
| 8 | 42103322 | rs7829991 | -2.41 | 1.159 | 2.158 | -0.156 | 0.017 | -0.04 | -0.04 | 0 | 0.09 | 0.45 | 0.07 | 0.01 | 0.04 | 0.03 |
| 8 | 42135178 | rs11986055 | -2.856 | 0.977 | 1.991 | -0.084 | -0.042 | -0.007 | -0.007 | 0.196 | 0.14 | 0.72 | 0.09 | 0.17 | 0.05 | 0.23 |
| 8 | 42140804 | rs4560769 | -3.178 | 1.001 | 2.003 | -0.095 | 0.007 | 0.005 | 0.005 | 2.409 | 0.17 | 0.83 | 0.12 | 0.16 | 0.09 | 0.26 |
| 8 | 42143739 | rs10099598 | 1.047 | 0.861 | 2.045 | -0.174 | -0.014 | -0.034 | -0.034 | 0.005 | 0.93 | 0.85 | 0.96 | 0.99 | 0.96 | 0.98 |
| 8 | 42147894 | rs2294097 | -2.806 | 0.883 | 1.784 | -0.162 | -0.04 | -0.006 | -0.006 | 1.669 | 0.14 | 0.72 | 0.09 | 0.16 | 0.05 | 0.23 |
| 8 | 42157902 | rs2272733 | 2.872 | 0.895 | 1.781 | -0.158 | -0.005 | -0.007 | -0.007 | 0 | 0.81 | 0.23 | 0.86 | 0.84 | 0.88 | 0.72 |
| 8 | 42180716 | rs10958713 | 1.015 | 0.546 | 1.371 | -0.176 | -0.102 | -0.176 | -0.176 | 0 | 0.34 | 0.03 | 0.51 | 0.54 | 0.35 | 0.45 |
| 8 | 42187154 | rs6474388 | 2.67 | 0.473 | 2.531 | -0.129 | -0.028 | -0.013 | -0.013 | 0 | 0.84 | 0.28 | 0.89 | 0.83 | 0.91 | 0.75 |
| 8 | 42201087 | rs2976239 | 1.877 | -0.3 | 2.72 | -0.089 | -0.005 | -0.037 | -0.037 | 0 | 0.91 | 0.61 | 0.95 | 0.99 | 0.96 | 0.98 |
| 8 | 42224155 | rs3136788 | -1.435 | -0.308 | 2.809 | -0.081 | 0.025 | 0.102 | 0.102 | 0 | 0.06 | 0.27 | 0.04 | 0.1 | 0.04 | 0.03 |
| 8 | 42236108 | rs10097867 | -3.051 | -0.311 | 2.64 | -0.086 | -0.036 | -0.017 | -0.017 | 0 | 0.16 | 0.7 | 0.1 | 0.17 | 0.09 | 0.25 |
| 8 | 42238574 | rs11993621 | -3.136 | -0.314 | 2.544 | -0.107 | -0.013 | -0.007 | -0.007 | 0.55 | 0.17 | 0.81 | 0.1 | 0.17 | 0.09 | 0.27 |
| 8 | 42240338 | rs12335053 | -3.236 | -0.28 | 2.704 | -0.001 | -0.047 | -0.009 | -0.009 | 0 | 0.15 | 0.67 | 0.08 | 0.17 | 0.07 | 0.21 |
| 8 | 42247345 | rs28510568 | -2.01 | -0.276 | 2.845 | -0.046 | 0.013 | 0.01 | 0.01 | 2.167 | 0.05 | 0.34 | 0.03 | 0 | 0 | 0.02 |
| 8 | 42259299 | rs3736649 | -3.141 | -0.272 | 2.759 | -0.043 | -0.04 | -0.017 | -0.017 | 0.225 | 0.15 | 0.66 | 0.08 | 0.16 | 0.07 | 0.21 |
| 8 | 42263157 | rs6773 | -3.141 | -0.272 | 2.759 | -0.043 | -0.04 | -0.017 | -0.017 | 0 | 0.15 | 0.66 | 0.08 | 0.16 | 0.07 | 0.22 |
| 8 | 42274960 | rs3763510 | -1.352 | -0.292 | 2.595 | -0.019 | -0.007 | 0.041 | 0.041 | 0 | 0.06 | 0.19 | 0.03 | 0.1 | 0.03 | 0.06 |
| 8 | 42286117 | rs6651364 | -2.652 | -0.312 | 1.889 | -0.039 | -0.043 | -0.019 | -0.019 | 0 | 0.16 | 0.69 | 0.08 | 0.15 | 0.07 | 0.22 |
| 8 | 42292952 | rs11993672 | 0.096 | -0.189 | 1.407 | -0.126 | -0.008 | -0.025 | -0.025 | 0 | 0.05 | 0.14 | 0.01 | 0 | 0.02 | 0 |
| 8 | 42301696 | rs7845666 | -1.442 | -0.216 | 1.328 | -0.14 | -0.014 | -0.039 | -0.039 | 0 | 0.09 | 0.42 | 0.04 | 0 | 0.02 | 0.01 |
| 8 | 42323980 | rs6999392 | -1.593 | -0.253 | 1.034 | -0.158 | -0.042 | -0.063 | -0.063 | 0 | 0.11 | 0.62 | 0.04 | 0 | 0.02 | 0.01 |
| 8 | 42324760 | rs2978457 | -0.949 | -0.465 | -0.481 | -0.313 | -0.016 | -0.038 | -0.038 | 0.645 | 0.52 | 0.9 | 0.71 | 0.49 | 0.49 | 0.47 |
| 8 | 42334926 | rs10808961 | 0.073 | -0.999 | -0.735 | -0.414 | -0.028 | -0.041 | -0.041 | 0 | 0.32 | 0.2 | 0.51 | 0.34 | 0.34 | 0.3 |
| 8 | 42340942 | rs4737046 | -0.873 | -1.236 | -0.962 | -0.397 | 0.014 | -0.02 | -0.02 | 0 | 0.47 | 0.85 | 0.55 | 0.44 | 0.37 | 0.43 |
| 8 | 42344238 | rs4368963 | 0.282 | -1.017 | -1.003 | -0.4 | -0.047 | -0.033 | -0.033 | 0.087 | 0.28 | 0.16 | 0.5 | 0.32 | 0.35 | 0.24 |
| 8 | 42351311 | rs6988483 | -0.757 | -1.817 | -0.896 | -0.494 | -0.013 | 0.004 | 0.004 | 0 | 0.57 | 0.44 | 0.52 | 0.41 | 0.51 | 0.49 |
| 8 | 42352007 | rs9643888 | 0.784 | -1.918 | -1.124 | -0.585 | -0.081 | -0.065 | -0.065 | 0.047 | 0.3 | 0.13 | 0.4 | 0.38 | 0.4 | 0.25 |
| 8 | 42354257 | rs7818789 | 0.172 | -1.938 | -1.122 | -0.6 | -0.042 | -0.028 | -0.028 | 0.562 | 0.36 | 0.23 | 0.41 | 0.4 | 0.41 | 0.38 |
| 8 | 42363980 | rs4436107 | 0.527 | -2.075 | -1.148 | -0.868 | 0.019 | 0.098 | 0.098 | 0 | 0.16 | 0.01 | 0.08 | 0 | 0.08 | 0.09 |
| 8 | 42374546 | rs13281070 | -0.658 | -2.182 | -0.753 | -0.926 | 0.011 | -0.004 | -0.004 | 0 | 0.26 | 0.21 | 0.11 | 0.03 | 0.11 | 0.23 |
| 8 | 42388301 | rs2923428 | -1.332 | -2.141 | -0.955 | -0.624 | -0.083 | -0.112 | -0.112 | 0 | 0.74 | 0.88 | 0.62 | 0.54 | 0.61 | 0.74 |
| 8 | 42399906 | rs13273204 | 2.023 | -2.354 | -1.033 | -0.777 | -0.117 | -0.118 | -0.118 | 0 | 0.28 | 0.02 | 0.32 | 0.43 | 0.4 | 0.4 |
| 8 | 42404822 | rs6990159 | 0.161 | -2.184 | -0.832 | -0.633 | -0.006 | 0.004 | 0.004 | 0.078 | 0.08 | 0.26 | 0.12 | 0.03 | 0.02 | 0.1 |
| 8 | 42413943 | rs2974354 | -0.491 | -2.199 | -0.711 | -0.642 | -0.029 | -0.039 | -0.039 | 0 | 0.65 | 0.68 | 0.51 | 0.51 | 0.59 | 0.62 |
| 8 | 42436651 | rs2974342 | -1.451 | -2.138 | -1.117 | -0.672 | -0.086 | -0.116 | -0.116 | 0.536 | 0.74 | 0.88 | 0.62 | 0.54 | 0.61 | 0.74 |
| 8 | 42446238 | rs2356610 | -1.435 | -2.189 | -1.107 | -0.66 | -0.09 | -0.072 | -0.072 | 0.725 | 0.74 | 0.88 | 0.64 | 0.6 | 0.61 | 0.74 |
| 8 | 42450441 | rs7813319 | -0.041 | -2.23 | -1.159 | -0.882 | -0.08 | -0.083 | -0.083 | 0 | 0.22 | 0.38 | 0.1 | 0.05 | 0.08 | 0.14 |
| 8 | 42455846 | rs2974312 | 0.537 | -1.697 | -1.721 | -0.405 | -0.114 | -0.118 | -0.118 | 0 | 0.4 | 0.07 | 0.38 | 0.52 | 0.5 | 0.47 |
| 8 | 42466416 | rs2923419 | 1.05 | -1.763 | -1.146 | -0.052 | 0.113 | 0.062 | 0.062 | 0.037 | 0.72 | 0.12 | 0.75 | 0.88 | 0.92 | 0.75 |
| 8 | 42468218 | rs13251378 | 0.972 | -0.831 | -0.995 | 0.249 | -0.11 | -0.154 | -0.154 | 0 | 0.25 | 0.02 | 0.32 | 0.48 | 0.39 | 0.34 |
| 8 | 42468348 | rs4737051 | -0.693 | -1.18 | 0.528 | 0.066 | -0.017 | 0.019 | 0.019 | 0 | 0.07 | 0.16 | 0.13 | 0.07 | 0.02 | 0.07 |

| | | | | | | | | | | | | | | | | |
|---|---|---|---|---|---|---|---|---|---|---|---|---|---|---|---|---|
| 8 | 42475077 | rs2974319 | 0.681 | -1.238 | 0.701 | -0.189 | -0.03 | -0.068 | -0.068 | 0 | 0.87 | 0.75 | 0.94 | 1 | 0.92 | 0.9 |
| 8 | 42476205 | rs11996441 | -0.929 | -1.327 | 1.411 | -0.064 | -0.008 | 0.051 | 0.051 | 0.049 | 0.08 | 0.31 | 0.13 | 0.09 | 0.02 | 0.09 |
| 8 | 42481880 | rs2974325 | 0.694 | -1.278 | 1.125 | -0.132 | -0.013 | -0.053 | -0.053 | 0 | 0.9 | 0.79 | 0.94 | 1 | 0.93 | 0.91 |
| 8 | 42483326 | rs4737055 | 0.674 | -1.265 | 1.132 | -0.126 | -0.011 | -0.051 | -0.051 | 0 | 0.9 | 0.79 | 0.95 | 1 | 0.95 | 0.91 |
| 8 | 42489842 | rs7844566 | -1.847 | -1.448 | 1.266 | 0.11 | -0.062 | 0.013 | 0.013 | 0.979 | 0.19 | 0.52 | 0.18 | 0.18 | 0.06 | 0.16 |
| 8 | 42501717 | rs7013926 | -1.626 | -1.42 | 1.359 | 0.081 | -0.051 | -0.006 | -0.006 | 0 | 0.16 | 0.34 | 0.15 | 0.15 | 0.06 | 0.16 |
| 8 | 42505466 | rs6989472 | 1.788 | -1.751 | 1.528 | -0.263 | -0.043 | -0.023 | -0.023 | 0 | 0.68 | 0.51 | 0.76 | 0.79 | 0.83 | 0.75 |
| 8 | 42507740 | rs11785591 | -1.489 | -1.77 | 1.786 | -0.256 | 0.002 | -0.006 | -0.006 | 0 | 0.1 | 0.14 | 0.09 | 0.06 | 0.08 | 0.08 |
| 8 | 42509597 | rs1376442 | -2.373 | -1.784 | 1.959 | -0.28 | -0.017 | 0.042 | 0.042 | 0.546 | 0.19 | 0.41 | 0.22 | 0.21 | 0.13 | 0.17 |
| 8 | 42515801 | rs1868859 | 1.435 | -1.711 | 1.937 | -0.419 | -0.033 | -0.087 | -0.087 | 0.997 | 0.81 | 0.69 | 0.94 | 1 | 0.93 | 0.9 |
| 8 | 42518723 | rs4305884 | 0.77 | -1.679 | 1.9 | -0.388 | -0.021 | -0.056 | -0.056 | 0 | 0.86 | 0.77 | 0.96 | 1 | 0.94 | 0.91 |
| 8 | 42521401 | rs13270610 | 0.046 | -1.641 | 2.219 | -0.281 | 0.016 | 0.165 | 0.165 | 0.554 | 0.93 | 0.79 | 0.88 | 0.86 | 0.96 | 0.92 |
| 8 | 42524584 | rs10958725 | 2.345 | -1.515 | 1.969 | -0.236 | 0.026 | 0.015 | 0.015 | 0.535 | 0.65 | 0.25 | 0.7 | 0.77 | 0.78 | 0.74 |
| 8 | 42531565 | rs16891530 | -1.044 | -1.525 | 2.029 | -0.317 | 0.021 | 0.012 | 0.012 | 0 | 0.13 | 0.1 | 0.02 | 0 | 0.04 | 0.06 |
| 8 | 42544017 | rs13273442 | 2.401 | -1.601 | 2.005 | -0.346 | 0.013 | 0.012 | 0.012 | 0.134 | 0.65 | 0.35 | 0.7 | 0.79 | 0.78 | 0.73 |
| 8 | 42548239 | rs1530847 | 1.747 | -1.607 | 2.173 | -0.35 | -0.005 | 0.046 | 0.046 | 0.465 | 0.8 | 0.6 | 0.74 | 0.8 | 0.84 | 0.79 |
| 8 | 42551064 | rs6474413 | 2.478 | -1.625 | 1.669 | -0.353 | 0.034 | 0.031 | 0.031 | 0.58 | 0.63 | 0.25 | 0.7 | 0.79 | 0.78 | 0.73 |
| 8 | 42552633 | rs4950 | 2.414 | -1.661 | 1.182 | -0.371 | 0.058 | 0.052 | 0.052 | 0.5 | 0.63 | 0.17 | 0.69 | 0.79 | 0.78 | 0.73 |
| 8 | 42552908 | rs1530848 | 2.493 | -1.659 | 1.167 | -0.37 | 0.064 | 0.058 | 0.058 | 0.78 | 0.63 | 0.12 | 0.69 | 0.79 | 0.77 | 0.72 |
| 8 | 42582583 | rs7815274 | -1.131 | -1.533 | 1.089 | -0.332 | 0.032 | 0.028 | 0.028 | 0 | 0.13 | 0.1 | 0.03 | 0 | 0.04 | 0.06 |
| 8 | 42587065 | rs4952 | -0.774 | -1.531 | 1.066 | -0.331 | 0.04 | 0.071 | 0.071 | 0 | 0.12 | 0.01 | 0.02 | 0 | 0.04 | 0.06 |
| 8 | 42604791 | rs6982753 | -2.091 | -1.65 | 1.004 | -0.279 | 0.015 | -0.01 | -0.01 | 0.439 | 0.28 | 0.84 | 0.28 | 0.23 | 0.2 | 0.21 |
| 8 | 42618286 | rs2196128 | -2.216 | -1.523 | 0.987 | -0.279 | 0.014 | -0.001 | -0.001 | 0 | 0.29 | 0.76 | 0.34 | 0.23 | 0.22 | 0.22 |
| 8 | 42618390 | rs10109429 | -1.348 | -1.585 | 1.743 | -0.31 | -0.009 | 0.217 | 0.217 | 0 | 0.12 | 0.41 | 0.2 | 0.23 | 0.05 | 0.18 |
| 8 | 42625663 | rs16891620 | -1.178 | -1.486 | 1.504 | -0.01 | 0.005 | -0.066 | -0.066 | 0 | 0.13 | 0.24 | 0.12 | 0 | 0.13 | 0.02 |
| 8 | 42629895 | rs10107450 | -2.917 | -1.68 | 1.047 | -0.074 | 0.048 | 0.045 | 0.045 | 0.668 | 0.36 | 0.88 | 0.35 | 0.21 | 0.23 | 0.29 |
| 8 | 42652859 | rs11986893 | -2.875 | -1.704 | 1.109 | -0.004 | 0.02 | 0.014 | 0.014 | 0.841 | 0.32 | 0.72 | 0.33 | 0.21 | 0.2 | 0.24 |
| 8 | 42659048 | rs7825128 | -2.034 | -1.613 | 1.648 | -0.072 | -0.029 | 0.021 | 0.021 | 0 | 0.14 | 0.3 | 0.22 | 0.14 | 0.07 | 0.16 |
| 8 | 42661847 | rs6985527 | -2.284 | -1.621 | 1.416 | -0.059 | 0.02 | 0.008 | 0.008 | 3.047 | 0.26 | 0.83 | 0.32 | 0.19 | 0.18 | 0.17 |
| 8 | 42674538 | rs7825957 | -0.888 | -1.659 | 1.698 | -0.04 | 0.001 | 0.056 | 0.056 | 0.402 | 0.08 | 0.32 | 0.22 | 0.08 | 0.03 | 0.07 |
| 8 | 42705702 | rs6474422 | -1.322 | -1.584 | 0.187 | -0.293 | -0.057 | -0.009 | -0.009 | 0 | 0.69 | 0.99 | 0.79 | 0.77 | 0.68 | 0.65 |
| 8 | 42711606 | rs7014595 | -1.219 | -1.028 | 0.303 | -0.381 | 0.025 | -0.054 | -0.054 | 0.548 | 0.09 | 0.3 | 0.07 | 0 | 0.08 | 0.08 |
| 8 | 47947701 | rs2287654 | 0.501 | -1.92 | -0.808 | -1.003 | 0.011 | 0 | 0 | 0 | 0.21 | 0.01 | 0.07 | 0.12 | 0.14 | 0.27 |
| 8 | 47986462 | rs1846504 | 0.336 | -1.497 | -0.309 | -0.783 | -0.003 | -0.004 | -0.004 | 0 | 0.53 | 0.48 | 0.63 | 0.73 | 0.71 | 0.39 |
| 8 | 48033959 | rs11136110 | 0.121 | -1.585 | -0.397 | -1.145 | -0.003 | -0.013 | -0.013 | 0 | 0.55 | 0.45 | 0.84 | 0.82 | 0.72 | 0.54 |
| 8 | 48089751 | rs10102598 | 0.427 | -1.733 | -0.178 | -1.04 | 0.029 | 0.03 | 0.03 | 1.559 | 0.49 | 0.38 | 0.56 | 0.62 | 0.64 | 0.32 |
| 8 | 48124855 | rs6558205 | 0.36 | -1.964 | -0.274 | -1.205 | 0.046 | 0.065 | 0.065 | 0 | 0.36 | 0.29 | 0.13 | 0.18 | 0.24 | 0.43 |
| 8 | 48129224 | rs9785104 | -0.536 | -1.961 | -0.024 | -1.161 | -0.04 | 0.03 | 0.03 | 0 | 0.08 | 0.26 | 0.22 | 0.08 | 0.01 | 0.19 |
| 8 | 48150346 | rs10095466 | -0.023 | -1.783 | 0.018 | -1.283 | 0.028 | 0.054 | 0.054 | 0.135 | 0.43 | 0.37 | 0.28 | 0.19 | 0.32 | 0.45 |
| 8 | 48172962 | rs35332377 | 0.107 | -1.591 | -0.008 | -1.156 | 0.062 | 0.112 | 0.112 | 0.146 | 0.42 | 0.28 | 0.15 | 0.2 | 0.28 | 0.44 |
| 8 | 48240805 | rs6473996 | -0.037 | -1.549 | 0.065 | -1.135 | -0.015 | -0.03 | -0.03 | 0 | 0.56 | 0.47 | 0.81 | 0.79 | 0.71 | 0.54 |
| 8 | 48281478 | rs10504060 | 2.159 | -0.625 | -0.44 | -1.382 | -0.062 | 0.063 | 0.063 | 1.415 | 0.11 | 0.01 | 0.1 | 0 | 0.16 | 0.07 |
| 8 | 48290688 | rs7839055 | 1.191 | -0.869 | -0.35 | -0.869 | 0.069 | 0.086 | 0.086 | 0.138 | 0.06 | 0.25 | 0.07 | 0.1 | 0.07 | 0.06 |
| 8 | 48353258 | rs2306975 | 1.053 | -0.614 | -1.213 | -0.449 | -0.096 | -0.091 | -0.091 | 3.721 | 0.14 | 0.01 | 0.21 | 0.32 | 0.3 | 0.17 |
| 8 | 48423371 | rs4873463 | 0.256 | -2.327 | -0.766 | -1.227 | 0.081 | 0.186 | 0.186 | 0.316 | 0.5 | 0.72 | 0.79 | 0.81 | 0.65 | 0.53 |
| 8 | 48436044 | rs16927307 | 1.269 | -2.27 | -0.645 | -1.143 | 0.096 | 0.024 | 0.024 | 0 | 0.05 | 0.24 | 0.07 | 0 | 0.07 | 0.05 |
| 8 | 48515523 | rs4608108 | 1.325 | -2.316 | -1.164 | -1.6 | -0.116 | -0.045 | -0.045 | 1.679 | 0.36 | 0.04 | 0.49 | 0.63 | 0.58 | 0.27 |
| 8 | 48559397 | rs10094128 | 0.307 | -2.44 | -0.995 | -1.661 | 0.067 | 0.17 | 0.17 | 0.577 | 0.5 | 0.69 | 0.77 | 0.8 | 0.65 | 0.5 |
| 8 | 48606556 | rs16930230 | 0.998 | -2.725 | -1.221 | -1.872 | 0.012 | 0.057 | 0.057 | 0 | 0.29 | 0.01 | 0.12 | 0.14 | 0.2 | 0.32 |
| 8 | 48654784 | rs10088698 | -0.638 | -2.243 | -0.343 | -1.501 | -0.024 | 0.036 | 0.036 | 0.292 | 0.12 | 0.51 | 0.28 | 0.16 | 0.09 | 0.2 |
| 8 | 48710955 | rs7830743 | 0.711 | -2.134 | -0.27 | -1.497 | 0.021 | 0.076 | 0.076 | 0 | 0.08 | 0.31 | 0.26 | 0.08 | 0.05 | 0.2 |
| 8 | 48722783 | rs7832898 | 0.211 | -2.164 | -0.281 | -1.501 | -0.001 | 0.133 | 0.133 | 0 | 0.09 | 0.35 | 0.27 | 0.18 | 0.06 | 0.22 |
| 8 | 48741570 | rs4521758 | -1.265 | -2.181 | -0.368 | -1.541 | -0.027 | -0.01 | -0.01 | 0 | 0.19 | 0.65 | 0.32 | 0.18 | 0.11 | 0.25 |
| 8 | 48770702 | rs7003908 | -0.223 | -2.234 | -0.588 | -1.755 | 0.059 | 0.117 | 0.117 | 0 | 0.51 | 0.68 | 0.71 | 0.76 | 0.64 | 0.56 |
| 8 | 48803675 | rs10109984 | -1.282 | -2.178 | -0.21 | -1.165 | -0.05 | 0.019 | 0.019 | 2.142 | 0.63 | 0.91 | 0.51 | 0.35 | 0.42 | 0.72 |
| 8 | 48816716 | rs2213178 | 0.316 | -2.197 | -0.242 | -1.225 | -0.027 | 0.174 | 0.174 | 0.427 | 0.39 | 0.02 | 0.17 | 0.12 | 0.3 | 0.46 |
| 8 | 48858933 | rs4873772 | -0.317 | -1.496 | 0.383 | -0.883 | 0.042 | 0.14 | 0.14 | 0.255 | 0.52 | 0.67 | 0.78 | 0.86 | 0.65 | 0.52 |
| 8 | 48873261 | rs1551655 | -2.126 | -0.652 | 1.468 | 0.015 | -0.055 | 0.027 | 0.027 | 0.424 | 0.13 | 0.63 | 0.29 | 0.19 | 0.06 | 0.24 |
| 8 | 48882899 | rs9657054 | -2.178 | -0.643 | 1.281 | 0.019 | -0.056 | 0.025 | 0.025 | 0.323 | 0.14 | 0.63 | 0.29 | 0.18 | 0.06 | 0.24 |
| 8 | 48885049 | rs1894311 | -0.936 | -0.616 | 1.301 | 0.037 | 0.013 | 0.152 | 0.152 | 0 | 0.08 | 0.33 | 0.26 | 0.19 | 0.06 | 0.21 |
| 8 | 48909724 | rs4873266 | -1.015 | -0.573 | 1.194 | 0.062 | 0.01 | 0.152 | 0.152 | 0 | 0.08 | 0.33 | 0.26 | 0.18 | 0.06 | 0.21 |
| 8 | 48936111 | rs28641816 | -1.929 | -0.573 | 0.974 | 0.046 | -0.056 | 0.027 | 0.027 | 0.418 | 0.14 | 0.63 | 0.29 | 0.16 | 0.06 | 0.25 |
| 8 | 48974616 | rs10808897 | -1.981 | -0.319 | 0.853 | 0.184 | -0.058 | 0.022 | 0.022 | 0.24 | 0.14 | 0.63 | 0.29 | 0.16 | 0.06 | 0.24 |
| 8 | 49003800 | rs11997106 | -1.83 | -0.552 | 0.235 | 0.275 | -0.046 | 0.024 | 0.024 | 0.378 | 0.14 | 0.64 | 0.29 | 0.18 | 0.07 | 0.22 |
| 8 | 49015537 | rs28457765 | -1.168 | -0.246 | 0.108 | 0.409 | 0.016 | 0.161 | 0.161 | 0 | 0.1 | 0.42 | 0.27 | 0.18 | 0.07 | 0.19 |
| 8 | 49039681 | rs6474040 | -0.545 | -0.54 | -1.343 | 0.132 | 0.052 | 0.002 | 0.002 | 0 | 0.43 | 0.82 | 0.41 | 0.41 | 0.3 | 0.55 |
| 8 | 49046707 | rs4433176 | -0.892 | -1.093 | -1.995 | -0.159 | 0.111 | 0.015 | 0.015 | 0 | 0.48 | 0.9 | 0.43 | 0.46 | 0.29 | 0.53 |
| 8 | 49047317 | rs7005646 | -0.378 | -1.64 | -1.798 | -0.329 | 0 | 0 | 0 | 0.05 | 0.14 | 0.13 | 0.07 | 0.15 | 0.12 | 0.2 |
| 8 | 49047878 | rs7004442 | -0.748 | -1.634 | -1.752 | -0.327 | 0.067 | 0.038 | 0.038 | 0 | 0.34 | 0.76 | 0.35 | 0.31 | 0.17 | 0.32 |

| | | | | | | | | | | | | | | | | |
|---|---|---|---|---|---|---|---|---|---|---|---|---|---|---|---|---|
| 8 | 49062168 | rs13281542 | -0.078 | -1.839 | -1.399 | -0.443 | -0.04 | 0 | 0 | 0 | 0.23 | 0.6 | 0.28 | 0.31 | 0.08 | 0.32 |
| 8 | 49064147 | rs10112655 | -0.266 | -1.866 | -1.44 | -0.456 | 0.035 | 0.002 | 0.002 | 0 | 0.24 | 0.72 | 0.29 | 0.31 | 0.09 | 0.32 |
| 8 | 49064827 | rs11781302 | -0.311 | -2.121 | -1.647 | -0.698 | 0.021 | 0.026 | 0.026 | 0 | 0.13 | 0.08 | 0.07 | 0 | 0.1 | 0.04 |
| 8 | 49089476 | rs4526384 | 0.959 | -1.554 | -1.293 | -0.414 | 0.008 | 0.001 | 0.001 | 0 | 0.62 | 0.38 | 0.65 | 0.54 | 0.75 | 0.5 |
| 8 | 49093329 | rs3934735 | 1.057 | -1.6 | -1.354 | -0.437 | 0.056 | 0.002 | 0.002 | 0 | 0.59 | 0.2 | 0.63 | 0.54 | 0.75 | 0.5 |
| 8 | 49093365 | rs3934732 | 0.115 | -1.616 | -1.331 | -0.415 | -0.025 | -0.007 | -0.007 | 0 | 0.75 | 0.41 | 0.71 | 0.71 | 0.87 | 0.67 |
| 8 | 49140396 | rs10808911 | 0.564 | -1.267 | -1.447 | -0.439 | -0.012 | -0.005 | -0.005 | 0 | 0.72 | 0.37 | 0.69 | 0.7 | 0.83 | 0.66 |
| 8 | 49156890 | rs10099462 | 0.05 | -1.346 | -1.035 | -0.458 | 0.003 | -0.002 | -0.002 | 0.057 | 0.8 | 0.81 | 0.73 | 0.74 | 0.84 | 0.68 |
| 8 | 49161509 | rs7832085 | -0.42 | -1.161 | -1.236 | -0.352 | -0.048 | -0.046 | -0.046 | 0.067 | 0.11 | 0.03 | 0.07 | 0.18 | 0.12 | 0.16 |
| 8 | 49170958 | rs4552930 | -0.529 | -1.164 | -1.238 | -0.355 | -0.048 | -0.047 | -0.047 | 0.032 | 0.12 | 0.03 | 0.07 | 0.18 | 0.12 | 0.16 |
| 8 | 49193586 | rs9650185 | 1.545 | -1.382 | -1.236 | -0.253 | 0.032 | -0.027 | -0.027 | 0.032 | 0.56 | 0.16 | 0.61 | 0.49 | 0.7 | 0.47 |
| 8 | 49194671 | rs4317605 | 0.235 | -1.298 | -1.184 | -0.337 | -0.005 | -0.054 | -0.054 | 0 | 0.91 | 0.99 | 0.97 | 0.78 | 0.94 | 0.85 |
| 8 | 49206743 | rs13280326 | 1.845 | -1.368 | -1.33 | -0.593 | -0.009 | 0.009 | 0.009 | 0 | 0.45 | 0.15 | 0.54 | 0.5 | 0.6 | 0.29 |
| 8 | 49209616 | rs12675189 | 0.504 | -1.421 | -1.255 | -0.626 | 0.001 | 0.001 | 0.001 | 0 | 0.69 | 0.36 | 0.7 | 0.71 | 0.83 | 0.64 |
| 8 | 49209652 | rs7008284 | 1.686 | -1.276 | -1.347 | -0.526 | -0.009 | 0.006 | 0.006 | 0 | 0.45 | 0.16 | 0.53 | 0.5 | 0.6 | 0.29 |
| 8 | 49213910 | rs1022140 | 0.247 | -1.248 | -0.475 | -0.114 | -0.014 | 0.005 | 0.005 | 0.039 | 0.83 | 0.74 | 0.76 | 0.72 | 0.91 | 0.7 |
| 8 | 49217135 | rs11782777 | 0.732 | -1.273 | -0.405 | -0.072 | -0.038 | 0.008 | 0.008 | 0 | 0.77 | 0.37 | 0.73 | 0.64 | 0.9 | 0.69 |
| 8 | 49220071 | rs17622439 | -0.852 | -1.219 | -0.56 | -0.203 | -0.003 | 0.005 | 0.005 | 0 | 0.18 | 0.16 | 0.15 | 0.13 | 0.19 | 0.26 |
| 8 | 49226169 | rs10441555 | -0.608 | -1.267 | -0.561 | -0.248 | -0.007 | -0.044 | -0.044 | 0 | 0.09 | 0.19 | 0.07 | 0.02 | 0.07 | 0.22 |
| 8 | 49237796 | rs11786911 | 0.82 | -0.909 | -0.614 | -0.333 | -0.03 | 0.003 | 0.003 | 0 | 0.75 | 0.49 | 0.75 | 0.66 | 0.82 | 0.66 |
| 8 | 49252776 | rs1828443 | -0.483 | -0.978 | -0.462 | -0.227 | -0.006 | -0.037 | -0.037 | 0 | 0.09 | 0.24 | 0.22 | 0.04 | 0.07 | 0.23 |
| 8 | 49252862 | rs907174 | 0.694 | -0.964 | -0.48 | -0.212 | -0.051 | 0.012 | 0.012 | 0 | 0.77 | 0.5 | 0.87 | 0.66 | 0.88 | 0.69 |
| 8 | 49253060 | rs907175 | 1.404 | -0.965 | -0.52 | -0.257 | -0.007 | 0.003 | 0.003 | 0 | 0.66 | 0.41 | 0.78 | 0.56 | 0.75 | 0.65 |
| 8 | 49283346 | rs12334834 | -2.24 | -0.517 | -0.209 | -0.003 | -0.002 | -0.007 | -0.007 | 0 | 0.3 | 0.45 | 0.21 | 0.29 | 0.27 | 0.42 |
| 8 | 49287012 | rs7832580 | -1.804 | -0.388 | -0.175 | -0.032 | -0.008 | -0.062 | -0.062 | 0 | 0.2 | 0.39 | 0.13 | 0.08 | 0.15 | 0.37 |
| 8 | 49287411 | rs13259359 | -2.26 | -0.368 | -0.303 | 0.026 | 0.007 | -0.01 | -0.01 | 0 | 0.32 | 0.5 | 0.23 | 0.29 | 0.29 | 0.47 |
| 8 | 49309167 | rs6471752 | -1.006 | -0.391 | -0.405 | 0.068 | -0.022 | -0.027 | -0.027 | 0 | 0.14 | 0.12 | 0.12 | 0.25 | 0.18 | 0.09 |
| 8 | 49323385 | rs6987448 | -0.057 | -0.515 | -0.342 | -0.009 | 0.011 | 0.019 | 0.019 | 1.041 | 0.08 | 0.02 | 0.03 | 0 | 0.05 | 0.14 |
| 8 | 49333212 | rs13273913 | -0.806 | -0.511 | -0.337 | 0.003 | 0.001 | 0 | 0 | 0 | 0.1 | 0.12 | 0.05 | 0.07 | 0.07 | 0.16 |
| 8 | 49339492 | rs13255660 | -2.056 | -0.583 | -0.26 | -0.049 | -0.051 | -0.08 | -0.08 | 0 | 0.21 | 0.48 | 0.14 | 0.12 | 0.12 | 0.4 |
| 8 | 49358454 | rs9298006 | -0.873 | -0.536 | -0.309 | 0.105 | -0.03 | -0.003 | -0.003 | 0 | 0.11 | 0.36 | 0.14 | 0.08 | 0.05 | 0.24 |
| 8 | 49361164 | rs10096760 | -2.055 | -0.343 | -0.39 | 0.126 | -0.001 | -0.003 | -0.003 | 0 | 0.32 | 0.46 | 0.27 | 0.31 | 0.28 | 0.46 |
| 8 | 49387740 | rs6995880 | 2.159 | -1.044 | -0.855 | -0.152 | -0.033 | -0.072 | -0.072 | 0 | 0.47 | 0.08 | 0.54 | 0.49 | 0.6 | 0.31 |
| 8 | 49404291 | rs11995181 | -1.19 | -0.571 | -0.301 | -0.029 | 0.006 | 0.025 | 0.025 | 0 | 0.22 | 0.39 | 0.26 | 0.24 | 0.2 | 0.29 |
| 8 | 49409564 | rs13250982 | -0.636 | -0.535 | -0.346 | -0.025 | 0.004 | 0.007 | 0.007 | 0.067 | 0.11 | 0.11 | 0.05 | 0.07 | 0.07 | 0.16 |
| 8 | 49441535 | rs2891639 | -1.162 | -0.531 | -0.208 | 0.172 | 0.009 | 0.028 | 0.028 | 0 | 0.21 | 0.4 | 0.25 | 0.24 | 0.19 | 0.29 |
| 8 | 49457532 | rs16937767 | -1.225 | -0.54 | -0.237 | 0.133 | 0.011 | 0.029 | 0.029 | 0 | 0.22 | 0.47 | 0.25 | 0.22 | 0.19 | 0.3 |
| 8 | 49469523 | rs6990833 | -2.021 | -0.57 | -0.356 | 0.105 | -0.008 | -0.002 | -0.002 | 0 | 0.32 | 0.54 | 0.31 | 0.3 | 0.26 | 0.47 |
| 8 | 49496154 | rs13250857 | 0.105 | -0.401 | -0.579 | 0.073 | -0.002 | 0.03 | 0.03 | 1.137 | 0.07 | 0 | 0.03 | 0.02 | 0.05 | 0.15 |
| 8 | 49509249 | rs13278035 | -0.295 | -0.648 | -0.575 | 0.053 | 0.024 | 0.031 | 0.031 | 0.162 | 0.09 | 0.02 | 0.06 | 0.02 | 0.06 | 0.17 |
| 8 | 49509863 | rs13251934 | -0.438 | -0.61 | -0.547 | 0.022 | 0.008 | 0.024 | 0.024 | 0 | 0.12 | 0.08 | 0.07 | 0.02 | 0.09 | 0.17 |
| 8 | 49514893 | rs16938118 | -1.589 | -0.712 | -0.246 | 0.149 | -0.014 | -0.004 | -0.004 | 0.707 | 0.3 | 0.43 | 0.3 | 0.27 | 0.25 | 0.47 |
| 8 | 49545585 | rs1905108 | 1.293 | -0.503 | 0.113 | 1.084 | -0.006 | 0.029 | 0.029 | 0 | 0.66 | 0.48 | 0.63 | 0.49 | 0.67 | 0.74 |
| 8 | 49552005 | rs10098811 | 0.265 | -0.286 | 0.284 | 1.387 | 0.001 | 0.013 | 0.013 | 0 | 0.11 | 0.11 | 0.21 | 0.21 | 0.15 | 0.07 |
| 8 | 49555905 | rs900240 | 2.021 | -0.576 | 0.008 | 1.411 | -0.02 | -0.06 | -0.06 | 0.245 | 0.58 | 0.06 | 0.5 | 0.39 | 0.63 | 0.45 |
| 8 | 49558124 | rs7822883 | -0.061 | -0.6 | 0.828 | 1.312 | -0.025 | 0.021 | 0.021 | 0 | 0.85 | 0.78 | 0.9 | 0.76 | 0.9 | 0.87 |
| 8 | 49567481 | rs6999198 | -1.467 | -0.592 | 1.179 | 1.477 | 0.064 | 0.159 | 0.159 | 0 | 0.2 | 0.48 | 0.27 | 0.38 | 0.26 | 0.15 |
| 8 | 49567497 | rs6999204 | -0.857 | -0.589 | 1.298 | 1.486 | 0.059 | 0.081 | 0.081 | 0.108 | 0.06 | 0.26 | 0.04 | 0.08 | 0.05 | 0.06 |
| 8 | 49567832 | rs11992055 | -0.587 | -0.579 | 1.286 | 1.501 | 0.02 | 0.055 | 0.055 | 0 | 0.14 | 0.22 | 0.22 | 0.3 | 0.21 | 0.09 |
| 8 | 49569508 | rs1384217 | -0.201 | -0.618 | 1.282 | 1.488 | -0.016 | -0.003 | -0.003 | 0 | 0.86 | 0.8 | 0.9 | 0.83 | 0.9 | 0.89 |
| 8 | 49572291 | rs6988001 | 0.472 | -0.602 | 1.201 | 1.503 | 0.002 | 0.024 | 0.024 | 0 | 0.1 | 0.13 | 0.2 | 0.24 | 0.15 | 0.07 |
| 8 | 49576282 | rs4242451 | 1.222 | -0.667 | 1.201 | 1.428 | -0.038 | 0.001 | 0.001 | 0 | 0.74 | 0.46 | 0.75 | 0.68 | 0.83 | 0.59 |
| 8 | 49581767 | rs878995 | -1.153 | -0.529 | 0.547 | 1.188 | -0.022 | 0.063 | 0.063 | 0 | 0.13 | 0.39 | 0.15 | 0.17 | 0.06 | 0.34 |
| 8 | 49588463 | rs1473441 | 0.949 | -0.334 | 0.529 | 1.101 | 0.007 | 0.088 | 0.088 | 0 | 0.72 | 0.49 | 0.66 | 0.54 | 0.7 | 0.81 |
| 8 | 49589585 | rs6984609 | -1.993 | -0.703 | 0.761 | 0.534 | -0.012 | -0.011 | -0.011 | 0 | 0.1 | 0.57 | 0.16 | 0.07 | 0.05 | 0.03 |
| 8 | 49589706 | rs6984786 | -2.152 | -1.118 | 0.541 | 0.125 | -0.021 | -0.041 | -0.041 | 1.179 | 0.18 | 0.71 | 0.15 | 0.09 | 0.11 | 0.42 |
| 8 | 49598167 | rs1482262 | 1.921 | -0.782 | -0.258 | 0.38 | -0.004 | -0.059 | -0.059 | 1.957 | 0.69 | 0.08 | 0.57 | 0.66 | 0.73 | 0.57 |
| 8 | 49604591 | rs10090044 | -0.501 | -0.675 | -0.097 | 0.425 | -0.001 | 0 | 0 | 0 | 0.06 | 0.08 | 0.05 | 0.08 | 0.08 | 0.04 |
| 8 | 49606500 | rs10097910 | -0.753 | -0.778 | -0.079 | 0.432 | -0.04 | -0.058 | -0.058 | 0 | 0.91 | 0.99 | 0.8 | 0.78 | 0.86 | 0.94 |
| 8 | 49607042 | rs11997496 | -1.169 | -0.807 | 0.262 | 0.398 | -0.023 | -0.004 | -0.004 | 10.743 | 0.06 | 0.13 | 0.08 | 0.05 | 0 | 0.28 |
| 8 | 49630150 | rs2891648 | 1.777 | -1.166 | -0.175 | -0.019 | 0.06 | 0.017 | 0.017 | 0 | 0.64 | 0.14 | 0.61 | 0.71 | 0.72 | 0.55 |
| 8 | 49634092 | rs11994469 | 0.067 | -0.993 | -0.07 | 0.047 | -0.015 | -0.025 | -0.025 | 0 | 0.06 | 0.12 | 0.06 | 0.01 | 0.03 | 0.06 |
| 8 | 49647435 | rs1126259 | -0.127 | -0.934 | -0.109 | 0.11 | -0.011 | -0.018 | -0.018 | 0 | 0.87 | 0.88 | 0.75 | 0.77 | 0.81 | 0.89 |
| 8 | 49656604 | rs11785222 | -0.976 | -1.069 | 0.578 | 0.051 | -0.034 | -0.024 | -0.024 | 0 | 0.1 | 0.25 | 0.05 | 0.04 | 0.05 | 0.17 |
| 8 | 49682858 | rs11779699 | -0.524 | -1.063 | 0.1 | 0.042 | 0.006 | 0.006 | 0.006 | 0.002 | 0.08 | 0.07 | 0.03 | 0.04 | 0.05 | 0.17 |
| 8 | 49684185 | rs16938743 | -0.773 | -1.085 | 0.275 | 0.121 | -0.005 | 0.022 | 0.022 | 1.523 | 0.07 | 0.49 | 0.05 | 0.03 | 0 | 0.11 |
| 8 | 49691705 | rs17712506 | 0.99 | -1.605 | -1.195 | -0.451 | -0.017 | 0.056 | 0.056 | 0 | 0.15 | 0.01 | 0.09 | 0.02 | 0.1 | 0.18 |
| 8 | 49695775 | rs12548278 | 1.419 | -1.827 | -1.25 | -0.587 | 0.043 | 0.018 | 0.018 | 0 | 0.58 | 0.12 | 0.56 | 0.67 | 0.64 | 0.49 |
| 8 | 49718452 | rs6472054 | -0.953 | -2.006 | -1.149 | -0.564 | 0.007 | 0.02 | 0.02 | 0 | 0.49 | 0.4 | 0.46 | 0.56 | 0.47 | 0.52 |

| | | | | | | | | | | | | | | | | |
|---|---|---|---|---|---|---|---|---|---|---|---|---|---|---|---|---|
| 8 | 49718692 | rs4873087 | -0.821 | -2.356 | -0.716 | -0.759 | -0.045 | 0.038 | 0.038 | 0 | 0.23 | 0.43 | 0.28 | 0.26 | 0.19 | 0.34 |
| 8 | 49729020 | rs4873290 | -1.12 | -2.167 | -0.881 | -0.757 | -0.043 | 0.016 | 0.016 | 0 | 0.24 | 0.54 | 0.26 | 0.26 | 0.14 | 0.31 |
| 8 | 49745347 | rs6472078 | -1.322 | -2.024 | -1.125 | -0.812 | 0.064 | -0.015 | -0.015 | 0 | 0.29 | 0.88 | 0.32 | 0.26 | 0.16 | 0.37 |
| 8 | 49769454 | rs10957281 | -2.014 | -2.564 | -1.514 | -1.179 | 0.068 | -0.05 | -0.05 | 0.452 | 0.49 | 0.93 | 0.46 | 0.57 | 0.34 | 0.55 |
| 8 | 49773584 | rs4075257 | 0.502 | -2.401 | -1.045 | -1.2 | 0.006 | -0.044 | -0.044 | 0 | 0.09 | 0.27 | 0.09 | 0.01 | 0.04 | 0.08 |
| 8 | 49780841 | rs16938996 | 0.082 | -2.178 | -0.654 | -0.967 | 0.009 | 0.01 | 0.01 | 0 | 0.07 | 0.26 | 0.05 | 0.04 | 0.04 | 0.21 |
| 8 | 49784645 | rs10808733 | 0.835 | -1.818 | -1.424 | -1.333 | -0.001 | -0.057 | -0.057 | 1.083 | 0.33 | 0.29 | 0.35 | 0.01 | 0.34 | 0.33 |
| 8 | 49789561 | rs6472114 | -0.707 | -1.941 | -2.064 | -1.364 | -0.058 | 0.261 | 0.261 | 0 | 0.56 | 0.84 | 0.47 | 0.05 | 0.48 | 0.68 |
| 8 | 49790942 | rs16939026 | 0.649 | -1.605 | -1.745 | -0.984 | 0.009 | 0.009 | 0.009 | 0 | 0.07 | 0.08 | 0.02 | 0.01 | 0.04 | 0.23 |
| 8 | 49816187 | rs894076 | -1.62 | -1.523 | -1.82 | -0.834 | -0.006 | -0.033 | -0.033 | 0.1 | 0.9 | 0.97 | 0.83 | 0.81 | 0.89 | 0.96 |
| 8 | 49822483 | rs12114565 | -0.56 | -1.715 | -1.516 | -1.003 | -0.04 | -0.041 | -0.041 | 0 | 0.2 | 0.48 | 0.1 | 0.15 | 0.12 | 0.46 |
| 8 | 49824013 | rs16939055 | 1.6 | -1.452 | -1.985 | -0.984 | -0.11 | -0.121 | -0.121 | 0.956 | 0.29 | 0.06 | 0.3 | 0.42 | 0.38 | 0.14 |
| 8 | 49824205 | rs1023768 | 0.617 | -1.124 | -1.646 | -1.169 | -0.005 | -0.003 | -0.003 | 0.021 | 0.31 | 0.24 | 0.34 | 0.01 | 0.34 | 0.28 |
| 8 | 49842246 | rs16939083 | 0.684 | -1.036 | -1.332 | 0.103 | 0.048 | -0.061 | -0.061 | 0.272 | 0.13 | 0 | 0.08 | 0.22 | 0.05 | 0.1 |
| 8 | 49861225 | rs870397 | 0.392 | -0.015 | -1.801 | -0.215 | -0.096 | -0.01 | -0.01 | 1.193 | 0.39 | 0.07 | 0.38 | 0.67 | 0.41 | 0.27 |
| 8 | 49880975 | rs11774804 | -0.278 | 0.028 | -1.372 | -0.192 | 0.015 | 0.098 | 0.098 | 0 | 0.53 | 0.32 | 0.45 | 0.8 | 0.54 | 0.58 |
| 8 | 49881987 | rs2203065 | -0.211 | -0.017 | -1.228 | -0.138 | -0.001 | 0.033 | 0.033 | 0 | 0.52 | 0.45 | 0.42 | 0.76 | 0.45 | 0.56 |
| 8 | 49900765 | rs17642290 | 0.369 | -0.738 | -2.195 | 0.167 | -0.01 | -0.009 | -0.009 | 0 | 0.44 | 0.17 | 0.46 | 0.4 | 0.53 | 0.37 |
| 8 | 49926047 | rs4873090 | 1.155 | -1.04 | -2.247 | 0.112 | -0.06 | -0.056 | -0.056 | 0 | 0.28 | 0.15 | 0.37 | 0.39 | 0.4 | 0.18 |
| 8 | 49930674 | rs12680366 | -0.463 | -1.373 | -1.493 | -0.387 | 0.008 | -0.04 | -0.04 | 1.29 | 0.63 | 0.6 | 0.67 | 0.82 | 0.58 | 0.58 |
| 8 | 49941703 | rs6982145 | 1.88 | -0.929 | -1.805 | -0.521 | -0.096 | 0.111 | 0.111 | 0 | 0.19 | 0.02 | 0.14 | 0 | 0.26 | 0.06 |
| 8 | 49957895 | rs7835336 | -0.37 | -1.931 | -1.366 | -0.5 | -0.011 | 0.077 | 0.077 | 0 | 0.59 | 0.31 | 0.64 | 0.81 | 0.57 | 0.58 |
| 8 | 49960530 | rs1467089 | 1.044 | -2.177 | -1.55 | -0.018 | 0.018 | -0.077 | -0.077 | 0.219 | 0.18 | 0.01 | 0.21 | 0.28 | 0.12 | 0.2 |
| 8 | 49966480 | rs9650217 | 0.27 | -2.029 | -1.477 | 0.084 | 0.049 | -0.09 | -0.09 | 0.225 | 0.27 | 0.01 | 0.24 | 0.34 | 0.17 | 0.38 |
| 8 | 49975746 | rs2385127 | -0.415 | -1.925 | -1.427 | -0.206 | 0.004 | 0.088 | 0.088 | 0 | 0.58 | 0.31 | 0.66 | 0.81 | 0.62 | 0.56 |
| 8 | 49978424 | rs341815 | 0.75 | -1.96 | -1.21 | -0.253 | -0.004 | 0.016 | 0.016 | 0 | 0.35 | 0.24 | 0.28 | 0.07 | 0.36 | 0.39 |
| 8 | 49978554 | rs10504070 | 1.306 | -1.754 | -1.147 | 0.013 | -0.018 | -0.01 | -0.01 | 0.111 | 0.3 | 0.27 | 0.29 | 0.32 | 0.39 | 0.17 |
| 8 | 49987230 | rs2292399 | 1.349 | -1.733 | -1.059 | 0.012 | -0.083 | -0.047 | -0.047 | 0 | 0.31 | 0.12 | 0.28 | 0.33 | 0.41 | 0.18 |
| 8 | 49988587 | rs10957357 | 1.26 | -1.732 | -1.066 | 0.013 | -0.023 | -0.011 | -0.011 | 0 | 0.31 | 0.27 | 0.29 | 0.33 | 0.41 | 0.18 |
| 8 | 49995284 | rs1876718 | 0.444 | -1.663 | -1.349 | 0.094 | 0.059 | -0.163 | -0.163 | 0 | 0.23 | 0.03 | 0.33 | 0.46 | 0.13 | 0.23 |
| 8 | 50001946 | rs341828 | 0.372 | -1.629 | -1.223 | -0.037 | -0.003 | 0.048 | 0.048 | 0 | 0.39 | 0.25 | 0.3 | 0.1 | 0.38 | 0.54 |
| 8 | 50014500 | rs9643395 | 0.992 | -1.433 | -0.934 | 0.172 | 0.017 | 0.005 | 0.005 | 0 | 0.33 | 0.39 | 0.32 | 0.33 | 0.42 | 0.17 |
| 8 | 50030513 | rs7819763 | -0.328 | -1.386 | -1.755 | 0.212 | -0.026 | 0.087 | 0.087 | 0.164 | 0.6 | 0.8 | 0.68 | 0.74 | 0.55 | 0.41 |
| 8 | 50037784 | rs341801 | 0.605 | -1.161 | -1.6 | 0.337 | 0.004 | -0.002 | -0.002 | 1.255 | 0.37 | 0.58 | 0.34 | 0.32 | 0.42 | 0.17 |
| 8 | 50044748 | rs1397797 | 0.344 | -1.068 | -1.515 | 0.457 | 0.009 | 0.036 | 0.036 | 0 | 0.95 | 0.9 | 0.93 | 0.86 | 0.93 | 0.94 |
| 8 | 50052500 | rs1606133 | 0.512 | -1.562 | -1.893 | 0.678 | 0.015 | -0.092 | -0.092 | 0.062 | 0.17 | 0.01 | 0.23 | 0.29 | 0.15 | 0.23 |
| 8 | 50054112 | rs1828494 | -0.883 | -1.88 | -2.027 | 0.178 | -0.018 | -0.018 | -0.018 | 0 | 0.59 | 0.7 | 0.62 | 0.52 | 0.52 | 0.75 |
| 8 | 50057792 | rs341821 | -0.02 | -1.919 | -2.017 | 0.188 | 0.02 | 0.073 | 0.073 | 0 | 0.94 | 0.83 | 0.92 | 0.86 | 0.93 | 0.94 |
| 8 | 50058707 | rs10081464 | 1.226 | -1.88 | -2.016 | 0.169 | -0.05 | -0.013 | -0.013 | 0.641 | 0.35 | 0.12 | 0.3 | 0.33 | 0.41 | 0.19 |
| 8 | 50058775 | rs17645042 | 0.269 | -1.597 | -1.841 | -0.1 | -0.041 | 0.087 | 0.087 | 0.18 | 0.33 | 0.14 | 0.33 | 0.13 | 0.36 | 0.51 |
| 8 | 50058986 | rs341822 | 0.619 | -1.211 | -1.224 | 0.399 | 0.024 | 0.057 | 0.057 | 0 | 0.95 | 0.83 | 0.95 | 0.91 | 0.93 | 0.95 |
| 8 | 50063992 | rs2667289 | 0.02 | -1.347 | -1.634 | 0.374 | 0.032 | -0.069 | -0.069 | 0 | 0.75 | 0.93 | 0.78 | 0.59 | 0.81 | 0.74 |
| 8 | 50070098 | rs13264833 | 0.18 | -1.385 | -1.603 | 0.229 | 0.015 | -0.071 | -0.071 | 0 | 0.7 | 0.91 | 0.76 | 0.56 | 0.77 | 0.67 |
| 8 | 50072010 | rs7841742 | 0.67 | -1.537 | -1.102 | 0.161 | 0.001 | -0.002 | -0.002 | 0 | 0.65 | 0.41 | 0.7 | 0.55 | 0.77 | 0.66 |
| 8 | 50080612 | rs11987140 | 0.504 | -1.681 | -1.455 | -0.114 | 0.005 | 0.024 | 0.024 | 0 | 0.31 | 0.23 | 0.31 | 0.16 | 0.28 | 0.4 |
| 8 | 50108120 | rs1480390 | -1.064 | -1.633 | -1.466 | -0.077 | 0 | 0 | 0 | 0.023 | 0.95 | 0.95 | 0.97 | 0.95 | 0.95 | 0.97 |
| 8 | 50108420 | rs1480391 | 0.355 | -1.707 | -1.462 | -0.105 | 0.007 | 0.023 | 0.023 | 0 | 0.33 | 0.3 | 0.32 | 0.18 | 0.28 | 0.4 |
| 8 | 50111746 | rs2385320 | 1.385 | -1.658 | -1.79 | -0.045 | -0.08 | -0.033 | -0.033 | 0.054 | 0.33 | 0.15 | 0.4 | 0.38 | 0.49 | 0.26 |
| 8 | 50115532 | rs7814408 | -0.146 | -1.636 | -1.81 | 0.002 | 0.017 | -0.064 | -0.064 | 0.01 | 0.7 | 0.91 | 0.76 | 0.58 | 0.79 | 0.68 |
| 8 | 50125078 | rs1124263 | 1.29 | -1.435 | -1.821 | -0.022 | -0.088 | -0.032 | -0.032 | 0 | 0.33 | 0.13 | 0.37 | 0.36 | 0.48 | 0.26 |
| 8 | 50145188 | rs985986 | 0.12 | -1.488 | -1.619 | 0.079 | 0.02 | -0.041 | -0.041 | 0 | 0.26 | 0.19 | 0.24 | 0.41 | 0.19 | 0.33 |
| 8 | 50165831 | rs1383273 | 0.413 | -1.624 | -1.358 | -0.009 | -0.014 | 0.014 | 0.014 | 0 | 0.66 | 0.48 | 0.71 | 0.55 | 0.77 | 0.67 |
| 8 | 50187902 | rs11995494 | 0.406 | -1.708 | -1.693 | -0.167 | 0.01 | 0.046 | 0.046 | 0 | 0.32 | 0.18 | 0.31 | 0.15 | 0.28 | 0.41 |
| 8 | 50195185 | rs17664823 | 0.464 | -1.689 | -1.782 | -0.174 | 0.003 | 0.049 | 0.049 | 0 | 0.31 | 0.13 | 0.31 | 0.15 | 0.28 | 0.41 |
| 8 | 50200680 | rs2385334 | -0.671 | -1.824 | -1.514 | -0.1 | -0.001 | -0.002 | -0.002 | 0 | 0.94 | 0.93 | 0.97 | 0.95 | 0.96 | 0.98 |
| 8 | 50210823 | rs16906613 | 0.211 | -2.003 | -1.804 | -0.047 | -0.007 | -0.124 | -0.124 | 0 | 0.24 | 0.02 | 0.2 | 0.4 | 0.19 | 0.28 |
| 8 | 50233005 | rs9650228 | 0.282 | -2.033 | -1.832 | -0.236 | -0.005 | -0.107 | -0.107 | 0.116 | 0.23 | 0.04 | 0.2 | 0.43 | 0.18 | 0.26 |
| 8 | 50235530 | rs16906822 | 0.819 | -2.353 | -1.948 | -0.55 | 0.001 | 0.001 | 0.001 | 0 | 0.17 | 0.14 | 0.22 | 0.12 | 0.13 | 0.32 |
| 8 | 50235564 | rs17665879 | 0.686 | -2.491 | -2.029 | -0.736 | 0.008 | 0.019 | 0.019 | 0 | 0.16 | 0.11 | 0.08 | 0.02 | 0.11 | 0.12 |
| 8 | 50236545 | rs1519833 | 1.055 | -2.647 | -2.031 | -0.809 | 0.058 | 0.009 | 0.009 | 0 | 0.5 | 0.31 | 0.63 | 0.51 | 0.66 | 0.57 |
| 8 | 50252962 | rs1519835 | 2.062 | -2.632 | -2.073 | -0.813 | -0.16 | -0.109 | -0.109 | 1.121 | 0.31 | 0.03 | 0.41 | 0.38 | 0.53 | 0.25 |
| 8 | 50253358 | rs1356760 | 0.02 | -2.474 | -1.48 | -0.548 | 0.022 | 0.008 | 0.008 | 0 | 0.91 | 0.49 | 0.92 | 0.95 | 0.96 | 0.96 |
| 8 | 50259111 | rs2139454 | 1.42 | -2.496 | -1.689 | -0.556 | 0.058 | -0.011 | -0.011 | 0 | 0.48 | 0.2 | 0.63 | 0.49 | 0.66 | 0.57 |
| 8 | 50261435 | rs10504079 | 0.522 | -2.449 | -1.713 | -0.538 | 0.009 | 0.026 | 0.026 | 0 | 0.15 | 0.08 | 0.08 | 0.02 | 0.11 | 0.12 |
| 8 | 50265730 | rs2222291 | -0.367 | -2.217 | -1.555 | -0.488 | 0.041 | -0.041 | -0.041 | 0 | 0.41 | 0.13 | 0.28 | 0.43 | 0.3 | 0.39 |
| 8 | 50275156 | rs13260210 | -0.412 | -2.223 | -1.543 | -0.492 | 0.048 | -0.037 | -0.037 | 0 | 0.42 | 0.12 | 0.29 | 0.43 | 0.3 | 0.4 |
| 8 | 50279613 | rs10808760 | 0.633 | -1.828 | -0.56 | -1.071 | -0.006 | -0.018 | -0.018 | 3.03 | 0.9 | 0.36 | 0.92 | 0.95 | 0.96 | 0.96 |
| 8 | 50283147 | rs7001086 | 0.299 | -1.752 | -0.724 | -1.179 | 0.019 | 0.022 | 0.022 | 0 | 0.81 | 0.88 | 0.85 | 0.91 | 0.89 | 0.75 |
| 8 | 50290167 | rs1401823 | 0.903 | -1.626 | -0.554 | -1.27 | 0.003 | 0.015 | 0.015 | 0 | 0.77 | 0.8 | 0.8 | 0.91 | 0.82 | 0.71 |

| | | | | | | | | | | | | | | | | |
|---|---|---|---|---|---|---|---|---|---|---|---|---|---|---|---|---|
| 8 | 50294735 | rs7828647 | 1.324 | -1.735 | 0.411 | -1.322 | -0.005 | -0.062 | -0.062 | 0.017 | 0.71 | 0.47 | 0.74 | 0.88 | 0.79 | 0.69 |
| 8 | 50306276 | rs1401817 | -1.262 | -1.536 | 0.396 | -1.242 | 0.102 | -0.024 | -0.024 | 0.218 | 0.06 | 0.18 | 0.09 | 0.01 | 0.1 | 0.06 |
| 8 | 50315105 | rs729592 | 1.268 | -1.817 | 0.422 | -1.291 | 0.032 | 0.051 | 0.051 | 0.281 | 0.72 | 0.19 | 0.77 | 0.88 | 0.85 | 0.74 |
| 8 | 50348450 | rs1674269 | -0.673 | -1.771 | 0.42 | -1.295 | 0.027 | 0.034 | 0.034 | 0 | 0.2 | 0.08 | 0.14 | 0.09 | 0.11 | 0.24 |
| 8 | 50392001 | rs1674258 | -0.728 | -1.744 | 0.108 | -1.266 | 0.018 | 0.019 | 0.019 | 0 | 0.2 | 0.12 | 0.15 | 0.11 | 0.11 | 0.25 |
| 8 | 50396852 | rs1715350 | -0.834 | -1.702 | 0.142 | -1.246 | -0.001 | 0.021 | 0.021 | 0 | 0.2 | 0.12 | 0.18 | 0.09 | 0.16 | 0.3 |
| 8 | 50403271 | rs1715347 | 1.491 | -1.464 | 0.546 | -1.148 | 0.023 | -0.035 | -0.035 | 0 | 0.92 | 0.35 | 0.92 | 0.99 | 0.96 | 0.98 |
| 8 | 50416266 | rs1715326 | 0.87 | -1.448 | 0.754 | -1.135 | 0.126 | 0.051 | 0.051 | 2.643 | 0.94 | 0.51 | 0.94 | 0.99 | 0.96 | 0.99 |
| 8 | 50430756 | rs9643612 | 1.098 | -2.131 | -0.051 | -1.683 | 0.033 | 0.214 | 0.214 | 1.89 | 0.54 | 0.08 | 0.61 | 0.81 | 0.66 | 0.55 |
| 8 | 50445456 | rs2956830 | -0.358 | -2.013 | 0.42 | -1.439 | -0.012 | 0.005 | 0.005 | 0 | 0.12 | 0.26 | 0.14 | 0.11 | 0.07 | 0.21 |
| 8 | 50462970 | rs2960488 | -0.292 | -2.139 | 0.367 | -1.501 | -0.008 | -0.004 | -0.004 | 0 | 0.13 | 0.15 | 0.14 | 0.08 | 0.07 | 0.21 |
| 8 | 50481632 | rs1585260 | 0.903 | -2.576 | 0.092 | -1.367 | 0.054 | -0.062 | -0.062 | 0.014 | 0.2 | 0.02 | 0.06 | 0.27 | 0.11 | 0.17 |
| 8 | 50505231 | rs7831417 | 0.48 | -1.466 | 1.21 | -1.401 | -0.015 | 0.001 | 0.001 | 0 | 0.87 | 0.73 | 0.86 | 0.89 | 0.93 | 0.79 |
| 8 | 50509258 | rs13251699 | 0.565 | -1.466 | 1.23 | -1.402 | -0.015 | 0.005 | 0.005 | 0 | 0.87 | 0.66 | 0.86 | 0.89 | 0.93 | 0.79 |
| 8 | 50514468 | rs6986382 | 0.543 | -1.466 | 1.23 | -1.402 | -0.014 | 0.006 | 0.006 | 0 | 0.87 | 0.66 | 0.86 | 0.89 | 0.93 | 0.79 |
| 8 | 50515677 | rs12678100 | -0.19 | -1.468 | 1.251 | -1.402 | -0.005 | 0.001 | 0.001 | 0 | 0.89 | 0.86 | 0.87 | 0.92 | 0.93 | 0.79 |
| 8 | 50516102 | rs10957531 | -0.19 | -1.468 | 1.251 | -1.402 | -0.005 | 0.001 | 0.001 | 0 | 0.89 | 0.86 | 0.87 | 0.92 | 0.93 | 0.79 |
| 8 | 50520852 | rs11779800 | 0.579 | -1.468 | 1.24 | -1.402 | -0.015 | 0.005 | 0.005 | 0 | 0.87 | 0.66 | 0.86 | 0.89 | 0.93 | 0.79 |
| 8 | 50523731 | rs2132526 | 0.579 | -1.468 | 1.24 | -1.402 | -0.015 | 0.005 | 0.005 | 0 | 0.87 | 0.66 | 0.86 | 0.89 | 0.93 | 0.79 |
| 8 | 50528355 | rs7830871 | -0.089 | -1.49 | 1.191 | -1.403 | -0.002 | -0.001 | -0.001 | 0.002 | 0.88 | 0.89 | 0.86 | 0.92 | 0.92 | 0.79 |
| 8 | 50532935 | rs1908709 | -0.96 | -1.431 | 1.306 | -1.387 | 0.155 | 0.089 | 0.089 | 1.341 | 0.05 | 0.46 | 0.06 | 0.04 | 0.04 | 0.01 |
| 8 | 50539763 | rs10104767 | -0.994 | -1.434 | 1.295 | -1.389 | 0.155 | 0.089 | 0.089 | 1.298 | 0.06 | 0.45 | 0.06 | 0.04 | 0.04 | 0.01 |
| 8 | 50560660 | rs10098245 | -1.492 | -1.442 | 1.267 | -1.391 | 0.127 | 0.076 | 0.076 | 2.801 | 0.06 | 0.46 | 0.06 | 0.07 | 0.04 | 0.01 |
| 8 | 50580952 | rs1587191 | -0.161 | -1.475 | 1.217 | -1.408 | 0 | 0 | 0 | 0 | 0.88 | 0.9 | 0.86 | 0.92 | 0.92 | 0.78 |
| 8 | 50581476 | rs7824171 | 1.554 | -1.471 | 1.204 | -1.422 | 0.006 | 0.009 | 0.009 | 1.367 | 0.79 | 0.19 | 0.79 | 0.85 | 0.88 | 0.78 |
| 8 | 50584283 | rs7812615 | -1.613 | -1.454 | 1.236 | -1.388 | 0.121 | 0.071 | 0.071 | 2.389 | 0.07 | 0.46 | 0.06 | 0.07 | 0.04 | 0.01 |
| 8 | 50609940 | rs7386188 | 0.958 | -1.482 | 1.569 | -1.381 | 0.15 | 0.085 | 0.085 | 1.21 | 0.94 | 0.55 | 0.94 | 0.96 | 0.96 | 0.99 |
| 8 | 50625113 | rs9643640 | 1.302 | -1.446 | 1.569 | -1.379 | 0.138 | 0.073 | 0.073 | 2.379 | 0.93 | 0.53 | 0.94 | 0.93 | 0.95 | 0.99 |
| 8 | 50632220 | rs1982925 | 1.286 | -1.608 | 1.308 | -1.398 | 0.006 | 0.008 | 0.008 | 1.404 | 0.8 | 0.19 | 0.79 | 0.84 | 0.88 | 0.78 |
| 8 | 50647262 | rs1390586 | -0.754 | -1.779 | 1.269 | -1.473 | -0.002 | -0.014 | -0.014 | 0 | 0.17 | 0.21 | 0.19 | 0.11 | 0.16 | 0.23 |
| 8 | 50653706 | rs7818806 | 1.705 | -1.847 | 1.335 | -1.509 | 0.013 | 0.038 | 0.038 | 0.887 | 0.75 | 0.16 | 0.75 | 0.84 | 0.82 | 0.76 |
| 8 | 50658801 | rs12546003 | 0.023 | -2.466 | 0.924 | -1.772 | 0.053 | 0.09 | 0.09 | 0 | 0.24 | 0.08 | 0.13 | 0.05 | 0.13 | 0.19 |
| 8 | 50660250 | rs1603400 | -0.647 | -2.082 | 1.641 | -1.54 | -0.008 | -0.015 | -0.015 | 0 | 0.17 | 0.22 | 0.17 | 0.11 | 0.15 | 0.24 |
| 8 | 50662483 | rs16912132 | -0.769 | -2.074 | 1.76 | -1.536 | -0.014 | -0.028 | -0.028 | 0 | 0.17 | 0.3 | 0.17 | 0.11 | 0.15 | 0.24 |
| 8 | 50662598 | rs13251494 | 0.635 | -2.058 | 1.804 | -1.536 | 0.098 | 0.049 | 0.049 | 0.285 | 0.95 | 0.65 | 0.96 | 0.99 | 0.98 | 0.99 |
| 8 | 50683876 | rs4873376 | 1.618 | -2.087 | 1.747 | -1.552 | 0.017 | 0.034 | 0.034 | 0.27 | 0.76 | 0.18 | 0.78 | 0.88 | 0.83 | 0.75 |
| 8 | 50685486 | rs12545000 | 0.733 | -1.911 | 1.866 | -1.472 | 0.103 | 0.051 | 0.051 | 0.695 | 0.94 | 0.58 | 0.96 | 0.99 | 0.98 | 0.99 |
| 8 | 50727379 | rs1979536 | 1.413 | -2.074 | 1.204 | -1.508 | 0.024 | 0.041 | 0.041 | 0.531 | 0.75 | 0.15 | 0.78 | 0.84 | 0.84 | 0.7 |
| 8 | 50732569 | rs17674246 | -0.178 | -2.024 | 1.138 | -1.549 | -0.01 | 0.01 | 0.01 | 0 | 0.13 | 0.06 | 0.07 | 0 | 0.15 | 0.06 |
| 8 | 50752038 | rs11786785 | 1.733 | -1.896 | 0.847 | -1.743 | 0.027 | 0.253 | 0.253 | 0.322 | 0.52 | 0.07 | 0.65 | 0.83 | 0.65 | 0.51 |
| 8 | 50758648 | rs17747567 | -0.412 | -1.593 | 1.19 | -1.528 | -0.002 | -0.025 | -0.025 | 0.721 | 0.15 | 0.14 | 0.09 | 0 | 0.17 | 0.07 |
| 8 | 50777802 | rs11785147 | 1.6 | -1.616 | 0.665 | -1.529 | 0.004 | 0.012 | 0.012 | 0.022 | 0.76 | 0.25 | 0.78 | 0.88 | 0.82 | 0.7 |
| 8 | 50781413 | rs13255117 | 1.127 | -1.507 | 0.817 | -1.26 | -0.016 | 0.08 | 0.08 | 0.25 | 0.92 | 0.43 | 0.88 | 0.88 | 0.99 | 0.77 |
| 8 | 50786230 | rs17676043 | 1.285 | -1.528 | -0.068 | -1.594 | 0.031 | 0.207 | 0.207 | 0.362 | 0.55 | 0.07 | 0.67 | 0.83 | 0.69 | 0.53 |
| 8 | 50794211 | rs7003721 | 0.474 | -1.636 | 0.997 | -1.615 | -0.034 | 0.017 | 0.017 | 1.034 | 0.88 | 0.38 | 0.87 | 0.88 | 0.97 | 0.76 |
| 8 | 50808334 | rs451034 | 1.016 | -1.68 | 1.06 | -1.606 | -0.047 | 0.003 | 0.003 | 2.491 | 0.86 | 0.28 | 0.85 | 0.86 | 0.97 | 0.76 |
| 8 | 50813174 | rs11782987 | 0.227 | -1.655 | 0.972 | -1.603 | -0.008 | 0.025 | 0.025 | 0 | 0.07 | 0.17 | 0.1 | 0.12 | 0.03 | 0.24 |
| 8 | 50815672 | rs12549517 | 0.222 | -1.729 | 0.651 | -1.723 | -0.001 | 0.118 | 0.118 | 0 | 0.22 | 0.02 | 0.11 | 0.03 | 0.17 | 0.2 |
| 8 | 50824431 | rs967116 | -0.244 | -1.682 | 0.657 | -1.791 | -0.015 | 0.009 | 0.009 | 0 | 0.15 | 0.09 | 0.08 | 0 | 0.17 | 0.07 |
| 8 | 50834340 | rs13271446 | 2.108 | -2.125 | 0.263 | -1.448 | -0.111 | -0.11 | -0.11 | 0 | 0.37 | 0.04 | 0.61 | 0.55 | 0.58 | 0.38 |
| 8 | 50875494 | rs971926 | 1.246 | -2.229 | 0.835 | -1.493 | 0.005 | 0.013 | 0.013 | 0 | 0.72 | 0.18 | 0.76 | 0.86 | 0.8 | 0.71 |
| 8 | 50878647 | rs1486247 | 0.961 | -2.398 | 0.731 | -1.398 | 0.035 | -0.041 | -0.041 | 0 | 0.13 | 0.02 | 0.04 | 0.27 | 0.04 | 0.11 |
| 8 | 50880234 | rs1344326 | -1.5 | -1.526 | 1.418 | -1.944 | -0.048 | 0.004 | 0.004 | 1.477 | 0.14 | 0.73 | 0.16 | 0.14 | 0.03 | 0.23 |
| 8 | 50882830 | rs9650160 | 0.185 | -1.599 | 1.201 | -2.033 | 0.018 | 0.108 | 0.108 | 0 | 0.21 | 0.02 | 0.1 | 0.03 | 0.16 | 0.2 |
| 8 | 50890197 | rs318903 | -0.205 | -1.548 | 1.211 | -2.036 | -0.023 | 0.026 | 0.026 | 0 | 0.87 | 0.95 | 0.92 | 1 | 0.84 | 0.94 |
| 8 | 50891046 | rs9643689 | 2.098 | -1.58 | 1.118 | -2.033 | 0.063 | 0.242 | 0.242 | 0 | 0.51 | 0.11 | 0.66 | 0.83 | 0.64 | 0.52 |
| 8 | 50894694 | rs13266009 | 2.119 | -1.582 | 1.439 | -2.034 | 0.063 | 0.242 | 0.242 | 0 | 0.51 | 0.1 | 0.66 | 0.82 | 0.64 | 0.51 |
| 8 | 50898214 | rs10112215 | 1.609 | -1.505 | 1.454 | -1.97 | 0.006 | 0.015 | 0.015 | 0 | 0.72 | 0.19 | 0.76 | 0.86 | 0.8 | 0.71 |
| 8 | 50901335 | rs4873121 | -0.401 | -1.464 | 1.377 | -1.911 | -0.012 | 0.01 | 0.01 | 0 | 0.09 | 0.2 | 0.11 | 0.13 | 0.03 | 0.23 |
| 8 | 50905788 | rs186454 | 0.918 | -1.456 | 1.38 | -1.883 | -0.003 | 0.011 | 0.011 | 0 | 0.79 | 0.77 | 0.81 | 0.84 | 0.81 | 0.71 |
| 8 | 50914632 | rs17685307 | 0.125 | -1.513 | 1.168 | -1.947 | 0.023 | 0.103 | 0.103 | 0 | 0.21 | 0.02 | 0.1 | 0.03 | 0.16 | 0.2 |
| 8 | 50916525 | rs1552316 | 1.901 | -1.587 | 1.36 | -1.98 | 0.062 | 0.175 | 0.175 | 0 | 0.52 | 0.16 | 0.66 | 0.8 | 0.65 | 0.51 |
| 8 | 50917230 | rs318915 | 0.775 | -1.508 | 1.073 | -1.965 | -0.008 | 0.027 | 0.027 | 0 | 0.8 | 0.84 | 0.89 | 0.95 | 0.81 | 0.88 |
| 8 | 50921942 | rs13270712 | 0.056 | -1.518 | 1.05 | -1.968 | 0.026 | 0.107 | 0.107 | 0 | 0.21 | 0.02 | 0.1 | 0.03 | 0.16 | 0.2 |
| 8 | 50927333 | rs1484813 | 0.624 | -1.145 | 0.3 | -1.774 | -0.014 | 0.035 | 0.035 | 0 | 0.81 | 0.89 | 0.9 | 0.96 | 0.81 | 0.88 |
| 8 | 50932336 | rs12541061 | 0.798 | -1.139 | 0.313 | -1.771 | -0.004 | 0.011 | 0.011 | 0 | 0.8 | 0.82 | 0.89 | 0.94 | 0.81 | 0.88 |
| 8 | 50936174 | rs1021202 | 0.795 | -1.135 | 0.214 | -1.779 | -0.008 | 0.027 | 0.027 | 0 | 0.8 | 0.84 | 0.89 | 0.95 | 0.81 | 0.88 |
| 8 | 50943748 | rs11781074 | -0.306 | -1.216 | -0.019 | -1.847 | -0.005 | 0.142 | 0.142 | 0 | 0.25 | 0.02 | 0.12 | 0.03 | 0.21 | 0.22 |
| 8 | 50945075 | rs12546435 | 1.104 | -1.448 | 0.422 | -1.934 | -0.011 | -0.017 | -0.017 | 2.32 | 0.79 | 0.26 | 0.78 | 0.88 | 0.84 | 0.74 |

| | | | | | | | | | | | | | | | | |
|---|---|---|---|---|---|---|---|---|---|---|---|---|---|---|---|---|
| 8 | 50946702 | rs17758599 | 1.665 | -1.482 | 0.417 | -1.951 | 0.065 | 0.185 | 0.185 | 0 | 0.53 | 0.16 | 0.65 | 0.84 | 0.64 | 0.52 |
| 8 | 50957454 | rs12545107 | -0.185 | -1.487 | 0.375 | -1.957 | -0.004 | 0.144 | 0.144 | 0 | 0.25 | 0.02 | 0.12 | 0.03 | 0.2 | 0.22 |
| 8 | 50967030 | rs203959 | 0.104 | -1.491 | 0.896 | -1.958 | 0 | -0.031 | -0.031 | 0.181 | 0.86 | 0.86 | 0.92 | 0.98 | 0.85 | 0.92 |
| 8 | 50976537 | rs203949 | 1.011 | -1.508 | 1.399 | -1.973 | -0.016 | -0.048 | -0.048 | 0 | 0.62 | 0.5 | 0.72 | 0.85 | 0.71 | 0.56 |
| 8 | 50977672 | rs203946 | 0.114 | -1.502 | 1.434 | -1.969 | 0.002 | -0.045 | -0.045 | 0.305 | 0.86 | 0.82 | 0.92 | 0.98 | 0.85 | 0.92 |
| 8 | 50980320 | rs1000638 | -0.649 | -1.52 | 1.574 | -1.978 | 0.01 | 0.014 | 0.014 | 0.193 | 0.29 | 0.29 | 0.24 | 0.14 | 0.2 | 0.39 |
| 8 | 50982597 | rs203940 | 1.416 | -1.52 | 1.574 | -1.978 | -0.001 | -0.018 | -0.018 | 0 | 0.57 | 0.52 | 0.67 | 0.84 | 0.64 | 0.52 |
| 8 | 50988837 | rs203930 | 1.916 | -1.544 | 1.584 | -1.983 | 0.06 | 0.238 | 0.238 | 0 | 0.51 | 0.07 | 0.64 | 0.81 | 0.64 | 0.47 |
| 8 | 50990861 | rs203927 | 0.334 | -1.564 | 1.521 | -2.035 | 0 | -0.044 | -0.044 | 0 | 0.85 | 0.82 | 0.91 | 0.98 | 0.85 | 0.92 |
| 8 | 51013729 | rs10096871 | -0.573 | -1.571 | 1.456 | -2.011 | 0.024 | 0.038 | 0.038 | 0.254 | 0.29 | 0.24 | 0.24 | 0.14 | 0.2 | 0.39 |
| 8 | 51022682 | rs203908 | 1.399 | -1.495 | 1.491 | -1.986 | -0.004 | -0.024 | -0.024 | 0.089 | 0.57 | 0.52 | 0.67 | 0.84 | 0.64 | 0.53 |
| 8 | 51029371 | rs1903311 | -0.481 | -1.491 | 1.355 | -1.807 | 0.026 | 0.061 | 0.061 | 0 | 0.27 | 0.13 | 0.22 | 0.14 | 0.2 | 0.38 |
| 8 | 51040022 | rs10095718 | -1.153 | -1.455 | 1.725 | -1.731 | 0.074 | 0.076 | 0.076 | 3.897 | 0.05 | 0.46 | 0.03 | 0 | 0 | 0.01 |
| 8 | 51067154 | rs10098057 | -1.417 | -1.499 | 1.593 | -1.842 | 0.058 | 0.138 | 0.138 | 0.947 | 0.4 | 0.88 | 0.33 | 0.16 | 0.29 | 0.46 |
| 8 | 51070193 | rs987356 | -0.99 | -1.48 | 1.636 | -1.834 | 0.013 | 0.056 | 0.056 | 0 | 0.33 | 0.23 | 0.27 | 0.15 | 0.28 | 0.41 |
| 8 | 51070979 | rs11986160 | -0.99 | -1.48 | 1.636 | -1.834 | 0.013 | 0.056 | 0.056 | 0 | 0.33 | 0.23 | 0.27 | 0.15 | 0.28 | 0.41 |
| 8 | 51084731 | rs4873128 | -1.139 | -1.522 | 1.593 | -1.854 | 0.008 | 0.018 | 0.018 | 0 | 0.34 | 0.31 | 0.28 | 0.15 | 0.28 | 0.45 |
| 8 | 51088892 | rs898520 | -1.159 | -1.522 | 1.592 | -1.854 | 0.005 | 0.01 | 0.01 | 0 | 0.34 | 0.35 | 0.28 | 0.15 | 0.28 | 0.45 |
| 8 | 51091268 | rs7002235 | -1.149 | -1.522 | 1.593 | -1.854 | 0.005 | 0.01 | 0.01 | 0 | 0.34 | 0.35 | 0.28 | 0.15 | 0.28 | 0.45 |
| 8 | 51125957 | rs4873428 | -1.553 | -1.538 | 1.279 | -1.873 | 0.066 | 0.148 | 0.148 | 0.761 | 0.41 | 0.88 | 0.32 | 0.16 | 0.28 | 0.45 |
| 8 | 51127196 | rs4873135 | -1.553 | -1.538 | 1.279 | -1.873 | 0.066 | 0.148 | 0.148 | 0.761 | 0.41 | 0.88 | 0.32 | 0.16 | 0.28 | 0.45 |
| 8 | 51142922 | rs12544631 | -1.364 | -1.635 | 1.371 | -1.861 | -0.022 | -0.05 | -0.05 | 0 | 0.33 | 0.49 | 0.29 | 0.18 | 0.26 | 0.43 |
| 8 | 51164313 | rs4383986 | 0.928 | -1.545 | 1.257 | -1.926 | -0.043 | -0.056 | -0.056 | 0 | 0.89 | 0.6 | 0.97 | 1 | 0.99 | 0.98 |
| 8 | 51170362 | rs9298295 | -0.693 | -1.291 | 0.751 | -1.796 | 0 | 0.039 | 0.039 | 0 | 0.27 | 0.15 | 0.15 | 0.06 | 0.26 | 0.18 |
| 8 | 51182810 | rs1375948 | 1.625 | -0.608 | 0.569 | -1.82 | 0.023 | 0.19 | 0.19 | 0 | 0.49 | 0.17 | 0.61 | 0.82 | 0.57 | 0.49 |
| 8 | 51186606 | rs6473039 | 0.929 | -0.613 | 1.001 | -1.92 | 0.038 | -0.121 | -0.121 | 0 | 0.83 | 0.54 | 0.88 | 0.99 | 0.84 | 0.95 |
| 8 | 51188675 | rs1450124 | 1.36 | -0.646 | 0.837 | -1.961 | 0.042 | 0.087 | 0.087 | 0 | 0.58 | 0.24 | 0.68 | 0.84 | 0.7 | 0.58 |
| 8 | 51203110 | rs17769846 | -0.068 | -0.691 | 0.82 | -1.946 | 0.005 | 0.104 | 0.104 | 0 | 0.24 | 0.03 | 0.12 | 0.05 | 0.21 | 0.17 |
| 8 | 51227228 | rs16914704 | -1.142 | -0.635 | 0.994 | -1.934 | 0.012 | -0.039 | -0.039 | 0 | 0.1 | 0.17 | 0.08 | 0.02 | 0.13 | 0.1 |
| 8 | 51227506 | rs1450136 | -0.68 | -0.69 | 0.948 | -1.977 | 0.019 | 0.08 | 0.08 | 0 | 0.29 | 0.11 | 0.17 | 0.08 | 0.27 | 0.29 |
| 8 | 51227777 | rs13273999 | 2.06 | -0.171 | 0.929 | -2.032 | -0.02 | 0.265 | 0.265 | 0.051 | 0.5 | 0.15 | 0.67 | 0.9 | 0.52 | 0.64 |
| 8 | 51230812 | rs12541772 | 1.661 | -0.738 | 0.94 | -2.007 | 0.027 | 0.107 | 0.107 | 0 | 0.54 | 0.25 | 0.64 | 0.83 | 0.62 | 0.56 |
| 8 | 51256438 | rs2100233 | 0.675 | -0.662 | 0.794 | -1.839 | 0.023 | -0.084 | -0.084 | 0.247 | 0.87 | 0.45 | 0.9 | 1 | 0.91 | 0.96 |
| 8 | 51256566 | rs2957602 | 1.81 | -0.653 | 0.785 | -1.843 | 0.006 | -0.082 | -0.082 | 0 | 0.78 | 0.34 | 0.83 | 0.97 | 0.79 | 0.88 |
| 8 | 51264624 | rs930671 | 0.063 | -0.631 | 0.487 | -1.706 | -0.005 | 0.065 | 0.065 | 0 | 0.21 | 0.04 | 0.12 | 0.06 | 0.21 | 0.18 |
| 8 | 51269514 | rs16914760 | -1.629 | -0.603 | 0.57 | -1.627 | 0.017 | 0.022 | 0.022 | 0.045 | 0.11 | 0.28 | 0.18 | 0.13 | 0.13 | 0.29 |
| 8 | 51284360 | rs1117213 | -0.554 | -0.627 | 0.576 | -1.647 | -0.021 | -0.018 | -0.018 | 0 | 0.27 | 0.35 | 0.23 | 0.17 | 0.21 | 0.38 |
| 8 | 51287482 | rs16914781 | -1.822 | -0.613 | 0.737 | -1.641 | 0.001 | 0.004 | 0.004 | 0 | 0.44 | 0.56 | 0.37 | 0.19 | 0.43 | 0.5 |
| 8 | 51306463 | rs4529480 | 1.422 | -0.64 | 0.155 | -1.597 | 0 | 0.002 | 0.002 | 0 | 0.58 | 0.6 | 0.66 | 0.81 | 0.59 | 0.5 |
| 8 | 51313795 | rs7005672 | 0.686 | -0.714 | 0.399 | -1.426 | 0.001 | 0.004 | 0.004 | 0 | 0.71 | 0.74 | 0.75 | 0.84 | 0.72 | 0.6 |
| 8 | 51324592 | rs13282203 | -0.2 | -0.71 | 0.242 | -1.251 | 0.009 | 0.047 | 0.047 | 0 | 0.22 | 0.11 | 0.12 | 0.06 | 0.21 | 0.18 |
| 8 | 51328497 | rs12545003 | 0.657 | -0.625 | 0.114 | -1.198 | -0.01 | -0.013 | -0.013 | 0.405 | 0.69 | 0.64 | 0.73 | 0.8 | 0.71 | 0.6 |
| 8 | 51336218 | rs10957910 | 1.131 | -1.466 | -0.825 | -1.4 | 0.074 | 0.067 | 0.067 | 1.543 | 0.62 | 0.12 | 0.69 | 0.79 | 0.75 | 0.62 |
| 8 | 51337724 | rs10081470 | -0.478 | -1.703 | 0.306 | -1.568 | -0.044 | -0.129 | -0.129 | 0 | 0.2 | 0.44 | 0.1 | 0.03 | 0.1 | 0.11 |
| 8 | 51337745 | rs7839067 | 0.584 | -1.678 | 0.452 | -1.517 | -0.023 | 0.012 | 0.012 | 0.001 | 0.83 | 0.61 | 0.79 | 0.86 | 0.85 | 0.73 |
| 8 | 51348531 | rs10957914 | 1.356 | -1.748 | 0.689 | -1.554 | -0.001 | 0 | 0 | 1.875 | 0.78 | 0.25 | 0.77 | 0.83 | 0.84 | 0.68 |
| 8 | 51348758 | rs10957915 | 0.268 | -1.757 | 0.734 | -1.553 | -0.009 | 0.018 | 0.018 | 0 | 0.86 | 0.71 | 0.82 | 0.88 | 0.87 | 0.82 |
| 8 | 51368483 | rs7009024 | 1.507 | -1.682 | 0.724 | -1.529 | -0.002 | -0.002 | -0.002 | 0 | 0.68 | 0.34 | 0.72 | 0.81 | 0.78 | 0.72 |
| 8 | 51375321 | rs11785209 | 2.02 | -2.867 | -0.506 | -1.434 | -0.09 | -0.149 | -0.149 | 1.922 | 0.31 | 0.03 | 0.49 | 0.5 | 0.57 | 0.29 |
| 8 | 51403548 | rs2068238 | -0.246 | -2.891 | 0.663 | -1.59 | -0.005 | 0.014 | 0.014 | 0 | 0.89 | 0.81 | 0.82 | 0.88 | 0.88 | 0.83 |
| 8 | 51408356 | rs2128000 | 1.148 | -2.916 | 0.749 | -1.607 | -0.037 | -0.03 | -0.03 | 0 | 0.71 | 0.43 | 0.77 | 0.85 | 0.82 | 0.73 |
| 8 | 51443564 | rs218876 | -1.696 | -3.214 | 0.506 | -1.722 | 0.015 | 0.014 | 0.014 | 0 | 0.41 | 0.61 | 0.27 | 0.23 | 0.23 | 0.48 |
| 8 | 51444645 | rs7008861 | -0.324 | -3.182 | 0.534 | -1.705 | -0.073 | -0.11 | -0.11 | 0.208 | 0.18 | 0.4 | 0.05 | 0.03 | 0.06 | 0.11 |
| 8 | 51453745 | rs1542614 | -1.658 | -3.004 | 0.308 | -1.607 | 0.122 | 0.096 | 0.096 | 0.119 | 0.43 | 0.78 | 0.3 | 0.23 | 0.24 | 0.48 |
| 8 | 51459757 | rs1483643 | -0.723 | -2.882 | 0.305 | -1.557 | -0.01 | 0.044 | 0.044 | 0.057 | 0.15 | 0.46 | 0.2 | 0.15 | 0.15 | 0.22 |
| 8 | 51461440 | rs996521 | -1.555 | -2.925 | 0.304 | -1.581 | 0.057 | 0.048 | 0.048 | 0 | 0.42 | 0.7 | 0.29 | 0.23 | 0.24 | 0.47 |
| 8 | 51461513 | rs2128001 | -0.667 | -2.835 | 0.23 | -1.401 | 0.021 | 0.019 | 0.019 | 0 | 0.26 | 0.24 | 0.09 | 0.08 | 0.09 | 0.25 |
| 8 | 51462627 | rs16915033 | -1.141 | -1.622 | 0.679 | -1.087 | 0.022 | -0.045 | -0.045 | 3.166 | 0.12 | 0.34 | 0.1 | 0.04 | 0.1 | 0.11 |
| 8 | 51480478 | rs13262914 | 0.594 | -2.3 | 0.36 | -1.474 | 0.03 | 0.047 | 0.047 | 0 | 0.71 | 0.79 | 0.88 | 0.93 | 0.8 | 0.71 |
| 8 | 51483551 | rs6987557 | 1.486 | -2.368 | 0.466 | -1.554 | 0.063 | 0.085 | 0.085 | 0.704 | 0.59 | 0.23 | 0.69 | 0.79 | 0.71 | 0.49 |
| 8 | 51488614 | rs1351754 | 0.292 | -2.502 | -0.361 | -1.634 | 0.041 | 0.083 | 0.083 | 0 | 0.76 | 0.9 | 0.92 | 0.93 | 0.83 | 0.75 |
| 8 | 51491428 | rs2016596 | -1.157 | -2.354 | -0.226 | -1.547 | 0.005 | 0.074 | 0.074 | 0.726 | 0.12 | 0.55 | 0.19 | 0.14 | 0.09 | 0.23 |
| 8 | 51494398 | rs6473235 | -1.762 | -2.352 | -0.334 | -1.549 | -0.01 | 0.005 | 0.005 | 1.272 | 0.17 | 0.68 | 0.22 | 0.15 | 0.12 | 0.26 |
| 8 | 51519246 | rs11785733 | 2.033 | -2.466 | -1.035 | -1.649 | -0.023 | 0.067 | 0.067 | 0.636 | 0.45 | 0.04 | 0.56 | 0.72 | 0.61 | 0.39 |
| 8 | 51520664 | rs11994238 | -1.778 | -2.447 | -0.68 | -1.553 | -0.024 | 0.008 | 0.008 | 2.113 | 0.14 | 0.72 | 0.2 | 0.13 | 0.05 | 0.22 |
| 8 | 51523824 | rs9298341 | 0.48 | -2.371 | -0.797 | -1.511 | -0.001 | 0.142 | 0.142 | 0.339 | 0.72 | 0.98 | 0.89 | 0.91 | 0.76 | 0.69 |
| 8 | 51525852 | rs9298342 | 0.493 | -2.372 | -0.797 | -1.531 | 0 | 0.147 | 0.147 | 0.323 | 0.72 | 0.97 | 0.89 | 0.91 | 0.76 | 0.69 |
| 8 | 51533503 | rs6999996 | 0.489 | -2.369 | -0.795 | -1.53 | 0 | 0.147 | 0.147 | 0.329 | 0.72 | 0.98 | 0.89 | 0.91 | 0.76 | 0.69 |
| 8 | 51540209 | rs10088550 | -1.189 | -2.066 | -0.274 | -1.223 | -0.019 | 0.111 | 0.111 | 2.245 | 0.08 | 0.61 | 0.16 | 0.12 | 0.02 | 0.18 |

| | | | | | | | | | | | | | | | | |
|---|---|---|---|---|---|---|---|---|---|---|---|---|---|---|---|---|
| 8 | 51546026 | rs906657 | -1.38 | -2.496 | -1.282 | -1.272 | 0.085 | 0.09 | 0.09 | 0.623 | 0.41 | 0.92 | 0.31 | 0.23 | 0.26 | 0.49 |
| 8 | 51550158 | rs13273663 | 1.77 | -2.499 | -0.976 | -1.144 | 0.047 | 0.075 | 0.075 | 0.119 | 0.47 | 0.18 | 0.58 | 0.67 | 0.63 | 0.4 |
| 8 | 51551385 | rs6993752 | -0.718 | -2.605 | -0.007 | -1.166 | -0.006 | 0.091 | 0.091 | 0 | 0.19 | 0.42 | 0.28 | 0.25 | 0.14 | 0.29 |
| 8 | 51558988 | rs4873152 | 0.629 | -2.792 | 0.161 | -1.32 | -0.021 | -0.103 | -0.103 | 0.034 | 0.73 | 0.57 | 0.88 | 0.93 | 0.81 | 0.71 |
| 8 | 51564759 | rs10111120 | -0.579 | -2.66 | 0.203 | -1.181 | -0.001 | 0.078 | 0.078 | 0 | 0.18 | 0.36 | 0.28 | 0.25 | 0.14 | 0.29 |
| 8 | 51569293 | rs2271204 | -0.532 | -2.654 | 0.171 | -1.178 | 0.007 | 0.097 | 0.097 | 0 | 0.17 | 0.36 | 0.28 | 0.25 | 0.14 | 0.28 |
| 8 | 51617003 | rs1567311 | 1.601 | -2.623 | 0.089 | -1.131 | 0.067 | 0.074 | 0.074 | 0 | 0.52 | 0.23 | 0.71 | 0.69 | 0.7 | 0.47 |
| 8 | 51622250 | rs7831359 | -1.124 | -2.703 | 0.644 | -1.118 | -0.006 | -0.009 | -0.009 | 0.168 | 0.33 | 0.31 | 0.2 | 0.2 | 0.19 | 0.29 |
| 8 | 51631199 | rs12375352 | -0.958 | -2.664 | 0.666 | -1.12 | 0.006 | 0.007 | 0.007 | 0 | 0.31 | 0.24 | 0.18 | 0.2 | 0.19 | 0.28 |
| 8 | 51642397 | rs906655 | -0.219 | -2.514 | 0.533 | -1.082 | -0.007 | 0.017 | 0.017 | 0.316 | 0.89 | 0.92 | 0.95 | 1 | 0.9 | 0.92 |
| 8 | 51642452 | rs906656 | 1.611 | -2.756 | 0.647 | -1.107 | 0.009 | 0.014 | 0.014 | 0 | 0.56 | 0.61 | 0.75 | 0.77 | 0.7 | 0.63 |
| 8 | 51666225 | rs12674534 | -1.693 | -2.971 | 0.753 | -1.277 | -0.046 | -0.063 | -0.063 | 0 | 0.23 | 0.51 | 0.12 | 0.09 | 0.13 | 0.18 |
| 8 | 51697092 | rs7831651 | -2.114 | -2.863 | 0.733 | -1.261 | 0.034 | 0.066 | 0.066 | 0 | 0.49 | 0.36 | 0.28 | 0.21 | 0.33 | 0.41 |
| 8 | 51744861 | rs723596 | -0.825 | -2.81 | 0.657 | -1.221 | -0.002 | -0.003 | -0.003 | 0 | 0.3 | 0.26 | 0.19 | 0.21 | 0.18 | 0.31 |
| 8 | 51750280 | rs7816312 | -2.702 | -2.837 | 0.592 | -1.263 | 0.036 | 0.065 | 0.065 | 0 | 0.55 | 0.77 | 0.32 | 0.3 | 0.32 | 0.51 |
| 8 | 51762848 | rs13258554 | -0.853 | -2.748 | 0.588 | -0.963 | -0.005 | -0.003 | -0.003 | 0 | 0.3 | 0.26 | 0.19 | 0.25 | 0.18 | 0.32 |
| 8 | 51769869 | rs2392742 | -0.853 | -2.725 | 0.588 | -0.941 | -0.004 | -0.003 | -0.003 | 0 | 0.3 | 0.26 | 0.2 | 0.25 | 0.18 | 0.32 |
| 8 | 51772697 | rs6473440 | -1.937 | -2.809 | 0.539 | -0.946 | 0.058 | 0.027 | 0.027 | 0 | 0.44 | 0.69 | 0.27 | 0.34 | 0.22 | 0.46 |
| 8 | 51780761 | rs13251151 | -0.874 | -2.797 | 0.566 | -0.945 | 0.015 | 0.01 | 0.01 | 0 | 0.3 | 0.19 | 0.19 | 0.25 | 0.18 | 0.32 |
| 8 | 51785729 | rs7837228 | -1.892 | -2.779 | 0.493 | -0.935 | 0.059 | 0.027 | 0.027 | 0 | 0.45 | 0.69 | 0.27 | 0.34 | 0.22 | 0.45 |
| 8 | 51796456 | rs6999023 | -1.892 | -2.774 | 0.495 | -0.932 | 0.058 | 0.027 | 0.027 | 0 | 0.44 | 0.69 | 0.26 | 0.34 | 0.22 | 0.46 |
| 8 | 51800598 | rs17787432 | 1.269 | -2.854 | 0.369 | -0.971 | -0.004 | -0.027 | -0.027 | 0 | 0.05 | 0 | 0.01 | 0.09 | 0.04 | 0.05 |
| 8 | 51802055 | rs10102094 | -0.932 | -2.887 | 0.73 | -0.984 | -0.054 | -0.001 | -0.001 | 0 | 0.13 | 0.34 | 0.06 | 0.09 | 0.04 | 0.12 |
| 8 | 51811511 | rs11785367 | 0.354 | -3.014 | 0.55 | -1.049 | -0.037 | -0.012 | -0.012 | 0.021 | 0.15 | 0 | 0.14 | 0.12 | 0.1 | 0.13 |
| 8 | 51813110 | rs11784136 | -2.409 | -2.974 | 0.737 | -1.026 | 0.044 | 0.033 | 0.033 | 0 | 0.49 | 0.64 | 0.29 | 0.34 | 0.29 | 0.5 |
| 8 | 51815533 | rs4873161 | -0.777 | -2.979 | 0.809 | -1.028 | 0.004 | 0.003 | 0.003 | 0 | 0.3 | 0.25 | 0.19 | 0.26 | 0.18 | 0.33 |
| 8 | 51816730 | rs6473485 | -2.832 | -2.999 | 0.682 | -0.985 | 0.038 | 0.029 | 0.029 | 0 | 0.55 | 0.77 | 0.33 | 0.35 | 0.32 | 0.52 |
| 8 | 51820294 | rs7000660 | -1.363 | -2.968 | 0.739 | -0.979 | -0.006 | 0 | 0 | 0.156 | 0.34 | 0.58 | 0.22 | 0.34 | 0.18 | 0.42 |
| 8 | 51820442 | rs6987528 | -2.073 | -2.952 | 0.688 | -0.966 | 0.051 | 0.033 | 0.033 | 0 | 0.46 | 0.66 | 0.28 | 0.34 | 0.28 | 0.47 |
| 8 | 51835145 | rs11780693 | -1.892 | -2.897 | 0.628 | -0.935 | 0.038 | 0.022 | 0.022 | 0 | 0.43 | 0.66 | 0.27 | 0.34 | 0.26 | 0.48 |
| 8 | 51846792 | rs7840842 | -1.698 | -2.894 | 0.684 | -0.945 | -0.002 | -0.005 | -0.005 | 0 | 0.42 | 0.51 | 0.26 | 0.26 | 0.26 | 0.4 |
| 8 | 51852690 | rs11985362 | -0.803 | -2.911 | 0.701 | -0.954 | 0.004 | 0.002 | 0.002 | 0 | 0.3 | 0.24 | 0.19 | 0.25 | 0.18 | 0.34 |
| 8 | 51853185 | rs6999096 | -0.803 | -2.911 | 0.701 | -0.954 | 0.004 | 0.002 | 0.002 | 0 | 0.3 | 0.24 | 0.19 | 0.26 | 0.18 | 0.34 |
| 8 | 51854351 | rs6985047 | -0.235 | -2.966 | 0.659 | -0.961 | 0 | -0.048 | -0.048 | 0 | 0.1 | 0.17 | 0.06 | 0 | 0.08 | 0.06 |
| 8 | 51855262 | rs13268263 | -2.217 | -2.709 | 0.623 | -0.996 | 0.022 | 0.036 | 0.036 | 0 | 0.47 | 0.59 | 0.29 | 0.25 | 0.32 | 0.41 |
| 8 | 51855787 | rs6473534 | -2.555 | -2.567 | 0.669 | -0.946 | 0.04 | 0.051 | 0.051 | 0 | 0.52 | 0.78 | 0.34 | 0.35 | 0.34 | 0.51 |
| 8 | 51862244 | rs6473539 | -1.83 | -2.588 | 0.654 | -0.862 | -0.005 | -0.009 | -0.009 | 0 | 0.39 | 0.53 | 0.28 | 0.26 | 0.26 | 0.4 |
| 8 | 51862991 | rs1484125 | -1.83 | -2.588 | 0.654 | -0.862 | -0.005 | -0.009 | -0.009 | 0.776 | 0.39 | 0.53 | 0.28 | 0.26 | 0.26 | 0.4 |
| 8 | 51879826 | rs4873485 | -1.618 | -2.6 | 0.66 | -0.868 | -0.01 | -0.019 | -0.019 | 0 | 0.38 | 0.41 | 0.27 | 0.26 | 0.26 | 0.4 |
| 8 | 51897268 | rs34896506 | -0.978 | -2.594 | 0.579 | -0.875 | 0.01 | 0.007 | 0.007 | 0 | 0.3 | 0.23 | 0.22 | 0.26 | 0.19 | 0.35 |
| 8 | 51907056 | rs7818008 | -0.176 | -2.567 | 0.565 | -0.869 | 0.008 | -0.049 | -0.049 | 0 | 0.09 | 0.19 | 0.07 | 0 | 0.08 | 0.06 |
| 8 | 51925820 | rs10958231 | -1.744 | -2.53 | 0.426 | -0.816 | -0.002 | -0.003 | -0.003 | 1.067 | 0.39 | 0.57 | 0.29 | 0.26 | 0.26 | 0.4 |
| 8 | 51933728 | rs11782445 | -0.9 | -2.521 | 0.115 | -0.756 | 0.009 | 0.004 | 0.004 | 0 | 0.29 | 0.22 | 0.21 | 0.25 | 0.18 | 0.34 |
| 8 | 51945593 | rs7834177 | -0.895 | -2.411 | 0.096 | -0.736 | 0.006 | 0.002 | 0.002 | 0 | 0.28 | 0.22 | 0.21 | 0.25 | 0.19 | 0.34 |
| 8 | 51949268 | rs2168330 | 2.05 | -2.589 | -0.095 | -0.804 | 0.072 | 0.053 | 0.053 | 0 | 0.49 | 0.18 | 0.65 | 0.64 | 0.68 | 0.51 |
| 8 | 51954441 | rs11775658 | -1.77 | -2.518 | -0.165 | -1.014 | 0.012 | 0.023 | 0.023 | 0 | 0.45 | 0.61 | 0.36 | 0.27 | 0.33 | 0.38 |
| 8 | 51955260 | rs16915742 | -0.699 | -2.344 | -0.671 | -0.86 | -0.038 | -0.037 | -0.037 | 0.044 | 0.18 | 0.03 | 0.17 | 0.18 | 0.18 | 0.23 |
| 8 | 51959112 | rs17727577 | 0.076 | -3.472 | -0.774 | -1.332 | 0.018 | 0.015 | 0.015 | 3.089 | 0.25 | 0.18 | 0.09 | 0.06 | 0.1 | 0.07 |
| 8 | 51959409 | rs7017034 | -0.138 | -3.144 | -1.505 | -1.47 | -0.079 | 0.053 | 0.053 | 0.184 | 0.24 | 0.06 | 0.23 | 0.12 | 0.29 | 0.29 |
| 8 | 51964292 | rs4873165 | -0.07 | -2.847 | -1.48 | -0.581 | -0.008 | -0.048 | -0.048 | 0.053 | 0.28 | 0.11 | 0.4 | 0.37 | 0.25 | 0.38 |
| 8 | 51970609 | rs6990018 | -0.397 | -3.01 | -1.274 | -0.704 | 0.001 | -0.004 | -0.004 | 0 | 0.3 | 0.69 | 0.14 | 0.08 | 0.1 | 0.11 |
| 8 | 51974368 | rs11990832 | -1.433 | -2.721 | -1.291 | -0.444 | -0.015 | -0.048 | -0.048 | 0 | 0.59 | 0.8 | 0.56 | 0.45 | 0.39 | 0.5 |
| 8 | 51980821 | rs2219211 | -0.507 | -2.978 | -0.999 | -0.118 | 0.016 | -0.073 | -0.073 | 0 | 0.32 | 0.11 | 0.4 | 0.45 | 0.25 | 0.38 |
| 8 | 51981777 | rs7813671 | -0.888 | -3.127 | -0.093 | -0.538 | 0.026 | 0.01 | 0.01 | 0 | 0.53 | 0.62 | 0.4 | 0.35 | 0.34 | 0.48 |
| 8 | 51982333 | rs4873496 | 0.064 | -3.373 | -0.347 | -0.812 | 0.054 | 0.068 | 0.068 | 0.463 | 0.36 | 0.13 | 0.28 | 0.21 | 0.2 | 0.4 |
| 8 | 51982432 | rs7833425 | -0.684 | -3.268 | 0.713 | -0.787 | -0.051 | -0.014 | -0.014 | 0.006 | 0.19 | 0.42 | 0.15 | 0.14 | 0.15 | 0.11 |
| 8 | 51988578 | rs10109996 | 0.184 | -3.324 | 0.733 | -0.801 | 0.017 | 0.047 | 0.047 | 0.454 | 0.33 | 0.12 | 0.29 | 0.21 | 0.24 | 0.4 |
| 8 | 51996075 | rs4873166 | -1.083 | -2.829 | 1.041 | -0.634 | 0.005 | 0.003 | 0.003 | 0 | 0.56 | 0.63 | 0.45 | 0.36 | 0.39 | 0.52 |
| 8 | 52005527 | rs7845086 | -1.29 | -2.998 | 0.984 | -0.616 | 0.019 | 0 | 0 | 0.192 | 0.59 | 0.76 | 0.45 | 0.36 | 0.39 | 0.56 |
| 8 | 52022731 | rs4294195 | 0.226 | -3.009 | 0.807 | -0.626 | 0.026 | 0.074 | 0.074 | 1.019 | 0.37 | 0.12 | 0.3 | 0.21 | 0.27 | 0.44 |
| 8 | 52027415 | rs4873176 | -1.107 | -3.143 | 0.982 | -0.599 | 0.032 | 0.008 | 0.008 | 0.761 | 0.57 | 0.69 | 0.42 | 0.36 | 0.35 | 0.55 |
| 8 | 52027473 | rs4873178 | -1.036 | -3.144 | 0.981 | -0.6 | 0.035 | 0.01 | 0.01 | 0.708 | 0.56 | 0.68 | 0.42 | 0.36 | 0.35 | 0.55 |
| 8 | 52029420 | rs10090890 | 0.025 | -3.116 | 0.733 | -0.618 | 0.072 | 0.103 | 0.103 | 0.896 | 0.42 | 0.15 | 0.32 | 0.24 | 0.28 | 0.5 |
| 8 | 52031261 | rs6473576 | -1.207 | -3 | 0.504 | -0.57 | 0.015 | -0.02 | -0.02 | 0 | 0.61 | 0.79 | 0.44 | 0.37 | 0.38 | 0.58 |
| 8 | 52033866 | rs4266664 | -1.247 | -2.915 | 0.516 | -0.626 | 0.01 | -0.017 | -0.017 | 0.648 | 0.61 | 0.79 | 0.44 | 0.37 | 0.38 | 0.58 |
| 8 | 52035023 | rs11991676 | 0.473 | -2.907 | 0.817 | -0.583 | -0.018 | 0.003 | 0.003 | 0 | 0.12 | 0.16 | 0.08 | 0.12 | 0.09 | 0.07 |
| 8 | 52042419 | rs2198700 | -1.287 | -2.765 | 0.395 | -0.538 | 0.022 | 0.007 | 0.007 | 2.488 | 0.6 | 0.75 | 0.43 | 0.36 | 0.37 | 0.56 |
| 8 | 52048415 | rs6987549 | -0.894 | -2.801 | 0.536 | -0.462 | -0.01 | -0.001 | -0.001 | 1.618 | 0.55 | 0.5 | 0.41 | 0.42 | 0.38 | 0.57 |
| 8 | 52050166 | rs17735532 | 0.284 | -2.951 | -0.058 | -0.514 | 0.045 | 0.059 | 0.059 | 1.243 | 0.35 | 0.07 | 0.27 | 0.21 | 0.23 | 0.42 |

| | | | | | | | | | | | | | | | | |
|---|---|---|---|---|---|---|---|---|---|---|---|---|---|---|---|---|
| 8 | 52054326 | rs10102468 | -0.901 | -2.458 | 0.263 | -0.349 | 0.017 | -0.003 | -0.003 | 0.179 | 0.57 | 0.61 | 0.42 | 0.42 | 0.38 | 0.57 |
| 8 | 52070410 | rs10104514 | -0.588 | -2.71 | 0.316 | -0.696 | 0.089 | 0.053 | 0.053 | 2.766 | 0.51 | 0.67 | 0.32 | 0.38 | 0.3 | 0.5 |
| 8 | 52075739 | rs7827396 | -0.585 | -2.733 | 0.353 | -0.68 | 0.109 | 0.063 | 0.063 | 2.811 | 0.51 | 0.7 | 0.31 | 0.38 | 0.29 | 0.49 |
| 8 | 52092897 | rs7462308 | 0.338 | -2.812 | 0.147 | -0.784 | 0.074 | 0.112 | 0.112 | 0.631 | 0.36 | 0.11 | 0.25 | 0.21 | 0.22 | 0.41 |
| 8 | 52097911 | rs2028157 | 0.021 | -2.728 | 0.392 | -0.768 | -0.013 | -0.023 | -0.023 | 0 | 0.41 | 0.57 | 0.29 | 0.21 | 0.24 | 0.42 |
| 8 | 52101444 | rs10504112 | 0.529 | -2.599 | -0.166 | -0.602 | 0.012 | -0.013 | -0.013 | 0 | 0.1 | 0.07 | 0.03 | 0.18 | 0.06 | 0.06 |
| 8 | 52120526 | rs1119176 | -1.46 | -2.435 | -0.446 | -0.791 | -0.116 | -0.101 | -0.101 | 0 | 0.65 | 0.96 | 0.45 | 0.4 | 0.4 | 0.54 |
| 8 | 52138162 | rs10104140 | 0.278 | -2.449 | 0.097 | -0.754 | -0.044 | 0.055 | 0.055 | 1.121 | 0.13 | 0.31 | 0.05 | 0.19 | 0.05 | 0.05 |
| 8 | 52140746 | rs4873514 | 0.362 | -2.638 | 0.313 | -0.878 | 0.045 | 0.062 | 0.062 | 0.117 | 0.37 | 0.25 | 0.27 | 0.21 | 0.23 | 0.42 |
| 8 | 52141931 | rs7846374 | -0.226 | -2.161 | 0.245 | -0.753 | 0.003 | 0.004 | 0.004 | 0.109 | 0.5 | 0.56 | 0.31 | 0.39 | 0.29 | 0.49 |
| 8 | 52147218 | rs7821095 | -1.336 | -2.256 | -0.078 | -0.757 | -0.114 | -0.142 | -0.142 | 2.757 | 0.66 | 0.96 | 0.46 | 0.44 | 0.41 | 0.56 |
| 8 | 52147655 | rs10504115 | -0.886 | -2.42 | 0.21 | -0.597 | 0.074 | -0.047 | -0.047 | 0 | 0.58 | 0.88 | 0.38 | 0.44 | 0.31 | 0.53 |
| 8 | 52155807 | rs9298438 | -0.445 | -1.97 | -0.554 | -0.371 | 0.035 | 0.011 | 0.011 | 0.005 | 0.51 | 0.64 | 0.33 | 0.43 | 0.29 | 0.5 |
| 8 | 52157579 | rs4873188 | -1.193 | -1.862 | -0.93 | -0.41 | 0.082 | 0.013 | 0.013 | 1.522 | 0.39 | 0.9 | 0.27 | 0.34 | 0.23 | 0.29 |
| 8 | 52157888 | rs4873189 | -1.158 | -1.744 | -0.12 | -0.527 | 0.006 | 0.004 | 0.004 | 0.371 | 0.38 | 0.66 | 0.25 | 0.32 | 0.28 | 0.29 |
| 8 | 52160675 | rs10101136 | -0.656 | -1.867 | 0.541 | -0.45 | -0.062 | 0.055 | 0.055 | 0.553 | 0.16 | 0.43 | 0.07 | 0.2 | 0.08 | 0.12 |
| 8 | 52164871 | rs4463411 | 0.108 | -1.856 | 0.7 | -0.434 | -0.021 | 0.025 | 0.025 | 0 | 0.1 | 0.16 | 0.03 | 0.19 | 0.05 | 0.08 |
| 8 | 52181920 | rs7844947 | -1.178 | -2.029 | 1.223 | -0.647 | -0.011 | -0.045 | -0.045 | 4.227 | 0.29 | 0.6 | 0.22 | 0.13 | 0.23 | 0.21 |
| 8 | 52213702 | rs4645549 | -0.97 | -2.082 | 1.478 | -0.673 | -0.013 | -0.073 | -0.073 | 0.35 | 0.27 | 0.52 | 0.21 | 0.13 | 0.23 | 0.22 |
| 8 | 52226018 | rs4471026 | -0.491 | -2.095 | 1.49 | -0.651 | -0.001 | -0.008 | -0.008 | 0 | 0.22 | 0.24 | 0.16 | 0.1 | 0.21 | 0.17 |
| 8 | 52231952 | rs4873544 | -1.291 | -2.179 | 1.811 | -0.692 | -0.008 | -0.031 | -0.031 | 0 | 0.3 | 0.64 | 0.23 | 0.14 | 0.24 | 0.23 |
| 8 | 52232624 | rs1370446 | -1.291 | -2.179 | 1.811 | -0.692 | -0.008 | -0.031 | -0.031 | 4.64 | 0.3 | 0.64 | 0.23 | 0.14 | 0.24 | 0.23 |
| 8 | 52233276 | rs7827306 | -1.252 | -2.179 | 1.819 | -0.692 | -0.009 | -0.035 | -0.035 | 0 | 0.3 | 0.63 | 0.23 | 0.14 | 0.24 | 0.23 |
| 8 | 52233408 | rs7827446 | -1.252 | -2.179 | 1.819 | -0.692 | -0.009 | -0.035 | -0.035 | 0 | 0.3 | 0.63 | 0.23 | 0.14 | 0.24 | 0.23 |
| 8 | 52237082 | rs4500070 | -1.853 | -2.305 | 1.587 | -0.728 | 0.078 | 0.067 | 0.067 | 0.658 | 0.42 | 0.87 | 0.26 | 0.3 | 0.28 | 0.33 |
| 8 | 52249319 | rs9298444 | -0.521 | -2.236 | 1.651 | -0.704 | -0.002 | -0.025 | -0.025 | 0 | 0.24 | 0.27 | 0.18 | 0.09 | 0.2 | 0.18 |
| 8 | 52253034 | rs16916015 | -0.384 | -2.271 | 1.544 | -0.721 | -0.02 | 0.031 | 0.031 | 0 | 0.12 | 0.15 | 0.06 | 0.24 | 0.07 | 0.14 |
| 8 | 52263301 | rs11782525 | -1.291 | -2.148 | 1.537 | -0.999 | 0.014 | -0.002 | -0.002 | 0.23 | 0.05 | 0.23 | 0.05 | 0 | 0.02 | 0.03 |
| 8 | 52267567 | rs17261657 | -0.53 | -2.249 | 1.188 | -1.106 | -0.031 | 0.133 | 0.133 | 0.131 | 0.23 | 0.01 | 0.17 | 0.05 | 0.21 | 0.2 |
| 8 | 52269392 | rs2915475 | 1.137 | -2.268 | 1.437 | -1.112 | -0.006 | -0.057 | -0.057 | 0 | 0.69 | 0.33 | 0.78 | 0.94 | 0.77 | 0.79 |
| 8 | 52279739 | rs2915495 | 0.628 | -2.555 | 1.405 | -1.095 | -0.037 | -0.109 | -0.109 | 0 | 0.72 | 0.58 | 0.84 | 0.95 | 0.83 | 0.81 |
| 8 | 52296100 | rs9298446 | -0.933 | -2.395 | 1.398 | -0.906 | -0.032 | 0.049 | 0.049 | 0.103 | 0.17 | 0.27 | 0.09 | 0.27 | 0.09 | 0.12 |
| 8 | 52297656 | rs10504117 | -1.481 | -2.278 | 1.427 | -0.685 | -0.049 | 0.095 | 0.095 | 0.817 | 0.21 | 0.48 | 0.11 | 0.34 | 0.11 | 0.21 |
| 8 | 52297865 | rs994127 | 1.967 | -2.401 | 1.082 | -0.638 | 0.114 | 0.012 | 0.012 | 0.507 | 0.53 | 0.12 | 0.73 | 0.59 | 0.73 | 0.6 |
| 8 | 52308575 | rs2977008 | 0.255 | -2.397 | 0.721 | -0.667 | 0.006 | -0.002 | -0.002 | 0 | 0.76 | 0.76 | 0.86 | 0.93 | 0.85 | 0.82 |
| 8 | 52310188 | rs11782014 | 0.212 | -2.38 | 0.701 | -0.641 | -0.001 | -0.001 | -0.001 | 1.257 | 0.05 | 0.04 | 0.04 | 0.06 | 0.05 | 0.04 |
| 8 | 52311233 | rs2977010 | 0.279 | -2.362 | 0.63 | -0.658 | 0.001 | 0 | 0 | 0 | 0.74 | 0.75 | 0.85 | 0.87 | 0.83 | 0.72 |
| 8 | 52315863 | rs2977016 | 1.911 | -2.367 | 0.542 | -0.664 | 0.114 | 0.01 | 0.01 | 0.591 | 0.52 | 0.12 | 0.72 | 0.59 | 0.72 | 0.6 |
| 8 | 52321243 | rs2977020 | 0.395 | -2.073 | 0.635 | -0.674 | -0.009 | -0.035 | -0.035 | 0 | 0.76 | 0.74 | 0.86 | 0.93 | 0.85 | 0.82 |
| 8 | 52331018 | rs10093611 | -0.789 | -1.926 | 0.883 | -0.293 | -0.019 | 0.08 | 0.08 | 0 | 0.16 | 0.31 | 0.09 | 0.28 | 0.1 | 0.12 |
| 8 | 52335459 | rs16916228 | -1.126 | -2.027 | 0.965 | -0.401 | -0.017 | -0.033 | -0.033 | 6.788 | 0.06 | 0.13 | 0.05 | 0 | 0.04 | 0.06 |
| 8 | 52338934 | rs2392900 | 1.757 | -2.265 | 1.542 | -0.258 | -0.045 | 0.07 | 0.07 | 0.004 | 0.76 | 0.49 | 0.83 | 0.63 | 0.82 | 0.75 |
| 8 | 52358680 | rs994588 | 2.007 | -2.802 | 1.011 | -0.668 | 0.127 | -0.051 | -0.051 | 0 | 0.53 | 0.1 | 0.7 | 0.57 | 0.7 | 0.59 |
| 8 | 52360543 | rs7846408 | 1.101 | -2.773 | 1.225 | -0.884 | -0.035 | 0.019 | 0.019 | 0 | 0.79 | 0.6 | 0.84 | 0.73 | 0.83 | 0.85 |
| 8 | 52366300 | rs7833909 | 0.01 | -2.721 | 1.268 | -0.869 | -0.017 | -0.039 | -0.039 | 0.944 | 0.21 | 0.24 | 0.12 | 0.06 | 0.12 | 0.19 |
| 8 | 52368218 | rs1367819 | -1.874 | -2.981 | 1.059 | -1.018 | 0.092 | 0.063 | 0.063 | 0 | 0.43 | 0.75 | 0.28 | 0.35 | 0.29 | 0.33 |
| 8 | 52382217 | rs7841317 | -1.061 | -2.721 | 1.141 | -1.064 | 0.032 | -0.038 | -0.038 | 0 | 0.1 | 0.2 | 0.22 | 0.03 | 0.15 | 0.08 |
| 8 | 52384670 | rs7839439 | -0.129 | -2.715 | 1.206 | -1.047 | -0.066 | -0.111 | -0.111 | 1.828 | 0.23 | 0.43 | 0.13 | 0.06 | 0.11 | 0.19 |
| 8 | 52384884 | rs10958268 | -0.435 | -2.714 | 1.048 | -1.046 | -0.06 | -0.098 | -0.098 | 0.289 | 0.24 | 0.56 | 0.14 | 0.06 | 0.11 | 0.19 |
| 8 | 52392445 | rs10504120 | -1.361 | -2.472 | 0.62 | -1.297 | 0.014 | -0.039 | -0.039 | 0.265 | 0.16 | 0.21 | 0.27 | 0.06 | 0.21 | 0.19 |
| 8 | 52392763 | rs16916394 | 0.262 | -2.571 | 0.407 | -1.285 | 0.034 | -0.041 | -0.041 | 0 | 0.11 | 0.03 | 0.03 | 0.26 | 0.05 | 0.1 |
| 8 | 52393084 | rs12675403 | 2.326 | -2.687 | 0.304 | -1.298 | 0.026 | 0.037 | 0.037 | 0.114 | 0.47 | 0.09 | 0.55 | 0.6 | 0.63 | 0.51 |
| 8 | 52395667 | rs12544226 | 0.227 | -2.629 | 0.647 | -1.348 | -0.014 | -0.055 | -0.055 | 0 | 0.22 | 0.28 | 0.23 | 0.06 | 0.14 | 0.21 |
| 8 | 52398025 | rs4873554 | 0.227 | -2.629 | 0.647 | -1.348 | -0.014 | -0.055 | -0.055 | 0 | 0.22 | 0.28 | 0.23 | 0.06 | 0.14 | 0.21 |
| 8 | 52398302 | rs9643782 | -0.172 | -2.673 | 0.604 | -1.285 | 0.044 | -0.05 | -0.05 | 0 | 0.13 | 0.07 | 0.03 | 0.28 | 0.05 | 0.11 |
| 8 | 52399179 | rs13271514 | 1.199 | -2.682 | 0.719 | -1.291 | 0.035 | 0.017 | 0.017 | 0.411 | 0.63 | 0.31 | 0.7 | 0.66 | 0.81 | 0.68 |
| 8 | 52402412 | rs4873195 | -0.935 | -2.547 | 0.677 | -1.264 | -0.022 | -0.103 | -0.103 | 0 | 0.31 | 0.49 | 0.32 | 0.08 | 0.26 | 0.28 |
| 8 | 52415420 | rs10755971 | -2.228 | -2.447 | 0.537 | -1.262 | 0.058 | 0.088 | 0.088 | 0 | 0.51 | 0.75 | 0.41 | 0.38 | 0.38 | 0.48 |
| 8 | 52416912 | rs12541541 | -1.401 | -2.483 | 0.52 | -1.214 | 0.01 | 0.008 | 0.008 | 0 | 0.39 | 0.6 | 0.34 | 0.34 | 0.28 | 0.33 |
| 8 | 52420536 | rs17181677 | 2.528 | -2.958 | -0.333 | -1.274 | 0.002 | -0.025 | -0.025 | 1.249 | 0.13 | 0.01 | 0.17 | 0.14 | 0.11 | 0.09 |
| 8 | 52421186 | rs2013700 | -1.84 | -2.545 | 0.279 | -1.226 | 0.058 | 0.05 | 0.05 | 0 | 0.44 | 0.7 | 0.35 | 0.35 | 0.3 | 0.36 |
| 8 | 52421372 | rs11780225 | -0.447 | -2.346 | 0.475 | -1.115 | 0.007 | -0.038 | -0.038 | 0.228 | 0.09 | 0.22 | 0.09 | 0.02 | 0.11 | 0.07 |
| 8 | 52424709 | rs9298450 | -1.377 | -2.045 | 0.326 | -1.106 | 0.009 | 0.009 | 0.009 | 0 | 0.4 | 0.6 | 0.37 | 0.33 | 0.3 | 0.32 |
| 8 | 52432763 | rs7841155 | 0.286 | -2.062 | 0.601 | -1.095 | 0.008 | -0.057 | -0.057 | 3.343 | 0.16 | 0.34 | 0.21 | 0.07 | 0.12 | 0.17 |
| 8 | 52433074 | rs3097708 | -1.894 | -2.169 | 0.515 | -1.115 | 0.016 | 0.014 | 0.014 | 0.222 | 0.36 | 0.68 | 0.33 | 0.33 | 0.26 | 0.34 |
| 8 | 52435947 | rs11786371 | -0.529 | -2.105 | 0.288 | -1.072 | -0.025 | 0.016 | 0.016 | 0 | 0.17 | 0.23 | 0.11 | 0.25 | 0.13 | 0.12 |
| 8 | 52436762 | rs4598242 | 2.195 | -2.22 | -0.191 | -1.142 | 0.086 | 0.13 | 0.13 | 0.033 | 0.5 | 0.2 | 0.62 | 0.63 | 0.65 | 0.54 |
| 8 | 52439270 | rs4501569 | -0.5 | -2.021 | 0.433 | -0.801 | -0.004 | -0.058 | -0.058 | 3.496 | 0.19 | 0.36 | 0.31 | 0.09 | 0.14 | 0.19 |
| 8 | 52439489 | rs7000275 | -0.693 | -1.899 | 0.387 | -0.521 | -0.025 | 0.015 | 0.015 | 0.471 | 0.15 | 0.18 | 0.08 | 0.25 | 0.1 | 0.12 |

| | | | | | | | | | | | | | | | | |
|---|---|---|---|---|---|---|---|---|---|---|---|---|---|---|---|---|
| 8 | 52442708 | rs7003517 | -0.695 | -1.781 | 0.797 | -0.044 | -0.044 | 0.129 | 0.129 | 2.824 | 0.11 | 0.27 | 0.03 | 0.25 | 0.03 | 0.06 |
| 8 | 52445595 | rs6473615 | -0.89 | -1.814 | 0.685 | -0.089 | -0.034 | 0.06 | 0.06 | 0.162 | 0.11 | 0.19 | 0.03 | 0.25 | 0.03 | 0.07 |
| 8 | 52454721 | rs7843354 | -2.112 | -2.131 | 0.841 | -0.213 | -0.041 | 0 | 0 | 0.07 | 0.28 | 0.34 | 0.13 | 0.3 | 0.13 | 0.17 |
| 8 | 52458791 | rs4407871 | -0.021 | -1.771 | 0.795 | -0.365 | 0.004 | -0.016 | -0.016 | 0 | 0.16 | 0.21 | 0.37 | 0.08 | 0.18 | 0.17 |
| 8 | 52467061 | rs4873562 | -1.793 | -1.919 | 0.785 | -0.45 | -0.007 | -0.06 | -0.06 | 0 | 0.33 | 0.58 | 0.47 | 0.13 | 0.28 | 0.29 |
| 8 | 52472051 | rs4574836 | -0.825 | -1.963 | 0.883 | -0.415 | -0.007 | -0.069 | -0.069 | 0 | 0.2 | 0.55 | 0.34 | 0.08 | 0.18 | 0.18 |
| 8 | 52486885 | rs6985833 | -2.445 | -2.014 | 1.069 | -0.372 | 0.047 | 0.017 | 0.017 | 0.271 | 0.42 | 0.71 | 0.44 | 0.38 | 0.3 | 0.37 |
| 8 | 52493581 | rs10958280 | -1.539 | -2.019 | 1.081 | -0.373 | -0.012 | 0.002 | 0.002 | 0 | 0.24 | 0.24 | 0.11 | 0.31 | 0.12 | 0.21 |
| 8 | 52496113 | rs4873198 | -2.756 | -2.02 | 0.751 | -0.373 | 0.042 | -0.017 | -0.017 | 0.142 | 0.47 | 0.94 | 0.47 | 0.39 | 0.31 | 0.39 |
| 8 | 52496221 | rs4242467 | 0.628 | -1.779 | 1.033 | -0.261 | -0.005 | 0.009 | 0.009 | 0 | 0.12 | 0.02 | 0.25 | 0.08 | 0.11 | 0.15 |
| 8 | 52497178 | rs6473628 | -2.407 | -1.515 | 0.302 | -0.75 | -0.154 | -0.092 | -0.092 | 0 | 0.63 | 0.95 | 0.59 | 0.39 | 0.54 | 0.44 |
| 8 | 52497762 | rs10107128 | -0.606 | -1.835 | 2.288 | 0.347 | 0.002 | -0.032 | -0.032 | 0 | 0.17 | 0.31 | 0.31 | 0.08 | 0.18 | 0.16 |
| 8 | 52516170 | rs11990982 | -1.917 | -2.326 | 1.546 | 0.468 | 0.015 | -0.03 | -0.03 | 0 | 0.1 | 0.19 | 0.1 | 0.07 | 0.1 | 0.14 |
| 8 | 52518723 | rs13263546 | -0.662 | -2.264 | 1.28 | 0.499 | -0.005 | 0.004 | 0.004 | 0 | 0.07 | 0.06 | 0.08 | 0.07 | 0.09 | 0.11 |
| 8 | 52522097 | rs6984427 | -2.917 | -2.297 | 1.109 | 0.839 | 0.012 | -0.124 | -0.124 | 1.19 | 0.39 | 0.92 | 0.52 | 0.46 | 0.29 | 0.38 |
| 8 | 52522352 | rs6998159 | -1.011 | -1.938 | 1.563 | 1.033 | -0.014 | 0.085 | 0.085 | 0 | 0.17 | 0.46 | 0.36 | 0.23 | 0.16 | 0.16 |
| 8 | 52525554 | rs16916598 | 1.965 | -2.164 | 0.448 | 1.143 | -0.028 | -0.12 | -0.12 | 0 | 0.51 | 0.04 | 0.41 | 0.51 | 0.64 | 0.58 |
| 8 | 52526732 | rs10105672 | -1.098 | -1.212 | 1.649 | 1.65 | -0.009 | -0.012 | -0.012 | 0.495 | 0.09 | 0.14 | 0.08 | 0.06 | 0.04 | 0.09 |
| 8 | 52529097 | rs12677741 | 2.37 | -1.279 | 2.308 | 1.707 | 0 | -0.012 | -0.012 | 0 | 0.64 | 0.31 | 0.46 | 0.53 | 0.65 | 0.61 |
| 8 | 52532601 | rs10958282 | -0.702 | -1.274 | 2.25 | 1.753 | 0.015 | 0.041 | 0.041 | 0 | 0.14 | 0.22 | 0.4 | 0.23 | 0.2 | 0.16 |
| 8 | 52534104 | rs4873566 | 2.745 | -1.25 | 2.045 | 1.722 | -0.055 | -0.149 | -0.149 | 0.289 | 0.61 | 0.05 | 0.45 | 0.55 | 0.64 | 0.61 |
| 8 | 52534759 | rs6473630 | -1.701 | -1.205 | 2.297 | 1.684 | 0.004 | 0.018 | 0.018 | 0 | 0.23 | 0.34 | 0.46 | 0.27 | 0.23 | 0.25 |
| 8 | 52535738 | rs12681754 | 1.405 | -0.539 | 2.52 | 1.656 | -0.001 | -0.004 | -0.004 | 0 | 0.89 | 0.89 | 0.95 | 0.85 | 0.89 | 0.86 |
| 8 | 52538439 | rs17276712 | -1.17 | -0.491 | 2.708 | 1.514 | -0.016 | -0.017 | -0.017 | 0 | 0.08 | 0.04 | 0.11 | 0.15 | 0.13 | 0.11 |
| 8 | 52538501 | rs7017902 | -1.682 | -0.533 | 2.862 | 1.703 | -0.02 | 0.03 | 0.03 | 0 | 0.15 | 0.26 | 0.17 | 0.23 | 0.07 | 0.15 |
| 8 | 52548843 | rs7831588 | -2.84 | -0.501 | 2.686 | 1.66 | 0.002 | -0.02 | -0.02 | 0 | 0.29 | 0.75 | 0.32 | 0.36 | 0.28 | 0.33 |
| 8 | 52557431 | rs4873568 | 2.023 | -0.525 | 2.753 | 1.682 | 0.009 | 0.075 | 0.075 | 0 | 0.74 | 0.55 | 0.71 | 0.61 | 0.73 | 0.67 |
| 8 | 52574610 | rs4291264 | 2.087 | -0.364 | 2.707 | 1.772 | 0.022 | 0.07 | 0.07 | 0 | 0.74 | 0.54 | 0.69 | 0.61 | 0.7 | 0.68 |
| 8 | 52585139 | rs7007741 | 1.835 | -0.331 | 2.124 | 1.128 | -0.001 | 0.002 | 0.002 | 0 | 0.8 | 0.8 | 0.82 | 0.84 | 0.77 | 0.79 |
| 8 | 52607916 | rs7011666 | 1.962 | -0.316 | 2.208 | 1.138 | 0.038 | 0.009 | 0.009 | 0 | 0.85 | 0.75 | 0.79 | 0.82 | 0.73 | 0.81 |
| 8 | 52616195 | rs10111115 | 2.352 | -0.403 | 2.557 | 1.094 | 0.015 | -0.027 | -0.027 | 0.987 | 0.75 | 0.46 | 0.75 | 0.81 | 0.71 | 0.75 |
| 8 | 52619866 | rs2976209 | 1.844 | -0.365 | 2.636 | 1.113 | 0.089 | 0.026 | 0.026 | 0 | 0.85 | 0.65 | 0.78 | 0.82 | 0.73 | 0.8 |
| 8 | 52655519 | rs4242472 | 2.32 | -0.485 | 2.299 | 1.021 | 0.002 | -0.033 | -0.033 | 2.321 | 0.73 | 0.42 | 0.75 | 0.85 | 0.7 | 0.75 |
| 8 | 52667603 | rs12674620 | 1.478 | -0.458 | 2.084 | 1.029 | 0.022 | -0.016 | -0.016 | 0 | 0.83 | 0.72 | 0.83 | 0.87 | 0.77 | 0.8 |
| 8 | 52668449 | rs12678680 | -0.462 | -0.484 | 1.858 | 1.022 | -0.013 | -0.02 | -0.02 | 0 | 0.12 | 0.03 | 0.1 | 0.13 | 0.15 | 0.14 |
| 8 | 52671514 | rs6473654 | 1.928 | -0.512 | 1.923 | 1.001 | -0.002 | 0.039 | 0.039 | 0 | 0.76 | 0.86 | 0.83 | 0.86 | 0.73 | 0.77 |
| 8 | 52680282 | rs4257997 | 0.635 | -0.505 | 2.148 | 0.939 | -0.035 | -0.053 | -0.053 | 0 | 0.9 | 0.71 | 0.96 | 1 | 0.98 | 0.95 |
| 8 | 52686100 | rs4873600 | -0.032 | -0.502 | 2.031 | 0.941 | -0.031 | -0.04 | -0.04 | 0.007 | 0.92 | 0.88 | 0.97 | 1 | 0.98 | 0.96 |
| 8 | 52687736 | rs4873601 | -0.112 | -0.502 | 1.887 | 0.941 | -0.015 | -0.022 | -0.022 | 0 | 0.92 | 0.9 | 0.98 | 1 | 0.98 | 0.96 |
| 8 | 52693141 | rs5003717 | 1.759 | -0.594 | 1.491 | 1.398 | 0.068 | -0.105 | -0.105 | 0.871 | 0.74 | 0.09 | 0.78 | 0.63 | 0.88 | 0.82 |
| 8 | 52701004 | rs4242474 | 2.75 | -0.599 | 1.471 | 1.503 | -0.016 | -0.085 | -0.085 | 0 | 0.58 | 0.05 | 0.62 | 0.48 | 0.63 | 0.63 |
| 8 | 52714788 | rs718250 | 1.949 | -0.605 | 1.598 | 1.502 | 0.041 | -0.046 | -0.046 | 0 | 0.75 | 0.2 | 0.78 | 0.63 | 0.88 | 0.82 |
| 8 | 52743043 | rs13250860 | 1.707 | -0.606 | 1.561 | 1.55 | 0.067 | -0.105 | -0.105 | 0 | 0.74 | 0.1 | 0.78 | 0.63 | 0.88 | 0.82 |
| 8 | 52746153 | rs9298462 | 2.738 | -0.593 | 1.089 | 1.555 | -0.015 | -0.084 | -0.084 | 0 | 0.58 | 0.05 | 0.63 | 0.48 | 0.63 | 0.63 |
| 8 | 52747571 | rs9643465 | 1.037 | -0.6 | 1.192 | 1.558 | -0.024 | -0.024 | -0.024 | 0 | 0.86 | 0.48 | 0.9 | 0.96 | 0.92 | 0.92 |
| 8 | 52757093 | rs10504130 | -0.324 | -0.514 | 0.137 | 1.6 | -0.013 | -0.041 | -0.041 | 0.167 | 0.12 | 0.03 | 0.1 | 0.15 | 0.15 | 0.13 |
| 8 | 52757270 | rs4587320 | 1.561 | -0.43 | 0.386 | 1.469 | 0.067 | -0.105 | -0.105 | 0 | 0.74 | 0.1 | 0.78 | 0.63 | 0.88 | 0.82 |
| 8 | 52791790 | rs7012946 | 1.436 | -0.46 | 0.398 | 1.47 | 0.049 | -0.03 | -0.03 | 0 | 0.77 | 0.19 | 0.8 | 0.67 | 0.94 | 0.75 |
| 8 | 52806682 | rs9298467 | 1.722 | -0.401 | 0.062 | 1.419 | 0.007 | -0.1 | -0.1 | 0 | 0.7 | 0.08 | 0.72 | 0.63 | 0.78 | 0.76 |
| 8 | 52818658 | rs4242479 | -1.733 | -0.509 | 1.148 | 1.55 | -0.002 | 0.065 | 0.065 | 0.442 | 0.28 | 0.41 | 0.27 | 0.49 | 0.29 | 0.29 |
| 8 | 52831142 | rs17212214 | -0.781 | -0.461 | 1.35 | 1.627 | -0.048 | -0.04 | -0.04 | 0.021 | 0.13 | 0.03 | 0.11 | 0.15 | 0.23 | 0.16 |
| 8 | 52838295 | rs6473666 | -1.983 | -0.597 | 1.934 | 1.557 | -0.001 | -0.001 | -0.001 | 0.55 | 0.26 | 0.35 | 0.18 | 0.19 | 0.29 | 0.25 |
| 8 | 52844411 | rs16916968 | -2.384 | -0.761 | 1.73 | 1.474 | -0.002 | -0.001 | -0.001 | 0.812 | 0.3 | 0.67 | 0.22 | 0.2 | 0.28 | 0.25 |
| 8 | 52865175 | rs2138297 | -1.561 | -0.806 | 1.533 | 1.49 | -0.05 | 0.077 | 0.077 | 0 | 0.17 | 0.46 | 0.14 | 0.19 | 0.1 | 0.16 |
| 8 | 52870877 | rs6993367 | -2.755 | -0.833 | 1.445 | 1.596 | 0.027 | -0.085 | -0.085 | 0.133 | 0.41 | 0.95 | 0.33 | 0.54 | 0.31 | 0.39 |
| 8 | 52873414 | rs7838307 | -2.038 | -0.723 | 1.557 | 1.576 | -0.045 | 0.005 | 0.005 | 0 | 0.26 | 0.47 | 0.18 | 0.21 | 0.2 | 0.22 |
| 8 | 52882218 | rs7817892 | -1.834 | -0.493 | 2.015 | 1.947 | -0.039 | 0.108 | 0.108 | 0.345 | 0.16 | 0.48 | 0.14 | 0.21 | 0.1 | 0.12 |
| 8 | 52887541 | rs1015213 | -1.125 | -0.473 | 2.176 | 1.874 | 0.003 | -0.052 | -0.052 | 0.397 | 0.12 | 0.34 | 0.05 | 0.01 | 0.1 | 0.11 |
| 8 | 52897480 | rs16917026 | -1.904 | -0.433 | 1.929 | 2.008 | 0.031 | 0.162 | 0.162 | 0 | 0.06 | 0.19 | 0.1 | 0.19 | 0.08 | 0.08 |
| 8 | 52898827 | rs12678795 | -1.904 | -0.433 | 1.929 | 2.008 | 0.031 | 0.162 | 0.162 | 0 | 0.06 | 0.19 | 0.1 | 0.19 | 0.08 | 0.08 |
| 8 | 52910266 | rs16917074 | -1.323 | -0.517 | 1.683 | 2.096 | -0.014 | 0.244 | 0.244 | 0.044 | 0.11 | 0.32 | 0.14 | 0.36 | 0.04 | 0.1 |
| 8 | 52913908 | rs1399857 | -2.57 | -0.433 | 1.372 | 1.866 | 0.01 | -0.01 | -0.01 | 0.239 | 0.24 | 0.83 | 0.27 | 0.22 | 0.21 | 0.19 |
| 8 | 52914736 | rs6982212 | -2.399 | -0.412 | 1.456 | 1.879 | 0.013 | -0.007 | -0.007 | 0 | 0.23 | 0.79 | 0.26 | 0.21 | 0.16 | 0.13 |
| 8 | 52917469 | rs2554465 | -2.14 | -0.559 | 1.372 | 1.66 | -0.024 | -0.009 | -0.009 | 3.366 | 0.25 | 0.66 | 0.26 | 0.17 | 0.18 | 0.14 |
| 8 | 52920335 | rs2280179 | -2.586 | -0.558 | 1.359 | 1.687 | 0.022 | 0.003 | 0.003 | 0.56 | 0.29 | 0.79 | 0.3 | 0.22 | 0.21 | 0.16 |
| 8 | 52923780 | rs10088713 | -1.856 | -0.605 | 1.403 | 1.895 | 0.001 | 0.194 | 0.194 | 0 | 0.09 | 0.24 | 0.15 | 0.38 | 0.05 | 0.12 |
| 8 | 52923809 | rs10104090 | -2.154 | -0.387 | 1.358 | 1.912 | 0.005 | 0.177 | 0.177 | 0 | 0.13 | 0.29 | 0.16 | 0.39 | 0.07 | 0.13 |
| 8 | 52924095 | rs16917103 | -1.191 | -0.538 | 1.646 | 1.898 | -0.025 | -0.094 | -0.094 | 0 | 0.15 | 0.49 | 0.16 | 0.01 | 0.09 | 0.06 |
| 8 | 52926773 | rs3886248 | -1.328 | -0.271 | 1.346 | 2.196 | -0.015 | 0.083 | 0.083 | 0 | 0.1 | 0.13 | 0.12 | 0.35 | 0.04 | 0.1 |

| | | | | | | | | | | | | | | | | |
|---|---|---|---|---|---|---|---|---|---|---|---|---|---|---|---|---|
| 8 | 52928389 | rs4873633 | -1.526 | -0.249 | 1.308 | 2.162 | 0.001 | 0.074 | 0.074 | 0 | 0.1 | 0.15 | 0.13 | 0.34 | 0.09 | 0.17 |
| 8 | 52929150 | rs268926 | -1.49 | 0.095 | 1.062 | 2.019 | 0.004 | 0.037 | 0.037 | 0 | 0.1 | 0.16 | 0.1 | 0.17 | 0.15 | 0.14 |
| 8 | 52930270 | rs10092108 | -1.527 | -0.176 | 0.725 | 2.081 | -0.014 | 0.073 | 0.073 | 0 | 0.19 | 0.29 | 0.15 | 0.42 | 0.14 | 0.15 |
| 8 | 52931576 | rs2450145 | -1.919 | 0.615 | 0.172 | 1.513 | -0.078 | -0.082 | -0.082 | 0.024 | 0.43 | 0.96 | 0.47 | 0.59 | 0.43 | 0.46 |
| 8 | 52932617 | rs268931 | -1.625 | 0.574 | 0.734 | 1.55 | -0.004 | -0.025 | -0.025 | 0 | 0.29 | 0.76 | 0.35 | 0.47 | 0.3 | 0.31 |
| 8 | 52932841 | rs7815671 | -1.248 | 0.572 | 0.729 | 1.548 | -0.003 | -0.013 | -0.013 | 0 | 0.13 | 0.19 | 0.12 | 0.09 | 0.12 | 0.16 |
| 8 | 52935478 | rs185596 | -2.301 | 0.585 | 0.616 | 1.554 | -0.069 | -0.092 | -0.092 | 0 | 0.42 | 0.95 | 0.46 | 0.55 | 0.41 | 0.47 |
| 8 | 52945956 | rs7817996 | -1.183 | 0.603 | 0.906 | 1.219 | -0.007 | -0.011 | -0.011 | 0 | 0.12 | 0.19 | 0.1 | 0.09 | 0.1 | 0.11 |
| 8 | 52948365 | rs2249242 | -2.45 | 0.861 | 1.895 | 1.409 | -0.006 | 0.054 | 0.054 | 0 | 0.31 | 0.48 | 0.27 | 0.55 | 0.3 | 0.39 |
| 8 | 52949557 | rs12541286 | -0.902 | 0.856 | 1.939 | 1.401 | 0.004 | 0.008 | 0.008 | 0 | 0.1 | 0.09 | 0.08 | 0.07 | 0.09 | 0.1 |
| 8 | 52950361 | rs2459164 | -2.875 | 0.909 | 2.397 | 1.369 | -0.002 | 0.028 | 0.028 | 0 | 0.36 | 0.56 | 0.3 | 0.55 | 0.36 | 0.44 |
| 8 | 52951973 | rs2252399 | -3.047 | 0.905 | 2.379 | 1.37 | 0.001 | 0.007 | 0.007 | 0 | 0.38 | 0.61 | 0.43 | 0.55 | 0.4 | 0.46 |
| 8 | 52961530 | rs3892650 | 0.047 | 0.898 | 2.239 | 1.332 | 0.005 | -0.021 | -0.021 | 0 | 0.05 | 0.08 | 0.02 | 0 | 0.06 | 0.05 |
| 8 | 52961970 | rs2433236 | -1.177 | 0.893 | 2.304 | 1.348 | 0.003 | 0.06 | 0.06 | 0 | 0.19 | 0.28 | 0.17 | 0.52 | 0.2 | 0.29 |
| 8 | 52964522 | rs9785150 | -0.159 | 0.803 | 2.493 | 1.115 | 0.022 | -0.021 | -0.021 | 0 | 0.05 | 0.18 | 0.02 | 0 | 0.06 | 0.05 |
| 8 | 52971672 | rs2450153 | -1.392 | 0.758 | 2.554 | 1.259 | -0.002 | 0.02 | 0.02 | 0 | 0.22 | 0.26 | 0.26 | 0.48 | 0.22 | 0.26 |
| 8 | 52975387 | rs3909673 | -1.12 | 0.807 | 2.451 | 1.215 | 0.023 | 0.024 | 0.024 | 0 | 0.07 | 0.19 | 0.1 | 0.09 | 0.08 | 0.1 |
| 8 | 52987396 | rs11987246 | 2.631 | 0.823 | 2.233 | 1.142 | -0.06 | -0.13 | -0.13 | 0.208 | 0.61 | 0.11 | 0.46 | 0.46 | 0.59 | 0.6 |
| 8 | 52998935 | rs11774243 | 2.687 | 0.51 | 2.362 | 1.066 | -0.038 | -0.129 | -0.129 | 0 | 0.61 | 0.08 | 0.45 | 0.45 | 0.61 | 0.6 |
| 8 | 52999459 | rs9650306 | 0.963 | 0.509 | 2.72 | 1.052 | 0.003 | 0.054 | 0.054 | 0 | 0.8 | 0.72 | 0.72 | 0.51 | 0.76 | 0.73 |
| 8 | 53001694 | rs7843694 | -1.005 | 0.572 | 3.001 | 0.598 | 0.057 | -0.02 | -0.02 | 0 | 0.07 | 0.32 | 0.17 | 0 | 0.05 | 0.1 |
| 8 | 53006766 | rs10090913 | -1.793 | 0.551 | 2.536 | 0.616 | -0.006 | -0.073 | -0.073 | 0 | 0.18 | 0.56 | 0.26 | 0.05 | 0.16 | 0.14 |
| 8 | 53007343 | rs6990453 | -1.655 | 0.564 | 2.411 | 0.622 | -0.003 | -0.068 | -0.068 | 0 | 0.17 | 0.58 | 0.26 | 0.05 | 0.16 | 0.14 |
| 8 | 53009554 | rs3843918 | 1.92 | 0.789 | 1.56 | 0.507 | -0.084 | -0.078 | -0.078 | 0 | 0.54 | 0.04 | 0.42 | 0.46 | 0.52 | 0.59 |
| 8 | 53009606 | rs3849810 | -0.825 | 1.257 | 2.056 | 0.69 | 0.02 | 0.1 | 0.1 | 0.059 | 0.15 | 0.25 | 0.22 | 0.49 | 0.2 | 0.26 |
| 8 | 53010530 | rs1895885 | 1.322 | 0.746 | 2.566 | -0.077 | 0.005 | -0.091 | -0.091 | 1.053 | 0.75 | 0.57 | 0.67 | 0.95 | 0.72 | 0.84 |
| 8 | 53011018 | rs4087846 | 1.232 | 0.778 | 2.144 | -0.107 | -0.001 | -0.122 | -0.122 | 0 | 0.73 | 0.44 | 0.66 | 0.94 | 0.72 | 0.85 |
| 8 | 53031607 | rs1346159 | -1.958 | 0.714 | 2.442 | 0.347 | 0.002 | 0.002 | 0.002 | 0.027 | 0.21 | 0.78 | 0.17 | 0.07 | 0.18 | 0.16 |
| 8 | 53036734 | rs7002709 | -0.915 | 1.146 | 2.532 | 0.815 | 0.04 | 0.112 | 0.112 | 0 | 0.11 | 0.23 | 0.12 | 0.41 | 0.18 | 0.24 |
| 8 | 53037349 | rs2114226 | -1.793 | 0.726 | 2.533 | 0.069 | -0.006 | -0.038 | -0.038 | 0 | 0.23 | 0.68 | 0.16 | 0.07 | 0.18 | 0.21 |
| 8 | 53046681 | rs11990195 | -1.345 | 0.699 | 2.839 | -0.031 | 0.007 | -0.13 | -0.13 | 0.001 | 0.18 | 0.42 | 0.12 | 0 | 0.17 | 0.17 |
| 8 | 53049037 | rs3815868 | -2.103 | 0.674 | 2.381 | -0.049 | -0.004 | -0.034 | -0.034 | 0 | 0.27 | 0.7 | 0.16 | 0.05 | 0.19 | 0.22 |
| 8 | 53055483 | rs7010548 | -1.898 | 0.741 | 2.243 | -0.018 | 0.001 | -0.018 | -0.018 | 0 | 0.25 | 0.74 | 0.16 | 0.05 | 0.18 | 0.2 |
| 8 | 53068920 | rs17290977 | -1.183 | 0.857 | 1.941 | -0.011 | 0.002 | -0.044 | -0.044 | 0 | 0.2 | 0.3 | 0.13 | 0.11 | 0.19 | 0.2 |
| 8 | 53076976 | rs9792154 | -1.667 | 0.618 | 1.451 | 0.724 | 0.007 | 0.019 | 0.019 | 0 | 0.16 | 0.68 | 0.22 | 0.22 | 0.16 | 0.19 |
| 8 | 53088124 | rs7834888 | -2.041 | 0.223 | 1.671 | 0.423 | 0.067 | 0.113 | 0.113 | 0 | 0.06 | 0.49 | 0.05 | 0.09 | 0.04 | 0.16 |
| 8 | 53091806 | rs7837165 | -2.706 | 0.104 | 1.612 | 0.365 | -0.025 | 0.003 | 0.003 | 0.048 | 0.1 | 0.6 | 0.07 | 0.11 | 0.04 | 0.17 |
| 8 | 53098200 | rs6992634 | -2.286 | -0.073 | 1.306 | 0.335 | -0.047 | -0.014 | -0.014 | 0 | 0.16 | 0.61 | 0.09 | 0.18 | 0.07 | 0.23 |
| 8 | 53100810 | rs6989604 | -2.073 | -0.079 | 1.854 | 0.331 | 0.029 | 0.07 | 0.07 | 0 | 0.07 | 0.4 | 0.04 | 0.09 | 0.04 | 0.17 |
| 8 | 53103663 | rs7838823 | -1.884 | 0.011 | 1.148 | 0.252 | -0.034 | -0.034 | -0.034 | 0.104 | 0.14 | 0.6 | 0.09 | 0.1 | 0.07 | 0.07 |
| 8 | 53110206 | rs16917430 | -0.825 | 0.079 | 0.046 | -0.089 | 0.003 | 0.028 | 0.028 | 0 | 0.27 | 0.2 | 0.14 | 0.15 | 0.21 | 0.26 |
| 8 | 53111052 | rs2360804 | -1.669 | 0.205 | 0.972 | 0.179 | 0.027 | 0.033 | 0.033 | 0 | 0.15 | 0.57 | 0.2 | 0.13 | 0.11 | 0.13 |
| 8 | 53112386 | rs6983307 | -1.864 | -0.011 | 0.847 | -0.175 | -0.002 | -0.008 | -0.008 | 0 | 0.22 | 0.69 | 0.27 | 0.13 | 0.17 | 0.14 |
| 8 | 53114431 | rs7840334 | -1.784 | -0.02 | 1.438 | -0.18 | 0.018 | -0.007 | -0.007 | 0 | 0.2 | 0.43 | 0.25 | 0.13 | 0.17 | 0.14 |
| 8 | 53114569 | rs2360805 | -1.519 | 0.236 | 0.071 | -0.557 | -0.073 | 0.098 | 0.098 | 0.188 | 0.46 | 0.91 | 0.52 | 0.27 | 0.49 | 0.28 |
| 8 | 53115983 | rs16917460 | -1.123 | -0.499 | 0.26 | -0.477 | -0.011 | 0.012 | 0.012 | 0 | 0.05 | 0.05 | 0.1 | 0.13 | 0.02 | 0.04 |
| 8 | 53118591 | rs16917470 | -0.237 | -0.448 | 0.202 | -0.618 | 0.023 | 0.017 | 0.017 | 0 | 0.21 | 0.13 | 0.18 | 0.13 | 0.2 | 0.09 |
| 8 | 53125900 | rs2360808 | -2.201 | -0.69 | 0.516 | -0.732 | -0.035 | 0.096 | 0.096 | 0.098 | 0.1 | 0.38 | 0.14 | 0.12 | 0.02 | 0.05 |
| 8 | 53126985 | rs7015251 | -0.484 | -0.664 | 0.548 | -0.735 | -0.034 | -0.049 | -0.049 | 0 | 0.22 | 0.31 | 0.1 | 0.08 | 0.2 | 0.09 |
| 8 | 53128122 | rs6992299 | -0.547 | -0.644 | 0.516 | -0.721 | -0.04 | -0.039 | -0.039 | 0 | 0.23 | 0.37 | 0.2 | 0.1 | 0.2 | 0.09 |
| 8 | 53129669 | rs12155844 | -0.399 | -0.647 | 0.511 | -0.839 | -0.054 | -0.071 | -0.071 | 0 | 0.26 | 0.46 | 0.23 | 0.1 | 0.22 | 0.12 |
| 8 | 53133166 | rs7830945 | -2.064 | -0.503 | 0.351 | -0.877 | -0.003 | 0.024 | 0.024 | 0.34 | 0.08 | 0.23 | 0.23 | 0.09 | 0.05 | 0.07 |
| 8 | 53137329 | rs4074067 | -0.446 | -0.839 | 0.374 | -0.845 | -0.037 | -0.05 | -0.05 | 0 | 0.25 | 0.38 | 0.23 | 0.09 | 0.22 | 0.13 |
| 8 | 53141715 | rs16917539 | -0.42 | -0.84 | 0.362 | -0.846 | -0.025 | -0.035 | -0.035 | 0.402 | 0.25 | 0.33 | 0.23 | 0.09 | 0.22 | 0.13 |
| 8 | 53172806 | rs4481619 | -0.411 | -1.004 | -0.206 | -0.626 | 0.018 | 0.047 | 0.047 | 0.732 | 0.35 | 0.24 | 0.17 | 0.18 | 0.3 | 0.12 |
| 8 | 53185718 | rs714048 | 0.706 | -1.397 | -0.885 | -0.745 | 0.02 | 0.006 | 0.006 | 0.224 | 0.12 | 0.01 | 0.05 | 0.07 | 0.06 | 0.19 |
| 8 | 53188287 | rs7463107 | 0.758 | -0.819 | -0.2 | -0.441 | -0.004 | -0.007 | -0.007 | 0 | 0.6 | 0.57 | 0.7 | 0.73 | 0.67 | 0.83 |
| 8 | 53193238 | rs4639533 | 0.153 | -0.476 | -0.63 | -0.516 | 0.004 | 0.106 | 0.106 | 0 | 0.27 | 0.08 | 0.24 | 0.08 | 0.26 | 0.14 |
| 8 | 53195088 | rs10958312 | 1.67 | -1.15 | -0.396 | 0.482 | 0.018 | -0.027 | -0.027 | 0 | 0.48 | 0.06 | 0.52 | 0.43 | 0.59 | 0.68 |
| 8 | 53199089 | rs17315018 | 0.143 | -0.688 | -0.138 | -0.28 | 0.038 | 0.092 | 0.092 | 0 | 0.26 | 0.08 | 0.22 | 0.09 | 0.22 | 0.14 |
| 8 | 53202655 | rs16917636 | -0.241 | -0.666 | -0.133 | -0.259 | -0.033 | -0.044 | -0.044 | 0 | 0.11 | 0.01 | 0.08 | 0.15 | 0.11 | 0.03 |
| 8 | 53227219 | rs7824696 | -2.047 | 0.347 | 1.321 | 1.069 | -0.008 | 0.025 | 0.025 | 0 | 0.18 | 0.33 | 0.22 | 0.24 | 0.16 | 0.08 |
| 8 | 53242775 | rs2885068 | -1.838 | 0.519 | 1.075 | 1.227 | 0.001 | 0.031 | 0.031 | 0 | 0.18 | 0.37 | 0.22 | 0.24 | 0.14 | 0.09 |
| 8 | 53248420 | rs13258413 | -1.384 | 0.619 | 0.593 | 1.07 | 0.017 | 0.02 | 0.02 | 0 | 0.15 | 0.24 | 0.26 | 0.23 | 0.15 | 0.12 |
| 8 | 53259979 | rs9643471 | -1.645 | -0.134 | 0.815 | 1.219 | -0.042 | 0.081 | 0.081 | 0 | 0.16 | 0.39 | 0.07 | 0.23 | 0.06 | 0.15 |
| 8 | 53283773 | rs2360791 | 1.549 | -0.235 | 1.402 | 1.286 | -0.059 | 0.089 | 0.089 | 0.022 | 0.83 | 0.45 | 0.92 | 0.76 | 0.93 | 0.85 |
| 8 | 53287893 | rs6981742 | -1.828 | -0.264 | 1.432 | 1.293 | -0.05 | 0.122 | 0.122 | 0 | 0.15 | 0.42 | 0.08 | 0.22 | 0.07 | 0.16 |
| 8 | 53294648 | rs5015618 | -1.284 | -0.302 | 0.72 | 1.344 | 0.002 | -0.035 | -0.035 | 0 | 0.06 | 0 | 0.03 | 0.14 | 0.05 | 0.1 |
| 8 | 53295154 | rs1965704 | 1.732 | -0.323 | 1.045 | 1.289 | -0.031 | 0.019 | 0.019 | 0.202 | 0.83 | 0.26 | 0.9 | 0.78 | 0.93 | 0.84 |

| | | | | | | | | | | | | | | | | |
|---|---|---|---|---|---|---|---|---|---|---|---|---|---|---|---|---|
| 8 | 53303885 | rs6473725 | 0.507 | -0.069 | 0.482 | 1.231 | 0.005 | -0.009 | -0.009 | 0 | 0.62 | 0.31 | 0.65 | 0.58 | 0.7 | 0.65 |
| 8 | 53305054 | rs16917805 | 0.966 | 0.066 | -0.137 | 1.279 | -0.061 | -0.027 | -0.027 | 0 | 0.51 | 0.06 | 0.55 | 0.39 | 0.6 | 0.54 |
| 8 | 53313496 | rs1427469 | 0.202 | -0.116 | 1.566 | 1.636 | 0 | -0.001 | -0.001 | 1.022 | 0.68 | 0.29 | 0.68 | 0.67 | 0.72 | 0.67 |
| 8 | 53317158 | rs4489337 | 0.167 | -0.154 | 1.334 | 1.604 | -0.018 | 0.003 | 0.003 | 0 | 0.25 | 0.15 | 0.27 | 0.24 | 0.25 | 0.32 |
| 8 | 53318644 | rs1317186 | 0.459 | -0.22 | 1.298 | 1.616 | -0.006 | 0.035 | 0.035 | 0 | 0.7 | 0.79 | 0.84 | 0.75 | 0.73 | 0.72 |
| 8 | 53320710 | rs12678991 | 1.36 | -0.244 | 1.127 | 1.557 | 0.016 | -0.01 | -0.01 | 0 | 0.54 | 0.19 | 0.6 | 0.42 | 0.66 | 0.6 |
| 8 | 53322749 | rs7841071 | -0.757 | 0.458 | 0.91 | 1.06 | 0.017 | -0.001 | -0.001 | 0 | 0.36 | 0.52 | 0.38 | 0.34 | 0.37 | 0.33 |
| 8 | 53323786 | rs3758126 | -0.571 | 1.038 | -0.344 | 0.813 | -0.093 | -0.107 | -0.107 | 0.014 | 0.23 | 0.04 | 0.3 | 0.38 | 0.25 | 0.31 |
| 8 | 53328331 | rs7840491 | -0.293 | 0.341 | -0.137 | -0.458 | -0.02 | -0.164 | -0.164 | 0 | 0.28 | 0.45 | 0.23 | 0.01 | 0.22 | 0.13 |
| 8 | 53334176 | rs890599 | -0.015 | 0.532 | -1.188 | -0.262 | -0.001 | 0.05 | 0.05 | 0 | 0.44 | 0.22 | 0.54 | 0.66 | 0.54 | 0.31 |
| 8 | 53335677 | rs716583 | -0.851 | -0.161 | -1.191 | -0.724 | -0.017 | -0.013 | -0.013 | 0 | 0.78 | 0.83 | 0.72 | 0.76 | 0.71 | 0.67 |
| 8 | 53338897 | rs4873211 | 1.507 | 0.048 | -1.573 | -1.038 | -0.136 | -0.09 | -0.09 | 0 | 0.29 | 0.03 | 0.38 | 0.66 | 0.34 | 0.25 |
| 8 | 53341622 | rs13279773 | 1.698 | 0.101 | -1.616 | -0.174 | -0.138 | -0.1 | -0.1 | 0 | 0.27 | 0.02 | 0.38 | 0.34 | 0.33 | 0.23 |
| 8 | 53342616 | rs1001932 | 1.698 | 0.101 | -1.616 | -0.172 | -0.138 | -0.103 | -0.103 | 0 | 0.27 | 0.02 | 0.38 | 0.35 | 0.33 | 0.23 |
| 8 | 53343969 | rs1660554 | -0.501 | -0.32 | -0.286 | -0.679 | 0.009 | -0.007 | -0.007 | 0 | 0.72 | 0.42 | 0.67 | 0.76 | 0.69 | 0.63 |
| 8 | 53353780 | rs2613243 | 0.471 | -0.441 | 0.32 | -0.792 | -0.071 | -0.113 | -0.113 | 0 | 0.32 | 0.41 | 0.16 | 0.1 | 0.16 | 0.35 |
| 8 | 53355625 | rs1440795 | 0.492 | -0.451 | 0.388 | -0.802 | -0.064 | -0.102 | -0.102 | 0 | 0.32 | 0.39 | 0.16 | 0.09 | 0.16 | 0.35 |
| 8 | 53355822 | rs17228239 | 0.718 | -0.427 | 0.302 | -0.743 | -0.005 | 0.06 | 0.06 | 0 | 0.46 | 0.17 | 0.53 | 0.67 | 0.54 | 0.3 |
| 8 | 53375405 | rs2556559 | 0.623 | -0.699 | 0.581 | -0.823 | -0.07 | -0.109 | -0.109 | 0 | 0.31 | 0.4 | 0.15 | 0.1 | 0.16 | 0.34 |
| 8 | 53376653 | rs2582635 | 1.044 | -0.487 | 0.49 | -0.794 | -0.006 | -0.006 | -0.006 | 0 | 0.22 | 0.23 | 0.28 | 0.24 | 0.27 | 0.35 |
| 8 | 53377236 | rs10958322 | 0.688 | -0.389 | -0.168 | -0.733 | 0.008 | 0.06 | 0.06 | 0 | 0.45 | 0.13 | 0.53 | 0.65 | 0.57 | 0.31 |
| 8 | 53380905 | rs7012920 | -0.534 | -0.296 | 0.435 | -0.646 | -0.006 | -0.01 | -0.01 | 0 | 0.74 | 0.37 | 0.67 | 0.75 | 0.73 | 0.64 |
| 8 | 53387502 | rs2613266 | -0.66 | -0.475 | 0.451 | -0.813 | 0.025 | 0.051 | 0.051 | 0 | 0.51 | 0.67 | 0.44 | 0.33 | 0.44 | 0.67 |
| 8 | 53388439 | rs1440788 | 1.151 | -0.621 | 0.518 | -0.912 | 0.034 | 0.047 | 0.047 | 0 | 0.26 | 0.12 | 0.12 | 0.09 | 0.16 | 0.32 |
| 8 | 53390116 | rs920504 | 1.015 | -0.615 | 0.542 | -0.91 | 0.003 | 0.079 | 0.079 | 0 | 0.46 | 0.11 | 0.54 | 0.66 | 0.56 | 0.33 |
| 8 | 53391097 | rs1837630 | -0.807 | -0.598 | 0.604 | -0.901 | 0.035 | 0.078 | 0.078 | 0 | 0.52 | 0.75 | 0.45 | 0.33 | 0.44 | 0.67 |
| 8 | 53392062 | rs6982235 | 0.026 | -0.613 | 0.583 | -0.876 | 0.01 | 0.01 | 0.01 | 0.026 | 0.34 | 0.64 | 0.25 | 0.22 | 0.22 | 0.41 |
| 8 | 53393301 | rs2613258 | -0.224 | -0.418 | 0.722 | -0.78 | 0.016 | 0.003 | 0.003 | 0 | 0.23 | 0.56 | 0.32 | 0.23 | 0.27 | 0.34 |
| 8 | 53394051 | rs2582629 | -0.25 | -0.414 | 0.594 | -0.78 | 0.009 | -0.001 | -0.001 | 0.195 | 0.24 | 0.62 | 0.32 | 0.23 | 0.27 | 0.34 |
| 8 | 53394342 | rs7845332 | 0.338 | -0.433 | 0.584 | -0.787 | -0.001 | -0.002 | -0.002 | 0 | 0.74 | 0.26 | 0.67 | 0.76 | 0.73 | 0.66 |
| 8 | 53402811 | rs11776384 | 1.551 | -0.184 | 0.124 | -1.121 | -0.023 | 0.037 | 0.037 | 0 | 0.14 | 0.06 | 0.14 | 0 | 0.22 | 0.05 |
| 8 | 53403318 | rs2613250 | 1.15 | -0.283 | 0.712 | -0.91 | 0.055 | 0.045 | 0.045 | 3.692 | 0.94 | 0.59 | 0.96 | 0.98 | 1 | 0.99 |
| 8 | 53406508 | rs2613246 | 0.951 | -0.096 | -0.058 | -0.892 | -0.035 | 0.024 | 0.024 | 0 | 0.16 | 0.1 | 0.2 | 0.11 | 0.2 | 0.24 |
| 8 | 53408434 | rs11776276 | 0.903 | -0.286 | -0.04 | -1.08 | 0.019 | 0.027 | 0.027 | 0 | 0.25 | 0.15 | 0.15 | 0.08 | 0.16 | 0.31 |
| 8 | 53418983 | rs12679115 | 1.617 | 0.299 | -0.449 | -1.181 | -0.019 | -0.047 | -0.047 | 0 | 0.28 | 0.11 | 0.3 | 0.65 | 0.22 | 0.22 |
| 8 | 53431744 | rs7836283 | 1.123 | -0.293 | 0.192 | -1.54 | -0.025 | -0.036 | -0.036 | 0 | 0.14 | 0.21 | 0.1 | 0.05 | 0.09 | 0.11 |
| 8 | 53433015 | rs11774072 | 1.886 | -0.215 | 0.404 | -1.758 | -0.003 | -0.019 | -0.019 | 0 | 0.17 | 0.12 | 0.16 | 0 | 0.24 | 0.06 |
| 8 | 53434613 | rs984309 | -0.478 | -0.222 | 0.588 | -1.769 | 0.014 | -0.06 | -0.06 | 0 | 0.65 | 0.61 | 0.61 | 0.89 | 0.56 | 0.7 |
| 8 | 53436131 | rs7820696 | 1.31 | -0.324 | 0.529 | -1.781 | -0.025 | -0.019 | -0.019 | 0 | 0.09 | 0.17 | 0.09 | 0.05 | 0.05 | 0.09 |
| 8 | 53444854 | rs10504139 | -0.374 | -0.36 | 0.5 | -1.767 | 0.007 | -0.052 | -0.052 | 0 | 0.6 | 0.6 | 0.58 | 0.88 | 0.49 | 0.68 |
| 8 | 53445338 | rs13282005 | 0.996 | -0.371 | 0.644 | -1.773 | -0.013 | -0.007 | -0.007 | 0 | 0.18 | 0.27 | 0.14 | 0.18 | 0.14 | 0.31 |
| 8 | 53447256 | rs1440798 | -0.722 | -0.374 | 0.469 | -1.785 | -0.024 | -0.038 | -0.038 | 0 | 0.33 | 0.53 | 0.29 | 0.23 | 0.27 | 0.46 |
| 8 | 53449548 | rs4873659 | 0.379 | -0.325 | 0.114 | -1.67 | -0.005 | -0.025 | -0.025 | 0.005 | 0.14 | 0.2 | 0.14 | 0.05 | 0.14 | 0.15 |
| 8 | 53455537 | rs12679091 | 2.154 | -0.129 | -0.232 | -0.742 | -0.054 | -0.13 | -0.13 | 1.959 | 0.25 | 0.03 | 0.29 | 0.35 | 0.2 | 0.2 |
| 8 | 53473205 | rs12335010 | -0.896 | -0.058 | 0.044 | -1.239 | 0.012 | -0.021 | -0.021 | 0 | 0.79 | 0.76 | 0.74 | 0.89 | 0.73 | 0.73 |
| 8 | 53488242 | rs7826532 | -0.496 | -0.125 | -0.278 | -1.25 | 0.014 | 0.2 | 0.2 | 0 | 0.51 | 0.76 | 0.45 | 0.2 | 0.51 | 0.53 |
| 8 | 53494271 | rs7843825 | -1.035 | 0.114 | -0.237 | -1.133 | -0.046 | 0.019 | 0.019 | 0 | 0.81 | 0.92 | 0.75 | 0.83 | 0.74 | 0.75 |
| 8 | 53501285 | rs2360279 | -0.949 | 0.114 | -0.234 | -1.127 | -0.057 | 0.014 | 0.014 | 0 | 0.8 | 0.93 | 0.74 | 0.82 | 0.74 | 0.75 |
| 8 | 53505755 | rs4319121 | 0.057 | -0.082 | 0.166 | -1.367 | -0.029 | -0.047 | -0.047 | 0 | 0.23 | 0.44 | 0.18 | 0.13 | 0.18 | 0.38 |
| 8 | 53509414 | rs13255007 | 0.281 | -0.137 | 0.144 | -1.508 | 0.003 | 0.001 | 0.001 | 0 | 0.5 | 0.58 | 0.52 | 0.79 | 0.42 | 0.61 |
| 8 | 53510650 | rs4873668 | 0.218 | -0.173 | 0.239 | -1.45 | -0.008 | 0.081 | 0.081 | 0 | 0.32 | 0.15 | 0.22 | 0.03 | 0.38 | 0.21 |
| 8 | 53553177 | rs12549201 | 0.716 | -0.283 | 0.548 | -1.438 | -0.013 | -0.01 | -0.01 | 1.107 | 0.16 | 0.24 | 0.14 | 0.13 | 0.12 | 0.31 |
| 8 | 53555069 | rs2305427 | 2.28 | -0.051 | -0.275 | -1.667 | -0.014 | -0.05 | -0.05 | 0.435 | 0.23 | 0.09 | 0.32 | 0.65 | 0.2 | 0.24 |
| 8 | 53557226 | rs2108 | 0.912 | 0.033 | -0.257 | -1.669 | -0.08 | 0.037 | 0.037 | 0 | 0.23 | 0.08 | 0.28 | 0.18 | 0.3 | 0.24 |
| 8 | 53602318 | rs16918093 | 1.157 | -0.166 | 0.43 | -1.481 | 0.015 | 0.02 | 0.02 | 0 | 0.22 | 0.14 | 0.12 | 0 | 0.21 | 0.06 |
| 8 | 53605442 | rs7002395 | 0.342 | -0.128 | 0.644 | -1.309 | -0.019 | -0.063 | -0.063 | 0 | 0.73 | 0.56 | 0.76 | 0.84 | 0.76 | 0.6 |
| 8 | 53669478 | rs10095687 | 0.518 | -0.113 | 0.61 | -1.238 | -0.008 | -0.043 | -0.043 | 2.386 | 0.71 | 0.6 | 0.74 | 0.83 | 0.71 | 0.54 |
| 8 | 53677171 | rs4873679 | 0.828 | -0.175 | 0.616 | -1.269 | -0.02 | -0.06 | -0.06 | 0 | 0.68 | 0.47 | 0.74 | 0.83 | 0.71 | 0.54 |
| 8 | 53680808 | rs11785226 | 0.89 | -0.135 | 0.463 | -1.359 | 0.014 | 0.049 | 0.049 | 0 | 0.49 | 0.39 | 0.63 | 0.83 | 0.5 | 0.48 |
| 8 | 53684861 | rs7822058 | -1.068 | -0.023 | 0.16 | -1.275 | 0.001 | -0.013 | -0.013 | 0 | 0.91 | 0.91 | 0.91 | 0.96 | 0.89 | 0.92 |
| 8 | 53687126 | rs17253031 | 1.478 | -0.021 | -0.317 | -1.256 | 0.005 | 0.036 | 0.036 | 0 | 0.19 | 0.08 | 0.12 | 0 | 0.21 | 0.06 |
| 8 | 53689112 | rs17338055 | 1.387 | 0.043 | -0.163 | -1.229 | 0.005 | 0.011 | 0.011 | 0 | 0.19 | 0.11 | 0.13 | 0 | 0.21 | 0.06 |
| 8 | 53690181 | rs12677803 | 1.988 | 0.087 | -0.412 | -0.288 | -0.025 | -0.009 | -0.009 | 0 | 0.07 | 0.01 | 0.25 | 0.32 | 0.05 | 0.11 |
| 8 | 53691001 | rs4873221 | -0.114 | -0.015 | 0.191 | -0.868 | 0.016 | -0.001 | -0.001 | 0 | 0.85 | 0.64 | 0.83 | 0.86 | 0.83 | 0.81 |
| 8 | 53693490 | rs7460309 | -0.045 | -0.005 | 0.196 | -0.862 | 0.019 | 0.002 | 0.002 | 0 | 0.85 | 0.64 | 0.83 | 0.86 | 0.83 | 0.81 |
| 8 | 53697552 | rs6999665 | 0.902 | -0.117 | 0.121 | -0.891 | 0.021 | 0.091 | 0.091 | 0 | 0.51 | 0.39 | 0.63 | 0.82 | 0.5 | 0.49 |
| 8 | 53703969 | rs7461729 | 0.527 | -0.115 | 0.158 | -0.884 | -0.015 | -0.054 | -0.054 | 0 | 0.71 | 0.56 | 0.76 | 0.83 | 0.71 | 0.54 |
| 8 | 53705808 | rs17253706 | 1.322 | -0.137 | 0.042 | -0.943 | 0.005 | 0.039 | 0.039 | 0 | 0.19 | 0.09 | 0.12 | 0 | 0.21 | 0.06 |
| 8 | 53711446 | rs6473743 | 0.732 | 0.029 | 0.019 | -0.759 | 0.018 | 0.111 | 0.111 | 0 | 0.52 | 0.39 | 0.66 | 0.81 | 0.52 | 0.53 |

| | | | | | | | | | | | | | | | | |
|---|---|---|---|---|---|---|---|---|---|---|---|---|---|---|---|---|
| 8 | 53714558 | rs10091593 | -0.179 | 0.19 | 0.114 | -0.731 | 0.028 | 0.009 | 0.009 | 0 | 0.14 | 0.38 | 0.17 | 0.13 | 0.16 | 0.16 |
| 8 | 53720790 | rs2360244 | 1.128 | 0.169 | -0.33 | -0.764 | -0.026 | 0.146 | 0.146 | 0.135 | 0.47 | 0.19 | 0.59 | 0.73 | 0.48 | 0.49 |
| 8 | 53727578 | rs13248877 | 0.583 | 0.332 | -0.293 | -0.679 | -0.006 | 0.015 | 0.015 | 0 | 0.15 | 0.06 | 0.1 | 0.12 | 0.15 | 0.28 |
| 8 | 53729375 | rs2360235 | -1.041 | 0.297 | -0.355 | -0.685 | 0.014 | 0.131 | 0.131 | 0 | 0.49 | 0.69 | 0.4 | 0.27 | 0.51 | 0.51 |
| 8 | 53731368 | rs722804 | -0.768 | 0.282 | -0.365 | -0.714 | -0.004 | -0.012 | -0.012 | 0.905 | 0.3 | 0.51 | 0.27 | 0.27 | 0.3 | 0.44 |
| 8 | 53731530 | rs7001365 | 1.054 | 0.373 | -0.491 | -0.565 | 0.003 | 0.043 | 0.043 | 0 | 0.19 | 0.11 | 0.12 | 0 | 0.21 | 0.07 |
| 8 | 53733925 | rs16918186 | -0.261 | 0.374 | 0.164 | -0.345 | 0.017 | 0.039 | 0.039 | 0 | 0.13 | 0.33 | 0.16 | 0.15 | 0.15 | 0.16 |
| 8 | 53738033 | rs11775217 | -0.392 | 0.287 | 0.107 | -0.578 | -0.011 | -0.068 | -0.068 | 0 | 0.32 | 0.44 | 0.29 | 0.15 | 0.36 | 0.23 |
| 8 | 53738140 | rs34504406 | 0.589 | 0.249 | -0.064 | -0.719 | -0.005 | 0.017 | 0.017 | 0 | 0.15 | 0.04 | 0.1 | 0.12 | 0.15 | 0.29 |
| 8 | 53739232 | rs10095724 | 0.688 | 0.432 | -0.46 | -0.887 | -0.005 | 0.123 | 0.123 | 0 | 0.48 | 0.19 | 0.46 | 0.7 | 0.39 | 0.55 |
| 8 | 53740565 | rs2360242 | -0.788 | 0.459 | -0.359 | -0.839 | 0.011 | -0.018 | -0.018 | 0 | 0.8 | 0.7 | 0.76 | 0.85 | 0.76 | 0.78 |
| 8 | 53745838 | rs2884946 | -1.358 | 0.478 | -0.316 | -0.828 | -0.011 | 0.005 | 0.005 | 0 | 0.84 | 0.85 | 0.77 | 0.85 | 0.76 | 0.78 |
| 8 | 53748776 | rs4342624 | 0.868 | 0.248 | -0.173 | -0.652 | -0.001 | 0.06 | 0.06 | 0 | 0.51 | 0.25 | 0.53 | 0.57 | 0.46 | 0.33 |
| 8 | 53750022 | rs10504148 | 0.064 | 0.403 | -0.083 | -0.393 | 0.019 | 0.037 | 0.037 | 0 | 0.13 | 0.33 | 0.16 | 0.15 | 0.15 | 0.15 |
| 8 | 53755161 | rs6991827 | 0.968 | 0.493 | -0.552 | -0.248 | 0.003 | -0.022 | -0.022 | 0.077 | 0.33 | 0.19 | 0.41 | 0.57 | 0.25 | 0.26 |
| 8 | 53766263 | rs11780727 | 0.016 | 0.585 | -0.01 | 0.135 | 0.011 | 0.012 | 0.012 | 0 | 0.63 | 0.55 | 0.58 | 0.53 | 0.55 | 0.53 |
| 8 | 53780436 | rs3909686 | -0.408 | 0.639 | 0.015 | 0.227 | 0.003 | -0.005 | -0.005 | 0 | 0.68 | 0.66 | 0.67 | 0.75 | 0.64 | 0.68 |
| 8 | 53791714 | rs7824907 | -0.315 | 0.607 | 0.152 | 0.179 | 0.025 | -0.006 | -0.006 | 0 | 0.67 | 0.53 | 0.62 | 0.72 | 0.64 | 0.65 |
| 8 | 53793920 | rs16918264 | 1.708 | 0.534 | -0.45 | 0.046 | 0.008 | -0.085 | -0.085 | 0 | 0.26 | 0.08 | 0.32 | 0.44 | 0.2 | 0.2 |
| 8 | 53796924 | rs7002054 | 0.769 | 0.475 | -0.626 | -0.116 | -0.001 | -0.017 | -0.017 | 0.57 | 0.1 | 0.07 | 0.05 | 0.02 | 0.08 | 0.24 |
| 8 | 53801085 | rs11998157 | -0.885 | 0.507 | -0.677 | -0.033 | -0.036 | -0.009 | -0.009 | 0.084 | 0.69 | 0.81 | 0.65 | 0.71 | 0.64 | 0.65 |
| 8 | 53810264 | rs3906476 | -0.901 | 0.472 | -0.729 | -0.088 | -0.034 | -0.009 | -0.009 | 0 | 0.7 | 0.8 | 0.67 | 0.72 | 0.64 | 0.65 |
| 8 | 53814090 | rs17340869 | 1.333 | 0.271 | -0.943 | -0.482 | 0.004 | 0.055 | 0.055 | 0 | 0.22 | 0.14 | 0.13 | 0 | 0.23 | 0.08 |
| 8 | 53823427 | rs11988557 | 1.447 | 0.256 | -1.033 | -0.28 | -0.009 | -0.114 | -0.114 | 1.075 | 0.27 | 0.05 | 0.32 | 0.43 | 0.2 | 0.19 |
| 8 | 53827232 | rs17256515 | 1.228 | 0.286 | -0.851 | -0.444 | 0.016 | 0.025 | 0.025 | 0 | 0.23 | 0.16 | 0.12 | 0 | 0.19 | 0.08 |
| 8 | 53827622 | rs16918308 | 0.443 | 0.293 | -0.691 | -0.394 | 0.032 | 0.033 | 0.033 | 0.163 | 0.52 | 0.25 | 0.44 | 0.44 | 0.43 | 0.31 |
| 8 | 53829203 | rs3860850 | -0.057 | 0.218 | -0.303 | -0.32 | 0.007 | 0.003 | 0.003 | 0 | 0.62 | 0.44 | 0.51 | 0.47 | 0.52 | 0.58 |
| 8 | 53830605 | rs487551 | -0.966 | 0.365 | -0.404 | -0.33 | -0.059 | -0.045 | -0.045 | 0 | 0.66 | 0.88 | 0.56 | 0.55 | 0.59 | 0.72 |
| 8 | 53834494 | rs11985185 | 1.139 | 0.106 | -0.012 | -0.57 | 0.002 | -0.033 | -0.033 | 0 | 0.21 | 0.25 | 0.09 | 0.01 | 0.13 | 0.11 |
| 8 | 53842413 | rs2728918 | 0.24 | 0.319 | 0.052 | -0.481 | 0.004 | 0.005 | 0.005 | 0 | 0.37 | 0.4 | 0.37 | 0.49 | 0.4 | 0.39 |
| 8 | 53844559 | rs471570 | -1.127 | 0.51 | -0.014 | -0.306 | -0.073 | -0.073 | -0.073 | 1.209 | 0.66 | 0.91 | 0.55 | 0.52 | 0.6 | 0.74 |
| 8 | 53851462 | rs545125 | -0.773 | 0.505 | 0.072 | -0.315 | -0.004 | -0.003 | -0.003 | 0 | 0.62 | 0.65 | 0.49 | 0.49 | 0.54 | 0.73 |
| 8 | 53855212 | rs34461441 | -0.056 | 0.485 | 0.392 | -0.269 | 0.066 | 0.043 | 0.043 | 0 | 0.05 | 0.26 | 0.06 | 0.04 | 0.06 | 0.01 |
| 8 | 53874753 | rs7843363 | 0.648 | -0.187 | -0.121 | -0.773 | -0.013 | -0.068 | -0.068 | 0 | 0.24 | 0.28 | 0.1 | 0.01 | 0.15 | 0.11 |
| 8 | 53875701 | rs527238 | 0.498 | -0.124 | -0.82 | -0.587 | -0.101 | -0.122 | -0.122 | 0 | 0.34 | 0.04 | 0.46 | 0.52 | 0.36 | 0.38 |
| 8 | 53877010 | rs7007453 | 0.306 | -0.126 | -0.558 | -0.591 | 0.023 | 0.011 | 0.011 | 0 | 0.1 | 0.33 | 0.09 | 0.07 | 0.11 | 0.22 |
| 8 | 53877882 | rs16918378 | 0.416 | 0.26 | -1.34 | -0.259 | 0.02 | 0.012 | 0.012 | 0 | 0.1 | 0.23 | 0.1 | 0.11 | 0.13 | 0.14 |
| 8 | 53880054 | rs533402 | 0.947 | 0.382 | -2.096 | -0.049 | -0.059 | -0.159 | -0.159 | 0 | 0.26 | 0.04 | 0.38 | 0.42 | 0.24 | 0.25 |
| 8 | 53884020 | rs2555185 | -0.786 | 0.59 | -1.612 | 0.028 | -0.045 | -0.037 | -0.037 | 1.496 | 0.72 | 0.88 | 0.72 | 0.6 | 0.64 | 0.75 |
| 8 | 53885043 | rs569706 | -0.154 | -0.407 | -2.025 | -0.22 | 0.034 | -0.011 | -0.011 | 0.267 | 0.37 | 0.16 | 0.47 | 0.68 | 0.33 | 0.49 |
| 8 | 53885194 | rs567971 | -0.639 | -0.912 | -1.117 | -0.848 | -0.006 | -0.058 | -0.058 | 0 | 0.77 | 0.7 | 0.88 | 0.99 | 0.84 | 0.86 |
| 8 | 53886458 | rs2081607 | 0.43 | -0.797 | -0.823 | -0.551 | -0.029 | -0.054 | -0.054 | 0 | 0.26 | 0.43 | 0.14 | 0.15 | 0.17 | 0.15 |
| 8 | 53887419 | rs11987577 | 0.898 | -0.694 | -0.539 | -0.547 | 0.008 | -0.024 | -0.024 | 0 | 0.13 | 0.22 | 0.09 | 0.09 | 0.14 | 0.15 |
| 8 | 53888073 | rs2588084 | -0.892 | -0.773 | -1.057 | -0.513 | 0.035 | 0.11 | 0.11 | 0.279 | 0.47 | 0.87 | 0.33 | 0.34 | 0.36 | 0.37 |
| 8 | 53892198 | rs2555186 | -1.534 | -0.539 | -1.247 | -0.117 | -0.028 | -0.18 | -0.18 | 5.48 | 0.74 | 0.99 | 0.66 | 0.57 | 0.75 | 0.65 |
| 8 | 53898634 | rs360968 | 0.061 | -0.396 | -0.9 | -0.475 | 0.018 | 0.09 | 0.09 | 0 | 0.24 | 0.1 | 0.11 | 0.01 | 0.17 | 0.12 |
| 8 | 53900796 | rs1822590 | -0.291 | 0.102 | -0.087 | -0.007 | -0.002 | -0.013 | -0.013 | 0 | 0.33 | 0.42 | 0.28 | 0.3 | 0.28 | 0.19 |
| 8 | 53901286 | rs360956 | -0.698 | -0.476 | -0.354 | -0.05 | -0.008 | 0.032 | 0.032 | 0 | 0.53 | 0.85 | 0.35 | 0.4 | 0.4 | 0.47 |
| 8 | 53902717 | rs12541632 | 1.008 | -0.502 | -0.057 | -0.242 | 0.001 | 0.007 | 0.007 | 0 | 0.18 | 0.14 | 0.11 | 0.08 | 0.16 | 0.27 |
| 8 | 53903947 | rs360958 | -0.283 | -0.353 | -0.003 | -0.041 | 0.02 | 0.075 | 0.075 | 0 | 0.5 | 0.71 | 0.32 | 0.3 | 0.4 | 0.44 |
| 8 | 53908875 | rs360974 | -0.265 | -0.404 | -0.35 | 0.005 | 0.043 | 0.12 | 0.12 | 0 | 0.5 | 0.78 | 0.29 | 0.31 | 0.39 | 0.42 |
| 8 | 53925553 | rs360988 | -0.737 | -0.459 | -0.517 | -0.112 | -0.07 | -0.017 | -0.017 | 0.053 | 0.61 | 0.86 | 0.37 | 0.37 | 0.42 | 0.48 |
| 8 | 53926047 | rs7824346 | -0.132 | -0.644 | 0.301 | -0.235 | 0.048 | 0.015 | 0.015 | 0.784 | 0.07 | 0.44 | 0.05 | 0 | 0.02 | 0.01 |
| 8 | 53926962 | rs1365394 | -0.223 | -0.789 | -0.27 | -0.395 | 0.043 | 0.045 | 0.045 | 0.091 | 0.3 | 0.82 | 0.24 | 0.19 | 0.2 | 0.33 |
| 8 | 53927134 | rs183117 | -1.021 | -0.791 | -0.255 | -0.398 | -0.06 | -0.01 | -0.01 | 0.01 | 0.61 | 0.85 | 0.36 | 0.37 | 0.42 | 0.48 |
| 8 | 53927581 | rs360990 | 0.483 | -0.755 | -0.267 | -0.384 | -0.011 | 0.054 | 0.054 | 0.039 | 0.31 | 0.04 | 0.13 | 0.18 | 0.22 | 0.16 |
| 8 | 53930770 | rs360993 | 0.461 | -0.755 | -0.264 | -0.384 | -0.007 | 0.059 | 0.059 | 0.063 | 0.31 | 0.05 | 0.13 | 0.18 | 0.22 | 0.16 |
| 8 | 53930798 | rs1469322 | 1.249 | -0.749 | 0.29 | -0.377 | 0.005 | -0.018 | -0.018 | 0 | 0.05 | 0.19 | 0.03 | 0 | 0.02 | 0.02 |
| 8 | 53935438 | rs4416837 | 0.973 | -0.775 | 0.236 | -0.393 | -0.008 | -0.02 | -0.02 | 0.751 | 0.2 | 0.26 | 0.12 | 0.09 | 0.16 | 0.28 |
| 8 | 53935694 | rs1426562 | 0.007 | -0.762 | -0.059 | -0.384 | 0.036 | 0.032 | 0.032 | 0.037 | 0.28 | 0.8 | 0.19 | 0.18 | 0.19 | 0.32 |
| 8 | 53939511 | rs6473783 | -0.005 | -0.756 | -0.058 | -0.381 | 0.037 | 0.033 | 0.033 | 0.149 | 0.28 | 0.8 | 0.19 | 0.18 | 0.19 | 0.32 |
| 8 | 53946170 | rs4873692 | 0.68 | -0.561 | 0.015 | -0.276 | -0.069 | -0.016 | -0.016 | 0 | 0.39 | 0.13 | 0.64 | 0.63 | 0.58 | 0.52 |
| 8 | 53966642 | rs1596499 | 0.699 | -0.429 | -0.009 | -0.178 | -0.077 | -0.022 | -0.022 | 0.406 | 0.39 | 0.13 | 0.62 | 0.63 | 0.58 | 0.51 |
| 8 | 53968277 | rs4873695 | 0.231 | -0.457 | 0.32 | -0.198 | 0.001 | 0.002 | 0.002 | 0.029 | 0.45 | 0.55 | 0.72 | 0.64 | 0.62 | 0.53 |
| 8 | 53969642 | rs1031144 | 1.545 | 0.161 | -0.256 | 0.089 | 0.003 | -0.122 | -0.122 | 0 | 0.27 | 0.04 | 0.37 | 0.39 | 0.23 | 0.32 |
| 8 | 53973180 | rs4873696 | 0.987 | 0.252 | -0.227 | 0.115 | -0.021 | -0.025 | -0.025 | 0 | 0.11 | 0.07 | 0.25 | 0.28 | 0.35 | 0.17 |
| 8 | 53975616 | rs12543603 | 0.682 | -0.116 | -0.299 | -0.039 | -0.025 | -0.023 | -0.023 | 0 | 0.25 | 0.35 | 0.15 | 0.16 | 0.17 | 0.32 |
| 8 | 53981711 | rs2120560 | 0.009 | 0.043 | -0.228 | 0.028 | 0.005 | 0.012 | 0.012 | 0 | 0.45 | 0.58 | 0.72 | 0.68 | 0.62 | 0.52 |
| 8 | 53996131 | rs977505 | -0.007 | 0.048 | -0.225 | 0.03 | 0.006 | 0.013 | 0.013 | 0 | 0.45 | 0.58 | 0.72 | 0.68 | 0.62 | 0.52 |

| | | | | | | | | | | | | | | | | |
|---|---|---|---|---|---|---|---|---|---|---|---|---|---|---|---|---|
| 8 | 54025977 | rs1982297 | -2.379 | -0.097 | -0.405 | 0.038 | 0.026 | -0.03 | -0.03 | 0.007 | 0.94 | 1 | 0.99 | 0.91 | 0.98 | 0.94 |
| 8 | 54031984 | rs193334 | -0.494 | -0.203 | -0.3 | -0.038 | 0.041 | 0.028 | 0.028 | 0.471 | 0.66 | 0.8 | 0.82 | 0.74 | 0.78 | 0.82 |
| 8 | 54032307 | rs360816 | 0.67 | -0.04 | -0.209 | 0.004 | 0.083 | 0.049 | 0.049 | 0 | 0.12 | 0.21 | 0.25 | 0.28 | 0.34 | 0.18 |
| 8 | 54034788 | rs17351880 | 1.42 | -0.085 | -0.506 | -0.013 | 0.012 | 0.047 | 0.047 | 0 | 0.2 | 0.09 | 0.1 | 0.06 | 0.16 | 0.29 |
| 8 | 54038037 | rs9298491 | 0.945 | 0.522 | 0.057 | 0.502 | 0.004 | -0.009 | -0.009 | 0 | 0.29 | 0.23 | 0.43 | 0.4 | 0.25 | 0.33 |
| 8 | 54038107 | rs2553909 | -0.835 | 0.339 | -0.036 | 0.349 | 0.033 | -0.024 | -0.024 | 0 | 0.91 | 1 | 0.99 | 0.91 | 0.98 | 0.9 |
| 8 | 54038721 | rs2553915 | -0.64 | 1.103 | 0.344 | 0.616 | 0.015 | 0.034 | 0.034 | 0 | 0.45 | 0.64 | 0.71 | 0.68 | 0.62 | 0.52 |
| 8 | 54040235 | rs2588079 | -1.235 | 2.05 | 0.57 | 0.71 | 0.173 | 0.082 | 0.082 | 0 | 0.16 | 0.41 | 0.31 | 0.27 | 0.38 | 0.2 |
| 8 | 54041206 | rs7017147 | -1.547 | 0.572 | 0.943 | 0.251 | 0.088 | 0.045 | 0.045 | 0 | 0.05 | 0.31 | 0.08 | 0.01 | 0.04 | 0.03 |
| 8 | 54048844 | rs1895832 | 0.514 | 0.499 | 0.369 | 0.115 | 0.003 | 0.075 | 0.075 | 0 | 0.52 | 0.31 | 0.29 | 0.15 | 0.52 | 0.37 |
| 8 | 54055583 | rs2553877 | -0.613 | 0.548 | 0.366 | 0.433 | -0.006 | -0.013 | -0.013 | 0 | 0.9 | 0.9 | 0.68 | 0.73 | 0.87 | 0.89 |
| 8 | 54056834 | rs2588049 | 0.471 | 0.473 | 0.568 | 0.463 | 0.006 | -0.022 | -0.022 | 0 | 0.57 | 0.52 | 0.33 | 0.23 | 0.55 | 0.41 |
| 8 | 54057346 | rs1373387 | 0.727 | 0.445 | 0.726 | 0.447 | -0.014 | 0.141 | 0.141 | 0 | 0.83 | 0.49 | 0.63 | 0.64 | 0.86 | 0.82 |
| 8 | 54058042 | rs1346115 | 1.023 | 0.368 | 0.794 | 0.415 | 0.004 | 0.164 | 0.164 | 0.22 | 0.49 | 0.21 | 0.28 | 0.15 | 0.51 | 0.35 |
| 8 | 54067378 | rs360811 | 0.102 | 0.574 | 0.725 | 0.504 | 0.022 | 0.098 | 0.098 | 0 | 0.87 | 0.66 | 0.65 | 0.73 | 0.85 | 0.86 |
| 8 | 54068071 | rs13273443 | 0.271 | 0.684 | 0.313 | 0.545 | -0.033 | -0.065 | -0.065 | 0.213 | 0.3 | 0.13 | 0.32 | 0.49 | 0.3 | 0.44 |
| 8 | 54090578 | rs2059546 | 0.126 | 0.654 | 0.265 | 0.541 | -0.102 | -0.142 | -0.142 | 0.388 | 0.31 | 0.04 | 0.31 | 0.57 | 0.32 | 0.47 |
| 8 | 54105576 | rs10958349 | 0.169 | 0.698 | -0.289 | 0.554 | -0.025 | -0.053 | -0.053 | 0 | 0.31 | 0.18 | 0.32 | 0.5 | 0.31 | 0.44 |
| 8 | 54105645 | rs2553914 | -0.996 | 0.689 | -0.224 | 0.567 | 0.013 | 0.144 | 0.144 | 0 | 0.17 | 0.5 | 0.39 | 0.36 | 0.17 | 0.15 |
| 8 | 54108225 | rs2717655 | -0.724 | 0.835 | -0.485 | 0.459 | -0.021 | 0.046 | 0.046 | 0.059 | 0.42 | 0.3 | 0.66 | 0.77 | 0.46 | 0.55 |
| 8 | 54113011 | rs1425892 | 0.01 | 1.1 | -1.812 | 0.732 | 0 | 0.015 | 0.015 | 0.111 | 0.11 | 0.13 | 0.38 | 0.33 | 0.16 | 0.11 |
| 8 | 54114761 | rs1425894 | -0.06 | 0.97 | -2.013 | 0.471 | -0.097 | -0.129 | -0.129 | 0 | 0.29 | 0.04 | 0.25 | 0.42 | 0.3 | 0.42 |
| 8 | 54115304 | rs1425898 | -0.025 | 0.479 | -1.261 | -0.183 | 0.002 | -0.018 | -0.018 | 0 | 0.64 | 0.59 | 0.7 | 0.75 | 0.61 | 0.79 |
| 8 | 54117053 | rs17355157 | -0.597 | -0.337 | -1.798 | -0.154 | 0.012 | 0.054 | 0.054 | 0 | 0.24 | 0.06 | 0.23 | 0.12 | 0.17 | 0.21 |
| 8 | 54119214 | rs1425902 | 0.818 | -1.122 | -2.081 | -0.224 | 0.117 | 0.091 | 0.091 | 0.128 | 0.53 | 0.16 | 0.59 | 0.71 | 0.69 | 0.52 |
| 8 | 54119394 | rs1425903 | -0.446 | -1.628 | -0.973 | -0.548 | -0.02 | -0.069 | -0.069 | 0.701 | 0.18 | 0.41 | 0.28 | 0.08 | 0.14 | 0.12 |
| 8 | 54130075 | rs2114105 | 1.368 | -1.624 | -1.109 | -0.319 | 0.115 | 0.1 | 0.1 | 0.117 | 0.54 | 0.17 | 0.59 | 0.75 | 0.69 | 0.53 |
| 8 | 54130678 | rs6473795 | -1.272 | -1.806 | -0.791 | -0.42 | 0.016 | 0.017 | 0.017 | 0.545 | 0.31 | 0.69 | 0.34 | 0.13 | 0.18 | 0.29 |
| 8 | 54132413 | rs966247 | -0.352 | -1.683 | -0.884 | -0.172 | -0.012 | -0.051 | -0.051 | 0 | 0.86 | 0.99 | 0.94 | 0.86 | 0.87 | 0.82 |
| 8 | 54137488 | rs10101110 | -1.239 | -1.84 | 0.016 | -0.314 | -0.049 | -0.048 | -0.048 | 0.035 | 0.28 | 0.5 | 0.3 | 0.13 | 0.15 | 0.27 |
| 8 | 54138618 | rs7832417 | -1.066 | -1.88 | 0.176 | -0.328 | -0.019 | -0.063 | -0.063 | 0 | 0.21 | 0.54 | 0.29 | 0.08 | 0.15 | 0.13 |
| 8 | 54138824 | rs16918842 | 0.169 | -1.894 | 0.113 | -0.327 | 0.047 | 0.013 | 0.013 | 0 | 0.12 | 0.04 | 0.03 | 0.05 | 0.04 | 0.18 |
| 8 | 54140789 | rs16918853 | -0.477 | -1.471 | 0.485 | -0.196 | -0.003 | -0.043 | -0.043 | 0 | 0.15 | 0.37 | 0.26 | 0.08 | 0.11 | 0.1 |
| 8 | 54140948 | rs3802279 | -0.477 | -1.51 | 0.485 | -0.196 | -0.007 | -0.043 | -0.043 | 1.34 | 0.15 | 0.37 | 0.26 | 0.08 | 0.11 | 0.1 |
| 8 | 54141163 | rs3802281 | -1.199 | -1.471 | 0.491 | -0.202 | -0.017 | -0.065 | -0.065 | 0.013 | 0.21 | 0.53 | 0.29 | 0.08 | 0.15 | 0.13 |
| 8 | 54141824 | rs963549 | -1.238 | -1.457 | 0.515 | -0.19 | -0.01 | -0.058 | -0.058 | 1.253 | 0.19 | 0.53 | 0.27 | 0.08 | 0.14 | 0.1 |
| 8 | 54142154 | rs16918875 | 0.47 | -1.458 | 0.545 | -0.19 | 0.004 | -0.028 | -0.028 | 0 | 0.05 | 0.16 | 0.03 | 0 | 0.04 | 0.03 |
| 8 | 54142157 | rs702764 | -1.289 | -1.457 | 0.541 | -0.192 | -0.014 | -0.058 | -0.058 | 1.5 | 0.19 | 0.54 | 0.27 | 0.08 | 0.14 | 0.11 |
| 8 | 54142815 | rs7817710 | -0.469 | -1.431 | 0.418 | -0.169 | 0.011 | -0.027 | -0.027 | 0.209 | 0.14 | 0.37 | 0.24 | 0.08 | 0.1 | 0.07 |
| 8 | 54152268 | rs6651353 | 0.352 | -1.415 | 0.406 | 0.076 | 0.013 | 0.035 | 0.035 | 0 | 0.09 | 0.26 | 0.06 | 0.07 | 0.07 | 0.06 |
| 8 | 54152378 | rs997917 | -1.82 | -1.811 | 0.829 | 0.077 | 0.054 | -0.006 | -0.006 | 0 | 0.43 | 0.75 | 0.38 | 0.34 | 0.28 | 0.48 |
| 8 | 54153105 | rs6473798 | -1.694 | -1.817 | 0.837 | 0.076 | 0.035 | -0.013 | -0.013 | 0.489 | 0.39 | 0.69 | 0.36 | 0.34 | 0.24 | 0.45 |
| 8 | 54154311 | rs16918928 | -0.494 | -1.759 | 1.116 | 0.131 | -0.012 | 0.026 | 0.026 | 0 | 0.12 | 0.23 | 0.07 | 0.15 | 0.08 | 0.19 |
| 8 | 54157071 | rs2303433 | -0.292 | -1.759 | 1.092 | 0.131 | 0.001 | -0.003 | -0.003 | 0 | 0.11 | 0.11 | 0.05 | 0.15 | 0.08 | 0.19 |
| 8 | 54157217 | rs2303432 | -0.292 | -1.759 | 1.103 | 0.131 | 0.001 | -0.002 | -0.002 | 0 | 0.11 | 0.11 | 0.05 | 0.15 | 0.08 | 0.19 |
| 8 | 54157539 | rs7836120 | -1.15 | -1.799 | 1.235 | 0.158 | -0.018 | 0.047 | 0.047 | 0 | 0.21 | 0.4 | 0.12 | 0.21 | 0.15 | 0.25 |
| 8 | 54160092 | rs10111937 | -1.968 | -1.891 | 1.058 | 0.17 | 0.088 | -0.031 | -0.031 | 0 | 0.46 | 0.86 | 0.4 | 0.34 | 0.29 | 0.5 |
| 8 | 54160702 | rs16918941 | 0.84 | -1.893 | 0.983 | 0.171 | 0.001 | -0.005 | -0.005 | 0 | 0.09 | 0.07 | 0.04 | 0.07 | 0.07 | 0.06 |
| 8 | 54161116 | rs6985606 | -1.643 | -1.589 | 0.102 | 0.157 | -0.187 | -0.093 | -0.093 | 0.122 | 0.66 | 0.97 | 0.65 | 0.59 | 0.51 | 0.62 |
| 8 | 54164802 | rs3808627 | 1.839 | -1.762 | 0.166 | 0.166 | -0.049 | -0.036 | -0.036 | 0 | 0.19 | 0.03 | 0.23 | 0.25 | 0.21 | 0.12 |
| 8 | 54180915 | rs6993747 | -1.525 | -1.739 | 0.891 | 0.229 | 0.002 | 0 | 0 | 0 | 0.43 | 0.59 | 0.35 | 0.32 | 0.26 | 0.5 |
| 8 | 54190761 | rs9792306 | -0.205 | -1.579 | 0.82 | 0.081 | -0.011 | -0.007 | -0.007 | 0 | 0.22 | 0.24 | 0.24 | 0.11 | 0.12 | 0.24 |
| 8 | 54193095 | rs2059547 | -0.22 | -1.573 | 0.853 | 0.115 | -0.016 | -0.01 | -0.01 | 0.144 | 0.21 | 0.24 | 0.24 | 0.11 | 0.11 | 0.23 |
| 8 | 54194253 | rs2376421 | -1.623 | -1.585 | 0.842 | 0.457 | 0.08 | -0.045 | -0.045 | 0 | 0.43 | 0.78 | 0.37 | 0.37 | 0.26 | 0.5 |
| 8 | 54194489 | rs2376419 | -0.214 | -1.525 | 0.642 | 0.483 | -0.038 | -0.025 | -0.025 | 0.484 | 0.22 | 0.32 | 0.25 | 0.11 | 0.11 | 0.23 |
| 8 | 54195705 | rs9772353 | -0.621 | -0.953 | 0.428 | 0.678 | -0.004 | 0.019 | 0.019 | 0 | 0.2 | 0.24 | 0.1 | 0.26 | 0.15 | 0.26 |
| 8 | 54224120 | rs7462232 | -0.071 | -1.32 | 0.594 | 0.275 | 0.015 | 0.011 | 0.011 | 0 | 0.2 | 0.18 | 0.25 | 0.12 | 0.11 | 0.23 |
| 8 | 54237672 | rs10111713 | -1.263 | -1.413 | 0.851 | 0.429 | 0.007 | 0.003 | 0.003 | 0 | 0.33 | 0.68 | 0.34 | 0.25 | 0.21 | 0.42 |
| 8 | 54240255 | rs10110650 | 0.902 | -1.343 | 0.802 | 0.443 | 0.049 | -0.006 | -0.006 | 0.1 | 0.06 | 0.27 | 0.04 | 0 | 0.05 | 0.05 |
| 8 | 54256580 | rs12676091 | 1.315 | -1.526 | 0.389 | 0.531 | -0.014 | -0.057 | -0.057 | 0.071 | 0.1 | 0 | 0.02 | 0.13 | 0.08 | 0.04 |
| 8 | 54267021 | rs7011944 | -1.628 | -1.493 | 0.43 | 0.4 | 0.065 | -0.03 | -0.03 | 0 | 0.47 | 0.83 | 0.41 | 0.4 | 0.33 | 0.48 |
| 8 | 54273057 | rs6981962 | -0.796 | -1.392 | 0.763 | 0.398 | -0.012 | 0.017 | 0.017 | 0 | 0.13 | 0.58 | 0.1 | 0.13 | 0.09 | 0.2 |
| 8 | 54277247 | rs7465458 | -0.173 | -0.896 | 0.494 | 0.335 | 0.004 | 0.011 | 0.011 | 0 | 0.8 | 0.82 | 0.72 | 0.9 | 0.84 | 0.76 |
| 8 | 54277280 | rs10958353 | 1.292 | -0.901 | 0.454 | 0.336 | 0.033 | 0.029 | 0.029 | 0 | 0.66 | 0.18 | 0.6 | 0.76 | 0.75 | 0.56 |
| 8 | 54280178 | rs6473814 | 1.006 | -0.671 | -0.164 | 0.438 | 0.003 | -0.03 | -0.03 | 0.468 | 0.06 | 0.13 | 0.03 | 0 | 0.05 | 0.05 |
| 8 | 54290967 | rs10113505 | 1.431 | -0.874 | -0.879 | 0.37 | -0.153 | -0.096 | -0.096 | 0.045 | 0.3 | 0.03 | 0.32 | 0.42 | 0.46 | 0.32 |
| 8 | 54295098 | rs6981644 | 0.356 | -0.809 | 0.125 | 0.226 | -0.001 | -0.026 | -0.026 | 0 | 0.06 | 0.11 | 0.03 | 0 | 0.05 | 0.05 |
| 8 | 54302434 | rs6473821 | 0.565 | -0.783 | -0.21 | 0.205 | 0.012 | 0.021 | 0.021 | 0.005 | 0.77 | 0.82 | 0.71 | 0.89 | 0.84 | 0.77 |
| 8 | 54304926 | rs7006592 | -0.599 | -0.282 | -0.393 | 0.205 | -0.001 | 0.037 | 0.037 | 0 | 0.13 | 0.37 | 0.09 | 0.15 | 0.11 | 0.32 |

| | | | | | | | | | | | | | | | | |
|---|---|---|---|---|---|---|---|---|---|---|---|---|---|---|---|---|
| 8 | 54305649 | rs7011444 | 0.694 | -0.306 | -0.289 | 0.206 | -0.002 | -0.013 | -0.013 | 0 | 0.72 | 0.62 | 0.82 | 0.81 | 0.75 | 0.5 |
| 8 | 54308719 | rs3887557 | 0.535 | -0.088 | -0.505 | 0.381 | 0.003 | 0.001 | 0.001 | 0 | 0.59 | 0.43 | 0.57 | 0.6 | 0.68 | 0.56 |
| 8 | 54309084 | rs10958356 | 0.193 | -0.157 | -0.263 | 0.387 | -0.019 | -0.017 | -0.017 | 0.165 | 0.7 | 0.64 | 0.66 | 0.74 | 0.79 | 0.67 |
| 8 | 54312886 | rs16919123 | 0.148 | 0.159 | -0.738 | 0.5 | -0.02 | -0.024 | -0.024 | 0.687 | 0.09 | 0.03 | 0.08 | 0.14 | 0.1 | 0.11 |
| 8 | 54317551 | rs2126800 | 0.956 | 0.28 | -0.757 | 0.447 | -0.002 | -0.001 | -0.001 | 0 | 0.4 | 0.37 | 0.5 | 0.43 | 0.47 | 0.31 |
| 8 | 54320473 | rs10958357 | 0.571 | -0.167 | -1.199 | 0.347 | 0.046 | 0.012 | 0.012 | 0 | 0.59 | 0.2 | 0.6 | 0.67 | 0.65 | 0.64 |
| 8 | 54322867 | rs7820883 | -0.695 | -0.174 | -0.949 | 0.359 | 0.039 | 0.014 | 0.014 | 0 | 0.4 | 0.76 | 0.39 | 0.33 | 0.35 | 0.36 |
| 8 | 54330515 | rs1477973 | -0.112 | -0.021 | -0.508 | 0.449 | -0.018 | -0.023 | -0.023 | 1.678 | 0.85 | 0.89 | 0.71 | 0.73 | 0.79 | 0.76 |
| 8 | 54330686 | rs16919143 | -0.737 | -0.116 | -0.194 | 0.217 | -0.004 | -0.011 | -0.011 | 0 | 0.15 | 0.21 | 0.09 | 0.1 | 0.1 | 0.12 |
| 8 | 54333688 | rs2008573 | -1.674 | -0.05 | -0.025 | -0.042 | -0.013 | -0.06 | -0.06 | 0 | 0.27 | 0.43 | 0.19 | 0.13 | 0.18 | 0.32 |
| 8 | 54335865 | rs1477972 | -1.777 | -0.142 | -0.068 | -0.09 | -0.018 | -0.068 | -0.068 | 0 | 0.28 | 0.46 | 0.19 | 0.13 | 0.18 | 0.32 |
| 8 | 54343357 | rs736586 | 0.205 | -0.067 | -0.449 | -0.054 | -0.029 | 0.049 | 0.049 | 1.901 | 0.1 | 0.01 | 0.06 | 0.01 | 0.08 | 0.19 |
| 8 | 54344100 | rs6988606 | 1.575 | -0.233 | 0.006 | 0.072 | 0.002 | 0.063 | 0.063 | 1.178 | 0.5 | 0.26 | 0.49 | 0.62 | 0.57 | 0.39 |
| 8 | 54348873 | rs7835893 | 0.648 | -0.227 | 0.075 | 0.052 | 0.001 | -0.017 | -0.017 | 0 | 0.65 | 0.59 | 0.61 | 0.74 | 0.67 | 0.51 |
| 8 | 54366491 | rs10429416 | -0.708 | -0.149 | 0.072 | 0.119 | -0.004 | -0.021 | -0.021 | 0 | 0.15 | 0.3 | 0.09 | 0.1 | 0.1 | 0.12 |
| 8 | 54382933 | rs13251975 | 0.872 | -0.408 | 0.026 | 0.137 | -0.001 | -0.003 | -0.003 | 0 | 0.62 | 0.42 | 0.56 | 0.71 | 0.67 | 0.51 |
| 8 | 54388267 | rs1600076 | -0.887 | -0.759 | 0.111 | 0.23 | -0.031 | -0.027 | -0.027 | 0 | 0.09 | 0.18 | 0.07 | 0.03 | 0.04 | 0.06 |
| 8 | 54393352 | rs11775527 | -0.688 | -0.42 | -0.158 | 0.391 | -0.004 | -0.002 | -0.002 | 0 | 0.11 | 0.16 | 0.03 | 0.09 | 0.06 | 0.11 |
| 8 | 54397221 | rs7010328 | -1.203 | -0.648 | -0.235 | 0.813 | -0.002 | 0.001 | 0.001 | 1.205 | 0.35 | 0.56 | 0.46 | 0.46 | 0.26 | 0.53 |
| 8 | 54399129 | rs2126794 | -0.555 | -0.555 | -0.303 | 0.909 | -0.015 | -0.014 | -0.014 | 0 | 0.14 | 0.11 | 0.26 | 0.24 | 0.18 | 0.29 |
| 8 | 54404955 | rs899106 | 0.741 | -0.772 | -0.193 | 0.876 | -0.029 | -0.019 | -0.019 | 2.262 | 0.69 | 0.62 | 0.7 | 0.78 | 0.78 | 0.72 |
| 8 | 54406745 | rs2889303 | -0.559 | -0.864 | -0.28 | 0.784 | -0.066 | 0.022 | 0.022 | 0 | 0.84 | 0.98 | 0.85 | 0.92 | 0.78 | 0.85 |
| 8 | 54410905 | rs16919215 | -0.49 | -0.884 | -0.228 | 0.797 | -0.113 | 0.005 | 0.005 | 0.47 | 0.17 | 0.37 | 0.17 | 0.14 | 0.03 | 0.14 |
| 8 | 54414255 | rs2068035 | 1.444 | -0.721 | -0.137 | 0.784 | 0.004 | 0.004 | 0.004 | 0 | 0.54 | 0.44 | 0.41 | 0.56 | 0.58 | 0.49 |
| 8 | 54421221 | rs1477958 | 0.033 | -0.782 | -0.62 | 0.621 | -0.015 | 0.031 | 0.031 | 0 | 0.8 | 0.88 | 0.84 | 0.9 | 0.76 | 0.8 |
| 8 | 54428996 | rs10504154 | 1.905 | -0.925 | -1.127 | 0.715 | -0.111 | -0.029 | -0.029 | 0 | 0.43 | 0.07 | 0.32 | 0.39 | 0.52 | 0.22 |
| 8 | 54432737 | rs9643478 | 0.429 | -0.539 | -0.753 | 0.799 | -0.024 | 0.054 | 0.054 | 0 | 0.76 | 0.9 | 0.8 | 0.9 | 0.73 | 0.81 |
| 8 | 54433888 | rs7829521 | 0.108 | -0.593 | -0.619 | 0.951 | -0.046 | 0.017 | 0.017 | 0 | 0.7 | 0.89 | 0.6 | 0.7 | 0.66 | 0.67 |
| 8 | 54439839 | rs11992458 | -1.242 | -1.065 | 0.775 | 1.116 | -0.053 | -0.031 | -0.031 | 0 | 0.25 | 0.42 | 0.23 | 0.15 | 0.14 | 0.26 |
| 8 | 54440206 | rs10090582 | -1.523 | -1.033 | 0.418 | 1.001 | -0.047 | 0.028 | 0.028 | 0 | 0.28 | 0.54 | 0.26 | 0.3 | 0.17 | 0.47 |
| 8 | 54449109 | rs16919915 | -1.076 | -0.049 | 0.19 | 0.665 | 0.004 | -0.033 | -0.033 | 3.09 | 0.07 | 0.14 | 0.07 | 0 | 0.07 | 0.12 |
| 8 | 54453130 | rs9693226 | -0.913 | -0.121 | 0.431 | 0.685 | 0.012 | -0.031 | -0.031 | 0 | 0.06 | 0.16 | 0.05 | 0 | 0.05 | 0.02 |
| 8 | 54454745 | rs2376142 | -2.397 | -0.454 | 1.446 | 1.424 | 0 | 0.214 | 0.214 | 0 | 0.12 | 0.49 | 0.21 | 0.23 | 0.07 | 0.25 |
| 8 | 54456660 | rs9693381 | -0.784 | -0.434 | 1.44 | 1.246 | 0.013 | -0.03 | -0.03 | 0 | 0.06 | 0.16 | 0.05 | 0 | 0.05 | 0.02 |
| 8 | 54457471 | rs6473847 | -1.043 | -0.434 | 1.443 | 1.246 | 0.025 | -0.028 | -0.028 | 0 | 0.06 | 0.23 | 0.06 | 0 | 0.05 | 0.02 |
| 8 | 54460873 | rs6473849 | -1.995 | -0.434 | 1.529 | 1.333 | 0.022 | 0.037 | 0.037 | 0 | 0.1 | 0.45 | 0.19 | 0.07 | 0.06 | 0.04 |
| 8 | 54462119 | rs1551903 | 0.299 | -0.208 | 0.474 | 1.135 | -0.064 | -0.008 | -0.008 | 0 | 0.78 | 0.98 | 0.81 | 0.8 | 0.76 | 0.8 |
| 8 | 54477983 | rs1382123 | 0.26 | -0.295 | 0.953 | 0.944 | -0.002 | 0.025 | 0.025 | 0 | 0.79 | 0.89 | 0.85 | 0.93 | 0.78 | 0.83 |
| 8 | 54479940 | rs2169692 | 0.796 | -0.33 | 1.281 | 0.909 | 0.003 | -0.096 | -0.096 | 0.02 | 0.72 | 0.63 | 0.78 | 0.93 | 0.73 | 0.81 |
| 8 | 54485315 | rs12549903 | 0.796 | -0.33 | 1.288 | 0.909 | 0.003 | -0.094 | -0.094 | 0 | 0.72 | 0.65 | 0.78 | 0.93 | 0.73 | 0.81 |
| 8 | 54488781 | rs6983438 | 0.796 | -0.338 | 1.398 | 0.905 | 0.003 | -0.094 | -0.094 | 0.2 | 0.72 | 0.65 | 0.78 | 0.93 | 0.73 | 0.81 |
| 8 | 54493000 | rs1551912 | 1.831 | -1.041 | 0.987 | 1.263 | 0.042 | 0.051 | 0.051 | 0.74 | 0.51 | 0.27 | 0.43 | 0.41 | 0.61 | 0.3 |
| 8 | 54496270 | rs4873263 | -0.616 | -1.145 | 1.277 | 1.454 | -0.036 | 0.086 | 0.086 | 0 | 0.15 | 0.26 | 0.31 | 0.32 | 0.07 | 0.26 |
| 8 | 54497467 | rs1551913 | -0.02 | -1.105 | 1.197 | 1.329 | -0.001 | -0.038 | -0.038 | 0 | 0.78 | 0.78 | 0.84 | 0.93 | 0.78 | 0.83 |
| 8 | 54502600 | rs6991825 | -0.099 | -1.115 | 1.176 | 1.324 | -0.001 | 0.018 | 0.018 | 0 | 0.78 | 0.88 | 0.85 | 0.93 | 0.78 | 0.83 |
| 8 | 54506372 | rs13268090 | 1.938 | -1.13 | 1.219 | 1.451 | 0.041 | 0.045 | 0.045 | 0.863 | 0.51 | 0.2 | 0.42 | 0.41 | 0.61 | 0.29 |
| 8 | 54510605 | rs10958377 | 0.136 | -1.102 | 1.403 | 1.339 | -0.002 | -0.03 | -0.03 | 0.698 | 0.77 | 0.77 | 0.83 | 0.93 | 0.77 | 0.83 |
| 8 | 54511637 | rs13275975 | 1.927 | -1.158 | 1.439 | 1.401 | 0.035 | 0.002 | 0.002 | 0.801 | 0.52 | 0.23 | 0.54 | 0.47 | 0.62 | 0.32 |
| 8 | 54515304 | rs6473863 | 0.114 | -1.151 | 1.383 | 1.398 | -0.003 | -0.053 | -0.053 | 0 | 0.77 | 0.74 | 0.83 | 0.93 | 0.77 | 0.83 |
| 8 | 54517208 | rs7462955 | 0.236 | -1.268 | 1.324 | 1.337 | -0.004 | -0.051 | -0.051 | 0.07 | 0.76 | 0.74 | 0.83 | 0.93 | 0.77 | 0.83 |
| 8 | 54518324 | rs7462399 | 0.192 | -1.229 | 1.384 | 1.351 | -0.005 | -0.071 | -0.071 | 0.531 | 0.76 | 0.69 | 0.82 | 0.93 | 0.77 | 0.83 |
| 8 | 54520350 | rs7387992 | 0.112 | -1.222 | 1.345 | 1.332 | -0.003 | -0.053 | -0.053 | 0 | 0.76 | 0.74 | 0.83 | 0.93 | 0.77 | 0.83 |
| 8 | 54521248 | rs12680875 | -1.356 | -1.275 | 1.87 | 1.365 | -0.034 | 0.138 | 0.138 | 0 | 0.18 | 0.33 | 0.22 | 0.45 | 0.1 | 0.47 |
| 8 | 54539644 | rs7462827 | -0.011 | -1.247 | 1.903 | 1.151 | 0.001 | 0.004 | 0.004 | 0 | 0.77 | 0.85 | 0.85 | 0.92 | 0.77 | 0.83 |
| 8 | 54540154 | rs7387498 | -1.377 | -1.259 | 1.697 | 1.672 | -0.035 | 0.097 | 0.097 | 0 | 0.19 | 0.33 | 0.22 | 0.56 | 0.1 | 0.47 |
| 8 | 54540531 | rs10958387 | 1.906 | -1.265 | 1.575 | 1.683 | 0.039 | 0.078 | 0.078 | 0.446 | 0.52 | 0.21 | 0.54 | 0.36 | 0.62 | 0.34 |
| 8 | 54548081 | rs7465143 | -1.447 | -1.218 | 1.526 | 1.77 | -0.038 | 0.116 | 0.116 | 0 | 0.19 | 0.37 | 0.22 | 0.56 | 0.1 | 0.47 |
| 8 | 54554299 | rs7388039 | -1.284 | -1.224 | 1.388 | 1.829 | -0.008 | 0.038 | 0.038 | 0 | 0.17 | 0.2 | 0.23 | 0.4 | 0.11 | 0.27 |
| 8 | 54559770 | rs7817444 | 1.244 | -1.284 | 1.265 | 1.683 | 0.014 | 0.007 | 0.007 | 0.023 | 0.59 | 0.37 | 0.59 | 0.37 | 0.69 | 0.37 |
| 8 | 54565013 | rs11776780 | 1.555 | -1.272 | 1.273 | 1.653 | 0.017 | 0.013 | 0.013 | 0.012 | 0.56 | 0.37 | 0.56 | 0.39 | 0.64 | 0.35 |
| 8 | 54568805 | rs7463851 | -0.639 | -1.204 | 1.357 | 1.828 | -0.006 | 0.062 | 0.062 | 0 | 0.07 | 0.14 | 0.03 | 0.15 | 0.04 | 0.21 |
| 8 | 54572995 | rs7464428 | 0.496 | -1.112 | 1.567 | 1.707 | -0.019 | 0.037 | 0.037 | 0 | 0.74 | 0.69 | 0.69 | 0.54 | 0.81 | 0.61 |
| 8 | 54573305 | rs7461045 | 0.971 | -1.161 | 1.618 | 1.657 | -0.029 | 0.054 | 0.054 | 0.352 | 0.71 | 0.5 | 0.64 | 0.54 | 0.77 | 0.59 |
| 8 | 54574395 | rs6473882 | -1.745 | -1.283 | 1.518 | 1.687 | 0.008 | 0.013 | 0.013 | 0 | 0.17 | 0.36 | 0.13 | 0.13 | 0.19 | 0.27 |
| 8 | 54578044 | rs10958390 | 1.965 | -1.507 | 1.439 | 1.623 | 0.024 | 0.008 | 0.008 | 0.177 | 0.56 | 0.34 | 0.56 | 0.41 | 0.63 | 0.34 |
| 8 | 54578332 | rs13269498 | 0.4 | -1.504 | 1.318 | 1.628 | 0.024 | -0.037 | -0.037 | 0 | 0.74 | 0.84 | 0.71 | 0.54 | 0.82 | 0.61 |
| 8 | 54579007 | rs7018412 | -1.749 | -1.316 | 1.316 | 1.599 | 0.008 | 0.012 | 0.012 | 0 | 0.17 | 0.35 | 0.12 | 0.13 | 0.19 | 0.27 |
| 8 | 54584082 | rs13259308 | 0.94 | -1.544 | 0.841 | 1.452 | 0.021 | -0.009 | -0.009 | 1.119 | 0.64 | 0.71 | 0.68 | 0.56 | 0.75 | 0.62 |
| 8 | 54584436 | rs6983168 | 1.168 | -1.603 | 1.281 | 1.417 | -0.038 | 0.006 | 0.006 | 0 | 0.62 | 0.51 | 0.64 | 0.56 | 0.75 | 0.62 |

| | | | | | | | | | | | | | | | | |
|---|---|---|---|---|---|---|---|---|---|---|---|---|---|---|---|---|
| 8 | 54589344 | rs13255905 | 1.889 | -1.622 | 1.288 | 1.479 | 0.021 | 0.011 | 0.011 | 0 | 0.52 | 0.41 | 0.56 | 0.42 | 0.62 | 0.37 |
| 8 | 54589977 | rs6473885 | 0.951 | -1.575 | 0.826 | 1.371 | 0.022 | -0.009 | -0.009 | 0.097 | 0.64 | 0.71 | 0.68 | 0.56 | 0.75 | 0.62 |
| 8 | 54628383 | rs6988082 | -1.431 | -1.649 | 0.773 | 1.515 | -0.024 | 0.065 | 0.065 | 0.094 | 0.13 | 0.22 | 0.13 | 0.39 | 0.06 | 0.19 |
| 8 | 54628736 | rs3735833 | 0.779 | -1.83 | 0.287 | 1.686 | -0.056 | -0.04 | -0.04 | 0 | 0.67 | 0.98 | 0.79 | 0.73 | 0.65 | 0.5 |
| 8 | 54640433 | rs16919508 | -0.492 | -1.828 | 0.381 | 1.706 | -0.002 | 0.042 | 0.042 | 0.027 | 0.1 | 0.12 | 0.12 | 0.4 | 0.05 | 0.18 |
| 8 | 54647078 | rs4737554 | -1.485 | -1.851 | 0.609 | 1.721 | -0.045 | 0.113 | 0.113 | 0.17 | 0.14 | 0.33 | 0.14 | 0.35 | 0.06 | 0.19 |
| 8 | 54651305 | rs2376011 | -1.788 | -1.934 | 0.922 | 1.677 | -0.058 | 0.113 | 0.113 | 0.353 | 0.16 | 0.4 | 0.15 | 0.35 | 0.06 | 0.2 |
| 8 | 54658155 | rs4738884 | -1.419 | -1.927 | 0.888 | 1.687 | -0.035 | 0.071 | 0.071 | 0.192 | 0.14 | 0.26 | 0.13 | 0.35 | 0.06 | 0.19 |
| 8 | 54704493 | rs13259921 | 0.116 | -2.021 | 0.63 | 1.522 | -0.024 | -0.036 | -0.036 | 0 | 0.06 | 0.01 | 0.05 | 0.14 | 0.11 | 0.28 |
| 8 | 54708747 | rs13275279 | 2.881 | -1.324 | -0.306 | 0.53 | -0.143 | 0.205 | 0.205 | 0.142 | 0.28 | 0.02 | 0.22 | 0 | 0.38 | 0.07 |
| 8 | 54739072 | rs4737974 | -0.632 | -1.436 | 0.052 | 1.003 | -0.017 | 0.063 | 0.063 | 0.056 | 0.12 | 0.18 | 0.13 | 0.43 | 0.05 | 0.24 |
| 8 | 54746486 | rs7016502 | 0.577 | -1.157 | -0.155 | 0.969 | -0.081 | -0.029 | -0.029 | 0 | 0.66 | 0.97 | 0.78 | 0.75 | 0.63 | 0.5 |
| 8 | 54751691 | rs16919588 | 1.057 | -1.036 | 0.408 | 1.129 | 0.001 | -0.028 | -0.028 | 0 | 0.05 | 0.01 | 0.06 | 0.15 | 0.03 | 0.04 |
| 8 | 54753912 | rs6473894 | -1.598 | -0.882 | 0.517 | 0.827 | -0.042 | 0.109 | 0.109 | 3.085 | 0.1 | 0.65 | 0.12 | 0.28 | 0.03 | 0.21 |
| 8 | 54770860 | rs11783925 | 2.189 | -0.439 | -0.668 | 0.063 | -0.054 | 0.089 | 0.089 | 0 | 0.16 | 0.02 | 0.13 | 0 | 0.21 | 0.03 |
| 8 | 54780174 | rs6997050 | -1.113 | -0.736 | 0.168 | 0.852 | -0.037 | 0.142 | 0.142 | 0.057 | 0.09 | 0.3 | 0.08 | 0.3 | 0.03 | 0.32 |
| 8 | 54780209 | rs1466525 | 0.075 | -0.796 | -0.223 | 0.743 | -0.04 | 0.035 | 0.035 | 0 | 0.77 | 0.98 | 0.85 | 0.85 | 0.78 | 0.74 |
| 8 | 54788251 | rs7827611 | -1.714 | -0.662 | 0.916 | 1.268 | -0.033 | 0.204 | 0.204 | 0.517 | 0.1 | 0.43 | 0.1 | 0.3 | 0.03 | 0.32 |
| 8 | 54812468 | rs16919658 | -1.941 | -0.449 | 1.334 | 1.062 | 0.028 | 0.14 | 0.14 | 5.74 | 0.06 | 0.63 | 0.06 | 0.09 | 0.01 | 0.02 |
| 8 | 54822319 | rs7824575 | -0.738 | -0.414 | 0.296 | 1.058 | 0.03 | -0.045 | -0.045 | 0 | 0.32 | 0.82 | 0.23 | 0.37 | 0.28 | 0.38 |
| 8 | 54835697 | rs12708158 | 1.881 | -0.122 | 0.327 | 0.45 | -0.034 | 0.052 | 0.052 | 0 | 0.09 | 0.01 | 0.07 | 0 | 0.12 | 0.02 |
| 8 | 54852599 | rs16919683 | -1 | -0.411 | 0.718 | 0.658 | 0.008 | 0.087 | 0.087 | 0 | 0.05 | 0.32 | 0.05 | 0.09 | 0.01 | 0.02 |
| 8 | 54852787 | rs16919692 | -1.039 | -0.411 | 0.555 | 0.84 | 0.015 | -0.012 | -0.012 | 1.793 | 0.38 | 0.73 | 0.34 | 0.53 | 0.37 | 0.69 |
| 8 | 54855680 | rs9298496 | -0.569 | -0.362 | 0.385 | 0.645 | -0.006 | 0.038 | 0.038 | 0.213 | 0.33 | 0.39 | 0.29 | 0.43 | 0.37 | 0.64 |
| 8 | 54858494 | rs11783652 | 0.024 | -0.392 | -0.189 | -0.444 | -0.002 | -0.007 | -0.007 | 0 | 0.41 | 0.42 | 0.49 | 0.71 | 0.34 | 0.52 |
| 8 | 54866148 | rs10958393 | -0.606 | -0.408 | 0.105 | -0.482 | -0.003 | -0.006 | -0.006 | 0 | 0.62 | 0.38 | 0.64 | 0.84 | 0.59 | 0.72 |
| 8 | 54866491 | rs6981243 | -0.604 | -0.4 | 0.12 | -0.482 | -0.005 | 0.002 | 0.002 | 0 | 0.62 | 0.37 | 0.63 | 0.84 | 0.59 | 0.71 |
| 8 | 54882796 | rs10095164 | -0.777 | -0.794 | 0.774 | 0.275 | 0.041 | 0.023 | 0.023 | 2.968 | 0.05 | 0.61 | 0.06 | 0 | 0.02 | 0 |
| 8 | 54890848 | rs11996666 | -0.961 | -1.289 | 0.367 | 0.069 | 0.045 | 0.011 | 0.011 | 0 | 0.25 | 0.16 | 0.34 | 0.73 | 0.16 | 0.52 |
| 8 | 54892988 | rs4515510 | -0.848 | -1.299 | 0.361 | 0.063 | 0.048 | 0.013 | 0.013 | 0 | 0.25 | 0.16 | 0.34 | 0.73 | 0.16 | 0.52 |
| 8 | 55040026 | rs952194 | -0.899 | -1.218 | 0.328 | 0.114 | -0.001 | 0.002 | 0.002 | 0 | 0.25 | 0.25 | 0.28 | 0.74 | 0.16 | 0.53 |
| 8 | 55049255 | rs2271567 | -0.96 | -1.151 | 0.094 | 0.122 | 0.037 | 0.012 | 0.012 | 0 | 0.24 | 0.17 | 0.28 | 0.72 | 0.16 | 0.55 |
| 8 | 55071319 | rs4737503 | -0.682 | -0.828 | -0.299 | 0.276 | -0.003 | -0.001 | -0.001 | 0 | 0.28 | 0.31 | 0.29 | 0.71 | 0.2 | 0.49 |
| 8 | 55074884 | rs13275258 | -0.859 | -0.67 | -0.218 | 1.03 | -0.015 | 0.019 | 0.019 | 0 | 0.29 | 0.37 | 0.23 | 0.46 | 0.19 | 0.15 |
| 8 | 55079018 | rs311396 | 0.159 | -0.252 | -1.29 | 0.415 | -0.058 | 0.122 | 0.122 | 0 | 0.33 | 0.07 | 0.27 | 0.16 | 0.41 | 0.25 |
| 8 | 55094784 | rs379532 | -1.922 | 0.02 | -1.029 | 0.84 | -0.011 | -0.008 | -0.008 | 0 | 0.7 | 0.85 | 0.51 | 0.67 | 0.72 | 0.54 |
| 8 | 55100391 | rs395449 | -1.5 | -0.186 | -0.906 | 0.897 | -0.034 | -0.01 | -0.01 | 0.649 | 0.65 | 0.81 | 0.48 | 0.61 | 0.63 | 0.5 |
| 8 | 55120092 | rs7015797 | -1.575 | -0.32 | -1.12 | 1.296 | -0.073 | -0.095 | -0.095 | 0 | 0.65 | 0.91 | 0.44 | 0.52 | 0.63 | 0.49 |
| 8 | 55120946 | rs219028 | -0.697 | -0.42 | -1.007 | 1.312 | 0.004 | -0.013 | -0.013 | 0 | 0.52 | 0.27 | 0.36 | 0.45 | 0.6 | 0.33 |
| 8 | 55124075 | rs311404 | 0.29 | 0.315 | -1.053 | 1.153 | -0.026 | 0.079 | 0.079 | 0 | 0.29 | 0.08 | 0.26 | 0.17 | 0.35 | 0.22 |
| 8 | 55126346 | rs184319 | -0.917 | 0.374 | -0.782 | 1.56 | -0.002 | -0.005 | -0.005 | 0.038 | 0.59 | 0.62 | 0.46 | 0.48 | 0.63 | 0.46 |
| 8 | 55136553 | rs312112 | -0.303 | 0.11 | -0.893 | 1.787 | -0.022 | -0.038 | -0.038 | 0 | 0.87 | 0.99 | 0.89 | 0.83 | 0.83 | 0.83 |
| 8 | 55143250 | rs312110 | 1.376 | 0.051 | -1.115 | 1.847 | -0.003 | 0.137 | 0.137 | 0.566 | 0.49 | 0.09 | 0.65 | 0.28 | 0.53 | 0.51 |
| 8 | 55155640 | rs312108 | 0.981 | -0.177 | -0.061 | 2.289 | 0.011 | 0.007 | 0.007 | 0.465 | 0.6 | 0.33 | 0.63 | 0.35 | 0.64 | 0.55 |
| 8 | 55158960 | rs7824834 | -1.014 | 0.279 | 0.423 | 2.541 | -0.009 | 0.144 | 0.144 | 0 | 0.06 | 0.18 | 0.1 | 0.15 | 0.02 | 0.16 |
| 8 | 55159106 | rs312104 | 1.685 | 0.375 | 0.246 | 2.707 | -0.019 | -0.004 | -0.004 | 0 | 0.59 | 0.12 | 0.56 | 0.37 | 0.57 | 0.53 |
| 8 | 55161461 | rs379593 | 1.679 | 0.377 | 0.247 | 2.708 | -0.019 | -0.005 | -0.005 | 0.072 | 0.59 | 0.12 | 0.56 | 0.37 | 0.57 | 0.53 |
| 8 | 55162863 | rs6473925 | -1.674 | 0.343 | 0.609 | 2.64 | -0.004 | 0.115 | 0.115 | 0 | 0.09 | 0.24 | 0.12 | 0.19 | 0.08 | 0.21 |
| 8 | 55166676 | rs312134 | 0.57 | 0.522 | 0.842 | 2.386 | -0.002 | 0.027 | 0.027 | 0 | 0.81 | 0.6 | 0.78 | 0.77 | 0.79 | 0.78 |
| 8 | 55173791 | rs16919928 | -1.281 | 0.602 | 0.425 | 2.205 | 0.013 | 0 | 0 | 0 | 0.07 | 0.16 | 0.05 | 0.04 | 0.08 | 0.08 |
| 8 | 55178575 | rs2594783 | -1.496 | 0.696 | 0.175 | 2.51 | -0.006 | -0.008 | -0.008 | 1.752 | 0.35 | 0.68 | 0.33 | 0.48 | 0.4 | 0.29 |
| 8 | 55182899 | rs1448452 | -1.81 | 0.738 | -0.425 | 2.495 | 0.007 | -0.04 | -0.04 | 0.794 | 0.34 | 0.89 | 0.34 | 0.37 | 0.32 | 0.35 |
| 8 | 55186953 | rs2656263 | -2.221 | 0.737 | -0.51 | 2.7 | -0.099 | 0.011 | 0.011 | 0 | 0.45 | 0.94 | 0.46 | 0.66 | 0.47 | 0.5 |
| 8 | 55194674 | rs16919938 | 0.04 | 1.17 | -0.095 | 2.504 | -0.051 | -0.036 | -0.036 | 0.648 | 0.09 | 0.02 | 0.05 | 0.16 | 0.15 | 0.03 |
| 8 | 55198858 | rs2719401 | -2.227 | 0.996 | -0.138 | 2.323 | -0.096 | -0.125 | -0.125 | 0 | 0.35 | 0.91 | 0.38 | 0.53 | 0.42 | 0.31 |
| 8 | 55201740 | rs2594803 | -1.369 | 1.017 | 0.935 | 2.29 | 0.005 | 0.073 | 0.073 | 0 | 0.19 | 0.55 | 0.26 | 0.32 | 0.19 | 0.18 |
| 8 | 55209704 | rs2922294 | -2.251 | 1.113 | 1.362 | 2.397 | 0.028 | 0.03 | 0.03 | 0 | 0.31 | 0.59 | 0.32 | 0.5 | 0.41 | 0.3 |
| 8 | 55210805 | rs2594813 | -1.207 | 1.071 | 1.603 | 2.32 | 0.026 | 0.086 | 0.086 | 0 | 0.17 | 0.51 | 0.25 | 0.29 | 0.2 | 0.19 |
| 8 | 55217420 | rs2656239 | -2.336 | 1.109 | 1.499 | 2.319 | 0.005 | 0.003 | 0.003 | 0 | 0.33 | 0.67 | 0.35 | 0.46 | 0.42 | 0.27 |
| 8 | 55231713 | rs6992754 | -1.124 | 1.239 | 1.34 | 1.959 | 0 | 0.005 | 0.005 | 0 | 0.16 | 0.18 | 0.1 | 0.13 | 0.18 | 0.09 |
| 8 | 55243761 | rs12542396 | -0.872 | 1.152 | 1.252 | 2.507 | 0.024 | 0.149 | 0.149 | 0 | 0.08 | 0.19 | 0.24 | 0.26 | 0.09 | 0.1 |
| 8 | 55243937 | rs2719353 | -1.337 | 1.164 | 1.507 | 2.6 | 0.013 | 0.05 | 0.05 | 0 | 0.16 | 0.26 | 0.2 | 0.31 | 0.16 | 0.18 |
| 8 | 55247102 | rs16920077 | -1.963 | 1.339 | 1.565 | 2.596 | 0.011 | 0.079 | 0.079 | 0 | 0.24 | 0.55 | 0.36 | 0.39 | 0.27 | 0.22 |
| 8 | 55257452 | rs10504164 | -1.067 | 1.249 | 0.901 | 2.463 | 0.024 | 0.159 | 0.159 | 0 | 0.08 | 0.23 | 0.24 | 0.26 | 0.09 | 0.11 |
| 8 | 55258998 | rs2719364 | -1.333 | 1.2 | 0.671 | 2.001 | 0.034 | 0.054 | 0.054 | 0 | 0.28 | 0.53 | 0.31 | 0.45 | 0.34 | 0.28 |
| 8 | 55268914 | rs895822 | -1.131 | 1.822 | 0.164 | 2.019 | 0.025 | 0.031 | 0.031 | 0 | 0.13 | 0.15 | 0.26 | 0.28 | 0.21 | 0.18 |
| 8 | 55269940 | rs10958403 | -0.567 | 1.088 | -0.616 | 1.116 | 0.004 | 0.018 | 0.018 | 0 | 0.28 | 0.23 | 0.18 | 0.05 | 0.23 | 0.19 |
| 8 | 55273800 | rs7006361 | -0.756 | 0.914 | -1.794 | 0.954 | -0.164 | -0.063 | -0.063 | 0.01 | 0.58 | 0.97 | 0.64 | 0.56 | 0.52 | 0.56 |
| 8 | 55274324 | rs2719370 | -1.239 | 1.302 | -1.187 | 1.242 | -0.046 | 0.007 | 0.007 | 0 | 0.12 | 0.06 | 0.09 | 0.04 | 0.14 | 0.11 |

| | | | | | | | | | | | | | | | | |
|---|---|---|---|---|---|---|---|---|---|---|---|---|---|---|---|---|
| 8 | 55276083 | rs1979915 | -1.823 | 1.075 | -0.452 | 1.408 | 0.001 | -0.003 | -0.003 | 1.99 | 0.4 | 0.42 | 0.54 | 0.69 | 0.43 | 0.5 |
| 8 | 55278457 | rs2594797 | -1.361 | 1.151 | -0.18 | 1.101 | -0.009 | 0.017 | 0.017 | 0.157 | 0.27 | 0.24 | 0.18 | 0.12 | 0.27 | 0.24 |
| 8 | 55281514 | rs2938702 | -1.021 | 0.328 | 0.201 | 1.519 | -0.013 | 0.038 | 0.038 | 0 | 0.21 | 0.4 | 0.18 | 0.34 | 0.16 | 0.16 |
| 8 | 55292539 | rs4737564 | -1.562 | 0.468 | 0.553 | 1.731 | 0.026 | 0.074 | 0.074 | 0.174 | 0.19 | 0.48 | 0.47 | 0.27 | 0.17 | 0.42 |
| 8 | 55302231 | rs4738837 | -1.308 | 0.486 | 0.911 | 1.808 | 0.031 | -0.022 | -0.022 | 0 | 0.13 | 0.31 | 0.25 | 0.08 | 0.14 | 0.15 |
| 8 | 55304600 | rs7840228 | -2.123 | 0.484 | 0.909 | 1.815 | 0.032 | 0.075 | 0.075 | 0.186 | 0.19 | 0.48 | 0.47 | 0.26 | 0.18 | 0.41 |
| 8 | 55308291 | rs16920160 | -0.326 | 0.538 | 0.614 | 1.839 | -0.023 | -0.02 | -0.02 | 0 | 0.05 | 0 | 0.04 | 0.06 | 0.05 | 0.02 |
| 8 | 55309191 | rs7005290 | -1.891 | 0.533 | 0.755 | 2.194 | 0.008 | 0.069 | 0.069 | 0 | 0.17 | 0.34 | 0.41 | 0.3 | 0.16 | 0.4 |
| 8 | 55309287 | rs7001445 | -0.569 | 0.591 | 0.902 | 2.154 | 0.012 | -0.01 | -0.01 | 0.216 | 0.08 | 0.11 | 0.16 | 0.03 | 0.12 | 0.1 |
| 8 | 55310831 | rs13270739 | -1.754 | 0.615 | 1.314 | 2.198 | 0.019 | 0.131 | 0.131 | 0 | 0.1 | 0.29 | 0.29 | 0.29 | 0.09 | 0.32 |
| 8 | 55317296 | rs12056424 | -1.858 | 0.609 | 1.326 | 2.161 | 0.048 | -0.02 | -0.02 | 0.135 | 0.13 | 0.32 | 0.25 | 0.08 | 0.17 | 0.16 |
| 8 | 55318176 | rs1119484 | -0.405 | 0.589 | 1.287 | 2.2 | 0 | 0.106 | 0.106 | 0 | 0.06 | 0.15 | 0.21 | 0.25 | 0.03 | 0.27 |
| 8 | 55318559 | rs7834421 | -0.438 | 0.585 | 1.285 | 2.197 | -0.001 | 0.103 | 0.103 | 0 | 0.06 | 0.15 | 0.21 | 0.25 | 0.03 | 0.28 |
| 8 | 55325943 | rs16920210 | -0.152 | 0.141 | 0.955 | 2.563 | 0.019 | -0.026 | -0.026 | 0.168 | 0.11 | 0.04 | 0.08 | 0.26 | 0.06 | 0.13 |
| 8 | 55326450 | rs12542212 | -2.468 | 0.152 | 1.078 | 2.326 | 0.034 | 0.099 | 0.099 | 0.055 | 0.19 | 0.45 | 0.46 | 0.33 | 0.21 | 0.43 |
| 8 | 55327091 | rs10958409 | -1.377 | 0.202 | 1.109 | 2.352 | 0.005 | 0.03 | 0.03 | 0.034 | 0.13 | 0.18 | 0.37 | 0.29 | 0.14 | 0.39 |
| 8 | 55327862 | rs6987174 | -1.653 | 0.201 | 1.114 | 2.351 | 0.02 | 0.084 | 0.084 | 0 | 0.14 | 0.29 | 0.38 | 0.29 | 0.15 | 0.39 |
| 8 | 55329422 | rs1995532 | -2.598 | 0.229 | 0.956 | 2.248 | 0.044 | 0.098 | 0.098 | 0 | 0.23 | 0.47 | 0.5 | 0.4 | 0.26 | 0.45 |
| 8 | 55330814 | rs13258851 | -0.967 | -0.179 | 0.802 | 2.336 | -0.007 | -0.006 | -0.006 | 2.432 | 0.1 | 0.09 | 0.25 | 0.32 | 0.09 | 0.3 |
| 8 | 55340313 | rs13272749 | -1.418 | -0.193 | 1.028 | 2.214 | -0.01 | -0.019 | -0.019 | 1.36 | 0.17 | 0.12 | 0.4 | 0.4 | 0.17 | 0.42 |
| 8 | 55344415 | rs12550470 | -1.126 | -0.129 | 0.642 | 2.24 | -0.059 | -0.107 | -0.107 | 0 | 0.15 | 0.02 | 0.34 | 0.37 | 0.14 | 0.4 |
| 8 | 55355086 | rs6986417 | -1.303 | -0.131 | 0.678 | 2.306 | -0.015 | -0.029 | -0.029 | 2.224 | 0.16 | 0.1 | 0.4 | 0.45 | 0.18 | 0.42 |
| 8 | 55355961 | rs4738890 | -1.218 | -0.289 | 0.925 | 2.489 | -0.051 | 0.16 | 0.16 | 0 | 0.17 | 0.52 | 0.18 | 0.37 | 0.11 | 0.1 |
| 8 | 55365190 | rs4523231 | -1.32 | 0.008 | 0.469 | 2.348 | -0.005 | -0.005 | -0.005 | 0.488 | 0.16 | 0.15 | 0.35 | 0.4 | 0.14 | 0.41 |
| 8 | 55368646 | rs10099341 | -1.362 | 0.038 | 0.416 | 2.341 | -0.01 | -0.027 | -0.027 | 0 | 0.16 | 0.13 | 0.36 | 0.41 | 0.15 | 0.41 |
| 8 | 55369597 | rs12234926 | -1.167 | 0.07 | 0.185 | 2.356 | -0.04 | -0.085 | -0.085 | 0 | 0.15 | 0.05 | 0.35 | 0.41 | 0.15 | 0.41 |
| 8 | 55382206 | rs10102604 | -0.229 | -0.776 | -0.035 | 1.359 | -0.025 | 0.002 | 0.002 | 0 | 0.29 | 0.38 | 0.19 | 0.3 | 0.22 | 0.15 |
| 8 | 55382848 | rs4738913 | -0.434 | -0.966 | -0.851 | 0.754 | -0.049 | 0.099 | 0.099 | 0 | 0.78 | 0.99 | 0.82 | 0.96 | 0.8 | 0.85 |
| 8 | 55384043 | rs4738915 | -0.198 | -1.015 | -0.676 | 0.767 | -0.004 | 0.073 | 0.073 | 0 | 0.77 | 0.89 | 0.82 | 0.93 | 0.8 | 0.84 |
| 8 | 55385228 | rs11987090 | 0.06 | -1.514 | -0.663 | 1.318 | -0.007 | 0.212 | 0.212 | 0 | 0.11 | 0.4 | 0.34 | 0.26 | 0.06 | 0.31 |
| 8 | 55391896 | rs6981891 | -0.234 | -1.448 | -1.483 | 1.287 | 0.008 | 0.005 | 0.005 | 0 | 0.73 | 0.9 | 0.78 | 0.76 | 0.76 | 0.8 |
| 8 | 55394269 | rs1079005 | 0.526 | -1.413 | -1.58 | 1.353 | 0.03 | -0.033 | -0.033 | 0 | 0.63 | 0.32 | 0.41 | 0.51 | 0.71 | 0.51 |
| 8 | 55399389 | rs11786806 | 1.399 | -1.313 | -2.53 | 0.808 | -0.159 | 0.06 | 0.06 | 0.114 | 0.32 | 0.04 | 0.25 | 0.18 | 0.49 | 0.34 |
| 8 | 55402535 | rs13277097 | 0.44 | -1.432 | -2.322 | 0.843 | 0.02 | -0.1 | -0.1 | 0.461 | 0.25 | 0.05 | 0.12 | 0.3 | 0.2 | 0.11 |
| 8 | 55402569 | rs4476998 | -0.12 | -1.048 | -2.094 | 0.366 | -0.017 | 0.111 | 0.111 | 0 | 0.74 | 0.98 | 0.81 | 0.94 | 0.8 | 0.82 |
| 8 | 55406254 | rs7016063 | -0.047 | -0.956 | -1.71 | 1.036 | 0.013 | -0.098 | -0.098 | 0.934 | 0.26 | 0.08 | 0.22 | 0.46 | 0.28 | 0.2 |
| 8 | 55414428 | rs1072739 | 1.104 | -1.49 | -0.981 | 1.129 | 0.004 | 0.003 | 0.003 | 1.189 | 0.52 | 0.48 | 0.51 | 0.38 | 0.6 | 0.57 |
| 8 | 55421064 | rs10085958 | 0.462 | -1.482 | -0.599 | 1.098 | -0.047 | 0.038 | 0.038 | 0 | 0.78 | 0.58 | 0.68 | 0.72 | 0.85 | 0.71 |
| 8 | 55423687 | rs6473938 | 0.509 | -1.503 | -0.319 | 1.087 | -0.053 | 0.041 | 0.041 | 0 | 0.77 | 0.51 | 0.67 | 0.72 | 0.85 | 0.71 |
| 8 | 55427589 | rs7817469 | 1.101 | -1.545 | -0.234 | 1.081 | 0.03 | 0.023 | 0.023 | 1.267 | 0.53 | 0.4 | 0.48 | 0.37 | 0.6 | 0.57 |
| 8 | 55435633 | rs7820949 | 1.161 | -1.602 | -0.134 | 1.21 | 0.019 | 0.014 | 0.014 | 1.042 | 0.52 | 0.45 | 0.48 | 0.37 | 0.6 | 0.57 |
| 8 | 55435931 | rs7840175 | 0.521 | -1.593 | -0.125 | 1.218 | -0.051 | 0.037 | 0.037 | 0 | 0.77 | 0.5 | 0.67 | 0.72 | 0.85 | 0.71 |
| 8 | 55437524 | rs9298506 | 0.19 | -0.463 | -0.169 | 0.922 | 0.027 | 0.024 | 0.024 | 0 | 0.11 | 0.17 | 0.14 | 0.21 | 0.17 | 0.23 |
| 8 | 55441025 | rs10108776 | 0.626 | -0.526 | -0.208 | 0.894 | 0.021 | 0.016 | 0.016 | 1.017 | 0.52 | 0.43 | 0.48 | 0.37 | 0.6 | 0.57 |
| 8 | 55449840 | rs7009744 | -0.002 | -0.626 | -0.445 | 0.82 | 0.027 | -0.045 | -0.045 | 0 | 0.36 | 0.1 | 0.25 | 0.35 | 0.34 | 0.17 |
| 8 | 55459476 | rs13276481 | 1.647 | -0.535 | -0.417 | 0.643 | -0.107 | 0.23 | 0.23 | 5.34 | 0.39 | 0.07 | 0.31 | 0.05 | 0.42 | 0.28 |
| 8 | 55460789 | rs7825993 | -0.365 | -0.588 | 0.436 | 0.968 | -0.024 | 0.167 | 0.167 | 0 | 0.89 | 0.65 | 0.74 | 0.73 | 0.94 | 0.74 |
| 8 | 55464983 | rs4737605 | -0.046 | -0.692 | 0.121 | 0.687 | 0.035 | -0.019 | -0.019 | 0 | 0.36 | 0.16 | 0.25 | 0.36 | 0.34 | 0.17 |
| 8 | 55470545 | rs11781152 | -0.329 | -0.694 | 0.227 | 0.751 | -0.023 | 0.255 | 0.255 | 0 | 0.88 | 0.63 | 0.73 | 0.61 | 0.93 | 0.67 |
| 8 | 55471051 | rs16920503 | -0.046 | -0.652 | 0.062 | 0.704 | 0.025 | -0.043 | -0.043 | 1.908 | 0.36 | 0.11 | 0.25 | 0.36 | 0.34 | 0.17 |
| 8 | 55471417 | rs10504167 | -0.234 | 0.039 | 0.646 | 0.997 | 0.029 | 0.049 | 0.049 | 0 | 0.09 | 0.18 | 0.12 | 0.19 | 0.11 | 0.17 |
| 8 | 55472515 | rs7827437 | 0.862 | 0.298 | 0.589 | 0.867 | -0.016 | -0.024 | -0.024 | 0 | 0.61 | 0.52 | 0.51 | 0.26 | 0.66 | 0.45 |
| 8 | 55476999 | rs6473944 | 0.325 | 0.9 | 0.794 | 1.104 | 0.02 | 0.189 | 0.189 | 0 | 0.87 | 0.64 | 0.78 | 0.63 | 0.91 | 0.57 |
| 8 | 55482675 | rs6997903 | -0.539 | 1.124 | 0.43 | 0.727 | 0.054 | -0.01 | -0.01 | 0 | 0.16 | 0.33 | 0.25 | 0.16 | 0.24 | 0.14 |
| 8 | 55483583 | rs10216467 | -0.461 | 1.032 | 0.336 | 0.752 | 0.014 | -0.003 | -0.003 | 0.494 | 0.16 | 0.29 | 0.18 | 0.13 | 0.16 | 0.13 |
| 8 | 55486408 | rs12674538 | -1.051 | 0.968 | 0.082 | 0.684 | 0 | -0.001 | -0.001 | 0 | 0.41 | 0.57 | 0.44 | 0.5 | 0.45 | 0.31 |
| 8 | 55489041 | rs7832057 | 0.349 | 1.31 | -0.658 | 0.852 | -0.019 | -0.098 | -0.098 | 0 | 0.22 | 0.06 | 0.22 | 0.37 | 0.28 | 0.13 |
| 8 | 55505788 | rs4739022 | -0.242 | 1.493 | -0.7 | 0.391 | -0.064 | -0.077 | -0.077 | 1.67 | 0.88 | 0.99 | 0.8 | 0.7 | 0.86 | 0.56 |
| 8 | 55506499 | rs4737619 | 1.043 | 0.996 | -0.146 | -0.514 | 0.005 | -0.036 | -0.036 | 0 | 0.45 | 0.36 | 0.33 | 0.03 | 0.39 | 0.26 |
| 8 | 55511676 | rs7823055 | 0.702 | 1.132 | -0.309 | 0.242 | -0.001 | -0.016 | -0.016 | 0 | 0.6 | 0.56 | 0.48 | 0.21 | 0.58 | 0.46 |
| 8 | 55513201 | rs4739035 | 0.146 | 1.362 | -0.951 | 0.243 | -0.015 | 0.034 | 0.034 | 0 | 0.7 | 0.89 | 0.62 | 0.17 | 0.68 | 0.5 |
| 8 | 55522880 | rs3910515 | -0.842 | 1.642 | -0.736 | 0.503 | -0.085 | -0.007 | -0.007 | 0.628 | 0.15 | 0.01 | 0.27 | 0.09 | 0.25 | 0.2 |
| 8 | 55524841 | rs10112903 | -0.824 | 0.795 | 0.202 | 1.303 | -0.059 | 0.082 | 0.082 | 0.462 | 0.23 | 0.46 | 0.21 | 0.44 | 0.13 | 0.15 |
| 8 | 55529073 | rs9643828 | 1.125 | 1.458 | 0.613 | 1.488 | 0.024 | 0.018 | 0.018 | 0 | 0.81 | 0.73 | 0.72 | 0.73 | 0.71 | 0.6 |
| 8 | 55529612 | rs145290 | -0.861 | 1.777 | 0.649 | 1.287 | 0.004 | -0.011 | -0.011 | 0.011 | 0.28 | 0.21 | 0.39 | 0.66 | 0.28 | 0.28 |
| 8 | 55537138 | rs429668 | -0.824 | 1.844 | 0.885 | 1.559 | 0.006 | -0.037 | -0.037 | 0 | 0.27 | 0.17 | 0.31 | 0.44 | 0.27 | 0.25 |
| 8 | 55539057 | rs444772 | -0.761 | 1.844 | 0.885 | 1.559 | 0.005 | -0.047 | -0.047 | 0.054 | 0.27 | 0.14 | 0.31 | 0.44 | 0.27 | 0.25 |
| 8 | 55541450 | rs446227 | -0.711 | 1.82 | 0.91 | 1.547 | -0.017 | -0.094 | -0.094 | 0.616 | 0.25 | 0.08 | 0.28 | 0.44 | 0.26 | 0.24 |
| 8 | 55541513 | rs414352 | -0.881 | 1.82 | 0.911 | 1.547 | 0.006 | -0.037 | -0.037 | 0 | 0.27 | 0.17 | 0.31 | 0.44 | 0.27 | 0.25 |

| | | | | | | | | | | | | | | | | |
|---|---|---|---|---|---|---|---|---|---|---|---|---|---|---|---|---|
| 8 | 55541617 | rs441800 | -0.632 | 1.906 | 0.689 | 1.59 | -0.036 | -0.121 | -0.121 | 0.262 | 0.24 | 0.06 | 0.27 | 0.44 | 0.26 | 0.24 |
| 8 | 55562071 | rs421469 | -0.754 | 1.907 | 0.689 | 1.58 | -0.017 | -0.096 | -0.096 | 0 | 0.25 | 0.08 | 0.28 | 0.45 | 0.26 | 0.24 |
| 8 | 55574960 | rs446222 | 0.86 | 1.874 | 0.945 | 1.589 | 0.024 | 0.071 | 0.071 | 0 | 0.82 | 0.78 | 0.6 | 0.51 | 0.69 | 0.59 |
| 8 | 55632762 | rs1437782 | -1.249 | 1.821 | 0.926 | 1.622 | 0.018 | 0.054 | 0.054 | 0 | 0.19 | 0.21 | 0.4 | 0.49 | 0.31 | 0.41 |
| 8 | 55649012 | rs1437785 | 0.229 | 1.92 | 0.722 | 1.648 | -0.033 | -0.116 | -0.116 | 1.511 | 0.76 | 0.95 | 0.72 | 0.55 | 0.73 | 0.75 |
| 8 | 55656582 | rs2043774 | -0.211 | 1.929 | 0.719 | 1.933 | -0.056 | -0.042 | -0.042 | 0.582 | 0.81 | 0.98 | 0.76 | 0.82 | 0.74 | 0.93 |
| 8 | 55688171 | rs13278605 | -1.119 | 2.15 | 1.203 | 1.885 | 0.011 | 0.01 | 0.01 | 0 | 0.17 | 0.18 | 0.27 | 0.3 | 0.31 | 0.38 |
| 8 | 55689820 | rs13270740 | -0.225 | 1.969 | 1.07 | 1.946 | 0.001 | 0.016 | 0.016 | 0.391 | 0.07 | 0.07 | 0.07 | 0.23 | 0.15 | 0.23 |
| 8 | 55733711 | rs10112190 | -1.412 | 2.215 | 1.124 | 1.76 | 0.077 | 0.038 | 0.038 | 0 | 0.21 | 0.36 | 0.29 | 0.28 | 0.37 | 0.38 |
| 8 | 55735722 | rs10105032 | -1.172 | 2.252 | 1.245 | 1.819 | 0.059 | 0.026 | 0.026 | 0 | 0.19 | 0.28 | 0.26 | 0.25 | 0.34 | 0.37 |
| 8 | 55737195 | rs13282412 | -1.115 | 2.255 | 1.263 | 1.755 | 0.002 | -0.001 | -0.001 | 0 | 0.19 | 0.16 | 0.25 | 0.29 | 0.34 | 0.38 |
| 8 | 55744792 | rs7822624 | 0.222 | 1.838 | 0.739 | 1.313 | -0.009 | -0.098 | -0.098 | 0 | 0.7 | 0.91 | 0.69 | 0.52 | 0.69 | 0.71 |
| 8 | 55754697 | rs1498175 | -0.844 | 1.826 | 0.732 | 1.377 | 0.021 | 0.012 | 0.012 | 0 | 0.19 | 0.19 | 0.26 | 0.29 | 0.34 | 0.37 |
| 8 | 55761124 | rs10958432 | 0.047 | 1.74 | 0.31 | 1.376 | -0.028 | -0.123 | -0.123 | 0.847 | 0.72 | 0.93 | 0.7 | 0.52 | 0.69 | 0.71 |
| 8 | 55763773 | rs4737736 | -1.302 | 1.813 | 0.645 | 1.307 | 0.065 | 0.031 | 0.031 | 0 | 0.21 | 0.33 | 0.29 | 0.29 | 0.37 | 0.38 |
| 8 | 55777564 | rs6990422 | 0.698 | 2.035 | 0.166 | 1.029 | -0.026 | 0.011 | 0.011 | 0 | 0.66 | 0.78 | 0.57 | 0.7 | 0.56 | 0.76 |
| 8 | 55779505 | rs1396894 | 0.69 | 2.029 | 0.167 | 1.111 | -0.032 | 0.053 | 0.053 | 0 | 0.66 | 0.8 | 0.57 | 0.78 | 0.56 | 0.76 |
| 8 | 55779565 | rs1396895 | 0.859 | 2.008 | 1.057 | 1.048 | 0.017 | -0.048 | -0.048 | 0 | 0.62 | 0.49 | 0.52 | 0.78 | 0.52 | 0.76 |
| 8 | 55783975 | rs9298514 | -0.725 | 1.727 | 1.336 | 1.433 | 0.024 | 0.19 | 0.19 | 0 | 0.13 | 0.29 | 0.16 | 0.45 | 0.15 | 0.38 |
| 8 | 55798736 | rs1603708 | -0.866 | 1.621 | 1.152 | 0.866 | 0.014 | 0.03 | 0.03 | 0 | 0.13 | 0.17 | 0.36 | 0.24 | 0.18 | 0.23 |
| 8 | 55803742 | rs7824078 | 0.628 | 1.193 | 1.409 | 0.14 | -0.003 | -0.032 | -0.032 | 0 | 0.67 | 0.38 | 0.71 | 0.82 | 0.66 | 0.86 |
| 8 | 55814206 | rs969183 | 0.354 | 1.148 | 1.302 | 0.181 | -0.011 | -0.064 | -0.064 | 0.079 | 0.69 | 0.48 | 0.74 | 0.82 | 0.7 | 0.87 |
| 8 | 55820605 | rs6473956 | -0.113 | 1.2 | 0.38 | 0.217 | -0.018 | 0.045 | 0.045 | 0 | 0.75 | 0.9 | 0.71 | 0.8 | 0.71 | 0.89 |
| 8 | 55824925 | rs7011709 | 0.084 | 0.629 | 0.551 | -0.299 | 0.002 | 0.011 | 0.011 | 0.041 | 0.31 | 0.28 | 0.19 | 0.1 | 0.29 | 0.13 |
| 8 | 55828934 | rs11996465 | -1.077 | 0.404 | 1.071 | 0.17 | -0.008 | 0.198 | 0.198 | 1.742 | 0.08 | 0.39 | 0.05 | 0.14 | 0.01 | 0.04 |
| 8 | 55829630 | rs12675799 | -0.777 | 0.454 | 1.058 | 0.195 | 0.007 | 0.219 | 0.219 | 0.326 | 0.06 | 0.35 | 0.05 | 0.14 | 0.01 | 0.04 |
| 8 | 55850104 | rs270794 | 0.312 | 0.804 | 0.549 | 0.173 | -0.001 | 0.004 | 0.004 | 0 | 0.52 | 0.41 | 0.49 | 0.38 | 0.45 | 0.47 |
| 8 | 55851641 | rs270795 | 0.485 | 1.151 | 0.153 | 0.015 | -0.012 | 0.011 | 0.011 | 0 | 0.08 | 0.04 | 0.16 | 0.03 | 0.16 | 0.12 |
| 8 | 55865884 | rs1493680 | 1.466 | 0.685 | -0.526 | -0.299 | -0.001 | 0.054 | 0.054 | 0.591 | 0.33 | 0.02 | 0.26 | 0.17 | 0.23 | 0.3 |
| 8 | 55877790 | rs4737805 | -0.176 | 0.089 | -0.533 | -0.544 | -0.009 | -0.023 | -0.023 | 0.027 | 0.33 | 0.45 | 0.27 | 0.21 | 0.28 | 0.29 |
| 8 | 55878305 | rs17422653 | 1.728 | 0.045 | -0.615 | -0.589 | -0.013 | -0.024 | -0.024 | 0.237 | 0.2 | 0.02 | 0.22 | 0.17 | 0.17 | 0.18 |
| 8 | 55879485 | rs16920971 | 1.326 | 0.033 | -0.538 | -0.586 | -0.009 | 0.005 | 0.005 | 0 | 0.05 | 0.1 | 0.03 | 0.04 | 0.04 | 0.01 |
| 8 | 55879686 | rs16920973 | 1.286 | 0.077 | -0.691 | -0.66 | 0.017 | 0.081 | 0.081 | 0 | 0.17 | 0.23 | 0.32 | 0.53 | 0.25 | 0.4 |
| 8 | 55884105 | rs4737806 | -0.564 | -0.309 | -0.432 | -0.389 | -0.001 | -0.002 | -0.002 | 0 | 0.35 | 0.63 | 0.31 | 0.21 | 0.28 | 0.29 |
| 8 | 55887631 | rs4737807 | 0.808 | -0.332 | -0.463 | -0.416 | -0.04 | 0.104 | 0.104 | 0 | 0.22 | 0.04 | 0.18 | 0.03 | 0.2 | 0.23 |
| 8 | 55907294 | rs17331902 | 1.102 | -0.017 | -1.077 | -0.204 | -0.013 | -0.024 | -0.024 | 0 | 0.2 | 0.02 | 0.22 | 0.17 | 0.17 | 0.18 |
| 8 | 55911546 | rs4737819 | -0.442 | -0.022 | -0.962 | -0.199 | -0.008 | -0.016 | -0.016 | 0 | 0.34 | 0.53 | 0.3 | 0.21 | 0.28 | 0.31 |
| 8 | 55912151 | rs16921013 | 0.111 | 0.38 | -1.765 | -0.173 | -0.001 | -0.009 | -0.009 | 0 | 0.28 | 0.34 | 0.25 | 0.17 | 0.24 | 0.3 |
| 8 | 55913547 | rs17332195 | 0.412 | 0.339 | -2.094 | 0.023 | -0.016 | -0.023 | -0.023 | 0 | 0.2 | 0.02 | 0.22 | 0.17 | 0.17 | 0.19 |
| 8 | 55914456 | rs3934874 | -0.24 | 0.273 | -1.681 | -0.176 | 0.009 | 0.034 | 0.034 | 0 | 0.25 | 0.14 | 0.14 | 0.1 | 0.19 | 0.32 |
| 8 | 55918430 | rs4428652 | -0.201 | 0.17 | -2.225 | -0.167 | -0.031 | 0.028 | 0.028 | 0 | 0.59 | 0.88 | 0.66 | 0.3 | 0.58 | 0.47 |
| 8 | 55918679 | rs4236947 | -0.706 | -0.057 | -2.541 | 0.398 | -0.054 | -0.066 | -0.066 | 0.04 | 0.81 | 0.92 | 0.71 | 0.6 | 0.69 | 0.58 |
| 8 | 55919502 | rs3924878 | 0.694 | -0.107 | -1.739 | -0.316 | 0.002 | -0.104 | -0.104 | 2.118 | 0.25 | 0.48 | 0.34 | 0.07 | 0.26 | 0.08 |
| 8 | 55920886 | rs7006973 | 0.802 | -0.25 | -1.442 | -0.548 | -0.037 | -0.101 | -0.101 | 0.111 | 0.16 | 0.32 | 0.14 | 0 | 0.12 | 0.02 |
| 8 | 55927212 | rs4737826 | 0.843 | -0.275 | -1.25 | -0.593 | 0.006 | -0.005 | -0.005 | 0 | 0.11 | 0.14 | 0.2 | 0.07 | 0.15 | 0.07 |
| 8 | 55930998 | rs7842680 | 0.244 | -0.549 | -1.243 | -0.789 | 0.002 | -0.021 | -0.021 | 1.225 | 0.17 | 0.2 | 0.13 | 0.02 | 0.14 | 0.03 |
| 8 | 55937602 | rs4634616 | 1.468 | -0.529 | -1.026 | -0.766 | 0.004 | -0.01 | -0.01 | 0.003 | 0.29 | 0.34 | 0.31 | 0.22 | 0.29 | 0.38 |
| 8 | 55948440 | rs1383452 | -0.831 | -1.014 | -0.779 | -0.82 | -0.005 | -0.066 | -0.066 | 0.001 | 0.29 | 0.39 | 0.34 | 0.09 | 0.29 | 0.1 |
| 8 | 55949530 | rs1904861 | 1.144 | -0.895 | -0.789 | -0.555 | 0.045 | -0.089 | -0.089 | 0 | 0.21 | 0.02 | 0.05 | 0.29 | 0.11 | 0.11 |
| 8 | 55981777 | rs16921098 | -0.087 | -0.479 | -0.411 | -0.498 | -0.008 | -0.022 | -0.022 | 0.432 | 0.07 | 0.2 | 0.04 | 0.02 | 0.02 | 0.02 |
| 8 | 56004352 | rs996825 | 0.691 | -0.53 | -0.167 | -0.556 | 0.009 | -0.103 | -0.103 | 0 | 0.78 | 0.47 | 0.73 | 0.93 | 0.81 | 0.92 |
| 8 | 56010193 | rs6473970 | 0.727 | -0.546 | -0.335 | -0.471 | 0.005 | -0.07 | -0.07 | 0.043 | 0.78 | 0.45 | 0.73 | 0.9 | 0.81 | 0.92 |
| 8 | 56023769 | rs2061998 | 0.645 | -0.76 | -0.121 | -0.439 | 0.024 | 0.024 | 0.024 | 5.847 | 0.49 | 0.34 | 0.6 | 0.62 | 0.62 | 0.78 |
| 8 | 56037030 | rs1480774 | -0.226 | -0.585 | -0.103 | -0.429 | 0.031 | -0.03 | -0.03 | 0.16 | 0.86 | 0.73 | 0.79 | 0.93 | 0.83 | 0.93 |
| 8 | 56037758 | rs12546351 | -0.55 | -0.58 | -0.062 | -0.426 | -0.002 | 0.001 | 0.001 | 0 | 0.87 | 0.87 | 0.8 | 0.93 | 0.83 | 0.93 |
| 8 | 56038222 | rs6983704 | -0.224 | -0.883 | -0.154 | -0.289 | -0.011 | -0.005 | -0.005 | 0.218 | 0.4 | 0.59 | 0.2 | 0.29 | 0.23 | 0.17 |
| 8 | 56045247 | rs16921196 | 0.372 | -0.561 | -1.765 | -0.11 | 0.006 | -0.045 | -0.045 | 0 | 0.27 | 0.01 | 0.1 | 0.26 | 0.18 | 0.16 |
| 8 | 56047072 | rs12682291 | 1.22 | -0.264 | -2.412 | -0.267 | -0.081 | -0.016 | -0.016 | 0 | 0.27 | 0.02 | 0.22 | 0.16 | 0.29 | 0.34 |
| 8 | 56048794 | rs9643833 | 0.488 | 0.22 | -2.193 | 0.008 | 0.022 | -0.054 | -0.054 | 0 | 0.29 | 0.05 | 0.11 | 0.29 | 0.2 | 0.17 |
| 8 | 56049488 | rs7812409 | -0.452 | 1.074 | -1.518 | 0.195 | 0.024 | -0.001 | -0.001 | 0 | 0.91 | 0.8 | 0.82 | 0.94 | 0.86 | 0.94 |
| 8 | 56054696 | rs11785942 | 0.911 | 0.763 | -1.795 | 0.355 | -0.052 | -0.062 | -0.062 | 0 | 0.16 | 0.07 | 0.27 | 0.39 | 0.29 | 0.38 |
| 8 | 56059868 | rs997855 | 0.219 | -0.184 | -1.439 | -0.363 | 0.053 | 0.068 | 0.068 | 0.625 | 0.52 | 0.67 | 0.42 | 0.26 | 0.36 | 0.38 |
| 8 | 56060025 | rs997854 | 0.236 | 0.223 | -1.153 | -0.327 | 0.01 | 0.013 | 0.013 | 0 | 0.49 | 0.43 | 0.35 | 0.29 | 0.41 | 0.4 |
| 8 | 56062921 | rs7000240 | 0.053 | 1.494 | -1.289 | 0.605 | -0.029 | -0.022 | -0.022 | 0 | 0.77 | 0.79 | 0.5 | 0.61 | 0.58 | 0.65 |
| 8 | 56063806 | rs5008177 | 0.21 | 1.233 | -1.754 | 0.238 | 0.049 | 0.046 | 0.046 | 0.321 | 0.53 | 0.75 | 0.37 | 0.38 | 0.38 | 0.39 |
| 8 | 56063814 | rs10958444 | 0.436 | 0.476 | -1.73 | 0.01 | 0.026 | 0.007 | 0.007 | 0.014 | 0.4 | 0.7 | 0.34 | 0.37 | 0.3 | 0.35 |
| 8 | 56064834 | rs7836267 | 0.517 | 0.266 | -1.501 | 0.034 | -0.008 | -0.001 | -0.001 | 0 | 0.35 | 0.62 | 0.32 | 0.37 | 0.26 | 0.25 |
| 8 | 56066078 | rs7012574 | 0.511 | 0.271 | -1.498 | 0.036 | -0.008 | -0.001 | -0.001 | 0.142 | 0.35 | 0.62 | 0.32 | 0.37 | 0.26 | 0.26 |
| 8 | 56080772 | rs9792156 | 1.5 | 0.024 | -1.433 | 0.016 | 0.016 | 0.003 | 0.003 | 1.835 | 0.25 | 0.13 | 0.24 | 0.22 | 0.24 | 0.22 |

| | | | | | | | | | | | | | | | | |
|---|---|---|---|---|---|---|---|---|---|---|---|---|---|---|---|---|
| 8 | 56081879 | rs2199921 | 0.943 | 0.062 | -1.725 | 0.028 | -0.011 | 0.106 | 0.106 | 1.16 | 0.34 | 0.03 | 0.16 | 0.17 | 0.26 | 0.27 |
| 8 | 56096714 | rs6997561 | 0.43 | 0.614 | -1.259 | 0.649 | 0.005 | 0.049 | 0.049 | 0 | 0.18 | 0.22 | 0.28 | 0.32 | 0.23 | 0.23 |
| 8 | 56106535 | rs4416803 | 1.099 | 0.369 | -1.208 | 0.748 | 0.024 | -0.035 | -0.035 | 0 | 0.27 | 0.11 | 0.27 | 0.26 | 0.24 | 0.25 |
| 8 | 56111322 | rs4737264 | -0.178 | 0.982 | -0.098 | 0.426 | 0.035 | 0.005 | 0.005 | 0 | 0.11 | 0.14 | 0.19 | 0.14 | 0.2 | 0.1 |
| 8 | 56112160 | rs12546778 | -0.92 | 0.947 | -0.356 | 0.404 | 0.098 | 0.086 | 0.086 | 0.029 | 0.15 | 0.49 | 0.25 | 0.26 | 0.21 | 0.16 |
| 8 | 56115441 | rs6982136 | -0.328 | 0.799 | -1.41 | -0.006 | -0.028 | 0.129 | 0.129 | 0.046 | 0.52 | 0.82 | 0.63 | 0.69 | 0.52 | 0.65 |
| 8 | 56119057 | rs16921393 | 0.156 | 0.363 | -0.956 | 0.12 | -0.004 | 0 | 0 | 0 | 0.38 | 0.4 | 0.34 | 0.52 | 0.27 | 0.32 |
| 8 | 56120404 | rs13248919 | 0.933 | 0.988 | -1.778 | -0.168 | -0.072 | 0.201 | 0.201 | 11.121 | 0.34 | 0.02 | 0.18 | 0.1 | 0.29 | 0.19 |
| 8 | 56120462 | rs8180956 | 0.182 | 1.429 | -0.62 | 0.533 | 0.026 | 0.07 | 0.07 | 0 | 0.13 | 0.22 | 0.19 | 0.27 | 0.22 | 0.27 |
| 8 | 56127083 | rs13249396 | -0.848 | 1.303 | -0.943 | 0.157 | -0.04 | 0.042 | 0.042 | 0 | 0.13 | 0.07 | 0.13 | 0.03 | 0.2 | 0.14 |
| 8 | 56133320 | rs4341145 | 0.099 | 1.111 | -0.107 | 0.339 | 0.032 | 0.118 | 0.118 | 0 | 0.9 | 0.64 | 0.78 | 0.81 | 0.91 | 0.84 |
| 8 | 56135727 | rs11784187 | 0.943 | 0.664 | -0.357 | 0.053 | -0.012 | 0.053 | 0.053 | 10.093 | 0.43 | 0.27 | 0.35 | 0.28 | 0.48 | 0.32 |
| 8 | 56148811 | rs6473978 | 0.582 | 0.397 | -0.079 | 0.14 | -0.017 | 0.068 | 0.068 | 0.128 | 0.86 | 0.38 | 0.69 | 0.77 | 0.91 | 0.82 |
| 8 | 56152930 | rs16921482 | 0.469 | 0.38 | -0.185 | 0.129 | 0.048 | -0.079 | -0.079 | 0.851 | 0.34 | 0.16 | 0.22 | 0.48 | 0.24 | 0.37 |
| 8 | 56159069 | rs11786326 | 0.532 | 0.331 | -0.206 | 0.065 | 0.106 | 0.146 | 0.146 | 9.054 | 0.5 | 0.15 | 0.47 | 0.29 | 0.67 | 0.45 |
| 8 | 56161282 | rs4475454 | 1.02 | 0.189 | -0.301 | 0.026 | -0.011 | -0.193 | -0.193 | 3.971 | 0.29 | 0.02 | 0.21 | 0.48 | 0.23 | 0.35 |
| 8 | 56165073 | rs13259631 | 0.95 | 0.419 | -0.27 | 0.042 | -0.029 | 0.028 | 0.028 | 0 | 0.07 | 0.02 | 0.12 | 0.01 | 0.18 | 0.13 |
| 8 | 56167882 | rs17430331 | 1.272 | 0.339 | -0.253 | 0.057 | -0.054 | 0.086 | 0.086 | 9.476 | 0.41 | 0.1 | 0.33 | 0.27 | 0.48 | 0.31 |
| 8 | 56168439 | rs13273951 | 0.318 | -0.039 | -0.02 | 0.115 | 0.006 | 0.13 | 0.13 | 0.046 | 0.87 | 0.47 | 0.7 | 0.78 | 0.91 | 0.81 |
| 8 | 56170306 | rs7002941 | 0.341 | -0.043 | -0.022 | 0.113 | 0.005 | 0.129 | 0.129 | 0.068 | 0.87 | 0.47 | 0.7 | 0.78 | 0.91 | 0.81 |
| 8 | 56171940 | rs10958453 | 0.914 | -0.06 | -0.335 | 0.102 | -0.021 | 0.017 | 0.017 | 0.147 | 0.82 | 0.3 | 0.68 | 0.77 | 0.89 | 0.8 |
| 8 | 56184390 | rs1013087 | 1.14 | 0.156 | -0.627 | 0.04 | -0.121 | 0.071 | 0.071 | 6.086 | 0.4 | 0.05 | 0.35 | 0.27 | 0.52 | 0.32 |
| 8 | 56192368 | rs6988288 | -0.614 | 0.322 | -0.003 | 0.159 | -0.001 | 0.001 | 0.001 | 0 | 0.93 | 0.93 | 0.83 | 0.91 | 0.95 | 0.91 |
| 8 | 56193282 | rs4236975 | 1.937 | 0.296 | 0.041 | 0.153 | 0 | 0.002 | 0.002 | 0 | 0.72 | 0.26 | 0.56 | 0.75 | 0.73 | 0.67 |
| 8 | 56194700 | rs16921542 | 0.255 | 0.305 | 0.088 | 0.148 | -0.011 | 0.01 | 0.01 | 0 | 0.08 | 0.06 | 0.11 | 0.02 | 0.16 | 0.13 |
| 8 | 56201307 | rs4738045 | 1.511 | 0.281 | 0.088 | 0.166 | -0.027 | 0.008 | 0.008 | 0 | 0.79 | 0.31 | 0.67 | 0.76 | 0.89 | 0.79 |
| 8 | 56201783 | rs11786627 | -0.005 | 0.357 | 0.091 | 0.214 | -0.007 | 0.029 | 0.029 | 0 | 0.91 | 0.8 | 0.74 | 0.86 | 0.95 | 0.89 |
| 8 | 56212760 | rs16921571 | -1.639 | 0.363 | -0.229 | 0.006 | -0.001 | 0.006 | 0.006 | 0 | 0.27 | 0.72 | 0.37 | 0.2 | 0.25 | 0.32 |
| 8 | 56220350 | rs4737297 | -0.054 | 0.167 | 0.289 | 0.169 | 0.048 | 0.114 | 0.114 | 0 | 0.06 | 0.32 | 0.05 | 0.09 | 0.05 | 0.07 |
| 8 | 56223857 | rs6998105 | -0.303 | 0.117 | 0.366 | 0.116 | 0.002 | 0.015 | 0.015 | 0 | 0.1 | 0.29 | 0.2 | 0.09 | 0.05 | 0.11 |
| 8 | 56231059 | rs4738073 | -0.046 | -0.166 | -0.054 | 0.046 | -0.02 | -0.004 | -0.004 | 0 | 0.39 | 0.58 | 0.34 | 0.61 | 0.28 | 0.43 |
| 8 | 56231209 | rs16921598 | 0.628 | -0.153 | -0.441 | 0.047 | 0.044 | -0.058 | -0.058 | 0.019 | 0.32 | 0.16 | 0.27 | 0.52 | 0.23 | 0.36 |
| 8 | 56232929 | rs4472490 | -0.656 | 0.008 | -0.857 | -0.073 | 0.129 | 0.143 | 0.143 | 4.169 | 0.5 | 0.87 | 0.54 | 0.71 | 0.33 | 0.53 |
| 8 | 56246342 | rs10089802 | 1.404 | 0.215 | -1.027 | -0.235 | -0.015 | -0.015 | -0.015 | 0.036 | 0.1 | 0.04 | 0.1 | 0.14 | 0.15 | 0.13 |
| 8 | 56247138 | rs2939678 | -1.011 | -0.245 | -1.385 | -0.336 | -0.021 | -0.13 | -0.13 | 0 | 0.77 | 0.96 | 0.7 | 0.45 | 0.78 | 0.72 |
| 8 | 56248483 | rs1119277 | -0.35 | -0.094 | -0.795 | -0.765 | 0.012 | -0.061 | -0.061 | 0.044 | 0.16 | 0.24 | 0.14 | 0.02 | 0.17 | 0.15 |
| 8 | 56253581 | rs1532899 | -0.314 | -0.156 | -0.77 | -0.377 | -0.025 | 0.028 | 0.028 | 1.633 | 0.68 | 0.47 | 0.64 | 0.53 | 0.76 | 0.72 |
| 8 | 56258246 | rs2036619 | 1.262 | 0.106 | -0.817 | -0.31 | -0.035 | -0.049 | -0.049 | 0 | 0.16 | 0.05 | 0.34 | 0.25 | 0.23 | 0.16 |
| 8 | 56271688 | rs16921659 | 0.024 | -0.249 | -0.09 | 0.052 | 0.017 | 0.009 | 0.009 | 0 | 0.06 | 0.28 | 0.06 | 0 | 0.01 | 0.01 |
| 8 | 56275507 | rs1506672 | 0.517 | -0.166 | -0.786 | 0.091 | 0.011 | -0.002 | -0.002 | 0 | 0.54 | 0.43 | 0.57 | 0.56 | 0.6 | 0.56 |
| 8 | 56281600 | rs2658913 | -0.224 | 0.303 | -1.507 | -0.293 | -0.016 | 0.037 | 0.037 | 0 | 0.11 | 0.07 | 0.05 | 0.02 | 0.1 | 0.12 |
| 8 | 56282834 | rs4738099 | 0.846 | 0.482 | -1.523 | -0.239 | -0.134 | 0.036 | 0.036 | 5.205 | 0.4 | 0.06 | 0.51 | 0.33 | 0.5 | 0.35 |
| 8 | 56290516 | rs2622554 | -0.977 | 0.604 | -0.245 | -0.301 | 0.032 | -0.069 | -0.069 | 0.315 | 0.13 | 0.22 | 0.09 | 0.01 | 0.14 | 0.12 |
| 8 | 56295448 | rs951256 | -0.639 | 0.302 | 0.692 | -0.357 | 0.098 | 0.039 | 0.039 | 0 | 0.07 | 0.4 | 0.07 | 0.01 | 0.04 | 0.05 |
| 8 | 56298109 | rs982330 | 0.159 | -0.104 | 0.094 | -0.162 | -0.038 | -0.085 | -0.085 | 0.616 | 0.67 | 0.89 | 0.53 | 0.57 | 0.65 | 0.65 |
| 8 | 56298676 | rs2622546 | 0.227 | -0.105 | 0.215 | -0.153 | -0.013 | -0.049 | -0.049 | 0 | 0.67 | 0.82 | 0.53 | 0.56 | 0.65 | 0.65 |
| 8 | 56313541 | rs1848085 | -1.733 | -0.337 | 0.61 | -0.197 | -0.04 | -0.093 | -0.093 | 1.049 | 0.15 | 0.56 | 0.09 | 0.01 | 0.06 | 0.09 |
| 8 | 56318630 | rs2939691 | -0.063 | -0.091 | 0.288 | -0.088 | 0.08 | 0.018 | 0.018 | 0 | 0.06 | 0.29 | 0.06 | 0 | 0.04 | 0.05 |
| 8 | 56330759 | rs2658897 | -1.305 | -0.374 | -1.044 | -0.164 | -0.013 | -0.081 | -0.081 | 0.441 | 0.52 | 0.93 | 0.57 | 0.54 | 0.43 | 0.49 |
| 8 | 56338058 | rs6995409 | -0.925 | -0.453 | -0.14 | 0.116 | -0.047 | -0.031 | -0.031 | 5.117 | 0.13 | 0.43 | 0.05 | 0.07 | 0.04 | 0.05 |
| 8 | 56339042 | rs2976003 | 0.146 | -0.355 | -0.536 | 0.206 | -0.047 | -0.088 | -0.088 | 0 | 0.26 | 0.06 | 0.42 | 0.37 | 0.3 | 0.31 |
| 8 | 56340947 | rs10095520 | -0.397 | -0.42 | 0.405 | 0.08 | 0.046 | 0.12 | 0.12 | 0.74 | 0.06 | 0.42 | 0.03 | 0.1 | 0.01 | 0.1 |
| 8 | 56344186 | rs13439780 | -1.367 | -0.486 | -0.229 | 0.045 | -0.025 | -0.003 | -0.003 | 0 | 0.15 | 0.77 | 0.09 | 0.17 | 0.04 | 0.12 |
| 8 | 56347498 | rs2658917 | -0.113 | 0.09 | -0.688 | -0.556 | 0 | -0.064 | -0.064 | 0 | 0.31 | 0.38 | 0.26 | 0.12 | 0.32 | 0.22 |
| 8 | 56348061 | rs16921822 | 0.35 | -0.016 | -0.966 | -1.029 | 0.002 | -0.055 | -0.055 | 1.054 | 0.18 | 0.39 | 0.14 | 0.09 | 0.19 | 0.15 |
| 8 | 56349420 | rs2658914 | -0.804 | 0.032 | -1.578 | -1 | -0.011 | 0.214 | 0.214 | 0 | 0.5 | 0.81 | 0.41 | 0.21 | 0.51 | 0.38 |
| 8 | 56350638 | rs2975997 | -1.532 | -0.109 | -2.189 | -0.509 | -0.075 | -0.131 | -0.131 | 0.01 | 0.68 | 0.96 | 0.73 | 0.6 | 0.65 | 0.63 |
| 8 | 56352233 | rs2622596 | 0.918 | -0.136 | -1.807 | -0.754 | -0.021 | 0.065 | 0.065 | 0 | 0.26 | 0.11 | 0.24 | 0.12 | 0.3 | 0.22 |
| 8 | 56356869 | rs2622592 | -0.681 | -0.087 | -2.068 | -0.722 | -0.05 | 0.169 | 0.169 | 0 | 0.54 | 0.84 | 0.41 | 0.21 | 0.52 | 0.43 |
| 8 | 56358274 | rs2622590 | 0.33 | -0.228 | -2.182 | -0.203 | -0.004 | -0.044 | -0.044 | 0 | 0.16 | 0.09 | 0.33 | 0.43 | 0.14 | 0.25 |
| 8 | 56358827 | rs1876230 | -1.682 | -0.226 | -2.191 | -0.214 | -0.071 | -0.105 | -0.105 | 0 | 0.7 | 0.95 | 0.73 | 0.64 | 0.66 | 0.68 |
| 8 | 56363158 | rs6473987 | 0.29 | -0.324 | -2.002 | -0.326 | -0.012 | -0.048 | -0.048 | 0 | 0.24 | 0.6 | 0.2 | 0.09 | 0.21 | 0.24 |
| 8 | 56363564 | rs2939632 | 0.212 | -0.16 | -1.935 | -0.162 | -0.003 | -0.033 | -0.033 | 0.037 | 0.16 | 0.09 | 0.33 | 0.43 | 0.14 | 0.24 |
| 8 | 56366735 | rs2929043 | -0.419 | 0.255 | -2.256 | -0.456 | -0.012 | -0.035 | -0.035 | 0 | 0.44 | 0.19 | 0.5 | 0.55 | 0.44 | 0.45 |
| 8 | 56371745 | rs2929032 | -1.085 | 0.173 | -2.272 | -0.551 | 0.006 | -0.003 | -0.003 | 1.009 | 0.52 | 0.34 | 0.6 | 0.55 | 0.53 | 0.53 |
| 8 | 56379772 | rs2975967 | 0.572 | 0.108 | -2.352 | -0.538 | -0.003 | 0.07 | 0.07 | 0 | 0.31 | 0.13 | 0.19 | 0.12 | 0.32 | 0.24 |
| 8 | 56380810 | rs2939654 | -0.26 | 0.164 | -2.178 | -0.466 | -0.006 | -0.026 | -0.026 | 0.58 | 0.44 | 0.23 | 0.51 | 0.55 | 0.44 | 0.45 |
| 8 | 56387829 | rs10088270 | -0.058 | 0.202 | -2.165 | -0.489 | 0.012 | -0.045 | -0.045 | 1.382 | 0.24 | 0.61 | 0.23 | 0.09 | 0.27 | 0.24 |
| 8 | 56388469 | rs2939661 | -1.996 | 0.161 | -2.29 | -0.474 | -0.035 | -0.106 | -0.106 | 0 | 0.72 | 0.95 | 0.75 | 0.63 | 0.72 | 0.71 |

| | | | | | | | | | | | | | | | | |
|---|---|---|---|---|---|---|---|---|---|---|---|---|---|---|---|---|
| 8 | 56396012 | rs10105874 | -0.342 | 0.319 | -1.761 | -0.397 | 0.005 | -0.031 | -0.031 | 1.892 | 0.25 | 0.67 | 0.23 | 0.09 | 0.27 | 0.25 |
| 8 | 56403385 | rs2975950 | -1.17 | -0.021 | -2.36 | -0.02 | -0.023 | -0.009 | -0.009 | 0 | 0.6 | 0.87 | 0.68 | 0.61 | 0.57 | 0.7 |
| 8 | 56404225 | rs2929019 | -0.215 | 0.475 | -1.513 | 0.051 | -0.011 | 0.002 | 0.002 | 0.034 | 0.11 | 0.1 | 0.13 | 0 | 0.17 | 0.1 |
| 8 | 56405514 | rs1395393 | -1.17 | 0.478 | -1.445 | 0.624 | -0.02 | -0.027 | -0.027 | 0.376 | 0.33 | 0.73 | 0.54 | 0.4 | 0.38 | 0.33 |
| 8 | 56405885 | rs2658911 | -0.588 | 0.466 | -1.336 | 0.81 | 0.024 | -0.019 | -0.019 | 0 | 0.28 | 0.62 | 0.28 | 0.14 | 0.37 | 0.21 |
| 8 | 56412075 | rs1395392 | -0.586 | 0.496 | -1.456 | 0.68 | -0.002 | 0.001 | 0.001 | 2.404 | 0.28 | 0.71 | 0.24 | 0.13 | 0.32 | 0.16 |
| 8 | 56420623 | rs16921942 | 0.075 | 0.49 | -1.72 | 0.58 | -0.007 | 0.033 | 0.033 | 0 | 0.16 | 0.06 | 0.11 | 0.07 | 0.15 | 0.19 |
| 8 | 56421026 | rs2622557 | 0.76 | 0.555 | -1.569 | 0.774 | -0.012 | -0.04 | -0.04 | 0 | 0.68 | 0.21 | 0.51 | 0.59 | 0.67 | 0.73 |
| 8 | 56425185 | rs2063026 | 0.25 | 0.55 | -1.114 | 0.762 | 0.035 | -0.038 | -0.038 | 0 | 0.73 | 0.47 | 0.77 | 0.87 | 0.68 | 0.84 |
| 8 | 56439158 | rs16921969 | 0.094 | 0.011 | -1.473 | 0.255 | 0.014 | 0.026 | 0.026 | 0 | 0.14 | 0.07 | 0.09 | 0.07 | 0.1 | 0.12 |
| 8 | 56441998 | rs7830792 | 0.2 | 0.365 | -0.842 | 0.568 | 0.046 | -0.013 | -0.013 | 0 | 0.73 | 0.54 | 0.74 | 0.8 | 0.67 | 0.79 |
| 8 | 56447358 | rs10504196 | 1.684 | 0.475 | -1.748 | 0.602 | -0.071 | -0.024 | -0.024 | 0 | 0.24 | 0.04 | 0.2 | 0.25 | 0.28 | 0.33 |
| 8 | 56448940 | rs6984950 | -0.12 | 0.377 | -1.338 | 0.841 | 0.036 | 0.031 | 0.031 | 0 | 0.43 | 0.25 | 0.33 | 0.34 | 0.36 | 0.36 |
| 8 | 56450064 | rs4641035 | -0.408 | 0.4 | -1.307 | 0.676 | 0.011 | -0.008 | -0.008 | 0.406 | 0.23 | 0.07 | 0.27 | 0.16 | 0.19 | 0.12 |
| 8 | 56450100 | rs4598235 | -0.265 | 0.353 | -0.443 | 0.644 | 0.038 | -0.014 | -0.014 | 0.06 | 0.83 | 0.69 | 0.82 | 0.88 | 0.79 | 0.85 |
| 8 | 56459785 | rs6988893 | -0.307 | 0.259 | -0.228 | 0.627 | 0.039 | -0.015 | -0.015 | 0 | 0.83 | 0.75 | 0.81 | 0.88 | 0.77 | 0.81 |
| 8 | 56460563 | rs10504197 | 2.081 | 0.007 | -0.678 | 0.627 | -0.057 | 0.028 | 0.028 | 0 | 0.29 | 0.05 | 0.23 | 0.26 | 0.32 | 0.3 |
| 8 | 56462049 | rs7001273 | 0.349 | -0.259 | -0.768 | 0.844 | 0.03 | -0.03 | -0.03 | 0.222 | 0.17 | 0.1 | 0.23 | 0.2 | 0.1 | 0.24 |
| 8 | 56462853 | rs4336589 | 0.986 | -0.326 | -0.501 | 0.893 | -0.001 | 0.028 | 0.028 | 0.242 | 0.26 | 0.36 | 0.28 | 0.31 | 0.25 | 0.25 |
| 8 | 56466441 | rs16922038 | -0.366 | -0.249 | -0.505 | 0.805 | -0.007 | 0.001 | 0.001 | 1.195 | 0.43 | 0.49 | 0.51 | 0.51 | 0.35 | 0.49 |
| 8 | 56467152 | rs10108855 | 0.274 | -0.097 | -0.62 | 0.845 | 0.005 | -0.003 | -0.003 | 0.013 | 0.11 | 0.11 | 0.06 | 0.1 | 0.11 | 0.03 |
| 8 | 56469668 | rs7000898 | -0.532 | 0.156 | -0.567 | 0.777 | 0.019 | -0.007 | -0.007 | 0 | 0.84 | 0.8 | 0.83 | 0.88 | 0.78 | 0.81 |
| 8 | 56491679 | rs11774480 | 0.046 | -0.018 | -0.883 | 1.08 | 0.02 | -0.047 | -0.047 | 0 | 0.17 | 0.1 | 0.25 | 0.28 | 0.12 | 0.23 |
| 8 | 56492754 | rs4236995 | -1.153 | -0.118 | -0.831 | 1.017 | 0.004 | 0.012 | 0.012 | 0 | 0.55 | 0.63 | 0.59 | 0.59 | 0.46 | 0.5 |
| 8 | 56493156 | rs4738279 | 0.462 | 0.548 | -0.313 | 1.511 | 0.051 | 0.114 | 0.114 | 0 | 0.79 | 0.58 | 0.61 | 0.56 | 0.71 | 0.68 |
| 8 | 56494044 | rs17438085 | 1.456 | 0.276 | -0.824 | 1.065 | -0.041 | 0.146 | 0.146 | 4.966 | 0.45 | 0.07 | 0.29 | 0.22 | 0.42 | 0.36 |
| 8 | 56506148 | rs9298536 | -0.006 | 0.397 | -0.524 | 1.192 | -0.01 | -0.023 | -0.023 | 0 | 0.07 | 0.04 | 0.19 | 0.16 | 0.06 | 0.15 |
| 8 | 56506851 | rs13280510 | 1.313 | 0.29 | -0.724 | 1.295 | -0.019 | -0.013 | -0.013 | 0 | 0.67 | 0.24 | 0.52 | 0.63 | 0.65 | 0.57 |
| 8 | 56506919 | rs6473999 | -0.382 | 0.298 | -0.559 | 1.161 | 0.088 | -0.002 | -0.002 | 0 | 0.84 | 0.55 | 0.8 | 0.87 | 0.78 | 0.81 |
| 8 | 56511944 | rs16922098 | -0.812 | 0.269 | -0.542 | 1.182 | -0.013 | 0.03 | 0.03 | 0.11 | 0.16 | 0.29 | 0.29 | 0.23 | 0.13 | 0.22 |
| 8 | 56514919 | rs10109697 | 0.749 | 0.295 | -1.347 | 1.203 | -0.019 | -0.001 | -0.001 | 0 | 0.68 | 0.26 | 0.51 | 0.66 | 0.65 | 0.6 |
| 8 | 56516943 | rs13258270 | 1.678 | 0.366 | -1.627 | 1.277 | -0.059 | -0.081 | -0.081 | 0.109 | 0.14 | 0.01 | 0.15 | 0.23 | 0.18 | 0.13 |
| 8 | 56517086 | rs17445056 | 1.847 | 0.24 | -1.557 | 1.187 | -0.032 | 0.083 | 0.083 | 2.415 | 0.38 | 0.06 | 0.27 | 0.22 | 0.37 | 0.32 |
| 8 | 56518537 | rs7007618 | -0.384 | 0.776 | -1.193 | 1.388 | -0.007 | -0.013 | -0.013 | 0 | 0.08 | 0.07 | 0.2 | 0.15 | 0.07 | 0.14 |
| 8 | 56521518 | rs6981643 | 1.463 | 0.723 | -1.331 | 1.411 | 0 | -0.058 | -0.058 | 0.106 | 0.45 | 0.07 | 0.31 | 0.44 | 0.39 | 0.36 |
| 8 | 56526439 | rs951167 | -0.555 | 0.782 | -0.837 | 1.051 | -0.001 | 0.003 | 0.003 | 0 | 0.08 | 0.11 | 0.2 | 0.15 | 0.07 | 0.14 |
| 8 | 56535193 | rs7003461 | 0.048 | 0.844 | -0.812 | 1.054 | -0.005 | -0.015 | -0.015 | 0 | 0.23 | 0.21 | 0.23 | 0.44 | 0.26 | 0.25 |
| 8 | 56535514 | rs6474006 | 0.975 | 0.813 | -0.838 | 0.612 | -0.021 | 0.021 | 0.021 | 1.105 | 0.61 | 0.29 | 0.47 | 0.67 | 0.56 | 0.49 |
| 8 | 56540235 | rs10107657 | -1.367 | 0.288 | -0.321 | 0.209 | -0.008 | -0.077 | -0.077 | 0 | 0.28 | 0.4 | 0.1 | 0.05 | 0.2 | 0.3 |
| 8 | 56547810 | rs13250111 | 0.166 | 0.247 | -0.361 | 0.198 | 0.001 | 0.017 | 0.017 | 0 | 0.12 | 0.1 | 0.04 | 0.04 | 0.08 | 0.16 |
| 8 | 56550203 | rs10108816 | -0.824 | 0.247 | -0.28 | 0.202 | -0.004 | -0.041 | -0.041 | 0 | 0.17 | 0.3 | 0.06 | 0.05 | 0.11 | 0.19 |
| 8 | 56567649 | rs4738382 | -1.866 | 0.233 | 0.042 | 0.499 | 0 | 0 | 0 | 0 | 0.32 | 0.72 | 0.13 | 0.3 | 0.22 | 0.39 |
| 8 | 56568582 | rs4075037 | -0.7 | 0.595 | -0.653 | 0.074 | 0.001 | -0.032 | -0.032 | 1.819 | 0.06 | 0.23 | 0.03 | 0 | 0.03 | 0.04 |
| 8 | 56589428 | rs7460004 | -0.93 | 0.761 | -0.975 | 0.225 | 0.002 | 0.006 | 0.006 | 0.016 | 0.45 | 0.44 | 0.33 | 0.39 | 0.38 | 0.45 |
| 8 | 56592291 | rs10086858 | -0.645 | 0.71 | -1.632 | 0.594 | -0.004 | -0.052 | -0.052 | 0 | 0.34 | 0.14 | 0.28 | 0.39 | 0.26 | 0.32 |
| 8 | 56593195 | rs7463339 | 0.018 | 0.699 | -1.546 | 0.895 | -0.005 | -0.091 | -0.091 | 0 | 0.23 | 0.06 | 0.24 | 0.34 | 0.18 | 0.17 |
| 8 | 56599548 | rs4737395 | 0.675 | 1.278 | -0.875 | 0.993 | 0.01 | 0.006 | 0.006 | 0 | 0.91 | 0.9 | 0.84 | 0.87 | 0.86 | 0.87 |
| 8 | 56599777 | rs4738393 | -0.094 | 1.345 | -0.963 | 0.925 | -0.002 | 0 | 0 | 0 | 0.26 | 0.29 | 0.37 | 0.48 | 0.3 | 0.47 |
| 8 | 56602838 | rs7011147 | -0.215 | 1.315 | -1.386 | 0.488 | -0.097 | 0.036 | 0.036 | 0 | 0.79 | 0.95 | 0.75 | 0.85 | 0.68 | 0.68 |
| 8 | 56606377 | rs6994234 | -0.203 | 0.396 | -0.587 | 0.506 | 0.001 | 0.001 | 0.001 | 0 | 0.48 | 0.5 | 0.33 | 0.41 | 0.41 | 0.35 |
| 8 | 56609388 | rs6982054 | 0.04 | 0.254 | -0.509 | 0.195 | 0.005 | -0.013 | -0.013 | 0 | 0.33 | 0.28 | 0.19 | 0.4 | 0.22 | 0.28 |
| 8 | 56611628 | rs3923066 | -0.837 | 0.093 | -0.005 | 0.189 | -0.026 | 0.004 | 0.004 | 0.12 | 0.15 | 0.44 | 0.08 | 0.14 | 0.09 | 0.14 |
| 8 | 56624934 | rs7830783 | -1.059 | 0.003 | 0.04 | 0.128 | -0.001 | 0.019 | 0.019 | 0.139 | 0.51 | 0.68 | 0.36 | 0.42 | 0.46 | 0.36 |
| 8 | 56639770 | rs10105904 | -0.955 | 0.13 | 0.249 | 0.227 | -0.016 | 0.009 | 0.009 | 0 | 0.14 | 0.33 | 0.07 | 0.14 | 0.1 | 0.13 |
| 8 | 56651746 | rs6997351 | 0.139 | 0.478 | -0.099 | 0.147 | -0.018 | 0.017 | 0.017 | 0 | 0.66 | 0.74 | 0.73 | 0.71 | 0.69 | 0.64 |
| 8 | 56656195 | rs7836019 | 0.301 | 0.466 | -0.143 | 0.109 | -0.004 | 0.004 | 0.004 | 0 | 0.9 | 0.92 | 0.89 | 0.99 | 0.89 | 0.84 |
| 8 | 56660557 | rs765158 | -0.025 | 0.359 | 0.224 | -0.086 | 0.002 | -0.004 | -0.004 | 1.018 | 0.33 | 0.31 | 0.27 | 0.27 | 0.37 | 0.24 |
| 8 | 56672303 | rs6474019 | 1.091 | 0.204 | 0.316 | -0.091 | -0.009 | -0.057 | -0.057 | 0 | 0.79 | 0.6 | 0.84 | 0.9 | 0.84 | 0.79 |
| 8 | 56678876 | rs13274924 | 0.32 | 0.098 | 0.34 | 0.427 | 0.005 | -0.068 | -0.068 | 0.104 | 0.88 | 0.72 | 0.88 | 0.99 | 0.89 | 0.85 |
| 8 | 56683297 | rs7016151 | 0.274 | 0.106 | 0.331 | 0.431 | 0.004 | -0.064 | -0.064 | 0 | 0.88 | 0.74 | 0.88 | 0.99 | 0.89 | 0.85 |
| 8 | 56684796 | rs13265995 | 1.583 | -0.081 | 0.177 | 0.632 | -0.094 | 0.022 | 0.022 | 0.13 | 0.41 | 0.06 | 0.35 | 0.27 | 0.46 | 0.38 |
| 8 | 56685933 | rs13276393 | 0.263 | 0.2 | 0.332 | -0.026 | 0.005 | -0.061 | -0.061 | 0.318 | 0.88 | 0.74 | 0.88 | 0.99 | 0.89 | 0.85 |
| 8 | 56686224 | rs1818 | 0.338 | 0.153 | 0.282 | -0.054 | 0.001 | -0.067 | -0.067 | 0 | 0.88 | 0.74 | 0.88 | 0.99 | 0.89 | 0.85 |
| 8 | 56696726 | rs7816281 | 1.199 | 0.1 | 0.348 | -0.014 | 0.003 | 0.013 | 0.013 | 0 | 0.45 | 0.38 | 0.57 | 0.61 | 0.47 | 0.57 |
| 8 | 56701983 | rs896126 | -1.238 | 0.115 | 0.093 | -0.089 | -0.06 | -0.068 | -0.068 | 1.405 | 0.78 | 0.97 | 0.85 | 0.7 | 0.8 | 0.8 |
| 8 | 56706994 | rs2316972 | 0.809 | 0.103 | 0.143 | -0.112 | -0.001 | 0.004 | 0.004 | 0 | 0.56 | 0.56 | 0.69 | 0.61 | 0.64 | 0.59 |
| 8 | 56708701 | rs10100659 | -0.6 | 0.105 | 0.18 | -0.121 | 0.002 | -0.071 | -0.071 | 0.017 | 0.13 | 0.28 | 0.12 | 0.01 | 0.11 | 0.15 |
| 8 | 56710778 | rs10091188 | -0.565 | 0.105 | 0.18 | -0.121 | -0.001 | -0.04 | -0.04 | 0.01 | 0.12 | 0.21 | 0.12 | 0.01 | 0.11 | 0.15 |
| 8 | 56722691 | rs16922271 | 0.274 | 0.103 | 0.152 | -0.121 | -0.005 | -0.001 | -0.001 | 0 | 0.09 | 0.12 | 0.03 | 0.09 | 0.05 | 0.06 |

| | | | | | | | | | | | | | | | | |
|---|---|---|---|---|---|---|---|---|---|---|---|---|---|---|---|---|
| 8 | 56723557 | rs7823773 | -0.657 | 0.187 | 0.044 | -0.099 | 0.002 | -0.071 | -0.071 | 0 | 0.13 | 0.28 | 0.12 | 0.01 | 0.11 | 0.15 |
| 8 | 56755804 | rs930679 | -1.075 | 0.218 | -0.358 | 0.08 | -0.046 | -0.056 | -0.056 | 0 | 0.78 | 0.96 | 0.85 | 0.7 | 0.8 | 0.8 |
| 8 | 56757808 | rs12155521 | 1.138 | 0.114 | -0.036 | -0.26 | 0 | -0.021 | -0.021 | 0 | 0.1 | 0.08 | 0.1 | 0 | 0.17 | 0.03 |
| 8 | 56774541 | rs7827790 | -0.001 | 0.059 | 0.092 | -0.164 | -0.001 | 0 | 0 | 0 | 0.1 | 0.21 | 0.06 | 0.09 | 0.08 | 0.06 |
| 8 | 56776963 | rs16922295 | -1.149 | -0.098 | -0.073 | -0.217 | 0.004 | 0.004 | 0.004 | 0 | 0.17 | 0.23 | 0.07 | 0.09 | 0.1 | 0.12 |
| 8 | 56785133 | rs6985703 | 0.502 | 0.283 | -0.078 | -0.332 | -0.002 | 0.025 | 0.025 | 0 | 0.55 | 0.64 | 0.61 | 0.77 | 0.48 | 0.62 |
| 8 | 56785685 | rs333613 | -0.084 | 0.113 | -0.511 | -0.488 | -0.078 | 0.132 | 0.132 | 0 | 0.64 | 0.91 | 0.73 | 0.8 | 0.61 | 0.76 |
| 8 | 56791244 | rs2271400 | -0.159 | 0.075 | -0.421 | -0.483 | 0.009 | -0.02 | -0.02 | 0 | 0.69 | 0.65 | 0.74 | 0.77 | 0.69 | 0.7 |
| 8 | 56793493 | rs6474026 | -0.09 | -0.022 | -0.517 | -0.2 | 0.002 | 0.003 | 0.003 | 0 | 0.2 | 0.18 | 0.15 | 0.13 | 0.2 | 0.25 |
| 8 | 56797440 | rs7834615 | 0.881 | 0.273 | -0.609 | -0.293 | 0 | 0 | 0 | 0 | 0.43 | 0.54 | 0.52 | 0.66 | 0.4 | 0.53 |
| 8 | 56800997 | rs333616 | -0.267 | 0.137 | -1.554 | -0.735 | -0.13 | 0.179 | 0.179 | 0 | 0.6 | 0.97 | 0.68 | 0.82 | 0.59 | 0.68 |
| 8 | 56803179 | rs1511723 | -0.569 | 0.421 | -0.846 | -0.645 | 0.006 | -0.02 | -0.02 | 0 | 0.92 | 0.87 | 0.89 | 0.98 | 0.92 | 0.87 |
| 8 | 56805477 | rs7820484 | -0.431 | 0.218 | -0.327 | -0.479 | -0.005 | 0.025 | 0.025 | 0 | 0.07 | 0.19 | 0.05 | 0.1 | 0.03 | 0.04 |
| 8 | 56814010 | rs1979128 | -0.121 | -0.204 | -0.462 | -1.165 | 0.03 | 0.073 | 0.073 | 0 | 0.5 | 0.58 | 0.34 | 0.22 | 0.39 | 0.31 |
| 8 | 56823191 | rs4551325 | 0.625 | -0.607 | -0.55 | -1.676 | 0.019 | 0.015 | 0.015 | 0 | 0.56 | 0.66 | 0.7 | 0.95 | 0.68 | 0.68 |
| 8 | 56823408 | rs4436100 | 0.679 | -0.633 | -0.564 | -1.692 | 0.02 | 0.018 | 0.018 | 0 | 0.56 | 0.66 | 0.7 | 0.95 | 0.68 | 0.68 |
| 8 | 56825385 | rs1397976 | 0.729 | -0.581 | -0.259 | -1.706 | 0.001 | -0.053 | -0.053 | 0 | 0.54 | 0.56 | 0.68 | 0.95 | 0.67 | 0.67 |
| 8 | 56826016 | rs6474033 | -0.679 | -0.607 | -0.279 | -1.549 | -0.002 | 0.034 | 0.034 | 0 | 0.87 | 0.96 | 0.87 | 0.97 | 0.89 | 0.83 |
| 8 | 56844876 | rs6985030 | 0.252 | -0.597 | -0.128 | -1.277 | 0 | -0.047 | -0.047 | 0 | 0.26 | 0.28 | 0.13 | 0 | 0.14 | 0.12 |
| 8 | 56846269 | rs16922425 | -0.56 | -0.08 | -0.126 | -0.897 | -0.001 | -0.003 | -0.003 | 0 | 0.53 | 0.55 | 0.38 | 0.23 | 0.52 | 0.32 |
| 8 | 56849386 | rs7829816 | -0.122 | -0.174 | -0.082 | -0.943 | -0.037 | -0.106 | -0.106 | 0.747 | 0.31 | 0.51 | 0.18 | 0.03 | 0.18 | 0.15 |
| 8 | 56854172 | rs16922459 | 0.536 | -0.113 | -0.639 | -0.839 | 0.012 | -0.016 | -0.016 | 1.646 | 0.23 | 0.17 | 0.12 | 0 | 0.15 | 0.12 |
| 8 | 56866780 | rs2292419 | 1.044 | 0.451 | -1.051 | -0.324 | -0.112 | -0.09 | -0.09 | 0.143 | 0.2 | 0.01 | 0.2 | 0.28 | 0.32 | 0.17 |
| 8 | 56867945 | rs7828258 | -0.71 | -0.027 | -0.495 | -0.766 | 0 | -0.009 | -0.009 | 0 | 0.54 | 0.53 | 0.37 | 0.26 | 0.52 | 0.32 |
| 8 | 56869605 | rs2668016 | 0.58 | -0.026 | -0.553 | -0.841 | -0.021 | -0.009 | -0.009 | 0.127 | 0.27 | 0.1 | 0.17 | 0.26 | 0.28 | 0.16 |
| 8 | 56872082 | rs2719268 | 0.704 | 0.018 | -0.56 | -0.834 | -0.039 | 0.003 | 0.003 | 0.892 | 0.26 | 0.06 | 0.16 | 0.24 | 0.28 | 0.16 |
| 8 | 56873547 | rs2668021 | 0.116 | 0.008 | -0.137 | -1.035 | -0.019 | -0.146 | -0.146 | 0 | 0.27 | 0.41 | 0.21 | 0.01 | 0.24 | 0.13 |
| 8 | 56874017 | rs2719267 | -0.752 | -0.053 | -0.037 | -0.969 | 0.004 | 0.016 | 0.016 | 0 | 0.51 | 0.43 | 0.33 | 0.27 | 0.46 | 0.26 |
| 8 | 56882471 | rs1027986 | -0.77 | 0.006 | -0.055 | -1.01 | -0.001 | -0.005 | -0.005 | 0 | 0.51 | 0.46 | 0.33 | 0.25 | 0.46 | 0.26 |
| 8 | 56883581 | rs2719244 | 0.375 | -0.071 | 0.034 | -1.088 | 0.011 | 0.016 | 0.016 | 0 | 0.26 | 0.18 | 0.21 | 0.01 | 0.24 | 0.13 |
| 8 | 56884089 | rs952784 | 0.442 | 0.016 | 0.002 | -0.948 | -0.001 | 0 | 0 | 0.532 | 0.25 | 0.25 | 0.13 | 0.24 | 0.22 | 0.13 |
| 8 | 56892230 | rs16922508 | -0.615 | -0.379 | 0.388 | -1.074 | -0.02 | -0.071 | -0.071 | 1.879 | 0.33 | 0.54 | 0.25 | 0.04 | 0.25 | 0.13 |
| 8 | 56897951 | rs2667978 | 0.521 | -0.485 | 0.138 | -1.136 | 0.014 | 0.048 | 0.048 | 0.093 | 0.24 | 0.14 | 0.15 | 0 | 0.21 | 0.09 |
| 8 | 56899136 | rs16920176 | -0.983 | -0.489 | 0.35 | -1.122 | -0.025 | 0.007 | 0.007 | 0.313 | 0.09 | 0.32 | 0.09 | 0.04 | 0.03 | 0.03 |
| 8 | 56903610 | rs9650314 | 0.763 | -0.419 | -0.082 | -1.13 | -0.021 | 0.148 | 0.148 | 0.394 | 0.22 | 0.02 | 0.15 | 0 | 0.21 | 0.09 |
| 8 | 56906068 | rs10282821 | 0.639 | -0.209 | -0.238 | -0.954 | -0.061 | -0.077 | -0.077 | 0 | 0.18 | 0.01 | 0.1 | 0.24 | 0.2 | 0.14 |
| 8 | 56910701 | rs4738466 | -1.476 | -0.229 | -0.056 | -1.261 | -0.009 | 0.008 | 0.008 | 0.558 | 0.9 | 0.94 | 0.88 | 0.97 | 0.87 | 0.81 |
| 8 | 56924362 | rs2719236 | -0.66 | -1.107 | 0.479 | -1.19 | -0.045 | -0.127 | -0.127 | 0.133 | 0.34 | 0.49 | 0.25 | 0.03 | 0.25 | 0.13 |
| 8 | 56941697 | rs2719260 | -0.556 | -1.191 | 0.422 | -1.258 | 0.025 | 0.022 | 0.022 | 0.021 | 0.33 | 0.22 | 0.22 | 0.03 | 0.26 | 0.14 |
| 8 | 56943139 | rs12674523 | 0.622 | -1.313 | 0.206 | -0.718 | 0.008 | -0.036 | -0.036 | 0.148 | 0.15 | 0.13 | 0.19 | 0.38 | 0.12 | 0.36 |
| 8 | 56945975 | rs6988853 | -0.545 | -0.705 | 0.474 | -0.91 | 0.024 | 0.022 | 0.022 | 0.072 | 0.33 | 0.22 | 0.22 | 0.03 | 0.26 | 0.14 |
| 8 | 56981194 | rs6474042 | -0.761 | -0.666 | 0.858 | -0.893 | 0.013 | -0.008 | -0.008 | 0 | 0.35 | 0.63 | 0.27 | 0.03 | 0.26 | 0.11 |
| 8 | 56995675 | rs4641033 | -0.833 | -0.508 | 0.862 | -0.702 | 0.019 | 0.021 | 0.021 | 0 | 0.07 | 0.31 | 0.09 | 0.03 | 0.05 | 0.04 |
| 8 | 56998131 | rs7825808 | -1.017 | -0.572 | 0.973 | -0.707 | 0.019 | 0.042 | 0.042 | 0.022 | 0.08 | 0.41 | 0.08 | 0.03 | 0.04 | 0.04 |
| 8 | 57012896 | rs2976020 | -0.734 | -0.707 | 0.939 | -0.912 | 0.009 | -0.016 | -0.016 | 0 | 0.35 | 0.61 | 0.27 | 0.03 | 0.26 | 0.11 |
| 8 | 57028107 | rs7015840 | -0.951 | -0.695 | 0.379 | -0.924 | -0.041 | -0.158 | -0.158 | 0 | 0.34 | 0.45 | 0.25 | 0.01 | 0.26 | 0.11 |
| 8 | 57035027 | rs4386955 | 1.309 | -0.58 | 0.078 | -0.313 | -0.029 | -0.006 | -0.006 | 0.003 | 0.05 | 0.02 | 0.16 | 0.21 | 0.1 | 0.21 |
| 8 | 57036929 | rs2976033 | -1.074 | -0.949 | 0.326 | -0.727 | -0.032 | -0.103 | -0.103 | 0 | 0.34 | 0.48 | 0.26 | 0.03 | 0.26 | 0.11 |
| 8 | 57054698 | rs4330666 | -0.787 | -0.134 | -0.168 | -0.272 | 0.001 | -0.002 | -0.002 | 0 | 0.42 | 0.4 | 0.52 | 0.38 | 0.42 | 0.31 |
| 8 | 57098250 | rs36077344 | 0.551 | 0.012 | -0.283 | -0.579 | 0.013 | 0.031 | 0.031 | 0 | 0.06 | 0.01 | 0.02 | 0 | 0.05 | 0.02 |
| 8 | 57100149 | rs7833986 | -0.42 | 0.087 | 0.257 | -0.536 | -0.017 | -0.066 | -0.066 | 0 | 0.22 | 0.34 | 0.17 | 0.08 | 0.2 | 0.15 |
| 8 | 57135889 | rs6984782 | -0.708 | -0.083 | 0.505 | -0.404 | -0.041 | -0.068 | -0.068 | 0 | 0.2 | 0.43 | 0.17 | 0.1 | 0.14 | 0.15 |
| 8 | 57138676 | rs36112366 | 0.349 | -0.006 | 0.038 | -0.35 | -0.009 | 0.015 | 0.015 | 0.434 | 0.14 | 0.01 | 0.1 | 0.1 | 0.12 | 0.14 |
| 8 | 57160043 | rs11994717 | -0.552 | 0.002 | 0.508 | -0.417 | -0.035 | -0.092 | -0.092 | 1.232 | 0.2 | 0.43 | 0.19 | 0.07 | 0.16 | 0.15 |
| 8 | 57179020 | rs7815788 | -0.267 | 0.002 | 0.7 | -0.417 | -0.005 | -0.02 | -0.02 | 0.067 | 0.19 | 0.23 | 0.17 | 0.06 | 0.15 | 0.15 |
| 8 | 57182507 | rs13270709 | -0.267 | 0.035 | 0.698 | -0.417 | -0.004 | -0.018 | -0.018 | 0.022 | 0.19 | 0.23 | 0.17 | 0.06 | 0.15 | 0.15 |
| 8 | 57193032 | rs13275320 | -0.58 | 0.025 | 0.744 | -0.423 | -0.031 | -0.09 | -0.09 | 0 | 0.2 | 0.43 | 0.18 | 0.06 | 0.15 | 0.15 |
| 8 | 57194486 | rs4075156 | -1.094 | 0.278 | 0.232 | -0.535 | -0.008 | -0.035 | -0.035 | 0.06 | 0.38 | 0.54 | 0.31 | 0.23 | 0.38 | 0.27 |
| 8 | 57195664 | rs7463906 | 0.611 | 0.341 | -0.158 | -0.496 | -0.01 | 0.005 | 0.005 | 1.897 | 0.17 | 0.1 | 0.13 | 0.17 | 0.23 | 0.12 |
| 8 | 57198214 | rs11990562 | 0.082 | 0.424 | -0.25 | -0.387 | 0.01 | 0.022 | 0.022 | 0.171 | 0.18 | 0.14 | 0.15 | 0.06 | 0.15 | 0.15 |
| 8 | 57209655 | rs6474057 | 0.24 | 0.405 | -0.209 | -0.369 | 0.003 | 0 | 0 | 0 | 0.17 | 0.16 | 0.15 | 0.06 | 0.14 | 0.14 |
| 8 | 57210606 | rs4077818 | 0.418 | 0.278 | -0.371 | -0.31 | -0.003 | -0.002 | -0.002 | 0.054 | 0.18 | 0.2 | 0.17 | 0.19 | 0.23 | 0.12 |
| 8 | 57211120 | rs13279861 | -0.706 | 0.302 | -0.142 | -0.264 | -0.016 | -0.03 | -0.03 | 0 | 0.36 | 0.58 | 0.34 | 0.25 | 0.36 | 0.25 |
| 8 | 57214372 | rs4409393 | 0.22 | 0.583 | -0.738 | 0.124 | -0.041 | -0.023 | -0.023 | 0 | 0.34 | 0.15 | 0.35 | 0.39 | 0.36 | 0.44 |
| 8 | 57214496 | rs12680392 | 0.211 | 0.173 | -0.18 | 0.329 | -0.011 | -0.012 | -0.012 | 0.491 | 0.2 | 0.24 | 0.17 | 0.15 | 0.16 | 0.17 |
| 8 | 57214729 | rs13282231 | 0.484 | 0.182 | -0.464 | 0.331 | 0.004 | 0.005 | 0.005 | 0 | 0.18 | 0.16 | 0.17 | 0.15 | 0.15 | 0.16 |
| 8 | 57216066 | rs4419802 | 0.223 | 0.373 | -0.248 | 0.378 | 0.005 | -0.012 | -0.012 | 0 | 0.24 | 0.45 | 0.2 | 0.2 | 0.27 | 0.15 |
| 8 | 57222136 | rs7011286 | -0.159 | 0.595 | -0.507 | 0.546 | 0.025 | 0.036 | 0.036 | 0 | 0.68 | 0.47 | 0.71 | 0.53 | 0.58 | 0.57 |
| 8 | 57231373 | rs11993636 | 0.452 | 0.427 | -0.603 | 0.517 | 0.002 | 0.022 | 0.022 | 1.135 | 0.19 | 0.13 | 0.15 | 0.12 | 0.18 | 0.09 |

| | | | | | | | | | | | | | | | | |
|---|---|---|---|---|---|---|---|---|---|---|---|---|---|---|---|---|
| 8 | 57274893 | rs4599805 | 0.659 | 0.448 | -0.543 | 0.517 | 0.003 | -0.005 | -0.005 | 0 | 0.4 | 0.27 | 0.48 | 0.37 | 0.33 | 0.37 |
| 8 | 57295200 | rs7002482 | -0.323 | 0.423 | -0.609 | 0.573 | 0.001 | 0.001 | 0.001 | 0.065 | 0.77 | 0.8 | 0.81 | 0.81 | 0.79 | 0.79 |
| 8 | 57298728 | rs16920516 | 0.257 | 0.625 | -0.454 | 0.571 | -0.064 | -0.047 | -0.047 | 0 | 0.35 | 0.17 | 0.3 | 0.43 | 0.46 | 0.42 |
| 8 | 57299548 | rs16920523 | 0.433 | 0.625 | -0.471 | 0.573 | -0.073 | -0.054 | -0.054 | 0.333 | 0.33 | 0.11 | 0.29 | 0.43 | 0.45 | 0.41 |
| 8 | 57303823 | rs4737430 | -0.036 | 0.466 | -0.418 | 0.495 | -0.001 | 0 | 0 | 0 | 0.6 | 0.43 | 0.65 | 0.57 | 0.54 | 0.56 |
| 8 | 57313632 | rs10108774 | 0.797 | 0.559 | -0.362 | 0.836 | -0.029 | -0.022 | -0.022 | 0 | 0.24 | 0.1 | 0.37 | 0.25 | 0.3 | 0.19 |
| 8 | 57313906 | rs12545109 | -0.081 | 0.326 | -0.213 | 0.843 | -0.004 | 0.006 | 0.006 | 0.147 | 0.72 | 0.62 | 0.73 | 0.7 | 0.77 | 0.8 |
| 8 | 57316247 | rs13253375 | 0.756 | 0.261 | -0.14 | 0.782 | 0.001 | 0.03 | 0.03 | 0 | 0.43 | 0.31 | 0.32 | 0.38 | 0.45 | 0.51 |
| 8 | 57334081 | rs17760660 | 0.929 | 0.341 | -0.171 | 0.758 | -0.022 | -0.014 | -0.014 | 1.352 | 0.24 | 0.12 | 0.36 | 0.24 | 0.3 | 0.2 |
| 8 | 57349056 | rs7007715 | 0.235 | 0.194 | 0.113 | 0.738 | 0.003 | 0.011 | 0.011 | 0 | 0.52 | 0.55 | 0.37 | 0.45 | 0.48 | 0.61 |
| 8 | 57352109 | rs13439178 | -0.724 | 0.017 | 0.329 | 0.664 | -0.034 | -0.011 | -0.011 | 0 | 0.09 | 0.27 | 0.04 | 0.06 | 0.03 | 0.11 |
| 8 | 57352136 | rs13262975 | 0.843 | 0.102 | 0.188 | 0.705 | -0.002 | 0.035 | 0.035 | 0 | 0.43 | 0.28 | 0.32 | 0.39 | 0.45 | 0.5 |
| 8 | 57356734 | rs1437277 | -0.727 | 0.051 | -0.01 | 0.83 | -0.05 | 0.022 | 0.022 | 0.083 | 0.8 | 0.95 | 0.82 | 0.89 | 0.78 | 0.85 |
| 8 | 57360485 | rs2609997 | 1.018 | -0.008 | -0.026 | 0.841 | -0.087 | -0.002 | -0.002 | 0 | 0.42 | 0.08 | 0.3 | 0.39 | 0.45 | 0.5 |
| 8 | 57365950 | rs2576581 | 0.752 | 0.036 | 0.318 | 0.875 | 0.002 | 0.033 | 0.033 | 0 | 0.46 | 0.27 | 0.34 | 0.45 | 0.47 | 0.59 |
| 8 | 57380064 | rs752388 | 0.855 | 0.055 | 0.224 | 0.842 | -0.007 | 0.044 | 0.044 | 0 | 0.44 | 0.22 | 0.32 | 0.39 | 0.45 | 0.5 |
| 8 | 57384360 | rs2438063 | -0.424 | 0.25 | 0.083 | 0.693 | -0.022 | 0.028 | 0.028 | 0.344 | 0.78 | 0.87 | 0.8 | 0.9 | 0.76 | 0.86 |
| 8 | 57385366 | rs2609983 | 0.298 | 0.23 | 0.153 | 0.734 | 0 | 0.003 | 0.003 | 0 | 0.52 | 0.59 | 0.43 | 0.66 | 0.49 | 0.64 |
| 8 | 57388410 | rs2576599 | -0.697 | 0.318 | -0.602 | 0.755 | -0.05 | 0.022 | 0.022 | 0.14 | 0.8 | 0.96 | 0.81 | 0.89 | 0.78 | 0.86 |
| 8 | 57392602 | rs2317370 | 1.028 | 0.295 | -0.554 | 0.753 | -0.014 | -0.015 | -0.015 | 0.075 | 0.25 | 0.11 | 0.35 | 0.23 | 0.28 | 0.21 |
| 8 | 57394467 | rs2609978 | -0.509 | 0.326 | -0.546 | 0.696 | -0.049 | 0.023 | 0.023 | 0.245 | 0.8 | 0.96 | 0.81 | 0.89 | 0.78 | 0.86 |
| 8 | 57413439 | rs2610045 | 0.397 | 0.126 | -0.018 | 0.993 | -0.003 | 0.014 | 0.014 | 0 | 0.71 | 0.68 | 0.72 | 0.57 | 0.75 | 0.7 |
| 8 | 57420256 | rs7011835 | 0.746 | 0.1 | -0.061 | 1.036 | -0.001 | -0.001 | -0.001 | 0 | 0.05 | 0.07 | 0.02 | 0.1 | 0.02 | 0.09 |
| 8 | 57423356 | rs9298554 | 1.053 | 0.119 | -0.114 | 0.97 | -0.01 | -0.011 | -0.011 | 0 | 0.24 | 0.15 | 0.35 | 0.23 | 0.28 | 0.21 |
| 8 | 57426778 | rs1877571 | 0.815 | 0.053 | -0.109 | 0.947 | -0.001 | 0.005 | 0.005 | 0 | 0.46 | 0.47 | 0.38 | 0.33 | 0.48 | 0.5 |
| 8 | 57429360 | rs1973628 | -0.981 | 0.179 | -0.382 | 0.823 | -0.057 | 0.007 | 0.007 | 0.075 | 0.82 | 0.98 | 0.85 | 0.89 | 0.81 | 0.89 |
| 8 | 57429516 | rs1012178 | 1.163 | 0.245 | -0.21 | 0.76 | 0 | 0.036 | 0.036 | 0 | 0.2 | 0.09 | 0.18 | 0.01 | 0.22 | 0.16 |
| 8 | 57431729 | rs1440757 | 0.704 | 0.293 | -0.176 | 0.89 | -0.013 | -0.002 | -0.002 | 0 | 0.24 | 0.34 | 0.25 | 0.22 | 0.23 | 0.21 |
| 8 | 57433697 | rs7018454 | 0.419 | 0.454 | -0.419 | 0.951 | -0.009 | -0.003 | -0.003 | 0 | 0.13 | 0.07 | 0.11 | 0.1 | 0.14 | 0.17 |
| 8 | 57435745 | rs6991165 | -0.413 | 0.368 | -0.397 | 0.836 | 0.007 | 0.14 | 0.14 | 0 | 0.2 | 0.43 | 0.32 | 0.34 | 0.18 | 0.23 |
| 8 | 57436577 | rs1866823 | 0.494 | 0.367 | -1.038 | 0.296 | 0.004 | 0.066 | 0.066 | 0 | 0.5 | 0.25 | 0.44 | 0.42 | 0.57 | 0.51 |
| 8 | 57440366 | rs2099573 | 0.452 | 0.237 | -0.821 | -0.027 | -0.006 | 0.032 | 0.032 | 0 | 0.36 | 0.26 | 0.47 | 0.21 | 0.42 | 0.33 |
| 8 | 57441010 | rs16920712 | 0.696 | 0.262 | -1.195 | 0.196 | -0.001 | 0.007 | 0.007 | 0 | 0.16 | 0.17 | 0.2 | 0.24 | 0.14 | 0.14 |
| 8 | 57450478 | rs7015773 | 0.436 | 0.236 | -1.08 | 0.165 | -0.001 | 0.032 | 0.032 | 0 | 0.18 | 0.28 | 0.23 | 0.24 | 0.14 | 0.14 |
| 8 | 57451312 | rs2610034 | 0.072 | 0.412 | -1.609 | 0.251 | -0.025 | -0.057 | -0.057 | 0.634 | 0.57 | 0.84 | 0.72 | 0.45 | 0.56 | 0.46 |
| 8 | 57461226 | rs2610027 | -0.249 | 0.74 | -1.506 | 0.093 | -0.04 | 0.053 | 0.053 | 0 | 0.85 | 0.99 | 0.9 | 0.93 | 0.87 | 0.82 |
| 8 | 57469913 | rs2011519 | 0.419 | 0.804 | -1.358 | 0.528 | -0.017 | -0.051 | -0.051 | 0 | 0.23 | 0.13 | 0.19 | 0.5 | 0.3 | 0.36 |
| 8 | 57477626 | rs2582413 | 0.905 | 0.212 | -1.615 | -0.197 | -0.022 | 0.087 | 0.087 | 0 | 0.41 | 0.17 | 0.49 | 0.19 | 0.47 | 0.34 |
| 8 | 57479269 | rs2610022 | -0.392 | 0.239 | -0.586 | -0.277 | 0.021 | -0.05 | -0.05 | 0 | 0.87 | 0.64 | 0.9 | 0.92 | 0.9 | 0.83 |
| 8 | 57480837 | rs2670040 | 0.832 | -0.14 | -0.426 | -0.344 | -0.001 | 0.033 | 0.033 | 0 | 0.43 | 0.33 | 0.5 | 0.19 | 0.47 | 0.29 |
| 8 | 57482100 | rs2610015 | -0.991 | -0.141 | -0.385 | -0.358 | -0.018 | 0.027 | 0.027 | 0 | 0.88 | 0.95 | 0.93 | 0.91 | 0.89 | 0.77 |
| 8 | 57484435 | rs6474070 | 1.342 | -0.011 | -0.355 | 0.01 | -0.007 | -0.021 | -0.021 | 0.004 | 0.24 | 0.14 | 0.19 | 0.4 | 0.28 | 0.36 |
| 8 | 57491459 | rs9298555 | 1.138 | 0.017 | -0.303 | 0.024 | -0.001 | -0.002 | -0.002 | 0.005 | 0.25 | 0.21 | 0.19 | 0.4 | 0.28 | 0.36 |
| 8 | 57496477 | rs2609982 | 0.518 | -0.017 | -0.678 | 0.081 | -0.001 | 0.014 | 0.014 | 0 | 0.66 | 0.55 | 0.69 | 0.59 | 0.74 | 0.64 |
| 8 | 57498784 | rs16920009 | 1.364 | 0.388 | -1.459 | 0.053 | -0.02 | -0.051 | -0.051 | 0.226 | 0.16 | 0.01 | 0.24 | 0.15 | 0.17 | 0.13 |
| 8 | 57504132 | rs10504206 | 1.155 | 0.596 | -0.907 | 0.376 | -0.007 | -0.013 | -0.013 | 0 | 0.13 | 0.1 | 0.17 | 0.33 | 0.13 | 0.08 |
| 8 | 57504505 | rs2670057 | -0.81 | 0.594 | -0.971 | 0.346 | -0.047 | 0.009 | 0.009 | 0 | 0.9 | 0.99 | 0.93 | 0.92 | 0.89 | 0.78 |
| 8 | 57505667 | rs2246935 | -0.076 | 0.533 | -0.795 | 0.325 | -0.017 | -0.028 | -0.028 | 0.043 | 0.76 | 0.8 | 0.76 | 0.58 | 0.76 | 0.7 |
| 8 | 57506561 | rs2582385 | 0.349 | 0.506 | -0.676 | -0.065 | 0.006 | 0.02 | 0.02 | 0 | 0.44 | 0.39 | 0.52 | 0.2 | 0.44 | 0.29 |
| 8 | 57507464 | rs10095976 | -0.116 | 0.563 | -0.617 | 0.136 | 0.004 | -0.01 | -0.01 | 0 | 0.4 | 0.45 | 0.41 | 0.66 | 0.42 | 0.45 |
| 8 | 57508107 | rs2582390 | 0.26 | 0.761 | -0.85 | 0.467 | -0.008 | -0.014 | -0.014 | 0.225 | 0.65 | 0.73 | 0.61 | 0.58 | 0.66 | 0.66 |
| 8 | 57508470 | rs12676353 | 1.028 | 0.549 | -1.365 | 0.88 | -0.022 | -0.075 | -0.075 | 1.814 | 0.12 | 0 | 0.05 | 0.25 | 0.1 | 0.2 |
| 8 | 57512904 | rs16920810 | 0.217 | 0.196 | -1.158 | 0.367 | 0.002 | 0.021 | 0.021 | 0 | 0.23 | 0.2 | 0.13 | 0.13 | 0.22 | 0.14 |
| 8 | 57512964 | rs1440752 | -0.738 | 0.231 | -1.359 | 0.396 | -0.05 | -0.084 | -0.084 | 0 | 0.86 | 0.98 | 0.73 | 0.72 | 0.84 | 0.78 |
| 8 | 57515963 | rs4737442 | 1.196 | -0.18 | -0.894 | 0.123 | -0.033 | -0.025 | -0.025 | 0 | 0.2 | 0.41 | 0.3 | 0.11 | 0.15 | 0.12 |
| 8 | 57520504 | rs10097622 | 0.355 | -0.198 | -0.292 | -0.026 | 0.002 | 0.008 | 0.008 | 0 | 0.56 | 0.44 | 0.4 | 0.67 | 0.57 | 0.65 |
| 8 | 57520600 | rs4738532 | 0.374 | -0.267 | -0.007 | -0.053 | -0.025 | -0.007 | -0.007 | 0.166 | 0.23 | 0.35 | 0.17 | 0.19 | 0.15 | 0.14 |
| 8 | 57521401 | rs16920843 | 2.241 | -0.247 | -0.08 | -0.056 | -0.011 | -0.036 | -0.036 | 0.208 | 0.08 | 0.15 | 0.03 | 0 | 0.05 | 0.03 |
| 8 | 57521816 | rs10958485 | 1.031 | -0.366 | 0.229 | -0.122 | -0.062 | -0.058 | -0.058 | 0 | 0.42 | 0.16 | 0.33 | 0.54 | 0.48 | 0.55 |
| 8 | 57524603 | rs1371938 | -0.01 | -0.348 | 0.22 | -0.109 | -0.018 | 0.003 | 0.003 | 0 | 0.57 | 0.7 | 0.47 | 0.61 | 0.53 | 0.59 |
| 8 | 57529208 | rs1606502 | 0.331 | -0.318 | 0.305 | -0.073 | 0.005 | 0.014 | 0.014 | 0 | 0.55 | 0.38 | 0.39 | 0.63 | 0.56 | 0.65 |
| 8 | 57533247 | rs4316149 | 0.859 | -0.233 | -0.118 | -0.09 | -0.009 | -0.029 | -0.029 | 0 | 0.25 | 0.33 | 0.26 | 0.09 | 0.2 | 0.12 |
| 8 | 57534550 | rs7004591 | -1.32 | -0.142 | -0.371 | 0.009 | -0.078 | -0.058 | -0.058 | 0 | 0.84 | 0.98 | 0.76 | 0.78 | 0.77 | 0.79 |
| 8 | 57540239 | rs1364958 | 0.433 | -0.309 | -0.085 | -0.049 | -0.034 | -0.005 | -0.005 | 0 | 0.31 | 0.47 | 0.35 | 0.28 | 0.23 | 0.21 |
| 8 | 57541988 | rs16920883 | 2.892 | -0.273 | -0.274 | -0.074 | -0.001 | -0.014 | -0.014 | 0 | 0.06 | 0.11 | 0.03 | 0 | 0.05 | 0.01 |
| 8 | 57555158 | rs1425590 | -0.549 | -0.273 | -0.213 | -0.076 | -0.005 | 0.004 | 0.004 | 0 | 0.67 | 0.7 | 0.53 | 0.69 | 0.63 | 0.69 |
| 8 | 57555815 | rs7002347 | 0.857 | -0.301 | -0.12 | 0.072 | -0.018 | -0.013 | -0.013 | 0.151 | 0.43 | 0.28 | 0.35 | 0.47 | 0.48 | 0.55 |
| 8 | 57555984 | rs828181 | -2.11 | -0.084 | -0.719 | -0.032 | -0.004 | -0.009 | -0.009 | 0 | 0.85 | 0.82 | 0.78 | 0.91 | 0.86 | 0.91 |
| 8 | 57557038 | rs10504207 | 1.947 | -0.031 | -0.683 | 0.126 | -0.006 | -0.021 | -0.021 | 0.017 | 0.11 | 0.03 | 0.05 | 0.13 | 0.09 | 0.1 |

| | | | | | | | | | | | | | | | | |
|---|---|---|---|---|---|---|---|---|---|---|---|---|---|---|---|---|
| 8 | 57559056 | rs10958490 | -0.05 | 0.023 | -0.469 | -0.055 | 0.001 | 0.013 | 0.013 | 0.065 | 0.58 | 0.57 | 0.5 | 0.67 | 0.57 | 0.67 |
| 8 | 57560790 | rs10096961 | 0.892 | 0.498 | -0.832 | -0.389 | -0.01 | 0.012 | 0.012 | 0 | 0.22 | 0.18 | 0.19 | 0.05 | 0.29 | 0.15 |
| 8 | 57561703 | rs1425593 | -0.388 | -0.011 | -0.823 | -0.168 | -0.005 | -0.013 | -0.013 | 0 | 0.63 | 0.54 | 0.51 | 0.68 | 0.67 | 0.58 |
| 8 | 57564454 | rs1425594 | 0.983 | 0.058 | -0.91 | 0.006 | 0.022 | 0.017 | 0.017 | 0 | 0.14 | 0.19 | 0.26 | 0.22 | 0.23 | 0.18 |
| 8 | 57565864 | rs12679072 | 1.166 | -0.383 | -1.328 | 0.054 | -0.045 | -0.075 | -0.075 | 0 | 0.3 | 0.06 | 0.17 | 0.38 | 0.34 | 0.31 |
| 8 | 57568699 | rs6474076 | -0.331 | -0.379 | -1.069 | -0.216 | -0.007 | 0.031 | 0.031 | 0 | 0.55 | 0.65 | 0.35 | 0.68 | 0.56 | 0.65 |
| 8 | 57568972 | rs16920907 | 1.117 | -0.343 | -1.122 | -0.031 | -0.013 | -0.019 | -0.019 | 0 | 0.08 | 0.05 | 0.09 | 0.24 | 0.12 | 0.17 |
| 8 | 57572882 | rs919022 | 0.257 | -0.39 | -0.924 | -0.337 | 0.004 | 0 | 0 | 1.536 | 0.47 | 0.55 | 0.25 | 0.44 | 0.44 | 0.48 |
| 8 | 57575455 | rs4574840 | -0.375 | -0.353 | -1.006 | -0.237 | -0.004 | 0.021 | 0.021 | 0 | 0.56 | 0.61 | 0.45 | 0.72 | 0.56 | 0.65 |
| 8 | 57575851 | rs1364956 | -2.059 | -0.27 | -1.486 | -0.325 | -0.069 | -0.004 | -0.004 | 0 | 0.84 | 0.98 | 0.69 | 0.91 | 0.8 | 0.91 |
| 8 | 57576173 | rs10095131 | 1.392 | -0.459 | -0.807 | -0.002 | -0.006 | -0.02 | -0.02 | 0.277 | 0.28 | 0.18 | 0.16 | 0.36 | 0.34 | 0.32 |
| 8 | 57585856 | rs2317630 | 1.351 | -0.461 | -0.808 | -0.004 | -0.002 | -0.007 | -0.007 | 0 | 0.28 | 0.23 | 0.16 | 0.36 | 0.34 | 0.32 |
| 8 | 57589763 | rs3930712 | -0.055 | -0.288 | -0.827 | -0.039 | -0.001 | 0.112 | 0.112 | 0 | 0.5 | 0.68 | 0.42 | 0.72 | 0.52 | 0.56 |
| 8 | 57599805 | rs10111407 | -0.371 | -0.092 | -0.793 | 0.091 | -0.02 | 0.047 | 0.047 | 0 | 0.57 | 0.7 | 0.44 | 0.69 | 0.54 | 0.61 |
| 8 | 57601930 | rs7005958 | -0.025 | -0.197 | -0.654 | -0.335 | 0.015 | 0.008 | 0.008 | 1.499 | 0.49 | 0.55 | 0.25 | 0.41 | 0.42 | 0.44 |
| 8 | 57609009 | rs16920917 | 0.781 | -0.272 | -0.318 | -0.16 | -0.047 | -0.036 | -0.036 | 0 | 0.13 | 0.26 | 0.06 | 0.05 | 0.03 | 0.12 |
| 8 | 57609632 | rs13255891 | 1.007 | 0.083 | -0.904 | 0.266 | -0.03 | -0.001 | -0.001 | 0.88 | 0.07 | 0.01 | 0.2 | 0.51 | 0.1 | 0.23 |
| 8 | 57609997 | rs6981525 | -0.896 | -0.133 | -1.13 | -0.692 | -0.025 | -0.01 | -0.01 | 0 | 0.56 | 0.77 | 0.49 | 0.58 | 0.44 | 0.66 |
| 8 | 57610626 | rs980698 | 0.201 | -0.043 | -0.244 | -0.537 | -0.001 | 0.037 | 0.037 | 0 | 0.26 | 0.27 | 0.3 | 0.62 | 0.22 | 0.36 |
| 8 | 57610697 | rs980699 | -0.65 | -0.128 | -0.404 | -1.024 | -0.028 | 0.059 | 0.059 | 0 | 0.61 | 0.69 | 0.49 | 0.89 | 0.55 | 0.71 |
| 8 | 57632308 | rs16920935 | 0.311 | 0.234 | -0.218 | -0.732 | -0.002 | -0.005 | -0.005 | 0 | 0.25 | 0.18 | 0.3 | 0.62 | 0.23 | 0.31 |
| 8 | 57633546 | rs7827783 | 1.096 | 0.112 | -0.299 | -0.881 | -0.012 | -0.011 | -0.011 | 0 | 0.28 | 0.4 | 0.16 | 0.27 | 0.3 | 0.33 |
| 8 | 57637233 | rs1834413 | -0.17 | 0.011 | -0.743 | -1.31 | -0.015 | 0.148 | 0.148 | 1.808 | 0.53 | 0.77 | 0.47 | 0.88 | 0.51 | 0.62 |
| 8 | 57644072 | rs1014086 | 0.439 | 0.535 | -0.17 | -0.678 | 0.046 | 0.254 | 0.254 | 0.05 | 0.07 | 0.17 | 0.23 | 0.53 | 0.1 | 0.22 |
| 8 | 57645857 | rs1972688 | -0.755 | 0.437 | -1.574 | -1.505 | -0.029 | 0.049 | 0.049 | 0 | 0.83 | 0.98 | 0.7 | 0.94 | 0.83 | 0.92 |
| 8 | 57650265 | rs353713 | -0.054 | 0.822 | -1.058 | -1.311 | -0.027 | 0.064 | 0.064 | 0 | 0.23 | 0.07 | 0.23 | 0 | 0.25 | 0.17 |
| 8 | 57655067 | rs10958491 | 1.317 | 0.91 | -0.84 | -0.699 | -0.035 | -0.033 | -0.033 | 0 | 0.08 | 0 | 0.06 | 0.15 | 0.09 | 0.17 |
| 8 | 57659644 | rs17785543 | 0.905 | 0.894 | -0.947 | -0.748 | -0.01 | 0.004 | 0.004 | 0 | 0.2 | 0.11 | 0.11 | 0.14 | 0.25 | 0.21 |
| 8 | 57659691 | rs1834414 | -0.362 | 0.712 | -0.506 | -0.917 | 0.002 | 0.01 | 0.01 | 0 | 0.38 | 0.64 | 0.38 | 0.73 | 0.36 | 0.46 |
| 8 | 57666538 | rs17785796 | 0.947 | 0.672 | -0.825 | -0.906 | 0.003 | 0.043 | 0.043 | 0 | 0.27 | 0.14 | 0.18 | 0.22 | 0.24 | 0.26 |
| 8 | 57667226 | rs918909 | 0.08 | 0.691 | -0.152 | -0.463 | 0.001 | 0.209 | 0.209 | 0 | 0.15 | 0.42 | 0.26 | 0.58 | 0.11 | 0.24 |
| 8 | 57676537 | rs353753 | -0.369 | 0.922 | -0.48 | -0.446 | 0.011 | 0.054 | 0.054 | 0 | 0.85 | 0.76 | 0.72 | 0.42 | 0.79 | 0.76 |
| 8 | 57683021 | rs353749 | -0.406 | 0.918 | -0.41 | -0.448 | 0.019 | 0.08 | 0.08 | 0 | 0.85 | 0.75 | 0.71 | 0.42 | 0.79 | 0.76 |
| 8 | 57683880 | rs9298556 | 0.373 | 0.711 | -0.888 | -0.976 | -0.012 | -0.037 | -0.037 | 1.177 | 0.38 | 0.51 | 0.2 | 0.14 | 0.36 | 0.33 |
| 8 | 57688486 | rs114898 | -0.537 | 0.967 | -0.89 | -0.3 | -0.025 | -0.036 | -0.036 | 0 | 0.87 | 0.92 | 0.72 | 0.42 | 0.79 | 0.76 |
| 8 | 57692649 | rs353721 | 0.356 | 0.97 | -1.014 | -0.754 | -0.072 | 0.058 | 0.058 | 0 | 0.18 | 0.03 | 0.32 | 0.06 | 0.19 | 0.18 |
| 8 | 57699282 | rs181333 | -0.405 | 0.678 | -0.27 | -0.859 | -0.022 | -0.117 | -0.117 | 0.022 | 0.24 | 0.48 | 0.37 | 0.06 | 0.17 | 0.21 |
| 8 | 57700914 | rs353731 | 0.286 | 0.699 | -0.576 | -0.848 | -0.017 | 0.072 | 0.072 | 0 | 0.19 | 0.07 | 0.33 | 0.06 | 0.16 | 0.21 |
| 8 | 57701557 | rs353734 | -0.019 | 0.578 | -0.675 | -0.543 | -0.04 | -0.075 | -0.075 | 1.549 | 0.81 | 0.96 | 0.72 | 0.42 | 0.74 | 0.72 |
| 8 | 57707934 | rs1820391 | -0.811 | 0.428 | -0.334 | -0.663 | -0.044 | -0.098 | -0.098 | 0.134 | 0.18 | 0.6 | 0.11 | 0 | 0.11 | 0.1 |
| 8 | 57709500 | rs16921065 | 0.78 | 0.427 | -0.099 | -0.664 | -0.014 | -0.058 | -0.058 | 0 | 0.11 | 0.23 | 0.06 | 0 | 0.09 | 0.07 |
| 8 | 57709567 | rs2432730 | -0.617 | 0.631 | -1.025 | -0.76 | -0.018 | 0.072 | 0.072 | 0 | 0.8 | 0.93 | 0.66 | 0.94 | 0.8 | 0.8 |
| 8 | 57712283 | rs7821860 | 1.349 | 0.801 | -1.045 | -0.464 | -0.015 | 0.042 | 0.042 | 0.573 | 0.52 | 0.25 | 0.29 | 0.39 | 0.49 | 0.48 |
| 8 | 57718944 | rs2911710 | -0.224 | 0.92 | -0.52 | 0.285 | -0.025 | 0.002 | 0.002 | 1.96 | 0.1 | 0.04 | 0.25 | 0.55 | 0.2 | 0.23 |
| 8 | 57731917 | rs7004092 | -1.144 | -0.034 | -0.206 | -0.358 | -0.062 | -0.078 | -0.078 | 0 | 0.14 | 0.62 | 0.1 | 0 | 0.05 | 0.05 |
| 8 | 57755644 | rs1834197 | -0.392 | 0.487 | -0.753 | 0.39 | -0.013 | -0.039 | -0.039 | 1.969 | 0.2 | 0.15 | 0.41 | 0.59 | 0.2 | 0.43 |
| 8 | 57764223 | rs11994433 | -1.542 | 0.554 | -0.053 | 0.01 | 0.011 | 0.016 | 0.016 | 0.148 | 0.33 | 0.45 | 0.5 | 0.66 | 0.3 | 0.58 |
| 8 | 57767866 | rs7812952 | -1.108 | 0.543 | 0.017 | -0.015 | 0.024 | -0.052 | -0.052 | 0.282 | 0.13 | 0.38 | 0.08 | 0.05 | 0.1 | 0.15 |
| 8 | 57769001 | rs889982 | -1.488 | 0.142 | -0.131 | -0.191 | -0.001 | 0.009 | 0.009 | 0 | 0.45 | 0.55 | 0.62 | 0.9 | 0.43 | 0.69 |
| 8 | 57769505 | rs745623 | -1.498 | 0.106 | -0.15 | -0.21 | -0.001 | 0.011 | 0.011 | 0 | 0.45 | 0.55 | 0.62 | 0.9 | 0.43 | 0.69 |
| 8 | 57773798 | rs12682379 | -0.116 | -0.141 | -0.178 | -0.207 | 0.002 | 0.012 | 0.012 | 0.339 | 0.3 | 0.36 | 0.51 | 0.63 | 0.29 | 0.4 |
| 8 | 57775012 | rs2873550 | 0.541 | 0.048 | -0.445 | -0.034 | -0.012 | -0.025 | -0.025 | 0 | 0.24 | 0.17 | 0.47 | 0.59 | 0.27 | 0.38 |
| 8 | 57775276 | rs16921224 | -0.28 | 0.09 | -0.422 | 0.117 | -0.003 | 0.006 | 0.006 | 0 | 0.19 | 0.21 | 0.15 | 0.3 | 0.19 | 0.24 |
| 8 | 57775837 | rs873374 | -0.774 | 0.097 | -0.522 | 0.2 | 0.002 | -0.01 | -0.01 | 0.062 | 0.24 | 0.2 | 0.13 | 0.41 | 0.24 | 0.36 |
| 8 | 57776217 | rs889979 | 0.049 | -0.113 | -1.152 | -0.991 | -0.006 | 0.09 | 0.09 | 0 | 0.43 | 0.17 | 0.49 | 0.65 | 0.4 | 0.45 |
| 8 | 57777258 | rs2279970 | -0.907 | -0.197 | -1.074 | -1.089 | 0.002 | -0.05 | -0.05 | 0.014 | 0.6 | 0.43 | 0.69 | 0.89 | 0.57 | 0.69 |
| 8 | 57777574 | rs889974 | 0.21 | -0.202 | -1.193 | -1.051 | -0.044 | 0.075 | 0.075 | 0 | 0.41 | 0.14 | 0.6 | 0.65 | 0.4 | 0.43 |
| 8 | 57790753 | rs13268762 | 0.211 | -0.283 | -1.296 | -1.164 | 0.003 | 0.005 | 0.005 | 0 | 0.12 | 0.1 | 0.07 | 0 | 0.13 | 0.07 |
| 8 | 57792222 | rs4304324 | 0.234 | -0.189 | -1.094 | -0.99 | -0.015 | -0.004 | -0.004 | 0 | 0.19 | 0.3 | 0.1 | 0.24 | 0.17 | 0.26 |
| 8 | 57799184 | rs13272502 | -1.433 | 0.542 | -1.046 | -1.18 | -0.013 | 0.02 | 0.02 | 0 | 0.83 | 0.87 | 0.84 | 0.91 | 0.77 | 0.86 |
| 8 | 57799364 | rs17789704 | 0.81 | -0.027 | -0.682 | -0.882 | -0.003 | -0.071 | -0.071 | 0 | 0.22 | 0.3 | 0.12 | 0.02 | 0.2 | 0.17 |
| 8 | 57804484 | rs1911694 | -0.1 | -0.271 | -0.624 | -0.332 | -0.001 | -0.004 | -0.004 | 0 | 0.51 | 0.45 | 0.55 | 0.66 | 0.49 | 0.45 |
| 8 | 57811629 | rs11786143 | 0.791 | -0.924 | -0.681 | 0.21 | 0.011 | -0.013 | -0.013 | 0 | 0.12 | 0.08 | 0.06 | 0.2 | 0.1 | 0.09 |
| 8 | 57817498 | rs1508086 | -0.812 | -1.062 | -0.479 | -0.007 | -0.017 | 0.049 | 0.049 | 0 | 0.6 | 0.68 | 0.6 | 0.81 | 0.55 | 0.48 |
| 8 | 57822303 | rs11991623 | 0.714 | -1.177 | -0.433 | 0.062 | 0.027 | 0.028 | 0.028 | 0 | 0.15 | 0.07 | 0.08 | 0.06 | 0.14 | 0.11 |
| 8 | 57822720 | rs11992437 | -0.346 | -1.23 | -0.358 | 0.088 | 0.016 | 0.006 | 0.006 | 0 | 0.21 | 0.19 | 0.1 | 0.21 | 0.17 | 0.14 |
| 8 | 57823833 | rs10096356 | 0.651 | -1.069 | -0.169 | 0.049 | -0.024 | -0.025 | -0.025 | 0 | 0.15 | 0.2 | 0.08 | 0.06 | 0.15 | 0.11 |
| 8 | 57826145 | rs7816057 | 1.369 | -0.816 | -0.197 | 0.229 | 0.018 | 0.026 | 0.026 | 1.322 | 0.26 | 0.28 | 0.44 | 0.4 | 0.32 | 0.25 |
| 8 | 57834357 | rs4738555 | -0.442 | -0.776 | -0.444 | 0.2 | -0.009 | 0.061 | 0.061 | 0 | 0.57 | 0.67 | 0.6 | 0.82 | 0.56 | 0.48 |

| | | | | | | | | | | | | | | | | |
|---|---|---|---|---|---|---|---|---|---|---|---|---|---|---|---|---|
| 8 | 57835489 | rs17187027 | 0.585 | -1.149 | -0.422 | 0.574 | 0.005 | 0.004 | 0.004 | 0 | 0.1 | 0.06 | 0.05 | 0.07 | 0.1 | 0.07 |
| 8 | 57838101 | rs6992067 | -1.225 | -1.067 | -0.425 | 0.779 | 0.003 | -0.002 | -0.002 | 0 | 0.16 | 0.18 | 0.08 | 0.38 | 0.14 | 0.14 |
| 8 | 57839087 | rs9643495 | 0.671 | -0.589 | -0.177 | 0.712 | 0 | 0 | 0 | 1.572 | 0.61 | 0.64 | 0.71 | 0.63 | 0.56 | 0.53 |
| 8 | 57841969 | rs7820756 | -0.042 | -0.99 | 0.244 | 0.602 | -0.038 | -0.014 | -0.014 | 0 | 0.17 | 0.42 | 0.1 | 0.14 | 0.12 | 0.15 |
| 8 | 57848219 | rs16921401 | 0.319 | -1.066 | 0.567 | 0.719 | -0.027 | 0.024 | 0.024 | 0.109 | 0.15 | 0.24 | 0.14 | 0.16 | 0.11 | 0.18 |
| 8 | 57853143 | rs7835887 | -0.239 | -1.169 | 0.452 | 0.845 | -0.047 | 0.016 | 0.016 | 0.037 | 0.18 | 0.36 | 0.16 | 0.16 | 0.12 | 0.19 |
| 8 | 57861021 | rs1508088 | -0.333 | -1.251 | 0.22 | 0.793 | -0.001 | 0.005 | 0.005 | 0 | 0.89 | 0.96 | 0.94 | 0.89 | 0.87 | 0.88 |
| 8 | 57866883 | rs6474091 | -0.997 | -1.157 | 0.43 | 0.614 | -0.002 | -0.05 | -0.05 | 0 | 0.38 | 0.48 | 0.3 | 0.19 | 0.37 | 0.39 |
| 8 | 57870451 | rs12550392 | 1.575 | -2.079 | 0.008 | 0.796 | 0.039 | -0.004 | -0.004 | 0 | 0.58 | 0.38 | 0.62 | 0.57 | 0.71 | 0.59 |
| 8 | 57870847 | rs8718 | 0.055 | -1.467 | -0.406 | 0.862 | -0.001 | -0.001 | -0.001 | 0 | 0.87 | 0.88 | 0.94 | 0.88 | 0.87 | 0.85 |
| 8 | 57871462 | rs1044731 | 0.461 | -1.474 | -0.306 | 0.947 | -0.028 | 0.023 | 0.023 | 0.02 | 0.14 | 0.19 | 0.13 | 0.18 | 0.1 | 0.17 |
| 8 | 57874064 | rs1396115 | -0.58 | -1.496 | -0.304 | 1.286 | -0.023 | 0.069 | 0.069 | 0 | 0.19 | 0.23 | 0.29 | 0.43 | 0.17 | 0.25 |
| 8 | 57875588 | rs10003 | 0.437 | -1.436 | -0.479 | 1.246 | -0.023 | 0.017 | 0.017 | 0.03 | 0.14 | 0.16 | 0.13 | 0.18 | 0.1 | 0.17 |
| 8 | 57887848 | rs13271029 | 1.987 | -1.081 | -1.069 | 1.097 | -0.124 | -0.021 | -0.021 | 0 | 0.41 | 0.08 | 0.45 | 0.33 | 0.47 | 0.39 |
| 8 | 57901469 | rs10088040 | 0.489 | -0.81 | -0.326 | 0.935 | 0 | -0.001 | -0.001 | 0.242 | 0.85 | 0.87 | 0.93 | 0.88 | 0.86 | 0.84 |
| 8 | 57917944 | rs4738559 | -1.283 | -1.153 | -0.466 | 1.244 | 0.014 | -0.039 | -0.039 | 0.012 | 0.37 | 0.23 | 0.4 | 0.5 | 0.37 | 0.35 |
| 8 | 57918262 | rs7844631 | -0.96 | -0.998 | -0.131 | 1.247 | -0.002 | -0.009 | -0.009 | 0 | 0.25 | 0.14 | 0.21 | 0.23 | 0.29 | 0.25 |
| 8 | 57926726 | rs2089515 | -1.297 | -0.824 | -0.124 | 0.916 | -0.015 | 0.016 | 0.016 | 0 | 0.26 | 0.12 | 0.3 | 0.21 | 0.33 | 0.25 |
| 8 | 57942358 | rs2119783 | -2.402 | -1.036 | 0.61 | 0.976 | 0.011 | -0.009 | -0.009 | 0 | 0.43 | 0.58 | 0.51 | 0.52 | 0.4 | 0.49 |
| 8 | 57943607 | rs1561785 | -2.138 | -1.085 | 0.78 | 0.959 | -0.005 | 0.02 | 0.02 | 0 | 0.35 | 0.51 | 0.48 | 0.46 | 0.37 | 0.31 |
| 8 | 57944932 | rs16920046 | 0.09 | -1.084 | 0.847 | 0.92 | -0.001 | -0.002 | -0.002 | 0.364 | 0.1 | 0.11 | 0.09 | 0.05 | 0.11 | 0.08 |
| 8 | 57947475 | rs10958518 | -2.504 | -1.086 | 0.871 | 0.922 | -0.005 | 0.023 | 0.023 | 0 | 0.35 | 0.49 | 0.48 | 0.46 | 0.37 | 0.31 |
| 8 | 57952770 | rs4738563 | -0.324 | -1.091 | 0.888 | 0.966 | -0.006 | 0.024 | 0.024 | 0 | 0.11 | 0.14 | 0.22 | 0.27 | 0.12 | 0.11 |
| 8 | 57955330 | rs1372506 | -2.52 | -1.108 | 0.876 | 0.737 | -0.006 | 0.01 | 0.01 | 0 | 0.35 | 0.51 | 0.35 | 0.42 | 0.37 | 0.31 |
| 8 | 57960820 | rs10092697 | -2.523 | -1.064 | 0.889 | 0.233 | -0.007 | 0.001 | 0.001 | 0 | 0.35 | 0.49 | 0.3 | 0.39 | 0.37 | 0.31 |
| 8 | 57962459 | rs6998250 | -2.944 | -1.086 | 0.934 | 0.222 | 0.015 | 0.011 | 0.011 | 0 | 0.43 | 0.61 | 0.33 | 0.44 | 0.4 | 0.49 |
| 8 | 57965003 | rs1442123 | -2.504 | -1.057 | 0.93 | 0.237 | -0.006 | 0.001 | 0.001 | 0 | 0.34 | 0.48 | 0.3 | 0.39 | 0.36 | 0.31 |
| 8 | 57971479 | rs16921525 | -0.24 | -0.941 | 0.968 | 0.234 | -0.01 | -0.014 | -0.014 | 2.127 | 0.11 | 0.16 | 0.09 | 0.05 | 0.11 | 0.06 |
| 8 | 57973339 | rs17793486 | -0.098 | -0.927 | 0.946 | 0.241 | -0.001 | -0.001 | -0.001 | 0 | 0.1 | 0.11 | 0.08 | 0.05 | 0.11 | 0.06 |
| 8 | 57976625 | rs1992461 | -2.165 | -0.953 | 1.255 | 0.186 | -0.038 | -0.046 | -0.046 | 2.314 | 0.17 | 0.28 | 0.09 | 0.09 | 0.1 | 0.26 |
| 8 | 57977246 | rs7819796 | -2.06 | -0.986 | 1.246 | 0.254 | -0.047 | 0.041 | 0.041 | 0 | 0.22 | 0.48 | 0.17 | 0.28 | 0.19 | 0.19 |
| 8 | 57977503 | rs6474097 | -2.604 | -0.906 | 1.18 | 0.267 | 0.008 | 0.001 | 0.001 | 0 | 0.4 | 0.61 | 0.33 | 0.46 | 0.39 | 0.44 |
| 8 | 57979314 | rs16921553 | -0.836 | -0.3 | 1.139 | 0.33 | -0.008 | 0.003 | 0.003 | 0 | 0.12 | 0.09 | 0.14 | 0.13 | 0.18 | 0.07 |
| 8 | 57979584 | rs7825430 | -2.443 | -0.36 | 1.2 | 0.321 | 0.008 | -0.036 | -0.036 | 0 | 0.25 | 0.36 | 0.23 | 0.2 | 0.28 | 0.32 |
| 8 | 57981470 | rs10504216 | -0.001 | 0.11 | 1.207 | 0.559 | -0.049 | -0.035 | -0.035 | 0 | 0.08 | 0.02 | 0.11 | 0.15 | 0.17 | 0.06 |
| 8 | 57990401 | rs4361758 | -2.181 | 0.075 | 1.304 | 0.549 | 0.023 | 0.06 | 0.06 | 0.023 | 0.22 | 0.34 | 0.33 | 0.37 | 0.26 | 0.23 |
| 8 | 57992193 | rs10111393 | -1.552 | -0.031 | 1.36 | 0.439 | 0.004 | -0.012 | -0.012 | 0 | 0.1 | 0.27 | 0.07 | 0.04 | 0.08 | 0.12 |
| 8 | 57994818 | rs10107655 | -1.458 | -0.039 | 1.375 | 0.435 | 0.003 | -0.013 | -0.013 | 0 | 0.1 | 0.27 | 0.07 | 0.04 | 0.08 | 0.12 |
| 8 | 57995439 | rs11990366 | -1.216 | 0.135 | 1.525 | 0.586 | 0.027 | 0.024 | 0.024 | 0 | 0.08 | 0.27 | 0.05 | 0.04 | 0.05 | 0.09 |
| 8 | 58005923 | rs7835839 | -0.866 | 0.143 | 1.431 | 0.59 | 0.016 | 0.014 | 0.014 | 0 | 0.07 | 0.2 | 0.03 | 0.04 | 0.05 | 0.09 |
| 8 | 58008281 | rs7833294 | -2.786 | 0.213 | 1.397 | 0.737 | -0.019 | 0.149 | 0.149 | 0.81 | 0.07 | 0.24 | 0.17 | 0.19 | 0.01 | 0.05 |
| 8 | 58018805 | rs6994162 | -2.815 | 0.221 | 1.401 | 0.741 | -0.02 | 0.146 | 0.146 | 0.941 | 0.07 | 0.24 | 0.17 | 0.18 | 0.01 | 0.05 |
| 8 | 58024168 | rs2317990 | -1.051 | 0.435 | 1.126 | 0.773 | -0.034 | -0.043 | -0.043 | 0.027 | 0.11 | 0.06 | 0.26 | 0.33 | 0.18 | 0.11 |
| 8 | 58031161 | rs6474100 | -1.654 | 0.433 | 1.355 | 0.94 | 0.033 | 0.067 | 0.067 | 0 | 0.13 | 0.19 | 0.28 | 0.33 | 0.19 | 0.11 |
| 8 | 58037522 | rs10504217 | -1.655 | 0.438 | 1.357 | 0.943 | 0.033 | 0.067 | 0.067 | 0 | 0.13 | 0.19 | 0.28 | 0.33 | 0.19 | 0.11 |
| 8 | 58051471 | rs10958522 | -1.948 | 0.316 | 1.2 | 0.839 | 0.008 | 0.015 | 0.015 | 0 | 0.22 | 0.31 | 0.36 | 0.35 | 0.29 | 0.18 |
| 8 | 58051891 | rs12679963 | -1.332 | 0.1 | 1.242 | 0.852 | -0.026 | 0.04 | 0.04 | 0 | 0.13 | 0.26 | 0.23 | 0.2 | 0.11 | 0.12 |
| 8 | 58057655 | rs16921695 | -1.578 | 0.341 | 0.968 | 0.1 | 0.014 | -0.031 | -0.031 | 0.141 | 0.25 | 0.58 | 0.24 | 0.18 | 0.28 | 0.2 |
| 8 | 58065790 | rs7017199 | -1.589 | 0.306 | 0.967 | 0.099 | 0.012 | -0.031 | -0.031 | 0 | 0.25 | 0.58 | 0.24 | 0.18 | 0.28 | 0.2 |
| 8 | 58077248 | rs7006508 | -1.739 | 0.353 | 1.143 | 0.089 | 0.011 | -0.035 | -0.035 | 0 | 0.24 | 0.53 | 0.23 | 0.18 | 0.26 | 0.2 |
| 8 | 58084006 | rs6995671 | -1.642 | 0.307 | 1.135 | 0.089 | 0.012 | -0.031 | -0.031 | 0 | 0.25 | 0.58 | 0.24 | 0.18 | 0.28 | 0.2 |
| 8 | 58087231 | rs7841086 | -1.629 | 0.307 | 1.135 | 0.089 | 0.012 | -0.031 | -0.031 | 0.003 | 0.25 | 0.58 | 0.24 | 0.18 | 0.28 | 0.2 |
| 8 | 58096278 | rs12682208 | -1.725 | 0.3 | 1.574 | 0.082 | 0.025 | -0.032 | -0.032 | 5.269 | 0.24 | 0.52 | 0.23 | 0.18 | 0.27 | 0.2 |
| 8 | 58103463 | rs10808915 | -1.725 | 0.314 | 1.574 | 0.082 | 0.022 | -0.032 | -0.032 | 0.397 | 0.24 | 0.52 | 0.23 | 0.18 | 0.27 | 0.2 |
| 8 | 58112053 | rs7463453 | -1.692 | 0.203 | 1.922 | 0.012 | 0.039 | 0 | 0 | 2.773 | 0.14 | 0.29 | 0.15 | 0.15 | 0.17 | 0.06 |
| 8 | 58114743 | rs2318144 | -0.579 | 0.196 | 1.923 | 0.008 | -0.008 | -0.006 | -0.006 | 0 | 0.09 | 0.06 | 0.11 | 0.15 | 0.16 | 0.06 |
| 8 | 58128557 | rs903702 | -1.903 | 0.172 | 1.925 | -0.004 | 0.037 | -0.002 | -0.002 | 0 | 0.14 | 0.3 | 0.15 | 0.15 | 0.17 | 0.06 |
| 8 | 58137742 | rs11774303 | -2.446 | 0.137 | 1.847 | -0.021 | -0.001 | -0.001 | -0.001 | 0 | 0.28 | 0.72 | 0.27 | 0.18 | 0.28 | 0.2 |
| 8 | 58143735 | rs10504224 | -0.689 | 0.203 | 1.809 | 0.014 | -0.01 | -0.006 | -0.006 | 0 | 0.1 | 0.06 | 0.11 | 0.15 | 0.17 | 0.06 |
| 8 | 58145010 | rs16921937 | -1.888 | 0.197 | 1.841 | 0.011 | 0.037 | -0.002 | -0.002 | 0.037 | 0.14 | 0.3 | 0.15 | 0.15 | 0.17 | 0.06 |
| 8 | 58147283 | rs6993531 | -1.989 | 0.302 | 1.896 | -0.002 | 0.047 | -0.003 | -0.003 | 0 | 0.14 | 0.33 | 0.15 | 0.15 | 0.19 | 0.06 |
| 8 | 58148061 | rs17215412 | -2.125 | 0.049 | 1.338 | -0.463 | -0.001 | -0.029 | -0.029 | 4.315 | 0.38 | 0.6 | 0.31 | 0.21 | 0.37 | 0.25 |
| 8 | 58151537 | rs17804365 | -0.516 | 0.512 | 1.551 | -0.013 | -0.029 | -0.01 | -0.01 | 2.547 | 0.09 | 0.05 | 0.11 | 0.14 | 0.17 | 0.08 |
| 8 | 58159585 | rs6999005 | -2.72 | 0.322 | 1.439 | 0.019 | -0.001 | 0.004 | 0.004 | 0.284 | 0.32 | 0.69 | 0.29 | 0.24 | 0.32 | 0.22 |
| 8 | 58161484 | rs17805146 | -0.467 | 0.321 | 1.264 | 0.041 | -0.03 | -0.01 | -0.01 | 0 | 0.09 | 0.05 | 0.11 | 0.14 | 0.17 | 0.08 |
| 8 | 58169798 | rs17805332 | -1.196 | 0.144 | 1.39 | 0.069 | -0.001 | 0.001 | 0.001 | 0 | 0.16 | 0.15 | 0.14 | 0.14 | 0.19 | 0.1 |
| 8 | 58173412 | rs6985942 | -1.953 | 0.108 | 1.563 | 0.063 | 0.021 | -0.048 | -0.048 | 0 | 0.21 | 0.45 | 0.18 | 0.14 | 0.19 | 0.1 |
| 8 | 58179299 | rs10089625 | -0.622 | 0.282 | 1.053 | 0.196 | -0.013 | 0.006 | 0.006 | 0 | 0.11 | 0.27 | 0.11 | 0.1 | 0.11 | 0.12 |
| 8 | 58191293 | rs3814487 | -0.6 | 0.25 | 1.056 | 0.2 | -0.016 | 0.007 | 0.007 | 0 | 0.11 | 0.27 | 0.11 | 0.1 | 0.11 | 0.12 |

| | | | | | | | | | | | | | | | | |
|---|---|---|---|---|---|---|---|---|---|---|---|---|---|---|---|---|
| 8 | 58196495 | rs10504229 | -1.585 | 0.638 | 0.5 | 0.345 | 0.002 | -0.002 | -0.002 | 0.363 | 0.16 | 0.18 | 0.15 | 0.14 | 0.19 | 0.1 |
| 8 | 58197150 | rs10504230 | -1.317 | 0.569 | 0.304 | 0.574 | -0.023 | -0.017 | -0.017 | 0 | 0.13 | 0.08 | 0.13 | 0.18 | 0.16 | 0.07 |
| 8 | 58200173 | rs16921974 | -1.732 | 0.012 | 0.552 | 0.374 | -0.002 | -0.008 | -0.008 | 0.803 | 0.13 | 0.11 | 0.07 | 0.05 | 0.09 | 0.12 |
| 8 | 58203365 | rs16921975 | -1.255 | 0.13 | 1.844 | 1.125 | 0.052 | 0.07 | 0.07 | 0.448 | 0.08 | 0.31 | 0.08 | 0.09 | 0.1 | 0.08 |
| 8 | 58211166 | rs10956999 | -2.796 | 0.082 | 2.167 | 1.03 | -0.011 | 0.024 | 0.024 | 0.077 | 0.23 | 0.6 | 0.21 | 0.31 | 0.19 | 0.18 |
| 8 | 58215017 | rs2318149 | -2.594 | 0.077 | 2.136 | 1.028 | -0.006 | 0.062 | 0.062 | 0 | 0.21 | 0.5 | 0.21 | 0.31 | 0.19 | 0.18 |
| 8 | 58222395 | rs16921987 | -2.415 | -0.017 | 2.338 | 0.968 | 0.032 | -0.009 | -0.009 | 0.02 | 0.12 | 0.52 | 0.1 | 0.05 | 0.11 | 0.08 |
| 8 | 58224353 | rs10504231 | -2.111 | -0.019 | 2.301 | 0.974 | -0.015 | 0.04 | 0.04 | 0 | 0.15 | 0.27 | 0.13 | 0.24 | 0.1 | 0.11 |
| 8 | 58226060 | rs7013994 | -2.74 | -0.01 | 1.894 | 0.963 | -0.012 | 0.024 | 0.024 | 0.2 | 0.24 | 0.6 | 0.22 | 0.31 | 0.2 | 0.19 |
| 8 | 58241429 | rs16922011 | -2.685 | -0.021 | 1.77 | 0.958 | 0.002 | -0.003 | -0.003 | 0 | 0.26 | 0.71 | 0.23 | 0.31 | 0.2 | 0.19 |
| 8 | 58242433 | rs12334334 | -2.771 | -0.087 | 1.779 | 0.925 | 0.024 | -0.025 | -0.025 | 2.951 | 0.27 | 0.83 | 0.24 | 0.31 | 0.2 | 0.19 |
| 8 | 58251655 | rs1316941 | -2.297 | -0.261 | 1.966 | 0.695 | -0.036 | -0.007 | -0.007 | 0 | 0.18 | 0.35 | 0.11 | 0.15 | 0.1 | 0.11 |
| 8 | 58257650 | rs7817881 | -2.536 | -0.312 | 1.911 | 0.661 | -0.023 | -0.026 | -0.026 | 0.012 | 0.08 | 0.34 | 0.03 | 0 | 0 | 0.01 |
| 8 | 58265798 | rs12679553 | -2.828 | 0.306 | 1.428 | 0.742 | 0.001 | 0.009 | 0.009 | 0.13 | 0.28 | 0.77 | 0.19 | 0.22 | 0.25 | 0.2 |
| 8 | 58276682 | rs6984314 | -1.546 | -0.494 | 0.942 | 0.177 | 0.02 | 0.034 | 0.034 | 0.015 | 0.3 | 0.2 | 0.12 | 0.17 | 0.21 | 0.25 |
| 8 | 58282023 | rs2325652 | -2.092 | -0.702 | 0.92 | 0.661 | 0.069 | 0.022 | 0.022 | 0.225 | 0.4 | 0.77 | 0.22 | 0.35 | 0.28 | 0.29 |
| 8 | 58295759 | rs7840862 | -2.559 | -0.635 | 1.545 | 0.654 | -0.011 | 0.074 | 0.074 | 0.106 | 0.17 | 0.48 | 0.18 | 0.19 | 0.14 | 0.12 |
| 8 | 58297248 | rs11990740 | -2.127 | -0.646 | 1.717 | 0.65 | 0.014 | 0.156 | 0.156 | 0.054 | 0.09 | 0.34 | 0.14 | 0.19 | 0.09 | 0.06 |
| 8 | 58298665 | rs7017317 | -0.737 | -0.757 | 1.56 | 0.602 | -0.002 | -0.001 | -0.001 | 0 | 0.07 | 0.06 | 0.06 | 0.17 | 0.11 | 0.16 |
| 8 | 58305969 | rs6651295 | -0.862 | -0.761 | 1.591 | 0.592 | -0.001 | -0.028 | -0.028 | 0.415 | 0.07 | 0.1 | 0.04 | 0 | 0.05 | 0.06 |
| 8 | 58308757 | rs13253851 | -2.918 | -0.818 | 1.756 | 0.531 | 0.006 | 0.001 | 0.001 | 0 | 0.38 | 0.63 | 0.28 | 0.36 | 0.29 | 0.3 |
| 8 | 58311929 | rs13248932 | -2.444 | -0.687 | 1.656 | 0.326 | 0.006 | -0.006 | -0.006 | 0.849 | 0.16 | 0.25 | 0.11 | 0.17 | 0.16 | 0.22 |
| 8 | 58316363 | rs16922120 | -1.345 | -0.678 | 1.532 | 0.41 | 0.012 | 0.012 | 0.012 | 0.805 | 0.21 | 0.14 | 0.09 | 0.18 | 0.15 | 0.17 |
| 8 | 58328635 | rs9297977 | -0.974 | -0.537 | 1.634 | 0.5 | 0.025 | 0.046 | 0.046 | 0.166 | 0.07 | 0.14 | 0.07 | 0.18 | 0.11 | 0.15 |
| 8 | 58336344 | rs1316350 | -2.001 | -0.634 | 1.701 | 0.607 | 0.004 | 0.019 | 0.019 | 0 | 0.11 | 0.16 | 0.07 | 0.21 | 0.12 | 0.19 |
| 8 | 58340503 | rs17819759 | -2.2 | -0.86 | 1.815 | 0.465 | -0.006 | -0.011 | -0.011 | 0 | 0.3 | 0.27 | 0.14 | 0.21 | 0.22 | 0.24 |
| 8 | 58351220 | rs7833343 | -2.387 | -0.135 | 1.551 | 0.652 | -0.035 | -0.031 | -0.031 | 0 | 0.31 | 0.57 | 0.15 | 0.22 | 0.17 | 0.2 |
| 8 | 58353280 | rs9643360 | -2.387 | -0.135 | 1.536 | 0.652 | -0.038 | -0.034 | -0.034 | 0.341 | 0.31 | 0.57 | 0.15 | 0.22 | 0.18 | 0.2 |
| 8 | 58354950 | rs16922177 | -0.801 | -0.091 | 1.174 | 0.474 | -0.03 | -0.048 | -0.048 | 0.223 | 0.18 | 0.18 | 0.05 | 0.01 | 0.05 | 0.03 |
| 8 | 58361959 | rs6989168 | -2.215 | 0.509 | 0.879 | 0.623 | 0.016 | 0.009 | 0.009 | 0.379 | 0.25 | 0.64 | 0.27 | 0.35 | 0.33 | 0.26 |
| 8 | 58364003 | rs1868841 | -1.76 | -0.446 | -0.661 | 0.458 | -0.131 | 0.021 | 0.021 | 0.018 | 0.59 | 0.95 | 0.54 | 0.72 | 0.49 | 0.66 |
| 8 | 58364300 | rs10957001 | -0.819 | 0.02 | -0.414 | 0.307 | -0.053 | -0.058 | -0.058 | 0.167 | 0.12 | 0.02 | 0.11 | 0.29 | 0.21 | 0.2 |
| 8 | 58364585 | rs2325743 | -0.069 | 0.089 | -0.167 | 0.541 | -0.033 | -0.031 | -0.031 | 0 | 0.06 | 0.01 | 0.05 | 0.29 | 0.14 | 0.18 |
| 8 | 58364888 | rs2325744 | -2.204 | -0.004 | 0.249 | 0.498 | 0.011 | -0.005 | -0.005 | 0 | 0.31 | 0.6 | 0.27 | 0.35 | 0.32 | 0.3 |
| 8 | 58374820 | rs7010138 | -1.481 | -0.062 | 0.308 | 0.409 | 0.009 | -0.078 | -0.078 | 0.236 | 0.21 | 0.47 | 0.21 | 0.06 | 0.19 | 0.12 |
| 8 | 58379630 | rs11985440 | -2.201 | -0.077 | 0.199 | 0.471 | 0.007 | -0.007 | -0.007 | 0.034 | 0.32 | 0.6 | 0.28 | 0.36 | 0.34 | 0.31 |
| 8 | 58390733 | rs6983236 | -0.397 | -0.036 | 0.109 | 0.462 | 0.002 | -0.011 | -0.011 | 0 | 0.13 | 0.2 | 0.17 | 0.06 | 0.14 | 0.08 |
| 8 | 58394712 | rs1451151 | -2.238 | -0.13 | -0.265 | 0.449 | -0.02 | 0.016 | 0.016 | 1.373 | 0.35 | 0.86 | 0.32 | 0.37 | 0.34 | 0.37 |
| 8 | 58401879 | rs6471618 | -1.008 | -0.026 | 0.338 | 0.433 | 0.002 | -0.065 | -0.065 | 0 | 0.11 | 0.33 | 0.06 | 0 | 0.05 | 0.05 |
| 8 | 58405076 | rs4737459 | 0.35 | -0.388 | -0.356 | 0.66 | 0.011 | -0.02 | -0.02 | 0 | 0.33 | 0.28 | 0.35 | 0.39 | 0.29 | 0.34 |
| 8 | 58405517 | rs7843806 | -1.143 | -0.396 | -0.587 | 0.668 | 0.001 | 0.075 | 0.075 | 0.622 | 0.52 | 0.88 | 0.49 | 0.7 | 0.44 | 0.6 |
| 8 | 58423936 | rs7845957 | -1.525 | -0.571 | -1.181 | 0.291 | -0.176 | 0.026 | 0.026 | 0 | 0.65 | 0.98 | 0.59 | 0.8 | 0.51 | 0.71 |
| 8 | 58438221 | rs7828768 | -1.702 | -0.44 | -0.159 | 0.419 | 0.001 | 0.006 | 0.006 | 0.092 | 0.2 | 0.59 | 0.13 | 0.28 | 0.17 | 0.24 |
| 8 | 58443123 | rs6471625 | -1.008 | -0.451 | 0.14 | 0.195 | -0.014 | -0.058 | -0.058 | 0.609 | 0.1 | 0.27 | 0.06 | 0 | 0.04 | 0.08 |
| 8 | 58443908 | rs1868842 | -1.773 | -0.31 | -0.589 | 0.44 | 0.005 | -0.025 | -0.025 | 0 | 0.36 | 0.77 | 0.27 | 0.48 | 0.33 | 0.35 |
| 8 | 58449229 | rs2875745 | 0.545 | -0.51 | -0.802 | 0.546 | 0.02 | 0.007 | 0.007 | 0.139 | 0.42 | 0.67 | 0.4 | 0.34 | 0.26 | 0.45 |
| 8 | 58450654 | rs896208 | -1.825 | -0.1 | -1.611 | 0.566 | -0.163 | -0.039 | -0.039 | 0 | 0.72 | 0.98 | 0.63 | 0.8 | 0.6 | 0.76 |
| 8 | 58452793 | rs16922274 | 0.894 | -0.085 | -1.541 | 0.773 | 0.011 | -0.038 | -0.038 | 0.951 | 0.15 | 0.07 | 0.13 | 0.28 | 0.15 | 0.12 |
| 8 | 58452850 | rs10092797 | 0.537 | 0.019 | -1.395 | 0.661 | -0.05 | -0.031 | -0.031 | 0 | 0.09 | 0.02 | 0.07 | 0.18 | 0.17 | 0.15 |
| 8 | 58453749 | rs1470733 | -0.507 | -0.084 | -1.455 | 0.633 | -0.061 | -0.082 | -0.082 | 0 | 0.56 | 0.89 | 0.49 | 0.52 | 0.45 | 0.64 |
| 8 | 58453913 | rs16922281 | 0.075 | 0.528 | -0.954 | 0.6 | -0.054 | -0.032 | -0.032 | 0 | 0.09 | 0.01 | 0.06 | 0.18 | 0.17 | 0.15 |
| 8 | 58457374 | rs10109157 | 1.195 | 0.042 | -1.054 | 0.409 | -0.036 | -0.021 | -0.021 | 0.247 | 0.11 | 0.01 | 0.1 | 0.16 | 0.13 | 0.14 |
| 8 | 58457398 | rs921420 | -0.007 | -0.196 | -0.506 | 0.296 | -0.01 | 0.018 | 0.018 | 0 | 0.6 | 0.69 | 0.54 | 0.66 | 0.49 | 0.63 |
| 8 | 58461249 | rs1451142 | 0.118 | -0.142 | -0.265 | 0.4 | -0.003 | 0.002 | 0.002 | 0 | 0.6 | 0.63 | 0.54 | 0.63 | 0.5 | 0.64 |
| 8 | 58462857 | rs2053897 | -1.178 | -0.145 | -0.275 | 0.045 | -0.015 | 0 | 0 | 1.251 | 0.74 | 0.79 | 0.65 | 0.79 | 0.63 | 0.78 |
| 8 | 58466473 | rs1451141 | -0.107 | 0.22 | -0.254 | 0.111 | -0.026 | 0.001 | 0.001 | 0 | 0.64 | 0.72 | 0.57 | 0.64 | 0.54 | 0.65 |
| 8 | 58477732 | rs16922314 | 0.698 | 0.353 | -0.423 | 0.076 | -0.034 | -0.016 | -0.016 | 0.376 | 0.15 | 0.01 | 0.1 | 0.16 | 0.15 | 0.15 |
| 8 | 58478850 | rs10099945 | 0.351 | 0.125 | 0.151 | -0.018 | -0.001 | -0.001 | -0.001 | 0 | 0.58 | 0.55 | 0.53 | 0.64 | 0.48 | 0.65 |
| 8 | 58478923 | rs9297980 | 0.591 | 0.055 | 0.615 | -0.053 | -0.005 | 0.015 | 0.015 | 0 | 0.56 | 0.3 | 0.51 | 0.64 | 0.48 | 0.65 |
| 8 | 58488370 | rs4738603 | -0.265 | 0.131 | 0.321 | -0.011 | -0.093 | -0.015 | -0.015 | 0.056 | 0.65 | 0.87 | 0.58 | 0.65 | 0.54 | 0.65 |
| 8 | 58491229 | rs7814032 | -1.152 | -0.164 | 0.868 | -0.063 | 0.018 | -0.035 | -0.035 | 0 | 0.09 | 0.46 | 0.06 | 0 | 0.06 | 0.01 |
| 8 | 58492268 | rs7005726 | 0.049 | -0.136 | 0.794 | -0.049 | 0.017 | -0.022 | -0.022 | 0 | 0.06 | 0.22 | 0.04 | 0 | 0.06 | 0.01 |
| 8 | 58499130 | rs10109559 | 0.527 | -0.105 | 0.734 | 0.019 | 0.004 | 0.003 | 0.003 | 0 | 0.54 | 0.22 | 0.49 | 0.62 | 0.47 | 0.65 |
| 8 | 58504239 | rs1451136 | 1.375 | -0.62 | -0.057 | -0.205 | -0.048 | -0.118 | -0.118 | 0 | 0.19 | 0.01 | 0.2 | 0.36 | 0.19 | 0.23 |
| 8 | 58504764 | rs7821347 | -1.664 | -0.275 | 0.832 | -0.013 | -0.031 | -0.066 | -0.066 | 1.3 | 0.11 | 0.68 | 0.08 | 0 | 0.06 | 0.01 |
| 8 | 58507102 | rs2167316 | -1.954 | -0.042 | 0.69 | 0.032 | -0.09 | -0.052 | -0.052 | 0.031 | 0.81 | 0.93 | 0.61 | 0.79 | 0.68 | 0.83 |
| 8 | 58514086 | rs2325777 | 0.74 | -0.116 | 0.895 | -0.117 | -0.04 | -0.054 | -0.054 | 0.072 | 0.16 | 0.02 | 0.04 | 0.18 | 0.15 | 0.17 |
| 8 | 58514862 | rs16922402 | -0.609 | -0.074 | 1.321 | -0.281 | 0.06 | 0.004 | 0.004 | 0.625 | 0.07 | 0.41 | 0.07 | 0 | 0.06 | 0.01 |
| 8 | 58521337 | rs6982367 | -1.426 | -0.086 | 1.262 | -0.288 | -0.006 | -0.052 | -0.052 | 0.759 | 0.1 | 0.57 | 0.07 | 0 | 0.06 | 0.01 |

| | | | | | | | | | | | | | | | | |
|---|---|---|---|---|---|---|---|---|---|---|---|---|---|---|---|---|
| 8 | 58539133 | rs7002096 | -1.107 | -0.102 | 1.296 | -0.324 | -0.002 | -0.012 | -0.012 | 0 | 0.28 | 0.68 | 0.14 | 0.14 | 0.23 | 0.2 |
| 8 | 58542880 | rs10087587 | 0.006 | -0.097 | 1.395 | -0.317 | -0.004 | -0.018 | -0.018 | 0.347 | 0.19 | 0.27 | 0.07 | 0.14 | 0.17 | 0.18 |
| 8 | 58553324 | rs976900 | -0.712 | 0.123 | 0.966 | -0.208 | 0.059 | 0.023 | 0.023 | 0 | 0.72 | 0.55 | 0.5 | 0.72 | 0.62 | 0.82 |
| 8 | 58562713 | rs6471635 | 0.459 | 0.154 | 0.742 | -0.193 | 0.031 | -0.009 | -0.009 | 0.034 | 0.06 | 0.23 | 0.05 | 0 | 0.06 | 0.02 |
| 8 | 58564501 | rs13280052 | 1.741 | 0.169 | 0.646 | -0.183 | -0.058 | -0.049 | -0.049 | 0 | 0.19 | 0.07 | 0.43 | 0.28 | 0.32 | 0.16 |
| 8 | 58572471 | rs4323454 | -0.371 | 0.038 | 0.759 | -0.108 | 0.004 | 0.005 | 0.005 | 0 | 0.45 | 0.63 | 0.56 | 0.42 | 0.55 | 0.37 |
| 8 | 58597142 | rs6992409 | -0.803 | -0.198 | 0.867 | -0.18 | 0.036 | -0.017 | -0.017 | 0 | 0.08 | 0.48 | 0.07 | 0 | 0.06 | 0.02 |
| 8 | 58618515 | rs4738608 | 0.121 | -0.252 | 0.047 | -0.281 | -0.002 | 0.013 | 0.013 | 0.068 | 0.39 | 0.27 | 0.25 | 0.31 | 0.4 | 0.37 |
| 8 | 58624978 | rs4598240 | -0.064 | -0.264 | 0.013 | -0.268 | -0.008 | -0.005 | -0.005 | 0 | 0.42 | 0.49 | 0.27 | 0.38 | 0.39 | 0.44 |
| 8 | 58629713 | rs10957014 | 0.123 | -0.248 | -0.101 | -0.26 | -0.005 | -0.002 | -0.002 | 0.001 | 0.41 | 0.5 | 0.28 | 0.38 | 0.39 | 0.44 |
| 8 | 58631351 | rs11777744 | 1.004 | -0.02 | -0.212 | -0.398 | 0.003 | -0.002 | -0.002 | 0 | 0.27 | 0.37 | 0.46 | 0.24 | 0.34 | 0.19 |
| 8 | 58635952 | rs10092909 | -0.52 | -0.098 | -0.527 | -0.286 | 0.01 | -0.029 | -0.029 | 0 | 0.67 | 0.85 | 0.73 | 0.61 | 0.74 | 0.65 |
| 8 | 58639198 | rs7000187 | 0.158 | 0.118 | -0.258 | -0.07 | 0.003 | 0.008 | 0.008 | 0 | 0.48 | 0.42 | 0.28 | 0.29 | 0.4 | 0.41 |
| 8 | 58640353 | rs4738610 | -0.083 | -0.104 | -0.188 | -0.012 | 0 | 0 | 0 | 0 | 0.4 | 0.63 | 0.54 | 0.33 | 0.43 | 0.3 |
| 8 | 58643606 | rs4554453 | 1.583 | -0.227 | -0.274 | -0.099 | 0.01 | -0.009 | -0.009 | 0 | 0.16 | 0.1 | 0.09 | 0.18 | 0.13 | 0.15 |
| 8 | 58646608 | rs10957020 | -0.607 | 0.421 | 0.066 | 0.454 | -0.001 | 0 | 0 | 0 | 0.4 | 0.63 | 0.54 | 0.33 | 0.44 | 0.31 |
| 8 | 58646783 | rs4546648 | 1.211 | 0.432 | 0.01 | 0.418 | -0.08 | -0.054 | -0.054 | 0 | 0.2 | 0.08 | 0.42 | 0.25 | 0.32 | 0.18 |
| 8 | 58653248 | rs4737472 | -1.058 | 0.467 | -0.025 | 0.388 | -0.041 | -0.066 | -0.066 | 0 | 0.85 | 0.96 | 0.81 | 0.6 | 0.82 | 0.7 |
| 8 | 58654706 | rs6471648 | -1.058 | 0.467 | -0.025 | 0.388 | -0.041 | -0.066 | -0.066 | 0 | 0.85 | 0.96 | 0.81 | 0.6 | 0.82 | 0.7 |
| 8 | 58658346 | rs4738617 | 1.282 | 0.467 | -0.008 | 0.318 | 0.015 | 0.033 | 0.033 | 0 | 0.43 | 0.29 | 0.26 | 0.28 | 0.38 | 0.39 |
| 8 | 58664164 | rs12542320 | 0.692 | 0.469 | 0 | 0.226 | -0.002 | 0.003 | 0.003 | 0 | 0.64 | 0.4 | 0.68 | 0.52 | 0.7 | 0.56 |
| 8 | 58664288 | rs9694511 | -0.294 | 0.413 | -0.698 | 0.438 | 0.01 | 0.056 | 0.056 | 0 | 0.81 | 0.76 | 0.79 | 0.53 | 0.82 | 0.62 |
| 8 | 58664971 | rs11987382 | 1.023 | 0.425 | -0.877 | 0.435 | -0.011 | -0.031 | -0.031 | 0 | 0.12 | 0.1 | 0.05 | 0 | 0.07 | 0.05 |
| 8 | 58669917 | rs3923581 | -0.334 | 0.848 | -1.014 | 0.808 | 0.006 | 0.009 | 0.009 | 0 | 0.74 | 0.64 | 0.62 | 0.72 | 0.69 | 0.73 |
| 8 | 58670012 | rs4291258 | -0.823 | 0.842 | -1.033 | 0.812 | -0.08 | -0.052 | -0.052 | 0.403 | 0.78 | 0.96 | 0.65 | 0.79 | 0.69 | 0.8 |
| 8 | 58670352 | rs10096565 | 0.405 | 0.894 | -0.965 | 0.776 | -0.01 | -0.059 | -0.059 | 0 | 0.29 | 0.13 | 0.25 | 0.4 | 0.26 | 0.35 |
| 8 | 58671521 | rs10090263 | 0.209 | 1.244 | -0.127 | 0.848 | 0.016 | 0.009 | 0.009 | 0 | 0.74 | 0.64 | 0.62 | 0.72 | 0.68 | 0.73 |
| 8 | 58678194 | rs13265778 | -1.067 | 0.974 | 0.037 | 0.932 | 0.01 | 0.075 | 0.075 | 0.06 | 0.2 | 0.54 | 0.18 | 0.35 | 0.18 | 0.2 |
| 8 | 58679518 | rs13249012 | -0.693 | 0.986 | 0.428 | 0.937 | 0 | 0.04 | 0.04 | 0 | 0.1 | 0.1 | 0.1 | 0.35 | 0.11 | 0.18 |
| 8 | 58692011 | rs6651297 | -0.874 | 0.901 | 1.013 | 0.942 | -0.009 | 0.187 | 0.187 | 0 | 0.09 | 0.24 | 0.08 | 0.35 | 0.06 | 0.19 |
| 8 | 58694111 | rs1561081 | -1.584 | 0.848 | 0.763 | 0.859 | -0.01 | 0.099 | 0.099 | 0.084 | 0.19 | 0.53 | 0.14 | 0.35 | 0.13 | 0.21 |
| 8 | 58694706 | rs1436001 | -1.393 | 0.757 | 0.107 | 0.781 | 0.006 | 0.015 | 0.015 | 0 | 0.39 | 0.64 | 0.45 | 0.75 | 0.34 | 0.59 |
| 8 | 58694902 | rs11780226 | 0.209 | 0.82 | -0.288 | 0.902 | -0.003 | -0.042 | -0.042 | 1.11 | 0.2 | 0.11 | 0.31 | 0.4 | 0.21 | 0.38 |
| 8 | 58702529 | rs12155847 | -0.829 | 0.725 | -1.08 | -0.12 | -0.054 | -0.017 | -0.017 | 0 | 0.7 | 0.85 | 0.65 | 0.75 | 0.63 | 0.78 |
| 8 | 58707002 | rs7844344 | -0.749 | 0.269 | -1.48 | -0.306 | -0.121 | -0.062 | -0.062 | 0 | 0.66 | 0.97 | 0.64 | 0.74 | 0.59 | 0.78 |
| 8 | 58711322 | rs7822434 | 0.134 | 0.273 | -0.191 | -0.336 | 0 | 0.004 | 0.004 | 0.065 | 0.28 | 0.5 | 0.25 | 0.33 | 0.23 | 0.37 |
| 8 | 58713160 | rs724514 | 0.448 | 0.388 | -0.127 | -0.516 | -0.009 | 0 | 0 | 0 | 0.17 | 0.12 | 0.17 | 0 | 0.26 | 0.11 |
| 8 | 58740354 | rs1436024 | -1.101 | 0.023 | 0.032 | 0.09 | 0.012 | 0.125 | 0.125 | 0 | 0.12 | 0.24 | 0.2 | 0.43 | 0.16 | 0.23 |
| 8 | 58750124 | rs6990718 | -1.068 | 0.037 | -0.065 | -0.603 | -0.001 | -0.056 | -0.056 | 0.156 | 0.14 | 0.18 | 0.07 | 0 | 0.12 | 0.1 |
| 8 | 58753067 | rs13256472 | 0.406 | 0.167 | -0.083 | -0.775 | 0.001 | -0.075 | -0.075 | 0 | 0.15 | 0.24 | 0.16 | 0 | 0.21 | 0.06 |
| 8 | 58753462 | rs16922621 | 0.529 | 0.18 | -0.621 | -0.564 | 0.008 | -0.036 | -0.036 | 0 | 0.19 | 0.08 | 0.16 | 0.31 | 0.12 | 0.37 |
| 8 | 58754827 | rs13275521 | -0.951 | 0.692 | -0.151 | -0.686 | 0.02 | -0.03 | -0.03 | 0 | 0.09 | 0.1 | 0.1 | 0 | 0.14 | 0.09 |
| 8 | 58756626 | rs10091209 | -0.808 | 0.45 | -0.15 | -0.902 | -0.004 | -0.144 | -0.144 | 0.676 | 0.28 | 0.39 | 0.23 | 0 | 0.33 | 0.15 |
| 8 | 58757734 | rs1436003 | -0.884 | 0.158 | -0.24 | -0.185 | -0.005 | 0.002 | 0.002 | 0.48 | 0.28 | 0.58 | 0.2 | 0.3 | 0.19 | 0.26 |
| 8 | 58760048 | rs6471656 | -1.279 | 0.565 | -1.096 | -0.313 | -0.136 | 0.079 | 0.079 | 3.403 | 0.6 | 0.97 | 0.57 | 0.74 | 0.54 | 0.69 |
| 8 | 58761153 | rs1436006 | -1.177 | 0.807 | 0.036 | -0.154 | 0.031 | 0 | 0 | 0.209 | 0.3 | 0.49 | 0.25 | 0.3 | 0.33 | 0.28 |
| 8 | 58762889 | rs7834994 | -0.224 | 0.532 | 0.518 | -0.207 | 0.006 | 0.028 | 0.028 | 0 | 0.21 | 0.29 | 0.17 | 0.3 | 0.18 | 0.21 |
| 8 | 58766472 | rs17826681 | -0.062 | 0.729 | 0.221 | -0.159 | -0.027 | 0.038 | 0.038 | 0 | 0.08 | 0.03 | 0.06 | 0 | 0.14 | 0.06 |
| 8 | 58766723 | rs11988959 | -0.132 | 0.684 | 0.257 | -0.131 | 0.022 | 0.058 | 0.058 | 0.043 | 0.21 | 0.37 | 0.17 | 0.3 | 0.18 | 0.21 |
| 8 | 58773222 | rs10093879 | -0.894 | 0.685 | -0.286 | -0.03 | 0.016 | -0.001 | -0.001 | 0 | 0.3 | 0.43 | 0.24 | 0.3 | 0.31 | 0.28 |
| 8 | 58778726 | rs13271149 | -0.371 | 0.897 | -0.501 | 0.055 | -0.001 | 0 | 0 | 0.626 | 0.48 | 0.59 | 0.61 | 0.45 | 0.51 | 0.48 |
| 8 | 58779172 | rs6997028 | 0.173 | 1.417 | -0.111 | 0.363 | 0.106 | 0.029 | 0.029 | 2.151 | 0.74 | 0.51 | 0.75 | 0.77 | 0.64 | 0.84 |
| 8 | 58779457 | rs10099734 | -0.995 | 0.052 | -0.25 | 0.504 | 0.021 | -0.016 | -0.016 | 2.3 | 0.4 | 0.69 | 0.49 | 0.53 | 0.31 | 0.44 |
| 8 | 58780481 | rs13255201 | -1.438 | 0.243 | 0.066 | 0.017 | 0.007 | 0.018 | 0.018 | 1.554 | 0.31 | 0.71 | 0.38 | 0.22 | 0.23 | 0.24 |
| 8 | 58784889 | rs7010864 | -1.453 | 0.275 | 0.179 | -0.004 | 0 | -0.004 | -0.004 | 0 | 0.25 | 0.74 | 0.3 | 0.22 | 0.22 | 0.23 |
| 8 | 58788078 | rs16922703 | -1.86 | 0.146 | 0.876 | -0.038 | -0.006 | -0.048 | -0.048 | 2.021 | 0.16 | 0.44 | 0.14 | 0.14 | 0.11 | 0.09 |
| 8 | 58789168 | rs10504248 | -1.484 | 0.117 | 0.94 | -0.123 | -0.035 | -0.086 | -0.086 | 2.148 | 0.13 | 0.35 | 0.06 | 0 | 0.05 | 0.05 |
| 8 | 58795132 | rs13273087 | -2.189 | -0.282 | 0.692 | -0.102 | -0.053 | -0.114 | -0.114 | 0.039 | 0.28 | 0.51 | 0.28 | 0.07 | 0.17 | 0.2 |
| 8 | 58795321 | rs2082911 | -1.734 | -0.287 | 0.693 | -0.104 | -0.03 | -0.092 | -0.092 | 0.523 | 0.21 | 0.5 | 0.26 | 0.07 | 0.13 | 0.16 |
| 8 | 58799068 | rs7464304 | -2.372 | -0.289 | 0.78 | 0.255 | -0.024 | -0.014 | -0.014 | 0.242 | 0.3 | 0.5 | 0.35 | 0.27 | 0.21 | 0.24 |
| 8 | 58801821 | rs13253940 | -1.676 | -0.219 | -0.035 | 0.279 | 0.022 | 0.045 | 0.045 | 0 | 0.2 | 0.13 | 0.22 | 0.07 | 0.14 | 0.18 |
| 8 | 58803166 | rs4738638 | -0.615 | -0.195 | -0.129 | 0.299 | -0.003 | 0.013 | 0.013 | 0 | 0.12 | 0.07 | 0.21 | 0.07 | 0.11 | 0.14 |
| 8 | 58810226 | rs17235986 | -2.065 | -0.271 | -0.156 | 0.299 | 0.002 | -0.024 | -0.024 | 0 | 0.23 | 0.13 | 0.31 | 0.26 | 0.2 | 0.23 |
| 8 | 58814505 | rs752573 | -0.893 | -0.252 | -0.138 | 0.314 | -0.002 | -0.02 | -0.02 | 0.178 | 0.11 | 0.06 | 0.1 | 0.19 | 0.1 | 0.09 |
| 8 | 58816749 | rs11787276 | -0.689 | -0.131 | -0.058 | 0.398 | -0.002 | 0.013 | 0.013 | 0 | 0.12 | 0.07 | 0.21 | 0.07 | 0.1 | 0.14 |
| 8 | 58817444 | rs7002316 | -0.575 | -0.062 | -0.443 | 0.397 | -0.024 | -0.052 | -0.052 | 0 | 0.61 | 0.78 | 0.76 | 0.47 | 0.54 | 0.44 |
| 8 | 58819650 | rs6988560 | -2.735 | -0.436 | 0.64 | 0.724 | 0.012 | -0.03 | -0.03 | 0 | 0.25 | 0.86 | 0.37 | 0.35 | 0.2 | 0.22 |
| 8 | 58822602 | rs16922784 | -1.08 | -0.325 | 0.797 | 0.783 | 0 | 0.014 | 0.014 | 0 | 0.08 | 0.09 | 0.09 | 0.19 | 0.08 | 0.06 |
| 8 | 58826865 | rs1982135 | -0.747 | -0.297 | 1.227 | 0.765 | 0.049 | 0.042 | 0.042 | 0 | 0.06 | 0.25 | 0.2 | 0.09 | 0.09 | 0.04 |

| | | | | | | | | | | | | | | | | |
|---|---|---|---|---|---|---|---|---|---|---|---|---|---|---|---|---|
| 8 | 58827592 | rs10086387 | -1.384 | -0.435 | 1.264 | 0.618 | 0.021 | 0.028 | 0.028 | 1.335 | 0.09 | 0.42 | 0.06 | 0.07 | 0.03 | 0.09 |
| 8 | 58835639 | rs930319 | -1.931 | -0.381 | 0.824 | 0.665 | -0.018 | 0.041 | 0.041 | 1.128 | 0.17 | 0.55 | 0.17 | 0.26 | 0.12 | 0.17 |
| 8 | 58837634 | rs10504247 | -1.692 | -0.191 | 0.829 | 0.796 | 0.001 | 0.062 | 0.062 | 0 | 0.13 | 0.31 | 0.27 | 0.21 | 0.09 | 0.08 |
| 8 | 58840924 | rs1992045 | -0.606 | -0.031 | 0.577 | 0.76 | -0.002 | -0.024 | -0.024 | 0 | 0.11 | 0.3 | 0.13 | 0.06 | 0.07 | 0.09 |
| 8 | 58843946 | rs7005278 | -1.244 | -0.057 | 0.655 | 0.759 | 0.002 | 0.048 | 0.048 | 0 | 0.17 | 0.36 | 0.34 | 0.28 | 0.13 | 0.2 |
| 8 | 58844022 | rs13250896 | 0.35 | -0.533 | -0.278 | 1.057 | 0.017 | -0.027 | -0.027 | 0 | 0.18 | 0.12 | 0.21 | 0.41 | 0.12 | 0.16 |
| 8 | 58859213 | rs10957028 | -0.437 | -0.126 | -0.411 | 0.123 | 0.004 | -0.006 | -0.006 | 0 | 0.58 | 0.55 | 0.41 | 0.6 | 0.47 | 0.54 |
| 8 | 58863591 | rs6471662 | -0.651 | -0.257 | -0.554 | 0.07 | -0.115 | -0.027 | -0.027 | 0 | 0.63 | 0.95 | 0.45 | 0.61 | 0.48 | 0.54 |
| 8 | 58864437 | rs17237877 | 1.038 | -0.716 | -1.027 | -0.308 | -0.043 | 0.084 | 0.084 | 0.329 | 0.24 | 0.07 | 0.14 | 0.13 | 0.25 | 0.31 |
| 8 | 58878786 | rs4737483 | -0.578 | -0.345 | -0.838 | -0.326 | 0.014 | -0.006 | -0.006 | 0.315 | 0.59 | 0.49 | 0.43 | 0.58 | 0.49 | 0.54 |
| 8 | 58882858 | rs7016189 | -1.081 | -0.287 | -0.86 | -0.326 | -0.136 | -0.06 | -0.06 | 0 | 0.65 | 0.9 | 0.48 | 0.59 | 0.5 | 0.53 |
| 8 | 58886451 | rs4237024 | -1.093 | 0.453 | -0.517 | -0.047 | -0.059 | 0.042 | 0.042 | 0.113 | 0.91 | 1 | 0.92 | 0.96 | 0.82 | 0.89 |
| 8 | 58887848 | rs4610731 | -0.539 | -0.572 | 0.104 | 0.35 | -0.024 | 0.041 | 0.041 | 3.371 | 0.31 | 0.4 | 0.26 | 0.47 | 0.22 | 0.2 |
| 8 | 58889557 | rs6987258 | -0.589 | -0.44 | -0.003 | 0.393 | 0.04 | 0.016 | 0.016 | 0.651 | 0.06 | 0.3 | 0.03 | 0 | 0.01 | 0.01 |
| 8 | 58892317 | rs4272363 | 0.152 | -0.344 | -0.653 | 0.594 | 0.005 | -0.133 | -0.133 | 0 | 0.21 | 0.05 | 0.2 | 0.45 | 0.17 | 0.14 |
| 8 | 58904865 | rs13280976 | 0.795 | -0.131 | -0.587 | 0.756 | -0.011 | -0.09 | -0.09 | 0 | 0.13 | 0.01 | 0.16 | 0.47 | 0.11 | 0.11 |
| 8 | 58908747 | rs6994303 | -0.395 | -0.096 | -0.157 | 0.75 | -0.011 | 0.035 | 0.035 | 0.104 | 0.3 | 0.37 | 0.27 | 0.51 | 0.24 | 0.24 |
| 8 | 58928607 | rs7006812 | -0.916 | -0.107 | 0.502 | 0.995 | -0.017 | 0.108 | 0.108 | 0 | 0.1 | 0.27 | 0.09 | 0.17 | 0.05 | 0.14 |
| 8 | 58929633 | rs7841308 | -1.989 | 0.044 | 0.734 | 0.913 | 0 | 0.081 | 0.081 | 0.345 | 0.15 | 0.39 | 0.14 | 0.21 | 0.11 | 0.18 |
| 8 | 58929688 | rs7841432 | -0.907 | -0.083 | 0.476 | 0.785 | -0.005 | 0.018 | 0.018 | 0 | 0.53 | 0.7 | 0.46 | 0.6 | 0.43 | 0.43 |
| 8 | 58937750 | rs1122328 | -1.278 | -0.096 | -0.384 | 0.748 | -0.01 | 0.021 | 0.021 | 0.382 | 0.87 | 0.96 | 0.74 | 0.93 | 0.83 | 0.76 |
| 8 | 58941463 | rs17239618 | 0.445 | 0.019 | -0.043 | 0.825 | 0.006 | 0.01 | 0.01 | 0.022 | 0.38 | 0.17 | 0.3 | 0.39 | 0.33 | 0.25 |
| 8 | 58952723 | rs11780747 | 0.021 | 0.11 | 0.177 | 0.772 | -0.005 | 0.022 | 0.022 | 0 | 0.08 | 0.11 | 0.07 | 0.17 | 0.04 | 0.13 |
| 8 | 58961400 | rs13263290 | 0.403 | 0.189 | 0.323 | 0.254 | 0.018 | 0.004 | 0.004 | 0 | 0.05 | 0.12 | 0.06 | 0.05 | 0.06 | 0.03 |
| 8 | 58961721 | rs2009981 | 1.137 | 0.084 | 0.198 | 0.18 | -0.068 | 0.028 | 0.028 | 0.444 | 0.35 | 0.15 | 0.26 | 0.27 | 0.39 | 0.28 |
| 8 | 58962794 | rs10100238 | -0.27 | 0.003 | 0.477 | 0.15 | 0 | 0.06 | 0.06 | 1.788 | 0.51 | 0.8 | 0.39 | 0.39 | 0.5 | 0.49 |
| 8 | 58963084 | rs11998721 | 0.565 | -0.171 | 0.547 | 0.19 | 0.001 | -0.006 | -0.006 | 0.661 | 0.07 | 0.07 | 0.02 | 0.01 | 0.04 | 0.12 |
| 8 | 58963460 | rs954172 | 1.091 | -0.152 | 0.607 | 0.205 | -0.058 | 0.003 | 0.003 | 0.421 | 0.35 | 0.16 | 0.26 | 0.33 | 0.39 | 0.31 |
| 8 | 58978322 | rs6988260 | 0.008 | -0.103 | 1.225 | -0.208 | 0.041 | -0.075 | -0.075 | 0.188 | 0.17 | 0.4 | 0.16 | 0.05 | 0.17 | 0.06 |
| 8 | 58978805 | rs10093645 | -1.29 | -0.206 | 1.024 | -0.227 | 0.008 | 0.087 | 0.087 | 0.505 | 0.3 | 0.83 | 0.21 | 0.13 | 0.22 | 0.22 |
| 8 | 58983976 | rs16922990 | -1.045 | -0.169 | 0.838 | -0.221 | -0.005 | -0.07 | -0.07 | 3.12 | 0.28 | 0.6 | 0.2 | 0.07 | 0.21 | 0.2 |
| 8 | 58984664 | rs10504245 | 0.974 | -0.063 | 0.463 | -0.006 | 0.027 | -0.002 | -0.002 | 0 | 0.22 | 0.04 | 0.09 | 0.21 | 0.19 | 0.25 |
| 8 | 58986367 | rs13268233 | -1.628 | 0.738 | 1.46 | 0.059 | 0.005 | 0.033 | 0.033 | 1.063 | 0.29 | 0.76 | 0.21 | 0.13 | 0.22 | 0.22 |
| 8 | 58986459 | rs13273891 | -1.431 | 0.745 | 1.469 | 0.023 | -0.003 | -0.114 | -0.114 | 3.553 | 0.27 | 0.49 | 0.18 | 0.07 | 0.21 | 0.19 |
| 8 | 58997283 | rs16923020 | -0.797 | 0.496 | 0.993 | 0.179 | -0.026 | -0.042 | -0.042 | 0.072 | 0.13 | 0.64 | 0.09 | 0.03 | 0.05 | 0.12 |
| 8 | 59001612 | rs6471680 | -1.06 | 0.629 | 0.44 | 0.25 | 0 | 0.001 | 0.001 | 0.48 | 0.28 | 0.69 | 0.19 | 0.14 | 0.21 | 0.21 |
| 8 | 59028860 | rs9792139 | -0.181 | 1.065 | -0.285 | 0.514 | -0.043 | -0.029 | -0.029 | 0 | 0.19 | 0.09 | 0.11 | 0.26 | 0.21 | 0.24 |
| 8 | 59034188 | rs11997240 | -0.499 | 1.071 | -1.454 | 0.576 | -0.02 | -0.007 | -0.007 | 0 | 0.29 | 0.82 | 0.2 | 0.29 | 0.25 | 0.26 |
| 8 | 59037323 | rs7010991 | 0.106 | 1.465 | -1.908 | 0.444 | -0.067 | -0.013 | -0.013 | 0.724 | 0.75 | 0.95 | 0.52 | 0.74 | 0.66 | 0.72 |
| 8 | 59050909 | rs17195982 | -0.075 | 1.284 | -1.48 | 0.204 | -0.009 | 0.07 | 0.07 | 0 | 0.25 | 0.12 | 0.24 | 0.1 | 0.25 | 0.31 |
| 8 | 59053700 | rs4738647 | -0.181 | 1.707 | -1.698 | 0.7 | -0.03 | -0.036 | -0.036 | 0 | 0.29 | 0.26 | 0.43 | 0.41 | 0.42 | 0.36 |
| 8 | 59057413 | rs4738648 | -0.048 | 1.786 | -1.774 | 0.503 | -0.031 | -0.016 | -0.016 | 0 | 0.28 | 0.22 | 0.43 | 0.35 | 0.41 | 0.33 |
| 8 | 59058018 | rs17255646 | 0.308 | 1.465 | -1.816 | 0.182 | 0.006 | 0.11 | 0.11 | 0.178 | 0.29 | 0.09 | 0.25 | 0.08 | 0.25 | 0.3 |
| 8 | 59060275 | rs4738649 | -0.252 | 1.549 | -1.44 | 0.49 | -0.01 | -0.007 | -0.007 | 0 | 0.28 | 0.29 | 0.36 | 0.36 | 0.4 | 0.33 |
| 8 | 59066247 | rs2939966 | -0.253 | 1.248 | -1.288 | 0.307 | -0.005 | 0.05 | 0.05 | 0 | 0.52 | 0.66 | 0.58 | 0.74 | 0.5 | 0.57 |
| 8 | 59070935 | rs13270507 | -0.993 | 1.252 | -1.609 | 0.545 | -0.023 | -0.042 | -0.042 | 0.816 | 0.11 | 0.01 | 0.09 | 0.12 | 0.11 | 0.24 |
| 8 | 59071190 | rs7844155 | -0.058 | 1.169 | -1.673 | 0.239 | -0.01 | 0.026 | 0.026 | 0 | 0.45 | 0.27 | 0.45 | 0.69 | 0.43 | 0.38 |
| 8 | 59075434 | rs11780161 | -0.66 | 1.441 | -2.129 | 0.299 | -0.042 | 0.02 | 0.02 | 0.233 | 0.23 | 0.07 | 0.35 | 0.19 | 0.33 | 0.13 |
| 8 | 59079038 | rs4738652 | -0.74 | 1.771 | -2.036 | 0.385 | -0.02 | 0.007 | 0.007 | 0.821 | 0.06 | 0.02 | 0.13 | 0.05 | 0.09 | 0.09 |
| 8 | 59079368 | rs979444 | -0.272 | 1.667 | -1.639 | 0.828 | 0.031 | -0.001 | -0.001 | 0 | 0.49 | 0.36 | 0.36 | 0.5 | 0.59 | 0.3 |
| 8 | 59080312 | rs2939975 | -1.219 | 1.134 | -1.614 | 0.806 | 0.071 | 0.08 | 0.08 | 1.459 | 0.67 | 0.85 | 0.71 | 0.87 | 0.8 | 0.67 |
| 8 | 59081949 | rs2939976 | 0.023 | 1.099 | -0.86 | 0.818 | 0.032 | 0.242 | 0.242 | 0 | 0.11 | 0.39 | 0.21 | 0.32 | 0.12 | 0.28 |
| 8 | 59082305 | rs1876379 | 0.306 | 1.579 | -0.671 | 0.899 | -0.01 | 0.007 | 0.007 | 0 | 0.47 | 0.21 | 0.33 | 0.47 | 0.55 | 0.29 |
| 8 | 59083439 | rs7013758 | 0.019 | 1.545 | -0.654 | 0.894 | 0.011 | 0.001 | 0.001 | 0.057 | 0.5 | 0.44 | 0.37 | 0.5 | 0.58 | 0.3 |
| 8 | 59087057 | rs2063103 | -0.286 | 1.51 | 0.266 | 0.859 | 0.003 | -0.027 | -0.027 | 0.609 | 0.1 | 0.15 | 0.17 | 0.04 | 0.11 | 0.2 |
| 8 | 59088925 | rs4373508 | -0.548 | 1.516 | 0.522 | 1.051 | -0.005 | 0.004 | 0.004 | 0.663 | 0.59 | 0.52 | 0.52 | 0.54 | 0.65 | 0.49 |
| 8 | 59092710 | rs2977387 | -0.606 | 1.471 | 0.608 | 0.792 | 0.019 | 0.026 | 0.026 | 1.258 | 0.61 | 0.72 | 0.58 | 0.83 | 0.71 | 0.61 |
| 8 | 59102996 | rs1605385 | -0.633 | 1.426 | 0.514 | 1.568 | -0.004 | 0.002 | 0.002 | 0.287 | 0.64 | 0.65 | 0.71 | 0.6 | 0.68 | 0.57 |
| 8 | 59107029 | rs11781890 | 0.254 | 1.708 | 0.547 | 1.77 | 0.003 | 0.017 | 0.017 | 0.091 | 0.5 | 0.38 | 0.31 | 0.48 | 0.51 | 0.43 |
| 8 | 59113981 | rs11994585 | -0.792 | 1.537 | 0.556 | 1.798 | -0.002 | -0.001 | -0.001 | 0 | 0.63 | 0.66 | 0.74 | 0.65 | 0.73 | 0.57 |
| 8 | 59120810 | rs1532936 | -0.194 | 1.568 | 0.473 | 1.636 | 0.03 | -0.015 | -0.015 | 3.159 | 0.54 | 0.33 | 0.37 | 0.52 | 0.62 | 0.42 |
| 8 | 59121122 | rs1354817 | 0.941 | 1.584 | 0.355 | 1.643 | -0.056 | 0.014 | 0.014 | 0 | 0.42 | 0.11 | 0.25 | 0.43 | 0.47 | 0.31 |
| 8 | 59121842 | rs723312 | 0.421 | 1.605 | 0.364 | 1.617 | 0 | -0.002 | -0.002 | 0 | 0.06 | 0.04 | 0.34 | 0.08 | 0.12 | 0.07 |
| 8 | 59123864 | rs6471689 | 0.453 | 1.561 | 0.262 | 1.662 | -0.034 | 0.009 | 0.009 | 0 | 0.46 | 0.17 | 0.3 | 0.48 | 0.52 | 0.37 |
| 8 | 59135137 | rs880056 | -1.453 | 1.5 | 0.265 | 1.578 | -0.017 | 0.004 | 0.004 | 0.695 | 0.69 | 0.67 | 0.78 | 0.65 | 0.8 | 0.63 |
| 8 | 59138364 | rs2977370 | -0.816 | 1.709 | -0.17 | 1.376 | -0.004 | 0.001 | 0.001 | 0 | 0.07 | 0.12 | 0.07 | 0.04 | 0.1 | 0.05 |
| 8 | 59139714 | rs2970751 | -1.42 | 1.714 | -0.177 | 1.422 | 0.003 | 0.003 | 0.003 | 0 | 0.12 | 0.18 | 0.12 | 0.11 | 0.16 | 0.11 |
| 8 | 59148256 | rs4448263 | -0.05 | 1.447 | -0.063 | 1.268 | -0.001 | -0.001 | -0.001 | 0.19 | 0.47 | 0.5 | 0.29 | 0.38 | 0.49 | 0.36 |
| 8 | 59150985 | rs17198545 | 0.303 | 1.451 | -0.163 | 1.105 | -0.014 | 0.042 | 0.042 | 0 | 0.39 | 0.26 | 0.23 | 0.31 | 0.43 | 0.29 |

| | | | | | | | | | | | | | | | | |
|---|---|---|---|---|---|---|---|---|---|---|---|---|---|---|---|---|
| 8 | 59169071 | rs4072993 | -0.7 | 1.36 | 0.121 | 1.4 | -0.02 | 0 | 0 | 0 | 0.87 | 0.82 | 0.94 | 0.87 | 0.93 | 0.77 |
| 8 | 59171724 | rs16923230 | 1.049 | 0.967 | -0.173 | 1.094 | -0.001 | 0 | 0 | 0.658 | 0.28 | 0.26 | 0.2 | 0.27 | 0.36 | 0.22 |
| 8 | 59171744 | rs4535729 | 1.571 | 1.014 | 0.026 | 1.057 | 0.015 | -0.016 | -0.016 | 0 | 0.36 | 0.27 | 0.3 | 0.45 | 0.29 | 0.43 |
| 8 | 59171944 | rs4237026 | 0.458 | 1.007 | 0.041 | 1.039 | -0.002 | 0.005 | 0.005 | 0 | 0.47 | 0.49 | 0.7 | 0.56 | 0.5 | 0.48 |
| 8 | 59176934 | rs4639499 | 0.513 | 1.136 | -0.232 | 0.945 | -0.005 | 0.032 | 0.032 | 0 | 0.37 | 0.28 | 0.22 | 0.21 | 0.41 | 0.2 |
| 8 | 59179240 | rs9643504 | 0.326 | 1.215 | -0.206 | 0.965 | 0.01 | 0.001 | 0.001 | 0 | 0.1 | 0.17 | 0.36 | 0.08 | 0.15 | 0.05 |
| 8 | 59182575 | rs13260327 | 0.283 | 1.163 | -0.169 | 0.957 | 0.003 | 0.017 | 0.017 | 0.46 | 0.49 | 0.48 | 0.71 | 0.67 | 0.52 | 0.57 |
| 8 | 59183432 | rs6471696 | 0.242 | 1.298 | -0.199 | 0.98 | 0.027 | 0.011 | 0.011 | 0 | 0.1 | 0.18 | 0.4 | 0.12 | 0.21 | 0.05 |
| 8 | 59183963 | rs7843843 | 1.414 | 1.187 | -0.24 | 0.915 | -0.001 | -0.001 | -0.001 | 0 | 0.24 | 0.23 | 0.19 | 0.17 | 0.34 | 0.13 |
| 8 | 59193836 | rs4602872 | 1.444 | 0.706 | -0.521 | 0.846 | 0.024 | 0.031 | 0.031 | 0 | 0.13 | 0.05 | 0.03 | 0.04 | 0.06 | 0.07 |
| 8 | 59194313 | rs13249339 | 0.428 | 1.077 | -0.527 | 0.705 | 0.02 | -0.003 | -0.003 | 0 | 0.33 | 0.38 | 0.55 | 0.28 | 0.51 | 0.18 |
| 8 | 59194917 | rs7007933 | 1.166 | 1.043 | -0.428 | 0.763 | 0.016 | -0.011 | -0.011 | 0 | 0.41 | 0.38 | 0.35 | 0.56 | 0.34 | 0.5 |
| 8 | 59195870 | rs10808692 | -1.222 | 1.03 | -0.446 | 0.692 | -0.016 | -0.003 | -0.003 | 0 | 0.87 | 0.8 | 0.94 | 0.88 | 0.92 | 0.75 |
| 8 | 59196091 | rs12115045 | -1.29 | 0.977 | -0.357 | 1.048 | 0.014 | 0.019 | 0.019 | 0 | 0.57 | 0.67 | 0.72 | 0.76 | 0.71 | 0.47 |
| 8 | 59215609 | rs6471703 | 1.406 | 1.369 | -0.284 | 1.098 | 0.032 | -0.012 | -0.012 | 0 | 0.24 | 0.32 | 0.19 | 0.21 | 0.35 | 0.13 |
| 8 | 59223668 | rs12678446 | -0.082 | 1.316 | -0.276 | 1.201 | 0.003 | 0.105 | 0.105 | 0 | 0.1 | 0.17 | 0.13 | 0.43 | 0.14 | 0.21 |
| 8 | 59227553 | rs1551546 | 0.254 | 1.25 | -0.387 | 0.996 | 0.033 | 0.015 | 0.015 | 0 | 0.34 | 0.5 | 0.32 | 0.64 | 0.48 | 0.34 |
| 8 | 59229282 | rs2970744 | 2.205 | 1.135 | -0.582 | 0.865 | 0.034 | 0.083 | 0.083 | 0 | 0.3 | 0.12 | 0.21 | 0.13 | 0.21 | 0.29 |
| 8 | 59229893 | rs7006132 | 0.047 | 1.167 | -0.456 | 0.969 | 0.001 | 0.021 | 0.021 | 0 | 0.23 | 0.18 | 0.39 | 0.12 | 0.21 | 0.12 |
| 8 | 59235124 | rs13269231 | 0.057 | 1.172 | -0.444 | 0.972 | 0.001 | 0.016 | 0.016 | 0 | 0.22 | 0.18 | 0.39 | 0.12 | 0.21 | 0.12 |
| 8 | 59243493 | rs2925659 | -0.908 | 1.18 | -0.267 | 0.979 | -0.004 | -0.006 | -0.006 | 0.603 | 0.86 | 0.73 | 0.92 | 0.87 | 0.9 | 0.75 |
| 8 | 59244395 | rs12542723 | -0.005 | 1.198 | -0.351 | 1.281 | -0.009 | -0.017 | -0.017 | 0.997 | 0.24 | 0.11 | 0.41 | 0.18 | 0.26 | 0.19 |
| 8 | 59249026 | rs17259490 | 1.447 | 1.288 | -0.221 | 1.325 | 0.032 | -0.013 | -0.013 | 0 | 0.24 | 0.31 | 0.19 | 0.2 | 0.35 | 0.12 |
| 8 | 59250140 | rs17259642 | 1.474 | 1.282 | -0.212 | 1.326 | 0.02 | -0.009 | -0.009 | 0.03 | 0.24 | 0.28 | 0.19 | 0.2 | 0.34 | 0.12 |
| 8 | 59251676 | rs2925662 | -1.064 | 1.256 | -0.204 | 1.423 | -0.004 | 0.014 | 0.014 | 0.082 | 0.86 | 0.73 | 0.92 | 0.84 | 0.9 | 0.75 |
| 8 | 59254904 | rs7820691 | -0.571 | 1.259 | -0.233 | 1.415 | 0.004 | 0.001 | 0.001 | 0 | 0.48 | 0.41 | 0.6 | 0.38 | 0.61 | 0.31 |
| 8 | 59256501 | rs1463437 | 0.034 | 1.211 | -0.302 | 1.41 | -0.004 | -0.01 | -0.01 | 0 | 0.24 | 0.13 | 0.41 | 0.18 | 0.26 | 0.19 |
| 8 | 59261256 | rs13251750 | 0.441 | 1.344 | -0.359 | 1.395 | -0.017 | -0.04 | -0.04 | 0 | 0.19 | 0.07 | 0.37 | 0.2 | 0.23 | 0.19 |
| 8 | 59261484 | rs747400 | 1.35 | 1.378 | -0.315 | 1.485 | 0.02 | 0.002 | 0.002 | 0.206 | 0.38 | 0.19 | 0.31 | 0.38 | 0.32 | 0.36 |
| 8 | 59261869 | rs2925663 | -0.314 | 1.355 | -0.205 | 1.105 | -0.009 | 0.001 | 0.001 | 0 | 0.81 | 0.58 | 0.88 | 0.77 | 0.88 | 0.67 |
| 8 | 59265352 | rs10808694 | -0.615 | 1.374 | -0.208 | 1.146 | 0.002 | 0.007 | 0.007 | 0 | 0.48 | 0.42 | 0.62 | 0.65 | 0.6 | 0.38 |
| 8 | 59271041 | rs7839217 | -0.856 | 1.333 | -0.291 | 1.075 | 0 | 0.008 | 0.008 | 0 | 0.29 | 0.34 | 0.48 | 0.55 | 0.29 | 0.34 |
| 8 | 59279292 | rs2970761 | -1.855 | 1.339 | -0.344 | 0.904 | 0.017 | 0.093 | 0.093 | 0 | 0.68 | 0.87 | 0.75 | 0.88 | 0.72 | 0.72 |
| 8 | 59292825 | rs2326074 | -0.42 | 1.396 | -0.217 | 1.043 | 0.001 | -0.007 | -0.007 | 0 | 0.41 | 0.56 | 0.64 | 0.7 | 0.56 | 0.4 |
| 8 | 59298600 | rs7812664 | 1.392 | 1.498 | -0.111 | 1.183 | 0.021 | -0.009 | -0.009 | 0 | 0.24 | 0.3 | 0.19 | 0.19 | 0.33 | 0.13 |
| 8 | 59299065 | rs1457046 | -1.243 | 1.466 | 0.088 | 1.255 | 0.017 | 0.099 | 0.099 | 0.264 | 0.18 | 0.37 | 0.45 | 0.52 | 0.23 | 0.27 |
| 8 | 59299146 | rs4379435 | -1.483 | 1.404 | -0.178 | 1.118 | 0.001 | 0.049 | 0.049 | 0 | 0.48 | 0.69 | 0.6 | 0.68 | 0.46 | 0.59 |
| 8 | 59301494 | rs6996798 | -1.405 | 1.516 | 0.316 | 1.455 | 0.033 | 0.174 | 0.174 | 0 | 0.18 | 0.4 | 0.52 | 0.48 | 0.22 | 0.29 |
| 8 | 59303159 | rs17260396 | 0.592 | 0.856 | -0.225 | 0.91 | 0.032 | 0.056 | 0.056 | 0.068 | 0.3 | 0.14 | 0.13 | 0.17 | 0.2 | 0.31 |
| 8 | 59303550 | rs10102916 | -0.511 | 0.852 | -0.156 | 0.865 | -0.001 | 0.001 | 0.001 | 0 | 0.4 | 0.6 | 0.69 | 0.65 | 0.54 | 0.4 |
| 8 | 59305199 | rs13268967 | -0.022 | 0.997 | -0.202 | 1.479 | 0.043 | 0.044 | 0.044 | 0 | 0.34 | 0.47 | 0.62 | 0.46 | 0.5 | 0.32 |
| 8 | 59305455 | rs1160132 | 0.362 | 1.013 | -0.223 | 1.681 | 0.061 | 0.015 | 0.015 | 0 | 0.3 | 0.46 | 0.6 | 0.35 | 0.46 | 0.25 |
| 8 | 59305974 | rs1457049 | -1.775 | 1.025 | -0.572 | 1.617 | 0.001 | 0.069 | 0.069 | 0 | 0.72 | 0.89 | 0.83 | 0.83 | 0.74 | 0.71 |
| 8 | 59306447 | rs12115131 | 1.76 | 1.487 | -0.623 | 1.407 | -0.06 | -0.031 | -0.031 | 0 | 0.12 | 0.01 | 0.08 | 0.14 | 0.18 | 0.07 |
| 8 | 59307338 | rs12546225 | 0.182 | 1.028 | -0.284 | 1.411 | 0.016 | 0.019 | 0.019 | 0 | 0.07 | 0.1 | 0.42 | 0.18 | 0.14 | 0.15 |
| 8 | 59308732 | rs7835920 | -0.216 | 0.073 | -0.165 | 1.326 | -0.029 | -0.007 | -0.007 | 0 | 0.38 | 0.47 | 0.21 | 0.36 | 0.24 | 0.38 |
| 8 | 59309897 | rs10957052 | -0.105 | 0.304 | -0.103 | 1.274 | -0.024 | 0.065 | 0.065 | 0 | 0.29 | 0.45 | 0.2 | 0.48 | 0.26 | 0.25 |
| 8 | 59310424 | rs12544703 | -0.259 | 0.856 | -0.264 | 1.177 | 0.01 | 0.002 | 0.002 | 0 | 0.08 | 0.09 | 0.47 | 0.18 | 0.19 | 0.17 |
| 8 | 59317531 | rs13255706 | 0.696 | -0.316 | 0.242 | 0.777 | -0.026 | -0.084 | -0.084 | 0.003 | 0.24 | 0.29 | 0.11 | 0.03 | 0.15 | 0.2 |
| 8 | 59319824 | rs13274906 | 0.954 | -0.324 | 0.398 | 0.756 | 0.004 | -0.01 | -0.01 | 0 | 0.22 | 0.18 | 0.1 | 0.03 | 0.15 | 0.2 |
| 8 | 59320207 | rs11776617 | 1.017 | -0.535 | 0.553 | 0.899 | 0.001 | 0.001 | 0.001 | 0 | 0.22 | 0.22 | 0.12 | 0.16 | 0.17 | 0.12 |
| 8 | 59321015 | rs10100741 | 0.97 | -0.488 | 0.555 | 0.907 | -0.02 | -0.076 | -0.076 | 0 | 0.23 | 0.27 | 0.11 | 0.03 | 0.15 | 0.2 |
| 8 | 59324162 | rs2859998 | -0.045 | -0.442 | 0.543 | 1.312 | -0.001 | 0.008 | 0.008 | 0 | 0.2 | 0.21 | 0.54 | 0.48 | 0.29 | 0.31 |
| 8 | 59329438 | rs13277646 | -0.07 | -0.438 | 0.34 | 1.209 | -0.031 | -0.016 | -0.016 | 0 | 0.2 | 0.14 | 0.51 | 0.24 | 0.29 | 0.23 |
| 8 | 59353534 | rs13277801 | -1.537 | -0.414 | 0.386 | 1.1 | -0.012 | 0.053 | 0.053 | 0 | 0.68 | 0.76 | 0.79 | 0.77 | 0.64 | 0.67 |
| 8 | 59355053 | rs7006025 | 0.854 | -0.363 | 0.413 | 1.127 | -0.006 | -0.006 | -0.006 | 0 | 0.21 | 0.24 | 0.12 | 0.16 | 0.17 | 0.12 |
| 8 | 59371725 | rs7826120 | -1.408 | -0.3 | 0.201 | 1.105 | -0.008 | 0.057 | 0.057 | 0 | 0.66 | 0.76 | 0.8 | 0.76 | 0.64 | 0.67 |
| 8 | 59372456 | rs4541896 | 0.851 | -0.115 | 0.452 | 1.178 | 0.002 | 0.003 | 0.003 | 0 | 0.21 | 0.22 | 0.13 | 0.16 | 0.18 | 0.13 |
| 8 | 59380675 | rs16923491 | 0.83 | -0.113 | 0.462 | 1.14 | 0.002 | 0.005 | 0.005 | 0 | 0.21 | 0.18 | 0.12 | 0.16 | 0.17 | 0.13 |
| 8 | 59384675 | rs6471720 | -0.153 | -0.083 | 0.68 | 1.272 | 0.021 | 0.005 | 0.005 | 0 | 0.18 | 0.28 | 0.5 | 0.23 | 0.26 | 0.22 |
| 8 | 59385919 | rs2326077 | -1.765 | -0.245 | 0.862 | 1.241 | -0.005 | 0.017 | 0.017 | 0 | 0.69 | 0.76 | 0.8 | 0.76 | 0.63 | 0.66 |
| 8 | 59386354 | rs17202769 | 1.195 | -0.322 | 0.925 | 1.272 | 0.033 | 0.071 | 0.071 | 0.975 | 0.27 | 0.12 | 0.11 | 0.04 | 0.17 | 0.18 |
| 8 | 59387670 | rs10087236 | 0.592 | -0.316 | 0.916 | 1.292 | -0.014 | 0.015 | 0.015 | 0.233 | 0.31 | 0.29 | 0.17 | 0.37 | 0.2 | 0.31 |
| 8 | 59388531 | rs10090994 | -0.469 | -0.274 | 0.841 | 1.292 | 0.002 | 0 | 0 | 0.455 | 0.51 | 0.48 | 0.3 | 0.5 | 0.38 | 0.44 |
| 8 | 59390540 | rs6471722 | -0.533 | -0.175 | 0.652 | 1.448 | -0.013 | 0.043 | 0.043 | 0.24 | 0.3 | 0.56 | 0.22 | 0.49 | 0.25 | 0.26 |
| 8 | 59398461 | rs10504255 | -1.549 | -0.232 | -0.402 | 0.996 | -0.078 | 0.005 | 0.005 | 0 | 0.72 | 0.97 | 0.8 | 0.76 | 0.64 | 0.68 |
| 8 | 59400186 | rs10957056 | 0.676 | -0.194 | 0.162 | 0.793 | 0.002 | 0.007 | 0.007 | 0 | 0.21 | 0.17 | 0.12 | 0.15 | 0.18 | 0.14 |
| 8 | 59400340 | rs10957057 | 0.66 | -0.181 | 0.108 | 0.817 | -0.001 | -0.001 | -0.001 | 0 | 0.12 | 0.12 | 0.06 | 0.15 | 0.1 | 0.09 |
| 8 | 59403576 | rs8192879 | 0.307 | -0.191 | 0.214 | 0.885 | -0.006 | 0.075 | 0.075 | 0 | 0.42 | 0.2 | 0.51 | 0.29 | 0.38 | 0.41 |

| | | | | | | | | | | | | | | | | |
|---|---|---|---|---|---|---|---|---|---|---|---|---|---|---|---|---|
| 8 | 59404441 | rs8192877 | 1.668 | -0.28 | 0.307 | 0.55 | 0.002 | 0.017 | 0.017 | 0 | 0.16 | 0.1 | 0.09 | 0.04 | 0.13 | 0.07 |
| 8 | 59405940 | rs11786580 | 1.004 | -0.289 | 0.628 | 0.576 | -0.001 | -0.016 | -0.016 | 1.624 | 0.21 | 0.19 | 0.12 | 0.07 | 0.19 | 0.11 |
| 8 | 59410439 | rs1457043 | -0.378 | -0.117 | 0.72 | 0.936 | 0.001 | 0.002 | 0.002 | 0 | 0.62 | 0.41 | 0.71 | 0.4 | 0.56 | 0.5 |
| 8 | 59411042 | rs2162459 | -0.358 | -0.131 | 1.133 | 0.929 | 0 | 0 | 0 | 0 | 0.62 | 0.39 | 0.7 | 0.4 | 0.56 | 0.5 |
| 8 | 59411394 | rs6996287 | -0.358 | -0.131 | 1.133 | 0.929 | 0 | 0 | 0 | 0 | 0.62 | 0.39 | 0.7 | 0.4 | 0.56 | 0.5 |
| 8 | 59411967 | rs8192872 | -0.294 | -0.176 | 1.457 | 1.18 | 0.006 | 0.198 | 0.198 | 0 | 0.07 | 0.18 | 0.07 | 0.35 | 0.07 | 0.17 |
| 8 | 59411971 | rs8192871 | -0.279 | -0.174 | 1.135 | 1.18 | 0 | 0 | 0 | 0 | 0.62 | 0.39 | 0.7 | 0.4 | 0.56 | 0.5 |
| 8 | 59412066 | rs8192870 | -1.421 | -0.228 | 1.225 | 1.051 | -0.049 | 0.013 | 0.013 | 0 | 0.72 | 0.86 | 0.8 | 0.74 | 0.63 | 0.67 |
| 8 | 59412924 | rs3808607 | -0.644 | -0.173 | 1.108 | 1.075 | 0.001 | 0.008 | 0.008 | 0 | 0.65 | 0.39 | 0.71 | 0.5 | 0.57 | 0.55 |
| 8 | 59413190 | rs3824260 | -0.665 | -0.163 | 1.049 | 1.081 | 0.005 | 0.024 | 0.024 | 0 | 0.65 | 0.49 | 0.71 | 0.5 | 0.57 | 0.55 |
| 8 | 59417753 | rs13251066 | 0.17 | -0.414 | 0.798 | 0.632 | -0.029 | 0.02 | 0.02 | 0.042 | 0.4 | 0.12 | 0.55 | 0.33 | 0.38 | 0.39 |
| 8 | 59418342 | rs3903445 | 0.107 | 0.252 | 1.62 | 1.239 | 0.006 | 0 | 0 | 0 | 0.64 | 0.41 | 0.7 | 0.36 | 0.56 | 0.49 |
| 8 | 59420418 | rs1023649 | 0.126 | 0.259 | 1.638 | 1.242 | 0.002 | 0 | 0 | 0 | 0.64 | 0.39 | 0.7 | 0.36 | 0.56 | 0.49 |
| 8 | 59437464 | rs1834733 | 0.051 | 0.242 | 1.813 | 1.234 | 0.01 | 0.003 | 0.003 | 0 | 0.66 | 0.61 | 0.71 | 0.36 | 0.56 | 0.49 |
| 8 | 59446454 | rs1582824 | 0.237 | 0.274 | 1.778 | 1.249 | -0.006 | 0 | 0 | 0 | 0.63 | 0.33 | 0.69 | 0.36 | 0.55 | 0.54 |
| 8 | 59459972 | rs11776003 | 0.301 | 0.341 | 1.703 | 1.221 | 0.025 | 0.039 | 0.039 | 0 | 0.27 | 0.37 | 0.23 | 0.6 | 0.37 | 0.42 |
| 8 | 59460058 | rs12375332 | 0.533 | 0.44 | 1.414 | 1.102 | -0.047 | -0.008 | -0.008 | 0 | 0.12 | 0.01 | 0.08 | 0.1 | 0.14 | 0.12 |
| 8 | 59464692 | rs10113750 | -0.09 | 0.76 | 2.091 | 1.507 | 0.037 | 0.06 | 0.06 | 0 | 0.28 | 0.44 | 0.23 | 0.6 | 0.37 | 0.42 |
| 8 | 59496052 | rs3739327 | -0.47 | 0.724 | 2.053 | 1.743 | 0.013 | 0.025 | 0.025 | 0 | 0.3 | 0.59 | 0.25 | 0.6 | 0.37 | 0.43 |
| 8 | 59496416 | rs1050504 | 0.331 | 0.77 | 1.984 | 1.738 | -0.054 | -0.044 | -0.044 | 0 | 0.23 | 0.08 | 0.17 | 0.32 | 0.35 | 0.36 |
| 8 | 59498190 | rs12677290 | -0.571 | 0.681 | 2.011 | 1.742 | 0.01 | 0.016 | 0.016 | 0 | 0.31 | 0.6 | 0.26 | 0.6 | 0.39 | 0.43 |
| 8 | 59498685 | rs12679938 | 0.612 | 0.701 | 1.75 | 1.772 | -0.064 | -0.014 | -0.014 | 0 | 0.22 | 0.05 | 0.18 | 0.26 | 0.35 | 0.31 |
| 8 | 59499389 | rs2634527 | 1.091 | 0.805 | 1.595 | 1.383 | -0.027 | 0.015 | 0.015 | 0.409 | 0.58 | 0.22 | 0.61 | 0.36 | 0.46 | 0.52 |
| 8 | 59508375 | rs2279460 | -0.23 | 0.556 | 1.387 | 1.507 | 0.042 | 0.062 | 0.062 | 0 | 0.28 | 0.46 | 0.25 | 0.62 | 0.37 | 0.41 |
| 8 | 59511246 | rs2680904 | -0.395 | 0.65 | 1.106 | 1.266 | -0.008 | -0.033 | -0.033 | 0 | 0.92 | 0.79 | 0.93 | 0.98 | 0.94 | 0.97 |
| 8 | 59512334 | rs2680903 | -0.616 | 0.664 | 0.926 | 1.266 | -0.002 | -0.016 | -0.016 | 0 | 0.92 | 0.86 | 0.94 | 0.98 | 0.93 | 0.97 |
| 8 | 59515756 | rs2255834 | 1.055 | 0.739 | 0.99 | 1.367 | -0.03 | -0.042 | -0.042 | 0.31 | 0.6 | 0.24 | 0.63 | 0.43 | 0.51 | 0.58 |
| 8 | 59520711 | rs17272151 | 0.416 | 0.704 | 1.039 | 1.527 | -0.034 | -0.018 | -0.018 | 0 | 0.23 | 0.09 | 0.18 | 0.32 | 0.35 | 0.32 |
| 8 | 59522034 | rs2680899 | 0.978 | 0.678 | 1.048 | 1.533 | -0.032 | -0.048 | -0.048 | 0 | 0.6 | 0.23 | 0.64 | 0.43 | 0.51 | 0.58 |
| 8 | 59523394 | rs7841618 | -1.114 | 0.673 | 1.248 | 1.592 | 0.073 | 0.368 | 0.368 | 5.727 | 0.06 | 0.42 | 0.09 | 0.23 | 0.02 | 0.07 |
| 8 | 59532707 | rs4598239 | -0.018 | 0.83 | 0.694 | 1.334 | -0.001 | -0.001 | -0.001 | 0 | 0.26 | 0.19 | 0.21 | 0.34 | 0.39 | 0.32 |
| 8 | 59544413 | rs10091952 | 0.169 | 0.985 | -0.044 | 1.393 | -0.032 | -0.02 | -0.02 | 0 | 0.25 | 0.09 | 0.18 | 0.34 | 0.34 | 0.3 |
| 8 | 59554706 | rs10113323 | 0.195 | 0.979 | -0.494 | 1.548 | -0.005 | -0.003 | -0.003 | 0 | 0.26 | 0.16 | 0.19 | 0.34 | 0.35 | 0.32 |
| 8 | 59562087 | rs6985643 | -0.75 | 0.979 | -1.112 | 0.835 | -0.001 | -0.001 | -0.001 | 0 | 0.39 | 0.58 | 0.32 | 0.61 | 0.46 | 0.4 |
| 8 | 59568175 | rs17273109 | -0.653 | 0.444 | -1.49 | 0.352 | 0.005 | 0.016 | 0.016 | 0 | 0.12 | 0.12 | 0.07 | 0.02 | 0.11 | 0.1 |
| 8 | 59571287 | rs973807 | 0.018 | 0.213 | -1.91 | 0.5 | 0.077 | 0.041 | 0.041 | 3.106 | 0.49 | 0.71 | 0.38 | 0.63 | 0.32 | 0.3 |
| 8 | 59575608 | rs7824249 | 0.101 | 0.637 | -2.355 | 0.898 | -0.004 | -0.028 | -0.028 | 0 | 0.8 | 0.9 | 0.78 | 0.58 | 0.8 | 0.78 |
| 8 | 59578916 | rs731685 | -0.407 | 1.032 | -3.113 | 0.125 | -0.123 | -0.022 | -0.022 | 0 | 0.33 | 0.02 | 0.3 | 0.23 | 0.38 | 0.48 |
| 8 | 59580354 | rs7818823 | 0.249 | 1.254 | -3.019 | 0.144 | -0.065 | -0.044 | -0.044 | 1.744 | 0.19 | 0.01 | 0.14 | 0.18 | 0.22 | 0.21 |
| 8 | 59586468 | rs4737508 | -0.365 | 1.15 | -2.562 | -0.082 | -0.011 | 0.054 | 0.054 | 0 | 0.87 | 0.97 | 0.88 | 1 | 0.87 | 0.87 |
| 8 | 59595411 | rs2130070 | 0.421 | 1.126 | -2.292 | 0.297 | -0.021 | -0.007 | -0.007 | 0 | 0.07 | 0.01 | 0.04 | 0.06 | 0.09 | 0.19 |
| 8 | 59596533 | rs13265615 | 0.506 | 0.713 | -2.176 | 0.102 | -0.019 | 0.094 | 0.094 | 3.079 | 0.39 | 0.21 | 0.36 | 0.14 | 0.42 | 0.26 |
| 8 | 59596999 | rs4737509 | -0.494 | 0.668 | -0.861 | 1.263 | 0.011 | 0.252 | 0.252 | 0 | 0.09 | 0.22 | 0.09 | 0.45 | 0.07 | 0.1 |
| 8 | 59599500 | rs4738700 | 0.019 | 0.298 | -0.626 | 0.743 | -0.015 | 0.017 | 0.017 | 0 | 0.38 | 0.47 | 0.3 | 0.52 | 0.33 | 0.33 |
| 8 | 59600895 | rs750961 | -0.3 | 0.274 | -0.379 | 0.688 | 0.003 | 0 | 0 | 0 | 0.52 | 0.55 | 0.43 | 0.53 | 0.5 | 0.44 |
| 8 | 59611753 | rs10109961 | -0.355 | 0.242 | -0.127 | 1.035 | 0.01 | -0.005 | -0.005 | 0 | 0.53 | 0.7 | 0.44 | 0.56 | 0.51 | 0.45 |
| 8 | 59613064 | rs10957064 | -0.619 | 0.414 | -0.148 | 0.895 | -0.044 | 0.01 | 0.01 | 0 | 0.58 | 0.8 | 0.54 | 0.67 | 0.56 | 0.49 |
| 8 | 59613227 | rs16923680 | 0.133 | 0.319 | 0.503 | 0.801 | 0.004 | 0.001 | 0.001 | 0 | 0.5 | 0.56 | 0.41 | 0.56 | 0.5 | 0.43 |
| 8 | 59614516 | rs13266837 | 0.133 | 0.257 | 0.827 | 0.77 | 0.004 | 0.001 | 0.001 | 0 | 0.5 | 0.56 | 0.41 | 0.56 | 0.5 | 0.43 |
| 8 | 59616462 | rs16923708 | -0.194 | 0.29 | 0.771 | 0.928 | -0.004 | 0.135 | 0.135 | 0 | 0.09 | 0.16 | 0.09 | 0.41 | 0.08 | 0.11 |
| 8 | 59623189 | rs10435594 | 0.353 | 0.746 | 0.4 | 0.45 | -0.001 | 0.114 | 0.114 | 0 | 0.51 | 0.3 | 0.37 | 0.21 | 0.54 | 0.47 |
| 8 | 59634740 | rs6471742 | -0.952 | 0.619 | 0.371 | 0.631 | -0.048 | -0.003 | -0.003 | 0 | 0.67 | 0.89 | 0.59 | 0.74 | 0.65 | 0.62 |
| 8 | 59646515 | rs6471748 | -0.867 | 0.585 | 0.41 | 0.611 | -0.044 | 0.006 | 0.006 | 0 | 0.66 | 0.87 | 0.58 | 0.74 | 0.62 | 0.62 |
| 8 | 59650994 | rs16923737 | 0.051 | 0.412 | 1.241 | 0.991 | 0.009 | 0.217 | 0.217 | 0 | 0.08 | 0.26 | 0.12 | 0.4 | 0.07 | 0.12 |
| 8 | 59651286 | rs7846161 | 0.109 | 0.281 | 0.772 | 0.728 | 0.034 | 0.051 | 0.051 | 0 | 0.46 | 0.8 | 0.43 | 0.69 | 0.39 | 0.39 |
| 8 | 59659127 | rs13255364 | 1.705 | 0.25 | 0.604 | 0.73 | -0.015 | 0.065 | 0.065 | 0 | 0.28 | 0.06 | 0.18 | 0.15 | 0.26 | 0.23 |
| 8 | 59663018 | rs1466633 | 0.586 | 0.291 | 0.765 | 0.702 | -0.015 | 0.019 | 0.019 | 0 | 0.39 | 0.54 | 0.3 | 0.55 | 0.33 | 0.35 |
| 8 | 59669125 | rs2634507 | 0.337 | 0.304 | 0.162 | 0.516 | 0 | 0 | 0 | 0 | 0.06 | 0.05 | 0.1 | 0.14 | 0.05 | 0.04 |
| 8 | 59673125 | rs2726584 | 0.077 | 0.25 | 0.107 | 0.74 | 0.033 | -0.029 | -0.029 | 0 | 0.46 | 0.85 | 0.43 | 0.59 | 0.39 | 0.38 |
| 8 | 59675214 | rs1565236 | 1.692 | 0.283 | -0.018 | 0.694 | -0.059 | 0.041 | 0.041 | 0 | 0.28 | 0.02 | 0.16 | 0.15 | 0.26 | 0.23 |
| 8 | 59677317 | rs1568953 | -0.861 | 0.414 | 0.532 | 0.731 | -0.003 | 0 | 0 | 0 | 0.57 | 0.46 | 0.52 | 0.51 | 0.61 | 0.62 |
| 8 | 59677977 | rs7825289 | 0.536 | 0.471 | 0.207 | 0.639 | -0.01 | -0.008 | -0.008 | 0 | 0.33 | 0.29 | 0.29 | 0.34 | 0.39 | 0.36 |
| 8 | 59680009 | rs2634515 | -0.3 | 0.582 | 0.062 | 0.559 | 0.003 | 0.027 | 0.027 | 0 | 0.23 | 0.12 | 0.23 | 0.15 | 0.22 | 0.25 |
| 8 | 59682414 | rs2634519 | 0.201 | 0.457 | 0.205 | 1.07 | -0.001 | 0.021 | 0.021 | 0 | 0.07 | 0.05 | 0.17 | 0.34 | 0.09 | 0.11 |
| 8 | 59682671 | rs7000903 | 0.697 | 0.524 | 0 | 0.92 | -0.102 | -0.109 | -0.109 | 0 | 0.29 | 0.09 | 0.29 | 0.38 | 0.41 | 0.35 |
| 8 | 59682942 | rs2726551 | -1.485 | 0.301 | 0.044 | 0.71 | 0.002 | 0.179 | 0.179 | 0 | 0.58 | 0.91 | 0.63 | 0.85 | 0.61 | 0.63 |
| 8 | 59684180 | rs2634523 | -1.327 | 0.187 | 0.396 | 0.654 | 0.028 | 0.118 | 0.118 | 0 | 0.57 | 0.78 | 0.62 | 0.85 | 0.61 | 0.63 |
| 8 | 59685441 | rs2634526 | -0.565 | 0.185 | 0.594 | 0.751 | 0.006 | -0.016 | -0.016 | 0.193 | 0.23 | 0.19 | 0.29 | 0.44 | 0.23 | 0.35 |
| 8 | 59687870 | rs6999741 | 0.374 | 0.23 | 0.946 | 0.868 | -0.089 | -0.103 | -0.103 | 0.346 | 0.3 | 0.09 | 0.29 | 0.38 | 0.37 | 0.27 |

| | | | | | | | | | | | | | | | | |
|---|---|---|---|---|---|---|---|---|---|---|---|---|---|---|---|---|
| 8 | 59692275 | rs7815285 | 0.185 | 0.232 | 1.302 | 0.854 | -0.022 | -0.027 | -0.027 | 0 | 0.31 | 0.23 | 0.31 | 0.41 | 0.39 | 0.28 |
| 8 | 59694660 | rs955103 | -0.131 | 0.25 | 1.146 | 0.863 | 0.004 | 0.006 | 0.006 | 0 | 0.35 | 0.58 | 0.34 | 0.41 | 0.38 | 0.27 |
| 8 | 59698371 | rs1526166 | 0.312 | 0.703 | 0.314 | 0.883 | -0.057 | -0.115 | -0.115 | 0 | 0.28 | 0.06 | 0.26 | 0.4 | 0.3 | 0.23 |
| 8 | 59698429 | rs16923878 | -0.575 | 0.809 | 0.596 | 0.876 | 0.069 | 0.004 | 0.004 | 0.124 | 0.06 | 0.17 | 0.04 | 0.03 | 0.1 | 0.05 |
| 8 | 59702031 | rs11781790 | 0.704 | 0.8 | 0.407 | 0.886 | -0.051 | -0.096 | -0.096 | 0 | 0.25 | 0.06 | 0.18 | 0.34 | 0.29 | 0.2 |
| 8 | 59703143 | rs2726588 | -0.876 | 0.809 | 0.568 | 0.899 | -0.003 | 0.004 | 0.004 | 0 | 0.47 | 0.58 | 0.49 | 0.52 | 0.5 | 0.47 |
| 8 | 59703560 | rs16923954 | -0.742 | 0.925 | 0.251 | 0.607 | -0.021 | -0.007 | -0.007 | 0 | 0.12 | 0.27 | 0.17 | 0.09 | 0.09 | 0.18 |
| 8 | 59708309 | rs960089 | -0.305 | 0.715 | -0.295 | 0.357 | -0.01 | -0.04 | -0.04 | 0.107 | 0.08 | 0.15 | 0.05 | 0.01 | 0.08 | 0.05 |
| 8 | 59710250 | rs2726595 | -1.22 | 0.779 | -0.486 | 0.718 | -0.005 | 0.068 | 0.068 | 0.454 | 0.22 | 0.42 | 0.2 | 0.28 | 0.21 | 0.17 |
| 8 | 59712593 | rs10957070 | 0.387 | 1.066 | -1.298 | 0.698 | -0.061 | -0.126 | -0.126 | 0 | 0.19 | 0.01 | 0.18 | 0.38 | 0.24 | 0.23 |
| 8 | 59717661 | rs16923988 | 0.554 | 0.879 | -0.764 | -0.162 | -0.009 | -0.091 | -0.091 | 0 | 0.31 | 0.45 | 0.18 | 0.13 | 0.3 | 0.13 |
| 8 | 59718696 | rs2726599 | -0.029 | 1.469 | -0.984 | 0.339 | -0.006 | 0.002 | 0.002 | 0.013 | 0.38 | 0.29 | 0.37 | 0.59 | 0.38 | 0.47 |
| 8 | 59720604 | rs2726600 | -0.78 | 1.635 | -1.267 | -0.058 | 0.002 | 0.171 | 0.171 | 2.079 | 0.6 | 0.85 | 0.6 | 0.93 | 0.63 | 0.75 |
| 8 | 59722405 | rs16924001 | -0.888 | 1.742 | -0.372 | 1.3 | -0.017 | -0.014 | -0.014 | 0 | 0.06 | 0.03 | 0.07 | 0.07 | 0.1 | 0.05 |
| 8 | 59725048 | rs2693430 | -0.89 | 1.641 | -0.085 | 1.302 | 0.015 | 0.072 | 0.072 | 0.583 | 0.26 | 0.57 | 0.3 | 0.39 | 0.26 | 0.43 |
| 8 | 59727453 | rs10957071 | -1.43 | 2.053 | 0.011 | 1.274 | 0.064 | 0.065 | 0.065 | 0 | 0.1 | 0.2 | 0.16 | 0.21 | 0.21 | 0.28 |
| 8 | 59735489 | rs3779998 | -0.716 | 1.12 | 0.275 | 0.806 | -0.022 | -0.015 | -0.015 | 0 | 0.17 | 0.22 | 0.1 | 0.15 | 0.1 | 0.1 |
| 8 | 59736227 | rs17292859 | -0.247 | 1.454 | -0.592 | 0.284 | -0.054 | 0.058 | 0.058 | 0.418 | 0.17 | 0.02 | 0.12 | 0.07 | 0.23 | 0.19 |
| 8 | 59736400 | rs826727 | 1.425 | 1.451 | -1.04 | 0.368 | -0.021 | -0.018 | -0.018 | 0 | 0.44 | 0.18 | 0.56 | 0.41 | 0.44 | 0.47 |
| 8 | 59737479 | rs826728 | -0.378 | 1.352 | -0.426 | -0.005 | -0.008 | 0.007 | 0.007 | 0 | 0.86 | 0.91 | 0.87 | 0.9 | 0.79 | 0.88 |
| 8 | 59746240 | rs7017464 | -1.161 | 0.944 | -0.324 | -0.077 | 0.051 | 0.122 | 0.122 | 0.04 | 0.45 | 0.8 | 0.31 | 0.34 | 0.37 | 0.45 |
| 8 | 59752708 | rs826733 | -0.423 | 1.082 | -0.378 | -0.099 | -0.014 | 0.009 | 0.009 | 0 | 0.9 | 0.97 | 0.91 | 0.91 | 0.85 | 0.89 |
| 8 | 59752981 | rs10103680 | 0.086 | 0.815 | -0.151 | -0.227 | -0.03 | -0.11 | -0.11 | 0.053 | 0.17 | 0.39 | 0.13 | 0.01 | 0.1 | 0.07 |
| 8 | 59753928 | rs4738721 | -1.516 | 0.811 | -0.273 | -0.228 | 0.046 | 0.124 | 0.124 | 0.016 | 0.45 | 0.81 | 0.31 | 0.34 | 0.37 | 0.42 |
| 8 | 59755177 | rs10504263 | 0.202 | 0.865 | -0.307 | -0.264 | -0.025 | 0.077 | 0.077 | 0.005 | 0.2 | 0.08 | 0.15 | 0.06 | 0.25 | 0.23 |
| 8 | 59756898 | rs3780000 | 0.393 | 0.831 | 0.468 | -0.26 | -0.005 | -0.033 | -0.033 | 0.097 | 0.16 | 0.16 | 0.11 | 0.01 | 0.1 | 0.07 |
| 8 | 59761734 | rs1349115 | -0.888 | 0.845 | 0.506 | -0.271 | 0.008 | 0.057 | 0.057 | 0 | 0.36 | 0.25 | 0.26 | 0.07 | 0.34 | 0.3 |
| 8 | 59763797 | rs6992462 | -0.38 | 0.773 | 0.736 | -0.19 | 0.015 | 0.183 | 0.183 | 0 | 0.07 | 0.3 | 0.03 | 0.26 | 0.02 | 0.14 |
| 8 | 59763910 | rs1075418 | 0.231 | 0.788 | 0.474 | -0.244 | -0.066 | 0.078 | 0.078 | 0.213 | 0.19 | 0.02 | 0.15 | 0.06 | 0.25 | 0.23 |
| 8 | 59766131 | rs1375187 | -0.873 | 0.764 | 0.952 | -0.275 | 0 | -0.027 | -0.027 | 0.659 | 0.35 | 0.31 | 0.26 | 0.09 | 0.34 | 0.29 |
| 8 | 59766309 | rs1375186 | 0.162 | 0.776 | 0.799 | -0.263 | -0.027 | 0.08 | 0.08 | 0.012 | 0.2 | 0.08 | 0.15 | 0.06 | 0.25 | 0.23 |
| 8 | 59769120 | rs2726552 | -2.002 | 0.996 | 0.718 | -0.301 | -0.079 | 0.045 | 0.045 | 0.437 | 0.57 | 0.94 | 0.46 | 0.43 | 0.54 | 0.53 |
| 8 | 59776637 | rs4738723 | -1.125 | 1.009 | 0.827 | -0.297 | 0.014 | 0.077 | 0.077 | 0.223 | 0.42 | 0.73 | 0.34 | 0.33 | 0.39 | 0.44 |
| 8 | 59781086 | rs7838499 | -0.843 | 1.006 | 0.907 | -0.318 | -0.002 | -0.026 | -0.026 | 0.004 | 0.4 | 0.41 | 0.28 | 0.22 | 0.36 | 0.43 |
| 8 | 59781882 | rs2726553 | -0.547 | 0.937 | 1.04 | 0.019 | 0.017 | 0.002 | 0.002 | 0 | 0.1 | 0.21 | 0.11 | 0.11 | 0.13 | 0.06 |
| 8 | 59784291 | rs2594959 | -1.331 | 0.914 | 1.216 | 0.038 | 0.047 | 0.078 | 0.078 | 0 | 0.16 | 0.48 | 0.17 | 0.23 | 0.18 | 0.1 |
| 8 | 59784736 | rs6992117 | -0.653 | 0.487 | 0.99 | -0.215 | -0.046 | -0.055 | -0.055 | 2.049 | 0.25 | 0.48 | 0.17 | 0.15 | 0.17 | 0.13 |
| 8 | 59785412 | rs2726555 | -0.358 | 0.519 | -0.134 | -0.473 | 0.007 | 0.023 | 0.023 | 0.077 | 0.48 | 0.57 | 0.35 | 0.39 | 0.43 | 0.37 |
| 8 | 59788913 | rs7003713 | -0.138 | 0.321 | -0.31 | -0.743 | -0.013 | -0.055 | -0.055 | 0 | 0.26 | 0.3 | 0.16 | 0.06 | 0.2 | 0.22 |
| 8 | 59792739 | rs17295487 | 1.29 | 0.187 | -0.781 | -1.13 | -0.046 | 0.119 | 0.119 | 1.18 | 0.23 | 0.04 | 0.13 | 0.05 | 0.27 | 0.11 |
| 8 | 59792778 | rs2726556 | -2.161 | 0.707 | -1.033 | -0.541 | -0.106 | -0.147 | -0.147 | 0 | 0.74 | 0.97 | 0.66 | 0.61 | 0.64 | 0.56 |
| 8 | 59792800 | rs2726557 | -1.278 | 0.35 | -0.139 | -0.105 | 0.004 | -0.018 | -0.018 | 0 | 0.36 | 0.75 | 0.43 | 0.47 | 0.29 | 0.28 |
| 8 | 59795176 | rs1448543 | -0.749 | 0.613 | -0.111 | 0.084 | 0.14 | 0.361 | 0.361 | 0.015 | 0.09 | 0.37 | 0.28 | 0.4 | 0.11 | 0.06 |
| 8 | 59795785 | rs6984917 | -0.397 | 0.169 | -0.576 | -0.538 | -0.028 | -0.085 | -0.085 | 0 | 0.27 | 0.4 | 0.16 | 0.06 | 0.18 | 0.22 |
| 8 | 59796616 | rs7837243 | -0.324 | -0.164 | -1.357 | -0.489 | 0.023 | -0.018 | -0.018 | 0.375 | 0.26 | 0.17 | 0.3 | 0.32 | 0.18 | 0.19 |
| 8 | 59798998 | rs10105293 | 0.066 | 0.349 | -1.932 | -1.071 | -0.086 | 0.139 | 0.139 | 0.014 | 0.29 | 0.11 | 0.23 | 0.1 | 0.43 | 0.21 |
| 8 | 59801932 | rs1448542 | 1.31 | 0.551 | -2.012 | -1.121 | -0.084 | 0.05 | 0.05 | 0.489 | 0.16 | 0.02 | 0.13 | 0.06 | 0.26 | 0.11 |
| 8 | 59803719 | rs7845800 | -1.74 | -0.088 | -1.987 | -0.683 | -0.018 | -0.042 | -0.042 | 1.16 | 0.69 | 0.8 | 0.58 | 0.61 | 0.62 | 0.53 |
| 8 | 59805738 | rs6471757 | -0.991 | 0.191 | -2.158 | -0.66 | 0.002 | 0.014 | 0.014 | 0 | 0.6 | 0.79 | 0.39 | 0.4 | 0.64 | 0.46 |
| 8 | 59806390 | rs17230460 | 1.251 | -0.423 | -2.195 | -0.727 | -0.096 | 0.147 | 0.147 | 0.004 | 0.3 | 0.1 | 0.23 | 0.11 | 0.43 | 0.22 |
| 8 | 59810308 | rs2044643 | -1.885 | -0.405 | -1.958 | -0.699 | -0.022 | -0.048 | -0.048 | 1.525 | 0.69 | 0.84 | 0.59 | 0.62 | 0.62 | 0.53 |
| 8 | 59813809 | rs13252116 | 2.508 | -0.756 | -1.681 | -0.899 | 0.022 | 0.012 | 0.012 | 0.184 | 0.16 | 0.1 | 0.04 | 0 | 0.06 | 0.09 |
| 8 | 59817068 | rs1562612 | -0.946 | -0.297 | -0.837 | -0.462 | -0.008 | -0.011 | -0.011 | 0.093 | 0.6 | 0.69 | 0.36 | 0.55 | 0.56 | 0.46 |
| 8 | 59823372 | rs4737526 | 2.6 | -0.462 | -0.44 | -0.493 | -0.004 | -0.002 | -0.002 | 0 | 0.21 | 0.18 | 0.32 | 0.38 | 0.32 | 0.42 |
| 8 | 59826620 | rs10504268 | 2.123 | -0.445 | -0.264 | -0.528 | 0 | -0.001 | -0.001 | 0 | 0.09 | 0.05 | 0.21 | 0.08 | 0.06 | 0.08 |
| 8 | 59837504 | rs1463142 | 1.157 | -0.683 | -0.388 | -0.567 | -0.064 | 0.038 | 0.038 | 0.044 | 0.33 | 0.14 | 0.27 | 0.27 | 0.42 | 0.24 |
| 8 | 59838228 | rs1015794 | 2.679 | -0.676 | -0.185 | -0.662 | 0.015 | 0.052 | 0.052 | 0.181 | 0.21 | 0.25 | 0.33 | 0.37 | 0.32 | 0.42 |
| 8 | 59840108 | rs7815098 | 2.04 | -0.934 | -0.417 | -0.616 | 0.054 | 0.057 | 0.057 | 0 | 0.23 | 0.12 | 0.07 | 0.08 | 0.09 | 0.2 |
| 8 | 59842670 | rs10504269 | -0.76 | -0.477 | -0.175 | -0.361 | 0.005 | -0.001 | -0.001 | 0 | 0.48 | 0.6 | 0.55 | 0.54 | 0.53 | 0.35 |
| 8 | 59848644 | rs13282806 | 0.748 | -0.176 | -0.328 | 0.043 | 0.016 | 0.056 | 0.056 | 0 | 0.12 | 0.17 | 0.26 | 0.3 | 0.11 | 0.11 |
| 8 | 59856665 | rs17233300 | 1.908 | -0.17 | -0.651 | -0.073 | -0.025 | -0.006 | -0.006 | 0 | 0.08 | 0.02 | 0.21 | 0.08 | 0.06 | 0.06 |
| 8 | 59858040 | rs2197803 | 2.156 | -0.271 | -0.275 | -0.349 | 0.019 | 0.064 | 0.064 | 0.177 | 0.23 | 0.32 | 0.33 | 0.37 | 0.32 | 0.42 |
| 8 | 59867483 | rs1448547 | 2.276 | -0.276 | -0.38 | -0.365 | -0.002 | 0.001 | 0.001 | 0.131 | 0.22 | 0.19 | 0.32 | 0.37 | 0.32 | 0.42 |
| 8 | 59869976 | rs6471762 | 0.315 | -0.714 | -0.658 | -0.947 | 0.043 | 0.028 | 0.028 | 0.143 | 0.54 | 0.34 | 0.58 | 0.61 | 0.72 | 0.65 |
| 8 | 59872177 | rs1947178 | 0.041 | -0.864 | -0.427 | -0.949 | 0.02 | 0.021 | 0.021 | 0 | 0.57 | 0.41 | 0.62 | 0.83 | 0.78 | 0.69 |
| 8 | 59873636 | rs1463140 | 1.492 | -0.678 | -0.751 | -1.035 | -0.031 | -0.018 | -0.018 | 0 | 0.37 | 0.17 | 0.45 | 0.56 | 0.57 | 0.52 |
| 8 | 59882421 | rs17300113 | 1.404 | -1.001 | -0.485 | -1.079 | -0.003 | 0.004 | 0.004 | 0 | 0.09 | 0.08 | 0.23 | 0.07 | 0.07 | 0.06 |
| 8 | 59884719 | rs10156249 | 1.472 | -0.975 | -0.523 | -1.114 | 0.004 | -0.001 | -0.001 | 1.151 | 0.17 | 0.13 | 0.15 | 0.18 | 0.24 | 0.1 |
| 8 | 59887499 | rs10108787 | 0.787 | -1.508 | -0.389 | -1.388 | 0.043 | 0.03 | 0.03 | 0 | 0.33 | 0.21 | 0.14 | 0.05 | 0.18 | 0.2 |

| | | | | | | | | | | | | | | | | |
|---|---|---|---|---|---|---|---|---|---|---|---|---|---|---|---|---|
| 8 | 59892486 | rs12676605 | 1.266 | -1.314 | -0.607 | -1.127 | 0.014 | -0.014 | -0.014 | 0 | 0.41 | 0.19 | 0.48 | 0.55 | 0.63 | 0.56 |
| 8 | 59898060 | rs7833731 | 0.406 | -0.996 | -0.259 | -1.071 | 0.01 | 0.05 | 0.05 | 0.278 | 0.76 | 0.53 | 0.64 | 0.68 | 0.8 | 0.87 |
| 8 | 59899670 | rs7842858 | 2.361 | -0.978 | -0.803 | -1.121 | -0.107 | -0.119 | -0.119 | 0 | 0.23 | 0.05 | 0.33 | 0.38 | 0.38 | 0.46 |
| 8 | 59900124 | rs10504272 | 2.167 | -1.752 | -0.701 | -1.304 | 0.039 | 0.04 | 0.04 | 0.716 | 0.15 | 0.05 | 0.08 | 0.05 | 0.08 | 0.08 |
| 8 | 59900712 | rs7813241 | -0.376 | -1.708 | -0.751 | -1.293 | -0.009 | 0.005 | 0.005 | 0.095 | 0.35 | 0.65 | 0.41 | 0.45 | 0.21 | 0.3 |
| 8 | 59911651 | rs407016 | 0.342 | -1.708 | -0.863 | -1.31 | -0.004 | -0.001 | -0.001 | 0 | 0.59 | 0.42 | 0.56 | 0.64 | 0.72 | 0.79 |
| 8 | 59913779 | rs10504274 | -1.601 | -1.233 | -1.07 | -1.037 | -0.001 | 0.02 | 0.02 | 0 | 0.92 | 0.96 | 0.95 | 1 | 0.93 | 0.98 |
| 8 | 59921413 | rs826595 | -0.76 | -1.253 | -1.035 | -1.029 | -0.003 | -0.003 | -0.003 | 0 | 0.89 | 0.83 | 0.93 | 0.92 | 0.93 | 0.87 |
| 8 | 59923386 | rs6985657 | 0.992 | -1.031 | -1.042 | -0.93 | -0.002 | 0.008 | 0.008 | 0 | 0.11 | 0.08 | 0.15 | 0.07 | 0.08 | 0.06 |
| 8 | 59925618 | rs310362 | 1.053 | -1.174 | -1.15 | -1.011 | -0.017 | -0.054 | -0.054 | 0 | 0.36 | 0.15 | 0.55 | 0.55 | 0.63 | 0.57 |
| 8 | 59929562 | rs367116 | 0.562 | -0.063 | -0.027 | -0.831 | -0.005 | -0.03 | -0.03 | 0 | 0.73 | 0.69 | 0.74 | 0.89 | 0.83 | 0.89 |
| 8 | 59929953 | rs420969 | -0.951 | -0.029 | -0.212 | -0.84 | 0.018 | 0.039 | 0.039 | 0 | 0.85 | 0.95 | 0.94 | 1 | 0.91 | 0.97 |
| 8 | 59930264 | rs10504277 | 1.206 | 0.329 | -0.322 | -0.793 | -0.011 | -0.002 | -0.002 | 1.645 | 0.12 | 0.1 | 0.14 | 0.17 | 0.23 | 0.1 |
| 8 | 59933381 | rs746867 | -0.038 | -0.397 | -0.032 | -0.95 | -0.006 | 0.023 | 0.023 | 0.846 | 0.23 | 0.3 | 0.25 | 0.32 | 0.17 | 0.2 |
| 8 | 59937643 | rs4737532 | 0.251 | -0.271 | 0.114 | -0.388 | -0.004 | 0.002 | 0.002 | 0 | 0.71 | 0.69 | 0.64 | 0.54 | 0.85 | 0.56 |
| 8 | 59938578 | rs12547374 | 0.407 | 0.378 | -0.003 | -0.362 | -0.037 | 0.002 | 0.002 | 2.273 | 0.16 | 0.1 | 0.21 | 0.18 | 0.38 | 0.12 |
| 8 | 59939182 | rs851800 | -0.004 | 0.154 | -0.049 | -0.688 | -0.015 | -0.068 | -0.068 | 0.149 | 0.78 | 0.72 | 0.91 | 1 | 0.88 | 0.94 |
| 8 | 59941575 | rs1366290 | 0.678 | 0.407 | -0.27 | -0.366 | 0.006 | 0.017 | 0.017 | 0.205 | 0.49 | 0.4 | 0.4 | 0.36 | 0.45 | 0.41 |
| 8 | 59944515 | rs9694192 | -1.034 | -0.08 | 0.236 | 0.212 | 0.105 | 0.131 | 0.131 | 3.315 | 0.13 | 0.42 | 0.16 | 0.23 | 0.21 | 0.06 |
| 8 | 59952090 | rs2914087 | -0.127 | -0.373 | -0.886 | -0.039 | -0.018 | 0.08 | 0.08 | 0 | 0.87 | 0.99 | 0.94 | 1 | 0.89 | 0.94 |
| 8 | 59953076 | rs16924491 | 0.635 | -0.076 | -0.194 | 0.473 | -0.026 | -0.038 | -0.038 | 0 | 0.07 | 0.13 | 0.02 | 0 | 0.02 | 0.01 |
| 8 | 59962804 | rs396720 | 1.266 | 0.749 | -0.446 | 1.224 | -0.018 | -0.02 | -0.02 | 0 | 0.83 | 0.84 | 0.81 | 0.67 | 0.71 | 0.89 |
| 8 | 59963737 | rs4737534 | 0.028 | 1.062 | -0.451 | 1.36 | -0.024 | 0.052 | 0.052 | 0.057 | 0.22 | 0.26 | 0.35 | 0.59 | 0.14 | 0.52 |
| 8 | 59966167 | rs10504281 | -0.884 | 2.631 | -0.811 | 2.379 | 0.01 | 0.017 | 0.017 | 0 | 0.07 | 0.06 | 0.13 | 0.21 | 0.21 | 0.08 |
| 8 | 59979252 | rs443245 | 1.135 | 0.259 | -0.798 | 1.107 | -0.003 | -0.027 | -0.027 | 0 | 0.93 | 0.98 | 0.97 | 0.88 | 0.94 | 0.97 |
| 8 | 59984665 | rs2008398 | -1.332 | -0.123 | -0.525 | 0.971 | -0.008 | 0.01 | 0.01 | 2.391 | 0.17 | 0.76 | 0.33 | 0.24 | 0.13 | 0.12 |
| 8 | 59985792 | rs9792192 | -1.358 | -0.097 | -0.356 | 0.965 | 0.001 | -0.003 | -0.003 | 0 | 0.17 | 0.79 | 0.34 | 0.23 | 0.14 | 0.12 |
| 8 | 59996064 | rs6471767 | -1.564 | -0.077 | -0.436 | 0.862 | 0.006 | -0.006 | -0.006 | 0 | 0.19 | 0.82 | 0.34 | 0.23 | 0.16 | 0.13 |
| 8 | 60001055 | rs7846649 | -0.818 | -0.114 | 0.54 | 0.833 | 0.041 | 0.129 | 0.129 | 0 | 0.08 | 0.22 | 0.27 | 0.22 | 0.13 | 0.09 |
| 8 | 60002542 | rs1010313 | -1.839 | -0.097 | -0.185 | 0.839 | -0.008 | 0.007 | 0.007 | 2.499 | 0.18 | 0.76 | 0.34 | 0.23 | 0.16 | 0.12 |
| 8 | 60010000 | rs10112035 | -0.893 | -0.862 | -0.716 | 0.231 | -0.021 | -0.056 | -0.056 | 0.152 | 0.2 | 0.21 | 0.07 | 0 | 0.08 | 0.03 |
| 8 | 60012601 | rs2561145 | 0.949 | -0.274 | -1.928 | 0.342 | -0.046 | 0.065 | 0.065 | 0.011 | 0.48 | 0.08 | 0.45 | 0.71 | 0.54 | 0.78 |
| 8 | 60018743 | rs448650 | -0.608 | -0.609 | -1.46 | 1.288 | 0.024 | -0.065 | -0.065 | 0.02 | 0.25 | 0.1 | 0.43 | 0.64 | 0.23 | 0.17 |
| 8 | 60021908 | rs16924609 | -0.078 | -0.697 | -1.548 | 1.981 | 0.032 | -0.064 | -0.064 | 0.223 | 0.18 | 0.02 | 0.16 | 0.25 | 0.14 | 0.08 |
| 8 | 60022497 | rs4509306 | -1.398 | -1.258 | -0.86 | 1.829 | -0.018 | 0.023 | 0.023 | 0 | 0.22 | 0.31 | 0.31 | 0.33 | 0.15 | 0.11 |
| 8 | 60024449 | rs392450 | -1.563 | -0.98 | -0.579 | 2.138 | 0.023 | 0.095 | 0.095 | 0.605 | 0.1 | 0.26 | 0.07 | 0.22 | 0.09 | 0.18 |
| 8 | 60032805 | rs2290526 | -0.582 | -1.063 | -0.78 | 2.241 | 0.002 | -0.076 | -0.076 | 0 | 0.15 | 0.03 | 0.28 | 0.3 | 0.13 | 0.1 |
| 8 | 60034329 | rs10957078 | -0.222 | -0.658 | 0.17 | 1.947 | -0.001 | 0.001 | 0.001 | 0 | 0.22 | 0.28 | 0.32 | 0.26 | 0.23 | 0.15 |
| 8 | 60035233 | rs16924622 | -0.361 | -0.642 | 0.13 | 1.95 | 0.002 | -0.076 | -0.076 | 2.361 | 0.15 | 0.03 | 0.28 | 0.3 | 0.13 | 0.1 |
| 8 | 60046126 | rs4568609 | -1.42 | -0.817 | 0.146 | 1.823 | 0.008 | -0.012 | -0.012 | 0 | 0.38 | 0.32 | 0.62 | 0.56 | 0.36 | 0.25 |
| 8 | 60049840 | rs3110150 | -2.342 | -0.622 | 0.593 | 1.604 | -0.024 | 0.113 | 0.113 | 0 | 0.53 | 0.78 | 0.75 | 0.79 | 0.52 | 0.43 |
| 8 | 60056229 | rs4738757 | -1.76 | -0.477 | 1.243 | 1.743 | -0.001 | 0.002 | 0.002 | 0 | 0.44 | 0.57 | 0.64 | 0.53 | 0.43 | 0.26 |
| 8 | 60060836 | rs7838240 | 0.012 | -0.534 | 1.363 | 1.818 | -0.057 | 0.009 | 0.009 | 0 | 0.21 | 0.39 | 0.19 | 0.24 | 0.17 | 0.1 |
| 8 | 60063190 | rs16924684 | -0.15 | -0.426 | 1.417 | 1.92 | -0.011 | -0.08 | -0.08 | 0.543 | 0.14 | 0.01 | 0.27 | 0.26 | 0.12 | 0.1 |
| 8 | 60093954 | rs3122694 | -1.058 | -0.731 | 1.561 | 1.812 | -0.008 | 0.081 | 0.081 | 0.146 | 0.11 | 0.24 | 0.08 | 0.26 | 0.08 | 0.23 |
| 8 | 60096451 | rs4737537 | -0.836 | -0.692 | 1.276 | 1.909 | -0.013 | -0.031 | -0.031 | 0 | 0.19 | 0.12 | 0.44 | 0.32 | 0.2 | 0.16 |
| 8 | 60097745 | rs12547861 | 0.729 | -0.626 | 1.239 | 1.928 | 0.009 | -0.042 | -0.042 | 0.24 | 0.17 | 0.01 | 0.17 | 0.2 | 0.17 | 0.08 |
| 8 | 60097984 | rs3110134 | -1.719 | -0.677 | 1.554 | 2.025 | 0.012 | -0.006 | -0.006 | 0 | 0.37 | 0.71 | 0.32 | 0.41 | 0.33 | 0.32 |
| 8 | 60098543 | rs3122691 | -1.834 | -0.554 | 1.671 | 2.019 | 0.012 | 0.077 | 0.077 | 0.817 | 0.16 | 0.39 | 0.13 | 0.24 | 0.14 | 0.24 |
| 8 | 60128658 | rs3122656 | -2.149 | -0.647 | 1.639 | 1.948 | 0.002 | -0.003 | -0.003 | 0 | 0.3 | 0.73 | 0.39 | 0.57 | 0.21 | 0.34 |
| 8 | 60130296 | rs3110127 | -2.306 | -0.721 | 1.609 | 1.909 | -0.001 | 0 | 0 | 0 | 0.36 | 0.61 | 0.59 | 0.61 | 0.34 | 0.39 |
| 8 | 60132194 | rs10113215 | -1.138 | -0.701 | 1.493 | 1.984 | 0.022 | -0.02 | -0.02 | 0 | 0.36 | 0.31 | 0.55 | 0.48 | 0.33 | 0.19 |
| 8 | 60148051 | rs3110131 | -0.614 | -0.644 | 1.633 | 2.11 | -0.003 | 0.161 | 0.161 | 0 | 0.08 | 0.19 | 0.07 | 0.3 | 0.05 | 0.22 |
| 8 | 60148508 | rs10110151 | -0.212 | -0.71 | 1.535 | 2.123 | 0.004 | 0.021 | 0.021 | 0 | 0.22 | 0.14 | 0.22 | 0.18 | 0.26 | 0.1 |
| 8 | 60153096 | rs10504290 | -0.323 | -0.754 | 1.569 | 2.151 | 0.009 | -0.042 | -0.042 | 0.04 | 0.17 | 0.06 | 0.37 | 0.3 | 0.14 | 0.11 |
| 8 | 60155314 | rs10755941 | -1.653 | -0.672 | 2.008 | 2.086 | 0.002 | -0.01 | -0.01 | 0 | 0.43 | 0.39 | 0.67 | 0.53 | 0.46 | 0.25 |
| 8 | 60161232 | rs7006620 | -0.94 | -0.754 | 2.087 | 2.073 | -0.001 | 0.182 | 0.182 | 0 | 0.09 | 0.26 | 0.08 | 0.27 | 0.06 | 0.22 |
| 8 | 60164180 | rs4571722 | -1.505 | -0.906 | 1.856 | 1.91 | 0.017 | -0.007 | -0.007 | 0 | 0.41 | 0.33 | 0.59 | 0.46 | 0.4 | 0.21 |
| 8 | 60179086 | rs7837791 | -2.111 | -0.584 | 1.555 | 1.73 | 0.006 | 0 | 0 | 0 | 0.48 | 0.65 | 0.71 | 0.46 | 0.48 | 0.23 |
| 8 | 60180904 | rs10097959 | -2.563 | -0.502 | 1.71 | 1.495 | 0.027 | -0.026 | -0.026 | 0 | 0.11 | 0.24 | 0.14 | 0.05 | 0.16 | 0.03 |
| 8 | 60187289 | rs9298008 | -0.314 | -0.72 | 1.457 | 1.449 | -0.016 | -0.015 | -0.015 | 0 | 0.19 | 0.29 | 0.31 | 0.13 | 0.2 | 0.08 |
| 8 | 60190231 | rs4305889 | -1.892 | -0.566 | 1.253 | 1.721 | 0.003 | -0.024 | -0.024 | 0 | 0.08 | 0.12 | 0.07 | 0 | 0.1 | 0.01 |
| 8 | 60195008 | rs6989377 | -1.438 | -0.736 | 1.35 | 2.154 | 0.013 | 0.361 | 0.361 | 1.15 | 0.1 | 0.44 | 0.2 | 0.34 | 0.06 | 0.19 |
| 8 | 60200732 | rs12675589 | -1.892 | -0.731 | 0.719 | 1.655 | 0 | 0.011 | 0.011 | 1.041 | 0.2 | 0.26 | 0.32 | 0.32 | 0.18 | 0.17 |
| 8 | 60208754 | rs4237031 | -1.858 | -0.758 | 0.979 | 1.833 | 0.005 | 0.178 | 0.178 | 2.402 | 0.18 | 0.59 | 0.3 | 0.47 | 0.14 | 0.25 |
| 8 | 60209436 | rs10464899 | -2.237 | -1.114 | 0.699 | 1.26 | 0.008 | 0.031 | 0.031 | 0 | 0.39 | 0.74 | 0.6 | 0.78 | 0.33 | 0.41 |
| 8 | 60214029 | rs6997052 | -2.269 | -1.117 | 0.915 | 1.262 | 0.01 | 0.031 | 0.031 | 0 | 0.39 | 0.74 | 0.6 | 0.78 | 0.33 | 0.42 |
| 8 | 60215990 | rs7007825 | -1.446 | -1.235 | 1.393 | 1.503 | -0.015 | 0.163 | 0.163 | 0 | 0.21 | 0.56 | 0.25 | 0.5 | 0.16 | 0.27 |
| 8 | 60229452 | rs4237032 | -0.759 | -1.276 | 1.446 | 1.524 | -0.008 | -0.024 | -0.024 | 0.095 | 0.11 | 0.05 | 0.18 | 0.21 | 0.1 | 0.1 |

| | | | | | | | | | | | | | | | | |
|---|---|---|---|---|---|---|---|---|---|---|---|---|---|---|---|---|
| 8 | 60229952 | rs4446726 | -1.283 | -1.277 | 1.439 | 1.51 | -0.032 | 0.229 | 0.229 | 0 | 0.18 | 0.53 | 0.2 | 0.5 | 0.09 | 0.26 |
| 8 | 60237300 | rs6987780 | -2.002 | -1.286 | 1.383 | 1.528 | -0.002 | 0.004 | 0.004 | 0 | 0.18 | 0.22 | 0.36 | 0.27 | 0.17 | 0.15 |
| 8 | 60241157 | rs6980657 | -2.002 | -1.286 | 1.39 | 1.528 | -0.002 | 0.004 | 0.004 | 0 | 0.18 | 0.22 | 0.36 | 0.27 | 0.17 | 0.15 |
| 8 | 60253104 | rs4386970 | -0.53 | -1.29 | 1.313 | 1.518 | -0.002 | -0.029 | -0.029 | 0 | 0.06 | 0.1 | 0.14 | 0 | 0.05 | 0.03 |
| 8 | 60254022 | rs12680662 | -1.194 | -1.305 | 1.432 | 1.595 | -0.012 | 0.208 | 0.208 | 0 | 0.2 | 0.52 | 0.24 | 0.5 | 0.16 | 0.26 |
| 8 | 60270825 | rs7822546 | -1.84 | -1.277 | 1.253 | 1.197 | 0.082 | 0.144 | 0.144 | 2.985 | 0.05 | 0.44 | 0.19 | 0.07 | 0.02 | 0.03 |
| 8 | 60273980 | rs4545083 | -1.845 | -1.238 | -0.062 | 0.978 | 0.016 | 0.095 | 0.095 | 0.183 | 0.43 | 0.88 | 0.61 | 0.77 | 0.34 | 0.42 |
| 8 | 60285226 | rs7830834 | 1.436 | -1.484 | 0.474 | 1.806 | -0.068 | 0.192 | 0.192 | 0 | 0.82 | 0.57 | 0.79 | 0.51 | 0.94 | 0.87 |
| 8 | 60291079 | rs6471783 | 0.387 | -1.519 | 0.63 | 2.063 | -0.025 | 0.208 | 0.208 | 0.233 | 0.9 | 0.78 | 0.82 | 0.63 | 0.96 | 0.9 |
| 8 | 60305327 | rs11775907 | 1.754 | -1.522 | 0.637 | 2.039 | -0.071 | 0.215 | 0.215 | 0 | 0.82 | 0.56 | 0.79 | 0.48 | 0.94 | 0.87 |
| 8 | 60308317 | rs13280933 | 1.323 | -2.039 | -0.259 | 1.206 | 0.06 | 0.142 | 0.142 | 1.401 | 0.51 | 0.05 | 0.51 | 0.21 | 0.64 | 0.63 |
| 8 | 60338539 | rs10102694 | -1.151 | -1.407 | 0.419 | 1.366 | -0.019 | 0.02 | 0.02 | 0 | 0.08 | 0.14 | 0.02 | 0.11 | 0.02 | 0.04 |
| 8 | 60346384 | rs10088757 | -1.116 | -1.497 | 0.381 | 1.409 | -0.019 | 0.059 | 0.059 | 0 | 0.05 | 0.22 | 0.04 | 0.11 | 0.01 | 0.02 |
| 8 | 60360286 | rs28647268 | -1.617 | -1.501 | 0.365 | 1.407 | -0.028 | 0.047 | 0.047 | 0 | 0.07 | 0.28 | 0.05 | 0.11 | 0.01 | 0.03 |
| 8 | 60383409 | rs12677067 | 0.567 | -2.085 | -0.069 | 1.158 | -0.01 | -0.021 | -0.021 | 0 | 0.36 | 0.08 | 0.26 | 0.36 | 0.37 | 0.32 |
| 8 | 60388601 | rs17250119 | 1.081 | -2.117 | -0.612 | 1.102 | -0.115 | 0.102 | 0.102 | 2.388 | 0.35 | 0.04 | 0.42 | 0.1 | 0.51 | 0.53 |
| 8 | 60398832 | rs7829285 | 1.693 | -1.787 | 0.829 | 1.852 | -0.037 | 0.055 | 0.055 | 0.286 | 0.75 | 0.63 | 0.78 | 0.46 | 0.88 | 0.86 |
| 8 | 60413073 | rs10103231 | 1.49 | -1.787 | 0.893 | 1.852 | -0.075 | 0.121 | 0.121 | 0.481 | 0.74 | 0.53 | 0.76 | 0.46 | 0.88 | 0.86 |
| 8 | 60430255 | rs2056264 | 1.423 | -1.706 | 0.231 | 1.888 | -0.047 | 0.069 | 0.069 | 0.449 | 0.75 | 0.62 | 0.78 | 0.46 | 0.88 | 0.87 |
| 8 | 60445734 | rs2139231 | 1.2 | -1.509 | -0.618 | 1.909 | 0.052 | -0.112 | -0.112 | 1.274 | 0.79 | 0.94 | 0.81 | 0.47 | 0.89 | 0.88 |
| 8 | 60458178 | rs2056265 | 0.32 | -0.826 | -1.064 | 1.964 | 0.007 | -0.099 | -0.099 | 0 | 0.86 | 0.98 | 0.83 | 0.57 | 0.9 | 0.91 |
| 8 | 60469380 | rs10957100 | -1.037 | -0.634 | -0.567 | 1.87 | -0.024 | 0.083 | 0.083 | 0 | 0.19 | 0.27 | 0.23 | 0.52 | 0.1 | 0.31 |
| 8 | 60470686 | rs4607594 | 0.867 | 0.126 | -1.089 | 1.574 | -0.044 | -0.068 | -0.068 | 13.524 | 0.89 | 0.99 | 0.73 | 0.74 | 0.87 | 0.91 |
| 8 | 60476673 | rs6982160 | -1.042 | -0.249 | -0.003 | 1.29 | 0.011 | 0.043 | 0.043 | 0 | 0.05 | 0.28 | 0.04 | 0.03 | 0 | 0.02 |
| 8 | 60478117 | rs7827918 | -0.301 | -0.256 | -0.928 | 0.968 | 0.039 | 0.035 | 0.035 | 0.149 | 0.54 | 0.84 | 0.39 | 0.65 | 0.42 | 0.67 |
| 8 | 60486037 | rs7832257 | -1.04 | -0.555 | -0.269 | 1.566 | -0.016 | 0.162 | 0.162 | 0.141 | 0.13 | 0.47 | 0.14 | 0.29 | 0.09 | 0.13 |
| 8 | 60505005 | rs4588843 | 0.283 | -0.544 | -0.302 | 1.556 | -0.004 | -0.006 | -0.006 | 0.38 | 0.9 | 0.9 | 0.72 | 0.77 | 0.89 | 0.93 |
| 8 | 60510787 | rs11986421 | -0.128 | -0.63 | 0.104 | 1.512 | 0.005 | 0.035 | 0.035 | 0 | 0.05 | 0.27 | 0.04 | 0.04 | 0 | 0.02 |
| 8 | 60514487 | rs12547681 | 0.199 | -0.826 | 0.361 | 1.415 | 0.02 | 0.053 | 0.053 | 0.67 | 0.46 | 0.16 | 0.28 | 0.33 | 0.43 | 0.56 |
| 8 | 60545664 | rs4737547 | -0.792 | -0.389 | 0.439 | 1.133 | -0.004 | 0.01 | 0.01 | 0.587 | 0.58 | 0.8 | 0.39 | 0.64 | 0.43 | 0.69 |
| 8 | 60549318 | rs6990562 | 0.393 | -0.959 | 0.797 | 0.927 | 0.016 | -0.03 | -0.03 | 0.556 | 0.21 | 0.14 | 0.09 | 0.36 | 0.12 | 0.4 |
| 8 | 60568606 | rs4501567 | -0.216 | -0.925 | 1.548 | 0.919 | -0.038 | 0.046 | 0.046 | 0 | 0.26 | 0.39 | 0.14 | 0.59 | 0.13 | 0.49 |
| 8 | 60574673 | rs7007604 | -2.352 | -0.841 | 1.671 | 0.917 | -0.021 | 0.124 | 0.124 | 1 | 0.11 | 0.58 | 0.09 | 0.26 | 0.04 | 0.12 |
| 8 | 60580660 | rs10504295 | -1.41 | -1.14 | 1.448 | 0.725 | -0.007 | 0.029 | 0.029 | 0 | 0.57 | 0.79 | 0.39 | 0.66 | 0.44 | 0.72 |
| 8 | 60585342 | rs4644228 | 1.001 | -1.167 | 1.42 | 0.719 | -0.009 | -0.117 | -0.117 | 1.24 | 0.18 | 0.01 | 0.08 | 0.36 | 0.13 | 0.39 |
| 8 | 60587632 | rs16925119 | 0.585 | -1.185 | 1.52 | 0.71 | 0.002 | -0.001 | -0.001 | 0 | 0.2 | 0.2 | 0.11 | 0.36 | 0.13 | 0.39 |
| 8 | 60600281 | rs6981960 | -0.63 | -1.188 | 1.537 | 0.674 | -0.003 | -0.004 | -0.004 | 0.063 | 0.38 | 0.59 | 0.28 | 0.29 | 0.31 | 0.33 |
| 8 | 60604193 | rs16925150 | -1.232 | -1.195 | 1.797 | 0.674 | -0.007 | 0.215 | 0.215 | 0.161 | 0.06 | 0.27 | 0.06 | 0.25 | 0.01 | 0.1 |
| 8 | 60623348 | rs12541902 | -0.138 | -1.212 | 1.769 | 0.629 | 0.002 | -0.01 | -0.01 | 0.443 | 0.69 | 0.73 | 0.78 | 0.95 | 0.7 | 0.74 |
| 8 | 60626099 | rs4237035 | -0.657 | -1.221 | 1.565 | 0.621 | -0.017 | 0.162 | 0.162 | 0.138 | 0.73 | 0.95 | 0.8 | 0.98 | 0.73 | 0.75 |
| 8 | 60629462 | rs7846430 | 1.423 | -1.243 | 1.539 | 0.656 | -0.039 | 0.009 | 0.009 | 0 | 0.42 | 0.14 | 0.61 | 0.36 | 0.56 | 0.28 |
| 8 | 60649504 | rs6992992 | -0.031 | -1.206 | 1.645 | 0.718 | 0.01 | 0.008 | 0.008 | 0 | 0.68 | 0.73 | 0.75 | 0.71 | 0.73 | 0.69 |
| 8 | 60656194 | rs1034518 | 1.3 | -1.139 | 1.244 | 0.811 | -0.006 | 0.031 | 0.031 | 0 | 0.43 | 0.19 | 0.61 | 0.35 | 0.57 | 0.27 |
| 8 | 60660043 | rs6471801 | -0.589 | -1.177 | 1.101 | 0.746 | 0.002 | 0.118 | 0.118 | 0.071 | 0.72 | 0.92 | 0.81 | 0.95 | 0.73 | 0.76 |
| 8 | 60667231 | rs6471803 | 0.214 | -1.21 | 1.194 | 0.835 | 0.023 | 0.014 | 0.014 | 0.014 | 0.55 | 0.86 | 0.73 | 0.62 | 0.6 | 0.38 |
| 8 | 60686330 | rs9643513 | 0.987 | -1.177 | 1.188 | 0.869 | 0.006 | -0.059 | -0.059 | 0 | 0.17 | 0.05 | 0.07 | 0.33 | 0.13 | 0.38 |
| 8 | 60691309 | rs12678056 | 0.409 | -1.181 | 1.191 | 0.895 | 0 | 0.011 | 0.011 | 0 | 0.08 | 0.07 | 0.3 | 0.25 | 0.11 | 0.07 |
| 8 | 60694647 | rs867743 | 0.158 | -1.232 | 1.208 | 0.852 | 0.008 | -0.001 | -0.001 | 0 | 0.56 | 0.87 | 0.73 | 0.62 | 0.6 | 0.38 |
| 8 | 60694900 | rs907211 | 0.441 | -1.229 | 1.292 | 0.85 | 0.014 | 0.024 | 0.024 | 0 | 0.52 | 0.72 | 0.71 | 0.62 | 0.57 | 0.37 |
| 8 | 60696581 | rs10957108 | -0.883 | -1.234 | 1.439 | 0.869 | -0.02 | 0.193 | 0.193 | 0.447 | 0.07 | 0.32 | 0.06 | 0.26 | 0.01 | 0.1 |
| 8 | 60705885 | rs7011502 | 0.273 | -1.22 | 0.56 | 0.891 | 0.007 | 0.026 | 0.026 | 0 | 0.54 | 0.79 | 0.71 | 0.61 | 0.57 | 0.37 |
| 8 | 60708881 | rs4440623 | 0.014 | -1.632 | 0.134 | 0.627 | -0.046 | -0.039 | -0.039 | 0 | 0.64 | 0.94 | 0.76 | 0.65 | 0.67 | 0.4 |
| 8 | 60709953 | rs6986814 | 2.268 | -1.597 | 0.089 | 0.595 | -0.18 | 0.085 | 0.085 | 0.06 | 0.34 | 0.04 | 0.3 | 0.11 | 0.45 | 0.21 |
| 8 | 60711404 | rs10092060 | 1.602 | -1.598 | 0.195 | 0.627 | -0.028 | 0.017 | 0.017 | 0 | 0.42 | 0.15 | 0.61 | 0.34 | 0.56 | 0.27 |
| 8 | 60728159 | rs10103912 | 2.25 | -1.587 | -0.045 | 0.563 | -0.18 | 0.084 | 0.084 | 0.036 | 0.34 | 0.04 | 0.31 | 0.1 | 0.45 | 0.19 |
| 8 | 60746997 | rs6471835 | -0.038 | -1.603 | -0.046 | 0.977 | -0.097 | 0.009 | 0.009 | 0.009 | 0.66 | 0.98 | 0.77 | 0.77 | 0.68 | 0.43 |
| 8 | 60753118 | rs7837720 | 1.045 | -1.287 | 0.344 | 0.91 | -0.014 | -0.011 | -0.011 | 0 | 0.11 | 0.16 | 0.04 | 0.07 | 0.1 | 0.04 |
| 8 | 60756918 | rs10105639 | 1.073 | -1.287 | 0.344 | 0.794 | -0.013 | -0.027 | -0.027 | 0 | 0.11 | 0.16 | 0.04 | 0.05 | 0.1 | 0.03 |
| 8 | 60760020 | rs10105068 | 0.744 | -1.372 | 0.326 | 0.803 | -0.038 | 0.017 | 0.017 | 1.591 | 0.12 | 0.19 | 0.04 | 0.16 | 0.07 | 0.05 |
| 8 | 60775727 | rs7822560 | 0.427 | -1.389 | 0.299 | 0.791 | -0.02 | 0.012 | 0.012 | 1.35 | 0.13 | 0.19 | 0.05 | 0.16 | 0.1 | 0.06 |
| 8 | 60806161 | rs7010234 | 1.679 | -1.383 | 0.194 | 0.78 | 0.027 | 0.019 | 0.019 | 1.684 | 0.09 | 0.03 | 0.03 | 0.05 | 0.07 | 0.02 |
| 8 | 60818649 | rs6471842 | -1.493 | -2.089 | -0.338 | 0.411 | -0.058 | -0.007 | -0.007 | 0.082 | 0.63 | 0.86 | 0.39 | 0.66 | 0.46 | 0.75 |
| 8 | 60819091 | rs1497039 | -1.115 | -1.944 | -0.448 | 0.458 | -0.023 | -0.046 | -0.046 | 0.76 | 0.92 | 0.99 | 0.71 | 0.76 | 0.89 | 0.93 |
| 8 | 60828697 | rs17342242 | 0.406 | -1.972 | -0.404 | 0.295 | -0.046 | 0.16 | 0.16 | 0 | 0.24 | 0.01 | 0.16 | 0 | 0.23 | 0.24 |
| 8 | 60855552 | rs17343814 | 0.95 | -1.44 | 0.356 | 0.851 | -0.009 | -0.018 | -0.018 | 0.278 | 0.1 | 0.13 | 0.04 | 0.04 | 0.1 | 0.03 |
| 8 | 60883570 | rs488934 | -0.985 | -1.625 | 0.33 | 0.892 | -0.013 | -0.026 | -0.026 | 0 | 0.59 | 0.74 | 0.38 | 0.54 | 0.46 | 0.72 |
| 8 | 60887218 | rs480945 | -0.018 | -1.617 | 0.575 | 0.87 | 0.004 | -0.001 | -0.001 | 0 | 0.46 | 0.48 | 0.31 | 0.5 | 0.34 | 0.69 |
| 8 | 60888414 | rs673973 | -1.053 | -1.555 | 0.297 | 0.877 | -0.02 | 0.005 | 0.005 | 0 | 0.61 | 0.81 | 0.37 | 0.66 | 0.45 | 0.75 |
| 8 | 60910643 | rs541902 | -0.403 | -1.619 | 0.26 | 0.949 | 0.025 | 0.018 | 0.018 | 0 | 0.49 | 0.66 | 0.33 | 0.62 | 0.35 | 0.72 |

| | | | | | | | | | | | | | | | | |
|---|---|---|---|---|---|---|---|---|---|---|---|---|---|---|---|---|
| 8 | 60938732 | rs1214492 | -0.606 | -1.517 | -0.424 | 0.719 | -0.001 | -0.018 | -0.018 | 0 | 0.89 | 0.94 | 0.69 | 0.77 | 0.86 | 0.91 |
| 8 | 60944902 | rs490409 | -0.629 | -1.559 | -0.309 | 0.583 | 0.002 | -0.003 | -0.003 | 3.437 | 0.56 | 0.62 | 0.35 | 0.54 | 0.41 | 0.71 |
| 8 | 60956071 | rs562271 | 0.028 | -1.508 | -0.315 | 0.716 | -0.002 | -0.001 | -0.001 | 0 | 0.55 | 0.49 | 0.69 | 0.53 | 0.67 | 0.37 |
| 8 | 60958891 | rs473097 | -0.025 | -2.239 | 0.535 | 0.97 | -0.023 | 0.007 | 0.007 | 0 | 0.62 | 0.58 | 0.79 | 0.6 | 0.72 | 0.56 |
| 8 | 60961821 | rs569688 | -0.243 | -1.903 | 0.367 | 1.051 | -0.015 | -0.041 | -0.041 | 0.11 | 0.17 | 0.1 | 0.46 | 0.36 | 0.23 | 0.16 |
| 8 | 60964107 | rs507200 | 0.191 | -2.307 | 0.747 | 0.894 | 0.043 | -0.006 | -0.006 | 0 | 0.55 | 0.36 | 0.75 | 0.48 | 0.71 | 0.5 |
| 8 | 60966679 | rs2688655 | -0.699 | -2.312 | 0.438 | 0.78 | 0.029 | 0.005 | 0.005 | 0 | 0.68 | 0.91 | 0.84 | 0.74 | 0.75 | 0.68 |
| 8 | 60977819 | rs577313 | -0.711 | -2.565 | 0.506 | 0.653 | 0.123 | -0.01 | -0.01 | 0 | 0.49 | 0.76 | 0.26 | 0.49 | 0.31 | 0.44 |
| 8 | 60997355 | rs1464327 | 2.482 | -2.481 | 0.289 | 0.622 | -0.175 | 0.076 | 0.076 | 0.024 | 0.28 | 0.02 | 0.51 | 0.12 | 0.45 | 0.21 |
| 8 | 60998730 | rs2611355 | 0.697 | -2.557 | 0.605 | 0.78 | 0.017 | -0.005 | -0.005 | 0 | 0.3 | 0.08 | 0.16 | 0.25 | 0.24 | 0.32 |
| 8 | 60999098 | rs1980238 | -0.111 | -2.194 | 1.322 | 0.782 | -0.053 | -0.005 | -0.005 | 0 | 0.13 | 0.24 | 0.05 | 0.12 | 0.06 | 0.09 |
| 8 | 61000576 | rs1467300 | -1.143 | -2.208 | 1.467 | 0.803 | -0.088 | 0.047 | 0.047 | 1.857 | 0.18 | 0.61 | 0.09 | 0.27 | 0.06 | 0.12 |
| 8 | 61000585 | rs1467299 | -0.779 | -2.036 | 0.758 | 0.558 | -0.02 | 0.018 | 0.018 | 0 | 0.06 | 0.12 | 0.02 | 0.12 | 0.01 | 0.08 |
| 8 | 61001507 | rs1402028 | 1.529 | -1.97 | 0.673 | 0.536 | -0.023 | -0.043 | -0.043 | 1.723 | 0.08 | 0.13 | 0.03 | 0 | 0.05 | 0 |
| 8 | 61015172 | rs9969570 | -0.54 | -2.09 | 0.299 | 0.399 | -0.02 | -0.019 | -0.019 | 0 | 0.23 | 0.29 | 0.1 | 0.12 | 0.14 | 0.22 |
| 8 | 61018346 | rs16925742 | 1.413 | -2.003 | -0.229 | 0.381 | 0.025 | 0.034 | 0.034 | 0.066 | 0.11 | 0.04 | 0.04 | 0 | 0.09 | 0.05 |
| 8 | 61027810 | rs16925766 | 0.752 | -1.923 | -0.255 | 0.498 | -0.015 | -0.035 | -0.035 | 0.401 | 0.07 | 0 | 0.05 | 0.13 | 0.06 | 0.17 |
| 8 | 61032228 | rs2611366 | -1.178 | -2.598 | -0.013 | 0.537 | 0.072 | -0.004 | -0.004 | 0 | 0.48 | 0.68 | 0.26 | 0.5 | 0.31 | 0.41 |
| 8 | 61032713 | rs939341 | 1.459 | -2.701 | -0.058 | 0.633 | -0.001 | -0.058 | -0.058 | 0 | 0.2 | 0.03 | 0.11 | 0.22 | 0.16 | 0.17 |
| 8 | 61035433 | rs11998573 | -0.435 | -2.622 | 0.576 | 0.35 | -0.006 | 0.049 | 0.049 | 0.542 | 0.12 | 0.4 | 0.07 | 0.13 | 0.06 | 0.18 |
| 8 | 61035873 | rs2681553 | -0.354 | -2.646 | 0.35 | 0.303 | -0.001 | 0.002 | 0.002 | 0 | 0.82 | 0.84 | 0.79 | 0.65 | 0.79 | 0.72 |
| 8 | 61036683 | rs2611367 | 0.926 | -2.578 | 0.356 | 0.378 | 0.015 | -0.025 | -0.025 | 0.05 | 0.24 | 0.17 | 0.15 | 0.37 | 0.17 | 0.19 |
| 8 | 61046885 | rs2611368 | -0.71 | -2.302 | 0.333 | 0.125 | 0.03 | 0 | 0 | 0.083 | 0.44 | 0.66 | 0.27 | 0.53 | 0.31 | 0.38 |
| 8 | 61048614 | rs2681549 | -0.703 | -1.992 | 0.807 | 0.45 | 0.031 | 0.014 | 0.014 | 0 | 0.53 | 0.63 | 0.3 | 0.55 | 0.38 | 0.41 |
| 8 | 61050200 | rs2681548 | 0.992 | -1.842 | 0.777 | 0.417 | 0.032 | -0.009 | -0.009 | 0 | 0.36 | 0.21 | 0.22 | 0.37 | 0.3 | 0.24 |
| 8 | 61050211 | rs2611370 | 0.992 | -1.842 | 0.777 | 0.397 | 0.032 | -0.007 | -0.007 | 0 | 0.36 | 0.21 | 0.22 | 0.37 | 0.3 | 0.24 |
| 8 | 61051128 | rs13263818 | 2.28 | -1.789 | 0.472 | 0.303 | -0.147 | 0.052 | 0.052 | 0 | 0.26 | 0.04 | 0.48 | 0.14 | 0.4 | 0.24 |
| 8 | 61052304 | rs6998241 | 1.245 | -1.791 | 0.667 | 0.303 | -0.001 | -0.008 | -0.008 | 0 | 0.34 | 0.39 | 0.51 | 0.14 | 0.42 | 0.24 |
| 8 | 61056493 | rs2681543 | 0.85 | -1.63 | 0.564 | 0.42 | -0.005 | 0.055 | 0.055 | 0.202 | 0.7 | 0.61 | 0.74 | 0.51 | 0.72 | 0.49 |
| 8 | 61057832 | rs2611339 | 0.85 | -1.63 | 0.564 | 0.42 | -0.005 | 0.055 | 0.055 | 0.202 | 0.7 | 0.61 | 0.74 | 0.51 | 0.72 | 0.49 |
| 8 | 61064200 | rs2176835 | 1.205 | -1.572 | 0.176 | 0.4 | -0.097 | 0.077 | 0.077 | 0 | 0.33 | 0.09 | 0.53 | 0.19 | 0.46 | 0.38 |
| 8 | 61064522 | rs2681539 | 0.031 | -1.447 | 0.612 | 0.526 | -0.002 | 0.03 | 0.03 | 1.37 | 0.79 | 0.75 | 0.79 | 0.56 | 0.8 | 0.64 |
| 8 | 61067170 | rs2611348 | -0.884 | -1.212 | 0.186 | 0.607 | -0.059 | -0.108 | -0.108 | 0 | 0.85 | 0.99 | 0.81 | 0.61 | 0.82 | 0.75 |
| 8 | 61081263 | rs28608109 | 0.686 | -1.292 | 0.222 | 0.584 | 0 | 0 | 0 | 0 | 0.08 | 0.05 | 0.05 | 0.05 | 0.05 | 0.13 |
| 8 | 61082900 | rs17811959 | 0.804 | -1.284 | 0.227 | 0.588 | 0 | 0 | 0 | 0.71 | 0.07 | 0.05 | 0.05 | 0.05 | 0.05 | 0.13 |
| 8 | 61085525 | rs2681536 | -0.834 | -1.272 | 0.286 | 0.613 | -0.011 | -0.04 | -0.04 | 2.318 | 0.84 | 0.92 | 0.8 | 0.62 | 0.82 | 0.75 |
| 8 | 61089580 | rs2681554 | -0.528 | -1.571 | 0.773 | 0.454 | 0.041 | 0.021 | 0.021 | 0 | 0.49 | 0.34 | 0.3 | 0.41 | 0.38 | 0.38 |
| 8 | 61090641 | rs1520269 | 0.35 | -1.338 | 0.77 | 0.552 | 0.047 | 0.004 | 0.004 | 0.481 | 0.4 | 0.22 | 0.24 | 0.36 | 0.34 | 0.25 |
| 8 | 61094650 | rs12335239 | -0.151 | -1.327 | 0.977 | 0.549 | 0.006 | 0.048 | 0.048 | 0.08 | 0.06 | 0.22 | 0.02 | 0.07 | 0.03 | 0.14 |
| 8 | 61096176 | rs720208 | -0.706 | -1.348 | 0.586 | 0.52 | 0.031 | 0.001 | 0.001 | 0 | 0.49 | 0.58 | 0.31 | 0.48 | 0.4 | 0.49 |
| 8 | 61101126 | rs2242155 | 0.502 | -1.295 | -0.005 | 0.548 | 0.027 | -0.005 | -0.005 | 0 | 0.37 | 0.21 | 0.24 | 0.36 | 0.32 | 0.24 |
| 8 | 61101162 | rs2242156 | 0.502 | -1.295 | -0.005 | 0.548 | 0.027 | -0.005 | -0.005 | 0 | 0.37 | 0.21 | 0.24 | 0.36 | 0.32 | 0.24 |
| 8 | 61105199 | rs10098647 | -0.613 | -0.837 | -0.681 | 0.694 | 0.054 | -0.011 | -0.011 | 0 | 0.53 | 0.7 | 0.32 | 0.48 | 0.4 | 0.51 |
| 8 | 61117903 | rs7841174 | -0.618 | -0.83 | -0.875 | 0.731 | 0.059 | -0.027 | -0.027 | 0 | 0.53 | 0.78 | 0.33 | 0.47 | 0.4 | 0.51 |
| 8 | 61120869 | rs12677216 | 0.945 | -0.315 | -0.426 | 0.695 | -0.025 | 0.026 | 0.026 | 0 | 0.32 | 0.22 | 0.48 | 0.21 | 0.42 | 0.27 |
| 8 | 61122903 | rs11991354 | 0.888 | -0.345 | -0.108 | 0.672 | -0.011 | 0.011 | 0.011 | 0 | 0.33 | 0.26 | 0.48 | 0.21 | 0.42 | 0.28 |
| 8 | 61125863 | rs7829561 | 0.231 | -0.959 | 0.066 | 0.75 | 0 | 0.006 | 0.006 | 0 | 0.49 | 0.45 | 0.57 | 0.39 | 0.52 | 0.45 |
| 8 | 61130839 | rs7462276 | 1.028 | -0.965 | 0.054 | 0.8 | 0.019 | -0.025 | -0.025 | 2.724 | 0.14 | 0.05 | 0.08 | 0.18 | 0.1 | 0.17 |
| 8 | 61132386 | rs10100314 | 1.088 | -1.11 | 0.209 | 1.009 | 0.003 | 0 | 0 | 0 | 0.29 | 0.23 | 0.25 | 0.28 | 0.3 | 0.25 |
| 8 | 61132637 | rs10957123 | 0.656 | -1.095 | 0.304 | 0.925 | 0.001 | -0.001 | -0.001 | 0.019 | 0.33 | 0.2 | 0.27 | 0.31 | 0.35 | 0.3 |
| 8 | 61134995 | rs6471849 | -0.73 | -0.943 | -0.113 | 1.177 | 0.006 | -0.053 | -0.053 | 0 | 0.82 | 0.94 | 0.89 | 0.73 | 0.85 | 0.77 |
| 8 | 61140958 | rs7463207 | 1.461 | -0.932 | -0.109 | 1.1 | -0.033 | 0.045 | 0.045 | 0.008 | 0.34 | 0.19 | 0.48 | 0.21 | 0.41 | 0.28 |
| 8 | 61145146 | rs11784742 | -0.583 | -1.012 | -0.139 | 1.231 | 0.002 | -0.008 | -0.008 | 0 | 0.92 | 0.98 | 0.94 | 0.73 | 0.94 | 0.82 |
| 8 | 61145264 | rs6986917 | -0.125 | -1.03 | -0.052 | 1.212 | 0.013 | -0.006 | -0.006 | 0 | 0.46 | 0.63 | 0.4 | 0.52 | 0.44 | 0.49 |
| 8 | 61145347 | rs6987086 | -0.386 | -1.01 | -0.197 | 1.225 | 0.003 | -0.008 | -0.008 | 0 | 0.92 | 0.97 | 0.94 | 0.73 | 0.94 | 0.82 |
| 8 | 61146834 | rs7465501 | -0.409 | -0.993 | -0.102 | 1.215 | 0.003 | 0.003 | 0.003 | 0 | 0.5 | 0.56 | 0.43 | 0.53 | 0.49 | 0.54 |
| 8 | 61149123 | rs7465573 | 0.294 | -0.688 | -0.263 | 1.302 | -0.001 | 0.026 | 0.026 | 0.75 | 0.84 | 0.82 | 0.85 | 0.52 | 0.87 | 0.64 |
| 8 | 61149731 | rs6984526 | -0.314 | -0.469 | 0.026 | 1.056 | -0.006 | 0.013 | 0.013 | 0 | 0.55 | 0.63 | 0.39 | 0.36 | 0.52 | 0.54 |
| 8 | 61150249 | rs10081509 | 0.955 | -0.903 | 0.513 | 1.023 | -0.038 | 0.087 | 0.087 | 0 | 0.85 | 0.59 | 0.88 | 0.77 | 0.93 | 0.81 |
| 8 | 61151150 | rs12708003 | 0.557 | -0.196 | 0.255 | 0.854 | 0.018 | 0.045 | 0.045 | 0.588 | 0.25 | 0.44 | 0.43 | 0.37 | 0.3 | 0.24 |
| 8 | 61151574 | rs8180936 | -0.46 | -0.698 | -0.273 | 0.281 | 0.003 | 0.004 | 0.004 | 0 | 0.66 | 0.87 | 0.69 | 0.63 | 0.68 | 0.62 |
| 8 | 61156208 | rs6999752 | 0.776 | -1.142 | -0.207 | -0.098 | -0.065 | -0.023 | -0.023 | 0 | 0.14 | 0.35 | 0.05 | 0.1 | 0.05 | 0.13 |
| 8 | 61162506 | rs6998008 | 0.635 | -1.22 | -0.422 | -0.191 | 0.01 | -0.02 | -0.02 | 0 | 0.32 | 0.48 | 0.26 | 0.16 | 0.34 | 0.28 |
| 8 | 61178574 | rs7464181 | -0.469 | -1.213 | -0.511 | -0.162 | 0.02 | 0.085 | 0.085 | 0.209 | 0.49 | 0.85 | 0.4 | 0.27 | 0.42 | 0.43 |
| 8 | 61185722 | rs10216440 | -1.25 | -1.372 | -0.785 | -0.247 | 0.004 | 0.01 | 0.01 | 0 | 0.82 | 0.9 | 0.85 | 0.84 | 0.81 | 0.85 |
| 8 | 61187018 | rs7843385 | -0.044 | -1.351 | -0.27 | -0.259 | 0 | 0.004 | 0.004 | 0.133 | 0.45 | 0.63 | 0.36 | 0.27 | 0.41 | 0.43 |
| 8 | 61191590 | rs7841372 | 0.268 | -1.327 | -0.368 | -0.239 | -0.004 | 0.006 | 0.006 | 0 | 0.39 | 0.4 | 0.33 | 0.27 | 0.4 | 0.41 |
| 8 | 61200304 | rs16925975 | 1.49 | -1.155 | -0.453 | -0.09 | 0.002 | -0.001 | -0.001 | 0 | 0.15 | 0.15 | 0.14 | 0.21 | 0.11 | 0.21 |
| 8 | 61205607 | rs6983247 | 0.704 | -1.448 | -0.327 | -0.216 | -0.01 | 0.006 | 0.006 | 0 | 0.33 | 0.28 | 0.33 | 0.18 | 0.37 | 0.31 |

| | | | | | | | | | | | | | | | | |
|---|---|---|---|---|---|---|---|---|---|---|---|---|---|---|---|---|
| 8 | 61209152 | rs7820639 | 2.033 | -1.472 | -0.451 | -0.204 | 0.004 | -0.008 | -0.008 | 0 | 0.13 | 0.05 | 0.11 | 0.21 | 0.1 | 0.2 |
| 8 | 61209459 | rs7842871 | -0.179 | -1.574 | -0.331 | -0.284 | 0.043 | 0.025 | 0.025 | 1.768 | 0.33 | 0.76 | 0.21 | 0.31 | 0.17 | 0.34 |
| 8 | 61225652 | rs11775891 | 0.94 | -1.603 | -0.543 | -0.337 | 0.094 | 0.048 | 0.048 | 0.004 | 0.48 | 0.12 | 0.63 | 0.53 | 0.65 | 0.51 |
| 8 | 61226540 | rs7001157 | -0.56 | -1.434 | -0.491 | -0.237 | -0.044 | -0.055 | -0.055 | 0 | 0.62 | 0.84 | 0.51 | 0.51 | 0.53 | 0.63 |
| 8 | 61230529 | rs7464602 | 1.328 | -1.387 | -0.067 | -0.241 | 0.03 | 0.067 | 0.067 | 0.385 | 0.25 | 0.47 | 0.38 | 0.35 | 0.29 | 0.24 |
| 8 | 61233350 | rs9298035 | -1.018 | -1.126 | 0.261 | -0.412 | 0.004 | 0.024 | 0.024 | 0 | 0.82 | 0.9 | 0.85 | 0.87 | 0.82 | 0.84 |
| 8 | 61238290 | rs17740762 | 1.4 | -0.827 | -0.159 | -0.319 | -0.018 | 0.042 | 0.042 | 0 | 0.08 | 0.01 | 0.08 | 0 | 0.11 | 0.06 |
| 8 | 61250319 | rs6982717 | -0.137 | -0.753 | 0.265 | -0.087 | 0.002 | 0.011 | 0.011 | 0 | 0.47 | 0.57 | 0.52 | 0.69 | 0.45 | 0.54 |
| 8 | 61276247 | rs11998308 | -0.285 | -0.498 | 0.095 | 0.132 | -0.003 | -0.008 | -0.008 | 0 | 0.61 | 0.66 | 0.56 | 0.53 | 0.61 | 0.6 |
| 8 | 61287045 | rs12677500 | 0.828 | -0.962 | -0.461 | 0.085 | 0.001 | -0.055 | -0.055 | 0.616 | 0.15 | 0.01 | 0.19 | 0.26 | 0.12 | 0.2 |
| 8 | 61294126 | rs11787444 | 0.967 | -0.941 | -0.493 | -0.196 | 0.019 | 0.024 | 0.024 | 0 | 0.21 | 0.13 | 0.13 | 0.11 | 0.14 | 0.15 |
| 8 | 61306938 | rs1552071 | -0.071 | -0.483 | -0.172 | 0.12 | -0.001 | 0.001 | 0.001 | 0 | 0.61 | 0.52 | 0.41 | 0.51 | 0.4 | 0.51 |
| 8 | 61307115 | rs10102945 | -1.333 | 0.397 | -0.548 | 0.173 | -0.108 | -0.091 | -0.091 | 0 | 0.8 | 0.97 | 0.62 | 0.68 | 0.6 | 0.73 |
| 8 | 61309122 | rs10504311 | 0.914 | 0.099 | -0.422 | 0.056 | 0.031 | 0.026 | 0.026 | 0 | 0.41 | 0.28 | 0.26 | 0.3 | 0.29 | 0.33 |
| 8 | 61309198 | rs10504312 | -1.693 | 0.13 | -0.492 | 0.066 | -0.134 | -0.113 | -0.113 | 0 | 0.8 | 0.99 | 0.62 | 0.67 | 0.6 | 0.73 |
| 8 | 61311510 | rs2086393 | -0.569 | -0.144 | -0.103 | 0.146 | -0.002 | -0.001 | -0.001 | 0 | 0.38 | 0.61 | 0.35 | 0.37 | 0.32 | 0.4 |
| 8 | 61319467 | rs7017556 | 0.958 | -0.226 | -0.088 | 0.13 | 0.007 | 0.004 | 0.004 | 0 | 0.38 | 0.34 | 0.25 | 0.31 | 0.23 | 0.33 |
| 8 | 61322563 | rs7826226 | 1.103 | -0.227 | -0.101 | 0.129 | 0.032 | 0.015 | 0.015 | 0 | 0.37 | 0.26 | 0.23 | 0.31 | 0.23 | 0.33 |
| 8 | 61330269 | rs13438846 | 0.839 | -0.163 | 0.162 | 0.159 | 0.018 | 0.008 | 0.008 | 3.428 | 0.12 | 0.16 | 0.12 | 0.08 | 0.15 | 0.1 |
| 8 | 61341711 | rs4354350 | 0.4 | -0.36 | 0.378 | 0.065 | -0.004 | 0.001 | 0.001 | 0 | 0.54 | 0.44 | 0.37 | 0.57 | 0.38 | 0.53 |
| 8 | 61368068 | rs7013829 | 0.747 | -0.424 | 0.513 | 0.031 | -0.044 | -0.05 | -0.05 | 0.33 | 0.09 | 0.25 | 0.03 | 0 | 0.02 | 0.08 |
| 8 | 61369006 | rs7829407 | 1.395 | -0.445 | 0.428 | 0.033 | 0.004 | -0.013 | -0.013 | 0.055 | 0.16 | 0.08 | 0.13 | 0.27 | 0.15 | 0.2 |
| 8 | 61373340 | rs6471880 | -1.312 | -0.33 | 0.416 | 0.061 | -0.074 | -0.049 | -0.049 | 0 | 0.75 | 0.84 | 0.52 | 0.65 | 0.54 | 0.71 |
| 8 | 61373597 | rs13254748 | 0.956 | -0.284 | 0.43 | 0.049 | 0.014 | 0.006 | 0.006 | 3.459 | 0.11 | 0.15 | 0.12 | 0.08 | 0.15 | 0.1 |
| 8 | 61396829 | rs2013112 | 0.173 | -0.711 | 0.59 | 0.096 | -0.006 | 0.001 | 0.001 | 0 | 0.55 | 0.51 | 0.39 | 0.59 | 0.38 | 0.55 |
| 8 | 61418431 | rs2326561 | 0.302 | -0.717 | 0.502 | 0.081 | 0.009 | -0.003 | -0.003 | 0 | 0.53 | 0.39 | 0.38 | 0.57 | 0.38 | 0.52 |
| 8 | 61428280 | rs2875967 | -0.736 | -0.791 | 0.357 | 0.04 | -0.113 | -0.165 | -0.165 | 0 | 0.67 | 0.96 | 0.43 | 0.59 | 0.4 | 0.63 |
| 8 | 61446150 | rs16926265 | 0.998 | -0.812 | 0.586 | 0.038 | -0.036 | -0.042 | -0.042 | 0.072 | 0.08 | 0.23 | 0.02 | 0 | 0.02 | 0.08 |
| 8 | 61452046 | rs2272620 | -0.602 | -0.825 | 0.564 | 0.033 | -0.065 | -0.117 | -0.117 | 2.222 | 0.65 | 0.9 | 0.43 | 0.59 | 0.4 | 0.63 |
| 8 | 61458899 | rs2007283 | 1.371 | -0.862 | 0.603 | 0.044 | 0.001 | -0.003 | -0.003 | 0 | 0.15 | 0.11 | 0.14 | 0.27 | 0.16 | 0.2 |
| 8 | 61462541 | rs11778888 | 1.281 | -0.866 | 0.624 | 0.055 | 0.05 | 0.023 | 0.023 | 0 | 0.37 | 0.21 | 0.23 | 0.3 | 0.22 | 0.33 |
| 8 | 61481614 | rs10098769 | 0.525 | -0.459 | 0.764 | 0.204 | 0.019 | -0.006 | -0.006 | 0.19 | 0.53 | 0.35 | 0.37 | 0.57 | 0.38 | 0.52 |
| 8 | 61512060 | rs10957150 | 0.113 | -0.472 | 0.762 | 0.198 | 0.013 | -0.011 | -0.011 | 0.847 | 0.57 | 0.66 | 0.41 | 0.59 | 0.38 | 0.55 |
| 8 | 61541273 | rs631408 | -0.104 | -0.206 | 0.154 | 0.324 | -0.001 | -0.003 | -0.003 | 0 | 0.62 | 0.59 | 0.39 | 0.57 | 0.4 | 0.6 |
| 8 | 61548494 | rs626913 | 0.373 | 0.104 | -0.303 | 0.05 | -0.004 | 0.01 | 0.01 | 0.831 | 0.56 | 0.35 | 0.46 | 0.65 | 0.52 | 0.52 |
| 8 | 61550764 | rs669960 | 0.585 | 0.048 | -0.559 | 0.246 | 0.011 | 0.008 | 0.008 | 0 | 0.46 | 0.46 | 0.28 | 0.33 | 0.25 | 0.34 |
| 8 | 61559253 | rs654534 | -0.772 | 0.762 | -1.274 | -0.061 | -0.169 | -0.045 | -0.045 | 1.134 | 0.75 | 0.98 | 0.53 | 0.73 | 0.53 | 0.7 |
| 8 | 61566655 | rs657718 | -0.57 | 1.046 | -1.267 | -0.142 | -0.158 | -0.059 | -0.059 | 0 | 0.67 | 0.97 | 0.5 | 0.67 | 0.53 | 0.61 |
| 8 | 61568210 | rs623917 | -0.29 | 0.597 | -0.814 | -0.09 | -0.032 | -0.003 | -0.003 | 0 | 0.65 | 0.75 | 0.45 | 0.65 | 0.47 | 0.59 |
| 8 | 61580676 | rs622755 | -0.599 | 0.559 | -0.997 | -0.054 | -0.106 | -0.011 | -0.011 | 0 | 0.65 | 0.9 | 0.48 | 0.67 | 0.49 | 0.61 |
| 8 | 61581424 | rs663747 | 0.382 | 0.263 | -0.868 | -0.483 | -0.007 | -0.048 | -0.048 | 0 | 0.35 | 0.34 | 0.19 | 0.06 | 0.2 | 0.26 |
| 8 | 61586261 | rs618190 | -0.16 | 0.691 | -1.036 | 0.124 | -0.028 | -0.002 | -0.002 | 3.063 | 0.62 | 0.68 | 0.46 | 0.64 | 0.49 | 0.57 |
| 8 | 61597447 | rs7013969 | -0.436 | -0.112 | 0.187 | -0.175 | -0.027 | -0.055 | -0.055 | 0.146 | 0.1 | 0.3 | 0.06 | 0.01 | 0.03 | 0.09 |
| 8 | 61599122 | rs900503 | 0.333 | 0.096 | -1.192 | -0.399 | -0.094 | 0.091 | 0.091 | 3.993 | 0.44 | 0.04 | 0.62 | 0.74 | 0.59 | 0.72 |
| 8 | 61605768 | rs11990117 | -0.134 | -0.193 | -0.814 | -0.808 | 0.014 | 0.019 | 0.019 | 0.188 | 0.64 | 0.35 | 0.75 | 0.94 | 0.73 | 0.85 |
| 8 | 61606762 | rs12680437 | 1.444 | 0.186 | -1.237 | -0.185 | -0.182 | -0.123 | -0.123 | 0 | 0.31 | 0.03 | 0.57 | 0.39 | 0.49 | 0.46 |
| 8 | 61615461 | rs6981399 | 1.045 | 0.065 | -0.611 | -0.175 | 0.005 | -0.002 | -0.002 | 0 | 0.18 | 0.13 | 0.12 | 0.21 | 0.13 | 0.12 |
| 8 | 61620613 | rs4738817 | 1.151 | 0.034 | -0.689 | -0.044 | -0.141 | -0.145 | -0.145 | 0 | 0.34 | 0.03 | 0.56 | 0.42 | 0.46 | 0.55 |
| 8 | 61634534 | rs11989002 | 0.822 | -0.298 | -0.04 | -0.071 | -0.024 | 0 | 0 | 0 | 0.2 | 0.29 | 0.12 | 0.22 | 0.08 | 0.12 |
| 8 | 61635991 | rs10957157 | -1.555 | -0.399 | 0.139 | 0.577 | 0.002 | 0.331 | 0.331 | 0 | 0.07 | 0.21 | 0.12 | 0.39 | 0.04 | 0.33 |
| 8 | 61644700 | rs16926440 | 0.994 | -0.353 | -0.121 | -0.177 | -0.017 | -0.024 | -0.024 | 0 | 0.07 | 0.08 | 0.02 | 0.01 | 0.01 | 0.07 |
| 8 | 61648515 | rs13255519 | -1.347 | -0.296 | -0.155 | -0.263 | 0 | -0.017 | -0.017 | 0 | 0.75 | 0.72 | 0.82 | 0.94 | 0.77 | 0.93 |
| 8 | 61648551 | rs13256023 | 0.042 | -0.179 | -0.299 | -0.24 | -0.017 | -0.04 | -0.04 | 0 | 0.28 | 0.56 | 0.2 | 0.16 | 0.2 | 0.14 |
| 8 | 61652144 | rs4288413 | 0.652 | -0.033 | -0.496 | -0.044 | -0.141 | 0.074 | 0.074 | 2.554 | 0.39 | 0.06 | 0.6 | 0.78 | 0.55 | 0.72 |
| 8 | 61686062 | rs16926466 | 1.028 | 0.171 | 0.193 | 0.28 | -0.013 | -0.013 | -0.013 | 0 | 0.06 | 0.06 | 0.01 | 0 | 0 | 0.05 |
| 8 | 61701057 | rs4237036 | 0.654 | -0.011 | -0.194 | 0.147 | -0.013 | 0.155 | 0.155 | 2.521 | 0.48 | 0.06 | 0.64 | 0.78 | 0.66 | 0.73 |
| 8 | 61709500 | rs4391471 | -0.418 | -0.07 | -0.059 | 0.091 | 0.004 | 0.018 | 0.018 | 0.035 | 0.71 | 0.74 | 0.83 | 0.94 | 0.77 | 0.89 |
| 8 | 61712370 | rs4289855 | -1.442 | 0.016 | -0.123 | 0.11 | -0.073 | 0.096 | 0.096 | 1.172 | 0.79 | 0.97 | 0.84 | 0.96 | 0.77 | 0.95 |
| 8 | 61722552 | rs6988835 | 0.395 | 0.095 | 0.117 | 0.173 | -0.016 | -0.012 | -0.012 | 0 | 0.22 | 0.65 | 0.19 | 0.17 | 0.11 | 0.15 |
| 8 | 61726879 | rs10092214 | 1.146 | 0.387 | -0.154 | 0.279 | -0.116 | -0.174 | -0.174 | 0 | 0.38 | 0.03 | 0.56 | 0.5 | 0.46 | 0.52 |
| 8 | 61764076 | rs4342642 | -0.051 | 0.151 | 0.165 | -0.51 | -0.017 | 0.097 | 0.097 | 0.406 | 0.55 | 0.77 | 0.71 | 0.78 | 0.61 | 0.76 |
| 8 | 61765273 | rs3763592 | -0.558 | 0.096 | 0.355 | -0.534 | -0.022 | -0.05 | -0.05 | 0.087 | 0.11 | 0.43 | 0.08 | 0.01 | 0.04 | 0.05 |
| 8 | 61765560 | rs2068096 | -0.543 | 0.109 | 0.375 | -0.527 | -0.023 | -0.047 | -0.047 | 0 | 0.1 | 0.36 | 0.07 | 0.01 | 0.04 | 0.05 |
| 8 | 61780289 | rs6985168 | 1.077 | 0.143 | -0.414 | -0.467 | -0.147 | 0.047 | 0.047 | 1.146 | 0.42 | 0.06 | 0.6 | 0.74 | 0.56 | 0.69 |
| 8 | 61782103 | rs7826377 | -0.186 | 0.168 | -0.009 | -0.475 | -0.016 | -0.046 | -0.046 | 0 | 0.09 | 0.28 | 0.06 | 0.01 | 0.04 | 0.05 |
| 8 | 61782304 | rs7844586 | -1.068 | 0.203 | -0.521 | -0.56 | -0.078 | 0.117 | 0.117 | 0.302 | 0.77 | 0.97 | 0.85 | 0.96 | 0.75 | 0.94 |
| 8 | 61783211 | rs13280121 | 0.598 | 0.563 | -0.168 | -0.157 | 0.03 | 0.041 | 0.041 | 0 | 0.42 | 0.67 | 0.57 | 0.24 | 0.37 | 0.33 |
| 8 | 61784348 | rs4738835 | -0.634 | 0.55 | -0.632 | -0.111 | -0.043 | 0.098 | 0.098 | 0.959 | 0.61 | 0.88 | 0.75 | 0.8 | 0.68 | 0.8 |
| 8 | 61785874 | rs4237042 | -0.02 | 0.728 | -0.523 | -0.102 | 0.013 | 0.126 | 0.126 | 0.14 | 0.48 | 0.86 | 0.64 | 0.27 | 0.44 | 0.38 |

| | | | | | | | | | | | | | | | | |
|---|---|---|---|---|---|---|---|---|---|---|---|---|---|---|---|---|
| 8 | 61788144 | rs13279104 | -0.989 | 0.716 | -0.583 | -0.077 | -0.015 | -0.052 | -0.052 | 0.043 | 0.17 | 0.58 | 0.14 | 0.05 | 0.11 | 0.1 |
| 8 | 61789152 | rs7005218 | 1.391 | 0.958 | -1.336 | 0.164 | -0.022 | -0.002 | -0.002 | 0 | 0.29 | 0.03 | 0.47 | 0.2 | 0.31 | 0.27 |
| 8 | 61790325 | rs2280916 | 0.568 | 1.138 | -1.31 | 0.342 | -0.079 | -0.121 | -0.121 | 0 | 0.15 | 0.02 | 0.13 | 0.55 | 0.24 | 0.43 |
| 8 | 61797890 | rs4738839 | 1.377 | 0.796 | -1.383 | 0.031 | -0.027 | 0.007 | 0.007 | 0 | 0.29 | 0.02 | 0.47 | 0.19 | 0.31 | 0.28 |
| 8 | 61802076 | rs7837493 | 0.799 | 0.852 | -1.082 | 0.014 | -0.038 | 0.087 | 0.087 | 0 | 0.37 | 0.12 | 0.58 | 0.21 | 0.43 | 0.28 |
| 8 | 61802242 | rs7837548 | -0.178 | 0.747 | -0.696 | -0.014 | -0.023 | -0.038 | -0.038 | 0.461 | 0.65 | 0.51 | 0.79 | 0.82 | 0.8 | 0.78 |
| 8 | 61804265 | rs16926555 | 0.641 | 1.022 | -1.165 | 0.155 | -0.014 | -0.016 | -0.016 | 0 | 0.16 | 0.1 | 0.16 | 0.57 | 0.27 | 0.46 |
| 8 | 61804729 | rs13251621 | 0.592 | 0.902 | -0.789 | -0.121 | 0.01 | -0.067 | -0.067 | 1.973 | 0.21 | 0.58 | 0.16 | 0.07 | 0.19 | 0.08 |
| 8 | 61804740 | rs4738840 | 0.298 | 0.949 | -1.609 | 0.192 | -0.024 | 0.124 | 0.124 | 0.408 | 0.47 | 0.16 | 0.68 | 0.76 | 0.62 | 0.7 |
| 8 | 61805324 | rs12546463 | 0.151 | 1.047 | -1.602 | 0.22 | 0.005 | 0.128 | 0.128 | 0.042 | 0.49 | 0.22 | 0.69 | 0.76 | 0.63 | 0.7 |
| 8 | 61809348 | rs11991058 | -0.109 | 1.205 | -1.212 | 0.879 | -0.008 | -0.023 | -0.023 | 0 | 0.21 | 0.15 | 0.16 | 0.39 | 0.28 | 0.43 |
| 8 | 61809918 | rs6984381 | -0.49 | 0.924 | -1.639 | -0.073 | 0.011 | 0.007 | 0.007 | 0 | 0.78 | 0.89 | 0.75 | 0.79 | 0.86 | 0.84 |
| 8 | 61809929 | rs6984384 | 0.356 | 1.096 | -1.524 | -0.537 | -0.012 | 0.219 | 0.219 | 0.082 | 0.45 | 0.18 | 0.39 | 0.08 | 0.46 | 0.22 |
| 8 | 61812553 | rs6982175 | 0.316 | 1.629 | -0.631 | 0.066 | 0.047 | -0.005 | -0.005 | 0.111 | 0.93 | 0.87 | 0.94 | 0.93 | 0.91 | 0.99 |
| 8 | 61813615 | rs4289856 | 1.182 | 0.79 | 0.323 | -0.376 | 0.008 | -0.072 | -0.072 | 0 | 0.82 | 0.44 | 0.89 | 0.92 | 0.88 | 0.94 |
| 8 | 61814443 | rs17827642 | 1.464 | 0.922 | -0.94 | -0.918 | -0.068 | 0.162 | 0.162 | 2.79 | 0.4 | 0.05 | 0.53 | 0.16 | 0.41 | 0.11 |
| 8 | 61815042 | rs16926569 | -0.017 | 1.441 | -0.555 | -0.014 | -0.001 | 0.004 | 0.004 | 0 | 0.08 | 0.07 | 0.08 | 0.16 | 0.11 | 0.11 |
| 8 | 61815886 | rs10957167 | 1.541 | 1.552 | 0.246 | 0.445 | 0.075 | 0.059 | 0.059 | 0 | 0.76 | 0.55 | 0.78 | 0.37 | 0.68 | 0.61 |
| 8 | 61824056 | rs6993285 | -1.316 | 1.516 | 0.279 | 0.492 | -0.003 | 0.075 | 0.075 | 0 | 0.26 | 0.47 | 0.21 | 0.55 | 0.26 | 0.43 |
| 8 | 61827676 | rs12680914 | 0.476 | 1.533 | 0.277 | 0.547 | 0.003 | 0.092 | 0.092 | 0.056 | 0.07 | 0.07 | 0.07 | 0.39 | 0.13 | 0.27 |
| 8 | 61829154 | rs10957168 | -0.798 | 1.386 | 0.39 | 0.314 | -0.031 | 0.01 | 0.01 | 0 | 0.19 | 0.41 | 0.14 | 0.16 | 0.14 | 0.17 |
| 8 | 61829264 | rs12549487 | -0.798 | 1.386 | 0.39 | 0.314 | -0.031 | 0.01 | 0.01 | 0 | 0.19 | 0.41 | 0.14 | 0.16 | 0.14 | 0.17 |
| 8 | 61831999 | rs7008176 | 0.398 | 1.513 | 0.36 | 0.248 | 0.028 | -0.015 | -0.015 | 0 | 0.75 | 0.65 | 0.77 | 0.8 | 0.69 | 0.64 |
| 8 | 61833637 | rs7846503 | 0.271 | 1.081 | 0.601 | 0.259 | 0.062 | 0.011 | 0.011 | 0.186 | 0.9 | 0.62 | 0.92 | 0.92 | 0.92 | 0.99 |
| 8 | 61840558 | rs13257990 | -0.612 | 0.599 | 0.596 | 0.427 | -0.025 | 0.024 | 0.024 | 0 | 0.14 | 0.28 | 0.11 | 0.17 | 0.08 | 0.12 |
| 8 | 61849086 | rs10087102 | -0.021 | 0.902 | 0.094 | 0.786 | -0.023 | -0.006 | -0.006 | 0 | 0.1 | 0.05 | 0.07 | 0.39 | 0.16 | 0.35 |
| 8 | 61849890 | rs1982638 | 0.01 | 0.857 | 0.112 | 0.453 | 0.002 | 0.008 | 0.008 | 0.352 | 0.2 | 0.21 | 0.18 | 0.09 | 0.2 | 0.28 |
| 8 | 61854841 | rs4554462 | -2.042 | 0.767 | 0.15 | 0.366 | -0.001 | 0 | 0 | 2.108 | 0.52 | 0.53 | 0.39 | 0.71 | 0.51 | 0.88 |
| 8 | 61855118 | rs4515540 | -0.656 | 0.409 | 0.245 | 0.083 | -0.001 | -0.01 | -0.01 | 21.266 | 0.3 | 0.34 | 0.22 | 0.14 | 0.26 | 0.42 |
| 8 | 61858213 | rs10957174 | -1.097 | 0.667 | 0.157 | 0.213 | 0.004 | 0.004 | 0.004 | 0.023 | 0.43 | 0.44 | 0.31 | 0.32 | 0.34 | 0.53 |
| 8 | 61860596 | rs10108543 | -1.128 | 0.601 | 0.241 | 0.262 | -0.005 | -0.005 | -0.005 | 0 | 0.46 | 0.46 | 0.42 | 0.55 | 0.36 | 0.66 |
| 8 | 61862346 | rs7001966 | -0.896 | 0.87 | 0.68 | 0.095 | 0.014 | -0.049 | -0.049 | 0 | 0.18 | 0.31 | 0.15 | 0.06 | 0.2 | 0.29 |
| 8 | 61862880 | rs11988513 | -1.086 | 0.813 | 0.895 | 0.093 | 0.012 | -0.061 | -0.061 | 0 | 0.19 | 0.36 | 0.16 | 0.06 | 0.2 | 0.31 |
| 8 | 61869649 | rs12335258 | -0.321 | 0.805 | 0.813 | 0.077 | -0.006 | 0.014 | 0.014 | 0 | 0.13 | 0.05 | 0.09 | 0.06 | 0.15 | 0.25 |
| 8 | 61871269 | rs13280897 | -1.758 | 0.981 | 0.663 | 0.585 | 0.009 | 0.049 | 0.049 | 0 | 0.33 | 0.44 | 0.28 | 0.46 | 0.37 | 0.58 |
| 8 | 61877623 | rs882229 | 0.101 | 1.007 | 0.565 | 0.582 | -0.001 | 0 | 0 | 0 | 0.15 | 0.16 | 0.16 | 0.13 | 0.16 | 0.16 |
| 8 | 61882751 | rs12678105 | 0.307 | 1.107 | 0.229 | 0.706 | -0.042 | -0.098 | -0.098 | 0 | 0.13 | 0.01 | 0.11 | 0.4 | 0.16 | 0.27 |
| 8 | 61885271 | rs6992440 | -1.828 | 1.131 | 0.491 | 0.528 | -0.004 | 0.046 | 0.046 | 0 | 0.49 | 0.65 | 0.45 | 0.59 | 0.52 | 0.73 |
| 8 | 61888469 | rs10113311 | -1.523 | 1.299 | 0.378 | 0.539 | -0.001 | 0.001 | 0.001 | 0 | 0.43 | 0.59 | 0.42 | 0.55 | 0.46 | 0.7 |
| 8 | 61890812 | rs7842384 | -1.706 | 1.469 | 0.305 | 0.522 | -0.007 | 0.012 | 0.012 | 0.157 | 0.47 | 0.64 | 0.46 | 0.55 | 0.52 | 0.75 |
| 8 | 61893034 | rs6471907 | -0.393 | 1.521 | 0.08 | 0.512 | 0.015 | 0.072 | 0.072 | 0 | 0.86 | 0.73 | 0.69 | 0.69 | 0.87 | 0.85 |
| 8 | 61898217 | rs2280813 | 0.349 | 1.353 | -0.2 | 0.771 | -0.033 | -0.099 | -0.099 | 0.063 | 0.13 | 0.01 | 0.1 | 0.39 | 0.14 | 0.33 |
| 8 | 61904948 | rs9692725 | -0.892 | 1.437 | 0.434 | 0.234 | 0.011 | -0.069 | -0.069 | 0 | 0.22 | 0.38 | 0.16 | 0.07 | 0.22 | 0.29 |
| 8 | 61909133 | rs16926675 | -0.361 | 1.31 | 0.387 | 0.277 | -0.01 | -0.035 | -0.035 | 0 | 0.18 | 0.25 | 0.12 | 0.06 | 0.14 | 0.25 |
| 8 | 61911070 | rs10957177 | -1.108 | 1.33 | 0.084 | 0.343 | 0.003 | -0.021 | -0.021 | 0.019 | 0.28 | 0.43 | 0.28 | 0.16 | 0.29 | 0.36 |
| 8 | 61920845 | rs7841986 | 0.185 | 1.717 | -0.437 | 0.596 | 0.018 | 0.002 | 0.002 | 0 | 0.88 | 0.87 | 0.79 | 0.9 | 0.78 | 0.81 |
| 8 | 61922459 | rs3864678 | 0.452 | 1.39 | -0.667 | 1.275 | -0.032 | -0.062 | -0.062 | 0 | 0.1 | 0.01 | 0.07 | 0.35 | 0.12 | 0.25 |
| 8 | 61926797 | rs6989470 | -1.146 | 0.325 | -1.069 | 0.505 | 0.082 | 0.029 | 0.029 | 0 | 0.48 | 0.86 | 0.54 | 0.42 | 0.34 | 0.56 |
| 8 | 61929772 | rs3864664 | -0.459 | -0.273 | -0.381 | 0.303 | -0.002 | 0.015 | 0.015 | 0 | 0.16 | 0.21 | 0.32 | 0.3 | 0.11 | 0.21 |
| 8 | 61932528 | rs10504318 | -0.081 | -0.54 | -0.051 | 0.23 | -0.101 | -0.118 | -0.118 | 2.317 | 0.22 | 0.49 | 0.13 | 0.02 | 0.09 | 0.13 |
| 8 | 61933710 | rs16926700 | -0.355 | -0.477 | -0.367 | 0.264 | 0 | 0.006 | 0.006 | 0 | 0.15 | 0.18 | 0.32 | 0.3 | 0.12 | 0.21 |
| 8 | 61937299 | rs6995339 | -1.243 | -0.339 | -0.469 | 0.336 | 0.078 | 0.044 | 0.044 | 0 | 0.39 | 0.8 | 0.45 | 0.33 | 0.21 | 0.35 |
| 8 | 61938852 | rs12542604 | -0.604 | 0.006 | -0.182 | 0.544 | -0.002 | -0.002 | -0.002 | 1.092 | 0.08 | 0.15 | 0.16 | 0.14 | 0.06 | 0.2 |
| 8 | 61944091 | rs6986434 | -0.107 | 0.181 | -0.02 | 0.673 | 0.023 | 0.022 | 0.022 | 0 | 0.84 | 0.72 | 0.68 | 0.82 | 0.78 | 0.66 |
| 8 | 61944202 | rs7004557 | -0.612 | 0.146 | 0.382 | 0.387 | 0.015 | -0.013 | -0.013 | 0 | 0.16 | 0.23 | 0.24 | 0.15 | 0.18 | 0.11 |
| 8 | 61947098 | rs7814400 | 0.149 | -0.21 | 0.156 | 0.331 | -0.001 | 0.007 | 0.007 | 0 | 0.68 | 0.66 | 0.57 | 0.6 | 0.67 | 0.58 |
| 8 | 61947573 | rs1962862 | 0.612 | -0.42 | -0.079 | 0.529 | -0.009 | -0.012 | -0.012 | 0 | 0.54 | 0.35 | 0.36 | 0.51 | 0.53 | 0.47 |
| 8 | 61948567 | rs1871217 | -0.035 | -0.58 | -0.266 | 0.373 | -0.02 | -0.027 | -0.027 | 0.089 | 0.14 | 0.24 | 0.08 | 0.04 | 0.09 | 0.19 |
| 8 | 61949234 | rs1465667 | 0.585 | -0.293 | -0.063 | 0.523 | -0.005 | -0.006 | -0.006 | 0 | 0.54 | 0.42 | 0.37 | 0.51 | 0.53 | 0.48 |
| 8 | 61951203 | rs11986500 | -0.71 | -1.338 | 0.33 | 0.518 | -0.009 | -0.01 | -0.01 | 0.061 | 0.15 | 0.25 | 0.12 | 0.1 | 0.11 | 0.17 |
| 8 | 61954484 | rs6471908 | -1.389 | -2.219 | 0.419 | 0.667 | 0.009 | -0.012 | -0.012 | 0 | 0.6 | 0.66 | 0.41 | 0.41 | 0.41 | 0.55 |
| 8 | 61954970 | rs7827230 | -0.791 | -2.313 | 0.782 | 0.712 | -0.051 | 0.024 | 0.024 | 0.091 | 0.18 | 0.46 | 0.19 | 0.17 | 0.12 | 0.2 |
| 8 | 61962417 | rs10097565 | -0.6 | -2.294 | 0.8 | 0.724 | -0.036 | 0.026 | 0.026 | 0.001 | 0.17 | 0.39 | 0.19 | 0.17 | 0.12 | 0.21 |
| 8 | 61963699 | rs4738858 | -1.445 | -2.485 | 0.724 | 1.001 | 0.025 | 0.042 | 0.042 | 0 | 0.54 | 0.68 | 0.51 | 0.64 | 0.4 | 0.57 |
| 8 | 61964013 | rs16926764 | -1.379 | -2.499 | 0.727 | 1.099 | -0.029 | 0.06 | 0.06 | 0.08 | 0.29 | 0.52 | 0.36 | 0.47 | 0.25 | 0.35 |
| 8 | 61966066 | rs11998473 | 0.643 | -2.414 | 0.721 | 0.984 | -0.016 | -0.002 | -0.002 | 0.01 | 0.11 | 0.21 | 0.07 | 0.07 | 0.07 | 0.06 |
| 8 | 61966420 | rs4545092 | -1.77 | -2.197 | 0.248 | 0.9 | -0.031 | 0.019 | 0.019 | 0 | 0.64 | 0.79 | 0.56 | 0.68 | 0.49 | 0.65 |
| 8 | 61967680 | rs12548301 | -1.333 | -2.455 | 0.292 | 0.895 | 0.005 | 0.004 | 0.004 | 0 | 0.57 | 0.64 | 0.45 | 0.57 | 0.42 | 0.49 |
| 8 | 61968256 | rs12548415 | -1.573 | -2.352 | 0.105 | 0.704 | -0.012 | 0.013 | 0.013 | 0 | 0.63 | 0.74 | 0.54 | 0.66 | 0.47 | 0.64 |

| | | | | | | | | | | | | | | | | |
|---|---|---|---|---|---|---|---|---|---|---|---|---|---|---|---|---|
| 8 | 61969128 | rs13257940 | 1.092 | -2.711 | -0.314 | 0.582 | 0.009 | 0.007 | 0.007 | 0 | 0.07 | 0.03 | 0.04 | 0.05 | 0.04 | 0.13 |
| 8 | 61969213 | rs16926786 | 0.091 | -2.851 | -0.195 | 0.826 | 0.037 | -0.029 | -0.029 | 0 | 0.28 | 0.11 | 0.24 | 0.29 | 0.21 | 0.28 |
| 8 | 61971140 | rs1808634 | -0.047 | -2.779 | -0.148 | 0.87 | 0.046 | -0.025 | -0.025 | 2.44 | 0.3 | 0.11 | 0.25 | 0.29 | 0.23 | 0.29 |
| 8 | 61979873 | rs4738861 | -1.164 | -2.714 | -0.465 | 0.641 | 0.119 | 0.087 | 0.087 | 0 | 0.49 | 0.87 | 0.48 | 0.65 | 0.32 | 0.38 |
| 8 | 61992256 | rs10097298 | -0.866 | -2.68 | -0.267 | 0.705 | 0.033 | 0.012 | 0.012 | 0 | 0.49 | 0.57 | 0.46 | 0.61 | 0.36 | 0.54 |
| 8 | 61993755 | rs11996331 | 0.342 | -2.65 | -1.426 | 0.732 | -0.004 | 0 | 0 | 0 | 0.12 | 0.12 | 0.06 | 0.08 | 0.06 | 0.07 |
| 8 | 61994781 | rs6471915 | -0.942 | -2.538 | -1.381 | 0.773 | 0.089 | 0.024 | 0.024 | 0 | 0.51 | 0.8 | 0.49 | 0.61 | 0.36 | 0.54 |
| 8 | 61998100 | rs11987923 | 0.008 | -2.405 | -1.324 | 0.841 | 0.047 | 0.029 | 0.029 | 0 | 0.27 | 0.14 | 0.12 | 0.13 | 0.16 | 0.29 |
| 8 | 62000239 | rs1560555 | -1.18 | -1.705 | -1.524 | 1.033 | -0.045 | 0.023 | 0.023 | 0 | 0.63 | 0.91 | 0.54 | 0.69 | 0.43 | 0.61 |
| 8 | 62002590 | rs3940483 | -0.229 | -0.972 | -0.52 | 1.276 | 0.003 | -0.004 | -0.004 | 0.368 | 0.2 | 0.14 | 0.15 | 0.21 | 0.15 | 0.15 |
| 8 | 62002751 | rs1985815 | -0.689 | -0.721 | -0.671 | 1.076 | -0.029 | 0.02 | 0.02 | 0 | 0.32 | 0.43 | 0.22 | 0.29 | 0.22 | 0.21 |
| 8 | 62004772 | rs2979419 | 0.296 | 0.305 | -1.07 | 0.919 | -0.009 | -0.024 | -0.024 | 0 | 0.65 | 0.73 | 0.63 | 0.59 | 0.59 | 0.61 |
| 8 | 62005820 | rs10089771 | -1.012 | -0.651 | -1.288 | 0.601 | 0.043 | 0.008 | 0.008 | 0 | 0.23 | 0.11 | 0.23 | 0.22 | 0.12 | 0.17 |
| 8 | 62006099 | rs7008414 | -1.241 | -0.839 | -0.487 | 0.573 | -0.037 | 0.008 | 0.008 | 0 | 0.22 | 0.35 | 0.2 | 0.19 | 0.16 | 0.41 |
| 8 | 62006248 | rs6988682 | 0.112 | -0.365 | -0.56 | -0.239 | -0.002 | -0.024 | -0.024 | 0 | 0.56 | 0.52 | 0.66 | 0.88 | 0.63 | 0.75 |
| 8 | 62011683 | rs6471921 | -0.604 | -0.826 | 0.025 | 0.054 | -0.02 | -0.013 | -0.013 | 0 | 0.23 | 0.33 | 0.38 | 0.16 | 0.18 | 0.18 |
| 8 | 62024167 | rs16926864 | -0.539 | -0.857 | 0.139 | 0.013 | 0.003 | 0.002 | 0.002 | 0 | 0.23 | 0.23 | 0.37 | 0.16 | 0.18 | 0.18 |
| 8 | 62027498 | rs16926871 | -0.598 | -0.896 | 0.559 | 0.854 | -0.002 | 0.066 | 0.066 | 0 | 0.06 | 0.13 | 0.07 | 0.27 | 0.04 | 0.12 |
| 8 | 62036793 | rs3864667 | -1.488 | -1.217 | 0.511 | 1.693 | -0.006 | 0.066 | 0.066 | 0.153 | 0.13 | 0.19 | 0.22 | 0.61 | 0.1 | 0.36 |
| 8 | 62043233 | rs4738868 | -1.004 | -0.799 | 0.521 | 1.02 | 0.009 | 0.004 | 0.004 | 0 | 0.42 | 0.8 | 0.52 | 0.36 | 0.38 | 0.47 |
| 8 | 62044066 | rs1367975 | -0.313 | -0.81 | 0.817 | 0.948 | -0.006 | -0.008 | -0.008 | 0 | 0.28 | 0.28 | 0.4 | 0.16 | 0.22 | 0.3 |
| 8 | 62044143 | rs10113823 | -0.244 | -0.613 | 1.147 | 1.046 | 0.07 | 0.106 | 0.106 | 0.885 | 0.14 | 0.43 | 0.11 | 0.2 | 0.16 | 0.17 |
| 8 | 62045799 | rs9298048 | -0.983 | -0.557 | 0.862 | 1.107 | 0.018 | 0.095 | 0.095 | 0 | 0.19 | 0.28 | 0.23 | 0.54 | 0.22 | 0.39 |
| 8 | 62049778 | rs12678909 | 0.019 | -0.734 | 0.385 | 1.08 | 0.004 | 0.037 | 0.037 | 0 | 0.07 | 0.06 | 0.16 | 0.34 | 0.06 | 0.22 |
| 8 | 62054618 | rs4738870 | -0.48 | -0.93 | 0.423 | 0.539 | 0.023 | 0.041 | 0.041 | 0 | 0.28 | 0.14 | 0.34 | 0.13 | 0.21 | 0.29 |
| 8 | 62055785 | rs7830295 | -1.202 | -0.738 | 0.913 | 1.139 | -0.015 | 0.023 | 0.023 | 0 | 0.3 | 0.47 | 0.5 | 0.67 | 0.25 | 0.48 |
| 8 | 62062476 | rs16926940 | -0.131 | -0.687 | 0.872 | 0.627 | -0.004 | 0.035 | 0.035 | 0 | 0.05 | 0.12 | 0.07 | 0.15 | 0.04 | 0.14 |
| 8 | 62065249 | rs2017819 | -0.298 | -0.719 | 0.941 | 0.635 | 0 | 0 | 0 | 0 | 0.23 | 0.21 | 0.32 | 0.13 | 0.21 | 0.22 |
| 8 | 62066813 | rs17831865 | -0.298 | -0.719 | 0.941 | 0.635 | 0 | 0 | 0 | 0 | 0.23 | 0.21 | 0.32 | 0.13 | 0.21 | 0.22 |
| 8 | 62073809 | rs2129348 | 1.142 | -0.687 | 0.925 | 0.427 | -0.003 | -0.028 | -0.028 | 1.755 | 0.63 | 0.51 | 0.41 | 0.16 | 0.63 | 0.39 |
| 8 | 62083655 | rs16926976 | -0.563 | -0.613 | 0.833 | 0.8 | -0.006 | 0.039 | 0.039 | 0 | 0.06 | 0.17 | 0.07 | 0.13 | 0.04 | 0.11 |
| 8 | 62099814 | rs11785638 | -1.948 | -0.677 | 1.291 | 0.855 | -0.008 | 0.097 | 0.097 | 0.164 | 0.11 | 0.42 | 0.11 | 0.15 | 0.08 | 0.14 |
| 8 | 62116679 | rs2978521 | -0.545 | -0.195 | 1.249 | 0.831 | 0.006 | 0.012 | 0.012 | 0 | 0.26 | 0.38 | 0.39 | 0.29 | 0.26 | 0.31 |
| 8 | 62120560 | rs2931329 | -0.119 | -0.152 | 0.862 | 0.823 | -0.004 | 0.005 | 0.005 | 0 | 0.2 | 0.19 | 0.31 | 0.14 | 0.2 | 0.2 |
| 8 | 62127463 | rs1808033 | -1.262 | -0.337 | 0.736 | 0.449 | -0.001 | 0.004 | 0.004 | 0 | 0.39 | 0.55 | 0.59 | 0.81 | 0.39 | 0.62 |
| 8 | 62128707 | rs17765747 | -0.912 | -0.488 | 0.832 | 0.528 | 0.002 | -0.017 | -0.017 | 0 | 0.05 | 0.25 | 0.04 | 0 | 0.03 | 0.04 |
| 8 | 62147973 | rs10504324 | -1.046 | -0.487 | 0.972 | 0.512 | 0.01 | -0.014 | -0.014 | 0.319 | 0.05 | 0.26 | 0.03 | 0 | 0.04 | 0.04 |
| 8 | 62148532 | rs2931352 | -1.214 | -0.52 | 0.809 | 0.637 | 0.004 | -0.003 | -0.003 | 0 | 0.33 | 0.4 | 0.45 | 0.29 | 0.4 | 0.35 |
| 8 | 62170423 | rs2919318 | -0.033 | -0.156 | 0.328 | 0.415 | -0.071 | 0.056 | 0.056 | 0 | 0.23 | 0.06 | 0.33 | 0.11 | 0.32 | 0.22 |
| 8 | 62171565 | rs3852343 | -0.525 | -1.478 | -0.13 | -0.453 | -0.086 | -0.06 | -0.06 | 0 | 0.29 | 0.49 | 0.29 | 0.86 | 0.12 | 0.62 |
| 8 | 62173034 | rs1812576 | -0.516 | -1.449 | -0.711 | -0.578 | -0.081 | -0.064 | -0.064 | 5.227 | 0.31 | 0.49 | 0.3 | 0.86 | 0.14 | 0.63 |
| 8 | 62176063 | rs11995189 | -1.077 | -1.468 | -0.727 | -0.609 | 0.059 | 0.023 | 0.023 | 0.171 | 0.06 | 0.39 | 0.09 | 0 | 0.05 | 0.04 |
| 8 | 62190532 | rs2978520 | 0.465 | -1.245 | -0.921 | -0.467 | -0.073 | 0.066 | 0.066 | 0 | 0.23 | 0.06 | 0.35 | 0.11 | 0.32 | 0.21 |
| 8 | 62195086 | rs2919308 | -1.706 | -1.273 | -0.895 | -0.493 | -0.053 | 0.203 | 0.203 | 0.259 | 0.58 | 0.81 | 0.69 | 0.97 | 0.48 | 0.85 |
| 8 | 62198366 | rs16927139 | 0.206 | -1.218 | -0.961 | 0.297 | 0.011 | 0.05 | 0.05 | 0 | 0.06 | 0.04 | 0.12 | 0.39 | 0.03 | 0.2 |
| 8 | 62214626 | rs2978558 | -1.37 | -1.066 | -0.921 | -0.486 | -0.006 | 0.019 | 0.019 | 0.157 | 0.55 | 0.59 | 0.65 | 0.97 | 0.47 | 0.86 |
| 8 | 62216733 | rs2931297 | -0.118 | -0.108 | -1.333 | 0.111 | -0.054 | 0.056 | 0.056 | 0 | 0.24 | 0.09 | 0.36 | 0.12 | 0.34 | 0.22 |
| 8 | 62220151 | rs870002 | -0.735 | 0.22 | -1.144 | 0.398 | -0.001 | 0.001 | 0.001 | 0 | 0.45 | 0.44 | 0.52 | 0.82 | 0.5 | 0.6 |
| 8 | 62240450 | rs4144412 | -0.24 | 0.245 | -1.856 | 0.596 | -0.164 | -0.049 | -0.049 | 0.998 | 0.36 | 0.04 | 0.46 | 0.63 | 0.49 | 0.43 |
| 8 | 62247290 | rs10094788 | 0.217 | 0.589 | -1.856 | 0.056 | -0.136 | -0.026 | -0.026 | 0 | 0.29 | 0.01 | 0.29 | 0.26 | 0.37 | 0.32 |
| 8 | 62252388 | rs4738883 | -0.383 | 0.172 | -1.299 | -0.772 | -0.006 | 0.022 | 0.022 | 0 | 0.44 | 0.41 | 0.51 | 0.81 | 0.49 | 0.59 |
| 8 | 62279368 | rs17775880 | 0.117 | -0.215 | -1.103 | -0.649 | -0.109 | -0.006 | -0.006 | 0 | 0.35 | 0.08 | 0.48 | 0.67 | 0.48 | 0.51 |
| 8 | 62289749 | rs10111649 | 0.421 | -0.596 | -0.167 | 0.141 | 0.011 | 0.006 | 0.006 | 0.715 | 0.6 | 0.38 | 0.67 | 0.67 | 0.76 | 0.56 |
| 8 | 62323556 | rs1873888 | 0.205 | -0.072 | -0.432 | 0.483 | -0.036 | -0.014 | -0.014 | 0.555 | 0.63 | 0.51 | 0.69 | 0.67 | 0.76 | 0.55 |
| 8 | 62345391 | rs7831797 | -1.155 | 0.434 | -0.147 | 0.563 | 0.014 | 0 | 0 | 0 | 0.1 | 0.28 | 0.1 | 0.07 | 0.11 | 0.08 |
| 8 | 62346192 | rs13258160 | -1.052 | 0.423 | -0.122 | 0.558 | 0.013 | 0.002 | 0.002 | 0 | 0.09 | 0.24 | 0.1 | 0.07 | 0.11 | 0.07 |
| 8 | 62365737 | rs6471951 | 0.352 | 0.275 | -0.269 | 0.795 | 0.002 | -0.007 | -0.007 | 0 | 0.81 | 0.82 | 0.74 | 0.76 | 0.84 | 0.64 |
| 8 | 62365912 | rs16927318 | -1.314 | -0.243 | 0.553 | 0.785 | -0.014 | 0.029 | 0.029 | 4.515 | 0.09 | 0.5 | 0.05 | 0.06 | 0.02 | 0.08 |
| 8 | 62367876 | rs949927 | 1.322 | -0.222 | 0.576 | 0.762 | 0.031 | -0.004 | -0.004 | 0 | 0.71 | 0.2 | 0.68 | 0.7 | 0.82 | 0.57 |
| 8 | 62368825 | rs11991141 | -1.621 | -0.215 | 0.702 | 0.764 | -0.041 | -0.014 | -0.014 | 0 | 0.1 | 0.61 | 0.06 | 0.06 | 0.02 | 0.08 |
| 8 | 62370511 | rs2350222 | 1.738 | -0.47 | 0.534 | 0.73 | 0.022 | -0.044 | -0.044 | 0 | 0.64 | 0.11 | 0.59 | 0.62 | 0.74 | 0.53 |
| 8 | 62371260 | rs2291607 | 2.156 | -0.369 | 0.656 | 0.822 | 0.001 | -0.011 | -0.011 | 3.09 | 0.76 | 0.18 | 0.71 | 0.75 | 0.78 | 0.76 |
| 8 | 62377604 | rs1994851 | 0.532 | -0.5 | 0.75 | 0.759 | 0.011 | 0.024 | 0.024 | 0 | 0.91 | 0.81 | 0.85 | 0.88 | 0.88 | 0.87 |
| 8 | 62380439 | rs10107557 | -2.212 | -0.463 | 0.759 | 0.731 | -0.021 | -0.015 | -0.015 | 0.236 | 0.18 | 0.64 | 0.15 | 0.13 | 0.11 | 0.12 |
| 8 | 62384161 | rs6994836 | 1.296 | -0.5 | 1.278 | 0.853 | 0.006 | 0.077 | 0.077 | 0.411 | 0.86 | 0.45 | 0.83 | 0.81 | 0.88 | 0.82 |
| 8 | 62405121 | rs2640237 | -1.78 | -0.327 | 1.235 | 0.767 | 0.007 | -0.012 | -0.012 | 0 | 0.12 | 0.25 | 0.13 | 0.07 | 0.15 | 0.11 |
| 8 | 62405397 | rs4738899 | 2.414 | -0.588 | 0.757 | 1.436 | 0.005 | -0.071 | -0.071 | 0 | 0.65 | 0.12 | 0.62 | 0.51 | 0.69 | 0.67 |
| 8 | 62411474 | rs17789756 | 2.006 | -0.223 | -0.186 | 1.08 | -0.058 | 0.013 | 0.013 | 0 | 0.31 | 0.06 | 0.34 | 0.22 | 0.39 | 0.09 |
| 8 | 62419120 | rs7013189 | -1.469 | -0.342 | 0.674 | 1.365 | -0.019 | 0.076 | 0.076 | 0.675 | 0.23 | 0.61 | 0.25 | 0.42 | 0.16 | 0.19 |

| | | | | | | | | | | | | | | | | |
|---|---|---|---|---|---|---|---|---|---|---|---|---|---|---|---|---|
| 8 | 62419877 | rs17790020 | 0.021 | -0.392 | 0.482 | 1.343 | -0.031 | -0.012 | -0.012 | 0 | 0.08 | 0.03 | 0.08 | 0.07 | 0.1 | 0.04 |
| 8 | 62422705 | rs4612335 | 2.277 | -0.392 | 0.503 | 1.34 | 0.018 | -0.062 | -0.062 | 0 | 0.63 | 0.15 | 0.62 | 0.51 | 0.69 | 0.7 |
| 8 | 62427809 | rs16927440 | -2.408 | -0.402 | 0.506 | 1.374 | 0.011 | -0.017 | -0.017 | 1.246 | 0.36 | 0.71 | 0.35 | 0.5 | 0.28 | 0.3 |
| 8 | 62428412 | rs16927444 | 2.327 | -0.401 | 0.456 | 1.375 | 0.013 | -0.076 | -0.076 | 1.444 | 0.62 | 0.11 | 0.61 | 0.51 | 0.69 | 0.69 |
| 8 | 62439981 | rs7839684 | -1.462 | -0.264 | 0.701 | 1.354 | -0.009 | 0.021 | 0.021 | 0.634 | 0.13 | 0.07 | 0.1 | 0.07 | 0.12 | 0.11 |
| 8 | 62458019 | rs16927470 | 1.834 | -0.326 | 0.731 | 1.411 | 0.018 | -0.038 | -0.038 | 0 | 0.72 | 0.22 | 0.71 | 0.57 | 0.79 | 0.74 |
| 8 | 62461256 | rs10808719 | 1.244 | -0.271 | 0.841 | 1.396 | 0.02 | -0.026 | -0.026 | 0 | 0.94 | 0.77 | 0.95 | 1 | 0.94 | 0.94 |
| 8 | 62462704 | rs7825172 | -1.914 | -0.263 | 0.843 | 1.422 | -0.03 | 0.075 | 0.075 | 0.092 | 0.24 | 0.59 | 0.25 | 0.43 | 0.16 | 0.26 |
| 8 | 62463594 | rs10097222 | 2.229 | -0.264 | 0.789 | 1.418 | 0.003 | -0.086 | -0.086 | 0 | 0.64 | 0.1 | 0.61 | 0.5 | 0.69 | 0.7 |
| 8 | 62465580 | rs4289805 | -1.997 | -0.249 | 0.816 | 1.502 | -0.012 | 0.024 | 0.024 | 3.101 | 0.24 | 0.67 | 0.22 | 0.36 | 0.18 | 0.15 |
| 8 | 62465795 | rs4291265 | -1.997 | -0.249 | 0.816 | 1.525 | -0.012 | 0.025 | 0.025 | 3.101 | 0.24 | 0.67 | 0.22 | 0.36 | 0.18 | 0.15 |
| 8 | 62465814 | rs4379440 | -0.97 | -0.203 | 0.792 | 1.542 | -0.04 | 0.153 | 0.153 | 0.994 | 0.14 | 0.54 | 0.12 | 0.28 | 0.07 | 0.1 |
| 8 | 62476530 | rs6471958 | 0.683 | -0.16 | 0.641 | 1.558 | -0.001 | 0.02 | 0.02 | 0 | 0.84 | 0.78 | 0.77 | 0.62 | 0.83 | 0.85 |
| 8 | 62479934 | rs10090754 | 2.249 | -0.131 | 0.395 | 1.497 | 0.006 | -0.027 | -0.027 | 0 | 0.65 | 0.25 | 0.62 | 0.51 | 0.68 | 0.7 |
| 8 | 62484574 | rs16927563 | -1.074 | -0.149 | 0.613 | 1.695 | -0.034 | 0.197 | 0.197 | 0.869 | 0.13 | 0.46 | 0.12 | 0.28 | 0.07 | 0.1 |
| 8 | 62487467 | rs4237045 | 0.091 | 0.236 | -0.111 | 1.498 | -0.002 | 0.001 | 0.001 | 0 | 0.93 | 0.91 | 0.86 | 0.86 | 0.89 | 0.9 |
| 8 | 62489334 | rs16927574 | -0.347 | 0.085 | 0.002 | 1.588 | -0.002 | 0.048 | 0.048 | 0 | 0.08 | 0.16 | 0.1 | 0.24 | 0.08 | 0.06 |
| 8 | 62493835 | rs1574196 | -1.078 | -0.634 | 0.241 | 1.148 | -0.034 | -0.017 | -0.017 | 2.196 | 0.3 | 0.57 | 0.24 | 0.31 | 0.2 | 0.34 |
| 8 | 62497607 | rs2350620 | 1.609 | -1.354 | -0.383 | 1.468 | 0.116 | 0.04 | 0.04 | 0.232 | 0.51 | 0.17 | 0.57 | 0.39 | 0.65 | 0.34 |
| 8 | 62498627 | rs7015848 | -0.359 | -1.386 | -0.508 | 1.459 | 0.011 | -0.034 | -0.034 | 0.982 | 0.87 | 0.94 | 0.84 | 0.7 | 0.87 | 0.74 |
| 8 | 62509154 | rs13250696 | 2.004 | -0.702 | -0.58 | 1.168 | -0.043 | 0.007 | 0.007 | 0.603 | 0.44 | 0.07 | 0.46 | 0.35 | 0.53 | 0.32 |
| 8 | 62517963 | rs6985734 | 1.559 | -0.248 | -0.104 | 1.181 | 0.061 | -0.021 | -0.021 | 0.006 | 0.52 | 0.22 | 0.55 | 0.48 | 0.63 | 0.34 |
| 8 | 62521732 | rs16927639 | 0.252 | -0.218 | -0.388 | 1.388 | -0.006 | -0.021 | -0.021 | 1.502 | 0.11 | 0.07 | 0.12 | 0.21 | 0.12 | 0.09 |
| 8 | 62521927 | rs13256118 | 0.052 | -0.3 | -0.266 | 1.308 | 0.001 | -0.019 | -0.019 | 0 | 0.84 | 0.91 | 0.82 | 0.78 | 0.84 | 0.72 |
| 8 | 62525896 | rs2882460 | -1.005 | -0.147 | 0.191 | 1.153 | 0.005 | 0.002 | 0.002 | 0.034 | 0.3 | 0.26 | 0.23 | 0.3 | 0.21 | 0.39 |
| 8 | 62532696 | rs6471964 | -0.91 | -0.207 | 0.33 | 1.111 | 0.013 | 0.02 | 0.02 | 0 | 0.24 | 0.16 | 0.18 | 0.21 | 0.19 | 0.38 |
| 8 | 62537309 | rs6549 | 2.47 | -0.224 | 0.11 | 1.28 | -0.065 | -0.067 | -0.067 | 0 | 0.46 | 0.07 | 0.47 | 0.47 | 0.51 | 0.3 |
| 8 | 62540586 | rs11997478 | -0.685 | -0.233 | 0.291 | 1.261 | 0.004 | 0.006 | 0.006 | 0 | 0.24 | 0.2 | 0.18 | 0.21 | 0.19 | 0.38 |
| 8 | 62547948 | rs2350621 | -0.383 | -0.259 | 0.598 | 1.282 | -0.002 | -0.012 | -0.012 | 0 | 0.85 | 0.89 | 0.81 | 0.76 | 0.83 | 0.72 |
| 8 | 62554412 | rs7812327 | -1.46 | -0.268 | 1.343 | 1.263 | -0.001 | -0.005 | -0.005 | 0 | 0.32 | 0.39 | 0.31 | 0.25 | 0.31 | 0.4 |
| 8 | 62565669 | rs12677072 | 1.209 | -0.364 | 1.2 | 1.243 | 0.031 | -0.027 | -0.027 | 0 | 0.08 | 0.01 | 0.07 | 0.09 | 0.02 | 0.23 |
| 8 | 62566588 | rs17793991 | 2.797 | -0.095 | 1.123 | 1.812 | -0.099 | 0.021 | 0.021 | 0.766 | 0.44 | 0.06 | 0.45 | 0.31 | 0.48 | 0.17 |
| 8 | 62599098 | rs7015043 | 2.455 | -0.218 | 1.174 | 1.627 | 0.003 | 0.017 | 0.017 | 0 | 0.48 | 0.29 | 0.51 | 0.52 | 0.51 | 0.33 |
| 8 | 62603795 | rs6985520 | 2.347 | -0.209 | 1.163 | 1.631 | 0.004 | 0.01 | 0.01 | 0 | 0.5 | 0.39 | 0.52 | 0.52 | 0.52 | 0.33 |
| 8 | 62605852 | rs13271527 | -0.384 | -0.181 | 1.088 | 1.568 | 0 | -0.002 | -0.002 | 0 | 0.85 | 0.86 | 0.81 | 0.76 | 0.83 | 0.72 |
| 8 | 62617027 | rs7836864 | 0.041 | -0.189 | 1.141 | 1.615 | -0.003 | 0.014 | 0.014 | 0 | 0.83 | 0.79 | 0.8 | 0.75 | 0.83 | 0.72 |
| 8 | 62619207 | rs4377960 | 2.025 | -0.522 | 1.039 | 1.505 | 0.005 | 0.007 | 0.007 | 0 | 0.53 | 0.43 | 0.56 | 0.57 | 0.58 | 0.36 |
| 8 | 62623162 | rs10107157 | 0.011 | -0.338 | 0.867 | 1.547 | -0.009 | -0.025 | -0.025 | 0 | 0.05 | 0.1 | 0.02 | 0 | 0.03 | 0.04 |
| 8 | 62625232 | rs2350920 | 0.097 | -0.323 | 0.692 | 1.568 | -0.016 | -0.026 | -0.026 | 0.002 | 0.83 | 0.89 | 0.74 | 0.74 | 0.77 | 0.74 |
| 8 | 62633848 | rs7003213 | 0.192 | -0.09 | 0.326 | 0.985 | 0.003 | -0.002 | -0.002 | 0 | 0.63 | 0.6 | 0.61 | 0.68 | 0.58 | 0.66 |
| 8 | 62636162 | rs10094729 | -0.191 | -0.242 | 0.158 | 0.983 | 0.002 | 0.002 | 0.002 | 0 | 0.2 | 0.21 | 0.12 | 0.16 | 0.15 | 0.32 |
| 8 | 62636679 | rs6471972 | 1.389 | -0.193 | 0.248 | 1.001 | -0.021 | -0.006 | -0.006 | 0 | 0.44 | 0.29 | 0.51 | 0.52 | 0.44 | 0.35 |
| 8 | 62650548 | rs6999120 | 0.029 | -0.049 | -0.066 | 1.136 | -0.043 | -0.059 | -0.059 | 1.317 | 0.63 | 0.84 | 0.6 | 0.51 | 0.55 | 0.5 |
| 8 | 62651000 | rs16927824 | 0.156 | -0.207 | 0.359 | 1.288 | -0.002 | -0.022 | -0.022 | 0.249 | 0.14 | 0.24 | 0.09 | 0.11 | 0.13 | 0.1 |
| 8 | 62658110 | rs16927830 | 0.228 | -0.511 | -0.649 | 1.149 | 0.06 | -0.025 | -0.025 | 0 | 0.59 | 0.18 | 0.6 | 0.54 | 0.69 | 0.58 |
| 8 | 62660401 | rs4269537 | 0.811 | -0.42 | -0.664 | 1.108 | 0.056 | 0.086 | 0.086 | 0 | 0.53 | 0.24 | 0.53 | 0.25 | 0.64 | 0.38 |
| 8 | 62661193 | rs4322003 | -1.567 | -0.555 | 0.135 | 1.626 | -0.015 | 0.076 | 0.076 | 0 | 0.85 | 0.73 | 0.76 | 0.73 | 0.87 | 0.88 |
| 8 | 62661247 | rs4130092 | 0.064 | -0.502 | 0.134 | 1.679 | -0.001 | 0.034 | 0.034 | 0 | 0.06 | 0.08 | 0.05 | 0.21 | 0.05 | 0.2 |
| 8 | 62661593 | rs4295658 | 0.501 | -0.498 | 0.1 | 1.68 | -0.004 | -0.007 | -0.007 | 0 | 0.05 | 0.02 | 0.05 | 0.21 | 0.05 | 0.2 |
| 8 | 62668787 | rs10504334 | -0.767 | -0.614 | 0.309 | 1.551 | -0.028 | 0.032 | 0.032 | 0 | 0.16 | 0.33 | 0.15 | 0.25 | 0.1 | 0.39 |
| 8 | 62669364 | rs17801505 | 1.141 | -0.503 | -0.31 | 1.453 | -0.023 | -0.038 | -0.038 | 0 | 0.19 | 0.06 | 0.24 | 0.27 | 0.28 | 0.24 |
| 8 | 62673021 | rs11783343 | -0.538 | -0.669 | -0.092 | 1.322 | -0.015 | 0.083 | 0.083 | 0 | 0.78 | 0.59 | 0.71 | 0.64 | 0.81 | 0.68 |
| 8 | 62674301 | rs13248406 | 0.78 | -0.719 | -0.09 | 1.167 | -0.006 | 0 | 0 | 0 | 0.1 | 0.11 | 0.08 | 0.12 | 0.05 | 0.2 |
| 8 | 62676146 | rs13266879 | -1.494 | -0.583 | -0.054 | 1.114 | -0.009 | 0.057 | 0.057 | 0 | 0.85 | 0.67 | 0.75 | 0.78 | 0.86 | 0.88 |
| 8 | 62683228 | rs13249337 | -0.229 | -0.519 | -0.223 | 0.576 | 0.006 | -0.001 | -0.001 | 0 | 0.1 | 0.09 | 0.12 | 0.13 | 0.04 | 0.21 |
| 8 | 62684067 | rs10091115 | 1.438 | -0.362 | -0.225 | 0.378 | -0.01 | 0.018 | 0.018 | 0 | 0.34 | 0.27 | 0.23 | 0.2 | 0.34 | 0.22 |
| 8 | 62689524 | rs13266168 | 0.018 | -0.266 | -0.634 | 0.441 | 0.007 | 0 | 0 | 0.652 | 0.11 | 0.09 | 0.12 | 0.13 | 0.04 | 0.21 |
| 8 | 62690015 | rs1015905 | -0.895 | -0.195 | -0.53 | 0.528 | 0.004 | 0.032 | 0.032 | 0 | 0.71 | 0.65 | 0.69 | 0.6 | 0.7 | 0.81 |
| 8 | 62690750 | rs1495003 | 1.454 | -0.167 | -0.896 | 0.499 | -0.024 | 0.049 | 0.049 | 0 | 0.36 | 0.25 | 0.27 | 0.2 | 0.38 | 0.21 |
| 8 | 62691101 | rs2015490 | 0.839 | -0.129 | -0.78 | 0.599 | -0.003 | 0.007 | 0.007 | 0 | 0.05 | 0.11 | 0.06 | 0.05 | 0.05 | 0.18 |
| 8 | 62692745 | rs7009135 | -0.546 | -0.098 | -0.884 | 0.62 | -0.013 | 0 | 0 | 0.133 | 0.16 | 0.19 | 0.18 | 0.18 | 0.09 | 0.39 |
| 8 | 62694154 | rs17802118 | 1.111 | -0.055 | -0.988 | 0.628 | -0.014 | -0.013 | -0.013 | 0.057 | 0.2 | 0.08 | 0.25 | 0.22 | 0.27 | 0.2 |
| 8 | 62696220 | rs13255569 | 0.408 | -0.046 | -0.78 | 0.178 | 0.015 | -0.007 | -0.007 | 0 | 0.08 | 0.05 | 0.11 | 0.13 | 0.04 | 0.21 |
| 8 | 62697358 | rs7824375 | -0.333 | 0.111 | -0.785 | 0.192 | 0 | 0.007 | 0.007 | 0.01 | 0.51 | 0.49 | 0.44 | 0.38 | 0.43 | 0.61 |
| 8 | 62698611 | rs7012553 | 0.333 | 0.836 | -1.289 | 0.017 | -0.02 | 0.045 | 0.045 | 0 | 0.34 | 0.24 | 0.23 | 0.19 | 0.34 | 0.21 |
| 8 | 62704349 | rs7002859 | -0.395 | 0.369 | -1.149 | 0.047 | 0.043 | 0.081 | 0.081 | 0.033 | 0.48 | 0.66 | 0.57 | 0.78 | 0.37 | 0.52 |
| 8 | 62705616 | rs7012161 | 0.18 | -0.135 | -1.211 | 0.732 | 0.025 | -0.021 | -0.021 | 0 | 0.3 | 0.1 | 0.32 | 0.3 | 0.27 | 0.45 |
| 8 | 62715255 | rs7015292 | -0.179 | 0.084 | -0.721 | 1.122 | -0.045 | 0.279 | 0.279 | 0.058 | 0.15 | 0.56 | 0.24 | 0.48 | 0.09 | 0.07 |
| 8 | 62716334 | rs1495008 | -0.051 | 0.214 | -1.509 | 0.865 | -0.022 | -0.073 | -0.073 | 0.668 | 0.22 | 0.02 | 0.25 | 0.27 | 0.21 | 0.25 |

| | | | | | | | | | | | | | | | | |
|---|---|---|---|---|---|---|---|---|---|---|---|---|---|---|---|---|
| 8 | 62723254 | rs13260889 | 0.804 | 1.663 | -0.376 | 1.129 | 0.055 | 0.085 | 0.085 | 0 | 0.77 | 0.46 | 0.61 | 0.43 | 0.68 | 0.79 |
| 8 | 62728138 | rs1948508 | 0.05 | 2.097 | -0.125 | 1.308 | 0.049 | 0.03 | 0.03 | 0 | 0.93 | 0.86 | 0.88 | 0.9 | 0.86 | 0.86 |
| 8 | 62728469 | rs1390028 | 1.018 | 2.152 | -0.501 | 0.521 | 0.014 | 0.014 | 0.014 | 0 | 0.56 | 0.28 | 0.43 | 0.42 | 0.44 | 0.57 |
| 8 | 62729423 | rs10957204 | -0.061 | 2.146 | -0.409 | 0.521 | 0.035 | 0.277 | 0.277 | 0.058 | 0.17 | 0.44 | 0.28 | 0.47 | 0.18 | 0.07 |
| 8 | 62731202 | rs12676768 | 1.084 | 2.042 | -0.538 | 0.524 | 0.045 | 0.071 | 0.071 | 0 | 0.53 | 0.19 | 0.41 | 0.36 | 0.41 | 0.5 |
| 8 | 62733267 | rs10096647 | 0.924 | 2.072 | -0.531 | 0.535 | 0.016 | 0.014 | 0.014 | 0 | 0.56 | 0.28 | 0.43 | 0.42 | 0.44 | 0.57 |
| 8 | 62734074 | rs7832140 | -0.674 | 2.272 | -0.578 | 0.549 | -0.029 | -0.019 | -0.019 | 0 | 0.95 | 0.99 | 0.89 | 0.95 | 0.9 | 0.89 |
| 8 | 62735692 | rs7824857 | -1.017 | 1.957 | -0.25 | 0.749 | -0.034 | 0.038 | 0.038 | 0 | 0.42 | 0.78 | 0.52 | 0.62 | 0.51 | 0.43 |
| 8 | 62736289 | rs4737595 | -0.932 | 1.995 | 0.271 | 0.757 | 0.004 | 0.004 | 0.004 | 0 | 0.39 | 0.58 | 0.46 | 0.52 | 0.45 | 0.37 |
| 8 | 62738734 | rs16927978 | -0.461 | 1.722 | 0.546 | 1.085 | 0.021 | 0.192 | 0.192 | 0.006 | 0.17 | 0.4 | 0.28 | 0.45 | 0.2 | 0.08 |
| 8 | 62764345 | rs13268279 | 0.418 | 1.694 | -0.201 | 1.562 | -0.031 | -0.053 | -0.053 | 0 | 0.1 | 0.03 | 0.14 | 0.26 | 0.12 | 0.05 |
| 8 | 62773778 | rs7000025 | -0.9 | 1.388 | 0.335 | 1.641 | 0 | 0.017 | 0.017 | 0 | 0.16 | 0.14 | 0.24 | 0.52 | 0.16 | 0.12 |
| 8 | 62776307 | rs11997057 | -1.061 | 1.263 | 0.887 | 1.464 | 0.101 | 0.056 | 0.056 | 6.1 | 0.07 | 0.58 | 0.09 | 0.04 | 0.07 | 0.04 |
| 8 | 62781264 | rs16928019 | -0.002 | 1.221 | 1.063 | 1.498 | -0.004 | 0.018 | 0.018 | 0 | 0.05 | 0.05 | 0.08 | 0.2 | 0.04 | 0.06 |
| 8 | 62793604 | rs6471976 | -0.221 | 1.781 | 0.873 | 0.772 | -0.005 | 0.003 | 0.003 | 0.825 | 0.27 | 0.74 | 0.28 | 0.11 | 0.3 | 0.29 |
| 8 | 62795087 | rs16928045 | -0.141 | 1.787 | 0.711 | 0.779 | 0.041 | -0.048 | -0.048 | 0 | 0.25 | 0.58 | 0.27 | 0.11 | 0.3 | 0.29 |
| 8 | 62803036 | rs7015098 | 0.681 | 1.746 | -0.104 | 0.706 | -0.017 | 0.022 | 0.022 | 0 | 0.19 | 0.12 | 0.19 | 0.06 | 0.23 | 0.25 |
| 8 | 62805280 | rs12541401 | -1.202 | 1.702 | -0.249 | 0.725 | -0.088 | 0.047 | 0.047 | 0 | 0.44 | 0.91 | 0.53 | 0.63 | 0.49 | 0.43 |
| 8 | 62816151 | rs16928064 | 0.97 | 1.139 | -1.011 | 0.285 | -0.054 | -0.02 | -0.02 | 0.279 | 0.2 | 0.01 | 0.19 | 0.22 | 0.19 | 0.1 |
| 8 | 62816900 | rs7003847 | -0.919 | 0.967 | -0.553 | 0.521 | -0.03 | 0.083 | 0.083 | 0 | 0.46 | 0.84 | 0.52 | 0.63 | 0.49 | 0.46 |
| 8 | 62821059 | rs10957207 | 0.565 | 1.21 | -0.562 | 0.426 | -0.03 | 0.056 | 0.056 | 0.476 | 0.22 | 0.1 | 0.19 | 0.07 | 0.25 | 0.29 |
| 8 | 62826149 | rs7817048 | -0.896 | 1.185 | 0.234 | 1.005 | 0.043 | 0.142 | 0.142 | 0.139 | 0.2 | 0.38 | 0.35 | 0.53 | 0.24 | 0.17 |
| 8 | 62838152 | rs7824027 | -0.822 | 0.817 | 0.424 | 0.705 | -0.002 | 0.005 | 0.005 | 0 | 0.48 | 0.51 | 0.5 | 0.54 | 0.48 | 0.58 |
| 8 | 62839354 | rs6981356 | 0.201 | 0.638 | 0.403 | 0.648 | -0.005 | 0.005 | 0.005 | 0 | 0.31 | 0.27 | 0.25 | 0.06 | 0.32 | 0.45 |
| 8 | 62840483 | rs7008487 | -0.122 | 0.451 | 0.598 | 1.021 | 0.013 | 0.01 | 0.01 | 0 | 0.41 | 0.5 | 0.42 | 0.36 | 0.5 | 0.58 |
| 8 | 62848331 | rs16918895 | -0.038 | 0.414 | 0.721 | 1.192 | -0.027 | 0.102 | 0.102 | 0 | 0.07 | 0.14 | 0.09 | 0.27 | 0.04 | 0.06 |
| 8 | 62849976 | rs6471977 | 0.348 | 0.087 | 0.246 | 0.938 | -0.045 | -0.005 | -0.005 | 0 | 0.27 | 0.58 | 0.18 | 0.26 | 0.16 | 0.15 |
| 8 | 62851130 | rs344301 | 0.128 | 0.082 | 0.201 | 0.932 | 0.01 | -0.011 | -0.011 | 0 | 0.28 | 0.7 | 0.24 | 0.4 | 0.22 | 0.2 |
| 8 | 62866541 | rs344290 | 0.155 | 0.457 | 0.2 | 0.908 | -0.009 | -0.005 | -0.005 | 7.86 | 0.38 | 0.39 | 0.45 | 0.45 | 0.5 | 0.6 |
| 8 | 62873803 | rs657284 | -0.31 | 0.281 | 0.441 | 0.805 | 0.005 | 0.009 | 0.009 | 0 | 0.56 | 0.65 | 0.54 | 0.52 | 0.62 | 0.68 |
| 8 | 62885312 | rs2052985 | 0.333 | 0.353 | 0.4 | 0.891 | -0.023 | -0.033 | -0.033 | 0 | 0.37 | 0.27 | 0.42 | 0.39 | 0.44 | 0.55 |
| 8 | 62888372 | rs1346993 | 0.533 | 0.384 | 0.378 | 0.924 | -0.055 | -0.045 | -0.045 | 0.195 | 0.34 | 0.1 | 0.34 | 0.3 | 0.39 | 0.53 |
| 8 | 62892296 | rs12545751 | 0.278 | 0.348 | 0.465 | 1.007 | -0.024 | -0.011 | -0.011 | 0 | 0.37 | 0.28 | 0.4 | 0.31 | 0.45 | 0.55 |
| 8 | 62896709 | rs4445219 | 0.119 | 0.31 | 0.71 | 1 | -0.003 | -0.003 | -0.003 | 0.527 | 0.38 | 0.38 | 0.43 | 0.39 | 0.45 | 0.56 |
| 8 | 62899559 | rs344336 | -0.897 | 0.274 | 0.371 | 0.859 | -0.065 | 0.094 | 0.094 | 0.422 | 0.62 | 0.95 | 0.62 | 0.71 | 0.66 | 0.71 |
| 8 | 62901250 | rs7814379 | 0.095 | 0.334 | 0.393 | 0.756 | -0.052 | -0.011 | -0.011 | 0 | 0.4 | 0.21 | 0.41 | 0.31 | 0.48 | 0.57 |
| 8 | 62914580 | rs7461068 | 1.148 | 0.205 | 0.818 | 0.801 | -0.016 | -0.023 | -0.023 | 0 | 0.19 | 0.25 | 0.14 | 0.09 | 0.17 | 0.11 |
| 8 | 62914742 | rs344222 | 0.568 | 0.214 | 1.005 | 0.809 | -0.016 | -0.027 | -0.027 | 0 | 0.23 | 0.63 | 0.18 | 0.14 | 0.22 | 0.15 |
| 8 | 62928715 | rs344229 | 1.041 | 0.349 | 0.68 | 0.901 | -0.008 | -0.019 | -0.019 | 0 | 0.2 | 0.4 | 0.17 | 0.14 | 0.22 | 0.15 |
| 8 | 62940286 | rs12546000 | 0.523 | 0.378 | -0.041 | 0.876 | -0.052 | -0.071 | -0.071 | 0 | 0.32 | 0.1 | 0.36 | 0.33 | 0.37 | 0.52 |
| 8 | 62944691 | rs16928270 | 0.618 | -0.017 | 0.135 | 0.593 | -0.01 | -0.015 | -0.015 | 4.366 | 0.24 | 0.25 | 0.14 | 0.08 | 0.19 | 0.15 |
| 8 | 62952067 | rs16928321 | 0.771 | -0.035 | 0.18 | 0.592 | 0.008 | 0.008 | 0.008 | 0 | 0.23 | 0.18 | 0.14 | 0.08 | 0.2 | 0.15 |
| 8 | 62967144 | rs344268 | -0.776 | 0.074 | 0.383 | 0.788 | -0.101 | 0.094 | 0.094 | 0 | 0.6 | 0.96 | 0.61 | 0.7 | 0.62 | 0.72 |
| 8 | 62969408 | rs10957210 | 0.4 | 0.02 | 0.512 | 0.762 | 0.004 | 0.002 | 0.002 | 0 | 0.33 | 0.34 | 0.43 | 0.57 | 0.37 | 0.53 |
| 8 | 63003117 | rs3921407 | 0.043 | 0.472 | 0.393 | 0.987 | 0.02 | 0.029 | 0.029 | 0.084 | 0.54 | 0.25 | 0.5 | 0.58 | 0.57 | 0.4 |
| 8 | 63014215 | rs1607131 | 0.597 | -0.069 | 0.607 | 0.742 | -0.034 | -0.064 | -0.064 | 0 | 0.26 | 0.62 | 0.16 | 0.1 | 0.2 | 0.21 |
| 8 | 63014589 | rs1607130 | 0.013 | -0.137 | 0.349 | 0.669 | -0.037 | -0.034 | -0.034 | 0 | 0.34 | 0.21 | 0.43 | 0.41 | 0.43 | 0.53 |
| 8 | 63014705 | rs1607129 | 0.003 | -0.137 | 0.36 | 0.683 | -0.035 | -0.039 | -0.039 | 0 | 0.34 | 0.22 | 0.43 | 0.4 | 0.43 | 0.54 |
| 8 | 63020728 | rs4400381 | 0.72 | -0.157 | 0.486 | 0.598 | -0.028 | -0.078 | -0.078 | 0 | 0.25 | 0.62 | 0.16 | 0.06 | 0.21 | 0.22 |
| 8 | 63023132 | rs6471996 | 1.463 | -0.292 | 0.627 | 0.513 | 0.002 | -0.074 | -0.074 | 0.404 | 0.17 | 0.38 | 0.12 | 0.03 | 0.18 | 0.18 |
| 8 | 63023681 | rs7821951 | 1.613 | -0.283 | 0.568 | 0.517 | 0.001 | -0.055 | -0.055 | 0.003 | 0.16 | 0.27 | 0.11 | 0.03 | 0.18 | 0.18 |
| 8 | 63031804 | rs10101368 | 0.684 | -0.231 | 0.323 | 0.615 | -0.033 | -0.09 | -0.09 | 0 | 0.25 | 0.61 | 0.16 | 0.06 | 0.21 | 0.22 |
| 8 | 63033016 | rs11782329 | 1.256 | -0.03 | -0.076 | 0.936 | -0.079 | 0 | 0 | 0 | 0.22 | 0.04 | 0.26 | 0.13 | 0.25 | 0.42 |
| 8 | 63039751 | rs7000539 | -0.905 | -0.219 | 0.004 | 0.907 | -0.004 | 0.011 | 0.011 | 0 | 0.58 | 0.84 | 0.58 | 0.46 | 0.62 | 0.75 |
| 8 | 63040410 | rs13279917 | 0.226 | -0.028 | -0.032 | 1.258 | -0.053 | -0.08 | -0.08 | 1.393 | 0.29 | 0.11 | 0.38 | 0.39 | 0.37 | 0.49 |
| 8 | 63044535 | rs4617145 | -0.411 | 0.007 | 0.536 | 1.392 | -0.005 | 0.112 | 0.112 | 0.372 | 0.88 | 0.71 | 0.85 | 0.71 | 0.87 | 0.89 |
| 8 | 63051531 | rs10110503 | 0.289 | -0.6 | 0.52 | 1.128 | 0.014 | 0.043 | 0.043 | 0 | 0.53 | 0.39 | 0.42 | 0.31 | 0.56 | 0.47 |
| 8 | 63055668 | rs900493 | 0.967 | -0.655 | 0.375 | 0.769 | -0.034 | 0.068 | 0.068 | 0.117 | 0.41 | 0.23 | 0.32 | 0.3 | 0.46 | 0.37 |
| 8 | 63056052 | rs4445217 | -0.337 | -0.63 | 0.667 | 1.003 | -0.012 | 0.19 | 0.19 | 1.343 | 0.86 | 0.58 | 0.76 | 0.65 | 0.87 | 0.87 |
| 8 | 63064585 | rs10093587 | 1.386 | -0.672 | 0.463 | 1.018 | -0.032 | 0.024 | 0.024 | 0 | 0.28 | 0.07 | 0.29 | 0.16 | 0.31 | 0.4 |
| 8 | 63066732 | rs1483179 | 0.89 | -0.435 | 0.375 | 0.942 | 0.002 | -0.007 | -0.007 | 0 | 0.32 | 0.2 | 0.34 | 0.34 | 0.31 | 0.4 |
| 8 | 63078778 | rs4562309 | 0.675 | -0.288 | -0.088 | 0.945 | -0.006 | 0.004 | 0.004 | 0 | 0.34 | 0.32 | 0.36 | 0.34 | 0.31 | 0.4 |
| 8 | 63099432 | rs7007064 | -0.504 | -0.249 | -0.274 | 0.876 | -0.013 | 0.061 | 0.061 | 0 | 0.56 | 0.69 | 0.63 | 0.7 | 0.49 | 0.59 |
| 8 | 63106536 | rs1026902 | -1.93 | 0.263 | -0.925 | 0.664 | -0.067 | 0.017 | 0.017 | 0.144 | 0.83 | 0.98 | 0.88 | 0.86 | 0.82 | 0.86 |
| 8 | 63112852 | rs12542544 | 1.657 | -0.026 | -0.723 | 0.992 | 0 | 0.005 | 0.005 | 0 | 0.08 | 0.1 | 0.12 | 0.29 | 0.1 | 0.08 |
| 8 | 63114121 | rs7002834 | -0.532 | -0.09 | -0.475 | 0.541 | -0.023 | 0.034 | 0.034 | 0.03 | 0.58 | 0.7 | 0.59 | 0.62 | 0.52 | 0.58 |
| 8 | 63142888 | rs13270035 | -0.16 | -0.086 | 0.137 | 1.195 | 0.001 | -0.016 | -0.016 | 0.179 | 0.19 | 0.2 | 0.19 | 0.44 | 0.17 | 0.17 |
| 8 | 63146398 | rs6472000 | -0.446 | -0.118 | 0.263 | 1.007 | -0.006 | 0.004 | 0.004 | 0 | 0.75 | 0.8 | 0.76 | 0.74 | 0.7 | 0.69 |
| 8 | 63148154 | rs6995731 | 1.124 | -0.266 | 0.334 | 0.822 | 0.013 | 0.035 | 0.035 | 0 | 0.49 | 0.34 | 0.43 | 0.23 | 0.49 | 0.48 |

| | | | | | | | | | | | | | | | | |
|---|---|---|---|---|---|---|---|---|---|---|---|---|---|---|---|---|
| 8 | 63156123 | rs1384681 | 0.534 | -0.151 | -0.052 | 0.872 | -0.025 | 0.008 | 0.008 | 0 | 0.58 | 0.69 | 0.58 | 0.3 | 0.55 | 0.55 |
| 8 | 63158693 | rs9298062 | 0.82 | -0.58 | 0.421 | 0.707 | -0.001 | -0.002 | -0.002 | 0 | 0.59 | 0.57 | 0.54 | 0.53 | 0.64 | 0.58 |
| 8 | 63161013 | rs12549042 | 0.982 | -0.599 | 0.552 | 0.698 | -0.001 | -0.002 | -0.002 | 0 | 0.58 | 0.44 | 0.52 | 0.53 | 0.64 | 0.58 |
| 8 | 63165114 | rs13262465 | 0.994 | -0.607 | 0.548 | 0.693 | -0.001 | -0.002 | -0.002 | 0 | 0.58 | 0.44 | 0.52 | 0.53 | 0.64 | 0.58 |
| 8 | 63169294 | rs2625413 | -0.343 | -0.717 | 0.529 | 0.283 | -0.013 | -0.007 | -0.007 | 1.066 | 0.74 | 0.82 | 0.67 | 0.65 | 0.7 | 0.67 |
| 8 | 63170100 | rs1384684 | 1.151 | -0.71 | 0.52 | 0.288 | -0.003 | -0.013 | -0.013 | 0.541 | 0.58 | 0.48 | 0.6 | 0.64 | 0.64 | 0.58 |
| 8 | 63171482 | rs1351702 | 1.067 | -0.708 | 0.515 | 0.289 | -0.003 | -0.019 | -0.019 | 0 | 0.58 | 0.53 | 0.61 | 0.64 | 0.64 | 0.58 |
| 8 | 63180008 | rs2625414 | 0.233 | -0.702 | 0.413 | 0.266 | 0.014 | 0.031 | 0.031 | 0 | 0.14 | 0.05 | 0.05 | 0 | 0.06 | 0.09 |
| 8 | 63184105 | rs7017975 | 1.171 | -0.705 | 0.433 | 0.315 | -0.002 | -0.011 | -0.011 | 0 | 0.57 | 0.47 | 0.6 | 0.64 | 0.64 | 0.58 |
| 8 | 63198386 | rs1384678 | -0.191 | -0.355 | 0.512 | 0.587 | -0.013 | -0.008 | -0.008 | 0.275 | 0.74 | 0.82 | 0.67 | 0.64 | 0.7 | 0.67 |
| 8 | 63214002 | rs7829743 | 0.541 | -0.117 | -0.118 | 0.587 | -0.016 | -0.004 | -0.004 | 0 | 0.21 | 0.12 | 0.21 | 0.29 | 0.25 | 0.28 |
| 8 | 63247352 | rs6995804 | -0.112 | -0.574 | 0.031 | 0.181 | 0.005 | -0.017 | -0.017 | 0 | 0.83 | 0.87 | 0.94 | 1 | 0.92 | 0.85 |
| 8 | 63249323 | rs12678077 | 0.806 | -0.553 | 0.573 | 0.415 | -0.021 | -0.015 | -0.015 | 0.158 | 0.89 | 0.72 | 0.85 | 0.94 | 0.93 | 0.93 |
| 8 | 63249610 | rs2351950 | 1.598 | -0.617 | 0.879 | 0.383 | -0.027 | -0.017 | -0.017 | 0.02 | 0.87 | 0.53 | 0.84 | 0.94 | 0.93 | 0.93 |
| 8 | 63249830 | rs2351949 | 0.6 | -0.492 | 0.444 | 0.382 | -0.009 | -0.04 | -0.04 | 0 | 0.9 | 0.87 | 0.92 | 1 | 0.93 | 0.93 |
| 8 | 63254413 | rs12114607 | -0.131 | -0.409 | 0.203 | 1 | -0.001 | 0.04 | 0.04 | 0 | 0.13 | 0.14 | 0.22 | 0.4 | 0.14 | 0.09 |
| 8 | 63271251 | rs16928742 | 0.448 | -0.29 | 0.343 | 0.753 | 0.01 | 0.025 | 0.025 | 0.026 | 0.06 | 0.18 | 0.13 | 0.06 | 0.05 | 0.05 |
| 8 | 63272213 | rs10097272 | 1.265 | -0.881 | 0.04 | 0.582 | 0.035 | 0.014 | 0.014 | 0 | 0.51 | 0.35 | 0.48 | 0.48 | 0.59 | 0.66 |
| 8 | 63273943 | rs16928749 | 0.381 | -0.741 | 0.185 | 0.894 | 0 | 0.011 | 0.011 | 0 | 0.1 | 0.08 | 0.15 | 0.38 | 0.11 | 0.08 |
| 8 | 63278209 | rs2351667 | 0.308 | -0.789 | 0.406 | 0.782 | 0.002 | 0.051 | 0.051 | 0 | 0.86 | 0.84 | 0.8 | 0.6 | 0.85 | 0.9 |
| 8 | 63287487 | rs11774856 | -0.068 | -0.752 | 0.152 | 0.479 | 0.003 | 0.007 | 0.007 | 0 | 0.58 | 0.38 | 0.64 | 0.65 | 0.69 | 0.61 |
| 8 | 63293407 | rs10504346 | 0.811 | -0.69 | 0.059 | 0.515 | -0.005 | 0.026 | 0.026 | 0 | 0.1 | 0.08 | 0.19 | 0.4 | 0.14 | 0.08 |
| 8 | 63300112 | rs2351660 | 1.848 | -0.763 | 0.013 | 0.397 | -0.08 | 0.046 | 0.046 | 0.046 | 0.32 | 0.08 | 0.37 | 0.21 | 0.43 | 0.46 |
| 8 | 63308955 | rs6988778 | 1.113 | -0.729 | 0.237 | 0.452 | -0.001 | 0.021 | 0.021 | 0 | 0.07 | 0.23 | 0.14 | 0.06 | 0.05 | 0.05 |
| 8 | 63321678 | rs2034605 | -0.533 | -0.809 | 0.285 | 0.411 | -0.002 | -0.009 | -0.009 | 0 | 0.64 | 0.62 | 0.78 | 0.7 | 0.74 | 0.67 |
| 8 | 63331062 | rs1451843 | 0.959 | -0.832 | 0.206 | 0.424 | -0.007 | 0.013 | 0.013 | 0 | 0.07 | 0.23 | 0.13 | 0.06 | 0.05 | 0.05 |
| 8 | 63331209 | rs1451844 | 0.247 | -0.801 | 0.276 | 0.408 | 0.022 | 0.047 | 0.047 | 0 | 0.56 | 0.21 | 0.63 | 0.64 | 0.68 | 0.61 |
| 8 | 63339639 | rs4604422 | 1.044 | -0.792 | 0.195 | 0.363 | -0.038 | 0.113 | 0.113 | 0.031 | 0.45 | 0.13 | 0.44 | 0.24 | 0.54 | 0.52 |
| 8 | 63358921 | rs13266120 | 1.076 | -0.85 | 0.413 | 0.335 | -0.04 | 0.109 | 0.109 | 0.013 | 0.45 | 0.15 | 0.44 | 0.24 | 0.54 | 0.52 |
| 8 | 63359627 | rs4738973 | -0.422 | -0.842 | 0.407 | 0.4 | -0.007 | -0.019 | -0.019 | 0 | 0.63 | 0.47 | 0.77 | 0.7 | 0.73 | 0.67 |
| 8 | 63362315 | rs17243191 | 0.609 | -0.821 | 0.286 | 0.425 | 0.002 | 0.009 | 0.009 | 0 | 0.33 | 0.28 | 0.21 | 0.27 | 0.25 | 0.29 |
| 8 | 63366972 | rs13275695 | 1.479 | -0.817 | -0.002 | 0.535 | 0.007 | 0.032 | 0.032 | 0.384 | 0.14 | 0 | 0.06 | 0.05 | 0.12 | 0.07 |
| 8 | 63372633 | rs4430089 | 1.26 | -0.855 | 0.104 | 0.606 | -0.007 | 0.004 | 0.004 | 0 | 0.07 | 0.14 | 0.13 | 0.06 | 0.05 | 0.05 |
| 8 | 63375340 | rs12115033 | 0.701 | -0.775 | 0.137 | 0.685 | 0.004 | 0.069 | 0.069 | 0 | 0.1 | 0.12 | 0.2 | 0.38 | 0.14 | 0.09 |
| 8 | 63376719 | rs4738975 | 0.277 | -0.792 | 0.038 | 0.428 | 0.007 | 0.006 | 0.006 | 0 | 0.21 | 0.21 | 0.2 | 0.12 | 0.17 | 0.12 |
| 8 | 63381369 | rs10504348 | 0.235 | -0.787 | 0.057 | 0.406 | 0.001 | 0.001 | 0.001 | 0 | 0.21 | 0.23 | 0.2 | 0.11 | 0.17 | 0.12 |
| 8 | 63389536 | rs2084799 | 1.474 | -0.691 | 0.191 | 0.414 | 0.001 | -0.002 | -0.002 | 0 | 0.34 | 0.34 | 0.37 | 0.23 | 0.43 | 0.46 |
| 8 | 63390820 | rs1376759 | -0.345 | -0.57 | 0.36 | 0.774 | 0.001 | 0.002 | 0.002 | 0 | 0.66 | 0.68 | 0.77 | 0.69 | 0.74 | 0.67 |
| 8 | 63391700 | rs16928991 | 0.264 | -0.177 | 0.489 | 1.228 | 0.006 | 0.07 | 0.07 | 0 | 0.1 | 0.13 | 0.2 | 0.34 | 0.13 | 0.09 |
| 8 | 63395559 | rs4500073 | 0.154 | -0.479 | -0.048 | 0.614 | -0.002 | 0.004 | 0.004 | 0 | 0.29 | 0.23 | 0.24 | 0.29 | 0.26 | 0.34 |
| 8 | 63402743 | rs6982729 | 0.539 | -0.76 | 0.032 | 0.955 | -0.068 | 0.084 | 0.084 | 0.248 | 0.2 | 0.01 | 0.13 | 0.05 | 0.2 | 0.21 |
| 8 | 63403161 | rs4551346 | 1.611 | -1.111 | 0.134 | 1.051 | 0.005 | -0.003 | -0.003 | 0 | 0.12 | 0.1 | 0.13 | 0.14 | 0.1 | 0.08 |
| 8 | 63409288 | rs13255137 | 1.947 | -1.123 | 0.074 | 1.075 | 0.009 | -0.003 | -0.003 | 0.347 | 0.11 | 0 | 0.05 | 0.07 | 0.1 | 0.07 |
| 8 | 63415156 | rs896487 | -0.288 | -1.077 | 0.158 | 1.232 | 0.003 | 0.026 | 0.026 | 0.017 | 0.9 | 0.89 | 0.83 | 0.77 | 0.88 | 0.9 |
| 8 | 63415467 | rs896486 | 0.45 | -0.922 | 0.224 | 0.931 | -0.03 | 0.078 | 0.078 | 0.209 | 0.7 | 0.56 | 0.63 | 0.54 | 0.73 | 0.8 |
| 8 | 63422329 | rs11782265 | 1.661 | -0.853 | -0.017 | 0.772 | -0.09 | 0.102 | 0.102 | 0.956 | 0.4 | 0.09 | 0.39 | 0.22 | 0.47 | 0.48 |
| 8 | 63423994 | rs13280307 | 0.342 | -0.783 | 0.436 | 0.937 | -0.004 | 0.02 | 0.02 | 0 | 0.07 | 0.15 | 0.09 | 0.1 | 0.08 | 0.02 |
| 8 | 63426826 | rs11783420 | 0.598 | -0.81 | 0.632 | 1.007 | -0.021 | 0.083 | 0.083 | 0 | 0.76 | 0.56 | 0.68 | 0.64 | 0.77 | 0.82 |
| 8 | 63428793 | rs1451859 | 1.43 | -0.684 | 0.693 | 1.166 | 0.004 | 0.172 | 0.172 | 0.759 | 0.87 | 0.48 | 0.75 | 0.72 | 0.91 | 0.93 |
| 8 | 63434639 | rs6993486 | 0.074 | -0.608 | 0.174 | 1.215 | -0.002 | 0.04 | 0.04 | 0 | 0.93 | 0.94 | 0.78 | 0.73 | 0.91 | 0.93 |
| 8 | 63436369 | rs1079157 | 0.788 | -0.594 | -0.444 | 0.959 | 0 | -0.034 | -0.034 | 0 | 0.58 | 0.23 | 0.47 | 0.47 | 0.59 | 0.55 |
| 8 | 63443941 | rs11775866 | 0.956 | -0.566 | -0.804 | 0.888 | -0.076 | 0.101 | 0.101 | 0 | 0.42 | 0.09 | 0.38 | 0.26 | 0.48 | 0.48 |
| 8 | 63444070 | rs1869018 | -0.585 | -0.581 | -0.765 | 1.025 | 0.055 | 0.024 | 0.024 | 0.706 | 0.51 | 0.85 | 0.4 | 0.46 | 0.43 | 0.45 |
| 8 | 63445919 | rs11781413 | 0.973 | -0.561 | -0.858 | 1.005 | -0.076 | 0.073 | 0.073 | 0 | 0.42 | 0.09 | 0.38 | 0.28 | 0.48 | 0.48 |
| 8 | 63446925 | rs896491 | -0.194 | -0.696 | -0.343 | 0.754 | 0.02 | 0.008 | 0.008 | 0 | 0.45 | 0.41 | 0.37 | 0.44 | 0.43 | 0.45 |
| 8 | 63449156 | rs12544945 | 0.818 | -0.626 | -1.039 | 0.496 | -0.078 | -0.105 | -0.105 | 0.114 | 0.37 | 0.09 | 0.55 | 0.5 | 0.5 | 0.44 |
| 8 | 63449366 | rs7825567 | 0.37 | -0.236 | -0.693 | 0.241 | 0.028 | 0.014 | 0.014 | 0.344 | 0.37 | 0.44 | 0.53 | 0.57 | 0.43 | 0.48 |
| 8 | 63455403 | rs13256147 | 1.061 | -0.647 | -0.516 | 0.006 | -0.045 | 0.056 | 0.056 | 0.016 | 0.24 | 0.05 | 0.14 | 0.1 | 0.27 | 0.12 |
| 8 | 63455590 | rs16929130 | 0.297 | -0.568 | 0.051 | 0.435 | 0.007 | 0.003 | 0.003 | 0 | 0.51 | 0.53 | 0.54 | 0.45 | 0.57 | 0.54 |
| 8 | 63461952 | rs1031399 | 0.08 | -0.741 | 0.195 | 0.435 | 0.002 | 0.013 | 0.013 | 0 | 0.3 | 0.51 | 0.37 | 0.36 | 0.3 | 0.32 |
| 8 | 63462352 | rs1383194 | 0.272 | -0.754 | 0.218 | 0.429 | 0.013 | 0.027 | 0.027 | 0 | 0.28 | 0.46 | 0.36 | 0.36 | 0.3 | 0.32 |
| 8 | 63463719 | rs2928389 | -1.473 | -0.784 | 0.176 | 0.435 | 0.006 | -0.001 | -0.001 | 0.041 | 0.79 | 0.84 | 0.9 | 0.8 | 0.88 | 0.85 |
| 8 | 63468810 | rs827710 | 1.052 | -0.746 | 0.157 | 0.322 | -0.009 | -0.02 | -0.02 | 0 | 0.19 | 0.15 | 0.27 | 0.33 | 0.2 | 0.26 |
| 8 | 63477014 | rs827702 | 0.936 | -0.736 | 0.175 | 0.205 | -0.001 | -0.001 | -0.001 | 0 | 0.2 | 0.22 | 0.28 | 0.35 | 0.19 | 0.27 |
| 8 | 63480302 | rs13262030 | 1.106 | -0.599 | 0.214 | 0.107 | 0.02 | -0.039 | -0.039 | 0 | 0.09 | 0.24 | 0.09 | 0 | 0.1 | 0.05 |
| 8 | 63481760 | rs11787352 | 1.117 | -0.579 | 0.108 | 0.164 | -0.045 | 0.111 | 0.111 | 0 | 0.43 | 0.22 | 0.41 | 0.24 | 0.49 | 0.46 |
| 8 | 63490414 | rs852286 | -1.051 | -0.596 | -0.03 | 0.23 | -0.019 | 0.049 | 0.049 | 0 | 0.74 | 0.65 | 0.77 | 0.55 | 0.79 | 0.75 |
| 8 | 63491015 | rs11784165 | 0.381 | -0.578 | -0.193 | 0.202 | 0.025 | 0.122 | 0.122 | 0.019 | 0.52 | 0.22 | 0.48 | 0.25 | 0.6 | 0.5 |
| 8 | 63495593 | rs2882923 | -0.197 | -0.597 | -0.041 | 0.171 | -0.032 | -0.078 | -0.078 | 0 | 0.15 | 0.29 | 0.1 | 0.01 | 0.11 | 0.06 |

| | | | | | | | | | | | | | | | | |
|---|---|---|---|---|---|---|---|---|---|---|---|---|---|---|---|---|
| 8 | 63496398 | rs827687 | 1.079 | -0.53 | -0.226 | 0.242 | -0.008 | -0.015 | -0.015 | 0 | 0.18 | 0.15 | 0.26 | 0.29 | 0.2 | 0.25 |
| 8 | 63500539 | rs827690 | -1.728 | -0.535 | -0.696 | 0.294 | 0.047 | -0.048 | -0.048 | 0 | 0.8 | 0.96 | 0.91 | 0.74 | 0.87 | 0.81 |
| 8 | 63515522 | rs1531658 | 0.279 | -0.482 | -0.528 | 0.323 | 0.015 | 0.007 | 0.007 | 0.434 | 0.32 | 0.27 | 0.18 | 0.27 | 0.23 | 0.23 |
| 8 | 63516962 | rs1383196 | 0.69 | -0.231 | -0.711 | 0.28 | 0.02 | 0.004 | 0.004 | 0 | 0.46 | 0.24 | 0.54 | 0.44 | 0.57 | 0.53 |
| 8 | 63517575 | rs4374986 | 0.748 | -0.204 | -0.871 | 0.294 | -0.001 | -0.023 | -0.023 | 0 | 0.45 | 0.15 | 0.52 | 0.44 | 0.57 | 0.53 |
| 8 | 63521095 | rs11988063 | 1.157 | -0.05 | -0.778 | 0.196 | -0.032 | -0.036 | -0.036 | 0.321 | 0.31 | 0.13 | 0.41 | 0.36 | 0.37 | 0.51 |
| 8 | 63530880 | rs936613 | -0.704 | -0.613 | -0.555 | 0.468 | 0.067 | 0.044 | 0.044 | 0.03 | 0.48 | 0.75 | 0.42 | 0.64 | 0.34 | 0.43 |
| 8 | 63531664 | rs16929298 | 0.45 | -0.797 | -0.269 | 0.567 | -0.011 | 0 | 0 | 0 | 0.28 | 0.26 | 0.13 | 0.31 | 0.15 | 0.21 |
| 8 | 63538623 | rs16929302 | 0.543 | -0.76 | -0.123 | 0.581 | 0.019 | 0.098 | 0.098 | 0.205 | 0.2 | 0.51 | 0.29 | 0.32 | 0.19 | 0.22 |
| 8 | 63546131 | rs16929319 | 1.201 | -0.812 | -0.231 | 0.535 | 0.041 | -0.032 | -0.032 | 0 | 0.22 | 0.06 | 0.09 | 0.31 | 0.13 | 0.16 |
| 8 | 63551554 | rs9969432 | 0.565 | -0.704 | -0.191 | 0.57 | -0.03 | -0.031 | -0.031 | 0 | 0.08 | 0.13 | 0.04 | 0 | 0.02 | 0.05 |
| 8 | 63554717 | rs961829 | 0.83 | -0.636 | -0.155 | 0.637 | 0.037 | 0.111 | 0.111 | 0 | 0.17 | 0.37 | 0.27 | 0.33 | 0.19 | 0.23 |
| 8 | 63577738 | rs1351361 | -1.208 | -0.446 | -0.472 | 0.646 | -0.011 | -0.013 | -0.013 | 0 | 0.92 | 0.94 | 0.84 | 0.82 | 0.85 | 0.92 |
| 8 | 63581067 | rs1480190 | -0.999 | -0.776 | -0.53 | 0.588 | 0.088 | 0.046 | 0.046 | 0.492 | 0.49 | 0.85 | 0.43 | 0.64 | 0.35 | 0.41 |
| 8 | 63586615 | rs1471516 | 1.339 | -1.336 | -0.358 | 0.726 | 0.027 | -0.019 | -0.019 | 1.22 | 0.18 | 0.09 | 0.09 | 0.25 | 0.12 | 0.13 |
| 8 | 63588151 | rs1471517 | -0.765 | -1.352 | -0.258 | 0.643 | 0.069 | 0.006 | 0.006 | 0.465 | 0.45 | 0.69 | 0.38 | 0.57 | 0.29 | 0.38 |
| 8 | 63600769 | rs11985179 | -0.371 | -1.396 | 0.065 | 0.399 | 0 | 0.003 | 0.003 | 1.073 | 0.33 | 0.63 | 0.16 | 0.29 | 0.16 | 0.22 |
| 8 | 63602018 | rs16929407 | 0.64 | -1.277 | 0.209 | 0.445 | -0.035 | -0.038 | -0.038 | 0.048 | 0.08 | 0.22 | 0.04 | 0 | 0.02 | 0.04 |
| 8 | 63605027 | rs11989310 | -0.213 | -1.224 | 0.151 | 0.529 | -0.047 | 0.105 | 0.105 | 0 | 0.24 | 0.45 | 0.27 | 0.38 | 0.17 | 0.27 |
| 8 | 63614581 | rs1904574 | -0.964 | -1.195 | -0.105 | 0.585 | -0.01 | 0 | 0 | 0.72 | 0.77 | 0.8 | 0.9 | 0.71 | 0.87 | 0.85 |
| 8 | 63635874 | rs6995644 | 0.31 | -1.245 | -0.135 | 0.755 | 0.078 | -0.017 | -0.017 | 2.899 | 0.63 | 0.28 | 0.7 | 0.42 | 0.85 | 0.69 |
| 8 | 63643215 | rs2127560 | -0.092 | -1.039 | 0.06 | 0.447 | -0.07 | -0.022 | -0.022 | 0.069 | 0.68 | 0.53 | 0.84 | 0.71 | 0.84 | 0.78 |
| 8 | 63644144 | rs2054523 | -1.298 | -0.909 | -0.223 | 0.515 | 0.028 | -0.015 | -0.015 | 0 | 0.79 | 0.89 | 0.9 | 0.71 | 0.87 | 0.85 |
| 8 | 63657332 | rs4737612 | -0.297 | -0.717 | -0.217 | 0.581 | -0.011 | 0.004 | 0.004 | 0 | 0.69 | 0.65 | 0.82 | 0.64 | 0.79 | 0.78 |
| 8 | 63659367 | rs975688 | 0.595 | -0.727 | -0.359 | 0.568 | 0.043 | 0.054 | 0.054 | 0 | 0.51 | 0.37 | 0.61 | 0.31 | 0.65 | 0.55 |
| 8 | 63659610 | rs1319275 | 0.83 | -0.768 | -0.714 | 0.547 | 0.074 | 0.103 | 0.103 | 0.011 | 0.48 | 0.19 | 0.58 | 0.31 | 0.64 | 0.55 |
| 8 | 63659635 | rs4739003 | 1.234 | -0.753 | -1.125 | 0.558 | -0.004 | 0.054 | 0.054 | 0 | 0.16 | 0.25 | 0.21 | 0.33 | 0.14 | 0.23 |
| 8 | 63669208 | rs1471512 | 1.226 | -0.751 | -1.124 | 0.491 | -0.004 | 0.043 | 0.043 | 0 | 0.17 | 0.25 | 0.21 | 0.29 | 0.14 | 0.23 |
| 8 | 63670777 | rs12546333 | 1.961 | -0.309 | -2.52 | 0.294 | -0.062 | 0.129 | 0.129 | 0 | 0.23 | 0.02 | 0.31 | 0.06 | 0.25 | 0.39 |
| 8 | 63675162 | rs6989572 | 1.149 | -1.134 | -1.767 | 1.018 | 0.003 | -0.001 | -0.001 | 0 | 0.07 | 0.05 | 0.14 | 0.22 | 0.08 | 0.07 |
| 8 | 63676101 | rs10957236 | 1.508 | -0.93 | -1.804 | 0.481 | -0.084 | 0.215 | 0.215 | 0.06 | 0.4 | 0.03 | 0.44 | 0.13 | 0.56 | 0.48 |
| 8 | 63691194 | rs4524781 | -0.334 | -0.466 | -0.736 | 1.075 | 0 | 0 | 0 | 1.271 | 0.1 | 0.09 | 0.08 | 0.08 | 0.09 | 0.08 |
| 8 | 63695794 | rs1480124 | -0.237 | -0.467 | -0.734 | 1.068 | 0.01 | -0.005 | -0.005 | 0.832 | 0.78 | 0.82 | 0.9 | 0.71 | 0.88 | 0.86 |
| 8 | 63696643 | rs725329 | -0.828 | 0.335 | -0.677 | 0.939 | 0 | 0 | 0 | 0 | 0.1 | 0.09 | 0.08 | 0.1 | 0.09 | 0.08 |
| 8 | 63726318 | rs4554459 | -0.803 | 0.36 | -0.674 | 0.871 | 0 | 0 | 0 | 0 | 0.1 | 0.09 | 0.08 | 0.08 | 0.09 | 0.09 |
| 8 | 63730748 | rs1351349 | -0.308 | -0.428 | -1.063 | 0.971 | 0.033 | -0.021 | -0.021 | 1.01 | 0.8 | 0.9 | 0.91 | 0.71 | 0.88 | 0.86 |
| 8 | 63733171 | rs1383155 | -0.477 | -0.564 | -0.63 | 1.147 | -0.005 | 0.039 | 0.039 | 0 | 0.26 | 0.34 | 0.29 | 0.36 | 0.23 | 0.31 |
| 8 | 63734876 | rs1383153 | -0.342 | -0.751 | -0.245 | 1.041 | -0.033 | 0.006 | 0.006 | 0 | 0.77 | 0.68 | 0.88 | 0.71 | 0.88 | 0.86 |
| 8 | 63737308 | rs899676 | 0.887 | -0.657 | -0.274 | 0.703 | 0.082 | 0.088 | 0.088 | 0 | 0.5 | 0.26 | 0.6 | 0.35 | 0.65 | 0.55 |
| 8 | 63738525 | rs17265947 | -1.502 | 0.575 | -0.242 | 0.547 | 0.002 | 0.002 | 0.002 | 0 | 0.1 | 0.12 | 0.16 | 0.11 | 0.18 | 0.06 |
| 8 | 63741602 | rs1007897 | 0.428 | -0.688 | -0.234 | 0.636 | 0.004 | -0.001 | -0.001 | 0.174 | 0.62 | 0.64 | 0.76 | 0.61 | 0.75 | 0.73 |
| 8 | 63743617 | rs9785076 | 0.09 | -0.99 | -0.479 | 0.604 | 0.034 | 0.03 | 0.03 | 0 | 0.17 | 0.05 | 0.04 | 0.09 | 0.11 | 0.14 |
| 8 | 63750069 | rs13266003 | 1.848 | -1.604 | -1.141 | 0.606 | -0.022 | 0.088 | 0.088 | 0.685 | 0.42 | 0.04 | 0.36 | 0.19 | 0.59 | 0.48 |
| 8 | 63757124 | rs1480116 | -1.329 | -1.593 | -1.022 | 0.774 | -0.005 | -0.069 | -0.069 | 0.695 | 0.84 | 0.96 | 0.82 | 0.7 | 0.86 | 0.87 |
| 8 | 63768015 | rs2017227 | -0.155 | -1.602 | -0.465 | 0.525 | -0.012 | 0.067 | 0.067 | 0 | 0.75 | 0.67 | 0.58 | 0.44 | 0.79 | 0.72 |
| 8 | 63768731 | rs899680 | 0.844 | -1.452 | -0.453 | 0.412 | -0.007 | -0.003 | -0.003 | 1.184 | 0.56 | 0.49 | 0.53 | 0.33 | 0.68 | 0.58 |
| 8 | 63776740 | rs13274287 | 0.767 | -1.301 | -0.835 | 0.092 | 0.002 | 0.003 | 0.003 | 0.463 | 0.5 | 0.49 | 0.42 | 0.33 | 0.58 | 0.54 |
| 8 | 63778353 | rs7842329 | 0.04 | -1.255 | -0.925 | 0.257 | 0.025 | -0.029 | -0.029 | 0 | 0.64 | 0.85 | 0.56 | 0.36 | 0.69 | 0.64 |
| 8 | 63778587 | rs7018025 | 0.04 | -1.255 | -0.925 | 0.257 | 0.025 | -0.029 | -0.029 | 0 | 0.64 | 0.86 | 0.56 | 0.36 | 0.69 | 0.64 |
| 8 | 63781563 | rs7826606 | -0.025 | -1.269 | -0.911 | 0.274 | 0.029 | -0.021 | -0.021 | 0 | 0.64 | 0.84 | 0.56 | 0.36 | 0.7 | 0.64 |
| 8 | 63789980 | rs10808726 | -0.458 | -1.309 | -0.336 | 0.304 | -0.071 | 0.016 | 0.016 | 0 | 0.15 | 0.4 | 0.14 | 0.16 | 0.09 | 0.19 |
| 8 | 63807649 | rs979843 | -0.393 | -1.095 | -1.127 | -0.017 | 0.028 | 0.055 | 0.055 | 0.557 | 0.41 | 0.79 | 0.37 | 0.29 | 0.36 | 0.49 |
| 8 | 63808400 | rs1869733 | 1.224 | -1.242 | -1.49 | 0.006 | 0.003 | -0.06 | -0.06 | 0 | 0.2 | 0.13 | 0.39 | 0.53 | 0.18 | 0.2 |
| 8 | 63809069 | rs4237055 | -2.25 | -1.278 | -1.656 | -0.096 | -0.097 | 0.08 | 0.08 | 0 | 0.77 | 0.93 | 0.81 | 0.93 | 0.68 | 0.86 |
| 8 | 63809801 | rs958694 | 0.054 | -1.446 | -1.557 | -0.272 | 0.018 | 0.003 | 0.003 | 0 | 0.35 | 0.17 | 0.44 | 0.68 | 0.31 | 0.37 |
| 8 | 63812686 | rs1455583 | 1.307 | -1.385 | -1.486 | -0.193 | -0.005 | -0.091 | -0.091 | 0.037 | 0.2 | 0.06 | 0.38 | 0.58 | 0.18 | 0.2 |
| 8 | 63812929 | rs13280971 | 0.68 | -1.385 | -0.662 | -0.212 | -0.039 | 0.014 | 0.014 | 0 | 0.12 | 0.23 | 0.13 | 0.14 | 0.09 | 0.14 |
| 8 | 63817743 | rs16929812 | 0.162 | -1.406 | -0.483 | -0.273 | -0.053 | 0.031 | 0.031 | 0 | 0.14 | 0.29 | 0.14 | 0.18 | 0.09 | 0.19 |
| 8 | 63817927 | rs4376493 | -2.423 | -1.315 | -0.6 | -0.393 | -0.078 | 0.093 | 0.093 | 0 | 0.78 | 0.91 | 0.81 | 0.97 | 0.68 | 0.86 |
| 8 | 63822440 | rs1455576 | 0.046 | -1.396 | -0.649 | -0.397 | 0 | -0.013 | -0.013 | 0 | 0.48 | 0.54 | 0.63 | 0.7 | 0.48 | 0.53 |
| 8 | 63829727 | rs4524780 | -1.117 | -1.416 | -0.664 | -0.437 | -0.051 | 0.113 | 0.113 | 0.186 | 0.64 | 0.82 | 0.78 | 0.89 | 0.57 | 0.71 |
| 8 | 63830366 | rs2061133 | 1.018 | -1.317 | -0.696 | -0.325 | 0.009 | -0.029 | -0.029 | 0 | 0.27 | 0.37 | 0.23 | 0.12 | 0.27 | 0.3 |
| 8 | 63838474 | rs1455573 | 1.536 | -1.308 | -0.866 | 0.059 | 0.014 | -0.075 | -0.075 | 0 | 0.18 | 0.05 | 0.18 | 0.32 | 0.12 | 0.1 |
| 8 | 63841046 | rs7821634 | -0.252 | -1.183 | -0.656 | 0.025 | -0.055 | -0.01 | -0.01 | 0 | 0.27 | 0.37 | 0.17 | 0.28 | 0.2 | 0.34 |
| 8 | 63848556 | rs1350059 | -0.352 | -0.875 | -0.651 | 0.175 | 0.001 | 0.025 | 0.025 | 0.135 | 0.52 | 0.62 | 0.47 | 0.29 | 0.45 | 0.49 |
| 8 | 63857960 | rs1002905 | 1.187 | -0.662 | -0.832 | 0.279 | -0.054 | -0.065 | -0.065 | 0 | 0.1 | 0.01 | 0.31 | 0.25 | 0.16 | 0.14 |
| 8 | 63862040 | rs16929920 | 0.818 | -0.89 | -0.375 | 0.272 | 0 | -0.121 | -0.121 | 0 | 0.3 | 0.43 | 0.23 | 0.03 | 0.28 | 0.39 |
| 8 | 63867010 | rs13261995 | 0.754 | -0.735 | -0.781 | 0.474 | 0.002 | -0.016 | -0.016 | 0 | 0.11 | 0.08 | 0.16 | 0.22 | 0.1 | 0.16 |
| 8 | 63867493 | rs7835051 | -0.401 | -0.869 | -0.778 | 0.439 | -0.023 | -0.02 | -0.02 | 0 | 0.59 | 0.76 | 0.44 | 0.5 | 0.49 | 0.58 |

| | | | | | | | | | | | | | | | | |
|---|---|---|---|---|---|---|---|---|---|---|---|---|---|---|---|---|
| 8 | 63870658 | rs9298070 | -1.168 | -0.847 | -0.977 | 0.405 | -0.074 | -0.001 | -0.001 | 0 | 0.67 | 0.85 | 0.67 | 0.75 | 0.58 | 0.71 |
| 8 | 63874975 | rs1350058 | 0.022 | -1.044 | -0.755 | 0.344 | 0.035 | 0.039 | 0.039 | 0.055 | 0.51 | 0.69 | 0.4 | 0.41 | 0.4 | 0.49 |
| 8 | 63875082 | rs1350057 | 1.306 | -1.08 | -0.729 | 0.612 | 0.036 | -0.036 | -0.036 | 0 | 0.17 | 0.08 | 0.16 | 0.26 | 0.11 | 0.09 |
| 8 | 63879566 | rs12543454 | -1.019 | -0.874 | -0.107 | 0.784 | -0.015 | 0.111 | 0.111 | 0 | 0.19 | 0.3 | 0.38 | 0.47 | 0.16 | 0.26 |
| 8 | 63882096 | rs12056419 | 0.016 | -0.727 | -0.344 | 0.756 | 0.008 | -0.025 | -0.025 | 0 | 0.19 | 0.15 | 0.2 | 0.31 | 0.19 | 0.24 |
| 8 | 63883593 | rs7834588 | 0.273 | -0.777 | 0.18 | 0.564 | 0.003 | 0.025 | 0.025 | 0 | 0.33 | 0.51 | 0.42 | 0.4 | 0.33 | 0.49 |
| 8 | 63892585 | rs16930023 | 1.187 | -0.901 | 0.373 | 0.462 | 0.004 | -0.014 | -0.014 | 0 | 0.26 | 0.29 | 0.21 | 0.16 | 0.28 | 0.39 |
| 8 | 63893072 | rs16930025 | 0.735 | -1.06 | 0.593 | 0.494 | -0.014 | -0.016 | -0.016 | 0 | 0.05 | 0.16 | 0.02 | 0 | 0 | 0 |
| 8 | 63898146 | rs4737623 | -0.873 | -1.222 | 0.486 | 0.525 | -0.05 | 0.067 | 0.067 | 0 | 0.23 | 0.45 | 0.26 | 0.36 | 0.17 | 0.25 |
| 8 | 63901488 | rs4425750 | 1.216 | -1.336 | 0.1 | 0.453 | 0.042 | -0.022 | -0.022 | 0 | 0.18 | 0.09 | 0.18 | 0.25 | 0.11 | 0.1 |
| 8 | 63902886 | rs17193714 | 0.929 | -1.256 | 0.206 | 0.46 | -0.011 | -0.016 | -0.016 | 0 | 0.13 | 0.17 | 0.05 | 0.09 | 0.09 | 0.15 |
| 8 | 63905750 | rs10957263 | 0.394 | -1.163 | 0.236 | 0.488 | -0.002 | 0.03 | 0.03 | 0 | 0.15 | 0.17 | 0.24 | 0.37 | 0.16 | 0.25 |
| 8 | 63905899 | rs10957264 | 0.394 | -1.163 | 0.232 | 0.49 | -0.002 | 0.03 | 0.03 | 0 | 0.15 | 0.18 | 0.24 | 0.37 | 0.16 | 0.25 |
| 8 | 63906122 | rs16930057 | -0.49 | -1.364 | 0.225 | 0.327 | -0.044 | -0.029 | -0.029 | 0 | 0.37 | 0.41 | 0.24 | 0.28 | 0.2 | 0.24 |
| 8 | 63906828 | rs1031553 | -0.425 | -1.169 | 0.255 | 0.487 | -0.017 | 0.051 | 0.051 | 0 | 0.2 | 0.23 | 0.25 | 0.4 | 0.16 | 0.26 |
| 8 | 63915482 | rs16930073 | 1.072 | -1.06 | 0.061 | 0.546 | -0.001 | 0.003 | 0.003 | 0 | 0.08 | 0.07 | 0.15 | 0.21 | 0.09 | 0.1 |
| 8 | 63923414 | rs12548933 | -0.09 | -1.124 | 0.073 | 0.476 | 0.002 | -0.003 | -0.003 | 0.224 | 0.18 | 0.15 | 0.24 | 0.41 | 0.17 | 0.25 |
| 8 | 63927746 | rs17279558 | 1.59 | -1.111 | 0.076 | 0.461 | 0.002 | 0.018 | 0.018 | 0 | 0.11 | 0.02 | 0.04 | 0.09 | 0.09 | 0.15 |
| 8 | 63930289 | rs12677953 | -0.777 | -1.033 | 0.074 | 0.457 | 0 | 0.048 | 0.048 | 1.459 | 0.08 | 0.11 | 0.22 | 0.32 | 0.1 | 0.14 |
| 8 | 63933190 | rs1031552 | -0.711 | -1.028 | 0.076 | 0.46 | 0 | 0.049 | 0.049 | 1.34 | 0.08 | 0.11 | 0.22 | 0.32 | 0.1 | 0.14 |
| 8 | 63938764 | rs11545078 | 1.538 | -1.01 | 0.046 | 0.468 | 0.006 | 0.012 | 0.012 | 0 | 0.11 | 0.06 | 0.04 | 0.09 | 0.09 | 0.15 |
| 8 | 63941139 | rs3780127 | 1.538 | -1.01 | 0.046 | 0.468 | 0.006 | 0.012 | 0.012 | 0 | 0.11 | 0.06 | 0.04 | 0.09 | 0.09 | 0.15 |
| 8 | 63942478 | rs11786957 | 1.345 | -1.019 | 0.139 | 0.363 | 0.007 | -0.027 | -0.027 | 0.106 | 0.26 | 0.31 | 0.21 | 0.14 | 0.28 | 0.38 |
| 8 | 63942717 | rs4617146 | 0.467 | -1.083 | -0.222 | 0.392 | 0.041 | -0.082 | -0.082 | 0 | 0.26 | 0.15 | 0.33 | 0.43 | 0.2 | 0.2 |
| 8 | 63949912 | rs3780126 | 0.375 | -0.754 | -0.185 | 0.021 | 0.009 | 0.015 | 0.015 | 0 | 0.48 | 0.6 | 0.42 | 0.37 | 0.39 | 0.49 |
| 8 | 63953149 | rs12544045 | 0.036 | 0.182 | -0.706 | 0.294 | -0.029 | -0.007 | -0.007 | 0 | 0.19 | 0.12 | 0.23 | 0.22 | 0.28 | 0.29 |
| 8 | 63963847 | rs1031551 | -0.077 | 0.171 | -0.695 | 0.288 | -0.018 | 0 | 0 | 0.052 | 0.21 | 0.18 | 0.23 | 0.22 | 0.28 | 0.29 |
| 8 | 63969320 | rs4739045 | 0.02 | 0.108 | -0.246 | 0.524 | 0.023 | 0.035 | 0.035 | 0 | 0.51 | 0.62 | 0.43 | 0.37 | 0.39 | 0.49 |
| 8 | 64001936 | rs6472073 | -0.457 | 0.126 | -0.166 | 0.572 | -0.004 | 0.019 | 0.019 | 0 | 0.57 | 0.64 | 0.64 | 0.76 | 0.48 | 0.63 |
| 8 | 64019832 | rs7836669 | -0.497 | 0.383 | -0.548 | 0.706 | -0.003 | -0.011 | -0.011 | 0 | 0.6 | 0.66 | 0.62 | 0.5 | 0.61 | 0.57 |
| 8 | 64026280 | rs4314646 | 1.249 | 0.168 | -1.299 | 0.614 | 0.022 | -0.042 | -0.042 | 0 | 0.16 | 0.02 | 0.17 | 0.23 | 0.1 | 0.1 |
| 8 | 64028399 | rs6472077 | -0.292 | 0.766 | -0.728 | 0.925 | -0.011 | -0.03 | -0.03 | 0.069 | 0.61 | 0.75 | 0.62 | 0.5 | 0.61 | 0.57 |
| 8 | 64029902 | rs2117695 | 0.197 | 0.829 | -0.757 | 0.269 | -0.011 | -0.011 | -0.011 | 3.179 | 0.08 | 0.08 | 0.18 | 0.13 | 0.19 | 0.05 |
| 8 | 64034747 | rs4739050 | -0.612 | 0.46 | -0.954 | 0.179 | -0.125 | 0.186 | 0.186 | 0 | 0.66 | 0.95 | 0.8 | 0.9 | 0.61 | 0.72 |
| 8 | 64035651 | rs1030788 | -0.61 | 0.526 | -0.906 | 0.213 | -0.006 | -0.013 | -0.013 | 0 | 0.65 | 0.71 | 0.79 | 0.9 | 0.6 | 0.73 |
| 8 | 64040373 | rs17195767 | -0.224 | 0.583 | -0.921 | 1.229 | -0.017 | -0.034 | -0.034 | 0 | 0.07 | 0.01 | 0.13 | 0.13 | 0.13 | 0.09 |
| 8 | 64050344 | rs4739053 | -0.796 | 0.613 | -0.791 | 1.827 | -0.005 | 0.061 | 0.061 | 0 | 0.07 | 0.08 | 0.2 | 0.36 | 0.08 | 0.05 |
| 8 | 64072039 | rs13250662 | -0.568 | 0.538 | -0.061 | 1.484 | -0.006 | 0.005 | 0.005 | 0 | 0.3 | 0.35 | 0.28 | 0.39 | 0.33 | 0.22 |
| 8 | 64080486 | rs2241753 | 0.031 | 0.106 | 1.105 | 1.761 | 0.002 | 0.103 | 0.103 | 0 | 0.05 | 0.07 | 0.18 | 0.33 | 0.07 | 0.04 |
| 8 | 64080833 | rs2241754 | -0.379 | -0.004 | 1.103 | 1.711 | -0.053 | 0.064 | 0.064 | 0 | 0.22 | 0.35 | 0.18 | 0.43 | 0.12 | 0.17 |
| 8 | 64084228 | rs1435457 | -1.154 | -0.229 | 1.273 | 1.579 | -0.035 | -0.013 | -0.013 | 0 | 0.3 | 0.55 | 0.36 | 0.75 | 0.19 | 0.21 |
| 8 | 64087276 | rs7000981 | -1.154 | -0.229 | 1.313 | 1.579 | -0.032 | -0.012 | -0.012 | 0 | 0.3 | 0.56 | 0.37 | 0.75 | 0.19 | 0.21 |
| 8 | 64095407 | rs7015644 | 0.943 | -0.282 | 0.532 | 0.682 | -0.054 | 0.152 | 0.152 | 0 | 0.28 | 0.05 | 0.33 | 0.04 | 0.31 | 0.36 |
| 8 | 64104092 | rs4739066 | -0.241 | -0.328 | 0.98 | 1.245 | 0.02 | 0.205 | 0.205 | 2.286 | 0.08 | 0.21 | 0.19 | 0.33 | 0.07 | 0.04 |
| 8 | 64114563 | rs10088859 | 0.154 | -0.374 | 0.955 | 1.206 | -0.016 | 0.104 | 0.104 | 0.119 | 0.08 | 0.21 | 0.12 | 0.17 | 0.03 | 0.06 |
| 8 | 64124956 | rs7464 | -1.257 | -0.462 | 1.132 | 1.14 | 0.011 | 0.006 | 0.006 | 0.174 | 0.31 | 0.68 | 0.38 | 0.75 | 0.19 | 0.21 |
| 8 | 64127717 | rs10957277 | -1.115 | -0.458 | 1.154 | 1.151 | -0.033 | -0.016 | -0.016 | 0.217 | 0.3 | 0.55 | 0.37 | 0.76 | 0.19 | 0.21 |
| 8 | 64137932 | rs10104382 | -0.196 | -0.412 | 1.071 | 0.972 | -0.02 | 0.124 | 0.124 | 0.354 | 0.08 | 0.26 | 0.12 | 0.17 | 0.03 | 0.06 |
| 8 | 64148307 | rs10108512 | -0.07 | -0.408 | 1.035 | 0.974 | -0.023 | 0.102 | 0.102 | 0.222 | 0.08 | 0.23 | 0.12 | 0.17 | 0.03 | 0.06 |
| 8 | 64168483 | rs6472086 | 0.357 | -0.185 | 0.725 | 0.895 | 0.019 | 0.012 | 0.012 | 0 | 0.52 | 0.38 | 0.55 | 0.52 | 0.54 | 0.71 |
| 8 | 64175348 | rs7017562 | 0.04 | -0.309 | 0.99 | 0.842 | -0.052 | 0.016 | 0.016 | 0 | 0.19 | 0.31 | 0.15 | 0.29 | 0.09 | 0.12 |
| 8 | 64176338 | rs3924926 | 0.475 | -0.159 | 0.805 | 0.834 | 0.011 | -0.025 | -0.025 | 0 | 0.3 | 0.4 | 0.31 | 0.19 | 0.37 | 0.17 |
| 8 | 64177713 | rs10102076 | 0.662 | -0.123 | 0.403 | 0.853 | -0.009 | 0.012 | 0.012 | 0 | 0.29 | 0.24 | 0.3 | 0.19 | 0.37 | 0.17 |
| 8 | 64183260 | rs4739074 | -0.366 | -0.208 | 0.381 | 0.9 | 0 | 0 | 0 | 0 | 0.45 | 0.44 | 0.44 | 0.49 | 0.46 | 0.29 |
| 8 | 64185560 | rs6991229 | 0.676 | -0.288 | 0.269 | 0.886 | -0.023 | 0.018 | 0.018 | 0 | 0.16 | 0.21 | 0.14 | 0.3 | 0.09 | 0.12 |
| 8 | 64191555 | rs13266641 | 1.248 | -0.105 | -0.407 | 0.619 | -0.114 | 0.149 | 0.149 | 0 | 0.24 | 0.03 | 0.21 | 0.02 | 0.38 | 0.12 |
| 8 | 64193072 | rs12114802 | 0.492 | 0.1 | -0.316 | 0.958 | -0.002 | -0.01 | -0.01 | 0.259 | 0.07 | 0.04 | 0.04 | 0.19 | 0.09 | 0.05 |
| 8 | 64194110 | rs10088840 | -0.498 | 0.143 | -0.152 | 0.734 | 0.009 | 0.051 | 0.051 | 0 | 0.53 | 0.69 | 0.52 | 0.71 | 0.59 | 0.31 |
| 8 | 64198447 | rs11994501 | -0.333 | -0.007 | -0.398 | 0.709 | -0.001 | -0.002 | -0.002 | 3.181 | 0.52 | 0.48 | 0.38 | 0.49 | 0.54 | 0.31 |
| 8 | 64207363 | rs13260204 | -0.292 | 0.004 | -0.599 | 0.281 | 0.002 | -0.001 | -0.001 | 0.297 | 0.5 | 0.53 | 0.5 | 0.47 | 0.59 | 0.41 |
| 8 | 64210187 | rs7465268 | 0.297 | 0.262 | -1.23 | -0.603 | 0.003 | 0.014 | 0.014 | 0 | 0.11 | 0.09 | 0.06 | 0.02 | 0.09 | 0.2 |
| 8 | 64210708 | rs12678487 | 0.568 | 0.151 | -1.337 | -0.583 | -0.017 | 0.059 | 0.059 | 0.053 | 0.34 | 0.16 | 0.42 | 0.16 | 0.4 | 0.16 |
| 8 | 64211673 | rs12156261 | 0.345 | 0.128 | -1.556 | -0.933 | -0.036 | 0.136 | 0.136 | 0.453 | 0.34 | 0.17 | 0.31 | 0.03 | 0.35 | 0.42 |
| 8 | 64214573 | rs7003951 | 0.086 | 0.326 | -1.385 | -0.879 | 0 | 0.021 | 0.021 | 0 | 0.53 | 0.6 | 0.46 | 0.22 | 0.52 | 0.47 |
| 8 | 64215873 | rs10091141 | 0.582 | -0.02 | -1.05 | -1.579 | 0.006 | -0.003 | -0.003 | 0.054 | 0.4 | 0.38 | 0.29 | 0.05 | 0.36 | 0.35 |
| 8 | 64216591 | rs7465087 | -1.065 | -0.119 | -1.623 | -1.947 | -0.145 | 0.134 | 0.134 | 0 | 0.74 | 0.98 | 0.77 | 0.93 | 0.6 | 0.89 |
| 8 | 64220654 | rs7837234 | 0.979 | -0.269 | -1.367 | -1.784 | 0.004 | -0.01 | -0.01 | 0 | 0.39 | 0.37 | 0.28 | 0.04 | 0.38 | 0.36 |
| 8 | 64221219 | rs6472089 | -0.013 | -0.079 | -1.335 | -0.786 | -0.014 | -0.005 | -0.005 | 0 | 0.59 | 0.78 | 0.6 | 0.59 | 0.49 | 0.73 |
| 8 | 64248108 | rs7822639 | 0.238 | -0.061 | -0.805 | -0.485 | -0.001 | 0.03 | 0.03 | 0 | 0.11 | 0.09 | 0.15 | 0.39 | 0.08 | 0.2 |

| | | | | | | | | | | | | | | | | |
|---|---|---|---|---|---|---|---|---|---|---|---|---|---|---|---|---|
| 8 | 64259215 | rs9987202 | 1.333 | -0.478 | -1.246 | -0.741 | -0.05 | 0.172 | 0.172 | 0.789 | 0.37 | 0.09 | 0.29 | 0.12 | 0.41 | 0.33 |
| 8 | 64263770 | rs4737639 | -1.071 | -0.522 | -1.128 | -0.78 | -0.053 | 0.09 | 0.09 | 0.838 | 0.71 | 0.91 | 0.77 | 0.91 | 0.6 | 0.85 |
| 8 | 64264499 | rs7460898 | -0.061 | -0.611 | -0.318 | -0.536 | -0.034 | 0.062 | 0.062 | 0.015 | 0.21 | 0.34 | 0.29 | 0.41 | 0.11 | 0.29 |
| 8 | 64276115 | rs7459881 | 1.341 | -0.27 | -0.309 | -0.274 | 0.002 | 0.013 | 0.013 | 0 | 0.28 | 0.39 | 0.26 | 0.33 | 0.31 | 0.25 |
| 8 | 64290407 | rs6472099 | 0.304 | -0.33 | -0.055 | -0.04 | 0.003 | 0.064 | 0.064 | 0 | 0.09 | 0.13 | 0.2 | 0.37 | 0.08 | 0.18 |
| 8 | 64290572 | rs6472100 | 0.516 | -0.265 | -0.01 | -0.174 | -0.01 | 0.034 | 0.034 | 0 | 0.45 | 0.7 | 0.52 | 0.71 | 0.46 | 0.45 |
| 8 | 64296883 | rs10112358 | -0.839 | -0.276 | -0.096 | -0.331 | 0.015 | -0.003 | -0.003 | 0.013 | 0.43 | 0.34 | 0.5 | 0.59 | 0.31 | 0.61 |
| 8 | 64315765 | rs9298076 | -0.279 | -0.094 | -0.283 | -0.537 | -0.01 | 0.003 | 0.003 | 0.962 | 0.71 | 0.78 | 0.76 | 0.87 | 0.63 | 0.75 |
| 8 | 64316879 | rs7841852 | 1.641 | -0.242 | -0.394 | -0.318 | -0.002 | -0.003 | -0.003 | 0.609 | 0.35 | 0.36 | 0.28 | 0.34 | 0.37 | 0.28 |
| 8 | 64321899 | rs10957284 | -0.993 | -0.405 | -0.17 | 0.031 | -0.028 | 0.041 | 0.041 | 1.487 | 0.79 | 0.91 | 0.81 | 0.93 | 0.69 | 0.89 |
| 8 | 64323922 | rs17211259 | 0.647 | -0.436 | -0.133 | 0.099 | -0.013 | 0.026 | 0.026 | 0.332 | 0.2 | 0.04 | 0.19 | 0.13 | 0.2 | 0.37 |
| 8 | 64335958 | rs10113822 | 0.547 | -0.486 | -0.138 | -0.012 | -0.017 | 0.034 | 0.034 | 0.749 | 0.19 | 0.04 | 0.18 | 0.12 | 0.21 | 0.37 |
| 8 | 64349203 | rs2043178 | -0.825 | -0.402 | -0.508 | 0.013 | -0.025 | 0.021 | 0.021 | 1.511 | 0.36 | 0.46 | 0.47 | 0.51 | 0.27 | 0.47 |
| 8 | 64354619 | rs1560506 | -0.541 | -0.385 | -0.558 | -0.013 | -0.009 | 0.007 | 0.007 | 1.705 | 0.76 | 0.84 | 0.81 | 0.87 | 0.69 | 0.79 |
| 8 | 64360059 | rs974555 | 1.569 | -0.341 | -0.791 | -0.014 | -0.006 | -0.001 | -0.001 | 0 | 0.4 | 0.37 | 0.34 | 0.35 | 0.42 | 0.32 |
| 8 | 64367858 | rs10957287 | -0.097 | -0.002 | -1.302 | -0.243 | 0.015 | -0.042 | -0.042 | 0 | 0.1 | 0.01 | 0.16 | 0.12 | 0.05 | 0.08 |
| 8 | 64369361 | rs4512415 | -0.386 | 0.402 | -0.928 | -0.545 | -0.018 | 0.025 | 0.025 | 2.705 | 0.79 | 0.9 | 0.82 | 0.93 | 0.7 | 0.91 |
| 8 | 64381431 | rs2060103 | 0.085 | 0.15 | -1.11 | -0.297 | -0.023 | 0.026 | 0.026 | 0.709 | 0.19 | 0.02 | 0.17 | 0.13 | 0.18 | 0.35 |
| 8 | 64386244 | rs12546503 | 1.08 | 0.062 | -0.558 | -0.29 | 0.002 | 0.005 | 0.005 | 0.156 | 0.54 | 0.43 | 0.6 | 0.5 | 0.58 | 0.45 |
| 8 | 64389880 | rs12115114 | -1.26 | 0.214 | -0.649 | -0.07 | -0.049 | 0.022 | 0.022 | 0 | 0.91 | 0.99 | 0.91 | 0.94 | 0.82 | 0.95 |
| 8 | 64390729 | rs1431586 | 0.06 | 0.091 | -0.202 | -0.015 | -0.007 | 0.066 | 0.066 | 0 | 0.08 | 0.14 | 0.09 | 0.26 | 0.04 | 0.04 |
| 8 | 64392460 | rs10504364 | -1.256 | 0.175 | -0.189 | -0.067 | -0.01 | 0.009 | 0.009 | 0 | 0.9 | 0.93 | 0.9 | 0.94 | 0.82 | 0.95 |
| 8 | 64405226 | rs7013832 | 0.471 | 0.005 | -0.321 | -0.371 | -0.001 | -0.027 | -0.027 | 0 | 0.19 | 0.11 | 0.34 | 0.5 | 0.19 | 0.3 |
| 8 | 64428514 | rs7000642 | 0.933 | -0.059 | 0.359 | -0.414 | 0 | 0.003 | 0.003 | 0 | 0.54 | 0.41 | 0.56 | 0.51 | 0.56 | 0.5 |
| 8 | 64431251 | rs720864 | 0.254 | -0.059 | 0.461 | -0.796 | 0.002 | 0.017 | 0.017 | 0.408 | 0.24 | 0.18 | 0.22 | 0.17 | 0.22 | 0.42 |
| 8 | 64434129 | rs11985469 | -0.487 | -0.094 | 0.593 | -0.826 | 0 | -0.005 | -0.005 | 0 | 0.31 | 0.26 | 0.25 | 0.17 | 0.27 | 0.45 |
| 8 | 64442682 | rs1431595 | -0.338 | -0.068 | 0.95 | -0.852 | 0.024 | -0.068 | -0.068 | 0 | 0.85 | 0.68 | 0.88 | 0.95 | 0.82 | 0.95 |
| 8 | 64445521 | rs7835225 | -1.237 | -0.056 | 0.626 | -0.845 | -0.009 | 0.006 | 0.006 | 0 | 0.89 | 0.91 | 0.9 | 0.95 | 0.82 | 0.95 |
| 8 | 64446416 | rs1452489 | 1.103 | -0.052 | 0.538 | -0.816 | -0.005 | 0.148 | 0.148 | 0 | 0.53 | 0.27 | 0.62 | 0.77 | 0.55 | 0.49 |
| 8 | 64448090 | rs10504365 | 0.048 | 0.01 | 0.399 | -0.531 | -0.001 | -0.009 | -0.009 | 0 | 0.24 | 0.22 | 0.22 | 0.17 | 0.22 | 0.43 |
| 8 | 64454446 | rs7833224 | -0.208 | 0.228 | 0.59 | 0.095 | 0.001 | 0.036 | 0.036 | 0.136 | 0.22 | 0.24 | 0.35 | 0.48 | 0.21 | 0.34 |
| 8 | 64457202 | rs10098166 | 1.443 | 0.156 | 0.458 | -0.271 | 0.006 | 0.036 | 0.036 | 0.174 | 0.51 | 0.32 | 0.44 | 0.41 | 0.52 | 0.44 |
| 8 | 64458106 | rs4739099 | -0.48 | 0.169 | 0.685 | -0.293 | 0.003 | 0.049 | 0.049 | 0 | 0.27 | 0.38 | 0.42 | 0.52 | 0.24 | 0.47 |
| 8 | 64460952 | rs7832904 | 1.512 | 0.141 | 0.306 | -0.292 | 0.006 | 0.032 | 0.032 | 0.06 | 0.51 | 0.32 | 0.44 | 0.42 | 0.52 | 0.44 |
| 8 | 64468578 | rs4739103 | 1.414 | 0.223 | 0.411 | 0.299 | -0.038 | -0.026 | -0.026 | 0 | 0.12 | 0.01 | 0.18 | 0.13 | 0.15 | 0.08 |
| 8 | 64469050 | rs4739104 | -0.828 | 0.197 | 0.655 | 0.18 | 0.004 | 0.077 | 0.077 | 0 | 0.27 | 0.49 | 0.43 | 0.51 | 0.24 | 0.47 |
| 8 | 64471089 | rs11775359 | -0.675 | 0.241 | 0.445 | -0.262 | -0.001 | -0.01 | -0.01 | 0 | 0.86 | 0.86 | 0.89 | 0.95 | 0.81 | 0.94 |
| 8 | 64475617 | rs12675996 | -0.047 | 0.307 | 0.195 | 0.176 | -0.023 | -0.06 | -0.06 | 0.009 | 0.2 | 0.09 | 0.39 | 0.49 | 0.22 | 0.34 |
| 8 | 64476692 | rs1377116 | 0.57 | 0.382 | 0.829 | -0.451 | 0.026 | -0.118 | -0.118 | 1.58 | 0.78 | 0.48 | 0.82 | 0.91 | 0.77 | 0.82 |
| 8 | 64482634 | rs993513 | 0.708 | 0.336 | 0.722 | -0.45 | 0.013 | -0.101 | -0.101 | 0 | 0.78 | 0.44 | 0.81 | 0.92 | 0.78 | 0.82 |
| 8 | 64498360 | rs6472116 | -0.584 | 0.137 | 0.845 | -0.499 | 0.004 | -0.03 | -0.03 | 0.314 | 0.85 | 0.82 | 0.89 | 0.96 | 0.8 | 0.9 |
| 8 | 64501700 | rs2354861 | -0.641 | -0.318 | 0.44 | -0.294 | 0.001 | 0 | 0 | 0.156 | 0.43 | 0.57 | 0.63 | 0.72 | 0.41 | 0.6 |
| 8 | 64505710 | rs13273817 | 1.231 | -0.485 | -0.647 | -0.052 | -0.031 | -0.041 | -0.041 | 3.823 | 0.15 | 0.01 | 0.21 | 0.16 | 0.17 | 0.09 |
| 8 | 64508587 | rs896619 | -0.521 | -0.248 | -0.395 | -0.39 | -0.014 | 0.073 | 0.073 | 0.53 | 0.4 | 0.2 | 0.61 | 0.74 | 0.44 | 0.6 |
| 8 | 64533015 | rs1562987 | 0.172 | -0.096 | -0.273 | 0.159 | -0.009 | 0.092 | 0.092 | 0.121 | 0.05 | 0.09 | 0.04 | 0.22 | 0.02 | 0.13 |
| 8 | 64539024 | rs10441559 | 0.415 | -0.247 | -0.716 | 0.074 | -0.016 | -0.015 | -0.015 | 0 | 0.15 | 0.02 | 0.1 | 0.15 | 0.15 | 0.2 |
| 8 | 64547318 | rs12334387 | -0.423 | -0.483 | -0.371 | -0.366 | 0.005 | 0.048 | 0.048 | 0.663 | 0.43 | 0.35 | 0.64 | 0.72 | 0.45 | 0.52 |
| 8 | 64553145 | rs1149458 | 1.827 | -0.356 | -1.059 | -0.947 | -0.069 | 0.228 | 0.228 | 0.202 | 0.31 | 0.02 | 0.15 | 0.01 | 0.28 | 0.15 |
| 8 | 64565990 | rs1149459 | 2.156 | -0.412 | -1.191 | -1.125 | -0.062 | 0.191 | 0.191 | 0.469 | 0.27 | 0.02 | 0.13 | 0 | 0.25 | 0.11 |
| 8 | 64591472 | rs1149467 | -0.308 | -0.201 | 0.335 | -0.512 | 0.061 | -0.137 | -0.137 | 0.122 | 0.81 | 0.54 | 0.84 | 0.97 | 0.75 | 0.82 |
| 8 | 64594355 | rs1217113 | -0.924 | -0.212 | 0.802 | -0.52 | 0.03 | -0.063 | -0.063 | 0.125 | 0.85 | 0.8 | 0.87 | 0.98 | 0.75 | 0.84 |
| 8 | 64602654 | rs812603 | 1.137 | -0.298 | 0.484 | -0.47 | 0.003 | 0.013 | 0.013 | 0 | 0.67 | 0.37 | 0.76 | 0.75 | 0.7 | 0.59 |
| 8 | 64621373 | rs790566 | -0.829 | 0.226 | 0.346 | -0.639 | 0.024 | -0.054 | -0.054 | 0.668 | 0.85 | 0.83 | 0.87 | 0.98 | 0.75 | 0.84 |
| 8 | 64625292 | rs1217103 | -0.921 | 0.225 | 0.492 | -0.647 | 0.01 | -0.033 | -0.033 | 0 | 0.86 | 0.87 | 0.87 | 0.98 | 0.75 | 0.84 |
| 8 | 64628120 | rs6992714 | 0.779 | 0.239 | 0.474 | -0.649 | -0.038 | 0.033 | 0.033 | 0 | 0.14 | 0.07 | 0.12 | 0.02 | 0.25 | 0.16 |
| 8 | 64636395 | rs11994082 | 0.59 | 0.226 | 0.489 | -0.656 | 0.032 | -0.059 | -0.059 | 0 | 0.15 | 0.18 | 0.14 | 0.02 | 0.25 | 0.15 |
| 8 | 64640696 | rs16930623 | 0.454 | -0.07 | 0.64 | -0.714 | -0.018 | -0.026 | -0.026 | 0 | 0.08 | 0.18 | 0.09 | 0.03 | 0.02 | 0.19 |
| 8 | 64642614 | rs16930628 | 1.084 | -0.183 | -0.055 | -0.492 | -0.014 | -0.093 | -0.093 | 0.138 | 0.13 | 0.01 | 0.06 | 0.33 | 0.16 | 0.19 |
| 8 | 64665734 | rs10957293 | -0.89 | -0.68 | 0.004 | -1.128 | -0.09 | -0.113 | -0.113 | 0 | 0.23 | 0.49 | 0.16 | 0.04 | 0.07 | 0.29 |
| 8 | 64682296 | rs6980548 | 1.167 | -0.489 | -0.061 | -1.076 | 0.015 | -0.028 | -0.028 | 0 | 0.14 | 0.15 | 0.19 | 0.04 | 0.25 | 0.16 |
| 8 | 64685833 | rs7001081 | -1.62 | -0.139 | -0.783 | -0.45 | -0.016 | 0.046 | 0.046 | 0.141 | 0.53 | 0.79 | 0.61 | 0.65 | 0.54 | 0.79 |
| 8 | 64692412 | rs6472121 | -0.583 | -0.065 | -0.334 | -0.627 | -0.001 | -0.004 | -0.004 | 0.692 | 0.31 | 0.65 | 0.33 | 0.07 | 0.29 | 0.37 |
| 8 | 64700949 | rs1520929 | -0.133 | -0.165 | -0.401 | -0.412 | 0.002 | 0.016 | 0.016 | 0.059 | 0.45 | 0.45 | 0.56 | 0.68 | 0.47 | 0.48 |
| 8 | 64705372 | rs1541750 | 1.159 | -0.461 | 0.309 | -0.59 | -0.026 | -0.045 | -0.045 | 4.925 | 0.09 | 0.18 | 0.05 | 0 | 0.03 | 0.08 |
| 8 | 64708102 | rs11997548 | 0.178 | -0.8 | 0.221 | -0.842 | -0.007 | 0 | 0 | 0 | 0.33 | 0.37 | 0.52 | 0.68 | 0.42 | 0.39 |
| 8 | 64717274 | rs2049131 | 1.462 | -0.825 | -0.555 | -0.611 | -0.014 | -0.017 | -0.017 | 0 | 0.2 | 0.2 | 0.46 | 0.35 | 0.29 | 0.26 |
| 8 | 64718119 | rs755223 | -0.962 | -1.091 | -0.912 | -0.879 | -0.048 | 0.123 | 0.123 | 0.763 | 0.5 | 0.89 | 0.68 | 0.71 | 0.49 | 0.56 |
| 8 | 64743305 | rs705988 | 0.574 | -0.972 | -0.006 | -0.455 | 0.034 | 0.098 | 0.098 | 0 | 0.27 | 0.49 | 0.56 | 0.49 | 0.32 | 0.35 |
| 8 | 64748166 | rs17217477 | 1.933 | -0.594 | -0.076 | -0.528 | 0.01 | -0.01 | -0.01 | 0 | 0.06 | 0.07 | 0.11 | 0.03 | 0.07 | 0.12 |

| | | | | | | | | | | | | | | | | |
|---|---|---|---|---|---|---|---|---|---|---|---|---|---|---|---|---|
| 8 | 64753633 | rs10504367 | 1.343 | -1.041 | -0.187 | -0.707 | -0.013 | 0.028 | 0.028 | 0 | 0.15 | 0.16 | 0.07 | 0.32 | 0.12 | 0.15 |
| 8 | 64771414 | rs12548723 | 1.744 | -1.02 | -0.217 | -0.715 | -0.008 | -0.01 | -0.01 | 0 | 0.2 | 0.21 | 0.46 | 0.36 | 0.29 | 0.22 |
| 8 | 64779013 | rs6982152 | -1.759 | -0.766 | -0.61 | -0.951 | -0.054 | 0.064 | 0.064 | 0 | 0.85 | 0.94 | 0.82 | 0.95 | 0.77 | 0.76 |
| 8 | 64779731 | rs2052695 | 1.532 | -0.978 | -0.83 | -1.032 | -0.047 | 0.078 | 0.078 | 1.273 | 0.31 | 0.03 | 0.14 | 0.13 | 0.28 | 0.13 |
| 8 | 64795941 | rs10957299 | -0.511 | -0.928 | 0.049 | -0.634 | 0.007 | 0.042 | 0.042 | 0.38 | 0.47 | 0.64 | 0.64 | 0.82 | 0.46 | 0.6 |
| 8 | 64825975 | rs10481236 | 1.559 | -0.911 | -0.132 | -0.55 | -0.014 | 0.044 | 0.044 | 0 | 0.12 | 0.16 | 0.05 | 0.32 | 0.11 | 0.16 |
| 8 | 64828623 | rs7824499 | -0.922 | -0.642 | -0.35 | -0.644 | -0.003 | 0.128 | 0.128 | 0.394 | 0.53 | 0.76 | 0.66 | 0.82 | 0.49 | 0.63 |
| 8 | 64847410 | rs919037 | -0.931 | -0.586 | -0.454 | -0.355 | -0.007 | 0.114 | 0.114 | 0.023 | 0.56 | 0.77 | 0.56 | 0.81 | 0.49 | 0.64 |
| 8 | 64861657 | rs2114018 | 1.237 | -0.184 | -0.651 | 0.137 | -0.004 | 0.045 | 0.045 | 0.052 | 0.05 | 0.06 | 0.19 | 0.24 | 0.08 | 0.18 |
| 8 | 64878574 | rs10217013 | 0.057 | -0.319 | -0.447 | -0.002 | 0.002 | -0.002 | -0.002 | 0 | 0.41 | 0.36 | 0.34 | 0.58 | 0.39 | 0.44 |
| 8 | 64882069 | rs10109510 | 0.465 | -0.229 | -0.692 | 0.104 | 0.008 | -0.033 | -0.033 | 0 | 0.36 | 0.18 | 0.26 | 0.47 | 0.39 | 0.31 |
| 8 | 64882680 | rs13280549 | 0.726 | -0.313 | -0.497 | 0.315 | 0.016 | -0.042 | -0.042 | 2.059 | 0.42 | 0.13 | 0.27 | 0.59 | 0.44 | 0.36 |
| 8 | 64891089 | rs2068480 | 0.755 | -0.283 | -0.668 | 0.171 | -0.031 | -0.138 | -0.138 | 0.433 | 0.4 | 0.02 | 0.31 | 0.52 | 0.42 | 0.32 |
| 8 | 64909013 | rs11786760 | -0.611 | -0.3 | -0.223 | 0.123 | -0.023 | 0.084 | 0.084 | 0 | 0.59 | 0.76 | 0.54 | 0.83 | 0.52 | 0.58 |
| 8 | 64931948 | rs6988624 | -1.551 | -0.214 | -0.494 | -0.051 | -0.07 | 0.157 | 0.157 | 0 | 0.76 | 0.98 | 0.78 | 0.97 | 0.74 | 0.8 |
| 8 | 64960855 | rs1838420 | -0.445 | -0.294 | 0.673 | 0.904 | 0.001 | 0.024 | 0.024 | 0 | 0.53 | 0.62 | 0.52 | 0.73 | 0.5 | 0.5 |
| 8 | 64980755 | rs13255793 | -0.143 | -0.296 | 0.619 | 0.932 | 0.01 | 0.089 | 0.089 | 0 | 0.49 | 0.31 | 0.46 | 0.73 | 0.5 | 0.5 |
| 8 | 64989407 | rs16930838 | 1.367 | -0.333 | 0.387 | 1.238 | 0.005 | -0.082 | -0.082 | 0 | 0.12 | 0 | 0.04 | 0.38 | 0.1 | 0.17 |
| 8 | 64992908 | rs16930842 | 0.07 | -0.27 | 0.575 | 1.239 | 0.008 | 0.071 | 0.071 | 0 | 0.08 | 0.18 | 0.14 | 0.21 | 0.06 | 0.18 |
| 8 | 65003854 | rs6997807 | -0.468 | -0.289 | 0.616 | 1.162 | -0.018 | 0.083 | 0.083 | 0.036 | 0.07 | 0.24 | 0.04 | 0.11 | 0.01 | 0.07 |
| 8 | 65008255 | rs1444511 | -0.856 | -0.225 | 0.484 | 1.106 | -0.016 | 0.05 | 0.05 | 0 | 0.61 | 0.72 | 0.55 | 0.84 | 0.57 | 0.61 |
| 8 | 65014043 | rs13251387 | 0.521 | -0.186 | 0.072 | 1.241 | -0.003 | 0.003 | 0.003 | 0 | 0.38 | 0.27 | 0.42 | 0.36 | 0.4 | 0.36 |
| 8 | 65045688 | rs13270019 | 1.153 | -0.113 | -0.112 | 1.139 | 0.002 | -0.024 | -0.024 | 0 | 0.14 | 0.08 | 0.08 | 0.38 | 0.15 | 0.19 |
| 8 | 65047353 | rs2629637 | -1.169 | -0.183 | -0.172 | 1.305 | -0.015 | -0.025 | -0.025 | 0 | 0.88 | 0.97 | 0.81 | 0.87 | 0.88 | 0.82 |
| 8 | 65055813 | rs2083998 | -0.248 | -0.319 | 0.099 | 1.628 | 0.001 | -0.003 | -0.003 | 0 | 0.43 | 0.38 | 0.44 | 0.46 | 0.43 | 0.42 |
| 8 | 65057965 | rs13271001 | 0.857 | -0.158 | -0.542 | 1.399 | -0.082 | 0.057 | 0.057 | 0 | 0.28 | 0.01 | 0.28 | 0.14 | 0.34 | 0.17 |
| 8 | 65069567 | rs12549779 | 0.933 | 0.17 | 0.079 | 1.686 | -0.018 | -0.039 | -0.039 | 0 | 0.08 | 0.01 | 0.11 | 0.11 | 0.08 | 0.11 |
| 8 | 65085905 | rs7824886 | -0.975 | 0.187 | 1.148 | 1.742 | -0.013 | -0.037 | -0.037 | 0 | 0.07 | 0.25 | 0.03 | 0 | 0.02 | 0 |
| 8 | 65088190 | rs13271334 | 0.455 | 0.198 | 0.642 | 1.781 | -0.084 | -0.008 | -0.008 | 0 | 0.29 | 0.01 | 0.29 | 0.19 | 0.36 | 0.23 |
| 8 | 65101330 | rs13277244 | -1.025 | -0.201 | 2.34 | 2.992 | -0.018 | 0.098 | 0.098 | 0.24 | 0.17 | 0.25 | 0.09 | 0.52 | 0.12 | 0.27 |
| 8 | 65107937 | rs10504373 | -0.462 | -0.195 | 1.492 | 2.927 | 0.009 | -0.045 | -0.045 | 0 | 0.14 | 0.06 | 0.06 | 0.38 | 0.11 | 0.22 |
| 8 | 65115733 | rs13268763 | 1.803 | -0.2 | 1.729 | 2.592 | 0.001 | 0 | 0 | 0.052 | 0.71 | 0.3 | 0.77 | 0.3 | 0.77 | 0.63 |
| 8 | 65128677 | rs10088800 | -0.402 | -0.223 | 1.743 | 2.839 | 0.005 | -0.007 | -0.007 | 0 | 0.13 | 0.08 | 0.07 | 0.23 | 0.11 | 0.23 |
| 8 | 65143126 | rs16930969 | -1.151 | -0.222 | 2.182 | 2.986 | 0.003 | 0.306 | 0.306 | 0.266 | 0.06 | 0.25 | 0.03 | 0.28 | 0.03 | 0.05 |
| 8 | 65144104 | rs6983423 | -1.049 | -0.113 | 1.765 | 2.524 | -0.015 | -0.002 | -0.002 | 0 | 0.17 | 0.27 | 0.09 | 0.23 | 0.12 | 0.24 |
| 8 | 65174630 | rs10110463 | -0.293 | -0.045 | 1.01 | 2.575 | -0.022 | -0.043 | -0.043 | 0 | 0.13 | 0 | 0.06 | 0.23 | 0.14 | 0.23 |
| 8 | 65186616 | rs2356665 | -2.575 | -0.001 | 1.762 | 2.733 | 0.003 | 0.189 | 0.189 | 2.12 | 0.15 | 0.67 | 0.18 | 0.43 | 0.13 | 0.15 |
| 8 | 65191812 | rs6994670 | -1.574 | -0.343 | 2.178 | 2.345 | 0.01 | 0.314 | 0.314 | 2.081 | 0.08 | 0.41 | 0.05 | 0.23 | 0.03 | 0.05 |
| 8 | 65197740 | rs1371685 | -1.461 | -0.797 | 0.499 | 2.004 | 0.091 | -0.112 | -0.112 | 0 | 0.34 | 0.86 | 0.22 | 0.47 | 0.21 | 0.4 |
| 8 | 65213478 | rs971236 | -0.629 | -0.769 | 0.585 | 1.901 | 0.007 | 0.047 | 0.047 | 1.114 | 0.23 | 0.06 | 0.11 | 0.17 | 0.21 | 0.33 |
| 8 | 65235506 | rs10957308 | -0.841 | -0.878 | 0.64 | 1.867 | -0.002 | -0.017 | -0.017 | 0 | 0.1 | 0.19 | 0.14 | 0.05 | 0.12 | 0.1 |
| 8 | 65242166 | rs13257322 | -1.72 | -0.875 | 0.413 | 1.776 | 0.004 | -0.005 | -0.005 | 0 | 0.38 | 0.57 | 0.31 | 0.49 | 0.31 | 0.48 |
| 8 | 65254172 | rs17315206 | -0.332 | -0.849 | -0.701 | 1.412 | -0.02 | 0.021 | 0.021 | 3.159 | 0.21 | 0 | 0.1 | 0.17 | 0.19 | 0.32 |
| 8 | 65272478 | rs298192 | 1.09 | -0.823 | -0.67 | 1.532 | 0.082 | -0.043 | -0.043 | 1.836 | 0.56 | 0.17 | 0.64 | 0.57 | 0.66 | 0.5 |
| 8 | 65283691 | rs190938 | 0.83 | -0.593 | -0.057 | 1.535 | -0.012 | 0.001 | 0.001 | 0 | 0.72 | 0.72 | 0.8 | 0.6 | 0.84 | 0.67 |
| 8 | 65295491 | rs4737191 | 0.064 | -1.067 | -0.439 | 1.327 | 0.004 | -0.014 | -0.014 | 0 | 0.09 | 0.01 | 0.1 | 0.1 | 0.07 | 0.07 |
| 8 | 65303114 | rs298181 | 1.373 | -1.337 | -0.087 | 1.271 | -0.005 | -0.002 | -0.002 | 0.063 | 0.67 | 0.72 | 0.8 | 0.68 | 0.82 | 0.67 |
| 8 | 65305203 | rs298185 | 0.389 | -1.392 | 0.086 | 1.241 | 0.011 | 0.009 | 0.009 | 0 | 0.79 | 0.86 | 0.92 | 0.78 | 0.9 | 0.76 |
| 8 | 65332897 | rs2600478 | -1.761 | -1.383 | 0.351 | 1.416 | -0.008 | -0.005 | -0.005 | 0 | 0.33 | 0.29 | 0.2 | 0.32 | 0.18 | 0.33 |
| 8 | 65333446 | rs2600480 | -0.563 | -1.341 | 0.256 | 1.406 | 0.011 | 0.009 | 0.009 | 0 | 0.21 | 0.14 | 0.08 | 0.22 | 0.1 | 0.24 |
| 8 | 65344089 | rs10957312 | -0.022 | -1.247 | 0.214 | 1.485 | -0.001 | -0.011 | -0.011 | 0 | 0.09 | 0.03 | 0.11 | 0.09 | 0.1 | 0.06 |
| 8 | 65347906 | rs2612600 | -0.544 | -1.158 | 0.323 | 1.485 | 0.012 | 0.009 | 0.009 | 0 | 0.2 | 0.12 | 0.07 | 0.22 | 0.1 | 0.24 |
| 8 | 65351833 | rs2577898 | -1.858 | -1.249 | 0.91 | 1.436 | -0.044 | -0.034 | -0.034 | 1.074 | 0.34 | 0.42 | 0.21 | 0.32 | 0.2 | 0.34 |
| 8 | 65357040 | rs2600468 | -1.917 | -1.335 | 0.929 | 1.399 | -0.029 | -0.019 | -0.019 | 0 | 0.34 | 0.39 | 0.2 | 0.32 | 0.2 | 0.34 |
| 8 | 65362019 | rs2612595 | -1.424 | -1.185 | 0.826 | 1.416 | 0.025 | 0.016 | 0.016 | 0 | 0.3 | 0.19 | 0.18 | 0.32 | 0.21 | 0.32 |
| 8 | 65364994 | rs1823667 | -0.498 | -1.187 | 0.802 | 1.416 | 0.027 | 0.012 | 0.012 | 0 | 0.19 | 0.04 | 0.06 | 0.22 | 0.11 | 0.26 |
| 8 | 65370653 | rs2600476 | -1.513 | -1.187 | 0.816 | 1.416 | 0.019 | 0.013 | 0.013 | 0 | 0.31 | 0.22 | 0.19 | 0.32 | 0.21 | 0.32 |
| 8 | 65387467 | rs7836768 | -1.92 | -0.797 | 0.972 | 1.661 | 0.006 | -0.007 | -0.007 | 0 | 0.36 | 0.7 | 0.37 | 0.44 | 0.39 | 0.38 |
| 8 | 65393079 | rs10504377 | -0.317 | -0.495 | 0.77 | 1.45 | -0.004 | 0.047 | 0.047 | 0 | 0.13 | 0.29 | 0.11 | 0.24 | 0.09 | 0.26 |
| 8 | 65401078 | rs13439473 | -0.69 | -0.816 | 0.195 | 1.577 | -0.001 | -0.085 | -0.085 | 0 | 0.22 | 0.03 | 0.18 | 0.26 | 0.15 | 0.32 |
| 8 | 65402084 | rs3862072 | -1.887 | -0.933 | 0.397 | 1.427 | 0.006 | -0.005 | -0.005 | 0 | 0.44 | 0.55 | 0.42 | 0.46 | 0.4 | 0.46 |
| 8 | 65412042 | rs2218914 | -0.911 | -1.367 | -0.008 | 0.941 | -0.011 | -0.01 | -0.01 | 0 | 0.18 | 0.2 | 0.11 | 0.09 | 0.13 | 0.11 |
| 8 | 65422363 | rs16931290 | -0.597 | -1.062 | -0.292 | 1.127 | 0.028 | -0.02 | -0.02 | 0 | 0.19 | 0.02 | 0.09 | 0.2 | 0.1 | 0.3 |
| 8 | 65422496 | rs896493 | -0.931 | -1.023 | -0.124 | 1.127 | 0.004 | -0.002 | -0.002 | 1.983 | 0.24 | 0.19 | 0.15 | 0.3 | 0.21 | 0.32 |
| 8 | 65430900 | rs10504380 | -0.886 | -0.574 | -0.428 | 0.595 | 0.003 | -0.007 | -0.007 | 3.442 | 0.12 | 0.12 | 0.06 | 0 | 0.09 | 0.1 |
| 8 | 65432449 | rs7017425 | -1.493 | -0.722 | 0.059 | 1.193 | -0.005 | 0.037 | 0.037 | 0 | 0.23 | 0.32 | 0.27 | 0.29 | 0.26 | 0.18 |
| 8 | 65436090 | rs1376771 | -0.823 | -0.756 | 0.216 | 0.879 | -0.024 | 0.002 | 0.002 | 0.011 | 0.17 | 0.27 | 0.1 | 0.21 | 0.08 | 0.15 |
| 8 | 65438121 | rs7814393 | -1.48 | -1.995 | -0.451 | 1.1 | 0.086 | -0.018 | -0.018 | 0 | 0.42 | 0.78 | 0.43 | 0.55 | 0.31 | 0.43 |
| 8 | 65447831 | rs1451863 | -1.09 | -2.252 | 0.346 | 1.186 | -0.007 | 0.011 | 0.011 | 0 | 0.28 | 0.34 | 0.34 | 0.35 | 0.26 | 0.25 |

| | | | | | | | | | | | | | | | | |
|---|---|---|---|---|---|---|---|---|---|---|---|---|---|---|---|---|
| 8 | 65450254 | rs6981465 | -0.379 | -2.313 | 0.443 | 1.138 | -0.009 | 0.028 | 0.028 | 0.146 | 0.19 | 0.24 | 0.26 | 0.35 | 0.13 | 0.23 |
| 8 | 65463442 | rs6996198 | -1.91 | -2.307 | 0.505 | 1.082 | -0.002 | -0.001 | -0.001 | 3.435 | 0.21 | 0.24 | 0.16 | 0.2 | 0.18 | 0.2 |
| 8 | 65467510 | rs10441561 | -0.972 | -2.261 | 0.61 | 1.152 | -0.005 | 0.009 | 0.009 | 0.605 | 0.22 | 0.72 | 0.3 | 0.35 | 0.14 | 0.24 |
| 8 | 65481050 | rs6997421 | 0.66 | -2.323 | 0.596 | 1.104 | -0.023 | -0.05 | -0.05 | 0.009 | 0.11 | 0.2 | 0.05 | 0 | 0.06 | 0.15 |
| 8 | 65483808 | rs7841976 | -0.375 | -2.333 | 0.792 | 1.116 | 0.01 | 0.238 | 0.238 | 0 | 0.09 | 0.26 | 0.23 | 0.35 | 0.07 | 0.06 |
| 8 | 65485318 | rs6981395 | 0.661 | -2.327 | 0.723 | 1.102 | -0.023 | -0.05 | -0.05 | 0 | 0.11 | 0.2 | 0.05 | 0 | 0.06 | 0.15 |
| 8 | 65492071 | rs4737675 | -2.067 | -1.55 | 0.512 | 0.895 | 0.025 | -0.032 | -0.032 | 0 | 0.3 | 0.75 | 0.37 | 0.37 | 0.29 | 0.26 |
| 8 | 65496059 | rs9650210 | -0.623 | -1.489 | 0.547 | 0.823 | -0.003 | -0.007 | -0.007 | 0.375 | 0.08 | 0.05 | 0.08 | 0 | 0.15 | 0.02 |
| 8 | 65499192 | rs2280870 | -2.416 | -1.477 | 0.524 | 0.893 | 0.065 | -0.024 | -0.024 | 0 | 0.41 | 0.77 | 0.46 | 0.55 | 0.33 | 0.45 |
| 8 | 65500797 | rs9298085 | 1.695 | -1.888 | 0.25 | 0.875 | 0.017 | -0.023 | -0.023 | 0.057 | 0.6 | 0.34 | 0.54 | 0.47 | 0.63 | 0.51 |
| 8 | 65506155 | rs13276281 | 0.054 | -1.937 | 0.13 | 0.603 | 0.023 | -0.022 | -0.022 | 0 | 0.87 | 1 | 0.93 | 0.84 | 0.95 | 0.81 |
| 8 | 65507326 | rs1451869 | -0.55 | -1.944 | 0.126 | 0.599 | 0.009 | -0.03 | -0.03 | 0 | 0.9 | 1 | 0.93 | 0.84 | 0.95 | 0.81 |
| 8 | 65507922 | rs13278548 | 0.107 | -1.957 | 0.125 | 0.594 | 0.027 | -0.018 | -0.018 | 0 | 0.87 | 1 | 0.93 | 0.84 | 0.95 | 0.81 |
| 8 | 65509694 | rs7822429 | -0.445 | -1.993 | 0.109 | 0.681 | 0.011 | -0.029 | -0.029 | 0.202 | 0.89 | 1 | 0.93 | 0.84 | 0.96 | 0.81 |
| 8 | 65510610 | rs8192906 | 1.493 | -1.98 | 0.333 | 0.687 | 0.008 | -0.019 | -0.019 | 0 | 0.65 | 0.76 | 0.62 | 0.51 | 0.64 | 0.51 |
| 8 | 65519166 | rs3779870 | 1.479 | -1.975 | 0.318 | 0.69 | 0.008 | -0.02 | -0.02 | 0 | 0.65 | 0.77 | 0.62 | 0.51 | 0.64 | 0.51 |
| 8 | 65519300 | rs3779871 | 1.818 | -2.046 | 0.235 | 0.657 | 0.009 | 0.044 | 0.044 | 0 | 0.08 | 0 | 0.04 | 0.02 | 0.06 | 0.15 |
| 8 | 65523104 | rs3779874 | 1.196 | -1.977 | 0.312 | 0.726 | 0.017 | -0.028 | -0.028 | 0 | 0.71 | 0.79 | 0.69 | 0.51 | 0.78 | 0.52 |
| 8 | 65524742 | rs8192899 | 1.732 | -1.946 | 0.19 | 0.778 | -0.001 | -0.001 | -0.001 | 2.362 | 0.61 | 0.44 | 0.59 | 0.51 | 0.63 | 0.51 |
| 8 | 65525439 | rs4465006 | 1.763 | -1.943 | 0.175 | 0.694 | -0.001 | -0.001 | -0.001 | 0 | 0.61 | 0.44 | 0.59 | 0.51 | 0.63 | 0.51 |
| 8 | 65535996 | rs3824188 | -0.459 | -1.756 | 0.031 | 0.568 | 0.032 | -0.009 | -0.009 | 0 | 0.14 | 0.03 | 0.08 | 0.16 | 0.05 | 0.2 |
| 8 | 65538103 | rs4452822 | 2.022 | -1.786 | 0.248 | 0.747 | 0.043 | -0.014 | -0.014 | 0 | 0.53 | 0.28 | 0.53 | 0.44 | 0.57 | 0.35 |
| 8 | 65538483 | rs16931350 | -0.199 | -1.788 | 0.246 | 0.746 | 0.022 | -0.032 | -0.032 | 0.361 | 0.13 | 0 | 0.08 | 0.16 | 0.05 | 0.2 |
| 8 | 65543401 | rs7008736 | 0.165 | -1.887 | 0.379 | 0.69 | -0.024 | -0.04 | -0.04 | 0.255 | 0.81 | 0.86 | 0.7 | 0.62 | 0.68 | 0.72 |
| 8 | 65544125 | rs10808738 | 2.254 | -1.888 | 0.378 | 0.689 | 0.044 | -0.014 | -0.014 | 0 | 0.52 | 0.28 | 0.53 | 0.44 | 0.57 | 0.35 |
| 8 | 65547656 | rs4327886 | 1.722 | -1.899 | 0.541 | 0.683 | -0.005 | 0.025 | 0.025 | 1.966 | 0.64 | 0.48 | 0.6 | 0.46 | 0.63 | 0.51 |
| 8 | 65550357 | rs9298087 | 2.245 | -1.897 | 0.491 | 0.684 | 0.044 | -0.014 | -0.014 | 0.035 | 0.52 | 0.27 | 0.53 | 0.44 | 0.57 | 0.35 |
| 8 | 65554969 | rs7005708 | 1.789 | -1.898 | 0.586 | 0.684 | 0.001 | -0.023 | -0.023 | 0 | 0.66 | 0.69 | 0.61 | 0.46 | 0.63 | 0.51 |
| 8 | 65561378 | rs13282866 | 0.74 | -1.904 | 0.539 | 0.645 | 0.003 | 0.012 | 0.012 | 1.282 | 0.8 | 0.74 | 0.69 | 0.62 | 0.68 | 0.72 |
| 8 | 65564090 | rs4527914 | -0.275 | -1.904 | 0.535 | 0.645 | 0.032 | -0.009 | -0.009 | 0.008 | 0.14 | 0.03 | 0.08 | 0.16 | 0.05 | 0.2 |
| 8 | 65564428 | rs4350019 | -0.921 | -1.901 | 0.541 | 0.646 | 0.007 | 0.077 | 0.077 | 2.779 | 0.92 | 0.87 | 0.81 | 0.63 | 0.88 | 0.73 |
| 8 | 65569387 | rs4395923 | 2.382 | -1.902 | 0.542 | 0.646 | 0.044 | -0.014 | -0.014 | 0 | 0.52 | 0.28 | 0.53 | 0.44 | 0.57 | 0.35 |
| 8 | 65569997 | rs4397440 | -0.289 | -1.859 | 0.498 | 0.673 | 0.032 | -0.009 | -0.009 | 0.055 | 0.14 | 0.03 | 0.08 | 0.16 | 0.05 | 0.2 |
| 8 | 65570867 | rs13260351 | -0.904 | -1.849 | 0.477 | 0.677 | 0.007 | 0.077 | 0.077 | 2.779 | 0.92 | 0.87 | 0.82 | 0.63 | 0.88 | 0.73 |
| 8 | 65571564 | rs13266877 | 0.671 | -1.848 | 0.473 | 0.671 | -0.035 | 0.064 | 0.064 | 0.323 | 0.11 | 0.01 | 0.11 | 0.01 | 0.2 | 0.02 |
| 8 | 65575601 | rs4236934 | 1.935 | -1.858 | 0.489 | 0.679 | 0.002 | -0.028 | -0.028 | 0 | 0.66 | 0.7 | 0.62 | 0.46 | 0.63 | 0.51 |
| 8 | 65576786 | rs1007219 | 0.415 | -1.817 | 0.44 | 0.695 | -0.024 | -0.04 | -0.04 | 0 | 0.81 | 0.86 | 0.71 | 0.62 | 0.68 | 0.72 |
| 8 | 65580990 | rs9298088 | 2.178 | -1.882 | 0.448 | 0.633 | 0.046 | -0.013 | -0.013 | 0.366 | 0.52 | 0.28 | 0.53 | 0.44 | 0.57 | 0.35 |
| 8 | 65582637 | rs4486246 | 1.771 | -1.89 | 0.567 | 0.629 | 0.001 | -0.019 | -0.019 | 0 | 0.66 | 0.66 | 0.61 | 0.46 | 0.63 | 0.51 |
| 8 | 65585409 | rs6999999 | 2.159 | -1.843 | 0.238 | 0.654 | 0.046 | -0.013 | -0.013 | 0 | 0.52 | 0.28 | 0.53 | 0.44 | 0.57 | 0.35 |
| 8 | 65586542 | rs16931371 | 0.109 | -1.479 | -0.579 | 0.166 | -0.045 | -0.063 | -0.063 | 0 | 0.69 | 0.77 | 0.47 | 0.49 | 0.56 | 0.59 |
| 8 | 65587228 | rs1986181 | 0.356 | -1.303 | -0.12 | 0.16 | -0.009 | -0.095 | -0.095 | 0 | 0.78 | 0.87 | 0.73 | 0.45 | 0.82 | 0.51 |
| 8 | 65587797 | rs4406437 | 2.023 | -1.367 | -0.017 | 0.101 | 0.048 | -0.01 | -0.01 | 0.045 | 0.51 | 0.27 | 0.52 | 0.42 | 0.56 | 0.34 |
| 8 | 65591720 | rs17293712 | 1.37 | -1.329 | -0.022 | 0.096 | -0.031 | 0.062 | 0.062 | 0.548 | 0.11 | 0.01 | 0.11 | 0.01 | 0.2 | 0.02 |
| 8 | 65593996 | rs11998077 | 2.038 | -1.343 | -0.017 | 0.098 | 0.048 | -0.01 | -0.01 | 0 | 0.51 | 0.26 | 0.52 | 0.42 | 0.56 | 0.34 |
| 8 | 65594917 | rs7817744 | 1.338 | -1.295 | 0.325 | 0.145 | -0.003 | 0.034 | 0.034 | 1.467 | 0.64 | 0.53 | 0.58 | 0.44 | 0.62 | 0.5 |
| 8 | 65595251 | rs7834854 | 0.527 | -1.258 | 0.358 | 0.13 | -0.009 | 0.102 | 0.102 | 0.07 | 0.75 | 0.61 | 0.7 | 0.45 | 0.82 | 0.51 |
| 8 | 65595527 | rs10090584 | 0.37 | -1.216 | -0.095 | 0.152 | -0.009 | -0.096 | -0.096 | 0 | 0.78 | 0.87 | 0.73 | 0.45 | 0.82 | 0.51 |
| 8 | 65604777 | rs13261981 | 1.239 | -1.216 | -0.113 | 0.154 | -0.031 | 0.064 | 0.064 | 0.361 | 0.11 | 0.01 | 0.11 | 0.01 | 0.2 | 0.02 |
| 8 | 65605719 | rs7827287 | 2.152 | -1.067 | -0.175 | -0.062 | -0.018 | -0.018 | -0.018 | 0.05 | 0.47 | 0.26 | 0.49 | 0.43 | 0.51 | 0.32 |
| 8 | 65606916 | rs7002672 | -0.519 | -1.066 | -0.147 | -0.041 | -0.02 | -0.022 | -0.022 | 0.014 | 0.84 | 0.86 | 0.76 | 0.63 | 0.79 | 0.65 |
| 8 | 65608240 | rs13277887 | 1.244 | -0.986 | -0.277 | -0.085 | -0.004 | 0.005 | 0.005 | 0.382 | 0.11 | 0.07 | 0.11 | 0.01 | 0.2 | 0.02 |
| 8 | 65608361 | rs7830315 | 1.951 | -0.941 | -0.316 | 0.035 | -0.018 | -0.018 | -0.018 | 0.012 | 0.47 | 0.26 | 0.49 | 0.43 | 0.51 | 0.32 |
| 8 | 65614426 | rs2356986 | -1.111 | -1.084 | 0.251 | -0.166 | -0.052 | -0.069 | -0.069 | 0 | 0.35 | 0.54 | 0.27 | 0.19 | 0.27 | 0.35 |
| 8 | 65614741 | rs2884074 | -1.109 | -1.089 | 0.249 | -0.167 | -0.052 | -0.069 | -0.069 | 0 | 0.35 | 0.54 | 0.27 | 0.19 | 0.27 | 0.35 |
| 8 | 65618273 | rs2884073 | -0.917 | -1.098 | 0.17 | -0.267 | -0.034 | -0.168 | -0.168 | 0.051 | 0.29 | 0.54 | 0.21 | 0.03 | 0.28 | 0.18 |
| 8 | 65619966 | rs716283 | -0.917 | -1.098 | 0.17 | -0.267 | -0.034 | -0.168 | -0.168 | 0.063 | 0.29 | 0.54 | 0.21 | 0.03 | 0.28 | 0.18 |
| 8 | 65628180 | rs9918790 | -0.946 | -1.109 | 0.182 | -0.274 | -0.034 | -0.168 | -0.168 | 0 | 0.3 | 0.54 | 0.21 | 0.03 | 0.28 | 0.18 |
| 8 | 65630771 | rs4737677 | 1.262 | -1.207 | 0.429 | -0.337 | -0.029 | -0.049 | -0.049 | 0.056 | 0.12 | 0.2 | 0.06 | 0.02 | 0.05 | 0.16 |
| 8 | 65634357 | rs4385495 | -1.247 | -1.228 | 0.539 | -0.35 | 0.004 | -0.039 | -0.039 | 0.347 | 0.31 | 0.7 | 0.22 | 0.03 | 0.28 | 0.18 |
| 8 | 65636139 | rs9650211 | 1.407 | -1.244 | 0.35 | -0.36 | -0.035 | 0.058 | 0.058 | 0.108 | 0.1 | 0.01 | 0.1 | 0.01 | 0.19 | 0.02 |
| 8 | 65640260 | rs10808739 | -0.535 | -1.234 | 0.516 | -0.35 | -0.032 | -0.113 | -0.113 | 0.371 | 0.27 | 0.39 | 0.19 | 0.03 | 0.24 | 0.18 |
| 8 | 65641261 | rs4269577 | -1.219 | -1.238 | 0.479 | -0.329 | 0.019 | -0.035 | -0.035 | 0.016 | 0.93 | 0.99 | 0.94 | 0.84 | 0.98 | 0.83 |
| 8 | 65650493 | rs10098783 | -1.219 | -1.236 | 0.46 | -0.327 | 0.016 | -0.039 | -0.039 | 0 | 0.93 | 1 | 0.94 | 0.84 | 0.98 | 0.83 |
| 8 | 65655434 | rs11984829 | -1.152 | -1.183 | 0.522 | -0.373 | -0.036 | -0.173 | -0.173 | 2.535 | 0.3 | 0.54 | 0.21 | 0.03 | 0.28 | 0.18 |
| 8 | 65659136 | rs7009078 | -1.152 | -1.181 | 0.495 | -0.372 | -0.036 | -0.173 | -0.173 | 0 | 0.3 | 0.54 | 0.21 | 0.03 | 0.28 | 0.18 |
| 8 | 65663801 | rs13272719 | -1.055 | -1.277 | 0.49 | -0.355 | -0.054 | -0.161 | -0.161 | 0.008 | 0.29 | 0.52 | 0.2 | 0.03 | 0.25 | 0.18 |
| 8 | 65668267 | rs11996540 | -1.055 | -1.277 | 0.49 | -0.355 | -0.054 | -0.161 | -0.161 | 0 | 0.29 | 0.52 | 0.2 | 0.03 | 0.25 | 0.18 |
| 8 | 65668770 | rs9650212 | 1.377 | -1.298 | 0.455 | -0.37 | -0.035 | 0.058 | 0.058 | 0.108 | 0.1 | 0.01 | 0.1 | 0.01 | 0.19 | 0.02 |

| | | | | | | | | | | | | | | | | |
|---|---|---|---|---|---|---|---|---|---|---|---|---|---|---|---|---|
| 8 | 65670364 | rs10957322 | -0.711 | -1.301 | 0.526 | -0.371 | -0.029 | -0.104 | -0.104 | 0 | 0.27 | 0.38 | 0.18 | 0.03 | 0.24 | 0.18 |
| 8 | 65675055 | rs4737680 | -1.109 | -1.289 | 0.345 | -0.365 | -0.055 | -0.163 | -0.163 | 1.722 | 0.29 | 0.53 | 0.2 | 0.03 | 0.25 | 0.18 |
| 8 | 65676069 | rs7826214 | -1.125 | -1.298 | 0.341 | -0.369 | -0.055 | -0.163 | -0.163 | 0 | 0.29 | 0.52 | 0.2 | 0.03 | 0.25 | 0.18 |
| 8 | 65676153 | rs6472152 | -1.201 | -1.353 | 0.304 | -0.329 | 0.02 | -0.031 | -0.031 | 0 | 0.93 | 0.99 | 0.93 | 0.84 | 0.98 | 0.83 |
| 8 | 65681616 | rs7812644 | -1.09 | -1.355 | 0.305 | -0.35 | -0.055 | -0.163 | -0.163 | 0 | 0.29 | 0.52 | 0.2 | 0.03 | 0.25 | 0.18 |
| 8 | 65685957 | rs4737682 | -1.201 | -1.35 | 0.314 | -0.318 | 0.019 | -0.034 | -0.034 | 0 | 0.93 | 0.99 | 0.93 | 0.84 | 0.98 | 0.83 |
| 8 | 65686539 | rs13255280 | 1.405 | -1.353 | 0.311 | -0.325 | -0.035 | 0.058 | 0.058 | 0.108 | 0.1 | 0.01 | 0.1 | 0.01 | 0.19 | 0.02 |
| 8 | 65689094 | rs4236936 | -1.202 | -1.353 | 0.312 | -0.324 | 0.016 | -0.039 | -0.039 | 0.74 | 0.93 | 1 | 0.93 | 0.84 | 0.98 | 0.84 |
| 8 | 65695385 | rs12674544 | 0.082 | -1.357 | 0.314 | -0.328 | -0.088 | -0.119 | -0.119 | 0 | 0.18 | 0.43 | 0.09 | 0.02 | 0.05 | 0.16 |
| 8 | 65696040 | rs11992601 | -1.102 | -1.276 | 0.313 | -0.326 | -0.052 | -0.163 | -0.163 | 0 | 0.29 | 0.52 | 0.2 | 0.03 | 0.25 | 0.18 |
| 8 | 65696352 | rs13259721 | 1.311 | -1.283 | -0.388 | -0.314 | -0.034 | 0.059 | 0.059 | 0.199 | 0.1 | 0.02 | 0.1 | 0.01 | 0.19 | 0.02 |
| 8 | 65704658 | rs13258738 | 1.399 | -1.279 | -0.385 | -0.312 | -0.035 | 0.058 | 0.058 | 0.108 | 0.1 | 0.01 | 0.1 | 0.01 | 0.19 | 0.02 |
| 8 | 65704945 | rs7821085 | 1.385 | -1.279 | -0.26 | -0.31 | 0.031 | 0.035 | 0.035 | 0 | 0.63 | 0.3 | 0.72 | 0.81 | 0.73 | 0.66 |
| 8 | 65709467 | rs6987861 | 1.394 | -1.27 | -0.279 | -0.31 | 0.033 | 0.038 | 0.038 | 0 | 0.63 | 0.3 | 0.72 | 0.81 | 0.72 | 0.66 |
| 8 | 65711945 | rs7829301 | -0.883 | -0.592 | -0.595 | 0.183 | 0.019 | -0.035 | -0.035 | 0 | 0.93 | 0.99 | 0.93 | 0.84 | 0.98 | 0.83 |
| 8 | 65720734 | rs6980478 | 0.748 | -0.266 | -0.486 | 0.139 | -0.007 | 0.002 | 0.002 | 0.194 | 0.11 | 0.1 | 0.11 | 0.01 | 0.2 | 0.02 |
| 8 | 65730207 | rs6472155 | -0.892 | -0.199 | -0.726 | 0.353 | 0 | 0.012 | 0.012 | 0.693 | 0.49 | 0.53 | 0.6 | 0.69 | 0.51 | 0.59 |
| 8 | 65762549 | rs7820880 | -0.104 | 0.397 | -0.833 | 0.347 | 0.001 | -0.008 | -0.008 | 0.874 | 0.6 | 0.61 | 0.54 | 0.62 | 0.56 | 0.68 |
| 8 | 65768237 | rs9298089 | 0.732 | 0.327 | -0.645 | 0.066 | 0.016 | 0.04 | 0.04 | 0 | 0.39 | 0.26 | 0.29 | 0.19 | 0.29 | 0.42 |
| 8 | 65770668 | rs4077788 | 0.347 | 0.363 | -0.259 | 0.453 | 0.039 | 0.069 | 0.069 | 0 | 0.21 | 0.35 | 0.25 | 0.43 | 0.27 | 0.27 |
| 8 | 65780215 | rs4401898 | -0.516 | 0.188 | -0.545 | -0.02 | 0 | -0.017 | -0.017 | 0 | 0.4 | 0.63 | 0.62 | 0.81 | 0.4 | 0.43 |
| 8 | 65780516 | rs12544578 | -0.516 | 0.188 | -0.545 | -0.02 | 0 | -0.017 | -0.017 | 0 | 0.4 | 0.63 | 0.62 | 0.81 | 0.4 | 0.43 |
| 8 | 65783019 | rs6985116 | -0.778 | 0.207 | -0.678 | 0.029 | -0.002 | 0.005 | 0.005 | 0.039 | 0.54 | 0.61 | 0.67 | 0.78 | 0.51 | 0.52 |
| 8 | 65786716 | rs4273884 | 0.533 | 0.018 | -0.543 | 0.107 | 0.001 | 0 | 0 | 0.062 | 0.33 | 0.33 | 0.32 | 0.44 | 0.4 | 0.29 |
| 8 | 65788461 | rs7819350 | -0.121 | -0.198 | -0.35 | 0.411 | -0.024 | 0.011 | 0.011 | 0 | 0.33 | 0.4 | 0.35 | 0.36 | 0.24 | 0.26 |
| 8 | 65796450 | rs7820286 | -0.254 | 0.165 | -0.275 | 0.307 | 0.006 | 0.007 | 0.007 | 0 | 0.37 | 0.49 | 0.48 | 0.44 | 0.39 | 0.33 |
| 8 | 65796677 | rs7840142 | 0.574 | 0.232 | -0.277 | 0.216 | -0.019 | -0.006 | -0.006 | 0 | 0.11 | 0.08 | 0.31 | 0.44 | 0.2 | 0.23 |
| 8 | 65799727 | rs4573321 | -0.943 | 0.126 | -0.499 | -0.264 | 0.001 | 0.081 | 0.081 | 1.177 | 0.65 | 0.79 | 0.74 | 0.91 | 0.65 | 0.61 |
| 8 | 65800457 | rs7833587 | 0.429 | 0.353 | -0.354 | 0.038 | -0.004 | 0.019 | 0.019 | 0.58 | 0.11 | 0.12 | 0.32 | 0.44 | 0.2 | 0.23 |
| 8 | 65807063 | rs6999082 | 0.392 | 0.049 | -0.163 | 0.037 | -0.002 | 0.027 | 0.027 | 0.132 | 0.27 | 0.32 | 0.26 | 0.48 | 0.26 | 0.28 |
| 8 | 65810061 | rs10102238 | 1.233 | -0.053 | -0.209 | -0.035 | -0.013 | 0.032 | 0.032 | 0 | 0.19 | 0.09 | 0.11 | 0.11 | 0.21 | 0.14 |
| 8 | 65818520 | rs4260923 | -0.722 | -0.884 | -0.815 | -0.207 | -0.094 | -0.095 | -0.095 | 0.043 | 0.63 | 0.88 | 0.52 | 0.46 | 0.46 | 0.6 |
| 8 | 65818743 | rs4263805 | 0.961 | -1.158 | -0.515 | -0.266 | -0.011 | -0.001 | -0.001 | 0 | 0.12 | 0.15 | 0.09 | 0.09 | 0.04 | 0.23 |
| 8 | 65827319 | rs3935037 | 1.485 | -1.258 | -0.363 | -0.382 | -0.016 | -0.085 | -0.085 | 1.628 | 0.21 | 0.26 | 0.14 | 0 | 0.17 | 0.09 |
| 8 | 65828731 | rs7823966 | -0.832 | -1.204 | -0.371 | -0.316 | -0.087 | -0.094 | -0.094 | 0.06 | 0.63 | 0.87 | 0.56 | 0.47 | 0.46 | 0.58 |
| 8 | 65854402 | rs4466436 | 1.185 | -1.177 | -0.345 | -0.378 | -0.083 | -0.121 | -0.121 | 0.568 | 0.23 | 0.03 | 0.3 | 0.37 | 0.25 | 0.25 |
| 8 | 65856143 | rs4078079 | -0.199 | -1.108 | -0.302 | -0.332 | 0.027 | 0.058 | 0.058 | 1.109 | 0.51 | 0.73 | 0.47 | 0.37 | 0.42 | 0.35 |
| 8 | 65859623 | rs4529498 | 1.171 | -1.089 | -0.383 | -0.501 | -0.017 | -0.086 | -0.086 | 0 | 0.22 | 0.26 | 0.14 | 0 | 0.17 | 0.09 |
| 8 | 65897418 | rs7840002 | -0.797 | -1.196 | -0.762 | -0.015 | -0.065 | 0.059 | 0.059 | 0.022 | 0.67 | 0.88 | 0.64 | 0.76 | 0.53 | 0.7 |
| 8 | 65898764 | rs2002700 | -0.647 | -1.515 | -0.361 | 0.075 | -0.022 | -0.032 | -0.032 | 0 | 0.64 | 0.75 | 0.53 | 0.47 | 0.45 | 0.57 |
| 8 | 65899664 | rs16931576 | 0.334 | -1.532 | -0.37 | -0.058 | 0.023 | 0.026 | 0.026 | 0.349 | 0.46 | 0.31 | 0.43 | 0.39 | 0.4 | 0.36 |
| 8 | 65907742 | rs7011697 | 0.596 | -1.084 | -0.028 | 0.429 | -0.011 | -0.009 | -0.009 | 0.487 | 0.11 | 0.13 | 0.07 | 0.04 | 0.04 | 0.21 |
| 8 | 65918785 | rs7004914 | 0.361 | -1.086 | -0.101 | 0.656 | -0.009 | 0.088 | 0.088 | 0 | 0.09 | 0.14 | 0.18 | 0.3 | 0.07 | 0.17 |
| 8 | 65922325 | rs6983551 | 1.575 | -0.933 | -0.468 | 0.241 | -0.018 | 0.028 | 0.028 | 0 | 0.18 | 0.09 | 0.17 | 0.1 | 0.22 | 0.1 |
| 8 | 65932063 | rs4478612 | 0.694 | -0.972 | -0.508 | 0.189 | 0.039 | 0.075 | 0.075 | 0.215 | 0.28 | 0.16 | 0.16 | 0.08 | 0.18 | 0.12 |
| 8 | 65940453 | rs10104122 | -0.966 | -0.826 | -0.726 | 0.44 | -0.123 | -0.117 | -0.117 | 0 | 0.7 | 0.91 | 0.61 | 0.51 | 0.51 | 0.61 |
| 8 | 65941449 | rs10957333 | 1.674 | -0.873 | -0.714 | 0.372 | -0.032 | 0.043 | 0.043 | 0 | 0.18 | 0.08 | 0.18 | 0.1 | 0.22 | 0.07 |
| 8 | 65942775 | rs4556118 | -0.851 | -1.709 | 0.445 | 1.005 | -0.01 | -0.002 | -0.002 | 0 | 0.47 | 0.48 | 0.41 | 0.39 | 0.29 | 0.51 |
| 8 | 65960510 | rs10957334 | -0.485 | -1.56 | -0.069 | 1.232 | 0.001 | -0.002 | -0.002 | 0 | 0.34 | 0.33 | 0.35 | 0.48 | 0.28 | 0.34 |
| 8 | 65962086 | rs4333639 | -0.402 | -1.047 | -0.57 | 1.353 | 0.027 | 0.143 | 0.143 | 0 | 0.07 | 0.15 | 0.19 | 0.33 | 0.12 | 0.23 |
| 8 | 65963602 | rs4527915 | -1.133 | -1.054 | -1.121 | 0.377 | -0.014 | 0.17 | 0.17 | 0 | 0.58 | 0.91 | 0.62 | 0.81 | 0.37 | 0.8 |
| 8 | 65989176 | rs2956320 | -1.256 | -0.598 | -0.824 | 0.623 | 0.023 | 0.173 | 0.173 | 0 | 0.53 | 0.89 | 0.6 | 0.82 | 0.4 | 0.68 |
| 8 | 65989486 | rs2956319 | 0.238 | -0.667 | -0.244 | 0.498 | 0.015 | 0.018 | 0.018 | 0 | 0.21 | 0.2 | 0.14 | 0.16 | 0.17 | 0.14 |
| 8 | 65991911 | rs2956313 | -0.188 | -0.226 | 0.324 | 0.76 | -0.042 | 0.004 | 0.004 | 0 | 0.37 | 0.54 | 0.34 | 0.42 | 0.22 | 0.28 |
| 8 | 65995595 | rs10504382 | -0.617 | -0.091 | 0.115 | 0.85 | 0.011 | 0.113 | 0.113 | 0 | 0.08 | 0.13 | 0.2 | 0.43 | 0.14 | 0.24 |
| 8 | 65997304 | rs2956310 | -0.644 | -0.14 | 0.233 | 0.558 | 0.027 | 0.08 | 0.08 | 0 | 0.46 | 0.73 | 0.55 | 0.84 | 0.36 | 0.52 |
| 8 | 66008068 | rs16931735 | -1.177 | 0.641 | 0.78 | 1.864 | 0.03 | 0.184 | 0.184 | 0 | 0.08 | 0.17 | 0.15 | 0.42 | 0.12 | 0.24 |
| 8 | 66020732 | rs7842129 | -1.111 | 0.204 | 0.793 | 0.568 | -0.007 | -0.045 | -0.045 | 2.505 | 0.1 | 0.29 | 0.08 | 0.03 | 0.03 | 0.19 |
| 8 | 66026395 | rs4737698 | -1.144 | 0.972 | -0.458 | 1.363 | -0.003 | 0.003 | 0.003 | 0 | 0.11 | 0.13 | 0.21 | 0.28 | 0.15 | 0.25 |
| 8 | 66060125 | rs4737703 | 0.348 | 0.306 | -0.641 | 0.202 | 0.008 | -0.009 | -0.009 | 0.105 | 0.78 | 0.84 | 0.88 | 1 | 0.78 | 0.9 |
| 8 | 66066748 | rs9298099 | 0.37 | 0.287 | -0.598 | 0.193 | 0.007 | -0.015 | -0.015 | 0 | 0.78 | 0.84 | 0.88 | 1 | 0.78 | 0.9 |
| 8 | 66073614 | rs10095326 | 0.958 | -0.085 | -1.816 | 0.356 | -0.132 | -0.066 | -0.066 | 0.467 | 0.41 | 0.04 | 0.41 | 0.4 | 0.49 | 0.39 |
| 8 | 66074310 | rs6998613 | 0.113 | -0.433 | -0.545 | 0.826 | -0.011 | 0.024 | 0.024 | 0.108 | 0.32 | 0.46 | 0.43 | 0.39 | 0.26 | 0.42 |
| 8 | 66078557 | rs7842485 | 0.086 | -0.638 | -0.643 | 0.726 | 0.017 | 0.046 | 0.046 | 0 | 0.19 | 0.08 | 0.08 | 0 | 0.17 | 0.09 |
| 8 | 66079805 | rs4582599 | 0.761 | -0.839 | -0.077 | 0.709 | -0.003 | -0.005 | -0.005 | 0 | 0.48 | 0.46 | 0.49 | 0.61 | 0.53 | 0.49 |
| 8 | 66089371 | rs4737709 | 1.161 | -1.041 | -0.531 | 0.834 | -0.03 | -0.121 | -0.121 | 0.651 | 0.45 | 0.04 | 0.45 | 0.44 | 0.61 | 0.42 |
| 8 | 66101488 | rs7833932 | 0.462 | -1.055 | -0.315 | 0.678 | 0.012 | 0.034 | 0.034 | 0 | 0.11 | 0.02 | 0.04 | 0 | 0.06 | 0.03 |
| 8 | 66102313 | rs13253048 | -0.959 | -1.327 | 0.353 | 0.9 | 0.004 | 0.003 | 0.003 | 0 | 0.46 | 0.52 | 0.47 | 0.24 | 0.37 | 0.48 |
| 8 | 66102770 | rs12544654 | -0.26 | -1.266 | -0.264 | 0.937 | 0.011 | 0.036 | 0.036 | 0 | 0.33 | 0.2 | 0.41 | 0.24 | 0.31 | 0.45 |

| | | | | | | | | | | | | | | | | |
|---|---|---|---|---|---|---|---|---|---|---|---|---|---|---|---|---|
| 8 | 66126854 | rs4404953 | -1.489 | -1.159 | -0.559 | 0.69 | -0.017 | -0.077 | -0.077 | 0.305 | 0.54 | 0.96 | 0.53 | 0.57 | 0.4 | 0.55 |
| 8 | 66127657 | rs4294223 | -1.333 | -0.842 | 0.445 | 0.35 | -0.052 | -0.014 | -0.014 | 0.009 | 0.19 | 0.63 | 0.11 | 0.17 | 0.09 | 0.08 |
| 8 | 66129133 | rs6988815 | -1.37 | -0.366 | 0.449 | 0.53 | -0.014 | 0.05 | 0.05 | 1.328 | 0.14 | 0.49 | 0.13 | 0.17 | 0.1 | 0.11 |
| 8 | 66129816 | rs4310240 | -1.02 | -0.557 | 0.077 | 0.38 | -0.005 | -0.006 | -0.006 | 3.536 | 0.28 | 0.3 | 0.16 | 0.22 | 0.22 | 0.22 |
| 8 | 66129990 | rs4737206 | 0.632 | -0.332 | 0.108 | 0.505 | -0.012 | 0.018 | 0.018 | 0 | 0.66 | 0.57 | 0.54 | 0.53 | 0.73 | 0.59 |
| 8 | 66130782 | rs10098238 | -1.377 | 0.376 | 0.254 | -0.04 | -0.003 | 0.013 | 0.013 | 0 | 0.14 | 0.1 | 0.16 | 0.05 | 0.18 | 0.14 |
| 8 | 66134803 | rs10099098 | 1.867 | 0.527 | 0.882 | -0.21 | 0.001 | -0.11 | -0.11 | 0.074 | 0.75 | 0.51 | 0.76 | 0.95 | 0.75 | 0.79 |
| 8 | 66148098 | rs4236939 | 0.993 | 0.55 | 0.157 | -0.068 | 0.001 | -0.029 | -0.029 | 0.346 | 0.88 | 0.89 | 0.92 | 1 | 0.87 | 0.92 |
| 8 | 66148463 | rs4236940 | 0.848 | 0.224 | -0.263 | -0.05 | -0.01 | 0.065 | 0.065 | 0.005 | 0.89 | 0.99 | 0.95 | 1 | 0.92 | 0.93 |
| 8 | 66151929 | rs7017178 | 0.868 | 0.224 | -0.263 | -0.05 | -0.017 | 0.065 | 0.065 | 0.006 | 0.89 | 0.99 | 0.95 | 1 | 0.91 | 0.93 |
| 8 | 66165333 | rs6472189 | 1.26 | 0.129 | 0.066 | 0.753 | 0.001 | -0.001 | -0.001 | 0 | 0.77 | 0.78 | 0.75 | 0.47 | 0.79 | 0.57 |
| 8 | 66180525 | rs7815138 | 1.525 | 0.118 | 0.173 | 0.692 | -0.002 | 0.006 | 0.006 | 0.062 | 0.73 | 0.7 | 0.72 | 0.47 | 0.77 | 0.56 |
| 8 | 66182076 | rs10104374 | -1.153 | 0.126 | 0.165 | 0.714 | 0.007 | -0.02 | -0.02 | 0.001 | 0.19 | 0.15 | 0.19 | 0.5 | 0.16 | 0.39 |
| 8 | 66183866 | rs11998282 | 1.506 | 0.131 | 0.161 | 0.69 | -0.001 | -0.001 | -0.001 | 0.426 | 0.73 | 0.71 | 0.71 | 0.27 | 0.78 | 0.56 |
| 8 | 66188263 | rs1877829 | 1.287 | -0.146 | 0.301 | 0.028 | -0.009 | -0.004 | -0.004 | 0 | 0.74 | 0.69 | 0.54 | 0.24 | 0.82 | 0.54 |
| 8 | 66191608 | rs9643572 | -1.003 | -0.072 | 0.121 | 0.071 | 0.025 | -0.03 | -0.03 | 0 | 0.21 | 0.04 | 0.37 | 0.74 | 0.14 | 0.42 |
| 8 | 66206231 | rs7817180 | -0.854 | 0.244 | 0.113 | -0.05 | 0.002 | -0.011 | -0.011 | 0 | 0.2 | 0.09 | 0.39 | 0.76 | 0.17 | 0.53 |
| 8 | 66222068 | rs925677 | -1.013 | 0.348 | 0.582 | -0.036 | -0.001 | 0 | 0 | 0.113 | 0.2 | 0.19 | 0.4 | 0.76 | 0.17 | 0.53 |
| 8 | 66235381 | rs7830803 | -0.354 | 0.153 | 0.843 | -0.407 | 0.01 | -0.017 | -0.017 | 0.52 | 0.12 | 0.4 | 0.13 | 0.05 | 0.07 | 0.13 |
| 8 | 66261909 | rs10216541 | -0.769 | 0.117 | -0.03 | -0.187 | 0.003 | -0.032 | -0.032 | 0.434 | 0.21 | 0.07 | 0.4 | 0.72 | 0.17 | 0.55 |
| 8 | 66282047 | rs10216812 | -0.899 | 0.121 | 0.109 | -0.263 | 0.009 | -0.011 | -0.011 | 0 | 0.22 | 0.15 | 0.4 | 0.73 | 0.17 | 0.55 |
| 8 | 66285716 | rs2357274 | -0.928 | 0.111 | 0.119 | -0.269 | 0.009 | -0.01 | -0.01 | 0 | 0.23 | 0.16 | 0.4 | 0.73 | 0.17 | 0.55 |
| 8 | 66287730 | rs6472195 | 1.053 | 0.119 | 0.41 | -0.286 | -0.001 | 0 | 0 | 0 | 0.73 | 0.71 | 0.54 | 0.22 | 0.78 | 0.42 |
| 8 | 66289587 | rs1554057 | 1.53 | 0.095 | 0.848 | -0.408 | -0.007 | -0.019 | -0.019 | 0 | 0.86 | 0.41 | 0.82 | 0.89 | 0.92 | 0.88 |
| 8 | 66299317 | rs12678756 | -1.254 | -0.031 | 0.322 | -1.307 | 0 | -0.076 | -0.076 | 0.001 | 0.49 | 0.47 | 0.64 | 0.94 | 0.42 | 0.74 |
| 8 | 66299820 | rs12681255 | -0.695 | -0.24 | -0.069 | -1.386 | 0.001 | 0.07 | 0.07 | 0 | 0.37 | 0.17 | 0.4 | 0.74 | 0.31 | 0.59 |
| 8 | 66302199 | rs1473603 | -1.225 | -0.273 | 0.331 | -1.565 | -0.009 | -0.025 | -0.025 | 0 | 0.26 | 0.41 | 0.29 | 0.19 | 0.21 | 0.17 |
| 8 | 66304887 | rs10096019 | -0.109 | -0.267 | 0.287 | -1.568 | 0.022 | 0.017 | 0.017 | 0 | 0.28 | 0.14 | 0.37 | 0.74 | 0.23 | 0.58 |
| 8 | 66307473 | rs4527917 | -1.96 | -0.222 | 0.322 | -1.594 | -0.001 | -0.077 | -0.077 | 0.006 | 0.54 | 0.55 | 0.66 | 0.93 | 0.44 | 0.76 |
| 8 | 66308987 | rs1355957 | 1.725 | -0.425 | 0.325 | -1.583 | -0.001 | -0.117 | -0.117 | 0.575 | 0.47 | 0.49 | 0.33 | 0 | 0.53 | 0.23 |
| 8 | 66317472 | rs16932054 | -0.652 | -0.372 | 0.866 | -1.033 | -0.006 | -0.009 | -0.009 | 0 | 0.12 | 0.15 | 0.17 | 0.05 | 0.06 | 0.08 |
| 8 | 66321181 | rs1514524 | 0.783 | -0.39 | 0.939 | -1.04 | 0.012 | -0.021 | -0.021 | 0.439 | 0.85 | 0.67 | 0.77 | 0.88 | 0.88 | 0.88 |
| 8 | 66324293 | rs4737217 | 0.883 | -0.402 | 0.927 | -1.046 | 0.007 | -0.024 | -0.024 | 6.355 | 0.84 | 0.66 | 0.77 | 0.88 | 0.88 | 0.88 |
| 8 | 66327344 | rs13282622 | 1.373 | -0.547 | 1.137 | -1.142 | -0.005 | -0.035 | -0.035 | 0.013 | 0.57 | 0.53 | 0.41 | 0.15 | 0.65 | 0.3 |
| 8 | 66333628 | rs4737732 | -0.174 | -0.47 | 0.817 | -1.123 | 0.015 | 0.005 | 0.005 | 0 | 0.27 | 0.14 | 0.35 | 0.73 | 0.23 | 0.56 |
| 8 | 66337113 | rs7009259 | 1.315 | -0.499 | 0.972 | -1.27 | -0.004 | -0.026 | -0.026 | 0 | 0.56 | 0.53 | 0.42 | 0.16 | 0.65 | 0.32 |
| 8 | 66347828 | rs13261786 | 1.335 | -0.503 | 0.953 | -1.265 | 0.013 | 0.028 | 0.028 | 0 | 0.54 | 0.39 | 0.41 | 0.16 | 0.66 | 0.32 |
| 8 | 66355062 | rs10105301 | 0.987 | -0.469 | 0.562 | -1.158 | 0.008 | 0.041 | 0.041 | 0 | 0.61 | 0.7 | 0.44 | 0.16 | 0.66 | 0.32 |
| 8 | 66360589 | rs10087929 | -0.505 | -0.802 | 0.211 | -0.999 | -0.033 | -0.029 | -0.029 | 0 | 0.23 | 0.02 | 0.38 | 0.79 | 0.26 | 0.52 |
| 8 | 66365374 | rs13259837 | 2.077 | -0.256 | 0.632 | -1.319 | 0.003 | 0.104 | 0.104 | 0.017 | 0.48 | 0.27 | 0.35 | 0.14 | 0.51 | 0.26 |
| 8 | 66371436 | rs4131754 | 1.426 | -0.278 | 1.091 | -0.139 | -0.002 | 0.016 | 0.016 | 0 | 0.68 | 0.64 | 0.66 | 0.5 | 0.68 | 0.5 |
| 8 | 66377675 | rs7460061 | 0.981 | -0.404 | 0.835 | -0.056 | -0.038 | 0.056 | 0.056 | 0 | 0.75 | 0.65 | 0.75 | 0.51 | 0.84 | 0.58 |
| 8 | 66377699 | rs16932129 | 0.284 | -0.305 | 0.564 | -0.015 | 0.007 | -0.043 | -0.043 | 0 | 0.17 | 0.09 | 0.27 | 0.37 | 0.16 | 0.22 |
| 8 | 66392336 | rs11782361 | 1.261 | -0.296 | 0.753 | -0.209 | 0.012 | 0.013 | 0.013 | 0 | 0.59 | 0.62 | 0.49 | 0.15 | 0.68 | 0.36 |
| 8 | 66396331 | rs13271536 | -0.391 | -0.068 | 0.802 | 0.023 | 0.083 | -0.038 | -0.038 | 0 | 0.08 | 0.24 | 0.12 | 0.02 | 0.16 | 0.07 |
| 8 | 66408755 | rs10089578 | 0.718 | -0.165 | 0.521 | 0.398 | 0 | 0.026 | 0.026 | 0 | 0.91 | 0.93 | 0.86 | 0.56 | 0.88 | 0.65 |
| 8 | 66410169 | rs13267781 | 0.647 | -0.163 | 0.454 | 0.4 | -0.009 | -0.022 | -0.022 | 0 | 0.91 | 0.98 | 0.86 | 0.56 | 0.88 | 0.65 |
| 8 | 66410546 | rs4737737 | -0.738 | -0.367 | 0.457 | 0.198 | -0.01 | 0.052 | 0.052 | 1.041 | 0.2 | 0.09 | 0.18 | 0.09 | 0.2 | 0.15 |
| 8 | 66436618 | rs12541486 | -0.174 | -0.937 | 0.947 | -0.033 | -0.057 | 0.118 | 0.118 | 0.615 | 0.21 | 0.38 | 0.35 | 0.49 | 0.14 | 0.27 |
| 8 | 66437645 | rs7831778 | 2.299 | -1.435 | 0.536 | -0.578 | 0.007 | 0.102 | 0.102 | 0 | 0.43 | 0.22 | 0.3 | 0 | 0.54 | 0.18 |
| 8 | 66438548 | rs16932169 | 0.801 | -1.511 | 0.252 | -0.29 | 0.016 | -0.132 | -0.132 | 0 | 0.16 | 0.02 | 0.24 | 0.4 | 0.13 | 0.25 |
| 8 | 66438842 | rs5001575 | -0.266 | -0.768 | 0.161 | -0.313 | -0.035 | -0.076 | -0.076 | 0 | 0.89 | 0.99 | 0.85 | 0.59 | 0.88 | 0.62 |
| 8 | 66443377 | rs7011890 | 2.335 | -0.732 | 0.287 | -0.365 | -0.021 | 0.074 | 0.074 | 0 | 0.43 | 0.26 | 0.29 | 0 | 0.51 | 0.18 |
| 8 | 66449457 | rs6981589 | 1.335 | -0.649 | 0.006 | -0.227 | 0.001 | 0.002 | 0.002 | 0 | 0.64 | 0.63 | 0.61 | 0.52 | 0.64 | 0.44 |
| 8 | 66456186 | rs4271000 | 1.115 | -1.094 | 0.128 | -0.087 | -0.008 | 0.007 | 0.007 | 0 | 0.7 | 0.68 | 0.73 | 0.56 | 0.8 | 0.5 |
| 8 | 66456252 | rs4562361 | 2.118 | -1.238 | -0.253 | -0.2 | 0.103 | 0.187 | 0.187 | 0.001 | 0.49 | 0.19 | 0.39 | 0.02 | 0.67 | 0.24 |
| 8 | 66456541 | rs4288416 | 0.982 | -0.963 | -0.761 | 0.031 | 0.025 | -0.041 | -0.041 | 0.78 | 0.72 | 0.84 | 0.75 | 0.56 | 0.81 | 0.53 |
| 8 | 66463478 | rs6981992 | 0.812 | -0.476 | -1.055 | -0.354 | -0.002 | 0.126 | 0.126 | 1.705 | 0.7 | 0.93 | 0.54 | 0.1 | 0.69 | 0.31 |
| 8 | 66470243 | rs16932223 | 0.853 | -0.372 | -0.964 | 0.308 | -0.005 | -0.088 | -0.088 | 0 | 0.12 | 0.02 | 0.18 | 0.33 | 0.13 | 0.22 |
| 8 | 66470962 | rs6991838 | 1.141 | -0.409 | -1.442 | -0.29 | -0.001 | 0.331 | 0.331 | 0 | 0.52 | 0.91 | 0.42 | 0.06 | 0.54 | 0.25 |
| 8 | 66474197 | rs6472207 | 0.992 | -0.567 | -1.031 | -0.061 | 0.033 | -0.03 | -0.03 | 0.19 | 0.65 | 0.82 | 0.61 | 0.35 | 0.74 | 0.47 |
| 8 | 66480563 | rs4294622 | -0.798 | -0.502 | -1.143 | 0.015 | 0.008 | -0.085 | -0.085 | 0.045 | 0.32 | 0.11 | 0.35 | 0.38 | 0.32 | 0.28 |
| 8 | 66499226 | rs4320587 | -1.212 | -0.542 | -0.641 | -0.034 | 0.005 | -0.041 | -0.041 | 0.681 | 0.38 | 0.26 | 0.42 | 0.44 | 0.39 | 0.32 |
| 8 | 66515865 | rs1053087 | -1.107 | -0.541 | -0.593 | -0.033 | -0.002 | -0.062 | -0.062 | 0.327 | 0.37 | 0.2 | 0.41 | 0.44 | 0.39 | 0.32 |
| 8 | 66519001 | rs7823840 | -1.107 | -0.541 | -0.593 | -0.033 | -0.002 | -0.062 | -0.062 | 0 | 0.37 | 0.2 | 0.41 | 0.44 | 0.39 | 0.32 |
| 8 | 66556187 | rs10504389 | 0.19 | -0.546 | -0.643 | -0.073 | 0.008 | 0.032 | 0.032 | 0 | 0.16 | 0.07 | 0.13 | 0.04 | 0.15 | 0.05 |
| 8 | 66556292 | rs7842162 | 0.987 | -0.567 | -0.567 | -0.055 | 0 | -0.044 | -0.044 | 0 | 0.63 | 0.79 | 0.59 | 0.6 | 0.62 | 0.68 |
| 8 | 66565403 | rs10102473 | -1.433 | -0.765 | -0.11 | -0.135 | -0.006 | -0.1 | -0.1 | 0 | 0.87 | 0.99 | 0.81 | 0.67 | 0.86 | 0.79 |
| 8 | 66590805 | rs7002171 | -0.498 | -0.854 | 0.643 | -0.186 | 0.007 | 0.042 | 0.042 | 0.032 | 0.19 | 0.15 | 0.16 | 0.07 | 0.17 | 0.08 |

| | | | | | | | | | | | | | | | | |
|---|---|---|---|---|---|---|---|---|---|---|---|---|---|---|---|---|
| 8 | 66596320 | rs6991669 | -0.93 | -0.836 | 0.635 | -0.194 | -0.012 | 0.066 | 0.066 | 0 | 0.23 | 0.15 | 0.2 | 0.07 | 0.24 | 0.12 |
| 8 | 66631972 | rs953593 | -1.504 | -0.963 | 0.544 | -0.265 | 0.003 | -0.002 | -0.002 | 0 | 0.41 | 0.44 | 0.46 | 0.63 | 0.32 | 0.57 |
| 8 | 66640090 | rs16932315 | -0.43 | -0.91 | 0.426 | -0.306 | 0.022 | -0.029 | -0.029 | 0 | 0.07 | 0.18 | 0.09 | 0.03 | 0.09 | 0.05 |
| 8 | 66644713 | rs10808746 | -1.828 | -0.702 | -0.557 | -0.805 | -0.17 | -0.104 | -0.104 | 0.88 | 0.61 | 0.96 | 0.61 | 0.66 | 0.48 | 0.63 |
| 8 | 66645549 | rs7833031 | -1.309 | -0.845 | -0.12 | -0.62 | -0.012 | 0.002 | 0.002 | 0.094 | 0.4 | 0.5 | 0.47 | 0.67 | 0.31 | 0.54 |
| 8 | 66652487 | rs16932321 | -0.837 | -0.735 | -0.246 | -0.102 | 0.001 | 0.085 | 0.085 | 0 | 0.08 | 0.16 | 0.07 | 0.27 | 0.07 | 0.17 |
| 8 | 66653081 | rs6995183 | -0.1 | -0.993 | -0.395 | -0.779 | 0.007 | -0.014 | -0.014 | 0 | 0.7 | 0.72 | 0.66 | 0.56 | 0.71 | 0.77 |
| 8 | 66654703 | rs17396080 | 1.498 | -1.006 | -0.476 | -1.215 | -0.003 | -0.073 | -0.073 | 0.37 | 0.46 | 0.5 | 0.33 | 0.04 | 0.55 | 0.24 |
| 8 | 66657050 | rs12677891 | -1.039 | -0.455 | -0.364 | -0.367 | 0.006 | -0.114 | -0.114 | 0 | 0.22 | 0.09 | 0.31 | 0.56 | 0.17 | 0.51 |
| 8 | 66663253 | rs12542226 | -0.149 | -0.477 | -0.336 | -0.299 | 0.034 | -0.134 | -0.134 | 0 | 0.17 | 0.02 | 0.22 | 0.55 | 0.08 | 0.45 |
| 8 | 66679221 | rs17304935 | -1.16 | -0.1 | -0.049 | -1.149 | 0.027 | 0.038 | 0.038 | 0.588 | 0.35 | 0.86 | 0.38 | 0.2 | 0.3 | 0.16 |
| 8 | 66686087 | rs2063685 | -1.484 | 0.143 | 0.323 | -1.94 | -0.003 | 0.126 | 0.126 | 1.532 | 0.38 | 0.82 | 0.34 | 0.1 | 0.35 | 0.2 |
| 8 | 66693623 | rs7828699 | -1.883 | 0.162 | 0.245 | -2.018 | -0.012 | 0.22 | 0.22 | 0.38 | 0.46 | 0.83 | 0.4 | 0.1 | 0.43 | 0.23 |
| 8 | 66711501 | rs11783534 | 0.725 | 0.184 | 0.546 | -2.035 | -0.004 | -0.004 | -0.004 | 0 | 0.15 | 0.15 | 0.09 | 0 | 0.15 | 0.06 |
| 8 | 66712108 | rs7001138 | 0.147 | 0.176 | 0.556 | -2.046 | -0.003 | 0.026 | 0.026 | 0.722 | 0.1 | 0.04 | 0.06 | 0 | 0.08 | 0.03 |
| 8 | 66724605 | rs11987993 | 0.274 | 0.17 | 1.104 | -2.044 | 0.004 | -0.096 | -0.096 | 0 | 0.17 | 0.3 | 0.11 | 0 | 0.15 | 0.06 |
| 8 | 66736695 | rs6415604 | -1.547 | 0.17 | 1.014 | -1.991 | -0.025 | -0.092 | -0.092 | 2.068 | 0.19 | 0.51 | 0.16 | 0.07 | 0.12 | 0.11 |
| 8 | 66738896 | rs6984402 | 0.254 | 0.229 | 0.856 | -1.873 | -0.027 | 0.03 | 0.03 | 0 | 0.11 | 0.03 | 0.13 | 0.03 | 0.16 | 0.06 |
| 8 | 66755558 | rs6472227 | -1.433 | 0.089 | 1.161 | -1.91 | -0.009 | -0.077 | -0.077 | 0.458 | 0.34 | 0.62 | 0.25 | 0.06 | 0.25 | 0.13 |
| 8 | 66836909 | rs11994535 | -1.099 | 0.054 | 1.107 | -1.523 | -0.011 | -0.076 | -0.076 | 0.045 | 0.18 | 0.33 | 0.16 | 0.03 | 0.16 | 0.04 |
| 8 | 66838470 | rs6999945 | -0.115 | 0.008 | 1.133 | -1.505 | 0.004 | -0.03 | -0.03 | 0.414 | 0.09 | 0.3 | 0.08 | 0.03 | 0.07 | 0.01 |
| 8 | 66851731 | rs4601292 | -2.035 | -0.133 | 0.647 | -1.736 | 0.023 | 0.125 | 0.125 | 2.342 | 0.36 | 0.8 | 0.3 | 0.1 | 0.29 | 0.19 |
| 8 | 66870013 | rs11987304 | -1.922 | -0.14 | 0.762 | -1.758 | -0.008 | -0.071 | -0.071 | 0 | 0.35 | 0.51 | 0.29 | 0.07 | 0.3 | 0.19 |
| 8 | 66878476 | rs16932382 | -1.943 | -0.125 | 0.762 | -1.75 | -0.005 | -0.054 | -0.054 | 0 | 0.36 | 0.52 | 0.3 | 0.07 | 0.3 | 0.2 |
| 8 | 66887496 | rs7000670 | -2.42 | -0.135 | 0.652 | -1.748 | 0.026 | 0.243 | 0.243 | 1.453 | 0.46 | 0.83 | 0.38 | 0.1 | 0.38 | 0.21 |
| 8 | 66893679 | rs7841695 | -1.376 | -0.086 | 0.815 | -1.666 | -0.01 | -0.092 | -0.092 | 0 | 0.26 | 0.42 | 0.23 | 0.07 | 0.22 | 0.17 |
| 8 | 66900280 | rs6472239 | -0.769 | -0.205 | 0.838 | -1.607 | -0.01 | -0.071 | -0.071 | 0 | 0.14 | 0.31 | 0.11 | 0.02 | 0.05 | 0.09 |
| 8 | 66903149 | rs16932389 | -0.038 | -0.146 | 0.933 | -1.603 | 0.004 | -0.014 | -0.014 | 0 | 0.1 | 0.3 | 0.07 | 0.05 | 0.07 | 0.01 |
| 8 | 66913287 | rs17397530 | -1.311 | -0.203 | 0.799 | -1.672 | 0.014 | 0.049 | 0.049 | 0.222 | 0.22 | 0.11 | 0.14 | 0.02 | 0.14 | 0.14 |
| 8 | 66917753 | rs950729 | -2.088 | -0.191 | 0.814 | -1.685 | -0.004 | -0.05 | -0.05 | 0.422 | 0.33 | 0.3 | 0.27 | 0.06 | 0.28 | 0.18 |
| 8 | 66933620 | rs953177 | -0.481 | 0.01 | 0.933 | -1.414 | 0.004 | -0.032 | -0.032 | 0 | 0.11 | 0.53 | 0.1 | 0.03 | 0.07 | 0.01 |
| 8 | 66935270 | rs7006334 | 0.049 | 0.023 | 0.667 | -1.42 | 0.007 | -0.023 | -0.023 | 0.18 | 0.1 | 0.17 | 0.12 | 0.03 | 0.14 | 0.05 |
| 8 | 66940998 | rs16932422 | 0.04 | -0.218 | -0.053 | -1.419 | 0.002 | -0.024 | -0.024 | 1.478 | 0.09 | 0.3 | 0.07 | 0.03 | 0.07 | 0.01 |
| 8 | 66946079 | rs17398646 | -0.386 | -0.143 | -1.412 | -1.434 | 0.008 | 0.038 | 0.038 | 0.396 | 0.13 | 0.05 | 0.09 | 0.03 | 0.06 | 0.13 |
| 8 | 66948194 | rs16932433 | -0.182 | -0.095 | -1.612 | -1.399 | -0.003 | -0.014 | -0.014 | 0.113 | 0.1 | 0.12 | 0.09 | 0.03 | 0.12 | 0.04 |
| 8 | 66949849 | rs4636189 | -0.547 | -0.014 | -2.241 | -1.331 | 0.019 | 0.011 | 0.011 | 7.441 | 0.16 | 0.8 | 0.14 | 0.01 | 0.08 | 0.04 |
| 8 | 66955437 | rs2357667 | 0.281 | 0.88 | -2.67 | -1.21 | -0.039 | -0.005 | -0.005 | 0 | 0.07 | 0.02 | 0.06 | 0.06 | 0.13 | 0.06 |
| 8 | 66955651 | rs2357668 | -0.841 | 0.72 | -2.688 | -1.217 | -0.099 | 0.062 | 0.062 | 2.306 | 0.26 | 0.03 | 0.2 | 0.11 | 0.28 | 0.19 |
| 8 | 66960324 | rs7011042 | -1.25 | 0.254 | -2.334 | -0.895 | 0.009 | 0.111 | 0.111 | 1.855 | 0.34 | 0.9 | 0.3 | 0.13 | 0.25 | 0.19 |
| 8 | 66960476 | rs6994799 | -1.127 | 0.334 | -2.355 | -0.684 | -0.008 | 0.139 | 0.139 | 0.82 | 0.43 | 0.91 | 0.39 | 0.17 | 0.36 | 0.19 |
| 8 | 66963989 | rs9298109 | -1.129 | -0.223 | -1.467 | 0.182 | 0.02 | -0.013 | -0.013 | 0.145 | 0.27 | 0.78 | 0.4 | 0.37 | 0.17 | 0.22 |
| 8 | 66971754 | rs4368961 | -1.159 | -0.062 | -2.351 | 0.054 | -0.093 | 0.018 | 0.018 | 3.734 | 0.6 | 0.94 | 0.62 | 0.69 | 0.49 | 0.63 |
| 8 | 66974252 | rs13279522 | -0.679 | -0.205 | -1.599 | 0.219 | -0.014 | 0.007 | 0.007 | 0.47 | 0.25 | 0.75 | 0.39 | 0.35 | 0.14 | 0.21 |
| 8 | 66976206 | rs6472244 | 0.339 | -0.374 | -0.713 | 0.071 | -0.022 | -0.048 | -0.048 | 0 | 0.09 | 0.2 | 0.04 | 0 | 0.05 | 0.03 |
| 8 | 66980584 | rs13257752 | -0.054 | -0.416 | 0.207 | 0.133 | -0.014 | 0.05 | 0.05 | 0 | 0.13 | 0.16 | 0.31 | 0.34 | 0.08 | 0.18 |
| 8 | 66988197 | rs16932465 | 0.37 | -0.428 | 0.62 | 0.095 | -0.008 | -0.021 | -0.021 | 0 | 0.07 | 0.21 | 0.04 | 0.02 | 0.04 | 0.01 |
| 8 | 66995147 | rs4308714 | -1.453 | -0.438 | 0.252 | -0.052 | -0.082 | 0.009 | 0.009 | 2.692 | 0.61 | 0.94 | 0.63 | 0.7 | 0.52 | 0.64 |
| 8 | 67007650 | rs6991418 | -0.32 | -0.456 | 0.508 | -0.011 | -0.01 | -0.051 | -0.051 | 0 | 0.11 | 0.29 | 0.08 | 0.01 | 0.09 | 0.03 |
| 8 | 67008429 | rs13264487 | -1.697 | -0.136 | 0.381 | 0.02 | 0.008 | -0.008 | -0.008 | 0 | 0.26 | 0.78 | 0.39 | 0.34 | 0.17 | 0.21 |
| 8 | 67037934 | rs6472251 | -1.345 | -0.124 | 0.56 | -0.075 | -0.018 | -0.09 | -0.09 | 0 | 0.19 | 0.67 | 0.12 | 0.01 | 0.13 | 0.04 |
| 8 | 67045076 | rs10092687 | -1.345 | -0.129 | 1.072 | -0.08 | -0.018 | -0.094 | -0.094 | 0.236 | 0.19 | 0.67 | 0.12 | 0.01 | 0.13 | 0.04 |
| 8 | 67052316 | rs9643575 | 0.089 | 0.018 | 0.664 | -0.042 | -0.112 | -0.052 | -0.052 | 0.051 | 0.4 | 0.04 | 0.26 | 0.35 | 0.38 | 0.43 |
| 8 | 67052569 | rs7835340 | -1.746 | -0.058 | 0.502 | -0.48 | -0.068 | 0.097 | 0.097 | 1.587 | 0.69 | 0.95 | 0.72 | 0.87 | 0.59 | 0.73 |
| 8 | 67055530 | rs10099957 | -0.541 | -0.374 | 1.231 | -0.017 | 0.115 | 0.001 | 0.001 | 1.348 | 0.08 | 0.48 | 0.07 | 0 | 0.09 | 0.03 |
| 8 | 67058975 | rs16932561 | 0.819 | -0.43 | 1.152 | -0.008 | -0.028 | -0.029 | -0.029 | 1.91 | 0.05 | 0.18 | 0.02 | 0 | 0.01 | 0 |
| 8 | 67059253 | rs16932564 | -0.427 | -0.424 | 1.234 | -0.016 | 0.096 | -0.003 | -0.003 | 0.459 | 0.07 | 0.43 | 0.07 | 0 | 0.09 | 0.03 |
| 8 | 67075545 | rs7820617 | -0.479 | -0.403 | 1.036 | -0.164 | 0.124 | 0.007 | 0.007 | 1.05 | 0.08 | 0.49 | 0.07 | 0 | 0.09 | 0.03 |
| 8 | 67084379 | rs7835214 | 0.501 | -0.5 | 0.732 | -0.062 | 0.004 | 0.029 | 0.029 | 0 | 0.07 | 0.05 | 0.27 | 0.34 | 0.05 | 0.17 |
| 8 | 67091379 | rs3176921 | -1.235 | -0.447 | 0.574 | -0.145 | -0.014 | -0.086 | -0.086 | 0.85 | 0.13 | 0.68 | 0.1 | 0 | 0.09 | 0.03 |
| 8 | 67092076 | rs6999780 | -0.402 | -0.441 | 0.673 | -0.142 | 0.068 | -0.02 | -0.02 | 0.552 | 0.08 | 0.39 | 0.06 | 0 | 0.09 | 0.03 |
| 8 | 67098737 | rs1870393 | -1.583 | -0.386 | -0.168 | -0.112 | -0.012 | 0.02 | 0.02 | 0.506 | 0.21 | 0.75 | 0.39 | 0.35 | 0.14 | 0.2 |
| 8 | 67101269 | rs4419797 | -1.574 | -0.383 | -0.167 | -0.11 | -0.023 | 0.018 | 0.018 | 0.5 | 0.21 | 0.76 | 0.39 | 0.34 | 0.11 | 0.2 |
| 8 | 67102818 | rs1870394 | -1.6 | -0.398 | -0.168 | -0.121 | -0.012 | 0.018 | 0.018 | 0.47 | 0.21 | 0.75 | 0.4 | 0.35 | 0.14 | 0.2 |
| 8 | 67114738 | rs11996294 | -0.89 | -0.293 | -0.185 | -0.743 | -0.002 | -0.077 | -0.077 | 0 | 0.35 | 0.69 | 0.19 | 0 | 0.26 | 0.11 |
| 8 | 67117808 | rs2469250 | 0.199 | 0.356 | 0.173 | 0.282 | 0.034 | 0.012 | 0.012 | 0 | 0.69 | 0.52 | 0.68 | 0.63 | 0.61 | 0.62 |
| 8 | 67118308 | rs2446432 | 0.489 | 0.191 | -0.089 | 0.198 | -0.022 | 0.007 | 0.007 | 0.018 | 0.57 | 0.34 | 0.4 | 0.35 | 0.51 | 0.42 |
| 8 | 67120935 | rs2446430 | 0.243 | 0.087 | 0.224 | 0.183 | 0.006 | -0.002 | -0.002 | 0 | 0.6 | 0.5 | 0.43 | 0.35 | 0.54 | 0.44 |
| 8 | 67121343 | rs2469251 | 0.605 | -0.091 | -0.121 | 0.15 | 0.02 | 0.037 | 0.037 | 0 | 0.51 | 0.37 | 0.36 | 0.35 | 0.47 | 0.43 |
| 8 | 67127559 | rs16932663 | -1.124 | -0.329 | 0.746 | 0.18 | -0.033 | 0.06 | 0.06 | 0.594 | 0.1 | 0.49 | 0.06 | 0.08 | 0.02 | 0.01 |

| | | | | | | | | | | | | | | | | |
|---|---|---|---|---|---|---|---|---|---|---|---|---|---|---|---|---|
| 8 | 67132592 | rs2446440 | -0.451 | -0.37 | -0.731 | 0.068 | -0.038 | -0.031 | -0.031 | 0.547 | 0.71 | 0.95 | 0.73 | 0.67 | 0.75 | 0.61 |
| 8 | 67133049 | rs2446439 | 0.295 | -0.414 | 0.029 | 0.052 | -0.024 | -0.002 | -0.002 | 0.301 | 0.62 | 0.51 | 0.68 | 0.59 | 0.73 | 0.6 |
| 8 | 67139307 | rs2469299 | 0.11 | -0.33 | -0.225 | 0.118 | -0.033 | -0.077 | -0.077 | 0 | 0.28 | 0.09 | 0.27 | 0.41 | 0.24 | 0.4 |
| 8 | 67139649 | rs2469298 | -0.16 | -0.303 | 0.123 | 0.108 | 0 | 0.002 | 0.002 | 0 | 0.3 | 0.3 | 0.28 | 0.41 | 0.24 | 0.4 |
| 8 | 67169917 | rs2469260 | 0.417 | -0.23 | 0.103 | 0.13 | -0.022 | -0.015 | -0.015 | 2.99 | 0.24 | 0.11 | 0.26 | 0.33 | 0.24 | 0.39 |
| 8 | 67171962 | rs936430 | -0.087 | -0.235 | 0.238 | 0.101 | -0.001 | -0.004 | -0.004 | 0 | 0.29 | 0.26 | 0.28 | 0.41 | 0.24 | 0.4 |
| 8 | 67177062 | rs2446448 | -0.781 | -0.35 | -0.627 | -0.519 | -0.054 | 0.168 | 0.168 | 0 | 0.69 | 0.94 | 0.73 | 0.94 | 0.64 | 0.82 |
| 8 | 67182109 | rs16932700 | -0.091 | -0.315 | 0.075 | 0.148 | -0.006 | 0.027 | 0.027 | 0 | 0.08 | 0.21 | 0.04 | 0.09 | 0.05 | 0.01 |
| 8 | 67185752 | rs4737769 | -0.501 | -0.343 | -0.006 | 0.4 | -0.001 | 0.023 | 0.023 | 0 | 0.35 | 0.4 | 0.35 | 0.47 | 0.29 | 0.4 |
| 8 | 67187945 | rs11782521 | -0.003 | -0.326 | -0.217 | 0.517 | -0.014 | -0.01 | -0.01 | 0 | 0.3 | 0.19 | 0.29 | 0.39 | 0.29 | 0.39 |
| 8 | 67192379 | rs4737771 | -0.325 | -0.121 | -0.145 | 0.616 | -0.009 | 0.151 | 0.151 | 0.099 | 0.11 | 0.4 | 0.11 | 0.14 | 0.06 | 0.03 |
| 8 | 67192629 | rs16932725 | 0.353 | 0.12 | -1.006 | 0.361 | -0.081 | -0.065 | -0.065 | 0 | 0.27 | 0.05 | 0.3 | 0.33 | 0.27 | 0.34 |
| 8 | 67193708 | rs6988640 | -0.195 | -0.566 | 0.112 | -0.221 | -0.009 | -0.078 | -0.078 | 0 | 0.13 | 0.33 | 0.12 | 0 | 0.1 | 0.12 |
| 8 | 67194171 | rs12334877 | -0.696 | -0.702 | 0.042 | 0.223 | -0.012 | 0.03 | 0.03 | 0.089 | 0.18 | 0.32 | 0.18 | 0.25 | 0.18 | 0.3 |
| 8 | 67199420 | rs7006364 | -1.341 | -1.042 | 0.398 | 0.28 | -0.009 | 0.017 | 0.017 | 0.134 | 0.33 | 0.59 | 0.37 | 0.45 | 0.29 | 0.44 |
| 8 | 67202787 | rs16932761 | -0.506 | -1.033 | 0.333 | 0.117 | -0.006 | -0.008 | -0.008 | 1.242 | 0.26 | 0.29 | 0.22 | 0.25 | 0.25 | 0.37 |
| 8 | 67204931 | rs9298111 | -1.141 | -1.033 | 0.35 | 0.114 | -0.014 | 0.034 | 0.034 | 0 | 0.31 | 0.55 | 0.34 | 0.42 | 0.27 | 0.4 |
| 8 | 67209954 | rs4737779 | -1.202 | -1.07 | 0.472 | 0.17 | 0.002 | -0.006 | -0.006 | 0 | 0.32 | 0.64 | 0.38 | 0.44 | 0.27 | 0.4 |
| 8 | 67211176 | rs12541132 | -1.413 | -1.591 | -0.354 | -0.428 | -0.08 | -0.043 | -0.043 | 0 | 0.64 | 0.89 | 0.56 | 0.59 | 0.5 | 0.65 |
| 8 | 67216422 | rs10106904 | 1.276 | -1.603 | 0.288 | -0.526 | -0.01 | -0.052 | -0.052 | 0 | 0.09 | 0.15 | 0.09 | 0 | 0.07 | 0.11 |
| 8 | 67218257 | rs4737782 | -0.407 | -1.616 | 0.504 | -0.519 | 0.017 | 0.016 | 0.016 | 0 | 0.47 | 0.39 | 0.4 | 0.39 | 0.4 | 0.43 |
| 8 | 67220641 | rs16932800 | 0.375 | -1.57 | 0.218 | -0.576 | 0.001 | 0.017 | 0.017 | 0.048 | 0.18 | 0.21 | 0.17 | 0.26 | 0.18 | 0.23 |
| 8 | 67221614 | rs11779466 | 0.551 | -1.595 | 0.21 | -0.638 | 0.041 | 0.08 | 0.08 | 0.636 | 0.3 | 0.13 | 0.23 | 0.13 | 0.22 | 0.21 |
| 8 | 67224071 | rs4562299 | 1.372 | -1.643 | 0.292 | -0.553 | -0.007 | 0.005 | 0.005 | 1.656 | 0.08 | 0.1 | 0.17 | 0.13 | 0.03 | 0.07 |
| 8 | 67224883 | rs949988 | 0.661 | -1.656 | 0.572 | -0.457 | -0.013 | 0.229 | 0.229 | 1.07 | 0.09 | 0.36 | 0.28 | 0.29 | 0.04 | 0.1 |
| 8 | 67226315 | rs7018347 | -0.783 | -1.379 | 0.24 | -0.673 | 0.03 | 0.044 | 0.044 | 0.083 | 0.5 | 0.55 | 0.44 | 0.14 | 0.4 | 0.38 |
| 8 | 67254723 | rs10092482 | -0.182 | -1.481 | 0.273 | -0.673 | -0.001 | -0.002 | -0.002 | 1.814 | 0.28 | 0.25 | 0.3 | 0.15 | 0.2 | 0.25 |
| 8 | 67257751 | rs2114487 | -0.136 | -1.622 | 0.409 | -1.099 | 0.004 | -0.085 | -0.085 | 0 | 0.2 | 0.24 | 0.15 | 0.01 | 0.23 | 0.12 |
| 8 | 67260472 | rs7819605 | -1.925 | -1.865 | 0.434 | -1.027 | -0.033 | -0.006 | -0.006 | 0 | 0.62 | 0.72 | 0.59 | 0.43 | 0.51 | 0.61 |
| 8 | 67276508 | rs7007961 | 0.772 | -1.93 | 0.404 | -1.088 | 0.004 | -0.003 | -0.003 | 0 | 0.19 | 0.11 | 0.13 | 0 | 0.16 | 0.16 |
| 8 | 67292716 | rs9693689 | -2.359 | -1.97 | 0.206 | -1.135 | -0.111 | -0.051 | -0.051 | 1.726 | 0.67 | 0.89 | 0.72 | 0.59 | 0.56 | 0.65 |
| 8 | 67312348 | rs12541622 | 1.092 | -2.039 | 0.54 | -1.149 | -0.007 | 0.014 | 0.014 | 0.803 | 0.11 | 0.13 | 0.11 | 0.26 | 0.06 | 0.2 |
| 8 | 67326892 | rs17405528 | 0.462 | -2.101 | 0.65 | -1.21 | -0.011 | 0.03 | 0.03 | 0 | 0.18 | 0.07 | 0.14 | 0 | 0.22 | 0.11 |
| 8 | 67329108 | rs10957373 | 0.382 | -2.167 | 0.796 | -1.211 | -0.007 | -0.001 | -0.001 | 0.809 | 0.25 | 0.29 | 0.26 | 0.16 | 0.2 | 0.19 |
| 8 | 67330132 | rs6991180 | 1.926 | -2.149 | 0.822 | -1.202 | 0.006 | 0.151 | 0.151 | 0 | 0.05 | 0.15 | 0.13 | 0.16 | 0.04 | 0.03 |
| 8 | 67334099 | rs16919047 | 0.504 | -2.167 | 0.802 | -1.21 | 0.009 | -0.017 | -0.017 | 0 | 0.16 | 0.11 | 0.22 | 0.42 | 0.1 | 0.23 |
| 8 | 67336382 | rs12541098 | -1.311 | -2.171 | 0.853 | -1.193 | 0.049 | 0.037 | 0.037 | 0.834 | 0.49 | 0.64 | 0.62 | 0.59 | 0.33 | 0.52 |
| 8 | 67341378 | rs3739334 | 1.667 | -2.116 | 0.78 | -1.167 | -0.019 | 0.03 | 0.03 | 1.036 | 0.08 | 0.14 | 0.22 | 0.15 | 0.04 | 0.09 |
| 8 | 67342077 | rs3739337 | 1.634 | -2.124 | 0.776 | -1.171 | -0.019 | 0.029 | 0.029 | 1.074 | 0.08 | 0.14 | 0.22 | 0.15 | 0.04 | 0.09 |
| 8 | 67344005 | rs35352031 | -0.409 | -2.112 | 0.618 | -1.173 | -0.038 | 0.05 | 0.05 | 2.425 | 0.23 | 0.36 | 0.36 | 0.44 | 0.14 | 0.35 |
| 8 | 67345621 | rs2555576 | 0.399 | -2.036 | 0.596 | -1.203 | -0.001 | -0.037 | -0.037 | 0.029 | 0.18 | 0.13 | 0.14 | 0 | 0.22 | 0.1 |
| 8 | 67346451 | rs10095182 | 0.892 | -2 | 0.415 | -1.175 | 0.004 | -0.002 | -0.002 | 0 | 0.19 | 0.12 | 0.13 | 0 | 0.16 | 0.16 |
| 8 | 67348540 | rs12675850 | -0.792 | -1.952 | 0.364 | -1.103 | 0.004 | 0.014 | 0.014 | 0.145 | 0.46 | 0.56 | 0.42 | 0.18 | 0.43 | 0.32 |
| 8 | 67348907 | rs4737229 | 1.249 | -1.782 | -0.19 | -0.952 | -0.004 | -0.058 | -0.058 | 0 | 0.09 | 0 | 0.1 | 0.26 | 0.06 | 0.19 |
| 8 | 67353258 | rs753082 | -2.102 | -1.797 | -0.175 | -1.028 | -0.099 | -0.042 | -0.042 | 0.125 | 0.67 | 0.86 | 0.76 | 0.59 | 0.55 | 0.62 |
| 8 | 67358921 | rs2718988 | -2.137 | -1.789 | -0.19 | -1.042 | -0.097 | -0.044 | -0.044 | 0.912 | 0.68 | 0.87 | 0.76 | 0.6 | 0.55 | 0.62 |
| 8 | 67361264 | rs3739338 | 1.047 | -1.853 | -0.06 | -1.098 | 0.001 | -0.001 | -0.001 | 0.506 | 0.11 | 0.1 | 0.11 | 0.25 | 0.07 | 0.18 |
| 8 | 67363241 | rs2433594 | 0.646 | -1.93 | -0.038 | -1.236 | -0.002 | -0.024 | -0.024 | 0 | 0.16 | 0.11 | 0.15 | 0 | 0.21 | 0.09 |
| 8 | 67363612 | rs2465983 | 0.646 | -1.93 | -0.038 | -1.236 | -0.002 | -0.024 | -0.024 | 0 | 0.16 | 0.11 | 0.15 | 0 | 0.21 | 0.09 |
| 8 | 67365860 | rs7014306 | 0.076 | -1.973 | 0.111 | -1.257 | 0.002 | -0.078 | -0.078 | 0.043 | 0.19 | 0.21 | 0.15 | 0 | 0.21 | 0.09 |
| 8 | 67369184 | rs2303519 | 0.545 | -2.142 | 0.18 | -1.351 | -0.011 | -0.012 | -0.012 | 0.04 | 0.21 | 0.27 | 0.33 | 0.14 | 0.17 | 0.24 |
| 8 | 67369355 | rs2555588 | 0.172 | -2.141 | 0.191 | -1.355 | 0.003 | -0.084 | -0.084 | 0 | 0.19 | 0.21 | 0.15 | 0 | 0.21 | 0.08 |
| 8 | 67373278 | rs12335128 | 0.323 | -1.822 | 0.106 | -1.331 | -0.008 | -0.015 | -0.015 | 0 | 0.23 | 0.29 | 0.34 | 0.14 | 0.2 | 0.24 |
| 8 | 67375519 | rs2555575 | -2.537 | -2.132 | 0.032 | -1.27 | -0.08 | -0.04 | -0.04 | 0 | 0.67 | 0.86 | 0.76 | 0.58 | 0.55 | 0.62 |
| 8 | 67379951 | rs2555573 | 0.452 | -2.164 | 0.058 | -1.269 | -0.002 | -0.024 | -0.024 | 0 | 0.16 | 0.12 | 0.15 | 0 | 0.21 | 0.09 |
| 8 | 67380677 | rs13682 | -1.9 | -2.078 | 0.102 | -1.116 | -0.027 | 0.001 | 0.001 | 0 | 0.63 | 0.75 | 0.66 | 0.38 | 0.5 | 0.58 |
| 8 | 67383784 | rs2719008 | -1.742 | -1.962 | -0.105 | -1.083 | 0.002 | -0.003 | -0.003 | 0.128 | 0.59 | 0.55 | 0.64 | 0.35 | 0.49 | 0.53 |
| 8 | 67385088 | rs1978566 | -1.04 | -1.746 | -0.474 | -1.524 | 0.001 | -0.006 | -0.006 | 0 | 0.45 | 0.39 | 0.49 | 0.13 | 0.42 | 0.35 |
| 8 | 67398541 | rs7001334 | 0.123 | -2.005 | -0.805 | -1.195 | 0.034 | -0.048 | -0.048 | 0.012 | 0.19 | 0.12 | 0.36 | 0.35 | 0.11 | 0.26 |
| 8 | 67400061 | rs2433596 | -0.641 | -1.116 | -0.539 | -1.696 | 0.001 | -0.035 | -0.035 | 0 | 0.21 | 0.16 | 0.13 | 0 | 0.18 | 0.14 |
| 8 | 67400347 | rs2555586 | -0.383 | -0.67 | -0.163 | -1.297 | -0.002 | -0.046 | -0.046 | 1.55 | 0.14 | 0.14 | 0.11 | 0 | 0.14 | 0.11 |
| 8 | 67407191 | rs9657084 | -0.283 | -0.599 | 0.187 | -1.016 | -0.011 | -0.098 | -0.098 | 0 | 0.17 | 0.33 | 0.13 | 0 | 0.15 | 0.14 |
| 8 | 67409841 | rs957655 | -0.795 | -0.568 | -0.381 | -1.003 | -0.025 | -0.159 | -0.159 | 0 | 0.29 | 0.51 | 0.25 | 0.02 | 0.26 | 0.3 |
| 8 | 67416087 | rs11778287 | -0.359 | -0.319 | -1.35 | -0.441 | -0.007 | -0.009 | -0.009 | 0 | 0.18 | 0.01 | 0.25 | 0.11 | 0.12 | 0.22 |
| 8 | 67419420 | rs9298112 | -0.243 | -0.114 | -0.668 | -0.593 | -0.002 | -0.04 | -0.04 | 1.508 | 0.16 | 0.22 | 0.14 | 0 | 0.15 | 0.17 |
| 8 | 67451184 | rs4463406 | -0.893 | 0.278 | -0.392 | -0.307 | 0.015 | 0.072 | 0.072 | 0 | 0.45 | 0.82 | 0.39 | 0.31 | 0.37 | 0.41 |
| 8 | 67486915 | rs10504398 | 0.234 | 0.335 | 0.487 | -0.042 | -0.049 | -0.061 | -0.061 | 0 | 0.1 | 0.01 | 0.12 | 0.23 | 0.14 | 0.21 |
| 8 | 67492691 | rs11990331 | -0.485 | 0.297 | 0.77 | -0.07 | 0.029 | 0.01 | 0.01 | 2.799 | 0.07 | 0.43 | 0.06 | 0.02 | 0.03 | 0.06 |
| 8 | 67558098 | rs11992252 | -0.009 | 1.335 | 0.28 | -0.859 | 0.041 | 0.016 | 0.016 | 3.399 | 0.06 | 0.43 | 0.06 | 0.02 | 0.03 | 0.06 |

| | | | | | | | | | | | | | | | | |
|---|---|---|---|---|---|---|---|---|---|---|---|---|---|---|---|---|
| 8 | 67567093 | rs10504393 | -0.009 | 1.324 | 0.28 | -0.859 | 0.035 | 0.016 | 0.016 | 3.399 | 0.06 | 0.43 | 0.06 | 0.02 | 0.03 | 0.06 |
| 8 | 67577670 | rs3808602 | -0.009 | 1.324 | 0.28 | -0.859 | 0.035 | 0.016 | 0.016 | 3.399 | 0.06 | 0.43 | 0.06 | 0.02 | 0.03 | 0.06 |
| 8 | 67590460 | rs6983214 | -0.459 | 0.964 | -0.237 | -1.498 | -0.054 | 0.018 | 0.018 | 0 | 0.3 | 0.05 | 0.23 | 0.24 | 0.29 | 0.34 |
| 8 | 67592152 | rs1057463 | -0.459 | 0.965 | -0.239 | -1.498 | -0.054 | 0.018 | 0.018 | 0 | 0.3 | 0.05 | 0.23 | 0.24 | 0.29 | 0.34 |
| 8 | 67609366 | rs6472282 | -1.464 | 0.903 | -0.679 | -1.945 | -0.046 | 0.191 | 0.191 | 0 | 0.5 | 0.94 | 0.41 | 0.26 | 0.43 | 0.43 |
| 8 | 67627906 | rs11996861 | -0.467 | 0.84 | 0.326 | -1.78 | 0.047 | 0.022 | 0.022 | 0.237 | 0.07 | 0.44 | 0.11 | 0.03 | 0.05 | 0.09 |
| 8 | 67638069 | rs11776708 | 0.776 | 0.71 | -0.03 | -2.016 | 0.005 | 0.05 | 0.05 | 0 | 0.12 | 0.03 | 0.06 | 0 | 0.09 | 0.06 |
| 8 | 67677588 | rs11997967 | -0.798 | 0.65 | 0.282 | -1.961 | 0.04 | 0.017 | 0.017 | 0.93 | 0.07 | 0.37 | 0.11 | 0.03 | 0.05 | 0.11 |
| 8 | 67694007 | rs6998777 | -0.722 | 0.654 | 0.337 | -1.959 | 0.046 | 0.022 | 0.022 | 0.448 | 0.07 | 0.38 | 0.11 | 0.03 | 0.05 | 0.11 |
| 8 | 67756985 | rs16933079 | -0.662 | 0.497 | 0.351 | -1.968 | 0.025 | 0.006 | 0.006 | 0 | 0.05 | 0.25 | 0.1 | 0.03 | 0.04 | 0.11 |
| 8 | 67765182 | rs11780700 | -1.541 | 0.183 | 0.322 | -1.583 | 0.015 | 0.016 | 0.016 | 0 | 0.44 | 0.85 | 0.45 | 0.38 | 0.39 | 0.4 |
| 8 | 67784825 | rs11990461 | -0.894 | 0.183 | 0.411 | -1.561 | 0.05 | 0.026 | 0.026 | 0.28 | 0.06 | 0.37 | 0.11 | 0.03 | 0.05 | 0.11 |
| 8 | 67808179 | rs16933097 | -0.779 | 0.185 | 0.433 | -1.56 | 0.075 | 0.045 | 0.045 | 0.659 | 0.05 | 0.36 | 0.1 | 0.01 | 0.03 | 0.05 |
| 8 | 67840116 | rs7837333 | -0.12 | 0.174 | 0.768 | -1.621 | -0.021 | -0.074 | -0.074 | 0 | 0.26 | 0.08 | 0.23 | 0.37 | 0.26 | 0.26 |
| 8 | 67848215 | rs6984650 | -1.014 | 0.228 | 0.86 | -1.622 | 0.059 | 0.029 | 0.029 | 0.314 | 0.06 | 0.34 | 0.11 | 0.03 | 0.05 | 0.11 |
| 8 | 67901607 | rs16933138 | -1.118 | 0.219 | 0.859 | -1.621 | 0.05 | 0.026 | 0.026 | 0.429 | 0.06 | 0.37 | 0.11 | 0.02 | 0.04 | 0.11 |
| 8 | 67933838 | rs16933139 | -1.082 | 0.194 | 1.656 | -1.634 | 0.056 | 0.031 | 0.031 | 0.444 | 0.06 | 0.37 | 0.11 | 0.02 | 0.04 | 0.11 |
| 8 | 67957130 | rs7840028 | 0.784 | 0.164 | 1.62 | -1.683 | 0.04 | -0.028 | -0.028 | 0 | 0.07 | 0.23 | 0.07 | 0 | 0.07 | 0.03 |
| 8 | 67970243 | rs6997490 | -2.153 | -0.63 | 0.554 | -1.314 | -0.17 | -0.07 | -0.07 | 0.43 | 0.6 | 0.97 | 0.54 | 0.64 | 0.51 | 0.59 |
| 8 | 67981275 | rs6990298 | -0.782 | -0.613 | 0.666 | -1.35 | 0.055 | 0.03 | 0.03 | 0.358 | 0.06 | 0.38 | 0.11 | 0.02 | 0.04 | 0.11 |
| 8 | 68041790 | rs12681862 | 1.458 | -0.597 | 0.54 | -1.226 | -0.028 | -0.079 | -0.079 | 0 | 0.11 | 0 | 0.13 | 0.37 | 0.12 | 0.18 |
| 8 | 68041907 | rs10957381 | -1.07 | -0.635 | 0.827 | -1.174 | 0.011 | 0.004 | 0.004 | 0 | 0.43 | 0.74 | 0.42 | 0.41 | 0.38 | 0.4 |
| 8 | 68074137 | rs16933182 | -0.82 | -0.544 | 0.634 | -1.118 | 0.05 | 0.026 | 0.026 | 0.429 | 0.06 | 0.36 | 0.11 | 0.02 | 0.04 | 0.11 |
| 8 | 68087279 | rs2290585 | -0.82 | -0.544 | 0.633 | -1.118 | 0.055 | 0.03 | 0.03 | 0.429 | 0.06 | 0.37 | 0.11 | 0.02 | 0.04 | 0.11 |
| 8 | 68118795 | rs7002622 | 0.369 | -0.676 | 0.411 | -1.174 | -0.042 | -0.099 | -0.099 | 0 | 0.26 | 0.05 | 0.23 | 0.38 | 0.26 | 0.26 |
| 8 | 68122913 | rs7006064 | 1.09 | -0.599 | 0.591 | -1.168 | 0.027 | -0.035 | -0.035 | 0 | 0.07 | 0.19 | 0.07 | 0 | 0.07 | 0.03 |
| 8 | 68170932 | rs10504399 | -0.543 | -1.98 | 0.837 | -0.87 | 0.064 | 0.033 | 0.033 | 0.552 | 0.06 | 0.38 | 0.11 | 0.02 | 0.05 | 0.11 |
| 8 | 68196817 | rs12678025 | -0.42 | -2 | 0.408 | -0.748 | -0.004 | 0.009 | 0.009 | 0 | 0.34 | 0.43 | 0.34 | 0.41 | 0.3 | 0.36 |
| 8 | 68201031 | rs10105795 | -1.279 | -2.004 | 0.066 | -0.752 | -0.009 | -0.015 | -0.015 | 0 | 0.46 | 0.92 | 0.45 | 0.41 | 0.39 | 0.4 |
| 8 | 68213906 | rs11779287 | -0.073 | -1.813 | 0.214 | -0.662 | 0 | 0.001 | 0.001 | 0 | 0.31 | 0.32 | 0.3 | 0.4 | 0.29 | 0.3 |
| 8 | 68219564 | rs10957384 | -0.073 | -1.813 | 0.165 | -0.662 | -0.001 | -0.002 | -0.002 | 0.799 | 0.31 | 0.31 | 0.3 | 0.4 | 0.29 | 0.3 |
| 8 | 68235165 | rs7012901 | 0.977 | -1.362 | -0.427 | -0.506 | 0.024 | -0.032 | -0.032 | 0 | 0.07 | 0.19 | 0.07 | 0 | 0.07 | 0.03 |
| 8 | 68250349 | rs6472296 | -1.275 | -1.275 | -1.219 | -0.16 | -0.173 | -0.095 | -0.095 | 0.414 | 0.61 | 0.97 | 0.46 | 0.66 | 0.5 | 0.58 |
| 8 | 68277883 | rs13277736 | -1.027 | -1.373 | 0.056 | 0.052 | 0.017 | -0.002 | -0.002 | 0.077 | 0.42 | 0.66 | 0.34 | 0.41 | 0.33 | 0.38 |
| 8 | 68300140 | rs10504401 | 0.728 | -1.193 | 0.384 | 0.125 | -0.012 | 0.003 | 0.003 | 0 | 0.6 | 0.53 | 0.68 | 0.59 | 0.66 | 0.61 |
| 8 | 68319668 | rs12677893 | 0.183 | -1.172 | 0.799 | 0.252 | -0.014 | 0.044 | 0.044 | 0 | 0.18 | 0.23 | 0.16 | 0.37 | 0.15 | 0.23 |
| 8 | 68345016 | rs13275199 | -0.62 | -0.963 | -0.34 | -0.361 | 0.012 | 0.067 | 0.067 | 0.041 | 0.28 | 0.82 | 0.22 | 0.11 | 0.21 | 0.24 |
| 8 | 68352280 | rs7008075 | -0.676 | -1.258 | -0.244 | 0.018 | 0.007 | -0.013 | -0.013 | 0 | 0.2 | 0.11 | 0.17 | 0.26 | 0.18 | 0.19 |
| 8 | 68363156 | rs7840914 | -0.94 | -1.284 | 0.373 | 0.03 | -0.026 | 0.034 | 0.034 | 0 | 0.23 | 0.41 | 0.21 | 0.28 | 0.19 | 0.27 |
| 8 | 68369098 | rs12682102 | -0.28 | -1.171 | 0.409 | 0.225 | -0.003 | 0.028 | 0.028 | 0 | 0.11 | 0.12 | 0.14 | 0.25 | 0.12 | 0.17 |
| 8 | 68389059 | rs7820882 | -0.416 | -1.456 | 0.097 | 0.094 | -0.003 | -0.008 | -0.008 | 0 | 0.46 | 0.52 | 0.35 | 0.34 | 0.36 | 0.32 |
| 8 | 68390435 | rs11779485 | -0.677 | -1.421 | 0.018 | 0.083 | 0.023 | 0.054 | 0.054 | 0 | 0.49 | 0.72 | 0.37 | 0.37 | 0.38 | 0.37 |
| 8 | 68396915 | rs17343819 | 1.371 | -1.709 | -0.305 | -0.16 | -0.005 | 0.131 | 0.131 | 0 | 0.23 | 0.06 | 0.12 | 0 | 0.17 | 0.07 |
| 8 | 68408438 | rs7827961 | 0.395 | -1.367 | 0.363 | 0.197 | 0.011 | 0.043 | 0.043 | 0 | 0.44 | 0.22 | 0.31 | 0.34 | 0.36 | 0.32 |
| 8 | 68420477 | rs4737838 | -0.582 | -1.357 | 0.374 | 0.211 | 0.021 | 0.061 | 0.061 | 0.01 | 0.51 | 0.8 | 0.38 | 0.38 | 0.39 | 0.37 |
| 8 | 68429770 | rs11994817 | -0.857 | -1.255 | 0.696 | 0.151 | 0.07 | 0.066 | 0.066 | 3.851 | 0.06 | 0.55 | 0.06 | 0.03 | 0.02 | 0.04 |
| 8 | 68433121 | rs6472314 | -0.669 | -1.236 | 0.331 | 0.161 | 0.104 | 0.1 | 0.1 | 2.862 | 0.05 | 0.44 | 0.06 | 0.03 | 0.02 | 0.04 |
| 8 | 68433597 | rs2089045 | -1.681 | -1.544 | -0.636 | 0.264 | -0.179 | -0.09 | -0.09 | 0.405 | 0.66 | 0.96 | 0.53 | 0.64 | 0.51 | 0.58 |
| 8 | 68435226 | rs1503366 | -0.3 | -1.142 | -0.235 | 0.024 | 0.018 | 0.052 | 0.052 | 0 | 0.48 | 0.36 | 0.34 | 0.34 | 0.37 | 0.32 |
| 8 | 68440281 | rs6472316 | -0.382 | -1.2 | 0.337 | 0.118 | 0.076 | 0.095 | 0.095 | 0 | 0.08 | 0.41 | 0.13 | 0.06 | 0.05 | 0.04 |
| 8 | 68441599 | rs7017897 | 0.03 | -1.748 | -0.1 | 0.064 | 0.025 | 0.033 | 0.033 | 0.016 | 0.43 | 0.38 | 0.28 | 0.23 | 0.29 | 0.42 |
| 8 | 68445739 | rs7831538 | -0.021 | -1.736 | -0.179 | 0.07 | -0.007 | -0.011 | -0.011 | 0.051 | 0.44 | 0.48 | 0.29 | 0.23 | 0.29 | 0.41 |
| 8 | 68454942 | rs4737845 | -0.338 | -1.523 | -1.066 | 0.15 | 0.089 | 0.073 | 0.073 | 0.966 | 0.51 | 0.91 | 0.41 | 0.28 | 0.31 | 0.46 |
| 8 | 68458267 | rs10957391 | 0.409 | -1.592 | -1.287 | -0.107 | 0.018 | 0.065 | 0.065 | 1.008 | 0.42 | 0.1 | 0.65 | 0.72 | 0.65 | 0.54 |
| 8 | 68464647 | rs924742 | 0.42 | -1.536 | -1.268 | -0.194 | 0.021 | 0.092 | 0.092 | 0.861 | 0.42 | 0.12 | 0.65 | 0.74 | 0.65 | 0.53 |
| 8 | 68467195 | rs13268227 | 0.644 | -1.044 | -1.076 | 0.278 | -0.014 | -0.013 | -0.013 | 3.872 | 0.36 | 0.29 | 0.37 | 0.48 | 0.59 | 0.45 |
| 8 | 68478337 | rs11997518 | 0.401 | -1.895 | -0.867 | -0.263 | -0.006 | -0.001 | -0.001 | 0 | 0.56 | 0.53 | 0.49 | 0.49 | 0.7 | 0.54 |
| 8 | 68479877 | rs392092 | 0.433 | -1.953 | -0.679 | -0.283 | 0.016 | 0.001 | 0.001 | 0.509 | 0.54 | 0.39 | 0.46 | 0.49 | 0.7 | 0.51 |
| 8 | 68496522 | rs12681574 | 0.338 | -2.156 | -0.566 | -0.281 | -0.027 | -0.01 | -0.01 | 0.143 | 0.3 | 0.36 | 0.18 | 0.26 | 0.19 | 0.35 |
| 8 | 68497841 | rs406796 | -0.831 | -2.159 | -0.62 | -0.268 | 0.02 | 0.005 | 0.005 | 0.214 | 0.47 | 0.61 | 0.54 | 0.52 | 0.31 | 0.48 |
| 8 | 68497939 | rs16933415 | 0.52 | -2.062 | -0.651 | -0.238 | -0.008 | -0.036 | -0.036 | 0.607 | 0.13 | 0.08 | 0.32 | 0.24 | 0.11 | 0.1 |
| 8 | 68497964 | rs17430478 | 1.814 | -2.033 | -0.612 | -0.408 | 0.009 | -0.009 | -0.009 | 0.513 | 0.17 | 0.11 | 0.09 | 0 | 0.11 | 0.06 |
| 8 | 68501719 | rs16933420 | 1.075 | -1.883 | -0.677 | -0.406 | -0.034 | -0.134 | -0.134 | 0.055 | 0.22 | 0.33 | 0.12 | 0.01 | 0.15 | 0.11 |
| 8 | 68508674 | rs4737846 | -1.42 | -1.872 | -0.805 | -0.295 | -0.1 | -0.094 | -0.094 | 0.21 | 0.67 | 0.89 | 0.64 | 0.53 | 0.44 | 0.56 |
| 8 | 68509800 | rs7005683 | 1.172 | -2.12 | -0.695 | -1.186 | -0.027 | -0.023 | -0.023 | 0 | 0.14 | 0.32 | 0.12 | 0.09 | 0.11 | 0.24 |
| 8 | 68516125 | rs719113 | 1.427 | -2.144 | -0.688 | -1.132 | -0.03 | -0.106 | -0.106 | 0 | 0.2 | 0.31 | 0.12 | 0 | 0.14 | 0.07 |
| 8 | 68516681 | rs10808755 | -1.268 | -2.098 | -0.978 | -1.118 | -0.067 | -0.097 | -0.097 | 0 | 0.65 | 0.89 | 0.62 | 0.52 | 0.42 | 0.53 |
| 8 | 68519685 | rs2128105 | -1.177 | -2.068 | -0.901 | -0.772 | 0.045 | 0.011 | 0.011 | 1.032 | 0.47 | 0.65 | 0.51 | 0.51 | 0.3 | 0.48 |
| 8 | 68521098 | rs13275970 | 0.841 | -1.988 | -0.947 | -0.781 | 0.034 | -0.01 | -0.01 | 0 | 0.15 | 0.03 | 0.04 | 0.15 | 0.08 | 0.11 |

| | | | | | | | | | | | | | | | | |
|---|---|---|---|---|---|---|---|---|---|---|---|---|---|---|---|---|
| 8 | 68524513 | rs1682076 | 0.678 | -1.764 | -0.777 | -1.07 | -0.014 | -0.039 | -0.039 | 0.213 | 0.17 | 0.35 | 0.13 | 0.09 | 0.14 | 0.26 |
| 8 | 68525283 | rs7814309 | -0.338 | -1.264 | -0.713 | -0.83 | -0.002 | 0 | 0 | 0 | 0.44 | 0.57 | 0.49 | 0.5 | 0.27 | 0.45 |
| 8 | 68527890 | rs900114 | 0.124 | -0.055 | -0.706 | -1.142 | -0.012 | 0.035 | 0.035 | 0 | 0.4 | 0.31 | 0.53 | 0.76 | 0.48 | 0.44 |
| 8 | 68528197 | rs900896 | 0.026 | -0.588 | -0.242 | -0.674 | -0.02 | -0.058 | -0.058 | 7.063 | 0.35 | 0.49 | 0.29 | 0.18 | 0.26 | 0.35 |
| 8 | 68528848 | rs1809437 | 0.48 | -0.386 | -0.374 | -0.684 | -0.013 | -0.015 | -0.015 | 0 | 0.22 | 0.14 | 0.3 | 0.28 | 0.26 | 0.08 |
| 8 | 68534788 | rs2114050 | 0.476 | -0.194 | -0.46 | -0.573 | -0.013 | -0.072 | -0.072 | 0.033 | 0.3 | 0.55 | 0.28 | 0.12 | 0.24 | 0.24 |
| 8 | 68536470 | rs10957393 | 0.931 | -0.178 | -0.451 | -0.571 | -0.009 | -0.062 | -0.062 | 0 | 0.26 | 0.34 | 0.24 | 0.09 | 0.22 | 0.23 |
| 8 | 68540692 | rs9643584 | 0.411 | -0.148 | -0.437 | -0.556 | -0.012 | -0.056 | -0.056 | 0.077 | 0.31 | 0.35 | 0.25 | 0.1 | 0.25 | 0.3 |
| 8 | 68541850 | rs6998443 | 0.516 | 0.372 | -0.769 | -0.272 | 0.001 | -0.002 | -0.002 | 0 | 0.37 | 0.64 | 0.31 | 0.19 | 0.27 | 0.36 |
| 8 | 68556828 | rs450738 | 0.533 | 0.674 | -0.897 | 0.151 | -0.008 | -0.025 | -0.025 | 0.013 | 0.43 | 0.51 | 0.32 | 0.19 | 0.34 | 0.48 |
| 8 | 68557693 | rs594006 | 0.086 | 0.831 | -1.508 | 0.682 | -0.067 | -0.101 | -0.101 | 0 | 0.66 | 0.86 | 0.65 | 0.48 | 0.56 | 0.56 |
| 8 | 68574359 | rs10112728 | -0.989 | 0.37 | 0.062 | 0.558 | -0.012 | 0.024 | 0.024 | 0 | 0.11 | 0.28 | 0.1 | 0.17 | 0.05 | 0.08 |
| 8 | 68576417 | rs4737240 | -0.737 | 0.205 | 0.074 | -0.066 | 0.01 | 0.026 | 0.026 | 0.124 | 0.42 | 0.71 | 0.35 | 0.3 | 0.31 | 0.46 |
| 8 | 68580393 | rs10087670 | -1.488 | 0.09 | 1.449 | -0.213 | 0.041 | 0.106 | 0.106 | 0.003 | 0.08 | 0.43 | 0.09 | 0.15 | 0.03 | 0.03 |
| 8 | 68583722 | rs13257028 | -0.975 | -0.143 | 1.223 | -0.506 | 0.028 | 0.097 | 0.097 | 0.776 | 0.43 | 0.88 | 0.37 | 0.32 | 0.34 | 0.42 |
| 8 | 68588027 | rs6980851 | -0.815 | -0.178 | 1.35 | -0.469 | 0.022 | 0.056 | 0.056 | 0.526 | 0.41 | 0.81 | 0.36 | 0.32 | 0.34 | 0.43 |
| 8 | 68589259 | rs6986791 | -1.307 | -0.232 | 1.045 | -0.775 | -0.007 | 0.15 | 0.15 | 2.866 | 0.5 | 0.94 | 0.41 | 0.32 | 0.41 | 0.48 |
| 8 | 68598136 | rs11985114 | -0.112 | -0.298 | 1.792 | -0.709 | 0.001 | 0.002 | 0.002 | 0.021 | 0.32 | 0.31 | 0.22 | 0.12 | 0.24 | 0.36 |
| 8 | 68600111 | rs11987166 | -0.019 | -0.306 | 1.83 | -0.713 | 0.003 | 0.038 | 0.038 | 0 | 0.31 | 0.25 | 0.22 | 0.12 | 0.24 | 0.36 |
| 8 | 68604866 | rs16933522 | -0.916 | -0.347 | 2.286 | -0.715 | -0.007 | 0.045 | 0.045 | 0 | 0.07 | 0.24 | 0.08 | 0.16 | 0.03 | 0.05 |
| 8 | 68607109 | rs2098874 | -0.249 | -0.316 | 2.409 | -0.715 | -0.001 | -0.01 | -0.01 | 0 | 0.33 | 0.37 | 0.26 | 0.17 | 0.3 | 0.38 |
| 8 | 68613771 | rs1871830 | -0.738 | -0.342 | 2.42 | -0.7 | 0.019 | 0.043 | 0.043 | 0.291 | 0.41 | 0.75 | 0.36 | 0.33 | 0.34 | 0.43 |
| 8 | 68635306 | rs971191 | 0.869 | -0.474 | 2.305 | -0.633 | 0.041 | 0.072 | 0.072 | 0.808 | 0.57 | 0.12 | 0.64 | 0.65 | 0.66 | 0.58 |
| 8 | 68635940 | rs10216497 | -1.339 | -0.303 | 2.278 | -0.718 | 0.024 | 0.012 | 0.012 | 0.067 | 0.1 | 0.19 | 0.15 | 0.15 | 0.12 | 0.11 |
| 8 | 68636704 | rs2623840 | 0.589 | -0.184 | 2.507 | -0.703 | -0.007 | -0.017 | -0.017 | 0 | 0.65 | 0.51 | 0.68 | 0.68 | 0.68 | 0.61 |
| 8 | 68642376 | rs2553676 | -0.155 | -0.515 | 2.342 | -0.902 | 0.074 | 0.087 | 0.087 | 0.112 | 0.77 | 0.93 | 0.94 | 1 | 0.91 | 0.89 |
| 8 | 68650640 | rs2553670 | 0.289 | -0.502 | 2.54 | -0.646 | 0.023 | 0.022 | 0.022 | 0 | 0.71 | 0.78 | 0.85 | 0.84 | 0.83 | 0.72 |
| 8 | 68659146 | rs2553662 | 0.153 | -0.566 | 2.756 | -0.586 | 0.019 | 0.003 | 0.003 | 0 | 0.72 | 0.78 | 0.85 | 0.78 | 0.84 | 0.73 |
| 8 | 68659183 | rs2553661 | 0.878 | -0.572 | 2.695 | -0.586 | 0.059 | 0.025 | 0.025 | 0.314 | 0.64 | 0.26 | 0.75 | 0.73 | 0.81 | 0.67 |
| 8 | 68659293 | rs13250665 | -0.68 | -0.38 | 2.464 | -0.587 | -0.007 | 0.005 | 0.005 | 0 | 0.07 | 0.06 | 0.09 | 0.01 | 0.09 | 0.07 |
| 8 | 68660147 | rs2162145 | 1.066 | -0.27 | 2.486 | -0.488 | 0.085 | 0.02 | 0.02 | 0.5 | 0.62 | 0.18 | 0.72 | 0.67 | 0.75 | 0.66 |
| 8 | 68662662 | rs2553658 | 0.171 | 0.069 | 1.952 | -0.642 | 0.027 | 0.003 | 0.003 | 0.05 | 0.71 | 0.86 | 0.83 | 0.76 | 0.79 | 0.74 |
| 8 | 68665932 | rs2553656 | 1.358 | 0.304 | 1.399 | -0.619 | 0.006 | 0.062 | 0.062 | 0.06 | 0.53 | 0.11 | 0.55 | 0.63 | 0.61 | 0.56 |
| 8 | 68666611 | rs2929535 | 1.021 | 0.218 | 1.527 | -0.459 | 0.038 | 0.028 | 0.028 | 0.022 | 0.6 | 0.19 | 0.68 | 0.64 | 0.69 | 0.66 |
| 8 | 68676568 | rs16933648 | -1.472 | 0.274 | 1.686 | -0.208 | 0.016 | 0.023 | 0.023 | 0 | 0.05 | 0.23 | 0.07 | 0.06 | 0.01 | 0.05 |
| 8 | 68677337 | rs2623859 | 1.14 | 0.355 | 0.922 | -0.174 | 0.001 | 0.011 | 0.011 | 0.017 | 0.94 | 0.86 | 0.94 | 0.91 | 0.91 | 0.86 |
| 8 | 68678318 | rs10106534 | -1.155 | -0.171 | 0.567 | -0.336 | -0.021 | -0.013 | -0.013 | 1.4 | 0.17 | 0.16 | 0.12 | 0.11 | 0.08 | 0.09 |
| 8 | 68679270 | rs2623849 | 0.77 | 0.386 | 0.35 | -0.577 | 0.006 | -0.026 | -0.026 | 1.117 | 0.68 | 0.6 | 0.62 | 0.82 | 0.69 | 0.8 |
| 8 | 68686779 | rs4737247 | 0.72 | 0.419 | 0.485 | -0.631 | -0.004 | 0.007 | 0.007 | 0 | 0.74 | 0.78 | 0.66 | 0.83 | 0.74 | 0.84 |
| 8 | 68687678 | rs13274503 | -0.541 | 0.754 | -0.568 | 0.154 | -0.039 | -0.073 | -0.073 | 0 | 0.11 | 0.01 | 0.18 | 0.26 | 0.12 | 0.09 |
| 8 | 68695814 | rs10504409 | -0.844 | 0.762 | -0.534 | -0.168 | 0 | -0.021 | -0.021 | 3.889 | 0.11 | 0.08 | 0.06 | 0.01 | 0.06 | 0.04 |
| 8 | 68696848 | rs17433963 | -1.448 | 1.096 | -0.622 | -0.24 | 0.002 | -0.016 | -0.016 | 0 | 0.13 | 0.08 | 0.09 | 0.01 | 0.09 | 0.04 |
| 8 | 68697834 | rs11781049 | 0.833 | 0.749 | -1.595 | -0.564 | -0.038 | 0.07 | 0.07 | 0.044 | 0.16 | 0.01 | 0.06 | 0.02 | 0.15 | 0.18 |
| 8 | 68703928 | rs6980766 | 0.061 | 0.61 | -0.737 | -0.225 | 0.091 | 0.219 | 0.219 | 1.498 | 0.06 | 0.46 | 0.11 | 0.14 | 0.08 | 0.02 |
| 8 | 68706223 | rs2623813 | -0.06 | 0.613 | -1.091 | -0.396 | -0.009 | -0.015 | -0.015 | 0 | 0.82 | 0.88 | 0.77 | 0.83 | 0.8 | 0.89 |
| 8 | 68708780 | rs2623812 | -0.33 | 0.699 | -1.135 | -0.42 | -0.017 | -0.037 | -0.037 | 0.343 | 0.84 | 0.96 | 0.78 | 0.83 | 0.82 | 0.88 |
| 8 | 68711093 | rs1834446 | -0.348 | 0.678 | -1.144 | -0.431 | -0.011 | -0.043 | -0.043 | 0 | 0.85 | 0.96 | 0.79 | 0.84 | 0.84 | 0.89 |
| 8 | 68711214 | rs1834447 | 1.777 | 0.715 | -1.057 | -0.597 | 0.004 | 0.022 | 0.022 | 1.352 | 0.53 | 0.6 | 0.62 | 0.81 | 0.57 | 0.65 |
| 8 | 68718260 | rs4430075 | 0.905 | 0.707 | -0.963 | -0.491 | 0.002 | 0.017 | 0.017 | 0.069 | 0.66 | 0.7 | 0.71 | 0.82 | 0.67 | 0.7 |
| 8 | 68721847 | rs13278432 | -1.415 | 0.809 | -0.999 | -0.632 | 0.008 | -0.109 | -0.109 | 0 | 0.19 | 0.28 | 0.13 | 0.01 | 0.18 | 0.11 |
| 8 | 68723609 | rs10957397 | 1.76 | 0.861 | -0.989 | -0.53 | -0.002 | 0.014 | 0.014 | 1.034 | 0.56 | 0.65 | 0.63 | 0.8 | 0.56 | 0.63 |
| 8 | 68727993 | rs10087150 | 1.452 | 0.875 | -0.886 | -0.479 | -0.004 | 0.019 | 0.019 | 0.423 | 0.6 | 0.7 | 0.64 | 0.8 | 0.6 | 0.65 |
| 8 | 68733176 | rs11775871 | 0.782 | 0.721 | -0.729 | -0.26 | 0.001 | -0.003 | -0.003 | 1.401 | 0.8 | 0.85 | 0.72 | 0.83 | 0.82 | 0.87 |
| 8 | 68734769 | rs4585742 | 1.231 | 0.704 | -0.356 | -0.317 | -0.002 | -0.011 | -0.011 | 1.678 | 0.69 | 0.67 | 0.65 | 0.81 | 0.72 | 0.73 |
| 8 | 68735313 | rs10087234 | 0.008 | 0.716 | -0.578 | -0.43 | 0 | -0.015 | -0.015 | 0 | 0.13 | 0.14 | 0.06 | 0.01 | 0.09 | 0.13 |
| 8 | 68735398 | rs9785137 | 1.064 | 0.563 | -0.371 | -0.357 | 0 | 0 | 0 | 0 | 0.81 | 0.84 | 0.72 | 0.83 | 0.81 | 0.87 |
| 8 | 68740862 | rs10808756 | 0.313 | 0.491 | -0.703 | -0.286 | -0.02 | -0.027 | -0.027 | 0.02 | 0.94 | 0.98 | 0.93 | 0.83 | 0.9 | 0.94 |
| 8 | 68742553 | rs4602870 | 0.264 | 0.493 | -0.703 | -0.285 | -0.024 | -0.03 | -0.03 | 0.133 | 0.94 | 0.99 | 0.93 | 0.83 | 0.9 | 0.94 |
| 8 | 68751611 | rs13279743 | 0.851 | 0.357 | -0.698 | -0.368 | 0 | -0.002 | -0.002 | 0 | 0.83 | 0.82 | 0.89 | 0.82 | 0.84 | 0.83 |
| 8 | 68762313 | rs12545592 | 1.242 | 0.275 | -0.383 | -0.409 | -0.004 | 0.002 | 0.002 | 0 | 0.81 | 0.69 | 0.86 | 0.82 | 0.84 | 0.83 |
| 8 | 68763665 | rs6472338 | -0.175 | 0.395 | -0.705 | -0.356 | -0.003 | 0.009 | 0.009 | 0.073 | 0.11 | 0.08 | 0.07 | 0.02 | 0.1 | 0.11 |
| 8 | 68763825 | rs13260831 | -0.332 | 0.654 | -0.843 | -0.617 | -0.016 | 0.033 | 0.033 | 0.347 | 0.15 | 0.06 | 0.11 | 0 | 0.2 | 0.11 |
| 8 | 68773513 | rs7827110 | 0.473 | 0.508 | -0.704 | -0.237 | -0.002 | -0.009 | -0.009 | 0 | 0.84 | 0.86 | 0.89 | 0.82 | 0.84 | 0.83 |
| 8 | 68777877 | rs4737859 | 1.056 | 0.545 | -1.834 | -0.088 | -0.042 | 0.048 | 0.048 | 0.255 | 0.48 | 0.13 | 0.39 | 0.38 | 0.49 | 0.56 |
| 8 | 68779307 | rs7826474 | -0.009 | 0.102 | -1.946 | -0.139 | 0.011 | -0.123 | -0.123 | 1.08 | 0.72 | 0.96 | 0.62 | 0.44 | 0.76 | 0.73 |
| 8 | 68781711 | rs7011558 | -0.567 | 0.211 | -0.912 | -0.546 | 0.041 | -0.041 | -0.041 | 6.629 | 0.11 | 0.58 | 0.13 | 0.04 | 0.15 | 0.06 |
| 8 | 68784876 | rs4621791 | 0.852 | -0.271 | -0.864 | -0.611 | 0.013 | -0.007 | -0.007 | 0 | 0.62 | 0.35 | 0.51 | 0.45 | 0.6 | 0.67 |
| 8 | 68791306 | rs13267802 | 0.076 | -0.186 | -1.207 | -0.911 | -0.005 | 0.001 | 0.001 | 0 | 0.14 | 0.11 | 0.12 | 0 | 0.17 | 0.1 |
| 8 | 68792563 | rs4556082 | -0.438 | -0.145 | -0.889 | -0.88 | -0.009 | -0.045 | -0.045 | 0 | 0.75 | 0.81 | 0.57 | 0.45 | 0.63 | 0.71 |

| | | | | | | | | | | | | | | | | |
|---|---|---|---|---|---|---|---|---|---|---|---|---|---|---|---|---|
| 8 | 68793432 | rs7834766 | 1.021 | -0.035 | -0.765 | -0.57 | -0.022 | 0.025 | 0.025 | 0.029 | 0.47 | 0.22 | 0.39 | 0.44 | 0.42 | 0.57 |
| 8 | 68805608 | rs4487746 | -1.065 | -0.193 | 0.266 | -0.939 | 0.017 | -0.079 | -0.079 | 0.074 | 0.15 | 0.56 | 0.14 | 0.04 | 0.12 | 0.1 |
| 8 | 68812838 | rs9969491 | -0.275 | -0.172 | 0.354 | -0.926 | 0.041 | -0.047 | -0.047 | 0 | 0.11 | 0.29 | 0.09 | 0.04 | 0.1 | 0.09 |
| 8 | 68813159 | rs6472353 | -0.275 | -0.172 | 0.354 | -0.926 | 0.041 | -0.047 | -0.047 | 0.302 | 0.11 | 0.29 | 0.09 | 0.04 | 0.1 | 0.09 |
| 8 | 68813274 | rs6472354 | -0.57 | -0.837 | -0.418 | -0.99 | 0.008 | -0.047 | -0.047 | 0 | 0.69 | 0.88 | 0.64 | 0.42 | 0.64 | 0.65 |
| 8 | 68819659 | rs10089660 | -0.227 | -0.731 | -0.12 | -1.362 | 0.015 | -0.012 | -0.012 | 0 | 0.07 | 0.3 | 0.05 | 0 | 0.02 | 0.01 |
| 8 | 68821800 | rs6472357 | -0.965 | -1.002 | -0.195 | -1.53 | -0.016 | -0.079 | -0.079 | 0 | 0.21 | 0.61 | 0.15 | 0.04 | 0.12 | 0.1 |
| 8 | 68827224 | rs4367520 | -0.039 | -0.985 | -0.264 | -1.518 | 0.001 | -0.078 | -0.078 | 0 | 0.15 | 0.31 | 0.1 | 0.04 | 0.09 | 0.09 |
| 8 | 68846925 | rs7000218 | -0.54 | -0.897 | -0.929 | -1.485 | -0.017 | -0.075 | -0.075 | 0.103 | 0.22 | 0.66 | 0.15 | 0.04 | 0.12 | 0.1 |
| 8 | 68851094 | rs10107290 | -0.54 | -0.897 | -0.942 | -1.485 | -0.016 | -0.073 | -0.073 | 0.057 | 0.22 | 0.66 | 0.15 | 0.04 | 0.12 | 0.1 |
| 8 | 68851492 | rs4437632 | -0.726 | -0.613 | -1.703 | -1.519 | 0.056 | 0.187 | 0.187 | 6.88 | 0.39 | 0.83 | 0.27 | 0.04 | 0.25 | 0.19 |
| 8 | 68853609 | rs4344055 | -0.51 | -0.262 | -1.301 | -0.658 | 0.007 | -0.099 | -0.099 | 0 | 0.69 | 0.88 | 0.7 | 0.56 | 0.65 | 0.64 |
| 8 | 68857719 | rs4737864 | 0.359 | 0.435 | -0.795 | -0.489 | 0.003 | 0.009 | 0.009 | 5.793 | 0.28 | 0.28 | 0.4 | 0.51 | 0.35 | 0.43 |
| 8 | 68863901 | rs10156241 | -0.868 | 0.013 | -0.869 | -0.843 | 0.009 | -0.039 | -0.039 | 0 | 0.6 | 0.81 | 0.64 | 0.55 | 0.58 | 0.63 |
| 8 | 68870215 | rs10957404 | -0.255 | 0.069 | -0.522 | -0.816 | -0.001 | -0.001 | -0.001 | 0 | 0.45 | 0.52 | 0.49 | 0.61 | 0.39 | 0.56 |
| 8 | 68876488 | rs10091922 | 0.271 | 0.208 | -0.346 | -0.794 | 0.02 | -0.016 | -0.016 | 0.235 | 0.12 | 0.26 | 0.09 | 0.06 | 0.1 | 0.12 |
| 8 | 68885634 | rs10504412 | 0.814 | 0.131 | -0.598 | -0.842 | -0.026 | -0.109 | -0.109 | 0 | 0.29 | 0.09 | 0.37 | 0.55 | 0.26 | 0.38 |
| 8 | 68895841 | rs11776718 | 0.429 | 0.157 | -0.397 | -0.822 | 0.005 | -0.01 | -0.01 | 0.779 | 0.11 | 0.18 | 0.08 | 0.06 | 0.1 | 0.12 |
| 8 | 68898599 | rs10088446 | 0.004 | -0.083 | -0.016 | -0.813 | -0.002 | 0.004 | 0.004 | 4.53 | 0.43 | 0.41 | 0.47 | 0.61 | 0.38 | 0.57 |
| 8 | 68904277 | rs7016022 | -0.576 | -0.059 | -0.41 | -0.874 | 0.006 | -0.003 | -0.003 | 0.442 | 0.59 | 0.52 | 0.59 | 0.61 | 0.5 | 0.59 |
| 8 | 68904848 | rs6472366 | -0.393 | 0.033 | -0.839 | -0.834 | -0.003 | 0.009 | 0.009 | 6.422 | 0.57 | 0.37 | 0.57 | 0.61 | 0.51 | 0.59 |
| 8 | 68906392 | rs17436597 | 0.221 | -0.292 | -1.271 | -0.774 | -0.006 | 0.094 | 0.094 | 0 | 0.16 | 0.05 | 0.12 | 0 | 0.14 | 0.09 |
| 8 | 68909760 | rs6999098 | -0.224 | 0.037 | -0.832 | -0.138 | 0.011 | -0.024 | -0.024 | 0 | 0.62 | 0.7 | 0.64 | 0.57 | 0.62 | 0.71 |
| 8 | 68911168 | rs4471020 | -0.45 | 0.113 | -0.749 | -0.153 | -0.017 | 0.006 | 0.006 | 0.289 | 0.73 | 0.57 | 0.71 | 0.73 | 0.71 | 0.83 |
| 8 | 68916796 | rs13274993 | -0.015 | 0.16 | -1.135 | -0.119 | -0.004 | -0.001 | -0.001 | 0 | 0.11 | 0.08 | 0.09 | 0.12 | 0.13 | 0.17 |
| 8 | 68917686 | rs7013096 | -0.564 | 0.498 | -1.667 | -0.261 | -0.094 | 0.002 | 0.002 | 0.532 | 0.69 | 0.94 | 0.7 | 0.74 | 0.64 | 0.84 |
| 8 | 68921628 | rs10109326 | -0.554 | -0.097 | -1.712 | -0.654 | 0.087 | 0.134 | 0.134 | 2.791 | 0.43 | 0.84 | 0.33 | 0.19 | 0.3 | 0.43 |
| 8 | 68922270 | rs13274293 | 0.404 | -0.314 | -1.029 | -0.771 | -0.013 | 0.002 | 0.002 | 0 | 0.12 | 0.22 | 0.07 | 0.12 | 0.09 | 0.12 |
| 8 | 68927592 | rs4512367 | -0.777 | -0.364 | -1.092 | -0.793 | 0.082 | 0.125 | 0.125 | 2.815 | 0.43 | 0.83 | 0.33 | 0.19 | 0.3 | 0.45 |
| 8 | 68931680 | rs6984225 | 0.063 | -0.08 | -0.774 | -0.607 | -0.015 | -0.004 | -0.004 | 0 | 0.13 | 0.24 | 0.09 | 0.12 | 0.12 | 0.13 |
| 8 | 68933526 | rs4270945 | 0.344 | -0.38 | -0.345 | -0.834 | -0.001 | -0.01 | -0.01 | 0 | 0.19 | 0.18 | 0.18 | 0.05 | 0.16 | 0.22 |
| 8 | 68939640 | rs10957414 | -0.205 | -0.437 | -0.156 | -0.68 | -0.031 | -0.014 | -0.014 | 0 | 0.16 | 0.35 | 0.11 | 0.13 | 0.12 | 0.14 |
| 8 | 68944070 | rs4503081 | -1.287 | -0.038 | -0.75 | -0.462 | -0.118 | -0.069 | -0.069 | 0 | 0.74 | 0.97 | 0.69 | 0.72 | 0.66 | 0.85 |
| 8 | 68945041 | rs7008059 | 0.102 | 0.504 | 0.065 | -0.436 | 0.052 | 0.021 | 0.021 | 0 | 0.05 | 0.14 | 0.07 | 0.06 | 0.09 | 0.21 |
| 8 | 68947310 | rs16934023 | 0.32 | -0.027 | -0.52 | -0.584 | 0.006 | -0.006 | -0.006 | 0 | 0.14 | 0.09 | 0.11 | 0 | 0.11 | 0.1 |
| 8 | 68949670 | rs7817030 | -1.286 | 0.825 | -0.507 | -0.553 | -0.025 | 0.041 | 0.041 | 2.219 | 0.4 | 0.69 | 0.41 | 0.24 | 0.52 | 0.51 |
| 8 | 68951905 | rs920004 | -0.472 | 1.112 | -0.047 | -0.619 | 0.055 | -0.063 | -0.063 | 0 | 0.22 | 0.37 | 0.22 | 0.07 | 0.3 | 0.26 |
| 8 | 68955398 | rs6995885 | 0.007 | 1.149 | -0.03 | -0.638 | 0.131 | 0.011 | 0.011 | 0 | 0.08 | 0.19 | 0.11 | 0.06 | 0.2 | 0.19 |
| 8 | 68962961 | rs4737886 | -1.625 | 0.545 | -1.049 | -0.213 | -0.053 | -0.063 | -0.063 | 0 | 0.63 | 0.96 | 0.65 | 0.63 | 0.65 | 0.77 |
| 8 | 68964940 | rs3812454 | -1.764 | 0.915 | -0.198 | -0.388 | -0.044 | 0.085 | 0.085 | 2.538 | 0.43 | 0.83 | 0.47 | 0.27 | 0.52 | 0.52 |
| 8 | 68967380 | rs6985261 | -1.737 | 0.916 | -0.197 | -0.388 | -0.045 | 0.083 | 0.083 | 2.51 | 0.43 | 0.83 | 0.47 | 0.27 | 0.52 | 0.52 |
| 8 | 68968166 | rs1434774 | -1.753 | 0.918 | -0.491 | -0.387 | -0.073 | 0.088 | 0.088 | 2.627 | 0.43 | 0.89 | 0.48 | 0.27 | 0.52 | 0.52 |
| 8 | 68968171 | rs1434775 | -1.753 | 0.918 | -0.491 | -0.387 | -0.073 | 0.088 | 0.088 | 2.627 | 0.43 | 0.89 | 0.48 | 0.27 | 0.52 | 0.52 |
| 8 | 68969174 | rs7005292 | -0.916 | 0.893 | -0.438 | -0.403 | 0.005 | 0.016 | 0.016 | 0 | 0.3 | 0.23 | 0.32 | 0.2 | 0.32 | 0.28 |
| 8 | 68969814 | rs7009954 | -0.812 | 0.903 | -0.436 | -0.373 | 0.085 | -0.012 | -0.012 | 0 | 0.13 | 0.66 | 0.15 | 0.07 | 0.21 | 0.24 |
| 8 | 68970014 | rs6472375 | 0.676 | 0.875 | -0.47 | -0.393 | 0.001 | -0.023 | -0.023 | 0 | 0.14 | 0.13 | 0.12 | 0 | 0.11 | 0.07 |
| 8 | 68970297 | rs13262967 | 0.296 | 0.895 | -0.55 | -0.372 | -0.038 | -0.047 | -0.047 | 0.037 | 0.14 | 0.02 | 0.2 | 0.2 | 0.21 | 0.22 |
| 8 | 68972143 | rs13249271 | -0.424 | 0.93 | -0.306 | -0.234 | 0.004 | 0.016 | 0.016 | 0 | 0.3 | 0.23 | 0.32 | 0.2 | 0.32 | 0.28 |
| 8 | 68973152 | rs6472376 | -1.062 | 1.033 | -0.993 | 0.012 | -0.075 | 0.021 | 0.021 | 2.537 | 0.43 | 0.89 | 0.54 | 0.33 | 0.52 | 0.52 |
| 8 | 68974103 | rs17445951 | -0.954 | 1.648 | -0.09 | 0.387 | 0.071 | 0.027 | 0.027 | 0 | 0.09 | 0.14 | 0.17 | 0.11 | 0.2 | 0.18 |
| 8 | 68977493 | rs976343 | -0.24 | 0.832 | -0.206 | -0.085 | -0.001 | -0.003 | -0.003 | 0 | 0.71 | 0.64 | 0.71 | 0.75 | 0.71 | 0.74 |
| 8 | 68979378 | rs4442128 | 0.298 | 0.529 | -0.219 | 0.116 | 0.003 | -0.001 | -0.001 | 0 | 0.53 | 0.49 | 0.45 | 0.47 | 0.57 | 0.57 |
| 8 | 68980919 | rs7002771 | 1.379 | 0.678 | -0.456 | -0.057 | -0.018 | -0.01 | -0.01 | 0.368 | 0.29 | 0.11 | 0.21 | 0.3 | 0.31 | 0.23 |
| 8 | 68995746 | rs4592016 | 0.082 | 0.615 | -0.279 | -0.089 | 0.015 | 0.002 | 0.002 | 2.837 | 0.51 | 0.52 | 0.45 | 0.47 | 0.57 | 0.57 |
| 8 | 68998773 | rs7004495 | -0.102 | 0.646 | -0.454 | -0.063 | -0.003 | -0.008 | -0.008 | 0 | 0.41 | 0.59 | 0.51 | 0.47 | 0.4 | 0.5 |
| 8 | 68998892 | rs7004943 | 0.533 | 0.598 | -0.015 | -0.107 | -0.033 | 0.044 | 0.044 | 10.582 | 0.22 | 0.35 | 0.28 | 0.28 | 0.15 | 0.19 |
| 8 | 68998925 | rs12546214 | 1.397 | 0.554 | -0.027 | -0.097 | 0.023 | 0.002 | 0.002 | 0 | 0.19 | 0.24 | 0.23 | 0.19 | 0.26 | 0.33 |
| 8 | 69001175 | rs17368359 | 1.567 | 0.502 | -0.223 | -0.143 | -0.071 | -0.05 | -0.05 | 2.987 | 0.28 | 0.03 | 0.21 | 0.28 | 0.31 | 0.22 |
| 8 | 69002965 | rs12544121 | -0.846 | 0.846 | 0.44 | 0.039 | -0.001 | -0.009 | -0.009 | 0 | 0.43 | 0.59 | 0.51 | 0.47 | 0.4 | 0.48 |
| 8 | 69004844 | rs2196235 | 0.44 | 0.642 | 0.521 | 0.141 | 0.007 | -0.01 | -0.01 | 0 | 0.19 | 0.17 | 0.2 | 0.28 | 0.11 | 0.13 |
| 8 | 69007195 | rs13261414 | 0.786 | 0.649 | 0.542 | 0.151 | 0.008 | -0.012 | -0.012 | 0 | 0.17 | 0.16 | 0.2 | 0.28 | 0.12 | 0.16 |
| 8 | 69009268 | rs3812458 | 1.009 | 0.749 | 0.455 | 0.142 | 0.039 | -0.001 | -0.001 | 0 | 0.21 | 0.27 | 0.24 | 0.19 | 0.28 | 0.34 |
| 8 | 69010226 | rs768252 | 0.244 | 0.857 | 0.596 | -0.557 | 0.002 | 0.008 | 0.008 | 2.961 | 0.4 | 0.51 | 0.45 | 0.47 | 0.39 | 0.5 |
| 8 | 69019294 | rs7018104 | -0.186 | 0.563 | 0.538 | -0.507 | 0.003 | -0.003 | -0.003 | 0 | 0.47 | 0.61 | 0.47 | 0.47 | 0.41 | 0.53 |
| 8 | 69020496 | rs4260880 | -0.033 | 0.57 | 0.413 | -0.503 | -0.001 | -0.001 | -0.001 | 0.059 | 0.45 | 0.53 | 0.46 | 0.47 | 0.41 | 0.52 |
| 8 | 69038959 | rs967258 | -0.065 | 0.551 | 0.087 | -0.464 | 0.012 | 0.003 | 0.003 | 0.97 | 0.49 | 0.63 | 0.49 | 0.47 | 0.54 | 0.54 |
| 8 | 69041728 | rs6472384 | -0.225 | 1.771 | 0.379 | -0.314 | 0.165 | -0.003 | -0.003 | 0.112 | 0.06 | 0.26 | 0.09 | 0 | 0.13 | 0.01 |
| 8 | 69042910 | rs1434755 | 0.318 | 0.861 | -0.209 | -0.46 | 0.003 | 0.009 | 0.009 | 0.327 | 0.66 | 0.72 | 0.6 | 0.68 | 0.65 | 0.56 |
| 8 | 69051551 | rs6994582 | 0.521 | 0.89 | -0.03 | -0.944 | 0 | -0.037 | -0.037 | 0.021 | 0.58 | 0.55 | 0.59 | 0.88 | 0.5 | 0.72 |

| | | | | | | | | | | | | | | | | |
|---|---|---|---|---|---|---|---|---|---|---|---|---|---|---|---|---|
| 8 | 69053437 | rs7831336 | 0.834 | 0.622 | 0.277 | -0.598 | -0.011 | 0.042 | 0.042 | 0 | 0.13 | 0.25 | 0.23 | 0.19 | 0.12 | 0.14 |
| 8 | 69057145 | rs13251311 | -0.704 | 0.706 | -0.091 | -0.8 | -0.015 | 0.019 | 0.019 | 0.423 | 0.41 | 0.65 | 0.53 | 0.31 | 0.48 | 0.37 |
| 8 | 69057972 | rs7839608 | -0.338 | 0.693 | -0.102 | -0.812 | 0.006 | -0.02 | -0.02 | 0 | 0.29 | 0.63 | 0.33 | 0.12 | 0.37 | 0.23 |
| 8 | 69058109 | rs16934231 | 1.372 | 0.625 | -0.194 | -0.765 | -0.014 | -0.063 | -0.063 | 0 | 0.12 | 0.01 | 0.2 | 0.19 | 0.12 | 0.14 |
| 8 | 69062328 | rs7825160 | 0.204 | 0.955 | 0.322 | -0.788 | 0.031 | -0.022 | -0.022 | 0.02 | 0.21 | 0.29 | 0.26 | 0.19 | 0.29 | 0.2 |
| 8 | 69066793 | rs1434781 | -0.668 | 0.714 | 0.114 | -0.786 | 0.021 | -0.027 | -0.027 | 0 | 0.37 | 0.47 | 0.35 | 0.33 | 0.43 | 0.42 |
| 8 | 69067352 | rs4582543 | 1.726 | 0.605 | -0.212 | -0.856 | 0.021 | -0.039 | -0.039 | 0 | 0.36 | 0.23 | 0.33 | 0.48 | 0.32 | 0.41 |
| 8 | 69077685 | rs6984885 | 0.801 | 0.722 | -0.257 | -0.901 | -0.004 | -0.002 | -0.002 | 0 | 0.14 | 0.2 | 0.08 | 0.14 | 0.12 | 0.2 |
| 8 | 69077751 | rs874805 | -0.194 | 0.499 | -0.702 | -0.616 | -0.015 | 0.018 | 0.018 | 0.279 | 0.39 | 0.65 | 0.37 | 0.39 | 0.43 | 0.39 |
| 8 | 69081080 | rs1545066 | 0.761 | 0.606 | -0.658 | -0.476 | -0.031 | 0.022 | 0.022 | 0 | 0.76 | 0.95 | 0.71 | 0.88 | 0.75 | 0.83 |
| 8 | 69081683 | rs2380448 | 0.697 | 0.638 | -0.707 | -0.461 | -0.025 | 0.023 | 0.023 | 0 | 0.76 | 0.93 | 0.71 | 0.88 | 0.75 | 0.83 |
| 8 | 69081942 | rs10504420 | 0.354 | 0.658 | -0.529 | -0.164 | 0.006 | -0.009 | -0.009 | 0 | 0.11 | 0.07 | 0.21 | 0.12 | 0.11 | 0.14 |
| 8 | 69085741 | rs17449325 | -0.831 | 1.04 | 0.104 | -0.244 | 0.006 | -0.059 | -0.059 | 0 | 0.1 | 0.24 | 0.07 | 0 | 0.09 | 0.06 |
| 8 | 69100070 | rs1155491 | 0.5 | 1.059 | -0.468 | -0.098 | -0.004 | -0.003 | -0.003 | 0 | 0.86 | 0.88 | 0.91 | 1 | 0.84 | 0.96 |
| 8 | 69102348 | rs2033581 | 0.896 | 1.022 | -0.37 | 0.364 | -0.001 | -0.007 | -0.007 | 0.504 | 0.64 | 0.63 | 0.78 | 0.72 | 0.68 | 0.7 |
| 8 | 69102803 | rs13254763 | -0.433 | 1.254 | -1.298 | 0.04 | -0.038 | 0.036 | 0.036 | 0 | 0.19 | 0.04 | 0.12 | 0.1 | 0.21 | 0.17 |
| 8 | 69105897 | rs1404603 | 0.193 | 0.705 | -0.79 | 0.346 | 0.002 | 0.024 | 0.024 | 0 | 0.87 | 0.92 | 0.93 | 1 | 0.89 | 0.96 |
| 8 | 69106451 | rs13439155 | -0.302 | 1.097 | -0.493 | 1.209 | -0.017 | -0.017 | -0.017 | 0.169 | 0.06 | 0.03 | 0.11 | 0.12 | 0.16 | 0.12 |
| 8 | 69119399 | rs1019995 | -1.177 | 0.322 | 0.133 | 1.884 | -0.03 | 0.124 | 0.124 | 0 | 0.15 | 0.25 | 0.22 | 0.33 | 0.09 | 0.36 |
| 8 | 69119721 | rs9792403 | -1.261 | 0.107 | -0.151 | 2.482 | 0.018 | -0.025 | -0.025 | 0.772 | 0.15 | 0.12 | 0.22 | 0.6 | 0.1 | 0.37 |
| 8 | 69123889 | rs16934321 | -1.206 | 0.328 | 0.396 | 2.557 | 0.022 | 0.267 | 0.267 | 0 | 0.05 | 0.21 | 0.05 | 0.31 | 0.04 | 0.03 |
| 8 | 69125469 | rs7014680 | -0.604 | 0.391 | 0.644 | 1.699 | 0.012 | -0.034 | -0.034 | 0 | 0.2 | 0.52 | 0.25 | 0.12 | 0.17 | 0.21 |
| 8 | 69132033 | rs1404598 | -1.251 | 0.225 | 0.136 | 1.691 | 0.008 | -0.01 | -0.01 | 2.682 | 0.36 | 0.73 | 0.47 | 0.55 | 0.26 | 0.59 |
| 8 | 69138454 | rs10957423 | -0.26 | 0.019 | 0.184 | 0.578 | 0.009 | 0.019 | 0.019 | 0.39 | 0.42 | 0.62 | 0.5 | 0.87 | 0.34 | 0.61 |
| 8 | 69140097 | rs12549980 | 0.415 | -0.127 | 0.392 | 1.016 | 0.004 | -0.052 | -0.052 | 0 | 0.32 | 0.13 | 0.37 | 0.38 | 0.3 | 0.48 |
| 8 | 69143589 | rs2280637 | 0.389 | -0.127 | 0.708 | 1.009 | 0.005 | -0.03 | -0.03 | 0 | 0.32 | 0.19 | 0.38 | 0.39 | 0.3 | 0.48 |
| 8 | 69146175 | rs12681348 | -0.358 | -0.145 | 1.331 | 1.003 | -0.006 | -0.044 | -0.044 | 0 | 0.4 | 0.43 | 0.48 | 0.86 | 0.34 | 0.54 |
| 8 | 69147931 | rs10504423 | -0.678 | -0.131 | 1.292 | 1.066 | 0.001 | 0.248 | 0.248 | 0.307 | 0.07 | 0.24 | 0.1 | 0.47 | 0.04 | 0.06 |
| 8 | 69150976 | rs7819467 | -0.407 | -0.133 | 1.648 | 0.764 | 0.015 | 0.043 | 0.043 | 0.406 | 0.43 | 0.68 | 0.5 | 0.87 | 0.34 | 0.61 |
| 8 | 69151500 | rs10808759 | -0.977 | -0.312 | 1.094 | 0.303 | 0.004 | 0.193 | 0.193 | 0.353 | 0.55 | 0.8 | 0.61 | 0.92 | 0.44 | 0.74 |
| 8 | 69155129 | rs1997035 | 0.914 | 0.437 | 0.437 | 0.798 | -0.032 | -0.057 | -0.057 | 0 | 0.37 | 0.13 | 0.46 | 0.47 | 0.43 | 0.47 |
| 8 | 69155563 | rs9650229 | -1.532 | 0.555 | 0.154 | 0.42 | -0.013 | 0.09 | 0.09 | 2.782 | 0.74 | 0.94 | 0.78 | 0.89 | 0.75 | 0.84 |
| 8 | 69156021 | rs4737888 | 0.601 | 0.528 | 0.59 | 0.605 | 0.007 | 0.029 | 0.029 | 0 | 0.06 | 0.12 | 0.09 | 0.07 | 0.05 | 0.11 |
| 8 | 69162031 | rs1608757 | -1.298 | 0.473 | 0.851 | 0.602 | -0.027 | 0.196 | 0.196 | 0.354 | 0.18 | 0.51 | 0.16 | 0.4 | 0.12 | 0.22 |
| 8 | 69162824 | rs7844684 | 1.25 | 0.541 | 0.296 | 0.705 | -0.071 | -0.068 | -0.068 | 0 | 0.36 | 0.06 | 0.44 | 0.35 | 0.43 | 0.46 |
| 8 | 69170495 | rs4570138 | 0.194 | 0.544 | 0.808 | 0.591 | 0.005 | 0.054 | 0.054 | 0.616 | 0.51 | 0.35 | 0.46 | 0.66 | 0.56 | 0.69 |
| 8 | 69178875 | rs1404595 | -0.245 | 0.593 | 0.72 | 0.936 | -0.005 | -0.02 | -0.02 | 0.004 | 0.13 | 0.09 | 0.14 | 0.36 | 0.14 | 0.2 |
| 8 | 69180842 | rs10110416 | -0.801 | 0.628 | 0.944 | 0.894 | 0.054 | 0.05 | 0.05 | 1.838 | 0.07 | 0.32 | 0.08 | 0.06 | 0.06 | 0.1 |
| 8 | 69181681 | rs1404593 | 0.641 | 0.711 | 0.866 | 0.817 | -0.001 | -0.008 | -0.008 | 0 | 0.06 | 0.05 | 0.08 | 0 | 0.06 | 0.01 |
| 8 | 69186276 | rs12681097 | -0.083 | 0.708 | 0.831 | 0.848 | -0.004 | -0.016 | -0.016 | 0.041 | 0.13 | 0.09 | 0.13 | 0.32 | 0.14 | 0.15 |
| 8 | 69193258 | rs13271781 | 1.376 | 0.678 | 0.921 | 0.754 | -0.015 | -0.025 | -0.025 | 0.613 | 0.37 | 0.15 | 0.33 | 0.36 | 0.42 | 0.51 |
| 8 | 69193594 | rs13279518 | 0.025 | 0.677 | 1.035 | 0.446 | -0.005 | -0.031 | -0.031 | 0 | 0.56 | 0.46 | 0.53 | 0.74 | 0.61 | 0.75 |
| 8 | 69194301 | rs2380454 | -0.721 | 0.649 | 1.221 | 0.454 | 0.019 | 0.086 | 0.086 | 0.551 | 0.64 | 0.79 | 0.64 | 0.84 | 0.67 | 0.77 |
| 8 | 69195836 | rs1949847 | 0.164 | 0.668 | 1.034 | 0.468 | 0.014 | 0.009 | 0.009 | 0 | 0.07 | 0.18 | 0.1 | 0.11 | 0.06 | 0.02 |
| 8 | 69196430 | rs12542353 | -0.558 | 0.279 | 1.013 | 0.413 | 0 | -0.017 | -0.017 | 2.063 | 0.63 | 0.64 | 0.63 | 0.84 | 0.67 | 0.77 |
| 8 | 69199198 | rs11998295 | 0.19 | 0.232 | 0.878 | 0.595 | -0.009 | 0.043 | 0.043 | 0.136 | 0.06 | 0.22 | 0.09 | 0.11 | 0.03 | 0.02 |
| 8 | 69199953 | rs6472394 | -0.059 | 0.283 | 0.894 | 0.333 | -0.012 | -0.035 | -0.035 | 0 | 0.57 | 0.46 | 0.54 | 0.73 | 0.64 | 0.75 |
| 8 | 69200163 | rs12674787 | 0.451 | 0.339 | 0.745 | 0.333 | 0.027 | 0.089 | 0.089 | 0.524 | 0.51 | 0.24 | 0.47 | 0.67 | 0.58 | 0.66 |
| 8 | 69201490 | rs7827778 | -0.593 | 0.349 | 0.643 | 0.231 | 0.004 | 0.009 | 0.009 | 0.509 | 0.64 | 0.68 | 0.63 | 0.84 | 0.66 | 0.77 |
| 8 | 69203721 | rs4737256 | -1.616 | 0.377 | 0.457 | 0.571 | 0 | 0.095 | 0.095 | 0 | 0.26 | 0.53 | 0.29 | 0.46 | 0.23 | 0.26 |
| 8 | 69204305 | rs7820659 | -1.616 | 0.377 | 0.457 | 0.571 | 0 | 0.095 | 0.095 | 0 | 0.26 | 0.54 | 0.29 | 0.46 | 0.23 | 0.26 |
| 8 | 69214029 | rs1581542 | 0.08 | 0.302 | 0.147 | 0.464 | 0.005 | -0.01 | -0.01 | 0 | 0.53 | 0.72 | 0.47 | 0.4 | 0.55 | 0.58 |
| 8 | 69221116 | rs2676643 | 0.05 | -0.389 | -0.408 | 0.447 | -0.05 | -0.008 | -0.008 | 0 | 0.52 | 0.91 | 0.52 | 0.51 | 0.52 | 0.49 |
| 8 | 69221210 | rs13251356 | 1.294 | -0.548 | -0.317 | 0.113 | -0.09 | -0.062 | -0.062 | 0 | 0.34 | 0.09 | 0.34 | 0.37 | 0.44 | 0.44 |
| 8 | 69229128 | rs2692027 | -0.076 | -0.539 | -0.408 | 0.099 | -0.049 | -0.047 | -0.047 | 0 | 0.52 | 0.91 | 0.52 | 0.48 | 0.52 | 0.49 |
| 8 | 69236839 | rs814439 | 0.028 | -0.448 | -0.036 | 0.1 | -0.011 | -0.04 | -0.04 | 0.449 | 0.08 | 0.2 | 0.06 | 0 | 0.07 | 0.03 |
| 8 | 69242537 | rs868546 | 1.171 | -0.397 | -0.32 | 0.179 | -0.122 | -0.088 | -0.088 | 0.387 | 0.34 | 0.07 | 0.34 | 0.37 | 0.44 | 0.44 |
| 8 | 69252417 | rs2138677 | 1.006 | -0.259 | 0.062 | 0.252 | -0.02 | -0.009 | -0.009 | 0.366 | 0.37 | 0.31 | 0.39 | 0.37 | 0.45 | 0.44 |
| 8 | 69271567 | rs1880535 | -0.029 | -0.622 | -0.324 | 0.112 | -0.028 | -0.049 | -0.049 | 0.221 | 0.57 | 0.87 | 0.54 | 0.48 | 0.56 | 0.55 |
| 8 | 69273453 | rs11777852 | 1.217 | -0.804 | -0.581 | 0.1 | -0.047 | 0.009 | 0.009 | 0.041 | 0.26 | 0.04 | 0.34 | 0.24 | 0.3 | 0.24 |
| 8 | 69280367 | rs814462 | -0.398 | -0.799 | -0.523 | -0.068 | -0.051 | -0.086 | -0.086 | 0.012 | 0.66 | 0.89 | 0.57 | 0.48 | 0.59 | 0.58 |
| 8 | 69295584 | rs7838935 | 1.518 | -1.116 | -0.032 | -0.051 | -0.013 | -0.006 | -0.006 | 0.678 | 0.39 | 0.34 | 0.41 | 0.39 | 0.48 | 0.44 |
| 8 | 69297184 | rs7845014 | -0.086 | -1.003 | -0.055 | -0.058 | -0.005 | -0.035 | -0.035 | 0.253 | 0.08 | 0.13 | 0.07 | 0 | 0.06 | 0.06 |
| 8 | 69298788 | rs2591019 | 0.544 | -1.089 | 0.188 | -0.074 | -0.003 | -0.002 | -0.002 | 0 | 0.12 | 0.15 | 0.09 | 0.1 | 0.1 | 0.1 |
| 8 | 69300002 | rs2676652 | -0.976 | -1.098 | 0.209 | -0.086 | -0.012 | -0.025 | -0.025 | 0.121 | 0.2 | 0.29 | 0.17 | 0.1 | 0.15 | 0.16 |
| 8 | 69305180 | rs1517124 | 0.31 | -1.129 | 0.39 | -0.006 | 0.007 | -0.001 | -0.001 | 0 | 0.57 | 0.62 | 0.52 | 0.37 | 0.58 | 0.53 |
| 8 | 69306811 | rs2676658 | -0.769 | -1.025 | 0.408 | 0.086 | -0.001 | -0.031 | -0.031 | 0 | 0.7 | 0.78 | 0.61 | 0.47 | 0.67 | 0.63 |
| 8 | 69307133 | rs16934570 | 0.211 | -1.153 | 0.437 | -0.057 | -0.014 | -0.061 | -0.061 | 0 | 0.11 | 0.17 | 0.07 | 0 | 0.07 | 0.07 |
| 8 | 69313151 | rs2591011 | 0.593 | -1.065 | 0.522 | 0.087 | 0.028 | 0.009 | 0.009 | 0.337 | 0.52 | 0.38 | 0.49 | 0.39 | 0.6 | 0.55 |

| | | | | | | | | | | | | | | | | |
|---|---|---|---|---|---|---|---|---|---|---|---|---|---|---|---|---|
| 8 | 69317537 | rs1867633 | -0.422 | -1.005 | 0.634 | 0.071 | -0.006 | 0.05 | 0.05 | 0.028 | 0.08 | 0.3 | 0.07 | 0.1 | 0.02 | 0.02 |
| 8 | 69322475 | rs7013706 | -0.033 | -0.893 | 0.397 | -0.003 | 0.004 | -0.023 | -0.023 | 0 | 0.59 | 0.72 | 0.5 | 0.47 | 0.55 | 0.57 |
| 8 | 69325848 | rs2591025 | -0.634 | -0.819 | 0.194 | -0.052 | -0.038 | -0.123 | -0.123 | 7.4 | 0.19 | 0.41 | 0.16 | 0 | 0.12 | 0.11 |
| 8 | 69332645 | rs2138675 | -0.137 | -0.551 | -0.266 | 0.507 | -0.017 | -0.037 | -0.037 | 0 | 0.93 | 0.99 | 0.92 | 0.77 | 0.87 | 0.97 |
| 8 | 69346829 | rs10104993 | 1.177 | -0.642 | 0.011 | 0.369 | -0.007 | -0.002 | -0.002 | 0.134 | 0.38 | 0.33 | 0.34 | 0.35 | 0.42 | 0.48 |
| 8 | 69347872 | rs2591002 | -0.971 | -0.663 | 0.316 | 0.323 | -0.001 | 0 | 0 | 1.725 | 0.56 | 0.62 | 0.62 | 0.4 | 0.51 | 0.48 |
| 8 | 69348019 | rs2890426 | 0.274 | -1.021 | 0.295 | 0.153 | 0.007 | 0.024 | 0.024 | 0.212 | 0.35 | 0.28 | 0.47 | 0.3 | 0.36 | 0.36 |
| 8 | 69348758 | rs16934606 | 0.417 | -1.114 | 0.334 | 0.024 | -0.024 | -0.049 | -0.049 | 0 | 0.1 | 0.18 | 0.04 | 0 | 0.04 | 0.03 |
| 8 | 69349596 | rs3812450 | -0.413 | -1.196 | 0.212 | -0.023 | -0.001 | 0 | 0 | 0 | 0.1 | 0.16 | 0.12 | 0.11 | 0.11 | 0.02 |
| 8 | 69350332 | rs16934610 | 0.419 | -1.104 | 0.209 | -0.009 | -0.016 | -0.037 | -0.037 | 0 | 0.1 | 0.15 | 0.04 | 0 | 0.04 | 0.03 |
| 8 | 69350425 | rs2591003 | 0.193 | -0.909 | 0.383 | 0.193 | 0.001 | -0.001 | -0.001 | 0 | 0.56 | 0.42 | 0.54 | 0.46 | 0.64 | 0.6 |
| 8 | 69354328 | rs2676637 | -1.231 | -0.676 | 0.055 | -0.125 | -0.02 | -0.004 | -0.004 | 0 | 0.59 | 0.74 | 0.61 | 0.42 | 0.48 | 0.52 |
| 8 | 69357500 | rs1517116 | 0.113 | -1.076 | 0.185 | -0.692 | 0.031 | 0.077 | 0.077 | 0.333 | 0.55 | 0.74 | 0.64 | 0.69 | 0.64 | 0.58 |
| 8 | 69358538 | rs16934632 | -0.007 | -1.045 | 0.268 | -0.781 | 0 | -0.016 | -0.016 | 0 | 0.08 | 0.16 | 0.14 | 0 | 0.12 | 0.03 |
| 8 | 69360144 | rs11786594 | 1.225 | -1.235 | -0.041 | -0.89 | -0.021 | 0.073 | 0.073 | 1.81 | 0.23 | 0.03 | 0.26 | 0.13 | 0.22 | 0.22 |
| 8 | 69366270 | rs1113405 | 1.428 | -1.116 | -0.102 | -0.709 | -0.038 | 0.017 | 0.017 | 0 | 0.4 | 0.22 | 0.42 | 0.58 | 0.47 | 0.52 |
| 8 | 69371911 | rs4737905 | 0.466 | -1.265 | -0.346 | -0.666 | 0.016 | 0.032 | 0.032 | 0.399 | 0.55 | 0.65 | 0.62 | 0.68 | 0.64 | 0.59 |
| 8 | 69380609 | rs2676638 | 0.298 | -1.188 | -0.772 | -0.674 | 0.013 | 0.009 | 0.009 | 0 | 0.11 | 0.05 | 0.05 | 0 | 0.09 | 0.06 |
| 8 | 69384598 | rs1517113 | 0.354 | -1.06 | -0.701 | -0.581 | 0.025 | 0.059 | 0.059 | 0.155 | 0.56 | 0.73 | 0.63 | 0.7 | 0.64 | 0.57 |
| 8 | 69398810 | rs4466385 | -0.588 | -0.851 | -0.219 | -0.918 | -0.005 | -0.067 | -0.067 | 0.003 | 0.46 | 0.43 | 0.34 | 0.12 | 0.36 | 0.31 |
| 8 | 69401509 | rs6472399 | -1.063 | -1.061 | -0.063 | -0.456 | 0.015 | 0.025 | 0.025 | 0 | 0.53 | 0.61 | 0.39 | 0.36 | 0.4 | 0.48 |
| 8 | 69406281 | rs10096415 | -1.21 | -1.02 | -0.084 | -0.378 | -0.004 | 0.012 | 0.012 | 0 | 0.57 | 0.75 | 0.46 | 0.43 | 0.43 | 0.48 |
| 8 | 69408783 | rs4588839 | -0.632 | -0.755 | -0.242 | -0.915 | 0.001 | -0.024 | -0.024 | 0 | 0.47 | 0.45 | 0.38 | 0.19 | 0.37 | 0.31 |
| 8 | 69410725 | rs4644227 | -0.388 | -0.196 | -1.022 | -0.786 | -0.055 | 0.202 | 0.202 | 0 | 0.41 | 0.06 | 0.35 | 0.17 | 0.42 | 0.31 |
| 8 | 69411716 | rs4737262 | -0.54 | -0.082 | -0.724 | -0.714 | -0.02 | 0.109 | 0.109 | 0 | 0.4 | 0.2 | 0.37 | 0.18 | 0.42 | 0.3 |
| 8 | 69419194 | rs7838662 | -0.975 | 0.303 | -0.388 | -0.24 | 0.004 | -0.003 | -0.003 | 0 | 0.14 | 0.13 | 0.1 | 0.17 | 0.12 | 0.12 |
| 8 | 69421515 | rs6986715 | -1.048 | 0.235 | 0.342 | -0.273 | -0.011 | 0.011 | 0.011 | 0 | 0.15 | 0.41 | 0.13 | 0.17 | 0.12 | 0.12 |
| 8 | 69428822 | rs7826853 | -1.189 | -0.177 | 0.822 | -0.554 | -0.047 | -0.02 | -0.02 | 1.547 | 0.21 | 0.63 | 0.16 | 0.17 | 0.11 | 0.16 |
| 8 | 69430368 | rs4415295 | -0.207 | -0.002 | 0.236 | -0.731 | -0.074 | 0.033 | 0.033 | 0.103 | 0.27 | 0.07 | 0.34 | 0.26 | 0.4 | 0.24 |
| 8 | 69434195 | rs2289831 | -0.679 | -0.282 | 1.051 | -0.696 | -0.031 | -0.049 | -0.049 | 0 | 0.2 | 0.41 | 0.15 | 0.11 | 0.14 | 0.15 |
| 8 | 69435996 | rs13271303 | -1.672 | 0.125 | 1.056 | -0.427 | 0 | -0.002 | -0.002 | 0.159 | 0.49 | 0.52 | 0.48 | 0.42 | 0.51 | 0.4 |
| 8 | 69444341 | rs11986007 | -1.757 | 0.232 | 0.988 | -0.458 | 0.007 | 0.066 | 0.066 | 0 | 0.5 | 0.74 | 0.5 | 0.41 | 0.52 | 0.4 |
| 8 | 69446828 | rs7819503 | -1.683 | 0.191 | 1.17 | -0.478 | 0.004 | 0.082 | 0.082 | 0 | 0.5 | 0.79 | 0.5 | 0.41 | 0.52 | 0.4 |
| 8 | 69451670 | rs2380478 | -0.689 | 0.146 | 1.207 | -0.53 | -0.017 | -0.03 | -0.03 | 0.492 | 0.2 | 0.32 | 0.13 | 0.11 | 0.14 | 0.15 |
| 8 | 69457045 | rs4321986 | 1.081 | 0.46 | 0.847 | -0.644 | -0.015 | 0.031 | 0.031 | 0 | 0.13 | 0.03 | 0.14 | 0 | 0.17 | 0.07 |
| 8 | 69460337 | rs7826286 | 0.442 | 0.063 | 1.046 | -0.582 | -0.009 | -0.001 | -0.001 | 0 | 0.09 | 0.24 | 0.09 | 0.11 | 0.06 | 0.09 |
| 8 | 69468914 | rs10108264 | -0.324 | 0.146 | 0.665 | -0.552 | -0.056 | -0.001 | -0.001 | 0 | 0.28 | 0.09 | 0.33 | 0.29 | 0.37 | 0.25 |
| 8 | 69482825 | rs16934715 | -0.774 | 0.172 | 0.964 | -0.481 | -0.032 | -0.058 | -0.058 | 0 | 0.21 | 0.61 | 0.16 | 0.12 | 0.15 | 0.15 |
| 8 | 69484133 | rs1897905 | -0.15 | 0.189 | 0.577 | -0.476 | -0.041 | -0.002 | -0.002 | 0 | 0.27 | 0.12 | 0.33 | 0.29 | 0.36 | 0.24 |
| 8 | 69497650 | rs1434940 | -0.917 | -0.201 | 1.005 | -0.536 | -0.041 | -0.068 | -0.068 | 0.099 | 0.21 | 0.59 | 0.16 | 0.12 | 0.15 | 0.15 |
| 8 | 69503062 | rs13273576 | -0.827 | 0.164 | 0.807 | -0.401 | -0.011 | -0.005 | -0.005 | 0.016 | 0.33 | 0.31 | 0.48 | 0.4 | 0.51 | 0.32 |
| 8 | 69503895 | rs10504427 | 1.104 | 0.135 | 0.794 | -0.205 | 0.001 | -0.006 | -0.006 | 0 | 0.07 | 0.06 | 0.11 | 0.15 | 0.08 | 0.05 |
| 8 | 69509456 | rs4268113 | -1.464 | -0.051 | 1.088 | -0.327 | 0.014 | 0.012 | 0.012 | 0 | 0.49 | 0.61 | 0.54 | 0.44 | 0.57 | 0.41 |
| 8 | 69510360 | rs1473266 | 0.229 | -0.131 | 1.366 | -0.287 | -0.005 | 0.025 | 0.025 | 0 | 0.09 | 0.28 | 0.14 | 0.11 | 0.08 | 0.08 |
| 8 | 69513803 | rs1369231 | -0.73 | -0.17 | 1.186 | -0.345 | -0.045 | 0.013 | 0.013 | 0 | 0.37 | 0.23 | 0.4 | 0.32 | 0.48 | 0.32 |
| 8 | 69516939 | rs6472409 | -0.992 | -0.853 | 0.954 | -0.877 | -0.035 | -0.033 | -0.033 | 0 | 0.27 | 0.6 | 0.21 | 0.12 | 0.15 | 0.17 |
| 8 | 69520538 | rs7845972 | -0.998 | -0.574 | 0.825 | -0.963 | -0.022 | -0.038 | -0.038 | 0 | 0.28 | 0.6 | 0.25 | 0.11 | 0.21 | 0.17 |
| 8 | 69520815 | rs2380483 | -1.707 | -0.514 | 0.816 | -0.753 | 0.031 | -0.014 | -0.014 | 0 | 0.56 | 0.76 | 0.57 | 0.43 | 0.64 | 0.43 |
| 8 | 69532253 | rs2060440 | -0.332 | -0.483 | 1.138 | -0.755 | -0.02 | -0.032 | -0.032 | 0 | 0.18 | 0.4 | 0.21 | 0.11 | 0.18 | 0.14 |
| 8 | 69534542 | rs4313142 | -1.781 | -0.618 | 0.784 | -1.107 | 0.025 | -0.019 | -0.019 | 0.022 | 0.59 | 0.72 | 0.62 | 0.44 | 0.65 | 0.43 |
| 8 | 69539842 | rs1434927 | 0.469 | -0.479 | 0.663 | -0.997 | -0.11 | -0.055 | -0.055 | 0 | 0.26 | 0.08 | 0.28 | 0.31 | 0.43 | 0.26 |
| 8 | 69541218 | rs920016 | 0.983 | -0.504 | 1.051 | -0.927 | 0.005 | 0.006 | 0.006 | 0 | 0.09 | 0.16 | 0.24 | 0.12 | 0.14 | 0.08 |
| 8 | 69551547 | rs984878 | 1.07 | -0.579 | 0.997 | -0.888 | -0.024 | -0.044 | -0.044 | 0.369 | 0.08 | 0.15 | 0.03 | 0 | 0.04 | 0.05 |
| 8 | 69557825 | rs11780358 | 1.811 | -1.07 | 0.203 | -1.316 | -0.006 | 0.062 | 0.062 | 0 | 0.16 | 0.21 | 0.23 | 0.55 | 0.16 | 0.39 |
| 8 | 69559835 | rs10504430 | 1.815 | -1.232 | -0.041 | -1.579 | 0.011 | 0.038 | 0.038 | 0 | 0.21 | 0.1 | 0.15 | 0 | 0.16 | 0.11 |
| 8 | 69560298 | rs2164197 | 0.167 | -1.287 | 0.139 | -1.601 | 0.039 | 0.042 | 0.042 | 0 | 0.5 | 0.36 | 0.45 | 0.31 | 0.59 | 0.38 |
| 8 | 69562029 | rs16934837 | 1.93 | -0.85 | 0.58 | -1.076 | 0 | 0.021 | 0.021 | 0 | 0.15 | 0.15 | 0.21 | 0.51 | 0.14 | 0.35 |
| 8 | 69563695 | rs7825271 | -1.715 | -0.94 | 0.439 | -1.553 | 0.004 | -0.002 | -0.002 | 0 | 0.87 | 0.79 | 0.77 | 0.87 | 0.85 | 0.85 |
| 8 | 69568102 | rs6987972 | 1.06 | -0.969 | 0.783 | -1.41 | -0.029 | -0.03 | -0.03 | 0.65 | 0.09 | 0.16 | 0.03 | 0.02 | 0.04 | 0.05 |
| 8 | 69569483 | rs961465 | 0.232 | -1.125 | 0.709 | -1.494 | -0.025 | 0.039 | 0.039 | 0 | 0.3 | 0.4 | 0.31 | 0.56 | 0.23 | 0.42 |
| 8 | 69570332 | rs17387279 | 1.614 | -1.08 | 0.244 | -1.539 | -0.028 | 0.108 | 0.108 | 0 | 0.18 | 0.01 | 0.13 | 0 | 0.15 | 0.11 |
| 8 | 69574272 | rs1865443 | -0.877 | -0.947 | 0.406 | -1.436 | 0.007 | -0.006 | -0.006 | 0 | 0.58 | 0.7 | 0.49 | 0.58 | 0.43 | 0.58 |
| 8 | 69576337 | rs2196255 | 1.478 | -0.755 | 0.358 | -1.2 | -0.012 | 0.071 | 0.071 | 0 | 0.16 | 0.19 | 0.24 | 0.56 | 0.15 | 0.37 |
| 8 | 69577442 | rs2164198 | 0.199 | -0.713 | 0.379 | -1.53 | -0.01 | -0.075 | -0.075 | 0.586 | 0.28 | 0.28 | 0.18 | 0 | 0.19 | 0.17 |
| 8 | 69578637 | rs16919100 | 1.197 | -0.672 | 0.265 | -1.529 | 0.004 | 0.001 | 0.001 | 0 | 0.19 | 0.12 | 0.15 | 0 | 0.16 | 0.12 |
| 8 | 69580675 | rs763845 | 1.414 | -0.706 | 0.064 | -1.618 | 0.003 | -0.011 | -0.011 | 3.41 | 0.13 | 0.13 | 0.06 | 0.01 | 0.08 | 0.05 |
| 8 | 69580951 | rs12681467 | 1.166 | -0.671 | 0.035 | -1.559 | -0.008 | 0.033 | 0.033 | 0 | 0.2 | 0.22 | 0.26 | 0.57 | 0.18 | 0.35 |
| 8 | 69581985 | rs1434937 | -0.383 | -0.575 | -0.138 | -1.495 | 0.021 | 0.007 | 0.007 | 0 | 0.48 | 0.51 | 0.44 | 0.57 | 0.37 | 0.52 |
| 8 | 69586034 | rs1116986 | -0.603 | -0.196 | -0.253 | -1.542 | 0.01 | 0.008 | 0.008 | 0 | 0.52 | 0.52 | 0.45 | 0.57 | 0.42 | 0.54 |

| | | | | | | | | | | | | | | | | |
|---|---|---|---|---|---|---|---|---|---|---|---|---|---|---|---|---|
| 8 | 69611629 | rs7016101 | 0.141 | -0.416 | -0.116 | -1.414 | -0.042 | 0.05 | 0.05 | 0 | 0.3 | 0.49 | 0.3 | 0.54 | 0.21 | 0.41 |
| 8 | 69613880 | rs7836513 | -0.781 | -0.221 | -0.146 | -1.375 | -0.005 | -0.016 | -0.016 | 0 | 0.61 | 0.69 | 0.49 | 0.55 | 0.44 | 0.59 |
| 8 | 69620696 | rs9987198 | 0.619 | -0.802 | 0.217 | -1.263 | -0.028 | -0.055 | -0.055 | 7.348 | 0.1 | 0.27 | 0.04 | 0 | 0.04 | 0.05 |
| 8 | 69623368 | rs1821567 | -1.402 | 0.086 | -0.346 | -0.977 | 0 | -0.008 | -0.008 | 0 | 0.73 | 0.76 | 0.62 | 0.67 | 0.68 | 0.73 |
| 8 | 69629087 | rs16919108 | 0.725 | 0.129 | -0.735 | -0.901 | 0.023 | 0.039 | 0.039 | 1.82 | 0.14 | 0.08 | 0.07 | 0.01 | 0.1 | 0.07 |
| 8 | 69642205 | rs6996725 | -0.345 | 0.528 | -0.248 | -0.384 | 0 | -0.035 | -0.035 | 0 | 0.34 | 0.41 | 0.21 | 0.23 | 0.27 | 0.21 |
| 8 | 69642540 | rs16934967 | 0.094 | -0.016 | -0.408 | -0.745 | -0.013 | -0.087 | -0.087 | 1.12 | 0.28 | 0.33 | 0.15 | 0 | 0.17 | 0.17 |
| 8 | 69647914 | rs4737915 | -0.042 | 0.056 | -0.833 | -0.151 | -0.007 | -0.025 | -0.025 | 0 | 0.63 | 0.71 | 0.61 | 0.52 | 0.6 | 0.49 |
| 8 | 69665550 | rs9886368 | -0.502 | -0.679 | -0.16 | -0.475 | -0.031 | -0.121 | -0.121 | 0 | 0.22 | 0.37 | 0.11 | 0 | 0.13 | 0.16 |
| 8 | 69666889 | rs7838605 | 0.368 | -0.513 | -0.428 | -0.347 | 0 | -0.015 | -0.015 | 0.121 | 0.1 | 0.1 | 0.05 | 0 | 0.06 | 0.09 |
| 8 | 69675453 | rs6990164 | -0.545 | -1.121 | -0.259 | -0.946 | -0.021 | -0.063 | -0.063 | 0 | 0.37 | 0.53 | 0.2 | 0.11 | 0.23 | 0.27 |
| 8 | 69676946 | rs7845019 | -0.814 | -1.448 | -0.417 | -0.013 | 0.02 | 0.006 | 0.006 | 0 | 0.45 | 0.64 | 0.31 | 0.59 | 0.28 | 0.47 |
| 8 | 69678500 | rs11987041 | -0.244 | -1.453 | -0.433 | -0.021 | -0.017 | -0.055 | -0.055 | 0 | 0.36 | 0.41 | 0.18 | 0.11 | 0.23 | 0.27 |
| 8 | 69681165 | rs6472420 | -1.328 | -1.123 | -1.253 | -0.066 | -0.006 | 0.008 | 0.008 | 0 | 0.57 | 0.77 | 0.39 | 0.59 | 0.42 | 0.57 |
| 8 | 69682388 | rs4278138 | 0.56 | -1.183 | -1.51 | -0.346 | 0.005 | 0.068 | 0.068 | 1.033 | 0.17 | 0.05 | 0.11 | 0.06 | 0.12 | 0.11 |
| 8 | 69687852 | rs4374973 | -0.731 | -1.003 | -1.429 | -0.007 | 0.019 | 0.151 | 0.151 | 1.116 | 0.54 | 0.83 | 0.32 | 0.18 | 0.41 | 0.33 |
| 8 | 69688844 | rs7833861 | -1.049 | -1.183 | -1.449 | 0.587 | -0.031 | -0.003 | -0.003 | 0.065 | 0.6 | 0.85 | 0.39 | 0.49 | 0.44 | 0.49 |
| 8 | 69691896 | rs16935038 | 0.78 | -1.198 | -1.432 | 0.752 | 0.004 | 0.012 | 0.012 | 0 | 0.07 | 0.02 | 0.08 | 0.29 | 0.05 | 0.16 |
| 8 | 69693543 | rs2953950 | -0.75 | -1.173 | -1.285 | 0.08 | 0.064 | 0.138 | 0.138 | 0.935 | 0.53 | 0.82 | 0.31 | 0.21 | 0.37 | 0.33 |
| 8 | 69699756 | rs16935065 | -0.284 | -1.084 | -0.285 | 0.165 | -0.012 | -0.016 | -0.016 | 0 | 0.21 | 0.28 | 0.1 | 0.12 | 0.15 | 0.13 |
| 8 | 69713852 | rs2953960 | -1.699 | -1.641 | -0.503 | 0.125 | 0.067 | 0.01 | 0.01 | 0 | 0.52 | 0.84 | 0.32 | 0.49 | 0.37 | 0.46 |
| 8 | 69715381 | rs10957452 | -1.007 | -1.606 | 0.025 | 0.131 | -0.02 | 0.021 | 0.021 | 0 | 0.31 | 0.34 | 0.18 | 0.47 | 0.19 | 0.34 |
| 8 | 69721955 | rs6982162 | 0.438 | -1.587 | 0.602 | 0.072 | -0.015 | -0.071 | -0.071 | 0 | 0.17 | 0.21 | 0.07 | 0 | 0.11 | 0.08 |
| 8 | 69722223 | rs2958321 | -1.728 | -1.692 | 0.503 | 0.066 | 0.019 | 0.003 | 0.003 | 0 | 0.46 | 0.54 | 0.28 | 0.47 | 0.33 | 0.45 |
| 8 | 69723939 | rs2958322 | -0.901 | -1.329 | 0.551 | -0.381 | -0.015 | -0.118 | -0.118 | 0.761 | 0.3 | 0.34 | 0.15 | 0 | 0.25 | 0.17 |
| 8 | 69726331 | rs16935095 | -0.69 | -1.109 | 0.594 | 0.517 | 0.001 | 0.008 | 0.008 | 0.065 | 0.14 | 0.12 | 0.14 | 0.43 | 0.13 | 0.29 |
| 8 | 69728484 | rs16935101 | -2.232 | -1.321 | 1.164 | 0.278 | -0.016 | 0.037 | 0.037 | 0.917 | 0.3 | 0.53 | 0.26 | 0.44 | 0.28 | 0.37 |
| 8 | 69729569 | rs2958326 | -1.574 | -0.525 | 1.022 | 0.525 | -0.024 | -0.025 | -0.025 | 0 | 0.89 | 0.95 | 0.81 | 0.74 | 0.83 | 0.79 |
| 8 | 69732058 | rs7837210 | 0.745 | -0.53 | 1.111 | 0.503 | -0.022 | 0.039 | 0.039 | 0 | 0.07 | 0.02 | 0.07 | 0 | 0.11 | 0.07 |
| 8 | 69732887 | rs10093275 | 0.106 | -0.681 | 1.305 | 0.479 | 0.009 | -0.005 | -0.005 | 0 | 0.79 | 0.72 | 0.72 | 0.74 | 0.72 | 0.72 |
| 8 | 69735094 | rs2958330 | 0.44 | -0.647 | 1.332 | 0.496 | 0.013 | -0.014 | -0.014 | 0 | 0.77 | 0.62 | 0.71 | 0.73 | 0.71 | 0.71 |
| 8 | 69742300 | rs9643598 | -0.598 | -0.685 | 1.051 | 0.538 | -0.005 | 0.084 | 0.084 | 0.049 | 0.15 | 0.21 | 0.13 | 0.43 | 0.13 | 0.28 |
| 8 | 69745066 | rs7010802 | -1.487 | -0.665 | 0.524 | 0.536 | 0.001 | -0.002 | -0.002 | 0 | 0.27 | 0.68 | 0.24 | 0.44 | 0.2 | 0.36 |
| 8 | 69747974 | rs2953968 | -0.217 | -0.632 | 0.387 | 0.496 | 0.018 | 0.001 | 0.001 | 0 | 0.81 | 0.75 | 0.72 | 0.74 | 0.72 | 0.71 |
| 8 | 69750527 | rs4737920 | -0.548 | -1.503 | 0.22 | 0.785 | 0.002 | 0.011 | 0.011 | 0.766 | 0.11 | 0.09 | 0.11 | 0.44 | 0.1 | 0.27 |
| 8 | 69751572 | rs12550697 | -0.407 | -1.561 | 0.01 | 0.151 | 0.014 | -0.035 | -0.035 | 0 | 0.65 | 0.47 | 0.48 | 0.72 | 0.48 | 0.58 |
| 8 | 69752193 | rs13280929 | 0.842 | -1.904 | -0.604 | -0.041 | 0.08 | -0.007 | -0.007 | 0.253 | 0.4 | 0.16 | 0.3 | 0.28 | 0.26 | 0.29 |
| 8 | 69755610 | rs10086214 | 0.444 | -1.582 | -0.116 | -0.271 | 0.059 | 0.082 | 0.082 | 1.063 | 0.5 | 0.31 | 0.41 | 0.7 | 0.36 | 0.57 |
| 8 | 69757635 | rs6472423 | -0.656 | -1.5 | 0.083 | -0.016 | -0.087 | 0.046 | 0.046 | 0.923 | 0.68 | 0.92 | 0.51 | 0.76 | 0.45 | 0.59 |
| 8 | 69764515 | rs4145295 | 0.176 | -1.618 | 0.29 | -0.282 | 0.004 | 0.028 | 0.028 | 1.199 | 0.58 | 0.29 | 0.45 | 0.74 | 0.43 | 0.57 |
| 8 | 69765125 | rs7827405 | -1.685 | -1.142 | 1.299 | -0.055 | -0.013 | 0.034 | 0.034 | 2.707 | 0.09 | 0.52 | 0.04 | 0.04 | 0.01 | 0.01 |
| 8 | 69776152 | rs10104452 | -0.804 | -1.259 | 1.044 | -0.318 | -0.044 | -0.105 | -0.105 | 0 | 0.17 | 0.49 | 0.1 | 0.01 | 0.09 | 0.08 |
| 8 | 69784983 | rs7838608 | -1.136 | -1.293 | 0.516 | -0.331 | -0.009 | -0.031 | -0.031 | 0 | 0.2 | 0.71 | 0.13 | 0.02 | 0.1 | 0.09 |
| 8 | 69799082 | rs10091347 | -1.003 | -1.215 | 0.474 | -0.31 | -0.04 | -0.085 | -0.085 | 1.218 | 0.18 | 0.54 | 0.11 | 0.01 | 0.09 | 0.08 |
| 8 | 69837546 | rs13259685 | -0.453 | -0.974 | -0.346 | 0.038 | 0.006 | 0.001 | 0.001 | 0.011 | 0.32 | 0.25 | 0.28 | 0.3 | 0.26 | 0.29 |
| 8 | 69838267 | rs6472429 | -0.821 | -1.018 | -1.156 | -0.245 | -0.138 | 0.001 | 0.001 | 5.831 | 0.65 | 0.91 | 0.56 | 0.73 | 0.49 | 0.64 |
| 8 | 69840033 | rs9643401 | -1.221 | -1.581 | -0.242 | -0.086 | 0.026 | 0.003 | 0.003 | 0 | 0.51 | 0.63 | 0.39 | 0.47 | 0.35 | 0.4 |
| 8 | 69843576 | rs7463208 | -0.332 | -1.515 | -0.839 | -0.373 | -0.001 | -0.062 | -0.062 | 1.364 | 0.31 | 0.29 | 0.17 | 0 | 0.24 | 0.15 |
| 8 | 69850612 | rs4737926 | -1.354 | -1.482 | -1.116 | -0.202 | 0.006 | 0 | 0 | 0 | 0.5 | 0.55 | 0.39 | 0.52 | 0.35 | 0.4 |
| 8 | 69852661 | rs5009754 | -0.53 | -1.193 | -1.801 | -0.143 | -0.005 | -0.055 | -0.055 | 0 | 0.33 | 0.36 | 0.18 | 0.03 | 0.24 | 0.17 |
| 8 | 69855507 | rs7001339 | 0.381 | -1.252 | -1.866 | -0.134 | -0.03 | -0.071 | -0.071 | 0 | 0.18 | 0.33 | 0.1 | 0.03 | 0.1 | 0.07 |
| 8 | 69856252 | rs4737263 | -0.25 | -0.671 | -2.406 | 0.649 | -0.01 | -0.086 | -0.086 | 0 | 0.14 | 0.04 | 0.18 | 0.5 | 0.11 | 0.24 |
| 8 | 69863032 | rs4347001 | -0.511 | -0.743 | -1.444 | 0.454 | -0.021 | -0.027 | -0.027 | 0.033 | 0.09 | 0.01 | 0.04 | 0.11 | 0.1 | 0.18 |
| 8 | 69872733 | rs12056548 | -0.685 | -1.265 | -1.506 | 0.81 | 0.024 | -0.025 | -0.025 | 0.442 | 0.1 | 0.04 | 0.06 | 0.24 | 0.03 | 0.15 |
| 8 | 69877518 | rs4342610 | -1.312 | -2.467 | -1.186 | -0.124 | 0.005 | 0.007 | 0.007 | 1.398 | 0.37 | 0.64 | 0.24 | 0.27 | 0.18 | 0.25 |
| 8 | 69878508 | rs7841533 | -0.691 | -2.448 | -0.542 | -0.362 | -0.015 | -0.059 | -0.059 | 0.141 | 0.18 | 0.26 | 0.1 | 0.03 | 0.12 | 0.11 |
| 8 | 69880186 | rs17613171 | -0.523 | -2.446 | -0.63 | -0.34 | 0.01 | 0.043 | 0.043 | 0 | 0.15 | 0.04 | 0.08 | 0.03 | 0.11 | 0.1 |
| 8 | 69883505 | rs4737930 | 0.936 | -3.214 | -0.768 | -0.073 | 0.037 | -0.007 | -0.007 | 0 | 0.57 | 0.69 | 0.72 | 0.51 | 0.75 | 0.5 |
| 8 | 69883644 | rs17613378 | -0.494 | -3.22 | -0.465 | -0.294 | 0.008 | -0.017 | -0.017 | 0 | 0.2 | 0.17 | 0.11 | 0 | 0.15 | 0.11 |
| 8 | 69883967 | rs10504434 | 1.695 | -3.894 | -0.304 | -0.239 | 0.002 | -0.031 | -0.031 | 0 | 0.37 | 0.29 | 0.56 | 0.46 | 0.6 | 0.35 |
| 8 | 69885011 | rs10104581 | -0.12 | -4.111 | -0.611 | -0.271 | 0.016 | -0.044 | -0.044 | 0 | 0.77 | 0.9 | 0.79 | 0.61 | 0.84 | 0.65 |
| 8 | 69893813 | rs10090892 | 0.267 | -4.14 | -0.582 | -0.349 | 0.009 | 0.008 | 0.008 | 0 | 0.18 | 0.16 | 0.08 | 0 | 0.15 | 0.14 |
| 8 | 69894554 | rs7824920 | 0.164 | -3.943 | -0.222 | 0.032 | -0.04 | 0.021 | 0.021 | 1.219 | 0.09 | 0.16 | 0.1 | 0.11 | 0.02 | 0.09 |
| 8 | 69895021 | rs7825569 | -1.557 | -3.536 | -0.65 | 0.085 | -0.061 | -0.089 | -0.089 | 0 | 0.63 | 0.89 | 0.49 | 0.49 | 0.42 | 0.63 |
| 8 | 69900903 | rs10504435 | -0.005 | -3.053 | -0.017 | 0.182 | -0.024 | 0.027 | 0.027 | 0 | 0.07 | 0.14 | 0.11 | 0.12 | 0.02 | 0.12 |
| 8 | 69901440 | rs10111517 | 1.778 | -3.034 | -0.14 | 0.429 | -0.068 | 0.023 | 0.023 | 0.084 | 0.35 | 0.16 | 0.37 | 0.26 | 0.54 | 0.26 |
| 8 | 69919382 | rs7001238 | 1.038 | -2.99 | 0.719 | 0.68 | 0.031 | 0 | 0 | 0 | 0.56 | 0.34 | 0.53 | 0.58 | 0.69 | 0.55 |
| 8 | 69923119 | rs12680675 | -0.862 | -3.026 | 0.414 | 0.597 | 0.009 | -0.033 | -0.033 | 0 | 0.83 | 0.93 | 0.78 | 0.73 | 0.88 | 0.81 |
| 8 | 69927346 | rs12544142 | -0.813 | -3.029 | 0.614 | 0.595 | 0.001 | -0.004 | -0.004 | 0 | 0.83 | 0.87 | 0.77 | 0.73 | 0.89 | 0.81 |

| | | | | | | | | | | | | | | | | |
|---|---|---|---|---|---|---|---|---|---|---|---|---|---|---|---|---|
| 8 | 69930650 | rs1863590 | -0.002 | -2.845 | 0.903 | 0.388 | -0.003 | -0.07 | -0.07 | 0 | 0.18 | 0.28 | 0.11 | 0 | 0.17 | 0.14 |
| 8 | 69940494 | rs10101263 | -0.44 | -2.822 | 0.315 | 0.339 | -0.012 | -0.086 | -0.086 | 0.02 | 0.2 | 0.7 | 0.12 | 0 | 0.15 | 0.11 |
| 8 | 69941508 | rs6990230 | -0.607 | -2.616 | -0.558 | 0.569 | -0.047 | -0.049 | -0.049 | 0 | 0.31 | 0.51 | 0.29 | 0.16 | 0.2 | 0.26 |
| 8 | 69942195 | rs933938 | 1.615 | -2.591 | -0.935 | 0.838 | -0.053 | 0.091 | 0.091 | 1.244 | 0.44 | 0.09 | 0.37 | 0.24 | 0.58 | 0.34 |
| 8 | 69943607 | rs16935371 | -0.269 | -1.765 | -0.188 | 1.419 | 0.001 | 0.047 | 0.047 | 0 | 0.11 | 0.15 | 0.11 | 0.3 | 0.11 | 0.24 |
| 8 | 69943681 | rs2912498 | 0.515 | -1.572 | -0.445 | 1.249 | -0.012 | 0.071 | 0.071 | 0.804 | 0.72 | 0.55 | 0.64 | 0.4 | 0.79 | 0.61 |
| 8 | 69944403 | rs2912500 | 0.515 | -1.572 | -0.445 | 1.232 | -0.012 | 0.069 | 0.069 | 0.804 | 0.72 | 0.55 | 0.64 | 0.39 | 0.79 | 0.61 |
| 8 | 69947175 | rs2912504 | 0.334 | -1.422 | -1.371 | 1.058 | 0.023 | -0.07 | -0.07 | 0 | 0.73 | 0.86 | 0.67 | 0.59 | 0.81 | 0.66 |
| 8 | 69959105 | rs2981060 | 0.503 | 0.547 | -1.993 | 0.102 | 0.025 | 0.025 | 0.025 | 0 | 0.44 | 0.2 | 0.44 | 0.58 | 0.36 | 0.56 |
| 8 | 69966936 | rs2162353 | -0.008 | 0.259 | -1.482 | -0.178 | 0.026 | -0.001 | -0.001 | 0 | 0.69 | 0.41 | 0.52 | 0.67 | 0.49 | 0.73 |
| 8 | 69970959 | rs1465514 | -0.75 | 0.605 | -1.355 | -0.322 | -0.041 | -0.04 | -0.04 | 0 | 0.84 | 0.95 | 0.67 | 0.68 | 0.72 | 0.77 |
| 8 | 69973433 | rs10504436 | 0.766 | 0.573 | -1.33 | -0.408 | 0.007 | 0.001 | 0.001 | 0.561 | 0.24 | 0.2 | 0.08 | 0.09 | 0.13 | 0.16 |
| 8 | 69975158 | rs13277901 | 0.822 | 0.765 | -0.522 | -0.294 | 0.045 | -0.111 | -0.111 | 0.012 | 0.16 | 0.38 | 0.13 | 0 | 0.25 | 0.05 |
| 8 | 69977857 | rs2912518 | 0.332 | 0.785 | -0.637 | -0.257 | -0.039 | -0.036 | -0.036 | 0 | 0.16 | 0.05 | 0.26 | 0.28 | 0.27 | 0.23 |
| 8 | 69979828 | rs1365415 | 0.703 | 0.718 | -0.438 | -0.25 | 0.064 | -0.131 | -0.131 | 0.004 | 0.16 | 0.5 | 0.15 | 0 | 0.25 | 0.05 |
| 8 | 69982901 | rs1344618 | 1.278 | 0.758 | -0.613 | -0.228 | -0.021 | -0.03 | -0.03 | 0 | 0.11 | 0.03 | 0.17 | 0.29 | 0.15 | 0.2 |
| 8 | 69986093 | rs13264794 | 0.792 | 0.537 | -0.541 | -0.356 | 0.035 | -0.098 | -0.098 | 0 | 0.16 | 0.32 | 0.13 | 0 | 0.25 | 0.05 |
| 8 | 69986578 | rs1820476 | 0.991 | -0.649 | -1.43 | -0.037 | 0.004 | -0.002 | -0.002 | 0 | 0.09 | 0.06 | 0.1 | 0.11 | 0.05 | 0.1 |
| 8 | 69990756 | rs578611 | 0.317 | -0.385 | -1.484 | -0.151 | 0.02 | 0.01 | 0.01 | 0.216 | 0.32 | 0.2 | 0.22 | 0.3 | 0.2 | 0.34 |
| 8 | 69992380 | rs2912522 | -0.268 | -0.318 | -1.432 | -0.075 | 0.028 | -0.032 | -0.032 | 0 | 0.65 | 0.77 | 0.61 | 0.59 | 0.72 | 0.62 |
| 8 | 69994670 | rs12675570 | -0.624 | -0.487 | -1.137 | -0.298 | 0.074 | -0.04 | -0.04 | 0 | 0.34 | 0.14 | 0.28 | 0.38 | 0.17 | 0.5 |
| 8 | 69995213 | rs1896135 | -0.869 | -0.493 | -1.118 | -0.41 | 0.069 | -0.006 | -0.006 | 0 | 0.4 | 0.21 | 0.33 | 0.38 | 0.24 | 0.57 |
| 8 | 69996372 | rs514802 | -1.537 | -0.342 | -0.675 | -0.497 | 0.028 | 0.025 | 0.025 | 0.26 | 0.49 | 0.67 | 0.46 | 0.38 | 0.37 | 0.59 |
| 8 | 69997500 | rs6472439 | -0.788 | -0.323 | -0.754 | -0.486 | 0.062 | -0.007 | -0.007 | 0.261 | 0.4 | 0.21 | 0.32 | 0.38 | 0.25 | 0.57 |
| 8 | 70001951 | rs2472130 | -1.14 | -0.439 | -0.841 | -0.006 | 0.003 | 0 | 0 | 0.452 | 0.42 | 0.59 | 0.42 | 0.38 | 0.29 | 0.52 |
| 8 | 70003448 | rs2042731 | -0.243 | -0.535 | -1.655 | 0.499 | 0.026 | -0.046 | -0.046 | 0.119 | 0.17 | 0.05 | 0.23 | 0.27 | 0.11 | 0.3 |
| 8 | 70004698 | rs16935510 | -0.575 | -0.354 | -1.108 | 0.247 | 0.004 | 0 | 0 | 2.936 | 0.18 | 0.17 | 0.06 | 0.13 | 0.06 | 0.22 |
| 8 | 70005353 | rs17695441 | -0.341 | 0.562 | -0.855 | -0.547 | 0.012 | -0.052 | -0.052 | 0 | 0.38 | 0.49 | 0.45 | 0.19 | 0.44 | 0.39 |
| 8 | 70006121 | rs2472135 | -0.609 | 1.575 | -0.469 | -1.156 | 0.022 | -0.077 | -0.077 | 0 | 0.26 | 0.39 | 0.3 | 0 | 0.43 | 0.14 |
| 8 | 70006868 | rs16935518 | -0.404 | 1.499 | 0.185 | -0.443 | 0.012 | 0.11 | 0.11 | 0 | 0.13 | 0.26 | 0.19 | 0.34 | 0.13 | 0.21 |
| 8 | 70007605 | rs2912523 | -0.251 | 1.349 | -0.111 | -0.747 | -0.008 | 0.019 | 0.019 | 2.237 | 0.69 | 0.63 | 0.68 | 0.58 | 0.77 | 0.77 |
| 8 | 70007938 | rs629535 | 0.749 | 1.459 | -0.222 | -0.873 | -0.045 | 0.054 | 0.054 | 3.527 | 0.14 | 0.03 | 0.18 | 0 | 0.26 | 0.07 |
| 8 | 70009214 | rs2912524 | -0.336 | 1.558 | 0.339 | -0.892 | 0.01 | -0.027 | -0.027 | 0 | 0.69 | 0.71 | 0.68 | 0.57 | 0.77 | 0.77 |
| 8 | 70009905 | rs17623055 | 0.654 | 1.562 | 0.408 | -1.029 | 0.01 | 0.01 | 0.01 | 2.529 | 0.5 | 0.34 | 0.42 | 0.53 | 0.42 | 0.56 |
| 8 | 70009988 | rs659430 | 0.508 | 1.579 | 0.463 | -0.89 | 0.017 | -0.055 | -0.055 | 0 | 0.15 | 0.18 | 0.21 | 0 | 0.32 | 0.07 |
| 8 | 70014344 | rs638853 | 0.044 | 1.229 | 0.334 | -0.881 | -0.001 | -0.05 | -0.05 | 0.6 | 0.15 | 0.1 | 0.19 | 0.39 | 0.15 | 0.34 |
| 8 | 70014495 | rs10504437 | 1.843 | 1.202 | 0.059 | -1.221 | -0.006 | 0.118 | 0.118 | 0.115 | 0.26 | 0.06 | 0.12 | 0.06 | 0.2 | 0.21 |
| 8 | 70025018 | rs590695 | -0.897 | 1.252 | 0.092 | -1.251 | -0.099 | 0.261 | 0.261 | 0.075 | 0.59 | 0.89 | 0.68 | 0.97 | 0.52 | 0.71 |
| 8 | 70031128 | rs2064033 | 0.087 | 1.235 | 0.225 | -1.156 | -0.005 | 0 | 0 | 0 | 0.43 | 0.5 | 0.45 | 0.54 | 0.39 | 0.38 |
| 8 | 70043260 | rs16935565 | 0.881 | 1.039 | 0.159 | -1.367 | 0 | -0.013 | -0.013 | 0 | 0.22 | 0.2 | 0.2 | 0.13 | 0.22 | 0.17 |
| 8 | 70049892 | rs668858 | 0.5 | 1.067 | 0.213 | -1.186 | 0.001 | -0.001 | -0.001 | 0 | 0.43 | 0.47 | 0.36 | 0.48 | 0.35 | 0.56 |
| 8 | 70049904 | rs10086816 | 1.737 | 1.094 | 0.207 | -1.202 | 0.002 | -0.007 | -0.007 | 0 | 0.28 | 0.23 | 0.16 | 0.03 | 0.21 | 0.22 |
| 8 | 70055382 | rs16935581 | 0.622 | 1.123 | 0.321 | -1.097 | -0.001 | -0.037 | -0.037 | 0 | 0.23 | 0.28 | 0.21 | 0.13 | 0.22 | 0.17 |
| 8 | 70055548 | rs16935585 | 0.478 | 1.127 | 0.431 | -1.101 | 0.003 | -0.074 | -0.074 | 0 | 0.24 | 0.44 | 0.22 | 0.13 | 0.22 | 0.17 |
| 8 | 70056441 | rs11786948 | 0.735 | 1.13 | 0.393 | -1.1 | 0 | -0.016 | -0.016 | 0 | 0.22 | 0.21 | 0.2 | 0.13 | 0.22 | 0.17 |
| 8 | 70065359 | rs485219 | 0.562 | 1.215 | 0.379 | -0.802 | -0.012 | 0.032 | 0.032 | 1.338 | 0.26 | 0.39 | 0.28 | 0.54 | 0.2 | 0.4 |
| 8 | 70067613 | rs4737952 | 1.275 | 1.185 | 0.283 | -0.743 | 0.004 | -0.01 | -0.01 | 0.199 | 0.2 | 0.17 | 0.24 | 0.4 | 0.15 | 0.23 |
| 8 | 70070757 | rs16935654 | 0.75 | 1.186 | -0.007 | -0.95 | 0.002 | -0.007 | -0.007 | 0 | 0.1 | 0.09 | 0.14 | 0.3 | 0.09 | 0.13 |
| 8 | 70071420 | rs1462027 | -0.225 | 1.183 | -0.041 | -0.999 | -0.001 | 0.009 | 0.009 | 2.415 | 0.3 | 0.34 | 0.36 | 0.43 | 0.3 | 0.28 |
| 8 | 70074265 | rs11775631 | 0.741 | 1.181 | -0.046 | -1.038 | 0.003 | -0.028 | -0.028 | 4.673 | 0.2 | 0.24 | 0.21 | 0.13 | 0.2 | 0.15 |
| 8 | 70084415 | rs12545578 | 1.442 | 1.193 | -0.328 | -1.093 | 0.005 | 0.087 | 0.087 | 0.074 | 0.3 | 0.14 | 0.17 | 0.03 | 0.24 | 0.25 |
| 8 | 70091974 | rs931001 | -0.147 | 1.215 | -0.229 | -1.026 | 0.002 | -0.001 | -0.001 | 0 | 0.6 | 0.55 | 0.52 | 0.46 | 0.53 | 0.54 |
| 8 | 70093514 | rs17704313 | 0.544 | 1.227 | -0.51 | -0.974 | 0.015 | 0.043 | 0.043 | 0 | 0.39 | 0.24 | 0.31 | 0.33 | 0.33 | 0.38 |
| 8 | 70100216 | rs10504441 | 0.51 | 1.334 | -0.35 | -0.888 | 0.012 | -0.061 | -0.061 | 0.041 | 0.21 | 0.4 | 0.21 | 0.13 | 0.2 | 0.16 |
| 8 | 70104164 | rs9298146 | 0.582 | 1.356 | -0.284 | -0.883 | 0.011 | -0.048 | -0.048 | 1.091 | 0.21 | 0.34 | 0.21 | 0.13 | 0.2 | 0.16 |
| 8 | 70109328 | rs10957471 | -0.904 | 1.402 | -0.533 | -0.913 | -0.004 | 0.007 | 0.007 | 0.471 | 0.45 | 0.57 | 0.54 | 0.43 | 0.55 | 0.35 |
| 8 | 70115083 | rs10957472 | -0.347 | 1.267 | -0.58 | -1.131 | 0.006 | -0.05 | -0.05 | 0 | 0.35 | 0.52 | 0.38 | 0.13 | 0.42 | 0.21 |
| 8 | 70123722 | rs831737 | 1.487 | 1.256 | -1.451 | -1.132 | -0.039 | 0.171 | 0.171 | 0 | 0.31 | 0.08 | 0.19 | 0.03 | 0.3 | 0.21 |
| 8 | 70124044 | rs10504442 | 0.934 | 1.216 | -1.444 | -0.945 | -0.015 | -0.059 | -0.059 | 0 | 0.09 | 0.01 | 0.13 | 0.31 | 0.09 | 0.13 |
| 8 | 70132590 | rs705998 | 0.058 | 1.014 | -1.276 | -0.841 | 0.031 | 0.098 | 0.098 | 0.11 | 0.53 | 0.24 | 0.52 | 0.33 | 0.62 | 0.45 |
| 8 | 70134082 | rs831726 | 0.158 | 1.02 | -1.345 | -0.839 | 0.034 | 0.118 | 0.118 | 0.492 | 0.52 | 0.18 | 0.5 | 0.33 | 0.62 | 0.45 |
| 8 | 70143771 | rs13275450 | 0.621 | 1.036 | -1.221 | -0.83 | -0.003 | -0.015 | -0.015 | 0 | 0.2 | 0.2 | 0.17 | 0.12 | 0.17 | 0.15 |
| 8 | 70145512 | rs2655167 | 0.825 | 1.098 | -0.664 | -0.166 | -0.004 | 0.119 | 0.119 | 4.026 | 0.51 | 0.29 | 0.36 | 0.15 | 0.47 | 0.37 |
| 8 | 70146075 | rs2010049 | 0.093 | 0.513 | -0.035 | 0.51 | -0.018 | 0.039 | 0.039 | 0.677 | 0.73 | 0.41 | 0.69 | 0.46 | 0.79 | 0.6 |
| 8 | 70154118 | rs1381495 | 0.008 | 0.612 | -0.309 | 0.487 | -0.02 | 0.054 | 0.054 | 0.362 | 0.74 | 0.48 | 0.68 | 0.44 | 0.79 | 0.61 |
| 8 | 70155917 | rs2655173 | -0.017 | 0.636 | -0.297 | 0.271 | -0.018 | 0.109 | 0.109 | 0.433 | 0.74 | 0.48 | 0.68 | 0.52 | 0.8 | 0.62 |
| 8 | 70180251 | rs283324 | -0.467 | 0.798 | -0.641 | -0.38 | 0 | -0.029 | -0.029 | 0.485 | 0.18 | 0.19 | 0.12 | 0 | 0.22 | 0.18 |
| 8 | 70186364 | rs10089248 | 0.304 | 1.021 | -0.742 | -0.123 | -0.001 | 0.001 | 0.001 | 0 | 0.31 | 0.41 | 0.34 | 0.5 | 0.31 | 0.32 |
| 8 | 70188775 | rs283347 | -0.129 | 0.981 | -0.715 | -0.346 | 0.001 | -0.008 | -0.008 | 0.059 | 0.17 | 0.15 | 0.12 | 0 | 0.22 | 0.18 |

| | | | | | | | | | | | | | | | | |
|---|---|---|---|---|---|---|---|---|---|---|---|---|---|---|---|---|
| 8 | 70198662 | rs6981344 | 0.418 | 1.289 | -0.696 | -0.389 | -0.001 | -0.017 | -0.017 | 0.21 | 0.09 | 0.03 | 0.05 | 0 | 0.11 | 0.01 |
| 8 | 70201705 | rs283327 | -0.429 | 1.279 | -0.494 | -0.197 | -0.001 | -0.001 | -0.001 | 0 | 0.75 | 0.8 | 0.72 | 0.69 | 0.74 | 0.71 |
| 8 | 70204571 | rs283320 | 0.462 | 1.065 | -0.133 | -0.553 | -0.001 | -0.046 | -0.046 | 0 | 0.16 | 0.15 | 0.09 | 0 | 0.2 | 0.1 |
| 8 | 70206163 | rs1462030 | 0.174 | 1.033 | 0.282 | -0.329 | 0 | -0.002 | -0.002 | 0 | 0.58 | 0.45 | 0.6 | 0.66 | 0.54 | 0.61 |
| 8 | 70208823 | rs831730 | -0.106 | 0.855 | 0.676 | -0.387 | -0.003 | -0.012 | -0.012 | 0.232 | 0.65 | 0.59 | 0.65 | 0.71 | 0.63 | 0.66 |
| 8 | 70215500 | rs10808768 | 0.371 | 0.74 | 0.354 | 0.178 | -0.011 | -0.02 | -0.02 | 0 | 0.27 | 0.18 | 0.46 | 0.43 | 0.34 | 0.37 |
| 8 | 70222636 | rs7843638 | 0.231 | 0.662 | 0.422 | 0.139 | -0.005 | -0.01 | -0.01 | 0.229 | 0.28 | 0.23 | 0.47 | 0.42 | 0.34 | 0.37 |
| 8 | 70229970 | rs1383730 | 0.288 | 0.65 | 0.521 | 0.133 | -0.005 | -0.009 | -0.009 | 2.589 | 0.28 | 0.23 | 0.47 | 0.42 | 0.34 | 0.37 |
| 8 | 70237924 | rs6983293 | 0.535 | 0.526 | 0.836 | -0.099 | 0.004 | 0.043 | 0.043 | 0.174 | 0.29 | 0.47 | 0.21 | 0.33 | 0.24 | 0.23 |
| 8 | 70240509 | rs2046164 | -0.493 | 0.436 | 0.53 | -0.242 | 0.019 | 0.093 | 0.093 | 0.025 | 0.58 | 0.74 | 0.7 | 0.76 | 0.58 | 0.6 |
| 8 | 70243940 | rs12542665 | -0.654 | 0.393 | 0.175 | -0.226 | -0.011 | 0.095 | 0.095 | 0.257 | 0.61 | 0.94 | 0.73 | 0.76 | 0.6 | 0.62 |
| 8 | 70251823 | rs1481271 | -0.491 | 0.218 | 0.651 | -0.316 | 0.003 | 0.001 | 0.001 | 0 | 0.54 | 0.57 | 0.67 | 0.77 | 0.59 | 0.6 |
| 8 | 70253769 | rs4737961 | -0.493 | 0.397 | 0.962 | -0.064 | -0.026 | -0.022 | -0.022 | 0.197 | 0.41 | 0.78 | 0.52 | 0.47 | 0.44 | 0.46 |
| 8 | 70254430 | rs1026832 | -0.07 | 0.472 | 0.923 | -0.07 | 0.031 | 0.051 | 0.051 | 0 | 0.32 | 0.48 | 0.45 | 0.47 | 0.38 | 0.37 |
| 8 | 70273197 | rs955436 | -0.174 | 0.38 | 0.703 | -0.125 | 0.023 | 0.041 | 0.041 | 0 | 0.33 | 0.48 | 0.46 | 0.47 | 0.4 | 0.38 |
| 8 | 70284940 | rs1383731 | -0.575 | 0.393 | 0.467 | -0.131 | -0.025 | -0.031 | -0.031 | 0 | 0.42 | 0.78 | 0.58 | 0.49 | 0.45 | 0.46 |
| 8 | 70305107 | rs4737965 | 0.187 | 0.64 | 0.486 | 0.108 | -0.067 | -0.097 | -0.097 | 0 | 0.3 | 0.1 | 0.46 | 0.49 | 0.39 | 0.37 |
| 8 | 70305798 | rs1481278 | 0.136 | 0.614 | 1.029 | 0.036 | -0.007 | -0.049 | -0.049 | 0 | 0.09 | 0.24 | 0.07 | 0 | 0.06 | 0.1 |
| 8 | 70337897 | rs7829527 | -0.506 | 0.67 | 1.395 | 0.189 | -0.014 | -0.018 | -0.018 | 0.121 | 0.39 | 0.36 | 0.53 | 0.5 | 0.45 | 0.46 |
| 8 | 70344345 | rs7824828 | -0.552 | 0.324 | 0.77 | -0.487 | 0.006 | 0.018 | 0.018 | 0 | 0.46 | 0.62 | 0.54 | 0.64 | 0.41 | 0.41 |
| 8 | 70349040 | rs10106358 | -0.552 | 0.38 | 0.773 | -0.487 | 0.002 | 0.008 | 0.008 | 0 | 0.46 | 0.59 | 0.54 | 0.64 | 0.41 | 0.41 |
| 8 | 70351739 | rs10957492 | 0.046 | 1.07 | 1.212 | -0.104 | -0.005 | 0.188 | 0.188 | 0.006 | 0.17 | 0.43 | 0.25 | 0.43 | 0.13 | 0.25 |
| 8 | 70353579 | rs12676187 | -1.083 | 1.149 | 0.434 | -0.187 | -0.002 | 0.019 | 0.019 | 0 | 0.45 | 0.64 | 0.61 | 0.65 | 0.45 | 0.45 |
| 8 | 70356990 | rs7017364 | -0.43 | 1.145 | 0.391 | -0.185 | -0.021 | 0.024 | 0.024 | 0 | 0.37 | 0.53 | 0.36 | 0.43 | 0.32 | 0.36 |
| 8 | 70360110 | rs17642309 | 0.725 | 0.994 | 0.07 | -0.335 | -0.002 | 0.001 | 0.001 | 0 | 0.05 | 0.05 | 0.11 | 0 | 0.06 | 0.05 |
| 8 | 70365909 | rs1372153 | -1.482 | 0.953 | 0.028 | -0.408 | -0.021 | 0.104 | 0.104 | 0 | 0.53 | 0.81 | 0.67 | 0.65 | 0.49 | 0.47 |
| 8 | 70369664 | rs7003377 | -0.753 | 0.958 | -0.003 | -0.294 | -0.001 | -0.008 | -0.008 | 0 | 0.4 | 0.6 | 0.52 | 0.64 | 0.39 | 0.42 |
| 8 | 70373110 | rs920530 | -1.289 | 0.576 | 0.286 | 0.656 | -0.026 | 0.073 | 0.073 | 0 | 0.3 | 0.57 | 0.38 | 0.42 | 0.21 | 0.33 |
| 8 | 70373263 | rs12545665 | -0.228 | 0.969 | -0.085 | 1.048 | -0.003 | -0.025 | -0.025 | 0 | 0.06 | 0.11 | 0.09 | 0 | 0.05 | 0.05 |
| 8 | 70373384 | rs16935893 | 0.074 | 0.955 | -0.205 | 1.143 | -0.004 | -0.049 | -0.049 | 0 | 0.12 | 0.04 | 0.21 | 0.4 | 0.11 | 0.24 |
| 8 | 70373728 | rs12546220 | -1.284 | 1.114 | 0.051 | 1.036 | 0.005 | 0.018 | 0.018 | 1.683 | 0.29 | 0.49 | 0.51 | 0.61 | 0.26 | 0.37 |
| 8 | 70373965 | rs16935901 | -0.985 | 1.364 | -0.33 | 0.811 | 0.039 | 0.072 | 0.072 | 0 | 0.06 | 0.12 | 0.13 | 0.19 | 0.09 | 0.07 |
| 8 | 70374961 | rs12546999 | -0.655 | 0.735 | -0.809 | 0.022 | -0.001 | 0.034 | 0.034 | 0 | 0.17 | 0.07 | 0.14 | 0 | 0.16 | 0.1 |
| 8 | 70376825 | rs2244817 | -0.771 | 0.541 | -1.08 | 0.347 | -0.008 | -0.006 | -0.006 | 0 | 0.39 | 0.8 | 0.49 | 0.63 | 0.4 | 0.38 |
| 8 | 70378496 | rs2623047 | -0.674 | 0.474 | -0.303 | 0.427 | -0.001 | -0.001 | -0.001 | 0.196 | 0.36 | 0.68 | 0.48 | 0.63 | 0.39 | 0.38 |
| 8 | 70380084 | rs2583086 | -0.332 | 0.408 | 0.573 | 0.931 | 0.016 | -0.016 | -0.016 | 0 | 0.23 | 0.34 | 0.26 | 0.15 | 0.28 | 0.17 |
| 8 | 70380779 | rs2583085 | -0.559 | 0.359 | 0.94 | 0.907 | 0.007 | -0.025 | -0.025 | 0 | 0.26 | 0.57 | 0.28 | 0.15 | 0.28 | 0.23 |
| 8 | 70381185 | rs2583084 | -0.548 | 0.364 | 0.943 | 0.909 | 0.008 | -0.025 | -0.025 | 0 | 0.26 | 0.57 | 0.28 | 0.15 | 0.28 | 0.23 |
| 8 | 70381561 | rs2099800 | -0.126 | 0.355 | 1.134 | 0.936 | 0.002 | 0.162 | 0.162 | 0 | 0.11 | 0.26 | 0.21 | 0.48 | 0.1 | 0.14 |
| 8 | 70382781 | rs2725089 | -1.295 | 0.321 | 1.068 | 0.886 | -0.003 | -0.004 | -0.004 | 0 | 0.35 | 0.64 | 0.47 | 0.63 | 0.38 | 0.38 |
| 8 | 70382815 | rs2725088 | -0.754 | 0.245 | 1.213 | 0.502 | -0.028 | -0.007 | -0.007 | 0 | 0.15 | 0.41 | 0.07 | 0.08 | 0.09 | 0.16 |
| 8 | 70398386 | rs13263859 | -0.471 | -0.193 | 0.409 | 0.652 | 0.012 | -0.031 | -0.031 | 0 | 0.22 | 0.15 | 0.26 | 0.53 | 0.18 | 0.27 |
| 8 | 70399367 | rs2704033 | -1.657 | -0.419 | 0.093 | 0.334 | 0.055 | 0.054 | 0.054 | 0.521 | 0.45 | 0.78 | 0.44 | 0.68 | 0.37 | 0.48 |
| 8 | 70401379 | rs2583092 | 0.1 | -0.32 | 0.213 | 0.32 | -0.008 | 0.014 | 0.014 | 0 | 0.13 | 0.24 | 0.11 | 0.15 | 0.1 | 0.11 |
| 8 | 70404947 | rs2704035 | -1.552 | -0.963 | -0.313 | -0.139 | 0.114 | 0.165 | 0.165 | 1.137 | 0.49 | 0.87 | 0.44 | 0.73 | 0.36 | 0.56 |
| 8 | 70405148 | rs2725092 | -0.824 | -1.058 | -0.333 | -0.173 | 0.045 | 0.022 | 0.022 | 0.378 | 0.35 | 0.27 | 0.34 | 0.72 | 0.22 | 0.49 |
| 8 | 70406403 | rs6998565 | 1.859 | -0.913 | -1.048 | -0.639 | 0.012 | 0.034 | 0.034 | 2.616 | 0.18 | 0.06 | 0.17 | 0.09 | 0.18 | 0.08 |
| 8 | 70408174 | rs1441204 | -1.308 | 0.206 | -0.52 | -0.363 | 0.002 | 0 | 0 | 0 | 0.77 | 0.72 | 0.79 | 0.86 | 0.72 | 0.67 |
| 8 | 70410550 | rs2028442 | -0.494 | 0.163 | -0.719 | -0.291 | 0.029 | -0.033 | -0.033 | 0 | 0.66 | 0.59 | 0.61 | 0.81 | 0.56 | 0.68 |
| 8 | 70410776 | rs2245781 | -1.674 | 0.177 | -0.694 | -0.293 | -0.021 | 0.006 | 0.006 | 0 | 0.79 | 0.87 | 0.81 | 0.86 | 0.74 | 0.75 |
| 8 | 70410813 | rs1001537 | 0.216 | -0.791 | -0.944 | 0.136 | 0.036 | -0.109 | -0.109 | 0 | 0.21 | 0.03 | 0.25 | 0.52 | 0.15 | 0.26 |
| 8 | 70410888 | rs1530241 | -0.659 | -0.175 | -0.558 | 0.128 | 0.029 | -0.033 | -0.033 | 0 | 0.66 | 0.59 | 0.61 | 0.81 | 0.56 | 0.68 |
| 8 | 70412450 | rs6982250 | -0.644 | -0.646 | -0.646 | 0.744 | -0.007 | 0.021 | 0.021 | 0 | 0.23 | 0.23 | 0.3 | 0.49 | 0.2 | 0.24 |
| 8 | 70414160 | rs13264163 | -0.039 | -0.469 | -1.423 | 0.403 | -0.048 | 0.008 | 0.008 | 0 | 0.27 | 0.01 | 0.18 | 0.16 | 0.26 | 0.3 |
| 8 | 70414309 | rs6992749 | 1.063 | -0.387 | -1.316 | -0.056 | -0.017 | 0.026 | 0.026 | 0 | 0.31 | 0.24 | 0.29 | 0.12 | 0.36 | 0.32 |
| 8 | 70416361 | rs7833667 | 0.95 | -0.296 | -1.018 | 0.404 | 0.036 | 0.033 | 0.033 | 0 | 0.07 | 0.12 | 0.18 | 0.21 | 0.15 | 0.17 |
| 8 | 70418446 | rs1946937 | -1.271 | -0.715 | -0.385 | 0.194 | 0.003 | 0.003 | 0.003 | 0.298 | 0.73 | 0.78 | 0.83 | 0.88 | 0.79 | 0.88 |
| 8 | 70418782 | rs17716998 | 1.131 | -0.665 | -0.58 | 0.193 | -0.021 | 0.043 | 0.043 | 0 | 0.33 | 0.24 | 0.28 | 0.12 | 0.37 | 0.35 |
| 8 | 70424271 | rs1946936 | 0.279 | -0.553 | -0.599 | 0.344 | 0.015 | 0.082 | 0.082 | 0 | 0.1 | 0.16 | 0.33 | 0.62 | 0.16 | 0.23 |
| 8 | 70425623 | rs7820571 | 0.039 | -0.628 | -0.544 | 0.282 | 0.003 | 0.008 | 0.008 | 0 | 0.61 | 0.36 | 0.46 | 0.27 | 0.62 | 0.65 |
| 8 | 70433289 | rs1031017 | -1.18 | -1.043 | -0.962 | 0.224 | 0.029 | 0.037 | 0.037 | 0.536 | 0.75 | 0.88 | 0.83 | 0.88 | 0.79 | 0.87 |
| 8 | 70437216 | rs1441193 | 0.569 | -1.023 | -0.932 | 0.444 | -0.052 | 0.003 | 0.003 | 0 | 0.26 | 0.01 | 0.18 | 0.16 | 0.25 | 0.3 |
| 8 | 70440879 | rs16936001 | 0.586 | -1.088 | -0.322 | 0.496 | -0.019 | -0.007 | -0.007 | 0 | 0.19 | 0.21 | 0.11 | 0.11 | 0.1 | 0.12 |
| 8 | 70442828 | rs7836170 | -0.8 | -1.639 | -0.054 | 0.659 | 0.004 | 0.004 | 0.004 | 0 | 0.51 | 0.52 | 0.37 | 0.65 | 0.35 | 0.48 |
| 8 | 70444822 | rs13275051 | 0.069 | -1.254 | -0.221 | 0.767 | 0.016 | 0.052 | 0.052 | 0 | 0.28 | 0.1 | 0.09 | 0.14 | 0.23 | 0.3 |
| 8 | 70445363 | rs10957495 | -0.474 | -0.981 | -0.216 | 1.032 | -0.021 | 0.207 | 0.207 | 0 | 0.06 | 0.17 | 0.15 | 0.4 | 0.03 | 0.1 |
| 8 | 70446384 | rs13251613 | 0.647 | -1.037 | -0.379 | 0.914 | -0.041 | -0.011 | -0.011 | 0.203 | 0.22 | 0.01 | 0.07 | 0.14 | 0.2 | 0.21 |
| 8 | 70447058 | rs2197030 | -0.824 | -0.963 | 0.184 | 1.071 | -0.008 | 0.34 | 0.34 | 0 | 0.06 | 0.26 | 0.15 | 0.4 | 0.03 | 0.1 |
| 8 | 70448627 | rs16936031 | -0.713 | -0.96 | 0.463 | 1.108 | 0 | 0.36 | 0.36 | 0 | 0.06 | 0.24 | 0.15 | 0.4 | 0.03 | 0.1 |

| | | | | | | | | | | | | | | | | |
|---|---|---|---|---|---|---|---|---|---|---|---|---|---|---|---|---|
| 8 | 70449375 | rs10957496 | -0.191 | -1.078 | 0.396 | 1.23 | -0.019 | 0.039 | 0.039 | 0 | 0.29 | 0.37 | 0.22 | 0.45 | 0.21 | 0.3 |
| 8 | 70450071 | rs6472463 | -0.212 | -0.177 | 0.55 | 1.092 | 0.041 | 0.013 | 0.013 | 0.109 | 0.82 | 0.67 | 0.69 | 0.77 | 0.74 | 0.8 |
| 8 | 70451675 | rs12674825 | -0.899 | -0.106 | 0.435 | 0.827 | -0.008 | 0.238 | 0.238 | 0 | 0.06 | 0.24 | 0.13 | 0.35 | 0.04 | 0.11 |
| 8 | 70456080 | rs4737283 | 0.125 | -0.212 | 0.24 | 0.111 | -0.002 | -0.006 | -0.006 | 0 | 0.59 | 0.46 | 0.56 | 0.65 | 0.61 | 0.62 |
| 8 | 70456195 | rs4737990 | 0.067 | -0.214 | 0.283 | 0.109 | -0.001 | -0.003 | -0.003 | 0 | 0.6 | 0.57 | 0.58 | 0.65 | 0.61 | 0.62 |
| 8 | 70457752 | rs4737991 | 0.21 | -0.207 | -0.148 | 0.132 | 0.001 | 0.003 | 0.003 | 0 | 0.58 | 0.37 | 0.55 | 0.64 | 0.61 | 0.62 |
| 8 | 70459208 | rs4737994 | -0.651 | 0.015 | -0.488 | -0.536 | 0.002 | 0.118 | 0.118 | 0.377 | 0.54 | 0.72 | 0.56 | 0.86 | 0.47 | 0.63 |
| 8 | 70459803 | rs16936059 | -0.336 | -0.324 | -0.372 | -0.081 | -0.002 | 0.003 | 0.003 | 0 | 0.19 | 0.23 | 0.31 | 0.3 | 0.16 | 0.21 |
| 8 | 70463628 | rs10504451 | -0.538 | 0.127 | -0.819 | -0.619 | -0.003 | 0.08 | 0.08 | 0 | 0.58 | 0.8 | 0.38 | 0.29 | 0.45 | 0.52 |
| 8 | 70466347 | rs10104141 | -1.349 | -0.574 | -1.029 | -0.449 | -0.192 | -0.184 | -0.184 | 3.971 | 0.71 | 0.97 | 0.61 | 0.53 | 0.53 | 0.68 |
| 8 | 70469428 | rs10095284 | 0.143 | -0.864 | 0.078 | -1.073 | -0.022 | -0.03 | -0.03 | 0.625 | 0.3 | 0.38 | 0.19 | 0.19 | 0.19 | 0.21 |
| 8 | 70470537 | rs16936099 | 0.185 | -0.778 | -0.44 | -1.03 | 0.006 | 0.007 | 0.007 | 0 | 0.29 | 0.25 | 0.17 | 0.19 | 0.19 | 0.21 |
| 8 | 70474026 | rs17646697 | 0.161 | -0.963 | 0.069 | -0.598 | -0.001 | -0.012 | -0.012 | 0 | 0.17 | 0.16 | 0.1 | 0.09 | 0.17 | 0.22 |
| 8 | 70475308 | rs7838730 | -0.232 | -0.788 | -0.157 | -0.505 | -0.016 | -0.067 | -0.067 | 0 | 0.24 | 0.34 | 0.16 | 0.09 | 0.21 | 0.28 |
| 8 | 70477764 | rs16936114 | -0.344 | -0.162 | -0.467 | 0.319 | -0.016 | -0.002 | -0.002 | 0.734 | 0.2 | 0.24 | 0.14 | 0.2 | 0.13 | 0.17 |
| 8 | 70481100 | rs7841611 | -0.118 | 0.176 | -0.138 | 0.475 | -0.021 | -0.008 | -0.008 | 0 | 0.3 | 0.44 | 0.37 | 0.3 | 0.25 | 0.32 |
| 8 | 70481484 | rs7845415 | -0.11 | 0.136 | -0.026 | 0.456 | -0.023 | -0.009 | -0.009 | 0 | 0.3 | 0.46 | 0.38 | 0.3 | 0.25 | 0.32 |
| 8 | 70484161 | rs17646998 | 0.257 | 0.368 | -0.605 | 0.327 | 0.036 | 0.095 | 0.095 | 0 | 0.39 | 0.17 | 0.27 | 0.2 | 0.35 | 0.25 |
| 8 | 70485402 | rs723290 | -0.479 | 0.593 | -0.094 | 0.584 | -0.047 | -0.052 | -0.052 | 0.108 | 0.69 | 0.81 | 0.65 | 0.51 | 0.56 | 0.6 |
| 8 | 70489669 | rs4737995 | 0.548 | 0.386 | 0.312 | 0.517 | -0.017 | -0.014 | -0.014 | 0 | 0.31 | 0.41 | 0.37 | 0.27 | 0.22 | 0.35 |
| 8 | 70490015 | rs10108002 | 0.434 | 0.357 | -0.02 | 0.335 | 0.015 | 0.054 | 0.054 | 0 | 0.32 | 0.1 | 0.22 | 0.22 | 0.3 | 0.23 |
| 8 | 70498501 | rs2241882 | 0.238 | 0.762 | 0.22 | 0.694 | -0.046 | -0.034 | -0.034 | 2.477 | 0.38 | 0.18 | 0.34 | 0.54 | 0.44 | 0.54 |
| 8 | 70500255 | rs9643604 | 0.124 | 0.8 | 0.934 | 0.929 | -0.009 | 0 | 0 | 0 | 0.46 | 0.24 | 0.37 | 0.46 | 0.52 | 0.47 |
| 8 | 70503919 | rs17720093 | 1.175 | 0.758 | 1.428 | 0.742 | 0.025 | 0.127 | 0.127 | 6.807 | 0.25 | 0.04 | 0.1 | 0.01 | 0.19 | 0.11 |
| 8 | 70504309 | rs920528 | -0.808 | 0.814 | 1.663 | 0.782 | 0.009 | 0.007 | 0.007 | 0 | 0.76 | 0.7 | 0.74 | 0.75 | 0.7 | 0.79 |
| 8 | 70505272 | rs4737996 | -1.091 | 0.791 | 1.943 | 0.806 | -0.008 | -0.014 | -0.014 | 0 | 0.78 | 0.81 | 0.76 | 0.75 | 0.73 | 0.87 |
| 8 | 70507943 | rs12545167 | 0.525 | 0.748 | 1.957 | 0.8 | -0.042 | -0.011 | -0.011 | 0 | 0.3 | 0.49 | 0.37 | 0.31 | 0.21 | 0.38 |
| 8 | 70512693 | rs1078494 | 0.34 | 1 | 1.958 | 0.785 | -0.041 | -0.017 | -0.017 | 0.123 | 0.32 | 0.48 | 0.38 | 0.3 | 0.22 | 0.39 |
| 8 | 70512729 | rs1078495 | 0.549 | 0.999 | 1.94 | 0.784 | 0.016 | 0.019 | 0.019 | 4.041 | 0.3 | 0.23 | 0.34 | 0.21 | 0.2 | 0.22 |
| 8 | 70513477 | rs752039 | 0.447 | 1.032 | 1.922 | 0.772 | 0.002 | 0.001 | 0.001 | 2.539 | 0.31 | 0.28 | 0.36 | 0.3 | 0.22 | 0.41 |
| 8 | 70513720 | rs752038 | 1.289 | 1.069 | 1.65 | 0.78 | -0.003 | 0.162 | 0.162 | 6.366 | 0.24 | 0.01 | 0.1 | 0.01 | 0.19 | 0.1 |
| 8 | 70513855 | rs752040 | -0.878 | 1.167 | 1.862 | 0.846 | 0.027 | 0.023 | 0.023 | 0 | 0.77 | 0.63 | 0.72 | 0.75 | 0.7 | 0.81 |
| 8 | 70514116 | rs1155965 | 0.35 | 1.173 | 1.645 | 0.861 | -0.014 | 0.007 | 0.007 | 0.478 | 0.46 | 0.3 | 0.36 | 0.44 | 0.48 | 0.38 |
| 8 | 70516680 | rs2119530 | -1 | 1.214 | 1.265 | 0.478 | -0.015 | -0.023 | -0.023 | 0 | 0.78 | 0.86 | 0.77 | 0.75 | 0.73 | 0.85 |
| 8 | 70516940 | rs4737998 | 0.343 | 1.878 | 1.382 | 0.398 | -0.006 | -0.003 | -0.003 | 0 | 0.46 | 0.54 | 0.6 | 0.64 | 0.42 | 0.5 |
| 8 | 70516969 | rs4737284 | 0.202 | 2.093 | 1.503 | 0.688 | 0.006 | -0.002 | -0.002 | 1.948 | 0.48 | 0.47 | 0.51 | 0.52 | 0.38 | 0.42 |
| 8 | 70517558 | rs6472468 | -1.074 | 1.762 | 0.978 | 0.41 | -0.023 | -0.033 | -0.033 | 0 | 0.79 | 0.89 | 0.77 | 0.75 | 0.72 | 0.85 |
| 8 | 70522119 | rs7005314 | -0.172 | 1.766 | 1.052 | 0.019 | -0.002 | -0.006 | -0.006 | 0 | 0.39 | 0.31 | 0.2 | 0.12 | 0.4 | 0.41 |
| 8 | 70526296 | rs1866897 | 0.713 | 1.618 | 0.958 | 0.088 | 0.003 | -0.001 | -0.001 | 0 | 0.39 | 0.41 | 0.56 | 0.64 | 0.34 | 0.34 |
| 8 | 70527022 | rs2028444 | -1.146 | 1.646 | 0.526 | 0.074 | -0.014 | -0.013 | -0.013 | 0.177 | 0.79 | 0.89 | 0.79 | 0.76 | 0.74 | 0.76 |
| 8 | 70531474 | rs6472471 | 0.457 | 1.555 | 0.513 | -0.134 | 0.02 | 0.177 | 0.177 | 2.404 | 0.33 | 0.06 | 0.15 | 0.01 | 0.29 | 0.16 |
| 8 | 70537507 | rs12549880 | -0.483 | 1.821 | 0.547 | 0.207 | -0.03 | -0.021 | -0.021 | 0 | 0.76 | 0.83 | 0.76 | 0.68 | 0.63 | 0.5 |
| 8 | 70539911 | rs4738002 | -0.952 | 1.798 | 0.448 | 0.212 | -0.017 | -0.008 | -0.008 | 0 | 0.79 | 0.9 | 0.79 | 0.78 | 0.73 | 0.75 |
| 8 | 70540216 | rs3802275 | -0.498 | 1.846 | 0.583 | 0.169 | -0.01 | -0.007 | -0.007 | 1.778 | 0.74 | 0.82 | 0.75 | 0.68 | 0.63 | 0.49 |
| 8 | 70540868 | rs4284050 | 0.246 | 1.791 | 0.355 | 0.12 | 0.023 | 0.003 | 0.003 | 0 | 0.66 | 0.62 | 0.7 | 0.67 | 0.56 | 0.46 |
| 8 | 70542298 | rs12543013 | 0.416 | 1.664 | -0.009 | 0.158 | 0.004 | -0.015 | -0.015 | 0.013 | 0.42 | 0.46 | 0.6 | 0.76 | 0.43 | 0.6 |
| 8 | 70542444 | rs10957500 | -0.669 | 1.888 | -0.349 | 0.767 | -0.032 | -0.036 | -0.036 | 0 | 0.79 | 0.92 | 0.78 | 0.74 | 0.73 | 0.74 |
| 8 | 70542552 | rs4738003 | 0.167 | 1.324 | -0.199 | 0.722 | -0.02 | -0.008 | -0.008 | 1.098 | 0.64 | 0.75 | 0.68 | 0.62 | 0.53 | 0.46 |
| 8 | 70546219 | rs7831688 | -0.037 | 0.465 | -0.362 | 0.37 | 0.002 | 0 | 0 | 1.23 | 0.57 | 0.39 | 0.56 | 0.53 | 0.64 | 0.47 |
| 8 | 70553688 | rs4738011 | -0.057 | -0.073 | -0.547 | 0.074 | -0.006 | 0.012 | 0.012 | 0.073 | 0.56 | 0.76 | 0.59 | 0.55 | 0.61 | 0.47 |
| 8 | 70554689 | rs7814130 | 0.102 | 0.009 | -0.367 | -0.331 | 0.001 | 0.007 | 0.007 | 0 | 0.45 | 0.61 | 0.48 | 0.54 | 0.4 | 0.54 |
| 8 | 70556547 | rs10106852 | 0.533 | -0.258 | 0.307 | -0.397 | 0.001 | 0.002 | 0.002 | 0.146 | 0.34 | 0.37 | 0.43 | 0.43 | 0.38 | 0.46 |
| 8 | 70556602 | rs17665471 | -0.069 | -0.343 | 0.062 | -0.66 | 0.006 | 0.011 | 0.011 | 0.032 | 0.18 | 0.1 | 0.09 | 0.01 | 0.18 | 0.14 |
| 8 | 70558315 | rs4279601 | -0.384 | -0.417 | 0.287 | -0.644 | 0.022 | 0.002 | 0.002 | 0 | 0.5 | 0.31 | 0.5 | 0.43 | 0.54 | 0.62 |
| 8 | 70564677 | rs7001561 | -0.641 | -0.354 | 0.979 | -0.523 | -0.002 | -0.122 | -0.122 | 0 | 0.2 | 0.39 | 0.14 | 0.03 | 0.19 | 0.25 |
| 8 | 70564701 | rs6472472 | -0.936 | -0.323 | 0.651 | -0.593 | -0.048 | -0.055 | -0.055 | 0 | 0.55 | 0.87 | 0.57 | 0.43 | 0.55 | 0.54 |
| 8 | 70569973 | rs11991158 | 0.065 | -0.336 | 1.043 | -0.384 | -0.004 | -0.03 | -0.03 | 0.016 | 0.08 | 0.3 | 0.04 | 0.01 | 0.03 | 0.05 |
| 8 | 70571313 | rs3802278 | 0.525 | -0.128 | 0.379 | -0.085 | -0.004 | 0.008 | 0.008 | 0 | 0.32 | 0.34 | 0.44 | 0.4 | 0.32 | 0.19 |
| 8 | 70571531 | rs6990375 | 0.863 | -0.031 | 0.069 | -0.037 | 0 | -0.004 | -0.004 | 4.057 | 0.29 | 0.3 | 0.42 | 0.4 | 0.32 | 0.17 |
| 8 | 70573004 | rs3087714 | 0.8 | -0.013 | -0.032 | -0.026 | 0.004 | -0.022 | -0.022 | 0.018 | 0.3 | 0.26 | 0.42 | 0.4 | 0.32 | 0.17 |
| 8 | 70576130 | rs4738017 | -0.137 | 0.064 | -0.244 | 0.366 | -0.012 | -0.013 | -0.013 | 0 | 0.65 | 0.77 | 0.74 | 0.6 | 0.65 | 0.64 |
| 8 | 70578217 | rs11779416 | 0.267 | -0.084 | -0.398 | 0.45 | 0.003 | -0.017 | -0.017 | 0 | 0.59 | 0.71 | 0.7 | 0.44 | 0.65 | 0.52 |
| 8 | 70580284 | rs10103251 | 0.534 | 0.138 | -0.496 | 0.451 | 0.002 | 0.014 | 0.014 | 0.413 | 0.9 | 0.87 | 0.9 | 0.8 | 0.89 | 0.77 |
| 8 | 70580782 | rs16936266 | 0.488 | 0.008 | -0.599 | 0.462 | -0.009 | -0.048 | -0.048 | 0 | 0.29 | 0.11 | 0.19 | 0.36 | 0.24 | 0.21 |
| 8 | 70582105 | rs16936267 | -0.237 | 0.006 | -0.554 | 0.528 | -0.014 | 0.028 | 0.028 | 0 | 0.06 | 0.09 | 0.06 | 0.18 | 0.01 | 0.11 |
| 8 | 70582510 | rs13438797 | 0.288 | 0.157 | -0.531 | 0.404 | -0.006 | -0.025 | -0.025 | 0 | 0.3 | 0.18 | 0.2 | 0.36 | 0.26 | 0.24 |
| 8 | 70583728 | rs10103580 | 1.262 | 0.04 | 0.394 | 0.351 | 0.013 | 0.033 | 0.033 | 3.834 | 0.82 | 0.68 | 0.8 | 0.76 | 0.86 | 0.63 |
| 8 | 70584378 | rs11989109 | 1.025 | -0.136 | 0.401 | 0.318 | -0.002 | 0.041 | 0.041 | 10.705 | 0.84 | 0.63 | 0.84 | 0.81 | 0.88 | 0.76 |
| 8 | 70584809 | rs16936279 | -0.625 | -0.67 | -0.244 | 0.281 | 0.002 | -0.002 | -0.002 | 0 | 0.26 | 0.28 | 0.2 | 0.25 | 0.15 | 0.21 |

| | | | | | | | | | | | | | | | | |
|---|---|---|---|---|---|---|---|---|---|---|---|---|---|---|---|---|
| 8 | 70588099 | rs16936280 | -0.189 | -0.084 | 0.007 | -0.051 | -0.001 | -0.002 | -0.002 | 0.026 | 0.43 | 0.47 | 0.35 | 0.44 | 0.42 | 0.4 |
| 8 | 70588878 | rs10448034 | 0.351 | 0.25 | -0.014 | -0.561 | 0.004 | -0.035 | -0.035 | 0.034 | 0.12 | 0.16 | 0.1 | 0.01 | 0.16 | 0.08 |
| 8 | 70590302 | rs9298151 | -1.052 | 0.111 | 0.27 | -0.926 | 0 | -0.059 | -0.059 | 0 | 0.21 | 0.28 | 0.1 | 0.03 | 0.16 | 0.28 |
| 8 | 70595221 | rs2933075 | -0.057 | -0.261 | 0.662 | -0.707 | -0.001 | 0 | 0 | 0 | 0.44 | 0.55 | 0.35 | 0.44 | 0.37 | 0.55 |
| 8 | 70597549 | rs10504456 | 0.966 | -0.389 | 0.395 | -0.915 | -0.04 | 0.072 | 0.072 | 0 | 0.12 | 0.02 | 0.09 | 0 | 0.17 | 0.08 |
| 8 | 70598660 | rs10504457 | 1.219 | -0.501 | 0.724 | -0.906 | -0.016 | -0.026 | -0.026 | 0.085 | 0.3 | 0.14 | 0.25 | 0.38 | 0.3 | 0.36 |
| 8 | 70598818 | rs2380570 | 0.105 | -0.497 | 0.751 | -0.901 | -0.001 | 0 | 0 | 0 | 0.43 | 0.52 | 0.35 | 0.44 | 0.37 | 0.56 |
| 8 | 70602595 | rs2933070 | -0.234 | -0.446 | 1.011 | -0.913 | -0.002 | -0.01 | -0.01 | 0 | 0.13 | 0.15 | 0.07 | 0.06 | 0.07 | 0.19 |
| 8 | 70602926 | rs2933069 | 0.029 | -0.396 | 1.203 | -0.888 | 0.002 | -0.023 | -0.023 | 2.799 | 0.12 | 0.18 | 0.08 | 0.06 | 0.07 | 0.19 |
| 8 | 70603742 | rs11995058 | 0.498 | -0.402 | 1.352 | -0.898 | 0 | -0.001 | -0.001 | 0.614 | 0.06 | 0.08 | 0.05 | 0.06 | 0.09 | 0.06 |
| 8 | 70605072 | rs2380563 | 0.112 | -0.464 | 1.408 | -0.852 | 0.005 | -0.021 | -0.021 | 0 | 0.11 | 0.17 | 0.08 | 0.06 | 0.07 | 0.19 |
| 8 | 70605196 | rs2380564 | -0.688 | -0.555 | 1.771 | -0.912 | 0.003 | -0.042 | -0.042 | 0 | 0.14 | 0.28 | 0.1 | 0.06 | 0.08 | 0.2 |
| 8 | 70607162 | rs10504458 | 1.023 | -0.59 | 1.715 | -0.897 | -0.007 | -0.005 | -0.005 | 0 | 0.33 | 0.2 | 0.28 | 0.39 | 0.33 | 0.36 |
| 8 | 70607929 | rs11990100 | 0.005 | -0.642 | 1.664 | -0.898 | 0 | 0.001 | 0.001 | 0 | 0.08 | 0.09 | 0.04 | 0.08 | 0.07 | 0.07 |
| 8 | 70608241 | rs2933060 | 0.173 | -0.611 | 1.665 | -0.881 | 0.005 | -0.021 | -0.021 | 0 | 0.11 | 0.17 | 0.08 | 0.06 | 0.07 | 0.19 |
| 8 | 70609438 | rs7000631 | 0.127 | -0.6 | 1.761 | -0.875 | -0.001 | -0.003 | -0.003 | 0.343 | 0.46 | 0.53 | 0.38 | 0.44 | 0.4 | 0.57 |
| 8 | 70611563 | rs2933056 | 0.199 | -0.599 | 1.753 | -0.874 | 0.001 | 0.007 | 0.007 | 0 | 0.45 | 0.41 | 0.36 | 0.44 | 0.4 | 0.57 |
| 8 | 70616481 | rs13252820 | 1.105 | -0.572 | 1.64 | -0.934 | -0.006 | -0.005 | -0.005 | 0 | 0.11 | 0.1 | 0.1 | 0 | 0.16 | 0.08 |
| 8 | 70617568 | rs2959578 | -1.635 | -0.628 | 1.65 | -0.884 | -0.048 | -0.125 | -0.125 | 0 | 0.7 | 0.94 | 0.55 | 0.54 | 0.66 | 0.74 |
| 8 | 70617675 | rs2380572 | -1.217 | -0.651 | 1.854 | -0.908 | -0.008 | -0.035 | -0.035 | 0 | 0.22 | 0.36 | 0.13 | 0.14 | 0.15 | 0.28 |
| 8 | 70618511 | rs2933053 | -0.728 | -0.646 | 1.874 | -0.905 | 0.001 | 0.001 | 0.001 | 1.121 | 0.59 | 0.61 | 0.48 | 0.46 | 0.58 | 0.66 |
| 8 | 70618849 | rs6472477 | 0.118 | -0.645 | 1.789 | -0.907 | -0.004 | 0.031 | 0.031 | 1.874 | 0.48 | 0.4 | 0.4 | 0.4 | 0.51 | 0.45 |
| 8 | 70619599 | rs2933052 | -1.226 | -0.597 | 1.802 | -0.858 | -0.002 | 0.007 | 0.007 | 2.076 | 0.66 | 0.66 | 0.51 | 0.54 | 0.64 | 0.73 |
| 8 | 70619769 | rs2959577 | -1.226 | -0.557 | 1.802 | -0.858 | -0.002 | 0.007 | 0.007 | 2.076 | 0.66 | 0.66 | 0.51 | 0.54 | 0.64 | 0.73 |
| 8 | 70620267 | rs2933051 | -1.215 | -0.626 | 1.673 | -0.858 | -0.003 | 0.034 | 0.034 | 2.064 | 0.66 | 0.58 | 0.5 | 0.54 | 0.64 | 0.73 |
| 8 | 70620413 | rs17728769 | -0.446 | -0.629 | 1.67 | -0.884 | -0.004 | 0.009 | 0.009 | 0 | 0.21 | 0.2 | 0.14 | 0.08 | 0.24 | 0.15 |
| 8 | 70621828 | rs2959574 | -1.133 | -0.475 | 1.572 | -0.763 | -0.003 | 0.032 | 0.032 | 2.064 | 0.66 | 0.58 | 0.5 | 0.53 | 0.64 | 0.72 |
| 8 | 70622201 | rs2933046 | -1.133 | -0.475 | 1.572 | -0.763 | -0.003 | 0.032 | 0.032 | 2.064 | 0.66 | 0.58 | 0.5 | 0.53 | 0.64 | 0.72 |
| 8 | 70622208 | rs2933045 | -1.133 | -0.475 | 1.572 | -0.763 | -0.003 | 0.032 | 0.032 | 2.064 | 0.66 | 0.58 | 0.5 | 0.54 | 0.64 | 0.72 |
| 8 | 70622642 | rs2959569 | -1.554 | -0.425 | 1.287 | -0.733 | -0.01 | -0.053 | -0.053 | 0 | 0.7 | 0.83 | 0.53 | 0.54 | 0.65 | 0.73 |
| 8 | 70623097 | rs2959566 | -0.742 | -0.429 | 1.083 | -0.714 | -0.01 | -0.007 | -0.007 | 0 | 0.58 | 0.67 | 0.43 | 0.54 | 0.49 | 0.65 |
| 8 | 70623154 | rs2933044 | -0.734 | -0.43 | 1.083 | -0.652 | -0.01 | -0.009 | -0.009 | 0 | 0.58 | 0.67 | 0.46 | 0.55 | 0.5 | 0.65 |
| 8 | 70624372 | rs16936356 | -0.67 | -0.424 | 0.957 | -0.698 | 0.041 | 0.025 | 0.025 | 0 | 0.11 | 0.31 | 0.09 | 0.1 | 0.11 | 0.17 |
| 8 | 70624586 | rs2380585 | -0.143 | -0.313 | 0.349 | -0.732 | 0.002 | -0.011 | -0.011 | 0 | 0.16 | 0.15 | 0.14 | 0.05 | 0.21 | 0.19 |
| 8 | 70624938 | rs2860461 | 0.574 | -0.316 | 0.295 | -0.733 | -0.041 | 0.074 | 0.074 | 0 | 0.13 | 0.03 | 0.1 | 0 | 0.17 | 0.09 |
| 8 | 70625574 | rs17729279 | -0.09 | -0.366 | 0.306 | -0.747 | -0.01 | 0.008 | 0.008 | 0.753 | 0.49 | 0.82 | 0.43 | 0.5 | 0.47 | 0.55 |
| 8 | 70626284 | rs2959559 | -1.474 | -0.223 | -0.253 | -0.639 | -0.04 | -0.119 | -0.119 | 0.175 | 0.69 | 0.94 | 0.59 | 0.51 | 0.68 | 0.73 |
| 8 | 70626590 | rs2933040 | -0.241 | 0.031 | 0.361 | -0.681 | 0.002 | -0.011 | -0.011 | 0 | 0.16 | 0.15 | 0.14 | 0.05 | 0.21 | 0.2 |
| 8 | 70626917 | rs2959558 | -0.241 | 0.031 | 0.361 | -0.681 | 0.002 | -0.011 | -0.011 | 0 | 0.16 | 0.15 | 0.14 | 0.05 | 0.21 | 0.2 |
| 8 | 70627652 | rs2380584 | -1.145 | -0.476 | 0.1 | -0.711 | -0.002 | -0.022 | -0.022 | 0 | 0.33 | 0.65 | 0.26 | 0.11 | 0.29 | 0.35 |
| 8 | 70627815 | rs2933039 | -1.2 | -0.556 | -0.097 | -0.685 | 0.011 | 0.037 | 0.037 | 0.383 | 0.33 | 0.72 | 0.27 | 0.12 | 0.3 | 0.34 |
| 8 | 70628283 | rs2380580 | -0.765 | -0.735 | -0.918 | -0.296 | 0.018 | 0.036 | 0.036 | 0.713 | 0.52 | 0.84 | 0.41 | 0.4 | 0.47 | 0.49 |
| 8 | 70628850 | rs2933035 | -0.633 | -0.905 | 0.012 | -0.314 | 0.016 | -0.004 | -0.004 | 0.048 | 0.63 | 0.53 | 0.37 | 0.34 | 0.59 | 0.59 |
| 8 | 70631004 | rs2933031 | -0.895 | -1.019 | 0.632 | -0.343 | 0.023 | 0.007 | 0.007 | 2.093 | 0.65 | 0.58 | 0.39 | 0.36 | 0.59 | 0.7 |
| 8 | 70632524 | rs2933029 | -0.241 | -1.248 | 0.481 | -0.563 | 0.003 | -0.001 | -0.001 | 0.079 | 0.1 | 0.13 | 0.04 | 0.01 | 0.07 | 0.09 |
| 8 | 70642018 | rs4507768 | 0.438 | -1.422 | 0.446 | -0.672 | -0.031 | 0.046 | 0.046 | 0.374 | 0.4 | 0.14 | 0.25 | 0.28 | 0.42 | 0.41 |
| 8 | 70660446 | rs10106297 | -0.686 | -0.827 | 0.362 | -0.334 | -0.022 | -0.041 | -0.041 | 0.646 | 0.09 | 0.22 | 0.05 | 0.01 | 0.05 | 0.05 |
| 8 | 70660859 | rs10096246 | -0.744 | -0.831 | 0.348 | -0.335 | -0.023 | -0.039 | -0.039 | 0.259 | 0.1 | 0.23 | 0.05 | 0.01 | 0.05 | 0.06 |
| 8 | 70661354 | rs10110141 | -0.997 | -0.848 | 0.416 | -0.343 | -0.032 | -0.05 | -0.05 | 0 | 0.11 | 0.32 | 0.06 | 0.01 | 0.06 | 0.06 |
| 8 | 70667129 | rs2380598 | 1.439 | -1.194 | -0.48 | -0.506 | -0.103 | -0.09 | -0.09 | 0.192 | 0.24 | 0.07 | 0.54 | 0.63 | 0.39 | 0.28 |
| 8 | 70680983 | rs10090896 | -1.063 | -1.389 | -0.164 | -0.833 | -0.054 | -0.081 | -0.081 | 0.756 | 0.25 | 0.56 | 0.15 | 0.09 | 0.16 | 0.26 |
| 8 | 70696034 | rs16936432 | -0.59 | -1.168 | -0.585 | -0.778 | -0.043 | -0.082 | -0.082 | 0 | 0.22 | 0.42 | 0.12 | 0.07 | 0.15 | 0.18 |
| 8 | 70703141 | rs4360298 | -0.999 | -0.207 | -1.487 | 0.473 | -0.121 | -0.123 | -0.123 | 0.009 | 0.75 | 0.96 | 0.56 | 0.55 | 0.63 | 0.65 |
| 8 | 70706585 | rs12550173 | 0.937 | -0.191 | -1.471 | 0.424 | -0.1 | -0.111 | -0.111 | 0 | 0.25 | 0.05 | 0.45 | 0.46 | 0.37 | 0.36 |
| 8 | 70715243 | rs12542317 | -0.729 | 0.207 | -1.289 | 0.37 | -0.066 | -0.01 | -0.01 | 0.051 | 0.66 | 0.9 | 0.65 | 0.64 | 0.6 | 0.66 |
| 8 | 70717139 | rs7833221 | 0.035 | -0.16 | -0.631 | 0.315 | 0.003 | 0.023 | 0.023 | 0 | 0.23 | 0.28 | 0.21 | 0.34 | 0.22 | 0.37 |
| 8 | 70717334 | rs2380601 | -0.468 | 0.185 | -0.212 | -0.131 | -0.012 | -0.038 | -0.038 | 0 | 0.38 | 0.51 | 0.32 | 0.23 | 0.36 | 0.26 |
| 8 | 70719382 | rs2380602 | -0.354 | -0.483 | 0.09 | 0.097 | -0.002 | -0.006 | -0.006 | 0.566 | 0.36 | 0.37 | 0.27 | 0.31 | 0.34 | 0.41 |
| 8 | 70724024 | rs10504460 | 0.663 | -0.547 | 0.153 | 0.087 | 0.016 | 0.004 | 0.004 | 0 | 0.14 | 0.09 | 0.06 | 0 | 0.1 | 0.08 |
| 8 | 70724754 | rs16936455 | 0.858 | -0.489 | 0.095 | 0.188 | -0.015 | -0.007 | -0.007 | 0 | 0.05 | 0.02 | 0.09 | 0.23 | 0.06 | 0.08 |
| 8 | 70724911 | rs1873547 | -1.727 | -0.662 | 0.242 | 0.093 | -0.039 | -0.049 | -0.049 | 0.014 | 0.55 | 0.85 | 0.48 | 0.56 | 0.5 | 0.58 |
| 8 | 70725287 | rs10504461 | 0.795 | -0.627 | 0.157 | 0.106 | -0.016 | -0.011 | -0.011 | 3.277 | 0.05 | 0.02 | 0.09 | 0.23 | 0.06 | 0.08 |
| 8 | 70731059 | rs2380603 | -0.217 | -0.641 | 0.454 | 0.095 | -0.002 | -0.008 | -0.008 | 0 | 0.33 | 0.41 | 0.28 | 0.31 | 0.33 | 0.39 |
| 8 | 70732654 | rs2380604 | -1.053 | -0.13 | 0.456 | 0.36 | 0.001 | -0.001 | -0.001 | 0 | 0.53 | 0.62 | 0.44 | 0.53 | 0.49 | 0.55 |
| 8 | 70739817 | rs7833228 | -1.003 | -0.101 | 0.459 | 0.366 | 0.001 | -0.001 | -0.001 | 0 | 0.52 | 0.59 | 0.42 | 0.53 | 0.49 | 0.55 |
| 8 | 70740896 | rs2380606 | -1.159 | -0.051 | -0.178 | 0.382 | -0.117 | -0.131 | -0.131 | 0 | 0.56 | 0.95 | 0.48 | 0.55 | 0.5 | 0.58 |
| 8 | 70744812 | rs3750266 | -0.764 | -0.008 | 0.121 | 0.165 | 0.044 | 0.088 | 0.088 | 0 | 0.49 | 0.73 | 0.36 | 0.32 | 0.43 | 0.5 |
| 8 | 70755309 | rs7846409 | -1.047 | -0.041 | 0.043 | 0.422 | -0.028 | -0.051 | -0.051 | 0 | 0.56 | 0.84 | 0.46 | 0.59 | 0.48 | 0.63 |
| 8 | 70761794 | rs4737291 | -0.113 | -0.233 | 0.562 | 0.19 | 0.019 | 0.019 | 0.019 | 0 | 0.38 | 0.31 | 0.26 | 0.34 | 0.3 | 0.49 |

| | | | | | | | | | | | | | | | | |
|---|---|---|---|---|---|---|---|---|---|---|---|---|---|---|---|---|
| 8 | 70762608 | rs3829049 | 0.05 | -0.229 | 0.566 | 0.211 | 0.015 | 0.017 | 0.017 | 0 | 0.37 | 0.31 | 0.26 | 0.32 | 0.29 | 0.48 |
| 8 | 70767051 | rs3885979 | 0.83 | -0.159 | 0.959 | 0.21 | -0.011 | -0.048 | -0.048 | 0 | 0.08 | 0.28 | 0.06 | 0 | 0.08 | 0.04 |
| 8 | 70770391 | rs10095440 | 0.112 | -0.025 | 0.436 | 0.349 | 0.016 | 0.054 | 0.054 | 0 | 0.09 | 0.13 | 0.11 | 0.25 | 0.1 | 0.1 |
| 8 | 70783974 | rs13251224 | 0.005 | -0.286 | 0.038 | 0.208 | 0.009 | 0.01 | 0.01 | 0 | 0.36 | 0.32 | 0.26 | 0.29 | 0.29 | 0.39 |
| 8 | 70788124 | rs10957505 | -0.782 | -0.023 | -0.706 | 0.358 | 0.034 | -0.026 | -0.026 | 0.549 | 0.5 | 0.88 | 0.42 | 0.54 | 0.44 | 0.53 |
| 8 | 70794547 | rs7015359 | -0.845 | 0.665 | 0.717 | 0.414 | 0.036 | 0.084 | 0.084 | 0 | 0.16 | 0.39 | 0.17 | 0.27 | 0.18 | 0.12 |
| 8 | 70804517 | rs10957507 | -0.477 | 0.755 | 0.23 | 0.274 | -0.003 | -0.004 | -0.004 | 0 | 0.74 | 0.76 | 0.74 | 0.69 | 0.73 | 0.75 |
| 8 | 70805563 | rs16936522 | -0.288 | 0.922 | 1.144 | 0.626 | 0.073 | 0.002 | 0.002 | 0 | 0.05 | 0.26 | 0.04 | 0 | 0.04 | 0.04 |
| 8 | 70809261 | rs12680109 | -0.958 | 1.089 | 0.262 | 0.813 | -0.014 | -0.032 | -0.032 | 0 | 0.33 | 0.26 | 0.49 | 0.53 | 0.42 | 0.45 |
| 8 | 70809973 | rs9643611 | -0.039 | 1.088 | -0.201 | 1.041 | -0.062 | -0.12 | -0.12 | 3.97 | 0.17 | 0.03 | 0.26 | 0.43 | 0.24 | 0.2 |
| 8 | 70812050 | rs6472494 | -1.186 | 0.433 | 0.413 | 0.1 | -0.022 | -0.053 | -0.053 | 0 | 0.35 | 0.42 | 0.32 | 0.22 | 0.33 | 0.37 |
| 8 | 70813658 | rs17670800 | -0.349 | 0.658 | 0.392 | -0.056 | -0.001 | 0.013 | 0.013 | 0 | 0.17 | 0.1 | 0.22 | 0.08 | 0.21 | 0.27 |
| 8 | 70828273 | rs896381 | -0.368 | 0.626 | 0.39 | -0.006 | -0.002 | 0.011 | 0.011 | 0 | 0.17 | 0.11 | 0.23 | 0.09 | 0.22 | 0.27 |
| 8 | 70834713 | rs4738050 | -1.53 | 0.575 | 0.698 | -0.01 | -0.006 | 0.022 | 0.022 | 0 | 0.55 | 0.6 | 0.73 | 0.74 | 0.56 | 0.69 |
| 8 | 70836655 | rs7013896 | -1.675 | 0.685 | 0.748 | 0.07 | -0.003 | 0.034 | 0.034 | 0.119 | 0.56 | 0.69 | 0.74 | 0.74 | 0.6 | 0.7 |
| 8 | 70837397 | rs7843437 | -1.061 | 0.63 | 0.379 | 0.132 | 0.015 | 0.037 | 0.037 | 0 | 0.36 | 0.27 | 0.37 | 0.18 | 0.33 | 0.46 |
| 8 | 70845801 | rs12335270 | -1.409 | 0.582 | 0.005 | 0.128 | -0.015 | -0.039 | -0.039 | 0 | 0.39 | 0.52 | 0.4 | 0.18 | 0.34 | 0.47 |
| 8 | 70848714 | rs6983097 | -1.876 | 0.583 | -0.125 | 0.139 | -0.017 | 0.097 | 0.097 | 0.143 | 0.57 | 0.78 | 0.74 | 0.78 | 0.59 | 0.7 |
| 8 | 70854206 | rs13265196 | 0.593 | 0.666 | -0.093 | 0.199 | 0.001 | 0.019 | 0.019 | 0.563 | 0.17 | 0.21 | 0.34 | 0.56 | 0.26 | 0.23 |
| 8 | 70855209 | rs17435904 | -0.13 | 0.561 | -0.146 | 0.162 | -0.001 | 0.012 | 0.012 | 0 | 0.17 | 0.11 | 0.22 | 0.1 | 0.2 | 0.27 |
| 8 | 70855759 | rs16922791 | -0.054 | 0.549 | -0.123 | 0.164 | 0.042 | 0.13 | 0.13 | 2.432 | 0.05 | 0.27 | 0.1 | 0.08 | 0.02 | 0.11 |
| 8 | 70856897 | rs17671808 | -0.13 | 0.569 | -0.12 | 0.159 | -0.001 | 0.012 | 0.012 | 0 | 0.17 | 0.11 | 0.22 | 0.1 | 0.2 | 0.27 |
| 8 | 70857871 | rs10095238 | -0.171 | 0.545 | -0.122 | 0.146 | -0.032 | -0.045 | -0.045 | 0.021 | 0.19 | 0.31 | 0.16 | 0.08 | 0.12 | 0.2 |
| 8 | 70859624 | rs7836791 | -1.238 | 0.577 | -0.116 | 0.137 | -0.022 | -0.029 | -0.029 | 0 | 0.36 | 0.42 | 0.37 | 0.22 | 0.32 | 0.47 |
| 8 | 70860030 | rs7008027 | -0.291 | 0.557 | 0.003 | 0.23 | -0.033 | -0.017 | -0.017 | 0 | 0.19 | 0.31 | 0.16 | 0.12 | 0.12 | 0.2 |
| 8 | 70863894 | rs10105638 | -0.754 | 1.051 | -0.751 | 0.234 | 0 | 0.001 | 0.001 | 0.021 | 0.33 | 0.35 | 0.43 | 0.59 | 0.38 | 0.31 |
| 8 | 70864617 | rs17671921 | -0.215 | 1.583 | -0.781 | 0.602 | -0.001 | -0.026 | -0.026 | 0.091 | 0.12 | 0.11 | 0.08 | 0 | 0.14 | 0.1 |
| 8 | 70865668 | rs1809104 | -0.775 | 1.921 | -0.897 | 1.242 | -0.013 | -0.016 | -0.016 | 0.207 | 0.22 | 0.23 | 0.5 | 0.67 | 0.33 | 0.4 |
| 8 | 70870856 | rs364641 | 0.761 | 2.126 | -0.747 | 1.454 | 0.01 | 0.025 | 0.025 | 0.668 | 0.87 | 0.86 | 0.52 | 0.33 | 0.79 | 0.77 |
| 8 | 70883360 | rs13274442 | -0.297 | 1.114 | -1.662 | 0.505 | 0.046 | 0.092 | 0.092 | 0.717 | 0.23 | 0.08 | 0.17 | 0.05 | 0.16 | 0.14 |
| 8 | 70887993 | rs12682062 | 1.024 | 0.886 | -2.903 | -0.465 | -0.014 | 0.048 | 0.048 | 0 | 0.28 | 0.03 | 0.1 | 0.15 | 0.2 | 0.32 |
| 8 | 70893868 | rs13259430 | -0.09 | 1.352 | -2.154 | 0.127 | -0.001 | 0.016 | 0.016 | 1.423 | 0.84 | 0.85 | 0.56 | 0.43 | 0.75 | 0.78 |
| 8 | 70895777 | rs6981483 | -0.389 | 1.389 | -2.144 | 0.061 | -0.028 | -0.031 | -0.031 | 0 | 0.86 | 0.93 | 0.6 | 0.38 | 0.78 | 0.79 |
| 8 | 70908001 | rs10808773 | -0.16 | 1.393 | -1.89 | 0.188 | -0.008 | 0.022 | 0.022 | 0 | 0.62 | 0.77 | 0.5 | 0.29 | 0.61 | 0.55 |
| 8 | 70912296 | rs7840667 | 1.208 | 1.36 | -1.412 | -0.315 | -0.008 | 0.014 | 0.014 | 0 | 0.14 | 0.16 | 0.17 | 0.22 | 0.13 | 0.19 |
| 8 | 70912685 | rs10957513 | 1.208 | 1.36 | -1.412 | -0.315 | -0.008 | 0.014 | 0.014 | 0 | 0.14 | 0.16 | 0.17 | 0.22 | 0.13 | 0.19 |
| 8 | 70914507 | rs13258578 | 0.764 | 1.448 | -0.971 | -0.409 | 0.025 | 0.057 | 0.057 | 0 | 0.24 | 0.13 | 0.16 | 0.05 | 0.18 | 0.13 |
| 8 | 70914878 | rs304601 | -0.298 | 1.875 | -0.935 | -0.204 | -0.002 | 0.023 | 0.023 | 0.802 | 0.86 | 0.88 | 0.65 | 0.45 | 0.82 | 0.78 |
| 8 | 70919503 | rs398593 | 0.826 | 2.322 | -1.061 | -0.571 | -0.027 | 0.017 | 0.017 | 0 | 0.11 | 0.05 | 0.1 | 0 | 0.2 | 0.1 |
| 8 | 70926678 | rs1371322 | -0.209 | 2.293 | -1.156 | -0.481 | -0.025 | -0.058 | -0.058 | 0 | 0.18 | 0.12 | 0.32 | 0.58 | 0.2 | 0.22 |
| 8 | 70927807 | rs410015 | 0.352 | 2.223 | -0.212 | -0.84 | 0.118 | -0.037 | -0.037 | 0 | 0.07 | 0.22 | 0.1 | 0 | 0.19 | 0.07 |
| 8 | 70931383 | rs13277769 | 1.033 | 1.869 | -0.319 | -1.133 | 0.014 | 0.047 | 0.047 | 0 | 0.18 | 0.11 | 0.12 | 0.02 | 0.15 | 0.12 |
| 8 | 70931521 | rs4737292 | -1.208 | 2.019 | -0.174 | -1.062 | 0 | -0.009 | -0.009 | 0 | 0.9 | 0.91 | 0.95 | 1 | 0.89 | 0.93 |
| 8 | 70933578 | rs365955 | 0.465 | 2.028 | -0.101 | -0.908 | 0.105 | -0.037 | -0.037 | 0.035 | 0.07 | 0.18 | 0.1 | 0 | 0.19 | 0.07 |
| 8 | 70936591 | rs2580204 | -0.878 | 1.347 | -0.381 | -1.014 | 0.008 | 0.024 | 0.024 | 0 | 0.46 | 0.4 | 0.72 | 0.82 | 0.56 | 0.61 |
| 8 | 70940654 | rs4737293 | -0.139 | 1.029 | -0.983 | -0.581 | -0.022 | 0.036 | 0.036 | 0 | 0.36 | 0.14 | 0.58 | 0.78 | 0.35 | 0.43 |
| 8 | 70942229 | rs10092351 | 0.84 | 0.728 | -0.151 | 0.024 | -0.016 | -0.013 | -0.013 | 0.485 | 0.4 | 0.41 | 0.34 | 0.37 | 0.31 | 0.36 |
| 8 | 70942408 | rs7011482 | -0.058 | 1.095 | 0.127 | 0.432 | 0.005 | 0.088 | 0.088 | 0 | 0.16 | 0.26 | 0.14 | 0.34 | 0.12 | 0.21 |
| 8 | 70946629 | rs10504464 | 1.562 | 0.844 | -0.125 | 0.002 | 0.055 | 0.056 | 0.056 | 0 | 0.33 | 0.25 | 0.15 | 0.12 | 0.21 | 0.25 |
| 8 | 70947278 | rs7000079 | -0.318 | 0.875 | 0.266 | 0.008 | -0.012 | 0.012 | 0.012 | 1.386 | 0.62 | 0.65 | 0.53 | 0.76 | 0.49 | 0.65 |
| 8 | 70950759 | rs2380622 | -0.74 | 1.112 | 0.303 | 0.205 | 0.017 | 0.055 | 0.055 | 0.523 | 0.29 | 0.37 | 0.39 | 0.63 | 0.28 | 0.4 |
| 8 | 70954657 | rs17745746 | 1.382 | 1.754 | -0.434 | 0.139 | -0.022 | 0.027 | 0.027 | 0 | 0.05 | 0 | 0.1 | 0 | 0.18 | 0.08 |
| 8 | 70956288 | rs12678469 | -1.193 | 1.111 | 0.055 | 0.218 | 0.025 | 0.019 | 0.019 | 0.446 | 0.37 | 0.44 | 0.48 | 0.63 | 0.45 | 0.48 |
| 8 | 70961429 | rs7822888 | -0.496 | 0.869 | -0.108 | 0.024 | -0.001 | -0.003 | -0.003 | 0 | 0.64 | 0.61 | 0.6 | 0.73 | 0.62 | 0.7 |
| 8 | 70961514 | rs7844675 | 0.498 | 1.02 | 0.07 | 0.123 | 0.011 | -0.004 | -0.004 | 0 | 0.45 | 0.43 | 0.27 | 0.43 | 0.32 | 0.45 |
| 8 | 70962698 | rs10113363 | -0.724 | 0.534 | 0.048 | 0.066 | -0.029 | 0.004 | 0.004 | 1.244 | 0.67 | 0.85 | 0.62 | 0.73 | 0.62 | 0.7 |
| 8 | 70964399 | rs10089937 | -0.493 | 0.646 | -0.2 | 0.122 | -0.001 | -0.003 | -0.003 | 1.104 | 0.64 | 0.58 | 0.6 | 0.73 | 0.61 | 0.7 |
| 8 | 70967727 | rs3750226 | -0.415 | 0.663 | -0.581 | 0.13 | -0.004 | -0.018 | -0.018 | 0 | 0.63 | 0.51 | 0.61 | 0.73 | 0.61 | 0.7 |
| 8 | 70970514 | rs16936659 | 1.441 | -0.146 | -0.733 | 0.406 | 0.061 | 0.059 | 0.059 | 0.546 | 0.27 | 0.13 | 0.13 | 0.09 | 0.17 | 0.24 |
| 8 | 70979433 | rs6994178 | -0.86 | 0.766 | 0.139 | 1.236 | 0.01 | 0.206 | 0.206 | 0.036 | 0.08 | 0.28 | 0.11 | 0.32 | 0.04 | 0.12 |
| 8 | 70980738 | rs3750228 | -0.082 | 0.891 | 0.071 | 1.308 | 0.003 | 0.044 | 0.044 | 0 | 0.13 | 0.16 | 0.25 | 0.3 | 0.14 | 0.18 |
| 8 | 70981311 | rs3750229 | -0.315 | 0.83 | -0.741 | 1.249 | 0.01 | 0.186 | 0.186 | 0 | 0.07 | 0.21 | 0.1 | 0.32 | 0.04 | 0.12 |
| 8 | 70987280 | rs16936692 | -0.323 | 0.843 | -1.235 | 1.237 | 0.001 | 0.022 | 0.022 | 0.19 | 0.1 | 0.07 | 0.09 | 0.35 | 0.07 | 0.17 |
| 8 | 70992506 | rs4737299 | -0.813 | 1.038 | -1.272 | 1.099 | -0.001 | -0.003 | -0.003 | 0.378 | 0.22 | 0.74 | 0.4 | 0.33 | 0.19 | 0.26 |
| 8 | 70998422 | rs28664277 | 0.778 | 0.98 | -1.734 | 0.481 | 0.02 | 0.065 | 0.065 | 0.192 | 0.52 | 0.17 | 0.36 | 0.3 | 0.46 | 0.47 |
| 8 | 71004987 | rs13278052 | 0.585 | 0.196 | -1.978 | 0.367 | 0.047 | -0.071 | -0.071 | 0 | 0.3 | 0.05 | 0.16 | 0.35 | 0.21 | 0.3 |
| 8 | 71005097 | rs4364664 | -0.642 | 0.334 | -1.465 | 0.163 | 0.027 | 0.027 | 0.027 | 0 | 0.52 | 0.66 | 0.48 | 0.67 | 0.4 | 0.58 |
| 8 | 71005911 | rs10504466 | 0.056 | 0.254 | -0.726 | 0.297 | -0.029 | 0.012 | 0.012 | 0 | 0.39 | 0.4 | 0.22 | 0.51 | 0.26 | 0.39 |
| 8 | 71011912 | rs3892265 | 0.133 | 0.652 | -0.358 | 0.516 | 0 | 0.01 | 0.01 | 0.164 | 0.07 | 0.09 | 0.04 | 0.17 | 0.06 | 0.09 |

| | | | | | | | | | | | | | | | | |
|---|---|---|---|---|---|---|---|---|---|---|---|---|---|---|---|---|
| 8 | 71025282 | rs1046013 | 0.138 | 0.645 | -0.287 | 0.513 | -0.001 | -0.006 | -0.006 | 0 | 0.06 | 0.04 | 0.07 | 0.17 | 0.06 | 0.09 |
| 8 | 71035488 | rs4236979 | -1.34 | 0.3 | 0.397 | 0.49 | -0.029 | 0.212 | 0.212 | 0.007 | 0.2 | 0.4 | 0.18 | 0.51 | 0.14 | 0.34 |
| 8 | 71038715 | rs4545135 | -1.944 | 0.867 | 0.412 | 0.543 | 0.009 | 0.044 | 0.044 | 0.119 | 0.27 | 0.5 | 0.36 | 0.62 | 0.26 | 0.42 |
| 8 | 71048959 | rs16936754 | 0.346 | 0.792 | 0.211 | 0.619 | -0.001 | -0.006 | -0.006 | 0 | 0.07 | 0.04 | 0.07 | 0.17 | 0.07 | 0.09 |
| 8 | 71053907 | rs12674920 | 0.515 | 0.774 | 0.245 | 0.627 | 0.012 | -0.033 | -0.033 | 0.059 | 0.11 | 0.04 | 0.08 | 0.34 | 0.08 | 0.24 |
| 8 | 71101719 | rs3812429 | -0.674 | 0.932 | 0.617 | 0.598 | 0.026 | 0.056 | 0.056 | 0.383 | 0.1 | 0.18 | 0.24 | 0.3 | 0.14 | 0.17 |
| 8 | 71122418 | rs4623463 | 1.515 | 1.171 | -0.035 | 0.085 | -0.031 | 0.037 | 0.037 | 0.887 | 0.54 | 0.1 | 0.51 | 0.3 | 0.56 | 0.37 |
| 8 | 71129813 | rs4738080 | -0.591 | 0.914 | -0.016 | 0.158 | 0.034 | -0.068 | -0.068 | 0 | 0.86 | 0.96 | 0.92 | 0.62 | 0.91 | 0.67 |
| 8 | 71141432 | rs16936837 | 1.132 | 1.077 | 0.26 | 0.019 | -0.003 | -0.009 | -0.009 | 0 | 0.07 | 0.12 | 0.04 | 0.01 | 0.05 | 0.1 |
| 8 | 71151197 | rs2926715 | 1.073 | 1.109 | 0.523 | 0.102 | -0.001 | 0.001 | 0.001 | 0.204 | 0.73 | 0.41 | 0.8 | 0.45 | 0.82 | 0.58 |
| 8 | 71163768 | rs1460680 | 0.672 | 1.045 | 0.557 | 0.331 | -0.002 | -0.006 | -0.006 | 0 | 0.06 | 0.04 | 0.07 | 0.17 | 0.06 | 0.09 |
| 8 | 71164887 | rs2958367 | 0.851 | 0.958 | 0.807 | 0.334 | -0.045 | 0.132 | 0.132 | 0.019 | 0.75 | 0.6 | 0.81 | 0.45 | 0.82 | 0.57 |
| 8 | 71167994 | rs2926702 | -0.232 | 1.036 | 0.749 | 0.342 | 0.007 | -0.08 | -0.08 | 0 | 0.84 | 0.96 | 0.9 | 0.62 | 0.85 | 0.65 |
| 8 | 71200109 | rs16936880 | -0.624 | 0.825 | 1.407 | 0.368 | -0.047 | 0.13 | 0.13 | 0 | 0.19 | 0.36 | 0.12 | 0.38 | 0.12 | 0.33 |
| 8 | 71334489 | rs7823221 | -0.611 | 0.695 | 0.764 | 0.083 | 0.003 | 0 | 0 | 0 | 0.48 | 0.47 | 0.3 | 0.51 | 0.36 | 0.6 |
| 8 | 71336196 | rs6472528 | -1.088 | 0.537 | 0.668 | 0.028 | -0.029 | 0.057 | 0.057 | 0.594 | 0.31 | 0.47 | 0.22 | 0.53 | 0.23 | 0.53 |
| 8 | 71338185 | rs10504477 | -0.982 | 0.669 | 0.603 | -0.201 | 0.016 | 0.054 | 0.054 | 0 | 0.54 | 0.73 | 0.47 | 0.7 | 0.44 | 0.67 |
| 8 | 71340989 | rs16937001 | -0.398 | 0.693 | 0.539 | 0.037 | 0 | 0.032 | 0.032 | 0 | 0.18 | 0.2 | 0.26 | 0.48 | 0.2 | 0.41 |
| 8 | 71342777 | rs10089804 | -0.758 | 0.624 | 0.346 | -0.07 | 0.006 | 0.04 | 0.04 | 0 | 0.25 | 0.31 | 0.28 | 0.49 | 0.27 | 0.42 |
| 8 | 71346171 | rs7817760 | 1.929 | -0.052 | -0.012 | -0.216 | 0.015 | -0.001 | -0.001 | 0 | 0.19 | 0.11 | 0.09 | 0 | 0.13 | 0.06 |
| 8 | 71346772 | rs7011266 | 0.197 | -0.056 | -0.113 | -0.212 | 0 | -0.003 | -0.003 | 0.134 | 0.33 | 0.34 | 0.19 | 0.22 | 0.24 | 0.26 |
| 8 | 71348562 | rs16937012 | -0.67 | 0.264 | 0.016 | 0.314 | -0.007 | 0.006 | 0.006 | 0.006 | 0.14 | 0.22 | 0.11 | 0.22 | 0.1 | 0.2 |
| 8 | 71351952 | rs13259522 | -0.777 | -0.108 | -0.204 | 0.47 | -0.016 | 0.012 | 0.012 | 0 | 0.26 | 0.31 | 0.13 | 0.36 | 0.17 | 0.41 |
| 8 | 71352047 | rs13259607 | -0.608 | -0.004 | 0.127 | 0.607 | -0.029 | 0.092 | 0.092 | 0 | 0.16 | 0.29 | 0.11 | 0.32 | 0.11 | 0.29 |
| 8 | 71352148 | rs13259910 | -0.576 | 0.34 | -0.099 | 0.501 | -0.008 | 0.037 | 0.037 | 0 | 0.18 | 0.22 | 0.12 | 0.33 | 0.16 | 0.32 |
| 8 | 71358464 | rs2380646 | -0.025 | 1.69 | -0.466 | 0.774 | 0.028 | 0.034 | 0.034 | 0 | 0.11 | 0.16 | 0.28 | 0.36 | 0.26 | 0.16 |
| 8 | 71366165 | rs13262850 | 0.247 | 1.858 | -0.477 | 0.757 | 0.008 | 0.005 | 0.005 | 0 | 0.09 | 0.09 | 0.28 | 0.36 | 0.28 | 0.16 |
| 8 | 71371902 | rs1481046 | -0.726 | 1.138 | -0.831 | 0.055 | 0.009 | 0.015 | 0.015 | 0 | 0.54 | 0.63 | 0.75 | 0.65 | 0.63 | 0.53 |
| 8 | 71385557 | rs2380651 | -0.873 | 1.695 | -1.158 | 0.516 | -0.053 | -0.036 | -0.036 | 2.411 | 0.26 | 0.19 | 0.29 | 0.65 | 0.48 | 0.35 |
| 8 | 71387645 | rs10087358 | -0.853 | 1.751 | -0.948 | 0.193 | 0.164 | 0.06 | 0.06 | 0.092 | 0.2 | 0.48 | 0.26 | 0.33 | 0.4 | 0.23 |
| 8 | 71405288 | rs10106461 | -0.611 | 0.925 | -2.067 | -0.452 | -0.013 | 0.239 | 0.239 | 0.759 | 0.42 | 0.12 | 0.62 | 0.93 | 0.63 | 0.56 |
| 8 | 71428717 | rs2622647 | -0.93 | 0.943 | -1.831 | -0.462 | 0.013 | 0.028 | 0.028 | 0.233 | 0.45 | 0.41 | 0.64 | 0.93 | 0.63 | 0.56 |
| 8 | 71432425 | rs10216576 | -0.78 | 1.082 | -2.41 | -0.382 | 0.014 | 0.142 | 0.142 | 0.67 | 0.43 | 0.25 | 0.63 | 0.93 | 0.63 | 0.55 |
| 8 | 71439320 | rs10101215 | 0.265 | 1.46 | -3.069 | 0.029 | -0.081 | -0.1 | -0.1 | 0 | 0.2 | 0.06 | 0.2 | 0.48 | 0.35 | 0.33 |
| 8 | 71445528 | rs12543533 | 0.033 | 1.302 | -2.483 | -0.157 | -0.06 | -0.088 | -0.088 | 0 | 0.2 | 0.03 | 0.41 | 0.25 | 0.26 | 0.18 |
| 8 | 71454171 | rs427330 | -0.223 | 0.79 | -1.918 | -0.753 | -0.046 | 0.099 | 0.099 | 0.018 | 0.84 | 0.98 | 0.91 | 1 | 0.83 | 0.91 |
| 8 | 71464387 | rs17758683 | 0.331 | 1.306 | -2.027 | -0.522 | -0.144 | 0.036 | 0.036 | 0 | 0.24 | 0.02 | 0.2 | 0.11 | 0.35 | 0.17 |
| 8 | 71468885 | rs268583 | -0.161 | -0.21 | -1.135 | -0.464 | -0.05 | 0.01 | 0.01 | 0 | 0.72 | 0.46 | 0.53 | 0.67 | 0.84 | 0.76 |
| 8 | 71477089 | rs268579 | -0.46 | -0.287 | -0.635 | -0.514 | -0.02 | 0.01 | 0.01 | 0 | 0.75 | 0.67 | 0.55 | 0.68 | 0.84 | 0.76 |
| 8 | 71485350 | rs2035196 | 0.832 | -0.268 | -0.717 | -0.746 | -0.118 | 0.152 | 0.152 | 0 | 0.32 | 0.11 | 0.24 | 0.09 | 0.44 | 0.16 |
| 8 | 71486085 | rs8993 | -0.201 | -0.271 | -0.651 | -0.775 | -0.049 | 0.013 | 0.013 | 0 | 0.72 | 0.46 | 0.54 | 0.68 | 0.85 | 0.75 |
| 8 | 71508905 | rs268624 | 0.638 | -0.308 | -0.336 | -0.915 | 0.012 | 0.001 | 0.001 | 0 | 0.37 | 0.24 | 0.24 | 0.6 | 0.33 | 0.58 |
| 8 | 71513130 | rs268594 | 0.251 | -0.31 | -0.296 | -0.918 | -0.013 | -0.003 | -0.003 | 0 | 0.67 | 0.38 | 0.49 | 0.68 | 0.77 | 0.73 |
| 8 | 71526043 | rs268574 | 0.629 | -0.329 | -0.289 | -0.939 | 0.005 | 0 | 0 | 0 | 0.37 | 0.26 | 0.24 | 0.6 | 0.33 | 0.58 |
| 8 | 71529266 | rs13271014 | 1.446 | -0.463 | -0.031 | -1.025 | -0.013 | -0.041 | -0.041 | 0 | 0.16 | 0.16 | 0.13 | 0.02 | 0.09 | 0.1 |
| 8 | 71538686 | rs13280965 | 0.232 | -0.472 | 0.038 | -1.039 | -0.058 | -0.158 | -0.158 | 0.492 | 0.24 | 0.43 | 0.19 | 0.02 | 0.17 | 0.14 |
| 8 | 71544990 | rs16937164 | 0.772 | -0.473 | 0.067 | -1.039 | -0.07 | -0.124 | -0.124 | 0 | 0.19 | 0.36 | 0.15 | 0.02 | 0.09 | 0.11 |
| 8 | 71548605 | rs268569 | 0.55 | -0.338 | 0.019 | -1.088 | 0.005 | 0.004 | 0.004 | 0 | 0.39 | 0.26 | 0.26 | 0.6 | 0.37 | 0.59 |
| 8 | 71548751 | rs13258368 | 1.358 | -0.787 | 0.121 | -0.969 | -0.114 | 0.143 | 0.143 | 0 | 0.31 | 0.12 | 0.25 | 0.08 | 0.44 | 0.15 |
| 8 | 71550832 | rs7830986 | 0.734 | -0.984 | 0.176 | -0.943 | -0.076 | -0.135 | -0.135 | 0 | 0.19 | 0.4 | 0.15 | 0.02 | 0.09 | 0.11 |
| 8 | 71560424 | rs268629 | -0.153 | -0.834 | 0.069 | -0.852 | 0 | 0.034 | 0.034 | 0.389 | 0.92 | 0.89 | 0.69 | 0.7 | 0.94 | 0.86 |
| 8 | 71564667 | rs441890 | -0.517 | -0.854 | 0.074 | -1.101 | 0.011 | 0.03 | 0.03 | 0 | 0.59 | 0.66 | 0.38 | 0.64 | 0.42 | 0.74 |
| 8 | 71578518 | rs11987501 | 1.367 | -0.707 | 0.06 | -1.104 | -0.093 | 0.123 | 0.123 | 0 | 0.3 | 0.13 | 0.24 | 0.07 | 0.44 | 0.14 |
| 8 | 71582591 | rs11991560 | 1.271 | -0.806 | 0.146 | -1.149 | -0.024 | -0.057 | -0.057 | 0 | 0.17 | 0.2 | 0.13 | 0.02 | 0.09 | 0.11 |
| 8 | 71592313 | rs268634 | 0.491 | -0.812 | 0.139 | -1.166 | -0.008 | -0.007 | -0.007 | 0 | 0.38 | 0.37 | 0.25 | 0.63 | 0.32 | 0.58 |
| 8 | 71593198 | rs12681420 | 1.39 | -0.793 | 0.045 | -1.171 | -0.069 | 0.098 | 0.098 | 0 | 0.3 | 0.18 | 0.25 | 0.07 | 0.43 | 0.14 |
| 8 | 71602009 | rs12675271 | 0.857 | -0.816 | 0.037 | -1.186 | -0.045 | 0.097 | 0.097 | 0 | 0.36 | 0.23 | 0.3 | 0.07 | 0.52 | 0.17 |
| 8 | 71615553 | rs1477945 | 1.378 | -0.769 | 0.058 | -1.162 | -0.069 | 0.098 | 0.098 | 0 | 0.3 | 0.18 | 0.25 | 0.07 | 0.43 | 0.14 |
| 8 | 71624632 | rs268647 | 0.623 | -0.778 | 0.066 | -1.16 | -0.016 | -0.029 | -0.029 | 0 | 0.08 | 0.03 | 0.32 | 0.29 | 0.07 | 0.13 |
| 8 | 71626364 | rs1373481 | -1.163 | -0.949 | -0.083 | -1.293 | -0.035 | 0.044 | 0.044 | 0 | 0.9 | 0.99 | 0.9 | 0.98 | 0.86 | 0.92 |
| 8 | 71646980 | rs2290702 | 1.519 | -0.904 | -0.06 | -1.262 | -0.069 | 0.098 | 0.098 | 0 | 0.3 | 0.18 | 0.25 | 0.07 | 0.43 | 0.14 |
| 8 | 71647605 | rs2218430 | -0.47 | -0.915 | -0.027 | -1.165 | 0.016 | 0.01 | 0.01 | 0 | 0.63 | 0.75 | 0.76 | 0.38 | 0.68 | 0.42 |
| 8 | 71663110 | rs4581087 | 1.033 | -0.942 | 0.037 | -1.18 | -0.047 | -0.091 | -0.091 | 0 | 0.18 | 0.28 | 0.14 | 0.02 | 0.09 | 0.11 |
| 8 | 71676339 | rs4738101 | 0.689 | -0.902 | -0.015 | -1.153 | -0.016 | -0.028 | -0.028 | 0 | 0.08 | 0.03 | 0.32 | 0.29 | 0.07 | 0.13 |
| 8 | 71685822 | rs7845516 | 0.038 | -0.938 | 0.028 | -1.395 | 0.002 | -0.031 | -0.031 | 0 | 0.54 | 0.58 | 0.44 | 0.09 | 0.61 | 0.28 |
| 8 | 71718185 | rs921956 | -1.919 | -0.791 | -0.079 | -1.394 | -0.006 | 0.03 | 0.03 | 0 | 0.94 | 0.99 | 0.94 | 1 | 0.94 | 0.99 |
| 8 | 71755038 | rs1156956 | 0.06 | -0.821 | 0.112 | -1.47 | 0.001 | -0.042 | -0.042 | 0 | 0.53 | 0.52 | 0.43 | 0.09 | 0.61 | 0.28 |
| 8 | 71765094 | rs13274381 | 0.929 | -0.798 | 0.012 | -1.46 | -0.046 | 0.09 | 0.09 | 0 | 0.35 | 0.23 | 0.3 | 0.07 | 0.52 | 0.17 |
| 8 | 71781849 | rs6472551 | 0.939 | -0.794 | 0.01 | -1.463 | -0.045 | 0.09 | 0.09 | 0 | 0.35 | 0.22 | 0.3 | 0.07 | 0.52 | 0.17 |

| | | | | | | | | | | | | | | | | |
|---|---|---|---|---|---|---|---|---|---|---|---|---|---|---|---|---|
| 8 | 71805132 | rs4629903 | 1.474 | -0.928 | 0.037 | -1.465 | -0.09 | 0.125 | 0.125 | 0 | 0.3 | 0.13 | 0.24 | 0.07 | 0.43 | 0.14 |
| 8 | 71834440 | rs1473522 | 0.581 | -0.935 | 0.162 | -1.433 | -0.022 | -0.11 | -0.11 | 1.573 | 0.22 | 0.29 | 0.18 | 0.02 | 0.18 | 0.15 |
| 8 | 71837936 | rs35545971 | 1.26 | -0.928 | 0.137 | -1.425 | -0.036 | -0.075 | -0.075 | 0.378 | 0.18 | 0.21 | 0.13 | 0.02 | 0.09 | 0.11 |
| 8 | 71846504 | rs1493199 | 1.547 | -0.914 | -0.083 | -1.454 | -0.11 | 0.138 | 0.138 | 0 | 0.29 | 0.1 | 0.23 | 0.07 | 0.43 | 0.14 |
| 8 | 71875128 | rs2639919 | 0.866 | -0.875 | 0.017 | -1.402 | -0.018 | 0.026 | 0.026 | 0 | 0.37 | 0.3 | 0.3 | 0.07 | 0.51 | 0.18 |
| 8 | 71901737 | rs4577985 | 1.354 | -0.877 | -0.038 | -1.373 | -0.006 | -0.03 | -0.03 | 0 | 0.17 | 0.14 | 0.13 | 0.02 | 0.09 | 0.11 |
| 8 | 71913174 | rs2639942 | 0.066 | -0.825 | -0.023 | -1.357 | 0.007 | 0.013 | 0.013 | 0 | 0.56 | 0.65 | 0.44 | 0.09 | 0.61 | 0.3 |
| 8 | 71923817 | rs3103852 | -1.442 | -0.809 | -0.172 | -1.349 | -0.035 | 0.044 | 0.044 | 0 | 0.9 | 0.99 | 0.9 | 0.98 | 0.87 | 0.91 |
| 8 | 71928298 | rs1389204 | 0.443 | -0.557 | 0.199 | -1.139 | -0.016 | 0.036 | 0.036 | 0 | 0.39 | 0.31 | 0.3 | 0.07 | 0.52 | 0.21 |
| 8 | 71932086 | rs11784872 | 0.701 | -0.559 | -0.122 | -0.896 | -0.017 | -0.028 | -0.028 | 0 | 0.08 | 0.03 | 0.32 | 0.3 | 0.08 | 0.13 |
| 8 | 71944895 | rs4532626 | -1.313 | -0.496 | -0.075 | -1.107 | -0.082 | 0.123 | 0.123 | 0 | 0.69 | 0.87 | 0.76 | 0.93 | 0.57 | 0.85 |
| 8 | 71954596 | rs11779364 | 0.246 | -0.479 | 0.04 | -1.087 | 0.007 | 0.041 | 0.041 | 0 | 0.44 | 0.3 | 0.56 | 0.91 | 0.4 | 0.7 |
| 8 | 71966458 | rs3887553 | -0.338 | -0.428 | -0.025 | -0.699 | 0.012 | 0.012 | 0.012 | 0 | 0.1 | 0.22 | 0.07 | 0.11 | 0.1 | 0.08 |
| 8 | 71969430 | rs1481804 | 0.31 | -0.489 | -0.072 | -0.705 | -0.012 | 0.034 | 0.034 | 0 | 0.39 | 0.33 | 0.31 | 0.19 | 0.53 | 0.23 |
| 8 | 71978814 | rs13268251 | -0.128 | -0.516 | 0.007 | -0.767 | 0.018 | 0.058 | 0.058 | 0 | 0.52 | 0.37 | 0.42 | 0.1 | 0.6 | 0.29 |
| 8 | 71981239 | rs11995646 | 1.196 | -0.466 | -0.012 | -0.746 | -0.027 | -0.058 | -0.058 | 0 | 0.16 | 0.2 | 0.12 | 0.02 | 0.08 | 0.09 |
| 8 | 71988962 | rs2086538 | -0.05 | -0.392 | -0.342 | -0.624 | -0.048 | -0.061 | -0.061 | 0 | 0.26 | 0.5 | 0.19 | 0.14 | 0.18 | 0.17 |
| 8 | 71990280 | rs12674838 | 1.227 | -0.404 | -0.371 | -0.479 | -0.08 | 0.089 | 0.089 | 0 | 0.3 | 0.13 | 0.25 | 0.15 | 0.44 | 0.16 |
| 8 | 71999126 | rs12155783 | 0.351 | 0.194 | -0.161 | 0.169 | 0.003 | -0.009 | -0.009 | 0.763 | 0.53 | 0.48 | 0.42 | 0.17 | 0.61 | 0.3 |
| 8 | 72001931 | rs13251367 | 0.184 | 0.008 | -0.352 | 0.24 | 0.001 | 0.002 | 0.002 | 0 | 0.59 | 0.63 | 0.44 | 0.3 | 0.64 | 0.36 |
| 8 | 72010065 | rs2219989 | 0.428 | 0.448 | -1.04 | 0.129 | -0.025 | 0.032 | 0.032 | 0.093 | 0.06 | 0.01 | 0.05 | 0 | 0.09 | 0.06 |
| 8 | 72012331 | rs1383978 | 0.025 | 0.063 | -0.718 | 0.587 | 0.003 | -0.009 | -0.009 | 0 | 0.32 | 0.25 | 0.23 | 0.48 | 0.28 | 0.53 |
| 8 | 72013005 | rs1481795 | 0.72 | 0.378 | -0.658 | 0.524 | -0.002 | 0.007 | 0.007 | 0.68 | 0.92 | 0.91 | 0.7 | 0.72 | 0.93 | 0.88 |
| 8 | 72015209 | rs10957535 | -0.694 | -0.246 | -0.194 | -0.252 | -0.024 | -0.013 | -0.013 | 1.518 | 0.15 | 0.32 | 0.28 | 0.11 | 0.14 | 0.16 |
| 8 | 72017177 | rs7018254 | -0.911 | -0.923 | -0.545 | -1.372 | 0.073 | 0.125 | 0.125 | 0 | 0.51 | 0.68 | 0.47 | 0.22 | 0.36 | 0.52 |
| 8 | 72018175 | rs7825744 | -0.247 | -0.914 | -0.468 | -1.746 | -0.002 | -0.082 | -0.082 | 7.136 | 0.17 | 0.34 | 0.11 | 0.07 | 0.13 | 0.08 |
| 8 | 72018996 | rs3919903 | -0.356 | -1 | -0.482 | -1.788 | -0.006 | -0.089 | -0.089 | 0.029 | 0.18 | 0.35 | 0.12 | 0.07 | 0.13 | 0.08 |
| 8 | 72020610 | rs12677424 | -0.493 | -1.695 | -0.907 | -2.089 | 0 | 0 | 0 | 0.467 | 0.3 | 0.27 | 0.32 | 0.17 | 0.22 | 0.28 |
| 8 | 72021712 | rs1351513 | -1.94 | -1.454 | -1.424 | -2.061 | -0.14 | -0.075 | -0.075 | 2.745 | 0.69 | 0.91 | 0.55 | 0.34 | 0.55 | 0.64 |
| 8 | 72027611 | rs6981565 | -1.13 | -1.789 | -1.424 | -2.206 | -0.006 | 0.016 | 0.016 | 0 | 0.56 | 0.86 | 0.52 | 0.34 | 0.42 | 0.6 |
| 8 | 72039958 | rs1481790 | 0.505 | -2.055 | -0.906 | -2.349 | -0.035 | -0.005 | -0.005 | 0 | 0.18 | 0.38 | 0.11 | 0.16 | 0.09 | 0.15 |
| 8 | 72040468 | rs1481789 | 0.241 | -2.161 | -0.744 | -2.42 | 0.019 | 0.009 | 0.009 | 0 | 0.24 | 0.17 | 0.33 | 0.17 | 0.2 | 0.29 |
| 8 | 72043312 | rs1481787 | 0.398 | -1.866 | -0.474 | -2.285 | -0.011 | -0.077 | -0.077 | 0 | 0.12 | 0.26 | 0.06 | 0 | 0.11 | 0.16 |
| 8 | 72043557 | rs1481786 | -2.096 | -1.866 | -0.664 | -2.372 | -0.128 | -0.051 | -0.051 | 0.082 | 0.68 | 0.88 | 0.58 | 0.34 | 0.55 | 0.64 |
| 8 | 72043751 | rs1481785 | -0.657 | -2.505 | -0.431 | -2.246 | -0.033 | -0.045 | -0.045 | 0.022 | 0.3 | 0.57 | 0.16 | 0.17 | 0.21 | 0.31 |
| 8 | 72044809 | rs10957537 | 0.526 | -2.527 | -0.485 | -2.221 | -0.006 | -0.04 | -0.04 | 0 | 0.12 | 0.16 | 0.05 | 0 | 0.11 | 0.14 |
| 8 | 72045541 | rs7840218 | -1.972 | -1.869 | -0.622 | -2.227 | -0.128 | -0.062 | -0.062 | 0 | 0.69 | 0.9 | 0.57 | 0.33 | 0.5 | 0.62 |
| 8 | 72047006 | rs2890501 | -1.395 | -2.009 | -0.043 | -1.823 | 0.031 | 0.023 | 0.023 | 0 | 0.52 | 0.6 | 0.45 | 0.3 | 0.36 | 0.54 |
| 8 | 72047354 | rs1075532 | -0.599 | -2.014 | -0.094 | -1.831 | -0.007 | 0.007 | 0.007 | 0 | 0.13 | 0.22 | 0.09 | 0.14 | 0.09 | 0.27 |
| 8 | 72049934 | rs9298161 | 0.518 | -2.005 | -0.097 | -1.854 | 0 | -0.007 | -0.007 | 0 | 0.09 | 0.08 | 0.04 | 0 | 0.09 | 0.12 |
| 8 | 72050405 | rs1481812 | 0.447 | -2.006 | -0.129 | -1.855 | 0 | 0.01 | 0.01 | 0 | 0.09 | 0.08 | 0.03 | 0 | 0.09 | 0.12 |
| 8 | 72054930 | rs17780422 | -0.193 | -2.058 | -0.14 | -1.892 | 0.075 | 0.094 | 0.094 | 0.598 | 0.36 | 0.16 | 0.33 | 0.12 | 0.25 | 0.25 |
| 8 | 72056169 | rs1481803 | -1.366 | -1.982 | -0.086 | -1.848 | 0.034 | 0.029 | 0.029 | 0 | 0.52 | 0.6 | 0.45 | 0.27 | 0.36 | 0.54 |
| 8 | 72058762 | rs1077243 | -1.075 | -0.694 | 0.459 | -1.006 | -0.011 | 0.009 | 0.009 | 0 | 0.13 | 0.2 | 0.08 | 0.14 | 0.08 | 0.27 |
| 8 | 72062096 | rs7843772 | -1.229 | -0.662 | 0.733 | -0.657 | -0.018 | 0.108 | 0.108 | 0 | 0.18 | 0.4 | 0.3 | 0.29 | 0.12 | 0.31 |
| 8 | 72062212 | rs10095316 | -1.197 | -0.676 | 0.903 | -0.798 | 0.009 | -0.013 | -0.013 | 0 | 0.17 | 0.28 | 0.25 | 0.11 | 0.15 | 0.2 |
| 8 | 72065480 | rs4452818 | -1.663 | -0.771 | 1.212 | -0.869 | -0.01 | 0.086 | 0.086 | 0 | 0.22 | 0.46 | 0.34 | 0.27 | 0.17 | 0.36 |
| 8 | 72069159 | rs1481799 | -2.01 | -0.856 | 1.271 | -0.822 | -0.031 | 0.08 | 0.08 | 0 | 0.2 | 0.56 | 0.33 | 0.27 | 0.12 | 0.31 |
| 8 | 72071716 | rs7001898 | -1.375 | -0.716 | 1.176 | -0.798 | -0.009 | 0.122 | 0.122 | 0.061 | 0.16 | 0.37 | 0.29 | 0.27 | 0.11 | 0.3 |
| 8 | 72077955 | rs16937456 | -0.99 | -0.544 | 1.294 | -0.748 | 0.008 | 0.079 | 0.079 | 0.273 | 0.13 | 0.3 | 0.23 | 0.19 | 0.12 | 0.14 |
| 8 | 72078648 | rs7823338 | -1.46 | -0.336 | 1.558 | -1.213 | 0.072 | -0.039 | -0.039 | 0.168 | 0.09 | 0.33 | 0.06 | 0 | 0.09 | 0.04 |
| 8 | 72081678 | rs7819928 | -1.882 | -0.515 | 1.443 | -1.628 | -0.007 | -0.037 | -0.037 | 0 | 0.25 | 0.65 | 0.18 | 0.08 | 0.16 | 0.34 |
| 8 | 72083227 | rs16937466 | -0.626 | -0.48 | 1.505 | -1.618 | -0.032 | -0.041 | -0.041 | 0 | 0.17 | 0.31 | 0.12 | 0.08 | 0.07 | 0.31 |
| 8 | 72083588 | rs6999672 | 0.344 | -0.488 | 0.997 | -1.908 | 0.002 | -0.045 | -0.045 | 0.026 | 0.14 | 0.24 | 0.09 | 0.05 | 0.12 | 0.06 |
| 8 | 72085317 | rs16937472 | -0.118 | -0.331 | 0.515 | -1.747 | 0.065 | -0.029 | -0.029 | 0 | 0.07 | 0.2 | 0.05 | 0 | 0.09 | 0.04 |
| 8 | 72088171 | rs737534 | -0.972 | -0.416 | 0.377 | -1.792 | 0.019 | 0.031 | 0.031 | 0.215 | 0.34 | 0.72 | 0.23 | 0.13 | 0.19 | 0.37 |
| 8 | 72089478 | rs16937477 | -1.083 | -0.223 | 0.723 | -1.631 | 0.097 | -0.033 | -0.033 | 0 | 0.09 | 0.35 | 0.07 | 0 | 0.09 | 0.04 |
| 8 | 72093841 | rs10504502 | -0.433 | -0.132 | 0.537 | -1.586 | 0.102 | -0.02 | -0.02 | 0 | 0.07 | 0.27 | 0.06 | 0 | 0.09 | 0.04 |
| 8 | 72094119 | rs980299 | -0.834 | 0.045 | -0.117 | -1.809 | 0.001 | -0.055 | -0.055 | 0.254 | 0.35 | 0.36 | 0.23 | 0.06 | 0.31 | 0.26 |
| 8 | 72095503 | rs7845690 | 0.047 | 0.399 | -0.82 | -1.244 | 0.045 | 0.047 | 0.047 | 0.772 | 0.34 | 0.16 | 0.46 | 0.77 | 0.25 | 0.44 |
| 8 | 72096237 | rs7828540 | -0.762 | 0.711 | -0.98 | -1.698 | 0.022 | 0.11 | 0.11 | 1.736 | 0.61 | 0.29 | 0.75 | 0.93 | 0.71 | 0.84 |
| 8 | 72101055 | rs4579563 | -0.666 | 0.738 | -0.815 | -1.277 | 0.021 | 0.174 | 0.174 | 0.129 | 0.5 | 0.32 | 0.6 | 0.9 | 0.49 | 0.71 |
| 8 | 72105951 | rs4738110 | 0.514 | 0.799 | -0.934 | -1.052 | -0.017 | -0.027 | -0.027 | 0 | 0.29 | 0.05 | 0.4 | 0.72 | 0.27 | 0.46 |
| 8 | 72108660 | rs4072400 | -0.685 | 1.066 | 0.028 | -0.492 | -0.001 | -0.058 | -0.058 | 0.039 | 0.19 | 0.36 | 0.19 | 0.08 | 0.19 | 0.13 |
| 8 | 72116724 | rs10957540 | -0.479 | 1.133 | -0.083 | -0.462 | 0.001 | -0.032 | -0.032 | 0 | 0.17 | 0.25 | 0.19 | 0.09 | 0.19 | 0.12 |
| 8 | 72122358 | rs10957541 | 0.375 | 1.141 | -0.607 | 0.177 | -0.002 | -0.061 | -0.061 | 0 | 0.11 | 0.04 | 0.27 | 0.58 | 0.12 | 0.31 |
| 8 | 72126639 | rs9298164 | -0.991 | 1.264 | -0.557 | 0.25 | 0.001 | -0.004 | -0.004 | 0 | 0.27 | 0.24 | 0.44 | 0.75 | 0.3 | 0.5 |
| 8 | 72129009 | rs4738118 | -0.249 | 1.242 | -0.38 | 0.706 | 0.005 | -0.005 | -0.005 | 0.364 | 0.12 | 0.1 | 0.27 | 0.67 | 0.12 | 0.38 |
| 8 | 72130973 | rs9643622 | 0.042 | 1.289 | -0.512 | 0.551 | 0.002 | -0.017 | -0.017 | 0.021 | 0.11 | 0.06 | 0.3 | 0.55 | 0.11 | 0.29 |

| | | | | | | | | | | | | | | | | |
|---|---|---|---|---|---|---|---|---|---|---|---|---|---|---|---|---|
| 8 | 72139309 | rs13265957 | -0.637 | 1.559 | 0.097 | -0.627 | -0.003 | -0.003 | -0.003 | 0 | 0.39 | 0.31 | 0.25 | 0.11 | 0.39 | 0.16 |
| 8 | 72143359 | rs1481791 | -0.683 | 1.697 | 0.209 | -0.63 | 0.001 | -0.035 | -0.035 | 0.068 | 0.39 | 0.37 | 0.24 | 0.1 | 0.36 | 0.15 |
| 8 | 72158135 | rs7839496 | -0.513 | 1.973 | 0.35 | -0.414 | 0.023 | -0.074 | -0.074 | 0 | 0.31 | 0.39 | 0.22 | 0.07 | 0.34 | 0.14 |
| 8 | 72180407 | rs17710245 | -0.955 | 2.871 | -0.277 | 0.35 | -0.04 | -0.012 | -0.012 | 0 | 0.16 | 0.08 | 0.17 | 0.19 | 0.31 | 0.25 |
| 8 | 72180481 | rs16937535 | -0.023 | 3.121 | -0.264 | 0.345 | -0.003 | 0.002 | 0.002 | 0 | 0.08 | 0.04 | 0.06 | 0.06 | 0.13 | 0.07 |
| 8 | 72184898 | rs11991562 | -0.38 | 3.029 | 0.126 | 0.357 | 0.01 | 0.023 | 0.023 | 0 | 0.21 | 0.26 | 0.34 | 0.26 | 0.27 | 0.3 |
| 8 | 72189038 | rs9298166 | -0.369 | 2.942 | 0.339 | 0.314 | -0.025 | -0.039 | -0.039 | 7.502 | 0.19 | 0.1 | 0.32 | 0.26 | 0.27 | 0.3 |
| 8 | 72192110 | rs7011671 | -1.008 | 2.828 | 0.393 | 0.079 | 0 | -0.001 | -0.001 | 0 | 0.41 | 0.52 | 0.53 | 0.38 | 0.42 | 0.57 |
| 8 | 72195392 | rs6990158 | 0.054 | 2.883 | 0.24 | 0.092 | -0.034 | -0.046 | -0.046 | 6.413 | 0.2 | 0.1 | 0.32 | 0.26 | 0.27 | 0.31 |
| 8 | 72212365 | rs9657099 | 0.679 | 2.436 | 0.563 | -0.025 | 0.002 | 0.003 | 0.003 | 0 | 0.13 | 0.08 | 0.13 | 0.12 | 0.1 | 0.22 |
| 8 | 72212692 | rs7822041 | -1.26 | 2.345 | 0.831 | -0.135 | -0.011 | 0.065 | 0.065 | 2.04 | 0.45 | 0.58 | 0.73 | 0.82 | 0.49 | 0.68 |
| 8 | 72218918 | rs10957545 | -1.048 | 1.843 | 0.91 | -0.35 | 0.009 | 0.024 | 0.024 | 0 | 0.47 | 0.41 | 0.66 | 0.7 | 0.44 | 0.56 |
| 8 | 72220079 | rs7831030 | -1.781 | 1.846 | 0.926 | -0.443 | 0.003 | -0.007 | -0.007 | 0 | 0.58 | 0.59 | 0.8 | 0.89 | 0.64 | 0.77 |
| 8 | 72224557 | rs1031178 | 1.043 | 1.79 | 0.786 | -0.434 | -0.002 | 0.006 | 0.006 | 0 | 0.16 | 0.01 | 0.14 | 0.13 | 0.11 | 0.23 |
| 8 | 72225252 | rs9298170 | 0.82 | 1.862 | 1.055 | -0.292 | 0.001 | 0.001 | 0.001 | 0 | 0.18 | 0.18 | 0.26 | 0.13 | 0.18 | 0.17 |
| 8 | 72225345 | rs9298171 | -1.228 | 1.912 | 0.879 | -0.298 | -0.005 | 0.006 | 0.006 | 0.117 | 0.41 | 0.58 | 0.66 | 0.74 | 0.53 | 0.53 |
| 8 | 72226102 | rs10095184 | 0.351 | 1.78 | 0.415 | -0.036 | -0.012 | -0.01 | -0.01 | 0 | 0.11 | 0.07 | 0.14 | 0.19 | 0.23 | 0.21 |
| 8 | 72233576 | rs6472581 | -1.387 | 0.77 | 0.693 | -0.194 | 0.029 | 0.077 | 0.077 | 0 | 0.48 | 0.71 | 0.67 | 0.76 | 0.41 | 0.63 |
| 8 | 72233744 | rs7008755 | -0.358 | 0.845 | 0.747 | -0.141 | 0 | 0 | 0 | 0 | 0.25 | 0.31 | 0.31 | 0.18 | 0.25 | 0.3 |
| 8 | 72237879 | rs12682640 | -0.643 | 1.171 | 0.455 | -0.056 | -0.009 | -0.021 | -0.021 | 0 | 0.46 | 0.51 | 0.64 | 0.7 | 0.37 | 0.52 |
| 8 | 72242009 | rs7817242 | -1.31 | 1.969 | 0.971 | -0.03 | 0.089 | 0.004 | 0.004 | 0 | 0.05 | 0.2 | 0.05 | 0.03 | 0.09 | 0.06 |
| 8 | 72242605 | rs12679412 | 1.284 | 2.617 | 0.778 | -0.606 | 0.019 | -0.064 | -0.064 | 0 | 0.77 | 0.63 | 0.82 | 0.96 | 0.68 | 0.78 |
| 8 | 72244775 | rs17712087 | -0.228 | 2.797 | 0.563 | -0.824 | -0.01 | 0.022 | 0.022 | 0 | 0.2 | 0.13 | 0.13 | 0.06 | 0.21 | 0.25 |
| 8 | 72245087 | rs4738128 | 1.516 | 2.726 | 1.565 | -0.256 | 0.084 | 0 | 0 | 0 | 0.85 | 0.48 | 0.83 | 0.85 | 0.85 | 0.85 |
| 8 | 72248174 | rs3779748 | -1.462 | 2.103 | 0.774 | -0.243 | -0.004 | 0.011 | 0.011 | 0.29 | 0.3 | 0.26 | 0.51 | 0.72 | 0.3 | 0.43 |
| 8 | 72248739 | rs7014725 | -0.919 | 2.024 | 1.156 | -0.135 | 0.032 | 0.078 | 0.078 | 0 | 0.07 | 0.23 | 0.09 | 0.15 | 0.04 | 0.05 |
| 8 | 72249217 | rs17785083 | -0.198 | 2.042 | 0.976 | -0.154 | -0.002 | -0.026 | -0.026 | 0 | 0.07 | 0.09 | 0.04 | 0 | 0.06 | 0.03 |
| 8 | 72249453 | rs16937594 | -0.758 | 2.093 | 1.046 | 0.092 | 0.017 | 0.054 | 0.054 | 0 | 0.06 | 0.17 | 0.09 | 0.15 | 0.04 | 0.05 |
| 8 | 72250298 | rs1445402 | -0.956 | 2.092 | 1.065 | 0.093 | 0.038 | 0.086 | 0.086 | 0.024 | 0.07 | 0.3 | 0.09 | 0.15 | 0.04 | 0.05 |
| 8 | 72250370 | rs1900077 | -0.396 | 2.205 | 0.148 | 0.156 | -0.012 | -0.019 | -0.019 | 0 | 0.16 | 0.09 | 0.44 | 0.71 | 0.2 | 0.34 |
| 8 | 72251025 | rs4738129 | -0.164 | 2.048 | 0.099 | 0.459 | -0.007 | -0.041 | -0.041 | 0 | 0.14 | 0.07 | 0.23 | 0.35 | 0.15 | 0.25 |
| 8 | 72253074 | rs4737316 | 0.063 | 1.92 | 0.106 | -0.513 | -0.044 | 0.025 | 0.025 | 0 | 0.33 | 0.1 | 0.52 | 0.7 | 0.33 | 0.48 |
| 8 | 72253783 | rs6983593 | 2.012 | 2.088 | 0.978 | 0.034 | 0.008 | -0.012 | -0.012 | 0 | 0.8 | 0.7 | 0.8 | 0.83 | 0.77 | 0.73 |
| 8 | 72260254 | rs1445403 | 1.825 | 2.156 | 0.991 | 0.027 | 0.006 | -0.024 | -0.024 | 0.397 | 0.74 | 0.29 | 0.74 | 0.84 | 0.7 | 0.7 |
| 8 | 72263956 | rs2218488 | 1.434 | 2.205 | 0.937 | 0.047 | 0.008 | 0.004 | 0.004 | 0 | 0.89 | 0.82 | 0.87 | 0.84 | 0.85 | 0.79 |
| 8 | 72268013 | rs1822917 | 1.711 | 2.152 | 0.913 | 0.064 | 0.019 | 0.002 | 0.002 | 0.004 | 0.85 | 0.77 | 0.83 | 0.83 | 0.81 | 0.77 |
| 8 | 72269374 | rs3808393 | -1.337 | 1.909 | 0.511 | 0.021 | -0.005 | -0.047 | -0.047 | 0 | 0.23 | 0.69 | 0.21 | 0.12 | 0.21 | 0.26 |
| 8 | 72277458 | rs4510897 | 1.38 | 1.931 | -0.738 | 0.872 | 0.006 | 0.014 | 0.014 | 0 | 0.91 | 0.85 | 0.88 | 0.81 | 0.87 | 0.84 |
| 8 | 72280705 | rs1016365 | -0.47 | 2.387 | -1.184 | 0.665 | -0.035 | -0.003 | -0.003 | 0.318 | 0.22 | 0.13 | 0.33 | 0.22 | 0.33 | 0.34 |
| 8 | 72284088 | rs4738130 | -0.094 | 1.716 | -1.805 | 0.636 | -0.015 | -0.113 | -0.113 | 0.816 | 0.8 | 0.98 | 0.69 | 0.41 | 0.84 | 0.8 |
| 8 | 72286799 | rs11775363 | -0.717 | 2.204 | -0.95 | 0.793 | 0.011 | -0.002 | -0.002 | 0 | 0.27 | 0.28 | 0.41 | 0.28 | 0.42 | 0.41 |
| 8 | 72290326 | rs4560821 | -1.212 | 1.951 | -1.216 | 1.504 | -0.017 | -0.03 | -0.03 | 0 | 0.14 | 0.09 | 0.27 | 0.41 | 0.21 | 0.34 |
| 8 | 72292696 | rs4413805 | -0.865 | 2.29 | -0.921 | 1.422 | 0.006 | 0.007 | 0.007 | 0.001 | 0.22 | 0.24 | 0.31 | 0.72 | 0.24 | 0.41 |
| 8 | 72294588 | rs10957550 | -1.241 | 1.711 | -1.264 | 0.943 | -0.012 | -0.001 | -0.001 | 0.18 | 0.3 | 0.13 | 0.32 | 0.72 | 0.28 | 0.42 |
| 8 | 72301132 | rs7012037 | 0.798 | 1.782 | -0.988 | 0.926 | 0.005 | 0 | 0 | 0.015 | 0.73 | 0.7 | 0.66 | 0.27 | 0.72 | 0.58 |
| 8 | 72314205 | rs9643625 | -0.059 | 1.224 | -2.206 | -0.323 | -0.043 | 0.079 | 0.079 | 0 | 0.39 | 0.81 | 0.46 | 0.23 | 0.43 | 0.46 |
| 8 | 72316250 | rs7819542 | 0.861 | 1.403 | -0.857 | -0.473 | 0.04 | -0.086 | -0.086 | 0 | 0.27 | 0.52 | 0.32 | 0.14 | 0.3 | 0.28 |
| 8 | 72316836 | rs11782216 | 0.517 | 1.654 | -1.061 | 0.597 | -0.047 | -0.038 | -0.038 | 0 | 0.08 | 0.01 | 0.21 | 0.36 | 0.14 | 0.18 |
| 8 | 72328044 | rs4379481 | 0.045 | 1.498 | -1.121 | -0.032 | -0.03 | -0.007 | -0.007 | 0 | 0.75 | 0.89 | 0.69 | 0.25 | 0.72 | 0.61 |
| 8 | 72331194 | rs10105943 | -0.207 | 1.166 | -0.965 | -0.222 | 0.001 | -0.005 | -0.005 | 1.856 | 0.34 | 0.37 | 0.4 | 0.22 | 0.42 | 0.43 |
| 8 | 72341815 | rs4344110 | 0.472 | 1.22 | -0.923 | -0.284 | 0 | -0.001 | -0.001 | 1.907 | 0.11 | 0.09 | 0.13 | 0.09 | 0.12 | 0.19 |
| 8 | 72343966 | rs7825729 | 0.447 | 1.163 | -0.768 | -0.096 | -0.005 | 0.016 | 0.016 | 0.475 | 0.73 | 0.83 | 0.59 | 0.16 | 0.73 | 0.55 |
| 8 | 72345405 | rs4434652 | 1.047 | 1.172 | -0.254 | -0.082 | 0.035 | 0.111 | 0.111 | 0.001 | 0.49 | 0.26 | 0.38 | 0.11 | 0.45 | 0.38 |
| 8 | 72346203 | rs4286979 | 1.024 | 1.17 | -0.253 | -0.099 | 0.033 | 0.099 | 0.099 | 0.034 | 0.49 | 0.27 | 0.39 | 0.1 | 0.45 | 0.37 |
| 8 | 72347264 | rs6993604 | 0.764 | 0.693 | -0.623 | 0.529 | -0.01 | 0.013 | 0.013 | 0 | 0.75 | 0.88 | 0.61 | 0.16 | 0.74 | 0.58 |
| 8 | 72348646 | rs13264641 | 0.623 | 0.727 | -0.761 | 0.548 | -0.011 | 0.02 | 0.02 | 0.331 | 0.74 | 0.87 | 0.6 | 0.16 | 0.74 | 0.58 |
| 8 | 72351824 | rs11785552 | -0.938 | 0.098 | -0.619 | 0.695 | 0.006 | -0.003 | -0.003 | 0 | 0.23 | 0.19 | 0.23 | 0.34 | 0.14 | 0.31 |
| 8 | 72358977 | rs4276716 | -0.85 | 0.609 | -0.512 | 0.863 | -0.008 | -0.012 | -0.012 | 0 | 0.08 | 0.04 | 0.09 | 0.19 | 0.1 | 0.13 |
| 8 | 72360490 | rs34049985 | -0.219 | 0.916 | -0.031 | 0.924 | 0.045 | 0.2 | 0.2 | 0.45 | 0.21 | 0.43 | 0.49 | 0.5 | 0.27 | 0.22 |
| 8 | 72375328 | rs12675477 | -0.618 | 0.655 | -0.88 | 0.434 | -0.019 | -0.024 | -0.024 | 0 | 0.29 | 0.1 | 0.23 | 0.31 | 0.27 | 0.41 |
| 8 | 72381735 | rs13280162 | 1.114 | 0.788 | -0.313 | 0.324 | -0.005 | -0.013 | -0.013 | 0 | 0.13 | 0.09 | 0.11 | 0 | 0.19 | 0.07 |
| 8 | 72382593 | rs10089563 | -0.569 | 0.763 | -0.349 | 0.38 | -0.014 | -0.015 | -0.015 | 0.883 | 0.3 | 0.13 | 0.24 | 0.31 | 0.27 | 0.41 |
| 8 | 72392687 | rs10093418 | 1.135 | 0.626 | -0.63 | 0.018 | -0.001 | -0.052 | -0.052 | 0 | 0.1 | 0.01 | 0.07 | 0.19 | 0.1 | 0.19 |
| 8 | 72399360 | rs7846030 | 1.082 | 0.76 | 0.033 | -0.097 | 0.018 | -0.066 | -0.066 | 0 | 0.13 | 0.19 | 0.11 | 0 | 0.17 | 0.09 |
| 8 | 72405673 | rs7828769 | -0.78 | 0.583 | 0.162 | 0.161 | 0.001 | -0.035 | -0.035 | 0 | 0.33 | 0.26 | 0.4 | 0.44 | 0.32 | 0.46 |
| 8 | 72408502 | rs10216347 | -0.797 | 0.655 | 0.338 | -0.01 | 0.002 | 0.012 | 0.012 | 0 | 0.34 | 0.26 | 0.56 | 0.79 | 0.31 | 0.48 |
| 8 | 72411302 | rs7844274 | -0.04 | 0.517 | 0.606 | -0.008 | 0.042 | 0.001 | 0.001 | 0.198 | 0.19 | 0.55 | 0.21 | 0.16 | 0.26 | 0.24 |
| 8 | 72439883 | rs3763548 | -0.118 | 0.355 | 0.463 | -0.113 | 0 | -0.009 | -0.009 | 0 | 0.26 | 0.41 | 0.27 | 0.23 | 0.26 | 0.24 |
| 8 | 72443097 | rs724066 | 0.249 | 0.567 | 0.257 | -0.082 | -0.005 | -0.004 | -0.004 | 0 | 0.18 | 0.21 | 0.24 | 0.23 | 0.22 | 0.18 |

| | | | | | | | | | | | | | | | | |
|---|---|---|---|---|---|---|---|---|---|---|---|---|---|---|---|---|
| 8 | 72456571 | rs7826437 | -1.642 | 0.507 | 0.141 | -0.131 | -0.029 | -0.034 | -0.034 | 0 | 0.56 | 0.82 | 0.5 | 0.53 | 0.55 | 0.64 |
| 8 | 72464866 | rs13249127 | 1.428 | 0.316 | 0.198 | -0.179 | -0.011 | -0.004 | -0.004 | 0.43 | 0.12 | 0.06 | 0.09 | 0 | 0.18 | 0.08 |
| 8 | 72468670 | rs16937647 | -0.421 | 0.055 | 0.111 | -0.057 | -0.003 | 0.113 | 0.113 | 2.788 | 0.09 | 0.12 | 0.31 | 0.42 | 0.08 | 0.16 |
| 8 | 72469742 | rs4738141 | -1.387 | -0.003 | 0.136 | -0.212 | 0.037 | 0.034 | 0.034 | 0.822 | 0.38 | 0.69 | 0.51 | 0.74 | 0.25 | 0.54 |
| 8 | 72480444 | rs16937662 | -0.348 | 0.088 | -0.089 | -0.082 | 0.003 | 0.167 | 0.167 | 3.039 | 0.09 | 0.17 | 0.32 | 0.43 | 0.09 | 0.17 |
| 8 | 72480460 | rs16937664 | -0.361 | 0.088 | -0.088 | -0.082 | 0.002 | 0.165 | 0.165 | 3.06 | 0.09 | 0.17 | 0.32 | 0.43 | 0.09 | 0.17 |
| 8 | 72483983 | rs1156154 | -1.749 | 0.097 | -0.203 | -0.18 | 0.01 | 0.182 | 0.182 | 0.521 | 0.5 | 0.83 | 0.61 | 0.74 | 0.43 | 0.64 |
| 8 | 72487950 | rs11777502 | 0.804 | 0.056 | -0.196 | -0.15 | -0.004 | -0.016 | -0.016 | 0 | 0.15 | 0.13 | 0.21 | 0.25 | 0.19 | 0.17 |
| 8 | 72488774 | rs972738 | -0.397 | -0.011 | -0.138 | -0.058 | 0 | 0.124 | 0.124 | 2.627 | 0.09 | 0.14 | 0.32 | 0.43 | 0.09 | 0.17 |
| 8 | 72506454 | rs1345976 | 0.275 | -0.138 | -0.552 | -0.164 | 0.059 | 0 | 0 | 0 | 0.21 | 0.02 | 0.06 | 0.18 | 0.1 | 0.21 |
| 8 | 72512620 | rs1017260 | -1.674 | -0.021 | -0.561 | -0.162 | 0.01 | 0.172 | 0.172 | 0.885 | 0.5 | 0.87 | 0.61 | 0.74 | 0.43 | 0.64 |
| 8 | 72517046 | rs1035666 | -2.222 | -0.242 | -0.565 | -0.044 | -0.114 | 0.244 | 0.244 | 3.185 | 0.65 | 0.96 | 0.81 | 0.99 | 0.61 | 0.81 |
| 8 | 72519174 | rs1544844 | -0.966 | -0.67 | 0.631 | 0.902 | 0.004 | -0.002 | -0.002 | 0.8 | 0.27 | 0.26 | 0.15 | 0.31 | 0.16 | 0.39 |
| 8 | 72525305 | rs12541255 | -1.185 | -0.65 | 0.932 | 0.921 | 0.048 | 0.463 | 0.463 | 4.038 | 0.1 | 0.41 | 0.35 | 0.43 | 0.08 | 0.16 |
| 8 | 72527240 | rs16937724 | -1.309 | -0.545 | 0.709 | 0.949 | -0.034 | 0.015 | 0.015 | 0 | 0.32 | 0.57 | 0.42 | 0.62 | 0.19 | 0.38 |
| 8 | 72530703 | rs1035669 | -0.174 | -0.414 | 0.601 | 0.931 | 0.039 | 0.441 | 0.441 | 3.599 | 0.05 | 0.2 | 0.31 | 0.44 | 0.04 | 0.15 |
| 8 | 72531550 | rs1559574 | -1.144 | -0.358 | 0.005 | 0.263 | -0.036 | -0.004 | -0.004 | 0 | 0.31 | 0.46 | 0.18 | 0.31 | 0.19 | 0.41 |
| 8 | 72532881 | rs16937732 | -0.348 | 0.319 | -0.314 | -0.014 | 0.003 | -0.012 | -0.012 | 2.683 | 0.13 | 0.18 | 0.05 | 0.1 | 0.12 | 0.12 |
| 8 | 72533079 | rs1424871 | -1.042 | -0.409 | -0.766 | 0.219 | 0.045 | 0.006 | 0.006 | 0.228 | 0.4 | 0.73 | 0.39 | 0.4 | 0.23 | 0.42 |
| 8 | 72533336 | rs4738147 | -0.59 | -0.266 | -0.911 | 0.4 | 0.008 | -0.02 | -0.02 | 0 | 0.11 | 0.05 | 0.15 | 0.22 | 0.1 | 0.18 |
| 8 | 72533577 | rs4738148 | 0.289 | 0.174 | -1.123 | -0.449 | -0.007 | 0.02 | 0.02 | 0 | 0.58 | 0.64 | 0.52 | 0.29 | 0.56 | 0.42 |
| 8 | 72533772 | rs7833628 | -0.199 | 0.751 | -1.424 | -0.254 | -0.059 | -0.054 | -0.054 | 0 | 0.27 | 0.1 | 0.41 | 0.67 | 0.32 | 0.44 |
| 8 | 72534921 | rs17788534 | 0.555 | -0.564 | -0.913 | -1.113 | 0.034 | 0.052 | 0.052 | 1.791 | 0.21 | 0.06 | 0.08 | 0 | 0.17 | 0.03 |
| 8 | 72535162 | rs11775188 | -0.778 | -0.473 | -0.252 | -1.09 | 0.011 | -0.049 | -0.049 | 0.706 | 0.14 | 0.51 | 0.13 | 0.06 | 0.11 | 0.11 |
| 8 | 72541728 | rs10504511 | 0.036 | -0.311 | -0.778 | -1.128 | 0.007 | 0.004 | 0.004 | 0.686 | 0.26 | 0.2 | 0.14 | 0.06 | 0.25 | 0.07 |
| 8 | 72553255 | rs2977752 | 0.221 | -1.661 | -0.673 | -0.76 | -0.019 | 0.002 | 0.002 | 0 | 0.67 | 0.61 | 0.63 | 0.65 | 0.81 | 0.61 |
| 8 | 72553680 | rs7016584 | 0.445 | -1.981 | -0.784 | -1.476 | 0.007 | 0.002 | 0.002 | 0 | 0.31 | 0.22 | 0.23 | 0.06 | 0.22 | 0.3 |
| 8 | 72556469 | rs17727266 | 1.056 | -2.348 | -0.856 | -1.68 | 0.016 | 0.006 | 0.006 | 0.794 | 0.13 | 0.05 | 0.08 | 0 | 0.08 | 0.07 |
| 8 | 72561742 | rs7834018 | -1.079 | -2.782 | -0.84 | -1.666 | -0.023 | 0.09 | 0.09 | 0 | 0.84 | 0.71 | 0.71 | 0.66 | 0.88 | 0.63 |
| 8 | 72562516 | rs16937759 | 0.184 | -2.785 | -0.947 | -2.033 | 0.009 | -0.012 | -0.012 | 0 | 0.26 | 0.2 | 0.17 | 0.05 | 0.21 | 0.15 |
| 8 | 72565496 | rs7818637 | -0.086 | -2.896 | -0.717 | -2.044 | -0.025 | -0.089 | -0.089 | 0 | 0.28 | 0.38 | 0.19 | 0.05 | 0.21 | 0.16 |
| 8 | 72566244 | rs1364616 | 2.192 | -2.92 | -0.695 | -2.065 | -0.023 | -0.023 | -0.023 | 0 | 0.22 | 0.14 | 0.31 | 0.55 | 0.47 | 0.25 |
| 8 | 72567811 | rs6472596 | -0.556 | -3.103 | -0.633 | -2.145 | -0.066 | 0.016 | 0.016 | 0 | 0.66 | 0.5 | 0.62 | 0.66 | 0.81 | 0.63 |
| 8 | 72568275 | rs7822050 | 0.942 | -2.873 | -0.685 | -2.046 | 0.025 | 0.038 | 0.038 | 0.463 | 0.17 | 0.06 | 0.11 | 0.04 | 0.13 | 0.12 |
| 8 | 72570234 | rs16937772 | 1.234 | -2.794 | -0.679 | -2.031 | -0.002 | -0.024 | -0.024 | 0 | 0.1 | 0.11 | 0.06 | 0.01 | 0.08 | 0.05 |
| 8 | 72571144 | rs6986887 | -0.687 | -2.98 | -0.705 | -2.053 | -0.03 | 0.003 | 0.003 | 0 | 0.64 | 0.6 | 0.63 | 0.66 | 0.8 | 0.63 |
| 8 | 72573676 | rs10504513 | 0.681 | -2.692 | -0.999 | -2.006 | 0.022 | 0.062 | 0.062 | 0.503 | 0.18 | 0.05 | 0.12 | 0.04 | 0.16 | 0.14 |
| 8 | 72574871 | rs17795408 | 1.71 | -2.531 | -0.921 | -1.896 | -0.003 | 0.005 | 0.005 | 0 | 0.14 | 0.14 | 0.15 | 0.06 | 0.13 | 0.2 |
| 8 | 72585569 | rs11783976 | 1.638 | -2.538 | -0.883 | -1.917 | -0.001 | -0.002 | -0.002 | 0.735 | 0.14 | 0.15 | 0.15 | 0.05 | 0.13 | 0.22 |
| 8 | 72585932 | rs17795839 | 0.81 | -2.63 | -0.998 | -1.981 | 0.029 | 0.06 | 0.06 | 0.328 | 0.17 | 0.05 | 0.11 | 0.04 | 0.13 | 0.11 |
| 8 | 72586187 | rs2911739 | -0.387 | -2.487 | -0.802 | -1.814 | -0.037 | 0.084 | 0.084 | 0 | 0.74 | 0.52 | 0.65 | 0.64 | 0.8 | 0.59 |
| 8 | 72586292 | rs2911738 | 1.602 | -1.866 | -1.049 | -1.463 | -0.029 | -0.028 | -0.028 | 0 | 0.22 | 0.15 | 0.31 | 0.55 | 0.46 | 0.25 |
| 8 | 72594691 | rs1593272 | 0.644 | -1.714 | -0.949 | -1.278 | -0.009 | -0.018 | -0.018 | 0 | 0.43 | 0.28 | 0.39 | 0.58 | 0.54 | 0.26 |
| 8 | 72597645 | rs7463937 | 0.635 | -1.815 | -0.771 | -1.262 | -0.007 | 0.03 | 0.03 | 0.065 | 0.44 | 0.28 | 0.47 | 0.64 | 0.53 | 0.3 |
| 8 | 72597686 | rs1424883 | 0.167 | -2.273 | -0.417 | -1.535 | 0.017 | 0.017 | 0.017 | 0.063 | 0.62 | 0.72 | 0.76 | 0.72 | 0.71 | 0.64 |
| 8 | 72598621 | rs16937783 | 0.285 | -2.54 | -0.459 | -1.726 | 0.011 | -0.002 | -0.002 | 0 | 0.25 | 0.16 | 0.17 | 0.03 | 0.2 | 0.15 |
| 8 | 72599069 | rs16937784 | 0.142 | -2.536 | -0.396 | -1.719 | -0.009 | -0.073 | -0.073 | 0 | 0.27 | 0.28 | 0.18 | 0.03 | 0.23 | 0.18 |
| 8 | 72600528 | rs2911756 | -0.978 | -2.406 | -0.325 | -1.443 | 0.027 | -0.039 | -0.039 | 1.826 | 0.89 | 0.97 | 0.94 | 0.75 | 0.93 | 0.82 |
| 8 | 72600782 | rs2911755 | 1.258 | -2.131 | -0.136 | -1.467 | -0.006 | 0.01 | 0.01 | 0 | 0.14 | 0.13 | 0.14 | 0.04 | 0.14 | 0.23 |
| 8 | 72602833 | rs1424863 | 0.66 | -2.053 | 0.014 | -1.371 | 0.071 | 0.062 | 0.062 | 0.172 | 0.05 | 0.38 | 0.16 | 0.03 | 0.03 | 0.11 |
| 8 | 72604670 | rs918939 | 0.247 | -2.474 | -0.461 | -1.617 | -0.004 | -0.017 | -0.017 | 0 | 0.66 | 0.42 | 0.62 | 0.68 | 0.7 | 0.51 |
| 8 | 72605379 | rs2068674 | 1.578 | -1.904 | -0.727 | -1.421 | -0.1 | -0.041 | -0.041 | 0 | 0.25 | 0.1 | 0.56 | 0.65 | 0.49 | 0.36 |
| 8 | 72606051 | rs7464949 | 0.347 | -2.179 | 0.071 | -1.304 | 0.003 | 0.002 | 0.002 | 0.04 | 0.54 | 0.4 | 0.7 | 0.61 | 0.66 | 0.39 |
| 8 | 72606942 | rs13277245 | 0.139 | -2.243 | -0.416 | -1.365 | 0.058 | -0.057 | -0.057 | 0 | 0.29 | 0.07 | 0.11 | 0.34 | 0.19 | 0.43 |
| 8 | 72610485 | rs6472600 | -1.417 | -1.224 | 0.486 | -1.097 | -0.003 | 0.001 | 0.001 | 0 | 0.88 | 0.92 | 0.86 | 0.96 | 0.86 | 0.83 |
| 8 | 72614650 | rs7350140 | -0.086 | -1.304 | 0.836 | -1.119 | 0.031 | -0.047 | -0.047 | 0 | 0.81 | 0.59 | 0.79 | 0.91 | 0.8 | 0.79 |
| 8 | 72614772 | rs6472602 | -0.086 | -1.278 | 0.898 | -1.195 | 0.056 | -0.09 | -0.09 | 0 | 0.81 | 0.51 | 0.78 | 0.96 | 0.78 | 0.83 |
| 8 | 72617637 | rs4737325 | -0.327 | -1.347 | 0.939 | -1.231 | 0.029 | -0.054 | -0.054 | 0.499 | 0.82 | 0.65 | 0.8 | 0.94 | 0.81 | 0.83 |
| 8 | 72619613 | rs10283242 | -0.341 | -1.345 | 0.966 | -1.231 | 0.036 | -0.054 | -0.054 | 0 | 0.83 | 0.62 | 0.8 | 0.94 | 0.81 | 0.83 |
| 8 | 72622172 | rs6472604 | 1.185 | -1.257 | 0.453 | -0.871 | -0.06 | -0.043 | -0.043 | 0 | 0.29 | 0.18 | 0.58 | 0.57 | 0.48 | 0.45 |
| 8 | 72622194 | rs6472605 | 0.084 | -1.296 | 0.586 | -0.878 | -0.002 | 0 | 0 | 0 | 0.41 | 0.48 | 0.69 | 0.59 | 0.61 | 0.48 |
| 8 | 72629959 | rs4076783 | 1.056 | -0.563 | 0.25 | -0.721 | -0.044 | -0.042 | -0.042 | 0 | 0.29 | 0.14 | 0.48 | 0.59 | 0.44 | 0.49 |
| 8 | 72632682 | rs10109406 | 1.186 | -1.313 | -0.436 | -1.127 | 0.04 | -0.083 | -0.083 | 0 | 0.24 | 0.06 | 0.15 | 0.37 | 0.15 | 0.24 |
| 8 | 72632742 | rs6996515 | 0.276 | -1.339 | 0.266 | -1.273 | 0.001 | -0.023 | -0.023 | 0 | 0.56 | 0.58 | 0.66 | 0.95 | 0.6 | 0.73 |
| 8 | 72636402 | rs4737326 | -0.603 | -1.478 | 0.044 | -1.38 | -0.025 | 0.063 | 0.063 | 0 | 0.75 | 0.87 | 0.79 | 0.96 | 0.77 | 0.85 |
| 8 | 72636606 | rs4406444 | 0.896 | -1.533 | 0.297 | -1.275 | 0.025 | -0.023 | -0.023 | 0.196 | 0.07 | 0.33 | 0.08 | 0 | 0.08 | 0.01 |
| 8 | 72640116 | rs3890710 | -0.342 | -1.658 | 0.189 | -1.277 | 0.035 | 0.014 | 0.014 | 0 | 0.44 | 0.41 | 0.36 | 0.37 | 0.35 | 0.37 |
| 8 | 72642493 | rs10957568 | 0.268 | -1.66 | -0.062 | -1.278 | 0.068 | -0.049 | -0.049 | 0.199 | 0.37 | 0.07 | 0.27 | 0.37 | 0.27 | 0.36 |
| 8 | 72643654 | rs10808779 | -0.379 | -1.574 | 0.263 | -1.091 | -0.015 | -0.002 | -0.002 | 0 | 0.45 | 0.52 | 0.36 | 0.42 | 0.35 | 0.37 |

| | | | | | | | | | | | | | | | | |
|---|---|---|---|---|---|---|---|---|---|---|---|---|---|---|---|---|
| 8 | 72646159 | rs6472614 | 0.65 | -1.253 | 0.172 | -1.418 | 0 | -0.1 | -0.1 | 0 | 0.58 | 0.56 | 0.74 | 0.96 | 0.6 | 0.73 |
| 8 | 72647799 | rs9694975 | -0.255 | -1.013 | 0.041 | -1.272 | 0.001 | -0.047 | -0.047 | 0.574 | 0.72 | 0.68 | 0.83 | 0.96 | 0.77 | 0.85 |
| 8 | 72648090 | rs13281719 | 0.473 | -0.939 | -0.605 | -1.269 | 0.004 | 0.002 | 0.002 | 1.211 | 0.14 | 0.1 | 0.06 | 0.01 | 0.09 | 0.06 |
| 8 | 72649635 | rs13267135 | -1.576 | -0.868 | -1.021 | -1.176 | -0.057 | 0.032 | 0.032 | 0 | 0.86 | 0.99 | 0.9 | 0.97 | 0.86 | 0.9 |
| 8 | 72649840 | rs11989824 | -0.172 | -1.236 | -0.9 | -1.03 | 0.046 | -0.007 | -0.007 | 0 | 0.45 | 0.62 | 0.44 | 0.47 | 0.3 | 0.34 |
| 8 | 72659540 | rs10957569 | 1.002 | -0.695 | -0.999 | -0.649 | -0.034 | -0.034 | -0.034 | 0 | 0.3 | 0.21 | 0.38 | 0.51 | 0.45 | 0.47 |
| 8 | 72665665 | rs13252179 | 0.549 | -0.626 | -1.113 | -0.511 | 0.019 | -0.039 | -0.039 | 0 | 0.28 | 0.12 | 0.31 | 0.43 | 0.22 | 0.3 |
| 8 | 72669805 | rs13274769 | 0.562 | -0.601 | -1.102 | -0.568 | 0.015 | -0.023 | -0.023 | 0.671 | 0.28 | 0.13 | 0.23 | 0.38 | 0.22 | 0.3 |
| 8 | 72676364 | rs13266504 | 0.955 | -0.485 | -1.638 | -0.624 | 0.013 | -0.065 | -0.065 | 0.942 | 0.25 | 0.06 | 0.2 | 0.33 | 0.2 | 0.2 |
| 8 | 72677349 | rs4300047 | 0.452 | 0.258 | -1.275 | -0.33 | -0.04 | -0.04 | -0.04 | 0.009 | 0.29 | 0.2 | 0.47 | 0.55 | 0.45 | 0.42 |
| 8 | 72684106 | rs4738157 | 0.197 | 1.153 | -1.331 | -0.121 | -0.032 | -0.065 | -0.065 | 0.6 | 0.3 | 0.19 | 0.44 | 0.57 | 0.31 | 0.34 |
| 8 | 72684401 | rs4738159 | 0.393 | 1.205 | -1.418 | -0.133 | -0.068 | -0.15 | -0.15 | 3.084 | 0.28 | 0.07 | 0.3 | 0.58 | 0.28 | 0.37 |
| 8 | 72685824 | rs13276508 | -0.325 | 1.523 | -0.833 | -0.294 | 0.024 | -0.004 | -0.004 | 0.084 | 0.94 | 0.82 | 0.92 | 0.98 | 0.92 | 0.89 |
| 8 | 72687043 | rs7004854 | 0.058 | 1.143 | -0.708 | -0.622 | -0.004 | -0.052 | -0.052 | 0 | 0.33 | 0.46 | 0.33 | 0.16 | 0.37 | 0.33 |
| 8 | 72691106 | rs4305943 | -0.496 | 1.081 | -0.551 | -0.091 | 0.006 | -0.002 | -0.002 | 0 | 0.94 | 0.82 | 0.93 | 0.98 | 0.95 | 0.9 |
| 8 | 72695597 | rs4266692 | 0.477 | 1.132 | -0.911 | -0.002 | -0.006 | -0.064 | -0.064 | 0.035 | 0.28 | 0.11 | 0.32 | 0.62 | 0.26 | 0.42 |
| 8 | 72695849 | rs9643632 | 0.744 | 1.239 | -0.501 | -0.17 | -0.001 | 0.007 | 0.007 | 0.774 | 0.29 | 0.54 | 0.32 | 0.28 | 0.28 | 0.23 |
| 8 | 72695995 | rs4598294 | -0.365 | 1.055 | -0.608 | -0.708 | 0.002 | 0.005 | 0.005 | 0 | 0.67 | 0.57 | 0.67 | 0.68 | 0.67 | 0.67 |
| 8 | 72700062 | rs4345608 | -0.903 | 0.892 | -1.356 | -0.637 | -0.012 | 0.011 | 0.011 | 0 | 0.68 | 0.93 | 0.71 | 0.77 | 0.68 | 0.72 |
| 8 | 72700611 | rs6984067 | -0.411 | 1.051 | -1.158 | -0.48 | 0 | 0 | 0 | 0 | 0.56 | 0.39 | 0.5 | 0.7 | 0.57 | 0.63 |
| 8 | 72701380 | rs10101539 | -0.411 | 1.051 | -1.158 | -0.48 | 0 | 0 | 0 | 0 | 0.56 | 0.39 | 0.5 | 0.7 | 0.57 | 0.63 |
| 8 | 72702271 | rs12681425 | 0.002 | 0.584 | -1.897 | -0.663 | -0.035 | -0.106 | -0.106 | 0.102 | 0.33 | 0.02 | 0.28 | 0.65 | 0.25 | 0.28 |
| 8 | 72707870 | rs13267141 | 0.098 | 1.04 | -0.628 | -1.511 | -0.014 | 0.009 | 0.009 | 0 | 0.18 | 0.09 | 0.12 | 0.02 | 0.21 | 0.21 |
| 8 | 72709739 | rs4345609 | -0.393 | 0.64 | -0.266 | -1.887 | -0.001 | -0.001 | -0.001 | 0 | 0.74 | 0.75 | 0.67 | 0.32 | 0.8 | 0.66 |
| 8 | 72710510 | rs6472619 | 0.95 | 0.537 | 0.121 | -2.135 | -0.003 | 0.087 | 0.087 | 0 | 0.44 | 0.2 | 0.39 | 0.29 | 0.49 | 0.43 |
| 8 | 72712373 | rs6998291 | 0.343 | 0.565 | -0.055 | -2.112 | -0.002 | 0.01 | 0.01 | 0 | 0.56 | 0.64 | 0.54 | 0.3 | 0.59 | 0.45 |
| 8 | 72718314 | rs7459644 | -1.214 | 0.486 | -0.228 | -2.106 | -0.011 | -0.036 | -0.036 | 1.508 | 0.86 | 0.94 | 0.76 | 0.33 | 0.87 | 0.68 |
| 8 | 72718581 | rs4133002 | 1.349 | 0.48 | -0.197 | -2.092 | -0.002 | 0.014 | 0.014 | 0 | 0.09 | 0.07 | 0.13 | 0.01 | 0.1 | 0.03 |
| 8 | 72721007 | rs11779798 | 1.234 | 0.465 | -0.08 | -2.177 | -0.003 | 0.09 | 0.09 | 0 | 0.4 | 0.22 | 0.33 | 0.24 | 0.42 | 0.37 |
| 8 | 72722475 | rs10089527 | 0.37 | 0.363 | 0.011 | -2.16 | -0.037 | -0.07 | -0.07 | 0.02 | 0.18 | 0.41 | 0.16 | 0.06 | 0.14 | 0.08 |
| 8 | 72722483 | rs13257039 | 0.877 | 0.451 | -0.09 | -2.098 | -0.017 | 0.014 | 0.014 | 0 | 0.17 | 0.09 | 0.13 | 0.02 | 0.21 | 0.2 |
| 8 | 72724089 | rs10108922 | 0.335 | 0.359 | -0.257 | -2.238 | 0.002 | 0.022 | 0.022 | 0 | 0.22 | 0.15 | 0.23 | 0.06 | 0.24 | 0.08 |
| 8 | 72726695 | rs4387021 | -1.485 | 0.483 | -0.271 | -2.184 | -0.012 | -0.036 | -0.036 | 0.491 | 0.87 | 0.94 | 0.76 | 0.34 | 0.86 | 0.69 |
| 8 | 72731410 | rs7816767 | 1.399 | 0.42 | -0.095 | -2.359 | -0.001 | 0.014 | 0.014 | 0 | 0.1 | 0.07 | 0.13 | 0.01 | 0.1 | 0.03 |
| 8 | 72731717 | rs7009696 | -0.549 | 0.381 | -0.005 | -2.418 | -0.001 | -0.109 | -0.109 | 0 | 0.41 | 0.56 | 0.37 | 0.04 | 0.38 | 0.27 |
| 8 | 72739271 | rs3923705 | -0.599 | 0.423 | -0.287 | -2.358 | 0.005 | -0.088 | -0.088 | 0 | 0.42 | 0.5 | 0.39 | 0.08 | 0.46 | 0.3 |
| 8 | 72741396 | rs10091707 | -0.857 | 0.875 | -0.717 | -2.034 | -0.007 | 0.091 | 0.091 | 0.142 | 0.55 | 0.71 | 0.48 | 0.09 | 0.53 | 0.33 |
| 8 | 72742860 | rs13282959 | -0.189 | 1.419 | -0.989 | -1.097 | -0.016 | 0.013 | 0.013 | 0 | 0.19 | 0.09 | 0.14 | 0.02 | 0.22 | 0.24 |
| 8 | 72742993 | rs10086032 | -0.846 | 1.431 | -1.019 | -1.111 | -0.02 | 0.035 | 0.035 | 0 | 0.29 | 0.16 | 0.27 | 0.03 | 0.32 | 0.26 |
| 8 | 72743597 | rs13254594 | -0.273 | 1.529 | -1.071 | -0.928 | -0.028 | 0.043 | 0.043 | 0 | 0.19 | 0.07 | 0.13 | 0.02 | 0.22 | 0.21 |
| 8 | 72745949 | rs7828865 | 0.607 | 1.141 | -1.057 | -0.753 | 0.02 | 0.125 | 0.125 | 3.821 | 0.49 | 0.75 | 0.39 | 0.24 | 0.5 | 0.37 |
| 8 | 72746291 | rs4078228 | -0.302 | 1.21 | -1.25 | 0.046 | -0.035 | 0.051 | 0.051 | 1.58 | 0.29 | 0.07 | 0.24 | 0.13 | 0.28 | 0.33 |
| 8 | 72746562 | rs4078227 | -0.558 | 1.063 | -0.064 | 0.821 | 0.048 | 0.072 | 0.072 | 0 | 0.07 | 0.3 | 0.1 | 0.11 | 0.06 | 0.04 |
| 8 | 72751905 | rs7002809 | 0.131 | 0.46 | -1.35 | 0.85 | 0.013 | -0.039 | -0.039 | 0.288 | 0.63 | 0.84 | 0.58 | 0.47 | 0.65 | 0.47 |
| 8 | 72752227 | rs1481852 | 0.145 | 0.256 | -0.738 | 1.19 | 0.008 | -0.018 | -0.018 | 0 | 0.64 | 0.7 | 0.59 | 0.54 | 0.68 | 0.54 |
| 8 | 72752882 | rs3919904 | -0.07 | 0.155 | -0.964 | 0.644 | -0.011 | 0.026 | 0.026 | 0 | 0.1 | 0.02 | 0.11 | 0.06 | 0.15 | 0.07 |
| 8 | 72753740 | rs3779757 | 0.327 | -0.524 | -0.993 | 0.51 | 0.012 | -0.003 | -0.003 | 0 | 0.36 | 0.25 | 0.29 | 0.32 | 0.29 | 0.25 |
| 8 | 72753770 | rs3779758 | 0.335 | -0.515 | -0.996 | 0.507 | 0.013 | -0.003 | -0.003 | 0 | 0.36 | 0.25 | 0.29 | 0.32 | 0.29 | 0.25 |
| 8 | 72754342 | rs900126 | -0.583 | -0.659 | -0.592 | 0.484 | -0.013 | -0.031 | -0.031 | 0 | 0.22 | 0.45 | 0.18 | 0.24 | 0.19 | 0.18 |
| 8 | 72754748 | rs3779761 | -0.686 | -0.727 | -0.399 | 0.449 | -0.018 | -0.036 | -0.036 | 1.051 | 0.22 | 0.53 | 0.19 | 0.24 | 0.19 | 0.17 |
| 8 | 72757443 | rs979451 | -0.726 | -0.911 | -0.15 | 0.453 | 0.007 | -0.009 | -0.009 | 0 | 0.57 | 0.68 | 0.47 | 0.57 | 0.47 | 0.44 |
| 8 | 72757932 | rs7845912 | -0.181 | -0.997 | 0.334 | 0.438 | -0.036 | -0.087 | -0.087 | 0 | 0.17 | 0.37 | 0.15 | 0.05 | 0.14 | 0.07 |
| 8 | 72761149 | rs16937848 | -0.118 | -1.053 | 0.8 | 0.367 | -0.011 | -0.001 | -0.001 | 0.072 | 0.16 | 0.18 | 0.14 | 0.23 | 0.15 | 0.18 |
| 8 | 72761190 | rs2035376 | 0.708 | -1.137 | 0.485 | 0.351 | 0.043 | -0.026 | -0.026 | 0 | 0.35 | 0.14 | 0.28 | 0.33 | 0.28 | 0.24 |
| 8 | 72764061 | rs6994325 | -0.945 | -1.148 | 0.817 | 0.339 | -0.049 | -0.026 | -0.026 | 1.669 | 0.22 | 0.6 | 0.19 | 0.23 | 0.15 | 0.2 |
| 8 | 72768125 | rs6472637 | 0.761 | -1.159 | 0.791 | 0.337 | 0.041 | -0.017 | -0.017 | 0 | 0.35 | 0.18 | 0.29 | 0.33 | 0.28 | 0.26 |
| 8 | 72774642 | rs12674658 | -0.715 | -1.144 | 1.021 | 0.326 | -0.067 | -0.028 | -0.028 | 0.933 | 0.21 | 0.48 | 0.17 | 0.24 | 0.15 | 0.18 |
| 8 | 72777556 | rs900124 | -0.695 | -0.989 | 0.724 | 0.359 | 0.025 | -0.018 | -0.018 | 0 | 0.57 | 0.74 | 0.47 | 0.57 | 0.44 | 0.46 |
| 8 | 72782083 | rs7842844 | -0.114 | -0.341 | 0.59 | 0.277 | -0.001 | 0 | 0 | 0 | 0.15 | 0.13 | 0.15 | 0.11 | 0.2 | 0.08 |
| 8 | 72784886 | rs1600797 | 0.847 | -1.045 | 0.244 | 0.027 | 0.01 | -0.015 | -0.015 | 0.579 | 0.3 | 0.11 | 0.24 | 0.26 | 0.3 | 0.24 |
| 8 | 72786073 | rs9298183 | 0.847 | -1.037 | 0.244 | 0.027 | 0.006 | -0.015 | -0.015 | 0 | 0.3 | 0.11 | 0.24 | 0.26 | 0.3 | 0.24 |
| 8 | 72787399 | rs1117685 | -0.768 | -1.234 | 0.667 | 0.117 | -0.002 | 0.001 | 0.001 | 0.446 | 0.58 | 0.58 | 0.47 | 0.61 | 0.58 | 0.55 |
| 8 | 72805585 | rs1947915 | -0.958 | -1.428 | 0.702 | 0.018 | -0.044 | -0.011 | -0.011 | 0 | 0.63 | 0.79 | 0.5 | 0.61 | 0.59 | 0.56 |
| 8 | 72805974 | rs7010214 | 1.536 | -1.594 | 0.836 | -0.22 | -0.026 | -0.058 | -0.058 | 8.989 | 0.1 | 0.21 | 0.05 | 0 | 0.05 | 0.03 |
| 8 | 72811579 | rs1074088 | -0.413 | -1.456 | 0.754 | -0.115 | 0 | -0.001 | -0.001 | 0.059 | 0.5 | 0.46 | 0.36 | 0.34 | 0.48 | 0.37 |
| 8 | 72817776 | rs10504517 | 0.129 | -1.467 | 0.386 | -0.049 | 0.009 | -0.001 | -0.001 | 2.565 | 0.42 | 0.34 | 0.29 | 0.34 | 0.35 | 0.34 |
| 8 | 72830625 | rs1020312 | -1.029 | -0.812 | -0.24 | -0.032 | -0.104 | -0.091 | -0.091 | 0.472 | 0.7 | 0.93 | 0.53 | 0.6 | 0.63 | 0.58 |
| 8 | 72833262 | rs10504520 | 1.66 | -0.774 | -0.391 | -0.236 | -0.034 | 0.059 | 0.059 | 0 | 0.1 | 0.01 | 0.07 | 0 | 0.14 | 0.05 |
| 8 | 72835774 | rs6472649 | 0.533 | -0.999 | 0.221 | -0.225 | -0.018 | 0.025 | 0.025 | 0 | 0.14 | 0.26 | 0.14 | 0.27 | 0.12 | 0.21 |

| | | | | | | | | | | | | | | | |
|---|---|---|---|---|---|---|---|---|---|---|---|---|---|---|---|
| 8 | 72836859 | rs13270508 | 1.479 | -1.132 | 0.333 | -0.336 | -0.028 | -0.058 | -0.058 | 0 | 0.11 | 0.22 | 0.06 | 0 | 0.05 | 0.04 |
| 8 | 72840154 | rs1373301 | 0.388 | -1.131 | 0.332 | -0.313 | 0 | 0 | 0 | 0 | 0.43 | 0.4 | 0.27 | 0.34 | 0.36 | 0.32 |
| 8 | 72852760 | rs7835557 | 0.524 | -1.049 | 0.004 | -0.291 | -0.019 | 0.024 | 0.024 | 0 | 0.15 | 0.3 | 0.14 | 0.25 | 0.13 | 0.19 |
| 8 | 72855904 | rs9298191 | 0.268 | -1.06 | -0.001 | -0.285 | 0.015 | 0.006 | 0.006 | 0.002 | 0.45 | 0.62 | 0.3 | 0.34 | 0.36 | 0.34 |
| 8 | 72858966 | rs12114832 | -0.615 | -0.912 | -0.603 | -0.248 | -0.079 | -0.035 | -0.035 | 0 | 0.6 | 0.91 | 0.44 | 0.6 | 0.49 | 0.53 |
| 8 | 72861260 | rs7843557 | -0.843 | -1.008 | -0.349 | -0.385 | 0.032 | 0.057 | 0.057 | 0.035 | 0.39 | 0.78 | 0.3 | 0.26 | 0.32 | 0.29 |
| 8 | 72871111 | rs1481853 | -0.257 | -0.721 | -0.14 | -0.024 | -0.009 | -0.004 | -0.004 | 0 | 0.56 | 0.48 | 0.69 | 0.63 | 0.65 | 0.64 |
| 8 | 72872544 | rs4738191 | 1.415 | -0.732 | -0.448 | -0.197 | 0.012 | 0 | 0 | 0 | 0.18 | 0.09 | 0.13 | 0.21 | 0.15 | 0.24 |
| 8 | 72872842 | rs10429397 | 0.13 | -0.771 | -0.443 | -0.222 | -0.021 | -0.018 | -0.018 | 2.323 | 0.32 | 0.4 | 0.2 | 0.23 | 0.21 | 0.34 |
| 8 | 72875478 | rs12541619 | 1.135 | -0.597 | -1.138 | -0.164 | -0.076 | -0.077 | -0.077 | 0.014 | 0.26 | 0.11 | 0.46 | 0.42 | 0.35 | 0.35 |
| 8 | 72875874 | rs11778494 | 1.593 | -0.677 | -0.944 | 0.201 | -0.02 | -0.058 | -0.058 | 0 | 0.26 | 0.04 | 0.18 | 0.3 | 0.26 | 0.2 |
| 8 | 72882582 | rs7016825 | 0.436 | -0.35 | -0.61 | 0.195 | 0.003 | 0.003 | 0.003 | 0 | 0.38 | 0.43 | 0.56 | 0.42 | 0.5 | 0.4 |
| 8 | 72890067 | rs13258140 | 0.542 | -0.324 | -0.975 | 0.296 | -0.02 | -0.023 | -0.023 | 6.4 | 0.13 | 0.24 | 0.08 | 0.04 | 0.09 | 0.09 |
| 8 | 72890597 | rs11996971 | 1.494 | -0.201 | -1.796 | 0.516 | 0.006 | -0.03 | -0.03 | 0 | 0.21 | 0.07 | 0.14 | 0.29 | 0.19 | 0.19 |
| 8 | 72890601 | rs17737248 | -0.076 | 0.036 | -1.712 | 0.29 | 0.037 | 0.038 | 0.038 | 1.016 | 0.24 | 0.06 | 0.12 | 0.18 | 0.17 | 0.29 |
| 8 | 72890680 | rs6472655 | 0.174 | 1.174 | -1.386 | 0.501 | 0.007 | 0.006 | 0.006 | 0 | 0.07 | 0.11 | 0.06 | 0.08 | 0.12 | 0.08 |
| 8 | 72891667 | rs4738193 | 0.049 | 0.583 | -1.712 | 0.341 | 0.017 | -0.033 | -0.033 | 0 | 0.24 | 0.15 | 0.41 | 0.29 | 0.2 | 0.3 |
| 8 | 72899566 | rs7842160 | -0.783 | 1.596 | -1.804 | 0.278 | -0.048 | 0.037 | 0.037 | 0.049 | 0.92 | 0.99 | 0.91 | 0.99 | 0.78 | 0.91 |
| 8 | 72904776 | rs1036963 | -0.82 | 0.47 | -1.652 | 0.446 | -0.015 | 0.016 | 0.016 | 1.833 | 0.55 | 0.31 | 0.62 | 0.56 | 0.53 | 0.62 |
| 8 | 72904930 | rs1036964 | 0.204 | 0.475 | -1.684 | 0.421 | 0.027 | -0.026 | -0.026 | 0 | 0.27 | 0.18 | 0.43 | 0.29 | 0.21 | 0.31 |
| 8 | 72907897 | rs958898 | -0.071 | 0.728 | -1.414 | 0.295 | -0.007 | 0.002 | 0.002 | 1.696 | 0.3 | 0.12 | 0.19 | 0.27 | 0.31 | 0.3 |
| 8 | 72917231 | rs1155489 | -0.277 | 0.489 | -0.574 | -0.028 | -0.02 | -0.024 | -0.024 | 0.049 | 0.3 | 0.43 | 0.47 | 0.22 | 0.25 | 0.43 |
| 8 | 72917890 | rs7844113 | 0.075 | 0.298 | -0.365 | -0.037 | 0.01 | 0.002 | 0.002 | 3.835 | 0.65 | 0.65 | 0.71 | 0.63 | 0.51 | 0.58 |
| 8 | 72919087 | rs10957577 | 1.166 | 0.024 | 0.021 | -0.131 | 0.012 | -0.025 | -0.025 | 0 | 0.36 | 0.3 | 0.32 | 0.46 | 0.33 | 0.3 |
| 8 | 72919626 | rs1531750 | -0.336 | 0.086 | 0.031 | -0.324 | -0.021 | 0 | 0 | 0 | 0.71 | 0.8 | 0.8 | 0.98 | 0.6 | 0.74 |
| 8 | 72920794 | rs9298195 | 0.288 | 0.135 | -0.108 | -0.27 | 0.005 | 0 | 0 | 0 | 0.62 | 0.65 | 0.71 | 0.63 | 0.5 | 0.58 |
| 8 | 72921140 | rs2054597 | 1.252 | 0.256 | -0.426 | -0.206 | 0.011 | -0.023 | -0.023 | 0 | 0.36 | 0.3 | 0.32 | 0.46 | 0.33 | 0.29 |
| 8 | 72925434 | rs894886 | -0.712 | 0.664 | -0.445 | -0.307 | -0.008 | 0.006 | 0.006 | 0 | 0.84 | 0.82 | 0.82 | 0.92 | 0.73 | 0.8 |
| 8 | 72925689 | rs4738199 | 0.469 | 0.169 | -0.095 | -0.317 | 0.016 | 0.016 | 0.016 | 0 | 0.27 | 0.23 | 0.38 | 0.13 | 0.18 | 0.32 |
| 8 | 72928115 | rs6472656 | -0.321 | 0.351 | 0.254 | -0.022 | -0.003 | 0.082 | 0.082 | 1.704 | 0.23 | 0.44 | 0.17 | 0.36 | 0.22 | 0.27 |
| 8 | 72933521 | rs6996413 | 0.506 | 0.364 | 0.391 | -0.227 | 0.018 | -0.114 | -0.114 | 0.032 | 0.16 | 0.4 | 0.12 | 0 | 0.13 | 0.11 |
| 8 | 72933632 | rs6996723 | -0.28 | 0.064 | 0.277 | -0.474 | -0.018 | -0.105 | -0.105 | 0 | 0.26 | 0.54 | 0.16 | 0.01 | 0.18 | 0.17 |
| 8 | 72934032 | rs7827617 | 0.464 | -0.128 | -0.233 | -0.646 | 0 | -0.039 | -0.039 | 0.431 | 0.22 | 0.14 | 0.1 | 0.35 | 0.2 | 0.2 |
| 8 | 72935839 | rs959974 | -0.154 | -0.171 | -0.257 | -0.786 | 0 | -0.002 | -0.002 | 1.491 | 0.53 | 0.46 | 0.39 | 0.76 | 0.49 | 0.41 |
| 8 | 72936145 | rs959976 | 0.652 | 0.032 | -0.674 | -0.649 | -0.037 | -0.107 | -0.107 | 0 | 0.21 | 0.01 | 0.08 | 0.34 | 0.2 | 0.19 |
| 8 | 72940861 | rs4738202 | -0.605 | 0.301 | -0.037 | -0.931 | 0.068 | -0.138 | -0.138 | 0 | 0.78 | 0.56 | 0.79 | 0.98 | 0.69 | 0.85 |
| 8 | 72943511 | rs10100108 | 0.922 | -0.147 | -0.16 | -0.876 | -0.009 | 0.019 | 0.019 | 0.471 | 0.32 | 0.39 | 0.57 | 0.56 | 0.31 | 0.39 |
| 8 | 72945037 | rs13259803 | 0.293 | -0.31 | -0.177 | -0.859 | 0.023 | 0.071 | 0.071 | 0.437 | 0.36 | 0.27 | 0.18 | 0.13 | 0.27 | 0.33 |
| 8 | 72947312 | rs12545839 | 1.965 | -0.421 | -0.211 | -0.609 | -0.003 | 0.004 | 0.004 | 0 | 0.12 | 0.12 | 0.29 | 0.22 | 0.08 | 0.26 |
| 8 | 72948588 | rs13280644 | 2.189 | -0.384 | -0.433 | -0.71 | -0.028 | 0.074 | 0.074 | 2.998 | 0.12 | 0 | 0.06 | 0 | 0.09 | 0.11 |
| 8 | 72953158 | rs1025926 | 2.154 | -0.424 | -0.252 | -0.71 | 0 | 0.008 | 0.008 | 1.189 | 0.19 | 0.21 | 0.28 | 0.34 | 0.23 | 0.13 |
| 8 | 72959152 | rs6982184 | 0.396 | -0.337 | -0.574 | -0.698 | 0.021 | 0.132 | 0.132 | 0.313 | 0.35 | 0.2 | 0.16 | 0.06 | 0.27 | 0.22 |
| 8 | 72961252 | rs2383844 | -0.4 | -0.315 | -0.518 | -0.68 | 0.03 | 0.01 | 0.01 | 0.015 | 0.68 | 0.61 | 0.73 | 0.62 | 0.57 | 0.61 |
| 8 | 72963135 | rs1025927 | 1.969 | -0.337 | -0.528 | -0.697 | 0 | -0.021 | -0.021 | 0 | 0.13 | 0.15 | 0.08 | 0 | 0.09 | 0.11 |
| 8 | 72963258 | rs1025928 | -0.67 | -0.335 | -0.518 | -0.694 | 0.014 | -0.002 | -0.002 | 0 | 0.69 | 0.68 | 0.75 | 0.69 | 0.57 | 0.71 |
| 8 | 72963296 | rs1025929 | -0.332 | -0.528 | -0.27 | -0.371 | 0.001 | 0.002 | 0.002 | 0.098 | 0.54 | 0.44 | 0.65 | 0.62 | 0.48 | 0.51 |
| 8 | 72965123 | rs10504525 | 1.041 | -0.537 | -0.285 | -0.377 | -0.004 | 0.077 | 0.077 | 0 | 0.22 | 0.03 | 0.07 | 0.06 | 0.17 | 0.12 |
| 8 | 72965973 | rs3735942 | 0.794 | -0.516 | -0.273 | -0.366 | -0.009 | 0.019 | 0.019 | 0.275 | 0.32 | 0.36 | 0.57 | 0.56 | 0.31 | 0.39 |
| 8 | 72966002 | rs3735943 | -0.294 | -0.466 | -0.518 | -0.466 | -0.002 | -0.014 | -0.014 | 0 | 0.55 | 0.51 | 0.67 | 0.69 | 0.48 | 0.61 |
| 8 | 72969877 | rs7825042 | 1.178 | -0.464 | -0.508 | -0.483 | 0.01 | 0.091 | 0.091 | 0 | 0.14 | 0.29 | 0.08 | 0.28 | 0.13 | 0.14 |
| 8 | 72972017 | rs10101155 | 0.698 | -0.475 | -0.569 | -0.493 | -0.019 | 0.013 | 0.013 | 0.078 | 0.33 | 0.43 | 0.59 | 0.63 | 0.31 | 0.49 |
| 8 | 72974329 | rs10109581 | -0.519 | -0.004 | -0.709 | -0.022 | -0.003 | 0 | 0 | 0 | 0.36 | 0.4 | 0.16 | 0.34 | 0.3 | 0.26 |
| 8 | 72976997 | rs16937961 | 1.557 | -0.069 | -0.668 | -0.009 | -0.002 | -0.038 | -0.038 | 0 | 0.19 | 0.12 | 0.27 | 0.34 | 0.23 | 0.13 |
| 8 | 72977703 | rs920829 | 0.356 | -0.078 | -0.39 | 0.039 | 0.011 | 0.097 | 0.097 | 0 | 0.14 | 0.3 | 0.08 | 0.28 | 0.13 | 0.14 |
| 8 | 72981499 | rs10091093 | -0.976 | -0.067 | -0.409 | 0.045 | -0.003 | 0 | 0 | 0.843 | 0.36 | 0.4 | 0.16 | 0.34 | 0.3 | 0.26 |
| 8 | 72987638 | rs13268757 | -0.228 | 0.027 | -1.156 | -0.305 | 0.016 | 0.076 | 0.076 | 0 | 0.22 | 0.08 | 0.07 | 0.06 | 0.17 | 0.11 |
| 8 | 72989828 | rs2587559 | -0.325 | 0.293 | -0.467 | 0.302 | -0.001 | 0.062 | 0.062 | 0 | 0.16 | 0.25 | 0.1 | 0.3 | 0.15 | 0.13 |
| 8 | 72991038 | rs2587562 | -0.63 | 0.656 | -0.409 | -0.733 | -0.003 | 0.006 | 0.006 | 0.995 | 0.45 | 0.46 | 0.29 | 0.35 | 0.47 | 0.41 |
| 8 | 72993631 | rs2587567 | -0.451 | 0.698 | -0.424 | -0.957 | -0.032 | 0 | 0 | 0.001 | 0.35 | 0.19 | 0.24 | 0.35 | 0.41 | 0.36 |
| 8 | 72994478 | rs10093536 | 1.12 | 0.486 | -0.654 | -0.948 | -0.002 | 0.002 | 0.002 | 0.006 | 0.19 | 0.23 | 0.26 | 0.21 | 0.2 | 0.15 |
| 8 | 72998638 | rs2701455 | -2.006 | 0.166 | -1.06 | -1.214 | 0.019 | 0.051 | 0.051 | 0 | 0.82 | 0.96 | 0.88 | 0.93 | 0.87 | 0.81 |
| 8 | 73000452 | rs2701454 | -0.217 | 0.188 | -1.018 | -1.196 | -0.003 | 0.007 | 0.007 | 1.14 | 0.46 | 0.46 | 0.29 | 0.35 | 0.47 | 0.41 |
| 8 | 73000881 | rs17741463 | 1.798 | 0.231 | -1.082 | -1.202 | -0.038 | 0.068 | 0.068 | 0.338 | 0.2 | 0.07 | 0.15 | 0.04 | 0.27 | 0.23 |
| 8 | 73001870 | rs1443948 | -1.258 | 0.058 | -1.028 | -1.211 | -0.001 | -0.001 | -0.001 | 0 | 0.68 | 0.69 | 0.45 | 0.42 | 0.62 | 0.61 |
| 8 | 73004294 | rs2383847 | 1.578 | 0.095 | -0.973 | -1.174 | -0.009 | -0.033 | -0.033 | 0 | 0.2 | 0.2 | 0.14 | 0.07 | 0.14 | 0.2 |
| 8 | 73004807 | rs2701453 | -0.761 | 0.047 | -1.018 | -1.193 | 0.006 | -0.009 | -0.009 | 0.102 | 0.63 | 0.76 | 0.53 | 0.55 | 0.66 | 0.5 |
| 8 | 73005474 | rs7835630 | -0.092 | 0.043 | -0.66 | -1.292 | 0.005 | -0.021 | -0.021 | 1.991 | 0.43 | 0.45 | 0.42 | 0.24 | 0.5 | 0.34 |
| 8 | 73006453 | rs1443946 | 0.041 | 0.056 | -0.706 | -1.266 | 0.002 | 0 | 0 | 0.328 | 0.39 | 0.45 | 0.25 | 0.38 | 0.29 | 0.36 |
| 8 | 73007576 | rs2587577 | 1.886 | -0.045 | -0.312 | -1.433 | -0.002 | 0.011 | 0.011 | 0 | 0.15 | 0.17 | 0.1 | 0.31 | 0.18 | 0.11 |

| | | | | | | | | | | | | | | | | |
|---|---|---|---|---|---|---|---|---|---|---|---|---|---|---|---|---|
| 8 | 73007895 | rs7008847 | 0.63 | -0.006 | -0.381 | -1.407 | -0.002 | -0.036 | -0.036 | 0 | 0.3 | 0.36 | 0.19 | 0.07 | 0.19 | 0.26 |
| 8 | 73008249 | rs2701448 | 2.725 | 0.098 | -0.541 | -1.209 | -0.007 | -0.015 | -0.015 | 2.93 | 0.09 | 0.05 | 0.05 | 0.31 | 0.11 | 0.11 |
| 8 | 73012834 | rs2701440 | -1.662 | 0.056 | -0.466 | -1.192 | 0.001 | -0.008 | -0.008 | 0.844 | 0.77 | 0.84 | 0.62 | 0.62 | 0.77 | 0.69 |
| 8 | 73015538 | rs7823587 | 1.744 | 0.299 | -0.526 | -1.218 | -0.003 | -0.003 | -0.003 | 0 | 0.14 | 0.13 | 0.2 | 0.21 | 0.15 | 0.14 |
| 8 | 73019578 | rs13253169 | 2.241 | 0.2 | -0.324 | -1.334 | 0.012 | -0.045 | -0.045 | 3.554 | 0.08 | 0.34 | 0.05 | 0 | 0.04 | 0.04 |
| 8 | 73020254 | rs16938029 | 2.413 | 0.072 | -0.334 | -1.439 | -0.005 | -0.024 | -0.024 | 0 | 0.1 | 0.09 | 0.07 | 0 | 0.09 | 0.01 |
| 8 | 73023295 | rs16938030 | 2.437 | 0.11 | -0.267 | -1.396 | -0.015 | -0.068 | -0.068 | 0.585 | 0.18 | 0.07 | 0.36 | 0.42 | 0.2 | 0.3 |
| 8 | 73025542 | rs16938031 | 2.103 | 0.092 | -0.18 | -1.56 | 0.007 | -0.043 | -0.043 | 0 | 0.08 | 0.29 | 0.05 | 0 | 0.04 | 0.04 |
| 8 | 73030939 | rs6472663 | 1.607 | 0.194 | -0.469 | -1.612 | -0.033 | 0.051 | 0.051 | 0.343 | 0.18 | 0.07 | 0.13 | 0.03 | 0.25 | 0.19 |
| 8 | 73035769 | rs6988179 | 0.739 | -0.078 | -0.409 | -1.567 | 0.002 | 0 | 0 | 1.11 | 0.34 | 0.32 | 0.58 | 0.63 | 0.36 | 0.44 |
| 8 | 73038310 | rs11775093 | 0.411 | 0.196 | -0.797 | -1.503 | -0.004 | 0.029 | 0.029 | 0.194 | 0.35 | 0.2 | 0.23 | 0.05 | 0.4 | 0.41 |
| 8 | 73040059 | rs13277178 | -0.33 | 0.549 | -0.432 | -1.009 | -0.006 | -0.015 | -0.015 | 0 | 0.76 | 0.77 | 0.82 | 0.65 | 0.74 | 0.72 |
| 8 | 73040431 | rs7000490 | -0.291 | 1.103 | -0.005 | -0.655 | 0.013 | -0.007 | -0.007 | 0.053 | 0.36 | 0.5 | 0.42 | 0.27 | 0.38 | 0.19 |
| 8 | 73041162 | rs7005180 | 1.247 | 0.521 | 0.144 | -0.576 | 0.001 | 0.003 | 0.003 | 0 | 0.42 | 0.3 | 0.4 | 0.35 | 0.42 | 0.44 |
| 8 | 73045055 | rs6988449 | 1.051 | 0.472 | 0.247 | -0.685 | -0.006 | -0.005 | -0.005 | 0 | 0.14 | 0.09 | 0.29 | 0.34 | 0.19 | 0.19 |
| 8 | 73045090 | rs6988473 | -2.332 | 0.495 | 0.231 | -0.695 | -0.007 | 0.019 | 0.019 | 0.03 | 0.94 | 0.96 | 0.9 | 0.71 | 0.92 | 0.9 |
| 8 | 73045557 | rs13265648 | -1.14 | 0.48 | 0.246 | -0.723 | -0.001 | 0.007 | 0.007 | 0 | 0.52 | 0.61 | 0.5 | 0.36 | 0.5 | 0.46 |
| 8 | 73049287 | rs16938067 | 1.872 | 0.321 | 0.455 | -1.015 | -0.014 | -0.043 | -0.043 | 0 | 0.13 | 0.17 | 0.08 | 0 | 0.1 | 0.04 |
| 8 | 73049791 | rs11777836 | 0.475 | 0.457 | 0.394 | -0.929 | -0.001 | -0.059 | -0.059 | 0.025 | 0.25 | 0.38 | 0.13 | 0.02 | 0.2 | 0.23 |
| 8 | 73050258 | rs4737340 | 0.542 | 0.387 | 0.266 | -0.767 | -0.001 | 0.002 | 0.002 | 0.021 | 0.54 | 0.47 | 0.48 | 0.35 | 0.52 | 0.48 |
| 8 | 73053079 | rs10504532 | 0.124 | 0.277 | -0.083 | -0.816 | -0.001 | -0.045 | -0.045 | 0 | 0.27 | 0.33 | 0.17 | 0.02 | 0.23 | 0.21 |
| 8 | 73054988 | rs16938076 | 0.591 | 0.227 | -0.049 | -0.587 | 0.001 | 0.001 | 0.001 | 0 | 0.17 | 0.16 | 0.1 | 0.13 | 0.13 | 0.12 |
| 8 | 73055368 | rs7842978 | 0.752 | 0.406 | 0.456 | -0.414 | 0.07 | 0.1 | 0.1 | 0 | 0.14 | 0.36 | 0.3 | 0.21 | 0.17 | 0.12 |
| 8 | 73066167 | rs4467977 | -0.293 | 0.531 | 0.387 | -0.265 | 0.073 | 0.231 | 0.231 | 0 | 0.81 | 0.59 | 0.6 | 0.5 | 0.75 | 0.78 |
| 8 | 73067682 | rs4363243 | -0.55 | 0.443 | 0.199 | -0.63 | -0.025 | 0.018 | 0.018 | 0.048 | 0.8 | 0.5 | 0.8 | 0.77 | 0.82 | 0.87 |
| 8 | 73070659 | rs4737343 | 1.272 | 0.356 | -0.134 | -0.959 | -0.043 | -0.044 | -0.044 | 0 | 0.41 | 0.12 | 0.39 | 0.42 | 0.44 | 0.45 |
| 8 | 73088793 | rs7017623 | -0.219 | 0.352 | 0.181 | -0.955 | 0.003 | 0.039 | 0.039 | 0 | 0.68 | 0.6 | 0.53 | 0.42 | 0.65 | 0.67 |
| 8 | 73091053 | rs7010938 | 1.012 | 0.308 | 0.259 | -1.072 | -0.004 | 0.003 | 0.003 | 0 | 0.45 | 0.32 | 0.42 | 0.41 | 0.45 | 0.45 |
| 8 | 73104407 | rs7014898 | 0.819 | 0.298 | 0.278 | -1.176 | 0 | -0.023 | -0.023 | 0 | 0.23 | 0.27 | 0.11 | 0.02 | 0.19 | 0.22 |
| 8 | 73106916 | rs1963982 | 0.958 | 0.336 | 0.11 | -0.988 | -0.005 | 0.008 | 0.008 | 0 | 0.45 | 0.32 | 0.42 | 0.4 | 0.45 | 0.45 |
| 8 | 73107739 | rs11784756 | 1.881 | 0.297 | -0.044 | -1.009 | 0 | -0.011 | -0.011 | 0 | 0.11 | 0.15 | 0.05 | 0 | 0.07 | 0.08 |
| 8 | 73112685 | rs2128199 | -2.345 | 0.372 | -0.306 | -0.844 | -0.004 | 0.01 | 0.01 | 0.095 | 0.94 | 0.95 | 0.9 | 0.78 | 0.92 | 0.9 |
| 8 | 73122967 | rs10112844 | 1.467 | 0.35 | -0.346 | -0.914 | -0.035 | -0.019 | -0.019 | 0 | 0.41 | 0.15 | 0.39 | 0.4 | 0.44 | 0.46 |
| 8 | 73123327 | rs7002197 | -2.304 | 0.505 | -0.397 | -0.782 | -0.002 | 0.017 | 0.017 | 0.095 | 0.94 | 0.95 | 0.9 | 0.78 | 0.92 | 0.9 |
| 8 | 73129923 | rs4645599 | 1.85 | 0.491 | -1.064 | -1.277 | 0.005 | 0.113 | 0.113 | 0 | 0.23 | 0.03 | 0.13 | 0.03 | 0.24 | 0.21 |
| 8 | 73130333 | rs17750497 | 2.048 | 0.38 | -0.368 | -0.968 | -0.016 | -0.037 | -0.037 | 0.071 | 0.12 | 0.13 | 0.07 | 0 | 0.08 | 0.04 |
| 8 | 73134321 | rs276649 | 0.159 | 0.374 | -0.277 | -0.906 | -0.001 | 0.003 | 0.003 | 0 | 0.68 | 0.41 | 0.52 | 0.4 | 0.66 | 0.67 |
| 8 | 73140966 | rs1384155 | -0.751 | 0.316 | -0.399 | -0.923 | -0.026 | 0.03 | 0.03 | 0.014 | 0.8 | 0.63 | 0.81 | 0.76 | 0.84 | 0.85 |
| 8 | 73151205 | rs11778612 | 1.289 | 0.744 | -1.147 | -0.699 | -0.009 | 0.06 | 0.06 | 0.08 | 0.14 | 0.04 | 0.07 | 0.02 | 0.13 | 0.18 |
| 8 | 73152314 | rs10109245 | 1.487 | 0.77 | -1.092 | -0.607 | 0.004 | 0.033 | 0.033 | 0 | 0.12 | 0.14 | 0.28 | 0.35 | 0.17 | 0.18 |
| 8 | 73154728 | rs922784 | -1.328 | 0.563 | -0.783 | -0.4 | 0.06 | 0.224 | 0.224 | 0.02 | 0.92 | 0.68 | 0.87 | 0.75 | 0.92 | 0.9 |
| 8 | 73155431 | rs17824132 | 1.503 | 0.372 | -1.286 | -1.177 | 0.003 | 0.019 | 0.019 | 0 | 0.09 | 0.02 | 0.05 | 0 | 0.07 | 0.03 |
| 8 | 73161144 | rs7837478 | 0.563 | 0.503 | -1.043 | -0.946 | -0.03 | -0.117 | -0.117 | 1.499 | 0.26 | 0.44 | 0.15 | 0.01 | 0.19 | 0.12 |
| 8 | 73162530 | rs4738225 | -0.057 | 1.356 | -1.447 | 0.015 | -0.058 | -0.02 | -0.02 | 0 | 0.39 | 0.07 | 0.37 | 0.38 | 0.42 | 0.42 |
| 8 | 73164823 | rs13259433 | -0.028 | 1.536 | -1.285 | 0.013 | -0.048 | 0.001 | 0.001 | 0 | 0.38 | 0.09 | 0.26 | 0.37 | 0.42 | 0.41 |
| 8 | 73169918 | rs7836885 | -0.645 | 2.106 | -0.072 | -0.286 | 0.037 | -0.147 | -0.147 | 0 | 0.24 | 0.45 | 0.15 | 0.01 | 0.27 | 0.14 |
| 8 | 73170864 | rs13266548 | -0.416 | 2.302 | 0.246 | -0.315 | 0.04 | -0.105 | -0.105 | 0.624 | 0.22 | 0.36 | 0.14 | 0.01 | 0.28 | 0.14 |
| 8 | 73171497 | rs4588902 | -0.429 | 2.294 | 0.242 | -0.319 | 0.039 | -0.105 | -0.105 | 0 | 0.22 | 0.37 | 0.14 | 0.01 | 0.29 | 0.14 |
| 8 | 73172443 | rs6981433 | -1.212 | 2.559 | -0.039 | 0.535 | 0.028 | 0.013 | 0.013 | 0.56 | 0.33 | 0.45 | 0.3 | 0.36 | 0.39 | 0.39 |
| 8 | 73175049 | rs11783296 | 1.019 | 2.789 | -0.112 | 0.491 | -0.024 | 0.031 | 0.031 | 1.902 | 0.08 | 0.03 | 0.07 | 0.01 | 0.11 | 0.05 |
| 8 | 73180176 | rs7844036 | 1.756 | 2.942 | 0.298 | 0.767 | -0.004 | 0.026 | 0.026 | 0.005 | 0.56 | 0.33 | 0.59 | 0.36 | 0.5 | 0.41 |
| 8 | 73184939 | rs4737352 | -0.604 | 2.93 | 0.358 | 0.887 | 0.001 | 0.022 | 0.022 | 0.007 | 0.92 | 0.92 | 0.91 | 0.78 | 0.89 | 0.83 |
| 8 | 73185998 | rs7820206 | -0.766 | 2.953 | 0.614 | 0.884 | 0.032 | 0.061 | 0.061 | 0 | 0.25 | 0.44 | 0.24 | 0.35 | 0.28 | 0.34 |
| 8 | 73193532 | rs6985078 | -0.525 | 2.985 | 0.464 | 0.885 | 0.011 | 0.023 | 0.023 | 0 | 0.24 | 0.29 | 0.23 | 0.36 | 0.28 | 0.34 |
| 8 | 73201039 | rs7825651 | 1.728 | 3.015 | 0.628 | 0.784 | -0.004 | 0.028 | 0.028 | 0 | 0.57 | 0.32 | 0.58 | 0.33 | 0.5 | 0.42 |
| 8 | 73202486 | rs919541 | -0.405 | 3.019 | 0.906 | 0.841 | 0.018 | 0.026 | 0.026 | 0 | 0.23 | 0.28 | 0.25 | 0.37 | 0.28 | 0.32 |
| 8 | 73212994 | rs1864583 | 0.23 | 3.144 | 0.738 | 0.701 | -0.002 | 0.001 | 0.001 | 0 | 0.83 | 0.84 | 0.85 | 0.7 | 0.78 | 0.77 |
| 8 | 73213843 | rs7844770 | 1.647 | 2.81 | 0.395 | 0.506 | 0.033 | 0.112 | 0.112 | 0 | 0.48 | 0.3 | 0.54 | 0.2 | 0.45 | 0.34 |
| 8 | 73215921 | rs11778198 | 1.896 | 2.656 | -0.135 | 0.266 | 0.051 | 0.091 | 0.091 | 0 | 0.38 | 0.21 | 0.24 | 0.16 | 0.31 | 0.23 |
| 8 | 73225081 | rs767415 | -0.22 | 2.795 | -0.182 | 0.649 | -0.039 | -0.058 | -0.058 | 0.198 | 0.86 | 0.97 | 0.88 | 0.78 | 0.82 | 0.79 |
| 8 | 73229560 | rs2082200 | -1.074 | 3.222 | -0.805 | 0.858 | 0.016 | 0.023 | 0.023 | 0 | 0.34 | 0.37 | 0.32 | 0.5 | 0.37 | 0.42 |
| 8 | 73229999 | rs1158327 | -0.409 | 3.301 | -0.993 | 0.973 | -0.018 | 0.007 | 0.007 | 0.017 | 0.95 | 0.98 | 0.94 | 0.79 | 0.9 | 0.84 |
| 8 | 73231564 | rs13275862 | 0.269 | 2.695 | -0.837 | 0.001 | 0.028 | -0.045 | -0.045 | 0 | 0.1 | 0.15 | 0.06 | 0 | 0.13 | 0.06 |
| 8 | 73239680 | rs10504537 | -0.162 | 1.766 | -1.029 | 0.29 | -0.028 | -0.008 | -0.008 | 0 | 0.12 | 0.03 | 0.07 | 0.14 | 0.14 | 0.14 |
| 8 | 73240030 | rs7825542 | 0.466 | 0.739 | -0.913 | 0.029 | 0.078 | 0.056 | 0.056 | 0.093 | 0.51 | 0.64 | 0.29 | 0.35 | 0.38 | 0.36 |
| 8 | 73242673 | rs346613 | 0.522 | 1.581 | -0.15 | -0.164 | 0.075 | -0.05 | -0.05 | 0.424 | 0.83 | 0.53 | 0.83 | 0.95 | 0.72 | 0.82 |
| 8 | 73242883 | rs7839922 | -0.56 | 0.548 | -0.52 | 0.086 | -0.012 | 0.041 | 0.041 | 1.351 | 0.33 | 0.44 | 0.33 | 0.47 | 0.28 | 0.37 |
| 8 | 73243541 | rs7827852 | -0.241 | 0.196 | -1.38 | -0.811 | 0.005 | 0.011 | 0.011 | 0.287 | 0.35 | 0.33 | 0.21 | 0.27 | 0.32 | 0.38 |
| 8 | 73244357 | rs346617 | 0.232 | -0.355 | -1.517 | -0.984 | 0.036 | 0.04 | 0.04 | 4.005 | 0.64 | 0.3 | 0.81 | 0.9 | 0.79 | 0.82 |

| | | | | | | | | | | | | | | | | |
|---|---|---|---|---|---|---|---|---|---|---|---|---|---|---|---|---|
| 8 | 73248631 | rs346621 | 0.723 | 0.442 | -1.471 | -0.37 | 0 | -0.005 | -0.005 | 0 | 0.94 | 0.96 | 0.94 | 1 | 0.95 | 0.99 |
| 8 | 73252878 | rs346610 | -0.358 | 0.777 | -2.096 | -0.174 | -0.034 | -0.045 | -0.045 | 0.002 | 0.67 | 0.87 | 0.76 | 0.37 | 0.63 | 0.7 |
| 8 | 73266227 | rs878927 | -0.018 | 0.786 | -2.633 | -0.642 | -0.003 | 0.082 | 0.082 | 0.455 | 0.54 | 0.85 | 0.47 | 0.27 | 0.45 | 0.55 |
| 8 | 73268693 | rs16938182 | -0.05 | 0.895 | -2.831 | -0.683 | -0.028 | -0.029 | -0.029 | 0 | 0.17 | 0.02 | 0.17 | 0.14 | 0.17 | 0.09 |
| 8 | 73270921 | rs1864581 | 0.168 | 0.861 | -2.483 | -0.892 | -0.064 | 0.077 | 0.077 | 0.039 | 0.77 | 0.99 | 0.77 | 0.93 | 0.74 | 0.81 |
| 8 | 73272191 | rs714482 | -0.293 | 0.607 | -1.006 | -0.439 | 0.005 | -0.038 | -0.038 | 0 | 0.17 | 0.53 | 0.22 | 0.07 | 0.19 | 0.19 |
| 8 | 73278582 | rs830480 | 0.5 | 0.861 | -0.666 | -0.332 | 0.012 | -0.033 | -0.033 | 0 | 0.75 | 0.49 | 0.68 | 0.87 | 0.76 | 0.57 |
| 8 | 73280829 | rs830483 | 0.174 | 0.012 | -1.167 | -0.083 | 0.017 | -0.021 | -0.021 | 0 | 0.66 | 0.71 | 0.57 | 0.37 | 0.74 | 0.64 |
| 8 | 73285538 | rs830489 | 0.237 | -0.292 | -1.302 | -0.496 | -0.022 | 0.045 | 0.045 | 0 | 0.91 | 0.99 | 0.91 | 1 | 0.91 | 0.9 |
| 8 | 73286537 | rs6992322 | 0.2 | -1.125 | -1.745 | -0.863 | 0.032 | 0.003 | 0.003 | 0.693 | 0.18 | 0.01 | 0.08 | 0.12 | 0.13 | 0.16 |
| 8 | 73294051 | rs10100487 | -0.025 | -0.516 | -1.648 | -1.443 | 0.049 | 0.12 | 0.12 | 0 | 0.42 | 0.17 | 0.2 | 0.19 | 0.36 | 0.39 |
| 8 | 73298128 | rs830492 | -0.802 | -0.466 | -1.282 | -1.419 | -0.001 | -0.012 | -0.012 | 0 | 0.91 | 0.93 | 0.91 | 1 | 0.91 | 0.9 |
| 8 | 73312929 | rs830500 | 1.094 | -1.02 | -0.956 | -1.524 | -0.048 | 0.049 | 0.049 | 0 | 0.37 | 0.22 | 0.5 | 0.7 | 0.49 | 0.45 |
| 8 | 73324657 | rs830510 | 1.168 | -0.668 | -1.077 | -1.533 | -0.023 | -0.021 | -0.021 | 0.61 | 0.08 | 0.01 | 0.16 | 0.11 | 0.07 | 0.06 |
| 8 | 73328620 | rs830517 | -0.801 | -0.631 | -0.83 | -1.504 | 0.005 | -0.033 | -0.033 | 0.242 | 0.91 | 0.87 | 0.88 | 1 | 0.89 | 0.89 |
| 8 | 73338613 | rs4738243 | 0.883 | -0.64 | -0.458 | -1.416 | -0.024 | 0.022 | 0.022 | 0 | 0.37 | 0.31 | 0.48 | 0.7 | 0.46 | 0.43 |
| 8 | 73351215 | rs7006111 | 1.418 | -0.685 | -0.354 | -1.339 | 0.003 | 0.009 | 0.009 | 0 | 0.2 | 0.13 | 0.11 | 0.09 | 0.2 | 0.18 |
| 8 | 73352367 | rs16938210 | 0.396 | -0.651 | -0.15 | -1.256 | -0.024 | -0.019 | -0.019 | 0 | 0.2 | 0.25 | 0.11 | 0.11 | 0.13 | 0.14 |
| 8 | 73354297 | rs348148 | -0.198 | -1.122 | -0.303 | -1.498 | 0.016 | 0.038 | 0.038 | 0.279 | 0.42 | 0.33 | 0.33 | 0.19 | 0.32 | 0.42 |
| 8 | 73356505 | rs348153 | -1.026 | -0.968 | 0.001 | -1.631 | 0.031 | -0.019 | -0.019 | 0 | 0.95 | 0.84 | 0.95 | 1 | 0.94 | 0.97 |
| 8 | 73360577 | rs1430836 | 0.413 | -1.214 | 0.217 | -0.631 | 0.033 | 0 | 0 | 0.019 | 0.51 | 0.37 | 0.67 | 0.46 | 0.63 | 0.37 |
| 8 | 73360653 | rs1430838 | 0.992 | -1.213 | 0.226 | -0.495 | -0.031 | 0.06 | 0.06 | 0.234 | 0.43 | 0.19 | 0.49 | 0.27 | 0.55 | 0.29 |
| 8 | 73362337 | rs16938216 | 0.444 | -1.158 | 0.157 | -0.469 | 0.018 | 0.068 | 0.068 | 0 | 0.08 | 0.18 | 0.18 | 0.19 | 0.08 | 0.07 |
| 8 | 73377453 | rs1431652 | 2.295 | -1.234 | -0.275 | -0.64 | -0.153 | 0.001 | 0.001 | 7.575 | 0.27 | 0.03 | 0.41 | 0.18 | 0.44 | 0.24 |
| 8 | 73384265 | rs1367824 | 0.701 | -1.084 | 0.921 | -0.128 | 0.015 | -0.002 | -0.002 | 0 | 0.52 | 0.45 | 0.67 | 0.46 | 0.63 | 0.37 |
| 8 | 73390785 | rs10100027 | -1.071 | -1.124 | 0.45 | -0.377 | 0.022 | 0.067 | 0.067 | 0.164 | 0.76 | 0.91 | 0.89 | 0.93 | 0.82 | 0.79 |
| 8 | 73402635 | rs1155593 | 0.734 | -0.794 | 0.718 | -0.209 | 0.002 | 0.018 | 0.018 | 0.192 | 0.47 | 0.58 | 0.55 | 0.27 | 0.54 | 0.32 |
| 8 | 73404584 | rs11782518 | 1.061 | -0.705 | 0.624 | -0.157 | -0.007 | 0.01 | 0.01 | 0.552 | 0.42 | 0.37 | 0.49 | 0.27 | 0.48 | 0.29 |
| 8 | 73405168 | rs7824218 | 1.484 | -0.702 | -0.27 | -0.157 | -0.159 | -0.003 | -0.003 | 0 | 0.39 | 0.05 | 0.46 | 0.27 | 0.48 | 0.29 |
| 8 | 73405632 | rs7827943 | -0.616 | -0.737 | -0.042 | -0.341 | 0.027 | 0.082 | 0.082 | 0.689 | 0.69 | 0.85 | 0.85 | 0.93 | 0.76 | 0.73 |
| 8 | 73406973 | rs1431642 | -0.91 | -0.812 | -0.324 | -0.349 | 0.02 | 0.101 | 0.101 | 0.282 | 0.75 | 0.98 | 0.89 | 0.93 | 0.82 | 0.75 |
| 8 | 73407824 | rs2037771 | 0.875 | -0.666 | 0.056 | 0.064 | 0.073 | -0.015 | -0.015 | 0.092 | 0.49 | 0.23 | 0.65 | 0.46 | 0.63 | 0.33 |
| 8 | 73408450 | rs2116351 | -0.655 | -0.661 | 0.394 | 0.029 | 0.019 | 0.021 | 0.021 | 0.242 | 0.67 | 0.74 | 0.83 | 0.93 | 0.78 | 0.66 |
| 8 | 73408980 | rs1019880 | -0.122 | -0.599 | -0.208 | 0.051 | 0.046 | 0.09 | 0.09 | 0.866 | 0.6 | 0.29 | 0.76 | 0.93 | 0.72 | 0.61 |
| 8 | 73409225 | rs17827756 | 0.793 | -0.569 | -0.347 | 0.363 | -0.009 | -0.014 | -0.014 | 0.038 | 0.1 | 0.08 | 0.19 | 0.2 | 0.14 | 0.06 |
| 8 | 73410434 | rs1431640 | -0.003 | -0.398 | -0.239 | 0.505 | 0.04 | 0.079 | 0.079 | 0.77 | 0.54 | 0.3 | 0.61 | 0.75 | 0.64 | 0.6 |
| 8 | 73415017 | rs1554781 | 0.924 | -0.332 | -0.518 | 0.534 | -0.019 | 0.065 | 0.065 | 0.303 | 0.44 | 0.2 | 0.5 | 0.28 | 0.54 | 0.32 |
| 8 | 73418260 | rs10957603 | -1.199 | -0.025 | 0.249 | 1.281 | -0.004 | 0.174 | 0.174 | 0 | 0.11 | 0.25 | 0.12 | 0.47 | 0.1 | 0.28 |
| 8 | 73429848 | rs6985810 | -1.143 | -0.246 | 0.097 | 0.468 | -0.02 | -0.039 | -0.039 | 0 | 0.07 | 0.21 | 0.06 | 0 | 0.02 | 0.1 |
| 8 | 73430830 | rs1431647 | -0.797 | -0.804 | 0.681 | 0.858 | -0.019 | 0.215 | 0.215 | 0.902 | 0.18 | 0.45 | 0.16 | 0.48 | 0.13 | 0.24 |
| 8 | 73436094 | rs12550268 | 1.636 | -0.985 | -0.449 | 0.315 | -0.117 | 0.148 | 0.148 | 2.065 | 0.43 | 0.04 | 0.61 | 0.2 | 0.57 | 0.33 |
| 8 | 73443576 | rs4738265 | -0.208 | -0.971 | -0.326 | 0.344 | -0.001 | 0.001 | 0.001 | 0 | 0.68 | 0.67 | 0.82 | 0.68 | 0.72 | 0.66 |
| 8 | 73444663 | rs349337 | 0.663 | -1.022 | -0.587 | 0.3 | -0.001 | 0.006 | 0.006 | 0 | 0.12 | 0.1 | 0.07 | 0.29 | 0.1 | 0.15 |
| 8 | 73452488 | rs7824350 | 1.354 | -1.061 | -0.495 | 0.185 | 0.021 | 0.166 | 0.166 | 1.017 | 0.51 | 0.23 | 0.67 | 0.21 | 0.59 | 0.43 |
| 8 | 73452949 | rs16938234 | 0.181 | -0.863 | -0.543 | 0.227 | -0.015 | -0.032 | -0.032 | 2.571 | 0.07 | 0.13 | 0.05 | 0 | 0.02 | 0.1 |
| 8 | 73457339 | rs349353 | -0.929 | -0.882 | -0.806 | 0.21 | -0.028 | 0.072 | 0.072 | 0 | 0.82 | 0.98 | 0.9 | 0.97 | 0.83 | 0.83 |
| 8 | 73460714 | rs349352 | 0.624 | -0.883 | -0.724 | 0.364 | -0.001 | 0.023 | 0.023 | 0 | 0.12 | 0.12 | 0.07 | 0.31 | 0.1 | 0.14 |
| 8 | 73462574 | rs421151 | 0.614 | -0.878 | -0.722 | 0.366 | -0.001 | 0.023 | 0.023 | 0 | 0.12 | 0.12 | 0.07 | 0.31 | 0.1 | 0.14 |
| 8 | 73471686 | rs16938237 | 0.464 | -0.926 | -0.371 | 0.315 | 0.013 | 0.179 | 0.179 | 0 | 0.1 | 0.17 | 0.12 | 0.47 | 0.1 | 0.18 |
| 8 | 73472871 | rs349346 | 0.683 | -0.962 | -0.4 | 0.241 | 0 | -0.006 | -0.006 | 2.157 | 0.67 | 0.7 | 0.73 | 0.5 | 0.71 | 0.54 |
| 8 | 73486092 | rs1431651 | 1.636 | -1.119 | -0.942 | 0.183 | -0.013 | 0.203 | 0.203 | 2.352 | 0.48 | 0.07 | 0.63 | 0.2 | 0.61 | 0.43 |
| 8 | 73487114 | rs6989807 | -0.107 | -1.26 | -1.05 | 0.098 | 0.024 | 0.094 | 0.094 | 0 | 0.17 | 0.01 | 0.03 | 0.04 | 0.08 | 0.17 |
| 8 | 73488971 | rs349335 | -0.95 | -1.174 | -0.99 | 0.161 | -0.005 | 0.07 | 0.07 | 0 | 0.82 | 0.98 | 0.92 | 0.97 | 0.85 | 0.83 |
| 8 | 73491177 | rs1431656 | 0.598 | -0.949 | -0.273 | 0.406 | 0.021 | 0.008 | 0.008 | 0.126 | 0.58 | 0.28 | 0.76 | 0.67 | 0.71 | 0.6 |
| 8 | 73498540 | rs16938243 | -0.184 | -0.455 | -0.143 | 0.968 | 0.002 | 0.081 | 0.081 | 0 | 0.1 | 0.09 | 0.12 | 0.47 | 0.1 | 0.18 |
| 8 | 73503743 | rs349359 | -0.332 | -0.908 | 0.015 | 0.708 | -0.011 | 0.02 | 0.02 | 0 | 0.19 | 0.25 | 0.11 | 0.25 | 0.13 | 0.18 |
| 8 | 73505068 | rs12547092 | -0.178 | -0.91 | 0 | 0.982 | 0 | 0.062 | 0.062 | 0 | 0.11 | 0.11 | 0.15 | 0.5 | 0.11 | 0.19 |
| 8 | 73515826 | rs4738267 | 0.153 | -0.914 | 0.001 | 1.224 | 0 | 0.055 | 0.055 | 0 | 0.11 | 0.1 | 0.15 | 0.48 | 0.11 | 0.19 |
| 8 | 73517286 | rs4738269 | -1.125 | -1.042 | -0.189 | 0.552 | 0.001 | 0.005 | 0.005 | 0 | 0.18 | 0.13 | 0.19 | 0.53 | 0.16 | 0.24 |
| 8 | 73519062 | rs7009604 | -1.636 | -1.273 | 0.424 | 0.397 | -0.016 | 0.104 | 0.104 | 0 | 0.23 | 0.39 | 0.25 | 0.55 | 0.19 | 0.32 |
| 8 | 73519816 | rs17762432 | 1.549 | -1.816 | -0.638 | -0.57 | -0.168 | 0.065 | 0.065 | 1.61 | 0.32 | 0.06 | 0.51 | 0.17 | 0.51 | 0.28 |
| 8 | 73535966 | rs4738270 | -0.192 | -1.714 | -0.441 | -0.594 | -0.005 | -0.005 | -0.005 | 0 | 0.56 | 0.52 | 0.77 | 0.72 | 0.68 | 0.6 |
| 8 | 73536762 | rs7815263 | -0.497 | -1.678 | -0.453 | -0.56 | -0.005 | 0.071 | 0.071 | 0 | 0.18 | 0.25 | 0.2 | 0.53 | 0.17 | 0.24 |
| 8 | 73537285 | rs7002471 | -0.948 | -1.67 | -0.464 | -0.562 | -0.016 | 0.104 | 0.104 | 0 | 0.23 | 0.38 | 0.25 | 0.55 | 0.19 | 0.32 |
| 8 | 73569185 | rs1822062 | -1.142 | -1.72 | -0.658 | -0.689 | 0.018 | 0.033 | 0.033 | 0.429 | 0.75 | 0.87 | 0.87 | 0.96 | 0.81 | 0.76 |
| 8 | 73582267 | rs2246547 | 0.642 | -1.637 | -0.64 | -0.55 | -0.017 | 0.02 | 0.02 | 0 | 0.21 | 0.67 | 0.13 | 0.24 | 0.12 | 0.17 |
| 8 | 73582738 | rs2246456 | -1.099 | -1.707 | -0.726 | -0.644 | 0.034 | 0.091 | 0.091 | 0.059 | 0.4 | 0.82 | 0.34 | 0.79 | 0.29 | 0.44 |
| 8 | 73586681 | rs2245973 | 0.506 | -1.634 | -0.637 | -0.52 | -0.007 | 0.004 | 0.004 | 0 | 0.21 | 0.74 | 0.15 | 0.23 | 0.12 | 0.16 |
| 8 | 73587678 | rs11787421 | 1.12 | -1.646 | -0.791 | -0.565 | -0.125 | 0.093 | 0.093 | 5.595 | 0.35 | 0.08 | 0.42 | 0.17 | 0.56 | 0.3 |

| | | | | | | | | | | | | | | | | |
|---|---|---|---|---|---|---|---|---|---|---|---|---|---|---|---|---|
| 8 | 73588474 | rs2439332 | 0.824 | -1.586 | -0.564 | -0.504 | -0.014 | 0.046 | 0.046 | 0 | 0.19 | 0.36 | 0.08 | 0.23 | 0.12 | 0.16 |
| 8 | 73602555 | rs1550467 | 0.729 | -1.593 | -0.903 | -0.491 | -0.086 | 0.109 | 0.109 | 2.648 | 0.39 | 0.11 | 0.45 | 0.19 | 0.56 | 0.33 |
| 8 | 73603360 | rs2252442 | 0.594 | -1.335 | -0.678 | -0.314 | -0.019 | 0.014 | 0.014 | 0 | 0.2 | 0.62 | 0.12 | 0.21 | 0.12 | 0.16 |
| 8 | 73604023 | rs17831704 | 1.457 | -0.605 | -1.654 | -0.593 | -0.102 | -0.048 | -0.048 | 0.795 | 0.14 | 0.01 | 0.25 | 0.13 | 0.34 | 0.22 |
| 8 | 73617600 | rs1899077 | -0.373 | -1.333 | -1.553 | -0.473 | 0.037 | -0.088 | -0.088 | 0 | 0.6 | 0.94 | 0.62 | 0.41 | 0.69 | 0.57 |
| 8 | 73623643 | rs2251387 | -0.05 | -1.467 | -1.441 | -0.45 | 0.087 | -0.002 | -0.002 | 0 | 0.53 | 0.83 | 0.57 | 0.4 | 0.67 | 0.48 |
| 8 | 73631168 | rs2247778 | -0.59 | -1.474 | -1.45 | -0.454 | 0.044 | -0.102 | -0.102 | 0 | 0.6 | 0.95 | 0.62 | 0.42 | 0.7 | 0.59 |
| 8 | 73633028 | rs2247572 | 0.605 | -1.547 | -1.133 | -0.536 | -0.018 | 0.041 | 0.041 | 0.006 | 0.22 | 0.45 | 0.13 | 0.23 | 0.14 | 0.24 |
| 8 | 73633305 | rs2247563 | -0.672 | -1.52 | -1.175 | -0.518 | 0.044 | -0.102 | -0.102 | 0.042 | 0.6 | 0.95 | 0.62 | 0.42 | 0.7 | 0.58 |
| 8 | 73633849 | rs2247463 | -0.672 | -1.52 | -1.175 | -0.518 | 0.044 | -0.102 | -0.102 | 5.658 | 0.6 | 0.95 | 0.62 | 0.42 | 0.7 | 0.58 |
| 8 | 73635954 | rs1470122 | -0.357 | -1.245 | -1.006 | -0.342 | 0.087 | -0.003 | -0.003 | 0 | 0.54 | 0.87 | 0.57 | 0.4 | 0.67 | 0.48 |
| 8 | 73645338 | rs2053422 | 0.421 | -0.902 | -0.919 | -0.123 | -0.004 | 0.003 | 0.003 | 0 | 0.36 | 0.36 | 0.46 | 0.19 | 0.55 | 0.32 |
| 8 | 73645962 | rs16938338 | 0.997 | -0.943 | -1.423 | -0.077 | -0.028 | -0.043 | -0.043 | 0 | 0.09 | 0.01 | 0.14 | 0.54 | 0.1 | 0.17 |
| 8 | 73649227 | rs4373559 | 0.337 | -0.848 | -1.253 | -0.231 | -0.062 | 0.085 | 0.085 | 0 | 0.36 | 0.18 | 0.45 | 0.19 | 0.56 | 0.32 |
| 8 | 73658605 | rs2053421 | 0.274 | -0.648 | -0.859 | -0.011 | 0.053 | -0.013 | -0.013 | 0.163 | 0.45 | 0.29 | 0.4 | 0.56 | 0.27 | 0.51 |
| 8 | 73660602 | rs2249942 | -0.15 | 0.492 | -0.979 | -0.071 | 0.046 | -0.021 | -0.021 | 0 | 0.29 | 0.43 | 0.25 | 0.21 | 0.38 | 0.2 |
| 8 | 73665740 | rs1822059 | 1.386 | 0.391 | -1.158 | 0.222 | -0.027 | 0.007 | 0.007 | 0 | 0.07 | 0.01 | 0.13 | 0.52 | 0.1 | 0.16 |
| 8 | 73677061 | rs1837544 | 0.844 | 0.788 | -0.901 | 0.059 | 0.037 | 0.03 | 0.03 | 0 | 0.09 | 0.1 | 0.15 | 0.16 | 0.21 | 0.06 |
| 8 | 73678362 | rs2245261 | 0.51 | 0.056 | -1.327 | 0.048 | 0.083 | -0.002 | -0.002 | 0.356 | 0.25 | 0.47 | 0.22 | 0.2 | 0.34 | 0.16 |
| 8 | 73692121 | rs2251898 | -1.209 | -0.149 | -1.959 | -0.274 | -0.023 | 0.005 | 0.005 | 0 | 0.74 | 0.84 | 0.65 | 0.78 | 0.63 | 0.67 |
| 8 | 73702832 | rs1866739 | -0.488 | -1.142 | -1.915 | -0.001 | 0.105 | 0.024 | 0.024 | 0 | 0.49 | 0.23 | 0.41 | 0.47 | 0.29 | 0.51 |
| 8 | 73717976 | rs4738281 | -1.224 | 0.007 | -0.94 | 0.861 | -0.003 | 0.053 | 0.053 | 0.079 | 0.14 | 0.18 | 0.2 | 0.44 | 0.14 | 0.21 |
| 8 | 73723640 | rs1440342 | -0.859 | 0.115 | -0.864 | 0.986 | -0.002 | 0.091 | 0.091 | 0 | 0.11 | 0.16 | 0.18 | 0.44 | 0.11 | 0.2 |
| 8 | 73730097 | rs17186996 | 0.619 | -0.335 | -1.387 | -0.029 | 0.052 | 0.076 | 0.076 | 0 | 0.19 | 0.05 | 0.07 | 0.02 | 0.11 | 0.21 |
| 8 | 73734459 | rs1440333 | -0.325 | 0.766 | -1.578 | 0.203 | -0.021 | 0.026 | 0.026 | 0 | 0.82 | 0.88 | 0.84 | 0.9 | 0.74 | 0.86 |
| 8 | 73735090 | rs1440332 | 0.444 | 0.403 | -1.408 | -0.251 | -0.004 | 0.012 | 0.012 | 0 | 0.42 | 0.65 | 0.47 | 0.72 | 0.44 | 0.44 |
| 8 | 73736455 | rs893891 | -0.158 | 0.353 | -1.088 | -0.2 | -0.01 | -0.012 | -0.012 | 0 | 0.76 | 0.72 | 0.65 | 0.39 | 0.66 | 0.68 |
| 8 | 73736766 | rs1107217 | 0.083 | -0.252 | -1.38 | -0.732 | 0.041 | 0.077 | 0.077 | 0 | 0.55 | 0.59 | 0.39 | 0.18 | 0.38 | 0.53 |
| 8 | 73739252 | rs2919408 | -0.255 | -0.347 | -1.061 | -0.85 | -0.002 | -0.032 | -0.032 | 0.127 | 0.64 | 0.67 | 0.71 | 0.9 | 0.64 | 0.74 |
| 8 | 73741823 | rs1822065 | 0.228 | -0.373 | -1.144 | -0.826 | 0.001 | 0 | 0 | 0 | 0.58 | 0.38 | 0.56 | 0.38 | 0.57 | 0.57 |
| 8 | 73745679 | rs1550461 | -1.273 | -0.373 | -1.169 | -0.839 | -0.08 | 0.036 | 0.036 | 0 | 0.82 | 0.99 | 0.86 | 0.9 | 0.74 | 0.86 |
| 8 | 73746579 | rs2958416 | 0.305 | -0.447 | -1.136 | -0.821 | -0.039 | 0.097 | 0.097 | 0 | 0.24 | 0.61 | 0.3 | 0.53 | 0.16 | 0.28 |
| 8 | 73746989 | rs6472704 | 1.366 | -0.447 | -1.19 | -0.866 | -0.07 | -0.027 | -0.027 | 0 | 0.2 | 0.1 | 0.25 | 0.2 | 0.29 | 0.15 |
| 8 | 73747106 | rs7817529 | 0.988 | -0.461 | -1.175 | -0.885 | 0.018 | 0.028 | 0.028 | 0 | 0.37 | 0.25 | 0.31 | 0.18 | 0.29 | 0.43 |
| 8 | 73748305 | rs10099670 | 1.221 | -0.49 | -1.087 | -0.829 | -0.065 | -0.111 | -0.111 | 0 | 0.17 | 0.31 | 0.09 | 0 | 0.09 | 0.11 |
| 8 | 73748817 | rs7005762 | 1.263 | -0.566 | -1.184 | -0.943 | -0.006 | -0.007 | -0.007 | 1.667 | 0.18 | 0.21 | 0.23 | 0.16 | 0.18 | 0.21 |
| 8 | 73749053 | rs7006486 | 0.2 | -0.535 | -1.143 | -0.893 | -0.009 | 0.039 | 0.039 | 0 | 0.56 | 0.29 | 0.45 | 0.27 | 0.56 | 0.57 |
| 8 | 73773663 | rs9886650 | -0.048 | -0.451 | -1.049 | -1.08 | 0.038 | 0.357 | 0.357 | 0 | 0.93 | 0.7 | 0.79 | 0.47 | 0.94 | 0.83 |
| 8 | 73776287 | rs10106279 | 1.128 | -0.493 | -0.985 | -1.305 | -0.064 | -0.114 | -0.114 | 0.849 | 0.16 | 0.32 | 0.09 | 0 | 0.09 | 0.12 |
| 8 | 73778642 | rs10504543 | -0.176 | -0.304 | -0.749 | -1.314 | -0.002 | 0.042 | 0.042 | 0 | 0.39 | 0.32 | 0.37 | 0.22 | 0.42 | 0.31 |
| 8 | 73780011 | rs7816119 | 0.521 | -0.538 | -0.597 | -1.315 | -0.018 | 0.045 | 0.045 | 0 | 0.19 | 0.12 | 0.14 | 0.07 | 0.24 | 0.13 |
| 8 | 73795256 | rs11780116 | 0.747 | -0.461 | -0.516 | -1.207 | -0.044 | -0.11 | -0.11 | 0 | 0.17 | 0.29 | 0.1 | 0.02 | 0.1 | 0.15 |
| 8 | 73798992 | rs17189856 | 1.476 | -0.286 | -0.905 | -1.976 | 0.025 | 0.094 | 0.094 | 0 | 0.2 | 0.03 | 0.07 | 0.02 | 0.12 | 0.23 |
| 8 | 73804390 | rs7341626 | -0.552 | 0.126 | 0.174 | -1.241 | 0.001 | -0.005 | -0.005 | 2.965 | 0.32 | 0.25 | 0.19 | 0.08 | 0.34 | 0.27 |
| 8 | 73809700 | rs13250827 | -1.178 | 0.813 | 0.69 | -0.669 | -0.012 | -0.086 | -0.086 | 0 | 0.32 | 0.6 | 0.25 | 0.09 | 0.32 | 0.25 |
| 8 | 73809941 | rs4385505 | 0.281 | 0.867 | 0.809 | -0.445 | 0.032 | 0.217 | 0.217 | 0 | 0.94 | 0.83 | 0.74 | 0.41 | 0.92 | 0.78 |
| 8 | 73812129 | rs7015692 | -1.127 | 0.829 | 0.765 | -0.532 | -0.011 | -0.081 | -0.081 | 0 | 0.32 | 0.6 | 0.25 | 0.09 | 0.32 | 0.25 |
| 8 | 73813300 | rs13279934 | -0.207 | 0.946 | 0.402 | -0.502 | 0 | -0.006 | -0.006 | 0 | 0.19 | 0.18 | 0.14 | 0.09 | 0.22 | 0.15 |
| 8 | 73815083 | rs6472714 | 1.093 | 1.018 | 0.384 | -0.526 | 0.002 | -0.008 | -0.008 | 0 | 0.64 | 0.56 | 0.52 | 0.32 | 0.59 | 0.52 |
| 8 | 73815971 | rs7465440 | 0.798 | 1.056 | 0.58 | 0.198 | 0.005 | 0.032 | 0.032 | 0 | 0.82 | 0.73 | 0.66 | 0.4 | 0.81 | 0.67 |
| 8 | 73819023 | rs10102173 | 0.567 | 0.956 | 0.253 | -0.629 | -0.005 | 0.011 | 0.011 | 0 | 0.71 | 0.77 | 0.46 | 0.1 | 0.66 | 0.6 |
| 8 | 73821073 | rs751822 | 0.157 | 0.955 | 0.139 | -0.456 | 0.001 | 0 | 0 | 0 | 0.16 | 0.18 | 0.26 | 0.22 | 0.21 | 0.17 |
| 8 | 73821809 | rs750574 | -0.204 | 0.942 | 0.149 | -0.411 | -0.012 | -0.037 | -0.037 | 0 | 0.88 | 0.96 | 0.72 | 0.35 | 0.88 | 0.77 |
| 8 | 73823745 | rs2929576 | -0.088 | 0.897 | 0.477 | -1.233 | 0.002 | -0.059 | -0.059 | 1.191 | 0.12 | 0.24 | 0.08 | 0 | 0.11 | 0.1 |
| 8 | 73830100 | rs2128158 | -0.009 | 0.897 | 0.453 | -1.233 | 0.003 | -0.058 | -0.058 | 0 | 0.12 | 0.22 | 0.08 | 0 | 0.11 | 0.1 |
| 8 | 73832023 | rs1482022 | 0.011 | 0.922 | 0.228 | -1.25 | -0.001 | -0.005 | -0.005 | 0.446 | 0.7 | 0.73 | 0.46 | 0.13 | 0.66 | 0.59 |
| 8 | 73832242 | rs1482024 | 0.418 | 0.927 | -0.555 | -0.984 | -0.003 | 0.006 | 0.006 | 0 | 0.57 | 0.37 | 0.37 | 0.13 | 0.54 | 0.5 |
| 8 | 73833641 | rs13276074 | 0.437 | 0.954 | -0.54 | -0.992 | -0.003 | 0.008 | 0.008 | 0 | 0.57 | 0.37 | 0.37 | 0.13 | 0.54 | 0.5 |
| 8 | 73834777 | rs13251382 | 0.447 | 0.947 | -0.619 | -0.996 | -0.006 | 0.034 | 0.034 | 0 | 0.57 | 0.36 | 0.36 | 0.13 | 0.55 | 0.5 |
| 8 | 73835919 | rs2981092 | -0.891 | 0.692 | -0.285 | -1.303 | -0.009 | -0.072 | -0.072 | 0 | 0.22 | 0.6 | 0.18 | 0.07 | 0.21 | 0.16 |
| 8 | 73839856 | rs1545828 | -0.269 | 0.582 | -0.955 | -2.108 | -0.035 | 0.088 | 0.088 | 0 | 0.4 | 0.21 | 0.59 | 0.89 | 0.48 | 0.54 |
| 8 | 73845937 | rs11786867 | -0.263 | 0.427 | -0.043 | -2.186 | 0 | -0.02 | -0.02 | 0.062 | 0.42 | 0.42 | 0.6 | 0.89 | 0.48 | 0.54 |
| 8 | 73847386 | rs11784364 | 0.186 | 0.279 | -0.403 | -2.203 | -0.102 | 0.103 | 0.103 | 0.119 | 0.27 | 0.05 | 0.23 | 0.1 | 0.41 | 0.17 |
| 8 | 73850203 | rs11782118 | -0.319 | 0.142 | -0.449 | -2.273 | -0.027 | 0.043 | 0.043 | 0.028 | 0.31 | 0.2 | 0.28 | 0.09 | 0.43 | 0.2 |
| 8 | 73852726 | rs2929566 | 0.583 | -0.668 | -0.658 | -3.115 | 0.006 | 0.003 | 0.003 | 0 | 0.22 | 0.21 | 0.14 | 0.05 | 0.14 | 0.27 |
| 8 | 73853561 | rs922772 | -1.022 | -0.742 | -0.455 | -3.163 | 0.001 | 0.011 | 0.011 | 0.111 | 0.59 | 0.38 | 0.7 | 0.93 | 0.6 | 0.78 |
| 8 | 73857539 | rs4738296 | 1.273 | -0.78 | -0.356 | -3.152 | 0.01 | -0.023 | -0.023 | 0 | 0.08 | 0.03 | 0.31 | 0.79 | 0.04 | 0.33 |
| 8 | 73857877 | rs3735830 | -0.22 | -0.764 | -0.393 | -3.143 | 0.001 | 0 | 0 | 0 | 0.31 | 0.32 | 0.46 | 0.84 | 0.18 | 0.61 |
| 8 | 73862691 | rs10504545 | 0.551 | -0.483 | -0.41 | -3.003 | -0.06 | 0.085 | 0.085 | 0.428 | 0.28 | 0.13 | 0.25 | 0.09 | 0.41 | 0.17 |

| | | | | | | | | | | | | | | | | |
|---|---|---|---|---|---|---|---|---|---|---|---|---|---|---|---|---|
| 8 | 73863995 | rs2981104 | 0.623 | -0.678 | -0.538 | -3.023 | 0.016 | 0.045 | 0.045 | 0.072 | 0.18 | 0.07 | 0.11 | 0.04 | 0.11 | 0.2 |
| 8 | 73867230 | rs2975836 | -1.103 | -0.312 | -0.043 | -2.957 | 0.011 | -0.081 | -0.081 | 0 | 0.69 | 0.62 | 0.8 | 0.94 | 0.65 | 0.82 |
| 8 | 73870170 | rs2929562 | -0.333 | 0.089 | -0.104 | -2.745 | 0.004 | -0.088 | -0.088 | 0.59 | 0.33 | 0.53 | 0.3 | 0.12 | 0.42 | 0.25 |
| 8 | 73878312 | rs3894367 | -0.055 | -0.31 | 0.055 | -2.207 | -0.071 | -0.095 | -0.095 | 0 | 0.26 | 0.52 | 0.18 | 0.1 | 0.18 | 0.17 |
| 8 | 73889320 | rs9298210 | -0.171 | 0.056 | -0.289 | -2.324 | -0.011 | 0.07 | 0.07 | 0 | 0.56 | 0.68 | 0.35 | 0.15 | 0.55 | 0.31 |
| 8 | 73893239 | rs10089002 | 0.792 | -0.133 | -0.068 | -2.52 | 0 | -0.02 | -0.02 | 0 | 0.33 | 0.32 | 0.2 | 0.07 | 0.37 | 0.19 |
| 8 | 73901272 | rs6983580 | 0.869 | -0.138 | -0.094 | -2.522 | -0.018 | 0.039 | 0.039 | 0 | 0.32 | 0.24 | 0.2 | 0.07 | 0.37 | 0.19 |
| 8 | 73901862 | rs10088657 | 0.687 | -0.087 | -0.203 | -2.319 | 0.028 | 0.073 | 0.073 | 0.063 | 0.2 | 0.06 | 0.11 | 0.06 | 0.17 | 0.11 |
| 8 | 73918164 | rs2010441 | 0.81 | 0.251 | -0.282 | -1.83 | -0.054 | -0.072 | -0.072 | 0 | 0.22 | 0.06 | 0.51 | 0.69 | 0.31 | 0.41 |
| 8 | 73923302 | rs1574690 | 0.403 | -0.105 | 0.173 | -2.087 | -0.002 | 0.059 | 0.059 | 0 | 0.44 | 0.3 | 0.27 | 0.15 | 0.44 | 0.33 |
| 8 | 73925591 | rs2975842 | 0.133 | -0.371 | 0.396 | -2.299 | 0.015 | 0.12 | 0.12 | 0.776 | 0.54 | 0.22 | 0.32 | 0.19 | 0.54 | 0.44 |
| 8 | 73929721 | rs10095169 | 1.667 | -0.385 | 0.525 | -2.298 | -0.053 | 0.083 | 0.083 | 0 | 0.25 | 0.09 | 0.16 | 0.05 | 0.32 | 0.19 |
| 8 | 73939623 | rs6982126 | 0.697 | -0.208 | 0.751 | -2.085 | -0.011 | -0.031 | -0.031 | 0 | 0.3 | 0.33 | 0.18 | 0.16 | 0.23 | 0.27 |
| 8 | 73940350 | rs10099824 | 0.421 | -0.212 | 0.746 | -2.088 | -0.002 | 0.065 | 0.065 | 0 | 0.44 | 0.3 | 0.27 | 0.15 | 0.44 | 0.33 |
| 8 | 73953688 | rs12334407 | -0.963 | -0.209 | 0.735 | -2.085 | -0.007 | -0.008 | -0.008 | 0 | 0.75 | 0.75 | 0.46 | 0.31 | 0.68 | 0.59 |
| 8 | 73957540 | rs1545827 | 0.481 | -0.219 | 0.786 | -2.235 | -0.015 | 0.131 | 0.131 | 0 | 0.43 | 0.21 | 0.26 | 0.15 | 0.44 | 0.33 |
| 8 | 73978819 | rs2291220 | 0.634 | 0.05 | -0.585 | -2.042 | -0.011 | -0.04 | -0.04 | 0.018 | 0.3 | 0.33 | 0.18 | 0.14 | 0.23 | 0.27 |
| 8 | 73982161 | rs2291219 | -0.32 | 0.433 | -1.006 | -1.173 | -0.008 | -0.006 | -0.006 | 0.261 | 0.75 | 0.74 | 0.47 | 0.29 | 0.68 | 0.61 |
| 8 | 73986292 | rs1007750 | -0.245 | 0.128 | -1.025 | -1.423 | 0.018 | 0.049 | 0.049 | 0.352 | 0.41 | 0.32 | 0.59 | 0.81 | 0.38 | 0.55 |
| 8 | 73987487 | rs7015175 | -0.303 | 0.294 | -1.528 | -1.742 | -0.051 | 0.119 | 0.119 | 0 | 0.66 | 0.86 | 0.77 | 0.88 | 0.62 | 0.67 |
| 8 | 73990808 | rs7011514 | 0.259 | -0.173 | -0.493 | -1.422 | 0.003 | 0 | 0 | 0 | 0.27 | 0.24 | 0.17 | 0.24 | 0.18 | 0.27 |
| 8 | 73991213 | rs4738301 | -0.236 | 0.168 | -0.35 | -1.359 | 0.004 | 0.018 | 0.018 | 0 | 0.39 | 0.68 | 0.48 | 0.2 | 0.35 | 0.23 |
| 8 | 73993200 | rs10087744 | -0.267 | 0.256 | -0.327 | -1.15 | 0.006 | 0.025 | 0.025 | 0 | 0.53 | 0.56 | 0.59 | 0.2 | 0.55 | 0.34 |
| 8 | 73997147 | rs7006404 | -0.299 | 0.001 | 0.363 | -1.344 | 0.006 | 0.005 | 0.005 | 0.326 | 0.41 | 0.65 | 0.48 | 0.18 | 0.33 | 0.16 |
| 8 | 73999080 | rs10957639 | 1.193 | 0.086 | 0.299 | -1.233 | -0.032 | -0.071 | -0.071 | 1.999 | 0.25 | 0.11 | 0.3 | 0.56 | 0.32 | 0.49 |
| 8 | 73999481 | rs9298215 | -0.736 | 0.112 | 0.274 | -1.219 | -0.002 | 0.003 | 0.003 | 0 | 0.66 | 0.72 | 0.78 | 0.74 | 0.65 | 0.65 |
| 8 | 73999575 | rs10283250 | 0.966 | -0.38 | 0.46 | -1.477 | 0.003 | -0.074 | -0.074 | 0 | 0.13 | 0.21 | 0.12 | 0 | 0.13 | 0.01 |
| 8 | 74007347 | rs12677663 | -1.024 | -0.393 | 0.02 | -1.475 | -0.065 | 0.08 | 0.08 | 0 | 0.67 | 0.82 | 0.58 | 0.85 | 0.62 | 0.7 |
| 8 | 74009472 | rs6980601 | 0.861 | -0.459 | 0.323 | -1.535 | -0.007 | 0.009 | 0.009 | 0.481 | 0.28 | 0.23 | 0.35 | 0.81 | 0.34 | 0.56 |
| 8 | 74015005 | rs7821093 | -0.26 | -0.483 | 0.416 | -1.48 | -0.006 | -0.058 | -0.058 | 0.549 | 0.39 | 0.53 | 0.22 | 0.05 | 0.28 | 0.14 |
| 8 | 74017231 | rs6472735 | -0.123 | -0.469 | 0.388 | -1.455 | -0.036 | -0.121 | -0.121 | 0 | 0.37 | 0.47 | 0.22 | 0.05 | 0.27 | 0.14 |
| 8 | 74018612 | rs6472738 | 0.15 | -0.284 | 0.45 | -0.964 | -0.033 | -0.091 | -0.091 | 1.297 | 0.29 | 0.4 | 0.14 | 0.05 | 0.2 | 0.11 |
| 8 | 74020191 | rs6415628 | -0.858 | -0.169 | -0.272 | -0.951 | -0.057 | 0.15 | 0.15 | 0 | 0.58 | 0.82 | 0.52 | 0.86 | 0.54 | 0.68 |
| 8 | 74022804 | rs6984102 | -0.011 | -0.252 | -0.295 | -0.918 | -0.042 | -0.108 | -0.108 | 0 | 0.31 | 0.44 | 0.17 | 0.05 | 0.21 | 0.11 |
| 8 | 74028940 | rs6988466 | 0.466 | 0.194 | -0.964 | -0.207 | -0.011 | -0.025 | -0.025 | 0 | 0.26 | 0.23 | 0.32 | 0.55 | 0.33 | 0.49 |
| 8 | 74030836 | rs6472746 | -0.131 | -0.226 | -0.772 | -0.124 | -0.042 | 0.011 | 0.011 | 0.212 | 0.24 | 0.57 | 0.16 | 0.27 | 0.13 | 0.12 |
| 8 | 74034680 | rs6472748 | -0.337 | 0.391 | -1.54 | -0.577 | 0.005 | 0.209 | 0.209 | 0.021 | 0.49 | 0.85 | 0.49 | 0.82 | 0.46 | 0.68 |
| 8 | 74041760 | rs4644236 | -0.047 | 0.659 | -1.093 | -0.029 | -0.005 | 0.003 | 0.003 | 0 | 0.34 | 0.27 | 0.58 | 0.67 | 0.36 | 0.49 |
| 8 | 74048541 | rs7830267 | -0.661 | 0.11 | -1.612 | -0.595 | -0.096 | 0.155 | 0.155 | 0.044 | 0.58 | 0.92 | 0.7 | 0.77 | 0.49 | 0.62 |
| 8 | 74049175 | rs2383886 | -0.487 | 0.015 | -0.539 | -0.643 | -0.013 | 0.068 | 0.068 | 0 | 0.56 | 0.69 | 0.68 | 0.77 | 0.49 | 0.62 |
| 8 | 74052613 | rs10504547 | 0.742 | -0.12 | -0.345 | -0.597 | 0.031 | 0.053 | 0.053 | 0 | 0.18 | 0.07 | 0.09 | 0.04 | 0.12 | 0.12 |
| 8 | 74058130 | rs2383880 | 0.226 | 0.106 | 0.432 | -0.621 | -0.012 | -0.03 | -0.03 | 0 | 0.2 | 0.4 | 0.15 | 0.1 | 0.17 | 0.07 |
| 8 | 74058147 | rs2383879 | 0.512 | 0.109 | 0.544 | -0.728 | 0 | 0.009 | 0.009 | 0 | 0.35 | 0.26 | 0.53 | 0.67 | 0.34 | 0.56 |
| 8 | 74058617 | rs4559234 | 0.503 | 0.13 | 0.652 | -0.732 | 0 | 0 | 0 | 0 | 0.36 | 0.33 | 0.54 | 0.67 | 0.33 | 0.56 |
| 8 | 74060655 | rs10808793 | -0.505 | 0.143 | 1.134 | -0.748 | -0.02 | 0.088 | 0.088 | 0 | 0.54 | 0.77 | 0.67 | 0.71 | 0.49 | 0.62 |
| 8 | 74061293 | rs7839918 | 0.467 | 0.193 | 1.205 | -0.672 | -0.003 | -0.009 | -0.009 | 0.122 | 0.36 | 0.38 | 0.54 | 0.67 | 0.33 | 0.56 |
| 8 | 74063433 | rs6472750 | 0.832 | 0.029 | 1.1 | 0.036 | 0.015 | -0.023 | -0.023 | 0 | 0.3 | 0.37 | 0.26 | 0.22 | 0.35 | 0.43 |
| 8 | 74065619 | rs2383878 | 0.601 | 0.146 | 0.924 | -0.002 | 0.003 | -0.02 | -0.02 | 0 | 0.44 | 0.44 | 0.31 | 0.24 | 0.48 | 0.53 |
| 8 | 74071023 | rs12545313 | -0.439 | 0.159 | 0.436 | 0.285 | -0.025 | 0.058 | 0.058 | 0 | 0.66 | 0.9 | 0.59 | 0.74 | 0.64 | 0.73 |
| 8 | 74074261 | rs7014577 | 0.153 | 0.047 | 0.999 | 0.766 | -0.001 | 0.005 | 0.005 | 0 | 0.2 | 0.22 | 0.23 | 0.33 | 0.14 | 0.17 |
| 8 | 74080735 | rs13249487 | 0.575 | 0.028 | 0.6 | 0.431 | 0.019 | 0.026 | 0.026 | 0.008 | 0.17 | 0.07 | 0.1 | 0 | 0.15 | 0.04 |
| 8 | 74083763 | rs3824268 | 0.274 | 0.025 | 0.758 | 0.567 | 0.007 | -0.013 | -0.013 | 0 | 0.2 | 0.17 | 0.23 | 0.34 | 0.14 | 0.2 |
| 8 | 74089904 | rs4560775 | -0.916 | 0.011 | 1.099 | 0.581 | -0.006 | 0.003 | 0.003 | 0.844 | 0.39 | 0.6 | 0.4 | 0.42 | 0.31 | 0.24 |
| 8 | 74093696 | rs4607600 | 0.365 | 0.112 | 0.594 | 0.603 | 0.012 | -0.023 | -0.023 | 0 | 0.19 | 0.14 | 0.22 | 0.34 | 0.14 | 0.17 |
| 8 | 74094510 | rs16938583 | 0.237 | 0.137 | 0.648 | 0.548 | -0.019 | -0.021 | -0.021 | 0 | 0.18 | 0.31 | 0.16 | 0.08 | 0.17 | 0.05 |
| 8 | 74098792 | rs16938589 | 0.891 | 0.109 | 0.554 | 0.783 | 0.005 | -0.012 | -0.012 | 0.379 | 0.17 | 0.16 | 0.11 | 0.24 | 0.14 | 0.1 |
| 8 | 74102571 | rs11781596 | -0.918 | 0.127 | 0.629 | 0.774 | -0.021 | 0.012 | 0.012 | 0.83 | 0.39 | 0.49 | 0.36 | 0.48 | 0.32 | 0.24 |
| 8 | 74102646 | rs16938592 | 0.956 | 0.133 | 0.6 | 0.769 | 0.005 | 0.01 | 0.01 | 0 | 0.17 | 0.16 | 0.08 | 0.08 | 0.14 | 0.09 |
| 8 | 74102874 | rs11781678 | -0.841 | 0.18 | 0.558 | 0.531 | -0.022 | 0.008 | 0.008 | 0.726 | 0.39 | 0.48 | 0.39 | 0.49 | 0.31 | 0.24 |
| 8 | 74104797 | rs11987488 | 0.174 | 0.236 | 0.278 | 0.738 | -0.008 | 0.028 | 0.028 | 0 | 0.2 | 0.28 | 0.23 | 0.48 | 0.14 | 0.2 |
| 8 | 74109865 | rs7017538 | -0.581 | 0.977 | 0.597 | 0.22 | 0 | -0.001 | -0.001 | 0 | 0.27 | 0.3 | 0.32 | 0.25 | 0.26 | 0.16 |
| 8 | 74112761 | rs888901 | 0.798 | 1.397 | 0.782 | 0.053 | 0 | -0.001 | -0.001 | 0 | 0.64 | 0.43 | 0.58 | 0.68 | 0.6 | 0.7 |
| 8 | 74114567 | rs1422503 | 0.803 | 1.511 | 0.341 | 0.112 | 0.009 | -0.005 | -0.005 | 0.149 | 0.66 | 0.64 | 0.6 | 0.68 | 0.6 | 0.7 |
| 8 | 74115879 | rs2605867 | 0.818 | 1.56 | 0.176 | 0.124 | 0.026 | -0.011 | -0.011 | 0 | 0.66 | 0.58 | 0.59 | 0.68 | 0.59 | 0.7 |
| 8 | 74116795 | rs10957647 | 0.293 | 1.706 | -1.01 | 0.337 | -0.003 | 0.034 | 0.034 | 0 | 0.06 | 0.05 | 0.22 | 0.29 | 0.1 | 0.1 |
| 8 | 74125952 | rs6472760 | 0.202 | 1.699 | -1.172 | -0.428 | -0.049 | 0.094 | 0.094 | 0 | 0.72 | 0.91 | 0.84 | 0.97 | 0.65 | 0.8 |
| 8 | 74138294 | rs6472762 | 0.165 | 1.625 | -1.23 | 0.017 | -0.053 | -0.014 | -0.014 | 0 | 0.59 | 0.77 | 0.57 | 0.64 | 0.52 | 0.65 |
| 8 | 74139097 | rs6994157 | -0.554 | 1.349 | -1.449 | -0.092 | -0.036 | 0.137 | 0.137 | 0.881 | 0.73 | 0.95 | 0.86 | 0.97 | 0.71 | 0.77 |
| 8 | 74142151 | rs12544429 | 0.623 | 1.509 | -1.489 | 0.043 | -0.011 | -0.015 | -0.015 | 2.735 | 0.11 | 0.07 | 0.23 | 0.32 | 0.12 | 0.12 |

| | | | | | | | | | | | | | | | | |
|---|---|---|---|---|---|---|---|---|---|---|---|---|---|---|---|---|
| 8 | 74142522 | rs7825059 | 0.172 | 1.319 | -0.661 | 0.033 | -0.001 | -0.001 | -0.001 | 0 | 0.58 | 0.46 | 0.56 | 0.64 | 0.59 | 0.63 |
| 8 | 74143490 | rs2605879 | 0.579 | 0.601 | -1.032 | -0.087 | 0.012 | -0.002 | -0.002 | 0.007 | 0.23 | 0.1 | 0.13 | 0.25 | 0.19 | 0.23 |
| 8 | 74143591 | rs2546229 | 0.141 | 1.033 | -0.411 | 0.266 | 0.002 | 0.014 | 0.014 | 0 | 0.1 | 0.13 | 0.2 | 0.18 | 0.12 | 0.27 |
| 8 | 74146766 | rs2605877 | 0.351 | 0.889 | -0.374 | 0.012 | -0.001 | -0.016 | -0.016 | 0 | 0.68 | 0.67 | 0.77 | 0.8 | 0.69 | 0.72 |
| 8 | 74168618 | rs2546220 | -1.098 | 0.744 | -0.045 | 0.391 | 0.001 | 0.016 | 0.016 | 0 | 0.14 | 0.23 | 0.13 | 0.19 | 0.14 | 0.15 |
| 8 | 74169337 | rs2546217 | -1.768 | 0.621 | 0.692 | 0.43 | -0.009 | 0.021 | 0.021 | 0.144 | 0.21 | 0.66 | 0.22 | 0.26 | 0.19 | 0.22 |
| 8 | 74171719 | rs2546216 | -0.138 | 0.689 | 0.446 | 0.442 | -0.025 | -0.028 | -0.028 | 2.035 | 0.06 | 0.01 | 0.06 | 0.15 | 0.07 | 0.07 |
| 8 | 74172878 | rs7005232 | 0.98 | 0.528 | 0.484 | 0.046 | -0.004 | -0.009 | -0.009 | 0 | 0.63 | 0.45 | 0.54 | 0.71 | 0.64 | 0.45 |
| 8 | 74175913 | rs1895259 | 1.098 | 0.526 | -0.009 | 0.77 | 0.004 | -0.041 | -0.041 | 1.458 | 0.6 | 0.2 | 0.49 | 0.47 | 0.61 | 0.38 |
| 8 | 74182557 | rs12681551 | -1.166 | 0.786 | -0.217 | 0.793 | 0 | -0.033 | -0.033 | 0.8 | 0.11 | 0.03 | 0.08 | 0.18 | 0.08 | 0.09 |
| 8 | 74183252 | rs11996978 | 0.624 | 0.417 | -0.328 | 0.654 | 0.001 | -0.037 | -0.037 | 0 | 0.66 | 0.82 | 0.6 | 0.56 | 0.68 | 0.49 |
| 8 | 74184632 | rs1988833 | -0.786 | 0.739 | -0.671 | 0.378 | 0 | 0.002 | 0.002 | 1.23 | 0.53 | 0.57 | 0.64 | 0.62 | 0.55 | 0.79 |
| 8 | 74189562 | rs6983289 | -0.102 | 0.572 | -0.367 | 0.422 | -0.006 | 0.022 | 0.022 | 0 | 0.39 | 0.48 | 0.5 | 0.53 | 0.36 | 0.63 |
| 8 | 74195243 | rs4349972 | -0.785 | 0.077 | -0.469 | 0.193 | -0.006 | 0.013 | 0.013 | 0 | 0.62 | 0.64 | 0.67 | 0.72 | 0.58 | 0.82 |
| 8 | 74197195 | rs7815444 | 0.992 | -0.04 | -0.431 | 0.133 | 0 | 0 | 0 | 0 | 0.18 | 0.17 | 0.18 | 0.19 | 0.23 | 0.2 |
| 8 | 74198922 | rs7833488 | -1.514 | 0.086 | -1.01 | -0.2 | 0.006 | 0.025 | 0.025 | 1.132 | 0.76 | 0.79 | 0.86 | 0.92 | 0.76 | 0.93 |
| 8 | 74200796 | rs4738315 | 1.624 | -0.041 | -1.185 | -0.075 | -0.013 | -0.106 | -0.106 | 0 | 0.31 | 0.02 | 0.36 | 0.37 | 0.24 | 0.51 |
| 8 | 74202188 | rs13278189 | -1.686 | 0.221 | -1.073 | -0.108 | 0.003 | 0.013 | 0.013 | 0 | 0.78 | 0.8 | 0.87 | 0.92 | 0.77 | 0.93 |
| 8 | 74204808 | rs4237000 | -1.686 | 0.221 | -1.073 | -0.108 | 0.003 | 0.013 | 0.013 | 0 | 0.78 | 0.79 | 0.87 | 0.92 | 0.77 | 0.93 |
| 8 | 74206582 | rs7819550 | 0.533 | 1.181 | -0.383 | 0.531 | 0.001 | 0.003 | 0.003 | 0 | 0.14 | 0.16 | 0.19 | 0.2 | 0.18 | 0.11 |
| 8 | 74219319 | rs7827088 | 0.641 | 0.723 | 0.034 | 0.093 | 0.01 | 0.009 | 0.009 | 0 | 0.18 | 0.21 | 0.18 | 0.19 | 0.22 | 0.19 |
| 8 | 74230223 | rs6989495 | -0.904 | 0.762 | -0.352 | 0.112 | 0.001 | 0.002 | 0.002 | 0.398 | 0.32 | 0.57 | 0.31 | 0.35 | 0.33 | 0.37 |
| 8 | 74230909 | rs16938613 | -0.27 | 0.935 | -0.616 | -0.035 | -0.001 | -0.002 | -0.002 | 0 | 0.09 | 0.1 | 0.1 | 0.16 | 0.11 | 0.16 |
| 8 | 74246331 | rs12541771 | -0.199 | 0.985 | -0.722 | -0.053 | -0.006 | -0.006 | -0.006 | 2.828 | 0.09 | 0.06 | 0.1 | 0.15 | 0.11 | 0.17 |
| 8 | 74261334 | rs7835634 | -0.239 | 1.383 | -0.81 | -0.248 | -0.012 | 0.005 | 0.005 | 0 | 0.09 | 0.07 | 0.08 | 0.1 | 0.14 | 0.1 |
| 8 | 74261484 | rs7823039 | 0.249 | 1.723 | 0.035 | -0.397 | 0.034 | -0.067 | -0.067 | 0 | 0.81 | 0.51 | 0.84 | 0.88 | 0.79 | 0.84 |
| 8 | 74262063 | rs2383899 | -0.511 | 1.671 | 0.038 | -0.407 | 0.038 | -0.05 | -0.05 | 0 | 0.17 | 0.48 | 0.15 | 0.12 | 0.18 | 0.11 |
| 8 | 74263630 | rs12334531 | 1.504 | 1.684 | -0.292 | -0.469 | 0.056 | 0.039 | 0.039 | 0.118 | 0.49 | 0.24 | 0.51 | 0.53 | 0.42 | 0.58 |
| 8 | 74265008 | rs11992844 | -0.458 | 1.869 | 0.262 | -0.802 | 0.013 | -0.004 | -0.004 | 1.694 | 0.16 | 0.6 | 0.14 | 0.16 | 0.15 | 0.13 |
| 8 | 74265395 | rs4738328 | 0.738 | 2.043 | 0.252 | -0.829 | 0.021 | -0.019 | -0.019 | 0.53 | 0.83 | 0.74 | 0.85 | 0.86 | 0.79 | 0.81 |
| 8 | 74267120 | rs2004484 | -0.187 | 1.817 | 0.037 | -0.807 | 0.002 | 0.004 | 0.004 | 1.36 | 0.91 | 0.94 | 0.95 | 0.96 | 0.93 | 0.91 |
| 8 | 74268575 | rs2383902 | 0.311 | 1.929 | 0.431 | -0.839 | 0.013 | -0.013 | -0.013 | 0 | 0.28 | 0.35 | 0.33 | 0.25 | 0.36 | 0.19 |
| 8 | 74268826 | rs4634634 | 0.123 | 1.747 | 0.601 | -0.766 | 0.034 | -0.002 | -0.002 | 0.023 | 0.12 | 0.31 | 0.11 | 0.13 | 0.14 | 0.1 |
| 8 | 74269768 | rs6472767 | 0.766 | 1.488 | 0.034 | -0.868 | -0.002 | -0.006 | -0.006 | 0.228 | 0.83 | 0.8 | 0.9 | 0.9 | 0.83 | 0.77 |
| 8 | 74271939 | rs17288270 | 0.396 | 1.662 | 0.069 | -0.828 | -0.004 | -0.001 | -0.001 | 0 | 0.08 | 0.07 | 0.07 | 0.11 | 0.1 | 0.1 |
| 8 | 74272171 | rs13277653 | -0.022 | 1.51 | 0.083 | -0.874 | -0.003 | -0.03 | -0.03 | 0 | 0.89 | 0.85 | 0.94 | 0.98 | 0.91 | 0.91 |
| 8 | 74275261 | rs4363185 | 1.319 | 1.827 | 0.135 | -0.966 | 0.015 | -0.021 | -0.021 | 0 | 0.75 | 0.65 | 0.8 | 0.81 | 0.68 | 0.75 |
| 8 | 74275323 | rs17216125 | 1.114 | 1.85 | 0.259 | -1.067 | 0.014 | -0.023 | -0.023 | 0 | 0.24 | 0.27 | 0.24 | 0.16 | 0.26 | 0.12 |
| 8 | 74275972 | rs11783921 | 0.944 | 1.824 | 0.23 | -1.118 | 0.007 | -0.023 | -0.023 | 0 | 0.06 | 0.2 | 0.06 | 0.03 | 0.05 | 0.07 |
| 8 | 74276211 | rs10098698 | 0.98 | 1.855 | 0.089 | -1.049 | 0.028 | -0.044 | -0.044 | 0 | 0.24 | 0.37 | 0.26 | 0.16 | 0.26 | 0.12 |
| 8 | 74276843 | rs16938625 | 0.375 | 2.209 | -0.309 | -0.785 | -0.002 | -0.001 | -0.001 | 0.448 | 0.07 | 0.08 | 0.09 | 0.11 | 0.14 | 0.1 |
| 8 | 74278256 | rs4738329 | 0.612 | 1.872 | -0.439 | -0.902 | -0.015 | 0.027 | 0.027 | 0.442 | 0.85 | 0.93 | 0.92 | 0.92 | 0.82 | 0.86 |
| 8 | 74283000 | rs11785540 | 0.697 | 1.684 | -0.224 | -0.83 | 0.002 | -0.022 | -0.022 | 0 | 0.27 | 0.31 | 0.23 | 0.22 | 0.28 | 0.16 |
| 8 | 74283845 | rs12156358 | 0.354 | 1.63 | -0.232 | -0.799 | 0.005 | -0.024 | -0.024 | 0.018 | 0.31 | 0.34 | 0.27 | 0.25 | 0.36 | 0.26 |
| 8 | 74284561 | rs10112555 | 0.884 | 0.911 | -0.893 | -0.507 | 0.094 | 0.027 | 0.027 | 2.041 | 0.45 | 0.13 | 0.51 | 0.55 | 0.31 | 0.48 |
| 8 | 74291106 | rs10088300 | -1.113 | 1.244 | -0.042 | -0.855 | 0.023 | -0.06 | -0.06 | 0.002 | 0.22 | 0.35 | 0.19 | 0.07 | 0.27 | 0.2 |
| 8 | 74298129 | rs6472769 | 0.912 | 0.959 | 0.071 | -0.74 | -0.018 | -0.007 | -0.007 | 0 | 0.42 | 0.55 | 0.44 | 0.63 | 0.27 | 0.47 |
| 8 | 74299041 | rs10216697 | -0.014 | 0.596 | -0.052 | -0.901 | -0.002 | 0.013 | 0.013 | 0.744 | 0.45 | 0.56 | 0.54 | 0.67 | 0.42 | 0.63 |
| 8 | 74302491 | rs7832526 | -0.652 | 0.865 | 0.117 | -1.062 | -0.01 | 0.065 | 0.065 | 0.02 | 0.82 | 0.91 | 0.9 | 0.96 | 0.79 | 0.81 |
| 8 | 74307417 | rs9298226 | 0.161 | 1.119 | 0.197 | -1.12 | 0 | -0.008 | -0.008 | 0.542 | 0.11 | 0.09 | 0.05 | 0.04 | 0.1 | 0.13 |
| 8 | 74309881 | rs13266042 | 0.116 | 0.929 | 0.137 | -1.133 | -0.01 | 0.073 | 0.073 | 0 | 0.26 | 0.08 | 0.13 | 0.08 | 0.28 | 0.26 |
| 8 | 74310019 | rs4496960 | 0.823 | 0.768 | 0.458 | -1.144 | 0.002 | -0.027 | -0.027 | 0 | 0.32 | 0.4 | 0.35 | 0.22 | 0.34 | 0.16 |
| 8 | 74312565 | rs4316144 | -0.63 | 0.969 | 0.163 | -1.075 | 0.004 | 0.001 | 0.001 | 0 | 0.94 | 0.92 | 0.91 | 0.95 | 0.88 | 0.91 |
| 8 | 74316068 | rs4738350 | 1.213 | 0.817 | 0.344 | -1.111 | -0.019 | -0.012 | -0.012 | 0 | 0.06 | 0.16 | 0.16 | 0.02 | 0.03 | 0.01 |
| 8 | 74316253 | rs10957663 | 1.814 | 0.754 | 0.441 | -1.193 | -0.024 | 0.013 | 0.013 | 0 | 0.25 | 0.09 | 0.29 | 0.21 | 0.3 | 0.11 |
| 8 | 74316810 | rs7840753 | -1.273 | 0.581 | 0.297 | -1.276 | -0.025 | -0.031 | -0.031 | 0 | 0.7 | 0.82 | 0.69 | 0.37 | 0.69 | 0.51 |
| 8 | 74319636 | rs13276705 | 1.075 | 1.004 | 0.447 | -0.852 | 0.015 | -0.092 | -0.092 | 0.167 | 0.32 | 0.14 | 0.24 | 0.45 | 0.26 | 0.52 |
| 8 | 74322142 | rs6993015 | 0.819 | 1.078 | 0.726 | -1.139 | -0.005 | 0.01 | 0.01 | 0 | 0.58 | 0.38 | 0.7 | 0.83 | 0.54 | 0.63 |
| 8 | 74323056 | rs4487756 | 0.971 | 0.96 | 0.509 | -1.102 | 0.017 | -0.067 | -0.067 | 0.775 | 0.33 | 0.17 | 0.29 | 0.54 | 0.28 | 0.53 |
| 8 | 74326231 | rs10108199 | 1.918 | 1.014 | 0.545 | -1.068 | -0.004 | -0.004 | -0.004 | 0 | 0.24 | 0.14 | 0.28 | 0.27 | 0.27 | 0.09 |
| 8 | 74327010 | rs4737380 | -0.22 | 0.86 | 0.713 | -1.166 | 0 | -0.004 | -0.004 | 0 | 0.31 | 0.28 | 0.3 | 0.13 | 0.32 | 0.26 |
| 8 | 74329796 | rs13256021 | -0.052 | 0.353 | 0.384 | -1.143 | -0.011 | 0.045 | 0.045 | 0 | 0.54 | 0.39 | 0.36 | 0.65 | 0.53 | 0.66 |
| 8 | 74330174 | rs13254848 | 1.342 | 0.255 | 0.181 | -1.121 | 0.04 | -0.061 | -0.061 | 0 | 0.29 | 0.19 | 0.19 | 0.53 | 0.21 | 0.44 |
| 8 | 74332207 | rs6998213 | 0.152 | 0.396 | 0.39 | -1.176 | 0.054 | 0.041 | 0.041 | 3.409 | 0.51 | 0.24 | 0.67 | 0.42 | 0.64 | 0.39 |
| 8 | 74335491 | rs10283372 | 0.125 | 0.026 | 0.61 | -1.319 | -0.003 | 0.017 | 0.017 | 0 | 0.4 | 0.57 | 0.58 | 0.31 | 0.43 | 0.27 |
| 8 | 74335653 | rs11787279 | 1.618 | 0.261 | 0.567 | -1.165 | -0.031 | -0.002 | -0.002 | 0 | 0.25 | 0.1 | 0.36 | 0.29 | 0.33 | 0.13 |
| 8 | 74335805 | rs16938650 | -0.191 | -0.068 | 0.919 | -1.014 | 0.004 | -0.044 | -0.044 | 0 | 0.1 | 0.31 | 0.09 | 0.02 | 0.07 | 0.1 |
| 8 | 74343250 | rs16938654 | 0.202 | -0.59 | 0.917 | -0.945 | -0.023 | -0.014 | -0.014 | 0.424 | 0.09 | 0.25 | 0.19 | 0.04 | 0.03 | 0.13 |
| 8 | 74343932 | rs11997512 | 0.36 | -0.749 | 0.819 | -0.857 | -0.046 | 0.036 | 0.036 | 0 | 0.33 | 0.5 | 0.32 | 0.51 | 0.21 | 0.51 |

| | | | | | | | | | | | | | | | | |
|---|---|---|---|---|---|---|---|---|---|---|---|---|---|---|---|---|
| 8 | 74348353 | rs2383893 | -0.154 | -0.432 | 0.311 | -1.047 | -0.076 | 0.094 | 0.094 | 0.012 | 0.7 | 0.92 | 0.7 | 0.88 | 0.62 | 0.69 |
| 8 | 74348656 | rs2383894 | -0.489 | -0.419 | 0.253 | -1.034 | -0.092 | 0.105 | 0.105 | 0.086 | 0.73 | 0.95 | 0.84 | 0.91 | 0.65 | 0.76 |
| 8 | 74349858 | rs10957665 | 0.447 | -0.371 | 0.802 | -1.04 | 0.001 | -0.001 | -0.001 | 0 | 0.38 | 0.31 | 0.62 | 0.39 | 0.48 | 0.24 |
| 8 | 74355018 | rs4389931 | 0.144 | -0.346 | 0.816 | -0.892 | -0.001 | 0.021 | 0.021 | 0 | 0.45 | 0.38 | 0.63 | 0.35 | 0.48 | 0.33 |
| 8 | 74358571 | rs2383905 | 0.146 | -0.352 | 0.813 | -0.895 | -0.001 | 0.021 | 0.021 | 0 | 0.45 | 0.38 | 0.63 | 0.35 | 0.48 | 0.33 |
| 8 | 74360148 | rs4738359 | -0.219 | -0.387 | 1.094 | -1.115 | 0.006 | 0.034 | 0.034 | 0 | 0.49 | 0.66 | 0.69 | 0.42 | 0.5 | 0.34 |
| 8 | 74362943 | rs4738362 | -0.788 | -0.39 | 0.65 | -1.25 | -0.033 | 0.041 | 0.041 | 0 | 0.76 | 0.84 | 0.84 | 0.96 | 0.69 | 0.79 |
| 8 | 74365367 | rs7819319 | 2.11 | -0.351 | 0.347 | -1.377 | -0.039 | -0.089 | -0.089 | 0 | 0.24 | 0.03 | 0.37 | 0.38 | 0.26 | 0.11 |
| 8 | 74367182 | rs6472775 | 1.45 | -0.727 | 0.893 | -0.996 | -0.037 | -0.011 | -0.011 | 0 | 0.33 | 0.13 | 0.47 | 0.38 | 0.42 | 0.17 |
| 8 | 74370726 | rs7839905 | 0.715 | -0.616 | 1.005 | -0.949 | 0.022 | -0.031 | -0.031 | 0 | 0.28 | 0.23 | 0.17 | 0.5 | 0.2 | 0.45 |
| 8 | 74371001 | rs6472777 | -0.566 | -0.527 | 0.731 | -0.986 | -0.033 | 0.039 | 0.039 | 0.005 | 0.76 | 0.82 | 0.83 | 0.96 | 0.69 | 0.81 |
| 8 | 74372696 | rs10112982 | 0.705 | -0.871 | 0.837 | -0.936 | 0.003 | -0.004 | -0.004 | 1.38 | 0.43 | 0.43 | 0.5 | 0.43 | 0.46 | 0.29 |
| 8 | 74376476 | rs7834413 | 0.128 | -0.843 | 0.735 | -0.975 | -0.002 | 0 | 0 | 0.032 | 0.32 | 0.35 | 0.33 | 0.55 | 0.24 | 0.52 |
| 8 | 74378595 | rs4738365 | -0.235 | -0.607 | 0.921 | -0.831 | 0.014 | -0.059 | -0.059 | 0 | 0.75 | 0.71 | 0.82 | 0.96 | 0.7 | 0.78 |
| 8 | 74389082 | rs10808795 | 0.603 | -0.604 | 0.98 | -0.719 | 0.001 | -0.035 | -0.035 | 0 | 0.63 | 0.42 | 0.78 | 0.9 | 0.64 | 0.75 |
| 8 | 74389238 | rs16938673 | -0.062 | -0.671 | 1.025 | -0.747 | -0.034 | -0.03 | -0.03 | 0.489 | 0.11 | 0.24 | 0.04 | 0.04 | 0.06 | 0.12 |
| 8 | 74391780 | rs4237005 | 0.559 | -0.852 | 0.925 | -0.693 | 0.005 | -0.008 | -0.008 | 0.506 | 0.25 | 0.26 | 0.27 | 0.53 | 0.16 | 0.4 |
| 8 | 74392490 | rs13268812 | 0.076 | -0.859 | 0.917 | -0.694 | -0.001 | -0.001 | -0.001 | 0 | 0.57 | 0.63 | 0.72 | 0.45 | 0.59 | 0.59 |
| 8 | 74392900 | rs4738367 | 0.465 | -0.859 | 1.023 | -0.729 | 0.014 | -0.082 | -0.082 | 0 | 0.66 | 0.56 | 0.81 | 0.9 | 0.65 | 0.76 |
| 8 | 74393504 | rs7831225 | 1.148 | -0.841 | 0.952 | -0.702 | 0.009 | -0.013 | -0.013 | 0 | 0.21 | 0.16 | 0.12 | 0.49 | 0.12 | 0.29 |
| 8 | 74393626 | rs4738371 | -0.736 | -0.714 | 0.791 | -0.68 | 0.001 | -0.015 | -0.015 | 0 | 0.78 | 0.82 | 0.85 | 0.94 | 0.71 | 0.88 |
| 8 | 74394066 | rs4738372 | 0.529 | -0.436 | 0.436 | -0.468 | -0.005 | 0.006 | 0.006 | 0 | 0.46 | 0.43 | 0.69 | 0.41 | 0.52 | 0.47 |
| 8 | 74397713 | rs16938682 | -0.978 | -0.169 | 0.484 | -0.435 | -0.009 | -0.011 | -0.011 | 0.159 | 0.07 | 0.26 | 0.19 | 0.04 | 0.04 | 0.12 |
| 8 | 74402277 | rs11783372 | 0.675 | -0.198 | 0.207 | -0.65 | -0.003 | 0.011 | 0.011 | 0 | 0.61 | 0.31 | 0.64 | 0.88 | 0.62 | 0.62 |
| 8 | 74406223 | rs7018252 | 1.143 | -0.535 | -0.032 | -0.443 | -0.026 | 0.013 | 0.013 | 0 | 0.41 | 0.23 | 0.53 | 0.4 | 0.49 | 0.33 |
| 8 | 74409780 | rs7823459 | 1.097 | -0.534 | -0.031 | -0.442 | -0.025 | 0.014 | 0.014 | 0 | 0.41 | 0.23 | 0.53 | 0.4 | 0.49 | 0.33 |
| 8 | 74409852 | rs4738376 | 0.288 | -0.422 | 0.471 | -0.32 | -0.051 | 0.088 | 0.088 | 0.258 | 0.25 | 0.4 | 0.29 | 0.5 | 0.15 | 0.4 |
| 8 | 74419237 | rs7823883 | 0.018 | -0.184 | 0.58 | -0.509 | 0.061 | -0.094 | -0.094 | 3.97 | 0.73 | 0.46 | 0.66 | 0.91 | 0.66 | 0.76 |
| 8 | 74424985 | rs7818404 | -0.165 | -0.062 | 0.053 | -0.402 | -0.062 | 0.059 | 0.059 | 0.244 | 0.76 | 0.9 | 0.81 | 0.9 | 0.66 | 0.88 |
| 8 | 74426588 | rs11997691 | 0.551 | -0.335 | 0.285 | -0.193 | 0 | 0.001 | 0.001 | 0.069 | 0.49 | 0.53 | 0.54 | 0.42 | 0.53 | 0.4 |
| 8 | 74450320 | rs10957671 | 1.12 | -0.343 | -0.376 | -0.165 | -0.061 | -0.045 | -0.045 | 0 | 0.39 | 0.09 | 0.46 | 0.36 | 0.45 | 0.34 |
| 8 | 74453600 | rs7812995 | 1.215 | -0.43 | -0.404 | -0.096 | -0.062 | 0.003 | 0.003 | 0 | 0.38 | 0.1 | 0.39 | 0.31 | 0.44 | 0.34 |
| 8 | 74455870 | rs12549456 | 0.114 | -0.44 | 0.182 | -0.22 | -0.014 | 0.008 | 0.008 | 1.094 | 0.64 | 0.58 | 0.7 | 0.47 | 0.77 | 0.45 |
| 8 | 74459295 | rs6991770 | 1.585 | -0.601 | 0.041 | -0.256 | 0.025 | -0.079 | -0.079 | 0 | 0.17 | 0.05 | 0.09 | 0.42 | 0.1 | 0.26 |
| 8 | 74466473 | rs10108786 | 0.26 | -0.232 | 0.299 | -0.13 | 0.008 | -0.006 | -0.006 | 0 | 0.24 | 0.26 | 0.28 | 0.16 | 0.34 | 0.11 |
| 8 | 74470142 | rs10087203 | 0.095 | -0.451 | 0.391 | -0.009 | -0.006 | 0.001 | 0.001 | 1.045 | 0.64 | 0.61 | 0.7 | 0.47 | 0.78 | 0.45 |
| 8 | 74478390 | rs4500072 | -0.036 | 0.248 | 0.348 | 0.035 | -0.005 | 0.006 | 0.006 | 0 | 0.94 | 0.91 | 0.97 | 0.91 | 0.94 | 0.94 |
| 8 | 74480705 | rs7014309 | -0.294 | 0.793 | 0.043 | -0.082 | 0.018 | -0.004 | -0.004 | 0.17 | 0.14 | 0.24 | 0.19 | 0.13 | 0.16 | 0.07 |
| 8 | 74487451 | rs6651282 | 0.879 | 1.044 | -0.362 | -0.123 | -0.03 | 0.027 | 0.027 | 0 | 0.11 | 0.03 | 0.08 | 0.03 | 0.19 | 0.06 |
| 8 | 74492747 | rs6994418 | -0.789 | -0.023 | 0.08 | -0.374 | -0.016 | -0.01 | -0.01 | 0 | 0.29 | 0.59 | 0.27 | 0.26 | 0.22 | 0.22 |
| 8 | 74499435 | rs13277065 | 1.001 | -0.383 | 0.061 | -0.605 | 0.02 | 0.018 | 0.018 | 0 | 0.64 | 0.25 | 0.71 | 0.73 | 0.74 | 0.74 |
| 8 | 74502909 | rs716009 | -0.714 | -1.015 | -0.451 | -0.213 | 0.032 | 0.013 | 0.013 | 0.341 | 0.47 | 0.59 | 0.34 | 0.63 | 0.31 | 0.52 |
| 8 | 74507018 | rs905539 | 0.966 | -0.811 | -0.47 | -0.167 | 0.021 | 0.02 | 0.02 | 0 | 0.61 | 0.25 | 0.53 | 0.71 | 0.7 | 0.62 |
| 8 | 74527996 | rs949493 | 0.184 | -0.788 | -0.361 | -0.323 | -0.009 | -0.018 | -0.018 | 0.509 | 0.9 | 0.78 | 0.79 | 0.97 | 0.92 | 0.83 |
| 8 | 74530344 | rs9657038 | 0.276 | -0.995 | 0.089 | -0.41 | -0.01 | -0.002 | -0.002 | 0.129 | 0.1 | 0.15 | 0.05 | 0.11 | 0.08 | 0.09 |
| 8 | 74534821 | rs16938706 | 0.119 | -1.17 | 0.672 | 0.014 | -0.019 | 0.13 | 0.13 | 0 | 0.21 | 0.44 | 0.24 | 0.38 | 0.14 | 0.29 |
| 8 | 74538698 | rs10086512 | 1.615 | -1.241 | 0.582 | -0.031 | 0.03 | 0.027 | 0.027 | 0 | 0.43 | 0.2 | 0.45 | 0.34 | 0.61 | 0.32 |
| 8 | 74545820 | rs6472795 | -0.766 | -1.033 | 0.844 | 0.094 | -0.015 | 0.037 | 0.037 | 0 | 0.3 | 0.37 | 0.25 | 0.47 | 0.22 | 0.37 |
| 8 | 74569540 | rs10808796 | 0.111 | -1.048 | 0.871 | 0.093 | -0.005 | 0.028 | 0.028 | 0 | 0.2 | 0.24 | 0.2 | 0.35 | 0.14 | 0.26 |
| 8 | 74575374 | rs6992006 | 0.201 | -1.106 | 0.889 | 0.059 | -0.012 | -0.034 | -0.034 | 0 | 0.9 | 0.76 | 0.78 | 0.98 | 0.92 | 0.83 |
| 8 | 74591268 | rs16938715 | 0.503 | -1.078 | 0.878 | 0.1 | -0.002 | -0.001 | -0.001 | 0 | 0.09 | 0.09 | 0.04 | 0.11 | 0.08 | 0.1 |
| 8 | 74596237 | rs12677119 | -0.744 | -1.053 | 0.834 | 0.084 | -0.017 | 0.036 | 0.036 | 0 | 0.3 | 0.37 | 0.25 | 0.47 | 0.22 | 0.37 |
| 8 | 74601700 | rs10106560 | -0.721 | -1.049 | 0.836 | 0.086 | -0.017 | 0.037 | 0.037 | 0 | 0.3 | 0.37 | 0.25 | 0.47 | 0.22 | 0.37 |
| 8 | 74603781 | rs10086435 | -0.891 | -1.092 | 0.682 | 0.055 | -0.025 | 0.046 | 0.046 | 0 | 0.31 | 0.55 | 0.28 | 0.49 | 0.22 | 0.38 |
| 8 | 74620139 | rs17223062 | 1.732 | -0.538 | 0.077 | -0.278 | -0.137 | -0.059 | -0.059 | 0 | 0.33 | 0.05 | 0.36 | 0.34 | 0.46 | 0.27 |
| 8 | 74636240 | rs6996826 | -0.802 | -0.754 | 0.838 | 0.045 | -0.03 | 0.072 | 0.072 | 0 | 0.3 | 0.47 | 0.26 | 0.49 | 0.22 | 0.38 |
| 8 | 74638267 | rs7017254 | 0.138 | -0.763 | 0.889 | 0.067 | -0.015 | 0.01 | 0.01 | 0 | 0.1 | 0.2 | 0.06 | 0.13 | 0.08 | 0.12 |
| 8 | 74648812 | rs7009095 | 0.061 | -0.715 | 0.946 | -0.053 | -0.007 | 0.047 | 0.047 | 0 | 0.21 | 0.29 | 0.21 | 0.37 | 0.15 | 0.26 |
| 8 | 74661778 | rs10086736 | 1.441 | -0.74 | 0.661 | -0.127 | 0.031 | -0.012 | -0.012 | 0 | 0.59 | 0.23 | 0.5 | 0.6 | 0.67 | 0.46 |
| 8 | 74667729 | rs7836815 | 0.346 | -0.577 | 0.872 | -0.37 | -0.006 | -0.025 | -0.025 | 1.162 | 0.09 | 0.11 | 0.04 | 0.01 | 0.07 | 0.07 |
| 8 | 74669409 | rs12545117 | 1.615 | -0.665 | 0.714 | -0.426 | 0 | -0.014 | -0.014 | 1.662 | 0.75 | 0.23 | 0.67 | 0.95 | 0.82 | 0.72 |
| 8 | 74669790 | rs6980645 | -0.008 | -0.59 | 0.666 | -0.381 | 0.005 | -0.009 | -0.009 | 0 | 0.94 | 0.9 | 0.79 | 0.98 | 0.9 | 0.83 |
| 8 | 74670445 | rs1481510 | -0.008 | -0.59 | 0.666 | -0.381 | 0.005 | -0.009 | -0.009 | 0 | 0.94 | 0.91 | 0.79 | 0.98 | 0.9 | 0.83 |
| 8 | 74674174 | rs4389932 | -1.039 | -0.636 | 0.76 | -0.437 | -0.059 | -0.138 | -0.138 | 0 | 0.19 | 0.51 | 0.11 | 0.01 | 0.11 | 0.08 |
| 8 | 74676017 | rs1481508 | 1.384 | -0.511 | 0.519 | -0.427 | -0.006 | -0.051 | -0.051 | 0 | 0.74 | 0.3 | 0.65 | 0.94 | 0.77 | 0.72 |
| 8 | 74685404 | rs2046180 | 0.315 | -0.864 | -0.23 | -0.046 | -0.003 | -0.108 | -0.108 | 0 | 0.79 | 0.94 | 0.67 | 0.59 | 0.83 | 0.58 |
| 8 | 74693749 | rs12679779 | 1.445 | -0.654 | -0.491 | -0.202 | -0.084 | 0.026 | 0.026 | 0 | 0.41 | 0.06 | 0.54 | 0.29 | 0.53 | 0.3 |
| 8 | 74699975 | rs1564531 | 1.529 | -0.018 | 1.549 | -0.031 | 0.005 | -0.056 | -0.056 | 0.038 | 0.78 | 0.35 | 0.8 | 0.93 | 0.78 | 0.78 |
| 8 | 74703133 | rs17224289 | -0.508 | -0.378 | 1.743 | -0.091 | 0 | -0.016 | -0.016 | 0 | 0.13 | 0.13 | 0.07 | 0 | 0.13 | 0.08 |

| | | | | | | | | | | | | | | | | |
|---|---|---|---|---|---|---|---|---|---|---|---|---|---|---|---|---|
| 8 | 74727081 | rs6992853 | -1.208 | -0.317 | 0.572 | -0.061 | -0.02 | -0.089 | -0.089 | 2.498 | 0.18 | 0.64 | 0.14 | 0.02 | 0.13 | 0.1 |
| 8 | 74804726 | rs16938741 | -1.199 | -0.318 | 0.479 | -0.061 | -0.023 | -0.097 | -0.097 | 0 | 0.18 | 0.63 | 0.14 | 0.02 | 0.13 | 0.1 |
| 8 | 74809210 | rs10217047 | -0.746 | 0 | 0.214 | 0.818 | 0.018 | 0.086 | 0.086 | 0.082 | 0.08 | 0.18 | 0.07 | 0.32 | 0.11 | 0.16 |
| 8 | 74810030 | rs10957689 | -1.015 | -0.707 | 0.257 | 0.973 | -0.01 | 0.019 | 0.019 | 0 | 0.21 | 0.24 | 0.18 | 0.34 | 0.17 | 0.33 |
| 8 | 74817097 | rs11777718 | 0.754 | -1.128 | -0.54 | 0.436 | 0.014 | 0.029 | 0.029 | 0.17 | 0.24 | 0.2 | 0.14 | 0.12 | 0.18 | 0.17 |
| 8 | 74818529 | rs1842665 | -1.44 | -0.956 | -0.736 | 0.482 | -0.035 | -0.004 | -0.004 | 0.004 | 0.61 | 0.8 | 0.45 | 0.63 | 0.47 | 0.58 |
| 8 | 74819410 | rs4737396 | -0.276 | -0.696 | -0.609 | 0.488 | 0.033 | -0.044 | -0.044 | 0 | 0.12 | 0.06 | 0.12 | 0.33 | 0.07 | 0.22 |
| 8 | 74847760 | rs11787158 | 0.189 | 0.249 | -0.505 | -0.576 | -0.009 | -0.002 | -0.002 | 0 | 0.23 | 0.22 | 0.31 | 0.21 | 0.34 | 0.3 |
| 8 | 74853706 | rs1038292 | 0.382 | -0.239 | -0.399 | -0.995 | 0.01 | 0.007 | 0.007 | 0 | 0.27 | 0.22 | 0.12 | 0.09 | 0.17 | 0.22 |
| 8 | 74861484 | rs7830335 | -0.288 | 0.052 | -0.711 | -0.981 | -0.008 | -0.003 | -0.003 | 0 | 0.26 | 0.25 | 0.42 | 0.24 | 0.31 | 0.36 |
| 8 | 74868305 | rs12676883 | 1.124 | -0.054 | -0.222 | -1.022 | -0.027 | -0.032 | -0.032 | 4.747 | 0.22 | 0.26 | 0.17 | 0.13 | 0.15 | 0.16 |
| 8 | 74875342 | rs10504551 | -0.139 | -0.047 | -0.307 | -1.018 | -0.01 | -0.005 | -0.005 | 0.339 | 0.26 | 0.23 | 0.42 | 0.24 | 0.31 | 0.36 |
| 8 | 74877351 | rs2433208 | 1.084 | -0.173 | -0.506 | -1.165 | -0.023 | 0.048 | 0.048 | 0.198 | 0.09 | 0.01 | 0.04 | 0 | 0.11 | 0.07 |
| 8 | 74878696 | rs12547746 | -1.666 | -0.471 | -0.1 | -0.992 | -0.052 | 0.004 | 0.004 | 0 | 0.56 | 0.83 | 0.64 | 0.37 | 0.48 | 0.52 |
| 8 | 74881674 | rs7013120 | 0.539 | 0.025 | -0.177 | -0.803 | -0.02 | -0.008 | -0.008 | 0.011 | 0.18 | 0.13 | 0.23 | 0.19 | 0.27 | 0.25 |
| 8 | 74884255 | rs4738406 | 0.906 | 0.001 | -0.202 | -0.811 | -0.044 | -0.024 | -0.024 | 0 | 0.16 | 0.07 | 0.22 | 0.18 | 0.27 | 0.25 |
| 8 | 74884530 | rs6990813 | -0.294 | -0.04 | -0.136 | -0.801 | -0.006 | -0.002 | -0.002 | 0.045 | 0.26 | 0.26 | 0.3 | 0.23 | 0.3 | 0.34 |
| 8 | 74888049 | rs12682285 | -0.988 | -0.039 | -0.281 | -0.8 | 0 | 0.011 | 0.011 | 0 | 0.49 | 0.52 | 0.6 | 0.34 | 0.46 | 0.5 |
| 8 | 74893821 | rs1053079 | 0.054 | -0.062 | -0.3 | -0.805 | -0.05 | -0.057 | -0.057 | 0 | 0.29 | 0.4 | 0.36 | 0.15 | 0.17 | 0.21 |
| 8 | 74893880 | rs7108 | -1.008 | -0.058 | -0.301 | -0.805 | 0 | 0.009 | 0.009 | 0 | 0.49 | 0.51 | 0.6 | 0.34 | 0.46 | 0.5 |
| 8 | 74894025 | rs1053054 | 0.052 | -0.045 | -0.294 | -0.799 | -0.05 | -0.057 | -0.057 | 0 | 0.29 | 0.4 | 0.36 | 0.15 | 0.17 | 0.21 |
| 8 | 74894748 | rs2291217 | -0.018 | 0.662 | -0.89 | -0.457 | -0.029 | -0.011 | -0.011 | 0.369 | 0.2 | 0.12 | 0.23 | 0.19 | 0.29 | 0.29 |
| 8 | 74899141 | rs11775193 | 0.089 | 0.7 | -0.959 | -0.479 | -0.022 | -0.007 | -0.007 | 0.389 | 0.19 | 0.16 | 0.23 | 0.19 | 0.27 | 0.31 |
| 8 | 74907558 | rs7821865 | 0.108 | 0.748 | -0.855 | -0.421 | -0.029 | -0.01 | -0.01 | 0 | 0.2 | 0.13 | 0.23 | 0.19 | 0.29 | 0.29 |
| 8 | 74919713 | rs10504554 | -0.286 | -0.089 | 0.24 | -0.708 | -0.027 | -0.075 | -0.075 | 0 | 0.27 | 0.54 | 0.22 | 0.11 | 0.19 | 0.14 |
| 8 | 74919912 | rs17226566 | 0.037 | 0.393 | -0.324 | -0.498 | -0.092 | -0.037 | -0.037 | 0 | 0.18 | 0.01 | 0.19 | 0.16 | 0.23 | 0.15 |
| 8 | 74928092 | rs4738414 | -0.951 | -0.491 | 1.128 | -0.561 | -0.043 | -0.074 | -0.074 | 0.417 | 0.28 | 0.41 | 0.32 | 0.14 | 0.18 | 0.16 |
| 8 | 74946880 | rs7001809 | -1.124 | -0.498 | 1.127 | -0.565 | -0.038 | -0.07 | -0.07 | 0 | 0.3 | 0.54 | 0.23 | 0.14 | 0.18 | 0.14 |
| 8 | 74954880 | rs12545427 | -1.105 | -0.437 | 0.935 | -0.488 | -0.013 | -0.028 | -0.028 | 0.085 | 0.29 | 0.62 | 0.34 | 0.17 | 0.21 | 0.14 |
| 8 | 74959000 | rs6982935 | -1.02 | -0.399 | 0.894 | -0.509 | -0.04 | -0.083 | -0.083 | 0 | 0.29 | 0.49 | 0.2 | 0.16 | 0.21 | 0.13 |
| 8 | 74973218 | rs10504557 | 0.133 | -0.269 | 0.626 | -0.392 | 0.002 | -0.007 | -0.007 | 0 | 0.06 | 0.04 | 0.02 | 0 | 0.04 | 0.03 |
| 8 | 74975532 | rs1392125 | -0.71 | -0.352 | 0.875 | -0.398 | -0.027 | -0.036 | -0.036 | 0 | 0.08 | 0.19 | 0.04 | 0.01 | 0.04 | 0.02 |
| 8 | 74991564 | rs3857970 | -0.724 | -0.336 | -0.22 | -0.475 | 0.011 | 0.003 | 0.003 | 2.175 | 0.28 | 0.7 | 0.3 | 0.12 | 0.19 | 0.12 |
| 8 | 75002407 | rs12676850 | 0.455 | -0.057 | -0.202 | 0.346 | -0.012 | 0.017 | 0.017 | 6.688 | 0.67 | 0.43 | 0.56 | 0.59 | 0.71 | 0.68 |
| 8 | 75004222 | rs2036273 | 0.686 | 0.516 | -0.697 | 0.01 | -0.014 | 0.011 | 0.011 | 0.192 | 0.45 | 0.25 | 0.45 | 0.6 | 0.48 | 0.42 |
| 8 | 75006830 | rs6472819 | -0.428 | 0.216 | -0.294 | -0.687 | 0.002 | -0.021 | -0.021 | 0 | 0.35 | 0.38 | 0.35 | 0.07 | 0.31 | 0.29 |
| 8 | 75008065 | rs2010344 | -0.572 | 0.02 | 0.072 | -0.881 | -0.007 | -0.055 | -0.055 | 0 | 0.23 | 0.28 | 0.13 | 0.01 | 0.19 | 0.12 |
| 8 | 75008270 | rs1827153 | 0.699 | -0.128 | 0.509 | -1.098 | -0.007 | -0.087 | -0.087 | 0 | 0.77 | 0.65 | 0.77 | 0.99 | 0.79 | 0.87 |
| 8 | 75014430 | rs13249908 | -0.571 | 0.174 | 0.052 | -0.917 | -0.005 | -0.003 | -0.003 | 0 | 0.54 | 0.59 | 0.45 | 0.38 | 0.6 | 0.45 |
| 8 | 75020890 | rs16938797 | 0.748 | 0.045 | -0.33 | -0.737 | 0.023 | 0.008 | 0.008 | 0.113 | 0.07 | 0.03 | 0.02 | 0.05 | 0.03 | 0.19 |
| 8 | 75021897 | rs4738417 | 0.616 | 0.855 | -0.623 | -0.572 | -0.074 | -0.081 | -0.081 | 5.147 | 0.15 | 0.01 | 0.16 | 0.22 | 0.24 | 0.07 |
| 8 | 75024816 | rs904596 | -0.229 | 0.505 | -0.255 | -0.844 | 0.014 | -0.002 | -0.002 | 0 | 0.48 | 0.39 | 0.37 | 0.44 | 0.4 | 0.32 |
| 8 | 75026885 | rs6987422 | -0.423 | 0.367 | -0.315 | -1.091 | -0.011 | -0.03 | -0.03 | 1.394 | 0.18 | 0.24 | 0.15 | 0.04 | 0.13 | 0.22 |
| 8 | 75026965 | rs9657104 | 0.362 | 0.775 | -0.396 | -0.924 | 0.002 | -0.014 | -0.014 | 2.979 | 0.08 | 0.11 | 0.05 | 0.01 | 0.11 | 0.13 |
| 8 | 75032879 | rs2173578 | 0.001 | -0.225 | -0.194 | -0.346 | -0.004 | -0.001 | -0.001 | 0 | 0.45 | 0.56 | 0.32 | 0.41 | 0.34 | 0.44 |
| 8 | 75033393 | rs16938799 | 0.964 | -0.149 | -0.688 | -0.186 | -0.02 | -0.062 | -0.062 | 0 | 0.15 | 0.01 | 0.12 | 0.17 | 0.14 | 0.07 |
| 8 | 75042238 | rs1909530 | -0.682 | -0.543 | -0.144 | -0.154 | 0 | 0 | 0 | 0 | 0.59 | 0.64 | 0.41 | 0.61 | 0.38 | 0.46 |
| 8 | 75044020 | rs10957694 | -0.11 | -0.882 | 0.009 | -0.272 | -0.016 | -0.008 | -0.008 | 0 | 0.24 | 0.35 | 0.24 | 0.15 | 0.16 | 0.09 |
| 8 | 75046612 | rs4489317 | 0.567 | -0.84 | -0.062 | -0.441 | 0.004 | -0.031 | -0.031 | 0 | 0.09 | 0.11 | 0.06 | 0 | 0.1 | 0.05 |
| 8 | 75048111 | rs4607599 | 0.15 | -0.755 | -0.138 | -0.359 | -0.007 | -0.003 | -0.003 | 0 | 0.23 | 0.31 | 0.23 | 0.15 | 0.17 | 0.09 |
| 8 | 75048599 | rs11783882 | 0.151 | -0.76 | -0.135 | -0.222 | 0.011 | -0.008 | -0.008 | 0 | 0.24 | 0.23 | 0.23 | 0.23 | 0.17 | 0.1 |
| 8 | 75050742 | rs6472822 | -1.533 | -0.761 | -0.202 | -0.262 | -0.036 | -0.018 | -0.018 | 0.213 | 0.72 | 0.83 | 0.49 | 0.61 | 0.5 | 0.56 |
| 8 | 75051696 | rs1808016 | -0.758 | -0.836 | -0.576 | 0.067 | 0.003 | -0.001 | -0.001 | 0 | 0.34 | 0.36 | 0.26 | 0.32 | 0.22 | 0.25 |
| 8 | 75052641 | rs2133552 | 1.492 | -0.88 | -0.712 | 0.085 | 0.027 | -0.021 | -0.021 | 0.034 | 0.25 | 0.11 | 0.13 | 0.28 | 0.17 | 0.21 |
| 8 | 75055005 | rs12547483 | -1.402 | -0.381 | -0.169 | 0.128 | -0.016 | -0.002 | -0.002 | 0 | 0.36 | 0.46 | 0.28 | 0.32 | 0.22 | 0.26 |
| 8 | 75055057 | rs17249164 | -0.336 | 0.016 | -0.755 | -0.249 | 0.018 | 0.027 | 0.027 | 8.937 | 0.18 | 0.13 | 0.08 | 0.08 | 0.12 | 0.07 |
| 8 | 75056939 | rs16938807 | -0.347 | 0.177 | -0.262 | -0.083 | -0.019 | -0.009 | -0.009 | 2.594 | 0.15 | 0.25 | 0.13 | 0.15 | 0.1 | 0.12 |
| 8 | 75057013 | rs17249304 | -0.632 | 0.457 | -0.374 | -0.423 | -0.002 | -0.014 | -0.014 | 0 | 0.11 | 0.1 | 0.14 | 0 | 0.13 | 0.09 |
| 8 | 75061350 | rs17249487 | 0.929 | 0.449 | -0.463 | -0.304 | -0.006 | -0.017 | -0.017 | 0.001 | 0.09 | 0 | 0.09 | 0.08 | 0.07 | 0.05 |
| 8 | 75061454 | rs2977049 | 1.172 | 0.527 | 0 | -0.542 | 0.023 | -0.09 | -0.09 | 0.184 | 0.8 | 0.58 | 0.81 | 0.93 | 0.82 | 0.71 |
| 8 | 75062690 | rs16938809 | 0.257 | 0.506 | 0.042 | -0.51 | -0.007 | -0.002 | -0.002 | 0.599 | 0.07 | 0.09 | 0.02 | 0.07 | 0.02 | 0.14 |
| 8 | 75074938 | rs2936676 | 1.282 | 0.503 | 0.035 | -0.483 | -0.005 | -0.055 | -0.055 | 2.929 | 0.71 | 0.54 | 0.72 | 0.84 | 0.75 | 0.67 |
| 8 | 75080275 | rs2977054 | 0.759 | 0.607 | 0.097 | -0.442 | 0.018 | -0.047 | -0.047 | 0 | 0.78 | 0.66 | 0.75 | 0.91 | 0.77 | 0.81 |
| 8 | 75087343 | rs1875067 | 0.689 | 0.574 | -0.48 | -0.379 | 0.031 | 0.001 | 0.001 | 0 | 0.48 | 0.37 | 0.3 | 0.48 | 0.38 | 0.48 |
| 8 | 75088702 | rs2936680 | -0.432 | 0.702 | -0.284 | -0.404 | 0.027 | -0.067 | -0.067 | 1.315 | 0.87 | 0.8 | 0.85 | 1 | 0.85 | 0.87 |
| 8 | 75090201 | rs2936689 | 0.997 | 0.152 | -0.952 | -0.545 | -0.039 | 0.037 | 0.037 | 0.815 | 0.18 | 0.01 | 0.1 | 0.08 | 0.17 | 0.32 |
| 8 | 75092581 | rs6984028 | 1.311 | 0.206 | -0.748 | -0.425 | 0.01 | -0.011 | -0.011 | 0.271 | 0.29 | 0.25 | 0.19 | 0.4 | 0.21 | 0.16 |
| 8 | 75096991 | rs4385459 | -0.187 | 0.239 | -0.811 | -0.519 | -0.001 | 0.003 | 0.003 | 0 | 0.43 | 0.64 | 0.61 | 0.42 | 0.56 | 0.45 |
| 8 | 75098003 | rs7001077 | 1.038 | -0.136 | -0.686 | -0.34 | 0.023 | -0.001 | -0.001 | 0 | 0.32 | 0.26 | 0.19 | 0.36 | 0.21 | 0.22 |

| | | | | | | | | | | | | | | | | |
|---|---|---|---|---|---|---|---|---|---|---|---|---|---|---|---|---|
| 8 | 75099011 | rs7835959 | -0.011 | 0.254 | -0.399 | -0.175 | 0.004 | -0.045 | -0.045 | 0 | 0.86 | 0.69 | 0.85 | 0.95 | 0.87 | 0.83 |
| 8 | 75101074 | rs2936665 | 0.021 | 0.402 | -0.943 | -0.018 | 0.012 | -0.001 | -0.001 | 0 | 0.28 | 0.21 | 0.2 | 0.24 | 0.23 | 0.3 |
| 8 | 75109785 | rs2956060 | -0.11 | 0.487 | -0.949 | 0.089 | 0.005 | -0.014 | -0.014 | 0.136 | 0.3 | 0.21 | 0.22 | 0.31 | 0.25 | 0.38 |
| 8 | 75110661 | rs12678640 | 1.215 | 1.139 | -1.468 | 0.005 | -0.085 | -0.094 | -0.094 | 0.281 | 0.14 | 0.01 | 0.33 | 0.32 | 0.24 | 0.15 |
| 8 | 75111953 | rs2956055 | -0.157 | 0.601 | -0.777 | -0.307 | 0.002 | -0.004 | -0.004 | 0 | 0.3 | 0.28 | 0.25 | 0.31 | 0.25 | 0.39 |
| 8 | 75114021 | rs16938822 | 1.367 | 0.614 | -0.892 | -0.295 | -0.001 | -0.013 | -0.013 | 0 | 0.1 | 0.06 | 0.1 | 0.11 | 0.08 | 0.05 |
| 8 | 75116209 | rs2936686 | 0.293 | 0.601 | -0.476 | -0.351 | 0 | -0.001 | -0.001 | 0 | 0.2 | 0.19 | 0.15 | 0.21 | 0.17 | 0.33 |
| 8 | 75119342 | rs6472827 | -0.754 | 0.6 | -0.563 | -0.353 | -0.006 | -0.024 | -0.024 | 0 | 0.82 | 0.94 | 0.89 | 0.79 | 0.83 | 0.67 |
| 8 | 75123241 | rs7832806 | 0.693 | 0.505 | -0.414 | -0.537 | -0.006 | -0.022 | -0.022 | 0 | 0.66 | 0.56 | 0.73 | 0.78 | 0.71 | 0.53 |
| 8 | 75126432 | rs12545808 | 0.768 | 0.249 | -0.47 | -0.719 | -0.005 | -0.014 | -0.014 | 0 | 0.58 | 0.53 | 0.69 | 0.73 | 0.63 | 0.46 |
| 8 | 75129728 | rs10092100 | -0.581 | 0.214 | -0.48 | -0.718 | 0.001 | 0.013 | 0.013 | 0 | 0.21 | 0.2 | 0.16 | 0.1 | 0.21 | 0.2 |
| 8 | 75131422 | rs7017837 | -0.553 | 0.229 | -0.348 | -0.675 | 0 | 0.01 | 0.01 | 0 | 0.19 | 0.17 | 0.14 | 0.1 | 0.19 | 0.17 |
| 8 | 75131473 | rs6983474 | -0.644 | 0.212 | -0.3 | -0.684 | -0.001 | -0.034 | -0.034 | 0 | 0.19 | 0.28 | 0.18 | 0.11 | 0.19 | 0.17 |
| 8 | 75132803 | rs11996194 | -0.617 | -0.029 | -0.309 | -0.709 | -0.005 | -0.024 | -0.024 | 0.131 | 0.19 | 0.23 | 0.16 | 0.09 | 0.15 | 0.17 |
| 8 | 75134110 | rs11990179 | -0.235 | 0.268 | -0.121 | -0.699 | 0.003 | -0.011 | -0.011 | 0 | 0.76 | 0.73 | 0.83 | 0.83 | 0.75 | 0.59 |
| 8 | 75134473 | rs6999501 | -0.554 | 0.261 | -0.104 | -0.702 | -0.017 | 0.013 | 0.013 | 3.258 | 0.78 | 0.9 | 0.85 | 0.83 | 0.76 | 0.61 |
| 8 | 75135150 | rs4737409 | 0.88 | 0.153 | 0.229 | -0.748 | -0.004 | -0.017 | -0.017 | 0 | 0.61 | 0.48 | 0.69 | 0.72 | 0.64 | 0.47 |
| 8 | 75135813 | rs7005773 | -0.245 | 0.211 | 0.311 | -0.729 | 0.003 | -0.018 | -0.018 | 0 | 0.76 | 0.7 | 0.81 | 0.84 | 0.75 | 0.59 |
| 8 | 75141913 | rs4416813 | -0.278 | 0.119 | 0.411 | -0.775 | 0.002 | -0.035 | -0.035 | 0 | 0.74 | 0.62 | 0.8 | 0.84 | 0.75 | 0.59 |
| 8 | 75143753 | rs10957702 | 0.693 | -0.08 | 0.076 | -0.782 | 0.007 | 0.021 | 0.021 | 1.575 | 0.58 | 0.38 | 0.65 | 0.73 | 0.64 | 0.47 |
| 8 | 75154867 | rs17337083 | 1.319 | -0.25 | -0.292 | -0.55 | -0.013 | -0.01 | -0.01 | 0 | 0.1 | 0 | 0.09 | 0.07 | 0.07 | 0.03 |
| 8 | 75156489 | rs4738438 | 0.379 | -0.373 | 0.171 | -0.465 | 0.007 | 0.003 | 0.003 | 3.585 | 0.67 | 0.28 | 0.72 | 0.7 | 0.71 | 0.57 |
| 8 | 75187365 | rs16938845 | 0.818 | -0.153 | 0.449 | -0.288 | 0.006 | 0.005 | 0.005 | 3.069 | 0.12 | 0.06 | 0.09 | 0.1 | 0.09 | 0.03 |
| 8 | 75192666 | rs926110 | -0.05 | -0.202 | 0.014 | -0.28 | -0.018 | 0.018 | 0.018 | 1.442 | 0.3 | 0.12 | 0.22 | 0.2 | 0.32 | 0.41 |
| 8 | 75193280 | rs16938849 | 1.336 | 0.389 | -0.317 | 0.23 | -0.067 | -0.069 | -0.069 | 0.149 | 0.1 | 0.01 | 0.19 | 0.27 | 0.19 | 0.1 |
| 8 | 75194157 | rs12334848 | -0.626 | 0.366 | -0.158 | 0.14 | 0.006 | 0.035 | 0.035 | 0.028 | 0.42 | 0.23 | 0.32 | 0.29 | 0.4 | 0.47 |
| 8 | 75196465 | rs7832145 | -0.793 | 0.071 | 0.188 | 0.116 | -0.001 | -0.006 | -0.006 | 1.59 | 0.35 | 0.37 | 0.35 | 0.25 | 0.31 | 0.2 |
| 8 | 75205295 | rs741844 | -0.771 | 0.08 | 0.387 | 0.262 | -0.004 | 0.005 | 0.005 | 0.266 | 0.36 | 0.55 | 0.37 | 0.41 | 0.32 | 0.21 |
| 8 | 75207653 | rs4738443 | -1.773 | -0.005 | 0.14 | 0.341 | -0.021 | 0.007 | 0.007 | 0 | 0.63 | 0.83 | 0.61 | 0.62 | 0.59 | 0.59 |
| 8 | 75210077 | rs2142182 | 0.027 | -0.155 | 0.372 | 0.33 | 0.002 | 0.007 | 0.007 | 2.012 | 0.29 | 0.2 | 0.28 | 0.23 | 0.24 | 0.13 |
| 8 | 75211071 | rs16938857 | 0.227 | -0.142 | 0.527 | 0.378 | 0.012 | 0.107 | 0.107 | 0 | 0.05 | 0.18 | 0.04 | 0.19 | 0.06 | 0.05 |
| 8 | 75211163 | rs1022430 | -2.022 | -0.029 | 0.494 | 0.348 | -0.004 | 0.003 | 0.003 | 0 | 0.65 | 0.84 | 0.64 | 0.63 | 0.63 | 0.63 |
| 8 | 75211424 | rs1022429 | 0.693 | -0.037 | 0.549 | 0.341 | -0.036 | -0.006 | -0.006 | 2.809 | 0.25 | 0.13 | 0.19 | 0.19 | 0.3 | 0.4 |
| 8 | 75212074 | rs6988935 | -0.34 | 0.055 | 0.54 | 0.41 | 0.003 | 0.018 | 0.018 | 5.756 | 0.36 | 0.62 | 0.35 | 0.4 | 0.39 | 0.5 |
| 8 | 75213345 | rs16938859 | -0.897 | 0.007 | 0.178 | 0.446 | -0.006 | 0.008 | 0.008 | 1.633 | 0.36 | 0.4 | 0.33 | 0.41 | 0.3 | 0.18 |
| 8 | 75217901 | rs760245 | -0.183 | 0.418 | -0.288 | 0.718 | -0.03 | -0.004 | -0.004 | 0 | 0.22 | 0.1 | 0.32 | 0.26 | 0.25 | 0.15 |
| 8 | 75218725 | rs1003406 | 0.429 | 0.514 | -0.208 | 0.755 | -0.036 | -0.033 | -0.033 | 0 | 0.16 | 0.08 | 0.31 | 0.29 | 0.22 | 0.11 |
| 8 | 75224861 | rs13263396 | -1.732 | 0.399 | 0.159 | 0.831 | 0.053 | 0.172 | 0.172 | 0.187 | 0.06 | 0.34 | 0.06 | 0.1 | 0.05 | 0.05 |
| 8 | 75225381 | rs13263313 | -0.401 | 0.482 | -0.45 | 0.856 | 0.005 | 0.013 | 0.013 | 0 | 0.21 | 0.25 | 0.35 | 0.39 | 0.27 | 0.16 |
| 8 | 75232955 | rs4738445 | 0.4 | 0.536 | -0.96 | 0.772 | -0.037 | -0.03 | -0.03 | 0 | 0.16 | 0.08 | 0.31 | 0.28 | 0.22 | 0.11 |
| 8 | 75239978 | rs10504572 | -0.671 | 0.383 | -1.164 | 0.77 | 0.01 | -0.005 | -0.005 | 0 | 0.33 | 0.22 | 0.21 | 0.27 | 0.3 | 0.4 |
| 8 | 75242411 | rs2975213 | -1.419 | 0.364 | -1.595 | 0.737 | 0.004 | 0.025 | 0.025 | 0.114 | 0.49 | 0.2 | 0.51 | 0.55 | 0.52 | 0.51 |
| 8 | 75245770 | rs6989476 | -0.878 | -0.218 | -1.26 | 0.606 | 0.001 | 0.011 | 0.011 | 0.302 | 0.3 | 0.08 | 0.18 | 0.19 | 0.25 | 0.37 |
| 8 | 75248900 | rs1997560 | -0.88 | -0.086 | -0.822 | 0.936 | 0.006 | -0.009 | -0.009 | 0 | 0.25 | 0.22 | 0.17 | 0.3 | 0.19 | 0.33 |
| 8 | 75249396 | rs760244 | -0.704 | -0.216 | -1.242 | 0.951 | 0.023 | -0.035 | -0.035 | 1.381 | 0.3 | 0.16 | 0.19 | 0.3 | 0.24 | 0.35 |
| 8 | 75254086 | rs7004754 | 0.078 | -0.097 | -0.578 | 1.072 | 0.008 | -0.007 | -0.007 | 0.028 | 0.24 | 0.19 | 0.41 | 0.28 | 0.19 | 0.24 |
| 8 | 75257155 | rs16938884 | 0.013 | -0.26 | 0.206 | 0.991 | -0.013 | 0.004 | 0.004 | 0 | 0.25 | 0.3 | 0.42 | 0.28 | 0.19 | 0.24 |
| 8 | 75265600 | rs16938888 | 0.388 | -0.137 | -0.152 | 0.989 | -0.024 | -0.033 | -0.033 | 1.276 | 0.28 | 0.14 | 0.18 | 0.3 | 0.36 | 0.19 |
| 8 | 75266189 | rs13269312 | 0.723 | -0.27 | 0.504 | 0.867 | -0.019 | 0.014 | 0.014 | 0.102 | 0.43 | 0.28 | 0.34 | 0.42 | 0.44 | 0.54 |
| 8 | 75271627 | rs10096726 | 0.571 | -0.274 | 0.752 | 0.865 | -0.003 | 0.004 | 0.004 | 0 | 0.46 | 0.44 | 0.38 | 0.42 | 0.45 | 0.55 |
| 8 | 75275911 | rs4464961 | 0.797 | -0.334 | 0.652 | 0.925 | 0 | -0.003 | -0.003 | 0.019 | 0.05 | 0.04 | 0.06 | 0.12 | 0.05 | 0.05 |
| 8 | 75279647 | rs7842662 | 0.943 | -0.359 | 0.672 | 0.926 | -0.023 | 0.023 | 0.023 | 0 | 0.38 | 0.22 | 0.28 | 0.3 | 0.4 | 0.5 |
| 8 | 75292492 | rs6996971 | 0.289 | -0.412 | 0.705 | 0.903 | -0.005 | 0.01 | 0.01 | 0 | 0.47 | 0.6 | 0.4 | 0.42 | 0.46 | 0.53 |
| 8 | 75294975 | rs4471028 | 1.007 | -0.468 | 0.785 | 0.865 | -0.043 | 0.023 | 0.023 | 0.019 | 0.37 | 0.22 | 0.3 | 0.3 | 0.4 | 0.5 |
| 8 | 75307831 | rs10283076 | 0.171 | -0.427 | 0.833 | 1.086 | -0.011 | 0.074 | 0.074 | 0 | 0.48 | 0.71 | 0.38 | 0.34 | 0.46 | 0.53 |
| 8 | 75332823 | rs6472849 | 1.038 | -0.458 | 0.742 | 0.748 | 0 | 0 | 0 | 0 | 0.33 | 0.37 | 0.29 | 0.32 | 0.3 | 0.46 |
| 8 | 75335908 | rs10112529 | 0.694 | -0.297 | 0.607 | 0.689 | 0.001 | 0 | 0 | 0 | 0.41 | 0.62 | 0.38 | 0.44 | 0.35 | 0.51 |
| 8 | 75339134 | rs9650249 | -0.002 | -0.285 | 0.253 | 0.572 | -0.025 | 0.002 | 0.002 | 1.935 | 0.34 | 0.19 | 0.22 | 0.29 | 0.47 | 0.21 |
| 8 | 75347706 | rs16938906 | 0.459 | -0.152 | -0.033 | 0.728 | -0.017 | -0.004 | -0.004 | 0 | 0.26 | 0.37 | 0.42 | 0.26 | 0.18 | 0.26 |
| 8 | 75356405 | rs7014530 | -0.407 | 0.342 | -0.458 | 0.82 | -0.005 | 0.002 | 0.002 | 2.499 | 0.36 | 0.3 | 0.26 | 0.29 | 0.48 | 0.18 |
| 8 | 75369545 | rs17350701 | 0.203 | 0.407 | -1.166 | 0.916 | -0.082 | -0.112 | -0.112 | 8.788 | 0.32 | 0.04 | 0.35 | 0.36 | 0.38 | 0.2 |
| 8 | 75377173 | rs10088057 | -0.159 | 0.258 | -0.496 | 0.84 | -0.002 | -0.016 | -0.016 | 0 | 0.66 | 0.72 | 0.62 | 0.63 | 0.62 | 0.78 |
| 8 | 75395361 | rs10104370 | -0.407 | -0.057 | -0.268 | 0.727 | 0.042 | 0 | 0 | 0.573 | 0.49 | 0.85 | 0.64 | 0.47 | 0.45 | 0.37 |
| 8 | 75397835 | rs6472855 | -0.619 | 0.169 | -0.572 | 0.727 | -0.073 | 0.001 | 0.001 | 0 | 0.83 | 0.97 | 0.89 | 0.88 | 0.75 | 0.88 |
| 8 | 75402095 | rs13274943 | -0.904 | -0.38 | 0.398 | 0.592 | -0.012 | 0.035 | 0.035 | 0 | 0.1 | 0.21 | 0.09 | 0.18 | 0.08 | 0.1 |
| 8 | 75405213 | rs7461712 | 0.388 | -0.398 | 0.541 | 0.52 | 0.006 | 0.012 | 0.012 | 0 | 0.37 | 0.25 | 0.48 | 0.29 | 0.37 | 0.26 |
| 8 | 75407809 | rs10098494 | -0.666 | -0.31 | 0.617 | 0.61 | -0.001 | 0.099 | 0.099 | 0 | 0.1 | 0.36 | 0.11 | 0.2 | 0.08 | 0.1 |
| 8 | 75408063 | rs7459665 | -0.645 | -0.365 | 0.415 | 0.594 | 0.043 | -0.015 | -0.015 | 0 | 0.5 | 0.83 | 0.53 | 0.48 | 0.45 | 0.36 |
| 8 | 75411977 | rs4735782 | -0.79 | -0.309 | 0.663 | 0.638 | 0 | 0.115 | 0.115 | 0.001 | 0.1 | 0.38 | 0.11 | 0.2 | 0.07 | 0.1 |

| | | | | | | | | | | | | | | | | |
|---|---|---|---|---|---|---|---|---|---|---|---|---|---|---|---|---|
| 8 | 75419366 | rs11986395 | 0.143 | -0.044 | 0.418 | 0.667 | -0.001 | 0.009 | 0.009 | 0 | 0.39 | 0.37 | 0.42 | 0.3 | 0.42 | 0.25 |
| 8 | 75424387 | rs4735683 | -0.631 | -0.022 | -0.298 | 0.501 | -0.009 | -0.01 | -0.01 | 0 | 0.52 | 0.8 | 0.58 | 0.49 | 0.5 | 0.35 |
| 8 | 75425389 | rs4545073 | -0.619 | -0.009 | -0.377 | 0.508 | -0.003 | -0.004 | -0.004 | 0.281 | 0.52 | 0.76 | 0.58 | 0.49 | 0.5 | 0.35 |
| 8 | 75431219 | rs12541981 | -0.21 | 0.686 | -0.908 | 1.011 | -0.004 | 0.043 | 0.043 | 0 | 0.39 | 0.27 | 0.41 | 0.28 | 0.43 | 0.25 |
| 8 | 75453034 | rs7015958 | -0.11 | 0.911 | -0.968 | 0.142 | -0.009 | -0.009 | -0.009 | 0 | 0.75 | 0.84 | 0.65 | 0.64 | 0.67 | 0.56 |
| 8 | 75457070 | rs7845621 | 0.468 | 0.634 | -0.855 | -0.965 | 0.001 | 0.006 | 0.006 | 0 | 0.35 | 0.31 | 0.34 | 0.63 | 0.35 | 0.31 |
| 8 | 75462525 | rs4440611 | 0.52 | 0.293 | -0.343 | -1.506 | 0 | -0.024 | -0.024 | 0 | 0.31 | 0.28 | 0.22 | 0.02 | 0.28 | 0.26 |
| 8 | 75472479 | rs1365036 | 1.018 | 0.356 | -0.287 | -1.274 | -0.017 | -0.056 | -0.056 | 4.205 | 0.29 | 0.14 | 0.23 | 0.56 | 0.3 | 0.18 |
| 8 | 75472647 | rs1946352 | -0.382 | 0.025 | 0.469 | -1.433 | -0.011 | -0.021 | -0.021 | 0 | 0.14 | 0.42 | 0.14 | 0.08 | 0.08 | 0.14 |
| 8 | 75474529 | rs746460 | -0.373 | 0.222 | 0.514 | -1.578 | -0.006 | 0.008 | 0.008 | 0 | 0.66 | 0.75 | 0.5 | 0.65 | 0.6 | 0.47 |
| 8 | 75479624 | rs13260300 | -0.521 | 0.159 | 0.285 | -1.609 | -0.049 | -0.005 | -0.005 | 2.601 | 0.68 | 0.9 | 0.52 | 0.65 | 0.6 | 0.47 |
| 8 | 75480503 | rs7015712 | -0.207 | 0.128 | 0.318 | -1.647 | 0.005 | 0.06 | 0.06 | 0 | 0.31 | 0.74 | 0.27 | 0.09 | 0.24 | 0.28 |
| 8 | 75485868 | rs11995376 | 0.024 | 0.201 | 0.901 | -1.651 | 0.042 | 0.011 | 0.011 | 0 | 0.11 | 0.36 | 0.2 | 0.07 | 0.1 | 0.22 |
| 8 | 75485927 | rs1425765 | -1.257 | 0.179 | 0.549 | -1.633 | -0.087 | -0.083 | -0.083 | 0 | 0.77 | 0.98 | 0.62 | 0.65 | 0.68 | 0.57 |
| 8 | 75492280 | rs959422 | 1.215 | 0.142 | 0.545 | -1.673 | -0.046 | 0.106 | 0.106 | 0 | 0.19 | 0.02 | 0.09 | 0.01 | 0.2 | 0.15 |
| 8 | 75494355 | rs4557676 | -1.03 | 0.162 | 0.63 | -1.654 | -0.053 | -0.037 | -0.037 | 0.02 | 0.74 | 0.93 | 0.6 | 0.65 | 0.66 | 0.57 |
| 8 | 75513831 | rs6472867 | 0.405 | 0.032 | 0.935 | -1.741 | -0.004 | -0.017 | -0.017 | 0 | 0.45 | 0.52 | 0.43 | 0.65 | 0.39 | 0.39 |
| 8 | 75522602 | rs7824069 | -1.04 | 0.086 | 0.601 | -1.678 | -0.032 | -0.029 | -0.029 | 0 | 0.75 | 0.91 | 0.6 | 0.66 | 0.67 | 0.57 |
| 8 | 75553100 | rs1365032 | 0.353 | -0.374 | 1.272 | -2.085 | -0.014 | 0.018 | 0.018 | 0.091 | 0.6 | 0.34 | 0.4 | 0.65 | 0.49 | 0.37 |
| 8 | 75555433 | rs9298241 | 0.961 | -0.51 | 1.079 | -2.198 | 0.06 | 0.038 | 0.038 | 0.284 | 0.39 | 0.11 | 0.27 | 0.65 | 0.29 | 0.22 |
| 8 | 75564547 | rs1365038 | 0.798 | -0.418 | 1.085 | -2.078 | -0.035 | 0.081 | 0.081 | 0 | 0.55 | 0.22 | 0.37 | 0.65 | 0.48 | 0.33 |
| 8 | 75565803 | rs2553713 | -0.35 | -0.004 | 1.285 | -1.865 | 0.002 | -0.003 | -0.003 | 0 | 0.72 | 0.74 | 0.58 | 0.76 | 0.59 | 0.51 |
| 8 | 75599357 | rs1593410 | 1.033 | -0.462 | 1.006 | -2.136 | 0.039 | 0.045 | 0.045 | 1.847 | 0.38 | 0.05 | 0.26 | 0.64 | 0.27 | 0.21 |
| 8 | 75610456 | rs1304474 | -0.69 | -0.269 | 0.96 | -2.187 | -0.118 | 0.042 | 0.042 | 0 | 0.73 | 0.94 | 0.56 | 0.76 | 0.55 | 0.54 |
| 8 | 75614614 | rs269198 | -0.159 | -0.197 | 1.201 | -2.126 | 0.001 | -0.009 | -0.009 | 0 | 0.33 | 0.66 | 0.29 | 0.11 | 0.27 | 0.33 |
| 8 | 75625511 | rs269188 | -1.283 | 0.165 | 0.747 | -1.943 | -0.136 | -0.017 | -0.017 | 0 | 0.81 | 0.98 | 0.58 | 0.75 | 0.61 | 0.63 |
| 8 | 75640056 | rs16938940 | -1.481 | 0.013 | 1.211 | -1.839 | 0.104 | 0.087 | 0.087 | 0.51 | 0.05 | 0.49 | 0.06 | 0.02 | 0.02 | 0.07 |
| 8 | 75657127 | rs13268614 | 0.762 | 0.051 | 0.567 | -1.893 | -0.003 | 0.021 | 0.021 | 0.08 | 0.23 | 0.16 | 0.19 | 0.06 | 0.24 | 0.24 |
| 8 | 75661534 | rs4735772 | 0.477 | 0.024 | 0.573 | -1.906 | 0 | 0.007 | 0.007 | 0 | 0.25 | 0.22 | 0.21 | 0.06 | 0.24 | 0.24 |
| 8 | 75674877 | rs991394 | -0.054 | 0.191 | 0.07 | -1.828 | 0.027 | 0.142 | 0.142 | 0 | 0.36 | 0.83 | 0.24 | 0.04 | 0.28 | 0.34 |
| 8 | 75678475 | rs10464886 | 0.663 | 0.175 | -0.268 | -1.848 | 0.021 | 0.033 | 0.033 | 1.687 | 0.4 | 0.03 | 0.21 | 0.63 | 0.3 | 0.21 |
| 8 | 75682535 | rs1392966 | -0.059 | 0.282 | 0.593 | -1.767 | 0 | -0.033 | -0.033 | 0.058 | 0.32 | 0.62 | 0.22 | 0.04 | 0.28 | 0.34 |
| 8 | 75685474 | rs11987589 | -1.298 | 0.407 | 1.246 | -1.492 | 0.04 | -0.029 | -0.029 | 1.221 | 0.09 | 0.33 | 0.08 | 0.03 | 0.08 | 0.05 |
| 8 | 75691260 | rs7812753 | -1.773 | 0.405 | 1.859 | -1.47 | 0.038 | -0.057 | -0.057 | 0 | 0.11 | 0.46 | 0.1 | 0.03 | 0.1 | 0.02 |
| 8 | 75707662 | rs7842758 | -1.578 | 0.531 | 1.956 | -1.453 | 0.056 | -0.01 | -0.01 | 0 | 0.11 | 0.45 | 0.14 | 0.06 | 0.13 | 0.04 |
| 8 | 75709529 | rs16938959 | -0.906 | 0.529 | 2.078 | -1.455 | 0.046 | 0.004 | 0.004 | 0 | 0.08 | 0.32 | 0.13 | 0.06 | 0.11 | 0.04 |
| 8 | 75722130 | rs6985430 | -1.34 | 0.504 | 1.759 | -1.458 | 0.064 | -0.009 | -0.009 | 0.119 | 0.12 | 0.49 | 0.15 | 0.06 | 0.13 | 0.04 |
| 8 | 75737818 | rs1991860 | -0.512 | 0.336 | 0.855 | -2.21 | 0 | -0.031 | -0.031 | 4.028 | 0.42 | 0.63 | 0.29 | 0.07 | 0.42 | 0.34 |
| 8 | 75740387 | rs11784963 | -0.431 | 0.339 | 0.645 | -2.209 | 0 | -0.088 | -0.088 | 0 | 0.41 | 0.56 | 0.27 | 0.07 | 0.42 | 0.34 |
| 8 | 75741263 | rs17358302 | 0.504 | 0.34 | 0.633 | -2.216 | -0.007 | 0.111 | 0.111 | 0 | 0.3 | 0.09 | 0.13 | 0.01 | 0.28 | 0.29 |
| 8 | 75749982 | rs1433470 | 0.486 | 0.33 | 0.629 | -2.221 | -0.006 | 0.112 | 0.112 | 0.004 | 0.3 | 0.09 | 0.13 | 0.01 | 0.28 | 0.29 |
| 8 | 75763075 | rs2732011 | -0.43 | 0.329 | 0.605 | -2.213 | 0 | -0.084 | -0.084 | 0.081 | 0.41 | 0.56 | 0.27 | 0.07 | 0.42 | 0.34 |
| 8 | 75764715 | rs2732018 | -0.43 | 0.329 | 0.605 | -2.213 | 0 | -0.084 | -0.084 | 0 | 0.41 | 0.56 | 0.27 | 0.07 | 0.42 | 0.34 |
| 8 | 75767414 | rs2956606 | 0.993 | 0.568 | -1.072 | -1.868 | 0.069 | -0.067 | -0.067 | 0.632 | 0.33 | 0.06 | 0.17 | 0.57 | 0.19 | 0.23 |
| 8 | 75772574 | rs2570180 | -0.731 | 0.68 | -0.469 | -1.761 | -0.009 | 0.168 | 0.168 | 0 | 0.44 | 0.82 | 0.36 | 0.12 | 0.44 | 0.37 |
| 8 | 75774987 | rs11774767 | 0.124 | 0.55 | -0.84 | -1.551 | -0.004 | 0.096 | 0.096 | 0 | 0.3 | 0.12 | 0.14 | 0.01 | 0.29 | 0.3 |
| 8 | 75792274 | rs2732014 | -0.21 | 0.554 | -0.609 | -0.825 | 0.009 | 0.009 | 0.009 | 2.188 | 0.6 | 0.4 | 0.64 | 0.39 | 0.65 | 0.73 |
| 8 | 75798282 | rs2916163 | -0.151 | 0.506 | 0.939 | -0.813 | 0.004 | 0.002 | 0.002 | 2.44 | 0.61 | 0.62 | 0.67 | 0.41 | 0.67 | 0.73 |
| 8 | 75802572 | rs7842487 | 0.553 | 0.618 | 0.423 | -1.338 | -0.001 | -0.024 | -0.024 | 0 | 0.29 | 0.28 | 0.15 | 0.05 | 0.28 | 0.27 |
| 8 | 75803536 | rs1560784 | 0.078 | 0.825 | 0.326 | -1.45 | 0 | -0.01 | -0.01 | 3.408 | 0.35 | 0.32 | 0.27 | 0.11 | 0.35 | 0.33 |
| 8 | 75807939 | rs1122317 | -0.471 | 0.831 | 0.038 | -1.417 | 0.016 | 0.023 | 0.023 | 1.839 | 0.54 | 0.4 | 0.53 | 0.36 | 0.6 | 0.67 |
| 8 | 75808298 | rs1978737 | 0.534 | 0.945 | -0.197 | -1.29 | -0.022 | 0.002 | 0.002 | 0 | 0.26 | 0.2 | 0.4 | 0.31 | 0.32 | 0.4 |
| 8 | 75810723 | rs2954873 | 0.534 | 0.939 | -0.195 | -1.287 | -0.022 | 0.002 | 0.002 | 0 | 0.26 | 0.2 | 0.4 | 0.31 | 0.32 | 0.38 |
| 8 | 75813778 | rs2570154 | -0.084 | 0.885 | -0.113 | -1.286 | -0.02 | 0.007 | 0.007 | 0 | 0.31 | 0.26 | 0.51 | 0.36 | 0.39 | 0.45 |
| 8 | 75818891 | rs2731995 | -0.433 | 0.567 | 0.025 | -1.313 | -0.002 | 0.124 | 0.124 | 0 | 0.42 | 0.74 | 0.35 | 0.18 | 0.4 | 0.36 |
| 8 | 75819800 | rs10504589 | -0.189 | 0.564 | 0.182 | -1.342 | 0.002 | 0.027 | 0.027 | 0 | 0.37 | 0.67 | 0.24 | 0.12 | 0.35 | 0.3 |
| 8 | 75821686 | rs2570159 | 0.598 | 0.511 | 0.635 | -0.983 | 0.018 | -0.001 | -0.001 | 0 | 0.05 | 0.15 | 0.13 | 0.06 | 0.05 | 0.05 |
| 8 | 75825202 | rs728025 | -0.428 | 0.512 | 1.166 | -1.002 | 0.087 | 0.057 | 0.057 | 0.931 | 0.05 | 0.34 | 0.09 | 0.06 | 0.06 | 0.02 |
| 8 | 75829655 | rs16939000 | -0.609 | 0.503 | 1.056 | -1.006 | 0.106 | 0.073 | 0.073 | 0.524 | 0.06 | 0.37 | 0.09 | 0.06 | 0.06 | 0.02 |
| 8 | 75832460 | rs2954870 | -1.356 | 0.486 | 0.464 | -1.179 | 0 | 0.065 | 0.065 | 0 | 0.31 | 0.83 | 0.36 | 0.19 | 0.32 | 0.3 |
| 8 | 75833964 | rs16939001 | 0.184 | 0.186 | 0.598 | -1.167 | 0.01 | 0.012 | 0.012 | 0.337 | 0.17 | 0.15 | 0.14 | 0.1 | 0.18 | 0.22 |
| 8 | 75838269 | rs7842291 | -0.89 | 0.554 | 1.377 | -0.641 | 0.157 | 0.126 | 0.126 | 0.022 | 0.06 | 0.36 | 0.1 | 0.08 | 0.1 | 0.03 |
| 8 | 75838927 | rs2570174 | -2.013 | 0.542 | 1.017 | -0.33 | 0.007 | 0.017 | 0.017 | 1.218 | 0.13 | 0.73 | 0.25 | 0.18 | 0.14 | 0.08 |
| 8 | 75854525 | rs2383926 | -1.169 | -0.062 | 0.781 | -0.615 | -0.018 | -0.038 | -0.038 | 0.532 | 0.23 | 0.54 | 0.2 | 0.14 | 0.2 | 0.17 |
| 8 | 75861772 | rs6981885 | -0.685 | -0.061 | 0.92 | -0.618 | -0.005 | -0.047 | -0.047 | 0.462 | 0.19 | 0.38 | 0.14 | 0.08 | 0.17 | 0.14 |
| 8 | 75866175 | rs16939003 | -1.217 | -0.076 | 0.87 | -0.64 | -0.015 | -0.034 | -0.034 | 0.039 | 0.23 | 0.58 | 0.2 | 0.14 | 0.2 | 0.17 |
| 8 | 75870915 | rs7002052 | -1.291 | -0.051 | 0.936 | -0.647 | -0.014 | -0.034 | -0.034 | 1.167 | 0.24 | 0.57 | 0.2 | 0.14 | 0.2 | 0.16 |
| 8 | 75878055 | rs10092310 | -1.761 | -0.491 | 0.459 | -0.691 | 0.061 | 0.084 | 0.084 | 0 | 0.39 | 0.84 | 0.27 | 0.21 | 0.35 | 0.3 |
| 8 | 75883054 | rs2956605 | -1.958 | -0.231 | -0.316 | 0.024 | -0.066 | -0.047 | -0.047 | 0 | 0.61 | 0.95 | 0.61 | 0.62 | 0.64 | 0.59 |

| | | | | | | | | | | | | | | | | |
|---|---|---|---|---|---|---|---|---|---|---|---|---|---|---|---|---|
| 8 | 75883589 | rs16939008 | -0.765 | -0.206 | -0.115 | -0.027 | -0.003 | -0.006 | -0.006 | 0 | 0.32 | 0.6 | 0.18 | 0.16 | 0.3 | 0.26 |
| 8 | 75886297 | rs2925155 | 0.713 | 0.04 | -0.213 | 0.381 | -0.128 | -0.107 | -0.107 | 0 | 0.21 | 0.02 | 0.33 | 0.3 | 0.29 | 0.29 |
| 8 | 75890354 | rs1378617 | -0.346 | -0.16 | 0.49 | 0.042 | -0.01 | -0.018 | -0.018 | 0.675 | 0.19 | 0.33 | 0.09 | 0.12 | 0.14 | 0.11 |
| 8 | 75894701 | rs7017893 | 1.29 | -0.3 | 0.507 | 0.036 | -0.008 | -0.009 | -0.009 | 2.336 | 0.12 | 0.13 | 0.06 | 0.04 | 0.13 | 0.11 |
| 8 | 75894894 | rs6472881 | 0.429 | -0.334 | 0.614 | 0.015 | -0.006 | 0.027 | 0.027 | 1.722 | 0.11 | 0.29 | 0.07 | 0.12 | 0.07 | 0.02 |
| 8 | 75899434 | rs10957744 | -0.049 | -0.436 | 0.737 | -0.015 | -0.028 | -0.021 | -0.021 | 0 | 0.18 | 0.28 | 0.13 | 0.14 | 0.14 | 0.15 |
| 8 | 75900388 | rs4472488 | -1.676 | -0.457 | 0.757 | -0.03 | 0.074 | 0.066 | 0.066 | 0.14 | 0.4 | 0.82 | 0.25 | 0.29 | 0.31 | 0.27 |
| 8 | 75902603 | rs10216950 | -1.063 | -0.46 | 0.816 | -0.034 | -0.05 | -0.069 | -0.069 | 0 | 0.29 | 0.53 | 0.18 | 0.17 | 0.23 | 0.25 |
| 8 | 75902945 | rs7841751 | 0.221 | -0.397 | 0.833 | -0.027 | -0.01 | 0.024 | 0.024 | 2.244 | 0.12 | 0.33 | 0.07 | 0.12 | 0.07 | 0.02 |
| 8 | 75907518 | rs7010624 | -1.483 | -0.408 | 0.814 | -0.037 | 0.022 | 0.023 | 0.023 | 0.08 | 0.34 | 0.71 | 0.18 | 0.19 | 0.28 | 0.23 |
| 8 | 75910104 | rs2383928 | 0.319 | -0.311 | 0.824 | 0.014 | 0.006 | -0.035 | -0.035 | 0 | 0.11 | 0.24 | 0.05 | 0.03 | 0.09 | 0.1 |
| 8 | 75916686 | rs16939020 | -1.232 | -0.375 | 0.757 | 0.025 | -0.027 | -0.024 | -0.024 | 0 | 0.23 | 0.58 | 0.12 | 0.15 | 0.15 | 0.11 |
| 8 | 75925763 | rs1455794 | 1.118 | -0.261 | 0.448 | 0.071 | 0 | -0.001 | -0.001 | 0.638 | 0.08 | 0.09 | 0.02 | 0.03 | 0.07 | 0.09 |
| 8 | 75939401 | rs10957748 | -1.027 | -0.257 | 0.321 | 0.041 | -0.022 | -0.026 | -0.026 | 0.874 | 0.36 | 0.56 | 0.21 | 0.21 | 0.28 | 0.25 |
| 8 | 75945850 | rs2085276 | -1.027 | -0.257 | 0.321 | 0.041 | -0.022 | -0.026 | -0.026 | 0 | 0.36 | 0.56 | 0.22 | 0.21 | 0.28 | 0.25 |
| 8 | 75946695 | rs13248650 | 0.539 | 0.086 | -0.343 | 0.368 | -0.01 | -0.01 | -0.01 | 0.291 | 0.2 | 0.17 | 0.37 | 0.35 | 0.34 | 0.36 |
| 8 | 75976170 | rs10104443 | 1.153 | 0.169 | -0.425 | 0.363 | -0.104 | -0.122 | -0.122 | 0 | 0.17 | 0.02 | 0.32 | 0.35 | 0.26 | 0.31 |
| 8 | 75984881 | rs7009332 | -2.069 | 0.086 | -0.492 | 0.368 | -0.056 | -0.023 | -0.023 | 0 | 0.59 | 0.95 | 0.61 | 0.64 | 0.64 | 0.61 |
| 8 | 75995961 | rs1455813 | -1.141 | -0.099 | -0.298 | 0.34 | 0.026 | 0.019 | 0.019 | 0.127 | 0.37 | 0.67 | 0.22 | 0.29 | 0.28 | 0.25 |
| 8 | 76000165 | rs1378613 | -1.166 | 0.263 | -0.264 | 0.424 | -0.004 | 0.06 | 0.06 | 0 | 0.5 | 0.78 | 0.51 | 0.61 | 0.53 | 0.53 |
| 8 | 76002787 | rs1543024 | -0.98 | 0.192 | 0.336 | 0.41 | -0.005 | -0.016 | -0.016 | 0 | 0.46 | 0.68 | 0.53 | 0.5 | 0.55 | 0.6 |
| 8 | 76004157 | rs985094 | -0.395 | 0.261 | 0.591 | 0.396 | 0.006 | 0.009 | 0.009 | 0.5 | 0.35 | 0.4 | 0.47 | 0.41 | 0.41 | 0.46 |
| 8 | 76009527 | rs6981492 | 0.015 | 0.25 | 0.649 | 0.357 | 0.062 | 0.001 | 0.001 | 0 | 0.1 | 0.29 | 0.07 | 0.09 | 0.13 | 0.13 |
| 8 | 76011518 | rs7841235 | -1.994 | 0.026 | 0.405 | 0.113 | 0.005 | 0.023 | 0.023 | 2.365 | 0.34 | 0.75 | 0.27 | 0.26 | 0.29 | 0.21 |
| 8 | 76012812 | rs17370342 | -0.932 | 0.007 | 0.194 | 0.185 | -0.007 | -0.012 | -0.012 | 0 | 0.22 | 0.28 | 0.15 | 0.15 | 0.16 | 0.08 |
| 8 | 76013093 | rs7812482 | -1.797 | 0.008 | 0.005 | 0.205 | -0.035 | -0.05 | -0.05 | 0 | 0.62 | 0.95 | 0.65 | 0.64 | 0.68 | 0.64 |
| 8 | 76013464 | rs10504590 | -0.864 | 0.254 | 0.768 | 0.46 | -0.013 | -0.007 | -0.007 | 0 | 0.18 | 0.24 | 0.11 | 0.15 | 0.13 | 0.05 |
| 8 | 76022121 | rs11987494 | -1.038 | 0.377 | 0.918 | 1.057 | -0.027 | 0.076 | 0.076 | 0.332 | 0.13 | 0.33 | 0.09 | 0.19 | 0.09 | 0.04 |
| 8 | 76025955 | rs11991447 | -0.206 | 0.982 | 0.762 | 1.636 | 0.08 | 0.127 | 0.127 | 0.118 | 0.19 | 0.37 | 0.42 | 0.34 | 0.27 | 0.33 |
| 8 | 76037567 | rs1455805 | -1.698 | 0.504 | -0.293 | 1.176 | -0.063 | -0.028 | -0.028 | 0 | 0.5 | 0.94 | 0.61 | 0.46 | 0.51 | 0.51 |
| 8 | 76039322 | rs10504592 | -0.686 | 0.673 | 0.292 | 1.167 | -0.013 | -0.019 | -0.019 | 0 | 0.24 | 0.29 | 0.15 | 0.21 | 0.22 | 0.2 |
| 8 | 76040583 | rs16939034 | 0.964 | 0.549 | 0.198 | 1.165 | -0.027 | 0.001 | 0.001 | 0.038 | 0.09 | 0.17 | 0.04 | 0.12 | 0.08 | 0.09 |
| 8 | 76040666 | rs1455801 | 0.399 | 0.741 | -0.059 | 1.007 | -0.023 | 0.013 | 0.013 | 0 | 0.13 | 0 | 0.08 | 0.09 | 0.11 | 0.11 |
| 8 | 76044206 | rs4487733 | 0.42 | 0.762 | 0.265 | 1.026 | 0.041 | 0.128 | 0.128 | 0 | 0.15 | 0.42 | 0.14 | 0.25 | 0.18 | 0.21 |
| 8 | 76054904 | rs1350100 | -2.129 | 0.908 | -0.769 | 0.995 | -0.075 | 0.05 | 0.05 | 0 | 0.53 | 0.96 | 0.66 | 0.67 | 0.57 | 0.57 |
| 8 | 76057713 | rs6994721 | -1.798 | 1.26 | -0.04 | 0.842 | -0.015 | 0.004 | 0.004 | 0 | 0.39 | 0.69 | 0.56 | 0.34 | 0.43 | 0.42 |
| 8 | 76064717 | rs7824974 | -1.092 | 0.991 | -1.168 | 0.51 | -0.028 | 0.01 | 0.01 | 0 | 0.24 | 0.11 | 0.19 | 0.19 | 0.29 | 0.18 |
| 8 | 76067224 | rs16939038 | -0.304 | 0.815 | -1.034 | 0.59 | -0.022 | -0.014 | -0.014 | 0 | 0.11 | 0.02 | 0.08 | 0.12 | 0.11 | 0.11 |
| 8 | 76072260 | rs1605940 | -0.6 | 0.732 | -0.821 | 0.6 | -0.003 | 0.018 | 0.018 | 0 | 0.26 | 0.29 | 0.18 | 0.51 | 0.23 | 0.4 |
| 8 | 76081612 | rs6990712 | 0.476 | 0.379 | -0.491 | 0.094 | -0.011 | -0.046 | -0.046 | 5.869 | 0.19 | 0.31 | 0.13 | 0.12 | 0.19 | 0.16 |
| 8 | 76083848 | rs1355255 | -0.169 | 0.362 | -0.396 | 0.092 | 0 | -0.001 | -0.001 | 0 | 0.26 | 0.25 | 0.17 | 0.51 | 0.23 | 0.4 |
| 8 | 76089347 | rs7001087 | 0.926 | 0.682 | -1.023 | -0.576 | 0.003 | -0.043 | -0.043 | 0.29 | 0.24 | 0.32 | 0.24 | 0.07 | 0.24 | 0.14 |
| 8 | 76089712 | rs13253586 | 1.203 | 0.727 | -0.898 | -0.651 | -0.001 | 0.006 | 0.006 | 0 | 0.2 | 0.17 | 0.12 | 0.04 | 0.19 | 0.09 |
| 8 | 76103975 | rs11786534 | 1.335 | 0.707 | -1.029 | -0.291 | -0.022 | -0.012 | -0.012 | 0 | 0.05 | 0 | 0.04 | 0.24 | 0.07 | 0.11 |
| 8 | 76107659 | rs6989358 | 1.534 | 0.645 | -0.955 | -0.373 | -0.015 | -0.084 | -0.084 | 1.29 | 0.16 | 0.28 | 0.11 | 0.02 | 0.16 | 0.13 |
| 8 | 76111050 | rs7006190 | -0.323 | 0.614 | -0.825 | -0.352 | -0.009 | -0.003 | -0.003 | 0 | 0.39 | 0.5 | 0.25 | 0.39 | 0.34 | 0.46 |
| 8 | 76112835 | rs13273502 | 0.554 | 0.502 | -0.759 | -0.324 | -0.003 | -0.005 | -0.005 | 0 | 0.26 | 0.28 | 0.17 | 0.28 | 0.27 | 0.2 |
| 8 | 76120341 | rs13255145 | 1.539 | 0.451 | -0.787 | -0.349 | -0.017 | 0.057 | 0.057 | 0 | 0.18 | 0.08 | 0.11 | 0.02 | 0.19 | 0.08 |
| 8 | 76121824 | rs12547658 | -0.58 | 0.446 | -0.913 | -0.483 | 0.015 | 0.008 | 0.008 | 0.076 | 0.44 | 0.6 | 0.37 | 0.42 | 0.41 | 0.55 |
| 8 | 76126385 | rs1351226 | -1.961 | 0.497 | -1.098 | -0.448 | -0.092 | -0.063 | -0.063 | 0 | 0.71 | 0.94 | 0.56 | 0.7 | 0.68 | 0.74 |
| 8 | 76134023 | rs1995957 | -0.921 | 0.295 | -1.059 | -0.581 | -0.051 | 0.088 | 0.088 | 0 | 0.57 | 0.82 | 0.46 | 0.3 | 0.57 | 0.39 |
| 8 | 76139901 | rs17300285 | -0.544 | 0.3 | -1.064 | -0.623 | -0.006 | 0.159 | 0.159 | 0 | 0.52 | 0.8 | 0.36 | 0.28 | 0.5 | 0.31 |
| 8 | 76140370 | rs16939045 | 0.188 | 0.276 | -0.983 | -0.642 | -0.012 | -0.025 | -0.025 | 0 | 0.3 | 0.36 | 0.19 | 0.24 | 0.3 | 0.22 |
| 8 | 76141132 | rs1351224 | -1.28 | 0.491 | -1.38 | -0.455 | -0.107 | -0.094 | -0.094 | 0 | 0.65 | 0.92 | 0.51 | 0.46 | 0.6 | 0.64 |
| 8 | 76148994 | rs16939047 | 0.616 | 0.178 | -1.082 | -0.23 | 0.003 | -0.021 | -0.021 | 3.7 | 0.11 | 0.09 | 0.14 | 0.19 | 0.08 | 0.3 |
| 8 | 76170249 | rs978598 | 0.755 | 0.517 | -0.97 | -0.267 | 0 | 0.013 | 0.013 | 0 | 0.47 | 0.36 | 0.64 | 0.35 | 0.49 | 0.45 |
| 8 | 76198239 | rs7018058 | 0.378 | 0.426 | -0.897 | -0.292 | -0.012 | 0.004 | 0.004 | 0 | 0.16 | 0.24 | 0.09 | 0.21 | 0.12 | 0.08 |
| 8 | 76200829 | rs7015436 | 0.337 | 0.459 | -0.96 | -0.291 | 0.002 | 0 | 0 | 0 | 0.57 | 0.43 | 0.7 | 0.53 | 0.55 | 0.7 |
| 8 | 76215097 | rs907290 | -0.784 | 0.54 | -1.022 | -0.381 | -0.005 | 0 | 0 | 0 | 0.73 | 0.73 | 0.79 | 0.74 | 0.67 | 0.78 |
| 8 | 76215616 | rs10957761 | 0.279 | 0.433 | -1.223 | -0.436 | 0 | 0.012 | 0.012 | 0 | 0.5 | 0.38 | 0.65 | 0.35 | 0.51 | 0.49 |
| 8 | 76224183 | rs1949109 | 1.097 | 0.084 | -1.048 | -0.471 | -0.001 | -0.006 | -0.006 | 0.029 | 0.3 | 0.16 | 0.44 | 0.3 | 0.28 | 0.26 |
| 8 | 76224630 | rs17301576 | 0.432 | -0.039 | -1.421 | -0.5 | -0.006 | 0.054 | 0.054 | 0 | 0.25 | 0.15 | 0.24 | 0.06 | 0.29 | 0.23 |
| 8 | 76226176 | rs6997167 | 0.561 | -0.276 | -1.249 | -0.07 | 0.001 | -0.007 | -0.007 | 0 | 0.57 | 0.3 | 0.7 | 0.43 | 0.61 | 0.45 |
| 8 | 76230301 | rs6472903 | -1.506 | -0.195 | -1.291 | -0.11 | -0.008 | 0.007 | 0.007 | 0 | 0.87 | 0.89 | 0.92 | 0.97 | 0.84 | 0.9 |
| 8 | 76231410 | rs7016243 | 1.571 | -0.362 | -1.729 | -0.28 | -0.013 | -0.099 | -0.099 | 0 | 0.19 | 0.02 | 0.2 | 0.3 | 0.13 | 0.12 |
| 8 | 76256222 | rs11990539 | -0.589 | -0.382 | -0.678 | 0.553 | -0.053 | 0.035 | 0.035 | 0 | 0.25 | 0.4 | 0.37 | 0.31 | 0.12 | 0.32 |
| 8 | 76257007 | rs16939058 | 0.627 | -0.387 | -0.868 | 0.676 | 0.026 | -0.039 | -0.039 | 0.14 | 0.11 | 0.05 | 0.05 | 0.16 | 0.04 | 0.21 |
| 8 | 76258307 | rs16939060 | 0.196 | -0.032 | -0.812 | 0.571 | -0.043 | -0.042 | -0.042 | 0 | 0.07 | 0 | 0.07 | 0.25 | 0.15 | 0.17 |
| 8 | 76260998 | rs2926615 | -0.18 | -0.616 | -0.484 | 0.424 | 0.027 | 0.013 | 0.013 | 0 | 0.73 | 0.59 | 0.81 | 0.64 | 0.64 | 0.66 |

| | | | | | | | | | | | | | | | | |
|---|---|---|---|---|---|---|---|---|---|---|---|---|---|---|---|---|
| 8 | 76262855 | rs2926619 | 0.372 | -0.236 | -1.276 | 0.186 | -0.004 | 0.071 | 0.071 | 0 | 0.51 | 0.22 | 0.45 | 0.32 | 0.54 | 0.38 |
| 8 | 76282062 | rs16939063 | -0.63 | -1.224 | -0.494 | 0.361 | 0.025 | 0.017 | 0.017 | 0.581 | 0.48 | 0.61 | 0.44 | 0.32 | 0.24 | 0.44 |
| 8 | 76282915 | rs7840945 | -0.584 | -1.241 | -0.455 | 0.353 | -0.007 | -0.001 | -0.001 | 0 | 0.47 | 0.58 | 0.44 | 0.32 | 0.24 | 0.44 |
| 8 | 76292539 | rs6998702 | -0.415 | -0.903 | -0.283 | 0.024 | -0.012 | -0.031 | -0.031 | 0 | 0.31 | 0.38 | 0.36 | 0.1 | 0.17 | 0.19 |
| 8 | 76296789 | rs2980231 | 0.421 | 0.075 | 0.256 | 1.038 | 0.018 | -0.02 | -0.02 | 0.022 | 0.67 | 0.51 | 0.44 | 0.29 | 0.6 | 0.54 |
| 8 | 76299138 | rs1812736 | -0.515 | 0.308 | 0.07 | 1.608 | -0.017 | -0.115 | -0.115 | 2.706 | 0.82 | 0.98 | 0.66 | 0.64 | 0.81 | 0.72 |
| 8 | 76299353 | rs2014238 | 1.351 | 0.242 | 0.379 | 1.167 | -0.003 | 0.008 | 0.008 | 1.173 | 0.58 | 0.41 | 0.37 | 0.29 | 0.5 | 0.5 |
| 8 | 76299919 | rs6472910 | -0.449 | 0.515 | 0.577 | 1.26 | 0.096 | 0.29 | 0.29 | 1.108 | 0.06 | 0.33 | 0.13 | 0.23 | 0.07 | 0.08 |
| 8 | 76306257 | rs2926609 | -0.081 | 0.484 | 0.386 | 1.256 | -0.015 | -0.103 | -0.103 | 0 | 0.78 | 0.92 | 0.54 | 0.45 | 0.72 | 0.69 |
| 8 | 76307676 | rs10100727 | -0.858 | 0.455 | 0.775 | 1.326 | 0.128 | 0.105 | 0.105 | 0 | 0.07 | 0.34 | 0.06 | 0.11 | 0.07 | 0.11 |
| 8 | 76312110 | rs2980240 | -0.703 | 0.446 | 0.641 | 1.461 | -0.023 | -0.133 | -0.133 | 0 | 0.82 | 0.98 | 0.6 | 0.59 | 0.8 | 0.71 |
| 8 | 76315205 | rs2980237 | 0.145 | 0.514 | 0.746 | 1.475 | 0.003 | 0.034 | 0.034 | 0.009 | 0.75 | 0.68 | 0.5 | 0.37 | 0.72 | 0.6 |
| 8 | 76318607 | rs3758054 | -1.846 | 0.462 | 0.991 | 1.411 | 0.032 | -0.011 | -0.011 | 0 | 0.21 | 0.31 | 0.18 | 0.17 | 0.29 | 0.19 |
| 8 | 76323329 | rs1037831 | 1.427 | 0.397 | 0.946 | 1.412 | -0.002 | 0.001 | 0.001 | 2.323 | 0.57 | 0.45 | 0.42 | 0.37 | 0.5 | 0.49 |
| 8 | 76327920 | rs2980248 | 1.452 | 0.47 | 0.89 | 1.363 | -0.007 | -0.011 | -0.011 | 0 | 0.57 | 0.41 | 0.39 | 0.48 | 0.51 | 0.5 |
| 8 | 76338381 | rs17303296 | -0.496 | 0.503 | 0.562 | 1.091 | -0.002 | 0.012 | 0.012 | 4.194 | 0.13 | 0.07 | 0.08 | 0.02 | 0.15 | 0.06 |
| 8 | 76352636 | rs830771 | 1.391 | 0.355 | 0.499 | 1.199 | -0.005 | 0.014 | 0.014 | 2.793 | 0.56 | 0.41 | 0.37 | 0.34 | 0.51 | 0.48 |
| 8 | 76367712 | rs16939073 | -1.692 | 0.481 | 0.652 | 1.033 | 0.029 | -0.014 | -0.014 | 0 | 0.1 | 0.23 | 0.14 | 0.04 | 0.15 | 0.09 |
| 8 | 76367775 | rs2172 | 0.343 | 0.816 | 0.612 | 0.835 | 0.029 | 0.066 | 0.066 | 0 | 0.09 | 0.29 | 0.13 | 0.12 | 0.12 | 0.25 |
| 8 | 76386803 | rs10092405 | -2.146 | 0.756 | 0.484 | 0.954 | -0.006 | -0.049 | -0.049 | 0.926 | 0.33 | 0.82 | 0.36 | 0.38 | 0.35 | 0.45 |
| 8 | 76389346 | rs2037095 | -1.743 | 0.749 | 0.502 | 0.958 | 0.006 | -0.037 | -0.037 | 0 | 0.29 | 0.78 | 0.33 | 0.37 | 0.28 | 0.42 |
| 8 | 76396971 | rs11998082 | -1.105 | 0.701 | 0.567 | 0.858 | -0.018 | 0.043 | 0.043 | 2.746 | 0.18 | 0.46 | 0.13 | 0.22 | 0.14 | 0.16 |
| 8 | 76404081 | rs7008442 | -1.105 | 0.701 | 0.564 | 0.858 | -0.018 | 0.034 | 0.034 | 2.746 | 0.18 | 0.47 | 0.13 | 0.21 | 0.14 | 0.15 |
| 8 | 76404776 | rs6995394 | -0.63 | 0.721 | 0.542 | 0.896 | 0.029 | 0.037 | 0.037 | 0 | 0.15 | 0.32 | 0.23 | 0.16 | 0.21 | 0.3 |
| 8 | 76409284 | rs2926583 | -1.433 | 0.723 | 0.541 | 0.883 | 0.002 | 0.006 | 0.006 | 4.232 | 0.21 | 0.53 | 0.16 | 0.22 | 0.19 | 0.2 |
| 8 | 76409479 | rs830781 | -1.069 | 1.21 | 0.101 | 0.812 | -0.059 | -0.116 | -0.116 | 0.103 | 0.79 | 0.98 | 0.63 | 0.64 | 0.75 | 0.72 |
| 8 | 76410861 | rs2926585 | 0.159 | 1.193 | 0.417 | 0.814 | -0.024 | -0.016 | -0.016 | 0.014 | 0.12 | 0.26 | 0.09 | 0.08 | 0.07 | 0.07 |
| 8 | 76415489 | rs1380692 | -1.451 | 1.193 | 0.409 | 0.814 | 0.003 | 0.006 | 0.006 | 4.514 | 0.21 | 0.53 | 0.16 | 0.22 | 0.19 | 0.19 |
| 8 | 76417937 | rs2943559 | 0.028 | 1.109 | 0.331 | 0.747 | -0.02 | -0.012 | -0.012 | 0 | 0.11 | 0.25 | 0.08 | 0.08 | 0.07 | 0.07 |
| 8 | 76419046 | rs2943568 | -0.686 | 1.112 | 0.232 | 0.748 | -0.031 | -0.021 | -0.021 | 0 | 0.13 | 0.41 | 0.1 | 0.08 | 0.07 | 0.07 |
| 8 | 76419145 | rs6994092 | -1.227 | 1.172 | 0.204 | 0.769 | 0.055 | 0.06 | 0.06 | 0 | 0.22 | 0.43 | 0.28 | 0.3 | 0.34 | 0.43 |
| 8 | 76424482 | rs2977912 | -1.928 | 1.182 | -0.033 | 0.769 | -0.037 | -0.048 | -0.048 | 0 | 0.36 | 0.85 | 0.38 | 0.38 | 0.41 | 0.5 |
| 8 | 76427548 | rs2977916 | -0.977 | 1.223 | -0.097 | 0.847 | -0.054 | -0.111 | -0.111 | 0 | 0.79 | 0.98 | 0.63 | 0.64 | 0.75 | 0.72 |
| 8 | 76440857 | rs2941450 | 0.59 | 1.084 | 0.421 | 0.537 | -0.001 | 0 | 0 | 0 | 0.59 | 0.55 | 0.35 | 0.35 | 0.47 | 0.36 |
| 8 | 76440918 | rs2943584 | -1.162 | 0.975 | 0.519 | 0.329 | 0.03 | -0.012 | -0.012 | 2.302 | 0.14 | 0.41 | 0.11 | 0.08 | 0.12 | 0.13 |
| 8 | 76441911 | rs722989 | -0.4 | 0.485 | 0.46 | -0.154 | -0.005 | -0.009 | -0.009 | 1.578 | 0.2 | 0.25 | 0.12 | 0.2 | 0.14 | 0.11 |
| 8 | 76445721 | rs16939086 | 2.255 | 0.867 | -0.549 | -0.799 | -0.048 | -0.057 | -0.057 | 0.363 | 0.15 | 0.01 | 0.13 | 0.21 | 0.15 | 0.16 |
| 8 | 76451098 | rs2943547 | -0.575 | 0.716 | -0.618 | -0.896 | -0.049 | 0.013 | 0.013 | 1.281 | 0.56 | 0.88 | 0.43 | 0.4 | 0.53 | 0.48 |
| 8 | 76452313 | rs2943549 | -0.928 | 0.711 | -0.705 | -0.879 | -0.092 | -0.067 | -0.067 | 0 | 0.61 | 0.95 | 0.48 | 0.39 | 0.58 | 0.5 |
| 8 | 76459726 | rs1800922 | -0.947 | 0.718 | -0.707 | -0.88 | -0.089 | -0.069 | -0.069 | 0 | 0.61 | 0.95 | 0.49 | 0.39 | 0.59 | 0.5 |
| 8 | 76462383 | rs1399563 | 1.376 | 0.535 | -0.731 | -1.144 | 0.006 | -0.015 | -0.015 | 0.007 | 0.16 | 0.24 | 0.17 | 0.04 | 0.23 | 0.1 |
| 8 | 76463571 | rs2272668 | 0.752 | 0.533 | -0.712 | -1.144 | -0.002 | 0.018 | 0.018 | 0.672 | 0.26 | 0.26 | 0.19 | 0.08 | 0.25 | 0.11 |
| 8 | 76465360 | rs1805099 | 1.505 | 0.557 | -0.545 | -1.031 | 0.003 | 0.042 | 0.042 | 0.014 | 0.07 | 0.2 | 0.05 | 0.08 | 0.05 | 0.09 |
| 8 | 76465823 | rs2977942 | 1.505 | 0.557 | -0.545 | -1.031 | 0.003 | 0.042 | 0.042 | 0.014 | 0.07 | 0.2 | 0.05 | 0.08 | 0.05 | 0.09 |
| 8 | 76468077 | rs16939097 | 1.587 | 0.591 | -0.754 | -1.012 | 0 | 0 | 0 | 0.165 | 0.16 | 0.17 | 0.14 | 0.2 | 0.16 | 0.21 |
| 8 | 76468282 | rs1805098 | -1.166 | 0.608 | -1.009 | -1.049 | -0.088 | -0.064 | -0.064 | 0 | 0.61 | 0.95 | 0.49 | 0.39 | 0.59 | 0.5 |
| 8 | 76473939 | rs2943611 | -2.088 | 0.629 | -1.044 | -1.026 | -0.031 | -0.15 | -0.15 | 0.765 | 0.78 | 0.99 | 0.64 | 0.6 | 0.78 | 0.72 |
| 8 | 76474058 | rs1464093 | 1.447 | 0.71 | -0.781 | -1.175 | 0.003 | -0.033 | -0.033 | 0 | 0.16 | 0.21 | 0.11 | 0.01 | 0.21 | 0.1 |
| 8 | 76476396 | rs1805100 | -0.558 | 0.637 | -0.717 | -1.172 | -0.03 | 0.08 | 0.08 | 0 | 0.45 | 0.77 | 0.38 | 0.32 | 0.47 | 0.41 |
| 8 | 76476457 | rs2272669 | -0.558 | 0.637 | -0.717 | -1.172 | -0.03 | 0.08 | 0.08 | 0 | 0.45 | 0.77 | 0.38 | 0.32 | 0.47 | 0.41 |
| 8 | 76476670 | rs2941477 | -0.558 | 0.637 | -0.717 | -1.172 | -0.03 | 0.08 | 0.08 | 0 | 0.45 | 0.77 | 0.38 | 0.32 | 0.47 | 0.41 |
| 8 | 76476959 | rs2941479 | -1.326 | 0.619 | -0.731 | -1.193 | -0.058 | 0.01 | 0.01 | 1.143 | 0.6 | 0.88 | 0.43 | 0.36 | 0.58 | 0.5 |
| 8 | 76477368 | rs2941481 | -0.549 | 0.628 | -0.723 | -1.184 | -0.03 | 0.08 | 0.08 | 0 | 0.45 | 0.77 | 0.38 | 0.32 | 0.47 | 0.41 |
| 8 | 76477995 | rs2941482 | 1.377 | 0.595 | -0.548 | -1.175 | 0.005 | 0.045 | 0.045 | 0.152 | 0.07 | 0.22 | 0.05 | 0.08 | 0.05 | 0.08 |
| 8 | 76478616 | rs2941483 | -1.196 | 0.639 | -0.832 | -1.159 | -0.022 | 0.031 | 0.031 | 0.88 | 0.59 | 0.83 | 0.42 | 0.36 | 0.58 | 0.5 |
| 8 | 76478768 | rs2941484 | -0.472 | 0.717 | -0.802 | -1.136 | -0.018 | 0.075 | 0.075 | 0 | 0.45 | 0.77 | 0.36 | 0.32 | 0.43 | 0.41 |
| 8 | 76482263 | rs2941489 | -1.122 | 0.675 | -0.859 | -1.022 | -0.005 | 0.016 | 0.016 | 1.247 | 0.61 | 0.84 | 0.43 | 0.37 | 0.62 | 0.55 |
| 8 | 76492185 | rs10957767 | -1.21 | 0.671 | -0.913 | -1.024 | -0.032 | -0.005 | -0.005 | 0 | 0.62 | 0.89 | 0.44 | 0.37 | 0.62 | 0.55 |
| 8 | 76498914 | rs1449549 | 1.747 | 0.788 | -1.111 | -1.115 | -0.003 | -0.006 | -0.006 | 0 | 0.15 | 0.17 | 0.1 | 0.08 | 0.1 | 0.2 |
| 8 | 76519834 | rs2941422 | -1.252 | 0.764 | -1.121 | -1.089 | -0.012 | 0.009 | 0.009 | 1.186 | 0.61 | 0.84 | 0.44 | 0.37 | 0.62 | 0.52 |
| 8 | 76521095 | rs10086261 | -0.318 | 0.929 | -0.537 | -0.991 | 0.027 | -0.021 | -0.021 | 0 | 0.15 | 0.28 | 0.16 | 0.09 | 0.2 | 0.18 |
| 8 | 76524363 | rs9298259 | 2.429 | 0.873 | -0.798 | -0.931 | -0.045 | -0.079 | -0.079 | 0 | 0.15 | 0.01 | 0.11 | 0.24 | 0.14 | 0.16 |
| 8 | 76527199 | rs6997571 | 1.168 | 0.857 | -0.402 | -1.021 | 0 | -0.002 | -0.002 | 0.619 | 0.18 | 0.19 | 0.12 | 0.08 | 0.14 | 0.23 |
| 8 | 76532408 | rs2941433 | 1.359 | 1.137 | -0.768 | -1.101 | 0.007 | -0.044 | -0.044 | 0 | 0.13 | 0.21 | 0.1 | 0.01 | 0.19 | 0.06 |
| 8 | 76555824 | rs7822447 | 0.809 | 1.022 | -1.115 | -0.946 | 0.025 | 0.026 | 0.026 | 0.017 | 0.08 | 0.21 | 0.08 | 0.08 | 0.11 | 0.13 |
| 8 | 76560454 | rs2167065 | -0.516 | 1.078 | -1.839 | -0.86 | -0.038 | 0.076 | 0.076 | 0.046 | 0.45 | 0.8 | 0.38 | 0.33 | 0.5 | 0.44 |
| 8 | 76565110 | rs2922767 | -0.728 | 1.069 | -2.052 | -0.623 | -0.156 | -0.085 | -0.085 | 0 | 0.61 | 0.97 | 0.46 | 0.35 | 0.52 | 0.54 |
| 8 | 76565553 | rs2922766 | -0.939 | 1.538 | -1.803 | -0.312 | -0.098 | -0.045 | -0.045 | 0 | 0.54 | 0.97 | 0.62 | 0.39 | 0.5 | 0.53 |
| 8 | 76566968 | rs16939122 | 0.693 | 1.476 | -1.339 | -0.455 | 0.004 | 0.013 | 0.013 | 0 | 0.06 | 0.07 | 0.13 | 0.07 | 0.06 | 0.02 |

| | | | | | | | | | | | | | | | | |
|---|---|---|---|---|---|---|---|---|---|---|---|---|---|---|---|---|
| 8 | 76569842 | rs7815096 | 0.324 | 1.53 | -0.805 | -0.561 | 0.04 | 0.007 | 0.007 | 0 | 0.09 | 0.22 | 0.09 | 0.08 | 0.14 | 0.19 |
| 8 | 76570511 | rs1449547 | 1.069 | 1.453 | -0.78 | -0.561 | -0.006 | 0.013 | 0.013 | 0 | 0.07 | 0.17 | 0.05 | 0.08 | 0.05 | 0.12 |
| 8 | 76573834 | rs16939127 | 0.115 | 1.376 | -0.659 | -0.731 | -0.007 | -0.007 | -0.007 | 0 | 0.31 | 0.65 | 0.32 | 0.16 | 0.17 | 0.25 |
| 8 | 76573860 | rs10156227 | 0.822 | 1.466 | -0.642 | -0.578 | 0.021 | 0.041 | 0.041 | 0 | 0.08 | 0.21 | 0.07 | 0.13 | 0.09 | 0.12 |
| 8 | 76597662 | rs1375639 | -0.407 | 1.527 | -0.711 | -0.614 | 0.044 | 0.05 | 0.05 | 0 | 0.14 | 0.28 | 0.22 | 0.21 | 0.2 | 0.18 |
| 8 | 76598346 | rs10092128 | -0.407 | 1.527 | -0.711 | -0.614 | 0.044 | 0.05 | 0.05 | 0 | 0.14 | 0.28 | 0.22 | 0.21 | 0.2 | 0.18 |
| 8 | 76598853 | rs4377937 | -0.268 | 1.456 | -0.709 | -0.611 | 0.017 | 0.057 | 0.057 | 0 | 0.13 | 0.28 | 0.2 | 0.21 | 0.15 | 0.14 |
| 8 | 76608407 | rs2588790 | 1.731 | 1.409 | -1.076 | -0.603 | -0.017 | -0.184 | -0.184 | 0 | 0.3 | 0.02 | 0.24 | 0.46 | 0.29 | 0.34 |
| 8 | 76615663 | rs4735692 | 0.389 | 1.524 | -0.907 | -0.6 | 0.003 | 0.003 | 0.003 | 0.019 | 0.56 | 0.61 | 0.58 | 0.63 | 0.41 | 0.59 |
| 8 | 76636537 | rs16939148 | 1.484 | 1.088 | -0.256 | -0.71 | 0.009 | -0.008 | -0.008 | 0 | 0.21 | 0.17 | 0.12 | 0.01 | 0.11 | 0.22 |
| 8 | 76639216 | rs16939150 | 1.323 | 1.079 | -0.186 | -0.692 | -0.004 | -0.009 | -0.009 | 0.797 | 0.22 | 0.22 | 0.13 | 0.09 | 0.11 | 0.22 |
| 8 | 76656945 | rs1823373 | -0.682 | 1.379 | -0.43 | -0.687 | -0.081 | 0.064 | 0.064 | 1.634 | 0.7 | 0.91 | 0.8 | 0.83 | 0.61 | 0.77 |
| 8 | 76687708 | rs7350068 | 0.28 | 1.104 | -0.215 | -0.469 | 0.002 | 0.054 | 0.054 | 0 | 0.12 | 0.22 | 0.18 | 0.2 | 0.12 | 0.12 |
| 8 | 76694662 | rs2977330 | 1.696 | 1.305 | -0.58 | -0.559 | -0.06 | -0.046 | -0.046 | 0 | 0.15 | 0.01 | 0.13 | 0.18 | 0.2 | 0.17 |
| 8 | 76695262 | rs6983696 | 0.98 | 0.801 | -0.115 | -0.505 | 0.01 | -0.007 | -0.007 | 0 | 0.21 | 0.17 | 0.12 | 0.01 | 0.11 | 0.21 |
| 8 | 76702548 | rs16939169 | -0.364 | 1.081 | 0.251 | 0.002 | 0.056 | 0.093 | 0.093 | 0 | 0.07 | 0.3 | 0.17 | 0.11 | 0.07 | 0.13 |
| 8 | 76705405 | rs2977327 | 0.705 | 1.551 | -0.146 | -0.269 | -0.042 | 0.037 | 0.037 | 0.247 | 0.26 | 0.08 | 0.17 | 0.17 | 0.32 | 0.24 |
| 8 | 76711112 | rs9693229 | 0.95 | 1.443 | -0.137 | -0.472 | -0.014 | -0.077 | -0.077 | 0 | 0.29 | 0.4 | 0.19 | 0.06 | 0.19 | 0.21 |
| 8 | 76713409 | rs2920917 | -0.315 | 1.7 | -0.187 | -0.48 | -0.003 | -0.024 | -0.024 | 0 | 0.56 | 0.52 | 0.38 | 0.23 | 0.58 | 0.49 |
| 8 | 76715314 | rs7826617 | -0.448 | 1.871 | -0.047 | -0.297 | 0.085 | 0.096 | 0.096 | 0.111 | 0.09 | 0.33 | 0.18 | 0.12 | 0.11 | 0.16 |
| 8 | 76721284 | rs2977322 | 0.828 | 1.804 | -0.17 | -0.271 | -0.075 | 0.026 | 0.026 | 0.256 | 0.26 | 0.05 | 0.16 | 0.17 | 0.33 | 0.24 |
| 8 | 76721772 | rs10087318 | -0.101 | 1.405 | 0.669 | 0.218 | 0.042 | 0.081 | 0.081 | 0 | 0.07 | 0.24 | 0.16 | 0.11 | 0.07 | 0.14 |
| 8 | 76735899 | rs4463396 | 0.444 | 1.403 | 0.793 | 0.224 | 0.007 | 0.003 | 0.003 | 0 | 0.06 | 0.09 | 0.14 | 0.04 | 0.07 | 0.02 |
| 8 | 76738118 | rs13271620 | -0.937 | 1.396 | 0.767 | 0.389 | 0.04 | 0.161 | 0.161 | 0 | 0.09 | 0.29 | 0.25 | 0.22 | 0.09 | 0.16 |
| 8 | 76776862 | rs969826 | -0.293 | 1.376 | 0.294 | 0.232 | 0.001 | -0.009 | -0.009 | 0 | 0.46 | 0.48 | 0.52 | 0.65 | 0.48 | 0.57 |
| 8 | 76797744 | rs16939186 | -0.656 | 1.367 | 0.563 | 0.389 | 0.018 | 0.102 | 0.102 | 0 | 0.08 | 0.3 | 0.18 | 0.16 | 0.06 | 0.14 |
| 8 | 76804524 | rs1462436 | -0.68 | 1.543 | 0.591 | 0.403 | 0.093 | 0.11 | 0.11 | 0 | 0.05 | 0.2 | 0.21 | 0.14 | 0.13 | 0.08 |
| 8 | 76810686 | rs12679314 | 1.059 | 1.458 | 0.318 | 0.305 | -0.048 | -0.192 | -0.192 | 0 | 0.31 | 0.03 | 0.27 | 0.48 | 0.31 | 0.34 |
| 8 | 76811708 | rs3911770 | 0.337 | 1.314 | 0.408 | 0.302 | -0.01 | 0.022 | 0.022 | 0 | 0.38 | 0.32 | 0.56 | 0.69 | 0.43 | 0.42 |
| 8 | 76813695 | rs12156270 | -0.511 | 1.332 | 0.435 | 0.278 | -0.004 | 0.081 | 0.081 | 0 | 0.48 | 0.64 | 0.68 | 0.78 | 0.5 | 0.58 |
| 8 | 76826648 | rs4735650 | -0.246 | 1.399 | 0.273 | 0.167 | -0.001 | -0.006 | -0.006 | 0.065 | 0.44 | 0.54 | 0.64 | 0.77 | 0.48 | 0.55 |
| 8 | 76827190 | rs6472942 | -0.324 | 0.991 | 0.356 | 0.163 | 0.001 | 0.006 | 0.006 | 0 | 0.45 | 0.55 | 0.63 | 0.77 | 0.43 | 0.52 |
| 8 | 76832442 | rs1542514 | -0.262 | 0.905 | 0.511 | 0.156 | 0.005 | 0.018 | 0.018 | 0 | 0.44 | 0.6 | 0.6 | 0.77 | 0.4 | 0.51 |
| 8 | 76859393 | rs12676826 | -0.381 | 0.895 | 0.796 | 0.257 | 0.012 | 0.069 | 0.069 | 0 | 0.46 | 0.64 | 0.62 | 0.77 | 0.42 | 0.52 |
| 8 | 76864320 | rs16939205 | 1.53 | 0.7 | 0.48 | 0.97 | -0.033 | -0.017 | -0.017 | 0 | 0.07 | 0 | 0.16 | 0.36 | 0.12 | 0.12 |
| 8 | 76868708 | rs4735653 | -0.426 | 0.611 | 0.703 | 0.379 | 0.016 | 0.1 | 0.1 | 0 | 0.47 | 0.7 | 0.63 | 0.77 | 0.42 | 0.53 |
| 8 | 76877492 | rs1870889 | 0.787 | 0.611 | 0.719 | 0.351 | -0.03 | -0.137 | -0.137 | 0.359 | 0.32 | 0.06 | 0.26 | 0.47 | 0.34 | 0.34 |
| 8 | 76881491 | rs10504601 | -0.185 | 0.643 | 0.771 | 0.321 | 0.008 | 0.01 | 0.01 | 0 | 0.06 | 0.17 | 0.13 | 0.03 | 0.05 | 0.02 |
| 8 | 76890782 | rs4735655 | 0.395 | 0.681 | 0.824 | 0.323 | -0.007 | -0.001 | -0.001 | 0 | 0.36 | 0.44 | 0.38 | 0.63 | 0.34 | 0.48 |
| 8 | 76898602 | rs1565153 | -0.167 | 0.624 | 0.822 | 0.32 | 0.006 | 0.012 | 0.012 | 0 | 0.43 | 0.61 | 0.52 | 0.67 | 0.39 | 0.5 |
| 8 | 76906851 | rs4735710 | 0.095 | 0.622 | 0.79 | 0.328 | -0.009 | -0.007 | -0.007 | 0 | 0.38 | 0.49 | 0.4 | 0.64 | 0.36 | 0.49 |
| 8 | 76918780 | rs12676679 | -0.589 | 0.733 | 0.494 | 0.173 | 0.007 | 0.077 | 0.077 | 0 | 0.51 | 0.69 | 0.56 | 0.67 | 0.46 | 0.53 |
| 8 | 76939927 | rs16939230 | 1.831 | 0.738 | 0.417 | 0.194 | -0.01 | 0.002 | 0.002 | 0.424 | 0.07 | 0.03 | 0.1 | 0.04 | 0.08 | 0.02 |
| 8 | 76944293 | rs11988715 | -0.099 | 0.733 | 0.596 | 0.137 | 0.005 | 0.005 | 0.005 | 0.008 | 0.13 | 0.25 | 0.24 | 0.11 | 0.11 | 0.04 |
| 8 | 76946010 | rs2467179 | -0.722 | 0.745 | 0.391 | 0.193 | 0.002 | -0.005 | -0.005 | 0 | 0.65 | 0.69 | 0.7 | 0.44 | 0.66 | 0.53 |
| 8 | 76975765 | rs4007365 | -0.302 | 0.741 | 0.485 | 0.263 | 0.013 | 0.084 | 0.084 | 0 | 0.07 | 0.2 | 0.19 | 0.11 | 0.04 | 0.06 |
| 8 | 76980861 | rs830463 | 0.259 | 0.689 | 0.238 | 0.129 | 0.023 | 0.09 | 0.09 | 0 | 0.51 | 0.3 | 0.46 | 0.32 | 0.57 | 0.48 |
| 8 | 76995840 | rs7003370 | -0.443 | 1.027 | 0.5 | 0.134 | 0.055 | -0.029 | -0.029 | 0.822 | 0.07 | 0.21 | 0.1 | 0 | 0.06 | 0.04 |
| 8 | 77000163 | rs1585325 | -0.116 | 0.861 | 0.4 | 0.133 | -0.006 | -0.016 | -0.016 | 0.584 | 0.56 | 0.51 | 0.52 | 0.34 | 0.59 | 0.51 |
| 8 | 77005300 | rs10086934 | 0.319 | 0.862 | 0.384 | 0.132 | 0 | 0.085 | 0.085 | 0 | 0.49 | 0.29 | 0.43 | 0.33 | 0.54 | 0.47 |
| 8 | 77006395 | rs10091057 | -0.018 | 0.723 | 0.209 | 0.294 | -0.005 | 0 | 0 | 0 | 0.56 | 0.51 | 0.65 | 0.46 | 0.6 | 0.52 |
| 8 | 77032017 | rs1993620 | -0.947 | 0.571 | -0.466 | 0.215 | -0.049 | -0.173 | -0.173 | 0 | 0.72 | 0.96 | 0.76 | 0.55 | 0.72 | 0.72 |
| 8 | 77035228 | rs16939239 | 0.216 | 0.777 | 0.256 | 0.108 | 0.009 | -0.067 | -0.067 | 0 | 0.11 | 0.25 | 0.09 | 0.01 | 0.12 | 0.04 |
| 8 | 77039923 | rs7831448 | -1 | 0.621 | -0.103 | 0.006 | -0.019 | -0.076 | -0.076 | 0 | 0.71 | 0.89 | 0.62 | 0.45 | 0.68 | 0.75 |
| 8 | 77046383 | rs10504610 | 0.938 | 0.691 | 0.432 | -0.069 | -0.004 | -0.001 | -0.001 | 0.024 | 0.1 | 0.08 | 0.06 | 0.01 | 0.12 | 0.04 |
| 8 | 77059481 | rs10102325 | 0.287 | 0.571 | 0.711 | -0.081 | -0.001 | 0.004 | 0.004 | 0.096 | 0.56 | 0.38 | 0.46 | 0.44 | 0.54 | 0.69 |
| 8 | 77059673 | rs10102558 | -0.122 | 0.569 | 0.641 | -0.077 | 0 | -0.007 | -0.007 | 0 | 0.59 | 0.71 | 0.54 | 0.44 | 0.57 | 0.72 |
| 8 | 77063605 | rs13259287 | 1.531 | -0.4 | 0.36 | -0.648 | 0.026 | 0.024 | 0.024 | 0.037 | 0.14 | 0.07 | 0.03 | 0 | 0.05 | 0.08 |
| 8 | 77077752 | rs1431018 | -0.707 | -0.314 | 0.298 | -0.589 | -0.018 | -0.03 | -0.03 | 3.301 | 0.61 | 0.82 | 0.56 | 0.44 | 0.57 | 0.72 |
| 8 | 77086454 | rs6981145 | 0.025 | -0.403 | 0.242 | -0.577 | 0.003 | 0.058 | 0.058 | 0 | 0.53 | 0.3 | 0.45 | 0.34 | 0.53 | 0.51 |
| 8 | 77097912 | rs10100261 | -0.091 | -0.367 | 0.225 | -0.56 | -0.001 | 0.006 | 0.006 | 0 | 0.55 | 0.37 | 0.46 | 0.44 | 0.54 | 0.68 |
| 8 | 77117377 | rs16939252 | 1.869 | -0.681 | -0.065 | -0.672 | 0.006 | -0.035 | -0.035 | 0 | 0.08 | 0 | 0.11 | 0.12 | 0.05 | 0.04 |
| 8 | 77118604 | rs1367563 | 0.416 | -0.605 | 0.238 | -0.688 | 0.028 | -0.026 | -0.026 | 0.473 | 0.33 | 0.37 | 0.37 | 0.22 | 0.46 | 0.41 |
| 8 | 77120007 | rs1864605 | 0.282 | -0.609 | 0.085 | -0.638 | -0.064 | -0.024 | -0.024 | 0 | 0.27 | 0.51 | 0.2 | 0.22 | 0.12 | 0.31 |
| 8 | 77123729 | rs9650162 | -1.907 | -0.495 | -0.436 | -0.482 | -0.065 | -0.155 | -0.155 | 0 | 0.73 | 0.98 | 0.75 | 0.56 | 0.69 | 0.78 |
| 8 | 77131580 | rs1431015 | -1.182 | -0.5 | -0.404 | -0.504 | -0.083 | -0.096 | -0.096 | 1.967 | 0.65 | 0.92 | 0.57 | 0.44 | 0.58 | 0.73 |
| 8 | 77136588 | rs10092489 | 0.352 | -0.452 | -0.141 | -0.5 | -0.032 | 0.031 | 0.031 | 0 | 0.34 | 0.17 | 0.34 | 0.22 | 0.45 | 0.41 |
| 8 | 77137533 | rs17424935 | 2.141 | -0.361 | -0.259 | -0.409 | -0.029 | -0.033 | -0.033 | 0 | 0.08 | 0.03 | 0.18 | 0.12 | 0.12 | 0.05 |
| 8 | 77145581 | rs919577 | -0.766 | -0.466 | -0.052 | -0.516 | -0.009 | -0.015 | -0.015 | 0 | 0.61 | 0.7 | 0.53 | 0.44 | 0.57 | 0.72 |

| | | | | | | | | | | | | | | | | |
|---|---|---|---|---|---|---|---|---|---|---|---|---|---|---|---|---|
| 8 | 77154066 | rs1821018 | 0.386 | -0.518 | 0.198 | -0.574 | -0.035 | -0.013 | -0.013 | 0.812 | 0.26 | 0.33 | 0.17 | 0.22 | 0.11 | 0.3 |
| 8 | 77159098 | rs7017651 | 0.373 | -0.52 | 0.197 | -0.575 | -0.035 | -0.013 | -0.013 | 0 | 0.27 | 0.34 | 0.17 | 0.22 | 0.11 | 0.3 |
| 8 | 77159766 | rs1821016 | 0.226 | -0.515 | 0.027 | -0.573 | -0.074 | -0.032 | -0.032 | 0.18 | 0.28 | 0.5 | 0.2 | 0.22 | 0.11 | 0.3 |
| 8 | 77162694 | rs10112085 | 0.711 | -0.432 | -0.089 | -0.587 | -0.126 | 0.011 | 0.011 | 0 | 0.3 | 0.03 | 0.29 | 0.22 | 0.42 | 0.38 |
| 8 | 77166498 | rs12678607 | 0.326 | -0.521 | -0.038 | -0.546 | -0.032 | 0.024 | 0.024 | 0 | 0.34 | 0.2 | 0.35 | 0.22 | 0.46 | 0.43 |
| 8 | 77176352 | rs10957794 | -0.679 | -0.303 | -0.222 | -0.383 | -0.004 | -0.013 | -0.013 | 0 | 0.62 | 0.72 | 0.55 | 0.44 | 0.58 | 0.73 |
| 8 | 77187010 | rs7845313 | -0.686 | -0.303 | -0.268 | -0.383 | -0.008 | -0.02 | -0.02 | 0 | 0.62 | 0.74 | 0.55 | 0.44 | 0.58 | 0.73 |
| 8 | 77187032 | rs7845323 | -0.686 | -0.303 | -0.268 | -0.383 | -0.008 | -0.02 | -0.02 | 0 | 0.62 | 0.74 | 0.55 | 0.44 | 0.58 | 0.73 |
| 8 | 77191302 | rs1405350 | 1.824 | -0.245 | -0.556 | -0.316 | -0.032 | -0.034 | -0.034 | 0 | 0.08 | 0.03 | 0.19 | 0.13 | 0.13 | 0.05 |
| 8 | 77214164 | rs1568986 | -1.829 | -0.317 | -0.563 | -0.356 | -0.033 | -0.131 | -0.131 | 0 | 0.74 | 0.96 | 0.77 | 0.56 | 0.72 | 0.79 |
| 8 | 77218065 | rs10504616 | 0.547 | -0.582 | -0.236 | -0.81 | -0.016 | -0.015 | -0.015 | 0.704 | 0.24 | 0.26 | 0.16 | 0.12 | 0.11 | 0.13 |
| 8 | 77218289 | rs9298276 | 0.732 | 0.046 | -0.567 | -0.911 | -0.114 | -0.02 | -0.02 | 0 | 0.24 | 0.03 | 0.27 | 0.21 | 0.37 | 0.33 |
| 8 | 77228222 | rs1405348 | -0.649 | -0.173 | -0.189 | -0.979 | 0.009 | 0.049 | 0.049 | 0 | 0.44 | 0.32 | 0.4 | 0.22 | 0.55 | 0.47 |
| 8 | 77234939 | rs10110701 | 0.022 | -0.224 | -0.173 | -1.09 | -0.105 | 0.065 | 0.065 | 0 | 0.35 | 0.11 | 0.34 | 0.22 | 0.46 | 0.39 |
| 8 | 77236201 | rs1405351 | 0.324 | -0.202 | -0.166 | -1.078 | 0.019 | -0.055 | -0.055 | 0 | 0.1 | 0.23 | 0.06 | 0 | 0.1 | 0.08 |
| 8 | 77238392 | rs10102172 | -0.719 | -0.319 | 0.057 | -1.015 | 0.008 | 0.024 | 0.024 | 0 | 0.47 | 0.58 | 0.44 | 0.22 | 0.55 | 0.48 |
| 8 | 77243452 | rs13275924 | 1.288 | 0.538 | -0.394 | -1.113 | -0.066 | -0.035 | -0.035 | 0 | 0.16 | 0.01 | 0.17 | 0.13 | 0.23 | 0.19 |
| 8 | 77246394 | rs10156347 | -0.667 | -0.013 | 0.027 | -0.919 | -0.004 | -0.018 | -0.018 | 0.143 | 0.46 | 0.49 | 0.43 | 0.22 | 0.55 | 0.48 |
| 8 | 77248455 | rs7844445 | -0.75 | -0.553 | 0.696 | -0.888 | -0.044 | -0.06 | -0.06 | 0.078 | 0.27 | 0.6 | 0.11 | 0.1 | 0.1 | 0.29 |
| 8 | 77254882 | rs10504618 | 0.04 | -0.514 | 1.095 | -0.894 | 0.033 | -0.017 | -0.017 | 0 | 0.06 | 0.23 | 0.03 | 0 | 0.04 | 0.06 |
| 8 | 77255695 | rs10095764 | -1.139 | -0.539 | 1.128 | -0.943 | 0.039 | 0.055 | 0.055 | 0.017 | 0.29 | 0.77 | 0.15 | 0.1 | 0.12 | 0.33 |
| 8 | 77259013 | rs13272268 | 0.828 | -0.575 | 1.319 | -0.807 | 0.012 | 0 | 0 | 0 | 0.15 | 0.09 | 0.03 | 0.01 | 0.06 | 0.09 |
| 8 | 77262140 | rs16939265 | -0.456 | -0.531 | 1.187 | -0.785 | -0.064 | -0.075 | -0.075 | 0 | 0.22 | 0.51 | 0.12 | 0.1 | 0.08 | 0.28 |
| 8 | 77268791 | rs10504619 | -0.18 | -0.607 | 0.758 | -0.737 | -0.052 | -0.046 | -0.046 | 0 | 0.2 | 0.35 | 0.07 | 0.1 | 0.06 | 0.24 |
| 8 | 77270727 | rs16939267 | -0.619 | -0.738 | 0.63 | -0.692 | -0.056 | -0.036 | -0.036 | 0 | 0.21 | 0.63 | 0.12 | 0.11 | 0.04 | 0.23 |
| 8 | 77285055 | rs6984322 | -0.621 | -0.359 | 0.259 | -0.239 | 0.002 | -0.001 | -0.001 | 0 | 0.75 | 0.59 | 0.75 | 0.7 | 0.74 | 0.63 |
| 8 | 77287631 | rs7838786 | 1.336 | -0.086 | 0.045 | -0.43 | 0.002 | -0.026 | -0.026 | 0 | 0.09 | 0.18 | 0.1 | 0.01 | 0.11 | 0.04 |
| 8 | 77297117 | rs12680480 | -0.157 | -0.046 | 0.09 | -0.305 | -0.007 | -0.003 | -0.003 | 0 | 0.7 | 0.41 | 0.71 | 0.65 | 0.72 | 0.51 |
| 8 | 77302352 | rs2383971 | 0.502 | -0.015 | -0.021 | -0.257 | -0.001 | 0.008 | 0.008 | 0.564 | 0.61 | 0.27 | 0.57 | 0.55 | 0.6 | 0.47 |
| 8 | 77313242 | rs4319064 | -0.626 | 0.235 | 0.267 | -0.33 | 0.003 | 0.022 | 0.022 | 0 | 0.75 | 0.65 | 0.69 | 0.64 | 0.74 | 0.62 |
| 8 | 77323802 | rs13249980 | 1.139 | 0.755 | -0.76 | -0.103 | -0.063 | -0.062 | -0.062 | 0 | 0.17 | 0.03 | 0.29 | 0.25 | 0.31 | 0.18 |
| 8 | 77325204 | rs4471009 | 0.966 | 0.644 | -0.555 | -0.548 | 0.009 | -0.051 | -0.051 | 2.369 | 0.1 | 0.27 | 0.1 | 0 | 0.11 | 0.03 |
| 8 | 77326204 | rs10441573 | 0.381 | 0.234 | -1.005 | -0.335 | -0.038 | -0.066 | -0.066 | 0 | 0.37 | 0.16 | 0.35 | 0.48 | 0.43 | 0.32 |
| 8 | 77331219 | rs10093934 | -0.196 | -0.198 | -0.416 | -0.541 | -0.008 | -0.017 | -0.017 | 0 | 0.32 | 0.31 | 0.21 | 0.14 | 0.2 | 0.26 |
| 8 | 77346155 | rs6472969 | -0.428 | 0.515 | -0.557 | -0.103 | -0.004 | -0.011 | -0.011 | 1.032 | 0.37 | 0.46 | 0.48 | 0.75 | 0.35 | 0.38 |
| 8 | 77346237 | rs4374963 | 0.487 | 0.257 | -0.753 | -0.876 | -0.009 | -0.011 | -0.011 | 0 | 0.34 | 0.34 | 0.26 | 0.15 | 0.23 | 0.26 |
| 8 | 77366608 | rs2872032 | 1.03 | 0.024 | -1.178 | -1.042 | 0.026 | 0.021 | 0.021 | 0 | 0.28 | 0.07 | 0.21 | 0.15 | 0.2 | 0.22 |
| 8 | 77366932 | rs7812747 | -1.261 | 0.397 | -1.124 | -1.041 | -0.067 | 0.071 | 0.071 | 0 | 0.74 | 0.91 | 0.75 | 0.95 | 0.56 | 0.7 |
| 8 | 77373304 | rs2312425 | 0.238 | 0.119 | -0.805 | -0.996 | -0.016 | -0.02 | -0.02 | 0 | 0.4 | 0.52 | 0.3 | 0.22 | 0.27 | 0.29 |
| 8 | 77374126 | rs11996335 | -0.172 | 0.226 | -1.181 | -0.379 | -0.032 | -0.009 | -0.009 | 0 | 0.63 | 0.77 | 0.57 | 0.35 | 0.47 | 0.49 |
| 8 | 77377945 | rs13281210 | -0.264 | 0.603 | -1.363 | -0.893 | -0.059 | 0.084 | 0.084 | 0 | 0.29 | 0.14 | 0.24 | 0.14 | 0.36 | 0.34 |
| 8 | 77397120 | rs7812499 | -0.95 | -0.977 | -1.009 | -0.532 | 0.077 | 0.02 | 0.02 | 0 | 0.52 | 0.82 | 0.53 | 0.56 | 0.33 | 0.43 |
| 8 | 77402241 | rs10096780 | 0.117 | -1.127 | -0.489 | -0.549 | 0.009 | -0.048 | -0.048 | 0 | 0.25 | 0.16 | 0.36 | 0.51 | 0.21 | 0.33 |
| 8 | 77417193 | rs16939284 | -0.177 | -1.084 | 0.157 | -0.848 | -0.044 | -0.086 | -0.086 | 0.005 | 0.24 | 0.43 | 0.14 | 0.05 | 0.12 | 0.11 |
| 8 | 77419768 | rs10216685 | 0.345 | -1.089 | 0.32 | -0.802 | -0.036 | -0.059 | -0.059 | 3.155 | 0.16 | 0.33 | 0.09 | 0.05 | 0.08 | 0.09 |
| 8 | 77426373 | rs10108974 | 0.84 | -0.973 | 0.072 | -0.744 | 0.001 | -0.002 | -0.002 | 1.069 | 0.13 | 0.14 | 0.07 | 0.05 | 0.08 | 0.08 |
| 8 | 77439202 | rs12681151 | 1.295 | -0.236 | -0.317 | -0.421 | -0.087 | -0.005 | -0.005 | 0 | 0.16 | 0.01 | 0.13 | 0.11 | 0.26 | 0.21 |
| 8 | 77448801 | rs17346463 | 1.029 | -1.052 | 0.301 | -0.063 | 0.001 | 0.008 | 0.008 | 0 | 0.11 | 0 | 0.03 | 0.07 | 0.08 | 0.09 |
| 8 | 77450209 | rs9298282 | -1.498 | -1.208 | 0.465 | 0.03 | 0.046 | 0.015 | 0.015 | 0 | 0.51 | 0.75 | 0.56 | 0.58 | 0.37 | 0.44 |
| 8 | 77453769 | rs4380909 | -1.708 | -1.21 | 0.318 | 0.027 | 0.048 | 0.008 | 0.008 | 0 | 0.52 | 0.83 | 0.57 | 0.58 | 0.37 | 0.44 |
| 8 | 77456729 | rs4574824 | -0.737 | -1.255 | 0.398 | 0.008 | 0.046 | -0.04 | -0.04 | 0 | 0.36 | 0.2 | 0.4 | 0.58 | 0.26 | 0.39 |
| 8 | 77459705 | rs4735733 | -1.591 | -1.126 | 0.07 | 0.08 | 0.047 | 0.01 | 0.01 | 0 | 0.52 | 0.81 | 0.57 | 0.58 | 0.37 | 0.44 |
| 8 | 77472461 | rs4735662 | -1.046 | -0.517 | 0.255 | 0.412 | -0.018 | 0.054 | 0.054 | 0 | 0.19 | 0.41 | 0.37 | 0.26 | 0.13 | 0.15 |
| 8 | 77473175 | rs12056645 | -0.871 | -0.498 | 0.271 | 0.413 | -0.01 | 0.029 | 0.029 | 0 | 0.18 | 0.26 | 0.35 | 0.26 | 0.13 | 0.15 |
| 8 | 77476499 | rs10101497 | -1.217 | -0.403 | -0.095 | 0.48 | -0.019 | -0.017 | -0.017 | 0 | 0.35 | 0.53 | 0.48 | 0.3 | 0.28 | 0.24 |
| 8 | 77478373 | rs4601282 | -0.435 | 0.296 | -0.322 | 1.438 | 0.002 | 0.051 | 0.051 | 0 | 0.1 | 0.15 | 0.29 | 0.26 | 0.09 | 0.13 |
| 8 | 77492915 | rs11997355 | 0.416 | 0.11 | -0.32 | 1.498 | -0.002 | 0.038 | 0.038 | 4.268 | 0.09 | 0.09 | 0.12 | 0.32 | 0.07 | 0.07 |
| 8 | 77504365 | rs13257282 | -0.645 | 0.544 | -0.388 | 0.474 | -0.001 | -0.017 | -0.017 | 0.105 | 0.12 | 0.1 | 0.08 | 0.01 | 0.15 | 0.11 |
| 8 | 77507613 | rs7846606 | -1.592 | 0.324 | -0.001 | 0.604 | 0.015 | 0.024 | 0.024 | 0.032 | 0.21 | 0.39 | 0.22 | 0.27 | 0.22 | 0.28 |
| 8 | 77508430 | rs13279286 | -0.936 | -0.183 | -0.396 | -0.33 | 0.007 | 0.032 | 0.032 | 0 | 0.28 | 0.18 | 0.13 | 0.09 | 0.24 | 0.21 |
| 8 | 77512647 | rs16939318 | -0.596 | -0.2 | -0.48 | -0.32 | -0.003 | -0.008 | -0.008 | 0 | 0.26 | 0.22 | 0.11 | 0.09 | 0.12 | 0.2 |
| 8 | 77514719 | rs13278854 | -0.42 | 0.244 | -0.627 | -0.027 | -0.019 | 0.018 | 0.018 | 0 | 0.06 | 0.05 | 0.04 | 0 | 0.11 | 0.03 |
| 8 | 77515395 | rs10957809 | -0.63 | 0.23 | -0.587 | -0.034 | 0.001 | -0.013 | -0.013 | 0 | 0.07 | 0.1 | 0.05 | 0 | 0.11 | 0.03 |
| 8 | 77517534 | rs10504629 | -1.093 | 0.294 | -1.005 | -0.9 | -0.057 | 0.089 | 0.089 | 0.478 | 0.68 | 0.84 | 0.78 | 0.97 | 0.54 | 0.62 |
| 8 | 77526501 | rs1433962 | -0.666 | -0.228 | -0.694 | -0.747 | 0.041 | 0.089 | 0.089 | 0.203 | 0.46 | 0.66 | 0.25 | 0.2 | 0.32 | 0.31 |
| 8 | 77529466 | rs1560846 | 0.045 | -0.7 | -0.006 | -1.046 | -0.004 | -0.01 | -0.01 | 0 | 0.24 | 0.27 | 0.15 | 0.19 | 0.19 | 0.22 |
| 8 | 77560474 | rs11784020 | -1.376 | -0.952 | 0.094 | -0.927 | 0.078 | 0.131 | 0.131 | 0.253 | 0.47 | 0.76 | 0.23 | 0.21 | 0.31 | 0.3 |
| 8 | 77566481 | rs6996667 | 0.054 | -1.061 | 0.184 | -0.993 | 0.006 | 0.004 | 0.004 | 0 | 0.14 | 0.11 | 0.06 | 0.01 | 0.12 | 0.07 |
| 8 | 77567148 | rs7015611 | 1.231 | -1.156 | 0.782 | -0.84 | -0.013 | -0.008 | -0.008 | 0 | 0.08 | 0.26 | 0.03 | 0.02 | 0.03 | 0.01 |

| | | | | | | | | | | | | | | | | |
|---|---|---|---|---|---|---|---|---|---|---|---|---|---|---|---|---|
| 8 | 77570299 | rs16939332 | 0.319 | -1.008 | 1.036 | -0.669 | 0.02 | 0.016 | 0.016 | 0 | 0.18 | 0.08 | 0.06 | 0.14 | 0.09 | 0.18 |
| 8 | 77575477 | rs1433955 | -1.411 | -0.974 | 1.107 | 0.11 | -0.008 | 0.001 | 0.001 | 0 | 0.38 | 0.52 | 0.24 | 0.41 | 0.23 | 0.32 |
| 8 | 77576939 | rs1560845 | -0.173 | -0.435 | 1.089 | 0.403 | 0.008 | -0.003 | -0.003 | 0 | 0.06 | 0.14 | 0.06 | 0.04 | 0.06 | 0.03 |
| 8 | 77588548 | rs10105863 | -1.501 | -0.507 | 0.991 | 0.392 | 0.024 | -0.025 | -0.025 | 0 | 0.33 | 0.74 | 0.25 | 0.43 | 0.21 | 0.28 |
| 8 | 77604121 | rs1368849 | -0.199 | -0.504 | 1.486 | 0.545 | -0.04 | -0.018 | -0.018 | 0 | 0.14 | 0.27 | 0.07 | 0.1 | 0.05 | 0.12 |
| 8 | 77604189 | rs16939334 | 1.371 | -0.707 | 1.44 | 1.23 | -0.009 | 0.052 | 0.052 | 6.284 | 0.06 | 0.07 | 0.12 | 0.22 | 0.03 | 0.04 |
| 8 | 77604704 | rs7011404 | -1.181 | -0.249 | 1.614 | 1.459 | -0.055 | 0.058 | 0.058 | 0 | 0.2 | 0.6 | 0.18 | 0.34 | 0.06 | 0.19 |
| 8 | 77611625 | rs4735738 | -1.583 | -0.173 | 1.617 | 1.648 | 0.006 | -0.013 | -0.013 | 0 | 0.27 | 0.74 | 0.24 | 0.44 | 0.16 | 0.23 |
| 8 | 77631295 | rs7838141 | -0.622 | -0.103 | 1.158 | 1.617 | -0.023 | 0.023 | 0.023 | 0 | 0.18 | 0.32 | 0.11 | 0.18 | 0.1 | 0.15 |
| 8 | 77631769 | rs7842199 | -0.859 | -0.021 | 1.179 | 1.636 | -0.013 | 0.012 | 0.012 | 0 | 0.2 | 0.35 | 0.13 | 0.18 | 0.15 | 0.16 |
| 8 | 77632017 | rs7842138 | -1.361 | -0.026 | 1.165 | 1.637 | -0.018 | 0.048 | 0.048 | 0.299 | 0.25 | 0.6 | 0.22 | 0.42 | 0.16 | 0.23 |
| 8 | 77663038 | rs7822914 | -1.162 | 0.026 | 0.982 | 1.561 | -0.016 | 0.118 | 0.118 | 0 | 0.22 | 0.51 | 0.2 | 0.42 | 0.16 | 0.16 |
| 8 | 77671877 | rs6472983 | -1.304 | 0.049 | 0.917 | 1.572 | -0.01 | 0.042 | 0.042 | 0 | 0.23 | 0.64 | 0.22 | 0.42 | 0.16 | 0.17 |
| 8 | 77686271 | rs2733714 | -0.885 | 0.675 | 0.615 | 1.49 | -0.058 | -0.077 | -0.077 | 0 | 0.84 | 0.98 | 0.65 | 0.67 | 0.79 | 0.76 |
| 8 | 77690337 | rs11782906 | -0.83 | 0.164 | 1.38 | 1.388 | -0.015 | -0.012 | -0.012 | 0.038 | 0.18 | 0.28 | 0.11 | 0.1 | 0.11 | 0.1 |
| 8 | 77690563 | rs16939357 | -1.088 | 0.168 | 1.148 | 1.378 | -0.012 | 0.065 | 0.065 | 0 | 0.23 | 0.6 | 0.21 | 0.39 | 0.17 | 0.18 |
| 8 | 77695388 | rs6985531 | -0.969 | 0.251 | 0.673 | 1.393 | -0.01 | 0.109 | 0.109 | 0 | 0.22 | 0.53 | 0.21 | 0.39 | 0.17 | 0.17 |
| 8 | 77697692 | rs2728436 | -0.311 | 1.011 | -0.855 | 1.044 | -0.042 | -0.035 | -0.035 | 0 | 0.63 | 0.77 | 0.43 | 0.43 | 0.56 | 0.63 |
| 8 | 77698990 | rs16939362 | 0.416 | 0.186 | -0.059 | 1.04 | -0.029 | 0.103 | 0.103 | 3.932 | 0.08 | 0.23 | 0.13 | 0.19 | 0.03 | 0.07 |
| 8 | 77705211 | rs2254772 | -0.254 | 0.211 | -0.006 | 0.903 | -0.002 | 0.001 | 0.001 | 0 | 0.07 | 0.07 | 0.04 | 0.06 | 0.06 | 0.08 |
| 8 | 77705701 | rs10095142 | -0.265 | 0.058 | 0.533 | 0.819 | -0.012 | 0.121 | 0.121 | 1.198 | 0.11 | 0.36 | 0.17 | 0.19 | 0.05 | 0.07 |
| 8 | 77707613 | rs2626353 | 0.61 | -0.429 | 0.626 | 0.144 | 0.008 | 0.019 | 0.019 | 0.134 | 0.24 | 0.16 | 0.13 | 0.05 | 0.22 | 0.28 |
| 8 | 77718819 | rs2054682 | 0.018 | 0.003 | 0.906 | 0.544 | -0.013 | 0.115 | 0.115 | 1.213 | 0.11 | 0.35 | 0.17 | 0.19 | 0.05 | 0.07 |
| 8 | 77720427 | rs2728457 | -1.877 | -0.041 | 0.758 | 0.765 | 0.002 | -0.001 | -0.001 | 0 | 0.32 | 0.72 | 0.42 | 0.61 | 0.23 | 0.26 |
| 8 | 77727516 | rs2733727 | -0.218 | 0.168 | 0.512 | 0.766 | 0.001 | 0.001 | 0.001 | 0 | 0.08 | 0.07 | 0.03 | 0.07 | 0.05 | 0.08 |
| 8 | 77749487 | rs2728451 | -1.798 | 0.088 | -0.098 | 0.78 | -0.003 | 0.102 | 0.102 | 1.557 | 0.49 | 0.94 | 0.52 | 0.65 | 0.45 | 0.51 |
| 8 | 77749820 | rs2733735 | -1.658 | 0.089 | 0.239 | 0.78 | 0.015 | 0.087 | 0.087 | 1.98 | 0.48 | 0.83 | 0.5 | 0.65 | 0.45 | 0.5 |
| 8 | 77752035 | rs6986186 | -1.04 | -0.1 | 0.829 | 1.144 | 0.04 | 0.22 | 0.22 | 1.075 | 0.08 | 0.24 | 0.2 | 0.33 | 0.1 | 0.07 |
| 8 | 77754326 | rs1825367 | -1.712 | -0.376 | 0.425 | 0.948 | 0.036 | 0.026 | 0.026 | 0.148 | 0.43 | 0.82 | 0.48 | 0.6 | 0.38 | 0.49 |
| 8 | 77754904 | rs1825366 | -1.32 | -0.495 | 0.64 | 0.803 | -0.05 | -0.075 | -0.075 | 0 | 0.19 | 0.65 | 0.11 | 0.05 | 0.07 | 0.09 |
| 8 | 77760070 | rs7841393 | -1.712 | -0.362 | 0.441 | 0.926 | 0.034 | -0.019 | -0.019 | 0.118 | 0.27 | 0.83 | 0.37 | 0.6 | 0.12 | 0.23 |
| 8 | 77760079 | rs11993441 | -1.5 | -0.377 | 0.447 | 1.033 | -0.003 | 0.01 | 0.01 | 0.593 | 0.2 | 0.77 | 0.25 | 0.4 | 0.09 | 0.16 |
| 8 | 77762439 | rs16939378 | 1.107 | -0.163 | 0.315 | 1.041 | 0.026 | -0.031 | -0.031 | 0 | 0.07 | 0.03 | 0.11 | 0.18 | 0.03 | 0.05 |
| 8 | 77765865 | rs2728455 | -0.179 | -0.222 | 0.198 | 0.877 | -0.04 | 0.061 | 0.061 | 0 | 0.17 | 0.04 | 0.13 | 0.03 | 0.26 | 0.28 |
| 8 | 77768390 | rs16939380 | 1.215 | -0.243 | 0.202 | 0.988 | 0.024 | -0.039 | -0.039 | 0.156 | 0.07 | 0.01 | 0.19 | 0.19 | 0.05 | 0.05 |
| 8 | 77769819 | rs2733711 | -0.707 | -0.096 | 0.606 | 1.006 | 0.013 | 0.05 | 0.05 | 0 | 0.2 | 0.23 | 0.23 | 0.44 | 0.27 | 0.35 |
| 8 | 77774263 | rs13249915 | -1.297 | -0.055 | 0.961 | 0.527 | -0.01 | -0.107 | -0.107 | 0.096 | 0.2 | 0.57 | 0.11 | 0.02 | 0.15 | 0.1 |
| 8 | 77781087 | rs1872138 | 1.658 | -0.214 | 1.507 | 0.448 | 0.001 | -0.096 | -0.096 | 0 | 0.66 | 0.41 | 0.78 | 0.95 | 0.67 | 0.67 |
| 8 | 77784096 | rs10090153 | 2.023 | -0.327 | 1.461 | 0.806 | 0.011 | 0.017 | 0.017 | 0.225 | 0.56 | 0.25 | 0.52 | 0.39 | 0.62 | 0.51 |
| 8 | 77785447 | rs16939382 | 1.852 | -0.384 | 1.824 | 0.935 | -0.01 | 0.002 | 0.002 | 0 | 0.61 | 0.46 | 0.56 | 0.4 | 0.65 | 0.55 |
| 8 | 77787110 | rs1872139 | -0.357 | -0.242 | 1.562 | 0.769 | -0.024 | 0.036 | 0.036 | 0 | 0.86 | 0.93 | 0.88 | 0.98 | 0.81 | 0.75 |
| 8 | 77791860 | rs13252111 | 2.042 | -0.355 | 1.591 | 0.901 | 0.009 | 0.004 | 0.004 | 0.164 | 0.57 | 0.29 | 0.48 | 0.37 | 0.63 | 0.51 |
| 8 | 77792343 | rs12676674 | -0.785 | -0.374 | 1.884 | 0.937 | 0.005 | 0.074 | 0.074 | 0 | 0.14 | 0.2 | 0.22 | 0.43 | 0.14 | 0.15 |
| 8 | 77796946 | rs12675423 | 0.948 | -0.422 | 1.568 | 0.871 | -0.009 | -0.016 | -0.016 | 0.111 | 0.74 | 0.6 | 0.71 | 0.81 | 0.77 | 0.68 |
| 8 | 77819767 | rs1843027 | 2.352 | -0.307 | 0.768 | 0.299 | 0.003 | -0.038 | -0.038 | 0.273 | 0.5 | 0.05 | 0.46 | 0.46 | 0.56 | 0.49 |
| 8 | 77819954 | rs1483801 | 0.239 | 0.241 | 1.194 | -0.138 | 0.009 | -0.051 | -0.051 | 0.304 | 0.83 | 0.77 | 0.86 | 0.98 | 0.81 | 0.78 |
| 8 | 77824032 | rs12549946 | 1.12 | 0.352 | 0.613 | 0.01 | 0.022 | -0.083 | -0.083 | 0.502 | 0.7 | 0.49 | 0.73 | 0.93 | 0.64 | 0.68 |
| 8 | 77824384 | rs12550004 | 1.131 | 0.354 | 0.59 | 0.011 | 0.02 | -0.081 | -0.081 | 0.286 | 0.7 | 0.47 | 0.73 | 0.93 | 0.64 | 0.68 |
| 8 | 77834985 | rs7819311 | -1.006 | 0.4 | 0.21 | 0.233 | 0.001 | -0.049 | -0.049 | 0 | 0.16 | 0.51 | 0.16 | 0.04 | 0.15 | 0.12 |
| 8 | 77837751 | rs7842547 | 0.418 | 0.143 | -0.243 | 0.344 | -0.001 | -0.002 | -0.002 | 0 | 0.67 | 0.66 | 0.55 | 0.45 | 0.67 | 0.63 |
| 8 | 77844519 | rs1483813 | 0.887 | 0.294 | -1.125 | 0.424 | -0.021 | 0.068 | 0.068 | 0 | 0.51 | 0.16 | 0.39 | 0.4 | 0.52 | 0.51 |
| 8 | 77873929 | rs6981577 | 0.137 | 0.275 | -0.523 | -0.105 | -0.001 | -0.006 | -0.006 | 0.414 | 0.56 | 0.55 | 0.6 | 0.66 | 0.59 | 0.54 |
| 8 | 77876363 | rs4278125 | -0.086 | 0.454 | -0.52 | 0.191 | -0.003 | -0.068 | -0.068 | 0 | 0.27 | 0.15 | 0.23 | 0.46 | 0.26 | 0.29 |
| 8 | 77878913 | rs4641034 | 1.072 | 0.151 | 0 | -0.194 | 0.039 | -0.002 | -0.002 | 0 | 0.54 | 0.25 | 0.63 | 0.52 | 0.61 | 0.61 |
| 8 | 77881800 | rs4496943 | -0.216 | 0.233 | 0.445 | -0.197 | -0.005 | -0.029 | -0.029 | 0 | 0.11 | 0.17 | 0.07 | 0.01 | 0.11 | 0.07 |
| 8 | 77891126 | rs4236857 | 1.134 | 0.202 | 0.576 | -0.2 | 0.051 | -0.004 | -0.004 | 0 | 0.53 | 0.18 | 0.62 | 0.53 | 0.61 | 0.61 |
| 8 | 77896472 | rs10957830 | 0.732 | 0.045 | 0.755 | -0.138 | 0.001 | 0.022 | 0.022 | 0 | 0.21 | 0.29 | 0.35 | 0.32 | 0.21 | 0.25 |
| 8 | 77896845 | rs10106087 | 1.113 | 0.583 | 0.637 | -0.196 | -0.041 | 0.138 | 0.138 | 0 | 0.41 | 0.09 | 0.32 | 0.22 | 0.4 | 0.41 |
| 8 | 77898932 | rs4735749 | 0.743 | 0.582 | 0.617 | -0.216 | 0.007 | -0.014 | -0.014 | 0.009 | 0.56 | 0.3 | 0.65 | 0.54 | 0.58 | 0.62 |
| 8 | 77901019 | rs10808814 | 0.847 | 0.639 | 0.703 | -0.152 | 0.001 | 0.024 | 0.024 | 0 | 0.21 | 0.29 | 0.35 | 0.32 | 0.21 | 0.25 |
| 8 | 77927021 | rs9643717 | -0.403 | 0.813 | 1.318 | 0.141 | -0.033 | -0.063 | -0.063 | 0 | 0.3 | 0.15 | 0.27 | 0.44 | 0.37 | 0.31 |
| 8 | 77929911 | rs7827658 | 0.438 | 0.782 | 1.086 | 0.233 | -0.047 | 0.048 | 0.048 | 1.545 | 0.19 | 0.01 | 0.11 | 0.06 | 0.23 | 0.09 |
| 8 | 77974559 | rs16939411 | 0.124 | 0.331 | 1.094 | 0.182 | -0.015 | -0.028 | -0.028 | 0 | 0.07 | 0.13 | 0.03 | 0 | 0.03 | 0.03 |
| 8 | 77981645 | rs12677622 | 1.313 | 0.342 | 0.921 | 0.186 | 0.006 | -0.009 | -0.009 | 0.159 | 0.56 | 0.35 | 0.66 | 0.54 | 0.58 | 0.62 |
| 8 | 77988917 | rs4735757 | 0.548 | 0.392 | 0.54 | 0.209 | -0.027 | -0.042 | -0.042 | 0 | 0.9 | 0.79 | 0.96 | 0.99 | 0.95 | 0.94 |
| 8 | 77989317 | rs10101739 | 0.472 | 0.588 | -0.017 | 0.303 | -0.024 | -0.057 | -0.057 | 0.033 | 0.68 | 0.83 | 0.73 | 0.55 | 0.61 | 0.65 |
| 8 | 77998659 | rs13259780 | -0.101 | 0.08 | -1.218 | 0.843 | -0.128 | -0.097 | -0.097 | 0 | 0.24 | 0.02 | 0.17 | 0.34 | 0.33 | 0.21 |
| 8 | 78012371 | rs10504644 | 0.365 | 0.6 | -0.762 | 0.659 | 0.008 | -0.019 | -0.019 | 0 | 0.56 | 0.3 | 0.64 | 0.54 | 0.61 | 0.64 |
| 8 | 78026317 | rs1545507 | -0.471 | -0.014 | -0.687 | -0.139 | 0.017 | 0.044 | 0.044 | 0.309 | 0.17 | 0.08 | 0.12 | 0 | 0.16 | 0.05 |

| | | | | | | | | | | | | | | | | |
|---|---|---|---|---|---|---|---|---|---|---|---|---|---|---|---|---|
| 8 | 78040531 | rs16939429 | 0.797 | -0.547 | -0.596 | 0.149 | 0.018 | -0.001 | -0.001 | 0.928 | 0.47 | 0.36 | 0.53 | 0.54 | 0.63 | 0.5 |
| 8 | 78042456 | rs11990859 | 0.22 | -1.128 | -0.796 | 0.403 | -0.045 | 0.15 | 0.15 | 0.01 | 0.13 | 0.39 | 0.15 | 0.24 | 0.05 | 0.2 |
| 8 | 78043790 | rs17449553 | 1.505 | -0.842 | -1.15 | 0.285 | -0.086 | -0.064 | -0.064 | 0 | 0.38 | 0.08 | 0.43 | 0.45 | 0.55 | 0.41 |
| 8 | 78054417 | rs16939433 | 0.708 | -0.902 | -1.029 | 0.376 | 0.023 | -0.033 | -0.033 | 0 | 0.09 | 0.04 | 0.11 | 0.24 | 0.04 | 0.18 |
| 8 | 78057807 | rs10504645 | 0.142 | -1.071 | -1.114 | 0.104 | -0.003 | -0.006 | -0.006 | 0 | 0.65 | 0.38 | 0.66 | 0.79 | 0.75 | 0.83 |
| 8 | 78060354 | rs13439699 | 0.888 | -1.087 | -1.086 | 0.095 | 0.051 | -0.003 | -0.003 | 0 | 0.54 | 0.33 | 0.55 | 0.55 | 0.71 | 0.61 |
| 8 | 78068366 | rs10282777 | 0.524 | -0.985 | -1.023 | 0.023 | 0.032 | 0.093 | 0.093 | 0.09 | 0.58 | 0.75 | 0.68 | 0.78 | 0.68 | 0.67 |
| 8 | 78095409 | rs6996553 | -0.396 | -0.775 | -0.521 | 0.115 | 0.072 | 0.068 | 0.068 | 3.819 | 0.05 | 0.48 | 0.06 | 0 | 0.01 | 0 |
| 8 | 78123983 | rs16939458 | 0.321 | -0.788 | -0.705 | 0.879 | 0.003 | 0.01 | 0.01 | 0 | 0.1 | 0.1 | 0.11 | 0.27 | 0.04 | 0.19 |
| 8 | 78124431 | rs17380635 | 0.542 | -0.722 | -0.89 | 0.51 | -0.044 | 0.059 | 0.059 | 0 | 0.11 | 0.01 | 0.03 | 0.01 | 0.09 | 0.07 |
| 8 | 78135814 | rs12156042 | 0.269 | 0.555 | -1.075 | 0.51 | 0.053 | 0.045 | 0.045 | 0.32 | 0.12 | 0.19 | 0.39 | 0.34 | 0.33 | 0.25 |
| 8 | 78141371 | rs1452826 | -0.58 | -0.059 | -1.417 | 0.023 | 0.033 | 0.086 | 0.086 | 0 | 0.75 | 0.91 | 0.86 | 1 | 0.79 | 0.88 |
| 8 | 78153487 | rs6989596 | 1.393 | -0.181 | -1.088 | 0.057 | 0.044 | 0.046 | 0.046 | 0 | 0.46 | 0.25 | 0.38 | 0.65 | 0.36 | 0.56 |
| 8 | 78160049 | rs7828279 | 0.681 | -0.119 | -1.085 | 0.022 | -0.018 | 0.029 | 0.029 | 0 | 0.57 | 0.25 | 0.41 | 0.66 | 0.45 | 0.63 |
| 8 | 78160179 | rs7846385 | 2.504 | -0.37 | -1.268 | -0.225 | 0.007 | 0.03 | 0.03 | 0 | 0.31 | 0.12 | 0.25 | 0.19 | 0.29 | 0.3 |
| 8 | 78161374 | rs10755962 | -0.576 | -0.389 | -1.271 | -0.28 | 0.028 | 0.106 | 0.106 | 0 | 0.75 | 0.92 | 0.86 | 1 | 0.79 | 0.88 |
| 8 | 78171773 | rs1452796 | 1.062 | -0.23 | -1.032 | -0.146 | -0.042 | 0.059 | 0.059 | 0 | 0.11 | 0.01 | 0.03 | 0.01 | 0.09 | 0.07 |
| 8 | 78178485 | rs6473015 | 2.288 | -0.192 | -0.965 | -0.097 | 0.004 | 0.03 | 0.03 | 0 | 0.31 | 0.12 | 0.25 | 0.19 | 0.29 | 0.3 |
| 8 | 78181070 | rs11997859 | -1.008 | 0.517 | -1.16 | -0.251 | 0.004 | -0.005 | -0.005 | 0 | 0.31 | 0.31 | 0.6 | 0.77 | 0.42 | 0.48 |
| 8 | 78186862 | rs6473016 | -1.712 | 0.161 | -1.437 | -0.335 | 0.014 | 0.068 | 0.068 | 0 | 0.64 | 0.86 | 0.74 | 0.8 | 0.69 | 0.62 |
| 8 | 78192352 | rs4298474 | 0.606 | 0.173 | -1.47 | -0.313 | -0.009 | -0.017 | -0.017 | 0.414 | 0.17 | 0.04 | 0.05 | 0.21 | 0.13 | 0.18 |
| 8 | 78198545 | rs4526336 | -0.815 | 0.053 | -1.46 | -0.449 | -0.012 | -0.005 | -0.005 | 0 | 0.36 | 0.69 | 0.61 | 0.59 | 0.44 | 0.39 |
| 8 | 78200774 | rs4073914 | 1.414 | 0.048 | -1.446 | -0.453 | 0.002 | -0.015 | -0.015 | 3.815 | 0.11 | 0.13 | 0.08 | 0 | 0.13 | 0.04 |
| 8 | 78202736 | rs7814259 | -1.087 | 0.188 | -1.483 | -0.349 | -0.013 | 0.036 | 0.036 | 0 | 0.47 | 0.82 | 0.69 | 0.59 | 0.56 | 0.44 |
| 8 | 78219154 | rs12549538 | -0.185 | 0.553 | -1.31 | -0.414 | -0.051 | -0.051 | -0.051 | 0 | 0.14 | 0.06 | 0.45 | 0.63 | 0.28 | 0.33 |
| 8 | 78233600 | rs7005777 | -2.872 | 0.382 | -1.62 | -0.722 | -0.048 | 0.039 | 0.039 | 0 | 0.73 | 0.98 | 0.76 | 0.83 | 0.71 | 0.72 |
| 8 | 78236020 | rs7016767 | -1.412 | 0.267 | -1.627 | -0.849 | -0.013 | 0.157 | 0.157 | 0 | 0.55 | 0.84 | 0.65 | 0.8 | 0.54 | 0.56 |
| 8 | 78245535 | rs12674542 | -0.376 | 0.185 | -1.18 | -0.592 | -0.001 | -0.001 | -0.001 | 0 | 0.28 | 0.25 | 0.17 | 0.43 | 0.14 | 0.29 |
| 8 | 78271372 | rs1154839 | -1.427 | 0.389 | -1.188 | -0.564 | -0.002 | 0.028 | 0.028 | 0 | 0.92 | 0.97 | 0.95 | 1 | 0.92 | 0.99 |
| 8 | 78271818 | rs2338958 | 1.481 | 0.346 | -1.119 | -0.407 | -0.007 | -0.024 | -0.024 | 0.389 | 0.1 | 0.04 | 0.03 | 0.16 | 0.09 | 0.14 |
| 8 | 78285699 | rs7832361 | -0.028 | 0.334 | -0.76 | -0.49 | 0.018 | 0.014 | 0.014 | 0 | 0.26 | 0.62 | 0.48 | 0.37 | 0.41 | 0.26 |
| 8 | 78322385 | rs7820260 | -1.713 | 0.081 | -1.281 | -0.631 | 0.026 | 0.11 | 0.11 | 0 | 0.59 | 0.84 | 0.69 | 0.8 | 0.64 | 0.58 |
| 8 | 78324364 | rs12542969 | 1.7 | 0.053 | -1.341 | -0.513 | 0.002 | -0.001 | -0.001 | 0 | 0.05 | 0 | 0.1 | 0.24 | 0.02 | 0.1 |
| 8 | 78326076 | rs17376283 | 2.914 | -0.064 | -1.468 | -0.687 | 0.01 | 0.015 | 0.015 | 0 | 0.27 | 0.11 | 0.24 | 0.2 | 0.23 | 0.34 |
| 8 | 78334407 | rs10102576 | -1.409 | -0.013 | -1.372 | -0.774 | 0.004 | 0.119 | 0.119 | 0 | 0.46 | 0.8 | 0.61 | 0.77 | 0.5 | 0.53 |
| 8 | 78373108 | rs264824 | 1.679 | 0.141 | -1.097 | -0.575 | -0.012 | -0.024 | -0.024 | 0 | 0.12 | 0.03 | 0.04 | 0.19 | 0.1 | 0.14 |
| 8 | 78376314 | rs6996354 | 3.37 | 0.089 | -1.117 | -0.62 | 0.011 | -0.019 | -0.019 | 0 | 0.23 | 0.06 | 0.2 | 0.22 | 0.2 | 0.27 |
| 8 | 78392770 | rs166444 | -1.887 | 0.633 | -0.593 | -0.459 | 0.032 | -0.051 | -0.051 | 0.366 | 0.55 | 0.87 | 0.6 | 0.54 | 0.61 | 0.47 |
| 8 | 78394149 | rs17450081 | 1.39 | 0.713 | -0.567 | -0.379 | 0.016 | 0.038 | 0.038 | 0 | 0.13 | 0.07 | 0.08 | 0 | 0.13 | 0.05 |
| 8 | 78405264 | rs10504653 | 0.933 | 0.799 | -0.575 | -0.358 | 0.008 | 0.028 | 0.028 | 0 | 0.1 | 0.05 | 0.02 | 0.02 | 0.05 | 0.03 |
| 8 | 78407296 | rs264815 | 0.077 | 1.007 | -0.292 | -0.118 | -0.01 | -0.018 | -0.018 | 0.297 | 0.28 | 0.37 | 0.14 | 0.19 | 0.23 | 0.19 |
| 8 | 78426530 | rs2338964 | 0.726 | 1.549 | -0.233 | 0.43 | -0.029 | -0.029 | -0.029 | 0 | 0.07 | 0.02 | 0.31 | 0.27 | 0.25 | 0.16 |
| 8 | 78448923 | rs264809 | 1.14 | 0.548 | -0.057 | 0.082 | -0.012 | -0.027 | -0.027 | 0.716 | 0.13 | 0.03 | 0.05 | 0.2 | 0.11 | 0.15 |
| 8 | 78452205 | rs364324 | 0.563 | 0.537 | 0.187 | 0.073 | -0.006 | 0.002 | 0.002 | 0 | 0.15 | 0.34 | 0.07 | 0.2 | 0.11 | 0.15 |
| 8 | 78510649 | rs446337 | -0.332 | 0.151 | -0.456 | -0.258 | 0.022 | 0.04 | 0.04 | 0.159 | 0.34 | 0.2 | 0.16 | 0.24 | 0.28 | 0.18 |
| 8 | 78511219 | rs395261 | -1.417 | 0.37 | -0.318 | -0.158 | 0.071 | -0.011 | -0.011 | 0 | 0.52 | 0.78 | 0.59 | 0.57 | 0.61 | 0.42 |
| 8 | 78523727 | rs17466207 | 2.783 | -0.045 | -0.745 | -0.183 | 0.024 | 0.003 | 0.003 | 0 | 0.24 | 0.06 | 0.19 | 0.19 | 0.2 | 0.26 |
| 8 | 78527566 | rs13253986 | 0.159 | 0.445 | -0.71 | 0.018 | 0.001 | -0.01 | -0.01 | 0 | 0.23 | 0.19 | 0.11 | 0.04 | 0.18 | 0.08 |
| 8 | 78530899 | rs883227 | 0.454 | 1.007 | -0.704 | 0.32 | -0.051 | -0.049 | -0.049 | 0 | 0.2 | 0.08 | 0.25 | 0.45 | 0.33 | 0.26 |
| 8 | 78550000 | rs1912391 | -1.439 | 0.379 | -0.904 | 0.182 | 0.004 | -0.064 | -0.064 | 0 | 0.89 | 0.98 | 0.88 | 0.78 | 0.9 | 0.89 |
| 8 | 78551702 | rs1039857 | -1.558 | 0.878 | -0.023 | -0.15 | -0.008 | 0.003 | 0.003 | 0 | 0.44 | 0.45 | 0.54 | 0.59 | 0.53 | 0.44 |
| 8 | 78575277 | rs7818862 | 0.24 | 1.299 | -0.183 | 0.043 | -0.149 | -0.144 | -0.144 | 0.07 | 0.21 | 0.04 | 0.39 | 0.5 | 0.37 | 0.28 |
| 8 | 78589461 | rs7839056 | -0.718 | 1.253 | 0.08 | -0.131 | 0.047 | -0.039 | -0.039 | 0 | 0.18 | 0.34 | 0.15 | 0.08 | 0.17 | 0.17 |
| 8 | 78593877 | rs16939539 | 1.009 | 0.999 | 0.165 | -0.327 | -0.003 | -0.03 | -0.03 | 0.109 | 0.13 | 0.2 | 0.1 | 0 | 0.13 | 0.05 |
| 8 | 78619510 | rs1116418 | -1.592 | 1.066 | 0.297 | -0.025 | 0.058 | -0.007 | -0.007 | 0 | 0.54 | 0.7 | 0.65 | 0.57 | 0.67 | 0.49 |
| 8 | 78629747 | rs16939542 | -0.626 | 2.146 | 0.799 | 0.043 | 0.027 | -0.025 | -0.025 | 2.203 | 0.11 | 0.17 | 0.12 | 0.04 | 0.13 | 0.15 |
| 8 | 78630367 | rs7016717 | -0.245 | 2.009 | 0.796 | 0.022 | 0.074 | 0.022 | 0.022 | 0 | 0.07 | 0.36 | 0.04 | 0.02 | 0.04 | 0.02 |
| 8 | 78631154 | rs10504658 | -0.711 | 1.955 | -0.141 | 0.195 | 0.008 | -0.014 | -0.014 | 2.76 | 0.11 | 0.14 | 0.11 | 0.04 | 0.12 | 0.16 |
| 8 | 78632893 | rs10957863 | -1.192 | 2.124 | -0.095 | 0.478 | 0.097 | 0.035 | 0.035 | 1.378 | 0.17 | 0.23 | 0.46 | 0.3 | 0.41 | 0.34 |
| 8 | 78633267 | rs7824975 | -0.584 | 0.255 | -0.024 | -0.324 | -0.071 | -0.114 | -0.114 | 0.441 | 0.19 | 0.43 | 0.1 | 0.01 | 0.08 | 0.03 |
| 8 | 78637869 | rs1022374 | 0.762 | 0.78 | -0.763 | -0.308 | -0.037 | 0.027 | 0.027 | 0 | 0.31 | 0.16 | 0.41 | 0.2 | 0.39 | 0.29 |
| 8 | 78638028 | rs16939546 | -0.104 | 0.026 | -1.201 | -0.122 | 0.087 | 0.01 | 0.01 | 0 | 0.37 | 0.16 | 0.21 | 0.35 | 0.26 | 0.3 |
| 8 | 78644605 | rs1519349 | -0.273 | 0.113 | -0.53 | 0.027 | 0.028 | -0.027 | -0.027 | 0 | 0.79 | 0.66 | 0.77 | 0.86 | 0.69 | 0.7 |
| 8 | 78646880 | rs16939552 | 0.886 | -0.452 | -0.747 | -0.039 | 0.046 | -0.044 | -0.044 | 0 | 0.2 | 0.09 | 0.12 | 0.3 | 0.1 | 0.14 |
| 8 | 78658127 | rs1607547 | 0.616 | -0.576 | -0.778 | -0.222 | -0.002 | -0.003 | -0.003 | 0.266 | 0.58 | 0.5 | 0.64 | 0.56 | 0.59 | 0.55 |
| 8 | 78663393 | rs4451298 | 1.029 | -0.646 | -0.602 | -0.197 | 0.006 | 0.039 | 0.039 | 0 | 0.49 | 0.37 | 0.5 | 0.25 | 0.54 | 0.45 |
| 8 | 78683460 | rs11984782 | 0.851 | -0.653 | -0.7 | -0.184 | 0.017 | -0.011 | -0.011 | 0 | 0.21 | 0.16 | 0.13 | 0.3 | 0.1 | 0.14 |
| 8 | 78719309 | rs10086879 | -0.203 | -0.659 | -0.63 | -0.229 | 0.044 | -0.043 | -0.043 | 0.24 | 0.78 | 0.58 | 0.76 | 0.86 | 0.7 | 0.7 |
| 8 | 78724669 | rs13270760 | -0.156 | -0.704 | -0.552 | -0.191 | -0.066 | 0.041 | 0.041 | 0 | 0.3 | 0.43 | 0.28 | 0.61 | 0.16 | 0.24 |

| | | | | | | | | | | | | | | | | |
|---|---|---|---|---|---|---|---|---|---|---|---|---|---|---|---|---|
| 8 | 78730284 | rs13269674 | 0.647 | -0.634 | -0.761 | -0.277 | -0.007 | 0.054 | 0.054 | 0 | 0.15 | 0.03 | 0.07 | 0.05 | 0.15 | 0.16 |
| 8 | 78733701 | rs2339076 | 0.172 | -0.744 | -0.663 | -0.264 | 0.027 | 0.059 | 0.059 | 0 | 0.7 | 0.51 | 0.63 | 0.54 | 0.65 | 0.6 |
| 8 | 78738002 | rs2222263 | 0.714 | -0.832 | -0.678 | -0.269 | -0.061 | 0.027 | 0.027 | 0 | 0.22 | 0.36 | 0.15 | 0.29 | 0.11 | 0.14 |
| 8 | 78749262 | rs4739127 | 0.14 | -0.767 | -0.7 | -0.274 | 0.014 | 0.036 | 0.036 | 0 | 0.7 | 0.45 | 0.61 | 0.54 | 0.65 | 0.6 |
| 8 | 78770472 | rs2879351 | 2.213 | -0.785 | -0.777 | -0.318 | -0.071 | 0.04 | 0.04 | 0 | 0.32 | 0.11 | 0.4 | 0.2 | 0.4 | 0.29 |
| 8 | 78774883 | rs11991423 | 0.88 | -0.747 | -0.673 | -0.202 | 0.014 | -0.012 | -0.012 | 0 | 0.21 | 0.16 | 0.13 | 0.31 | 0.11 | 0.14 |
| 8 | 78778013 | rs4739175 | 0.873 | -0.64 | -0.517 | -0.257 | 0.015 | 0.08 | 0.08 | 0 | 0.49 | 0.34 | 0.49 | 0.25 | 0.55 | 0.45 |
| 8 | 78785042 | rs4739176 | 0.917 | -0.634 | -0.538 | -0.254 | 0.014 | 0.097 | 0.097 | 0 | 0.49 | 0.28 | 0.49 | 0.25 | 0.55 | 0.45 |
| 8 | 78812885 | rs13272656 | 0.438 | -0.378 | -0.434 | -0.155 | -0.009 | 0.049 | 0.049 | 0 | 0.16 | 0.05 | 0.07 | 0.05 | 0.15 | 0.16 |
| 8 | 78815409 | rs1948087 | 0.036 | -0.329 | -0.246 | -0.2 | 0.073 | -0.06 | -0.06 | 0 | 0.2 | 0.43 | 0.25 | 0.13 | 0.29 | 0.31 |
| 8 | 78821562 | rs17388515 | 0.488 | 0.038 | -0.67 | -0.255 | -0.007 | 0.049 | 0.049 | 0 | 0.16 | 0.03 | 0.06 | 0.05 | 0.15 | 0.16 |
| 8 | 78822971 | rs10957875 | 1.006 | 0.052 | -0.561 | -0.167 | -0.01 | -0.014 | -0.014 | 0.013 | 0.55 | 0.33 | 0.54 | 0.51 | 0.5 | 0.45 |
| 8 | 78835777 | rs10156234 | 1.077 | -0.236 | -0.52 | -0.134 | 0.027 | -0.016 | -0.016 | 0 | 0.57 | 0.33 | 0.59 | 0.51 | 0.64 | 0.46 |
| 8 | 78863442 | rs11998680 | -1.261 | -0.338 | -0.67 | -0.422 | 0.049 | 0.12 | 0.12 | 0.091 | 0.39 | 0.8 | 0.31 | 0.18 | 0.33 | 0.44 |
| 8 | 78864058 | rs959439 | 0.309 | -0.294 | -0.653 | -0.314 | -0.014 | -0.008 | -0.008 | 0 | 0.13 | 0.04 | 0.15 | 0.12 | 0.14 | 0.27 |
| 8 | 78870858 | rs7016182 | -0.326 | -0.323 | -0.335 | -0.391 | -0.022 | -0.072 | -0.072 | 0 | 0.18 | 0.59 | 0.16 | 0.05 | 0.16 | 0.13 |
| 8 | 78873842 | rs13269418 | 0.566 | -0.335 | 0.058 | -0.394 | -0.009 | -0.039 | -0.039 | 0.026 | 0.14 | 0.19 | 0.09 | 0.05 | 0.14 | 0.12 |
| 8 | 78879189 | rs4739185 | 1.405 | -0.707 | -0.337 | -0.419 | 0.052 | -0.031 | -0.031 | 0 | 0.55 | 0.25 | 0.54 | 0.49 | 0.62 | 0.47 |
| 8 | 78882007 | rs17481178 | 2.239 | -0.773 | -0.517 | -0.668 | -0.083 | 0.048 | 0.048 | 0 | 0.34 | 0.07 | 0.39 | 0.2 | 0.4 | 0.25 |
| 8 | 78885976 | rs7823046 | -1.214 | -0.974 | -0.214 | -0.706 | 0.002 | 0.002 | 0.002 | 0 | 0.33 | 0.65 | 0.3 | 0.18 | 0.29 | 0.42 |
| 8 | 78897147 | rs880091 | 0.209 | -0.922 | -0.367 | -0.66 | -0.013 | -0.001 | -0.001 | 0 | 0.15 | 0.04 | 0.14 | 0.13 | 0.13 | 0.29 |
| 8 | 78906904 | rs13261986 | -0.701 | -0.898 | -0.093 | -0.645 | -0.006 | -0.017 | -0.017 | 0.031 | 0.3 | 0.31 | 0.24 | 0.17 | 0.27 | 0.42 |
| 8 | 78941331 | rs7004112 | -1.055 | -0.977 | -0.067 | -0.435 | 0.012 | -0.008 | -0.008 | 0 | 0.43 | 0.33 | 0.38 | 0.47 | 0.36 | 0.53 |
| 8 | 78946230 | rs9643732 | -1.055 | -1.022 | -0.207 | -0.434 | 0.04 | -0.028 | -0.028 | 0 | 0.43 | 0.26 | 0.38 | 0.47 | 0.36 | 0.53 |
| 8 | 78951278 | rs2368508 | -1.072 | -1.023 | -0.064 | -0.382 | 0.03 | -0.016 | -0.016 | 0.061 | 0.43 | 0.3 | 0.38 | 0.48 | 0.36 | 0.53 |
| 8 | 78966585 | rs13281300 | -1.021 | -1.091 | -0.269 | -0.396 | 0.051 | -0.052 | -0.052 | 0 | 0.42 | 0.19 | 0.37 | 0.48 | 0.36 | 0.53 |
| 8 | 78974274 | rs11989865 | 0.664 | -1.333 | -0.166 | -0.368 | -0.015 | -0.007 | -0.007 | 1.789 | 0.18 | 0.18 | 0.06 | 0.2 | 0.1 | 0.2 |
| 8 | 78975396 | rs13276389 | 0.949 | -1.185 | -0.114 | -0.267 | 0.003 | 0.009 | 0.009 | 0 | 0.13 | 0.09 | 0.08 | 0.04 | 0.14 | 0.14 |
| 8 | 78990829 | rs2200015 | -1.21 | -1.094 | -0.439 | -0.288 | 0.015 | -0.01 | -0.01 | 0.336 | 0.42 | 0.66 | 0.41 | 0.48 | 0.32 | 0.47 |
| 8 | 78995713 | rs10957882 | -2.218 | -0.823 | -0.887 | -0.282 | -0.104 | -0.02 | -0.02 | 0 | 0.64 | 0.92 | 0.53 | 0.69 | 0.58 | 0.71 |
| 8 | 79000607 | rs2448962 | 1.096 | -0.526 | -0.487 | 0.142 | 0 | 0.034 | 0.034 | 0 | 0.08 | 0.06 | 0.13 | 0.29 | 0.05 | 0.09 |
| 8 | 79001991 | rs11780588 | 0.232 | -0.444 | -0.673 | 0.071 | -0.007 | -0.027 | -0.027 | 0 | 0.13 | 0.04 | 0.12 | 0.14 | 0.09 | 0.22 |
| 8 | 79003368 | rs2448961 | -0.399 | -0.459 | -0.116 | 0.092 | -0.022 | 0.043 | 0.043 | 0 | 0.22 | 0.39 | 0.23 | 0.34 | 0.19 | 0.23 |
| 8 | 79004739 | rs6997505 | 0.377 | 0.022 | -0.516 | -0.173 | -0.024 | 0.024 | 0.024 | 0 | 0.06 | 0.02 | 0.05 | 0 | 0.15 | 0.08 |
| 8 | 79008608 | rs1481513 | -0.892 | -0.397 | -0.202 | -0.353 | -0.007 | -0.054 | -0.054 | 0 | 0.38 | 0.52 | 0.28 | 0.13 | 0.42 | 0.4 |
| 8 | 79009590 | rs6473088 | -1.204 | -0.405 | -0.166 | -0.136 | 0.003 | 0.009 | 0.009 | 0 | 0.45 | 0.64 | 0.32 | 0.35 | 0.46 | 0.52 |
| 8 | 79013694 | rs16939608 | 0.809 | -0.367 | -0.266 | 0.03 | -0.015 | -0.028 | -0.028 | 0 | 0.05 | 0 | 0.03 | 0.22 | 0.04 | 0.12 |
| 8 | 79024940 | rs2887265 | -0.549 | -0.598 | -0.187 | -0.232 | -0.028 | -0.05 | -0.05 | 0 | 0.37 | 0.49 | 0.23 | 0.13 | 0.3 | 0.38 |
| 8 | 79031935 | rs13278866 | -0.502 | -0.66 | -0.045 | -0.263 | 0.043 | 0.081 | 0.081 | 0 | 0.34 | 0.22 | 0.21 | 0.13 | 0.3 | 0.38 |
| 8 | 79039530 | rs12677810 | 0.84 | -0.515 | -0.077 | -0.282 | 0.022 | 0.009 | 0.009 | 0 | 0.14 | 0.08 | 0.1 | 0.08 | 0.08 | 0.21 |
| 8 | 79042331 | rs10504670 | -0.381 | -0.516 | 0.136 | -0.228 | 0.003 | 0.06 | 0.06 | 0 | 0.06 | 0.15 | 0.06 | 0.24 | 0.04 | 0.14 |
| 8 | 79045302 | rs13251222 | 0.134 | -0.498 | 0.046 | -0.34 | -0.031 | 0.094 | 0.094 | 0 | 0.19 | 0.02 | 0.08 | 0.03 | 0.2 | 0.16 |
| 8 | 79047096 | rs6996222 | -0.508 | -0.422 | 0.249 | -0.392 | 0.009 | -0.092 | -0.092 | 0 | 0.25 | 0.35 | 0.16 | 0.03 | 0.34 | 0.17 |
| 8 | 79059629 | rs1481512 | -1.633 | -0.424 | 0.204 | -0.219 | 0.001 | 0.003 | 0.003 | 0 | 0.44 | 0.63 | 0.33 | 0.36 | 0.47 | 0.51 |
| 8 | 79074228 | rs11782332 | -0.914 | -0.511 | 0.476 | -0.252 | 0.001 | -0.006 | -0.006 | 3.827 | 0.36 | 0.38 | 0.25 | 0.12 | 0.39 | 0.32 |
| 8 | 79100584 | rs11990002 | -1.736 | -0.437 | 0.459 | -0.181 | 0.003 | 0.009 | 0.009 | 0 | 0.44 | 0.69 | 0.36 | 0.36 | 0.44 | 0.51 |
| 8 | 79108318 | rs16919529 | -0.83 | -0.479 | 0.284 | -0.117 | 0.029 | 0.128 | 0.128 | 0 | 0.08 | 0.26 | 0.1 | 0.26 | 0.05 | 0.2 |
| 8 | 79129583 | rs1978660 | 1.373 | -0.527 | 0.17 | -0.129 | 0.015 | -0.002 | -0.002 | 0 | 0.12 | 0.04 | 0.08 | 0.08 | 0.09 | 0.16 |
| 8 | 79136599 | rs16905551 | -0.458 | -0.444 | 0.266 | -0.106 | 0.007 | 0.09 | 0.09 | 0 | 0.06 | 0.17 | 0.08 | 0.26 | 0.04 | 0.19 |
| 8 | 79144277 | rs17485419 | -0.266 | -0.389 | 0.071 | -0.201 | -0.026 | 0.073 | 0.073 | 0 | 0.2 | 0.11 | 0.13 | 0.02 | 0.29 | 0.15 |
| 8 | 79171435 | rs2195688 | 1.163 | -0.56 | -0.313 | 0.436 | -0.044 | -0.077 | -0.077 | 0 | 0.11 | 0 | 0.17 | 0.31 | 0.17 | 0.13 |
| 8 | 79185324 | rs6980983 | -1.365 | -0.854 | 0.328 | 0.334 | 0.02 | 0.007 | 0.007 | 0 | 0.42 | 0.72 | 0.31 | 0.38 | 0.28 | 0.49 |
| 8 | 79196677 | rs7016181 | 0.017 | -0.745 | 0.207 | 0.065 | -0.001 | -0.015 | -0.015 | 0 | 0.18 | 0.2 | 0.1 | 0.05 | 0.15 | 0.14 |
| 8 | 79198642 | rs2369121 | -1.068 | -0.892 | 0.463 | -0.023 | -0.023 | -0.047 | -0.047 | 1.333 | 0.35 | 0.47 | 0.21 | 0.11 | 0.25 | 0.28 |
| 8 | 79214863 | rs11780328 | 0.042 | -0.826 | 0.22 | 0.004 | 0.002 | -0.012 | -0.012 | 0 | 0.18 | 0.18 | 0.1 | 0.03 | 0.17 | 0.14 |
| 8 | 79222482 | rs12679798 | 1.062 | -0.841 | 0.207 | 0.045 | 0.025 | 0.011 | 0.011 | 0 | 0.14 | 0.08 | 0.08 | 0.08 | 0.09 | 0.14 |
| 8 | 79247094 | rs13262405 | -0.346 | -0.765 | 0.447 | 0.018 | -0.01 | -0.056 | -0.056 | 2.583 | 0.2 | 0.3 | 0.12 | 0.03 | 0.17 | 0.14 |
| 8 | 79256864 | rs7016193 | 0.125 | -0.739 | 0.363 | 0.031 | 0.002 | -0.009 | -0.009 | 0 | 0.18 | 0.17 | 0.1 | 0.03 | 0.17 | 0.14 |
| 8 | 79262741 | rs7823679 | -0.615 | -0.587 | 0.758 | 0.387 | -0.005 | 0.01 | 0.01 | 0 | 0.23 | 0.37 | 0.19 | 0.25 | 0.21 | 0.33 |
| 8 | 79270433 | rs7003771 | -0.656 | -0.589 | 0.771 | 0.387 | -0.006 | 0.013 | 0.013 | 0 | 0.23 | 0.41 | 0.19 | 0.25 | 0.21 | 0.33 |
| 8 | 79276063 | rs2195687 | -0.954 | -0.56 | 0.765 | 0.321 | 0.02 | 0.013 | 0.013 | 0 | 0.41 | 0.71 | 0.29 | 0.3 | 0.3 | 0.43 |
| 8 | 79284507 | rs7016026 | -1.558 | -0.156 | 0.034 | 0.296 | -0.059 | -0.038 | -0.038 | 0.166 | 0.67 | 0.97 | 0.64 | 0.66 | 0.71 | 0.64 |
| 8 | 79302840 | rs7018296 | -0.264 | -0.534 | 0.609 | 0.259 | 0.017 | 0.047 | 0.047 | 0 | 0.29 | 0.2 | 0.16 | 0.11 | 0.27 | 0.28 |
| 8 | 79324247 | rs11990902 | -0.475 | -0.514 | 0.615 | 0.26 | 0.011 | 0.021 | 0.021 | 0 | 0.31 | 0.28 | 0.17 | 0.11 | 0.27 | 0.28 |
| 8 | 79334466 | rs7010579 | -1.216 | -0.549 | 0.672 | 0.333 | -0.003 | -0.001 | -0.001 | 0 | 0.39 | 0.66 | 0.28 | 0.3 | 0.3 | 0.43 |
| 8 | 79343367 | rs13254983 | 0.004 | -0.835 | 0.697 | 0.672 | 0.023 | -0.011 | -0.011 | 0 | 0.23 | 0.16 | 0.13 | 0.25 | 0.19 | 0.25 |
| 8 | 79347559 | rs7833623 | -0.866 | -0.872 | 0.94 | 0.713 | 0.02 | 0.156 | 0.156 | 0 | 0.09 | 0.34 | 0.13 | 0.21 | 0.06 | 0.16 |
| 8 | 79350458 | rs13278903 | -1.482 | -0.806 | 1.134 | 0.696 | 0.052 | -0.051 | -0.051 | 0.016 | 0.37 | 0.82 | 0.31 | 0.46 | 0.28 | 0.43 |
| 8 | 79364896 | rs13282611 | 0.21 | -0.759 | 0.97 | 0.697 | 0.018 | -0.013 | -0.013 | 0 | 0.22 | 0.16 | 0.13 | 0.25 | 0.2 | 0.26 |

| | | | | | | | | | | | | | | | | |
|---|---|---|---|---|---|---|---|---|---|---|---|---|---|---|---|---|
| 8 | 79374229 | rs10216378 | -0.694 | -0.705 | 0.977 | 0.755 | 0.019 | 0.159 | 0.159 | 0 | 0.09 | 0.33 | 0.12 | 0.22 | 0.06 | 0.15 |
| 8 | 79394667 | rs6984124 | -1.465 | -0.86 | 0.797 | 0.549 | 0.04 | -0.021 | -0.021 | 2.667 | 0.41 | 0.74 | 0.28 | 0.46 | 0.3 | 0.46 |
| 8 | 79399752 | rs4145315 | -1.289 | -0.701 | 1.162 | 0.824 | 0.043 | 0.185 | 0.185 | 0 | 0.1 | 0.4 | 0.13 | 0.21 | 0.08 | 0.15 |
| 8 | 79401038 | rs1520333 | -1.236 | -0.696 | 0.782 | 0.799 | 0.019 | -0.02 | -0.02 | 0 | 0.35 | 0.74 | 0.28 | 0.46 | 0.28 | 0.4 |
| 8 | 79402089 | rs16905701 | -0.926 | -0.098 | 0.534 | 0.938 | 0.017 | 0.147 | 0.147 | 0 | 0.08 | 0.27 | 0.11 | 0.22 | 0.06 | 0.15 |
| 8 | 79413343 | rs4294171 | -0.282 | -0.331 | 0.095 | 0.501 | -0.012 | 0.011 | 0.011 | 0 | 0.13 | 0.18 | 0.1 | 0.15 | 0.13 | 0.14 |
| 8 | 79420199 | rs7013822 | -0.45 | -0.847 | -0.17 | -0.321 | -0.017 | -0.032 | -0.032 | 0 | 0.25 | 0.35 | 0.14 | 0.09 | 0.18 | 0.16 |
| 8 | 79433432 | rs13275966 | 0.819 | -1.024 | -0.223 | -0.392 | 0.036 | 0.031 | 0.031 | 0 | 0.15 | 0.05 | 0.07 | 0.09 | 0.11 | 0.07 |
| 8 | 79438103 | rs9693457 | -1.147 | -1.103 | -0.145 | -0.527 | -0.028 | -0.057 | -0.057 | 0 | 0.3 | 0.42 | 0.15 | 0.09 | 0.2 | 0.18 |
| 8 | 79447414 | rs7837755 | -0.145 | -0.92 | -0.156 | -0.424 | 0.033 | 0.045 | 0.045 | 0 | 0.2 | 0.14 | 0.09 | 0.09 | 0.14 | 0.1 |
| 8 | 79451370 | rs10504676 | 0.561 | -0.884 | -0.118 | -0.379 | 0.024 | 0.022 | 0.022 | 0 | 0.16 | 0.16 | 0.09 | 0.09 | 0.11 | 0.08 |
| 8 | 79461182 | rs10504677 | 0.546 | -0.915 | -0.103 | -0.38 | 0.01 | 0.01 | 0.01 | 0 | 0.16 | 0.19 | 0.09 | 0.09 | 0.11 | 0.08 |
| 8 | 79475862 | rs7824063 | -0.155 | -0.859 | -0.178 | -0.44 | -0.045 | -0.042 | -0.042 | 0 | 0.2 | 0.36 | 0.12 | 0.09 | 0.11 | 0.09 |
| 8 | 79483835 | rs6473106 | -0.88 | -0.909 | -0.185 | -0.4 | -0.03 | -0.045 | -0.045 | 0 | 0.28 | 0.43 | 0.15 | 0.11 | 0.2 | 0.14 |
| 8 | 79497899 | rs967426 | -1.687 | -1.063 | -0.348 | -0.063 | 0.085 | -0.015 | -0.015 | 0.858 | 0.47 | 0.94 | 0.36 | 0.48 | 0.38 | 0.42 |
| 8 | 79501241 | rs16905824 | 0 | -1.047 | 0.218 | -0.708 | -0.012 | -0.006 | -0.006 | 0 | 0.18 | 0.23 | 0.1 | 0.11 | 0.11 | 0.1 |
| 8 | 79510525 | rs6990845 | -0.528 | -1.189 | 0.324 | -0.621 | -0.003 | -0.055 | -0.055 | 2.501 | 0.1 | 0.21 | 0.06 | 0 | 0.09 | 0.04 |
| 8 | 79516759 | rs1062248 | -1.075 | -1.233 | 0.33 | -0.636 | -0.009 | -0.016 | -0.016 | 0 | 0.27 | 0.36 | 0.14 | 0.11 | 0.2 | 0.15 |
| 8 | 79518376 | rs10504678 | 0.277 | -1.125 | 0.21 | -0.645 | -0.001 | -0.013 | -0.013 | 0 | 0.08 | 0.08 | 0.03 | 0.01 | 0.07 | 0.04 |
| 8 | 79523704 | rs16919538 | 0.31 | -1.089 | 0.244 | -0.627 | -0.002 | -0.025 | -0.025 | 0 | 0.08 | 0.11 | 0.04 | 0.01 | 0.07 | 0.05 |
| 8 | 79526456 | rs6473109 | -0.846 | -1.163 | 0.266 | -0.629 | -0.032 | -0.042 | -0.042 | 0 | 0.25 | 0.4 | 0.13 | 0.1 | 0.18 | 0.16 |
| 8 | 79532730 | rs2369294 | 0.511 | -1.1 | 0.032 | -0.569 | 0.032 | 0.009 | 0.009 | 0 | 0.15 | 0.05 | 0.07 | 0.09 | 0.1 | 0.08 |
| 8 | 79551511 | rs16905872 | -0.514 | -0.942 | 0.321 | -0.453 | -0.002 | -0.048 | -0.048 | 0 | 0.1 | 0.33 | 0.06 | 0.01 | 0.07 | 0.07 |
| 8 | 79553027 | rs12155808 | 0.192 | -1.228 | 0.328 | -0.448 | 0.016 | 0.017 | 0.017 | 0.046 | 0.75 | 0.77 | 0.89 | 0.9 | 0.83 | 0.87 |
| 8 | 79556148 | rs4739134 | 0.615 | -1.168 | 0.676 | -0.487 | -0.009 | -0.022 | -0.022 | 0 | 0.65 | 0.61 | 0.81 | 0.9 | 0.71 | 0.78 |
| 8 | 79556891 | rs1966115 | 0.714 | -1.178 | 0.824 | -0.492 | -0.036 | -0.064 | -0.064 | 0.16 | 0.64 | 0.45 | 0.8 | 0.9 | 0.71 | 0.78 |
| 8 | 79564187 | rs11782836 | 0.368 | -1.143 | 0.87 | -0.483 | 0.003 | -0.038 | -0.038 | 0 | 0.07 | 0.21 | 0.06 | 0 | 0.08 | 0.02 |
| 8 | 79576159 | rs979564 | 0.233 | -1.238 | 0.737 | -0.514 | -0.014 | -0.016 | -0.016 | 0.19 | 0.7 | 0.68 | 0.86 | 0.9 | 0.79 | 0.79 |
| 8 | 79577726 | rs3808619 | 0.796 | -1.252 | 1.013 | -0.522 | -0.041 | -0.07 | -0.07 | 0 | 0.63 | 0.5 | 0.8 | 0.9 | 0.71 | 0.77 |
| 8 | 79582496 | rs4641056 | 0.303 | -1.223 | 1.035 | -0.484 | -0.061 | -0.07 | -0.07 | 0 | 0.69 | 0.52 | 0.84 | 0.89 | 0.79 | 0.79 |
| 8 | 79610710 | rs1054283 | 0.766 | -1.236 | 1.047 | -0.519 | -0.041 | -0.072 | -0.072 | 0.303 | 0.63 | 0.5 | 0.8 | 0.9 | 0.72 | 0.77 |
| 8 | 79622363 | rs2717540 | 0.26 | -1.247 | 1.093 | -0.528 | -0.014 | -0.018 | -0.018 | 0 | 0.7 | 0.68 | 0.86 | 0.9 | 0.79 | 0.79 |
| 8 | 79632010 | rs1026920 | 0.763 | -1.234 | 1.04 | -0.411 | -0.041 | -0.072 | -0.072 | 1.338 | 0.63 | 0.5 | 0.8 | 0.9 | 0.71 | 0.77 |
| 8 | 79635864 | rs3739330 | 0.747 | -0.979 | 0.501 | -0.388 | -0.056 | -0.016 | -0.016 | 0.15 | 0.13 | 0 | 0.15 | 0.11 | 0.15 | 0.12 |
| 8 | 79643639 | rs12544041 | -0.264 | -1.248 | 0.824 | -0.346 | -0.011 | -0.014 | -0.014 | 0 | 0.29 | 0.28 | 0.14 | 0.1 | 0.21 | 0.21 |
| 8 | 79654145 | rs6993386 | -1.316 | -1.223 | 0.844 | 0.067 | 0.064 | 0.021 | 0.021 | 3.113 | 0.44 | 0.75 | 0.38 | 0.38 | 0.34 | 0.3 |
| 8 | 79659801 | rs16906063 | -1.048 | -0.8 | 1.178 | 0.297 | 0.004 | 0.1 | 0.1 | 0.429 | 0.15 | 0.46 | 0.24 | 0.28 | 0.13 | 0.11 |
| 8 | 79663408 | rs2583763 | -1.394 | -0.907 | 1.166 | 0.33 | -0.003 | 0.161 | 0.161 | 1.043 | 0.15 | 0.43 | 0.21 | 0.31 | 0.09 | 0.13 |
| 8 | 79664483 | rs1119642 | -1.732 | -0.913 | 1.162 | 0.317 | -0.014 | 0.079 | 0.079 | 1.855 | 0.2 | 0.43 | 0.24 | 0.31 | 0.16 | 0.15 |
| 8 | 79670387 | rs2953474 | -1.719 | -0.915 | 1.238 | 0.316 | -0.019 | 0.074 | 0.074 | 2.216 | 0.21 | 0.55 | 0.26 | 0.31 | 0.16 | 0.15 |
| 8 | 79671067 | rs2717548 | -1.741 | -0.894 | 1.226 | 0.316 | -0.012 | 0.078 | 0.078 | 1.67 | 0.2 | 0.43 | 0.24 | 0.31 | 0.16 | 0.15 |
| 8 | 79686236 | rs11776228 | 0.944 | -0.803 | 0.911 | 0.313 | -0.001 | 0.008 | 0.008 | 0.321 | 0.06 | 0.08 | 0.04 | 0 | 0.06 | 0.02 |
| 8 | 79688026 | rs10110519 | -1 | -0.892 | 1.037 | 0.336 | 0.011 | 0.202 | 0.202 | 0.808 | 0.12 | 0.45 | 0.19 | 0.29 | 0.07 | 0.09 |
| 8 | 79700362 | rs11990466 | 0.705 | -0.835 | 0.695 | 0.31 | -0.003 | -0.031 | -0.031 | 0 | 0.06 | 0.14 | 0.05 | 0 | 0.06 | 0.02 |
| 8 | 79703113 | rs11775538 | 0.705 | -0.835 | 0.695 | 0.31 | -0.003 | -0.031 | -0.031 | 1.742 | 0.06 | 0.14 | 0.05 | 0 | 0.06 | 0.02 |
| 8 | 79709926 | rs4739139 | 0.317 | -0.83 | 0.765 | 0.4 | 0.006 | 0.083 | 0.083 | 0 | 0.07 | 0.13 | 0.15 | 0.27 | 0.06 | 0.08 |
| 8 | 79712998 | rs16906115 | -0.265 | -0.819 | 0.552 | 0.369 | 0.006 | 0.17 | 0.17 | 1.311 | 0.12 | 0.4 | 0.19 | 0.29 | 0.07 | 0.09 |
| 8 | 79716536 | rs3888020 | -1.552 | -0.988 | -0.431 | -0.05 | 0.043 | -0.043 | -0.043 | 1.421 | 0.52 | 0.86 | 0.52 | 0.45 | 0.4 | 0.49 |
| 8 | 79718005 | rs6997891 | -0.067 | -0.128 | -0.206 | 0.541 | 0.001 | -0.032 | -0.032 | 0 | 0.11 | 0.13 | 0.07 | 0 | 0.11 | 0.06 |
| 8 | 79728046 | rs3885022 | -0.66 | -0.151 | -0.301 | 0.532 | -0.001 | -0.007 | -0.007 | 0 | 0.13 | 0.14 | 0.07 | 0.02 | 0.13 | 0.09 |
| 8 | 79743689 | rs2581738 | -0.901 | -0.184 | -0.177 | 1.01 | -0.009 | 0.129 | 0.129 | 0.062 | 0.17 | 0.39 | 0.31 | 0.36 | 0.13 | 0.25 |
| 8 | 79747724 | rs12677936 | -0.193 | 0.21 | -1.064 | 0.701 | -0.018 | -0.035 | -0.035 | 0.035 | 0.09 | 0 | 0.15 | 0.14 | 0.1 | 0.08 |
| 8 | 79750564 | rs2587155 | -0.868 | -0.136 | -0.611 | 0.446 | -0.015 | 0.003 | 0.003 | 2.703 | 0.37 | 0.58 | 0.43 | 0.36 | 0.31 | 0.4 |
| 8 | 79750813 | rs17506603 | -0.994 | -0.09 | -0.799 | -0.042 | 0.008 | 0.017 | 0.017 | 0 | 0.16 | 0.09 | 0.09 | 0.06 | 0.17 | 0.21 |
| 8 | 79753010 | rs4739140 | -1.443 | 0.233 | -1.686 | -0.58 | -0.098 | 0.21 | 0.21 | 2.173 | 0.62 | 0.98 | 0.72 | 0.9 | 0.63 | 0.57 |
| 8 | 79756515 | rs2369537 | -0.701 | 0.253 | -0.758 | -0.492 | -0.004 | 0.007 | 0.007 | 0 | 0.4 | 0.59 | 0.66 | 0.82 | 0.55 | 0.35 |
| 8 | 79758241 | rs4739111 | -0.85 | 0.181 | -0.301 | -0.102 | 0.018 | -0.016 | -0.016 | 1.008 | 0.35 | 0.53 | 0.61 | 0.72 | 0.45 | 0.28 |
| 8 | 79759499 | rs2010128 | -1.215 | 0.176 | -0.285 | 0.118 | 0.007 | 0.028 | 0.028 | 0.373 | 0.39 | 0.7 | 0.38 | 0.22 | 0.41 | 0.35 |
| 8 | 79766499 | rs11992182 | -0.674 | -0.153 | 0.064 | -0.125 | 0 | 0.001 | 0.001 | 0.611 | 0.25 | 0.22 | 0.22 | 0.15 | 0.29 | 0.18 |
| 8 | 79779026 | rs1596434 | -1.74 | -0.131 | 0 | 0.028 | 0.012 | 0.143 | 0.143 | 3.417 | 0.46 | 0.91 | 0.52 | 0.77 | 0.45 | 0.36 |
| 8 | 79790796 | rs1858120 | -1.542 | -0.182 | 0.875 | 0.121 | -0.006 | -0.019 | -0.019 | 1.556 | 0.39 | 0.5 | 0.47 | 0.7 | 0.43 | 0.34 |
| 8 | 79794041 | rs10808826 | -1.519 | -0.183 | 0.827 | 0.12 | -0.005 | -0.018 | -0.018 | 0 | 0.39 | 0.46 | 0.47 | 0.7 | 0.43 | 0.33 |
| 8 | 79802070 | rs6987092 | -1.017 | -0.21 | 1.109 | 0.129 | -0.002 | 0.118 | 0.118 | 0 | 0.2 | 0.39 | 0.31 | 0.55 | 0.22 | 0.17 |
| 8 | 79804155 | rs1443898 | -2.19 | -0.244 | 1.064 | 0.096 | 0.014 | 0.145 | 0.145 | 3.143 | 0.46 | 0.91 | 0.52 | 0.77 | 0.46 | 0.36 |
| 8 | 79805651 | rs12548956 | -0.727 | -0.241 | 1.095 | 0.098 | 0 | 0.054 | 0.054 | 0 | 0.18 | 0.24 | 0.29 | 0.55 | 0.22 | 0.17 |
| 8 | 79806660 | rs6993385 | -2.727 | -0.538 | 0.621 | -0.297 | -0.159 | 0.21 | 0.21 | 1.184 | 0.62 | 0.96 | 0.71 | 0.93 | 0.57 | 0.6 |
| 8 | 79808576 | rs1015028 | -2.038 | -0.789 | 1.123 | 0.035 | 0.019 | 0.086 | 0.086 | 2.379 | 0.44 | 0.85 | 0.52 | 0.7 | 0.46 | 0.34 |
| 8 | 79811685 | rs11996659 | -0.55 | -0.792 | 0.919 | -0.03 | -0.035 | -0.111 | -0.111 | 1.519 | 0.15 | 0.47 | 0.09 | 0 | 0.12 | 0.02 |
| 8 | 79812354 | rs10106730 | -0.741 | -0.292 | 0.866 | 0.375 | -0.011 | -0.015 | -0.015 | 0 | 0.21 | 0.31 | 0.18 | 0.15 | 0.21 | 0.17 |

| | | | | | | | | | | | | | | | | |
|---|---|---|---|---|---|---|---|---|---|---|---|---|---|---|---|---|
| 8 | 79820821 | rs9283993 | -1.33 | -0.09 | 0.516 | 0.532 | 0.005 | 0.03 | 0.03 | 0 | 0.17 | 0.24 | 0.31 | 0.65 | 0.18 | 0.3 |
| 8 | 79821298 | rs11782492 | -0.455 | -0.049 | -0.207 | 0.443 | 0.01 | 0.025 | 0.025 | 0 | 0.21 | 0.1 | 0.1 | 0.14 | 0.2 | 0.15 |
| 8 | 79821526 | rs10448016 | 0.147 | 0.004 | -0.346 | 0.501 | -0.037 | -0.016 | -0.016 | 0 | 0.08 | 0.01 | 0.23 | 0.55 | 0.12 | 0.16 |
| 8 | 79825547 | rs7000123 | -0.638 | -0.048 | -0.122 | 0.161 | -0.009 | -0.019 | -0.019 | 0.423 | 0.22 | 0.26 | 0.13 | 0.12 | 0.2 | 0.15 |
| 8 | 79831262 | rs6999801 | -1.236 | 0.181 | -0.43 | 0.146 | 0.01 | 0.046 | 0.046 | 0 | 0.38 | 0.73 | 0.46 | 0.77 | 0.36 | 0.42 |
| 8 | 79844006 | rs2120410 | -1.21 | 0.08 | -0.704 | 0.599 | 0.032 | -0.017 | -0.017 | 0 | 0.41 | 0.8 | 0.35 | 0.38 | 0.42 | 0.33 |
| 8 | 79844078 | rs4739119 | -0.723 | 0.079 | -0.585 | 0.799 | -0.004 | 0.01 | 0.01 | 0 | 0.07 | 0.1 | 0.12 | 0.19 | 0.1 | 0.05 |
| 8 | 79844115 | rs2067696 | -0.628 | -0.107 | 0.477 | 0.746 | 0.003 | 0.102 | 0.102 | 0 | 0.1 | 0.43 | 0.07 | 0.14 | 0.06 | 0.09 |
| 8 | 79845147 | rs6990279 | -0.706 | -0.16 | 0.979 | 0.752 | -0.001 | -0.011 | -0.011 | 0 | 0.12 | 0.16 | 0.06 | 0.07 | 0.17 | 0.07 |
| 8 | 79846184 | rs4739148 | -2.002 | -0.242 | 1.501 | 0.751 | -0.012 | 0.105 | 0.105 | 0 | 0.22 | 0.51 | 0.25 | 0.35 | 0.21 | 0.3 |
| 8 | 79846344 | rs11775912 | -0.578 | -0.248 | 1.343 | 0.46 | -0.013 | -0.018 | -0.018 | 0 | 0.12 | 0.19 | 0.1 | 0.09 | 0.11 | 0.2 |
| 8 | 79847288 | rs10098207 | -1.296 | -0.271 | 1.369 | 0.449 | -0.024 | -0.03 | -0.03 | 0.002 | 0.16 | 0.22 | 0.11 | 0.09 | 0.12 | 0.19 |
| 8 | 79847855 | rs925246 | -1.721 | -0.142 | 1.306 | 0.506 | -0.018 | -0.039 | -0.039 | 0.034 | 0.24 | 0.4 | 0.17 | 0.16 | 0.28 | 0.27 |
| 8 | 79856369 | rs6983483 | -1.825 | -0.175 | 1.229 | 0.825 | -0.004 | 0.005 | 0.005 | 0 | 0.26 | 0.4 | 0.23 | 0.27 | 0.24 | 0.27 |
| 8 | 79857156 | rs11781378 | -2.334 | -0.036 | 1.5 | 0.891 | -0.001 | -0.003 | -0.003 | 0 | 0.36 | 0.57 | 0.36 | 0.34 | 0.4 | 0.33 |
| 8 | 79860421 | rs17418058 | -2.071 | -0.056 | 1.595 | 0.908 | -0.003 | 0.007 | 0.007 | 0 | 0.29 | 0.25 | 0.21 | 0.17 | 0.35 | 0.3 |
| 8 | 79871922 | rs7003181 | -2.03 | -0.074 | 1.656 | 0.903 | -0.021 | 0.026 | 0.026 | 0 | 0.23 | 0.43 | 0.22 | 0.27 | 0.2 | 0.26 |
| 8 | 79873935 | rs7842502 | -2.679 | -0.232 | 0.782 | 0.246 | -0.042 | 0.17 | 0.17 | 0.044 | 0.59 | 0.82 | 0.59 | 0.94 | 0.57 | 0.54 |
| 8 | 79875067 | rs12680350 | 0.764 | 0.02 | 0.651 | 0.496 | 0 | 0.028 | 0.028 | 0 | 0.16 | 0.14 | 0.2 | 0.6 | 0.14 | 0.2 |
| 8 | 79875221 | rs12682560 | 1.681 | 0.026 | 0.541 | 1.174 | -0.016 | -0.063 | -0.063 | 0.469 | 0.11 | 0 | 0.04 | 0.2 | 0.07 | 0.12 |
| 8 | 79877645 | rs6988545 | 0.291 | 0.029 | 0.592 | 0.794 | 0.004 | -0.007 | -0.007 | 0 | 0.1 | 0.16 | 0.14 | 0.07 | 0.16 | 0.06 |
| 8 | 79881736 | rs17515867 | 0.459 | 0.011 | 0.525 | 0.805 | -0.008 | 0.019 | 0.019 | 0 | 0.12 | 0.05 | 0.05 | 0.05 | 0.14 | 0.07 |
| 8 | 79883523 | rs10464926 | 0.458 | 0.01 | 0.598 | 0.846 | 0 | 0.044 | 0.044 | 0 | 0.16 | 0.17 | 0.21 | 0.6 | 0.14 | 0.2 |
| 8 | 79883695 | rs10464927 | 0.458 | 0.01 | 0.598 | 0.846 | 0 | 0.044 | 0.044 | 0 | 0.16 | 0.17 | 0.21 | 0.6 | 0.14 | 0.2 |
| 8 | 79885746 | rs12056707 | 0.498 | 0.011 | 0.5 | 0.848 | 0 | 0.029 | 0.029 | 0 | 0.16 | 0.14 | 0.2 | 0.6 | 0.14 | 0.2 |
| 8 | 79888451 | rs7812820 | -1.918 | -0.158 | 0.441 | 0.661 | -0.001 | -0.002 | -0.002 | 0 | 0.47 | 0.4 | 0.43 | 0.77 | 0.52 | 0.53 |
| 8 | 79888705 | rs1834743 | -1.417 | -0.242 | 0.333 | 0.781 | -0.002 | 0.004 | 0.004 | 0.082 | 0.37 | 0.27 | 0.31 | 0.69 | 0.37 | 0.47 |
| 8 | 79892197 | rs10808828 | -0.29 | -0.103 | 0.417 | 1.016 | 0 | 0.018 | 0.018 | 0 | 0.21 | 0.18 | 0.23 | 0.59 | 0.19 | 0.24 |
| 8 | 79900591 | rs7845577 | 0.092 | -0.133 | 0.781 | 0.824 | 0.024 | -0.053 | -0.053 | 0 | 0.1 | 0.17 | 0.08 | 0 | 0.14 | 0.02 |
| 8 | 79906572 | rs2175085 | 1.191 | 0.775 | 0.091 | 0.341 | -0.082 | 0.046 | 0.046 | 0 | 0.13 | 0.01 | 0.16 | 0.04 | 0.24 | 0.12 |
| 8 | 79916893 | rs539254 | -1.058 | -0.745 | 1.401 | 1.135 | -0.01 | 0.039 | 0.039 | 0 | 0.28 | 0.36 | 0.26 | 0.55 | 0.24 | 0.22 |
| 8 | 79918459 | rs11987519 | -0.197 | -1.005 | 1.464 | 1.103 | -0.022 | 0.021 | 0.021 | 0 | 0.08 | 0.23 | 0.07 | 0.07 | 0.03 | 0.04 |
| 8 | 79924472 | rs683929 | -0.169 | -0.992 | 1.471 | 1.11 | -0.019 | 0.051 | 0.051 | 0 | 0.2 | 0.21 | 0.19 | 0.55 | 0.11 | 0.21 |
| 8 | 79954121 | rs503431 | -2.582 | -1.186 | 1.095 | 0.666 | -0.051 | 0.225 | 0.225 | 0.001 | 0.53 | 0.98 | 0.55 | 0.86 | 0.44 | 0.46 |
| 8 | 79971168 | rs675510 | -0.258 | -1.157 | 1.329 | 0.684 | -0.01 | -0.032 | -0.032 | 2.077 | 0.25 | 0.36 | 0.17 | 0.15 | 0.23 | 0.17 |
| 8 | 79973576 | rs17517571 | 0.444 | -1.179 | 1.303 | 0.695 | 0 | -0.005 | -0.005 | 0 | 0.11 | 0.09 | 0.1 | 0.12 | 0.12 | 0.06 |
| 8 | 79978105 | rs558228 | -1.735 | -1.209 | 1.504 | 0.709 | 0.028 | 0.031 | 0.031 | 0 | 0.42 | 0.66 | 0.44 | 0.74 | 0.33 | 0.42 |
| 8 | 79984111 | rs519121 | -1.796 | -1.207 | 1.506 | 0.675 | 0.042 | 0.08 | 0.08 | 0 | 0.42 | 0.7 | 0.44 | 0.82 | 0.32 | 0.44 |
| 8 | 79985408 | rs10504690 | 0.611 | -1.153 | 1.465 | 0.672 | -0.004 | 0.018 | 0.018 | 0.061 | 0.1 | 0.06 | 0.1 | 0.04 | 0.12 | 0.03 |
| 8 | 79991509 | rs16906486 | -0.841 | -1.155 | 1.594 | 0.671 | -0.014 | -0.044 | -0.044 | 0.257 | 0.18 | 0.25 | 0.16 | 0.04 | 0.15 | 0.05 |
| 8 | 80018703 | rs16906512 | -0.527 | -1.1 | 1.37 | 0.743 | -0.003 | 0.02 | 0.02 | 0.014 | 0.05 | 0.09 | 0.02 | 0.11 | 0.05 | 0.15 |
| 8 | 80022146 | rs5003440 | -2.468 | -1.157 | 1.15 | 0.667 | 0.03 | 0.178 | 0.178 | 0 | 0.52 | 0.77 | 0.55 | 0.87 | 0.45 | 0.48 |
| 8 | 80045426 | rs2222361 | -0.268 | -1.144 | 0.933 | 0.714 | 0.013 | 0.04 | 0.04 | 0.617 | 0.15 | 0.07 | 0.13 | 0.04 | 0.13 | 0.05 |
| 8 | 80058894 | rs12682020 | 0.315 | -1.218 | 1.275 | 1.137 | -0.02 | 0.006 | 0.006 | 0 | 0.18 | 0.22 | 0.1 | 0.2 | 0.1 | 0.16 |
| 8 | 80060941 | rs10504696 | -1.401 | -1.193 | 0.747 | 1.069 | 0.076 | -0.006 | -0.006 | 0 | 0.33 | 0.83 | 0.22 | 0.31 | 0.18 | 0.33 |
| 8 | 80062983 | rs7000109 | -2.87 | -1.121 | 0.599 | 0.86 | -0.074 | 0.207 | 0.207 | 0.185 | 0.54 | 0.97 | 0.55 | 0.86 | 0.45 | 0.47 |
| 8 | 80077927 | rs16906568 | -1.059 | -1.081 | 1.004 | 1.071 | 0 | 0.005 | 0.005 | 0 | 0.22 | 0.22 | 0.34 | 0.62 | 0.27 | 0.15 |
| 8 | 80082078 | rs16906578 | -0.078 | -1.099 | 1.1 | 1.19 | 0.007 | 0.068 | 0.068 | 0 | 0.13 | 0.15 | 0.25 | 0.56 | 0.16 | 0.11 |
| 8 | 80095928 | rs10108874 | -0.458 | -1.267 | 1.349 | 0.926 | -0.073 | 0.026 | 0.026 | 0 | 0.22 | 0.41 | 0.13 | 0.23 | 0.09 | 0.16 |
| 8 | 80097636 | rs1551712 | -1.195 | -1.215 | 0.721 | 0.935 | -0.017 | -0.003 | -0.003 | 1.866 | 0.29 | 0.66 | 0.19 | 0.26 | 0.17 | 0.31 |
| 8 | 80111894 | rs16906594 | -1.403 | -0.679 | 0.521 | 1.392 | 0.001 | 0.013 | 0.013 | 0 | 0.2 | 0.17 | 0.22 | 0.49 | 0.19 | 0.21 |
| 8 | 80123611 | rs6473142 | -0.525 | -0.747 | 0.558 | 0.852 | -0.005 | -0.039 | -0.039 | 0.002 | 0.11 | 0.1 | 0.07 | 0.01 | 0.09 | 0.02 |
| 8 | 80131103 | rs1227646 | -1.854 | -1.246 | 0.288 | 0.763 | 0.068 | 0.099 | 0.099 | 0.66 | 0.43 | 0.85 | 0.48 | 0.78 | 0.34 | 0.41 |
| 8 | 80141625 | rs1227630 | -2.287 | -1.174 | 0.106 | 0.618 | -0.041 | 0.133 | 0.133 | 1.771 | 0.5 | 0.97 | 0.55 | 0.78 | 0.43 | 0.41 |
| 8 | 80145056 | rs1227666 | -1.166 | -1.422 | 1.161 | 0.886 | -0.004 | 0.029 | 0.029 | 0 | 0.29 | 0.34 | 0.3 | 0.61 | 0.27 | 0.34 |
| 8 | 80155734 | rs1227632 | -1.154 | -1.42 | 1.181 | 0.887 | -0.004 | 0.032 | 0.032 | 0 | 0.28 | 0.34 | 0.3 | 0.61 | 0.27 | 0.34 |
| 8 | 80156474 | rs1227634 | -1.71 | -1.455 | 1.823 | 0.895 | -0.017 | -0.001 | -0.001 | 0 | 0.36 | 0.58 | 0.34 | 0.69 | 0.3 | 0.38 |
| 8 | 80161352 | rs1915554 | 0.405 | -1.449 | 1.737 | 0.897 | -0.034 | 0.027 | 0.027 | 0.849 | 0.08 | 0.23 | 0.04 | 0.08 | 0.03 | 0.05 |
| 8 | 80166841 | rs1234900 | -0.073 | -1.484 | 1.824 | 1.178 | 0.001 | -0.003 | -0.003 | 0.227 | 0.17 | 0.08 | 0.12 | 0.21 | 0.18 | 0.16 |
| 8 | 80174864 | rs1227655 | -1.165 | -1.473 | 1.919 | 1.115 | 0.002 | -0.033 | -0.033 | 0 | 0.28 | 0.17 | 0.28 | 0.61 | 0.27 | 0.34 |
| 8 | 80186148 | rs16906665 | 0.554 | -1.444 | 1.931 | 1.118 | -0.001 | 0.005 | 0.005 | 0 | 0.05 | 0.06 | 0.12 | 0.09 | 0.05 | 0.03 |
| 8 | 80186751 | rs1227649 | -2.427 | -1.489 | 1.871 | 1.108 | 0.056 | 0.102 | 0.102 | 0.56 | 0.44 | 0.85 | 0.48 | 0.78 | 0.35 | 0.41 |
| 8 | 80197250 | rs1870575 | -1.314 | -1.496 | 1.91 | 1.114 | -0.003 | 0.025 | 0.025 | 0 | 0.29 | 0.33 | 0.29 | 0.61 | 0.27 | 0.34 |
| 8 | 80199903 | rs7817332 | -2.445 | -1.487 | 1.816 | 1.092 | 0.056 | 0.105 | 0.105 | 0.637 | 0.44 | 0.86 | 0.48 | 0.79 | 0.35 | 0.41 |
| 8 | 80202384 | rs10504700 | 0.233 | -1.497 | 1.892 | 1.083 | -0.035 | 0.032 | 0.032 | 0.716 | 0.08 | 0.24 | 0.05 | 0.09 | 0.03 | 0.05 |
| 8 | 80211249 | rs10504702 | -0.505 | -1.514 | 2.168 | 0.938 | -0.01 | 0.092 | 0.092 | 0 | 0.12 | 0.22 | 0.16 | 0.41 | 0.08 | 0.21 |
| 8 | 80214806 | rs7829769 | -0.43 | -1.526 | 2.277 | 0.938 | 0.036 | 0.092 | 0.092 | 0 | 0.07 | 0.27 | 0.14 | 0.09 | 0.05 | 0.03 |
| 8 | 80220861 | rs1531346 | -1.143 | -1.468 | 2.243 | 0.939 | -0.025 | 0.057 | 0.057 | 0 | 0.17 | 0.31 | 0.2 | 0.41 | 0.1 | 0.22 |
| 8 | 80227400 | rs1460162 | -0.527 | -1.426 | 2.118 | 0.961 | -0.01 | 0.094 | 0.094 | 0 | 0.11 | 0.21 | 0.16 | 0.41 | 0.08 | 0.21 |

| | | | | | | | | | | | | | | | | |
|---|---|---|---|---|---|---|---|---|---|---|---|---|---|---|---|---|
| 8 | 80235462 | rs10504705 | -0.932 | -1.607 | 1.982 | 0.812 | -0.019 | -0.001 | -0.001 | 0 | 0.27 | 0.37 | 0.17 | 0.28 | 0.21 | 0.21 |
| 8 | 80237893 | rs1380634 | -1.785 | -1.581 | 1.802 | 0.807 | -0.021 | -0.002 | -0.002 | 0 | 0.38 | 0.53 | 0.33 | 0.69 | 0.29 | 0.42 |
| 8 | 80239298 | rs1350681 | -1.782 | -1.574 | 1.811 | 0.815 | -0.022 | -0.001 | -0.001 | 0 | 0.38 | 0.53 | 0.33 | 0.69 | 0.29 | 0.42 |
| 8 | 80241359 | rs16906746 | 0.144 | -1.576 | 1.962 | 0.861 | -0.005 | 0.074 | 0.074 | 0 | 0.1 | 0.15 | 0.16 | 0.41 | 0.08 | 0.21 |
| 8 | 80243314 | rs1460168 | -1.628 | -1.516 | 1.74 | 0.796 | -0.022 | -0.001 | -0.001 | 0 | 0.38 | 0.53 | 0.33 | 0.69 | 0.29 | 0.42 |
| 8 | 80246590 | rs16906775 | 1.569 | -1.423 | 1.443 | 0.724 | 0.016 | -0.033 | -0.033 | 0 | 0.06 | 0.13 | 0.07 | 0 | 0.08 | 0.01 |
| 8 | 80257277 | rs4074500 | -1.925 | -1.447 | 1.164 | 0.771 | 0.04 | 0.026 | 0.026 | 0 | 0.45 | 0.8 | 0.41 | 0.68 | 0.38 | 0.43 |
| 8 | 80259189 | rs11778785 | -0.708 | -1.457 | 0.868 | 0.821 | 0.004 | -0.004 | -0.004 | 0 | 0.23 | 0.15 | 0.13 | 0.27 | 0.19 | 0.2 |
| 8 | 80263178 | rs4739607 | -1.885 | -1.496 | 0.513 | 0.81 | 0.018 | -0.004 | -0.004 | 0.001 | 0.43 | 0.88 | 0.41 | 0.68 | 0.36 | 0.41 |
| 8 | 80264615 | rs4074079 | 0.318 | -1.686 | 1.085 | 1.08 | 0.005 | 0.117 | 0.117 | 0 | 0.08 | 0.15 | 0.16 | 0.41 | 0.08 | 0.21 |
| 8 | 80266553 | rs7462051 | -1.079 | -1.791 | 0.706 | 0.992 | 0.011 | -0.021 | -0.021 | 0 | 0.27 | 0.2 | 0.25 | 0.37 | 0.24 | 0.23 |
| 8 | 80272544 | rs6999948 | -2.2 | -1.748 | 0.567 | 0.953 | 0.072 | 0.123 | 0.123 | 0.252 | 0.47 | 0.89 | 0.51 | 0.77 | 0.37 | 0.45 |
| 8 | 80274461 | rs9643438 | -1.127 | -1.57 | 0.866 | 1.022 | -0.009 | 0.151 | 0.151 | 0.418 | 0.14 | 0.59 | 0.25 | 0.41 | 0.13 | 0.19 |
| 8 | 80278205 | rs10957934 | -0.901 | -1.49 | 0.385 | 1.002 | -0.001 | 0.009 | 0.009 | 0 | 0.24 | 0.24 | 0.25 | 0.36 | 0.24 | 0.24 |
| 8 | 80279515 | rs11985579 | -0.096 | -1.477 | 0.069 | 0.888 | -0.003 | -0.021 | -0.021 | 0 | 0.18 | 0.09 | 0.13 | 0.27 | 0.19 | 0.21 |
| 8 | 80290562 | rs12678425 | -1.666 | -1.637 | -0.09 | 0.554 | 0.07 | 0.086 | 0.086 | 0 | 0.46 | 0.85 | 0.5 | 0.76 | 0.37 | 0.44 |
| 8 | 80297006 | rs7462145 | -0.995 | -1.736 | -0.084 | 0.482 | -0.011 | -0.015 | -0.015 | 0 | 0.37 | 0.39 | 0.42 | 0.76 | 0.34 | 0.48 |
| 8 | 80297198 | rs12545192 | 0.054 | -1.519 | 0.197 | 0.855 | 0 | 0.044 | 0.044 | 0 | 0.1 | 0.09 | 0.17 | 0.42 | 0.12 | 0.21 |
| 8 | 80305930 | rs6998419 | -0.805 | -1.627 | 0.355 | 0.028 | -0.066 | -0.14 | -0.14 | 0.009 | 0.19 | 0.55 | 0.17 | 0 | 0.1 | 0.02 |
| 8 | 80308664 | rs10086740 | 1.309 | -1.461 | -0.948 | -0.012 | -0.026 | -0.056 | -0.056 | 0 | 0.23 | 0.01 | 0.28 | 0.23 | 0.21 | 0.34 |
| 8 | 80320484 | rs11991397 | -0.839 | -1.801 | 0.422 | -0.032 | -0.089 | 0.002 | 0.002 | 0.019 | 0.24 | 0.5 | 0.13 | 0.21 | 0.13 | 0.15 |
| 8 | 80328475 | rs7465277 | 0.754 | -1.984 | 0.53 | -0.194 | -0.01 | -0.023 | -0.023 | 0.002 | 0.13 | 0.18 | 0.1 | 0.05 | 0.08 | 0.11 |
| 8 | 80331745 | rs7465051 | 1.036 | -1.981 | 0.856 | -0.142 | -0.019 | -0.011 | -0.011 | 0 | 0.09 | 0.14 | 0.04 | 0.07 | 0.04 | 0.09 |
| 8 | 80349311 | rs7461233 | -0.004 | -1.542 | 0.834 | -0.192 | 0.02 | 0.079 | 0.079 | 0 | 0.06 | 0.13 | 0.07 | 0.14 | 0.08 | 0.05 |
| 8 | 80351787 | rs7836491 | 0.794 | -1.634 | 0.404 | -0.18 | 0.003 | 0.005 | 0.005 | 0 | 0.12 | 0.08 | 0.06 | 0.11 | 0.12 | 0.2 |
| 8 | 80352815 | rs7012426 | -1.134 | -1.282 | 0.998 | 0.069 | -0.082 | -0.001 | -0.001 | 0 | 0.24 | 0.54 | 0.14 | 0.21 | 0.13 | 0.15 |
| 8 | 80358313 | rs7460206 | -1.805 | -1.289 | 0.828 | 0.073 | 0.067 | 0.034 | 0.034 | 0.822 | 0.39 | 0.82 | 0.28 | 0.26 | 0.26 | 0.27 |
| 8 | 80364186 | rs9650266 | 0.985 | -1.142 | 0.761 | 0.115 | 0.001 | 0.047 | 0.047 | 1.562 | 0.09 | 0.01 | 0.1 | 0 | 0.08 | 0.02 |
| 8 | 80364499 | rs6473158 | -0.957 | -1.069 | 0.956 | 0.164 | -0.033 | -0.001 | -0.001 | 0 | 0.26 | 0.54 | 0.23 | 0.21 | 0.2 | 0.16 |
| 8 | 80380011 | rs10089772 | -0.945 | -1.297 | 0.57 | 0.306 | 0.001 | 0.001 | 0.001 | 0.144 | 0.36 | 0.61 | 0.24 | 0.28 | 0.2 | 0.28 |
| 8 | 80386461 | rs7836445 | -1.166 | -1.037 | 0.721 | 0.353 | -0.038 | -0.008 | -0.008 | 0 | 0.32 | 0.51 | 0.26 | 0.27 | 0.26 | 0.33 |
| 8 | 80387861 | rs11777412 | -0.03 | -0.395 | 0.757 | 0.466 | 0.006 | 0.002 | 0.002 | 0 | 0.09 | 0.19 | 0.09 | 0.11 | 0.11 | 0.14 |
| 8 | 80423136 | rs10092479 | -0.975 | -0.213 | 0.128 | 0.357 | -0.012 | -0.045 | -0.045 | 0 | 0.57 | 0.82 | 0.48 | 0.46 | 0.48 | 0.59 |
| 8 | 80430937 | rs903583 | -1.131 | 0.136 | 0.135 | 0.286 | -0.051 | -0.043 | -0.043 | 0.211 | 0.59 | 0.79 | 0.49 | 0.46 | 0.56 | 0.6 |
| 8 | 80431800 | rs6992970 | 0.284 | 0.114 | 0.358 | 0.257 | 0.007 | -0.006 | -0.006 | 0 | 0.11 | 0.15 | 0.09 | 0.08 | 0.19 | 0.18 |
| 8 | 80437943 | rs16906865 | 0.968 | -0.09 | 0.766 | 0.362 | -0.018 | 0.011 | 0.011 | 0 | 0.08 | 0.17 | 0.1 | 0.12 | 0.05 | 0.1 |
| 8 | 80459969 | rs16906894 | 1.178 | -0.085 | 0.687 | 0.365 | -0.006 | 0.001 | 0.001 | 0 | 0.08 | 0.08 | 0.09 | 0.12 | 0.05 | 0.1 |
| 8 | 80498998 | rs1157435 | -0.536 | -0.05 | 0.67 | 0.638 | 0.005 | 0.013 | 0.013 | 0 | 0.51 | 0.79 | 0.5 | 0.39 | 0.52 | 0.49 |
| 8 | 80502285 | rs6473170 | -1.655 | 0.508 | 0.265 | 0.952 | -0.055 | 0.026 | 0.026 | 3.093 | 0.69 | 0.92 | 0.76 | 0.73 | 0.67 | 0.8 |
| 8 | 80505098 | rs7007698 | -0.291 | 0.515 | 0.334 | 0.92 | 0.005 | 0.013 | 0.013 | 0 | 0.51 | 0.79 | 0.5 | 0.39 | 0.53 | 0.49 |
| 8 | 80510544 | rs17451666 | 1.237 | 0.389 | -0.252 | 0.581 | -0.033 | 0.073 | 0.073 | 0 | 0.13 | 0.01 | 0.07 | 0 | 0.14 | 0.07 |
| 8 | 80519541 | rs12681591 | 0.568 | 0.269 | -0.158 | 0.653 | -0.003 | 0.058 | 0.058 | 0.392 | 0.33 | 0.17 | 0.31 | 0.17 | 0.36 | 0.4 |
| 8 | 80519993 | rs4739846 | -0.487 | 0.211 | 0.605 | 0.749 | 0.003 | 0.004 | 0.004 | 0 | 0.48 | 0.48 | 0.47 | 0.39 | 0.52 | 0.49 |
| 8 | 80521603 | rs3812459 | -1.094 | 0.175 | 0.662 | 0.718 | -0.02 | -0.009 | -0.009 | 0 | 0.58 | 0.73 | 0.57 | 0.52 | 0.57 | 0.6 |
| 8 | 80522605 | rs3812460 | 0.422 | -0.018 | 0.515 | 0.822 | -0.002 | 0.028 | 0.028 | 0.511 | 0.33 | 0.28 | 0.34 | 0.19 | 0.37 | 0.41 |
| 8 | 80523536 | rs3829028 | 0.801 | 0.319 | 0.172 | 1.064 | -0.03 | -0.049 | -0.049 | 0 | 0.1 | 0.04 | 0.17 | 0.22 | 0.09 | 0.19 |
| 8 | 80528973 | rs7013510 | -0.295 | 0.187 | 0.299 | 0.868 | 0.005 | 0.035 | 0.035 | 0.028 | 0.48 | 0.42 | 0.4 | 0.18 | 0.5 | 0.49 |
| 8 | 80534144 | rs11787508 | -1.797 | 0.154 | 0.309 | 0.871 | -0.041 | 0.053 | 0.053 | 2.413 | 0.68 | 0.86 | 0.67 | 0.74 | 0.63 | 0.8 |
| 8 | 80535407 | rs4739857 | 0.44 | 0.294 | 0.358 | 0.723 | 0 | 0 | 0 | 0.352 | 0.34 | 0.33 | 0.32 | 0.18 | 0.35 | 0.42 |
| 8 | 80536776 | rs7846029 | 0.584 | 0.528 | 0.216 | 0.555 | -0.004 | 0.008 | 0.008 | 0 | 0.16 | 0.11 | 0.09 | 0 | 0.17 | 0.11 |
| 8 | 80538524 | rs1390248 | -0.878 | 0.496 | 0.504 | 0.906 | 0.014 | 0.07 | 0.07 | 0.558 | 0.26 | 0.43 | 0.33 | 0.42 | 0.26 | 0.28 |
| 8 | 80539224 | rs10112708 | -1.306 | 0.495 | 0.501 | 0.964 | -0.013 | 0.01 | 0.01 | 1.387 | 0.61 | 0.81 | 0.66 | 0.57 | 0.61 | 0.7 |
| 8 | 80544154 | rs16907010 | -0.218 | 0.552 | 0.422 | 1.004 | 0.026 | 0.204 | 0.204 | 2.607 | 0.12 | 0.31 | 0.24 | 0.42 | 0.11 | 0.2 |
| 8 | 80546219 | rs6473173 | -1.263 | 0.502 | 0.171 | 0.954 | -0.004 | 0.007 | 0.007 | 1.311 | 0.6 | 0.72 | 0.65 | 0.56 | 0.61 | 0.7 |
| 8 | 80553068 | rs2029818 | -0.469 | 0.596 | 0.073 | 0.504 | 0.006 | 0.07 | 0.07 | 0.123 | 0.48 | 0.34 | 0.39 | 0.15 | 0.51 | 0.5 |
| 8 | 80553916 | rs16907025 | -0.889 | 0.989 | 0.03 | 0.796 | 0.001 | -0.006 | -0.006 | 0.462 | 0.26 | 0.33 | 0.27 | 0.19 | 0.23 | 0.27 |
| 8 | 80562259 | rs2979715 | 0.677 | 1.359 | -0.026 | 1.174 | 0.016 | 0.106 | 0.106 | 0 | 0.86 | 0.75 | 0.65 | 0.62 | 0.81 | 0.74 |
| 8 | 80566038 | rs12676762 | 0.844 | 1.182 | -0.654 | 0.656 | 0.012 | -0.013 | -0.013 | 0 | 0.22 | 0.03 | 0.18 | 0.13 | 0.16 | 0.23 |
| 8 | 80568365 | rs13255802 | -0.099 | 1.442 | -0.505 | 0.599 | -0.007 | 0.006 | 0.006 | 0 | 0.12 | 0.07 | 0.09 | 0.08 | 0.15 | 0.14 |
| 8 | 80569874 | rs3793382 | 0.968 | 1.783 | -0.398 | 0.607 | -0.003 | -0.001 | -0.001 | 0 | 0.06 | 0.05 | 0.03 | 0.03 | 0.1 | 0.15 |
| 8 | 80571630 | rs7838627 | -1.212 | 1.897 | -0.079 | 0.648 | 0.019 | -0.017 | -0.017 | 0 | 0.19 | 0.3 | 0.13 | 0.13 | 0.21 | 0.17 |
| 8 | 80571728 | rs4739895 | 0.376 | 1.855 | -0.535 | 0.878 | -0.001 | 0.002 | 0.002 | 0 | 0.06 | 0.07 | 0.16 | 0.12 | 0.04 | 0.16 |
| 8 | 80572425 | rs4739651 | 0.068 | 2.062 | -0.702 | 0.767 | 0.01 | 0.09 | 0.09 | 0 | 0.86 | 0.8 | 0.62 | 0.43 | 0.81 | 0.85 |
| 8 | 80574314 | rs2120723 | 0.94 | 1.951 | -0.965 | 0.306 | -0.034 | 0.203 | 0.203 | 2.016 | 0.45 | 0.19 | 0.3 | 0.11 | 0.48 | 0.3 |
| 8 | 80582149 | rs1867496 | 0.383 | 1.96 | -1.026 | 0.341 | -0.026 | 0.037 | 0.037 | 0 | 0.07 | 0.01 | 0.06 | 0 | 0.12 | 0.04 |
| 8 | 80589219 | rs4739907 | 0.241 | 1.941 | -0.783 | 0.429 | -0.016 | -0.001 | -0.001 | 0 | 0.35 | 0.47 | 0.27 | 0.29 | 0.28 | 0.48 |
| 8 | 80590137 | rs2461035 | 1.29 | 1.854 | -1.177 | 0.33 | -0.014 | 0.234 | 0.234 | 0.487 | 0.34 | 0.05 | 0.19 | 0.01 | 0.31 | 0.18 |
| 8 | 80597253 | rs2467753 | -0.038 | 2.036 | -1.011 | 0.773 | 0.015 | 0.067 | 0.067 | 0 | 0.93 | 0.9 | 0.82 | 0.73 | 0.84 | 0.88 |
| 8 | 80597708 | rs2467754 | 1.074 | 1.44 | -1.169 | 0.28 | 0.02 | 0.166 | 0.166 | 0.535 | 0.35 | 0.12 | 0.2 | 0.01 | 0.31 | 0.18 |

| | | | | | | | | | | | | | | | | |
|---|---|---|---|---|---|---|---|---|---|---|---|---|---|---|---|---|
| 8 | 80600394 | rs1444554 | -0.412 | 1.548 | -0.372 | 1.484 | 0.006 | 0.209 | 0.209 | 0.083 | 0.09 | 0.17 | 0.28 | 0.55 | 0.07 | 0.07 |
| 8 | 80601680 | rs2956253 | -1.443 | 1.787 | -0.028 | 1.196 | 0.02 | 0.092 | 0.092 | 0.351 | 0.19 | 0.38 | 0.35 | 0.4 | 0.2 | 0.12 |
| 8 | 80610888 | rs1879561 | -0.472 | 1.736 | -0.395 | 1.083 | -0.01 | -0.004 | -0.004 | 0 | 0.89 | 0.92 | 0.88 | 0.86 | 0.85 | 0.73 |
| 8 | 80612714 | rs2920942 | -0.492 | 2.135 | -0.208 | 0.96 | 0.054 | 0.05 | 0.05 | 0 | 0.24 | 0.39 | 0.45 | 0.67 | 0.33 | 0.27 |
| 8 | 80614400 | rs2440629 | 1.69 | 1.86 | -0.293 | 0.76 | 0.013 | 0.071 | 0.071 | 0 | 0.51 | 0.33 | 0.29 | 0.17 | 0.43 | 0.38 |
| 8 | 80619089 | rs2947540 | 0.967 | 1.755 | 0.176 | 1.408 | -0.011 | 0.084 | 0.084 | 0.18 | 0.07 | 0.09 | 0.21 | 0.35 | 0.04 | 0.02 |
| 8 | 80627249 | rs16907106 | 1.056 | 1.78 | 0.086 | 1.229 | -0.002 | 0.026 | 0.026 | 0.019 | 0.06 | 0.08 | 0.03 | 0.11 | 0.04 | 0.11 |
| 8 | 80630850 | rs1348558 | 0.084 | 1.94 | -0.054 | 1.096 | 0.059 | 0.084 | 0.084 | 0 | 0.17 | 0.31 | 0.22 | 0.3 | 0.28 | 0.24 |
| 8 | 80633975 | rs2920947 | -1.496 | 1.437 | -0.42 | 1.204 | -0.019 | 0.063 | 0.063 | 0 | 0.28 | 0.4 | 0.35 | 0.53 | 0.21 | 0.23 |
| 8 | 80639886 | rs1348559 | -0.082 | 1.722 | -0.135 | 1.314 | 0.001 | 0.001 | 0.001 | 0 | 0.72 | 0.58 | 0.75 | 0.77 | 0.67 | 0.63 |
| 8 | 80640248 | rs1444549 | 0.324 | 1.74 | -0.819 | 1.326 | 0.008 | -0.001 | -0.001 | 0.144 | 0.68 | 0.46 | 0.72 | 0.73 | 0.61 | 0.59 |
| 8 | 80643439 | rs4433192 | 0.317 | 1.832 | -0.87 | 1.24 | 0.005 | -0.001 | -0.001 | 0 | 0.06 | 0.06 | 0.09 | 0.08 | 0.15 | 0.08 |
| 8 | 80649173 | rs2979708 | 1.299 | 1.638 | -0.957 | 1.304 | -0.008 | 0.09 | 0.09 | 0.998 | 0.51 | 0.12 | 0.32 | 0.32 | 0.4 | 0.46 |
| 8 | 80652442 | rs2979710 | -0.757 | 1.439 | -0.405 | 1.652 | -0.005 | 0.042 | 0.042 | 0.019 | 0.15 | 0.2 | 0.31 | 0.37 | 0.13 | 0.1 |
| 8 | 80656206 | rs962695 | -1.095 | 1.249 | -0.476 | 1.56 | 0.011 | 0.042 | 0.042 | 0 | 0.24 | 0.33 | 0.42 | 0.47 | 0.28 | 0.2 |
| 8 | 80658272 | rs7830221 | -0.611 | 0.968 | -0.779 | 1.698 | -0.001 | 0.005 | 0.005 | 0 | 0.11 | 0.07 | 0.06 | 0.25 | 0.12 | 0.1 |
| 8 | 80658578 | rs2467781 | -1.026 | 0.333 | -0.748 | 1.576 | 0.002 | 0.002 | 0.002 | 0.366 | 0.38 | 0.33 | 0.39 | 0.71 | 0.33 | 0.21 |
| 8 | 80660497 | rs2467778 | -0.107 | 0.33 | -0.68 | 1.612 | 0.027 | -0.038 | -0.038 | 0 | 0.26 | 0.18 | 0.31 | 0.47 | 0.2 | 0.12 |
| 8 | 80663308 | rs2461043 | -0.968 | -0.226 | -0.622 | 0.669 | -0.001 | 0 | 0 | 0.362 | 0.61 | 0.58 | 0.53 | 0.71 | 0.51 | 0.52 |
| 8 | 80664160 | rs6473176 | 0.743 | -0.527 | -0.476 | 0.38 | -0.021 | -0.096 | -0.096 | 0.131 | 0.18 | 0.33 | 0.13 | 0 | 0.17 | 0.26 |
| 8 | 80669115 | rs1444547 | -1.516 | -0.736 | -0.83 | 0.161 | 0.02 | -0.014 | -0.014 | 0 | 0.71 | 0.66 | 0.59 | 0.84 | 0.6 | 0.68 |
| 8 | 80669734 | rs7843886 | 0.882 | -0.93 | -0.912 | 0.108 | -0.029 | -0.107 | -0.107 | 0 | 0.2 | 0.36 | 0.15 | 0.01 | 0.17 | 0.3 |
| 8 | 80674165 | rs6473177 | 1.151 | -0.894 | -1.323 | 0.441 | 0.003 | -0.006 | -0.006 | 0 | 0.09 | 0.1 | 0.06 | 0.19 | 0.09 | 0.14 |
| 8 | 80674788 | rs7004035 | 0.633 | -0.817 | -1.526 | 0.618 | -0.035 | -0.069 | -0.069 | 0 | 0.14 | 0.01 | 0.13 | 0.23 | 0.18 | 0.13 |
| 8 | 80675246 | rs2467774 | 1.179 | -0.949 | -1.508 | 0.628 | 0.014 | -0.102 | -0.102 | 1.071 | 0.24 | 0.02 | 0.25 | 0.41 | 0.19 | 0.1 |
| 8 | 80676529 | rs1046472 | -0.816 | -0.363 | -1.885 | 0.095 | -0.086 | -0.073 | -0.073 | 0 | 0.77 | 0.99 | 0.84 | 0.78 | 0.72 | 0.77 |
| 8 | 80678865 | rs960978 | 0.817 | 0.092 | -1.539 | -0.042 | 0.048 | -0.007 | -0.007 | 0.463 | 0.22 | 0.35 | 0.38 | 0.2 | 0.28 | 0.23 |
| 8 | 80683321 | rs2467779 | 1.292 | 0.33 | -1.534 | 0.297 | -0.004 | 0.005 | 0.005 | 0 | 0.21 | 0.03 | 0.18 | 0.22 | 0.2 | 0.08 |
| 8 | 80683922 | rs2920950 | 0.977 | 1.158 | -1.066 | 0.368 | 0.132 | 0.016 | 0.016 | 0 | 0.11 | 0.29 | 0.22 | 0.11 | 0.21 | 0.13 |
| 8 | 80691801 | rs2467777 | -0.267 | 0.461 | -1.801 | 0.547 | -0.055 | -0.076 | -0.076 | 0 | 0.73 | 0.97 | 0.82 | 0.68 | 0.71 | 0.8 |
| 8 | 80699369 | rs4739987 | -0.084 | 0.512 | -1.181 | -0.251 | -0.007 | -0.085 | -0.085 | 0 | 0.38 | 0.52 | 0.47 | 0.1 | 0.36 | 0.52 |
| 8 | 80702099 | rs7819714 | -0.376 | 0.34 | -0.756 | -0.093 | 0.001 | 0 | 0 | 0 | 0.31 | 0.65 | 0.48 | 0.27 | 0.31 | 0.37 |
| 8 | 80705936 | rs7838496 | 0.715 | 0.325 | -0.72 | -0.078 | 0.01 | 0.079 | 0.079 | 0 | 0.14 | 0.34 | 0.15 | 0.18 | 0.12 | 0.18 |
| 8 | 80706953 | rs2920949 | 0.5 | 0.521 | -0.939 | 0.023 | 0.001 | -0.004 | -0.004 | 0 | 0.39 | 0.41 | 0.49 | 0.37 | 0.38 | 0.25 |
| 8 | 80708610 | rs2467796 | -0.292 | 0.608 | -0.577 | 0.061 | 0.013 | 0.001 | 0.001 | 0 | 0.6 | 0.53 | 0.51 | 0.57 | 0.56 | 0.66 |
| 8 | 80710881 | rs2440655 | -1.781 | 0.53 | -1.107 | -0.061 | -0.053 | -0.109 | -0.109 | 0 | 0.82 | 0.99 | 0.87 | 0.67 | 0.8 | 0.85 |
| 8 | 80711103 | rs2467798 | 0.903 | 0.487 | -1.066 | -0.088 | 0.039 | 0.05 | 0.05 | 0 | 0.42 | 0.29 | 0.3 | 0.27 | 0.34 | 0.39 |
| 8 | 80712063 | rs2467799 | 2.15 | 0.05 | -1.066 | -0.021 | 0.007 | -0.013 | -0.013 | 0.454 | 0.21 | 0.09 | 0.17 | 0.27 | 0.19 | 0.06 |
| 8 | 80712375 | rs13270957 | 0.963 | -0.015 | -0.901 | 0.02 | -0.001 | 0.003 | 0.003 | 0 | 0.13 | 0.17 | 0.14 | 0.18 | 0.11 | 0.18 |
| 8 | 80714713 | rs2247008 | -0.489 | -0.002 | -0.81 | -0.093 | -0.024 | -0.053 | -0.053 | 0 | 0.72 | 0.87 | 0.77 | 0.55 | 0.66 | 0.76 |
| 8 | 80714935 | rs2278667 | 0.071 | 0.042 | -0.267 | -0.246 | -0.012 | 0.102 | 0.102 | 0 | 0.13 | 0.35 | 0.14 | 0.18 | 0.09 | 0.16 |
| 8 | 80719847 | rs4329312 | -0.196 | 0.103 | -0.484 | -0.35 | 0.009 | 0.032 | 0.032 | 1.807 | 0.46 | 0.71 | 0.58 | 0.32 | 0.45 | 0.65 |
| 8 | 80728052 | rs1838440 | -0.796 | 0.04 | -0.481 | -0.281 | -0.016 | -0.046 | -0.046 | 0 | 0.55 | 0.89 | 0.67 | 0.45 | 0.57 | 0.74 |
| 8 | 80729805 | rs12547549 | 0.919 | 0.056 | -0.498 | -0.272 | 0.038 | 0.04 | 0.04 | 0 | 0.08 | 0.11 | 0.09 | 0.14 | 0.13 | 0.09 |
| 8 | 80733475 | rs2457400 | 1.134 | -0.059 | -0.835 | -0.247 | -0.025 | 0.087 | 0.087 | 2.119 | 0.33 | 0.01 | 0.14 | 0.14 | 0.27 | 0.11 |
| 8 | 80733696 | rs10110615 | 0.834 | -0.173 | -0.781 | -0.358 | 0.015 | -0.026 | -0.026 | 0 | 0.21 | 0.3 | 0.33 | 0.15 | 0.26 | 0.29 |
| 8 | 80735073 | rs1396856 | -0.682 | 0.036 | -1.156 | -0.168 | -0.005 | -0.034 | -0.034 | 0 | 0.55 | 0.85 | 0.65 | 0.45 | 0.57 | 0.74 |
| 8 | 80739345 | rs2467769 | -1.227 | 0.594 | -0.678 | 0.426 | -0.043 | -0.077 | -0.077 | 0 | 0.86 | 0.97 | 0.8 | 0.59 | 0.79 | 0.84 |
| 8 | 80740809 | rs2467768 | -0.335 | 0.478 | -0.105 | 0.385 | 0.065 | 0.153 | 0.153 | 0 | 0.8 | 0.65 | 0.73 | 0.45 | 0.72 | 0.76 |
| 8 | 80743270 | rs1396858 | 0.198 | -0.005 | -0.262 | 0.065 | -0.017 | 0 | 0 | 2.193 | 0.24 | 0.35 | 0.23 | 0.2 | 0.17 | 0.35 |
| 8 | 80744549 | rs17534643 | 1.376 | 0.011 | -0.412 | -0.333 | 0.015 | 0.025 | 0.025 | 0 | 0.15 | 0.1 | 0.09 | 0 | 0.13 | 0.21 |
| 8 | 80745070 | rs2457398 | -1.568 | 0.21 | -0.552 | -0.135 | -0.024 | -0.065 | -0.065 | 0.13 | 0.85 | 0.95 | 0.82 | 0.59 | 0.81 | 0.83 |
| 8 | 80745281 | rs2467764 | -0.931 | 0.05 | -0.13 | -0.276 | 0.012 | 0.075 | 0.075 | 0 | 0.8 | 0.69 | 0.69 | 0.52 | 0.79 | 0.74 |
| 8 | 80745551 | rs10109063 | -0.231 | -0.128 | -0.316 | -0.665 | 0.02 | 0.021 | 0.021 | 0 | 0.31 | 0.37 | 0.41 | 0.38 | 0.39 | 0.43 |
| 8 | 80746904 | rs2467762 | -1.106 | -0.02 | -0.452 | -0.528 | 0.011 | 0.074 | 0.074 | 0 | 0.8 | 0.69 | 0.69 | 0.52 | 0.79 | 0.73 |
| 8 | 80756095 | rs2467787 | -0.187 | 0.216 | -0.679 | -0.361 | 0.005 | 0.003 | 0.003 | 1.577 | 0.59 | 0.43 | 0.36 | 0.37 | 0.53 | 0.44 |
| 8 | 80762327 | rs176085 | 0.956 | 0.179 | -1.389 | -1.239 | -0.046 | 0.075 | 0.075 | 3.18 | 0.31 | 0.01 | 0.15 | 0.14 | 0.27 | 0.1 |
| 8 | 80764370 | rs378052 | 0.256 | 0.099 | -0.476 | -0.787 | -0.001 | -0.008 | -0.008 | 0 | 0.56 | 0.54 | 0.56 | 0.3 | 0.56 | 0.36 |
| 8 | 80766354 | rs390691 | 0.145 | 0.55 | -0.677 | -0.238 | -0.001 | -0.017 | -0.017 | 0 | 0.57 | 0.47 | 0.56 | 0.29 | 0.62 | 0.55 |
| 8 | 80774955 | rs4740029 | 0.077 | 0.394 | -0.52 | 0.066 | 0 | 0 | 0 | 0 | 0.41 | 0.46 | 0.48 | 0.59 | 0.39 | 0.51 |
| 8 | 80782703 | rs6473185 | -1.558 | 0.218 | 0.748 | -0.006 | -0.005 | 0.076 | 0.076 | 1.343 | 0.11 | 0.5 | 0.15 | 0.15 | 0.05 | 0.13 |
| 8 | 80783998 | rs17456596 | 0.9 | 0.322 | 0.177 | -0.434 | -0.01 | 0.035 | 0.035 | 1.319 | 0.57 | 0.3 | 0.49 | 0.65 | 0.57 | 0.53 |
| 8 | 80785374 | rs2123270 | -0.341 | 0.44 | 0.07 | -0.515 | -0.118 | 0.029 | 0.029 | 3.676 | 0.76 | 0.98 | 0.69 | 0.85 | 0.68 | 0.75 |
| 8 | 80785667 | rs16907251 | -0.535 | 0.423 | 0.834 | -0.504 | 0.003 | 0.005 | 0.005 | 0.481 | 0.06 | 0.21 | 0.03 | 0.04 | 0.04 | 0.1 |
| 8 | 80786648 | rs6999925 | 0.791 | 0.403 | 0.618 | -0.491 | -0.002 | 0.007 | 0.007 | 2.218 | 0.65 | 0.34 | 0.61 | 0.8 | 0.64 | 0.65 |
| 8 | 80799281 | rs166872 | 0.87 | 0.668 | 0.304 | -0.785 | -0.07 | 0.028 | 0.028 | 1.441 | 0.16 | 0.02 | 0.28 | 0.08 | 0.27 | 0.12 |
| 8 | 80799543 | rs7825524 | -1.91 | 0.506 | 0.855 | -0.753 | -0.008 | 0.039 | 0.039 | 0 | 0.13 | 0.5 | 0.17 | 0.12 | 0.08 | 0.1 |
| 8 | 80812152 | rs2001721 | 1.14 | 0.605 | 0.604 | -0.54 | -0.01 | 0.017 | 0.017 | 2.802 | 0.62 | 0.24 | 0.5 | 0.65 | 0.62 | 0.59 |
| 8 | 80815412 | rs6473190 | -0.106 | 0.718 | 0.425 | -0.455 | -0.11 | -0.027 | -0.027 | 0 | 0.77 | 0.99 | 0.69 | 0.79 | 0.69 | 0.8 |

| | | | | | | | | | | | | | | | | |
|---|---|---|---|---|---|---|---|---|---|---|---|---|---|---|---|---|
| 8 | 80821153 | rs13439520 | -1.827 | 0.747 | 1.133 | -0.472 | 0.071 | 0.194 | 0.194 | 0 | 0.05 | 0.4 | 0.13 | 0.12 | 0.02 | 0.1 |
| 8 | 80830068 | rs12549933 | 0.626 | 0.805 | 0.821 | -0.649 | 0.008 | -0.016 | -0.016 | 0 | 0.7 | 0.4 | 0.53 | 0.74 | 0.66 | 0.76 |
| 8 | 80831713 | rs1274931 | 0.47 | 1.076 | -0.046 | -1.151 | -0.007 | 0.008 | 0.008 | 0.01 | 0.63 | 0.38 | 0.66 | 0.86 | 0.62 | 0.67 |
| 8 | 80841972 | rs2100346 | -0.479 | 0.981 | -0.002 | -1.267 | -0.039 | -0.006 | -0.006 | 0 | 0.79 | 0.92 | 0.7 | 0.81 | 0.74 | 0.81 |
| 8 | 80845186 | rs9298332 | -0.454 | 1.196 | 0.292 | -1.337 | -0.03 | 0.03 | 0.03 | 0.357 | 0.8 | 0.9 | 0.72 | 0.87 | 0.74 | 0.85 |
| 8 | 80853574 | rs720302 | -1.777 | 0.611 | 0.637 | -1.229 | -0.003 | 0.043 | 0.043 | 0 | 0.07 | 0.18 | 0.04 | 0.09 | 0.04 | 0.07 |
| 8 | 80856693 | rs2123269 | -0.67 | 0.582 | 1.047 | -1.305 | -0.001 | -0.001 | -0.001 | 0 | 0.76 | 0.81 | 0.68 | 0.84 | 0.71 | 0.76 |
| 8 | 80867359 | rs6987013 | 0.582 | 0.344 | 0.824 | -1.497 | -0.015 | 0.012 | 0.012 | 1.356 | 0.6 | 0.16 | 0.38 | 0.65 | 0.64 | 0.64 |
| 8 | 80870910 | rs2053962 | -0.499 | 0.413 | 1.428 | -1.472 | 0.017 | -0.046 | -0.046 | 0 | 0.72 | 0.57 | 0.64 | 0.84 | 0.7 | 0.76 |
| 8 | 80873478 | rs10094992 | -0.511 | 0.413 | 1.428 | -1.472 | 0.017 | -0.045 | -0.045 | 0 | 0.72 | 0.57 | 0.64 | 0.84 | 0.7 | 0.77 |
| 8 | 80882568 | rs2044968 | 0.041 | 0.479 | 1.424 | -1.427 | -0.01 | 0.009 | 0.009 | 3.422 | 0.69 | 0.2 | 0.58 | 0.77 | 0.7 | 0.73 |
| 8 | 80903607 | rs1562986 | -1.033 | 0.476 | 1.573 | -1.517 | -0.001 | -0.001 | -0.001 | 0 | 0.77 | 0.8 | 0.68 | 0.85 | 0.72 | 0.77 |
| 8 | 80909229 | rs7014295 | -0.002 | 0.498 | 1.389 | -1.504 | -0.004 | 0.014 | 0.014 | 3.369 | 0.69 | 0.23 | 0.59 | 0.78 | 0.7 | 0.73 |
| 8 | 80924116 | rs17458102 | 1.405 | 1.01 | 1.479 | -1.387 | -0.037 | 0.036 | 0.036 | 0 | 0.07 | 0.01 | 0.06 | 0 | 0.15 | 0.03 |
| 8 | 80939650 | rs12156383 | 0.486 | -0.257 | 1.193 | -1.576 | 0.018 | 0.187 | 0.187 | 0.304 | 0.3 | 0.05 | 0.14 | 0.01 | 0.26 | 0.29 |
| 8 | 80952186 | rs7827979 | -2.473 | 0.285 | 2.16 | -0.749 | 0.053 | 0.156 | 0.156 | 1.443 | 0.08 | 0.52 | 0.09 | 0.09 | 0.04 | 0.07 |
| 8 | 80956545 | rs10957954 | -2.326 | 0.29 | 1.993 | -0.734 | -0.01 | 0.071 | 0.071 | 1.509 | 0.08 | 0.67 | 0.1 | 0.08 | 0.03 | 0.07 |
| 8 | 80965830 | rs2303447 | -0.143 | 0.181 | 1.245 | -0.882 | -0.001 | -0.005 | -0.005 | 0.016 | 0.61 | 0.4 | 0.5 | 0.69 | 0.64 | 0.61 |
| 8 | 80970355 | rs10107002 | -0.884 | 0.177 | 0.228 | -0.923 | -0.055 | 0.051 | 0.051 | 6.595 | 0.68 | 0.93 | 0.59 | 0.76 | 0.65 | 0.68 |
| 8 | 80972451 | rs7837150 | -0.881 | 0.189 | 0.275 | -0.903 | -0.065 | 0.06 | 0.06 | 5.67 | 0.67 | 0.93 | 0.59 | 0.78 | 0.63 | 0.68 |
| 8 | 80980662 | rs11984461 | -0.152 | 0.062 | 0.852 | -0.902 | -0.004 | 0.016 | 0.016 | 7.984 | 0.59 | 0.31 | 0.5 | 0.71 | 0.59 | 0.59 |
| 8 | 80981317 | rs1025187 | -0.181 | -0.362 | 1.223 | -1.199 | -0.057 | -0.066 | -0.066 | 0.196 | 0.28 | 0.5 | 0.16 | 0.09 | 0.19 | 0.34 |
| 8 | 80986578 | rs7845122 | 0.594 | -0.377 | 1.596 | -1.171 | -0.033 | 0.009 | 0.009 | 8.281 | 0.38 | 0.2 | 0.4 | 0.7 | 0.44 | 0.32 |
| 8 | 80990280 | rs6473198 | -1.073 | -0.416 | 1.602 | -1.151 | 0.009 | -0.009 | -0.009 | 0 | 0.65 | 0.54 | 0.55 | 0.72 | 0.62 | 0.61 |
| 8 | 80993478 | rs16907441 | 0.073 | -0.471 | 1.204 | -1.116 | -0.018 | -0.019 | -0.019 | 0.018 | 0.26 | 0.29 | 0.14 | 0.09 | 0.17 | 0.32 |
| 8 | 81003013 | rs4740098 | -1.057 | -0.298 | 0.658 | -1.114 | -0.036 | 0.065 | 0.065 | 5.712 | 0.66 | 0.88 | 0.66 | 0.8 | 0.64 | 0.62 |
| 8 | 81005170 | rs2217063 | -0.4 | -0.91 | 1.095 | -1.113 | 0 | 0 | 0 | 0 | 0.49 | 0.5 | 0.49 | 0.71 | 0.52 | 0.47 |
| 8 | 81005880 | rs4740099 | -1.161 | -0.581 | 0.846 | -1.051 | -0.107 | 0.049 | 0.049 | 6.201 | 0.65 | 0.91 | 0.63 | 0.79 | 0.58 | 0.64 |
| 8 | 81014719 | rs6473199 | -1.16 | -0.543 | 0.916 | -0.975 | -0.085 | 0.062 | 0.062 | 5.578 | 0.65 | 0.89 | 0.63 | 0.79 | 0.58 | 0.62 |
| 8 | 81019703 | rs1074786 | 0.217 | -0.775 | 1.132 | -0.952 | -0.045 | 0.027 | 0.027 | 7.901 | 0.4 | 0.18 | 0.39 | 0.72 | 0.43 | 0.33 |
| 8 | 81024974 | rs12679499 | -0.157 | -0.767 | 1.496 | -0.957 | -0.004 | 0.044 | 0.044 | 8.253 | 0.46 | 0.34 | 0.49 | 0.79 | 0.46 | 0.41 |
| 8 | 81035358 | rs9643749 | -0.209 | -0.763 | 1.304 | -0.97 | -0.01 | 0.079 | 0.079 | 8.014 | 0.45 | 0.28 | 0.48 | 0.78 | 0.46 | 0.41 |
| 8 | 81035634 | rs9643750 | 0.573 | -0.75 | 1.131 | -0.966 | -0.076 | 0.001 | 0.001 | 8.369 | 0.38 | 0.08 | 0.37 | 0.71 | 0.42 | 0.33 |
| 8 | 81036045 | rs6473202 | 0.768 | -0.829 | 1.438 | -1.013 | -0.023 | -0.063 | -0.063 | 0 | 0.15 | 0.22 | 0.07 | 0 | 0.09 | 0.21 |
| 8 | 81045600 | rs13261075 | -1.219 | -0.769 | 1.342 | -0.994 | -0.05 | 0.065 | 0.065 | 5.664 | 0.64 | 0.81 | 0.6 | 0.79 | 0.54 | 0.62 |
| 8 | 81058344 | rs7840008 | -0.311 | -0.726 | 1.518 | -0.971 | -0.003 | 0.073 | 0.073 | 7.257 | 0.47 | 0.6 | 0.53 | 0.79 | 0.46 | 0.41 |
| 8 | 81059514 | rs1863439 | 0.214 | -0.573 | 1.171 | -0.875 | -0.025 | 0.019 | 0.019 | 8.278 | 0.39 | 0.25 | 0.4 | 0.71 | 0.43 | 0.33 |
| 8 | 81059558 | rs1863440 | -1.095 | -0.525 | 0.897 | -0.86 | -0.094 | 0.074 | 0.074 | 5.465 | 0.62 | 0.87 | 0.61 | 0.77 | 0.55 | 0.61 |
| 8 | 81065580 | rs7014791 | 0.08 | -0.471 | 0.74 | -0.836 | 0.007 | -0.016 | -0.016 | 0 | 0.42 | 0.49 | 0.43 | 0.73 | 0.43 | 0.33 |
| 8 | 81068774 | rs4740122 | 0.395 | -0.742 | 0.818 | -0.824 | -0.06 | -0.047 | -0.047 | 0.907 | 0.2 | 0.38 | 0.18 | 0.07 | 0.11 | 0.29 |
| 8 | 81077969 | rs10755964 | -0.42 | -0.189 | 0.586 | -0.26 | -0.001 | -0.005 | -0.005 | 0 | 0.4 | 0.44 | 0.42 | 0.69 | 0.38 | 0.32 |
| 8 | 81082118 | rs4739721 | -0.832 | -0.216 | 0.496 | -0.188 | -0.011 | -0.002 | -0.002 | 0.652 | 0.13 | 0.4 | 0.15 | 0.05 | 0.07 | 0.2 |
| 8 | 81090148 | rs6473211 | -0.819 | -0.112 | -0.581 | 0.184 | 0.001 | 0.072 | 0.072 | 3.812 | 0.51 | 0.85 | 0.54 | 0.65 | 0.42 | 0.48 |
| 8 | 81094701 | rs4740131 | -1.62 | -0.562 | 0.31 | 0.459 | 0.029 | 0 | 0 | 0 | 0.45 | 0.74 | 0.48 | 0.46 | 0.37 | 0.42 |
| 8 | 81095371 | rs7839986 | 0.509 | -0.607 | 0.265 | 0.429 | -0.017 | 0.057 | 0.057 | 0 | 0.11 | 0.03 | 0.06 | 0 | 0.12 | 0.04 |
| 8 | 81110456 | rs7830310 | -0.969 | -0.729 | 0.858 | 0.353 | -0.01 | -0.088 | -0.088 | 0.402 | 0.28 | 0.12 | 0.33 | 0.46 | 0.27 | 0.27 |
| 8 | 81113769 | rs7845298 | -0.914 | -0.716 | 1.612 | 0.369 | -0.023 | 0.14 | 0.14 | 0 | 0.22 | 0.53 | 0.35 | 0.46 | 0.16 | 0.23 |
| 8 | 81120189 | rs4415363 | -1.616 | -0.659 | 1.611 | 0.358 | -0.003 | 0.002 | 0.002 | 0 | 0.42 | 0.43 | 0.47 | 0.52 | 0.36 | 0.43 |
| 8 | 81137030 | rs7007995 | -1.975 | -0.494 | 1.17 | 0.397 | 0.034 | 0.015 | 0.015 | 0 | 0.48 | 0.77 | 0.53 | 0.53 | 0.38 | 0.43 |
| 8 | 81142630 | rs7011326 | -2.139 | -0.768 | 1.279 | 0.515 | 0.067 | 0.018 | 0.018 | 0 | 0.48 | 0.78 | 0.53 | 0.52 | 0.36 | 0.43 |
| 8 | 81143345 | rs4740144 | -1.141 | -0.774 | 1.399 | 0.49 | -0.002 | -0.078 | -0.078 | 0 | 0.31 | 0.13 | 0.4 | 0.51 | 0.28 | 0.29 |
| 8 | 81149186 | rs4739729 | -1.48 | -0.647 | 1.78 | 0.448 | -0.002 | -0.019 | -0.019 | 0 | 0.33 | 0.26 | 0.45 | 0.51 | 0.34 | 0.3 |
| 8 | 81161499 | rs11785423 | 1.074 | -1.136 | 1.965 | 0.505 | 0.009 | -0.043 | -0.043 | 0 | 0.68 | 0.83 | 0.59 | 0.5 | 0.71 | 0.69 |
| 8 | 81175152 | rs10108355 | 1.69 | -0.776 | 2.803 | 0.756 | -0.002 | -0.001 | -0.001 | 0 | 0.56 | 0.42 | 0.5 | 0.5 | 0.62 | 0.56 |
| 8 | 81186078 | rs11784890 | 1.359 | -0.769 | 2.679 | 0.76 | 0.011 | -0.005 | -0.005 | 0 | 0.59 | 0.69 | 0.53 | 0.51 | 0.62 | 0.56 |
| 8 | 81195187 | rs4740149 | 1.856 | -0.732 | 2.52 | 0.991 | 0.053 | -0.013 | -0.013 | 0 | 0.53 | 0.18 | 0.46 | 0.52 | 0.61 | 0.57 |
| 8 | 81197858 | rs6473217 | 0.738 | -0.472 | 2.585 | 1.241 | 0 | 0 | 0 | 0.508 | 0.81 | 0.83 | 0.64 | 0.51 | 0.82 | 0.74 |
| 8 | 81236791 | rs16907577 | -0.643 | -0.684 | 2.465 | 0.651 | -0.019 | -0.047 | -0.047 | 0 | 0.08 | 0.19 | 0.04 | 0 | 0.02 | 0.01 |
| 8 | 81245003 | rs6473219 | -1.457 | -0.575 | 1.818 | 0.734 | 0.011 | 0.019 | 0.019 | 1.808 | 0.39 | 0.66 | 0.34 | 0.24 | 0.32 | 0.25 |
| 8 | 81246699 | rs7822342 | 1.671 | -0.493 | 1.707 | 1.191 | 0.05 | 0.038 | 0.038 | 2.446 | 0.51 | 0.23 | 0.42 | 0.44 | 0.58 | 0.52 |
| 8 | 81256542 | rs13275909 | 1.584 | -0.568 | 1.921 | 1.117 | 0.047 | 0.042 | 0.042 | 2.35 | 0.52 | 0.25 | 0.43 | 0.42 | 0.6 | 0.51 |
| 8 | 81259826 | rs13275219 | 0.859 | -0.669 | 2.306 | 1.178 | -0.009 | 0.006 | 0.006 | 0 | 0.63 | 0.55 | 0.5 | 0.42 | 0.65 | 0.55 |
| 8 | 81273883 | rs6987042 | 1.537 | -0.534 | 1.92 | 1.164 | 0.035 | 0.03 | 0.03 | 0.958 | 0.55 | 0.28 | 0.44 | 0.41 | 0.62 | 0.53 |
| 8 | 81275860 | rs11786704 | 1.467 | -0.536 | 1.919 | 1.163 | 0.041 | 0.036 | 0.036 | 0.937 | 0.55 | 0.24 | 0.44 | 0.41 | 0.62 | 0.52 |
| 8 | 81281007 | rs6473225 | 1.588 | -0.531 | 1.756 | 1.165 | 0.049 | 0.041 | 0.041 | 1.133 | 0.53 | 0.18 | 0.44 | 0.41 | 0.62 | 0.53 |
| 8 | 81285759 | rs6473226 | 1.456 | -0.357 | 1.882 | 1.183 | 0.018 | 0.013 | 0.013 | 1.56 | 0.57 | 0.31 | 0.53 | 0.43 | 0.63 | 0.54 |
| 8 | 81288634 | rs952559 | 1.028 | -0.357 | 2.018 | 1.18 | -0.005 | 0.002 | 0.002 | 0 | 0.63 | 0.46 | 0.56 | 0.43 | 0.65 | 0.54 |
| 8 | 81295224 | rs1543857 | 1.56 | -0.237 | 1.286 | 1.084 | 0.017 | 0.006 | 0.006 | 1.129 | 0.55 | 0.26 | 0.52 | 0.45 | 0.61 | 0.54 |
| 8 | 81299603 | rs7003011 | 1.353 | -0.166 | 1.208 | 1.12 | 0.006 | 0 | 0 | 0 | 0.57 | 0.37 | 0.53 | 0.45 | 0.62 | 0.54 |

| | | | | | | | | | | | | | | | | |
|---|---|---|---|---|---|---|---|---|---|---|---|---|---|---|---|---|
| 8 | 81307007 | rs10808842 | -1.027 | 0.673 | 1.106 | 1.501 | -0.002 | -0.021 | -0.021 | 0 | 0.29 | 0.2 | 0.33 | 0.34 | 0.31 | 0.41 |
| 8 | 81312277 | rs11784307 | -1.357 | 0.708 | 0.656 | 1.349 | 0.012 | 0.022 | 0.022 | 1.321 | 0.35 | 0.51 | 0.42 | 0.55 | 0.38 | 0.44 |
| 8 | 81318892 | rs10091870 | -0.56 | 0.904 | -0.256 | 1.185 | -0.04 | -0.057 | -0.057 | 0 | 0.09 | 0.01 | 0.06 | 0.2 | 0.12 | 0.18 |
| 8 | 81319427 | rs4739748 | -1.04 | 0.97 | 0.036 | 1.151 | 0.025 | 0.177 | 0.177 | 0 | 0.12 | 0.24 | 0.25 | 0.46 | 0.13 | 0.25 |
| 8 | 81320578 | rs16907659 | -0.483 | 0.659 | -0.743 | 0.591 | 0.003 | 0.086 | 0.086 | 0 | 0.1 | 0.15 | 0.23 | 0.32 | 0.08 | 0.2 |
| 8 | 81330198 | rs484681 | -0.277 | 0.643 | -0.864 | 0.121 | -0.005 | 0.002 | 0.002 | 0.617 | 0.17 | 0.14 | 0.1 | 0 | 0.21 | 0.15 |
| 8 | 81331943 | rs16907682 | -0.874 | 0.489 | -0.756 | 0.77 | -0.002 | 0.014 | 0.014 | 1.275 | 0.16 | 0.16 | 0.24 | 0.3 | 0.1 | 0.19 |
| 8 | 81332842 | rs17554231 | -0.495 | 0.545 | -1.115 | -0.274 | 0 | -0.002 | -0.002 | 0.316 | 0.15 | 0.12 | 0.09 | 0.06 | 0.12 | 0.11 |
| 8 | 81335062 | rs272604 | 0.364 | -0.328 | -1.039 | -0.768 | -0.006 | -0.016 | -0.016 | 0.114 | 0.8 | 0.77 | 0.92 | 0.94 | 0.9 | 0.73 |
| 8 | 81338686 | rs367273 | 0.554 | 0.337 | -1.457 | -1.067 | 0.008 | 0.098 | 0.098 | 0.259 | 0.58 | 0.27 | 0.6 | 0.81 | 0.62 | 0.62 |
| 8 | 81353654 | rs9298338 | -0.995 | 0.305 | -1.473 | -0.595 | 0.001 | -0.031 | -0.031 | 0 | 0.09 | 0.03 | 0.05 | 0.2 | 0.08 | 0.11 |
| 8 | 81358791 | rs4610800 | -1.439 | -0.209 | -1.015 | -1.083 | -0.005 | -0.014 | -0.014 | 0 | 0.26 | 0.71 | 0.2 | 0.17 | 0.19 | 0.17 |
| 8 | 81364205 | rs6998967 | -0.785 | -0.641 | -0.561 | -1.52 | -0.043 | -0.133 | -0.133 | 0.052 | 0.27 | 0.46 | 0.15 | 0.05 | 0.19 | 0.2 |
| 8 | 81381871 | rs7830160 | -1.686 | -0.78 | -0.543 | -1.591 | 0.071 | 0.091 | 0.091 | 0.406 | 0.38 | 0.8 | 0.26 | 0.27 | 0.28 | 0.3 |
| 8 | 81389791 | rs11993814 | -1.656 | -0.784 | -0.546 | -1.594 | 0.068 | 0.088 | 0.088 | 0.457 | 0.38 | 0.81 | 0.26 | 0.27 | 0.28 | 0.29 |
| 8 | 81397268 | rs10095139 | 0.483 | -0.735 | -0.507 | -1.601 | 0.003 | -0.008 | -0.008 | 0 | 0.05 | 0.02 | 0.03 | 0.23 | 0.05 | 0.07 |
| 8 | 81407333 | rs4587368 | 0.291 | -1.082 | -0.834 | -2.019 | -0.003 | -0.024 | -0.024 | 0.591 | 0.09 | 0.1 | 0.05 | 0 | 0.06 | 0.02 |
| 8 | 81407796 | rs562210 | -2.146 | -1.139 | -0.927 | -2.075 | 0.109 | 0.212 | 0.212 | 0.041 | 0.48 | 0.93 | 0.31 | 0.28 | 0.34 | 0.32 |
| 8 | 81407856 | rs3863246 | -1.195 | -1.033 | -0.74 | -1.924 | -0.041 | -0.044 | -0.044 | 0.374 | 0.33 | 0.49 | 0.18 | 0.28 | 0.24 | 0.26 |
| 8 | 81415556 | rs16907771 | 0.601 | -0.994 | -0.744 | -1.876 | -0.001 | -0.027 | -0.027 | 1.013 | 0.05 | 0 | 0.02 | 0.25 | 0.05 | 0.07 |
| 8 | 81429892 | rs10504722 | -1.181 | -0.964 | -0.858 | -1.843 | -0.043 | -0.045 | -0.045 | 0 | 0.33 | 0.49 | 0.18 | 0.28 | 0.24 | 0.26 |
| 8 | 81438310 | rs1051919 | -0.584 | -0.806 | -0.758 | -1.914 | -0.047 | -0.156 | -0.156 | 1.71 | 0.27 | 0.47 | 0.16 | 0.03 | 0.19 | 0.19 |
| 8 | 81443654 | rs272612 | -1.095 | 0.35 | -1.804 | -1.872 | -0.078 | 0.08 | 0.08 | 0.029 | 0.74 | 0.96 | 0.68 | 0.89 | 0.68 | 0.82 |
| 8 | 81449141 | rs1863593 | -0.988 | -0.499 | -0.692 | -1.222 | -0.045 | 0.056 | 0.056 | 1.331 | 0.12 | 0.64 | 0.27 | 0.17 | 0.07 | 0.07 |
| 8 | 81450564 | rs693015 | -1.813 | -0.145 | -0.782 | -0.842 | -0.007 | 0.073 | 0.073 | 0.734 | 0.17 | 0.66 | 0.31 | 0.4 | 0.19 | 0.19 |
| 8 | 81451995 | rs381349 | -0.719 | -0.134 | -0.765 | -0.901 | 0.021 | 0.043 | 0.043 | 0 | 0.06 | 0.07 | 0.04 | 0.22 | 0.11 | 0.12 |
| 8 | 81452333 | rs446027 | -1.842 | -0.596 | -1.332 | -1.489 | 0.05 | 0.033 | 0.033 | 0.047 | 0.43 | 0.91 | 0.44 | 0.38 | 0.36 | 0.36 |
| 8 | 81455763 | rs10957982 | -1.34 | -1.167 | -1.047 | -1.706 | 0.011 | 0.011 | 0.011 | 0 | 0.32 | 0.74 | 0.36 | 0.16 | 0.22 | 0.22 |
| 8 | 81461974 | rs440837 | -1.771 | -1.187 | -1.178 | -1.72 | 0.099 | 0.115 | 0.115 | 0 | 0.38 | 0.86 | 0.39 | 0.16 | 0.25 | 0.24 |
| 8 | 81462608 | rs382796 | -1.649 | -1.028 | -0.193 | -1.448 | -0.062 | 0.019 | 0.019 | 0.669 | 0.16 | 0.45 | 0.22 | 0.16 | 0.11 | 0.1 |
| 8 | 81464082 | rs10109619 | -1.425 | -0.842 | -0.213 | -1.381 | 0.033 | 0.056 | 0.056 | 0 | 0.08 | 0.26 | 0.1 | 0.1 | 0.06 | 0.15 |
| 8 | 81464148 | rs575452 | -2.271 | -1.012 | -0.405 | -1.513 | 0.052 | 0.028 | 0.028 | 1.721 | 0.27 | 0.91 | 0.31 | 0.22 | 0.16 | 0.23 |
| 8 | 81469704 | rs272592 | -2.228 | -1.028 | -0.601 | -1.518 | 0.05 | 0.024 | 0.024 | 1.907 | 0.26 | 0.91 | 0.31 | 0.22 | 0.16 | 0.23 |
| 8 | 81470120 | rs272594 | -1.617 | -0.66 | -0.472 | -1.365 | 0.015 | -0.01 | -0.01 | 0.203 | 0.16 | 0.43 | 0.16 | 0.1 | 0.12 | 0.16 |
| 8 | 81476229 | rs17471946 | -1.154 | -0.661 | -1.056 | -1.329 | 0.008 | -0.015 | -0.015 | 0 | 0.1 | 0.01 | 0.14 | 0.13 | 0.09 | 0.1 |
| 8 | 81476365 | rs17471953 | -1.363 | -0.706 | -1.026 | -1.337 | 0.022 | 0.002 | 0.002 | 0 | 0.1 | 0.05 | 0.15 | 0.13 | 0.09 | 0.1 |
| 8 | 81478754 | rs4324961 | 0.198 | -0.863 | -1.261 | -1.492 | -0.005 | 0.015 | 0.015 | 0.08 | 0.15 | 0.01 | 0.1 | 0.08 | 0.11 | 0.25 |
| 8 | 81482337 | rs272610 | -1.312 | -0.762 | -0.965 | -1.114 | 0.007 | 0.01 | 0.01 | 0 | 0.5 | 0.41 | 0.52 | 0.36 | 0.42 | 0.54 |
| 8 | 81490424 | rs7464898 | 0.241 | -0.531 | -1.13 | -0.921 | -0.003 | -0.022 | -0.022 | 0.245 | 0.83 | 0.92 | 0.67 | 0.76 | 0.83 | 0.72 |
| 8 | 81497016 | rs2033129 | 0.135 | -0.926 | -0.599 | -0.867 | -0.033 | -0.044 | -0.044 | 0 | 0.19 | 0.35 | 0.13 | 0.09 | 0.11 | 0.21 |
| 8 | 81500656 | rs7005189 | -0.335 | -0.676 | -1.272 | -0.64 | -0.011 | -0.084 | -0.084 | 0.008 | 0.17 | 0.01 | 0.32 | 0.25 | 0.14 | 0.29 |
| 8 | 81501052 | rs11988829 | 1.378 | -0.464 | -1.213 | -0.898 | -0.002 | -0.016 | -0.016 | 0 | 0.1 | 0.08 | 0.1 | 0 | 0.15 | 0.02 |
| 8 | 81503192 | rs4739755 | -0.404 | -0.568 | -1.3 | -0.751 | -0.021 | -0.105 | -0.105 | 0 | 0.19 | 0.01 | 0.33 | 0.29 | 0.16 | 0.31 |
| 8 | 81505848 | rs11989997 | 0.595 | -0.814 | -0.966 | -1.346 | 0.001 | 0.02 | 0.02 | 0.175 | 0.16 | 0.07 | 0.09 | 0 | 0.17 | 0.02 |
| 8 | 81514063 | rs11992883 | -0.399 | -0.838 | -1.037 | -1.514 | 0.005 | 0.059 | 0.059 | 0.666 | 0.25 | 0.09 | 0.12 | 0 | 0.22 | 0.04 |
| 8 | 81518320 | rs1026804 | -0.843 | -1.163 | -0.217 | -1.288 | -0.043 | -0.105 | -0.105 | 0.538 | 0.3 | 0.5 | 0.2 | 0.08 | 0.17 | 0.26 |
| 8 | 81518364 | rs1026805 | 0.146 | -0.9 | -0.375 | -1.044 | 0.01 | 0.012 | 0.012 | 0.744 | 0.19 | 0.18 | 0.12 | 0.08 | 0.11 | 0.25 |
| 8 | 81528234 | rs16907900 | -1.196 | -0.818 | 0.424 | -0.982 | -0.002 | -0.077 | -0.077 | 0 | 0.12 | 0.54 | 0.11 | 0 | 0.08 | 0.02 |
| 8 | 81533311 | rs6473236 | -1.085 | -0.816 | 0.744 | -0.998 | 0.033 | -0.059 | -0.059 | 4.21 | 0.11 | 0.49 | 0.1 | 0 | 0.08 | 0.02 |
| 8 | 81545195 | rs1045667 | 0.937 | -0.457 | 0.016 | -0.664 | -0.109 | -0.102 | -0.102 | 0 | 0.29 | 0.01 | 0.37 | 0.41 | 0.37 | 0.3 |
| 8 | 81551652 | rs10755966 | 1.024 | -0.409 | 0.249 | -0.185 | -0.112 | -0.106 | -0.106 | 3.209 | 0.29 | 0.01 | 0.37 | 0.41 | 0.37 | 0.28 |
| 8 | 81569563 | rs7018189 | -0.371 | -0.587 | 0.959 | -0.456 | 0.004 | -0.057 | -0.057 | 0.011 | 0.1 | 0.24 | 0.06 | 0 | 0.07 | 0.02 |
| 8 | 81570177 | rs6998930 | -0.532 | -0.589 | 0.906 | -0.457 | 0.016 | -0.061 | -0.061 | 0 | 0.11 | 0.39 | 0.08 | 0 | 0.07 | 0.02 |
| 8 | 81576965 | rs7018449 | -0.485 | -0.567 | 0.926 | -0.446 | 0.034 | -0.052 | -0.052 | 0 | 0.1 | 0.44 | 0.09 | 0 | 0.07 | 0.02 |
| 8 | 81596608 | rs6992359 | -0.802 | -0.612 | 0.999 | -0.469 | 0.015 | -0.072 | -0.072 | 1.104 | 0.12 | 0.46 | 0.09 | 0 | 0.08 | 0.02 |
| 8 | 81677834 | rs1489582 | -0.927 | -0.308 | 1.003 | -0.67 | 0.023 | -0.11 | -0.11 | 0 | 0.16 | 0.51 | 0.18 | 0 | 0.15 | 0.05 |
| 8 | 81704158 | rs11987529 | -1.251 | -0.633 | 0.291 | -1.124 | -0.003 | -0.046 | -0.046 | 0 | 0.35 | 0.59 | 0.26 | 0.09 | 0.27 | 0.39 |
| 8 | 81705396 | rs7006546 | -0.432 | -0.647 | 0.157 | -1.149 | 0.046 | -0.103 | -0.103 | 0.512 | 0.14 | 0.48 | 0.17 | 0 | 0.15 | 0.05 |
| 8 | 81709470 | rs12548319 | 0.646 | -0.705 | -0.278 | -1.155 | 0.013 | 0.059 | 0.059 | 0 | 0.19 | 0 | 0.08 | 0.09 | 0.12 | 0.38 |
| 8 | 81710837 | rs4308687 | -1.378 | -0.802 | -0.626 | -1.18 | -0.005 | 0.105 | 0.105 | 0 | 0.57 | 0.83 | 0.35 | 0.29 | 0.4 | 0.55 |
| 8 | 81751247 | rs1157139 | 0.878 | -0.577 | -0.761 | -0.75 | 0.048 | -0.012 | -0.012 | 0 | 0.23 | 0.03 | 0.07 | 0.2 | 0.13 | 0.15 |
| 8 | 81753399 | rs7837508 | -0.03 | 0.582 | 0.151 | -0.47 | 0.238 | 0.046 | 0.046 | 0 | 0.06 | 0.38 | 0.13 | 0 | 0.09 | 0.02 |
| 8 | 81757711 | rs16908087 | 0.78 | -0.334 | -0.229 | -0.573 | 0.001 | -0.003 | -0.003 | 0 | 0.07 | 0.04 | 0.04 | 0 | 0.08 | 0.02 |
| 8 | 81771461 | rs1199028 | 1.087 | -0.111 | -0.776 | -0.141 | -0.052 | -0.097 | -0.097 | 0 | 0.15 | 0.03 | 0.14 | 0.29 | 0.22 | 0.11 |
| 8 | 81807098 | rs7825743 | 0.231 | -0.087 | -0.413 | -0.238 | 0.007 | -0.004 | -0.004 | 0 | 0.36 | 0.3 | 0.64 | 0.37 | 0.51 | 0.36 |
| 8 | 81811905 | rs2084088 | 0.57 | -0.056 | -0.336 | -0.227 | -0.018 | 0.002 | 0.002 | 0 | 0.31 | 0.22 | 0.55 | 0.37 | 0.44 | 0.34 |
| 8 | 81816313 | rs2895689 | 1.633 | -0.039 | -0.595 | -0.194 | -0.122 | -0.091 | -0.091 | 0.006 | 0.23 | 0.02 | 0.49 | 0.37 | 0.36 | 0.31 |
| 8 | 81820942 | rs12542283 | 0.149 | -0.089 | -0.248 | -0.229 | 0.032 | -0.019 | -0.019 | 0 | 0.37 | 0.41 | 0.65 | 0.37 | 0.51 | 0.36 |
| 8 | 81825137 | rs17475818 | -0.135 | -0.455 | -0.245 | -0.04 | 0.039 | -0.07 | -0.07 | 0 | 0.12 | 0.19 | 0.13 | 0 | 0.15 | 0.05 |

| | | | | | | | | | | | | | | | | |
|---|---|---|---|---|---|---|---|---|---|---|---|---|---|---|---|---|
| 8 | 81828477 | rs1866276 | 0.561 | -0.876 | -1.055 | -0.112 | -0.086 | -0.018 | -0.018 | 0 | 0.4 | 0.1 | 0.56 | 0.47 | 0.47 | 0.62 |
| 8 | 81829801 | rs11997228 | 0.784 | -1.68 | -1.002 | -0.018 | 0.004 | 0.006 | 0.006 | 0 | 0.12 | 0.08 | 0.06 | 0.1 | 0.09 | 0.3 |
| 8 | 81830223 | rs1822971 | 0.106 | -1.586 | -0.895 | 0.059 | 0.072 | 0.051 | 0.051 | 0.216 | 0.49 | 0.17 | 0.66 | 0.48 | 0.61 | 0.66 |
| 8 | 81839692 | rs1445570 | 1.318 | -1.816 | -0.957 | 0.206 | -0.119 | -0.039 | -0.039 | 0 | 0.32 | 0.09 | 0.61 | 0.39 | 0.44 | 0.38 |
| 8 | 81840875 | rs7007683 | -0.66 | -2.294 | -0.51 | 0.354 | 0.014 | 0.076 | 0.076 | 0.371 | 0.82 | 0.63 | 0.85 | 0.74 | 0.78 | 0.88 |
| 8 | 81843186 | rs6980848 | 0.726 | -2.428 | -0.387 | 0.287 | -0.007 | -0.012 | -0.012 | 0 | 0.12 | 0.14 | 0.05 | 0.09 | 0.09 | 0.26 |
| 8 | 81843869 | rs7003901 | 1.145 | -2.629 | -0.369 | 0.207 | -0.012 | 0.012 | 0.012 | 0.108 | 0.39 | 0.23 | 0.68 | 0.38 | 0.53 | 0.39 |
| 8 | 81845260 | rs1445565 | 0.413 | -2.597 | -0.304 | 0.24 | -0.003 | 0.028 | 0.028 | 0 | 0.69 | 0.39 | 0.79 | 0.65 | 0.68 | 0.63 |
| 8 | 81846682 | rs10504727 | 0.866 | -2.623 | -0.493 | 0.196 | 0.081 | 0.028 | 0.028 | 2.927 | 0.29 | 0.1 | 0.1 | 0.28 | 0.16 | 0.23 |
| 8 | 81847103 | rs12548193 | 1.148 | -2.033 | -0.055 | 0.312 | 0.001 | -0.008 | -0.008 | 0 | 0.62 | 0.33 | 0.65 | 0.61 | 0.61 | 0.58 |
| 8 | 81847757 | rs6996716 | 1.428 | -1.921 | 0.279 | 0.803 | 0.001 | 0.01 | 0.01 | 0 | 0.68 | 0.33 | 0.67 | 0.61 | 0.64 | 0.6 |
| 8 | 81855797 | rs1445559 | 0.037 | -1.823 | 0.365 | 0.908 | 0.054 | 0.134 | 0.134 | 0 | 0.87 | 0.6 | 0.83 | 0.79 | 0.81 | 0.88 |
| 8 | 81860373 | rs1445558 | 1.144 | -1.789 | 0.374 | 0.887 | 0.034 | 0.042 | 0.042 | 0.735 | 0.79 | 0.39 | 0.74 | 0.81 | 0.73 | 0.85 |
| 8 | 81862346 | rs895275 | 0.866 | -1.918 | 0.341 | 0.943 | 0.026 | 0.081 | 0.081 | 0 | 0.76 | 0.49 | 0.74 | 0.7 | 0.72 | 0.64 |
| 8 | 81869780 | rs2028222 | 1.435 | -1.917 | 0.063 | 0.885 | -0.01 | -0.012 | -0.012 | 0 | 0.68 | 0.17 | 0.67 | 0.74 | 0.66 | 0.62 |
| 8 | 81870851 | rs2028814 | -0.132 | -1.706 | 0.422 | 1.123 | 0.062 | 0.108 | 0.108 | 0 | 0.89 | 0.64 | 0.83 | 0.82 | 0.81 | 0.88 |
| 8 | 81871222 | rs4739609 | -0.058 | -1.707 | 0.427 | 1.122 | 0.065 | 0.113 | 0.113 | 0 | 0.89 | 0.61 | 0.83 | 0.82 | 0.81 | 0.88 |
| 8 | 81873308 | rs877418 | 1.612 | -1.972 | -0.34 | 0.84 | -0.037 | 0.011 | 0.011 | 0.162 | 0.4 | 0.17 | 0.54 | 0.36 | 0.48 | 0.35 |
| 8 | 81878654 | rs1866274 | -0.224 | -1.888 | -0.574 | 0.279 | -0.012 | -0.019 | -0.019 | 0 | 0.28 | 0.3 | 0.19 | 0.16 | 0.16 | 0.26 |
| 8 | 81878670 | rs920983 | 0.894 | -1.869 | -0.542 | 0.398 | 0.008 | 0.011 | 0.011 | 2.535 | 0.57 | 0.64 | 0.65 | 0.66 | 0.58 | 0.62 |
| 8 | 81880096 | rs11743 | 0.087 | -1.482 | -0.105 | 0.147 | 0.023 | -0.016 | -0.016 | 0 | 0.11 | 0.03 | 0.09 | 0.14 | 0.06 | 0.22 |
| 8 | 81881160 | rs10088860 | -1.054 | -1.498 | 0.094 | 0.127 | -0.051 | -0.029 | -0.029 | 0.547 | 0.18 | 0.37 | 0.16 | 0.14 | 0.11 | 0.24 |
| 8 | 81882850 | rs3739342 | -1.071 | -1.499 | 0.095 | 0.127 | -0.055 | -0.034 | -0.034 | 0 | 0.18 | 0.42 | 0.16 | 0.13 | 0.11 | 0.24 |
| 8 | 81883905 | rs6984342 | -0.027 | -1.513 | -0.163 | 0.105 | -0.005 | -0.001 | -0.001 | 0 | 0.12 | 0.14 | 0.09 | 0.13 | 0.06 | 0.23 |
| 8 | 81885077 | rs1445560 | -0.068 | -1.579 | -0.324 | 0.131 | -0.047 | -0.06 | -0.06 | 0 | 0.66 | 0.95 | 0.77 | 0.41 | 0.64 | 0.57 |
| 8 | 81886515 | rs3816314 | -1.551 | -1.395 | 0.663 | 0.445 | -0.058 | -0.035 | -0.035 | 0 | 0.18 | 0.37 | 0.15 | 0.13 | 0.1 | 0.24 |
| 8 | 81886686 | rs7017984 | -1.345 | -1.322 | -0.415 | 0.482 | -0.021 | -0.013 | -0.013 | 0 | 0.17 | 0.22 | 0.14 | 0.13 | 0.1 | 0.24 |
| 8 | 81887806 | rs16908229 | -0.237 | -0.762 | -0.599 | 0.483 | -0.009 | -0.025 | -0.025 | 0.182 | 0.05 | 0.08 | 0.04 | 0 | 0.04 | 0.01 |
| 8 | 81891232 | rs7012200 | -0.85 | -1.218 | -1.184 | 0.543 | 0.062 | 0.039 | 0.039 | 0 | 0.43 | 0.83 | 0.33 | 0.59 | 0.35 | 0.53 |
| 8 | 81891362 | rs6473257 | -0.6 | -1.609 | -0.213 | 0.841 | -0.019 | 0.045 | 0.045 | 0.886 | 0.32 | 0.58 | 0.25 | 0.45 | 0.27 | 0.28 |
| 8 | 81894604 | rs9650270 | -0.643 | -2.112 | -0.757 | 0.793 | -0.016 | -0.133 | -0.133 | 0 | 0.19 | 0.03 | 0.16 | 0.53 | 0.21 | 0.29 |
| 8 | 81895868 | rs1530050 | -1.654 | -2.424 | -1.197 | 0.41 | 0.044 | 0.014 | 0.014 | 0.33 | 0.55 | 0.97 | 0.35 | 0.67 | 0.37 | 0.62 |
| 8 | 81900844 | rs6998543 | 0.169 | -2.556 | -0.11 | 0.193 | -0.077 | -0.15 | -0.15 | 0 | 0.27 | 0.56 | 0.12 | 0 | 0.1 | 0.16 |
| 8 | 81904003 | rs2289830 | -1.018 | -2.519 | -0.027 | 0.336 | -0.019 | -0.002 | -0.002 | 1.95 | 0.46 | 0.5 | 0.28 | 0.52 | 0.32 | 0.44 |
| 8 | 81905294 | rs895270 | -0.521 | -2.291 | -0.269 | 0.48 | 0.003 | -0.013 | -0.013 | 0.051 | 0.2 | 0.18 | 0.17 | 0.5 | 0.21 | 0.26 |
| 8 | 81905397 | rs2016465 | -1.119 | -2.104 | -0.239 | 0.597 | 0.126 | 0.035 | 0.035 | 0 | 0.5 | 0.85 | 0.32 | 0.53 | 0.33 | 0.43 |
| 8 | 81905533 | rs895267 | -1.056 | -2.122 | -0.169 | 0.529 | 0.11 | 0.004 | 0.004 | 0 | 0.49 | 0.77 | 0.31 | 0.48 | 0.3 | 0.42 |
| 8 | 81911211 | rs7818013 | 0.228 | -1.999 | -0.055 | 1.051 | 0.025 | -0.055 | -0.055 | 0 | 0.82 | 0.94 | 0.91 | 0.61 | 0.86 | 0.76 |
| 8 | 81920610 | rs2400340 | 0.975 | -2.478 | -0.196 | 0.502 | 0.097 | -0.033 | -0.033 | 0 | 0.47 | 0.21 | 0.73 | 0.48 | 0.67 | 0.5 |
| 8 | 81930070 | rs1347756 | 1.104 | -2.56 | 0.519 | 0.57 | 0.053 | -0.026 | -0.026 | 0 | 0.45 | 0.1 | 0.67 | 0.42 | 0.62 | 0.48 |
| 8 | 81936438 | rs2705495 | 0.605 | -2.666 | 0.969 | 0.447 | 0.005 | -0.039 | -0.039 | 0.815 | 0.28 | 0.3 | 0.11 | 0 | 0.14 | 0.14 |
| 8 | 81944983 | rs7007766 | 0.505 | -2.433 | 0.822 | 0.64 | 0.01 | 0 | 0 | 0 | 0.51 | 0.45 | 0.76 | 0.48 | 0.7 | 0.51 |
| 8 | 81948444 | rs4739611 | 0.968 | -2.343 | 0.2 | 0.609 | 0.05 | -0.013 | -0.013 | 0 | 0.43 | 0.15 | 0.67 | 0.41 | 0.62 | 0.46 |
| 8 | 81951523 | rs2705496 | 0.13 | -2.643 | 0.683 | 0.677 | -0.043 | 0.082 | 0.082 | 0.22 | 0.79 | 0.65 | 0.83 | 0.53 | 0.86 | 0.79 |
| 8 | 81954342 | rs2623026 | 0.486 | -2.856 | -0.011 | 0.305 | 0.036 | 0.036 | 0.036 | 0 | 0.3 | 0.19 | 0.11 | 0.07 | 0.19 | 0.16 |
| 8 | 81954377 | rs6989890 | 0.798 | -2.691 | -0.278 | 0.532 | 0.102 | -0.02 | -0.02 | 0.006 | 0.44 | 0.15 | 0.68 | 0.39 | 0.65 | 0.51 |
| 8 | 81956152 | rs2705499 | 0.02 | -2.342 | -0.148 | -0.226 | 0.022 | 0.019 | 0.019 | 0 | 0.33 | 0.28 | 0.17 | 0.1 | 0.24 | 0.28 |
| 8 | 81956395 | rs7000832 | 0.638 | -1.18 | -1.121 | 0.157 | -0.175 | -0.13 | -0.13 | 0.988 | 0.28 | 0.02 | 0.32 | 0.4 | 0.4 | 0.45 |
| 8 | 81958305 | rs1026830 | 1 | -2.41 | 0.114 | -0.735 | -0.07 | -0.013 | -0.013 | 0 | 0.67 | 0.42 | 0.77 | 0.74 | 0.83 | 0.66 |
| 8 | 81962644 | rs4585701 | -0.46 | -2.201 | 0.331 | -0.672 | -0.032 | -0.04 | -0.04 | 0 | 0.12 | 0.35 | 0.05 | 0.03 | 0.06 | 0.13 |
| 8 | 81962952 | rs16908481 | 0.008 | -1.729 | -0.242 | -0.357 | 0.051 | -0.067 | -0.067 | 3.257 | 0.15 | 0.03 | 0.14 | 0.22 | 0.09 | 0.18 |
| 8 | 81971745 | rs7815826 | -0.581 | -1.143 | 0.023 | -1.111 | -0.004 | 0.017 | 0.017 | 0 | 0.53 | 0.6 | 0.76 | 0.86 | 0.51 | 0.69 |
| 8 | 81973288 | rs4265143 | -0.635 | -1.176 | 0.026 | -1.113 | -0.038 | 0.199 | 0.199 | 0 | 0.54 | 0.79 | 0.75 | 0.86 | 0.52 | 0.7 |
| 8 | 81974075 | rs11986552 | -0.557 | -1.19 | 0.848 | -1.105 | -0.005 | 0.022 | 0.022 | 0 | 0.53 | 0.6 | 0.73 | 0.86 | 0.52 | 0.7 |
| 8 | 81976972 | rs11778741 | 0.634 | -1.518 | 1.086 | -1.17 | -0.023 | 0.018 | 0.018 | 0 | 0.31 | 0.26 | 0.58 | 0.76 | 0.4 | 0.46 |
| 8 | 81978867 | rs10957998 | 0.624 | -1.596 | 1.308 | -1.236 | -0.14 | -0.023 | -0.023 | 0 | 0.32 | 0.06 | 0.53 | 0.68 | 0.44 | 0.49 |
| 8 | 81981261 | rs12676670 | 0.595 | -1.583 | 1.574 | -1.229 | -0.063 | 0.006 | 0.006 | 0 | 0.33 | 0.16 | 0.55 | 0.68 | 0.44 | 0.46 |
| 8 | 81982892 | rs10108428 | -0.541 | -1.266 | 1.885 | -1.237 | -0.008 | 0.131 | 0.131 | 0 | 0.55 | 0.74 | 0.77 | 0.86 | 0.55 | 0.71 |
| 8 | 81991349 | rs4515498 | -0.479 | -1.271 | 1.165 | -1.145 | -0.033 | 0.081 | 0.081 | 0 | 0.55 | 0.85 | 0.67 | 0.67 | 0.55 | 0.59 |
| 8 | 81997301 | rs7834107 | -0.06 | -1.336 | 1.414 | -1.177 | 0 | 0.122 | 0.122 | 0 | 0.49 | 0.14 | 0.58 | 0.67 | 0.55 | 0.6 |
| 8 | 82002872 | rs7007992 | -1.662 | -1.025 | 1.806 | -1.316 | 0.001 | -0.008 | -0.008 | 0 | 0.08 | 0.56 | 0.08 | 0 | 0.02 | 0.05 |
| 8 | 82004857 | rs7018273 | -2.12 | -1.039 | 0.876 | -1.322 | -0.039 | -0.045 | -0.045 | 0 | 0.1 | 0.8 | 0.1 | 0 | 0.02 | 0.05 |
| 8 | 82011187 | rs4237093 | 0.11 | -0.926 | 0.287 | -1.531 | 0.004 | 0.02 | 0.02 | 0 | 0.49 | 0.55 | 0.63 | 0.75 | 0.49 | 0.67 |
| 8 | 82013498 | rs9987382 | -0.2 | -0.72 | -0.638 | -1.45 | 0.013 | -0.031 | -0.031 | 0 | 0.07 | 0.23 | 0.05 | 0 | 0.04 | 0.04 |
| 8 | 82015265 | rs7835961 | -0.157 | -0.548 | -1.79 | -1.305 | -0.047 | 0.12 | 0.12 | 0 | 0.55 | 0.87 | 0.67 | 0.75 | 0.49 | 0.7 |
| 8 | 82016264 | rs10504729 | -0.464 | -0.985 | -1.566 | -1.376 | 0.06 | 0.076 | 0.076 | 0 | 0.42 | 0.73 | 0.25 | 0.15 | 0.33 | 0.37 |
| 8 | 82017372 | rs10504730 | -0.226 | -0.666 | -2.032 | -0.917 | 0.02 | -0.043 | -0.043 | 0.216 | 0.19 | 0.03 | 0.07 | 0.19 | 0.16 | 0.19 |
| 8 | 82032183 | rs4480065 | 0.823 | -0.459 | -2.914 | -2.128 | -0.006 | 0.184 | 0.184 | 3.382 | 0.41 | 0.09 | 0.58 | 0.85 | 0.35 | 0.6 |
| 8 | 82033862 | rs10504732 | 0.768 | 0.423 | -2.086 | -2.588 | 0.031 | -0.107 | -0.107 | 0.244 | 0.21 | 0.32 | 0.2 | 0 | 0.31 | 0.11 |

| | | | | | | | | | | | | | | | | |
|---|---|---|---|---|---|---|---|---|---|---|---|---|---|---|---|---|
| 8 | 82036542 | rs12677218 | 0.486 | 0.304 | -2.085 | -2.705 | -0.065 | 0.18 | 0.18 | 1.358 | 0.62 | 0.17 | 0.66 | 1 | 0.5 | 0.78 |
| 8 | 82039618 | rs16908663 | -0.337 | 0.322 | -1.955 | -2.539 | -0.002 | 0.048 | 0.048 | 0.18 | 0.4 | 0.3 | 0.27 | 0.17 | 0.46 | 0.29 |
| 8 | 82042292 | rs7839273 | 0.433 | 0.766 | -1.599 | -2.658 | -0.045 | 0.059 | 0.059 | 0.065 | 0.21 | 0.1 | 0.16 | 0 | 0.32 | 0.12 |
| 8 | 82048061 | rs4598215 | 0.243 | 0.7 | -0.902 | -1.966 | -0.039 | 0.035 | 0.035 | 0 | 0.91 | 0.98 | 0.92 | 1 | 0.83 | 0.89 |
| 8 | 82056340 | rs2895702 | 1.12 | 0.62 | -0.874 | -1.986 | -0.012 | -0.005 | -0.005 | 0 | 0.82 | 0.17 | 0.81 | 1 | 0.79 | 0.86 |
| 8 | 82061692 | rs17494473 | -0.332 | 0.433 | -1.109 | -2.172 | 0.009 | 0.019 | 0.019 | 0.014 | 0.1 | 0.05 | 0.04 | 0 | 0.06 | 0.03 |
| 8 | 82063205 | rs4739795 | 1.652 | 0.419 | -1.051 | -2.235 | -0.029 | 0.095 | 0.095 | 1.806 | 0.7 | 0.13 | 0.77 | 1 | 0.72 | 0.81 |
| 8 | 82063993 | rs6986622 | 1.317 | 0.421 | -0.877 | -2.234 | 0.001 | -0.064 | -0.064 | 0 | 0.72 | 0.34 | 0.8 | 1 | 0.72 | 0.81 |
| 8 | 82073833 | rs13266020 | 1.663 | 0.391 | -1.118 | -2.25 | -0.041 | 0.141 | 0.141 | 1.868 | 0.7 | 0.09 | 0.76 | 1 | 0.72 | 0.81 |
| 8 | 82096911 | rs1896216 | 0.75 | 0.243 | 0.48 | -2.05 | 0.041 | -0.085 | -0.085 | 0.227 | 0.86 | 0.69 | 0.9 | 1 | 0.83 | 0.83 |
| 8 | 82119503 | rs6473264 | 1.32 | -0.144 | -0.24 | -2.328 | 0.012 | 0.025 | 0.025 | 1.979 | 0.69 | 0.21 | 0.81 | 1 | 0.75 | 0.79 |
| 8 | 82149338 | rs4739619 | 1.314 | 0 | -0.72 | -2.088 | 0.003 | 0.152 | 0.152 | 3.08 | 0.69 | 0.1 | 0.8 | 0.96 | 0.74 | 0.78 |
| 8 | 82154061 | rs16908834 | -0.706 | 0.002 | -0.339 | -0.521 | -0.001 | 0.095 | 0.095 | 0 | 0.06 | 0.03 | 0.08 | 0.46 | 0.09 | 0.12 |
| 8 | 82156582 | rs10504737 | -0.345 | -0.4 | -0.258 | -1.481 | -0.023 | 0.007 | 0.007 | 0 | 0.21 | 0.23 | 0.27 | 0.76 | 0.14 | 0.44 |
| 8 | 82158027 | rs10090760 | 1.329 | 0.641 | 0.386 | -1.58 | -0.003 | -0.038 | -0.038 | 0 | 0.65 | 0.38 | 0.56 | 0.83 | 0.67 | 0.77 |
| 8 | 82159397 | rs4739801 | -0.788 | 0.745 | 0.004 | -1.552 | 0.001 | 0.001 | 0.001 | 0 | 0.14 | 0.2 | 0.32 | 0.17 | 0.14 | 0.09 |
| 8 | 82165658 | rs11782731 | 1.345 | 0.912 | 0.06 | -1.466 | -0.002 | 0.014 | 0.014 | 1.64 | 0.74 | 0.2 | 0.58 | 0.83 | 0.77 | 0.83 |
| 8 | 82166028 | rs4338057 | 0.179 | 0.577 | 0.691 | -1.501 | 0.024 | -0.061 | -0.061 | 0 | 0.9 | 0.69 | 0.94 | 1 | 0.91 | 0.92 |
| 8 | 82172031 | rs7828212 | 1.387 | 0.588 | 0.624 | -1.515 | 0 | -0.048 | -0.048 | 0.172 | 0.74 | 0.23 | 0.85 | 1 | 0.82 | 0.86 |
| 8 | 82183000 | rs12549270 | 0.036 | 0.64 | 0.629 | -1.477 | 0.032 | -0.07 | -0.07 | 0 | 0.89 | 0.66 | 0.95 | 1 | 0.91 | 0.92 |
| 8 | 82199438 | rs482745 | 1.387 | 0.653 | 0.504 | -1.469 | -0.005 | -0.067 | -0.067 | 0 | 0.73 | 0.25 | 0.84 | 1 | 0.8 | 0.84 |
| 8 | 82200518 | rs202275 | 0.309 | 1.18 | 0.763 | -0.826 | 0.024 | -0.077 | -0.077 | 0 | 0.88 | 0.69 | 0.93 | 1 | 0.89 | 0.91 |
| 8 | 82201519 | rs202277 | 0.62 | 1.191 | 0.144 | -0.821 | 0.024 | -0.1 | -0.1 | 0 | 0.86 | 0.51 | 0.9 | 1 | 0.89 | 0.91 |
| 8 | 82202118 | rs202279 | 0.62 | 1.191 | 0.144 | -0.821 | 0.024 | -0.1 | -0.1 | 0.006 | 0.86 | 0.51 | 0.9 | 1 | 0.89 | 0.91 |
| 8 | 82225697 | rs202296 | 0.633 | 0.7 | -0.229 | -0.483 | 0.012 | -0.107 | -0.107 | 0 | 0.85 | 0.46 | 0.88 | 1 | 0.86 | 0.89 |
| 8 | 82232665 | rs10100646 | -0.074 | 0.858 | -1.514 | -0.114 | -0.025 | 0.014 | 0.014 | 0.016 | 0.36 | 0.12 | 0.26 | 0.25 | 0.38 | 0.51 |
| 8 | 82233820 | rs890555 | 0.596 | 0.609 | -1.345 | -0.434 | -0.001 | -0.002 | -0.002 | 0.013 | 0.65 | 0.31 | 0.68 | 0.76 | 0.68 | 0.79 |
| 8 | 82234543 | rs7841723 | 0.394 | 0.477 | -0.443 | -0.499 | 0.01 | -0.028 | -0.028 | 0 | 0.7 | 0.62 | 0.72 | 0.76 | 0.68 | 0.79 |
| 8 | 82236582 | rs13251566 | -1.026 | -0.112 | -0.525 | -0.746 | 0.011 | 0.014 | 0.014 | 0 | 0.26 | 0.24 | 0.23 | 0.23 | 0.19 | 0.14 |
| 8 | 82236841 | rs2081678 | -1.413 | 0.199 | 0.155 | -0.362 | 0.022 | 0.006 | 0.006 | 0.228 | 0.21 | 0.55 | 0.28 | 0.27 | 0.2 | 0.13 |
| 8 | 82237403 | rs2102113 | -1.365 | -0.346 | -0.232 | 0.514 | 0.003 | -0.007 | -0.007 | 0.087 | 0.25 | 0.72 | 0.29 | 0.57 | 0.18 | 0.18 |
| 8 | 82244169 | rs10958013 | -1.459 | -0.365 | -0.322 | 0.466 | 0.022 | -0.051 | -0.051 | 0.115 | 0.27 | 0.8 | 0.34 | 0.55 | 0.2 | 0.19 |
| 8 | 82245226 | rs1566558 | -1.126 | -0.508 | -1.267 | -0.092 | -0.034 | 0.155 | 0.155 | 5.862 | 0.69 | 0.99 | 0.77 | 0.92 | 0.78 | 0.73 |
| 8 | 82245451 | rs1909936 | -0.322 | -0.623 | 0.234 | 0.844 | 0.028 | 0.252 | 0.252 | 0.489 | 0.07 | 0.43 | 0.07 | 0.25 | 0.02 | 0.07 |
| 8 | 82246092 | rs12155567 | -0.67 | -0.565 | -0.473 | 0.709 | 0.004 | 0 | 0 | 0 | 0.17 | 0.21 | 0.22 | 0.18 | 0.18 | 0.08 |
| 8 | 82249548 | rs10435626 | -1.208 | -0.17 | -1.218 | 1.002 | -0.004 | 0.023 | 0.023 | 2.29 | 0.24 | 0.7 | 0.33 | 0.59 | 0.2 | 0.2 |
| 8 | 82251964 | rs1346291 | -0.098 | 0.696 | -1.498 | 0.731 | 0 | 0 | 0 | 0 | 0.73 | 0.75 | 0.79 | 0.72 | 0.79 | 0.86 |
| 8 | 82256186 | rs13266574 | -0.45 | 1.608 | -1.528 | 1.119 | -0.08 | -0.061 | -0.061 | 0 | 0.17 | 0.01 | 0.14 | 0.17 | 0.29 | 0.14 |
| 8 | 82263832 | rs10958016 | -0.454 | 0.884 | -0.81 | 1.17 | -0.015 | 0.034 | 0.034 | 0.211 | 0.33 | 0.48 | 0.42 | 0.42 | 0.32 | 0.44 |
| 8 | 82266889 | rs12234970 | -0.684 | 0.477 | -0.379 | 1.74 | -0.013 | 0.018 | 0.018 | 0 | 0.25 | 0.35 | 0.34 | 0.7 | 0.19 | 0.26 |
| 8 | 82277240 | rs11774397 | 0.39 | 0.915 | 0.301 | 1.832 | 0.005 | -0.032 | -0.032 | 0.336 | 0.08 | 0.14 | 0.08 | 0 | 0.1 | 0.03 |
| 8 | 82277361 | rs7816915 | -0.247 | 0.838 | 0.62 | 1.839 | 0.015 | 0.094 | 0.094 | 0 | 0.1 | 0.3 | 0.25 | 0.28 | 0.1 | 0.07 |
| 8 | 82284413 | rs17582647 | -0.161 | 0.615 | 0.554 | 1.305 | 0.02 | 0.021 | 0.021 | 0 | 0.19 | 0.09 | 0.08 | 0 | 0.15 | 0.14 |
| 8 | 82289838 | rs1450790 | -1.555 | 0.452 | 0.669 | 1.178 | -0.003 | -0.001 | -0.001 | 0 | 0.38 | 0.67 | 0.45 | 0.69 | 0.32 | 0.31 |
| 8 | 82307171 | rs10087852 | -0.701 | 1.294 | -0.063 | 0.851 | -0.051 | -0.036 | -0.036 | 0 | 0.92 | 0.99 | 0.76 | 0.92 | 0.81 | 0.82 |
| 8 | 82321682 | rs975353 | 0.072 | 0.584 | 0.127 | 0.741 | 0.023 | 0.028 | 0.028 | 0 | 0.19 | 0.07 | 0.07 | 0 | 0.15 | 0.14 |
| 8 | 82334430 | rs10958022 | -0.867 | 0.548 | 0.698 | 0.903 | -0.006 | -0.024 | -0.024 | 0 | 0.29 | 0.33 | 0.22 | 0.11 | 0.24 | 0.21 |
| 8 | 82335354 | rs7012897 | -1.554 | 0.438 | 1.143 | 1.097 | 0.017 | 0.01 | 0.01 | 0 | 0.39 | 0.71 | 0.45 | 0.74 | 0.3 | 0.31 |
| 8 | 82336076 | rs16909095 | -0.872 | 0.237 | 1.517 | 1.019 | -0.012 | -0.041 | -0.041 | 0 | 0.3 | 0.4 | 0.22 | 0.08 | 0.22 | 0.19 |
| 8 | 82336285 | rs10097139 | -2.105 | 0.254 | 0.885 | 0.444 | -0.132 | 0.265 | 0.265 | 0 | 0.59 | 0.97 | 0.73 | 0.94 | 0.51 | 0.58 |
| 8 | 82337084 | rs6984711 | -1.196 | -0.31 | 1.503 | 0.823 | -0.012 | 0.043 | 0.043 | 0 | 0.2 | 0.36 | 0.31 | 0.74 | 0.14 | 0.17 |
| 8 | 82352251 | rs7821877 | -1.255 | -0.214 | 1.489 | 0.858 | -0.004 | 0.066 | 0.066 | 0 | 0.19 | 0.36 | 0.33 | 0.75 | 0.14 | 0.17 |
| 8 | 82353078 | rs2167255 | -1.516 | -0.236 | 1.53 | 0.746 | 0.03 | 0.028 | 0.028 | 0 | 0.41 | 0.75 | 0.46 | 0.75 | 0.31 | 0.31 |
| 8 | 82353503 | rs10112249 | -1.188 | -0.234 | 1.509 | 0.748 | -0.004 | 0.066 | 0.066 | 0 | 0.19 | 0.36 | 0.33 | 0.75 | 0.14 | 0.17 |
| 8 | 82353594 | rs10097844 | -1.374 | -0.284 | 1.484 | 0.722 | -0.008 | 0.058 | 0.058 | 0 | 0.2 | 0.36 | 0.33 | 0.75 | 0.14 | 0.17 |
| 8 | 82355621 | rs6473276 | -1.524 | -0.215 | 1.554 | 0.584 | 0.032 | 0.03 | 0.03 | 0 | 0.41 | 0.76 | 0.46 | 0.75 | 0.31 | 0.31 |
| 8 | 82357112 | rs2229015 | -1.235 | -0.14 | 1.495 | 0.672 | -0.004 | 0.066 | 0.066 | 0 | 0.19 | 0.36 | 0.33 | 0.75 | 0.14 | 0.17 |
| 8 | 82361129 | rs3824088 | 0.113 | -0.131 | 1.525 | 0.768 | 0.044 | 0.316 | 0.316 | 0 | 0.08 | 0.27 | 0.21 | 0.63 | 0.08 | 0.1 |
| 8 | 82363342 | rs1156770 | -1.105 | -0.134 | 1.509 | 0.489 | -0.002 | 0.077 | 0.077 | 0 | 0.18 | 0.33 | 0.33 | 0.73 | 0.14 | 0.15 |
| 8 | 82383934 | rs4739815 | -1.546 | 0.007 | 1.218 | 0.61 | 0.02 | 0.004 | 0.004 | 0 | 0.35 | 0.75 | 0.43 | 0.74 | 0.26 | 0.3 |
| 8 | 82390760 | rs1054135 | -0.205 | 0.128 | 1.06 | 0.625 | 0.044 | 0.217 | 0.217 | 0 | 0.09 | 0.28 | 0.22 | 0.68 | 0.09 | 0.13 |
| 8 | 82394421 | rs7018409 | -0.512 | -0.094 | 0.563 | 0.188 | 0.006 | 0.004 | 0.004 | 0 | 0.22 | 0.16 | 0.17 | 0.07 | 0.16 | 0.17 |
| 8 | 82409004 | rs6473282 | -0.543 | -0.176 | 0.269 | 0.146 | 0.019 | 0.053 | 0.053 | 0 | 0.22 | 0.08 | 0.16 | 0.06 | 0.16 | 0.14 |
| 8 | 82414063 | rs6992709 | -0.151 | -0.101 | 0.302 | 0.17 | 0.006 | 0.011 | 0.011 | 0.13 | 0.16 | 0.06 | 0.08 | 0 | 0.15 | 0.1 |
| 8 | 82419512 | rs1485999 | -1.081 | -0.476 | 0.375 | -0.293 | 0.034 | 0.042 | 0.042 | 0 | 0.4 | 0.67 | 0.43 | 0.14 | 0.3 | 0.4 |
| 8 | 82425347 | rs1486004 | -1.106 | -0.501 | 0.174 | -0.308 | 0.084 | 0.122 | 0.122 | 0 | 0.41 | 0.81 | 0.44 | 0.14 | 0.3 | 0.4 |
| 8 | 82426412 | rs7821748 | -1.533 | -0.326 | 0.044 | -0.318 | 0.036 | 0.189 | 0.189 | 0 | 0.49 | 0.94 | 0.65 | 0.78 | 0.42 | 0.52 |
| 8 | 82427790 | rs10504740 | -2.368 | -0.088 | 1.411 | 0.163 | 0.049 | 0.125 | 0.125 | 2.611 | 0.06 | 0.55 | 0.14 | 0.06 | 0.01 | 0.03 |
| 8 | 82435447 | rs11996913 | -0.439 | -0.004 | 1.408 | 0.04 | 0.031 | -0.016 | -0.016 | 1.94 | 0.06 | 0.27 | 0.04 | 0 | 0.03 | 0.05 |

| | | | | | | | | | | | | | | | | |
|---|---|---|---|---|---|---|---|---|---|---|---|---|---|---|---|---|
| 8 | 82435456 | rs16909302 | -0.673 | -0.045 | 1.5 | -0.129 | -0.017 | -0.084 | -0.084 | 0.458 | 0.14 | 0.3 | 0.09 | 0 | 0.12 | 0.06 |
| 8 | 82437457 | rs11994340 | -0.413 | -0.061 | 1.497 | -0.009 | 0.028 | -0.018 | -0.018 | 0 | 0.06 | 0.26 | 0.04 | 0 | 0.03 | 0.05 |
| 8 | 82439221 | rs16909306 | -0.575 | 0.082 | 1.35 | -0.007 | -0.015 | -0.078 | -0.078 | 0.593 | 0.13 | 0.29 | 0.08 | 0 | 0.11 | 0.05 |
| 8 | 82441040 | rs9298367 | -1.408 | 0.179 | 1.425 | 0.298 | -0.007 | 0.024 | 0.024 | 0.218 | 0.21 | 0.59 | 0.14 | 0.23 | 0.16 | 0.18 |
| 8 | 82443387 | rs11985337 | -1.657 | 0.049 | 0.985 | 0.187 | 0.006 | 0.006 | 0.006 | 0.179 | 0.3 | 0.78 | 0.18 | 0.23 | 0.2 | 0.23 |
| 8 | 82444396 | rs6992664 | 0.483 | -0.019 | 0.774 | 0.188 | -0.023 | -0.028 | -0.028 | 0 | 0.07 | 0.1 | 0.01 | 0 | 0.01 | 0.02 |
| 8 | 82463838 | rs10095951 | 0.696 | 0.418 | -0.01 | 0.882 | -0.024 | -0.05 | -0.05 | 0 | 0.12 | 0.04 | 0.14 | 0.26 | 0.14 | 0.05 |
| 8 | 82464906 | rs10046697 | -0.861 | 0.284 | -0.223 | 1.057 | -0.03 | -0.093 | -0.093 | 0 | 0.2 | 0.06 | 0.17 | 0.35 | 0.21 | 0.17 |
| 8 | 82466485 | rs7824133 | -0.195 | 0.297 | -0.217 | 1.058 | -0.009 | -0.032 | -0.032 | 0 | 0.07 | 0.01 | 0.03 | 0.09 | 0.07 | 0.11 |
| 8 | 82471545 | rs396738 | -1.768 | 0.119 | 0.404 | 1.108 | 0 | -0.001 | -0.001 | 0.568 | 0.37 | 0.68 | 0.28 | 0.49 | 0.29 | 0.31 |
| 8 | 82483592 | rs7010208 | 1.457 | 0.052 | 0.921 | 1.078 | -0.014 | 0.009 | 0.009 | 0.029 | 0.66 | 0.54 | 0.75 | 0.65 | 0.75 | 0.76 |
| 8 | 82493729 | rs2556575 | 1.297 | 0.033 | 0.985 | 1.083 | -0.027 | 0.02 | 0.02 | 0.114 | 0.68 | 0.46 | 0.75 | 0.65 | 0.79 | 0.77 |
| 8 | 82495971 | rs1593714 | 1.051 | -0.051 | 1.001 | 0.98 | -0.062 | -0.047 | -0.047 | 0.067 | 0.81 | 0.55 | 0.89 | 0.92 | 0.93 | 0.82 |
| 8 | 82499131 | rs2114695 | 1.587 | 0.179 | 0.847 | 0.961 | 0.001 | -0.005 | -0.005 | 4.119 | 0.65 | 0.29 | 0.72 | 0.51 | 0.73 | 0.7 |
| 8 | 82503999 | rs2581529 | 1.801 | -0.219 | 0.347 | -0.016 | -0.037 | 0.16 | 0.16 | 0.239 | 0.45 | 0.18 | 0.55 | 0.1 | 0.56 | 0.56 |
| 8 | 82517136 | rs10958032 | 1.008 | 0.111 | 0.343 | 0.291 | 0.005 | -0.017 | -0.017 | 0 | 0.13 | 0.16 | 0.09 | 0.06 | 0.16 | 0.06 |
| 8 | 82523326 | rs2195099 | -2.056 | 0.181 | -0.03 | -0.146 | -0.002 | 0.055 | 0.055 | 0 | 0.6 | 0.76 | 0.49 | 0.88 | 0.56 | 0.5 |
| 8 | 82545437 | rs890601 | 0.693 | 0.504 | 0.062 | 0.474 | -0.047 | -0.057 | -0.057 | 0 | 0.1 | 0.01 | 0.15 | 0.58 | 0.16 | 0.09 |
| 8 | 82545970 | rs2021174 | 1.117 | 0.45 | 0.031 | 0.488 | 0.012 | -0.025 | -0.025 | 0 | 0.05 | 0.01 | 0.03 | 0.09 | 0.03 | 0.1 |
| 8 | 82549124 | rs2957018 | 0.713 | 0.457 | 0.033 | 0.49 | -0.047 | -0.057 | -0.057 | 0.241 | 0.1 | 0.01 | 0.15 | 0.58 | 0.16 | 0.09 |
| 8 | 82556653 | rs2912806 | 0.713 | 0.457 | 0.033 | 0.49 | -0.047 | -0.057 | -0.057 | 0.007 | 0.1 | 0.01 | 0.15 | 0.58 | 0.16 | 0.09 |
| 8 | 82570631 | rs1058401 | -0.791 | 0.24 | 0.194 | 0.267 | 0 | -0.001 | -0.001 | 2.647 | 0.3 | 0.35 | 0.29 | 0.74 | 0.32 | 0.24 |
| 8 | 82571434 | rs2955014 | 0.817 | 0.238 | 0.192 | 0.276 | -0.047 | -0.058 | -0.058 | 0 | 0.1 | 0.01 | 0.15 | 0.59 | 0.15 | 0.09 |
| 8 | 82582057 | rs7016327 | 0.838 | 0.168 | 0.157 | 0.197 | 0.014 | 0.029 | 0.029 | 0 | 0.15 | 0.06 | 0.07 | 0 | 0.13 | 0.12 |
| 8 | 82583771 | rs1967328 | -0.851 | 0.431 | 0.133 | 0.541 | 0 | -0.001 | -0.001 | 7.86 | 0.3 | 0.35 | 0.29 | 0.74 | 0.32 | 0.24 |
| 8 | 82587368 | rs2268431 | 0.571 | 0.521 | 0.132 | 0.562 | -0.047 | -0.056 | -0.056 | 0.103 | 0.1 | 0.01 | 0.15 | 0.59 | 0.15 | 0.09 |
| 8 | 82598908 | rs2912821 | -1.438 | 0.53 | 0.336 | 0.488 | -0.003 | -0.015 | -0.015 | 0 | 0.41 | 0.51 | 0.34 | 0.74 | 0.37 | 0.3 |
| 8 | 82601846 | rs2955008 | -1.482 | 0.584 | -0.007 | 0.457 | -0.002 | -0.016 | -0.016 | 0 | 0.41 | 0.39 | 0.36 | 0.87 | 0.41 | 0.37 |
| 8 | 82614003 | rs2955015 | 0.573 | 0.625 | -0.049 | 0.595 | -0.047 | -0.056 | -0.056 | 0.009 | 0.1 | 0.01 | 0.15 | 0.59 | 0.15 | 0.09 |
| 8 | 82614590 | rs6415649 | -1.527 | 0.561 | 0.005 | 0.586 | 0.001 | 0.017 | 0.017 | 0.189 | 0.86 | 0.92 | 0.93 | 0.94 | 0.86 | 0.94 |
| 8 | 82617029 | rs16909509 | 0.77 | 0.531 | 0.039 | 0.57 | -0.016 | 0.022 | 0.022 | 0 | 0.06 | 0.14 | 0.05 | 0.08 | 0.03 | 0.1 |
| 8 | 82628098 | rs11985309 | 1.269 | 0.551 | -0.02 | 0.517 | 0.004 | -0.063 | -0.063 | 0.389 | 0.58 | 0.53 | 0.63 | 0.13 | 0.59 | 0.63 |
| 8 | 82632694 | rs7826125 | 1.269 | 0.551 | -0.025 | 0.517 | 0.005 | -0.069 | -0.069 | 1.027 | 0.58 | 0.53 | 0.62 | 0.13 | 0.59 | 0.63 |
| 8 | 82638489 | rs1844358 | -0.505 | 0.426 | -0.759 | 0.716 | -0.015 | -0.059 | -0.059 | 0 | 0.2 | 0.1 | 0.2 | 0.59 | 0.2 | 0.15 |
| 8 | 82646469 | rs16909527 | 0.413 | 0.383 | -0.647 | 0.622 | -0.01 | 0.005 | 0.005 | 0 | 0.06 | 0.09 | 0.04 | 0.08 | 0.03 | 0.1 |
| 8 | 82651241 | rs7828550 | 1.211 | 0.447 | -0.733 | 0.599 | -0.001 | 0.001 | 0.001 | 0.025 | 0.43 | 0.41 | 0.6 | 0.35 | 0.53 | 0.6 |
| 8 | 82653644 | rs11782652 | 0.333 | 0.564 | -1.225 | 0.043 | 0.003 | 0.007 | 0.007 | 0 | 0.11 | 0.07 | 0.05 | 0 | 0.08 | 0.07 |
| 8 | 82657693 | rs13276421 | -0.106 | 0.396 | -1.3 | 0.143 | 0.023 | 0.066 | 0.066 | 0.35 | 0.7 | 0.84 | 0.84 | 0.88 | 0.76 | 0.84 |
| 8 | 82657970 | rs11987442 | 0.636 | 0.539 | -1.301 | 0.554 | -0.001 | 0 | 0 | 0 | 0.11 | 0.09 | 0.14 | 0.24 | 0.11 | 0.13 |
| 8 | 82664723 | rs4537255 | 0.427 | 0.463 | -1.23 | 0.478 | 0.014 | 0.019 | 0.019 | 0 | 0.6 | 0.71 | 0.7 | 0.65 | 0.64 | 0.71 |
| 8 | 82668074 | rs17633897 | 0.109 | 0.604 | -1.45 | 0.58 | 0.029 | 0.062 | 0.062 | 0 | 0.14 | 0.01 | 0.06 | 0 | 0.1 | 0.12 |
| 8 | 82669563 | rs16909563 | 0.121 | 1.667 | -1.272 | 1.362 | -0.023 | 0.006 | 0.006 | 0 | 0.08 | 0.04 | 0.19 | 0.37 | 0.17 | 0.26 |
| 8 | 82680267 | rs1387551 | -0.118 | 0.782 | -1.113 | 0.687 | -0.01 | 0.006 | 0.006 | 0 | 0.13 | 0.17 | 0.19 | 0.18 | 0.09 | 0.2 |
| 8 | 82691708 | rs10112819 | 1.544 | 0.625 | -1.474 | -0.23 | -0.002 | -0.066 | -0.066 | 0 | 0.31 | 0.64 | 0.29 | 0.02 | 0.32 | 0.21 |
| 8 | 82691809 | rs7815272 | -0.766 | 0.761 | -1.762 | 0.113 | -0.055 | -0.096 | -0.096 | 2.952 | 0.75 | 0.93 | 0.71 | 0.45 | 0.74 | 0.81 |
| 8 | 82694974 | rs7846264 | 0.869 | 0.537 | -1.925 | -0.117 | 0.023 | 0.092 | 0.092 | 0.327 | 0.5 | 0.68 | 0.47 | 0.27 | 0.59 | 0.53 |
| 8 | 82695743 | rs10100803 | -0.213 | 0.512 | -1.043 | -0.137 | -0.058 | -0.033 | -0.033 | 0.187 | 0.27 | 0.53 | 0.28 | 0.18 | 0.16 | 0.28 |
| 8 | 82696813 | rs7461058 | 0.695 | 0.726 | -1.449 | -0.142 | -0.064 | -0.067 | -0.067 | 1.023 | 0.14 | 0.02 | 0.15 | 0.24 | 0.2 | 0.27 |
| 8 | 82698328 | rs7836040 | -0.844 | 0.653 | -1.361 | -0.048 | -0.045 | -0.105 | -0.105 | 0 | 0.75 | 0.94 | 0.71 | 0.45 | 0.75 | 0.81 |
| 8 | 82702122 | rs4551296 | 0.951 | 0.675 | -1.182 | 0.088 | -0.006 | 0.028 | 0.028 | 0 | 0.1 | 0.18 | 0.18 | 0.18 | 0.07 | 0.15 |
| 8 | 82705464 | rs10091077 | 0.167 | 0.758 | -1.537 | 0.056 | 0.028 | -0.027 | -0.027 | 0 | 0.59 | 0.84 | 0.64 | 0.45 | 0.66 | 0.68 |
| 8 | 82712929 | rs11776932 | 1.716 | 0.873 | -1.121 | -0.265 | 0.002 | -0.127 | -0.127 | 0 | 0.36 | 0.52 | 0.29 | 0.03 | 0.41 | 0.29 |
| 8 | 82720700 | rs2600604 | 1.732 | 0.882 | -1.123 | -0.278 | 0.003 | -0.13 | -0.13 | 0 | 0.36 | 0.52 | 0.28 | 0.03 | 0.41 | 0.27 |
| 8 | 82722994 | rs10958045 | 0.121 | 0.881 | -1.144 | -0.136 | 0.013 | -0.001 | -0.001 | 0 | 0.59 | 0.66 | 0.61 | 0.4 | 0.67 | 0.71 |
| 8 | 82723869 | rs774469 | 1.843 | 0.887 | -1.175 | -0.133 | 0.002 | -0.127 | -0.127 | 0.116 | 0.36 | 0.52 | 0.29 | 0.03 | 0.41 | 0.29 |
| 8 | 82727515 | rs2635482 | -0.403 | 1.037 | -1.188 | 0.072 | -0.036 | -0.068 | -0.068 | 1.526 | 0.23 | 0.1 | 0.32 | 0.38 | 0.27 | 0.41 |
| 8 | 82752291 | rs12056632 | -0.403 | 1.037 | -1.188 | 0.072 | -0.036 | -0.068 | -0.068 | 0.611 | 0.23 | 0.1 | 0.32 | 0.38 | 0.27 | 0.41 |
| 8 | 82754023 | rs12542216 | 0.613 | 1.068 | -1.225 | 0.123 | -0.064 | -0.059 | -0.059 | 0 | 0.14 | 0.01 | 0.15 | 0.2 | 0.2 | 0.27 |
| 8 | 82760263 | rs1010504 | 0.997 | 0.997 | -1.187 | 0.088 | 0.001 | -0.004 | -0.004 | 0.361 | 0.09 | 0.07 | 0.17 | 0.18 | 0.07 | 0.14 |
| 8 | 82769370 | rs17713368 | 1.178 | 1.222 | -1.213 | 0.073 | -0.038 | -0.026 | -0.026 | 0 | 0.09 | 0.01 | 0.1 | 0.08 | 0.12 | 0.17 |
| 8 | 82791846 | rs10111831 | -1.721 | 1.367 | -1.111 | 0.142 | -0.002 | -0.153 | -0.153 | 0.006 | 0.63 | 0.53 | 0.71 | 0.98 | 0.6 | 0.69 |
| 8 | 82796272 | rs10481273 | -1.012 | 1.24 | -1.334 | 0.066 | 0.023 | 0.177 | 0.177 | 0 | 0.48 | 0.23 | 0.61 | 0.98 | 0.52 | 0.61 |
| 8 | 82825514 | rs4610705 | 0.967 | 1.254 | -1.336 | 0.487 | -0.041 | -0.034 | -0.034 | 0 | 0.1 | 0.01 | 0.1 | 0.1 | 0.13 | 0.18 |
| 8 | 82833013 | rs1966850 | 1.368 | 1.181 | -1.138 | 0.034 | 0.001 | -0.114 | -0.114 | 0 | 0.4 | 0.5 | 0.32 | 0.02 | 0.42 | 0.3 |
| 8 | 82849452 | rs10103353 | -0.482 | 1.202 | -1.432 | 0.285 | -0.016 | 0.068 | 0.068 | 0.051 | 0.44 | 0.12 | 0.43 | 0.69 | 0.43 | 0.45 |
| 8 | 82849658 | rs10103624 | 0.673 | 1.527 | -0.703 | 1.244 | -0.02 | -0.029 | -0.029 | 0.002 | 0.16 | 0 | 0.07 | 0.16 | 0.14 | 0.11 |
| 8 | 82862754 | rs1451991 | 0.358 | 1.518 | -0.248 | 1.182 | 0.031 | 0.111 | 0.111 | 0 | 0.84 | 0.59 | 0.74 | 0.7 | 0.84 | 0.74 |
| 8 | 82879546 | rs2400541 | 0.288 | 1.696 | -1.095 | 1.057 | -0.078 | -0.143 | -0.143 | 0.623 | 0.72 | 0.97 | 0.69 | 0.54 | 0.68 | 0.63 |
| 8 | 82892518 | rs1376842 | -0.314 | 1.468 | -0.224 | 0.721 | 0.009 | 0.011 | 0.011 | 0 | 0.41 | 0.49 | 0.44 | 0.45 | 0.54 | 0.32 |

| | | | | | | | | | | | | | | | | |
|---|---|---|---|---|---|---|---|---|---|---|---|---|---|---|---|---|
| 8 | 82896400 | rs10808847 | -1.437 | 1.542 | -0.43 | 0.47 | 0.01 | 0.076 | 0.076 | 0.247 | 0.66 | 0.85 | 0.77 | 0.82 | 0.74 | 0.62 |
| 8 | 82904399 | rs16909786 | -0.154 | 1.622 | 0.168 | 1.176 | -0.004 | 0.005 | 0.005 | 0 | 0.22 | 0.23 | 0.3 | 0.31 | 0.17 | 0.26 |
| 8 | 82904748 | rs16909787 | 1.452 | 1.582 | -0.048 | 1.208 | 0.005 | -0.035 | -0.035 | 0 | 0.09 | 0 | 0.21 | 0.14 | 0.06 | 0.08 |
| 8 | 82916953 | rs7830040 | -0.468 | 1.689 | 0.321 | 1.235 | -0.011 | -0.01 | -0.01 | 0.003 | 0.39 | 0.34 | 0.55 | 0.51 | 0.49 | 0.32 |
| 8 | 82918392 | rs10095650 | -1.646 | 1.708 | 0.133 | 1.178 | 0.031 | 0.09 | 0.09 | 0 | 0.58 | 0.75 | 0.73 | 0.82 | 0.66 | 0.56 |
| 8 | 82921787 | rs13274868 | 0.499 | 1.739 | 0.208 | 1.33 | -0.099 | 0.079 | 0.079 | 0 | 0.24 | 0.01 | 0.18 | 0.1 | 0.34 | 0.17 |
| 8 | 82921979 | rs4529421 | -0.043 | 1.599 | 0.382 | 1.176 | -0.088 | -0.097 | -0.097 | 0.662 | 0.29 | 0.1 | 0.32 | 0.37 | 0.43 | 0.25 |
| 8 | 82930326 | rs7826017 | -1.473 | 1.563 | 0.548 | 1.12 | 0.018 | 0.047 | 0.047 | 0 | 0.52 | 0.63 | 0.66 | 0.7 | 0.6 | 0.52 |
| 8 | 82937296 | rs16909815 | 0.614 | 1.609 | 0.662 | 1.161 | 0.003 | 0.003 | 0.003 | 0 | 0.12 | 0.13 | 0.09 | 0.17 | 0.11 | 0.18 |
| 8 | 82939088 | rs10216960 | -0.705 | 1.576 | 0.789 | 1.116 | -0.001 | -0.001 | -0.001 | 0 | 0.42 | 0.41 | 0.43 | 0.53 | 0.53 | 0.43 |
| 8 | 82942569 | rs12682120 | -1.361 | 1.501 | 0.81 | 1.049 | 0.001 | 0.002 | 0.002 | 0 | 0.52 | 0.48 | 0.64 | 0.69 | 0.6 | 0.51 |
| 8 | 82945765 | rs7824502 | -1.101 | 1.709 | 0.593 | 0.967 | 0.038 | 0.135 | 0.135 | 0.048 | 0.25 | 0.45 | 0.33 | 0.48 | 0.3 | 0.33 |
| 8 | 82947791 | rs1112855 | 0.984 | 1.307 | 0.153 | 1.101 | 0.006 | -0.001 | -0.001 | 0 | 0.09 | 0.05 | 0.22 | 0.13 | 0.06 | 0.07 |
| 8 | 82947950 | rs1112854 | 0.807 | 1.3 | 0.193 | 1.097 | -0.007 | 0 | 0 | 0 | 0.1 | 0.1 | 0.22 | 0.13 | 0.06 | 0.07 |
| 8 | 82950636 | rs955311 | 0.807 | 1.3 | 0.193 | 1.097 | -0.007 | 0 | 0 | 2.076 | 0.1 | 0.1 | 0.22 | 0.13 | 0.06 | 0.07 |
| 8 | 82953371 | rs1507090 | -0.834 | 1.404 | -0.084 | 1.073 | -0.001 | -0.001 | -0.001 | 0 | 0.42 | 0.58 | 0.59 | 0.51 | 0.5 | 0.32 |
| 8 | 82960488 | rs9298382 | -0.372 | 1.599 | -0.896 | 1.176 | -0.055 | -0.079 | -0.079 | 0 | 0.31 | 0.2 | 0.3 | 0.38 | 0.43 | 0.24 |
| 8 | 82961957 | rs998249 | 0.181 | 1.004 | -1.088 | 0.738 | -0.076 | 0.01 | 0.01 | 0 | 0.26 | 0.05 | 0.21 | 0.18 | 0.37 | 0.17 |
| 8 | 82966648 | rs13251139 | -0.627 | 0.424 | -0.17 | 1.106 | 0 | 0 | 0 | 0 | 0.15 | 0.21 | 0.11 | 0.23 | 0.13 | 0.18 |
| 8 | 82970410 | rs7824840 | 0.545 | 0.393 | -0.484 | 1.047 | 0.008 | -0.017 | -0.017 | 0 | 0.09 | 0 | 0.07 | 0.1 | 0.06 | 0.08 |
| 8 | 82994362 | rs1507088 | -1.161 | 0.302 | 0.35 | 1.282 | -0.015 | 0.062 | 0.062 | 0 | 0.26 | 0.5 | 0.22 | 0.41 | 0.21 | 0.3 |
| 8 | 83007997 | rs10102998 | -1.218 | 0.36 | 0.712 | 1.284 | -0.002 | 0.058 | 0.058 | 0 | 0.26 | 0.43 | 0.22 | 0.4 | 0.25 | 0.3 |
| 8 | 83036411 | rs4739638 | -0.582 | 0.382 | 0.349 | 1.299 | -0.026 | -0.063 | -0.063 | 0 | 0.14 | 0.06 | 0.09 | 0.31 | 0.16 | 0.21 |
| 8 | 83054058 | rs7007666 | -0.78 | 0.035 | 0.586 | 1.181 | -0.044 | -0.034 | -0.034 | 0 | 0.17 | 0.53 | 0.19 | 0.12 | 0.1 | 0.13 |
| 8 | 83072460 | rs6987297 | -0.725 | 0.046 | 0.591 | 1.185 | -0.043 | -0.032 | -0.032 | 0.251 | 0.16 | 0.53 | 0.19 | 0.12 | 0.1 | 0.13 |
| 8 | 83088553 | rs6473329 | -1.77 | -0.166 | 0.128 | 1.085 | 0.021 | -0.018 | -0.018 | 0.59 | 0.35 | 0.69 | 0.3 | 0.45 | 0.28 | 0.34 |
| 8 | 83098845 | rs1454829 | -1.227 | 0.015 | -0.453 | 0.807 | 0.013 | 0.048 | 0.048 | 0.706 | 0.4 | 0.78 | 0.48 | 0.68 | 0.36 | 0.49 |
| 8 | 83115706 | rs4357283 | -0.861 | -0.087 | 0.606 | 0.929 | 0.056 | 0.115 | 0.115 | 1.95 | 0.06 | 0.47 | 0.05 | 0.08 | 0.01 | 0.11 |
| 8 | 83152911 | rs1563244 | -1.023 | -0.046 | 0.783 | 1.266 | -0.001 | 0.013 | 0.013 | 1.288 | 0.28 | 0.69 | 0.4 | 0.5 | 0.27 | 0.4 |
| 8 | 83163527 | rs1378211 | -0.706 | -0.055 | 0.816 | 1.254 | 0.042 | 0.095 | 0.095 | 1.006 | 0.06 | 0.41 | 0.05 | 0.08 | 0.01 | 0.1 |
| 8 | 83176155 | rs7001569 | 0.44 | -0.108 | 0.974 | 1.206 | -0.002 | 0.003 | 0.003 | 0 | 0.12 | 0.09 | 0.08 | 0.18 | 0.08 | 0.08 |
| 8 | 83179915 | rs1242391 | -1.402 | -0.474 | 0.474 | 0.912 | 0.003 | -0.033 | -0.033 | 0.725 | 0.91 | 1 | 0.94 | 0.93 | 0.95 | 0.87 |
| 8 | 83183692 | rs1904275 | 0.869 | 0.072 | 0.827 | 1.157 | -0.005 | -0.058 | -0.058 | 0 | 0.1 | 0.01 | 0.07 | 0.18 | 0.08 | 0.09 |
| 8 | 83219272 | rs11993752 | -0.582 | 0.133 | 1.086 | 1.146 | 0.022 | 0.133 | 0.133 | 0.017 | 0.23 | 0.44 | 0.37 | 0.44 | 0.26 | 0.3 |
| 8 | 83222777 | rs7845376 | -0.014 | 0.222 | 0.892 | 1.668 | -0.031 | -0.038 | -0.038 | 1.418 | 0.1 | 0.04 | 0.3 | 0.33 | 0.18 | 0.07 |
| 8 | 83236695 | rs6987592 | -0.048 | -0.039 | 1.343 | 1.693 | -0.002 | 0.065 | 0.065 | 0.031 | 0.12 | 0.36 | 0.07 | 0.1 | 0.08 | 0.23 |
| 8 | 83252586 | rs10504754 | -1.978 | -0.083 | 0.437 | 1.269 | 0.045 | 0.147 | 0.147 | 0.805 | 0.5 | 0.81 | 0.52 | 0.75 | 0.41 | 0.62 |
| 8 | 83258347 | rs4739861 | -0.859 | -0.051 | 0.982 | 1.466 | -0.004 | 0.059 | 0.059 | 0 | 0.27 | 0.41 | 0.38 | 0.58 | 0.25 | 0.29 |
| 8 | 83262803 | rs17720196 | -2.192 | -0.057 | 1.114 | 1.453 | 0.04 | 0.128 | 0.128 | 0.691 | 0.49 | 0.78 | 0.51 | 0.75 | 0.41 | 0.62 |
| 8 | 83265571 | rs10092418 | -1.42 | -0.013 | 0.923 | 1.486 | -0.004 | 0.007 | 0.007 | 0 | 0.36 | 0.44 | 0.41 | 0.62 | 0.33 | 0.51 |
| 8 | 83271395 | rs16910284 | -0.997 | 0.082 | 1.239 | 1.885 | -0.008 | 0.101 | 0.101 | 1.262 | 0.1 | 0.33 | 0.08 | 0.25 | 0.09 | 0.2 |
| 8 | 83271680 | rs12679925 | -0.997 | 0.082 | 1.239 | 1.885 | -0.008 | 0.101 | 0.101 | 1.262 | 0.1 | 0.33 | 0.09 | 0.25 | 0.09 | 0.28 |
| 8 | 83273087 | rs1031874 | -0.975 | 0.082 | 1.238 | 1.9 | -0.008 | 0.106 | 0.106 | 1.173 | 0.1 | 0.33 | 0.08 | 0.25 | 0.09 | 0.19 |
| 8 | 83273096 | rs17720831 | -0.659 | 0.055 | 1.186 | 1.9 | 0.007 | -0.02 | -0.02 | 0 | 0.2 | 0.04 | 0.1 | 0.17 | 0.16 | 0.33 |
| 8 | 83281436 | rs7828444 | -1.606 | 0.076 | 1.174 | 1.878 | -0.008 | 0.011 | 0.011 | 0 | 0.3 | 0.37 | 0.18 | 0.42 | 0.25 | 0.52 |
| 8 | 83292295 | rs1471464 | 0.716 | 0.225 | 0.898 | 1.755 | -0.003 | -0.043 | -0.043 | 0 | 0.1 | 0.01 | 0.07 | 0.13 | 0.08 | 0.11 |
| 8 | 83293872 | rs1031877 | -1.313 | 0.416 | 1.335 | 2.142 | -0.008 | 0.044 | 0.044 | 0 | 0.22 | 0.34 | 0.33 | 0.45 | 0.16 | 0.42 |
| 8 | 83298113 | rs1904276 | -1.377 | 0.47 | 1.764 | 2.082 | 0.005 | 0.057 | 0.057 | 0 | 0.22 | 0.43 | 0.38 | 0.38 | 0.18 | 0.21 |
| 8 | 83299022 | rs16910330 | 0.239 | 0.383 | 1.696 | 2.107 | -0.047 | -0.07 | -0.07 | 0 | 0.11 | 0.01 | 0.29 | 0.32 | 0.11 | 0.1 |
| 8 | 83299295 | rs13277937 | -2.394 | 0.347 | 1.906 | 2.083 | 0.005 | 0.016 | 0.016 | 0.034 | 0.39 | 0.67 | 0.49 | 0.75 | 0.34 | 0.61 |
| 8 | 83303150 | rs4739876 | -1.114 | 0.292 | 1.745 | 1.901 | 0.002 | 0.042 | 0.042 | 0 | 0.2 | 0.3 | 0.36 | 0.38 | 0.17 | 0.21 |
| 8 | 83312232 | rs7833204 | -2.048 | 0.865 | 1.949 | 1.822 | 0.004 | 0.009 | 0.009 | 0 | 0.33 | 0.6 | 0.47 | 0.41 | 0.33 | 0.45 |
| 8 | 83314667 | rs10958097 | -1.741 | 0.859 | 2.045 | 1.819 | 0 | 0.001 | 0.001 | 0 | 0.3 | 0.32 | 0.44 | 0.41 | 0.32 | 0.44 |
| 8 | 83315548 | rs10504758 | 0.044 | 0.88 | 2.032 | 1.761 | 0.005 | -0.005 | -0.005 | 3.027 | 0.09 | 0.11 | 0.09 | 0.06 | 0.14 | 0.14 |
| 8 | 83319341 | rs11993414 | -1.94 | 0.992 | 1.779 | 1.648 | -0.002 | -0.006 | -0.006 | 0.05 | 0.43 | 0.65 | 0.54 | 0.58 | 0.41 | 0.5 |
| 8 | 83326590 | rs7016611 | -1.089 | 0.988 | 1.871 | 1.607 | -0.007 | -0.046 | -0.046 | 0.057 | 0.3 | 0.2 | 0.42 | 0.53 | 0.26 | 0.36 |
| 8 | 83326784 | rs7016525 | -0.225 | 0.965 | 1.873 | 1.617 | 0 | -0.001 | -0.001 | 0 | 0.18 | 0.17 | 0.12 | 0.22 | 0.15 | 0.26 |
| 8 | 83329505 | rs10504759 | 0.134 | 0.955 | 2.009 | 1.521 | 0.003 | -0.007 | -0.007 | 0 | 0.1 | 0.17 | 0.1 | 0.07 | 0.14 | 0.13 |
| 8 | 83334060 | rs7819753 | -2.223 | 0.957 | 1.848 | 1.546 | -0.019 | 0.071 | 0.071 | 0 | 0.48 | 0.74 | 0.57 | 0.72 | 0.48 | 0.59 |
| 8 | 83334905 | rs7013518 | -2.123 | 0.965 | 1.837 | 1.55 | -0.009 | 0.037 | 0.037 | 0 | 0.47 | 0.64 | 0.56 | 0.72 | 0.48 | 0.59 |
| 8 | 83344525 | rs925673 | -1.991 | 1.01 | 1.845 | 1.574 | -0.002 | 0.005 | 0.005 | 0 | 0.46 | 0.53 | 0.55 | 0.72 | 0.47 | 0.59 |
| 8 | 83371624 | rs16910467 | -0.744 | 1.545 | 1.81 | 1.454 | 0.043 | 0.03 | 0.03 | 0 | 0.14 | 0.26 | 0.14 | 0.21 | 0.25 | 0.22 |
| 8 | 83384151 | rs979908 | -1.625 | 1.591 | 1.819 | 1.51 | 0.02 | 0.046 | 0.046 | 0 | 0.27 | 0.6 | 0.45 | 0.5 | 0.35 | 0.33 |
| 8 | 83386881 | rs9643764 | -0.794 | 1.558 | 1.879 | 1.485 | 0.036 | 0.188 | 0.188 | 0.12 | 0.18 | 0.49 | 0.36 | 0.43 | 0.17 | 0.2 |
| 8 | 83408662 | rs12056325 | 0.048 | 1.524 | 1.59 | 1.473 | 0.03 | 0.12 | 0.12 | 0 | 0.13 | 0.34 | 0.33 | 0.29 | 0.11 | 0.11 |
| 8 | 83412270 | rs950947 | 0.099 | 1.531 | 1.247 | 1.477 | 0.007 | 0.047 | 0.047 | 0 | 0.13 | 0.24 | 0.32 | 0.29 | 0.11 | 0.11 |
| 8 | 83416246 | rs17667124 | 0.16 | 1.616 | 1.058 | 1.424 | 0.005 | -0.003 | -0.003 | 0 | 0.1 | 0.12 | 0.1 | 0.07 | 0.18 | 0.14 |
| 8 | 83416323 | rs17667153 | 0.16 | 1.616 | 1.038 | 1.424 | 0.002 | -0.002 | -0.002 | 0 | 0.1 | 0.12 | 0.1 | 0.07 | 0.18 | 0.14 |
| 8 | 83424234 | rs7829925 | -0.536 | 0.785 | 1.133 | 1.583 | -0.005 | 0.077 | 0.077 | 0 | 0.2 | 0.27 | 0.18 | 0.43 | 0.15 | 0.32 |

| | | | | | | | | | | | | | | | | |
|---|---|---|---|---|---|---|---|---|---|---|---|---|---|---|---|---|
| 8 | 83424302 | rs7830076 | -0.479 | 0.73 | 1.417 | 1.61 | 0.049 | 0.118 | 0.118 | 0.701 | 0.06 | 0.3 | 0.06 | 0.13 | 0.06 | 0.08 |
| 8 | 83427215 | rs12679968 | -0.256 | 0.836 | 0.614 | 1.409 | 0.026 | 0.092 | 0.092 | 0 | 0.13 | 0.28 | 0.34 | 0.3 | 0.12 | 0.11 |
| 8 | 83432030 | rs7013595 | -1.474 | 0.596 | -0.5 | 1.008 | -0.079 | 0.098 | 0.098 | 0 | 0.52 | 0.93 | 0.63 | 0.72 | 0.47 | 0.66 |
| 8 | 83432358 | rs7842921 | -0.206 | 0.5 | 0.512 | 1.171 | -0.019 | 0.034 | 0.034 | 0 | 0.21 | 0.38 | 0.31 | 0.28 | 0.13 | 0.15 |
| 8 | 83439477 | rs1514487 | -0.094 | 0.462 | 0.582 | 1.317 | -0.008 | 0.008 | 0.008 | 0 | 0.21 | 0.14 | 0.13 | 0.16 | 0.21 | 0.27 |
| 8 | 83439540 | rs7009509 | -0.742 | 0.426 | 1.041 | 1.498 | -0.006 | 0.043 | 0.043 | 0.001 | 0.24 | 0.64 | 0.42 | 0.48 | 0.18 | 0.21 |
| 8 | 83449849 | rs4316152 | -1.144 | 0.407 | 1.08 | 1.521 | 0 | 0 | 0 | 0 | 0.28 | 0.71 | 0.45 | 0.55 | 0.2 | 0.34 |
| 8 | 83476458 | rs7000227 | -0.345 | 0.57 | 1.22 | 1.393 | -0.012 | 0.024 | 0.024 | 0 | 0.2 | 0.31 | 0.31 | 0.29 | 0.13 | 0.15 |
| 8 | 83489460 | rs9650274 | -0.189 | 0.337 | 0.595 | 1.038 | -0.001 | 0 | 0 | 8.108 | 0.2 | 0.18 | 0.13 | 0.16 | 0.21 | 0.28 |
| 8 | 83504664 | rs11778345 | -0.374 | 0.194 | 0.384 | 1.135 | -0.001 | 0 | 0 | 0.026 | 0.2 | 0.19 | 0.29 | 0.29 | 0.13 | 0.15 |
| 8 | 83508855 | rs13281657 | 0.498 | 0.009 | 0.038 | 0.001 | 0.009 | 0.072 | 0.072 | 0 | 0.46 | 0.3 | 0.37 | 0.15 | 0.53 | 0.37 |
| 8 | 83515171 | rs11987007 | -0.247 | -0.055 | 0.533 | 0.546 | 0.011 | 0.273 | 0.273 | 0.116 | 0.09 | 0.31 | 0.14 | 0.41 | 0.07 | 0.19 |
| 8 | 83516216 | rs9298389 | -1.559 | 0.023 | 0.085 | 0.077 | -0.11 | 0.173 | 0.173 | 0 | 0.69 | 0.94 | 0.73 | 0.93 | 0.63 | 0.76 |
| 8 | 83524772 | rs7002811 | -0.469 | -0.007 | 0.233 | 0.173 | 0.007 | 0.039 | 0.039 | 0 | 0.53 | 0.73 | 0.58 | 0.57 | 0.49 | 0.51 |
| 8 | 83542128 | rs10111398 | -0.021 | -0.215 | 0.623 | 0.005 | 0 | 0 | 0 | 0 | 0.45 | 0.4 | 0.52 | 0.57 | 0.44 | 0.47 |
| 8 | 83548428 | rs16910695 | 0.551 | -0.145 | 0.673 | 0.027 | -0.01 | -0.043 | -0.043 | 0.013 | 0.08 | 0.24 | 0.05 | 0 | 0.04 | 0.04 |
| 8 | 83552280 | rs7007229 | -0.726 | -0.275 | 0.653 | -0.033 | 0 | -0.003 | -0.003 | 0 | 0.58 | 0.53 | 0.56 | 0.65 | 0.52 | 0.67 |
| 8 | 83556325 | rs7008791 | -1.241 | -0.269 | 0.949 | -0.019 | -0.032 | -0.057 | -0.057 | 0.252 | 0.19 | 0.45 | 0.1 | 0.1 | 0.11 | 0.21 |
| 8 | 83567219 | rs16910716 | -0.459 | -0.149 | 0.972 | -0.036 | -0.003 | -0.017 | -0.017 | 0 | 0.16 | 0.2 | 0.07 | 0.09 | 0.12 | 0.22 |
| 8 | 83573982 | rs6473376 | -0.663 | -0.16 | 0.954 | -0.036 | -0.007 | -0.039 | -0.039 | 0.308 | 0.17 | 0.34 | 0.09 | 0.09 | 0.12 | 0.22 |
| 8 | 83574432 | rs7833831 | -0.097 | -0.285 | 0.542 | -0.18 | -0.001 | 0.018 | 0.018 | 0 | 0.55 | 0.63 | 0.39 | 0.16 | 0.5 | 0.47 |
| 8 | 83585352 | rs4323467 | -0.19 | -0.181 | 0.532 | -0.094 | 0.009 | 0.015 | 0.015 | 0 | 0.56 | 0.66 | 0.49 | 0.38 | 0.57 | 0.52 |
| 8 | 83588608 | rs7819717 | -1.435 | -0.061 | 0.792 | 0.149 | 0.032 | 0.09 | 0.09 | 1.457 | 0.18 | 0.56 | 0.21 | 0.3 | 0.18 | 0.27 |
| 8 | 83591879 | rs4739899 | -1.222 | -0.058 | 0.44 | 0.268 | 0.004 | -0.001 | -0.001 | 0 | 0.47 | 0.55 | 0.44 | 0.5 | 0.35 | 0.64 |
| 8 | 83599801 | rs12155922 | 1.204 | -0.097 | 0.337 | -0.121 | -0.003 | -0.004 | -0.004 | 0 | 0.06 | 0.04 | 0.04 | 0 | 0.07 | 0.06 |
| 8 | 83601965 | rs11996822 | -0.95 | -0.215 | 0.588 | -0.01 | -0.006 | -0.001 | -0.001 | 0.009 | 0.14 | 0.26 | 0.09 | 0.12 | 0.11 | 0.2 |
| 8 | 83615856 | rs17731245 | -0.134 | -0.289 | 0.476 | -0.261 | 0.006 | -0.075 | -0.075 | 0 | 0.17 | 0.2 | 0.12 | 0.01 | 0.15 | 0.22 |
| 8 | 83616309 | rs16910767 | 1.252 | -0.301 | 0.456 | -0.24 | 0 | 0.001 | 0.001 | 0 | 0.06 | 0.06 | 0.03 | 0.08 | 0.03 | 0.14 |
| 8 | 83620515 | rs4623417 | -0.684 | -0.403 | 0.531 | -0.38 | 0.014 | -0.004 | -0.004 | 0 | 0.64 | 0.75 | 0.6 | 0.41 | 0.68 | 0.68 |
| 8 | 83620829 | rs16910782 | -0.048 | -0.325 | 0.555 | -0.263 | 0 | 0.061 | 0.061 | 0 | 0.07 | 0.24 | 0.05 | 0.11 | 0.04 | 0.14 |
| 8 | 83621901 | rs12543690 | 0.362 | -0.245 | 0.32 | -0.275 | 0.011 | 0.029 | 0.029 | 0 | 0.47 | 0.39 | 0.47 | 0.39 | 0.52 | 0.45 |
| 8 | 83629042 | rs4270965 | 1.16 | -0.096 | 0.153 | -0.214 | -0.062 | 0.037 | 0.037 | 0 | 0.36 | 0.17 | 0.39 | 0.28 | 0.46 | 0.3 |
| 8 | 83636018 | rs2177732 | 0.988 | 0.009 | -0.295 | -0.121 | -0.09 | -0.033 | -0.033 | 0 | 0.35 | 0.14 | 0.37 | 0.41 | 0.45 | 0.43 |
| 8 | 83640539 | rs7822192 | -0.606 | -0.164 | -0.052 | -0.404 | -0.005 | -0.094 | -0.094 | 0.263 | 0.23 | 0.66 | 0.18 | 0.01 | 0.18 | 0.25 |
| 8 | 83650902 | rs10958126 | -0.352 | -0.136 | -0.53 | -0.137 | 0.036 | 0.056 | 0.056 | 0 | 0.58 | 0.77 | 0.54 | 0.33 | 0.6 | 0.67 |
| 8 | 83658408 | rs1404768 | 0.754 | -0.24 | -0.321 | -0.034 | -0.026 | -0.014 | -0.014 | 0 | 0.38 | 0.28 | 0.4 | 0.44 | 0.47 | 0.45 |
| 8 | 83668549 | rs4739902 | 0.756 | -0.276 | -0.27 | -0.05 | -0.021 | -0.011 | -0.011 | 0 | 0.38 | 0.29 | 0.4 | 0.43 | 0.48 | 0.44 |
| 8 | 83669120 | rs6473383 | 0.199 | -0.438 | -0.26 | -0.208 | 0.028 | -0.105 | -0.105 | 0.149 | 0.16 | 0.36 | 0.11 | 0.01 | 0.14 | 0.23 |
| 8 | 83670807 | rs9643449 | -0.399 | -0.586 | -0.303 | -0.305 | 0.024 | -0.006 | -0.006 | 0 | 0.59 | 0.67 | 0.55 | 0.45 | 0.66 | 0.69 |
| 8 | 83672742 | rs11997468 | 0.197 | -0.59 | -0.298 | -0.314 | 0.012 | -0.129 | -0.129 | 1.12 | 0.21 | 0.37 | 0.14 | 0.01 | 0.18 | 0.26 |
| 8 | 83678803 | rs1917633 | 0.843 | -0.453 | -0.246 | -0.218 | -0.031 | -0.016 | -0.016 | 0 | 0.38 | 0.28 | 0.41 | 0.44 | 0.48 | 0.43 |
| 8 | 83687961 | rs4541922 | 0.871 | -0.78 | -0.619 | 0.11 | -0.016 | 0.071 | 0.071 | 0 | 0.14 | 0.23 | 0.26 | 0.24 | 0.1 | 0.08 |
| 8 | 83690134 | rs1404756 | -1.216 | -0.733 | -1.031 | 0.047 | 0.019 | -0.037 | -0.037 | 0 | 0.77 | 0.91 | 0.82 | 0.69 | 0.76 | 0.8 |
| 8 | 83691079 | rs7465344 | 2.066 | -0.397 | -1.033 | 0.041 | -0.125 | -0.014 | -0.014 | 0 | 0.27 | 0.04 | 0.31 | 0.23 | 0.39 | 0.17 |
| 8 | 83697485 | rs2204418 | 1.368 | -0.508 | -0.746 | -0.017 | -0.014 | 0.01 | 0.01 | 0 | 0.33 | 0.25 | 0.29 | 0.08 | 0.4 | 0.26 |
| 8 | 83701808 | rs999464 | 0.767 | -1.15 | -0.579 | 0.443 | -0.011 | 0.043 | 0.043 | 0 | 0.13 | 0.2 | 0.26 | 0.24 | 0.1 | 0.08 |
| 8 | 83702826 | rs1525335 | -1.186 | -1.073 | -1.162 | 0.482 | -0.022 | -0.081 | -0.081 | 0.792 | 0.78 | 0.99 | 0.83 | 0.69 | 0.76 | 0.8 |
| 8 | 83705468 | rs1997055 | -0.356 | -0.447 | -0.469 | 0.069 | -0.013 | 0.02 | 0.02 | 0 | 0.24 | 0.25 | 0.21 | 0.37 | 0.18 | 0.33 |
| 8 | 83706036 | rs16910965 | 0.091 | -0.445 | -0.471 | 0.053 | -0.019 | 0.008 | 0.008 | 0 | 0.2 | 0.22 | 0.15 | 0.23 | 0.13 | 0.22 |
| 8 | 83717114 | rs10504767 | 1.161 | -0.424 | -0.53 | 0.021 | -0.112 | 0.152 | 0.152 | 0 | 0.33 | 0.06 | 0.32 | 0.09 | 0.46 | 0.39 |
| 8 | 83719927 | rs7839678 | -0.51 | -0.302 | -0.36 | 0.098 | 0 | -0.001 | -0.001 | 0 | 0.6 | 0.6 | 0.55 | 0.46 | 0.64 | 0.72 |
| 8 | 83720591 | rs12547546 | 0.925 | -0.206 | -0.33 | -0.313 | -0.069 | 0.131 | 0.131 | 0 | 0.35 | 0.12 | 0.33 | 0.1 | 0.46 | 0.39 |
| 8 | 83723890 | rs13268825 | 0.862 | -0.101 | -0.577 | -0.258 | -0.012 | 0.017 | 0.017 | 0 | 0.36 | 0.34 | 0.34 | 0.09 | 0.46 | 0.39 |
| 8 | 83724668 | rs7829384 | 0.126 | 0.195 | -0.641 | 0.304 | 0 | 0 | 0 | 0 | 0.57 | 0.56 | 0.49 | 0.33 | 0.59 | 0.6 |
| 8 | 83734479 | rs9886528 | -0.347 | 0.949 | -0.43 | 0.346 | 0.027 | 0.043 | 0.043 | 0.077 | 0.31 | 0.34 | 0.34 | 0.48 | 0.41 | 0.44 |
| 8 | 83734562 | rs6473386 | -0.87 | 0.191 | -0.82 | -0.584 | 0.02 | 0.02 | 0.02 | 0 | 0.74 | 0.83 | 0.84 | 0.99 | 0.78 | 0.81 |
| 8 | 83738448 | rs7842063 | 1.413 | 0.067 | -0.949 | -0.49 | 0.034 | 0.022 | 0.022 | 0 | 0.27 | 0.14 | 0.19 | 0.2 | 0.19 | 0.21 |
| 8 | 83748196 | rs10111901 | 1.107 | 0.556 | -0.633 | -0.856 | 0.009 | -0.05 | -0.05 | 0 | 0.12 | 0.16 | 0.05 | 0 | 0.11 | 0.09 |
| 8 | 83753850 | rs1525339 | 0.765 | 0.887 | 0.157 | -0.474 | 0.044 | 0.139 | 0.139 | 0 | 0.19 | 0.33 | 0.31 | 0.48 | 0.29 | 0.35 |
| 8 | 83754263 | rs1997056 | -0.238 | 0.852 | 0.27 | -0.462 | 0.016 | -0.007 | -0.007 | 0.97 | 0.32 | 0.36 | 0.48 | 0.76 | 0.48 | 0.53 |
| 8 | 83755658 | rs4739913 | -1.16 | 0.801 | 0.41 | -0.509 | 0.021 | 0.04 | 0.04 | 0 | 0.74 | 0.88 | 0.84 | 0.99 | 0.77 | 0.82 |
| 8 | 83759424 | rs2895840 | 0.943 | 0.808 | 0.417 | 0.047 | 0.041 | 0.039 | 0.039 | 0 | 0.18 | 0.33 | 0.27 | 0.32 | 0.29 | 0.23 |
| 8 | 83761875 | rs1014032 | 0.831 | 0.77 | 0.208 | 0.038 | -0.01 | -0.011 | -0.011 | 0 | 0.15 | 0.11 | 0.18 | 0.27 | 0.19 | 0.17 |
| 8 | 83762709 | rs16911067 | 1.02 | 0 | 0.218 | -0.336 | -0.002 | 0.047 | 0.047 | 0 | 0.06 | 0.12 | 0.08 | 0.16 | 0.05 | 0.12 |
| 8 | 83762822 | rs10102377 | 1.283 | -0.032 | 0.181 | -0.375 | 0.02 | 0.028 | 0.028 | 0 | 0.34 | 0.17 | 0.29 | 0.23 | 0.25 | 0.28 |
| 8 | 83770383 | rs1997054 | -0.983 | 0.292 | 0.146 | -0.716 | 0.024 | 0.087 | 0.087 | 0 | 0.74 | 0.94 | 0.84 | 0.99 | 0.77 | 0.82 |
| 8 | 83776723 | rs4285487 | 0.463 | 0.33 | 0.184 | -0.232 | -0.019 | 0.008 | 0.008 | 0 | 0.41 | 0.5 | 0.4 | 0.4 | 0.3 | 0.41 |
| 8 | 83789467 | rs7008434 | 0.058 | 0.788 | 0.631 | -0.462 | 0.003 | 0.006 | 0.006 | 0 | 0.84 | 0.84 | 0.82 | 0.73 | 0.79 | 0.82 |
| 8 | 83794841 | rs12682090 | 1.275 | 0.348 | 1.132 | -1.054 | 0 | 0.003 | 0.003 | 0 | 0.59 | 0.39 | 0.64 | 0.72 | 0.63 | 0.62 |

| | | | | | | | | | | | | | | | | |
|---|---|---|---|---|---|---|---|---|---|---|---|---|---|---|---|---|
| 8 | 83826701 | rs6473393 | 1.217 | 0.326 | 1.265 | -1.288 | -0.021 | 0.094 | 0.094 | 0 | 0.58 | 0.79 | 0.68 | 0.73 | 0.58 | 0.58 |
| 8 | 83827104 | rs4445223 | 1.618 | 0.305 | 1.404 | -1.298 | 0.007 | 0.111 | 0.111 | 0.022 | 0.53 | 0.22 | 0.59 | 0.73 | 0.58 | 0.58 |
| 8 | 83852033 | rs1584545 | -1.861 | 0.354 | 1.768 | -1.479 | 0.001 | 0.028 | 0.028 | 0 | 0.48 | 0.67 | 0.4 | 0.18 | 0.45 | 0.42 |
| 8 | 83855731 | rs11780716 | -1.495 | 0.321 | 1.848 | -1.496 | -0.024 | 0.115 | 0.115 | 0 | 0.43 | 0.22 | 0.33 | 0.18 | 0.45 | 0.42 |
| 8 | 83856081 | rs11784473 | -0.144 | 0.331 | 1.773 | -1.551 | -0.038 | -0.011 | -0.011 | 0.168 | 0.17 | 0.03 | 0.19 | 0.17 | 0.25 | 0.21 |
| 8 | 83857175 | rs10105219 | -2.048 | 0.354 | 1.845 | -1.526 | 0.002 | 0.099 | 0.099 | 0.021 | 0.48 | 0.73 | 0.4 | 0.18 | 0.45 | 0.42 |
| 8 | 83858931 | rs13281508 | -1.825 | 0.403 | 1.601 | -1.492 | -0.003 | 0.041 | 0.041 | 0 | 0.45 | 0.41 | 0.36 | 0.18 | 0.45 | 0.42 |
| 8 | 83865438 | rs10109906 | -1.525 | 0.251 | 1.219 | -1.419 | -0.017 | 0.114 | 0.114 | 0 | 0.41 | 0.2 | 0.33 | 0.18 | 0.41 | 0.4 |
| 8 | 83865622 | rs1037293 | -0.735 | 0.168 | 1.332 | -1.46 | 0.01 | 0.053 | 0.053 | 0 | 0.24 | 0.15 | 0.14 | 0.01 | 0.19 | 0.19 |
| 8 | 83888761 | rs2167107 | -1.221 | -0.186 | 1.711 | -1.45 | -0.012 | -0.088 | -0.088 | 1.053 | 0.27 | 0.64 | 0.18 | 0.01 | 0.2 | 0.19 |
| 8 | 83892519 | rs2400926 | -0.008 | -0.183 | 1.283 | -1.433 | -0.004 | -0.008 | -0.008 | 0 | 0.11 | 0.06 | 0.07 | 0.01 | 0.13 | 0.13 |
| 8 | 83910616 | rs1597382 | 0 | -0.45 | 1.165 | -1.384 | -0.023 | -0.01 | -0.01 | 0 | 0.15 | 0.02 | 0.16 | 0.13 | 0.16 | 0.17 |
| 8 | 83920104 | rs959284 | 1.624 | -0.361 | 1.276 | -1.519 | -0.005 | 0.06 | 0.06 | 0.113 | 0.61 | 0.76 | 0.69 | 0.86 | 0.61 | 0.63 |
| 8 | 83928705 | rs754497 | 0.581 | -0.37 | 1.192 | -1.533 | -0.014 | 0.046 | 0.046 | 0 | 0.8 | 0.93 | 0.88 | 0.99 | 0.8 | 0.8 |
| 8 | 83932784 | rs1449829 | -1.341 | -0.871 | 1.614 | -1.167 | 0.005 | 0.015 | 0.015 | 0 | 0.28 | 0.21 | 0.25 | 0.15 | 0.25 | 0.32 |
| 8 | 83936116 | rs10103661 | -2.026 | -1.019 | 1.888 | -1.075 | 0.012 | 0.017 | 0.017 | 0.038 | 0.32 | 0.72 | 0.31 | 0.15 | 0.22 | 0.32 |
| 8 | 83943924 | rs10086141 | -0.746 | -1.002 | 1.68 | -0.999 | -0.028 | -0.081 | -0.081 | 0 | 0.14 | 0.31 | 0.09 | 0.01 | 0.08 | 0.16 |
| 8 | 83968656 | rs1823437 | -0.222 | -0.891 | 0.926 | -0.888 | -0.009 | -0.026 | -0.026 | 2.87 | 0.87 | 0.98 | 0.86 | 0.86 | 0.89 | 0.85 |
| 8 | 83972553 | rs1375781 | 2.149 | -1.204 | 1.218 | -1.115 | 0.048 | 0.065 | 0.065 | 1.47 | 0.65 | 0.16 | 0.68 | 0.85 | 0.78 | 0.68 |
| 8 | 83974398 | rs11786610 | -0.332 | -1.15 | 1.553 | -1.088 | -0.01 | -0.038 | -0.038 | 0 | 0.13 | 0.18 | 0.08 | 0 | 0.08 | 0.06 |
| 8 | 83981662 | rs1449814 | 1.879 | -1.181 | 1.6 | -1.109 | 0.034 | 0.045 | 0.045 | 1.044 | 0.65 | 0.23 | 0.67 | 0.85 | 0.77 | 0.68 |
| 8 | 83984810 | rs896010 | 0.268 | -1.119 | 1.901 | -1.055 | -0.002 | -0.001 | -0.001 | 0 | 0.83 | 0.82 | 0.83 | 0.86 | 0.89 | 0.85 |
| 8 | 83986887 | rs993429 | -0.042 | -1.12 | 1.953 | -1.056 | 0.005 | -0.002 | -0.002 | 0.066 | 0.85 | 0.91 | 0.84 | 0.86 | 0.89 | 0.85 |
| 8 | 83988800 | rs10958141 | 0.41 | -1.156 | 1.988 | -1.066 | -0.009 | -0.004 | -0.004 | 0 | 0.82 | 0.78 | 0.82 | 0.85 | 0.89 | 0.76 |
| 8 | 83991231 | rs1375785 | 0.246 | -1.113 | 1.71 | -1.045 | -0.009 | -0.003 | -0.003 | 0 | 0.83 | 0.78 | 0.82 | 0.85 | 0.89 | 0.76 |
| 8 | 83991555 | rs1375786 | 1.498 | -1.056 | 0.941 | -1.017 | -0.011 | -0.016 | -0.016 | 0 | 0.69 | 0.31 | 0.71 | 0.85 | 0.81 | 0.69 |
| 8 | 83991660 | rs10504776 | 0.208 | -1.035 | 0.755 | -1.007 | -0.003 | 0.05 | 0.05 | 0 | 0.1 | 0.01 | 0.06 | 0 | 0.08 | 0.06 |
| 8 | 83992851 | rs6473412 | 0.91 | -0.913 | 0.883 | -0.916 | -0.03 | -0.041 | -0.041 | 0 | 0.69 | 0.41 | 0.72 | 0.85 | 0.81 | 0.69 |
| 8 | 83997827 | rs726723 | 0.924 | -1.544 | 0.046 | -1.248 | 0.145 | 0.2 | 0.2 | 2.652 | 0.54 | 0.16 | 0.65 | 0.86 | 0.71 | 0.62 |
| 8 | 84000103 | rs4739920 | 0.574 | -0.961 | 0.704 | -0.838 | -0.001 | 0.062 | 0.062 | 0 | 0.57 | 0.74 | 0.58 | 0.75 | 0.63 | 0.59 |
| 8 | 84009539 | rs12680160 | 0.185 | -0.878 | 0.797 | -0.256 | -0.002 | 0.076 | 0.076 | 0.225 | 0.06 | 0.14 | 0.06 | 0.12 | 0.07 | 0.07 |
| 8 | 84019357 | rs7000740 | 1.366 | -0.898 | 0.45 | -0.305 | -0.086 | -0.03 | -0.03 | 0 | 0.45 | 0.04 | 0.45 | 0.62 | 0.55 | 0.5 |
| 8 | 84025426 | rs1449802 | 1.252 | -0.438 | 1.801 | 0.272 | -0.001 | -0.001 | -0.001 | 0 | 0.61 | 0.58 | 0.62 | 0.67 | 0.66 | 0.54 |
| 8 | 84025585 | rs6998971 | 0.407 | -0.475 | 1.788 | 0.26 | -0.024 | 0.021 | 0.021 | 0 | 0.72 | 0.94 | 0.71 | 0.8 | 0.74 | 0.6 |
| 8 | 84037048 | rs13265895 | 0.404 | -0.451 | 1.834 | 0.272 | -0.025 | 0.021 | 0.021 | 0 | 0.72 | 0.94 | 0.71 | 0.8 | 0.74 | 0.59 |
| 8 | 84059341 | rs6981282 | 1.451 | -0.482 | 2.61 | 0.263 | 0 | 0 | 0 | 0.056 | 0.59 | 0.45 | 0.61 | 0.67 | 0.67 | 0.53 |
| 8 | 84077412 | rs6991732 | 1.476 | -0.498 | 2.412 | 0.29 | 0.001 | 0 | 0 | 0 | 0.6 | 0.42 | 0.61 | 0.67 | 0.68 | 0.54 |
| 8 | 84091688 | rs7826679 | -1.712 | -0.498 | 2.321 | 0.349 | -0.014 | 0.091 | 0.091 | 1.646 | 0.12 | 0.41 | 0.11 | 0.13 | 0.09 | 0.07 |
| 8 | 84097208 | rs12680888 | 0.578 | -0.305 | 1.583 | 0.342 | -0.03 | 0.024 | 0.024 | 0 | 0.7 | 0.94 | 0.7 | 0.8 | 0.71 | 0.6 |
| 8 | 84113893 | rs7829784 | -1.053 | 0.001 | 2.35 | 0.686 | 0.007 | -0.003 | -0.003 | 0.011 | 0.27 | 0.41 | 0.25 | 0.21 | 0.27 | 0.36 |
| 8 | 84114931 | rs1568528 | 1.002 | -0.054 | 2.596 | 0.691 | 0.014 | 0.12 | 0.12 | 0 | 0.94 | 0.75 | 0.84 | 0.87 | 0.97 | 0.92 |
| 8 | 84153041 | rs4739669 | -0.71 | -0.009 | 2.607 | 0.647 | 0.009 | -0.055 | -0.055 | 0.061 | 0.23 | 0.36 | 0.18 | 0.08 | 0.2 | 0.3 |
| 8 | 84155214 | rs10106717 | -0.438 | 0.014 | 1.976 | 0.595 | -0.014 | 0.032 | 0.032 | 0 | 0.2 | 0.15 | 0.1 | 0.02 | 0.19 | 0.12 |
| 8 | 84173698 | rs16911725 | 1.19 | 0.004 | 1.659 | 0.679 | 0 | -0.002 | -0.002 | 0 | 0.67 | 0.37 | 0.6 | 0.67 | 0.7 | 0.56 |
| 8 | 84211031 | rs1915806 | 1.409 | 0.12 | 0.401 | 0.798 | -0.003 | -0.064 | -0.064 | 0 | 0.59 | 0.06 | 0.54 | 0.61 | 0.66 | 0.52 |
| 8 | 84211567 | rs6985259 | -0.163 | 1.042 | 0.215 | 1.441 | -0.093 | -0.101 | -0.101 | 0 | 0.14 | 0.01 | 0.24 | 0.38 | 0.27 | 0.25 |
| 8 | 84216690 | rs2128002 | 1.155 | 0.392 | 1.013 | 1.346 | -0.003 | -0.007 | -0.007 | 0.522 | 0.67 | 0.47 | 0.62 | 0.67 | 0.73 | 0.59 |
| 8 | 84217036 | rs969299 | -0.476 | 0.351 | 1.473 | 1.317 | -0.008 | 0 | 0 | 0 | 0.22 | 0.16 | 0.12 | 0.02 | 0.21 | 0.15 |
| 8 | 84217328 | rs1402849 | 0.928 | 0.331 | 1.662 | 1.333 | -0.008 | 0.179 | 0.179 | 0 | 0.94 | 0.82 | 0.78 | 0.68 | 0.96 | 0.73 |
| 8 | 84219847 | rs785964 | -0.313 | 0.381 | 1.328 | 1.191 | -0.074 | 0.098 | 0.098 | 0 | 0.19 | 0 | 0.08 | 0.02 | 0.17 | 0.14 |
| 8 | 84275516 | rs12375340 | -1.503 | 0.14 | 2.187 | 1.699 | -0.013 | 0.082 | 0.082 | 0 | 0.07 | 0.21 | 0.16 | 0.13 | 0.03 | 0.09 |
| 8 | 84279884 | rs952017 | -1.55 | 0.105 | 2.288 | 1.773 | 0.001 | 0.281 | 0.281 | 0.077 | 0.09 | 0.31 | 0.15 | 0.38 | 0.08 | 0.24 |
| 8 | 84281027 | rs1481486 | 1.977 | 0.031 | 1.751 | 1.612 | 0.011 | -0.107 | -0.107 | 0.47 | 0.6 | 0.1 | 0.53 | 0.48 | 0.68 | 0.52 |
| 8 | 84319445 | rs10107680 | 0.23 | 0.102 | 2.369 | 1.496 | -0.062 | 0.106 | 0.106 | 0 | 0.8 | 0.98 | 0.89 | 0.98 | 0.81 | 0.88 |
| 8 | 84323391 | rs7828142 | -1.129 | 0.132 | 2.695 | 1.683 | -0.016 | 0.058 | 0.058 | 0 | 0.07 | 0.09 | 0.07 | 0.21 | 0.07 | 0.06 |
| 8 | 84334585 | rs13265809 | -0.747 | 0.682 | 2.501 | 2.042 | 0.033 | 0.053 | 0.053 | 0 | 0.17 | 0.3 | 0.31 | 0.36 | 0.26 | 0.39 |
| 8 | 84337288 | rs7817496 | -1.547 | 0.64 | 2.34 | 1.832 | -0.021 | 0.129 | 0.129 | 0.293 | 0.09 | 0.25 | 0.09 | 0.21 | 0.07 | 0.06 |
| 8 | 84373550 | rs8180969 | 0.615 | 0.679 | 2.25 | 1.608 | -0.002 | -0.022 | -0.022 | 0 | 0.77 | 0.79 | 0.87 | 0.98 | 0.81 | 0.86 |
| 8 | 84374321 | rs2053744 | -0.513 | 0.747 | 2.176 | 2.17 | -0.012 | -0.003 | -0.003 | 0 | 0.14 | 0.13 | 0.29 | 0.44 | 0.26 | 0.38 |
| 8 | 84378819 | rs7004708 | -1.155 | 0.702 | 2.344 | 2.31 | -0.017 | 0.1 | 0.1 | 0.145 | 0.07 | 0.18 | 0.08 | 0.21 | 0.07 | 0.06 |
| 8 | 84389871 | rs1552548 | 1.816 | 0.694 | 2.519 | 2.265 | -0.008 | -0.007 | -0.007 | 0.204 | 0.57 | 0.32 | 0.4 | 0.42 | 0.52 | 0.39 |
| 8 | 84394525 | rs2121621 | -0.671 | 1.067 | 2.291 | 2.475 | -0.012 | -0.003 | -0.003 | 0 | 0.14 | 0.13 | 0.29 | 0.44 | 0.26 | 0.38 |
| 8 | 84428411 | rs1447259 | -0.813 | 1.31 | 1.999 | 2.394 | -0.046 | -0.043 | -0.043 | 0 | 0.12 | 0.09 | 0.19 | 0.22 | 0.24 | 0.19 |
| 8 | 84442025 | rs1374606 | 1.443 | 0.989 | 2.439 | 2.401 | -0.005 | 0.022 | 0.022 | 0.148 | 0.72 | 0.45 | 0.7 | 0.62 | 0.79 | 0.62 |
| 8 | 84452541 | rs1447255 | 1.045 | 1.057 | 1.998 | 2.388 | -0.009 | 0 | 0 | 0 | 0.77 | 0.87 | 0.87 | 0.76 | 0.81 | 0.7 |
| 8 | 84477913 | rs6997745 | 1.295 | 1.026 | 2.166 | 2.298 | -0.002 | -0.006 | -0.006 | 0 | 0.73 | 0.75 | 0.68 | 0.42 | 0.73 | 0.56 |
| 8 | 84492059 | rs901958 | 0.21 | 1.094 | 1.99 | 2.299 | -0.062 | -0.051 | -0.051 | 0 | 0.86 | 0.97 | 0.91 | 0.79 | 0.82 | 0.85 |
| 8 | 84492735 | rs12547052 | -1.383 | 1.008 | 2.064 | 2.255 | 0.005 | 0.003 | 0.003 | 0.018 | 0.13 | 0.1 | 0.23 | 0.37 | 0.09 | 0.29 |
| 8 | 84493827 | rs10958156 | -1.305 | 1.409 | 2.488 | 2.078 | 0.101 | 0.085 | 0.085 | 0.028 | 0.16 | 0.53 | 0.29 | 0.2 | 0.23 | 0.19 |

| | | | | | | | | | | | | | | | | |
|---|---|---|---|---|---|---|---|---|---|---|---|---|---|---|---|---|
| 8 | 84499541 | rs2220456 | -1.305 | 1.409 | 2.5 | 2.089 | 0.104 | 0.092 | 0.092 | 0.028 | 0.16 | 0.49 | 0.28 | 0.2 | 0.23 | 0.19 |
| 8 | 84511490 | rs4612338 | 0.406 | 1.429 | 2.644 | 1.869 | 0.066 | -0.102 | -0.102 | 1.416 | 0.86 | 0.58 | 0.9 | 1 | 0.85 | 0.92 |
| 8 | 84547452 | rs10958161 | -1.655 | 1.367 | 2.69 | 2.297 | 0.002 | 0.021 | 0.021 | 0 | 0.14 | 0.14 | 0.15 | 0.56 | 0.12 | 0.35 |
| 8 | 84559621 | rs13265681 | -2.286 | 1.393 | 2.763 | 2.264 | -0.007 | 0.002 | 0.002 | 0 | 0.26 | 0.26 | 0.5 | 0.77 | 0.35 | 0.52 |
| 8 | 84561877 | rs4639509 | -0.634 | 1.486 | 2.426 | 2.127 | -0.06 | -0.057 | -0.057 | 0.968 | 0.12 | 0.03 | 0.34 | 0.21 | 0.23 | 0.17 |
| 8 | 84570892 | rs4623415 | -0.746 | 1.631 | 2.455 | 2.092 | -0.06 | -0.047 | -0.047 | 0.454 | 0.11 | 0.03 | 0.34 | 0.19 | 0.24 | 0.17 |
| 8 | 84572978 | rs10958163 | -0.855 | 1.699 | 2.485 | 2.378 | 0.001 | -0.014 | -0.014 | 0 | 0.1 | 0.06 | 0.16 | 0.38 | 0.11 | 0.29 |
| 8 | 84573177 | rs12682163 | -0.695 | 1.644 | 2.834 | 2.36 | 0.023 | -0.011 | -0.011 | 0 | 0.15 | 0.28 | 0.11 | 0.12 | 0.17 | 0.14 |
| 8 | 84576995 | rs4739962 | -2.116 | 1.681 | 2.811 | 2.347 | 0.017 | 0.037 | 0.037 | 0 | 0.23 | 0.3 | 0.52 | 0.57 | 0.35 | 0.46 |
| 8 | 84581698 | rs7845183 | -2.735 | 1.676 | 2.821 | 2.308 | 0.003 | 0 | 0 | 0 | 0.4 | 0.55 | 0.63 | 0.65 | 0.52 | 0.59 |
| 8 | 84592640 | rs4265180 | -1.916 | 1.679 | 2.585 | 2.352 | -0.029 | 0.121 | 0.121 | 0.395 | 0.16 | 0.63 | 0.22 | 0.44 | 0.11 | 0.26 |
| 8 | 84595022 | rs7814353 | -2.296 | 1.72 | 2.371 | 2.352 | -0.01 | -0.061 | -0.061 | 0.71 | 0.31 | 0.83 | 0.31 | 0.49 | 0.28 | 0.37 |
| 8 | 84615691 | rs4574844 | -0.696 | 1.882 | 2.077 | 2.447 | -0.047 | -0.041 | -0.041 | 0 | 0.12 | 0.07 | 0.36 | 0.21 | 0.25 | 0.17 |
| 8 | 84616700 | rs4379441 | -2.694 | 1.831 | 2.414 | 2.416 | -0.001 | 0.002 | 0.002 | 0.012 | 0.41 | 0.62 | 0.64 | 0.7 | 0.53 | 0.54 |
| 8 | 84618780 | rs10958166 | -1.611 | 1.855 | 2.257 | 2.066 | -0.022 | -0.035 | -0.035 | 0 | 0.24 | 0.1 | 0.22 | 0.39 | 0.25 | 0.32 |
| 8 | 84624128 | rs4388459 | -0.807 | 2.043 | 2.202 | 2.134 | -0.029 | -0.022 | -0.022 | 0 | 0.12 | 0.11 | 0.37 | 0.38 | 0.25 | 0.17 |
| 8 | 84626294 | rs4739969 | -0.799 | 2.041 | 2.209 | 2.133 | -0.025 | -0.014 | -0.014 | 0 | 0.12 | 0.12 | 0.37 | 0.38 | 0.25 | 0.17 |
| 8 | 84633804 | rs12550806 | -2.338 | 1.726 | 2.266 | 1.819 | 0.002 | 0.015 | 0.015 | 0 | 0.35 | 0.39 | 0.31 | 0.49 | 0.35 | 0.38 |
| 8 | 84652949 | rs4412368 | -1.374 | 1.918 | 2.221 | 1.627 | 0.021 | -0.096 | -0.096 | 0 | 0.22 | 0.34 | 0.14 | 0.08 | 0.26 | 0.14 |
| 8 | 84658822 | rs7818253 | -2.333 | 1.936 | 2.147 | 1.632 | 0.017 | 0.019 | 0.019 | 0 | 0.36 | 0.46 | 0.52 | 0.45 | 0.51 | 0.34 |
| 8 | 84661820 | rs4272373 | -1.879 | 1.953 | 1.869 | 1.745 | 0.006 | 0.029 | 0.029 | 0 | 0.22 | 0.28 | 0.43 | 0.45 | 0.33 | 0.25 |
| 8 | 84671396 | rs13251395 | 0.1 | 1.739 | 1.728 | 1.675 | 0.004 | 0.003 | 0.003 | 0 | 0.07 | 0.02 | 0.04 | 0.08 | 0.08 | 0.06 |
| 8 | 84671732 | rs4626593 | -0.778 | 2.006 | 1.704 | 1.728 | -0.027 | -0.036 | -0.036 | 0 | 0.12 | 0.09 | 0.37 | 0.37 | 0.22 | 0.2 |
| 8 | 84680146 | rs9650286 | -0.419 | 2.009 | 1.841 | 1.834 | -0.068 | 0.08 | 0.08 | 0 | 0.13 | 0.02 | 0.08 | 0 | 0.17 | 0.08 |
| 8 | 84683145 | rs7003433 | -1.917 | 2.148 | 2.031 | 1.84 | 0.012 | 0.121 | 0.121 | 0 | 0.15 | 0.3 | 0.31 | 0.44 | 0.23 | 0.24 |
| 8 | 84695925 | rs4739974 | -0.907 | 2.148 | 2.102 | 1.844 | 0.016 | 0.14 | 0.14 | 0 | 0.06 | 0.11 | 0.25 | 0.37 | 0.14 | 0.18 |
| 8 | 84701890 | rs6473474 | -2.254 | 2.152 | 2.124 | 1.826 | 0.01 | 0.015 | 0.015 | 0 | 0.28 | 0.34 | 0.39 | 0.44 | 0.4 | 0.31 |
| 8 | 84705521 | rs7010758 | -0.492 | 2.133 | 1.985 | 1.755 | -0.01 | -0.016 | -0.016 | 0 | 0.09 | 0.15 | 0.06 | 0.07 | 0.08 | 0.06 |
| 8 | 84732202 | rs4132243 | -0.731 | 2.242 | 2.063 | 1.822 | 0.053 | -0.087 | -0.087 | 0 | 0.14 | 0.29 | 0.1 | 0 | 0.17 | 0.08 |
| 8 | 84736781 | rs4739981 | -1.6 | 1.836 | 1.495 | 2.244 | 0.037 | 0.219 | 0.219 | 0 | 0.08 | 0.23 | 0.27 | 0.36 | 0.14 | 0.22 |
| 8 | 84739785 | rs7004331 | 1.157 | 1.756 | 1.256 | 1.855 | 0 | 0.075 | 0.075 | 0 | 0.73 | 0.5 | 0.52 | 0.43 | 0.69 | 0.74 |
| 8 | 84740129 | rs6473482 | -2.498 | 2.394 | 1.424 | 1.848 | 0.078 | 0.2 | 0.2 | 2.311 | 0.12 | 0.55 | 0.14 | 0.29 | 0.14 | 0.19 |
| 8 | 84748939 | rs11781410 | -1.956 | 2.944 | 1.262 | 1.571 | -0.011 | -0.014 | -0.014 | 0.61 | 0.23 | 0.21 | 0.21 | 0.37 | 0.32 | 0.3 |
| 8 | 84751766 | rs12682602 | -2.605 | 2.947 | 1.08 | 1.543 | -0.012 | -0.014 | -0.014 | 0 | 0.29 | 0.71 | 0.27 | 0.37 | 0.34 | 0.31 |
| 8 | 84754153 | rs12547395 | -2.692 | 2.932 | 0.967 | 2.084 | -0.029 | 0.018 | 0.018 | 0.018 | 0.42 | 0.79 | 0.49 | 0.57 | 0.53 | 0.44 |
| 8 | 84758943 | rs10087833 | -2.312 | 3.029 | 0.99 | 2.007 | -0.025 | 0.014 | 0.014 | 0 | 0.35 | 0.74 | 0.42 | 0.29 | 0.43 | 0.25 |
| 8 | 84764257 | rs11786147 | -1.932 | 2.958 | 1.219 | 2.013 | -0.015 | -0.002 | -0.002 | 1.304 | 0.3 | 0.22 | 0.36 | 0.3 | 0.41 | 0.25 |
| 8 | 84767563 | rs11781052 | -1.94 | 2.959 | 1.219 | 2.013 | -0.015 | -0.001 | -0.001 | 0.463 | 0.3 | 0.22 | 0.36 | 0.29 | 0.41 | 0.25 |
| 8 | 84771480 | rs7819156 | 0.613 | 3.01 | 1.205 | 2.132 | -0.012 | 0.029 | 0.029 | 0 | 0.94 | 0.96 | 0.92 | 0.71 | 0.9 | 0.81 |
| 8 | 84772442 | rs4590440 | 2.162 | 3.019 | 1.217 | 2.131 | -0.031 | -0.004 | -0.004 | 0 | 0.63 | 0.73 | 0.57 | 0.4 | 0.49 | 0.57 |
| 8 | 84776101 | rs12541971 | -0.267 | 3.065 | 1.2 | 2.102 | -0.016 | -0.029 | -0.029 | 0 | 0.13 | 0.05 | 0.21 | 0.21 | 0.18 | 0.12 |
| 8 | 84777289 | rs4237105 | -1.866 | 3.038 | 1.209 | 2.095 | -0.016 | -0.002 | -0.002 | 0 | 0.3 | 0.22 | 0.36 | 0.29 | 0.41 | 0.25 |
| 8 | 84799266 | rs12675334 | 2.419 | 2.804 | 1.132 | 1.791 | -0.05 | 0.01 | 0.01 | 0.103 | 0.58 | 0.16 | 0.5 | 0.4 | 0.48 | 0.55 |
| 8 | 84806193 | rs4492374 | 1.663 | 2.908 | 1.477 | 1.651 | 0.012 | 0.013 | 0.013 | 0 | 0.91 | 0.9 | 0.87 | 0.78 | 0.81 | 0.77 |
| 8 | 84810178 | rs7003447 | 2.739 | 2.89 | 2.075 | 1.67 | -0.009 | -0.006 | -0.006 | 0 | 0.76 | 0.24 | 0.71 | 0.68 | 0.66 | 0.68 |
| 8 | 84823255 | rs7820174 | 2.141 | 2.936 | 1.887 | 1.709 | 0.024 | 0.02 | 0.02 | 0 | 0.8 | 0.72 | 0.78 | 0.68 | 0.69 | 0.69 |
| 8 | 84823691 | rs1866984 | -0.151 | 2.918 | 1.685 | 1.295 | -0.014 | -0.006 | -0.006 | 0 | 0.91 | 0.99 | 0.94 | 0.91 | 0.88 | 0.92 |
| 8 | 84823737 | rs6473514 | 0.149 | 2.914 | 1.69 | 1.298 | 0 | -0.003 | -0.003 | 0.213 | 0.89 | 0.87 | 0.92 | 0.91 | 0.87 | 0.92 |
| 8 | 84828638 | rs6473515 | 1.652 | 2.836 | 1.702 | 1.406 | 0.006 | 0 | 0 | 0 | 0.81 | 0.78 | 0.82 | 0.75 | 0.69 | 0.69 |
| 8 | 84832969 | rs6473516 | 1.985 | 1.935 | 1.262 | 0.649 | 0.01 | -0.05 | -0.05 | 0.563 | 0.66 | 0.47 | 0.72 | 0.77 | 0.62 | 0.61 |
| 8 | 84834888 | rs7834265 | 0.976 | 1.99 | 0.744 | 0.607 | -0.022 | 0.051 | 0.051 | 0 | 0.78 | 0.9 | 0.82 | 0.92 | 0.71 | 0.8 |
| 8 | 84859944 | rs1442036 | 1.082 | 1.844 | 1.543 | 0.467 | -0.023 | 0.063 | 0.063 | 0 | 0.76 | 0.91 | 0.82 | 0.92 | 0.71 | 0.8 |
| 8 | 84864301 | rs731956 | -1.509 | 1.998 | 1.768 | 0.672 | 0.129 | 0.135 | 0.135 | 0 | 0.07 | 0.38 | 0.11 | 0.15 | 0.09 | 0.2 |
| 8 | 84866246 | rs10100059 | -2.307 | 2.016 | 2.063 | 0.669 | 0.074 | 0.077 | 0.077 | 0.461 | 0.12 | 0.52 | 0.15 | 0.18 | 0.13 | 0.25 |
| 8 | 84879603 | rs1971841 | -1.994 | 2.017 | 1.883 | 0.666 | 0.005 | 0.031 | 0.031 | 0.038 | 0.11 | 0.32 | 0.11 | 0.18 | 0.11 | 0.24 |
| 8 | 84890104 | rs7820311 | -2.282 | 2.024 | 1.576 | 0.673 | 0.053 | 0.056 | 0.056 | 1.288 | 0.13 | 0.58 | 0.16 | 0.18 | 0.13 | 0.25 |
| 8 | 84911664 | rs3847186 | 0.865 | 2.011 | 0.47 | 0.53 | -0.03 | 0.027 | 0.027 | 0 | 0.79 | 0.9 | 0.82 | 0.89 | 0.69 | 0.82 |
| 8 | 84914149 | rs4739995 | 2.114 | 1.798 | -0.245 | 0.324 | -0.008 | 0.202 | 0.202 | 0 | 0.44 | 0.05 | 0.4 | 0.16 | 0.33 | 0.36 |
| 8 | 84917254 | rs6473520 | 1.569 | 2.277 | 0.518 | 0.383 | -0.015 | 0.023 | 0.023 | 0 | 0.72 | 0.25 | 0.71 | 0.86 | 0.64 | 0.75 |
| 8 | 84919577 | rs11778058 | -0.912 | 2.361 | 0.264 | 0.368 | -0.013 | 0.009 | 0.009 | 0 | 0.16 | 0.08 | 0.13 | 0.16 | 0.14 | 0.22 |
| 8 | 84922270 | rs16912373 | -1.965 | 2.353 | 1.101 | 0.305 | -0.021 | -0.034 | -0.034 | 0 | 0.28 | 0.48 | 0.23 | 0.24 | 0.22 | 0.27 |
| 8 | 84925709 | rs16912403 | -2.036 | 2.472 | 1.729 | 0.861 | 0.009 | 0.05 | 0.05 | 0 | 0.25 | 0.37 | 0.26 | 0.59 | 0.24 | 0.39 |
| 8 | 84933435 | rs16912418 | -1.566 | 2.453 | 1.64 | 0.818 | -0.004 | -0.006 | -0.006 | 0.334 | 0.22 | 0.29 | 0.18 | 0.24 | 0.18 | 0.27 |
| 8 | 84944712 | rs1596134 | -1.074 | 2.485 | 1.802 | 0.82 | 0.003 | 0.015 | 0.015 | 0 | 0.17 | 0.25 | 0.21 | 0.25 | 0.16 | 0.25 |
| 8 | 84948131 | rs965417 | 0.386 | 2.528 | 1.626 | 0.826 | 0.019 | -0.016 | -0.016 | 0 | 0.93 | 0.92 | 0.92 | 1 | 0.84 | 0.91 |
| 8 | 84953839 | rs6991736 | -0.763 | 2.408 | 1.556 | 0.818 | 0.002 | 0.116 | 0.116 | 0 | 0.1 | 0.19 | 0.13 | 0.34 | 0.11 | 0.13 |
| 8 | 84966400 | rs4298483 | 0.783 | 2.385 | 1.192 | 0.378 | 0.105 | -0.035 | -0.035 | 0.172 | 0.92 | 0.78 | 0.9 | 1 | 0.84 | 0.91 |
| 8 | 84974524 | rs7000681 | -1.995 | 2.137 | 0.744 | 0.23 | -0.025 | 0.011 | 0.011 | 0 | 0.45 | 0.76 | 0.47 | 0.58 | 0.5 | 0.47 |
| 8 | 84986353 | rs13268001 | -0.938 | 2.492 | 0.885 | 0.245 | 0.017 | -0.048 | -0.048 | 0 | 0.14 | 0.18 | 0.13 | 0 | 0.22 | 0.06 |

| | | | | | | | | | | | | | | | | |
|---|---|---|---|---|---|---|---|---|---|---|---|---|---|---|---|---|
| 8 | 84987529 | rs16922863 | -0.365 | 2.487 | 0.967 | 0.337 | -0.004 | 0.006 | 0.006 | 0 | 0.16 | 0.11 | 0.13 | 0.14 | 0.14 | 0.27 |
| 8 | 84996380 | rs13281887 | -0.954 | 2.366 | 0.94 | 0.165 | 0.02 | -0.049 | -0.049 | 2.063 | 0.14 | 0.18 | 0.13 | 0 | 0.22 | 0.06 |
| 8 | 85000872 | rs4285486 | -1.274 | 1.882 | 0.982 | 0.375 | 0.001 | 0.025 | 0.025 | 0 | 0.27 | 0.36 | 0.32 | 0.58 | 0.27 | 0.42 |
| 8 | 85003092 | rs11989306 | 0.003 | 1.357 | 1.037 | -0.098 | -0.017 | -0.059 | -0.059 | 0 | 0.12 | 0.18 | 0.06 | 0 | 0.08 | 0.03 |
| 8 | 85013772 | rs10958203 | 1.345 | 0.841 | 0.605 | -0.408 | 0.006 | 0.022 | 0.022 | 0.061 | 0.34 | 0.22 | 0.49 | 0.28 | 0.31 | 0.38 |
| 8 | 85032004 | rs10808857 | -0.184 | 0.806 | 0.216 | -0.376 | -0.084 | -0.012 | -0.012 | 1.077 | 0.36 | 0.15 | 0.28 | 0.34 | 0.48 | 0.39 |
| 8 | 85032101 | rs10738143 | -0.075 | 0.89 | -0.077 | -0.115 | 0.012 | -0.023 | -0.023 | 0.382 | 0.75 | 0.82 | 0.74 | 0.62 | 0.8 | 0.77 |
| 8 | 85038334 | rs7820224 | -0.166 | 0.952 | -0.304 | -0.125 | 0.013 | -0.024 | -0.024 | 0.339 | 0.74 | 0.79 | 0.73 | 0.61 | 0.8 | 0.77 |
| 8 | 85053510 | rs16912483 | -0.064 | 0.148 | -0.216 | -0.388 | -0.01 | -0.006 | -0.006 | 0.112 | 0.37 | 0.33 | 0.26 | 0.37 | 0.38 | 0.39 |
| 8 | 85054405 | rs6990992 | 1.22 | 0.111 | -0.317 | -0.406 | 0.009 | 0.051 | 0.051 | 0.39 | 0.37 | 0.19 | 0.43 | 0.22 | 0.37 | 0.41 |
| 8 | 85065107 | rs7841440 | 1.203 | 0.284 | -0.772 | -0.288 | -0.008 | -0.035 | -0.035 | 0 | 0.37 | 0.5 | 0.47 | 0.22 | 0.36 | 0.38 |
| 8 | 85070206 | rs6473525 | 0.212 | 0.268 | -1.111 | 0.357 | -0.023 | -0.093 | -0.093 | 2.069 | 0.75 | 0.94 | 0.74 | 0.59 | 0.74 | 0.77 |
| 8 | 85077271 | rs893282 | 0.951 | 0.26 | -0.914 | 0.291 | -0.021 | -0.036 | -0.036 | 0 | 0.4 | 0.47 | 0.44 | 0.22 | 0.32 | 0.41 |
| 8 | 85079709 | rs2033732 | 0.805 | 0.39 | -0.961 | 0.736 | -0.009 | -0.072 | -0.072 | 0 | 0.77 | 0.91 | 0.77 | 0.59 | 0.77 | 0.76 |
| 8 | 85101510 | rs10099120 | -0.986 | -0.588 | 0.025 | 0.409 | -0.08 | 0.014 | 0.014 | 0 | 0.22 | 0.53 | 0.2 | 0.26 | 0.11 | 0.11 |
| 8 | 85107214 | rs7010829 | -0.958 | -0.594 | 0.185 | 0.406 | -0.069 | 0.012 | 0.012 | 0 | 0.21 | 0.42 | 0.18 | 0.26 | 0.11 | 0.11 |
| 8 | 85110976 | rs4427165 | -0.24 | -1.886 | -1.405 | -0.91 | 0.129 | 0.226 | 0.226 | 0 | 0.39 | 0.07 | 0.21 | 0.17 | 0.2 | 0.34 |
| 8 | 85112377 | rs9298419 | -0.291 | -2.147 | -1.803 | -1.604 | 0.112 | 0.192 | 0.192 | 0 | 0.45 | 0.15 | 0.28 | 0.19 | 0.29 | 0.4 |
| 8 | 85113675 | rs1550209 | -0.368 | -2.304 | -0.954 | -1.721 | 0.051 | 0.062 | 0.062 | 0 | 0.48 | 0.39 | 0.31 | 0.19 | 0.3 | 0.4 |
| 8 | 85117456 | rs6473531 | -0.872 | -0.583 | -1.225 | -1.256 | -0.004 | 0.007 | 0.007 | 0.823 | 0.23 | 0.27 | 0.17 | 0.11 | 0.26 | 0.19 |
| 8 | 85121515 | rs1550210 | 0.886 | 0.295 | -1.326 | -0.493 | -0.006 | -0.008 | -0.008 | 0.076 | 0.61 | 0.69 | 0.43 | 0.5 | 0.4 | 0.49 |
| 8 | 85122058 | rs12375429 | -0.397 | -0.102 | -1.072 | -0.493 | -0.008 | 0.005 | 0.005 | 0.78 | 0.2 | 0.22 | 0.16 | 0.12 | 0.25 | 0.17 |
| 8 | 85130404 | rs1978828 | -0.728 | -0.25 | 0.009 | -0.565 | 0.01 | -0.027 | -0.027 | 0 | 0.23 | 0.48 | 0.19 | 0.12 | 0.25 | 0.16 |
| 8 | 85132271 | rs10099886 | 0.728 | -0.64 | 0.073 | -0.785 | 0.016 | 0.04 | 0.04 | 0 | 0.41 | 0.26 | 0.26 | 0.24 | 0.29 | 0.29 |
| 8 | 85134247 | rs9298423 | 0.171 | -0.539 | 0.318 | -0.797 | 0.01 | 0.022 | 0.022 | 0 | 0.51 | 0.38 | 0.33 | 0.23 | 0.38 | 0.37 |
| 8 | 85134333 | rs10108169 | -0.937 | -0.167 | 0.088 | -0.163 | -0.047 | -0.027 | -0.027 | 0 | 0.71 | 0.83 | 0.66 | 0.64 | 0.6 | 0.54 |
| 8 | 85135902 | rs2053244 | -0.979 | -0.188 | 0.039 | -0.129 | -0.047 | -0.027 | -0.027 | 0 | 0.72 | 0.84 | 0.66 | 0.64 | 0.6 | 0.58 |
| 8 | 85137818 | rs16912559 | -0.858 | -0.068 | 0.541 | 0.317 | -0.004 | 0.137 | 0.137 | 0 | 0.2 | 0.41 | 0.33 | 0.43 | 0.22 | 0.21 |
| 8 | 85138070 | rs7007401 | -0.877 | -0.07 | 0.439 | 0.316 | -0.001 | 0.199 | 0.199 | 0 | 0.2 | 0.51 | 0.34 | 0.43 | 0.22 | 0.21 |
| 8 | 85141824 | rs4740003 | 0.817 | 0.377 | -0.144 | 0.58 | -0.088 | -0.058 | -0.058 | 0 | 0.26 | 0.07 | 0.3 | 0.35 | 0.36 | 0.4 |
| 8 | 85143228 | rs7014531 | -1.207 | -0.08 | 0.219 | 0.388 | 0.036 | 0.065 | 0.065 | 0 | 0.55 | 0.69 | 0.71 | 0.75 | 0.66 | 0.61 |
| 8 | 85143822 | rs1529989 | 0.18 | -0.075 | 0.083 | 0.534 | -0.089 | -0.053 | -0.053 | 0 | 0.33 | 0.12 | 0.52 | 0.59 | 0.47 | 0.51 |
| 8 | 85146868 | rs16912577 | -0.838 | 0.108 | 0.31 | 0.644 | -0.008 | 0.006 | 0.006 | 0.216 | 0.17 | 0.27 | 0.14 | 0.16 | 0.15 | 0.1 |
| 8 | 85148907 | rs12676075 | -0.208 | -0.074 | 0.352 | 0.681 | -0.011 | 0.05 | 0.05 | 2.397 | 0.08 | 0.21 | 0.07 | 0.16 | 0.07 | 0.06 |
| 8 | 85149183 | rs10958215 | -1.391 | -0.09 | -0.07 | 0.405 | 0.058 | 0.108 | 0.108 | 0.012 | 0.54 | 0.8 | 0.72 | 0.75 | 0.66 | 0.61 |
| 8 | 85149583 | rs10504794 | 0.125 | 0.352 | -0.379 | 0.374 | -0.066 | -0.014 | -0.014 | 0.038 | 0.31 | 0.13 | 0.36 | 0.35 | 0.45 | 0.4 |
| 8 | 85151258 | rs13261079 | 0.782 | 0.461 | -0.466 | 0.531 | -0.072 | -0.041 | -0.041 | 1.313 | 0.28 | 0.09 | 0.31 | 0.35 | 0.36 | 0.43 |
| 8 | 85151612 | rs7825715 | -0.957 | -0.128 | 0.439 | 0.506 | 0.011 | -0.009 | -0.009 | 0 | 0.54 | 0.58 | 0.5 | 0.47 | 0.55 | 0.65 |
| 8 | 85152853 | rs10112823 | -0.45 | -0.216 | 0.847 | 0.455 | -0.001 | -0.004 | -0.004 | 0 | 0.49 | 0.46 | 0.47 | 0.34 | 0.52 | 0.61 |
| 8 | 85155391 | rs13268314 | -1.013 | -0.211 | 0.81 | 0.467 | 0.012 | -0.009 | -0.009 | 0 | 0.54 | 0.58 | 0.51 | 0.47 | 0.55 | 0.64 |
| 8 | 85160418 | rs16912601 | 0.345 | -0.213 | 0.777 | 0.489 | -0.001 | 0.012 | 0.012 | 0 | 0.05 | 0.03 | 0.03 | 0.13 | 0.03 | 0.06 |
| 8 | 85165783 | rs10094547 | -0.234 | -0.245 | 1.061 | 0.414 | 0.018 | -0.05 | -0.05 | 0 | 0.1 | 0.28 | 0.1 | 0.01 | 0.1 | 0.04 |
| 8 | 85176983 | rs4392892 | -1.014 | -0.263 | 0.952 | 0.401 | -0.076 | -0.012 | -0.012 | 0.832 | 0.67 | 0.91 | 0.67 | 0.67 | 0.61 | 0.44 |
| 8 | 85183072 | rs12542046 | 1.031 | -0.452 | 1.135 | 0.3 | 0.006 | -0.018 | -0.018 | 1.34 | 0.43 | 0.37 | 0.49 | 0.53 | 0.44 | 0.33 |
| 8 | 85185158 | rs12680668 | -0.504 | -0.407 | 1.126 | 0.305 | -0.014 | 0.008 | 0.008 | 0 | 0.11 | 0.14 | 0.06 | 0.13 | 0.05 | 0.09 |
| 8 | 85229266 | rs7002872 | -1.109 | -0.395 | 0.712 | 0.29 | -0.021 | -0.002 | -0.002 | 0.109 | 0.67 | 0.79 | 0.66 | 0.66 | 0.61 | 0.46 |
| 8 | 85239703 | rs10504792 | 0.284 | -0.287 | 0.196 | 0.513 | 0.009 | -0.01 | -0.01 | 0.227 | 0.08 | 0.03 | 0.04 | 0.14 | 0.04 | 0.04 |
| 8 | 85257540 | rs1907549 | 0.78 | -0.192 | 0.171 | 0.443 | -0.006 | -0.051 | -0.051 | 0 | 0.43 | 0.23 | 0.47 | 0.52 | 0.45 | 0.32 |
| 8 | 85272419 | rs4639510 | 0.982 | -0.231 | -0.043 | 0.452 | -0.029 | -0.099 | -0.099 | 0 | 0.4 | 0.17 | 0.45 | 0.52 | 0.43 | 0.33 |
| 8 | 85276246 | rs6982764 | -0.575 | -0.088 | 0.281 | 0.397 | -0.004 | -0.024 | -0.024 | 0 | 0.06 | 0.1 | 0.04 | 0 | 0.04 | 0.02 |
| 8 | 85284531 | rs11998502 | -1.677 | -0.174 | 0.933 | 0.462 | -0.024 | 0.029 | 0.029 | 1.33 | 0.16 | 0.41 | 0.17 | 0.14 | 0.11 | 0.05 |
| 8 | 85314840 | rs2061866 | 0.699 | 0.003 | 0.873 | 0.418 | -0.006 | 0.004 | 0.004 | 0.463 | 0.39 | 0.26 | 0.38 | 0.36 | 0.38 | 0.6 |
| 8 | 85328171 | rs317952 | -0.188 | -0.02 | 1.016 | 0.4 | 0.004 | 0.022 | 0.022 | 0 | 0.5 | 0.56 | 0.51 | 0.36 | 0.5 | 0.63 |
| 8 | 85330924 | rs7010781 | 0.678 | -0.02 | 0.817 | 0.44 | -0.006 | 0.007 | 0.007 | 0.369 | 0.39 | 0.25 | 0.38 | 0.36 | 0.38 | 0.61 |
| 8 | 85352136 | rs783508 | 0.018 | -0.063 | 0.728 | 0.493 | -0.007 | 0.035 | 0.035 | 0 | 0.47 | 0.64 | 0.52 | 0.36 | 0.5 | 0.56 |
| 8 | 85366469 | rs711012 | -0.812 | -0.142 | 0.742 | 0.581 | 0 | -0.017 | -0.017 | 0 | 0.58 | 0.62 | 0.61 | 0.47 | 0.55 | 0.63 |
| 8 | 85372283 | rs9298424 | 0.724 | -0.1 | 0.605 | 0.518 | 0.012 | -0.011 | -0.011 | 0 | 0.38 | 0.47 | 0.43 | 0.32 | 0.39 | 0.54 |
| 8 | 85383821 | rs7013087 | 1.053 | -0.09 | 0.546 | 0.513 | -0.009 | 0.002 | 0.002 | 0 | 0.34 | 0.25 | 0.38 | 0.32 | 0.36 | 0.53 |
| 8 | 85396742 | rs16912908 | 0.21 | -0.066 | 0.161 | 0.539 | 0.005 | -0.007 | -0.007 | 0 | 0.1 | 0.05 | 0.09 | 0.15 | 0.06 | 0.09 |
| 8 | 85403656 | rs783783 | 1.223 | 0.047 | -0.286 | 0.757 | -0.063 | -0.046 | -0.046 | 0.052 | 0.33 | 0.03 | 0.16 | 0.35 | 0.33 | 0.24 |
| 8 | 85417425 | rs711037 | -0.378 | 0.453 | 0.1 | 0.697 | 0 | -0.007 | -0.007 | 0 | 0.91 | 0.91 | 0.91 | 1 | 0.9 | 0.99 |
| 8 | 85424315 | rs711041 | -0.141 | 0.444 | 0.259 | 0.71 | 0.002 | -0.007 | -0.007 | 0.376 | 0.56 | 0.59 | 0.53 | 0.68 | 0.54 | 0.47 |
| 8 | 85452289 | rs7820792 | 0.621 | 0.462 | 0.418 | 0.253 | 0.006 | -0.004 | -0.004 | 0 | 0.35 | 0.34 | 0.38 | 0.32 | 0.36 | 0.52 |
| 8 | 85464739 | rs1393990 | 0.243 | 0.509 | 0.07 | 0.23 | 0.007 | -0.016 | -0.016 | 0 | 0.39 | 0.52 | 0.43 | 0.31 | 0.38 | 0.52 |
| 8 | 85470451 | rs1252202 | 0.616 | 1.265 | 0.193 | 0.345 | 0.006 | 0.003 | 0.003 | 0 | 0.86 | 0.84 | 0.81 | 0.85 | 0.77 | 0.85 |
| 8 | 85474803 | rs1252217 | -0.816 | 1.262 | 0.144 | 0.344 | -0.008 | -0.006 | -0.006 | 0.029 | 0.14 | 0.09 | 0.18 | 0.15 | 0.23 | 0.13 |
| 8 | 85500162 | rs1252207 | -0.424 | 0.869 | 0.163 | -0.043 | 0.002 | -0.002 | -0.002 | 0 | 0.59 | 0.71 | 0.63 | 0.38 | 0.6 | 0.7 |
| 8 | 85509648 | rs1681355 | -0.395 | 0.277 | 0.239 | 0.132 | -0.001 | -0.001 | -0.001 | 0 | 0.14 | 0.09 | 0.18 | 0.14 | 0.23 | 0.13 |
| 8 | 85526604 | rs7842614 | -1.163 | 0.139 | 1.022 | 0.03 | 0.037 | 0.009 | 0.009 | 1.028 | 0.19 | 0.44 | 0.24 | 0.15 | 0.26 | 0.17 |

| | | | | | | | | | | | | | | | | |
|---|---|---|---|---|---|---|---|---|---|---|---|---|---|---|---|---|
| 8 | 85653792 | rs9886620 | -1.138 | 0.765 | 1.01 | -0.027 | 0.057 | -0.004 | -0.004 | 0 | 0.22 | 0.35 | 0.28 | 0.15 | 0.4 | 0.16 |
| 8 | 85657553 | rs10112163 | -1.009 | 0.712 | 1.047 | 0.004 | 0.02 | 0.002 | 0.002 | 1.05 | 0.19 | 0.36 | 0.23 | 0.13 | 0.26 | 0.16 |
| 8 | 85667846 | rs6988126 | -1.722 | 0.466 | 1.766 | 0.071 | 0.012 | -0.009 | -0.009 | 0 | 0.22 | 0.59 | 0.25 | 0.13 | 0.27 | 0.15 |
| 8 | 85683502 | rs2718972 | -1.12 | 0.528 | 1.715 | 0.105 | 0.003 | 0.001 | 0.001 | 0.405 | 0.18 | 0.19 | 0.22 | 0.14 | 0.26 | 0.16 |
| 8 | 85706462 | rs2467212 | -1.378 | 0.535 | 1.23 | 0.143 | 0.043 | 0.024 | 0.024 | 0.929 | 0.19 | 0.41 | 0.21 | 0.14 | 0.26 | 0.15 |
| 8 | 85711775 | rs6996709 | 0.782 | 0.561 | 0.634 | 0.17 | -0.001 | -0.001 | -0.001 | 0 | 0.82 | 0.82 | 0.8 | 0.86 | 0.74 | 0.85 |
| 8 | 87159653 | rs2128930 | -0.262 | 0.426 | 1.438 | -0.297 | -0.007 | -0.004 | -0.004 | 0 | 0.87 | 0.92 | 0.75 | 0.86 | 0.87 | 0.8 |
| 8 | 87166396 | rs6983781 | -0.901 | 0.472 | 1.518 | -0.305 | 0.002 | -0.001 | -0.001 | 0 | 0.18 | 0.16 | 0.34 | 0.17 | 0.28 | 0.32 |
| 8 | 87214088 | rs12550469 | 0.172 | 0.314 | 1.427 | -0.248 | 0.002 | 0 | 0 | 0 | 0.12 | 0.1 | 0.25 | 0.15 | 0.13 | 0.2 |
| 8 | 87216600 | rs11784259 | -0.612 | 0.293 | 1.655 | -0.259 | 0.022 | 0.01 | 0.01 | 0 | 0.16 | 0.34 | 0.28 | 0.17 | 0.15 | 0.23 |
| 8 | 87226700 | rs9656982 | 0.071 | -0.26 | 1.474 | -0.215 | -0.012 | 0.001 | 0.001 | 0 | 0.12 | 0.2 | 0.24 | 0.14 | 0.08 | 0.2 |
| 8 | 87227153 | rs10956518 | -0.005 | -0.264 | 1.466 | -0.217 | -0.01 | 0 | 0 | 0 | 0.12 | 0.21 | 0.24 | 0.14 | 0.08 | 0.2 |
| 8 | 87245253 | rs4310184 | 0.267 | -0.065 | -0.349 | 0.381 | 0.013 | -0.024 | -0.024 | 0 | 0.31 | 0.18 | 0.46 | 0.48 | 0.25 | 0.4 |
| 8 | 87249971 | rs4406372 | -0.862 | -0.13 | -0.153 | 0.631 | -0.001 | 0.074 | 0.074 | 0 | 0.06 | 0.16 | 0.08 | 0.13 | 0.04 | 0.09 |
| 8 | 87293797 | rs13278499 | 0.489 | -0.005 | -0.626 | 0.555 | 0.008 | 0.027 | 0.027 | 0.287 | 0.8 | 0.45 | 0.81 | 0.82 | 0.79 | 0.83 |
| 8 | 87314758 | rs2953525 | 1.209 | 0.017 | -0.635 | 0.111 | 0 | -0.002 | -0.002 | 0 | 0.76 | 0.2 | 0.82 | 0.89 | 0.78 | 0.76 |
| 8 | 87330713 | rs7003059 | 0.037 | 0.053 | -0.67 | 0.112 | 0.003 | -0.005 | -0.005 | 0 | 0.35 | 0.29 | 0.52 | 0.54 | 0.29 | 0.48 |
| 8 | 87338202 | rs16905416 | -0.234 | 0.21 | -0.952 | -0.015 | 0.021 | -0.025 | -0.025 | 0 | 0.06 | 0.13 | 0.05 | 0 | 0.09 | 0.08 |
| 8 | 87374654 | rs6471296 | 0.009 | 0.13 | -0.853 | -0.036 | 0.068 | -0.01 | -0.01 | 0 | 0.58 | 0.25 | 0.41 | 0.46 | 0.7 | 0.58 |
| 8 | 87381607 | rs17681826 | 0.178 | 0.154 | -0.834 | 0.011 | -0.009 | 0.011 | 0.011 | 0 | 0.14 | 0.07 | 0.08 | 0.07 | 0.19 | 0.16 |
| 8 | 87400459 | rs6471314 | -0.899 | 0.12 | -0.893 | -0.131 | 0.021 | -0.045 | -0.045 | 0 | 0.69 | 0.85 | 0.5 | 0.47 | 0.73 | 0.59 |
| 8 | 87434132 | rs12155981 | 0.284 | 0.187 | -0.902 | -0.158 | -0.009 | 0.015 | 0.015 | 0 | 0.14 | 0.07 | 0.08 | 0.05 | 0.19 | 0.14 |
| 8 | 87442135 | rs10092168 | -0.855 | 0.082 | -0.908 | -0.12 | -0.002 | -0.061 | -0.061 | 0.039 | 0.27 | 0.62 | 0.2 | 0.05 | 0.28 | 0.17 |
| 8 | 87482244 | rs10100368 | -0.848 | 0.037 | -0.902 | -0.144 | -0.002 | -0.061 | -0.061 | 0.12 | 0.27 | 0.62 | 0.2 | 0.05 | 0.28 | 0.17 |
| 8 | 87484913 | rs13042 | 0.208 | 0.154 | -0.88 | -0.042 | 0.005 | -0.006 | -0.006 | 0 | 0.35 | 0.33 | 0.53 | 0.53 | 0.31 | 0.48 |
| 8 | 87486168 | rs4961193 | 0.553 | 0.161 | -0.985 | -0.016 | 0.017 | -0.04 | -0.04 | 0 | 0.31 | 0.17 | 0.51 | 0.53 | 0.28 | 0.41 |
| 8 | 87502855 | rs7820697 | -0.788 | 0.219 | -0.882 | -0.029 | -0.001 | -0.001 | -0.001 | 0 | 0.52 | 0.48 | 0.66 | 0.58 | 0.55 | 0.63 |
| 8 | 87518985 | rs10107571 | -0.34 | 0.591 | -0.841 | 0.303 | 0.016 | -0.026 | -0.026 | 0 | 0.17 | 0.17 | 0.13 | 0.05 | 0.25 | 0.15 |
| 8 | 87519315 | rs6980476 | -0.34 | 0.591 | -0.841 | 0.303 | 0.016 | -0.026 | -0.026 | 0 | 0.17 | 0.17 | 0.13 | 0.05 | 0.25 | 0.15 |
| 8 | 87526439 | rs6985066 | 0.228 | 0.64 | -0.855 | 0.349 | 0.025 | -0.049 | -0.049 | 0 | 0.31 | 0.15 | 0.52 | 0.52 | 0.27 | 0.41 |
| 8 | 87529297 | rs11781016 | 0.85 | 0.668 | -0.88 | 0.361 | 0.001 | 0.013 | 0.013 | 0 | 0.51 | 0.62 | 0.34 | 0.43 | 0.48 | 0.43 |
| 8 | 87540872 | rs10956871 | -0.402 | 0.711 | -1.007 | 0.378 | 0.021 | -0.033 | -0.033 | 0 | 0.17 | 0.21 | 0.14 | 0.05 | 0.26 | 0.15 |
| 8 | 87544174 | rs10087788 | -0.008 | 0.379 | -1.105 | 0.933 | 0.031 | -0.055 | -0.055 | 0 | 0.68 | 0.85 | 0.49 | 0.48 | 0.73 | 0.58 |
| 8 | 87562330 | rs7823951 | -0.817 | 0.884 | -0.628 | 0.742 | 0.03 | 0.003 | 0.003 | 0 | 0.07 | 0.1 | 0.05 | 0.05 | 0.12 | 0.09 |
| 8 | 87564586 | rs1542081 | -0.51 | 0.274 | -1.179 | 1.328 | 0.019 | -0.08 | -0.08 | 0 | 0.16 | 0.04 | 0.07 | 0.33 | 0.11 | 0.2 |
| 8 | 87567193 | rs2304789 | -0.077 | 0.047 | -1.105 | 0.079 | 0.001 | -0.062 | -0.062 | 0 | 0.27 | 0.28 | 0.17 | 0.01 | 0.33 | 0.23 |
| 8 | 87567305 | rs3816240 | 0.321 | -0.414 | -0.861 | -0.04 | -0.014 | -0.014 | -0.014 | 0 | 0.63 | 0.57 | 0.43 | 0.3 | 0.74 | 0.58 |
| 8 | 87567848 | rs10504825 | 1.432 | -0.573 | -0.81 | -0.175 | -0.096 | 0.06 | 0.06 | 0 | 0.33 | 0.12 | 0.24 | 0.26 | 0.38 | 0.29 |
| 8 | 87568644 | rs2304787 | 0.222 | -0.602 | -0.821 | -0.161 | -0.014 | -0.014 | -0.014 | 4.332 | 0.63 | 0.57 | 0.43 | 0.3 | 0.74 | 0.58 |
| 8 | 87570665 | rs1062850 | 0.804 | -0.808 | -1.095 | -0.454 | 0.066 | 0.11 | 0.11 | 0 | 0.51 | 0.31 | 0.34 | 0.27 | 0.62 | 0.49 |
| 8 | 87570728 | rs1062851 | 0.805 | -0.817 | -1.099 | -0.458 | 0.066 | 0.11 | 0.11 | 0 | 0.51 | 0.31 | 0.34 | 0.27 | 0.62 | 0.49 |
| 8 | 87571049 | rs1044719 | 0.831 | -0.817 | -1.099 | -0.458 | 0.068 | 0.112 | 0.112 | 0 | 0.51 | 0.3 | 0.34 | 0.27 | 0.62 | 0.49 |
| 8 | 87571252 | rs1062852 | 0.67 | -1.076 | -0.863 | -0.084 | -0.001 | 0.013 | 0.013 | 0.092 | 0.18 | 0.16 | 0.47 | 0.38 | 0.16 | 0.24 |
| 8 | 87571822 | rs1062853 | 0.826 | -1.084 | -0.79 | -0.094 | 0.003 | -0.045 | -0.045 | 0 | 0.17 | 0.08 | 0.47 | 0.35 | 0.16 | 0.21 |
| 8 | 87572663 | rs1044728 | -0.398 | -1.146 | -0.342 | -0.125 | -0.014 | 0.021 | 0.021 | 0.009 | 0.3 | 0.33 | 0.56 | 0.41 | 0.26 | 0.29 |
| 8 | 87573120 | rs1044730 | 1.006 | -1.162 | -0.1 | -0.166 | -0.004 | -0.007 | -0.007 | 0.569 | 0.17 | 0.18 | 0.11 | 0.01 | 0.24 | 0.2 |
| 8 | 87574385 | rs4961198 | -0.169 | -1.17 | 0.264 | -0.131 | -0.004 | 0.008 | 0.008 | 0 | 0.28 | 0.24 | 0.54 | 0.4 | 0.26 | 0.27 |
| 8 | 87578048 | rs6471396 | -1.343 | -1.069 | 0.183 | -0.058 | 0.018 | -0.095 | -0.095 | 0 | 0.84 | 0.98 | 0.93 | 0.7 | 0.9 | 0.8 |
| 8 | 87585119 | rs6471400 | 0.859 | -1.133 | 0.122 | -0.057 | 0.033 | -0.033 | -0.033 | 0 | 0.62 | 0.28 | 0.72 | 0.5 | 0.69 | 0.63 |
| 8 | 87585863 | rs13268560 | -0.504 | -1.149 | 0.56 | -0.051 | 0.016 | 0.007 | 0.007 | 0.06 | 0.08 | 0.26 | 0.11 | 0.11 | 0.09 | 0.11 |
| 8 | 87586151 | rs990193 | -1.57 | -1.157 | 0.22 | -0.051 | 0.006 | -0.099 | -0.099 | 0 | 0.85 | 0.98 | 0.93 | 0.67 | 0.88 | 0.8 |
| 8 | 87586393 | rs990192 | 0.828 | -1.103 | 0.23 | -0.03 | 0.033 | -0.033 | -0.033 | 0 | 0.62 | 0.28 | 0.72 | 0.5 | 0.69 | 0.63 |
| 8 | 87588198 | rs3735972 | 0.742 | -0.895 | 0.256 | 0.153 | 0.006 | 0.002 | 0.002 | 0 | 0.05 | 0.07 | 0.09 | 0.1 | 0.09 | 0.07 |
| 8 | 87591778 | rs958401 | 0.567 | -0.87 | 0.247 | 0.143 | 0.005 | 0.001 | 0.001 | 0 | 0.06 | 0.07 | 0.09 | 0.1 | 0.11 | 0.07 |
| 8 | 87614243 | rs1441249 | 1.399 | -0.931 | -0.021 | 0.105 | -0.138 | 0.128 | 0.128 | 0.115 | 0.43 | 0.03 | 0.29 | 0.26 | 0.48 | 0.32 |
| 8 | 87615964 | rs4961202 | -0.134 | -0.863 | 0.205 | 0.179 | 0.002 | -0.006 | -0.006 | 0 | 0.29 | 0.19 | 0.48 | 0.24 | 0.28 | 0.33 |
| 8 | 87628371 | rs10093381 | 0.394 | -0.798 | 0.212 | 0.414 | 0.003 | -0.005 | -0.005 | 0.181 | 0.23 | 0.24 | 0.09 | 0.32 | 0.18 | 0.23 |
| 8 | 87629659 | rs7823042 | -0.242 | -0.797 | 0.252 | 0.363 | -0.023 | 0.033 | 0.033 | 0 | 0.29 | 0.6 | 0.2 | 0.41 | 0.27 | 0.28 |
| 8 | 87629825 | rs13249366 | -0.241 | -0.451 | 0.036 | 0.4 | -0.002 | 0.006 | 0.006 | 0.091 | 0.21 | 0.22 | 0.43 | 0.27 | 0.16 | 0.34 |
| 8 | 87630789 | rs12056692 | 0.321 | -0.379 | -0.234 | 0.44 | 0.001 | 0.024 | 0.024 | 0.151 | 0.78 | 0.55 | 0.53 | 0.77 | 0.81 | 0.71 |
| 8 | 87631110 | rs3779802 | 0.097 | 0.809 | -1.327 | 0.053 | -0.062 | -0.016 | -0.016 | 0 | 0.23 | 0.09 | 0.25 | 0.34 | 0.37 | 0.3 |
| 8 | 87638453 | rs13258590 | 0.428 | 1.128 | -2.104 | 0.211 | -0.054 | 0.056 | 0.056 | 0.099 | 0.49 | 0.07 | 0.3 | 0.38 | 0.45 | 0.41 |
| 8 | 87642150 | rs3779800 | -0.704 | 1.257 | -0.931 | 0.26 | 0.041 | -0.024 | -0.024 | 0 | 0.25 | 0.55 | 0.37 | 0.22 | 0.3 | 0.23 |
| 8 | 87652115 | rs6991672 | -0.401 | 1.27 | -0.383 | 0.277 | -0.001 | 0 | 0 | 0 | 0.06 | 0.04 | 0.06 | 0.08 | 0.07 | 0.11 |
| 8 | 87654215 | rs4961205 | -0.411 | 1.39 | 0.372 | 0.166 | 0.001 | 0.005 | 0.005 | 0 | 0.17 | 0.23 | 0.3 | 0.15 | 0.16 | 0.1 |
| 8 | 87660100 | rs13265557 | -0.266 | 1.408 | 0.365 | 0.208 | -0.002 | 0.001 | 0.001 | 0 | 0.06 | 0.04 | 0.06 | 0.08 | 0.07 | 0.11 |
| 8 | 87664244 | rs9297943 | 1.594 | 1.383 | 0.555 | 0.018 | 0.003 | -0.007 | -0.007 | 2.359 | 0.71 | 0.38 | 0.56 | 0.78 | 0.64 | 0.76 |
| 8 | 87669792 | rs2013536 | -1.119 | 1.379 | 0.375 | 0.187 | 0.003 | -0.005 | -0.005 | 0 | 0.37 | 0.25 | 0.54 | 0.54 | 0.38 | 0.52 |
| 8 | 87671396 | rs3779794 | 0.591 | 1.349 | 0.339 | 0.209 | 0 | -0.091 | -0.091 | 0 | 0.21 | 0.07 | 0.25 | 0.39 | 0.21 | 0.4 |

| | | | | | | | | | | | | | | | | |
|---|---|---|---|---|---|---|---|---|---|---|---|---|---|---|---|---|
| 8 | 87673633 | rs7017987 | 0.205 | 1.353 | 0.429 | 0.16 | 0.002 | 0 | 0 | 0 | 0.15 | 0.18 | 0.29 | 0.13 | 0.15 | 0.1 |
| 8 | 87675854 | rs3779792 | -0.27 | 1.263 | 0.831 | 0.15 | 0.064 | 0.02 | 0.02 | 0.059 | 0.06 | 0.3 | 0.07 | 0.08 | 0.07 | 0.1 |
| 8 | 87676710 | rs3779791 | 0.705 | 1.348 | 0.807 | 0.097 | 0.084 | -0.067 | -0.067 | 0 | 0.86 | 0.59 | 0.84 | 0.92 | 0.79 | 0.87 |
| 8 | 87677217 | rs4960986 | -0.507 | 1.34 | 0.541 | 0.097 | -0.004 | -0.009 | -0.009 | 0 | 0.92 | 0.93 | 0.93 | 1 | 0.86 | 0.97 |
| 8 | 87679098 | rs3735967 | 1.824 | 1.394 | 0.377 | 0.165 | 0.023 | 0.064 | 0.064 | 0.383 | 0.5 | 0.32 | 0.32 | 0.37 | 0.43 | 0.37 |
| 8 | 87679463 | rs3735966 | -0.247 | 1.358 | 0.165 | 0.219 | 0.049 | 0.01 | 0.01 | 0 | 0.06 | 0.23 | 0.06 | 0.08 | 0.07 | 0.1 |
| 8 | 87690019 | rs10504826 | 1.249 | 1.107 | -0.379 | 0.287 | -0.012 | 0.049 | 0.049 | 1.313 | 0.72 | 0.12 | 0.58 | 0.86 | 0.72 | 0.86 |
| 8 | 87692895 | rs12548374 | 0.354 | 1.117 | -0.142 | 0.432 | 0.005 | -0.004 | -0.004 | 0 | 0.11 | 0.06 | 0.26 | 0.11 | 0.11 | 0.07 |
| 8 | 87694905 | rs7842410 | 1.061 | 1.08 | -0.503 | 0.297 | -0.049 | 0.022 | 0.022 | 0.41 | 0.51 | 0.1 | 0.34 | 0.47 | 0.48 | 0.48 |
| 8 | 87713975 | rs2957792 | 1.402 | 0.812 | -0.972 | -0.37 | 0.026 | -0.023 | -0.023 | 0 | 0.32 | 0.12 | 0.23 | 0.36 | 0.23 | 0.17 |
| 8 | 87716283 | rs1458118 | 1.246 | 0.885 | 0.344 | -0.415 | -0.01 | 0.051 | 0.051 | 0 | 0.25 | 0.42 | 0.29 | 0.37 | 0.24 | 0.42 |
| 8 | 87720350 | rs11987550 | 1.148 | 0.87 | 0.351 | -0.45 | -0.015 | 0.034 | 0.034 | 0 | 0.07 | 0.14 | 0.08 | 0.12 | 0.04 | 0.04 |
| 8 | 87720419 | rs13278423 | -0.572 | 0.933 | 0.303 | -0.516 | -0.012 | -0.014 | -0.014 | 0.012 | 0.52 | 0.73 | 0.58 | 0.51 | 0.51 | 0.76 |
| 8 | 87721349 | rs7814749 | -1.436 | 1.038 | 0.223 | -0.552 | -0.033 | -0.006 | -0.006 | 0 | 0.63 | 0.87 | 0.69 | 0.64 | 0.63 | 0.82 |
| 8 | 87721437 | rs4961212 | 0.286 | 0.957 | 0.334 | -0.429 | -0.027 | 0.04 | 0.04 | 0 | 0.32 | 0.54 | 0.37 | 0.49 | 0.28 | 0.46 |
| 8 | 87754395 | rs1458119 | -1.436 | 0.796 | 0.21 | -0.137 | 0.002 | -0.011 | -0.011 | 0 | 0.66 | 0.75 | 0.72 | 0.57 | 0.7 | 0.82 |
| 8 | 87759861 | rs6990710 | 1.097 | 0.722 | 0.424 | 0.071 | -0.002 | 0.014 | 0.014 | 0 | 0.24 | 0.35 | 0.3 | 0.32 | 0.25 | 0.43 |
| 8 | 87761007 | rs4961216 | 0.524 | 0.738 | 0.583 | 0.04 | -0.02 | 0.025 | 0.025 | 0 | 0.11 | 0.24 | 0.11 | 0.12 | 0.07 | 0.04 |
| 8 | 87768331 | rs10956972 | -0.609 | 0.803 | 0.416 | 0.014 | -0.003 | -0.018 | -0.018 | 0.925 | 0.45 | 0.72 | 0.49 | 0.47 | 0.49 | 0.53 |
| 8 | 87775582 | rs16917400 | 1.182 | 0.813 | 0.335 | 0.037 | -0.001 | 0.015 | 0.015 | 0 | 0.25 | 0.35 | 0.3 | 0.32 | 0.26 | 0.43 |
| 8 | 87776019 | rs1982563 | -0.536 | 0.84 | -0.416 | 0.058 | -0.011 | -0.036 | -0.036 | 0.032 | 0.46 | 0.79 | 0.49 | 0.47 | 0.49 | 0.53 |
| 8 | 87777391 | rs10085929 | 0.008 | 0.851 | -0.371 | 0.05 | 0.001 | -0.001 | -0.001 | 0 | 0.35 | 0.65 | 0.39 | 0.36 | 0.42 | 0.49 |
| 8 | 87780862 | rs11786561 | -0.339 | 0.631 | -0.14 | 0.293 | -0.001 | 0.001 | 0.001 | 0 | 0.16 | 0.12 | 0.18 | 0.11 | 0.21 | 0.06 |
| 8 | 87784959 | rs10955046 | -0.856 | 0.132 | -0.399 | 0.533 | 0.026 | -0.038 | -0.038 | 0 | 0.7 | 0.82 | 0.8 | 0.64 | 0.82 | 0.84 |
| 8 | 87786254 | rs1598888 | -1.401 | 0.099 | -0.626 | 0.436 | -0.001 | -0.03 | -0.03 | 0 | 0.86 | 0.93 | 0.88 | 0.73 | 0.89 | 0.9 |
| 8 | 87787661 | rs13256575 | -0.617 | -0.092 | 0.182 | 0.331 | -0.019 | 0.055 | 0.055 | 1.348 | 0.81 | 0.61 | 0.82 | 0.73 | 0.89 | 0.88 |
| 8 | 87797573 | rs13257199 | 0.62 | -0.23 | -0.021 | 0.32 | 0.022 | -0.022 | -0.022 | 0 | 0.61 | 0.3 | 0.66 | 0.52 | 0.71 | 0.7 |
| 8 | 87799663 | rs7830108 | -0.001 | 0.017 | 0.222 | -0.074 | -0.01 | -0.025 | -0.025 | 0 | 0.72 | 0.6 | 0.73 | 0.78 | 0.75 | 0.72 |
| 8 | 87804309 | rs1379736 | 0.007 | 0.029 | 0.352 | -0.097 | -0.01 | -0.029 | -0.029 | 0 | 0.72 | 0.57 | 0.73 | 0.78 | 0.76 | 0.72 |
| 8 | 87806604 | rs7834685 | -0.697 | 0.176 | 0.391 | -0.27 | 0 | -0.006 | -0.006 | 0 | 0.7 | 0.74 | 0.76 | 0.59 | 0.74 | 0.73 |
| 8 | 87807203 | rs11786304 | 1.252 | -0.05 | 0.114 | -0.56 | -0.077 | -0.056 | -0.056 | 0.002 | 0.35 | 0.15 | 0.49 | 0.56 | 0.46 | 0.5 |
| 8 | 87813527 | rs17693567 | -0.329 | -0.074 | 0.115 | -0.805 | 0.007 | 0.01 | 0.01 | 0 | 0.28 | 0.14 | 0.22 | 0 | 0.27 | 0.17 |
| 8 | 87813597 | rs7018237 | -0.813 | -0.081 | 0.241 | -0.626 | -0.002 | 0.031 | 0.031 | 1.139 | 0.69 | 0.62 | 0.75 | 0.58 | 0.74 | 0.73 |
| 8 | 87833711 | rs10808349 | -0.567 | 0.246 | 0.262 | -0.425 | -0.006 | -0.005 | -0.005 | 0 | 0.58 | 0.75 | 0.71 | 0.57 | 0.63 | 0.7 |
| 8 | 87844003 | rs7846655 | -0.101 | 0.292 | 0.276 | -0.457 | -0.002 | -0.058 | -0.058 | 0.116 | 0.17 | 0.18 | 0.17 | 0 | 0.2 | 0.16 |
| 8 | 87846941 | rs10955072 | 1.61 | 0.468 | -0.133 | -0.23 | -0.074 | -0.067 | -0.067 | 0 | 0.12 | 0.01 | 0.14 | 0.19 | 0.21 | 0.26 |
| 8 | 87853542 | rs4320564 | 0.591 | 0.223 | 0.521 | -0.563 | 0.001 | 0.003 | 0.003 | 0 | 0.39 | 0.51 | 0.52 | 0.57 | 0.42 | 0.53 |
| 8 | 87862663 | rs16894253 | 0.174 | 0.623 | 0.393 | -0.411 | -0.017 | -0.053 | -0.053 | 0 | 0.09 | 0.28 | 0.08 | 0 | 0.07 | 0.05 |
| 8 | 87867347 | rs2045506 | 0.928 | 0.607 | 0.22 | -0.439 | -0.007 | -0.018 | -0.018 | 0 | 0.41 | 0.25 | 0.53 | 0.54 | 0.45 | 0.52 |
| 8 | 87876114 | rs2934950 | -0.512 | 0.83 | 0.068 | -0.595 | -0.003 | -0.005 | -0.005 | 0 | 0.64 | 0.72 | 0.73 | 0.59 | 0.65 | 0.72 |
| 8 | 87878846 | rs2453451 | -0.824 | 0.666 | -0.006 | -0.624 | 0.001 | -0.016 | -0.016 | 0 | 0.2 | 0.22 | 0.18 | 0.02 | 0.2 | 0.19 |
| 8 | 87881132 | rs2446120 | -0.502 | 0.527 | 0.09 | -0.549 | -0.026 | -0.03 | -0.03 | 0 | 0.65 | 0.83 | 0.72 | 0.59 | 0.64 | 0.72 |
| 8 | 87890899 | rs13248329 | 0.223 | 0.45 | 0.206 | -0.471 | -0.001 | -0.002 | -0.002 | 0.175 | 0.1 | 0.09 | 0.07 | 0.03 | 0.07 | 0.09 |
| 8 | 87895691 | rs13269825 | 0.966 | 0.44 | 0.358 | -0.532 | 0.001 | 0.005 | 0.005 | 0 | 0.43 | 0.52 | 0.55 | 0.56 | 0.46 | 0.55 |
| 8 | 87900893 | rs6468523 | 0.104 | 0.399 | 0.388 | -0.569 | -0.019 | 0.001 | 0.001 | 0.006 | 0.55 | 0.74 | 0.64 | 0.58 | 0.53 | 0.66 |
| 8 | 87905053 | rs7831076 | -0.462 | 0.327 | 0.531 | -0.277 | -0.026 | -0.034 | -0.034 | 0 | 0.12 | 0.22 | 0.09 | 0.03 | 0.07 | 0.11 |
| 8 | 87913985 | rs13248854 | 0.957 | 0.573 | 0.719 | -0.255 | -0.001 | -0.002 | -0.002 | 0 | 0.52 | 0.43 | 0.57 | 0.55 | 0.53 | 0.61 |
| 8 | 87914099 | rs4504655 | -0.125 | 0.602 | 0.818 | -0.31 | 0.002 | 0.011 | 0.011 | 0 | 0.34 | 0.37 | 0.33 | 0.42 | 0.33 | 0.26 |
| 8 | 87918128 | rs4373537 | -1.05 | 0.633 | 0.456 | -0.283 | -0.009 | 0.034 | 0.034 | 0 | 0.93 | 0.98 | 0.95 | 1 | 0.92 | 0.94 |
| 8 | 87935589 | rs7016229 | -0.013 | 0.596 | 0.51 | -0.259 | -0.005 | -0.015 | -0.015 | 0 | 0.33 | 0.22 | 0.25 | 0.4 | 0.33 | 0.27 |
| 8 | 87936056 | rs10808353 | 0.708 | 0.608 | 0.479 | -0.193 | 0 | -0.001 | -0.001 | 0 | 0.54 | 0.59 | 0.64 | 0.57 | 0.54 | 0.61 |
| 8 | 87958064 | rs7835558 | 0.218 | 0.495 | 0.231 | -0.167 | -0.002 | 0.006 | 0.006 | 0 | 0.63 | 0.57 | 0.7 | 0.6 | 0.64 | 0.64 |
| 8 | 87958212 | rs7822655 | 0.753 | 0.43 | 0.033 | -0.213 | -0.003 | -0.001 | -0.001 | 0.013 | 0.55 | 0.48 | 0.65 | 0.58 | 0.56 | 0.56 |
| 8 | 87974669 | rs6996849 | 0.962 | 0.731 | -0.533 | -0.203 | 0.004 | 0.02 | 0.02 | 0 | 0.51 | 0.26 | 0.6 | 0.57 | 0.51 | 0.54 |
| 8 | 87979777 | rs11778879 | 0.85 | 0.883 | -0.388 | -0.337 | -0.005 | 0.053 | 0.053 | 0 | 0.59 | 0.26 | 0.63 | 0.77 | 0.56 | 0.69 |
| 8 | 87982674 | rs6468587 | 0.857 | 0.711 | -0.009 | -0.349 | -0.003 | -0.134 | -0.134 | 0 | 0.81 | 0.48 | 0.83 | 0.97 | 0.8 | 0.88 |
| 8 | 88006771 | rs1524834 | -0.159 | 0.827 | -0.763 | -0.094 | -0.005 | 0.028 | 0.028 | 0 | 0.43 | 0.63 | 0.31 | 0.26 | 0.4 | 0.36 |
| 8 | 88007114 | rs1852027 | -0.019 | 1.003 | -0.65 | -0.148 | 0 | 0 | 0 | 0 | 0.54 | 0.48 | 0.59 | 0.64 | 0.54 | 0.61 |
| 8 | 88009165 | rs6468613 | -0.282 | 0.576 | -0.824 | -0.044 | -0.003 | -0.006 | -0.006 | 0 | 0.54 | 0.64 | 0.47 | 0.44 | 0.52 | 0.52 |
| 8 | 88010106 | rs1880473 | 0.11 | 0.3 | -0.931 | 0.452 | 0.004 | -0.007 | -0.007 | 0 | 0.19 | 0.17 | 0.08 | 0.31 | 0.15 | 0.25 |
| 8 | 88032231 | rs2453446 | -0.403 | 0.34 | -0.6 | 0.289 | 0.007 | 0.007 | 0.007 | 0.301 | 0.36 | 0.55 | 0.29 | 0.6 | 0.38 | 0.41 |
| 8 | 88037170 | rs2453449 | -0.451 | 0.335 | -0.631 | 0.287 | 0.001 | 0.001 | 0.001 | 0.553 | 0.36 | 0.61 | 0.3 | 0.6 | 0.39 | 0.42 |
| 8 | 88043523 | rs17623133 | 1.179 | 0.15 | -0.899 | 0.085 | -0.003 | 0.01 | 0.01 | 0 | 0.3 | 0.25 | 0.47 | 0.23 | 0.35 | 0.19 |
| 8 | 88043829 | rs11786526 | 1.514 | 0.244 | -1.155 | 0.358 | 0.035 | 0.019 | 0.019 | 0 | 0.14 | 0.19 | 0.11 | 0.19 | 0.22 | 0.14 |
| 8 | 88051932 | rs2949554 | 0.311 | -0.002 | -1.882 | 0.426 | 0.025 | 0.054 | 0.054 | 1.667 | 0.4 | 0.14 | 0.29 | 0.27 | 0.34 | 0.4 |
| 8 | 88052397 | rs2943177 | -0.873 | 0.054 | -1.667 | 0.341 | -0.019 | -0.115 | -0.115 | 0 | 0.83 | 0.99 | 0.85 | 0.6 | 0.84 | 0.75 |
| 8 | 88055465 | rs4596672 | 0.57 | -0.087 | -1.529 | 0.008 | 0.023 | 0.02 | 0.02 | 0.232 | 0.3 | 0.09 | 0.28 | 0.22 | 0.25 | 0.34 |
| 8 | 88064750 | rs2943174 | 0.778 | 0.274 | -1.718 | -0.301 | -0.071 | -0.003 | -0.003 | 1.841 | 0.2 | 0.06 | 0.37 | 0.19 | 0.32 | 0.19 |
| 8 | 88073155 | rs11986813 | -0.325 | -0.263 | -0.66 | -0.45 | -0.017 | -0.06 | -0.06 | 0 | 0.11 | 0.47 | 0.08 | 0.01 | 0.06 | 0.05 |

| | | | | | | | | | | | | | | | | |
|---|---|---|---|---|---|---|---|---|---|---|---|---|---|---|---|---|
| 8 | 88075043 | rs2974279 | -0.313 | -0.345 | -1.444 | -0.408 | 0.064 | 0.044 | 0.044 | 0.437 | 0.54 | 0.86 | 0.6 | 0.36 | 0.63 | 0.42 |
| 8 | 88075198 | rs2974278 | -1.11 | -0.387 | -1.496 | -0.253 | 0.006 | -0.131 | -0.131 | 0 | 0.82 | 0.99 | 0.85 | 0.57 | 0.87 | 0.73 |
| 8 | 88088357 | rs6997613 | -0.026 | -0.041 | -1.19 | -0.648 | -0.005 | 0.072 | 0.072 | 0.806 | 0.47 | 0.88 | 0.5 | 0.3 | 0.55 | 0.4 |
| 8 | 88090264 | rs35096718 | 0.797 | 0.001 | -0.789 | -0.738 | -0.004 | -0.03 | -0.03 | 0 | 0.06 | 0.12 | 0.04 | 0 | 0.04 | 0.05 |
| 8 | 88109732 | rs6468651 | 0.437 | 0.19 | -0.661 | -0.84 | 0.077 | 0.056 | 0.056 | 0.747 | 0.12 | 0.38 | 0.26 | 0.15 | 0.21 | 0.12 |
| 8 | 88115158 | rs2974277 | 0.096 | -0.056 | -1.287 | -1.23 | 0.003 | 0 | 0 | 0 | 0.19 | 0.15 | 0.21 | 0.07 | 0.17 | 0.18 |
| 8 | 88115347 | rs2974276 | 0.613 | -0.111 | -1.447 | -1.276 | -0.007 | 0.029 | 0.029 | 0 | 0.13 | 0 | 0.14 | 0.07 | 0.1 | 0.17 |
| 8 | 88117846 | rs17623943 | 0.975 | -0.121 | -1.559 | -1.291 | -0.077 | -0.042 | -0.042 | 0 | 0.15 | 0.03 | 0.27 | 0.16 | 0.27 | 0.12 |
| 8 | 88121700 | rs6994542 | 0.268 | 0.135 | -1.785 | -1.387 | -0.021 | 0.073 | 0.073 | 1.13 | 0.47 | 0.84 | 0.45 | 0.31 | 0.49 | 0.34 |
| 8 | 88125316 | rs2943179 | 0.088 | -0.183 | -1.708 | -1.403 | 0 | -0.007 | -0.007 | 0 | 0.2 | 0.15 | 0.21 | 0.04 | 0.18 | 0.17 |
| 8 | 88132717 | rs7844876 | 0.094 | -0.08 | -1.214 | -1.256 | 0.027 | 0.043 | 0.043 | 0 | 0.5 | 0.63 | 0.47 | 0.35 | 0.56 | 0.4 |
| 8 | 88133822 | rs6415434 | -0.098 | -0.06 | -1.874 | -1.252 | 0.033 | 0.069 | 0.069 | 0.061 | 0.52 | 0.84 | 0.49 | 0.35 | 0.56 | 0.38 |
| 8 | 88153131 | rs1431761 | 0.211 | -0.085 | -1.756 | -1.174 | -0.039 | 0.045 | 0.045 | 0.263 | 0.45 | 0.83 | 0.44 | 0.36 | 0.5 | 0.38 |
| 8 | 88156383 | rs17624357 | 1.897 | -0.053 | -1.782 | -1.222 | -0.052 | 0.043 | 0.043 | 0.014 | 0.25 | 0.08 | 0.13 | 0.15 | 0.26 | 0.22 |
| 8 | 88166193 | rs1431758 | 1.049 | -0.311 | -1.177 | -0.969 | -0.008 | 0.004 | 0.004 | 0 | 0.41 | 0.39 | 0.4 | 0.35 | 0.5 | 0.38 |
| 8 | 88176198 | rs2943161 | 0.58 | 0.086 | -2.09 | -0.711 | -0.008 | 0.074 | 0.074 | 0.071 | 0.49 | 0.84 | 0.46 | 0.35 | 0.52 | 0.41 |
| 8 | 88190551 | rs997597 | -0.101 | 0.238 | -1.573 | -0.577 | -0.003 | -0.005 | -0.005 | 0 | 0.2 | 0.15 | 0.23 | 0.04 | 0.21 | 0.14 |
| 8 | 88191970 | rs2878863 | -0.934 | -0.04 | -1.621 | 0.175 | 0.026 | -0.097 | -0.097 | 0 | 0.87 | 0.99 | 0.88 | 0.59 | 0.91 | 0.79 |
| 8 | 88235341 | rs13273720 | 0.456 | 0.358 | -1.415 | -0.522 | -0.076 | 0.089 | 0.089 | 0 | 0.39 | 0.14 | 0.49 | 0.19 | 0.46 | 0.2 |
| 8 | 88244548 | rs7844935 | -1.139 | 0.209 | -0.051 | -0.318 | 0.046 | -0.029 | -0.029 | 0.705 | 0.09 | 0.55 | 0.09 | 0 | 0.07 | 0.07 |
| 8 | 88253171 | rs4961016 | 0.185 | 0.185 | -0.072 | -0.147 | -0.024 | 0.007 | 0.007 | 0 | 0.64 | 0.54 | 0.62 | 0.39 | 0.68 | 0.46 |
| 8 | 88260285 | rs7462260 | 0.016 | 0.683 | 0.155 | -0.134 | 0 | 0 | 0 | 0.029 | 0.66 | 0.69 | 0.52 | 0.4 | 0.63 | 0.53 |
| 8 | 88261791 | rs7833568 | -0.189 | 0.688 | 0.208 | -0.137 | 0.002 | -0.007 | -0.007 | 0 | 0.69 | 0.79 | 0.57 | 0.4 | 0.7 | 0.54 |
| 8 | 88289035 | rs7844806 | 0.555 | 0.732 | 0.134 | -0.425 | -0.006 | -0.071 | -0.071 | 0.112 | 0.26 | 0.43 | 0.17 | 0.15 | 0.25 | 0.3 |
| 8 | 88292520 | rs4961020 | 0.607 | 0.976 | -0.196 | -0.178 | -0.066 | 0.051 | 0.051 | 0 | 0.39 | 0.13 | 0.38 | 0.25 | 0.45 | 0.25 |
| 8 | 88321459 | rs12547621 | -0.851 | 1.056 | 0.634 | 0.077 | 0.105 | 0.082 | 0.082 | 0.017 | 0.08 | 0.42 | 0.11 | 0.08 | 0.1 | 0.11 |
| 8 | 88331031 | rs13282346 | -0.141 | 0.822 | 0.49 | -0.048 | -0.005 | 0.1 | 0.1 | 0 | 0.49 | 0.81 | 0.4 | 0.26 | 0.49 | 0.35 |
| 8 | 88341350 | rs7819936 | 0.675 | 0.804 | 0.481 | -0.065 | -0.007 | 0.041 | 0.041 | 0 | 0.4 | 0.35 | 0.29 | 0.19 | 0.39 | 0.22 |
| 8 | 88356270 | rs10094122 | 1.081 | 0.905 | 0.493 | -0.107 | -0.006 | 0.064 | 0.064 | 0 | 0.36 | 0.2 | 0.25 | 0.18 | 0.33 | 0.21 |
| 8 | 88358049 | rs7829334 | -0.66 | 0.907 | 0.264 | -0.182 | -0.003 | -0.065 | -0.065 | 0 | 0.64 | 0.94 | 0.59 | 0.42 | 0.71 | 0.59 |
| 8 | 88361114 | rs10098438 | 1.384 | 0.812 | 0.359 | -0.205 | -0.023 | 0.073 | 0.073 | 0 | 0.36 | 0.11 | 0.25 | 0.18 | 0.33 | 0.21 |
| 8 | 88379205 | rs10504830 | -0.66 | 0.818 | 0.692 | -0.192 | 0.068 | 0.042 | 0.042 | 0 | 0.09 | 0.4 | 0.11 | 0.08 | 0.1 | 0.14 |
| 8 | 88387183 | rs4344102 | 0.947 | 0.821 | 0.646 | -0.209 | -0.011 | 0.059 | 0.059 | 0 | 0.4 | 0.3 | 0.29 | 0.19 | 0.39 | 0.23 |
| 8 | 88407275 | rs10101259 | 0.087 | 0.816 | 0.569 | -0.2 | -0.001 | 0.108 | 0.108 | 0 | 0.5 | 0.81 | 0.4 | 0.26 | 0.49 | 0.37 |
| 8 | 88410905 | rs4415337 | 1.03 | 0.617 | 0.298 | 0.071 | -0.032 | 0.007 | 0.007 | 0 | 0.16 | 0.05 | 0.26 | 0.15 | 0.18 | 0.15 |
| 8 | 88411051 | rs10087899 | 1.413 | 0.924 | 0.577 | 0.097 | -0.045 | 0.07 | 0.07 | 0 | 0.36 | 0.07 | 0.24 | 0.18 | 0.33 | 0.2 |
| 8 | 88411745 | rs10955278 | 1.183 | 0.927 | 0.639 | 0.113 | 0 | -0.001 | -0.001 | 0 | 0.15 | 0.11 | 0.18 | 0.15 | 0.22 | 0.22 |
| 8 | 88414581 | rs4961033 | -0.15 | 0.928 | 0.695 | 0.083 | 0.001 | 0.083 | 0.083 | 0 | 0.49 | 0.75 | 0.4 | 0.26 | 0.49 | 0.37 |
| 8 | 88439189 | rs4961034 | 1.186 | 0.98 | 0.537 | 0.105 | -0.019 | 0.081 | 0.081 | 0 | 0.37 | 0.11 | 0.25 | 0.18 | 0.33 | 0.2 |
| 8 | 88441967 | rs7843243 | -0.336 | 1.148 | 1.211 | 0.272 | 0.014 | 0.005 | 0.005 | 0 | 0.07 | 0.1 | 0.08 | 0.08 | 0.1 | 0.11 |
| 8 | 88450505 | rs7007340 | -1.207 | 1.161 | 0.871 | 0.295 | 0.108 | 0.02 | 0.02 | 0 | 0.11 | 0.46 | 0.13 | 0.08 | 0.16 | 0.16 |
| 8 | 88463304 | rs2905419 | 0.689 | 1.075 | -0.034 | 0.427 | -0.01 | 0.001 | 0.001 | 0 | 0.15 | 0.09 | 0.19 | 0.22 | 0.23 | 0.31 |
| 8 | 88465534 | rs2905418 | 0.431 | 1.167 | -0.091 | 0.324 | -0.015 | 0.013 | 0.013 | 0 | 0.18 | 0.1 | 0.15 | 0.19 | 0.26 | 0.31 |
| 8 | 88472853 | rs12114848 | 0.008 | 1.363 | 0.163 | 0.38 | 0.027 | 0.012 | 0.012 | 0 | 0.06 | 0.13 | 0.05 | 0.05 | 0.11 | 0.11 |
| 8 | 88472924 | rs404469 | 0.716 | 1.605 | 0.022 | 0.45 | -0.014 | -0.014 | -0.014 | 0 | 0.09 | 0.05 | 0.28 | 0.19 | 0.13 | 0.12 |
| 8 | 88480902 | rs428838 | -1.373 | 1.275 | -0.266 | 0.276 | 0.022 | -0.106 | -0.106 | 0.147 | 0.73 | 0.92 | 0.74 | 0.61 | 0.79 | 0.78 |
| 8 | 88481760 | rs2977762 | -0.455 | 1.019 | 0.4 | 0.177 | 0.013 | -0.02 | -0.02 | 0 | 0.28 | 0.56 | 0.23 | 0.24 | 0.38 | 0.45 |
| 8 | 88484377 | rs443440 | 0.61 | 1.075 | 0.352 | 0.574 | -0.019 | -0.011 | -0.011 | 0.091 | 0.46 | 0.24 | 0.61 | 0.48 | 0.47 | 0.35 |
| 8 | 88484882 | rs1160280 | 0.249 | 1.362 | 0.604 | 0.553 | 0.022 | -0.031 | -0.031 | 0 | 0.2 | 0.33 | 0.17 | 0.19 | 0.27 | 0.34 |
| 8 | 88488238 | rs455071 | 0.531 | 1.437 | 0.151 | 0.849 | -0.016 | -0.019 | -0.019 | 0 | 0.09 | 0.05 | 0.29 | 0.2 | 0.13 | 0.12 |
| 8 | 88495559 | rs7000330 | -0.506 | 1.733 | 0.457 | 0.78 | 0.046 | 0.016 | 0.016 | 0 | 0.07 | 0.16 | 0.09 | 0.09 | 0.15 | 0.12 |
| 8 | 88500608 | rs412932 | -1.527 | 1.511 | 0.006 | 1.009 | -0.002 | -0.075 | -0.075 | 0 | 0.76 | 0.99 | 0.86 | 0.76 | 0.83 | 0.81 |
| 8 | 88505366 | rs452325 | -1.154 | 1.259 | 0.751 | 0.961 | 0.018 | 0.007 | 0.007 | 0.312 | 0.33 | 0.57 | 0.52 | 0.44 | 0.48 | 0.54 |
| 8 | 88512286 | rs391916 | -1.181 | 1.251 | 0.782 | 0.957 | 0.01 | 0.004 | 0.004 | 0.384 | 0.33 | 0.61 | 0.52 | 0.44 | 0.48 | 0.54 |
| 8 | 88513775 | rs463060 | -0.653 | 1.298 | 0.583 | 1.029 | 0.036 | 0.036 | 0.036 | 0.705 | 0.16 | 0.25 | 0.35 | 0.26 | 0.25 | 0.24 |
| 8 | 88515393 | rs459578 | -1.041 | 1.616 | 0.312 | 0.984 | -0.047 | -0.084 | -0.084 | 0.925 | 0.68 | 0.92 | 0.73 | 0.63 | 0.68 | 0.72 |
| 8 | 88519408 | rs462993 | 0.224 | 2.038 | 0.857 | 0.974 | -0.002 | -0.001 | -0.001 | 0 | 0.1 | 0.09 | 0.26 | 0.17 | 0.13 | 0.12 |
| 8 | 88534855 | rs428068 | -1.129 | 1.958 | 0.98 | 0.798 | 0.079 | 0.141 | 0.141 | 0.451 | 0.17 | 0.39 | 0.45 | 0.37 | 0.25 | 0.24 |
| 8 | 88540932 | rs2337086 | -0.197 | 1.62 | 0.831 | 0.509 | 0.032 | -0.048 | -0.048 | 0 | 0.19 | 0.45 | 0.18 | 0.17 | 0.25 | 0.2 |
| 8 | 88541187 | rs10086700 | 0.111 | 1.61 | 0.959 | 0.504 | 0.028 | -0.033 | -0.033 | 0.101 | 0.18 | 0.36 | 0.17 | 0.17 | 0.25 | 0.2 |
| 8 | 88548180 | rs17712764 | 1.098 | 1.593 | 0.42 | 0.512 | -0.019 | 0.026 | 0.026 | 0 | 0.32 | 0.09 | 0.16 | 0.21 | 0.29 | 0.37 |
| 8 | 88549744 | rs2142195 | -0.08 | 1.601 | 0.483 | 0.505 | -0.003 | 0.067 | 0.067 | 0.126 | 0.52 | 0.25 | 0.31 | 0.38 | 0.54 | 0.59 |
| 8 | 88555634 | rs13260754 | 0.781 | 1.277 | 0.409 | 0.534 | 0.004 | 0.045 | 0.045 | 0.023 | 0.35 | 0.14 | 0.16 | 0.21 | 0.29 | 0.39 |
| 8 | 88574333 | rs2056862 | -0.421 | 1.375 | 0.053 | 0.523 | -0.002 | -0.005 | -0.005 | 0 | 0.58 | 0.57 | 0.39 | 0.38 | 0.63 | 0.62 |
| 8 | 88578142 | rs741868 | 0.319 | 1.434 | -0.214 | 0.467 | -0.06 | 0.032 | 0.032 | 0 | 0.19 | 0.07 | 0.2 | 0.17 | 0.35 | 0.2 |
| 8 | 88579766 | rs11994666 | -0.154 | 1.316 | 0.095 | 0.61 | -0.001 | 0.005 | 0.005 | 0 | 0.57 | 0.63 | 0.39 | 0.38 | 0.6 | 0.62 |
| 8 | 88584919 | rs2337088 | -0.231 | 1.297 | -0.839 | 0.743 | 0.015 | 0.02 | 0.02 | 0.061 | 0.58 | 0.26 | 0.38 | 0.38 | 0.65 | 0.67 |
| 8 | 88590643 | rs10101821 | -0.952 | 1.448 | -1.336 | 0.656 | 0.014 | 0.008 | 0.008 | 0.19 | 0.68 | 0.94 | 0.79 | 0.79 | 0.79 | 0.86 |
| 8 | 88593519 | rs10955331 | -0.675 | 1.213 | -1.103 | 0.622 | -0.009 | -0.002 | -0.002 | 0 | 0.66 | 0.82 | 0.7 | 0.72 | 0.68 | 0.74 |

| | | | | | | | | | | | | | | | | |
|---|---|---|---|---|---|---|---|---|---|---|---|---|---|---|---|---|
| 8 | 88595985 | rs13265907 | 0.487 | 1.13 | -1.368 | 0.445 | -0.007 | 0.067 | 0.067 | 0.033 | 0.38 | 0.13 | 0.17 | 0.21 | 0.32 | 0.44 |
| 8 | 88596505 | rs16870389 | -0.739 | 1.084 | -0.739 | 0.715 | -0.028 | 0.051 | 0.051 | 0.346 | 0.23 | 0.61 | 0.41 | 0.45 | 0.2 | 0.29 |
| 8 | 88596553 | rs17713950 | -0.011 | 1.425 | -1.148 | 0.391 | -0.024 | 0.039 | 0.039 | 0.024 | 0.14 | 0.05 | 0.13 | 0.1 | 0.23 | 0.2 |
| 8 | 88600593 | rs10504836 | 0.028 | 1.535 | -1.213 | 0.438 | -0.055 | 0.031 | 0.031 | 0 | 0.17 | 0.06 | 0.19 | 0.18 | 0.27 | 0.24 |
| 8 | 88647577 | rs10088650 | -0.637 | 1.156 | -1.316 | 0.386 | -0.03 | -0.008 | -0.008 | 1.049 | 0.4 | 0.24 | 0.6 | 0.58 | 0.48 | 0.45 |
| 8 | 88651037 | rs11782013 | 0.768 | 0.98 | -1.248 | 0.445 | -0.01 | -0.055 | -0.055 | 0 | 0.2 | 0.1 | 0.42 | 0.42 | 0.22 | 0.21 |
| 8 | 88652017 | rs967023 | 0.683 | 0.984 | -1.238 | 0.447 | -0.004 | -0.027 | -0.027 | 1.254 | 0.21 | 0.15 | 0.42 | 0.42 | 0.22 | 0.21 |
| 8 | 88670142 | rs7844301 | -0.828 | 0.994 | -1.301 | 0.452 | -0.006 | -0.039 | -0.039 | 0.113 | 0.79 | 0.86 | 0.58 | 0.58 | 0.78 | 0.8 |
| 8 | 88696839 | rs10103568 | 0.427 | 1.305 | -1.203 | 0.351 | -0.032 | -0.016 | -0.016 | 4.689 | 0.61 | 0.77 | 0.38 | 0.48 | 0.49 | 0.58 |
| 8 | 88706004 | rs12542080 | 0.356 | 1.352 | -1.415 | 0.351 | -0.097 | 0.061 | 0.061 | 0.027 | 0.18 | 0.02 | 0.19 | 0.11 | 0.29 | 0.22 |
| 8 | 88706202 | rs13264571 | 2.047 | 0.728 | -1.933 | -0.059 | -0.041 | -0.049 | -0.049 | 0 | 0.25 | 0.03 | 0.14 | 0.25 | 0.24 | 0.33 |
| 8 | 88712279 | rs9642798 | 1.056 | 0.418 | -1.702 | -0.113 | 0.018 | -0.017 | -0.017 | 2.403 | 0.07 | 0.03 | 0.08 | 0.17 | 0.03 | 0.1 |
| 8 | 88730843 | rs7833080 | 0.711 | 0.488 | -0.934 | -0.214 | -0.011 | -0.095 | -0.095 | 0.974 | 0.25 | 0.39 | 0.13 | 0.06 | 0.21 | 0.15 |
| 8 | 88743769 | rs10090077 | -1.515 | 0.517 | -1.464 | 0.589 | 0.024 | -0.032 | -0.032 | 0 | 0.84 | 0.9 | 0.74 | 0.77 | 0.93 | 0.93 |
| 8 | 88778152 | rs7003865 | -1.919 | 0.485 | -1.368 | 0.259 | -0.001 | 0 | 0 | 0 | 0.79 | 0.79 | 0.86 | 0.78 | 0.82 | 0.73 |
| 8 | 88784195 | rs16872607 | 0.677 | 0.594 | -1.373 | 0.603 | -0.008 | -0.054 | -0.054 | 2.08 | 0.19 | 0.1 | 0.41 | 0.41 | 0.2 | 0.14 |
| 8 | 88792344 | rs12678345 | -0.506 | 0.436 | -1.474 | 0.284 | 0.019 | -0.006 | -0.006 | 0 | 0.32 | 0.21 | 0.53 | 0.66 | 0.27 | 0.34 |
| 8 | 88795281 | rs13248173 | 1.296 | 0.398 | -1.707 | -0.072 | -0.007 | 0.006 | 0.006 | 0 | 0.1 | 0.09 | 0.03 | 0.05 | 0.11 | 0.1 |
| 8 | 88799185 | rs4960945 | 1.267 | 0.388 | -1.695 | -0.021 | 0.006 | -0.06 | -0.06 | 0 | 0.17 | 0.1 | 0.39 | 0.41 | 0.18 | 0.13 |
| 8 | 88799819 | rs7813504 | 0.662 | 0.318 | -1.672 | -0.379 | -0.032 | -0.115 | -0.115 | 1.688 | 0.23 | 0.45 | 0.14 | 0.06 | 0.19 | 0.13 |
| 8 | 88803575 | rs7818882 | -1.592 | 0.42 | -1.805 | -0.214 | -0.039 | 0.034 | 0.034 | 0.088 | 0.64 | 0.8 | 0.71 | 0.77 | 0.55 | 0.56 |
| 8 | 88804137 | rs11776728 | 0.196 | 0.546 | -1.63 | 0.082 | 0.002 | -0.032 | -0.032 | 0.4 | 0.24 | 0.18 | 0.45 | 0.51 | 0.23 | 0.24 |
| 8 | 88827729 | rs1995727 | 1.748 | 0.536 | -1.574 | 0.484 | 0.042 | -0.048 | -0.048 | 0 | 0.13 | 0.06 | 0.23 | 0.23 | 0.05 | 0.06 |
| 8 | 88828202 | rs13270057 | -0.045 | 0.538 | -1.622 | 0.267 | 0.005 | -0.066 | -0.066 | 0.065 | 0.31 | 0.52 | 0.2 | 0.16 | 0.3 | 0.22 |
| 8 | 88831715 | rs7819401 | -0.506 | 0.386 | -1.504 | 0.564 | 0.029 | 0.079 | 0.079 | 0 | 0.08 | 0.16 | 0.2 | 0.32 | 0.14 | 0.17 |
| 8 | 88846956 | rs7825784 | -0.095 | 0.176 | -1.432 | 0.507 | 0.004 | -0.006 | -0.006 | 2.114 | 0.25 | 0.31 | 0.46 | 0.55 | 0.19 | 0.2 |
| 8 | 88852543 | rs6995710 | 0.054 | 0.188 | -1.466 | 0.513 | 0.001 | -0.002 | -0.002 | 0 | 0.25 | 0.25 | 0.46 | 0.55 | 0.2 | 0.2 |
| 8 | 88856948 | rs1507883 | -0.392 | 0.528 | -1.328 | 0.34 | 0.003 | 0.064 | 0.064 | 0 | 0.12 | 0.18 | 0.27 | 0.36 | 0.15 | 0.15 |
| 8 | 88868340 | rs12543318 | -0.882 | 0.723 | -1.477 | -0.055 | 0.055 | 0.055 | 0.055 | 0 | 0.51 | 0.81 | 0.56 | 0.42 | 0.65 | 0.52 |
| 8 | 88881431 | rs10429299 | -0.173 | 0.643 | -1.292 | -0.331 | -0.014 | -0.036 | -0.036 | 0 | 0.29 | 0.63 | 0.23 | 0.11 | 0.2 | 0.17 |
| 8 | 88891871 | rs10100405 | -1.98 | 0.578 | -1.711 | -0.165 | -0.034 | -0.025 | -0.025 | 0 | 0.73 | 0.99 | 0.85 | 0.83 | 0.77 | 0.69 |
| 8 | 88896480 | rs1911665 | -1.161 | 0.57 | -1.626 | -0.23 | -0.008 | 0.022 | 0.022 | 0 | 0.64 | 0.83 | 0.78 | 0.7 | 0.68 | 0.58 |
| 8 | 88896639 | rs17644659 | 1.07 | 0.867 | -1.647 | -0.359 | -0.038 | 0.033 | 0.033 | 0 | 0.14 | 0.06 | 0.17 | 0 | 0.28 | 0.19 |
| 8 | 88901809 | rs11985946 | 0.609 | 0.566 | -0.882 | -0.436 | 0.019 | -0.01 | -0.01 | 0.253 | 0.07 | 0.36 | 0.05 | 0 | 0.03 | 0.01 |
| 8 | 88918331 | rs1034832 | -1.014 | 0.289 | -1.038 | -0.44 | 0 | -0.001 | -0.001 | 0.812 | 0.43 | 0.56 | 0.45 | 0.4 | 0.53 | 0.5 |
| 8 | 88930612 | rs11989278 | 0.435 | 0.312 | -0.952 | -0.393 | -0.007 | -0.046 | -0.046 | 5.526 | 0.09 | 0.3 | 0.06 | 0 | 0.05 | 0.02 |
| 8 | 88934441 | rs1006364 | -2.422 | 0.286 | -1.39 | -0.315 | -0.033 | -0.018 | -0.018 | 0 | 0.75 | 0.99 | 0.85 | 0.85 | 0.79 | 0.74 |
| 8 | 88937177 | rs11782427 | -0.095 | 0.572 | -1.235 | -0.49 | -0.013 | -0.041 | -0.041 | 0 | 0.22 | 0.11 | 0.26 | 0.39 | 0.25 | 0.3 |
| 8 | 88956612 | rs4375050 | 1.305 | 0.619 | -1.297 | -0.48 | -0.013 | -0.02 | -0.02 | 0 | 0.12 | 0.01 | 0.09 | 0.11 | 0.1 | 0.15 |
| 8 | 88970682 | rs4400429 | -1.624 | 0.39 | -1.211 | -0.431 | -0.008 | 0.007 | 0.007 | 0 | 0.64 | 0.87 | 0.79 | 0.69 | 0.69 | 0.62 |
| 8 | 88981540 | rs1477915 | -1.023 | 0.456 | -1.053 | -0.538 | -0.005 | 0.008 | 0.008 | 0 | 0.43 | 0.6 | 0.46 | 0.36 | 0.54 | 0.53 |
| 8 | 88998663 | rs2169655 | 0.142 | 0.577 | -0.873 | -0.53 | 0.004 | 0.008 | 0.008 | 0 | 0.33 | 0.48 | 0.42 | 0.31 | 0.38 | 0.29 |
| 8 | 89006607 | rs730316 | -0.031 | 0.516 | -0.917 | -0.556 | -0.008 | -0.043 | -0.043 | 0 | 0.21 | 0.09 | 0.25 | 0.4 | 0.21 | 0.3 |
| 8 | 89007711 | rs4961073 | 1.343 | 0.498 | -0.943 | -0.624 | 0.012 | 0.006 | 0.006 | 0 | 0.12 | 0 | 0.09 | 0.08 | 0.07 | 0.14 |
| 8 | 89011448 | rs6469158 | -0.889 | 0.494 | -0.872 | -0.682 | 0.006 | 0.019 | 0.019 | 0 | 0.51 | 0.6 | 0.66 | 0.67 | 0.57 | 0.56 |
| 8 | 89013301 | rs17646041 | 1.404 | 0.475 | -0.875 | -0.737 | -0.038 | 0.033 | 0.033 | 0.009 | 0.13 | 0.05 | 0.15 | 0 | 0.29 | 0.21 |
| 8 | 89014455 | rs2664363 | 1.218 | 0.293 | -0.786 | -0.687 | -0.055 | 0.078 | 0.078 | 0.736 | 0.19 | 0.39 | 0.27 | 0.29 | 0.08 | 0.07 |
| 8 | 89017568 | rs2664364 | -1.699 | 0.324 | -0.81 | -0.753 | 0.006 | 0.007 | 0.007 | 0 | 0.64 | 0.81 | 0.78 | 0.69 | 0.7 | 0.62 |
| 8 | 89018125 | rs2616510 | 0.298 | 0.345 | -0.688 | -0.716 | -0.029 | 0.011 | 0.011 | 0.55 | 0.29 | 0.62 | 0.38 | 0.29 | 0.19 | 0.11 |
| 8 | 89021717 | rs10089030 | 0.776 | 0.464 | -0.573 | -0.818 | -0.002 | -0.077 | -0.077 | 1.828 | 0.13 | 0.31 | 0.13 | 0.01 | 0.12 | 0.03 |
| 8 | 89030490 | rs17718917 | 0.241 | 0.631 | -0.889 | -0.77 | -0.025 | 0.015 | 0.015 | 0 | 0.08 | 0 | 0.02 | 0.03 | 0.08 | 0.12 |
| 8 | 89036579 | rs2616483 | -0.367 | 0.432 | -0.296 | -0.398 | -0.001 | 0.003 | 0.003 | 0 | 0.41 | 0.63 | 0.53 | 0.32 | 0.43 | 0.34 |
| 8 | 89041070 | rs2664367 | -0.573 | 0.73 | -0.017 | -0.427 | 0.001 | 0.005 | 0.005 | 0 | 0.49 | 0.55 | 0.53 | 0.37 | 0.48 | 0.47 |
| 8 | 89041596 | rs2616500 | 0.278 | 0.725 | 0.085 | -0.576 | -0.034 | 0.012 | 0.012 | 0.33 | 0.36 | 0.48 | 0.46 | 0.39 | 0.2 | 0.25 |
| 8 | 89042888 | rs2616499 | -1.043 | 0.698 | -0.11 | -0.514 | 0.006 | 0.007 | 0.007 | 0 | 0.57 | 0.64 | 0.68 | 0.67 | 0.61 | 0.58 |
| 8 | 89046119 | rs10106275 | -0.451 | 0.707 | -0.119 | -0.523 | -0.028 | 0.007 | 0.007 | 0 | 0.27 | 0.2 | 0.25 | 0.28 | 0.47 | 0.36 |
| 8 | 89049531 | rs2664369 | -0.851 | 0.724 | -0.175 | -0.485 | -0.008 | -0.009 | -0.009 | 0 | 0.55 | 0.52 | 0.67 | 0.67 | 0.61 | 0.58 |
| 8 | 89051297 | rs2664370 | -0.817 | 0.846 | 0.026 | -0.136 | 0.004 | 0.005 | 0.005 | 0 | 0.5 | 0.55 | 0.66 | 0.67 | 0.57 | 0.55 |
| 8 | 89056569 | rs10097366 | 0.489 | 0.653 | -0.223 | -0.093 | 0 | -0.001 | -0.001 | 0 | 0.2 | 0.19 | 0.19 | 0.29 | 0.26 | 0.19 |
| 8 | 89061298 | rs2616496 | -0.764 | 0.635 | -0.693 | -0.127 | 0.026 | 0.046 | 0.046 | 0 | 0.52 | 0.69 | 0.63 | 0.68 | 0.41 | 0.41 |
| 8 | 89069654 | rs16877270 | 0.076 | 0.366 | -0.151 | -0.068 | 0.056 | -0.078 | -0.078 | 0 | 0.15 | 0.27 | 0.17 | 0.01 | 0.3 | 0.23 |
| 8 | 89073084 | rs1477908 | -0.467 | -0.605 | -0.64 | 0.688 | -0.007 | 0.005 | 0.005 | 0 | 0.29 | 0.31 | 0.43 | 0.41 | 0.15 | 0.21 |
| 8 | 89083979 | rs2616488 | -0.591 | -0.393 | -0.919 | 0.7 | 0.02 | -0.028 | -0.028 | 0 | 0.31 | 0.25 | 0.38 | 0.39 | 0.22 | 0.24 |
| 8 | 89084837 | rs7826929 | -1.05 | -0.744 | -0.543 | 0.272 | -0.043 | -0.022 | -0.022 | 0 | 0.19 | 0.26 | 0.31 | 0.1 | 0.12 | 0.29 |
| 8 | 89087367 | rs2054414 | -0.884 | -1.143 | -1.156 | 0.194 | 0.047 | 0.021 | 0.021 | 0.17 | 0.22 | 0.08 | 0.13 | 0.21 | 0.21 | 0.23 |
| 8 | 89088009 | rs2616506 | -0.975 | -1.464 | -0.755 | 0.467 | -0.002 | -0.001 | -0.001 | 0 | 0.43 | 0.53 | 0.41 | 0.64 | 0.38 | 0.45 |
| 8 | 89091272 | rs17663841 | 0.285 | -1.515 | -1.314 | 0 | -0.015 | 0.033 | 0.033 | 0 | 0.21 | 0.06 | 0.16 | 0.13 | 0.24 | 0.12 |
| 8 | 89092101 | rs977231 | -0.536 | -1.196 | -1.543 | -0.324 | 0.003 | 0.002 | 0.002 | 0 | 0.43 | 0.55 | 0.43 | 0.64 | 0.39 | 0.49 |
| 8 | 89093298 | rs2664352 | -0.696 | -1.16 | -1.499 | -0.3 | 0.005 | 0.011 | 0.011 | 0 | 0.49 | 0.57 | 0.5 | 0.58 | 0.47 | 0.4 |

| | | | | | | | | | | | | | | | | |
|---|---|---|---|---|---|---|---|---|---|---|---|---|---|---|---|---|
| 8 | 89095141 | rs11782395 | 2.018 | -0.04 | -1.739 | -0.587 | -0.03 | 0.001 | 0.001 | 0 | 0.19 | 0.09 | 0.19 | 0.18 | 0.22 | 0.2 |
| 8 | 89098320 | rs2664357 | 0.455 | -0.055 | -1.478 | -0.483 | -0.009 | 0.096 | 0.096 | 0 | 0.28 | 0.45 | 0.36 | 0.46 | 0.27 | 0.3 |
| 8 | 89110193 | rs9771895 | -0.768 | -0.391 | -0.945 | -0.86 | 0.003 | 0.045 | 0.045 | 0.003 | 0.49 | 0.43 | 0.45 | 0.18 | 0.48 | 0.43 |
| 8 | 89115041 | rs4961076 | 0.478 | -0.869 | -0.809 | 0.108 | -0.024 | -0.025 | -0.025 | 0 | 0.05 | 0 | 0.2 | 0.16 | 0.09 | 0.1 |
| 8 | 89119305 | rs10089111 | -0.964 | -1.13 | -0.341 | -0.138 | 0.003 | 0.003 | 0.003 | 0.005 | 0.46 | 0.56 | 0.4 | 0.35 | 0.4 | 0.43 |
| 8 | 89120562 | rs9297422 | 0.308 | -1.289 | -0.335 | -0.344 | 0.001 | 0.011 | 0.011 | 0 | 0.23 | 0.18 | 0.19 | 0.14 | 0.25 | 0.13 |
| 8 | 89125990 | rs16878625 | 0.188 | -1.109 | -0.652 | -0.251 | -0.002 | 0.024 | 0.024 | 0 | 0.08 | 0.15 | 0.05 | 0.19 | 0.09 | 0.09 |
| 8 | 89129953 | rs1477917 | 0.853 | -1.332 | -0.662 | -0.27 | -0.005 | 0.022 | 0.022 | 0 | 0.06 | 0.1 | 0.03 | 0.18 | 0.07 | 0.1 |
| 8 | 89131809 | rs2664361 | 1.852 | -1.563 | -1.028 | -0.504 | -0.071 | -0.063 | -0.063 | 3.93 | 0.38 | 0.15 | 0.34 | 0.48 | 0.44 | 0.37 |
| 8 | 89136754 | rs16878818 | 1.001 | -1.611 | -1.038 | -0.59 | 0.009 | 0.008 | 0.008 | 0.204 | 0.1 | 0.08 | 0.08 | 0.01 | 0.08 | 0.11 |
| 8 | 89147717 | rs10099888 | 0.556 | -1.632 | -1.126 | -0.578 | -0.014 | 0.029 | 0.029 | 0 | 0.23 | 0.06 | 0.19 | 0.14 | 0.26 | 0.13 |
| 8 | 89157919 | rs1996637 | -1.751 | -1.583 | -1 | -0.599 | -0.06 | -0.055 | -0.055 | 0 | 0.61 | 0.82 | 0.66 | 0.52 | 0.55 | 0.62 |
| 8 | 89176613 | rs1519938 | -0.799 | -1.351 | -0.547 | -0.276 | -0.002 | 0.001 | 0.001 | 0 | 0.42 | 0.46 | 0.45 | 0.51 | 0.45 | 0.42 |
| 8 | 89188454 | rs6981717 | -1.085 | -2.003 | -0.094 | -0.733 | 0.007 | -0.006 | -0.006 | 0.144 | 0.35 | 0.33 | 0.22 | 0.05 | 0.33 | 0.26 |
| 8 | 89207265 | rs2176771 | 1.166 | -1.738 | 0.546 | 0.048 | -0.009 | 0.005 | 0.005 | 0 | 0.7 | 0.38 | 0.48 | 0.42 | 0.77 | 0.63 |
| 8 | 89209114 | rs1519941 | -0.445 | -1.724 | 0.86 | 0.064 | 0.005 | -0.021 | -0.021 | 0.212 | 0.93 | 0.8 | 0.93 | 0.99 | 0.94 | 0.93 |
| 8 | 89209945 | rs12546847 | -0.477 | -1.748 | 0.881 | 0.083 | -0.033 | 0.127 | 0.127 | 0 | 0.21 | 0.4 | 0.44 | 0.54 | 0.14 | 0.25 |
| 8 | 89209965 | rs17722347 | 0.534 | -1.716 | 0.774 | 0.097 | 0.004 | 0.014 | 0.014 | 0 | 0.06 | 0.03 | 0.05 | 0.01 | 0.06 | 0.07 |
| 8 | 89211196 | rs4961080 | 0.743 | -1.763 | 0.865 | 0.13 | 0.008 | -0.02 | -0.02 | 0 | 0.12 | 0.07 | 0.4 | 0.5 | 0.11 | 0.21 |
| 8 | 89218210 | rs10504847 | 0.2 | -1.726 | 0.794 | 0.103 | -0.004 | -0.013 | -0.013 | 0 | 0.07 | 0.09 | 0.07 | 0.01 | 0.06 | 0.07 |
| 8 | 89220059 | rs13261974 | -1.267 | -1.708 | 0.842 | 0.126 | -0.043 | 0.067 | 0.067 | 0 | 0.27 | 0.5 | 0.5 | 0.54 | 0.2 | 0.33 |
| 8 | 89222426 | rs6469298 | 0.195 | -1.923 | 0.446 | -0.409 | 0.002 | -0.014 | -0.014 | 2.639 | 0.77 | 0.76 | 0.88 | 0.97 | 0.81 | 0.8 |
| 8 | 89224646 | rs17666351 | 0.466 | -1.837 | 0.313 | -0.35 | -0.003 | -0.013 | -0.013 | 7.167 | 0.07 | 0.07 | 0.07 | 0.01 | 0.06 | 0.07 |
| 8 | 89225076 | rs13261169 | 0.055 | -1.933 | 0.29 | -0.429 | 0.016 | 0.046 | 0.046 | 0 | 0.21 | 0.15 | 0.1 | 0.04 | 0.19 | 0.19 |
| 8 | 89226636 | rs1401861 | 1.343 | -1.926 | 0.168 | -0.215 | -0.018 | -0.05 | -0.05 | 0 | 0.1 | 0.01 | 0.39 | 0.41 | 0.11 | 0.11 |
| 8 | 89227970 | rs13256568 | 0.169 | -1.944 | 0.214 | -0.347 | 0.007 | -0.004 | -0.004 | 0 | 0.2 | 0.17 | 0.11 | 0.01 | 0.19 | 0.19 |
| 8 | 89232797 | rs1879201 | 0.984 | -1.956 | 0.655 | -0.339 | -0.053 | -0.168 | -0.168 | 0 | 0.66 | 0.49 | 0.8 | 0.99 | 0.74 | 0.73 |
| 8 | 89246009 | rs16880099 | 1.219 | -1.677 | 0.959 | 0.067 | -0.034 | 0.086 | 0.086 | 0 | 0.76 | 0.43 | 0.55 | 0.48 | 0.86 | 0.73 |
| 8 | 89248590 | rs7826477 | -0.309 | -1.502 | 0.551 | 0.152 | -0.005 | 0.103 | 0.103 | 0 | 0.17 | 0.24 | 0.42 | 0.52 | 0.13 | 0.25 |
| 8 | 89258247 | rs16880226 | -1.039 | -1.355 | 0.62 | -0.452 | -0.008 | 0.008 | 0.008 | 0.417 | 0.1 | 0.4 | 0.05 | 0.09 | 0.04 | 0.14 |
| 8 | 89264537 | rs16880322 | -1.012 | -1.349 | 0.577 | -0.449 | -0.008 | 0.007 | 0.007 | 0.18 | 0.1 | 0.38 | 0.04 | 0.09 | 0.04 | 0.14 |
| 8 | 89268543 | rs6994019 | 1.155 | -1.661 | -0.141 | -0.814 | 0.074 | 0.118 | 0.118 | 0.175 | 0.59 | 0.23 | 0.75 | 0.9 | 0.72 | 0.6 |
| 8 | 89272062 | rs1467251 | -0.59 | -1.671 | 0.731 | -0.822 | -0.016 | -0.076 | -0.076 | 0 | 0.2 | 0.37 | 0.13 | 0.01 | 0.18 | 0.15 |
| 8 | 89273678 | rs10955542 | 1.737 | -1.774 | 1.116 | -0.755 | -0.006 | -0.025 | -0.025 | 0 | 0.66 | 0.34 | 0.69 | 0.88 | 0.74 | 0.67 |
| 8 | 89279580 | rs2222294 | -0.256 | -1.744 | 1.295 | -0.763 | -0.012 | -0.073 | -0.073 | 0.122 | 0.85 | 0.68 | 0.9 | 0.99 | 0.85 | 0.91 |
| 8 | 89287763 | rs16880580 | 0.473 | -1.751 | 1.304 | -0.766 | 0.002 | 0.019 | 0.019 | 0 | 0.12 | 0.1 | 0.08 | 0.01 | 0.14 | 0.08 |
| 8 | 89292303 | rs7817382 | 1.031 | -1.676 | 1.209 | -0.908 | -0.005 | -0.079 | -0.079 | 0.185 | 0.75 | 0.57 | 0.85 | 0.94 | 0.75 | 0.75 |
| 8 | 89310568 | rs10100297 | 2.062 | -1.679 | 1.111 | -0.928 | 0.026 | 0.07 | 0.07 | 0 | 0.6 | 0.26 | 0.65 | 0.83 | 0.67 | 0.56 |
| 8 | 89311785 | rs7834743 | -0.224 | -1.655 | 1.261 | -0.894 | 0.008 | -0.012 | -0.012 | 0 | 0.91 | 0.84 | 0.96 | 0.95 | 0.92 | 0.88 |
| 8 | 89322905 | rs3851539 | 2.04 | -1.781 | 0.801 | -0.345 | 0.042 | 0.042 | 0.042 | 0 | 0.49 | 0.23 | 0.37 | 0.44 | 0.56 | 0.42 |
| 8 | 89327738 | rs10504846 | -0.408 | -1.431 | 1.098 | -0.511 | -0.008 | -0.007 | -0.007 | 0.288 | 0.11 | 0.12 | 0.17 | 0.11 | 0.07 | 0.17 |
| 8 | 89338155 | rs7835845 | -1.697 | -1.569 | 1.297 | -0.388 | -0.022 | -0.056 | -0.056 | 0.095 | 0.31 | 0.55 | 0.28 | 0.16 | 0.28 | 0.39 |
| 8 | 89368594 | rs7006944 | 1.505 | -1.587 | 1.394 | -0.431 | -0.026 | -0.078 | -0.078 | 0 | 0.71 | 0.48 | 0.8 | 0.9 | 0.74 | 0.7 |
| 8 | 89374660 | rs4483202 | 1.94 | -1.601 | 1.538 | -0.441 | 0.003 | 0.002 | 0.002 | 0 | 0.64 | 0.34 | 0.67 | 0.84 | 0.68 | 0.55 |
| 8 | 89376806 | rs16881745 | 0.43 | -1.574 | 1.519 | -0.413 | 0.008 | -0.008 | -0.008 | 2.7 | 0.07 | 0.14 | 0.13 | 0.06 | 0.06 | 0.15 |
| 8 | 89383727 | rs1504801 | -1.053 | -1.579 | 1.605 | -0.416 | 0.005 | 0.011 | 0.011 | 0 | 0.21 | 0.2 | 0.13 | 0.1 | 0.22 | 0.21 |
| 8 | 89390692 | rs7002751 | -1.257 | -1.578 | 1.628 | -0.416 | -0.017 | -0.04 | -0.04 | 0 | 0.23 | 0.4 | 0.15 | 0.1 | 0.22 | 0.21 |
| 8 | 89411565 | rs10093191 | 0.071 | -1.519 | 1.521 | -0.382 | 0.007 | -0.011 | -0.011 | 0 | 0.08 | 0.22 | 0.14 | 0.06 | 0.06 | 0.19 |
| 8 | 89415843 | rs13278603 | -0.171 | -1.549 | 1.447 | -0.415 | -0.011 | 0.046 | 0.046 | 0 | 0.15 | 0.07 | 0.11 | 0.04 | 0.18 | 0.11 |
| 8 | 89426999 | rs10504840 | -1.484 | -1.658 | 1.542 | -0.48 | -0.015 | -0.052 | -0.052 | 0 | 0.3 | 0.37 | 0.19 | 0.09 | 0.27 | 0.27 |
| 8 | 89434405 | rs1504797 | -2.034 | -1.56 | 1.548 | -0.206 | 0.004 | 0.01 | 0.01 | 0 | 0.37 | 0.67 | 0.32 | 0.21 | 0.33 | 0.45 |
| 8 | 89452923 | rs7819898 | 0.32 | -1.49 | 1.459 | -0.294 | 0.001 | -0.009 | -0.009 | 0.308 | 0.07 | 0.13 | 0.13 | 0.06 | 0.06 | 0.16 |
| 8 | 89462321 | rs1601817 | -1.885 | -1.513 | 1.388 | -0.304 | -0.008 | -0.041 | -0.041 | 2.003 | 0.36 | 0.55 | 0.31 | 0.15 | 0.33 | 0.45 |
| 8 | 89479917 | rs821118 | -1.652 | -1.851 | 1.548 | -0.316 | -0.008 | -0.022 | -0.022 | 0 | 0.33 | 0.37 | 0.27 | 0.15 | 0.31 | 0.42 |
| 8 | 89486032 | rs1685550 | -0.845 | -1.885 | 1.415 | 0.126 | 0.004 | 0.001 | 0.001 | 0.112 | 0.87 | 0.91 | 0.71 | 0.58 | 0.89 | 0.86 |
| 8 | 89486513 | rs1685549 | -1.776 | -1.856 | 1.431 | -0.117 | -0.019 | -0.052 | -0.052 | 2.985 | 0.34 | 0.46 | 0.28 | 0.15 | 0.31 | 0.42 |
| 8 | 89499611 | rs821109 | 0.347 | -1.96 | 1.305 | -0.103 | 0.006 | 0.009 | 0.009 | 0 | 0.79 | 0.81 | 0.89 | 0.91 | 0.79 | 0.8 |
| 8 | 89513786 | rs16883803 | 0.364 | -2.036 | 1.248 | -0.087 | 0.011 | 0.003 | 0.003 | 0 | 0.11 | 0.05 | 0.02 | 0.07 | 0.07 | 0.11 |
| 8 | 89514775 | rs6469448 | -0.279 | -1.923 | 1.333 | -0.015 | 0.01 | 0.014 | 0.014 | 0 | 0.21 | 0.17 | 0.11 | 0.1 | 0.21 | 0.2 |
| 8 | 89517922 | rs7013582 | -1.62 | -1.93 | 1.56 | -0.046 | -0.014 | -0.031 | -0.031 | 0 | 0.34 | 0.57 | 0.3 | 0.17 | 0.31 | 0.44 |
| 8 | 89520084 | rs4961099 | 0.592 | -1.968 | 1.49 | 0.166 | 0.004 | -0.009 | -0.009 | 0 | 0.13 | 0.1 | 0.28 | 0.35 | 0.11 | 0.15 |
| 8 | 89524792 | rs4524843 | 1.575 | -1.712 | 1.724 | -0.395 | 0.009 | 0.014 | 0.014 | 0.005 | 0.63 | 0.32 | 0.7 | 0.84 | 0.69 | 0.55 |
| 8 | 89529961 | rs1872922 | -0.967 | -1.662 | 1.669 | -0.36 | -0.002 | -0.025 | -0.025 | 0 | 0.12 | 0.27 | 0.18 | 0.08 | 0.1 | 0.27 |
| 8 | 89530078 | rs11989655 | -1.724 | -1.729 | 1.819 | -0.398 | -0.005 | -0.011 | -0.011 | 0.58 | 0.35 | 0.6 | 0.3 | 0.17 | 0.29 | 0.44 |
| 8 | 89536805 | rs11988146 | -0.817 | -1.691 | 1.854 | -0.354 | -0.002 | -0.028 | -0.028 | 0 | 0.12 | 0.22 | 0.17 | 0.07 | 0.1 | 0.27 |
| 8 | 89573045 | rs7814164 | -0.337 | -1.683 | 1.684 | -0.318 | 0.007 | 0.007 | 0.007 | 0 | 0.21 | 0.18 | 0.11 | 0.1 | 0.2 | 0.17 |
| 8 | 89579649 | rs16884419 | -1.242 | -1.701 | 1.779 | -0.331 | -0.028 | -0.032 | -0.032 | 0 | 0.29 | 0.43 | 0.25 | 0.17 | 0.26 | 0.35 |
| 8 | 89585048 | rs7838490 | -0.367 | -1.739 | 1.78 | -0.355 | -0.011 | -0.014 | -0.014 | 0 | 0.21 | 0.26 | 0.12 | 0.1 | 0.2 | 0.2 |
| 8 | 89599194 | rs12680992 | 0.401 | -1.718 | 1.462 | -0.342 | 0.001 | 0.002 | 0.002 | 0 | 0.07 | 0.04 | 0.12 | 0.07 | 0.06 | 0.15 |

| | | | | | | | | | | | | | | | | |
|---|---|---|---|---|---|---|---|---|---|---|---|---|---|---|---|---|
| 8 | 89600751 | rs13274015 | -1.789 | -1.76 | 1.468 | -0.329 | 0.026 | 0.041 | 0.041 | 0.121 | 0.37 | 0.74 | 0.32 | 0.22 | 0.3 | 0.46 |
| 8 | 89605999 | rs9969514 | -0.484 | -1.753 | 1.597 | -0.303 | -0.01 | -0.022 | -0.022 | 0 | 0.21 | 0.27 | 0.13 | 0.09 | 0.18 | 0.2 |
| 8 | 89610780 | rs11994353 | -1.687 | -1.943 | 1.488 | -0.365 | 0.005 | 0.005 | 0.005 | 0 | 0.36 | 0.64 | 0.3 | 0.16 | 0.28 | 0.44 |
| 8 | 89615855 | rs13272330 | 0.003 | -1.938 | 1.488 | -0.364 | 0.004 | 0.024 | 0.024 | 0 | 0.18 | 0.08 | 0.11 | 0.08 | 0.18 | 0.17 |
| 8 | 89616587 | rs10086545 | 1.64 | -1.917 | 1.491 | -0.35 | 0.031 | 0.044 | 0.044 | 0.003 | 0.61 | 0.28 | 0.69 | 0.82 | 0.7 | 0.55 |
| 8 | 89640705 | rs4961102 | 1.825 | -2.626 | 0.526 | -0.765 | 0.087 | 0.148 | 0.148 | 0.402 | 0.48 | 0.06 | 0.62 | 0.78 | 0.66 | 0.5 |
| 8 | 89646080 | rs12334864 | -0.382 | -2.757 | 0.882 | -0.829 | -0.039 | -0.03 | -0.03 | 0.239 | 0.24 | 0.43 | 0.24 | 0.15 | 0.18 | 0.26 |
| 8 | 89649177 | rs11995369 | -1.167 | -2.808 | 0.977 | -0.858 | 0.011 | 0.011 | 0.011 | 0 | 0.35 | 0.68 | 0.28 | 0.15 | 0.22 | 0.27 |
| 8 | 89665567 | rs7816883 | 1.472 | -2.807 | 0.878 | -0.893 | 0.016 | 0.012 | 0.012 | 0 | 0.09 | 0.03 | 0.04 | 0 | 0.04 | 0.01 |
| 8 | 89668921 | rs4961103 | 2.099 | -2.84 | 0.778 | -0.909 | 0.102 | 0.194 | 0.194 | 0.147 | 0.48 | 0.09 | 0.63 | 0.82 | 0.66 | 0.5 |
| 8 | 89670067 | rs7011535 | -0.18 | -2.82 | 0.894 | -0.912 | -0.014 | -0.042 | -0.042 | 0.542 | 0.14 | 0.2 | 0.08 | 0.05 | 0.1 | 0.18 |
| 8 | 89677161 | rs13257525 | -0.091 | -2.72 | 0.74 | -0.852 | -0.008 | -0.008 | -0.008 | 1.478 | 0.22 | 0.27 | 0.22 | 0.13 | 0.18 | 0.24 |
| 8 | 89679613 | rs4573324 | -1.823 | -2.695 | 0.603 | -0.842 | 0.134 | 0.224 | 0.224 | 0.072 | 0.49 | 0.86 | 0.35 | 0.16 | 0.31 | 0.45 |
| 8 | 89721771 | rs7833665 | -0.194 | -2.539 | 0.752 | -0.528 | -0.01 | -0.012 | -0.012 | 0 | 0.23 | 0.28 | 0.22 | 0.13 | 0.18 | 0.24 |
| 8 | 89729597 | rs1603793 | -1.875 | -2.423 | -0.058 | -0.53 | 0.065 | 0.119 | 0.119 | 0.072 | 0.52 | 0.91 | 0.45 | 0.24 | 0.35 | 0.47 |
| 8 | 89745416 | rs1391636 | -1.67 | -2.429 | -0.033 | -0.552 | 0.116 | 0.239 | 0.239 | 0.112 | 0.5 | 0.88 | 0.37 | 0.2 | 0.33 | 0.44 |
| 8 | 89753984 | rs11997183 | 0.125 | -2.387 | 0.03 | -0.555 | 0.018 | 0.026 | 0.026 | 0 | 0.2 | 0.11 | 0.2 | 0.09 | 0.17 | 0.24 |
| 8 | 89754060 | rs11997212 | -1.485 | -2.455 | 0.198 | -0.602 | -0.009 | -0.031 | -0.031 | 0 | 0.46 | 0.47 | 0.34 | 0.15 | 0.33 | 0.43 |
| 8 | 89760311 | rs7004633 | -0.6 | -2.449 | 0.109 | -0.401 | -0.042 | -0.004 | -0.004 | 0 | 0.27 | 0.48 | 0.31 | 0.19 | 0.21 | 0.34 |
| 8 | 89761913 | rs10090518 | -0.937 | -2.187 | 0.119 | -0.429 | -0.045 | -0.07 | -0.07 | 0.015 | 0.25 | 0.6 | 0.11 | 0.05 | 0.1 | 0.17 |
| 8 | 89763009 | rs10109897 | 0.942 | -1.864 | -0.27 | 0.303 | -0.012 | 0.022 | 0.022 | 0 | 0.64 | 0.5 | 0.58 | 0.56 | 0.7 | 0.56 |
| 8 | 89773651 | rs1353739 | 0.61 | -1.844 | -0.332 | 0.6 | -0.007 | 0.074 | 0.074 | 0 | 0.8 | 0.64 | 0.77 | 0.69 | 0.84 | 0.74 |
| 8 | 89774351 | rs968030 | -1.006 | -1.701 | -0.403 | 0.739 | -0.001 | -0.038 | -0.038 | 0 | 0.91 | 0.98 | 0.92 | 0.76 | 0.92 | 0.88 |
| 8 | 89794519 | rs1498523 | 1.315 | -1.654 | -0.716 | 0.627 | -0.01 | 0.007 | 0.007 | 0 | 0.39 | 0.34 | 0.37 | 0.34 | 0.53 | 0.33 |
| 8 | 89795446 | rs3942951 | 0.637 | -1.646 | -1.285 | 0.593 | -0.006 | -0.008 | -0.008 | 0.039 | 0.07 | 0.04 | 0.12 | 0.08 | 0.08 | 0.15 |
| 8 | 89799474 | rs1909270 | 1.635 | -1.636 | -1.412 | 0.559 | -0.01 | -0.032 | -0.032 | 0 | 0.1 | 0.14 | 0.04 | 0 | 0.05 | 0.01 |
| 8 | 89799516 | rs1909271 | 0.691 | -1.662 | -1.523 | 0.59 | 0.001 | 0.001 | 0.001 | 0 | 0.5 | 0.5 | 0.42 | 0.41 | 0.58 | 0.35 |
| 8 | 89800500 | rs12679056 | 1.905 | -1.416 | -1.751 | 0.523 | -0.197 | -0.095 | -0.095 | 0.015 | 0.32 | 0.03 | 0.33 | 0.34 | 0.48 | 0.27 |
| 8 | 89800771 | rs10481132 | 1.531 | -1.298 | -1.42 | 0.507 | -0.001 | -0.019 | -0.019 | 0 | 0.1 | 0.12 | 0.04 | 0 | 0.05 | 0.01 |
| 8 | 89809252 | rs11781588 | 1.092 | -0.807 | -1.429 | 1.006 | 0 | -0.001 | -0.001 | 0.874 | 0.49 | 0.51 | 0.42 | 0.37 | 0.58 | 0.35 |
| 8 | 89818717 | rs9297551 | -0.288 | -0.077 | -0.542 | 1.282 | -0.003 | -0.038 | -0.038 | 0.268 | 0.91 | 0.98 | 0.92 | 0.76 | 0.91 | 0.89 |
| 8 | 89843820 | rs1498539 | 1.358 | -0.096 | -0.513 | 1.207 | -0.008 | 0.09 | 0.09 | 0 | 0.79 | 0.64 | 0.75 | 0.64 | 0.83 | 0.73 |
| 8 | 89845702 | rs13262829 | 0.056 | 0.12 | -1.428 | 1.316 | -0.003 | -0.037 | -0.037 | 0.071 | 0.91 | 0.98 | 0.91 | 0.72 | 0.91 | 0.89 |
| 8 | 89864220 | rs10086779 | 0.242 | 0.249 | -1.546 | 0.581 | 0.016 | 0.023 | 0.023 | 0 | 0.32 | 0.17 | 0.24 | 0.24 | 0.27 | 0.36 |
| 8 | 89875994 | rs963994 | -0.844 | 0.878 | -2.162 | 0.852 | -0.079 | -0.079 | -0.079 | 0 | 0.84 | 0.98 | 0.8 | 0.77 | 0.74 | 0.81 |
| 8 | 89888548 | rs2454331 | -0.028 | 0.787 | -2.239 | 0.665 | -0.018 | -0.056 | -0.056 | 0 | 0.51 | 0.92 | 0.55 | 0.56 | 0.43 | 0.45 |
| 8 | 89891698 | rs1391629 | 0.265 | 0.757 | -1.871 | 0.478 | 0.002 | -0.011 | -0.011 | 0 | 0.12 | 0.14 | 0.19 | 0.04 | 0.14 | 0.2 |
| 8 | 89894046 | rs10091936 | 0.704 | 0.399 | -1.919 | 0.113 | -0.01 | -0.066 | -0.066 | 0.186 | 0.26 | 0.15 | 0.3 | 0.52 | 0.27 | 0.24 |
| 8 | 89903393 | rs10110062 | 0.745 | 0.392 | -1.973 | 0.131 | -0.033 | -0.063 | -0.063 | 0 | 0.5 | 0.08 | 0.47 | 0.45 | 0.57 | 0.56 |
| 8 | 89903532 | rs12545320 | 0.761 | 0.35 | -1.864 | 0.084 | -0.005 | 0.004 | 0.004 | 0.003 | 0.1 | 0.06 | 0.18 | 0.04 | 0.14 | 0.19 |
| 8 | 89906765 | rs10091322 | 0.194 | 0.302 | -1.812 | 0.099 | 0.025 | -0.045 | -0.045 | 1.959 | 0.59 | 0.13 | 0.64 | 0.49 | 0.69 | 0.73 |
| 8 | 89907014 | rs11783597 | 1.08 | 0.366 | -2.021 | 0.179 | -0.054 | -0.034 | -0.034 | 0 | 0.18 | 0.04 | 0.21 | 0.18 | 0.25 | 0.19 |
| 8 | 89917982 | rs7005417 | -0.262 | 0.281 | -0.64 | 0.134 | 0.001 | 0.006 | 0.006 | 1.576 | 0.38 | 0.62 | 0.5 | 0.56 | 0.39 | 0.43 |
| 8 | 89923420 | rs10504867 | 1.739 | 0.199 | -0.729 | 0.115 | -0.062 | -0.054 | -0.054 | 1.412 | 0.31 | 0.01 | 0.24 | 0.26 | 0.28 | 0.36 |
| 8 | 89923973 | rs6984367 | 0.92 | 0.224 | -0.528 | 0.134 | -0.013 | -0.065 | -0.065 | 0.315 | 0.26 | 0.13 | 0.29 | 0.51 | 0.23 | 0.25 |
| 8 | 89931641 | rs6986236 | -0.006 | 0.221 | -0.409 | 0.094 | 0.051 | 0.004 | 0.004 | 0 | 0.12 | 0.5 | 0.2 | 0.04 | 0.13 | 0.18 |
| 8 | 89940072 | rs1548121 | -0.197 | 0.072 | 0.002 | 0.191 | 0.004 | 0.014 | 0.014 | 0 | 0.38 | 0.59 | 0.49 | 0.49 | 0.37 | 0.45 |
| 8 | 89941029 | rs12550183 | 0.679 | 0.13 | 0.033 | 0.232 | 0.011 | -0.007 | -0.007 | 0.011 | 0.09 | 0.17 | 0.16 | 0.04 | 0.12 | 0.18 |
| 8 | 89941889 | rs13439196 | 0.958 | 0.115 | -0.174 | 0.215 | -0.036 | -0.017 | -0.017 | 1.836 | 0.2 | 0.06 | 0.21 | 0.19 | 0.25 | 0.17 |
| 8 | 89946032 | rs11775804 | 1.732 | 0.104 | -0.225 | 0.235 | -0.064 | -0.084 | -0.084 | 0 | 0.3 | 0.02 | 0.25 | 0.32 | 0.29 | 0.36 |
| 8 | 89954658 | rs1038620 | -0.315 | 0.254 | 0.733 | 0.454 | -0.006 | 0.006 | 0.006 | 0 | 0.37 | 0.5 | 0.35 | 0.45 | 0.3 | 0.29 |
| 8 | 89968423 | rs7812413 | 0.508 | 0.572 | 0.096 | 0.625 | 0 | 0 | 0 | 0 | 0.08 | 0.09 | 0.16 | 0.04 | 0.12 | 0.18 |
| 8 | 89982095 | rs13271290 | 0.165 | 0.517 | 0.048 | 0.668 | -0.007 | -0.032 | -0.032 | 0 | 0.28 | 0.16 | 0.31 | 0.45 | 0.25 | 0.28 |
| 8 | 89996790 | rs1872892 | 0.49 | 0.108 | 0.605 | 0.402 | 0.005 | -0.081 | -0.081 | 0 | 0.83 | 0.6 | 0.89 | 0.97 | 0.84 | 0.83 |
| 8 | 89998242 | rs1586475 | 0.282 | 0.006 | 0.125 | 0.603 | -0.106 | -0.114 | -0.114 | 1.75 | 0.41 | 0.07 | 0.35 | 0.54 | 0.44 | 0.36 |
| 8 | 90002125 | rs1565148 | 0.151 | 0.073 | 0.478 | 0.688 | 0.062 | 0.011 | 0.011 | 0 | 0.9 | 0.65 | 0.92 | 0.97 | 0.89 | 0.85 |
| 8 | 90007392 | rs2046675 | 0.958 | -0.046 | 0.71 | 0.835 | -0.001 | 0.013 | 0.013 | 1.878 | 0.82 | 0.28 | 0.68 | 0.83 | 0.83 | 0.78 |
| 8 | 90010448 | rs1619456 | 1.211 | -0.062 | 0.858 | 0.812 | -0.003 | -0.005 | -0.005 | 0 | 0.77 | 0.35 | 0.68 | 0.85 | 0.78 | 0.76 |
| 8 | 90015212 | rs1487791 | 1.512 | -0.066 | 0.724 | 0.793 | 0.002 | -0.03 | -0.03 | 0 | 0.73 | 0.15 | 0.51 | 0.74 | 0.73 | 0.73 |
| 8 | 90017135 | rs1240140 | 0.996 | -0.017 | 0.857 | 0.764 | -0.001 | 0.003 | 0.003 | 2.055 | 0.82 | 0.28 | 0.69 | 0.86 | 0.83 | 0.79 |
| 8 | 90018397 | rs12548743 | -0.139 | 0.053 | 0.803 | 0.792 | -0.011 | -0.013 | -0.013 | 0 | 0.11 | 0.08 | 0.25 | 0.13 | 0.16 | 0.21 |
| 8 | 90019419 | rs13260549 | -0.583 | 0.058 | 0.893 | 0.795 | 0.047 | 0.058 | 0.058 | 0 | 0.13 | 0.37 | 0.29 | 0.13 | 0.17 | 0.21 |
| 8 | 90028155 | rs17678109 | 0.898 | 0.078 | 0.51 | 0.585 | -0.067 | 0.056 | 0.056 | 0 | 0.21 | 0.04 | 0.2 | 0.12 | 0.3 | 0.16 |
| 8 | 90031688 | rs7004307 | 0.016 | -0.27 | 0.972 | 0.776 | 0 | 0.03 | 0.03 | 0 | 0.89 | 0.62 | 0.79 | 0.88 | 0.9 | 0.95 |
| 8 | 90047329 | rs1240118 | -0.579 | -0.36 | 0.669 | 0.672 | 0.014 | 0.025 | 0.025 | 0 | 0.58 | 0.58 | 0.79 | 0.8 | 0.66 | 0.6 |
| 8 | 90048522 | rs10955882 | 0.507 | -0.344 | 0.328 | 0.68 | 0.009 | 0.126 | 0.126 | 0.072 | 0.4 | 0.09 | 0.18 | 0.2 | 0.35 | 0.4 |
| 8 | 90066677 | rs13254288 | 1.119 | -0.371 | 0.314 | 0.666 | -0.006 | 0.064 | 0.064 | 0 | 0.33 | 0.08 | 0.17 | 0.2 | 0.29 | 0.38 |
| 8 | 90067523 | rs4391478 | 0.667 | -0.319 | 0.271 | 0.683 | -0.05 | 0.076 | 0.076 | 0.052 | 0.38 | 0.09 | 0.21 | 0.22 | 0.39 | 0.52 |
| 8 | 90072115 | rs922913 | 0.521 | -0.657 | 0.65 | 0.649 | 0.031 | 0.087 | 0.087 | 0 | 0.41 | 0.3 | 0.21 | 0.16 | 0.35 | 0.39 |

| | | | | | | | | | | | | | | | | |
|---|---|---|---|---|---|---|---|---|---|---|---|---|---|---|---|---|
| 8 | 90074006 | rs1531846 | -0.308 | -0.658 | 0.976 | 0.812 | -0.009 | -0.012 | -0.012 | 0 | 0.56 | 0.43 | 0.6 | 0.68 | 0.61 | 0.56 |
| 8 | 90074757 | rs1240049 | 0.54 | -0.654 | 0.878 | 0.77 | -0.015 | -0.039 | -0.039 | 1.429 | 0.09 | 0.22 | 0.03 | 0 | 0.06 | 0.02 |
| 8 | 90078197 | rs10955896 | 1.236 | -0.646 | 0.812 | 0.79 | -0.017 | 0.113 | 0.113 | 0.058 | 0.31 | 0.04 | 0.18 | 0.14 | 0.29 | 0.37 |
| 8 | 90094210 | rs1681444 | -0.773 | -0.694 | 1.043 | 0.823 | 0.005 | -0.019 | -0.019 | 0 | 0.91 | 0.93 | 0.83 | 0.84 | 0.9 | 0.93 |
| 8 | 90104975 | rs13274815 | 0.973 | -0.718 | 1.165 | 0.791 | 0.019 | 0.057 | 0.057 | 0 | 0.34 | 0.25 | 0.32 | 0.15 | 0.29 | 0.37 |
| 8 | 90105157 | rs1240069 | -0.484 | -0.714 | 1.163 | 0.787 | 0.031 | 0.054 | 0.054 | 0 | 0.58 | 0.68 | 0.65 | 0.7 | 0.62 | 0.57 |
| 8 | 90114345 | rs1483382 | 0.23 | -0.596 | 1.286 | 1.05 | 0.02 | 0.001 | 0.001 | 0 | 0.47 | 0.17 | 0.4 | 0.57 | 0.54 | 0.49 |
| 8 | 90116102 | rs2706820 | -0.366 | -0.526 | 1.317 | 1.057 | -0.006 | 0.008 | 0.008 | 0 | 0.9 | 0.83 | 0.83 | 0.84 | 0.92 | 0.94 |
| 8 | 90119525 | rs10441513 | 0.133 | -0.529 | 1.346 | 1.063 | -0.017 | 0.081 | 0.081 | 0 | 0.88 | 0.57 | 0.8 | 0.82 | 0.92 | 0.93 |
| 8 | 90132402 | rs7462065 | -0.102 | -0.554 | 1.181 | 1.143 | 0.065 | 0.069 | 0.069 | 0 | 0.51 | 0.22 | 0.59 | 0.66 | 0.61 | 0.54 |
| 8 | 90137645 | rs1483359 | -0.124 | -0.565 | 1.249 | 1.202 | -0.013 | 0.049 | 0.049 | 0 | 0.9 | 0.71 | 0.82 | 0.82 | 0.92 | 0.93 |
| 8 | 90137908 | rs7014508 | 0.838 | -0.579 | 1.055 | 1.197 | -0.04 | 0.072 | 0.072 | 0.441 | 0.85 | 0.38 | 0.79 | 0.8 | 0.9 | 0.91 |
| 8 | 90158554 | rs11779424 | -0.139 | -0.61 | 0.818 | 1.177 | 0.06 | 0.062 | 0.062 | 0 | 0.52 | 0.23 | 0.59 | 0.66 | 0.61 | 0.51 |
| 8 | 90161392 | rs4960963 | 0.433 | -0.771 | 0.635 | 1.44 | 0.023 | -0.009 | -0.009 | 0 | 0.47 | 0.15 | 0.4 | 0.49 | 0.55 | 0.45 |
| 8 | 90185423 | rs4961118 | 0.137 | -0.868 | 0.704 | 1.328 | 0.069 | 0.017 | 0.017 | 0 | 0.52 | 0.21 | 0.59 | 0.58 | 0.63 | 0.51 |
| 8 | 90186443 | rs1483373 | 1.463 | -0.874 | 0.859 | 1.33 | 0.008 | 0.295 | 0.295 | 6.771 | 0.91 | 0.45 | 0.81 | 0.74 | 0.96 | 0.94 |
| 8 | 90186798 | rs1483374 | 0.051 | -0.885 | 0.85 | 1.324 | 0.064 | 0.014 | 0.014 | 0 | 0.53 | 0.21 | 0.61 | 0.58 | 0.64 | 0.56 |
| 8 | 90190241 | rs10955936 | 0.115 | -0.938 | 0.851 | 1.434 | 0.07 | 0.004 | 0.004 | 0 | 0.53 | 0.21 | 0.59 | 0.47 | 0.63 | 0.53 |
| 8 | 90193247 | rs7842444 | 1.771 | -0.761 | 1.185 | 1.188 | -0.008 | 0.282 | 0.282 | 7.232 | 0.91 | 0.45 | 0.81 | 0.73 | 0.96 | 0.91 |
| 8 | 90201937 | rs1351726 | 1.129 | -0.739 | 0.355 | 1.19 | -0.048 | 0.057 | 0.057 | 3.987 | 0.82 | 0.29 | 0.77 | 0.7 | 0.9 | 0.88 |
| 8 | 90211013 | rs2046316 | 0.242 | -0.397 | 0.069 | 1.196 | -0.011 | -0.025 | -0.025 | 2.438 | 0.44 | 0.23 | 0.53 | 0.52 | 0.49 | 0.4 |
| 8 | 90211100 | rs2046315 | 1.264 | -0.638 | 0.669 | 1.392 | -0.03 | 0.065 | 0.065 | 2.914 | 0.81 | 0.35 | 0.73 | 0.66 | 0.86 | 0.76 |
| 8 | 90220200 | rs318258 | -1.753 | -1.036 | 1.372 | 1.366 | -0.082 | 0.194 | 0.194 | 2.544 | 0.14 | 0.57 | 0.22 | 0.34 | 0.05 | 0.22 |
| 8 | 90236993 | rs318283 | -1.958 | -1.014 | 1.705 | 1.291 | 0.039 | 0.101 | 0.101 | 4.247 | 0.06 | 0.49 | 0.04 | 0.05 | 0.01 | 0.02 |
| 8 | 90238321 | rs318285 | -2.022 | -1.058 | 1.641 | 1.35 | -0.075 | 0.151 | 0.151 | 2.738 | 0.15 | 0.61 | 0.22 | 0.34 | 0.06 | 0.22 |
| 8 | 90241246 | rs6469919 | -0.409 | -1.171 | 1.489 | 0.984 | 0.117 | 0.001 | 0.001 | 0 | 0.51 | 0.83 | 0.43 | 0.44 | 0.37 | 0.55 |
| 8 | 90246015 | rs6469928 | -1.192 | -1.127 | 1.709 | 1.065 | -0.045 | 0.188 | 0.188 | 0.202 | 0.12 | 0.42 | 0.2 | 0.28 | 0.05 | 0.2 |
| 8 | 90252801 | rs16893311 | -1.975 | -1.158 | 1.701 | 1.056 | -0.069 | 0.156 | 0.156 | 2.636 | 0.14 | 0.64 | 0.22 | 0.34 | 0.06 | 0.22 |
| 8 | 90259170 | rs13266695 | 1.074 | -1.224 | 1.359 | 0.975 | 0.019 | 0.155 | 0.155 | 0.302 | 0.34 | 0.07 | 0.18 | 0.1 | 0.3 | 0.31 |
| 8 | 90263457 | rs6999851 | 0.093 | -1.203 | 1.377 | 1.011 | -0.002 | 0.027 | 0.027 | 0 | 0.08 | 0.11 | 0.18 | 0.28 | 0.05 | 0.18 |
| 8 | 90270072 | rs7813237 | -1.514 | -1.209 | 1.314 | 0.997 | -0.061 | 0.24 | 0.24 | 0.633 | 0.13 | 0.48 | 0.21 | 0.34 | 0.06 | 0.22 |
| 8 | 90272202 | rs7835780 | -0.058 | -0.72 | 0.926 | 1.099 | 0.127 | 0.008 | 0.008 | 0 | 0.5 | 0.83 | 0.43 | 0.44 | 0.37 | 0.53 |
| 8 | 90283685 | rs7818636 | 0.453 | -0.735 | 0.95 | 1.062 | 0.072 | 0.016 | 0.016 | 0 | 0.43 | 0.18 | 0.37 | 0.38 | 0.36 | 0.52 |
| 8 | 90289326 | rs10102049 | 0.453 | -0.735 | 0.958 | 1.062 | 0.072 | 0.017 | 0.017 | 0 | 0.43 | 0.18 | 0.37 | 0.38 | 0.36 | 0.51 |
| 8 | 90297599 | rs6469964 | 0.404 | -0.753 | 1.248 | 1.054 | 0.039 | 0.018 | 0.018 | 0 | 0.45 | 0.35 | 0.39 | 0.38 | 0.36 | 0.51 |
| 8 | 90300401 | rs13274960 | -0.338 | -0.73 | 0.951 | 1.273 | 0.06 | -0.013 | -0.013 | 0.372 | 0.53 | 0.83 | 0.6 | 0.53 | 0.42 | 0.63 |
| 8 | 90307568 | rs1479901 | -0.788 | -0.726 | 1.117 | 1.211 | 0.058 | 0.013 | 0.013 | 0 | 0.58 | 0.81 | 0.63 | 0.63 | 0.64 | 0.5 |
| 8 | 90307652 | rs1479902 | -0.469 | -0.739 | 1.396 | 1.218 | 0.005 | 0.002 | 0.002 | 0 | 0.54 | 0.53 | 0.61 | 0.58 | 0.64 | 0.48 |
| 8 | 90314239 | rs2879240 | -0.137 | -0.754 | 1.165 | 1.211 | 0.126 | 0.037 | 0.037 | 0 | 0.51 | 0.19 | 0.58 | 0.58 | 0.65 | 0.47 |
| 8 | 90315707 | rs11780858 | -0.63 | -0.677 | 0.926 | 1.225 | 0.067 | 0.019 | 0.019 | 0 | 0.57 | 0.81 | 0.63 | 0.63 | 0.64 | 0.49 |
| 8 | 90331518 | rs10087030 | -0.535 | -0.684 | 1.116 | 1.417 | 0.006 | 0.001 | 0.001 | 0 | 0.54 | 0.46 | 0.59 | 0.55 | 0.64 | 0.45 |
| 8 | 90333189 | rs10104777 | -0.202 | -0.794 | 1.112 | 1.362 | 0.092 | 0.02 | 0.02 | 0 | 0.51 | 0.28 | 0.57 | 0.55 | 0.64 | 0.44 |
| 8 | 90335076 | rs16894311 | 0.012 | -1.296 | 1.079 | 1.228 | -0.01 | -0.002 | -0.002 | 0.008 | 0.45 | 0.47 | 0.43 | 0.45 | 0.58 | 0.37 |
| 8 | 90337941 | rs1038160 | 0.985 | -1.208 | 0.632 | 1.3 | 0.06 | 0.006 | 0.006 | 0 | 0.42 | 0.19 | 0.38 | 0.38 | 0.36 | 0.53 |
| 8 | 90339068 | rs9650082 | 0.245 | -1.185 | 0.497 | 1.174 | 0.011 | 0 | 0 | 0 | 0.42 | 0.36 | 0.41 | 0.43 | 0.58 | 0.36 |
| 8 | 90348104 | rs10086910 | 1.129 | -0.334 | -0.757 | 1.229 | -0.137 | 0.032 | 0.032 | 0 | 0.24 | 0.03 | 0.23 | 0.12 | 0.37 | 0.16 |
| 8 | 90354767 | rs10087480 | 0.284 | -1.083 | -0.357 | 1.465 | -0.043 | 0.125 | 0.125 | 0.514 | 0.84 | 0.69 | 0.67 | 0.58 | 0.91 | 0.83 |
| 8 | 90355884 | rs13261143 | 0.357 | -0.85 | -0.554 | 1.307 | -0.008 | 0.235 | 0.235 | 0.484 | 0.94 | 0.77 | 0.83 | 0.68 | 0.97 | 0.91 |
| 8 | 90357820 | rs1564402 | 0.447 | -0.446 | -1.419 | 0.465 | 0.07 | 0.121 | 0.121 | 0 | 0.39 | 0.14 | 0.21 | 0.14 | 0.31 | 0.46 |
| 8 | 90366827 | rs2086343 | -0.431 | -0.479 | -1.216 | 0.998 | 0.008 | -0.056 | -0.056 | 0.088 | 0.29 | 0.23 | 0.51 | 0.44 | 0.26 | 0.25 |
| 8 | 90369994 | rs6470016 | -0.483 | -0.361 | -1.028 | 0.442 | -0.028 | 0.057 | 0.057 | 0.002 | 0.62 | 0.76 | 0.73 | 0.81 | 0.52 | 0.52 |
| 8 | 90372173 | rs983898 | 0.548 | -0.927 | -0.783 | 0.717 | 0.053 | -0.028 | -0.028 | 0.002 | 0.36 | 0.19 | 0.17 | 0.39 | 0.27 | 0.27 |
| 8 | 90372447 | rs983899 | -0.266 | -0.783 | -0.729 | 0.643 | -0.023 | -0.009 | -0.009 | 0 | 0.59 | 0.3 | 0.65 | 0.55 | 0.54 | 0.47 |
| 8 | 90374239 | rs4314690 | -0.87 | -0.765 | -0.808 | 0.64 | -0.045 | 0.014 | 0.014 | 0 | 0.67 | 0.81 | 0.73 | 0.7 | 0.62 | 0.6 |
| 8 | 90385087 | rs4458917 | 0.558 | -1.12 | -0.214 | 0.561 | -0.029 | -0.023 | -0.023 | 0 | 0.31 | 0.38 | 0.12 | 0.17 | 0.19 | 0.24 |
| 8 | 90406301 | rs715934 | 0.924 | -1.305 | -0.114 | 0.842 | -0.02 | 0.107 | 0.107 | 0.202 | 0.13 | 0.27 | 0.12 | 0.33 | 0.13 | 0.16 |
| 8 | 90407910 | rs12544520 | 1.052 | -1.247 | -0.783 | 0.712 | -0.085 | -0.007 | -0.007 | 0.001 | 0.19 | 0.02 | 0.44 | 0.15 | 0.26 | 0.21 |
| 8 | 90409007 | rs11786321 | 1.326 | -1.143 | -0.596 | 0.681 | -0.053 | -0.015 | -0.015 | 0 | 0.31 | 0.16 | 0.26 | 0.36 | 0.38 | 0.39 |
| 8 | 90449190 | rs2029276 | 1.51 | -1.186 | -0.417 | 0.64 | 0.036 | 0.115 | 0.115 | 0.359 | 0.25 | 0.07 | 0.1 | 0.06 | 0.17 | 0.12 |
| 8 | 90476925 | rs10216900 | 1.072 | -1.253 | -0.384 | 0.758 | -0.047 | 0.006 | 0.006 | 0 | 0.36 | 0.21 | 0.29 | 0.33 | 0.4 | 0.38 |
| 8 | 90478557 | rs10956094 | 0.944 | -1.226 | -0.261 | 0.759 | -0.029 | 0.046 | 0.046 | 0 | 0.37 | 0.26 | 0.23 | 0.22 | 0.39 | 0.39 |
| 8 | 90488877 | rs1487669 | 1.749 | -1.235 | -0.376 | 0.766 | 0.017 | 0.002 | 0.002 | 0.062 | 0.22 | 0.02 | 0.14 | 0.15 | 0.16 | 0.12 |
| 8 | 90490180 | rs1155439 | 1.002 | -1.141 | -0.275 | 0.883 | -0.042 | 0.051 | 0.051 | 0 | 0.36 | 0.19 | 0.22 | 0.24 | 0.39 | 0.39 |
| 8 | 90520649 | rs10103760 | -1.11 | -1.093 | 0.186 | 0.875 | -0.018 | 0.116 | 0.116 | 0.09 | 0.87 | 0.75 | 0.88 | 0.68 | 0.88 | 0.84 |
| 8 | 90528845 | rs7846085 | 1.038 | -1.194 | 0.091 | 0.81 | -0.041 | 0.015 | 0.015 | 0 | 0.35 | 0.2 | 0.36 | 0.31 | 0.38 | 0.34 |
| 8 | 90539951 | rs1386648 | 0.611 | -1.179 | 0.156 | 0.826 | -0.006 | -0.007 | -0.007 | 0 | 0.39 | 0.34 | 0.37 | 0.45 | 0.39 | 0.48 |
| 8 | 90541496 | rs1386646 | 1.74 | -1.171 | -0.043 | 0.461 | 0.016 | -0.002 | -0.002 | 0 | 0.23 | 0.02 | 0.14 | 0.17 | 0.17 | 0.12 |
| 8 | 90541694 | rs10504879 | 0.92 | -1.067 | 0.017 | 0.451 | -0.034 | 0.068 | 0.068 | 0.02 | 0.2 | 0.09 | 0.31 | 0.08 | 0.27 | 0.21 |
| 8 | 90545792 | rs1546745 | 1.069 | -1.117 | 0.087 | 0.484 | -0.015 | 0.007 | 0.007 | 0 | 0.35 | 0.28 | 0.37 | 0.32 | 0.38 | 0.33 |

| | | | | | | | | | | | | | | | | |
|---|---|---|---|---|---|---|---|---|---|---|---|---|---|---|---|---|
| 8 | 90546979 | rs7833686 | 1.738 | -1.125 | -0.072 | 0.458 | 0.015 | -0.003 | -0.003 | 0 | 0.23 | 0.02 | 0.14 | 0.17 | 0.16 | 0.12 |
| 8 | 90564114 | rs17768420 | 0.897 | -0.936 | 0.101 | 0.624 | -0.083 | 0.065 | 0.065 | 0.154 | 0.19 | 0.02 | 0.3 | 0.09 | 0.27 | 0.21 |
| 8 | 90571218 | rs10504880 | 0.854 | -0.931 | 0.124 | 0.626 | -0.036 | 0.061 | 0.061 | 0 | 0.2 | 0.09 | 0.3 | 0.09 | 0.27 | 0.21 |
| 8 | 90571660 | rs7830141 | 1.08 | -0.809 | 0.268 | 0.868 | -0.013 | 0.115 | 0.115 | 0 | 0.12 | 0.2 | 0.11 | 0.33 | 0.12 | 0.18 |
| 8 | 90574472 | rs10096970 | -0.257 | -0.833 | 0.07 | 0.614 | 0.006 | 0.051 | 0.051 | 0 | 0.52 | 0.59 | 0.5 | 0.75 | 0.51 | 0.67 |
| 8 | 90576635 | rs1510465 | 0.419 | -0.712 | -0.049 | 0.674 | -0.006 | 0.002 | 0.002 | 0 | 0.38 | 0.35 | 0.3 | 0.38 | 0.39 | 0.48 |
| 8 | 90576918 | rs10081423 | 0.777 | -0.735 | -0.058 | 0.689 | -0.004 | 0.003 | 0.003 | 0 | 0.35 | 0.33 | 0.29 | 0.27 | 0.38 | 0.33 |
| 8 | 90578495 | rs1510466 | -0.093 | -0.761 | -0.023 | 0.716 | 0.006 | 0.021 | 0.021 | 0 | 0.49 | 0.58 | 0.4 | 0.62 | 0.51 | 0.51 |
| 8 | 90578560 | rs13269643 | 0.822 | -0.804 | 0.012 | 0.724 | -0.015 | 0.115 | 0.115 | 0.03 | 0.14 | 0.2 | 0.11 | 0.34 | 0.13 | 0.18 |
| 8 | 90579197 | rs7842612 | 1.55 | -0.695 | -0.362 | 0.546 | 0 | 0.007 | 0.007 | 0 | 0.1 | 0.11 | 0.14 | 0.22 | 0.11 | 0.21 |
| 8 | 90584054 | rs10092477 | 0.644 | -0.862 | -0.088 | 0.803 | 0.004 | -0.002 | -0.002 | 0 | 0.39 | 0.41 | 0.3 | 0.38 | 0.39 | 0.49 |
| 8 | 90597465 | rs9297680 | -0.046 | -0.767 | -0.687 | 0.848 | 0.005 | 0.046 | 0.046 | 0 | 0.52 | 0.59 | 0.41 | 0.73 | 0.51 | 0.66 |
| 8 | 90597595 | rs2132347 | 0.469 | -0.321 | -1.787 | 0.606 | -0.088 | 0.03 | 0.03 | 0 | 0.19 | 0.02 | 0.38 | 0.1 | 0.27 | 0.21 |
| 8 | 90603149 | rs311638 | 0.151 | 0.006 | -1.132 | 1.012 | -0.035 | -0.042 | -0.042 | 0.806 | 0.37 | 0.19 | 0.34 | 0.49 | 0.39 | 0.49 |
| 8 | 90607469 | rs311667 | -0.592 | 0.029 | -0.227 | 1.076 | -0.02 | 0.072 | 0.072 | 1.495 | 0.23 | 0.6 | 0.19 | 0.39 | 0.24 | 0.25 |
| 8 | 90608086 | rs7835902 | 0.109 | 0.784 | -0.996 | 0.573 | -0.101 | 0.038 | 0.038 | 0.643 | 0.17 | 0.02 | 0.29 | 0.08 | 0.3 | 0.19 |
| 8 | 90608246 | rs1510470 | -0.158 | 0.434 | -0.811 | 0.317 | 0.002 | 0 | 0 | 0 | 0.44 | 0.63 | 0.45 | 0.58 | 0.39 | 0.54 |
| 8 | 90613278 | rs827725 | 0.024 | 0.292 | -0.921 | -0.049 | 0.018 | 0.032 | 0.032 | 0 | 0.31 | 0.38 | 0.43 | 0.41 | 0.38 | 0.35 |
| 8 | 90615624 | rs557807 | -0.323 | 0.404 | -1.211 | -0.375 | 0.009 | 0.001 | 0.001 | 0.263 | 0.51 | 0.4 | 0.53 | 0.46 | 0.56 | 0.47 |
| 8 | 90615793 | rs606565 | 1.581 | 0.416 | -0.992 | -0.258 | -0.005 | 0.1 | 0.1 | 0 | 0.12 | 0.18 | 0.13 | 0.32 | 0.09 | 0.09 |
| 8 | 90616054 | rs13282828 | 1.363 | 0.284 | -0.998 | -0.069 | -0.008 | -0.033 | -0.033 | 0 | 0.13 | 0.05 | 0.23 | 0.26 | 0.14 | 0.22 |
| 8 | 90617010 | rs1510469 | 0.805 | 0.139 | -0.596 | -0.163 | -0.005 | -0.005 | -0.005 | 2.463 | 0.24 | 0.2 | 0.16 | 0.3 | 0.24 | 0.29 |
| 8 | 90620856 | rs17696708 | 1.174 | 0.203 | -0.578 | -0.208 | -0.022 | 0.048 | 0.048 | 0 | 0.2 | 0.11 | 0.13 | 0.04 | 0.2 | 0.13 |
| 8 | 90623228 | rs521592 | -1.668 | 0.167 | -0.822 | -0.232 | 0.019 | 0.17 | 0.17 | 0.003 | 0.72 | 0.97 | 0.85 | 0.98 | 0.76 | 0.86 |
| 8 | 90624692 | rs7824985 | 0.787 | 0.287 | -0.734 | -0.132 | -0.004 | -0.005 | -0.005 | 0 | 0.26 | 0.2 | 0.16 | 0.31 | 0.25 | 0.31 |
| 8 | 90626359 | rs593072 | -0.328 | 0.28 | -0.603 | -0.127 | -0.004 | 0.026 | 0.026 | 0.053 | 0.46 | 0.6 | 0.68 | 0.68 | 0.52 | 0.56 |
| 8 | 90627436 | rs2644012 | -1.85 | 0.246 | -0.633 | -0.195 | 0.008 | 0.155 | 0.155 | 0.001 | 0.73 | 0.97 | 0.86 | 0.99 | 0.77 | 0.87 |
| 8 | 90628008 | rs17696834 | 1.42 | 0.629 | -0.506 | 0.06 | -0.058 | 0.064 | 0.064 | 0 | 0.19 | 0.02 | 0.09 | 0.07 | 0.18 | 0.14 |
| 8 | 90629430 | rs656153 | -1.04 | 0.415 | -0.269 | 0.206 | 0.001 | 0.001 | 0.001 | 0 | 0.65 | 0.63 | 0.78 | 0.72 | 0.7 | 0.69 |
| 8 | 90632741 | rs218921 | 0.558 | 0.876 | -0.069 | 1.069 | -0.004 | 0.104 | 0.104 | 0 | 0.12 | 0.18 | 0.13 | 0.31 | 0.1 | 0.09 |
| 8 | 90635791 | rs4562370 | -0.28 | 1.096 | -0.493 | 0.606 | 0 | 0.006 | 0.006 | 0 | 0.14 | 0.17 | 0.24 | 0.27 | 0.14 | 0.21 |
| 8 | 90637625 | rs160414 | 0.019 | 0.909 | -0.438 | -0.399 | -0.006 | -0.11 | -0.11 | 0.393 | 0.41 | 0.49 | 0.25 | 0.05 | 0.35 | 0.26 |
| 8 | 90638201 | rs2665636 | 0.98 | 1.774 | -0.466 | 0.971 | 0.001 | 0.011 | 0.011 | 0.189 | 0.89 | 0.86 | 0.79 | 0.71 | 0.74 | 0.7 |
| 8 | 90648728 | rs160410 | -0.086 | 0.717 | -0.8 | -0.609 | -0.05 | 0.126 | 0.126 | 0.139 | 0.69 | 0.89 | 0.68 | 0.95 | 0.64 | 0.82 |
| 8 | 90660515 | rs160449 | 0.379 | -0.123 | -0.171 | -0.46 | -0.002 | 0.025 | 0.025 | 0 | 0.57 | 0.34 | 0.59 | 0.76 | 0.59 | 0.64 |
| 8 | 90662941 | rs160451 | 0.253 | -0.157 | 0.293 | -0.431 | 0.003 | -0.024 | -0.024 | 0 | 0.58 | 0.47 | 0.6 | 0.75 | 0.59 | 0.64 |
| 8 | 90671024 | rs160426 | 0.446 | -0.209 | 0.504 | -0.249 | -0.018 | 0.001 | 0.001 | 0 | 0.06 | 0.1 | 0.03 | 0.06 | 0.03 | 0.02 |
| 8 | 90678123 | rs12216864 | 0.792 | -0.265 | 0.471 | -0.281 | 0 | 0.119 | 0.119 | 0 | 0.5 | 0.15 | 0.56 | 0.7 | 0.56 | 0.63 |
| 8 | 90681455 | rs160436 | -0.306 | -0.276 | 0.59 | -0.152 | -0.028 | 0.014 | 0.014 | 0.321 | 0.08 | 0.28 | 0.03 | 0.06 | 0.03 | 0.02 |
| 8 | 90708428 | rs6981616 | 0.276 | -0.318 | 1.015 | -0.182 | -0.019 | 0.091 | 0.091 | 1.016 | 0.1 | 0.23 | 0.04 | 0.19 | 0.03 | 0.06 |
| 8 | 90726598 | rs218896 | 0.433 | -0.189 | 0.595 | -0.35 | -0.015 | -0.004 | -0.004 | 0.056 | 0.3 | 0.26 | 0.35 | 0.05 | 0.38 | 0.31 |
| 8 | 90729507 | rs218899 | -1.025 | -0.698 | -0.292 | -0.563 | -0.08 | -0.016 | -0.016 | 0.354 | 0.63 | 0.93 | 0.51 | 0.32 | 0.6 | 0.46 |
| 8 | 90736846 | rs218906 | 0.294 | -0.321 | 0.436 | -0.379 | 0.008 | 0.029 | 0.029 | 0 | 0.1 | 0.21 | 0.06 | 0.14 | 0.07 | 0.06 |
| 8 | 90737258 | rs7841459 | 0.8 | -0.555 | 0.303 | -0.617 | 0.008 | 0.012 | 0.012 | 3.317 | 0.14 | 0.09 | 0.07 | 0.01 | 0.14 | 0.02 |
| 8 | 90741114 | rs218908 | -1.089 | -0.517 | 0.23 | -0.679 | -0.054 | 0.004 | 0.004 | 0.028 | 0.63 | 0.9 | 0.48 | 0.3 | 0.61 | 0.43 |
| 8 | 90758337 | rs13274635 | 0.231 | -0.674 | 0.498 | -0.841 | -0.024 | 0.092 | 0.092 | 0.599 | 0.43 | 0.27 | 0.37 | 0.06 | 0.52 | 0.35 |
| 8 | 90776033 | rs428484 | 0.239 | -0.717 | 0.564 | 0.107 | -0.018 | 0.051 | 0.051 | 0.51 | 0.43 | 0.31 | 0.36 | 0.33 | 0.51 | 0.36 |
| 8 | 90776293 | rs16900429 | -0.908 | -0.816 | 0.525 | 0.11 | -0.032 | 0.017 | 0.017 | 3.915 | 0.21 | 0.63 | 0.13 | 0.24 | 0.1 | 0.09 |
| 8 | 90776812 | rs39502 | 0.475 | -0.887 | 0.503 | 0.048 | -0.046 | 0.141 | 0.141 | 0.563 | 0.39 | 0.17 | 0.34 | 0.05 | 0.49 | 0.21 |
| 8 | 90778513 | rs42490 | -0.992 | -0.888 | 0.869 | 0.062 | -0.006 | -0.012 | -0.012 | 0 | 0.61 | 0.72 | 0.46 | 0.49 | 0.61 | 0.44 |
| 8 | 90795302 | rs39508 | 0.249 | -0.893 | 0.885 | 0.063 | -0.016 | 0.049 | 0.049 | 0.545 | 0.43 | 0.31 | 0.37 | 0.33 | 0.51 | 0.36 |
| 8 | 90795768 | rs10097279 | -0.649 | -0.885 | 0.88 | 0.047 | -0.035 | 0.004 | 0.004 | 0.535 | 0.19 | 0.41 | 0.1 | 0.16 | 0.1 | 0.08 |
| 8 | 90802995 | rs16900627 | -0.47 | -0.996 | 0.903 | 0.07 | -0.072 | 0.013 | 0.013 | 1.257 | 0.18 | 0.41 | 0.08 | 0.16 | 0.07 | 0.07 |
| 8 | 90803724 | rs39765 | 1.244 | -1.016 | 0.445 | -0.024 | -0.096 | 0.188 | 0.188 | 0.416 | 0.29 | 0.04 | 0.28 | 0.04 | 0.36 | 0.2 |
| 8 | 90804165 | rs39509 | -1.133 | -0.902 | 0.57 | 0.152 | -0.131 | -0.132 | -0.132 | 0.779 | 0.64 | 0.95 | 0.48 | 0.57 | 0.58 | 0.44 |
| 8 | 90808212 | rs42343 | 0.13 | -0.661 | 0.863 | 0.196 | -0.001 | 0 | 0 | 0.123 | 0.44 | 0.44 | 0.38 | 0.41 | 0.51 | 0.37 |
| 8 | 90823687 | rs40457 | -0.267 | -0.635 | 0 | 0.109 | -0.028 | 0.028 | 0.028 | 0 | 0.59 | 0.76 | 0.45 | 0.3 | 0.59 | 0.33 |
| 8 | 90827701 | rs12679456 | 0.337 | -0.745 | -0.081 | 0.357 | -0.02 | 0.026 | 0.026 | 0 | 0.08 | 0.11 | 0.01 | 0.23 | 0.02 | 0.14 |
| 8 | 90828052 | rs40638 | -0.946 | -0.712 | -0.164 | 0.331 | -0.133 | -0.156 | -0.156 | 0.043 | 0.68 | 0.96 | 0.48 | 0.54 | 0.61 | 0.47 |
| 8 | 90837630 | rs374957 | 0.114 | -0.996 | 0.629 | 0.272 | -0.006 | -0.016 | -0.016 | 1.318 | 0.61 | 0.5 | 0.43 | 0.3 | 0.67 | 0.34 |
| 8 | 90837747 | rs16900888 | 0.161 | -1.261 | 0.712 | 0.016 | 0.001 | 0 | 0 | 2.091 | 0.19 | 0.18 | 0.1 | 0.17 | 0.19 | 0.09 |
| 8 | 90840996 | rs40383 | 1.035 | -1.283 | 0.691 | -0.062 | -0.039 | 0.118 | 0.118 | 0.02 | 0.39 | 0.16 | 0.3 | 0.07 | 0.46 | 0.23 |
| 8 | 90846042 | rs39519 | -0.551 | -1.29 | 0.601 | -0.028 | -0.009 | 0.006 | 0.006 | 0 | 0.64 | 0.82 | 0.47 | 0.32 | 0.68 | 0.35 |
| 8 | 90849305 | rs10094579 | 0.165 | -1.299 | 0.537 | -0.027 | -0.003 | 0.006 | 0.006 | 1.868 | 0.19 | 0.21 | 0.11 | 0.25 | 0.19 | 0.1 |
| 8 | 90857989 | rs12679902 | -0.434 | -1.267 | 0.174 | 0.187 | -0.023 | 0.093 | 0.093 | 0.105 | 0.27 | 0.5 | 0.27 | 0.48 | 0.26 | 0.3 |
| 8 | 90863793 | rs381948 | 1.013 | -0.981 | -0.776 | 0.122 | -0.06 | 0.197 | 0.197 | 0.062 | 0.38 | 0.09 | 0.22 | 0.05 | 0.43 | 0.22 |
| 8 | 90864625 | rs425140 | -0.726 | -0.454 | -0.592 | 0.701 | -0.084 | -0.147 | -0.147 | 0 | 0.79 | 0.96 | 0.56 | 0.53 | 0.74 | 0.54 |
| 8 | 90881645 | rs16901238 | -0.174 | -0.739 | -0.148 | 0.738 | -0.004 | 0 | 0 | 0 | 0.17 | 0.16 | 0.18 | 0.16 | 0.08 | 0.08 |
| 8 | 90881910 | rs11782542 | -0.943 | -0.718 | -0.212 | 0.294 | 0.006 | 0.01 | 0.01 | 0 | 0.42 | 0.35 | 0.56 | 0.93 | 0.33 | 0.61 |

| | | | | | | | | | | | | | | | | |
|---|---|---|---|---|---|---|---|---|---|---|---|---|---|---|---|---|
| 8 | 90886015 | rs10098655 | -1.159 | -0.55 | 0.233 | 1.484 | -0.004 | 0.002 | 0.002 | 1.812 | 0.28 | 0.74 | 0.4 | 0.41 | 0.19 | 0.28 |
| 8 | 90894988 | rs16901369 | -0.184 | -0.296 | 0.222 | 1.393 | -0.005 | 0.005 | 0.005 | 0 | 0.05 | 0.07 | 0.04 | 0.07 | 0.03 | 0.1 |
| 8 | 90895010 | rs2697668 | -1.591 | -0.394 | 0.129 | 1.624 | 0.066 | -0.101 | -0.101 | 0.116 | 0.36 | 0.9 | 0.47 | 0.51 | 0.23 | 0.44 |
| 8 | 90897511 | rs989384 | -1.668 | -0.378 | 0.265 | 1.774 | 0.05 | 0.016 | 0.016 | 3.692 | 0.34 | 0.82 | 0.33 | 0.32 | 0.21 | 0.38 |
| 8 | 90902053 | rs2735376 | -1.631 | -0.332 | 0.235 | 1.626 | 0.072 | -0.075 | -0.075 | 0.716 | 0.42 | 0.88 | 0.5 | 0.5 | 0.28 | 0.45 |
| 8 | 90914272 | rs2250670 | -1.582 | -0.373 | 1.074 | 1.587 | 0.054 | -0.034 | -0.034 | 0 | 0.42 | 0.81 | 0.48 | 0.5 | 0.27 | 0.45 |
| 8 | 90917706 | rs2697671 | -0.693 | -0.405 | 1.459 | 1.597 | 0.002 | 0.002 | 0.002 | 0 | 0.3 | 0.28 | 0.28 | 0.26 | 0.22 | 0.26 |
| 8 | 90918220 | rs6983345 | -1.995 | -0.513 | 1.522 | 1.487 | 0.007 | 0.207 | 0.207 | 3.522 | 0.11 | 0.53 | 0.2 | 0.24 | 0.06 | 0.18 |
| 8 | 90923519 | rs9656966 | -1.75 | -0.53 | 1.601 | 1.472 | -0.019 | 0.017 | 0.017 | 3.249 | 0.11 | 0.53 | 0.07 | 0.07 | 0.03 | 0.12 |
| 8 | 90925344 | rs2284133 | -2.094 | -0.524 | 1.6 | 1.477 | -0.012 | 0.152 | 0.152 | 3.421 | 0.12 | 0.57 | 0.2 | 0.24 | 0.06 | 0.18 |
| 8 | 90930785 | rs1881467 | -0.65 | -0.103 | 1.525 | 1.823 | -0.002 | -0.014 | -0.014 | 0.201 | 0.3 | 0.25 | 0.31 | 0.42 | 0.3 | 0.31 |
| 8 | 90931334 | rs2697674 | -1.912 | -0.108 | 1.508 | 1.797 | 0.01 | -0.003 | -0.003 | 0 | 0.43 | 0.91 | 0.53 | 0.66 | 0.35 | 0.49 |
| 8 | 90945833 | rs14448 | -1.728 | 0.043 | 1.691 | 1.925 | 0.012 | 0.226 | 0.226 | 1.598 | 0.11 | 0.51 | 0.2 | 0.24 | 0.06 | 0.18 |
| 8 | 90946056 | rs9995 | -0.911 | 0.02 | 1.302 | 1.865 | 0.005 | -0.004 | -0.004 | 0.011 | 0.34 | 0.3 | 0.32 | 0.42 | 0.3 | 0.35 |
| 8 | 90947537 | rs1063053 | -0.871 | 0.054 | 0.412 | 1.771 | 0.008 | -0.009 | -0.009 | 0 | 0.33 | 0.25 | 0.31 | 0.42 | 0.29 | 0.35 |
| 8 | 90949425 | rs13312969 | -0.366 | 0.179 | 0.312 | 1.509 | -0.003 | 0 | 0 | 0 | 0.05 | 0.04 | 0.03 | 0.06 | 0.03 | 0.1 |
| 8 | 90954481 | rs6470522 | -0.752 | 0.319 | 0.009 | 1.735 | -0.018 | -0.026 | -0.026 | 0 | 0.9 | 0.97 | 0.84 | 0.69 | 0.81 | 0.83 |
| 8 | 90958422 | rs1061302 | -1.115 | -0.471 | 0.901 | 1.816 | 0.027 | -0.031 | -0.031 | 0.414 | 0.38 | 0.22 | 0.38 | 0.47 | 0.31 | 0.45 |
| 8 | 90967711 | rs709816 | -1.763 | -0.349 | 1.589 | 1.867 | 0.058 | 0.064 | 0.064 | 0.834 | 0.47 | 0.85 | 0.55 | 0.66 | 0.34 | 0.53 |
| 8 | 90970862 | rs1805818 | -1.114 | -0.349 | 1.546 | 1.854 | 0.014 | -0.009 | -0.009 | 0 | 0.38 | 0.3 | 0.38 | 0.47 | 0.31 | 0.45 |
| 8 | 90970935 | rs2234744 | -1.056 | -0.349 | 1.544 | 1.854 | 0.028 | -0.029 | -0.029 | 0.235 | 0.38 | 0.22 | 0.38 | 0.47 | 0.31 | 0.45 |
| 8 | 90995019 | rs1063045 | -1.133 | -0.356 | 1.937 | 1.864 | 0.012 | -0.008 | -0.008 | 0.571 | 0.39 | 0.31 | 0.38 | 0.47 | 0.31 | 0.45 |
| 8 | 91002586 | rs1805801 | -1.849 | -0.371 | 1.906 | 1.88 | 0.047 | 0.061 | 0.061 | 1.128 | 0.47 | 0.95 | 0.56 | 0.66 | 0.34 | 0.53 |
| 8 | 91007806 | rs4961166 | -1.062 | -0.333 | 1.851 | 1.913 | 0.028 | -0.033 | -0.033 | 0 | 0.38 | 0.22 | 0.38 | 0.47 | 0.31 | 0.45 |
| 8 | 91019060 | rs1805808 | -1.297 | -0.326 | 1.851 | 1.898 | -0.012 | 0.004 | 0.004 | 0 | 0.42 | 0.61 | 0.4 | 0.47 | 0.31 | 0.47 |
| 8 | 91020834 | rs1011683 | -1.242 | -0.322 | 1.823 | 1.882 | -0.003 | 0.001 | 0.001 | 0 | 0.41 | 0.53 | 0.4 | 0.47 | 0.31 | 0.46 |
| 8 | 91022867 | rs1805860 | -1.588 | -0.236 | 1.828 | 2.081 | 0.013 | 0.157 | 0.157 | 0.127 | 0.08 | 0.19 | 0.19 | 0.25 | 0.05 | 0.16 |
| 8 | 91035913 | rs7000324 | -2.468 | -0.25 | 1.855 | 2.079 | -0.02 | 0.147 | 0.147 | 5.368 | 0.13 | 0.61 | 0.22 | 0.26 | 0.06 | 0.17 |
| 8 | 91067028 | rs12679577 | -0.382 | -0.416 | 0.806 | 1.94 | 0.011 | -0.045 | -0.045 | 0.111 | 0.3 | 0.15 | 0.32 | 0.4 | 0.26 | 0.3 |
| 8 | 91067602 | rs1805870 | -0.411 | -0.437 | 0.796 | 1.93 | 0.013 | -0.044 | -0.044 | 0 | 0.3 | 0.15 | 0.32 | 0.4 | 0.26 | 0.3 |
| 8 | 91069944 | rs12549239 | 0.138 | -0.468 | 0.744 | 1.952 | 0.023 | -0.034 | -0.034 | 0 | 0.25 | 0.02 | 0.26 | 0.24 | 0.17 | 0.24 |
| 8 | 91081580 | rs1805873 | -0.202 | -0.536 | 0.978 | 1.783 | -0.006 | 0.014 | 0.014 | 0 | 0.05 | 0.07 | 0.07 | 0.1 | 0.03 | 0.07 |
| 8 | 91082062 | rs1805874 | 0.903 | -0.526 | 1.009 | 1.868 | 0.025 | 0.019 | 0.019 | 0 | 0.62 | 0.75 | 0.5 | 0.29 | 0.69 | 0.53 |
| 8 | 91094086 | rs7815279 | 0.198 | -0.53 | 0.961 | 1.717 | 0.018 | -0.021 | -0.021 | 0 | 0.69 | 0.8 | 0.68 | 0.59 | 0.74 | 0.7 |
| 8 | 91098934 | rs12171665 | 1.085 | -0.595 | 1.277 | 1.712 | 0.042 | -0.024 | -0.024 | 0.004 | 0.59 | 0.23 | 0.6 | 0.49 | 0.71 | 0.62 |
| 8 | 91101529 | rs10104321 | 0.153 | -0.579 | 1.198 | 1.709 | 0.017 | -0.033 | -0.033 | 0 | 0.7 | 0.83 | 0.68 | 0.59 | 0.74 | 0.69 |
| 8 | 91111193 | rs6988013 | 0.516 | -0.541 | 1.374 | 1.707 | -0.01 | 0.004 | 0.004 | 0 | 0.67 | 0.54 | 0.66 | 0.58 | 0.74 | 0.69 |
| 8 | 91114536 | rs12541465 | -1.03 | -0.546 | 1.358 | 1.689 | 0.005 | 0.081 | 0.081 | 0.836 | 0.07 | 0.26 | 0.06 | 0.1 | 0.03 | 0.07 |
| 8 | 91127680 | rs6470708 | 0.948 | -0.467 | 0.926 | 1.632 | 0.047 | -0.031 | -0.031 | 0 | 0.59 | 0.21 | 0.59 | 0.49 | 0.71 | 0.62 |
| 8 | 91130683 | rs10100369 | 1.053 | -0.341 | 0.993 | 1.638 | 0.01 | 0.01 | 0.01 | 0.101 | 0.58 | 0.38 | 0.46 | 0.29 | 0.68 | 0.53 |
| 8 | 91132141 | rs6470713 | 0.127 | -0.441 | 1.061 | 1.828 | 0.002 | -0.055 | -0.055 | 0 | 0.7 | 0.89 | 0.69 | 0.59 | 0.74 | 0.7 |
| 8 | 91133678 | rs6983707 | 0.127 | -0.441 | 1.061 | 1.828 | 0.002 | -0.055 | -0.055 | 0 | 0.7 | 0.89 | 0.69 | 0.59 | 0.74 | 0.7 |
| 8 | 91136078 | rs2205108 | 1.289 | -0.442 | 1.233 | 1.7 | 0.034 | 0.033 | 0.033 | 0.278 | 0.58 | 0.3 | 0.46 | 0.29 | 0.68 | 0.53 |
| 8 | 91142473 | rs2049815 | 0.6 | -0.421 | 0.991 | 1.676 | 0.01 | -0.02 | -0.02 | 0 | 0.66 | 0.74 | 0.53 | 0.39 | 0.71 | 0.6 |
| 8 | 91151614 | rs7014961 | 0.6 | -0.449 | 1.044 | 1.676 | 0.014 | -0.028 | -0.028 | 0 | 0.66 | 0.77 | 0.54 | 0.39 | 0.71 | 0.6 |
| 8 | 91153308 | rs10113507 | 1.069 | -0.456 | 1.2 | 1.676 | -0.01 | -0.009 | -0.009 | 2.604 | 0.6 | 0.47 | 0.48 | 0.29 | 0.69 | 0.53 |
| 8 | 91158208 | rs978953 | 0.511 | -0.155 | 0.605 | 1.833 | -0.001 | 0.004 | 0.004 | 0 | 0.69 | 0.64 | 0.52 | 0.39 | 0.72 | 0.61 |
| 8 | 91160946 | rs10088826 | 0.391 | -0.145 | 0.41 | 1.838 | 0.003 | -0.019 | -0.019 | 0 | 0.69 | 0.74 | 0.54 | 0.39 | 0.72 | 0.61 |
| 8 | 91163167 | rs16904010 | -0.504 | -0.142 | 0.4 | 1.81 | -0.01 | 0.025 | 0.025 | 0 | 0.05 | 0.11 | 0.04 | 0.1 | 0.02 | 0.07 |
| 8 | 91171735 | rs2189261 | 0.965 | -0.047 | 0.225 | 1.669 | 0 | 0 | 0 | 0 | 0.65 | 0.61 | 0.5 | 0.3 | 0.7 | 0.54 |
| 8 | 91175874 | rs10106310 | 0.15 | 0.425 | -0.443 | 1.714 | -0.049 | -0.102 | -0.102 | 7.518 | 0.73 | 0.93 | 0.68 | 0.56 | 0.75 | 0.72 |
| 8 | 91176950 | rs7011930 | -0.112 | -0.016 | -0.329 | 1.795 | -0.017 | -0.102 | -0.102 | 0 | 0.75 | 0.93 | 0.69 | 0.57 | 0.79 | 0.71 |
| 8 | 91182886 | rs988851 | 0.351 | -0.026 | -0.245 | 1.632 | 0.007 | -0.035 | -0.035 | 0 | 0.71 | 0.8 | 0.55 | 0.37 | 0.76 | 0.62 |
| 8 | 91186718 | rs10956500 | 0.365 | -0.06 | -0.254 | 1.605 | 0.009 | -0.048 | -0.048 | 0 | 0.71 | 0.8 | 0.66 | 0.45 | 0.76 | 0.63 |
| 8 | 91190214 | rs2214417 | -0.601 | -0.337 | 0.004 | 1.334 | -0.012 | 0.007 | 0.007 | 0 | 0.4 | 0.49 | 0.5 | 0.42 | 0.3 | 0.38 |
| 8 | 91191414 | rs6998900 | -0.456 | -0.262 | 0.007 | 1.47 | -0.007 | -0.003 | -0.003 | 0 | 0.37 | 0.46 | 0.27 | 0.34 | 0.3 | 0.3 |
| 8 | 91194822 | rs12677020 | -0.239 | -0.268 | 0.06 | 1.472 | 0.007 | -0.026 | -0.026 | 0.04 | 0.33 | 0.16 | 0.23 | 0.35 | 0.27 | 0.29 |
| 8 | 91200676 | rs7826142 | -1.483 | -0.326 | 0.929 | 1.806 | 0.035 | 0.348 | 0.348 | 0 | 0.06 | 0.2 | 0.36 | 0.3 | 0.05 | 0.22 |
| 8 | 91201889 | rs10504892 | -0.639 | -0.412 | 0.774 | 1.479 | 0.03 | 0.044 | 0.044 | 0 | 0.38 | 0.23 | 0.55 | 0.74 | 0.27 | 0.46 |
| 8 | 91205664 | rs7823556 | -0.508 | -0.485 | 0.714 | 1.465 | 0.044 | 0.081 | 0.081 | 0 | 0.36 | 0.15 | 0.54 | 0.75 | 0.27 | 0.45 |
| 8 | 91209785 | rs4961178 | -0.804 | -0.538 | 1.522 | 1.411 | -0.01 | -0.038 | -0.038 | 0.06 | 0.39 | 0.57 | 0.6 | 0.77 | 0.31 | 0.47 |
| 8 | 91212707 | rs1235385 | 1.001 | -0.619 | 0.993 | 0.783 | -0.008 | 0.017 | 0.017 | 0 | 0.14 | 0.03 | 0.1 | 0.11 | 0.2 | 0.15 |
| 8 | 91214786 | rs7841791 | -0.038 | -0.547 | 1.192 | 0.897 | 0.04 | 0.062 | 0.062 | 0 | 0.34 | 0.18 | 0.53 | 0.76 | 0.26 | 0.44 |
| 8 | 91222654 | rs11997469 | -0.357 | -0.537 | 1.41 | 0.941 | 0.008 | 0.023 | 0.023 | 0 | 0.41 | 0.65 | 0.62 | 0.77 | 0.33 | 0.47 |
| 8 | 91229131 | rs6470808 | -1.832 | -0.513 | 1.596 | 1.269 | 0.039 | 0.459 | 0.459 | 0.068 | 0.07 | 0.25 | 0.34 | 0.41 | 0.05 | 0.22 |
| 8 | 91235642 | rs10111402 | -0.301 | -0.562 | 1.449 | 1.214 | -0.009 | -0.04 | -0.04 | 0 | 0.4 | 0.54 | 0.57 | 0.77 | 0.33 | 0.46 |
| 8 | 91238566 | rs12682044 | 0.182 | -0.612 | 1.337 | 1.221 | 0.016 | -0.004 | -0.004 | 0 | 0.33 | 0.24 | 0.19 | 0.36 | 0.22 | 0.22 |
| 8 | 91238763 | rs12679936 | 0.346 | -0.567 | 1.313 | 1.204 | 0.045 | -0.042 | -0.042 | 0 | 0.32 | 0.08 | 0.19 | 0.36 | 0.23 | 0.24 |
| 8 | 91241853 | rs1235387 | -0.697 | -0.855 | 0.957 | 0.835 | 0.011 | 0.054 | 0.054 | 0 | 0.78 | 0.97 | 0.84 | 0.87 | 0.77 | 0.81 |

| | | | | | | | | | | | | | | | | |
|---|---|---|---|---|---|---|---|---|---|---|---|---|---|---|---|---|
| 8 | 91249844 | rs9642892 | 0.08 | -0.907 | 1.704 | 1.687 | 0.044 | -0.024 | -0.024 | 0 | 0.33 | 0.16 | 0.19 | 0.36 | 0.23 | 0.24 |
| 8 | 91252346 | rs2021728 | -0.467 | -0.89 | 1.684 | 1.641 | -0.005 | -0.03 | -0.03 | 0 | 0.4 | 0.45 | 0.55 | 0.78 | 0.33 | 0.49 |
| 8 | 91258967 | rs16904195 | -1.845 | -0.763 | 1.245 | 1.786 | 0.053 | 0.266 | 0.266 | 0 | 0.07 | 0.23 | 0.36 | 0.41 | 0.1 | 0.25 |
| 8 | 91267588 | rs10956539 | -0.285 | -0.944 | 1.312 | 1.722 | -0.011 | 0.002 | 0.002 | 0 | 0.37 | 0.51 | 0.27 | 0.38 | 0.31 | 0.29 |
| 8 | 91291686 | rs2107071 | -1.05 | -1.003 | 0.735 | 1.145 | 0.046 | 0.311 | 0.311 | 0.279 | 0.51 | 0.93 | 0.63 | 0.86 | 0.37 | 0.57 |
| 8 | 91293849 | rs1882851 | -1.874 | -1.365 | 1.538 | 1.388 | 0.083 | 0.079 | 0.079 | 3.445 | 0.05 | 0.5 | 0.04 | 0.01 | 0 | 0 |
| 8 | 91299951 | rs2339111 | -0.961 | -1.504 | 1.038 | 1.337 | 0.115 | 0.192 | 0.192 | 0.027 | 0.45 | 0.86 | 0.59 | 0.77 | 0.31 | 0.46 |
| 8 | 91301523 | rs9643260 | -0.312 | -1.455 | 1.34 | 1.874 | 0.043 | -0.038 | -0.038 | 0 | 0.35 | 0.13 | 0.28 | 0.41 | 0.25 | 0.3 |
| 8 | 91302177 | rs16904279 | -0.555 | -1.065 | 2.331 | 2.081 | -0.033 | 0.023 | 0.023 | 0 | 0.37 | 0.54 | 0.34 | 0.41 | 0.28 | 0.28 |
| 8 | 91303944 | rs2214416 | -0.994 | -1.076 | 2.263 | 1.782 | 0.078 | 0.145 | 0.145 | 0.503 | 0.43 | 0.84 | 0.61 | 0.77 | 0.32 | 0.45 |
| 8 | 91305844 | rs12550585 | -0.636 | -1.071 | 2.702 | 1.792 | -0.003 | -0.01 | -0.01 | 0 | 0.38 | 0.38 | 0.56 | 0.76 | 0.32 | 0.44 |
| 8 | 91324589 | rs1989936 | -0.863 | -0.93 | 2.29 | 1.764 | 0.038 | 0.076 | 0.076 | 0 | 0.45 | 0.73 | 0.64 | 0.76 | 0.35 | 0.46 |
| 8 | 91336887 | rs10956590 | -0.868 | -1.003 | 2.642 | 2.209 | 0.049 | 0.068 | 0.068 | 0 | 0.45 | 0.75 | 0.65 | 0.66 | 0.38 | 0.45 |
| 8 | 91339579 | rs1859179 | -0.274 | -0.997 | 2.641 | 2.32 | 0.027 | -0.058 | -0.058 | 4.711 | 0.38 | 0.13 | 0.35 | 0.42 | 0.34 | 0.32 |
| 8 | 91345282 | rs4734445 | -0.385 | -0.988 | 2.645 | 2.306 | 0.019 | -0.017 | -0.017 | 0 | 0.38 | 0.33 | 0.35 | 0.44 | 0.3 | 0.29 |
| 8 | 91350756 | rs13265958 | -0.35 | -0.988 | 2.524 | 2.309 | 0.023 | -0.023 | -0.023 | 0 | 0.38 | 0.27 | 0.34 | 0.43 | 0.3 | 0.3 |
| 8 | 91357816 | rs4551412 | -2.036 | -0.973 | 2.881 | 2.35 | 0.045 | 0.314 | 0.314 | 1.64 | 0.06 | 0.37 | 0.27 | 0.23 | 0.04 | 0.14 |
| 8 | 91361833 | rs4510907 | -0.67 | -1.032 | 2.997 | 2.338 | -0.017 | -0.019 | -0.019 | 0 | 0.43 | 0.46 | 0.59 | 0.68 | 0.34 | 0.43 |
| 8 | 91366482 | rs16904410 | -0.197 | -1.03 | 2.863 | 2.338 | 0.034 | -0.048 | -0.048 | 0.365 | 0.37 | 0.15 | 0.33 | 0.45 | 0.3 | 0.3 |
| 8 | 91374909 | rs7827909 | -0.68 | -0.983 | 2.624 | 1.892 | 0.005 | 0.006 | 0.006 | 0 | 0.43 | 0.52 | 0.6 | 0.68 | 0.34 | 0.44 |
| 8 | 91375302 | rs6470995 | -0.716 | -0.989 | 2.724 | 1.889 | 0.033 | 0.038 | 0.038 | 0 | 0.43 | 0.58 | 0.6 | 0.68 | 0.34 | 0.44 |
| 8 | 91377153 | rs4576466 | -1.048 | -1.001 | 2.715 | 1.873 | 0.089 | 0.119 | 0.119 | 0.096 | 0.48 | 0.82 | 0.65 | 0.68 | 0.38 | 0.45 |
| 8 | 91381801 | rs6470999 | -0.841 | -1.023 | 2.461 | 1.919 | 0.059 | 0.067 | 0.067 | 0 | 0.45 | 0.72 | 0.62 | 0.67 | 0.34 | 0.44 |
| 8 | 91397407 | rs6999291 | -0.677 | -0.797 | 2.501 | 1.907 | -0.004 | -0.007 | -0.007 | 0 | 0.43 | 0.39 | 0.6 | 0.65 | 0.36 | 0.45 |
| 8 | 91411745 | rs4403455 | -1.109 | -1.032 | 2.552 | 2.174 | 0.05 | 0 | 0 | 0.002 | 0.47 | 0.69 | 0.48 | 0.51 | 0.38 | 0.38 |
| 8 | 91416005 | rs2339149 | -0.25 | -1.078 | 2.47 | 2.137 | 0.019 | -0.039 | -0.039 | 0 | 0.36 | 0.19 | 0.4 | 0.46 | 0.34 | 0.33 |
| 8 | 91434040 | rs730679 | -0.952 | -0.973 | 2.455 | 2.244 | 0.012 | -0.011 | -0.011 | 0.348 | 0.41 | 0.63 | 0.46 | 0.43 | 0.35 | 0.31 |
| 8 | 91443772 | rs2103445 | -0.112 | -0.834 | 1.586 | 2.335 | 0.007 | -0.073 | -0.073 | 0 | 0.31 | 0.08 | 0.35 | 0.37 | 0.29 | 0.25 |
| 8 | 91453910 | rs7826818 | -1.419 | -0.701 | 1.804 | 1.717 | 0.012 | -0.008 | -0.008 | 0 | 0.05 | 0.28 | 0.04 | 0 | 0.03 | 0.01 |
| 8 | 91460670 | rs2142145 | -1.031 | -0.47 | 1.264 | 1.997 | 0.026 | 0.007 | 0.007 | 0 | 0.47 | 0.71 | 0.4 | 0.44 | 0.39 | 0.38 |
| 8 | 91463449 | rs7829073 | -1.012 | -0.3 | 1.413 | 2.047 | 0.01 | 0 | 0 | 0 | 0.35 | 0.73 | 0.28 | 0.35 | 0.28 | 0.33 |
| 8 | 91472515 | rs2205158 | -0.626 | -0.352 | 2.242 | 2.013 | 0.004 | -0.007 | -0.007 | 0 | 0.37 | 0.32 | 0.35 | 0.41 | 0.36 | 0.33 |
| 8 | 91475724 | rs9642910 | -1.048 | -0.389 | 2.548 | 1.987 | 0.011 | -0.01 | -0.01 | 0.024 | 0.43 | 0.65 | 0.38 | 0.49 | 0.36 | 0.36 |
| 8 | 91483618 | rs2205152 | -0.7 | -0.357 | 2.8 | 1.948 | -0.008 | 0.013 | 0.013 | 0 | 0.37 | 0.49 | 0.39 | 0.46 | 0.35 | 0.33 |
| 8 | 91491152 | rs10111060 | -1.219 | -0.358 | 2.505 | 1.927 | 0.039 | -0.031 | -0.031 | 0 | 0.46 | 0.85 | 0.45 | 0.52 | 0.38 | 0.37 |
| 8 | 91526934 | rs2187983 | -1.52 | 0.009 | 2.091 | 1.698 | -0.053 | 0.23 | 0.23 | 2.547 | 0.5 | 0.98 | 0.64 | 0.76 | 0.43 | 0.53 |
| 8 | 91528313 | rs4735201 | -0.59 | 0.115 | 2.395 | 1.75 | 0.019 | -0.043 | -0.043 | 0 | 0.38 | 0.26 | 0.5 | 0.5 | 0.34 | 0.33 |
| 8 | 91555603 | rs1476428 | -0.957 | 0.037 | 2.558 | 0.961 | 0.003 | 0.007 | 0.007 | 0.462 | 0.43 | 0.61 | 0.62 | 0.7 | 0.37 | 0.34 |
| 8 | 91556317 | rs2187982 | -0.675 | 0.022 | 2.674 | 0.948 | 0.003 | 0.008 | 0.008 | 0 | 0.39 | 0.36 | 0.58 | 0.7 | 0.36 | 0.34 |
| 8 | 91563137 | rs7013893 | -0.659 | 0.035 | 2.689 | 0.966 | 0.009 | 0.016 | 0.016 | 0 | 0.39 | 0.33 | 0.57 | 0.7 | 0.34 | 0.34 |
| 8 | 91569144 | rs4734247 | -0.929 | 0.064 | 2.618 | 0.978 | 0.011 | 0.017 | 0.017 | 0.369 | 0.43 | 0.65 | 0.62 | 0.7 | 0.36 | 0.34 |
| 8 | 91592086 | rs11988668 | -1.268 | 0.162 | 2.626 | 0.962 | 0.025 | 0.017 | 0.017 | 0 | 0.06 | 0.37 | 0.07 | 0.01 | 0.04 | 0.06 |
| 8 | 91609175 | rs6471183 | -2.023 | 0.148 | 2.723 | 0.965 | 0.032 | 0.016 | 0.016 | 1.015 | 0.09 | 0.5 | 0.09 | 0.01 | 0.06 | 0.06 |
| 8 | 91635731 | rs13262354 | -0.743 | 0.148 | 2.365 | 0.692 | 0.003 | 0.005 | 0.005 | 0 | 0.38 | 0.37 | 0.57 | 0.71 | 0.34 | 0.34 |
| 8 | 91656468 | rs4571698 | -2.324 | 0.092 | 2.292 | 0.69 | 0.029 | 0.03 | 0.03 | 3.474 | 0.08 | 0.56 | 0.08 | 0.03 | 0.06 | 0.04 |
| 8 | 91682497 | rs4623371 | -0.855 | 0.143 | 1.866 | 0.424 | 0.007 | -0.011 | -0.011 | 0 | 0.39 | 0.45 | 0.62 | 0.74 | 0.48 | 0.47 |
| 8 | 91704043 | rs6998736 | -1.193 | 0.201 | 2.174 | 0.237 | 0.002 | 0.058 | 0.058 | 0.836 | 0.48 | 0.67 | 0.67 | 0.75 | 0.54 | 0.5 |
| 8 | 91710052 | rs6997009 | -1.127 | 0.178 | 2.394 | 0.231 | 0 | 0.019 | 0.019 | 0.614 | 0.47 | 0.54 | 0.65 | 0.75 | 0.54 | 0.5 |
| 8 | 91734482 | rs2339228 | -1.288 | 0.075 | 2.371 | 0.169 | -0.008 | 0.136 | 0.136 | 2.119 | 0.49 | 0.79 | 0.68 | 0.75 | 0.54 | 0.47 |
| 8 | 91738830 | rs2339230 | -1.143 | 0.161 | 2.308 | 0.204 | 0 | -0.008 | -0.008 | 2.125 | 0.47 | 0.54 | 0.65 | 0.74 | 0.53 | 0.5 |
| 8 | 91776478 | rs1394503 | -1.127 | 0.105 | 2.159 | 0.203 | 0 | 0.029 | 0.029 | 0.645 | 0.47 | 0.44 | 0.64 | 0.78 | 0.53 | 0.51 |
| 8 | 91781241 | rs1976495 | -1.22 | 0.116 | 1.426 | 0.2 | 0.002 | 0.027 | 0.027 | 1.021 | 0.48 | 0.62 | 0.66 | 0.76 | 0.53 | 0.5 |
| 8 | 91804805 | rs9643314 | -1.376 | -0.382 | 1.491 | 0.302 | -0.002 | -0.007 | -0.007 | 0.001 | 0.29 | 0.32 | 0.51 | 0.82 | 0.29 | 0.4 |
| 8 | 91817642 | rs1818193 | -1.587 | -0.502 | 1.757 | 0.248 | -0.021 | -0.053 | -0.053 | 0 | 0.31 | 0.49 | 0.55 | 0.79 | 0.31 | 0.39 |
| 8 | 91839576 | rs2062882 | -1.889 | -1.117 | 1.431 | -0.243 | 0.058 | 0.186 | 0.186 | 0.437 | 0.46 | 0.84 | 0.65 | 0.84 | 0.42 | 0.45 |
| 8 | 91849608 | rs11997330 | -1.581 | -0.862 | 1.99 | 0.152 | 0.11 | 0.112 | 0.112 | 0.324 | 0.05 | 0.38 | 0.07 | 0.05 | 0.06 | 0.05 |
| 8 | 91855808 | rs6994971 | -1.218 | -0.903 | 1.698 | 0.465 | -0.013 | 0.011 | 0.011 | 0.78 | 0.23 | 0.4 | 0.48 | 0.72 | 0.23 | 0.28 |
| 8 | 91857581 | rs10956781 | -1.245 | -0.904 | 1.659 | 0.464 | -0.016 | 0.017 | 0.017 | 0.929 | 0.23 | 0.43 | 0.49 | 0.72 | 0.23 | 0.28 |
| 8 | 91931256 | rs1003996 | -0.985 | -0.906 | 1.405 | 0.84 | 0 | 0.01 | 0.01 | 0 | 0.21 | 0.23 | 0.48 | 0.55 | 0.28 | 0.3 |
| 8 | 91937320 | rs2339289 | -0.985 | -0.906 | 1.405 | 0.84 | 0 | 0.01 | 0.01 | 0 | 0.21 | 0.23 | 0.48 | 0.55 | 0.28 | 0.3 |
| 8 | 91953138 | rs7018395 | -1.008 | -0.845 | 1.357 | 0.855 | -0.006 | -0.033 | -0.033 | 0 | 0.19 | 0.12 | 0.47 | 0.55 | 0.28 | 0.3 |
| 8 | 91964558 | rs10490979 | -0.332 | -1.087 | 0.983 | 0.526 | 0.007 | 0.028 | 0.028 | 0.63 | 0.14 | 0.02 | 0.04 | 0.05 | 0.07 | 0.07 |
| 8 | 91968527 | rs1060583 | -0.49 | -1.126 | 1.669 | 0.506 | -0.019 | -0.028 | -0.028 | 0.001 | 0.16 | 0.22 | 0.05 | 0.05 | 0.07 | 0.07 |
| 8 | 92016879 | rs10112439 | 0.564 | -1.135 | 1.644 | 0.623 | -0.006 | -0.023 | -0.023 | 0.954 | 0.81 | 0.88 | 0.52 | 0.45 | 0.72 | 0.7 |
| 8 | 92044528 | rs7846412 | 1.836 | -1.14 | 1.352 | 0.528 | 0.024 | 0.099 | 0.099 | 3.541 | 0.58 | 0.17 | 0.41 | 0.33 | 0.62 | 0.67 |
| 8 | 92045726 | rs12375263 | -0.31 | -1.178 | 1.634 | 0.472 | -0.03 | -0.077 | -0.077 | 0 | 0.14 | 0.28 | 0.05 | 0.01 | 0.07 | 0.04 |
| 8 | 92049496 | rs10106732 | 1.186 | -1.112 | 1.686 | 0.539 | 0.003 | -0.003 | -0.003 | 0 | 0.72 | 0.49 | 0.47 | 0.34 | 0.69 | 0.71 |
| 8 | 92050774 | rs7016813 | 1.108 | -1.115 | 1.704 | 0.537 | 0.006 | 0.001 | 0.001 | 1.579 | 0.73 | 0.51 | 0.47 | 0.34 | 0.69 | 0.71 |
| 8 | 92055426 | rs7820781 | -0.923 | -1.076 | 2.003 | 0.418 | -0.047 | -0.074 | -0.074 | 0 | 0.18 | 0.44 | 0.09 | 0.05 | 0.09 | 0.05 |

| | | | | | | | | | | | | | | | | |
|---|---|---|---|---|---|---|---|---|---|---|---|---|---|---|---|---|
| 8 | 92056288 | rs4260862 | -0.839 | -1.077 | 2.004 | 0.418 | -0.043 | -0.067 | -0.067 | 0 | 0.17 | 0.39 | 0.08 | 0.05 | 0.09 | 0.05 |
| 8 | 92060665 | rs6999873 | -0.275 | -0.96 | 1.629 | 0.42 | -0.022 | -0.056 | -0.056 | 0.973 | 0.14 | 0.2 | 0.05 | 0.01 | 0.07 | 0.04 |
| 8 | 92073814 | rs10110267 | 1.3 | -0.922 | 1.698 | 0.882 | 0 | 0.001 | 0.001 | 0 | 0.65 | 0.71 | 0.47 | 0.63 | 0.63 | 0.69 |
| 8 | 92107070 | rs12542178 | 1.314 | -0.779 | 1.684 | 1.008 | 0 | 0.004 | 0.004 | 0 | 0.64 | 0.7 | 0.47 | 0.63 | 0.62 | 0.69 |
| 8 | 92128840 | rs10956794 | 1.784 | -0.993 | 1.362 | 0.745 | 0.007 | -0.028 | -0.028 | 0 | 0.53 | 0.2 | 0.38 | 0.58 | 0.57 | 0.66 |
| 8 | 92136728 | rs11785003 | 1.784 | -0.993 | 1.374 | 0.745 | 0.008 | -0.026 | -0.026 | 0 | 0.53 | 0.21 | 0.38 | 0.58 | 0.57 | 0.66 |
| 8 | 92137446 | rs11778758 | 1.784 | -0.993 | 1.374 | 0.745 | 0.008 | -0.026 | -0.026 | 0 | 0.53 | 0.21 | 0.38 | 0.58 | 0.57 | 0.66 |
| 8 | 92140589 | rs12546650 | 1.888 | -0.996 | 1.356 | 0.814 | -0.019 | 0.112 | 0.112 | 0.317 | 0.52 | 0.13 | 0.36 | 0.35 | 0.57 | 0.65 |
| 8 | 92148195 | rs7846033 | 0.687 | -1.022 | 1.801 | 0.696 | 0.006 | 0.017 | 0.017 | 0 | 0.74 | 0.75 | 0.5 | 0.62 | 0.67 | 0.71 |
| 8 | 92149429 | rs7009170 | 0.515 | -1.073 | 1.805 | 0.669 | -0.066 | -0.101 | -0.101 | 0 | 0.77 | 0.95 | 0.51 | 0.63 | 0.67 | 0.73 |
| 8 | 92155114 | rs10956795 | 1.723 | -1.084 | 1.859 | 0.664 | -0.01 | -0.056 | -0.056 | 0.187 | 0.67 | 0.22 | 0.42 | 0.58 | 0.65 | 0.71 |
| 8 | 92156541 | rs4128119 | 1.723 | -1.084 | 1.86 | 0.664 | -0.009 | -0.054 | -0.054 | 0 | 0.67 | 0.23 | 0.42 | 0.59 | 0.65 | 0.71 |
| 8 | 92159279 | rs4360257 | 0.799 | -1.056 | 1.861 | 0.688 | 0.006 | 0.017 | 0.017 | 0 | 0.74 | 0.76 | 0.5 | 0.62 | 0.67 | 0.71 |
| 8 | 92179965 | rs6471268 | 1.935 | -1.105 | 1.936 | 0.627 | 0 | -0.02 | -0.02 | 0 | 0.57 | 0.36 | 0.39 | 0.58 | 0.57 | 0.66 |
| 8 | 92181214 | rs7015294 | 0.725 | -1.041 | 1.701 | 0.665 | 0.001 | 0.004 | 0.004 | 0 | 0.75 | 0.79 | 0.5 | 0.61 | 0.67 | 0.72 |
| 8 | 92183658 | rs7842678 | 0.725 | -1.041 | 1.701 | 0.665 | 0.001 | 0.004 | 0.004 | 0 | 0.75 | 0.79 | 0.5 | 0.61 | 0.67 | 0.72 |
| 8 | 92193710 | rs7844122 | 0.708 | -1.045 | 1.68 | 0.669 | 0.001 | 0.005 | 0.005 | 0 | 0.75 | 0.8 | 0.5 | 0.6 | 0.66 | 0.72 |
| 8 | 92198211 | rs7832313 | 1.001 | -1.372 | 1.738 | 0.559 | 0.002 | 0.01 | 0.01 | 0 | 0.73 | 0.77 | 0.5 | 0.62 | 0.67 | 0.67 |
| 8 | 92203762 | rs3923174 | 2.015 | -1.427 | 1.515 | 0.54 | 0.02 | -0.008 | -0.008 | 0 | 0.54 | 0.24 | 0.38 | 0.59 | 0.57 | 0.62 |
| 8 | 92222283 | rs4339600 | 0.489 | -1.045 | 1.141 | 0.694 | -0.053 | -0.104 | -0.104 | 0 | 0.78 | 0.96 | 0.54 | 0.63 | 0.72 | 0.73 |
| 8 | 92227075 | rs11784415 | 2.371 | -1.246 | 0.765 | 0.627 | -0.146 | 0.121 | 0.121 | 0.751 | 0.43 | 0.03 | 0.28 | 0.25 | 0.51 | 0.59 |
| 8 | 92229281 | rs10093786 | 1.096 | -0.44 | 1.844 | 1.004 | 0.01 | 0.028 | 0.028 | 0 | 0.75 | 0.75 | 0.51 | 0.63 | 0.68 | 0.71 |
| 8 | 92241460 | rs4476968 | 1.39 | -0.418 | 1.837 | 1.068 | 0.013 | 0.036 | 0.036 | 0 | 0.72 | 0.7 | 0.48 | 0.6 | 0.65 | 0.7 |
| 8 | 92242762 | rs4431560 | 1.911 | -0.571 | 1.266 | 0.852 | -0.016 | 0.023 | 0.023 | 0.345 | 0.57 | 0.13 | 0.35 | 0.39 | 0.6 | 0.68 |
| 8 | 92243681 | rs10108620 | -1.132 | -0.251 | 1.878 | 0.38 | -0.006 | 0.01 | 0.01 | 0 | 0.05 | 0.19 | 0.05 | 0.04 | 0.03 | 0.01 |
| 8 | 92271863 | rs7814787 | -1.246 | -0.284 | 2.264 | 0.061 | -0.022 | -0.119 | -0.119 | 1.598 | 0.24 | 0.38 | 0.16 | 0.02 | 0.16 | 0.08 |
| 8 | 92271995 | rs10102839 | -0.689 | -0.281 | 2.494 | 0.05 | 0.02 | 0.019 | 0.019 | 0.108 | 0.2 | 0.14 | 0.08 | 0 | 0.1 | 0.07 |
| 8 | 92275768 | rs4492333 | 1.996 | -0.283 | 2.576 | 0.299 | 0.005 | -0.046 | -0.046 | 0 | 0.62 | 0.17 | 0.46 | 0.65 | 0.68 | 0.69 |
| 8 | 92285722 | rs7812726 | -1.256 | -0.26 | 2.519 | 0.299 | -0.023 | -0.12 | -0.12 | 0 | 0.24 | 0.38 | 0.16 | 0.02 | 0.15 | 0.07 |
| 8 | 92295114 | rs4330648 | 1.418 | -0.264 | 2.205 | 0.306 | -0.013 | 0.025 | 0.025 | 0.029 | 0.7 | 0.59 | 0.59 | 0.68 | 0.77 | 0.72 |
| 8 | 92299861 | rs4289769 | 0.462 | -0.194 | 2.07 | 0.353 | 0.023 | 0.157 | 0.157 | 0 | 0.88 | 0.78 | 0.66 | 0.68 | 0.85 | 0.79 |
| 8 | 92300107 | rs10097220 | 1.8 | -0.202 | 1.382 | 0.354 | -0.002 | -0.055 | -0.055 | 0 | 0.62 | 0.15 | 0.46 | 0.65 | 0.67 | 0.69 |
| 8 | 92318522 | rs935633 | -0.699 | -0.446 | 1.342 | -0.226 | 0.002 | -0.031 | -0.031 | 0 | 0.21 | 0.23 | 0.08 | 0 | 0.1 | 0.07 |
| 8 | 92327826 | rs980904 | -0.699 | -0.446 | 1.342 | -0.226 | 0.002 | -0.031 | -0.031 | 0 | 0.21 | 0.23 | 0.09 | 0 | 0.1 | 0.07 |
| 8 | 92335402 | rs7827280 | -0.926 | -0.42 | 1.243 | -0.216 | -0.029 | -0.118 | -0.118 | 0 | 0.25 | 0.47 | 0.17 | 0.03 | 0.15 | 0.1 |
| 8 | 92345747 | rs7459480 | 1.159 | 0.627 | 0.538 | 0.764 | 0.004 | 0.01 | 0.01 | 0 | 0.88 | 0.87 | 0.81 | 0.81 | 0.79 | 0.8 |
| 8 | 92347533 | rs12334455 | 0.954 | 0.22 | 1.399 | 0.547 | -0.016 | 0.225 | 0.225 | 1.895 | 0.85 | 0.57 | 0.71 | 0.69 | 0.87 | 0.82 |
| 8 | 92348647 | rs6471275 | 0.641 | 0.215 | 1.195 | 0.568 | -0.022 | 0.104 | 0.104 | 0.86 | 0.87 | 0.72 | 0.75 | 0.7 | 0.91 | 0.82 |
| 8 | 92360193 | rs6471277 | -1.184 | 0.323 | 0.769 | -0.476 | -0.004 | -0.022 | -0.022 | 0 | 0.33 | 0.7 | 0.3 | 0.02 | 0.34 | 0.28 |
| 8 | 92360396 | rs921231 | -1.255 | 0.455 | 1.462 | -0.49 | 0.136 | 0.017 | 0.017 | 0.465 | 0.08 | 0.41 | 0.14 | 0.03 | 0.13 | 0.1 |
| 8 | 92360576 | rs921232 | -1.215 | 0.429 | 1.488 | -0.547 | 0.008 | -0.091 | -0.091 | 0.031 | 0.27 | 0.56 | 0.25 | 0.03 | 0.3 | 0.26 |
| 8 | 92364100 | rs10109254 | -1.211 | 0.428 | 1.726 | -0.467 | 0.005 | -0.067 | -0.067 | 0 | 0.27 | 0.32 | 0.22 | 0.04 | 0.28 | 0.21 |
| 8 | 92374753 | rs17722765 | -0.771 | 0.436 | 1.415 | -0.46 | -0.016 | 0.095 | 0.095 | 0 | 0.23 | 0.09 | 0.15 | 0.01 | 0.22 | 0.19 |
| 8 | 92391870 | rs7818318 | -1.69 | 0.464 | 1.681 | -0.213 | 0.043 | -0.038 | -0.038 | 1.333 | 0.12 | 0.33 | 0.14 | 0.03 | 0.12 | 0.1 |
| 8 | 92398290 | rs10102673 | -2.067 | 0.588 | 1.859 | -0.242 | 0.056 | -0.063 | -0.063 | 0 | 0.14 | 0.43 | 0.17 | 0.03 | 0.16 | 0.11 |
| 8 | 92402207 | rs10504904 | -0.556 | 0.304 | -0.479 | -0.004 | -0.001 | -0.068 | -0.068 | 0.03 | 0.41 | 0.09 | 0.46 | 0.36 | 0.35 | 0.42 |
| 8 | 92407524 | rs9297895 | -1.776 | 0.566 | 1.102 | -0.148 | 0.058 | -0.013 | -0.013 | 0 | 0.08 | 0.25 | 0.11 | 0.04 | 0.08 | 0.05 |
| 8 | 92419127 | rs9283962 | -1.822 | 0.672 | 1.114 | -0.152 | 0.054 | -0.022 | -0.022 | 0 | 0.08 | 0.25 | 0.11 | 0.04 | 0.09 | 0.05 |
| 8 | 92479701 | rs1505219 | 0.59 | 0.294 | 0.28 | -0.393 | -0.021 | -0.002 | -0.002 | 0 | 0.42 | 0.25 | 0.39 | 0.62 | 0.48 | 0.46 |
| 8 | 92513770 | rs6471285 | -1.659 | 0.102 | 0.928 | -0.684 | 0.012 | -0.041 | -0.041 | 0.475 | 0.1 | 0.63 | 0.12 | 0.05 | 0.1 | 0.05 |
| 8 | 92546836 | rs843094 | 0.57 | -0.291 | -0.232 | -1.138 | -0.049 | 0.165 | 0.165 | 0 | 0.25 | 0.01 | 0.18 | 0.01 | 0.25 | 0.24 |
| 8 | 92560661 | rs1457056 | -0.876 | -0.34 | -0.172 | -0.669 | -0.084 | -0.005 | -0.005 | 0 | 0.63 | 0.89 | 0.5 | 0.39 | 0.55 | 0.62 |
| 8 | 92578051 | rs2607106 | -1.154 | -0.648 | -0.681 | -0.928 | -0.1 | -0.048 | -0.048 | 2.744 | 0.71 | 0.92 | 0.51 | 0.4 | 0.6 | 0.71 |
| 8 | 92580131 | rs6471286 | -1.442 | -0.707 | 0.362 | -0.893 | 0.014 | -0.004 | -0.004 | 0.538 | 0.08 | 0.66 | 0.09 | 0.05 | 0.04 | 0.02 |
| 8 | 92599661 | rs843092 | -0.56 | -0.785 | 0.013 | -0.944 | -0.093 | 0.001 | 0.001 | 0 | 0.61 | 0.9 | 0.48 | 0.4 | 0.54 | 0.61 |
| 8 | 92618959 | rs843083 | -0.255 | -0.772 | 0.692 | -0.57 | -0.004 | 0 | 0 | 0 | 0.58 | 0.38 | 0.44 | 0.46 | 0.55 | 0.61 |
| 8 | 92621938 | rs865214 | -0.407 | -0.787 | 1.373 | -0.577 | -0.006 | -0.003 | -0.003 | 1.336 | 0.61 | 0.63 | 0.47 | 0.46 | 0.55 | 0.61 |
| 8 | 92629886 | rs2657191 | 0.015 | -0.801 | 1.008 | -0.613 | -0.034 | 0.019 | 0.019 | 2.384 | 0.55 | 0.2 | 0.41 | 0.42 | 0.51 | 0.6 |
| 8 | 92642528 | rs10956804 | 0.599 | -0.797 | 0.847 | -0.598 | -0.079 | -0.057 | -0.057 | 0.188 | 0.36 | 0.12 | 0.51 | 0.54 | 0.45 | 0.38 |
| 8 | 92667587 | rs11985424 | 0.395 | -0.863 | 1.049 | -0.632 | -0.073 | -0.001 | -0.001 | 0.035 | 0.37 | 0.2 | 0.54 | 0.61 | 0.48 | 0.39 |
| 8 | 92678041 | rs6471291 | 0.062 | -0.828 | 1.675 | -0.653 | 0.017 | -0.008 | -0.008 | 0 | 0.4 | 0.53 | 0.57 | 0.61 | 0.48 | 0.39 |
| 8 | 92681583 | rs4452760 | -0.774 | -1.181 | 1.385 | -0.888 | 0.012 | 0.108 | 0.108 | 0.743 | 0.54 | 0.9 | 0.64 | 0.72 | 0.59 | 0.54 |
| 8 | 92687716 | rs4734795 | 0.337 | -1.125 | 1.544 | -0.831 | 0.007 | -0.006 | -0.006 | 8.353 | 0.41 | 0.57 | 0.54 | 0.63 | 0.48 | 0.39 |
| 8 | 92688278 | rs12545507 | 0.48 | -0.977 | 1.169 | -0.791 | 0.011 | 0.003 | 0.003 | 0 | 0.38 | 0.46 | 0.54 | 0.55 | 0.44 | 0.38 |
| 8 | 92698851 | rs4304298 | 0.48 | -0.978 | 1.166 | -0.791 | 0.011 | 0.004 | 0.004 | 0 | 0.38 | 0.46 | 0.55 | 0.55 | 0.44 | 0.38 |
| 8 | 92721954 | rs10283141 | -2.029 | -0.926 | 1.225 | -0.736 | 0.112 | 0.12 | 0.12 | 2.096 | 0.05 | 0.41 | 0.07 | 0.07 | 0.04 | 0.02 |
| 8 | 92730829 | rs11787346 | -0.25 | -1.299 | 0.01 | -0.979 | 0.014 | 0.048 | 0.048 | 0 | 0.57 | 0.12 | 0.57 | 0.67 | 0.61 | 0.65 |
| 8 | 92736906 | rs13253001 | 0.767 | -0.99 | 0.283 | -0.83 | -0.061 | -0.037 | -0.037 | 0 | 0.35 | 0.13 | 0.5 | 0.57 | 0.43 | 0.38 |
| 8 | 92746400 | rs10956807 | 0.44 | -1.163 | 0.392 | -0.86 | -0.043 | 0.024 | 0.024 | 0 | 0.37 | 0.25 | 0.56 | 0.66 | 0.47 | 0.39 |

| | | | | | | | | | | | | | | | | |
|---|---|---|---|---|---|---|---|---|---|---|---|---|---|---|---|---|
| 8 | 92756492 | rs4734843 | -0.08 | -1.351 | 0.596 | -0.967 | 0.01 | 0.067 | 0.067 | 0 | 0.53 | 0.35 | 0.6 | 0.69 | 0.55 | 0.53 |
| 8 | 92780504 | rs11992146 | -0.421 | -1.033 | 0.331 | -0.963 | 0.001 | 0.024 | 0.024 | 0 | 0.51 | 0.46 | 0.57 | 0.7 | 0.5 | 0.52 |
| 8 | 92786474 | rs10956810 | 0.451 | -1.077 | 0.054 | -0.767 | -0.016 | 0.004 | 0.004 | 0 | 0.14 | 0.26 | 0.15 | 0.08 | 0.09 | 0.13 |
| 8 | 92793786 | rs4734868 | 1.348 | -0.969 | -0.397 | -0.649 | -0.016 | -0.022 | -0.022 | 0.479 | 0.21 | 0.18 | 0.35 | 0.5 | 0.33 | 0.25 |
| 8 | 92799912 | rs10156332 | -0.634 | -0.833 | -0.791 | -0.755 | -0.014 | 0.069 | 0.069 | 0 | 0.56 | 0.83 | 0.61 | 0.67 | 0.52 | 0.52 |
| 8 | 92814026 | rs12681007 | -0.319 | -0.701 | -0.613 | -0.605 | -0.035 | 0.055 | 0.055 | 1.553 | 0.42 | 0.81 | 0.57 | 0.67 | 0.46 | 0.4 |
| 8 | 92817138 | rs4734890 | -0.757 | -0.351 | -0.626 | -0.476 | 0.019 | -0.004 | -0.004 | 0 | 0.85 | 0.79 | 0.83 | 0.92 | 0.76 | 0.96 |
| 8 | 92820523 | rs10956814 | 0.522 | -0.455 | -0.372 | -0.373 | 0.015 | 0.001 | 0.001 | 0 | 0.71 | 0.54 | 0.68 | 0.82 | 0.67 | 0.78 |
| 8 | 92823082 | rs12546664 | 0.614 | -0.465 | -0.293 | -0.371 | 0.002 | -0.002 | -0.002 | 0.16 | 0.13 | 0.12 | 0.13 | 0.1 | 0.09 | 0.18 |
| 8 | 92823859 | rs4323435 | 0.461 | -0.521 | -0.313 | -0.371 | 0.015 | -0.001 | -0.001 | 0 | 0.71 | 0.53 | 0.68 | 0.82 | 0.67 | 0.78 |
| 8 | 92835327 | rs4509277 | 0.212 | 0.014 | -0.528 | -0.402 | 0.025 | -0.009 | -0.009 | 7.077 | 0.72 | 0.52 | 0.62 | 0.84 | 0.67 | 0.78 |
| 8 | 92838655 | rs7017906 | 1.369 | 0.013 | -0.416 | 0.105 | -0.008 | -0.004 | -0.004 | 2.558 | 0.25 | 0.22 | 0.27 | 0.29 | 0.34 | 0.21 |
| 8 | 92838980 | rs17732285 | 1.376 | -0.575 | -0.56 | 0.122 | -0.021 | -0.021 | -0.021 | 0 | 0.07 | 0 | 0.03 | 0.08 | 0.08 | 0.17 |
| 8 | 92845510 | rs4435678 | -0.4 | -0.073 | -0.059 | 0.029 | -0.008 | 0.005 | 0.005 | 0 | 0.25 | 0.48 | 0.31 | 0.16 | 0.23 | 0.19 |
| 8 | 92856096 | rs13250505 | 0.81 | -0.012 | -0.536 | 0.023 | -0.014 | -0.01 | -0.01 | 0 | 0.13 | 0.21 | 0.1 | 0.07 | 0.12 | 0.03 |
| 8 | 92866188 | rs12679380 | 1.095 | 0.105 | -0.777 | 0.08 | -0.001 | -0.003 | -0.003 | 1.074 | 0.11 | 0.09 | 0.18 | 0.09 | 0.09 | 0.16 |
| 8 | 92867682 | rs16914603 | 1.017 | 0.255 | -1.017 | 0.249 | -0.007 | -0.006 | -0.006 | 0.098 | 0.12 | 0.17 | 0.09 | 0.07 | 0.12 | 0.03 |
| 8 | 92867821 | rs1373169 | -0.722 | 0.524 | -1.481 | 0.487 | 0 | 0 | 0 | 0 | 0.72 | 0.72 | 0.66 | 0.73 | 0.54 | 0.76 |
| 8 | 92869618 | rs10094421 | -0.093 | -0.502 | -1.654 | 0.69 | 0.04 | -0.049 | -0.049 | 0 | 0.34 | 0.2 | 0.32 | 0.58 | 0.26 | 0.44 |
| 8 | 92870759 | rs10095605 | -0.591 | 0.194 | -1.306 | 0.863 | -0.018 | -0.028 | -0.028 | 0 | 0.35 | 0.21 | 0.4 | 0.39 | 0.4 | 0.29 |
| 8 | 92873618 | rs4734909 | -0.274 | -0.198 | -0.911 | 0.659 | -0.002 | 0.023 | 0.023 | 0 | 0.19 | 0.09 | 0.27 | 0.16 | 0.19 | 0.22 |
| 8 | 92873970 | rs16914618 | -0.141 | -0.076 | -1.03 | 0.789 | -0.009 | -0.022 | -0.022 | 0 | 0.07 | 0.02 | 0.13 | 0.13 | 0.11 | 0.1 |
| 8 | 92874936 | rs16914621 | -0.498 | -0.948 | -1.08 | 0.765 | 0.032 | -0.002 | -0.002 | 0.009 | 0.57 | 0.79 | 0.55 | 0.62 | 0.38 | 0.68 |
| 8 | 92874959 | rs10096445 | -0.093 | -1.257 | -0.176 | 0.677 | 0.015 | -0.002 | -0.002 | 0 | 0.48 | 0.53 | 0.49 | 0.53 | 0.35 | 0.58 |
| 8 | 92880676 | rs1867307 | -0.556 | -0.989 | -0.695 | 0.557 | 0.024 | 0.075 | 0.075 | 0 | 0.57 | 0.83 | 0.6 | 0.74 | 0.38 | 0.65 |
| 8 | 92895301 | rs1443556 | -1.051 | -0.826 | -0.432 | -0.226 | -0.108 | 0.106 | 0.106 | 0 | 0.67 | 0.89 | 0.72 | 0.87 | 0.49 | 0.81 |
| 8 | 92896022 | rs10090064 | 0.103 | -1.12 | -0.296 | -0.291 | -0.001 | -0.008 | -0.008 | 0.023 | 0.22 | 0.21 | 0.27 | 0.15 | 0.21 | 0.24 |
| 8 | 92896491 | rs4734923 | -0.406 | -1.204 | 0.089 | -0.095 | 0.003 | 0.004 | 0.004 | 0 | 0.56 | 0.62 | 0.63 | 0.61 | 0.45 | 0.66 |
| 8 | 92909705 | rs10103170 | -0.245 | -1.196 | 0.02 | -0.094 | -0.002 | -0.002 | -0.002 | 1.842 | 0.52 | 0.47 | 0.62 | 0.59 | 0.44 | 0.65 |
| 8 | 92916704 | rs6992905 | 0.602 | -1.225 | -0.179 | -0.094 | 0.011 | -0.011 | -0.011 | 0 | 0.11 | 0.07 | 0.08 | 0.27 | 0.04 | 0.12 |
| 8 | 92921867 | rs7825946 | -0.484 | -1.209 | -0.23 | -0.117 | 0.005 | 0.006 | 0.006 | 0 | 0.55 | 0.65 | 0.63 | 0.57 | 0.46 | 0.67 |
| 8 | 92928669 | rs1443561 | -0.954 | -1.011 | -0.663 | -0.495 | -0.17 | 0.108 | 0.108 | 0.245 | 0.66 | 0.97 | 0.74 | 0.87 | 0.49 | 0.81 |
| 8 | 92928753 | rs1443562 | 0.122 | -0.992 | -0.59 | -0.486 | 0.02 | 0.025 | 0.025 | 0 | 0.44 | 0.63 | 0.47 | 0.72 | 0.3 | 0.59 |
| 8 | 92933434 | rs4734934 | -1.102 | -0.856 | -1.142 | -0.429 | -0.17 | 0.094 | 0.094 | 0.374 | 0.67 | 0.97 | 0.74 | 0.87 | 0.48 | 0.79 |
| 8 | 92942911 | rs1443564 | -0.107 | -0.259 | -0.875 | -0.176 | 0.001 | 0.036 | 0.036 | 0 | 0.2 | 0.08 | 0.25 | 0.17 | 0.19 | 0.23 |
| 8 | 92965409 | rs2218367 | 0.051 | -0.095 | -0.464 | 0.141 | -0.01 | 0.017 | 0.017 | 0 | 0.32 | 0.37 | 0.35 | 0.33 | 0.28 | 0.48 |
| 8 | 92967910 | rs10098853 | -0.283 | 0.029 | 0.211 | -0.602 | 0.008 | -0.02 | -0.02 | 0 | 0.17 | 0.25 | 0.28 | 0.16 | 0.16 | 0.16 |
| 8 | 92970153 | rs4500123 | -0.281 | -0.142 | 0.554 | -0.759 | 0.003 | -0.01 | -0.01 | 1.088 | 0.1 | 0.26 | 0.09 | 0.09 | 0.08 | 0.11 |
| 8 | 92972780 | rs2977676 | -0.694 | -0.751 | -0.266 | -1.242 | -0.096 | 0.119 | 0.119 | 0.186 | 0.66 | 0.91 | 0.68 | 0.89 | 0.48 | 0.79 |
| 8 | 92976563 | rs2976501 | 0.377 | -0.61 | 0.731 | -1.858 | -0.012 | -0.114 | -0.114 | 0 | 0.71 | 0.48 | 0.8 | 0.99 | 0.76 | 0.85 |
| 8 | 92980734 | rs2976502 | -0.058 | -0.649 | 0.868 | -1.878 | 0.001 | -0.013 | -0.013 | 0 | 0.75 | 0.78 | 0.82 | 0.99 | 0.76 | 0.85 |
| 8 | 93009484 | rs16914707 | 0.299 | -0.861 | 0.723 | -1.971 | 0.005 | -0.028 | -0.028 | 1.696 | 0.13 | 0 | 0.1 | 0.15 | 0.13 | 0.08 |
| 8 | 93051254 | rs4734170 | -0.599 | -0.466 | 1.533 | -1.934 | -0.013 | -0.153 | -0.153 | 1.114 | 0.28 | 0.45 | 0.2 | 0.01 | 0.24 | 0.15 |
| 8 | 93060896 | rs10089969 | -0.342 | -0.423 | 1.204 | -1.913 | -0.002 | -0.057 | -0.057 | 0.567 | 0.26 | 0.28 | 0.19 | 0.01 | 0.24 | 0.15 |
| 8 | 93098924 | rs720444 | 0.107 | -0.446 | 1.094 | -1.898 | 0.001 | -0.002 | -0.002 | 0 | 0.76 | 0.78 | 0.83 | 0.99 | 0.78 | 0.86 |
| 8 | 93103921 | rs17747233 | 0.481 | -0.396 | 0.933 | -2.015 | 0.001 | -0.013 | -0.013 | 0 | 0.05 | 0.05 | 0.07 | 0 | 0.06 | 0.08 |
| 8 | 93104805 | rs1552314 | 0.742 | -0.483 | 0.916 | -1.984 | 0.027 | 0.133 | 0.133 | 1.908 | 0.69 | 0.13 | 0.78 | 0.99 | 0.78 | 0.86 |
| 8 | 93141987 | rs12056584 | 0.307 | -0.188 | 0.848 | -1.831 | -0.01 | -0.117 | -0.117 | 3.087 | 0.73 | 0.61 | 0.81 | 0.99 | 0.78 | 0.86 |
| 8 | 93160730 | rs993790 | 0.025 | -0.013 | 0.217 | -1.739 | -0.009 | 0.062 | 0.062 | 0 | 0.77 | 0.91 | 0.84 | 0.99 | 0.78 | 0.86 |
| 8 | 93162375 | rs6471322 | 0.821 | 0.551 | -0.096 | -2.024 | -0.003 | -0.032 | -0.032 | 0 | 0.62 | 0.66 | 0.76 | 0.99 | 0.64 | 0.79 |
| 8 | 93167342 | rs7006836 | -0.119 | 0.401 | -0.354 | -1.98 | -0.004 | 0.041 | 0.041 | 0 | 0.77 | 0.84 | 0.84 | 0.99 | 0.77 | 0.85 |
| 8 | 93173000 | rs990747 | -0.101 | 0.416 | -0.366 | -1.972 | -0.001 | 0.008 | 0.008 | 0 | 0.77 | 0.81 | 0.83 | 0.99 | 0.77 | 0.85 |
| 8 | 93179976 | rs6471325 | 0.331 | 0.569 | -0.408 | -1.677 | -0.001 | -0.068 | -0.068 | 0 | 0.61 | 0.61 | 0.76 | 0.99 | 0.64 | 0.78 |
| 8 | 93184065 | rs3857914 | -0.226 | 0.955 | -0.347 | -1.096 | 0.006 | -0.032 | -0.032 | 0 | 0.27 | 0.33 | 0.22 | 0.16 | 0.32 | 0.18 |
| 8 | 93187902 | rs10956824 | 0.666 | 0.681 | -0.455 | -0.691 | -0.004 | 0.001 | 0.001 | 0.097 | 0.36 | 0.27 | 0.41 | 0.34 | 0.37 | 0.47 |
| 8 | 93189631 | rs3857915 | 1.307 | 0.452 | -0.668 | -0.241 | -0.002 | -0.048 | -0.048 | 2.723 | 0.2 | 0.07 | 0.12 | 0.25 | 0.16 | 0.18 |
| 8 | 93191681 | rs12680455 | 2.034 | 0.228 | -0.878 | -0.388 | 0 | -0.048 | -0.048 | 0 | 0.08 | 0.01 | 0.14 | 0.25 | 0.04 | 0.1 |
| 8 | 93196771 | rs1444499 | 1.831 | 0.214 | -0.775 | -0.395 | 0.011 | -0.027 | -0.027 | 0 | 0.08 | 0.04 | 0.14 | 0.25 | 0.04 | 0.1 |
| 8 | 93201255 | rs1373529 | 0.574 | 0.158 | -0.532 | -0.487 | 0 | -0.001 | -0.001 | 0 | 0.46 | 0.54 | 0.58 | 0.59 | 0.41 | 0.57 |
| 8 | 93202128 | rs2218439 | 0.009 | 0 | -0.071 | -0.552 | 0.007 | -0.053 | -0.053 | 0 | 0.65 | 0.58 | 0.7 | 0.83 | 0.57 | 0.75 |
| 8 | 93202710 | rs1552313 | 0.009 | -0.03 | -0.074 | -0.287 | 0.004 | -0.023 | -0.023 | 0.146 | 0.65 | 0.59 | 0.58 | 0.76 | 0.57 | 0.75 |
| 8 | 93209764 | rs6471329 | -0.992 | -0.155 | 0.18 | -0.306 | -0.001 | -0.023 | -0.023 | 0.014 | 0.9 | 0.92 | 0.9 | 1 | 0.88 | 0.94 |
| 8 | 93210301 | rs7814997 | -0.992 | -0.155 | 0.18 | -0.312 | -0.001 | -0.026 | -0.026 | 0 | 0.9 | 0.92 | 0.9 | 1 | 0.88 | 0.94 |
| 8 | 93213010 | rs2046395 | 0.603 | -0.142 | 0.275 | -0.23 | 0.011 | 0.02 | 0.02 | 0 | 0.55 | 0.3 | 0.42 | 0.55 | 0.54 | 0.69 |
| 8 | 93213535 | rs13279070 | 0.566 | -0.154 | 0.248 | -0.688 | -0.011 | -0.012 | -0.012 | 0.451 | 0.3 | 0.37 | 0.33 | 0.22 | 0.21 | 0.27 |
| 8 | 93213672 | rs7008512 | -1.522 | -0.192 | 0.487 | -0.728 | 0.002 | -0.05 | -0.05 | 0 | 0.9 | 0.83 | 0.9 | 1 | 0.88 | 0.94 |
| 8 | 93213730 | rs13271152 | 0.586 | -0.148 | 0.422 | -0.67 | -0.011 | -0.016 | -0.016 | 0.738 | 0.3 | 0.37 | 0.21 | 0.21 | 0.21 | 0.27 |
| 8 | 93221658 | rs6471333 | 0.43 | -0.083 | 0.002 | -0.674 | -0.013 | -0.024 | -0.024 | 0 | 0.33 | 0.35 | 0.25 | 0.22 | 0.26 | 0.28 |
| 8 | 93237876 | rs4286925 | 1.227 | 0.073 | -0.572 | -0.558 | -0.129 | -0.127 | -0.127 | 0.382 | 0.32 | 0.03 | 0.23 | 0.51 | 0.36 | 0.47 |

| | | | | | | | | | | | | | | | | |
|---|---|---|---|---|---|---|---|---|---|---|---|---|---|---|---|---|
| 8 | 93247362 | rs6471335 | 1.231 | 0.023 | -0.301 | -0.516 | 0.002 | 0.002 | 0.002 | 0 | 0.21 | 0.28 | 0.39 | 0.26 | 0.26 | 0.2 |
| 8 | 93262882 | rs1444506 | 1.042 | 0.1 | -0.75 | -0.42 | -0.121 | -0.114 | -0.114 | 0.105 | 0.33 | 0.04 | 0.23 | 0.48 | 0.38 | 0.47 |
| 8 | 93264453 | rs4734221 | -0.423 | -0.252 | -0.17 | -0.409 | -0.002 | -0.005 | -0.005 | 1.139 | 0.68 | 0.68 | 0.74 | 0.79 | 0.75 | 0.74 |
| 8 | 93268649 | rs10755939 | 1.3 | 0.055 | -0.864 | -0.347 | -0.026 | -0.008 | -0.008 | 0 | 0.21 | 0.09 | 0.17 | 0.25 | 0.3 | 0.31 |
| 8 | 93272783 | rs9657011 | 0.059 | -0.087 | -1.07 | -0.4 | 0.029 | -0.019 | -0.019 | 0 | 0.43 | 0.82 | 0.59 | 0.52 | 0.39 | 0.4 |
| 8 | 93279099 | rs967226 | 2.039 | -0.076 | -0.939 | -0.571 | 0.021 | -0.044 | -0.044 | 0 | 0.1 | 0.23 | 0.07 | 0.01 | 0.16 | 0.07 |
| 8 | 93283799 | rs6993870 | 1.026 | -0.136 | -1.495 | -0.458 | -0.056 | -0.072 | -0.072 | 0.006 | 0.31 | 0.03 | 0.34 | 0.25 | 0.27 | 0.29 |
| 8 | 93284681 | rs17748153 | 1.557 | -0.024 | -1.273 | -0.334 | -0.077 | -0.047 | -0.047 | 0 | 0.21 | 0.04 | 0.16 | 0.25 | 0.3 | 0.33 |
| 8 | 93287501 | rs16914874 | 2.038 | -0.261 | -1.124 | -0.307 | 0.021 | -0.048 | -0.048 | 1.974 | 0.08 | 0.01 | 0.03 | 0.23 | 0.02 | 0.06 |
| 8 | 93294854 | rs1992631 | -0.911 | -0.358 | -0.595 | -0.401 | 0.002 | -0.002 | -0.002 | 1.688 | 0.75 | 0.76 | 0.63 | 0.73 | 0.75 | 0.71 |
| 8 | 93298074 | rs12681205 | 0.074 | -0.393 | -0.612 | -0.517 | -0.005 | 0.044 | 0.044 | 0 | 0.62 | 0.16 | 0.52 | 0.72 | 0.59 | 0.66 |
| 8 | 93305142 | rs2199926 | 0.45 | -0.66 | -0.82 | -0.484 | 0.008 | -0.103 | -0.103 | 0 | 0.41 | 0.09 | 0.36 | 0.47 | 0.28 | 0.32 |
| 8 | 93329382 | rs10097299 | 2.237 | -0.727 | -0.852 | -0.602 | -0.012 | 0.051 | 0.051 | 0 | 0.15 | 0.05 | 0.2 | 0 | 0.18 | 0.18 |
| 8 | 93330786 | rs2928914 | 0.51 | -0.672 | -0.901 | -0.557 | 0.016 | -0.097 | -0.097 | 0 | 0.41 | 0.1 | 0.36 | 0.48 | 0.28 | 0.32 |
| 8 | 93334453 | rs1480890 | -0.91 | -0.92 | -0.695 | -0.52 | -0.002 | 0 | 0 | 0 | 0.75 | 0.74 | 0.62 | 0.73 | 0.75 | 0.71 |
| 8 | 93355038 | rs1480889 | 0.63 | -0.633 | -0.651 | -0.433 | 0.013 | -0.014 | -0.014 | 0.025 | 0.34 | 0.41 | 0.26 | 0.25 | 0.49 | 0.41 |
| 8 | 93357620 | rs1480895 | 0.935 | -0.858 | -0.889 | -0.527 | 0.028 | -0.095 | -0.095 | 0.2 | 0.29 | 0.1 | 0.26 | 0.49 | 0.17 | 0.24 |
| 8 | 93358703 | rs12675587 | 2.17 | -0.954 | -0.952 | -0.493 | 0.014 | -0.026 | -0.026 | 0 | 0.09 | 0.07 | 0.04 | 0.24 | 0.03 | 0.05 |
| 8 | 93359043 | rs10956833 | 0.577 | -0.964 | -0.843 | -0.504 | 0.002 | -0.005 | -0.005 | 0.15 | 0.31 | 0.28 | 0.28 | 0.49 | 0.17 | 0.24 |
| 8 | 93360383 | rs12682312 | 0.734 | -0.915 | -1.228 | -0.477 | 0.028 | -0.065 | -0.065 | 0 | 0.3 | 0.16 | 0.27 | 0.49 | 0.17 | 0.24 |
| 8 | 93363999 | rs16914926 | 2.405 | -0.819 | -1.382 | -0.429 | 0.002 | -0.049 | -0.049 | 0 | 0.08 | 0 | 0.03 | 0.24 | 0.03 | 0.05 |
| 8 | 93369299 | rs16914930 | 1.636 | -0.776 | -1.328 | -0.481 | 0.005 | -0.07 | -0.07 | 0.057 | 0.2 | 0.03 | 0.23 | 0.25 | 0.14 | 0.19 |
| 8 | 93369304 | rs10956836 | -0.007 | -1.056 | -0.724 | -0.335 | 0.002 | 0.002 | 0.002 | 0 | 0.65 | 0.69 | 0.54 | 0.74 | 0.66 | 0.65 |
| 8 | 93393331 | rs10808648 | 1.134 | -0.473 | -0.676 | 0.262 | -0.044 | -0.077 | -0.077 | 0 | 0.23 | 0.07 | 0.3 | 0.54 | 0.31 | 0.46 |
| 8 | 93420729 | rs10106912 | 1.629 | -0.602 | -0.917 | 0.072 | -0.011 | 0.051 | 0.051 | 0 | 0.25 | 0.07 | 0.25 | 0.19 | 0.22 | 0.25 |
| 8 | 93424621 | rs9297909 | 0.358 | -0.494 | -0.821 | 0.122 | 0.001 | 0.101 | 0.101 | 0 | 0.5 | 0.29 | 0.55 | 0.74 | 0.53 | 0.71 |
| 8 | 93426059 | rs4735174 | 1.472 | -0.525 | -0.811 | 0.081 | 0.002 | 0.006 | 0.006 | 0 | 0.25 | 0.23 | 0.25 | 0.19 | 0.22 | 0.24 |
| 8 | 93438176 | rs10107525 | 0.204 | -0.615 | -0.768 | 0.356 | -0.003 | -0.001 | -0.001 | 0 | 0.76 | 0.76 | 0.73 | 0.74 | 0.79 | 0.83 |
| 8 | 93443393 | rs12155836 | 0.661 | -0.222 | -0.763 | 0.81 | -0.013 | -0.038 | -0.038 | 0 | 0.26 | 0.16 | 0.31 | 0.52 | 0.32 | 0.48 |
| 8 | 93448756 | rs16914980 | 1.271 | -0.122 | -0.935 | 0.661 | -0.068 | -0.063 | -0.063 | 0 | 0.21 | 0.04 | 0.16 | 0.28 | 0.29 | 0.29 |
| 8 | 93450212 | rs1072934 | 0.997 | -0.084 | -0.732 | 0.143 | -0.034 | -0.075 | -0.075 | 0.083 | 0.27 | 0.1 | 0.32 | 0.49 | 0.34 | 0.53 |
| 8 | 93452462 | rs559606 | -0.177 | -0.388 | -0.364 | -0.087 | -0.013 | 0.04 | 0.04 | 0 | 0.9 | 0.83 | 0.9 | 0.75 | 0.91 | 0.89 |
| 8 | 93454053 | rs16914984 | 1.179 | -0.404 | -0.326 | -0.43 | 0.022 | -0.072 | -0.072 | 0 | 0.24 | 0.34 | 0.2 | 0.03 | 0.28 | 0.11 |
| 8 | 93454352 | rs1111435 | 0.489 | -0.722 | -0.073 | -0.092 | 0.016 | 0.014 | 0.014 | 0.793 | 0.54 | 0.43 | 0.63 | 0.54 | 0.65 | 0.67 |
| 8 | 93459258 | rs7818242 | 0.939 | -0.524 | -0.8 | -0.299 | 0.01 | 0.056 | 0.056 | 0.794 | 0.25 | 0.1 | 0.24 | 0.2 | 0.2 | 0.18 |
| 8 | 93470840 | rs491546 | 0.352 | -0.27 | -0.401 | -0.321 | -0.01 | -0.034 | -0.034 | 0.927 | 0.09 | 0.1 | 0.03 | 0 | 0.06 | 0.04 |
| 8 | 93475137 | rs3735912 | -1.026 | 0.045 | 0.203 | 0.057 | 0.005 | 0.051 | 0.051 | 0 | 0.12 | 0.25 | 0.1 | 0.25 | 0.1 | 0.12 |
| 8 | 93482380 | rs2046130 | 0.781 | -0.158 | -0.117 | -0.334 | -0.005 | -0.07 | -0.07 | 0.013 | 0.26 | 0.32 | 0.25 | 0 | 0.2 | 0.1 |
| 8 | 93482694 | rs13249669 | -0.505 | -0.172 | -0.651 | -0.268 | 0.011 | 0.024 | 0.024 | 1.314 | 0.38 | 0.64 | 0.36 | 0.25 | 0.3 | 0.22 |
| 8 | 93493264 | rs574048 | 0.61 | 0.126 | -1.56 | -0.172 | -0.029 | 0.1 | 0.1 | 2.102 | 0.39 | 0.07 | 0.45 | 0.74 | 0.41 | 0.64 |
| 8 | 93493424 | rs12545226 | 0.809 | 0.101 | -1.517 | -0.512 | -0.007 | 0.136 | 0.136 | 0.052 | 0.26 | 0.1 | 0.25 | 0 | 0.22 | 0.13 |
| 8 | 93493736 | rs519138 | 0.788 | 0.205 | -1.354 | 0.383 | -0.046 | -0.085 | -0.085 | 0 | 0.37 | 0.05 | 0.31 | 0.42 | 0.4 | 0.5 |
| 8 | 93499817 | rs1597301 | 0.584 | 0.406 | -0.312 | 0.907 | -0.018 | -0.016 | -0.016 | 0 | 0.6 | 0.33 | 0.54 | 0.46 | 0.58 | 0.62 |
| 8 | 93508014 | rs561608 | 0.125 | 0.523 | 0.248 | 0.894 | 0.03 | 0.052 | 0.052 | 0 | 0.7 | 0.53 | 0.5 | 0.65 | 0.66 | 0.61 |
| 8 | 93514906 | rs482619 | 0.326 | 0.015 | 0.262 | 0.993 | -0.018 | 0.038 | 0.038 | 0.093 | 0.71 | 0.45 | 0.53 | 0.63 | 0.73 | 0.6 |
| 8 | 93518426 | rs538100 | -0.283 | 0.033 | 0.042 | 1.01 | -0.003 | 0.026 | 0.026 | 0 | 0.11 | 0.15 | 0.07 | 0.34 | 0.1 | 0.1 |
| 8 | 93521765 | rs543027 | 0.019 | -0.362 | -0.013 | 0.915 | 0.012 | -0.009 | -0.009 | 0 | 0.2 | 0.11 | 0.07 | 0.31 | 0.1 | 0.13 |
| 8 | 93524692 | rs522576 | -0.692 | -0.152 | -0.09 | 0.621 | -0.001 | -0.004 | -0.004 | 1.834 | 0.85 | 0.85 | 0.83 | 0.96 | 0.78 | 0.9 |
| 8 | 93534196 | rs4535720 | 0.21 | -0.117 | -0.257 | 0.672 | 0.014 | 0.011 | 0.011 | 0.014 | 0.65 | 0.41 | 0.6 | 0.63 | 0.59 | 0.71 |
| 8 | 93535437 | rs6989429 | -0.038 | -0.186 | -0.905 | 0.749 | 0.031 | 0.009 | 0.009 | 0.36 | 0.36 | 0.23 | 0.19 | 0.33 | 0.26 | 0.24 |
| 8 | 93540332 | rs7828100 | 0.752 | -0.395 | -1.467 | 0.479 | -0.101 | -0.006 | -0.006 | 0 | 0.38 | 0.09 | 0.51 | 0.36 | 0.47 | 0.49 |
| 8 | 93543383 | rs6989464 | 0.59 | -0.277 | -1.347 | 0.533 | -0.059 | 0.039 | 0.039 | 0.568 | 0.42 | 0.17 | 0.53 | 0.35 | 0.48 | 0.53 |
| 8 | 93553650 | rs1449232 | -0.121 | 0.474 | -1.261 | 0.221 | -0.15 | 0.186 | 0.186 | 0 | 0.68 | 0.95 | 0.6 | 0.95 | 0.52 | 0.8 |
| 8 | 93553737 | rs7844026 | -0.064 | 0.466 | -0.991 | 0.216 | -0.078 | 0.113 | 0.113 | 0 | 0.67 | 0.88 | 0.6 | 0.95 | 0.52 | 0.8 |
| 8 | 93554257 | rs10956850 | 0.315 | 0.146 | -0.829 | 0.924 | 0.023 | 0.03 | 0.03 | 0.285 | 0.49 | 0.26 | 0.47 | 0.62 | 0.46 | 0.61 |
| 8 | 93563752 | rs2197890 | -1.356 | 0.201 | 0.356 | 1.946 | -0.007 | 0.154 | 0.154 | 0.028 | 0.13 | 0.25 | 0.25 | 0.55 | 0.1 | 0.32 |
| 8 | 93564449 | rs13258527 | -1.421 | 0.16 | 0.54 | 1.939 | -0.009 | 0.178 | 0.178 | 0.022 | 0.14 | 0.28 | 0.25 | 0.55 | 0.1 | 0.32 |
| 8 | 93567028 | rs2618137 | -0.861 | 0.186 | 1.395 | 1.681 | 0.04 | -0.031 | -0.031 | 0 | 0.08 | 0.31 | 0.11 | 0 | 0.07 | 0.04 |
| 8 | 93567187 | rs13250219 | -1.088 | 0.18 | 1.468 | 1.685 | 0.002 | 0.013 | 0.013 | 0 | 0.13 | 0.12 | 0.14 | 0.55 | 0.1 | 0.32 |
| 8 | 93568065 | rs2256182 | -0.401 | 0.758 | 1.067 | 1.142 | -0.048 | 0.062 | 0.062 | 0 | 0.86 | 0.94 | 0.8 | 0.99 | 0.75 | 0.92 |
| 8 | 93568943 | rs4735183 | 0.863 | 0.609 | 1.198 | 1.111 | 0.027 | -0.127 | -0.127 | 0.019 | 0.76 | 0.54 | 0.68 | 0.99 | 0.68 | 0.89 |
| 8 | 93575263 | rs12680247 | -0.397 | 0.675 | 0.575 | 1.692 | -0.008 | -0.035 | -0.035 | 0.263 | 0.1 | 0.04 | 0.05 | 0.15 | 0.09 | 0.17 |
| 8 | 93577910 | rs2291016 | 1.319 | 0.774 | 0.746 | 1.729 | -0.016 | -0.036 | -0.036 | 0 | 0.62 | 0.22 | 0.51 | 0.48 | 0.58 | 0.57 |
| 8 | 93584392 | rs6993453 | -2.08 | 0.447 | 1.387 | 1.873 | -0.01 | 0.042 | 0.042 | 0.635 | 0.2 | 0.65 | 0.15 | 0.3 | 0.14 | 0.25 |
| 8 | 93584776 | rs10111610 | -2.08 | 0.447 | 1.387 | 1.873 | -0.01 | 0.042 | 0.042 | 0.635 | 0.2 | 0.65 | 0.15 | 0.3 | 0.14 | 0.25 |
| 8 | 93585934 | rs17757201 | -0.767 | 0.449 | 1.529 | 1.874 | 0.006 | -0.001 | -0.001 | 0 | 0.07 | 0.09 | 0.06 | 0.05 | 0.09 | 0.16 |
| 8 | 93586221 | rs16915067 | -1.34 | 0.165 | 1.925 | 1.914 | 0.005 | 0.11 | 0.11 | 0.272 | 0.05 | 0.27 | 0.05 | 0.08 | 0.01 | 0.03 |
| 8 | 93592320 | rs2618131 | -1.269 | 0.516 | 0.949 | 1.721 | 0.01 | 0.052 | 0.052 | 3.499 | 0.13 | 0.25 | 0.11 | 0.18 | 0.11 | 0.1 |
| 8 | 93593396 | rs11993886 | -1.575 | 0.453 | 0.905 | 2.453 | 0.009 | 0.337 | 0.337 | 0.035 | 0.07 | 0.2 | 0.14 | 0.51 | 0.07 | 0.2 |

| | | | | | | | | | | | | | | | | |
|---|---|---|---|---|---|---|---|---|---|---|---|---|---|---|---|---|
| 8 | 93605944 | rs7012115 | -0.641 | 0.297 | 0.415 | 1.302 | -0.002 | -0.01 | -0.01 | 0.211 | 0.31 | 0.32 | 0.45 | 0.25 | 0.3 | 0.34 |
| 8 | 93608054 | rs7832130 | 0.269 | 0.387 | -0.203 | 1.033 | -0.012 | 0.08 | 0.08 | 2.367 | 0.28 | 0.49 | 0.28 | 0.52 | 0.27 | 0.34 |
| 8 | 93613214 | rs1463184 | -1.181 | 0.363 | -0.354 | 0.749 | -0.043 | 0.091 | 0.091 | 0 | 0.6 | 0.82 | 0.74 | 0.76 | 0.59 | 0.7 |
| 8 | 93615407 | rs10097667 | 0.241 | 0.543 | -0.255 | 1.337 | 0.008 | 0.075 | 0.075 | 0.619 | 0.27 | 0.52 | 0.3 | 0.32 | 0.3 | 0.2 |
| 8 | 93618272 | rs748801 | 0.029 | 0.534 | -0.336 | 1.285 | -0.001 | -0.001 | -0.001 | 0 | 0.3 | 0.27 | 0.44 | 0.49 | 0.31 | 0.5 |
| 8 | 93618825 | rs2167029 | 0.404 | 0.647 | -0.859 | 1.4 | -0.01 | -0.011 | -0.011 | 0 | 0.07 | 0.02 | 0.19 | 0.39 | 0.07 | 0.23 |
| 8 | 93620945 | rs1449241 | -0.821 | 0.569 | -0.629 | 0.519 | -0.01 | 0.079 | 0.079 | 0 | 0.58 | 0.77 | 0.73 | 0.81 | 0.6 | 0.7 |
| 8 | 93626295 | rs1551029 | -0.859 | 0.531 | -1.08 | 0.07 | -0.064 | 0.067 | 0.067 | 0 | 0.63 | 0.87 | 0.68 | 0.8 | 0.55 | 0.7 |
| 8 | 93631538 | rs2618126 | -0.864 | 0.458 | -1.099 | 0.036 | -0.062 | 0.067 | 0.067 | 0 | 0.63 | 0.86 | 0.68 | 0.8 | 0.55 | 0.7 |
| 8 | 93637168 | rs2618121 | -0.147 | -0.087 | -0.453 | 0.149 | -0.014 | -0.009 | -0.009 | 0.058 | 0.28 | 0.33 | 0.36 | 0.33 | 0.23 | 0.44 |
| 8 | 93638037 | rs935710 | -1.464 | -0.087 | -0.417 | 0.201 | -0.055 | 0.063 | 0.063 | 0 | 0.64 | 0.84 | 0.67 | 0.8 | 0.56 | 0.7 |
| 8 | 93641556 | rs1868470 | -0.361 | -0.692 | 0.452 | 0.726 | -0.009 | 0.088 | 0.088 | 0 | 0.09 | 0.24 | 0.05 | 0.27 | 0.06 | 0.22 |
| 8 | 93644497 | rs12547140 | 0.216 | -0.896 | 0.452 | 0.431 | 0.001 | -0.016 | -0.016 | 0 | 0.19 | 0.17 | 0.27 | 0.06 | 0.16 | 0.23 |
| 8 | 93645236 | rs10097386 | -1.046 | -0.944 | 0.364 | 0.179 | 0.003 | 0 | 0 | 0 | 0.45 | 0.57 | 0.51 | 0.53 | 0.37 | 0.46 |
| 8 | 93649154 | rs16915148 | 0.554 | -0.999 | 0.361 | -0.014 | 0.002 | -0.028 | -0.028 | 0 | 0.15 | 0.22 | 0.12 | 0.03 | 0.16 | 0.09 |
| 8 | 93650885 | rs1449240 | -1.92 | -0.958 | 0.219 | 0.033 | 0.025 | 0.113 | 0.113 | 0 | 0.54 | 0.83 | 0.57 | 0.79 | 0.43 | 0.67 |
| 8 | 93658348 | rs2595599 | -0.995 | -0.986 | 0.64 | 0.239 | -0.02 | -0.003 | -0.003 | 5.578 | 0.32 | 0.6 | 0.38 | 0.76 | 0.21 | 0.52 |
| 8 | 93667572 | rs7018123 | -0.897 | -0.958 | 0.805 | 0.525 | -0.009 | 0.012 | 0.012 | 0.631 | 0.35 | 0.48 | 0.44 | 0.53 | 0.3 | 0.37 |
| 8 | 93671662 | rs2339657 | 0.778 | -0.944 | 0.78 | 0.493 | 0.007 | 0.036 | 0.036 | 0 | 0.16 | 0.07 | 0.14 | 0.03 | 0.14 | 0.18 |
| 8 | 93672476 | rs9297912 | -0.287 | -0.958 | 0.9 | 0.591 | -0.004 | 0.134 | 0.134 | 0.046 | 0.18 | 0.36 | 0.25 | 0.49 | 0.16 | 0.18 |
| 8 | 93676347 | rs2447955 | 0.256 | -0.956 | 0.937 | 0.337 | -0.024 | 0.048 | 0.048 | 0 | 0.89 | 0.79 | 0.92 | 0.73 | 0.93 | 0.78 |
| 8 | 93677234 | rs16915185 | 0.042 | -0.882 | 0.874 | 0.259 | 0.034 | -0.015 | -0.015 | 0 | 0.05 | 0.19 | 0.04 | 0 | 0.05 | 0.05 |
| 8 | 93681178 | rs2461429 | 1.092 | -0.831 | 0.703 | 0.438 | -0.003 | -0.003 | -0.003 | 0 | 0.58 | 0.44 | 0.67 | 0.61 | 0.66 | 0.52 |
| 8 | 93683272 | rs2461430 | 1.001 | -0.82 | 0.688 | 0.448 | -0.003 | -0.002 | -0.002 | 0 | 0.59 | 0.55 | 0.67 | 0.61 | 0.65 | 0.51 |
| 8 | 93696461 | rs7817246 | -0.802 | -0.798 | 0.321 | 0.699 | -0.002 | 0 | 0 | 0 | 0.23 | 0.23 | 0.23 | 0.35 | 0.2 | 0.38 |
| 8 | 93698134 | rs2879537 | -2.082 | -0.828 | -0.256 | 1.002 | 0.056 | 0.031 | 0.031 | 0 | 0.43 | 0.91 | 0.49 | 0.72 | 0.31 | 0.57 |
| 8 | 93739076 | rs16915219 | -0.255 | -0.588 | 0 | 1.114 | -0.001 | -0.001 | -0.001 | 1.049 | 0.18 | 0.12 | 0.17 | 0.04 | 0.18 | 0.21 |
| 8 | 93791210 | rs12541611 | -0.235 | -0.562 | 0.14 | 1 | 0.003 | 0 | 0 | 1.3 | 0.18 | 0.1 | 0.16 | 0.03 | 0.16 | 0.17 |
| 8 | 93831343 | rs7009343 | 0.751 | -0.475 | 0.127 | 1.167 | 0.004 | 0.022 | 0.022 | 0 | 0.95 | 0.96 | 0.98 | 0.69 | 0.97 | 0.78 |
| 8 | 93871997 | rs1109399 | -1.017 | -0.116 | 0.27 | 0.599 | -0.008 | -0.052 | -0.052 | 0 | 0.3 | 0.3 | 0.24 | 0.07 | 0.21 | 0.27 |
| 8 | 93908488 | rs13249658 | 0.939 | 0.716 | -0.088 | 0.873 | 0.015 | 0.018 | 0.018 | 0.049 | 0.79 | 0.8 | 0.8 | 0.6 | 0.72 | 0.62 |
| 8 | 93932701 | rs7463093 | 1.658 | 0.516 | -0.195 | 0.73 | 0 | 0 | 0 | 0.298 | 0.66 | 0.72 | 0.81 | 0.55 | 0.66 | 0.46 |
| 8 | 93939545 | rs13439759 | 0.19 | 0.632 | -0.431 | 0.905 | -0.003 | 0.059 | 0.059 | 0 | 0.2 | 0.42 | 0.2 | 0.29 | 0.17 | 0.24 |
| 8 | 93942913 | rs4735193 | 1.918 | 0.676 | -0.599 | 0.773 | 0.027 | 0.017 | 0.017 | 0 | 0.49 | 0.27 | 0.61 | 0.34 | 0.46 | 0.24 |
| 8 | 93977130 | rs7016705 | 0.847 | 0.867 | -0.259 | 0.65 | 0.087 | 0.03 | 0.03 | 0.489 | 0.05 | 0.33 | 0.05 | 0 | 0.04 | 0 |
| 8 | 93985173 | rs1847552 | 0.853 | 0.828 | -0.716 | 0.692 | -0.001 | -0.01 | -0.01 | 0.526 | 0.14 | 0.1 | 0.09 | 0.2 | 0.13 | 0.19 |
| 8 | 93995489 | rs172160 | -1.894 | 0.419 | -1.465 | 0.412 | 0.013 | -0.015 | -0.015 | 0 | 0.63 | 0.75 | 0.53 | 0.65 | 0.66 | 0.71 |
| 8 | 93996721 | rs1875882 | 1.057 | 0.661 | -1.425 | 0.403 | -0.021 | -0.007 | -0.007 | 0 | 0.11 | 0.04 | 0.11 | 0.1 | 0.23 | 0.2 |
| 8 | 94017751 | rs6471356 | -0.197 | 0.565 | -1.339 | 0.309 | 0.019 | -0.028 | -0.028 | 0 | 0.24 | 0.48 | 0.22 | 0.2 | 0.21 | 0.24 |
| 8 | 94023146 | rs279955 | -1.994 | 0.252 | -1.478 | 0.396 | 0.008 | -0.004 | -0.004 | 0 | 0.64 | 0.77 | 0.53 | 0.69 | 0.66 | 0.8 |
| 8 | 94032622 | rs279978 | -1.796 | 0.604 | -1.222 | 0.483 | 0.016 | -0.006 | -0.006 | 0 | 0.58 | 0.7 | 0.46 | 0.62 | 0.63 | 0.74 |
| 8 | 94032666 | rs279979 | -0.844 | 0.623 | -1.223 | 0.451 | -0.033 | -0.028 | -0.028 | 0 | 0.39 | 0.24 | 0.32 | 0.49 | 0.45 | 0.56 |
| 8 | 94032743 | rs2339774 | 0.117 | 0.704 | -0.912 | 0.691 | 0.014 | -0.047 | -0.047 | 0.01 | 0.19 | 0.47 | 0.14 | 0.13 | 0.17 | 0.18 |
| 8 | 94034543 | rs279981 | 0.353 | 0.318 | -1.616 | 0.81 | 0.026 | -0.076 | -0.076 | 0.299 | 0.19 | 0.03 | 0.18 | 0.35 | 0.18 | 0.22 |
| 8 | 94034920 | rs172159 | -0.872 | 0.678 | -1.174 | 0.505 | 0.019 | 0.003 | 0.003 | 0 | 0.55 | 0.67 | 0.45 | 0.62 | 0.6 | 0.72 |
| 8 | 94038491 | rs279985 | 0.68 | 1.192 | -0.903 | 0.5 | -0.008 | 0.02 | 0.02 | 0 | 0.62 | 0.29 | 0.56 | 0.29 | 0.57 | 0.6 |
| 8 | 94039383 | rs4735195 | 1.32 | 1.206 | -1.083 | 1.232 | -0.004 | -0.07 | -0.07 | 0 | 0.11 | 0 | 0.06 | 0.35 | 0.09 | 0.19 |
| 8 | 94041758 | rs2553698 | 0.49 | 1.302 | -1.006 | 1.28 | -0.005 | 0.057 | 0.057 | 0 | 0.78 | 0.35 | 0.83 | 0.51 | 0.81 | 0.68 |
| 8 | 94041928 | rs6987915 | -0.266 | 1.367 | -0.243 | 1.144 | 0.103 | 0.079 | 0.079 | 0.104 | 0.1 | 0.49 | 0.1 | 0.12 | 0.1 | 0.12 |
| 8 | 94046456 | rs12550679 | 0.206 | 1.207 | -0.741 | 0.993 | -0.04 | -0.034 | -0.034 | 0 | 0.17 | 0.08 | 0.27 | 0.24 | 0.23 | 0.08 |
| 8 | 94060339 | rs279968 | -1.245 | 1.424 | -0.488 | 0.648 | -0.016 | 0.017 | 0.017 | 0 | 0.42 | 0.72 | 0.46 | 0.64 | 0.48 | 0.46 |
| 8 | 94061810 | rs279971 | -1.211 | 1.906 | -0.134 | 0.75 | 0.068 | 0.028 | 0.028 | 0 | 0.18 | 0.62 | 0.3 | 0.27 | 0.24 | 0.21 |
| 8 | 94062885 | rs279973 | -0.731 | 1.805 | 0.315 | 0.575 | 0.132 | -0.023 | -0.023 | 0.162 | 0.1 | 0.48 | 0.07 | 0.04 | 0.1 | 0.17 |
| 8 | 94068426 | rs7015011 | 0.662 | 1.827 | 0.375 | 0.594 | 0.025 | -0.011 | -0.011 | 0 | 0.05 | 0.07 | 0.04 | 0.04 | 0.08 | 0.15 |
| 8 | 94080831 | rs1472748 | 1.109 | 1.189 | 0.09 | 0.697 | 0.008 | -0.014 | -0.014 | 0 | 0.17 | 0.13 | 0.14 | 0.31 | 0.09 | 0.2 |
| 8 | 94081629 | rs279468 | 0.02 | 1.324 | 0.133 | 0.695 | -0.007 | 0.03 | 0.03 | 2.367 | 0.26 | 0.58 | 0.2 | 0.35 | 0.16 | 0.36 |
| 8 | 94086267 | rs279461 | -1.132 | 1.241 | -0.731 | 0.052 | 0.017 | 0.035 | 0.035 | 0 | 0.59 | 0.83 | 0.62 | 0.72 | 0.59 | 0.7 |
| 8 | 94086392 | rs279460 | -0.179 | 1.412 | -0.027 | 0.294 | 0.072 | -0.014 | -0.014 | 0.008 | 0.07 | 0.23 | 0.04 | 0.04 | 0.07 | 0.16 |
| 8 | 94092222 | rs10808656 | -0.105 | 1.239 | -0.166 | 0.221 | 0 | 0 | 0 | 0 | 0.38 | 0.38 | 0.43 | 0.63 | 0.4 | 0.5 |
| 8 | 94092846 | rs10100582 | 0.398 | 1.011 | -0.911 | 0.227 | -0.007 | 0.005 | 0.005 | 0 | 0.1 | 0.17 | 0.05 | 0.09 | 0.08 | 0.04 |
| 8 | 94110848 | rs279479 | -0.509 | 1.017 | -0.98 | 0.011 | -0.003 | 0.058 | 0.058 | 0.41 | 0.51 | 0.64 | 0.49 | 0.73 | 0.49 | 0.57 |
| 8 | 94123271 | rs279472 | 0.415 | 1.527 | -1.008 | 0.534 | -0.015 | -0.02 | -0.02 | 0.017 | 0.17 | 0.12 | 0.24 | 0.29 | 0.28 | 0.32 |
| 8 | 94128737 | rs1483463 | 0.487 | 0.855 | -1.343 | 0.142 | 0.031 | 0.051 | 0.051 | 4.456 | 0.36 | 0.13 | 0.32 | 0.23 | 0.32 | 0.21 |
| 8 | 94128797 | rs1483462 | -0.189 | 0.786 | -1.254 | 0.131 | 0.007 | 0.015 | 0.015 | 0.674 | 0.44 | 0.42 | 0.4 | 0.29 | 0.39 | 0.36 |
| 8 | 94134191 | rs11776167 | -0.165 | 0.773 | -1.605 | 0.158 | 0.012 | 0.043 | 0.043 | 0.78 | 0.43 | 0.35 | 0.39 | 0.29 | 0.4 | 0.36 |
| 8 | 94139765 | rs1847554 | 0.146 | 0.495 | -1.612 | 0.226 | 0.016 | 0.029 | 0.029 | 2.635 | 0.38 | 0.32 | 0.35 | 0.23 | 0.34 | 0.21 |
| 8 | 94141855 | rs16915660 | 1.056 | 0.655 | -2.373 | 0.487 | 0.001 | -0.035 | -0.035 | 0 | 0.1 | 0.02 | 0.16 | 0.19 | 0.08 | 0.12 |
| 8 | 94143207 | rs1483460 | 0.191 | 0.716 | -2.057 | 0.132 | 0.012 | 0.048 | 0.048 | 4.592 | 0.35 | 0.13 | 0.32 | 0.23 | 0.34 | 0.2 |
| 8 | 94158704 | rs733330 | 0.355 | 0.782 | -2.211 | -0.062 | -0.015 | 0.095 | 0.095 | 1.057 | 0.38 | 0.17 | 0.3 | 0.25 | 0.35 | 0.35 |

| | | | | | | | | | | | | | | | | |
|---|---|---|---|---|---|---|---|---|---|---|---|---|---|---|---|---|
| 8 | 94159583 | rs278578 | 0.381 | 0.858 | -1.672 | 0.025 | -0.009 | 0.083 | 0.083 | 0.412 | 0.45 | 0.29 | 0.43 | 0.23 | 0.44 | 0.43 |
| 8 | 94162412 | rs11991822 | 0.276 | 1.55 | -0.929 | 0.723 | 0.051 | 0.124 | 0.124 | 0.022 | 0.24 | 0.45 | 0.31 | 0.43 | 0.33 | 0.35 |
| 8 | 94167493 | rs7818649 | 1.241 | 1.578 | -1.072 | 0.553 | 0.001 | 0.028 | 0.028 | 0.098 | 0.11 | 0.03 | 0.05 | 0.05 | 0.1 | 0.14 |
| 8 | 94168001 | rs17697619 | 1.241 | 1.578 | -1.072 | 0.553 | 0.001 | 0.028 | 0.028 | 0.098 | 0.11 | 0.03 | 0.05 | 0.05 | 0.1 | 0.14 |
| 8 | 94168620 | rs278565 | -1.346 | 1.606 | -1.002 | 0.764 | -0.013 | -0.025 | -0.025 | 0 | 0.81 | 0.89 | 0.68 | 0.71 | 0.75 | 0.83 |
| 8 | 94170598 | rs2623301 | 0.414 | 1.411 | -0.781 | 0.759 | -0.002 | 0.005 | 0.005 | 0 | 0.08 | 0.06 | 0.04 | 0 | 0.07 | 0.07 |
| 8 | 94174117 | rs278572 | 0.731 | 1.43 | -0.575 | 0.778 | 0.042 | 0.119 | 0.119 | 1.287 | 0.77 | 0.6 | 0.75 | 0.57 | 0.73 | 0.59 |
| 8 | 94177968 | rs278595 | -0.586 | 1.364 | -0.705 | 0.538 | 0 | -0.028 | -0.028 | 0.343 | 0.82 | 0.78 | 0.86 | 1 | 0.81 | 0.9 |
| 8 | 94192318 | rs1374629 | 0.634 | 1.466 | -0.53 | 0.85 | 0.067 | 0.237 | 0.237 | 1.071 | 0.15 | 0.33 | 0.21 | 0.41 | 0.2 | 0.33 |
| 8 | 94195920 | rs278587 | -0.115 | 1.447 | -0.577 | 0.846 | 0 | -0.001 | -0.001 | 0 | 0.92 | 0.9 | 0.89 | 0.92 | 0.93 | 0.86 |
| 8 | 94201147 | rs2449607 | 0.473 | 1.506 | -0.592 | 0.903 | 0.031 | 0.014 | 0.014 | 0.356 | 0.68 | 0.55 | 0.72 | 0.68 | 0.61 | 0.65 |
| 8 | 94210422 | rs278530 | 0.367 | 1.486 | -0.552 | 0.827 | 0.046 | 0.107 | 0.107 | 1.992 | 0.74 | 0.56 | 0.73 | 0.59 | 0.68 | 0.62 |
| 8 | 94218476 | rs7845022 | 2.052 | 1.482 | -0.687 | 0.803 | -0.001 | -0.013 | -0.013 | 0 | 0.1 | 0.1 | 0.09 | 0 | 0.11 | 0.03 |
| 8 | 94237286 | rs1531848 | 0.678 | 1.457 | -0.794 | 0.832 | 0.008 | 0.014 | 0.014 | 0.621 | 0.63 | 0.46 | 0.63 | 0.58 | 0.57 | 0.62 |
| 8 | 94242350 | rs278562 | 0.715 | 1.439 | -0.79 | 0.657 | -0.004 | -0.005 | -0.005 | 0.796 | 0.63 | 0.37 | 0.62 | 0.61 | 0.57 | 0.62 |
| 8 | 94245892 | rs1447314 | -0.008 | 1.312 | -0.985 | 0.628 | -0.003 | -0.013 | -0.013 | 0 | 0.62 | 0.64 | 0.66 | 0.54 | 0.62 | 0.63 |
| 8 | 94250101 | rs1471690 | 0.183 | 1.336 | -1.148 | 0.904 | 0.059 | 0.129 | 0.129 | 0.863 | 0.17 | 0.31 | 0.2 | 0.37 | 0.24 | 0.29 |
| 8 | 94250295 | rs961544 | 1.358 | 1.255 | -1.423 | 0.655 | -0.003 | 0.098 | 0.098 | 0 | 0.31 | 0.05 | 0.34 | 0.15 | 0.27 | 0.31 |
| 8 | 94254642 | rs17699161 | 0.399 | 1.289 | -1.161 | 0.763 | -0.002 | 0.002 | 0.002 | 0 | 0.48 | 0.41 | 0.47 | 0.43 | 0.5 | 0.54 |
| 8 | 94256127 | rs43870 | 0.271 | 0.723 | -1.555 | 0.6 | 0.005 | 0.019 | 0.019 | 0 | 0.66 | 0.6 | 0.7 | 0.54 | 0.63 | 0.55 |
| 8 | 94259105 | rs17771939 | 1.016 | -0.406 | -1.427 | 0.839 | 0.001 | 0.131 | 0.131 | 1.51 | 0.32 | 0.05 | 0.34 | 0.17 | 0.27 | 0.28 |
| 8 | 94259971 | rs16915813 | -0.605 | -0.221 | -0.319 | 1.868 | -0.012 | 0.031 | 0.031 | 1.774 | 0.2 | 0.26 | 0.14 | 0.29 | 0.16 | 0.33 |
| 8 | 94260527 | rs19441 | -0.224 | -0.254 | -0.263 | 1.933 | -0.024 | 0.081 | 0.081 | 0.257 | 0.17 | 0.44 | 0.23 | 0.25 | 0.14 | 0.16 |
| 8 | 94260987 | rs278536 | -0.386 | -0.191 | -0.414 | 1.71 | -0.018 | 0.148 | 0.148 | 0 | 0.18 | 0.39 | 0.24 | 0.4 | 0.15 | 0.23 |
| 8 | 94262608 | rs278539 | -0.908 | -0.008 | -1.028 | 0.991 | 0.03 | 0.029 | 0.029 | 1.545 | 0.37 | 0.69 | 0.38 | 0.72 | 0.29 | 0.51 |
| 8 | 94264684 | rs1374636 | -0.967 | 0.162 | -0.848 | 1.093 | 0.025 | 0.026 | 0.026 | 1.598 | 0.38 | 0.69 | 0.38 | 0.66 | 0.3 | 0.57 |
| 8 | 94273434 | rs16915825 | -0.736 | 0.818 | -0.675 | 1.591 | 0 | 0.024 | 0.024 | 0 | 0.13 | 0.12 | 0.22 | 0.35 | 0.12 | 0.16 |
| 8 | 94276132 | rs17699529 | -0.226 | 0.621 | -0.76 | 0.971 | -0.029 | 0.033 | 0.033 | 0.262 | 0.1 | 0.05 | 0.16 | 0 | 0.18 | 0.18 |
| 8 | 94278345 | rs7002977 | 0.746 | 0.05 | -0.387 | 1.055 | -0.019 | 0.01 | 0.01 | 1.629 | 0.25 | 0.29 | 0.25 | 0.34 | 0.18 | 0.24 |
| 8 | 94281440 | rs6995488 | -1.035 | 0.503 | 0.08 | 1.85 | -0.009 | 0.034 | 0.034 | 0.637 | 0.25 | 0.3 | 0.28 | 0.41 | 0.19 | 0.44 |
| 8 | 94285944 | rs16915833 | -0.193 | -0.113 | 0.048 | 1.907 | 0.001 | 0.011 | 0.011 | 0.103 | 0.14 | 0.11 | 0.2 | 0.34 | 0.07 | 0.2 |
| 8 | 94286289 | rs10106229 | -1.547 | 0.289 | 0.183 | 1.212 | 0.005 | 0.012 | 0.012 | 0 | 0.25 | 0.2 | 0.25 | 0.18 | 0.27 | 0.46 |
| 8 | 94286326 | rs10091769 | -1.546 | 0.288 | 0.2 | 1.211 | 0.005 | 0.01 | 0.01 | 0 | 0.25 | 0.2 | 0.25 | 0.18 | 0.27 | 0.46 |
| 8 | 94296841 | rs10103267 | -0.523 | 0.352 | 0.093 | 1.128 | 0.002 | 0.002 | 0.002 | 0 | 0.14 | 0.11 | 0.11 | 0.12 | 0.11 | 0.23 |
| 8 | 94298184 | rs4735211 | -0.652 | 0.468 | -0.155 | 1.176 | -0.009 | 0.004 | 0.004 | 0 | 0.26 | 0.35 | 0.27 | 0.28 | 0.27 | 0.49 |
| 8 | 94305178 | rs1550883 | -1.164 | 1.018 | 0.424 | 1.631 | 0.015 | 0.015 | 0.015 | 0 | 0.09 | 0.17 | 0.12 | 0.12 | 0.13 | 0.15 |
| 8 | 94310204 | rs4978306 | -1.119 | 1 | 0.417 | 1.624 | 0.001 | 0.035 | 0.035 | 0 | 0.07 | 0.17 | 0.11 | 0.12 | 0.07 | 0.14 |
| 8 | 94313865 | rs7824191 | 0.496 | 0.491 | 1.197 | 0.479 | -0.004 | -0.001 | -0.001 | 0 | 0.67 | 0.42 | 0.72 | 0.72 | 0.72 | 0.71 |
| 8 | 94316905 | rs10808658 | -0.168 | 0.491 | 1.245 | 0.47 | 0.001 | 0.001 | 0.001 | 1.173 | 0.75 | 0.8 | 0.85 | 0.82 | 0.79 | 0.85 |
| 8 | 94327562 | rs368998 | -0.237 | 0.454 | 1.397 | 0.781 | -0.006 | 0.002 | 0.002 | 0.947 | 0.75 | 0.68 | 0.78 | 0.81 | 0.8 | 0.85 |
| 8 | 94330970 | rs277781 | -0.129 | 0.44 | 1.265 | 0.771 | 0.003 | 0.004 | 0.004 | 0.966 | 0.75 | 0.82 | 0.8 | 0.83 | 0.8 | 0.85 |
| 8 | 94339212 | rs277789 | -0.206 | 0.294 | 1.193 | 0.702 | 0.006 | -0.003 | -0.003 | 0 | 0.74 | 0.82 | 0.79 | 0.79 | 0.8 | 0.85 |
| 8 | 94355021 | rs277776 | -0.233 | 0.189 | 1.147 | 0.66 | 0.002 | 0.004 | 0.004 | 1.496 | 0.77 | 0.81 | 0.81 | 0.81 | 0.8 | 0.87 |
| 8 | 94355601 | rs7817416 | -0.692 | 0.349 | 1.113 | -0.074 | -0.004 | 0.031 | 0.031 | 0.9 | 0.52 | 0.59 | 0.48 | 0.38 | 0.55 | 0.54 |
| 8 | 94362144 | rs2339866 | -1.321 | 0.6 | 0.192 | -0.772 | -0.083 | 0.094 | 0.094 | 0 | 0.85 | 0.99 | 0.87 | 1 | 0.75 | 0.92 |
| 8 | 94368227 | rs16915883 | 0.395 | 0.566 | 0.093 | -0.013 | -0.04 | -0.001 | -0.001 | 0.423 | 0.07 | 0.01 | 0.27 | 0.44 | 0.1 | 0.18 |
| 8 | 94370510 | rs13263830 | 0.829 | 0.145 | 0.379 | -0.299 | 0.004 | 0.011 | 0.011 | 1.555 | 0.31 | 0.23 | 0.26 | 0.22 | 0.27 | 0.24 |
| 8 | 94370855 | rs1367793 | 1.17 | 0.062 | 0.67 | -0.358 | 0.001 | -0.016 | -0.016 | 6.315 | 0.25 | 0.31 | 0.25 | 0.16 | 0.3 | 0.3 |
| 8 | 94375844 | rs16915889 | 0.887 | 0.063 | 0.662 | -0.362 | 0 | -0.024 | -0.024 | 0.026 | 0.27 | 0.49 | 0.29 | 0.21 | 0.3 | 0.3 |
| 8 | 94376545 | rs7387715 | -0.153 | 0.174 | 0.503 | -0.307 | -0.032 | 0.003 | 0.003 | 1.751 | 0.58 | 0.8 | 0.55 | 0.43 | 0.56 | 0.54 |
| 8 | 94384501 | rs6991797 | 0.465 | 0.182 | 0.696 | -0.69 | 0.045 | -0.088 | -0.088 | 0 | 0.18 | 0.49 | 0.21 | 0.11 | 0.19 | 0.27 |
| 8 | 94385560 | rs1864673 | -0.308 | 0.204 | 0.636 | -0.723 | 0.008 | -0.039 | -0.039 | 0 | 0.23 | 0.62 | 0.28 | 0.18 | 0.24 | 0.39 |
| 8 | 94391484 | rs12334722 | -0.063 | 0.202 | 0.389 | -0.667 | -0.003 | -0.029 | -0.029 | 0 | 0.06 | 0.12 | 0.07 | 0.03 | 0.05 | 0.12 |
| 8 | 94405276 | rs6471376 | -0.063 | 0.315 | 0.039 | -0.777 | 0.019 | 0.027 | 0.027 | 0.053 | 0.52 | 0.3 | 0.31 | 0.41 | 0.45 | 0.36 |
| 8 | 94407069 | rs6983017 | 1.889 | 0.282 | 0.21 | -0.883 | -0.005 | -0.054 | -0.054 | 0.182 | 0.2 | 0.22 | 0.13 | 0 | 0.18 | 0.04 |
| 8 | 94407211 | rs7006918 | -0.136 | 0.437 | -0.015 | -0.6 | -0.029 | -0.015 | -0.015 | 0 | 0.9 | 0.97 | 0.67 | 0.53 | 0.87 | 0.73 |
| 8 | 94428704 | rs1367794 | -0.854 | 0.021 | -0.266 | -0.881 | -0.025 | 0.022 | 0.022 | 1.01 | 0.83 | 0.99 | 0.86 | 0.92 | 0.86 | 0.94 |
| 8 | 94436215 | rs3857899 | -2.225 | 0.024 | -0.025 | -1.004 | 0.009 | 0.009 | 0.009 | 4.721 | 0.82 | 0.88 | 0.9 | 0.99 | 0.83 | 0.96 |
| 8 | 94439189 | rs902758 | -2.162 | 0.009 | 0.079 | -1.011 | 0.01 | 0.011 | 0.011 | 4.216 | 0.82 | 0.87 | 0.9 | 0.99 | 0.83 | 0.96 |
| 8 | 94468378 | rs7837480 | 0.145 | -0.047 | 0.142 | -0.998 | -0.016 | 0.02 | 0.02 | 0.627 | 0.55 | 0.33 | 0.31 | 0.37 | 0.5 | 0.36 |
| 8 | 94471469 | rs1491947 | -0.158 | -0.067 | -0.014 | -0.77 | -0.009 | 0.008 | 0.008 | 0 | 0.09 | 0.03 | 0.37 | 0.51 | 0.1 | 0.26 |
| 8 | 94504160 | rs1491948 | -1.788 | -0.099 | 0.045 | -0.947 | 0.005 | -0.007 | -0.007 | 1.107 | 0.8 | 0.85 | 0.88 | 0.99 | 0.81 | 0.95 |
| 8 | 94534211 | rs7003293 | 0.016 | -0.28 | -0.595 | -0.802 | 0.016 | -0.013 | -0.013 | 0 | 0.62 | 0.64 | 0.49 | 0.46 | 0.69 | 0.58 |
| 8 | 94537065 | rs4336580 | 0.312 | 0.003 | -0.582 | -0.895 | 0.022 | -0.025 | -0.025 | 0 | 0.61 | 0.66 | 0.49 | 0.51 | 0.67 | 0.6 |
| 8 | 94541853 | rs4487724 | -0.675 | 0.12 | -0.592 | -1.031 | 0.003 | -0.028 | -0.028 | 0.33 | 0.72 | 0.69 | 0.86 | 0.98 | 0.76 | 0.85 |
| 8 | 94547806 | rs10092702 | -1.007 | 0.294 | -0.677 | -0.821 | 0.003 | -0.05 | -0.05 | 0 | 0.75 | 0.85 | 0.57 | 0.52 | 0.74 | 0.78 |
| 8 | 94573456 | rs7005045 | -1.405 | 0.591 | -0.707 | -0.779 | 0.003 | -0.081 | -0.081 | 0.003 | 0.78 | 0.9 | 0.6 | 0.52 | 0.76 | 0.79 |
| 8 | 94583653 | rs4236810 | -2.633 | 0.593 | -0.662 | -0.823 | 0.004 | -0.001 | -0.001 | 0 | 0.83 | 0.89 | 0.89 | 1 | 0.81 | 0.96 |
| 8 | 94587666 | rs4617121 | 0.972 | 0.932 | -1.027 | -1.054 | -0.039 | 0.056 | 0.056 | 0.012 | 0.25 | 0.07 | 0.26 | 0.19 | 0.29 | 0.37 |

| | | | | | | | | | | | | | | | | |
|---|---|---|---|---|---|---|---|---|---|---|---|---|---|---|---|---|
| 8 | 94627082 | rs10099602 | 0.493 | 0.839 | -0.322 | -0.87 | -0.001 | 0.003 | 0.003 | 0.007 | 0.39 | 0.35 | 0.27 | 0.32 | 0.38 | 0.3 |
| 8 | 94633233 | rs4567005 | 0.44 | 0.91 | -0.662 | -0.9 | -0.013 | -0.008 | -0.008 | 0.652 | 0.39 | 0.53 | 0.25 | 0.33 | 0.29 | 0.29 |
| 8 | 94639779 | rs11783982 | -1.979 | 1.123 | -0.909 | -0.704 | -0.024 | 0.03 | 0.03 | 0 | 0.94 | 0.99 | 0.69 | 0.52 | 0.93 | 0.82 |
| 8 | 94648740 | rs13274827 | 0.188 | 0.932 | -0.776 | -0.934 | 0.023 | 0.021 | 0.021 | 0.56 | 0.44 | 0.64 | 0.31 | 0.32 | 0.3 | 0.37 |
| 8 | 94654207 | rs12056774 | 0.473 | 0.6 | -0.879 | -1.014 | -0.006 | -0.001 | -0.001 | 0 | 0.35 | 0.63 | 0.26 | 0.32 | 0.22 | 0.31 |
| 8 | 94662817 | rs13281056 | 0.132 | 1.094 | -1.003 | -1.168 | -0.009 | 0.068 | 0.068 | 0.37 | 0.36 | 0.22 | 0.33 | 0.18 | 0.37 | 0.47 |
| 8 | 94667346 | rs11995221 | 0.943 | 1.182 | -0.864 | -1.104 | -0.008 | 0.046 | 0.046 | 0.265 | 0.52 | 0.68 | 0.41 | 0.34 | 0.51 | 0.31 |
| 8 | 94671580 | rs7814823 | 0.212 | 1.223 | -0.89 | -1.103 | -0.01 | 0.05 | 0.05 | 0.124 | 0.35 | 0.23 | 0.33 | 0.18 | 0.37 | 0.41 |
| 8 | 94671771 | rs2976369 | 2.34 | 1.487 | -0.852 | -0.816 | -0.03 | 0.034 | 0.034 | 0.432 | 0.18 | 0.08 | 0.14 | 0 | 0.28 | 0.05 |
| 8 | 94672947 | rs2976370 | -1.6 | 1.354 | -0.742 | -0.754 | -0.013 | -0.023 | -0.023 | 0 | 0.75 | 0.81 | 0.6 | 0.53 | 0.65 | 0.78 |
| 8 | 94675612 | rs11786927 | 1.8 | 1.06 | -0.805 | -0.641 | -0.001 | -0.007 | -0.007 | 0 | 0.05 | 0.07 | 0.02 | 0.02 | 0.05 | 0.05 |
| 8 | 94690198 | rs2914960 | 1.027 | 1.539 | -0.881 | -0.986 | -0.01 | 0.048 | 0.048 | 0.767 | 0.37 | 0.22 | 0.33 | 0.13 | 0.45 | 0.3 |
| 8 | 94692527 | rs11781676 | 0.543 | 1.284 | -0.681 | -0.752 | 0.039 | 0.017 | 0.017 | 0 | 0.39 | 0.17 | 0.24 | 0.34 | 0.3 | 0.32 |
| 8 | 94700401 | rs17710303 | 0.54 | 1.265 | -0.65 | -0.684 | -0.016 | 0.029 | 0.029 | 0.002 | 0.06 | 0.01 | 0.06 | 0.04 | 0.05 | 0.15 |
| 8 | 94711093 | rs10105040 | 0.584 | 1.037 | -0.739 | -0.815 | 0.048 | 0.008 | 0.008 | 0 | 0.38 | 0.17 | 0.23 | 0.33 | 0.29 | 0.3 |
| 8 | 94715291 | rs6471393 | -1.563 | 1.386 | -0.424 | -0.838 | -0.022 | -0.042 | -0.042 | 0 | 0.77 | 0.79 | 0.63 | 0.51 | 0.69 | 0.74 |
| 8 | 94721057 | rs2976366 | -1.128 | 1.291 | 0.191 | -0.857 | 0.021 | 0.079 | 0.079 | 0.009 | 0.73 | 0.57 | 0.62 | 0.51 | 0.68 | 0.69 |
| 8 | 94739345 | rs2914959 | 2.001 | 1.286 | 0.25 | -0.861 | -0.103 | -0.16 | -0.16 | 2.049 | 0.19 | 0.02 | 0.33 | 0.48 | 0.3 | 0.24 |
| 8 | 94742350 | rs2914953 | 0.856 | 1.248 | 0.534 | -0.818 | 0.016 | 0.074 | 0.074 | 0.05 | 0.29 | 0.55 | 0.39 | 0.5 | 0.32 | 0.31 |
| 8 | 94744813 | rs10504930 | 1.089 | 1.248 | 0.536 | -0.818 | 0.021 | 0.078 | 0.078 | 0.016 | 0.27 | 0.43 | 0.38 | 0.5 | 0.32 | 0.32 |
| 8 | 94744960 | rs10504931 | 2.506 | 1.201 | 0.102 | -1.112 | -0.078 | 0.032 | 0.032 | 0.099 | 0.19 | 0.01 | 0.18 | 0.12 | 0.23 | 0.23 |
| 8 | 94745050 | rs11549490 | 0.979 | 0.972 | 0.209 | -1.044 | 0.014 | 0.068 | 0.068 | 0.04 | 0.29 | 0.56 | 0.39 | 0.5 | 0.32 | 0.31 |
| 8 | 94745386 | rs16916186 | 0.992 | 0.972 | 0.209 | -1.044 | 0.015 | 0.069 | 0.069 | 0.021 | 0.29 | 0.56 | 0.39 | 0.5 | 0.32 | 0.31 |
| 8 | 94749565 | rs16916191 | 0.968 | 0.974 | 0.209 | -1.043 | 0.014 | 0.068 | 0.068 | 0.049 | 0.29 | 0.56 | 0.39 | 0.5 | 0.32 | 0.31 |
| 8 | 94755226 | rs3134012 | -1.626 | 1.24 | 0.39 | -0.634 | -0.022 | 0.031 | 0.031 | 0.008 | 0.78 | 0.86 | 0.74 | 0.85 | 0.73 | 0.71 |
| 8 | 94760940 | rs3097422 | -0.408 | 1.129 | 0.437 | -0.64 | -0.021 | 0.046 | 0.046 | 0.094 | 0.65 | 0.75 | 0.63 | 0.83 | 0.6 | 0.63 |
| 8 | 94761486 | rs3097423 | -0.371 | 1.125 | 0.435 | -0.642 | -0.02 | 0.048 | 0.048 | 0.078 | 0.65 | 0.75 | 0.63 | 0.83 | 0.6 | 0.63 |
| 8 | 94808165 | rs3134031 | -0.343 | 1.204 | 0.597 | -0.033 | -0.021 | 0.018 | 0.018 | 0 | 0.65 | 0.76 | 0.67 | 0.73 | 0.61 | 0.66 |
| 8 | 94831346 | rs3133973 | -0.41 | 1.025 | 0.474 | -0.661 | -0.036 | 0.077 | 0.077 | 0.145 | 0.65 | 0.81 | 0.65 | 0.83 | 0.6 | 0.65 |
| 8 | 94837810 | rs12674712 | -0.01 | 1.034 | 0.488 | -0.695 | 0.007 | -0.057 | -0.057 | 0 | 0.61 | 0.61 | 0.64 | 0.84 | 0.59 | 0.59 |
| 8 | 94838313 | rs12675188 | 0.033 | 0.978 | 0.328 | -0.698 | -0.003 | 0.016 | 0.016 | 0 | 0.59 | 0.4 | 0.61 | 0.84 | 0.61 | 0.59 |
| 8 | 94856652 | rs4735243 | -0.041 | 0.281 | 0.327 | -0.628 | 0.006 | -0.059 | -0.059 | 0.75 | 0.6 | 0.53 | 0.59 | 0.85 | 0.5 | 0.6 |
| 8 | 94868279 | rs10100603 | 1.093 | 0.422 | -0.265 | -0.548 | -0.001 | -0.021 | -0.021 | 0 | 0.11 | 0.1 | 0.11 | 0 | 0.13 | 0.04 |
| 8 | 94873128 | rs4236814 | -0.108 | -0.022 | -0.369 | -0.41 | 0.006 | 0.036 | 0.036 | 0.081 | 0.57 | 0.67 | 0.59 | 0.68 | 0.47 | 0.57 |
| 8 | 94882313 | rs7826194 | -0.355 | -0.529 | -0.679 | -0.305 | -0.006 | 0.081 | 0.081 | 0.584 | 0.59 | 0.81 | 0.62 | 0.72 | 0.45 | 0.58 |
| 8 | 94892120 | rs4236815 | -1.778 | -0.459 | -0.975 | -0.344 | -0.158 | 0.094 | 0.094 | 0.035 | 0.76 | 0.97 | 0.75 | 0.89 | 0.59 | 0.75 |
| 8 | 94895433 | rs4735246 | 0.066 | -2.479 | -0.011 | 0.012 | 0.008 | 0.015 | 0.015 | 0.002 | 0.52 | 0.46 | 0.55 | 0.68 | 0.43 | 0.49 |
| 8 | 94898694 | rs10956888 | 0.023 | -2.417 | 0.035 | 0.009 | 0.003 | 0.003 | 0.003 | 0.006 | 0.52 | 0.49 | 0.54 | 0.69 | 0.43 | 0.48 |
| 8 | 94908250 | rs4993569 | -0.528 | -2.289 | 0.139 | 0.069 | 0 | 0.01 | 0.01 | 0.091 | 0.58 | 0.63 | 0.59 | 0.7 | 0.44 | 0.57 |
| 8 | 94909097 | rs4735255 | 0.917 | -2.737 | -0.043 | 0.401 | -0.004 | 0.049 | 0.049 | 0 | 0.25 | 0.34 | 0.24 | 0.44 | 0.24 | 0.28 |
| 8 | 94909829 | rs7837769 | -0.35 | -2.779 | -0.18 | -0.752 | 0.002 | 0.001 | 0.001 | 0.244 | 0.34 | 0.35 | 0.38 | 0.26 | 0.21 | 0.31 |
| 8 | 94922985 | rs7830919 | -0.638 | -2.502 | 0.067 | -1.06 | 0.008 | 0.012 | 0.012 | 0.265 | 0.54 | 0.37 | 0.57 | 0.66 | 0.41 | 0.5 |
| 8 | 94928911 | rs28507397 | -0.695 | -2.498 | 0.108 | -1.058 | 0.002 | 0.003 | 0.003 | 0.151 | 0.54 | 0.4 | 0.57 | 0.66 | 0.41 | 0.5 |
| 8 | 94938346 | rs10099485 | -0.638 | -2.577 | 0.189 | -1.05 | -0.001 | -0.002 | -0.002 | 0 | 0.55 | 0.49 | 0.62 | 0.68 | 0.41 | 0.5 |
| 8 | 94942750 | rs3115899 | -0.089 | -2.631 | -0.197 | -1.065 | 0.054 | 0.021 | 0.021 | 0.416 | 0.46 | 0.25 | 0.5 | 0.62 | 0.33 | 0.47 |
| 8 | 94944609 | rs3133976 | -0.174 | -2.612 | -0.226 | -1.057 | 0.043 | 0.024 | 0.024 | 0.411 | 0.47 | 0.3 | 0.55 | 0.64 | 0.33 | 0.47 |
| 8 | 94948378 | rs3133980 | -0.926 | -2.329 | 0.191 | -0.448 | 0 | -0.007 | -0.007 | 0 | 0.08 | 0.2 | 0.13 | 0.07 | 0.08 | 0.04 |
| 8 | 94948595 | rs3115904 | -0.86 | -2.322 | 0.154 | -0.461 | 0.001 | -0.014 | -0.014 | 0.006 | 0.08 | 0.18 | 0.09 | 0.05 | 0.08 | 0.03 |
| 8 | 94954056 | rs4236821 | -0.536 | -2.717 | 0.029 | -0.379 | -0.003 | -0.001 | -0.001 | 0 | 0.33 | 0.31 | 0.42 | 0.63 | 0.18 | 0.41 |
| 8 | 94968525 | rs3134003 | -0.972 | -1.966 | 0.456 | -0.096 | -0.011 | 0.004 | 0.004 | 0 | 0.06 | 0.18 | 0.05 | 0.05 | 0.02 | 0.03 |
| 8 | 94969154 | rs17785173 | 0.199 | -2.036 | 0.301 | -0.267 | -0.01 | -0.006 | -0.006 | 0.095 | 0.13 | 0.07 | 0.14 | 0.19 | 0.18 | 0.11 |
| 8 | 94973212 | rs3134010 | -0.872 | -2.152 | 0.303 | -0.344 | -0.013 | -0.007 | -0.007 | 0 | 0.29 | 0.66 | 0.28 | 0.29 | 0.24 | 0.23 |
| 8 | 94974328 | rs755600 | 0.027 | -2.105 | 0.584 | -0.314 | -0.008 | -0.01 | -0.01 | 1.73 | 0.23 | 0.33 | 0.18 | 0.21 | 0.22 | 0.19 |
| 8 | 94987448 | rs10956893 | -0.841 | -2.171 | 0.275 | -0.118 | -0.001 | -0.002 | -0.002 | 0 | 0.45 | 0.57 | 0.47 | 0.7 | 0.38 | 0.42 |
| 8 | 94991734 | rs10808664 | -1.069 | -2.008 | -0.042 | -0.145 | 0.049 | 0.094 | 0.094 | 0 | 0.48 | 0.76 | 0.48 | 0.71 | 0.38 | 0.42 |
| 8 | 94992958 | rs16916331 | 0.052 | -1.572 | -0.383 | -0.019 | 0.008 | -0.032 | -0.032 | 0 | 0.23 | 0.14 | 0.24 | 0.47 | 0.18 | 0.22 |
| 8 | 94998256 | rs12681770 | 0.186 | -1.017 | -0.439 | -0.638 | 0.006 | 0.009 | 0.009 | 0 | 0.42 | 0.34 | 0.46 | 0.66 | 0.39 | 0.4 |
| 8 | 94999197 | rs4579497 | -0.244 | -0.38 | -0.87 | -0.522 | 0.027 | 0.057 | 0.057 | 0 | 0.53 | 0.71 | 0.52 | 0.71 | 0.42 | 0.49 |
| 8 | 95002352 | rs7002680 | -0.194 | -0.129 | -1.219 | -0.399 | 0 | -0.001 | -0.001 | 0.182 | 0.53 | 0.55 | 0.54 | 0.71 | 0.46 | 0.52 |
| 8 | 95006099 | rs2978129 | 0.184 | -0.439 | -1.803 | -0.672 | 0.035 | 0.138 | 0.138 | 0 | 0.63 | 0.86 | 0.78 | 0.91 | 0.66 | 0.67 |
| 8 | 95007871 | rs2978130 | 0.131 | -0.506 | -1.683 | 0.061 | -0.005 | -0.025 | -0.025 | 0 | 0.68 | 0.81 | 0.71 | 0.49 | 0.66 | 0.65 |
| 8 | 95007941 | rs3018850 | 0.817 | -1.532 | -2.667 | -0.488 | -0.156 | 0.116 | 0.116 | 0.643 | 0.37 | 0.04 | 0.36 | 0.17 | 0.45 | 0.33 |
| 8 | 95008144 | rs16916361 | -0.337 | -2.148 | -2.186 | 0.563 | 0.047 | -0.058 | -0.058 | 0 | 0.24 | 0.1 | 0.17 | 0.41 | 0.16 | 0.34 |
| 8 | 95009020 | rs876324 | -0.993 | -2.434 | -1.436 | 0.381 | -0.002 | 0.001 | 0.001 | 0.127 | 0.41 | 0.61 | 0.49 | 0.74 | 0.24 | 0.54 |
| 8 | 95009811 | rs877979 | -0.534 | -2.323 | -0.418 | 0.324 | -0.022 | 0.031 | 0.031 | 0.191 | 0.18 | 0.34 | 0.34 | 0.32 | 0.12 | 0.2 |
| 8 | 95011132 | rs7016698 | -1.244 | -2.357 | -0.521 | 0.489 | -0.026 | -0.049 | -0.049 | 0 | 0.78 | 0.91 | 0.75 | 0.73 | 0.74 | 0.79 |
| 8 | 95011920 | rs3018851 | -1.266 | -2.62 | -0.109 | 0.511 | 0.066 | 0.036 | 0.036 | 1.652 | 0.46 | 0.7 | 0.52 | 0.71 | 0.29 | 0.56 |
| 8 | 95012376 | rs7008341 | 0.17 | -2.726 | 0.158 | 0.5 | 0 | -0.001 | -0.001 | 0.011 | 0.22 | 0.22 | 0.16 | 0.13 | 0.2 | 0.29 |
| 8 | 95023498 | rs2978169 | 0.282 | -2.571 | 0.254 | 0.698 | -0.013 | 0.058 | 0.058 | 0 | 0.13 | 0.24 | 0.29 | 0.32 | 0.07 | 0.19 |

| | | | | | | | | | | | | | | | | |
|---|---|---|---|---|---|---|---|---|---|---|---|---|---|---|---|---|
| 8 | 95027506 | rs4276654 | 1.49 | -2.601 | -0.127 | 0.764 | -0.038 | -0.081 | -0.081 | 0.748 | 0.16 | 0.01 | 0.2 | 0.22 | 0.19 | 0.12 |
| 8 | 95030552 | rs3018860 | -1.416 | -2.871 | 0.432 | 0.627 | 0.072 | 0.019 | 0.019 | 1.737 | 0.41 | 0.7 | 0.47 | 0.72 | 0.23 | 0.51 |
| 8 | 95032386 | rs10089749 | -0.047 | -2.879 | 0.652 | 0.638 | -0.032 | -0.018 | -0.018 | 0 | 0.21 | 0.3 | 0.15 | 0.15 | 0.15 | 0.25 |
| 8 | 95034705 | rs3018862 | 0.285 | -2.672 | 0.611 | 0.784 | -0.019 | 0.086 | 0.086 | 0 | 0.12 | 0.22 | 0.29 | 0.36 | 0.07 | 0.19 |
| 8 | 95037331 | rs6995326 | -0.599 | -2.558 | 0.412 | 0.486 | -0.026 | 0.012 | 0.012 | 0.062 | 0.25 | 0.3 | 0.13 | 0.36 | 0.12 | 0.3 |
| 8 | 95042062 | rs4449754 | 0.499 | -2.24 | 0.401 | 0.046 | -0.003 | -0.035 | -0.035 | 0 | 0.59 | 0.45 | 0.4 | 0.12 | 0.66 | 0.42 |
| 8 | 95044718 | rs2978142 | 0.318 | -2.07 | 0.446 | 0.521 | -0.015 | 0.001 | 0.001 | 0.02 | 0.67 | 0.43 | 0.51 | 0.37 | 0.72 | 0.54 |
| 8 | 95046638 | rs4735264 | -0.817 | -2.109 | 1.169 | 1.125 | -0.026 | 0.058 | 0.058 | 0 | 0.14 | 0.37 | 0.27 | 0.21 | 0.1 | 0.21 |
| 8 | 95047412 | rs4735265 | -1.271 | -2.34 | 1.054 | 0.959 | -0.033 | -0.004 | -0.004 | 1.262 | 0.26 | 0.33 | 0.34 | 0.29 | 0.16 | 0.37 |
| 8 | 95050753 | rs2978138 | -2.063 | -2.485 | 0.943 | 0.875 | 0.028 | -0.017 | -0.017 | 0 | 0.36 | 0.68 | 0.47 | 0.63 | 0.2 | 0.48 |
| 8 | 95065264 | rs10086104 | -0.695 | -2.317 | 0.562 | 0.642 | 0.019 | 0.064 | 0.064 | 0.341 | 0.19 | 0.07 | 0.15 | 0.11 | 0.17 | 0.15 |
| 8 | 95065747 | rs10105107 | -0.656 | -2.329 | 0.752 | 0.636 | 0.019 | 0.04 | 0.04 | 0.112 | 0.2 | 0.12 | 0.15 | 0.11 | 0.17 | 0.15 |
| 8 | 95067942 | rs7013178 | -1.283 | -2.532 | 1.591 | 0.825 | -0.038 | 0.095 | 0.095 | 1.464 | 0.14 | 0.54 | 0.25 | 0.19 | 0.05 | 0.21 |
| 8 | 95068998 | rs3018892 | -1.625 | -2.646 | 1.888 | 0.918 | -0.04 | 0.145 | 0.145 | 0 | 0.17 | 0.32 | 0.32 | 0.51 | 0.08 | 0.32 |
| 8 | 95071420 | rs3018897 | -2.351 | -2.684 | 1.585 | 0.905 | 0.072 | -0.024 | -0.024 | 0 | 0.4 | 0.76 | 0.51 | 0.62 | 0.25 | 0.47 |
| 8 | 95073377 | rs3018898 | 0.024 | -2.437 | 1.577 | 1.16 | -0.011 | 0.178 | 0.178 | 0 | 0.06 | 0.16 | 0.11 | 0.34 | 0.03 | 0.11 |
| 8 | 95074595 | rs3018899 | -2.228 | -2.548 | 1.874 | 1.167 | -0.071 | 0.167 | 0.167 | 0 | 0.2 | 0.41 | 0.33 | 0.55 | 0.08 | 0.3 |
| 8 | 95078169 | rs3018901 | -0.449 | -2.498 | 1.886 | 1.314 | -0.008 | 0.269 | 0.269 | 0 | 0.07 | 0.31 | 0.13 | 0.35 | 0.03 | 0.12 |
| 8 | 95078190 | rs3018902 | -1.738 | -2.384 | 1.61 | 1.178 | -0.048 | 0.08 | 0.08 | 0 | 0.26 | 0.54 | 0.28 | 0.47 | 0.19 | 0.25 |
| 8 | 95078963 | rs2978151 | -2.761 | -2.443 | 1.612 | 1.181 | 0.017 | -0.003 | -0.003 | 0 | 0.39 | 0.69 | 0.5 | 0.67 | 0.25 | 0.44 |
| 8 | 95083587 | rs10096362 | -0.236 | -2.148 | 0.926 | 0.935 | 0.008 | 0.034 | 0.034 | 0.067 | 0.16 | 0.05 | 0.14 | 0.12 | 0.16 | 0.14 |
| 8 | 95084753 | rs4734267 | -0.391 | -1.931 | 0.998 | 1.073 | -0.025 | 0.017 | 0.017 | 0 | 0.08 | 0.11 | 0.18 | 0.19 | 0.02 | 0.16 |
| 8 | 95086899 | rs3018843 | -0.482 | -1.222 | 0.38 | 0.096 | -0.001 | -0.006 | -0.006 | 0 | 0.45 | 0.35 | 0.5 | 0.63 | 0.39 | 0.54 |
| 8 | 95086921 | rs2978156 | -0.246 | -1.348 | 0.469 | 0.036 | -0.009 | -0.019 | -0.019 | 0 | 0.43 | 0.23 | 0.48 | 0.63 | 0.38 | 0.55 |
| 8 | 95087055 | rs3018845 | -0.827 | -1.267 | 0.613 | -0.152 | 0.002 | -0.003 | -0.003 | 0 | 0.44 | 0.53 | 0.49 | 0.42 | 0.49 | 0.44 |
| 8 | 95087796 | rs2978157 | 0.081 | -1.333 | 0.774 | -0.092 | -0.004 | -0.004 | -0.004 | 0 | 0.31 | 0.25 | 0.32 | 0.42 | 0.32 | 0.38 |
| 8 | 95090785 | rs16916461 | 0.208 | -1.921 | 0.836 | -0.002 | -0.002 | 0 | 0 | 0 | 0.17 | 0.16 | 0.18 | 0.24 | 0.09 | 0.2 |
| 8 | 95093852 | rs2978140 | -2.143 | -2.012 | 0.579 | -0.086 | 0.085 | -0.013 | -0.013 | 0 | 0.47 | 0.83 | 0.5 | 0.64 | 0.31 | 0.5 |
| 8 | 95095966 | rs7013751 | 0.368 | -2.046 | 1.001 | -0.126 | 0.032 | -0.069 | -0.069 | 0 | 0.12 | 0.46 | 0.15 | 0 | 0.11 | 0.05 |
| 8 | 95097669 | rs2513755 | 0.278 | -1.99 | 0.528 | -0.08 | -0.02 | 0.024 | 0.024 | 0 | 0.84 | 0.75 | 0.81 | 0.61 | 0.95 | 0.79 |
| 8 | 95104497 | rs2513759 | -0.571 | -2.021 | 0.445 | -0.124 | -0.015 | -0.017 | -0.017 | 0 | 0.27 | 0.6 | 0.3 | 0.25 | 0.22 | 0.25 |
| 8 | 95107477 | rs2513764 | 0.404 | -2.11 | 0.416 | -0.073 | -0.039 | 0.049 | 0.049 | 0 | 0.84 | 0.69 | 0.8 | 0.61 | 0.95 | 0.79 |
| 8 | 95114655 | rs10100317 | 1.182 | -1.952 | -0.024 | -0.103 | 0.031 | -0.014 | -0.014 | 0 | 0.59 | 0.35 | 0.56 | 0.39 | 0.76 | 0.52 |
| 8 | 95124272 | rs2513782 | 1.134 | -1.299 | -0.248 | -0.223 | -0.007 | 0.023 | 0.023 | 0 | 0.07 | 0.03 | 0.08 | 0 | 0.1 | 0.05 |
| 8 | 95125708 | rs2446826 | -0.846 | -1.769 | -0.323 | -0.514 | 0.001 | 0 | 0 | 0.258 | 0.33 | 0.66 | 0.3 | 0.18 | 0.24 | 0.29 |
| 8 | 95126478 | rs2513784 | -0.025 | -1.774 | -0.494 | -0.67 | 0.01 | -0.001 | -0.001 | 0 | 0.17 | 0.13 | 0.11 | 0 | 0.15 | 0.07 |
| 8 | 95126706 | rs12546486 | 1.221 | -1.822 | -1.001 | -0.58 | -0.041 | 0.015 | 0.015 | 0.003 | 0.37 | 0.14 | 0.38 | 0.29 | 0.55 | 0.43 |
| 8 | 95127612 | rs2446823 | 0.799 | -1.888 | -1.105 | -0.584 | -0.007 | -0.042 | -0.042 | 0 | 0.15 | 0.18 | 0.1 | 0 | 0.12 | 0.06 |
| 8 | 95130632 | rs2340019 | 0.012 | -1.97 | -0.8 | -0.462 | -0.01 | -0.012 | -0.012 | 0 | 0.58 | 0.54 | 0.67 | 0.69 | 0.69 | 0.68 |
| 8 | 95131183 | rs2249967 | -1.352 | -1.99 | -0.74 | -0.461 | 0.006 | -0.022 | -0.022 | 0 | 0.75 | 0.86 | 0.79 | 0.69 | 0.81 | 0.76 |
| 8 | 95132887 | rs10956902 | 2.324 | -1.781 | -1.177 | -0.768 | 0.016 | -0.04 | -0.04 | 0 | 0.22 | 0.07 | 0.19 | 0.29 | 0.2 | 0.33 |
| 8 | 95139766 | rs9417 | -1.597 | -1.637 | -1.249 | -1.44 | 0.004 | -0.018 | -0.018 | 0 | 0.74 | 0.85 | 0.77 | 0.69 | 0.81 | 0.74 |
| 8 | 95143138 | rs2513797 | 1.289 | -1.737 | -1.214 | -1.472 | 0.01 | 0.001 | 0.001 | 0.016 | 0.14 | 0.13 | 0.1 | 0 | 0.12 | 0.06 |
| 8 | 95143172 | rs1051624 | 0.662 | -1.811 | -1.294 | -1.362 | -0.009 | -0.011 | -0.011 | 0 | 0.41 | 0.28 | 0.39 | 0.53 | 0.57 | 0.48 |
| 8 | 95150036 | rs6471432 | 0.914 | -1.698 | -1.43 | -1.339 | -0.014 | -0.018 | -0.018 | 0.445 | 0.2 | 0.32 | 0.28 | 0.16 | 0.14 | 0.21 |
| 8 | 95153257 | rs2340022 | 2.037 | -1.644 | -1.447 | -1.33 | -0.082 | 0.068 | 0.068 | 0 | 0.14 | 0.04 | 0.19 | 0 | 0.35 | 0.08 |
| 8 | 95158259 | rs2251734 | -0.34 | -1.618 | -1.418 | -1.302 | -0.016 | -0.039 | -0.039 | 0 | 0.34 | 0.45 | 0.38 | 0.16 | 0.26 | 0.28 |
| 8 | 95158765 | rs7837503 | 2.575 | -1.723 | -1.674 | -1.217 | 0.004 | 0.029 | 0.029 | 0 | 0.15 | 0 | 0.06 | 0.06 | 0.1 | 0.22 |
| 8 | 95159158 | rs2514801 | -1.767 | -1.215 | -1.394 | -1.363 | 0.007 | -0.013 | -0.013 | 1.55 | 0.76 | 0.81 | 0.81 | 0.47 | 0.8 | 0.61 |
| 8 | 95166730 | rs2514804 | -1.34 | -1.172 | -1.024 | -0.851 | -0.024 | -0.018 | -0.018 | 0 | 0.65 | 0.73 | 0.64 | 0.47 | 0.49 | 0.57 |
| 8 | 95170675 | rs2446869 | -0.62 | -0.389 | -1.479 | -1.179 | 0.009 | 0.014 | 0.014 | 0.43 | 0.46 | 0.4 | 0.37 | 0.27 | 0.36 | 0.38 |
| 8 | 95171183 | rs3779687 | -0.26 | 0.412 | -1.148 | -0.466 | -0.001 | -0.001 | -0.001 | 0 | 0.32 | 0.29 | 0.26 | 0.26 | 0.24 | 0.31 |
| 8 | 95173672 | rs4446705 | -0.316 | 0.596 | -1.685 | -0.552 | -0.028 | 0.054 | 0.054 | 0.235 | 0.34 | 0.15 | 0.22 | 0.2 | 0.33 | 0.24 |
| 8 | 95175109 | rs13256327 | -0.12 | 0.585 | -1.661 | -1.065 | -0.025 | 0.11 | 0.11 | 0 | 0.31 | 0.13 | 0.19 | 0.1 | 0.3 | 0.21 |
| 8 | 95176015 | rs11776675 | 0.531 | 0.452 | -2.16 | -1.597 | -0.024 | 0.212 | 0.212 | 0.369 | 0.4 | 0.04 | 0.2 | 0.13 | 0.33 | 0.37 |
| 8 | 95176118 | rs11786144 | 0.214 | 0.416 | -1.22 | -1.443 | 0.052 | 0.075 | 0.075 | 0 | 0.49 | 0.62 | 0.28 | 0.29 | 0.33 | 0.47 |
| 8 | 95176404 | rs7842215 | 0.117 | 0.291 | -1.211 | -1.339 | 0.057 | 0.119 | 0.119 | 0 | 0.52 | 0.78 | 0.32 | 0.32 | 0.37 | 0.51 |
| 8 | 95181246 | rs2514809 | -0.501 | 0.541 | -1.643 | -1.522 | -0.014 | -0.009 | -0.009 | 4.165 | 0.69 | 0.94 | 0.58 | 0.34 | 0.7 | 0.6 |
| 8 | 95183490 | rs2340029 | 1.136 | 0.934 | -1.353 | -1.62 | -0.014 | 0.113 | 0.113 | 0.048 | 0.45 | 0.21 | 0.27 | 0.32 | 0.43 | 0.48 |
| 8 | 95188583 | rs16912 | 0.014 | 1.204 | -1.167 | -1.554 | -0.015 | -0.002 | -0.002 | 0 | 0.14 | 0.09 | 0.22 | 0 | 0.26 | 0.06 |
| 8 | 95191326 | rs2340031 | -0.009 | 1.204 | -1.231 | -1.554 | -0.043 | 0.036 | 0.036 | 0 | 0.14 | 0.07 | 0.21 | 0 | 0.26 | 0.06 |
| 8 | 95195548 | rs1600765 | -0.802 | 1.172 | -1.285 | -1.525 | 0.028 | -0.019 | -0.019 | 0.674 | 0.72 | 0.94 | 0.6 | 0.37 | 0.76 | 0.62 |
| 8 | 95196794 | rs10098632 | 0.405 | 1.355 | 0.133 | -1.551 | 0.113 | 0.023 | 0.023 | 0 | 0.06 | 0.28 | 0.05 | 0.03 | 0.05 | 0.02 |
| 8 | 95199215 | rs2613217 | 0.727 | 1.186 | -0.456 | -1.521 | 0.003 | 0.118 | 0.118 | 0 | 0.52 | 0.26 | 0.32 | 0.31 | 0.45 | 0.52 |
| 8 | 95205117 | rs10956904 | 0.921 | 1.139 | -0.75 | -1.554 | -0.003 | 0.183 | 0.183 | 0 | 0.49 | 0.14 | 0.29 | 0.32 | 0.45 | 0.46 |
| 8 | 95218579 | rs971398 | -0.175 | 1.522 | -0.573 | -1.538 | 0.001 | -0.022 | -0.022 | 0 | 0.12 | 0.11 | 0.19 | 0 | 0.19 | 0.05 |
| 8 | 95225173 | rs6989467 | -0.285 | 1.088 | -0.451 | -1.507 | 0.022 | 0.022 | 0.022 | 0 | 0.7 | 0.86 | 0.62 | 0.35 | 0.7 | 0.59 |
| 8 | 95227768 | rs2607058 | 0.451 | 1.193 | -0.212 | -1.492 | -0.009 | 0.034 | 0.034 | 0.395 | 0.58 | 0.64 | 0.39 | 0.34 | 0.51 | 0.53 |
| 8 | 95232449 | rs2613225 | 0.939 | 0.821 | -0.326 | -1.612 | -0.01 | 0.172 | 0.172 | 0.029 | 0.51 | 0.11 | 0.31 | 0.34 | 0.52 | 0.44 |

| | | | | | | | | | | | | | | | | |
|---|---|---|---|---|---|---|---|---|---|---|---|---|---|---|---|---|
| 8 | 95233336 | rs2607060 | -1.749 | 0.549 | 0.128 | -1.614 | -0.026 | -0.052 | -0.052 | 0 | 0.23 | 0.68 | 0.29 | 0.07 | 0.14 | 0.15 |
| 8 | 95234716 | rs2445704 | -0.832 | 0.58 | 0.12 | -1.591 | -0.03 | -0.023 | -0.023 | 0 | 0.75 | 0.87 | 0.6 | 0.41 | 0.66 | 0.59 |
| 8 | 95235933 | rs2446861 | -0.139 | 0.545 | 0.943 | -1.624 | 0.027 | -0.018 | -0.018 | 0 | 0.69 | 0.51 | 0.5 | 0.35 | 0.61 | 0.51 |
| 8 | 95236826 | rs10504935 | -0.048 | 0.486 | 0.985 | -1.647 | 0.004 | 0.004 | 0.004 | 0 | 0.08 | 0.08 | 0.14 | 0.01 | 0.06 | 0.04 |
| 8 | 95238312 | rs2100602 | -0.405 | 0.56 | 1.139 | -1.686 | -0.015 | -0.041 | -0.041 | 0 | 0.08 | 0.2 | 0.04 | 0 | 0.03 | 0.03 |
| 8 | 95239328 | rs4735277 | 0.354 | 0.622 | 1.463 | -1.662 | 0.026 | 0.012 | 0.012 | 0.488 | 0.06 | 0.2 | 0.08 | 0.06 | 0.05 | 0.07 |
| 8 | 95240271 | rs2446856 | 0.261 | 0.602 | 1.288 | -1.568 | 0.003 | -0.03 | -0.03 | 1.92 | 0.63 | 0.47 | 0.38 | 0.37 | 0.59 | 0.49 |
| 8 | 95254478 | rs2607070 | -1.122 | 0.099 | 1.107 | -1.912 | -0.058 | -0.057 | -0.057 | 0 | 0.28 | 0.49 | 0.22 | 0.19 | 0.19 | 0.17 |
| 8 | 95256726 | rs2607073 | -0.941 | 0.075 | 1.353 | -1.922 | 0.021 | 0.029 | 0.029 | 0 | 0.25 | 0.19 | 0.17 | 0.14 | 0.18 | 0.14 |
| 8 | 95258065 | rs2445709 | -0.704 | -0.162 | 1.328 | -2.178 | 0 | 0.01 | 0.01 | 0 | 0.68 | 0.78 | 0.6 | 0.34 | 0.69 | 0.52 |
| 8 | 95259559 | rs2515114 | -0.468 | -0.258 | 1.497 | -2.232 | -0.004 | -0.032 | -0.032 | 6.913 | 0.66 | 0.63 | 0.55 | 0.29 | 0.69 | 0.49 |
| 8 | 95259942 | rs930869 | 1.294 | -0.274 | 1.571 | -2.254 | -0.063 | 0.149 | 0.149 | 0 | 0.38 | 0.16 | 0.35 | 0.15 | 0.5 | 0.35 |
| 8 | 95261955 | rs17036 | -0.802 | -0.574 | 1.713 | -2.274 | 0 | -0.023 | -0.023 | 0 | 0.69 | 0.66 | 0.58 | 0.29 | 0.7 | 0.49 |
| 8 | 95262253 | rs1050616 | 0.266 | -0.591 | 1.472 | -2.268 | 0.02 | 0.08 | 0.08 | 0.002 | 0.56 | 0.31 | 0.41 | 0.29 | 0.62 | 0.46 |
| 8 | 95263273 | rs2623166 | 0.068 | -0.64 | 1.507 | -2.247 | 0.014 | 0.009 | 0.009 | 0 | 0.18 | 0.15 | 0.06 | 0.15 | 0.12 | 0.11 |
| 8 | 95267327 | rs10216999 | -1.038 | -0.156 | 1.579 | -2.013 | -0.012 | -0.022 | -0.022 | 0.034 | 0.11 | 0.31 | 0.18 | 0.05 | 0.08 | 0.07 |
| 8 | 95274016 | rs2445715 | 0.096 | 0.159 | 1.616 | -2.051 | -0.001 | -0.029 | -0.029 | 0 | 0.6 | 0.63 | 0.45 | 0.28 | 0.61 | 0.48 |
| 8 | 95274357 | rs2250208 | 0.114 | 0.154 | 1.343 | -2.054 | 0.005 | 0.046 | 0.046 | 0 | 0.59 | 0.37 | 0.43 | 0.28 | 0.61 | 0.48 |
| 8 | 95275933 | rs2514791 | 0.292 | 0.152 | 1.329 | -2.044 | 0.009 | 0.088 | 0.088 | 0 | 0.57 | 0.3 | 0.42 | 0.28 | 0.61 | 0.47 |
| 8 | 95278160 | rs16916672 | -0.025 | 0.187 | 1.318 | -2.046 | 0.036 | -0.023 | -0.023 | 3.319 | 0.06 | 0.16 | 0.11 | 0 | 0.07 | 0.04 |
| 8 | 95279199 | rs923313 | 0.006 | 0.208 | 0.489 | -2.003 | -0.002 | -0.029 | -0.029 | 0.531 | 0.59 | 0.52 | 0.44 | 0.29 | 0.61 | 0.48 |
| 8 | 95279760 | rs977274 | 1.187 | 0.406 | -0.771 | -1.973 | -0.048 | 0.162 | 0.162 | 0 | 0.4 | 0.18 | 0.36 | 0.14 | 0.48 | 0.33 |
| 8 | 95281278 | rs17735424 | -0.231 | 0.097 | -0.673 | -1.413 | -0.004 | 0.013 | 0.013 | 0 | 0.16 | 0.08 | 0.06 | 0.14 | 0.13 | 0.14 |
| 8 | 95282557 | rs2514792 | 0.119 | 0.387 | -0.296 | -1.305 | -0.009 | 0.086 | 0.086 | 0 | 0.55 | 0.68 | 0.49 | 0.14 | 0.54 | 0.4 |
| 8 | 95287395 | rs2445720 | 0.877 | -0.362 | 0.512 | -0.112 | -0.016 | 0.114 | 0.114 | 0.035 | 0.89 | 0.62 | 0.88 | 0.79 | 0.94 | 0.87 |
| 8 | 95289326 | rs2515144 | 0.235 | -0.484 | 0.208 | -0.3 | -0.042 | -0.024 | -0.024 | 0.493 | 0.73 | 0.52 | 0.8 | 0.78 | 0.78 | 0.82 |
| 8 | 95303456 | rs2124856 | 1.198 | -0.455 | -1.314 | 0.107 | 0.011 | 0.153 | 0.153 | 0 | 0.49 | 0.08 | 0.45 | 0.31 | 0.56 | 0.38 |
| 8 | 95314221 | rs12548050 | -0.339 | 0.064 | -0.72 | 0.497 | -0.001 | -0.027 | -0.027 | 0 | 0.21 | 0.16 | 0.33 | 0.51 | 0.22 | 0.42 |
| 8 | 95316256 | rs2515137 | -0.048 | -0.101 | -0.32 | 0.099 | -0.014 | -0.03 | -0.03 | 0 | 0.79 | 0.7 | 0.85 | 0.91 | 0.8 | 0.88 |
| 8 | 95324332 | rs2450564 | 0.485 | -0.439 | -0.169 | 1.004 | -0.021 | 0.21 | 0.21 | 0 | 0.92 | 0.79 | 0.94 | 0.67 | 0.97 | 0.84 |
| 8 | 95345128 | rs2515131 | 1.417 | -1.185 | 0.344 | 0.553 | -0.025 | 0.03 | 0.03 | 5.584 | 0.73 | 0.38 | 0.66 | 0.57 | 0.87 | 0.63 |
| 8 | 95365165 | rs2515164 | 0.1 | -1.14 | 0.438 | 0.541 | -0.022 | 0.214 | 0.214 | 0 | 0.92 | 0.77 | 0.94 | 0.67 | 0.97 | 0.84 |
| 8 | 95365885 | rs6997115 | -0.599 | -1.048 | -0.202 | 0.287 | 0.009 | -0.007 | -0.007 | 0 | 0.42 | 0.36 | 0.55 | 0.46 | 0.35 | 0.42 |
| 8 | 95366370 | rs2470728 | 1.286 | -1.198 | 0.089 | 0.211 | 0.059 | 0.207 | 0.207 | 0.036 | 0.48 | 0.2 | 0.37 | 0.21 | 0.61 | 0.42 |
| 8 | 95383875 | rs10956911 | -0.495 | -1.177 | 0.005 | 0.173 | 0.006 | -0.006 | -0.006 | 0 | 0.38 | 0.36 | 0.51 | 0.46 | 0.33 | 0.4 |
| 8 | 95401265 | rs12681366 | -0.502 | -1.087 | 0.011 | 0.151 | 0.007 | -0.008 | -0.008 | 0 | 0.39 | 0.35 | 0.51 | 0.46 | 0.35 | 0.39 |
| 8 | 95410986 | rs7817230 | 0.041 | -1.169 | 0.27 | 0.103 | -0.031 | -0.029 | -0.029 | 0 | 0.09 | 0.25 | 0.14 | 0.03 | 0.02 | 0.1 |
| 8 | 95419698 | rs2291439 | -0.662 | -1.122 | 0.068 | -0.034 | 0.008 | -0.008 | -0.008 | 0 | 0.41 | 0.35 | 0.54 | 0.47 | 0.38 | 0.42 |
| 8 | 95450289 | rs3019281 | 0.443 | -1.008 | -0.507 | -0.004 | -0.018 | -0.1 | -0.1 | 1.86 | 0.28 | 0.08 | 0.35 | 0.43 | 0.34 | 0.29 |
| 8 | 95461946 | rs935052 | -1.418 | -1.231 | -0.558 | -0.161 | 0.018 | 0.205 | 0.205 | 0.828 | 0.55 | 0.84 | 0.64 | 0.83 | 0.43 | 0.6 |
| 8 | 95484581 | rs10095298 | -0.067 | -1.011 | -0.296 | -0.02 | -0.003 | -0.052 | -0.052 | 0 | 0.33 | 0.16 | 0.4 | 0.43 | 0.36 | 0.32 |
| 8 | 95497989 | rs3102854 | 0.045 | -1.243 | -0.616 | -0.843 | 0.035 | 0.012 | 0.012 | 0 | 0.4 | 0.32 | 0.26 | 0.32 | 0.22 | 0.44 |
| 8 | 95499295 | rs3098726 | 0.231 | -1.299 | -0.692 | -0.921 | -0.01 | 0.136 | 0.136 | 0 | 0.1 | 0.27 | 0.06 | 0.24 | 0.05 | 0.16 |
| 8 | 95502554 | rs3762053 | 0.955 | -1.333 | -0.41 | -1.001 | 0.035 | 0.146 | 0.146 | 0 | 0.3 | 0.03 | 0.2 | 0.07 | 0.17 | 0.28 |
| 8 | 95505296 | rs3099407 | -1.475 | -1.275 | -0.487 | -1.152 | -0.062 | 0.023 | 0.023 | 0 | 0.79 | 0.87 | 0.78 | 0.83 | 0.58 | 0.8 |
| 8 | 95508721 | rs2289455 | 1.097 | -1.202 | -0.563 | -1.077 | 0.035 | 0.146 | 0.146 | 0 | 0.3 | 0.03 | 0.2 | 0.08 | 0.17 | 0.28 |
| 8 | 95516508 | rs3098705 | -0.155 | -1.179 | -0.512 | -1.059 | 0.018 | 0.012 | 0.012 | 0 | 0.44 | 0.37 | 0.3 | 0.32 | 0.25 | 0.46 |
| 8 | 95531419 | rs1866844 | 0.096 | -1.177 | -0.48 | -1.054 | 0.024 | 0.008 | 0.008 | 0 | 0.39 | 0.33 | 0.26 | 0.32 | 0.22 | 0.44 |
| 8 | 95536943 | rs3133648 | -0.17 | -1.299 | -0.118 | -1.133 | 0.02 | -0.036 | -0.036 | 0 | 0.66 | 0.6 | 0.7 | 0.81 | 0.53 | 0.75 |
| 8 | 95538468 | rs2890827 | -0.228 | -1.323 | 0.014 | -1.146 | -0.014 | -0.009 | -0.009 | 0 | 0.46 | 0.46 | 0.32 | 0.32 | 0.25 | 0.46 |
| 8 | 95538708 | rs7835534 | 0.839 | -1.338 | -0.03 | -1.153 | 0.001 | -0.016 | -0.016 | 0.145 | 0.05 | 0.03 | 0.05 | 0 | 0.04 | 0.02 |
| 8 | 95541223 | rs957447 | 1.956 | -1.348 | 0.005 | -1.154 | -0.028 | -0.127 | -0.127 | 0 | 0.23 | 0.06 | 0.41 | 0.49 | 0.31 | 0.31 |
| 8 | 95541302 | rs957448 | 1.478 | -1.348 | 0.004 | -1.154 | -0.001 | 0.007 | 0.007 | 0.584 | 0.27 | 0.27 | 0.44 | 0.49 | 0.31 | 0.31 |
| 8 | 95558694 | rs1992370 | 1.896 | -1.348 | -0.016 | -1.154 | -0.035 | -0.142 | -0.142 | 0.041 | 0.24 | 0.05 | 0.41 | 0.49 | 0.31 | 0.31 |
| 8 | 95559092 | rs3133651 | -0.593 | -1.348 | 0.023 | -1.162 | 0.036 | -0.038 | -0.038 | 8.933 | 0.7 | 0.58 | 0.73 | 0.81 | 0.56 | 0.77 |
| 8 | 95564340 | rs17716313 | 0.658 | -1.372 | -0.039 | -1.17 | 0.047 | 0.157 | 0.157 | 0 | 0.34 | 0.08 | 0.24 | 0.08 | 0.2 | 0.3 |
| 8 | 95564533 | rs12679345 | 1.918 | -1.251 | 0.002 | -1.132 | -0.036 | -0.141 | -0.141 | 1.031 | 0.23 | 0.05 | 0.41 | 0.49 | 0.31 | 0.31 |
| 8 | 95565273 | rs3133657 | -0.555 | -1.239 | 0.111 | -1.126 | 0.031 | -0.032 | -0.032 | 0 | 0.7 | 0.6 | 0.74 | 0.81 | 0.56 | 0.77 |
| 8 | 95565762 | rs3133659 | -0.121 | -1.23 | 0.016 | -0.978 | 0.014 | 0.009 | 0.009 | 0 | 0.44 | 0.36 | 0.3 | 0.32 | 0.25 | 0.46 |
| 8 | 95582606 | rs12680842 | 1.593 | -0.125 | -0.38 | -0.78 | -0.031 | -0.092 | -0.092 | 0 | 0.27 | 0.11 | 0.44 | 0.48 | 0.38 | 0.37 |
| 8 | 95591774 | rs3099420 | 0.078 | -0.171 | -0.297 | -0.847 | -0.016 | -0.006 | -0.006 | 0 | 0.42 | 0.49 | 0.24 | 0.32 | 0.23 | 0.4 |
| 8 | 95604859 | rs2114212 | 1.079 | 0.399 | -0.119 | -0.679 | -0.004 | -0.004 | -0.004 | 0 | 0.63 | 0.36 | 0.59 | 0.43 | 0.58 | 0.57 |
| 8 | 95616980 | rs4735302 | 1.748 | 0.366 | -0.217 | -0.703 | -0.01 | -0.035 | -0.035 | 0 | 0.28 | 0.19 | 0.45 | 0.42 | 0.37 | 0.37 |
| 8 | 95628254 | rs3096604 | -0.206 | 0.414 | 0.242 | -0.684 | 0.007 | 0.007 | 0.007 | 0 | 0.46 | 0.41 | 0.29 | 0.33 | 0.33 | 0.45 |
| 8 | 95628814 | rs4734283 | 1.157 | 0.539 | 0.673 | -0.521 | 0.026 | 0.04 | 0.04 | 6.185 | 0.7 | 0.41 | 0.67 | 0.5 | 0.64 | 0.67 |
| 8 | 95629254 | rs3096605 | -0.065 | 0.513 | 0.716 | -0.521 | 0.005 | 0.186 | 0.186 | 0 | 0.09 | 0.32 | 0.08 | 0.25 | 0.06 | 0.15 |
| 8 | 95630829 | rs3133622 | 0.377 | 0.701 | 0.506 | -0.537 | 0.035 | -0.007 | -0.007 | 0 | 0.79 | 0.68 | 0.73 | 0.75 | 0.7 | 0.81 |
| 8 | 95632185 | rs4735305 | 0.914 | 0.769 | 0.294 | -0.145 | 0.004 | 0.007 | 0.007 | 6.535 | 0.72 | 0.67 | 0.6 | 0.5 | 0.68 | 0.7 |
| 8 | 95634950 | rs756474 | 0.106 | 0.633 | 0.268 | -0.413 | -0.005 | -0.026 | -0.026 | 0 | 0.37 | 0.39 | 0.23 | 0.1 | 0.27 | 0.3 |

| | | | | | | | | | | | | | | | | |
|---|---|---|---|---|---|---|---|---|---|---|---|---|---|---|---|---|
| 8 | 95636000 | rs17716607 | 1.042 | 0.628 | -0.456 | -0.591 | -0.013 | 0.135 | 0.135 | 0 | 0.22 | 0.03 | 0.11 | 0 | 0.16 | 0.12 |
| 8 | 95636052 | rs2890805 | 0.403 | 1.366 | 0.103 | -0.262 | 0.05 | 0.019 | 0.019 | 0.607 | 0.84 | 0.76 | 0.67 | 0.71 | 0.71 | 0.74 |
| 8 | 95638496 | rs10504938 | 1.528 | 1.357 | -0.041 | -0.168 | 0.026 | -0.07 | -0.07 | 0.662 | 0.38 | 0.14 | 0.36 | 0.43 | 0.37 | 0.38 |
| 8 | 95641609 | rs2052716 | 0.715 | 1.583 | 0.186 | -0.232 | 0.001 | 0.007 | 0.007 | 0.006 | 0.49 | 0.56 | 0.54 | 0.59 | 0.56 | 0.5 |
| 8 | 95643315 | rs10156335 | -0.538 | 1.714 | 0.521 | -0.1 | -0.003 | -0.004 | -0.004 | 0 | 0.13 | 0.09 | 0.11 | 0.22 | 0.14 | 0.07 |
| 8 | 95645917 | rs3133630 | 1.548 | 1.921 | 0.717 | -0.33 | 0.015 | 0.002 | 0.002 | 0 | 0.73 | 0.45 | 0.78 | 0.67 | 0.75 | 0.75 |
| 8 | 95647792 | rs13262758 | -0.675 | 1.889 | 0.498 | -0.498 | -0.001 | 0.004 | 0.004 | 0 | 0.12 | 0.1 | 0.1 | 0.05 | 0.13 | 0.06 |
| 8 | 95649072 | rs4735307 | -1.213 | 2.028 | 0.644 | -0.689 | 0 | -0.008 | -0.008 | 0 | 0.49 | 0.43 | 0.46 | 0.29 | 0.45 | 0.39 |
| 8 | 95654628 | rs7826830 | 0.562 | 1.414 | 0.144 | -0.681 | 0.014 | 0.124 | 0.124 | 5.986 | 0.27 | 0.06 | 0.12 | 0.02 | 0.17 | 0.2 |
| 8 | 95655050 | rs7386115 | 0.849 | 1.477 | 1.185 | -0.384 | 0.026 | -0.028 | -0.028 | 0 | 0.87 | 0.4 | 0.88 | 0.95 | 0.86 | 0.94 |
| 8 | 95657956 | rs3808381 | 1.75 | 1.557 | 0.814 | -0.144 | -0.018 | -0.011 | -0.011 | 0.153 | 0.77 | 0.18 | 0.72 | 0.77 | 0.72 | 0.82 |
| 8 | 95660772 | rs6990716 | 1.366 | 1.357 | 1.235 | -0.098 | 0.052 | 0.092 | 0.092 | 0.004 | 0.87 | 0.49 | 0.76 | 0.77 | 0.87 | 0.88 |
| 8 | 95666621 | rs10956922 | 0.096 | 1.281 | 1.176 | -0.725 | -0.004 | -0.068 | -0.068 | 0.408 | 0.6 | 0.58 | 0.76 | 0.95 | 0.65 | 0.75 |
| 8 | 95675372 | rs1895874 | 0.959 | 1.328 | 0.991 | -0.63 | -0.009 | 0.094 | 0.094 | 0 | 0.49 | 0.25 | 0.55 | 0.75 | 0.54 | 0.64 |
| 8 | 95677132 | rs1426171 | 1.647 | 1.358 | 1.057 | -0.649 | 0.028 | -0.005 | -0.005 | 0 | 0.77 | 0.6 | 0.74 | 0.76 | 0.74 | 0.8 |
| 8 | 95685147 | rs11786992 | -0.371 | 1.324 | 0.968 | -0.608 | 0 | -0.017 | -0.017 | 6.245 | 0.39 | 0.45 | 0.36 | 0.22 | 0.35 | 0.29 |
| 8 | 95685587 | rs1365205 | 1.876 | 1.342 | 0.749 | -0.628 | 0.01 | -0.059 | -0.059 | 0.003 | 0.39 | 0.13 | 0.39 | 0.54 | 0.39 | 0.51 |
| 8 | 95686850 | rs17717431 | 1.073 | 1.475 | 0.47 | -0.611 | -0.041 | 0.158 | 0.158 | 3.858 | 0.26 | 0.04 | 0.14 | 0 | 0.23 | 0.18 |
| 8 | 95687335 | rs2033074 | 1.949 | 1.483 | 1.167 | -0.233 | 0.051 | 0.079 | 0.079 | 0 | 0.71 | 0.54 | 0.57 | 0.54 | 0.63 | 0.69 |
| 8 | 95700688 | rs2554399 | 0.669 | 1.307 | 0.679 | -0.202 | 0.022 | 0.097 | 0.097 | 0 | 0.88 | 0.71 | 0.8 | 0.79 | 0.88 | 0.89 |
| 8 | 95702969 | rs2722886 | 0.468 | 1.289 | 0.691 | -0.204 | 0.018 | 0.092 | 0.092 | 0 | 0.89 | 0.71 | 0.8 | 0.81 | 0.9 | 0.9 |
| 8 | 95707468 | rs11787473 | -0.868 | 1.267 | 0.253 | -0.231 | -0.002 | 0.001 | 0.001 | 0 | 0.13 | 0.11 | 0.08 | 0.1 | 0.14 | 0.13 |
| 8 | 95732766 | rs10109430 | -0.041 | 1.201 | 0.449 | -0.101 | 0.013 | 0.09 | 0.09 | 0 | 0.09 | 0.18 | 0.17 | 0.18 | 0.09 | 0.07 |
| 8 | 95734714 | rs6471480 | -0.849 | 1.033 | 0.466 | -0.308 | 0.032 | 0.042 | 0.042 | 1.321 | 0.53 | 0.65 | 0.56 | 0.32 | 0.55 | 0.42 |
| 8 | 95740643 | rs7461991 | -1.5 | 0.912 | 0.089 | -0.283 | -0.041 | -0.174 | -0.174 | 0 | 0.63 | 0.98 | 0.66 | 0.47 | 0.63 | 0.51 |
| 8 | 95744466 | rs6988635 | -0.888 | 1.032 | 0.465 | -0.507 | 0.005 | 0.007 | 0.007 | 0.458 | 0.54 | 0.66 | 0.47 | 0.38 | 0.52 | 0.38 |
| 8 | 95758766 | rs12234995 | -2.117 | 0.702 | 1.165 | -0.868 | -0.002 | 0.013 | 0.013 | 0 | 0.35 | 0.71 | 0.37 | 0.22 | 0.34 | 0.26 |
| 8 | 95760323 | rs7837249 | -1.546 | 0.768 | 1.362 | -0.911 | -0.017 | -0.048 | -0.048 | 0 | 0.22 | 0.64 | 0.15 | 0.07 | 0.17 | 0.05 |
| 8 | 95770914 | rs10101523 | -2.49 | 0.739 | 1.673 | -0.943 | -0.002 | 0.024 | 0.024 | 0 | 0.36 | 0.76 | 0.38 | 0.22 | 0.35 | 0.26 |
| 8 | 95773014 | rs7830822 | -1.917 | 0.767 | 1.564 | -0.933 | 0.008 | -0.031 | -0.031 | 0 | 0.23 | 0.34 | 0.27 | 0.15 | 0.23 | 0.22 |
| 8 | 95774806 | rs4524755 | -1.233 | 1.307 | 1.029 | -0.668 | -0.025 | -0.003 | -0.003 | 0 | 0.13 | 0.06 | 0.22 | 0.15 | 0.18 | 0.21 |
| 8 | 95789654 | rs7017149 | -1.216 | 1.143 | 1.288 | -0.57 | 0.015 | -0.037 | -0.037 | 0 | 0.13 | 0.31 | 0.1 | 0.05 | 0.13 | 0.04 |
| 8 | 95806454 | rs6415570 | 1.042 | 1.119 | 1.047 | -0.581 | 0.01 | -0.044 | -0.044 | 0 | 0.86 | 0.63 | 0.9 | 0.95 | 0.87 | 0.96 |
| 8 | 95809165 | rs12541990 | 1.385 | 1.302 | 0.146 | -0.468 | -0.024 | 0.002 | 0.002 | 1.741 | 0.77 | 0.8 | 0.71 | 0.75 | 0.71 | 0.73 |
| 8 | 95817874 | rs6986659 | 0.844 | 1.543 | 0.585 | -0.446 | 0.06 | 0.055 | 0.055 | 0.325 | 0.89 | 0.77 | 0.77 | 0.8 | 0.8 | 0.77 |
| 8 | 95820969 | rs11786021 | -0.132 | 1.291 | 0.386 | -1.332 | 0.007 | 0.006 | 0.006 | 2.383 | 0.27 | 0.22 | 0.11 | 0.05 | 0.19 | 0.14 |
| 8 | 95838869 | rs12677346 | 0.577 | 1.251 | 0.369 | -1.207 | 0.036 | 0.033 | 0.033 | 0 | 0.87 | 0.76 | 0.75 | 0.79 | 0.76 | 0.77 |
| 8 | 95853708 | rs11996455 | 1.654 | 0.906 | 0.534 | -1.49 | -0.005 | 0.025 | 0.025 | 1.519 | 0.48 | 0.38 | 0.56 | 0.65 | 0.5 | 0.59 |
| 8 | 95875676 | rs6991067 | 1.712 | 0.825 | 0.931 | -1.526 | -0.012 | 0.048 | 0.048 | 1.792 | 0.47 | 0.29 | 0.56 | 0.65 | 0.5 | 0.59 |
| 8 | 95879035 | rs16917102 | -0.138 | 0.856 | 1.048 | -1.584 | -0.001 | -0.028 | -0.028 | 0 | 0.29 | 0.28 | 0.12 | 0.05 | 0.21 | 0.14 |
| 8 | 95883232 | rs7844932 | -0.5 | 0.83 | 1.124 | -1.572 | 0.008 | 0.012 | 0.012 | 2.695 | 0.32 | 0.31 | 0.13 | 0.14 | 0.22 | 0.16 |
| 8 | 95886551 | rs10956928 | 0.727 | 0.842 | 1.218 | -1.607 | 0.048 | 0.027 | 0.027 | 0 | 0.84 | 0.7 | 0.72 | 0.8 | 0.72 | 0.77 |
| 8 | 95888307 | rs1129152 | 2.025 | 1.192 | 1.095 | -1.727 | -0.005 | 0.039 | 0.039 | 2.059 | 0.44 | 0.25 | 0.53 | 0.65 | 0.46 | 0.55 |
| 8 | 95888981 | rs716733 | -0.119 | 1.21 | 1.028 | -1.906 | 0.004 | -0.011 | -0.011 | 0 | 0.29 | 0.26 | 0.11 | 0.05 | 0.2 | 0.13 |
| 8 | 95891113 | rs6987256 | 0.687 | 1.236 | 0.949 | -1.834 | 0.047 | 0.026 | 0.026 | 0 | 0.84 | 0.7 | 0.72 | 0.8 | 0.72 | 0.77 |
| 8 | 95909498 | rs16917117 | 0.38 | 0.873 | 0.895 | -1.858 | 0.016 | 0.022 | 0.022 | 3.083 | 0.24 | 0.14 | 0.07 | 0.05 | 0.16 | 0.12 |
| 8 | 95913425 | rs2956216 | 0.596 | 1.086 | 1.306 | -1.762 | 0.027 | 0.023 | 0.023 | 0 | 0.85 | 0.77 | 0.75 | 0.8 | 0.76 | 0.78 |
| 8 | 95923032 | rs477653 | 1.254 | 1.12 | 1.41 | -1.779 | 0.039 | 0.026 | 0.026 | 0 | 0.82 | 0.66 | 0.72 | 0.79 | 0.75 | 0.77 |
| 8 | 95925490 | rs551270 | 0.169 | 1.136 | 1.41 | -1.773 | 0.009 | 0.007 | 0.007 | 0 | 0.87 | 0.86 | 0.76 | 0.8 | 0.76 | 0.78 |
| 8 | 95927702 | rs13264488 | 0.646 | 1.19 | 1.026 | -2.023 | -0.031 | 0.043 | 0.043 | 0 | 0.08 | 0.01 | 0.05 | 0 | 0.12 | 0.07 |
| 8 | 95931782 | rs2123647 | 1.921 | 0.787 | 1.381 | -1.828 | 0.009 | -0.012 | -0.012 | 0 | 0.72 | 0.54 | 0.65 | 0.78 | 0.69 | 0.73 |
| 8 | 95935102 | rs2467674 | 1.963 | 0.82 | 1.24 | -1.812 | -0.044 | -0.031 | -0.031 | 0.662 | 0.81 | 0.46 | 0.86 | 0.89 | 0.87 | 0.88 |
| 8 | 95937502 | rs7845219 | 1.982 | 0.729 | 1.215 | -1.887 | 0.014 | 0.094 | 0.094 | 0.128 | 0.49 | 0.28 | 0.56 | 0.74 | 0.53 | 0.61 |
| 8 | 95938422 | rs7760 | 1.024 | 0.823 | 1.281 | -1.753 | -0.004 | 0.036 | 0.036 | 0 | 0.88 | 0.59 | 0.9 | 0.89 | 0.9 | 0.9 |
| 8 | 95939594 | rs2945554 | 2.106 | 0.829 | 1.232 | -1.751 | 0.022 | -0.008 | -0.008 | 0.527 | 0.74 | 0.51 | 0.66 | 0.78 | 0.69 | 0.73 |
| 8 | 95942244 | rs896849 | 1.826 | 0.754 | 1.284 | -1.736 | -0.038 | -0.023 | -0.023 | 2.142 | 0.82 | 0.51 | 0.87 | 0.89 | 0.87 | 0.88 |
| 8 | 95951273 | rs1011796 | 1.901 | 0.75 | 1.381 | -1.627 | 0.015 | 0.015 | 0.015 | 0 | 0.76 | 0.43 | 0.67 | 0.78 | 0.7 | 0.73 |
| 8 | 95952477 | rs2956210 | 1.473 | 0.771 | 1.405 | -1.61 | 0.04 | 0.046 | 0.046 | 0 | 0.79 | 0.52 | 0.69 | 0.78 | 0.72 | 0.74 |
| 8 | 95965135 | rs493506 | 1.077 | 0.821 | 1.342 | -1.584 | 0.07 | 0.172 | 0.172 | 1.751 | 0.92 | 0.55 | 0.92 | 0.89 | 0.92 | 0.92 |
| 8 | 95965743 | rs2515226 | -0.799 | 0.74 | 1.2 | -1.737 | 0.001 | 0.067 | 0.067 | 0 | 0.37 | 0.2 | 0.34 | 0.16 | 0.36 | 0.3 |
| 8 | 95973465 | rs4735337 | 1.372 | 0.636 | 1.045 | -1.678 | 0.011 | 0.038 | 0.038 | 0 | 0.53 | 0.37 | 0.58 | 0.73 | 0.56 | 0.59 |
| 8 | 95986340 | rs7827478 | 1.257 | 0.654 | 1.185 | -1.683 | 0.024 | 0.023 | 0.023 | 0 | 0.76 | 0.5 | 0.71 | 0.78 | 0.73 | 0.71 |
| 8 | 96000919 | rs4734295 | 1.679 | 0.789 | 0.78 | -1.767 | -0.015 | 0.05 | 0.05 | 0 | 0.46 | 0.31 | 0.52 | 0.71 | 0.48 | 0.57 |
| 8 | 96020400 | rs7812312 | 0.16 | 0.674 | 0.53 | -1.753 | 0.042 | 0.079 | 0.079 | 1.19 | 0.24 | 0.12 | 0.13 | 0.05 | 0.13 | 0.13 |
| 8 | 96027392 | rs7002940 | 0.176 | 0.762 | 0.777 | -1.751 | 0.029 | 0.042 | 0.042 | 1.276 | 0.24 | 0.14 | 0.15 | 0.05 | 0.13 | 0.13 |
| 8 | 96029002 | rs7826196 | 0.176 | 0.762 | 0.777 | -1.751 | 0.029 | 0.042 | 0.042 | 1.276 | 0.24 | 0.14 | 0.14 | 0.05 | 0.13 | 0.13 |
| 8 | 96036422 | rs10092003 | 0.277 | 0.769 | 0.623 | -1.748 | 0.04 | 0.091 | 0.091 | 1.129 | 0.23 | 0.1 | 0.13 | 0.05 | 0.13 | 0.13 |
| 8 | 96036741 | rs11776189 | 0.277 | 0.769 | 0.621 | -1.748 | 0.04 | 0.094 | 0.094 | 1.129 | 0.23 | 0.1 | 0.13 | 0.05 | 0.13 | 0.13 |
| 8 | 96043829 | rs11780931 | 0.198 | 0.724 | 0.857 | -1.77 | 0.018 | 0.018 | 0.018 | 1.307 | 0.24 | 0.15 | 0.14 | 0.05 | 0.13 | 0.13 |

| | | | | | | | | | | | | | | | | |
|---|---|---|---|---|---|---|---|---|---|---|---|---|---|---|---|---|
| 8 | 96049339 | rs2678827 | -1.304 | 1.213 | 1.042 | -1.432 | -0.008 | -0.006 | -0.006 | 0 | 0.24 | 0.19 | 0.26 | 0.26 | 0.36 | 0.3 |
| 8 | 96062601 | rs2678822 | -1.117 | 1.287 | 0.988 | -1.386 | 0.003 | 0.003 | 0.003 | 0 | 0.22 | 0.22 | 0.26 | 0.26 | 0.34 | 0.3 |
| 8 | 96081402 | rs877350 | 1.426 | 1.146 | 1.006 | -1.69 | 0.077 | -0.008 | -0.008 | 0 | 0.92 | 0.63 | 0.89 | 1 | 0.9 | 0.98 |
| 8 | 96083924 | rs2678825 | 0.99 | 1.006 | 0.501 | -2.168 | -0.002 | -0.051 | -0.051 | 0 | 0.57 | 0.53 | 0.6 | 0.83 | 0.54 | 0.73 |
| 8 | 96087025 | rs2599706 | -0.025 | 1.172 | 0.311 | -2.14 | 0.018 | -0.006 | -0.006 | 0.048 | 0.95 | 0.95 | 0.9 | 1 | 0.8 | 0.96 |
| 8 | 96088038 | rs6982086 | -0.632 | 1.045 | 0.462 | -2.096 | 0.007 | 0.008 | 0.008 | 0.216 | 0.35 | 0.26 | 0.27 | 0.24 | 0.27 | 0.24 |
| 8 | 96090259 | rs2599702 | 0.883 | 1.023 | 0.441 | -2.103 | -0.001 | -0.022 | -0.022 | 0.007 | 0.59 | 0.61 | 0.61 | 0.76 | 0.54 | 0.73 |
| 8 | 96099666 | rs10087402 | 0.836 | 0.813 | -0.092 | -2.225 | 0.031 | 0.043 | 0.043 | 0 | 0.16 | 0.03 | 0.17 | 0.14 | 0.09 | 0.12 |
| 8 | 96101786 | rs4734298 | 1.419 | 0.868 | 0.168 | -2.387 | -0.023 | 0.062 | 0.062 | 0.015 | 0.71 | 0.81 | 0.81 | 0.88 | 0.62 | 0.85 |
| 8 | 96102659 | rs11781447 | -0.627 | 0.744 | 0.151 | -2.324 | 0.005 | 0.028 | 0.028 | 0 | 0.18 | 0.06 | 0.07 | 0.12 | 0.18 | 0.12 |
| 8 | 96106037 | rs4590408 | -0.217 | 0.506 | 0.208 | -2.412 | 0 | -0.005 | -0.005 | 0.881 | 0.1 | 0.13 | 0.04 | 0.04 | 0.08 | 0.09 |
| 8 | 96109135 | rs16917325 | -0.901 | 0.067 | 0.651 | -2.531 | -0.049 | -0.037 | -0.037 | 0.219 | 0.2 | 0.4 | 0.08 | 0.14 | 0.14 | 0.11 |
| 8 | 96109370 | rs1000286 | -1.09 | -0.024 | 0.64 | -2.64 | -0.068 | -0.058 | -0.058 | 0 | 0.26 | 0.41 | 0.1 | 0.14 | 0.15 | 0.13 |
| 8 | 96115476 | rs7833057 | -1.234 | -0.016 | 0.17 | -3.008 | -0.042 | -0.151 | -0.151 | 0 | 0.32 | 0.5 | 0.2 | 0 | 0.19 | 0.13 |
| 8 | 96120072 | rs4392868 | -0.102 | 0.276 | -0.209 | -3.069 | 0.036 | -0.019 | -0.019 | 0 | 0.93 | 0.9 | 0.88 | 1 | 0.76 | 0.96 |
| 8 | 96120302 | rs10808675 | 1.645 | 0.22 | -0.323 | -3.057 | -0.01 | 0.056 | 0.056 | 0 | 0.6 | 0.28 | 0.67 | 1 | 0.57 | 0.83 |
| 8 | 96123381 | rs2044035 | 1.908 | 0.231 | -0.387 | -3.111 | -0.016 | 0.137 | 0.137 | 0 | 0.54 | 0.27 | 0.62 | 0.98 | 0.52 | 0.77 |
| 8 | 96127132 | rs16917360 | 0.23 | 0.473 | -0.671 | -2.736 | -0.015 | -0.001 | -0.001 | 0 | 0.06 | 0.01 | 0.05 | 0.02 | 0.05 | 0.07 |
| 8 | 96132104 | rs212551 | 0.678 | 0.387 | -0.381 | -2.785 | -0.016 | 0.002 | 0.002 | 0 | 0.79 | 0.84 | 0.82 | 1 | 0.71 | 0.9 |
| 8 | 96139617 | rs1470769 | -1.597 | 0.077 | -0.412 | -2.606 | -0.032 | -0.04 | -0.04 | 0 | 0.38 | 0.51 | 0.23 | 0.29 | 0.3 | 0.2 |
| 8 | 96139702 | rs16917364 | -1.21 | -0.165 | -0.078 | -2.835 | -0.068 | -0.153 | -0.153 | 0 | 0.3 | 0.51 | 0.18 | 0 | 0.13 | 0.11 |
| 8 | 96141011 | rs7839455 | -1.755 | 0.043 | -0.897 | -2.843 | 0.038 | 0.344 | 0.344 | 0.609 | 0.5 | 0.84 | 0.34 | 0 | 0.39 | 0.2 |
| 8 | 96147295 | rs7842334 | -0.308 | 1.108 | -0.067 | -1.824 | -0.006 | -0.03 | -0.03 | 0 | 0.14 | 0.12 | 0.07 | 0 | 0.05 | 0.06 |
| 8 | 96160914 | rs10504941 | -0.612 | 1.228 | -0.044 | -1.813 | 0.039 | -0.016 | -0.016 | 0 | 0.06 | 0.05 | 0.1 | 0 | 0.2 | 0.04 |
| 8 | 96162964 | rs13263827 | -0.705 | 1.174 | -0.213 | -1.798 | 0.096 | -0.034 | -0.034 | 0 | 0.06 | 0.14 | 0.1 | 0 | 0.2 | 0.04 |
| 8 | 96164922 | rs11985227 | -0.455 | -0.004 | -0.76 | -1.519 | -0.009 | -0.033 | -0.033 | 0 | 0.14 | 0.12 | 0.07 | 0 | 0.05 | 0.06 |
| 8 | 96179683 | rs297563 | -0.688 | 0.431 | -1.237 | -1.508 | -0.022 | 0.118 | 0.118 | 0 | 0.6 | 0.79 | 0.41 | 0.17 | 0.62 | 0.24 |
| 8 | 96186655 | rs297573 | -0.037 | 0.317 | -0.505 | -1.586 | 0 | -0.006 | -0.006 | 0.039 | 0.32 | 0.36 | 0.21 | 0.27 | 0.34 | 0.19 |
| 8 | 96187103 | rs297574 | -0.368 | 0.203 | -0.1 | -1.625 | -0.002 | -0.013 | -0.013 | 0 | 0.35 | 0.61 | 0.22 | 0.27 | 0.34 | 0.19 |
| 8 | 96193895 | rs10093480 | -0.559 | 0.036 | 0.144 | -1.59 | -0.006 | -0.053 | -0.053 | 0 | 0.13 | 0.17 | 0.11 | 0 | 0.1 | 0.07 |
| 8 | 96193976 | rs6982829 | 0.147 | -0.307 | 0.28 | -1.712 | -0.009 | -0.004 | -0.004 | 0 | 0.18 | 0.23 | 0.07 | 0.16 | 0.12 | 0.07 |
| 8 | 96194737 | rs10956941 | -1.259 | -0.191 | 0.002 | -1.392 | -0.017 | -0.011 | -0.011 | 0 | 0.32 | 0.41 | 0.19 | 0.3 | 0.23 | 0.22 |
| 8 | 96195171 | rs7009196 | -1.203 | -0.246 | -0.057 | -1.469 | -0.033 | -0.038 | -0.038 | 0.529 | 0.32 | 0.45 | 0.19 | 0.2 | 0.22 | 0.14 |
| 8 | 96195723 | rs297551 | 0.768 | 0.168 | -0.477 | -1.339 | -0.018 | 0.024 | 0.024 | 0 | 0.23 | 0.13 | 0.23 | 0.09 | 0.31 | 0.18 |
| 8 | 96199585 | rs17730693 | -0.249 | -0.075 | -0.445 | -1.058 | -0.004 | 0.05 | 0.05 | 0 | 0.11 | 0.03 | 0.09 | 0 | 0.1 | 0.07 |
| 8 | 96200507 | rs7002762 | 0.064 | -0.207 | -0.229 | -0.938 | -0.001 | -0.012 | -0.012 | 0.017 | 0.5 | 0.46 | 0.35 | 0.23 | 0.56 | 0.19 |
| 8 | 96202284 | rs11784608 | -0.95 | 0.012 | -0.306 | -0.61 | -0.022 | -0.066 | -0.066 | 0.135 | 0.15 | 0.27 | 0.11 | 0 | 0.09 | 0.08 |
| 8 | 96204142 | rs11787209 | -0.145 | 0.269 | -0.751 | -0.219 | 0.012 | 0.024 | 0.024 | 0 | 0.48 | 0.38 | 0.37 | 0.4 | 0.58 | 0.32 |
| 8 | 96205920 | rs998980 | 0.139 | 0.425 | -0.792 | -0.05 | 0.01 | 0.009 | 0.009 | 0.686 | 0.58 | 0.5 | 0.61 | 0.52 | 0.51 | 0.55 |
| 8 | 96206553 | rs2439641 | -0.573 | 0.024 | -0.654 | -0.092 | -0.026 | 0 | 0 | 0 | 0.17 | 0.22 | 0.12 | 0.15 | 0.11 | 0.22 |
| 8 | 96206789 | rs2439640 | -0.607 | -0.286 | -1.138 | -0.174 | 0.001 | -0.001 | -0.001 | 0 | 0.3 | 0.27 | 0.39 | 0.68 | 0.22 | 0.43 |
| 8 | 96206803 | rs16917410 | 0.856 | -0.117 | -1.358 | -0.005 | 0.022 | -0.05 | -0.05 | 0.69 | 0.11 | 0.04 | 0.25 | 0.53 | 0.06 | 0.23 |
| 8 | 96209033 | rs4734300 | 0.92 | -0.173 | -1.464 | -0.864 | -0.013 | 0.084 | 0.084 | 0.87 | 0.34 | 0.17 | 0.21 | 0 | 0.34 | 0.13 |
| 8 | 96211820 | rs12545256 | 0.179 | 0.263 | -1.452 | -1.177 | -0.005 | 0.099 | 0.099 | 1.562 | 0.48 | 0.28 | 0.34 | 0.04 | 0.48 | 0.29 |
| 8 | 96212069 | rs2439636 | -0.777 | 0.367 | -1.291 | -1.043 | -0.001 | -0.006 | -0.006 | 3.777 | 0.44 | 0.51 | 0.42 | 0.32 | 0.45 | 0.53 |
| 8 | 96214292 | rs12155945 | 0.832 | 0.444 | -0.85 | -0.97 | 0.012 | -0.012 | -0.012 | 0 | 0.08 | 0.08 | 0.1 | 0.04 | 0.12 | 0.12 |
| 8 | 96217043 | rs2514565 | -0.701 | 0.204 | -0.842 | -1.055 | 0.003 | 0.019 | 0.019 | 0 | 0.5 | 0.56 | 0.6 | 0.73 | 0.47 | 0.67 |
| 8 | 96217634 | rs6995298 | -0.611 | 0.18 | -0.87 | -0.994 | -0.006 | -0.023 | -0.023 | 0 | 0.79 | 0.82 | 0.65 | 0.31 | 0.77 | 0.66 |
| 8 | 96218944 | rs2439632 | -0.504 | 0.255 | -0.418 | -0.608 | -0.009 | -0.028 | -0.028 | 0 | 0.32 | 0.42 | 0.27 | 0.17 | 0.28 | 0.31 |
| 8 | 96221015 | rs880641 | 0.026 | 0.744 | -0.394 | -0.349 | -0.001 | -0.001 | -0.001 | 0 | 0.84 | 0.89 | 0.83 | 0.72 | 0.79 | 0.76 |
| 8 | 96226340 | rs7824657 | 1.489 | 0.961 | -0.643 | -0.484 | 0.001 | -0.005 | -0.005 | 0.047 | 0.66 | 0.45 | 0.68 | 0.68 | 0.66 | 0.6 |
| 8 | 96229760 | rs13273059 | 0.142 | 1.383 | -1.346 | -0.121 | -0.032 | -0.045 | -0.045 | 0.417 | 0.87 | 0.95 | 0.85 | 0.72 | 0.81 | 0.76 |
| 8 | 96230428 | rs730820 | 0.995 | 1.219 | -1.231 | -0.165 | -0.015 | -0.009 | -0.009 | 0 | 0.76 | 0.84 | 0.76 | 0.68 | 0.69 | 0.64 |
| 8 | 96232747 | rs894153 | 0.404 | 0.966 | -2.124 | -0.128 | -0.007 | -0.148 | -0.148 | 2.338 | 0.36 | 0.07 | 0.34 | 0.54 | 0.33 | 0.43 |
| 8 | 96234075 | rs13278698 | 0.143 | 1.728 | -1.452 | -0.162 | -0.015 | 0.012 | 0.012 | 6.71 | 0.27 | 0.21 | 0.3 | 0.14 | 0.34 | 0.21 |
| 8 | 96237529 | rs4308702 | 0.135 | 1.61 | -1.414 | 0.015 | -0.002 | -0.002 | -0.002 | 0.154 | 0.12 | 0.08 | 0.13 | 0.13 | 0.16 | 0.2 |
| 8 | 96237596 | rs4509290 | -0.876 | 1.792 | -1.89 | -0.432 | -0.034 | 0.038 | 0.038 | 0.681 | 0.3 | 0.16 | 0.26 | 0.21 | 0.38 | 0.49 |
| 8 | 96239113 | rs879999 | -1.216 | 2.055 | -1.889 | -0.169 | -0.043 | 0.039 | 0.039 | 1.205 | 0.4 | 0.22 | 0.32 | 0.29 | 0.49 | 0.54 |
| 8 | 96239849 | rs13258855 | -0.807 | 2.021 | -1.496 | -0.156 | -0.017 | 0.002 | 0.002 | 0.354 | 0.32 | 0.2 | 0.26 | 0.29 | 0.38 | 0.49 |
| 8 | 96242237 | rs4734301 | 1.679 | 1.712 | -0.694 | -0.362 | -0.002 | 0 | 0 | 1.021 | 0.43 | 0.46 | 0.51 | 0.6 | 0.35 | 0.41 |
| 8 | 96245252 | rs2439630 | -1.747 | 1.624 | -0.388 | -0.403 | 0 | -0.003 | -0.003 | 0.358 | 0.54 | 0.51 | 0.45 | 0.29 | 0.58 | 0.57 |
| 8 | 96246687 | rs13276926 | 2.259 | 1.662 | -0.278 | -0.4 | -0.005 | 0.003 | 0.003 | 1.525 | 0.08 | 0.07 | 0.06 | 0 | 0.1 | 0.05 |
| 8 | 96247268 | rs10086865 | 2.272 | 1.851 | 0.043 | -0.274 | 0.015 | -0.014 | -0.014 | 0 | 0.36 | 0.3 | 0.38 | 0.46 | 0.3 | 0.31 |
| 8 | 96248443 | rs1441393 | 0.276 | 1.892 | -0.363 | -0.53 | -0.001 | 0.01 | 0.01 | 0 | 0.18 | 0.13 | 0.14 | 0.09 | 0.2 | 0.24 |
| 8 | 96251039 | rs17731290 | 0.811 | 1.779 | -0.394 | -0.149 | -0.052 | -0.035 | -0.035 | 0 | 0.13 | 0.02 | 0.11 | 0.18 | 0.18 | 0.25 |
| 8 | 96256866 | rs2514554 | 1.439 | 1.701 | -0.042 | 0.001 | 0.01 | 0.003 | 0.003 | 0 | 0.51 | 0.43 | 0.56 | 0.54 | 0.46 | 0.47 |
| 8 | 96263842 | rs13267037 | 1.993 | 1.573 | -0.1 | -0.068 | -0.017 | 0.034 | 0.034 | 1.866 | 0.09 | 0.04 | 0.06 | 0 | 0.1 | 0.05 |
| 8 | 96263909 | rs2165498 | 1.135 | 1.438 | 0.06 | -0.091 | -0.021 | -0.026 | -0.026 | 0 | 0.07 | 0.15 | 0.11 | 0.01 | 0.04 | 0.02 |
| 8 | 96273255 | rs13270004 | -0.711 | 1.729 | -0.003 | -0.11 | -0.009 | 0.037 | 0.037 | 0 | 0.33 | 0.2 | 0.24 | 0.14 | 0.34 | 0.32 |

| | | | | | | | | | | | | | | | | |
|---|---|---|---|---|---|---|---|---|---|---|---|---|---|---|---|---|
| 8 | 96276292 | rs7835921 | -1.407 | 1.703 | 0.005 | -0.049 | -0.001 | 0.003 | 0.003 | 0 | 0.46 | 0.49 | 0.39 | 0.29 | 0.47 | 0.5 |
| 8 | 96282926 | rs10956944 | 0.723 | 1.725 | -0.033 | -0.04 | -0.019 | 0.004 | 0.004 | 0.191 | 0.15 | 0.05 | 0.13 | 0.14 | 0.17 | 0.23 |
| 8 | 96283161 | rs2514557 | -1.604 | 1.652 | -0.021 | 0.066 | 0.008 | 0.001 | 0.001 | 0 | 0.51 | 0.58 | 0.44 | 0.46 | 0.54 | 0.52 |
| 8 | 96301075 | rs10504946 | 0.157 | 1.121 | -0.068 | 0.056 | -0.026 | -0.056 | -0.056 | 0 | 0.12 | 0.27 | 0.12 | 0.01 | 0.1 | 0.05 |
| 8 | 96301273 | rs7814035 | 0.675 | 1.146 | -0.277 | 0.091 | -0.024 | -0.044 | -0.044 | 0.385 | 0.1 | 0.23 | 0.12 | 0.01 | 0.07 | 0.03 |
| 8 | 96302648 | rs16917453 | 1.54 | 1.449 | -0.726 | 0.456 | -0.013 | -0.032 | -0.032 | 0.316 | 0.05 | 0.01 | 0.12 | 0.1 | 0.04 | 0.1 |
| 8 | 96308144 | rs17669929 | -0.745 | 1.522 | -0.597 | 0.036 | 0.004 | 0.002 | 0.002 | 0.166 | 0.27 | 0.24 | 0.22 | 0.29 | 0.32 | 0.46 |
| 8 | 96308367 | rs17669960 | 0.351 | 0.859 | -0.775 | -0.591 | 0.004 | 0.008 | 0.008 | 0 | 0.09 | 0.04 | 0.04 | 0 | 0.07 | 0.04 |
| 8 | 96314064 | rs7812444 | -0.729 | 0.641 | -1.019 | -1.052 | -0.002 | 0 | 0 | 2.16 | 0.53 | 0.44 | 0.35 | 0.16 | 0.56 | 0.2 |
| 8 | 96320757 | rs6989157 | 0.967 | 0.473 | -1.194 | -1.156 | -0.003 | -0.043 | -0.043 | 0 | 0.14 | 0.15 | 0.09 | 0 | 0.13 | 0.05 |
| 8 | 96321503 | rs13277271 | -0.156 | 0.309 | -1.544 | -1.211 | -0.024 | 0.09 | 0.09 | 0 | 0.3 | 0.1 | 0.23 | 0.11 | 0.33 | 0.19 |
| 8 | 96322185 | rs13268795 | -0.186 | 0.303 | -1.558 | -1.214 | -0.024 | 0.093 | 0.093 | 0 | 0.3 | 0.1 | 0.23 | 0.11 | 0.33 | 0.19 |
| 8 | 96329180 | rs4521728 | 1.301 | 0.282 | -1.584 | -1.248 | -0.013 | 0.046 | 0.046 | 0 | 0.15 | 0.07 | 0.1 | 0.01 | 0.19 | 0.06 |
| 8 | 96331853 | rs10956946 | 0.338 | 0.194 | -1.598 | -1.315 | -0.001 | 0.074 | 0.074 | 0 | 0.23 | 0.12 | 0.14 | 0.01 | 0.24 | 0.06 |
| 8 | 96332304 | rs1388953 | 0.497 | 0.152 | -1.584 | -1.35 | 0.007 | 0.008 | 0.008 | 0 | 0.15 | 0.08 | 0.13 | 0.11 | 0.12 | 0.14 |
| 8 | 96332994 | rs744382 | -0.569 | 0.259 | -1.504 | -1.233 | 0.002 | 0.14 | 0.14 | 0 | 0.38 | 0.15 | 0.27 | 0.12 | 0.36 | 0.2 |
| 8 | 96334914 | rs4735360 | -0.997 | 0.379 | -1.191 | -0.983 | -0.003 | 0.018 | 0.018 | 0.214 | 0.54 | 0.43 | 0.4 | 0.26 | 0.52 | 0.26 |
| 8 | 96344427 | rs4734305 | 0.415 | 0.303 | -1.311 | -0.592 | 0.01 | 0.012 | 0.012 | 0 | 0.62 | 0.73 | 0.76 | 0.71 | 0.66 | 0.79 |
| 8 | 96356624 | rs7842046 | -0.559 | 0.274 | -1.011 | -0.354 | 0.011 | -0.019 | -0.019 | 1.599 | 0.37 | 0.72 | 0.32 | 0.56 | 0.32 | 0.43 |
| 8 | 96359448 | rs4358776 | 0.358 | 0.173 | -0.029 | -0.321 | 0.017 | -0.099 | -0.099 | 1.322 | 0.22 | 0.46 | 0.14 | 0.13 | 0.24 | 0.25 |
| 8 | 96361869 | rs10099677 | 0.451 | 0.131 | 0.308 | -0.364 | 0.004 | -0.028 | -0.028 | 0 | 0.2 | 0.21 | 0.11 | 0.13 | 0.21 | 0.25 |
| 8 | 96363243 | rs16917486 | 1.254 | 0.214 | 0.033 | -0.459 | -0.003 | -0.005 | -0.005 | 0 | 0.21 | 0.19 | 0.35 | 0.4 | 0.24 | 0.36 |
| 8 | 96364688 | rs875062 | -0.124 | 0.338 | 0.088 | -0.254 | -0.001 | 0.027 | 0.027 | 0 | 0.25 | 0.47 | 0.16 | 0.33 | 0.23 | 0.27 |
| 8 | 96366766 | rs17732667 | 0.724 | 0.617 | -0.415 | -0.783 | -0.004 | 0.003 | 0.003 | 0 | 0.18 | 0.14 | 0.11 | 0.12 | 0.21 | 0.26 |
| 8 | 96370246 | rs10089336 | -0.556 | 0.615 | -0.699 | -0.795 | -0.001 | 0.033 | 0.033 | 0.005 | 0.31 | 0.71 | 0.2 | 0.12 | 0.3 | 0.31 |
| 8 | 96378214 | rs2318550 | 1.171 | 1.157 | -1.343 | -0.388 | -0.076 | -0.131 | -0.131 | 0 | 0.17 | 0.02 | 0.33 | 0.41 | 0.22 | 0.27 |
| 8 | 96378601 | rs10099630 | -0.329 | 0.868 | -0.445 | -1.117 | 0.009 | -0.039 | -0.039 | 3.546 | 0.22 | 0.26 | 0.14 | 0.1 | 0.28 | 0.3 |
| 8 | 96393670 | rs3103764 | 0.14 | 0.863 | -1.098 | -0.807 | -0.027 | 0.044 | 0.044 | 0.691 | 0.86 | 0.92 | 0.9 | 1 | 0.81 | 0.91 |
| 8 | 96404286 | rs1108144 | 0.632 | 0.181 | -1.056 | -1.444 | -0.005 | -0.055 | -0.055 | 0 | 0.6 | 0.55 | 0.72 | 0.89 | 0.56 | 0.8 |
| 8 | 96405931 | rs1352989 | 0.299 | 0.093 | -0.911 | -1.516 | 0.014 | 0.053 | 0.053 | 0.004 | 0.25 | 0.1 | 0.14 | 0.11 | 0.27 | 0.12 |
| 8 | 96406796 | rs3133749 | 0.127 | 0.406 | -0.72 | -1.374 | 0.015 | -0.109 | -0.109 | 0 | 0.83 | 0.65 | 0.86 | 1 | 0.82 | 0.91 |
| 8 | 96407350 | rs13268209 | -1.01 | -0.172 | -1 | -1.561 | 0.013 | 0.01 | 0.01 | 0.073 | 0.5 | 0.43 | 0.31 | 0.11 | 0.46 | 0.19 |
| 8 | 96414012 | rs12543022 | -0.066 | -0.06 | -1.19 | -1.289 | 0.006 | 0.026 | 0.026 | 0.958 | 0.13 | 0.06 | 0.06 | 0.04 | 0.09 | 0.04 |
| 8 | 96415129 | rs7827971 | 0.405 | -0.219 | -0.921 | -1.223 | -0.009 | -0.024 | -0.024 | 0 | 0.06 | 0.08 | 0.04 | 0 | 0.04 | 0.01 |
| 8 | 96415683 | rs10504943 | -0.63 | -0.746 | -0.679 | -1.322 | 0.008 | 0.003 | 0.003 | 0.709 | 0.27 | 0.24 | 0.14 | 0.05 | 0.18 | 0.11 |
| 8 | 96416544 | rs4132840 | -0.709 | -0.763 | -0.571 | -1.319 | 0.013 | 0.013 | 0.013 | 0.296 | 0.18 | 0.14 | 0.09 | 0.06 | 0.09 | 0.1 |
| 8 | 96417185 | rs7825742 | -0.234 | -0.705 | -0.577 | -1.291 | 0.008 | 0.008 | 0.008 | 0 | 0.21 | 0.19 | 0.15 | 0.06 | 0.17 | 0.1 |
| 8 | 96418497 | rs6997004 | -0.708 | -0.837 | -0.066 | -1.371 | -0.062 | -0.089 | -0.089 | 0.147 | 0.23 | 0.43 | 0.17 | 0.05 | 0.16 | 0.09 |
| 8 | 96419021 | rs7001679 | 1.04 | -0.795 | -0.242 | -1.323 | 0.009 | 0 | 0 | 0.699 | 0.1 | 0.06 | 0.03 | 0 | 0.05 | 0.03 |
| 8 | 96423887 | rs872529 | -0.022 | -0.022 | -0.554 | -0.71 | 0.026 | 0.043 | 0.043 | 0 | 0.1 | 0.16 | 0.13 | 0.21 | 0.21 | 0.12 |
| 8 | 96425734 | rs1027266 | -1.069 | -0.145 | -0.595 | -0.778 | -0.003 | -0.006 | -0.006 | 0.038 | 0.3 | 0.23 | 0.25 | 0.25 | 0.38 | 0.22 |
| 8 | 96428448 | rs2087791 | -1.336 | -0.104 | -0.403 | -0.616 | -0.011 | -0.006 | -0.006 | 0 | 0.4 | 0.44 | 0.28 | 0.24 | 0.36 | 0.24 |
| 8 | 96430728 | rs7829745 | -0.23 | 0.614 | -0.416 | -0.012 | 0.017 | 0.052 | 0.052 | 0 | 0.08 | 0.15 | 0.1 | 0.2 | 0.16 | 0.12 |
| 8 | 96435186 | rs2055055 | 0.316 | -0.395 | -0.476 | -0.394 | -0.011 | -0.014 | -0.014 | 0.283 | 0.72 | 0.69 | 0.88 | 0.86 | 0.81 | 0.9 |
| 8 | 96435634 | rs2035892 | -0.336 | -0.128 | 0.242 | -0.094 | 0.035 | 0.155 | 0.155 | 0 | 0.08 | 0.23 | 0.12 | 0.18 | 0.09 | 0.15 |
| 8 | 96435774 | rs2035893 | -0.885 | -0.225 | 0.321 | -0.185 | 0.015 | 0.077 | 0.077 | 0 | 0.15 | 0.31 | 0.15 | 0.18 | 0.15 | 0.18 |
| 8 | 96435888 | rs7463005 | -1.47 | -0.687 | 0.418 | -0.66 | 0.004 | 0.005 | 0.005 | 0 | 0.4 | 0.54 | 0.34 | 0.25 | 0.37 | 0.35 |
| 8 | 96436087 | rs10956949 | -0.201 | -0.637 | 0.127 | -0.905 | -0.005 | -0.028 | -0.028 | 0 | 0.25 | 0.22 | 0.2 | 0.06 | 0.22 | 0.17 |
| 8 | 96439026 | rs7824060 | -1.004 | -0.727 | -0.192 | -0.905 | -0.009 | -0.047 | -0.047 | 0 | 0.35 | 0.32 | 0.23 | 0.05 | 0.28 | 0.17 |
| 8 | 96442676 | rs4060029 | -0.613 | -0.164 | -0.119 | -0.666 | 0 | -0.013 | -0.013 | 0 | 0.23 | 0.21 | 0.2 | 0.05 | 0.24 | 0.22 |
| 8 | 96442763 | rs4060030 | -1.53 | -0.614 | -0.062 | -0.31 | 0.008 | 0.002 | 0.002 | 0 | 0.35 | 0.59 | 0.29 | 0.22 | 0.26 | 0.33 |
| 8 | 96444628 | rs3133755 | -0.671 | -0.665 | -0.336 | -0.475 | -0.003 | -0.021 | -0.021 | 0 | 0.42 | 0.61 | 0.32 | 0.2 | 0.31 | 0.25 |
| 8 | 96446676 | rs6471530 | 0.386 | -0.489 | -0.79 | -0.089 | -0.024 | -0.07 | -0.07 | 0 | 0.79 | 0.91 | 0.8 | 0.72 | 0.73 | 0.71 |
| 8 | 96446999 | rs1873745 | 0.174 | -0.888 | -0.383 | 0.284 | 0.019 | 0.034 | 0.034 | 0 | 0.2 | 0.34 | 0.27 | 0.31 | 0.2 | 0.17 |
| 8 | 96447726 | rs7009057 | 1.942 | -1.259 | -0.272 | -0.276 | -0.006 | 0.007 | 0.007 | 0 | 0.42 | 0.31 | 0.51 | 0.41 | 0.5 | 0.5 |
| 8 | 96451828 | rs16917667 | 2.136 | -1.315 | -0.173 | -0.281 | -0.015 | -0.028 | -0.028 | 0 | 0.07 | 0.11 | 0.02 | 0 | 0.02 | 0 |
| 8 | 96452047 | rs7823934 | -0.075 | -1.305 | -0.158 | -0.202 | 0.028 | 0.097 | 0.097 | 0 | 0.8 | 0.66 | 0.73 | 0.55 | 0.77 | 0.7 |
| 8 | 96452309 | rs12542369 | -0.06 | -1.306 | -0.159 | -0.203 | 0.028 | 0.096 | 0.096 | 0 | 0.79 | 0.66 | 0.73 | 0.55 | 0.77 | 0.7 |
| 8 | 96452712 | rs2221222 | -0.032 | -1.323 | 0.088 | -0.212 | 0.045 | 0.144 | 0.144 | 0 | 0.79 | 0.62 | 0.73 | 0.55 | 0.77 | 0.71 |
| 8 | 96454743 | rs7001207 | 0.717 | -1.384 | 0.041 | -0.251 | 0.016 | 0.019 | 0.019 | 0 | 0.67 | 0.55 | 0.6 | 0.47 | 0.64 | 0.61 |
| 8 | 96455453 | rs3133764 | 2.198 | -1.831 | -0.051 | -0.792 | -0.032 | 0.072 | 0.072 | 0 | 0.39 | 0.19 | 0.34 | 0.23 | 0.45 | 0.35 |
| 8 | 96456072 | rs3133767 | -0.759 | -1.607 | 0.027 | -0.739 | -0.015 | -0.01 | -0.01 | 0 | 0.28 | 0.34 | 0.21 | 0.18 | 0.22 | 0.25 |
| 8 | 96456588 | rs3133768 | -1.077 | -1.534 | 0.178 | -0.684 | -0.002 | -0.018 | -0.018 | 0 | 0.39 | 0.62 | 0.3 | 0.18 | 0.34 | 0.31 |
| 8 | 96462961 | rs6999006 | -0.345 | -1.256 | 0.332 | -0.512 | 0.002 | 0.003 | 0.003 | 0 | 0.34 | 0.36 | 0.23 | 0.08 | 0.26 | 0.2 |
| 8 | 96467671 | rs1532658 | -0.938 | -1.041 | 0.227 | -0.337 | 0.012 | 0.035 | 0.035 | 0 | 0.5 | 0.62 | 0.33 | 0.18 | 0.39 | 0.32 |
| 8 | 96470300 | rs6471533 | 0.203 | -0.97 | 0.216 | -0.205 | 0 | 0.002 | 0.002 | 0 | 0.63 | 0.48 | 0.55 | 0.4 | 0.68 | 0.57 |
| 8 | 96470675 | rs4236834 | -1.281 | -1.386 | -0.177 | -0.375 | -0.012 | 0.011 | 0.011 | 0 | 0.61 | 0.74 | 0.6 | 0.67 | 0.46 | 0.55 |
| 8 | 96472090 | rs12675383 | -0.275 | -1.138 | 0.365 | 0.496 | 0 | -0.013 | -0.013 | 0 | 0.16 | 0.1 | 0.16 | 0.28 | 0.18 | 0.24 |
| 8 | 96472423 | rs7835982 | -0.286 | -1.235 | 0.649 | 0.774 | 0 | 0 | 0 | 0 | 0.32 | 0.39 | 0.3 | 0.28 | 0.23 | 0.3 |

| | | | | | | | | | | | | | | | | |
|---|---|---|---|---|---|---|---|---|---|---|---|---|---|---|---|---|
| 8 | 96479988 | rs3133723 | -1.961 | -1.28 | 0.642 | 0.577 | 0.005 | -0.005 | -0.005 | 0.858 | 0.33 | 0.6 | 0.36 | 0.57 | 0.23 | 0.43 |
| 8 | 96483451 | rs10504950 | 0.9 | -1.065 | 0.236 | 0.687 | -0.001 | -0.015 | -0.015 | 0 | 0.08 | 0.02 | 0.1 | 0.08 | 0.08 | 0.11 |
| 8 | 96484147 | rs12545121 | -0.206 | -0.992 | 0.363 | 0.786 | -0.004 | 0.083 | 0.083 | 0 | 0.12 | 0.16 | 0.2 | 0.47 | 0.12 | 0.3 |
| 8 | 96489189 | rs2133947 | -1.481 | -0.873 | 0.067 | 0.46 | -0.031 | 0.001 | 0.001 | 0 | 0.54 | 0.81 | 0.51 | 0.56 | 0.44 | 0.55 |
| 8 | 96499699 | rs10956953 | -1.132 | -0.827 | 0.292 | 0.332 | 0.005 | 0.012 | 0.012 | 0 | 0.49 | 0.66 | 0.48 | 0.56 | 0.42 | 0.55 |
| 8 | 96507982 | rs3133732 | -1.399 | -1.277 | 0.391 | 0.227 | 0.004 | -0.004 | -0.004 | 0 | 0.33 | 0.6 | 0.37 | 0.6 | 0.25 | 0.43 |
| 8 | 96515888 | rs1909881 | -0.479 | -1.19 | 0.539 | 0.468 | 0.004 | 0.015 | 0.015 | 0 | 0.13 | 0.17 | 0.1 | 0.19 | 0.13 | 0.14 |
| 8 | 96516607 | rs6984692 | -0.611 | -1.168 | 0.396 | 0.383 | -0.002 | -0.021 | -0.021 | 0 | 0.42 | 0.45 | 0.26 | 0.19 | 0.33 | 0.26 |
| 8 | 96517216 | rs6993445 | 0.503 | -1.076 | -0.202 | 0.443 | 0.003 | 0.018 | 0.018 | 0.014 | 0.08 | 0.08 | 0.07 | 0.19 | 0.11 | 0.13 |
| 8 | 96518204 | rs12676175 | -0.138 | -0.99 | -0.389 | 0.611 | -0.027 | 0.115 | 0.115 | 0 | 0.14 | 0.26 | 0.24 | 0.42 | 0.1 | 0.29 |
| 8 | 96518612 | rs3104898 | -1.124 | -0.455 | -1.009 | 0.524 | -0.054 | -0.019 | -0.019 | 0 | 0.59 | 0.86 | 0.51 | 0.61 | 0.43 | 0.55 |
| 8 | 96521988 | rs1566516 | -1.005 | -0.626 | -1.044 | 0.523 | -0.031 | 0.002 | 0.002 | 0 | 0.6 | 0.85 | 0.52 | 0.63 | 0.43 | 0.58 |
| 8 | 96522225 | rs1009699 | -0.588 | -1.362 | -0.394 | 0.903 | -0.025 | 0.022 | 0.022 | 0 | 0.36 | 0.56 | 0.42 | 0.51 | 0.24 | 0.34 |
| 8 | 96522595 | rs904925 | -1.223 | -1.09 | -0.059 | 1.352 | -0.009 | 0.015 | 0.015 | 0.74 | 0.2 | 0.25 | 0.28 | 0.24 | 0.19 | 0.33 |
| 8 | 96524316 | rs3104891 | -0.106 | -0.82 | -0.036 | 1.244 | 0.032 | 0.107 | 0.107 | 0.15 | 0.09 | 0.19 | 0.16 | 0.23 | 0.09 | 0.18 |
| 8 | 96528845 | rs6471541 | -0.367 | -0.863 | -0.182 | 0.551 | -0.026 | -0.035 | -0.035 | 0 | 0.31 | 0.5 | 0.15 | 0.17 | 0.22 | 0.21 |
| 8 | 96529410 | rs1001095 | 0.643 | -0.982 | -0.31 | 0.428 | -0.007 | 0.014 | 0.014 | 1.639 | 0.19 | 0.18 | 0.09 | 0 | 0.18 | 0.07 |
| 8 | 96530572 | rs904923 | -0.33 | -0.88 | -0.096 | 0.541 | -0.058 | 0.091 | 0.091 | 0.108 | 0.13 | 0.37 | 0.06 | 0.17 | 0.03 | 0.14 |
| 8 | 96530648 | rs904924 | -0.975 | -0.758 | -0.678 | 0.651 | -0.001 | 0.001 | 0.001 | 0.242 | 0.4 | 0.41 | 0.48 | 0.52 | 0.37 | 0.49 |
| 8 | 96530971 | rs3104884 | -0.699 | -0.574 | -0.247 | 1.563 | 0 | 0.107 | 0.107 | 1.095 | 0.18 | 0.24 | 0.36 | 0.51 | 0.18 | 0.39 |
| 8 | 96533604 | rs2202056 | -1.071 | -0.447 | -0.539 | 0.817 | -0.027 | 0.015 | 0.015 | 1.104 | 0.29 | 0.42 | 0.42 | 0.68 | 0.24 | 0.54 |
| 8 | 96534806 | rs3133745 | -0.878 | -0.881 | -1.259 | 0.453 | 0.061 | 0.033 | 0.033 | 3.913 | 0.31 | 0.82 | 0.28 | 0.21 | 0.16 | 0.29 |
| 8 | 96534917 | rs3133746 | -0.629 | -0.777 | -1.118 | 0.218 | -0.004 | -0.004 | -0.004 | 0 | 0.54 | 0.52 | 0.52 | 0.63 | 0.4 | 0.47 |
| 8 | 96536023 | rs11776832 | 0.497 | -0.417 | -1.726 | -0.696 | -0.05 | 0.153 | 0.153 | 0 | 0.31 | 0.11 | 0.17 | 0.02 | 0.32 | 0.12 |
| 8 | 96537844 | rs1472560 | -0.527 | -0.541 | -1.231 | -0.894 | 0.003 | 0.068 | 0.068 | 0.773 | 0.5 | 0.38 | 0.45 | 0.06 | 0.46 | 0.39 |
| 8 | 96543556 | rs3134233 | -0.63 | -0.388 | -0.765 | 0.595 | -0.001 | 0.005 | 0.005 | 0.665 | 0.21 | 0.17 | 0.32 | 0.36 | 0.18 | 0.45 |
| 8 | 96549915 | rs3105001 | -0.876 | -0.416 | -0.665 | 0.589 | -0.014 | 0.036 | 0.036 | 0.275 | 0.23 | 0.3 | 0.35 | 0.38 | 0.2 | 0.44 |
| 8 | 96551323 | rs10808678 | 0.159 | -0.381 | -0.815 | -0.082 | 0.013 | 0.021 | 0.021 | 0 | 0.45 | 0.53 | 0.27 | 0.21 | 0.33 | 0.26 |
| 8 | 96551404 | rs7011044 | 0.639 | -0.975 | -0.794 | 0.058 | -0.042 | -0.007 | -0.007 | 0 | 0.65 | 0.52 | 0.53 | 0.32 | 0.79 | 0.52 |
| 8 | 96551874 | rs3105000 | 0.297 | 0.275 | -0.824 | -0.153 | -0.057 | 0.01 | 0.01 | 0 | 0.32 | 0.23 | 0.46 | 0.24 | 0.46 | 0.33 |
| 8 | 96552742 | rs7835538 | 0.864 | 0.01 | -0.441 | -0.514 | -0.001 | -0.028 | -0.028 | 0 | 0.19 | 0.16 | 0.12 | 0.01 | 0.2 | 0.08 |
| 8 | 96553127 | rs3134227 | 0.345 | 0.087 | -0.726 | -0.736 | -0.074 | -0.002 | -0.002 | 0.955 | 0.32 | 0.12 | 0.41 | 0.25 | 0.43 | 0.37 |
| 8 | 96555479 | rs3104999 | -0.629 | -0.281 | 0.014 | -0.557 | -0.004 | 0.003 | 0.003 | 0.191 | 0.57 | 0.48 | 0.57 | 0.47 | 0.65 | 0.68 |
| 8 | 96555664 | rs3134216 | -0.253 | -0.362 | 0.039 | -0.609 | -0.011 | -0.013 | -0.013 | 0 | 0.42 | 0.3 | 0.45 | 0.46 | 0.46 | 0.54 |
| 8 | 96560359 | rs11776205 | 0.546 | -0.709 | -0.018 | -0.732 | 0.001 | 0.017 | 0.017 | 0 | 0.29 | 0.33 | 0.3 | 0.6 | 0.27 | 0.48 |
| 8 | 96563049 | rs3134212 | -0.792 | -0.238 | -0.356 | -0.767 | -0.01 | 0.068 | 0.068 | 0.032 | 0.56 | 0.72 | 0.66 | 0.87 | 0.43 | 0.69 |
| 8 | 96564170 | rs17749656 | 1.428 | 0.04 | -0.748 | -0.453 | -0.053 | 0.057 | 0.057 | 0.311 | 0.24 | 0.09 | 0.22 | 0.13 | 0.34 | 0.22 |
| 8 | 96572791 | rs3104992 | -0.64 | -0.166 | -0.216 | -0.335 | 0.001 | -0.018 | -0.018 | 0 | 0.4 | 0.49 | 0.51 | 0.71 | 0.36 | 0.62 |
| 8 | 96572941 | rs7816064 | 1.289 | 0.048 | -0.512 | -0.071 | -0.006 | -0.015 | -0.015 | 0 | 0.18 | 0.18 | 0.23 | 0.39 | 0.18 | 0.22 |
| 8 | 96573283 | rs13271214 | 0.089 | -0.123 | -0.375 | -0.156 | -0.002 | -0.022 | -0.022 | 0 | 0.31 | 0.28 | 0.34 | 0.64 | 0.29 | 0.37 |
| 8 | 96573370 | rs3134240 | -0.38 | 0.58 | -0.299 | -0.223 | 0.01 | 0.006 | 0.006 | 0.25 | 0.9 | 0.78 | 0.94 | 0.87 | 0.91 | 0.79 |
| 8 | 96574975 | rs3134238 | 0.192 | 0.441 | -0.538 | -0.572 | -0.004 | 0 | 0 | 0.776 | 0.36 | 0.37 | 0.48 | 0.32 | 0.48 | 0.42 |
| 8 | 96576196 | rs3104988 | 0.192 | 0.441 | -0.516 | -0.572 | -0.013 | 0 | 0 | 0.106 | 0.36 | 0.33 | 0.48 | 0.32 | 0.48 | 0.42 |
| 8 | 96582595 | rs1604259 | -0.008 | 0.123 | -0.469 | -0.817 | 0.007 | 0.005 | 0.005 | 0 | 0.45 | 0.43 | 0.39 | 0.67 | 0.32 | 0.51 |
| 8 | 96583551 | rs2318878 | -1.789 | 0.017 | -0.683 | -0.872 | 0 | -0.024 | -0.024 | 1.16 | 0.8 | 0.84 | 0.87 | 0.99 | 0.8 | 0.93 |
| 8 | 96586615 | rs9297962 | 1.703 | -0.015 | -1.005 | -0.553 | -0.077 | 0.044 | 0.044 | 1.056 | 0.2 | 0.04 | 0.19 | 0.07 | 0.31 | 0.2 |
| 8 | 96592937 | rs4532570 | 0.131 | -0.698 | -0.344 | -0.171 | -0.063 | -0.013 | -0.013 | 0 | 0.23 | 0.35 | 0.11 | 0.22 | 0.07 | 0.27 |
| 8 | 96593539 | rs2318876 | 0.307 | -0.466 | -0.489 | -0.162 | -0.014 | -0.006 | -0.006 | 0 | 0.56 | 0.48 | 0.63 | 0.33 | 0.72 | 0.49 |
| 8 | 96593716 | rs16917906 | 0.299 | 0.029 | -0.997 | 0.127 | -0.008 | -0.066 | -0.066 | 0 | 0.25 | 0.14 | 0.25 | 0.45 | 0.22 | 0.37 |
| 8 | 96594034 | rs3104966 | 0.081 | 0.29 | -0.74 | -0.429 | -0.017 | 0.034 | 0.034 | 0 | 0.41 | 0.36 | 0.44 | 0.19 | 0.48 | 0.38 |
| 8 | 96594137 | rs3104965 | 0.177 | -0.013 | -0.117 | -0.458 | -0.048 | -0.017 | -0.017 | 0 | 0.76 | 0.57 | 0.8 | 0.77 | 0.89 | 0.8 |
| 8 | 96594239 | rs4735376 | -0.172 | 0.162 | -0.39 | -0.519 | 0.004 | 0 | 0 | 0.011 | 0.37 | 0.28 | 0.4 | 0.74 | 0.3 | 0.54 |
| 8 | 96594769 | rs2873824 | -0.457 | 0.683 | -0.048 | -0.36 | 0.042 | 0.105 | 0.105 | 0.373 | 0.86 | 0.74 | 0.75 | 0.66 | 0.8 | 0.76 |
| 8 | 96596354 | rs904921 | 0.677 | 0.188 | -0.188 | -0.704 | 0.017 | 0.01 | 0.01 | 0 | 0.49 | 0.4 | 0.4 | 0.47 | 0.4 | 0.39 |
| 8 | 96596749 | rs3134245 | -0.114 | 0.329 | -0.577 | -0.772 | -0.038 | -0.009 | -0.009 | 0 | 0.67 | 0.79 | 0.61 | 0.62 | 0.53 | 0.54 |
| 8 | 96620404 | rs3104899 | 0.044 | 0.045 | -0.538 | -0.906 | 0.036 | 0.017 | 0.017 | 0 | 0.46 | 0.65 | 0.51 | 0.61 | 0.3 | 0.45 |
| 8 | 96625550 | rs3134211 | 0.297 | 0.107 | -0.483 | -0.953 | -0.002 | 0.047 | 0.047 | 0 | 0.09 | 0.21 | 0.15 | 0.14 | 0.09 | 0.1 |
| 8 | 96628523 | rs12549245 | 2.263 | 0.488 | -0.727 | -0.892 | -0.047 | -0.036 | -0.036 | 1.064 | 0.08 | 0.01 | 0.15 | 0.11 | 0.17 | 0.13 |
| 8 | 96630632 | rs11776030 | 0.295 | 0.237 | -0.671 | -0.994 | 0.09 | 0.209 | 0.209 | 0.151 | 0.48 | 0.16 | 0.41 | 0.15 | 0.67 | 0.32 |
| 8 | 96636912 | rs12155746 | 1.475 | 0.478 | -0.614 | -1.038 | -0.087 | 0.062 | 0.062 | 0 | 0.15 | 0.02 | 0.12 | 0.04 | 0.25 | 0.11 |
| 8 | 96637260 | rs4366053 | 0.063 | 0.254 | -0.354 | -1.042 | -0.024 | 0.016 | 0.016 | 0 | 0.17 | 0.31 | 0.21 | 0.15 | 0.13 | 0.2 |
| 8 | 96649711 | rs5028874 | 1.374 | 0.392 | -0.582 | -1.113 | -0.044 | 0.048 | 0.048 | 0 | 0.15 | 0.06 | 0.12 | 0.04 | 0.25 | 0.19 |
| 8 | 96651700 | rs9649979 | 1.418 | 0.397 | -0.586 | -1.121 | -0.047 | 0.051 | 0.051 | 0 | 0.15 | 0.05 | 0.12 | 0.04 | 0.25 | 0.19 |
| 8 | 96659417 | rs1484593 | 1.393 | 0.157 | -0.525 | -1.143 | -0.01 | -0.023 | -0.023 | 2.908 | 0.1 | 0.14 | 0.05 | 0 | 0.04 | 0.02 |
| 8 | 96671587 | rs997257 | 1.027 | 0.285 | 0.003 | -1.183 | 0.036 | -0.064 | -0.064 | 0.265 | 0.17 | 0.25 | 0.15 | 0.04 | 0.25 | 0.19 |
| 8 | 96672404 | rs1385237 | 1.312 | -0.116 | -0.161 | -1.104 | -0.005 | 0.008 | 0.008 | 0 | 0.29 | 0.31 | 0.3 | 0.48 | 0.18 | 0.39 |
| 8 | 96675932 | rs6471554 | 1.729 | -0.072 | -0.532 | -1.203 | -0.023 | 0.056 | 0.056 | 0.137 | 0.21 | 0.08 | 0.13 | 0.01 | 0.24 | 0.08 |
| 8 | 96685295 | rs7013599 | -0.125 | -0.074 | -0.447 | -1.044 | -0.013 | 0.058 | 0.058 | 0 | 0.46 | 0.28 | 0.41 | 0.34 | 0.49 | 0.32 |
| 8 | 96694204 | rs1470389 | 1.434 | -0.058 | -0.346 | -1.056 | -0.028 | -0.046 | -0.046 | 0 | 0.1 | 0.21 | 0.05 | 0 | 0.04 | 0.02 |

| | | | | | | | | | | | | | | | | |
|---|---|---|---|---|---|---|---|---|---|---|---|---|---|---|---|---|
| 8 | 96694458 | rs3134204 | 1.036 | -0.004 | -0.597 | -1.046 | -0.056 | -0.023 | -0.023 | 0.105 | 0.21 | 0.09 | 0.25 | 0.18 | 0.33 | 0.21 |
| 8 | 96700246 | rs1470386 | -0.167 | 0.096 | -0.456 | -0.994 | 0.001 | -0.03 | -0.03 | 0 | 0.35 | 0.55 | 0.33 | 0.19 | 0.38 | 0.23 |
| 8 | 96701836 | rs3099088 | -0.216 | 0.498 | -0.546 | -1.518 | -0.002 | -0.053 | -0.053 | 0 | 0.56 | 0.58 | 0.39 | 0.08 | 0.61 | 0.3 |
| 8 | 96703930 | rs3099087 | 0.33 | 0.041 | -0.578 | -1.309 | 0.003 | -0.026 | -0.026 | 0 | 0.34 | 0.35 | 0.21 | 0.04 | 0.28 | 0.1 |
| 8 | 96705190 | rs3102484 | -0.519 | 0.01 | -0.516 | -1.185 | 0 | -0.001 | -0.001 | 0 | 0.62 | 0.66 | 0.51 | 0.49 | 0.46 | 0.57 |
| 8 | 96706331 | rs4735382 | 0.098 | -0.026 | -0.556 | -0.625 | -0.004 | 0.007 | 0.007 | 0 | 0.29 | 0.31 | 0.29 | 0.48 | 0.18 | 0.47 |
| 8 | 96710585 | rs13280459 | -0.813 | -0.016 | -0.67 | -1.786 | 0.02 | 0.043 | 0.043 | 0 | 0.42 | 0.66 | 0.29 | 0.04 | 0.34 | 0.31 |
| 8 | 96713178 | rs10102578 | -0.732 | -0.246 | -0.455 | -1.827 | -0.015 | -0.112 | -0.112 | 0 | 0.32 | 0.55 | 0.21 | 0.03 | 0.25 | 0.28 |
| 8 | 96717277 | rs13261503 | 1.446 | -0.078 | -0.776 | -1.671 | 0.003 | -0.001 | -0.001 | 0 | 0.06 | 0.03 | 0.02 | 0 | 0.03 | 0.03 |
| 8 | 96720443 | rs10089847 | -0.755 | -0.232 | -0.622 | -1.8 | -0.011 | 0.076 | 0.076 | 0 | 0.27 | 0.11 | 0.15 | 0.03 | 0.26 | 0.31 |
| 8 | 96723902 | rs7012001 | -0.522 | -0.223 | -0.372 | -1.785 | -0.002 | -0.053 | -0.053 | 0 | 0.22 | 0.24 | 0.15 | 0.03 | 0.22 | 0.3 |
| 8 | 96725932 | rs4236837 | -1.188 | 0.213 | -1.32 | -1.628 | -0.103 | -0.009 | -0.009 | 0.369 | 0.76 | 0.94 | 0.75 | 0.82 | 0.6 | 0.81 |
| 8 | 96726063 | rs1545898 | -0.745 | 0.06 | -0.887 | -1.655 | 0 | 0.055 | 0.055 | 0 | 0.29 | 0.14 | 0.17 | 0.04 | 0.27 | 0.31 |
| 8 | 96730999 | rs6998540 | -0.548 | 0.283 | -0.692 | -1.407 | -0.001 | -0.017 | -0.017 | 0 | 0.19 | 0.14 | 0.13 | 0.04 | 0.24 | 0.21 |
| 8 | 96732452 | rs6990507 | 0.491 | -0.33 | -0.391 | -1.807 | 0.041 | 0.073 | 0.073 | 0 | 0.51 | 0.39 | 0.56 | 0.88 | 0.35 | 0.54 |
| 8 | 96733694 | rs2008555 | -0.516 | -0.068 | 0.003 | -1.168 | 0.02 | 0.002 | 0.002 | 0 | 0.18 | 0.31 | 0.21 | 0.14 | 0.25 | 0.22 |
| 8 | 96734320 | rs1825443 | 0.86 | -1.04 | -0.072 | -1.574 | -0.004 | -0.045 | -0.045 | 0.006 | 0.21 | 0.22 | 0.12 | 0.01 | 0.14 | 0.17 |
| 8 | 96736624 | rs11984568 | -1.09 | -0.856 | -0.377 | -1.579 | -0.067 | 0.069 | 0.069 | 0 | 0.72 | 0.82 | 0.71 | 0.94 | 0.53 | 0.77 |
| 8 | 96738171 | rs3102480 | 0.769 | -1.174 | -0.703 | -1.52 | 0.034 | 0.031 | 0.031 | 0 | 0.33 | 0.16 | 0.38 | 0.8 | 0.21 | 0.34 |
| 8 | 96739885 | rs4270930 | -0.282 | -1.019 | -0.689 | -1.446 | -0.002 | -0.006 | -0.006 | 0 | 0.34 | 0.3 | 0.27 | 0.14 | 0.3 | 0.4 |
| 8 | 96742830 | rs6471558 | -0.445 | 0.061 | -0.455 | -0.719 | 0.04 | -0.051 | -0.051 | 0 | 0.69 | 0.59 | 0.67 | 0.87 | 0.51 | 0.74 |
| 8 | 96744403 | rs2016731 | -0.583 | 0.282 | -0.84 | 0.442 | 0.014 | 0.056 | 0.056 | 0 | 0.11 | 0.14 | 0.21 | 0.25 | 0.17 | 0.24 |
| 8 | 96746100 | rs3102477 | -0.833 | 0.129 | -1.01 | 1.136 | -0.002 | 0.129 | 0.129 | 0 | 0.07 | 0.15 | 0.16 | 0.31 | 0.05 | 0.21 |
| 8 | 96746613 | rs11780919 | 0.026 | -0.041 | -1.817 | 1.258 | 0.011 | -0.057 | -0.057 | 0 | 0.14 | 0.03 | 0.17 | 0.24 | 0.12 | 0.17 |
| 8 | 96747238 | rs6471559 | 0.061 | -0.226 | -2.188 | 0.818 | 0 | 0.008 | 0.008 | 0.282 | 0.58 | 0.9 | 0.58 | 0.62 | 0.39 | 0.57 |
| 8 | 96749765 | rs3134496 | -0.73 | -0.225 | -1.782 | 1.38 | -0.038 | -0.079 | -0.079 | 0 | 0.88 | 0.99 | 0.77 | 0.72 | 0.86 | 0.87 |
| 8 | 96750281 | rs6982567 | -0.48 | -0.865 | -1.061 | 0.906 | -0.022 | 0.004 | 0.004 | 0 | 0.25 | 0.35 | 0.26 | 0.27 | 0.19 | 0.2 |
| 8 | 96751374 | rs3134497 | -0.359 | -1.08 | -1.27 | 1.023 | -0.019 | 0.016 | 0.016 | 0 | 0.3 | 0.39 | 0.3 | 0.55 | 0.2 | 0.25 |
| 8 | 96751803 | rs7819896 | -0.236 | -0.275 | -2.041 | 0.797 | -0.075 | -0.068 | -0.068 | 0 | 0.78 | 0.99 | 0.79 | 0.78 | 0.74 | 0.75 |
| 8 | 96752955 | rs13275914 | -0.354 | -0.401 | -2.083 | 0.459 | -0.019 | -0.039 | -0.039 | 0.296 | 0.58 | 0.21 | 0.57 | 0.53 | 0.56 | 0.57 |
| 8 | 96753076 | rs13268649 | 0.063 | -0.96 | -1.044 | 0.193 | -0.054 | 0.146 | 0.146 | 0.215 | 0.2 | 0.5 | 0.32 | 0.46 | 0.12 | 0.26 |
| 8 | 96757443 | rs11995026 | 0.773 | -0.913 | -0.996 | 0.403 | -0.016 | 0.04 | 0.04 | 0 | 0.16 | 0.24 | 0.28 | 0.39 | 0.12 | 0.27 |
| 8 | 96758008 | rs4236838 | -2.135 | -0.996 | -1.135 | 0.417 | 0.004 | -0.001 | -0.001 | 0 | 0.92 | 0.94 | 0.96 | 0.93 | 0.93 | 0.91 |
| 8 | 96758878 | rs3102476 | 0.724 | -1.021 | -0.971 | 0.705 | -0.002 | 0.041 | 0.041 | 0 | 0.1 | 0.15 | 0.19 | 0.26 | 0.08 | 0.12 |
| 8 | 96762253 | rs1484584 | -0.007 | -1.078 | -1.134 | 0.404 | 0.036 | -0.016 | -0.016 | 0 | 0.57 | 0.73 | 0.56 | 0.48 | 0.67 | 0.56 |
| 8 | 96762545 | rs3107060 | 0.285 | -0.902 | -1.353 | 0.291 | -0.013 | -0.019 | -0.019 | 0 | 0.32 | 0.16 | 0.34 | 0.3 | 0.34 | 0.33 |
| 8 | 96764258 | rs7817825 | -0.847 | -0.681 | -1.396 | 0.319 | 0.003 | 0.058 | 0.058 | 0 | 0.73 | 0.89 | 0.83 | 0.85 | 0.79 | 0.82 |
| 8 | 96770906 | rs7830018 | -0.552 | -0.61 | -0.325 | 0.767 | 0.008 | -0.005 | -0.005 | 0 | 0.57 | 0.56 | 0.6 | 0.58 | 0.49 | 0.51 |
| 8 | 96771076 | rs13253287 | 1.737 | -0.497 | -0.196 | 0.675 | 0.025 | -0.008 | -0.008 | 0 | 0.21 | 0.22 | 0.17 | 0.17 | 0.33 | 0.24 |
| 8 | 96773262 | rs4263730 | -0.083 | -0.889 | -0.321 | 0.498 | -0.014 | -0.01 | -0.01 | 0.98 | 0.45 | 0.7 | 0.45 | 0.48 | 0.5 | 0.52 |
| 8 | 96773469 | rs12155779 | 0.711 | -0.891 | -0.402 | 0.523 | -0.001 | 0.012 | 0.012 | 0 | 0.34 | 0.2 | 0.33 | 0.33 | 0.31 | 0.21 |
| 8 | 96784526 | rs12676394 | 0.507 | -0.961 | -0.535 | 0.39 | 0.002 | 0.01 | 0.01 | 0 | 0.3 | 0.22 | 0.31 | 0.27 | 0.3 | 0.29 |
| 8 | 96784636 | rs12678569 | -1.669 | -1.005 | -0.598 | 0.391 | -0.012 | -0.01 | -0.01 | 0 | 0.8 | 0.97 | 0.8 | 0.82 | 0.83 | 0.73 |
| 8 | 96784717 | rs3107047 | 0.915 | -1.085 | -0.1 | 0.347 | 0.042 | 0.064 | 0.064 | 0 | 0.31 | 0.46 | 0.36 | 0.46 | 0.42 | 0.37 |
| 8 | 96786148 | rs10106907 | 0.035 | -1.077 | -0.239 | 0.347 | 0.012 | 0.064 | 0.064 | 0 | 0.44 | 0.2 | 0.39 | 0.35 | 0.4 | 0.35 |
| 8 | 96787822 | rs1817551 | 0.071 | -1.011 | -0.116 | 0.392 | 0.003 | 0.021 | 0.021 | 0.137 | 0.49 | 0.7 | 0.46 | 0.55 | 0.51 | 0.44 |
| 8 | 96788760 | rs3099105 | 0.714 | -0.822 | 0.011 | 0.519 | 0.037 | 0.062 | 0.062 | 0 | 0.31 | 0.47 | 0.36 | 0.47 | 0.41 | 0.38 |
| 8 | 96790061 | rs3134517 | 0.854 | -0.79 | -0.014 | 0.533 | 0.036 | 0.066 | 0.066 | 0 | 0.3 | 0.43 | 0.35 | 0.47 | 0.4 | 0.38 |
| 8 | 96794835 | rs3099096 | 0.35 | -0.881 | -0.378 | 0.542 | 0.028 | -0.003 | -0.003 | 0.018 | 0.49 | 0.43 | 0.47 | 0.48 | 0.64 | 0.56 |
| 8 | 96797851 | rs3107041 | -0.006 | -0.553 | -0.073 | 0.76 | -0.005 | 0.009 | 0.009 | 0 | 0.68 | 0.66 | 0.58 | 0.57 | 0.74 | 0.63 |
| 8 | 96801202 | rs990530 | -1.021 | -0.29 | -0.536 | 0.69 | 0.013 | 0.016 | 0.016 | 0.407 | 0.79 | 0.88 | 0.79 | 0.82 | 0.83 | 0.76 |
| 8 | 96804065 | rs3107038 | 0.447 | -0.235 | -0.407 | 0.76 | 0.02 | -0.002 | -0.002 | 0 | 0.51 | 0.34 | 0.31 | 0.46 | 0.56 | 0.55 |
| 8 | 96804762 | rs1465992 | 1.789 | -0.069 | -0.583 | 0.803 | -0.035 | -0.014 | -0.014 | 0 | 0.19 | 0.12 | 0.14 | 0.21 | 0.31 | 0.27 |
| 8 | 96812747 | rs3107035 | 1.841 | -0.092 | -0.537 | 0.81 | -0.025 | -0.009 | -0.009 | 0 | 0.19 | 0.13 | 0.14 | 0.21 | 0.3 | 0.27 |
| 8 | 96814338 | rs3107034 | -0.268 | -0.243 | -0.504 | 0.769 | -0.012 | 0.028 | 0.028 | 0 | 0.56 | 0.74 | 0.65 | 0.61 | 0.52 | 0.48 |
| 8 | 96815218 | rs4236839 | -1.246 | -0.272 | -0.726 | 0.777 | 0.003 | 0.019 | 0.019 | 0.46 | 0.78 | 0.97 | 0.8 | 0.83 | 0.83 | 0.75 |
| 8 | 96819401 | rs2873955 | 0.271 | -0.42 | -0.365 | 0.842 | -0.005 | -0.011 | -0.011 | 0 | 0.47 | 0.33 | 0.58 | 0.52 | 0.47 | 0.39 |
| 8 | 96823305 | rs3134498 | -0.802 | -0.374 | -0.36 | 0.806 | -0.028 | 0.012 | 0.012 | 0 | 0.59 | 0.79 | 0.65 | 0.62 | 0.53 | 0.5 |
| 8 | 96835380 | rs568815 | -0.596 | -0.221 | -0.216 | 0.765 | 0.011 | 0.009 | 0.009 | 0.005 | 0.7 | 0.41 | 0.8 | 0.7 | 0.64 | 0.65 |
| 8 | 96846278 | rs559167 | -1.317 | -0.366 | -0.537 | 0.535 | -0.022 | 0.017 | 0.017 | 0 | 0.76 | 0.8 | 0.83 | 0.81 | 0.67 | 0.72 |
| 8 | 96846973 | rs506767 | 0.417 | -0.362 | -0.396 | 0.605 | -0.017 | -0.026 | -0.026 | 0.788 | 0.27 | 0.36 | 0.34 | 0.2 | 0.23 | 0.25 |
| 8 | 96848744 | rs472920 | -0.237 | -0.435 | -0.795 | 0.972 | -0.009 | -0.01 | -0.01 | 0.628 | 0.61 | 0.31 | 0.71 | 0.59 | 0.56 | 0.53 |
| 8 | 96852520 | rs515782 | 0.981 | -0.624 | -1.06 | 0.785 | 0.013 | 0.003 | 0.003 | 0 | 0.21 | 0.06 | 0.3 | 0.2 | 0.18 | 0.23 |
| 8 | 96853415 | rs16918205 | 0.271 | -0.651 | -1.042 | 0.624 | -0.002 | -0.019 | -0.019 | 0 | 0.08 | 0.08 | 0.03 | 0 | 0.05 | 0.05 |
| 8 | 96855256 | rs551363 | -0.317 | -0.374 | -1.617 | 1.032 | 0.024 | -0.03 | -0.03 | 0 | 0.35 | 0.24 | 0.38 | 0.47 | 0.25 | 0.44 |
| 8 | 96858955 | rs4735401 | 0.994 | -0.23 | -2.32 | 0.374 | -0.005 | 0.031 | 0.031 | 0 | 0.22 | 0.02 | 0.11 | 0.12 | 0.16 | 0.19 |
| 8 | 96859623 | rs17688634 | 0.474 | -0.044 | -1.717 | 0.577 | -0.01 | 0.019 | 0.019 | 0 | 0.38 | 0.25 | 0.41 | 0.35 | 0.39 | 0.29 |
| 8 | 96863057 | rs577904 | -0.59 | -0.009 | -1.452 | 0.578 | 0.015 | -0.003 | -0.003 | 0 | 0.21 | 0.04 | 0.3 | 0.21 | 0.18 | 0.25 |
| 8 | 96864355 | rs561330 | -0.409 | -0.016 | -1.337 | 0.601 | 0.014 | -0.006 | -0.006 | 0 | 0.17 | 0.04 | 0.23 | 0.19 | 0.15 | 0.24 |

| | | | | | | | | | | | | | | | | |
|---|---|---|---|---|---|---|---|---|---|---|---|---|---|---|---|---|
| 8 | 96870639 | rs3864656 | -0.359 | 0.009 | -1.214 | 0.657 | 0.022 | -0.022 | -0.022 | 0 | 0.19 | 0.11 | 0.26 | 0.29 | 0.15 | 0.17 |
| 8 | 96871771 | rs550254 | -0.859 | 0.134 | -0.199 | 0.375 | 0.082 | -0.018 | -0.018 | 0 | 0.09 | 0.47 | 0.09 | 0 | 0.11 | 0.05 |
| 8 | 96874831 | rs564500 | -0.522 | 0.38 | -1.309 | 0.355 | -0.023 | -0.082 | -0.082 | 0 | 0.63 | 0.85 | 0.57 | 0.5 | 0.63 | 0.55 |
| 8 | 96881703 | rs554014 | -0.306 | 0.278 | -1.252 | 0.305 | 0.002 | -0.042 | -0.042 | 0 | 0.61 | 0.77 | 0.56 | 0.5 | 0.62 | 0.54 |
| 8 | 96885413 | rs552100 | -0.453 | 0.302 | -1.438 | 0.339 | -0.023 | -0.087 | -0.087 | 0.165 | 0.62 | 0.86 | 0.58 | 0.5 | 0.62 | 0.54 |
| 8 | 96893825 | rs980160 | 0.734 | 0.572 | -0.717 | 0.137 | -0.005 | 0.003 | 0.003 | 0 | 0.31 | 0.23 | 0.23 | 0.2 | 0.35 | 0.33 |
| 8 | 96896514 | rs994741 | 0.675 | 0.574 | -0.5 | 0.114 | -0.014 | 0.014 | 0.014 | 0 | 0.32 | 0.26 | 0.25 | 0.2 | 0.4 | 0.34 |
| 8 | 96896725 | rs1994222 | 0.393 | 0.67 | -0.687 | 0.309 | -0.006 | -0.005 | -0.005 | 0 | 0.53 | 0.58 | 0.51 | 0.48 | 0.49 | 0.55 |
| 8 | 96896913 | rs11995162 | 0.363 | 0.975 | -1.674 | 0.634 | 0.018 | -0.03 | -0.03 | 0 | 0.25 | 0.09 | 0.17 | 0.36 | 0.2 | 0.2 |
| 8 | 96897432 | rs753896 | -0.205 | 1.613 | -1.096 | 0.522 | -0.017 | -0.019 | -0.019 | 1.055 | 0.13 | 0.08 | 0.22 | 0.15 | 0.17 | 0.17 |
| 8 | 96898466 | rs678287 | -0.906 | 1.794 | -0.877 | 0.233 | -0.02 | 0.027 | 0.027 | 0 | 0.05 | 0.01 | 0.07 | 0 | 0.17 | 0.06 |
| 8 | 96900502 | rs2319361 | 0.844 | 0.988 | -0.787 | -0.083 | 0.001 | -0.011 | -0.011 | 0 | 0.86 | 0.85 | 0.9 | 0.99 | 0.85 | 0.92 |
| 8 | 96900622 | rs2873957 | -0.26 | 0.662 | -0.452 | 0.199 | -0.01 | 0.072 | 0.072 | 0.286 | 0.3 | 0.52 | 0.43 | 0.47 | 0.27 | 0.36 |
| 8 | 96900721 | rs2319362 | -0.286 | 0.652 | -0.182 | 0.19 | -0.009 | 0.058 | 0.058 | 0.243 | 0.3 | 0.49 | 0.43 | 0.47 | 0.26 | 0.36 |
| 8 | 96901876 | rs920200 | 1.658 | 0.768 | -0.97 | -0.21 | -0.073 | 0.066 | 0.066 | 0.423 | 0.29 | 0.06 | 0.22 | 0.18 | 0.35 | 0.29 |
| 8 | 96903504 | rs983072 | 0.244 | 0.473 | -0.334 | -0.174 | 0.004 | -0.059 | -0.059 | 0.115 | 0.9 | 0.72 | 0.92 | 0.99 | 0.89 | 0.93 |
| 8 | 96904970 | rs13274725 | 0.72 | 0.354 | 0.332 | 0.032 | 0.003 | -0.001 | -0.001 | 0 | 0.63 | 0.4 | 0.54 | 0.83 | 0.7 | 0.69 |
| 8 | 96905699 | rs10111313 | -0.458 | 0.357 | 0.583 | 0.08 | 0.024 | 0.134 | 0.134 | 1 | 0.15 | 0.37 | 0.13 | 0.31 | 0.16 | 0.14 |
| 8 | 96909360 | rs17768143 | 0.972 | 0.366 | 0.323 | 0.072 | 0 | -0.075 | -0.075 | 0 | 0.59 | 0.1 | 0.46 | 0.52 | 0.62 | 0.58 |
| 8 | 96913121 | rs10093358 | 1.076 | 0.365 | 0.569 | 0.099 | 0.002 | 0.057 | 0.057 | 0.153 | 0.07 | 0.05 | 0.08 | 0.31 | 0.12 | 0.12 |
| 8 | 96922072 | rs874123 | 0.74 | 0.236 | 0.763 | 0.024 | 0.013 | 0.017 | 0.017 | 0 | 0.19 | 0.1 | 0.31 | 0.16 | 0.14 | 0.23 |
| 8 | 96924546 | rs10955052 | 1.477 | 0.219 | 0.68 | 0.008 | 0 | -0.077 | -0.077 | 0 | 0.54 | 0.08 | 0.44 | 0.52 | 0.59 | 0.56 |
| 8 | 96925157 | rs969314 | 1.327 | 0.194 | 0.89 | -0.011 | 0.017 | -0.02 | -0.02 | 0 | 0.57 | 0.24 | 0.46 | 0.52 | 0.61 | 0.58 |
| 8 | 96930815 | rs6983038 | -0.091 | 0.186 | 0.87 | 0.072 | -0.029 | -0.031 | -0.031 | 0 | 0.26 | 0.34 | 0.35 | 0.18 | 0.19 | 0.24 |
| 8 | 96935794 | rs1025667 | 1.367 | 0.194 | 0.37 | 0.188 | 0.032 | -0.106 | -0.106 | 0 | 0.61 | 0.09 | 0.47 | 0.52 | 0.67 | 0.62 |
| 8 | 96959819 | rs10504958 | 1.337 | 0.47 | 0.835 | 0.343 | 0.018 | 0.074 | 0.074 | 0.997 | 0.81 | 0.45 | 0.81 | 0.7 | 0.77 | 0.8 |
| 8 | 96965786 | rs6468480 | 0.019 | 0.527 | 0.187 | 0.208 | -0.015 | 0.032 | 0.032 | 0 | 0.93 | 0.99 | 0.95 | 0.99 | 0.92 | 0.93 |
| 8 | 96969356 | rs11987725 | 0.711 | 0.504 | 0.278 | 0.176 | -0.001 | -0.01 | -0.01 | 0 | 0.92 | 0.93 | 0.93 | 0.99 | 0.91 | 0.93 |
| 8 | 96977010 | rs2033720 | -0.516 | 0.359 | 0.7 | 0.224 | -0.003 | 0.025 | 0.025 | 0 | 0.28 | 0.38 | 0.41 | 0.47 | 0.26 | 0.37 |
| 8 | 96978748 | rs893225 | 0.591 | 0.363 | 0.284 | 0.277 | 0.008 | 0.013 | 0.013 | 0 | 0.18 | 0.1 | 0.31 | 0.15 | 0.14 | 0.24 |
| 8 | 96979871 | rs10955055 | 1.09 | 0.307 | 0.004 | 0.188 | -0.034 | -0.104 | -0.104 | 0.426 | 0.59 | 0.08 | 0.47 | 0.53 | 0.62 | 0.56 |
| 8 | 96980358 | rs17769517 | 0.634 | 0.024 | -0.021 | 0.199 | -0.067 | -0.133 | -0.133 | 0 | 0.28 | 0.01 | 0.24 | 0.35 | 0.29 | 0.25 |
| 8 | 96985292 | rs10100514 | 0.934 | 0.04 | 0.327 | 0.27 | -0.011 | 0.031 | 0.031 | 0 | 0.91 | 0.95 | 0.94 | 1 | 0.88 | 0.93 |
| 8 | 96988158 | rs9642934 | 1.631 | -0.127 | -0.028 | 0.203 | -0.056 | 0.157 | 0.157 | 0 | 0.42 | 0.15 | 0.44 | 0.19 | 0.44 | 0.37 |
| 8 | 96988274 | rs11780335 | 0.292 | 0.192 | -0.082 | 0.922 | -0.016 | -0.079 | -0.079 | 0 | 0.19 | 0.03 | 0.16 | 0.28 | 0.18 | 0.23 |
| 8 | 97004203 | rs1421221 | 1.175 | -0.368 | 0.365 | 0.709 | 0.05 | 0.047 | 0.047 | 0 | 0.5 | 0.24 | 0.62 | 0.63 | 0.56 | 0.47 |
| 8 | 97022165 | rs261572 | 1.756 | -0.1 | 1.125 | 0.648 | -0.003 | -0.01 | -0.01 | 0 | 0.87 | 0.3 | 0.86 | 0.89 | 0.88 | 0.91 |
| 8 | 97026378 | rs261579 | -0.577 | -0.316 | 0.873 | 0.75 | 0.023 | 0.013 | 0.013 | 0 | 0.36 | 0.14 | 0.26 | 0.3 | 0.31 | 0.42 |
| 8 | 97030042 | rs10110393 | 1.089 | -0.323 | 1.233 | 0.794 | 0.013 | 0.011 | 0.011 | 0 | 0.53 | 0.44 | 0.64 | 0.6 | 0.58 | 0.49 |
| 8 | 97035898 | rs261552 | 1.062 | -0.159 | 1.307 | 0.545 | 0.002 | 0.009 | 0.009 | 0.05 | 0.54 | 0.43 | 0.65 | 0.63 | 0.54 | 0.46 |
| 8 | 97036115 | rs261553 | 0.267 | -0.22 | 1.507 | 0.494 | -0.006 | 0.007 | 0.007 | 0 | 0.72 | 0.64 | 0.74 | 0.76 | 0.73 | 0.7 |
| 8 | 97041113 | rs893222 | 1.757 | -0.436 | 0.628 | 0.324 | -0.054 | -0.061 | -0.061 | 0 | 0.46 | 0.1 | 0.6 | 0.56 | 0.53 | 0.42 |
| 8 | 97043469 | rs261558 | -0.869 | -0.164 | 1.542 | 0.014 | 0 | -0.08 | -0.08 | 0.035 | 0.21 | 0.35 | 0.1 | 0.11 | 0.17 | 0.24 |
| 8 | 97051579 | rs4734328 | 1.22 | -0.489 | 1.893 | 0.1 | -0.007 | -0.003 | -0.003 | 0 | 0.57 | 0.55 | 0.7 | 0.59 | 0.63 | 0.51 |
| 8 | 97052453 | rs16894219 | 0.044 | -0.47 | 1.9 | 0.075 | -0.019 | -0.021 | -0.021 | 0 | 0.14 | 0.14 | 0.03 | 0.11 | 0.07 | 0.14 |
| 8 | 97053086 | rs497805 | -0.334 | -0.473 | 1.701 | 0.05 | -0.002 | -0.003 | -0.003 | 0.15 | 0.26 | 0.22 | 0.26 | 0.3 | 0.3 | 0.35 |
| 8 | 97053344 | rs11780042 | -0.563 | -0.46 | 1.798 | 0.03 | -0.027 | -0.066 | -0.066 | 0 | 0.17 | 0.41 | 0.08 | 0.11 | 0.12 | 0.2 |
| 8 | 97055706 | rs1421223 | -0.534 | -0.443 | 1.45 | 0.039 | -0.027 | -0.063 | -0.063 | 0.549 | 0.17 | 0.4 | 0.09 | 0.11 | 0.11 | 0.2 |
| 8 | 97059472 | rs13265857 | 1.472 | -0.448 | 0.439 | 0.101 | 0.015 | 0.019 | 0.019 | 0 | 0.53 | 0.2 | 0.64 | 0.59 | 0.59 | 0.45 |
| 8 | 97060715 | rs573233 | -0.634 | -0.146 | 0.756 | 0.125 | 0.005 | 0.001 | 0.001 | 0 | 0.27 | 0.24 | 0.2 | 0.26 | 0.32 | 0.35 |
| 8 | 97061202 | rs7003742 | 1.245 | 0.082 | 0.162 | 0.053 | -0.031 | 0.031 | 0.031 | 0.404 | 0.65 | 0.16 | 0.69 | 0.8 | 0.64 | 0.62 |
| 8 | 97063942 | rs532255 | -0.854 | -0.01 | 0.469 | 0.234 | 0.005 | 0.009 | 0.009 | 0.004 | 0.23 | 0.31 | 0.2 | 0.26 | 0.24 | 0.38 |
| 8 | 97069446 | rs935190 | -1.072 | 0.089 | 0.464 | 0.177 | -0.002 | -0.003 | -0.003 | 0 | 0.2 | 0.22 | 0.18 | 0.3 | 0.32 | 0.42 |
| 8 | 97071558 | rs519577 | 1.206 | -0.698 | 1.012 | -0.385 | -0.053 | -0.1 | -0.1 | 0.798 | 0.69 | 0.57 | 0.87 | 0.88 | 0.81 | 0.77 |
| 8 | 97072309 | rs526146 | 1.046 | -0.657 | 1.234 | -0.36 | 0.03 | -0.001 | -0.001 | 0 | 0.89 | 0.77 | 0.92 | 0.88 | 0.88 | 0.87 |
| 8 | 97081252 | rs580234 | 0.205 | -0.592 | 0.797 | -0.241 | -0.019 | -0.013 | -0.013 | 2.571 | 0.13 | 0.07 | 0.12 | 0.2 | 0.2 | 0.3 |
| 8 | 97081921 | rs1421217 | 2.022 | -1.057 | 0.486 | -0.348 | 0.031 | 0.086 | 0.086 | 0 | 0.5 | 0.24 | 0.71 | 0.66 | 0.6 | 0.47 |
| 8 | 97097254 | rs2543090 | 1.212 | -1.08 | 0.596 | -0.342 | 0.016 | 0.015 | 0.015 | 0.33 | 0.65 | 0.32 | 0.81 | 0.86 | 0.8 | 0.77 |
| 8 | 97100460 | rs7017245 | 1.948 | -0.996 | 0.104 | -0.537 | -0.011 | 0.057 | 0.057 | 0 | 0.45 | 0.24 | 0.68 | 0.66 | 0.55 | 0.44 |
| 8 | 97102537 | rs10096979 | 2.3 | -1.432 | -0.507 | -0.238 | -0.162 | -0.02 | -0.02 | 1.645 | 0.33 | 0.04 | 0.54 | 0.24 | 0.48 | 0.23 |
| 8 | 97102896 | rs16894295 | 0.121 | -1.546 | 0.211 | -0.879 | 0.002 | 0.003 | 0.003 | 3.227 | 0.1 | 0.1 | 0.04 | 0.07 | 0.05 | 0.05 |
| 8 | 97108727 | rs542253 | -0.193 | -1.543 | 0.337 | -0.878 | -0.027 | -0.005 | -0.005 | 0 | 0.89 | 0.85 | 0.96 | 0.92 | 0.95 | 0.95 |
| 8 | 97108910 | rs7015987 | 1.174 | -1.688 | 0.335 | -0.962 | 0.011 | 0.003 | 0.003 | 0 | 0.12 | 0.11 | 0.04 | 0 | 0.05 | 0.09 |
| 8 | 97110026 | rs4735413 | 1.724 | -1.376 | 0.158 | -0.9 | 0.047 | 0.078 | 0.078 | 0 | 0.49 | 0.26 | 0.7 | 0.66 | 0.6 | 0.45 |
| 8 | 97112032 | rs546474 | 0.233 | -0.953 | 0 | -0.914 | -0.045 | -0.008 | -0.008 | 0 | 0.88 | 0.79 | 0.95 | 0.92 | 0.95 | 0.95 |
| 8 | 97122652 | rs2587000 | -0.277 | -0.554 | -0.584 | -1.035 | -0.07 | -0.016 | -0.016 | 0 | 0.18 | 0.02 | 0.13 | 0.14 | 0.21 | 0.32 |
| 8 | 97125655 | rs478101 | 0.427 | -0.531 | 0.207 | -1.474 | 0.033 | 0.016 | 0.016 | 0 | 0.92 | 0.75 | 0.94 | 0.94 | 0.91 | 0.87 |
| 8 | 97125876 | rs4734332 | 1.491 | -1.14 | 0.13 | -2.018 | 0.024 | 0.071 | 0.071 | 0 | 0.47 | 0.32 | 0.69 | 0.84 | 0.62 | 0.51 |
| 8 | 97125941 | rs13266786 | 1.479 | -1.154 | 0.13 | -2.024 | 0.025 | 0.072 | 0.072 | 0 | 0.48 | 0.32 | 0.69 | 0.84 | 0.62 | 0.51 |

| | | | | | | | | | | | | | | | | |
|---|---|---|---|---|---|---|---|---|---|---|---|---|---|---|---|---|
| 8 | 97126738 | rs2052380 | 0.844 | -0.929 | 0.339 | -1.868 | -0.012 | -0.05 | -0.05 | 0 | 0.59 | 0.49 | 0.76 | 0.87 | 0.68 | 0.61 |
| 8 | 97136462 | rs512030 | -0.195 | -0.765 | 0.046 | -1.535 | -0.04 | -0.019 | -0.019 | 0.444 | 0.17 | 0.04 | 0.13 | 0.17 | 0.2 | 0.24 |
| 8 | 97139953 | rs477593 | -0.088 | -0.441 | -0.375 | -2.023 | -0.077 | 0.049 | 0.049 | 0 | 0.3 | 0.09 | 0.23 | 0.18 | 0.37 | 0.32 |
| 8 | 97150045 | rs2247792 | -0.695 | -0.919 | 0.18 | -2.262 | -0.003 | -0.029 | -0.029 | 0.317 | 0.45 | 0.49 | 0.27 | 0.18 | 0.4 | 0.4 |
| 8 | 97153758 | rs2440198 | -0.066 | -0.505 | -0.53 | -2.072 | -0.064 | 0.053 | 0.053 | 0 | 0.31 | 0.11 | 0.24 | 0.17 | 0.37 | 0.33 |
| 8 | 97164904 | rs2255182 | -0.26 | -0.807 | -0.283 | -2.36 | -0.019 | 0.054 | 0.054 | 0 | 0.43 | 0.34 | 0.38 | 0.24 | 0.49 | 0.48 |
| 8 | 97169327 | rs2514527 | -1.473 | -1.168 | -0.387 | -2.721 | -0.021 | 0.022 | 0.022 | 0 | 0.69 | 0.78 | 0.44 | 0.22 | 0.6 | 0.61 |
| 8 | 97174637 | rs2514530 | 0.244 | -1.21 | -0.545 | -2.701 | -0.153 | 0.106 | 0.106 | 0.323 | 0.39 | 0.05 | 0.29 | 0.18 | 0.46 | 0.47 |
| 8 | 97179713 | rs6986442 | 0.03 | -1.407 | 0.1 | -2.816 | -0.085 | -0.178 | -0.178 | 4.345 | 0.27 | 0.4 | 0.13 | 0 | 0.14 | 0.14 |
| 8 | 97181050 | rs2245832 | -1.169 | -1.023 | 0.2 | -2.473 | -0.013 | 0.012 | 0.012 | 0 | 0.69 | 0.69 | 0.45 | 0.24 | 0.6 | 0.61 |
| 8 | 97182976 | rs2514537 | -1.371 | -0.98 | -0.149 | -2.452 | -0.047 | 0.018 | 0.018 | 0 | 0.71 | 0.82 | 0.46 | 0.24 | 0.6 | 0.61 |
| 8 | 97191039 | rs2514542 | 0.218 | -1.041 | -0.33 | -2.421 | -0.061 | 0.064 | 0.064 | 1.626 | 0.38 | 0.1 | 0.34 | 0.24 | 0.43 | 0.48 |
| 8 | 97194671 | rs10216734 | 0.467 | -1.27 | 0.36 | -2.459 | -0.035 | -0.082 | -0.082 | 0 | 0.2 | 0.27 | 0.08 | 0 | 0.1 | 0.14 |
| 8 | 97201453 | rs2440212 | 0.21 | -0.948 | -0.072 | -2.705 | -0.076 | 0.097 | 0.097 | 1.957 | 0.42 | 0.1 | 0.35 | 0.24 | 0.47 | 0.49 |
| 8 | 97201855 | rs11776025 | 0.351 | -0.937 | -0.046 | -2.691 | -0.017 | -0.059 | -0.059 | 0.153 | 0.22 | 0.23 | 0.1 | 0 | 0.13 | 0.15 |
| 8 | 97206319 | rs2440213 | -1.324 | -0.756 | 0.224 | -2.568 | 0.017 | -0.021 | -0.021 | 0.212 | 0.67 | 0.53 | 0.47 | 0.24 | 0.6 | 0.64 |
| 8 | 97207924 | rs2514549 | -1.541 | -0.741 | 0.066 | -2.56 | 0 | 0 | 0 | 0 | 0.69 | 0.67 | 0.47 | 0.24 | 0.6 | 0.64 |
| 8 | 97215912 | rs6985783 | 0.159 | -1.343 | -0.205 | -2.696 | 0.043 | 0.062 | 0.062 | 0 | 0.23 | 0.07 | 0.08 | 0 | 0.12 | 0.11 |
| 8 | 97219690 | rs11991704 | 0.509 | -0.9 | -0.386 | -2.623 | -0.012 | 0.029 | 0.029 | 0.052 | 0.32 | 0.18 | 0.27 | 0.22 | 0.34 | 0.39 |
| 8 | 97221895 | rs11992876 | -0.958 | -0.107 | -0.146 | -2.102 | 0.002 | -0.003 | -0.003 | 2.118 | 0.66 | 0.67 | 0.43 | 0.22 | 0.53 | 0.58 |
| 8 | 97232444 | rs11776157 | 0.404 | -0.203 | -0.491 | -1.912 | -0.008 | 0.102 | 0.102 | 0 | 0.35 | 0.13 | 0.26 | 0.15 | 0.36 | 0.28 |
| 8 | 97234255 | rs2643342 | -1.115 | -0.126 | -0.405 | -1.754 | -0.05 | 0.055 | 0.055 | 0.309 | 0.6 | 0.83 | 0.4 | 0.25 | 0.48 | 0.45 |
| 8 | 97252713 | rs10105238 | -0.369 | -0.147 | 0.392 | -1.702 | 0.001 | 0.01 | 0.01 | 0 | 0.1 | 0.07 | 0.06 | 0 | 0.08 | 0.04 |
| 8 | 97253788 | rs10955069 | -0.717 | -0.27 | 0.625 | -1.818 | 0.006 | 0.022 | 0.022 | 0 | 0.49 | 0.54 | 0.35 | 0.18 | 0.44 | 0.33 |
| 8 | 97254372 | rs10112328 | -1.093 | -0.271 | 0.595 | -1.818 | 0.003 | 0.118 | 0.118 | 0.031 | 0.54 | 0.78 | 0.37 | 0.21 | 0.44 | 0.4 |
| 8 | 97268077 | rs7838479 | 0.405 | -0.261 | 0.733 | -1.879 | -0.015 | 0.075 | 0.075 | 0 | 0.36 | 0.11 | 0.26 | 0.18 | 0.36 | 0.29 |
| 8 | 97269738 | rs10095443 | -0.736 | -0.27 | 0.744 | -1.886 | 0.005 | 0.021 | 0.021 | 0 | 0.49 | 0.54 | 0.35 | 0.18 | 0.44 | 0.33 |
| 8 | 97273543 | rs3808386 | 0.365 | -0.17 | -0.027 | -1.819 | -0.009 | 0.071 | 0.071 | 0 | 0.36 | 0.13 | 0.26 | 0.18 | 0.36 | 0.29 |
| 8 | 97295677 | rs6987316 | -1.645 | -0.578 | -0.087 | -1.766 | -0.028 | -0.088 | -0.088 | 0 | 0.18 | 0.66 | 0.1 | 0.01 | 0.09 | 0.12 |
| 8 | 97312442 | rs17707686 | -0.3 | -0.146 | -0.33 | -1.554 | 0.001 | -0.005 | -0.005 | 0.429 | 0.11 | 0.09 | 0.06 | 0 | 0.08 | 0.04 |
| 8 | 97317409 | rs3808480 | -0.182 | 0.058 | -0.393 | -1.025 | 0.019 | 0.018 | 0.018 | 0 | 0.38 | 0.27 | 0.27 | 0.25 | 0.32 | 0.32 |
| 8 | 97327229 | rs11786602 | -0.608 | 0.427 | 0.105 | -1.049 | 0.005 | 0.002 | 0.002 | 0.144 | 0.48 | 0.53 | 0.36 | 0.09 | 0.48 | 0.33 |
| 8 | 97328308 | rs2643341 | 0.118 | 0.218 | 0.094 | -0.461 | -0.002 | 0.033 | 0.033 | 0 | 0.94 | 0.91 | 0.86 | 0.71 | 0.94 | 0.87 |
| 8 | 97329366 | rs13255914 | -0.449 | 0.219 | 0.467 | -0.514 | 0 | -0.029 | -0.029 | 0 | 0.49 | 0.45 | 0.37 | 0.1 | 0.5 | 0.35 |
| 8 | 97337513 | rs4734338 | 0.98 | 0.145 | 0.447 | -0.537 | 0.022 | 0.022 | 0.022 | 0.562 | 0.42 | 0.13 | 0.44 | 0.61 | 0.43 | 0.51 |
| 8 | 97341390 | rs11784595 | 0.808 | 0.167 | 0.551 | -0.542 | 0.03 | 0.042 | 0.042 | 0.619 | 0.45 | 0.21 | 0.48 | 0.61 | 0.47 | 0.51 |
| 8 | 97342090 | rs2280264 | 0.8 | 0.167 | 0.618 | -0.542 | 0.03 | 0.04 | 0.04 | 0.556 | 0.45 | 0.23 | 0.48 | 0.61 | 0.46 | 0.51 |
| 8 | 97342582 | rs7001397 | -0.521 | 0.159 | 0.796 | -0.56 | 0.005 | 0.006 | 0.006 | 0.185 | 0.48 | 0.5 | 0.36 | 0.1 | 0.48 | 0.34 |
| 8 | 97345590 | rs3735986 | 0.362 | 0.754 | 1.06 | -0.809 | 0.021 | -0.001 | -0.001 | 4.308 | 0.76 | 0.68 | 0.65 | 0.7 | 0.69 | 0.77 |
| 8 | 97352390 | rs7819735 | -0.509 | 0.349 | 1.563 | -1.065 | 0.017 | -0.043 | -0.043 | 0 | 0.1 | 0.24 | 0.1 | 0.05 | 0.12 | 0.12 |
| 8 | 97352536 | rs10103493 | 0.143 | 0.127 | 1.669 | -1.191 | -0.051 | -0.034 | -0.034 | 0.402 | 0.23 | 0.3 | 0.08 | 0.09 | 0.11 | 0.17 |
| 8 | 97365321 | rs16894509 | -0.655 | 0.001 | 1.97 | -1.264 | -0.023 | -0.035 | -0.035 | 0 | 0.29 | 0.36 | 0.17 | 0.15 | 0.23 | 0.3 |
| 8 | 97366006 | rs11997560 | -1.728 | 0.097 | 2.324 | -1.432 | 0.03 | 0.088 | 0.088 | 0.149 | 0.53 | 0.69 | 0.42 | 0.15 | 0.55 | 0.41 |
| 8 | 97366701 | rs6997395 | 0.356 | 0.105 | 2.353 | -1.429 | -0.033 | 0.008 | 0.008 | 0.621 | 0.21 | 0.16 | 0.22 | 0 | 0.31 | 0.11 |
| 8 | 97369332 | rs7013150 | -1.401 | 0.075 | 2.392 | -1.411 | 0.009 | 0.004 | 0.004 | 0 | 0.5 | 0.5 | 0.38 | 0.15 | 0.55 | 0.41 |
| 8 | 97369634 | rs2640806 | -1.054 | 0.054 | 2.337 | -1.427 | 0 | -0.032 | -0.032 | 0 | 0.44 | 0.46 | 0.29 | 0.09 | 0.42 | 0.28 |
| 8 | 97371385 | rs7835633 | -0.36 | 0.074 | 2.348 | -1.34 | 0.026 | -0.039 | -0.039 | 0 | 0.1 | 0.32 | 0.12 | 0.06 | 0.13 | 0.13 |
| 8 | 97376980 | rs2678332 | -1.053 | 0.115 | 2.134 | -1.375 | 0 | -0.052 | -0.052 | 0 | 0.36 | 0.65 | 0.37 | 0.06 | 0.46 | 0.23 |
| 8 | 97384973 | rs10504962 | 0.42 | 0.016 | 2.093 | -1.368 | -0.011 | -0.002 | -0.002 | 1.614 | 0.19 | 0.17 | 0.05 | 0.09 | 0.09 | 0.17 |
| 8 | 97386841 | rs2244711 | -1.594 | 0.08 | 1.829 | -1.426 | 0.039 | 0.108 | 0.108 | 0.881 | 0.55 | 0.73 | 0.43 | 0.15 | 0.57 | 0.42 |
| 8 | 97397154 | rs784803 | 1.679 | -0.01 | 1.799 | -1.321 | 0.036 | 0.086 | 0.086 | 0.634 | 0.43 | 0.21 | 0.55 | 0.69 | 0.43 | 0.52 |
| 8 | 97402724 | rs750500 | 0.281 | -0.056 | 1.862 | -1.581 | -0.09 | 0.074 | 0.074 | 1.354 | 0.23 | 0.08 | 0.23 | 0.01 | 0.34 | 0.12 |
| 8 | 97404339 | rs811480 | -1.96 | -0.243 | 1.74 | -1.594 | 0.023 | 0.112 | 0.112 | 2.104 | 0.59 | 0.8 | 0.45 | 0.22 | 0.57 | 0.5 |
| 8 | 97404721 | rs13251611 | -1.679 | -0.234 | 2.085 | -1.609 | 0.014 | 0.028 | 0.028 | 1.652 | 0.56 | 0.62 | 0.41 | 0.13 | 0.57 | 0.42 |
| 8 | 97408966 | rs7840314 | -0.169 | -0.238 | 1.742 | -1.569 | -0.001 | -0.019 | -0.019 | 0.257 | 0.11 | 0.16 | 0.1 | 0.07 | 0.13 | 0.14 |
| 8 | 97414597 | rs17710204 | 0.777 | -0.368 | 1.662 | -1.514 | 0.049 | 0.047 | 0.047 | 0 | 0.17 | 0.03 | 0.04 | 0.06 | 0.09 | 0.16 |
| 8 | 97425295 | rs7832352 | -0.866 | 0.097 | 1.908 | -1.287 | -0.009 | 0.018 | 0.018 | 1.723 | 0.35 | 0.3 | 0.44 | 0.12 | 0.48 | 0.27 |
| 8 | 97432021 | rs746736 | -1.851 | -0.173 | 1.468 | -1.391 | 0.054 | 0.135 | 0.135 | 2.481 | 0.56 | 0.93 | 0.57 | 0.24 | 0.6 | 0.49 |
| 8 | 97454130 | rs12550085 | -0.769 | -0.17 | 1.808 | -1.358 | -0.005 | 0.012 | 0.012 | 1.062 | 0.35 | 0.29 | 0.44 | 0.12 | 0.51 | 0.27 |
| 8 | 97457533 | rs1020344 | -0.693 | -0.203 | 1.98 | -1.141 | -0.051 | -0.023 | -0.023 | 0 | 0.2 | 0.62 | 0.12 | 0.14 | 0.09 | 0.21 |
| 8 | 97467696 | rs1444569 | -1.796 | -0.305 | 1.086 | -1.346 | 0.012 | 0.146 | 0.146 | 0.32 | 0.55 | 0.82 | 0.55 | 0.24 | 0.53 | 0.51 |
| 8 | 97468774 | rs974272 | -0.823 | -0.184 | 1.615 | -1.377 | 0.024 | -0.077 | -0.077 | 1.113 | 0.37 | 0.41 | 0.33 | 0.15 | 0.44 | 0.35 |
| 8 | 97474401 | rs12675232 | -0.682 | -0.217 | 2.003 | -0.992 | -0.022 | 0.033 | 0.033 | 0.282 | 0.11 | 0.34 | 0.09 | 0.15 | 0.06 | 0.2 |
| 8 | 97476956 | rs6468507 | -1.381 | -0.223 | 1.629 | -0.976 | -0.043 | 0.004 | 0.004 | 1.711 | 0.16 | 0.62 | 0.26 | 0.19 | 0.07 | 0.21 |
| 8 | 97478729 | rs16894605 | -1.256 | -0.185 | 1.245 | -0.952 | -0.051 | 0.015 | 0.015 | 1.604 | 0.16 | 0.57 | 0.24 | 0.19 | 0.07 | 0.21 |
| 8 | 97482056 | rs16894608 | -0.706 | 0.771 | 0.676 | -0.661 | -0.007 | -0.011 | -0.011 | 2.369 | 0.31 | 0.36 | 0.19 | 0.23 | 0.24 | 0.41 |
| 8 | 97483767 | rs1992087 | -0.203 | 1.705 | 0.358 | 0.601 | -0.031 | -0.032 | -0.032 | 0 | 0.15 | 0.11 | 0.32 | 0.27 | 0.26 | 0.25 |
| 8 | 97489329 | rs11990608 | -1.349 | 0.954 | 0.52 | 0.169 | 0.115 | 0.055 | 0.055 | 3.255 | 0.06 | 0.36 | 0.08 | 0.04 | 0.07 | 0.03 |
| 8 | 97492799 | rs2640820 | -0.109 | 0.786 | -0.403 | 0.32 | 0.001 | -0.007 | -0.007 | 0 | 0.12 | 0.09 | 0.07 | 0.13 | 0.07 | 0.11 |

| | | | | | | | | | | | | | | | | |
|---|---|---|---|---|---|---|---|---|---|---|---|---|---|---|---|---|
| 8 | 97494338 | rs1971855 | -0.646 | 0.912 | -0.952 | 0.095 | -0.001 | 0.02 | 0.02 | 0 | 0.61 | 0.79 | 0.62 | 0.63 | 0.57 | 0.64 |
| 8 | 97494356 | rs12545081 | -0.787 | 1.072 | -0.375 | 0.368 | 0.032 | 0.035 | 0.035 | 0 | 0.07 | 0.2 | 0.09 | 0.11 | 0.09 | 0.15 |
| 8 | 97494572 | rs11786386 | -0.481 | 1.208 | -0.928 | 0.065 | -0.02 | 0.012 | 0.012 | 0.093 | 0.21 | 0.15 | 0.17 | 0.1 | 0.31 | 0.17 |
| 8 | 97495996 | rs10808350 | -0.627 | 0.815 | -0.968 | 0.304 | -0.008 | -0.03 | -0.03 | 0 | 0.15 | 0.12 | 0.32 | 0.6 | 0.16 | 0.37 |
| 8 | 97496509 | rs2452992 | -1.175 | 0.907 | -1.22 | -0.323 | -0.009 | -0.019 | -0.019 | 0.52 | 0.76 | 0.81 | 0.73 | 0.66 | 0.69 | 0.75 |
| 8 | 97499703 | rs16894633 | 0.265 | 0.841 | -1.239 | -0.057 | -0.003 | 0.014 | 0.014 | 0 | 0.12 | 0.1 | 0.3 | 0.44 | 0.11 | 0.28 |
| 8 | 97505545 | rs2437770 | -1.385 | 0.871 | -1.235 | 0.035 | -0.031 | -0.033 | -0.033 | 0 | 0.77 | 0.92 | 0.78 | 0.74 | 0.7 | 0.78 |
| 8 | 97507658 | rs2437771 | -1.303 | 0.91 | -1.245 | 0.048 | -0.01 | -0.012 | -0.012 | 0 | 0.77 | 0.84 | 0.76 | 0.73 | 0.7 | 0.78 |
| 8 | 97508361 | rs1348563 | -0.184 | 0.449 | -1.051 | -0.492 | -0.005 | 0.002 | 0.002 | 0 | 0.46 | 0.43 | 0.56 | 0.48 | 0.4 | 0.5 |
| 8 | 97508555 | rs895033 | 1.204 | 0.218 | -1.104 | -0.955 | 0.051 | -0.044 | -0.044 | 4.893 | 0.16 | 0.29 | 0.17 | 0.05 | 0.23 | 0.15 |
| 8 | 97510132 | rs13270556 | 0.918 | -0.486 | -1.069 | -0.984 | -0.041 | 0.024 | 0.024 | 0 | 0.27 | 0.13 | 0.17 | 0.24 | 0.3 | 0.27 |
| 8 | 97512221 | rs16894649 | 0.862 | -0.461 | -1.164 | -0.824 | -0.007 | -0.007 | -0.007 | 3.746 | 0.12 | 0.09 | 0.3 | 0.5 | 0.11 | 0.26 |
| 8 | 97512977 | rs2437772 | -1.534 | 0.296 | -0.682 | -0.516 | -0.002 | 0 | 0 | 0.302 | 0.76 | 0.75 | 0.74 | 0.79 | 0.67 | 0.78 |
| 8 | 97516984 | rs2582814 | 0.909 | 0.304 | -0.674 | -0.584 | 0.055 | -0.076 | -0.076 | 0 | 0.3 | 0.17 | 0.38 | 0.51 | 0.17 | 0.36 |
| 8 | 97517389 | rs2439518 | -1.03 | 0.437 | -0.152 | -0.64 | -0.012 | 0.052 | 0.052 | 0 | 0.46 | 0.63 | 0.37 | 0.29 | 0.5 | 0.41 |
| 8 | 97518087 | rs2008026 | 0.288 | 0.443 | -0.124 | -0.642 | -0.015 | 0.046 | 0.046 | 0 | 0.25 | 0.17 | 0.18 | 0.08 | 0.26 | 0.14 |
| 8 | 97518630 | rs2439519 | -1.121 | -0.296 | 0.469 | 0.198 | 0.003 | 0.013 | 0.013 | 0 | 0.21 | 0.46 | 0.19 | 0.21 | 0.24 | 0.27 |
| 8 | 97522509 | rs2589183 | -0.485 | -0.202 | 0.562 | 0.219 | 0 | -0.045 | -0.045 | 0 | 0.16 | 0.28 | 0.15 | 0.05 | 0.19 | 0.15 |
| 8 | 97534651 | rs2575735 | 0.079 | -1.022 | -0.212 | 0.065 | 0.002 | 0 | 0 | 1.662 | 0.57 | 0.6 | 0.48 | 0.2 | 0.69 | 0.4 |
| 8 | 97534816 | rs2582822 | -0.123 | -0.99 | 0.068 | -0.529 | -0.01 | -0.09 | -0.09 | 0.033 | 0.19 | 0.3 | 0.1 | 0 | 0.18 | 0.12 |
| 8 | 97537041 | rs2439525 | 0.381 | -1.88 | -0.257 | 0.109 | 0.015 | -0.041 | -0.041 | 0.736 | 0.79 | 0.81 | 0.61 | 0.43 | 0.82 | 0.58 |
| 8 | 97537952 | rs7003874 | 1.414 | -2.354 | 0.038 | -0.168 | -0.043 | 0.032 | 0.032 | 0 | 0.32 | 0.17 | 0.3 | 0.19 | 0.48 | 0.27 |
| 8 | 97538205 | rs2575734 | 0.451 | -2.655 | 0.839 | -0.264 | -0.022 | -0.027 | -0.027 | 1.684 | 0.21 | 0.37 | 0.14 | 0.15 | 0.18 | 0.13 |
| 8 | 97539802 | rs2589213 | 1.144 | -2.782 | 0.902 | -0.375 | -0.003 | -0.042 | -0.042 | 0 | 0.23 | 0.23 | 0.15 | 0 | 0.14 | 0.15 |
| 8 | 97542143 | rs874643 | -0.152 | -2.764 | 0.701 | -0.264 | 0.013 | -0.093 | -0.093 | 0.102 | 0.8 | 0.92 | 0.62 | 0.43 | 0.82 | 0.56 |
| 8 | 97543390 | rs895034 | 0.24 | -2.813 | 1.065 | -0.312 | -0.023 | -0.027 | -0.027 | 0 | 0.22 | 0.36 | 0.14 | 0.15 | 0.18 | 0.13 |
| 8 | 97544581 | rs2440681 | -0.252 | -2.721 | 0.994 | -0.275 | 0.038 | 0.05 | 0.05 | 0 | 0.45 | 0.32 | 0.27 | 0.23 | 0.34 | 0.32 |
| 8 | 97547130 | rs2437774 | 0.025 | -2.544 | 0.921 | 0.423 | 0.027 | -0.062 | -0.062 | 0 | 0.82 | 0.89 | 0.7 | 0.46 | 0.88 | 0.59 |
| 8 | 97549700 | rs2589203 | 0.719 | -2.737 | 1.273 | 0.368 | 0.056 | 0.003 | 0.003 | 0 | 0.54 | 0.31 | 0.5 | 0.41 | 0.71 | 0.4 |
| 8 | 97551112 | rs2589201 | 0.395 | -2.748 | 1.26 | 0.948 | 0.022 | 0.015 | 0.015 | 0 | 0.6 | 0.63 | 0.68 | 0.76 | 0.75 | 0.6 |
| 8 | 97556565 | rs2589198 | 0.568 | -2.754 | 1.414 | 1.15 | -0.035 | 0.018 | 0.018 | 0 | 0.81 | 0.69 | 0.79 | 0.77 | 0.9 | 0.73 |
| 8 | 97558840 | rs2582840 | 0.321 | -2.78 | 1.296 | 1.159 | -0.011 | 0.007 | 0.007 | 0 | 0.82 | 0.81 | 0.82 | 0.77 | 0.91 | 0.73 |
| 8 | 97559162 | rs10100191 | 0.265 | -3.114 | 1.388 | 1.11 | -0.039 | 0.029 | 0.029 | 0 | 0.21 | 0.32 | 0.26 | 0.36 | 0.14 | 0.24 |
| 8 | 97559665 | rs16894717 | -0.33 | -3.03 | 1.135 | 1.028 | -0.022 | -0.026 | -0.026 | 0 | 0.06 | 0.12 | 0.02 | 0 | 0.01 | 0.01 |
| 8 | 97563644 | rs2582842 | 1.011 | -3.065 | 0.988 | 1.161 | 0.119 | 0.04 | 0.04 | 0.207 | 0.54 | 0.2 | 0.51 | 0.33 | 0.75 | 0.48 |
| 8 | 97564661 | rs1444573 | 1.223 | -2.587 | 0.753 | 1.02 | 0.109 | 0.134 | 0.134 | 0.295 | 0.51 | 0.16 | 0.38 | 0.25 | 0.65 | 0.43 |
| 8 | 97564882 | rs1444572 | -1.146 | -2.395 | 1.031 | 1.236 | 0.048 | 0.183 | 0.183 | 0 | 0.08 | 0.25 | 0.22 | 0.43 | 0.14 | 0.22 |
| 8 | 97566797 | rs2582843 | 0.434 | -2.492 | 1.034 | 1.223 | -0.071 | -0.013 | -0.013 | 0.027 | 0.63 | 0.5 | 0.69 | 0.73 | 0.81 | 0.68 |
| 8 | 97567106 | rs1984456 | 0.896 | -2.399 | 0.339 | 1.046 | 0.075 | 0.055 | 0.055 | 0.018 | 0.55 | 0.21 | 0.46 | 0.33 | 0.68 | 0.46 |
| 8 | 97574582 | rs2118060 | 0.066 | -2.289 | 0.201 | 1.189 | 0.021 | -0.027 | -0.027 | 0 | 0.88 | 0.95 | 0.79 | 0.72 | 0.94 | 0.79 |
| 8 | 97578113 | rs2016529 | 0.776 | -2.578 | 0.327 | 1.332 | 0.05 | -0.022 | -0.022 | 0.181 | 0.62 | 0.21 | 0.6 | 0.53 | 0.79 | 0.57 |
| 8 | 97578672 | rs16894747 | -1.05 | -1.196 | 0.465 | 1.329 | 0.021 | 0.13 | 0.13 | 0 | 0.05 | 0.18 | 0.1 | 0.16 | 0.05 | 0.15 |
| 8 | 97583489 | rs7831863 | 0.68 | -1.238 | -0.197 | 0.436 | 0.009 | 0.042 | 0.042 | 0 | 0.6 | 0.3 | 0.52 | 0.12 | 0.66 | 0.52 |
| 8 | 97596782 | rs2439510 | 0.859 | -1.668 | 0.205 | 1.874 | -0.024 | 0.081 | 0.081 | 0 | 0.83 | 0.71 | 0.68 | 0.48 | 0.86 | 0.78 |
| 8 | 97604112 | rs16894821 | -0.675 | -1.685 | 0.528 | 1.989 | -0.025 | 0.047 | 0.047 | 0 | 0.11 | 0.19 | 0.17 | 0.21 | 0.09 | 0.13 |
| 8 | 97605800 | rs1126681 | -0.003 | -1.921 | 0.213 | 1.73 | -0.06 | -0.073 | -0.073 | 1.13 | 0.25 | 0.05 | 0.16 | 0.36 | 0.22 | 0.38 |
| 8 | 97607422 | rs2704256 | 0.127 | -1.923 | 0.213 | 1.719 | -0.062 | -0.067 | -0.067 | 0 | 0.25 | 0.05 | 0.16 | 0.35 | 0.23 | 0.38 |
| 8 | 97607515 | rs2250503 | -0.961 | -1.898 | 0.251 | 1.707 | 0.006 | -0.023 | -0.023 | 1.731 | 0.36 | 0.24 | 0.33 | 0.59 | 0.32 | 0.51 |
| 8 | 97610088 | rs16894859 | 0.098 | -0.494 | 0.352 | 1.087 | 0.023 | -0.041 | -0.041 | 2.048 | 0.16 | 0.37 | 0.13 | 0.08 | 0.14 | 0.12 |
| 8 | 97614625 | rs3816208 | 0.428 | -0.504 | 0.263 | 1.161 | 0.016 | -0.044 | -0.044 | 0 | 0.08 | 0.03 | 0.14 | 0.21 | 0.07 | 0.12 |
| 8 | 97614661 | rs1042381 | -0.175 | -0.355 | 0.175 | 0.97 | 0.023 | 0.021 | 0.021 | 0.373 | 0.18 | 0.39 | 0.16 | 0.23 | 0.17 | 0.16 |
| 8 | 97615235 | rs1561158 | 0.575 | -0.184 | -0.454 | 0.293 | -0.004 | 0.02 | 0.02 | 0 | 0.23 | 0.11 | 0.15 | 0.2 | 0.17 | 0.34 |
| 8 | 97618681 | rs2704240 | 0.994 | -0.096 | -0.623 | 0.066 | -0.012 | 0.123 | 0.123 | 0.604 | 0.45 | 0.25 | 0.39 | 0.06 | 0.54 | 0.29 |
| 8 | 97625450 | rs2651458 | 0.364 | -0.464 | -0.758 | 0.56 | 0.079 | 0.045 | 0.045 | 2.007 | 0.58 | 0.17 | 0.66 | 0.34 | 0.78 | 0.51 |
| 8 | 97629906 | rs10955078 | -0.568 | -0.43 | -1.053 | 0.551 | 0.001 | -0.004 | -0.004 | 0.915 | 0.75 | 0.82 | 0.69 | 0.68 | 0.76 | 0.74 |
| 8 | 97630529 | rs10090346 | 0.389 | -0.648 | -0.258 | 0.344 | -0.024 | -0.004 | -0.004 | 0 | 0.33 | 0.55 | 0.2 | 0.34 | 0.21 | 0.38 |
| 8 | 97631133 | rs11986468 | 0.227 | -0.753 | -0.087 | 0.291 | -0.001 | 0 | 0 | 0 | 0.34 | 0.66 | 0.21 | 0.34 | 0.21 | 0.38 |
| 8 | 97631573 | rs714046 | 0.928 | -0.891 | 0.283 | 0.24 | 0 | -0.001 | -0.001 | 0.448 | 0.28 | 0.27 | 0.14 | 0.32 | 0.18 | 0.37 |
| 8 | 97634019 | rs11776740 | 0.187 | -0.914 | -0.013 | 0.241 | 0.018 | 0.007 | 0.007 | 1.732 | 0.36 | 0.73 | 0.22 | 0.33 | 0.21 | 0.4 |
| 8 | 97643212 | rs750341 | 0.806 | -0.853 | -0.716 | 0.005 | -0.094 | 0.005 | 0.005 | 0 | 0.35 | 0.11 | 0.52 | 0.34 | 0.57 | 0.37 |
| 8 | 97643454 | rs750343 | 0.754 | -0.846 | -0.73 | 0.008 | -0.105 | -0.003 | -0.003 | 0 | 0.35 | 0.08 | 0.52 | 0.34 | 0.57 | 0.37 |
| 8 | 97651358 | rs2704253 | -0.229 | -1.235 | -0.325 | -0.043 | -0.061 | 0.04 | 0.04 | 0 | 0.73 | 0.61 | 0.77 | 0.68 | 0.87 | 0.79 |
| 8 | 97655880 | rs10086882 | -0.135 | -0.616 | -0.509 | -0.406 | -0.004 | 0.06 | 0.06 | 0 | 0.51 | 0.32 | 0.29 | 0.34 | 0.43 | 0.56 |
| 8 | 97659298 | rs2704247 | 0.038 | -0.216 | -0.241 | -0.17 | 0.013 | -0.012 | -0.012 | 0.976 | 0.14 | 0.25 | 0.18 | 0.11 | 0.16 | 0.19 |
| 8 | 97662068 | rs2651447 | -1.909 | -0.399 | -0.179 | 0.053 | -0.055 | 0.106 | 0.106 | 0 | 0.85 | 0.76 | 0.85 | 0.68 | 0.94 | 0.89 |
| 8 | 97664572 | rs7813101 | 0.84 | -0.32 | -0.307 | -0.26 | 0.03 | -0.033 | -0.033 | 0.19 | 0.62 | 0.25 | 0.58 | 0.56 | 0.72 | 0.64 |
| 8 | 97682211 | rs2651482 | 1.044 | -0.39 | -0.565 | -0.697 | 0.015 | 0.06 | 0.06 | 0 | 0.56 | 0.24 | 0.57 | 0.29 | 0.64 | 0.48 |
| 8 | 97684364 | rs2704245 | 0.553 | -0.558 | -0.261 | -0.428 | -0.025 | 0.03 | 0.03 | 0 | 0.1 | 0.22 | 0.05 | 0.1 | 0.03 | 0.08 |
| 8 | 97685211 | rs2455052 | 0.199 | -0.473 | -0.47 | -0.378 | 0.004 | 0.02 | 0.02 | 0 | 0.68 | 0.61 | 0.64 | 0.39 | 0.67 | 0.56 |

| | | | | | | | | | | | | | | | | |
|---|---|---|---|---|---|---|---|---|---|---|---|---|---|---|---|---|
| 8 | 97685591 | rs12675345 | 2.026 | -0.646 | -0.812 | -0.652 | -0.052 | 0.104 | 0.104 | 0.403 | 0.46 | 0.12 | 0.46 | 0.29 | 0.5 | 0.39 |
| 8 | 97688582 | rs10447975 | 2.649 | -0.285 | -0.503 | -0.309 | 0.009 | 0.049 | 0.049 | 0 | 0.11 | 0.03 | 0.03 | 0 | 0.07 | 0.05 |
| 8 | 97693790 | rs2464483 | -0.064 | -0.082 | -0.13 | -0.085 | 0.017 | 0.061 | 0.061 | 0.354 | 0.79 | 0.7 | 0.7 | 0.4 | 0.75 | 0.61 |
| 8 | 97735490 | rs2319918 | -0.667 | -0.176 | -0.112 | -0.278 | 0.013 | 0.041 | 0.041 | 0.013 | 0.32 | 0.47 | 0.38 | 0.57 | 0.35 | 0.39 |
| 8 | 97741971 | rs3857894 | 0.455 | -0.849 | -0.311 | -1.256 | 0.001 | 0.169 | 0.169 | 0.219 | 0.54 | 0.25 | 0.68 | 0.86 | 0.56 | 0.66 |
| 8 | 97761434 | rs16895064 | 1.938 | -0.772 | -0.16 | -1.265 | 0.002 | -0.057 | -0.057 | 0.014 | 0.1 | 0.21 | 0.05 | 0 | 0.08 | 0.04 |
| 8 | 97772970 | rs2319924 | -0.221 | -0.772 | -0.182 | -1.283 | 0.003 | -0.079 | -0.079 | 0.028 | 0.65 | 0.5 | 0.74 | 0.86 | 0.64 | 0.7 |
| 8 | 97780226 | rs1075479 | -1.294 | -0.838 | -0.22 | -1.407 | -0.001 | 0.031 | 0.031 | 0 | 0.77 | 0.84 | 0.82 | 0.95 | 0.78 | 0.81 |
| 8 | 97795719 | rs2319927 | 0.677 | -0.867 | -0.248 | -1.405 | -0.015 | 0.218 | 0.218 | 0.411 | 0.53 | 0.21 | 0.68 | 0.88 | 0.56 | 0.67 |
| 8 | 97796731 | rs3763558 | -0.299 | -0.727 | -0.142 | -1.218 | 0.047 | -0.016 | -0.016 | 0 | 0.11 | 0.31 | 0.07 | 0.07 | 0.13 | 0.1 |
| 8 | 97833936 | rs11779519 | 0.056 | -0.63 | -0.659 | -1.101 | 0.004 | 0.057 | 0.057 | 0 | 0.61 | 0.32 | 0.66 | 0.88 | 0.61 | 0.69 |
| 8 | 97847358 | rs1864384 | 0.066 | -0.629 | -0.662 | -1.101 | 0.004 | 0.054 | 0.054 | 0 | 0.61 | 0.32 | 0.66 | 0.88 | 0.61 | 0.69 |
| 8 | 97847774 | rs2319931 | 0.432 | -0.611 | -0.668 | -1.099 | 0.001 | 0.136 | 0.136 | 0.335 | 0.53 | 0.29 | 0.63 | 0.88 | 0.53 | 0.65 |
| 8 | 97851907 | rs1982002 | 0.043 | -0.403 | -0.592 | -0.839 | -0.001 | -0.034 | -0.034 | 0.947 | 0.63 | 0.44 | 0.67 | 0.88 | 0.61 | 0.69 |
| 8 | 97861325 | rs4734351 | 1.129 | -0.433 | -0.339 | -0.397 | 0.01 | 0.004 | 0.004 | 0 | 0.28 | 0.25 | 0.44 | 0.38 | 0.43 | 0.24 |
| 8 | 97864038 | rs13268611 | 0.073 | -1.164 | 0.02 | -0.743 | -0.006 | 0.006 | 0.006 | 0 | 0.42 | 0.52 | 0.47 | 0.36 | 0.45 | 0.3 |
| 8 | 97868090 | rs4735441 | 0.565 | -1.521 | -0.393 | -0.492 | 0.013 | -0.047 | -0.047 | 0 | 0.13 | 0.02 | 0.15 | 0.18 | 0.11 | 0.19 |
| 8 | 97871757 | rs2319936 | 0.047 | -1.363 | -0.129 | -0.563 | -0.015 | 0.042 | 0.042 | 0 | 0.45 | 0.64 | 0.49 | 0.34 | 0.48 | 0.33 |
| 8 | 97878729 | rs4469424 | -2.229 | -1.295 | 0.127 | -0.579 | -0.045 | -0.047 | -0.047 | 0 | 0.76 | 0.92 | 0.6 | 0.63 | 0.67 | 0.77 |
| 8 | 97883333 | rs10108534 | 0.888 | -1.404 | 0.277 | -0.637 | 0.004 | -0.002 | -0.002 | 1.191 | 0.37 | 0.3 | 0.44 | 0.33 | 0.45 | 0.28 |
| 8 | 97885269 | rs28698145 | 0.14 | -1.319 | -0.041 | -0.591 | -0.015 | 0.04 | 0.04 | 0 | 0.45 | 0.64 | 0.49 | 0.34 | 0.48 | 0.34 |
| 8 | 97920157 | rs4584121 | 0.281 | -1.263 | -0.009 | -0.679 | -0.009 | -0.083 | -0.083 | 0.701 | 0.34 | 0.47 | 0.18 | 0.11 | 0.25 | 0.24 |
| 8 | 97932044 | rs6983702 | -0.672 | -1.143 | -0.629 | -0.494 | 0.005 | -0.023 | -0.023 | 0 | 0.81 | 0.81 | 0.79 | 0.48 | 0.85 | 0.65 |
| 8 | 97944133 | rs4644202 | -1.522 | -1.122 | -0.932 | -0.496 | -0.158 | -0.17 | -0.17 | 0 | 0.68 | 0.98 | 0.57 | 0.46 | 0.55 | 0.63 |
| 8 | 97944696 | rs13278940 | -0.381 | -1.286 | -0.677 | -0.491 | 0.009 | -0.007 | -0.007 | 0.367 | 0.35 | 0.26 | 0.39 | 0.35 | 0.34 | 0.47 |
| 8 | 97950759 | rs10447959 | -0.997 | -1.408 | -0.319 | -0.57 | 0.001 | 0.001 | 0.001 | 0 | 0.45 | 0.52 | 0.43 | 0.35 | 0.36 | 0.49 |
| 8 | 97952965 | rs2853259 | -0.528 | -1.391 | -0.37 | -0.53 | 0.004 | 0.002 | 0.002 | 0 | 0.37 | 0.39 | 0.4 | 0.34 | 0.34 | 0.47 |
| 8 | 97966545 | rs2853255 | -1.388 | -1.145 | -0.157 | -0.414 | 0.039 | 0.021 | 0.021 | 1.121 | 0.55 | 0.83 | 0.5 | 0.41 | 0.41 | 0.54 |
| 8 | 98009019 | rs2853275 | -0.993 | -0.52 | -0.444 | -1.014 | 0.036 | 0.036 | 0.036 | 1.466 | 0.45 | 0.81 | 0.44 | 0.38 | 0.39 | 0.46 |
| 8 | 98018264 | rs10441498 | 0.964 | -0.466 | -0.725 | -1.02 | -0.002 | 0.047 | 0.047 | 0.406 | 0.14 | 0.05 | 0.19 | 0 | 0.14 | 0.11 |
| 8 | 98025685 | rs6468527 | 0.283 | 0.057 | -0.331 | -0.606 | 0.011 | 0.013 | 0.013 | 1.002 | 0.56 | 0.3 | 0.58 | 0.61 | 0.61 | 0.54 |
| 8 | 98035986 | rs10955094 | -0.332 | 0.088 | -0.788 | -0.522 | 0.007 | -0.01 | -0.01 | 1.491 | 0.64 | 0.74 | 0.66 | 0.62 | 0.66 | 0.56 |
| 8 | 98050511 | rs2513402 | 0.393 | -0.504 | -0.549 | -0.519 | 0.069 | 0.037 | 0.037 | 1.36 | 0.54 | 0.17 | 0.55 | 0.62 | 0.66 | 0.54 |
| 8 | 98073236 | rs2513341 | -0.545 | -0.296 | -0.64 | -0.581 | 0.046 | 0.006 | 0.006 | 0 | 0.67 | 0.78 | 0.73 | 0.69 | 0.78 | 0.6 |
| 8 | 98078587 | rs2513345 | -0.341 | -0.492 | -0.635 | -0.565 | 0.029 | 0.014 | 0.014 | 0 | 0.64 | 0.77 | 0.65 | 0.69 | 0.7 | 0.59 |
| 8 | 98080225 | rs2513399 | 1.67 | -0.571 | -0.933 | -0.734 | -0.156 | -0.184 | -0.184 | 0.061 | 0.25 | 0.02 | 0.42 | 0.5 | 0.4 | 0.27 |
| 8 | 98081414 | rs2513349 | 0.224 | -0.622 | -0.016 | -0.366 | 0.023 | 0.014 | 0.014 | 0 | 0.63 | 0.69 | 0.63 | 0.69 | 0.7 | 0.58 |
| 8 | 98090586 | rs2513405 | 0.869 | -0.368 | -0.053 | -0.159 | 0.025 | 0.029 | 0.029 | 0 | 0.48 | 0.34 | 0.52 | 0.6 | 0.54 | 0.43 |
| 8 | 98104633 | rs11992953 | -0.559 | -0.174 | -0.148 | -0.001 | 0.028 | 0.029 | 0.029 | 0.68 | 0.09 | 0.29 | 0.14 | 0.07 | 0.06 | 0.04 |
| 8 | 98106445 | rs10098932 | -0.559 | -0.174 | -0.165 | -0.001 | 0.028 | 0.029 | 0.029 | 0.68 | 0.09 | 0.29 | 0.14 | 0.07 | 0.06 | 0.04 |
| 8 | 98109070 | rs2513361 | 0.302 | 0.124 | -1.025 | 0.152 | 0.022 | 0.021 | 0.021 | 0 | 0.61 | 0.73 | 0.74 | 0.68 | 0.64 | 0.49 |
| 8 | 98122722 | rs13264693 | -0.296 | 0.177 | -1.028 | 0.252 | 0.021 | 0.007 | 0.007 | 0.165 | 0.37 | 0.25 | 0.27 | 0.33 | 0.35 | 0.51 |
| 8 | 98130180 | rs7844514 | -0.746 | 0.267 | -0.118 | 0.288 | 0.03 | 0.031 | 0.031 | 0 | 0.06 | 0.15 | 0.12 | 0.07 | 0.06 | 0.05 |
| 8 | 98134018 | rs1431890 | -0.84 | -0.158 | 0.183 | 0.491 | -0.013 | 0.097 | 0.097 | 0 | 0.22 | 0.39 | 0.27 | 0.63 | 0.19 | 0.32 |
| 8 | 98140470 | rs7817272 | -0.867 | 0.154 | 0.17 | 0.542 | 0.002 | 0.107 | 0.107 | 0 | 0.22 | 0.39 | 0.27 | 0.62 | 0.21 | 0.32 |
| 8 | 98147539 | rs920576 | -1.006 | 0.047 | 0.457 | 0.175 | -0.002 | 0.06 | 0.06 | 0 | 0.24 | 0.32 | 0.28 | 0.63 | 0.22 | 0.33 |
| 8 | 98150967 | rs2436051 | 1.142 | -0.637 | 0.027 | -0.072 | -0.021 | -0.03 | -0.03 | 0 | 0.68 | 0.6 | 0.66 | 0.22 | 0.75 | 0.57 |
| 8 | 98151238 | rs16895256 | 0.585 | -1.277 | -0.687 | -0.657 | 0.002 | -0.028 | -0.028 | 0.823 | 0.13 | 0 | 0.09 | 0.12 | 0.12 | 0.24 |
| 8 | 98151295 | rs16895257 | 0.397 | -1.309 | -0.167 | -0.803 | -0.004 | -0.036 | -0.036 | 0 | 0.07 | 0.18 | 0.14 | 0 | 0.06 | 0.06 |
| 8 | 98153281 | rs4556066 | -0.534 | -1.22 | -0.227 | -0.621 | -0.025 | 0.045 | 0.045 | 0 | 0.27 | 0.43 | 0.29 | 0.63 | 0.2 | 0.33 |
| 8 | 98163017 | rs2436046 | -0.063 | -2.032 | -0.364 | -0.649 | 0.008 | -0.011 | -0.011 | 0 | 0.31 | 0.26 | 0.2 | 0.57 | 0.19 | 0.41 |
| 8 | 98165873 | rs2436047 | 0.255 | -2.707 | -0.281 | -1.066 | 0.002 | 0.002 | 0.002 | 0 | 0.68 | 0.69 | 0.66 | 0.85 | 0.8 | 0.69 |
| 8 | 98166122 | rs16895264 | 0.351 | -2.624 | -0.204 | -0.98 | 0.01 | 0.044 | 0.044 | 0 | 0.07 | 0.19 | 0.09 | 0.1 | 0.07 | 0.08 |
| 8 | 98166913 | rs1835740 | -0.188 | -2.731 | -0.645 | -1.035 | 0.021 | -0.013 | -0.013 | 0 | 0.66 | 0.75 | 0.78 | 0.38 | 0.78 | 0.57 |
| 8 | 98168703 | rs982502 | 0.284 | -2.637 | -0.571 | -0.828 | 0.02 | -0.024 | -0.024 | 0 | 0.33 | 0.24 | 0.23 | 0.54 | 0.2 | 0.42 |
| 8 | 98169720 | rs16895268 | -0.783 | -2.58 | -0.327 | -0.941 | -0.039 | -0.025 | -0.025 | 0.181 | 0.29 | 0.55 | 0.37 | 0.21 | 0.21 | 0.31 |
| 8 | 98176043 | rs2044048 | -0.204 | -2.556 | -0.716 | -0.866 | 0.047 | -0.017 | -0.017 | 0 | 0.63 | 0.78 | 0.75 | 0.38 | 0.79 | 0.55 |
| 8 | 98177363 | rs1155199 | 0.808 | -2.015 | -0.517 | -0.84 | -0.015 | 0.038 | 0.038 | 0 | 0.37 | 0.17 | 0.36 | 0.3 | 0.59 | 0.3 |
| 8 | 98177956 | rs7845920 | 1.625 | -1.474 | -0.632 | -0.86 | -0.168 | 0.051 | 0.051 | 0 | 0.28 | 0.04 | 0.22 | 0.16 | 0.45 | 0.2 |
| 8 | 98178978 | rs1431884 | -0.392 | -1.48 | 0.844 | -0.293 | 0.077 | 0.152 | 0.152 | 1.46 | 0.06 | 0.27 | 0.08 | 0.15 | 0.06 | 0.09 |
| 8 | 98180213 | rs6990629 | 0.167 | -1.468 | 0.614 | -0.742 | 0.007 | -0.044 | -0.044 | 0.011 | 0.09 | 0.27 | 0.24 | 0 | 0.09 | 0.06 |
| 8 | 98187086 | rs1835742 | 1.205 | -1.821 | 0.087 | -0.477 | 0.082 | 0.158 | 0.158 | 0 | 0.5 | 0.19 | 0.45 | 0.21 | 0.62 | 0.29 |
| 8 | 98193889 | rs11783877 | -0.149 | -1.416 | 0.123 | 0.165 | 0.012 | -0.008 | -0.008 | 0.095 | 0.17 | 0.11 | 0.13 | 0.18 | 0.12 | 0.24 |
| 8 | 98196280 | rs4734357 | -0.343 | -1.489 | 0.301 | 0.091 | 0 | 0.012 | 0.012 | 0 | 0.82 | 0.85 | 0.85 | 0.4 | 0.84 | 0.64 |
| 8 | 98204790 | rs7838062 | -0.147 | -1.061 | 0.084 | -0.462 | 0.019 | 0.011 | 0.011 | 3.344 | 0.14 | 0.09 | 0.09 | 0.12 | 0.1 | 0.22 |
| 8 | 98225875 | rs12549505 | 0.287 | -0.715 | -0.484 | -0.406 | 0.013 | 0.031 | 0.031 | 0 | 0.11 | 0.05 | 0.22 | 0.04 | 0.06 | 0.09 |
| 8 | 98226375 | rs10738132 | 0.454 | -0.754 | -0.274 | -0.79 | 0.005 | 0.004 | 0.004 | 0 | 0.47 | 0.49 | 0.47 | 0.17 | 0.62 | 0.3 |
| 8 | 98227007 | rs10110404 | -0.129 | -0.831 | -0.27 | -0.655 | -0.002 | -0.004 | -0.004 | 8.933 | 0.58 | 0.59 | 0.52 | 0.29 | 0.68 | 0.51 |
| 8 | 98231744 | rs11784000 | -0.291 | -0.467 | -0.464 | -0.256 | -0.002 | -0.001 | -0.001 | 0 | 0.53 | 0.47 | 0.4 | 0.42 | 0.57 | 0.52 |

| | | | | | | | | | | | | | | | | |
|---|---|---|---|---|---|---|---|---|---|---|---|---|---|---|---|---|
| 8 | 98232135 | rs16895317 | 0.682 | -0.246 | -0.697 | -0.087 | 0.008 | 0 | 0 | 0.962 | 0.11 | 0.05 | 0.05 | 0.12 | 0.06 | 0.22 |
| 8 | 98237755 | rs10101931 | 0.489 | -0.097 | -0.675 | -0.201 | -0.007 | 0.002 | 0.002 | 0 | 0.37 | 0.32 | 0.32 | 0.3 | 0.47 | 0.24 |
| 8 | 98239350 | rs16895344 | 0.085 | -0.194 | -0.648 | -0.231 | 0.01 | 0.007 | 0.007 | 0.921 | 0.11 | 0.05 | 0.05 | 0.08 | 0.06 | 0.23 |
| 8 | 98240326 | rs7831704 | -0.379 | 0.31 | -0.582 | -0.225 | 0.013 | 0.034 | 0.034 | 0 | 0.49 | 0.47 | 0.35 | 0.25 | 0.44 | 0.41 |
| 8 | 98246924 | rs7010112 | -0.564 | 0.398 | -0.243 | -0.616 | -0.002 | 0.069 | 0.069 | 0 | 0.55 | 0.77 | 0.45 | 0.22 | 0.44 | 0.35 |
| 8 | 98248330 | rs6998596 | 0.053 | -0.152 | -0.149 | -0.962 | -0.026 | -0.073 | -0.073 | 0.672 | 0.31 | 0.54 | 0.22 | 0.09 | 0.24 | 0.19 |
| 8 | 98260909 | rs17739295 | 0.426 | 0.195 | -0.911 | -0.688 | -0.015 | -0.006 | -0.006 | 1.552 | 0.15 | 0.06 | 0.17 | 0.13 | 0.15 | 0.14 |
| 8 | 98262162 | rs2436042 | -0.204 | -0.102 | -0.815 | -0.779 | 0.039 | 0.076 | 0.076 | 0 | 0.4 | 0.79 | 0.34 | 0.23 | 0.33 | 0.32 |
| 8 | 98262244 | rs2116422 | -0.076 | -0.202 | -0.197 | -0.859 | 0.041 | 0.032 | 0.032 | 0 | 0.11 | 0.37 | 0.14 | 0.1 | 0.11 | 0.12 |
| 8 | 98271030 | rs2513380 | -0.656 | -0.183 | -0.563 | -0.95 | 0.03 | 0.112 | 0.112 | 0 | 0.43 | 0.77 | 0.37 | 0.2 | 0.38 | 0.3 |
| 8 | 98280132 | rs6468543 | 0.345 | -0.286 | -0.626 | -1.109 | -0.022 | 0.037 | 0.037 | 0 | 0.16 | 0.05 | 0.13 | 0 | 0.24 | 0.07 |
| 8 | 98284737 | rs2583503 | -0.855 | -0.382 | -0.247 | -1.117 | -0.007 | -0.035 | -0.035 | 0 | 0.27 | 0.56 | 0.25 | 0.12 | 0.26 | 0.2 |
| 8 | 98285120 | rs2567771 | -1.152 | -0.378 | -0.306 | -1.112 | 0 | -0.01 | -0.01 | 0 | 0.3 | 0.59 | 0.28 | 0.12 | 0.32 | 0.21 |
| 8 | 98288164 | rs2289496 | 1.618 | -0.981 | -1.148 | -0.727 | -0.127 | -0.115 | -0.115 | 0.261 | 0.32 | 0.02 | 0.38 | 0.37 | 0.39 | 0.4 |
| 8 | 98289714 | rs2635164 | -0.727 | -0.959 | -0.116 | -0.712 | 0.01 | -0.027 | -0.027 | 1.729 | 0.15 | 0.51 | 0.18 | 0.09 | 0.14 | 0.15 |
| 8 | 98296209 | rs2635165 | -0.539 | -0.835 | 0.251 | -0.571 | 0.014 | 0.046 | 0.046 | 0 | 0.12 | 0.37 | 0.14 | 0.14 | 0.09 | 0.21 |
| 8 | 98301557 | rs2635139 | -0.666 | -0.832 | 0.217 | -0.587 | 0.012 | 0.034 | 0.034 | 0.056 | 0.13 | 0.36 | 0.14 | 0.14 | 0.1 | 0.21 |
| 8 | 98302185 | rs2635140 | -1.006 | -0.833 | -0.378 | -0.589 | 0 | -0.007 | -0.007 | 0.022 | 0.34 | 0.63 | 0.31 | 0.16 | 0.34 | 0.3 |
| 8 | 98324440 | rs2635143 | -0.446 | -1.21 | -0.462 | -0.276 | -0.007 | -0.004 | -0.004 | 0 | 0.47 | 0.53 | 0.34 | 0.43 | 0.36 | 0.42 |
| 8 | 98325837 | rs4400351 | -0.577 | -1.81 | 0.548 | -0.411 | 0.003 | -0.052 | -0.052 | 0 | 0.36 | 0.36 | 0.19 | 0.01 | 0.31 | 0.15 |
| 8 | 98326152 | rs4593512 | -0.624 | -1.818 | 0.699 | -0.377 | -0.009 | -0.095 | -0.095 | 0 | 0.37 | 0.44 | 0.21 | 0.01 | 0.31 | 0.15 |
| 8 | 98327059 | rs16895514 | -0.612 | -1.818 | 0.637 | -0.377 | -0.002 | -0.066 | -0.066 | 0.334 | 0.36 | 0.42 | 0.21 | 0.01 | 0.31 | 0.15 |
| 8 | 98331852 | rs7825243 | -0.471 | -1.808 | 0.634 | 0.547 | -0.013 | 0.183 | 0.183 | 0 | 0.08 | 0.24 | 0.07 | 0.3 | 0.04 | 0.15 |
| 8 | 98350880 | rs6468549 | -1.053 | -1.753 | 0.304 | 0.679 | 0.001 | 0 | 0 | 0.148 | 0.45 | 0.61 | 0.28 | 0.38 | 0.34 | 0.32 |
| 8 | 98357081 | rs4520131 | 1.735 | -1.844 | -0.39 | 0.842 | -0.099 | -0.068 | -0.068 | 0.174 | 0.39 | 0.04 | 0.61 | 0.38 | 0.58 | 0.46 |
| 8 | 98357382 | rs11786542 | -0.071 | -1.761 | 0.175 | 1.254 | -0.012 | 0.089 | 0.089 | 0 | 0.07 | 0.1 | 0.07 | 0.32 | 0.04 | 0.16 |
| 8 | 98360795 | rs2247105 | 1.239 | -1.998 | 0.714 | 1.045 | -0.038 | 0.041 | 0.041 | 0.889 | 0.74 | 0.39 | 0.78 | 0.4 | 0.87 | 0.61 |
| 8 | 98362363 | rs17741775 | 0.95 | -1.886 | 0.519 | 0.473 | 0 | -0.016 | -0.016 | 0 | 0.13 | 0.12 | 0.08 | 0.01 | 0.12 | 0.05 |
| 8 | 98364076 | rs16895579 | 0.229 | -1.849 | 0.454 | 0.823 | -0.002 | 0.016 | 0.016 | 0 | 0.06 | 0.07 | 0.06 | 0.24 | 0.04 | 0.16 |
| 8 | 98364203 | rs2679765 | 1.011 | -1.996 | 0.851 | 0.649 | -0.018 | -0.001 | -0.001 | 0 | 0.68 | 0.6 | 0.84 | 0.72 | 0.82 | 0.72 |
| 8 | 98364250 | rs2853327 | -0.4 | -1.745 | 0.83 | 0.776 | 0 | 0.006 | 0.006 | 0.191 | 0.94 | 0.89 | 0.92 | 0.87 | 0.95 | 0.95 |
| 8 | 98364628 | rs2679764 | 0.732 | -1.587 | 1.038 | 0.617 | -0.001 | 0.022 | 0.022 | 1.889 | 0.73 | 0.41 | 0.74 | 0.64 | 0.77 | 0.74 |
| 8 | 98372762 | rs16895640 | -0.227 | -1.656 | 1.204 | 0.436 | -0.042 | -0.048 | -0.048 | 0.514 | 0.1 | 0.27 | 0.04 | 0.01 | 0.04 | 0.06 |
| 8 | 98377549 | rs6468553 | 1.619 | -1.894 | 0.913 | 0.665 | 0.01 | 0.002 | 0.002 | 3.72 | 0.55 | 0.33 | 0.7 | 0.39 | 0.7 | 0.6 |
| 8 | 98387588 | rs10955117 | 0.651 | -1.698 | 0.92 | 0.808 | 0.003 | -0.004 | -0.004 | 1.138 | 0.79 | 0.81 | 0.83 | 0.44 | 0.86 | 0.69 |
| 8 | 98387725 | rs2679778 | -1.254 | -1.791 | 1.212 | 0.704 | 0.004 | 0.005 | 0.005 | 1.899 | 0.31 | 0.71 | 0.19 | 0.16 | 0.18 | 0.2 |
| 8 | 98388568 | rs2853286 | -0.71 | -1.697 | 1.938 | 1.086 | -0.027 | 0.016 | 0.016 | 0 | 0.14 | 0.37 | 0.1 | 0.15 | 0.06 | 0.17 |
| 8 | 98389164 | rs2853288 | -1.38 | -1.906 | 1.888 | 1.042 | -0.058 | -0.011 | -0.011 | 0 | 0.27 | 0.46 | 0.13 | 0.22 | 0.12 | 0.25 |
| 8 | 98395246 | rs10100673 | 2.119 | -1.784 | 1.738 | 0.92 | 0.089 | 0.011 | 0.011 | 0 | 0.53 | 0.2 | 0.67 | 0.37 | 0.7 | 0.48 |
| 8 | 98395951 | rs4735463 | 2.068 | -1.781 | 1.898 | 0.921 | 0.062 | 0.01 | 0.01 | 0 | 0.53 | 0.28 | 0.68 | 0.37 | 0.7 | 0.48 |
| 8 | 98399988 | rs7840010 | 1.942 | -1.794 | 1.941 | 0.912 | 0.024 | -0.001 | -0.001 | 1.459 | 0.57 | 0.31 | 0.7 | 0.37 | 0.72 | 0.56 |
| 8 | 98413480 | rs16895730 | -0.28 | -1.82 | 2.02 | 0.899 | -0.032 | -0.014 | -0.014 | 3.106 | 0.14 | 0.19 | 0.05 | 0.07 | 0.06 | 0.07 |
| 8 | 98423535 | rs16892200 | -0.126 | -1.804 | 2.101 | 0.907 | -0.032 | -0.014 | -0.014 | 1.167 | 0.14 | 0.19 | 0.05 | 0.07 | 0.06 | 0.07 |
| 8 | 98427531 | rs2513387 | 0.114 | -1.793 | 1.991 | 0.83 | -0.019 | -0.073 | -0.073 | 0.216 | 0.85 | 0.97 | 0.85 | 0.57 | 0.84 | 0.82 |
| 8 | 98428688 | rs4368946 | -0.94 | -1.856 | 2.24 | 0.776 | -0.068 | -0.087 | -0.087 | 0 | 0.19 | 0.36 | 0.08 | 0.05 | 0.08 | 0.18 |
| 8 | 98428772 | rs2331465 | 2.457 | -1.883 | 1.846 | 0.821 | 0.116 | 0.023 | 0.023 | 0 | 0.51 | 0.14 | 0.67 | 0.37 | 0.7 | 0.47 |
| 8 | 98431651 | rs2253976 | 1.311 | -1.851 | 1.555 | 0.84 | -0.036 | 0.038 | 0.038 | 0 | 0.68 | 0.59 | 0.77 | 0.51 | 0.77 | 0.71 |
| 8 | 98435107 | rs2853305 | -0.783 | -1.532 | 1.108 | 0.914 | -0.023 | -0.003 | -0.003 | 0 | 0.15 | 0.28 | 0.09 | 0.14 | 0.07 | 0.25 |
| 8 | 98444949 | rs2513390 | 0.326 | -1.502 | 0.915 | 0.938 | 0.011 | -0.011 | -0.011 | 0 | 0.1 | 0.09 | 0.06 | 0.14 | 0.05 | 0.16 |
| 8 | 98446184 | rs13281649 | 1.491 | -1.504 | 1.067 | 1.056 | 0.001 | -0.003 | -0.003 | 0 | 0.69 | 0.67 | 0.71 | 0.34 | 0.77 | 0.57 |
| 8 | 98447831 | rs2460228 | -0.228 | -1.343 | 0.864 | 1.094 | -0.01 | 0.017 | 0.017 | 0 | 0.11 | 0.17 | 0.13 | 0.2 | 0.09 | 0.17 |
| 8 | 98451430 | rs12549368 | 1.927 | -1.197 | 0.278 | 1.061 | 0.048 | -0.085 | -0.085 | 0 | 0.58 | 0.03 | 0.62 | 0.35 | 0.73 | 0.56 |
| 8 | 98452499 | rs572175 | -0.72 | -0.83 | 1.317 | 1.427 | -0.025 | 0.025 | 0.025 | 0 | 0.13 | 0.23 | 0.13 | 0.2 | 0.08 | 0.17 |
| 8 | 98453225 | rs2460241 | 0.915 | -0.736 | 0.882 | 1.463 | -0.001 | 0.04 | 0.04 | 0 | 0.92 | 0.91 | 0.86 | 0.6 | 0.91 | 0.9 |
| 8 | 98454136 | rs2448166 | 0.915 | -0.736 | 0.882 | 1.463 | -0.001 | 0.04 | 0.04 | 0 | 0.92 | 0.91 | 0.86 | 0.6 | 0.91 | 0.9 |
| 8 | 98456632 | rs2513393 | 1.838 | -0.768 | 1.17 | 1.441 | -0.005 | 0.184 | 0.184 | 1.068 | 0.86 | 0.57 | 0.82 | 0.59 | 0.89 | 0.83 |
| 8 | 98458835 | rs663366 | -1.145 | -0.413 | 0.812 | 1.486 | -0.019 | 0.031 | 0.031 | 0.155 | 0.19 | 0.34 | 0.15 | 0.24 | 0.11 | 0.25 |
| 8 | 98459872 | rs2448191 | 0.858 | -0.153 | 0.268 | 1.541 | 0 | 0.068 | 0.068 | 0 | 0.94 | 0.92 | 0.88 | 0.61 | 0.92 | 0.91 |
| 8 | 98466755 | rs2460245 | -0.72 | 0.118 | 0.996 | 0.73 | 0.031 | -0.07 | -0.07 | 0 | 0.14 | 0.27 | 0.13 | 0.01 | 0.21 | 0.04 |
| 8 | 98466902 | rs2448149 | 0.08 | -0.442 | 0.301 | 0.732 | 0.042 | -0.056 | -0.056 | 0.338 | 0.71 | 0.85 | 0.76 | 0.54 | 0.82 | 0.7 |
| 8 | 98471716 | rs586380 | -0.17 | -0.293 | 0.267 | 0.575 | 0.031 | -0.07 | -0.07 | 0 | 0.14 | 0.28 | 0.13 | 0.01 | 0.21 | 0.04 |
| 8 | 98472201 | rs499177 | 1.545 | -0.583 | -0.529 | 0.519 | -0.104 | -0.082 | -0.082 | 1.401 | 0.43 | 0.05 | 0.53 | 0.4 | 0.55 | 0.55 |
| 8 | 98474916 | rs6995779 | -1.268 | -0.824 | -0.242 | 0.547 | 0.046 | -0.013 | -0.013 | 0.023 | 0.31 | 0.77 | 0.22 | 0.34 | 0.17 | 0.35 |
| 8 | 98477111 | rs591736 | -1.241 | -0.827 | 0.154 | 0.546 | 0.005 | -0.002 | -0.002 | 1.04 | 0.31 | 0.69 | 0.21 | 0.34 | 0.17 | 0.35 |
| 8 | 98485861 | rs10808356 | 0.75 | -0.71 | 0.093 | -0.018 | 0.017 | 0.018 | 0.018 | 1.782 | 0.28 | 0.17 | 0.24 | 0.19 | 0.23 | 0.24 |
| 8 | 98519919 | rs11784202 | -0.125 | -0.842 | -0.343 | 0.255 | 0.063 | 0.022 | 0.022 | 1.222 | 0.49 | 0.19 | 0.4 | 0.58 | 0.38 | 0.39 |
| 8 | 98526298 | rs597018 | 0.061 | -0.806 | -0.128 | 0.413 | -0.025 | 0.03 | 0.03 | 0.608 | 0.87 | 0.39 | 0.9 | 0.84 | 0.94 | 0.86 |
| 8 | 98553065 | rs16895910 | 0.915 | -0.661 | -0.561 | 0.465 | -0.018 | -0.121 | -0.121 | 0 | 0.2 | 0.01 | 0.16 | 0.38 | 0.16 | 0.19 |
| 8 | 98572450 | rs17750806 | 0.574 | -0.064 | -0.7 | 0.333 | -0.124 | -0.019 | -0.019 | 0 | 0.34 | 0.06 | 0.47 | 0.26 | 0.52 | 0.46 |

| | | | | | | | | | | | | | | | | |
|---|---|---|---|---|---|---|---|---|---|---|---|---|---|---|---|---|
| 8 | 98578620 | rs1871311 | 0.629 | 0.001 | 0.403 | 0.679 | 0.023 | 0.084 | 0.084 | 0 | 0.89 | 0.52 | 0.89 | 0.84 | 0.91 | 0.85 |
| 8 | 98580680 | rs7815922 | 0.922 | 0.117 | -0.353 | 0.723 | -0.002 | -0.005 | -0.005 | 0.582 | 0.9 | 0.91 | 0.94 | 0.82 | 0.89 | 0.82 |
| 8 | 98581814 | rs1351205 | 0.346 | -0.001 | -0.651 | 1.102 | -0.005 | -0.049 | -0.049 | 0.067 | 0.19 | 0.05 | 0.16 | 0.35 | 0.15 | 0.18 |
| 8 | 98581957 | rs7015322 | -0.781 | -0.281 | -0.336 | 1.197 | -0.015 | 0.03 | 0.03 | 0 | 0.15 | 0.25 | 0.4 | 0.22 | 0.12 | 0.24 |
| 8 | 98582216 | rs1382606 | -1.856 | -0.131 | -1.183 | 0.807 | -0.14 | 0.083 | 0.083 | 4.338 | 0.57 | 0.96 | 0.74 | 0.76 | 0.51 | 0.57 |
| 8 | 98582908 | rs1871313 | -1.429 | -0.042 | -1.085 | 0.885 | 0.049 | 0.081 | 0.081 | 2.869 | 0.49 | 0.92 | 0.68 | 0.67 | 0.38 | 0.49 |
| 8 | 98585277 | rs7013802 | 0.624 | -0.212 | -0.218 | 0.701 | 0.022 | -0.08 | -0.08 | 0 | 0.12 | 0.34 | 0.09 | 0 | 0.11 | 0.02 |
| 8 | 98586824 | rs1471463 | -1.215 | -0.072 | -0.214 | 0.898 | -0.034 | 0.145 | 0.145 | 0.358 | 0.18 | 0.46 | 0.44 | 0.32 | 0.14 | 0.25 |
| 8 | 98594829 | rs10808357 | -0.65 | -0.21 | -0.758 | 0.902 | 0.025 | -0.032 | -0.032 | 0 | 0.14 | 0.14 | 0.38 | 0.28 | 0.1 | 0.24 |
| 8 | 98600607 | rs2448142 | -1.535 | 0.006 | -1.117 | 0.568 | -0.125 | 0.104 | 0.104 | 3.272 | 0.56 | 0.96 | 0.74 | 0.77 | 0.5 | 0.56 |
| 8 | 98600657 | rs4735480 | -1.19 | -0.192 | -0.334 | 0.798 | -0.02 | 0.225 | 0.225 | 0.335 | 0.12 | 0.35 | 0.45 | 0.37 | 0.1 | 0.18 |
| 8 | 98600885 | rs2512416 | 1.084 | -0.157 | -0.336 | 0.766 | -0.01 | 0.019 | 0.019 | 0 | 0.07 | 0.03 | 0.02 | 0 | 0.08 | 0.02 |
| 8 | 98600977 | rs2448140 | -0.819 | -0.161 | -0.256 | 0.81 | -0.001 | -0.002 | -0.002 | 0 | 0.44 | 0.54 | 0.29 | 0.4 | 0.4 | 0.38 |
| 8 | 98615175 | rs7823849 | -0.796 | -0.064 | -0.166 | 0.848 | -0.01 | 0.126 | 0.126 | 0 | 0.22 | 0.43 | 0.5 | 0.48 | 0.22 | 0.31 |
| 8 | 98621340 | rs9694091 | 0.713 | 0.082 | -0.245 | 0.36 | 0.001 | -0.018 | -0.018 | 0.585 | 0.12 | 0.14 | 0.09 | 0.04 | 0.15 | 0.04 |
| 8 | 98625615 | rs2512426 | -0.484 | -0.161 | 0.22 | 0.598 | 0 | -0.014 | -0.014 | 0.456 | 0.32 | 0.36 | 0.22 | 0.25 | 0.3 | 0.21 |
| 8 | 98628286 | rs2512430 | -0.539 | -0.648 | 0.223 | 0.546 | -0.005 | -0.006 | -0.006 | 0 | 0.33 | 0.34 | 0.19 | 0.26 | 0.22 | 0.27 |
| 8 | 98635877 | rs2512436 | -1.696 | -1.052 | -0.259 | 0.533 | 0.12 | 0.025 | 0.025 | 0 | 0.5 | 0.87 | 0.5 | 0.58 | 0.32 | 0.49 |
| 8 | 98645779 | rs2512442 | -1.132 | -0.804 | 0.051 | 0.693 | 0.014 | 0.003 | 0.003 | 0.366 | 0.39 | 0.67 | 0.43 | 0.62 | 0.26 | 0.5 |
| 8 | 98647447 | rs2448193 | 0.942 | -0.668 | -0.241 | 0.694 | 0.003 | 0.025 | 0.025 | 0 | 0.08 | 0.04 | 0.03 | 0 | 0.06 | 0.05 |
| 8 | 98648478 | rs2512444 | -0.186 | -0.675 | -0.208 | 0.776 | 0.013 | -0.029 | -0.029 | 0 | 0.13 | 0.11 | 0.23 | 0.3 | 0.09 | 0.19 |
| 8 | 98655698 | rs2468028 | 0.341 | -0.653 | -0.251 | 0.804 | 0.007 | -0.045 | -0.045 | 0 | 0.1 | 0.05 | 0.22 | 0.29 | 0.09 | 0.18 |
| 8 | 98655937 | rs16896059 | 0.552 | -0.742 | -0.18 | 0.724 | 0.011 | -0.009 | -0.009 | 0 | 0.22 | 0.18 | 0.15 | 0.28 | 0.16 | 0.21 |
| 8 | 98659491 | rs2516278 | -1.809 | -0.933 | -0.137 | 0.507 | 0.099 | 0.052 | 0.052 | 0.01 | 0.48 | 0.8 | 0.48 | 0.62 | 0.32 | 0.58 |
| 8 | 98664528 | rs4448244 | 1.254 | -1.01 | -0.205 | 0.566 | -0.027 | -0.093 | -0.093 | 0 | 0.17 | 0.01 | 0.12 | 0.33 | 0.15 | 0.18 |
| 8 | 98673250 | rs16896067 | 0.057 | -0.765 | 0.3 | 0.711 | 0.032 | -0.037 | -0.037 | 1.312 | 0.3 | 0.11 | 0.17 | 0.38 | 0.22 | 0.34 |
| 8 | 98676847 | rs2438215 | -1.268 | -0.712 | 0.546 | 0.718 | 0.02 | 0.012 | 0.012 | 0 | 0.46 | 0.58 | 0.45 | 0.62 | 0.32 | 0.57 |
| 8 | 98710543 | rs2513973 | -0.447 | -0.344 | 0.688 | 0.778 | -0.015 | 0.221 | 0.221 | 0 | 0.13 | 0.38 | 0.28 | 0.29 | 0.09 | 0.23 |
| 8 | 98737749 | rs2438211 | 0.19 | -0.217 | -0.391 | 0.645 | 0.01 | -0.021 | -0.021 | 0.888 | 0.11 | 0.07 | 0.24 | 0.29 | 0.09 | 0.23 |
| 8 | 98737909 | rs2449512 | 0.243 | -0.203 | -0.384 | 0.652 | 0.009 | -0.019 | -0.019 | 0.52 | 0.11 | 0.06 | 0.23 | 0.29 | 0.09 | 0.23 |
| 8 | 98738361 | rs1311 | 0.243 | -0.203 | -0.384 | 0.652 | 0.009 | -0.019 | -0.019 | 0.749 | 0.11 | 0.06 | 0.23 | 0.29 | 0.09 | 0.23 |
| 8 | 98762432 | rs2513980 | -1.295 | 0.031 | -0.696 | 0.245 | 0.012 | 0.046 | 0.046 | 0 | 0.52 | 0.65 | 0.5 | 0.65 | 0.42 | 0.62 |
| 8 | 98766434 | rs7009821 | -0.503 | -0.192 | -0.923 | -0.074 | 0.001 | 0.001 | 0.001 | 0 | 0.34 | 0.56 | 0.23 | 0.35 | 0.24 | 0.3 |
| 8 | 98776879 | rs7815283 | 0.461 | 0.304 | -1.881 | -0.174 | -0.011 | -0.04 | -0.04 | 0 | 0.3 | 0.05 | 0.16 | 0.34 | 0.29 | 0.19 |
| 8 | 98785523 | rs2449529 | -0.161 | 0.598 | -1.444 | -0.069 | 0.022 | 0.016 | 0.016 | 0 | 0.51 | 0.36 | 0.28 | 0.46 | 0.44 | 0.47 |
| 8 | 98787795 | rs2002450 | 0.164 | 1.01 | -1.505 | -0.224 | -0.016 | 0.026 | 0.026 | 0 | 0.08 | 0.03 | 0.03 | 0.01 | 0.07 | 0.05 |
| 8 | 98803729 | rs6991424 | 0.135 | 0.56 | -1.552 | -0.214 | -0.01 | 0.015 | 0.015 | 0.093 | 0.42 | 0.37 | 0.28 | 0.38 | 0.46 | 0.27 |
| 8 | 98808157 | rs10955132 | -0.263 | 0.374 | -1.016 | -0.277 | 0.003 | 0.012 | 0.012 | 0 | 0.52 | 0.58 | 0.32 | 0.38 | 0.46 | 0.27 |
| 8 | 98811697 | rs2468013 | 0.447 | 0.247 | -1.397 | 0.159 | -0.164 | 0.062 | 0.062 | 0 | 0.4 | 0.06 | 0.42 | 0.21 | 0.49 | 0.45 |
| 8 | 98816199 | rs2917959 | 0.325 | 0.142 | -1.419 | 0.48 | -0.162 | -0.09 | -0.09 | 0 | 0.42 | 0.05 | 0.43 | 0.41 | 0.49 | 0.5 |
| 8 | 98823826 | rs4735498 | 0.187 | -0.161 | -0.362 | 0.446 | -0.012 | -0.012 | -0.012 | 3.192 | 0.46 | 0.34 | 0.43 | 0.55 | 0.46 | 0.4 |
| 8 | 98827531 | rs7818775 | 0.171 | -0.161 | -0.371 | 0.454 | -0.008 | -0.005 | -0.005 | 0 | 0.46 | 0.36 | 0.44 | 0.55 | 0.46 | 0.41 |
| 8 | 98841651 | rs2449535 | 0.056 | -0.128 | -0.487 | 0.472 | -0.002 | -0.003 | -0.003 | 0 | 0.47 | 0.39 | 0.45 | 0.55 | 0.46 | 0.43 |
| 8 | 98846219 | rs2449545 | -0.299 | -0.095 | -0.186 | 0.944 | -0.048 | -0.029 | -0.029 | 0 | 0.18 | 0.03 | 0.15 | 0.15 | 0.19 | 0.17 |
| 8 | 98847255 | rs2449539 | 0.917 | -0.407 | 0.198 | 1.224 | 0.039 | -0.046 | -0.046 | 0 | 0.12 | 0.07 | 0.22 | 0.35 | 0.03 | 0.23 |
| 8 | 98849303 | rs2512073 | 0.426 | -0.387 | 0.251 | 1.209 | -0.095 | 0.096 | 0.096 | 0.343 | 0.79 | 0.54 | 0.66 | 0.61 | 0.93 | 0.71 |
| 8 | 98851098 | rs2449520 | -0.799 | 0.017 | 0.126 | 0.856 | -0.011 | 0.001 | 0.001 | 0 | 0.18 | 0.28 | 0.16 | 0.19 | 0.17 | 0.15 |
| 8 | 98856699 | rs11991498 | 0.227 | 0.177 | 0.091 | 1.019 | -0.001 | 0.004 | 0.004 | 0 | 0.17 | 0.2 | 0.15 | 0.23 | 0.15 | 0.14 |
| 8 | 98856976 | rs2513970 | -1.832 | 0.297 | -0.305 | 1.008 | 0.005 | 0 | 0 | 0 | 0.62 | 0.92 | 0.64 | 0.66 | 0.68 | 0.6 |
| 8 | 98859437 | rs7017753 | -1.341 | 1.431 | 0.511 | 0.98 | 0.002 | -0.002 | -0.002 | 2.54 | 0.4 | 0.51 | 0.39 | 0.37 | 0.55 | 0.28 |
| 8 | 98861844 | rs11781043 | -0.644 | 1.894 | 0.089 | 1.066 | -0.038 | -0.037 | -0.037 | 0 | 0.26 | 0.18 | 0.34 | 0.36 | 0.36 | 0.2 |
| 8 | 98862550 | rs4501557 | -0.278 | 0.898 | 0.273 | 0.637 | 0.099 | -0.036 | -0.036 | 2.864 | 0.24 | 0.44 | 0.27 | 0.16 | 0.4 | 0.15 |
| 8 | 98864794 | rs3412 | -1.124 | 0.894 | -0.53 | 0.832 | 0.028 | 0.034 | 0.034 | 0 | 0.47 | 0.72 | 0.47 | 0.36 | 0.6 | 0.32 |
| 8 | 98866057 | rs10504983 | -0.485 | 0.599 | -0.17 | 0.931 | 0.006 | 0.045 | 0.045 | 0.1 | 0.18 | 0.44 | 0.16 | 0.2 | 0.16 | 0.13 |
| 8 | 98871690 | rs6996243 | 0.309 | 0.995 | -1.03 | 1.072 | -0.041 | -0.049 | -0.049 | 0.295 | 0.68 | 0.74 | 0.61 | 0.48 | 0.57 | 0.55 |
| 8 | 98874693 | rs10099863 | 0.585 | -0.459 | -0.524 | 1.058 | -0.072 | 0.041 | 0.041 | 0.234 | 0.69 | 0.49 | 0.69 | 0.41 | 0.85 | 0.59 |
| 8 | 98876595 | rs4631429 | -0.332 | -0.372 | -1.508 | 1.001 | -0.064 | -0.016 | -0.016 | 0.471 | 0.32 | 0.07 | 0.22 | 0.31 | 0.36 | 0.31 |
| 8 | 98877537 | rs6468601 | -0.603 | -0.427 | -1.104 | 1.084 | -0.005 | 0.022 | 0.022 | 0 | 0.82 | 0.81 | 0.76 | 0.66 | 0.87 | 0.71 |
| 8 | 98880097 | rs35148174 | 1.518 | -0.724 | -0.705 | 0.647 | -0.044 | 0.037 | 0.037 | 0 | 0.36 | 0.19 | 0.43 | 0.3 | 0.45 | 0.36 |
| 8 | 98880839 | rs28366014 | -0.351 | -0.708 | -0.377 | 0.635 | 0.003 | 0.057 | 0.057 | 3.62 | 0.09 | 0.52 | 0.08 | 0.05 | 0.02 | 0.02 |
| 8 | 98889153 | rs4360265 | 1.55 | -0.658 | -0.819 | 0.627 | -0.079 | 0.043 | 0.043 | 0 | 0.34 | 0.13 | 0.43 | 0.25 | 0.45 | 0.32 |
| 8 | 98893864 | rs4077920 | -0.438 | -0.904 | -0.054 | 0.659 | -0.049 | 0.071 | 0.071 | 0 | 0.81 | 0.7 | 0.76 | 0.66 | 0.92 | 0.77 |
| 8 | 98894110 | rs4559216 | 0.281 | -0.913 | 0.046 | 0.613 | -0.047 | 0.074 | 0.074 | 0 | 0.76 | 0.6 | 0.71 | 0.65 | 0.86 | 0.69 |
| 8 | 98897461 | rs7816792 | -0.379 | -0.946 | 0.017 | 0.71 | -0.049 | 0.105 | 0.105 | 0 | 0.81 | 0.67 | 0.76 | 0.61 | 0.91 | 0.77 |
| 8 | 98902730 | rs7815790 | 0.507 | -0.559 | -0.154 | 0.499 | 0.013 | 0.009 | 0.009 | 0 | 0.51 | 0.45 | 0.56 | 0.45 | 0.58 | 0.45 |
| 8 | 98908198 | rs2444889 | 0.687 | -0.267 | -0.814 | 0.241 | -0.037 | 0.032 | 0.032 | 0 | 0.46 | 0.14 | 0.56 | 0.44 | 0.51 | 0.4 |
| 8 | 98910287 | rs11996075 | -1.012 | -0.397 | -0.486 | -0.145 | 0.005 | 0.005 | 0.005 | 3.423 | 0.16 | 0.8 | 0.14 | 0.11 | 0.12 | 0.18 |
| 8 | 98917811 | rs4735508 | 0.82 | -0.215 | -0.1 | -0.493 | 0.007 | -0.033 | -0.033 | 0 | 0.1 | 0.2 | 0.3 | 0.06 | 0.1 | 0.16 |
| 8 | 98925477 | rs1869612 | -1.04 | -0.482 | -0.685 | -1.019 | -0.001 | -0.015 | -0.015 | 0 | 0.36 | 0.57 | 0.51 | 0.15 | 0.38 | 0.35 |

| | | | | | | | | | | | | | | | | |
|---|---|---|---|---|---|---|---|---|---|---|---|---|---|---|---|---|
| 8 | 98927699 | rs12676720 | 0.126 | -0.154 | -1.4 | -1.147 | 0.004 | 0.055 | 0.055 | 0 | 0.42 | 0.21 | 0.31 | 0.73 | 0.36 | 0.52 |
| 8 | 98933711 | rs2513834 | 0.605 | 0.09 | -1.27 | -1.242 | -0.004 | 0.045 | 0.045 | 0 | 0.38 | 0.25 | 0.31 | 0.75 | 0.34 | 0.51 |
| 8 | 98934748 | rs2444907 | 0.268 | 0.224 | -1.184 | -1.267 | -0.003 | 0.065 | 0.065 | 0 | 0.43 | 0.27 | 0.31 | 0.76 | 0.39 | 0.56 |
| 8 | 98936178 | rs10087445 | 1.282 | 0.348 | -1.387 | -1.393 | -0.047 | -0.032 | -0.032 | 0 | 0.11 | 0.01 | 0.13 | 0.08 | 0.18 | 0.09 |
| 8 | 98938149 | rs6468603 | -2.133 | 0.238 | -1.068 | -1.61 | -0.001 | -0.01 | -0.01 | 0 | 0.77 | 0.77 | 0.8 | 0.93 | 0.71 | 0.88 |
| 8 | 98938388 | rs1378124 | -1.059 | 0.166 | -0.75 | -1.671 | -0.017 | -0.131 | -0.131 | 0 | 0.66 | 0.49 | 0.75 | 0.93 | 0.67 | 0.84 |
| 8 | 98941602 | rs2124086 | 0.214 | 0.226 | -1.307 | -1.639 | 0 | 0.184 | 0.184 | 0 | 0.49 | 0.25 | 0.55 | 0.85 | 0.43 | 0.67 |
| 8 | 98942017 | rs1037529 | 0.249 | 0.194 | -1.324 | -1.655 | -0.005 | 0.174 | 0.174 | 0 | 0.48 | 0.25 | 0.55 | 0.85 | 0.43 | 0.67 |
| 8 | 98943545 | rs2290471 | -1.081 | 0.304 | -0.808 | -1.034 | 0.002 | -0.069 | -0.069 | 3.112 | 0.28 | 0.5 | 0.26 | 0.08 | 0.29 | 0.18 |
| 8 | 98943598 | rs2290472 | 0.622 | 0.798 | -1.389 | -0.543 | -0.051 | -0.031 | -0.031 | 0 | 0.11 | 0.01 | 0.13 | 0.08 | 0.18 | 0.09 |
| 8 | 98944327 | rs930817 | 1.647 | 0.687 | -1.044 | -0.458 | -0.022 | 0.003 | 0.003 | 0 | 0.08 | 0 | 0.26 | 0.07 | 0.09 | 0.19 |
| 8 | 98948973 | rs9297280 | -0.233 | 0.533 | -0.881 | -0.116 | -0.005 | 0.003 | 0.003 | 0 | 0.43 | 0.41 | 0.3 | 0.36 | 0.35 | 0.3 |
| 8 | 98949206 | rs2513843 | 1.022 | 0.6 | -1.327 | -0.053 | -0.052 | -0.018 | -0.018 | 0 | 0.4 | 0.2 | 0.31 | 0.42 | 0.46 | 0.4 |
| 8 | 98952826 | rs2512024 | -0.667 | 0.485 | -0.708 | 0.557 | 0.008 | 0.059 | 0.059 | 0 | 0.93 | 0.88 | 0.86 | 0.84 | 0.91 | 0.92 |
| 8 | 98953076 | rs2085144 | 0.57 | 0.391 | -1.356 | 0.337 | -0.036 | -0.073 | -0.073 | 1.958 | 0.28 | 0.01 | 0.17 | 0.26 | 0.2 | 0.11 |
| 8 | 98960813 | rs2512028 | -0.662 | 0.579 | -0.622 | 0.037 | -0.009 | 0.022 | 0.022 | 0 | 0.55 | 0.57 | 0.38 | 0.68 | 0.45 | 0.55 |
| 8 | 98963817 | rs897189 | -0.844 | 0.618 | -0.239 | -0.083 | -0.001 | -0.047 | -0.047 | 0.336 | 0.59 | 0.47 | 0.47 | 0.77 | 0.56 | 0.62 |
| 8 | 98964771 | rs2444882 | 0.13 | 0.581 | -0.812 | 0.216 | -0.086 | -0.131 | -0.131 | 0 | 0.39 | 0.07 | 0.33 | 0.44 | 0.41 | 0.24 |
| 8 | 98969239 | rs2512032 | -1.02 | 0.546 | -0.436 | 0.014 | 0 | -0.024 | -0.024 | 0 | 0.6 | 0.43 | 0.46 | 0.78 | 0.56 | 0.62 |
| 8 | 98970479 | rs2512033 | 0.11 | 0.604 | -0.758 | 0.319 | -0.093 | -0.115 | -0.115 | 2.499 | 0.4 | 0.04 | 0.32 | 0.41 | 0.4 | 0.23 |
| 8 | 98980663 | rs2444871 | -0.144 | 0.596 | 0.568 | 0.459 | -0.002 | 0.022 | 0.022 | 0.019 | 0.8 | 0.75 | 0.87 | 0.57 | 0.82 | 0.62 |
| 8 | 98981529 | rs2512040 | 1.234 | 0.461 | 0.056 | 0.101 | -0.039 | 0.104 | 0.104 | 0.302 | 0.43 | 0.1 | 0.3 | 0.21 | 0.44 | 0.17 |
| 8 | 98984840 | rs4735516 | 1.721 | 0.417 | 0.349 | 0.03 | -0.001 | -0.005 | -0.005 | 0 | 0.09 | 0.13 | 0.26 | 0.11 | 0.09 | 0.17 |
| 8 | 98987643 | rs11779647 | 2.118 | 0.324 | 0.237 | -0.149 | -0.009 | 0.023 | 0.023 | 0 | 0.25 | 0.15 | 0.18 | 0 | 0.31 | 0.09 |
| 8 | 98987918 | rs2512043 | 0.185 | 0.345 | 0.434 | -0.081 | 0 | -0.004 | -0.004 | 0 | 0.43 | 0.33 | 0.35 | 0.42 | 0.38 | 0.24 |
| 8 | 98990218 | rs2513849 | 0.614 | 0.376 | 0.613 | 0.337 | 0.002 | -0.032 | -0.032 | 0 | 0.68 | 0.7 | 0.56 | 0.44 | 0.69 | 0.34 |
| 8 | 98991223 | rs1869608 | -0.187 | 0.373 | 0.946 | -0.042 | -0.001 | 0.008 | 0.008 | 0 | 0.15 | 0.33 | 0.32 | 0.17 | 0.13 | 0.28 |
| 8 | 98991719 | rs4735518 | 0.27 | 0.372 | 0.788 | -0.042 | -0.001 | -0.002 | -0.002 | 0 | 0.13 | 0.13 | 0.31 | 0.16 | 0.13 | 0.27 |
| 8 | 98994154 | rs7835120 | -1.508 | 0.312 | 1.021 | -0.084 | 0.017 | 0.018 | 0.018 | 4.922 | 0.09 | 0.53 | 0.1 | 0.04 | 0.04 | 0.09 |
| 8 | 99006736 | rs11559202 | 1.045 | 0.501 | 0.341 | -0.2 | -0.003 | 0.022 | 0.022 | 0 | 0.17 | 0.15 | 0.11 | 0.08 | 0.15 | 0.11 |
| 8 | 99006748 | rs11559201 | 1.045 | 0.501 | 0.341 | -0.2 | -0.003 | 0.022 | 0.022 | 0 | 0.17 | 0.15 | 0.11 | 0.08 | 0.15 | 0.11 |
| 8 | 99008015 | rs4735521 | -0.097 | 0.443 | 0.627 | -0.231 | -0.006 | -0.031 | -0.031 | 0 | 0.29 | 0.54 | 0.4 | 0.21 | 0.24 | 0.31 |
| 8 | 99009024 | rs16896518 | 1.092 | 0.456 | 0.689 | -0.15 | -0.002 | 0.001 | 0.001 | 0 | 0.1 | 0.11 | 0.26 | 0.13 | 0.09 | 0.19 |
| 8 | 99009191 | rs10098159 | 0.757 | 0.513 | 0.619 | -0.585 | 0.004 | -0.075 | -0.075 | 0 | 0.45 | 0.5 | 0.31 | 0.08 | 0.51 | 0.21 |
| 8 | 99014550 | rs10955140 | 0.29 | 0.29 | 0.607 | -0.477 | 0.025 | 0.112 | 0.112 | 0 | 0.56 | 0.74 | 0.62 | 0.21 | 0.63 | 0.42 |
| 8 | 99016188 | rs2513806 | 0.812 | 0.278 | 0.463 | -0.067 | 0.029 | -0.013 | -0.013 | 0.365 | 0.22 | 0.14 | 0.15 | 0.22 | 0.14 | 0.1 |
| 8 | 99016723 | rs2444891 | -0.891 | 0.543 | 0.493 | 0.311 | -0.003 | -0.008 | -0.008 | 1.591 | 0.9 | 0.93 | 0.85 | 0.84 | 0.87 | 0.78 |
| 8 | 99020949 | rs2444895 | 0.008 | 0.604 | 0.406 | 0.225 | 0.003 | -0.011 | -0.011 | 0.176 | 0.7 | 0.45 | 0.71 | 0.76 | 0.7 | 0.65 |
| 8 | 99022009 | rs2444896 | 0.023 | 0.422 | 0.075 | 0.274 | -0.003 | -0.015 | -0.015 | 0 | 0.8 | 0.83 | 0.61 | 0.7 | 0.78 | 0.59 |
| 8 | 99030280 | rs2248014 | -0.151 | 0.294 | -0.267 | 0.031 | 0.001 | -0.054 | -0.054 | 0 | 0.71 | 0.84 | 0.74 | 0.43 | 0.72 | 0.51 |
| 8 | 99030718 | rs11991161 | 1.183 | 0.298 | -0.156 | 0.02 | -0.001 | -0.012 | -0.012 | 0 | 0.18 | 0.18 | 0.12 | 0.08 | 0.15 | 0.11 |
| 8 | 99044528 | rs2255317 | 1.219 | 0.302 | -0.551 | 0.02 | -0.005 | 0.011 | 0.011 | 3.094 | 0.14 | 0.12 | 0.12 | 0.22 | 0.09 | 0.08 |
| 8 | 99045978 | rs2255464 | -0.442 | 0.422 | -1.015 | -0.083 | 0.029 | 0.006 | 0.006 | 0.001 | 0.45 | 0.73 | 0.53 | 0.43 | 0.33 | 0.4 |
| 8 | 99046040 | rs2279120 | 1.693 | 0.411 | -1.031 | -0.092 | -0.049 | 0.022 | 0.022 | 0 | 0.16 | 0.01 | 0.1 | 0.08 | 0.15 | 0.11 |
| 8 | 99048329 | rs2615 | -0.43 | 0.336 | -1.065 | -0.129 | 0.029 | 0.006 | 0.006 | 0.001 | 0.45 | 0.73 | 0.53 | 0.43 | 0.33 | 0.4 |
| 8 | 99052895 | rs2444860 | -0.41 | 0.731 | -1.116 | 0.102 | -0.012 | -0.017 | -0.017 | 0 | 0.87 | 0.93 | 0.81 | 0.71 | 0.8 | 0.74 |
| 8 | 99055222 | rs2877453 | 1.498 | 0.865 | -1.125 | -0.719 | -0.048 | 0.074 | 0.074 | 0 | 0.28 | 0.12 | 0.22 | 0 | 0.41 | 0.12 |
| 8 | 99057903 | rs13255477 | 0.726 | 0.841 | -0.964 | -0.546 | -0.02 | 0.219 | 0.219 | 0.184 | 0.44 | 0.13 | 0.32 | 0.08 | 0.56 | 0.23 |
| 8 | 99088251 | rs2959644 | -1.237 | 1.297 | -0.276 | 0.422 | 0.003 | 0.005 | 0.005 | 0 | 0.54 | 0.4 | 0.65 | 0.72 | 0.58 | 0.71 |
| 8 | 99088581 | rs13278732 | 0.478 | 1.604 | -1.077 | 0.458 | -0.004 | 0.006 | 0.006 | 0 | 0.26 | 0.25 | 0.17 | 0.07 | 0.24 | 0.22 |
| 8 | 99088913 | rs2513829 | -1.315 | 1.625 | -0.585 | 1.658 | -0.004 | -0.004 | -0.004 | 0 | 0.29 | 0.27 | 0.5 | 0.64 | 0.34 | 0.49 |
| 8 | 99089452 | rs2447513 | -1.318 | 1.626 | -0.584 | 1.624 | -0.004 | -0.004 | -0.004 | 0.001 | 0.29 | 0.27 | 0.49 | 0.63 | 0.34 | 0.48 |
| 8 | 99091103 | rs2045806 | 1.265 | 1.813 | -0.42 | 1.258 | 0.031 | 0.037 | 0.037 | 0 | 0.78 | 0.61 | 0.63 | 0.69 | 0.69 | 0.6 |
| 8 | 99091245 | rs2045805 | -0.857 | 1.552 | -1.348 | 0.997 | -0.04 | -0.032 | -0.032 | 1.493 | 0.1 | 0 | 0.27 | 0.13 | 0.13 | 0.14 |
| 8 | 99094948 | rs2514335 | 0.378 | 1.48 | -1.118 | 0.981 | -0.011 | -0.007 | -0.007 | 0.091 | 0.66 | 0.31 | 0.56 | 0.42 | 0.61 | 0.59 |
| 8 | 99100559 | rs16896628 | -0.671 | 1.107 | -1.027 | 1.07 | 0 | -0.002 | -0.002 | 0.094 | 0.25 | 0.23 | 0.43 | 0.39 | 0.29 | 0.24 |
| 8 | 99100816 | rs6998768 | -0.72 | 1.09 | -0.331 | 0.575 | 0.014 | -0.047 | -0.047 | 0 | 0.11 | 0.27 | 0.09 | 0.02 | 0.1 | 0.1 |
| 8 | 99100932 | rs2444861 | -0.091 | 1.046 | -0.639 | 0.855 | -0.001 | -0.016 | -0.016 | 0.401 | 0.27 | 0.21 | 0.4 | 0.4 | 0.3 | 0.37 |
| 8 | 99101139 | rs2514344 | -0.081 | 0.733 | -0.88 | 0.643 | 0.01 | -0.029 | -0.029 | 0 | 0.4 | 0.28 | 0.4 | 0.5 | 0.36 | 0.3 |
| 8 | 99102403 | rs768420 | 0.11 | 0.612 | -1.137 | 0.905 | -0.008 | -0.023 | -0.023 | 0 | 0.22 | 0.11 | 0.21 | 0.25 | 0.2 | 0.2 |
| 8 | 99105726 | rs2306129 | 0.77 | 0.661 | -0.962 | 0.258 | 0.031 | -0.002 | -0.002 | 1.695 | 0.47 | 0.32 | 0.38 | 0.46 | 0.42 | 0.45 |
| 8 | 99109800 | rs1462975 | 1.371 | 0.795 | -1.071 | 0.371 | -0.013 | -0.001 | -0.001 | 0 | 0.07 | 0.03 | 0.14 | 0.26 | 0.1 | 0.16 |
| 8 | 99110123 | rs2514332 | -0.673 | 0.833 | -0.444 | 0.391 | 0.026 | 0.118 | 0.118 | 0 | 0.95 | 0.87 | 0.94 | 0.8 | 0.94 | 0.97 |
| 8 | 99114970 | rs8218 | 1.257 | 0.878 | -0.27 | 0.178 | 0.023 | -0.017 | -0.017 | 0 | 0.11 | 0.12 | 0.25 | 0.04 | 0.22 | 0.23 |
| 8 | 99115082 | rs11537947 | 1.154 | 0.676 | -0.356 | 0.085 | 0.014 | 0.035 | 0.035 | 0 | 0.16 | 0.09 | 0.06 | 0 | 0.1 | 0.04 |
| 8 | 99115359 | rs1462977 | 1.504 | 0.476 | -0.725 | 0.404 | -0.021 | -0.037 | -0.037 | 0.114 | 0.26 | 0.06 | 0.22 | 0.29 | 0.29 | 0.28 |
| 8 | 99116201 | rs2045800 | 0.744 | 0.932 | -0.377 | 0.136 | -0.012 | -0.009 | -0.009 | 0 | 0.35 | 0.22 | 0.37 | 0.56 | 0.4 | 0.46 |
| 8 | 99121408 | rs2447500 | -0.355 | 0.698 | -0.215 | 0.054 | -0.007 | -0.009 | -0.009 | 0.24 | 0.57 | 0.41 | 0.72 | 0.66 | 0.68 | 0.79 |
| 8 | 99125841 | rs2514337 | -0.757 | 1.133 | 0.252 | 0.439 | 0.066 | 0.058 | 0.058 | 2.155 | 0.29 | 0.49 | 0.54 | 0.41 | 0.4 | 0.53 |

| | | | | | | | | | | | | | | | | |
|---|---|---|---|---|---|---|---|---|---|---|---|---|---|---|---|---|
| 8 | 99128077 | rs10088925 | -0.381 | 0.955 | 0.319 | 0.034 | 0.06 | -0.049 | -0.049 | 0 | 0.2 | 0.48 | 0.33 | 0.12 | 0.25 | 0.34 |
| 8 | 99129507 | rs2071598 | 0.618 | 1.301 | 0.378 | 0.21 | 0.102 | 0.005 | 0.005 | 0.135 | 0.08 | 0.16 | 0.23 | 0.1 | 0.21 | 0.24 |
| 8 | 99133678 | rs4734384 | 0.095 | 0.392 | -0.302 | -0.116 | 0.031 | 0.072 | 0.072 | 0.778 | 0.4 | 0.12 | 0.31 | 0.25 | 0.35 | 0.36 |
| 8 | 99137153 | rs10808358 | -0.384 | 0.447 | -0.156 | -0.105 | -0.001 | -0.01 | -0.01 | 0 | 0.59 | 0.45 | 0.43 | 0.29 | 0.43 | 0.49 |
| 8 | 99139610 | rs4735529 | -0.289 | 0.601 | -0.129 | -0.1 | 0.008 | -0.01 | -0.01 | 0 | 0.6 | 0.45 | 0.47 | 0.3 | 0.53 | 0.49 |
| 8 | 99140208 | rs3802195 | -0.818 | 0.425 | -0.235 | 0.18 | -0.016 | 0.004 | 0.004 | 0.741 | 0.78 | 0.58 | 0.76 | 0.76 | 0.81 | 0.87 |
| 8 | 99140379 | rs3802196 | -1.247 | 0.538 | -0.749 | 0.258 | -0.001 | -0.021 | -0.021 | 0 | 0.8 | 0.93 | 0.79 | 0.79 | 0.82 | 0.91 |
| 8 | 99141814 | rs7840318 | -0.023 | 0.42 | -0.829 | 0.202 | 0.004 | 0.013 | 0.013 | 2.691 | 0.53 | 0.4 | 0.56 | 0.7 | 0.57 | 0.78 |
| 8 | 99146605 | rs16896659 | 0.935 | 0.416 | -1.229 | 0.803 | -0.022 | -0.066 | -0.066 | 0 | 0.11 | 0.01 | 0.11 | 0.25 | 0.13 | 0.16 |
| 8 | 99150174 | rs717230 | -0.505 | -0.076 | -0.693 | 0.329 | -0.015 | 0.003 | 0.003 | 0.64 | 0.78 | 0.7 | 0.85 | 0.76 | 0.83 | 0.87 |
| 8 | 99153088 | rs17184326 | 0.167 | 0.06 | -0.519 | -0.281 | 0 | -0.049 | -0.049 | 0 | 0.16 | 0.16 | 0.1 | 0 | 0.14 | 0.09 |
| 8 | 99153160 | rs17184361 | 0.321 | 0.105 | -0.594 | -0.352 | 0 | -0.034 | -0.034 | 0.296 | 0.15 | 0.13 | 0.09 | 0 | 0.12 | 0.06 |
| 8 | 99154285 | rs16896680 | -0.016 | 0.064 | -0.511 | -0.358 | 0 | -0.036 | -0.036 | 0.121 | 0.17 | 0.16 | 0.1 | 0 | 0.15 | 0.08 |
| 8 | 99155281 | rs17765430 | -0.251 | 0.124 | -0.531 | -0.355 | 0.015 | -0.021 | -0.021 | 0 | 0.05 | 0.06 | 0.05 | 0 | 0.1 | 0.05 |
| 8 | 99155532 | rs16896685 | -0.01 | 0.229 | 0.022 | -0.35 | 0.028 | 0.027 | 0.027 | 0 | 0.12 | 0.34 | 0.23 | 0.15 | 0.12 | 0.18 |
| 8 | 99159594 | rs11996826 | -1.25 | -0.149 | -0.379 | -0.491 | 0.007 | 0.082 | 0.082 | 0 | 0.39 | 0.79 | 0.39 | 0.2 | 0.31 | 0.38 |
| 8 | 99162888 | rs4735530 | -1.296 | -0.154 | -1.341 | -0.413 | -0.087 | -0.179 | -0.179 | 4.169 | 0.63 | 0.98 | 0.65 | 0.46 | 0.62 | 0.68 |
| 8 | 99166896 | rs9297283 | 0.374 | -0.219 | -0.659 | -0.458 | -0.01 | -0.014 | -0.014 | 0.516 | 0.21 | 0.3 | 0.28 | 0.16 | 0.17 | 0.2 |
| 8 | 99168383 | rs16896695 | 0.92 | -0.466 | -1.007 | -0.493 | -0.022 | -0.032 | -0.032 | 1.529 | 0.09 | 0.13 | 0.03 | 0.02 | 0.04 | 0.12 |
| 8 | 99175119 | rs2514342 | -0.51 | -0.225 | -1.249 | -0.315 | 0.014 | 0.062 | 0.062 | 0 | 0.45 | 0.69 | 0.43 | 0.23 | 0.36 | 0.43 |
| 8 | 99178531 | rs753316 | 1.055 | 0.022 | -1.521 | -0.183 | 0.003 | 0.007 | 0.007 | 0 | 0.19 | 0.22 | 0.25 | 0.33 | 0.29 | 0.26 |
| 8 | 99182032 | rs2447498 | -0.868 | 0.056 | -1.785 | -0.22 | -0.038 | -0.114 | -0.114 | 0 | 0.65 | 0.97 | 0.68 | 0.56 | 0.65 | 0.71 |
| 8 | 99189845 | rs2443573 | -0.346 | 0.533 | -0.919 | -0.063 | 0.019 | 0.037 | 0.037 | 0 | 0.72 | 0.58 | 0.7 | 0.42 | 0.69 | 0.63 |
| 8 | 99191257 | rs2447494 | 0.225 | -0.068 | -1.318 | -0.061 | 0.003 | 0.016 | 0.016 | 0 | 0.21 | 0.13 | 0.27 | 0.16 | 0.18 | 0.26 |
| 8 | 99192115 | rs6990795 | 0.261 | -0.106 | -0.868 | -0.108 | 0.01 | 0.011 | 0.011 | 0 | 0.35 | 0.26 | 0.31 | 0.26 | 0.33 | 0.3 |
| 8 | 99195204 | rs736191 | 0.258 | 0.542 | -0.456 | 0.208 | -0.001 | -0.005 | -0.005 | 0 | 0.56 | 0.5 | 0.57 | 0.44 | 0.47 | 0.67 |
| 8 | 99199066 | rs896416 | 0.393 | 0.805 | -0.194 | 0.445 | -0.001 | 0.003 | 0.003 | 0 | 0.41 | 0.5 | 0.57 | 0.58 | 0.43 | 0.62 |
| 8 | 99201718 | rs16896768 | 0.752 | 0.707 | 0.204 | 0.42 | 0.015 | -0.011 | -0.011 | 0.386 | 0.36 | 0.24 | 0.48 | 0.43 | 0.34 | 0.53 |
| 8 | 99203508 | rs2053934 | 0.568 | 0.82 | 0.38 | 0.236 | 0.011 | 0.03 | 0.03 | 0.035 | 0.34 | 0.26 | 0.22 | 0.08 | 0.28 | 0.22 |
| 8 | 99204723 | rs4734392 | 0.721 | 0.835 | 0.308 | 0.262 | -0.009 | -0.016 | -0.016 | 0.185 | 0.4 | 0.25 | 0.56 | 0.58 | 0.44 | 0.62 |
| 8 | 99205111 | rs3735885 | 0.138 | 0.981 | 0.17 | 0.18 | 0 | 0.003 | 0.003 | 0.082 | 0.51 | 0.54 | 0.65 | 0.59 | 0.5 | 0.67 |
| 8 | 99208244 | rs7010905 | 0.786 | 0.92 | 0.33 | 0.149 | 0.016 | 0.061 | 0.061 | 0.037 | 0.33 | 0.17 | 0.21 | 0.08 | 0.27 | 0.22 |
| 8 | 99209248 | rs10095887 | 1.114 | 0.753 | 0.423 | 0.244 | 0.013 | -0.009 | -0.009 | 0 | 0.36 | 0.29 | 0.48 | 0.43 | 0.35 | 0.53 |
| 8 | 99211688 | rs7838769 | 0.025 | 0.8 | 0.211 | 0.276 | 0 | 0.002 | 0.002 | 0.13 | 0.52 | 0.54 | 0.65 | 0.59 | 0.51 | 0.67 |
| 8 | 99225337 | rs6468620 | 0.923 | 0.794 | 0.302 | 0.192 | -0.005 | -0.001 | -0.001 | 0.688 | 0.38 | 0.39 | 0.5 | 0.41 | 0.35 | 0.55 |
| 8 | 99226821 | rs10093856 | 0.824 | 0.782 | -0.013 | 0.186 | -0.011 | -0.002 | -0.002 | 0 | 0.39 | 0.55 | 0.52 | 0.41 | 0.35 | 0.55 |
| 8 | 99227157 | rs10955157 | -0.236 | 0.235 | -0.463 | -0.057 | -0.009 | -0.003 | -0.003 | 0.06 | 0.17 | 0.09 | 0.14 | 0.15 | 0.16 | 0.11 |
| 8 | 99235179 | rs1993111 | 0.531 | 0.218 | -0.635 | -0.22 | 0.02 | 0.049 | 0.049 | 0 | 0.12 | 0.04 | 0.07 | 0 | 0.08 | 0.03 |
| 8 | 99243178 | rs6468624 | 0.852 | 0.349 | -0.447 | -0.011 | -0.012 | 0.02 | 0.02 | 0 | 0.66 | 0.42 | 0.71 | 0.46 | 0.74 | 0.76 |
| 8 | 99245008 | rs12550682 | 0.698 | 0.726 | -1.247 | 0.336 | -0.055 | -0.018 | -0.018 | 0 | 0.35 | 0.13 | 0.5 | 0.38 | 0.47 | 0.54 |
| 8 | 99253639 | rs2443566 | 0.444 | 0.402 | -0.667 | 1.027 | -0.001 | 0.04 | 0.04 | 0 | 0.88 | 0.82 | 0.9 | 0.58 | 0.91 | 0.89 |
| 8 | 99258187 | rs17288024 | -0.166 | 0.412 | -0.792 | -0.163 | -0.001 | 0.031 | 0.031 | 0 | 0.37 | 0.24 | 0.35 | 0.13 | 0.43 | 0.24 |
| 8 | 99259641 | rs10105510 | -0.098 | 0.383 | -1.073 | -0.135 | -0.037 | 0.029 | 0.029 | 0.929 | 0.27 | 0.09 | 0.42 | 0.2 | 0.32 | 0.33 |
| 8 | 99262445 | rs7016352 | 1.29 | -0.08 | -1.058 | 0.021 | -0.005 | -0.02 | -0.02 | 0 | 0.47 | 0.21 | 0.36 | 0.48 | 0.55 | 0.46 |
| 8 | 99270705 | rs1993113 | -0.246 | -0.434 | -0.679 | -0.021 | -0.012 | 0.052 | 0.052 | 0.362 | 0.83 | 0.81 | 0.84 | 0.64 | 0.89 | 0.79 |
| 8 | 99297875 | rs6982941 | -0.394 | -0.122 | -0.4 | -0.729 | -0.012 | -0.068 | -0.068 | 0.846 | 0.22 | 0.31 | 0.11 | 0 | 0.15 | 0.16 |
| 8 | 99314591 | rs4735551 | -0.621 | 0.416 | -0.409 | -0.558 | -0.033 | 0.045 | 0.045 | 0.174 | 0.37 | 0.33 | 0.51 | 0.18 | 0.49 | 0.38 |
| 8 | 99315582 | rs7812348 | -0.949 | 0.095 | 0.35 | -0.787 | -0.004 | 0.006 | 0.006 | 0.874 | 0.43 | 0.63 | 0.56 | 0.19 | 0.5 | 0.48 |
| 8 | 99318081 | rs4388427 | 0.674 | 0.65 | 0.324 | -0.741 | -0.02 | -0.019 | -0.019 | 0 | 0.09 | 0.04 | 0.32 | 0.18 | 0.18 | 0.13 |
| 8 | 99321817 | rs11775839 | 0.286 | 0.569 | 0.334 | -0.846 | -0.032 | 0.069 | 0.069 | 0 | 0.22 | 0.12 | 0.15 | 0 | 0.29 | 0.15 |
| 8 | 99324856 | rs12546215 | 0.652 | -0.048 | 0.188 | -0.428 | -0.002 | 0.008 | 0.008 | 1.898 | 0.13 | 0.13 | 0.12 | 0.24 | 0.09 | 0.06 |
| 8 | 99329605 | rs7820378 | 0.107 | -0.078 | 0.593 | -0.444 | -0.034 | 0.072 | 0.072 | 2.198 | 0.16 | 0.33 | 0.15 | 0.24 | 0.09 | 0.06 |
| 8 | 99336995 | rs4644205 | -1.34 | -0.099 | 0.206 | -0.541 | 0.016 | 0.003 | 0.003 | 0 | 0.53 | 0.84 | 0.67 | 0.43 | 0.57 | 0.44 |
| 8 | 99352637 | rs4346964 | -0.205 | 0.207 | 0.268 | -0.265 | -0.034 | -0.087 | -0.087 | 0 | 0.23 | 0.11 | 0.44 | 0.43 | 0.28 | 0.19 |
| 8 | 99355819 | rs4734399 | -1.267 | 0.065 | 0.35 | -0.426 | 0.017 | 0.018 | 0.018 | 0.02 | 0.49 | 0.71 | 0.66 | 0.43 | 0.57 | 0.43 |
| 8 | 99357924 | rs11784029 | 0.423 | 0.125 | 0.577 | -0.361 | 0.026 | -0.14 | -0.14 | 0 | 0.22 | 0.43 | 0.18 | 0 | 0.28 | 0.14 |
| 8 | 99360067 | rs6468632 | -0.965 | 0.065 | 0.298 | -0.129 | 0.022 | 0.023 | 0.023 | 0.009 | 0.5 | 0.82 | 0.65 | 0.42 | 0.56 | 0.47 |
| 8 | 99366439 | rs7828548 | -0.804 | -0.219 | 0.788 | 0.383 | 0.009 | 0.035 | 0.035 | 0 | 0.31 | 0.47 | 0.45 | 0.42 | 0.3 | 0.29 |
| 8 | 99366466 | rs7828561 | -0.075 | -0.021 | 0.872 | 0.587 | 0.012 | 0.023 | 0.023 | 0 | 0.18 | 0.33 | 0.34 | 0.23 | 0.18 | 0.18 |
| 8 | 99368788 | rs4487725 | -0.263 | -0.283 | 0.197 | 0.959 | 0.02 | -0.004 | -0.004 | 0.002 | 0.43 | 0.34 | 0.32 | 0.47 | 0.31 | 0.42 |
| 8 | 99373167 | rs6989037 | 0.489 | -0.056 | 0.088 | 1.015 | -0.002 | 0.001 | 0.001 | 0.348 | 0.12 | 0.12 | 0.11 | 0.19 | 0.08 | 0.06 |
| 8 | 99374659 | rs4476980 | 0.189 | 0.01 | 0.353 | 1.025 | 0.003 | 0 | 0 | 0 | 0.49 | 0.58 | 0.59 | 0.5 | 0.56 | 0.44 |
| 8 | 99374877 | rs10955167 | 0.622 | -0.004 | 0.19 | 1.076 | 0.025 | 0.008 | 0.008 | 0.046 | 0.32 | 0.19 | 0.22 | 0.27 | 0.23 | 0.34 |
| 8 | 99379259 | rs7831552 | 0.509 | 0.718 | 0.032 | 1.084 | 0.008 | -0.006 | -0.006 | 0 | 0.38 | 0.47 | 0.5 | 0.32 | 0.47 | 0.35 |
| 8 | 99379585 | rs11988085 | 0.314 | 1.487 | -0.017 | 1.475 | -0.012 | -0.015 | -0.015 | 0 | 0.83 | 0.88 | 0.84 | 0.78 | 0.79 | 0.77 |
| 8 | 99384890 | rs4565432 | 0.824 | 1.581 | 0.601 | 1.834 | 0.031 | 0.039 | 0.039 | 0 | 0.72 | 0.6 | 0.65 | 0.39 | 0.69 | 0.68 |
| 8 | 99385200 | rs4628231 | 0.927 | 1.631 | 0.663 | 1.777 | 0.03 | 0.021 | 0.021 | 0 | 0.7 | 0.59 | 0.64 | 0.39 | 0.69 | 0.69 |
| 8 | 99386594 | rs6984462 | -0.503 | 1.651 | 0.884 | 2.276 | -0.017 | 0.081 | 0.081 | 0 | 0.19 | 0.25 | 0.25 | 0.35 | 0.14 | 0.18 |
| 8 | 99386716 | rs7834444 | 1.174 | 1.5 | 0.891 | 2.38 | 0.03 | -0.031 | -0.031 | 0.123 | 0.71 | 0.57 | 0.47 | 0.23 | 0.7 | 0.64 |

| | | | | | | | | | | | | | | | | |
|---|---|---|---|---|---|---|---|---|---|---|---|---|---|---|---|---|
| 8 | 99389419 | rs7000370 | 1.187 | 1.501 | 0.886 | 2.38 | 0.028 | -0.031 | -0.031 | 0.053 | 0.71 | 0.56 | 0.47 | 0.22 | 0.7 | 0.62 |
| 8 | 99392416 | rs4735557 | 0.816 | 1.271 | 0.551 | 2.409 | 0.009 | 0.065 | 0.065 | 0.322 | 0.91 | 0.87 | 0.93 | 0.74 | 0.91 | 0.87 |
| 8 | 99402038 | rs10093553 | -0.543 | 1.226 | 0.355 | 1.464 | 0.027 | 0.017 | 0.017 | 0.233 | 0.37 | 0.53 | 0.46 | 0.6 | 0.49 | 0.51 |
| 8 | 99404946 | rs6468636 | 0.77 | 1.03 | 0.602 | 1.926 | -0.013 | 0.031 | 0.031 | 0 | 0.91 | 0.85 | 0.96 | 0.85 | 0.94 | 0.79 |
| 8 | 99413330 | rs6987962 | 0.851 | 1.097 | 0.67 | 1.935 | -0.011 | 0.04 | 0.04 | 0 | 0.91 | 0.82 | 0.96 | 0.85 | 0.94 | 0.79 |
| 8 | 99414574 | rs4735558 | 0.146 | 0.586 | -0.093 | 2.139 | -0.014 | -0.13 | -0.13 | 0 | 0.16 | 0.01 | 0.44 | 0.45 | 0.16 | 0.12 |
| 8 | 99414902 | rs6997978 | 0.622 | 0.593 | -0.087 | 2.218 | 0.021 | -0.028 | -0.028 | 2.995 | 0.75 | 0.8 | 0.52 | 0.42 | 0.78 | 0.66 |
| 8 | 99415437 | rs17310732 | -0.268 | 1.229 | 0.241 | 1.671 | 0.014 | -0.003 | -0.003 | 0.016 | 0.09 | 0.13 | 0.08 | 0.09 | 0.13 | 0.05 |
| 8 | 99420915 | rs4130404 | -0.956 | 1.09 | 1.019 | 1.341 | 0.044 | -0.056 | -0.056 | 3.185 | 0.16 | 0.38 | 0.11 | 0.07 | 0.17 | 0.12 |
| 8 | 99441904 | rs3802197 | 1.315 | 1.037 | 0.668 | 1.43 | 0 | 0.014 | 0.014 | 0.811 | 0.74 | 0.4 | 0.84 | 0.77 | 0.77 | 0.66 |
| 8 | 99445122 | rs11783149 | -0.97 | 1.303 | 0.314 | 1.192 | 0.027 | -0.025 | -0.025 | 0 | 0.14 | 0.26 | 0.1 | 0.08 | 0.16 | 0.1 |
| 8 | 99450646 | rs12546344 | -0.496 | 1.745 | -0.619 | 1.326 | -0.096 | -0.158 | -0.158 | 0 | 0.31 | 0.05 | 0.49 | 0.45 | 0.37 | 0.16 |
| 8 | 99452962 | rs10955174 | -1.074 | 1.522 | 0.123 | 1.08 | 0 | 0 | 0 | 0 | 0.43 | 0.48 | 0.56 | 0.51 | 0.46 | 0.26 |
| 8 | 99455311 | rs11774066 | 0.018 | 1.595 | -0.17 | 1.053 | -0.022 | -0.142 | -0.142 | 0 | 0.25 | 0.05 | 0.43 | 0.43 | 0.26 | 0.11 |
| 8 | 99461041 | rs16897007 | -0.493 | 1.556 | 0.675 | 1.022 | 0.039 | 0.004 | 0.004 | 0 | 0.11 | 0.19 | 0.13 | 0.08 | 0.17 | 0.08 |
| 8 | 99467856 | rs10955176 | -1.074 | 1.518 | 0.589 | 1.019 | -0.002 | -0.021 | -0.021 | 0 | 0.39 | 0.36 | 0.55 | 0.51 | 0.44 | 0.21 |
| 8 | 99477666 | rs4995528 | -1.2 | 1.449 | 0.662 | 1.211 | -0.001 | 0.003 | 0.003 | 0 | 0.43 | 0.46 | 0.58 | 0.56 | 0.47 | 0.27 |
| 8 | 99483391 | rs16897031 | 0.421 | 1.75 | 0.541 | 1.315 | -0.003 | -0.018 | -0.018 | 0.034 | 0.07 | 0.04 | 0.06 | 0.14 | 0.05 | 0.19 |
| 8 | 99497671 | rs4370499 | -0.676 | 1.789 | 0.603 | 1.331 | -0.023 | -0.098 | -0.098 | 2.775 | 0.35 | 0.17 | 0.54 | 0.48 | 0.41 | 0.18 |
| 8 | 99500187 | rs4348458 | -1.31 | 1.665 | 0.75 | 1.198 | 0.001 | 0.011 | 0.011 | 0 | 0.47 | 0.55 | 0.62 | 0.65 | 0.49 | 0.4 |
| 8 | 99525658 | rs4735564 | -0.88 | 1.658 | 0.578 | 1.215 | 0.001 | -0.024 | -0.024 | 1.889 | 0.39 | 0.32 | 0.55 | 0.51 | 0.42 | 0.21 |
| 8 | 99547555 | rs10955184 | -0.758 | 1.506 | 0.548 | 1.175 | 0.001 | -0.002 | -0.002 | 0 | 0.4 | 0.33 | 0.56 | 0.55 | 0.44 | 0.22 |
| 8 | 99552527 | rs1893884 | -1.086 | 1.522 | 0.584 | 1.233 | 0.005 | -0.006 | -0.006 | 0.079 | 0.44 | 0.45 | 0.56 | 0.5 | 0.43 | 0.26 |
| 8 | 99553244 | rs6991043 | -0.79 | 1.459 | 0.421 | 1.309 | -0.001 | -0.019 | -0.019 | 1.39 | 0.35 | 0.28 | 0.51 | 0.55 | 0.38 | 0.18 |
| 8 | 99553256 | rs7011216 | -0.513 | 1.738 | 0.325 | 1.338 | -0.008 | -0.021 | -0.021 | 1.479 | 0.93 | 0.96 | 0.95 | 0.88 | 0.95 | 0.87 |
| 8 | 99554231 | rs7016626 | -0.884 | 1.767 | 0.211 | 1.299 | -0.008 | -0.048 | -0.048 | 1.056 | 0.39 | 0.22 | 0.52 | 0.5 | 0.41 | 0.21 |
| 8 | 99558354 | rs16897076 | -0.627 | 1.824 | 0.301 | 1.38 | -0.043 | -0.099 | -0.099 | 0.05 | 0.36 | 0.14 | 0.52 | 0.45 | 0.4 | 0.21 |
| 8 | 99625091 | rs4735568 | -1.217 | 1.758 | 0.389 | 1.996 | -0.008 | -0.036 | -0.036 | 0 | 0.22 | 0.21 | 0.43 | 0.48 | 0.29 | 0.14 |
| 8 | 99661377 | rs4734410 | -1.607 | 1.729 | 0.386 | 1.819 | -0.016 | -0.047 | -0.047 | 0 | 0.25 | 0.22 | 0.44 | 0.44 | 0.32 | 0.19 |
| 8 | 99686213 | rs10096329 | -2.165 | 1.536 | 0.557 | 1.473 | -0.007 | 0.014 | 0.014 | 2.224 | 0.41 | 0.74 | 0.59 | 0.58 | 0.44 | 0.38 |
| 8 | 99730651 | rs6988087 | -1.921 | 1.758 | 0.75 | 1.562 | 0.029 | 0.062 | 0.062 | 0.653 | 0.29 | 0.52 | 0.51 | 0.52 | 0.36 | 0.21 |
| 8 | 99734384 | rs4131873 | -1.457 | 1.773 | 0.144 | 1.56 | -0.05 | -0.108 | -0.108 | 0 | 0.24 | 0.11 | 0.43 | 0.45 | 0.32 | 0.2 |
| 8 | 99737278 | rs4735574 | -1.452 | 1.772 | 0.144 | 1.63 | -0.046 | -0.1 | -0.1 | 0 | 0.24 | 0.11 | 0.43 | 0.44 | 0.3 | 0.19 |
| 8 | 99748557 | rs4549741 | -1.565 | 1.765 | 0.213 | 1.611 | -0.015 | -0.046 | -0.046 | 0 | 0.24 | 0.2 | 0.43 | 0.45 | 0.3 | 0.2 |
| 8 | 99762654 | rs35798019 | -1.793 | 2.323 | -0.127 | 1.499 | -0.008 | -0.014 | -0.014 | 0 | 0.28 | 0.3 | 0.46 | 0.45 | 0.35 | 0.2 |
| 8 | 99766557 | rs11778767 | -1.556 | 2.325 | -0.219 | 1.506 | -0.068 | -0.116 | -0.116 | 0 | 0.27 | 0.09 | 0.44 | 0.45 | 0.34 | 0.2 |
| 8 | 99767861 | rs34366206 | -0.682 | 2.288 | -0.249 | 1.526 | -0.091 | -0.151 | -0.151 | 0 | 0.21 | 0.04 | 0.43 | 0.47 | 0.31 | 0.15 |
| 8 | 99794459 | rs4255105 | 1.03 | 2.235 | -0.179 | 1.533 | -0.055 | -0.124 | -0.124 | 0.076 | 0.77 | 0.92 | 0.58 | 0.52 | 0.7 | 0.85 |
| 8 | 99799101 | rs7845271 | 1.684 | 2.287 | -0.035 | 1.562 | -0.008 | -0.011 | -0.011 | 0 | 0.69 | 0.64 | 0.51 | 0.46 | 0.63 | 0.77 |
| 8 | 99824050 | rs2515219 | 1.322 | 2.25 | -0.047 | 1.554 | -0.011 | -0.007 | -0.007 | 0.718 | 0.72 | 0.75 | 0.74 | 0.73 | 0.67 | 0.8 |
| 8 | 99846972 | rs3019293 | -1.671 | 2.322 | -0.044 | 0.962 | -0.017 | -0.055 | -0.055 | 0 | 0.29 | 0.23 | 0.44 | 0.48 | 0.33 | 0.21 |
| 8 | 99874520 | rs3019290 | -1.077 | 2.423 | 0.052 | 0.989 | -0.042 | -0.093 | -0.093 | 0.065 | 0.23 | 0.1 | 0.43 | 0.47 | 0.31 | 0.15 |
| 8 | 99895405 | rs3019286 | -1.765 | 2.411 | 0.077 | 0.965 | -0.027 | -0.039 | -0.039 | 0.69 | 0.3 | 0.24 | 0.47 | 0.47 | 0.38 | 0.24 |
| 8 | 99938470 | rs11987297 | -1.566 | 2.099 | 0.517 | 0.536 | 0.022 | 0.003 | 0.003 | 0.461 | 0.1 | 0.23 | 0.11 | 0.06 | 0.12 | 0.12 |
| 8 | 99955574 | rs985794 | -1.387 | 2.334 | 0.076 | 0.271 | -0.033 | 0.013 | 0.013 | 0 | 0.95 | 0.99 | 0.97 | 0.97 | 0.91 | 0.98 |
| 8 | 99999226 | rs4735592 | -0.602 | 2.347 | 0.507 | 0.251 | 0.011 | 0.109 | 0.109 | 0.789 | 0.18 | 0.31 | 0.42 | 0.47 | 0.21 | 0.13 |
| 8 | 100024447 | rs4510823 | -0.593 | 2.334 | 0.668 | 0.244 | 0.011 | 0.109 | 0.109 | 0.789 | 0.18 | 0.31 | 0.42 | 0.47 | 0.21 | 0.12 |
| 8 | 100032844 | rs13253769 | 2.341 | 2.072 | 1.177 | 0.118 | -0.015 | 0 | 0 | 0.17 | 0.6 | 0.27 | 0.44 | 0.41 | 0.6 | 0.67 |
| 8 | 100136131 | rs7459833 | 0.786 | 2.064 | 1.355 | 0.134 | 0.004 | 0.121 | 0.121 | 1.329 | 0.79 | 0.6 | 0.58 | 0.53 | 0.79 | 0.88 |
| 8 | 100141089 | rs7461275 | -0.53 | 2.102 | 1.335 | 0.133 | -0.018 | -0.037 | -0.037 | 0 | 0.86 | 0.91 | 0.64 | 0.59 | 0.8 | 0.9 |
| 8 | 100180871 | rs4442113 | -1.797 | 2.001 | 1.376 | 0.108 | 0.001 | 0.007 | 0.007 | 0.745 | 0.3 | 0.68 | 0.52 | 0.56 | 0.3 | 0.29 |
| 8 | 100222436 | rs4735607 | -0.359 | 1.997 | 1.15 | 0.124 | 0.003 | 0.05 | 0.05 | 0 | 0.85 | 0.83 | 0.64 | 0.59 | 0.83 | 0.9 |
| 8 | 100252754 | rs13263759 | 0.291 | 1.955 | 1.307 | 0.096 | -0.003 | 0.002 | 0.002 | 0.011 | 0.08 | 0.05 | 0.09 | 0.03 | 0.13 | 0.05 |
| 8 | 100293807 | rs4446702 | -0.671 | 1.987 | 1.201 | -0.258 | -0.021 | -0.054 | -0.054 | 0 | 0.87 | 0.94 | 0.65 | 0.59 | 0.83 | 0.9 |
| 8 | 100336535 | rs2291547 | 2.959 | 1.584 | 0.599 | -0.75 | -0.034 | -0.053 | -0.053 | 0 | 0.34 | 0.03 | 0.26 | 0.36 | 0.34 | 0.48 |
| 8 | 100341366 | rs3103731 | -0.08 | 1.51 | 0.701 | -0.66 | 0.011 | 0.109 | 0.109 | 0.577 | 0.82 | 0.71 | 0.58 | 0.53 | 0.79 | 0.88 |
| 8 | 100350083 | rs11995956 | -0.431 | 1.421 | 0.674 | -0.695 | -0.007 | 0.006 | 0.006 | 1.735 | 0.06 | 0.12 | 0.08 | 0.06 | 0.04 | 0.11 |
| 8 | 100369465 | rs3110405 | -0.753 | 1.48 | 0.716 | -0.667 | 0.003 | 0.051 | 0.051 | 0 | 0.85 | 0.83 | 0.64 | 0.59 | 0.83 | 0.9 |
| 8 | 100387299 | rs957539 | -0.555 | 1.479 | 0.733 | -0.673 | 0.008 | 0.131 | 0.131 | 0.978 | 0.2 | 0.4 | 0.42 | 0.47 | 0.21 | 0.12 |
| 8 | 100432678 | rs10504989 | 2.517 | 1.475 | 0.628 | -0.714 | -0.061 | -0.014 | -0.014 | 0 | 0.39 | 0.03 | 0.3 | 0.35 | 0.42 | 0.5 |
| 8 | 100432976 | rs3103705 | -1.918 | 1.825 | 1.237 | -0.273 | 0.002 | -0.014 | -0.014 | 0 | 0.34 | 0.79 | 0.53 | 0.56 | 0.31 | 0.3 |
| 8 | 100464671 | rs1492076 | -0.281 | 2.034 | 0.558 | -0.042 | -0.024 | 0.027 | 0.027 | 0 | 0.92 | 0.98 | 0.96 | 0.97 | 0.9 | 0.96 |
| 8 | 100484151 | rs3134160 | -1.011 | 2.158 | 0.54 | 0.102 | 0.008 | 0.109 | 0.109 | 0.152 | 0.19 | 0.32 | 0.42 | 0.48 | 0.21 | 0.12 |
| 8 | 100539155 | rs12542214 | -1.273 | 1.248 | 0.569 | 0.275 | 0.003 | 0.116 | 0.116 | 1.682 | 0.21 | 0.4 | 0.43 | 0.48 | 0.21 | 0.12 |
| 8 | 100569248 | rs965657 | -1.115 | 1.305 | 0.406 | 0.304 | 0.006 | 0.087 | 0.087 | 0.096 | 0.19 | 0.3 | 0.42 | 0.48 | 0.21 | 0.12 |
| 8 | 100583364 | rs12548482 | -0.167 | 1.485 | -0.073 | 0.39 | -0.013 | -0.033 | -0.033 | 0 | 0.13 | 0.09 | 0.35 | 0.41 | 0.17 | 0.1 |
| 8 | 100635818 | rs7008019 | 0.132 | 1.665 | 0.061 | 0.557 | -0.012 | -0.006 | -0.006 | 0 | 0.11 | 0.06 | 0.33 | 0.41 | 0.13 | 0.09 |
| 8 | 100635915 | rs4735627 | -0.57 | 1.65 | 0.076 | 0.547 | -0.004 | 0.026 | 0.026 | 0.009 | 0.13 | 0.14 | 0.39 | 0.48 | 0.17 | 0.12 |
| 8 | 100638489 | rs6985574 | -0.386 | 1.662 | 0.086 | 0.553 | -0.006 | 0.022 | 0.022 | 0 | 0.13 | 0.13 | 0.38 | 0.48 | 0.16 | 0.12 |

| | | | | | | | | | | | | | | | | |
|---|---|---|---|---|---|---|---|---|---|---|---|---|---|---|---|---|
| 8 | 100645287 | rs16897517 | -0.19 | 1.657 | 0.296 | 0.54 | 0.011 | 0.091 | 0.091 | 0 | 0.12 | 0.16 | 0.33 | 0.41 | 0.13 | 0.09 |
| 8 | 100650223 | rs1386374 | -1.817 | 1.658 | 0.526 | 0.414 | 0.04 | 0.154 | 0.154 | 0.05 | 0.23 | 0.45 | 0.46 | 0.48 | 0.24 | 0.22 |
| 8 | 100708595 | rs4735633 | -1.835 | 1.582 | -0.216 | 0.407 | 0.011 | 0.014 | 0.014 | 0.441 | 0.07 | 0.16 | 0.09 | 0.06 | 0.09 | 0.07 |
| 8 | 100758980 | rs1788148 | 0.361 | 1.391 | -0.287 | 0.474 | -0.02 | -0.007 | -0.007 | 0 | 0.09 | 0.04 | 0.33 | 0.43 | 0.13 | 0.09 |
| 8 | 100812300 | rs10092463 | 0.882 | 0.741 | -0.477 | 0.06 | -0.014 | -0.031 | -0.031 | 0.629 | 0.37 | 0.03 | 0.28 | 0.35 | 0.34 | 0.58 |
| 8 | 100824550 | rs921313 | -2.143 | 1.101 | 0.88 | 0.375 | 0.033 | 0.056 | 0.056 | 4.685 | 0.06 | 0.27 | 0.07 | 0.05 | 0.04 | 0.02 |
| 8 | 100829138 | rs2442756 | -1.617 | 1.071 | 0.807 | 0.367 | 0.005 | 0.015 | 0.015 | 0.194 | 0.33 | 0.34 | 0.49 | 0.49 | 0.36 | 0.26 |
| 8 | 100848480 | rs12674812 | 0.177 | 1.151 | 0.661 | 0.452 | -0.059 | -0.092 | -0.092 | 0.039 | 0.14 | 0.01 | 0.34 | 0.36 | 0.17 | 0.15 |
| 8 | 100850010 | rs7844422 | -1.9 | 1.12 | 1.155 | 0.389 | 0.01 | 0.028 | 0.028 | 0.035 | 0.33 | 0.55 | 0.52 | 0.5 | 0.36 | 0.28 |
| 8 | 100850412 | rs12680071 | 0.252 | 1.097 | 1.275 | 0.431 | -0.026 | -0.051 | -0.051 | 0 | 0.14 | 0.05 | 0.34 | 0.35 | 0.18 | 0.13 |
| 8 | 100860657 | rs4434590 | -2.35 | 0.516 | 2 | -0.209 | 0.013 | 0.001 | 0.001 | 2.062 | 0.18 | 0.5 | 0.17 | 0.14 | 0.18 | 0.11 |
| 8 | 100865836 | rs6468694 | -1.498 | 0.546 | 2.002 | -0.235 | 0.002 | 0.002 | 0.002 | 0.349 | 0.12 | 0.11 | 0.11 | 0.12 | 0.13 | 0.08 |
| 8 | 100870382 | rs6468697 | 2.56 | 0.513 | 1.968 | -0.329 | 0.008 | -0.037 | -0.037 | 0 | 0.77 | 0.52 | 0.78 | 0.87 | 0.77 | 0.81 |
| 8 | 100882147 | rs12546882 | -2.657 | 0.541 | 2.189 | -0.036 | 0.021 | 0 | 0 | 2.47 | 0.18 | 0.47 | 0.16 | 0.13 | 0.17 | 0.11 |
| 8 | 100889987 | rs4409375 | -2.108 | 0.574 | 2.163 | -0.019 | 0.011 | 0.01 | 0.01 | 0.689 | 0.15 | 0.28 | 0.14 | 0.11 | 0.14 | 0.09 |
| 8 | 100890542 | rs4626565 | -2.681 | 0.573 | 1.929 | -0.019 | 0.018 | -0.005 | -0.005 | 0 | 0.19 | 0.47 | 0.17 | 0.13 | 0.18 | 0.12 |
| 8 | 100890795 | rs7844439 | -2.652 | 0.591 | 1.942 | -0.01 | 0.021 | -0.003 | -0.003 | 0.123 | 0.18 | 0.46 | 0.17 | 0.14 | 0.18 | 0.12 |
| 8 | 100891210 | rs4510829 | -2.652 | 0.591 | 1.942 | -0.01 | 0.02 | -0.003 | -0.003 | 0 | 0.18 | 0.47 | 0.17 | 0.13 | 0.18 | 0.12 |
| 8 | 100891630 | rs1135382 | -2.635 | 0.591 | 1.935 | -0.01 | 0.024 | -0.002 | -0.002 | 0 | 0.18 | 0.44 | 0.17 | 0.13 | 0.18 | 0.12 |
| 8 | 100892645 | rs7828241 | -2.66 | 0.581 | 1.91 | -0.015 | 0.022 | -0.003 | -0.003 | 0 | 0.19 | 0.45 | 0.17 | 0.13 | 0.18 | 0.12 |
| 8 | 100897383 | rs13253073 | -1.926 | 0.584 | 1.521 | -0.008 | 0 | 0 | 0 | 0.412 | 0.14 | 0.11 | 0.12 | 0.12 | 0.14 | 0.09 |
| 8 | 100903890 | rs12544943 | -1.776 | 0.676 | 1.545 | 0.011 | 0.001 | 0.001 | 0.001 | 0.473 | 0.13 | 0.11 | 0.12 | 0.11 | 0.14 | 0.09 |
| 8 | 100916952 | rs13277184 | -2.188 | 0.42 | 1.261 | 0.104 | 0.001 | 0.004 | 0.004 | 2.804 | 0.17 | 0.5 | 0.16 | 0.13 | 0.17 | 0.11 |
| 8 | 100924271 | rs7837596 | -1.73 | 0.545 | 1.361 | 0.197 | 0.063 | 0.099 | 0.099 | 0.067 | 0.11 | 0.45 | 0.15 | 0.11 | 0.12 | 0.05 |
| 8 | 100926037 | rs7830618 | -2.153 | -0.138 | 1.003 | 0.147 | 0.06 | 0.004 | 0.004 | 1.867 | 0.24 | 0.85 | 0.19 | 0.17 | 0.15 | 0.19 |
| 8 | 100970952 | rs10107791 | -2.023 | 0.096 | 1.396 | 0.08 | -0.001 | 0.015 | 0.015 | 0.302 | 0.14 | 0.48 | 0.14 | 0.1 | 0.12 | 0.14 |
| 8 | 100993013 | rs16897831 | -1.106 | 0.053 | 1.033 | 0.105 | -0.023 | 0.015 | 0.015 | 0 | 0.1 | 0.14 | 0.09 | 0.08 | 0.06 | 0.11 |
| 8 | 100993147 | rs7830936 | -1.155 | 0.169 | 0.452 | 0.185 | 0.073 | 0.059 | 0.059 | 0.011 | 0.07 | 0.27 | 0.06 | 0.06 | 0.08 | 0.08 |
| 8 | 100994389 | rs6468704 | -1.269 | 0.087 | -0.853 | 0.156 | 0.037 | 0.013 | 0.013 | 3.146 | 0.27 | 0.85 | 0.21 | 0.17 | 0.21 | 0.21 |
| 8 | 100997714 | rs6985209 | -0.562 | 0.275 | -1.275 | -0.122 | -0.007 | 0.039 | 0.039 | 0 | 0.21 | 0.07 | 0.26 | 0.17 | 0.22 | 0.27 |
| 8 | 101009423 | rs3133704 | -0.166 | 0.074 | -2.175 | -0.204 | -0.044 | 0.019 | 0.019 | 0.596 | 0.6 | 0.92 | 0.7 | 0.64 | 0.6 | 0.62 |
| 8 | 101012421 | rs4734443 | 0.779 | -0.494 | -2.325 | -0.258 | -0.079 | -0.085 | -0.085 | 0 | 0.32 | 0.11 | 0.44 | 0.47 | 0.42 | 0.33 |
| 8 | 101024505 | rs7006527 | -0.178 | -0.905 | -1.24 | -0.457 | -0.02 | -0.025 | -0.025 | 0 | 0.19 | 0.29 | 0.15 | 0.07 | 0.16 | 0.24 |
| 8 | 101026725 | rs13276275 | 1.66 | -0.591 | -1.626 | -0.576 | -0.122 | -0.163 | -0.163 | 0 | 0.23 | 0.01 | 0.41 | 0.44 | 0.33 | 0.25 |
| 8 | 101041644 | rs3133692 | -1.184 | -0.718 | -1.173 | -0.583 | -0.013 | 0.033 | 0.033 | 0.137 | 0.59 | 0.88 | 0.7 | 0.64 | 0.6 | 0.61 |
| 8 | 101043237 | rs11779445 | -0.9 | -0.697 | -0.541 | -0.574 | -0.002 | -0.001 | -0.001 | 0 | 0.54 | 0.54 | 0.63 | 0.55 | 0.6 | 0.57 |
| 8 | 101074593 | rs2446925 | -0.533 | -1.391 | -0.818 | -0.863 | -0.007 | -0.041 | -0.041 | 4.412 | 0.53 | 0.13 | 0.57 | 0.48 | 0.57 | 0.56 |
| 8 | 101085778 | rs1380994 | 0.527 | -1.186 | -0.595 | -0.778 | -0.034 | -0.027 | -0.027 | 0.254 | 0.35 | 0.24 | 0.51 | 0.57 | 0.43 | 0.37 |
| 8 | 101095141 | rs2514701 | -0.79 | -1.191 | -0.625 | -0.795 | -0.001 | -0.01 | -0.01 | 0 | 0.54 | 0.54 | 0.66 | 0.64 | 0.56 | 0.6 |
| 8 | 101095559 | rs2446909 | 0.679 | -0.744 | -0.813 | -0.607 | -0.093 | -0.095 | -0.095 | 0 | 0.32 | 0.09 | 0.44 | 0.47 | 0.43 | 0.34 |
| 8 | 101102768 | rs3133719 | -1.072 | -0.716 | 0.364 | -0.592 | 0.024 | 0.058 | 0.058 | 2.574 | 0.13 | 0.54 | 0.13 | 0.14 | 0.09 | 0.1 |
| 8 | 101104014 | rs3133722 | -0.767 | -0.717 | 0.69 | -0.592 | 0.016 | 0.041 | 0.041 | 0.973 | 0.12 | 0.34 | 0.1 | 0.14 | 0.09 | 0.1 |
| 8 | 101106663 | rs11780823 | -0.198 | -0.972 | 0.482 | -0.981 | 0.027 | 0.034 | 0.034 | 0 | 0.22 | 0.18 | 0.14 | 0.07 | 0.13 | 0.22 |
| 8 | 101110024 | rs1993707 | -1.426 | -1.095 | 0.031 | -0.909 | -0.051 | 0.005 | 0.005 | 0.059 | 0.66 | 0.9 | 0.71 | 0.67 | 0.57 | 0.62 |
| 8 | 101114794 | rs1460930 | 1.297 | -0.948 | -0.208 | -0.803 | -0.121 | -0.173 | -0.173 | 0 | 0.3 | 0.02 | 0.43 | 0.52 | 0.38 | 0.28 |
| 8 | 101117554 | rs1471294 | 1.269 | -0.94 | -0.194 | -0.73 | -0.116 | -0.173 | -0.173 | 0.001 | 0.3 | 0.02 | 0.43 | 0.51 | 0.38 | 0.28 |
| 8 | 101117734 | rs1471293 | -1.177 | -0.989 | -0.29 | -0.877 | -0.081 | 0.03 | 0.03 | 0.019 | 0.64 | 0.88 | 0.73 | 0.7 | 0.56 | 0.56 |
| 8 | 101119037 | rs3735976 | 0.441 | -1.103 | -0.341 | -0.944 | 0.036 | 0.089 | 0.089 | 0 | 0.21 | 0.02 | 0.18 | 0.08 | 0.16 | 0.23 |
| 8 | 101137897 | rs2453626 | -0.919 | -0.947 | 0.184 | -0.904 | -0.024 | 0.011 | 0.011 | 0 | 0.61 | 0.71 | 0.69 | 0.65 | 0.56 | 0.53 |
| 8 | 101145709 | rs2446917 | 1.03 | -0.694 | 0.385 | -0.897 | 0.006 | 0.03 | 0.03 | 0 | 0.31 | 0.37 | 0.43 | 0.52 | 0.36 | 0.29 |
| 8 | 101150355 | rs11775440 | -0.173 | -0.723 | 0.394 | -0.939 | 0.03 | 0.061 | 0.061 | 0 | 0.25 | 0.12 | 0.25 | 0.12 | 0.21 | 0.25 |
| 8 | 101153024 | rs1460934 | -0.621 | -0.662 | 0.465 | -0.852 | -0.011 | 0.092 | 0.092 | 0 | 0.53 | 0.73 | 0.69 | 0.73 | 0.56 | 0.57 |
| 8 | 101166455 | rs2453643 | 0.94 | -0.554 | 0.397 | -0.771 | 0.011 | 0.048 | 0.048 | 0 | 0.31 | 0.4 | 0.44 | 0.52 | 0.36 | 0.29 |
| 8 | 101167395 | rs2514690 | 1.23 | -0.553 | 0.222 | -0.77 | -0.023 | -0.06 | -0.06 | 0 | 0.3 | 0.13 | 0.4 | 0.52 | 0.36 | 0.29 |
| 8 | 101167695 | rs2008612 | 0.969 | -0.524 | 0.139 | -0.757 | 0.002 | 0.01 | 0.01 | 0 | 0.31 | 0.3 | 0.43 | 0.52 | 0.36 | 0.29 |
| 8 | 101174426 | rs16898015 | -0.244 | -0.58 | 0.094 | -0.771 | -0.009 | -0.012 | -0.012 | 0.023 | 0.2 | 0.31 | 0.25 | 0.16 | 0.19 | 0.27 |
| 8 | 101180268 | rs11775395 | 0.106 | -0.611 | -0.063 | -0.865 | 0.027 | 0.079 | 0.079 | 0 | 0.24 | 0.13 | 0.24 | 0.07 | 0.21 | 0.24 |
| 8 | 101183535 | rs10464841 | 0.266 | -0.66 | -0.189 | -0.892 | 0.008 | 0.116 | 0.116 | 0 | 0.23 | 0.02 | 0.24 | 0.07 | 0.21 | 0.24 |
| 8 | 101203737 | rs3802149 | 0.325 | -0.669 | -0.108 | -0.893 | 0.022 | 0.1 | 0.1 | 0 | 0.23 | 0.05 | 0.24 | 0.07 | 0.21 | 0.24 |
| 8 | 101206391 | rs17335870 | 0.325 | -0.669 | -0.108 | -0.893 | 0.022 | 0.1 | 0.1 | 0 | 0.23 | 0.05 | 0.24 | 0.07 | 0.21 | 0.24 |
| 8 | 101210471 | rs7816209 | 0.325 | -0.669 | 0.03 | -0.893 | 0.024 | 0.068 | 0.068 | 0 | 0.23 | 0.07 | 0.24 | 0.07 | 0.21 | 0.24 |
| 8 | 101222499 | rs2514680 | 0.566 | -0.65 | 0.192 | -0.925 | 0 | -0.001 | -0.001 | 0 | 0.36 | 0.67 | 0.48 | 0.61 | 0.36 | 0.34 |
| 8 | 101226443 | rs1866788 | 1.121 | -0.641 | 0.295 | -0.952 | -0.012 | -0.038 | -0.038 | 0 | 0.31 | 0.18 | 0.41 | 0.52 | 0.36 | 0.3 |
| 8 | 101232701 | rs2919471 | -1.092 | -0.318 | -0.058 | -1.133 | -0.052 | -0.025 | -0.025 | 0.285 | 0.65 | 0.86 | 0.72 | 0.6 | 0.62 | 0.6 |
| 8 | 101243347 | rs7839147 | -0.283 | -0.381 | 0.51 | -1.123 | 0.035 | 0.093 | 0.093 | 1.641 | 0.06 | 0.38 | 0.07 | 0.06 | 0 | 0.05 |
| 8 | 101249762 | rs1788189 | -0.974 | -0.445 | 0.687 | -1.19 | -0.006 | -0.006 | -0.006 | 0.268 | 0.63 | 0.72 | 0.69 | 0.53 | 0.62 | 0.54 |
| 8 | 101252680 | rs6511 | -0.972 | -0.394 | 0.673 | -1.204 | -0.007 | -0.006 | -0.006 | 0 | 0.64 | 0.72 | 0.72 | 0.53 | 0.62 | 0.54 |
| 8 | 101253184 | rs1788190 | 1.26 | -0.469 | 0.408 | -1.354 | -0.107 | -0.157 | -0.157 | 0.01 | 0.31 | 0.03 | 0.4 | 0.51 | 0.37 | 0.3 |
| 8 | 101262063 | rs17421246 | 0.591 | -0.481 | 0.515 | -1.387 | 0.013 | 0.025 | 0.025 | 0 | 0.23 | 0.12 | 0.18 | 0.02 | 0.21 | 0.2 |

| | | | | | | | | | | | | | | | | |
|---|---|---|---|---|---|---|---|---|---|---|---|---|---|---|---|---|
| 8 | 101268237 | rs1660350 | 1.285 | -0.472 | 0.433 | -1.378 | -0.108 | -0.157 | -0.157 | 0 | 0.31 | 0.03 | 0.4 | 0.51 | 0.37 | 0.3 |
| 8 | 101277640 | rs1660322 | 1.882 | -0.415 | 0.218 | -1.558 | -0.037 | -0.043 | -0.043 | 1.544 | 0.27 | 0.1 | 0.21 | 0.34 | 0.31 | 0.36 |
| 8 | 101283085 | rs7837658 | -0.334 | -0.48 | 0.306 | -1.49 | 0.033 | 0.095 | 0.095 | 2.191 | 0.06 | 0.49 | 0.08 | 0.06 | 0 | 0.06 |
| 8 | 101309456 | rs1660331 | -1.16 | -0.214 | -0.85 | -1.513 | -0.082 | -0.131 | -0.131 | 0 | 0.68 | 0.92 | 0.6 | 0.48 | 0.63 | 0.7 |
| 8 | 101338257 | rs16898244 | 0.02 | -0.633 | -0.413 | -1.853 | -0.001 | -0.046 | -0.046 | 0 | 0.26 | 0.28 | 0.23 | 0.02 | 0.23 | 0.24 |
| 8 | 101342599 | rs6985608 | 0.507 | -0.358 | -0.775 | -1.703 | 0.058 | 0.033 | 0.033 | 0.947 | 0.34 | 0.77 | 0.23 | 0.25 | 0.18 | 0.37 |
| 8 | 101349662 | rs7816786 | -0.31 | -0.387 | -0.917 | -1.905 | 0.057 | 0.151 | 0.151 | 0.152 | 0.54 | 0.89 | 0.39 | 0.25 | 0.38 | 0.55 |
| 8 | 101354153 | rs6468712 | 0.602 | -0.213 | -0.411 | -1.893 | -0.014 | 0.009 | 0.009 | 0.309 | 0.36 | 0.45 | 0.29 | 0.42 | 0.27 | 0.36 |
| 8 | 101361955 | rs7015060 | -1.083 | -0.045 | -0.373 | -1.837 | -0.03 | -0.012 | -0.012 | 3.077 | 0.62 | 0.71 | 0.5 | 0.43 | 0.5 | 0.59 |
| 8 | 101365738 | rs6994382 | 0.998 | 0.068 | -0.31 | -1.755 | 0.02 | -0.005 | -0.005 | 0.234 | 0.11 | 0.18 | 0.09 | 0.11 | 0.14 | 0.05 |
| 8 | 101369801 | rs17423888 | 1.312 | -0.152 | -0.39 | -1.902 | 0.012 | -0.004 | -0.004 | 0 | 0.18 | 0.1 | 0.13 | 0.01 | 0.12 | 0.22 |
| 8 | 101371613 | rs16898286 | -0.386 | -0.178 | 0.148 | -1.945 | 0.05 | -0.074 | -0.074 | 0 | 0.16 | 0.41 | 0.16 | 0.07 | 0.2 | 0.05 |
| 8 | 101372374 | rs7003064 | -1.251 | -0.205 | 0.049 | -1.992 | -0.005 | -0.096 | -0.096 | 0 | 0.38 | 0.5 | 0.32 | 0.13 | 0.36 | 0.28 |
| 8 | 101382999 | rs9297295 | -1.349 | -0.39 | -0.014 | -1.986 | 0.006 | 0 | 0 | 0 | 0.4 | 0.64 | 0.32 | 0.08 | 0.36 | 0.27 |
| 8 | 101388283 | rs11779535 | 0.678 | -0.307 | -0.242 | -1.926 | 0.005 | -0.026 | -0.026 | 0.057 | 0.22 | 0.16 | 0.15 | 0.01 | 0.16 | 0.22 |
| 8 | 101389567 | rs4455801 | -0.816 | 0.198 | -0.325 | -1.561 | -0.127 | -0.082 | -0.082 | 0 | 0.65 | 0.92 | 0.52 | 0.43 | 0.5 | 0.59 |
| 8 | 101392669 | rs9886554 | 1.073 | 0.584 | -0.395 | -1.735 | -0.048 | 0.017 | 0.017 | 0 | 0.08 | 0 | 0.06 | 0.06 | 0.14 | 0.04 |
| 8 | 101394041 | rs3019087 | 0.82 | 0.256 | -0.321 | -1.85 | 0.003 | -0.026 | -0.026 | 0.114 | 0.23 | 0.26 | 0.19 | 0.05 | 0.29 | 0.19 |
| 8 | 101394661 | rs13278341 | -0.438 | -0.407 | -0.492 | -1.689 | -0.003 | -0.039 | -0.039 | 1.974 | 0.49 | 0.54 | 0.38 | 0.13 | 0.61 | 0.34 |
| 8 | 101397135 | rs16898307 | 0.005 | 0.18 | -0.066 | -1.042 | -0.002 | -0.034 | -0.034 | 0.922 | 0.08 | 0.2 | 0.09 | 0 | 0.07 | 0.01 |
| 8 | 101400303 | rs16898310 | 1.526 | 0.033 | -0.9 | -0.805 | 0.046 | -0.05 | -0.05 | 0 | 0.23 | 0.07 | 0.23 | 0.29 | 0.16 | 0.32 |
| 8 | 101403617 | rs17425247 | 0.23 | 0.365 | -0.458 | -0.587 | -0.013 | -0.038 | -0.038 | 0.103 | 0.15 | 0.14 | 0.08 | 0.04 | 0.09 | 0.13 |
| 8 | 101404950 | rs1075394 | 0.512 | 0.634 | -0.77 | -0.541 | 0.006 | 0.016 | 0.016 | 0 | 0.5 | 0.67 | 0.52 | 0.4 | 0.51 | 0.45 |
| 8 | 101408041 | rs2979002 | -0.09 | 0.403 | -1.464 | -0.318 | -0.109 | -0.092 | -0.092 | 0 | 0.66 | 0.93 | 0.6 | 0.43 | 0.6 | 0.59 |
| 8 | 101408103 | rs3019091 | -0.604 | -0.449 | -0.734 | -0.102 | 0.007 | -0.021 | -0.021 | 0 | 0.78 | 0.83 | 0.71 | 0.65 | 0.83 | 0.69 |
| 8 | 101421798 | rs6994682 | -1.349 | -0.44 | -0.531 | -0.88 | 0.019 | 0.012 | 0.012 | 2.668 | 0.21 | 0.84 | 0.23 | 0.04 | 0.12 | 0.13 |
| 8 | 101427609 | rs16898354 | -0.211 | -0.202 | -0.116 | -0.608 | -0.011 | -0.034 | -0.034 | 0.114 | 0.08 | 0.22 | 0.06 | 0.04 | 0.05 | 0.09 |
| 8 | 101430680 | rs4075553 | -0.834 | 0.129 | -0.505 | -0.711 | -0.003 | 0.007 | 0.007 | 0 | 0.41 | 0.59 | 0.4 | 0.22 | 0.45 | 0.26 |
| 8 | 101448416 | rs2000479 | -0.547 | 0.489 | -0.956 | -0.214 | -0.011 | 0.029 | 0.029 | 0 | 0.33 | 0.3 | 0.31 | 0.19 | 0.4 | 0.16 |
| 8 | 101459442 | rs7008598 | 0.329 | -0.156 | -0.786 | 0.754 | 0.012 | -0.02 | -0.02 | 0 | 0.8 | 0.88 | 0.79 | 0.51 | 0.85 | 0.45 |
| 8 | 101461664 | rs6999599 | -0.554 | -0.244 | -0.151 | 0.284 | 0.005 | -0.051 | -0.051 | 0 | 0.1 | 0.29 | 0.07 | 0.03 | 0.07 | 0.03 |
| 8 | 101470503 | rs6468719 | 0.895 | -0.33 | 0.366 | 0.306 | -0.011 | 0.018 | 0.018 | 0.774 | 0.7 | 0.65 | 0.72 | 0.48 | 0.78 | 0.42 |
| 8 | 101472510 | rs7001419 | 0.969 | -0.463 | 0.85 | 0.24 | -0.037 | 0.06 | 0.06 | 1.439 | 0.69 | 0.55 | 0.71 | 0.48 | 0.78 | 0.42 |
| 8 | 101473019 | rs1892736 | 1.116 | -0.46 | 0.589 | 0.241 | -0.031 | 0.041 | 0.041 | 0.815 | 0.68 | 0.44 | 0.7 | 0.48 | 0.78 | 0.42 |
| 8 | 101474782 | rs736178 | 0.111 | 0.117 | -0.24 | -0.892 | 0.011 | -0.066 | -0.066 | 0.114 | 0.22 | 0.3 | 0.25 | 0.01 | 0.35 | 0.16 |
| 8 | 101475221 | rs745346 | 1.401 | -0.029 | -0.83 | -0.565 | -0.119 | -0.017 | -0.017 | 0 | 0.4 | 0.03 | 0.38 | 0.33 | 0.4 | 0.19 |
| 8 | 101478499 | rs7015865 | 0.966 | 0.052 | -0.397 | -0.354 | 0.002 | 0.028 | 0.028 | 2.145 | 0.47 | 0.55 | 0.46 | 0.26 | 0.42 | 0.19 |
| 8 | 101479723 | rs12155948 | -0.155 | 0.377 | -1.05 | -0.463 | -0.078 | 0.062 | 0.062 | 0.193 | 0.17 | 0.01 | 0.14 | 0.04 | 0.25 | 0.21 |
| 8 | 101486501 | rs2154637 | 1.037 | 0.129 | -1.195 | -0.254 | -0.103 | 0.013 | 0.013 | 2.558 | 0.45 | 0.08 | 0.38 | 0.33 | 0.48 | 0.22 |
| 8 | 101492376 | rs11781587 | -0.94 | 0.767 | -0.127 | -0.472 | 0.037 | -0.119 | -0.119 | 0.827 | 0.23 | 0.49 | 0.19 | 0.03 | 0.27 | 0.22 |
| 8 | 101494012 | rs6997292 | -0.93 | 0.775 | -0.123 | -0.468 | 0.038 | -0.119 | -0.119 | 0.775 | 0.23 | 0.48 | 0.18 | 0.03 | 0.27 | 0.22 |
| 8 | 101498052 | rs7001302 | -0.823 | 0.7 | -1.286 | -0.15 | -0.018 | 0.056 | 0.056 | 1.769 | 0.36 | 0.9 | 0.31 | 0.26 | 0.34 | 0.29 |
| 8 | 101503820 | rs16898428 | -0.733 | 0.803 | -0.103 | -0.204 | 0.038 | -0.019 | -0.019 | 0.879 | 0.09 | 0.29 | 0.14 | 0.04 | 0.09 | 0.07 |
| 8 | 101504357 | rs4734457 | -0.911 | 1.03 | 0.134 | 0.484 | 0.01 | 0.075 | 0.075 | 0.031 | 0.07 | 0.16 | 0.27 | 0.24 | 0.11 | 0.03 |
| 8 | 101504803 | rs2022916 | 0.326 | 1.173 | -0.697 | -0.529 | -0.001 | -0.003 | -0.003 | 0.012 | 0.5 | 0.49 | 0.47 | 0.26 | 0.48 | 0.28 |
| 8 | 101505135 | rs4734459 | 0.402 | 1.356 | -0.606 | -0.553 | 0 | -0.001 | -0.001 | 0 | 0.5 | 0.48 | 0.48 | 0.26 | 0.51 | 0.3 |
| 8 | 101506305 | rs7826305 | 0.116 | 1.001 | -0.864 | -0.97 | 0.005 | 0.052 | 0.052 | 0 | 0.62 | 0.7 | 0.4 | 0.15 | 0.63 | 0.46 |
| 8 | 101520098 | rs6468721 | -0.129 | 1.572 | -0.869 | -1.12 | -0.014 | -0.004 | -0.004 | 0.419 | 0.2 | 0.14 | 0.16 | 0.01 | 0.25 | 0.21 |
| 8 | 101520351 | rs719870 | -0.225 | 1.206 | -1.177 | -1.43 | 0.033 | 0.133 | 0.133 | 0 | 0.5 | 0.28 | 0.36 | 0.16 | 0.62 | 0.41 |
| 8 | 101525085 | rs7832617 | -0.972 | 0.966 | -1.113 | -0.575 | -0.008 | -0.085 | -0.085 | 0 | 0.85 | 0.95 | 0.72 | 0.45 | 0.85 | 0.65 |
| 8 | 101531518 | rs17349841 | 0.579 | 1.126 | -1.258 | -0.938 | 0.01 | 0.086 | 0.086 | 0 | 0.34 | 0.21 | 0.25 | 0.13 | 0.36 | 0.2 |
| 8 | 101533868 | rs15989 | 0.336 | 1.331 | -0.426 | -0.942 | -0.001 | -0.001 | -0.001 | 0 | 0.41 | 0.45 | 0.36 | 0.28 | 0.4 | 0.24 |
| 8 | 101542205 | rs2469652 | -0.436 | 1.189 | -0.116 | -1.017 | -0.004 | 0.063 | 0.063 | 0.002 | 0.52 | 0.87 | 0.5 | 0.33 | 0.49 | 0.38 |
| 8 | 101545455 | rs12545394 | 0.143 | 1.129 | 0.015 | -0.963 | 0.005 | 0.139 | 0.139 | 0.039 | 0.08 | 0.39 | 0.13 | 0.15 | 0.04 | 0.04 |
| 8 | 101552505 | rs2000480 | 0 | 1.187 | -0.449 | -1.446 | 0.013 | 0.098 | 0.098 | 0 | 0.5 | 0.31 | 0.4 | 0.18 | 0.49 | 0.39 |
| 8 | 101554481 | rs17428846 | 1.874 | 1.007 | -0.62 | -1.363 | 0.018 | 0.036 | 0.036 | 0 | 0.22 | 0.08 | 0.15 | 0.11 | 0.24 | 0.18 |
| 8 | 101554696 | rs2469649 | 0.033 | 1.021 | 0.116 | -0.918 | -0.013 | 0.05 | 0.05 | 0 | 0.39 | 0.22 | 0.33 | 0.18 | 0.39 | 0.37 |
| 8 | 101555649 | rs17428950 | 0.829 | 0.817 | -0.121 | -0.72 | 0.002 | -0.012 | -0.012 | 0.759 | 0.05 | 0.04 | 0.07 | 0 | 0.1 | 0.02 |
| 8 | 101573820 | rs7835153 | 0.71 | 0.442 | -0.228 | -0.79 | -0.028 | 0.107 | 0.107 | 0 | 0.37 | 0.2 | 0.28 | 0.14 | 0.43 | 0.22 |
| 8 | 101575211 | rs2469654 | -0.113 | 0.095 | 0.665 | -0.643 | -0.01 | 0.016 | 0.016 | 0 | 0.78 | 0.78 | 0.62 | 0.28 | 0.81 | 0.59 |
| 8 | 101577980 | rs11785829 | 1.257 | 0.101 | 0.357 | -0.829 | -0.007 | 0.08 | 0.08 | 0 | 0.41 | 0.1 | 0.32 | 0.24 | 0.42 | 0.26 |
| 8 | 101580997 | rs2105508 | -0.006 | 0.154 | 1.328 | -0.958 | -0.034 | 0.074 | 0.074 | 0 | 0.29 | 0.13 | 0.22 | 0.07 | 0.33 | 0.31 |
| 8 | 101584521 | rs2469653 | -0.673 | 0.05 | 1.758 | -0.923 | 0.016 | -0.06 | -0.06 | 0.506 | 0.35 | 0.4 | 0.26 | 0.11 | 0.4 | 0.34 |
| 8 | 101585249 | rs1609 | -0.33 | 0.063 | 1.623 | -0.918 | -0.013 | 0.011 | 0.011 | 0 | 0.32 | 0.26 | 0.25 | 0.11 | 0.39 | 0.33 |
| 8 | 101586133 | rs2022923 | -0.864 | 0.112 | 1.291 | -0.715 | 0 | -0.003 | -0.003 | 0 | 0.42 | 0.55 | 0.37 | 0.28 | 0.44 | 0.37 |
| 8 | 101594707 | rs4548151 | -0.756 | 0.482 | 0.724 | -0.47 | -0.014 | 0.017 | 0.017 | 0 | 0.32 | 0.3 | 0.27 | 0.11 | 0.38 | 0.31 |
| 8 | 101595906 | rs2507778 | -0.965 | 0.323 | 0.689 | -0.579 | 0.008 | -0.015 | -0.015 | 0 | 0.33 | 0.54 | 0.44 | 0.25 | 0.34 | 0.37 |
| 8 | 101597286 | rs16898600 | -0.357 | 0.605 | 0.356 | -0.385 | 0.003 | -0.003 | -0.003 | 0 | 0.06 | 0.04 | 0.19 | 0.15 | 0.02 | 0.1 |
| 8 | 101597298 | rs16898602 | -1.107 | 0.841 | 0.276 | -0.399 | 0.004 | -0.028 | -0.028 | 0 | 0.07 | 0.12 | 0.04 | 0 | 0.07 | 0.05 |

| | | | | | | | | | | | | | | | | |
|---|---|---|---|---|---|---|---|---|---|---|---|---|---|---|---|---|
| 8 | 101611200 | rs2096548 | -0.723 | 0.523 | -0.305 | -0.283 | 0.007 | -0.046 | -0.046 | 2.738 | 0.33 | 0.46 | 0.43 | 0.18 | 0.33 | 0.36 |
| 8 | 101620374 | rs2469666 | -0.374 | 0.529 | -1.055 | 0.484 | -0.011 | -0.033 | -0.033 | 0.832 | 0.39 | 0.15 | 0.42 | 0.49 | 0.37 | 0.47 |
| 8 | 101625336 | rs7001509 | -0.985 | 0.113 | -0.269 | 0.654 | -0.021 | 0.047 | 0.047 | 0 | 0.11 | 0.25 | 0.09 | 0.17 | 0.07 | 0.15 |
| 8 | 101626241 | rs752036 | -0.55 | 0.257 | -0.194 | 0.17 | 0.007 | -0.022 | -0.022 | 0 | 0.37 | 0.43 | 0.5 | 0.24 | 0.4 | 0.47 |
| 8 | 101628449 | rs7017809 | -1.143 | -0.293 | 0.338 | 0.01 | -0.018 | 0.007 | 0.007 | 0.373 | 0.36 | 0.51 | 0.47 | 0.48 | 0.29 | 0.55 |
| 8 | 101637840 | rs7004022 | -0.861 | -0.247 | 0.799 | 0.037 | -0.019 | 0.024 | 0.024 | 0.242 | 0.14 | 0.31 | 0.2 | 0.25 | 0.09 | 0.27 |
| 8 | 101637901 | rs7004470 | -1.715 | -0.181 | 0.958 | -0.129 | -0.016 | -0.01 | -0.01 | 0.11 | 0.24 | 0.46 | 0.26 | 0.27 | 0.19 | 0.32 |
| 8 | 101644172 | rs7821477 | -2.172 | -0.154 | 0.878 | -0.2 | -0.001 | -0.004 | -0.004 | 0 | 0.33 | 0.67 | 0.33 | 0.27 | 0.26 | 0.4 |
| 8 | 101648271 | rs2155882 | -0.454 | -0.107 | 0.706 | -0.169 | 0.018 | -0.049 | -0.049 | 0 | 0.1 | 0.29 | 0.08 | 0.01 | 0.08 | 0.07 |
| 8 | 101648698 | rs7823425 | -1.574 | -0.12 | 0.753 | -0.157 | -0.016 | -0.013 | -0.013 | 0.032 | 0.23 | 0.53 | 0.28 | 0.26 | 0.17 | 0.35 |
| 8 | 101658926 | rs7815631 | 0.226 | 0.088 | 0.341 | -0.143 | -0.034 | 0.021 | 0.021 | 0 | 0.08 | 0.01 | 0.04 | 0.02 | 0.09 | 0.05 |
| 8 | 101667422 | rs3115892 | -0.786 | 0.137 | 0.706 | -0.166 | 0.044 | -0.043 | -0.043 | 0 | 0.1 | 0.29 | 0.09 | 0.01 | 0.09 | 0.07 |
| 8 | 101668920 | rs1693586 | -2.227 | 0.361 | -0.378 | -0.076 | -0.105 | -0.142 | -0.142 | 0.363 | 0.55 | 0.98 | 0.68 | 0.57 | 0.51 | 0.57 |
| 8 | 101669027 | rs1693587 | -0.43 | 0.352 | -0.071 | -0.099 | -0.02 | -0.015 | -0.015 | 0.153 | 0.2 | 0.11 | 0.32 | 0.26 | 0.23 | 0.24 |
| 8 | 101669273 | rs3016889 | -1.607 | 0.302 | 0.522 | -0.127 | -0.007 | -0.011 | -0.011 | 0 | 0.31 | 0.65 | 0.33 | 0.3 | 0.26 | 0.32 |
| 8 | 101669793 | rs3016888 | -0.994 | 0.354 | 0.469 | -0.116 | 0.002 | -0.003 | -0.003 | 0 | 0.17 | 0.25 | 0.16 | 0.17 | 0.18 | 0.23 |
| 8 | 101670653 | rs1786349 | -2.413 | 0.116 | 0.051 | -0.059 | -0.13 | -0.13 | -0.13 | 0 | 0.53 | 0.98 | 0.64 | 0.56 | 0.46 | 0.57 |
| 8 | 101670866 | rs1693590 | -2.174 | 0.177 | 0.322 | 0.072 | -0.005 | -0.028 | -0.028 | 0 | 0.49 | 0.83 | 0.61 | 0.56 | 0.44 | 0.57 |
| 8 | 101671221 | rs7002825 | -1.379 | 0.115 | 0.672 | 0.209 | -0.009 | 0.025 | 0.025 | 0.109 | 0.27 | 0.52 | 0.35 | 0.4 | 0.2 | 0.26 |
| 8 | 101675584 | rs1693551 | -1.978 | -0.061 | -0.599 | -0.184 | -0.118 | -0.12 | -0.12 | 0.046 | 0.56 | 0.98 | 0.64 | 0.56 | 0.47 | 0.55 |
| 8 | 101675872 | rs2935545 | -1.146 | -0.2 | 0.099 | -0.271 | 0.008 | 0.012 | 0.012 | 0 | 0.41 | 0.63 | 0.27 | 0.3 | 0.32 | 0.45 |
| 8 | 101676200 | rs10097948 | 0.226 | -0.36 | -0.03 | -0.37 | -0.029 | 0.02 | 0.02 | 0.02 | 0.2 | 0.07 | 0.14 | 0.16 | 0.23 | 0.19 |
| 8 | 101676363 | rs1786342 | -1.934 | -0.455 | 0.276 | -0.407 | 0.013 | 0.033 | 0.033 | 0 | 0.44 | 0.78 | 0.58 | 0.42 | 0.41 | 0.35 |
| 8 | 101676572 | rs7831729 | -1.496 | -0.437 | 0.55 | -0.187 | -0.031 | 0.033 | 0.033 | 0 | 0.26 | 0.5 | 0.42 | 0.4 | 0.2 | 0.3 |
| 8 | 101676969 | rs1693566 | -1.984 | -0.373 | -0.031 | -0.155 | 0.068 | 0.003 | 0.003 | 0.234 | 0.36 | 0.89 | 0.5 | 0.41 | 0.23 | 0.36 |
| 8 | 101677774 | rs1693569 | -0.39 | -0.346 | 0.091 | -0.166 | 0.003 | 0.028 | 0.028 | 0 | 0.12 | 0.2 | 0.35 | 0.25 | 0.14 | 0.1 |
| 8 | 101680355 | rs2935548 | -1.468 | -0.345 | 0.234 | -0.208 | -0.017 | -0.015 | -0.015 | 3.294 | 0.24 | 0.7 | 0.15 | 0.15 | 0.1 | 0.26 |
| 8 | 101685694 | rs879391 | -1.234 | 0.108 | -0.894 | -0.133 | 0.016 | 0.059 | 0.059 | 0 | 0.45 | 0.84 | 0.55 | 0.41 | 0.39 | 0.33 |
| 8 | 101691690 | rs1786331 | -0.435 | 0.828 | -1.299 | 0.21 | -0.114 | -0.059 | -0.059 | 0.545 | 0.23 | 0.02 | 0.19 | 0.25 | 0.33 | 0.36 |
| 8 | 101709538 | rs1693543 | -1.876 | -0.264 | -1.901 | 0.183 | -0.02 | -0.049 | -0.049 | 0.492 | 0.5 | 0.89 | 0.48 | 0.52 | 0.46 | 0.5 |
| 8 | 101717689 | rs1693552 | -0.169 | -0.252 | -1.161 | 0.165 | -0.016 | 0.003 | 0.003 | 0 | 0.27 | 0.2 | 0.21 | 0.25 | 0.34 | 0.36 |
| 8 | 101722366 | rs1039153 | -0.069 | -0.239 | -1.156 | 0.172 | -0.017 | -0.001 | -0.001 | 0 | 0.25 | 0.19 | 0.2 | 0.25 | 0.34 | 0.37 |
| 8 | 101737445 | rs2029997 | -0.652 | -0.426 | 0.071 | 0.14 | -0.019 | 0.016 | 0.016 | 0 | 0.17 | 0.24 | 0.23 | 0.27 | 0.1 | 0.14 |
| 8 | 101760349 | rs3115893 | 1.384 | -0.55 | 0.382 | 0.061 | -0.005 | 0 | 0 | 0 | 0.74 | 0.29 | 0.69 | 0.73 | 0.88 | 0.86 |
| 8 | 101763118 | rs2978107 | -0.006 | 0.094 | -0.092 | 0.043 | 0.002 | 0.004 | 0.004 | 0.01 | 0.85 | 0.83 | 0.75 | 0.77 | 0.84 | 0.86 |
| 8 | 101766580 | rs2978087 | 0.876 | 0.109 | 0.145 | 0.242 | -0.004 | 0.048 | 0.048 | 0 | 0.77 | 0.61 | 0.68 | 0.66 | 0.8 | 0.8 |
| 8 | 101770400 | rs2978094 | 0.834 | 0.201 | -0.473 | 0.27 | 0 | 0.004 | 0.004 | 0 | 0.76 | 0.78 | 0.71 | 0.65 | 0.78 | 0.8 |
| 8 | 101774134 | rs2935555 | 0.343 | 0.654 | -1.352 | 0.172 | -0.045 | 0.025 | 0.025 | 0 | 0.23 | 0.1 | 0.18 | 0.18 | 0.33 | 0.33 |
| 8 | 101793314 | rs12056810 | 1.718 | 0.573 | -1.58 | 0.415 | -0.076 | -0.097 | -0.097 | 1.5 | 0.45 | 0.02 | 0.46 | 0.44 | 0.46 | 0.47 |
| 8 | 101796638 | rs1982970 | -0.242 | 0.75 | -0.822 | 0.353 | -0.017 | 0.023 | 0.023 | 0 | 0.26 | 0.17 | 0.19 | 0.18 | 0.3 | 0.34 |
| 8 | 101803629 | rs3105429 | -0.133 | 0.842 | -0.96 | 0.246 | -0.06 | 0.01 | 0.01 | 0.444 | 0.78 | 0.92 | 0.82 | 0.81 | 0.7 | 0.68 |
| 8 | 101806147 | rs3108908 | 0.853 | 0.257 | -0.144 | 0.521 | 0.004 | 0.019 | 0.019 | 0.475 | 0.65 | 0.43 | 0.6 | 0.53 | 0.64 | 0.62 |
| 8 | 101806532 | rs3105433 | -0.985 | 0.034 | -0.252 | 0.532 | -0.029 | -0.027 | -0.027 | 1.089 | 0.84 | 0.98 | 0.86 | 0.82 | 0.82 | 0.71 |
| 8 | 101810846 | rs10808368 | 0.086 | 0.016 | -0.01 | 0.567 | -0.003 | 0.036 | 0.036 | 0 | 0.09 | 0.11 | 0.19 | 0.29 | 0.05 | 0.07 |
| 8 | 101812693 | rs3108910 | 0.917 | -0.029 | 0.097 | 0.568 | -0.023 | 0.058 | 0.058 | 0.318 | 0.7 | 0.51 | 0.64 | 0.53 | 0.76 | 0.64 |
| 8 | 101813156 | rs3847153 | -0.287 | -0.525 | 0.278 | 0.686 | -0.009 | 0.093 | 0.093 | 0 | 0.1 | 0.24 | 0.2 | 0.29 | 0.05 | 0.07 |
| 8 | 101816789 | rs7844324 | -0.065 | -0.918 | -0.212 | 0.277 | -0.017 | -0.003 | -0.003 | 0 | 0.59 | 0.66 | 0.5 | 0.5 | 0.51 | 0.49 |
| 8 | 101817561 | rs16898766 | 1.144 | -1.781 | -0.132 | -0.084 | -0.001 | 0.004 | 0.004 | 0.027 | 0.34 | 0.25 | 0.31 | 0.37 | 0.33 | 0.38 |
| 8 | 101817926 | rs3893206 | -0.543 | -1.751 | -0.239 | -0.114 | -0.031 | -0.015 | -0.015 | 0 | 0.6 | 0.73 | 0.5 | 0.52 | 0.5 | 0.5 |
| 8 | 101821033 | rs7003310 | 0.655 | -1.654 | -0.152 | -0.445 | -0.013 | 0.025 | 0.025 | 0 | 0.45 | 0.27 | 0.51 | 0.67 | 0.48 | 0.57 |
| 8 | 101821313 | rs2173214 | -0.218 | -1.656 | -0.092 | -0.478 | -0.054 | -0.048 | -0.048 | 0 | 0.26 | 0.43 | 0.17 | 0.14 | 0.15 | 0.1 |
| 8 | 101821743 | rs7004284 | -0.964 | -1.343 | -0.464 | -0.355 | -0.029 | 0.021 | 0.021 | 0 | 0.69 | 0.77 | 0.68 | 0.81 | 0.58 | 0.67 |
| 8 | 101825384 | rs2380244 | -0.037 | -1.712 | -0.639 | -0.151 | 0.019 | 0.055 | 0.055 | 0 | 0.51 | 0.68 | 0.56 | 0.67 | 0.45 | 0.57 |
| 8 | 101833267 | rs3105438 | -0.606 | -1.178 | -0.553 | -0.336 | -0.005 | -0.004 | -0.004 | 0 | 0.19 | 0.76 | 0.17 | 0.09 | 0.12 | 0.03 |
| 8 | 101835049 | rs3105439 | -0.759 | -0.462 | -0.271 | -0.356 | 0.007 | -0.026 | -0.026 | 0.093 | 0.14 | 0.75 | 0.13 | 0.08 | 0.15 | 0.03 |
| 8 | 101841731 | rs12680064 | -0.039 | -1.092 | -0.389 | 0.022 | -0.04 | 0.088 | 0.088 | 0 | 0.25 | 0.41 | 0.32 | 0.36 | 0.18 | 0.27 |
| 8 | 101844126 | rs4734479 | -0.007 | -0.975 | -0.755 | 0.08 | -0.016 | 0.041 | 0.041 | 0 | 0.24 | 0.3 | 0.31 | 0.36 | 0.18 | 0.27 |
| 8 | 101849796 | rs3105453 | 0.063 | 0.525 | -0.907 | -0.531 | -0.104 | 0.099 | 0.099 | 0 | 0.65 | 0.98 | 0.64 | 0.79 | 0.45 | 0.65 |
| 8 | 101859831 | rs7816400 | -0.546 | 1.113 | 0.046 | 0.095 | 0.002 | -0.003 | -0.003 | 0 | 0.3 | 0.66 | 0.26 | 0.28 | 0.32 | 0.28 |
| 8 | 101862078 | rs4734484 | 0.214 | 1.059 | 0.323 | -0.014 | 0.001 | -0.002 | -0.002 | 0 | 0.12 | 0.13 | 0.16 | 0.05 | 0.24 | 0.18 |
| 8 | 101862313 | rs4734487 | -1.486 | 2.068 | 0.66 | -0.207 | 0.019 | -0.012 | -0.012 | 0 | 0.35 | 0.57 | 0.33 | 0.29 | 0.46 | 0.31 |
| 8 | 101863176 | rs7840123 | 0.225 | 1.956 | 0.922 | -0.142 | -0.022 | 0.035 | 0.035 | 0.276 | 0.16 | 0.26 | 0.11 | 0.22 | 0.11 | 0.1 |
| 8 | 101873079 | rs4734489 | 0.082 | 1.602 | 0.294 | -0.347 | -0.027 | 0.013 | 0.013 | 0 | 0.28 | 0.22 | 0.24 | 0.27 | 0.37 | 0.3 |
| 8 | 101873671 | rs11994674 | -1.164 | 0.757 | -0.393 | -0.477 | 0.08 | -0.06 | -0.06 | 0 | 0.64 | 0.89 | 0.6 | 0.49 | 0.71 | 0.62 |
| 8 | 101875635 | rs7009643 | -0.378 | 0.856 | -0.195 | -0.515 | -0.017 | 0.044 | 0.044 | 0.572 | 0.39 | 0.79 | 0.36 | 0.3 | 0.49 | 0.33 |
| 8 | 101878357 | rs9642967 | -0.852 | 1.152 | 0.177 | -0.359 | -0.002 | 0.003 | 0.003 | 0 | 0.35 | 0.64 | 0.36 | 0.18 | 0.33 | 0.32 |
| 8 | 101879598 | rs7823558 | -0.29 | 0.49 | -0.347 | 0.055 | -0.003 | -0.006 | -0.006 | 0 | 0.26 | 0.31 | 0.26 | 0.19 | 0.22 | 0.29 |
| 8 | 101887030 | rs2135121 | -0.525 | 0.352 | -0.742 | 0.364 | 0.012 | -0.018 | -0.018 | 0.097 | 0.19 | 0.1 | 0.27 | 0.2 | 0.2 | 0.19 |
| 8 | 101892985 | rs2135126 | -0.495 | 0.381 | -0.799 | 0.182 | -0.02 | -0.001 | -0.001 | 0 | 0.59 | 0.68 | 0.56 | 0.56 | 0.55 | 0.64 |

| | | | | | | | | | | | | | | | | |
|---|---|---|---|---|---|---|---|---|---|---|---|---|---|---|---|---|
| 8 | 101898707 | rs6993937 | 1.096 | 0.014 | -0.936 | 0.192 | -0.044 | 0.02 | 0.02 | 0.061 | 0.25 | 0.07 | 0.2 | 0.2 | 0.25 | 0.24 |
| 8 | 101901901 | rs4734492 | 0.393 | -0.49 | -0.24 | -0.133 | 0 | -0.016 | -0.016 | 0 | 0.33 | 0.32 | 0.27 | 0.23 | 0.28 | 0.28 |
| 8 | 101905680 | rs3134370 | 0.3 | -0.603 | -0.08 | -0.093 | -0.041 | 0.047 | 0.047 | 0.675 | 0.76 | 0.55 | 0.72 | 0.68 | 0.8 | 0.67 |
| 8 | 101908844 | rs4734014 | 0.527 | -0.573 | -0.082 | -0.182 | 0 | -0.016 | -0.016 | 0 | 0.33 | 0.35 | 0.27 | 0.22 | 0.28 | 0.28 |
| 8 | 101913022 | rs4236763 | 1.569 | -0.576 | -0.056 | -0.34 | -0.004 | 0.003 | 0.003 | 0.013 | 0.24 | 0.2 | 0.21 | 0.14 | 0.25 | 0.25 |
| 8 | 101915310 | rs3134373 | 0.751 | -0.565 | -0.166 | -0.198 | -0.065 | -0.029 | -0.029 | 0 | 0.42 | 0.16 | 0.43 | 0.49 | 0.5 | 0.4 |
| 8 | 101920724 | rs2102291 | 0.588 | 0.078 | -0.19 | -0.269 | -0.01 | 0.03 | 0.03 | 0 | 0.69 | 0.52 | 0.67 | 0.65 | 0.69 | 0.59 |
| 8 | 101927695 | rs13254653 | 0.416 | 0.424 | -0.053 | -0.148 | 0.009 | -0.044 | -0.044 | 0 | 0.37 | 0.47 | 0.29 | 0.15 | 0.35 | 0.3 |
| 8 | 101937057 | rs10505003 | 0.77 | 1.102 | 0.241 | 0.299 | -0.027 | 0.015 | 0.015 | 0.458 | 0.05 | 0.01 | 0.02 | 0.02 | 0.07 | 0.04 |
| 8 | 101937374 | rs3105460 | 0.52 | 1.174 | 0.108 | 0.126 | -0.051 | -0.059 | -0.059 | 0 | 0.72 | 0.85 | 0.63 | 0.62 | 0.59 | 0.67 |
| 8 | 101937455 | rs964917 | 1.369 | 1.139 | 0.387 | 0.106 | 0.009 | 0.012 | 0.012 | 0.595 | 0.65 | 0.47 | 0.58 | 0.63 | 0.57 | 0.66 |
| 8 | 101943404 | rs17365661 | -0.054 | 0.68 | 0.46 | 0.236 | 0.008 | -0.018 | -0.018 | 0.18 | 0.06 | 0.09 | 0.03 | 0.02 | 0.07 | 0.04 |
| 8 | 101957311 | rs17366009 | 0.369 | 0.721 | 0.333 | 0.257 | -0.004 | 0.002 | 0.002 | 0.8 | 0.05 | 0.03 | 0.02 | 0.02 | 0.07 | 0.04 |
| 8 | 101975479 | rs2386922 | -0.861 | 0.732 | 0.36 | 0.33 | -0.01 | -0.015 | -0.015 | 0 | 0.28 | 0.3 | 0.39 | 0.4 | 0.41 | 0.32 |
| 8 | 101986864 | rs2923774 | -0.12 | 1.004 | 0.275 | 0.136 | -0.032 | -0.004 | -0.004 | 0 | 0.19 | 0.14 | 0.19 | 0.21 | 0.36 | 0.22 |
| 8 | 101988226 | rs4002957 | -1.605 | 0.965 | 0.326 | 0.108 | -0.003 | 0.002 | 0.002 | 0.182 | 0.33 | 0.67 | 0.28 | 0.21 | 0.45 | 0.28 |
| 8 | 101992193 | rs11986776 | -1.64 | 0.764 | 0.795 | 0 | 0.011 | -0.022 | -0.022 | 0.047 | 0.18 | 0.51 | 0.26 | 0.13 | 0.17 | 0.08 |
| 8 | 101992759 | rs17464037 | 0.175 | 0.795 | 0.605 | -0.004 | -0.029 | 0.025 | 0.025 | 0.698 | 0.05 | 0.01 | 0.02 | 0 | 0.07 | 0.04 |
| 8 | 101994075 | rs10955246 | -0.817 | 0.373 | 0.918 | 0.209 | 0.002 | 0.036 | 0.036 | 2.076 | 0.12 | 0.34 | 0.23 | 0.12 | 0.09 | 0.04 |
| 8 | 101994414 | rs12335167 | -1.032 | 0.376 | 0.759 | 0.21 | -0.004 | 0.033 | 0.033 | 2.352 | 0.13 | 0.47 | 0.24 | 0.12 | 0.09 | 0.04 |
| 8 | 102001227 | rs16898881 | -0.723 | 0.145 | 0.389 | 0.268 | -0.035 | 0.042 | 0.042 | 2.092 | 0.12 | 0.32 | 0.22 | 0.12 | 0.06 | 0.04 |
| 8 | 102002608 | rs3019270 | -0.563 | 0.36 | -0.27 | -0.137 | 0.002 | 0.127 | 0.127 | 0 | 0.48 | 0.26 | 0.61 | 0.8 | 0.53 | 0.52 |
| 8 | 102003418 | rs2923772 | -0.431 | 0.361 | 0.224 | 0.461 | -0.001 | 0.005 | 0.005 | 0 | 0.27 | 0.27 | 0.32 | 0.55 | 0.32 | 0.37 |
| 8 | 102006405 | rs6468747 | -1.382 | 0.274 | 1.319 | 0.104 | 0.08 | 0.104 | 0.104 | 1.861 | 0.07 | 0.49 | 0.13 | 0.11 | 0.07 | 0.12 |
| 8 | 102007701 | rs1026228 | 0.521 | 0.502 | 0.872 | 0.012 | -0.032 | -0.003 | -0.003 | 0 | 0.61 | 0.21 | 0.64 | 0.59 | 0.53 | 0.49 |
| 8 | 102008828 | rs6468749 | -0.258 | 0.513 | 0.767 | -0.115 | -0.057 | -0.021 | -0.021 | 0 | 0.69 | 0.84 | 0.78 | 0.67 | 0.6 | 0.61 |
| 8 | 102012805 | rs11785425 | -0.076 | -0.126 | 1.099 | -0.661 | 0.05 | 0.134 | 0.134 | 0 | 0.38 | 0.15 | 0.32 | 0.05 | 0.24 | 0.27 |
| 8 | 102028996 | rs16898906 | -1.607 | 0.274 | 1.362 | -0.262 | 0.009 | 0.004 | 0.004 | 0 | 0.07 | 0.2 | 0.09 | 0.1 | 0.06 | 0.13 |
| 8 | 102050512 | rs6994935 | -1.8 | -0.137 | 1.405 | -0.395 | -0.039 | 0.004 | 0.004 | 8.996 | 0.1 | 0.68 | 0.09 | 0.11 | 0.02 | 0.05 |
| 8 | 102053606 | rs1451640 | -0.065 | 0.017 | 0.603 | -1.002 | 0.001 | -0.023 | -0.023 | 0.266 | 0.35 | 0.4 | 0.35 | 0.06 | 0.28 | 0.37 |
| 8 | 102058350 | rs4734509 | -0.195 | 0.289 | -0.163 | -0.847 | -0.006 | -0.065 | -0.065 | 0 | 0.42 | 0.59 | 0.39 | 0.09 | 0.3 | 0.44 |
| 8 | 102060415 | rs4734512 | -0.354 | 0.144 | -1.308 | -0.761 | 0.074 | 0.17 | 0.17 | 0.218 | 0.44 | 0.77 | 0.41 | 0.09 | 0.29 | 0.43 |
| 8 | 102068160 | rs4734021 | -0.549 | 0.721 | -1.366 | 0.187 | -0.012 | -0.04 | -0.04 | 0 | 0.33 | 0.18 | 0.32 | 0.41 | 0.41 | 0.39 |
| 8 | 102068621 | rs6996970 | -0.329 | 1.025 | -1.508 | 0.437 | 0.001 | -0.006 | -0.006 | 0 | 0.24 | 0.17 | 0.23 | 0.3 | 0.24 | 0.28 |
| 8 | 102068804 | rs3019260 | -0.093 | 0.208 | -1.106 | -0.021 | -0.005 | 0.011 | 0.011 | 0 | 0.35 | 0.36 | 0.5 | 0.47 | 0.33 | 0.36 |
| 8 | 102073521 | rs6468757 | -0.881 | 0.334 | -0.07 | -0.411 | 0.01 | -0.087 | -0.087 | 0.251 | 0.15 | 0.54 | 0.14 | 0.11 | 0.14 | 0.15 |
| 8 | 102084543 | rs13279158 | -0.611 | 0.305 | 0.09 | -0.352 | 0.009 | -0.043 | -0.043 | 0 | 0.12 | 0.17 | 0.11 | 0.14 | 0.14 | 0.14 |
| 8 | 102086007 | rs13259960 | -0.611 | 0.305 | 0.09 | -0.352 | 0.009 | -0.043 | -0.043 | 0 | 0.12 | 0.17 | 0.11 | 0.14 | 0.14 | 0.14 |
| 8 | 102087549 | rs12679969 | -0.235 | 0.386 | -0.161 | 0.323 | -0.007 | 0.031 | 0.031 | 0.039 | 0.07 | 0.03 | 0.15 | 0.34 | 0.11 | 0.29 |
| 8 | 102091906 | rs776272 | -0.466 | 0.311 | -0.292 | -0.483 | -0.027 | -0.021 | -0.021 | 0 | 0.73 | 0.51 | 0.67 | 0.72 | 0.78 | 0.81 |
| 8 | 102092673 | rs2514616 | 0.219 | 0.296 | -0.789 | -1.064 | -0.017 | 0.011 | 0.011 | 0 | 0.35 | 0.27 | 0.49 | 0.68 | 0.47 | 0.52 |
| 8 | 102100773 | rs703811 | -0.889 | -0.32 | -0.418 | -0.951 | -0.011 | -0.006 | -0.006 | 0 | 0.73 | 0.64 | 0.68 | 0.72 | 0.77 | 0.81 |
| 8 | 102101619 | rs776263 | -1.425 | -0.365 | -0.66 | -0.989 | -0.016 | -0.025 | -0.025 | 1.973 | 0.78 | 0.96 | 0.72 | 0.81 | 0.79 | 0.82 |
| 8 | 102103227 | rs776260 | -1.41 | -0.384 | -0.613 | -0.983 | -0.015 | -0.01 | -0.01 | 0.05 | 0.78 | 0.94 | 0.71 | 0.8 | 0.77 | 0.82 |
| 8 | 102109037 | rs166995 | 0.277 | -0.135 | -0.376 | -1.027 | -0.025 | 0.017 | 0.017 | 0 | 0.36 | 0.24 | 0.49 | 0.69 | 0.48 | 0.51 |
| 8 | 102109143 | rs2599528 | -0.443 | -0.066 | 0.101 | -0.461 | -0.018 | -0.016 | -0.016 | 0.029 | 0.73 | 0.55 | 0.66 | 0.73 | 0.77 | 0.8 |
| 8 | 102110257 | rs192604 | -0.647 | 0.038 | -0.299 | -0.423 | -0.004 | -0.001 | -0.001 | 0 | 0.75 | 0.71 | 0.67 | 0.72 | 0.77 | 0.8 |
| 8 | 102118012 | rs4734517 | -0.647 | -0.495 | -0.683 | -0.109 | 0.082 | 0.057 | 0.057 | 0.62 | 0.44 | 0.84 | 0.3 | 0.34 | 0.3 | 0.54 |
| 8 | 102120338 | rs11996948 | -0.291 | -0.628 | 0.605 | -0.319 | -0.055 | -0.129 | -0.129 | 2.226 | 0.33 | 0.48 | 0.11 | 0.05 | 0.19 | 0.26 |
| 8 | 102123897 | rs7005641 | -1.094 | -0.535 | 1.259 | -0.23 | 0.027 | 0.019 | 0.019 | 3.229 | 0.06 | 0.35 | 0.02 | 0.01 | 0.01 | 0.01 |
| 8 | 102127338 | rs1374498 | 0.256 | -0.483 | 0.723 | -0.073 | -0.019 | 0.017 | 0.017 | 0 | 0.05 | 0.02 | 0.15 | 0.29 | 0.1 | 0.25 |
| 8 | 102128719 | rs872433 | 1.452 | -0.321 | 0.076 | -0.24 | -0.042 | -0.005 | -0.005 | 0 | 0.33 | 0.02 | 0.33 | 0.18 | 0.34 | 0.23 |
| 8 | 102130291 | rs752047 | 0.287 | -0.253 | 0.515 | -0.338 | -0.019 | 0.018 | 0.018 | 0 | 0.26 | 0.15 | 0.22 | 0.14 | 0.35 | 0.2 |
| 8 | 102131740 | rs13253653 | 0.268 | -0.349 | 0.583 | -0.581 | 0.02 | 0.017 | 0.017 | 0 | 0.27 | 0.18 | 0.08 | 0.04 | 0.17 | 0.26 |
| 8 | 102132835 | rs895602 | -0.383 | -0.048 | 1.014 | -0.242 | 0.035 | -0.004 | -0.004 | 0 | 0.34 | 0.48 | 0.4 | 0.37 | 0.46 | 0.42 |
| 8 | 102133008 | rs746769 | 0.492 | -0.012 | 0.734 | -0.176 | -0.013 | 0.065 | 0.065 | 0.029 | 0.55 | 0.22 | 0.43 | 0.24 | 0.5 | 0.49 |
| 8 | 102133238 | rs2441704 | -0.319 | 0.344 | 0.626 | 0.204 | 0.03 | 0.264 | 0.264 | 0 | 0.95 | 0.87 | 0.82 | 0.59 | 0.95 | 0.92 |
| 8 | 102133330 | rs4734518 | 0.479 | 0.235 | 0.386 | -0.227 | 0 | -0.003 | -0.003 | 0 | 0.43 | 0.3 | 0.48 | 0.41 | 0.47 | 0.47 |
| 8 | 102133555 | rs2100097 | -0.591 | 0.174 | 0.843 | 0.245 | 0.058 | 0.152 | 0.152 | 0.287 | 0.09 | 0.29 | 0.21 | 0.28 | 0.12 | 0.24 |
| 8 | 102137549 | rs7821200 | -0.521 | -0.381 | 0.579 | -0.286 | -0.025 | -0.072 | -0.072 | 0 | 0.17 | 0.34 | 0.09 | 0.02 | 0.12 | 0.12 |
| 8 | 102140698 | rs2053741 | -0.284 | -0.585 | 0.474 | -0.524 | -0.079 | -0.155 | -0.155 | 0 | 0.26 | 0.49 | 0.14 | 0.01 | 0.14 | 0.13 |
| 8 | 102142243 | rs2441709 | 0.584 | -0.134 | -0.506 | -0.353 | -0.001 | 0 | 0 | 0 | 0.45 | 0.38 | 0.57 | 0.46 | 0.53 | 0.5 |
| 8 | 102145463 | rs2441703 | 0.183 | 0.538 | -0.552 | 0.263 | 0.011 | 0.068 | 0.068 | 0 | 0.69 | 0.49 | 0.66 | 0.5 | 0.7 | 0.64 |
| 8 | 102150552 | rs1814492 | 0.188 | 1.211 | -1.829 | 0.632 | -0.171 | -0.186 | -0.186 | 0 | 0.71 | 0.98 | 0.67 | 0.48 | 0.52 | 0.58 |
| 8 | 102150937 | rs2004474 | 0.254 | 1.154 | -1.667 | 0.578 | -0.124 | -0.133 | -0.133 | 0.56 | 0.71 | 0.88 | 0.65 | 0.48 | 0.51 | 0.58 |
| 8 | 102155413 | rs1470517 | 0.541 | 0.402 | -1.783 | 0.022 | 0.003 | -0.017 | -0.017 | 0.205 | 0.25 | 0.2 | 0.24 | 0.37 | 0.16 | 0.22 |
| 8 | 102159018 | rs17381828 | -0.531 | 0.333 | -2.046 | 0.38 | -0.04 | -0.035 | -0.035 | 0.537 | 0.76 | 0.82 | 0.68 | 0.58 | 0.53 | 0.65 |
| 8 | 102162717 | rs10112963 | 0.357 | 0.07 | -2.04 | 0.105 | 0.103 | 0.158 | 0.158 | 0.862 | 0.51 | 0.8 | 0.4 | 0.25 | 0.35 | 0.43 |
| 8 | 102164667 | rs10092729 | 0.663 | -0.827 | -1.288 | -0.645 | -0.024 | -0.09 | -0.09 | 0.657 | 0.3 | 0.4 | 0.15 | 0.03 | 0.21 | 0.12 |

| | | | | | | | | | | | | | | | | |
|---|---|---|---|---|---|---|---|---|---|---|---|---|---|---|---|---|
| 8 | 102169000 | rs16899055 | 0.492 | -0.729 | -0.891 | -0.659 | -0.006 | -0.027 | -0.027 | 1.115 | 0.09 | 0.15 | 0.05 | 0.02 | 0.08 | 0.05 |
| 8 | 102170103 | rs2441701 | -0.531 | -0.366 | -1.401 | -0.662 | -0.109 | -0.114 | -0.114 | 0 | 0.68 | 0.95 | 0.65 | 0.44 | 0.55 | 0.57 |
| 8 | 102180058 | rs11996854 | 0.494 | -0.66 | -0.713 | -0.67 | -0.018 | -0.032 | -0.032 | 0 | 0.42 | 0.5 | 0.3 | 0.24 | 0.27 | 0.36 |
| 8 | 102181249 | rs7828323 | 0.425 | -0.663 | -0.658 | -0.671 | 0.014 | 0.016 | 0.016 | 0.279 | 0.42 | 0.59 | 0.31 | 0.24 | 0.27 | 0.36 |
| 8 | 102183601 | rs6992385 | 0.409 | -0.67 | -0.661 | -0.654 | 0.014 | 0.017 | 0.017 | 0.3 | 0.42 | 0.59 | 0.31 | 0.24 | 0.27 | 0.36 |
| 8 | 102192178 | rs7011529 | 0.662 | -0.605 | -0.604 | -0.734 | -0.017 | -0.061 | -0.061 | 1.77 | 0.39 | 0.53 | 0.28 | 0.1 | 0.26 | 0.3 |
| 8 | 102200403 | rs11987160 | -0.157 | -0.559 | -0.343 | -0.643 | 0.063 | 0.14 | 0.14 | 0.746 | 0.52 | 0.8 | 0.36 | 0.24 | 0.36 | 0.43 |
| 8 | 102203690 | rs2441706 | 0.935 | -0.543 | -0.482 | -0.721 | -0.069 | -0.098 | -0.098 | 0.185 | 0.25 | 0.02 | 0.32 | 0.33 | 0.25 | 0.22 |
| 8 | 102205158 | rs16899096 | 0.452 | -0.845 | -0.105 | -0.881 | -0.013 | -0.061 | -0.061 | 0.285 | 0.43 | 0.57 | 0.32 | 0.11 | 0.34 | 0.36 |
| 8 | 102207849 | rs10093422 | -0.232 | -0.847 | -0.151 | -0.871 | 0.069 | 0.154 | 0.154 | 0.762 | 0.52 | 0.8 | 0.36 | 0.23 | 0.35 | 0.43 |
| 8 | 102208863 | rs7018179 | -0.247 | -0.841 | -0.148 | -0.867 | 0.068 | 0.152 | 0.152 | 0.896 | 0.52 | 0.8 | 0.36 | 0.23 | 0.35 | 0.43 |
| 8 | 102213785 | rs2212702 | -1.192 | -0.928 | -0.01 | -0.89 | -0.083 | -0.033 | -0.033 | 0.112 | 0.15 | 0.4 | 0.05 | 0.13 | 0.03 | 0.08 |
| 8 | 102215628 | rs10086418 | 2.102 | -0.851 | 0.029 | -0.852 | -0.023 | -0.012 | -0.012 | 3.352 | 0.06 | 0.13 | 0.01 | 0.02 | 0.01 | 0.01 |
| 8 | 102220016 | rs1447013 | -0.24 | -0.86 | 0.043 | -0.88 | 0.084 | 0.176 | 0.176 | 0.641 | 0.51 | 0.82 | 0.36 | 0.23 | 0.35 | 0.43 |
| 8 | 102227923 | rs1264202 | -1.478 | -0.881 | 0.014 | -0.919 | -0.051 | -0.07 | -0.07 | 0.495 | 0.75 | 0.87 | 0.67 | 0.52 | 0.6 | 0.65 |
| 8 | 102230186 | rs6468762 | -0.415 | -1.063 | 0.418 | -1.279 | -0.019 | 0.013 | 0.013 | 0 | 0.07 | 0.22 | 0.04 | 0.11 | 0.02 | 0.07 |
| 8 | 102236994 | rs1447016 | 0.543 | -1.059 | 0.34 | -1.289 | -0.003 | -0.033 | -0.033 | 0.095 | 0.37 | 0.4 | 0.3 | 0.09 | 0.32 | 0.35 |
| 8 | 102240854 | rs6990686 | 1.34 | -0.901 | 0.232 | -1.093 | -0.025 | -0.017 | -0.017 | 0 | 0.06 | 0.14 | 0.01 | 0.02 | 0.01 | 0.01 |
| 8 | 102247516 | rs10093920 | 1.202 | -0.91 | 0.319 | -1.147 | -0.03 | -0.021 | -0.021 | 0 | 0.07 | 0.14 | 0.01 | 0.02 | 0.01 | 0.01 |
| 8 | 102247938 | rs17468203 | 0.604 | -1.204 | 0.376 | -1.394 | 0.011 | 0.114 | 0.114 | 0 | 0.34 | 0.16 | 0.27 | 0.09 | 0.29 | 0.33 |
| 8 | 102248181 | rs7846554 | -0.547 | -1.232 | 0.342 | -1.417 | 0.082 | 0.152 | 0.152 | 0.56 | 0.48 | 0.8 | 0.33 | 0.11 | 0.31 | 0.34 |
| 8 | 102252632 | rs1947124 | 0.086 | -1.229 | 0.417 | -1.415 | -0.023 | -0.088 | -0.088 | 0 | 0.39 | 0.55 | 0.31 | 0.1 | 0.3 | 0.34 |
| 8 | 102257415 | rs11998374 | -1.274 | -1.214 | 0.675 | -0.805 | -0.08 | -0.069 | -0.069 | 0 | 0.13 | 0.42 | 0.04 | 0.02 | 0.02 | 0.01 |
| 8 | 102258242 | rs746920 | -0.157 | -0.904 | 0.133 | -0.651 | -0.038 | -0.027 | -0.027 | 0 | 0.08 | 0.25 | 0.02 | 0.02 | 0.01 | 0.01 |
| 8 | 102261691 | rs1529605 | 0.469 | -0.428 | -0.892 | -0.992 | -0.027 | 0.129 | 0.129 | 0 | 0.44 | 0.27 | 0.38 | 0.2 | 0.45 | 0.56 |
| 8 | 102271186 | rs506389 | -0.289 | -0.293 | -1.113 | -0.258 | 0.006 | -0.022 | -0.022 | 0 | 0.77 | 0.82 | 0.78 | 0.52 | 0.76 | 0.78 |
| 8 | 102272195 | rs475207 | 0.174 | 0.03 | -0.688 | 0.025 | -0.003 | -0.015 | -0.015 | 0.945 | 0.27 | 0.16 | 0.35 | 0.34 | 0.3 | 0.22 |
| 8 | 102279321 | rs6985943 | 0.62 | 0.069 | -0.987 | -0.356 | -0.06 | 0.069 | 0.069 | 0 | 0.34 | 0.13 | 0.26 | 0.23 | 0.37 | 0.44 |
| 8 | 102279538 | rs614961 | 0.176 | -0.179 | -0.536 | -0.031 | -0.002 | -0.003 | -0.003 | 0.016 | 0.29 | 0.25 | 0.29 | 0.38 | 0.35 | 0.24 |
| 8 | 102281301 | rs474304 | -0.406 | -0.555 | -0.003 | 0.053 | 0.01 | -0.01 | -0.01 | 0 | 0.72 | 0.76 | 0.7 | 0.64 | 0.8 | 0.72 |
| 8 | 102281315 | rs4734522 | 1.131 | -0.446 | -0.313 | -0.02 | -0.045 | 0.119 | 0.119 | 0 | 0.41 | 0.16 | 0.31 | 0.22 | 0.48 | 0.47 |
| 8 | 102281374 | rs643556 | 0.104 | -0.545 | -0.166 | -0.004 | 0.026 | -0.004 | -0.004 | 0.445 | 0.64 | 0.34 | 0.64 | 0.64 | 0.78 | 0.69 |
| 8 | 102293420 | rs622435 | -0.108 | -0.28 | -0.597 | 0.157 | 0 | 0.02 | 0.02 | 0 | 0.31 | 0.56 | 0.37 | 0.42 | 0.3 | 0.24 |
| 8 | 102295395 | rs668179 | -0.25 | -0.253 | -0.722 | 0.131 | 0.013 | 0.012 | 0.012 | 0 | 0.32 | 0.57 | 0.41 | 0.42 | 0.39 | 0.25 |
| 8 | 102295995 | rs681304 | -0.214 | -0.217 | -0.714 | -0.355 | 0.013 | -0.018 | -0.018 | 0 | 0.32 | 0.57 | 0.36 | 0.23 | 0.38 | 0.24 |
| 8 | 102298428 | rs4546625 | 0.393 | 0 | -0.967 | -0.751 | -0.015 | 0.079 | 0.079 | 0 | 0.41 | 0.22 | 0.28 | 0.23 | 0.39 | 0.46 |
| 8 | 102301122 | rs6468764 | 0.205 | 0.168 | -1.045 | -0.663 | -0.008 | 0.093 | 0.093 | 0 | 0.43 | 0.25 | 0.28 | 0.24 | 0.39 | 0.44 |
| 8 | 102307638 | rs10102355 | -0.958 | -0.287 | -0.325 | -0.685 | -0.028 | -0.077 | -0.077 | 0 | 0.14 | 0.32 | 0.04 | 0.03 | 0.05 | 0.05 |
| 8 | 102308173 | rs9297302 | -0.13 | -0.078 | -1.328 | -0.788 | -0.063 | 0.052 | 0.052 | 0.292 | 0.59 | 0.92 | 0.43 | 0.3 | 0.48 | 0.57 |
| 8 | 102315465 | rs7001476 | -1.079 | -0.439 | -0.176 | -0.992 | -0.026 | -0.08 | -0.08 | 0 | 0.14 | 0.57 | 0.09 | 0.04 | 0.08 | 0.13 |
| 8 | 102317954 | rs4734525 | -0.418 | -0.463 | -0.921 | -1.272 | -0.109 | 0.029 | 0.029 | 0.815 | 0.64 | 0.98 | 0.42 | 0.28 | 0.54 | 0.58 |
| 8 | 102323874 | rs7814017 | -0.716 | -0.557 | 0.214 | -1.442 | -0.018 | -0.103 | -0.103 | 0 | 0.2 | 0.35 | 0.11 | 0.05 | 0.13 | 0.16 |
| 8 | 102330393 | rs10505007 | 1.387 | -0.794 | 0.114 | -1.533 | -0.009 | 0.015 | 0.015 | 0 | 0.37 | 0.33 | 0.29 | 0.24 | 0.4 | 0.41 |
| 8 | 102335433 | rs10094106 | -1.539 | -0.922 | 0.281 | -1.646 | -0.015 | -0.134 | -0.134 | 0 | 0.3 | 0.54 | 0.15 | 0.05 | 0.23 | 0.19 |
| 8 | 102335896 | rs13276224 | -1.02 | -0.873 | 0.07 | -1.623 | 0 | 0.005 | 0.005 | 0 | 0.26 | 0.2 | 0.14 | 0.05 | 0.22 | 0.19 |
| 8 | 102346409 | rs477351 | -1.392 | -0.931 | -0.348 | -0.795 | 0.008 | -0.126 | -0.126 | 3.749 | 0.79 | 0.99 | 0.8 | 0.68 | 0.84 | 0.82 |
| 8 | 102358215 | rs13250690 | -0.632 | -0.502 | 0.338 | -1.248 | -0.001 | -0.014 | -0.014 | 0 | 0.15 | 0.15 | 0.1 | 0.04 | 0.16 | 0.14 |
| 8 | 102358879 | rs16899294 | -1.755 | -0.551 | 0.637 | -1.267 | -0.01 | -0.127 | -0.127 | 0.167 | 0.3 | 0.53 | 0.16 | 0.05 | 0.23 | 0.19 |
| 8 | 102375267 | rs16867579 | -0.878 | -0.54 | 0.977 | -1.057 | -0.037 | -0.063 | -0.063 | 0 | 0.11 | 0.42 | 0.03 | 0.01 | 0.01 | 0.03 |
| 8 | 102379297 | rs10100433 | -1.639 | -0.507 | 0.895 | -1.038 | 0.033 | 0.019 | 0.019 | 3.36 | 0.06 | 0.36 | 0.02 | 0.01 | 0 | 0.03 |
| 8 | 102387329 | rs10106346 | -2.139 | -0.749 | 0.629 | -1.249 | -0.022 | -0.105 | -0.105 | 0.016 | 0.33 | 0.56 | 0.2 | 0.06 | 0.22 | 0.21 |
| 8 | 102392085 | rs13258434 | -1.509 | -0.751 | 0.498 | -1.266 | -0.003 | -0.054 | -0.054 | 0.725 | 0.28 | 0.32 | 0.17 | 0.05 | 0.22 | 0.19 |
| 8 | 102394180 | rs10098190 | -1.278 | -0.759 | 0.37 | -1.269 | 0.005 | 0.03 | 0.03 | 0 | 0.26 | 0.2 | 0.17 | 0.05 | 0.22 | 0.19 |
| 8 | 102397915 | rs7002389 | -2.215 | -0.788 | 0.41 | -1.301 | -0.028 | -0.152 | -0.152 | 0.29 | 0.34 | 0.48 | 0.2 | 0.05 | 0.24 | 0.22 |
| 8 | 102398444 | rs9297304 | -1.32 | -0.468 | -0.251 | -1.11 | 0.004 | 0.029 | 0.029 | 0 | 0.26 | 0.21 | 0.17 | 0.05 | 0.22 | 0.19 |
| 8 | 102398676 | rs10086534 | -0.773 | -0.489 | -0.262 | -1.122 | -0.003 | -0.001 | -0.001 | 0 | 0.2 | 0.17 | 0.17 | 0.05 | 0.21 | 0.19 |
| 8 | 102401046 | rs10097780 | -1.221 | -0.444 | -0.237 | -1.096 | 0.004 | 0.029 | 0.029 | 0 | 0.26 | 0.21 | 0.17 | 0.05 | 0.22 | 0.19 |
| 8 | 102402505 | rs930148 | -1.221 | -0.444 | -0.237 | -1.096 | 0.004 | 0.029 | 0.029 | 0 | 0.26 | 0.2 | 0.17 | 0.05 | 0.22 | 0.19 |
| 8 | 102408547 | rs16867610 | -0.494 | -0.354 | 0.106 | -1.004 | 0.025 | -0.068 | -0.068 | 0 | 0.14 | 0.34 | 0.16 | 0.04 | 0.16 | 0.18 |
| 8 | 102409702 | rs17393430 | 1.769 | -0.311 | -0.967 | -1.134 | -0.12 | 0.006 | 0.006 | 0 | 0.32 | 0.01 | 0.23 | 0.2 | 0.37 | 0.36 |
| 8 | 102413837 | rs16867625 | 0.454 | -0.172 | -0.262 | -0.935 | 0.002 | -0.012 | -0.012 | 0 | 0.1 | 0.1 | 0.03 | 0 | 0.04 | 0.02 |
| 8 | 102414789 | rs899296 | -0.224 | -0.125 | 0.024 | -0.698 | 0.031 | -0.015 | -0.015 | 0 | 0.14 | 0.29 | 0.16 | 0.12 | 0.18 | 0.17 |
| 8 | 102419151 | rs10098995 | -1.571 | -0.193 | 0.105 | -0.743 | -0.006 | -0.068 | -0.068 | 0 | 0.25 | 0.44 | 0.18 | 0.12 | 0.22 | 0.18 |
| 8 | 102419210 | rs10098920 | -1.608 | -0.361 | 0.363 | -0.843 | -0.013 | -0.048 | -0.048 | 0.58 | 0.25 | 0.61 | 0.2 | 0.13 | 0.21 | 0.2 |
| 8 | 102424918 | rs1264478 | 1.754 | -0.103 | 0.224 | -0.205 | -0.002 | 0.01 | 0.01 | 0 | 0.6 | 0.34 | 0.55 | 0.57 | 0.57 | 0.55 |
| 8 | 102424950 | rs1264477 | 1.755 | -0.1 | 0.222 | -0.228 | -0.002 | 0.013 | 0.013 | 0 | 0.6 | 0.34 | 0.55 | 0.58 | 0.57 | 0.55 |
| 8 | 102425991 | rs1264466 | 0.775 | -0.047 | 0.086 | -0.168 | -0.003 | -0.003 | -0.003 | 0 | 0.74 | 0.69 | 0.71 | 0.7 | 0.75 | 0.73 |
| 8 | 102428258 | rs1264471 | -0.193 | 0.278 | 0.239 | 0.141 | 0.014 | 0.032 | 0.032 | 0 | 0.86 | 0.8 | 0.74 | 0.7 | 0.8 | 0.75 |
| 8 | 102430331 | rs16867692 | -0.505 | 0.336 | -0.069 | -0.112 | 0.037 | -0.01 | -0.01 | 0 | 0.14 | 0.33 | 0.16 | 0.12 | 0.18 | 0.18 |

| | | | | | | | | | | | | | | | | |
|---|---|---|---|---|---|---|---|---|---|---|---|---|---|---|---|---|
| 8 | 102447003 | rs1264461 | -0.055 | 0.273 | -0.687 | 0.143 | -0.007 | -0.002 | -0.002 | 0.069 | 0.86 | 0.9 | 0.77 | 0.81 | 0.8 | 0.75 |
| 8 | 102447087 | rs1148489 | 0.405 | -0.213 | -0.965 | 0.399 | -0.018 | -0.17 | -0.17 | 0 | 0.21 | 0.01 | 0.29 | 0.5 | 0.17 | 0.17 |
| 8 | 102450006 | rs12335148 | -0.312 | -0.141 | -0.24 | -0.238 | 0.024 | -0.064 | -0.064 | 3.536 | 0.18 | 0.28 | 0.18 | 0.03 | 0.25 | 0.2 |
| 8 | 102462292 | rs507201 | 1.124 | -0.41 | -0.335 | -0.097 | 0.002 | 0.001 | 0.001 | 0.736 | 0.68 | 0.63 | 0.54 | 0.28 | 0.75 | 0.56 |
| 8 | 102464584 | rs554232 | 0.407 | -0.457 | -0.467 | -0.149 | -0.026 | -0.055 | -0.055 | 0 | 0.82 | 0.91 | 0.6 | 0.28 | 0.83 | 0.62 |
| 8 | 102465300 | rs660416 | 1.999 | -0.483 | -0.189 | -0.203 | 0.003 | 0.048 | 0.048 | 0.962 | 0.5 | 0.35 | 0.34 | 0.18 | 0.52 | 0.35 |
| 8 | 102465428 | rs16867710 | 1.267 | -0.522 | -0.15 | -0.227 | -0.003 | -0.02 | -0.02 | 0 | 0.06 | 0.08 | 0.03 | 0 | 0.04 | 0.01 |
| 8 | 102466767 | rs633142 | 1.931 | -0.536 | -0.152 | -0.177 | 0.003 | 0.061 | 0.061 | 1.024 | 0.49 | 0.34 | 0.34 | 0.18 | 0.52 | 0.35 |
| 8 | 102467380 | rs575661 | -0.664 | -0.494 | -0.397 | 0.335 | -0.008 | -0.022 | -0.022 | 0.009 | 0.92 | 1 | 0.87 | 0.55 | 0.94 | 0.84 |
| 8 | 102467706 | rs618854 | 1.814 | -0.44 | -0.214 | -0.121 | 0.001 | 0.017 | 0.017 | 0.821 | 0.51 | 0.42 | 0.35 | 0.24 | 0.52 | 0.36 |
| 8 | 102471246 | rs7005727 | -0.365 | -0.649 | 0.154 | -0.088 | 0.022 | -0.054 | -0.054 | 0 | 0.17 | 0.25 | 0.18 | 0.04 | 0.24 | 0.21 |
| 8 | 102472926 | rs474949 | -0.732 | -1.091 | 0.2 | 0.319 | 0 | -0.001 | -0.001 | 0.227 | 0.49 | 0.51 | 0.49 | 0.41 | 0.41 | 0.46 |
| 8 | 102477742 | rs670006 | 1.24 | -2.124 | 0.755 | 0.802 | -0.053 | 0.151 | 0.151 | 0 | 0.85 | 0.69 | 0.77 | 0.41 | 0.95 | 0.8 |
| 8 | 102480148 | rs10102933 | 0.464 | -1.769 | 0.087 | 0.858 | -0.032 | -0.029 | -0.029 | 0 | 0.06 | 0 | 0.16 | 0.27 | 0.09 | 0.18 |
| 8 | 102482839 | rs615581 | 0.973 | -1.917 | 0.286 | 0.943 | 0.009 | -0.073 | -0.073 | 0 | 0.85 | 0.94 | 0.77 | 0.41 | 0.91 | 0.8 |
| 8 | 102484486 | rs539674 | 1.694 | -2.043 | 0.412 | 0.79 | -0.01 | 0.001 | 0.001 | 0.015 | 0.63 | 0.41 | 0.65 | 0.4 | 0.79 | 0.68 |
| 8 | 102485762 | rs572589 | 1.092 | -1.879 | 0.034 | 1.388 | 0.029 | -0.046 | -0.046 | 0.019 | 0.84 | 0.91 | 0.78 | 0.39 | 0.92 | 0.79 |
| 8 | 102488478 | rs619777 | 1.093 | -1.919 | 0.631 | 1.494 | -0.048 | 0.135 | 0.135 | 0 | 0.87 | 0.76 | 0.78 | 0.44 | 0.97 | 0.85 |
| 8 | 102489035 | rs508148 | -1.195 | -2.399 | 0.925 | 1.007 | -0.069 | -0.091 | -0.091 | 0 | 0.2 | 0.4 | 0.11 | 0.02 | 0.1 | 0.11 |
| 8 | 102489704 | rs2470622 | 1.576 | -2.261 | 0.701 | 1.239 | -0.054 | 0.032 | 0.032 | 0.009 | 0.76 | 0.68 | 0.74 | 0.41 | 0.9 | 0.83 |
| 8 | 102490380 | rs2509765 | 1.511 | -1.629 | 0.837 | 1.423 | -0.039 | 0.054 | 0.054 | 0.192 | 0.76 | 0.62 | 0.59 | 0.37 | 0.83 | 0.74 |
| 8 | 102494523 | rs6999959 | -1.596 | -1.394 | 0.775 | 1.027 | -0.014 | -0.017 | -0.017 | 0 | 0.13 | 0.21 | 0.09 | 0.07 | 0.09 | 0.1 |
| 8 | 102495233 | rs518898 | 0.742 | -1.379 | 1.164 | 1.582 | 0.015 | 0.083 | 0.083 | 0 | 0.76 | 0.61 | 0.6 | 0.5 | 0.73 | 0.68 |
| 8 | 102495593 | rs10435531 | -0.032 | -1.594 | 0.764 | 1.88 | 0.016 | -0.005 | -0.005 | 3.009 | 0.07 | 0.03 | 0.06 | 0.22 | 0.04 | 0.06 |
| 8 | 102497411 | rs566942 | -1.926 | -1.718 | 1.563 | 1.81 | 0.007 | 0.009 | 0.009 | 0.514 | 0.39 | 0.65 | 0.41 | 0.64 | 0.34 | 0.43 |
| 8 | 102502448 | rs570630 | -1.403 | -1.917 | 1.796 | 1.753 | -0.004 | 0.051 | 0.051 | 0.147 | 0.3 | 0.53 | 0.35 | 0.44 | 0.3 | 0.38 |
| 8 | 102506764 | rs674788 | 1.376 | -2.234 | 1.998 | 1.737 | -0.107 | 0.1 | 0.1 | 0 | 0.8 | 0.53 | 0.79 | 0.72 | 0.93 | 0.81 |
| 8 | 102506831 | rs567113 | 1.376 | -2.234 | 1.998 | 1.737 | -0.107 | 0.1 | 0.1 | 0 | 0.8 | 0.53 | 0.79 | 0.72 | 0.93 | 0.81 |
| 8 | 102509319 | rs502216 | 0.912 | -2.243 | 2.031 | 1.734 | -0.063 | 0.086 | 0.086 | 0 | 0.85 | 0.7 | 0.81 | 0.71 | 0.95 | 0.85 |
| 8 | 102509799 | rs476321 | -1.038 | -2.146 | 1.709 | 1.794 | -0.001 | 0.041 | 0.041 | 0 | 0.24 | 0.31 | 0.3 | 0.42 | 0.25 | 0.33 |
| 8 | 102511718 | rs608337 | 0.903 | -2.157 | 1.668 | 1.801 | -0.094 | 0.089 | 0.089 | 0 | 0.81 | 0.58 | 0.79 | 0.72 | 0.94 | 0.83 |
| 8 | 102514213 | rs533968 | -0.016 | -2.008 | 1.409 | 1.665 | -0.001 | -0.02 | -0.02 | 0.194 | 0.09 | 0.09 | 0.04 | 0 | 0.07 | 0.04 |
| 8 | 102514354 | rs642074 | 2.39 | -2.012 | 0.873 | 1.597 | 0.072 | 0.205 | 0.205 | 0.047 | 0.49 | 0.09 | 0.44 | 0.24 | 0.63 | 0.46 |
| 8 | 102516385 | rs671557 | -1.416 | -1.52 | 1.246 | 1.58 | -0.019 | 0.096 | 0.096 | 0 | 0.18 | 0.47 | 0.26 | 0.33 | 0.16 | 0.2 |
| 8 | 102520853 | rs751603 | 0.241 | -1.383 | 0.767 | 1.755 | -0.016 | -0.029 | -0.029 | 0 | 0.06 | 0 | 0.18 | 0.22 | 0.08 | 0.15 |
| 8 | 102520975 | rs515743 | 1.097 | -1.278 | 0.907 | 1.553 | 0.001 | -0.001 | -0.001 | 0 | 0.66 | 0.37 | 0.59 | 0.54 | 0.79 | 0.7 |
| 8 | 102527840 | rs471757 | 0.992 | -0.666 | -0.197 | 1.249 | 0.012 | 0.003 | 0.003 | 0 | 0.51 | 0.32 | 0.48 | 0.42 | 0.56 | 0.56 |
| 8 | 102529633 | rs488780 | -1.092 | -0.599 | -0.235 | 1.166 | 0.006 | -0.005 | -0.005 | 0 | 0.47 | 0.65 | 0.42 | 0.49 | 0.44 | 0.4 |
| 8 | 102532638 | rs16867810 | -1 | -0.788 | -0.131 | 1.184 | -0.007 | 0.019 | 0.019 | 0 | 0.19 | 0.23 | 0.21 | 0.32 | 0.15 | 0.21 |
| 8 | 102536405 | rs10955255 | 0.737 | -0.558 | -1.284 | 0.125 | -0.045 | 0.12 | 0.12 | 0 | 0.39 | 0.14 | 0.48 | 0.24 | 0.59 | 0.27 |
| 8 | 102542441 | rs2127035 | 1.008 | -0.708 | -0.559 | -0.029 | -0.017 | 0.021 | 0.021 | 0 | 0.36 | 0.26 | 0.45 | 0.23 | 0.53 | 0.25 |
| 8 | 102544575 | rs922466 | 0.656 | -1.022 | 0.065 | 0.15 | 0.009 | -0.026 | -0.026 | 0 | 0.42 | 0.47 | 0.49 | 0.29 | 0.55 | 0.27 |
| 8 | 102550842 | rs17476242 | 1.124 | -1.029 | -0.063 | 0.06 | -0.074 | 0.118 | 0.118 | 0 | 0.37 | 0.13 | 0.47 | 0.23 | 0.58 | 0.25 |
| 8 | 102552499 | rs10955257 | 0.514 | -1.128 | -0.007 | 0.005 | 0.005 | 0.024 | 0.024 | 0 | 0.46 | 0.38 | 0.52 | 0.28 | 0.61 | 0.27 |
| 8 | 102552777 | rs1981361 | 0.871 | -1.059 | -0.014 | 0.036 | -0.007 | 0.023 | 0.023 | 0 | 0.4 | 0.31 | 0.5 | 0.28 | 0.6 | 0.26 |
| 8 | 102554217 | rs17398001 | -0.299 | -0.978 | -0.064 | -0.378 | 0 | -0.033 | -0.033 | 0 | 0.21 | 0.24 | 0.07 | 0.01 | 0.15 | 0.21 |
| 8 | 102555060 | rs470801 | -0.143 | -1.749 | -0.321 | 0.498 | -0.001 | 0.004 | 0.004 | 0 | 0.19 | 0.12 | 0.17 | 0.14 | 0.16 | 0.24 |
| 8 | 102557218 | rs666026 | 0.852 | -2.413 | -0.148 | 1.247 | -0.003 | 0 | 0 | 1.768 | 0.86 | 0.86 | 0.86 | 0.69 | 0.95 | 0.92 |
| 8 | 102567412 | rs608799 | 1.141 | -2.77 | 0.022 | 1.182 | 0.03 | -0.007 | -0.007 | 1.268 | 0.62 | 0.29 | 0.55 | 0.43 | 0.77 | 0.59 |
| 8 | 102571852 | rs2509767 | 1.264 | -2.603 | -0.006 | 1.074 | 0.02 | -0.004 | -0.004 | 0.187 | 0.61 | 0.34 | 0.51 | 0.43 | 0.68 | 0.59 |
| 8 | 102575106 | rs16867827 | -0.594 | -2.844 | 0.519 | 1.669 | -0.025 | 0.051 | 0.051 | 0.022 | 0.07 | 0.14 | 0.1 | 0.18 | 0.02 | 0.03 |
| 8 | 102578341 | rs523352 | 0.369 | -2.887 | 0.283 | 1.461 | 0.015 | -0.02 | -0.02 | 0 | 0.93 | 0.97 | 0.95 | 0.88 | 0.97 | 0.94 |
| 8 | 102586349 | rs16867839 | 0.381 | -2.955 | 0.179 | 1.547 | -0.042 | -0.011 | -0.011 | 0 | 0.16 | 0 | 0.22 | 0.11 | 0.16 | 0.29 |
| 8 | 102587259 | rs1474412 | -1.208 | -3.008 | 0.593 | 1.528 | 0.013 | -0.015 | -0.015 | 0.642 | 0.33 | 0.31 | 0.38 | 0.43 | 0.24 | 0.36 |
| 8 | 102590235 | rs496887 | 1.314 | -3.099 | 0.69 | 1.285 | -0.094 | 0.127 | 0.127 | 0.5 | 0.81 | 0.44 | 0.87 | 0.7 | 0.95 | 0.83 |
| 8 | 102592085 | rs10481099 | -0.964 | -2.842 | 0.665 | 0.86 | -0.054 | 0.014 | 0.014 | 0.123 | 0.13 | 0.43 | 0.09 | 0.11 | 0.04 | 0.13 |
| 8 | 102593321 | rs10955258 | -0.589 | -2.628 | 0.052 | 0.666 | -0.001 | -0.004 | -0.004 | 0 | 0.28 | 0.26 | 0.3 | 0.2 | 0.23 | 0.35 |
| 8 | 102595267 | rs479690 | 1.306 | -2.824 | 0.223 | 0.891 | -0.055 | 0.012 | 0.012 | 0.003 | 0.74 | 0.37 | 0.86 | 0.73 | 0.89 | 0.82 |
| 8 | 102595587 | rs667737 | 1.477 | -2.982 | 0.389 | 0.81 | -0.041 | -0.012 | -0.012 | 1.463 | 0.72 | 0.38 | 0.86 | 0.79 | 0.85 | 0.8 |
| 8 | 102596210 | rs6988306 | -1.017 | -2.956 | 0.6 | 0.701 | 0.028 | 0.035 | 0.035 | 0 | 0.19 | 0.11 | 0.08 | 0.04 | 0.13 | 0.15 |
| 8 | 102599384 | rs560667 | 1.323 | -2.924 | 0.968 | 0.854 | -0.015 | 0.118 | 0.118 | 0.359 | 0.93 | 0.8 | 0.94 | 0.8 | 0.97 | 0.95 |
| 8 | 102600608 | rs7823289 | 1.696 | -2.831 | -0.012 | 0.935 | 0.019 | 0.002 | 0.002 | 0 | 0.46 | 0.29 | 0.48 | 0.44 | 0.62 | 0.37 |
| 8 | 102601618 | rs9297306 | 1.075 | -2.614 | 0.06 | 1.013 | 0.023 | -0.007 | -0.007 | 0 | 0.57 | 0.35 | 0.56 | 0.53 | 0.73 | 0.57 |
| 8 | 102604306 | rs9642972 | 2.197 | -2.825 | -0.702 | 1.18 | -0.068 | 0.002 | 0.002 | 0 | 0.35 | 0.1 | 0.44 | 0.31 | 0.6 | 0.31 |
| 8 | 102607311 | rs6980991 | -0.862 | -3.016 | 0.13 | 0.885 | -0.016 | -0.03 | -0.03 | 0 | 0.19 | 0.41 | 0.1 | 0.08 | 0.15 | 0.2 |
| 8 | 102609456 | rs16867971 | -1.546 | -3.112 | -0.26 | 0.967 | 0.018 | 0.093 | 0.093 | 0.475 | 0.52 | 0.86 | 0.49 | 0.33 | 0.38 | 0.52 |
| 8 | 102612520 | rs6468782 | -2.214 | -2.963 | -0.079 | 1.143 | -0.019 | -0.01 | -0.01 | 0.009 | 0.6 | 0.82 | 0.54 | 0.42 | 0.39 | 0.66 |
| 8 | 102614570 | rs4734546 | -2.068 | -2.792 | 0.018 | 1.268 | -0.009 | 0.017 | 0.017 | 0 | 0.58 | 0.8 | 0.52 | 0.37 | 0.39 | 0.66 |
| 8 | 102622590 | rs4734560 | -1.914 | -2.842 | 0.093 | 1.185 | 0.002 | 0.008 | 0.008 | 0 | 0.56 | 0.78 | 0.53 | 0.42 | 0.39 | 0.66 |

| | | | | | | | | | | | | | | | | |
|---|---|---|---|---|---|---|---|---|---|---|---|---|---|---|---|---|
| 8 | 102627955 | rs682769 | -1.013 | -2.816 | 0.337 | 0.989 | -0.026 | -0.055 | -0.055 | 0.054 | 0.36 | 0.44 | 0.38 | 0.2 | 0.23 | 0.35 |
| 8 | 102645085 | rs2387625 | 2.28 | -2.76 | -0.262 | 1.27 | -0.084 | 0.01 | 0.01 | 0 | 0.37 | 0.11 | 0.41 | 0.3 | 0.55 | 0.27 |
| 8 | 102647858 | rs7010796 | -1.177 | -2.848 | 0.751 | 1.386 | 0.012 | 0.091 | 0.091 | 1.392 | 0.08 | 0.36 | 0.05 | 0.09 | 0.03 | 0.03 |
| 8 | 102653718 | rs4734030 | -1.296 | -3.025 | 0.451 | 1.513 | -0.001 | 0.005 | 0.005 | 0.651 | 0.35 | 0.59 | 0.41 | 0.38 | 0.27 | 0.41 |
| 8 | 102654182 | rs2226399 | 2.512 | -2.883 | 0.275 | 1.426 | -0.089 | 0.002 | 0.002 | 0.025 | 0.37 | 0.11 | 0.41 | 0.31 | 0.56 | 0.28 |
| 8 | 102660160 | rs1892756 | -1.406 | -2.943 | 0.738 | 1.779 | -0.003 | 0.004 | 0.004 | 0.847 | 0.34 | 0.62 | 0.44 | 0.38 | 0.25 | 0.52 |
| 8 | 102660776 | rs1892754 | -1.422 | -2.925 | 1.602 | 2.147 | -0.007 | -0.014 | -0.014 | 0.05 | 0.3 | 0.54 | 0.42 | 0.28 | 0.2 | 0.39 |
| 8 | 102660827 | rs1892753 | -1.888 | -2.923 | 1.55 | 2.095 | -0.007 | 0.008 | 0.008 | 2.267 | 0.34 | 0.61 | 0.46 | 0.42 | 0.21 | 0.52 |
| 8 | 102666132 | rs10808374 | -1.917 | -2.736 | 1.351 | 2.048 | 0.037 | 0.02 | 0.02 | 1.259 | 0.4 | 0.74 | 0.46 | 0.32 | 0.25 | 0.4 |
| 8 | 102675287 | rs4734038 | -2.342 | -2.84 | 1.746 | 2.121 | 0.003 | 0.007 | 0.007 | 0 | 0.51 | 0.56 | 0.49 | 0.35 | 0.38 | 0.61 |
| 8 | 102678146 | rs12549642 | -1.907 | -2.891 | 1.889 | 2.13 | -0.002 | 0 | 0 | 0.31 | 0.42 | 0.48 | 0.46 | 0.41 | 0.34 | 0.6 |
| 8 | 102678972 | rs3735713 | -1.823 | -2.889 | 1.892 | 2.13 | 0 | 0 | 0 | 0.685 | 0.41 | 0.45 | 0.44 | 0.36 | 0.33 | 0.6 |
| 8 | 102679216 | rs3824090 | -0.328 | -2.998 | 1.973 | 2.118 | 0.002 | -0.001 | -0.001 | 0 | 0.22 | 0.29 | 0.34 | 0.23 | 0.18 | 0.33 |
| 8 | 102679217 | rs3824091 | -0.844 | -3.005 | 1.976 | 2.115 | -0.004 | -0.009 | -0.009 | 0.551 | 0.25 | 0.37 | 0.34 | 0.24 | 0.18 | 0.36 |
| 8 | 102679341 | rs16868177 | -0.068 | -3.004 | 1.965 | 2.11 | -0.007 | -0.034 | -0.034 | 0 | 0.09 | 0.1 | 0.04 | 0 | 0.05 | 0.02 |
| 8 | 102679909 | rs3735715 | 3.126 | -3.048 | 1.938 | 2.135 | -0.092 | -0.083 | -0.083 | 0 | 0.41 | 0.05 | 0.45 | 0.42 | 0.6 | 0.35 |
| 8 | 102680168 | rs16868178 | -0.509 | -3.029 | 2.11 | 2.152 | 0.007 | 0.018 | 0.018 | 0 | 0.16 | 0.1 | 0.1 | 0.12 | 0.15 | 0.22 |
| 8 | 102680405 | rs3735717 | 3.08 | -3.016 | 2.125 | 2.137 | -0.048 | -0.041 | -0.041 | 0 | 0.42 | 0.09 | 0.45 | 0.42 | 0.6 | 0.36 |
| 8 | 102681337 | rs6989650 | -0.342 | -2.959 | 2.103 | 2.111 | 0.002 | -0.001 | -0.001 | 1.744 | 0.22 | 0.29 | 0.34 | 0.23 | 0.18 | 0.34 |
| 8 | 102681760 | rs6468783 | -0.075 | -2.935 | 2.012 | 2.122 | -0.007 | -0.034 | -0.034 | 0 | 0.09 | 0.1 | 0.04 | 0 | 0.05 | 0.02 |
| 8 | 102690179 | rs11994981 | -0.575 | -2.973 | 1.987 | 2.104 | 0.005 | -0.001 | -0.001 | 0.028 | 0.23 | 0.39 | 0.34 | 0.24 | 0.17 | 0.37 |
| 8 | 102690192 | rs10099482 | -0.508 | -2.957 | 1.751 | 2.108 | -0.006 | 0.018 | 0.018 | 0 | 0.16 | 0.08 | 0.1 | 0.12 | 0.15 | 0.22 |
| 8 | 102698284 | rs16868198 | -1.483 | -2.858 | 1.928 | 2.209 | -0.021 | 0.067 | 0.067 | 1.703 | 0.28 | 0.49 | 0.4 | 0.39 | 0.19 | 0.46 |
| 8 | 102700295 | rs1131863 | -1.138 | -2.764 | 1.331 | 2.128 | -0.011 | 0.018 | 0.018 | 0.132 | 0.28 | 0.11 | 0.35 | 0.25 | 0.23 | 0.35 |
| 8 | 102703451 | rs7826700 | 1.916 | -2.688 | 1.343 | 2.264 | 0.045 | -0.009 | -0.009 | 0 | 0.54 | 0.19 | 0.42 | 0.54 | 0.66 | 0.46 |
| 8 | 102704506 | rs13281022 | 2.001 | -2.712 | 1.332 | 2.305 | 0.046 | 0.012 | 0.012 | 1.585 | 0.53 | 0.19 | 0.38 | 0.41 | 0.65 | 0.37 |
| 8 | 102708371 | rs2387620 | -2.206 | -2.691 | 1.477 | 2.382 | 0.051 | 0.02 | 0.02 | 1.357 | 0.47 | 0.69 | 0.51 | 0.46 | 0.32 | 0.62 |
| 8 | 102712894 | rs6468788 | 0.045 | -2.491 | 1.529 | 2.485 | -0.003 | 0.008 | 0.008 | 3.388 | 0.92 | 0.89 | 0.87 | 0.87 | 0.95 | 0.98 |
| 8 | 102718362 | rs11782150 | 2.742 | -2.508 | 1.258 | 2.203 | -0.022 | 0.045 | 0.045 | 0.048 | 0.44 | 0.12 | 0.33 | 0.36 | 0.6 | 0.36 |
| 8 | 102729403 | rs7816488 | -0.855 | -2.445 | 1.526 | 2.124 | -0.002 | -0.005 | -0.005 | 0 | 0.19 | 0.21 | 0.07 | 0.09 | 0.15 | 0.18 |
| 8 | 102740499 | rs16868250 | -0.524 | -2.448 | 1.749 | 2.283 | 0.001 | 0.006 | 0.006 | 0.622 | 0.18 | 0.2 | 0.33 | 0.25 | 0.15 | 0.31 |
| 8 | 102741806 | rs9987401 | -2.992 | -2.48 | 1.34 | 2.083 | -0.031 | -0.006 | -0.006 | 0 | 0.56 | 0.94 | 0.63 | 0.62 | 0.38 | 0.64 |
| 8 | 102743031 | rs16868262 | -0.665 | -2.415 | 1.616 | 2.073 | 0.003 | 0.037 | 0.037 | 0.027 | 0.2 | 0.24 | 0.5 | 0.49 | 0.19 | 0.42 |
| 8 | 102744231 | rs1080599 | -2.279 | -2.69 | 1.732 | 2.046 | 0.007 | 0.006 | 0.006 | 0 | 0.37 | 0.66 | 0.18 | 0.22 | 0.22 | 0.23 |
| 8 | 102745291 | rs10107483 | -1.465 | -2.186 | 1.788 | 2.457 | -0.043 | -0.026 | -0.026 | 0 | 0.25 | 0.55 | 0.12 | 0.17 | 0.15 | 0.21 |
| 8 | 102745421 | rs6993923 | -0.934 | -1.652 | 1.453 | 2.641 | 0 | 0.009 | 0.009 | 0 | 0.17 | 0.2 | 0.36 | 0.37 | 0.16 | 0.39 |
| 8 | 102746223 | rs12545512 | -0.701 | -1.541 | 1.429 | 2.7 | -0.002 | 0 | 0 | 0 | 0.16 | 0.15 | 0.36 | 0.37 | 0.16 | 0.39 |
| 8 | 102747613 | rs16868290 | -1.991 | -1.567 | 1.946 | 2.199 | -0.029 | 0.111 | 0.111 | 0 | 0.13 | 0.43 | 0.07 | 0.18 | 0.08 | 0.13 |
| 8 | 102751489 | rs1573285 | 1.099 | -0.769 | 1.76 | 1.958 | 0.045 | 0.039 | 0.039 | 0.245 | 0.85 | 0.73 | 0.69 | 0.73 | 0.83 | 0.63 |
| 8 | 102754874 | rs1125334 | 2.125 | -0.949 | 2.162 | 2.008 | -0.002 | -0.004 | -0.004 | 3.513 | 0.73 | 0.26 | 0.55 | 0.64 | 0.76 | 0.6 |
| 8 | 102756851 | rs6468792 | 2.02 | -0.953 | 2.218 | 2.006 | 0.002 | 0.005 | 0.005 | 4.337 | 0.74 | 0.3 | 0.55 | 0.65 | 0.76 | 0.6 |
| 8 | 102761332 | rs1073373 | 1.134 | -0.958 | 2.242 | 2.037 | 0.034 | -0.013 | -0.013 | 0 | 0.83 | 0.76 | 0.76 | 0.84 | 0.79 | 0.73 |
| 8 | 102762074 | rs2387618 | 2.213 | -0.964 | 2.413 | 2.044 | 0.023 | 0.021 | 0.021 | 4.272 | 0.76 | 0.39 | 0.58 | 0.68 | 0.78 | 0.61 |
| 8 | 102767252 | rs12545537 | 2.246 | -0.956 | 2.56 | 2.033 | 0.025 | -0.008 | -0.008 | 0.278 | 0.76 | 0.47 | 0.64 | 0.76 | 0.76 | 0.72 |
| 8 | 102774580 | rs4734593 | 1.375 | -0.94 | 2.592 | 2.013 | 0.071 | -0.022 | -0.022 | 0 | 0.82 | 0.66 | 0.76 | 0.84 | 0.78 | 0.73 |
| 8 | 102781449 | rs4734594 | 2.384 | -0.952 | 2.528 | 2.004 | 0.009 | 0.004 | 0.004 | 0.615 | 0.75 | 0.37 | 0.62 | 0.7 | 0.77 | 0.64 |
| 8 | 102781496 | rs4734595 | 0.706 | -0.929 | 2.471 | 1.999 | 0.005 | 0 | 0 | 0 | 0.86 | 0.86 | 0.78 | 0.84 | 0.79 | 0.75 |
| 8 | 102798565 | rs1465167 | 2.218 | -1.016 | 1.646 | 2.028 | 0.001 | 0.007 | 0.007 | 1.233 | 0.73 | 0.32 | 0.56 | 0.64 | 0.77 | 0.63 |
| 8 | 102806358 | rs6468795 | 1.172 | -0.953 | 1.196 | 1.912 | -0.012 | -0.004 | -0.004 | 0 | 0.83 | 0.86 | 0.68 | 0.79 | 0.78 | 0.75 |
| 8 | 102811002 | rs6468799 | 0.717 | -0.931 | 1.202 | 1.923 | -0.004 | -0.004 | -0.004 | 0 | 0.86 | 0.88 | 0.71 | 0.79 | 0.79 | 0.74 |
| 8 | 102836613 | rs10105774 | 2.273 | -1.249 | 0.802 | 2.121 | 0.014 | 0.052 | 0.052 | 0.094 | 0.52 | 0.03 | 0.41 | 0.31 | 0.64 | 0.51 |
| 8 | 102840673 | rs6468801 | 1.132 | -0.181 | 1.539 | 2.056 | 0.041 | 0.056 | 0.056 | 0 | 0.83 | 0.69 | 0.63 | 0.67 | 0.79 | 0.65 |
| 8 | 102842872 | rs7829693 | 1.307 | -0.403 | 1.118 | 1.808 | 0.045 | 0.052 | 0.052 | 0 | 0.78 | 0.52 | 0.61 | 0.66 | 0.75 | 0.66 |
| 8 | 102847202 | rs2155939 | -1.789 | -0.407 | 0.875 | 1.543 | 0 | 0 | 0 | 0 | 0.26 | 0.72 | 0.38 | 0.28 | 0.26 | 0.34 |
| 8 | 102859307 | rs1264866 | -2.115 | -0.569 | 0.903 | 1.572 | -0.002 | 0.01 | 0.01 | 0.587 | 0.28 | 0.79 | 0.4 | 0.29 | 0.25 | 0.34 |
| 8 | 102863119 | rs472963 | -1.563 | -0.547 | 0.93 | 1.579 | 0.018 | 0.006 | 0.006 | 0 | 0.23 | 0.53 | 0.32 | 0.27 | 0.22 | 0.34 |
| 8 | 102867399 | rs16868555 | 0.01 | -0.526 | 0.617 | 1.607 | -0.064 | -0.062 | -0.062 | 0 | 0.1 | 0.01 | 0.2 | 0.26 | 0.17 | 0.22 |
| 8 | 102873333 | rs486196 | -0.413 | -0.445 | 0.681 | 1.422 | 0.033 | 0.069 | 0.069 | 0 | 0.21 | 0.39 | 0.32 | 0.39 | 0.23 | 0.34 |
| 8 | 102889127 | rs478917 | -0.301 | -0.407 | 0.688 | 1.4 | 0.028 | 0.069 | 0.069 | 0 | 0.21 | 0.37 | 0.31 | 0.38 | 0.21 | 0.34 |
| 8 | 102899450 | rs508386 | -1.349 | -0.452 | 0.575 | 1.398 | -0.001 | -0.007 | -0.007 | 0 | 0.3 | 0.8 | 0.41 | 0.38 | 0.26 | 0.36 |
| 8 | 102899476 | rs513159 | -0.874 | -0.391 | 0.484 | 1.416 | 0 | 0.001 | 0.001 | 0.075 | 0.26 | 0.73 | 0.36 | 0.35 | 0.23 | 0.35 |
| 8 | 102914320 | rs550281 | -0.805 | -0.248 | 0.527 | 1.364 | 0.011 | 0.04 | 0.04 | 0 | 0.24 | 0.61 | 0.35 | 0.39 | 0.23 | 0.34 |
| 8 | 102920114 | rs7828198 | -0.864 | -0.454 | 0.307 | 0.951 | -0.053 | 0.014 | 0.014 | 1.155 | 0.2 | 0.46 | 0.14 | 0.22 | 0.13 | 0.12 |
| 8 | 102942620 | rs561252 | 0.229 | 0.352 | -0.863 | 0.824 | 0 | -0.024 | -0.024 | 0.038 | 0.94 | 1 | 0.92 | 0.92 | 0.96 | 0.94 |
| 8 | 102944863 | rs517811 | 0.347 | 0.428 | -0.901 | 0.917 | 0.002 | -0.005 | -0.005 | 0 | 0.6 | 0.63 | 0.6 | 0.47 | 0.61 | 0.65 |
| 8 | 102955639 | rs7843596 | -1.557 | -0.111 | 0.248 | 1.245 | 0.004 | 0.133 | 0.133 | 0 | 0.06 | 0.26 | 0.1 | 0.1 | 0 | 0.01 |
| 8 | 102965754 | rs2387333 | -1.291 | -0.935 | 0.427 | 0.966 | -0.058 | 0.012 | 0.012 | 1.234 | 0.24 | 0.6 | 0.28 | 0.26 | 0.14 | 0.18 |
| 8 | 102970640 | rs6991019 | 0.476 | -0.823 | 0.082 | 0.818 | 0.003 | 0.006 | 0.006 | 0 | 0.46 | 0.66 | 0.52 | 0.42 | 0.39 | 0.48 |
| 8 | 102972179 | rs7839884 | 1.31 | -0.514 | 0.007 | 0.815 | -0.011 | 0.009 | 0.009 | 0.165 | 0.36 | 0.26 | 0.4 | 0.32 | 0.36 | 0.44 |

| | | | | | | | | | | | | | | | | |
|---|---|---|---|---|---|---|---|---|---|---|---|---|---|---|---|---|
| 8 | 102976004 | rs718262 | 0.142 | -0.543 | 0.087 | 0.786 | -0.002 | 0.006 | 0.006 | 0 | 0.52 | 0.54 | 0.64 | 0.6 | 0.49 | 0.6 |
| 8 | 102984091 | rs6982001 | 1.034 | -0.644 | 0.184 | 0.693 | -0.002 | 0.008 | 0.008 | 0 | 0.38 | 0.32 | 0.43 | 0.33 | 0.36 | 0.45 |
| 8 | 102984279 | rs6982161 | -0.038 | -0.569 | 0.423 | 0.636 | 0.001 | 0 | 0 | 0 | 0.55 | 0.49 | 0.61 | 0.47 | 0.53 | 0.6 |
| 8 | 102984312 | rs6982566 | 0.222 | -0.56 | 0.226 | 0.63 | 0.002 | 0.005 | 0.005 | 0 | 0.52 | 0.38 | 0.51 | 0.47 | 0.47 | 0.56 |
| 8 | 102986395 | rs12544769 | -0.269 | -0.516 | 0.002 | 0.579 | 0.006 | -0.024 | -0.024 | 0 | 0.14 | 0.07 | 0.16 | 0.17 | 0.14 | 0.12 |
| 8 | 102996980 | rs16868855 | -1.56 | -0.651 | -0.548 | 0.095 | 0.067 | 0.06 | 0.06 | 1.345 | 0.34 | 0.82 | 0.26 | 0.22 | 0.17 | 0.29 |
| 8 | 103000553 | rs4734049 | 0.151 | -0.701 | -0.272 | -0.013 | 0.002 | -0.038 | -0.038 | 0 | 0.15 | 0.31 | 0.15 | 0.1 | 0.09 | 0.17 |
| 8 | 103001164 | rs10282818 | 1.811 | -0.718 | -0.537 | 0.137 | -0.083 | -0.093 | -0.093 | 1.293 | 0.25 | 0.02 | 0.35 | 0.27 | 0.29 | 0.26 |
| 8 | 103001914 | rs7814951 | -0.278 | -0.41 | -0.27 | 0.516 | 0.003 | 0.033 | 0.033 | 0 | 0.17 | 0.34 | 0.22 | 0.21 | 0.12 | 0.24 |
| 8 | 103002579 | rs4734609 | -0.538 | -0.393 | -0.268 | 0.513 | -0.011 | 0.021 | 0.021 | 0 | 0.19 | 0.35 | 0.22 | 0.21 | 0.12 | 0.22 |
| 8 | 103004858 | rs7829974 | -0.478 | -0.286 | -0.422 | 0.651 | -0.055 | 0.019 | 0.019 | 0 | 0.61 | 0.87 | 0.77 | 0.67 | 0.51 | 0.62 |
| 8 | 103018338 | rs10505016 | -0.507 | -0.297 | -0.454 | 0.636 | -0.074 | 0.003 | 0.003 | 0 | 0.61 | 0.88 | 0.7 | 0.66 | 0.5 | 0.6 |
| 8 | 103021475 | rs11783903 | 0.487 | -0.282 | -0.273 | 0.679 | 0 | -0.001 | -0.001 | 0 | 0.45 | 0.52 | 0.53 | 0.45 | 0.4 | 0.4 |
| 8 | 103028289 | rs10105250 | -0.531 | 0.173 | -0.378 | 0.768 | -0.008 | -0.031 | -0.031 | 0 | 0.81 | 0.86 | 0.87 | 0.67 | 0.77 | 0.69 |
| 8 | 103032394 | rs13248449 | 0.922 | 0.262 | -0.491 | 0.778 | 0 | 0.017 | 0.017 | 0 | 0.63 | 0.57 | 0.64 | 0.45 | 0.66 | 0.48 |
| 8 | 103035545 | rs1269715 | -0.46 | 0.312 | -0.552 | 0.522 | -0.037 | -0.062 | -0.062 | 0 | 0.8 | 0.93 | 0.79 | 0.66 | 0.74 | 0.67 |
| 8 | 103039917 | rs1269711 | 0.116 | 0.276 | 0.016 | 0.206 | 0.005 | -0.019 | -0.019 | 0 | 0.86 | 0.69 | 0.84 | 0.88 | 0.91 | 0.84 |
| 8 | 103044676 | rs1269707 | 0.018 | 0.512 | -0.614 | 0.159 | -0.005 | 0.005 | 0.005 | 0 | 0.91 | 0.93 | 0.9 | 0.92 | 0.85 | 0.96 |
| 8 | 103044932 | rs1269705 | 0.197 | 0.614 | -0.59 | -0.957 | -0.004 | -0.075 | -0.075 | 0 | 0.6 | 0.49 | 0.71 | 0.86 | 0.55 | 0.89 |
| 8 | 103044961 | rs6987078 | 0.554 | 0.578 | -0.83 | -0.379 | -0.012 | -0.001 | -0.001 | 0 | 0.29 | 0.34 | 0.23 | 0.32 | 0.21 | 0.48 |
| 8 | 103047139 | rs1623102 | -1.547 | 0.792 | -0.488 | -0.477 | 0.001 | -0.005 | -0.005 | 0.001 | 0.92 | 0.9 | 0.93 | 0.92 | 0.88 | 0.97 |
| 8 | 103048003 | rs16868908 | 0.783 | 0.532 | -0.364 | -0.846 | -0.009 | -0.017 | -0.017 | 0.201 | 0.09 | 0.17 | 0.06 | 0.05 | 0.07 | 0.04 |
| 8 | 103048115 | rs11777589 | -0.311 | 0.606 | 0.04 | -1.654 | -0.01 | -0.103 | -0.103 | 0 | 0.34 | 0.44 | 0.19 | 0.05 | 0.33 | 0.12 |
| 8 | 103050664 | rs7014685 | -1.08 | 0.756 | -0.168 | -1.983 | 0.013 | -0.071 | -0.071 | 0 | 0.28 | 0.43 | 0.21 | 0.14 | 0.29 | 0.23 |
| 8 | 103054099 | rs7007251 | 0.171 | 0.702 | -0.081 | -1.967 | -0.029 | -0.066 | -0.066 | 0 | 0.15 | 0.37 | 0.1 | 0.05 | 0.11 | 0.06 |
| 8 | 103054644 | rs1893533 | -1.36 | 0.849 | -0.165 | -1.287 | 0.017 | -0.011 | -0.011 | 0.696 | 0.31 | 0.41 | 0.31 | 0.26 | 0.37 | 0.27 |
| 8 | 103056517 | rs11776754 | 1.832 | 0.839 | -0.249 | -1.281 | 0.006 | 0.02 | 0.02 | 0 | 0.08 | 0.03 | 0.04 | 0.05 | 0.07 | 0.04 |
| 8 | 103056645 | rs12334499 | 1.31 | 0.866 | -0.258 | -1.287 | -0.06 | 0.028 | 0.028 | 0 | 0.13 | 0.04 | 0.11 | 0.08 | 0.17 | 0.16 |
| 8 | 103058601 | rs6991144 | 1.415 | 0.861 | -0.211 | -1.298 | -0.007 | -0.016 | -0.016 | 0 | 0.11 | 0.17 | 0.06 | 0.05 | 0.08 | 0.04 |
| 8 | 103059848 | rs7000754 | -1.276 | 0.813 | -0.148 | -1.319 | -0.003 | 0.001 | 0.001 | 0.077 | 0.31 | 0.29 | 0.29 | 0.27 | 0.36 | 0.27 |
| 8 | 103063862 | rs1573312 | -1.157 | 0.739 | -0.023 | -1.4 | -0.001 | -0.002 | -0.002 | 0 | 0.46 | 0.59 | 0.37 | 0.27 | 0.5 | 0.3 |
| 8 | 103067199 | rs1269701 | 1.008 | -0.271 | -0.307 | -1.716 | 0.033 | 0.151 | 0.151 | 0 | 0.47 | 0.29 | 0.66 | 0.77 | 0.54 | 0.75 |
| 8 | 103067627 | rs2155235 | 2.409 | -0.222 | -0.384 | -1.637 | 0.003 | -0.008 | -0.008 | 0 | 0.14 | 0.13 | 0.06 | 0 | 0.13 | 0.02 |
| 8 | 103067638 | rs17414701 | 2.312 | -0.142 | -0.588 | -1.504 | -0.009 | -0.006 | -0.006 | 0 | 0.11 | 0.01 | 0.09 | 0.1 | 0.07 | 0.13 |
| 8 | 103070048 | rs1110079 | -0.383 | -0.199 | -0.29 | -1.59 | -0.027 | -0.099 | -0.099 | 0.092 | 0.65 | 0.52 | 0.81 | 0.87 | 0.68 | 0.92 |
| 8 | 103070301 | rs17414857 | 0.442 | -0.159 | -0.323 | -1.334 | -0.013 | -0.021 | -0.021 | 0 | 0.17 | 0.21 | 0.12 | 0.09 | 0.11 | 0.15 |
| 8 | 103071053 | rs1269695 | 0.266 | -0.116 | -0.263 | -1.258 | -0.008 | -0.028 | -0.028 | 0 | 0.11 | 0.22 | 0.06 | 0.02 | 0.07 | 0.1 |
| 8 | 103071366 | rs1893535 | 0.482 | -0.083 | -0.522 | -1.526 | -0.013 | 0.154 | 0.154 | 0 | 0.47 | 0.25 | 0.64 | 0.82 | 0.45 | 0.68 |
| 8 | 103072851 | rs11775854 | 0.855 | -0.127 | -0.277 | -1.31 | -0.006 | -0.014 | -0.014 | 0 | 0.15 | 0.2 | 0.1 | 0.08 | 0.17 | 0.06 |
| 8 | 103073460 | rs1269693 | 0.211 | -0.156 | -0.962 | -1.309 | -0.004 | 0.002 | 0.002 | 0 | 0.56 | 0.5 | 0.49 | 0.5 | 0.6 | 0.61 |
| 8 | 103076379 | rs11785772 | 0.036 | -0.256 | -0.878 | -1.659 | 0.049 | 0.177 | 0.177 | 0.45 | 0.36 | 0.88 | 0.25 | 0.03 | 0.34 | 0.1 |
| 8 | 103077667 | rs10110783 | 0.575 | -0.271 | -0.718 | -1.637 | -0.013 | -0.101 | -0.101 | 0.209 | 0.18 | 0.46 | 0.14 | 0.03 | 0.19 | 0.05 |
| 8 | 103077849 | rs635670 | 0.394 | 0.362 | -0.822 | -1.634 | 0.001 | 0.003 | 0.003 | 0 | 0.5 | 0.38 | 0.44 | 0.57 | 0.52 | 0.56 |
| 8 | 103079505 | rs10090764 | 1.213 | 0.377 | -0.838 | -1.565 | -0.008 | -0.053 | -0.053 | 0 | 0.15 | 0.28 | 0.11 | 0.03 | 0.17 | 0.05 |
| 8 | 103085156 | rs11784616 | 0.414 | 0.409 | -0.849 | -1.768 | 0.007 | -0.048 | -0.048 | 0 | 0.19 | 0.28 | 0.15 | 0.03 | 0.26 | 0.12 |
| 8 | 103085612 | rs610125 | -2.362 | 0.463 | -0.991 | -1.751 | -0.033 | -0.133 | -0.133 | 0 | 0.82 | 0.99 | 0.72 | 0.61 | 0.85 | 0.73 |
| 8 | 103087806 | rs687542 | 1.279 | 0.617 | -0.934 | -1.653 | -0.086 | -0.093 | -0.093 | 0 | 0.28 | 0.04 | 0.26 | 0.42 | 0.34 | 0.4 |
| 8 | 103090257 | rs10955279 | 1.373 | 1.039 | -0.521 | -1.475 | -0.005 | -0.036 | -0.036 | 0 | 0.14 | 0.18 | 0.11 | 0.04 | 0.17 | 0.05 |
| 8 | 103093093 | rs7830150 | -0.737 | 1.06 | -0.61 | -1.467 | 0.032 | 0.037 | 0.037 | 0 | 0.12 | 0.31 | 0.15 | 0.15 | 0.13 | 0.19 |
| 8 | 103094185 | rs688431 | -2.498 | 1.023 | -0.723 | -1.497 | -0.032 | -0.126 | -0.126 | 0.269 | 0.83 | 0.99 | 0.73 | 0.61 | 0.86 | 0.74 |
| 8 | 103097606 | rs11777456 | -0.474 | 1.057 | -0.553 | -1.439 | 0.004 | -0.03 | -0.03 | 0.169 | 0.26 | 0.6 | 0.26 | 0.18 | 0.3 | 0.24 |
| 8 | 103106554 | rs556388 | 0.253 | 1.015 | -0.543 | -1.469 | -0.005 | 0.017 | 0.017 | 0 | 0.48 | 0.34 | 0.38 | 0.45 | 0.53 | 0.47 |
| 8 | 103107967 | rs501493 | 1.711 | 1.331 | -0.682 | -1.625 | -0.019 | -0.079 | -0.079 | 0 | 0.26 | 0.05 | 0.22 | 0.43 | 0.26 | 0.38 |
| 8 | 103109921 | rs652008 | -0.729 | 1.276 | -0.609 | -1.614 | 0.002 | -0.001 | -0.001 | 0 | 0.59 | 0.64 | 0.52 | 0.61 | 0.66 | 0.67 |
| 8 | 103111461 | rs6992873 | 0.295 | 1.344 | -0.673 | -1.658 | 0.003 | -0.042 | -0.042 | 0.028 | 0.22 | 0.28 | 0.15 | 0.02 | 0.28 | 0.09 |
| 8 | 103114573 | rs501344 | 0.835 | 1.367 | -0.462 | -1.573 | -0.025 | -0.034 | -0.034 | 0.356 | 0.31 | 0.15 | 0.27 | 0.43 | 0.34 | 0.44 |
| 8 | 103114802 | rs13278975 | 2.89 | 1.356 | -0.126 | -1.576 | -0.003 | -0.045 | -0.045 | 0 | 0.12 | 0.13 | 0.08 | 0 | 0.13 | 0.02 |
| 8 | 103116959 | rs7826367 | -0.425 | 1.761 | 0.589 | -0.998 | 0.045 | -0.062 | -0.062 | 0.295 | 0.1 | 0.28 | 0.08 | 0 | 0.13 | 0.05 |
| 8 | 103120655 | rs7013304 | -0.749 | 1.792 | 0.519 | -1.019 | 0.007 | -0.135 | -0.135 | 0 | 0.29 | 0.52 | 0.28 | 0.03 | 0.32 | 0.18 |
| 8 | 103121989 | rs9649919 | 1.274 | 1.786 | 0.539 | -1.004 | 0 | -0.005 | -0.005 | 0 | 0.08 | 0.05 | 0.11 | 0.01 | 0.07 | 0.06 |
| 8 | 103124212 | rs6987653 | -0.33 | 1.831 | 0.587 | -0.985 | 0.004 | -0.079 | -0.079 | 0 | 0.23 | 0.39 | 0.17 | 0.02 | 0.28 | 0.09 |
| 8 | 103127270 | rs7833656 | -1.115 | 1.679 | 0.561 | -0.981 | 0.019 | -0.126 | -0.126 | 0 | 0.35 | 0.52 | 0.32 | 0.04 | 0.43 | 0.23 |
| 8 | 103129949 | rs10111484 | 0.435 | 1.994 | 0.54 | -0.722 | -0.03 | -0.083 | -0.083 | 0.009 | 0.17 | 0.4 | 0.13 | 0.02 | 0.18 | 0.05 |
| 8 | 103130360 | rs503342 | 0.001 | 2.166 | 0.616 | -0.528 | -0.037 | -0.032 | -0.032 | 0 | 0.35 | 0.19 | 0.26 | 0.44 | 0.37 | 0.42 |
| 8 | 103132854 | rs527421 | -0.522 | 2.071 | 0.356 | -0.451 | -0.006 | 0 | 0 | 0 | 0.44 | 0.59 | 0.5 | 0.61 | 0.43 | 0.62 |
| 8 | 103138045 | rs648119 | -0.423 | 1.563 | 0.571 | -0.724 | -0.001 | -0.002 | -0.002 | 0.348 | 0.44 | 0.42 | 0.37 | 0.44 | 0.52 | 0.5 |
| 8 | 103139683 | rs6468813 | -0.106 | 1.081 | 0.326 | -0.883 | 0.009 | 0.012 | 0.012 | 0 | 0.19 | 0.21 | 0.12 | 0.07 | 0.18 | 0.16 |
| 8 | 103144592 | rs6980591 | -1.628 | 1.61 | 0.932 | -0.503 | 0.077 | -0.08 | -0.08 | 0 | 0.17 | 0.46 | 0.27 | 0.04 | 0.21 | 0.22 |
| 8 | 103159452 | rs13268773 | -1.183 | 1.583 | 0.252 | -0.424 | -0.049 | -0.139 | -0.139 | 0.023 | 0.65 | 0.98 | 0.64 | 0.5 | 0.68 | 0.75 |

| | | | | | | | | | | | | | | | | |
|---|---|---|---|---|---|---|---|---|---|---|---|---|---|---|---|---|
| 8 | 103164438 | rs2051274 | -0.436 | 1.537 | 0.634 | -0.435 | -0.005 | 0.017 | 0.017 | 0.014 | 0.54 | 0.68 | 0.54 | 0.45 | 0.56 | 0.54 |
| 8 | 103165042 | rs1111912 | 0.142 | 1.58 | 0.676 | -0.403 | 0.001 | 0.011 | 0.011 | 0 | 0.47 | 0.47 | 0.34 | 0.41 | 0.49 | 0.49 |
| 8 | 103176946 | rs10955282 | 0.275 | 1.511 | 0.404 | -0.386 | 0.001 | -0.001 | -0.001 | 2.814 | 0.44 | 0.6 | 0.47 | 0.48 | 0.46 | 0.48 |
| 8 | 103184794 | rs1265119 | -0.294 | 1.236 | -0.33 | -0.26 | -0.02 | 0.003 | 0.003 | 0 | 0.53 | 0.76 | 0.55 | 0.48 | 0.53 | 0.53 |
| 8 | 103187804 | rs2067029 | -0.288 | 1.156 | -0.092 | 0.23 | 0.063 | 0.082 | 0.082 | 0 | 0.07 | 0.32 | 0.2 | 0.07 | 0.05 | 0.04 |
| 8 | 103190530 | rs1265122 | -0.253 | 1.302 | -1.487 | 0.266 | -0.015 | -0.06 | -0.06 | 0 | 0.6 | 0.89 | 0.61 | 0.51 | 0.62 | 0.71 |
| 8 | 103193781 | rs1265125 | -1.075 | 1.251 | -0.768 | 0 | 0.007 | -0.034 | -0.034 | 0 | 0.07 | 0.11 | 0.04 | 0 | 0.06 | 0.07 |
| 8 | 103195988 | rs10094658 | 0.228 | 1.601 | -1.219 | -0.548 | -0.004 | 0.099 | 0.099 | 0 | 0.4 | 0.27 | 0.29 | 0.06 | 0.47 | 0.28 |
| 8 | 103198203 | rs747822 | -0.358 | 0.478 | -0.744 | 0.369 | -0.001 | -0.001 | -0.001 | 0 | 0.61 | 0.62 | 0.63 | 0.47 | 0.7 | 0.69 |
| 8 | 103200036 | rs1265138 | -1.125 | 0.648 | -0.645 | 1.048 | -0.001 | 0.069 | 0.069 | 0 | 0.08 | 0.13 | 0.08 | 0.31 | 0.05 | 0.26 |
| 8 | 103200321 | rs1265139 | -1.414 | 0.836 | -0.88 | 0.322 | 0 | -0.011 | -0.011 | 0 | 0.11 | 0.11 | 0.07 | 0.01 | 0.11 | 0.08 |
| 8 | 103200931 | rs10505018 | -0.694 | 0.117 | -1.223 | 0.749 | 0.021 | -0.027 | -0.027 | 0 | 0.22 | 0.13 | 0.38 | 0.3 | 0.14 | 0.3 |
| 8 | 103204972 | rs11786397 | -0.971 | 0.006 | -1.308 | -0.182 | 0.01 | 0.019 | 0.019 | 0 | 0.17 | 0.14 | 0.17 | 0.03 | 0.13 | 0.12 |
| 8 | 103208028 | rs1265143 | -0.025 | 0.374 | -2.019 | -0.83 | -0.049 | 0.07 | 0.07 | 0 | 0.13 | 0.01 | 0.06 | 0.01 | 0.17 | 0.12 |
| 8 | 103209964 | rs3931977 | 1.223 | -0.037 | -1.562 | -0.534 | 0 | 0 | 0 | 0 | 0.47 | 0.38 | 0.51 | 0.52 | 0.46 | 0.57 |
| 8 | 103210211 | rs16869246 | 0.246 | -0.07 | -1.584 | -0.493 | 0.001 | 0.003 | 0.003 | 0 | 0.08 | 0.09 | 0.15 | 0 | 0.07 | 0.1 |
| 8 | 103213460 | rs1265151 | -1.542 | -0.354 | -1.212 | -0.641 | -0.03 | 0.089 | 0.089 | 0 | 0.88 | 0.84 | 0.85 | 0.58 | 0.94 | 0.83 |
| 8 | 103216898 | rs1052069 | 0.965 | -0.177 | -1.279 | -0.734 | 0.005 | 0.017 | 0.017 | 0 | 0.08 | 0.04 | 0.04 | 0.03 | 0.08 | 0.02 |
| 8 | 103217482 | rs3204695 | 0.796 | -0.189 | -1.299 | -0.752 | -0.011 | 0.037 | 0.037 | 0 | 0.07 | 0 | 0.12 | 0 | 0.06 | 0.1 |
| 8 | 103218688 | rs16918482 | -0.459 | -0.195 | -1.23 | -0.757 | 0.008 | 0.04 | 0.04 | 0 | 0.15 | 0.08 | 0.17 | 0.03 | 0.13 | 0.12 |
| 8 | 103219240 | rs4102401 | 0.799 | -0.195 | -1.231 | -0.757 | -0.011 | 0.037 | 0.037 | 0 | 0.07 | 0 | 0.12 | 0 | 0.06 | 0.1 |
| 8 | 103219363 | rs3907099 | 0.768 | -0.195 | -1.23 | -0.756 | 0.003 | 0.003 | 0.003 | 0 | 0.08 | 0.08 | 0.05 | 0.03 | 0.08 | 0.02 |
| 8 | 103219629 | rs16869269 | -0.458 | -0.195 | -1.23 | -0.757 | 0.008 | 0.04 | 0.04 | 0 | 0.15 | 0.08 | 0.17 | 0.03 | 0.13 | 0.12 |
| 8 | 103244599 | rs16869295 | 0.674 | 0.029 | -1.147 | -0.681 | 0.004 | 0.008 | 0.008 | 0 | 0.08 | 0.07 | 0.05 | 0.02 | 0.08 | 0.02 |
| 8 | 103247134 | rs2853229 | -1.487 | -0.244 | -0.58 | -0.893 | 0.015 | 0.006 | 0.006 | 0 | 0.5 | 0.72 | 0.49 | 0.5 | 0.51 | 0.38 |
| 8 | 103249111 | rs11777942 | 0.951 | -0.061 | -0.619 | -0.809 | 0.005 | 0.018 | 0.018 | 0 | 0.08 | 0.04 | 0.04 | 0.02 | 0.08 | 0.02 |
| 8 | 103268783 | rs2607662 | -1.389 | -0.431 | -0.704 | -1.078 | -0.018 | -0.023 | -0.023 | 0.007 | 0.55 | 0.77 | 0.51 | 0.51 | 0.55 | 0.38 |
| 8 | 103274006 | rs2853236 | -2.302 | 0.082 | -1.222 | -1.592 | -0.008 | -0.068 | -0.068 | 0 | 0.77 | 0.88 | 0.81 | 0.52 | 0.76 | 0.63 |
| 8 | 103276753 | rs12707772 | 1.22 | 0.12 | -1.135 | -1.601 | -0.02 | 0.02 | 0.02 | 0.026 | 0.08 | 0.06 | 0.06 | 0.01 | 0.17 | 0.03 |
| 8 | 103277847 | rs2978432 | 0.401 | 0.013 | -1.106 | -1.724 | 0.008 | 0.02 | 0.02 | 0 | 0.13 | 0.11 | 0.15 | 0.03 | 0.12 | 0.07 |
| 8 | 103279922 | rs13261195 | 2.278 | 0.018 | -1.115 | -1.718 | 0 | 0.033 | 0.033 | 0 | 0.07 | 0 | 0.02 | 0.02 | 0.06 | 0.02 |
| 8 | 103282518 | rs2978431 | 1.472 | -0.241 | -0.136 | -1.914 | 0.001 | -0.039 | -0.039 | 0 | 0.12 | 0.14 | 0.05 | 0.01 | 0.12 | 0.14 |
| 8 | 103288343 | rs2853238 | -1.666 | -0.121 | -0.157 | -1.91 | -0.017 | -0.017 | -0.017 | 0 | 0.55 | 0.75 | 0.53 | 0.48 | 0.55 | 0.39 |
| 8 | 103291591 | rs2512408 | -1.494 | -0.164 | -0.007 | -1.928 | 0 | -0.001 | -0.001 | 0 | 0.53 | 0.61 | 0.5 | 0.48 | 0.54 | 0.38 |
| 8 | 103297707 | rs2293982 | 0.8 | -0.108 | -0.154 | -1.839 | -0.019 | 0.02 | 0.02 | 0 | 0.1 | 0.06 | 0.07 | 0.02 | 0.18 | 0.03 |
| 8 | 103301555 | rs2168689 | -0.361 | -0.134 | -0.166 | -1.855 | -0.036 | 0.074 | 0.074 | 0 | 0.21 | 0.09 | 0.21 | 0.04 | 0.27 | 0.09 |
| 8 | 103306033 | rs16869352 | 2.197 | -0.181 | -0.122 | -1.854 | 0.001 | 0.035 | 0.035 | 0 | 0.07 | 0 | 0.02 | 0.02 | 0.06 | 0.02 |
| 8 | 103326735 | rs2512411 | 1.875 | -0.163 | -0.244 | -1.885 | -0.006 | 0.012 | 0.012 | 0 | 0.11 | 0.05 | 0.05 | 0.01 | 0.12 | 0.14 |
| 8 | 103339881 | rs2445948 | -1.38 | -0.229 | -0.208 | -1.966 | 0.007 | 0.073 | 0.073 | 0 | 0.5 | 0.61 | 0.4 | 0.07 | 0.52 | 0.3 |
| 8 | 103346407 | rs10505025 | 2.204 | -0.241 | -0.489 | -1.945 | 0.032 | 0.051 | 0.051 | 0.544 | 0.13 | 0.01 | 0.08 | 0.01 | 0.11 | 0.05 |
| 8 | 103356112 | rs10481053 | -0.759 | -0.235 | -0.437 | -1.94 | -0.001 | 0.088 | 0.088 | 0 | 0.34 | 0.2 | 0.31 | 0.05 | 0.39 | 0.16 |
| 8 | 103373304 | rs10090707 | 0.313 | -0.195 | -0.418 | -1.89 | -0.026 | 0.05 | 0.05 | 0 | 0.17 | 0.08 | 0.1 | 0.04 | 0.24 | 0.05 |
| 8 | 103397353 | rs10955285 | 1.644 | 0.047 | -0.251 | -1.659 | 0.009 | 0.008 | 0.008 | 0.085 | 0.14 | 0.11 | 0.1 | 0.01 | 0.12 | 0.07 |
| 8 | 103407126 | rs10505029 | 1.587 | 0.792 | -0.585 | -1.767 | -0.034 | 0.05 | 0.05 | 1.856 | 0.13 | 0.03 | 0.26 | 0.03 | 0.12 | 0.17 |
| 8 | 103428008 | rs13248227 | -1.476 | 0.761 | -0.021 | -1.295 | 0.008 | -0.009 | -0.009 | 0 | 0.47 | 0.7 | 0.57 | 0.51 | 0.49 | 0.37 |
| 8 | 103430407 | rs13269273 | -0.031 | 1.344 | -0.043 | -0.786 | 0.041 | 0.186 | 0.186 | 0 | 0.1 | 0.19 | 0.25 | 0.45 | 0.17 | 0.1 |
| 8 | 103440763 | rs4609174 | -1.23 | 1.243 | -0.126 | -0.916 | 0.039 | 0.091 | 0.091 | 0 | 0.27 | 0.43 | 0.42 | 0.48 | 0.33 | 0.28 |
| 8 | 103444927 | rs4734621 | -1.629 | 1.579 | -0.159 | -0.835 | 0.077 | 0.09 | 0.09 | 0 | 0.29 | 0.48 | 0.45 | 0.49 | 0.41 | 0.34 |
| 8 | 103456352 | rs4269511 | -1.082 | 1.632 | -0.058 | -0.723 | 0.052 | 0.169 | 0.169 | 0 | 0.15 | 0.33 | 0.28 | 0.45 | 0.19 | 0.14 |
| 8 | 103469290 | rs3923790 | -0.675 | 1.829 | -0.298 | -0.544 | 0.055 | 0.149 | 0.149 | 0.107 | 0.19 | 0.44 | 0.2 | 0.37 | 0.22 | 0.29 |
| 8 | 103483466 | rs7836225 | -0.817 | 1.423 | -0.671 | -0.731 | -0.001 | -0.002 | -0.002 | 1.337 | 0.39 | 0.57 | 0.44 | 0.44 | 0.45 | 0.38 |
| 8 | 103490914 | rs7010489 | -1.024 | 1.382 | -0.686 | -0.728 | -0.002 | -0.004 | -0.004 | 1.788 | 0.45 | 0.58 | 0.48 | 0.45 | 0.57 | 0.39 |
| 8 | 103498788 | rs13249355 | 0.675 | 1.661 | -0.831 | -0.553 | -0.029 | 0.018 | 0.018 | 0 | 0.06 | 0.01 | 0.05 | 0.01 | 0.13 | 0.02 |
| 8 | 103515898 | rs2436949 | -0.648 | 1.582 | -0.74 | -1.169 | 0.001 | -0.014 | -0.014 | 0 | 0.47 | 0.62 | 0.43 | 0.07 | 0.49 | 0.36 |
| 8 | 103517702 | rs4317537 | -1.274 | 1.601 | -0.624 | -0.891 | 0.023 | 0.016 | 0.016 | 0 | 0.25 | 0.47 | 0.34 | 0.28 | 0.33 | 0.27 |
| 8 | 103520099 | rs1106867 | 0.835 | 1.576 | -0.682 | -0.904 | 0.006 | 0.014 | 0.014 | 0 | 0.26 | 0.17 | 0.25 | 0.03 | 0.19 | 0.26 |
| 8 | 103524176 | rs4734058 | -0.739 | 1.766 | -0.254 | -0.413 | 0.017 | 0.058 | 0.058 | 0 | 0.15 | 0.31 | 0.26 | 0.26 | 0.18 | 0.19 |
| 8 | 103524461 | rs6468833 | -1.223 | 1.691 | -0.406 | -0.478 | -0.004 | 0.016 | 0.016 | 0.096 | 0.47 | 0.66 | 0.48 | 0.3 | 0.49 | 0.44 |
| 8 | 103529405 | rs10089843 | -1.87 | 2.057 | -0.202 | 0.248 | 0.044 | 0.059 | 0.059 | 0.117 | 0.29 | 0.53 | 0.5 | 0.48 | 0.41 | 0.32 |
| 8 | 103532303 | rs16869493 | -0.005 | 1.76 | -0.272 | 0.13 | 0.026 | -0.033 | -0.033 | 0 | 0.08 | 0.01 | 0.16 | 0.2 | 0.04 | 0.15 |
| 8 | 103534797 | rs7007146 | -0.628 | 1.77 | -0.284 | 0.044 | 0.001 | -0.027 | -0.027 | 0 | 0.45 | 0.56 | 0.41 | 0.28 | 0.5 | 0.32 |
| 8 | 103541717 | rs16869497 | -0.698 | 1.486 | -0.531 | 0.234 | -0.033 | -0.068 | -0.068 | 0.208 | 0.12 | 0.03 | 0.33 | 0.4 | 0.14 | 0.23 |
| 8 | 103542538 | rs4734636 | -1.263 | 1.499 | -0.456 | -0.029 | 0.013 | -0.003 | -0.003 | 0.261 | 0.57 | 0.39 | 0.56 | 0.45 | 0.62 | 0.49 |
| 8 | 103548562 | rs1347322 | 0.136 | 1.52 | -0.535 | -0.174 | 0.002 | -0.043 | -0.043 | 0 | 0.15 | 0.22 | 0.09 | 0.01 | 0.2 | 0.04 |
| 8 | 103549754 | rs16869515 | 0.477 | 1.517 | -0.525 | -0.12 | -0.016 | 0.102 | 0.102 | 0 | 0.31 | 0.16 | 0.15 | 0.06 | 0.29 | 0.22 |
| 8 | 103550014 | rs892503 | -1.255 | 1.455 | -0.273 | 0.045 | 0.012 | 0.026 | 0.026 | 0 | 0.31 | 0.49 | 0.45 | 0.43 | 0.36 | 0.27 |
| 8 | 103550517 | rs16869518 | -0.665 | 0.813 | -0.135 | 0.111 | 0.014 | -0.056 | -0.056 | 0 | 0.1 | 0.27 | 0.06 | 0 | 0.12 | 0.03 |
| 8 | 103551385 | rs8180987 | -0.309 | 0.966 | -0.678 | 0.575 | 0.002 | -0.034 | -0.034 | 2.307 | 0.06 | 0.01 | 0.09 | 0.2 | 0.04 | 0.17 |
| 8 | 103553857 | rs13274948 | 0.769 | 0.855 | -0.629 | 0.174 | 0.018 | 0.073 | 0.073 | 0 | 0.22 | 0.07 | 0.09 | 0.04 | 0.14 | 0.18 |

| | | | | | | | | | | | | | | | | |
|---|---|---|---|---|---|---|---|---|---|---|---|---|---|---|---|---|
| 8 | 103555913 | rs1000672 | -1.537 | 1.193 | 0.227 | 1.039 | -0.001 | 0 | 0 | 0 | 0.13 | 0.19 | 0.13 | 0.18 | 0.14 | 0.19 |
| 8 | 103557043 | rs2436908 | 1.813 | 1.331 | 0.554 | 1.539 | 0.032 | 0.145 | 0.145 | 0 | 0.8 | 0.56 | 0.66 | 0.43 | 0.77 | 0.73 |
| 8 | 103557944 | rs2511758 | 0.696 | 1.443 | 0.439 | 1.674 | 0.064 | 0.287 | 0.287 | 0 | 0.94 | 0.8 | 0.79 | 0.62 | 0.9 | 0.92 |
| 8 | 103558857 | rs6986714 | -1.421 | 1.14 | 0.405 | 1.032 | -0.003 | -0.003 | -0.003 | 0 | 0.16 | 0.25 | 0.15 | 0.16 | 0.14 | 0.18 |
| 8 | 103559001 | rs16869546 | -0.784 | 1.08 | 0.291 | 1.157 | -0.003 | 0.011 | 0.011 | 0 | 0.05 | 0.07 | 0.09 | 0.15 | 0.04 | 0.15 |
| 8 | 103560356 | rs2511733 | 0.457 | 1.39 | 0.45 | 0.842 | 0.038 | -0.014 | -0.014 | 1.347 | 0.67 | 0.52 | 0.5 | 0.31 | 0.59 | 0.53 |
| 8 | 103562249 | rs16869559 | -0.642 | 1.185 | 0.43 | 1.069 | -0.014 | 0.037 | 0.037 | 0 | 0.06 | 0.1 | 0.08 | 0.17 | 0.03 | 0.22 |
| 8 | 103566428 | rs998201 | 0.087 | 1.546 | 0.552 | 1.402 | 0.003 | 0.009 | 0.009 | 0 | 0.22 | 0.26 | 0.4 | 0.34 | 0.26 | 0.32 |
| 8 | 103566787 | rs919953 | 1.487 | 1.091 | 0.808 | 1.65 | 0.005 | 0.087 | 0.087 | 0.008 | 0.89 | 0.86 | 0.86 | 0.51 | 0.88 | 0.69 |
| 8 | 103572247 | rs4734641 | 0.599 | 0.944 | 0.705 | 1.182 | 0.008 | 0.067 | 0.067 | 0 | 0.83 | 0.75 | 0.89 | 0.63 | 0.83 | 0.87 |
| 8 | 103576066 | rs2436891 | -0.785 | 1.043 | 0.314 | 0.117 | 0.004 | -0.046 | -0.046 | 0.72 | 0.21 | 0.26 | 0.23 | 0.01 | 0.27 | 0.09 |
| 8 | 103576658 | rs2436889 | -0.785 | 1.043 | 0.314 | 0.117 | 0.004 | -0.046 | -0.046 | 0 | 0.21 | 0.26 | 0.23 | 0.01 | 0.27 | 0.09 |
| 8 | 103578377 | rs12681882 | 1.153 | 0.044 | -0.65 | -0.306 | 0.048 | 0.054 | 0.054 | 0 | 0.25 | 0.06 | 0.11 | 0.13 | 0.12 | 0.17 |
| 8 | 103581213 | rs13263248 | -0.005 | 0.056 | -0.5 | 0.021 | -0.001 | -0.001 | -0.001 | 1.209 | 0.11 | 0.08 | 0.07 | 0.15 | 0.1 | 0.2 |
| 8 | 103584064 | rs2012106 | 0.377 | 0.386 | 0.086 | 0.508 | 0.034 | 0.018 | 0.018 | 0 | 0.74 | 0.53 | 0.67 | 0.69 | 0.68 | 0.81 |
| 8 | 103584390 | rs2511745 | -0.159 | 0.456 | 0.088 | 0.429 | 0.007 | -0.036 | -0.036 | 0 | 0.2 | 0.19 | 0.23 | 0.01 | 0.26 | 0.1 |
| 8 | 103584651 | rs4276657 | 0.788 | 0.2 | -0.198 | 0.762 | -0.06 | -0.051 | -0.051 | 0 | 0.36 | 0.2 | 0.49 | 0.38 | 0.43 | 0.45 |
| 8 | 103585093 | rs2511748 | -0.669 | 0.201 | -0.188 | 0.754 | -0.048 | 0.027 | 0.027 | 0 | 0.59 | 0.74 | 0.44 | 0.34 | 0.51 | 0.46 |
| 8 | 103591184 | rs2436915 | -0.209 | 0.095 | -0.092 | 0.398 | 0.007 | 0.02 | 0.02 | 0 | 0.51 | 0.49 | 0.29 | 0.3 | 0.42 | 0.37 |
| 8 | 103591237 | rs2436916 | -0.208 | 0.019 | -0.788 | 0.356 | -0.011 | -0.003 | -0.003 | 0 | 0.23 | 0.15 | 0.18 | 0.22 | 0.29 | 0.15 |
| 8 | 103591361 | rs4734646 | 1.256 | -0.306 | -0.903 | 0.396 | 0.055 | 0.071 | 0.071 | 0 | 0.26 | 0.08 | 0.1 | 0.08 | 0.13 | 0.21 |
| 8 | 103594122 | rs2436919 | -0.775 | 0.567 | -0.69 | 0.726 | -0.015 | -0.032 | -0.032 | 0 | 0.11 | 0.04 | 0.1 | 0.2 | 0.11 | 0.16 |
| 8 | 103597492 | rs965660 | -1.309 | 0.733 | -0.262 | 1.532 | 0.01 | 0.134 | 0.134 | 0.51 | 0.09 | 0.16 | 0.13 | 0.48 | 0.1 | 0.11 |
| 8 | 103599696 | rs10505031 | 0.508 | 0.935 | -0.816 | 1.057 | -0.015 | 0.016 | 0.016 | 1.907 | 0.15 | 0.03 | 0.22 | 0.09 | 0.14 | 0.08 |
| 8 | 103602649 | rs2511751 | -1.271 | 0.916 | -0.593 | 0.521 | -0.002 | 0.047 | 0.047 | 0 | 0.48 | 0.61 | 0.71 | 0.72 | 0.5 | 0.47 |
| 8 | 103602767 | rs6468837 | 0.089 | 0.815 | -0.141 | 0.515 | -0.003 | -0.004 | -0.004 | 0 | 0.26 | 0.21 | 0.43 | 0.63 | 0.33 | 0.36 |
| 8 | 103616695 | rs2436930 | -0.623 | 0.841 | -0.067 | 0.593 | 0.002 | 0.021 | 0.021 | 0.018 | 0.24 | 0.27 | 0.35 | 0.55 | 0.25 | 0.19 |
| 8 | 103619902 | rs2436932 | 0.729 | 0.855 | -0.091 | 0.764 | -0.026 | -0.01 | -0.01 | 7.432 | 0.16 | 0.03 | 0.25 | 0.13 | 0.17 | 0.12 |
| 8 | 103620164 | rs2511679 | 0.124 | 1.082 | 0.063 | 0.549 | -0.001 | 0.005 | 0.005 | 0 | 0.68 | 0.58 | 0.59 | 0.65 | 0.65 | 0.52 |
| 8 | 103620332 | rs13250522 | 1.615 | 1.082 | 0.124 | 0.494 | 0.01 | 0.07 | 0.07 | 0 | 0.45 | 0.3 | 0.23 | 0.1 | 0.39 | 0.33 |
| 8 | 103624715 | rs2511674 | 0.457 | 1.119 | 0.145 | 0.586 | -0.011 | 0.014 | 0.014 | 0.355 | 0.66 | 0.71 | 0.51 | 0.23 | 0.57 | 0.45 |
| 8 | 103630028 | rs17186961 | 0.541 | 1.126 | 0.348 | 0.604 | -0.007 | -0.011 | -0.011 | 0 | 0.24 | 0.28 | 0.34 | 0.16 | 0.22 | 0.28 |
| 8 | 103630839 | rs2436952 | -0.49 | 1.215 | 0.458 | 0.784 | -0.009 | -0.015 | -0.015 | 0 | 0.92 | 0.91 | 0.93 | 0.82 | 0.88 | 0.8 |
| 8 | 103635994 | rs4734648 | 0.897 | 1.216 | 0.426 | 0.718 | -0.006 | -0.003 | -0.003 | 0.02 | 0.2 | 0.27 | 0.18 | 0.18 | 0.18 | 0.22 |
| 8 | 103639941 | rs6998277 | 0.529 | 1.219 | 0.226 | 0.684 | -0.019 | -0.023 | -0.023 | 0 | 0.22 | 0.45 | 0.2 | 0.17 | 0.18 | 0.22 |
| 8 | 103640065 | rs7844766 | 1.842 | 1.296 | -0.068 | 0.48 | 0.011 | 0.047 | 0.047 | 0 | 0.39 | 0.25 | 0.19 | 0.08 | 0.31 | 0.3 |
| 8 | 103640433 | rs2511662 | 0.563 | 1.309 | -0.038 | 0.429 | -0.005 | 0.009 | 0.009 | 0 | 0.21 | 0.13 | 0.41 | 0.15 | 0.22 | 0.17 |
| 8 | 103640721 | rs1978752 | 0.905 | 1.322 | 0.134 | 0.683 | 0.001 | -0.005 | -0.005 | 3.669 | 0.61 | 0.38 | 0.6 | 0.23 | 0.53 | 0.47 |
| 8 | 103649935 | rs2436964 | -0.286 | 1.302 | 0.365 | 1.691 | -0.017 | -0.024 | -0.024 | 0 | 0.93 | 0.96 | 0.93 | 0.82 | 0.88 | 0.8 |
| 8 | 103656997 | rs16869676 | -0.404 | 1.316 | 0.461 | 1.767 | 0.023 | 0.209 | 0.209 | 0 | 0.05 | 0.11 | 0.1 | 0.42 | 0.08 | 0.07 |
| 8 | 103658711 | rs6981411 | 0.789 | 1.265 | 0.519 | 1.623 | 0.008 | 0.02 | 0.02 | 0 | 0.87 | 0.79 | 0.88 | 0.82 | 0.85 | 0.81 |
| 8 | 103658945 | rs2511718 | -0.784 | 1.246 | 0.302 | 1.777 | 0.012 | 0.103 | 0.103 | 0 | 0.09 | 0.11 | 0.24 | 0.43 | 0.12 | 0.11 |
| 8 | 103662219 | rs3191333 | 1.881 | 1.148 | -0.183 | 1.285 | 0.015 | 0.123 | 0.123 | 0 | 0.44 | 0.22 | 0.3 | 0.17 | 0.35 | 0.32 |
| 8 | 103663912 | rs11552577 | 1.089 | 1.132 | -0.269 | 1.317 | -0.029 | -0.036 | -0.036 | 0.028 | 0.16 | 0.02 | 0.27 | 0.15 | 0.19 | 0.14 |
| 8 | 103669991 | rs4734654 | 1.578 | 1.065 | 0.014 | 1.315 | 0.019 | 0.109 | 0.109 | 0 | 0.44 | 0.25 | 0.29 | 0.17 | 0.35 | 0.32 |
| 8 | 103676345 | rs1076030 | 1.229 | 1.078 | 0.204 | 1.35 | 0.005 | 0.012 | 0.012 | 0 | 0.49 | 0.39 | 0.34 | 0.25 | 0.38 | 0.46 |
| 8 | 103680416 | rs2099408 | 1.27 | 1.116 | 0.223 | 1.214 | 0.003 | 0.005 | 0.005 | 0 | 0.48 | 0.39 | 0.34 | 0.18 | 0.38 | 0.37 |
| 8 | 103683191 | rs930266 | 0.787 | 1.199 | 0.118 | 1.29 | -0.001 | 0 | 0 | 0 | 0.56 | 0.57 | 0.56 | 0.62 | 0.47 | 0.48 |
| 8 | 103688381 | rs10106298 | 1.154 | 1.121 | -0.74 | 0.645 | 0.019 | 0.12 | 0.12 | 0 | 0.45 | 0.26 | 0.3 | 0.18 | 0.35 | 0.37 |
| 8 | 103693821 | rs9886429 | 1.187 | 1.024 | -0.692 | 0.409 | -0.009 | 0.03 | 0.03 | 0 | 0.45 | 0.33 | 0.31 | 0.02 | 0.49 | 0.2 |
| 8 | 103694272 | rs2511692 | -0.002 | 0.568 | -0.846 | 0.914 | 0.014 | -0.026 | -0.026 | 0 | 0.27 | 0.18 | 0.18 | 0.32 | 0.24 | 0.49 |
| 8 | 103694457 | rs1434277 | 1.142 | 0.917 | -1.12 | 0.752 | -0.03 | 0.108 | 0.108 | 0 | 0.39 | 0.2 | 0.27 | 0.02 | 0.39 | 0.17 |
| 8 | 103697783 | rs11986282 | -0.927 | 0.501 | -0.324 | 1.08 | 0 | -0.077 | -0.077 | 0 | 0.13 | 0.22 | 0.06 | 0 | 0.12 | 0.12 |
| 8 | 103711907 | rs2511698 | -1.22 | 0.485 | -1.079 | 1.799 | -0.011 | -0.036 | -0.036 | 0.105 | 0.24 | 0.15 | 0.46 | 0.6 | 0.28 | 0.41 |
| 8 | 103714346 | rs2513929 | -1.183 | 0.45 | -0.737 | 2.181 | -0.001 | 0.032 | 0.032 | 0 | 0.2 | 0.29 | 0.3 | 0.37 | 0.2 | 0.29 |
| 8 | 103715226 | rs2511694 | -1.471 | 0.047 | -0.905 | 1.933 | 0.005 | -0.002 | -0.002 | 0 | 0.34 | 0.61 | 0.27 | 0.61 | 0.33 | 0.62 |
| 8 | 103718551 | rs4734660 | -0.778 | 0.262 | -1.062 | 2.026 | -0.001 | 0.026 | 0.026 | 0 | 0.07 | 0.11 | 0.1 | 0.18 | 0.07 | 0.32 |
| 8 | 103720185 | rs13249179 | 1.111 | 0.128 | -0.934 | 2.315 | -0.011 | 0.07 | 0.07 | 0 | 0.85 | 0.79 | 0.86 | 0.6 | 0.9 | 0.84 |
| 8 | 103724898 | rs13278114 | 0.584 | -0.169 | -1.674 | 1.3 | 0.055 | 0.024 | 0.024 | 0 | 0.54 | 0.68 | 0.64 | 0.36 | 0.66 | 0.43 |
| 8 | 103726919 | rs1574480 | -0.129 | 0.462 | -1.955 | 1.354 | -0.012 | -0.016 | -0.016 | 0 | 0.1 | 0.05 | 0.32 | 0.29 | 0.16 | 0.1 |
| 8 | 103727429 | rs2513931 | 0.37 | -0.118 | -1.686 | 1.387 | -0.015 | 0.009 | 0.009 | 0 | 0.91 | 0.82 | 0.91 | 0.83 | 0.94 | 0.66 |
| 8 | 103729146 | rs2511690 | 0.875 | -0.119 | -1.736 | 1.501 | -0.002 | 0.002 | 0.002 | 0 | 0.84 | 0.78 | 0.8 | 0.71 | 0.85 | 0.56 |
| 8 | 103730189 | rs2511688 | 0.094 | 0.231 | -1.89 | 1.39 | -0.004 | -0.113 | -0.113 | 0 | 0.27 | 0.04 | 0.18 | 0.38 | 0.2 | 0.13 |
| 8 | 103730879 | rs11786065 | -0.847 | 0.783 | -1.116 | 1.041 | -0.016 | -0.039 | -0.039 | 0 | 0.07 | 0.01 | 0.11 | 0.13 | 0.08 | 0.1 |
| 8 | 103732048 | rs4734663 | -1.201 | 0.942 | -0.272 | 0.9 | 0.012 | 0.019 | 0.019 | 0 | 0.21 | 0.38 | 0.47 | 0.21 | 0.23 | 0.18 |
| 8 | 103732069 | rs4734664 | -1.606 | 1.205 | 0.284 | 0.897 | 0.027 | 0.022 | 0.022 | 0 | 0.25 | 0.54 | 0.51 | 0.24 | 0.3 | 0.22 |
| 8 | 103732321 | rs7010198 | -1.216 | 1.062 | 0.991 | 0.916 | 0.001 | 0.048 | 0.048 | 0 | 0.19 | 0.36 | 0.22 | 0.2 | 0.22 | 0.22 |
| 8 | 103733632 | rs9642793 | -1.24 | 1.143 | 0.95 | 0.703 | 0 | -0.007 | -0.007 | 0 | 0.1 | 0.14 | 0.29 | 0.1 | 0.11 | 0.17 |
| 8 | 103734106 | rs9642982 | -0.427 | 1.112 | 0.943 | 0.699 | 0.002 | 0.004 | 0.004 | 0 | 0.07 | 0.11 | 0.28 | 0.1 | 0.09 | 0.04 |

| | | | | | | | | | | | | | | | | |
|---|---|---|---|---|---|---|---|---|---|---|---|---|---|---|---|---|
| 8 | 103735201 | rs2513940 | -1.319 | 1.16 | 0.739 | -0.28 | 0.029 | 0.025 | 0.025 | 2.028 | 0.55 | 0.71 | 0.77 | 0.34 | 0.68 | 0.62 |
| 8 | 103735424 | rs2513941 | 1.129 | 1.291 | 0.911 | -0.532 | -0.003 | 0.006 | 0.006 | 1.703 | 0.66 | 0.43 | 0.42 | 0.67 | 0.63 | 0.57 |
| 8 | 103738201 | rs2436863 | -0.639 | 0.819 | 0.582 | -0.879 | -0.003 | -0.008 | -0.008 | 0.45 | 0.35 | 0.31 | 0.47 | 0.14 | 0.43 | 0.28 |
| 8 | 103741756 | rs871561 | 0.933 | 0.91 | 0.307 | -0.862 | -0.052 | 0.064 | 0.064 | 0 | 0.2 | 0.08 | 0.17 | 0.02 | 0.31 | 0.19 |
| 8 | 103742658 | rs13274698 | -0.481 | 0.87 | 0.351 | -0.858 | -0.004 | 0.003 | 0.003 | 1.575 | 0.27 | 0.43 | 0.31 | 0.22 | 0.28 | 0.38 |
| 8 | 103744282 | rs1469764 | 0.899 | 0.981 | 0.024 | -0.876 | -0.018 | 0.005 | 0.005 | 0 | 0.21 | 0.14 | 0.17 | 0.02 | 0.3 | 0.2 |
| 8 | 103748538 | rs17188664 | 0.107 | 0.885 | -0.222 | -0.666 | 0.001 | 0.008 | 0.008 | 0 | 0.12 | 0.04 | 0.28 | 0.12 | 0.11 | 0.09 |
| 8 | 103750385 | rs11988192 | -0.504 | 0.82 | 0.138 | -0.465 | 0.008 | 0.021 | 0.021 | 1.067 | 0.24 | 0.38 | 0.33 | 0.24 | 0.29 | 0.25 |
| 8 | 103754111 | rs10955290 | -0.003 | 1.054 | -0.155 | -0.1 | -0.003 | -0.008 | -0.008 | 0 | 0.26 | 0.25 | 0.28 | 0.34 | 0.3 | 0.26 |
| 8 | 103755301 | rs2513892 | -0.808 | 0.717 | -0.384 | 0.239 | 0.003 | -0.014 | -0.014 | 0 | 0.65 | 0.74 | 0.77 | 0.48 | 0.7 | 0.73 |
| 8 | 103769678 | rs2513896 | -0.836 | 0.973 | -0.335 | 0.371 | -0.001 | -0.001 | -0.001 | 0 | 0.34 | 0.66 | 0.48 | 0.39 | 0.36 | 0.43 |
| 8 | 103770205 | rs871187 | -0.578 | 1.09 | 0.481 | 0.514 | 0.022 | 0.033 | 0.033 | 0.265 | 0.19 | 0.36 | 0.17 | 0.24 | 0.2 | 0.26 |
| 8 | 103775175 | rs10505035 | -0.801 | 0.927 | 0.312 | 0.239 | 0.001 | 0.008 | 0.008 | 0.332 | 0.24 | 0.19 | 0.43 | 0.21 | 0.24 | 0.35 |
| 8 | 103778809 | rs7841186 | 0.026 | 0.923 | 0.581 | 0.321 | 0.014 | 0.028 | 0.028 | 0.464 | 0.06 | 0.15 | 0.05 | 0.13 | 0.09 | 0.01 |
| 8 | 103780586 | rs10100732 | -1.79 | 0.625 | 1.007 | 0.056 | 0.002 | -0.007 | -0.007 | 0 | 0.13 | 0.57 | 0.08 | 0.13 | 0.11 | 0.02 |
| 8 | 103782719 | rs1529796 | -1.512 | 0.61 | 0.623 | 0.075 | 0.015 | 0.016 | 0.016 | 0 | 0.35 | 0.79 | 0.48 | 0.34 | 0.32 | 0.37 |
| 8 | 103787072 | rs2513897 | -1.323 | 0.238 | -0.335 | -0.643 | 0.013 | 0.096 | 0.096 | 0 | 0.68 | 0.9 | 0.81 | 0.91 | 0.66 | 0.69 |
| 8 | 103788755 | rs7009612 | -0.309 | 0.332 | 0.275 | -0.399 | 0.021 | 0.051 | 0.051 | 1.142 | 0.08 | 0.38 | 0.07 | 0.12 | 0.07 | 0.02 |
| 8 | 103793370 | rs1865336 | 0.14 | 0.442 | -0.379 | -0.202 | -0.011 | -0.028 | -0.028 | 0 | 0.08 | 0.02 | 0.34 | 0.22 | 0.11 | 0.25 |
| 8 | 103794864 | rs1991924 | -0.324 | 0.337 | -0.211 | -0.367 | -0.003 | 0.003 | 0.003 | 0 | 0.32 | 0.38 | 0.46 | 0.39 | 0.32 | 0.4 |
| 8 | 103798301 | rs2513899 | -1.313 | 0.348 | -0.715 | -0.62 | 0.011 | 0.074 | 0.074 | 0 | 0.69 | 0.88 | 0.83 | 0.9 | 0.69 | 0.76 |
| 8 | 103801557 | rs768878 | 0.94 | 0.79 | -0.791 | -0.186 | -0.001 | -0.005 | -0.005 | 0 | 0.26 | 0.31 | 0.21 | 0.14 | 0.23 | 0.16 |
| 8 | 103804097 | rs12680146 | -0.375 | 0.847 | -1.052 | 0.311 | -0.021 | -0.021 | -0.021 | 0 | 0.12 | 0.07 | 0.2 | 0.48 | 0.22 | 0.2 |
| 8 | 103805097 | rs2513905 | -0.018 | 0.712 | -0.915 | 0.146 | 0 | -0.012 | -0.012 | 0 | 0.57 | 0.71 | 0.66 | 0.44 | 0.52 | 0.58 |
| 8 | 103806914 | rs2436868 | -0.862 | -0.007 | -0.787 | -0.745 | 0.013 | 0.025 | 0.025 | 0 | 0.69 | 0.78 | 0.85 | 0.91 | 0.74 | 0.78 |
| 8 | 103809995 | rs10955293 | 0.118 | -0.292 | -1.279 | -0.645 | 0.024 | 0.008 | 0.008 | 0.538 | 0.33 | 0.14 | 0.23 | 0.32 | 0.23 | 0.25 |
| 8 | 103811092 | rs2436867 | -0.603 | -0.176 | -0.738 | -0.681 | 0.017 | 0 | 0 | 0 | 0.46 | 0.63 | 0.57 | 0.46 | 0.42 | 0.52 |
| 8 | 103826037 | rs2436854 | -0.1 | -0.29 | -0.202 | -0.867 | 0.002 | -0.001 | -0.001 | 0.645 | 0.39 | 0.4 | 0.25 | 0.39 | 0.28 | 0.43 |
| 8 | 103830692 | rs2513920 | 0.631 | -0.503 | -0.305 | -1.183 | -0.019 | 0.051 | 0.051 | 0.197 | 0.29 | 0.11 | 0.16 | 0.16 | 0.26 | 0.4 |
| 8 | 103838004 | rs10112564 | 0.684 | -0.501 | -0.232 | -1.149 | -0.034 | 0.075 | 0.075 | 0.018 | 0.08 | 0.28 | 0.04 | 0.21 | 0.01 | 0.03 |
| 8 | 103845372 | rs2304347 | 2.297 | -0.513 | -0.263 | -1.149 | 0.024 | -0.027 | -0.027 | 0 | 0.05 | 0 | 0.02 | 0.21 | 0.01 | 0.03 |
| 8 | 103845526 | rs2304348 | 0.79 | -0.571 | -0.407 | -1.267 | -0.027 | 0.003 | 0.003 | 0 | 0.07 | 0.01 | 0.27 | 0.04 | 0.12 | 0.07 |
| 8 | 103851052 | rs1062048 | 0.803 | -0.585 | -0.41 | -1.27 | -0.024 | 0.048 | 0.048 | 0.345 | 0.29 | 0.06 | 0.16 | 0.15 | 0.26 | 0.4 |
| 8 | 103855691 | rs2304344 | -0.236 | -0.431 | -0.348 | -1.309 | -0.006 | 0.005 | 0.005 | 0 | 0.09 | 0.1 | 0.31 | 0.07 | 0.17 | 0.08 |
| 8 | 103857316 | rs2570940 | -1.429 | -0.366 | -0.571 | -1.899 | -0.032 | -0.056 | -0.056 | 0 | 0.7 | 0.88 | 0.63 | 0.43 | 0.62 | 0.64 |
| 8 | 103857774 | rs6991188 | 1.865 | -0.336 | -0.561 | -1.892 | 0.002 | -0.14 | -0.14 | 0 | 0.22 | 0.38 | 0.11 | 0 | 0.17 | 0.13 |
| 8 | 103864271 | rs1019976 | 0.995 | -0.212 | -0.665 | -1.597 | -0.023 | 0.048 | 0.048 | 0.345 | 0.29 | 0.06 | 0.16 | 0.15 | 0.26 | 0.4 |
| 8 | 103871206 | rs2679758 | -0.739 | -0.23 | -0.561 | -1.597 | 0.003 | 0.005 | 0.005 | 0 | 0.47 | 0.47 | 0.52 | 0.43 | 0.45 | 0.51 |
| 8 | 103876325 | rs2679745 | 1.018 | -0.43 | -0.558 | -1.546 | 0.01 | 0.029 | 0.029 | 0.58 | 0.26 | 0.06 | 0.12 | 0.15 | 0.2 | 0.36 |
| 8 | 103877434 | rs1055376 | -0.608 | -0.426 | -0.527 | -1.552 | -0.011 | -0.034 | -0.034 | 0 | 0.4 | 0.49 | 0.47 | 0.22 | 0.38 | 0.44 |
| 8 | 103885884 | rs2436856 | 0.663 | -0.445 | -0.523 | -1.555 | 0.022 | 0.036 | 0.036 | 0.275 | 0.29 | 0.17 | 0.13 | 0.15 | 0.2 | 0.36 |
| 8 | 103894326 | rs4734674 | 1.742 | -0.413 | -0.54 | -1.546 | -0.019 | 0.021 | 0.021 | 0.52 | 0.05 | 0 | 0.26 | 0 | 0.08 | 0.02 |
| 8 | 103912514 | rs678839 | -0.09 | -0.404 | -0.537 | -1.536 | -0.012 | -0.022 | -0.022 | 0 | 0.34 | 0.41 | 0.2 | 0.22 | 0.3 | 0.43 |
| 8 | 103914746 | rs601588 | -0.773 | -0.397 | -0.595 | -1.507 | -0.007 | -0.003 | -0.003 | 0.047 | 0.46 | 0.5 | 0.49 | 0.43 | 0.39 | 0.46 |
| 8 | 103916392 | rs10505039 | 1.563 | -0.225 | -1.33 | -1.422 | -0.022 | 0.084 | 0.084 | 0.698 | 0.17 | 0.01 | 0.05 | 0.02 | 0.12 | 0.07 |
| 8 | 103926036 | rs581610 | 0.814 | -0.064 | -1.097 | -1.232 | 0.019 | 0.036 | 0.036 | 0.37 | 0.28 | 0.13 | 0.12 | 0.15 | 0.2 | 0.36 |
| 8 | 103928940 | rs608236 | -1.145 | 0.007 | -1.35 | -1.295 | -0.136 | -0.132 | -0.132 | 1.021 | 0.7 | 0.97 | 0.6 | 0.44 | 0.56 | 0.6 |
| 8 | 103940088 | rs529802 | 0.866 | -0.229 | -1.081 | -1.341 | 0.005 | 0.009 | 0.009 | 0 | 0.28 | 0.23 | 0.13 | 0.19 | 0.24 | 0.37 |
| 8 | 103940219 | rs6468847 | 0.927 | -0.368 | -0.706 | -1.506 | -0.021 | -0.056 | -0.056 | 0.111 | 0.3 | 0.49 | 0.14 | 0.19 | 0.18 | 0.17 |
| 8 | 103940336 | rs587946 | -1.224 | -0.086 | -0.972 | -1.597 | -0.02 | -0.036 | -0.036 | 1.339 | 0.72 | 0.85 | 0.63 | 0.42 | 0.62 | 0.62 |
| 8 | 103943414 | rs6998032 | -0.107 | -0.015 | -1.507 | -1.554 | -0.053 | 0.039 | 0.039 | 0 | 0.13 | 0.01 | 0.35 | 0.05 | 0.2 | 0.09 |
| 8 | 103945860 | rs7841955 | -0.31 | -0.069 | -1.428 | -1.579 | 0.009 | 0.071 | 0.071 | 0 | 0.44 | 0.68 | 0.51 | 0.23 | 0.39 | 0.25 |
| 8 | 103950997 | rs571241 | 0.912 | -0.135 | -1.499 | -1.46 | 0.008 | 0.017 | 0.017 | 0 | 0.28 | 0.18 | 0.13 | 0.19 | 0.24 | 0.37 |
| 8 | 103953319 | rs1533182 | 0.141 | -0.111 | -1.484 | -1.404 | -0.045 | 0.058 | 0.058 | 0.207 | 0.11 | 0.01 | 0.33 | 0.01 | 0.17 | 0.02 |
| 8 | 103953803 | rs650884 | 0.069 | -0.205 | -1.363 | -1.311 | -0.008 | 0.076 | 0.076 | 0.445 | 0.41 | 0.21 | 0.49 | 0.23 | 0.43 | 0.46 |
| 8 | 103956525 | rs2668880 | -0.216 | -0.044 | -1.24 | -1.231 | -0.004 | -0.019 | -0.019 | 0 | 0.46 | 0.46 | 0.52 | 0.23 | 0.43 | 0.48 |
| 8 | 103960710 | rs563201 | 0.64 | -0.308 | -0.681 | -0.456 | -0.003 | 0.069 | 0.069 | 0.002 | 0.49 | 0.31 | 0.51 | 0.24 | 0.44 | 0.53 |
| 8 | 103963878 | rs2458285 | 0.312 | -0.356 | -0.662 | -0.383 | -0.007 | 0.023 | 0.023 | 0 | 0.55 | 0.68 | 0.35 | 0.38 | 0.48 | 0.56 |
| 8 | 103967694 | rs4075572 | 0.339 | 0.395 | -0.852 | 0.062 | 0.002 | 0.03 | 0.03 | 0 | 0.52 | 0.36 | 0.54 | 0.26 | 0.49 | 0.53 |
| 8 | 103972485 | rs2460221 | -0.339 | -0.203 | -0.826 | 0.368 | -0.021 | -0.005 | -0.005 | 0.167 | 0.32 | 0.63 | 0.22 | 0.25 | 0.19 | 0.24 |
| 8 | 103972511 | rs2460222 | 0.153 | 0.376 | -1.027 | 0.916 | -0.019 | -0.03 | -0.03 | 0.194 | 0.88 | 0.95 | 0.78 | 0.45 | 0.77 | 0.78 |
| 8 | 103978699 | rs2458286 | 0.585 | -0.158 | -1.072 | 0.48 | 0.063 | 0.007 | 0.007 | 0 | 0.52 | 0.23 | 0.72 | 0.56 | 0.63 | 0.62 |
| 8 | 103979714 | rs2458287 | -0.37 | -0.196 | -1.259 | 0.605 | -0.02 | -0.073 | -0.073 | 0 | 0.83 | 0.94 | 0.88 | 0.61 | 0.79 | 0.77 |
| 8 | 103980407 | rs7827655 | -0.306 | -0.349 | -0.286 | 0.008 | -0.055 | -0.112 | -0.112 | 0 | 0.29 | 0.52 | 0.16 | 0.05 | 0.16 | 0.15 |
| 8 | 103992590 | rs556059 | -0.9 | 0.659 | -0.234 | 0.743 | 0.029 | 0.074 | 0.074 | 0 | 0.06 | 0.13 | 0.05 | 0.19 | 0.12 | 0.05 |
| 8 | 103993578 | rs525817 | -0.289 | 0.371 | -1.366 | -0.021 | -0.025 | 0.078 | 0.078 | 0 | 0.51 | 0.89 | 0.39 | 0.33 | 0.43 | 0.4 |
| 8 | 104003232 | rs11781860 | 0.876 | -0.059 | 0.065 | -0.341 | -0.005 | -0.019 | -0.019 | 0 | 0.3 | 0.32 | 0.21 | 0.13 | 0.21 | 0.27 |
| 8 | 104008681 | rs628793 | -0.872 | -0.031 | 0.123 | -0.314 | 0.055 | 0.134 | 0.134 | 0.756 | 0.11 | 0.45 | 0.09 | 0.16 | 0.14 | 0.06 |
| 8 | 104009564 | rs515024 | -0.802 | -0.132 | -0.257 | -0.4 | -0.047 | 0.05 | 0.05 | 0.06 | 0.53 | 0.9 | 0.4 | 0.34 | 0.45 | 0.4 |

| | | | | | | | | | | | | | | | | |
|---|---|---|---|---|---|---|---|---|---|---|---|---|---|---|---|---|
| 8 | 104012949 | rs543736 | 0.141 | -0.164 | -0.108 | -0.312 | -0.005 | -0.006 | -0.006 | 0 | 0.38 | 0.55 | 0.28 | 0.3 | 0.34 | 0.31 |
| 8 | 104032954 | rs11985451 | 0.102 | -0.267 | 0.039 | -0.31 | -0.009 | -0.01 | -0.01 | 1.912 | 0.38 | 0.47 | 0.25 | 0.3 | 0.33 | 0.32 |
| 8 | 104036135 | rs2048549 | -0.068 | -0.844 | -0.284 | -0.554 | -0.008 | -0.008 | -0.008 | 0 | 0.43 | 0.51 | 0.26 | 0.3 | 0.34 | 0.32 |
| 8 | 104037558 | rs11775734 | 0.548 | -0.797 | -0.282 | -0.557 | -0.01 | -0.032 | -0.032 | 0 | 0.34 | 0.43 | 0.22 | 0.14 | 0.25 | 0.27 |
| 8 | 104038155 | rs924211 | 1.294 | -0.215 | -0.172 | -0.486 | 0 | -0.003 | -0.003 | 0 | 0.05 | 0.06 | 0.02 | 0.03 | 0.04 | 0.02 |
| 8 | 104039314 | rs3808551 | 1.134 | -0.161 | -0.317 | -0.575 | -0.043 | 0.012 | 0.012 | 0 | 0.08 | 0.01 | 0.07 | 0.04 | 0.1 | 0.09 |
| 8 | 104040970 | rs7013362 | -0.204 | -0.203 | -0.26 | -0.585 | 0 | -0.004 | -0.004 | 0 | 0.47 | 0.54 | 0.33 | 0.34 | 0.43 | 0.41 |
| 8 | 104057065 | rs2458283 | -0.97 | -0.295 | -0.593 | -0.278 | 0.015 | -0.045 | -0.045 | 5.532 | 0.74 | 0.82 | 0.86 | 0.6 | 0.77 | 0.73 |
| 8 | 104062733 | rs2458290 | -0.236 | -0.324 | -0.622 | -0.337 | 0.025 | -0.016 | -0.016 | 0 | 0.38 | 0.44 | 0.62 | 0.33 | 0.46 | 0.41 |
| 8 | 104067005 | rs1398270 | 0.854 | -0.32 | -0.847 | -0.354 | -0.017 | 0.027 | 0.027 | 0 | 0.14 | 0.07 | 0.1 | 0.07 | 0.14 | 0.11 |
| 8 | 104069151 | rs2458291 | -1.017 | -0.349 | -0.847 | -0.339 | 0.021 | -0.051 | -0.051 | 0 | 0.74 | 0.81 | 0.86 | 0.6 | 0.78 | 0.73 |
| 8 | 104070145 | rs2515200 | -0.277 | -0.169 | -0.564 | 0.062 | -0.056 | 0.013 | 0.013 | 0.009 | 0.36 | 0.17 | 0.58 | 0.33 | 0.46 | 0.41 |
| 8 | 104070255 | rs2454043 | -1.675 | -0.466 | -0.918 | -0.271 | -0.036 | -0.041 | -0.041 | 0 | 0.59 | 0.9 | 0.8 | 0.6 | 0.61 | 0.6 |
| 8 | 104081996 | rs1158921 | -0.788 | 0.489 | -0.404 | 0.17 | 0.023 | 0.007 | 0.007 | 0 | 0.22 | 0.47 | 0.17 | 0.21 | 0.22 | 0.16 |
| 8 | 104082411 | rs4734684 | 0.773 | 0.485 | -0.425 | 0.168 | -0.036 | -0.021 | -0.021 | 2.053 | 0.22 | 0.08 | 0.48 | 0.26 | 0.32 | 0.3 |
| 8 | 104083651 | rs2453994 | -1.234 | 0.414 | 0.015 | 0.342 | 0.022 | 0.013 | 0.013 | 0 | 0.22 | 0.47 | 0.17 | 0.23 | 0.22 | 0.16 |
| 8 | 104083786 | rs16870033 | -0.285 | 0.459 | -0.141 | 0.367 | 0.06 | 0.131 | 0.131 | 0.635 | 0.07 | 0.32 | 0.07 | 0.12 | 0.08 | 0.05 |
| 8 | 104085444 | rs2453997 | -0.306 | 0.605 | -1.006 | 0.203 | -0.014 | 0.014 | 0.014 | 0.028 | 0.15 | 0.08 | 0.1 | 0.11 | 0.14 | 0.11 |
| 8 | 104088562 | rs2515206 | 0.071 | 0.896 | -0.664 | 0.531 | 0.015 | 0.03 | 0.03 | 0 | 0.83 | 0.75 | 0.62 | 0.75 | 0.75 | 0.7 |
| 8 | 104089482 | rs16870038 | 0.13 | 0.811 | -1.308 | 0.263 | -0.018 | -0.057 | -0.057 | 1.056 | 0.27 | 0.13 | 0.24 | 0.57 | 0.28 | 0.39 |
| 8 | 104090901 | rs7829540 | 0.821 | 0.798 | -1.367 | 0.71 | -0.014 | -0.034 | -0.034 | 0 | 0.15 | 0.1 | 0.16 | 0.46 | 0.16 | 0.3 |
| 8 | 104094370 | rs7819080 | 0.816 | 0.636 | -1.114 | 1.138 | -0.014 | -0.033 | -0.033 | 0.104 | 0.15 | 0.1 | 0.12 | 0.37 | 0.16 | 0.3 |
| 8 | 104097396 | rs2047375 | 0.884 | -0.024 | -1.041 | -0.714 | 0.002 | 0.003 | 0.003 | 0 | 0.33 | 0.3 | 0.49 | 0.23 | 0.3 | 0.44 |
| 8 | 104098515 | rs16870060 | 1.039 | 0.127 | -1.169 | -0.562 | -0.024 | -0.045 | -0.045 | 0.001 | 0.07 | 0.01 | 0.07 | 0.18 | 0.1 | 0.09 |
| 8 | 104098620 | rs1110115 | -0.332 | 0.095 | -1.066 | -0.964 | 0.009 | 0.008 | 0.008 | 0 | 0.58 | 0.45 | 0.69 | 0.49 | 0.59 | 0.56 |
| 8 | 104099607 | rs17201443 | 1.28 | 0.09 | -1.1 | -0.811 | -0.046 | -0.055 | -0.055 | 0 | 0.14 | 0.01 | 0.13 | 0.15 | 0.17 | 0.04 |
| 8 | 104099952 | rs13278305 | 1.768 | 0.048 | -1.043 | -1.037 | -0.008 | -0.062 | -0.062 | 0 | 0.28 | 0.26 | 0.15 | 0.01 | 0.16 | 0.19 |
| 8 | 104102910 | rs2454039 | -0.599 | 0.208 | -0.817 | -0.984 | 0.022 | -0.019 | -0.019 | 0.563 | 0.88 | 0.54 | 0.93 | 0.94 | 0.9 | 0.92 |
| 8 | 104108882 | rs2454015 | 1.644 | 0.211 | -1.181 | -1.035 | -0.027 | -0.039 | -0.039 | 0 | 0.19 | 0.1 | 0.29 | 0.44 | 0.27 | 0.24 |
| 8 | 104109811 | rs2515210 | 0.217 | -0.347 | -0.888 | -0.997 | 0.016 | -0.012 | -0.012 | 0 | 0.4 | 0.47 | 0.39 | 0.65 | 0.44 | 0.39 |
| 8 | 104110575 | rs13258732 | 2.077 | -0.433 | -1.406 | -1.05 | 0.017 | 0.156 | 0.156 | 0 | 0.23 | 0 | 0.06 | 0 | 0.12 | 0.16 |
| 8 | 104111500 | rs1982974 | 0.776 | 0.072 | -0.961 | -0.583 | -0.043 | -0.038 | -0.038 | 0 | 0.15 | 0.01 | 0.13 | 0.13 | 0.17 | 0.03 |
| 8 | 104113219 | rs7825005 | 0.459 | 0.043 | -0.79 | -0.559 | 0.001 | 0.001 | 0.001 | 0 | 0.16 | 0.17 | 0.13 | 0.13 | 0.16 | 0.05 |
| 8 | 104114304 | rs872586 | 0.836 | 0.072 | -0.926 | -0.522 | -0.001 | -0.002 | -0.002 | 0 | 0.14 | 0.1 | 0.13 | 0.13 | 0.15 | 0.03 |
| 8 | 104116112 | rs2454025 | 1.026 | 0.211 | -1.139 | -0.529 | -0.004 | -0.005 | -0.005 | 0 | 0.25 | 0.26 | 0.31 | 0.42 | 0.32 | 0.26 |
| 8 | 104117312 | rs4734686 | 1.156 | 0.225 | -1.188 | -0.534 | -0.013 | -0.017 | -0.017 | 0 | 0.07 | 0.03 | 0.27 | 0.21 | 0.13 | 0.32 |
| 8 | 104117437 | rs979671 | -0.052 | 0.346 | -1.116 | -0.502 | -0.011 | -0.004 | -0.004 | 0 | 0.4 | 0.35 | 0.45 | 0.55 | 0.49 | 0.29 |
| 8 | 104118184 | rs6983249 | 0.434 | 0.414 | -1.228 | -0.436 | -0.009 | -0.002 | -0.002 | 0 | 0.16 | 0.09 | 0.14 | 0.13 | 0.18 | 0.04 |
| 8 | 104121049 | rs16870106 | 0.729 | -0.005 | -1.14 | -0.544 | 0.001 | -0.002 | -0.002 | 0 | 0.15 | 0.15 | 0.04 | 0.17 | 0.1 | 0.12 |
| 8 | 104121210 | rs10111377 | 0.158 | -0.194 | -1.366 | -0.757 | 0.025 | 0.028 | 0.028 | 0 | 0.5 | 0.55 | 0.5 | 0.39 | 0.39 | 0.64 |
| 8 | 104121274 | rs2515173 | -0.324 | 0.39 | -1.403 | -0.608 | 0.059 | 0.168 | 0.168 | 0.711 | 0.5 | 0.18 | 0.39 | 0.73 | 0.57 | 0.4 |
| 8 | 104121862 | rs2047377 | -0.839 | 0.697 | -1.224 | -0.326 | -0.015 | 0.075 | 0.075 | 0 | 0.68 | 0.81 | 0.8 | 0.91 | 0.66 | 0.83 |
| 8 | 104122683 | rs964166 | 0.155 | 1.119 | -0.582 | 0.634 | 0.026 | 0.053 | 0.053 | 0 | 0.23 | 0.24 | 0.31 | 0.44 | 0.3 | 0.27 |
| 8 | 104122872 | rs1318739 | -0.344 | 0.329 | -0.72 | 0.377 | -0.013 | -0.001 | -0.001 | 0 | 0.2 | 0.28 | 0.33 | 0.23 | 0.15 | 0.38 |
| 8 | 104124629 | rs983674 | -0.164 | 0.705 | -1.176 | 0.275 | -0.015 | 0.061 | 0.061 | 0.042 | 0.53 | 0.79 | 0.54 | 0.63 | 0.46 | 0.45 |
| 8 | 104125084 | rs12677471 | 0.842 | 0.525 | -1.327 | -0.07 | 0.003 | 0.044 | 0.044 | 0 | 0.23 | 0.17 | 0.1 | 0.05 | 0.21 | 0.15 |
| 8 | 104125807 | rs1497455 | -0.639 | 0.512 | -1.129 | -0.063 | 0.002 | 0.025 | 0.025 | 0 | 0.49 | 0.6 | 0.52 | 0.63 | 0.43 | 0.41 |
| 8 | 104126852 | rs7832497 | 0.045 | 0.493 | -0.227 | 0.205 | 0.001 | 0.018 | 0.018 | 0 | 0.28 | 0.38 | 0.43 | 0.58 | 0.25 | 0.31 |
| 8 | 104127468 | rs9297330 | -0.507 | 0.588 | -0.638 | 0.511 | 0.017 | -0.063 | -0.063 | 1.411 | 0.29 | 0.83 | 0.41 | 0.43 | 0.24 | 0.24 |
| 8 | 104129007 | rs1391117 | 0.408 | 0.813 | -0.45 | 0.222 | 0 | -0.002 | -0.002 | 0 | 0.46 | 0.55 | 0.47 | 0.44 | 0.48 | 0.56 |
| 8 | 104133226 | rs3019159 | -1.246 | 0.645 | -0.889 | 0.399 | -0.01 | -0.071 | -0.071 | 0 | 0.86 | 0.95 | 0.73 | 0.6 | 0.88 | 0.86 |
| 8 | 104135844 | rs2454000 | -0.327 | 0.91 | -0.392 | 0.241 | 0.003 | 0.024 | 0.024 | 0 | 0.8 | 0.72 | 0.68 | 0.59 | 0.79 | 0.85 |
| 8 | 104135949 | rs7004923 | 0.017 | 0.845 | -0.013 | 0.059 | -0.004 | 0.026 | 0.026 | 0 | 0.32 | 0.19 | 0.22 | 0.16 | 0.34 | 0.28 |
| 8 | 104137518 | rs2454007 | 1.402 | 0.83 | -0.113 | -0.241 | -0.001 | -0.008 | -0.008 | 0 | 0.3 | 0.29 | 0.32 | 0.13 | 0.25 | 0.2 |
| 8 | 104138586 | rs2388493 | -0.31 | 1.116 | -0.491 | -0.288 | 0.005 | -0.001 | -0.001 | 0.053 | 0.47 | 0.57 | 0.35 | 0.49 | 0.5 | 0.65 |
| 8 | 104141425 | rs963599 | -0.174 | 1.176 | -0.659 | -0.246 | -0.018 | 0.015 | 0.015 | 0.081 | 0.67 | 0.71 | 0.56 | 0.28 | 0.61 | 0.52 |
| 8 | 104141937 | rs10955309 | 1.28 | 1.314 | -0.463 | 0.039 | 0.005 | 0.008 | 0.008 | 0 | 0.14 | 0.21 | 0.14 | 0.23 | 0.2 | 0.33 |
| 8 | 104143919 | rs1473541 | 0.005 | 0.994 | -0.509 | -0.258 | -0.002 | 0.002 | 0.002 | 0 | 0.64 | 0.68 | 0.6 | 0.37 | 0.7 | 0.69 |
| 8 | 104144178 | rs7829807 | 0.757 | 1.04 | -0.785 | -0.432 | -0.004 | -0.007 | -0.007 | 0.806 | 0.24 | 0.24 | 0.2 | 0.31 | 0.33 | 0.39 |
| 8 | 104146731 | rs3736042 | 1.763 | 1.139 | -0.76 | -0.396 | 0.014 | 0.06 | 0.06 | 0 | 0.12 | 0.21 | 0.14 | 0.3 | 0.2 | 0.27 |
| 8 | 104151423 | rs6999622 | 0.302 | 0.804 | -0.94 | -1.205 | -0.025 | 0.039 | 0.039 | 0 | 0.15 | 0.09 | 0.1 | 0 | 0.18 | 0.08 |
| 8 | 104154713 | rs2454013 | 0.892 | 0.675 | -0.683 | -1.3 | 0 | -0.001 | -0.001 | 0.446 | 0.45 | 0.5 | 0.53 | 0.56 | 0.51 | 0.55 |
| 8 | 104154965 | rs2454014 | 0.88 | 0.682 | -0.611 | -1.297 | 0.001 | -0.002 | -0.002 | 0.096 | 0.46 | 0.55 | 0.53 | 0.56 | 0.51 | 0.55 |
| 8 | 104156603 | rs2515190 | -0.646 | 0.567 | -0.447 | -1.223 | 0.034 | -0.029 | -0.029 | 0 | 0.67 | 0.79 | 0.67 | 0.58 | 0.77 | 0.65 |
| 8 | 104156874 | rs10113728 | 1.053 | 0.44 | -0.532 | -1.349 | -0.008 | -0.047 | -0.047 | 0.216 | 0.22 | 0.21 | 0.08 | 0 | 0.15 | 0.19 |
| 8 | 104159118 | rs1874091 | 1.828 | 0.553 | -0.429 | -1.246 | -0.017 | -0.05 | -0.05 | 0.854 | 0.3 | 0.4 | 0.34 | 0.15 | 0.25 | 0.23 |
| 8 | 104168865 | rs1845430 | -0.406 | 0.613 | -0.593 | -1.434 | -0.004 | -0.001 | -0.001 | 1.258 | 0.45 | 0.41 | 0.41 | 0.43 | 0.39 | 0.37 |
| 8 | 104171886 | rs6468861 | 0.417 | 0.313 | -1.023 | -1.695 | -0.019 | -0.029 | -0.029 | 0 | 0.33 | 0.26 | 0.28 | 0.43 | 0.44 | 0.43 |
| 8 | 104175576 | rs6468862 | -0.731 | 0.251 | -1.046 | -2.208 | -0.001 | -0.079 | -0.079 | 0 | 0.46 | 0.39 | 0.24 | 0.03 | 0.44 | 0.31 |

| | | | | | | | | | | | | | | | | |
|---|---|---|---|---|---|---|---|---|---|---|---|---|---|---|---|---|
| 8 | 104175861 | rs7009527 | 2.602 | 0.375 | -1.13 | -2.055 | -0.034 | 0.019 | 0.019 | 0.432 | 0.08 | 0 | 0.05 | 0.02 | 0.11 | 0.03 |
| 8 | 104176568 | rs6468864 | -2.239 | 0.345 | -1.036 | -2.003 | -0.002 | -0.019 | -0.019 | 0 | 0.79 | 0.69 | 0.7 | 0.87 | 0.83 | 0.81 |
| 8 | 104178577 | rs10955311 | 2.162 | 0.216 | -1.305 | -2.047 | -0.045 | 0.046 | 0.046 | 0 | 0.08 | 0.01 | 0.06 | 0 | 0.13 | 0.01 |
| 8 | 104180743 | rs7843452 | 2.133 | -0.012 | -0.998 | -2.096 | -0.041 | -0.054 | -0.054 | 4.152 | 0.1 | 0.17 | 0.03 | 0 | 0.02 | 0.03 |
| 8 | 104180922 | rs10481102 | 3.151 | 0.101 | -1.088 | -2.095 | -0.032 | 0.052 | 0.052 | 0 | 0.1 | 0 | 0.04 | 0.01 | 0.13 | 0.14 |
| 8 | 104181000 | rs1497456 | 2.129 | 0.228 | -0.679 | -1.809 | 0.007 | 0.037 | 0.037 | 0 | 0.19 | 0.23 | 0.37 | 0.49 | 0.29 | 0.29 |
| 8 | 104182696 | rs13258575 | -0.556 | 0.499 | -0.508 | -1.541 | 0.004 | -0.029 | -0.029 | 0 | 0.37 | 0.31 | 0.17 | 0.03 | 0.31 | 0.28 |
| 8 | 104183205 | rs10955312 | -1.819 | 0.403 | -0.558 | -1.526 | 0.006 | 0.014 | 0.014 | 0 | 0.62 | 0.49 | 0.47 | 0.43 | 0.56 | 0.58 |
| 8 | 104186211 | rs1000461 | 0.789 | 0.615 | -0.131 | -1.166 | -0.022 | 0.103 | 0.103 | 0.047 | 0.19 | 0.29 | 0.25 | 0.39 | 0.12 | 0.29 |
| 8 | 104187968 | rs954782 | 0.105 | 0.759 | -0.182 | -1.22 | 0.002 | 0.064 | 0.064 | 0 | 0.27 | 0.37 | 0.31 | 0.39 | 0.25 | 0.31 |
| 8 | 104189378 | rs12707776 | -0.661 | 0.688 | -0.409 | -1.254 | -0.049 | 0.131 | 0.131 | 0 | 0.33 | 0.13 | 0.19 | 0.03 | 0.4 | 0.26 |
| 8 | 104191982 | rs16870228 | 0.679 | 0.206 | -0.166 | -0.857 | 0 | 0.005 | 0.005 | 0.249 | 0.18 | 0.15 | 0.25 | 0.39 | 0.15 | 0.29 |
| 8 | 104194973 | rs1494272 | -1.235 | 0.282 | -0.023 | -0.953 | -0.006 | 0.019 | 0.019 | 0.105 | 0.3 | 0.57 | 0.33 | 0.4 | 0.24 | 0.39 |
| 8 | 104196139 | rs3956319 | 1.399 | 0.319 | 0.018 | -1.03 | 0.001 | 0.001 | 0.001 | 0 | 0.15 | 0.1 | 0.23 | 0.32 | 0.11 | 0.18 |
| 8 | 104196509 | rs4645535 | -0.25 | 0.406 | 0.256 | -0.91 | 0.064 | -0.044 | -0.044 | 0 | 0.09 | 0.36 | 0.07 | 0.01 | 0.09 | 0.08 |
| 8 | 104196713 | rs713368 | -1.323 | 0.522 | -0.317 | -0.915 | -0.001 | 0.013 | 0.013 | 0 | 0.51 | 0.6 | 0.46 | 0.4 | 0.49 | 0.55 |
| 8 | 104199801 | rs1032469 | -1.06 | 0.756 | -0.042 | -1.016 | 0.066 | -0.015 | -0.015 | 0 | 0.11 | 0.35 | 0.07 | 0.05 | 0.12 | 0.15 |
| 8 | 104200005 | rs16870240 | 0.583 | 0.696 | -0.047 | -0.99 | -0.005 | 0.005 | 0.005 | 0 | 0.05 | 0.11 | 0.02 | 0.05 | 0.04 | 0.13 |
| 8 | 104201401 | rs12548629 | 0.066 | 0.522 | -0.434 | -1.142 | 0.007 | -0.004 | -0.004 | 0 | 0.3 | 0.13 | 0.32 | 0.29 | 0.29 | 0.32 |
| 8 | 104205431 | rs16870249 | 1.169 | 0.49 | -0.633 | -1.219 | -0.028 | -0.094 | -0.094 | 1.824 | 0.16 | 0.01 | 0.22 | 0.28 | 0.13 | 0.18 |
| 8 | 104208330 | rs13270175 | -1.714 | 0.344 | -0.517 | -1.366 | -0.001 | 0.007 | 0.007 | 0.325 | 0.49 | 0.42 | 0.44 | 0.37 | 0.49 | 0.56 |
| 8 | 104213889 | rs7017690 | 2.419 | 0.274 | -0.739 | -1.584 | 0 | 0.005 | 0.005 | 0 | 0.11 | 0.07 | 0.04 | 0 | 0.08 | 0.02 |
| 8 | 104215466 | rs1157213 | -2.097 | 0.192 | -0.635 | -1.599 | 0.001 | 0.001 | 0.001 | 0.004 | 0.62 | 0.55 | 0.49 | 0.37 | 0.57 | 0.57 |
| 8 | 104217953 | rs13439041 | 1.126 | 0.595 | -0.612 | -1.426 | -0.012 | 0.068 | 0.068 | 0 | 0.14 | 0.02 | 0.08 | 0.01 | 0.16 | 0.16 |
| 8 | 104222025 | rs2201369 | 0.468 | 0.654 | -0.636 | -1.426 | 0.003 | 0.032 | 0.032 | 0 | 0.16 | 0.04 | 0.07 | 0.07 | 0.17 | 0.26 |
| 8 | 104226946 | rs7001084 | -1.331 | 0.883 | -0.445 | -1.388 | -0.001 | 0.001 | 0.001 | 1.978 | 0.47 | 0.41 | 0.44 | 0.36 | 0.5 | 0.55 |
| 8 | 104229793 | rs11782632 | 0.665 | 0.56 | -0.61 | -1.353 | -0.001 | 0.025 | 0.025 | 0.019 | 0.15 | 0.05 | 0.07 | 0.07 | 0.17 | 0.25 |
| 8 | 104231944 | rs9656787 | 0.244 | 0.649 | -0.49 | -1.339 | -0.003 | -0.005 | -0.005 | 0 | 0.23 | 0.18 | 0.28 | 0.29 | 0.21 | 0.28 |
| 8 | 104238182 | rs10107615 | -0.072 | -0.045 | -0.54 | -1.381 | 0.009 | 0.06 | 0.06 | 0 | 0.27 | 0.16 | 0.14 | 0.07 | 0.26 | 0.28 |
| 8 | 104244309 | rs12543038 | 1.258 | 0.015 | -0.669 | -1.348 | -0.021 | -0.096 | -0.096 | 0 | 0.16 | 0.01 | 0.23 | 0.28 | 0.12 | 0.18 |
| 8 | 104249839 | rs11785583 | 0.537 | 0.051 | -0.638 | -1.359 | -0.007 | 0.093 | 0.093 | 0 | 0.22 | 0.09 | 0.11 | 0.01 | 0.22 | 0.15 |
| 8 | 104256725 | rs4734695 | -0.736 | 0.027 | -0.316 | -1.351 | 0.005 | 0.01 | 0.01 | 0 | 0.45 | 0.35 | 0.37 | 0.27 | 0.38 | 0.35 |
| 8 | 104278419 | rs3104033 | -1.543 | 0.181 | -0.443 | -1.275 | -0.011 | -0.001 | -0.001 | 0.19 | 0.6 | 0.61 | 0.47 | 0.34 | 0.52 | 0.56 |
| 8 | 104281357 | rs2220826 | -0.406 | 0.173 | -0.791 | -1.385 | 0.004 | 0.066 | 0.066 | 0.673 | 0.35 | 0.23 | 0.35 | 0.17 | 0.34 | 0.3 |
| 8 | 104287845 | rs1522707 | 0.526 | 0.393 | -0.75 | -1.257 | 0.002 | 0 | 0 | 0 | 0.2 | 0.11 | 0.25 | 0.2 | 0.16 | 0.21 |
| 8 | 104290148 | rs6990501 | 1.364 | 0.491 | -0.861 | -1.1 | -0.006 | 0.019 | 0.019 | 0 | 0.07 | 0.03 | 0.04 | 0.01 | 0.1 | 0.1 |
| 8 | 104292965 | rs827541 | -1.179 | 0.526 | -0.574 | -1.054 | -0.016 | 0.024 | 0.024 | 0.228 | 0.6 | 0.65 | 0.47 | 0.27 | 0.53 | 0.56 |
| 8 | 104299426 | rs827540 | -0.177 | 0.398 | -0.624 | -1.267 | 0.005 | 0.046 | 0.046 | 0 | 0.33 | 0.24 | 0.17 | 0.07 | 0.31 | 0.3 |
| 8 | 104299936 | rs1095713 | -1.074 | 0.31 | -0.339 | -1.063 | -0.006 | 0.001 | 0.001 | 0.144 | 0.56 | 0.58 | 0.44 | 0.36 | 0.48 | 0.59 |
| 8 | 104312432 | rs827528 | 0.187 | 0.172 | -0.19 | -1.189 | 0.005 | 0.043 | 0.043 | 0 | 0.3 | 0.22 | 0.17 | 0.01 | 0.27 | 0.17 |
| 8 | 104322866 | rs16870322 | 0.882 | 0.197 | -0.148 | -1.219 | -0.009 | 0.006 | 0.006 | 0 | 0.05 | 0.11 | 0.02 | 0.06 | 0.04 | 0.13 |
| 8 | 104323026 | rs827536 | -1.254 | 0.245 | -0.168 | -1.215 | -0.015 | 0.002 | 0.002 | 0.266 | 0.57 | 0.64 | 0.44 | 0.36 | 0.48 | 0.58 |
| 8 | 104328309 | rs2131860 | 0.599 | 0.265 | -0.353 | -1.168 | -0.014 | -0.077 | -0.077 | 1.849 | 0.19 | 0.06 | 0.24 | 0.29 | 0.17 | 0.28 |
| 8 | 104328590 | rs2131858 | 1.623 | 0.157 | -0.661 | -1.256 | -0.089 | -0.127 | -0.127 | 0 | 0.25 | 0.1 | 0.28 | 0.56 | 0.36 | 0.31 |
| 8 | 104333838 | rs703810 | 1.502 | 0.461 | -0.26 | -1.198 | 0.001 | 0.073 | 0.073 | 0 | 0.16 | 0.05 | 0.07 | 0 | 0.12 | 0.05 |
| 8 | 104335413 | rs2948451 | 1.485 | 0.459 | -0.254 | -1.199 | 0.003 | 0.069 | 0.069 | 0 | 0.16 | 0.06 | 0.07 | 0 | 0.12 | 0.05 |
| 8 | 104337096 | rs3808554 | 1.259 | 0.442 | 0.068 | -0.984 | -0.026 | -0.012 | -0.012 | 1.555 | 0.39 | 0.3 | 0.51 | 0.64 | 0.46 | 0.41 |
| 8 | 104337367 | rs3808553 | 1.277 | 0.444 | 0.07 | -0.98 | -0.026 | -0.012 | -0.012 | 0 | 0.39 | 0.3 | 0.51 | 0.64 | 0.46 | 0.4 |
| 8 | 104343686 | rs1053917 | 1.356 | 0.418 | 0.108 | -0.992 | -0.032 | -0.017 | -0.017 | 0.263 | 0.38 | 0.28 | 0.5 | 0.63 | 0.45 | 0.39 |
| 8 | 104344620 | rs7827146 | 0.383 | 0.518 | 0.277 | -0.923 | 0.013 | 0.037 | 0.037 | 0 | 0.06 | 0.21 | 0.03 | 0.06 | 0.04 | 0.13 |
| 8 | 104345778 | rs1403294 | 0.072 | 0.642 | 0.029 | -0.818 | 0.002 | 0.047 | 0.047 | 0 | 0.32 | 0.24 | 0.17 | 0.11 | 0.3 | 0.27 |
| 8 | 104353333 | rs951826 | -1.178 | 0.633 | -0.185 | -0.844 | -0.005 | 0.074 | 0.074 | 0 | 0.76 | 0.88 | 0.87 | 0.99 | 0.77 | 0.85 |
| 8 | 104356037 | rs827577 | -0.181 | 0.64 | -0.198 | -0.835 | -0.048 | 0.084 | 0.084 | 0 | 0.63 | 0.84 | 0.66 | 0.8 | 0.63 | 0.66 |
| 8 | 104358312 | rs827584 | 0.95 | 1.08 | -0.165 | -0.533 | -0.02 | -0.012 | -0.012 | 0.727 | 0.38 | 0.31 | 0.5 | 0.63 | 0.44 | 0.39 |
| 8 | 104363338 | rs3133836 | 0.497 | 1.736 | 0.379 | -0.189 | -0.021 | -0.033 | -0.033 | 0.012 | 0.09 | 0.01 | 0.2 | 0.21 | 0.1 | 0.19 |
| 8 | 104365116 | rs16870364 | -0.31 | 1.549 | 0.635 | -0.399 | -0.002 | -0.014 | -0.014 | 0 | 0.35 | 0.59 | 0.21 | 0.13 | 0.31 | 0.22 |
| 8 | 104365623 | rs844137 | 1.309 | 2.049 | 0.321 | -0.131 | 0.008 | -0.018 | -0.018 | 0 | 0.45 | 0.29 | 0.47 | 0.57 | 0.42 | 0.57 |
| 8 | 104367673 | rs2575700 | 0.426 | 1.91 | 0.921 | 0.031 | 0.014 | 0.037 | 0.037 | 1.818 | 0.86 | 0.8 | 0.75 | 0.77 | 0.82 | 0.83 |
| 8 | 104369485 | rs719842 | 0.157 | 1.906 | 1.087 | 0.03 | 0.018 | 0.078 | 0.078 | 0.064 | 0.1 | 0.18 | 0.2 | 0.23 | 0.11 | 0.16 |
| 8 | 104373506 | rs10105450 | 0.382 | 1.899 | 1.045 | -0.181 | 0.002 | -0.018 | -0.018 | 0.076 | 0.26 | 0.25 | 0.16 | 0.01 | 0.24 | 0.12 |
| 8 | 104388574 | rs7845210 | 0.145 | 1.94 | 0.777 | -0.267 | -0.018 | -0.158 | -0.158 | 0 | 0.28 | 0.46 | 0.18 | 0.01 | 0.24 | 0.12 |
| 8 | 104394744 | rs3098233 | 1.083 | 2.003 | 0.182 | -0.152 | 0.002 | 0.017 | 0.017 | 3.484 | 0.81 | 0.79 | 0.73 | 0.67 | 0.77 | 0.74 |
| 8 | 104400808 | rs3133803 | -0.483 | 2.028 | 0.165 | -0.126 | -0.006 | -0.004 | -0.004 | 0 | 0.13 | 0.07 | 0.23 | 0.14 | 0.15 | 0.18 |
| 8 | 104404392 | rs4512347 | -0.17 | 2.028 | 0.171 | -0.115 | 0.009 | 0.073 | 0.073 | 0 | 0.05 | 0.09 | 0.04 | 0.19 | 0.08 | 0.08 |
| 8 | 104408510 | rs17228538 | 0.459 | 2.02 | 0.18 | -0.177 | -0.002 | -0.053 | -0.053 | 0.488 | 0.27 | 0.29 | 0.16 | 0.01 | 0.24 | 0.12 |
| 8 | 104410928 | rs1061196 | 0.459 | 2.02 | 0.18 | -0.177 | -0.002 | -0.053 | -0.053 | 0 | 0.27 | 0.29 | 0.16 | 0.01 | 0.24 | 0.12 |
| 8 | 104412388 | rs2241777 | -1.158 | 2.031 | 0.158 | -0.172 | 0 | -0.003 | -0.003 | 0 | 0.46 | 0.5 | 0.43 | 0.35 | 0.47 | 0.37 |
| 8 | 104415118 | rs3133810 | 0.673 | 2.027 | 0.138 | -0.17 | -0.029 | -0.042 | -0.042 | 0 | 0.08 | 0 | 0.18 | 0.14 | 0.09 | 0.16 |
| 8 | 104423486 | rs3134287 | -1.252 | 1.87 | 0.233 | -0.469 | -0.004 | 0.023 | 0.023 | 2.276 | 0.48 | 0.68 | 0.45 | 0.35 | 0.48 | 0.38 |

| | | | | | | | | | | | | | | | | |
|---|---|---|---|---|---|---|---|---|---|---|---|---|---|---|---|---|
| 8 | 104425304 | rs3098241 | -0.06 | 2.102 | -0.748 | -0.358 | -0.009 | -0.01 | -0.01 | 0 | 0.14 | 0.12 | 0.22 | 0.33 | 0.17 | 0.24 |
| 8 | 104427359 | rs3134295 | -1.14 | 2.102 | -0.771 | -0.358 | 0 | -0.003 | -0.003 | 0.385 | 0.46 | 0.5 | 0.43 | 0.35 | 0.47 | 0.37 |
| 8 | 104427541 | rs3134296 | -0.198 | 2.102 | -0.769 | -0.352 | 0.001 | 0.028 | 0.028 | 0.003 | 0.14 | 0.16 | 0.23 | 0.33 | 0.17 | 0.24 |
| 8 | 104428530 | rs750606 | 0.119 | 2.119 | -0.822 | -0.344 | -0.025 | -0.045 | -0.045 | 0 | 0.13 | 0.07 | 0.22 | 0.33 | 0.17 | 0.24 |
| 8 | 104452199 | rs3098218 | -1.012 | 2.014 | -0.803 | -0.263 | -0.001 | -0.002 | -0.002 | 0 | 0.47 | 0.5 | 0.45 | 0.34 | 0.52 | 0.38 |
| 8 | 104452281 | rs3098217 | -0.938 | 1.895 | -0.923 | 0.881 | 0 | 0.011 | 0.011 | 3.357 | 0.19 | 0.2 | 0.27 | 0.33 | 0.23 | 0.25 |
| 8 | 104467990 | rs7821398 | -0.433 | 2.399 | -1.155 | 0.243 | -0.046 | 0.039 | 0.039 | 0 | 0.07 | 0.01 | 0.09 | 0 | 0.17 | 0.05 |
| 8 | 104471749 | rs1898254 | -0.291 | 2.556 | -1.671 | -0.179 | -0.007 | 0.024 | 0.024 | 2.426 | 0.22 | 0.11 | 0.13 | 0.13 | 0.21 | 0.19 |
| 8 | 104487840 | rs2437769 | -1.659 | 1.671 | -0.718 | -0.788 | -0.014 | -0.006 | -0.006 | 0 | 0.84 | 0.99 | 0.89 | 0.88 | 0.82 | 0.7 |
| 8 | 104490503 | rs1347603 | -0.766 | 1.172 | -0.846 | -1.614 | 0.02 | 0.087 | 0.087 | 0 | 0.69 | 0.91 | 0.84 | 0.88 | 0.71 | 0.64 |
| 8 | 104572771 | rs2449911 | 1.131 | 0.995 | 0.022 | -1.798 | 0.006 | 0.028 | 0.028 | 0.16 | 0.34 | 0.21 | 0.56 | 0.76 | 0.32 | 0.39 |
| 8 | 104602790 | rs1365287 | -0.146 | 1.033 | 0.216 | -1.809 | -0.001 | -0.011 | -0.011 | 0.02 | 0.59 | 0.62 | 0.73 | 0.87 | 0.54 | 0.58 |
| 8 | 104607119 | rs7813485 | 1.115 | 1.032 | 0.258 | -1.807 | -0.01 | 0.022 | 0.022 | 2.478 | 0.22 | 0.13 | 0.14 | 0.11 | 0.22 | 0.2 |
| 8 | 104614003 | rs2469970 | 1.461 | 0.713 | 0.19 | -1.482 | -0.007 | -0.063 | -0.063 | 1.24 | 0.29 | 0.12 | 0.43 | 0.62 | 0.27 | 0.31 |
| 8 | 104634925 | rs17234011 | 1.265 | 0.898 | 0.076 | -1.734 | -0.02 | -0.003 | -0.003 | 0 | 0.06 | 0.02 | 0.04 | 0.05 | 0.09 | 0.08 |
| 8 | 104670180 | rs4734725 | 0.932 | 0.873 | 0.261 | -1.743 | 0.008 | 0.02 | 0.02 | 0 | 0.35 | 0.28 | 0.58 | 0.76 | 0.32 | 0.38 |
| 8 | 104672044 | rs10097250 | -0.353 | 0.88 | 0.177 | -1.757 | 0.021 | -0.041 | -0.041 | 0 | 0.32 | 0.41 | 0.25 | 0.12 | 0.39 | 0.25 |
| 8 | 104690593 | rs7834353 | 0.944 | 0.884 | 0.132 | -1.752 | 0.064 | -0.055 | -0.055 | 0 | 0.1 | 0.25 | 0.11 | 0 | 0.17 | 0.06 |
| 8 | 104723816 | rs1176675 | -1.491 | 0.946 | 0.179 | -1.782 | 0.038 | 0.101 | 0.101 | 0 | 0.7 | 0.96 | 0.85 | 0.88 | 0.74 | 0.64 |
| 8 | 104726802 | rs1521059 | 1.817 | 0.915 | 0.195 | -1.8 | 0.01 | 0.02 | 0.02 | 2.54 | 0.16 | 0.09 | 0.1 | 0.06 | 0.13 | 0.11 |
| 8 | 104729301 | rs17234986 | 1.319 | 0.931 | 0.19 | -1.796 | -0.02 | -0.003 | -0.003 | 0.027 | 0.06 | 0.02 | 0.04 | 0.05 | 0.09 | 0.08 |
| 8 | 104737956 | rs1593501 | -0.996 | 0.902 | 0.183 | -1.817 | -0.051 | 0.134 | 0.134 | 0 | 0.66 | 0.97 | 0.82 | 0.87 | 0.65 | 0.61 |
| 8 | 104745597 | rs1607910 | 0.155 | 0.855 | 0.192 | -1.826 | 0.007 | -0.031 | -0.031 | 0 | 0.28 | 0.36 | 0.22 | 0.12 | 0.31 | 0.23 |
| 8 | 104825219 | rs3110468 | 1.111 | 0.844 | 0.104 | -1.873 | 0.006 | 0.023 | 0.023 | 0.138 | 0.33 | 0.14 | 0.54 | 0.76 | 0.3 | 0.39 |
| 8 | 104834911 | rs4734731 | -0.202 | 0.859 | 0.081 | -1.873 | 0.064 | -0.068 | -0.068 | 0.002 | 0.15 | 0.26 | 0.15 | 0 | 0.28 | 0.1 |
| 8 | 104840500 | rs2388784 | 2.09 | 0.602 | -0.545 | -2.328 | -0.012 | -0.084 | -0.084 | 0.013 | 0.17 | 0.04 | 0.42 | 0.61 | 0.14 | 0.23 |
| 8 | 104882403 | rs924165 | 1.5 | 0.348 | -0.517 | -2.384 | 0.042 | 0.067 | 0.067 | 0 | 0.32 | 0.16 | 0.16 | 0.12 | 0.26 | 0.35 |
| 8 | 104901578 | rs10955332 | 1.642 | 0.158 | -0.521 | -2.503 | 0.003 | 0.005 | 0.005 | 2.457 | 0.2 | 0.16 | 0.09 | 0.11 | 0.17 | 0.17 |
| 8 | 104908955 | rs924164 | -1.994 | 0.369 | -0.679 | -2.481 | -0.051 | -0.12 | -0.12 | 11.775 | 0.82 | 0.98 | 0.55 | 0.4 | 0.81 | 0.75 |
| 8 | 104913292 | rs2132643 | 1.63 | 0.344 | -0.468 | -2.597 | -0.053 | 0.046 | 0.046 | 0 | 0.08 | 0.01 | 0.09 | 0 | 0.19 | 0.06 |
| 8 | 104919431 | rs10093057 | -0.298 | 0.349 | -0.307 | -2.625 | 0.018 | -0.042 | -0.042 | 0 | 0.36 | 0.42 | 0.23 | 0.11 | 0.44 | 0.25 |
| 8 | 104925755 | rs13273017 | -0.45 | 0.428 | -0.192 | -2.574 | 0.018 | 0.028 | 0.028 | 0.053 | 0.66 | 0.82 | 0.41 | 0.24 | 0.7 | 0.58 |
| 8 | 104927491 | rs16870826 | -0.879 | 0.409 | -0.249 | -2.546 | 0.032 | -0.016 | -0.016 | 0 | 0.11 | 0.33 | 0.06 | 0.05 | 0.09 | 0.08 |
| 8 | 104936546 | rs13259052 | 1.339 | -0.076 | -0.6 | -2.69 | 0.034 | 0.131 | 0.131 | 0 | 0.34 | 0.03 | 0.16 | 0.13 | 0.28 | 0.35 |
| 8 | 104950095 | rs17817357 | -0.039 | -0.004 | -0.169 | -2.694 | 0.021 | -0.019 | -0.019 | 0 | 0.28 | 0.32 | 0.3 | 0.11 | 0.43 | 0.23 |
| 8 | 104956705 | rs6983436 | -0.414 | -0.107 | -0.15 | -2.566 | 0.013 | 0.015 | 0.015 | 0 | 0.65 | 0.73 | 0.49 | 0.24 | 0.71 | 0.58 |
| 8 | 104985516 | rs10104167 | 0.611 | 0.28 | -0.373 | -2.249 | 0.005 | -0.013 | -0.013 | 0.827 | 0.19 | 0.24 | 0.21 | 0.06 | 0.24 | 0.13 |
| 8 | 105002995 | rs17239114 | 0.983 | 0.415 | -0.655 | -2.181 | -0.026 | 0.026 | 0.026 | 1.749 | 0.17 | 0.13 | 0.21 | 0.07 | 0.24 | 0.13 |
| 8 | 105023672 | rs12678653 | 1.762 | 0.239 | -0.619 | -2.025 | 0.02 | 0.003 | 0.003 | 0 | 0.07 | 0.01 | 0 | 0.05 | 0.01 | 0.05 |
| 8 | 105053383 | rs10086223 | 0.987 | 0.2 | -0.397 | -2.318 | -0.002 | -0.003 | -0.003 | 1.258 | 0.34 | 0.39 | 0.27 | 0.15 | 0.42 | 0.4 |
| 8 | 105065091 | rs4549737 | -0.424 | -0.266 | -0.159 | -2.31 | -0.003 | -0.004 | -0.004 | 0 | 0.69 | 0.66 | 0.52 | 0.26 | 0.7 | 0.62 |
| 8 | 105078342 | rs4734087 | -0.724 | -0.2 | 0.028 | -1.955 | 0 | 0.008 | 0.008 | 0 | 0.74 | 0.73 | 0.58 | 0.39 | 0.74 | 0.71 |
| 8 | 105078706 | rs13252107 | 1.469 | -0.109 | 0.225 | -1.87 | 0.001 | -0.001 | -0.001 | 0.928 | 0.28 | 0.34 | 0.17 | 0.12 | 0.23 | 0.37 |
| 8 | 105122037 | rs16870937 | 2.016 | -0.16 | 0.183 | -1.706 | 0.01 | 0.026 | 0.026 | 0 | 0.23 | 0.2 | 0.15 | 0.12 | 0.21 | 0.32 |
| 8 | 105122201 | rs7461951 | 0.235 | 0.122 | 0.283 | -1.732 | 0.005 | 0.05 | 0.05 | 0 | 0.17 | 0.22 | 0.33 | 0.59 | 0.19 | 0.25 |
| 8 | 105130066 | rs7000750 | 1.189 | 0.468 | 0.431 | -1.739 | -0.021 | -0.025 | -0.025 | 1.485 | 0.38 | 0.51 | 0.2 | 0.19 | 0.25 | 0.39 |
| 8 | 105140610 | rs11996084 | -0.191 | 0.702 | -0.126 | -1.01 | -0.082 | 0.209 | 0.209 | 0.653 | 0.6 | 0.89 | 0.63 | 0.92 | 0.47 | 0.77 |
| 8 | 105143482 | rs6468904 | 0.409 | 0.497 | 0.057 | -0.808 | 0.016 | 0.135 | 0.135 | 0 | 0.13 | 0.24 | 0.31 | 0.58 | 0.16 | 0.24 |
| 8 | 105153213 | rs3133284 | 0.265 | 0.497 | -0.093 | -0.632 | 0.028 | 0.172 | 0.172 | 0.155 | 0.55 | 0.77 | 0.59 | 0.92 | 0.41 | 0.75 |
| 8 | 105159724 | rs2441863 | 0.44 | 0.615 | 0.292 | -0.075 | 0.016 | 0.249 | 0.249 | 0 | 0.1 | 0.27 | 0.27 | 0.59 | 0.08 | 0.21 |
| 8 | 105160350 | rs10955341 | 0.601 | 0.879 | 0.144 | -0.427 | 0.03 | 0.032 | 0.032 | 0 | 0.67 | 0.54 | 0.54 | 0.42 | 0.6 | 0.65 |
| 8 | 105164011 | rs12549681 | 0.641 | 0.746 | -0.075 | -0.704 | -0.008 | 0.015 | 0.015 | 0.407 | 0.26 | 0.26 | 0.26 | 0.1 | 0.31 | 0.13 |
| 8 | 105168074 | rs13270802 | 1.547 | 0.656 | -0.712 | -0.859 | 0.044 | 0.063 | 0.063 | 0 | 0.37 | 0.2 | 0.2 | 0.17 | 0.31 | 0.44 |
| 8 | 105170013 | rs1596975 | -0.4 | 0.733 | -0.859 | -0.825 | -0.06 | 0.088 | 0.088 | 0 | 0.78 | 0.94 | 0.84 | 1 | 0.72 | 0.88 |
| 8 | 105174941 | rs1545370 | 1.436 | 0.444 | -0.782 | -0.659 | 0.048 | 0.057 | 0.057 | 0 | 0.38 | 0.16 | 0.23 | 0.22 | 0.33 | 0.45 |
| 8 | 105178819 | rs2511636 | -0.79 | 0.632 | -0.609 | -0.647 | -0.017 | 0.017 | 0.017 | 0 | 0.82 | 0.9 | 0.86 | 1 | 0.75 | 0.88 |
| 8 | 105185621 | rs4236772 | -0.263 | 0.574 | -0.018 | -0.181 | 0.008 | 0.002 | 0.002 | 0.006 | 0.35 | 0.56 | 0.5 | 0.64 | 0.37 | 0.34 |
| 8 | 105197239 | rs2511642 | 0.913 | 0.343 | -0.31 | -0.305 | 0.016 | 0.064 | 0.064 | 0 | 0.45 | 0.32 | 0.25 | 0.19 | 0.42 | 0.47 |
| 8 | 105211090 | rs2441905 | -1.804 | -0.233 | -0.458 | -0.469 | -0.03 | 0.001 | 0.001 | 0.155 | 0.67 | 0.92 | 0.71 | 0.72 | 0.64 | 0.53 |
| 8 | 105216321 | rs2511549 | -0.635 | -0.304 | -0.124 | -0.504 | 0.008 | 0.009 | 0.009 | 0 | 0.51 | 0.53 | 0.66 | 0.66 | 0.58 | 0.44 |
| 8 | 105217958 | rs17819924 | 1.264 | -0.303 | -0.043 | -0.495 | -0.021 | 0.028 | 0.028 | 0 | 0.19 | 0.09 | 0.25 | 0.11 | 0.25 | 0.1 |
| 8 | 105221758 | rs7018164 | 0.887 | -0.318 | -0.095 | -0.491 | -0.031 | -0.014 | -0.014 | 0.112 | 0.14 | 0.29 | 0.05 | 0.05 | 0.07 | 0.08 |
| 8 | 105226492 | rs2511558 | -0.805 | -0.307 | -0.123 | -0.479 | -0.001 | -0.002 | -0.002 | 0 | 0.35 | 0.36 | 0.25 | 0.22 | 0.29 | 0.27 |
| 8 | 105227985 | rs6468909 | 0.421 | -0.342 | -0.001 | -0.492 | -0.042 | 0.013 | 0.013 | 0 | 0.17 | 0.26 | 0.12 | 0.19 | 0.07 | 0.17 |
| 8 | 105229023 | rs895625 | -0.666 | -0.327 | -0.026 | -0.486 | -0.009 | -0.013 | -0.013 | 0 | 0.35 | 0.41 | 0.26 | 0.22 | 0.3 | 0.27 |
| 8 | 105232453 | rs2441919 | -0.666 | -0.327 | -0.026 | -0.486 | -0.009 | -0.013 | -0.013 | 0 | 0.35 | 0.41 | 0.26 | 0.22 | 0.3 | 0.27 |
| 8 | 105234363 | rs2441921 | -0.72 | -0.304 | -0.023 | -0.489 | -0.019 | -0.023 | -0.023 | 0 | 0.36 | 0.46 | 0.26 | 0.22 | 0.3 | 0.27 |
| 8 | 105268769 | rs1868009 | 1.065 | -0.289 | -0.003 | -0.485 | -0.023 | -0.018 | -0.018 | 1.282 | 0.13 | 0.19 | 0.04 | 0.04 | 0.07 | 0.09 |
| 8 | 105278802 | rs4734754 | -0.223 | -0.155 | -0.187 | -0.432 | 0.016 | 0.03 | 0.03 | 0 | 0.34 | 0.48 | 0.53 | 0.64 | 0.35 | 0.33 |

| | | | | | | | | | | | | | | | | |
|---|---|---|---|---|---|---|---|---|---|---|---|---|---|---|---|---|
| 8 | 105281322 | rs2441822 | 0.565 | -0.117 | -0.789 | -0.397 | -0.003 | 0.016 | 0.016 | 0 | 0.21 | 0.11 | 0.21 | 0.16 | 0.23 | 0.19 |
| 8 | 105292163 | rs10106504 | -0.468 | 0.167 | -0.573 | -0.298 | -0.006 | 0.001 | 0.001 | 0 | 0.44 | 0.59 | 0.62 | 0.65 | 0.47 | 0.41 |
| 8 | 105292565 | rs16871199 | 0.919 | 0.134 | -0.285 | -0.263 | 0.007 | 0.122 | 0.122 | 0 | 0.2 | 0.36 | 0.33 | 0.54 | 0.18 | 0.29 |
| 8 | 105292598 | rs17821383 | 1.096 | 0.2 | -0.372 | -0.348 | 0 | 0 | 0 | 0 | 0.19 | 0.14 | 0.24 | 0.11 | 0.22 | 0.1 |
| 8 | 105293427 | rs2511605 | -1.966 | 0.059 | -0.731 | -0.283 | -0.088 | 0.055 | 0.055 | 0.247 | 0.7 | 0.97 | 0.79 | 0.88 | 0.65 | 0.61 |
| 8 | 105295346 | rs4734761 | -0.76 | 0.154 | -0.575 | -0.332 | -0.002 | 0.015 | 0.015 | 0 | 0.56 | 0.59 | 0.65 | 0.69 | 0.51 | 0.5 |
| 8 | 105304314 | rs2441845 | 1.822 | 0.16 | -0.848 | -0.284 | 0.004 | -0.019 | -0.019 | 0 | 0.07 | 0.07 | 0.07 | 0.01 | 0.13 | 0.02 |
| 8 | 105307981 | rs1374520 | 1.588 | 0.126 | -0.867 | -0.302 | 0.001 | -0.018 | -0.018 | 0 | 0.08 | 0.07 | 0.07 | 0.01 | 0.13 | 0.02 |
| 8 | 105311074 | rs2511628 | 2.137 | 0.174 | -0.853 | -0.287 | -0.011 | 0.026 | 0.026 | 0.047 | 0.07 | 0.01 | 0.04 | 0 | 0.1 | 0.02 |
| 8 | 105312256 | rs7818735 | -0.484 | -0.164 | -1.024 | -0.266 | -0.02 | 0.006 | 0.006 | 0 | 0.47 | 0.77 | 0.64 | 0.62 | 0.45 | 0.41 |
| 8 | 105328181 | rs1374526 | 1.987 | -0.505 | -0.813 | -0.154 | -0.001 | -0.019 | -0.019 | 1.107 | 0.07 | 0.06 | 0.05 | 0.01 | 0.09 | 0.02 |
| 8 | 105328529 | rs2166646 | 0.546 | -0.48 | -0.617 | -0.117 | 0.023 | 0.076 | 0.076 | 0 | 0.2 | 0.29 | 0.38 | 0.52 | 0.22 | 0.3 |
| 8 | 105330622 | rs2441864 | 2.121 | -0.459 | -0.861 | -0.187 | 0.003 | 0.027 | 0.027 | 0.046 | 0.07 | 0.02 | 0.04 | 0 | 0.09 | 0.02 |
| 8 | 105332817 | rs2458422 | -1.824 | -0.722 | -1.308 | -0.366 | -0.138 | 0.097 | 0.097 | 0.054 | 0.67 | 0.96 | 0.77 | 0.89 | 0.59 | 0.63 |
| 8 | 105334224 | rs1533242 | -1.159 | -0.747 | -1.241 | -0.236 | -0.081 | 0.012 | 0.012 | 0 | 0.58 | 0.9 | 0.53 | 0.68 | 0.48 | 0.51 |
| 8 | 105336038 | rs2458424 | -0.795 | -0.804 | -1.416 | -0.296 | -0.067 | -0.038 | -0.038 | 0 | 0.5 | 0.93 | 0.67 | 0.63 | 0.51 | 0.46 |
| 8 | 105337871 | rs2514658 | -1.835 | -0.63 | -1.378 | -0.23 | -0.143 | 0.082 | 0.082 | 0.152 | 0.68 | 0.99 | 0.77 | 0.88 | 0.58 | 0.63 |
| 8 | 105338752 | rs4299971 | -0.479 | -0.53 | -0.714 | -0.188 | 0.01 | 0.017 | 0.017 | 0 | 0.4 | 0.5 | 0.46 | 0.6 | 0.39 | 0.42 |
| 8 | 105343889 | rs1812215 | -1.167 | -0.829 | -1.031 | -0.13 | -0.053 | 0.125 | 0.125 | 0.31 | 0.54 | 0.9 | 0.57 | 0.74 | 0.44 | 0.53 |
| 8 | 105351943 | rs2514666 | -0.401 | -0.338 | -0.335 | 0.035 | 0.021 | 0.053 | 0.053 | 0 | 0.79 | 0.66 | 0.76 | 0.56 | 0.76 | 0.7 |
| 8 | 105351998 | rs2458415 | -0.389 | -0.345 | -0.338 | 0.033 | 0.02 | 0.05 | 0.05 | 0 | 0.78 | 0.66 | 0.76 | 0.57 | 0.76 | 0.7 |
| 8 | 105362656 | rs2514668 | -0.263 | -0.532 | -0.989 | 0.032 | 0.019 | 0.039 | 0.039 | 0 | 0.74 | 0.4 | 0.72 | 0.55 | 0.74 | 0.69 |
| 8 | 105367096 | rs2458431 | -0.254 | -0.556 | -0.994 | 0.04 | 0.016 | 0.036 | 0.036 | 0 | 0.74 | 0.39 | 0.72 | 0.55 | 0.73 | 0.68 |
| 8 | 105368408 | rs4734780 | 0.38 | -1.202 | -1.097 | -0.11 | 0.002 | 0.004 | 0.004 | 2.359 | 0.14 | 0.13 | 0.03 | 0.07 | 0.05 | 0.08 |
| 8 | 105371276 | rs16871341 | 0.26 | -0.376 | -1.143 | -0.075 | 0.003 | 0.029 | 0.029 | 0 | 0.14 | 0.06 | 0.19 | 0.14 | 0.13 | 0.11 |
| 8 | 105374885 | rs4734783 | 0.376 | -0.643 | -0.794 | 0.215 | 0 | 0 | 0 | 0.065 | 0.77 | 0.74 | 0.84 | 0.82 | 0.79 | 0.7 |
| 8 | 105374981 | rs10808389 | 0.799 | -0.801 | -1.279 | 0.083 | 0.011 | 0.064 | 0.064 | 0.419 | 0.45 | 0.26 | 0.47 | 0.26 | 0.55 | 0.45 |
| 8 | 105377050 | rs2853177 | 0.767 | -0.72 | -1.511 | 0.424 | 0.049 | 0.144 | 0.144 | 0.733 | 0.52 | 0.06 | 0.49 | 0.26 | 0.64 | 0.42 |
| 8 | 105386546 | rs2853143 | -0.531 | -0.538 | -0.817 | 0.773 | -0.003 | -0.014 | -0.014 | 0 | 0.91 | 0.94 | 0.9 | 0.82 | 0.88 | 0.73 |
| 8 | 105387457 | rs1317221 | 0.167 | -0.705 | -0.407 | 0.861 | -0.05 | 0.071 | 0.071 | 0.287 | 0.73 | 0.48 | 0.65 | 0.54 | 0.83 | 0.52 |
| 8 | 105393021 | rs2853149 | 0.338 | 0.252 | -1.706 | 0.18 | -0.139 | 0.034 | 0.034 | 0 | 0.38 | 0.04 | 0.44 | 0.26 | 0.48 | 0.28 |
| 8 | 105393687 | rs2253336 | 0.637 | -0.207 | -0.776 | 0.691 | -0.023 | 0.077 | 0.077 | 1.176 | 0.85 | 0.38 | 0.68 | 0.7 | 0.9 | 0.79 |
| 8 | 105393876 | rs16871361 | -0.939 | -0.25 | -0.035 | 0.588 | 0.013 | -0.013 | -0.013 | 0 | 0.07 | 0.23 | 0.06 | 0.02 | 0.07 | 0.01 |
| 8 | 105394245 | rs16871363 | -0.275 | -0.333 | 0.137 | 0.237 | -0.013 | 0.02 | 0.02 | 0.759 | 0.29 | 0.67 | 0.21 | 0.43 | 0.2 | 0.33 |
| 8 | 105394423 | rs16871364 | -0.468 | -0.333 | 0.175 | 0.361 | -0.005 | 0.006 | 0.006 | 1.825 | 0.3 | 0.71 | 0.24 | 0.51 | 0.21 | 0.33 |
| 8 | 105394466 | rs16871365 | -0.942 | -0.381 | 0.06 | 0.432 | 0.029 | -0.053 | -0.053 | 0.062 | 0.34 | 0.82 | 0.28 | 0.51 | 0.28 | 0.34 |
| 8 | 105395476 | rs2853150 | -0.741 | -0.36 | 0.128 | 0.569 | -0.01 | 0.12 | 0.12 | 0 | 0.95 | 0.9 | 0.78 | 0.73 | 0.98 | 0.83 |
| 8 | 105396792 | rs2853151 | -0.755 | -0.36 | 0.128 | 0.59 | -0.01 | 0.12 | 0.12 | 0 | 0.95 | 0.9 | 0.78 | 0.74 | 0.98 | 0.83 |
| 8 | 105399186 | rs3793358 | 0.798 | -0.563 | 0.675 | 0.833 | 0.008 | 0.234 | 0.234 | 0 | 0.12 | 0.34 | 0.12 | 0.38 | 0.12 | 0.27 |
| 8 | 105412148 | rs3793357 | 1.615 | -0.617 | 0.286 | 0.671 | -0.014 | 0.006 | 0.006 | 2.259 | 0.1 | 0.01 | 0.05 | 0.04 | 0.06 | 0.04 |
| 8 | 105414175 | rs7014456 | -0.46 | -0.66 | 1.221 | 0.637 | 0.024 | 0.056 | 0.056 | 1.304 | 0.06 | 0.33 | 0.06 | 0.09 | 0.03 | 0.02 |
| 8 | 105418862 | rs1962267 | 1.381 | -0.121 | -0.178 | -0.084 | -0.12 | 0.166 | 0.166 | 0 | 0.34 | 0.02 | 0.31 | 0.12 | 0.41 | 0.26 |
| 8 | 105422209 | rs2853160 | 0.256 | -0.076 | 0.252 | 0.028 | 0 | -0.022 | -0.022 | 0 | 0.53 | 0.52 | 0.44 | 0.23 | 0.5 | 0.33 |
| 8 | 105423221 | rs3793356 | -0.638 | -0.027 | -0.12 | 0.148 | -0.008 | 0.168 | 0.168 | 0.032 | 0.94 | 0.83 | 0.77 | 0.73 | 0.98 | 0.82 |
| 8 | 105423590 | rs3793355 | 1.025 | 0.136 | -0.448 | 0.11 | -0.087 | 0.075 | 0.075 | 0.167 | 0.26 | 0.01 | 0.18 | 0.08 | 0.29 | 0.22 |
| 8 | 105426922 | rs2333874 | -1.325 | 0.295 | -0.788 | 0.262 | -0.013 | -0.022 | -0.022 | 0 | 0.7 | 0.83 | 0.54 | 0.6 | 0.69 | 0.62 |
| 8 | 105431528 | rs10505058 | -1.304 | 0.294 | -0.801 | 0.259 | -0.012 | -0.023 | -0.023 | 0 | 0.7 | 0.83 | 0.54 | 0.59 | 0.69 | 0.62 |
| 8 | 105440637 | rs2280010 | 0.966 | 0.266 | -1.186 | 0.824 | -0.048 | -0.056 | -0.056 | 0.004 | 0.25 | 0.04 | 0.19 | 0.25 | 0.24 | 0.31 |
| 8 | 105445959 | rs2333871 | 0.861 | 0.265 | -1.189 | 0.824 | -0.062 | -0.069 | -0.069 | 0.001 | 0.25 | 0.01 | 0.18 | 0.24 | 0.25 | 0.31 |
| 8 | 105449151 | rs2250601 | 0.065 | 0.19 | -0.826 | 0.73 | 0 | -0.001 | -0.001 | 0.312 | 0.65 | 0.38 | 0.54 | 0.6 | 0.69 | 0.67 |
| 8 | 105451737 | rs2853172 | 0.383 | 0.38 | -0.447 | 0.579 | -0.003 | 0.002 | 0.002 | 0 | 0.4 | 0.39 | 0.36 | 0.36 | 0.44 | 0.36 |
| 8 | 105463690 | rs2669429 | 0.407 | 0.298 | -0.792 | 0.519 | 0.001 | 0.036 | 0.036 | 0 | 0.43 | 0.26 | 0.42 | 0.36 | 0.55 | 0.35 |
| 8 | 105468464 | rs13264598 | 0.521 | 0.315 | -0.699 | 0.359 | 0.005 | 0.021 | 0.021 | 0 | 0.11 | 0.07 | 0.07 | 0.04 | 0.1 | 0.08 |
| 8 | 105469973 | rs11783979 | -0.808 | 0.329 | -1.25 | 0.615 | -0.036 | -0.01 | -0.01 | 0 | 0.63 | 0.79 | 0.5 | 0.58 | 0.58 | 0.61 |
| 8 | 105471913 | rs1355691 | 1.073 | 0.048 | -0.533 | 0.528 | -0.01 | -0.016 | -0.016 | 0 | 0.2 | 0.12 | 0.17 | 0.26 | 0.24 | 0.26 |
| 8 | 105474935 | rs3133278 | -0.044 | -0.272 | -0.307 | 0.308 | 0.01 | 0.006 | 0.006 | 0 | 0.35 | 0.36 | 0.26 | 0.3 | 0.28 | 0.26 |
| 8 | 105478933 | rs2298840 | 0.804 | -0.43 | -0.182 | 0.341 | -0.009 | 0.003 | 0.003 | 5.463 | 0.25 | 0.29 | 0.21 | 0.31 | 0.18 | 0.15 |
| 8 | 105480180 | rs2959025 | 0.054 | -0.007 | -0.522 | 0.535 | -0.083 | -0.104 | -0.104 | 0.406 | 0.28 | 0.09 | 0.21 | 0.41 | 0.36 | 0.37 |
| 8 | 105481705 | rs2942570 | -1.482 | -0.16 | -0.031 | 0.413 | -0.002 | 0.005 | 0.005 | 0 | 0.63 | 0.67 | 0.51 | 0.72 | 0.59 | 0.63 |
| 8 | 105482475 | rs3750187 | 0.999 | -0.285 | 0.1 | 0.391 | -0.007 | 0 | 0 | 0.001 | 0.28 | 0.31 | 0.23 | 0.29 | 0.2 | 0.17 |
| 8 | 105486661 | rs10101743 | 0.659 | -0.351 | 0.023 | 0.378 | 0.002 | -0.003 | -0.003 | 0.329 | 0.09 | 0.07 | 0.23 | 0.17 | 0.05 | 0.21 |
| 8 | 105495023 | rs2853132 | 0.228 | -0.299 | -0.032 | 0.285 | 0.053 | 0.037 | 0.037 | 0.122 | 0.53 | 0.31 | 0.65 | 0.67 | 0.66 | 0.67 |
| 8 | 105501813 | rs13279448 | 1.585 | -0.507 | -0.055 | 0.158 | 0.011 | 0.015 | 0.015 | 0 | 0.11 | 0.07 | 0.04 | 0.03 | 0.07 | 0.06 |
| 8 | 105508391 | rs3018437 | -3.672 | -0.375 | -0.399 | -0.275 | -0.022 | 0.072 | 0.072 | 0.011 | 0.88 | 1 | 0.93 | 1 | 0.85 | 0.95 |
| 8 | 105508810 | rs12546843 | 1.539 | -0.3 | 0.003 | 0.056 | -0.005 | -0.001 | -0.001 | 0 | 0.28 | 0.3 | 0.24 | 0.28 | 0.22 | 0.17 |
| 8 | 105509049 | rs3018438 | 0.293 | -0.077 | 0.049 | 0.128 | -0.035 | -0.054 | -0.054 | 0 | 0.18 | 0.38 | 0.11 | 0.05 | 0.12 | 0.16 |
| 8 | 105605417 | rs7461146 | -2.426 | 0.254 | -0.469 | 0.295 | -0.032 | -0.031 | -0.031 | 0 | 0.95 | 0.99 | 0.95 | 0.89 | 0.92 | 0.92 |
| 8 | 105623638 | rs13268122 | 1.58 | -0.477 | -0.446 | -0.331 | 0.011 | 0.019 | 0.019 | 0 | 0.1 | 0.06 | 0.04 | 0.03 | 0.07 | 0.07 |
| 8 | 105635003 | rs12681417 | 1.23 | -0.671 | -0.513 | -0.41 | 0.019 | 0.004 | 0.004 | 3.9 | 0.3 | 0.26 | 0.24 | 0.28 | 0.24 | 0.18 |

| | | | | | | | | | | | | | | | | |
|---|---|---|---|---|---|---|---|---|---|---|---|---|---|---|---|---|
| 8 | 105641421 | rs2918072 | 1.925 | -0.656 | -0.692 | -0.485 | -0.041 | 0.061 | 0.061 | 0 | 0.28 | 0.11 | 0.23 | 0.13 | 0.37 | 0.21 |
| 8 | 105641573 | rs2880074 | 1.536 | -0.416 | -0.763 | -0.305 | 0.003 | 0.004 | 0.004 | 0 | 0.1 | 0.1 | 0.04 | 0.03 | 0.07 | 0.07 |
| 8 | 105647803 | rs2918073 | -1.666 | -0.231 | -0.815 | -0.116 | -0.093 | -0.108 | -0.108 | 0.016 | 0.85 | 0.99 | 0.85 | 0.63 | 0.77 | 0.73 |
| 8 | 105666107 | rs16871642 | 1.666 | -0.542 | -0.404 | -0.099 | 0.013 | -0.018 | -0.018 | 0 | 0.06 | 0.04 | 0.23 | 0.19 | 0.03 | 0.18 |
| 8 | 105672475 | rs1438353 | -0.8 | -0.35 | 0.012 | -0.41 | 0.032 | 0.026 | 0.026 | 0.358 | 0.48 | 0.62 | 0.35 | 0.31 | 0.36 | 0.31 |
| 8 | 105672856 | rs7834249 | 0.624 | -0.261 | -0.391 | -0.349 | -0.001 | 0 | 0 | 0 | 0.3 | 0.31 | 0.25 | 0.25 | 0.24 | 0.17 |
| 8 | 105674943 | rs12679911 | -0.296 | -0.32 | -0.511 | -0.33 | -0.014 | 0.135 | 0.135 | 0.392 | 0.45 | 0.25 | 0.28 | 0.11 | 0.45 | 0.36 |
| 8 | 105675462 | rs11990797 | 0.003 | -0.362 | 0.042 | -0.123 | -0.029 | 0.014 | 0.014 | 0 | 0.21 | 0.45 | 0.17 | 0.19 | 0.16 | 0.17 |
| 8 | 105684804 | rs11775279 | 0.621 | -0.208 | -0.368 | -0.488 | 0.004 | 0.03 | 0.03 | 0.216 | 0.17 | 0.08 | 0.11 | 0.01 | 0.15 | 0.18 |
| 8 | 105684996 | rs987474 | -0.689 | -0.288 | -0.232 | -0.686 | 0.001 | 0.106 | 0.106 | 0.039 | 0.5 | 0.7 | 0.36 | 0.22 | 0.48 | 0.41 |
| 8 | 105692826 | rs7006961 | 0.525 | -0.136 | -0.09 | -0.485 | 0.011 | -0.032 | -0.032 | 1.821 | 0.31 | 0.52 | 0.23 | 0.21 | 0.33 | 0.23 |
| 8 | 105696848 | rs4734804 | 0.825 | 0.194 | -0.79 | -0.051 | -0.056 | -0.08 | -0.08 | 0 | 0.16 | 0.04 | 0.17 | 0.41 | 0.21 | 0.26 |
| 8 | 105698914 | rs6985113 | -0.729 | -0.676 | -0.472 | -0.415 | -0.029 | 0.035 | 0.035 | 0 | 0.67 | 0.61 | 0.5 | 0.5 | 0.72 | 0.63 |
| 8 | 105700583 | rs4734805 | 0.856 | 0.161 | -0.848 | -0.499 | -0.013 | 0.06 | 0.06 | 2.964 | 0.21 | 0.08 | 0.12 | 0.01 | 0.19 | 0.17 |
| 8 | 105711531 | rs6984008 | -0.132 | -0.065 | -0.742 | -0.814 | -0.019 | 0.084 | 0.084 | 1.087 | 0.42 | 0.29 | 0.3 | 0.09 | 0.47 | 0.35 |
| 8 | 105713065 | rs1866313 | 1.12 | 0.045 | -0.773 | -0.806 | -0.048 | 0.039 | 0.039 | 0 | 0.19 | 0.13 | 0.16 | 0.08 | 0.27 | 0.19 |
| 8 | 105715533 | rs1000421 | 1.101 | -0.001 | -0.701 | -0.558 | -0.005 | 0.009 | 0.009 | 0.013 | 0.14 | 0.17 | 0.18 | 0.42 | 0.22 | 0.24 |
| 8 | 105716380 | rs13279815 | 1.065 | 0.142 | -0.627 | -0.561 | 0.019 | 0.075 | 0.075 | 0 | 0.15 | 0.27 | 0.19 | 0.41 | 0.22 | 0.23 |
| 8 | 105717183 | rs1438342 | 1.449 | -0.311 | -0.988 | -0.526 | 0.012 | -0.046 | -0.046 | 0 | 0.14 | 0.01 | 0.12 | 0.21 | 0.1 | 0.21 |
| 8 | 105723307 | rs7821869 | -1.961 | -0.43 | -0.939 | -0.523 | 0.052 | 0.054 | 0.054 | 4.167 | 0.72 | 0.94 | 0.8 | 0.86 | 0.77 | 0.8 |
| 8 | 105724847 | rs1438349 | -0.295 | 0.917 | -0.344 | -0.022 | -0.033 | 0.041 | 0.041 | 1.274 | 0.32 | 0.23 | 0.24 | 0.09 | 0.4 | 0.32 |
| 8 | 105732953 | rs718635 | 0.978 | 0.942 | -0.566 | 0.015 | -0.042 | -0.051 | -0.051 | 0 | 0.13 | 0.05 | 0.17 | 0.41 | 0.22 | 0.24 |
| 8 | 105744498 | rs1992214 | -1.225 | 1.469 | -0.891 | -0.07 | 0.021 | -0.001 | -0.001 | 0 | 0.48 | 0.5 | 0.44 | 0.5 | 0.64 | 0.56 |
| 8 | 105751982 | rs9297349 | 0.987 | 1.517 | -1.375 | -0.696 | -0.04 | 0.005 | 0.005 | 0.001 | 0.11 | 0.08 | 0.11 | 0.08 | 0.23 | 0.17 |
| 8 | 105769530 | rs13263724 | -0.094 | 1.053 | -1.255 | -0.628 | -0.022 | 0.08 | 0.08 | 3.474 | 0.2 | 0.04 | 0.11 | 0.01 | 0.17 | 0.15 |
| 8 | 105778689 | rs13267303 | -0.465 | 1.029 | -0.856 | -0.514 | 0.005 | -0.014 | -0.014 | 0.014 | 0.31 | 0.32 | 0.39 | 0.13 | 0.42 | 0.33 |
| 8 | 105780263 | rs10808393 | -0.128 | -0.368 | -0.869 | -0.54 | -0.012 | 0.015 | 0.015 | 0 | 0.35 | 0.39 | 0.28 | 0.49 | 0.27 | 0.29 |
| 8 | 105784723 | rs3847147 | 1.329 | 0.135 | -1.068 | -0.954 | -0.004 | 0.002 | 0.002 | 0 | 0.15 | 0.13 | 0.15 | 0.07 | 0.29 | 0.12 |
| 8 | 105785762 | rs1438346 | 0.353 | -0.513 | -1.171 | -1.407 | -0.001 | -0.034 | -0.034 | 0 | 0.3 | 0.31 | 0.22 | 0.07 | 0.36 | 0.18 |
| 8 | 105787284 | rs930383 | -1.547 | -0.761 | -1.591 | -1.391 | -0.032 | -0.072 | -0.072 | 0 | 0.84 | 0.96 | 0.78 | 0.72 | 0.8 | 0.88 |
| 8 | 105792850 | rs16871828 | 0.573 | -0.865 | -1.388 | -1.271 | -0.03 | 0.011 | 0.011 | 0 | 0.38 | 0.47 | 0.4 | 0.52 | 0.29 | 0.29 |
| 8 | 105797936 | rs1123406 | 0.556 | -0.868 | -1.39 | -1.27 | -0.03 | 0.007 | 0.007 | 0 | 0.38 | 0.47 | 0.4 | 0.54 | 0.29 | 0.29 |
| 8 | 105805344 | rs1347779 | 0.458 | -0.707 | -1.337 | -1.236 | -0.009 | -0.009 | -0.009 | 0 | 0.46 | 0.48 | 0.48 | 0.65 | 0.32 | 0.55 |
| 8 | 105805489 | rs10505066 | 0.805 | -0.366 | -0.579 | -0.86 | 0.111 | 0.424 | 0.424 | 0 | 0.06 | 0.28 | 0.1 | 0.3 | 0.09 | 0.11 |
| 8 | 105808348 | rs6468957 | 0.001 | -0.758 | -1.383 | -1.06 | 0.018 | 0.14 | 0.14 | 0.13 | 0.2 | 0.46 | 0.17 | 0.3 | 0.2 | 0.19 |
| 8 | 105810397 | rs12547291 | 0.117 | -1.173 | -1.709 | -1.321 | 0.023 | 0.085 | 0.085 | 0 | 0.53 | 0.34 | 0.51 | 0.26 | 0.54 | 0.61 |
| 8 | 105815145 | rs10505067 | 1.693 | -1.253 | -1.834 | -1.366 | 0.031 | 0.022 | 0.022 | 0.024 | 0.23 | 0.07 | 0.28 | 0.17 | 0.18 | 0.42 |
| 8 | 105817209 | rs17265772 | 0.432 | -1.236 | -1.831 | -1.401 | 0.055 | 0.075 | 0.075 | 0.704 | 0.38 | 0.23 | 0.36 | 0.18 | 0.24 | 0.49 |
| 8 | 105822000 | rs13280230 | 1.051 | -1.245 | -1.659 | -1.382 | 0.011 | -0.065 | -0.065 | 0 | 0.28 | 0.1 | 0.32 | 0.43 | 0.28 | 0.21 |
| 8 | 105823389 | rs1460238 | -0.407 | -1.071 | -1.443 | -1.24 | -0.001 | 0.138 | 0.138 | 0.508 | 0.8 | 0.52 | 0.82 | 0.69 | 0.81 | 0.82 |
| 8 | 105823681 | rs1870590 | 0.435 | -1.131 | -1.705 | -1.289 | 0.054 | 0.002 | 0.002 | 2.212 | 0.41 | 0.21 | 0.39 | 0.44 | 0.33 | 0.27 |
| 8 | 105827218 | rs9297351 | -0.312 | -1.187 | -1.59 | -1.345 | 0.001 | 0.128 | 0.128 | 0.613 | 0.79 | 0.45 | 0.72 | 0.65 | 0.79 | 0.82 |
| 8 | 105832227 | rs13439166 | 1.312 | -0.97 | -2 | -0.964 | 0.027 | 0 | 0 | 1.5 | 0.23 | 0.07 | 0.28 | 0.19 | 0.18 | 0.43 |
| 8 | 105851222 | rs10955359 | 0.478 | -0.619 | -1.787 | -0.79 | 0.028 | -0.049 | -0.049 | 0 | 0.28 | 0.07 | 0.23 | 0.39 | 0.27 | 0.21 |
| 8 | 105857776 | rs2085769 | 0.435 | -0.109 | -1.503 | -0.75 | -0.016 | 0.103 | 0.103 | 0.376 | 0.46 | 0.28 | 0.45 | 0.24 | 0.46 | 0.57 |
| 8 | 105863172 | rs10093584 | 0.443 | -0.106 | -1.504 | -0.749 | -0.014 | 0.1 | 0.1 | 0.377 | 0.46 | 0.28 | 0.45 | 0.25 | 0.46 | 0.57 |
| 8 | 105863417 | rs1962961 | 0.989 | -0.24 | -1.466 | -0.713 | 0.011 | -0.044 | -0.044 | 1.489 | 0.23 | 0.09 | 0.22 | 0.39 | 0.23 | 0.2 |
| 8 | 105864131 | rs1020826 | -0.051 | -0.253 | -1.425 | -0.79 | 0.04 | 0.268 | 0.268 | 0.928 | 0.86 | 0.43 | 0.74 | 0.64 | 0.85 | 0.85 |
| 8 | 105872179 | rs1031720 | 1.624 | -0.338 | -1.631 | -1.062 | 0.012 | 0.01 | 0.01 | 0 | 0.24 | 0.18 | 0.29 | 0.16 | 0.17 | 0.44 |
| 8 | 105873020 | rs11787091 | -0.265 | -0.252 | -1.628 | -1.091 | -0.002 | -0.005 | -0.005 | 0 | 0.65 | 0.39 | 0.55 | 0.25 | 0.67 | 0.67 |
| 8 | 105873463 | rs4734824 | 1.591 | -0.228 | -1.643 | -0.962 | 0.011 | 0.01 | 0.01 | 0 | 0.24 | 0.18 | 0.29 | 0.16 | 0.17 | 0.44 |
| 8 | 105876738 | rs728378 | 0.766 | 0.652 | -1.165 | -0.628 | -0.056 | 0.08 | 0.08 | 0 | 0.22 | 0.1 | 0.16 | 0.09 | 0.29 | 0.14 |
| 8 | 105878647 | rs4734825 | 0.299 | 0.579 | 0.05 | 0.248 | 0.063 | 0.319 | 0.319 | 0.296 | 0.87 | 0.54 | 0.75 | 0.64 | 0.85 | 0.85 |
| 8 | 105878832 | rs10094493 | 0.484 | 0.59 | -0.489 | 0.344 | 0.003 | 0.021 | 0.021 | 0.198 | 0.7 | 0.37 | 0.67 | 0.64 | 0.69 | 0.77 |
| 8 | 105883980 | rs7004002 | -0.9 | 0.707 | 0.082 | -0.021 | 0.021 | -0.098 | -0.098 | 0 | 0.16 | 0.37 | 0.11 | 0 | 0.19 | 0.17 |
| 8 | 105884484 | rs7004837 | -1.26 | 0.357 | 0.803 | 0.923 | -0.001 | 0.098 | 0.098 | 0 | 0.17 | 0.33 | 0.22 | 0.31 | 0.16 | 0.15 |
| 8 | 105884605 | rs6468964 | -0.385 | 0.361 | 0.097 | 0.623 | -0.002 | 0.001 | 0.001 | 0 | 0.45 | 0.42 | 0.6 | 0.5 | 0.5 | 0.51 |
| 8 | 105887854 | rs6468966 | 0.601 | 0.365 | 0.186 | 1.012 | 0.005 | 0.006 | 0.006 | 0 | 0.09 | 0.14 | 0.11 | 0.09 | 0.13 | 0.11 |
| 8 | 105890188 | rs1460235 | 0.532 | 0.144 | 0.768 | 0.908 | -0.001 | -0.001 | -0.001 | 0 | 0.33 | 0.31 | 0.5 | 0.42 | 0.36 | 0.37 |
| 8 | 105890308 | rs13275783 | 1.114 | -0.207 | 0.041 | 0.335 | 0.029 | -0.089 | -0.089 | 0 | 0.28 | 0.05 | 0.21 | 0.44 | 0.23 | 0.33 |
| 8 | 105894884 | rs2169164 | -0.406 | -0.084 | 0.214 | 0.317 | -0.001 | 0 | 0 | 0 | 0.45 | 0.44 | 0.36 | 0.52 | 0.41 | 0.46 |
| 8 | 105895914 | rs2125917 | -0.581 | -0.111 | 0.276 | 0.258 | 0.009 | 0.019 | 0.019 | 0 | 0.47 | 0.6 | 0.47 | 0.75 | 0.43 | 0.47 |
| 8 | 105905155 | rs10100692 | 1.618 | -0.083 | 0.129 | 0.256 | -0.026 | 0.004 | 0.004 | 0 | 0.3 | 0.07 | 0.36 | 0.2 | 0.31 | 0.36 |
| 8 | 105906441 | rs10955366 | -0.196 | -0.081 | 0.22 | 0.261 | 0.004 | -0.005 | -0.005 | 0 | 0.15 | 0.32 | 0.14 | 0.09 | 0.15 | 0.14 |
| 8 | 105912844 | rs12545436 | -0.224 | -0.098 | 0.33 | 0.313 | 0.004 | -0.011 | -0.011 | 0 | 0.71 | 0.37 | 0.7 | 0.77 | 0.71 | 0.83 |
| 8 | 105915256 | rs1545699 | 1.769 | -0.088 | 0.096 | 0.37 | -0.029 | -0.077 | -0.077 | 0 | 0.29 | 0.07 | 0.38 | 0.3 | 0.3 | 0.35 |
| 8 | 105930209 | rs2169166 | -1.016 | -0.072 | 0.259 | 0.363 | -0.02 | 0.023 | 0.023 | 0 | 0.77 | 0.84 | 0.84 | 0.94 | 0.73 | 0.84 |
| 8 | 105930490 | rs1809286 | 0.959 | -0.083 | 0.15 | 0.951 | -0.055 | -0.038 | -0.038 | 0.357 | 0.57 | 0.13 | 0.57 | 0.51 | 0.56 | 0.66 |
| 8 | 105936645 | rs284500 | -1.012 | -0.014 | 0.242 | 0.983 | -0.017 | -0.009 | -0.009 | 0 | 0.78 | 0.81 | 0.8 | 0.68 | 0.73 | 0.8 |

| | | | | | | | | | | | | | | | | |
|---|---|---|---|---|---|---|---|---|---|---|---|---|---|---|---|---|
| 8 | 105942054 | rs284476 | -0.153 | -0.065 | 0.185 | 0.815 | 0.001 | -0.079 | -0.079 | 0.015 | 0.14 | 0.28 | 0.09 | 0 | 0.15 | 0.14 |
| 8 | 105943784 | rs13278098 | 0.558 | 0.047 | -0.19 | 1.088 | 0.016 | 0.029 | 0.029 | 0 | 0.3 | 0.18 | 0.2 | 0.22 | 0.26 | 0.3 |
| 8 | 105946333 | rs284486 | 0.871 | 0.244 | -0.596 | 1.146 | -0.044 | -0.034 | -0.034 | 0 | 0.36 | 0.1 | 0.44 | 0.29 | 0.4 | 0.38 |
| 8 | 105957540 | rs16872085 | 0.479 | 0.14 | -0.146 | 1.732 | 0.008 | -0.009 | -0.009 | 0 | 0.09 | 0.05 | 0.07 | 0.27 | 0.09 | 0.1 |
| 8 | 105958020 | rs284489 | -1.076 | 0.117 | 0.098 | 1.394 | 0.002 | -0.005 | -0.005 | 0.019 | 0.36 | 0.68 | 0.42 | 0.41 | 0.34 | 0.32 |
| 8 | 105965285 | rs284494 | -1.02 | 0.009 | 0.264 | 1.28 | -0.013 | 0.051 | 0.051 | 3.652 | 0.16 | 0.6 | 0.27 | 0.24 | 0.15 | 0.1 |
| 8 | 105965737 | rs284496 | -1.244 | 0.032 | 0.186 | 1.366 | 0.002 | -0.006 | -0.006 | 0.002 | 0.36 | 0.68 | 0.42 | 0.43 | 0.34 | 0.27 |
| 8 | 105981846 | rs6991677 | -0.218 | 0.374 | -0.415 | 1.095 | -0.007 | 0.002 | 0.002 | 0 | 0.2 | 0.11 | 0.15 | 0.19 | 0.19 | 0.18 |
| 8 | 105987231 | rs4358773 | -0.201 | 0.648 | -0.045 | 0.811 | 0.002 | -0.002 | -0.002 | 0 | 0.24 | 0.21 | 0.17 | 0.2 | 0.3 | 0.29 |
| 8 | 105989161 | rs6991780 | -1.09 | 0.201 | -0.301 | 0.88 | 0.013 | 0.065 | 0.065 | 0 | 0.51 | 0.79 | 0.43 | 0.66 | 0.55 | 0.59 |
| 8 | 105989262 | rs12677340 | 0.351 | 0.177 | -0.039 | 1.013 | 0 | 0.007 | 0.007 | 0 | 0.19 | 0.19 | 0.16 | 0.41 | 0.13 | 0.21 |
| 8 | 105992696 | rs977396 | -0.701 | 0.061 | -0.398 | 0.874 | -0.019 | -0.045 | -0.045 | 3.146 | 0.87 | 0.99 | 0.8 | 0.82 | 0.86 | 0.94 |
| 8 | 106004969 | rs12543306 | 0.459 | 0.258 | -0.084 | 1.008 | 0.02 | -0.045 | -0.045 | 0 | 0.19 | 0.12 | 0.14 | 0.4 | 0.13 | 0.21 |
| 8 | 106010200 | rs1879688 | -1.341 | 0.265 | -0.068 | 0.95 | 0 | 0.062 | 0.062 | 0 | 0.51 | 0.85 | 0.44 | 0.66 | 0.55 | 0.59 |
| 8 | 106021780 | rs9297354 | 0.219 | 0.327 | 0.208 | 0.989 | 0.037 | 0.07 | 0.07 | 1.246 | 0.06 | 0.34 | 0.07 | 0.06 | 0.05 | 0.08 |
| 8 | 106024987 | rs16872183 | 0.305 | 0.302 | -0.518 | 1.11 | 0.012 | -0.021 | -0.021 | 0 | 0.19 | 0.14 | 0.14 | 0.4 | 0.13 | 0.21 |
| 8 | 106032537 | rs1402820 | -0.097 | 0.935 | -0.135 | 1.029 | 0 | 0 | 0 | 0.376 | 0.22 | 0.19 | 0.18 | 0.22 | 0.3 | 0.27 |
| 8 | 106037189 | rs3107546 | 0.297 | 0.925 | 0.083 | 0.872 | 0.048 | -0.014 | -0.014 | 0 | 0.88 | 0.73 | 0.86 | 0.92 | 0.82 | 0.85 |
| 8 | 106037345 | rs3110295 | -0.381 | 0.587 | -0.683 | 1.121 | -0.051 | -0.057 | -0.057 | 0 | 0.33 | 0.12 | 0.32 | 0.42 | 0.38 | 0.33 |
| 8 | 106040818 | rs16872216 | 1.101 | 0.434 | -0.684 | 1.353 | -0.001 | 0.066 | 0.066 | 0 | 0.05 | 0.07 | 0.06 | 0.3 | 0.02 | 0.08 |
| 8 | 106041827 | rs1521776 | 1.687 | 0.186 | -0.91 | 1.131 | -0.054 | 0.126 | 0.126 | 0.117 | 0.44 | 0.06 | 0.43 | 0.2 | 0.41 | 0.43 |
| 8 | 106052343 | rs3101274 | -0.394 | 0.385 | -0.64 | 1.149 | -0.035 | -0.03 | -0.03 | 0 | 0.33 | 0.16 | 0.31 | 0.41 | 0.38 | 0.34 |
| 8 | 106075248 | rs285787 | 0.743 | 0.557 | 0.038 | 0.938 | 0.057 | -0.007 | -0.007 | 0.003 | 0.87 | 0.68 | 0.84 | 0.9 | 0.81 | 0.85 |
| 8 | 106083175 | rs285798 | 1.645 | -0.167 | -0.388 | 0.846 | -0.028 | 0.109 | 0.109 | 0 | 0.39 | 0.16 | 0.42 | 0.19 | 0.39 | 0.39 |
| 8 | 106086536 | rs3107547 | 0.914 | 0.12 | -0.204 | 0.989 | -0.001 | 0.001 | 0.001 | 0.292 | 0.1 | 0.12 | 0.15 | 0.18 | 0.1 | 0.03 |
| 8 | 106091472 | rs285839 | 1.076 | 0.23 | -0.115 | 0.984 | -0.003 | -0.018 | -0.018 | 0 | 0.67 | 0.69 | 0.68 | 0.57 | 0.69 | 0.74 |
| 8 | 106095659 | rs16872304 | 1.176 | 0.194 | -0.336 | 0.987 | 0.024 | -0.03 | -0.03 | 2.057 | 0.12 | 0.07 | 0.04 | 0.24 | 0.05 | 0.16 |
| 8 | 106102180 | rs285823 | 1.484 | 0.063 | -0.218 | 0.953 | -0.01 | 0.012 | 0.012 | 1.935 | 0.69 | 0.25 | 0.63 | 0.72 | 0.67 | 0.82 |
| 8 | 106102889 | rs285822 | 0.555 | 0.232 | 0 | 0.806 | 0.016 | -0.015 | -0.015 | 0 | 0.84 | 0.81 | 0.82 | 0.9 | 0.78 | 0.86 |
| 8 | 106105268 | rs3107548 | 0.259 | 0.15 | 0.058 | 0.87 | 0.005 | -0.007 | -0.007 | 0 | 0.22 | 0.25 | 0.19 | 0.41 | 0.14 | 0.19 |
| 8 | 106105811 | rs2045672 | -1.36 | 0.268 | -0.228 | 0.858 | 0.001 | -0.007 | -0.007 | 0 | 0.5 | 0.49 | 0.46 | 0.81 | 0.46 | 0.54 |
| 8 | 106106177 | rs285818 | -0.737 | 0.286 | -0.381 | 0.86 | -0.015 | 0.009 | 0.009 | 0 | 0.92 | 0.94 | 0.91 | 1 | 0.87 | 0.94 |
| 8 | 106106951 | rs17192765 | 0.257 | 0.314 | -0.474 | 0.899 | -0.031 | -0.042 | -0.042 | 0 | 0.22 | 0.09 | 0.21 | 0.32 | 0.27 | 0.29 |
| 8 | 106108593 | rs1460583 | 0.065 | 0.262 | -0.432 | 0.751 | -0.035 | -0.046 | -0.046 | 0 | 0.24 | 0.09 | 0.24 | 0.38 | 0.31 | 0.35 |
| 8 | 106109017 | rs627350 | 1.816 | 0.221 | -0.491 | 0.806 | -0.058 | 0.104 | 0.104 | 0 | 0.39 | 0.11 | 0.42 | 0.19 | 0.4 | 0.39 |
| 8 | 106109149 | rs2685536 | 0.947 | 0.573 | -0.123 | 0.793 | 0.028 | -0.007 | -0.007 | 0 | 0.64 | 0.53 | 0.64 | 0.62 | 0.55 | 0.58 |
| 8 | 106111662 | rs10505072 | -0.438 | 0.828 | -0.505 | 0.989 | -0.034 | -0.05 | -0.05 | 0 | 0.23 | 0.1 | 0.24 | 0.42 | 0.31 | 0.4 |
| 8 | 106113818 | rs17278434 | -0.395 | 0.622 | -0.208 | 0.643 | 0 | -0.005 | -0.005 | 0 | 0.23 | 0.29 | 0.3 | 0.23 | 0.28 | 0.2 |
| 8 | 106115172 | rs7819593 | -0.401 | 0.184 | -0.296 | 0.497 | -0.049 | 0.034 | 0.034 | 0 | 0.36 | 0.49 | 0.3 | 0.46 | 0.27 | 0.42 |
| 8 | 106115255 | rs285774 | 0.839 | 0.333 | -0.656 | 0.263 | 0.052 | -0.032 | -0.032 | 0 | 0.54 | 0.16 | 0.59 | 0.54 | 0.58 | 0.52 |
| 8 | 106120450 | rs285802 | -0.167 | 0.444 | -0.355 | 0.317 | 0.017 | -0.038 | -0.038 | 0 | 0.91 | 0.84 | 0.91 | 1 | 0.86 | 0.94 |
| 8 | 106124371 | rs1460586 | -0.407 | 0.318 | -0.457 | 0.361 | -0.017 | 0.006 | 0.006 | 0 | 0.4 | 0.52 | 0.34 | 0.45 | 0.31 | 0.42 |
| 8 | 106125471 | rs285809 | 1.241 | 0.653 | -1.401 | -0.249 | -0.122 | 0.092 | 0.092 | 0.012 | 0.4 | 0.01 | 0.42 | 0.19 | 0.39 | 0.32 |
| 8 | 106128825 | rs285811 | 0.373 | 1.025 | -0.128 | -0.034 | 0.065 | -0.053 | -0.053 | 8.478 | 0.91 | 0.73 | 0.88 | 1 | 0.83 | 0.94 |
| 8 | 106130379 | rs2613122 | 0.475 | 1.024 | -0.092 | -0.056 | 0.079 | -0.056 | -0.056 | 0 | 0.9 | 0.66 | 0.88 | 1 | 0.83 | 0.95 |
| 8 | 106136318 | rs6986587 | -1.025 | 0.802 | -0.89 | -0.537 | 0.002 | -0.005 | -0.005 | 0 | 0.73 | 0.76 | 0.67 | 0.77 | 0.73 | 0.73 |
| 8 | 106142109 | rs16872439 | 0.859 | 0.343 | -1.881 | -0.393 | 0.016 | -0.029 | -0.029 | 0.051 | 0.17 | 0.06 | 0.05 | 0.2 | 0.07 | 0.3 |
| 8 | 106142576 | rs6990385 | -1.974 | 0.365 | -2.334 | -0.481 | -0.06 | -0.033 | -0.033 | 0 | 0.79 | 0.97 | 0.7 | 0.77 | 0.76 | 0.76 |
| 8 | 106147155 | rs3884501 | 0.417 | 0.056 | -2.266 | -0.576 | -0.007 | -0.026 | -0.026 | 0 | 0.42 | 0.2 | 0.49 | 0.57 | 0.48 | 0.39 |
| 8 | 106152318 | rs898560 | -0.983 | -0.059 | -2.895 | -0.79 | 0.007 | 0.004 | 0.004 | 0.076 | 0.7 | 0.97 | 0.8 | 0.75 | 0.82 | 0.68 |
| 8 | 106156492 | rs2045673 | 0.647 | 0.7 | -2.7 | -0.476 | -0.011 | -0.09 | -0.09 | 0 | 0.31 | 0.08 | 0.35 | 0.55 | 0.37 | 0.36 |
| 8 | 106162551 | rs13249350 | 1.407 | 1.143 | -2.286 | -0.341 | -0.088 | -0.066 | -0.066 | 0.367 | 0.15 | 0 | 0.25 | 0.2 | 0.22 | 0.22 |
| 8 | 106175867 | rs11785817 | 0.526 | 1.085 | -1.924 | -0.415 | 0.012 | -0.03 | -0.03 | 0 | 0.09 | 0.09 | 0.1 | 0 | 0.17 | 0.1 |
| 8 | 106197292 | rs768388 | 0.777 | 1.007 | -2.558 | -0.433 | -0.022 | -0.08 | -0.08 | 0 | 0.27 | 0.08 | 0.52 | 0.59 | 0.36 | 0.31 |
| 8 | 106207444 | rs13262572 | 1.39 | 0.922 | -2.519 | -0.372 | -0.087 | -0.061 | -0.061 | 0.014 | 0.16 | 0.01 | 0.12 | 0.19 | 0.22 | 0.22 |
| 8 | 106224061 | rs2342460 | -0.214 | 1.006 | -2.199 | -0.146 | -0.083 | -0.064 | -0.064 | 0 | 0.35 | 0.1 | 0.61 | 0.59 | 0.52 | 0.4 |
| 8 | 106231172 | rs1350723 | -0.445 | 0.887 | -1.816 | -0.066 | -0.008 | -0.03 | -0.03 | 0 | 0.76 | 0.83 | 0.48 | 0.37 | 0.64 | 0.73 |
| 8 | 106233193 | rs12156003 | 0.042 | 0.248 | -1.43 | -0.668 | 0.001 | 0.043 | 0.043 | 0.002 | 0.42 | 0.3 | 0.28 | 0.2 | 0.37 | 0.32 |
| 8 | 106234642 | rs6468988 | 1.13 | 0.255 | -1.409 | -0.619 | -0.005 | -0.068 | -0.068 | 0 | 0.32 | 0.52 | 0.2 | 0.13 | 0.27 | 0.38 |
| 8 | 106247695 | rs2613130 | -1.354 | 0.305 | -1.801 | -0.564 | -0.039 | -0.077 | -0.077 | 0 | 0.79 | 0.93 | 0.48 | 0.38 | 0.64 | 0.74 |
| 8 | 106248394 | rs1903335 | 0.162 | 0.133 | -1.639 | -0.734 | -0.023 | 0.12 | 0.12 | 0.093 | 0.41 | 0.13 | 0.26 | 0.19 | 0.37 | 0.32 |
| 8 | 106254571 | rs2957470 | 1.606 | 0.342 | -1.682 | 0.18 | -0.062 | -0.075 | -0.075 | 0 | 0.16 | 0.02 | 0.43 | 0.25 | 0.26 | 0.15 |
| 8 | 106255824 | rs7841616 | 1.208 | 0.134 | -1.15 | 0.034 | -0.001 | -0.027 | -0.027 | 0 | 0.36 | 0.42 | 0.21 | 0.17 | 0.32 | 0.42 |
| 8 | 106256501 | rs10110132 | 1.369 | 0.096 | -0.944 | 0.027 | 0 | -0.02 | -0.02 | 0 | 0.35 | 0.37 | 0.19 | 0.12 | 0.31 | 0.4 |
| 8 | 106257210 | rs9642799 | 0.404 | 0.37 | -1.551 | 0.017 | 0.02 | -0.065 | -0.065 | 0.07 | 0.33 | 0.11 | 0.23 | 0.58 | 0.27 | 0.34 |
| 8 | 106257855 | rs12155577 | -0.328 | 0.507 | -1.374 | -0.21 | -0.063 | 0.029 | 0.029 | 0 | 0.72 | 0.85 | 0.53 | 0.82 | 0.58 | 0.7 |
| 8 | 106257921 | rs2941647 | 0.132 | 0.505 | -1.308 | -0.042 | -0.015 | -0.038 | -0.038 | 0 | 0.84 | 0.85 | 0.73 | 0.43 | 0.75 | 0.79 |
| 8 | 106259387 | rs10955383 | 0.609 | 0.315 | -1.329 | -0.679 | 0.005 | -0.021 | -0.021 | 0.589 | 0.28 | 0.15 | 0.28 | 0.7 | 0.26 | 0.27 |
| 8 | 106260115 | rs12678274 | 0.424 | 0.332 | -1.244 | -0.705 | -0.001 | 0 | 0 | 0 | 0.3 | 0.27 | 0.31 | 0.72 | 0.27 | 0.28 |

| | | | | | | | | | | | | | | | | |
|---|---|---|---|---|---|---|---|---|---|---|---|---|---|---|---|---|
| 8 | 106269323 | rs2342785 | -0.016 | 0.271 | -1.148 | -0.821 | -0.005 | 0.06 | 0.06 | 0 | 0.46 | 0.29 | 0.63 | 0.83 | 0.52 | 0.43 |
| 8 | 106282960 | rs4734853 | 1.264 | 0.164 | -1.112 | -1.058 | -0.01 | 0.085 | 0.085 | 0.511 | 0.33 | 0.15 | 0.18 | 0.1 | 0.32 | 0.3 |
| 8 | 106283030 | rs2941627 | -0.616 | 0.137 | -0.681 | -0.99 | 0.03 | -0.042 | -0.042 | 0 | 0.89 | 0.73 | 0.89 | 0.97 | 0.88 | 0.89 |
| 8 | 106296090 | rs17209630 | 1.206 | -0.269 | -1.081 | -1.081 | -0.018 | 0.044 | 0.044 | 0 | 0.24 | 0.07 | 0.4 | 0.11 | 0.28 | 0.25 |
| 8 | 106296720 | rs6468994 | -0.345 | 0.166 | -1.014 | -1.171 | -0.005 | 0.003 | 0.003 | 0 | 0.65 | 0.7 | 0.47 | 0.87 | 0.56 | 0.63 |
| 8 | 106297815 | rs4078084 | 1.03 | -0.015 | -1.225 | -1.304 | -0.011 | -0.017 | -0.017 | 0.016 | 0.27 | 0.11 | 0.28 | 0.75 | 0.27 | 0.21 |
| 8 | 106299441 | rs2957437 | 1.486 | -0.135 | -1.221 | -1.257 | -0.051 | 0.025 | 0.025 | 0 | 0.23 | 0.02 | 0.39 | 0.11 | 0.28 | 0.25 |
| 8 | 106302875 | rs13250684 | 1.798 | -0.313 | -1.014 | -1.262 | 0.001 | 0.044 | 0.044 | 0 | 0.27 | 0.15 | 0.14 | 0.11 | 0.27 | 0.22 |
| 8 | 106307524 | rs7819944 | 0.64 | -0.229 | -1.144 | -1.226 | 0.01 | 0.011 | 0.011 | 0.062 | 0.39 | 0.65 | 0.21 | 0.11 | 0.3 | 0.43 |
| 8 | 106310577 | rs2001569 | 1.012 | 0 | -1.309 | -0.951 | 0.001 | 0.046 | 0.046 | 0 | 0.24 | 0.13 | 0.4 | 0.11 | 0.29 | 0.23 |
| 8 | 106319151 | rs10101839 | -0.48 | 0.006 | -1.137 | -0.935 | 0.008 | 0.017 | 0.017 | 0 | 0.41 | 0.62 | 0.37 | 0.77 | 0.32 | 0.49 |
| 8 | 106326772 | rs2069199 | -0.082 | -0.001 | -0.927 | -0.765 | 0 | -0.003 | -0.003 | 0 | 0.45 | 0.44 | 0.62 | 0.75 | 0.43 | 0.48 |
| 8 | 106337464 | rs10103822 | 0.329 | 0.066 | -0.812 | 0.07 | -0.043 | 0.237 | 0.237 | 0.458 | 0.15 | 0.36 | 0.17 | 0.51 | 0.07 | 0.25 |
| 8 | 106338217 | rs1113601 | -0.49 | 0.281 | -1.011 | -0.166 | 0.023 | -0.038 | -0.038 | 0 | 0.87 | 0.78 | 0.86 | 0.97 | 0.84 | 0.87 |
| 8 | 106377941 | rs17211035 | 1.231 | 0.3 | -1.36 | -0.118 | -0.04 | 0.001 | 0.001 | 0 | 0.12 | 0.03 | 0.11 | 0.11 | 0.16 | 0.16 |
| 8 | 106381365 | rs4734858 | 0.78 | 0.404 | -1.161 | -0.116 | 0.004 | -0.008 | -0.008 | 0 | 0.13 | 0.09 | 0.18 | 0.5 | 0.12 | 0.27 |
| 8 | 106385049 | rs17283482 | 1.038 | 0.441 | -1.283 | -0.19 | -0.074 | -0.018 | -0.018 | 0 | 0.12 | 0.01 | 0.09 | 0.12 | 0.16 | 0.16 |
| 8 | 106386119 | rs2957457 | 1.036 | -0.243 | -1.535 | -0.266 | -0.017 | -0.066 | -0.066 | 0 | 0.31 | 0.02 | 0.44 | 0.23 | 0.32 | 0.26 |
| 8 | 106386944 | rs2929062 | 0.889 | 0.531 | -0.776 | 0.093 | -0.024 | -0.012 | -0.012 | 0 | 0.6 | 0.26 | 0.56 | 0.34 | 0.56 | 0.46 |
| 8 | 106397116 | rs6999452 | 0.89 | 0.639 | -0.805 | 0.149 | 0.002 | 0.001 | 0.001 | 0 | 0.61 | 0.6 | 0.55 | 0.34 | 0.56 | 0.44 |
| 8 | 106401887 | rs2054577 | 0.876 | 0.682 | -0.904 | 0.172 | 0.007 | 0.003 | 0.003 | 0 | 0.61 | 0.56 | 0.59 | 0.35 | 0.56 | 0.46 |
| 8 | 106406847 | rs988260 | 0.911 | 0.749 | -1.1 | 0.183 | 0 | 0.001 | 0.001 | 0 | 0.61 | 0.59 | 0.6 | 0.36 | 0.54 | 0.43 |
| 8 | 106417183 | rs17211906 | 0.078 | 0.813 | -1.449 | -0.008 | 0.001 | 0.004 | 0.004 | 0 | 0.41 | 0.64 | 0.43 | 0.28 | 0.38 | 0.41 |
| 8 | 106419754 | rs10464844 | 0.892 | 0.71 | -1.31 | 0.022 | -0.02 | -0.02 | -0.02 | 0 | 0.29 | 0.63 | 0.35 | 0.18 | 0.23 | 0.26 |
| 8 | 106440076 | rs2035364 | 1.452 | 0.82 | -1.609 | 0.156 | -0.045 | -0.034 | -0.034 | 4.085 | 0.08 | 0 | 0.05 | 0.13 | 0.12 | 0.16 |
| 8 | 106457633 | rs1871745 | 0.918 | 0.807 | -1.657 | 0.058 | -0.022 | -0.017 | -0.017 | 2.163 | 0.29 | 0.61 | 0.35 | 0.19 | 0.23 | 0.26 |
| 8 | 106461141 | rs11777142 | 0.85 | 0.336 | -1.624 | 0.225 | 0.005 | -0.007 | -0.007 | 0 | 0.33 | 0.02 | 0.25 | 0.2 | 0.3 | 0.18 |
| 8 | 106468396 | rs7822940 | -0.202 | 0.672 | -1.311 | 0.177 | -0.013 | 0.015 | 0.015 | 0.557 | 0.55 | 0.34 | 0.53 | 0.65 | 0.51 | 0.48 |
| 8 | 106468592 | rs7822979 | -0.12 | 0.685 | -1.349 | 0.178 | 0 | -0.001 | -0.001 | 0.089 | 0.56 | 0.44 | 0.54 | 0.65 | 0.51 | 0.48 |
| 8 | 106482105 | rs6983550 | -0.767 | 0.746 | -1.811 | 0.185 | -0.002 | 0.008 | 0.008 | 0 | 0.65 | 0.65 | 0.6 | 0.79 | 0.63 | 0.63 |
| 8 | 106492618 | rs7009208 | 0.338 | 0.506 | -1.662 | 0.298 | 0.062 | 0.074 | 0.074 | 0.015 | 0.11 | 0.34 | 0.06 | 0.15 | 0.12 | 0.16 |
| 8 | 106503845 | rs2122444 | -0.097 | 0.001 | -2.931 | 0.146 | 0.007 | 0.018 | 0.018 | 1.515 | 0.35 | 0.02 | 0.25 | 0.2 | 0.33 | 0.18 |
| 8 | 106505971 | rs2622637 | -0.576 | 0.89 | -2.295 | 0.544 | -0.005 | -0.022 | -0.022 | 0 | 0.37 | 0.25 | 0.48 | 0.5 | 0.38 | 0.43 |
| 8 | 106509975 | rs2622633 | -0.046 | 1.075 | -2.053 | 0.273 | 0.003 | 0.006 | 0.006 | 0 | 0.25 | 0.2 | 0.26 | 0.09 | 0.23 | 0.31 |
| 8 | 106510518 | rs1825074 | -0.055 | 1.692 | -2.194 | 0.829 | -0.106 | -0.059 | -0.059 | 0 | 0.78 | 0.96 | 0.85 | 0.77 | 0.67 | 0.77 |
| 8 | 106511690 | rs302954 | -0.521 | 1.34 | -1.314 | 0.875 | 0.026 | 0.037 | 0.037 | 0 | 0.31 | 0.44 | 0.43 | 0.46 | 0.36 | 0.21 |
| 8 | 106512481 | rs302953 | 0.099 | 1.039 | -2.612 | 0.519 | -0.027 | -0.004 | -0.004 | 0.877 | 0.5 | 0.09 | 0.42 | 0.43 | 0.57 | 0.53 |
| 8 | 106513301 | rs6988664 | -0.039 | 1.113 | -2.06 | 0.617 | 0.03 | 0.012 | 0.012 | 0.828 | 0.38 | 0.67 | 0.32 | 0.33 | 0.29 | 0.38 |
| 8 | 106515179 | rs12680351 | 0.727 | 0.663 | -1.711 | 0.352 | 0 | -0.009 | -0.009 | 0 | 0.22 | 0.14 | 0.23 | 0.26 | 0.21 | 0.17 |
| 8 | 106517641 | rs302959 | -0.453 | 0.96 | -2.588 | 0.002 | -0.146 | 0.087 | 0.087 | 0.093 | 0.71 | 0.95 | 0.73 | 0.85 | 0.6 | 0.69 |
| 8 | 106517677 | rs302960 | 1.72 | 0.867 | -2.659 | 0.081 | -0.044 | -0.156 | -0.156 | 0.12 | 0.26 | 0.05 | 0.36 | 0.4 | 0.25 | 0.23 |
| 8 | 106518366 | rs302963 | 1.476 | 0.787 | -2.586 | 0.064 | 0.022 | -0.074 | -0.074 | 0.766 | 0.28 | 0.18 | 0.37 | 0.4 | 0.23 | 0.25 |
| 8 | 106518412 | rs302964 | 0.1 | 0.797 | -2.664 | 0.061 | -0.096 | 0.051 | 0.051 | 0 | 0.66 | 0.86 | 0.68 | 0.7 | 0.52 | 0.62 |
| 8 | 106520051 | rs4734871 | -1.864 | 1.004 | -2.67 | 0.125 | -0.046 | -0.131 | -0.131 | 0.035 | 0.74 | 0.96 | 0.64 | 0.65 | 0.73 | 0.79 |
| 8 | 106521845 | rs7007968 | 0.115 | 0.506 | -2.475 | -0.198 | 0.049 | 0.048 | 0.048 | 1.334 | 0.4 | 0.79 | 0.34 | 0.27 | 0.3 | 0.39 |
| 8 | 106523185 | rs10094510 | 0.387 | 0.396 | -1.678 | -0.245 | -0.009 | -0.01 | -0.01 | 0 | 0.34 | 0.39 | 0.29 | 0.26 | 0.29 | 0.37 |
| 8 | 106527975 | rs16873287 | -0.083 | 0.412 | -1.39 | -0.24 | 0.045 | 0.052 | 0.052 | 1.241 | 0.4 | 0.76 | 0.34 | 0.27 | 0.3 | 0.39 |
| 8 | 106529045 | rs12675041 | -0.412 | 0.414 | -1.419 | -0.268 | 0.053 | -0.023 | -0.023 | 0 | 0.45 | 0.84 | 0.37 | 0.45 | 0.35 | 0.47 |
| 8 | 106532477 | rs6996138 | 0.165 | 0.524 | -1.306 | -0.17 | -0.004 | -0.004 | -0.004 | 0.462 | 0.37 | 0.54 | 0.31 | 0.29 | 0.29 | 0.37 |
| 8 | 106536253 | rs2343298 | -1.593 | 1.008 | -1.434 | -0.181 | -0.019 | -0.103 | -0.103 | 0.368 | 0.72 | 0.89 | 0.64 | 0.59 | 0.71 | 0.73 |
| 8 | 106537490 | rs13250492 | 0.645 | 0.725 | -1.473 | -0.329 | -0.006 | 0.007 | 0.007 | 0 | 0.17 | 0.08 | 0.09 | 0.16 | 0.16 | 0.13 |
| 8 | 106537714 | rs996488 | 0.115 | 0.661 | -1.388 | -0.53 | -0.001 | 0.006 | 0.006 | 0.147 | 0.42 | 0.59 | 0.48 | 0.4 | 0.43 | 0.52 |
| 8 | 106538051 | rs1450164 | 0.697 | 0.796 | -1.118 | -0.497 | 0.02 | -0.006 | -0.006 | 0 | 0.13 | 0.21 | 0.2 | 0.14 | 0.19 | 0.2 |
| 8 | 106543197 | rs6988352 | -0.396 | 0.272 | -1.11 | -0.528 | -0.013 | -0.015 | -0.015 | 0.635 | 0.25 | 0.3 | 0.11 | 0.19 | 0.19 | 0.18 |
| 8 | 106545190 | rs11786961 | 0.833 | 0.233 | -1.116 | -0.524 | 0.007 | 0.003 | 0.003 | 0 | 0.16 | 0.09 | 0.07 | 0.16 | 0.13 | 0.13 |
| 8 | 106545794 | rs13280496 | -1.013 | 0.135 | -1.142 | -0.452 | -0.005 | -0.018 | -0.018 | 0 | 0.37 | 0.53 | 0.3 | 0.34 | 0.35 | 0.39 |
| 8 | 106547983 | rs1375961 | 1.011 | 0.186 | -1.232 | -0.535 | 0 | 0 | 0 | 0 | 0.12 | 0.15 | 0.17 | 0.14 | 0.14 | 0.19 |
| 8 | 106550669 | rs11778194 | 0.661 | 0.175 | -1.356 | -0.5 | -0.004 | 0.004 | 0.004 | 0 | 0.17 | 0.08 | 0.09 | 0.17 | 0.16 | 0.14 |
| 8 | 106558235 | rs16873365 | 1.734 | -0.146 | -1.553 | -0.756 | -0.044 | -0.07 | -0.07 | 0 | 0.15 | 0.01 | 0.17 | 0.26 | 0.16 | 0.14 |
| 8 | 106561277 | rs1037306 | -0.429 | 0.273 | -0.962 | -0.575 | -0.004 | -0.017 | -0.017 | 0 | 0.2 | 0.32 | 0.21 | 0.16 | 0.19 | 0.24 |
| 8 | 106564068 | rs13254292 | 0.418 | 0.404 | -1.005 | -0.414 | -0.005 | 0.029 | 0.029 | 0 | 0.17 | 0.08 | 0.09 | 0.12 | 0.16 | 0.14 |
| 8 | 106573578 | rs3735953 | -1.417 | 0.36 | -1.066 | -0.66 | -0.106 | -0.152 | -0.152 | 0.023 | 0.71 | 0.96 | 0.59 | 0.62 | 0.6 | 0.72 |
| 8 | 106582169 | rs13279114 | 0.779 | 0.434 | -0.857 | -0.658 | 0.01 | -0.001 | -0.001 | 0 | 0.11 | 0.2 | 0.17 | 0.13 | 0.14 | 0.18 |
| 8 | 106583124 | rs4734879 | 0.887 | 0.426 | -0.912 | -0.638 | -0.009 | 0.002 | 0.002 | 0 | 0.36 | 0.42 | 0.31 | 0.4 | 0.29 | 0.36 |
| 8 | 106583872 | rs1470684 | -0.59 | 0.705 | -1.66 | -0.443 | -0.019 | -0.042 | -0.042 | 0 | 0.64 | 0.74 | 0.56 | 0.59 | 0.58 | 0.67 |
| 8 | 106588140 | rs13270854 | 0.264 | 1.015 | -1.849 | -0.369 | -0.018 | 0.062 | 0.062 | 0 | 0.16 | 0.03 | 0.07 | 0.04 | 0.15 | 0.12 |
| 8 | 106591802 | rs16873415 | -0.439 | 1.011 | -1.591 | -0.034 | -0.01 | -0.035 | -0.035 | 0.174 | 0.55 | 0.82 | 0.5 | 0.56 | 0.46 | 0.61 |
| 8 | 106592145 | rs16873418 | 0.034 | 0.51 | -1.399 | -0.136 | 0.032 | 0.005 | 0.005 | 0 | 0.43 | 0.7 | 0.33 | 0.44 | 0.32 | 0.43 |
| 8 | 106593662 | rs6995272 | 0.452 | 1.287 | -1.079 | 0.468 | 0.006 | 0.008 | 0.008 | 0 | 0.7 | 0.62 | 0.45 | 0.53 | 0.54 | 0.51 |

| | | | | | | | | | | | | | | | | |
|---|---|---|---|---|---|---|---|---|---|---|---|---|---|---|---|---|
| 8 | 106606008 | rs7015353 | -0.554 | 0.788 | -1.453 | -0.075 | 0.006 | 0.052 | 0.052 | 0 | 0.33 | 0.19 | 0.26 | 0.18 | 0.34 | 0.42 |
| 8 | 106610368 | rs10111812 | 1.134 | 0.947 | -1.226 | -0.987 | 0.01 | -0.009 | -0.009 | 0 | 0.4 | 0.33 | 0.22 | 0.01 | 0.32 | 0.18 |
| 8 | 106610810 | rs6982399 | 0.824 | 1.118 | -1.694 | -0.789 | -0.001 | -0.002 | -0.002 | 0 | 0.09 | 0.1 | 0.1 | 0.14 | 0.14 | 0.24 |
| 8 | 106624958 | rs1375960 | 0.668 | 0.975 | -1.788 | -0.914 | 0.024 | 0.132 | 0.132 | 0 | 0.21 | 0.02 | 0.07 | 0.01 | 0.14 | 0.15 |
| 8 | 106625030 | rs16873442 | 1.062 | 1.246 | -1.83 | -0.058 | -0.012 | 0.008 | 0.008 | 0 | 0.07 | 0.03 | 0.22 | 0.36 | 0.08 | 0.06 |
| 8 | 106627197 | rs1037304 | -0.808 | 1.014 | -1.537 | -0.368 | 0.005 | 0.035 | 0.035 | 0 | 0.33 | 0.26 | 0.26 | 0.18 | 0.34 | 0.42 |
| 8 | 106628965 | rs1817023 | -0.221 | 1.387 | -1.196 | 0.139 | 0.01 | 0.008 | 0.008 | 0 | 0.12 | 0.15 | 0.18 | 0.17 | 0.2 | 0.27 |
| 8 | 106651423 | rs13263514 | 0.334 | 1.394 | -1.228 | 0.132 | 0.007 | 0.003 | 0.003 | 0 | 0.09 | 0.13 | 0.09 | 0.13 | 0.13 | 0.24 |
| 8 | 106654054 | rs9656866 | -0.393 | 1.373 | -1.058 | 0.121 | 0.016 | 0.011 | 0.011 | 0 | 0.12 | 0.18 | 0.18 | 0.16 | 0.19 | 0.27 |
| 8 | 106664873 | rs10105946 | -0.487 | 1.507 | -1.198 | 0.831 | -0.005 | 0.01 | 0.01 | 0.035 | 0.9 | 0.94 | 0.77 | 0.72 | 0.8 | 0.71 |
| 8 | 106665439 | rs11985933 | 0.106 | 0.57 | -1.473 | 0.378 | -0.005 | -0.005 | -0.005 | 0 | 0.39 | 0.58 | 0.44 | 0.46 | 0.42 | 0.31 |
| 8 | 106672991 | rs6986134 | 0.117 | 0.212 | -1.725 | 0.12 | -0.001 | -0.004 | -0.004 | 2.324 | 0.64 | 0.6 | 0.6 | 0.66 | 0.64 | 0.54 |
| 8 | 106673800 | rs1442330 | -0.123 | -0.003 | -1.174 | 0.193 | -0.003 | -0.002 | -0.002 | 0.016 | 0.39 | 0.56 | 0.44 | 0.46 | 0.42 | 0.31 |
| 8 | 106674812 | rs2218279 | -0.316 | -0.089 | -1.181 | 0.288 | 0.004 | 0.003 | 0.003 | 0 | 0.42 | 0.7 | 0.51 | 0.47 | 0.43 | 0.33 |
| 8 | 106675898 | rs1372618 | -1.161 | -0.403 | -0.539 | 0.098 | -0.016 | -0.021 | -0.021 | 0 | 0.1 | 0.2 | 0.08 | 0.04 | 0.07 | 0.1 |
| 8 | 106692498 | rs4734889 | -1.307 | -0.812 | -0.255 | -0.185 | -0.059 | -0.09 | -0.09 | 0.075 | 0.3 | 0.53 | 0.17 | 0.08 | 0.2 | 0.23 |
| 8 | 106696670 | rs10104574 | -0.74 | -0.201 | -0.814 | 0.328 | -0.017 | -0.018 | -0.018 | 0.005 | 0.21 | 0.07 | 0.27 | 0.22 | 0.21 | 0.33 |
| 8 | 106704846 | rs7008088 | -0.455 | -0.308 | -0.343 | 0.21 | -0.006 | -0.013 | -0.013 | 0.073 | 0.16 | 0.16 | 0.06 | 0.03 | 0.09 | 0.12 |
| 8 | 106707739 | rs6469010 | -0.943 | 0.045 | 0.243 | 0.612 | 0.011 | 0.033 | 0.033 | 0 | 0.14 | 0.22 | 0.22 | 0.14 | 0.13 | 0.33 |
| 8 | 106715359 | rs7826613 | -0.496 | -0.266 | 0.482 | 0.394 | 0.033 | 0.035 | 0.035 | 0.108 | 0.16 | 0.08 | 0.06 | 0.02 | 0.08 | 0.09 |
| 8 | 106715767 | rs7845449 | -0.444 | -0.367 | 0.363 | 0.878 | -0.014 | 0.228 | 0.228 | 0.382 | 0.17 | 0.34 | 0.12 | 0.5 | 0.16 | 0.06 |
| 8 | 106719773 | rs6980689 | -1.089 | -0.287 | 0.209 | 0.464 | -0.011 | 0.009 | 0.009 | 0 | 0.29 | 0.37 | 0.34 | 0.63 | 0.27 | 0.38 |
| 8 | 106720087 | rs1442335 | -1.568 | -0.058 | -0.119 | 0.477 | 0.045 | 0.099 | 0.099 | 0.449 | 0.49 | 0.66 | 0.46 | 0.69 | 0.42 | 0.51 |
| 8 | 106724370 | rs10808407 | -1.284 | 0.414 | 0.332 | 0.706 | -0.002 | -0.026 | -0.026 | 0 | 0.32 | 0.58 | 0.26 | 0.16 | 0.34 | 0.16 |
| 8 | 106725186 | rs7003445 | -1.726 | 0.463 | 0.441 | 1.071 | 0.004 | 0.015 | 0.015 | 0 | 0.42 | 0.67 | 0.37 | 0.33 | 0.4 | 0.25 |
| 8 | 106729039 | rs16873564 | -1.177 | 0.303 | 1.182 | 0.973 | -0.004 | -0.08 | -0.08 | 0.803 | 0.29 | 0.4 | 0.21 | 0.04 | 0.26 | 0.14 |
| 8 | 106730235 | rs7823089 | -1.876 | 0.294 | 1.178 | 1.206 | -0.009 | 0.132 | 0.132 | 0 | 0.13 | 0.27 | 0.24 | 0.31 | 0.11 | 0.11 |
| 8 | 106746435 | rs6469013 | -2.095 | 0.371 | 1.108 | 1.271 | -0.009 | 0.003 | 0.003 | 0 | 0.34 | 0.57 | 0.31 | 0.33 | 0.3 | 0.22 |
| 8 | 106755759 | rs1442326 | -1.839 | 0.447 | 0.881 | 1.388 | -0.011 | -0.051 | -0.051 | 4.67 | 0.28 | 0.15 | 0.37 | 0.35 | 0.29 | 0.2 |
| 8 | 106763729 | rs2343547 | -1.833 | 0.5 | 1.02 | 1.672 | -0.001 | 0.028 | 0.028 | 0 | 0.1 | 0.1 | 0.1 | 0.33 | 0.1 | 0.1 |
| 8 | 106773803 | rs7817128 | -1.565 | 0.509 | 1.356 | 1.773 | 0.03 | 0.051 | 0.051 | 0 | 0.11 | 0.47 | 0.16 | 0.09 | 0.08 | 0.03 |
| 8 | 106778064 | rs6469014 | -2.572 | 0.514 | 1.332 | 1.724 | -0.002 | 0.209 | 0.209 | 0 | 0.18 | 0.53 | 0.26 | 0.41 | 0.16 | 0.13 |
| 8 | 106780628 | rs10505082 | -2.572 | 0.511 | 1.332 | 1.724 | 0.002 | 0.209 | 0.209 | 0 | 0.18 | 0.53 | 0.26 | 0.41 | 0.16 | 0.13 |
| 8 | 106781720 | rs10105425 | -2.564 | 0.511 | 1.328 | 1.713 | 0.003 | 0.22 | 0.22 | 0 | 0.18 | 0.52 | 0.26 | 0.41 | 0.16 | 0.13 |
| 8 | 106787615 | rs10505084 | -2.258 | 0.505 | 1.278 | 1.736 | 0.047 | 0.123 | 0.123 | 2.666 | 0.08 | 0.45 | 0.06 | 0.1 | 0.07 | 0.06 |
| 8 | 106787935 | rs10505085 | -0.615 | 0.438 | 1.291 | 1.568 | 0.001 | -0.025 | -0.025 | 0 | 0.06 | 0.09 | 0.05 | 0 | 0.06 | 0.02 |
| 8 | 106792591 | rs1442324 | -2.248 | 0.941 | 0.581 | 1.102 | -0.096 | -0.068 | -0.068 | 0 | 0.5 | 0.96 | 0.6 | 0.46 | 0.53 | 0.31 |
| 8 | 106812527 | rs6990997 | -2.001 | 0.911 | 1.021 | 0.608 | 0.103 | -0.01 | -0.01 | 0 | 0.08 | 0.43 | 0.09 | 0.01 | 0.09 | 0.03 |
| 8 | 106813672 | rs920628 | -1.964 | 0.959 | 1.011 | 0.59 | 0.054 | 0.023 | 0.023 | 4.222 | 0.06 | 0.39 | 0.05 | 0.01 | 0.03 | 0.05 |
| 8 | 106815517 | rs11995760 | -1.929 | 0.96 | 0.984 | 0.59 | 0.056 | 0.025 | 0.025 | 5.53 | 0.06 | 0.39 | 0.05 | 0.01 | 0.03 | 0.05 |
| 8 | 106816281 | rs6469016 | -1.961 | 0.96 | 0.984 | 0.59 | 0.054 | 0.023 | 0.023 | 4.222 | 0.06 | 0.39 | 0.05 | 0.01 | 0.03 | 0.05 |
| 8 | 106837240 | rs2958705 | -2.547 | 0.992 | 0.55 | 0.696 | -0.1 | -0.044 | -0.044 | 0.077 | 0.5 | 0.97 | 0.5 | 0.38 | 0.52 | 0.36 |
| 8 | 106840940 | rs2028553 | -1.581 | 0.843 | 0.719 | 0.782 | 0.009 | -0.053 | -0.053 | 0 | 0.28 | 0.45 | 0.22 | 0.13 | 0.3 | 0.17 |
| 8 | 106853152 | rs2920051 | -2.722 | 1.077 | 0.749 | 0.808 | 0.061 | 0.038 | 0.038 | 1.592 | 0.12 | 0.44 | 0.13 | 0.13 | 0.16 | 0.11 |
| 8 | 106853639 | rs1372614 | -2.141 | 0.914 | 0.779 | 0.851 | 0.044 | 0.082 | 0.082 | 2.768 | 0.09 | 0.4 | 0.1 | 0.13 | 0.1 | 0.08 |
| 8 | 106876334 | rs11996743 | -2.168 | 0.871 | 0.213 | 0.7 | -0.027 | -0.024 | -0.024 | 0.012 | 0.45 | 0.88 | 0.46 | 0.38 | 0.44 | 0.4 |
| 8 | 106880649 | rs11986537 | -2.138 | 0.816 | 0.454 | 0.866 | 0.014 | -0.091 | -0.091 | 0 | 0.26 | 0.89 | 0.32 | 0.36 | 0.23 | 0.31 |
| 8 | 106890347 | rs11986214 | -2.238 | 0.63 | 1.28 | 0.785 | 0.035 | 0.031 | 0.031 | 2.126 | 0.06 | 0.39 | 0.04 | 0.01 | 0 | 0.03 |
| 8 | 106899886 | rs7829063 | -1.345 | 0.704 | 1.224 | 0.898 | -0.005 | 0.101 | 0.101 | 0 | 0.13 | 0.28 | 0.19 | 0.23 | 0.12 | 0.24 |
| 8 | 106900146 | rs16873901 | -0.679 | 0.768 | 0.617 | 0.892 | 0.001 | 0.001 | 0.001 | 0 | 0.08 | 0.14 | 0.08 | 0.13 | 0.13 | 0.04 |
| 8 | 106905987 | rs7002447 | -2.434 | 0.749 | 0.446 | 0.843 | 0.009 | -0.09 | -0.09 | 0.391 | 0.27 | 0.89 | 0.32 | 0.37 | 0.25 | 0.31 |
| 8 | 106916390 | rs16873925 | -2.405 | 0.739 | 0.459 | 0.806 | 0.005 | -0.081 | -0.081 | 0.804 | 0.27 | 0.89 | 0.3 | 0.34 | 0.25 | 0.3 |
| 8 | 106916907 | rs16873927 | -0.209 | 0.75 | 0.662 | 0.896 | 0.004 | 0.02 | 0.02 | 0.689 | 0.07 | 0.15 | 0.13 | 0.16 | 0.11 | 0.04 |
| 8 | 106917162 | rs6995395 | -1.509 | 0.703 | 1.292 | 0.822 | -0.018 | 0.07 | 0.07 | 0 | 0.15 | 0.4 | 0.12 | 0.17 | 0.11 | 0.23 |
| 8 | 106925995 | rs4543516 | -1.975 | 0.697 | 1.791 | 0.658 | 0.024 | -0.041 | -0.041 | 0.811 | 0.11 | 0.48 | 0.08 | 0.02 | 0.07 | 0.1 |
| 8 | 106926096 | rs12542647 | -1.007 | 0.702 | 1.749 | 0.858 | -0.006 | -0.039 | -0.039 | 0 | 0.12 | 0.05 | 0.26 | 0.32 | 0.14 | 0.2 |
| 8 | 106926778 | rs10955406 | -2.35 | 0.353 | 1.134 | 0.42 | -0.043 | -0.075 | -0.075 | 0 | 0.46 | 0.96 | 0.47 | 0.4 | 0.4 | 0.36 |
| 8 | 106942537 | rs7007134 | -1.842 | 0.381 | 1.531 | 0.456 | 0.011 | -0.056 | -0.056 | 0 | 0.12 | 0.46 | 0.08 | 0.02 | 0.09 | 0.1 |
| 8 | 106943245 | rs1353449 | -2.594 | 0.383 | 1.505 | 0.495 | 0.012 | -0.021 | -0.021 | 0.57 | 0.26 | 0.81 | 0.37 | 0.34 | 0.23 | 0.3 |
| 8 | 106952331 | rs7824634 | -1.671 | 0.474 | 1.757 | 0.383 | 0.055 | 0.023 | 0.023 | 0.312 | 0.1 | 0.51 | 0.09 | 0.07 | 0.09 | 0.07 |
| 8 | 106962618 | rs10098367 | -2.929 | 0.457 | 1.669 | 0.561 | 0.008 | -0.048 | -0.048 | 1.085 | 0.29 | 0.84 | 0.43 | 0.43 | 0.28 | 0.33 |
| 8 | 106965511 | rs6997487 | -2.689 | 0.41 | 1.352 | 0.22 | -0.035 | 0.027 | 0.027 | 0.015 | 0.53 | 0.75 | 0.56 | 0.36 | 0.52 | 0.43 |
| 8 | 106974182 | rs16873995 | -0.47 | 0.417 | 1.253 | 0.223 | -0.001 | 0.048 | 0.048 | 0 | 0.27 | 0.12 | 0.15 | 0 | 0.26 | 0.12 |
| 8 | 106989558 | rs6469023 | -1.666 | 0.332 | 1.199 | 0.225 | -0.005 | 0.013 | 0.013 | 0.416 | 0.43 | 0.18 | 0.46 | 0.35 | 0.44 | 0.35 |
| 8 | 107024505 | rs1496182 | -1.853 | 0.338 | 1.216 | 0.214 | 0.002 | 0.01 | 0.01 | 0.138 | 0.44 | 0.33 | 0.48 | 0.36 | 0.44 | 0.38 |
| 8 | 107031319 | rs1603363 | -1.825 | 0.336 | 1.203 | 0.213 | 0.002 | 0.013 | 0.013 | 0.047 | 0.44 | 0.31 | 0.48 | 0.36 | 0.44 | 0.38 |
| 8 | 107042333 | rs7005320 | -0.667 | 0.362 | 1.078 | 0.215 | 0.037 | -0.023 | -0.023 | 0 | 0.08 | 0.27 | 0.06 | 0.01 | 0.08 | 0.05 |
| 8 | 107043797 | rs6415444 | -2.539 | 0.364 | 0.742 | 0.23 | -0.114 | -0.017 | -0.017 | 0.322 | 0.55 | 0.92 | 0.57 | 0.36 | 0.52 | 0.43 |
| 8 | 107090000 | rs12541635 | 1.824 | -0.078 | 0.797 | -0.085 | 0.003 | 0.001 | 0.001 | 0.278 | 0.52 | 0.54 | 0.42 | 0.61 | 0.53 | 0.63 |

| | | | | | | | | | | | | | | | | |
|---|---|---|---|---|---|---|---|---|---|---|---|---|---|---|---|---|
| 8 | 107091636 | rs10955413 | 1.125 | 0.009 | 0.535 | 0.093 | -0.001 | 0.022 | 0.022 | 0.466 | 0.83 | 0.83 | 0.58 | 0.61 | 0.82 | 0.76 |
| 8 | 107093203 | rs6469035 | -0.438 | 0.189 | 0.389 | 0.193 | -0.038 | -0.044 | -0.044 | 0.052 | 0.93 | 0.99 | 0.83 | 0.85 | 0.88 | 0.96 |
| 8 | 107096458 | rs4734130 | 1.407 | 0.029 | 0.598 | -0.024 | 0.01 | 0.007 | 0.007 | 0.057 | 0.53 | 0.33 | 0.42 | 0.61 | 0.57 | 0.62 |
| 8 | 107104545 | rs6989082 | 0.39 | -0.013 | 0.356 | -0.405 | -0.037 | -0.064 | -0.064 | 0.425 | 0.69 | 0.9 | 0.54 | 0.68 | 0.67 | 0.67 |
| 8 | 107111290 | rs6469040 | -2.332 | -0.066 | 0.406 | -0.473 | 0.003 | 0.1 | 0.1 | 8.05 | 0.24 | 0.93 | 0.2 | 0.08 | 0.2 | 0.14 |
| 8 | 107115261 | rs7831526 | -0.755 | -0.072 | 0.785 | -0.481 | -0.005 | -0.097 | -0.097 | 0 | 0.13 | 0.46 | 0.11 | 0 | 0.1 | 0.04 |
| 8 | 107125612 | rs7463839 | 0.157 | 0.071 | 0.718 | -0.596 | 0.022 | 0.025 | 0.025 | 0.009 | 0.9 | 0.76 | 0.81 | 0.89 | 0.84 | 0.88 |
| 8 | 107162795 | rs11776357 | 0.233 | 0.027 | 0.595 | -0.725 | -0.006 | -0.051 | -0.051 | 0.307 | 0.1 | 0.14 | 0.07 | 0 | 0.1 | 0.02 |
| 8 | 107163978 | rs1366963 | 1.803 | 0.038 | 0.004 | -0.725 | -0.007 | 0.068 | 0.068 | 7.294 | 0.73 | 0.06 | 0.69 | 0.88 | 0.74 | 0.83 |
| 8 | 107169324 | rs2914875 | 1.786 | 0.014 | 0.108 | -0.732 | 0.001 | 0.003 | 0.003 | 1.956 | 0.75 | 0.22 | 0.71 | 0.88 | 0.74 | 0.84 |
| 8 | 107170543 | rs13282530 | 0.425 | 0.003 | -0.566 | -1.193 | -0.056 | -0.001 | -0.001 | 0 | 0.15 | 0.01 | 0.11 | 0.1 | 0.16 | 0.07 |
| 8 | 107175621 | rs2914860 | 0.049 | 0.006 | -0.277 | -0.923 | -0.1 | -0.029 | -0.029 | 0 | 0.36 | 0.03 | 0.41 | 0.27 | 0.37 | 0.35 |
| 8 | 107187838 | rs6995297 | -1.779 | -0.12 | 1.141 | -0.673 | -0.016 | -0.061 | -0.061 | 0.467 | 0.17 | 0.72 | 0.12 | 0.01 | 0.1 | 0.04 |
| 8 | 107210629 | rs1503565 | 0.004 | 0.043 | 0.854 | -0.491 | 0 | 0.002 | 0.002 | 0 | 0.85 | 0.83 | 0.71 | 0.84 | 0.8 | 0.87 |
| 8 | 107214049 | rs7824769 | -1.811 | -0.124 | 1.399 | -0.542 | 0.023 | 0.025 | 0.025 | 7.65 | 0.06 | 0.36 | 0.03 | 0.01 | 0 | 0.03 |
| 8 | 107219431 | rs1503567 | 1.17 | -0.033 | 1.294 | -0.63 | -0.001 | -0.013 | -0.013 | 0 | 0.77 | 0.56 | 0.64 | 0.84 | 0.74 | 0.85 |
| 8 | 107225803 | rs2062826 | 1.653 | -0.008 | 0.718 | -0.713 | 0.013 | 0.032 | 0.032 | 0.461 | 0.68 | 0.23 | 0.65 | 0.83 | 0.67 | 0.82 |
| 8 | 107230594 | rs11993836 | 1.486 | 0.006 | 0.965 | -0.696 | -0.008 | -0.021 | -0.021 | 0 | 0.69 | 0.43 | 0.66 | 0.83 | 0.68 | 0.82 |
| 8 | 107243508 | rs968817 | -0.062 | 0.037 | 0.784 | -0.723 | 0.003 | 0.006 | 0.006 | 0 | 0.85 | 0.81 | 0.74 | 0.84 | 0.78 | 0.87 |
| 8 | 107249559 | rs958630 | 1.654 | -0.068 | 0.62 | -0.792 | 0.017 | 0.051 | 0.051 | 1.172 | 0.67 | 0.2 | 0.63 | 0.83 | 0.67 | 0.83 |
| 8 | 107250284 | rs3897874 | -0.081 | -0.076 | 0.754 | -0.799 | -0.001 | -0.044 | -0.044 | 0.055 | 0.1 | 0.37 | 0.06 | 0.01 | 0.07 | 0.02 |
| 8 | 107254937 | rs1393976 | 1.368 | -0.059 | -0.277 | -0.769 | 0.017 | 0.067 | 0.067 | 3.649 | 0.65 | 0.11 | 0.62 | 0.83 | 0.67 | 0.82 |
| 8 | 107276200 | rs1018145 | -0.988 | 0.005 | -0.134 | -0.785 | 0.017 | 0.016 | 0.016 | 3.718 | 0.27 | 0.77 | 0.3 | 0.16 | 0.2 | 0.18 |
| 8 | 107278277 | rs2938305 | -0.975 | 0.013 | -0.132 | -0.781 | 0.016 | 0.016 | 0.016 | 3.597 | 0.27 | 0.77 | 0.3 | 0.16 | 0.2 | 0.18 |
| 8 | 107280050 | rs2938304 | -0.298 | 0.817 | -0.807 | -1.001 | -0.101 | -0.028 | -0.028 | 0 | 0.8 | 0.99 | 0.77 | 0.84 | 0.73 | 0.79 |
| 8 | 107298011 | rs1503559 | -0.276 | 0.287 | 0.491 | -0.924 | -0.026 | -0.12 | -0.12 | 0 | 0.19 | 0.48 | 0.13 | 0.02 | 0.13 | 0.07 |
| 8 | 107300650 | rs3019308 | 0.938 | 0.238 | -0.096 | -0.956 | 0.016 | -0.025 | -0.025 | 0 | 0.34 | 0.23 | 0.31 | 0.58 | 0.32 | 0.41 |
| 8 | 107312556 | rs1503573 | -0.234 | 0.267 | -0.114 | -1.001 | 0.02 | 0.027 | 0.027 | 0 | 0.52 | 0.73 | 0.45 | 0.6 | 0.47 | 0.48 |
| 8 | 107323253 | rs1587737 | 0.279 | 0.236 | 0.072 | -0.962 | 0.034 | 0.032 | 0.032 | 0.129 | 0.05 | 0.14 | 0.07 | 0.08 | 0.06 | 0.03 |
| 8 | 107326839 | rs7013285 | 0.505 | 0.213 | 0.239 | -0.973 | -0.015 | -0.09 | -0.09 | 0.482 | 0.18 | 0.46 | 0.12 | 0.02 | 0.12 | 0.07 |
| 8 | 107326900 | rs7013315 | -0.649 | 0.061 | -0.141 | -1.055 | -0.003 | -0.014 | -0.014 | 0 | 0.27 | 0.68 | 0.22 | 0.1 | 0.2 | 0.21 |
| 8 | 107328133 | rs1472698 | -0.963 | 0.408 | -0.17 | -0.993 | -0.002 | -0.017 | -0.017 | 0 | 0.63 | 0.76 | 0.64 | 0.42 | 0.58 | 0.55 |
| 8 | 107338553 | rs6469058 | -0.422 | 0.46 | -0.115 | -1.081 | -0.005 | -0.034 | -0.034 | 0.508 | 0.26 | 0.65 | 0.23 | 0.1 | 0.2 | 0.12 |
| 8 | 107344358 | rs2345799 | -1.063 | 0.414 | -0.237 | -1.108 | -0.002 | -0.015 | -0.015 | 0 | 0.63 | 0.76 | 0.64 | 0.42 | 0.58 | 0.55 |
| 8 | 107346131 | rs7814702 | 0.068 | 0.353 | 0.171 | -1.103 | -0.021 | -0.118 | -0.118 | 0.184 | 0.21 | 0.5 | 0.15 | 0.02 | 0.14 | 0.07 |
| 8 | 107355371 | rs905557 | -0.383 | 0.196 | -0.047 | -1.249 | -0.026 | -0.104 | -0.104 | 0 | 0.23 | 0.62 | 0.14 | 0 | 0.14 | 0.17 |
| 8 | 107372260 | rs13277825 | -0.213 | 0.439 | 0.083 | -1.004 | -0.001 | 0.004 | 0.004 | 0 | 0.46 | 0.58 | 0.52 | 0.4 | 0.43 | 0.5 |
| 8 | 107375446 | rs16874440 | 0.277 | 0.307 | 0.102 | -1.042 | -0.001 | 0.027 | 0.027 | 0 | 0.39 | 0.22 | 0.45 | 0.32 | 0.36 | 0.46 |
| 8 | 107376277 | rs7833003 | 0.082 | 0.352 | -0.14 | -1.017 | -0.006 | -0.023 | -0.023 | 0.043 | 0.42 | 0.49 | 0.46 | 0.32 | 0.37 | 0.48 |
| 8 | 107386281 | rs17337119 | 0.853 | 0.288 | -0.752 | -0.605 | -0.079 | -0.064 | -0.064 | 0 | 0.38 | 0.07 | 0.46 | 0.38 | 0.46 | 0.36 |
| 8 | 107386941 | rs7836368 | 0.081 | 0.175 | 0.173 | -0.752 | 0.044 | -0.017 | -0.017 | 0 | 0.07 | 0.23 | 0.05 | 0 | 0.05 | 0.01 |
| 8 | 107388668 | rs1954756 | -0.504 | 0.116 | 0.336 | -0.841 | -0.009 | -0.136 | -0.136 | 0 | 0.23 | 0.5 | 0.14 | 0 | 0.16 | 0.05 |
| 8 | 107392965 | rs7839683 | -1.134 | 0.037 | -0.274 | -0.711 | 0.026 | 0.103 | 0.103 | 0.571 | 0.29 | 0.85 | 0.21 | 0.08 | 0.17 | 0.09 |
| 8 | 107395988 | rs1954755 | 0.671 | 0.236 | -0.358 | -0.587 | -0.005 | 0.049 | 0.049 | 0 | 0.15 | 0.07 | 0.07 | 0 | 0.1 | 0.04 |
| 8 | 107398704 | rs2022979 | 0.723 | 0.379 | -0.186 | -0.389 | -0.023 | -0.049 | -0.049 | 0 | 0.41 | 0.19 | 0.52 | 0.48 | 0.5 | 0.42 |
| 8 | 107401813 | rs2022976 | 0.691 | 0.188 | 0.106 | 0.753 | -0.034 | -0.022 | -0.022 | 1.725 | 0.05 | 0 | 0.14 | 0.21 | 0.12 | 0.18 |
| 8 | 107405714 | rs2211924 | 0.066 | -0.086 | 0.386 | 0.343 | -0.006 | 0.025 | 0.025 | 0 | 0.62 | 0.47 | 0.62 | 0.49 | 0.67 | 0.47 |
| 8 | 107408180 | rs10087808 | -0.052 | -0.009 | 0.042 | 0.381 | 0.002 | -0.016 | -0.016 | 0 | 0.63 | 0.62 | 0.64 | 0.49 | 0.67 | 0.47 |
| 8 | 107410067 | rs2022970 | 0.138 | 0.08 | -0.448 | 0.455 | -0.003 | 0.01 | 0.01 | 0 | 0.6 | 0.44 | 0.6 | 0.49 | 0.64 | 0.44 |
| 8 | 107411853 | rs7827856 | -0.691 | -1.01 | -0.524 | 0.361 | 0.015 | 0.009 | 0.009 | 0 | 0.39 | 0.35 | 0.21 | 0.36 | 0.26 | 0.36 |
| 8 | 107411992 | rs10110475 | 0.41 | -0.962 | 0.127 | 0.878 | -0.011 | 0.181 | 0.181 | 0.312 | 0.15 | 0.36 | 0.22 | 0.42 | 0.12 | 0.16 |
| 8 | 107413542 | rs10955421 | -0.214 | -1.224 | -0.17 | 0.675 | 0.027 | -0.001 | -0.001 | 0 | 0.32 | 0.23 | 0.2 | 0.36 | 0.26 | 0.35 |
| 8 | 107419779 | rs9297382 | 0.324 | -1.248 | 0.155 | 0.694 | -0.015 | 0.265 | 0.265 | 0.773 | 0.16 | 0.48 | 0.23 | 0.42 | 0.11 | 0.16 |
| 8 | 107424062 | rs10109171 | -0.803 | -1.478 | -0.04 | 0.223 | -0.013 | -0.005 | -0.005 | 0 | 0.37 | 0.57 | 0.34 | 0.68 | 0.26 | 0.49 |
| 8 | 107462505 | rs1621663 | -1.417 | -1.558 | -0.095 | 0.164 | 0.073 | 0.141 | 0.141 | 1.781 | 0.48 | 0.84 | 0.45 | 0.78 | 0.36 | 0.52 |
| 8 | 107474139 | rs1670387 | -1.417 | -1.616 | -0.422 | 0.135 | 0.073 | 0.141 | 0.141 | 1.781 | 0.48 | 0.84 | 0.45 | 0.78 | 0.36 | 0.52 |
| 8 | 107477037 | rs7817974 | 0.038 | -1.583 | -0.141 | 0.741 | -0.019 | 0.114 | 0.114 | 1.242 | 0.18 | 0.55 | 0.24 | 0.37 | 0.13 | 0.17 |
| 8 | 107477527 | rs12548886 | -0.024 | -1.305 | -0.757 | 0.827 | 0.017 | -0.061 | -0.061 | 0 | 0.19 | 0.1 | 0.11 | 0.41 | 0.13 | 0.33 |
| 8 | 107486045 | rs2510817 | -0.227 | -1.132 | -0.667 | 0.748 | 0.004 | -0.028 | -0.028 | 0 | 0.2 | 0.12 | 0.28 | 0.26 | 0.21 | 0.17 |
| 8 | 107502130 | rs2444322 | -0.053 | -1.125 | -0.467 | 0.743 | -0.001 | -0.001 | -0.001 | 0 | 0.1 | 0.15 | 0.08 | 0.09 | 0.11 | 0.03 |
| 8 | 107503845 | rs1453226 | 0.469 | -1.941 | -0.197 | 0.189 | -0.008 | -0.003 | -0.003 | 0 | 0.6 | 0.5 | 0.67 | 0.34 | 0.73 | 0.52 |
| 8 | 107503974 | rs1453224 | -0.394 | -2.027 | -0.371 | 0.136 | -0.011 | -0.012 | -0.012 | 0.223 | 0.14 | 0.27 | 0.11 | 0.12 | 0.15 | 0.05 |
| 8 | 107506132 | rs2510831 | -1.594 | -2.236 | -0.558 | 0.282 | -0.01 | -0.057 | -0.057 | 0 | 0.9 | 0.98 | 0.96 | 0.69 | 0.92 | 0.87 |
| 8 | 107509019 | rs16874768 | 1.529 | -2.16 | -0.563 | 0.219 | -0.03 | 0.111 | 0.111 | 0.383 | 0.45 | 0.06 | 0.54 | 0.22 | 0.57 | 0.48 |
| 8 | 107511717 | rs7459717 | 1.888 | -2.03 | -1.084 | -0.213 | -0.185 | 0.009 | 0.009 | 0.276 | 0.34 | 0.03 | 0.43 | 0.22 | 0.46 | 0.39 |
| 8 | 107515351 | rs1789983 | -0.388 | -1.85 | -0.098 | 0.076 | 0.026 | -0.055 | -0.055 | 0 | 0.76 | 0.9 | 0.9 | 0.62 | 0.84 | 0.68 |
| 8 | 107519504 | rs2023127 | 0.914 | -1.833 | -0.08 | 0.115 | 0.072 | 0.045 | 0.045 | 0.018 | 0.56 | 0.23 | 0.63 | 0.32 | 0.69 | 0.51 |
| 8 | 107524424 | rs10108020 | 1.591 | -1.667 | -0.119 | 0.244 | -0.005 | -0.038 | -0.038 | 1.017 | 0.11 | 0.05 | 0.2 | 0.28 | 0.11 | 0.15 |
| 8 | 107524985 | rs2444306 | 0.615 | -1.641 | -0.011 | 0.149 | 0.019 | 0.005 | 0.005 | 0 | 0.59 | 0.39 | 0.64 | 0.36 | 0.7 | 0.52 |

| | | | | | | | | | | | | | | | | |
|---|---|---|---|---|---|---|---|---|---|---|---|---|---|---|---|---|
| 8 | 107529963 | rs7819358 | -0.616 | -0.883 | 0.178 | 0.241 | -0.001 | 0.003 | 0.003 | 1.411 | 0.26 | 0.65 | 0.32 | 0.4 | 0.25 | 0.21 |
| 8 | 107539741 | rs1901783 | 0.379 | -0.858 | 0.248 | 0.021 | -0.006 | 0.003 | 0.003 | 0 | 0.65 | 0.56 | 0.78 | 0.65 | 0.66 | 0.68 |
| 8 | 107544383 | rs1453227 | 1.283 | -0.831 | 0.211 | 0.052 | -0.003 | 0.026 | 0.026 | 1.367 | 0.16 | 0.19 | 0.23 | 0.25 | 0.14 | 0.16 |
| 8 | 107545590 | rs6469074 | 1.369 | -0.816 | 0.218 | 0.059 | -0.001 | 0.029 | 0.029 | 1.088 | 0.15 | 0.19 | 0.23 | 0.25 | 0.14 | 0.16 |
| 8 | 107546489 | rs1620490 | 1.102 | -0.81 | -0.13 | -0.111 | 0.001 | -0.009 | -0.009 | 0.221 | 0.08 | 0.09 | 0.08 | 0 | 0.09 | 0.02 |
| 8 | 107546500 | rs4734924 | 1.243 | -0.76 | -0.119 | -0.08 | -0.053 | 0.038 | 0.038 | 0.077 | 0.45 | 0.13 | 0.51 | 0.37 | 0.51 | 0.48 |
| 8 | 107562052 | rs1789968 | 0.696 | -0.557 | -0.102 | -0.19 | 0.002 | 0.005 | 0.005 | 0 | 0.09 | 0.08 | 0.07 | 0 | 0.1 | 0.02 |
| 8 | 107562507 | rs1901778 | 0.271 | -0.281 | -0.278 | -0.141 | -0.003 | -0.012 | -0.012 | 0 | 0.68 | 0.79 | 0.81 | 0.65 | 0.67 | 0.69 |
| 8 | 107567329 | rs4734925 | 1.35 | -0.479 | 0.057 | -0.096 | 0 | 0.024 | 0.024 | 0.22 | 0.43 | 0.37 | 0.49 | 0.23 | 0.43 | 0.46 |
| 8 | 107572320 | rs1453213 | 0.011 | -0.466 | 0.256 | -0.073 | -0.003 | 0.006 | 0.006 | 0.359 | 0.7 | 0.66 | 0.66 | 0.62 | 0.72 | 0.81 |
| 8 | 107579392 | rs2851394 | 0.596 | -0.469 | 0.227 | -0.077 | -0.005 | 0.006 | 0.006 | 0.015 | 0.65 | 0.49 | 0.61 | 0.58 | 0.66 | 0.76 |
| 8 | 107582421 | rs1453212 | 0.79 | -0.361 | 0.079 | 0.065 | -0.001 | -0.007 | -0.007 | 0 | 0.48 | 0.54 | 0.54 | 0.26 | 0.49 | 0.51 |
| 8 | 107585062 | rs7823515 | 0.897 | -0.131 | -0.172 | 0.248 | 0 | 0.007 | 0.007 | 0.55 | 0.18 | 0.14 | 0.25 | 0.34 | 0.17 | 0.17 |
| 8 | 107600050 | rs3110776 | -0.847 | 0.112 | -0.79 | -0.246 | 0 | 0.056 | 0.056 | 0 | 0.52 | 0.73 | 0.32 | 0.35 | 0.47 | 0.38 |
| 8 | 107606729 | rs10092734 | 0.471 | -0.037 | -0.963 | -0.228 | 0.003 | -0.013 | -0.013 | 0 | 0.23 | 0.17 | 0.24 | 0.36 | 0.18 | 0.22 |
| 8 | 107612591 | rs3101545 | -1.422 | -0.06 | -0.972 | -0.238 | -0.036 | 0.023 | 0.023 | 0.02 | 0.64 | 0.8 | 0.49 | 0.67 | 0.54 | 0.51 |
| 8 | 107613017 | rs13277536 | 0.891 | -0.004 | -0.673 | -0.15 | -0.005 | 0.034 | 0.034 | 0 | 0.21 | 0.24 | 0.25 | 0.4 | 0.18 | 0.15 |
| 8 | 107614824 | rs3110432 | -1.148 | 0.101 | -0.77 | -0.177 | 0.012 | -0.012 | -0.012 | 0 | 0.93 | 0.94 | 0.92 | 0.97 | 0.86 | 0.92 |
| 8 | 107615482 | rs9297384 | 0.895 | 0.038 | -0.851 | -0.169 | 0.001 | -0.037 | -0.037 | 0 | 0.2 | 0.14 | 0.24 | 0.4 | 0.18 | 0.16 |
| 8 | 107617360 | rs10505093 | 2.499 | -0.005 | -0.989 | -0.179 | -0.096 | -0.069 | -0.069 | 0 | 0.27 | 0.02 | 0.41 | 0.29 | 0.31 | 0.42 |
| 8 | 107623453 | rs3101525 | 0.021 | -0.098 | -0.5 | -0.155 | -0.011 | -0.018 | -0.018 | 0 | 0.38 | 0.46 | 0.22 | 0.28 | 0.34 | 0.27 |
| 8 | 107634635 | rs13277585 | 0.087 | -0.072 | -0.536 | -0.102 | -0.009 | 0.017 | 0.017 | 0 | 0.06 | 0.03 | 0.06 | 0.03 | 0.05 | 0.08 |
| 8 | 107635026 | rs12114857 | 0.777 | -0.057 | -0.396 | -0.136 | -0.019 | 0.08 | 0.08 | 0.148 | 0.2 | 0.34 | 0.21 | 0.37 | 0.16 | 0.15 |
| 8 | 107637500 | rs10109609 | 1.809 | -0.037 | -0.524 | -0.118 | 0.003 | -0.063 | -0.063 | 0 | 0.15 | 0.05 | 0.17 | 0.34 | 0.13 | 0.14 |
| 8 | 107641107 | rs10096751 | 1.21 | -0.028 | -0.483 | -0.099 | -0.002 | 0.013 | 0.013 | 0 | 0.18 | 0.17 | 0.2 | 0.37 | 0.16 | 0.15 |
| 8 | 107658187 | rs3101528 | -1.177 | 0.004 | -0.483 | -0.259 | -0.015 | 0.014 | 0.014 | 0 | 0.62 | 0.72 | 0.49 | 0.68 | 0.53 | 0.5 |
| 8 | 107667793 | rs2126582 | 0.009 | 0.469 | -0.405 | -0.45 | -0.003 | -0.016 | -0.016 | 0 | 0.44 | 0.54 | 0.26 | 0.21 | 0.38 | 0.27 |
| 8 | 107668532 | rs3101531 | 2.18 | 0.71 | -1.061 | -0.653 | -0.023 | -0.053 | -0.053 | 0 | 0.12 | 0.01 | 0.04 | 0.16 | 0.1 | 0.12 |
| 8 | 107669928 | rs2282509 | -1.719 | 0.739 | -1.02 | -0.435 | -0.04 | -0.022 | -0.022 | 0 | 0.67 | 0.83 | 0.48 | 0.61 | 0.54 | 0.51 |
| 8 | 107682407 | rs28921387 | 1.915 | 0.486 | -0.56 | -0.667 | -0.009 | -0.031 | -0.031 | 0.018 | 0.07 | 0.09 | 0.06 | 0 | 0.06 | 0.02 |
| 8 | 107695648 | rs3102433 | -0.567 | 0.565 | -0.676 | -0.763 | 0.002 | 0.002 | 0.002 | 0.466 | 0.3 | 0.68 | 0.21 | 0.27 | 0.21 | 0.16 |
| 8 | 107698907 | rs3102434 | -0.711 | 0.558 | -0.677 | -0.882 | -0.009 | -0.016 | -0.016 | 0 | 0.32 | 0.61 | 0.17 | 0.2 | 0.22 | 0.16 |
| 8 | 107708167 | rs12548149 | 2.279 | 0.524 | -0.834 | -0.835 | -0.011 | -0.101 | -0.101 | 0.255 | 0.18 | 0.03 | 0.13 | 0.33 | 0.13 | 0.21 |
| 8 | 107709845 | rs3134127 | 2.694 | 0.5 | -0.835 | -0.855 | -0.038 | -0.056 | -0.056 | 0.374 | 0.12 | 0.01 | 0.04 | 0.2 | 0.11 | 0.13 |
| 8 | 107710800 | rs1580429 | 2.054 | 0.573 | -0.799 | -0.859 | -0.016 | 0.025 | 0.025 | 0 | 0.15 | 0.07 | 0.12 | 0.09 | 0.19 | 0.13 |
| 8 | 107730433 | rs776959 | 0.748 | 0.521 | -0.808 | -0.839 | -0.028 | -0.072 | -0.072 | 0 | 0.33 | 0.09 | 0.28 | 0.4 | 0.31 | 0.34 |
| 8 | 107750110 | rs776938 | -2.06 | 0.538 | -0.771 | -0.906 | -0.025 | -0.015 | -0.015 | 0 | 0.67 | 0.79 | 0.48 | 0.64 | 0.55 | 0.49 |
| 8 | 107751849 | rs3016494 | 1.499 | 0.73 | -0.716 | -0.942 | 0.005 | -0.006 | -0.006 | 1.161 | 0.08 | 0.07 | 0.09 | 0.04 | 0.16 | 0.08 |
| 8 | 107759557 | rs28924694 | -0.138 | 0.364 | -0.032 | -0.848 | -0.035 | -0.118 | -0.118 | 0 | 0.18 | 0.35 | 0.11 | 0 | 0.11 | 0.03 |
| 8 | 107782065 | rs13259080 | 1.763 | 0.505 | -0.635 | -0.581 | 0.002 | -0.079 | -0.079 | 0.026 | 0.19 | 0.06 | 0.19 | 0.36 | 0.14 | 0.22 |
| 8 | 107782335 | rs11996457 | -0.424 | 0.421 | -0.419 | -0.757 | -0.022 | -0.045 | -0.045 | 0.251 | 0.31 | 0.54 | 0.17 | 0.19 | 0.22 | 0.16 |
| 8 | 107782395 | rs11996466 | -0.824 | 0.477 | -0.383 | -0.733 | 0.013 | 0.002 | 0.002 | 0 | 0.49 | 0.56 | 0.3 | 0.52 | 0.36 | 0.38 |
| 8 | 107789571 | rs1784486 | 1.03 | 0.682 | -0.413 | -0.773 | -0.009 | 0.122 | 0.122 | 0 | 0.53 | 0.21 | 0.65 | 0.7 | 0.56 | 0.71 |
| 8 | 107790707 | rs11990338 | 0.295 | 0.347 | 0.08 | -0.652 | 0 | -0.031 | -0.031 | 0 | 0.26 | 0.38 | 0.15 | 0.19 | 0.23 | 0.16 |
| 8 | 107791095 | rs16875079 | 0.837 | 0.394 | 0.228 | -0.527 | 0.007 | 0.005 | 0.005 | 0 | 0.19 | 0.37 | 0.12 | 0.19 | 0.15 | 0.13 |
| 8 | 107794157 | rs2981240 | 1.482 | 0.382 | -0.345 | -1.304 | 0.012 | 0.058 | 0.058 | 1.48 | 0.22 | 0.07 | 0.15 | 0.08 | 0.21 | 0.19 |
| 8 | 107796783 | rs1385157 | -0.7 | 0.38 | -0.233 | -1.322 | 0.007 | 0.194 | 0.194 | 0 | 0.47 | 0.81 | 0.36 | 0.17 | 0.43 | 0.33 |
| 8 | 107807716 | rs2930483 | 1.549 | 0.404 | -0.245 | -1.299 | 0.012 | 0.059 | 0.059 | 1.881 | 0.22 | 0.07 | 0.14 | 0.08 | 0.21 | 0.19 |
| 8 | 107810891 | rs1872283 | -2.42 | 0.517 | -0.549 | -1.368 | -0.028 | -0.023 | -0.023 | 0 | 0.75 | 0.85 | 0.62 | 0.67 | 0.65 | 0.59 |
| 8 | 107814129 | rs11987857 | 1.166 | 0.444 | -0.049 | -1.381 | 0.039 | -0.093 | -0.093 | 0.004 | 0.2 | 0.4 | 0.17 | 0.06 | 0.21 | 0.13 |
| 8 | 107816174 | rs3015946 | -0.451 | 0.413 | -0.052 | -1.381 | 0 | 0.056 | 0.056 | 0 | 0.44 | 0.6 | 0.34 | 0.14 | 0.42 | 0.32 |
| 8 | 107819358 | rs11989160 | 0.88 | 0.461 | -0.072 | -1.208 | 0.034 | -0.11 | -0.11 | 0 | 0.21 | 0.45 | 0.17 | 0.06 | 0.21 | 0.13 |
| 8 | 107823148 | rs2449837 | 0.819 | 0.41 | -0.609 | -1.026 | -0.03 | -0.169 | -0.169 | 0 | 0.27 | 0.03 | 0.28 | 0.49 | 0.26 | 0.26 |
| 8 | 107825657 | rs16875124 | 0.638 | 0.327 | -0.227 | -1.218 | -0.002 | -0.085 | -0.085 | 0.099 | 0.35 | 0.37 | 0.22 | 0.06 | 0.29 | 0.22 |
| 8 | 107825988 | rs2513858 | -2.136 | 0.337 | -0.335 | -1.216 | -0.021 | -0.025 | -0.025 | 0 | 0.73 | 0.85 | 0.73 | 0.67 | 0.67 | 0.6 |
| 8 | 107827299 | rs2930488 | 1.253 | 0.401 | 0.212 | -1.211 | 0.006 | 0.002 | 0.002 | 0 | 0.06 | 0.09 | 0.07 | 0.1 | 0.11 | 0.09 |
| 8 | 107830861 | rs2981225 | -0.668 | 0.341 | 0.202 | -1.26 | 0.009 | 0.107 | 0.107 | 0 | 0.45 | 0.76 | 0.31 | 0.18 | 0.4 | 0.32 |
| 8 | 107839322 | rs16875160 | 0.407 | 0.462 | 0.016 | -1.222 | 0 | -0.013 | -0.013 | 0 | 0.33 | 0.26 | 0.2 | 0.06 | 0.28 | 0.21 |
| 8 | 107840533 | rs1484415 | -0.23 | 0.469 | 0.088 | -1.225 | 0 | -0.056 | -0.056 | 0.089 | 0.38 | 0.39 | 0.26 | 0.11 | 0.36 | 0.29 |
| 8 | 107840797 | rs3018923 | 1.51 | 0.6 | -0.204 | -0.875 | 0.008 | 0.002 | 0.002 | 0 | 0.05 | 0.12 | 0.06 | 0.05 | 0.08 | 0.08 |
| 8 | 107842295 | rs2930492 | 1.12 | 0.477 | -0.209 | -0.952 | -0.001 | -0.005 | -0.005 | 0.03 | 0.19 | 0.18 | 0.18 | 0.08 | 0.23 | 0.18 |
| 8 | 107842816 | rs2930493 | -0.703 | 0.141 | -0.116 | -1.04 | 0.025 | 0.129 | 0.129 | 0 | 0.48 | 0.82 | 0.33 | 0.23 | 0.4 | 0.33 |
| 8 | 107842970 | rs3018925 | -0.293 | 0.121 | 0.019 | -1.054 | 0 | -0.004 | -0.004 | 0.41 | 0.42 | 0.58 | 0.29 | 0.18 | 0.37 | 0.3 |
| 8 | 107843050 | rs2449838 | 1.462 | -0.147 | -0.63 | -1.015 | -0.046 | -0.125 | -0.125 | 0.1 | 0.2 | 0.02 | 0.36 | 0.42 | 0.25 | 0.17 |
| 8 | 107845898 | rs16875176 | 0.754 | -0.09 | 0.402 | -1.157 | 0 | -0.002 | -0.002 | 0 | 0.22 | 0.23 | 0.1 | 0.09 | 0.14 | 0.11 |
| 8 | 107860362 | rs10504087 | -0.757 | -0.067 | 0.833 | -0.757 | -0.015 | 0.007 | 0.007 | 0 | 0.13 | 0.31 | 0.09 | 0.11 | 0.09 | 0.09 |
| 8 | 107861997 | rs3018929 | 0.336 | -0.129 | 0.895 | -1.06 | -0.007 | -0.002 | -0.002 | 0.034 | 0.28 | 0.19 | 0.23 | 0.02 | 0.31 | 0.23 |
| 8 | 107867258 | rs3018932 | -1.243 | -0.151 | 1.031 | -1.075 | 0.004 | 0.093 | 0.093 | 0.532 | 0.44 | 0.76 | 0.37 | 0.17 | 0.43 | 0.32 |
| 8 | 107870353 | rs2981258 | -0.894 | -0.049 | 0.775 | -1.035 | -0.001 | 0.02 | 0.02 | 0.031 | 0.42 | 0.66 | 0.35 | 0.13 | 0.41 | 0.32 |

| | | | | | | | | | | | | | | | | |
|---|---|---|---|---|---|---|---|---|---|---|---|---|---|---|---|---|
| 8 | 107870380 | rs16875192 | 0.501 | 0.076 | 0.208 | -0.854 | 0 | -0.005 | -0.005 | 0 | 0.08 | 0.06 | 0.06 | 0.11 | 0.09 | 0.09 |
| 8 | 107874800 | rs2981248 | 0.623 | 0.19 | 0.087 | -0.918 | -0.043 | 0.068 | 0.068 | 0 | 0.24 | 0.11 | 0.24 | 0.02 | 0.32 | 0.2 |
| 8 | 107880864 | rs16875208 | -0.535 | -0.01 | 0.566 | -0.501 | -0.013 | 0.004 | 0.004 | 0.014 | 0.13 | 0.23 | 0.1 | 0.15 | 0.1 | 0.09 |
| 8 | 107881453 | rs16875210 | -1.029 | -0.028 | 0.701 | -0.514 | -0.023 | 0.003 | 0.003 | 0.481 | 0.15 | 0.36 | 0.11 | 0.15 | 0.11 | 0.09 |
| 8 | 107884189 | rs16875214 | -0.357 | -0.153 | 0.606 | -0.476 | -0.026 | 0.009 | 0.009 | 0.014 | 0.12 | 0.23 | 0.08 | 0.15 | 0.07 | 0.08 |
| 8 | 107893477 | rs2981256 | -1.061 | -0.266 | 0.027 | -0.458 | -0.079 | -0.053 | -0.053 | 0.425 | 0.63 | 0.88 | 0.51 | 0.56 | 0.51 | 0.65 |
| 8 | 107898709 | rs3018907 | 1.093 | -0.462 | -0.036 | -0.68 | -0.063 | 0.108 | 0.108 | 0.082 | 0.27 | 0.08 | 0.24 | 0.02 | 0.35 | 0.27 |
| 8 | 107907395 | rs4236779 | -1.263 | -0.459 | 0.056 | -0.668 | -0.057 | -0.033 | -0.033 | 0 | 0.63 | 0.85 | 0.51 | 0.56 | 0.51 | 0.64 |
| 8 | 107908032 | rs6469084 | -0.946 | -0.67 | 0.647 | -0.083 | 0.061 | -0.034 | -0.034 | 0 | 0.36 | 0.78 | 0.27 | 0.54 | 0.16 | 0.4 |
| 8 | 107908221 | rs16875259 | -0.069 | -0.491 | 0.594 | 0.077 | -0.027 | 0.011 | 0.011 | 0 | 0.11 | 0.22 | 0.07 | 0.11 | 0.05 | 0.09 |
| 8 | 107909009 | rs4269515 | 0.551 | -0.121 | 0.49 | 0.185 | -0.014 | 0.042 | 0.042 | 0 | 0.18 | 0.19 | 0.15 | 0.39 | 0.09 | 0.28 |
| 8 | 107916446 | rs11992975 | 0.488 | -0.12 | 0.465 | 0.186 | -0.028 | 0.084 | 0.084 | 0 | 0.19 | 0.25 | 0.15 | 0.39 | 0.09 | 0.28 |
| 8 | 107921739 | rs3922732 | 0.685 | -0.119 | 0.083 | -0.037 | 0.018 | -0.03 | -0.03 | 0 | 0.18 | 0.12 | 0.14 | 0.44 | 0.09 | 0.27 |
| 8 | 107924334 | rs4270934 | -0.195 | 0.034 | 0.049 | -0.46 | 0.003 | -0.002 | -0.002 | 0.135 | 0.49 | 0.31 | 0.35 | 0.49 | 0.44 | 0.55 |
| 8 | 107925727 | rs4734141 | -0.263 | 0.083 | 0.163 | -0.635 | -0.007 | 0.124 | 0.124 | 0.068 | 0.74 | 0.5 | 0.52 | 0.5 | 0.71 | 0.7 |
| 8 | 107928113 | rs7836820 | -1.798 | 0.227 | 0.017 | -0.562 | -0.043 | -0.093 | -0.093 | 0.205 | 0.87 | 0.99 | 0.63 | 0.61 | 0.79 | 0.79 |
| 8 | 107930586 | rs17227435 | 1.609 | 0.221 | 0.055 | -0.702 | -0.025 | 0.064 | 0.064 | 0 | 0.24 | 0.08 | 0.16 | 0.01 | 0.28 | 0.16 |
| 8 | 107932600 | rs4339615 | -1.299 | 0.283 | 0.489 | -0.502 | 0.005 | 0.013 | 0.013 | 0 | 0.84 | 0.78 | 0.67 | 0.78 | 0.77 | 0.79 |
| 8 | 107939696 | rs16875331 | 0.491 | 0.237 | 0.386 | -0.554 | -0.001 | 0.001 | 0.001 | 0 | 0.25 | 0.26 | 0.2 | 0.53 | 0.13 | 0.32 |
| 8 | 107940668 | rs2447243 | 0.04 | 0.398 | 0.523 | -0.449 | -0.004 | 0.025 | 0.025 | 0 | 0.54 | 0.38 | 0.44 | 0.74 | 0.46 | 0.61 |
| 8 | 107940792 | rs2447241 | -1 | 0.303 | 0.578 | -0.45 | 0.012 | 0.022 | 0.022 | 0 | 0.83 | 0.67 | 0.65 | 0.78 | 0.76 | 0.79 |
| 8 | 107944965 | rs1002901 | 0.861 | 0.326 | 0.373 | -0.443 | 0 | -0.036 | -0.036 | 0 | 0.29 | 0.28 | 0.19 | 0.04 | 0.31 | 0.18 |
| 8 | 107945064 | rs2447234 | 0.033 | 0.329 | 0.384 | -0.441 | -0.003 | 0.024 | 0.024 | 0 | 0.54 | 0.39 | 0.46 | 0.74 | 0.46 | 0.6 |
| 8 | 107946720 | rs2447232 | -0.431 | 0.414 | 0.536 | 0.452 | 0.013 | 0.041 | 0.041 | 0 | 0.83 | 0.67 | 0.65 | 0.76 | 0.77 | 0.79 |
| 8 | 107954511 | rs1381484 | -0.404 | 0.326 | 0.578 | 0.747 | -0.004 | 0.019 | 0.019 | 0.043 | 0.1 | 0.18 | 0.08 | 0.13 | 0.08 | 0.12 |
| 8 | 107955559 | rs1462003 | 0.937 | 0.063 | 0.801 | 0.738 | -0.02 | -0.002 | -0.002 | 0 | 0.67 | 0.57 | 0.7 | 0.36 | 0.78 | 0.57 |
| 8 | 107957800 | rs9297387 | 0.763 | 0.053 | 0.65 | 0.753 | 0.009 | -0.004 | -0.004 | 0 | 0.69 | 0.72 | 0.74 | 0.38 | 0.78 | 0.57 |
| 8 | 107961779 | rs2927110 | -0.074 | 0.149 | 0.517 | 0.786 | 0.012 | -0.054 | -0.054 | 1.357 | 0.78 | 0.89 | 0.81 | 0.5 | 0.84 | 0.67 |
| 8 | 107962124 | rs2927109 | -0.074 | 0.122 | 0.507 | 0.786 | 0.014 | -0.058 | -0.058 | 1.88 | 0.78 | 0.89 | 0.81 | 0.5 | 0.85 | 0.67 |
| 8 | 107967119 | rs1381485 | 1.558 | 0.32 | 0.12 | 0.382 | -0.057 | 0.142 | 0.142 | 1.549 | 0.41 | 0.08 | 0.29 | 0.21 | 0.43 | 0.22 |
| 8 | 107979407 | rs7818910 | 0.894 | 0.277 | 0.568 | 0.258 | 0.033 | 0.006 | 0.006 | 0.075 | 0.61 | 0.27 | 0.65 | 0.38 | 0.72 | 0.52 |
| 8 | 107986935 | rs3019943 | 0.13 | 0.319 | 0.734 | 0.301 | -0.012 | 0.011 | 0.011 | 0 | 0.09 | 0.18 | 0.07 | 0.1 | 0.07 | 0.11 |
| 8 | 108001216 | rs7004403 | 1.599 | 0.401 | 0.598 | 0.223 | -0.052 | 0.121 | 0.121 | 1.598 | 0.42 | 0.09 | 0.29 | 0.22 | 0.44 | 0.22 |
| 8 | 108008913 | rs1462014 | 0.439 | 0.434 | 0.628 | 0.204 | -0.009 | 0.005 | 0.005 | 0.153 | 0.2 | 0.18 | 0.36 | 0.15 | 0.28 | 0.3 |
| 8 | 108012065 | rs10101809 | 0.761 | 0.341 | 0.755 | 0.396 | -0.006 | 0 | 0 | 0 | 0.63 | 0.43 | 0.67 | 0.4 | 0.72 | 0.54 |
| 8 | 108035477 | rs1462011 | 0.779 | 0.328 | 0.803 | 0.385 | 0.004 | 0 | 0 | 0.095 | 0.63 | 0.39 | 0.66 | 0.38 | 0.72 | 0.52 |
| 8 | 108039422 | rs2927155 | -0.004 | 0.258 | 1.119 | 0.465 | -0.013 | 0.064 | 0.064 | 0 | 0.73 | 0.64 | 0.75 | 0.51 | 0.79 | 0.67 |
| 8 | 108047032 | rs6469090 | 1.083 | 0.236 | 1.02 | 0.453 | 0.028 | 0.004 | 0.004 | 0.004 | 0.61 | 0.28 | 0.65 | 0.38 | 0.72 | 0.52 |
| 8 | 108051016 | rs1564018 | 0.303 | 0.257 | 0.92 | 0.461 | -0.002 | 0.001 | 0.001 | 0 | 0.08 | 0.08 | 0.06 | 0.11 | 0.07 | 0.11 |
| 8 | 108058541 | rs2920097 | 0.046 | 0.29 | 0.943 | 0.457 | 0.001 | -0.005 | -0.005 | 0 | 0.73 | 0.78 | 0.75 | 0.51 | 0.79 | 0.65 |
| 8 | 108062735 | rs2584375 | 0.404 | 0.438 | 0.66 | 0.293 | -0.022 | 0.013 | 0.013 | 0.107 | 0.2 | 0.15 | 0.35 | 0.16 | 0.29 | 0.3 |
| 8 | 108065427 | rs1462000 | -0.338 | 0.35 | 0.314 | 0.16 | -0.005 | -0.096 | -0.096 | 0.298 | 0.77 | 0.91 | 0.78 | 0.51 | 0.79 | 0.67 |
| 8 | 108066634 | rs10955435 | 1.662 | 0.341 | 0.227 | 0.174 | -0.029 | 0.118 | 0.118 | 1.18 | 0.42 | 0.11 | 0.29 | 0.23 | 0.43 | 0.23 |
| 8 | 108067119 | rs2920086 | 0.03 | 0.534 | 0.505 | 0.41 | 0 | 0 | 0 | 0 | 0.76 | 0.8 | 0.76 | 0.51 | 0.79 | 0.67 |
| 8 | 108068283 | rs3019920 | 0.196 | 0.556 | 0.492 | 0.42 | -0.013 | 0.073 | 0.073 | 0 | 0.73 | 0.63 | 0.75 | 0.51 | 0.79 | 0.65 |
| 8 | 108080619 | rs7822406 | -0.45 | 0.626 | 0.339 | 0.571 | -0.011 | 0.079 | 0.079 | 0 | 0.18 | 0.38 | 0.12 | 0.3 | 0.13 | 0.17 |
| 8 | 108084988 | rs1461993 | -0.957 | 1.078 | -0.318 | 0.407 | 0.027 | -0.007 | -0.007 | 0 | 0.28 | 0.48 | 0.43 | 0.28 | 0.34 | 0.43 |
| 8 | 108086462 | rs13255596 | -0.049 | 1.039 | -1.194 | 0.144 | -0.007 | 0.012 | 0.012 | 0.311 | 0.14 | 0.08 | 0.08 | 0.01 | 0.15 | 0.15 |
| 8 | 108100495 | rs13272391 | -0.422 | 0.822 | -1.407 | 0.489 | -0.008 | -0.011 | -0.011 | 0 | 0.19 | 0.1 | 0.13 | 0.26 | 0.18 | 0.17 |
| 8 | 108119964 | rs3015943 | -0.368 | 0.731 | -1.64 | 0.354 | 0.012 | -0.039 | -0.039 | 0.019 | 0.63 | 0.78 | 0.67 | 0.55 | 0.71 | 0.61 |
| 8 | 108122180 | rs2920102 | -0.678 | 0.463 | -2.112 | 0.776 | 0.015 | -0.084 | -0.084 | 0.015 | 0.83 | 1 | 0.84 | 0.79 | 0.9 | 0.74 |
| 8 | 108124063 | rs1954728 | 0.628 | 0.854 | -2.224 | 0.199 | -0.041 | 0.028 | 0.028 | 0 | 0.24 | 0.11 | 0.39 | 0.16 | 0.39 | 0.22 |
| 8 | 108151432 | rs10108503 | 0.918 | 1.356 | -2.4 | 0.041 | -0.033 | 0.049 | 0.049 | 0 | 0.18 | 0.06 | 0.13 | 0.01 | 0.28 | 0.06 |
| 8 | 108155732 | rs1954720 | 1.022 | 1.467 | -2.308 | 0.313 | -0.029 | -0.029 | -0.029 | 0 | 0.05 | 0 | 0.26 | 0.17 | 0.11 | 0.16 |
| 8 | 108158383 | rs2584363 | -1.002 | 0.999 | -2.448 | 0.246 | -0.004 | -0.005 | -0.005 | 0 | 0.79 | 0.94 | 0.87 | 0.74 | 0.81 | 0.82 |
| 8 | 108158452 | rs2584364 | 0.635 | 0.793 | -1.955 | 0.223 | -0.001 | -0.001 | -0.001 | 0.064 | 0.1 | 0.09 | 0.09 | 0.11 | 0.12 | 0.2 |
| 8 | 108175323 | rs721872 | 0.911 | 0.407 | -1.42 | 0.147 | 0.014 | -0.015 | -0.015 | 0 | 0.38 | 0.27 | 0.32 | 0.43 | 0.26 | 0.37 |
| 8 | 108175983 | rs2584370 | -0.449 | 0.71 | -1.8 | 0.12 | -0.011 | -0.038 | -0.038 | 0 | 0.71 | 0.89 | 0.79 | 0.62 | 0.74 | 0.67 |
| 8 | 108185425 | rs7829806 | 0.291 | 0.472 | -0.876 | 0.296 | 0.007 | 0.006 | 0.006 | 0.476 | 0.46 | 0.59 | 0.42 | 0.56 | 0.34 | 0.48 |
| 8 | 108185786 | rs7830253 | -1.105 | 0.596 | -1.192 | 0.362 | -0.034 | -0.056 | -0.056 | 0 | 0.14 | 0.04 | 0.1 | 0.16 | 0.15 | 0.18 |
| 8 | 108192360 | rs10808412 | 0.068 | 0.95 | -0.635 | 0.242 | -0.003 | -0.019 | -0.019 | 0 | 0.57 | 0.48 | 0.69 | 0.76 | 0.61 | 0.67 |
| 8 | 108193330 | rs2022962 | 0.722 | 1.208 | -0.444 | 0.828 | 0.003 | -0.003 | -0.003 | 0 | 0.47 | 0.37 | 0.45 | 0.47 | 0.45 | 0.45 |
| 8 | 108193562 | rs12547442 | 0.214 | 1.124 | -0.452 | 0.828 | -0.031 | -0.043 | -0.043 | 0 | 0.09 | 0.03 | 0.24 | 0.29 | 0.15 | 0.22 |
| 8 | 108200124 | rs2022958 | 0.488 | 1.414 | -0.312 | 1.209 | 0.004 | 0.003 | 0.003 | 0.294 | 0.49 | 0.38 | 0.59 | 0.52 | 0.49 | 0.5 |
| 8 | 108202969 | rs1954741 | 1.555 | 1.33 | 0.029 | 1.178 | 0.015 | 0.037 | 0.037 | 2.048 | 0.37 | 0.18 | 0.33 | 0.24 | 0.33 | 0.28 |
| 8 | 108207576 | rs2584382 | 0.838 | 1.258 | 0.91 | 1.151 | 0.018 | 0.095 | 0.095 | 0.874 | 0.79 | 0.66 | 0.62 | 0.49 | 0.76 | 0.58 |
| 8 | 108212339 | rs7833208 | 0.253 | 1.283 | 0.702 | 1.167 | -0.002 | 0.006 | 0.006 | 0.068 | 0.1 | 0.08 | 0.24 | 0.29 | 0.15 | 0.22 |
| 8 | 108218786 | rs2844270 | -0.436 | 1.271 | 0.716 | 1.161 | 0 | 0.008 | 0.008 | 0.003 | 0.91 | 0.92 | 0.88 | 0.8 | 0.91 | 0.8 |
| 8 | 108224765 | rs4305876 | 0.66 | 1.29 | 0.682 | 1.167 | -0.003 | 0.002 | 0.002 | 1.217 | 0.81 | 0.83 | 0.64 | 0.51 | 0.76 | 0.58 |

| | | | | | | | | | | | | | | | | |
|---|---|---|---|---|---|---|---|---|---|---|---|---|---|---|---|---|
| 8 | 108226328 | rs7012260 | -0.342 | 1.254 | 0.589 | 1.108 | -0.001 | 0.002 | 0.002 | 0 | 0.91 | 0.91 | 0.88 | 0.8 | 0.91 | 0.8 |
| 8 | 108226806 | rs9643029 | -0.433 | 1.37 | 0.524 | 1.093 | -0.021 | -0.04 | -0.04 | 0.527 | 0.11 | 0.04 | 0.12 | 0.23 | 0.14 | 0.23 |
| 8 | 108234516 | rs2022950 | 0.661 | 1.215 | 0.566 | 0.976 | -0.003 | 0.091 | 0.091 | 0 | 0.31 | 0.14 | 0.16 | 0.02 | 0.31 | 0.13 |
| 8 | 108238820 | rs7464479 | 0.107 | 1.232 | 0.537 | 1.123 | -0.013 | -0.001 | -0.001 | 0 | 0.07 | 0.04 | 0.09 | 0.24 | 0.11 | 0.17 |
| 8 | 108240629 | rs10099794 | 0.124 | 1.103 | 0.734 | 1.073 | 0.002 | 0.056 | 0.056 | 0 | 0.88 | 0.73 | 0.85 | 0.78 | 0.9 | 0.8 |
| 8 | 108247892 | rs10094991 | 1.941 | 1.062 | 0.714 | 0.978 | 0.01 | 0.033 | 0.033 | 2.415 | 0.36 | 0.18 | 0.32 | 0.24 | 0.32 | 0.32 |
| 8 | 108247913 | rs1954732 | -0.414 | 1.131 | 0.806 | 1.038 | 0 | 0.009 | 0.009 | 0 | 0.91 | 0.9 | 0.86 | 0.8 | 0.91 | 0.81 |
| 8 | 108248735 | rs16875907 | -0.046 | 1.113 | 0.957 | 0.819 | 0.006 | 0.038 | 0.038 | 0 | 0.07 | 0.13 | 0.06 | 0.16 | 0.09 | 0.17 |
| 8 | 108252238 | rs6469101 | 0.082 | 0.949 | 1.039 | 0.648 | 0.009 | 0.029 | 0.029 | 0 | 0.11 | 0.21 | 0.24 | 0.32 | 0.16 | 0.18 |
| 8 | 108260740 | rs1892765 | 0.98 | 0.914 | 0.898 | 0.604 | 0.006 | 0.003 | 0.003 | 0 | 0.47 | 0.53 | 0.42 | 0.43 | 0.42 | 0.49 |
| 8 | 108266983 | rs10100390 | 0.372 | 0.832 | 0.707 | 0.473 | -0.003 | -0.073 | -0.073 | 5.435 | 0.34 | 0.3 | 0.18 | 0.02 | 0.31 | 0.13 |
| 8 | 108270037 | rs2507802 | -0.223 | 0.836 | -0.049 | 0.654 | 0.003 | 0.044 | 0.044 | 0.035 | 0.89 | 0.86 | 0.82 | 0.77 | 0.87 | 0.8 |
| 8 | 108273016 | rs2514878 | 1.12 | 0.728 | -0.44 | 0.212 | -0.03 | -0.038 | -0.038 | 0 | 0.41 | 0.1 | 0.38 | 0.4 | 0.4 | 0.49 |
| 8 | 108279912 | rs11777978 | -1.292 | 0.597 | -0.202 | 0.443 | 0.007 | 0.058 | 0.058 | 2.72 | 0.31 | 0.48 | 0.47 | 0.55 | 0.32 | 0.43 |
| 8 | 108287577 | rs9297393 | 0.84 | 0.399 | -0.91 | 0.104 | -0.031 | 0.014 | 0.014 | 0 | 0.43 | 0.11 | 0.34 | 0.37 | 0.4 | 0.36 |
| 8 | 108287597 | rs10090742 | 0.301 | 0.101 | -0.398 | 0.086 | -0.002 | 0.005 | 0.005 | 0 | 0.54 | 0.49 | 0.58 | 0.59 | 0.48 | 0.55 |
| 8 | 108292664 | rs13257393 | 0.264 | 0.058 | -0.482 | 0.391 | 0 | 0.023 | 0.023 | 0 | 0.1 | 0.1 | 0.17 | 0.33 | 0.11 | 0.25 |
| 8 | 108299501 | rs4595097 | 0.425 | 0.197 | -0.544 | 0.401 | -0.008 | -0.015 | -0.015 | 0 | 0.84 | 0.87 | 0.77 | 0.5 | 0.79 | 0.63 |
| 8 | 108301798 | rs4324901 | 1.701 | 0.224 | -1.066 | 0.108 | -0.116 | -0.055 | -0.055 | 0 | 0.35 | 0.02 | 0.41 | 0.35 | 0.38 | 0.34 |
| 8 | 108307058 | rs6982586 | 0.253 | 0.286 | -0.286 | -0.043 | -0.014 | -0.091 | -0.091 | 0 | 0.33 | 0.34 | 0.14 | 0.01 | 0.21 | 0.13 |
| 8 | 108307337 | rs10505105 | 0.775 | 0.339 | -0.406 | 0.04 | -0.044 | -0.048 | -0.048 | 0.061 | 0.09 | 0.02 | 0.17 | 0.13 | 0.19 | 0.15 |
| 8 | 108314971 | rs4133395 | -0.652 | 0.207 | -0.043 | 0.075 | 0.005 | 0.012 | 0.012 | 0 | 0.46 | 0.51 | 0.33 | 0.15 | 0.41 | 0.29 |
| 8 | 108315277 | rs4472483 | -1.233 | 0.226 | -0.248 | 0.163 | -0.059 | -0.052 | -0.052 | 0.049 | 0.56 | 0.83 | 0.52 | 0.48 | 0.52 | 0.53 |
| 8 | 108327348 | rs16876018 | 0.658 | -0.023 | 0.007 | 0.189 | -0.038 | -0.04 | -0.04 | 0.03 | 0.07 | 0.17 | 0.02 | 0.01 | 0.01 | 0.04 |
| 8 | 108329796 | rs4397378 | -0.483 | 0.351 | -0.777 | 0.39 | -0.023 | -0.029 | -0.029 | 14.722 | 0.94 | 0.97 | 0.95 | 0.85 | 0.91 | 0.88 |
| 8 | 108332485 | rs4341142 | -0.083 | 0.181 | -0.696 | 0.31 | 0.004 | 0.08 | 0.08 | 0.64 | 0.13 | 0.28 | 0.19 | 0.34 | 0.13 | 0.24 |
| 8 | 108363596 | rs16876090 | 0.718 | 0.243 | -1.368 | 0.019 | -0.021 | -0.017 | -0.017 | 0.885 | 0.06 | 0.02 | 0.05 | 0.11 | 0.09 | 0.12 |
| 8 | 108366032 | rs1283651 | -0.719 | 0.053 | -1.442 | -0.364 | -0.02 | -0.115 | -0.115 | 0 | 0.68 | 0.63 | 0.87 | 0.98 | 0.8 | 0.86 |
| 8 | 108366127 | rs1433195 | 0.73 | 0.494 | -1.389 | -0.018 | -0.026 | -0.029 | -0.029 | 0 | 0.1 | 0.03 | 0.31 | 0.14 | 0.22 | 0.16 |
| 8 | 108372446 | rs2436556 | 0.564 | 0.387 | -1.283 | 0.012 | -0.002 | 0.031 | 0.031 | 0.079 | 0.1 | 0.09 | 0.18 | 0.38 | 0.11 | 0.24 |
| 8 | 108377347 | rs1283658 | -0.547 | 0.244 | -1.239 | -0.071 | -0.049 | -0.11 | -0.11 | 0 | 0.81 | 0.64 | 0.92 | 0.98 | 0.91 | 0.89 |
| 8 | 108377901 | rs2163870 | 0.408 | -0.114 | -1.691 | -0.343 | 0.006 | -0.055 | -0.055 | 0 | 0.41 | 0.17 | 0.3 | 0.49 | 0.35 | 0.41 |
| 8 | 108378311 | rs1672149 | 0.447 | 0.085 | -1.927 | -0.11 | 0.003 | 0.005 | 0.005 | 0.099 | 0.4 | 0.47 | 0.62 | 0.49 | 0.56 | 0.48 |
| 8 | 108378623 | rs2514857 | 0.646 | 0.09 | -2.342 | -0.112 | -0.109 | -0.119 | -0.119 | 0.015 | 0.37 | 0.08 | 0.59 | 0.49 | 0.55 | 0.48 |
| 8 | 108383325 | rs2445365 | 0.849 | 0.085 | -2.563 | -0.092 | -0.013 | -0.112 | -0.112 | 0 | 0.32 | 0.04 | 0.24 | 0.38 | 0.24 | 0.29 |
| 8 | 108383641 | rs1672195 | -0.231 | 0.359 | -2.07 | -0.104 | -0.028 | -0.086 | -0.086 | 1.763 | 0.73 | 0.53 | 0.86 | 0.87 | 0.81 | 0.77 |
| 8 | 108386442 | rs1347183 | 0.181 | 0.48 | -2.13 | -0.041 | -0.002 | -0.003 | -0.003 | 0 | 0.47 | 0.51 | 0.54 | 0.52 | 0.47 | 0.45 |
| 8 | 108389120 | rs2436557 | 0.807 | 1.166 | -2.665 | 0.538 | -0.119 | -0.088 | -0.088 | 0 | 0.26 | 0.03 | 0.32 | 0.35 | 0.34 | 0.32 |
| 8 | 108389187 | rs2514862 | 0.818 | 1.172 | -2.662 | 0.541 | -0.119 | -0.088 | -0.088 | 0 | 0.26 | 0.03 | 0.32 | 0.35 | 0.34 | 0.32 |
| 8 | 108392928 | rs1433176 | 0.107 | 0.77 | -2.637 | 0.555 | -0.06 | -0.064 | -0.064 | 0 | 0.49 | 0.05 | 0.59 | 0.47 | 0.61 | 0.53 |
| 8 | 108395025 | rs16876199 | -0.585 | 0.79 | -1.495 | 1.409 | -0.002 | 0.243 | 0.243 | 0 | 0.07 | 0.14 | 0.13 | 0.47 | 0.05 | 0.11 |
| 8 | 108398406 | rs2916083 | 0.784 | 0.515 | -0.938 | 0.717 | -0.013 | -0.045 | -0.045 | 0 | 0.76 | 0.7 | 0.72 | 0.92 | 0.82 | 0.75 |
| 8 | 108400823 | rs1283706 | 1.944 | 0.876 | -1.317 | 0.639 | -0.031 | 0.029 | 0.029 | 2.277 | 0.55 | 0.09 | 0.46 | 0.33 | 0.6 | 0.49 |
| 8 | 108403482 | rs1283703 | 0.014 | 0.754 | -1.108 | 0.592 | 0.004 | 0.03 | 0.03 | 0 | 0.84 | 0.9 | 0.78 | 0.94 | 0.88 | 0.76 |
| 8 | 108409423 | rs11785702 | -0.978 | 0.816 | -0.473 | 0.646 | -0.004 | 0.203 | 0.203 | 0 | 0.12 | 0.26 | 0.19 | 0.55 | 0.09 | 0.22 |
| 8 | 108416439 | rs1283687 | -0.236 | 1.575 | -0.968 | 0.278 | -0.049 | -0.01 | -0.01 | 0 | 0.12 | 0.05 | 0.11 | 0.12 | 0.19 | 0.22 |
| 8 | 108422946 | rs892248 | 1.549 | 0.964 | 0.013 | 0.429 | 0.012 | 0.004 | 0.004 | 0.232 | 0.68 | 0.29 | 0.63 | 0.76 | 0.78 | 0.64 |
| 8 | 108424451 | rs1654751 | 0.994 | 0.951 | 0.588 | 0.39 | -0.043 | -0.036 | -0.036 | 0 | 0.73 | 0.56 | 0.72 | 0.85 | 0.82 | 0.75 |
| 8 | 108427493 | rs1283679 | 0.607 | 0.97 | 0.464 | 0.404 | 0.009 | 0.005 | 0.005 | 0.006 | 0.78 | 0.86 | 0.75 | 0.85 | 0.82 | 0.75 |
| 8 | 108431161 | rs1283673 | 1.698 | 1.423 | 0.81 | 0.436 | 0.007 | -0.005 | -0.005 | 0.251 | 0.63 | 0.5 | 0.61 | 0.71 | 0.61 | 0.54 |
| 8 | 108434677 | rs1283670 | -0.445 | 1.582 | 0.635 | 0.373 | -0.01 | 0.004 | 0.004 | 0 | 0.09 | 0.07 | 0.1 | 0.06 | 0.13 | 0.17 |
| 8 | 108437679 | rs1283701 | -2.134 | 1.503 | 0.866 | 0.313 | 0.062 | -0.036 | -0.036 | 0 | 0.21 | 0.6 | 0.21 | 0.11 | 0.31 | 0.26 |
| 8 | 108438906 | rs1283699 | 0.727 | 1.383 | 0.756 | 0.38 | 0.005 | 0.001 | 0.001 | 0 | 0.78 | 0.82 | 0.75 | 0.85 | 0.82 | 0.73 |
| 8 | 108439898 | rs1004460 | -0.17 | 1.541 | 0.871 | 0.365 | 0.026 | -0.002 | -0.002 | 0 | 0.07 | 0.07 | 0.06 | 0.05 | 0.17 | 0.09 |
| 8 | 108440057 | rs1004459 | -0.019 | 1.599 | 0.838 | 0.336 | -0.007 | 0 | 0 | 7.014 | 0.06 | 0.05 | 0.08 | 0.06 | 0.09 | 0.16 |
| 8 | 108445788 | rs11786476 | -1.936 | 1.737 | 0.809 | 0.44 | 0 | 0.009 | 0.009 | 0 | 0.21 | 0.22 | 0.28 | 0.59 | 0.23 | 0.24 |
| 8 | 108445984 | rs1283681 | 1.015 | 1.735 | 0.855 | 0.413 | -0.004 | 0.002 | 0.002 | 1.002 | 0.75 | 0.8 | 0.73 | 0.84 | 0.71 | 0.64 |
| 8 | 108446403 | rs13280030 | -0.279 | 1.72 | 0.809 | 0.505 | -0.018 | -0.015 | -0.015 | 0 | 0.1 | 0.03 | 0.17 | 0.45 | 0.12 | 0.13 |
| 8 | 108452794 | rs1433184 | -1.127 | 1.848 | 0.908 | 0.385 | 0.006 | 0 | 0 | 0 | 0.13 | 0.12 | 0.14 | 0.1 | 0.24 | 0.24 |
| 8 | 108454681 | rs13273578 | -1.813 | 1.833 | 0.947 | 0.386 | 0 | 0.01 | 0.01 | 0 | 0.21 | 0.22 | 0.28 | 0.59 | 0.23 | 0.24 |
| 8 | 108456805 | rs4114169 | 2.622 | 1.84 | 0.949 | 0.34 | 0.002 | -0.004 | -0.004 | 0.597 | 0.5 | 0.51 | 0.38 | 0.2 | 0.43 | 0.25 |
| 8 | 108457353 | rs1654680 | 0.065 | 1.655 | 0.863 | 0.409 | 0 | 0 | 0 | 0 | 0.82 | 0.84 | 0.78 | 0.89 | 0.86 | 0.72 |
| 8 | 108459674 | rs10102601 | -1.483 | 1.69 | 1.246 | 0.384 | 0.097 | -0.038 | -0.038 | 0 | 0.12 | 0.35 | 0.1 | 0.04 | 0.19 | 0.09 |
| 8 | 108460133 | rs1283722 | 0.058 | 1.432 | 1.054 | 0.478 | -0.006 | 0 | 0 | 0 | 0.06 | 0.04 | 0.08 | 0.06 | 0.09 | 0.15 |
| 8 | 108462641 | rs1433166 | 2.306 | 1.342 | 1.212 | 0.427 | 0 | 0.013 | 0.013 | 0.375 | 0.56 | 0.45 | 0.43 | 0.24 | 0.58 | 0.34 |
| 8 | 108463603 | rs13274462 | -1.916 | 1.288 | 1.178 | 0.555 | -0.008 | 0.085 | 0.085 | 0 | 0.23 | 0.33 | 0.29 | 0.61 | 0.21 | 0.26 |
| 8 | 108472328 | rs1283720 | -0.619 | 1.186 | 0.816 | 0.264 | -0.006 | 0.013 | 0.013 | 0 | 0.13 | 0.09 | 0.11 | 0.06 | 0.15 | 0.18 |
| 8 | 108476951 | rs1433168 | 2.756 | 0.841 | 0.502 | 0.012 | -0.005 | 0.026 | 0.026 | 0.975 | 0.46 | 0.17 | 0.47 | 0.65 | 0.47 | 0.38 |
| 8 | 108485912 | rs2195989 | 0.144 | 0.84 | 0.712 | -0.032 | 0.026 | -0.046 | -0.046 | 0 | 0.15 | 0.28 | 0.09 | 0.04 | 0.18 | 0.09 |

| | | | | | | | | | | | | | | | |
|---|---|---|---|---|---|---|---|---|---|---|---|---|---|---|---|
| 8 | 108489996 | rs951759 | 1.658 | 0.844 | 0.691 | -0.015 | -0.005 | -0.008 | -0.008 | 0 | 0.63 | 0.54 | 0.57 | 0.7 | 0.65 | 0.46 |
| 8 | 108490998 | rs1157411 | 1.692 | 0.839 | 0.701 | -0.018 | -0.006 | -0.009 | -0.009 | 0 | 0.62 | 0.53 | 0.57 | 0.7 | 0.65 | 0.46 |
| 8 | 108499883 | rs13257142 | 2.466 | 0.824 | 0.641 | 0.075 | -0.013 | 0.116 | 0.116 | 0.768 | 0.5 | 0.17 | 0.53 | 0.75 | 0.5 | 0.49 |
| 8 | 108502688 | rs6469116 | 1.285 | 0.812 | 0.599 | 0.06 | 0.002 | 0.02 | 0.02 | 0 | 0.63 | 0.71 | 0.63 | 0.79 | 0.65 | 0.57 |
| 8 | 108503702 | rs13272688 | 0.464 | 0.922 | 0.452 | 0.223 | -0.006 | 0.047 | 0.047 | 0 | 0.08 | 0.04 | 0.14 | 0.45 | 0.08 | 0.11 |
| 8 | 108515373 | rs9297395 | -1.615 | 0.976 | 0.697 | -0.085 | 0.048 | -0.063 | -0.063 | 0 | 0.16 | 0.58 | 0.16 | 0.06 | 0.18 | 0.13 |
| 8 | 108518274 | rs1433174 | 2.958 | 0.933 | 0.094 | -0.081 | -0.035 | 0.041 | 0.041 | 0.368 | 0.41 | 0.07 | 0.37 | 0.31 | 0.43 | 0.33 |
| 8 | 108522376 | rs6469117 | -1.65 | 1.068 | 0.728 | 0.158 | 0.045 | 0.075 | 0.075 | 0.092 | 0.22 | 0.38 | 0.5 | 0.58 | 0.26 | 0.36 |
| 8 | 108522424 | rs7003565 | 2.443 | 1.019 | 0.247 | 0.07 | 0.015 | 0.136 | 0.136 | 0.018 | 0.49 | 0.18 | 0.32 | 0.3 | 0.49 | 0.43 |
| 8 | 108532275 | rs16876379 | -0.875 | 1.519 | 0.814 | 0.614 | 0.021 | 0.012 | 0.012 | 0 | 0.11 | 0.24 | 0.21 | 0.11 | 0.16 | 0.21 |
| 8 | 108533278 | rs1368493 | -0.426 | 1.575 | 0.51 | 0.801 | -0.003 | 0.081 | 0.081 | 0.019 | 0.07 | 0.03 | 0.25 | 0.48 | 0.1 | 0.13 |
| 8 | 108537188 | rs10955457 | -1.385 | 1.466 | 0.459 | 0.766 | 0.036 | 0.154 | 0.154 | 0.862 | 0.2 | 0.39 | 0.38 | 0.49 | 0.21 | 0.22 |
| 8 | 108543427 | rs281712 | 0.416 | 0.825 | -0.262 | 0.784 | -0.053 | -0.056 | -0.056 | 0 | 0.16 | 0.08 | 0.35 | 0.4 | 0.26 | 0.18 |
| 8 | 108543767 | rs4734976 | 0.776 | 0.593 | -0.442 | 0.783 | -0.003 | -0.001 | -0.001 | 0 | 0.22 | 0.25 | 0.21 | 0.28 | 0.13 | 0.33 |
| 8 | 108544206 | rs10103715 | 1.57 | 0.27 | -0.849 | 0.548 | -0.038 | 0.117 | 0.117 | 0 | 0.37 | 0.11 | 0.27 | 0.12 | 0.43 | 0.17 |
| 8 | 108545055 | rs281713 | 0.078 | 0.322 | -0.811 | 0.616 | 0.004 | 0.011 | 0.011 | 0 | 0.26 | 0.31 | 0.41 | 0.46 | 0.32 | 0.27 |
| 8 | 108547201 | rs16876408 | 0.555 | 0.33 | -0.709 | 0.765 | 0 | 0.002 | 0.002 | 0.033 | 0.07 | 0.09 | 0.06 | 0.1 | 0.07 | 0.01 |
| 8 | 108575039 | rs6993722 | 1.279 | 0.708 | -0.401 | 0.72 | -0.011 | 0.092 | 0.092 | 0 | 0.46 | 0.19 | 0.32 | 0.19 | 0.48 | 0.26 |
| 8 | 108582653 | rs394569 | -0.063 | 1.556 | -0.488 | 0.775 | 0.031 | 0.036 | 0.036 | 0 | 0.19 | 0.29 | 0.37 | 0.39 | 0.27 | 0.19 |
| 8 | 108592204 | rs281664 | -0.543 | 1.613 | -0.674 | 0.783 | 0.029 | 0.05 | 0.05 | 0 | 0.25 | 0.54 | 0.4 | 0.41 | 0.28 | 0.25 |
| 8 | 108603343 | rs281687 | -0.389 | 1.512 | -0.795 | 0.873 | -0.033 | -0.038 | -0.038 | 0 | 0.92 | 0.98 | 0.92 | 0.88 | 0.9 | 0.83 |
| 8 | 108611644 | rs2014769 | -0.618 | 1.235 | -0.496 | 0.705 | 0.032 | 0.049 | 0.049 | 0 | 0.23 | 0.43 | 0.39 | 0.41 | 0.28 | 0.25 |
| 8 | 108623345 | rs281721 | -0.595 | 1.179 | -0.499 | 0.671 | 0.034 | 0.035 | 0.035 | 0 | 0.25 | 0.55 | 0.41 | 0.4 | 0.3 | 0.25 |
| 8 | 108626828 | rs190735 | -0.112 | 1.074 | -0.251 | 0.648 | 0.019 | 0.016 | 0.016 | 0 | 0.18 | 0.26 | 0.37 | 0.38 | 0.29 | 0.18 |
| 8 | 108632019 | rs10505110 | 1.108 | 1.331 | -0.544 | 0.861 | 0.033 | 0.051 | 0.051 | 0 | 0.68 | 0.45 | 0.51 | 0.47 | 0.6 | 0.59 |
| 8 | 108635344 | rs16876511 | -0.25 | 1.313 | -0.26 | 0.374 | 0.003 | 0.02 | 0.02 | 0 | 0.05 | 0.24 | 0.03 | 0.02 | 0.01 | 0.07 |
| 8 | 108637820 | rs9297397 | 1.172 | 0.961 | -0.986 | -0.272 | -0.019 | 0.117 | 0.117 | 0.511 | 0.32 | 0.12 | 0.21 | 0.02 | 0.34 | 0.16 |
| 8 | 108639617 | rs281747 | -0.056 | 0.869 | -0.975 | -0.134 | -0.002 | -0.002 | -0.002 | 3.076 | 0.44 | 0.42 | 0.39 | 0.58 | 0.36 | 0.59 |
| 8 | 108644707 | rs1969215 | 1.104 | 0.724 | -1.092 | 0.038 | -0.001 | -0.026 | -0.026 | 0 | 0.07 | 0 | 0.03 | 0.07 | 0.06 | 0.1 |
| 8 | 108646193 | rs366575 | 0.118 | 1.158 | -0.855 | -0.119 | 0.001 | 0.002 | 0.002 | 0 | 0.45 | 0.41 | 0.48 | 0.58 | 0.39 | 0.62 |
| 8 | 108646968 | rs412080 | 0.041 | 1.356 | -0.788 | -0.376 | 0.02 | -0.021 | -0.021 | 0 | 0.8 | 0.68 | 0.73 | 0.83 | 0.73 | 0.86 |
| 8 | 108647423 | rs9297398 | -0.163 | 0.898 | -0.923 | -0.5 | -0.006 | -0.047 | -0.047 | 0 | 0.43 | 0.46 | 0.53 | 0.79 | 0.38 | 0.63 |
| 8 | 108649072 | rs903489 | -0.057 | 1.421 | -1.005 | -0.533 | -0.012 | 0.009 | 0.009 | 0 | 0.83 | 0.87 | 0.75 | 0.95 | 0.72 | 0.89 |
| 8 | 108650188 | rs1494971 | -0.081 | 0.052 | -1.086 | -0.361 | 0.021 | 0.01 | 0.01 | 0 | 0.25 | 0.18 | 0.25 | 0.16 | 0.16 | 0.27 |
| 8 | 108654627 | rs6994899 | 0.214 | -0.433 | -0.292 | 0.239 | 0.031 | 0.023 | 0.023 | 2.641 | 0.49 | 0.63 | 0.38 | 0.29 | 0.32 | 0.44 |
| 8 | 108660612 | rs7820162 | 0.021 | -0.48 | 0.096 | 0.652 | 0 | 0.001 | 0.001 | 0 | 0.07 | 0.09 | 0.06 | 0.15 | 0.08 | 0.06 |
| 8 | 108669676 | rs2036032 | -0.111 | -0.639 | -0.034 | 0.926 | 0.012 | -0.01 | -0.01 | 0 | 0.57 | 0.65 | 0.34 | 0.51 | 0.41 | 0.5 |
| 8 | 108669727 | rs2036033 | 0.856 | -0.637 | -0.228 | 0.904 | 0.048 | 0.071 | 0.071 | 4.254 | 0.44 | 0.31 | 0.22 | 0.14 | 0.31 | 0.41 |
| 8 | 108673901 | rs7006253 | -0.358 | -0.593 | -0.339 | 0.982 | -0.017 | 0.006 | 0.006 | 2.065 | 0.61 | 0.91 | 0.49 | 0.65 | 0.42 | 0.53 |
| 8 | 108679029 | rs6469128 | -0.313 | -0.527 | -0.246 | 0.622 | 0.011 | -0.021 | -0.021 | 0 | 0.59 | 0.74 | 0.36 | 0.51 | 0.41 | 0.52 |
| 8 | 108683736 | rs2860020 | 0.489 | -0.554 | -0.424 | -0.019 | 0.042 | 0.057 | 0.057 | 3.278 | 0.44 | 0.35 | 0.23 | 0.14 | 0.31 | 0.4 |
| 8 | 108685788 | rs16876583 | -0.499 | -0.255 | 0.384 | 0.301 | 0.019 | 0.016 | 0.016 | 1.715 | 0.06 | 0.33 | 0.04 | 0 | 0.01 | 0.02 |
| 8 | 108685880 | rs6995417 | -0.129 | -0.361 | -0.033 | 0.026 | 0.002 | 0.005 | 0.005 | 0.504 | 0.44 | 0.54 | 0.25 | 0.27 | 0.39 | 0.55 |
| 8 | 108686103 | rs7016097 | 0.358 | -0.208 | -0.093 | 0.041 | 0.028 | 0.018 | 0.018 | 0 | 0.06 | 0.15 | 0.06 | 0.13 | 0.08 | 0.05 |
| 8 | 108686594 | rs1353308 | -0.312 | -0.578 | -0.084 | 0.756 | 0 | 0 | 0 | 0 | 0.47 | 0.58 | 0.37 | 0.65 | 0.41 | 0.57 |
| 8 | 108689222 | rs4734156 | -0.683 | -0.482 | -0.116 | 1.097 | -0.029 | 0.318 | 0.318 | 1.397 | 0.09 | 0.38 | 0.16 | 0.38 | 0.02 | 0.05 |
| 8 | 108692786 | rs16876591 | 0.103 | 0.177 | -0.818 | 0.818 | -0.026 | -0.03 | -0.03 | 0 | 0.14 | 0.04 | 0.22 | 0.25 | 0.2 | 0.23 |
| 8 | 108698283 | rs6469131 | -1.391 | -0.096 | -0.581 | 1.058 | -0.002 | -0.018 | -0.018 | 0 | 0.32 | 0.71 | 0.41 | 0.43 | 0.33 | 0.38 |
| 8 | 108702663 | rs2118254 | -1.555 | -0.214 | -1.081 | 0.697 | -0.12 | 0.187 | 0.187 | 0.155 | 0.68 | 0.95 | 0.66 | 0.95 | 0.58 | 0.79 |
| 8 | 108709307 | rs13253953 | 1.376 | -0.228 | -0.881 | 1.054 | 0.03 | 0.044 | 0.044 | 1.092 | 0.3 | 0.06 | 0.16 | 0.24 | 0.23 | 0.37 |
| 8 | 108716270 | rs10505111 | 0.027 | 0.116 | 0.262 | 1.104 | 0.057 | -0.018 | -0.018 | 0.013 | 0.08 | 0.27 | 0.07 | 0.02 | 0.09 | 0.06 |
| 8 | 108723445 | rs4734159 | -1.783 | 0.136 | -0.299 | 1.12 | -0.014 | -0.087 | -0.087 | 0 | 0.34 | 0.82 | 0.42 | 0.43 | 0.33 | 0.38 |
| 8 | 108730393 | rs7004951 | -0.244 | 0.499 | -0.396 | 0.517 | -0.026 | -0.036 | -0.036 | 0 | 0.18 | 0.05 | 0.26 | 0.25 | 0.23 | 0.28 |
| 8 | 108744809 | rs7817374 | -1.056 | 0.402 | -0.515 | 0.562 | 0.024 | 0.002 | 0.002 | 0.205 | 0.28 | 0.51 | 0.35 | 0.27 | 0.32 | 0.35 |
| 8 | 108752542 | rs13278956 | 0.118 | 0.165 | -1.51 | 0.082 | -0.025 | 0.2 | 0.2 | 0.178 | 0.4 | 0.07 | 0.3 | 0.11 | 0.38 | 0.52 |
| 8 | 108753147 | rs7834863 | -1.342 | 0.092 | -0.311 | 1.344 | 0.001 | 0.204 | 0.204 | 0.097 | 0.07 | 0.24 | 0.1 | 0.36 | 0.04 | 0.08 |
| 8 | 108757215 | rs6469142 | 0.456 | 0.189 | -0.799 | 1.073 | -0.004 | -0.014 | -0.014 | 0 | 0.44 | 0.23 | 0.31 | 0.47 | 0.43 | 0.33 |
| 8 | 108759874 | rs9297399 | -0.507 | -0.001 | -0.246 | 1.1 | -0.006 | 0.093 | 0.093 | 0 | 0.12 | 0.26 | 0.21 | 0.29 | 0.1 | 0.2 |
| 8 | 108762559 | rs10955465 | 1.443 | -0.182 | -0.754 | 0.936 | -0.087 | 0.19 | 0.19 | 2.304 | 0.35 | 0.05 | 0.24 | 0.06 | 0.4 | 0.23 |
| 8 | 108764653 | rs7828691 | -0.252 | -0.133 | -0.463 | 0.978 | -0.009 | 0.04 | 0.04 | 0.16 | 0.58 | 0.68 | 0.54 | 0.77 | 0.53 | 0.53 |
| 8 | 108768057 | rs7001159 | -0.328 | 0.178 | -0.481 | 0.987 | 0 | -0.001 | -0.001 | 0 | 0.06 | 0.08 | 0.03 | 0.08 | 0.04 | 0.02 |
| 8 | 108768347 | rs10098451 | 1.084 | 0.427 | -0.927 | 0.617 | -0.08 | 0.183 | 0.183 | 2.662 | 0.35 | 0.05 | 0.22 | 0.07 | 0.36 | 0.25 |
| 8 | 108772269 | rs4735009 | -0.351 | 0.393 | -0.637 | 0.727 | 0 | -0.005 | -0.005 | 4.986 | 0.77 | 0.79 | 0.8 | 0.63 | 0.81 | 0.87 |
| 8 | 108787464 | rs1866015 | 0.692 | 0.462 | -0.459 | 0.95 | -0.005 | 0.015 | 0.015 | 1.719 | 0.67 | 0.43 | 0.61 | 0.4 | 0.76 | 0.77 |
| 8 | 108787967 | rs893227 | 0.754 | 0.544 | -0.802 | 0.934 | 0.063 | 0.15 | 0.15 | 0 | 0.85 | 0.62 | 0.66 | 0.63 | 0.85 | 0.89 |
| 8 | 108788603 | rs10087176 | 0.974 | 0.261 | -1.748 | 0.698 | -0.111 | 0.029 | 0.029 | 1.878 | 0.42 | 0.06 | 0.3 | 0.24 | 0.46 | 0.38 |
| 8 | 108792197 | rs16892437 | 0.9 | 0.457 | -1.275 | 1.05 | -0.027 | -0.038 | -0.038 | 0.007 | 0.07 | 0 | 0.08 | 0.16 | 0.08 | 0.12 |
| 8 | 108795106 | rs1437253 | -0.411 | 0.282 | -0.912 | 0.662 | -0.005 | 0.014 | 0.014 | 1.931 | 0.27 | 0.68 | 0.26 | 0.53 | 0.16 | 0.24 |
| 8 | 108801017 | rs2348081 | 0.49 | 0.06 | -0.98 | 0.312 | 0.002 | 0.012 | 0.012 | 0.087 | 0.52 | 0.31 | 0.33 | 0.43 | 0.47 | 0.43 |

| | | | | | | | | | | | | | | | | |
|---|---|---|---|---|---|---|---|---|---|---|---|---|---|---|---|---|
| 8 | 108809266 | rs1353303 | 0.222 | -0.015 | -1.015 | 0.408 | -0.052 | 0.031 | 0.031 | 4.949 | 0.2 | 0.4 | 0.07 | 0.24 | 0.09 | 0.12 |
| 8 | 108813231 | rs7843427 | -0.305 | 1.021 | -1.722 | 0.124 | -0.012 | -0.006 | -0.006 | 0.206 | 0.73 | 0.77 | 0.4 | 0.66 | 0.55 | 0.55 |
| 8 | 108814617 | rs2934941 | -0.158 | 0.576 | -2.099 | -0.112 | -0.034 | 0.049 | 0.049 | 4.474 | 0.32 | 0.04 | 0.19 | 0.18 | 0.29 | 0.28 |
| 8 | 108818909 | rs1299342 | 0.308 | 0.887 | -1.491 | -0.428 | -0.02 | 0.182 | 0.182 | 0.094 | 0.55 | 0.21 | 0.34 | 0.17 | 0.58 | 0.55 |
| 8 | 108839030 | rs12543466 | -1.049 | 0.429 | 0.468 | 0.545 | 0.055 | 0.298 | 0.298 | 6.061 | 0.06 | 0.42 | 0.06 | 0.2 | 0.01 | 0.07 |
| 8 | 108841064 | rs11997086 | -1.421 | 0.32 | 0.984 | 0.108 | -0.008 | 0.003 | 0.003 | 4.861 | 0.12 | 0.46 | 0.08 | 0.09 | 0.06 | 0.03 |
| 8 | 108841582 | rs13269271 | -0.103 | 0.834 | 0.688 | -0.603 | 0.001 | -0.011 | -0.011 | 0 | 0.29 | 0.29 | 0.28 | 0.12 | 0.36 | 0.27 |
| 8 | 108850819 | rs10109988 | -1.295 | 0.537 | 0.405 | -0.464 | 0.019 | -0.043 | -0.043 | 0 | 0.69 | 0.81 | 0.68 | 0.64 | 0.73 | 0.66 |
| 8 | 108856341 | rs11987403 | -0.49 | 0.507 | 1.171 | -0.945 | -0.01 | -0.027 | -0.027 | 0 | 0.11 | 0.33 | 0.06 | 0.07 | 0.05 | 0.18 |
| 8 | 108857416 | rs12548708 | 0.755 | 0.644 | 1.346 | -0.915 | -0.024 | -0.039 | -0.039 | 0 | 0.28 | 0.16 | 0.39 | 0.63 | 0.32 | 0.24 |
| 8 | 108858606 | rs10092441 | -0.336 | 0.957 | 1.634 | -0.814 | -0.008 | -0.016 | -0.016 | 0 | 0.1 | 0.19 | 0.04 | 0.07 | 0.05 | 0.18 |
| 8 | 108866822 | rs17309585 | 0.765 | 0.814 | 1.083 | -1.108 | -0.05 | 0.201 | 0.201 | 0.033 | 0.34 | 0.05 | 0.21 | 0.03 | 0.38 | 0.26 |
| 8 | 108869762 | rs1494917 | -0.633 | 0.824 | 1.271 | -1.074 | 0.008 | 0 | 0 | 0 | 0.32 | 0.62 | 0.42 | 0.71 | 0.34 | 0.4 |
| 8 | 108870037 | rs4734161 | -1.399 | 0.861 | 0.767 | -1.051 | 0.017 | 0.008 | 0.008 | 0 | 0.69 | 0.79 | 0.68 | 0.74 | 0.73 | 0.66 |
| 8 | 108873492 | rs7015214 | -0.002 | 0.969 | 0.513 | -0.876 | -0.001 | -0.001 | -0.001 | 0.156 | 0.24 | 0.28 | 0.38 | 0.71 | 0.3 | 0.27 |
| 8 | 108876210 | rs1603145 | -0.193 | 0.789 | 0.57 | -0.797 | -0.002 | -0.001 | -0.001 | 0 | 0.3 | 0.35 | 0.4 | 0.71 | 0.33 | 0.29 |
| 8 | 108899563 | rs1874203 | -1.103 | 0.652 | 0.085 | -0.975 | -0.019 | -0.073 | -0.073 | 0.123 | 0.7 | 0.42 | 0.78 | 0.89 | 0.77 | 0.82 |
| 8 | 108901118 | rs6987451 | -0.581 | 0.685 | 0.053 | -0.961 | -0.003 | -0.004 | -0.004 | 0 | 0.37 | 0.36 | 0.54 | 0.86 | 0.38 | 0.57 |
| 8 | 108909499 | rs16876914 | 0.387 | 0.667 | 0.021 | -0.778 | 0.003 | 0.019 | 0.019 | 0 | 0.08 | 0.08 | 0.15 | 0.14 | 0.1 | 0.17 |
| 8 | 108910484 | rs10955473 | 0.237 | 0.783 | -0.068 | -0.897 | -0.007 | -0.019 | -0.019 | 0 | 0.26 | 0.18 | 0.32 | 0.72 | 0.25 | 0.38 |
| 8 | 108910785 | rs10955474 | 0.159 | 0.75 | -0.048 | -0.901 | -0.004 | -0.013 | -0.013 | 0 | 0.26 | 0.2 | 0.33 | 0.72 | 0.25 | 0.38 |
| 8 | 108912166 | rs555008 | -1.023 | -0.196 | 0.184 | -0.847 | -0.024 | -0.089 | -0.089 | 0 | 0.69 | 0.42 | 0.78 | 0.91 | 0.77 | 0.82 |
| 8 | 108917591 | rs4735020 | 0.339 | -0.088 | -0.682 | -0.88 | -0.002 | -0.009 | -0.009 | 0 | 0.32 | 0.08 | 0.29 | 0.25 | 0.29 | 0.34 |
| 8 | 108922953 | rs4735022 | 0.459 | -0.007 | -0.696 | -1.031 | -0.018 | 0.024 | 0.024 | 0 | 0.31 | 0.07 | 0.25 | 0.17 | 0.29 | 0.33 |
| 8 | 108924771 | rs17311083 | 1.231 | -0.015 | -0.729 | -1.061 | -0.017 | 0.121 | 0.121 | 0.26 | 0.24 | 0.02 | 0.1 | 0.03 | 0.19 | 0.16 |
| 8 | 108926546 | rs681606 | 0.624 | -0.001 | -0.698 | -1.028 | -0.107 | -0.057 | -0.057 | 0 | 0.31 | 0.09 | 0.5 | 0.64 | 0.47 | 0.33 |
| 8 | 108937395 | rs2453417 | 0.536 | 0.146 | -0.636 | -0.967 | -0.104 | -0.053 | -0.053 | 0 | 0.32 | 0.09 | 0.5 | 0.64 | 0.47 | 0.33 |
| 8 | 108942260 | rs10505116 | 0.488 | 0.224 | 0.097 | -0.341 | -0.005 | 0.005 | 0.005 | 0 | 0.07 | 0.03 | 0.15 | 0.18 | 0.1 | 0.18 |
| 8 | 108943013 | rs668941 | 0.206 | 0.182 | 0.133 | -0.433 | 0.021 | 0.02 | 0.02 | 0 | 0.61 | 0.24 | 0.61 | 0.69 | 0.67 | 0.64 |
| 8 | 108953881 | rs1826700 | -0.478 | 0.198 | 0.236 | -0.549 | -0.002 | -0.031 | -0.031 | 0.23 | 0.68 | 0.37 | 0.77 | 0.94 | 0.77 | 0.82 |
| 8 | 108970367 | rs601558 | 0.558 | 0.188 | -0.392 | -0.534 | 0.012 | 0.044 | 0.044 | 0.049 | 0.56 | 0.08 | 0.6 | 0.69 | 0.66 | 0.6 |
| 8 | 108972645 | rs541730 | 0.56 | 0.201 | -0.389 | -0.527 | 0.029 | 0.058 | 0.058 | 0.226 | 0.56 | 0.1 | 0.6 | 0.69 | 0.66 | 0.6 |
| 8 | 108974658 | rs656624 | 0.324 | 0.52 | -0.227 | -0.383 | 0.02 | 0.027 | 0.027 | 0.037 | 0.61 | 0.25 | 0.6 | 0.7 | 0.66 | 0.64 |
| 8 | 108989834 | rs684786 | -0.766 | 0.878 | -0.766 | -0.185 | 0.009 | 0.048 | 0.048 | 0 | 0.74 | 0.82 | 0.81 | 0.92 | 0.77 | 0.82 |
| 8 | 108992158 | rs651858 | -0.223 | 0.799 | -0.668 | -0.098 | 0 | 0.003 | 0.003 | 0.004 | 0.66 | 0.72 | 0.64 | 0.7 | 0.67 | 0.63 |
| 8 | 109012769 | rs1443930 | 0.112 | 0.855 | -0.745 | -0.003 | -0.001 | -0.001 | -0.001 | 0 | 0.69 | 0.74 | 0.5 | 0.44 | 0.66 | 0.75 |
| 8 | 109015020 | rs13282910 | 0.099 | 0.433 | -0.864 | -1.215 | -0.004 | -0.083 | -0.083 | 0 | 0.35 | 0.34 | 0.24 | 0.02 | 0.32 | 0.26 |
| 8 | 109015798 | rs2514830 | 0.913 | 0.384 | -1.096 | -1.754 | -0.005 | -0.099 | -0.099 | 0.023 | 0.25 | 0.41 | 0.24 | 0.08 | 0.25 | 0.22 |
| 8 | 109020400 | rs989291 | -0.467 | 0.336 | -1.714 | -1.724 | -0.177 | 0.172 | 0.172 | 0.017 | 0.68 | 0.96 | 0.51 | 0.11 | 0.6 | 0.54 |
| 8 | 109028325 | rs2443786 | -1.004 | 0.381 | -1.807 | -1.59 | -0.106 | -0.148 | -0.148 | 0 | 0.76 | 0.99 | 0.59 | 0.42 | 0.73 | 0.78 |
| 8 | 109039417 | rs1452018 | 1.519 | 0.132 | -1.49 | -1.785 | -0.009 | 0.041 | 0.041 | 0.115 | 0.21 | 0.11 | 0.21 | 0.1 | 0.24 | 0.22 |
| 8 | 109043781 | rs2443782 | -0.063 | -0.207 | -1.174 | -2.038 | 0.018 | 0.022 | 0.022 | 2.009 | 0.42 | 0.7 | 0.32 | 0.11 | 0.33 | 0.4 |
| 8 | 109046727 | rs2443779 | -1.215 | -0.168 | -1.766 | -1.99 | -0.102 | -0.143 | -0.143 | 0 | 0.74 | 0.98 | 0.59 | 0.42 | 0.73 | 0.76 |
| 8 | 109052656 | rs4735036 | 1.47 | -0.09 | -1.553 | -1.825 | -0.033 | 0.172 | 0.172 | 3.965 | 0.25 | 0.01 | 0.18 | 0.01 | 0.26 | 0.13 |
| 8 | 109054668 | rs16877095 | 1.29 | 0.183 | -0.904 | -1.233 | -0.04 | -0.009 | -0.009 | 0 | 0.07 | 0.01 | 0.07 | 0.29 | 0.13 | 0.23 |
| 8 | 109060280 | rs988886 | 0.856 | 0.012 | -0.993 | -1.549 | -0.021 | 0.026 | 0.026 | 0 | 0.17 | 0.1 | 0.21 | 0.11 | 0.24 | 0.21 |
| 8 | 109060572 | rs2514840 | -0.439 | -0.518 | -0.655 | -1.903 | -0.008 | -0.131 | -0.131 | 0.221 | 0.33 | 0.51 | 0.27 | 0.08 | 0.31 | 0.35 |
| 8 | 109060667 | rs2514841 | -0.674 | -1.08 | -1.141 | -2.052 | 0.009 | 0.031 | 0.031 | 0 | 0.49 | 0.62 | 0.36 | 0.41 | 0.48 | 0.48 |
| 8 | 109062481 | rs2443775 | -1.161 | -0.83 | -1.745 | -1.648 | -0.145 | -0.137 | -0.137 | 0.031 | 0.6 | 0.96 | 0.43 | 0.5 | 0.52 | 0.62 |
| 8 | 109084909 | rs11783906 | -0.202 | -0.259 | -1.075 | -1.108 | 0.005 | -0.045 | -0.045 | 0.111 | 0.25 | 0.16 | 0.13 | 0.4 | 0.23 | 0.28 |
| 8 | 109090197 | rs16877149 | -0.003 | -0.221 | -1.206 | -1.088 | 0 | -0.074 | -0.074 | 0 | 0.24 | 0.09 | 0.12 | 0.4 | 0.23 | 0.29 |
| 8 | 109094747 | rs10955475 | 0.342 | -0.073 | -1.346 | -0.924 | -0.018 | -0.129 | -0.129 | 0.295 | 0.22 | 0.03 | 0.12 | 0.38 | 0.21 | 0.26 |
| 8 | 109096029 | rs374810 | -0.782 | -0.308 | -1.301 | -1.265 | -0.002 | -0.028 | -0.028 | 0 | 0.47 | 0.18 | 0.35 | 0.49 | 0.48 | 0.54 |
| 8 | 109114261 | rs389329 | 0.909 | -0.299 | -1.404 | -1.274 | -0.024 | 0.063 | 0.063 | 0 | 0.2 | 0.04 | 0.2 | 0.11 | 0.25 | 0.2 |
| 8 | 109127521 | rs10505119 | 0.485 | -0.382 | -1.259 | -1.1 | -0.018 | -0.129 | -0.129 | 0 | 0.22 | 0.03 | 0.12 | 0.38 | 0.21 | 0.26 |
| 8 | 109145537 | rs1450596 | -0.923 | -0.661 | -0.358 | -1.256 | -0.018 | 0.017 | 0.017 | 0.388 | 0.34 | 0.62 | 0.19 | 0.38 | 0.24 | 0.34 |
| 8 | 109146371 | rs10103270 | 0.304 | -0.546 | -0.295 | -1.178 | 0.007 | -0.058 | -0.058 | 0.044 | 0.23 | 0.15 | 0.12 | 0.38 | 0.21 | 0.27 |
| 8 | 109164182 | rs672161 | -0.243 | -0.435 | -0.095 | -1.331 | -0.015 | -0.047 | -0.047 | 1.234 | 0.4 | 0.19 | 0.33 | 0.49 | 0.46 | 0.46 |
| 8 | 109207275 | rs13250911 | -0.014 | -0.449 | 0.173 | -1.351 | -0.012 | -0.027 | -0.027 | 0 | 0.05 | 0.12 | 0.02 | 0 | 0.02 | 0.08 |
| 8 | 109213088 | rs661036 | -1.109 | -0.59 | 0.58 | -1.427 | 0.007 | -0.002 | -0.002 | 0 | 0.52 | 0.59 | 0.39 | 0.5 | 0.48 | 0.54 |
| 8 | 109228008 | rs611744 | -1.234 | -0.595 | 0.567 | -1.429 | 0.007 | -0.011 | -0.011 | 0.511 | 0.53 | 0.69 | 0.39 | 0.5 | 0.48 | 0.54 |
| 8 | 109248046 | rs16877533 | 1.113 | -0.071 | 0.139 | -1.548 | -0.068 | -0.078 | -0.078 | 0 | 0.12 | 0.01 | 0.16 | 0.3 | 0.26 | 0.22 |
| 8 | 109272006 | rs2298605 | -0.332 | -0.117 | 0.269 | -1.55 | -0.051 | -0.057 | -0.057 | 0 | 0.46 | 0.11 | 0.4 | 0.5 | 0.57 | 0.48 |
| 8 | 109280487 | rs16877589 | 0.62 | -0.17 | 0.385 | -1.591 | -0.001 | -0.025 | -0.025 | 0.784 | 0.22 | 0.23 | 0.22 | 0.11 | 0.24 | 0.2 |
| 8 | 109288399 | rs6469168 | -1.229 | -0.152 | 0.01 | -1.059 | -0.001 | -0.001 | -0.001 | 0.055 | 0.57 | 0.58 | 0.5 | 0.54 | 0.6 | 0.57 |
| 8 | 109288442 | rs7826090 | 0.119 | -0.151 | -0.003 | -1.049 | -0.025 | -0.094 | -0.094 | 0.148 | 0.28 | 0.46 | 0.26 | 0.15 | 0.25 | 0.21 |
| 8 | 109291233 | rs1975804 | 2.165 | -0.202 | -0.658 | -1.44 | -0.075 | -0.124 | -0.124 | 0 | 0.18 | 0.03 | 0.35 | 0.46 | 0.23 | 0.25 |
| 8 | 109298794 | rs7815105 | -1.477 | -0.444 | -0.624 | -1.428 | -0.088 | -0.117 | -0.117 | 0 | 0.68 | 0.94 | 0.55 | 0.53 | 0.64 | 0.69 |
| 8 | 109339587 | rs13267980 | -0.065 | -0.358 | -0.504 | -1.398 | -0.023 | -0.092 | -0.092 | 0.744 | 0.32 | 0.53 | 0.28 | 0.15 | 0.27 | 0.28 |

| | | | | | | | | | | | | | | | | |
|---|---|---|---|---|---|---|---|---|---|---|---|---|---|---|---|---|
| 8 | 109343201 | rs13253916 | 0.765 | -0.007 | -0.938 | -1.085 | -0.084 | -0.139 | -0.139 | 0 | 0.26 | 0.04 | 0.22 | 0.39 | 0.34 | 0.29 |
| 8 | 109348150 | rs6994837 | -0.061 | -0.055 | -0.772 | -1.053 | -0.002 | 0.004 | 0.004 | 0.206 | 0.52 | 0.29 | 0.45 | 0.55 | 0.58 | 0.53 |
| 8 | 109354102 | rs4734174 | -1.193 | -0.141 | -0.815 | -1.19 | -0.065 | -0.096 | -0.096 | 0 | 0.69 | 0.9 | 0.55 | 0.55 | 0.67 | 0.72 |
| 8 | 109357798 | rs7844203 | -0.106 | -0.121 | -0.31 | -1.171 | -0.023 | -0.108 | -0.108 | 0 | 0.14 | 0.52 | 0.09 | 0 | 0.08 | 0.19 |
| 8 | 109360772 | rs1375021 | -0.372 | -0.122 | -0.472 | -1.171 | -0.039 | -0.103 | -0.103 | 0 | 0.16 | 0.61 | 0.1 | 0 | 0.08 | 0.19 |
| 8 | 109379543 | rs12056624 | 1.93 | 0.171 | -1.027 | -0.813 | -0.063 | -0.109 | -0.109 | 0 | 0.15 | 0.01 | 0.3 | 0.35 | 0.17 | 0.15 |
| 8 | 109392461 | rs9650035 | 1.015 | 0.057 | -1.514 | -1.291 | -0.074 | -0.163 | -0.163 | 0 | 0.25 | 0.03 | 0.2 | 0.39 | 0.3 | 0.3 |
| 8 | 109395713 | rs16877774 | 0.781 | -0.074 | -1.232 | -1.45 | -0.003 | -0.088 | -0.088 | 3.833 | 0.28 | 0.4 | 0.27 | 0.12 | 0.3 | 0.31 |
| 8 | 109396626 | rs11781805 | -2.254 | 0.074 | -1.227 | -1.294 | -0.06 | -0.105 | -0.105 | 0 | 0.84 | 0.97 | 0.66 | 0.56 | 0.8 | 0.83 |
| 8 | 109410157 | rs1893885 | 0.625 | -0.034 | -0.754 | -1.252 | -0.017 | -0.07 | -0.07 | 0 | 0.13 | 0.48 | 0.08 | 0.04 | 0.07 | 0.16 |
| 8 | 109415550 | rs16877821 | 1.293 | 0.007 | -1.166 | -1.27 | -0.074 | -0.163 | -0.163 | 2.07 | 0.25 | 0.03 | 0.2 | 0.39 | 0.3 | 0.3 |
| 8 | 109420357 | rs16877824 | 0.752 | -0.064 | -0.902 | -1.349 | -0.01 | -0.065 | -0.065 | 0.951 | 0.13 | 0.42 | 0.08 | 0.04 | 0.07 | 0.16 |
| 8 | 109433315 | rs10111457 | -0.251 | -0.063 | -1.277 | -1.457 | 0.022 | 0.191 | 0.191 | 0.321 | 0.41 | 0.83 | 0.35 | 0.17 | 0.37 | 0.47 |
| 8 | 109455234 | rs10100704 | -0.086 | 0.154 | -1.078 | -1.303 | -0.002 | -0.002 | -0.002 | 0 | 0.53 | 0.42 | 0.45 | 0.51 | 0.59 | 0.64 |
| 8 | 109455863 | rs3737502 | 1.2 | 0.058 | -1.152 | -1.371 | -0.033 | -0.118 | -0.118 | 0.112 | 0.27 | 0.08 | 0.21 | 0.38 | 0.31 | 0.32 |
| 8 | 109498656 | rs10111934 | -0.943 | -0.025 | -0.826 | -1.182 | -0.044 | -0.064 | -0.064 | 0 | 0.65 | 0.85 | 0.54 | 0.55 | 0.66 | 0.79 |
| 8 | 109500113 | rs7009774 | 0.697 | -0.043 | -0.764 | -1.199 | -0.015 | -0.065 | -0.065 | 0.048 | 0.13 | 0.43 | 0.08 | 0.05 | 0.07 | 0.15 |
| 8 | 109509609 | rs11985955 | -0.99 | -0.039 | -0.762 | -1.197 | -0.044 | -0.065 | -0.065 | 0 | 0.66 | 0.85 | 0.54 | 0.55 | 0.66 | 0.79 |
| 8 | 109520281 | rs7002520 | -0.086 | -0.04 | -0.741 | -1.197 | -0.002 | -0.002 | -0.002 | 0.09 | 0.52 | 0.39 | 0.46 | 0.51 | 0.59 | 0.64 |
| 8 | 109520708 | rs7833253 | 1.291 | -0.059 | -0.754 | -1.219 | 0.001 | -0.028 | -0.028 | 0 | 0.1 | 0.39 | 0.07 | 0.05 | 0.05 | 0.12 |
| 8 | 109546289 | rs1480558 | -1.077 | -0.057 | -0.866 | -1.197 | -0.055 | -0.075 | -0.075 | 0 | 0.66 | 0.88 | 0.55 | 0.55 | 0.66 | 0.79 |
| 8 | 109548766 | rs6992425 | 0.243 | 0.007 | -0.473 | -1.108 | -0.011 | -0.07 | -0.07 | 2.119 | 0.14 | 0.41 | 0.08 | 0.05 | 0.08 | 0.15 |
| 8 | 109567733 | rs16878137 | 0.743 | -0.059 | -0.468 | -1.037 | -0.018 | -0.047 | -0.047 | 0.173 | 0.12 | 0.42 | 0.07 | 0.05 | 0.05 | 0.12 |
| 8 | 109568589 | rs1383354 | -0.902 | -0.083 | -0.816 | -1.054 | -0.04 | -0.041 | -0.041 | 0 | 0.64 | 0.82 | 0.54 | 0.56 | 0.64 | 0.76 |
| 8 | 109571545 | rs1871686 | -0.376 | 0.178 | -0.912 | -0.826 | -0.001 | 0.003 | 0.003 | 0.302 | 0.53 | 0.37 | 0.47 | 0.51 | 0.59 | 0.64 |
| 8 | 109577887 | rs13261211 | 0.361 | 0.342 | -1.205 | -0.308 | -0.032 | -0.114 | -0.114 | 0.116 | 0.27 | 0.09 | 0.21 | 0.38 | 0.31 | 0.32 |
| 8 | 109587652 | rs16878162 | 0.515 | 0.44 | -1.327 | -0.136 | -0.068 | -0.14 | -0.14 | 0.052 | 0.24 | 0.02 | 0.21 | 0.36 | 0.31 | 0.32 |
| 8 | 109595712 | rs1979443 | -0.708 | 0.095 | -0.375 | -0.131 | -0.005 | -0.027 | -0.027 | 0 | 0.37 | 0.55 | 0.47 | 0.71 | 0.35 | 0.42 |
| 8 | 109611346 | rs2882102 | 0.855 | 0.213 | -0.285 | 0.449 | 0 | 0.059 | 0.059 | 0 | 0.1 | 0.09 | 0.23 | 0.42 | 0.11 | 0.22 |
| 8 | 109611747 | rs2349200 | 0.306 | 0.227 | -0.08 | 0.432 | 0.003 | 0.157 | 0.157 | 0 | 0.12 | 0.25 | 0.26 | 0.43 | 0.13 | 0.26 |
| 8 | 109614094 | rs2928826 | 0.096 | 0.008 | -0.044 | 0.294 | -0.001 | -0.001 | -0.001 | 0 | 0.24 | 0.22 | 0.2 | 0.28 | 0.22 | 0.16 |
| 8 | 109639362 | rs1497639 | -1.267 | 0.086 | -0.686 | 0.085 | 0.014 | 0.054 | 0.054 | 1.588 | 0.47 | 0.7 | 0.52 | 0.71 | 0.39 | 0.39 |
| 8 | 109654860 | rs1039167 | 0.815 | 0.064 | -0.46 | 0.608 | 0 | 0.065 | 0.065 | 0 | 0.1 | 0.09 | 0.23 | 0.41 | 0.11 | 0.22 |
| 8 | 109660089 | rs2935762 | -0.791 | -0.052 | -0.754 | 0.118 | 0.017 | 0.031 | 0.031 | 1.552 | 0.4 | 0.7 | 0.48 | 0.71 | 0.32 | 0.37 |
| 8 | 109664109 | rs2935755 | 0.779 | -0.064 | -0.758 | 0.273 | 0.008 | -0.017 | -0.017 | 0.001 | 0.16 | 0.12 | 0.17 | 0.28 | 0.12 | 0.11 |
| 8 | 109664146 | rs2935754 | 0.313 | 0.05 | -0.753 | 0.413 | 0.003 | -0.025 | -0.025 | 0 | 0.22 | 0.13 | 0.21 | 0.29 | 0.18 | 0.12 |
| 8 | 109667084 | rs1497628 | 0.005 | 0.239 | -0.322 | 0.197 | 0.003 | 0.09 | 0.09 | 0 | 0.22 | 0.34 | 0.45 | 0.4 | 0.2 | 0.3 |
| 8 | 109680768 | rs1497630 | 0.374 | 0.254 | -0.505 | -0.488 | 0 | -0.014 | -0.014 | 1.084 | 0.42 | 0.53 | 0.43 | 0.69 | 0.4 | 0.61 |
| 8 | 109682418 | rs3019409 | 0.713 | 0.185 | -0.53 | -0.746 | -0.001 | -0.001 | -0.001 | 0.154 | 0.27 | 0.21 | 0.22 | 0.29 | 0.28 | 0.15 |
| 8 | 109698889 | rs3019415 | 1.501 | 0.18 | -0.593 | -0.764 | -0.023 | -0.038 | -0.038 | 0 | 0.2 | 0.09 | 0.2 | 0.29 | 0.22 | 0.13 |
| 8 | 109700663 | rs3019418 | -1.225 | 0.123 | -0.569 | -0.765 | -0.018 | 0.082 | 0.082 | 0 | 0.73 | 0.88 | 0.67 | 0.98 | 0.7 | 0.76 |
| 8 | 109701780 | rs10093487 | 0.978 | 0.239 | -0.108 | -0.459 | -0.01 | 0.106 | 0.106 | 0 | 0.16 | 0.27 | 0.25 | 0.41 | 0.12 | 0.26 |
| 8 | 109703295 | rs1391201 | 0.116 | 0.258 | -0.137 | -0.204 | 0.001 | 0.008 | 0.008 | 0 | 0.45 | 0.62 | 0.44 | 0.56 | 0.41 | 0.56 |
| 8 | 109703425 | rs1391203 | -0.734 | 0.299 | -0.21 | -0.162 | 0.001 | 0.082 | 0.082 | 0 | 0.66 | 0.83 | 0.65 | 0.85 | 0.65 | 0.69 |
| 8 | 109705054 | rs11774262 | 1.573 | 0.302 | -0.213 | -0.138 | -0.008 | -0.001 | -0.001 | 0 | 0.25 | 0.19 | 0.18 | 0.27 | 0.28 | 0.35 |
| 8 | 109710863 | rs10096148 | 0.704 | 0.018 | -0.078 | 0.303 | -0.007 | 0.058 | 0.058 | 0 | 0.15 | 0.22 | 0.24 | 0.29 | 0.11 | 0.22 |
| 8 | 109717628 | rs3019335 | 0.489 | 0.058 | -0.146 | 0.28 | -0.005 | -0.008 | -0.008 | 0 | 0.24 | 0.2 | 0.21 | 0.29 | 0.26 | 0.14 |
| 8 | 109722443 | rs3019343 | -0.402 | 0.095 | -0.301 | 0.033 | 0 | 0 | 0 | 1.022 | 0.39 | 0.4 | 0.45 | 0.58 | 0.36 | 0.34 |
| 8 | 109725508 | rs1027479 | -0.933 | -0.011 | -0.412 | 0.018 | 0.02 | 0.151 | 0.151 | 0.736 | 0.48 | 0.81 | 0.49 | 0.78 | 0.4 | 0.42 |
| 8 | 109726503 | rs1391200 | -0.468 | 0.05 | -0.62 | 0.149 | -0.003 | -0.001 | -0.001 | 1.464 | 0.39 | 0.44 | 0.44 | 0.58 | 0.33 | 0.36 |
| 8 | 109730646 | rs13256244 | -0.102 | 0.09 | -0.664 | 0.035 | 0.004 | -0.005 | -0.005 | 0 | 0.42 | 0.42 | 0.49 | 0.51 | 0.31 | 0.41 |
| 8 | 109733537 | rs2935791 | 0.217 | 0.378 | -1.121 | -0.127 | -0.028 | 0.044 | 0.044 | 0.122 | 0.31 | 0.11 | 0.29 | 0.26 | 0.36 | 0.16 |
| 8 | 109735830 | rs7820557 | -0.043 | -0.058 | -0.703 | 0.157 | 0.031 | -0.027 | -0.027 | 0 | 0.41 | 0.75 | 0.39 | 0.47 | 0.29 | 0.44 |
| 8 | 109748009 | rs2935788 | -1.434 | 0.207 | -0.91 | -0.193 | -0.098 | 0.033 | 0.033 | 0 | 0.73 | 0.95 | 0.68 | 0.86 | 0.66 | 0.64 |
| 8 | 109749049 | rs6993219 | 0.299 | -0.032 | -0.524 | -0.087 | -0.003 | 0.009 | 0.009 | 0 | 0.36 | 0.53 | 0.36 | 0.44 | 0.29 | 0.43 |
| 8 | 109754941 | rs3019359 | -1.278 | -0.117 | -0.064 | -0.041 | -0.002 | -0.001 | -0.001 | 0 | 0.66 | 0.69 | 0.65 | 0.85 | 0.61 | 0.63 |
| 8 | 109770234 | rs3019901 | -0.777 | -0.303 | 0.688 | 0.328 | 0 | -0.002 | -0.002 | 0.137 | 0.33 | 0.32 | 0.32 | 0.4 | 0.35 | 0.22 |
| 8 | 109774461 | rs1039164 | -0.52 | -0.304 | 0.723 | 0.328 | 0 | -0.009 | -0.009 | 0 | 0.3 | 0.27 | 0.31 | 0.4 | 0.35 | 0.2 |
| 8 | 109776218 | rs2935792 | -1.086 | -0.313 | 0.904 | 0.308 | -0.01 | 0.01 | 0.01 | 0 | 0.35 | 0.51 | 0.32 | 0.4 | 0.35 | 0.22 |
| 8 | 109776684 | rs16878801 | 1.1 | -0.315 | 0.851 | 0.372 | 0.002 | 0.082 | 0.082 | 9.48 | 0.05 | 0.14 | 0.03 | 0.12 | 0.04 | 0.07 |
| 8 | 109782511 | rs6469203 | -1.136 | -0.369 | 1.044 | 0.348 | -0.003 | 0 | 0 | 0 | 0.36 | 0.61 | 0.34 | 0.4 | 0.36 | 0.25 |
| 8 | 109787856 | rs7816300 | 0.006 | -0.492 | 0.679 | 0.359 | -0.028 | -0.003 | -0.003 | 0.066 | 0.25 | 0.09 | 0.28 | 0.27 | 0.31 | 0.16 |
| 8 | 109792588 | rs7845404 | -0.165 | -0.485 | 0.687 | 0.362 | -0.012 | 0.009 | 0.009 | 0.711 | 0.27 | 0.15 | 0.28 | 0.27 | 0.31 | 0.16 |
| 8 | 109800541 | rs11995185 | -0.793 | -0.199 | 1.044 | 0.421 | 0 | 0.123 | 0.123 | 7.664 | 0.1 | 0.5 | 0.06 | 0.12 | 0.05 | 0.1 |
| 8 | 109831197 | rs7819657 | -0.423 | 0.034 | 0.216 | 0.453 | -0.002 | 0 | 0 | 0 | 0.23 | 0.25 | 0.25 | 0.24 | 0.3 | 0.17 |
| 8 | 109855813 | rs12542619 | -0.634 | -0.146 | 0.138 | 0.452 | 0.004 | 0.018 | 0.018 | 0 | 0.24 | 0.27 | 0.26 | 0.39 | 0.31 | 0.23 |
| 8 | 109857400 | rs13253194 | -0.594 | -0.066 | 0.14 | 0.473 | 0.044 | 0.146 | 0.146 | 0 | 0.13 | 0.27 | 0.18 | 0.39 | 0.21 | 0.25 |
| 8 | 109875397 | rs11996192 | -1.012 | -0.538 | -0.155 | 0.409 | -0.032 | 0.011 | 0.011 | 0 | 0.35 | 0.52 | 0.29 | 0.4 | 0.32 | 0.32 |
| 8 | 109887539 | rs4617129 | 0.238 | -0.129 | -0.752 | 0.196 | -0.009 | 0.027 | 0.027 | 0.181 | 0.7 | 0.83 | 0.5 | 0.71 | 0.68 | 0.82 |

| | | | | | | | | | | | | | | | | |
|---|---|---|---|---|---|---|---|---|---|---|---|---|---|---|---|---|
| 8 | 109888569 | rs7815781 | 0.825 | -0.389 | -0.553 | 0.257 | -0.016 | -0.018 | -0.018 | 0.088 | 0.58 | 0.48 | 0.44 | 0.62 | 0.64 | 0.78 |
| 8 | 109907526 | rs6469211 | -0.123 | 0.037 | -1.435 | 0.389 | -0.079 | -0.059 | -0.059 | 0 | 0.74 | 0.99 | 0.5 | 0.69 | 0.66 | 0.84 |
| 8 | 109912830 | rs4734184 | 0.077 | 0.605 | -1.583 | 0.758 | -0.053 | -0.058 | -0.058 | 0 | 0.15 | 0.04 | 0.11 | 0.23 | 0.19 | 0.35 |
| 8 | 109917023 | rs12547648 | -0.094 | -0.186 | -1.291 | 0.329 | -0.009 | -0.027 | -0.027 | 0 | 0.29 | 0.12 | 0.5 | 0.37 | 0.32 | 0.21 |
| 8 | 109918080 | rs6983279 | -0.836 | -0.69 | -0.657 | -0.032 | -0.004 | -0.004 | -0.004 | 0 | 0.43 | 0.59 | 0.54 | 0.37 | 0.34 | 0.24 |
| 8 | 109919044 | rs6988622 | -0.101 | -0.461 | -0.861 | 0.218 | -0.011 | -0.059 | -0.059 | 0.211 | 0.31 | 0.14 | 0.53 | 0.46 | 0.36 | 0.29 |
| 8 | 109921525 | rs7000420 | -0.182 | -0.169 | -0.382 | 0.398 | 0.006 | 0.007 | 0.007 | 2.133 | 0.4 | 0.39 | 0.54 | 0.37 | 0.36 | 0.39 |
| 8 | 109923059 | rs4735078 | -0.669 | 0.037 | -0.721 | 0.4 | 0.007 | 0.006 | 0.006 | 0 | 0.52 | 0.64 | 0.62 | 0.47 | 0.48 | 0.57 |
| 8 | 109924310 | rs12681052 | -0.198 | -0.364 | -0.833 | 0.193 | 0.018 | 0.017 | 0.017 | 0 | 0.27 | 0.21 | 0.36 | 0.19 | 0.18 | 0.24 |
| 8 | 109928145 | rs4735079 | -0.3 | -0.517 | -0.957 | 0.05 | 0.033 | 0.07 | 0.07 | 1.855 | 0.29 | 0.12 | 0.39 | 0.13 | 0.2 | 0.23 |
| 8 | 109940377 | rs4336584 | -1.217 | -0.488 | -0.5 | 0.089 | 0.026 | 0.059 | 0.059 | 0 | 0.47 | 0.76 | 0.51 | 0.28 | 0.34 | 0.54 |
| 8 | 109961231 | rs13439408 | -0.478 | -0.614 | 0.134 | 0.061 | -0.023 | -0.071 | -0.071 | 0 | 0.11 | 0.35 | 0.05 | 0 | 0.04 | 0.16 |
| 8 | 109965998 | rs12543660 | -1.087 | -0.677 | 0.202 | 0.034 | -0.021 | -0.018 | -0.018 | 0 | 0.4 | 0.5 | 0.46 | 0.28 | 0.27 | 0.49 |
| 8 | 109966356 | rs12543745 | 0.401 | -0.658 | 0.029 | 0.026 | 0.016 | 0.068 | 0.068 | 0 | 0.24 | 0.09 | 0.37 | 0.08 | 0.19 | 0.16 |
| 8 | 109973032 | rs12677646 | 0.081 | -0.592 | 0.05 | -0.038 | 0.011 | -0.013 | -0.013 | 0 | 0.28 | 0.15 | 0.47 | 0.28 | 0.23 | 0.32 |
| 8 | 109978529 | rs10086780 | -0.976 | -0.593 | 0.123 | -0.022 | 0.018 | 0.016 | 0.016 | 0.007 | 0.41 | 0.65 | 0.54 | 0.29 | 0.27 | 0.49 |
| 8 | 109979163 | rs12681212 | -1.531 | -0.345 | -0.595 | 0.077 | -0.137 | -0.127 | -0.127 | 0 | 0.62 | 0.98 | 0.67 | 0.55 | 0.48 | 0.75 |
| 8 | 109979852 | rs7824283 | -0.376 | -0.283 | 0.44 | -0.063 | -0.006 | -0.03 | -0.03 | 0 | 0.08 | 0.15 | 0.03 | 0 | 0.04 | 0.16 |
| 8 | 109987530 | rs7015029 | -0.961 | -0.217 | 0.769 | 0.073 | 0.047 | 0.043 | 0.043 | 6.212 | 0.06 | 0.41 | 0.05 | 0 | 0 | 0 |
| 8 | 109991637 | rs4236795 | -1.199 | 0.05 | -0.3 | 0.147 | -0.047 | -0.043 | -0.043 | 5.132 | 0.56 | 0.82 | 0.58 | 0.51 | 0.47 | 0.63 |
| 8 | 109991668 | rs4620259 | 0.461 | -0.117 | -0.303 | -0.729 | 0.029 | 0.084 | 0.084 | 0 | 0.26 | 0.16 | 0.38 | 0.07 | 0.18 | 0.16 |
| 8 | 110002080 | rs17520 | -1.746 | -0.145 | -0.114 | -0.448 | -0.036 | 0.059 | 0.059 | 0 | 0.58 | 0.75 | 0.64 | 0.71 | 0.49 | 0.66 |
| 8 | 110007151 | rs7838433 | -1.157 | -0.141 | -0.075 | -0.407 | -0.001 | -0.001 | -0.001 | 0 | 0.32 | 0.57 | 0.26 | 0.63 | 0.29 | 0.5 |
| 8 | 110010886 | rs10429304 | -0.41 | -0.138 | 0.48 | -0.395 | 0.002 | 0.005 | 0.005 | 0 | 0.22 | 0.21 | 0.2 | 0.63 | 0.25 | 0.37 |
| 8 | 110020230 | rs4735081 | -0.711 | -0.11 | 0.615 | -0.381 | 0.015 | 0.036 | 0.036 | 0 | 0.26 | 0.46 | 0.23 | 0.63 | 0.26 | 0.37 |
| 8 | 110026577 | rs9643043 | -1.214 | 0.012 | 0.536 | -0.386 | 0.006 | 0.088 | 0.088 | 0 | 0.5 | 0.7 | 0.69 | 0.72 | 0.44 | 0.59 |
| 8 | 110030691 | rs6469215 | -1.539 | 0.277 | 0.097 | -0.591 | -0.033 | 0.088 | 0.088 | 0 | 0.58 | 0.85 | 0.78 | 0.72 | 0.52 | 0.6 |
| 8 | 110033532 | rs6988172 | 0.166 | 0.379 | 0.393 | -0.363 | 0.053 | 0.21 | 0.21 | 0 | 0.14 | 0.31 | 0.21 | 0.55 | 0.18 | 0.25 |
| 8 | 110059051 | rs11986219 | -0.261 | 0.357 | -0.189 | -0.423 | 0.019 | 0.096 | 0.096 | 0 | 0.35 | 0.19 | 0.44 | 0.16 | 0.28 | 0.23 |
| 8 | 110075358 | rs4469429 | 0.39 | 0.609 | 0.076 | -0.033 | 0.032 | 0.131 | 0.131 | 0 | 0.12 | 0.2 | 0.19 | 0.55 | 0.18 | 0.25 |
| 8 | 110085818 | rs4506180 | -1.062 | 0.47 | -0.427 | -0.257 | -0.033 | 0 | 0 | 0 | 0.53 | 0.8 | 0.64 | 0.54 | 0.49 | 0.66 |
| 8 | 110088241 | rs6469232 | 0.117 | 0.273 | 0.006 | -0.645 | -0.006 | -0.031 | -0.031 | 0 | 0.14 | 0.22 | 0.11 | 0.07 | 0.1 | 0.12 |
| 8 | 110090458 | rs3110052 | 0.726 | 0.293 | -0.448 | -1.014 | -0.026 | 0.097 | 0.097 | 0 | 0.2 | 0.03 | 0.36 | 0.05 | 0.19 | 0.1 |
| 8 | 110093871 | rs3110043 | -1.049 | 0.175 | 0.4 | -1.103 | 0.014 | 0.022 | 0.022 | 0 | 0.44 | 0.63 | 0.54 | 0.13 | 0.33 | 0.41 |
| 8 | 110095633 | rs3110040 | -1.71 | -0.003 | 0.405 | -0.738 | -0.004 | -0.023 | -0.023 | 0.293 | 0.6 | 0.8 | 0.76 | 0.56 | 0.57 | 0.74 |
| 8 | 110104203 | rs3134108 | -1.457 | 0.006 | 0.449 | -0.733 | -0.003 | 0.002 | 0.002 | 0 | 0.58 | 0.57 | 0.74 | 0.56 | 0.57 | 0.74 |
| 8 | 110109255 | rs16879095 | -0.602 | -0.034 | 0.898 | -0.673 | 0.04 | 0.036 | 0.036 | 4.785 | 0.06 | 0.48 | 0.04 | 0 | 0 | 0 |
| 8 | 110115676 | rs7832552 | -0.234 | -0.215 | -0.003 | -1.074 | -0.026 | -0.067 | -0.067 | 1.692 | 0.26 | 0.16 | 0.48 | 0.46 | 0.27 | 0.42 |
| 8 | 110122139 | rs4129682 | -1.966 | -0.549 | -0.387 | -1.359 | -0.024 | -0.033 | -0.033 | 0 | 0.53 | 0.84 | 0.67 | 0.51 | 0.41 | 0.62 |
| 8 | 110125762 | rs4546626 | -1.165 | -0.514 | -0.02 | -1.328 | -0.001 | 0.001 | 0.001 | 1.994 | 0.37 | 0.42 | 0.54 | 0.46 | 0.32 | 0.51 |
| 8 | 110126270 | rs4446710 | -1.165 | -0.514 | -0.02 | -1.328 | -0.001 | 0.001 | 0.001 | 1.994 | 0.37 | 0.42 | 0.54 | 0.46 | 0.32 | 0.51 |
| 8 | 110133437 | rs5784 | 0.907 | -0.352 | -0.384 | -1.206 | 0.008 | 0.049 | 0.049 | 0.074 | 0.09 | 0.03 | 0.04 | 0 | 0.05 | 0.05 |
| 8 | 110150671 | rs4734194 | -1.684 | -0.152 | -0.44 | -1.291 | -0.065 | -0.049 | -0.049 | 0.545 | 0.57 | 0.85 | 0.68 | 0.51 | 0.45 | 0.64 |
| 8 | 110174335 | rs9969620 | -0.12 | 0.531 | -0.089 | -1.096 | -0.01 | -0.015 | -0.015 | 0 | 0.24 | 0.19 | 0.48 | 0.46 | 0.29 | 0.45 |
| 8 | 110202230 | rs10955512 | 0.717 | 0.618 | 0.042 | -1.001 | 0.037 | 0.086 | 0.086 | 0.28 | 0.14 | 0.18 | 0.46 | 0.46 | 0.24 | 0.29 |
| 8 | 110217266 | rs7823896 | -0.825 | -0.018 | 0.134 | -1.24 | 0.001 | -0.01 | -0.01 | 0 | 0.31 | 0.71 | 0.2 | 0.06 | 0.13 | 0.19 |
| 8 | 110238631 | rs17364037 | -0.147 | 0.081 | -0.264 | -1.507 | -0.019 | -0.017 | -0.017 | 0.36 | 0.32 | 0.27 | 0.53 | 0.46 | 0.41 | 0.52 |
| 8 | 110249729 | rs2980635 | -1.216 | -0.147 | -0.049 | -1.504 | -0.006 | -0.012 | -0.012 | 0 | 0.46 | 0.77 | 0.59 | 0.46 | 0.39 | 0.59 |
| 8 | 110250441 | rs2926216 | -0.255 | -0.135 | 0.07 | -1.511 | 0.019 | 0.025 | 0.025 | 0.433 | 0.31 | 0.35 | 0.51 | 0.46 | 0.36 | 0.51 |
| 8 | 110254895 | rs1548082 | 0.171 | -0.106 | -0.358 | -1.48 | -0.026 | -0.03 | -0.03 | 0 | 0.28 | 0.18 | 0.49 | 0.46 | 0.36 | 0.52 |
| 8 | 110256090 | rs1531298 | -2.054 | -0.107 | -0.397 | -1.485 | -0.06 | -0.07 | -0.07 | 0.026 | 0.59 | 0.89 | 0.69 | 0.52 | 0.49 | 0.7 |
| 8 | 110281480 | rs2125513 | -2.045 | -0.142 | -0.354 | -1.487 | -0.044 | -0.05 | -0.05 | 0 | 0.59 | 0.86 | 0.68 | 0.52 | 0.48 | 0.7 |
| 8 | 110300579 | rs2980614 | -2.549 | -0.181 | -0.747 | -1.594 | -0.048 | -0.1 | -0.1 | 0 | 0.66 | 0.94 | 0.77 | 0.58 | 0.61 | 0.77 |
| 8 | 110301998 | rs2980618 | 0.174 | -0.352 | -0.489 | -1.506 | -0.026 | -0.03 | -0.03 | 1.36 | 0.28 | 0.18 | 0.49 | 0.46 | 0.36 | 0.51 |
| 8 | 110302047 | rs2980619 | -1.6 | 0.218 | -0.87 | -1.154 | -0.066 | -0.073 | -0.073 | 0.419 | 0.59 | 0.89 | 0.69 | 0.52 | 0.48 | 0.7 |
| 8 | 110333207 | rs16879302 | 0.67 | 0.164 | -0.534 | -1.027 | 0.005 | 0.01 | 0.01 | 0 | 0.13 | 0.1 | 0.1 | 0.06 | 0.09 | 0.11 |
| 8 | 110345261 | rs2926244 | -1.17 | 0.443 | -1.132 | -0.801 | -0.076 | -0.072 | -0.072 | 0.121 | 0.59 | 0.9 | 0.69 | 0.5 | 0.47 | 0.7 |
| 8 | 110351258 | rs1458919 | -1.175 | 0.044 | -0.013 | -0.684 | 0.03 | 0.084 | 0.084 | 0 | 0.27 | 0.47 | 0.53 | 0.44 | 0.34 | 0.36 |
| 8 | 110358672 | rs2926239 | -0.15 | 0.147 | 0.025 | -0.532 | 0.005 | 0.036 | 0.036 | 0.007 | 0.13 | 0.14 | 0.4 | 0.36 | 0.16 | 0.24 |
| 8 | 110360958 | rs2926236 | -2.124 | -0.394 | -0.737 | -1.04 | -0.105 | -0.147 | -0.147 | 1.813 | 0.66 | 0.94 | 0.73 | 0.5 | 0.57 | 0.75 |
| 8 | 110372477 | rs6982826 | -0.11 | -0.386 | 0.359 | -1.291 | -0.006 | -0.169 | -0.169 | 0.079 | 0.3 | 0.43 | 0.15 | 0 | 0.2 | 0.29 |
| 8 | 110376861 | rs7008687 | -0.102 | -0.386 | 0.359 | -1.291 | -0.006 | -0.169 | -0.169 | 0 | 0.3 | 0.43 | 0.15 | 0 | 0.2 | 0.29 |
| 8 | 110392129 | rs921960 | -1.608 | -0.429 | 0.395 | -1.103 | 0.008 | 0.015 | 0.015 | 0 | 0.51 | 0.55 | 0.64 | 0.39 | 0.52 | 0.66 |
| 8 | 110408527 | rs17367052 | 0.547 | -0.325 | 0.245 | -1.055 | -0.049 | -0.006 | -0.006 | 0 | 0.1 | 0.01 | 0.11 | 0.08 | 0.19 | 0.11 |
| 8 | 110416813 | rs1457286 | -1.763 | -0.448 | 0.59 | -1.12 | 0.008 | 0.018 | 0.018 | 0 | 0.53 | 0.69 | 0.66 | 0.4 | 0.52 | 0.71 |
| 8 | 110418922 | rs1037695 | -1.612 | -0.488 | 0.913 | -1.14 | 0.011 | 0.017 | 0.017 | 0 | 0.51 | 0.61 | 0.65 | 0.4 | 0.52 | 0.71 |
| 8 | 110420158 | rs1379365 | -0.35 | -0.496 | 0.977 | -1.189 | 0.005 | 0.021 | 0.021 | 0 | 0.17 | 0.16 | 0.47 | 0.39 | 0.31 | 0.34 |
| 8 | 110420384 | rs13254245 | -0.35 | -0.496 | 0.977 | -1.189 | 0.005 | 0.021 | 0.021 | 0 | 0.17 | 0.16 | 0.47 | 0.39 | 0.31 | 0.34 |
| 8 | 110420429 | rs13252234 | -0.35 | -0.496 | 0.977 | -1.189 | 0.005 | 0.021 | 0.021 | 0 | 0.17 | 0.16 | 0.47 | 0.39 | 0.31 | 0.34 |

| | | | | | | | | | | | | | | | | |
|---|---|---|---|---|---|---|---|---|---|---|---|---|---|---|---|---|
| 8 | 110425552 | rs10100944 | 0.218 | -0.499 | 1.037 | -1.27 | 0 | -0.099 | -0.099 | 0.51 | 0.28 | 0.31 | 0.14 | 0 | 0.2 | 0.3 |
| 8 | 110442726 | rs11987659 | -1.326 | -0.698 | 1.097 | -1.273 | -0.066 | -0.08 | -0.08 | 0 | 0.13 | 0.55 | 0.06 | 0 | 0.01 | 0.07 |
| 8 | 110448635 | rs10093885 | -0.516 | -0.624 | 1.209 | -1.299 | -0.007 | -0.109 | -0.109 | 0 | 0.33 | 0.35 | 0.15 | 0 | 0.21 | 0.32 |
| 8 | 110455321 | rs1673408 | -1.806 | -0.607 | 1.368 | -1.241 | 0.007 | 0.008 | 0.008 | 0 | 0.52 | 0.55 | 0.66 | 0.4 | 0.51 | 0.72 |
| 8 | 110458819 | rs17368044 | 0.188 | -0.627 | 1.401 | -1.256 | 0 | -0.044 | -0.044 | 0 | 0.28 | 0.23 | 0.14 | 0 | 0.2 | 0.29 |
| 8 | 110460488 | rs1673407 | -2.081 | -0.617 | 1.222 | -1.249 | 0.007 | 0.008 | 0.008 | 0 | 0.54 | 0.75 | 0.67 | 0.4 | 0.51 | 0.72 |
| 8 | 110465481 | rs11778756 | 0.648 | -0.597 | 0.956 | -1.274 | -0.031 | 0.116 | 0.116 | 0 | 0.25 | 0.07 | 0.13 | 0 | 0.2 | 0.28 |
| 8 | 110473907 | rs1026439 | -2.268 | -0.646 | 0.969 | -1.234 | -0.045 | -0.07 | -0.07 | 0 | 0.59 | 0.89 | 0.7 | 0.41 | 0.53 | 0.72 |
| 8 | 110475991 | rs1783144 | -1.719 | -0.543 | 1.094 | -1.152 | 0.009 | 0.005 | 0.005 | 0 | 0.52 | 0.57 | 0.66 | 0.41 | 0.51 | 0.72 |
| 8 | 110476993 | rs1673398 | -1.044 | -0.525 | 1.081 | -1.151 | 0 | -0.001 | -0.001 | 0 | 0.44 | 0.39 | 0.63 | 0.41 | 0.5 | 0.71 |
| 8 | 110491877 | rs1630942 | -0.773 | -0.325 | 1.316 | -0.973 | 0.005 | 0 | 0 | 0 | 0.36 | 0.38 | 0.52 | 0.33 | 0.34 | 0.61 |
| 8 | 110510993 | rs9774677 | 0.666 | -0.305 | 1.353 | -0.899 | -0.028 | -0.041 | -0.041 | 1.416 | 0.07 | 0.16 | 0.03 | 0 | 0.01 | 0 |
| 8 | 110522965 | rs2607624 | -0.929 | -0.171 | 1.308 | -0.817 | 0.053 | 0.237 | 0.237 | 0.088 | 0.11 | 0.35 | 0.4 | 0.32 | 0.14 | 0.23 |
| 8 | 110528149 | rs1673411 | -1.32 | -0.267 | 1.319 | -0.91 | 0.002 | 0.009 | 0.009 | 0 | 0.44 | 0.6 | 0.55 | 0.33 | 0.34 | 0.61 |
| 8 | 110539186 | rs1783174 | -1.023 | -0.238 | 1.31 | -0.619 | 0.053 | 0.237 | 0.237 | 0.088 | 0.11 | 0.35 | 0.4 | 0.32 | 0.14 | 0.23 |
| 8 | 110545510 | rs2022935 | 0.093 | -0.183 | 0.762 | -0.588 | 0.03 | 0.151 | 0.151 | 0 | 0.92 | 0.84 | 0.63 | 0.68 | 0.87 | 0.83 |
| 8 | 110551134 | rs3808380 | -0.791 | -0.286 | 0.46 | -0.62 | -0.05 | -0.052 | -0.052 | 0 | 0.91 | 0.99 | 0.84 | 0.71 | 0.85 | 0.82 |
| 8 | 110552577 | rs1044230 | -0.093 | -0.285 | 0.616 | -0.667 | -0.003 | 0.01 | 0.01 | 0 | 0.87 | 0.9 | 0.72 | 0.68 | 0.8 | 0.75 |
| 8 | 110558622 | rs721076 | -0.863 | -0.48 | 0.953 | -0.781 | -0.038 | -0.091 | -0.091 | 0.573 | 0.13 | 0.46 | 0.07 | 0 | 0.04 | 0.02 |
| 8 | 110563924 | rs2703386 | -1.681 | -0.541 | 0.23 | -0.897 | 0.018 | 0.047 | 0.047 | 0.149 | 0.47 | 0.79 | 0.56 | 0.34 | 0.36 | 0.62 |
| 8 | 110567774 | rs2022934 | -1.762 | -0.545 | 0.241 | -0.899 | 0.014 | 0.05 | 0.05 | 0.498 | 0.48 | 0.85 | 0.57 | 0.34 | 0.36 | 0.62 |
| 8 | 110570060 | rs2844232 | -2.55 | -0.587 | 0.058 | -0.921 | -0.082 | -0.146 | -0.146 | 0.075 | 0.68 | 0.93 | 0.73 | 0.44 | 0.61 | 0.78 |
| 8 | 110577755 | rs17450206 | -1.483 | -0.278 | 1.14 | -0.269 | 0.013 | -0.032 | -0.032 | 0 | 0.34 | 0.68 | 0.28 | 0 | 0.16 | 0.33 |
| 8 | 110584426 | rs6988341 | 0.376 | -0.161 | 0.848 | -0.194 | 0.007 | 0.028 | 0.028 | 0 | 0.18 | 0.1 | 0.21 | 0 | 0.12 | 0.3 |
| 8 | 110585288 | rs7824177 | -1.349 | -0.262 | 1.009 | -0.24 | 0.019 | -0.023 | -0.023 | 0 | 0.36 | 0.68 | 0.28 | 0 | 0.16 | 0.33 |
| 8 | 110594148 | rs2844235 | -0.575 | -0.185 | 0.654 | 0.196 | 0.007 | 0.053 | 0.053 | 0 | 0.08 | 0.08 | 0.25 | 0.33 | 0.17 | 0.22 |
| 8 | 110594818 | rs2844236 | -0.588 | -0.194 | 0.657 | 0.199 | -0.011 | -0.01 | -0.01 | 0 | 0.13 | 0.08 | 0.26 | 0.33 | 0.19 | 0.24 |
| 8 | 110613000 | rs6982592 | -0.275 | -0.427 | 0.737 | 0.124 | 0.02 | 0.058 | 0.058 | 0 | 0.23 | 0.1 | 0.22 | 0 | 0.14 | 0.31 |
| 8 | 110616994 | rs1892759 | -1.533 | -0.478 | 1.242 | 0.515 | 0.006 | -0.006 | -0.006 | 0 | 0.36 | 0.68 | 0.4 | 0.36 | 0.25 | 0.47 |
| 8 | 110622831 | rs1954714 | -1.464 | -0.341 | 1.464 | 0.525 | -0.011 | 0.011 | 0.011 | 0.274 | 0.32 | 0.58 | 0.27 | 0.41 | 0.21 | 0.42 |
| 8 | 110627005 | rs2844245 | -1.247 | -0.324 | 1.494 | 0.534 | -0.027 | 0.039 | 0.039 | 0.378 | 0.31 | 0.47 | 0.26 | 0.41 | 0.21 | 0.42 |
| 8 | 110630182 | rs2022930 | -0.77 | -0.339 | 1.622 | 0.518 | 0.009 | 0.152 | 0.152 | 0.379 | 0.11 | 0.21 | 0.16 | 0.41 | 0.12 | 0.17 |
| 8 | 110630664 | rs11989290 | 0.168 | -0.36 | 1.644 | 0.505 | 0.021 | 0.049 | 0.049 | 0 | 0.18 | 0.08 | 0.09 | 0 | 0.09 | 0.25 |
| 8 | 110643295 | rs6469270 | -1.109 | -0.339 | 1.752 | 0.538 | -0.015 | 0.028 | 0.028 | 0.404 | 0.3 | 0.4 | 0.26 | 0.41 | 0.21 | 0.42 |
| 8 | 110648272 | rs16879844 | -1.034 | -0.325 | 1.55 | 0.545 | -0.005 | 0.009 | 0.009 | 0.467 | 0.29 | 0.32 | 0.25 | 0.41 | 0.21 | 0.42 |
| 8 | 110651323 | rs10481055 | -0.541 | -0.183 | 1.328 | 0.423 | 0 | 0 | 0 | 0 | 0.24 | 0.23 | 0.17 | 0.19 | 0.2 | 0.3 |
| 8 | 110655337 | rs11998387 | -1.809 | -0.302 | 1.176 | 0.523 | 0.035 | -0.03 | -0.03 | 0 | 0.42 | 0.77 | 0.35 | 0.51 | 0.32 | 0.47 |
| 8 | 110660908 | rs7841022 | -0.758 | 0.346 | 1.771 | 1.007 | 0.01 | 0.069 | 0.069 | 0.708 | 0.09 | 0.42 | 0.05 | 0.11 | 0.01 | 0.02 |
| 8 | 110666155 | rs10107091 | 0.483 | 0.344 | 1.823 | 1.006 | -0.001 | -0.021 | -0.021 | 0.516 | 0.06 | 0.19 | 0.03 | 0 | 0.01 | 0 |
| 8 | 110670303 | rs13268243 | -2.498 | 0.335 | 1.547 | 1.038 | 0.015 | 0.146 | 0.146 | 0 | 0.21 | 0.55 | 0.25 | 0.52 | 0.22 | 0.22 |
| 8 | 110676044 | rs16879903 | -0.952 | 0.312 | 1.57 | 1.011 | 0.003 | 0.061 | 0.061 | 0.394 | 0.09 | 0.45 | 0.05 | 0.11 | 0.01 | 0.02 |
| 8 | 110689378 | rs3133926 | -1.979 | 0.408 | 1.228 | 0.916 | 0.027 | 0.151 | 0.151 | 0 | 0.2 | 0.37 | 0.24 | 0.52 | 0.22 | 0.22 |
| 8 | 110701498 | rs3935694 | -1.351 | 0.42 | 0.21 | 0.391 | 0.011 | -0.003 | -0.003 | 4.403 | 0.11 | 0.2 | 0.08 | 0.12 | 0.12 | 0.06 |
| 8 | 110714321 | rs10102228 | -0.564 | 0.361 | -0.463 | 0.45 | -0.001 | -0.002 | -0.002 | 0 | 0.08 | 0.05 | 0.07 | 0.12 | 0.08 | 0.04 |
| 8 | 110721705 | rs3133942 | -0.657 | 0.254 | -0.68 | 0.386 | 0.009 | 0.001 | 0.001 | 0.836 | 0.48 | 0.71 | 0.59 | 0.53 | 0.46 | 0.58 |
| 8 | 110722139 | rs3108844 | 1.164 | -0.018 | -1.228 | 0.106 | -0.105 | -0.056 | -0.056 | 0 | 0.37 | 0.07 | 0.32 | 0.39 | 0.45 | 0.28 |
| 8 | 110724604 | rs12547163 | 0.582 | -0.175 | -1.147 | 0.027 | 0 | 0.145 | 0.145 | 0 | 0.28 | 0.07 | 0.35 | 0.06 | 0.24 | 0.38 |
| 8 | 110725294 | rs3133945 | 0.395 | -0.177 | -1.076 | 0.094 | -0.017 | 0.037 | 0.037 | 0 | 0.07 | 0.03 | 0.13 | 0.34 | 0.13 | 0.18 |
| 8 | 110742458 | rs12547355 | -0.138 | -0.24 | -0.704 | 0.068 | -0.005 | -0.047 | -0.047 | 0 | 0.36 | 0.23 | 0.49 | 0.4 | 0.37 | 0.56 |
| 8 | 110742889 | rs3108875 | 0.167 | -0.176 | -0.716 | 0.095 | 0.009 | 0.104 | 0.104 | 0 | 0.08 | 0.14 | 0.15 | 0.34 | 0.15 | 0.19 |
| 8 | 110747110 | rs3108879 | -0.325 | -0.219 | -0.314 | 0.023 | 0 | -0.001 | -0.001 | 0 | 0.38 | 0.39 | 0.51 | 0.42 | 0.37 | 0.59 |
| 8 | 110749551 | rs3133965 | -0.253 | -0.026 | -0.363 | 0.104 | 0.049 | 0.265 | 0.265 | 0 | 0.1 | 0.3 | 0.17 | 0.36 | 0.16 | 0.22 |
| 8 | 110753672 | rs7463805 | 0.671 | -0.221 | -0.575 | -0.114 | 0.008 | 0 | 0 | 0 | 0.48 | 0.36 | 0.4 | 0.52 | 0.53 | 0.31 |
| 8 | 110758319 | rs16879981 | -0.412 | 0.063 | -0.698 | -0.242 | -0.002 | -0.005 | -0.005 | 0 | 0.12 | 0.11 | 0.08 | 0.06 | 0.09 | 0.1 |
| 8 | 110764919 | rs8180959 | -0.002 | 0.71 | -1.072 | 0.197 | -0.037 | -0.089 | -0.089 | 0 | 0.2 | 0.07 | 0.21 | 0.43 | 0.29 | 0.15 |
| 8 | 110765282 | rs2350101 | 0.037 | 0.734 | -1.087 | 0.205 | -0.046 | -0.103 | -0.103 | 0 | 0.19 | 0.04 | 0.2 | 0.42 | 0.27 | 0.15 |
| 8 | 110770293 | rs1601181 | -0.947 | 0.393 | -0.623 | -0.147 | 0.002 | -0.024 | -0.024 | 0.054 | 0.4 | 0.44 | 0.48 | 0.8 | 0.47 | 0.55 |
| 8 | 110782414 | rs3133916 | -0.067 | -0.159 | -0.42 | -0.003 | 0.009 | 0.142 | 0.142 | 0.08 | 0.18 | 0.38 | 0.28 | 0.38 | 0.19 | 0.4 |
| 8 | 110791959 | rs3102242 | -0.206 | -0.318 | -0.142 | -0.134 | -0.01 | 0.129 | 0.129 | 0 | 0.19 | 0.38 | 0.27 | 0.38 | 0.18 | 0.39 |
| 8 | 110808944 | rs1552246 | -0.691 | 0.389 | -0.47 | -0.266 | -0.132 | -0.096 | -0.096 | 0 | 0.74 | 0.95 | 0.67 | 0.59 | 0.56 | 0.85 |
| 8 | 110821596 | rs12548341 | -0.015 | -0.169 | -0.007 | -0.151 | -0.019 | 0.105 | 0.105 | 0 | 0.22 | 0.4 | 0.27 | 0.37 | 0.21 | 0.45 |
| 8 | 110843613 | rs2129035 | -0.692 | -0.201 | 0.061 | -0.609 | -0.004 | 0.001 | 0.001 | 0 | 0.32 | 0.65 | 0.34 | 0.56 | 0.28 | 0.52 |
| 8 | 110856481 | rs7813186 | -0.846 | -0.361 | -0.266 | -0.926 | 0.019 | 0.019 | 0.019 | 0.09 | 0.57 | 0.81 | 0.45 | 0.59 | 0.41 | 0.73 |
| 8 | 110866734 | rs10104270 | -0.843 | 0.037 | 0.416 | 0.029 | -0.013 | 0.01 | 0.01 | 0.016 | 0.32 | 0.6 | 0.35 | 0.59 | 0.29 | 0.52 |
| 8 | 110870227 | rs7820832 | -0.245 | 0.01 | 0.55 | 0.009 | 0.018 | -0.009 | -0.009 | 0 | 0.07 | 0.26 | 0.05 | 0 | 0.03 | 0.03 |
| 8 | 110888220 | rs1505652 | -0.714 | 0.15 | -0.162 | 0.1 | -0.012 | 0.008 | 0.008 | 0.02 | 0.32 | 0.6 | 0.35 | 0.59 | 0.29 | 0.52 |
| 8 | 110934142 | rs10107566 | -0.792 | 0.446 | 0.407 | -0.401 | -0.048 | -0.035 | -0.035 | 0 | 0.21 | 0.6 | 0.11 | 0.15 | 0.1 | 0.07 |
| 8 | 110948848 | rs7007649 | -0.447 | 0.419 | 0.691 | -0.415 | -0.029 | -0.008 | -0.008 | 0 | 0.16 | 0.44 | 0.1 | 0.15 | 0.1 | 0.06 |
| 8 | 110975199 | rs1605128 | 0.144 | 0.318 | 0.431 | -0.341 | 0 | 0.008 | 0.008 | 0.21 | 0.22 | 0.22 | 0.39 | 0.43 | 0.22 | 0.49 |

| | | | | | | | | | | | | | | | | |
|---|---|---|---|---|---|---|---|---|---|---|---|---|---|---|---|---|
| 8 | 110978142 | rs6469290 | -0.176 | 0.43 | 0.356 | -0.263 | -0.036 | -0.023 | -0.023 | 0 | 0.16 | 0.45 | 0.1 | 0.15 | 0.1 | 0.06 |
| 8 | 110981060 | rs6980482 | 0.295 | 0.428 | 0.327 | -0.264 | -0.004 | -0.002 | -0.002 | 0 | 0.13 | 0.13 | 0.06 | 0.15 | 0.08 | 0.05 |
| 8 | 110982522 | rs4735145 | -0.203 | 0.464 | 0.007 | -0.246 | -0.044 | -0.033 | -0.033 | 0.04 | 0.18 | 0.52 | 0.1 | 0.15 | 0.1 | 0.06 |
| 8 | 110990156 | rs10092842 | -0.898 | 0.655 | -0.611 | -0.622 | -0.06 | -0.028 | -0.028 | 0 | 0.66 | 0.86 | 0.64 | 0.58 | 0.53 | 0.79 |
| 8 | 111028667 | rs11992891 | -0.275 | 0.566 | -0.789 | 0.279 | -0.004 | 0.028 | 0.028 | 0 | 0.21 | 0.27 | 0.37 | 0.38 | 0.21 | 0.47 |
| 8 | 111047461 | rs971850 | -0.387 | 0.829 | -1.772 | -0.042 | -0.071 | 0.024 | 0.024 | 0 | 0.57 | 0.9 | 0.63 | 0.63 | 0.52 | 0.66 |
| 8 | 111047932 | rs7462775 | -0.028 | 0.887 | -2.003 | 0.106 | -0.019 | -0.071 | -0.071 | 0.52 | 0.25 | 0.16 | 0.43 | 0.47 | 0.3 | 0.47 |
| 8 | 111051403 | rs10505135 | -0.405 | 0.798 | -1.795 | -0.598 | 0.002 | 0.002 | 0.002 | 0.011 | 0.58 | 0.4 | 0.73 | 0.6 | 0.68 | 0.65 |
| 8 | 111055485 | rs1450757 | 0.9 | 0.752 | -1.681 | -0.604 | -0.033 | 0.063 | 0.063 | 0.87 | 0.31 | 0.16 | 0.3 | 0.13 | 0.39 | 0.16 |
| 8 | 111055739 | rs1562820 | -1.443 | 0.55 | -1.817 | -0.652 | -0.001 | -0.022 | -0.022 | 1.737 | 0.81 | 0.96 | 0.87 | 0.78 | 0.84 | 0.78 |
| 8 | 111056336 | rs973815 | 0.559 | 0.531 | -1.788 | -0.633 | -0.002 | -0.008 | -0.008 | 0 | 0.26 | 0.24 | 0.43 | 0.47 | 0.28 | 0.49 |
| 8 | 111061186 | rs996249 | -0.132 | 1.048 | -2.022 | -0.661 | 0.002 | 0.004 | 0.004 | 0.019 | 0.57 | 0.39 | 0.71 | 0.61 | 0.62 | 0.64 |
| 8 | 111064847 | rs16880318 | 1.558 | 0.829 | -2.106 | -0.883 | 0.012 | 0.016 | 0.016 | 0 | 0.11 | 0.06 | 0.04 | 0.1 | 0.08 | 0.07 |
| 8 | 111074894 | rs13248987 | -1.445 | 0.837 | -1.87 | -0.891 | -0.016 | -0.009 | -0.009 | 0 | 0.8 | 0.9 | 0.86 | 0.78 | 0.79 | 0.76 |
| 8 | 111077863 | rs4032759 | 0.877 | 0.934 | -1.868 | -0.867 | -0.013 | -0.08 | -0.08 | 0 | 0.25 | 0.15 | 0.42 | 0.47 | 0.28 | 0.48 |
| 8 | 111083330 | rs16880352 | 0.221 | 0.545 | -0.916 | -0.885 | -0.015 | -0.033 | -0.033 | 0 | 0.22 | 0.37 | 0.13 | 0.17 | 0.18 | 0.14 |
| 8 | 111087094 | rs10107946 | 1.034 | 0.274 | -1.169 | -0.902 | -0.056 | 0.133 | 0.133 | 1.738 | 0.4 | 0.12 | 0.35 | 0.19 | 0.42 | 0.28 |
| 8 | 111088894 | rs16880357 | 0.449 | 0.113 | -1.091 | -0.974 | -0.013 | 0.015 | 0.015 | 0.11 | 0.15 | 0.4 | 0.1 | 0.15 | 0.11 | 0.11 |
| 8 | 111093796 | rs12335311 | 1.13 | -0.045 | -1.247 | -0.895 | -0.069 | 0.135 | 0.135 | 1.713 | 0.4 | 0.09 | 0.34 | 0.19 | 0.42 | 0.27 |
| 8 | 111101361 | rs17476650 | 1.607 | -0.019 | -0.434 | -0.828 | 0.006 | -0.006 | -0.006 | 0 | 0.09 | 0.12 | 0.07 | 0.08 | 0.1 | 0.07 |
| 8 | 111102673 | rs6469295 | -1.517 | -0.043 | -0.609 | -0.865 | -0.04 | -0.014 | -0.014 | 0 | 0.8 | 0.97 | 0.79 | 0.77 | 0.77 | 0.75 |
| 8 | 111113707 | rs7011544 | 0.014 | -0.051 | -0.298 | -0.826 | -0.004 | -0.005 | -0.005 | 0.584 | 0.36 | 0.52 | 0.31 | 0.39 | 0.34 | 0.35 |
| 8 | 111116148 | rs13257183 | 0.643 | -0.092 | -0.149 | -0.88 | -0.001 | 0.005 | 0.005 | 0 | 0.22 | 0.23 | 0.33 | 0.43 | 0.24 | 0.37 |
| 8 | 111116619 | rs16880373 | 1.2 | -0.067 | -0.184 | -0.846 | -0.021 | 0.004 | 0.004 | 0 | 0.23 | 0.06 | 0.2 | 0.24 | 0.23 | 0.23 |
| 8 | 111117738 | rs4734220 | -0.02 | -0.067 | -0.176 | -0.847 | -0.002 | -0.004 | -0.004 | 0 | 0.36 | 0.52 | 0.31 | 0.39 | 0.35 | 0.36 |
| 8 | 111122904 | rs10097037 | 1.146 | -0.184 | -0.333 | -0.816 | -0.02 | 0.004 | 0.004 | 0 | 0.23 | 0.06 | 0.21 | 0.24 | 0.23 | 0.25 |
| 8 | 111138684 | rs1823506 | -0.614 | -0.479 | 0.037 | -0.778 | -0.001 | 0.006 | 0.006 | 0 | 0.35 | 0.43 | 0.42 | 0.45 | 0.31 | 0.5 |
| 8 | 111149619 | rs1982401 | 0.693 | -0.404 | -0.794 | -0.768 | -0.073 | -0.133 | -0.133 | 1.093 | 0.19 | 0.04 | 0.35 | 0.43 | 0.26 | 0.37 |
| 8 | 111158893 | rs715731 | -0.731 | -0.477 | -0.634 | -0.777 | 0 | -0.006 | -0.006 | 6.639 | 0.34 | 0.35 | 0.39 | 0.44 | 0.29 | 0.5 |
| 8 | 111168599 | rs13266032 | -1.088 | -0.44 | -0.994 | -0.778 | 0.017 | -0.01 | -0.01 | 0 | 0.4 | 0.8 | 0.55 | 0.55 | 0.31 | 0.55 |
| 8 | 111169622 | rs7465667 | -1.597 | -0.529 | -1.302 | -1.06 | -0.054 | 0.052 | 0.052 | 0.021 | 0.53 | 0.93 | 0.64 | 0.63 | 0.45 | 0.62 |
| 8 | 111174133 | rs13272142 | 0.629 | -0.034 | -1.09 | -0.577 | -0.08 | -0.128 | -0.128 | 0.279 | 0.18 | 0.04 | 0.35 | 0.46 | 0.27 | 0.33 |
| 8 | 111176772 | rs13260817 | -0.852 | -0.189 | -0.837 | -0.622 | 0.01 | 0.048 | 0.048 | 0 | 0.29 | 0.54 | 0.51 | 0.55 | 0.28 | 0.37 |
| 8 | 111181385 | rs4147310 | -0.884 | -0.377 | -0.658 | -0.707 | 0.002 | 0.017 | 0.017 | 0 | 0.42 | 0.67 | 0.59 | 0.61 | 0.41 | 0.45 |
| 8 | 111181468 | rs13272100 | 0.089 | -0.369 | -0.714 | -0.697 | -0.021 | -0.034 | -0.034 | 0 | 0.31 | 0.18 | 0.42 | 0.52 | 0.39 | 0.4 |
| 8 | 111182246 | rs11984657 | -0.562 | -0.547 | -0.089 | -0.761 | -0.037 | 0.038 | 0.038 | 4.687 | 0.11 | 0.44 | 0.16 | 0.1 | 0.02 | 0.05 |
| 8 | 111187061 | rs4371977 | -1.098 | 0.334 | -1.086 | -0.758 | -0.035 | 0.03 | 0.03 | 0 | 0.94 | 1 | 0.93 | 1 | 0.88 | 0.99 |
| 8 | 111208985 | rs13439825 | -0.2 | -0.61 | -0.739 | -0.588 | -0.05 | -0.016 | -0.016 | 0.059 | 0.19 | 0.29 | 0.29 | 0.15 | 0.07 | 0.18 |
| 8 | 111218857 | rs10086143 | -0.342 | -0.431 | -1.014 | -0.925 | 0.005 | 0.087 | 0.087 | 0 | 0.69 | 0.9 | 0.62 | 0.85 | 0.7 | 0.75 |
| 8 | 111221549 | rs12375273 | 0.771 | -1.392 | -0.878 | -1.042 | 0.061 | 0.043 | 0.043 | 1.348 | 0.47 | 0.26 | 0.49 | 0.65 | 0.63 | 0.56 |
| 8 | 111244015 | rs16880528 | 0.43 | -1.371 | -0.68 | -0.991 | -0.064 | 0.042 | 0.042 | 1.758 | 0.17 | 0.39 | 0.12 | 0.21 | 0.07 | 0.19 |
| 8 | 111255089 | rs7006763 | -0.9 | -1.083 | -0.694 | -1.078 | -0.025 | 0.086 | 0.086 | 0 | 0.68 | 0.9 | 0.6 | 0.84 | 0.67 | 0.74 |
| 8 | 111272209 | rs7003392 | -0.684 | -0.999 | -0.696 | -1.086 | -0.004 | 0.074 | 0.074 | 0 | 0.65 | 0.84 | 0.58 | 0.84 | 0.65 | 0.72 |
| 8 | 111280576 | rs10505143 | 1.15 | -1.004 | -1.152 | -1.013 | -0.016 | 0.039 | 0.039 | 0.456 | 0.12 | 0.2 | 0.09 | 0.21 | 0.07 | 0.12 |
| 8 | 111290187 | rs7007400 | -0.851 | -0.725 | -1.879 | -0.884 | -0.084 | 0.083 | 0.083 | 0 | 0.67 | 0.97 | 0.59 | 0.84 | 0.65 | 0.73 |
| 8 | 111298543 | rs2948998 | 0.282 | -0.347 | -1.618 | -0.722 | 0.004 | 0.017 | 0.017 | 0.909 | 0.44 | 0.29 | 0.46 | 0.63 | 0.58 | 0.57 |
| 8 | 111302825 | rs13274029 | 0.054 | 0.148 | -1.443 | -0.563 | -0.024 | 0.007 | 0.007 | 0.224 | 0.13 | 0.05 | 0.22 | 0.13 | 0.14 | 0.25 |
| 8 | 111316902 | rs6469302 | -0.9 | -0.326 | -1.62 | -0.765 | 0.069 | 0.057 | 0.057 | 0 | 0.42 | 0.85 | 0.37 | 0.34 | 0.26 | 0.47 |
| 8 | 111327488 | rs1949758 | -0.184 | -0.557 | -1.309 | -0.791 | -0.001 | -0.006 | -0.006 | 0 | 0.2 | 0.22 | 0.27 | 0.13 | 0.17 | 0.3 |
| 8 | 111342240 | rs7832047 | -1.273 | -0.342 | -1.745 | -0.826 | 0.07 | 0.045 | 0.045 | 0 | 0.48 | 0.95 | 0.43 | 0.36 | 0.33 | 0.51 |
| 8 | 111354022 | rs17480167 | 1.009 | 0.734 | -1.3 | -0.812 | -0.03 | 0.029 | 0.029 | 0 | 0.06 | 0.01 | 0.08 | 0 | 0.17 | 0.02 |
| 8 | 111380551 | rs6469303 | -0.023 | -0.79 | -0.341 | -0.739 | -0.071 | 0.064 | 0.064 | 1.341 | 0.17 | 0.52 | 0.11 | 0.23 | 0.06 | 0.14 |
| 8 | 111421582 | rs11993192 | 0.131 | -0.996 | -0.393 | -0.856 | -0.053 | -0.059 | -0.059 | 0 | 0.11 | 0.28 | 0.02 | 0 | 0.02 | 0.1 |
| 8 | 111427941 | rs11991389 | 1.034 | -0.939 | -0.595 | -0.846 | -0.024 | -0.037 | -0.037 | 0 | 0.07 | 0.15 | 0.02 | 0 | 0.02 | 0.06 |
| 8 | 111432405 | rs4500059 | -0.737 | -0.805 | -1.123 | -0.791 | 0.091 | 0.019 | 0.019 | 3.336 | 0.29 | 0.84 | 0.14 | 0.24 | 0.09 | 0.23 |
| 8 | 111438655 | rs7008113 | -1.007 | -0.691 | -1.427 | -1.207 | 0.125 | 0.203 | 0.203 | 2.6 | 0.37 | 0.94 | 0.18 | 0.09 | 0.18 | 0.26 |
| 8 | 111457175 | rs12542501 | -0.731 | -0.209 | -0.675 | -0.985 | 0.003 | -0.013 | -0.013 | 0 | 0.13 | 0.19 | 0.14 | 0.08 | 0.15 | 0.17 |
| 8 | 111463358 | rs12155933 | 0.614 | -0.283 | -0.849 | -1.319 | -0.014 | 0.022 | 0.022 | 0 | 0.08 | 0.03 | 0.03 | 0 | 0.09 | 0.02 |
| 8 | 111468172 | rs10103911 | -0.92 | -0.303 | -0.27 | -1.176 | 0.015 | 0 | 0 | 0.391 | 0.14 | 0.31 | 0.18 | 0.09 | 0.16 | 0.17 |
| 8 | 111478321 | rs13267290 | -0.847 | -0.366 | -0.354 | -1.165 | 0.004 | 0.004 | 0.004 | 0.043 | 0.16 | 0.22 | 0.18 | 0.19 | 0.19 | 0.19 |
| 8 | 111493342 | rs13264696 | -0.424 | -0.3 | 0.215 | -1.502 | 0.004 | -0.1 | -0.1 | 0.524 | 0.19 | 0.29 | 0.09 | 0 | 0.18 | 0.11 |
| 8 | 111498521 | rs4735157 | -1.359 | -0.364 | 0.524 | -1.533 | 0.003 | 0.014 | 0.014 | 2.083 | 0.3 | 0.71 | 0.27 | 0.19 | 0.28 | 0.31 |
| 8 | 111509813 | rs13264567 | -1.651 | -0.359 | 0.522 | -1.538 | -0.005 | 0.062 | 0.062 | 0.611 | 0.37 | 0.75 | 0.29 | 0.19 | 0.37 | 0.33 |
| 8 | 111518434 | rs13265247 | 0.468 | -0.321 | 0.665 | -1.369 | -0.007 | -0.079 | -0.079 | 0 | 0.13 | 0.26 | 0.07 | 0 | 0.09 | 0.11 |
| 8 | 111539679 | rs11993254 | -1.735 | -0.408 | 0.535 | -1.526 | 0.006 | 0.093 | 0.093 | 2.148 | 0.34 | 0.87 | 0.28 | 0.2 | 0.3 | 0.31 |
| 8 | 111558924 | rs13250442 | 0.393 | -0.456 | 0.715 | -1.437 | -0.001 | -0.085 | -0.085 | 0.288 | 0.13 | 0.28 | 0.08 | 0 | 0.1 | 0.11 |
| 8 | 111580170 | rs1494232 | -0.093 | -0.518 | 0.682 | -1.335 | -0.012 | -0.121 | -0.121 | 0 | 0.16 | 0.45 | 0.09 | 0 | 0.1 | 0.11 |
| 8 | 111592072 | rs7460087 | -1.469 | -0.526 | 0.474 | -1.311 | 0.019 | 0.048 | 0.048 | 2.027 | 0.31 | 0.83 | 0.24 | 0.19 | 0.24 | 0.3 |
| 8 | 111593882 | rs7825323 | -0.909 | -0.549 | 0.658 | -1.326 | -0.001 | -0.013 | -0.013 | 0 | 0.25 | 0.68 | 0.22 | 0.19 | 0.24 | 0.26 |

| | | | | | | | | | | | | | | | | |
|---|---|---|---|---|---|---|---|---|---|---|---|---|---|---|---|---|
| 8 | 111607634 | rs13276609 | -0.086 | -0.484 | 0.453 | -1.291 | 0.03 | 0.05 | 0.05 | 0 | 0.11 | 0.29 | 0.14 | 0.18 | 0.14 | 0.15 |
| 8 | 111621137 | rs1552880 | 1.919 | -0.767 | -0.838 | -1.415 | -0.14 | -0.044 | -0.044 | 0.061 | 0.26 | 0.01 | 0.25 | 0.18 | 0.39 | 0.17 |
| 8 | 111626655 | rs7006232 | -1.109 | -0.925 | -0.364 | -1.126 | 0.008 | 0.037 | 0.037 | 1.442 | 0.32 | 0.8 | 0.28 | 0.18 | 0.27 | 0.29 |
| 8 | 111628625 | rs1353208 | -1.109 | -0.925 | -0.364 | -1.126 | 0.008 | 0.037 | 0.037 | 1.442 | 0.32 | 0.8 | 0.28 | 0.18 | 0.27 | 0.29 |
| 8 | 111635765 | rs7008570 | -1.15 | -0.962 | -0.479 | -1.145 | 0.013 | 0.076 | 0.076 | 1.438 | 0.32 | 0.85 | 0.29 | 0.15 | 0.27 | 0.28 |
| 8 | 111642122 | rs12681997 | 0.618 | -0.957 | -0.501 | -1.143 | -0.005 | -0.009 | -0.009 | 0 | 0.09 | 0.09 | 0.1 | 0.14 | 0.1 | 0.1 |
| 8 | 111645187 | rs10505145 | 0.236 | -0.958 | -0.425 | -1.143 | 0.007 | 0.016 | 0.016 | 0.004 | 0.1 | 0.21 | 0.11 | 0.14 | 0.1 | 0.1 |
| 8 | 111646626 | rs11987479 | -1.151 | -0.942 | -0.636 | -1.121 | 0.01 | 0.089 | 0.089 | 1.665 | 0.32 | 0.87 | 0.29 | 0.15 | 0.27 | 0.28 |
| 8 | 111649375 | rs2351182 | -1.357 | -1.23 | -0.837 | -1.246 | 0.028 | 0.137 | 0.137 | 4.805 | 0.38 | 0.92 | 0.29 | 0.14 | 0.29 | 0.29 |
| 8 | 111649680 | rs6996931 | 1.818 | -1.494 | -1.048 | -1.466 | -0.097 | 0.06 | 0.06 | 0 | 0.39 | 0.09 | 0.32 | 0.23 | 0.51 | 0.22 |
| 8 | 111652574 | rs6984552 | 0.624 | -1.977 | -0.099 | -1.408 | -0.012 | -0.092 | -0.092 | 0.193 | 0.17 | 0.28 | 0.11 | 0 | 0.13 | 0.12 |
| 8 | 111653713 | rs16880943 | -0.088 | -1.982 | 0.023 | -1.41 | -0.001 | 0 | 0 | 0 | 0.14 | 0.16 | 0.11 | 0.14 | 0.12 | 0.11 |
| 8 | 111665105 | rs10505149 | -0.115 | -1.565 | 0.461 | -1.199 | -0.03 | -0.13 | -0.13 | 0 | 0.2 | 0.58 | 0.14 | 0 | 0.13 | 0.12 |
| 8 | 111673039 | rs6983816 | -0.647 | -1.566 | 0.464 | -1.2 | -0.005 | -0.052 | -0.052 | 0.038 | 0.23 | 0.73 | 0.18 | 0.01 | 0.17 | 0.18 |
| 8 | 111681660 | rs1587014 | -0.075 | -1.549 | 0.085 | -1.159 | -0.031 | -0.147 | -0.147 | 0.109 | 0.2 | 0.47 | 0.13 | 0 | 0.13 | 0.12 |
| 8 | 111693132 | rs7819148 | -1.098 | -1.818 | -0.393 | -0.422 | 0.03 | -0.06 | -0.06 | 0 | 0.9 | 1 | 0.75 | 0.56 | 0.96 | 0.76 |
| 8 | 111697594 | rs16881028 | 0.071 | -1.693 | -0.104 | -1.055 | -0.025 | -0.107 | -0.107 | 0 | 0.19 | 0.32 | 0.11 | 0 | 0.13 | 0.12 |
| 8 | 111737050 | rs13273439 | 0.057 | -1.726 | -0.152 | -1.073 | 0.01 | 0.001 | 0.001 | 0.087 | 0.18 | 0.15 | 0.1 | 0 | 0.13 | 0.12 |
| 8 | 111747867 | rs10103670 | 1.586 | -2.021 | -0.388 | -1.186 | 0.119 | -0.014 | -0.014 | 0 | 0.51 | 0.08 | 0.48 | 0.42 | 0.69 | 0.43 |
| 8 | 111752010 | rs13281768 | 0.122 | -1.768 | -0.169 | -1.206 | 0.005 | -0.017 | -0.017 | 0 | 0.18 | 0.19 | 0.1 | 0 | 0.13 | 0.12 |
| 8 | 111787113 | rs11987524 | 0.449 | -2.102 | -0.295 | -1.17 | 0.047 | 0.055 | 0.055 | 0.641 | 0.17 | 0.07 | 0.06 | 0 | 0.07 | 0.11 |
| 8 | 111803586 | rs7010569 | 1.232 | -2.243 | -0.182 | -1.147 | 0.052 | 0.003 | 0.003 | 0.038 | 0.54 | 0.32 | 0.53 | 0.43 | 0.75 | 0.43 |
| 8 | 111811966 | rs10105924 | 1.488 | -2.262 | -0.301 | -1.157 | 0.19 | -0.028 | -0.028 | 0 | 0.51 | 0.08 | 0.5 | 0.43 | 0.75 | 0.43 |
| 8 | 111826308 | rs4573245 | -0.39 | -2.38 | -0.205 | -1.215 | 0.015 | -0.142 | -0.142 | 0.471 | 0.78 | 0.96 | 0.66 | 0.44 | 0.85 | 0.62 |
| 8 | 111826826 | rs1489356 | 1.326 | -2.418 | 0.21 | -1.241 | 0 | -0.001 | -0.001 | 0 | 0.56 | 0.52 | 0.54 | 0.43 | 0.75 | 0.43 |
| 8 | 111830080 | rs16881074 | -0.566 | -2.351 | 0.33 | -1.253 | -0.092 | -0.165 | -0.165 | 0.436 | 0.24 | 0.49 | 0.12 | 0.01 | 0.1 | 0.2 |
| 8 | 111842653 | rs2130185 | 2.232 | -2.04 | 0.013 | -1.497 | -0.152 | 0.009 | 0.009 | 0 | 0.34 | 0.01 | 0.27 | 0.24 | 0.56 | 0.2 |
| 8 | 111846111 | rs1489365 | 0.049 | -2.412 | 0.798 | -1.17 | -0.011 | 0.018 | 0.018 | 0 | 0.76 | 0.72 | 0.63 | 0.43 | 0.84 | 0.62 |
| 8 | 111851695 | rs1387412 | 1.623 | -2.413 | 0.579 | -1.169 | 0.184 | 0 | 0 | 0.412 | 0.51 | 0.12 | 0.51 | 0.43 | 0.74 | 0.42 |
| 8 | 111852200 | rs1489360 | -0.165 | -2.167 | 0.754 | -1.082 | -0.062 | -0.112 | -0.112 | 0 | 0.2 | 0.33 | 0.07 | 0 | 0.07 | 0.12 |
| 8 | 111869446 | rs7004409 | -0.073 | -2.161 | 0.622 | -1.078 | -0.064 | -0.114 | -0.114 | 0 | 0.2 | 0.33 | 0.07 | 0 | 0.07 | 0.11 |
| 8 | 111869916 | rs7009162 | 0.39 | -2.176 | 0.668 | -1.086 | -0.012 | -0.043 | -0.043 | 0 | 0.17 | 0.18 | 0.06 | 0 | 0.07 | 0.11 |
| 8 | 111885336 | rs16881302 | 0.058 | -2.111 | 0.311 | -1.053 | -0.065 | -0.114 | -0.114 | 0 | 0.2 | 0.33 | 0.07 | 0 | 0.07 | 0.11 |
| 8 | 111894555 | rs2171707 | -1.656 | -2.174 | 0.085 | -1.04 | 0.167 | -0.001 | -0.001 | 0.764 | 0.48 | 0.85 | 0.49 | 0.58 | 0.25 | 0.59 |
| 8 | 111910956 | rs6469330 | -1.637 | -2.166 | 0 | -1.036 | 0.167 | -0.005 | -0.005 | 0 | 0.47 | 0.87 | 0.49 | 0.58 | 0.25 | 0.57 |
| 8 | 111929209 | rs1546013 | -0.161 | -2.084 | 0 | -0.964 | -0.065 | -0.114 | -0.114 | 0.543 | 0.8 | 0.65 | 0.93 | 1 | 0.93 | 0.89 |
| 8 | 111932394 | rs1489354 | -0.161 | -2.084 | -0.026 | -0.964 | -0.064 | -0.112 | -0.112 | 0 | 0.8 | 0.65 | 0.93 | 1 | 0.93 | 0.89 |
| 8 | 111943344 | rs10955548 | 0.41 | -1.893 | -0.7 | -0.733 | 0.033 | -0.123 | -0.123 | 0 | 0.18 | 0.04 | 0.3 | 0.57 | 0.11 | 0.36 |
| 8 | 111952230 | rs4289849 | -0.018 | -1.697 | -0.59 | -0.677 | -0.033 | 0.085 | 0.085 | 0 | 0.24 | 0.39 | 0.39 | 0.58 | 0.16 | 0.45 |
| 8 | 111991956 | rs13268040 | 0.176 | -1.119 | -0.361 | -1.007 | -0.077 | -0.136 | -0.136 | 0 | 0.81 | 0.55 | 0.93 | 1 | 0.93 | 0.88 |
| 8 | 112021630 | rs4493937 | 0.532 | -0.127 | -1.559 | -0.734 | -0.088 | 0.069 | 0.069 | 1.065 | 0.4 | 0.07 | 0.28 | 0.25 | 0.5 | 0.27 |
| 8 | 112028234 | rs4876879 | 0.459 | -0.965 | -0.512 | -0.903 | -0.033 | -0.016 | -0.016 | 1.743 | 0.7 | 0.38 | 0.76 | 0.77 | 0.84 | 0.73 |
| 8 | 112032097 | rs1427188 | -0.834 | -1.064 | -0.46 | -0.987 | 0.014 | 0.022 | 0.022 | 0 | 0.9 | 0.93 | 0.92 | 1 | 0.94 | 0.87 |
| 8 | 112033379 | rs12546516 | 0.806 | -1.081 | -0.322 | -0.954 | -0.019 | -0.009 | -0.009 | 0.351 | 0.68 | 0.36 | 0.76 | 0.77 | 0.84 | 0.73 |
| 8 | 112060308 | rs13278392 | 1.169 | -1.22 | -0.168 | -1.024 | 0.07 | 0.042 | 0.042 | 0 | 0.65 | 0.18 | 0.74 | 0.77 | 0.84 | 0.73 |
| 8 | 112062191 | rs10099529 | 0.731 | -1.191 | -0.179 | -1.009 | -0.027 | -0.013 | -0.013 | 0 | 0.69 | 0.39 | 0.76 | 0.77 | 0.85 | 0.73 |
| 8 | 112079211 | rs1896238 | 0.667 | -0.862 | 0.38 | -1.028 | -0.016 | -0.018 | -0.018 | 0 | 0.69 | 0.59 | 0.75 | 0.77 | 0.77 | 0.73 |
| 8 | 112132555 | rs1019707 | 0.587 | -1.099 | 0.421 | -1.092 | -0.05 | -0.027 | -0.027 | 0.346 | 0.7 | 0.45 | 0.77 | 0.79 | 0.84 | 0.73 |
| 8 | 112145558 | rs1593667 | 0.76 | -1.082 | 0.296 | -1.084 | -0.028 | -0.015 | -0.015 | 0.309 | 0.68 | 0.36 | 0.76 | 0.79 | 0.84 | 0.73 |
| 8 | 112191432 | rs1427181 | 0.895 | -1.208 | 0.061 | -1.239 | 0.083 | 0.071 | 0.071 | 0.073 | 0.65 | 0.18 | 0.73 | 0.8 | 0.83 | 0.73 |
| 8 | 112198405 | rs17516585 | 2.143 | -0.87 | -0.792 | -1.566 | -0.149 | 0.114 | 0.114 | 0 | 0.33 | 0.01 | 0.2 | 0.11 | 0.43 | 0.19 |
| 8 | 112226560 | rs4074825 | 0.297 | -1.366 | 0.077 | -1.35 | -0.037 | -0.091 | -0.091 | 0 | 0.14 | 0.35 | 0.1 | 0 | 0.06 | 0.12 |
| 8 | 112238390 | rs4332164 | -1.315 | -1.29 | -0.586 | -1.187 | -0.015 | -0.111 | -0.111 | 0 | 0.84 | 0.99 | 0.7 | 0.41 | 0.87 | 0.59 |
| 8 | 112270826 | rs13250058 | 1.102 | -1.056 | -0.387 | -1.103 | 0.129 | 0.194 | 0.194 | 0 | 0.49 | 0.13 | 0.43 | 0.2 | 0.7 | 0.33 |
| 8 | 112271971 | rs16881844 | 0.616 | -1.007 | -0.424 | -1.001 | 0.011 | -0.041 | -0.041 | 0 | 0.15 | 0.01 | 0.14 | 0.2 | 0.1 | 0.15 |
| 8 | 112274634 | rs9643057 | 0.575 | -0.963 | -0.423 | -1.027 | -0.003 | -0.033 | -0.033 | 0 | 0.15 | 0.01 | 0.14 | 0.18 | 0.11 | 0.16 |
| 8 | 112276970 | rs4504665 | -0.268 | -1.062 | -0.067 | -1.104 | -0.031 | 0.025 | 0.025 | 0 | 0.71 | 0.6 | 0.58 | 0.38 | 0.81 | 0.48 |
| 8 | 112286614 | rs10088186 | 1.143 | -1.032 | -0.04 | -1.122 | 0.111 | 0.182 | 0.182 | 0 | 0.48 | 0.15 | 0.42 | 0.2 | 0.7 | 0.32 |
| 8 | 112290740 | rs10104559 | 1.193 | -1.032 | -0.051 | -1.122 | 0.105 | 0.188 | 0.188 | 0 | 0.48 | 0.1 | 0.41 | 0.2 | 0.7 | 0.33 |
| 8 | 112325064 | rs10102107 | 0.711 | -1.011 | 0.017 | -1.129 | -0.011 | -0.024 | -0.024 | 0 | 0.53 | 0.46 | 0.43 | 0.16 | 0.7 | 0.32 |
| 8 | 112335650 | rs10955574 | 0.01 | -0.918 | -0.144 | -1.055 | 0.013 | 0.004 | 0.004 | 0 | 0.19 | 0.13 | 0.17 | 0.18 | 0.15 | 0.17 |
| 8 | 112340385 | rs13280188 | 1.032 | -0.906 | -0.125 | -1.217 | 0.05 | 0.085 | 0.085 | 0 | 0.48 | 0.3 | 0.4 | 0.16 | 0.66 | 0.3 |
| 8 | 112359706 | rs10955576 | 0.36 | -0.909 | -0.344 | -0.984 | 0.007 | 0.002 | 0.002 | 0 | 0.16 | 0.12 | 0.14 | 0.15 | 0.09 | 0.17 |
| 8 | 112370516 | rs13280988 | 1.167 | -0.477 | -0.835 | -1.316 | -0.031 | 0.23 | 0.23 | 0 | 0.43 | 0.07 | 0.29 | 0.07 | 0.62 | 0.22 |
| 8 | 112378482 | rs13274435 | 0.695 | 0.043 | -0.541 | -1.202 | 0.006 | -0.006 | -0.006 | 0 | 0.06 | 0.08 | 0.06 | 0 | 0.14 | 0.02 |
| 8 | 112379434 | rs6983489 | -0.414 | -0.358 | -0.068 | -1.212 | -0.011 | 0.008 | 0.008 | 0 | 0.74 | 0.7 | 0.59 | 0.3 | 0.78 | 0.5 |
| 8 | 112404802 | rs1573937 | 1.162 | -0.319 | 0.024 | -1.24 | 0.008 | 0.133 | 0.133 | 0 | 0.46 | 0.2 | 0.38 | 0.16 | 0.56 | 0.29 |
| 8 | 112417948 | rs10094094 | 1.219 | -0.214 | 0.049 | -1.225 | -0.031 | 0.025 | 0.025 | 0 | 0.06 | 0.01 | 0.06 | 0 | 0.15 | 0.02 |
| 8 | 112432365 | rs13267927 | 0.975 | -0.217 | 0.085 | -1.227 | -0.012 | 0.012 | 0.012 | 0 | 0.06 | 0.05 | 0.06 | 0 | 0.14 | 0.02 |

| | | | | | | | | | | | | | | | | |
|---|---|---|---|---|---|---|---|---|---|---|---|---|---|---|---|---|
| 8 | 112454301 | rs2606203 | 0.141 | -0.506 | 0.363 | -1.387 | -0.031 | -0.089 | -0.089 | 0 | 0.14 | 0.31 | 0.08 | 0 | 0.07 | 0.12 |
| 8 | 112455664 | rs7005948 | 0.333 | -0.528 | 0.123 | -1.104 | -0.026 | 0.031 | 0.031 | 0 | 0.14 | 0.35 | 0.15 | 0.17 | 0.09 | 0.18 |
| 8 | 112488720 | rs10108623 | 0.601 | -0.336 | -0.045 | -1.097 | 0.016 | -0.017 | -0.017 | 0 | 0.07 | 0.12 | 0.07 | 0 | 0.15 | 0.02 |
| 8 | 112488761 | rs10093778 | -0.032 | -0.697 | 0.151 | -1.111 | 0.007 | -0.002 | -0.002 | 0.703 | 0.68 | 0.7 | 0.62 | 0.31 | 0.81 | 0.49 |
| 8 | 112489722 | rs6469354 | 0.367 | -0.664 | 0.31 | -1.086 | -0.028 | 0.011 | 0.011 | 0.043 | 0.14 | 0.27 | 0.16 | 0.15 | 0.09 | 0.18 |
| 8 | 112491668 | rs13264220 | -0.169 | -0.644 | -0.209 | -1.086 | 0.006 | -0.003 | -0.003 | 0 | 0.7 | 0.72 | 0.64 | 0.29 | 0.82 | 0.51 |
| 8 | 112533575 | rs2717181 | -0.801 | -0.307 | -0.491 | -0.911 | -0.004 | 0.002 | 0.002 | 0 | 0.88 | 0.9 | 0.86 | 0.84 | 0.92 | 0.82 |
| 8 | 112548351 | rs2717188 | -0.595 | -0.335 | -0.462 | -0.924 | -0.011 | 0.004 | 0.004 | 0.02 | 0.87 | 0.86 | 0.85 | 0.84 | 0.92 | 0.82 |
| 8 | 112552852 | rs2606205 | 1.092 | -0.47 | -1.135 | -1.048 | 0.005 | -0.083 | -0.083 | 0 | 0.26 | 0.06 | 0.43 | 0.7 | 0.21 | 0.46 |
| 8 | 112556471 | rs4876699 | 0.527 | -0.281 | -0.851 | -1.167 | 0.058 | 0.176 | 0.176 | 0 | 0.47 | 0.11 | 0.32 | 0.14 | 0.65 | 0.31 |
| 8 | 112565069 | rs4876704 | 1.238 | -0.298 | -0.882 | -1.131 | -0.142 | 0.137 | 0.137 | 0 | 0.34 | 0.01 | 0.23 | 0.07 | 0.43 | 0.2 |
| 8 | 112579771 | rs2352165 | 0.741 | -0.145 | -0.493 | -0.99 | -0.069 | 0.173 | 0.173 | 0 | 0.42 | 0.17 | 0.27 | 0.08 | 0.5 | 0.27 |
| 8 | 112589344 | rs13273084 | 0.768 | -0.144 | -0.715 | -0.359 | -0.06 | -0.125 | -0.125 | 0 | 0.31 | 0.04 | 0.43 | 0.43 | 0.31 | 0.33 |
| 8 | 112591382 | rs1807420 | 0.705 | -0.138 | -0.754 | -0.347 | -0.003 | 0.016 | 0.016 | 0 | 0.46 | 0.59 | 0.29 | 0.13 | 0.5 | 0.29 |
| 8 | 112592247 | rs6986024 | 0.575 | -0.314 | -0.476 | 0.072 | -0.002 | 0.021 | 0.021 | 0 | 0.78 | 0.78 | 0.73 | 0.56 | 0.8 | 0.62 |
| 8 | 112598361 | rs7828470 | -0.766 | -0.22 | -0.93 | 0.101 | -0.015 | -0.048 | -0.048 | 2.891 | 0.88 | 0.96 | 0.8 | 0.57 | 0.87 | 0.7 |
| 8 | 112604086 | rs7002030 | -0.094 | -0.213 | -0.844 | 0.255 | -0.007 | -0.048 | -0.048 | 0 | 0.8 | 0.89 | 0.73 | 0.59 | 0.81 | 0.64 |
| 8 | 112607588 | rs13279316 | 0.74 | 0.198 | -1.544 | -0.039 | 0.03 | 0.045 | 0.045 | 0 | 0.33 | 0.1 | 0.14 | 0.16 | 0.27 | 0.27 |
| 8 | 112608658 | rs10088527 | 0.006 | 0.401 | -0.908 | 0.048 | 0 | -0.001 | -0.001 | 0 | 0.45 | 0.46 | 0.57 | 0.43 | 0.54 | 0.37 |
| 8 | 112616415 | rs4876731 | 0.382 | 0.439 | -1.525 | -0.07 | -0.014 | -0.008 | -0.008 | 0.221 | 0.24 | 0.06 | 0.12 | 0.2 | 0.22 | 0.25 |
| 8 | 112622355 | rs13250751 | 0.422 | 0.416 | -1.683 | 0.021 | -0.057 | 0.013 | 0.013 | 0.084 | 0.42 | 0.18 | 0.38 | 0.37 | 0.5 | 0.45 |
| 8 | 112624201 | rs7842968 | -0.395 | 0.245 | -0.861 | 0.057 | -0.005 | 0.014 | 0.014 | 0 | 0.69 | 0.43 | 0.51 | 0.6 | 0.72 | 0.69 |
| 8 | 112630334 | rs7357550 | -0.867 | 0.026 | -0.947 | -0.039 | -0.043 | -0.092 | -0.092 | 0.359 | 0.75 | 0.93 | 0.55 | 0.6 | 0.73 | 0.69 |
| 8 | 112633913 | rs4142767 | 0.953 | 0.014 | -0.818 | -0.173 | -0.003 | -0.011 | -0.011 | 0 | 0.08 | 0.1 | 0.06 | 0.06 | 0.06 | 0.09 |
| 8 | 112649830 | rs13281532 | 1.513 | 0.383 | -1.287 | -0.467 | -0.03 | -0.082 | -0.082 | 0 | 0.27 | 0.03 | 0.24 | 0.3 | 0.25 | 0.32 |
| 8 | 112651697 | rs4876786 | 1.688 | 0.013 | -1.208 | -0.327 | -0.017 | 0.012 | 0.012 | 0 | 0.2 | 0 | 0.09 | 0.1 | 0.17 | 0.18 |
| 8 | 112656042 | rs9694019 | 1.173 | 0.04 | -1.092 | -0.321 | -0.001 | -0.01 | -0.01 | 0 | 0.26 | 0.09 | 0.12 | 0.23 | 0.22 | 0.24 |
| 8 | 112661852 | rs13260133 | 0.821 | 0.064 | -0.311 | -0.32 | -0.074 | -0.009 | -0.009 | 0 | 0.35 | 0.13 | 0.32 | 0.3 | 0.43 | 0.36 |
| 8 | 112673991 | rs12675078 | -0.832 | -0.056 | 0.427 | -0.307 | -0.001 | -0.003 | -0.003 | 0 | 0.72 | 0.73 | 0.54 | 0.59 | 0.72 | 0.7 |
| 8 | 112675571 | rs12677714 | -0.915 | -0.058 | 0.512 | -0.308 | -0.002 | -0.013 | -0.013 | 0 | 0.72 | 0.75 | 0.55 | 0.59 | 0.72 | 0.7 |
| 8 | 112681391 | rs13259383 | 0.955 | -0.042 | 0.289 | -0.335 | -0.12 | -0.041 | -0.041 | 0 | 0.33 | 0.05 | 0.34 | 0.3 | 0.45 | 0.36 |
| 8 | 112692491 | rs13282733 | 0.765 | -0.011 | 0.611 | -0.32 | -0.02 | 0.004 | 0.004 | 0 | 0.36 | 0.3 | 0.36 | 0.3 | 0.44 | 0.36 |
| 8 | 112693175 | rs7837084 | 0.277 | 0.07 | 0.442 | -0.261 | 0.005 | 0.004 | 0.004 | 0 | 0.35 | 0.35 | 0.18 | 0.29 | 0.28 | 0.34 |
| 8 | 112712038 | rs3749413 | 0.714 | 0.482 | 0.628 | -0.098 | 0.008 | -0.033 | -0.033 | 0.042 | 0.08 | 0.19 | 0.07 | 0.03 | 0.06 | 0.11 |
| 8 | 112713528 | rs1479468 | -0.589 | 0.472 | 0.196 | 0.043 | 0.002 | 0.01 | 0.01 | 0.071 | 0.75 | 0.72 | 0.56 | 0.6 | 0.72 | 0.7 |
| 8 | 112738096 | rs7814781 | 0.59 | 0.386 | -0.151 | -0.058 | -0.046 | 0.019 | 0.019 | 0 | 0.37 | 0.15 | 0.37 | 0.26 | 0.45 | 0.43 |
| 8 | 112740051 | rs7842868 | 0.307 | 0.007 | -0.127 | -0.139 | 0.029 | 0.033 | 0.033 | 0.244 | 0.34 | 0.69 | 0.19 | 0.18 | 0.24 | 0.19 |
| 8 | 112746822 | rs4876874 | -0.816 | 0.002 | 0.27 | -0.062 | -0.005 | -0.014 | -0.014 | 0 | 0.73 | 0.78 | 0.56 | 0.61 | 0.74 | 0.69 |
| 8 | 112752080 | rs1904355 | 0.974 | 0.641 | 0.063 | -0.105 | -0.059 | -0.011 | -0.011 | 0 | 0.34 | 0.1 | 0.31 | 0.25 | 0.38 | 0.39 |
| 8 | 112752967 | rs7836329 | 0.004 | 0.645 | 0.376 | -0.11 | 0.095 | 0.058 | 0.058 | 0.005 | 0.06 | 0.3 | 0.06 | 0.07 | 0.06 | 0.06 |
| 8 | 112753418 | rs9642811 | -0.501 | 0.626 | 0.294 | -0.11 | -0.022 | -0.026 | -0.026 | 0 | 0.08 | 0.06 | 0.07 | 0.19 | 0.12 | 0.12 |
| 8 | 112758125 | rs7826225 | 0.232 | 0.614 | 0.329 | -0.128 | 0.001 | -0.002 | -0.002 | 0 | 0.4 | 0.4 | 0.36 | 0.32 | 0.45 | 0.45 |
| 8 | 112764621 | rs17524343 | -0.93 | 0.602 | -0.066 | -0.081 | 0.01 | 0.02 | 0.02 | 0 | 0.09 | 0.18 | 0.08 | 0.19 | 0.12 | 0.12 |
| 8 | 112767456 | rs13275382 | 1.08 | 0.601 | -0.114 | -0.09 | -0.064 | -0.013 | -0.013 | 0 | 0.33 | 0.07 | 0.3 | 0.25 | 0.38 | 0.38 |
| 8 | 112781860 | rs13263479 | -1.156 | 0.759 | 0.221 | 0.116 | 0.011 | 0.021 | 0.021 | 0 | 0.09 | 0.18 | 0.08 | 0.19 | 0.12 | 0.12 |
| 8 | 112788583 | rs13257287 | -0.319 | 0.551 | 0.143 | 0.195 | -0.001 | -0.003 | -0.003 | 0 | 0.71 | 0.65 | 0.56 | 0.39 | 0.72 | 0.6 |
| 8 | 112803730 | rs992953 | -0.942 | 0.603 | -0.104 | 0.18 | -0.077 | -0.116 | -0.116 | 0 | 0.77 | 0.95 | 0.58 | 0.6 | 0.74 | 0.71 |
| 8 | 112809411 | rs13252457 | -0.028 | 0.246 | 0.239 | -0.171 | 0 | -0.001 | -0.001 | 0 | 0.46 | 0.57 | 0.41 | 0.45 | 0.49 | 0.49 |
| 8 | 112819969 | rs1397364 | 0.176 | 0.261 | 0.712 | -0.14 | 0.006 | 0.018 | 0.018 | 0.425 | 0.66 | 0.55 | 0.51 | 0.33 | 0.66 | 0.54 |
| 8 | 112821478 | rs13249789 | 0.54 | 0.221 | 0.732 | -0.164 | -0.036 | 0.047 | 0.047 | 0.251 | 0.42 | 0.23 | 0.38 | 0.24 | 0.48 | 0.42 |
| 8 | 112823931 | rs7009072 | -1.27 | 0.177 | 0.651 | -0.135 | -0.074 | -0.118 | -0.118 | 0.114 | 0.77 | 0.95 | 0.58 | 0.6 | 0.74 | 0.71 |
| 8 | 112825223 | rs10955602 | -1.023 | 0.17 | 0.774 | -0.111 | 0.015 | 0.129 | 0.129 | 0.162 | 0.11 | 0.34 | 0.07 | 0.27 | 0.08 | 0.16 |
| 8 | 112855963 | rs5016753 | 1.589 | 0.149 | 0.702 | -0.221 | 0.044 | 0.049 | 0.049 | 0.011 | 0.23 | 0.07 | 0.11 | 0.08 | 0.18 | 0.12 |
| 8 | 112886310 | rs16882789 | 0.047 | 0.216 | 0.796 | -0.221 | 0.076 | 0.041 | 0.041 | 0.015 | 0.06 | 0.29 | 0.05 | 0.07 | 0.06 | 0.06 |
| 8 | 112888825 | rs10087987 | 1.581 | 0.206 | 0.412 | -0.351 | 0.044 | 0.049 | 0.049 | 0.011 | 0.23 | 0.08 | 0.12 | 0.08 | 0.18 | 0.12 |
| 8 | 112898361 | rs408383 | 0.489 | 0.156 | 0.383 | -0.317 | -0.053 | 0.062 | 0.062 | 0.425 | 0.41 | 0.17 | 0.37 | 0.24 | 0.48 | 0.42 |
| 8 | 112931908 | rs13248419 | 1.255 | 0.36 | 0.026 | -0.528 | -0.075 | 0.062 | 0.062 | 0 | 0.11 | 0.02 | 0.14 | 0 | 0.32 | 0.07 |
| 8 | 112974855 | rs1585436 | 1.519 | -0.087 | 0.063 | -0.449 | 0.011 | 0.059 | 0.059 | 0.044 | 0.25 | 0.03 | 0.13 | 0.08 | 0.2 | 0.13 |
| 8 | 112977188 | rs1460592 | 0.799 | -0.158 | 0.462 | -0.347 | -0.001 | 0.059 | 0.059 | 0.304 | 0.25 | 0.44 | 0.35 | 0.37 | 0.24 | 0.3 |
| 8 | 112997877 | rs10282859 | 0.745 | -0.275 | 0.591 | -0.397 | -0.022 | -0.021 | -0.021 | 0 | 0.32 | 0.43 | 0.2 | 0.26 | 0.27 | 0.2 |
| 8 | 113074557 | rs4876276 | -1.444 | -0.112 | 0.379 | -0.334 | -0.053 | -0.11 | -0.11 | 0 | 0.79 | 0.95 | 0.67 | 0.63 | 0.76 | 0.7 |
| 8 | 113092014 | rs4436153 | 0.149 | -0.203 | 0.428 | -0.374 | 0.044 | 0.007 | 0.007 | 2.06 | 0.38 | 0.73 | 0.29 | 0.39 | 0.28 | 0.27 |
| 8 | 113099309 | rs4391464 | -1.381 | -0.037 | 0.108 | -0.273 | -0.053 | -0.109 | -0.109 | 2.267 | 0.79 | 0.95 | 0.67 | 0.63 | 0.76 | 0.69 |
| 8 | 113110939 | rs6469403 | -1.338 | -0.203 | 0.272 | -0.315 | 0.03 | 0.055 | 0.055 | 0.188 | 0.11 | 0.47 | 0.08 | 0.14 | 0.09 | 0.08 |
| 8 | 113114520 | rs9297463 | 0.266 | -0.201 | 0.098 | -0.341 | 0.001 | 0.025 | 0.025 | 0.375 | 0.67 | 0.4 | 0.58 | 0.5 | 0.67 | 0.6 |
| 8 | 113115858 | rs13260685 | 0.82 | -0.153 | -0.444 | -0.412 | -0.163 | -0.011 | -0.011 | 0 | 0.34 | 0.05 | 0.41 | 0.24 | 0.44 | 0.42 |
| 8 | 113162234 | rs7831133 | -1.537 | -0.209 | -0.581 | -0.401 | -0.061 | -0.115 | -0.115 | 0 | 0.81 | 0.96 | 0.66 | 0.63 | 0.76 | 0.7 |
| 8 | 113167988 | rs13266472 | 0.782 | -0.226 | -0.594 | -0.404 | -0.168 | 0.016 | 0.016 | 0.002 | 0.35 | 0.05 | 0.42 | 0.23 | 0.45 | 0.42 |
| 8 | 113171397 | rs2353261 | -0.131 | -0.35 | -0.43 | -0.542 | -0.007 | -0.011 | -0.011 | 0.042 | 0.44 | 0.44 | 0.25 | 0.24 | 0.35 | 0.26 |

| | | | | | | | | | | | | | | | | |
|---|---|---|---|---|---|---|---|---|---|---|---|---|---|---|---|---|
| 8 | 113181316 | rs17627747 | 1.607 | -0.36 | -0.467 | -0.48 | 0.013 | 0.064 | 0.064 | 0 | 0.26 | 0.03 | 0.13 | 0.09 | 0.22 | 0.13 |
| 8 | 113181722 | rs2248571 | -1.441 | -0.29 | -0.352 | -0.429 | -0.005 | -0.012 | -0.012 | 0.148 | 0.8 | 0.83 | 0.66 | 0.63 | 0.76 | 0.7 |
| 8 | 113194305 | rs1548988 | -0.631 | -0.271 | -0.369 | -0.451 | -0.067 | 0.069 | 0.069 | 0.098 | 0.61 | 0.89 | 0.54 | 0.77 | 0.49 | 0.56 |
| 8 | 113206719 | rs10283383 | 0.498 | -0.198 | -0.071 | -0.282 | 0.01 | 0.132 | 0.132 | 0.921 | 0.06 | 0.35 | 0.03 | 0.18 | 0 | 0.05 |
| 8 | 113213815 | rs10955616 | -0.702 | -0.233 | -1.205 | -0.444 | 0.029 | -0.004 | -0.004 | 0 | 0.4 | 0.85 | 0.45 | 0.69 | 0.33 | 0.45 |
| 8 | 113215758 | rs1420855 | 0.277 | -0.303 | -1.231 | -0.495 | 0.002 | -0.024 | -0.024 | 0 | 0.46 | 0.22 | 0.48 | 0.45 | 0.46 | 0.42 |
| 8 | 113219539 | rs10505186 | 1.445 | -0.28 | -1.303 | -0.53 | 0.021 | 0.049 | 0.049 | 0.449 | 0.26 | 0.12 | 0.14 | 0.09 | 0.22 | 0.12 |
| 8 | 113234556 | rs10505185 | 1.098 | -0.261 | -1.309 | -0.467 | -0.001 | 0.001 | 0.001 | 0.49 | 0.21 | 0.18 | 0.35 | 0.37 | 0.24 | 0.33 |
| 8 | 113237045 | rs4876458 | 2.69 | -0.566 | -1.527 | -0.709 | 0.001 | 0.091 | 0.091 | 0 | 0.16 | 0 | 0.09 | 0 | 0.13 | 0.07 |
| 8 | 113239131 | rs7845263 | -0.031 | -0.278 | -1.313 | -0.587 | 0.092 | 0.037 | 0.037 | 1.239 | 0.46 | 0.77 | 0.24 | 0.41 | 0.3 | 0.26 |
| 8 | 113241088 | rs1592624 | -0.829 | -0.172 | -1.354 | -0.358 | -0.138 | 0.002 | 0.002 | 0 | 0.66 | 0.94 | 0.59 | 0.76 | 0.57 | 0.59 |
| 8 | 113256222 | rs6995441 | 0.045 | -0.109 | -1.189 | -0.296 | 0.071 | 0.031 | 0.031 | 0.463 | 0.45 | 0.72 | 0.25 | 0.42 | 0.32 | 0.26 |
| 8 | 113264578 | rs929684 | -0.785 | -0.136 | -1.282 | -0.322 | -0.122 | 0.019 | 0.019 | 0 | 0.65 | 0.94 | 0.58 | 0.76 | 0.56 | 0.58 |
| 8 | 113265869 | rs17630545 | 1.408 | 0.374 | -0.883 | -0.109 | 0.001 | 0.059 | 0.059 | 0.084 | 0.25 | 0.06 | 0.14 | 0.09 | 0.23 | 0.12 |
| 8 | 113276635 | rs16883325 | 0.76 | 0.201 | -0.845 | -0.04 | -0.053 | -0.094 | -0.094 | 0 | 0.19 | 0.08 | 0.32 | 0.36 | 0.25 | 0.31 |
| 8 | 113302023 | rs929686 | 0.597 | 0.224 | -0.774 | -0.02 | 0 | 0.005 | 0.005 | 0.416 | 0.2 | 0.21 | 0.32 | 0.35 | 0.25 | 0.31 |
| 8 | 113322357 | rs11778532 | 0.841 | 0.222 | -0.859 | -0.008 | -0.069 | -0.115 | -0.115 | 0 | 0.19 | 0.05 | 0.31 | 0.35 | 0.25 | 0.31 |
| 8 | 113323471 | rs17637578 | 1.494 | 0.333 | -0.943 | -0.093 | -0.017 | 0.065 | 0.065 | 0.093 | 0.25 | 0.07 | 0.15 | 0.1 | 0.23 | 0.13 |
| 8 | 113333060 | rs12155729 | 0.263 | 0.329 | -0.819 | -0.077 | -0.015 | -0.012 | -0.012 | 1.85 | 0.45 | 0.29 | 0.48 | 0.48 | 0.48 | 0.44 |
| 8 | 113337756 | rs12155552 | 1.511 | 0.337 | -0.856 | -0.078 | -0.022 | 0.064 | 0.064 | 0.047 | 0.25 | 0.06 | 0.15 | 0.1 | 0.23 | 0.13 |
| 8 | 113357431 | rs7845794 | -0.068 | 0.285 | -0.716 | -0.059 | 0.002 | 0 | 0 | 0 | 0.49 | 0.41 | 0.52 | 0.48 | 0.52 | 0.45 |
| 8 | 113364696 | rs11778209 | 0.882 | 0.329 | -0.963 | -0.015 | -0.052 | -0.103 | -0.103 | 0 | 0.19 | 0.07 | 0.32 | 0.39 | 0.24 | 0.31 |
| 8 | 113382392 | rs12676507 | 0.249 | 0.329 | -0.951 | -0.03 | -0.071 | -0.077 | -0.077 | 0.458 | 0.45 | 0.14 | 0.46 | 0.47 | 0.48 | 0.44 |
| 8 | 113387038 | rs11778443 | 0.613 | 0.661 | -0.745 | 0.187 | -0.05 | -0.103 | -0.103 | 0 | 0.19 | 0.06 | 0.32 | 0.38 | 0.24 | 0.31 |
| 8 | 113387333 | rs11785419 | 0.63 | 0.661 | -0.745 | 0.187 | -0.05 | -0.103 | -0.103 | 0 | 0.19 | 0.07 | 0.32 | 0.38 | 0.24 | 0.31 |
| 8 | 113390874 | rs13263903 | 0.858 | 0.881 | -0.325 | -0.097 | 0.064 | -0.011 | -0.011 | 0.464 | 0.15 | 0.35 | 0.11 | 0.15 | 0.24 | 0.12 |
| 8 | 113404924 | rs9649933 | 0.433 | 0.716 | -0.955 | -0.029 | -0.006 | 0.005 | 0.005 | 1.302 | 0.51 | 0.33 | 0.51 | 0.48 | 0.51 | 0.48 |
| 8 | 113405543 | rs7461456 | 0.875 | 0.926 | -0.752 | -0.225 | 0.068 | -0.001 | -0.001 | 0 | 0.16 | 0.4 | 0.1 | 0.17 | 0.21 | 0.12 |
| 8 | 113430184 | rs7824071 | 0.58 | 0.921 | -0.814 | -0.241 | 0.073 | -0.016 | -0.016 | 0.121 | 0.17 | 0.5 | 0.12 | 0.17 | 0.23 | 0.12 |
| 8 | 113441607 | rs2883427 | 0.516 | 0.81 | -1.012 | -0.204 | -0.001 | -0.007 | -0.007 | 0 | 0.51 | 0.31 | 0.5 | 0.46 | 0.5 | 0.46 |
| 8 | 113460930 | rs11779960 | 0.714 | 0.981 | -0.821 | -0.107 | -0.011 | -0.051 | -0.051 | 0 | 0.21 | 0.15 | 0.35 | 0.37 | 0.26 | 0.3 |
| 8 | 113469872 | rs16883621 | 0.802 | 1.015 | -0.7 | -0.133 | 0.11 | -0.003 | -0.003 | 0.224 | 0.16 | 0.43 | 0.12 | 0.17 | 0.23 | 0.12 |
| 8 | 113484935 | rs12155677 | 0.97 | 0.707 | -0.789 | -0.052 | -0.014 | 0.058 | 0.058 | 1.159 | 0.1 | 0.01 | 0.05 | 0 | 0.07 | 0.05 |
| 8 | 113485645 | rs10505174 | 0.106 | 0.917 | -0.413 | 0.291 | 0 | 0.012 | 0.012 | 0 | 0.23 | 0.24 | 0.36 | 0.37 | 0.25 | 0.3 |
| 8 | 113486216 | rs10505173 | -0.231 | 0.817 | -0.54 | 0.341 | 0.005 | -0.053 | -0.053 | 5.018 | 0.28 | 0.22 | 0.38 | 0.5 | 0.3 | 0.44 |
| 8 | 113510766 | rs10097465 | -0.465 | 0.86 | -0.686 | 0.331 | 0 | 0.049 | 0.049 | 0.043 | 0.3 | 0.46 | 0.41 | 0.49 | 0.33 | 0.44 |
| 8 | 113518467 | rs896760 | 0.094 | 0.942 | -0.698 | 0.206 | 0 | 0.012 | 0.012 | 0 | 0.25 | 0.28 | 0.34 | 0.45 | 0.25 | 0.28 |
| 8 | 113520770 | rs7835439 | -0.011 | 1.051 | -0.348 | 0.175 | 0.007 | 0.026 | 0.026 | 0 | 0.26 | 0.35 | 0.38 | 0.45 | 0.3 | 0.27 |
| 8 | 113522557 | rs4876287 | 1.617 | 0.93 | -0.902 | -0.288 | -0.02 | 0.059 | 0.059 | 1.651 | 0.25 | 0.03 | 0.13 | 0.11 | 0.23 | 0.15 |
| 8 | 113527268 | rs2251983 | 0.292 | 0.908 | -0.936 | -0.38 | 0 | 0 | 0 | 0 | 0.52 | 0.55 | 0.42 | 0.46 | 0.51 | 0.44 |
| 8 | 113535887 | rs1377371 | -1.323 | 0.805 | -1.19 | -0.378 | -0.002 | 0.051 | 0.051 | 8.842 | 0.8 | 0.95 | 0.81 | 0.91 | 0.81 | 0.7 |
| 8 | 113565333 | rs10105193 | 0.928 | 0.575 | -1.03 | -0.246 | -0.005 | -0.024 | -0.024 | 0 | 0.22 | 0.19 | 0.36 | 0.45 | 0.26 | 0.27 |
| 8 | 113567276 | rs10112543 | 0.793 | 0.57 | -0.994 | -0.249 | 0 | 0.005 | 0.005 | 0 | 0.23 | 0.24 | 0.36 | 0.45 | 0.26 | 0.27 |
| 8 | 113569549 | rs9650045 | 1.671 | 0.668 | -1.135 | -0.268 | -0.03 | 0.059 | 0.059 | 1.388 | 0.25 | 0.04 | 0.13 | 0.11 | 0.24 | 0.15 |
| 8 | 113587644 | rs12676958 | 1.038 | 0.439 | -0.502 | -0.131 | -0.028 | -0.008 | -0.008 | 0.13 | 0.46 | 0.18 | 0.48 | 0.56 | 0.48 | 0.42 |
| 8 | 113590548 | rs12056653 | 0.587 | 0.842 | -0.301 | 0.075 | -0.01 | 0.021 | 0.021 | 0 | 0.21 | 0.42 | 0.26 | 0.18 | 0.18 | 0.13 |
| 8 | 113618413 | rs7823392 | -1.143 | 0.804 | -0.383 | 0.075 | -0.002 | 0.009 | 0.009 | 1.673 | 0.52 | 0.7 | 0.52 | 0.45 | 0.51 | 0.58 |
| 8 | 113626453 | rs12679012 | 0.164 | 1.027 | -0.539 | 0.183 | -0.013 | -0.023 | -0.023 | 1.108 | 0.05 | 0.02 | 0.05 | 0.17 | 0.07 | 0.15 |
| 8 | 113636554 | rs4876288 | 0.645 | 0.995 | -0.483 | 0.112 | -0.01 | 0.015 | 0.015 | 0 | 0.2 | 0.37 | 0.25 | 0.18 | 0.19 | 0.14 |
| 8 | 113638880 | rs7822475 | -0.408 | 1.001 | -0.751 | 0.246 | 0 | 0.016 | 0.016 | 0.863 | 0.44 | 0.62 | 0.46 | 0.27 | 0.44 | 0.43 |
| 8 | 113647451 | rs4876289 | -0.776 | 1.096 | -0.678 | 0.384 | 0.003 | 0.014 | 0.014 | 0 | 0.06 | 0.09 | 0.06 | 0.14 | 0.07 | 0.15 |
| 8 | 113652365 | rs16883889 | -0.213 | 1.164 | -0.681 | -0.055 | 0.002 | -0.019 | -0.019 | 0 | 0.23 | 0.24 | 0.22 | 0.09 | 0.25 | 0.26 |
| 8 | 113653100 | rs10091592 | -0.2 | 0.73 | -1.128 | -0.032 | 0.034 | 0.014 | 0.014 | 0.086 | 0.47 | 0.27 | 0.59 | 0.44 | 0.6 | 0.52 |
| 8 | 113655596 | rs10505169 | 0.307 | -0.196 | -0.687 | -0.204 | 0.005 | 0.091 | 0.091 | 0 | 0.27 | 0.46 | 0.3 | 0.44 | 0.28 | 0.29 |
| 8 | 113662583 | rs17652171 | 0.52 | -0.555 | -0.814 | -0.557 | 0.048 | 0.059 | 0.059 | 0.05 | 0.3 | 0.18 | 0.12 | 0.08 | 0.15 | 0.15 |
| 8 | 113664153 | rs16883931 | 0.673 | -0.58 | -0.336 | -0.358 | 0.011 | 0.053 | 0.053 | 0 | 0.24 | 0.34 | 0.29 | 0.37 | 0.28 | 0.29 |
| 8 | 113671290 | rs7814711 | -1.072 | -0.006 | -1.078 | -0.533 | -0.023 | -0.073 | -0.073 | 0.165 | 0.82 | 0.91 | 0.57 | 0.59 | 0.74 | 0.68 |
| 8 | 113673291 | rs6985219 | -0.347 | -0.141 | -1.101 | -0.658 | -0.01 | -0.02 | -0.02 | 0.075 | 0.59 | 0.75 | 0.43 | 0.53 | 0.44 | 0.47 |
| 8 | 113681944 | rs13271162 | -1.161 | -0.157 | -0.935 | -0.658 | -0.005 | -0.04 | -0.04 | 0.024 | 0.82 | 0.89 | 0.59 | 0.61 | 0.76 | 0.67 |
| 8 | 113683162 | rs6469434 | 1.919 | -0.299 | -1.154 | -0.838 | -0.036 | 0.043 | 0.043 | 0 | 0.12 | 0.05 | 0.09 | 0 | 0.2 | 0.08 |
| 8 | 113687481 | rs10955631 | -0.544 | -0.243 | -1.21 | -0.806 | -0.011 | -0.014 | -0.014 | 0 | 0.58 | 0.78 | 0.45 | 0.54 | 0.45 | 0.47 |
| 8 | 113748216 | rs12679365 | 0.373 | -0.263 | -1.049 | -0.794 | 0.002 | 0.081 | 0.081 | 0 | 0.25 | 0.55 | 0.29 | 0.36 | 0.25 | 0.28 |
| 8 | 113789291 | rs2355821 | -1.19 | -0.365 | -1.161 | -0.886 | 0.05 | 0.093 | 0.093 | 0 | 0.7 | 0.86 | 0.85 | 0.92 | 0.81 | 0.84 |
| 8 | 113797994 | rs2355822 | 0.514 | -0.334 | -1.252 | -0.867 | -0.013 | -0.012 | -0.012 | 0 | 0.42 | 0.22 | 0.55 | 0.46 | 0.55 | 0.53 |
| 8 | 113801484 | rs9297479 | -1.004 | -0.172 | -1.229 | -0.912 | 0.053 | 0.113 | 0.113 | 0 | 0.71 | 0.93 | 0.86 | 0.92 | 0.82 | 0.84 |
| 8 | 113813757 | rs6999481 | -0.855 | -0.099 | -0.984 | -0.916 | 0.02 | 0.027 | 0.027 | 0 | 0.69 | 0.75 | 0.83 | 0.91 | 0.79 | 0.87 |
| 8 | 113815714 | rs1991138 | -0.023 | -0.014 | -1.032 | -0.955 | 0 | 0.005 | 0.005 | 0 | 0.42 | 0.58 | 0.68 | 0.74 | 0.47 | 0.63 |
| 8 | 113824046 | rs13261160 | 1.152 | 0.005 | -1.101 | -0.951 | -0.096 | 0.033 | 0.033 | 0.215 | 0.26 | 0.06 | 0.13 | 0.17 | 0.33 | 0.24 |
| 8 | 113825058 | rs2432742 | 1.44 | -0.035 | -1.121 | -1.005 | 0.057 | 0.091 | 0.091 | 0 | 0.26 | 0.03 | 0.13 | 0.09 | 0.15 | 0.13 |

| | | | | | | | | | | | | | | | | |
|---|---|---|---|---|---|---|---|---|---|---|---|---|---|---|---|---|
| 8 | 113841720 | rs373880 | 0.85 | 0.003 | -0.947 | -0.975 | 0.02 | -0.045 | -0.045 | 0 | 0.28 | 0.36 | 0.16 | 0.07 | 0.33 | 0.22 |
| 8 | 113852201 | rs231316 | -0.733 | 0.194 | -1.179 | -0.748 | 0.03 | 0.065 | 0.065 | 0 | 0.7 | 0.9 | 0.84 | 0.82 | 0.77 | 0.85 |
| 8 | 113859094 | rs2432736 | 0.896 | 0.102 | -1.226 | -0.563 | 0.056 | 0.059 | 0.059 | 0 | 0.28 | 0.06 | 0.14 | 0.12 | 0.18 | 0.15 |
| 8 | 113861073 | rs231312 | 1.44 | 0.837 | -1.192 | -0.428 | -0.095 | 0.054 | 0.054 | 0 | 0.2 | 0.04 | 0.14 | 0.07 | 0.3 | 0.2 |
| 8 | 113878100 | rs1458999 | -1.009 | 0.792 | -1.159 | -0.471 | -0.066 | 0.059 | 0.059 | 0 | 0.76 | 0.9 | 0.85 | 0.93 | 0.66 | 0.78 |
| 8 | 113894331 | rs1026488 | -1.532 | 0.794 | -1.231 | -0.439 | -0.095 | 0.054 | 0.054 | 0 | 0.8 | 0.96 | 0.86 | 0.93 | 0.7 | 0.8 |
| 8 | 113917259 | rs10097475 | -0.166 | 0.032 | -0.947 | -0.232 | 0.003 | 0.002 | 0.002 | 0 | 0.31 | 0.41 | 0.41 | 0.41 | 0.36 | 0.39 |
| 8 | 113929201 | rs1459000 | -1.677 | 0.607 | -0.997 | -0.346 | -0.02 | 0.048 | 0.048 | 0.019 | 0.79 | 0.91 | 0.9 | 0.93 | 0.79 | 0.82 |
| 8 | 113942738 | rs1380134 | -1.702 | 0.586 | -0.695 | -0.207 | -0.006 | 0.019 | 0.019 | 0.06 | 0.79 | 0.9 | 0.9 | 0.93 | 0.79 | 0.82 |
| 8 | 113946062 | rs1870405 | 0.214 | 0.564 | -0.652 | -0.182 | 0.003 | 0.009 | 0.009 | 0 | 0.29 | 0.42 | 0.4 | 0.41 | 0.32 | 0.36 |
| 8 | 113955195 | rs13270310 | -0.011 | 0.363 | -0.403 | 0.032 | 0.002 | 0.001 | 0.001 | 0 | 0.48 | 0.44 | 0.49 | 0.52 | 0.43 | 0.44 |
| 8 | 113960534 | rs7833953 | -0.303 | 0.232 | 0.528 | 0.958 | -0.011 | 0.379 | 0.379 | 0 | 0.11 | 0.42 | 0.31 | 0.42 | 0.06 | 0.23 |
| 8 | 113961958 | rs10505196 | -0.355 | 0.4 | 0.383 | -0.041 | 0.01 | 0.037 | 0.037 | 0.682 | 0.46 | 0.35 | 0.68 | 0.76 | 0.42 | 0.5 |
| 8 | 113995218 | rs6993468 | -1.171 | 0.434 | 0.503 | -0.149 | -0.032 | 0.172 | 0.172 | 0.809 | 0.56 | 0.78 | 0.76 | 0.95 | 0.49 | 0.71 |
| 8 | 114009227 | rs1857718 | -1.557 | 0.564 | 0.474 | -0.183 | -0.069 | 0.226 | 0.226 | 1.605 | 0.59 | 0.89 | 0.78 | 0.95 | 0.55 | 0.74 |
| 8 | 114025995 | rs1969112 | -1.189 | 0.552 | 0.495 | -0.202 | -0.041 | 0.193 | 0.193 | 0.993 | 0.56 | 0.8 | 0.76 | 0.95 | 0.49 | 0.71 |
| 8 | 114029421 | rs964764 | -1.189 | 0.552 | 0.495 | -0.202 | -0.041 | 0.193 | 0.193 | 0.993 | 0.56 | 0.8 | 0.76 | 0.95 | 0.49 | 0.71 |
| 8 | 114082617 | rs17661766 | 0.744 | 0.494 | 0.315 | -0.24 | -0.013 | 0.023 | 0.023 | 0.218 | 0.15 | 0.06 | 0.09 | 0.08 | 0.16 | 0.12 |
| 8 | 114130275 | rs1480442 | 0.157 | 0.452 | 0.408 | -0.182 | -0.012 | 0.013 | 0.013 | 0 | 0.32 | 0.39 | 0.61 | 0.67 | 0.27 | 0.39 |
| 8 | 114154128 | rs12543781 | -0.113 | 0.47 | 0.402 | -0.196 | -0.002 | 0.006 | 0.006 | 0 | 0.34 | 0.4 | 0.63 | 0.67 | 0.34 | 0.44 |
| 8 | 114160622 | rs10955648 | 0.1 | 0.46 | 0.49 | -0.195 | -0.002 | 0.165 | 0.165 | 0.016 | 0.09 | 0.34 | 0.05 | 0.21 | 0.05 | 0.17 |
| 8 | 114162023 | rs9642819 | 0.569 | 0.43 | 0.166 | -0.211 | 0.007 | -0.03 | -0.03 | 2.167 | 0.29 | 0.16 | 0.58 | 0.67 | 0.27 | 0.39 |
| 8 | 114186003 | rs2219898 | 0.048 | 0.44 | 0.179 | -0.23 | -0.019 | 0.013 | 0.013 | 0 | 0.33 | 0.44 | 0.61 | 0.68 | 0.27 | 0.39 |
| 8 | 114194727 | rs17680534 | 0.758 | 0.466 | 0.091 | -0.292 | -0.046 | 0.013 | 0.013 | 0.3 | 0.15 | 0.01 | 0.08 | 0.07 | 0.17 | 0.12 |
| 8 | 114195858 | rs1017829 | -1.647 | 0.456 | 0.122 | -0.296 | -0.061 | 0.221 | 0.221 | 1.434 | 0.59 | 0.88 | 0.78 | 0.95 | 0.55 | 0.73 |
| 8 | 114198799 | rs1480459 | -1.359 | 0.388 | -0.076 | -0.184 | -0.004 | -0.001 | -0.001 | 0 | 0.31 | 0.58 | 0.17 | 0.27 | 0.29 | 0.35 |
| 8 | 114199799 | rs12675523 | 0.344 | 0.497 | 0.114 | -0.069 | 0.013 | 0.169 | 0.169 | 0 | 0.09 | 0.32 | 0.04 | 0.2 | 0.04 | 0.19 |
| 8 | 114218991 | rs2177112 | -1.488 | 0.557 | 0.015 | -0.116 | 0.001 | -0.001 | -0.001 | 0 | 0.32 | 0.55 | 0.19 | 0.27 | 0.33 | 0.37 |
| 8 | 114227020 | rs12682363 | -0.729 | 0.561 | 0.019 | -0.111 | -0.021 | 0.135 | 0.135 | 0.266 | 0.13 | 0.47 | 0.06 | 0.2 | 0.05 | 0.21 |
| 8 | 114265455 | rs12547550 | 0.649 | 0.553 | -0.019 | -0.143 | -0.014 | 0.017 | 0.017 | 0 | 0.28 | 0.36 | 0.57 | 0.68 | 0.22 | 0.39 |
| 8 | 114270739 | rs7834149 | 0.223 | 0.585 | -0.015 | -0.141 | -0.008 | 0.026 | 0.026 | 0.041 | 0.18 | 0.1 | 0.12 | 0.07 | 0.2 | 0.12 |
| 8 | 114274275 | rs6996697 | 0.769 | 0.585 | -0.017 | -0.143 | -0.013 | 0.027 | 0.027 | 0.215 | 0.15 | 0.04 | 0.09 | 0.07 | 0.17 | 0.12 |
| 8 | 114277500 | rs11986420 | -0.827 | 0.546 | -0.023 | -0.139 | 0.019 | 0.112 | 0.112 | 0.685 | 0.14 | 0.45 | 0.08 | 0.2 | 0.12 | 0.25 |
| 8 | 114285601 | rs7001577 | -1.177 | 0.553 | -0.058 | -0.132 | -0.001 | -0.001 | -0.001 | 0 | 0.32 | 0.55 | 0.2 | 0.27 | 0.33 | 0.34 |
| 8 | 114296513 | rs6993801 | 0.193 | 0.557 | -0.019 | -0.07 | 0.016 | 0.198 | 0.198 | 0 | 0.09 | 0.37 | 0.06 | 0.2 | 0.05 | 0.17 |
| 8 | 114321829 | rs1904217 | 1.449 | 0.615 | -0.661 | -0.124 | -0.099 | 0.242 | 0.242 | 0.82 | 0.41 | 0.07 | 0.25 | 0.05 | 0.51 | 0.28 |
| 8 | 114371899 | rs720405 | 1.144 | 0.798 | 0.355 | 0.674 | -0.002 | 0 | 0 | 0.314 | 0.7 | 0.35 | 0.81 | 0.73 | 0.71 | 0.66 |
| 8 | 114389690 | rs10505203 | -0.777 | 1.002 | 0.378 | 1.124 | 0.022 | 0.186 | 0.186 | 0.027 | 0.1 | 0.4 | 0.07 | 0.2 | 0.06 | 0.17 |
| 8 | 114417130 | rs4531082 | -0.048 | 2.288 | -1.174 | 0.091 | -0.065 | 0.044 | 0.044 | 0.003 | 0.83 | 0.92 | 0.85 | 0.95 | 0.67 | 0.87 |
| 8 | 114440178 | rs16884651 | -0.113 | 1.774 | -1.184 | 0.641 | 0.001 | -0.023 | -0.023 | 0.373 | 0.23 | 0.12 | 0.54 | 0.44 | 0.24 | 0.37 |
| 8 | 114445055 | rs16884680 | 0.616 | 1.204 | -1.018 | 0.613 | -0.006 | 0.01 | 0.01 | 0 | 0.13 | 0.16 | 0.08 | 0.2 | 0.11 | 0.18 |
| 8 | 114445215 | rs1055168 | -1.008 | 1.202 | -0.233 | 0.581 | -0.002 | -0.004 | -0.004 | 0.066 | 0.43 | 0.45 | 0.71 | 0.5 | 0.57 | 0.51 |
| 8 | 114447079 | rs2927868 | -0.368 | 1.133 | 0.003 | 0.458 | 0.011 | -0.024 | -0.024 | 0 | 0.23 | 0.26 | 0.2 | 0.05 | 0.37 | 0.14 |
| 8 | 114457117 | rs1513520 | -0.912 | 1.135 | -0.131 | 0.612 | -0.021 | -0.021 | -0.021 | 0 | 0.39 | 0.37 | 0.67 | 0.52 | 0.52 | 0.51 |
| 8 | 114460679 | rs16892653 | 0.461 | 1.132 | -0.149 | 0.616 | -0.004 | -0.066 | -0.066 | 0.844 | 0.19 | 0.05 | 0.5 | 0.47 | 0.19 | 0.37 |
| 8 | 114461585 | rs2927874 | 0.217 | 1.144 | -0.106 | 0.603 | -0.013 | 0.01 | 0.01 | 0 | 0.19 | 0.23 | 0.16 | 0.05 | 0.33 | 0.14 |
| 8 | 114461787 | rs16884721 | 0.899 | 0.477 | 0.419 | 0.536 | -0.033 | 0.092 | 0.092 | 0 | 0.08 | 0.19 | 0.03 | 0.19 | 0.01 | 0.06 |
| 8 | 114485653 | rs12545448 | -0.06 | 0.597 | 0.38 | 0.497 | 0 | -0.001 | -0.001 | 0.776 | 0.24 | 0.18 | 0.55 | 0.47 | 0.23 | 0.37 |
| 8 | 114509172 | rs10107411 | 0.666 | 0.532 | 0.277 | 0.469 | -0.015 | -0.026 | -0.026 | 2.028 | 0.59 | 0.76 | 0.3 | 0.48 | 0.44 | 0.49 |
| 8 | 114509631 | rs4565485 | 0.693 | 0.533 | 0.261 | 0.469 | -0.01 | -0.018 | -0.018 | 0 | 0.58 | 0.69 | 0.3 | 0.48 | 0.44 | 0.49 |
| 8 | 114520181 | rs12548727 | 0.551 | 0.534 | 0.188 | 0.5 | -0.003 | -0.072 | -0.072 | 0 | 0.19 | 0.05 | 0.5 | 0.47 | 0.19 | 0.37 |
| 8 | 114528445 | rs9886503 | 0.882 | 0.543 | 0.289 | 0.462 | -0.002 | -0.004 | -0.004 | 3.022 | 0.57 | 0.55 | 0.29 | 0.48 | 0.43 | 0.49 |
| 8 | 114532488 | rs11786058 | -0.893 | 0.613 | 0.166 | 0.446 | -0.065 | 0.048 | 0.048 | 0.01 | 0.83 | 0.92 | 0.85 | 0.95 | 0.67 | 0.87 |
| 8 | 114545719 | rs10110326 | -0.874 | 0.614 | 0.174 | 0.446 | -0.035 | 0.029 | 0.029 | 0.015 | 0.83 | 0.89 | 0.84 | 0.95 | 0.67 | 0.87 |
| 8 | 114564791 | rs10505205 | 0.539 | 0.554 | 0.138 | 0.513 | -0.003 | -0.072 | -0.072 | 0 | 0.19 | 0.06 | 0.5 | 0.47 | 0.19 | 0.37 |
| 8 | 114569143 | rs10081457 | -0.322 | 0.517 | 0.29 | 0.473 | 0.003 | -0.028 | -0.028 | 0 | 0.06 | 0.25 | 0.05 | 0 | 0.04 | 0 |
| 8 | 114572537 | rs17612497 | 1.426 | 0.385 | 0.427 | 0.481 | 0.011 | 0.052 | 0.052 | 0.198 | 0.48 | 0.34 | 0.26 | 0.27 | 0.42 | 0.42 |
| 8 | 114581966 | rs16884874 | 0.558 | 0.473 | 0.202 | 0.496 | -0.002 | -0.07 | -0.07 | 0 | 0.2 | 0.08 | 0.5 | 0.47 | 0.19 | 0.38 |
| 8 | 114584737 | rs7461229 | -0.978 | 0.464 | 0.196 | 0.476 | -0.064 | 0.045 | 0.045 | 0 | 0.83 | 0.92 | 0.85 | 0.95 | 0.67 | 0.87 |
| 8 | 114589007 | rs12680097 | 0.497 | -0.062 | 0.238 | 0.589 | -0.003 | -0.074 | -0.074 | 0.856 | 0.19 | 0.05 | 0.5 | 0.47 | 0.19 | 0.38 |
| 8 | 114599447 | rs4492406 | 0.68 | -0.116 | 0.15 | 0.624 | 0.006 | -0.032 | -0.032 | 4.388 | 0.15 | 0.07 | 0.4 | 0.49 | 0.13 | 0.3 |
| 8 | 114604322 | rs4486239 | -1.377 | 0.524 | 0.334 | 0.451 | -0.006 | -0.007 | -0.007 | 0.035 | 0.45 | 0.43 | 0.64 | 0.73 | 0.36 | 0.5 |
| 8 | 114621134 | rs12549169 | -0.913 | 0.585 | 0.335 | 0.429 | 0 | 0 | 0 | 0 | 0.36 | 0.39 | 0.51 | 0.53 | 0.32 | 0.42 |
| 8 | 114625436 | rs10099644 | 0.724 | 0.598 | 0.2 | 0.488 | 0.006 | -0.031 | -0.031 | 0 | 0.15 | 0.07 | 0.4 | 0.48 | 0.13 | 0.3 |
| 8 | 114645013 | rs1348382 | -0.999 | 0.615 | 0.339 | 0.455 | 0.003 | -0.008 | -0.008 | 0 | 0.4 | 0.35 | 0.63 | 0.54 | 0.38 | 0.47 |
| 8 | 114647140 | rs10088331 | -0.957 | 0.611 | 0.419 | 0.453 | -0.002 | 0.002 | 0.002 | 0.069 | 0.41 | 0.44 | 0.64 | 0.54 | 0.38 | 0.48 |
| 8 | 114664356 | rs7832929 | -1.543 | 0.581 | 0.374 | 0.385 | 0.046 | 0.066 | 0.066 | 0 | 0.51 | 0.68 | 0.66 | 0.72 | 0.39 | 0.54 |
| 8 | 114673166 | rs17613558 | 0.808 | -0.064 | -0.223 | 0.102 | 0.015 | 0.054 | 0.054 | 0 | 0.17 | 0.07 | 0.08 | 0.05 | 0.14 | 0.07 |
| 8 | 114676890 | rs10102855 | -1.419 | 0.157 | 0.294 | -0.069 | 0.043 | 0.069 | 0.069 | 0 | 0.5 | 0.64 | 0.65 | 0.73 | 0.39 | 0.5 |

| | | | | | | | | | | | | | | | | |
|---|---|---|---|---|---|---|---|---|---|---|---|---|---|---|---|---|
| 8 | 114677210 | rs10088339 | 0.527 | 0.112 | 0.571 | 0.096 | 0.003 | -0.022 | -0.022 | 0 | 0.21 | 0.14 | 0.51 | 0.5 | 0.19 | 0.38 |
| 8 | 114683788 | rs1443344 | -1.486 | 0.117 | 0.53 | 0.086 | 0.043 | 0.069 | 0.069 | 0 | 0.5 | 0.64 | 0.65 | 0.73 | 0.39 | 0.5 |
| 8 | 114696874 | rs16884991 | 0.877 | 0.203 | 0.376 | 0.056 | 0.021 | 0.056 | 0.056 | 0 | 0.17 | 0.07 | 0.08 | 0.05 | 0.14 | 0.07 |
| 8 | 114698419 | rs12386964 | -0.318 | 0.211 | 0.56 | 0.336 | 0.003 | 0.059 | 0.059 | 0 | 0.27 | 0.34 | 0.55 | 0.5 | 0.24 | 0.41 |
| 8 | 114703295 | rs7826446 | -0.705 | -0.034 | 0.448 | 0.214 | -0.009 | 0.047 | 0.047 | 0.471 | 0.3 | 0.41 | 0.55 | 0.5 | 0.24 | 0.41 |
| 8 | 114719996 | rs12680503 | 0.021 | 0.095 | 0.516 | 0.305 | -0.004 | 0.06 | 0.06 | 0.398 | 0.23 | 0.34 | 0.53 | 0.5 | 0.2 | 0.38 |
| 8 | 114725187 | rs9297501 | 0.383 | 0.097 | 0.396 | 0.317 | 0.001 | -0.008 | -0.008 | 0 | 0.21 | 0.16 | 0.51 | 0.5 | 0.19 | 0.38 |
| 8 | 114729848 | rs9297502 | -1.077 | 0.439 | 0.375 | 0.14 | -0.001 | 0 | 0 | 0 | 0.48 | 0.49 | 0.68 | 0.55 | 0.46 | 0.49 |
| 8 | 114731745 | rs1257826 | 0.648 | 0.533 | 0.171 | -0.046 | -0.004 | -0.001 | -0.001 | 0 | 0.18 | 0.19 | 0.15 | 0.15 | 0.3 | 0.24 |
| 8 | 114738009 | rs10087099 | -0.065 | 0.253 | 0.194 | 0.191 | 0.004 | 0.026 | 0.026 | 0.287 | 0.26 | 0.28 | 0.57 | 0.5 | 0.28 | 0.39 |
| 8 | 114740737 | rs10109202 | -0.001 | 0.253 | 0.194 | 0.209 | 0.004 | 0.029 | 0.029 | 0.213 | 0.26 | 0.28 | 0.57 | 0.5 | 0.28 | 0.39 |
| 8 | 114743324 | rs7814760 | 0.11 | 0.243 | 0.229 | 0.19 | -0.016 | -0.059 | -0.059 | 0 | 0.24 | 0.11 | 0.55 | 0.5 | 0.27 | 0.39 |
| 8 | 114747352 | rs1470304 | -0.159 | 0.123 | 0.311 | 0.154 | -0.009 | -0.001 | -0.001 | 0 | 0.23 | 0.25 | 0.1 | 0.23 | 0.14 | 0.1 |
| 8 | 114749009 | rs4876313 | -1.687 | 0.134 | 0.331 | 0.15 | 0.001 | 0.037 | 0.037 | 0 | 0.53 | 0.63 | 0.69 | 0.73 | 0.46 | 0.52 |
| 8 | 114758168 | rs1452757 | 0.022 | 0.151 | 0.344 | 0.177 | -0.007 | -0.033 | -0.033 | 0.746 | 0.25 | 0.17 | 0.56 | 0.5 | 0.27 | 0.39 |
| 8 | 114758912 | rs16885086 | 1.266 | 0.147 | 0.221 | 0.182 | -0.019 | 0.113 | 0.113 | 0.058 | 0.06 | 0.18 | 0.02 | 0.17 | 0.01 | 0.03 |
| 8 | 114761849 | rs11996253 | -0.483 | 0.129 | 0.202 | 0.17 | -0.023 | -0.008 | -0.008 | 0.244 | 0.27 | 0.35 | 0.12 | 0.23 | 0.17 | 0.19 |
| 8 | 114763795 | rs10096263 | -1.396 | 0.54 | 0.525 | 0.734 | -0.001 | 0.001 | 0.001 | 0 | 0.36 | 0.55 | 0.61 | 0.68 | 0.33 | 0.45 |
| 8 | 114766393 | rs10107672 | -0.38 | 0.941 | -0.135 | 0.549 | -0.009 | -0.03 | -0.03 | 0 | 0.23 | 0.14 | 0.55 | 0.5 | 0.27 | 0.39 |
| 8 | 114776941 | rs6986020 | -0.771 | 0.775 | -0.192 | 0.702 | 0.009 | 0.028 | 0.028 | 0.02 | 0.31 | 0.38 | 0.59 | 0.5 | 0.32 | 0.41 |
| 8 | 114788424 | rs9297508 | 1.084 | 0.407 | -1.156 | -0.006 | -0.091 | 0.182 | 0.182 | 0 | 0.33 | 0.05 | 0.19 | 0.06 | 0.41 | 0.3 |
| 8 | 114789982 | rs9297509 | 0.533 | 0.347 | -0.637 | 0.334 | 0.021 | 0.116 | 0.116 | 0.035 | 0.51 | 0.23 | 0.3 | 0.23 | 0.58 | 0.47 |
| 8 | 114791261 | rs1901696 | -0.754 | 0.24 | -0.495 | 1.027 | 0 | 0.127 | 0.127 | 0.019 | 0.94 | 0.95 | 0.64 | 0.53 | 0.92 | 0.86 |
| 8 | 114793032 | rs7005303 | -0.117 | 0.003 | 0.149 | 0.972 | -0.002 | -0.032 | -0.032 | 0 | 0.35 | 0.41 | 0.3 | 0.14 | 0.3 | 0.32 |
| 8 | 114795014 | rs9649939 | 0.731 | 0.001 | 0.144 | 1.007 | -0.016 | 0.168 | 0.168 | 1.073 | 0.86 | 0.64 | 0.62 | 0.38 | 0.88 | 0.78 |
| 8 | 114795407 | rs13273350 | 0.081 | -0.051 | 0.398 | 0.947 | -0.002 | 0.01 | 0.01 | 0 | 0.33 | 0.27 | 0.29 | 0.17 | 0.3 | 0.31 |
| 8 | 114799224 | rs2044999 | 0.187 | -0.091 | 0.5 | 1.031 | 0 | -0.001 | -0.001 | 0.548 | 0.63 | 0.63 | 0.34 | 0.38 | 0.63 | 0.64 |
| 8 | 114800664 | rs7006134 | -1.225 | -0.114 | 0.528 | 1.045 | -0.047 | 0.062 | 0.062 | 3.655 | 0.12 | 0.38 | 0.04 | 0.17 | 0.05 | 0.16 |
| 8 | 114801383 | rs896680 | 0.285 | -0.082 | 0.442 | 0.964 | 0.007 | 0.081 | 0.081 | 0 | 0.85 | 0.83 | 0.52 | 0.45 | 0.83 | 0.77 |
| 8 | 114805297 | rs7833659 | 1.049 | -0.064 | 0.35 | 0.91 | -0.002 | 0.034 | 0.034 | 0 | 0.21 | 0.12 | 0.17 | 0.08 | 0.19 | 0.12 |
| 8 | 114810353 | rs1947374 | 0.184 | -0.074 | 0.516 | 0.991 | -0.001 | 0.149 | 0.149 | 0.344 | 0.92 | 0.84 | 0.63 | 0.53 | 0.92 | 0.84 |
| 8 | 114812254 | rs7012827 | 2.375 | -0.103 | 0.312 | 0.822 | -0.097 | 0.194 | 0.194 | 0 | 0.34 | 0.05 | 0.19 | 0.05 | 0.42 | 0.29 |
| 8 | 114814714 | rs10101872 | 1.282 | -0.159 | 0.517 | 0.927 | 0.004 | 0.027 | 0.027 | 0 | 0.55 | 0.38 | 0.32 | 0.23 | 0.59 | 0.57 |
| 8 | 114814755 | rs11777753 | -0.129 | -0.141 | 0.274 | 0.869 | -0.001 | -0.002 | -0.002 | 0 | 0.3 | 0.32 | 0.21 | 0.23 | 0.27 | 0.24 |
| 8 | 114822319 | rs11998298 | 0.249 | -0.118 | 0.256 | 0.951 | -0.008 | 0.025 | 0.025 | 0.05 | 0.09 | 0.15 | 0.03 | 0.16 | 0.07 | 0.11 |
| 8 | 114824947 | rs13274148 | 0.792 | -0.025 | 0.252 | 1.05 | 0 | 0.006 | 0.006 | 0 | 0.84 | 0.81 | 0.55 | 0.54 | 0.86 | 0.82 |
| 8 | 114825281 | rs10808463 | -0.76 | 0.035 | 0.315 | 1.104 | 0.002 | 0.161 | 0.161 | 0 | 0.94 | 0.84 | 0.63 | 0.55 | 0.93 | 0.94 |
| 8 | 114828429 | rs13267919 | 0.959 | -0.015 | 0.19 | 0.704 | 0.001 | 0.017 | 0.017 | 0 | 0.58 | 0.61 | 0.33 | 0.22 | 0.59 | 0.57 |
| 8 | 114832814 | rs10103525 | 2.114 | 0.071 | -0.15 | 0.657 | -0.103 | 0.195 | 0.195 | 0.008 | 0.34 | 0.05 | 0.19 | 0.07 | 0.42 | 0.29 |
| 8 | 114833306 | rs4436151 | 1.052 | -0.111 | -0.049 | 0.775 | -0.001 | -0.002 | -0.002 | 0 | 0.79 | 0.79 | 0.51 | 0.3 | 0.79 | 0.73 |
| 8 | 114836998 | rs13268405 | -0.111 | -0.063 | -0.051 | 0.853 | 0.002 | 0.093 | 0.093 | 0.101 | 0.88 | 0.83 | 0.61 | 0.4 | 0.86 | 0.84 |
| 8 | 114839446 | rs13254570 | 2.034 | -0.018 | -0.301 | 0.588 | -0.073 | 0.175 | 0.175 | 0 | 0.35 | 0.09 | 0.2 | 0.06 | 0.43 | 0.31 |
| 8 | 114839718 | rs7819200 | 0.992 | -0.06 | -0.184 | 0.762 | -0.002 | 0.034 | 0.034 | 0 | 0.79 | 0.7 | 0.53 | 0.41 | 0.82 | 0.76 |
| 8 | 114841349 | rs12156378 | 0.588 | -0.052 | 0.175 | 0.746 | 0.002 | 0.003 | 0.003 | 0 | 0.2 | 0.19 | 0.13 | 0.14 | 0.17 | 0.19 |
| 8 | 114841751 | rs12682240 | 1.922 | -0.005 | -0.052 | 0.764 | -0.073 | 0.175 | 0.175 | 0 | 0.35 | 0.09 | 0.2 | 0.06 | 0.43 | 0.31 |
| 8 | 114842242 | rs11997248 | 0.477 | 0.087 | -0.065 | 0.901 | 0.004 | 0.129 | 0.129 | 0 | 0.85 | 0.71 | 0.57 | 0.51 | 0.85 | 0.85 |
| 8 | 114848963 | rs7011798 | -1.059 | 0.058 | 0.095 | 0.617 | -0.002 | -0.003 | -0.003 | 0.992 | 0.52 | 0.51 | 0.4 | 0.41 | 0.44 | 0.57 |
| 8 | 114849850 | rs9650053 | 1.865 | 0.076 | 0.097 | 0.832 | -0.02 | 0.045 | 0.045 | 0 | 0.35 | 0.19 | 0.29 | 0.14 | 0.43 | 0.31 |
| 8 | 114850678 | rs16885210 | 0.367 | 0.022 | 0.194 | 0.832 | -0.027 | -0.046 | -0.046 | 2.764 | 0.21 | 0.35 | 0.14 | 0.08 | 0.15 | 0.16 |
| 8 | 114850899 | rs13248967 | 0.263 | 0.307 | -0.197 | 1.081 | 0.01 | 0.134 | 0.134 | 0.012 | 0.87 | 0.71 | 0.69 | 0.56 | 0.87 | 0.88 |
| 8 | 114864148 | rs2357037 | -0.241 | 0.252 | -0.704 | 0.698 | -0.011 | -0.014 | -0.014 | 0 | 0.31 | 0.19 | 0.29 | 0.34 | 0.3 | 0.41 |
| 8 | 114867123 | rs16892681 | -0.712 | 0.117 | -0.177 | 0.326 | -0.03 | -0.081 | -0.081 | 0 | 0.37 | 0.53 | 0.22 | 0.12 | 0.25 | 0.3 |
| 8 | 114869929 | rs6991624 | -1.304 | 0.414 | -0.042 | 0.643 | -0.002 | 0.016 | 0.016 | 0 | 0.52 | 0.73 | 0.48 | 0.44 | 0.47 | 0.69 |
| 8 | 114871075 | rs13275149 | -0.519 | 0.202 | 0.234 | 0.497 | -0.003 | -0.005 | -0.005 | 0 | 0.31 | 0.25 | 0.34 | 0.33 | 0.28 | 0.51 |
| 8 | 114873380 | rs1912816 | -1.054 | 0.12 | 0.216 | 0.574 | -0.016 | -0.016 | -0.016 | 1.14 | 0.33 | 0.67 | 0.3 | 0.25 | 0.27 | 0.34 |
| 8 | 114874435 | rs13251159 | 0.166 | 0.247 | -0.045 | 0.663 | -0.013 | 0.002 | 0.002 | 0 | 0.19 | 0.08 | 0.17 | 0.19 | 0.2 | 0.35 |
| 8 | 114876759 | rs1397965 | -0.178 | 0.459 | 0.306 | 1.321 | -0.005 | -0.001 | -0.001 | 0 | 0.12 | 0.2 | 0.11 | 0.12 | 0.08 | 0.15 |
| 8 | 114882527 | rs1397961 | -0.197 | 0.507 | 0.902 | 1.515 | -0.024 | 0.046 | 0.046 | 0 | 0.17 | 0.42 | 0.11 | 0.15 | 0.14 | 0.12 |
| 8 | 114892901 | rs7833314 | -0.631 | 0.28 | 0.297 | 0.79 | -0.005 | 0 | 0 | 0.08 | 0.23 | 0.7 | 0.15 | 0.21 | 0.19 | 0.19 |
| 8 | 114908475 | rs2956648 | 0.95 | 0.862 | -0.846 | 0.06 | -0.037 | 0.203 | 0.203 | 0 | 0.45 | 0.15 | 0.27 | 0.11 | 0.48 | 0.35 |
| 8 | 114909016 | rs7834762 | 0.764 | 0.802 | -0.155 | 0.149 | 0.011 | 0.01 | 0.01 | 0 | 0.13 | 0.2 | 0.09 | 0.1 | 0.16 | 0.04 |
| 8 | 114913979 | rs12548343 | -0.888 | 0.57 | -0.587 | -0.228 | 0.03 | 0.072 | 0.072 | 0 | 0.49 | 0.61 | 0.65 | 0.76 | 0.4 | 0.54 |
| 8 | 114915810 | rs966210 | 1.147 | 0.694 | -0.51 | -0.309 | -0.008 | 0.111 | 0.111 | 0 | 0.46 | 0.19 | 0.32 | 0.24 | 0.55 | 0.41 |
| 8 | 114918152 | rs2458071 | 1.495 | 0.659 | -0.582 | 0.058 | -0.088 | 0.089 | 0.089 | 0 | 0.39 | 0.1 | 0.28 | 0.19 | 0.49 | 0.35 |
| 8 | 114924899 | rs7842617 | 0.163 | 0.472 | -0.391 | -0.012 | -0.012 | -0.003 | -0.003 | 0.055 | 0.27 | 0.32 | 0.18 | 0.22 | 0.24 | 0.15 |
| 8 | 114924989 | rs869476 | -0.307 | 0.476 | -0.337 | -0.018 | 0.005 | 0.003 | 0.003 | 0 | 0.42 | 0.38 | 0.27 | 0.34 | 0.34 | 0.41 |
| 8 | 114932476 | rs1912817 | 0.483 | 0.502 | -0.342 | 0.067 | 0.021 | 0.199 | 0.199 | 0 | 0.91 | 0.74 | 0.61 | 0.58 | 0.9 | 0.89 |
| 8 | 114933739 | rs2468871 | 0.833 | 0.506 | -0.575 | -0.321 | 0.01 | 0.051 | 0.051 | 0 | 0.49 | 0.36 | 0.34 | 0.24 | 0.56 | 0.48 |
| 8 | 114942214 | rs13439749 | 0.975 | 0.53 | -0.606 | -0.329 | 0 | -0.002 | -0.002 | 0.145 | 0.07 | 0.08 | 0.03 | 0.05 | 0.06 | 0.06 |

| | | | | | | | | | | | | | | | | |
|---|---|---|---|---|---|---|---|---|---|---|---|---|---|---|---|---|
| 8 | 114944774 | rs960956 | 0.837 | 0.527 | -0.585 | -0.312 | 0.011 | 0.047 | 0.047 | 0 | 0.49 | 0.37 | 0.35 | 0.24 | 0.56 | 0.48 |
| 8 | 114954678 | rs16885345 | 0.924 | 0.527 | -0.515 | -0.24 | -0.009 | -0.015 | -0.015 | 0 | 0.19 | 0.22 | 0.12 | 0.12 | 0.12 | 0.32 |
| 8 | 114956591 | rs2458064 | 0.739 | 0.502 | -0.774 | -0.686 | 0.005 | 0.068 | 0.068 | 0.043 | 0.53 | 0.16 | 0.37 | 0.31 | 0.57 | 0.47 |
| 8 | 114959293 | rs1511704 | 1.148 | 0.44 | -0.841 | -0.437 | -0.038 | -0.152 | -0.152 | 0.585 | 0.19 | 0.01 | 0.4 | 0.52 | 0.16 | 0.32 |
| 8 | 114963014 | rs1511706 | 0.028 | 0.318 | -0.584 | -0.604 | -0.024 | -0.044 | -0.044 | 0.063 | 0.8 | 0.59 | 0.87 | 0.87 | 0.86 | 0.83 |
| 8 | 114965111 | rs990895 | 1.432 | 0.337 | -0.644 | -0.599 | -0.038 | -0.147 | -0.147 | 0.371 | 0.18 | 0.01 | 0.39 | 0.52 | 0.15 | 0.32 |
| 8 | 114965243 | rs10112981 | 0.447 | 0.259 | -0.584 | -0.626 | -0.012 | -0.005 | -0.005 | 0.04 | 0.62 | 0.49 | 0.48 | 0.35 | 0.7 | 0.51 |
| 8 | 114966953 | rs11782400 | 0.566 | 0.26 | -0.722 | -0.618 | 0.023 | 0.008 | 0.008 | 0 | 0.15 | 0.06 | 0.09 | 0.12 | 0.09 | 0.12 |
| 8 | 114978005 | rs10505210 | 1.256 | 0.432 | -0.683 | -0.4 | -0.006 | -0.084 | -0.084 | 0.499 | 0.18 | 0.08 | 0.41 | 0.51 | 0.16 | 0.32 |
| 8 | 114978390 | rs7835123 | 0.979 | 0.56 | -0.437 | -0.546 | 0 | -0.032 | -0.032 | 0 | 0.44 | 0.41 | 0.31 | 0.19 | 0.46 | 0.33 |
| 8 | 114979407 | rs1511700 | 0.445 | 0.576 | -0.436 | -0.526 | -0.001 | -0.016 | -0.016 | 0 | 0.62 | 0.49 | 0.72 | 0.7 | 0.63 | 0.65 |
| 8 | 114979484 | rs13263301 | 1.4 | 0.561 | -0.455 | -0.552 | -0.009 | -0.008 | -0.008 | 0 | 0.05 | 0.03 | 0.06 | 0.11 | 0.09 | 0.08 |
| 8 | 114979501 | rs1511699 | 0.499 | 0.65 | -0.449 | -0.555 | 0 | -0.019 | -0.019 | 0 | 0.62 | 0.46 | 0.71 | 0.7 | 0.63 | 0.65 |
| 8 | 114980457 | rs7016357 | 1.175 | 0.616 | -0.601 | -0.599 | -0.029 | 0.125 | 0.125 | 0 | 0.42 | 0.19 | 0.29 | 0.18 | 0.46 | 0.32 |
| 8 | 114985828 | rs7823309 | 0.113 | 0.583 | -0.522 | -0.563 | -0.013 | -0.038 | -0.038 | 0 | 0.67 | 0.52 | 0.79 | 0.81 | 0.71 | 0.74 |
| 8 | 114991022 | rs4147331 | -0.288 | 0.533 | -1.103 | -0.699 | -0.002 | -0.013 | -0.013 | 0.033 | 0.38 | 0.44 | 0.26 | 0.32 | 0.38 | 0.35 |
| 8 | 114992222 | rs7826990 | 1.455 | 0.83 | -1.556 | -1.064 | 0.007 | 0.031 | 0.031 | 0 | 0.26 | 0.13 | 0.19 | 0.13 | 0.24 | 0.21 |
| 8 | 115002551 | rs4364660 | 0.294 | 0.902 | -1.52 | -1.008 | -0.01 | 0.109 | 0.109 | 0 | 0.43 | 0.24 | 0.29 | 0.18 | 0.46 | 0.33 |
| 8 | 115003742 | rs7821078 | 0.193 | 0.901 | -0.996 | -1.227 | 0.008 | 0.029 | 0.029 | 0.058 | 0.6 | 0.25 | 0.72 | 0.68 | 0.64 | 0.65 |
| 8 | 115003926 | rs7824832 | 0.007 | 0.895 | -0.959 | -1.238 | -0.003 | -0.008 | -0.008 | 0.805 | 0.35 | 0.44 | 0.25 | 0.32 | 0.34 | 0.34 |
| 8 | 115004205 | rs7825278 | 1.014 | 1.032 | -1.8 | -1.284 | -0.002 | -0.007 | -0.007 | 0 | 0.15 | 0.16 | 0.11 | 0.04 | 0.2 | 0.08 |
| 8 | 115005081 | rs10098737 | 0.463 | 0.687 | -1.147 | -0.798 | 0.009 | 0.03 | 0.03 | 0.2 | 0.6 | 0.24 | 0.71 | 0.68 | 0.64 | 0.65 |
| 8 | 115007395 | rs2175484 | 0.136 | 0.652 | -1.202 | -0.867 | 0.005 | 0.037 | 0.037 | 0.595 | 0.67 | 0.27 | 0.78 | 0.81 | 0.72 | 0.74 |
| 8 | 115010308 | rs13257991 | 0.601 | 0.799 | -1.144 | -0.769 | 0.008 | 0.054 | 0.054 | 0.222 | 0.56 | 0.25 | 0.7 | 0.68 | 0.6 | 0.64 |
| 8 | 115011589 | rs7825514 | 0.102 | 0.852 | -0.184 | -0.669 | 0.045 | 0.057 | 0.057 | 0 | 0.91 | 0.74 | 0.89 | 0.86 | 0.89 | 0.95 |
| 8 | 115012364 | rs3920209 | -0.27 | 0.407 | -0.559 | -1.339 | 0.002 | 0.002 | 0.002 | 0 | 0.37 | 0.37 | 0.24 | 0.19 | 0.34 | 0.2 |
| 8 | 115018622 | rs10086932 | 0.768 | 0.194 | -0.474 | -1.843 | 0 | -0.012 | -0.012 | 0 | 0.56 | 0.55 | 0.7 | 0.78 | 0.6 | 0.8 |
| 8 | 115022709 | rs10109903 | -1.228 | 0.121 | -0.169 | -1.757 | 0.02 | -0.027 | -0.027 | 0 | 0.9 | 0.86 | 0.91 | 0.98 | 0.83 | 0.94 |
| 8 | 115027676 | rs1397970 | 0.342 | -0.226 | -0.293 | -1.901 | 0.004 | 0.011 | 0.011 | 0.001 | 0.37 | 0.35 | 0.53 | 0.75 | 0.34 | 0.5 |
| 8 | 115029201 | rs4552933 | -0.165 | -0.285 | -0.28 | -1.934 | -0.002 | -0.061 | -0.061 | 0 | 0.79 | 0.67 | 0.89 | 0.94 | 0.81 | 0.9 |
| 8 | 115034776 | rs4445256 | 1.115 | -0.225 | -0.394 | -1.917 | -0.011 | 0.033 | 0.033 | 0 | 0.42 | 0.32 | 0.36 | 0.19 | 0.48 | 0.4 |
| 8 | 115039038 | rs6983475 | 0.033 | -0.369 | -0.517 | -1.974 | 0.002 | 0.003 | 0.003 | 0 | 0.41 | 0.32 | 0.44 | 0.62 | 0.41 | 0.53 |
| 8 | 115041135 | rs6993921 | 0.231 | -0.079 | -0.544 | -1.964 | 0.003 | -0.036 | -0.036 | 0 | 0.2 | 0.22 | 0.12 | 0.03 | 0.24 | 0.11 |
| 8 | 115046511 | rs6993039 | 1.443 | -0.018 | -0.582 | -1.909 | 0.01 | -0.021 | -0.021 | 0 | 0.11 | 0.13 | 0.08 | 0.02 | 0.17 | 0.06 |
| 8 | 115060120 | rs9297521 | 1.524 | -0.015 | -0.581 | -1.888 | 0.011 | -0.019 | -0.019 | 0 | 0.1 | 0.13 | 0.08 | 0.02 | 0.17 | 0.06 |
| 8 | 115063598 | rs10955698 | 1.551 | -0.031 | -0.829 | -1.995 | -0.15 | 0.045 | 0.045 | 0.865 | 0.33 | 0.02 | 0.31 | 0.17 | 0.42 | 0.37 |
| 8 | 115070619 | rs4329300 | 2.125 | -0.167 | -0.83 | -1.926 | -0.012 | -0.126 | -0.126 | 0 | 0.17 | 0.01 | 0.3 | 0.55 | 0.15 | 0.27 |
| 8 | 115095516 | rs13266408 | 1.378 | -0.059 | -0.506 | -1.953 | -0.008 | 0.02 | 0.02 | 0.383 | 0.34 | 0.25 | 0.31 | 0.17 | 0.39 | 0.37 |
| 8 | 115097202 | rs4538916 | -2.018 | -0.116 | -0.442 | -2.018 | -0.002 | -0.003 | -0.003 | 0 | 0.87 | 0.88 | 0.83 | 0.83 | 0.85 | 0.95 |
| 8 | 115101787 | rs7006291 | 0.406 | -0.154 | -0.333 | -2.065 | 0.022 | -0.083 | -0.083 | 0.361 | 0.21 | 0.32 | 0.15 | 0.03 | 0.25 | 0.12 |
| 8 | 115104817 | rs11779159 | 0.479 | -0.149 | -0.324 | -2.038 | 0.028 | -0.075 | -0.075 | 0 | 0.2 | 0.31 | 0.15 | 0.04 | 0.24 | 0.12 |
| 8 | 115113221 | rs12549856 | 1.91 | -0.199 | -0.486 | -1.969 | 0.013 | -0.064 | -0.064 | 0 | 0.21 | 0.09 | 0.32 | 0.59 | 0.18 | 0.42 |
| 8 | 115119439 | rs12675001 | 1.104 | 0.229 | -0.354 | -1.863 | -0.008 | 0.022 | 0.022 | 0.411 | 0.34 | 0.25 | 0.31 | 0.17 | 0.39 | 0.37 |
| 8 | 115123578 | rs7000905 | 1.085 | 0.225 | -0.329 | -1.895 | -0.007 | 0.018 | 0.018 | 0.338 | 0.34 | 0.26 | 0.31 | 0.17 | 0.39 | 0.37 |
| 8 | 115129791 | rs12545023 | 1.041 | 0.071 | -0.369 | -1.922 | 0.011 | -0.007 | -0.007 | 0.248 | 0.31 | 0.26 | 0.36 | 0.63 | 0.2 | 0.46 |
| 8 | 115130148 | rs13277450 | -0.451 | 0.313 | -0.26 | -1.773 | -0.013 | 0.007 | 0.007 | 0 | 0.77 | 0.78 | 0.75 | 0.81 | 0.68 | 0.9 |
| 8 | 115162781 | rs7003100 | 0.473 | 0.252 | -0.769 | -1.79 | 0.015 | 0.005 | 0.005 | 0.885 | 0.4 | 0.61 | 0.44 | 0.65 | 0.28 | 0.51 |
| 8 | 115172410 | rs13257428 | 1.238 | 0.453 | -1.028 | -1.757 | -0.126 | 0.051 | 0.051 | 1.074 | 0.33 | 0.02 | 0.29 | 0.17 | 0.39 | 0.38 |
| 8 | 115184664 | rs1515685 | 0.063 | 0.73 | -0.379 | -1.366 | 0.009 | -0.007 | -0.007 | 0.194 | 0.76 | 0.71 | 0.75 | 0.82 | 0.68 | 0.89 |
| 8 | 115195537 | rs7009818 | 0.143 | 0.577 | 0.056 | -1.128 | 0.061 | 0.038 | 0.038 | 0.087 | 0.13 | 0.3 | 0.16 | 0.18 | 0.18 | 0.08 |
| 8 | 115200723 | rs7812989 | -0.342 | 0.392 | 0.106 | -1.725 | 0.004 | 0.003 | 0.003 | 0 | 0.47 | 0.45 | 0.41 | 0.65 | 0.32 | 0.51 |
| 8 | 115206234 | rs1606891 | -1.443 | 0.34 | -0.155 | -1.908 | -0.149 | 0.093 | 0.093 | 0.829 | 0.63 | 0.98 | 0.64 | 0.83 | 0.52 | 0.62 |
| 8 | 115213656 | rs7011881 | -1.44 | 0.387 | -0.154 | -1.908 | -0.145 | 0.089 | 0.089 | 0.852 | 0.63 | 0.98 | 0.64 | 0.83 | 0.53 | 0.61 |
| 8 | 115220918 | rs7015999 | -0.095 | 0.446 | 0.49 | -1.858 | 0.174 | 0.196 | 0.196 | 0.003 | 0.05 | 0.32 | 0.14 | 0.16 | 0.1 | 0.03 |
| 8 | 115228215 | rs7836915 | 1.698 | 0.073 | 0.264 | -1.904 | 0 | 0.003 | 0.003 | 0 | 0.07 | 0.09 | 0.09 | 0.06 | 0.06 | 0.03 |
| 8 | 115229759 | rs16885580 | 1.289 | 0.013 | 0.308 | -1.927 | -0.008 | -0.037 | -0.037 | 0.059 | 0.09 | 0.13 | 0.04 | 0.02 | 0.08 | 0.05 |
| 8 | 115235429 | rs12680086 | 0.292 | 0.173 | 0.292 | -1.824 | 0.005 | 0.002 | 0.002 | 0 | 0.39 | 0.35 | 0.37 | 0.65 | 0.27 | 0.48 |
| 8 | 115237434 | rs7007631 | 1.694 | 0.2 | 0.183 | -1.808 | -0.001 | 0.002 | 0.002 | 0 | 0.08 | 0.05 | 0.06 | 0.02 | 0.09 | 0.08 |
| 8 | 115242543 | rs17632129 | 1.755 | 0.2 | 0.173 | -1.808 | -0.003 | 0.011 | 0.011 | 0 | 0.08 | 0.03 | 0.06 | 0.02 | 0.09 | 0.08 |
| 8 | 115247555 | rs1515687 | 0.298 | 0.141 | 0.27 | -1.802 | 0.02 | 0.005 | 0.005 | 0 | 0.39 | 0.3 | 0.37 | 0.65 | 0.27 | 0.48 |
| 8 | 115261134 | rs2176118 | 0.375 | 0.388 | 0.482 | -1.777 | -0.002 | -0.05 | -0.05 | 1.504 | 0.15 | 0.21 | 0.09 | 0.01 | 0.13 | 0.08 |
| 8 | 115262197 | rs16885615 | 0.354 | 0.119 | 0.343 | -1.927 | 0.051 | 0.03 | 0.03 | 0 | 0.46 | 0.29 | 0.4 | 0.66 | 0.3 | 0.5 |
| 8 | 115268116 | rs4876562 | 1.313 | 0.147 | 0.295 | -1.798 | 0 | -0.005 | -0.005 | 0 | 0.08 | 0.09 | 0.04 | 0 | 0.05 | 0.02 |
| 8 | 115285808 | rs1022314 | 1.207 | 0.142 | 0.412 | -1.8 | 0.002 | -0.037 | -0.037 | 1.509 | 0.08 | 0.16 | 0.05 | 0 | 0.05 | 0.02 |
| 8 | 115291932 | rs17706527 | 1.963 | 0.044 | 0.424 | -1.73 | 0 | 0.006 | 0.006 | 0 | 0.07 | 0.05 | 0.04 | 0.01 | 0.07 | 0.06 |
| 8 | 115320834 | rs10808467 | -0.764 | 0.324 | 0.089 | -1.872 | -0.106 | 0.118 | 0.118 | 0.254 | 0.61 | 0.89 | 0.62 | 0.83 | 0.47 | 0.6 |
| 8 | 115321191 | rs12677904 | 1.739 | -0.144 | -0.165 | -2.042 | 0.038 | -0.097 | -0.097 | 0 | 0.25 | 0.05 | 0.38 | 0.58 | 0.17 | 0.26 |
| 8 | 115321807 | rs6997580 | -0.76 | 0.231 | 0.008 | -1.857 | -0.07 | 0.111 | 0.111 | 0.123 | 0.61 | 0.84 | 0.61 | 0.83 | 0.47 | 0.6 |
| 8 | 115342194 | rs11991902 | -0.304 | 0.224 | 0.048 | -1.847 | 0.031 | 0.152 | 0.152 | 0 | 0.52 | 0.74 | 0.56 | 0.84 | 0.38 | 0.55 |

| | | | | | | | | | | | | | | | | |
|---|---|---|---|---|---|---|---|---|---|---|---|---|---|---|---|---|
| 8 | 115345232 | rs10113591 | -0.445 | 0.266 | 0.006 | -1.837 | 0.019 | 0.179 | 0.179 | 0 | 0.54 | 0.79 | 0.58 | 0.84 | 0.4 | 0.55 |
| 8 | 115351857 | rs1497194 | -0.75 | 0.205 | -0.138 | -1.824 | -0.084 | 0.153 | 0.153 | 0.218 | 0.61 | 0.87 | 0.62 | 0.86 | 0.47 | 0.6 |
| 8 | 115360395 | rs971445 | -0.253 | 0.078 | 0.057 | -1.834 | 0.075 | 0.049 | 0.049 | 0.043 | 0.52 | 0.7 | 0.52 | 0.66 | 0.3 | 0.51 |
| 8 | 115396892 | rs182353 | -0.747 | 0.145 | 0.004 | -1.914 | -0.066 | 0.137 | 0.137 | 0.008 | 0.61 | 0.83 | 0.61 | 0.86 | 0.47 | 0.6 |
| 8 | 115404627 | rs10808469 | 0.849 | 0.321 | -0.016 | -1.75 | 0.024 | 0.015 | 0.015 | 0 | 0.1 | 0.12 | 0.09 | 0.18 | 0.19 | 0.09 |
| 8 | 115445405 | rs11787235 | 1.146 | 0.329 | -0.06 | -1.739 | -0.019 | -0.019 | -0.019 | 0 | 0.09 | 0.06 | 0.08 | 0.18 | 0.17 | 0.09 |
| 8 | 115447503 | rs290631 | 0.28 | -0.005 | 0.232 | -1.854 | 0.065 | 0.044 | 0.044 | 0 | 0.46 | 0.26 | 0.48 | 0.67 | 0.28 | 0.51 |
| 8 | 115455967 | rs2590903 | -0.389 | -0.435 | 0.264 | -1.997 | -0.013 | -0.018 | -0.018 | 0 | 0.59 | 0.46 | 0.54 | 0.73 | 0.35 | 0.61 |
| 8 | 115458963 | rs13257128 | 1.11 | -0.237 | -0.244 | -2.045 | -0.081 | 0.099 | 0.099 | 0.014 | 0.29 | 0.14 | 0.33 | 0.09 | 0.46 | 0.29 |
| 8 | 115461491 | rs1373897 | 1.752 | -2.305 | -0.609 | -1.97 | 0.059 | -0.114 | -0.114 | 0.081 | 0.26 | 0.03 | 0.34 | 0.6 | 0.15 | 0.26 |
| 8 | 115464957 | rs16885867 | 0.274 | -2.044 | -0.276 | -1.977 | -0.009 | 0 | 0 | 0 | 0.07 | 0.08 | 0.04 | 0.07 | 0.02 | 0.07 |
| 8 | 115465477 | rs11996955 | -0.38 | -1.819 | -0.337 | -1.884 | 0.023 | 0.05 | 0.05 | 0 | 0.79 | 0.9 | 0.91 | 0.94 | 0.85 | 0.72 |
| 8 | 115474758 | rs7014060 | 0.858 | -1.406 | -0.119 | -1.824 | -0.022 | 0.025 | 0.025 | 0.018 | 0.31 | 0.25 | 0.35 | 0.09 | 0.52 | 0.35 |
| 8 | 115475287 | rs4242562 | 1.096 | -1.516 | -0.364 | -1.156 | 0.008 | -0.005 | -0.005 | 0 | 0.14 | 0.11 | 0.13 | 0.21 | 0.08 | 0.2 |
| 8 | 115477600 | rs6991398 | 0.047 | -1.829 | -0.458 | -1.221 | -0.011 | -0.034 | -0.034 | 3.163 | 0.76 | 0.86 | 0.66 | 0.36 | 0.77 | 0.67 |
| 8 | 115492603 | rs4354332 | -0.887 | -1.85 | -0.448 | -1.059 | -0.01 | -0.004 | -0.004 | 0 | 0.84 | 0.89 | 0.89 | 0.82 | 0.83 | 0.89 |
| 8 | 115498999 | rs6469525 | 0.576 | -2.108 | -0.093 | -1.27 | 0 | 0.001 | 0.001 | 1.507 | 0.39 | 0.36 | 0.24 | 0.27 | 0.25 | 0.31 |
| 8 | 115501579 | rs13276268 | 1.296 | -2.082 | 0.032 | -1.279 | 0.007 | -0.027 | -0.027 | 0.101 | 0.32 | 0.33 | 0.36 | 0.09 | 0.53 | 0.35 |
| 8 | 115504434 | rs7836298 | 1.318 | -2.181 | 0.123 | -1.283 | -0.009 | 0.004 | 0.004 | 0 | 0.14 | 0.16 | 0.14 | 0.21 | 0.1 | 0.2 |
| 8 | 115506170 | rs1822958 | -0.009 | -2.232 | 0.091 | -1.318 | -0.005 | -0.035 | -0.035 | 0 | 0.76 | 0.86 | 0.67 | 0.36 | 0.79 | 0.67 |
| 8 | 115506842 | rs10955711 | 0.285 | -2.406 | 0.329 | -1.19 | -0.024 | -0.019 | -0.019 | 0 | 0.43 | 0.44 | 0.3 | 0.27 | 0.26 | 0.32 |
| 8 | 115509819 | rs4876329 | 1.285 | -2.754 | 0.476 | -0.677 | 0.016 | -0.012 | -0.012 | 1.623 | 0.14 | 0.07 | 0.12 | 0.21 | 0.1 | 0.2 |
| 8 | 115511772 | rs16886032 | 0.064 | -2.7 | 0.369 | -0.653 | 0.005 | -0.026 | -0.026 | 2.129 | 0.18 | 0.14 | 0.3 | 0.63 | 0.13 | 0.29 |
| 8 | 115532721 | rs1390808 | 1.732 | -2.775 | 0.447 | -0.659 | 0.048 | 0.079 | 0.079 | 3.601 | 0.24 | 0.08 | 0.1 | 0.07 | 0.15 | 0.11 |
| 8 | 115543198 | rs16886058 | 1.145 | -2.765 | 0.407 | -0.673 | -0.003 | 0.001 | 0.001 | 0 | 0.16 | 0.16 | 0.16 | 0.21 | 0.12 | 0.2 |
| 8 | 115543245 | rs10505222 | 1.721 | -2.805 | 0.41 | -0.697 | 0.011 | 0.007 | 0.007 | 2.505 | 0.25 | 0.19 | 0.1 | 0.06 | 0.15 | 0.11 |
| 8 | 115546011 | rs10505223 | 0.304 | -2.812 | 0.32 | -0.695 | 0.011 | -0.023 | -0.023 | 0 | 0.17 | 0.13 | 0.28 | 0.64 | 0.11 | 0.29 |
| 8 | 115548047 | rs7825159 | -1.454 | -2.779 | 0.307 | -0.691 | -0.014 | 0.133 | 0.133 | 0.826 | 0.64 | 0.85 | 0.62 | 0.91 | 0.39 | 0.61 |
| 8 | 115553747 | rs16886089 | 1.101 | -2.676 | 0.061 | -0.528 | 0.008 | -0.011 | -0.011 | 0 | 0.08 | 0 | 0.22 | 0.46 | 0.04 | 0.22 |
| 8 | 115562322 | rs10108622 | 0.137 | -2.633 | 0.274 | -0.656 | -0.038 | -0.033 | -0.033 | 0 | 0.39 | 0.48 | 0.21 | 0.24 | 0.24 | 0.19 |
| 8 | 115571779 | rs12676786 | -1.118 | -1.971 | 0.122 | -0.324 | -0.012 | -0.02 | -0.02 | 0.104 | 0.37 | 0.4 | 0.5 | 0.85 | 0.24 | 0.5 |
| 8 | 115578307 | rs7014900 | -0.546 | -1.796 | 0.157 | -0.202 | -0.034 | 0.011 | 0.011 | 0 | 0.3 | 0.43 | 0.27 | 0.38 | 0.2 | 0.28 |
| 8 | 115589625 | rs11987649 | 0.147 | -2.138 | 0.175 | -0.419 | -0.026 | -0.025 | -0.025 | 1.786 | 0.39 | 0.52 | 0.28 | 0.24 | 0.24 | 0.25 |
| 8 | 115603659 | rs7000026 | -2.113 | -2.022 | -0.187 | -0.645 | -0.034 | 0.049 | 0.049 | 0 | 0.91 | 0.99 | 0.92 | 1 | 0.87 | 0.96 |
| 8 | 115611596 | rs7816178 | -0.098 | -2.264 | 0.53 | -0.188 | -0.015 | -0.014 | -0.014 | 0 | 0.39 | 0.41 | 0.27 | 0.24 | 0.24 | 0.25 |
| 8 | 115620846 | rs990064 | 1.727 | -2.301 | 0.023 | -0.266 | 0.024 | 0.034 | 0.034 | 1.285 | 0.21 | 0.14 | 0.09 | 0.06 | 0.13 | 0.09 |
| 8 | 115626102 | rs10955722 | -0.828 | -2.27 | -0.029 | -0.131 | 0.091 | 0.105 | 0.105 | 0.002 | 0.52 | 0.76 | 0.57 | 0.74 | 0.31 | 0.49 |
| 8 | 115650409 | rs6469536 | -0.942 | -2.134 | -0.134 | -0.012 | 0.083 | 0.043 | 0.043 | 0.109 | 0.53 | 0.8 | 0.37 | 0.4 | 0.33 | 0.37 |
| 8 | 115674664 | rs6992289 | 0.177 | -2.288 | 0.166 | -0.047 | 0.024 | 0.018 | 0.018 | 1.605 | 0.33 | 0.24 | 0.13 | 0.23 | 0.19 | 0.19 |
| 8 | 115683736 | rs6985846 | -0.197 | -2.19 | 0.098 | 0.005 | 0.013 | 0.012 | 0.012 | 1.117 | 0.38 | 0.65 | 0.21 | 0.23 | 0.21 | 0.21 |
| 8 | 115688964 | rs4610784 | -0.187 | -2.189 | 0.12 | 0.005 | 0.009 | 0.009 | 0.009 | 1.097 | 0.38 | 0.63 | 0.21 | 0.23 | 0.21 | 0.21 |
| 8 | 115699735 | rs1454796 | -0.348 | -2.258 | 0.079 | -0.007 | -0.008 | 0.003 | 0.003 | 0 | 0.26 | 0.24 | 0.21 | 0.33 | 0.2 | 0.28 |
| 8 | 115702679 | rs10091896 | 0.872 | -2.271 | 0.076 | 0.742 | -0.008 | 0.086 | 0.086 | 0 | 0.11 | 0.1 | 0.09 | 0.42 | 0.07 | 0.11 |
| 8 | 115711436 | rs16886291 | 0.916 | -2.26 | 0.094 | -0.202 | 0.006 | -0.015 | -0.015 | 0.043 | 0.15 | 0.05 | 0.15 | 0.17 | 0.12 | 0.17 |
| 8 | 115712241 | rs7827613 | 1.163 | -2.27 | 0.108 | -0.215 | -0.069 | -0.104 | -0.104 | 0.005 | 0.26 | 0.52 | 0.16 | 0.07 | 0.14 | 0.1 |
| 8 | 115720329 | rs9297531 | -0.301 | -2.186 | -0.084 | -0.169 | 0.014 | -0.011 | -0.011 | 1.324 | 0.26 | 0.14 | 0.2 | 0.33 | 0.2 | 0.28 |
| 8 | 115721304 | rs1378177 | 0.844 | -1.985 | 0.173 | -0.06 | -0.008 | 0.008 | 0.008 | 0 | 0.1 | 0.09 | 0.05 | 0.16 | 0.07 | 0.11 |
| 8 | 115726321 | rs7006212 | -0.194 | -1.813 | -0.149 | 0.011 | 0.005 | 0.005 | 0.005 | 1.094 | 0.38 | 0.62 | 0.21 | 0.23 | 0.21 | 0.2 |
| 8 | 115732048 | rs16886335 | 0.659 | -1.444 | -0.509 | 0.476 | 0.004 | 0.011 | 0.011 | 0 | 0.07 | 0 | 0.24 | 0.51 | 0.05 | 0.23 |
| 8 | 115732418 | rs7839457 | -0.138 | -2.084 | 0.105 | 0.07 | -0.031 | 0.005 | 0.005 | 0.098 | 0.36 | 0.45 | 0.4 | 0.58 | 0.19 | 0.33 |
| 8 | 115735288 | rs16886337 | -0.214 | -2.281 | 0.008 | -0.209 | -0.032 | 0.022 | 0.022 | 0 | 0.26 | 0.4 | 0.22 | 0.33 | 0.19 | 0.27 |
| 8 | 115738188 | rs16886345 | 0.398 | -1.156 | 0.516 | 0.269 | 0.004 | 0.073 | 0.073 | 0 | 0.07 | 0.05 | 0.25 | 0.51 | 0.05 | 0.23 |
| 8 | 115741063 | rs6981925 | -0.705 | -1.199 | 0.555 | 0.193 | 0.013 | 0.013 | 0.013 | 0 | 0.5 | 0.49 | 0.54 | 0.75 | 0.3 | 0.49 |
| 8 | 115745774 | rs7842799 | -0.678 | -1.2 | 0.554 | 0.192 | 0.012 | 0.012 | 0.012 | 0 | 0.49 | 0.49 | 0.54 | 0.75 | 0.3 | 0.49 |
| 8 | 115760498 | rs6989633 | -0.485 | -1.316 | 1.082 | 0.124 | 0.016 | 0.048 | 0.048 | 0.373 | 0.1 | 0.42 | 0.12 | 0.08 | 0.07 | 0.07 |
| 8 | 115765911 | rs7823356 | -1.142 | -1.414 | 1.185 | 0.173 | -0.033 | 0.244 | 0.244 | 1.502 | 0.13 | 0.36 | 0.36 | 0.54 | 0.07 | 0.25 |
| 8 | 115769026 | rs10090535 | -0.992 | -1.503 | 1.165 | 0.086 | 0.094 | 0.116 | 0.116 | 0 | 0.5 | 0.72 | 0.61 | 0.77 | 0.29 | 0.51 |
| 8 | 115769507 | rs1454781 | 0.281 | -1.444 | 1.03 | 0.12 | 0.007 | 0.006 | 0.006 | 0.39 | 0.36 | 0.37 | 0.25 | 0.24 | 0.22 | 0.26 |
| 8 | 115774366 | rs10104793 | 0.007 | -1.341 | 0.804 | 0.235 | -0.014 | 0.136 | 0.136 | 0 | 0.09 | 0.15 | 0.31 | 0.54 | 0.05 | 0.22 |
| 8 | 115785829 | rs4332167 | 0.903 | -1.345 | 0.528 | -0.036 | 0.071 | 0.087 | 0.087 | 2.599 | 0.29 | 0.06 | 0.17 | 0.15 | 0.17 | 0.2 |
| 8 | 115801693 | rs16886401 | 0.283 | -1.409 | 0.981 | -0.045 | -0.018 | 0.02 | 0.02 | 0 | 0.06 | 0.07 | 0.03 | 0.15 | 0.02 | 0.07 |
| 8 | 115810238 | rs1470968 | -1.138 | -0.806 | 1.062 | -0.116 | 0.004 | 0.004 | 0.004 | 0 | 0.57 | 0.59 | 0.6 | 0.86 | 0.38 | 0.57 |
| 8 | 115810385 | rs1470969 | -0.866 | -0.826 | 1.154 | -0.126 | 0.031 | 0.097 | 0.097 | 0 | 0.53 | 0.3 | 0.57 | 0.86 | 0.38 | 0.57 |
| 8 | 115821293 | rs13252246 | 1.439 | -0.921 | 0.81 | -0.209 | -0.023 | 0.022 | 0.022 | 0 | 0.08 | 0.02 | 0.04 | 0.02 | 0.11 | 0.06 |
| 8 | 115821351 | rs2357704 | -0.506 | -0.927 | 0.821 | -0.142 | 0.074 | 0.113 | 0.113 | 0 | 0.48 | 0.21 | 0.5 | 0.78 | 0.33 | 0.51 |
| 8 | 115839310 | rs10505228 | 1.404 | -0.79 | 0.058 | -0.117 | 0.055 | 0.052 | 0.052 | 3.805 | 0.24 | 0.02 | 0.16 | 0.13 | 0.14 | 0.15 |
| 8 | 115839385 | rs10505229 | -1.144 | -0.808 | 0.324 | -0.117 | -0.007 | 0.005 | 0.005 | 0 | 0.23 | 0.6 | 0.17 | 0.25 | 0.18 | 0.19 |
| 8 | 115843061 | rs9297534 | -0.654 | -0.782 | 0.342 | -0.103 | 0.037 | 0.027 | 0.027 | 0 | 0.48 | 0.62 | 0.33 | 0.38 | 0.32 | 0.34 |
| 8 | 115850338 | rs4242563 | -0.449 | -0.779 | 0.387 | -0.129 | -0.008 | -0.006 | -0.006 | 0 | 0.46 | 0.53 | 0.32 | 0.38 | 0.32 | 0.34 |

| | | | | | | | | | | | | | | | | |
|---|---|---|---|---|---|---|---|---|---|---|---|---|---|---|---|---|
| 8 | 115851352 | rs7004742 | -0.435 | -0.775 | 0.39 | -0.131 | -0.009 | -0.008 | -0.008 | 1.263 | 0.46 | 0.53 | 0.32 | 0.36 | 0.32 | 0.34 |
| 8 | 115853096 | rs6986502 | 0.013 | -0.637 | 0.322 | -0.368 | -0.007 | -0.019 | -0.019 | 0 | 0.39 | 0.46 | 0.29 | 0.23 | 0.3 | 0.26 |
| 8 | 115858519 | rs7842690 | -0.586 | 0.209 | 0.68 | -0.284 | 0.041 | -0.04 | -0.04 | 0 | 0.17 | 0.54 | 0.14 | 0.1 | 0.16 | 0.11 |
| 8 | 115863500 | rs10093354 | -0.469 | 0.798 | 0.193 | 0.038 | -0.043 | 0.07 | 0.07 | 0 | 0.62 | 0.84 | 0.7 | 0.81 | 0.51 | 0.61 |
| 8 | 115866116 | rs2884258 | -2.239 | 1.288 | 1.356 | 0.841 | 0 | 0.007 | 0.007 | 1.968 | 0.2 | 0.76 | 0.19 | 0.27 | 0.21 | 0.25 |
| 8 | 115878730 | rs7820058 | -1.434 | 1.024 | 1.404 | 0.749 | -0.006 | -0.003 | -0.003 | 0 | 0.13 | 0.08 | 0.05 | 0.12 | 0.11 | 0.17 |
| 8 | 115884048 | rs11988351 | -3.137 | 1.387 | 1.767 | 0.66 | -0.032 | -0.005 | -0.005 | 0.411 | 0.35 | 0.78 | 0.32 | 0.3 | 0.36 | 0.38 |
| 8 | 115892114 | rs10097028 | -2.101 | 1.125 | 2.431 | 0.648 | 0.036 | 0.014 | 0.014 | 0 | 0.21 | 0.38 | 0.23 | 0.17 | 0.24 | 0.21 |
| 8 | 115908822 | rs10099099 | -0.496 | 1.139 | 2.365 | 0.635 | -0.004 | 0.009 | 0.009 | 0 | 0.07 | 0.06 | 0.12 | 0.01 | 0.15 | 0.04 |
| 8 | 115909561 | rs13438839 | -1.882 | 1.092 | 2.72 | 0.627 | 0.086 | 0.072 | 0.072 | 0 | 0.16 | 0.43 | 0.19 | 0.17 | 0.19 | 0.1 |
| 8 | 115913210 | rs7828196 | -0.542 | 1.184 | 2.518 | 0.571 | -0.009 | -0.029 | -0.029 | 0.101 | 0.05 | 0.1 | 0.05 | 0 | 0.04 | 0.11 |
| 8 | 115923992 | rs11989244 | -1.921 | 1.351 | 2.68 | 0.752 | 0.101 | 0.049 | 0.049 | 0 | 0.16 | 0.35 | 0.2 | 0.16 | 0.23 | 0.12 |
| 8 | 115929098 | rs16886637 | -1.806 | 1.025 | 2.757 | 0.748 | -0.028 | 0.046 | 0.046 | 1.162 | 0.13 | 0.49 | 0.06 | 0.12 | 0.05 | 0.14 |
| 8 | 115934365 | rs4515577 | -1.438 | 1.029 | 2.703 | 0.75 | -0.014 | 0.055 | 0.055 | 0.032 | 0.11 | 0.41 | 0.05 | 0.12 | 0.05 | 0.14 |
| 8 | 115963398 | rs10091791 | -0.881 | 1.099 | 2.695 | 0.805 | 0.003 | 0.027 | 0.027 | 0 | 0.09 | 0.13 | 0.08 | 0.16 | 0.09 | 0.08 |
| 8 | 115964638 | rs1872780 | -2.452 | 1.06 | 2.827 | 0.805 | -0.015 | 0.093 | 0.093 | 0 | 0.2 | 0.46 | 0.12 | 0.28 | 0.14 | 0.23 |
| 8 | 115971723 | rs16886751 | -0.954 | 1.075 | 2.624 | 0.797 | -0.009 | 0.007 | 0.007 | 0 | 0.09 | 0.15 | 0.03 | 0.12 | 0.05 | 0.14 |
| 8 | 115976488 | rs16886754 | -0.896 | 1.075 | 2.579 | 0.796 | 0.008 | -0.008 | -0.008 | 0 | 0.09 | 0.07 | 0.02 | 0.12 | 0.05 | 0.15 |
| 8 | 115983884 | rs673380 | -2.08 | 1.089 | 2.709 | 0.814 | -0.006 | 0.047 | 0.047 | 0 | 0.18 | 0.31 | 0.11 | 0.28 | 0.14 | 0.23 |
| 8 | 115986109 | rs666095 | -2.929 | 1.26 | 2.854 | 0.777 | 0.054 | 0.054 | 0.054 | 0 | 0.25 | 0.52 | 0.23 | 0.29 | 0.28 | 0.27 |
| 8 | 115988349 | rs10505243 | -1.894 | 1.189 | 2.936 | 0.756 | 0.076 | 0.012 | 0.012 | 0 | 0.15 | 0.4 | 0.16 | 0.13 | 0.19 | 0.18 |
| 8 | 115989490 | rs2631813 | -2.714 | 1.188 | 2.754 | 0.768 | 0.033 | 0.037 | 0.037 | 0 | 0.25 | 0.62 | 0.25 | 0.29 | 0.28 | 0.27 |
| 8 | 115996511 | rs17733014 | -0.637 | 1.236 | 2.279 | 0.78 | 0.071 | -0.017 | -0.017 | 0 | 0.06 | 0.16 | 0.13 | 0.01 | 0.14 | 0.04 |
| 8 | 115999185 | rs11994126 | 0.058 | 1.013 | 1.788 | 1.132 | -0.008 | 0.124 | 0.124 | 0.084 | 0.08 | 0.13 | 0.33 | 0.48 | 0.06 | 0.16 |
| 8 | 116003403 | rs12546832 | 0.058 | 1.013 | 1.788 | 1.132 | -0.008 | 0.124 | 0.124 | 0.084 | 0.08 | 0.13 | 0.33 | 0.48 | 0.06 | 0.16 |
| 8 | 116009906 | rs1684978 | -0.894 | 1.003 | 1.491 | 0.677 | 0.005 | 0.046 | 0.046 | 0 | 0.09 | 0.14 | 0.08 | 0.16 | 0.08 | 0.08 |
| 8 | 116027062 | rs11784785 | -1.401 | 1.084 | 1.548 | 0.673 | 0.063 | -0.001 | -0.001 | 0 | 0.17 | 0.41 | 0.16 | 0.13 | 0.18 | 0.08 |
| 8 | 116027759 | rs4876590 | 0.459 | 1.001 | 0.414 | 0.709 | 0.008 | 0.05 | 0.05 | 0 | 0.92 | 0.88 | 0.91 | 0.84 | 0.92 | 0.91 |
| 8 | 116031304 | rs10955738 | 1.746 | 0.777 | -0.297 | 0.57 | -0.12 | -0.072 | -0.072 | 0.115 | 0.62 | 0.04 | 0.65 | 0.68 | 0.62 | 0.55 |
| 8 | 116046422 | rs7012500 | -1.88 | 0.007 | -0.047 | 0.754 | 0.002 | -0.034 | -0.034 | 0 | 0.38 | 0.91 | 0.27 | 0.33 | 0.31 | 0.46 |
| 8 | 116053820 | rs10464853 | -0.45 | -0.63 | 0.027 | 1.019 | 0.003 | -0.005 | -0.005 | 0 | 0.14 | 0.03 | 0.06 | 0.12 | 0.11 | 0.18 |
| 8 | 116054292 | rs7838203 | -0.669 | -0.606 | 0.358 | 0.933 | -0.028 | -0.04 | -0.04 | 0 | 0.08 | 0.32 | 0.03 | 0.01 | 0.02 | 0.04 |
| 8 | 116054890 | rs6993473 | -1.752 | -0.489 | -0.537 | 1.191 | 0.064 | 0.079 | 0.079 | 1.447 | 0.39 | 0.76 | 0.49 | 0.81 | 0.24 | 0.47 |
| 8 | 116081696 | rs2036014 | -0.714 | 0.387 | -0.405 | 2.007 | -0.004 | -0.032 | -0.032 | 0 | 0.09 | 0 | 0.3 | 0.48 | 0.08 | 0.18 |
| 8 | 116095815 | rs7013769 | -1.154 | 0.575 | 0.437 | 1.584 | 0.051 | 0.193 | 0.193 | 0 | 0.08 | 0.29 | 0.1 | 0.19 | 0.07 | 0.08 |
| 8 | 116110725 | rs218361 | 0.31 | 0.018 | -0.168 | 1.478 | 0.012 | 0.004 | 0.004 | 0.506 | 0.55 | 0.27 | 0.6 | 0.57 | 0.58 | 0.4 |
| 8 | 116112960 | rs16886947 | -0.637 | -0.072 | 0.393 | 1.419 | -0.021 | 0.13 | 0.13 | 1.575 | 0.07 | 0.18 | 0.06 | 0.26 | 0.03 | 0.13 |
| 8 | 116116608 | rs309610 | -0.546 | -0.228 | 0.121 | 1.215 | -0.075 | 0.154 | 0.154 | 0 | 0.74 | 0.97 | 0.81 | 0.98 | 0.68 | 0.71 |
| 8 | 116118689 | rs309607 | 0.356 | -0.188 | 0.386 | 1.226 | 0.002 | 0.005 | 0.005 | 0.062 | 0.62 | 0.37 | 0.7 | 0.72 | 0.63 | 0.49 |
| 8 | 116129344 | rs309605 | 0.287 | -0.149 | 0.208 | 1.213 | 0 | 0.001 | 0.001 | 0 | 0.62 | 0.39 | 0.7 | 0.74 | 0.64 | 0.49 |
| 8 | 116146327 | rs2132025 | -0.361 | -0.079 | -0.264 | 1.173 | 0.002 | 0.015 | 0.015 | 0 | 0.37 | 0.68 | 0.31 | 0.28 | 0.35 | 0.51 |
| 8 | 116157283 | rs4559271 | -0.773 | -0.033 | -1.467 | 1.159 | -0.15 | -0.132 | -0.132 | 0.576 | 0.68 | 0.98 | 0.55 | 0.61 | 0.6 | 0.72 |
| 8 | 116161903 | rs1494754 | 0.301 | -0.495 | -0.726 | 1.21 | 0 | 0.008 | 0.008 | 0 | 0.44 | 0.41 | 0.48 | 0.34 | 0.42 | 0.33 |
| 8 | 116162223 | rs1389913 | 0.349 | -0.52 | -0.889 | 1.873 | -0.02 | -0.011 | -0.011 | 0 | 0.06 | 0.01 | 0.05 | 0.25 | 0.08 | 0.1 |
| 8 | 116168410 | rs7016195 | 0.173 | -0.932 | -0.458 | 1.4 | -0.018 | 0.013 | 0.013 | 0 | 0.88 | 0.86 | 0.89 | 0.79 | 0.95 | 0.85 |
| 8 | 116170577 | rs7836816 | 0.727 | -1.127 | -0.667 | 1.297 | -0.103 | -0.04 | -0.04 | 0 | 0.45 | 0.05 | 0.45 | 0.58 | 0.49 | 0.42 |
| 8 | 116173246 | rs16887070 | -0.568 | -1.075 | -0.016 | 1.195 | -0.011 | 0.015 | 0.015 | 0.476 | 0.05 | 0.25 | 0.04 | 0.04 | 0 | 0.17 |
| 8 | 116176124 | rs777801 | 1.178 | -1.085 | 0.043 | 1.188 | -0.076 | 0.053 | 0.053 | 0 | 0.82 | 0.51 | 0.84 | 0.75 | 0.94 | 0.68 |
| 8 | 116182549 | rs809001 | 0.663 | -0.998 | 0.262 | 1.028 | -0.026 | 0.029 | 0.029 | 0 | 0.31 | 0.23 | 0.3 | 0.1 | 0.44 | 0.23 |
| 8 | 116188663 | rs1519084 | 0.612 | -1.036 | 0.334 | 1.138 | 0.001 | -0.012 | -0.012 | 0.008 | 0.32 | 0.32 | 0.32 | 0.1 | 0.45 | 0.24 |
| 8 | 116188975 | rs1546139 | 0.414 | -1.362 | 0.173 | 1.422 | -0.002 | -0.001 | -0.001 | 0.57 | 0.88 | 0.89 | 0.89 | 0.79 | 0.95 | 0.85 |
| 8 | 116210888 | rs777814 | 1.168 | -1.615 | 0.451 | 1.802 | 0.011 | -0.03 | -0.03 | 0 | 0.83 | 0.91 | 0.82 | 0.59 | 0.86 | 0.78 |
| 8 | 116210958 | rs777815 | 1.072 | -1.436 | 0.255 | 1.224 | -0.091 | 0.071 | 0.071 | 0 | 0.27 | 0.11 | 0.27 | 0.09 | 0.4 | 0.24 |
| 8 | 116211228 | rs16887143 | -0.34 | -1.555 | 0.901 | 1.257 | -0.018 | -0.003 | -0.003 | 0 | 0.1 | 0.33 | 0.12 | 0.06 | 0.03 | 0.05 |
| 8 | 116222460 | rs777825 | 0.738 | -1.454 | 0.486 | 1.44 | -0.008 | 0 | 0 | 0.343 | 0.9 | 0.91 | 0.92 | 0.86 | 0.94 | 0.89 |
| 8 | 116232225 | rs13277974 | 0.402 | -1.456 | 0.714 | 1.392 | -0.014 | -0.041 | -0.041 | 0.091 | 0.07 | 0.18 | 0.05 | 0 | 0.03 | 0.03 |
| 8 | 116232937 | rs777831 | -1.14 | -1.49 | 0.564 | 1.422 | -0.021 | 0.009 | 0.009 | 0 | 0.1 | 0.13 | 0.08 | 0.18 | 0.05 | 0.23 |
| 8 | 116235058 | rs777835 | -1.021 | -1.557 | 0.529 | 1.313 | -0.002 | -0.005 | -0.005 | 1.708 | 0.71 | 0.67 | 0.7 | 0.91 | 0.58 | 0.74 |
| 8 | 116236561 | rs13248611 | 1.144 | -1.466 | -0.115 | 1.448 | -0.047 | -0.021 | -0.021 | 0 | 0.39 | 0.05 | 0.36 | 0.39 | 0.4 | 0.31 |
| 8 | 116244283 | rs2634839 | -1.572 | -1.371 | 0.808 | 2.164 | -0.028 | 0.08 | 0.08 | 0 | 0.11 | 0.18 | 0.08 | 0.35 | 0.05 | 0.21 |
| 8 | 116246667 | rs16887197 | -0.308 | -1.183 | 0.875 | 2.082 | 0 | 0.075 | 0.075 | 0 | 0.08 | 0.15 | 0.14 | 0.27 | 0.08 | 0.1 |
| 8 | 116248980 | rs2680386 | -1.028 | -1.197 | 0.647 | 1.406 | -0.009 | 0.078 | 0.078 | 0 | 0.07 | 0.16 | 0.04 | 0.21 | 0.03 | 0.18 |
| 8 | 116253370 | rs16887231 | -1.443 | -1.319 | 0.825 | 1.322 | -0.066 | 0.058 | 0.058 | 0 | 0.19 | 0.43 | 0.24 | 0.21 | 0.08 | 0.33 |
| 8 | 116253683 | rs1589196 | 0.988 | -1.188 | 0.448 | 1.115 | -0.002 | 0.001 | 0.001 | 0 | 0.31 | 0.31 | 0.25 | 0.28 | 0.38 | 0.21 |
| 8 | 116255153 | rs11996430 | -0.595 | -1.231 | 0.356 | 1.145 | -0.048 | -0.032 | -0.032 | 0 | 0.52 | 0.88 | 0.51 | 0.48 | 0.46 | 0.54 |
| 8 | 116259336 | rs4242564 | -1.072 | -1.529 | 0.145 | 1.11 | -0.126 | -0.096 | -0.096 | 0 | 0.62 | 0.95 | 0.58 | 0.61 | 0.53 | 0.62 |
| 8 | 116262065 | rs17657793 | 0.394 | -1.445 | 0.629 | 1.169 | -0.006 | 0.011 | 0.011 | 0 | 0.41 | 0.73 | 0.41 | 0.39 | 0.4 | 0.35 |
| 8 | 116270215 | rs7005063 | 0.52 | -1.549 | 0.841 | 1.159 | -0.002 | 0.001 | 0.001 | 0 | 0.4 | 0.61 | 0.41 | 0.39 | 0.41 | 0.35 |
| 8 | 116270987 | rs12235003 | 0.094 | -1.552 | 0.788 | 1.175 | -0.018 | 0.077 | 0.077 | 0 | 0.06 | 0.18 | 0.15 | 0.11 | 0.03 | 0.11 |

| | | | | | | | | | | | | | | | | |
|---|---|---|---|---|---|---|---|---|---|---|---|---|---|---|---|---|
| 8 | 116276618 | rs2033995 | -0.676 | -1.613 | 0.698 | 1.166 | -0.088 | -0.044 | -0.044 | 0 | 0.55 | 0.91 | 0.56 | 0.6 | 0.48 | 0.56 |
| 8 | 116281087 | rs10096834 | 0.959 | -1.459 | 0.809 | 1.147 | 0.011 | -0.016 | -0.016 | 0 | 0.34 | 0.42 | 0.27 | 0.28 | 0.39 | 0.24 |
| 8 | 116281423 | rs17714308 | -0.037 | -1.45 | 0.62 | 1.166 | -0.004 | -0.012 | -0.012 | 0 | 0.46 | 0.79 | 0.48 | 0.48 | 0.43 | 0.47 |
| 8 | 116285104 | rs10113076 | 0.526 | -1.341 | 0.659 | 1.155 | 0.009 | -0.023 | -0.023 | 0 | 0.39 | 0.55 | 0.3 | 0.28 | 0.43 | 0.29 |
| 8 | 116295809 | rs2946337 | 0.729 | -1.313 | 0.264 | 0.977 | -0.041 | 0.004 | 0.004 | 0.118 | 0.24 | 0.04 | 0.21 | 0.19 | 0.27 | 0.18 |
| 8 | 116298537 | rs16887279 | -0.485 | -1.267 | 1.361 | 1.14 | -0.012 | 0.034 | 0.034 | 0 | 0.05 | 0.15 | 0.07 | 0.08 | 0.01 | 0.12 |
| 8 | 116299406 | rs7010965 | -2.353 | -1.444 | 1.176 | 1.17 | 0 | -0.006 | -0.006 | 0 | 0.23 | 0.8 | 0.31 | 0.29 | 0.08 | 0.3 |
| 8 | 116304944 | rs17800694 | -1.044 | -1.42 | 1.291 | 1.103 | -0.02 | 0.007 | 0.007 | 0 | 0.11 | 0.14 | 0.15 | 0.12 | 0.05 | 0.16 |
| 8 | 116305599 | rs10955747 | -1.648 | -1.157 | 1.366 | 0.941 | -0.05 | 0.07 | 0.07 | 3.565 | 0.12 | 0.64 | 0.15 | 0.16 | 0.03 | 0.14 |
| 8 | 116327775 | rs9969656 | -1.681 | -1.022 | 0.581 | 0.284 | -0.06 | -0.072 | -0.072 | 0.673 | 0.21 | 0.71 | 0.19 | 0.05 | 0.07 | 0.18 |
| 8 | 116333231 | rs16887312 | -1.156 | -0.166 | 0.229 | 1.026 | -0.016 | 0.057 | 0.057 | 0 | 0.08 | 0.24 | 0.12 | 0.13 | 0.05 | 0.12 |
| 8 | 116340191 | rs7833220 | -1.156 | -0.166 | 0.229 | 0.589 | -0.016 | -0.014 | -0.014 | 0.041 | 0.08 | 0.24 | 0.12 | 0.05 | 0.05 | 0.12 |
| 8 | 116359584 | rs7011253 | -0.449 | 0.544 | -0.31 | 0.706 | 0.007 | 0.002 | 0.002 | 0.554 | 0.37 | 0.38 | 0.48 | 0.43 | 0.49 | 0.52 |
| 8 | 116367545 | rs984713 | -1.221 | 0.91 | -0.064 | 0.741 | -0.001 | -0.005 | -0.005 | 0 | 0.23 | 0.66 | 0.24 | 0.25 | 0.26 | 0.21 |
| 8 | 116387977 | rs7016277 | -0.332 | 0.837 | 0.916 | 0.729 | 0.021 | 0.036 | 0.036 | 0 | 0.09 | 0.28 | 0.08 | 0.12 | 0.09 | 0.07 |
| 8 | 116394075 | rs17658378 | 1.42 | 0.18 | -0.018 | -0.143 | 0.002 | 0.06 | 0.06 | 0 | 0.13 | 0.01 | 0.11 | 0 | 0.12 | 0.04 |
| 8 | 116396326 | rs13265665 | 1.381 | -0.313 | -0.392 | -0.36 | 0.025 | -0.05 | -0.05 | 0 | 0.46 | 0.06 | 0.3 | 0.41 | 0.35 | 0.31 |
| 8 | 116401505 | rs231166 | 0.355 | -0.083 | -0.001 | -0.296 | 0.012 | 0.001 | 0.001 | 0 | 0.13 | 0.35 | 0.16 | 0.13 | 0.18 | 0.14 |
| 8 | 116405191 | rs231173 | -0.226 | 0.101 | -0.109 | -0.047 | -0.019 | -0.003 | -0.003 | 0 | 0.72 | 0.82 | 0.62 | 0.65 | 0.62 | 0.52 |
| 8 | 116417563 | rs231147 | -0.711 | 0.257 | 0.165 | 0.519 | -0.004 | 0.042 | 0.042 | 0.015 | 0.44 | 0.61 | 0.57 | 0.77 | 0.48 | 0.57 |
| 8 | 116421936 | rs800897 | -0.852 | 0.277 | -0.066 | 1.304 | -0.028 | 0.002 | 0.002 | 0.063 | 0.45 | 0.83 | 0.55 | 0.57 | 0.48 | 0.54 |
| 8 | 116425860 | rs800899 | -0.364 | 0.353 | 0.286 | 1.208 | -0.009 | 0.021 | 0.021 | 0.001 | 0.39 | 0.35 | 0.6 | 0.8 | 0.48 | 0.57 |
| 8 | 116429998 | rs808994 | 0.305 | 0.241 | -0.193 | 1.09 | 0.002 | 0.01 | 0.01 | 0 | 0.85 | 0.9 | 0.84 | 0.9 | 0.81 | 0.86 |
| 8 | 116430861 | rs1180628 | 0.646 | 0.364 | -0.284 | 1.252 | -0.012 | -0.069 | -0.069 | 0.217 | 0.84 | 0.9 | 0.7 | 0.64 | 0.79 | 0.83 |
| 8 | 116437154 | rs800892 | 0.015 | 0.485 | -0.198 | 1.018 | -0.027 | -0.021 | -0.021 | 0 | 0.37 | 0.26 | 0.5 | 0.55 | 0.46 | 0.53 |
| 8 | 116439639 | rs800890 | 0.594 | 0.528 | -0.198 | 0.857 | -0.015 | 0.006 | 0.006 | 0 | 0.85 | 0.93 | 0.84 | 0.88 | 0.79 | 0.84 |
| 8 | 116449599 | rs3808407 | 1.147 | 0.369 | 0.054 | 0.824 | 0.045 | 0.055 | 0.055 | 0 | 0.45 | 0.27 | 0.25 | 0.33 | 0.37 | 0.43 |
| 8 | 116472478 | rs1180647 | -0.481 | 1.199 | 1.148 | 0.612 | -0.001 | -0.012 | -0.012 | 0 | 0.39 | 0.35 | 0.32 | 0.16 | 0.44 | 0.26 |
| 8 | 116480753 | rs1483582 | -0.036 | 1.176 | 1.13 | 0.592 | -0.031 | 0.066 | 0.066 | 0 | 0.65 | 0.78 | 0.71 | 0.84 | 0.56 | 0.7 |
| 8 | 116522025 | rs2293888 | -1.759 | 1.938 | 1.126 | 0.877 | 0.024 | 0.087 | 0.087 | 0 | 0.16 | 0.26 | 0.47 | 0.43 | 0.21 | 0.3 |
| 8 | 116537833 | rs2884365 | 1.952 | 1.843 | 0.689 | 0.85 | 0.033 | -0.029 | -0.029 | 1.376 | 0.42 | 0.07 | 0.21 | 0.4 | 0.36 | 0.35 |
| 8 | 116539067 | rs12549788 | 0.131 | 1.962 | 0.95 | 0.105 | -0.007 | 0.01 | 0.01 | 0.011 | 0.64 | 0.67 | 0.72 | 0.84 | 0.56 | 0.63 |
| 8 | 116570630 | rs6469597 | 0.127 | 2.016 | 0.112 | 0.203 | -0.03 | 0.076 | 0.076 | 0 | 0.66 | 0.86 | 0.73 | 0.83 | 0.58 | 0.68 |
| 8 | 116592355 | rs6469601 | 1.884 | 1.459 | -0.074 | 0.468 | 0.038 | -0.075 | -0.075 | 1.428 | 0.46 | 0.07 | 0.33 | 0.49 | 0.39 | 0.37 |
| 8 | 116594538 | rs17659542 | -0.025 | 2.045 | 0.036 | 0.571 | -0.036 | -0.031 | -0.031 | 0.184 | 0.06 | 0.01 | 0.21 | 0.08 | 0.12 | 0.07 |
| 8 | 116602459 | rs12679749 | 0.236 | 1.72 | 0.102 | 0.079 | -0.091 | 0.118 | 0.118 | 0.455 | 0.66 | 0.96 | 0.75 | 0.82 | 0.58 | 0.67 |
| 8 | 116629732 | rs1569364 | 0.499 | 1.971 | 0.59 | 0.37 | -0.01 | 0.017 | 0.017 | 0.012 | 0.64 | 0.32 | 0.69 | 0.85 | 0.57 | 0.66 |
| 8 | 116630311 | rs2737217 | -0.019 | 1.946 | 0.534 | 0.36 | -0.135 | 0.099 | 0.099 | 0.45 | 0.7 | 0.97 | 0.75 | 0.85 | 0.58 | 0.72 |
| 8 | 116631278 | rs2737218 | -1.488 | 2.005 | 0.659 | 0.661 | 0.015 | 0.045 | 0.045 | 0 | 0.14 | 0.21 | 0.34 | 0.32 | 0.18 | 0.26 |
| 8 | 116635688 | rs2737219 | 0.628 | 1.836 | 0.636 | 0.044 | -0.013 | 0.019 | 0.019 | 0.086 | 0.67 | 0.32 | 0.7 | 0.86 | 0.61 | 0.66 |
| 8 | 116635942 | rs2721939 | 0.398 | 1.926 | 0.187 | 0.089 | 0.028 | -0.055 | -0.055 | 0 | 0.69 | 0.62 | 0.72 | 0.86 | 0.61 | 0.66 |
| 8 | 116650491 | rs727581 | -2.402 | 1.286 | 0.545 | 0.966 | -0.018 | -0.028 | -0.028 | 0.459 | 0.33 | 0.9 | 0.5 | 0.34 | 0.32 | 0.46 |
| 8 | 116660365 | rs2737247 | -2.311 | 1.227 | 1.054 | 1.078 | -0.003 | -0.012 | -0.012 | 0 | 0.31 | 0.77 | 0.49 | 0.36 | 0.32 | 0.4 |
| 8 | 116662038 | rs2721965 | -2.073 | 1.017 | 1.554 | 1.011 | -0.018 | -0.026 | -0.026 | 0 | 0.22 | 0.13 | 0.41 | 0.36 | 0.31 | 0.33 |
| 8 | 116670347 | rs3808477 | -1.88 | 1.334 | 1.804 | 1.29 | -0.069 | -0.07 | -0.07 | 0.368 | 0.11 | 0.01 | 0.32 | 0.3 | 0.24 | 0.2 |
| 8 | 116670428 | rs7828416 | -2.317 | 1.025 | 2.132 | 0.981 | -0.02 | -0.028 | -0.028 | 0 | 0.08 | 0.62 | 0.07 | 0 | 0.01 | 0.06 |
| 8 | 116695112 | rs179456 | -1.867 | 1.28 | 2.03 | 1.268 | 0.01 | 0.036 | 0.036 | 0 | 0.09 | 0.09 | 0.28 | 0.35 | 0.21 | 0.2 |
| 8 | 116700847 | rs7842702 | -2.692 | 1.255 | 3.349 | 1.245 | 0.043 | 0.039 | 0.039 | 2.263 | 0.06 | 0.42 | 0.05 | 0 | 0 | 0.02 |
| 8 | 116701130 | rs12682033 | 2.987 | 1.262 | 3.24 | 1.25 | 0.099 | 0.202 | 0.202 | 0.122 | 0.85 | 0.49 | 0.67 | 0.64 | 0.79 | 0.78 |
| 8 | 116746477 | rs1880814 | 2.845 | 1.286 | 3.454 | 1.215 | 0.087 | 0.144 | 0.144 | 0 | 0.84 | 0.48 | 0.67 | 0.67 | 0.77 | 0.78 |
| 8 | 116760997 | rs16887723 | -3.632 | 1.112 | 3.26 | 1.087 | -0.05 | -0.052 | -0.052 | 0 | 0.1 | 0.71 | 0.07 | 0 | 0 | 0.03 |
| 8 | 116801780 | rs800518 | 1.975 | 1.374 | 2.926 | 1.405 | 0.002 | 0.041 | 0.041 | 0 | 0.9 | 0.89 | 0.57 | 0.57 | 0.78 | 0.76 |
| 8 | 116811437 | rs10106649 | -2.39 | 1.28 | 3.305 | 1.15 | 0.032 | 0.033 | 0.033 | 2.251 | 0.07 | 0.49 | 0.15 | 0.03 | 0.01 | 0.07 |
| 8 | 116816841 | rs16887791 | -1.824 | 1.314 | 3.102 | 1.152 | 0.054 | 0.051 | 0.051 | 0.387 | 0.06 | 0.4 | 0.14 | 0.03 | 0.01 | 0.07 |
| 8 | 116822134 | rs9918880 | -2.236 | 1.279 | 3.003 | 1.152 | 0.035 | 0.026 | 0.026 | 2.095 | 0.06 | 0.39 | 0.03 | 0 | 0 | 0.02 |
| 8 | 116824700 | rs2254401 | 1.41 | 1.522 | 2.106 | 1.34 | -0.049 | -0.023 | -0.023 | 0 | 0.92 | 0.98 | 0.69 | 0.63 | 0.79 | 0.82 |
| 8 | 116825690 | rs1474518 | 1.886 | 1.505 | 2.239 | 1.283 | 0.033 | 0.09 | 0.09 | 0 | 0.91 | 0.85 | 0.68 | 0.63 | 0.79 | 0.82 |
| 8 | 116837897 | rs7842594 | -2.752 | 1.274 | 1.973 | 1.017 | -0.044 | -0.042 | -0.042 | 0 | 0.11 | 0.77 | 0.1 | 0.03 | 0.01 | 0.08 |
| 8 | 116845729 | rs800526 | 1.961 | 1.75 | 1.628 | 1.253 | 0.032 | 0.09 | 0.09 | 0 | 0.92 | 0.86 | 0.51 | 0.63 | 0.8 | 0.81 |
| 8 | 116849719 | rs800523 | 1.748 | 1.728 | 1.361 | 1.263 | -0.016 | 0.016 | 0.016 | 0 | 0.92 | 0.94 | 0.63 | 0.61 | 0.8 | 0.79 |
| 8 | 116853988 | rs702770 | 1.748 | 1.728 | 1.361 | 1.263 | -0.013 | 0.021 | 0.021 | 0 | 0.92 | 0.94 | 0.63 | 0.62 | 0.8 | 0.79 |
| 8 | 116872821 | rs17717380 | 0.335 | 1.119 | 1.102 | 0.367 | -0.017 | 0.043 | 0.043 | 0 | 0.08 | 0 | 0.07 | 0 | 0.1 | 0.02 |
| 8 | 116873609 | rs7841676 | -1.925 | 0.988 | 2.529 | 0.626 | 0.001 | 0.002 | 0.002 | 0 | 0.11 | 0.45 | 0.13 | 0.05 | 0.04 | 0.16 |
| 8 | 116883864 | rs12545387 | -2.463 | 0.905 | 3.031 | 0.675 | -0.017 | -0.017 | -0.017 | 0 | 0.13 | 0.35 | 0.3 | 0.09 | 0.07 | 0.21 |
| 8 | 116902341 | rs7812637 | 1.404 | 1.046 | 3.174 | 0.69 | 0.043 | -0.039 | -0.039 | 0.056 | 0.88 | 0.65 | 0.86 | 0.98 | 0.84 | 0.91 |
| 8 | 116902470 | rs7812812 | -2.265 | 1.038 | 3.216 | 0.686 | -0.017 | -0.012 | -0.012 | 0 | 0.12 | 0.33 | 0.3 | 0.09 | 0.06 | 0.19 |
| 8 | 116902508 | rs10097735 | 2.921 | 1.069 | 3.18 | 0.686 | -0.001 | -0.009 | -0.009 | 0 | 0.76 | 0.25 | 0.56 | 0.89 | 0.78 | 0.71 |
| 8 | 116904676 | rs2205258 | 0.909 | 1.059 | 3.065 | 0.68 | -0.038 | 0.045 | 0.045 | 0 | 0.91 | 0.98 | 0.89 | 1 | 0.84 | 0.92 |
| 8 | 116934158 | rs6991030 | 1.409 | 1.064 | 3.086 | 0.682 | 0.039 | -0.037 | -0.037 | 0 | 0.89 | 0.64 | 0.86 | 0.98 | 0.83 | 0.9 |

| | | | | | | | | | | | | | | | | |
|---|---|---|---|---|---|---|---|---|---|---|---|---|---|---|---|---|
| 8 | 116935231 | rs11786434 | 3.009 | 1.025 | 3.102 | 0.785 | -0.003 | -0.01 | -0.01 | 0 | 0.75 | 0.24 | 0.54 | 0.86 | 0.77 | 0.71 |
| 8 | 116938631 | rs926135 | -2.452 | 0.96 | 3.015 | 0.795 | -0.02 | 0.02 | 0.02 | 0 | 0.13 | 0.4 | 0.33 | 0.12 | 0.06 | 0.19 |
| 8 | 116947667 | rs7825602 | -2.26 | 0.952 | 2.737 | 0.763 | -0.034 | -0.015 | -0.015 | 0 | 0.16 | 0.3 | 0.33 | 0.12 | 0.07 | 0.3 |
| 8 | 116948169 | rs760297 | 1.416 | 1.215 | 2.717 | 0.731 | 0.035 | -0.032 | -0.032 | 0.004 | 0.89 | 0.65 | 0.86 | 0.97 | 0.84 | 0.9 |
| 8 | 116954592 | rs4140855 | 1.808 | 1.186 | 2.67 | 0.686 | 0.022 | -0.042 | -0.042 | 0 | 0.87 | 0.65 | 0.86 | 0.97 | 0.83 | 0.89 |
| 8 | 116959520 | rs932267 | 1.438 | 1.204 | 2.503 | 0.694 | -0.004 | 0.003 | 0.003 | 0 | 0.88 | 0.9 | 0.88 | 0.97 | 0.83 | 0.89 |
| 8 | 116960729 | rs7828742 | 1.824 | 1.533 | 2.303 | 0.916 | -0.007 | -0.007 | -0.007 | 1.213 | 0.77 | 0.75 | 0.75 | 0.68 | 0.6 | 0.81 |
| 8 | 116967356 | rs10216416 | 2.709 | 1.361 | 1.999 | 0.75 | -0.164 | -0.097 | -0.097 | 0.013 | 0.58 | 0.03 | 0.48 | 0.61 | 0.53 | 0.56 |
| 8 | 116979531 | rs11784903 | 2.553 | 1.334 | 2.024 | 0.712 | -0.162 | -0.118 | -0.118 | 0 | 0.59 | 0.03 | 0.49 | 0.6 | 0.53 | 0.57 |
| 8 | 116980396 | rs11993108 | -0.694 | 1.324 | 2.431 | 0.723 | 0.004 | -0.027 | -0.027 | 0 | 0.09 | 0.24 | 0.05 | 0.04 | 0.04 | 0.07 |
| 8 | 116980929 | rs17741331 | -1.392 | 1.419 | 2.454 | 0.726 | -0.013 | -0.026 | -0.026 | 0.012 | 0.19 | 0.13 | 0.29 | 0.36 | 0.28 | 0.23 |
| 8 | 116981938 | rs9297548 | 2.771 | 1.427 | 2.428 | 0.718 | -0.162 | -0.106 | -0.106 | 0 | 0.57 | 0.03 | 0.49 | 0.6 | 0.53 | 0.56 |
| 8 | 116988247 | rs10110738 | 1.93 | 1.682 | 2.97 | 1.106 | -0.047 | -0.045 | -0.045 | 0.007 | 0.73 | 0.84 | 0.61 | 0.64 | 0.58 | 0.7 |
| 8 | 116988947 | rs10086506 | 1.93 | 1.682 | 2.983 | 1.106 | -0.043 | -0.041 | -0.041 | 0 | 0.73 | 0.84 | 0.61 | 0.64 | 0.58 | 0.7 |
| 8 | 116997615 | rs12156416 | 0.675 | 1.705 | 2.919 | 1.008 | -0.041 | 0.035 | 0.035 | 0 | 0.92 | 0.99 | 0.89 | 0.98 | 0.85 | 0.93 |
| 8 | 117007850 | rs2064009 | 1.977 | 1.721 | 3.043 | 1.049 | -0.04 | -0.042 | -0.042 | 0.016 | 0.72 | 0.82 | 0.6 | 0.62 | 0.56 | 0.69 |
| 8 | 117011944 | rs2205264 | 2.121 | 1.74 | 3.217 | 1.091 | 0.016 | 0.018 | 0.018 | 0 | 0.71 | 0.65 | 0.59 | 0.62 | 0.56 | 0.69 |
| 8 | 117014143 | rs7014419 | 0.616 | 1.7 | 3.168 | 1.108 | -0.034 | 0.03 | 0.03 | 0 | 0.93 | 0.98 | 0.9 | 0.98 | 0.85 | 0.93 |
| 8 | 117017561 | rs1079800 | 1.269 | 1.369 | 3.214 | 1.134 | -0.031 | -0.032 | -0.032 | 0.094 | 0.74 | 0.81 | 0.61 | 0.63 | 0.6 | 0.68 |
| 8 | 117022324 | rs2223065 | -0.7 | 1.349 | 3.214 | 1.217 | 0.006 | -0.008 | -0.008 | 0 | 0.05 | 0.03 | 0.18 | 0.17 | 0.02 | 0.16 |
| 8 | 117038175 | rs6469617 | 0.712 | 1.338 | 3.169 | 0.919 | -0.035 | -0.036 | -0.036 | 0 | 0.74 | 0.82 | 0.61 | 0.63 | 0.6 | 0.69 |
| 8 | 117049546 | rs13252360 | 0.768 | 1.339 | 3.127 | 0.919 | -0.032 | -0.033 | -0.033 | 0 | 0.74 | 0.81 | 0.61 | 0.63 | 0.6 | 0.69 |
| 8 | 117050089 | rs2223063 | 0.744 | 1.339 | 3.126 | 0.919 | -0.031 | -0.032 | -0.032 | 0 | 0.74 | 0.81 | 0.61 | 0.63 | 0.6 | 0.69 |
| 8 | 117061748 | rs2223064 | 0.819 | 1.342 | 3.171 | 0.921 | 0.005 | 0.006 | 0.006 | 0 | 0.73 | 0.73 | 0.6 | 0.63 | 0.6 | 0.69 |
| 8 | 117069149 | rs13272758 | 0.895 | 1.324 | 2.999 | 0.903 | 0.004 | 0.005 | 0.005 | 0 | 0.73 | 0.72 | 0.6 | 0.63 | 0.6 | 0.67 |
| 8 | 117077882 | rs10109846 | 1.012 | 1.383 | 2.631 | 0.934 | -0.004 | -0.004 | -0.004 | 0.344 | 0.72 | 0.7 | 0.61 | 0.63 | 0.6 | 0.68 |
| 8 | 117087613 | rs2049841 | 1.048 | 1.424 | 2.723 | 0.953 | 0.02 | 0.023 | 0.023 | 0 | 0.72 | 0.64 | 0.6 | 0.63 | 0.6 | 0.69 |
| 8 | 117113215 | rs4567068 | 0.712 | 1.246 | 2.032 | 0.945 | -0.001 | -0.001 | -0.001 | 0 | 0.74 | 0.72 | 0.6 | 0.61 | 0.6 | 0.68 |
| 8 | 117122598 | rs1574542 | 1.358 | 1.283 | 1.06 | 0.974 | -0.027 | -0.028 | -0.028 | 0.753 | 0.67 | 0.19 | 0.56 | 0.61 | 0.6 | 0.62 |
| 8 | 117128202 | rs9792120 | 1.27 | 1.259 | 0.751 | 1.054 | -0.046 | -0.054 | -0.054 | 0 | 0.68 | 0.14 | 0.55 | 0.61 | 0.63 | 0.62 |
| 8 | 117129410 | rs2223059 | 1.08 | 1.185 | 0.635 | 1.153 | -0.001 | -0.001 | -0.001 | 0 | 0.7 | 0.29 | 0.56 | 0.6 | 0.63 | 0.64 |
| 8 | 117132729 | rs972395 | 0.719 | 1.223 | 0.486 | 1.272 | 0.044 | 0.065 | 0.065 | 0 | 0.74 | 0.56 | 0.6 | 0.6 | 0.65 | 0.68 |
| 8 | 117135496 | rs6993240 | -1.676 | 0.193 | 0.961 | 1.975 | -0.045 | 0.133 | 0.133 | 0 | 0.19 | 0.58 | 0.26 | 0.48 | 0.11 | 0.23 |
| 8 | 117136227 | rs6469633 | -1.385 | -0.638 | 1.114 | 1.776 | -0.032 | -0.005 | -0.005 | 1.447 | 0.31 | 0.58 | 0.27 | 0.32 | 0.23 | 0.34 |
| 8 | 117146997 | rs13262647 | -0.937 | -0.602 | 0.934 | 1.667 | -0.004 | 0.002 | 0.002 | 0 | 0.1 | 0.12 | 0.12 | 0.08 | 0.11 | 0.05 |
| 8 | 117154992 | rs4573309 | 1.264 | -1.673 | 1.216 | 1.962 | -0.079 | 0.057 | 0.057 | 0.118 | 0.71 | 0.61 | 0.59 | 0.5 | 0.88 | 0.49 |
| 8 | 117167843 | rs1005502 | 2.18 | -1.77 | 0.559 | 1.872 | 0.097 | -0.101 | -0.101 | 0.485 | 0.56 | 0.03 | 0.4 | 0.43 | 0.75 | 0.38 |
| 8 | 117168712 | rs1022440 | 1.974 | -1.573 | 1.629 | 1.608 | 0.036 | 0.012 | 0.012 | 0.135 | 0.93 | 0.64 | 0.91 | 1 | 0.95 | 0.94 |
| 8 | 117190718 | rs17662374 | -1.268 | -1.508 | 1.333 | 1.658 | -0.026 | 0.006 | 0.006 | 0 | 0.08 | 0.02 | 0.03 | 0.05 | 0.08 | 0.02 |
| 8 | 117236486 | rs799865 | 2.339 | -1.812 | 2.222 | 1.394 | -0.024 | -0.102 | -0.102 | 0.875 | 0.76 | 0.49 | 0.86 | 0.94 | 0.82 | 0.79 |
| 8 | 117243066 | rs799871 | 1.67 | -1.754 | 2.159 | 1.42 | -0.003 | -0.025 | -0.025 | 0 | 0.81 | 0.8 | 0.92 | 0.94 | 0.84 | 0.8 |
| 8 | 117245476 | rs799875 | 1.566 | -1.504 | 2.024 | 1.423 | -0.002 | 0.011 | 0.011 | 0 | 0.82 | 0.86 | 0.91 | 0.94 | 0.82 | 0.79 |
| 8 | 117245789 | rs811645 | 1.566 | -1.504 | 2.024 | 1.423 | -0.002 | 0.009 | 0.009 | 0 | 0.82 | 0.86 | 0.91 | 0.94 | 0.82 | 0.79 |
| 8 | 117256951 | rs799885 | -1.814 | -1.686 | 1.757 | 1.93 | -0.045 | 0.049 | 0.049 | 0 | 0.3 | 0.5 | 0.52 | 0.51 | 0.17 | 0.42 |
| 8 | 117264451 | rs799860 | -0.898 | -1.766 | 1.301 | 2.289 | -0.003 | -0.031 | -0.031 | 0 | 0.12 | 0 | 0.08 | 0.12 | 0.08 | 0.1 |
| 8 | 117265106 | rs799855 | -2.42 | -1.769 | 1.418 | 2.08 | 0.061 | -0.041 | -0.041 | 3.105 | 0.33 | 0.82 | 0.55 | 0.52 | 0.18 | 0.42 |
| 8 | 117278641 | rs7015270 | -2.697 | -1.654 | 1.457 | 2.635 | 0.041 | 0.372 | 0.372 | 2.292 | 0.06 | 0.47 | 0.06 | 0.27 | 0.02 | 0.01 |
| 8 | 117293884 | rs8181015 | 2.373 | -1.781 | 1.041 | 2.039 | 0.032 | -0.076 | -0.076 | 0.007 | 0.49 | 0.03 | 0.43 | 0.48 | 0.67 | 0.38 |
| 8 | 117294102 | rs16888254 | -2.958 | -1.467 | 1.833 | 1.973 | 0.034 | 0.132 | 0.132 | 2.441 | 0.06 | 0.44 | 0.06 | 0.1 | 0.02 | 0.01 |
| 8 | 117300171 | rs7822185 | -3.017 | -1.495 | 1.793 | 1.77 | 0.009 | 0.061 | 0.061 | 1.521 | 0.07 | 0.43 | 0.1 | 0.06 | 0.01 | 0.09 |
| 8 | 117305398 | rs1918719 | -1.991 | -1.492 | 1.809 | 2.032 | -0.012 | 0.034 | 0.034 | 0.198 | 0.24 | 0.29 | 0.35 | 0.43 | 0.2 | 0.26 |
| 8 | 117306264 | rs4876641 | -2.086 | -1.451 | 1.919 | 2.029 | -0.022 | 0.054 | 0.054 | 0.503 | 0.26 | 0.42 | 0.36 | 0.43 | 0.2 | 0.26 |
| 8 | 117306606 | rs16888276 | -1.348 | -1.472 | 1.768 | 2.041 | -0.007 | -0.002 | -0.002 | 0 | 0.15 | 0.18 | 0.06 | 0.16 | 0.12 | 0.08 |
| 8 | 117329669 | rs1527937 | -0.224 | -2.462 | 1.194 | 1.258 | -0.025 | -0.02 | -0.02 | 0 | 0.21 | 0.23 | 0.11 | 0.11 | 0.1 | 0.26 |
| 8 | 117332218 | rs12547047 | -2.128 | -2.391 | 0.772 | 1.318 | 0.085 | 0.016 | 0.016 | 0.714 | 0.5 | 0.71 | 0.51 | 0.54 | 0.32 | 0.51 |
| 8 | 117334573 | rs1881129 | -0.83 | -1.734 | 0.898 | 1.777 | 0.003 | 0.007 | 0.007 | 0 | 0.09 | 0.08 | 0.3 | 0.27 | 0.07 | 0.17 |
| 8 | 117336860 | rs6469638 | -1.54 | -1.575 | 1.036 | 0.755 | 0.003 | 0 | 0 | 0.049 | 0.14 | 0.16 | 0.08 | 0.15 | 0.11 | 0.07 |
| 8 | 117351973 | rs2316692 | -1.68 | -1.585 | 1.076 | 0.728 | -0.022 | -0.007 | -0.007 | 0 | 0.16 | 0.27 | 0.09 | 0.15 | 0.11 | 0.07 |
| 8 | 117360816 | rs6991997 | -1.963 | -1.52 | 0.79 | 0.623 | 0.005 | 0.007 | 0.007 | 3.804 | 0.2 | 0.22 | 0.1 | 0.15 | 0.16 | 0.1 |
| 8 | 117365414 | rs11778307 | -2.102 | -1.734 | 0.548 | 0.51 | 0.053 | 0.068 | 0.068 | 3.036 | 0.37 | 0.8 | 0.24 | 0.22 | 0.25 | 0.29 |
| 8 | 117380851 | rs10090183 | -1.207 | -1.499 | 0.445 | 0.515 | -0.041 | -0.003 | -0.003 | 0 | 0.28 | 0.39 | 0.43 | 0.69 | 0.18 | 0.47 |
| 8 | 117385888 | rs13439023 | -1.154 | -1.493 | 0.435 | 0.519 | -0.039 | -0.002 | -0.002 | 0 | 0.27 | 0.4 | 0.43 | 0.69 | 0.18 | 0.47 |
| 8 | 117392156 | rs1527934 | -0.872 | -1.348 | -0.607 | 0.606 | -0.009 | 0.006 | 0.006 | 10.361 | 0.12 | 0.19 | 0.06 | 0.18 | 0.08 | 0.03 |
| 8 | 117394185 | rs6469640 | -1.895 | -0.644 | -1.11 | 0.15 | 0.021 | 0.315 | 0.315 | 1.365 | 0.5 | 0.96 | 0.56 | 0.86 | 0.34 | 0.56 |
| 8 | 117410485 | rs11776115 | 1.425 | -0.629 | -0.852 | 0.333 | 0.055 | 0.26 | 0.26 | 2.072 | 0.54 | 0.09 | 0.45 | 0.13 | 0.69 | 0.47 |
| 8 | 117415285 | rs11986435 | -1.274 | -0.482 | -0.008 | 1.379 | -0.022 | 0.107 | 0.107 | 2.574 | 0.23 | 0.46 | 0.36 | 0.52 | 0.19 | 0.3 |
| 8 | 117418375 | rs7822344 | 0.945 | -0.537 | -0.304 | 1.284 | 0.01 | -0.012 | -0.012 | 0 | 0.67 | 0.71 | 0.61 | 0.44 | 0.77 | 0.43 |
| 8 | 117418945 | rs10955775 | -0.646 | -0.331 | 0.561 | 1.053 | 0.099 | 0.038 | 0.038 | 1.448 | 0.06 | 0.39 | 0.07 | 0 | 0.05 | 0.01 |
| 8 | 117429757 | rs4512403 | 0.81 | -0.551 | 0.235 | 1.269 | 0.017 | -0.086 | -0.086 | 0 | 0.75 | 0.89 | 0.67 | 0.52 | 0.81 | 0.49 |

| | | | | | | | | | | | | | | | | |
|---|---|---|---|---|---|---|---|---|---|---|---|---|---|---|---|---|
| 8 | 117429845 | rs4515584 | -1.264 | 0.107 | 0.48 | 0.987 | 0.086 | -0.027 | -0.027 | 2.209 | 0.08 | 0.52 | 0.11 | 0 | 0.1 | 0.03 |
| 8 | 117445358 | rs4242565 | 0.408 | -0.01 | 0.122 | 1.148 | -0.025 | 0 | 0 | 0 | 0.91 | 0.99 | 0.75 | 0.89 | 0.92 | 0.79 |
| 8 | 117457413 | rs4466427 | 1.18 | 0.209 | -0.09 | 1.017 | 0.022 | -0.091 | -0.091 | 0.084 | 0.72 | 0.88 | 0.66 | 0.53 | 0.8 | 0.47 |
| 8 | 117462506 | rs13263375 | 2.299 | -0.066 | -0.314 | 1.294 | 0.007 | 0.115 | 0.115 | 0 | 0.54 | 0.23 | 0.39 | 0.3 | 0.58 | 0.36 |
| 8 | 117462992 | rs13270180 | 2.47 | -0.062 | -0.333 | 1.296 | -0.006 | 0.135 | 0.135 | 0 | 0.52 | 0.08 | 0.38 | 0.3 | 0.58 | 0.36 |
| 8 | 117467008 | rs4317609 | 1.886 | -0.111 | -0.148 | 1.132 | 0.019 | -0.028 | -0.028 | 0 | 0.6 | 0.35 | 0.56 | 0.53 | 0.67 | 0.42 |
| 8 | 117468999 | rs11774963 | -0.572 | 0.008 | 0.648 | 0.824 | 0.092 | -0.016 | -0.016 | 0.014 | 0.07 | 0.31 | 0.09 | 0 | 0.11 | 0.02 |
| 8 | 117475196 | rs4345586 | 1.135 | -0.337 | 0.132 | 1.266 | 0.032 | -0.002 | -0.002 | 0 | 0.81 | 0.86 | 0.73 | 0.79 | 0.91 | 0.76 |
| 8 | 117475597 | rs4129295 | 1.807 | -0.307 | 0.569 | 1.475 | -0.01 | 0.004 | 0.004 | 0 | 0.61 | 0.42 | 0.63 | 0.51 | 0.74 | 0.45 |
| 8 | 117475871 | rs3925055 | -1.584 | 0.171 | 0.535 | 0.32 | 0.003 | -0.075 | -0.075 | 1.088 | 0.28 | 0.57 | 0.19 | 0.11 | 0.3 | 0.17 |
| 8 | 117478072 | rs7839897 | -0.043 | -0.109 | -0.161 | 0.406 | 0.008 | -0.052 | -0.052 | 1.169 | 0.78 | 0.84 | 0.61 | 0.52 | 0.83 | 0.72 |
| 8 | 117480311 | rs7001801 | -0.043 | -0.109 | -0.161 | 0.392 | 0.008 | -0.054 | -0.054 | 0 | 0.78 | 0.84 | 0.61 | 0.51 | 0.83 | 0.72 |
| 8 | 117482886 | rs13278681 | -1.357 | -0.004 | -0.397 | -0.198 | 0.001 | -0.008 | -0.008 | 0.041 | 0.35 | 0.25 | 0.25 | 0.35 | 0.38 | 0.39 |
| 8 | 117483379 | rs6983244 | 2.107 | 0.023 | -0.14 | -0.284 | -0.002 | -0.008 | -0.008 | 0.001 | 0.4 | 0.38 | 0.34 | 0.2 | 0.46 | 0.34 |
| 8 | 117484739 | rs4458908 | -0.878 | 0.073 | -0.124 | -0.3 | 0.046 | -0.081 | -0.081 | 0 | 0.13 | 0.61 | 0.15 | 0 | 0.15 | 0.03 |
| 8 | 117487056 | rs13248744 | 0.479 | -0.049 | -0.38 | -0.248 | -0.023 | -0.111 | -0.111 | 0 | 0.16 | 0 | 0.1 | 0.35 | 0.17 | 0.28 |
| 8 | 117487677 | rs7005255 | 0.679 | -0.178 | -0.441 | -0.333 | 0.034 | -0.059 | -0.059 | 2.162 | 0.67 | 0.81 | 0.57 | 0.52 | 0.77 | 0.63 |
| 8 | 117491474 | rs4268149 | -0.873 | -0.123 | -0.17 | -0.274 | 0.017 | 0.035 | 0.035 | 0 | 0.26 | 0.36 | 0.23 | 0.35 | 0.31 | 0.3 |
| 8 | 117505270 | rs13252528 | 2.018 | -0.297 | -0.541 | -0.416 | 0.011 | 0.001 | 0.001 | 0.585 | 0.5 | 0.35 | 0.41 | 0.52 | 0.55 | 0.54 |
| 8 | 117519662 | rs4128874 | 0.317 | -0.426 | -0.777 | -0.381 | 0.003 | -0.005 | -0.005 | 0 | 0.69 | 0.68 | 0.58 | 0.52 | 0.77 | 0.62 |
| 8 | 117523035 | rs4361786 | -0.31 | -0.431 | -0.924 | -0.375 | 0.007 | -0.017 | -0.017 | 0 | 0.76 | 0.79 | 0.61 | 0.52 | 0.83 | 0.68 |
| 8 | 117534096 | rs4876359 | -0.117 | -0.255 | -0.971 | -0.665 | 0.013 | -0.034 | -0.034 | 0 | 0.73 | 0.79 | 0.6 | 0.52 | 0.8 | 0.67 |
| 8 | 117541607 | rs13279799 | 0.456 | -0.3 | -1.484 | -0.535 | -0.002 | -0.056 | -0.056 | 0.027 | 0.17 | 0.06 | 0.1 | 0.35 | 0.18 | 0.26 |
| 8 | 117553553 | rs16888508 | -0.387 | -0.193 | -1.256 | -0.838 | -0.03 | -0.055 | -0.055 | 0 | 0.12 | 0.23 | 0.05 | 0 | 0.06 | 0.07 |
| 8 | 117564278 | rs16888518 | -0.037 | 0.026 | -1.582 | -0.448 | 0.001 | -0.009 | -0.009 | 0.059 | 0.07 | 0.06 | 0.03 | 0 | 0.06 | 0.06 |
| 8 | 117571728 | rs12676238 | 2.534 | 0.182 | -1.66 | -0.331 | -0.117 | 0.077 | 0.077 | 0 | 0.28 | 0.03 | 0.36 | 0.14 | 0.36 | 0.24 |
| 8 | 117578764 | rs16888536 | 0.355 | 0.099 | -0.158 | 0.73 | 0.009 | -0.038 | -0.038 | 0 | 0.07 | 0.15 | 0.07 | 0 | 0.07 | 0.01 |
| 8 | 117584829 | rs2033691 | 0.473 | -0.464 | -0.254 | 1.3 | 0.011 | -0.028 | -0.028 | 1.552 | 0.75 | 0.84 | 0.86 | 0.64 | 0.78 | 0.67 |
| 8 | 117592429 | rs6469650 | -2.123 | -1.061 | -0.231 | 0.974 | -0.041 | 0.148 | 0.148 | 0 | 0.59 | 0.84 | 0.59 | 0.85 | 0.48 | 0.59 |
| 8 | 117592926 | rs1436764 | -1.025 | -1.416 | 0.443 | 1.731 | 0.004 | -0.025 | -0.025 | 0.764 | 0.37 | 0.29 | 0.43 | 0.51 | 0.37 | 0.46 |
| 8 | 117594864 | rs13439805 | 2.423 | -1.315 | 0.274 | 1.352 | -0.042 | 0.157 | 0.157 | 0 | 0.42 | 0.16 | 0.33 | 0.14 | 0.52 | 0.44 |
| 8 | 117596952 | rs12679048 | -0.906 | -1.612 | 1.026 | 1.816 | -0.011 | 0.033 | 0.033 | 0 | 0.09 | 0.2 | 0.13 | 0.17 | 0.09 | 0.05 |
| 8 | 117599967 | rs2317215 | -0.377 | -1.704 | 0.704 | 2.16 | -0.013 | -0.118 | -0.118 | 0.428 | 0.23 | 0.06 | 0.27 | 0.49 | 0.21 | 0.28 |
| 8 | 117600602 | rs7006843 | 1.869 | -1.687 | 0.759 | 2.153 | 0.028 | -0.013 | -0.013 | 0 | 0.68 | 0.26 | 0.59 | 0.66 | 0.76 | 0.75 |
| 8 | 117605106 | rs4876665 | 2.416 | -1.626 | 0.64 | 1.921 | -0.006 | 0.16 | 0.16 | 0 | 0.44 | 0.17 | 0.32 | 0.16 | 0.55 | 0.46 |
| 8 | 117616701 | rs1317644 | 0.7 | -1.533 | 1.077 | 2.045 | -0.004 | 0 | 0 | 0 | 0.82 | 0.77 | 0.75 | 0.84 | 0.84 | 0.8 |
| 8 | 117616931 | rs7814028 | -0.866 | -1.94 | 1.131 | 2.348 | -0.002 | 0.008 | 0.008 | 0 | 0.09 | 0.07 | 0.13 | 0.18 | 0.09 | 0.06 |
| 8 | 117617135 | rs1106331 | -0.5 | -1.936 | 0.95 | 2.217 | -0.028 | -0.144 | -0.144 | 0 | 0.23 | 0.03 | 0.24 | 0.45 | 0.21 | 0.24 |
| 8 | 117621592 | rs2511653 | -0.766 | -2.032 | 1.163 | 2.214 | 0.012 | -0.024 | -0.024 | 0 | 0.25 | 0.18 | 0.25 | 0.46 | 0.19 | 0.23 |
| 8 | 117623719 | rs2450114 | 0.723 | -1.774 | 1.652 | 2.38 | -0.01 | 0.041 | 0.041 | 0 | 0.84 | 0.47 | 0.77 | 0.85 | 0.88 | 0.8 |
| 8 | 117624483 | rs1529943 | -1.053 | -1.626 | 0.941 | 2.362 | 0.004 | -0.033 | -0.033 | 0 | 0.09 | 0.03 | 0.13 | 0.23 | 0.1 | 0.06 |
| 8 | 117630683 | rs16892766 | -0.338 | -1.653 | 1.123 | 2.101 | -0.007 | -0.042 | -0.042 | 0 | 0.15 | 0.15 | 0.06 | 0 | 0.09 | 0.08 |
| 8 | 117631964 | rs6469653 | 2.137 | -1.695 | 0.867 | 2.154 | 0.066 | 0.222 | 0.222 | 0 | 0.49 | 0.13 | 0.39 | 0.17 | 0.61 | 0.52 |
| 8 | 117635577 | rs1074584 | 1.955 | -1.697 | 1.173 | 2.159 | 0.064 | 0.088 | 0.088 | 0 | 0.52 | 0.36 | 0.56 | 0.24 | 0.66 | 0.63 |
| 8 | 117636863 | rs10094824 | 1.534 | -1.729 | 1.262 | 2.503 | -0.031 | 0.006 | 0.006 | 0 | 0.61 | 0.54 | 0.53 | 0.41 | 0.71 | 0.63 |
| 8 | 117639532 | rs2015069 | -0.111 | -1.725 | 0.947 | 2.504 | 0.024 | -0.115 | -0.115 | 0 | 0.82 | 0.98 | 0.8 | 0.58 | 0.86 | 0.84 |
| 8 | 117640315 | rs11986063 | -0.577 | -1.639 | 1.168 | 1.76 | -0.019 | -0.039 | -0.039 | 1.906 | 0.17 | 0.22 | 0.07 | 0.05 | 0.1 | 0.14 |
| 8 | 117640939 | rs11987050 | -0.289 | -1.557 | 1.032 | 2.123 | -0.01 | 0.069 | 0.069 | 0 | 0.05 | 0.03 | 0.14 | 0.32 | 0.09 | 0.05 |
| 8 | 117641610 | rs11987235 | 1.158 | -1.668 | 1.078 | 2.025 | -0.019 | 0.048 | 0.048 | 1.703 | 0.75 | 0.34 | 0.56 | 0.47 | 0.78 | 0.73 |
| 8 | 117642345 | rs12542425 | 2.414 | -1.638 | 0.263 | 1.829 | 0.033 | 0.309 | 0.309 | 0.423 | 0.48 | 0.05 | 0.36 | 0.11 | 0.63 | 0.55 |
| 8 | 117643773 | rs9650068 | 0.549 | -1.305 | 0.697 | 2.084 | -0.01 | 0.098 | 0.098 | 0.403 | 0.82 | 0.65 | 0.62 | 0.48 | 0.81 | 0.77 |
| 8 | 117644145 | rs17743419 | 0.092 | -1.325 | 0.158 | 2.105 | 0.008 | -0.06 | -0.06 | 0 | 0.13 | 0.04 | 0.15 | 0.32 | 0.1 | 0.07 |
| 8 | 117645687 | rs7839361 | -0.065 | -1.221 | 0.162 | 1.779 | -0.025 | -0.072 | -0.072 | 0.06 | 0.86 | 0.98 | 0.8 | 0.67 | 0.83 | 0.85 |
| 8 | 117647033 | rs4610785 | 0.334 | -1.128 | 0.373 | 1.721 | 0.008 | -0.02 | -0.02 | 0.001 | 0.13 | 0.06 | 0.13 | 0.34 | 0.09 | 0.06 |
| 8 | 117648454 | rs4406425 | 2.285 | -1.397 | 0.167 | 1.564 | 0.03 | 0.288 | 0.288 | 0.426 | 0.48 | 0.05 | 0.37 | 0.12 | 0.65 | 0.55 |
| 8 | 117648719 | rs6988452 | 0.081 | -1.348 | 0.415 | 1.571 | -0.013 | 0.045 | 0.045 | 0.028 | 0.15 | 0.2 | 0.14 | 0.34 | 0.09 | 0.06 |
| 8 | 117649905 | rs4565487 | -1.57 | -1.364 | 0.834 | 1.452 | -0.024 | 0.011 | 0.011 | 0.41 | 0.33 | 0.56 | 0.22 | 0.39 | 0.2 | 0.23 |
| 8 | 117652242 | rs17743495 | 0.443 | -1.339 | 0.773 | 1.409 | 0.007 | -0.018 | -0.018 | 0 | 0.14 | 0.07 | 0.13 | 0.39 | 0.1 | 0.06 |
| 8 | 117656781 | rs1347626 | -0.681 | -1.353 | 0.778 | 1.456 | 0.011 | -0.055 | -0.055 | 0 | 0.88 | 0.97 | 0.8 | 0.67 | 0.89 | 0.85 |
| 8 | 117669202 | rs10808483 | 2.342 | -1.361 | 0.765 | 1.408 | 0.029 | 0.299 | 0.299 | 0.405 | 0.48 | 0.05 | 0.37 | 0.12 | 0.65 | 0.55 |
| 8 | 117670802 | rs4602925 | 0.479 | -1.318 | 0.998 | 1.423 | -0.02 | -0.057 | -0.057 | 0 | 0.12 | 0.14 | 0.05 | 0 | 0.06 | 0.09 |
| 8 | 117681031 | rs7833054 | -0.719 | -1.273 | 0.953 | 1.462 | 0.01 | -0.052 | -0.052 | 0.217 | 0.88 | 0.97 | 0.8 | 0.67 | 0.89 | 0.84 |
| 8 | 117703509 | rs7823271 | -0.913 | -1.234 | 1.014 | 1.434 | -0.025 | -0.121 | -0.121 | 3.12 | 0.19 | 0.45 | 0.08 | 0.01 | 0.11 | 0.17 |
| 8 | 117706218 | rs12675038 | 0.468 | -1.226 | 0.947 | 1.457 | 0.007 | -0.017 | -0.017 | 0 | 0.14 | 0.07 | 0.13 | 0.39 | 0.1 | 0.06 |
| 8 | 117722555 | rs7846316 | -0.905 | -1.226 | 0.942 | 1.486 | 0.016 | -0.044 | -0.044 | 0.056 | 0.88 | 0.97 | 0.93 | 0.73 | 0.91 | 0.86 |
| 8 | 117725175 | rs7825662 | -1.041 | -1.24 | 0.991 | 1.103 | -0.013 | -0.003 | -0.003 | 0 | 0.21 | 0.46 | 0.26 | 0.17 | 0.15 | 0.22 |
| 8 | 117735209 | rs16888699 | -0.663 | -1.243 | 1.045 | 1.099 | -0.016 | -0.088 | -0.088 | 0.018 | 0.18 | 0.26 | 0.07 | 0.01 | 0.11 | 0.17 |
| 8 | 117741913 | rs17811450 | -1.805 | -1.239 | 0.623 | 1.104 | -0.019 | 0.021 | 0.021 | 1.793 | 0.35 | 0.53 | 0.39 | 0.56 | 0.24 | 0.29 |
| 8 | 117763873 | rs4876679 | 0.402 | -1.209 | 0.572 | 1.025 | 0.013 | -0.009 | -0.009 | 1.751 | 0.74 | 0.81 | 0.66 | 0.28 | 0.79 | 0.79 |

| | | | | | | | | | | | | | | | | |
|---|---|---|---|---|---|---|---|---|---|---|---|---|---|---|---|---|
| 8 | 117771775 | rs4372028 | 1.929 | -1.214 | 0.576 | 0.904 | 0.067 | 0.097 | 0.097 | 0.197 | 0.53 | 0.27 | 0.57 | 0.26 | 0.68 | 0.61 |
| 8 | 117778570 | rs10283122 | 1.929 | -1.214 | 0.576 | 0.904 | 0.067 | 0.097 | 0.097 | 0.197 | 0.53 | 0.27 | 0.57 | 0.26 | 0.68 | 0.61 |
| 8 | 117778673 | rs10283129 | 0.519 | -1.082 | 0.597 | 0.972 | 0.01 | -0.003 | -0.003 | 0 | 0.73 | 0.78 | 0.66 | 0.28 | 0.79 | 0.79 |
| 8 | 117783914 | rs1133950 | 0.324 | -1.074 | -0.112 | 0.954 | -0.017 | -0.057 | -0.057 | 0 | 0.13 | 0.14 | 0.05 | 0 | 0.06 | 0.09 |
| 8 | 117783975 | rs16888728 | -0.185 | -0.923 | -0.84 | 1.058 | -0.022 | -0.125 | -0.125 | 0 | 0.19 | 0.46 | 0.08 | 0.01 | 0.11 | 0.17 |
| 8 | 117795600 | rs17664079 | 1.627 | -0.973 | -0.943 | 1.067 | 0.094 | 0.146 | 0.146 | 0.571 | 0.51 | 0.14 | 0.55 | 0.27 | 0.68 | 0.61 |
| 8 | 117797719 | rs6469660 | -1.319 | -0.514 | -0.835 | 1.676 | -0.027 | -0.002 | -0.002 | 0 | 0.29 | 0.36 | 0.36 | 0.72 | 0.21 | 0.22 |
| 8 | 117809105 | rs13279543 | -0.41 | 0.08 | -1.768 | 1.395 | 0.002 | -0.052 | -0.052 | 0 | 0.12 | 0.04 | 0.21 | 0.33 | 0.12 | 0.14 |
| 8 | 117817256 | rs4876683 | 0.539 | 0.231 | -2.386 | 0.145 | -0.039 | 0.253 | 0.253 | 0.66 | 0.47 | 0.05 | 0.32 | 0.11 | 0.56 | 0.51 |
| 8 | 117819007 | rs16888856 | -0.573 | -0.397 | -1.148 | 0.024 | -0.013 | -0.018 | -0.018 | 2.959 | 0.19 | 0.47 | 0.23 | 0.12 | 0.19 | 0.43 |
| 8 | 117825068 | rs6991815 | 0.736 | -1.142 | -1.985 | 0.179 | 0.007 | -0.056 | -0.056 | 0.038 | 0.16 | 0 | 0.21 | 0.19 | 0.1 | 0.14 |
| 8 | 117830986 | rs10101691 | -1.389 | -1.108 | -1.918 | -0.29 | -0.084 | 0.086 | 0.086 | 0 | 0.69 | 0.92 | 0.71 | 0.85 | 0.58 | 0.78 |
| 8 | 117838623 | rs10109496 | 0.786 | -1.253 | -1.531 | 0.139 | -0.009 | -0.026 | -0.026 | 0 | 0.17 | 0.09 | 0.23 | 0.27 | 0.21 | 0.44 |
| 8 | 117841222 | rs10081574 | 0.774 | -1.263 | -1.565 | 0.095 | 0.002 | -0.004 | -0.004 | 3.151 | 0.17 | 0.21 | 0.21 | 0.12 | 0.19 | 0.43 |
| 8 | 117841774 | rs10108200 | 0.744 | -1.26 | -1.564 | 0.097 | 0.001 | -0.005 | -0.005 | 0 | 0.17 | 0.21 | 0.21 | 0.12 | 0.19 | 0.43 |
| 8 | 117842055 | rs10955793 | -0.151 | -1.449 | -1.342 | -0.092 | 0.023 | 0.25 | 0.25 | 0 | 0.17 | 0.52 | 0.22 | 0.56 | 0.19 | 0.13 |
| 8 | 117850062 | rs7824338 | -1.048 | -1.565 | -1.599 | -0.493 | -0.098 | 0.169 | 0.169 | 0 | 0.62 | 0.92 | 0.69 | 0.88 | 0.55 | 0.78 |
| 8 | 117864217 | rs1050838 | 1.444 | -1.547 | -1.433 | -0.222 | -0.021 | -0.056 | -0.056 | 0.031 | 0.18 | 0.01 | 0.25 | 0.17 | 0.15 | 0.16 |
| 8 | 117869498 | rs2921787 | 0.116 | -1.692 | -0.958 | -0.246 | -0.056 | 0.006 | 0.006 | 0.187 | 0.12 | 0.35 | 0.08 | 0.17 | 0.02 | 0.04 |
| 8 | 117873811 | rs2289937 | 0.116 | -1.692 | -0.958 | 0.012 | -0.056 | 0.055 | 0.055 | 0.187 | 0.12 | 0.35 | 0.08 | 0.22 | 0.02 | 0.04 |
| 8 | 117884270 | rs11786746 | 0.734 | -1.759 | -0.451 | -0.151 | 0.018 | 0.196 | 0.196 | 0 | 0.11 | 0.28 | 0.14 | 0.38 | 0.13 | 0.09 |
| 8 | 117899211 | rs959692 | -0.987 | -1.797 | 0.072 | -0.183 | -0.002 | -0.002 | -0.002 | 0.361 | 0.6 | 0.6 | 0.55 | 0.47 | 0.47 | 0.74 |
| 8 | 117902254 | rs2921771 | 0.344 | -1.784 | -0.405 | -0.152 | -0.054 | 0.002 | 0.002 | 0.069 | 0.11 | 0.3 | 0.07 | 0.17 | 0.02 | 0.04 |
| 8 | 117914100 | rs6997684 | -0.038 | -1.696 | -0.653 | -0.141 | 0.004 | 0.034 | 0.034 | 0 | 0.45 | 0.16 | 0.45 | 0.31 | 0.42 | 0.69 |
| 8 | 117915762 | rs1548064 | -0.299 | -1.731 | 0.023 | 0.248 | -0.012 | 0.003 | 0.003 | 0 | 0.35 | 0.56 | 0.43 | 0.54 | 0.29 | 0.26 |
| 8 | 117915914 | rs1548063 | -0.428 | -1.754 | 0.009 | 0.224 | 0.019 | -0.012 | -0.012 | 1.067 | 0.37 | 0.66 | 0.44 | 0.48 | 0.29 | 0.26 |
| 8 | 117916235 | rs720132 | -1.283 | -1.671 | -0.145 | 0.079 | -0.01 | 0.021 | 0.021 | 0 | 0.61 | 0.7 | 0.7 | 0.87 | 0.59 | 0.79 |
| 8 | 117916281 | rs720133 | -0.157 | -1.055 | -0.238 | 0.668 | -0.002 | -0.011 | -0.011 | 0.189 | 0.28 | 0.23 | 0.27 | 0.38 | 0.29 | 0.53 |
| 8 | 117917627 | rs12682047 | -0.199 | -1.559 | 0.623 | 1.182 | 0.004 | 0.094 | 0.094 | 0 | 0.1 | 0.15 | 0.13 | 0.31 | 0.12 | 0.08 |
| 8 | 117926775 | rs3020164 | -1.154 | -1.54 | 0.308 | 1.111 | 0 | 0.001 | 0.001 | 0 | 0.44 | 0.48 | 0.49 | 0.63 | 0.34 | 0.32 |
| 8 | 117927937 | rs1584220 | 0.381 | -1.5 | -0.352 | 1.171 | 0.003 | 0.061 | 0.061 | 0 | 0.09 | 0.07 | 0.13 | 0.3 | 0.12 | 0.07 |
| 8 | 117928825 | rs3020167 | 0.126 | -1.299 | -0.941 | 1.271 | 0.027 | -0.006 | -0.006 | 0 | 0.28 | 0.23 | 0.35 | 0.32 | 0.22 | 0.22 |
| 8 | 117929458 | rs2921739 | -1.007 | -1.239 | -0.946 | 0.759 | 0.049 | 0.021 | 0.021 | 0 | 0.46 | 0.64 | 0.51 | 0.63 | 0.34 | 0.32 |
| 8 | 117936482 | rs16889265 | -0.378 | -0.264 | -1.245 | 0.249 | 0.025 | 0.016 | 0.016 | 0 | 0.27 | 0.22 | 0.3 | 0.17 | 0.21 | 0.22 |
| 8 | 117948841 | rs1348794 | 0.299 | 0.64 | -1.145 | 0.31 | -0.003 | 0.046 | 0.046 | 0.197 | 0.61 | 0.74 | 0.78 | 0.82 | 0.61 | 0.72 |
| 8 | 117951521 | rs6998067 | -0.579 | 0.164 | -0.773 | 1.208 | 0.032 | 0 | 0 | 0 | 0.36 | 0.74 | 0.47 | 0.62 | 0.28 | 0.32 |
| 8 | 117952142 | rs16889293 | -0.205 | 0.305 | -0.864 | 0.878 | 0.013 | 0.021 | 0.021 | 0.016 | 0.31 | 0.22 | 0.32 | 0.18 | 0.28 | 0.3 |
| 8 | 117957039 | rs1446534 | 0.113 | 0.128 | -1.202 | 1.289 | -0.012 | -0.114 | -0.114 | 1.559 | 0.25 | 0.01 | 0.29 | 0.32 | 0.25 | 0.26 |
| 8 | 117958555 | rs3020106 | 0.291 | 0.511 | -2.191 | 0.548 | -0.062 | 0.072 | 0.072 | 0 | 0.23 | 0.06 | 0.12 | 0.12 | 0.33 | 0.34 |
| 8 | 117960147 | rs2921722 | -0.642 | -0.023 | -1.614 | 0.23 | -0.007 | -0.004 | -0.004 | 0.42 | 0.55 | 0.42 | 0.45 | 0.45 | 0.6 | 0.6 |
| 8 | 117974268 | rs3020115 | -0.762 | -0.188 | -0.756 | 0.11 | 0 | 0 | 0 | 0.622 | 0.57 | 0.6 | 0.51 | 0.45 | 0.6 | 0.6 |
| 8 | 118001867 | rs4269571 | 0.715 | -0.626 | 0.219 | 0.445 | -0.047 | -0.001 | -0.001 | 0 | 0.45 | 0.16 | 0.51 | 0.46 | 0.47 | 0.52 |
| 8 | 118002807 | rs6469667 | -0.352 | -0.631 | 0.305 | 0.437 | -0.012 | -0.001 | -0.001 | 0 | 0.48 | 0.31 | 0.38 | 0.44 | 0.52 | 0.47 |
| 8 | 118006571 | rs3020119 | 0.759 | -0.552 | -0.311 | 0.645 | -0.156 | 0.086 | 0.086 | 0 | 0.44 | 0.02 | 0.45 | 0.29 | 0.47 | 0.51 |
| 8 | 118022485 | rs6469668 | -0.304 | -0.557 | -0.245 | 0.406 | 0.039 | 0.009 | 0.009 | 0 | 0.45 | 0.74 | 0.38 | 0.54 | 0.39 | 0.48 |
| 8 | 118024548 | rs2938860 | -0.17 | -0.569 | -0.101 | 0.394 | 0.004 | 0.005 | 0.005 | 0 | 0.87 | 0.88 | 0.88 | 0.83 | 0.84 | 0.97 |
| 8 | 118025645 | rs3019885 | 0.901 | -0.558 | -0.066 | 0.396 | -0.07 | 0.114 | 0.114 | 0.013 | 0.43 | 0.1 | 0.55 | 0.29 | 0.46 | 0.52 |
| 8 | 118028297 | rs11781136 | 0.558 | -0.421 | 0.133 | 0.546 | -0.015 | -0.017 | -0.017 | 0 | 0.09 | 0.03 | 0.06 | 0.18 | 0.13 | 0.03 |
| 8 | 118029019 | rs7009891 | -0.868 | -0.653 | -0.194 | 0.078 | -0.059 | 0.087 | 0.087 | 0.05 | 0.63 | 0.85 | 0.47 | 0.73 | 0.55 | 0.57 |
| 8 | 118034120 | rs1001646 | -0.889 | -0.666 | -0.193 | 0.079 | -0.057 | 0.085 | 0.085 | 0.034 | 0.63 | 0.86 | 0.47 | 0.73 | 0.55 | 0.57 |
| 8 | 118040071 | rs2047962 | 0.187 | -0.768 | -0.185 | -0.01 | 0.004 | 0.001 | 0.001 | 0 | 0.43 | 0.61 | 0.36 | 0.55 | 0.39 | 0.46 |
| 8 | 118044282 | rs11990353 | 0.153 | -0.329 | 0.096 | 0.123 | -0.012 | -0.03 | -0.03 | 0 | 0.09 | 0.1 | 0.04 | 0.01 | 0.04 | 0.08 |
| 8 | 118052405 | rs7011057 | -1.348 | -0.069 | 0.297 | 0.305 | 0.004 | 0.007 | 0.007 | 0.021 | 0.22 | 0.71 | 0.22 | 0.23 | 0.23 | 0.16 |
| 8 | 118057538 | rs11780215 | -0.023 | 0.178 | -0.251 | 0.387 | -0.048 | -0.047 | -0.047 | 0 | 0.08 | 0.01 | 0.08 | 0.14 | 0.17 | 0.02 |
| 8 | 118060748 | rs7016437 | -0.195 | -0.052 | -0.103 | 0.374 | 0.003 | 0.01 | 0.01 | 0 | 0.09 | 0.09 | 0.06 | 0.14 | 0.14 | 0.02 |
| 8 | 118076115 | rs1505521 | 0.159 | -0.22 | 0.508 | 0.277 | -0.003 | -0.043 | -0.043 | 0 | 0.1 | 0.27 | 0.09 | 0.01 | 0.04 | 0.07 |
| 8 | 118078696 | rs2062947 | 0.904 | -0.642 | -0.632 | 0.169 | 0.018 | 0.009 | 0.009 | 0 | 0.29 | 0.01 | 0.09 | 0.21 | 0.19 | 0.31 |
| 8 | 118078751 | rs6980503 | -1.328 | -0.069 | 0.776 | 0.409 | 0.019 | 0.019 | 0.019 | 0 | 0.21 | 0.47 | 0.17 | 0.16 | 0.18 | 0.16 |
| 8 | 118088055 | rs12679997 | 0.249 | -0.14 | 0.734 | 0.52 | 0.002 | -0.009 | -0.009 | 0 | 0.06 | 0.03 | 0.03 | 0.12 | 0.06 | 0.13 |
| 8 | 118113304 | rs16889402 | -1.851 | -0.127 | 0.794 | 0.51 | 0.002 | 0.027 | 0.027 | 0 | 0.27 | 0.54 | 0.21 | 0.28 | 0.25 | 0.29 |
| 8 | 118121273 | rs10107643 | -1.831 | -0.168 | 0.968 | 0.529 | 0.022 | 0.105 | 0.105 | 0 | 0.18 | 0.36 | 0.15 | 0.32 | 0.23 | 0.23 |
| 8 | 118121734 | rs4876702 | -0.549 | -0.166 | 0.951 | 0.538 | 0.023 | 0.064 | 0.064 | 0 | 0.11 | 0.2 | 0.09 | 0.2 | 0.15 | 0.1 |
| 8 | 118123205 | rs7014190 | -2.451 | -0.203 | 1.043 | 0.511 | 0.007 | -0.025 | -0.025 | 0 | 0.3 | 0.78 | 0.25 | 0.34 | 0.26 | 0.3 |
| 8 | 118137545 | rs12676721 | -1.06 | -0.215 | 0.709 | 0.55 | -0.042 | -0.07 | -0.07 | 0 | 0.14 | 0.04 | 0.08 | 0.27 | 0.21 | 0.15 |
| 8 | 118143743 | rs6469674 | 0.094 | -0.221 | 0.725 | 0.509 | -0.012 | -0.05 | -0.05 | 0.437 | 0.11 | 0.29 | 0.09 | 0.01 | 0.04 | 0.07 |
| 8 | 118150993 | rs11989843 | -1.203 | 0.011 | 0.916 | 0.711 | 0.005 | 0.004 | 0.004 | 0 | 0.21 | 0.31 | 0.15 | 0.16 | 0.19 | 0.09 |
| 8 | 118170261 | rs10505312 | -1.207 | 0.294 | 0.755 | 0.82 | 0.018 | 0.102 | 0.102 | 0 | 0.18 | 0.34 | 0.13 | 0.33 | 0.21 | 0.24 |
| 8 | 118174635 | rs10505310 | -1.283 | 0.31 | 0.636 | 0.826 | 0.024 | 0.129 | 0.129 | 0 | 0.19 | 0.43 | 0.15 | 0.33 | 0.21 | 0.24 |
| 8 | 118174941 | rs4876370 | -0.897 | 0.443 | -0.356 | 0.793 | 0.003 | 0.018 | 0.018 | 0 | 0.18 | 0.24 | 0.11 | 0.27 | 0.21 | 0.16 |

| | | | | | | | | | | | | | | | |
|---|---|---|---|---|---|---|---|---|---|---|---|---|---|---|---|
| 8 | 118181090 | rs10955804 | -1.194 | 0.327 | -0.752 | 0.661 | 0.005 | -0.015 | -0.015 | 0 | 0.3 | 0.74 | 0.23 | 0.34 | 0.25 | 0.3 |
| 8 | 118183551 | rs2464592 | 0.384 | 0.332 | -1.235 | -0.358 | 0.031 | 0.1 | 0.1 | 0 | 0.59 | 0.22 | 0.69 | 0.82 | 0.69 | 0.69 |
| 8 | 118183587 | rs2466299 | 0.23 | 0.254 | -1.294 | -0.594 | 0.016 | 0.02 | 0.02 | 0 | 0.68 | 0.26 | 0.75 | 0.82 | 0.8 | 0.77 |
| 8 | 118184783 | rs13266634 | 1.258 | 0.971 | -2.067 | -0.146 | -0.029 | -0.058 | -0.058 | 0 | 0.17 | 0.07 | 0.27 | 0.46 | 0.28 | 0.25 |
| 8 | 118185025 | rs3802177 | 1.212 | 0.852 | -1.663 | -0.177 | -0.029 | -0.06 | -0.06 | 0 | 0.17 | 0.08 | 0.27 | 0.46 | 0.28 | 0.25 |
| 8 | 118185041 | rs2466295 | 0.371 | 1.141 | -1.931 | -0.46 | 0.014 | 0.153 | 0.153 | 0 | 0.54 | 0.21 | 0.65 | 0.82 | 0.61 | 0.65 |
| 8 | 118185733 | rs11558471 | 1.169 | 1.139 | -2.001 | -0.513 | -0.033 | -0.07 | -0.07 | 0 | 0.18 | 0.08 | 0.27 | 0.46 | 0.29 | 0.26 |
| 8 | 118185938 | rs2466293 | 0.788 | 1.058 | -2.033 | -0.573 | -0.027 | -0.007 | -0.007 | 0 | 0.41 | 0.12 | 0.4 | 0.36 | 0.37 | 0.43 |
| 8 | 118187826 | rs10282940 | 0.632 | 0.609 | -0.914 | -0.898 | -0.009 | -0.067 | -0.067 | 1.293 | 0.15 | 0.25 | 0.13 | 0.01 | 0.09 | 0.11 |
| 8 | 118189566 | rs7813920 | 0.541 | 0.595 | -0.264 | -0.905 | -0.012 | -0.092 | -0.092 | 0.05 | 0.16 | 0.32 | 0.14 | 0.01 | 0.1 | 0.12 |
| 8 | 118190393 | rs2466291 | 0.968 | 0.959 | -0.501 | -1.065 | 0.038 | 0.191 | 0.191 | 0.065 | 0.49 | 0.19 | 0.6 | 0.82 | 0.48 | 0.63 |
| 8 | 118200429 | rs4617186 | 2.124 | -0.104 | -0.887 | -0.87 | 0.029 | -0.062 | -0.062 | 0.067 | 0.23 | 0.01 | 0.23 | 0.16 | 0.16 | 0.22 |
| 8 | 118214520 | rs2464582 | -1.523 | 0.24 | -0.254 | -1.032 | -0.01 | 0.015 | 0.015 | 0 | 0.83 | 0.94 | 0.89 | 0.89 | 0.81 | 0.85 |
| 8 | 118215301 | rs6998120 | 0.301 | 0.253 | -0.256 | -1.036 | 0 | -0.013 | -0.013 | 0 | 0.11 | 0.12 | 0.07 | 0.07 | 0.11 | 0.08 |
| 8 | 118217271 | rs10105491 | -0.364 | 0.324 | -0.421 | -0.982 | -0.003 | -0.11 | -0.11 | 0 | 0.26 | 0.41 | 0.2 | 0.08 | 0.2 | 0.2 |
| 8 | 118218521 | rs2464577 | -0.877 | 0.15 | -0.619 | -0.563 | -0.001 | -0.004 | -0.004 | 0 | 0.82 | 0.77 | 0.88 | 0.9 | 0.82 | 0.84 |
| 8 | 118229762 | rs1995222 | 0.381 | 0.564 | -0.543 | -0.108 | 0.017 | 0.033 | 0.033 | 1.19 | 0.27 | 0.34 | 0.41 | 0.47 | 0.44 | 0.2 |
| 8 | 118232629 | rs12545006 | 0.345 | 0.562 | -0.524 | -0.089 | 0.01 | 0.021 | 0.021 | 0.358 | 0.28 | 0.27 | 0.4 | 0.47 | 0.45 | 0.2 |
| 8 | 118234918 | rs10955806 | -0.873 | 0.41 | -1.56 | -0.353 | -0.017 | -0.049 | -0.049 | 0 | 0.58 | 0.88 | 0.77 | 0.52 | 0.62 | 0.38 |
| 8 | 118236041 | rs7828267 | -0.501 | 0.416 | -0.854 | -0.53 | -0.002 | -0.061 | -0.061 | 0 | 0.16 | 0.19 | 0.12 | 0.01 | 0.12 | 0.1 |
| 8 | 118237006 | rs10505309 | -0.478 | 0.172 | -1.611 | -0.336 | 0.017 | 0.023 | 0.023 | 0 | 0.15 | 0.09 | 0.09 | 0.02 | 0.07 | 0.12 |
| 8 | 118238117 | rs1499430 | -0.428 | 0.877 | -2.063 | 0.436 | -0.075 | -0.071 | -0.071 | 0 | 0.73 | 0.97 | 0.85 | 0.71 | 0.75 | 0.48 |
| 8 | 118240824 | rs1392044 | 0.289 | 0.896 | -2.224 | 0.149 | -0.044 | -0.005 | -0.005 | 0.214 | 0.32 | 0.15 | 0.34 | 0.38 | 0.41 | 0.18 |
| 8 | 118241220 | rs2649102 | -0.552 | 0.518 | -1.234 | 0.185 | -0.013 | -0.016 | -0.016 | 0 | 0.72 | 0.53 | 0.75 | 0.79 | 0.75 | 0.54 |
| 8 | 118245643 | rs1499433 | 0.895 | 1.086 | -1.79 | 0.174 | -0.057 | -0.073 | -0.073 | 0 | 0.23 | 0.05 | 0.28 | 0.35 | 0.29 | 0.16 |
| 8 | 118248838 | rs7831051 | 0.086 | 0.783 | -1.173 | 0.107 | -0.008 | 0.01 | 0.01 | 0.472 | 0.29 | 0.58 | 0.26 | 0.32 | 0.27 | 0.21 |
| 8 | 118249754 | rs1392037 | -1.692 | 0.685 | -1.448 | -0.018 | -0.017 | 0.026 | 0.026 | 2.773 | 0.75 | 0.96 | 0.87 | 0.83 | 0.77 | 0.56 |
| 8 | 118250328 | rs904544 | 0.066 | 0.675 | -1.193 | -0.037 | -0.012 | -0.002 | -0.002 | 0 | 0.44 | 0.31 | 0.5 | 0.48 | 0.46 | 0.33 |
| 8 | 118250570 | rs2009769 | 1.915 | 0.759 | -1.353 | -0.109 | -0.041 | -0.043 | -0.043 | 0 | 0.13 | 0.01 | 0.18 | 0.13 | 0.12 | 0.17 |
| 8 | 118251913 | rs1793733 | 0.029 | 0.882 | -0.861 | -0.047 | -0.009 | -0.025 | -0.025 | 0.229 | 0.44 | 0.26 | 0.57 | 0.51 | 0.45 | 0.34 |
| 8 | 118254700 | rs2464594 | 0.597 | 1.137 | -0.224 | 0.303 | -0.019 | -0.014 | -0.014 | 0 | 0.26 | 0.22 | 0.31 | 0.36 | 0.33 | 0.15 |
| 8 | 118257185 | rs1499417 | -0.211 | 1.154 | 0.197 | 0.255 | 0.005 | 0.014 | 0.014 | 0 | 0.27 | 0.28 | 0.33 | 0.33 | 0.3 | 0.22 |
| 8 | 118260487 | rs1793729 | 0.291 | 1.491 | 0.536 | 0.021 | -0.025 | -0.029 | -0.029 | 0 | 0.25 | 0.15 | 0.28 | 0.37 | 0.32 | 0.16 |
| 8 | 118265880 | rs1793723 | 0.018 | 1.461 | 0.699 | -0.094 | 0 | 0 | 0 | 0 | 0.27 | 0.33 | 0.3 | 0.36 | 0.32 | 0.16 |
| 8 | 118272454 | rs1695687 | -0.353 | 1.53 | 0.832 | -0.098 | 0.025 | 0.034 | 0.034 | 0 | 0.3 | 0.45 | 0.46 | 0.39 | 0.36 | 0.17 |
| 8 | 118276623 | rs1695688 | -0.28 | 1.541 | 0.834 | -0.09 | 0.015 | 0.019 | 0.019 | 0 | 0.29 | 0.35 | 0.45 | 0.4 | 0.37 | 0.18 |
| 8 | 118276821 | rs2705254 | 0.294 | 1.531 | 0.746 | -0.226 | 0.004 | -0.066 | -0.066 | 3.117 | 0.15 | 0.25 | 0.16 | 0.01 | 0.19 | 0.07 |
| 8 | 118284380 | rs1499439 | -1.371 | 1.528 | 0.646 | -0.218 | -0.004 | 0.015 | 0.015 | 1.016 | 0.25 | 0.82 | 0.38 | 0.05 | 0.28 | 0.09 |
| 8 | 118288747 | rs1499437 | -0.268 | 1.177 | 0.569 | -0.118 | 0.023 | 0.029 | 0.029 | 0 | 0.31 | 0.45 | 0.52 | 0.39 | 0.37 | 0.18 |
| 8 | 118289276 | rs1695725 | -1.331 | 1.174 | 0.449 | -0.132 | -0.018 | 0.016 | 0.016 | 0 | 0.47 | 0.85 | 0.64 | 0.4 | 0.56 | 0.25 |
| 8 | 118290724 | rs1695724 | -0.074 | 1.408 | 0.419 | -0.003 | 0.002 | 0.004 | 0.004 | 0 | 0.28 | 0.32 | 0.45 | 0.39 | 0.36 | 0.17 |
| 8 | 118291899 | rs16889547 | 0.086 | 1.292 | 0.026 | 0.051 | -0.003 | -0.071 | -0.071 | 0 | 0.15 | 0.27 | 0.15 | 0.01 | 0.17 | 0.07 |
| 8 | 118293451 | rs7011815 | -0.528 | 1.461 | -0.291 | 0.149 | 0.006 | -0.033 | -0.033 | 0 | 0.18 | 0.28 | 0.31 | 0.04 | 0.24 | 0.09 |
| 8 | 118296603 | rs1793709 | -0.208 | 1.026 | -0.897 | 0.269 | -0.005 | 0.004 | 0.004 | 0 | 0.42 | 0.62 | 0.48 | 0.37 | 0.49 | 0.23 |
| 8 | 118300497 | rs1793705 | -0.156 | 0.499 | -1.174 | 0.253 | -0.024 | 0.053 | 0.053 | 0 | 0.57 | 0.76 | 0.66 | 0.31 | 0.57 | 0.59 |
| 8 | 118302950 | rs2511543 | -0.329 | 0.574 | -1.214 | 0.505 | -0.044 | -0.009 | -0.009 | 0.042 | 0.6 | 0.88 | 0.81 | 0.37 | 0.59 | 0.6 |
| 8 | 118306363 | rs1472487 | -0.306 | -0.526 | -0.113 | -0.28 | -0.089 | -0.133 | -0.133 | 0 | 0.25 | 0.45 | 0.1 | 0.03 | 0.09 | 0.24 |
| 8 | 118307005 | rs1793704 | -1.078 | -0.679 | -0.316 | -0.325 | -0.031 | 0.016 | 0.016 | 0 | 0.64 | 0.88 | 0.72 | 0.29 | 0.6 | 0.57 |
| 8 | 118307354 | rs1793703 | 0.271 | -0.685 | -0.18 | -0.258 | -0.011 | 0.023 | 0.023 | 0 | 0.4 | 0.32 | 0.52 | 0.24 | 0.5 | 0.33 |
| 8 | 118308214 | rs16889584 | -0.316 | -0.77 | 0.079 | -0.334 | -0.104 | -0.096 | -0.096 | 0 | 0.25 | 0.5 | 0.23 | 0.09 | 0.08 | 0.23 |
| 8 | 118309831 | rs924386 | -0.364 | -0.592 | 0.107 | 0.556 | 0.039 | 0.266 | 0.266 | 0.15 | 0.1 | 0.31 | 0.09 | 0.33 | 0.1 | 0.13 |
| 8 | 118311155 | rs1695695 | 1.356 | -0.219 | -0.453 | 0.09 | -0.052 | 0.057 | 0.057 | 0 | 0.21 | 0.08 | 0.35 | 0.09 | 0.32 | 0.2 |
| 8 | 118311980 | rs7017745 | 0.433 | 0.033 | -0.142 | 0.388 | 0 | 0.124 | 0.124 | 0 | 0.08 | 0.12 | 0.13 | 0.34 | 0.08 | 0.12 |
| 8 | 118313693 | rs989511 | 1.138 | -0.15 | 0.432 | 0.182 | -0.001 | 0 | 0 | 0 | 0.34 | 0.31 | 0.19 | 0.39 | 0.35 | 0.31 |
| 8 | 118318971 | rs13277475 | 0.477 | -0.098 | -0.14 | 0.321 | -0.034 | -0.026 | -0.026 | 0 | 0.12 | 0.03 | 0.16 | 0.14 | 0.15 | 0.09 |
| 8 | 118320851 | rs4876717 | -0.472 | -0.453 | -0.024 | 0.548 | 0.003 | -0.005 | -0.005 | 0 | 0.25 | 0.73 | 0.29 | 0.38 | 0.14 | 0.37 |
| 8 | 118321722 | rs11996643 | -0.591 | -0.457 | 0.016 | 0.536 | -0.042 | 0.038 | 0.038 | 2.509 | 0.25 | 0.62 | 0.28 | 0.37 | 0.14 | 0.37 |
| 8 | 118325941 | rs1695693 | -1.135 | -0.385 | -0.339 | 0.471 | -0.051 | 0.051 | 0.051 | 0.36 | 0.71 | 0.89 | 0.67 | 0.86 | 0.66 | 0.79 |
| 8 | 118332179 | rs1695690 | -0.007 | -0.467 | -0.081 | 0.427 | 0.006 | 0.017 | 0.017 | 0 | 0.53 | 0.43 | 0.62 | 0.75 | 0.59 | 0.54 |
| 8 | 118336206 | rs7843021 | 0.378 | -0.318 | -0.408 | 0.423 | 0 | 0.006 | 0.006 | 0 | 0.08 | 0.05 | 0.13 | 0.28 | 0.08 | 0.11 |
| 8 | 118339749 | rs992385 | 0.683 | -0.163 | -0.508 | 0.463 | -0.065 | -0.008 | -0.008 | 0 | 0.4 | 0.16 | 0.34 | 0.42 | 0.48 | 0.42 |
| 8 | 118340002 | rs1793687 | -0.611 | 0.34 | -0.668 | 0.263 | -0.053 | 0.028 | 0.028 | 0 | 0.69 | 0.91 | 0.67 | 0.79 | 0.65 | 0.79 |
| 8 | 118347262 | rs7836463 | -0.494 | -0.015 | -0.256 | 0.27 | -0.028 | -0.079 | -0.079 | 0 | 0.29 | 0.42 | 0.2 | 0.11 | 0.23 | 0.3 |
| 8 | 118348077 | rs16889608 | 0.377 | 0.016 | -0.69 | 0.263 | 0.023 | 0.03 | 0.03 | 0 | 0.21 | 0.11 | 0.11 | 0.03 | 0.15 | 0.26 |
| 8 | 118352994 | rs13262016 | 1.079 | 0.43 | -0.85 | 0.418 | -0.004 | 0 | 0 | 0.117 | 0.36 | 0.37 | 0.29 | 0.38 | 0.29 | 0.35 |
| 8 | 118356334 | rs1603942 | -0.626 | 0.572 | -0.79 | 0.588 | -0.071 | 0.03 | 0.03 | 0.154 | 0.69 | 0.83 | 0.57 | 0.78 | 0.54 | 0.74 |
| 8 | 118359997 | rs1603941 | -0.584 | 0.604 | -0.841 | 0.604 | -0.042 | 0.025 | 0.025 | 0.168 | 0.69 | 0.77 | 0.56 | 0.78 | 0.54 | 0.74 |
| 8 | 118366317 | rs7843259 | -0.766 | 0.481 | -0.684 | 0.851 | -0.019 | 0.024 | 0.024 | 0 | 0.31 | 0.39 | 0.27 | 0.39 | 0.23 | 0.4 |
| 8 | 118367827 | rs2853190 | -0.731 | 0.714 | -1.527 | 0.815 | -0.014 | -0.076 | -0.076 | 0 | 0.32 | 0.12 | 0.37 | 0.4 | 0.37 | 0.44 |

| | | | | | | | | | | | | | | | |
|---|---|---|---|---|---|---|---|---|---|---|---|---|---|---|---|
| 8 | 118376314 | rs3018441 | -0.324 | 0.931 | -1.686 | 1.027 | -0.009 | -0.055 | -0.055 | 0 | 0.8 | 0.91 | 0.82 | 0.59 | 0.8 | 0.74 |
| 8 | 118381545 | rs2649097 | -0.445 | 0.534 | -1.564 | 0.916 | -0.001 | -0.001 | -0.001 | 0 | 0.1 | 0.19 | 0.22 | 0.12 | 0.13 | 0.12 |
| 8 | 118388429 | rs1793750 | 0.388 | 1.046 | -1.741 | 0.99 | -0.082 | -0.11 | -0.11 | 0 | 0.22 | 0.02 | 0.12 | 0.37 | 0.26 | 0.23 |
| 8 | 118390198 | rs11777279 | -0.819 | 1.58 | -2.051 | 0.889 | -0.011 | 0.003 | 0.003 | 0 | 0.15 | 0.09 | 0.2 | 0.15 | 0.23 | 0.21 |
| 8 | 118391516 | rs7820779 | 0.214 | 1.59 | -2.044 | 0.867 | -0.048 | -0.005 | -0.005 | 0 | 0.09 | 0.01 | 0.12 | 0.06 | 0.18 | 0.1 |
| 8 | 118398295 | rs1352561 | -0.127 | 1.19 | -1.955 | 0.789 | -0.008 | -0.001 | -0.001 | 0 | 0.08 | 0.08 | 0.08 | 0.09 | 0.05 | 0.12 |
| 8 | 118410695 | rs10505301 | 1.662 | 0.724 | -2.824 | 0.432 | -0.056 | -0.057 | -0.057 | 0.177 | 0.17 | 0.03 | 0.12 | 0.24 | 0.22 | 0.24 |
| 8 | 118418008 | rs1586601 | 0.138 | 0.856 | -2.276 | 0.427 | -0.04 | -0.051 | -0.051 | 0 | 0.33 | 0.21 | 0.33 | 0.48 | 0.44 | 0.39 |
| 8 | 118431887 | rs969592 | -0.962 | 0.361 | -1.315 | 0.192 | 0 | 0 | 0 | 0 | 0.61 | 0.61 | 0.74 | 0.69 | 0.64 | 0.69 |
| 8 | 118442583 | rs11987213 | 0.33 | 0.361 | -1.276 | -0.252 | 0.005 | 0.003 | 0.003 | 0 | 0.15 | 0.11 | 0.3 | 0.13 | 0.12 | 0.18 |
| 8 | 118457084 | rs1548124 | -0.381 | 0.71 | -1.051 | 0.021 | 0.009 | -0.017 | -0.017 | 0 | 0.59 | 0.2 | 0.51 | 0.41 | 0.66 | 0.48 |
| 8 | 118476814 | rs10099281 | 0.075 | 0.79 | -0.557 | -0.007 | 0.001 | 0.014 | 0.014 | 0 | 0.53 | 0.42 | 0.4 | 0.27 | 0.55 | 0.37 |
| 8 | 118478847 | rs7015627 | -0.122 | 0.283 | -0.288 | -0.065 | -0.014 | 0.008 | 0.008 | 0 | 0.08 | 0.09 | 0.12 | 0.13 | 0.06 | 0.12 |
| 8 | 118481119 | rs17740228 | -0.41 | 0.311 | -0.44 | -0.056 | 0.021 | -0.022 | -0.022 | 0.216 | 0.09 | 0.03 | 0.12 | 0.14 | 0.07 | 0.12 |
| 8 | 118489697 | rs1489407 | 0.024 | 0.429 | 0.097 | -0.252 | -0.001 | 0.006 | 0.006 | 0 | 0.53 | 0.42 | 0.4 | 0.23 | 0.48 | 0.37 |
| 8 | 118511339 | rs1489404 | -0.697 | 0.552 | 0.598 | -0.257 | -0.006 | 0 | 0 | 0 | 0.63 | 0.61 | 0.54 | 0.36 | 0.54 | 0.48 |
| 8 | 118515249 | rs2873645 | -1.444 | 0.558 | 0.51 | -0.441 | -0.002 | 0.027 | 0.027 | 0 | 0.22 | 0.49 | 0.33 | 0.28 | 0.26 | 0.23 |
| 8 | 118524413 | rs1871583 | -0.555 | 0.672 | 0.591 | -0.416 | 0.011 | -0.012 | -0.012 | 0 | 0.61 | 0.52 | 0.48 | 0.33 | 0.55 | 0.47 |
| 8 | 118532964 | rs6469696 | -0.871 | 0.723 | 0.577 | -0.536 | 0.004 | -0.007 | -0.007 | 0 | 0.63 | 0.6 | 0.48 | 0.24 | 0.53 | 0.47 |
| 8 | 118536441 | rs6993408 | -0.537 | 1.079 | 0.708 | -0.315 | 0.001 | -0.002 | -0.002 | 0.103 | 0.09 | 0.06 | 0.08 | 0.04 | 0.07 | 0.12 |
| 8 | 118542735 | rs10505295 | -0.395 | 1.434 | 0.996 | -0.171 | 0.027 | -0.005 | -0.005 | 0 | 0.1 | 0.15 | 0.17 | 0.06 | 0.17 | 0.08 |
| 8 | 118558257 | rs16889867 | -1.027 | 1.429 | 1.033 | -0.188 | 0.044 | 0.006 | 0.006 | 0 | 0.12 | 0.25 | 0.25 | 0.12 | 0.2 | 0.09 |
| 8 | 118560491 | rs922587 | -1.249 | 1.892 | 0.904 | -0.277 | 0.012 | -0.006 | -0.006 | 0 | 0.14 | 0.14 | 0.21 | 0.06 | 0.24 | 0.14 |
| 8 | 118567694 | rs11994257 | -0.677 | 1.497 | 1.196 | -0.348 | -0.016 | 0.024 | 0.024 | 0 | 0.63 | 0.73 | 0.51 | 0.27 | 0.54 | 0.45 |
| 8 | 118571202 | rs16889881 | -2.08 | 1.59 | 2.024 | 0.168 | 0.08 | -0.065 | -0.065 | 0.739 | 0.2 | 0.43 | 0.26 | 0.08 | 0.27 | 0.15 |
| 8 | 118573949 | rs755982 | -0.605 | 0.87 | 1.657 | 0.351 | -0.034 | 0.009 | 0.009 | 0 | 0.12 | 0.04 | 0.18 | 0.06 | 0.16 | 0.08 |
| 8 | 118574630 | rs749492 | -0.598 | 0.871 | 1.666 | 0.351 | -0.03 | 0.01 | 0.01 | 0 | 0.12 | 0.04 | 0.18 | 0.06 | 0.16 | 0.08 |
| 8 | 118602983 | rs732095 | -1.968 | 0.863 | 2.028 | 0.49 | 0.011 | -0.009 | -0.009 | 0 | 0.21 | 0.3 | 0.27 | 0.14 | 0.26 | 0.16 |
| 8 | 118608973 | rs6986953 | -1.701 | 0.903 | 2.032 | 0.483 | 0.081 | 0.075 | 0.075 | 0 | 0.1 | 0.45 | 0.14 | 0.09 | 0.11 | 0.08 |
| 8 | 118611634 | rs17746363 | -1.261 | 0.85 | 1.881 | 0.458 | -0.001 | -0.001 | -0.001 | 0.607 | 0.14 | 0.13 | 0.17 | 0.06 | 0.19 | 0.08 |
| 8 | 118613782 | rs11991406 | -2.416 | 0.834 | 1.622 | 0.52 | 0.013 | -0.002 | -0.002 | 0 | 0.24 | 0.54 | 0.36 | 0.2 | 0.28 | 0.17 |
| 8 | 118617032 | rs1396347 | -1.2 | 0.835 | 0.309 | 0.388 | -0.002 | 0.002 | 0.002 | 0 | 0.14 | 0.15 | 0.17 | 0.06 | 0.17 | 0.08 |
| 8 | 118626279 | rs10100465 | -0.873 | 0.71 | 0.445 | 0.722 | 0.005 | 0.004 | 0.004 | 2.964 | 0.21 | 0.31 | 0.17 | 0.24 | 0.22 | 0.25 |
| 8 | 118632412 | rs1396354 | -1.185 | 0.549 | 1.249 | 0.725 | 0.03 | 0.027 | 0.027 | 3.944 | 0.23 | 0.51 | 0.25 | 0.27 | 0.23 | 0.28 |
| 8 | 118638878 | rs10089009 | -0.964 | 0.552 | 1.418 | 0.726 | 0.014 | 0.009 | 0.009 | 2.806 | 0.21 | 0.34 | 0.18 | 0.24 | 0.22 | 0.28 |
| 8 | 118641568 | rs1160120 | -1.693 | 0.233 | 1.611 | 0.792 | 0.003 | -0.014 | -0.014 | 0 | 0.29 | 0.82 | 0.34 | 0.37 | 0.24 | 0.33 |
| 8 | 118671467 | rs4604472 | 1.1 | 0.119 | 1.809 | 0.951 | -0.001 | -0.001 | -0.001 | 1.723 | 0.74 | 0.27 | 0.73 | 0.75 | 0.77 | 0.71 |
| 8 | 118676625 | rs10955823 | 0.804 | 0.194 | 1.609 | 0.979 | 0.01 | 0.023 | 0.023 | 3.221 | 0.78 | 0.56 | 0.77 | 0.75 | 0.79 | 0.72 |
| 8 | 118687239 | rs1508558 | 1.634 | 0.086 | 0.218 | 1.31 | -0.002 | 0.026 | 0.026 | 3.986 | 0.58 | 0.12 | 0.61 | 0.37 | 0.67 | 0.42 |
| 8 | 118706776 | rs218017 | 0.799 | 0.058 | 0.308 | 1.347 | -0.047 | -0.045 | -0.045 | 0 | 0.77 | 0.91 | 0.77 | 0.71 | 0.72 | 0.72 |
| 8 | 118707556 | rs218019 | 1.784 | -0.096 | 0.303 | 1.421 | 0.004 | -0.028 | -0.028 | 0 | 0.6 | 0.17 | 0.66 | 0.45 | 0.66 | 0.49 |
| 8 | 118707726 | rs3115775 | 0.075 | -0.155 | 0.276 | 1.443 | -0.013 | -0.027 | -0.027 | 0 | 0.08 | 0.02 | 0.04 | 0.29 | 0.07 | 0.25 |
| 8 | 118708824 | rs218020 | 1.893 | -0.226 | 0.322 | 1.372 | 0.018 | -0.014 | -0.014 | 0 | 0.6 | 0.22 | 0.67 | 0.45 | 0.68 | 0.45 |
| 8 | 118710620 | rs218023 | 0.684 | -0.088 | 0.239 | 1.392 | -0.032 | -0.013 | -0.013 | 2.324 | 0.78 | 0.88 | 0.82 | 0.79 | 0.75 | 0.7 |
| 8 | 118713826 | rs13249969 | -0.038 | 0 | 0.209 | 1.48 | -0.047 | 0.001 | 0.001 | 0.427 | 0.1 | 0.01 | 0.13 | 0.07 | 0.13 | 0.17 |
| 8 | 118714803 | rs218011 | 0.526 | -0.525 | -0.116 | 1.184 | -0.022 | 0.015 | 0.015 | 0 | 0.71 | 0.88 | 0.8 | 0.79 | 0.71 | 0.69 |
| 8 | 118716319 | rs3134388 | 0.058 | -0.395 | 0.102 | 1.356 | -0.001 | 0.044 | 0.044 | 0 | 0.07 | 0.07 | 0.09 | 0.33 | 0.07 | 0.22 |
| 8 | 118717588 | rs3134389 | 0.067 | -0.396 | 0.096 | 1.381 | 0 | 0.021 | 0.021 | 0 | 0.07 | 0.05 | 0.09 | 0.34 | 0.07 | 0.22 |
| 8 | 118718840 | rs3105771 | -1.152 | -0.518 | 0.333 | 1.348 | -0.017 | 0.135 | 0.135 | 0.013 | 0.13 | 0.27 | 0.17 | 0.42 | 0.09 | 0.27 |
| 8 | 118718959 | rs926148 | 1.912 | -0.753 | 0.048 | 1.073 | -0.009 | 0.039 | 0.039 | 0.166 | 0.46 | 0.1 | 0.57 | 0.37 | 0.59 | 0.36 |
| 8 | 118719250 | rs734649 | -1.014 | -0.688 | 0.166 | 1.062 | -0.017 | 0.135 | 0.135 | 0.013 | 0.13 | 0.27 | 0.17 | 0.42 | 0.09 | 0.27 |
| 8 | 118726160 | rs3134395 | -1.917 | -0.793 | -0.45 | 0.972 | 0.015 | -0.043 | -0.043 | 0 | 0.54 | 0.88 | 0.42 | 0.51 | 0.39 | 0.65 |
| 8 | 118729267 | rs3105760 | 0.407 | -0.811 | -0.483 | 0.954 | -0.001 | -0.023 | -0.023 | 0 | 0.08 | 0.03 | 0.04 | 0.28 | 0.06 | 0.24 |
| 8 | 118729295 | rs1567303 | -0.861 | -0.944 | -0.193 | 0.792 | 0.006 | 0.059 | 0.059 | 0 | 0.34 | 0.23 | 0.22 | 0.09 | 0.3 | 0.37 |
| 8 | 118731317 | rs1396349 | -1.725 | -0.832 | -0.18 | 0.859 | 0.038 | 0.006 | 0.006 | 0.057 | 0.52 | 0.81 | 0.37 | 0.46 | 0.38 | 0.64 |
| 8 | 118738651 | rs11776828 | 0.022 | -0.372 | -0.532 | 0.45 | -0.009 | 0.025 | 0.025 | 0 | 0.22 | 0.16 | 0.22 | 0.07 | 0.23 | 0.19 |
| 8 | 118751200 | rs4876736 | -0.436 | -0.35 | 0.213 | 1.065 | -0.003 | 0.015 | 0.015 | 0.102 | 0.12 | 0.22 | 0.11 | 0.17 | 0.1 | 0.15 |
| 8 | 118752520 | rs3115767 | -0.047 | -0.393 | 0.118 | 1.326 | 0 | -0.016 | -0.016 | 0 | 0.09 | 0.04 | 0.04 | 0.33 | 0.08 | 0.24 |
| 8 | 118756256 | rs4876738 | -0.517 | -0.245 | 0.373 | 1.153 | -0.014 | 0.015 | 0.015 | 0.093 | 0.11 | 0.22 | 0.11 | 0.17 | 0.09 | 0.15 |
| 8 | 118759092 | rs12682151 | -1.147 | 0.057 | -0.491 | 0.821 | -0.007 | -0.052 | -0.052 | 0 | 0.38 | 0.78 | 0.44 | 0.52 | 0.38 | 0.42 |
| 8 | 118761500 | rs1396353 | 0.25 | 0.666 | -0.702 | 1.13 | 0.006 | 0.011 | 0.011 | 0 | 0.19 | 0.19 | 0.29 | 0.35 | 0.28 | 0.27 |
| 8 | 118764706 | rs741922 | -0.64 | 0.297 | -0.733 | 0.825 | -0.001 | 0.002 | 0.002 | 0.018 | 0.11 | 0.13 | 0.1 | 0.17 | 0.1 | 0.15 |
| 8 | 118770442 | rs7011355 | -0.105 | 0.218 | -1.361 | 0.47 | 0.012 | 0.039 | 0.039 | 0 | 0.64 | 0.81 | 0.65 | 0.72 | 0.69 | 0.55 |
| 8 | 118774997 | rs720896 | 0.184 | 0.361 | -1.185 | 0.85 | -0.011 | -0.009 | -0.009 | 0 | 0.48 | 0.67 | 0.37 | 0.52 | 0.46 | 0.41 |
| 8 | 118776037 | rs13267691 | 0.321 | 0.237 | -0.246 | 0.788 | -0.002 | -0.003 | -0.003 | 0 | 0.44 | 0.42 | 0.35 | 0.51 | 0.46 | 0.41 |
| 8 | 118784316 | rs6981439 | -0.732 | -0.194 | -0.161 | 0.528 | -0.009 | 0.029 | 0.029 | 2.643 | 0.23 | 0.61 | 0.31 | 0.28 | 0.24 | 0.36 |
| 8 | 118794474 | rs7836738 | 1.52 | -0.318 | -0.942 | 0.783 | -0.127 | -0.019 | -0.019 | 0 | 0.29 | 0.03 | 0.27 | 0.25 | 0.36 | 0.29 |
| 8 | 118805719 | rs12548168 | 0.109 | -0.144 | -0.651 | 0.889 | 0 | 0.023 | 0.023 | 0.126 | 0.19 | 0.29 | 0.29 | 0.28 | 0.24 | 0.35 |
| 8 | 118806423 | rs4876754 | -1.015 | -0.236 | -0.542 | 0.749 | 0.005 | 0.015 | 0.015 | 0 | 0.64 | 0.74 | 0.64 | 0.79 | 0.7 | 0.77 |

| | | | | | | | | | | | | | | | | |
|---|---|---|---|---|---|---|---|---|---|---|---|---|---|---|---|---|
| 8 | 118808566 | rs4876390 | -0.978 | -0.016 | -0.598 | 0.862 | 0.002 | -0.001 | -0.001 | 0.267 | 0.66 | 0.66 | 0.69 | 0.67 | 0.64 | 0.69 |
| 8 | 118808893 | rs10113582 | 0.825 | 0.079 | -0.874 | 1.024 | -0.014 | -0.027 | -0.027 | 0 | 0.06 | 0.01 | 0.04 | 0.19 | 0.07 | 0.07 |
| 8 | 118814084 | rs4242575 | -0.173 | 0.252 | -1.034 | 0.639 | -0.009 | 0.02 | 0.02 | 0 | 0.53 | 0.3 | 0.66 | 0.55 | 0.53 | 0.4 |
| 8 | 118819578 | rs7837891 | 0.451 | 0.475 | -1.254 | 0.754 | 0.043 | -0.001 | -0.001 | 0 | 0.44 | 0.16 | 0.55 | 0.38 | 0.42 | 0.3 |
| 8 | 118820371 | rs17429936 | 1.035 | 0.237 | -1.686 | 0.421 | 0.028 | 0.11 | 0.11 | 0 | 0.26 | 0.05 | 0.36 | 0.07 | 0.23 | 0.19 |
| 8 | 118820586 | rs12675718 | 0.384 | 0.631 | -1.327 | 0.71 | -0.039 | -0.044 | -0.044 | 0 | 0.08 | 0.01 | 0.1 | 0.17 | 0.11 | 0.1 |
| 8 | 118823167 | rs17439576 | 0.306 | 0.627 | -1.316 | 0.708 | -0.022 | -0.026 | -0.026 | 0 | 0.08 | 0.04 | 0.1 | 0.17 | 0.11 | 0.1 |
| 8 | 118827839 | rs11989122 | -0.782 | 0.638 | -0.694 | 0.276 | 0.002 | 0.003 | 0.003 | 0 | 0.66 | 0.64 | 0.77 | 0.6 | 0.69 | 0.61 |
| 8 | 118828761 | rs10955836 | 0.638 | 0.441 | -0.726 | 0.545 | -0.002 | -0.001 | -0.001 | 0.326 | 0.83 | 0.81 | 0.75 | 0.35 | 0.75 | 0.59 |
| 8 | 118830036 | rs7000499 | 0.366 | 0.415 | -0.712 | 0.702 | -0.001 | 0.041 | 0.041 | 0 | 0.85 | 0.81 | 0.87 | 0.46 | 0.8 | 0.57 |
| 8 | 118830320 | rs6982912 | 0.283 | 0.758 | -0.278 | 0.2 | -0.006 | 0.042 | 0.042 | 0 | 0.53 | 0.36 | 0.62 | 0.24 | 0.57 | 0.29 |
| 8 | 118830820 | rs10955837 | 0.268 | 0.753 | -0.281 | 0.197 | -0.007 | 0.041 | 0.041 | 0 | 0.53 | 0.36 | 0.62 | 0.24 | 0.57 | 0.29 |
| 8 | 118830894 | rs4876757 | 0.9 | 0.827 | -0.043 | 0.225 | 0.001 | -0.007 | -0.007 | 0.144 | 0.32 | 0.35 | 0.24 | 0.22 | 0.23 | 0.27 |
| 8 | 118835137 | rs4366105 | 1.227 | 0.728 | -0.464 | -0.243 | 0.001 | 0.114 | 0.114 | 0 | 0.29 | 0.05 | 0.3 | 0.09 | 0.27 | 0.19 |
| 8 | 118838295 | rs17439784 | 0.608 | 0.759 | -0.366 | 0.049 | -0.026 | -0.028 | -0.028 | 0 | 0.1 | 0.04 | 0.1 | 0.4 | 0.19 | 0.3 |
| 8 | 118844039 | rs12682457 | 0.538 | 0.885 | -0.561 | 0.055 | -0.017 | -0.003 | -0.003 | 0.029 | 0.1 | 0.06 | 0.1 | 0.4 | 0.19 | 0.3 |
| 8 | 118846051 | rs11989463 | 1.067 | 0.676 | -0.342 | -0.162 | -0.007 | 0.037 | 0.037 | 0 | 0.2 | 0.09 | 0.19 | 0.04 | 0.25 | 0.11 |
| 8 | 118853829 | rs10103003 | 0.06 | 0.735 | -0.555 | -0.33 | -0.001 | -0.001 | -0.001 | 0 | 0.46 | 0.45 | 0.55 | 0.62 | 0.51 | 0.57 |
| 8 | 118856307 | rs17430287 | 0.953 | 0.593 | -0.801 | -0.364 | -0.053 | 0.113 | 0.113 | 0 | 0.19 | 0.02 | 0.18 | 0.02 | 0.25 | 0.11 |
| 8 | 118871476 | rs4463453 | 0.748 | 0.893 | -0.887 | -0.539 | -0.029 | 0.114 | 0.114 | 0.714 | 0.34 | 0.12 | 0.34 | 0.09 | 0.36 | 0.21 |
| 8 | 118873577 | rs4478606 | -0.785 | 0.929 | -0.386 | -0.088 | -0.013 | 0.008 | 0.008 | 1.239 | 0.7 | 0.59 | 0.74 | 0.63 | 0.8 | 0.71 |
| 8 | 118879130 | rs12677136 | 0.037 | 1.414 | -0.092 | 0.491 | -0.015 | -0.008 | -0.008 | 0 | 0.1 | 0.05 | 0.09 | 0.4 | 0.16 | 0.29 |
| 8 | 118881446 | rs4876765 | 0.777 | 1.048 | 0.041 | -0.397 | 0.005 | 0.112 | 0.112 | 0.014 | 0.55 | 0.74 | 0.42 | 0.15 | 0.43 | 0.32 |
| 8 | 118883490 | rs17503900 | -0.3 | 0.698 | 0.715 | 0.196 | -0.001 | -0.02 | -0.02 | 0 | 0.3 | 0.23 | 0.28 | 0.48 | 0.35 | 0.43 |
| 8 | 118885642 | rs12548703 | 0.688 | 0.816 | 0.626 | 0.204 | 0.006 | 0.007 | 0.007 | 0.524 | 0.2 | 0.15 | 0.14 | 0.07 | 0.12 | 0.15 |
| 8 | 118887266 | rs11986974 | -1.456 | 0.728 | 1.133 | 0.214 | 0.043 | 0.231 | 0.231 | 0.068 | 0.17 | 0.43 | 0.17 | 0.43 | 0.25 | 0.29 |
| 8 | 118891896 | rs11785986 | -0.609 | 0.788 | 1.143 | 0.248 | 0.031 | 0.155 | 0.155 | 0 | 0.13 | 0.23 | 0.16 | 0.41 | 0.23 | 0.27 |
| 8 | 118895712 | rs17450772 | 0.7 | 0.134 | 0.768 | 0.014 | -0.002 | -0.008 | -0.008 | 0 | 0.2 | 0.19 | 0.15 | 0.08 | 0.12 | 0.16 |
| 8 | 118901585 | rs4480148 | 0.776 | 0.545 | 0.686 | 0.114 | 0.007 | 0.026 | 0.026 | 0.119 | 0.79 | 0.76 | 0.8 | 0.5 | 0.69 | 0.67 |
| 8 | 118903024 | rs9692869 | -0.827 | 0.487 | 1.011 | 0.188 | -0.006 | -0.023 | -0.023 | 1 | 0.37 | 0.33 | 0.31 | 0.47 | 0.41 | 0.47 |
| 8 | 118905711 | rs17450835 | -0.405 | 0.474 | 0.818 | 0.19 | -0.06 | -0.094 | -0.094 | 0.459 | 0.17 | 0.08 | 0.16 | 0.39 | 0.29 | 0.29 |
| 8 | 118906580 | rs17430533 | 0.873 | 0.376 | 0.986 | 0.138 | 0.026 | 0.053 | 0.053 | 0.645 | 0.19 | 0.06 | 0.13 | 0.07 | 0.12 | 0.16 |
| 8 | 118919249 | rs4515578 | -0.527 | 0.354 | 1.253 | 0.092 | -0.009 | -0.004 | -0.004 | 0.016 | 0.77 | 0.79 | 0.8 | 0.83 | 0.84 | 0.82 |
| 8 | 118919779 | rs7007139 | -0.222 | 0.355 | 1.193 | 0.092 | -0.052 | -0.03 | -0.03 | 0 | 0.75 | 0.57 | 0.78 | 0.83 | 0.84 | 0.81 |
| 8 | 118919973 | rs13265834 | -0.216 | 0.339 | 1.172 | 0.085 | -0.043 | -0.025 | -0.025 | 0.516 | 0.75 | 0.62 | 0.79 | 0.83 | 0.84 | 0.81 |
| 8 | 118921721 | rs13275942 | 1.072 | 0.347 | 0.997 | 0.074 | 0.005 | 0.021 | 0.021 | 0 | 0.56 | 0.26 | 0.6 | 0.43 | 0.55 | 0.52 |
| 8 | 118924792 | rs11781245 | -0.27 | 0.694 | 1.239 | 0.402 | -0.071 | -0.094 | -0.094 | 0 | 0.15 | 0.06 | 0.15 | 0.39 | 0.29 | 0.28 |
| 8 | 118934480 | rs3860883 | 1.525 | 0.522 | 1.214 | 0.216 | -0.008 | -0.023 | -0.023 | 0 | 0.73 | 0.25 | 0.7 | 0.51 | 0.65 | 0.67 |
| 8 | 118938994 | rs1542519 | 1.229 | 0.445 | 1.266 | 0.277 | 0.013 | 0.043 | 0.043 | 0.285 | 0.77 | 0.68 | 0.82 | 0.61 | 0.69 | 0.69 |
| 8 | 118940928 | rs2447541 | 0.686 | 0.473 | 1.299 | 0.318 | -0.029 | -0.051 | -0.051 | 0.244 | 0.8 | 0.86 | 0.83 | 0.61 | 0.69 | 0.7 |
| 8 | 118940981 | rs7840309 | -0.866 | 0.445 | 1.378 | 0.54 | -0.033 | 0.046 | 0.046 | 0 | 0.95 | 1 | 0.92 | 0.62 | 0.85 | 0.84 |
| 8 | 118953887 | rs4876395 | 0.169 | 0.186 | 1.484 | 0.052 | 0.004 | 0.115 | 0.115 | 0 | 0.07 | 0.2 | 0.1 | 0.19 | 0.05 | 0.16 |
| 8 | 118958519 | rs10955842 | -0.578 | 0.183 | 1.502 | 0.046 | 0.008 | 0.13 | 0.13 | 0 | 0.08 | 0.3 | 0.12 | 0.18 | 0.05 | 0.16 |
| 8 | 118961385 | rs11562687 | -1.514 | 0.367 | 1.475 | -0.017 | 0.052 | 0.033 | 0.033 | 1.555 | 0.05 | 0.33 | 0.03 | 0 | 0.02 | 0.02 |
| 8 | 118989097 | rs10087446 | -0.284 | 0.346 | 0.867 | -0.103 | -0.033 | -0.07 | -0.07 | 0 | 0.79 | 0.91 | 0.82 | 0.61 | 0.68 | 0.71 |
| 8 | 118992150 | rs10955844 | 0.385 | 0.027 | 0.708 | -0.378 | -0.001 | -0.001 | -0.001 | 0 | 0.33 | 0.35 | 0.3 | 0.2 | 0.23 | 0.34 |
| 8 | 118997146 | rs12543161 | 0.375 | 0.031 | 0.708 | -0.351 | 0 | 0 | 0 | 0.246 | 0.33 | 0.35 | 0.31 | 0.29 | 0.24 | 0.35 |
| 8 | 119007276 | rs7006696 | 0.673 | 0.128 | 0.385 | -0.41 | -0.009 | 0.017 | 0.017 | 0 | 0.42 | 0.36 | 0.49 | 0.32 | 0.41 | 0.35 |
| 8 | 119012781 | rs13255959 | 0.739 | 0.129 | 0.223 | -0.428 | -0.021 | 0.024 | 0.024 | 0 | 0.4 | 0.27 | 0.49 | 0.33 | 0.42 | 0.35 |
| 8 | 119017973 | rs4876785 | 0.439 | 0.095 | 0.518 | -0.294 | -0.001 | 0 | 0 | 0 | 0.45 | 0.45 | 0.5 | 0.33 | 0.44 | 0.37 |
| 8 | 119018671 | rs11562702 | 0.601 | 0.65 | -0.469 | -0.402 | -0.03 | 0.026 | 0.026 | 0 | 0.05 | 0.01 | 0.12 | 0 | 0.08 | 0.13 |
| 8 | 119018957 | rs6986291 | 0.27 | 0.635 | -0.285 | -0.299 | -0.002 | 0.006 | 0.006 | 0 | 0.47 | 0.44 | 0.52 | 0.42 | 0.44 | 0.37 |
| 8 | 119023819 | rs2126630 | 0.055 | 0.202 | -0.687 | -0.53 | -0.049 | 0.056 | 0.056 | 0.595 | 0.33 | 0.06 | 0.39 | 0.19 | 0.31 | 0.42 |
| 8 | 119028915 | rs1037693 | -0.555 | 0.026 | -0.287 | -0.618 | 0.027 | 0.086 | 0.086 | 0 | 0.45 | 0.75 | 0.48 | 0.29 | 0.37 | 0.47 |
| 8 | 119045329 | rs4876399 | -0.269 | -0.009 | 0.23 | -0.552 | 0 | 0.046 | 0.046 | 1.88 | 0.52 | 0.6 | 0.53 | 0.27 | 0.47 | 0.4 |
| 8 | 119048686 | rs7829981 | -0.147 | -0.018 | 0.621 | -0.56 | 0 | 0.007 | 0.007 | 2.208 | 0.51 | 0.51 | 0.51 | 0.27 | 0.46 | 0.39 |
| 8 | 119050918 | rs10955850 | 0.08 | -0.113 | 1.145 | -0.608 | -0.003 | 0.105 | 0.105 | 1.707 | 0.48 | 0.25 | 0.49 | 0.27 | 0.46 | 0.39 |
| 8 | 119057067 | rs13277610 | 0.25 | -0.164 | 1.201 | -0.613 | -0.025 | 0.159 | 0.159 | 6.369 | 0.46 | 0.16 | 0.48 | 0.22 | 0.45 | 0.35 |
| 8 | 119061121 | rs9649944 | -0.407 | 0 | 0.59 | -0.628 | -0.008 | 0.09 | 0.09 | 3.03 | 0.54 | 0.7 | 0.53 | 0.26 | 0.47 | 0.4 |
| 8 | 119079196 | rs2451157 | -0.724 | 0.319 | 0.568 | 0.225 | 0.012 | 0.204 | 0.204 | 0.017 | 0.2 | 0.46 | 0.14 | 0.48 | 0.19 | 0.24 |
| 8 | 119081993 | rs11562721 | 0.491 | -0.027 | -0.144 | -0.153 | -0.007 | -0.024 | -0.024 | 0.012 | 0.16 | 0.2 | 0.05 | 0.01 | 0.08 | 0.13 |
| 8 | 119082077 | rs2255931 | -1.124 | -0.052 | 0.07 | -0.156 | -0.02 | 0.04 | 0.04 | 0.02 | 0.33 | 0.55 | 0.17 | 0.5 | 0.27 | 0.37 |
| 8 | 119088127 | rs17431728 | 0.164 | 0.238 | -0.696 | -0.318 | -0.006 | -0.024 | -0.024 | 0.443 | 0.16 | 0.2 | 0.05 | 0.01 | 0.08 | 0.13 |
| 8 | 119091168 | rs9297575 | 0.106 | 1.046 | -1.166 | 0.005 | -0.079 | 0.027 | 0.027 | 0 | 0.35 | 0.15 | 0.48 | 0.31 | 0.41 | 0.37 |
| 8 | 119093994 | rs6983199 | -1.184 | 2.018 | -0.096 | 0.467 | 0.114 | -0.019 | -0.019 | 0 | 0.25 | 0.38 | 0.43 | 0.2 | 0.43 | 0.34 |
| 8 | 119095507 | rs2445904 | -1.437 | 1.24 | -0.562 | 0.934 | 0.001 | 0.052 | 0.052 | 1.136 | 0.4 | 0.81 | 0.4 | 0.67 | 0.34 | 0.42 |
| 8 | 119098527 | rs10955854 | -0.332 | 1.436 | -0.228 | 0.898 | -0.007 | -0.004 | -0.004 | 0 | 0.13 | 0.11 | 0.26 | 0.13 | 0.19 | 0.13 |
| 8 | 119100285 | rs7826515 | -0.527 | 1.284 | 0.566 | 1.484 | -0.039 | 0.057 | 0.057 | 5.51 | 0.19 | 0.26 | 0.07 | 0.35 | 0.09 | 0.17 |
| 8 | 119103081 | rs2468151 | 0.932 | 1.356 | 0.611 | 1.494 | -0.045 | 0.043 | 0.043 | 4.595 | 0.76 | 0.41 | 0.88 | 0.43 | 0.89 | 0.81 |

| | | | | | | | | | | | | | | | | |
|---|---|---|---|---|---|---|---|---|---|---|---|---|---|---|---|---|
| 8 | 119104646 | rs2445905 | 1.466 | 1.388 | 0.297 | 1.399 | 0.013 | 0.038 | 0.038 | 0.976 | 0.63 | 0.29 | 0.61 | 0.3 | 0.69 | 0.68 |
| 8 | 119106346 | rs2445906 | 1.685 | 1.367 | 0.173 | 1.42 | -0.004 | 0.068 | 0.068 | 0.842 | 0.6 | 0.21 | 0.59 | 0.29 | 0.65 | 0.59 |
| 8 | 119107051 | rs11992017 | -0.487 | 1.004 | 0.352 | 1.475 | 0.004 | -0.003 | -0.003 | 0 | 0.16 | 0.12 | 0.09 | 0.35 | 0.12 | 0.25 |
| 8 | 119111527 | rs10505327 | 2.036 | 1.156 | -0.316 | 0.723 | -0.107 | 0.238 | 0.238 | 0 | 0.45 | 0.1 | 0.34 | 0.19 | 0.54 | 0.31 |
| 8 | 119113078 | rs2468139 | 0.467 | 1.346 | 0.426 | 0.668 | -0.013 | -0.02 | -0.02 | 0.017 | 0.79 | 0.56 | 0.9 | 0.81 | 0.83 | 0.87 |
| 8 | 119113236 | rs10755915 | 1.711 | 1.323 | 0.435 | 1.095 | -0.038 | -0.069 | -0.069 | 0.395 | 0.62 | 0.22 | 0.61 | 0.59 | 0.63 | 0.59 |
| 8 | 119113979 | rs2468137 | -0.912 | 1.333 | 0.463 | 1.098 | 0.035 | 0.069 | 0.069 | 0.173 | 0.16 | 0.31 | 0.29 | 0.22 | 0.2 | 0.28 |
| 8 | 119121535 | rs7014053 | 1.495 | 1.332 | 0.467 | 1.09 | -0.008 | -0.014 | -0.014 | 0 | 0.66 | 0.43 | 0.64 | 0.6 | 0.63 | 0.58 |
| 8 | 119126897 | rs2514759 | -0.729 | 1.336 | 0.465 | 1.13 | 0.027 | 0.048 | 0.048 | 0 | 0.14 | 0.21 | 0.24 | 0.21 | 0.2 | 0.29 |
| 8 | 119127950 | rs2514760 | 0.555 | 1.301 | 0.747 | 1.111 | -0.013 | -0.021 | -0.021 | 0 | 0.78 | 0.55 | 0.86 | 0.81 | 0.83 | 0.87 |
| 8 | 119128860 | rs1563564 | 0.748 | 1.294 | 0.648 | 1.108 | -0.018 | -0.024 | -0.024 | 0 | 0.77 | 0.47 | 0.85 | 0.81 | 0.83 | 0.87 |
| 8 | 119133417 | rs4147528 | 2.284 | 1.303 | 0.052 | 0.854 | -0.041 | -0.049 | -0.049 | 0 | 0.43 | 0.15 | 0.35 | 0.51 | 0.49 | 0.39 |
| 8 | 119133623 | rs2445922 | 0.165 | 1.696 | 0.112 | 0.756 | -0.031 | -0.04 | -0.04 | 0 | 0.08 | 0.02 | 0.08 | 0.13 | 0.12 | 0.13 |
| 8 | 119146587 | rs2514730 | -0.719 | 1.364 | 0.239 | 0.66 | -0.023 | -0.018 | -0.018 | 0 | 0.27 | 0.13 | 0.29 | 0.25 | 0.31 | 0.36 |
| 8 | 119154296 | rs726960 | -0.699 | 0.96 | 0.38 | 0.42 | -0.002 | -0.075 | -0.075 | 1.748 | 0.19 | 0.23 | 0.08 | 0 | 0.16 | 0.11 |
| 8 | 119157032 | rs2514731 | 0.904 | 0.92 | 0.258 | 1.066 | -0.001 | 0.012 | 0.012 | 0 | 0.68 | 0.4 | 0.43 | 0.51 | 0.68 | 0.53 |
| 8 | 119163143 | rs4876796 | 0.16 | 0.993 | 0.02 | 1.158 | -0.001 | 0 | 0 | 0.202 | 0.83 | 0.8 | 0.83 | 0.81 | 0.86 | 0.84 |
| 8 | 119164828 | rs11785842 | 0.114 | 0.964 | 0.034 | 1.168 | -0.001 | 0.001 | 0.001 | 0.117 | 0.83 | 0.8 | 0.83 | 0.81 | 0.86 | 0.84 |
| 8 | 119167509 | rs6469726 | -0.679 | 1.034 | -0.174 | 1.168 | -0.001 | -0.002 | -0.002 | 0 | 0.27 | 0.3 | 0.33 | 0.23 | 0.26 | 0.21 |
| 8 | 119167532 | rs7005385 | -0.679 | 1.034 | -0.174 | 1.158 | -0.001 | -0.003 | -0.003 | 1.811 | 0.27 | 0.3 | 0.33 | 0.23 | 0.26 | 0.21 |
| 8 | 119167602 | rs7005426 | -0.551 | 1.15 | -0.082 | 1.226 | -0.004 | -0.004 | -0.004 | 2.395 | 0.24 | 0.17 | 0.31 | 0.22 | 0.26 | 0.21 |
| 8 | 119171312 | rs11774887 | 1.106 | 1.077 | 0.328 | 1.122 | -0.004 | 0.013 | 0.013 | 0 | 0.76 | 0.46 | 0.68 | 0.73 | 0.81 | 0.59 |
| 8 | 119179060 | rs9297578 | 1.247 | 1.131 | -0.153 | 1.149 | -0.011 | 0.041 | 0.041 | 0 | 0.55 | 0.31 | 0.47 | 0.37 | 0.63 | 0.45 |
| 8 | 119179464 | rs7832658 | -1.42 | 0.466 | 0.592 | 1.048 | -0.038 | -0.043 | -0.043 | 0 | 0.17 | 0.31 | 0.05 | 0.03 | 0.07 | 0.13 |
| 8 | 119192723 | rs7820728 | -0.308 | 0.473 | 0.931 | 1.057 | -0.005 | -0.013 | -0.013 | 0 | 0.06 | 0.06 | 0.01 | 0 | 0.02 | 0.07 |
| 8 | 119192998 | rs2445920 | 0.688 | 0.794 | 1.1 | 1.641 | -0.004 | -0.005 | -0.005 | 2.752 | 0.74 | 0.73 | 0.63 | 0.62 | 0.74 | 0.73 |
| 8 | 119194911 | rs2514734 | 2.008 | 0.532 | 1.198 | 2.017 | 0.033 | 0.011 | 0.011 | 0 | 0.57 | 0.22 | 0.48 | 0.37 | 0.69 | 0.54 |
| 8 | 119204861 | rs3802184 | 0.782 | 1.129 | 1.259 | 2.236 | 0.021 | 0.022 | 0.022 | 0 | 0.74 | 0.67 | 0.53 | 0.39 | 0.59 | 0.59 |
| 8 | 119205873 | rs2451153 | 1.062 | 0.393 | 1.137 | 2.064 | -0.007 | -0.01 | -0.01 | 0 | 0.71 | 0.73 | 0.63 | 0.62 | 0.72 | 0.66 |
| 8 | 119207029 | rs10955863 | -1.583 | 0.302 | 1.621 | 1.919 | -0.029 | 0.104 | 0.104 | 0 | 0.1 | 0.3 | 0.12 | 0.15 | 0.02 | 0.11 |
| 8 | 119207847 | rs2514981 | 1.895 | 0.319 | 1.653 | 1.911 | 0.008 | 0.003 | 0.003 | 0 | 0.58 | 0.38 | 0.5 | 0.37 | 0.69 | 0.54 |
| 8 | 119207938 | rs2514742 | 1.837 | 0.332 | 1.592 | 1.918 | 0.001 | 0 | 0 | 0 | 0.59 | 0.41 | 0.5 | 0.37 | 0.69 | 0.54 |
| 8 | 119208131 | rs9886406 | -0.019 | 0.569 | 1.293 | 1.729 | -0.009 | -0.016 | -0.016 | 0.37 | 0.06 | 0.1 | 0.01 | 0 | 0.02 | 0.07 |
| 8 | 119209773 | rs2514743 | 1.267 | 0.727 | 1.071 | 1.857 | -0.003 | -0.019 | -0.019 | 0 | 0.67 | 0.71 | 0.62 | 0.52 | 0.7 | 0.65 |
| 8 | 119212592 | rs11562802 | -1.324 | 0.723 | 1.034 | 1.435 | 0.02 | -0.007 | -0.007 | 0 | 0.09 | 0.17 | 0.16 | 0.04 | 0.08 | 0.2 |
| 8 | 119213727 | rs10955864 | 1.748 | 0.571 | 1.277 | 1.791 | -0.001 | 0 | 0 | 0 | 0.6 | 0.36 | 0.4 | 0.33 | 0.67 | 0.31 |
| 8 | 119217328 | rs17451949 | -0.368 | 0.621 | 1.115 | 1.513 | 0.021 | -0.001 | -0.001 | 0 | 0.12 | 0.22 | 0.12 | 0.09 | 0.17 | 0.04 |
| 8 | 119226785 | rs10092413 | -1.612 | 0.503 | 1.507 | 1.547 | 0.036 | -0.019 | -0.019 | 0 | 0.26 | 0.45 | 0.29 | 0.17 | 0.26 | 0.22 |
| 8 | 119229093 | rs7819465 | 0.563 | 0.412 | 1.466 | 1.707 | 0.012 | -0.008 | -0.008 | 0.08 | 0.87 | 0.91 | 0.66 | 0.36 | 0.92 | 0.59 |
| 8 | 119234570 | rs2514966 | 2.092 | 0.408 | 1.535 | 1.67 | 0.003 | -0.042 | -0.042 | 7.551 | 0.61 | 0.49 | 0.51 | 0.19 | 0.67 | 0.37 |
| 8 | 119238582 | rs4876805 | -1.76 | 0.417 | 1.346 | 1.651 | -0.018 | -0.022 | -0.022 | 0 | 0.29 | 0.56 | 0.16 | 0.17 | 0.19 | 0.21 |
| 8 | 119238894 | rs2451136 | 1.179 | 0.678 | 1.138 | 1.824 | -0.015 | 0.046 | 0.046 | 0 | 0.85 | 0.79 | 0.65 | 0.34 | 0.92 | 0.53 |
| 8 | 119248227 | rs11562709 | -0.885 | 0.34 | 1.012 | 1.462 | 0.008 | -0.07 | -0.07 | 0 | 0.12 | 0.21 | 0.06 | 0 | 0.07 | 0.17 |
| 8 | 119251587 | rs2450218 | -1.364 | 0.154 | 0.574 | 2.191 | 0.002 | 0.021 | 0.021 | 0 | 0.13 | 0.11 | 0.22 | 0.49 | 0.09 | 0.37 |
| 8 | 119260904 | rs2450206 | -1.367 | 0.232 | 0.389 | 2.151 | 0.008 | -0.043 | -0.043 | 0 | 0.16 | 0.08 | 0.27 | 0.51 | 0.13 | 0.39 |
| 8 | 119264856 | rs2450213 | -2.12 | -0.233 | 1.057 | 2.1 | -0.074 | 0.153 | 0.153 | 2.336 | 0.21 | 0.54 | 0.23 | 0.58 | 0.09 | 0.4 |
| 8 | 119269786 | rs2450216 | -1.126 | -0.244 | 0.988 | 2.104 | 0 | 0.029 | 0.029 | 0 | 0.14 | 0.11 | 0.22 | 0.57 | 0.09 | 0.33 |
| 8 | 119275733 | rs7846380 | -1.379 | -0.244 | 1.319 | 1.964 | -0.006 | 0.117 | 0.117 | 4.084 | 0.07 | 0.37 | 0.14 | 0.14 | 0.01 | 0.02 |
| 8 | 119276240 | rs7834609 | -1.786 | -0.521 | 1.091 | 2.102 | -0.001 | 0.001 | 0.001 | 0.137 | 0.31 | 0.66 | 0.32 | 0.51 | 0.23 | 0.33 |
| 8 | 119279232 | rs2450217 | -0.175 | -0.161 | 0.707 | 1.922 | -0.05 | -0.059 | -0.059 | 0.042 | 0.14 | 0.03 | 0.24 | 0.22 | 0.13 | 0.28 |
| 8 | 119280694 | rs6989069 | -1.863 | -0.483 | 0.765 | 1.766 | 0.04 | -0.039 | -0.039 | 0 | 0.37 | 0.76 | 0.34 | 0.55 | 0.25 | 0.42 |
| 8 | 119293911 | rs4637872 | -1.567 | -0.668 | 0.817 | 1.799 | 0.008 | -0.003 | -0.003 | 0 | 0.32 | 0.71 | 0.29 | 0.33 | 0.2 | 0.21 |
| 8 | 119294282 | rs10110578 | -0.959 | -0.672 | 0.963 | 1.839 | 0.013 | 0.032 | 0.032 | 0 | 0.25 | 0.19 | 0.12 | 0.08 | 0.18 | 0.13 |
| 8 | 119296422 | rs721816 | -0.898 | -0.741 | 1.202 | 1.876 | -0.034 | 0.159 | 0.159 | 0 | 0.11 | 0.29 | 0.18 | 0.34 | 0.04 | 0.22 |
| 8 | 119296910 | rs3763564 | -0.973 | -0.713 | 1.3 | 1.875 | -0.038 | 0.219 | 0.219 | 0 | 0.11 | 0.36 | 0.18 | 0.39 | 0.04 | 0.28 |
| 8 | 119298134 | rs9297582 | -0.989 | -0.79 | 0.606 | 1.759 | 0.004 | 0.004 | 0.004 | 0 | 0.25 | 0.22 | 0.13 | 0.15 | 0.19 | 0.12 |
| 8 | 119303457 | rs2450219 | -2.292 | -1.36 | -0.51 | 1.047 | -0.06 | 0.161 | 0.161 | 0.004 | 0.53 | 0.92 | 0.69 | 0.78 | 0.38 | 0.7 |
| 8 | 119304622 | rs2325945 | 0.267 | -1.392 | -0.471 | 1.134 | -0.057 | -0.082 | -0.082 | 0.44 | 0.15 | 0.04 | 0.26 | 0.3 | 0.16 | 0.28 |
| 8 | 119309329 | rs3849851 | -1.459 | -1.457 | -0.159 | 0.984 | 0.046 | -0.016 | -0.016 | 0 | 0.37 | 0.74 | 0.33 | 0.43 | 0.21 | 0.36 |
| 8 | 119312168 | rs7829943 | -0.28 | -1.163 | 0.086 | 0.647 | 0.008 | 0.004 | 0.004 | 0 | 0.19 | 0.32 | 0.13 | 0.15 | 0.19 | 0.06 |
| 8 | 119319468 | rs16890815 | -0.388 | -1.402 | 0.127 | 0.665 | -0.041 | 0.012 | 0.012 | 0 | 0.09 | 0.19 | 0.14 | 0.13 | 0.01 | 0.07 |
| 8 | 119327908 | rs1385806 | -1.481 | -0.489 | -0.282 | 0.374 | 0.005 | 0.025 | 0.025 | 0.103 | 0.4 | 0.86 | 0.36 | 0.34 | 0.34 | 0.22 |
| 8 | 119332036 | rs6469733 | -1.561 | -0.548 | -0.238 | 0.353 | 0.009 | 0.031 | 0.031 | 0.114 | 0.42 | 0.84 | 0.37 | 0.34 | 0.36 | 0.22 |
| 8 | 119338870 | rs2450221 | -2.156 | -0.68 | -0.748 | 0.132 | -0.043 | 0.107 | 0.107 | 0.021 | 0.59 | 0.92 | 0.63 | 0.79 | 0.53 | 0.71 |
| 8 | 119343495 | rs7005520 | -1.005 | -0.633 | 0.307 | 0.026 | -0.033 | -0.041 | -0.041 | 0 | 0.21 | 0.46 | 0.21 | 0.18 | 0.15 | 0.16 |
| 8 | 119344964 | rs10110149 | -1.488 | -0.57 | -0.122 | 0.008 | 0.007 | 0.047 | 0.047 | 0.115 | 0.42 | 0.86 | 0.37 | 0.29 | 0.37 | 0.23 |
| 8 | 119348323 | rs6993964 | -1.459 | -0.53 | 0.012 | 0.028 | 0.003 | 0.021 | 0.021 | 0 | 0.4 | 0.76 | 0.35 | 0.29 | 0.37 | 0.22 |
| 8 | 119351241 | rs2450316 | -1.723 | -0.294 | -0.553 | -0.461 | 0.008 | 0.072 | 0.072 | 0 | 0.6 | 0.79 | 0.75 | 0.85 | 0.59 | 0.7 |
| 8 | 119351765 | rs16890858 | 0.503 | -0.304 | -0.086 | -0.71 | 0.02 | -0.096 | -0.096 | 1.798 | 0.16 | 0.35 | 0.12 | 0 | 0.2 | 0.04 |

| | | | | | | | | | | | | | | | | |
|---|---|---|---|---|---|---|---|---|---|---|---|---|---|---|---|---|
| 8 | 119351791 | rs10955872 | 0.157 | -0.562 | -0.278 | -0.595 | -0.058 | 0.008 | 0.008 | 0 | 0.12 | 0.28 | 0.15 | 0.16 | 0.02 | 0.15 |
| 8 | 119352278 | rs2450203 | -1.21 | -0.461 | -0.356 | -0.561 | -0.005 | -0.013 | -0.013 | 0 | 0.47 | 0.51 | 0.6 | 0.68 | 0.58 | 0.55 |
| 8 | 119353654 | rs2450324 | -1.226 | -0.482 | -0.356 | -0.564 | -0.005 | -0.013 | -0.013 | 0 | 0.47 | 0.51 | 0.6 | 0.68 | 0.58 | 0.55 |
| 8 | 119355129 | rs2450322 | 0.356 | -0.292 | -0.854 | 0.092 | -0.036 | -0.037 | -0.037 | 0.127 | 0.15 | 0.06 | 0.13 | 0.18 | 0.23 | 0.09 |
| 8 | 119356466 | rs2515029 | -1.136 | -0.311 | -0.778 | -0.203 | -0.007 | 0.079 | 0.079 | 0 | 0.42 | 0.33 | 0.62 | 0.85 | 0.39 | 0.66 |
| 8 | 119374066 | rs1825629 | -0.913 | 0.134 | -1.153 | 0.031 | -0.072 | -0.141 | -0.141 | 0 | 0.77 | 0.98 | 0.6 | 0.54 | 0.77 | 0.55 |
| 8 | 119376778 | rs6983935 | -0.013 | 0.553 | -1.072 | -0.235 | -0.009 | 0.006 | 0.006 | 0 | 0.21 | 0.22 | 0.25 | 0.18 | 0.26 | 0.16 |
| 8 | 119385578 | rs11786582 | 2.474 | 0.382 | -1.182 | -0.416 | 0.02 | 0.047 | 0.047 | 0 | 0.27 | 0.09 | 0.15 | 0.14 | 0.21 | 0.26 |
| 8 | 119388668 | rs4876817 | -0.481 | 0.312 | -1.089 | -0.492 | -0.002 | -0.006 | -0.006 | 0 | 0.68 | 0.88 | 0.53 | 0.32 | 0.69 | 0.48 |
| 8 | 119389854 | rs10086119 | 0.266 | 0.306 | -0.979 | -0.575 | 0.019 | 0.074 | 0.074 | 0 | 0.56 | 0.67 | 0.38 | 0.14 | 0.6 | 0.36 |
| 8 | 119390889 | rs12541679 | 0.399 | 0.281 | -0.968 | -0.54 | 0 | 0 | 0 | 0 | 0.19 | 0.21 | 0.23 | 0.18 | 0.24 | 0.14 |
| 8 | 119391791 | rs5020517 | -0.429 | 0.188 | -0.668 | -0.403 | -0.001 | -0.005 | -0.005 | 0 | 0.68 | 0.87 | 0.53 | 0.32 | 0.69 | 0.48 |
| 8 | 119391866 | rs12171673 | 1.32 | 0.176 | -0.622 | -0.442 | -0.042 | 0.063 | 0.063 | 0 | 0.13 | 0.03 | 0.09 | 0 | 0.2 | 0.03 |
| 8 | 119400402 | rs2450209 | 0.618 | 0.254 | -0.507 | -0.224 | 0 | 0.016 | 0.016 | 0 | 0.09 | 0.09 | 0.06 | 0.21 | 0.08 | 0.07 |
| 8 | 119400473 | rs9297584 | 0.668 | 0.277 | -0.354 | -0.292 | -0.017 | 0.015 | 0.015 | 0 | 0.17 | 0.12 | 0.11 | 0 | 0.24 | 0.06 |
| 8 | 119403582 | rs6469744 | 0.377 | 0.275 | -0.241 | -0.294 | 0.022 | -0.103 | -0.103 | 0 | 0.19 | 0.32 | 0.13 | 0 | 0.24 | 0.06 |
| 8 | 119403784 | rs2515062 | 0.514 | 0.067 | -0.965 | -0.055 | 0.001 | 0.014 | 0.014 | 0 | 0.09 | 0.09 | 0.06 | 0.22 | 0.08 | 0.07 |
| 8 | 119405522 | rs2515063 | -1.108 | 0.349 | -1.056 | -0.307 | -0.053 | -0.126 | -0.126 | 0.092 | 0.76 | 0.96 | 0.59 | 0.54 | 0.77 | 0.55 |
| 8 | 119411462 | rs2450330 | -1.06 | 0.342 | -1.045 | -0.311 | -0.037 | -0.106 | -0.106 | 1.839 | 0.76 | 0.95 | 0.59 | 0.54 | 0.77 | 0.55 |
| 8 | 119413545 | rs2450212 | -1.197 | 0.377 | -1.049 | -0.278 | -0.056 | -0.074 | -0.074 | 0 | 0.78 | 0.96 | 0.6 | 0.65 | 0.77 | 0.56 |
| 8 | 119416748 | rs7832475 | -1.003 | 0.498 | -0.733 | -0.274 | -0.01 | 0.04 | 0.04 | 0 | 0.41 | 0.75 | 0.38 | 0.29 | 0.45 | 0.24 |
| 8 | 119416966 | rs7817780 | 0.315 | 0.531 | -0.379 | -0.463 | -0.003 | -0.001 | -0.001 | 0 | 0.18 | 0.14 | 0.1 | 0 | 0.21 | 0.06 |
| 8 | 119417628 | rs2450214 | 1.356 | 0.469 | -0.652 | -0.404 | 0.012 | -0.026 | -0.026 | 0 | 0.32 | 0.19 | 0.21 | 0.36 | 0.29 | 0.31 |
| 8 | 119418276 | rs17749504 | -0.212 | 0.837 | -0.615 | -0.666 | -0.002 | 0.044 | 0.044 | 0 | 0.18 | 0.12 | 0.09 | 0 | 0.18 | 0.06 |
| 8 | 119420259 | rs4876823 | -0.818 | 0.745 | -0.44 | -0.609 | 0.02 | -0.125 | -0.125 | 0 | 0.28 | 0.45 | 0.22 | 0.01 | 0.35 | 0.11 |
| 8 | 119427347 | rs9297586 | 1.293 | 0.257 | -1.089 | 0.224 | -0.141 | 0.015 | 0.015 | 0.106 | 0.27 | 0.02 | 0.21 | 0.15 | 0.39 | 0.3 |
| 8 | 119427392 | rs4876412 | 0.588 | 0.361 | -0.723 | 0.389 | -0.022 | 0.093 | 0.093 | 0.026 | 0.94 | 0.8 | 0.96 | 0.78 | 0.98 | 0.78 |
| 8 | 119428764 | rs10505334 | -0.05 | 0.237 | -1.099 | 0.41 | 0.001 | 0.003 | 0.003 | 0.448 | 0.38 | 0.17 | 0.27 | 0.35 | 0.35 | 0.2 |
| 8 | 119428944 | rs4077747 | 0.355 | 0.331 | -0.783 | 0.452 | 0.001 | 0.008 | 0.008 | 2.255 | 0.22 | 0.24 | 0.45 | 0.29 | 0.23 | 0.28 |
| 8 | 119432108 | rs2514988 | 0.175 | 0.381 | -0.754 | 0.626 | -0.005 | -0.053 | -0.053 | 0.555 | 0.25 | 0.15 | 0.18 | 0.39 | 0.25 | 0.21 |
| 8 | 119432132 | rs2514989 | 0.225 | 0.382 | -0.753 | 0.622 | -0.004 | -0.034 | -0.034 | 0 | 0.24 | 0.15 | 0.18 | 0.33 | 0.24 | 0.21 |
| 8 | 119432245 | rs2514990 | -0.01 | 0.338 | -0.809 | 0.526 | 0.012 | 0.001 | 0.001 | 0 | 0.82 | 0.91 | 0.88 | 0.81 | 0.86 | 0.75 |
| 8 | 119433803 | rs6469746 | 0.606 | 0.389 | -0.384 | 0.383 | 0.015 | -0.011 | -0.011 | 0 | 0.55 | 0.62 | 0.68 | 0.47 | 0.61 | 0.55 |
| 8 | 119437946 | rs13251354 | 0.082 | 0.501 | -0.199 | 0.356 | 0.004 | 0.031 | 0.031 | 0.099 | 0.67 | 0.42 | 0.7 | 0.5 | 0.66 | 0.6 |
| 8 | 119448453 | rs12675551 | 0.779 | 0.633 | -0.766 | 0.178 | -0.057 | -0.043 | -0.043 | 0 | 0.52 | 0.09 | 0.6 | 0.5 | 0.57 | 0.57 |
| 8 | 119452526 | rs10955883 | 0.326 | 0.474 | -0.143 | 0.11 | -0.003 | -0.002 | -0.002 | 0 | 0.59 | 0.37 | 0.64 | 0.5 | 0.57 | 0.58 |
| 8 | 119454157 | rs10108907 | -0.522 | 0.398 | 0.255 | 0.018 | -0.007 | -0.013 | -0.013 | 0 | 0.73 | 0.73 | 0.75 | 0.52 | 0.68 | 0.67 |
| 8 | 119455953 | rs13256814 | 1.754 | 0.542 | -0.472 | -0.141 | -0.018 | 0.108 | 0.108 | 0.101 | 0.35 | 0.02 | 0.4 | 0.14 | 0.32 | 0.33 |
| 8 | 119463191 | rs4876841 | 0.494 | 0.429 | -0.407 | -0.16 | -0.051 | -0.038 | -0.038 | 0 | 0.57 | 0.15 | 0.61 | 0.5 | 0.57 | 0.58 |
| 8 | 119464353 | rs4876843 | 0.443 | 0.141 | -0.116 | -0.009 | -0.042 | -0.038 | -0.038 | 0.033 | 0.11 | 0.01 | 0.11 | 0.13 | 0.16 | 0.11 |
| 8 | 119470337 | rs10095856 | -0.792 | 0.028 | 0.36 | -0.086 | -0.026 | -0.135 | -0.135 | 0 | 0.17 | 0.46 | 0.11 | 0 | 0.1 | 0.04 |
| 8 | 119480896 | rs10104506 | -0.864 | 0.207 | 0.235 | -0.082 | 0.001 | -0.105 | -0.105 | 0 | 0.17 | 0.55 | 0.13 | 0.02 | 0.11 | 0.06 |
| 8 | 119486034 | rs2515000 | 0.065 | -0.168 | -0.65 | 0.405 | -0.019 | -0.073 | -0.073 | 0.803 | 0.8 | 0.95 | 0.84 | 0.64 | 0.79 | 0.7 |
| 8 | 119492296 | rs10955889 | 1.662 | -0.252 | -1.345 | 0.366 | -0.095 | -0.094 | -0.094 | 1.282 | 0.42 | 0.05 | 0.54 | 0.4 | 0.53 | 0.53 |
| 8 | 119492578 | rs2515011 | 0.601 | -0.394 | -1.024 | 0.669 | 0.006 | -0.041 | -0.041 | 0 | 0.18 | 0.05 | 0.16 | 0.24 | 0.17 | 0.12 |
| 8 | 119494300 | rs2515032 | -0.641 | -0.379 | -0.63 | 0.776 | -0.001 | 0.001 | 0.001 | 0 | 0.32 | 0.34 | 0.3 | 0.39 | 0.35 | 0.28 |
| 8 | 119495145 | rs2515033 | 0.303 | -0.061 | -0.52 | 0.945 | -0.016 | 0.007 | 0.007 | 0 | 0.87 | 0.84 | 0.94 | 0.8 | 0.95 | 0.82 |
| 8 | 119496990 | rs2515036 | 0.421 | 0.264 | -0.517 | 0.917 | -0.001 | -0.01 | -0.01 | 0.185 | 0.8 | 0.83 | 0.84 | 0.71 | 0.79 | 0.71 |
| 8 | 119498112 | rs2515037 | -0.7 | 0.162 | -0.632 | 0.631 | 0.009 | 0.022 | 0.022 | 0 | 0.4 | 0.68 | 0.34 | 0.69 | 0.34 | 0.65 |
| 8 | 119499751 | rs4876847 | 0.242 | 0.279 | -0.549 | 1.027 | -0.006 | 0.086 | 0.086 | 0 | 0.07 | 0.11 | 0.06 | 0.27 | 0.08 | 0.1 |
| 8 | 119501453 | rs2515041 | -0.442 | 0.274 | -1.38 | 0.57 | -0.017 | 0.066 | 0.066 | 0.089 | 0.51 | 0.82 | 0.41 | 0.68 | 0.49 | 0.49 |
| 8 | 119505000 | rs2515049 | 1.008 | -0.066 | -1.512 | 0.466 | -0.029 | -0.051 | -0.051 | 0 | 0.29 | 0.07 | 0.25 | 0.36 | 0.31 | 0.23 |
| 8 | 119506499 | rs2515050 | -0.343 | 0.186 | -1.083 | -0.045 | 0.045 | 0.028 | 0.028 | 0.084 | 0.49 | 0.75 | 0.38 | 0.63 | 0.44 | 0.36 |
| 8 | 119511032 | rs2515055 | -0.941 | 0.225 | -1.196 | -0.07 | -0.057 | -0.001 | -0.001 | 0.391 | 0.67 | 0.87 | 0.53 | 0.75 | 0.58 | 0.77 |
| 8 | 119513852 | rs7836391 | 0.65 | 0.4 | -0.311 | -0.206 | 0.012 | -0.005 | -0.005 | 0.542 | 0.31 | 0.44 | 0.47 | 0.26 | 0.37 | 0.22 |
| 8 | 119525962 | rs4559272 | 0.526 | 0.364 | -0.161 | -0.195 | 0.006 | 0.003 | 0.003 | 0 | 0.32 | 0.52 | 0.49 | 0.28 | 0.37 | 0.24 |
| 8 | 119526617 | rs4637874 | -0.439 | 0.271 | -0.341 | -0.157 | -0.001 | 0 | 0 | 0 | 0.57 | 0.48 | 0.67 | 0.39 | 0.62 | 0.71 |
| 8 | 119534455 | rs13280603 | 0.76 | 0.275 | -0.307 | -0.15 | -0.004 | 0.006 | 0.006 | 0 | 0.36 | 0.33 | 0.49 | 0.26 | 0.43 | 0.24 |
| 8 | 119539512 | rs4876418 | -0.444 | 0.255 | -0.842 | -0.137 | -0.001 | -0.001 | -0.001 | 1.104 | 0.57 | 0.47 | 0.67 | 0.38 | 0.62 | 0.71 |
| 8 | 119542307 | rs4876852 | 0.37 | 0.325 | -0.967 | -0.132 | 0.002 | -0.005 | -0.005 | 0 | 0.22 | 0.23 | 0.18 | 0.13 | 0.2 | 0.47 |
| 8 | 119549041 | rs7844282 | -0.626 | -0.113 | -1.566 | 0.084 | -0.018 | -0.097 | -0.097 | 0.019 | 0.63 | 0.95 | 0.7 | 0.39 | 0.62 | 0.71 |
| 8 | 119552699 | rs12541178 | 0.066 | 0.404 | -1.183 | 0.174 | -0.026 | 0.019 | 0.019 | 0 | 0.39 | 0.31 | 0.5 | 0.27 | 0.48 | 0.39 |
| 8 | 119552755 | rs12548521 | 0.072 | 0.41 | -1.18 | 0.177 | -0.026 | 0.018 | 0.018 | 0 | 0.39 | 0.31 | 0.5 | 0.27 | 0.48 | 0.39 |
| 8 | 119556430 | rs6469756 | -0.258 | -0.14 | -1.485 | 0.639 | 0.006 | -0.089 | -0.089 | 0 | 0.72 | 0.92 | 0.88 | 0.53 | 0.78 | 0.8 |
| 8 | 119567424 | rs4421378 | 0.7 | 0.68 | -0.248 | 0.729 | -0.006 | -0.035 | -0.035 | 0 | 0.68 | 0.8 | 0.8 | 0.44 | 0.7 | 0.56 |
| 8 | 119567719 | rs4641102 | -0.088 | 0.509 | -0.538 | 0.381 | 0 | -0.028 | -0.028 | 0.165 | 0.17 | 0.16 | 0.12 | 0 | 0.16 | 0.1 |
| 8 | 119568066 | rs12550842 | 1.694 | 0.291 | -0.778 | 0.246 | -0.086 | -0.106 | -0.106 | 0 | 0.46 | 0.03 | 0.62 | 0.43 | 0.5 | 0.44 |
| 8 | 119576978 | rs3936020 | 1.429 | 0.271 | -1.017 | 0.27 | -0.105 | -0.082 | -0.082 | 0 | 0.51 | 0.03 | 0.66 | 0.47 | 0.57 | 0.46 |
| 8 | 119579349 | rs4512404 | -0.884 | 0.313 | 0.053 | 0.389 | -0.003 | -0.05 | -0.05 | 0.24 | 0.19 | 0.2 | 0.12 | 0 | 0.17 | 0.1 |

| | | | | | | | | | | | | | | | | |
|---|---|---|---|---|---|---|---|---|---|---|---|---|---|---|---|---|
| 8 | 119580807 | rs10505343 | -1.039 | 0.39 | 0.314 | 0.653 | -0.006 | -0.007 | -0.007 | 0 | 0.27 | 0.28 | 0.15 | 0.23 | 0.22 | 0.18 |
| 8 | 119580973 | rs17756861 | -0.978 | 0.421 | -0.032 | 0.648 | 0.009 | 0.01 | 0.01 | 0 | 0.26 | 0.21 | 0.14 | 0.23 | 0.22 | 0.18 |
| 8 | 119596377 | rs2514582 | 1.582 | 0.918 | 0.345 | 1.007 | -0.005 | 0.026 | 0.026 | 0.211 | 0.6 | 0.22 | 0.69 | 0.36 | 0.66 | 0.49 |
| 8 | 119598072 | rs2460962 | 1.143 | 0.921 | 1.226 | 1.094 | -0.002 | 0.033 | 0.033 | 0 | 0.7 | 0.53 | 0.76 | 0.41 | 0.73 | 0.64 |
| 8 | 119602579 | rs1607925 | -0.83 | 0.926 | 1.235 | 1.067 | 0.014 | -0.012 | -0.012 | 3.669 | 0.09 | 0.18 | 0.05 | 0.06 | 0.07 | 0.14 |
| 8 | 119602607 | rs1607924 | 1.185 | 0.947 | 1.247 | 1.086 | -0.004 | 0.02 | 0.02 | 0 | 0.7 | 0.43 | 0.75 | 0.42 | 0.73 | 0.64 |
| 8 | 119604236 | rs12676135 | -0.849 | 1.167 | 0.917 | 0.778 | 0.014 | -0.009 | -0.009 | 0 | 0.08 | 0.17 | 0.05 | 0.06 | 0.07 | 0.14 |
| 8 | 119605555 | rs10505342 | -0.752 | 1.178 | 0.867 | 0.783 | 0.011 | -0.008 | -0.008 | 0 | 0.08 | 0.13 | 0.05 | 0.06 | 0.07 | 0.15 |
| 8 | 119605603 | rs1915541 | 0.844 | 0.877 | 0.792 | 1.079 | -0.003 | 0 | 0 | 0.511 | 0.8 | 0.79 | 0.81 | 0.82 | 0.77 | 0.73 |
| 8 | 119607281 | rs10505340 | -0.873 | 0.891 | 1.082 | 1.164 | 0.003 | -0.031 | -0.031 | 0 | 0.07 | 0.18 | 0.05 | 0 | 0.08 | 0.04 |
| 8 | 119611476 | rs2514591 | 1.472 | 1.088 | 1.184 | 1.41 | 0.071 | 0.105 | 0.105 | 0 | 0.94 | 0.84 | 0.88 | 0.8 | 0.89 | 0.88 |
| 8 | 119627671 | rs17750541 | -1.027 | 0.936 | 0.714 | 1.282 | -0.033 | 0.031 | 0.031 | 0 | 0.15 | 0.03 | 0.09 | 0.08 | 0.14 | 0.25 |
| 8 | 119628230 | rs2460968 | 1.472 | 0.871 | 1.575 | 1.239 | 0.076 | 0.143 | 0.143 | 0.874 | 0.89 | 0.55 | 0.85 | 0.8 | 0.9 | 0.86 |
| 8 | 119641169 | rs1521154 | -1.797 | 0.834 | 1.445 | 1.455 | 0.007 | 0.042 | 0.042 | 0 | 0.26 | 0.33 | 0.17 | 0.5 | 0.23 | 0.34 |
| 8 | 119651801 | rs7006088 | -1.802 | 0.797 | 1.406 | 1.431 | 0.005 | 0.034 | 0.034 | 0.002 | 0.27 | 0.34 | 0.18 | 0.5 | 0.24 | 0.34 |
| 8 | 119663714 | rs10453061 | -1.577 | 0.79 | 1.545 | 1.576 | -0.002 | 0.001 | 0.001 | 0 | 0.24 | 0.24 | 0.17 | 0.14 | 0.23 | 0.31 |
| 8 | 119676755 | rs7845199 | -1.573 | 0.859 | 0.907 | 1.066 | -0.002 | -0.001 | -0.001 | 0 | 0.28 | 0.3 | 0.18 | 0.5 | 0.25 | 0.34 |
| 8 | 119681282 | rs7834374 | -1.4 | 0.695 | 0.981 | 0.455 | 0.058 | -0.047 | -0.047 | 0 | 0.1 | 0.41 | 0.08 | 0 | 0.07 | 0.04 |
| 8 | 119683126 | rs7459445 | -1.091 | 0.732 | -0.194 | 0.912 | 0.004 | -0.016 | -0.016 | 0 | 0.07 | 0.02 | 0.13 | 0.15 | 0.05 | 0.15 |
| 8 | 119685652 | rs13438904 | -0.853 | 0.914 | 0.277 | 0.677 | 0.016 | 0.025 | 0.025 | 0 | 0.16 | 0.24 | 0.08 | 0.25 | 0.18 | 0.15 |
| 8 | 119696794 | rs10100960 | -1.179 | 0.4 | 0.304 | -0.081 | 0.003 | 0.002 | 0.002 | 1.333 | 0.25 | 0.81 | 0.14 | 0.1 | 0.15 | 0.13 |
| 8 | 119697571 | rs16891361 | -0.813 | 0.409 | 1.049 | -0.115 | 0.008 | -0.067 | -0.067 | 0 | 0.12 | 0.27 | 0.05 | 0 | 0.08 | 0.04 |
| 8 | 119698024 | rs4876424 | -0.271 | 0.435 | 1.179 | 0.043 | -0.009 | 0.002 | 0.002 | 0 | 0.1 | 0.19 | 0.06 | 0.1 | 0.06 | 0.09 |
| 8 | 119704124 | rs13273785 | -0.04 | 0.197 | 0.391 | -0.897 | -0.001 | -0.064 | -0.064 | 0 | 0.36 | 0.44 | 0.36 | 0.06 | 0.35 | 0.19 |
| 8 | 119722725 | rs2102978 | 0.903 | 0.075 | 0.072 | -1.078 | -0.05 | 0.006 | 0.006 | 0.277 | 0.09 | 0.02 | 0.24 | 0.06 | 0.18 | 0.11 |
| 8 | 119723897 | rs10104977 | -0.888 | -0.091 | 0.321 | -1.002 | 0.005 | 0.249 | 0.249 | 0 | 0.49 | 0.88 | 0.45 | 0.17 | 0.44 | 0.28 |
| 8 | 119725997 | rs7815129 | -0.089 | 0.044 | 0.24 | -0.941 | -0.001 | -0.074 | -0.074 | 0 | 0.35 | 0.44 | 0.36 | 0.06 | 0.35 | 0.19 |
| 8 | 119726242 | rs10113233 | -0.059 | 0.011 | 0.243 | -0.938 | -0.003 | -0.075 | -0.075 | 0 | 0.35 | 0.44 | 0.36 | 0.06 | 0.34 | 0.19 |
| 8 | 119742870 | rs3103985 | -0.902 | -0.297 | -0.596 | 0.335 | 0.017 | -0.108 | -0.108 | 0.333 | 0.82 | 0.98 | 0.86 | 0.66 | 0.88 | 0.73 |
| 8 | 119754459 | rs16891419 | -1.124 | 0.301 | 0.111 | -0.102 | 0.05 | -0.033 | -0.033 | 0 | 0.06 | 0.18 | 0.04 | 0 | 0.07 | 0.04 |
| 8 | 119758541 | rs4876856 | 0.145 | -0.254 | -0.051 | -0.145 | 0.004 | 0.003 | 0.003 | 0 | 0.1 | 0.07 | 0.05 | 0.07 | 0.07 | 0.08 |
| 8 | 119760559 | rs13251655 | 0.719 | -0.875 | 0.124 | -0.014 | 0.065 | 0.03 | 0.03 | 0 | 0.55 | 0.29 | 0.68 | 0.41 | 0.69 | 0.47 |
| 8 | 119766194 | rs11988997 | 0.54 | -0.865 | 0.187 | 0.096 | 0.004 | 0.003 | 0.003 | 0 | 0.1 | 0.07 | 0.05 | 0.07 | 0.07 | 0.08 |
| 8 | 119768611 | rs7000310 | -0.577 | -0.747 | 0.296 | 0.196 | 0.03 | -0.077 | -0.077 | 0 | 0.74 | 0.9 | 0.78 | 0.48 | 0.78 | 0.57 |
| 8 | 119769809 | rs13253934 | -0.067 | -0.669 | 0.362 | 0.235 | -0.029 | 0.025 | 0.025 | 0 | 0.7 | 0.64 | 0.77 | 0.48 | 0.77 | 0.55 |
| 8 | 119777586 | rs1493937 | -0.888 | -0.251 | -0.022 | 0.539 | -0.013 | 0.017 | 0.017 | 2.035 | 0.3 | 0.63 | 0.41 | 0.56 | 0.2 | 0.43 |
| 8 | 119778259 | rs959454 | 0.191 | -0.241 | 0.085 | 0.818 | 0.003 | 0.019 | 0.019 | 0 | 0.11 | 0.08 | 0.3 | 0.45 | 0.08 | 0.18 |
| 8 | 119780061 | rs4384011 | -1.379 | -0.46 | -0.032 | 0.501 | 0.002 | 0.003 | 0.003 | 0 | 0.48 | 0.54 | 0.5 | 0.68 | 0.38 | 0.61 |
| 8 | 119784816 | rs11787376 | -0.059 | -0.609 | 0.375 | 0.433 | -0.001 | -0.021 | -0.021 | 0.408 | 0.31 | 0.36 | 0.17 | 0.15 | 0.25 | 0.39 |
| 8 | 119786804 | rs16891482 | 1.01 | -0.631 | 0.236 | 0.935 | 0 | -0.002 | -0.002 | 0 | 0.09 | 0.07 | 0.05 | 0.11 | 0.04 | 0.06 |
| 8 | 119790734 | rs3133593 | 1.757 | 0.179 | -0.828 | 0.327 | -0.087 | 0.024 | 0.024 | 0 | 0.16 | 0.01 | 0.24 | 0.07 | 0.29 | 0.11 |
| 8 | 119793583 | rs10955898 | 0.185 | -0.386 | -0.5 | 0.462 | 0.012 | 0.115 | 0.115 | 0 | 0.15 | 0.25 | 0.31 | 0.37 | 0.12 | 0.34 |
| 8 | 119796643 | rs4876860 | -1.315 | -0.546 | -0.16 | 0.248 | 0.036 | 0.09 | 0.09 | 0 | 0.51 | 0.78 | 0.52 | 0.68 | 0.38 | 0.63 |
| 8 | 119800914 | rs1874031 | 1.458 | -0.862 | -0.842 | -0.218 | -0.113 | 0.1 | 0.1 | 0 | 0.31 | 0.05 | 0.39 | 0.13 | 0.47 | 0.23 |
| 8 | 119800929 | rs1874030 | 0.476 | -1.207 | -0.59 | -0.069 | -0.01 | 0.015 | 0.015 | 0 | 0.05 | 0.09 | 0.04 | 0.1 | 0.05 | 0.17 |
| 8 | 119801271 | rs1908222 | -0.025 | -1.158 | -0.644 | 0.187 | -0.005 | 0.01 | 0.01 | 0.056 | 0.07 | 0.09 | 0.12 | 0.13 | 0.06 | 0.18 |
| 8 | 119806441 | rs4563918 | -0.999 | -1.299 | -0.405 | 0 | -0.004 | -0.018 | -0.018 | 0 | 0.7 | 0.69 | 0.84 | 0.79 | 0.75 | 0.79 |
| 8 | 119806880 | rs4876429 | -0.36 | -1.425 | -0.214 | -0.053 | -0.019 | 0.06 | 0.06 | 0 | 0.2 | 0.29 | 0.34 | 0.56 | 0.14 | 0.38 |
| 8 | 119814670 | rs3103975 | -1.918 | -1.404 | -0.487 | -0.046 | -0.007 | -0.006 | -0.006 | 0.305 | 0.79 | 0.77 | 0.87 | 0.82 | 0.82 | 0.85 |
| 8 | 119815147 | rs11781292 | 1.96 | -1.4 | -0.59 | -0.243 | -0.043 | 0.012 | 0.012 | 0 | 0.15 | 0.01 | 0.09 | 0.11 | 0.16 | 0.16 |
| 8 | 119819044 | rs3134098 | -0.82 | -1.398 | -0.466 | -0.193 | 0 | 0.012 | 0.012 | 0 | 0.68 | 0.67 | 0.68 | 0.48 | 0.71 | 0.67 |
| 8 | 119821583 | rs3103980 | -0.624 | -1.185 | -0.036 | -0.115 | -0.077 | 0.033 | 0.033 | 0.231 | 0.2 | 0.48 | 0.19 | 0.25 | 0.1 | 0.26 |
| 8 | 119821860 | rs3134036 | 1.078 | -1.095 | -0.457 | -0.217 | -0.054 | 0.085 | 0.085 | 0 | 0.33 | 0.19 | 0.4 | 0.13 | 0.46 | 0.25 |
| 8 | 119822843 | rs3134047 | -0.343 | -1.474 | 0.364 | -0.155 | -0.044 | 0.042 | 0.042 | 0.521 | 0.17 | 0.34 | 0.17 | 0.25 | 0.09 | 0.22 |
| 8 | 119829141 | rs11776031 | 0.362 | -1.3 | 0.314 | -0.235 | 0.012 | -0.044 | -0.044 | 1.759 | 0.12 | 0.23 | 0.06 | 0.03 | 0.1 | 0.08 |
| 8 | 119830953 | rs903613 | 0.386 | -1.296 | 0.316 | -0.233 | 0.012 | -0.044 | -0.044 | 0.063 | 0.12 | 0.23 | 0.06 | 0.03 | 0.1 | 0.08 |
| 8 | 119836572 | rs1948467 | 1.536 | -1.387 | -0.485 | 0.026 | -0.136 | -0.008 | -0.008 | 0 | 0.33 | 0.06 | 0.44 | 0.25 | 0.54 | 0.26 |
| 8 | 119848956 | rs1493935 | -0.184 | -1.506 | -0.328 | 0.051 | 0.021 | -0.014 | -0.014 | 0 | 0.53 | 0.73 | 0.63 | 0.49 | 0.64 | 0.53 |
| 8 | 119856209 | rs3134086 | -1.211 | -1.773 | -0.334 | 0.111 | 0.043 | 0.008 | 0.008 | 2.381 | 0.7 | 0.96 | 0.86 | 0.72 | 0.78 | 0.72 |
| 8 | 119859803 | rs3103969 | 0.888 | -1.686 | 0.444 | 0.063 | 0.003 | -0.012 | -0.012 | 0 | 0.42 | 0.55 | 0.64 | 0.25 | 0.59 | 0.32 |
| 8 | 119872439 | rs2035979 | 1.305 | -1.771 | 0.1 | -0.005 | -0.176 | -0.038 | -0.038 | 0 | 0.35 | 0.03 | 0.58 | 0.25 | 0.55 | 0.3 |
| 8 | 119880918 | rs7829526 | 1.371 | -1.924 | 0.427 | -0.128 | -0.003 | -0.055 | -0.055 | 0 | 0.11 | 0.17 | 0.08 | 0 | 0.1 | 0.06 |
| 8 | 119883396 | rs13340559 | 0.106 | -2.126 | 0.641 | -0.07 | 0.024 | 0.02 | 0.02 | 0 | 0.34 | 0.22 | 0.12 | 0.32 | 0.22 | 0.44 |
| 8 | 119886091 | rs3103991 | -0.621 | -2.386 | 0.975 | 0.104 | -0.005 | 0.006 | 0.006 | 0 | 0.72 | 0.74 | 0.78 | 0.41 | 0.8 | 0.69 |
| 8 | 119886101 | rs3134095 | -1.264 | -2.015 | 1.226 | 0.258 | 0.001 | 0.025 | 0.025 | 0 | 0.95 | 0.93 | 0.89 | 0.88 | 0.95 | 0.96 |
| 8 | 119886889 | rs16891617 | 1.699 | -2.022 | 1.276 | 0.198 | -0.001 | -0.024 | -0.024 | 0 | 0.1 | 0.1 | 0.07 | 0 | 0.1 | 0.07 |
| 8 | 119897215 | rs4242590 | 0.578 | -1.782 | 1.511 | 0.394 | 0.022 | 0.194 | 0.194 | 0.015 | 0.9 | 0.75 | 0.84 | 0.7 | 0.93 | 0.9 |
| 8 | 119897611 | rs4242591 | 0.791 | -1.807 | 1.52 | 0.382 | 0.016 | 0.22 | 0.22 | 1.823 | 0.88 | 0.4 | 0.81 | 0.7 | 0.92 | 0.9 |
| 8 | 119898843 | rs11986290 | 0.961 | -1.92 | 1.217 | 0.19 | 0.014 | 0.015 | 0.015 | 0 | 0.19 | 0.07 | 0.13 | 0.14 | 0.14 | 0.14 |

| | | | | | | | | | | | | | | | | |
|---|---|---|---|---|---|---|---|---|---|---|---|---|---|---|---|---|
| 8 | 119909351 | rs13279492 | 0.694 | -1.854 | 1.078 | 0.005 | -0.004 | -0.006 | -0.006 | 0 | 0.14 | 0.1 | 0.08 | 0.01 | 0.14 | 0.17 |
| 8 | 119909831 | rs6987473 | 1.241 | -1.915 | 0.983 | 0.081 | 0.031 | -0.003 | -0.003 | 0 | 0.27 | 0.13 | 0.1 | 0.33 | 0.19 | 0.28 |
| 8 | 119910605 | rs4593581 | -0.677 | -1.765 | 1.167 | 0.119 | 0.009 | 0.058 | 0.058 | 0 | 0.93 | 0.8 | 0.88 | 0.88 | 0.95 | 0.96 |
| 8 | 119912178 | rs4377989 | 0.332 | -1.768 | 1.127 | 0.117 | 0.017 | 0.087 | 0.087 | 0.709 | 0.9 | 0.51 | 0.85 | 0.88 | 0.95 | 0.96 |
| 8 | 119922993 | rs13277230 | 2.153 | -1.711 | 0.78 | 0.088 | -0.163 | -0.082 | -0.082 | 0 | 0.28 | 0.03 | 0.49 | 0.24 | 0.45 | 0.3 |
| 8 | 119934031 | rs11573962 | -1.047 | -1.674 | 1.235 | 0.046 | 0.046 | 0.042 | 0.042 | 6.813 | 0.05 | 0.48 | 0.04 | 0 | 0 | 0 |
| 8 | 119937741 | rs10955911 | 0.728 | -1.566 | 0.907 | 0.117 | 0.001 | 0.001 | 0.001 | 0 | 0.22 | 0.15 | 0.14 | 0.14 | 0.16 | 0.16 |
| 8 | 119945166 | rs1564858 | 0.826 | -1.363 | 0.361 | 0.169 | -0.013 | 0.027 | 0.027 | 0.103 | 0.12 | 0.05 | 0.07 | 0.01 | 0.12 | 0.16 |
| 8 | 119946807 | rs3102724 | -0.5 | -1.651 | 0.578 | 0.262 | 0.041 | 0.022 | 0.022 | 0 | 0.52 | 0.56 | 0.28 | 0.46 | 0.36 | 0.45 |
| 8 | 119949768 | rs2875845 | 0.93 | -1.739 | 0.602 | 0.31 | 0.029 | 0.039 | 0.039 | 0 | 0.22 | 0.08 | 0.13 | 0.13 | 0.16 | 0.16 |
| 8 | 119951692 | rs1905786 | -1.319 | -1.683 | 0.398 | 0.379 | -0.041 | 0.018 | 0.018 | 0.195 | 0.64 | 0.74 | 0.45 | 0.76 | 0.47 | 0.57 |
| 8 | 119951773 | rs1032128 | 0.32 | -1.733 | 0.713 | 0.358 | -0.005 | 0.004 | 0.004 | 0 | 0.34 | 0.33 | 0.28 | 0.63 | 0.28 | 0.39 |
| 8 | 119953468 | rs3134057 | -0.827 | -1.692 | 0.746 | 0.359 | 0.005 | 0.003 | 0.003 | 0.423 | 0.57 | 0.52 | 0.29 | 0.46 | 0.39 | 0.48 |
| 8 | 119954108 | rs3134058 | -1.095 | -1.694 | 0.701 | 0.335 | -0.003 | -0.018 | -0.018 | 0 | 0.6 | 0.74 | 0.34 | 0.64 | 0.42 | 0.53 |
| 8 | 119954995 | rs11573856 | 2.176 | -1.888 | 0.554 | 0.339 | 0.019 | -0.026 | -0.026 | 0.193 | 0.11 | 0.01 | 0.04 | 0.18 | 0.08 | 0.08 |
| 8 | 119959657 | rs3134063 | -1.321 | -1.141 | 0.882 | 0.305 | -0.063 | -0.045 | -0.045 | 0.008 | 0.66 | 0.79 | 0.39 | 0.65 | 0.51 | 0.65 |
| 8 | 119963843 | rs10505346 | -0.17 | -1.172 | 0.986 | 0.268 | -0.026 | -0.049 | -0.049 | 0 | 0.3 | 0.33 | 0.15 | 0.13 | 0.19 | 0.18 |
| 8 | 119965070 | rs3102735 | 0.09 | -1.007 | 0.909 | 0.356 | 0.008 | 0.002 | 0.002 | 0 | 0.13 | 0.23 | 0.17 | 0.13 | 0.13 | 0.15 |
| 8 | 119968733 | rs13270766 | 1.638 | -0.851 | 0.613 | 0.4 | -0.003 | -0.002 | -0.002 | 0.032 | 0.06 | 0.04 | 0.05 | 0.01 | 0.09 | 0.12 |
| 8 | 119975589 | rs1564859 | 0.543 | -0.947 | 0.761 | 0.404 | -0.023 | 0.017 | 0.017 | 0.206 | 0.36 | 0.53 | 0.19 | 0.51 | 0.26 | 0.33 |
| 8 | 119975670 | rs1564860 | -0.425 | -0.954 | 0.721 | 0.386 | -0.015 | -0.026 | -0.026 | 0 | 0.36 | 0.45 | 0.32 | 0.26 | 0.3 | 0.35 |
| 8 | 119976144 | rs1485302 | -0.796 | -0.887 | 0.572 | 0.484 | -0.009 | -0.025 | -0.025 | 0.069 | 0.62 | 0.62 | 0.4 | 0.58 | 0.44 | 0.54 |
| 8 | 119976542 | rs7839059 | -0.91 | -0.891 | 0.6 | 0.444 | 0.006 | 0.011 | 0.011 | 0 | 0.43 | 0.57 | 0.33 | 0.26 | 0.33 | 0.36 |
| 8 | 119977337 | rs12679857 | 0.305 | -0.935 | 0.712 | 0.546 | -0.012 | 0.013 | 0.013 | 0 | 0.37 | 0.49 | 0.25 | 0.52 | 0.32 | 0.44 |
| 8 | 119979097 | rs6987559 | 1.431 | -0.905 | 0.45 | 0.583 | 0.02 | -0.001 | -0.001 | 1.063 | 0.27 | 0.19 | 0.13 | 0.33 | 0.19 | 0.3 |
| 8 | 119979249 | rs7837123 | 0.482 | -0.326 | 0.2 | 0.692 | 0.041 | 0.064 | 0.064 | 0 | 0.09 | 0.28 | 0.11 | 0.19 | 0.12 | 0.14 |
| 8 | 119986235 | rs4876876 | -0.765 | -0.28 | -0.142 | 0.795 | -0.071 | 0.004 | 0.004 | 0 | 0.66 | 0.84 | 0.49 | 0.76 | 0.52 | 0.65 |
| 8 | 120008371 | rs6469792 | -0.959 | -0.007 | -0.938 | 0.922 | -0.106 | -0.108 | -0.108 | 0.452 | 0.7 | 0.84 | 0.46 | 0.59 | 0.53 | 0.68 |
| 8 | 120013090 | rs13264791 | -1.092 | -0.149 | -0.988 | 0.758 | -0.16 | -0.031 | -0.031 | 0 | 0.71 | 0.94 | 0.51 | 0.79 | 0.54 | 0.71 |
| 8 | 120015963 | rs1485284 | 1.101 | -0.581 | -0.708 | 0.689 | 0.005 | -0.002 | -0.002 | 5.611 | 0.3 | 0.26 | 0.13 | 0.38 | 0.21 | 0.3 |
| 8 | 120016060 | rs4354338 | -1.45 | -0.545 | -0.877 | 0.687 | -0.088 | 0 | 0 | 0.248 | 0.72 | 0.86 | 0.52 | 0.8 | 0.58 | 0.72 |
| 8 | 120016099 | rs6469794 | -1.529 | -0.543 | -0.877 | 0.686 | -0.114 | -0.012 | -0.012 | 0 | 0.72 | 0.92 | 0.53 | 0.8 | 0.58 | 0.72 |
| 8 | 120017079 | rs9650075 | -1.014 | -0.561 | -0.802 | 0.708 | -0.067 | -0.039 | -0.039 | 0 | 0.68 | 0.75 | 0.46 | 0.63 | 0.52 | 0.69 |
| 8 | 120018042 | rs6469796 | -1.491 | -0.545 | -0.823 | 0.718 | -0.114 | -0.012 | -0.012 | 0.045 | 0.72 | 0.92 | 0.52 | 0.8 | 0.58 | 0.72 |
| 8 | 120025341 | rs1074162 | 1.054 | -0.579 | -0.717 | 0.719 | 0.016 | -0.005 | -0.005 | 0 | 0.31 | 0.24 | 0.14 | 0.38 | 0.22 | 0.3 |
| 8 | 120027489 | rs1586275 | 0.786 | -0.117 | -1.325 | 0.858 | 0.01 | -0.014 | -0.014 | 0.034 | 0.1 | 0.02 | 0.03 | 0.13 | 0.05 | 0.14 |
| 8 | 120039462 | rs1485295 | -0.881 | 0.295 | -1.18 | 0.612 | -0.09 | 0 | 0 | 0.248 | 0.72 | 0.86 | 0.53 | 0.8 | 0.58 | 0.72 |
| 8 | 120047880 | rs17771512 | 0.786 | 0.256 | -1.126 | 0.546 | -0.009 | 0.014 | 0.014 | 0 | 0.07 | 0.04 | 0.06 | 0.01 | 0.1 | 0.14 |
| 8 | 120048149 | rs6469805 | -0.516 | 0.398 | -1.039 | 0.883 | -0.072 | -0.049 | -0.049 | 4.397 | 0.7 | 0.78 | 0.48 | 0.63 | 0.55 | 0.7 |
| 8 | 120052238 | rs6993813 | -0.516 | 0.398 | -1.039 | 0.883 | -0.075 | -0.051 | -0.051 | 0.054 | 0.7 | 0.79 | 0.48 | 0.63 | 0.55 | 0.7 |
| 8 | 120071898 | rs1485315 | 0.063 | -0.059 | -0.315 | 0.991 | -0.005 | 0 | 0 | 2.78 | 0.3 | 0.33 | 0.21 | 0.69 | 0.25 | 0.32 |
| 8 | 120071939 | rs1385511 | 0.017 | -0.033 | -0.281 | 0.997 | -0.006 | 0 | 0 | 0 | 0.31 | 0.34 | 0.21 | 0.69 | 0.25 | 0.32 |
| 8 | 120072087 | rs1485316 | 0.213 | -0.08 | -0.264 | 1.243 | 0.001 | -0.003 | -0.003 | 0.055 | 0.28 | 0.23 | 0.17 | 0.49 | 0.2 | 0.29 |
| 8 | 120076702 | rs16891935 | -0.209 | 0.11 | -0.079 | 1.255 | -0.003 | 0.058 | 0.058 | 0 | 0.09 | 0.2 | 0.08 | 0.26 | 0.05 | 0.09 |
| 8 | 120080115 | rs1385498 | 0.418 | 0.082 | -0.376 | 0.756 | -0.007 | 0.003 | 0.003 | 0 | 0.43 | 0.34 | 0.35 | 0.65 | 0.47 | 0.34 |
| 8 | 120080814 | rs1872424 | 0.319 | -0.086 | -0.334 | 0.669 | -0.001 | 0.009 | 0.009 | 0 | 0.46 | 0.39 | 0.36 | 0.65 | 0.47 | 0.34 |
| 8 | 120097576 | rs2450063 | -0.517 | 0.337 | -0.336 | 0.709 | -0.067 | -0.038 | -0.038 | 0 | 0.73 | 0.94 | 0.53 | 0.73 | 0.68 | 0.61 |
| 8 | 120100059 | rs11993915 | 0.242 | 0.128 | 0.354 | 0.474 | 0.002 | -0.082 | -0.082 | 0 | 0.14 | 0.26 | 0.09 | 0 | 0.12 | 0.06 |
| 8 | 120107625 | rs13254750 | 0.193 | -0.005 | 0.431 | 0.411 | -0.007 | 0.039 | 0.039 | 0 | 0.13 | 0.05 | 0.1 | 0.02 | 0.14 | 0.16 |
| 8 | 120109987 | rs16891997 | 0.73 | -0.154 | 0.456 | 0.404 | 0.013 | -0.001 | -0.001 | 2.201 | 0.1 | 0.06 | 0.02 | 0.06 | 0.02 | 0.09 |
| 8 | 120111411 | rs17179583 | 0.2 | -0.056 | 0.437 | 0.402 | -0.01 | 0.04 | 0.04 | 0 | 0.13 | 0.05 | 0.11 | 0.02 | 0.16 | 0.17 |
| 8 | 120112982 | rs1351953 | 0.919 | 0.134 | 0.176 | 0.646 | -0.022 | -0.021 | -0.021 | 0 | 0.09 | 0.03 | 0.09 | 0.14 | 0.16 | 0.19 |
| 8 | 120120901 | rs7825193 | -0.933 | 0.338 | -0.033 | 0.524 | 0.042 | 0.074 | 0.074 | 2.515 | 0.12 | 0.52 | 0.12 | 0.2 | 0.12 | 0.07 |
| 8 | 120121084 | rs2450076 | 1.253 | 0.333 | -0.024 | 0.575 | -0.007 | -0.008 | -0.008 | 0 | 0.43 | 0.26 | 0.33 | 0.46 | 0.46 | 0.33 |
| 8 | 120127586 | rs11994775 | -0.115 | 0.345 | -0.073 | 0.571 | 0.002 | 0.004 | 0.004 | 0 | 0.66 | 0.39 | 0.45 | 0.54 | 0.62 | 0.59 |
| 8 | 120128070 | rs6469812 | 0.142 | 0.366 | -0.158 | 0.559 | -0.071 | 0.025 | 0.025 | 0 | 0.61 | 0.86 | 0.43 | 0.71 | 0.54 | 0.45 |
| 8 | 120133272 | rs9650076 | -0.914 | 0.409 | -0.126 | 0.596 | -0.052 | -0.032 | -0.032 | 0 | 0.73 | 0.91 | 0.53 | 0.73 | 0.68 | 0.62 |
| 8 | 120135125 | rs13269842 | -0.305 | 0.461 | -0.157 | 0.578 | -0.059 | 0.02 | 0.02 | 0 | 0.22 | 0.06 | 0.2 | 0.16 | 0.32 | 0.35 |
| 8 | 120143152 | rs6989728 | -1.374 | 0.441 | 0.29 | 0.518 | -0.019 | -0.006 | -0.006 | 0.062 | 0.28 | 0.61 | 0.2 | 0.28 | 0.23 | 0.29 |
| 8 | 120144574 | rs16892098 | -0.634 | 0.429 | 0.322 | 0.547 | -0.056 | 0.074 | 0.074 | 5.984 | 0.17 | 0.55 | 0.1 | 0.26 | 0.08 | 0.11 |
| 8 | 120163391 | rs13248274 | 1.375 | 0.471 | 0.119 | 0.623 | 0.005 | 0.014 | 0.014 | 0 | 0.46 | 0.3 | 0.34 | 0.44 | 0.46 | 0.33 |
| 8 | 120167195 | rs17785868 | 1.935 | 0.927 | -0.047 | 0.511 | -0.044 | -0.032 | -0.032 | 0 | 0.07 | 0.01 | 0.31 | 0.09 | 0.18 | 0.13 |
| 8 | 120180777 | rs6990281 | -1.13 | 0.354 | 0.208 | 0.458 | -0.032 | -0.069 | -0.069 | 0.339 | 0.26 | 0.55 | 0.16 | 0.08 | 0.2 | 0.26 |
| 8 | 120184525 | rs1003465 | -1.13 | 0.354 | 0.208 | 0.458 | -0.032 | -0.069 | -0.069 | 0 | 0.26 | 0.55 | 0.16 | 0.08 | 0.2 | 0.26 |
| 8 | 120187354 | rs2468173 | -0.6 | 0.854 | -0.449 | 0.821 | -0.059 | -0.055 | -0.055 | 7.176 | 0.74 | 0.91 | 0.54 | 0.71 | 0.68 | 0.6 |
| 8 | 120187709 | rs2468175 | 0.282 | 0.8 | -0.409 | 0.779 | -0.017 | -0.041 | -0.041 | 0.945 | 0.64 | 0.81 | 0.52 | 0.65 | 0.66 | 0.52 |
| 8 | 120190485 | rs16892211 | 0.473 | 0.354 | -0.31 | 0.731 | -0.001 | 0.002 | 0.002 | 0.149 | 0.1 | 0.09 | 0.02 | 0.06 | 0.02 | 0.09 |
| 8 | 120201893 | rs16892216 | -0.097 | 0.498 | -0.225 | 0.857 | 0.009 | -0.002 | -0.002 | 0.023 | 0.08 | 0.1 | 0.11 | 0.05 | 0.14 | 0.16 |
| 8 | 120205486 | rs13265548 | -0.993 | 0.57 | -0.071 | 0.869 | 0.012 | 0.001 | 0.001 | 0 | 0.48 | 0.42 | 0.64 | 0.4 | 0.56 | 0.66 |

| | | | | | | | | | | | | | | | | |
|---|---|---|---|---|---|---|---|---|---|---|---|---|---|---|---|---|
| 8 | 120206785 | rs4871372 | -1.507 | 0.564 | -0.514 | 0.87 | 0.013 | -0.016 | -0.016 | 0 | 0.55 | 0.93 | 0.73 | 0.56 | 0.61 | 0.69 |
| 8 | 120210057 | rs6469818 | -0.845 | 1.305 | -0.194 | 0.745 | -0.001 | 0 | 0 | 0.478 | 0.41 | 0.37 | 0.62 | 0.34 | 0.53 | 0.61 |
| 8 | 120215613 | rs7000127 | -0.317 | 1.335 | -0.31 | 0.748 | -0.023 | -0.004 | -0.004 | 0 | 0.32 | 0.22 | 0.51 | 0.29 | 0.39 | 0.43 |
| 8 | 120215645 | rs7000278 | -0.236 | 1.407 | -0.856 | 0.784 | -0.064 | -0.02 | -0.02 | 0.272 | 0.32 | 0.12 | 0.5 | 0.29 | 0.39 | 0.43 |
| 8 | 120216957 | rs16892234 | -0.891 | 1.394 | -0.532 | 0.279 | 0.021 | -0.008 | -0.008 | 0 | 0.09 | 0.18 | 0.1 | 0.05 | 0.14 | 0.16 |
| 8 | 120217356 | rs10091367 | -0.565 | 1.268 | -0.665 | 0.35 | -0.004 | -0.001 | -0.001 | 0.388 | 0.35 | 0.33 | 0.55 | 0.4 | 0.48 | 0.54 |
| 8 | 120218302 | rs9297594 | -0.455 | 1.028 | -0.738 | -0.772 | 0 | 0.038 | 0.038 | 0 | 0.52 | 0.62 | 0.68 | 0.9 | 0.49 | 0.64 |
| 8 | 120219129 | rs3812491 | -0.563 | 0.934 | -0.076 | -0.717 | 0.045 | 0.038 | 0.038 | 0.543 | 0.05 | 0.29 | 0.06 | 0.03 | 0.06 | 0.09 |
| 8 | 120222541 | rs12677274 | 0.496 | 1.254 | -0.722 | -0.326 | -0.074 | -0.115 | -0.115 | 1.398 | 0.19 | 0.02 | 0.26 | 0.57 | 0.22 | 0.2 |
| 8 | 120229785 | rs1364704 | -0.122 | 0.19 | -0.259 | -0.888 | -0.057 | -0.037 | -0.037 | 1.92 | 0.87 | 0.68 | 0.93 | 0.96 | 0.98 | 0.85 |
| 8 | 120234269 | rs2468191 | 0.833 | -0.075 | -0.125 | -1.298 | 0.024 | 0.023 | 0.023 | 0.012 | 0.58 | 0.3 | 0.53 | 0.64 | 0.67 | 0.47 |
| 8 | 120235752 | rs3824331 | 0.809 | -0.221 | 0.067 | -1.388 | -0.004 | -0.021 | -0.021 | 1 | 0.21 | 0.08 | 0.34 | 0.32 | 0.21 | 0.35 |
| 8 | 120239647 | rs1425068 | -0.211 | -0.175 | 0.115 | -1.507 | -0.065 | -0.052 | -0.052 | 0 | 0.86 | 0.62 | 0.93 | 0.97 | 0.98 | 0.85 |
| 8 | 120258996 | rs2326224 | 0.813 | -0.199 | 0.021 | -1.519 | -0.004 | -0.012 | -0.012 | 2.068 | 0.2 | 0.08 | 0.33 | 0.28 | 0.2 | 0.36 |
| 8 | 120259928 | rs1016698 | 1.174 | -0.349 | 0.06 | -1.595 | 0.029 | 0.036 | 0.036 | 0 | 0.57 | 0.3 | 0.55 | 0.68 | 0.68 | 0.45 |
| 8 | 120271776 | rs2252251 | -0.65 | -0.355 | 0.128 | -1.569 | 0.007 | 0 | 0 | 0.96 | 0.88 | 0.89 | 0.96 | 0.97 | 0.98 | 0.94 |
| 8 | 120275270 | rs2251880 | -1.212 | -0.344 | 0.127 | -1.563 | 0.007 | 0.003 | 0.003 | 0 | 0.91 | 0.93 | 0.96 | 0.98 | 0.98 | 0.94 |
| 8 | 120277153 | rs16892267 | -0.862 | -0.327 | 0.162 | -1.551 | 0.007 | 0.057 | 0.057 | 3.502 | 0.07 | 0.42 | 0.04 | 0.04 | 0 | 0.09 |
| 8 | 120286224 | rs1118144 | -0.283 | -0.381 | 0.239 | -1.599 | -0.022 | -0.048 | -0.048 | 0 | 0.87 | 0.79 | 0.95 | 0.98 | 0.96 | 0.93 |
| 8 | 120294297 | rs2432969 | -1.365 | -0.397 | 0.621 | -1.607 | 0.005 | 0.001 | 0.001 | 0 | 0.91 | 0.93 | 0.96 | 0.98 | 0.98 | 0.94 |
| 8 | 120295792 | rs2033042 | 1.48 | -0.473 | 0.073 | -2.337 | -0.047 | 0.023 | 0.023 | 0 | 0.35 | 0.14 | 0.34 | 0.65 | 0.4 | 0.26 |
| 8 | 120303034 | rs1425058 | 0.269 | -0.549 | 0.516 | -1.715 | -0.07 | -0.07 | -0.07 | 0 | 0.83 | 0.53 | 0.92 | 0.94 | 0.96 | 0.85 |
| 8 | 120327008 | rs1425065 | 0.992 | -0.525 | 0.341 | -1.719 | 0.026 | 0.031 | 0.031 | 0 | 0.57 | 0.32 | 0.55 | 0.66 | 0.68 | 0.46 |
| 8 | 120334254 | rs4870752 | -0.593 | -0.383 | 0.037 | -1.772 | -0.015 | -0.005 | -0.005 | 0 | 0.77 | 0.92 | 0.62 | 0.71 | 0.72 | 0.59 |
| 8 | 120334539 | rs13253204 | 1.296 | -0.435 | 0.112 | -1.802 | 0.035 | 0.065 | 0.065 | 0 | 0.52 | 0.28 | 0.48 | 0.65 | 0.59 | 0.4 |
| 8 | 120338208 | rs2447188 | -0.345 | -0.37 | -0.106 | -1.766 | -0.001 | 0.003 | 0.003 | 0 | 0.75 | 0.83 | 0.61 | 0.73 | 0.72 | 0.59 |
| 8 | 120339493 | rs1381339 | -0.902 | -0.485 | 0.189 | -1.6 | -0.035 | -0.074 | -0.074 | 0 | 0.22 | 0.41 | 0.12 | 0.07 | 0.13 | 0.19 |
| 8 | 120345252 | rs2447187 | -0.415 | -0.347 | -0.132 | -1.448 | -0.006 | -0.005 | -0.005 | 0 | 0.31 | 0.3 | 0.38 | 0.3 | 0.22 | 0.43 |
| 8 | 120346710 | rs2447186 | -0.319 | -0.322 | -0.293 | -1.436 | 0.008 | 0.007 | 0.007 | 0 | 0.3 | 0.23 | 0.37 | 0.3 | 0.22 | 0.43 |
| 8 | 120350167 | rs1026594 | 0.441 | -0.223 | -0.416 | -1.363 | -0.003 | -0.006 | -0.006 | 0 | 0.19 | 0.09 | 0.32 | 0.25 | 0.2 | 0.33 |
| 8 | 120350348 | rs972393 | 1.026 | -0.127 | -0.259 | -1.687 | 0.02 | 0.083 | 0.083 | 0 | 0.51 | 0.29 | 0.47 | 0.69 | 0.54 | 0.36 |
| 8 | 120356892 | rs2447176 | -0.32 | -0.534 | -0.015 | -2.021 | -0.005 | -0.045 | -0.045 | 0 | 0.81 | 0.79 | 0.87 | 1 | 0.85 | 0.89 |
| 8 | 120357114 | rs2447175 | 0.094 | -0.474 | 0.222 | -1.919 | -0.013 | -0.021 | -0.021 | 0 | 0.78 | 0.74 | 0.87 | 0.9 | 0.84 | 0.82 |
| 8 | 120365319 | rs2470027 | 1.077 | -0.547 | 0.164 | -1.538 | 0 | 0.023 | 0.023 | 0 | 0.56 | 0.37 | 0.47 | 0.69 | 0.54 | 0.33 |
| 8 | 120367602 | rs1903512 | 2.264 | -0.504 | -1.071 | -2.057 | -0.05 | 0.109 | 0.109 | 0.206 | 0.17 | 0.01 | 0.1 | 0 | 0.2 | 0.04 |
| 8 | 120376789 | rs2326244 | 0.886 | 0.079 | -0.381 | -1.576 | -0.015 | -0.006 | -0.006 | 0 | 0.17 | 0.09 | 0.36 | 0.21 | 0.28 | 0.45 |
| 8 | 120382592 | rs2447154 | -0.004 | -0.018 | -0.413 | -1.65 | 0.002 | 0.002 | 0.002 | 0 | 0.78 | 0.78 | 0.87 | 0.9 | 0.84 | 0.82 |
| 8 | 120414548 | rs13261466 | 1.294 | 0.118 | -0.977 | -1.527 | -0.078 | -0.07 | -0.07 | 0 | 0.2 | 0.02 | 0.28 | 0.69 | 0.22 | 0.19 |
| 8 | 120431506 | rs11538929 | 0.963 | -0.413 | -0.184 | -0.778 | -0.019 | 0.048 | 0.048 | 0 | 0.11 | 0.02 | 0.04 | 0 | 0.11 | 0.07 |
| 8 | 120432799 | rs17187503 | -0.352 | -0.401 | 0.698 | -0.248 | 0.004 | -0.019 | -0.019 | 0 | 0.2 | 0.16 | 0.14 | 0 | 0.18 | 0.11 |
| 8 | 120433976 | rs7816205 | -0.349 | -0.404 | 0.697 | -0.249 | 0.003 | -0.019 | -0.019 | 0 | 0.2 | 0.16 | 0.14 | 0 | 0.18 | 0.11 |
| 8 | 120434684 | rs17187655 | -0.365 | -0.38 | 1.028 | -0.239 | 0.004 | -0.019 | -0.019 | 0 | 0.2 | 0.16 | 0.14 | 0 | 0.18 | 0.11 |
| 8 | 120435705 | rs11785549 | -0.333 | -0.38 | 1.028 | -0.239 | 0.003 | -0.02 | -0.02 | 0 | 0.2 | 0.16 | 0.14 | 0 | 0.18 | 0.11 |
| 8 | 120435812 | rs2071518 | -1.658 | -0.432 | 1.336 | -0.182 | -0.02 | -0.027 | -0.027 | 0.056 | 0.31 | 0.59 | 0.19 | 0.16 | 0.22 | 0.31 |
| 8 | 120435908 | rs2071519 | -0.954 | -0.361 | 1.18 | -0.143 | -0.011 | -0.02 | -0.02 | 0.885 | 0.24 | 0.29 | 0.16 | 0.13 | 0.21 | 0.26 |
| 8 | 120436541 | rs11526 | -0.339 | -0.361 | 1.18 | -0.143 | 0.004 | -0.019 | -0.019 | 0 | 0.2 | 0.16 | 0.14 | 0 | 0.18 | 0.11 |
| 8 | 120444158 | rs1461693 | -0.888 | -0.338 | 0.791 | -0.141 | -0.009 | -0.012 | -0.012 | 0 | 0.25 | 0.32 | 0.17 | 0.16 | 0.21 | 0.26 |
| 8 | 120459480 | rs16892630 | -1.436 | -0.078 | 0.35 | 0.204 | -0.008 | 0.05 | 0.05 | 0.257 | 0.07 | 0.17 | 0.03 | 0.16 | 0.04 | 0.2 |
| 8 | 120467179 | rs4871221 | -1.057 | -0.147 | 0.377 | 0.245 | -0.032 | 0.011 | 0.011 | 0 | 0.24 | 0.44 | 0.19 | 0.26 | 0.15 | 0.26 |
| 8 | 120492161 | rs4354341 | -0.646 | 0.082 | 0.131 | 0.463 | -0.009 | 0.032 | 0.032 | 0 | 0.37 | 0.21 | 0.48 | 0.27 | 0.42 | 0.62 |
| 8 | 120500278 | rs2326263 | -0.646 | 0.082 | 0.131 | 0.463 | -0.009 | 0.032 | 0.032 | 0 | 0.37 | 0.21 | 0.48 | 0.27 | 0.42 | 0.62 |
| 8 | 120501413 | rs6996111 | -0.955 | -0.001 | 0.607 | 0.415 | -0.002 | -0.021 | -0.021 | 0 | 0.42 | 0.54 | 0.51 | 0.27 | 0.43 | 0.64 |
| 8 | 120516889 | rs10086941 | -0.651 | 0.143 | -0.078 | 0.424 | -0.016 | 0.014 | 0.014 | 0 | 0.36 | 0.17 | 0.47 | 0.32 | 0.43 | 0.63 |
| 8 | 120544948 | rs6993464 | -0.688 | -0.059 | -0.152 | 0.486 | -0.019 | 0.021 | 0.021 | 0.078 | 0.41 | 0.16 | 0.51 | 0.34 | 0.45 | 0.62 |
| 8 | 120550360 | rs1446071 | -0.868 | -0.136 | 0.538 | 0.446 | -0.001 | -0.007 | -0.007 | 0.293 | 0.43 | 0.44 | 0.53 | 0.34 | 0.46 | 0.62 |
| 8 | 120558078 | rs16892729 | -0.486 | -0.423 | 0.918 | -0.055 | -0.017 | -0.062 | -0.062 | 0 | 0.27 | 0.38 | 0.18 | 0.05 | 0.2 | 0.16 |
| 8 | 120567358 | rs4871352 | -0.243 | -0.12 | 0.486 | 0.207 | 0.002 | 0 | 0 | 0.041 | 0.2 | 0.19 | 0.41 | 0.34 | 0.3 | 0.43 |
| 8 | 120569877 | rs2305129 | -1.109 | -0.32 | 0.534 | 0.419 | -0.014 | 0.042 | 0.042 | 0 | 0.09 | 0.23 | 0.12 | 0.14 | 0.04 | 0.14 |
| 8 | 120573798 | rs10808504 | -2.229 | -0.291 | 0.663 | 0.39 | -0.004 | -0.036 | -0.036 | 0 | 0.57 | 0.81 | 0.69 | 0.5 | 0.55 | 0.77 |
| 8 | 120575184 | rs1058913 | 0.319 | -0.355 | 0.49 | 0.262 | -0.025 | 0.08 | 0.08 | 0 | 0.22 | 0.02 | 0.13 | 0.04 | 0.19 | 0.12 |
| 8 | 120577030 | rs3816330 | -1.432 | -0.202 | 0.867 | 0.501 | -0.014 | 0.037 | 0.037 | 0 | 0.09 | 0.21 | 0.12 | 0.13 | 0.04 | 0.16 |
| 8 | 120582871 | rs16892786 | -0.373 | -0.376 | 0.775 | 0.183 | -0.019 | -0.013 | -0.013 | 0 | 0.14 | 0.17 | 0.07 | 0.06 | 0.09 | 0.16 |
| 8 | 120583148 | rs2305125 | -1.923 | -0.774 | 0.573 | 0.159 | -0.013 | 0.003 | 0.003 | 0.003 | 0.58 | 0.64 | 0.68 | 0.62 | 0.51 | 0.8 |
| 8 | 120584983 | rs7012314 | -0.105 | -0.694 | 0.58 | 0.167 | -0.013 | -0.009 | -0.009 | 0 | 0.14 | 0.16 | 0.08 | 0.06 | 0.1 | 0.18 |
| 8 | 120586411 | rs3793375 | -0.34 | -0.696 | 0.605 | 0.166 | -0.045 | -0.036 | -0.036 | 0 | 0.16 | 0.28 | 0.08 | 0.06 | 0.1 | 0.18 |
| 8 | 120592482 | rs2289887 | -0.575 | -0.438 | 0.815 | 0.146 | -0.001 | 0.02 | 0.02 | 0 | 0.07 | 0.14 | 0.13 | 0.12 | 0.05 | 0.13 |
| 8 | 120592815 | rs7832704 | -0.01 | -0.43 | 0.836 | 0.166 | 0.07 | 0.078 | 0.078 | 0 | 0.18 | 0.39 | 0.4 | 0.29 | 0.27 | 0.32 |
| 8 | 120594802 | rs4871364 | -0.544 | -0.436 | 0.856 | 0.087 | -0.031 | -0.076 | -0.076 | 1.893 | 0.26 | 0.52 | 0.15 | 0.14 | 0.17 | 0.22 |
| 8 | 120597417 | rs1992721 | -0.753 | -0.477 | 0.23 | 0.046 | -0.009 | -0.024 | -0.024 | 0 | 0.29 | 0.18 | 0.52 | 0.42 | 0.32 | 0.52 |

| | | | | | | | | | | | | | | | | |
|---|---|---|---|---|---|---|---|---|---|---|---|---|---|---|---|---|
| 8 | 120600965 | rs16892827 | 0.508 | -0.541 | 0.044 | 0.015 | 0.041 | 0.023 | 0.023 | 0 | 0.17 | 0.06 | 0.06 | 0.13 | 0.09 | 0.18 |
| 8 | 120613747 | rs2289888 | -1.581 | -0.357 | 0.156 | -0.069 | -0.006 | -0.007 | -0.007 | 0 | 0.54 | 0.65 | 0.52 | 0.53 | 0.47 | 0.74 |
| 8 | 120618120 | rs4871401 | -1.551 | -0.327 | 0.142 | -0.075 | -0.004 | -0.005 | -0.005 | 0 | 0.54 | 0.6 | 0.52 | 0.53 | 0.47 | 0.74 |
| 8 | 120636850 | rs6469835 | -0.912 | -0.001 | 0.081 | 0.367 | 0.009 | 0.061 | 0.061 | 0 | 0.27 | 0.45 | 0.28 | 0.42 | 0.28 | 0.4 |
| 8 | 120638194 | rs6982863 | -0.842 | 0.088 | -0.195 | 0.274 | 0.005 | 0.023 | 0.023 | 0 | 0.29 | 0.33 | 0.28 | 0.42 | 0.3 | 0.44 |
| 8 | 120639217 | rs6469836 | -0.711 | 0.453 | -0.333 | 0.094 | 0.015 | 0.041 | 0.041 | 0 | 0.25 | 0.29 | 0.31 | 0.45 | 0.27 | 0.45 |
| 8 | 120641067 | rs11786983 | -1.274 | -0.228 | -0.649 | -0.904 | -0.021 | -0.085 | -0.085 | 1.054 | 0.4 | 0.44 | 0.17 | 0.05 | 0.22 | 0.36 |
| 8 | 120644433 | rs16892900 | 0.417 | -0.153 | -0.539 | -0.872 | -0.025 | -0.064 | -0.064 | 0 | 0.16 | 0.29 | 0.09 | 0.06 | 0.08 | 0.15 |
| 8 | 120653842 | rs7840386 | -0.87 | -0.06 | 0.087 | -0.81 | 0.038 | 0.04 | 0.04 | 0.299 | 0.42 | 0.65 | 0.19 | 0.08 | 0.22 | 0.37 |
| 8 | 120670531 | rs13267849 | -0.811 | 0.811 | 0.014 | -0.754 | 0.035 | 0.056 | 0.056 | 0.021 | 0.23 | 0.33 | 0.31 | 0.49 | 0.34 | 0.49 |
| 8 | 120679258 | rs10505371 | 0.148 | 0.597 | -0.318 | -0.906 | -0.022 | 0.062 | 0.062 | 0 | 0.13 | 0.01 | 0.05 | 0.02 | 0.12 | 0.21 |
| 8 | 120679572 | rs10505372 | -0.683 | 0.621 | -0.15 | -0.825 | 0.007 | 0.014 | 0.014 | 0.095 | 0.23 | 0.32 | 0.3 | 0.49 | 0.34 | 0.49 |
| 8 | 120701433 | rs17216215 | 0.687 | 0.631 | -0.336 | -0.864 | -0.05 | 0.027 | 0.027 | 0 | 0.11 | 0.04 | 0.07 | 0.06 | 0.17 | 0.16 |
| 8 | 120701853 | rs6986529 | -1.091 | 0.28 | -0.352 | -1.329 | -0.002 | 0 | 0 | 0 | 0.53 | 0.53 | 0.42 | 0.57 | 0.48 | 0.66 |
| 8 | 120705817 | rs7004630 | 0.183 | 0.043 | -0.213 | -1.221 | -0.006 | -0.043 | -0.043 | 3.931 | 0.31 | 0.27 | 0.12 | 0.04 | 0.13 | 0.18 |
| 8 | 120706244 | rs1433948 | -1.846 | 0.098 | -1.061 | -1.081 | -0.023 | -0.07 | -0.07 | 0 | 0.67 | 0.91 | 0.5 | 0.58 | 0.57 | 0.83 |
| 8 | 120707435 | rs11997615 | -1.831 | 0.079 | -1.067 | -1.121 | -0.011 | -0.048 | -0.048 | 0.008 | 0.67 | 0.86 | 0.49 | 0.59 | 0.57 | 0.83 |
| 8 | 120707600 | rs9297602 | -1.165 | 0.78 | -0.992 | -0.523 | -0.019 | -0.02 | -0.02 | 0.079 | 0.39 | 0.36 | 0.37 | 0.55 | 0.48 | 0.7 |
| 8 | 120715175 | rs4871530 | -0.474 | 0.262 | -1.05 | -1.078 | 0.043 | 0.134 | 0.134 | 0 | 0.38 | 0.16 | 0.17 | 0.05 | 0.27 | 0.38 |
| 8 | 120718441 | rs6989251 | 0.416 | 0.695 | -0.27 | -1.257 | 0.081 | -0.011 | -0.011 | 0 | 0.07 | 0.25 | 0.05 | 0 | 0.06 | 0.03 |
| 8 | 120719691 | rs7012791 | 1.068 | 0.992 | -0.483 | -1.017 | -0.001 | 0.007 | 0.007 | 0 | 0.41 | 0.61 | 0.55 | 0.4 | 0.41 | 0.27 |
| 8 | 120738163 | rs2164021 | -0.324 | 0.797 | -0.352 | -1.079 | 0.042 | -0.051 | -0.051 | 0.123 | 0.1 | 0.51 | 0.07 | 0.01 | 0.06 | 0.16 |
| 8 | 120742400 | rs4380974 | -0.52 | 0.829 | -0.363 | -1.076 | -0.008 | -0.01 | -0.01 | 2.039 | 0.79 | 0.76 | 0.72 | 0.45 | 0.67 | 0.65 |
| 8 | 120744349 | rs956748 | -0.314 | 0.795 | -0.278 | -1.063 | 0.053 | -0.047 | -0.047 | 0 | 0.1 | 0.48 | 0.07 | 0.02 | 0.06 | 0.16 |
| 8 | 120744399 | rs956749 | 0.875 | 0.793 | -0.179 | -1.063 | 0.077 | -0.012 | -0.012 | 2.901 | 0.07 | 0.23 | 0.04 | 0 | 0.06 | 0.03 |
| 8 | 120749166 | rs10092747 | -1.607 | 0.992 | -0.391 | -1.111 | -0.048 | -0.071 | -0.071 | 0 | 0.89 | 0.99 | 0.82 | 0.48 | 0.82 | 0.75 |
| 8 | 120757008 | rs2289011 | 2.222 | 0.977 | -0.415 | -1.142 | -0.058 | -0.076 | -0.076 | 0 | 0.3 | 0.05 | 0.48 | 0.39 | 0.35 | 0.12 |
| 8 | 120762233 | rs7818587 | -1.923 | 0.978 | -0.421 | -0.662 | -0.049 | -0.061 | -0.061 | 0 | 0.9 | 0.99 | 0.84 | 0.59 | 0.84 | 0.78 |
| 8 | 120772055 | rs9693207 | 1.059 | 0.915 | -0.284 | -1.051 | 0 | -0.001 | -0.001 | 0 | 0.4 | 0.53 | 0.56 | 0.4 | 0.41 | 0.28 |
| 8 | 120792195 | rs6469845 | -0.163 | 0.897 | -0.473 | -1.231 | -0.005 | -0.059 | -0.059 | 0 | 0.38 | 0.51 | 0.17 | 0.04 | 0.21 | 0.28 |
| 8 | 120793296 | rs7002501 | -0.163 | 0.897 | -0.473 | -1.231 | -0.005 | -0.059 | -0.059 | 0 | 0.38 | 0.51 | 0.17 | 0.04 | 0.21 | 0.28 |
| 8 | 120795607 | rs13268676 | 0.501 | 0.931 | -0.483 | -1.182 | -0.014 | -0.011 | -0.011 | 0.83 | 0.69 | 0.68 | 0.66 | 0.43 | 0.55 | 0.4 |
| 8 | 120803654 | rs11996390 | 0.28 | 0.914 | -0.378 | -1.18 | -0.014 | -0.077 | -0.077 | 0 | 0.35 | 0.31 | 0.14 | 0.03 | 0.2 | 0.15 |
| 8 | 120841323 | rs10108071 | -0.47 | 1.02 | -0.719 | -1.129 | -0.008 | -0.01 | -0.01 | 0.255 | 0.79 | 0.77 | 0.73 | 0.45 | 0.68 | 0.65 |
| 8 | 120845035 | rs16893214 | 0.236 | 0.987 | -0.63 | -1.175 | -0.032 | -0.126 | -0.126 | 0.071 | 0.36 | 0.43 | 0.15 | 0.03 | 0.2 | 0.15 |
| 8 | 120846589 | rs14027 | 1.779 | 1.056 | -0.96 | -1.038 | -0.018 | -0.03 | -0.03 | 0 | 0.32 | 0.15 | 0.5 | 0.4 | 0.35 | 0.25 |
| 8 | 120847188 | rs1055130 | -0.24 | 1.013 | -0.674 | -0.701 | -0.009 | -0.013 | -0.013 | 0 | 0.79 | 0.77 | 0.73 | 0.45 | 0.68 | 0.66 |
| 8 | 120862080 | rs6983932 | -1.608 | 0.818 | -0.979 | -0.739 | -0.033 | -0.025 | -0.025 | 0.121 | 0.66 | 0.86 | 0.49 | 0.6 | 0.61 | 0.75 |
| 8 | 120866948 | rs1509347 | 0.212 | 0.653 | -0.559 | -1.304 | -0.002 | 0 | 0 | 0 | 0.08 | 0.05 | 0.06 | 0.02 | 0.13 | 0.25 |
| 8 | 120869022 | rs7815122 | 0.044 | 0.281 | -0.41 | -1.607 | -0.014 | -0.043 | -0.043 | 0 | 0.3 | 0.31 | 0.12 | 0.04 | 0.13 | 0.28 |
| 8 | 120873342 | rs7843326 | -1.09 | 0.351 | -0.266 | -1.693 | 0.051 | 0.188 | 0.188 | 0 | 0.44 | 0.73 | 0.24 | 0.06 | 0.31 | 0.56 |
| 8 | 120873916 | rs6981251 | 0.568 | 0.214 | -0.165 | -1.633 | 0.019 | 0.008 | 0.008 | 0.224 | 0.26 | 0.2 | 0.1 | 0.02 | 0.11 | 0.12 |
| 8 | 120876023 | rs7823109 | 0.32 | 0.501 | 0.163 | -1.363 | 0.184 | 0.06 | 0.06 | 0.292 | 0.05 | 0.29 | 0.06 | 0 | 0.07 | 0.03 |
| 8 | 120884499 | rs4870988 | -1.588 | 0.321 | -0.292 | -1.581 | -0.002 | -0.017 | -0.017 | 0 | 0.29 | 0.66 | 0.22 | 0.06 | 0.32 | 0.54 |
| 8 | 120884536 | rs4870991 | -1.37 | 0.357 | -0.219 | -1.564 | 0.019 | -0.111 | -0.111 | 0.508 | 0.27 | 0.47 | 0.2 | 0.06 | 0.32 | 0.54 |
| 8 | 120896235 | rs6469858 | -0.793 | 0.249 | -0.492 | -1.514 | 0.042 | 0.065 | 0.065 | 2.74 | 0.77 | 0.92 | 0.92 | 0.98 | 0.88 | 0.93 |
| 8 | 120909236 | rs7814126 | -0.603 | 0.666 | 0.02 | -1.374 | 0.104 | -0.055 | -0.055 | 0.329 | 0.13 | 0.33 | 0.11 | 0.02 | 0.2 | 0.27 |
| 8 | 120910380 | rs6987580 | 1.262 | 0.449 | -0.076 | -1.63 | 0.015 | 0.222 | 0.222 | 0 | 0.48 | 0.26 | 0.7 | 0.92 | 0.56 | 0.39 |
| 8 | 120913535 | rs9649948 | -0.391 | 0.358 | -0.003 | -1.669 | 0.025 | 0.034 | 0.034 | 0.363 | 0.68 | 0.79 | 0.85 | 0.96 | 0.78 | 0.85 |
| 8 | 120916234 | rs12679478 | 0.266 | 0.356 | -0.045 | -1.671 | -0.002 | -0.044 | -0.044 | 0.258 | 0.61 | 0.6 | 0.81 | 0.94 | 0.76 | 0.66 |
| 8 | 120922341 | rs7015470 | 1.343 | 0.378 | -0.118 | -1.66 | 0.016 | 0.223 | 0.223 | 0 | 0.48 | 0.26 | 0.7 | 0.92 | 0.56 | 0.39 |
| 8 | 120924030 | rs10088028 | 0.315 | 0.183 | -0.055 | -1.656 | -0.015 | -0.026 | -0.026 | 0 | 0.06 | 0.09 | 0.03 | 0.01 | 0.02 | 0.19 |
| 8 | 120936177 | rs1519810 | -0.878 | 0.313 | 0.031 | -1.636 | 0.062 | -0.067 | -0.067 | 0.606 | 0.14 | 0.36 | 0.11 | 0.03 | 0.19 | 0.26 |
| 8 | 120936411 | rs4871786 | -0.367 | 0.164 | -0.054 | -1.642 | -0.006 | -0.075 | -0.075 | 0 | 0.38 | 0.38 | 0.19 | 0.04 | 0.24 | 0.34 |
| 8 | 120948797 | rs10955944 | 0.413 | 0.337 | -0.1 | -1.545 | -0.001 | -0.001 | -0.001 | 0 | 0.08 | 0.06 | 0.05 | 0.03 | 0.12 | 0.23 |
| 8 | 120951602 | rs9918814 | -0.538 | 0.237 | -0.016 | -1.629 | 0.012 | 0.018 | 0.018 | 0 | 0.4 | 0.31 | 0.21 | 0.05 | 0.34 | 0.4 |
| 8 | 120958270 | rs12541789 | 1.62 | 0.669 | -0.288 | -0.729 | -0.047 | -0.079 | -0.079 | 0.033 | 0.38 | 0.17 | 0.52 | 0.51 | 0.41 | 0.17 |
| 8 | 120960461 | rs1554376 | 0.594 | 0.652 | -0.12 | -0.746 | 0.025 | 0.053 | 0.053 | 0 | 0.72 | 0.67 | 0.7 | 0.57 | 0.64 | 0.52 |
| 8 | 120962526 | rs3930200 | -0.688 | 0.673 | 0.189 | -0.649 | 0.029 | 0.098 | 0.098 | 0 | 0.84 | 0.8 | 0.76 | 0.59 | 0.75 | 0.75 |
| 8 | 120964825 | rs1464277 | 1.198 | 0.721 | -0.001 | -0.67 | -0.023 | -0.028 | -0.028 | 0 | 0.43 | 0.31 | 0.56 | 0.52 | 0.44 | 0.19 |
| 8 | 120990068 | rs908945 | -0.268 | 0.42 | 0.446 | -0.484 | 0.033 | 0.116 | 0.116 | 0 | 0.83 | 0.74 | 0.73 | 0.6 | 0.76 | 0.76 |
| 8 | 120993134 | rs4871819 | 0.844 | 0.431 | 0.567 | -0.711 | 0.025 | 0.054 | 0.054 | 0 | 0.72 | 0.53 | 0.66 | 0.6 | 0.67 | 0.55 |
| 8 | 120994394 | rs11984766 | 1.056 | 0.158 | 0.472 | -0.99 | -0.009 | -0.014 | -0.014 | 0 | 0.43 | 0.3 | 0.53 | 0.5 | 0.48 | 0.21 |
| 8 | 121009824 | rs17822044 | -0.047 | 0.096 | 0.267 | -1.243 | 0.021 | 0.071 | 0.071 | 0 | 0.34 | 0.22 | 0.22 | 0.13 | 0.32 | 0.48 |
| 8 | 121013770 | rs2271900 | 0.541 | 0.116 | 0.432 | -1.15 | 0.006 | 0.019 | 0.019 | 0 | 0.73 | 0.47 | 0.72 | 0.71 | 0.73 | 0.65 |
| 8 | 121018318 | rs7000361 | -0.746 | 0.138 | 0.478 | -1.2 | 0.037 | -0.042 | -0.042 | 2.961 | 0.1 | 0.39 | 0.07 | 0 | 0.08 | 0.06 |
| 8 | 121019737 | rs2037344 | 0.655 | 0.163 | 0.452 | -1.185 | 0.012 | 0.028 | 0.028 | 0 | 0.73 | 0.54 | 0.73 | 0.71 | 0.73 | 0.65 |
| 8 | 121019769 | rs2037343 | 0.711 | 0.163 | 0.433 | -1.187 | 0.008 | 0.02 | 0.02 | 0 | 0.73 | 0.48 | 0.72 | 0.71 | 0.73 | 0.65 |
| 8 | 121023967 | rs7833689 | 1.496 | 0.149 | 0.374 | -1.22 | -0.021 | -0.01 | -0.01 | 0 | 0.39 | 0.25 | 0.5 | 0.58 | 0.41 | 0.17 |

| | | | | | | | | | | | | | | | | |
|---|---|---|---|---|---|---|---|---|---|---|---|---|---|---|---|---|
| 8 | 121026021 | rs6469890 | 0.762 | 0.142 | 0.525 | -1.231 | -0.002 | 0.017 | 0.017 | 0 | 0.48 | 0.61 | 0.57 | 0.58 | 0.49 | 0.23 |
| 8 | 121027419 | rs7386139 | -0.373 | 0.05 | 0.535 | -1.229 | -0.004 | -0.009 | -0.009 | 0 | 0.82 | 0.82 | 0.79 | 0.71 | 0.8 | 0.71 |
| 8 | 121034642 | rs4871825 | -0.522 | 0.016 | 0.534 | -1.265 | -0.004 | -0.039 | -0.039 | 1.485 | 0.43 | 0.54 | 0.28 | 0.13 | 0.39 | 0.57 |
| 8 | 121037192 | rs2176751 | 0.79 | 0.144 | 0.459 | -1.129 | 0.011 | 0.026 | 0.026 | 0 | 0.72 | 0.46 | 0.71 | 0.7 | 0.71 | 0.67 |
| 8 | 121045227 | rs1519812 | -0.541 | -0.017 | -0.147 | -1.278 | 0.009 | -0.002 | -0.002 | 0 | 0.43 | 0.58 | 0.26 | 0.13 | 0.37 | 0.57 |
| 8 | 121058175 | rs7836934 | 0.273 | 0.11 | -0.006 | -1.106 | 0.082 | 0.008 | 0.008 | 0 | 0.05 | 0.23 | 0.05 | 0 | 0.07 | 0.03 |
| 8 | 121059203 | rs11778912 | 2.068 | 0.33 | -0.973 | -1.695 | -0.068 | 0.084 | 0.084 | 0 | 0.14 | 0 | 0.09 | 0 | 0.17 | 0.03 |
| 8 | 121062077 | rs1051534 | 1.456 | -0.557 | -0.692 | -1.286 | -0.057 | -0.063 | -0.063 | 0 | 0.32 | 0.13 | 0.49 | 0.56 | 0.4 | 0.14 |
| 8 | 121062335 | rs10112515 | 0.2 | -1.018 | -0.721 | -1.539 | -0.023 | -0.01 | -0.01 | 0 | 0.08 | 0.12 | 0.03 | 0.05 | 0.01 | 0.26 |
| 8 | 121066470 | rs10808506 | 0.343 | -1.149 | -0.794 | -1.68 | 0.017 | 0.095 | 0.095 | 0 | 0.32 | 0.16 | 0.18 | 0.11 | 0.29 | 0.27 |
| 8 | 121071231 | rs10955949 | -0.156 | -0.858 | -0.222 | -1.062 | -0.035 | -0.021 | -0.021 | 0 | 0.08 | 0.17 | 0.03 | 0.05 | 0.01 | 0.26 |
| 8 | 121074228 | rs6998748 | -0.609 | -0.881 | -0.197 | -1.074 | -0.043 | -0.028 | -0.028 | 0.109 | 0.1 | 0.2 | 0.03 | 0.05 | 0.01 | 0.27 |
| 8 | 121081936 | rs4871830 | -0.631 | -0.598 | -0.29 | -0.926 | -0.029 | -0.026 | -0.026 | 0 | 0.95 | 1 | 0.95 | 0.93 | 0.91 | 0.88 |
| 8 | 121101436 | rs16893529 | -0.773 | -0.796 | 0.061 | -1.1 | 0.004 | -0.119 | -0.119 | 0 | 0.16 | 0.39 | 0.11 | 0 | 0.17 | 0.15 |
| 8 | 121101522 | rs876084 | 0.794 | -1.147 | -0.596 | -1.117 | 0.013 | 0.03 | 0.03 | 0.584 | 0.57 | 0.36 | 0.67 | 0.7 | 0.61 | 0.36 |
| 8 | 121103751 | rs16893536 | -0.237 | -1.054 | -0.256 | -0.997 | -0.002 | -0.019 | -0.019 | 0 | 0.35 | 0.63 | 0.26 | 0.12 | 0.33 | 0.35 |
| 8 | 121104385 | rs10104973 | -0.44 | -1.188 | -0.268 | -0.981 | 0.002 | 0.002 | 0.002 | 0.201 | 0.73 | 0.76 | 0.8 | 0.75 | 0.77 | 0.72 |
| 8 | 121104786 | rs9969584 | 1.182 | -1.163 | -0.474 | -0.966 | -0.006 | 0.004 | 0.004 | 0 | 0.41 | 0.36 | 0.55 | 0.63 | 0.44 | 0.37 |
| 8 | 121123494 | rs12678023 | -0.48 | -1.02 | -0.688 | -0.891 | 0.001 | 0.002 | 0.002 | 0.31 | 0.74 | 0.78 | 0.8 | 0.76 | 0.77 | 0.72 |
| 8 | 121130164 | rs7827219 | 1.328 | -0.632 | -0.849 | -0.704 | -0.072 | -0.047 | -0.047 | 0 | 0.35 | 0.12 | 0.49 | 0.56 | 0.44 | 0.18 |
| 8 | 121140963 | rs7820004 | -0.132 | -0.55 | 0.22 | -0.198 | -0.005 | 0.047 | 0.047 | 0.513 | 0.07 | 0.34 | 0.07 | 0.07 | 0 | 0.18 |
| 8 | 121177524 | rs4601361 | 0.607 | -0.294 | -0.996 | -0.124 | 0.04 | 0.028 | 0.028 | 0 | 0.62 | 0.24 | 0.69 | 0.69 | 0.76 | 0.5 |
| 8 | 121178772 | rs13276675 | -0.599 | 0.809 | -0.65 | 0.263 | 0.006 | 0.005 | 0.005 | 0 | 0.19 | 0.27 | 0.21 | 0.24 | 0.25 | 0.27 |
| 8 | 121179997 | rs2124471 | 0.039 | 1.265 | -0.207 | -0.097 | 0.04 | -0.032 | -0.032 | 0 | 0.86 | 0.71 | 0.88 | 0.92 | 0.83 | 0.88 |
| 8 | 121180667 | rs2198751 | 0.42 | 0.895 | -1.085 | -0.134 | -0.014 | -0.078 | -0.078 | 0 | 0.37 | 0.17 | 0.47 | 0.47 | 0.39 | 0.24 |
| 8 | 121183650 | rs12541735 | -0.1 | 0.659 | -0.725 | -0.416 | -0.018 | 0.002 | 0.002 | 0 | 0.67 | 0.73 | 0.68 | 0.66 | 0.59 | 0.56 |
| 8 | 121190297 | rs13257307 | 0.657 | 0.322 | -0.15 | -0.452 | -0.026 | -0.042 | -0.042 | 0.747 | 0.28 | 0.36 | 0.2 | 0.18 | 0.19 | 0.31 |
| 8 | 121194900 | rs7005932 | 0.304 | 0.299 | -0.299 | -0.463 | -0.023 | -0.034 | -0.034 | 0 | 0.31 | 0.55 | 0.21 | 0.18 | 0.2 | 0.31 |
| 8 | 121209000 | rs2305597 | 0.465 | 0.447 | -0.231 | -0.561 | -0.028 | -0.048 | -0.048 | 0 | 0.32 | 0.48 | 0.23 | 0.19 | 0.26 | 0.35 |
| 8 | 121210069 | rs2305598 | -0.536 | 0.364 | -0.744 | -0.617 | -0.021 | 0.029 | 0.029 | 0 | 0.6 | 0.82 | 0.6 | 0.66 | 0.46 | 0.56 |
| 8 | 121210250 | rs2305599 | 0.8 | 0.394 | -0.703 | -0.6 | 0.013 | -0.049 | -0.049 | 0 | 0.25 | 0.19 | 0.35 | 0.47 | 0.2 | 0.21 |
| 8 | 121215991 | rs2305600 | -0.042 | 0.631 | -0.334 | -0.2 | -0.02 | 0.024 | 0.024 | 0 | 0.6 | 0.8 | 0.59 | 0.67 | 0.46 | 0.59 |
| 8 | 121228679 | rs4870723 | 0.064 | 0.62 | -0.257 | -0.202 | -0.005 | 0.01 | 0.01 | 0 | 0.58 | 0.63 | 0.57 | 0.67 | 0.47 | 0.59 |
| 8 | 121234756 | rs6989074 | -0.152 | 0.627 | -0.273 | -0.198 | -0.015 | 0.016 | 0.016 | 0 | 0.6 | 0.73 | 0.59 | 0.67 | 0.47 | 0.59 |
| 8 | 121240646 | rs1993392 | -0.636 | 0.839 | -0.358 | -0.252 | -0.005 | 0.004 | 0.004 | 0 | 0.72 | 0.79 | 0.7 | 0.85 | 0.61 | 0.67 |
| 8 | 121240812 | rs2305602 | 0.305 | 0.416 | -0.628 | -0.029 | -0.021 | 0.11 | 0.11 | 0 | 0.11 | 0.19 | 0.23 | 0.47 | 0.03 | 0.18 |
| 8 | 121256150 | rs2305603 | 0.547 | 0.435 | -0.588 | 0.061 | -0.013 | 0.077 | 0.077 | 0 | 0.1 | 0.14 | 0.22 | 0.41 | 0.03 | 0.17 |
| 8 | 121257231 | rs1010830 | -0.231 | 0.49 | -0.764 | -0.355 | -0.042 | 0.086 | 0.086 | 0 | 0.6 | 0.81 | 0.63 | 0.8 | 0.5 | 0.59 |
| 8 | 121259270 | rs13258640 | 0.922 | 0.487 | -0.594 | -0.385 | -0.003 | -0.008 | -0.008 | 0 | 0.46 | 0.49 | 0.36 | 0.39 | 0.43 | 0.41 |
| 8 | 121259843 | rs2305604 | 0.981 | 0.479 | -0.641 | -0.389 | 0.002 | 0.007 | 0.007 | 0 | 0.45 | 0.4 | 0.35 | 0.39 | 0.43 | 0.41 |
| 8 | 121259862 | rs2305605 | 0.824 | 0.533 | -1.196 | -0.21 | -0.002 | 0.06 | 0.06 | 0 | 0.09 | 0.08 | 0.21 | 0.41 | 0.03 | 0.17 |
| 8 | 121262775 | rs2305607 | 0.053 | 0.53 | -1.186 | -0.218 | -0.005 | 0.051 | 0.051 | 0 | 0.12 | 0.16 | 0.24 | 0.41 | 0.07 | 0.18 |
| 8 | 121266654 | rs10111291 | 0.577 | 0.719 | -1.275 | -0.307 | -0.003 | 0.017 | 0.017 | 0 | 0.15 | 0.08 | 0.11 | 0.05 | 0.16 | 0.09 |
| 8 | 121267736 | rs3750255 | 0.49 | 0.827 | -1.208 | 0.039 | -0.005 | 0.071 | 0.071 | 0 | 0.1 | 0.14 | 0.23 | 0.4 | 0.05 | 0.17 |
| 8 | 121268269 | rs1563400 | -0.217 | 1.014 | -1.235 | -0.162 | -0.028 | -0.019 | -0.019 | 0 | 0.65 | 0.81 | 0.51 | 0.43 | 0.61 | 0.5 |
| 8 | 121269836 | rs7813088 | 0.413 | 1.447 | -0.937 | -0.228 | -0.004 | 0.002 | 0.002 | 0 | 0.69 | 0.62 | 0.58 | 0.73 | 0.63 | 0.68 |
| 8 | 121270124 | rs7813506 | -0.164 | 0.943 | -1.173 | -0.536 | -0.035 | -0.044 | -0.044 | 0.005 | 0.65 | 0.78 | 0.49 | 0.54 | 0.48 | 0.6 |
| 8 | 121270355 | rs7814222 | -0.986 | 0.935 | -0.94 | -0.787 | -0.024 | -0.03 | -0.03 | 0 | 0.75 | 0.78 | 0.62 | 0.58 | 0.61 | 0.62 |
| 8 | 121275761 | rs959370 | -0.338 | 0.602 | -0.617 | -0.738 | -0.028 | -0.029 | -0.029 | 0 | 0.62 | 0.73 | 0.6 | 0.59 | 0.58 | 0.54 |
| 8 | 121276075 | rs17833457 | 0.801 | 0.582 | -0.537 | -0.76 | 0.003 | -0.013 | -0.013 | 2.943 | 0.1 | 0.12 | 0.1 | 0.03 | 0.16 | 0.08 |
| 8 | 121280052 | rs11781559 | 0.321 | 0.58 | -0.358 | -0.75 | 0.004 | 0 | 0 | 0 | 0.51 | 0.41 | 0.49 | 0.55 | 0.42 | 0.47 |
| 8 | 121292407 | rs1869831 | -0.591 | 0.594 | -1.033 | -0.78 | -0.078 | -0.086 | -0.086 | 0.29 | 0.64 | 0.88 | 0.6 | 0.58 | 0.58 | 0.54 |
| 8 | 121301474 | rs1563389 | 0.11 | 0.543 | -0.963 | -0.756 | 0.025 | -0.01 | -0.01 | 0.087 | 0.53 | 0.69 | 0.51 | 0.55 | 0.42 | 0.47 |
| 8 | 121312336 | rs1531131 | 0.178 | 0.546 | -0.977 | -0.755 | 0.022 | -0.004 | -0.004 | 0 | 0.52 | 0.6 | 0.5 | 0.55 | 0.42 | 0.47 |
| 8 | 121320682 | rs7006555 | 0.274 | 0.455 | -0.942 | -0.709 | 0.024 | -0.002 | -0.002 | 0 | 0.49 | 0.54 | 0.49 | 0.55 | 0.4 | 0.45 |
| 8 | 121325031 | rs7004099 | 0.095 | 0.506 | -1.547 | -0.665 | 0.057 | -0.023 | -0.023 | 0 | 0.52 | 0.8 | 0.51 | 0.55 | 0.4 | 0.45 |
| 8 | 121325670 | rs6469915 | -0.561 | 0.595 | -1.636 | -0.677 | -0.138 | -0.122 | -0.122 | 0 | 0.63 | 0.93 | 0.6 | 0.58 | 0.55 | 0.52 |
| 8 | 121327673 | rs7844334 | -0.135 | 0.25 | -0.943 | -0.132 | 0.058 | -0.001 | -0.001 | 0 | 0.5 | 0.7 | 0.5 | 0.55 | 0.39 | 0.45 |
| 8 | 121333937 | rs11988416 | 0.307 | 0.559 | -0.218 | 0 | 0.001 | -0.029 | -0.029 | 0 | 0.11 | 0.1 | 0.06 | 0 | 0.11 | 0.09 |
| 8 | 121336940 | rs7001937 | -1.12 | 0.401 | 0.09 | -0.059 | 0.001 | -0.022 | -0.022 | 0.892 | 0.39 | 0.57 | 0.31 | 0.19 | 0.42 | 0.36 |
| 8 | 121343774 | rs2034843 | -1.153 | 0.595 | 0.709 | 0.475 | -0.001 | 0.002 | 0.002 | 0 | 0.39 | 0.56 | 0.44 | 0.44 | 0.4 | 0.49 |
| 8 | 121353416 | rs1563390 | -1.055 | 0.507 | 0.784 | 0.447 | -0.004 | 0.009 | 0.009 | 0 | 0.4 | 0.54 | 0.44 | 0.44 | 0.4 | 0.49 |
| 8 | 121353651 | rs1563391 | -1.299 | 0.429 | 0.528 | 0.452 | 0 | -0.013 | -0.013 | 2.909 | 0.43 | 0.67 | 0.46 | 0.44 | 0.43 | 0.51 |
| 8 | 121355875 | rs4871060 | -1.249 | 0.449 | 0.438 | 0.625 | -0.001 | 0.002 | 0.002 | 0 | 0.43 | 0.65 | 0.59 | 0.53 | 0.44 | 0.51 |
| 8 | 121377780 | rs7821312 | -0.778 | 0.641 | 0.456 | 0.13 | -0.018 | 0.022 | 0.022 | 0 | 0.31 | 0.23 | 0.22 | 0.21 | 0.38 | 0.32 |
| 8 | 121383900 | rs1129353 | 0.811 | 0.303 | 1.3 | 0.642 | -0.038 | 0.06 | 0.06 | 0.135 | 0.81 | 0.45 | 0.73 | 0.8 | 0.9 | 0.77 |
| 8 | 121384163 | rs2429 | 0.84 | 0.299 | 1.298 | 0.64 | -0.04 | 0.059 | 0.059 | 0.261 | 0.8 | 0.45 | 0.73 | 0.8 | 0.9 | 0.77 |
| 8 | 121386925 | rs4870728 | -0.486 | 0.402 | 1.207 | 0.571 | 0 | -0.001 | -0.001 | 3.086 | 0.21 | 0.21 | 0.12 | 0.21 | 0.2 | 0.25 |
| 8 | 121391011 | rs6991421 | 1.52 | 0.427 | 1.216 | 0.567 | 0.009 | -0.023 | -0.023 | 0 | 0.6 | 0.25 | 0.61 | 0.58 | 0.69 | 0.52 |

| | | | | | | | | | | | | | | | | |
|---|---|---|---|---|---|---|---|---|---|---|---|---|---|---|---|---|
| 8 | 121408195 | rs6650 | 1.628 | 0.416 | 1.091 | 0.551 | 0.005 | -0.054 | -0.054 | 0.001 | 0.59 | 0.13 | 0.61 | 0.58 | 0.69 | 0.52 |
| 8 | 121430599 | rs13263962 | 0.688 | 0.395 | 1.339 | 0.551 | -0.035 | 0.077 | 0.077 | 0.098 | 0.81 | 0.5 | 0.74 | 0.8 | 0.9 | 0.77 |
| 8 | 121458163 | rs7812590 | -0.232 | 0.406 | 0.926 | 0.551 | -0.014 | 0.042 | 0.042 | 0 | 0.88 | 0.82 | 0.77 | 0.8 | 0.93 | 0.81 |
| 8 | 121498963 | rs4545141 | 0.792 | 0.495 | 0.137 | 0.229 | -0.003 | 0.023 | 0.023 | 0 | 0.66 | 0.47 | 0.64 | 0.58 | 0.72 | 0.56 |
| 8 | 121517348 | rs4279642 | 0.798 | 0.568 | -0.719 | 0.345 | 0.008 | -0.031 | -0.031 | 0 | 0.6 | 0.21 | 0.62 | 0.58 | 0.69 | 0.52 |
| 8 | 121526747 | rs12155569 | 0.525 | 0.616 | -0.595 | 0.471 | 0.006 | -0.031 | -0.031 | 0 | 0.6 | 0.21 | 0.62 | 0.58 | 0.69 | 0.52 |
| 8 | 121530061 | rs13258768 | 0.58 | 0.568 | -0.618 | 0.485 | 0.008 | -0.042 | -0.042 | 0 | 0.59 | 0.18 | 0.61 | 0.58 | 0.69 | 0.52 |
| 8 | 121549677 | rs3924784 | -0.654 | 0.565 | -0.831 | 0.489 | 0.011 | -0.025 | -0.025 | 0 | 0.4 | 0.77 | 0.39 | 0.42 | 0.3 | 0.48 |
| 8 | 121550719 | rs7003420 | -0.698 | 0.636 | -1.334 | 0.525 | 0.007 | -0.054 | -0.054 | 0 | 0.41 | 0.88 | 0.4 | 0.42 | 0.31 | 0.48 |
| 8 | 121555764 | rs3922897 | 0.16 | 0.957 | -1.052 | 0.612 | -0.022 | -0.004 | -0.004 | 0.232 | 0.66 | 0.78 | 0.6 | 0.62 | 0.58 | 0.61 |
| 8 | 121561369 | rs7015410 | 0.378 | 0.018 | -0.553 | 0.793 | -0.011 | 0.057 | 0.057 | 0 | 0.74 | 0.56 | 0.62 | 0.41 | 0.79 | 0.65 |
| 8 | 121562418 | rs7839488 | 1.096 | -0.851 | -1.155 | -0.036 | -0.013 | 0.092 | 0.092 | 0 | 0.41 | 0.15 | 0.43 | 0.23 | 0.61 | 0.39 |
| 8 | 121564211 | rs7814292 | -0.263 | -1.075 | -0.737 | -0.198 | 0.008 | 0.007 | 0.007 | 0.003 | 0.37 | 0.69 | 0.24 | 0.18 | 0.22 | 0.28 |
| 8 | 121568694 | rs4409432 | 0.168 | -1.175 | -0.289 | 0.238 | 0.014 | 0.031 | 0.031 | 0 | 0.92 | 0.77 | 0.95 | 0.82 | 0.92 | 0.8 |
| 8 | 121568921 | rs4395927 | 1.464 | -1.304 | -0.435 | 0.158 | -0.041 | 0.053 | 0.053 | 0 | 0.38 | 0.15 | 0.38 | 0.24 | 0.56 | 0.38 |
| 8 | 121571266 | rs4297067 | -0.034 | -1.433 | 0.379 | 0.886 | -0.019 | 0.056 | 0.056 | 0 | 0.21 | 0.32 | 0.31 | 0.34 | 0.14 | 0.16 |
| 8 | 121575255 | rs7842050 | 0.789 | -1.461 | 0.331 | 0.554 | -0.017 | 0.06 | 0.06 | 0 | 0.72 | 0.44 | 0.64 | 0.37 | 0.78 | 0.65 |
| 8 | 121580362 | rs4871071 | 1.376 | -1.661 | -0.343 | 0.228 | -0.013 | 0.043 | 0.043 | 0 | 0.09 | 0.02 | 0.04 | 0 | 0.08 | 0.06 |
| 8 | 121587165 | rs6984991 | 1.409 | -1.791 | -0.401 | 0.407 | -0.018 | 0.051 | 0.051 | 0 | 0.41 | 0.12 | 0.41 | 0.25 | 0.61 | 0.39 |
| 8 | 121593187 | rs4871072 | 0.029 | -1.966 | -0.037 | 0.39 | -0.029 | -0.029 | -0.029 | 0.047 | 0.63 | 0.53 | 0.74 | 0.69 | 0.75 | 0.55 |
| 8 | 121597725 | rs7823782 | 1.036 | -1.98 | 0.125 | 0.427 | -0.025 | 0.057 | 0.057 | 0 | 0.72 | 0.6 | 0.65 | 0.38 | 0.8 | 0.64 |
| 8 | 121603802 | rs7015089 | -0.253 | -2.472 | 0.114 | 0.455 | -0.02 | 0.051 | 0.051 | 0 | 0.2 | 0.25 | 0.32 | 0.44 | 0.12 | 0.16 |
| 8 | 121607294 | rs7827807 | 0.489 | -2.448 | 0.153 | 0.367 | 0 | -0.001 | -0.001 | 0 | 0.3 | 0.23 | 0.22 | 0.13 | 0.18 | 0.25 |
| 8 | 121609298 | rs7008488 | 0.301 | -2.536 | -0.225 | 0.338 | -0.013 | -0.009 | -0.009 | 0 | 0.64 | 0.42 | 0.7 | 0.69 | 0.74 | 0.56 |
| 8 | 121616345 | rs4870736 | -0.58 | -2.501 | -0.236 | 0.353 | -0.001 | 0.019 | 0.019 | 0 | 0.94 | 0.9 | 0.97 | 0.82 | 0.94 | 0.84 |
| 8 | 121616369 | rs7017780 | 0.37 | -2.494 | -0.3 | 0.306 | -0.023 | -0.029 | -0.029 | 0 | 0.28 | 0.31 | 0.26 | 0.14 | 0.2 | 0.28 |
| 8 | 121626094 | rs6986444 | 0.969 | -2.458 | -1.016 | 0.478 | -0.029 | 0.068 | 0.068 | 0.019 | 0.82 | 0.69 | 0.7 | 0.38 | 0.91 | 0.68 |
| 8 | 121629752 | rs10099784 | -0.296 | -2.509 | -1.153 | 0.302 | 0.077 | 0.114 | 0.114 | 0 | 0.48 | 0.67 | 0.33 | 0.14 | 0.29 | 0.34 |
| 8 | 121630257 | rs6987986 | -0.454 | -1.994 | -0.587 | 1.113 | -0.015 | 0.061 | 0.061 | 0 | 0.11 | 0.13 | 0.26 | 0.44 | 0.03 | 0.16 |
| 8 | 121639595 | rs13258818 | 1.264 | -1.852 | -1.298 | 0.762 | 0.009 | 0.059 | 0.059 | 0.162 | 0.42 | 0.06 | 0.39 | 0.25 | 0.62 | 0.45 |
| 8 | 121639620 | rs7838111 | 0.187 | -1.796 | -0.425 | 0.424 | -0.003 | -0.014 | -0.014 | 0 | 0.7 | 0.69 | 0.78 | 0.88 | 0.75 | 0.9 |
| 8 | 121642058 | rs10108624 | 0.617 | -1.863 | -0.235 | 0.317 | -0.008 | -0.021 | -0.021 | 0 | 0.65 | 0.6 | 0.79 | 0.76 | 0.69 | 0.74 |
| 8 | 121643025 | rs4324958 | 0.617 | -1.818 | -0.235 | 0.317 | -0.007 | -0.021 | -0.021 | 0 | 0.65 | 0.59 | 0.79 | 0.76 | 0.68 | 0.74 |
| 8 | 121643985 | rs4433191 | 0.048 | -1.745 | -0.307 | 0.569 | 0.001 | 0.013 | 0.013 | 0 | 0.06 | 0.08 | 0.03 | 0.15 | 0.06 | 0.16 |
| 8 | 121653902 | rs13265370 | 1.601 | -1.322 | -0.696 | 0.562 | -0.087 | 0.081 | 0.081 | 0.698 | 0.44 | 0.07 | 0.34 | 0.26 | 0.5 | 0.46 |
| 8 | 121654755 | rs7825018 | 0.661 | -1.163 | -0.04 | 0.189 | 0.05 | 0.135 | 0.135 | 0 | 0.91 | 0.76 | 0.84 | 0.74 | 0.89 | 0.85 |
| 8 | 121666841 | rs10955984 | -0.308 | -1.288 | -0.09 | 0.039 | -0.008 | 0.088 | 0.088 | 0 | 0.27 | 0.15 | 0.16 | 0.07 | 0.27 | 0.11 |
| 8 | 121669415 | rs7821158 | 1.035 | -1.24 | 0.149 | 0.091 | 0.006 | -0.012 | -0.012 | 0 | 0.64 | 0.52 | 0.63 | 0.69 | 0.61 | 0.74 |
| 8 | 121679674 | rs11779540 | -0.463 | -1.408 | -0.294 | 0.02 | -0.033 | 0.099 | 0.099 | 0 | 0.27 | 0.14 | 0.23 | 0.06 | 0.32 | 0.13 |
| 8 | 121680857 | rs7818703 | 0.673 | -1.162 | 0.151 | 0.413 | 0.066 | 0.23 | 0.23 | 0 | 0.94 | 0.72 | 0.83 | 0.78 | 0.94 | 0.87 |
| 8 | 121686293 | rs4360333 | -0.86 | -1.365 | -0.733 | 0.131 | -0.089 | -0.023 | -0.023 | 0 | 0.32 | 0.07 | 0.37 | 0.28 | 0.38 | 0.28 |
| 8 | 121686691 | rs7828909 | -2.016 | -1.956 | -0.928 | -0.15 | -0.019 | 0.049 | 0.049 | 0.406 | 0.55 | 0.92 | 0.49 | 0.34 | 0.44 | 0.43 |
| 8 | 121690802 | rs4387018 | -0.284 | -2.037 | 0.053 | -0.202 | -0.05 | -0.045 | -0.045 | 0 | 0.14 | 0.44 | 0.06 | 0.07 | 0.06 | 0.15 |
| 8 | 121694650 | rs11989782 | 0.089 | -2.101 | -0.551 | -0.195 | -0.068 | 0.006 | 0.006 | 0 | 0.13 | 0.01 | 0.17 | 0.06 | 0.25 | 0.09 |
| 8 | 121699811 | rs10283394 | 0.136 | -2.908 | -0.051 | 0.529 | -0.008 | 0.141 | 0.141 | 0.112 | 0.06 | 0.08 | 0.18 | 0.41 | 0.02 | 0.1 |
| 8 | 121701132 | rs4242317 | -0.936 | -2.997 | 0.104 | 0.344 | 0.021 | 0.001 | 0.001 | 0 | 0.27 | 0.16 | 0.29 | 0.26 | 0.23 | 0.23 |
| 8 | 121707556 | rs9642852 | 0.11 | -2.717 | 0.289 | 0.42 | -0.005 | -0.006 | -0.006 | 0 | 0.08 | 0.11 | 0.13 | 0.07 | 0.09 | 0.05 |
| 8 | 121710675 | rs7841636 | 0.187 | -2.837 | 0.567 | 0.344 | -0.039 | -0.015 | -0.015 | 2.621 | 0.1 | 0.33 | 0.05 | 0.07 | 0.03 | 0.16 |
| 8 | 121713417 | rs4871099 | -1.752 | -2.976 | -0.189 | 0.188 | -0.011 | -0.004 | -0.004 | 0 | 0.4 | 0.64 | 0.4 | 0.69 | 0.19 | 0.48 |
| 8 | 121715079 | rs4871100 | -1.554 | -2.97 | -0.146 | 0.191 | 0.055 | 0.037 | 0.037 | 0.432 | 0.4 | 0.72 | 0.41 | 0.69 | 0.19 | 0.48 |
| 8 | 121718491 | rs4242319 | -1.956 | -2.766 | -0.951 | 0.262 | 0.151 | 0.198 | 0.198 | 2.565 | 0.51 | 0.98 | 0.55 | 0.75 | 0.29 | 0.53 |
| 8 | 121732670 | rs4304359 | -0.733 | -1.801 | -0.008 | 1.035 | 0.001 | -0.002 | -0.002 | 0 | 0.18 | 0.15 | 0.17 | 0.2 | 0.13 | 0.19 |
| 8 | 121739448 | rs4478618 | -0.199 | -1.651 | 0.116 | 1.32 | 0.009 | 0.022 | 0.022 | 0 | 0.11 | 0.08 | 0.17 | 0.44 | 0.03 | 0.13 |
| 8 | 121752559 | rs4871109 | -0.985 | -0.936 | 0.459 | 0.679 | -0.017 | 0.033 | 0.033 | 0 | 0.14 | 0.25 | 0.16 | 0.17 | 0.09 | 0.15 |
| 8 | 121763223 | rs4871122 | -0.923 | -0.921 | 0.398 | 0.666 | -0.016 | 0.03 | 0.03 | 0 | 0.14 | 0.25 | 0.16 | 0.16 | 0.09 | 0.15 |
| 8 | 121772783 | rs10108016 | -0.493 | -1.732 | -0.813 | 0.208 | 0.12 | 0.02 | 0.02 | 0.202 | 0.47 | 0.89 | 0.5 | 0.52 | 0.29 | 0.43 |
| 8 | 121793178 | rs4345604 | -0.843 | -1.879 | -0.374 | -0.169 | -0.019 | -0.063 | -0.063 | 0 | 0.13 | 0.25 | 0.07 | 0 | 0.08 | 0.13 |
| 8 | 121796183 | rs4871131 | 0.693 | -2.193 | -0.658 | -0.107 | -0.04 | 0.056 | 0.056 | 0 | 0.21 | 0.03 | 0.22 | 0.1 | 0.24 | 0.16 |
| 8 | 121798339 | rs6993049 | 0.278 | -2.137 | -0.62 | -0.069 | 0.013 | -0.028 | -0.028 | 0 | 0.06 | 0.15 | 0.05 | 0 | 0.05 | 0.03 |
| 8 | 121805251 | rs6469953 | -1.385 | -2.168 | -0.752 | 0.215 | 0.093 | 0.012 | 0.012 | 0.017 | 0.51 | 0.88 | 0.51 | 0.48 | 0.35 | 0.46 |
| 8 | 121810549 | rs10090787 | -0.683 | -2.222 | -0.559 | 0.151 | -0.009 | -0.056 | -0.056 | 0 | 0.36 | 0.47 | 0.32 | 0.11 | 0.32 | 0.29 |
| 8 | 121814601 | rs10955990 | -1.231 | -2.021 | -1.461 | 0.224 | 0.087 | 0 | 0 | 0.098 | 0.51 | 0.89 | 0.52 | 0.48 | 0.34 | 0.46 |
| 8 | 121818295 | rs13438834 | -0.157 | -1.934 | -1.391 | 0.2 | -0.006 | 0.058 | 0.058 | 0 | 0.27 | 0.13 | 0.27 | 0.14 | 0.29 | 0.2 |
| 8 | 121818671 | rs6985848 | -1.13 | -2.253 | -2.062 | 0.176 | 0.009 | -0.068 | -0.068 | 1.953 | 0.58 | 0.93 | 0.52 | 0.52 | 0.36 | 0.42 |
| 8 | 121827840 | rs7822849 | -0.661 | -2.245 | -2.322 | 0.424 | -0.059 | -0.114 | -0.114 | 0.762 | 0.76 | 0.98 | 0.74 | 0.64 | 0.72 | 0.73 |
| 8 | 121830295 | rs13282943 | 1.471 | -2.122 | -2.047 | -0.022 | -0.041 | 0.04 | 0.04 | 0 | 0.3 | 0.19 | 0.29 | 0.15 | 0.47 | 0.32 |
| 8 | 121832965 | rs4129740 | -0.408 | -2.396 | -1.342 | 0.217 | -0.008 | 0.011 | 0.011 | 0 | 0.28 | 0.34 | 0.34 | 0.4 | 0.19 | 0.3 |
| 8 | 121834372 | rs4335166 | 1.034 | -2.451 | -1.532 | 0.636 | -0.035 | 0.032 | 0.032 | 0 | 0.14 | 0.2 | 0.18 | 0.23 | 0.02 | 0.13 |
| 8 | 121840359 | rs10090646 | -0.307 | -2.276 | -1.828 | 0.708 | -0.014 | -0.046 | -0.046 | 0 | 0.15 | 0.05 | 0.22 | 0.41 | 0.16 | 0.29 |

| | | | | | | | | | | | | | | | | |
|---|---|---|---|---|---|---|---|---|---|---|---|---|---|---|---|---|
| 8 | 121840706 | rs10091004 | 0.12 | -2.293 | -1.84 | 0.709 | -0.015 | -0.027 | -0.027 | 0 | 0.12 | 0.05 | 0.2 | 0.39 | 0.15 | 0.28 |
| 8 | 121841059 | rs7013980 | 0.133 | -2.669 | -1.49 | 0.532 | -0.025 | -0.015 | -0.015 | 0 | 0.14 | 0.22 | 0.14 | 0.09 | 0.06 | 0.13 |
| 8 | 121841341 | rs7014184 | 0.133 | -2.669 | -1.458 | 0.532 | -0.026 | -0.016 | -0.016 | 0 | 0.14 | 0.23 | 0.14 | 0.09 | 0.06 | 0.13 |
| 8 | 121845280 | rs7007426 | 1.805 | -2.76 | -1.363 | 0.532 | 0.059 | 0.021 | 0.021 | 0 | 0.31 | 0.44 | 0.29 | 0.26 | 0.42 | 0.37 |
| 8 | 121845554 | rs12546614 | 1.31 | -2.853 | -1.484 | 0.547 | 0.017 | -0.006 | -0.006 | 0.726 | 0.15 | 0.12 | 0.2 | 0.19 | 0.03 | 0.11 |
| 8 | 121846538 | rs4642694 | 0.17 | -2.481 | -1.276 | 0.693 | -0.032 | -0.021 | -0.021 | 0 | 0.15 | 0.27 | 0.15 | 0.09 | 0.06 | 0.12 |
| 8 | 121847643 | rs4369032 | -0.735 | -2.484 | -1.314 | 0.686 | 0.038 | 0.1 | 0.1 | 0.279 | 0.6 | 0.85 | 0.65 | 0.74 | 0.63 | 0.78 |
| 8 | 121852680 | rs7017161 | 1.21 | -2.121 | -0.978 | 0.418 | 0.044 | 0.014 | 0.014 | 0 | 0.32 | 0.48 | 0.29 | 0.26 | 0.42 | 0.37 |
| 8 | 121853405 | rs4871134 | -0.431 | -2.279 | -0.753 | 0.303 | -0.031 | -0.019 | -0.019 | 0 | 0.15 | 0.25 | 0.15 | 0.09 | 0.06 | 0.13 |
| 8 | 121855464 | rs6994902 | -0.328 | -2.266 | -0.853 | 0.31 | -0.026 | -0.016 | -0.016 | 0 | 0.14 | 0.23 | 0.14 | 0.09 | 0.06 | 0.13 |
| 8 | 121861465 | rs10104069 | -0.68 | -2.078 | -1.698 | 0.13 | -0.003 | 0.023 | 0.023 | 0 | 0.73 | 0.86 | 0.8 | 0.87 | 0.69 | 0.81 |
| 8 | 121868551 | rs4871137 | 0.674 | -2.622 | -1.522 | 0.187 | 0.082 | 0.042 | 0.042 | 0.207 | 0.55 | 0.22 | 0.69 | 0.71 | 0.63 | 0.73 |
| 8 | 121873345 | rs6996149 | -0.722 | -2.35 | -1.476 | 0.064 | 0.005 | 0.021 | 0.021 | 0.197 | 0.72 | 0.83 | 0.79 | 0.87 | 0.68 | 0.81 |
| 8 | 121875083 | rs7836397 | 1.449 | -1.443 | -1.876 | -0.016 | -0.183 | 0.054 | 0.054 | 0 | 0.28 | 0.01 | 0.3 | 0.13 | 0.42 | 0.22 |
| 8 | 121886409 | rs7840108 | -0.37 | -1.814 | -0.447 | -0.253 | -0.004 | -0.03 | -0.03 | 0.789 | 0.2 | 0.21 | 0.13 | 0.04 | 0.15 | 0.16 |
| 8 | 121891108 | rs13279396 | -1.33 | -2.143 | -0.029 | -0.361 | -0.001 | -0.018 | -0.018 | 0 | 0.75 | 0.7 | 0.75 | 0.9 | 0.71 | 0.85 |
| 8 | 121891873 | rs7014203 | 0.303 | -2.222 | 0.136 | -0.198 | -0.012 | 0.147 | 0.147 | 0 | 0.06 | 0.21 | 0.05 | 0.14 | 0.01 | 0.11 |
| 8 | 121893978 | rs12681019 | 1.36 | -2.274 | 0.216 | -0.23 | 0.037 | 0.05 | 0.05 | 0 | 0.47 | 0.21 | 0.56 | 0.72 | 0.55 | 0.59 |
| 8 | 121896825 | rs10955995 | 1.36 | -2.274 | 0.216 | -0.23 | 0.037 | 0.05 | 0.05 | 0 | 0.47 | 0.21 | 0.56 | 0.72 | 0.55 | 0.59 |
| 8 | 121897899 | rs11985559 | -0.1 | -2.268 | 0.25 | -0.229 | -0.016 | -0.06 | -0.06 | 0 | 0.21 | 0.26 | 0.14 | 0.04 | 0.15 | 0.16 |
| 8 | 121904711 | rs7842191 | -1.105 | -2.27 | 0.242 | -0.224 | -0.058 | -0.018 | -0.018 | 0 | 0.27 | 0.48 | 0.19 | 0.18 | 0.16 | 0.26 |
| 8 | 121907688 | rs6469961 | -0.833 | -2.219 | 0.12 | -0.219 | -0.02 | 0.011 | 0.011 | 0 | 0.69 | 0.5 | 0.7 | 0.76 | 0.69 | 0.74 |
| 8 | 121914565 | rs7830584 | -1.424 | -2.17 | -0.065 | -0.201 | 0.037 | 0.048 | 0.048 | 0 | 0.52 | 0.76 | 0.44 | 0.28 | 0.45 | 0.41 |
| 8 | 121917686 | rs10096496 | 0.61 | -1.927 | -0.05 | -0.047 | -0.015 | -0.022 | -0.022 | 0 | 0.29 | 0.44 | 0.3 | 0.16 | 0.3 | 0.2 |
| 8 | 121918368 | rs3922773 | 0.46 | -1.935 | -0.17 | 0.312 | -0.002 | 0.002 | 0.002 | 0 | 0.06 | 0.09 | 0.14 | 0.23 | 0.1 | 0.12 |
| 8 | 121921290 | rs2891944 | 0.656 | -1.96 | -0.208 | 0.274 | -0.015 | -0.021 | -0.021 | 0 | 0.29 | 0.45 | 0.3 | 0.15 | 0.3 | 0.2 |
| 8 | 121926125 | rs10110670 | 0.365 | -2.046 | -0.408 | -0.015 | 0.004 | 0.018 | 0.018 | 0.355 | 0.36 | 0.29 | 0.24 | 0.3 | 0.37 | 0.26 |
| 8 | 121926352 | rs10110895 | 0.361 | -2.041 | -0.393 | -0.006 | 0.004 | 0.014 | 0.014 | 0.338 | 0.36 | 0.31 | 0.24 | 0.3 | 0.37 | 0.26 |
| 8 | 121928254 | rs6990987 | -1.349 | -1.746 | -0.269 | 0.022 | -0.002 | -0.002 | -0.002 | 0 | 0.59 | 0.59 | 0.36 | 0.48 | 0.5 | 0.55 |
| 8 | 121928369 | rs10505382 | -0.151 | -1.635 | -0.34 | -0.421 | 0 | -0.002 | -0.002 | 0 | 0.18 | 0.15 | 0.09 | 0.04 | 0.14 | 0.16 |
| 8 | 121928562 | rs10505383 | 0.965 | -1.486 | -0.605 | -0.387 | -0.002 | -0.009 | -0.009 | 0 | 0.11 | 0.08 | 0.04 | 0 | 0.09 | 0.13 |
| 8 | 121937245 | rs10105297 | 0.18 | -1.727 | -0.276 | 0.089 | -0.02 | -0.027 | -0.027 | 0 | 0.07 | 0.04 | 0.05 | 0.22 | 0.1 | 0.1 |
| 8 | 121938422 | rs7826823 | -0.925 | -1.888 | -0.245 | 0.067 | 0.019 | 0.012 | 0.012 | 0.567 | 0.5 | 0.58 | 0.33 | 0.45 | 0.41 | 0.52 |
| 8 | 121946555 | rs939141 | -0.044 | -1.802 | -0.523 | 0.039 | -0.009 | -0.032 | -0.032 | 0 | 0.1 | 0 | 0.1 | 0.08 | 0.08 | 0.09 |
| 8 | 121947406 | rs12680036 | -0.514 | -1.552 | 0.057 | -0.185 | 0.012 | 0.031 | 0.031 | 0 | 0.09 | 0.18 | 0.06 | 0.22 | 0.1 | 0.1 |
| 8 | 121948544 | rs13439619 | -0.386 | -1.595 | 0.065 | -0.17 | 0.017 | 0.015 | 0.015 | 0.839 | 0.4 | 0.34 | 0.21 | 0.36 | 0.33 | 0.43 |
| 8 | 121950167 | rs16894449 | -0.247 | -1.572 | 0.065 | -0.16 | 0.004 | 0.003 | 0.003 | 0.728 | 0.4 | 0.4 | 0.22 | 0.36 | 0.33 | 0.43 |
| 8 | 121967381 | rs7821732 | 0.55 | -0.511 | -0.556 | 0.526 | 0.003 | -0.012 | -0.012 | 0.088 | 0.32 | 0.19 | 0.3 | 0.34 | 0.31 | 0.41 |
| 8 | 121971039 | rs7824584 | -0.69 | -0.922 | -0.385 | 0.479 | 0.005 | 0 | 0 | 0 | 0.67 | 0.63 | 0.63 | 0.67 | 0.54 | 0.71 |
| 8 | 121977328 | rs4871144 | -0.612 | -1.056 | -0.246 | 0.491 | 0.013 | -0.004 | -0.004 | 0 | 0.66 | 0.55 | 0.61 | 0.68 | 0.54 | 0.73 |
| 8 | 121982031 | rs10505385 | 1.232 | -0.855 | -0.593 | 0.664 | 0.005 | -0.059 | -0.059 | 0 | 0.12 | 0.04 | 0.22 | 0.32 | 0.07 | 0.14 |
| 8 | 121984068 | rs7828438 | 1.041 | -0.82 | -0.473 | 0.633 | -0.012 | -0.014 | -0.014 | 0 | 0.31 | 0.23 | 0.3 | 0.35 | 0.34 | 0.43 |
| 8 | 121984358 | rs17196271 | 0.071 | -0.771 | -0.43 | 0.039 | 0.001 | 0.014 | 0.014 | 0 | 0.47 | 0.4 | 0.54 | 0.67 | 0.42 | 0.57 |
| 8 | 121988612 | rs2386232 | -0.546 | -1.018 | -0.571 | 0.126 | 0.008 | -0.009 | -0.009 | 0 | 0.59 | 0.47 | 0.58 | 0.62 | 0.49 | 0.69 |
| 8 | 121993791 | rs1655015 | 0.073 | -0.849 | -0.647 | 0.385 | 0.001 | 0.007 | 0.007 | 0 | 0.46 | 0.42 | 0.48 | 0.42 | 0.4 | 0.29 |
| 8 | 121994660 | rs1560774 | 0.409 | -0.936 | -0.712 | 0.333 | -0.016 | 0.002 | 0.002 | 0 | 0.42 | 0.24 | 0.45 | 0.43 | 0.4 | 0.22 |
| 8 | 122000602 | rs1433396 | 1.488 | -0.805 | -0.744 | 0.238 | -0.099 | 0.166 | 0.166 | 0 | 0.27 | 0.04 | 0.16 | 0 | 0.32 | 0.08 |
| 8 | 122025559 | rs9969592 | -0.205 | -0.798 | -0.686 | 0.326 | -0.009 | 0.08 | 0.08 | 0.167 | 0.07 | 0.17 | 0.11 | 0.19 | 0.04 | 0.36 |
| 8 | 122035990 | rs1714646 | 1.011 | -0.773 | -0.802 | 0.387 | -0.121 | 0.075 | 0.075 | 0 | 0.31 | 0.04 | 0.18 | 0.13 | 0.36 | 0.22 |
| 8 | 122045814 | rs1433401 | 1.106 | -0.841 | -0.547 | 0.414 | -0.034 | 0.064 | 0.064 | 0 | 0.28 | 0.13 | 0.18 | 0 | 0.32 | 0.08 |
| 8 | 122049626 | rs12548955 | -1.513 | -1.343 | 0.541 | 1.681 | -0.04 | 0.042 | 0.042 | 0 | 0.24 | 0.41 | 0.4 | 0.63 | 0.12 | 0.5 |
| 8 | 122074926 | rs17199454 | 0.04 | -1.034 | 0.566 | 1.433 | 0.026 | 0.033 | 0.033 | 0.826 | 0.15 | 0 | 0.03 | 0.05 | 0.07 | 0.12 |
| 8 | 122078334 | rs4871157 | -1.398 | -0.966 | 0.614 | 1.494 | -0.011 | 0.079 | 0.079 | 0.237 | 0.08 | 0.17 | 0.11 | 0.19 | 0.04 | 0.36 |
| 8 | 122088488 | rs13277349 | -0.432 | -0.954 | 0.319 | 1.525 | 0.011 | 0.009 | 0.009 | 0.139 | 0.18 | 0.08 | 0.19 | 0.13 | 0.14 | 0.17 |
| 8 | 122096261 | rs2043375 | -2.683 | 0.111 | 0.733 | 2.196 | 0.006 | 0.202 | 0.202 | 0.233 | 0.13 | 0.45 | 0.29 | 0.27 | 0.11 | 0.41 |
| 8 | 122099514 | rs4871158 | -1.732 | -0.072 | -0.388 | 2.008 | -0.001 | 0.019 | 0.019 | 0.12 | 0.07 | 0.08 | 0.1 | 0.19 | 0.03 | 0.36 |
| 8 | 122108725 | rs4870757 | -1.445 | -0.038 | -0.459 | 1.843 | 0.004 | 0.001 | 0.001 | 0.052 | 0.06 | 0.06 | 0.09 | 0.16 | 0.03 | 0.35 |
| 8 | 122111031 | rs1394425 | -0.436 | -0.106 | -0.643 | 2.194 | 0.001 | -0.052 | -0.052 | 0.058 | 0.12 | 0.03 | 0.21 | 0.4 | 0.07 | 0.18 |
| 8 | 122111867 | rs7005496 | -1.979 | -0.244 | -0.035 | 2.212 | -0.03 | 0.057 | 0.057 | 0 | 0.25 | 0.47 | 0.42 | 0.58 | 0.15 | 0.54 |
| 8 | 122111938 | rs16894660 | -1.142 | -0.095 | -0.37 | 2.452 | -0.009 | 0.126 | 0.126 | 0 | 0.06 | 0.18 | 0.11 | 0.18 | 0.03 | 0.28 |
| 8 | 122117842 | rs13275987 | -0.61 | 0.215 | -0.891 | 2.03 | 0 | -0.055 | -0.055 | 0 | 0.12 | 0.03 | 0.21 | 0.4 | 0.07 | 0.18 |
| 8 | 122128261 | rs10216393 | -0.555 | 0.219 | -0.771 | 1.783 | 0.001 | -0.005 | -0.005 | 3.018 | 0.29 | 0.29 | 0.27 | 0.57 | 0.28 | 0.41 |
| 8 | 122129577 | rs4871162 | -1.153 | 0.352 | -0.842 | 1.192 | 0 | 0.063 | 0.063 | 0.371 | 0.53 | 0.36 | 0.65 | 0.83 | 0.51 | 0.71 |
| 8 | 122131996 | rs6997963 | -0.714 | -0.081 | -0.497 | 0.997 | 0.045 | 0.042 | 0.042 | 1.357 | 0.05 | 0.59 | 0.06 | 0 | 0 | 0 |
| 8 | 122139552 | rs2060319 | 0.013 | 0.014 | -0.7 | 1.375 | 0.006 | -0.036 | -0.036 | 0 | 0.17 | 0.11 | 0.38 | 0.37 | 0.13 | 0.29 |
| 8 | 122141692 | rs2163920 | -0.445 | -0.024 | -0.521 | 1.514 | -0.005 | 0.013 | 0.013 | 0 | 0.32 | 0.27 | 0.24 | 0.13 | 0.34 | 0.25 |
| 8 | 122144267 | rs1989696 | -0.377 | -0.121 | -0.602 | 1.501 | -0.006 | 0.025 | 0.025 | 0 | 0.2 | 0.22 | 0.13 | 0.46 | 0.15 | 0.26 |
| 8 | 122148511 | rs1433394 | -0.683 | -0.251 | -0.705 | 1.392 | -0.011 | -0.084 | -0.084 | 0 | 0.24 | 0.08 | 0.38 | 0.38 | 0.18 | 0.41 |
| 8 | 122149252 | rs1036265 | -1.757 | -0.337 | -0.498 | 1.168 | 0 | 0.003 | 0.003 | 0 | 0.42 | 0.59 | 0.55 | 0.72 | 0.36 | 0.63 |

| | | | | | | | | | | | | | | | | |
|---|---|---|---|---|---|---|---|---|---|---|---|---|---|---|---|---|
| 8 | 122156253 | rs1433393 | -2.305 | -0.184 | -0.116 | 1.152 | -0.056 | 0.176 | 0.176 | 0 | 0.57 | 0.89 | 0.66 | 0.84 | 0.5 | 0.73 |
| 8 | 122156901 | rs10111630 | -2.296 | -0.184 | -0.116 | 1.152 | -0.062 | 0.176 | 0.176 | 0 | 0.56 | 0.88 | 0.64 | 0.84 | 0.48 | 0.73 |
| 8 | 122166846 | rs7006781 | 0.564 | -0.161 | 1.571 | 1.161 | -0.008 | -0.001 | -0.001 | 0.012 | 0.14 | 0.27 | 0.12 | 0.12 | 0.14 | 0.09 |
| 8 | 122174537 | rs7017023 | -1.841 | -0.173 | 1.575 | 1.178 | 0.001 | 0.008 | 0.008 | 0 | 0.43 | 0.62 | 0.51 | 0.72 | 0.37 | 0.66 |
| 8 | 122177081 | rs1368605 | -1.604 | -0.173 | 1.515 | 1.232 | -0.023 | -0.017 | -0.017 | 0 | 0.34 | 0.5 | 0.44 | 0.72 | 0.22 | 0.61 |
| 8 | 122182342 | rs1604898 | -2.274 | -0.126 | 1.329 | 1.138 | -0.047 | 0.167 | 0.167 | 0 | 0.57 | 0.85 | 0.63 | 0.84 | 0.51 | 0.8 |
| 8 | 122182732 | rs4871174 | -0.886 | -0.173 | 1.501 | 1.306 | 0.012 | -0.051 | -0.051 | 1.267 | 0.27 | 0.19 | 0.4 | 0.39 | 0.19 | 0.43 |
| 8 | 122188915 | rs11987945 | -0.305 | -0.215 | 1.466 | 1.698 | -0.007 | 0.003 | 0.003 | 0.114 | 0.15 | 0.19 | 0.1 | 0.18 | 0.05 | 0.07 |
| 8 | 122191388 | rs1827656 | -0.305 | -0.215 | 1.466 | 1.698 | -0.007 | 0.003 | 0.003 | 0.114 | 0.15 | 0.19 | 0.1 | 0.18 | 0.05 | 0.07 |
| 8 | 122191502 | rs1847547 | -0.359 | -0.226 | 1.502 | 1.722 | -0.02 | 0.01 | 0.01 | 0.213 | 0.16 | 0.25 | 0.11 | 0.18 | 0.05 | 0.07 |
| 8 | 122192062 | rs1354506 | -1.514 | -0.227 | 1.799 | 1.694 | 0 | -0.001 | -0.001 | 0.082 | 0.41 | 0.4 | 0.51 | 0.66 | 0.43 | 0.73 |
| 8 | 122193547 | rs17203449 | -0.633 | -0.217 | 1.775 | 1.729 | -0.028 | -0.06 | -0.06 | 0 | 0.27 | 0.09 | 0.38 | 0.34 | 0.26 | 0.51 |
| 8 | 122198250 | rs7004241 | 0.238 | -0.03 | 1.909 | 1.669 | 0.023 | -0.046 | -0.046 | 0.28 | 0.09 | 0.18 | 0.08 | 0 | 0.14 | 0.05 |
| 8 | 122201164 | rs1504658 | -2.543 | -0.03 | 1.74 | 1.428 | -0.101 | 0.182 | 0.182 | 0.064 | 0.58 | 0.92 | 0.64 | 0.84 | 0.49 | 0.8 |
| 8 | 122223650 | rs1553504 | -1.718 | -0.095 | 2.424 | 1.67 | 0.009 | 0.004 | 0.004 | 0.926 | 0.38 | 0.48 | 0.52 | 0.6 | 0.44 | 0.68 |
| 8 | 122226775 | rs1002898 | -0.377 | -0.17 | 2.472 | 1.581 | -0.023 | 0.08 | 0.08 | 0 | 0.15 | 0.36 | 0.13 | 0.24 | 0.08 | 0.1 |
| 8 | 122235345 | rs1504645 | -0.347 | -0.164 | 2.45 | 1.58 | -0.019 | 0.068 | 0.068 | 0 | 0.15 | 0.29 | 0.12 | 0.24 | 0.08 | 0.1 |
| 8 | 122236020 | rs12542986 | -0.724 | -0.205 | 2.307 | 1.881 | -0.032 | -0.063 | -0.063 | 0 | 0.25 | 0.13 | 0.4 | 0.39 | 0.3 | 0.46 |
| 8 | 122238834 | rs9297638 | -0.736 | -0.099 | 2.068 | 1.88 | -0.004 | 0.166 | 0.166 | 0.535 | 0.07 | 0.22 | 0.06 | 0.28 | 0.03 | 0.17 |
| 8 | 122245126 | rs4871178 | -1.929 | -0.087 | 1.903 | 1.719 | 0.005 | -0.006 | -0.006 | 0 | 0.4 | 0.48 | 0.52 | 0.67 | 0.45 | 0.68 |
| 8 | 122249282 | rs17272172 | 0.32 | -0.085 | 1.315 | 1.622 | -0.013 | 0.02 | 0.02 | 0 | 0.07 | 0.05 | 0.05 | 0 | 0.12 | 0.04 |
| 8 | 122257320 | rs2656118 | 1.713 | -0.52 | 1.393 | 1.456 | 0.024 | 0.095 | 0.095 | 0 | 0.54 | 0.34 | 0.4 | 0.08 | 0.61 | 0.34 |
| 8 | 122259074 | rs4871180 | -0.851 | -0.571 | 1.471 | 1.727 | -0.003 | -0.041 | -0.041 | 0 | 0.24 | 0.17 | 0.44 | 0.39 | 0.27 | 0.39 |
| 8 | 122259282 | rs2656117 | 1.375 | -0.644 | 1.467 | 2.024 | 0.007 | 0.195 | 0.195 | 0.83 | 0.85 | 0.53 | 0.86 | 0.61 | 0.9 | 0.76 |
| 8 | 122263639 | rs1354505 | -0.465 | -0.614 | 1.722 | 1.941 | 0.06 | 0.355 | 0.355 | 0 | 0.06 | 0.32 | 0.06 | 0.29 | 0.03 | 0.15 |
| 8 | 122264777 | rs16894980 | -0.651 | -0.609 | 1.233 | 1.265 | -0.023 | 0.024 | 0.024 | 0 | 0.06 | 0.19 | 0.1 | 0.08 | 0.01 | 0.12 |
| 8 | 122267244 | rs2174352 | -1.482 | -0.916 | 0.908 | 1.003 | 0.018 | -0.033 | -0.033 | 1.897 | 0.34 | 0.18 | 0.45 | 0.4 | 0.29 | 0.45 |
| 8 | 122268345 | rs2251503 | 2.831 | -1.011 | 0.815 | 0.766 | -0.009 | 0.235 | 0.235 | 0.043 | 0.43 | 0.12 | 0.32 | 0.01 | 0.61 | 0.23 |
| 8 | 122272740 | rs1354504 | -1.623 | -0.996 | 0.796 | 1.021 | -0.001 | 0 | 0 | 0 | 0.34 | 0.4 | 0.48 | 0.5 | 0.24 | 0.53 |
| 8 | 122276120 | rs10956011 | -1.082 | -0.871 | 0.426 | 1.016 | 0.009 | -0.041 | -0.041 | 0.015 | 0.25 | 0.18 | 0.4 | 0.43 | 0.19 | 0.36 |
| 8 | 122280341 | rs987231 | 1.297 | -0.519 | 0.681 | 1.179 | 0.027 | -0.023 | -0.023 | 0 | 0.69 | 0.26 | 0.72 | 0.45 | 0.82 | 0.58 |
| 8 | 122294605 | rs2256123 | 1.681 | -0.563 | 0.521 | 1.146 | 0.088 | -0.039 | -0.039 | 0 | 0.63 | 0.21 | 0.71 | 0.45 | 0.8 | 0.56 |
| 8 | 122301156 | rs10505398 | -1.251 | 0.5 | 0.65 | 1.141 | -0.114 | -0.121 | -0.121 | 1.567 | 0.24 | 0.08 | 0.37 | 0.45 | 0.34 | 0.4 |
| 8 | 122301168 | rs13278708 | -1.338 | 0.508 | 0.71 | 1.146 | 0.007 | 0.009 | 0.009 | 1.181 | 0.24 | 0.26 | 0.39 | 0.45 | 0.34 | 0.4 |
| 8 | 122302309 | rs4871182 | -2.069 | 0.487 | 0.714 | 1.167 | 0.013 | 0.021 | 0.021 | 0 | 0.37 | 0.56 | 0.56 | 0.59 | 0.38 | 0.5 |
| 8 | 122316171 | rs13252333 | -1.201 | 0.449 | 0.591 | 1.397 | -0.001 | 0.022 | 0.022 | 0 | 0.15 | 0.18 | 0.38 | 0.4 | 0.14 | 0.32 |
| 8 | 122322450 | rs13257834 | -1.035 | 0.462 | 0.598 | 0.868 | -0.001 | 0.003 | 0.003 | 0 | 0.14 | 0.17 | 0.22 | 0.19 | 0.12 | 0.29 |
| 8 | 122325332 | rs10956023 | -1.619 | 0.904 | 0.113 | 1.177 | -0.034 | -0.04 | -0.04 | 0 | 0.23 | 0.12 | 0.39 | 0.69 | 0.3 | 0.38 |
| 8 | 122329172 | rs13256476 | -2.025 | 0.387 | 0.825 | 0.427 | -0.015 | -0.015 | -0.015 | 0 | 0.13 | 0.4 | 0.1 | 0.08 | 0.08 | 0.07 |
| 8 | 122329373 | rs7007558 | -2.056 | 0.371 | 1.471 | 0.33 | 0.006 | -0.009 | -0.009 | 0 | 0.1 | 0.33 | 0.09 | 0.05 | 0.06 | 0.08 |
| 8 | 122330471 | rs13267240 | -0.916 | 1.001 | 1.064 | 0.844 | 0.002 | 0.029 | 0.029 | 0 | 0.17 | 0.22 | 0.27 | 0.47 | 0.22 | 0.29 |
| 8 | 122330565 | rs12547273 | -0.393 | 1.333 | 0.427 | 0.486 | -0.036 | -0.03 | -0.03 | 0.75 | 0.2 | 0.07 | 0.2 | 0.25 | 0.23 | 0.27 |
| 8 | 122337780 | rs7834611 | -1.267 | 1.352 | 0.766 | 0.237 | 0.03 | 0.078 | 0.078 | 0.706 | 0.13 | 0.41 | 0.29 | 0.25 | 0.14 | 0.1 |
| 8 | 122340986 | rs17276050 | -0.36 | 1.49 | -0.099 | 0.342 | -0.059 | -0.062 | -0.062 | 0.004 | 0.21 | 0.04 | 0.2 | 0.27 | 0.25 | 0.33 |
| 8 | 122352055 | rs975037 | -1.333 | 1.313 | 0.41 | 0.617 | 0.002 | -0.001 | -0.001 | 1.228 | 0.28 | 0.31 | 0.52 | 0.73 | 0.34 | 0.39 |
| 8 | 122355850 | rs2129536 | 0.324 | 0.742 | 0.691 | 0.754 | 0.03 | 0.056 | 0.056 | 0.128 | 0.84 | 0.45 | 0.67 | 0.7 | 0.82 | 0.93 |
| 8 | 122358307 | rs1386443 | -1.448 | 0.215 | 0.933 | 0.333 | 0.001 | -0.046 | -0.046 | 0.05 | 0.48 | 0.57 | 0.53 | 0.96 | 0.39 | 0.6 |
| 8 | 122358919 | rs13257411 | -0.745 | 0.029 | 0.605 | 1.05 | 0.034 | -0.038 | -0.038 | 0 | 0.24 | 0.16 | 0.15 | 0.39 | 0.12 | 0.24 |
| 8 | 122359138 | rs13255877 | -0.66 | 0.032 | 0.725 | 1.051 | 0.018 | -0.018 | -0.018 | 0.101 | 0.25 | 0.22 | 0.16 | 0.39 | 0.12 | 0.24 |
| 8 | 122367142 | rs10505400 | 0.232 | 0.179 | 0.348 | 1.144 | -0.089 | -0.04 | -0.04 | 0.005 | 0.18 | 0.04 | 0.13 | 0.19 | 0.29 | 0.26 |
| 8 | 122375165 | rs6415475 | 2.005 | 0.179 | 0.476 | 1.131 | -0.014 | 0.15 | 0.15 | 0 | 0.48 | 0.22 | 0.48 | 0.13 | 0.55 | 0.42 |
| 8 | 122377814 | rs16895201 | 0.251 | 0.189 | 0.617 | 1.13 | -0.044 | -0.012 | -0.012 | 0 | 0.18 | 0.1 | 0.14 | 0.19 | 0.29 | 0.29 |
| 8 | 122385445 | rs1487164 | 1.357 | 0.203 | 0.736 | 1.162 | -0.013 | 0.005 | 0.005 | 0.006 | 0.67 | 0.37 | 0.62 | 0.33 | 0.83 | 0.69 |
| 8 | 122385614 | rs1487163 | 2.034 | 0.214 | 0.765 | 1.154 | -0.025 | 0.154 | 0.154 | 0 | 0.47 | 0.18 | 0.48 | 0.13 | 0.54 | 0.41 |
| 8 | 122392117 | rs7006267 | -0.247 | -0.098 | 1.171 | 1.388 | 0.027 | 0.105 | 0.105 | 0 | 0.95 | 0.88 | 0.82 | 0.72 | 0.94 | 0.93 |
| 8 | 122393258 | rs7824421 | -0.238 | -0.36 | 1.17 | 1.388 | 0.015 | 0.105 | 0.105 | 0 | 0.95 | 0.88 | 0.82 | 0.72 | 0.95 | 0.93 |
| 8 | 122394987 | rs7833779 | -1.034 | -0.383 | 1.161 | 1.355 | -0.078 | 0.062 | 0.062 | 0 | 0.28 | 0.5 | 0.2 | 0.4 | 0.11 | 0.26 |
| 8 | 122415207 | rs6469999 | 1.554 | -0.356 | 1.037 | 1.318 | 0.01 | -0.006 | -0.006 | 0 | 0.66 | 0.32 | 0.62 | 0.34 | 0.84 | 0.69 |
| 8 | 122426082 | rs13258852 | -0.856 | 0.112 | 1.105 | 1.56 | 0.027 | -0.05 | -0.05 | 0.033 | 0.22 | 0.14 | 0.15 | 0.4 | 0.12 | 0.24 |
| 8 | 122438189 | rs1386442 | 0.346 | 0.531 | 0.683 | 1.469 | 0.015 | 0.089 | 0.089 | 0 | 0.94 | 0.87 | 0.8 | 0.73 | 0.95 | 0.93 |
| 8 | 122458999 | rs7005145 | 0.736 | 0.787 | 0.123 | 1.086 | -0.033 | 0.031 | 0.031 | 0.047 | 0.71 | 0.81 | 0.82 | 0.8 | 0.64 | 0.65 |
| 8 | 122462734 | rs11989957 | -0.178 | 0.201 | 0.058 | 1.595 | 0.007 | -0.017 | -0.017 | 0 | 0.13 | 0.08 | 0.13 | 0.19 | 0.08 | 0.05 |
| 8 | 122462932 | rs11990704 | -0.178 | 0.201 | 0.058 | 1.595 | 0.007 | -0.017 | -0.017 | 0 | 0.13 | 0.08 | 0.13 | 0.19 | 0.08 | 0.05 |
| 8 | 122464572 | rs1482189 | -1.388 | -0.051 | 0.647 | 1.808 | -0.026 | 0.171 | 0.171 | 0.335 | 0.19 | 0.46 | 0.25 | 0.48 | 0.1 | 0.23 |
| 8 | 122480008 | rs1482190 | -0.097 | -0.209 | 0.552 | 1.238 | 0.006 | -0.014 | -0.014 | 0 | 0.25 | 0.29 | 0.18 | 0.17 | 0.31 | 0.36 |
| 8 | 122482236 | rs13274955 | 0.227 | -0.28 | 0.458 | 1.489 | 0.015 | -0.019 | -0.019 | 0.367 | 0.1 | 0.08 | 0.04 | 0.18 | 0.04 | 0.13 |
| 8 | 122483648 | rs10956045 | -0.529 | -0.069 | 0.675 | 1.415 | -0.001 | 0 | 0 | 0 | 0.36 | 0.48 | 0.22 | 0.37 | 0.36 | 0.51 |
| 8 | 122484460 | rs1482188 | -0.647 | -0.041 | 0.082 | 1.327 | 0.001 | 0.001 | 0.001 | 0.147 | 0.37 | 0.62 | 0.24 | 0.36 | 0.33 | 0.51 |
| 8 | 122489509 | rs13274659 | -1.244 | 0.092 | -0.71 | 1.012 | -0.152 | 0.128 | 0.128 | 0.094 | 0.57 | 0.96 | 0.56 | 0.85 | 0.46 | 0.62 |

| | | | | | | | | | | | | | | | | |
|---|---|---|---|---|---|---|---|---|---|---|---|---|---|---|---|---|
| 8 | 122496474 | rs16895496 | -2.271 | 0.038 | 0.857 | 1.517 | -0.021 | 0.015 | 0.015 | 1.188 | 0.14 | 0.62 | 0.13 | 0.16 | 0.07 | 0.04 |
| 8 | 122497385 | rs7842523 | -0.758 | 0.167 | 0.312 | 1.62 | -0.001 | 0.021 | 0.021 | 0 | 0.06 | 0.09 | 0.05 | 0.16 | 0.05 | 0.04 |
| 8 | 122501989 | rs4870766 | -1.794 | 0.174 | 0.308 | 1.651 | -0.002 | 0.082 | 0.082 | 1.595 | 0.16 | 0.68 | 0.3 | 0.43 | 0.11 | 0.11 |
| 8 | 122506731 | rs16895527 | -1.443 | 0.074 | 1.33 | 1.105 | 0.033 | -0.045 | -0.045 | 0 | 0.09 | 0.23 | 0.06 | 0 | 0.06 | 0.01 |
| 8 | 122513645 | rs6985125 | -2.184 | -0.324 | 1.462 | 0.84 | -0.024 | -0.082 | -0.082 | 2.982 | 0.2 | 0.58 | 0.12 | 0.02 | 0.09 | 0.12 |
| 8 | 122514171 | rs13261248 | -1.97 | -0.71 | 0.287 | 0.181 | 0.01 | 0.074 | 0.074 | 0.336 | 0.44 | 0.95 | 0.4 | 0.29 | 0.31 | 0.43 |
| 8 | 122516874 | rs1542682 | -1.203 | -0.692 | 0.882 | -0.106 | -0.003 | -0.077 | -0.077 | 0 | 0.17 | 0.26 | 0.09 | 0 | 0.14 | 0.15 |
| 8 | 122518935 | rs16895584 | -0.683 | -0.758 | 1.342 | -0.057 | -0.011 | -0.062 | -0.062 | 0 | 0.11 | 0.17 | 0.04 | 0 | 0.05 | 0.05 |
| 8 | 122524093 | rs900212 | -2.816 | -1.036 | 1.226 | 0.014 | 0.039 | 0.05 | 0.05 | 1.728 | 0.37 | 0.84 | 0.31 | 0.24 | 0.24 | 0.26 |
| 8 | 122532443 | rs11986236 | -2.865 | -1.038 | 1.203 | 0.01 | 0.038 | 0.047 | 0.047 | 1.331 | 0.37 | 0.91 | 0.32 | 0.24 | 0.24 | 0.26 |
| 8 | 122535739 | rs1482184 | -2.08 | -1.04 | 1.408 | 0.011 | -0.026 | 0.027 | 0.027 | 0.571 | 0.22 | 0.4 | 0.22 | 0.24 | 0.16 | 0.21 |
| 8 | 122539123 | rs1564569 | -2.778 | -1.035 | 1.441 | 0.01 | 0.031 | 0.037 | 0.037 | 1.56 | 0.36 | 0.77 | 0.29 | 0.23 | 0.24 | 0.26 |
| 8 | 122546725 | rs6470004 | -1.244 | -1.029 | 1.303 | 0.012 | -0.001 | -0.089 | -0.089 | 1.894 | 0.14 | 0.34 | 0.07 | 0 | 0.09 | 0.07 |
| 8 | 122548940 | rs1482183 | -1.965 | -0.976 | 1.349 | 0.024 | -0.02 | 0.013 | 0.013 | 0.189 | 0.2 | 0.3 | 0.18 | 0.21 | 0.15 | 0.19 |
| 8 | 122551370 | rs6470005 | -2.712 | -1.014 | 1.257 | 0.02 | -0.01 | -0.01 | -0.01 | 0 | 0.33 | 0.6 | 0.25 | 0.2 | 0.22 | 0.24 |
| 8 | 122561919 | rs4871204 | -1.628 | -0.707 | 1.245 | -0.168 | -0.003 | -0.057 | -0.057 | 0 | 0.15 | 0.27 | 0.07 | 0.04 | 0.1 | 0.11 |
| 8 | 122562447 | rs2128207 | -2.589 | -0.802 | 1.058 | -0.255 | 0.031 | 0.118 | 0.118 | 0.454 | 0.28 | 0.88 | 0.21 | 0.04 | 0.16 | 0.18 |
| 8 | 122563815 | rs7846194 | -1.221 | -0.978 | 1.265 | -0.216 | -0.012 | -0.077 | -0.077 | 0 | 0.13 | 0.28 | 0.06 | 0 | 0.07 | 0.06 |
| 8 | 122577017 | rs4871207 | -2.575 | -1.163 | 0.942 | -0.427 | 0.076 | 0.134 | 0.134 | 0 | 0.32 | 0.87 | 0.2 | 0.03 | 0.16 | 0.19 |
| 8 | 122580578 | rs4871208 | -0.933 | -1.203 | 1.304 | -0.429 | 0.002 | -0.05 | -0.05 | 0 | 0.09 | 0.31 | 0.05 | 0 | 0.04 | 0.03 |
| 8 | 122581626 | rs7839029 | -2.261 | -1.142 | 1.375 | -0.444 | 0.019 | 0.016 | 0.016 | 1.128 | 0.25 | 0.74 | 0.17 | 0.03 | 0.11 | 0.1 |
| 8 | 122587014 | rs6986092 | -2.813 | -1.465 | 0.877 | -0.027 | 0.094 | -0.035 | -0.035 | 0 | 0.39 | 0.93 | 0.31 | 0.29 | 0.22 | 0.32 |
| 8 | 122598274 | rs4870775 | -2.036 | -1.452 | 1.401 | -0.043 | -0.017 | 0.006 | 0.006 | 0.007 | 0.26 | 0.72 | 0.23 | 0.26 | 0.15 | 0.31 |
| 8 | 122598324 | rs4870776 | -1.909 | -1.393 | 1.39 | -0.013 | -0.046 | 0.048 | 0.048 | 0 | 0.24 | 0.52 | 0.2 | 0.26 | 0.15 | 0.31 |
| 8 | 122601353 | rs7826054 | -1.084 | -0.85 | 1.118 | -1.06 | -0.007 | -0.069 | -0.069 | 0 | 0.15 | 0.24 | 0.07 | 0 | 0.1 | 0.09 |
| 8 | 122606303 | rs1384184 | -1.738 | -0.953 | 0.462 | -1.863 | -0.014 | 0.082 | 0.082 | 0 | 0.59 | 0.74 | 0.65 | 0.97 | 0.44 | 0.72 |
| 8 | 122608079 | rs2016573 | 0.258 | -1.078 | 0.56 | -1.605 | 0.002 | 0.007 | 0.007 | 0 | 0.27 | 0.17 | 0.47 | 0.84 | 0.24 | 0.5 |
| 8 | 122615461 | rs12056571 | -1.059 | -0.572 | 0.844 | -1.879 | 0.004 | 0.006 | 0.006 | 0 | 0.56 | 0.61 | 0.6 | 0.96 | 0.41 | 0.65 |
| 8 | 122615781 | rs16895753 | -1.059 | -0.572 | 0.844 | -1.879 | 0.004 | 0.006 | 0.006 | 0 | 0.56 | 0.61 | 0.6 | 0.96 | 0.41 | 0.65 |
| 8 | 122619283 | rs1384186 | -0.056 | -0.241 | 0.497 | -1.921 | 0.006 | 0.044 | 0.044 | 0 | 0.43 | 0.25 | 0.56 | 0.92 | 0.43 | 0.59 |
| 8 | 122621909 | rs11995252 | 0.002 | -0.229 | 0.473 | -1.915 | -0.015 | -0.062 | -0.062 | 2.149 | 0.11 | 0.26 | 0.05 | 0 | 0.06 | 0.1 |
| 8 | 122622697 | rs11991792 | -1.151 | -0.229 | 0.566 | -1.928 | -0.006 | -0.019 | -0.019 | 0.15 | 0.58 | 0.6 | 0.63 | 0.93 | 0.51 | 0.7 |
| 8 | 122623115 | rs10956050 | -0.541 | 0.024 | 0.166 | -1.471 | 0.003 | 0.004 | 0.004 | 0 | 0.44 | 0.34 | 0.56 | 0.92 | 0.43 | 0.59 |
| 8 | 122623421 | rs10505408 | -0.425 | 0.085 | 0 | -1.441 | 0.002 | 0.037 | 0.037 | 0 | 0.43 | 0.28 | 0.56 | 0.92 | 0.43 | 0.59 |
| 8 | 122632611 | rs11989271 | -0.562 | 0.021 | 0.657 | -1.015 | 0 | -0.044 | -0.044 | 0 | 0.09 | 0.24 | 0.07 | 0.01 | 0.06 | 0.06 |
| 8 | 122634104 | rs4612347 | -0.341 | -0.046 | 0.51 | -0.818 | 0 | -0.02 | -0.02 | 0 | 0.16 | 0.11 | 0.36 | 0.71 | 0.21 | 0.3 |
| 8 | 122638262 | rs6984484 | -0.713 | -0.469 | 0.761 | -1.49 | -0.028 | -0.089 | -0.089 | 0.455 | 0.35 | 0.44 | 0.27 | 0.11 | 0.25 | 0.21 |
| 8 | 122639443 | rs11992999 | -0.578 | -0.468 | 0.742 | -1.489 | -0.012 | -0.049 | -0.049 | 0.253 | 0.34 | 0.38 | 0.27 | 0.11 | 0.25 | 0.21 |
| 8 | 122642363 | rs11986885 | 0.606 | -0.441 | 0.606 | -1.477 | 0 | -0.007 | -0.007 | 0 | 0.08 | 0.07 | 0.06 | 0.01 | 0.06 | 0.06 |
| 8 | 122650052 | rs7831954 | -1.564 | -0.613 | 0.889 | -1.57 | 0.009 | 0.21 | 0.21 | 0.147 | 0.54 | 0.83 | 0.65 | 0.85 | 0.47 | 0.49 |
| 8 | 122653247 | rs1057308 | -0.842 | -0.584 | 1.246 | -1.525 | -0.014 | -0.001 | -0.001 | 0.724 | 0.26 | 0.49 | 0.44 | 0.73 | 0.28 | 0.3 |
| 8 | 122660248 | rs10089677 | -1.457 | -0.538 | 0.824 | -1.615 | -0.017 | 0.222 | 0.222 | 0 | 0.56 | 0.86 | 0.65 | 0.85 | 0.49 | 0.5 |
| 8 | 122674244 | rs7008033 | -0.789 | -0.621 | 1.443 | -1.626 | 0.013 | 0.009 | 0.009 | 0 | 0.46 | 0.54 | 0.57 | 0.84 | 0.42 | 0.48 |
| 8 | 122679098 | rs6470026 | -0.603 | -0.708 | 1.315 | -1.484 | 0.002 | 0.003 | 0.003 | 0.029 | 0.4 | 0.62 | 0.43 | 0.66 | 0.26 | 0.43 |
| 8 | 122683440 | rs9297647 | -0.799 | -0.743 | 1.442 | -1.654 | 0.001 | -0.004 | -0.004 | 0 | 0.4 | 0.58 | 0.36 | 0.14 | 0.37 | 0.2 |
| 8 | 122684194 | rs16892997 | -0.297 | -0.744 | 1.481 | -1.656 | -0.015 | -0.054 | -0.054 | 0.789 | 0.33 | 0.5 | 0.29 | 0.13 | 0.3 | 0.18 |
| 8 | 122686768 | rs10110528 | -1.87 | -0.571 | 0.914 | -1.687 | -0.042 | 0.185 | 0.185 | 0 | 0.57 | 0.83 | 0.69 | 0.83 | 0.54 | 0.5 |
| 8 | 122686888 | rs17299925 | 0.037 | -0.309 | 0.902 | -1.404 | -0.008 | -0.008 | -0.008 | 0 | 0.09 | 0.05 | 0.18 | 0.13 | 0.18 | 0.07 |
| 8 | 122687025 | rs9642857 | 0.217 | -0.427 | 0.855 | -1.328 | 0.002 | -0.003 | -0.003 | 0 | 0.17 | 0.15 | 0.33 | 0.69 | 0.18 | 0.31 |
| 8 | 122688397 | rs1008786 | -0.335 | -0.289 | 1.104 | -1.36 | 0.02 | -0.053 | -0.053 | 0.697 | 0.11 | 0.41 | 0.1 | 0.01 | 0.08 | 0.02 |
| 8 | 122688842 | rs17299960 | 0.488 | -0.156 | 0.441 | -1.321 | -0.014 | -0.019 | -0.019 | 0 | 0.08 | 0.04 | 0.08 | 0.13 | 0.16 | 0.05 |
| 8 | 122692580 | rs10113295 | -1.599 | -0.547 | 0.134 | -1.492 | -0.096 | 0.161 | 0.161 | 0 | 0.6 | 0.93 | 0.71 | 0.78 | 0.55 | 0.5 |
| 8 | 122696798 | rs9297650 | 0.744 | -0.762 | 0.711 | -1.299 | 0.021 | 0.037 | 0.037 | 0 | 0.23 | 0.11 | 0.09 | 0 | 0.15 | 0.12 |
| 8 | 122701377 | rs16895861 | -1.134 | -0.76 | 1.344 | -1.205 | 0.079 | 0.075 | 0.075 | 3.036 | 0.05 | 0.51 | 0.07 | 0 | 0.01 | 0 |
| 8 | 122701402 | rs3903303 | 0.203 | -0.618 | 0.784 | -1.071 | -0.004 | 0.013 | 0.013 | 0 | 0.16 | 0.14 | 0.4 | 0.64 | 0.18 | 0.31 |
| 8 | 122706453 | rs9642859 | -0.718 | -0.608 | 1.075 | -1.205 | -0.014 | -0.015 | -0.015 | 0 | 0.42 | 0.45 | 0.54 | 0.66 | 0.38 | 0.43 |
| 8 | 122708178 | rs6982790 | 0.391 | -0.526 | 0.644 | -1.365 | 0.012 | 0.02 | 0.02 | 0 | 0.25 | 0.15 | 0.13 | 0 | 0.2 | 0.12 |
| 8 | 122761187 | rs7011279 | -1.507 | -0.052 | 0.361 | -1.467 | -0.021 | 0.221 | 0.221 | 0.006 | 0.53 | 0.8 | 0.66 | 0.85 | 0.55 | 0.53 |
| 8 | 122766075 | rs6991370 | -1.602 | -0.068 | 0.345 | -1.474 | -0.04 | 0.245 | 0.245 | 0.054 | 0.54 | 0.86 | 0.7 | 0.85 | 0.56 | 0.53 |
| 8 | 122771217 | rs11779972 | 0.438 | -0.203 | 0.51 | -0.947 | -0.004 | -0.037 | -0.037 | 0 | 0.15 | 0.08 | 0.31 | 0.57 | 0.18 | 0.12 |
| 8 | 122776605 | rs17243236 | 0.461 | -0.234 | 1.452 | -0.957 | 0.018 | 0.028 | 0.028 | 0 | 0.26 | 0.19 | 0.23 | 0.14 | 0.21 | 0.32 |
| 8 | 122781931 | rs10106505 | -0.729 | -0.226 | 0.953 | -0.907 | 0.031 | 0.037 | 0.037 | 0 | 0.37 | 0.74 | 0.33 | 0.27 | 0.32 | 0.4 |
| 8 | 122782960 | rs17243536 | 0.508 | -0.311 | 1.14 | -0.941 | 0.004 | 0.004 | 0.004 | 0 | 0.24 | 0.25 | 0.22 | 0.15 | 0.15 | 0.32 |
| 8 | 122788769 | rs279621 | 1.198 | 0.212 | 0.447 | -1.233 | -0.032 | 0.034 | 0.034 | 0 | 0.07 | 0 | 0.04 | 0.01 | 0.08 | 0.14 |
| 8 | 122790388 | rs7016012 | -0.329 | 0.216 | 0.688 | -1.151 | -0.005 | -0.011 | -0.011 | 0 | 0.37 | 0.61 | 0.34 | 0.3 | 0.35 | 0.41 |
| 8 | 122793637 | rs279629 | -1.704 | 0.269 | 0.414 | -1.245 | -0.111 | 0.217 | 0.217 | 0 | 0.61 | 0.94 | 0.72 | 0.87 | 0.58 | 0.68 |
| 8 | 122801313 | rs279640 | -0.677 | 0.26 | 0.573 | -1.178 | 0.007 | 0.018 | 0.018 | 0.201 | 0.44 | 0.66 | 0.39 | 0.31 | 0.4 | 0.55 |
| 8 | 122815018 | rs1030980 | 0.997 | -0.247 | 1.373 | -0.786 | -0.027 | 0.078 | 0.078 | 0 | 0.75 | 0.55 | 0.63 | 0.45 | 0.78 | 0.78 |
| 8 | 122818932 | rs1371718 | -0.286 | -0.163 | 1.374 | -0.754 | 0.001 | -0.027 | -0.027 | 0 | 0.93 | 0.87 | 0.95 | 0.99 | 0.95 | 0.9 |

| | | | | | | | | | | | | | | | | |
|---|---|---|---|---|---|---|---|---|---|---|---|---|---|---|---|---|
| 8 | 122838008 | rs6999162 | -0.572 | -0.123 | 1.096 | -0.846 | 0.005 | 0.021 | 0.021 | 0.183 | 0.16 | 0.26 | 0.17 | 0.3 | 0.2 | 0.33 |
| 8 | 122842642 | rs4474034 | 0.896 | -0.076 | 1 | -0.87 | -0.001 | -0.001 | -0.001 | 0 | 0.7 | 0.67 | 0.74 | 0.71 | 0.72 | 0.57 |
| 8 | 122848829 | rs10091098 | 1.641 | -0.345 | 0.818 | -1.04 | -0.003 | -0.047 | -0.047 | 0 | 0.56 | 0.43 | 0.47 | 0.13 | 0.58 | 0.44 |
| 8 | 122851357 | rs4582571 | -0.801 | -0.359 | 0.63 | -0.986 | 0 | -0.006 | -0.006 | 1.935 | 0.94 | 0.96 | 0.95 | 0.99 | 0.95 | 0.9 |
| 8 | 122864050 | rs16896250 | -1.407 | -0.294 | 0.515 | -0.99 | -0.008 | -0.068 | -0.068 | 0.143 | 0.38 | 0.57 | 0.5 | 0.85 | 0.38 | 0.45 |
| 8 | 122870120 | rs1397347 | 1.551 | -0.295 | 0.515 | -0.998 | -0.002 | -0.055 | -0.055 | 0.558 | 0.57 | 0.43 | 0.46 | 0.13 | 0.57 | 0.46 |
| 8 | 122875056 | rs6988392 | -0.64 | -0.015 | -0.012 | -0.726 | -0.001 | 0 | 0 | 0 | 0.22 | 0.26 | 0.21 | 0.28 | 0.24 | 0.34 |
| 8 | 122876201 | rs4341166 | 0.803 | -0.031 | 0.004 | -0.708 | -0.003 | 0 | 0 | 0 | 0.72 | 0.72 | 0.75 | 0.71 | 0.71 | 0.57 |
| 8 | 122888939 | rs1567461 | -0.534 | -0.238 | -0.173 | -0.564 | -0.002 | 0 | 0 | 0.583 | 0.24 | 0.24 | 0.17 | 0.28 | 0.3 | 0.25 |
| 8 | 122889159 | rs11781753 | 0.938 | -0.403 | -0.321 | -0.59 | -0.006 | -0.032 | -0.032 | 0 | 0.9 | 0.9 | 0.95 | 0.99 | 0.93 | 0.9 |
| 8 | 122890163 | rs11775964 | -0.601 | -0.682 | 0.029 | -0.515 | -0.005 | -0.02 | -0.02 | 0 | 0.26 | 0.48 | 0.23 | 0.26 | 0.25 | 0.38 |
| 8 | 122890522 | rs16896361 | 0.549 | -0.95 | -0.292 | -0.403 | 0.009 | -0.018 | -0.018 | 0.222 | 0.21 | 0.16 | 0.27 | 0.57 | 0.14 | 0.18 |
| 8 | 122892013 | rs1397346 | 1.436 | -0.97 | -0.292 | -0.452 | -0.004 | 0.172 | 0.172 | 0 | 0.47 | 0.23 | 0.45 | 0.16 | 0.55 | 0.4 |
| 8 | 122894115 | rs16896375 | 0.744 | -1.461 | -0.181 | -0.303 | -0.013 | -0.02 | -0.02 | 0 | 0.05 | 0.01 | 0.02 | 0.13 | 0.04 | 0.24 |
| 8 | 122895259 | rs7012540 | -0.982 | -1.433 | -0.185 | -0.282 | 0.003 | -0.019 | -0.019 | 0 | 0.94 | 0.93 | 0.95 | 0.99 | 0.94 | 0.96 |
| 8 | 122897272 | rs7816916 | 1.358 | -1.401 | -0.337 | -0.282 | 0.04 | 0.187 | 0.187 | 0 | 0.48 | 0.23 | 0.45 | 0.16 | 0.58 | 0.4 |
| 8 | 122906770 | rs9643167 | 0.446 | -1.715 | -0.347 | -0.129 | 0.009 | -0.011 | -0.011 | 0 | 0.2 | 0.14 | 0.26 | 0.57 | 0.11 | 0.16 |
| 8 | 122909687 | rs956225 | 0.786 | -1.719 | -0.459 | -0.239 | -0.025 | -0.023 | -0.023 | 0 | 0.15 | 0.01 | 0.1 | 0.13 | 0.15 | 0.14 |
| 8 | 122917182 | rs6470049 | -0.692 | -1.495 | -0.152 | -0.161 | 0.002 | 0.001 | 0.001 | 0 | 0.32 | 0.62 | 0.34 | 0.31 | 0.27 | 0.46 |
| 8 | 122926401 | rs7830188 | 0.25 | -1.252 | -0.024 | -0.04 | -0.034 | -0.047 | -0.047 | 0 | 0.29 | 0.44 | 0.24 | 0.18 | 0.21 | 0.42 |
| 8 | 122926477 | rs2037211 | -0.501 | -1.25 | -0.129 | 0.263 | -0.01 | -0.007 | -0.007 | 0 | 0.46 | 0.55 | 0.52 | 0.74 | 0.27 | 0.57 |
| 8 | 122927021 | rs4871214 | -0.612 | -0.841 | -0.344 | 0.514 | 0.051 | 0.063 | 0.063 | 0 | 0.52 | 0.62 | 0.57 | 0.76 | 0.34 | 0.62 |
| 8 | 122929553 | rs13271824 | 0.296 | -0.157 | -0.772 | 1.241 | 0.05 | -0.11 | -0.11 | 0.545 | 0.15 | 0.02 | 0.22 | 0.45 | 0.06 | 0.12 |
| 8 | 122944987 | rs1980018 | 0.048 | 1.123 | -0.534 | 0.166 | -0.001 | 0.013 | 0.013 | 0 | 0.59 | 0.6 | 0.6 | 0.75 | 0.43 | 0.61 |
| 8 | 122955670 | rs977771 | 0.085 | 1.011 | -0.528 | 0.241 | 0.001 | -0.004 | -0.004 | 0.046 | 0.61 | 0.58 | 0.61 | 0.75 | 0.47 | 0.66 |
| 8 | 122961848 | rs13280903 | -0.01 | 0.843 | -0.934 | 0.254 | 0.001 | 0.048 | 0.048 | 0 | 0.6 | 0.7 | 0.59 | 0.76 | 0.41 | 0.66 |
| 8 | 122970273 | rs16896441 | -0.733 | 1.176 | 0.069 | 0.645 | 0.007 | 0.011 | 0.011 | 0 | 0.34 | 0.4 | 0.33 | 0.43 | 0.41 | 0.52 |
| 8 | 122976795 | rs10956063 | -0.298 | 1.484 | -0.022 | 1.017 | -0.031 | -0.03 | -0.03 | 0.387 | 0.08 | 0.02 | 0.06 | 0.26 | 0.13 | 0.13 |
| 8 | 122982390 | rs6989617 | -0.298 | 1.47 | 0.067 | 0.866 | 0.012 | 0.049 | 0.049 | 0.318 | 0.21 | 0.09 | 0.17 | 0.14 | 0.2 | 0.17 |
| 8 | 122985747 | rs1158702 | -0.104 | 1.419 | 0.067 | 1.323 | 0.01 | 0 | 0 | 0 | 0.19 | 0.08 | 0.16 | 0.19 | 0.2 | 0.17 |
| 8 | 122989850 | rs16896479 | -0.992 | 1.396 | 0.137 | 1.298 | -0.012 | -0.015 | -0.015 | 0 | 0.28 | 0.39 | 0.28 | 0.25 | 0.28 | 0.4 |
| 8 | 122990776 | rs16896489 | -0.395 | 1.378 | 0.153 | 1.532 | -0.031 | -0.02 | -0.02 | 0 | 0.09 | 0.02 | 0.06 | 0.32 | 0.13 | 0.14 |
| 8 | 123004940 | rs6651164 | -1.575 | 1.206 | 0.994 | 1.432 | 0.067 | 0.152 | 0.152 | 0.019 | 0.14 | 0.44 | 0.16 | 0.25 | 0.15 | 0.16 |
| 8 | 123019652 | rs7820645 | 0.182 | 1.173 | 0.465 | 0.469 | -0.001 | 0.022 | 0.022 | 1.79 | 0.22 | 0.26 | 0.43 | 0.49 | 0.19 | 0.32 |
| 8 | 123022065 | rs7014335 | -0.842 | 1.092 | 0.586 | 0.163 | 0.009 | -0.06 | -0.06 | 0.93 | 0.34 | 0.51 | 0.51 | 0.8 | 0.32 | 0.45 |
| 8 | 123031994 | rs2017155 | -0.351 | 0.149 | -0.128 | 0.034 | 0.031 | -0.007 | -0.007 | 0 | 0.39 | 0.21 | 0.59 | 0.54 | 0.33 | 0.4 |
| 8 | 123043110 | rs4262329 | 0.654 | 0.623 | -0.774 | -0.762 | -0.002 | 0.015 | 0.015 | 0 | 0.32 | 0.27 | 0.18 | 0.09 | 0.32 | 0.25 |
| 8 | 123043443 | rs17317732 | 0.64 | 0.598 | -0.773 | -0.762 | -0.001 | 0.016 | 0.016 | 0 | 0.33 | 0.27 | 0.18 | 0.1 | 0.31 | 0.25 |
| 8 | 123046516 | rs3902292 | 0.17 | 0.413 | -1.166 | -0.456 | -0.013 | 0.208 | 0.208 | 0 | 0.5 | 0.19 | 0.39 | 0.2 | 0.49 | 0.43 |
| 8 | 123075591 | rs3914097 | -0.636 | -0.316 | 0.397 | 0.05 | -0.041 | -0.028 | -0.028 | 0.207 | 0.23 | 0.6 | 0.24 | 0.25 | 0.14 | 0.18 |
| 8 | 123088145 | rs1995863 | -1.276 | 0.115 | -1.346 | -0.368 | -0.078 | 0.208 | 0.208 | 0.277 | 0.61 | 0.98 | 0.71 | 0.91 | 0.58 | 0.8 |
| 8 | 123093555 | rs16896682 | 0.169 | 1.053 | -0.776 | 0.817 | -0.147 | -0.152 | -0.152 | 0 | 0.28 | 0.03 | 0.32 | 0.47 | 0.37 | 0.43 |
| 8 | 123101743 | rs12541121 | 0.084 | 0.984 | -0.774 | 0.709 | -0.152 | -0.104 | -0.104 | 0.728 | 0.28 | 0.05 | 0.33 | 0.56 | 0.38 | 0.53 |
| 8 | 123132855 | rs1605945 | -0.699 | 0.6 | -0.159 | 0.951 | 0.014 | 0.006 | 0.006 | 0.295 | 0.41 | 0.75 | 0.54 | 0.64 | 0.35 | 0.6 |
| 8 | 123135434 | rs16896731 | -0.891 | 0.679 | 0.203 | 1.013 | 0.048 | 0.12 | 0.12 | 0 | 0.11 | 0.28 | 0.14 | 0.24 | 0.15 | 0.17 |
| 8 | 123140360 | rs2136635 | -0.026 | 0.452 | -0.017 | 0.55 | 0.001 | -0.002 | -0.002 | 0.454 | 0.11 | 0.1 | 0.14 | 0.25 | 0.1 | 0.21 |
| 8 | 123159934 | rs1397094 | -0.868 | 0.464 | 0.601 | 0.647 | 0.017 | 0.028 | 0.028 | 5.061 | 0.1 | 0.55 | 0.08 | 0.1 | 0.06 | 0.13 |
| 8 | 123165047 | rs16896806 | -1.246 | 0.428 | 0.02 | 0.628 | -0.015 | -0.012 | -0.012 | 0.003 | 0.12 | 0.77 | 0.1 | 0.1 | 0.06 | 0.13 |
| 8 | 123170223 | rs7816590 | 0.38 | 0.583 | -0.355 | 0.707 | -0.052 | -0.113 | -0.113 | 0 | 0.24 | 0.04 | 0.34 | 0.52 | 0.24 | 0.39 |
| 8 | 123179784 | rs12544067 | 0.347 | 0.624 | -0.039 | 0.752 | -0.005 | -0.034 | -0.034 | 0 | 0.24 | 0.18 | 0.35 | 0.49 | 0.23 | 0.38 |
| 8 | 123185248 | rs4410914 | -1.137 | 0.562 | 0.283 | 0.489 | 0.005 | -0.031 | -0.031 | 0 | 0.24 | 0.53 | 0.28 | 0.22 | 0.24 | 0.33 |
| 8 | 123192373 | rs13262033 | 0.481 | 0.562 | 0.111 | 0.542 | -0.005 | -0.018 | -0.018 | 1.09 | 0.24 | 0.2 | 0.35 | 0.53 | 0.24 | 0.4 |
| 8 | 123219109 | rs4642655 | -1.328 | 0.525 | -0.196 | 0.013 | -0.006 | 0.071 | 0.071 | 0 | 0.62 | 0.72 | 0.77 | 0.84 | 0.59 | 0.79 |
| 8 | 123226553 | rs4375006 | -0.191 | 0.528 | 0.213 | 0.074 | 0.102 | 0.176 | 0.176 | 3.015 | 0.06 | 0.47 | 0.13 | 0.15 | 0.04 | 0.13 |
| 8 | 123236745 | rs2023116 | -0.123 | 0.483 | -0.051 | 0.019 | -0.003 | 0.016 | 0.016 | 0 | 0.16 | 0.14 | 0.22 | 0.1 | 0.17 | 0.2 |
| 8 | 123250735 | rs10090889 | 0.129 | 0.462 | -0.216 | 0.047 | -0.064 | 0.021 | 0.021 | 0 | 0.39 | 0.07 | 0.41 | 0.58 | 0.37 | 0.45 |
| 8 | 123271871 | rs2023113 | 0.204 | 0.438 | -0.174 | 0.006 | -0.003 | 0.095 | 0.095 | 0 | 0.4 | 0.18 | 0.42 | 0.69 | 0.37 | 0.51 |
| 8 | 123278754 | rs10216683 | 0.176 | 0.449 | -0.253 | 0.015 | -0.072 | 0.023 | 0.023 | 0 | 0.39 | 0.04 | 0.41 | 0.6 | 0.37 | 0.47 |
| 8 | 123281374 | rs4380946 | -0.619 | 0.493 | 0.041 | -0.071 | -0.001 | -0.044 | -0.044 | 0.08 | 0.18 | 0.29 | 0.23 | 0.09 | 0.17 | 0.2 |
| 8 | 123282439 | rs12682632 | -0.512 | 0.481 | 0.22 | 0.157 | 0.095 | 0.168 | 0.168 | 2.749 | 0.07 | 0.44 | 0.13 | 0.15 | 0.04 | 0.13 |
| 8 | 123297701 | rs4871240 | -0.602 | 0.379 | -0.779 | 0.094 | -0.019 | 0.23 | 0.23 | 0.015 | 0.52 | 0.12 | 0.6 | 0.78 | 0.51 | 0.7 |
| 8 | 123302128 | rs13270547 | -0.519 | 0.369 | -0.747 | 0.12 | 0 | 0.206 | 0.206 | 0.17 | 0.51 | 0.16 | 0.6 | 0.77 | 0.51 | 0.7 |
| 8 | 123304604 | rs17320342 | -1.278 | 0.406 | -0.826 | 0.033 | -0.088 | 0.237 | 0.237 | 0.611 | 0.63 | 0.94 | 0.77 | 0.92 | 0.56 | 0.84 |
| 8 | 123314768 | rs7013943 | -0.525 | 0.713 | -0.667 | 1.346 | -0.021 | -0.118 | -0.118 | 0.236 | 0.28 | 0.09 | 0.48 | 0.47 | 0.27 | 0.3 |
| 8 | 123321180 | rs16896907 | -0.673 | 0.815 | -0.49 | 1.539 | -0.003 | -0.002 | -0.002 | 0.075 | 0.07 | 0.07 | 0.04 | 0.16 | 0.08 | 0.13 |
| 8 | 123325745 | rs7006321 | 0.217 | 0.936 | -0.513 | 1.543 | -0.025 | -0.102 | -0.102 | 0.58 | 0.76 | 0.91 | 0.53 | 0.5 | 0.74 | 0.76 |
| 8 | 123325803 | rs7006639 | 1.019 | 0.942 | -0.41 | 1.505 | -0.015 | 0.002 | 0.002 | 0 | 0.68 | 0.79 | 0.48 | 0.35 | 0.66 | 0.64 |
| 8 | 123329358 | rs17268347 | 0.298 | 0.918 | -0.387 | 1.545 | -0.026 | -0.029 | -0.029 | 0 | 0.05 | 0 | 0.02 | 0.11 | 0.07 | 0.11 |
| 8 | 123330083 | rs4871243 | 0.314 | 0.928 | -0.419 | 1.839 | -0.004 | -0.046 | -0.046 | 2.678 | 0.78 | 0.85 | 0.78 | 0.51 | 0.79 | 0.76 |

| | | | | | | | | | | | | | | | | |
|---|---|---|---|---|---|---|---|---|---|---|---|---|---|---|---|---|
| 8 | 123330779 | rs16896924 | -0.703 | 0.934 | -0.357 | 1.734 | -0.002 | 0 | 0 | 0.017 | 0.07 | 0.06 | 0.04 | 0.16 | 0.08 | 0.12 |
| 8 | 123345261 | rs4606068 | -1.1 | 0.893 | 0.079 | 1.773 | 0.011 | 0.097 | 0.097 | 0 | 0.09 | 0.18 | 0.15 | 0.23 | 0.08 | 0.13 |
| 8 | 123352919 | rs6415477 | -0.424 | 1.062 | -0.11 | 1.908 | -0.037 | -0.038 | -0.038 | 0.262 | 0.92 | 0.99 | 0.72 | 0.85 | 0.9 | 0.9 |
| 8 | 123369269 | rs7826716 | 0.679 | 0.831 | -0.034 | 1.884 | 0.002 | -0.049 | -0.049 | 0 | 0.14 | 0.05 | 0.08 | 0.2 | 0.12 | 0.1 |
| 8 | 123374679 | rs2023118 | -0.206 | 0.828 | 0.17 | 1.454 | 0.003 | 0.001 | 0.001 | 0 | 0.66 | 0.64 | 0.55 | 0.6 | 0.63 | 0.54 |
| 8 | 123384530 | rs4355771 | 0.91 | 0.741 | 0.292 | 1.201 | 0.018 | 0.144 | 0.144 | 0 | 0.46 | 0.28 | 0.35 | 0.17 | 0.45 | 0.36 |
| 8 | 123401184 | rs4871253 | 0.932 | 0.697 | 0.304 | 1.195 | 0.006 | 0.195 | 0.195 | 0.029 | 0.45 | 0.22 | 0.26 | 0.06 | 0.45 | 0.26 |
| 8 | 123409560 | rs4492387 | 0.58 | 0.729 | 0.619 | 1.439 | 0.002 | 0.012 | 0.012 | 0 | 0.5 | 0.47 | 0.47 | 0.4 | 0.51 | 0.43 |
| 8 | 123410777 | rs13279923 | -0.267 | 0.734 | 0.762 | 1.596 | 0.005 | 0.222 | 0.222 | 0 | 0.2 | 0.45 | 0.22 | 0.43 | 0.18 | 0.16 |
| 8 | 123434634 | rs4397408 | -0.836 | 0.872 | 0.958 | 1.677 | 0.127 | 0.373 | 0.373 | 1.466 | 0.06 | 0.42 | 0.14 | 0.23 | 0.06 | 0.08 |
| 8 | 123443180 | rs17270266 | -0.089 | 1.166 | 0.826 | 1.668 | -0.005 | -0.003 | -0.003 | 1.26 | 0.67 | 0.75 | 0.59 | 0.65 | 0.67 | 0.55 |
| 8 | 123445244 | rs4549775 | 0.411 | 1.161 | -0.545 | 1.251 | -0.013 | -0.005 | -0.005 | 0 | 0.6 | 0.28 | 0.44 | 0.42 | 0.61 | 0.46 |
| 8 | 123448480 | rs7014937 | -0.082 | 1.027 | -1.147 | 0.332 | -0.001 | 0.12 | 0.12 | 0.336 | 0.47 | 0.82 | 0.54 | 0.75 | 0.51 | 0.56 |
| 8 | 123448950 | rs7813226 | 0.364 | 1.134 | -1.01 | 0.182 | -0.003 | 0.002 | 0.002 | 0 | 0.17 | 0.14 | 0.11 | 0.07 | 0.21 | 0.04 |
| 8 | 123448989 | rs17270918 | 0.732 | 0.783 | -0.809 | 0.251 | 0 | 0 | 0 | 0 | 0.14 | 0.12 | 0.09 | 0.1 | 0.12 | 0.04 |
| 8 | 123449128 | rs4259418 | -0.166 | 0.791 | -0.393 | 0.43 | -0.002 | 0.004 | 0.004 | 0 | 0.64 | 0.52 | 0.66 | 0.62 | 0.67 | 0.72 |
| 8 | 123449477 | rs10098937 | 0.431 | 0.49 | -0.69 | -0.176 | 0.009 | 0.053 | 0.053 | 0 | 0.23 | 0.12 | 0.1 | 0.05 | 0.18 | 0.24 |
| 8 | 123450875 | rs4412375 | 1.092 | 0.806 | -0.087 | 0.451 | 0.004 | 0.032 | 0.032 | 0 | 0.23 | 0.31 | 0.45 | 0.32 | 0.28 | 0.35 |
| 8 | 123451096 | rs4570163 | 0.541 | 0.877 | -0.025 | 0.541 | 0.007 | 0.044 | 0.044 | 0 | 0.28 | 0.61 | 0.49 | 0.38 | 0.32 | 0.47 |
| 8 | 123451833 | rs4415320 | 0.153 | 0.993 | -0.365 | 0.189 | 0.004 | -0.024 | -0.024 | 0 | 0.27 | 0.31 | 0.2 | 0.1 | 0.33 | 0.17 |
| 8 | 123453832 | rs4870795 | 1.276 | 1.074 | -0.407 | 0.176 | -0.001 | -0.033 | -0.033 | 0 | 0.17 | 0.16 | 0.12 | 0 | 0.2 | 0.04 |
| 8 | 123455152 | rs7003395 | 0.65 | 0.797 | -0.225 | 0.211 | -0.021 | 0.016 | 0.016 | 0 | 0.35 | 0.53 | 0.56 | 0.37 | 0.32 | 0.5 |
| 8 | 123456890 | rs7464984 | 0.926 | 0.785 | -0.148 | 0.179 | -0.006 | 0.001 | 0.001 | 0 | 0.33 | 0.36 | 0.54 | 0.31 | 0.32 | 0.4 |
| 8 | 123459231 | rs17271627 | 0.933 | 0.807 | -0.192 | 0.183 | -0.002 | 0.001 | 0.001 | 0 | 0.1 | 0.08 | 0.06 | 0.04 | 0.12 | 0.12 |
| 8 | 123463044 | rs7846137 | -0.585 | 0.82 | -0.18 | 0.189 | 0.007 | -0.011 | -0.011 | 0 | 0.61 | 0.75 | 0.75 | 0.42 | 0.66 | 0.66 |
| 8 | 123464305 | rs4316158 | 0.056 | 0.692 | -0.323 | 0.213 | -0.002 | 0.009 | 0.009 | 0 | 0.51 | 0.58 | 0.67 | 0.38 | 0.53 | 0.54 |
| 8 | 123467576 | rs2981073 | 0.687 | 0.606 | -0.431 | 0.248 | -0.009 | 0.007 | 0.007 | 0 | 0.34 | 0.41 | 0.55 | 0.38 | 0.32 | 0.5 |
| 8 | 123470712 | rs6980713 | 0.3 | 0.45 | -0.724 | 0.581 | 0.002 | -0.012 | -0.012 | 0 | 0.37 | 0.31 | 0.27 | 0.48 | 0.34 | 0.32 |
| 8 | 123471833 | rs6999334 | -0.748 | 0.439 | -0.683 | 0.557 | 0.009 | -0.034 | -0.034 | 1.443 | 0.83 | 0.76 | 0.87 | 0.94 | 0.79 | 0.96 |
| 8 | 123487582 | rs9656942 | 0.344 | -0.24 | -1.431 | 1.197 | 0.016 | -0.014 | -0.014 | 1.545 | 0.27 | 0.18 | 0.22 | 0.44 | 0.16 | 0.21 |
| 8 | 123495037 | rs4506214 | -0.063 | 0.543 | -1.646 | 1.13 | -0.04 | -0.004 | -0.004 | 0 | 0.59 | 0.81 | 0.42 | 0.63 | 0.46 | 0.57 |
| 8 | 123496567 | rs6990139 | -0.102 | 0.588 | -1.973 | 0.445 | -0.064 | 0.092 | 0.092 | 0 | 0.59 | 0.87 | 0.53 | 0.79 | 0.46 | 0.61 |
| 8 | 123505014 | rs7820388 | -0.154 | -0.234 | -1.386 | 0.768 | 0.019 | -0.017 | -0.017 | 0 | 0.12 | 0.05 | 0.07 | 0.2 | 0.06 | 0.11 |
| 8 | 123507509 | rs4518674 | -0.268 | -0.455 | -1.541 | 0.638 | -0.025 | -0.02 | -0.02 | 0.268 | 0.62 | 0.73 | 0.48 | 0.52 | 0.47 | 0.53 |
| 8 | 123511247 | rs11784208 | 0.462 | -0.496 | -0.966 | 0.709 | 0.003 | 0.009 | 0.009 | 0 | 0.47 | 0.61 | 0.36 | 0.21 | 0.41 | 0.36 |
| 8 | 123511652 | rs12543592 | 0.185 | -0.438 | -0.994 | 1.21 | 0.028 | 0.012 | 0.012 | 0 | 0.5 | 0.7 | 0.52 | 0.47 | 0.41 | 0.43 |
| 8 | 123522322 | rs4506215 | 0.748 | -0.855 | -1.201 | 0.618 | 0.002 | 0.011 | 0.011 | 0 | 0.27 | 0.2 | 0.13 | 0.23 | 0.22 | 0.25 |
| 8 | 123525406 | rs4637834 | 0.64 | -0.971 | -0.314 | 0.37 | -0.017 | -0.012 | -0.012 | 0 | 0.21 | 0.31 | 0.32 | 0.17 | 0.18 | 0.16 |
| 8 | 123527159 | rs4871268 | -0.401 | -0.726 | -0.829 | -0.215 | 0.069 | 0.09 | 0.09 | 0.133 | 0.5 | 0.66 | 0.43 | 0.27 | 0.34 | 0.35 |
| 8 | 123536333 | rs4242336 | -0.777 | -0.388 | -0.179 | -0.646 | -0.018 | -0.005 | -0.005 | 0 | 0.88 | 0.99 | 0.96 | 0.91 | 0.92 | 0.88 |
| 8 | 123536419 | rs4870799 | 0.388 | 0.26 | -0.147 | -0.615 | -0.032 | -0.03 | -0.03 | 0 | 0.18 | 0.14 | 0.5 | 0.52 | 0.42 | 0.42 |
| 8 | 123536456 | rs4242337 | 0.044 | -0.166 | 0.294 | -1.038 | -0.002 | -0.021 | -0.021 | 0 | 0.73 | 0.66 | 0.87 | 0.91 | 0.81 | 0.76 |
| 8 | 123537015 | rs4368984 | 0.852 | -0.27 | 0.444 | -1.108 | 0.007 | 0.007 | 0.007 | 0 | 0.53 | 0.42 | 0.36 | 0.39 | 0.39 | 0.34 |
| 8 | 123537857 | rs6986952 | 1.203 | -0.686 | 0.349 | -1.389 | -0.038 | -0.005 | -0.005 | 1.444 | 0.34 | 0.36 | 0.26 | 0.36 | 0.21 | 0.29 |
| 8 | 123537948 | rs7834917 | 0.688 | 0.148 | 0.22 | -1.031 | 0.002 | 0.01 | 0.01 | 0 | 0.14 | 0.11 | 0.06 | 0 | 0.11 | 0.11 |
| 8 | 123538147 | rs7005211 | -0.557 | 0.185 | 0.239 | -1.009 | 0 | 0 | 0 | 0 | 0.39 | 0.36 | 0.61 | 0.55 | 0.6 | 0.48 |
| 8 | 123543258 | rs10086964 | 0.77 | 0.519 | -1.029 | -0.846 | -0.043 | -0.041 | -0.041 | 0 | 0.16 | 0.08 | 0.48 | 0.52 | 0.39 | 0.39 |
| 8 | 123544259 | rs1523217 | -1.046 | 0.004 | -0.752 | -1.768 | -0.009 | -0.112 | -0.112 | 0 | 0.39 | 0.49 | 0.21 | 0.03 | 0.32 | 0.2 |
| 8 | 123545374 | rs16893023 | -0.36 | 0.435 | -1.172 | -0.963 | 0 | -0.036 | -0.036 | 0.725 | 0.22 | 0.23 | 0.14 | 0.01 | 0.2 | 0.07 |
| 8 | 123552740 | rs1055142 | 0.046 | 0.785 | -1.97 | -0.878 | -0.074 | 0.102 | 0.102 | 1.934 | 0.63 | 0.88 | 0.7 | 0.79 | 0.52 | 0.66 |
| 8 | 123552796 | rs16886 | 0.396 | -0.102 | -2.737 | -1.341 | -0.034 | -0.017 | -0.017 | 0 | 0.29 | 0.05 | 0.46 | 0.69 | 0.32 | 0.34 |
| 8 | 123555635 | rs2948640 | 0.368 | -0.08 | -2.995 | -1.803 | -0.068 | 0.079 | 0.079 | 0.052 | 0.32 | 0.15 | 0.24 | 0.18 | 0.38 | 0.24 |
| 8 | 123556500 | rs4871275 | -0.195 | -0.005 | -2.84 | -1.65 | -0.047 | 0.137 | 0.137 | 2.352 | 0.57 | 0.83 | 0.7 | 0.79 | 0.53 | 0.62 |
| 8 | 123569427 | rs10091964 | -0.116 | -0.112 | -1.98 | -1.725 | -0.008 | 0.042 | 0.042 | 2.623 | 0.56 | 0.64 | 0.69 | 0.82 | 0.53 | 0.62 |
| 8 | 123570236 | rs2140327 | 0.542 | -0.366 | -2.168 | -1.315 | -0.066 | -0.053 | -0.053 | 0 | 0.26 | 0.02 | 0.44 | 0.74 | 0.32 | 0.38 |
| 8 | 123603473 | rs1880014 | -0.259 | -0.889 | -2.081 | -1.103 | 0.033 | 0.037 | 0.037 | 0.047 | 0.42 | 0.74 | 0.58 | 0.7 | 0.38 | 0.58 |
| 8 | 123615949 | rs7846496 | -0.42 | -1.143 | -1.651 | -0.993 | 0.028 | 0.027 | 0.027 | 1.901 | 0.22 | 0.81 | 0.17 | 0.08 | 0.08 | 0.22 |
| 8 | 123616097 | rs12547950 | -0.392 | -1.127 | -1.288 | -0.985 | -0.019 | -0.012 | -0.012 | 0.604 | 0.22 | 0.75 | 0.17 | 0.08 | 0.08 | 0.22 |
| 8 | 123626804 | rs4871293 | -0.335 | 0.021 | -1.698 | -1.104 | -0.013 | 0.054 | 0.054 | 0.037 | 0.38 | 0.21 | 0.31 | 0.23 | 0.38 | 0.24 |
| 8 | 123633185 | rs13270758 | -0.251 | 0.216 | -1.587 | -0.94 | -0.032 | 0.022 | 0.022 | 0 | 0.3 | 0.15 | 0.27 | 0.21 | 0.34 | 0.17 |
| 8 | 123646965 | rs4870806 | 0.524 | 0.009 | -1.172 | -0.747 | -0.006 | 0.041 | 0.041 | 0.979 | 0.61 | 0.23 | 0.49 | 0.74 | 0.61 | 0.81 |
| 8 | 123647245 | rs1568684 | 0.368 | 0.036 | -0.317 | -0.72 | 0.006 | -0.008 | -0.008 | 0 | 0.67 | 0.61 | 0.52 | 0.73 | 0.61 | 0.81 |
| 8 | 123648234 | rs6990818 | 0.631 | 0.29 | -0.352 | -0.195 | -0.017 | -0.012 | -0.012 | 0.718 | 0.54 | 0.22 | 0.38 | 0.5 | 0.5 | 0.57 |
| 8 | 123649642 | rs2140330 | 1.09 | 0.244 | -0.374 | 0.374 | 0.028 | -0.025 | -0.025 | 0 | 0.46 | 0.21 | 0.32 | 0.47 | 0.43 | 0.52 |
| 8 | 123651639 | rs9692955 | 0.257 | 0.359 | 0.186 | 0.35 | 0.022 | -0.022 | -0.022 | 0 | 0.1 | 0.24 | 0.1 | 0.06 | 0.09 | 0.17 |
| 8 | 123651925 | rs1357478 | 0.811 | 0.287 | 0.208 | 0.369 | 0.025 | 0.01 | 0.01 | 0 | 0.66 | 0.41 | 0.43 | 0.6 | 0.57 | 0.65 |
| 8 | 123651942 | rs1357477 | 1.039 | 0.43 | 0.265 | 0.4 | -0.006 | -0.008 | -0.008 | 0 | 0.2 | 0.22 | 0.11 | 0.14 | 0.14 | 0.14 |
| 8 | 123657902 | rs940123 | -0.462 | 0.475 | -0.2 | 0.427 | -0.022 | -0.043 | -0.043 | 0 | 0.32 | 0.57 | 0.22 | 0.2 | 0.23 | 0.29 |
| 8 | 123658247 | rs17654120 | 0.665 | 0.488 | -0.107 | 0.436 | 0.007 | -0.013 | -0.013 | 0 | 0.09 | 0.11 | 0.09 | 0.06 | 0.09 | 0.15 |

| | | | | | | | | | | | | | | | | |
|---|---|---|---|---|---|---|---|---|---|---|---|---|---|---|---|---|
| 8 | 123658340 | rs4545104 | -0.787 | 0.508 | -0.205 | 0.478 | 0.009 | 0.006 | 0.006 | 0 | 0.79 | 0.78 | 0.54 | 0.66 | 0.66 | 0.81 |
| 8 | 123658457 | rs10505425 | -0.306 | 0.511 | -0.144 | 0.477 | -0.027 | -0.054 | -0.054 | 0 | 0.31 | 0.49 | 0.21 | 0.21 | 0.23 | 0.29 |
| 8 | 123661005 | rs4458884 | 0.104 | 1.375 | -0.193 | 1.048 | -0.003 | -0.003 | -0.003 | 3.354 | 0.77 | 0.78 | 0.53 | 0.65 | 0.63 | 0.81 |
| 8 | 123668493 | rs1916317 | -1.008 | 0.518 | 0 | 0.623 | 0.001 | -0.024 | -0.024 | 0.505 | 0.11 | 0.23 | 0.11 | 0.07 | 0.09 | 0.15 |
| 8 | 123673402 | rs6984347 | -0.222 | -0.226 | 0.21 | 0.986 | -0.045 | -0.021 | -0.021 | 0 | 0.18 | 0.3 | 0.06 | 0.13 | 0.05 | 0.1 |
| 8 | 123676513 | rs4595130 | -0.987 | -0.776 | 0.487 | 1.304 | -0.087 | 0.033 | 0.033 | 0 | 0.3 | 0.44 | 0.16 | 0.34 | 0.12 | 0.4 |
| 8 | 123676967 | rs6984873 | -1.47 | -0.004 | 0.626 | 0.924 | 0.004 | 0.075 | 0.075 | 0 | 0.49 | 0.75 | 0.45 | 0.65 | 0.54 | 0.58 |
| 8 | 123679458 | rs7827027 | 0.178 | 0.021 | 0.782 | 0.933 | 0.01 | 0.001 | 0.001 | 0 | 0.18 | 0.2 | 0.27 | 0.31 | 0.41 | 0.18 |
| 8 | 123680920 | rs16897280 | 0.581 | 0.004 | 0.815 | 0.939 | -0.013 | 0.028 | 0.028 | 0 | 0.11 | 0.14 | 0.1 | 0.21 | 0.07 | 0.3 |
| 8 | 123684295 | rs7836045 | 0.69 | 0.004 | 0.834 | 0.795 | -0.007 | 0.004 | 0.004 | 0 | 0.11 | 0.11 | 0.1 | 0.15 | 0.07 | 0.3 |
| 8 | 123689670 | rs4871296 | -1.01 | -0.05 | 0.925 | 0.761 | -0.016 | -0.018 | -0.018 | 0 | 0.39 | 0.68 | 0.34 | 0.49 | 0.46 | 0.31 |
| 8 | 123689919 | rs4870809 | -0.038 | 0.012 | 0.921 | 0.625 | 0.008 | 0.005 | 0.005 | 0 | 0.19 | 0.21 | 0.27 | 0.35 | 0.41 | 0.22 |
| 8 | 123691458 | rs16897288 | 0.451 | 0.011 | 1.039 | -0.058 | -0.06 | -0.07 | -0.07 | 0 | 0.18 | 0.35 | 0.07 | 0.06 | 0.07 | 0.1 |
| 8 | 123692892 | rs16897290 | 0.481 | 0.015 | 0.996 | -0.055 | -0.046 | -0.057 | -0.057 | 0 | 0.18 | 0.29 | 0.07 | 0.06 | 0.06 | 0.1 |
| 8 | 123696613 | rs11774246 | -0.536 | 0.324 | 0.905 | 0.103 | -0.039 | -0.027 | -0.027 | 0.353 | 0.28 | 0.18 | 0.34 | 0.57 | 0.48 | 0.49 |
| 8 | 123698785 | rs876742 | -0.591 | 0.284 | 0.878 | 0.107 | -0.015 | -0.011 | -0.011 | 1.692 | 0.29 | 0.24 | 0.34 | 0.56 | 0.47 | 0.46 |
| 8 | 123711143 | rs7018013 | 0.099 | -0.167 | 0.714 | -0.325 | -0.029 | -0.041 | -0.041 | 0 | 0.24 | 0.28 | 0.06 | 0.06 | 0.08 | 0.11 |
| 8 | 123718989 | rs11997170 | 0.012 | -0.152 | 0.702 | -0.32 | -0.085 | -0.118 | -0.118 | 0 | 0.25 | 0.45 | 0.09 | 0.06 | 0.09 | 0.14 |
| 8 | 123730877 | rs7822857 | -0.535 | 0.24 | 0.611 | -0.317 | -0.006 | -0.058 | -0.058 | 0 | 0.34 | 0.39 | 0.17 | 0.07 | 0.29 | 0.22 |
| 8 | 123744737 | rs11781433 | 0.168 | 0.328 | 0.689 | -0.248 | -0.001 | -0.025 | -0.025 | 0 | 0.15 | 0.17 | 0.12 | 0 | 0.22 | 0.1 |
| 8 | 123744784 | rs17356990 | 0.649 | 0.442 | 0.499 | -0.209 | -0.028 | -0.016 | -0.016 | 0 | 0.1 | 0.01 | 0.06 | 0.08 | 0.12 | 0.21 |
| 8 | 123748284 | rs4291275 | -0.648 | 0.408 | 1.025 | -0.15 | -0.011 | 0.01 | 0.01 | 0.887 | 0.37 | 0.64 | 0.6 | 0.29 | 0.48 | 0.38 |
| 8 | 123749087 | rs4486207 | 0.675 | 0.413 | 1.132 | -0.138 | 0.009 | 0.018 | 0.018 | 0.008 | 0.2 | 0.26 | 0.46 | 0.29 | 0.25 | 0.28 |
| 8 | 123755268 | rs11779978 | 0.151 | 0.503 | 1.054 | -0.132 | 0.022 | -0.082 | -0.082 | 0.066 | 0.16 | 0.28 | 0.13 | 0 | 0.22 | 0.1 |
| 8 | 123766714 | rs12544595 | 0.953 | 0.546 | 0.588 | -0.057 | -0.001 | -0.001 | -0.001 | 0 | 0.17 | 0.14 | 0.45 | 0.29 | 0.25 | 0.19 |
| 8 | 123768374 | rs7002192 | -0.261 | 0.541 | 0.638 | -0.153 | 0.04 | -0.133 | -0.133 | 0.015 | 0.18 | 0.51 | 0.17 | 0 | 0.22 | 0.1 |
| 8 | 123769076 | rs4871302 | -0.526 | 0.394 | -0.04 | -0.204 | -0.01 | -0.082 | -0.082 | 5.307 | 0.35 | 0.43 | 0.16 | 0.07 | 0.29 | 0.19 |
| 8 | 123769451 | rs10089735 | -0.774 | 1.628 | -0.586 | 0.535 | 0.017 | -0.004 | -0.004 | 0 | 0.33 | 0.38 | 0.55 | 0.29 | 0.47 | 0.3 |
| 8 | 123775124 | rs9918775 | -0.365 | 0.68 | -1.301 | 0.79 | 0 | -0.001 | -0.001 | 0 | 0.24 | 0.7 | 0.52 | 0.32 | 0.25 | 0.22 |
| 8 | 123779111 | rs13267440 | -0.516 | 0.677 | -1.584 | 0.363 | -0.019 | 0.036 | 0.036 | 0 | 0.16 | 0.03 | 0.07 | 0.11 | 0.18 | 0.25 |
| 8 | 123783276 | rs1562180 | 0.584 | 0.557 | -1.209 | 0.804 | -0.003 | 0.009 | 0.009 | 0 | 0.89 | 0.87 | 0.82 | 0.84 | 0.9 | 0.87 |
| 8 | 123786515 | rs13271342 | -0.155 | 1.216 | -2.72 | 1.186 | -0.07 | 0.053 | 0.053 | 0.045 | 0.27 | 0.02 | 0.19 | 0.19 | 0.3 | 0.24 |
| 8 | 123787061 | rs6980880 | 1.06 | 0.689 | -1.698 | 1.195 | 0.014 | 0.021 | 0.021 | 0 | 0.56 | 0.32 | 0.53 | 0.6 | 0.59 | 0.6 |
| 8 | 123788289 | rs6986574 | 1.047 | 0.22 | -1.757 | 1.276 | 0.036 | 0.012 | 0.012 | 0 | 0.54 | 0.24 | 0.59 | 0.54 | 0.6 | 0.58 |
| 8 | 123793166 | rs11777561 | -0.197 | 0.234 | -1.982 | 1.158 | -0.057 | 0.074 | 0.074 | 0.618 | 0.29 | 0.02 | 0.24 | 0.18 | 0.3 | 0.25 |
| 8 | 123793984 | rs13261590 | -0.312 | 0.481 | -1.345 | 1.364 | 0.002 | 0.012 | 0.012 | 0 | 0.92 | 0.87 | 0.92 | 0.87 | 0.91 | 0.86 |
| 8 | 123801083 | rs7009772 | 0.25 | 0.257 | -1.544 | 1.483 | -0.001 | 0 | 0 | 0 | 0.67 | 0.66 | 0.57 | 0.47 | 0.64 | 0.53 |
| 8 | 123802686 | rs4871305 | 0.084 | 0.42 | -1.448 | 1.504 | 0.014 | 0.037 | 0.037 | 0 | 0.87 | 0.82 | 0.89 | 0.77 | 0.85 | 0.77 |
| 8 | 123805336 | rs10956100 | 0.349 | 0.442 | -2.047 | 0.988 | -0.01 | -0.044 | -0.044 | 0 | 0.23 | 0.03 | 0.26 | 0.29 | 0.23 | 0.21 |
| 8 | 123806196 | rs1445169 | -0.355 | 0.314 | -1.137 | 0.734 | -0.016 | -0.044 | -0.044 | 0 | 0.36 | 0.49 | 0.29 | 0.27 | 0.35 | 0.25 |
| 8 | 123806240 | rs2384844 | -0.033 | 0.297 | -0.955 | 0.727 | -0.023 | 0.069 | 0.069 | 4.093 | 0.32 | 0.08 | 0.22 | 0.23 | 0.34 | 0.22 |
| 8 | 123807378 | rs11775943 | -0.105 | 0.298 | 0.109 | 0.727 | 0.003 | 0.059 | 0.059 | 3.557 | 0.33 | 0.16 | 0.24 | 0.23 | 0.34 | 0.22 |
| 8 | 123808821 | rs11777048 | -1.104 | 0.118 | 0.304 | 0.727 | -0.009 | 0.003 | 0.003 | 0.003 | 0.59 | 0.66 | 0.58 | 0.43 | 0.55 | 0.53 |
| 8 | 123814825 | rs748856 | -0.81 | 0.186 | 0.517 | 0.763 | -0.003 | -0.003 | -0.003 | 0.012 | 0.54 | 0.48 | 0.55 | 0.48 | 0.49 | 0.51 |
| 8 | 123816210 | rs4870814 | -0.415 | 0.027 | 0.399 | 0.727 | -0.009 | 0.019 | 0.019 | 0 | 0.77 | 0.9 | 0.91 | 0.82 | 0.8 | 0.8 |
| 8 | 123817194 | rs11783697 | 1.463 | 0.108 | 0.555 | 0.882 | -0.001 | 0.008 | 0.008 | 0.09 | 0.36 | 0.46 | 0.38 | 0.37 | 0.33 | 0.26 |
| 8 | 123818170 | rs4870815 | 0.762 | -0.319 | 0.386 | 0.769 | 0.008 | 0.03 | 0.03 | 0 | 0.06 | 0.11 | 0.26 | 0.16 | 0.08 | 0.14 |
| 8 | 123819255 | rs6991737 | -1.063 | -0.295 | 0.334 | 0.611 | 0 | 0 | 0 | 0 | 0.57 | 0.43 | 0.56 | 0.53 | 0.54 | 0.6 |
| 8 | 123821456 | rs9642865 | -1.724 | -0.086 | 0.169 | 0.821 | 0.009 | -0.002 | -0.002 | 0 | 0.52 | 0.73 | 0.57 | 0.63 | 0.5 | 0.69 |
| 8 | 123822629 | rs7830373 | 0.285 | -0.001 | 0.116 | 0.931 | -0.004 | 0.012 | 0.012 | 0.23 | 0.21 | 0.37 | 0.21 | 0.29 | 0.22 | 0.2 |
| 8 | 123823987 | rs12544338 | 1.089 | 0.139 | -0.815 | 0.956 | -0.02 | -0.004 | -0.004 | 0.001 | 0.05 | 0.03 | 0.26 | 0.22 | 0.08 | 0.14 |
| 8 | 123827049 | rs16897499 | 0.352 | -0.193 | -0.745 | 0.547 | 0.005 | -0.001 | -0.001 | 0.111 | 0.2 | 0.2 | 0.07 | 0.07 | 0.11 | 0.17 |
| 8 | 123830795 | rs11777569 | 0.157 | 0.405 | -0.705 | 1.08 | -0.006 | -0.022 | -0.022 | 0 | 0.18 | 0.1 | 0.17 | 0.3 | 0.22 | 0.2 |
| 8 | 123834815 | rs13273011 | -0.658 | 0.323 | -0.309 | 1.15 | 0.039 | 0.171 | 0.171 | 1.077 | 0.11 | 0.43 | 0.3 | 0.26 | 0.13 | 0.21 |
| 8 | 123840945 | rs12675789 | 0.583 | 0.592 | -1.757 | 0.573 | -0.02 | 0.053 | 0.053 | 3.284 | 0.43 | 0.06 | 0.35 | 0.24 | 0.4 | 0.27 |
| 8 | 123853184 | rs10100547 | -0.149 | 0.749 | -0.77 | 0.582 | -0.005 | 0.013 | 0.013 | 0 | 0.48 | 0.66 | 0.69 | 0.54 | 0.52 | 0.49 |
| 8 | 123854444 | rs11778542 | 0.755 | 0.626 | -0.327 | 0.418 | 0.01 | 0.005 | 0.005 | 0.037 | 0.27 | 0.2 | 0.25 | 0.22 | 0.25 | 0.21 |
| 8 | 123856788 | rs7002043 | -0.532 | 1.093 | -0.333 | 0.479 | -0.033 | -0.05 | -0.05 | 0.051 | 0.76 | 0.85 | 0.52 | 0.61 | 0.68 | 0.67 |
| 8 | 123858679 | rs7012131 | -0.654 | 0.856 | 0.363 | -0.061 | 0.084 | 0.01 | 0.01 | 0.136 | 0.08 | 0.36 | 0.08 | 0.02 | 0.09 | 0.05 |
| 8 | 123864916 | rs6982380 | 0.628 | 0.517 | 0.087 | 0.073 | 0 | -0.001 | -0.001 | 0 | 0.34 | 0.36 | 0.23 | 0.38 | 0.32 | 0.41 |
| 8 | 123866583 | rs12549062 | 0.833 | 0.559 | -0.243 | 0.073 | -0.002 | -0.017 | -0.017 | 6.538 | 0.32 | 0.2 | 0.21 | 0.38 | 0.32 | 0.41 |
| 8 | 123870345 | rs4870820 | 0.078 | 0.316 | -0.304 | -0.136 | -0.029 | -0.037 | -0.037 | 0.306 | 0.36 | 0.43 | 0.3 | 0.2 | 0.27 | 0.23 |
| 8 | 123877155 | rs11776799 | 0.159 | 0.7 | -1.488 | 0.568 | 0.023 | -0.034 | -0.034 | 0 | 0.18 | 0.05 | 0.18 | 0.23 | 0.17 | 0.18 |
| 8 | 123880227 | rs10109650 | -0.079 | 0.922 | -1.236 | 0.459 | 0 | -0.001 | -0.001 | 0 | 0.36 | 0.35 | 0.51 | 0.37 | 0.38 | 0.44 |
| 8 | 123881190 | rs4870821 | -0.043 | 1.074 | -1.227 | 0.213 | -0.01 | 0.033 | 0.033 | 0.094 | 0.49 | 0.81 | 0.36 | 0.41 | 0.48 | 0.47 |
| 8 | 123881218 | rs6470120 | 0.337 | 0.871 | -1.227 | 0.451 | -0.001 | -0.001 | -0.001 | 0 | 0.4 | 0.58 | 0.28 | 0.37 | 0.36 | 0.35 |
| 8 | 123890211 | rs4870822 | -0.438 | 0.862 | -1.543 | 0.354 | -0.013 | 0.028 | 0.028 | 0 | 0.22 | 0.07 | 0.2 | 0.17 | 0.23 | 0.15 |
| 8 | 123892069 | rs4871320 | -0.236 | 0.204 | -1.669 | 0.476 | 0.023 | -0.001 | -0.001 | 0 | 0.39 | 0.69 | 0.28 | 0.4 | 0.3 | 0.28 |
| 8 | 123894892 | rs17367785 | 0.365 | 0.376 | -1.226 | 0.546 | -0.007 | -0.002 | -0.002 | 0.055 | 0.57 | 0.42 | 0.55 | 0.54 | 0.5 | 0.64 |

| | | | | | | | | | | | | | | | | |
|---|---|---|---|---|---|---|---|---|---|---|---|---|---|---|---|---|
| 8 | 123897867 | rs4871323 | 0.175 | 0.379 | -0.909 | 0.501 | 0 | 0 | 0 | 0 | 0.61 | 0.43 | 0.58 | 0.54 | 0.55 | 0.64 |
| 8 | 123915944 | rs13264706 | 0.271 | 0.235 | 0.024 | 0.536 | -0.022 | -0.016 | -0.016 | 1.705 | 0.63 | 0.77 | 0.65 | 0.55 | 0.53 | 0.67 |
| 8 | 123925998 | rs7812750 | 0.921 | 0.166 | -0.17 | 0.507 | 0.02 | 0.029 | 0.029 | 0 | 0.47 | 0.33 | 0.53 | 0.39 | 0.49 | 0.56 |
| 8 | 123927562 | rs7463533 | -0.438 | 0.577 | -0.317 | 0.801 | -0.031 | -0.038 | -0.038 | 0 | 0.76 | 0.86 | 0.77 | 0.58 | 0.67 | 0.8 |
| 8 | 123928859 | rs7843271 | -0.928 | 0.531 | -0.101 | 0.747 | -0.055 | -0.07 | -0.07 | 0.826 | 0.87 | 0.97 | 0.9 | 0.71 | 0.84 | 0.83 |
| 8 | 123930626 | rs10099926 | -0.277 | 0.518 | 0.002 | 0.693 | -0.01 | -0.012 | -0.012 | 0 | 0.83 | 0.9 | 0.87 | 0.72 | 0.81 | 0.82 |
| 8 | 123931864 | rs16897667 | -0.291 | 0.08 | 0.239 | 0.363 | -0.021 | -0.01 | -0.01 | 0 | 0.27 | 0.36 | 0.23 | 0.23 | 0.22 | 0.27 |
| 8 | 123932061 | rs7832429 | -0.111 | 0.108 | -0.035 | 0.398 | -0.001 | -0.002 | -0.002 | 0.196 | 0.27 | 0.22 | 0.19 | 0.18 | 0.22 | 0.25 |
| 8 | 123938787 | rs7828669 | -0.613 | 0.105 | -0.383 | 0.317 | -0.035 | 0.004 | 0.004 | 0 | 0.31 | 0.51 | 0.29 | 0.33 | 0.23 | 0.28 |
| 8 | 123941078 | rs11775763 | -0.23 | 0.243 | -0.688 | 0.339 | -0.017 | 0 | 0 | 0 | 0.27 | 0.32 | 0.18 | 0.28 | 0.22 | 0.25 |
| 8 | 123946096 | rs4870828 | 0.621 | 0.285 | -0.712 | 0.439 | -0.001 | -0.001 | -0.001 | 0 | 0.61 | 0.68 | 0.72 | 0.55 | 0.61 | 0.58 |
| 8 | 123948459 | rs10087621 | 0.83 | 0.136 | -0.416 | -0.304 | -0.001 | -0.027 | -0.027 | 0 | 0.68 | 0.67 | 0.8 | 0.81 | 0.71 | 0.77 |
| 8 | 123950590 | rs7826199 | 1.039 | 0.067 | -0.307 | -0.293 | -0.011 | -0.035 | -0.035 | 0.043 | 0.64 | 0.56 | 0.73 | 0.77 | 0.69 | 0.75 |
| 8 | 123952169 | rs10956109 | 0.875 | 0.185 | -0.73 | -0.341 | 0.001 | -0.024 | -0.024 | 0 | 0.67 | 0.67 | 0.8 | 0.81 | 0.68 | 0.76 |
| 8 | 123952293 | rs7015548 | -0.257 | 0.37 | -1.63 | -0.194 | -0.001 | -0.002 | -0.002 | 0.078 | 0.85 | 0.88 | 0.89 | 0.81 | 0.84 | 0.8 |
| 8 | 123954718 | rs6989244 | 0.626 | 0.282 | -1.889 | -0.165 | 0.011 | 0.031 | 0.031 | 0.793 | 0.65 | 0.27 | 0.68 | 0.77 | 0.7 | 0.8 |
| 8 | 123961799 | rs4870831 | -0.058 | 0.764 | -2.383 | -0.18 | -0.036 | 0.076 | 0.076 | 0 | 0.76 | 0.92 | 0.81 | 0.87 | 0.74 | 0.83 |
| 8 | 123964431 | rs3204141 | 0.154 | 0.925 | -2.799 | 0.009 | -0.072 | 0.074 | 0.074 | 0.794 | 0.29 | 0.02 | 0.33 | 0.13 | 0.35 | 0.43 |
| 8 | 123964953 | rs3808600 | 0.156 | 0.936 | -2.734 | 0.031 | -0.076 | 0.073 | 0.073 | 0.467 | 0.29 | 0.01 | 0.33 | 0.13 | 0.36 | 0.43 |
| 8 | 123967175 | rs7011777 | 0.053 | 0.676 | -2.697 | 0.162 | -0.041 | 0.075 | 0.075 | 0.718 | 0.26 | 0.01 | 0.29 | 0.1 | 0.24 | 0.45 |
| 8 | 123969205 | rs7002339 | -0.272 | 0.89 | -1.993 | 0.427 | -0.006 | 0.003 | 0.003 | 0 | 0.73 | 0.82 | 0.76 | 0.75 | 0.66 | 0.78 |
| 8 | 123971081 | rs7841763 | 0.085 | 0.49 | -1.533 | 0.21 | -0.004 | -0.018 | -0.018 | 0 | 0.08 | 0.17 | 0.06 | 0.02 | 0.1 | 0.15 |
| 8 | 123973068 | rs10108684 | 0.391 | 0.742 | -1.899 | 0.768 | -0.052 | -0.08 | -0.08 | 0.003 | 0.11 | 0.01 | 0.11 | 0.46 | 0.14 | 0.22 |
| 8 | 123973561 | rs10108984 | -0.377 | 0.784 | -1.809 | 0.683 | -0.063 | 0.005 | 0.005 | 0 | 0.7 | 0.93 | 0.69 | 0.74 | 0.56 | 0.62 |
| 8 | 123980551 | rs11779459 | 1.097 | 0.516 | -1.352 | 0.499 | 0.004 | 0.013 | 0.013 | 0 | 0.47 | 0.56 | 0.45 | 0.24 | 0.37 | 0.37 |
| 8 | 123980697 | rs7844465 | -0.923 | 0.58 | -1.454 | 0.535 | -0.016 | 0.007 | 0.007 | 0 | 0.74 | 0.83 | 0.72 | 0.77 | 0.66 | 0.78 |
| 8 | 123983143 | rs16897813 | 1.224 | 0.446 | -1.66 | 0.379 | 0.004 | -0.011 | -0.011 | 0 | 0.13 | 0.06 | 0.2 | 0.13 | 0.11 | 0.19 |
| 8 | 123983434 | rs17370716 | 1.295 | 0.507 | -1.653 | 0.489 | 0.002 | 0 | 0 | 2.076 | 0.07 | 0 | 0.1 | 0.05 | 0.05 | 0.04 |
| 8 | 123985708 | rs3802266 | 0.058 | 0.542 | -0.668 | 0.536 | -0.003 | 0.02 | 0.02 | 0 | 0.54 | 0.64 | 0.5 | 0.26 | 0.47 | 0.51 |
| 8 | 123986649 | rs2833 | 0.038 | 0.664 | -1.028 | 0.576 | -0.009 | 0.033 | 0.033 | 0 | 0.56 | 0.72 | 0.51 | 0.25 | 0.47 | 0.51 |
| 8 | 123988424 | rs13248065 | 0.51 | 0.525 | -0.946 | 0.927 | 0.002 | 0.031 | 0.031 | 0.399 | 0.32 | 0.27 | 0.24 | 0.09 | 0.26 | 0.17 |
| 8 | 123989445 | rs10216833 | -0.16 | 0.323 | -0.828 | 1.291 | -0.052 | -0.078 | -0.078 | 0 | 0.11 | 0.01 | 0.11 | 0.46 | 0.14 | 0.23 |
| 8 | 123993765 | rs11994870 | -0.08 | 0.262 | -0.625 | 1.233 | 0 | 0.066 | 0.066 | 0 | 0.1 | 0.09 | 0.09 | 0.48 | 0.08 | 0.22 |
| 8 | 123995186 | rs10106512 | 0.24 | 0.061 | -0.551 | 0.126 | 0 | 0.008 | 0.008 | 0.083 | 0.3 | 0.26 | 0.23 | 0.08 | 0.27 | 0.15 |
| 8 | 124001089 | rs12674855 | -0.749 | -0.847 | 0.544 | 1.297 | -0.042 | 0.126 | 0.126 | 0 | 0.23 | 0.51 | 0.19 | 0.46 | 0.16 | 0.28 |
| 8 | 124004186 | rs11993361 | 0.022 | -0.9 | 0.618 | 1.339 | -0.001 | 0.011 | 0.011 | 0 | 0.07 | 0.15 | 0.06 | 0.08 | 0.07 | 0.08 |
| 8 | 124008339 | rs4510872 | -0.894 | -0.882 | 0.627 | 1.314 | -0.029 | 0.095 | 0.095 | 0 | 0.22 | 0.43 | 0.18 | 0.46 | 0.16 | 0.27 |
| 8 | 124014634 | rs6982326 | -1.251 | -1.303 | 0.745 | 0.83 | -0.008 | -0.004 | -0.004 | 0.156 | 0.31 | 0.3 | 0.19 | 0.25 | 0.23 | 0.22 |
| 8 | 124021543 | rs12235095 | 0.673 | -1.152 | 1.051 | 0.69 | 0.012 | 0.065 | 0.065 | 0 | 0.88 | 0.69 | 0.79 | 0.78 | 0.86 | 0.86 |
| 8 | 124023374 | rs16897861 | 1.55 | -1.463 | 1.133 | 0.919 | -0.001 | 0.001 | 0.001 | 0 | 0.65 | 0.36 | 0.54 | 0.47 | 0.64 | 0.54 |
| 8 | 124023538 | rs11780596 | 1.742 | -2.041 | 0.335 | 0.342 | -0.015 | -0.066 | -0.066 | 0 | 0.54 | 0.06 | 0.46 | 0.47 | 0.57 | 0.52 |
| 8 | 124023621 | rs6982014 | -0.517 | -1.965 | 0.558 | 0.372 | -0.003 | 0.016 | 0.016 | 0.024 | 0.85 | 0.86 | 0.83 | 0.77 | 0.85 | 0.76 |
| 8 | 124024823 | rs12549382 | 1.426 | -2.156 | 0.23 | 0.368 | 0.032 | 0.11 | 0.11 | 1.205 | 0.51 | 0.18 | 0.53 | 0.34 | 0.55 | 0.44 |
| 8 | 124026900 | rs1059588 | 1.34 | -2.174 | 0.392 | 0.391 | 0.007 | -0.037 | -0.037 | 0.105 | 0.13 | 0.02 | 0.06 | 0.17 | 0.1 | 0.11 |
| 8 | 124027301 | rs2294086 | -0.33 | -1.875 | 0.633 | 0.299 | -0.013 | 0.058 | 0.058 | 0 | 0.14 | 0.36 | 0.14 | 0.21 | 0.1 | 0.12 |
| 8 | 124027508 | rs11691 | 0.585 | -1.879 | 0.633 | 0.297 | -0.01 | -0.044 | -0.044 | 0 | 0.09 | 0.15 | 0.07 | 0.01 | 0.06 | 0.02 |
| 8 | 124027562 | rs7812 | -0.317 | -1.879 | 0.626 | 0.298 | -0.012 | 0.06 | 0.06 | 0 | 0.14 | 0.36 | 0.14 | 0.21 | 0.1 | 0.12 |
| 8 | 124027628 | rs7159 | 0.883 | -1.884 | 0.577 | 0.25 | -0.006 | 0.009 | 0.009 | 1.038 | 0.69 | 0.5 | 0.65 | 0.7 | 0.67 | 0.7 |
| 8 | 124028479 | rs11781842 | 2.424 | -2.617 | -0.208 | 0.041 | -0.192 | 0.077 | 0.077 | 0.796 | 0.29 | 0.02 | 0.33 | 0.12 | 0.44 | 0.33 |
| 8 | 124028631 | rs10090419 | 0.448 | -1.356 | 0.73 | 0.554 | 0.001 | -0.001 | -0.001 | 0 | 0.11 | 0.14 | 0.19 | 0.08 | 0.15 | 0.1 |
| 8 | 124031541 | rs2272722 | 0.412 | -1.347 | 0.559 | 0.564 | -0.004 | -0.027 | -0.027 | 0.574 | 0.08 | 0.1 | 0.07 | 0.01 | 0.06 | 0.02 |
| 8 | 124038045 | rs12677755 | 1.259 | -1.276 | 0.545 | 0.653 | -0.007 | 0.002 | 0.002 | 2.68 | 0.68 | 0.47 | 0.65 | 0.7 | 0.67 | 0.7 |
| 8 | 124042733 | rs2891672 | -0.844 | -1.197 | 0.001 | 0.624 | -0.029 | -0.06 | -0.06 | 1.931 | 0.91 | 0.98 | 0.92 | 0.79 | 0.9 | 0.82 |
| 8 | 124049057 | rs13255713 | 1.195 | -1.108 | 0.153 | 0.68 | 0.009 | -0.02 | -0.02 | 0.001 | 0.63 | 0.32 | 0.61 | 0.5 | 0.64 | 0.61 |
| 8 | 124056914 | rs4871340 | -1.042 | -0.733 | 0.041 | 0.714 | -0.005 | 0.034 | 0.034 | 0.295 | 0.24 | 0.58 | 0.33 | 0.3 | 0.25 | 0.22 |
| 8 | 124068128 | rs7826050 | 0.342 | -0.499 | -0.296 | 0.684 | -0.001 | 0.011 | 0.011 | 0 | 0.74 | 0.74 | 0.71 | 0.67 | 0.72 | 0.65 |
| 8 | 124069629 | rs6470135 | 0.918 | 0.284 | -0.79 | -0.001 | -0.01 | 0.043 | 0.043 | 0 | 0.39 | 0.28 | 0.25 | 0.22 | 0.38 | 0.36 |
| 8 | 124070432 | rs17313372 | -0.624 | 0.412 | -1.045 | -0.098 | -0.023 | 0.084 | 0.084 | 0 | 0.22 | 0.06 | 0.24 | 0.06 | 0.24 | 0.25 |
| 8 | 124076688 | rs6992733 | -0.268 | 1.227 | -0.516 | 0.093 | 0.041 | -0.025 | -0.025 | 0 | 0.09 | 0.17 | 0.23 | 0.03 | 0.16 | 0.1 |
| 8 | 124080340 | rs4871342 | -0.892 | 0.862 | -0.926 | 0.38 | -0.001 | 0.012 | 0.012 | 0.305 | 0.56 | 0.68 | 0.68 | 0.62 | 0.56 | 0.61 |
| 8 | 124084685 | rs2280913 | 0.006 | 0.838 | -1.24 | 0.389 | -0.031 | -0.002 | -0.002 | 0.072 | 0.15 | 0.03 | 0.2 | 0.1 | 0.19 | 0.19 |
| 8 | 124089878 | rs2385161 | -0.761 | 1.038 | -1.126 | 0.491 | 0.021 | -0.014 | -0.014 | 0 | 0.61 | 0.81 | 0.73 | 0.57 | 0.67 | 0.62 |
| 8 | 124096227 | rs4871345 | -0.756 | 1.309 | -0.225 | 0.055 | 0.002 | -0.011 | -0.011 | 0 | 0.29 | 0.68 | 0.51 | 0.15 | 0.34 | 0.31 |
| 8 | 124107382 | rs11785137 | -0.135 | 1.512 | 1.122 | -0.004 | -0.006 | -0.008 | -0.008 | 1.221 | 0.21 | 0.24 | 0.24 | 0.15 | 0.19 | 0.24 |
| 8 | 124114147 | rs11786727 | -0.839 | 1.504 | 1.357 | -0.021 | 0.013 | -0.037 | -0.037 | 2.258 | 0.24 | 0.45 | 0.36 | 0.15 | 0.29 | 0.26 |
| 8 | 124115146 | rs10956118 | 0.649 | 1.431 | 1.25 | 0.102 | 0.001 | -0.004 | -0.004 | 0 | 0.2 | 0.25 | 0.16 | 0.17 | 0.19 | 0.21 |
| 8 | 124120097 | rs10505431 | 0.146 | 1.388 | 1.363 | 0.099 | 0.001 | 0.001 | 0.001 | 0 | 0.19 | 0.14 | 0.23 | 0.15 | 0.18 | 0.24 |
| 8 | 124121798 | rs16897969 | -0.456 | 1.417 | 1.343 | 0.104 | 0.008 | -0.03 | -0.03 | 0 | 0.24 | 0.39 | 0.36 | 0.15 | 0.29 | 0.26 |
| 8 | 124127595 | rs952656 | -0.371 | 1.439 | 1.353 | 0.107 | 0.008 | -0.022 | -0.022 | 0 | 0.23 | 0.33 | 0.35 | 0.15 | 0.29 | 0.26 |

| | | | | | | | | | | | | | | | | |
|---|---|---|---|---|---|---|---|---|---|---|---|---|---|---|---|---|
| 8 | 124128838 | rs4871351 | -1.192 | 1.179 | 1.122 | 0.001 | -0.001 | -0.002 | -0.002 | 0 | 0.48 | 0.5 | 0.61 | 0.36 | 0.43 | 0.53 |
| 8 | 124133518 | rs13252935 | -0.454 | 1.644 | 1.017 | 0.067 | 0.004 | -0.01 | -0.01 | 0.366 | 0.1 | 0.12 | 0.27 | 0.05 | 0.14 | 0.1 |
| 8 | 124135054 | rs13271524 | -0.477 | 1.63 | 1.063 | 0.06 | 0.017 | -0.026 | -0.026 | 0.182 | 0.1 | 0.17 | 0.28 | 0.05 | 0.14 | 0.1 |
| 8 | 124137984 | rs7824293 | 0.038 | 1.582 | 0.866 | 0.038 | -0.003 | 0.014 | 0.014 | 0.159 | 0.08 | 0.05 | 0.2 | 0.05 | 0.06 | 0.09 |
| 8 | 124138891 | rs16898013 | 0.166 | 1.609 | 0.968 | 0.051 | -0.001 | 0 | 0 | 0.385 | 0.08 | 0.08 | 0.2 | 0.05 | 0.06 | 0.09 |
| 8 | 124140634 | rs2385167 | 0.299 | 1.61 | 0.921 | 0.052 | -0.009 | 0.018 | 0.018 | 0.402 | 0.08 | 0.04 | 0.2 | 0.05 | 0.06 | 0.09 |
| 8 | 124145469 | rs10505429 | 0.308 | 1.595 | 1.019 | 0.064 | -0.003 | 0.006 | 0.006 | 1.406 | 0.2 | 0.15 | 0.15 | 0.14 | 0.19 | 0.2 |
| 8 | 124152245 | rs6470144 | -1.279 | 1.57 | 1.211 | 0.211 | 0.007 | -0.015 | -0.015 | 0 | 0.32 | 0.61 | 0.47 | 0.23 | 0.34 | 0.3 |
| 8 | 124154697 | rs10101626 | 0.295 | 1.539 | 1.228 | 0.298 | 0.001 | 0 | 0 | 0.962 | 0.2 | 0.25 | 0.16 | 0.18 | 0.19 | 0.2 |
| 8 | 124170826 | rs1713736 | 0.022 | 1.743 | -0.17 | 0.412 | -0.037 | -0.017 | -0.017 | 0 | 0.74 | 0.93 | 0.68 | 0.77 | 0.74 | 0.72 |
| 8 | 124172209 | rs1670189 | 0.319 | 1.797 | 0.494 | 0.229 | 0.028 | -0.025 | -0.025 | 2.072 | 0.73 | 0.63 | 0.69 | 0.83 | 0.68 | 0.76 |
| 8 | 124176245 | rs7008277 | 0.322 | 1.804 | -0.061 | 0.35 | -0.022 | -0.007 | -0.007 | 0.229 | 0.2 | 0.04 | 0.08 | 0.17 | 0.18 | 0.19 |
| 8 | 124177926 | rs3107193 | -0.164 | 1.825 | -0.088 | 0.378 | -0.044 | -0.005 | -0.005 | 0.161 | 0.78 | 0.95 | 0.79 | 0.82 | 0.76 | 0.76 |
| 8 | 124179301 | rs17323102 | 0.467 | 1.415 | -0.746 | -0.199 | 0.01 | 0.022 | 0.022 | 0 | 0.16 | 0.02 | 0.06 | 0.09 | 0.12 | 0.18 |
| 8 | 124183551 | rs7822076 | 0.01 | 1.604 | 0.224 | -0.201 | 0.014 | -0.011 | -0.011 | 0.802 | 0.23 | 0.27 | 0.14 | 0.17 | 0.25 | 0.2 |
| 8 | 124184067 | rs3107194 | 0.145 | 1.584 | 0.414 | -0.204 | 0.04 | -0.027 | -0.027 | 0 | 0.74 | 0.61 | 0.71 | 0.83 | 0.69 | 0.78 |
| 8 | 124190715 | rs1621182 | 1.41 | 1.068 | -0.773 | -0.839 | -0.082 | 0.135 | 0.135 | 1.234 | 0.28 | 0.01 | 0.22 | 0.04 | 0.33 | 0.25 |
| 8 | 124194968 | rs3739291 | 0.34 | 1.243 | -0.969 | -0.18 | -0.04 | -0.024 | -0.024 | 0.017 | 0.2 | 0.02 | 0.09 | 0.18 | 0.18 | 0.18 |
| 8 | 124199440 | rs1670171 | 0.077 | 1.217 | -0.815 | -0.154 | -0.013 | 0.004 | 0.004 | 0 | 0.76 | 0.85 | 0.84 | 0.79 | 0.72 | 0.8 |
| 8 | 124203215 | rs7833092 | 0.059 | 1.042 | -0.954 | -0.011 | -0.019 | -0.005 | -0.005 | 0.064 | 0.22 | 0.07 | 0.21 | 0.2 | 0.2 | 0.2 |
| 8 | 124206324 | rs7813708 | 0.128 | 1.045 | -0.969 | 0.115 | -0.047 | -0.045 | -0.045 | 0.033 | 0.22 | 0.02 | 0.2 | 0.21 | 0.2 | 0.2 |
| 8 | 124209157 | rs6997530 | -0.221 | 0.912 | 0.114 | 0.042 | -0.001 | -0.005 | -0.005 | 1.632 | 0.26 | 0.38 | 0.25 | 0.21 | 0.21 | 0.21 |
| 8 | 124213844 | rs11786273 | -0.194 | 0.903 | 0.143 | -0.028 | -0.001 | -0.043 | -0.043 | 0 | 0.26 | 0.32 | 0.25 | 0.09 | 0.21 | 0.2 |
| 8 | 124214051 | rs11779458 | -0.215 | 0.912 | 0.17 | -0.031 | 0 | -0.026 | -0.026 | 0 | 0.25 | 0.29 | 0.24 | 0.09 | 0.21 | 0.21 |
| 8 | 124219044 | rs1713730 | 0.668 | 1.026 | -0.227 | 0.309 | -0.04 | -0.024 | -0.024 | 0 | 0.39 | 0.11 | 0.49 | 0.64 | 0.4 | 0.47 |
| 8 | 124254528 | rs13248008 | -0.196 | 1.179 | 1.023 | -0.134 | -0.024 | 0.025 | 0.025 | 0.429 | 0.31 | 0.26 | 0.3 | 0.14 | 0.45 | 0.26 |
| 8 | 124264201 | rs1670177 | 1.649 | 0.796 | 0.641 | 0.088 | -0.08 | -0.015 | -0.015 | 0.597 | 0.3 | 0.01 | 0.19 | 0.19 | 0.31 | 0.34 |
| 8 | 124264927 | rs1044714 | 0.621 | 1.197 | 1.332 | -0.173 | -0.01 | 0.024 | 0.024 | 0.479 | 0.21 | 0.13 | 0.22 | 0.1 | 0.24 | 0.16 |
| 8 | 124300658 | rs11782961 | 0.688 | 1.385 | 1.348 | 0.191 | -0.024 | 0.011 | 0.011 | 0.598 | 0.2 | 0.06 | 0.22 | 0.16 | 0.24 | 0.17 |
| 8 | 124332900 | rs9021 | -0.62 | 1.324 | 1.747 | 0.78 | -0.001 | -0.039 | -0.039 | 0 | 0.6 | 0.76 | 0.52 | 0.52 | 0.64 | 0.52 |
| 8 | 124411678 | rs34573020 | 1.017 | 1.264 | -0.19 | 0.85 | -0.002 | -0.067 | -0.067 | 0 | 0.33 | 0.18 | 0.46 | 0.49 | 0.29 | 0.42 |
| 8 | 124416004 | rs6470146 | 0.551 | 1.586 | 0.641 | 0.849 | 0.033 | -0.063 | -0.063 | 5.563 | 0.81 | 0.72 | 0.83 | 0.97 | 0.62 | 0.86 |
| 8 | 124421397 | rs4307325 | -0.082 | 1.493 | 0.745 | 0.958 | 0.022 | -0.017 | -0.017 | 0.131 | 0.4 | 0.34 | 0.39 | 0.57 | 0.3 | 0.42 |
| 8 | 124426096 | rs3857957 | 0.831 | 1.504 | 0.731 | 0.871 | 0.085 | -0.095 | -0.095 | 1.895 | 0.79 | 0.52 | 0.81 | 0.92 | 0.62 | 0.86 |
| 8 | 124426145 | rs7821700 | 1.209 | 1.402 | 0.551 | 0.946 | 0.029 | -0.055 | -0.055 | 0 | 0.22 | 0.01 | 0.21 | 0.23 | 0.11 | 0.2 |
| 8 | 124429923 | rs3779989 | 0.936 | 1.984 | 1.212 | 0.44 | 0.076 | -0.077 | -0.077 | 0 | 0.8 | 0.65 | 0.82 | 0.93 | 0.63 | 0.86 |
| 8 | 124440174 | rs6999234 | 0.566 | 1.967 | 1.137 | 0.511 | 0.014 | -0.009 | -0.009 | 0.017 | 0.4 | 0.34 | 0.39 | 0.59 | 0.31 | 0.43 |
| 8 | 124441504 | rs7005105 | 0.413 | 2.066 | 1.242 | 0.548 | 0.024 | -0.027 | -0.027 | 0 | 0.93 | 0.84 | 0.94 | 0.99 | 0.89 | 0.96 |
| 8 | 124448689 | rs9297669 | 0.413 | 2.066 | 1.242 | 0.548 | 0.024 | -0.027 | -0.027 | 0.091 | 0.93 | 0.84 | 0.94 | 0.99 | 0.89 | 0.96 |
| 8 | 124448736 | rs7014678 | 0.575 | 2.066 | 1.205 | 0.551 | 0.014 | -0.009 | -0.009 | 0 | 0.4 | 0.34 | 0.39 | 0.59 | 0.31 | 0.43 |
| 8 | 124449466 | rs3824250 | 0.575 | 2.066 | 1.205 | 0.551 | 0.014 | -0.009 | -0.009 | 0 | 0.4 | 0.34 | 0.39 | 0.59 | 0.31 | 0.43 |
| 8 | 124453662 | rs13269287 | 0.575 | 2.066 | 1.205 | 0.551 | 0.014 | -0.009 | -0.009 | 0 | 0.4 | 0.34 | 0.39 | 0.59 | 0.31 | 0.43 |
| 8 | 124456727 | rs7822061 | 0.584 | 2.141 | 1.28 | 0.59 | 0.014 | -0.009 | -0.009 | 3.633 | 0.4 | 0.34 | 0.39 | 0.59 | 0.31 | 0.43 |
| 8 | 124460546 | rs13265801 | 0.581 | 2.153 | 1.286 | 0.596 | 0.014 | -0.009 | -0.009 | 0.229 | 0.4 | 0.34 | 0.39 | 0.59 | 0.31 | 0.43 |
| 8 | 124461256 | rs3824254 | 0.739 | 2.254 | 0.373 | 0.645 | -0.007 | -0.03 | -0.03 | 0 | 0.39 | 0.17 | 0.42 | 0.34 | 0.34 | 0.43 |
| 8 | 124473238 | rs3824252 | -0.662 | 2.009 | -0.086 | 0.851 | 0 | 0.002 | 0.002 | 0 | 0.54 | 0.58 | 0.51 | 0.65 | 0.55 | 0.52 |
| 8 | 124473894 | rs3901852 | -1.035 | 2.804 | -0.052 | 1.155 | -0.034 | 0.027 | 0.027 | 0 | 0.12 | 0.08 | 0.1 | 0.07 | 0.24 | 0.09 |
| 8 | 124490937 | rs2385273 | 1.507 | 1.944 | 0.489 | 1.673 | 0.017 | 0.064 | 0.064 | 0 | 0.94 | 0.8 | 0.81 | 0.83 | 0.91 | 0.88 |
| 8 | 124498734 | rs12548263 | 1.757 | 2.096 | 0.737 | 2.121 | -0.001 | 0.005 | 0.005 | 0.231 | 0.76 | 0.3 | 0.5 | 0.58 | 0.74 | 0.76 |
| 8 | 124510274 | rs2280915 | -1.524 | 1.787 | 0.719 | 2.453 | -0.001 | 0.065 | 0.065 | 0 | 0.05 | 0.08 | 0.21 | 0.25 | 0.06 | 0.07 |
| 8 | 124515854 | rs4870855 | -2.353 | 1.664 | 1.53 | 1.744 | 0.036 | 0.112 | 0.112 | 2.42 | 0.09 | 0.4 | 0.21 | 0.15 | 0.09 | 0.11 |
| 8 | 124517475 | rs10089389 | -1.347 | 1.705 | 0.909 | 1.597 | -0.001 | -0.001 | -0.001 | 0 | 0.06 | 0.09 | 0.04 | 0.05 | 0.06 | 0.05 |
| 8 | 124520424 | rs16898473 | -1.264 | 2.207 | 1.08 | 1.509 | 0.046 | -0.029 | -0.029 | 2.33 | 0.06 | 0.17 | 0.05 | 0.01 | 0.11 | 0.06 |
| 8 | 124525377 | rs3739288 | -1.747 | 2.258 | 1.003 | 2.451 | 0.031 | 0.235 | 0.235 | 0 | 0.08 | 0.21 | 0.19 | 0.48 | 0.1 | 0.11 |
| 8 | 124530373 | rs16898497 | -1.186 | 1.944 | 0.282 | 2.529 | 0 | 0.01 | 0.01 | 0.094 | 0.07 | 0.11 | 0.25 | 0.15 | 0.09 | 0.07 |
| 8 | 124535292 | rs16898508 | -1.575 | 1.969 | 0.412 | 2.215 | 0.05 | 0.186 | 0.186 | 0 | 0.11 | 0.28 | 0.2 | 0.42 | 0.16 | 0.12 |
| 8 | 124540644 | rs4242341 | -0.086 | 1.743 | -1.135 | 1.464 | -0.062 | -0.075 | -0.075 | 0.072 | 0.61 | 0.92 | 0.59 | 0.56 | 0.6 | 0.47 |
| 8 | 124542485 | rs7830622 | -0.302 | 1.632 | -1.035 | 1.506 | 0.001 | 0 | 0 | 0.383 | 0.06 | 0.06 | 0.02 | 0.15 | 0.04 | 0.07 |
| 8 | 124545147 | rs2294090 | -0.686 | 2.051 | -0.47 | 0.842 | 0.032 | -0.01 | -0.01 | 3.467 | 0.32 | 0.6 | 0.42 | 0.25 | 0.48 | 0.36 |
| 8 | 124552303 | rs7835298 | -0.719 | 2.433 | 0.259 | 0.831 | 0.05 | 0.018 | 0.018 | 0.998 | 0.29 | 0.41 | 0.39 | 0.36 | 0.44 | 0.31 |
| 8 | 124555235 | rs6470154 | -1.204 | 1.807 | -0.345 | 0.449 | 0.038 | -0.014 | -0.014 | 0.122 | 0.55 | 0.9 | 0.59 | 0.5 | 0.63 | 0.46 |
| 8 | 124555833 | rs2891779 | 0.641 | 1.515 | -0.207 | 0.461 | 0.018 | 0.027 | 0.027 | 0.24 | 0.19 | 0.12 | 0.13 | 0.15 | 0.14 | 0.09 |
| 8 | 124556006 | rs16898553 | -0.428 | 2.006 | -0.01 | 0.534 | -0.025 | -0.041 | -0.041 | 0 | 0.2 | 0.1 | 0.15 | 0.33 | 0.24 | 0.28 |
| 8 | 124565203 | rs6990663 | -0.857 | 1.804 | -0.192 | 0.288 | 0.005 | -0.005 | -0.005 | 0 | 0.43 | 0.62 | 0.53 | 0.43 | 0.59 | 0.58 |
| 8 | 124565311 | rs16898561 | 1.245 | 1.785 | -0.177 | 0.373 | 0.004 | 0.027 | 0.027 | 0 | 0.11 | 0.12 | 0.08 | 0.18 | 0.1 | 0.25 |
| 8 | 124566356 | rs6996389 | -0.325 | 2.164 | 0.149 | 0.309 | -0.024 | 0.008 | 0.008 | 0 | 0.25 | 0.19 | 0.24 | 0.23 | 0.32 | 0.3 |
| 8 | 124571581 | rs12541094 | -0.847 | 1.989 | 0.37 | 0.307 | 0.004 | 0.003 | 0.003 | 0 | 0.37 | 0.41 | 0.32 | 0.36 | 0.41 | 0.53 |
| 8 | 124574279 | rs4388446 | -1.507 | 1.878 | 0.525 | 0.127 | 0.036 | 0.001 | 0.001 | 0.462 | 0.48 | 0.63 | 0.58 | 0.39 | 0.64 | 0.6 |
| 8 | 124580116 | rs7816339 | -0.291 | 0.784 | -0.129 | -0.47 | 0.004 | 0.003 | 0.003 | 0.129 | 0.57 | 0.66 | 0.57 | 0.6 | 0.58 | 0.63 |

| | | | | | | | | | | | | | | | | |
|---|---|---|---|---|---|---|---|---|---|---|---|---|---|---|---|---|
| 8 | 124581970 | rs7844912 | -0.621 | 0.634 | -0.108 | -0.453 | 0.013 | 0 | 0 | 0.477 | 0.62 | 0.68 | 0.76 | 0.62 | 0.75 | 0.66 |
| 8 | 124583930 | rs16898583 | 1.631 | 0.797 | -0.063 | -0.347 | 0.004 | 0.051 | 0.051 | 0 | 0.12 | 0.2 | 0.08 | 0.18 | 0.09 | 0.23 |
| 8 | 124586305 | rs2385296 | 0.397 | 0.946 | 0.029 | -0.508 | -0.001 | -0.002 | -0.002 | 1.621 | 0.45 | 0.44 | 0.47 | 0.6 | 0.46 | 0.53 |
| 8 | 124587044 | rs16898590 | 1.702 | 0.398 | 0.141 | -0.243 | 0.007 | 0.055 | 0.055 | 0 | 0.11 | 0.16 | 0.08 | 0.22 | 0.09 | 0.26 |
| 8 | 124591674 | rs16898594 | 1.296 | 0.052 | 0.102 | -0.273 | 0 | 0.041 | 0.041 | 0 | 0.12 | 0.16 | 0.04 | 0.19 | 0.1 | 0.25 |
| 8 | 124591925 | rs17253999 | 0.625 | 0.137 | -0.339 | -0.253 | -0.087 | -0.014 | -0.014 | 0 | 0.24 | 0.1 | 0.37 | 0.24 | 0.42 | 0.3 |
| 8 | 124592374 | rs4870857 | 0.911 | -0.389 | -0.189 | -0.545 | 0.011 | 0.005 | 0.005 | 0 | 0.48 | 0.44 | 0.52 | 0.43 | 0.38 | 0.35 |
| 8 | 124593007 | rs2385456 | 0.876 | -0.317 | -0.729 | -0.538 | -0.088 | -0.013 | -0.013 | 0 | 0.24 | 0.12 | 0.37 | 0.24 | 0.42 | 0.3 |
| 8 | 124593490 | rs4382457 | -1.892 | -0.785 | -0.708 | -0.434 | -0.016 | -0.002 | -0.002 | 0 | 0.87 | 0.83 | 0.95 | 0.86 | 0.95 | 0.93 |
| 8 | 124600993 | rs4870858 | 1.162 | -0.47 | -0.7 | -0.374 | 0.058 | -0.013 | -0.013 | 0.183 | 0.41 | 0.19 | 0.44 | 0.43 | 0.31 | 0.27 |
| 8 | 124601535 | rs12547541 | 1.629 | -0.275 | -0.662 | -0.154 | 0.025 | -0.045 | -0.045 | 0 | 0.28 | 0.15 | 0.37 | 0.43 | 0.2 | 0.23 |
| 8 | 124604759 | rs9650084 | 1.177 | 0.221 | -0.664 | -0.441 | -0.002 | -0.005 | -0.005 | 0.505 | 0.06 | 0.07 | 0.06 | 0 | 0.07 | 0.09 |
| 8 | 124608162 | rs2891781 | 0.641 | 0.145 | -0.604 | -0.342 | 0.007 | 0.014 | 0.014 | 0.148 | 0.48 | 0.5 | 0.47 | 0.62 | 0.38 | 0.57 |
| 8 | 124614367 | rs9656946 | -0.219 | -0.506 | -0.696 | -0.513 | 0.032 | 0.033 | 0.033 | 2.793 | 0.75 | 0.82 | 0.89 | 0.88 | 0.85 | 0.92 |
| 8 | 124616796 | rs6470161 | -0.285 | -0.409 | -0.278 | -0.73 | -0.021 | -0.032 | -0.032 | 0 | 0.75 | 0.65 | 0.86 | 0.9 | 0.81 | 0.93 |
| 8 | 124618652 | rs2385295 | 0.869 | -0.002 | -0.293 | -0.572 | 0.012 | 0.029 | 0.029 | 0.076 | 0.57 | 0.28 | 0.56 | 0.68 | 0.62 | 0.58 |
| 8 | 124624056 | rs12542467 | 1.042 | 0.201 | -0.38 | -0.695 | -0.019 | 0.061 | 0.061 | 0 | 0.53 | 0.21 | 0.53 | 0.68 | 0.55 | 0.55 |
| 8 | 124626983 | rs4871396 | -1.145 | 0.371 | -0.532 | -0.748 | -0.028 | 0 | 0 | 0.204 | 0.84 | 0.91 | 0.9 | 0.89 | 0.83 | 0.87 |
| 8 | 124630352 | rs7821932 | 0.549 | 0.256 | 0.273 | -0.85 | -0.007 | -0.073 | -0.073 | 0 | 0.13 | 0.26 | 0.07 | 0 | 0.1 | 0.06 |
| 8 | 124631864 | rs2385441 | -0.05 | 0.297 | -0.296 | -0.815 | -0.003 | -0.046 | -0.046 | 0 | 0.73 | 0.89 | 0.82 | 0.72 | 0.78 | 0.68 |
| 8 | 124637959 | rs16898635 | 0.645 | 0.403 | -0.026 | -0.719 | -0.013 | -0.046 | -0.046 | 1.005 | 0.12 | 0.01 | 0.09 | 0.17 | 0.1 | 0.19 |
| 8 | 124641220 | rs7001678 | -0.279 | 0.225 | 0.488 | -0.831 | 0.025 | -0.091 | -0.091 | 0 | 0.17 | 0.48 | 0.25 | 0.05 | 0.18 | 0.13 |
| 8 | 124646022 | rs12545556 | 0.192 | -0.001 | 0.528 | -0.781 | 0.019 | -0.051 | -0.051 | 0.213 | 0.13 | 0.45 | 0.22 | 0.04 | 0.12 | 0.09 |
| 8 | 124647861 | rs4871398 | 0.227 | -0.034 | 0.067 | -0.806 | -0.004 | -0.002 | -0.002 | 0 | 0.7 | 0.75 | 0.78 | 0.72 | 0.72 | 0.6 |
| 8 | 124648032 | rs2385463 | 0.67 | -0.154 | 0.209 | -0.786 | -0.002 | 0.002 | 0.002 | 0.006 | 0.65 | 0.64 | 0.74 | 0.73 | 0.66 | 0.57 |
| 8 | 124648269 | rs1985958 | -1.381 | -0.185 | -0.246 | -0.254 | -0.033 | 0.031 | 0.031 | 0 | 0.47 | 0.85 | 0.5 | 0.57 | 0.47 | 0.6 |
| 8 | 124649459 | rs10956139 | 0.377 | 0.259 | 0.012 | 0.026 | -0.009 | 0.002 | 0.002 | 0 | 0.61 | 0.68 | 0.59 | 0.63 | 0.6 | 0.56 |
| 8 | 124650289 | rs16898657 | -0.768 | 0.329 | 0.652 | 0.308 | 0.075 | 0.142 | 0.142 | 0.284 | 0.1 | 0.35 | 0.11 | 0.15 | 0.11 | 0.09 |
| 8 | 124654560 | rs7388387 | 0.815 | 0.187 | 0.326 | 0.252 | 0 | 0.053 | 0.053 | 0 | 0.78 | 0.7 | 0.67 | 0.58 | 0.78 | 0.65 |
| 8 | 124656137 | rs16898668 | 1.234 | 0.145 | 0.047 | 0.291 | 0.007 | -0.004 | -0.004 | 5.988 | 0.08 | 0.05 | 0.07 | 0.22 | 0.03 | 0.2 |
| 8 | 124658210 | rs16898671 | 0.741 | 0.038 | 0.174 | -0.034 | 0.012 | 0.04 | 0.04 | 0 | 0.12 | 0.21 | 0.08 | 0.16 | 0.1 | 0.2 |
| 8 | 124659121 | rs1122389 | 1.132 | -0.099 | -0.235 | -0.324 | -0.014 | 0.025 | 0.025 | 1.064 | 0.23 | 0.18 | 0.15 | 0.1 | 0.26 | 0.18 |
| 8 | 124659329 | rs11991744 | 0.131 | -0.366 | 0.273 | 0.022 | -0.012 | 0.016 | 0.016 | 0 | 0.84 | 0.82 | 0.77 | 0.72 | 0.89 | 0.73 |
| 8 | 124661974 | rs4871404 | 0.272 | -0.12 | 0.155 | -0.119 | 0.02 | 0.016 | 0.016 | 0 | 0.19 | 0.33 | 0.11 | 0.17 | 0.17 | 0.26 |
| 8 | 124662772 | rs4871405 | -0.55 | -0.1 | -0.161 | -0.295 | -0.001 | 0 | 0 | 0 | 0.93 | 0.89 | 0.84 | 0.93 | 0.93 | 0.94 |
| 8 | 124663895 | rs16898689 | -0.329 | -0.313 | -0.184 | -0.358 | -0.036 | 0.022 | 0.022 | 0 | 0.15 | 0.23 | 0.13 | 0.22 | 0.07 | 0.24 |
| 8 | 124664130 | rs11780509 | 0.847 | -0.238 | -0.326 | -0.379 | -0.023 | 0.006 | 0.006 | 0 | 0.75 | 0.66 | 0.69 | 0.71 | 0.8 | 0.68 |
| 8 | 124664167 | rs11784192 | 0.661 | -0.23 | -0.322 | -0.375 | -0.042 | 0.024 | 0.024 | 0 | 0.78 | 0.66 | 0.71 | 0.71 | 0.86 | 0.7 |
| 8 | 124664792 | rs7387544 | 0.585 | -0.206 | -0.277 | -0.363 | -0.023 | 0.007 | 0.007 | 0 | 0.75 | 0.67 | 0.69 | 0.71 | 0.8 | 0.68 |
| 8 | 124666429 | rs4367519 | 0.749 | -0.065 | -0.587 | -0.127 | 0.019 | -0.023 | -0.023 | 0 | 0.09 | 0.04 | 0.08 | 0.23 | 0.05 | 0.21 |
| 8 | 124675680 | rs11997295 | 0.086 | 0.246 | -0.372 | -0.157 | 0.018 | 0.002 | 0.002 | 0.277 | 0.2 | 0.43 | 0.12 | 0.14 | 0.17 | 0.27 |
| 8 | 124683734 | rs10429365 | 0.405 | 0.442 | -1.209 | -0.07 | 0.001 | -0.019 | -0.019 | 0 | 0.55 | 0.23 | 0.55 | 0.52 | 0.57 | 0.45 |
| 8 | 124685509 | rs4871408 | -0.351 | 0.356 | -1.143 | -0.337 | 0 | 0 | 0 | 0 | 0.34 | 0.65 | 0.5 | 0.32 | 0.31 | 0.33 |
| 8 | 124688771 | rs9643184 | 0.651 | 0.3 | -1.789 | -0.396 | 0.019 | -0.002 | -0.002 | 0 | 0.29 | 0.07 | 0.18 | 0.28 | 0.22 | 0.29 |
| 8 | 124689847 | rs4870864 | -0.536 | 1.305 | -1.605 | 0.132 | -0.037 | -0.113 | -0.113 | 0 | 0.74 | 0.92 | 0.73 | 0.48 | 0.69 | 0.72 |
| 8 | 124690960 | rs4242345 | 0.332 | 1.217 | -1.033 | -0.008 | 0.012 | -0.01 | -0.01 | 0 | 0.58 | 0.53 | 0.4 | 0.38 | 0.5 | 0.62 |
| 8 | 124693444 | rs3739283 | -0.276 | 1.303 | -1.531 | 0.276 | -0.016 | -0.044 | -0.044 | 0 | 0.75 | 0.85 | 0.72 | 0.44 | 0.69 | 0.71 |
| 8 | 124693641 | rs4314634 | 0.419 | 1.183 | -1.703 | -0.157 | -0.003 | -0.001 | -0.001 | 0.154 | 0.25 | 0.28 | 0.34 | 0.08 | 0.28 | 0.26 |
| 8 | 124695748 | rs7459561 | -0.456 | 1.251 | -2.031 | 0.485 | -0.03 | -0.134 | -0.134 | 0.015 | 0.78 | 0.96 | 0.69 | 0.46 | 0.76 | 0.77 |
| 8 | 124696867 | rs2294015 | 0.58 | 1.233 | -2.663 | 0.508 | -0.109 | -0.123 | -0.123 | 0 | 0.66 | 0.94 | 0.66 | 0.5 | 0.56 | 0.74 |
| 8 | 124699175 | rs17256536 | 0.021 | 0.464 | -2.721 | -0.105 | 0.018 | 0.049 | 0.049 | 2.287 | 0.43 | 0.24 | 0.35 | 0.3 | 0.4 | 0.32 |
| 8 | 124700878 | rs3739282 | 0.264 | 0.48 | -2.367 | -0.428 | 0.002 | 0.007 | 0.007 | 0 | 0.32 | 0.26 | 0.41 | 0.15 | 0.3 | 0.36 |
| 8 | 124704361 | rs7000525 | 0.49 | 0.204 | -2.345 | -0.526 | -0.001 | 0.039 | 0.039 | 0.174 | 0.31 | 0.12 | 0.16 | 0.21 | 0.3 | 0.17 |
| 8 | 124708875 | rs6988728 | 1.873 | -0.363 | -2.708 | -0.638 | -0.107 | -0.085 | -0.085 | 0.493 | 0.21 | 0.01 | 0.29 | 0.26 | 0.26 | 0.45 |
| 8 | 124709501 | rs3779980 | -0.229 | -0.703 | -2.359 | -1.407 | 0.022 | 0.064 | 0.064 | 0 | 0.48 | 0.69 | 0.63 | 0.3 | 0.52 | 0.69 |
| 8 | 124709857 | rs7386945 | -1.47 | -0.778 | -2.631 | -1.461 | -0.038 | -0.069 | -0.069 | 0 | 0.89 | 0.99 | 0.86 | 0.5 | 0.87 | 0.92 |
| 8 | 124710664 | rs6995099 | 1.611 | -1.01 | -2.15 | -1.378 | -0.001 | -0.018 | -0.018 | 0 | 0.07 | 0.06 | 0.06 | 0.01 | 0.08 | 0.04 |
| 8 | 124710729 | rs2294013 | 0.885 | -1.109 | -2.069 | -1.459 | 0.028 | 0.066 | 0.066 | 0.133 | 0.37 | 0.21 | 0.21 | 0.19 | 0.34 | 0.21 |
| 8 | 124711210 | rs1080109 | -0.667 | -0.965 | -1.181 | -1.384 | -0.004 | -0.12 | -0.12 | 0 | 0.81 | 0.95 | 0.79 | 0.47 | 0.82 | 0.88 |
| 8 | 124711414 | rs4601291 | -0.373 | -1.12 | -0.761 | -1.421 | 0.01 | -0.074 | -0.074 | 1.315 | 0.25 | 0.47 | 0.32 | 0.1 | 0.26 | 0.24 |
| 8 | 124712699 | rs4581037 | 0.502 | -1.066 | -0.366 | -1.366 | 0 | -0.031 | -0.031 | 0 | 0.38 | 0.48 | 0.39 | 0.22 | 0.36 | 0.53 |
| 8 | 124712992 | rs6470175 | 0.176 | -0.976 | -0.597 | -1.346 | -0.002 | 0.015 | 0.015 | 0 | 0.43 | 0.57 | 0.45 | 0.22 | 0.43 | 0.54 |
| 8 | 124715120 | rs17257150 | 0.922 | -0.778 | -0.777 | -1.245 | -0.01 | 0.005 | 0.005 | 0 | 0.13 | 0.12 | 0.25 | 0.07 | 0.18 | 0.17 |
| 8 | 124716201 | rs11786766 | 1.207 | -0.902 | -0.703 | -1.288 | 0.011 | 0.029 | 0.029 | 0.015 | 0.28 | 0.17 | 0.13 | 0.18 | 0.26 | 0.19 |
| 8 | 124717641 | rs11774189 | 0.477 | -0.793 | -0.494 | -1.146 | 0.016 | 0.033 | 0.033 | 0 | 0.36 | 0.28 | 0.21 | 0.19 | 0.34 | 0.22 |
| 8 | 124718333 | rs16898757 | -0.318 | -0.19 | -0.649 | -0.851 | -0.002 | -0.039 | -0.039 | 1.308 | 0.08 | 0.22 | 0.04 | 0.01 | 0.03 | 0.03 |
| 8 | 124725328 | rs13277433 | 1.027 | 0.009 | -1.601 | -0.54 | -0.101 | -0.056 | -0.056 | 0 | 0.26 | 0.01 | 0.31 | 0.24 | 0.28 | 0.43 |
| 8 | 124732388 | rs2010301 | 1.207 | 0.327 | -0.874 | -0.386 | -0.011 | 0.052 | 0.052 | 1.963 | 0.09 | 0.01 | 0.03 | 0 | 0.09 | 0.01 |
| 8 | 124733999 | rs11998151 | -0.585 | 0.156 | -0.617 | -0.529 | -0.001 | 0.029 | 0.029 | 0 | 0.36 | 0.31 | 0.26 | 0.18 | 0.37 | 0.25 |

| | | | | | | | | | | | | | | | | |
|---|---|---|---|---|---|---|---|---|---|---|---|---|---|---|---|---|
| 8 | 124734801 | rs3898733 | 0.47 | 0.211 | -0.468 | -0.563 | -0.002 | 0.003 | 0.003 | 0 | 0.26 | 0.27 | 0.2 | 0.16 | 0.28 | 0.2 |
| 8 | 124741583 | rs17258233 | 0.53 | 0.419 | -0.386 | -1.038 | 0.001 | -0.014 | -0.014 | 0 | 0.24 | 0.26 | 0.26 | 0.08 | 0.25 | 0.17 |
| 8 | 124743796 | rs4006384 | 0.35 | -0.165 | -0.048 | -0.91 | -0.001 | -0.028 | -0.028 | 0.719 | 0.13 | 0.12 | 0.1 | 0 | 0.14 | 0.06 |
| 8 | 124744639 | rs2385507 | -0.288 | -0.305 | -0.068 | -0.937 | 0.003 | 0.061 | 0.061 | 0.81 | 0.52 | 0.34 | 0.4 | 0.26 | 0.5 | 0.56 |
| 8 | 124749610 | rs6470179 | -1.114 | -0.304 | -0.066 | -0.965 | 0.007 | 0.012 | 0.012 | 3.117 | 0.88 | 0.94 | 0.94 | 1 | 0.9 | 0.95 |
| 8 | 124749704 | rs10956146 | 0.366 | -0.324 | 0.003 | -0.959 | -0.009 | 0.053 | 0.053 | 0.01 | 0.44 | 0.33 | 0.57 | 0.72 | 0.5 | 0.42 |
| 8 | 124750521 | rs13279803 | 0.337 | -0.173 | -0.53 | -0.882 | 0.012 | 0.05 | 0.05 | 0 | 0.42 | 0.28 | 0.34 | 0.26 | 0.4 | 0.5 |
| 8 | 124754410 | rs1030590 | 0.22 | 0.149 | -1.343 | -0.628 | -0.006 | 0.042 | 0.042 | 0.012 | 0.44 | 0.35 | 0.57 | 0.72 | 0.5 | 0.42 |
| 8 | 124757661 | rs17350191 | 0.049 | 0.359 | -1.927 | -0.378 | -0.043 | 0.087 | 0.087 | 0 | 0.36 | 0.03 | 0.28 | 0.2 | 0.33 | 0.47 |
| 8 | 124772451 | rs1978681 | 0.846 | 0.569 | -0.451 | -0.172 | 0.035 | -0.014 | -0.014 | 0.04 | 0.95 | 0.8 | 0.96 | 1 | 0.91 | 0.92 |
| 8 | 124776245 | rs12550235 | -0.832 | 0.494 | 0.43 | 0.268 | 0.019 | 0.141 | 0.141 | 0.618 | 0.14 | 0.3 | 0.33 | 0.29 | 0.17 | 0.24 |
| 8 | 124793043 | rs2163534 | 1.127 | 0.274 | -0.33 | -0.336 | -0.091 | -0.066 | -0.066 | 0 | 0.41 | 0.06 | 0.39 | 0.43 | 0.42 | 0.49 |
| 8 | 124804669 | rs7828192 | -1.559 | -0.216 | 0.516 | -0.173 | 0.005 | -0.006 | -0.006 | 0.768 | 0.25 | 0.76 | 0.4 | 0.33 | 0.23 | 0.27 |
| 8 | 124806084 | rs16898871 | -0.295 | -0.224 | 0.607 | -0.165 | 0.009 | 0.116 | 0.116 | 1.379 | 0.14 | 0.26 | 0.31 | 0.32 | 0.17 | 0.24 |
| 8 | 124814272 | rs17260111 | 1.337 | -0.288 | 0.323 | -0.352 | -0.059 | 0.087 | 0.087 | 0.01 | 0.15 | 0.01 | 0.08 | 0.01 | 0.16 | 0.05 |
| 8 | 124814481 | rs4871415 | -0.951 | -0.638 | 1.205 | 0.468 | 0.013 | 0.146 | 0.146 | 2.34 | 0.15 | 0.34 | 0.32 | 0.32 | 0.16 | 0.24 |
| 8 | 124817145 | rs11991217 | -0.746 | -0.648 | 1.287 | 0.454 | -0.012 | -0.045 | -0.045 | 0.965 | 0.09 | 0.31 | 0.07 | 0.01 | 0.07 | 0.03 |
| 8 | 124824581 | rs2060112 | -1.093 | -0.59 | 1.235 | 0.499 | 0.005 | 0.217 | 0.217 | 3.71 | 0.15 | 0.45 | 0.24 | 0.31 | 0.14 | 0.21 |
| 8 | 124829395 | rs2001966 | 0.313 | -1.105 | 1.223 | 0.883 | -0.045 | -0.085 | -0.085 | 0.141 | 0.13 | 0.01 | 0.15 | 0.29 | 0.15 | 0.2 |
| 8 | 124831459 | rs10481071 | -2.302 | -1.255 | 2.096 | 1.011 | -0.014 | 0.171 | 0.171 | 3.497 | 0.13 | 0.53 | 0.27 | 0.31 | 0.09 | 0.22 |
| 8 | 124831834 | rs10481151 | -2.267 | -1.24 | 2.104 | 1.019 | -0.012 | 0.175 | 0.175 | 3.499 | 0.13 | 0.53 | 0.27 | 0.31 | 0.09 | 0.22 |
| 8 | 124833311 | rs3889940 | -1.921 | -1.003 | 1.932 | 0.892 | 0.003 | -0.005 | -0.005 | 0 | 0.06 | 0.27 | 0.03 | 0.01 | 0.03 | 0.03 |
| 8 | 124834384 | rs11997795 | -1.998 | -1.017 | 1.579 | 0.873 | -0.023 | -0.068 | -0.068 | 1.38 | 0.11 | 0.52 | 0.09 | 0.01 | 0.07 | 0.04 |
| 8 | 124835357 | rs964059 | 2.032 | -0.76 | 1.694 | 1.01 | 0.007 | -0.069 | -0.069 | 1.25 | 0.88 | 0.54 | 0.91 | 0.99 | 0.87 | 0.87 |
| 8 | 124837723 | rs9792207 | -0.441 | -0.776 | 1.255 | 1.214 | 0.006 | -0.066 | -0.066 | 0 | 0.12 | 0.01 | 0.15 | 0.18 | 0.06 | 0.08 |
| 8 | 124845645 | rs1431668 | 2.286 | 0.08 | 2.029 | 0.993 | -0.007 | -0.09 | -0.09 | 0 | 0.86 | 0.52 | 0.89 | 0.99 | 0.87 | 0.83 |
| 8 | 124846425 | rs6470188 | 1.734 | 0.21 | 1.299 | 1.093 | -0.005 | -0.023 | -0.023 | 0 | 0.92 | 0.91 | 0.96 | 0.99 | 0.91 | 0.84 |
| 8 | 124850733 | rs6470189 | 1.676 | 0.322 | 1.369 | 1.374 | 0.003 | 0.197 | 0.197 | 0 | 0.86 | 0.68 | 0.75 | 0.61 | 0.83 | 0.68 |
| 8 | 124855776 | rs1431671 | 1.039 | 0.625 | 0.898 | 1.457 | 0.01 | 0.095 | 0.095 | 0 | 0.88 | 0.81 | 0.74 | 0.66 | 0.83 | 0.67 |
| 8 | 124857300 | rs6470190 | 1.049 | 0.645 | 0.493 | 1.467 | 0.04 | 0.233 | 0.233 | 0.023 | 0.86 | 0.58 | 0.73 | 0.66 | 0.83 | 0.7 |
| 8 | 124861459 | rs12677352 | -0.975 | 0.645 | -0.313 | 1.088 | -0.001 | 0.005 | 0.005 | 0 | 0.1 | 0.09 | 0.04 | 0.18 | 0.08 | 0.12 |
| 8 | 124862503 | rs6470192 | -0.054 | 1.067 | -0.598 | 0.335 | 0 | 0.014 | 0.014 | 0 | 0.51 | 0.5 | 0.32 | 0.29 | 0.49 | 0.36 |
| 8 | 124862604 | rs7006880 | -0.369 | 0.96 | -0.701 | 0.711 | 0.003 | 0.019 | 0.019 | 0 | 0.76 | 0.64 | 0.68 | 0.67 | 0.73 | 0.69 |
| 8 | 124864404 | rs1897318 | 0.589 | 0.858 | -0.815 | -0.036 | -0.007 | 0.015 | 0.015 | 0 | 0.4 | 0.33 | 0.28 | 0.14 | 0.45 | 0.22 |
| 8 | 124864868 | rs17352760 | -1.081 | 0.995 | -0.889 | 0.006 | -0.005 | 0.004 | 0.004 | 0 | 0.07 | 0.03 | 0.1 | 0.02 | 0.07 | 0.07 |
| 8 | 124865882 | rs17352788 | -0.481 | 0.665 | -1.3 | -0.626 | 0.001 | 0.061 | 0.061 | 0 | 0.22 | 0.14 | 0.13 | 0.03 | 0.26 | 0.1 |
| 8 | 124866062 | rs2385522 | -0.452 | 0.515 | -0.991 | -0.962 | -0.007 | -0.023 | -0.023 | 0 | 0.55 | 0.55 | 0.57 | 0.75 | 0.65 | 0.53 |
| 8 | 124867374 | rs16893043 | 0.047 | 0.188 | -0.441 | -0.9 | -0.006 | -0.013 | -0.013 | 0 | 0.14 | 0.19 | 0.1 | 0.06 | 0.13 | 0.12 |
| 8 | 124882382 | rs7823205 | 0.948 | -0.277 | -0.657 | -1.288 | 0 | 0.172 | 0.172 | 0.962 | 0.4 | 0.11 | 0.29 | 0.15 | 0.35 | 0.36 |
| 8 | 124882439 | rs6470196 | -0.264 | -0.326 | -0.096 | -1.226 | -0.035 | 0.039 | 0.039 | 0 | 0.63 | 0.69 | 0.46 | 0.22 | 0.5 | 0.49 |
| 8 | 124895809 | rs7843017 | -0.799 | -0.051 | -0.141 | -0.775 | -0.01 | -0.005 | -0.005 | 0.058 | 0.77 | 0.82 | 0.67 | 0.71 | 0.73 | 0.85 |
| 8 | 124896389 | rs6980985 | 0.361 | -0.144 | 0.823 | -0.851 | -0.003 | -0.006 | -0.006 | 0 | 0.59 | 0.54 | 0.53 | 0.63 | 0.6 | 0.73 |
| 8 | 124898469 | rs4871417 | 0.867 | -0.073 | 0.832 | -0.772 | 0.004 | 0.035 | 0.035 | 0 | 0.15 | 0.19 | 0.2 | 0.46 | 0.22 | 0.36 |
| 8 | 124907620 | rs7012454 | 0.788 | -0.084 | 0.756 | -0.82 | -0.008 | -0.012 | -0.012 | 0 | 0.14 | 0.18 | 0.13 | 0.08 | 0.13 | 0.13 |
| 8 | 124908497 | rs10111524 | -0.022 | -0.101 | 0.938 | -0.753 | -0.001 | -0.001 | -0.001 | 2.003 | 0.64 | 0.63 | 0.61 | 0.59 | 0.61 | 0.73 |
| 8 | 124919648 | rs7823126 | 0.311 | -0.112 | 0.99 | -0.76 | -0.005 | -0.004 | -0.004 | 0 | 0.59 | 0.46 | 0.58 | 0.59 | 0.6 | 0.72 |
| 8 | 124922448 | rs16899029 | 0.67 | -0.034 | 1.024 | -0.664 | 0.039 | 0.12 | 0.12 | 0 | 0.16 | 0.3 | 0.27 | 0.44 | 0.22 | 0.35 |
| 8 | 124922736 | rs11779980 | 1.073 | -0.062 | 0.512 | -0.743 | 0.033 | 0.145 | 0.145 | 0.781 | 0.46 | 0.18 | 0.31 | 0.15 | 0.38 | 0.36 |
| 8 | 124924961 | rs4388448 | -0.171 | -0.217 | 0.936 | -0.675 | 0.001 | 0.008 | 0.008 | 0.104 | 0.57 | 0.65 | 0.57 | 0.59 | 0.54 | 0.69 |
| 8 | 124927711 | rs4355755 | -0.199 | -0.1 | 1.19 | -0.694 | 0.052 | 0.168 | 0.168 | 0 | 0.21 | 0.43 | 0.33 | 0.44 | 0.26 | 0.42 |
| 8 | 124930887 | rs12541518 | 0.286 | -0.265 | 1.187 | -0.729 | 0.016 | 0.048 | 0.048 | 0 | 0.51 | 0.62 | 0.38 | 0.22 | 0.44 | 0.42 |
| 8 | 124932905 | rs4242349 | -1.15 | -0.228 | 1.158 | -0.476 | 0.003 | 0.006 | 0.006 | 0 | 0.36 | 0.57 | 0.45 | 0.52 | 0.39 | 0.56 |
| 8 | 124934558 | rs746747 | 0.158 | 0.067 | 1.138 | -0.102 | -0.017 | -0.017 | -0.017 | 0.81 | 0.14 | 0.13 | 0.27 | 0.42 | 0.21 | 0.4 |
| 8 | 124938412 | rs4871429 | -1.168 | -0.152 | 1.283 | -0.421 | 0.004 | 0.011 | 0.011 | 0 | 0.32 | 0.33 | 0.43 | 0.45 | 0.4 | 0.52 |
| 8 | 124939645 | rs16899060 | 0.558 | -0.163 | 1.41 | -0.436 | -0.002 | -0.062 | -0.062 | 0 | 0.14 | 0.19 | 0.12 | 0 | 0.15 | 0.07 |
| 8 | 124945090 | rs4537289 | 0.246 | -0.09 | 0.84 | -0.326 | -0.043 | -0.071 | -0.071 | 0 | 0.15 | 0.08 | 0.26 | 0.44 | 0.21 | 0.4 |
| 8 | 124946585 | rs7842762 | -1.293 | -0.113 | 0.811 | -0.349 | 0.007 | 0.028 | 0.028 | 0.128 | 0.37 | 0.49 | 0.45 | 0.45 | 0.41 | 0.51 |
| 8 | 124946823 | rs7018193 | 0.28 | -0.053 | 0.647 | -0.461 | -0.002 | -0.043 | -0.043 | 0 | 0.17 | 0.19 | 0.15 | 0.01 | 0.19 | 0.12 |
| 8 | 124948042 | rs11774078 | 0.682 | 0.263 | 0.37 | -0.047 | -0.003 | -0.01 | -0.01 | 2.025 | 0.42 | 0.57 | 0.35 | 0.26 | 0.36 | 0.44 |
| 8 | 124948479 | rs16899111 | 0.466 | 0.506 | 0.088 | 0.183 | -0.005 | -0.013 | -0.013 | 0 | 0.47 | 0.53 | 0.36 | 0.29 | 0.37 | 0.44 |
| 8 | 124951527 | rs4870880 | -1.101 | 0.763 | -0.565 | 0.873 | -0.003 | -0.006 | -0.006 | 0 | 0.23 | 0.24 | 0.32 | 0.4 | 0.26 | 0.34 |
| 8 | 124970848 | rs10956157 | 0.286 | 0.981 | -0.849 | 0.862 | -0.029 | 0.011 | 0.011 | 0.201 | 0.34 | 0.2 | 0.26 | 0.29 | 0.41 | 0.2 |
| 8 | 124971455 | rs11993947 | -0.069 | 0.934 | -0.822 | 0.837 | -0.012 | 0.019 | 0.019 | 0 | 0.38 | 0.3 | 0.28 | 0.29 | 0.42 | 0.2 |
| 8 | 124980708 | rs17270769 | 0.397 | 1.014 | -1.038 | 0.819 | -0.051 | 0.005 | 0.005 | 0.369 | 0.33 | 0.13 | 0.25 | 0.31 | 0.41 | 0.19 |
| 8 | 124987478 | rs7465584 | -0.117 | 0.716 | -0.931 | 1.068 | -0.009 | 0.002 | 0.002 | 0.255 | 0.42 | 0.4 | 0.36 | 0.44 | 0.49 | 0.28 |
| 8 | 124989360 | rs16899163 | -1.047 | 1.022 | -0.909 | 1.778 | -0.048 | -0.081 | -0.081 | 0 | 0.14 | 0.04 | 0.22 | 0.34 | 0.19 | 0.34 |
| 8 | 125006489 | rs7818302 | -1.276 | 0.49 | -0.5 | 0.916 | -0.017 | 0.012 | 0.012 | 0 | 0.05 | 0.18 | 0.03 | 0.06 | 0.01 | 0.01 |
| 8 | 125022733 | rs7844981 | -1.08 | 0.789 | -0.819 | 1.056 | -0.001 | 0 | 0 | 0 | 0.12 | 0.11 | 0.08 | 0.26 | 0.09 | 0.3 |
| 8 | 125027300 | rs1074934 | -0.507 | 1.055 | -0.643 | 0.245 | 0.009 | -0.062 | -0.062 | 0 | 0.17 | 0.19 | 0.13 | 0.01 | 0.24 | 0.08 |

| | | | | | | | | | | | | | | | | |
|---|---|---|---|---|---|---|---|---|---|---|---|---|---|---|---|---|
| 8 | 125031615 | rs4460354 | -0.412 | 0.918 | -0.59 | 0.572 | -0.002 | -0.012 | -0.012 | 0 | 0.62 | 0.68 | 0.4 | 0.51 | 0.59 | 0.48 |
| 8 | 125032732 | rs7003861 | -0.392 | 1.543 | -0.637 | 0.543 | -0.017 | -0.012 | -0.012 | 0.127 | 0.61 | 0.68 | 0.36 | 0.52 | 0.53 | 0.48 |
| 8 | 125033678 | rs7820272 | -0.077 | 0.91 | -0.346 | 0.738 | 0.003 | -0.001 | -0.001 | 0 | 0.55 | 0.33 | 0.34 | 0.43 | 0.57 | 0.43 |
| 8 | 125040676 | rs4631446 | 0.01 | 0.673 | -0.099 | 1.229 | -0.021 | 0.132 | 0.132 | 0 | 0.85 | 0.7 | 0.75 | 0.61 | 0.86 | 0.89 |
| 8 | 125044118 | rs4373503 | 0.178 | 0.508 | -0.434 | 0.796 | -0.01 | 0.025 | 0.025 | 0 | 0.28 | 0.21 | 0.33 | 0.17 | 0.35 | 0.35 |
| 8 | 125050657 | rs4871453 | -0.048 | 0.235 | -0.287 | 1.039 | -0.005 | 0.036 | 0.036 | 0.001 | 0.84 | 0.76 | 0.73 | 0.63 | 0.87 | 0.81 |
| 8 | 125051716 | rs4242353 | -0.309 | 0.256 | -0.376 | 1.107 | 0.002 | -0.03 | -0.03 | 0 | 0.86 | 0.91 | 0.75 | 0.64 | 0.87 | 0.82 |
| 8 | 125058587 | rs4871456 | 0.515 | 0.208 | -0.029 | 0.998 | -0.001 | 0 | 0 | 0 | 0.55 | 0.51 | 0.45 | 0.52 | 0.52 | 0.45 |
| 8 | 125058717 | rs4871457 | -0.253 | 0.421 | 0.111 | 1.112 | 0.002 | -0.015 | -0.015 | 1.113 | 0.85 | 0.88 | 0.74 | 0.61 | 0.87 | 0.79 |
| 8 | 125058981 | rs4871458 | -0.088 | 0.414 | 0.146 | 1.111 | -0.003 | 0.033 | 0.033 | 0 | 0.84 | 0.79 | 0.73 | 0.59 | 0.87 | 0.79 |
| 8 | 125060558 | rs7001300 | -0.123 | 0.422 | 0.141 | 1.108 | -0.004 | 0.033 | 0.033 | 0.011 | 0.84 | 0.79 | 0.73 | 0.62 | 0.87 | 0.79 |
| 8 | 125072426 | rs4483140 | -0.246 | 0.537 | 0.136 | 1.063 | 0 | -0.011 | -0.011 | 0.021 | 0.84 | 0.87 | 0.74 | 0.69 | 0.87 | 0.83 |
| 8 | 125074916 | rs9987271 | 0.36 | 0.368 | 0.471 | 1.15 | 0.043 | 0.034 | 0.034 | 0 | 0.25 | 0.37 | 0.34 | 0.35 | 0.38 | 0.28 |
| 8 | 125075728 | rs10102566 | -0.405 | 0.327 | 0.454 | 1.156 | 0 | -0.011 | -0.011 | 0 | 0.84 | 0.87 | 0.72 | 0.66 | 0.87 | 0.82 |
| 8 | 125076589 | rs7012186 | -0.283 | 0.312 | 0.531 | 1.133 | -0.003 | 0.044 | 0.044 | 0.005 | 0.83 | 0.77 | 0.71 | 0.67 | 0.87 | 0.82 |
| 8 | 125081858 | rs7006395 | 1.461 | -0.006 | -0.147 | 0.57 | 0.002 | 0.014 | 0.014 | 0 | 0.26 | 0.2 | 0.13 | 0.01 | 0.24 | 0.18 |
| 8 | 125083483 | rs12542273 | 0.893 | 0.453 | 0.631 | 1.281 | -0.036 | 0.023 | 0.023 | 0 | 0.2 | 0.09 | 0.28 | 0.15 | 0.31 | 0.26 |
| 8 | 125095089 | rs13261229 | -0.319 | 0.108 | 0.471 | 1.186 | 0.001 | -0.002 | -0.002 | 0 | 0.86 | 0.87 | 0.74 | 0.69 | 0.9 | 0.81 |
| 8 | 125107201 | rs6470219 | 0.241 | 0.062 | 0.489 | 1.102 | -0.012 | 0.097 | 0.097 | 0.408 | 0.81 | 0.61 | 0.68 | 0.64 | 0.87 | 0.82 |
| 8 | 125108977 | rs4870888 | 0.615 | 0.138 | -0.066 | 0.946 | -0.013 | 0.06 | 0.06 | 0.864 | 0.54 | 0.22 | 0.33 | 0.32 | 0.49 | 0.54 |
| 8 | 125117277 | rs4430074 | 0.661 | 0.028 | 0.276 | 1.256 | -0.008 | 0.033 | 0.033 | 0.032 | 0.82 | 0.74 | 0.7 | 0.68 | 0.87 | 0.82 |
| 8 | 125132234 | rs13278430 | 1.008 | 0.164 | 0.07 | 0.942 | -0.019 | 0.022 | 0.022 | 0 | 0.65 | 0.52 | 0.51 | 0.38 | 0.74 | 0.58 |
| 8 | 125133870 | rs4363179 | 0.907 | 0.173 | 0.048 | 0.852 | -0.036 | 0.032 | 0.032 | 0 | 0.67 | 0.52 | 0.53 | 0.38 | 0.79 | 0.57 |
| 8 | 125150123 | rs4413755 | 1.085 | 0.115 | 0.339 | 0.87 | -0.002 | 0.001 | 0.001 | 0 | 0.66 | 0.61 | 0.52 | 0.38 | 0.75 | 0.6 |
| 8 | 125172349 | rs4242357 | 0.981 | 0.066 | 0.364 | 0.906 | -0.009 | 0.009 | 0.009 | 0 | 0.67 | 0.6 | 0.55 | 0.36 | 0.78 | 0.6 |
| 8 | 125180387 | rs11775425 | 0.771 | 0.073 | 0.146 | 0.928 | 0.013 | -0.045 | -0.045 | 0.004 | 0.71 | 0.82 | 0.56 | 0.37 | 0.79 | 0.59 |
| 8 | 125199582 | rs7823009 | -0.962 | 0.215 | 0.798 | 1.414 | 0.035 | 0.129 | 0.129 | 0 | 0.12 | 0.33 | 0.2 | 0.31 | 0.12 | 0.15 |
| 8 | 125215606 | rs7815027 | -0.322 | 0.173 | 0.367 | 0.808 | -0.001 | 0 | 0 | 0 | 0.67 | 0.69 | 0.69 | 0.68 | 0.68 | 0.83 |
| 8 | 125216329 | rs6470235 | -1.008 | 0.371 | 0.421 | 0.756 | -0.002 | 0.042 | 0.042 | 0 | 0.3 | 0.43 | 0.36 | 0.5 | 0.25 | 0.4 |
| 8 | 125221563 | rs11994044 | -0.231 | 0.311 | 0.365 | 0.624 | -0.001 | -0.003 | -0.003 | 0 | 0.19 | 0.19 | 0.17 | 0.18 | 0.15 | 0.25 |
| 8 | 125222043 | rs9643190 | 0.316 | 0.598 | 0.351 | 0.877 | 0.002 | 0.042 | 0.042 | 0 | 0.09 | 0.12 | 0.18 | 0.31 | 0.1 | 0.16 |
| 8 | 125224719 | rs12548675 | -1.169 | 0.816 | 0.06 | 0.24 | -0.013 | 0.017 | 0.017 | 0 | 0.81 | 0.85 | 0.82 | 0.94 | 0.77 | 0.86 |
| 8 | 125225255 | rs7837873 | -1.464 | 0.276 | 0.077 | 0.438 | 0.043 | 0.141 | 0.141 | 0 | 0.5 | 0.81 | 0.55 | 0.81 | 0.4 | 0.5 |
| 8 | 125232152 | rs10096791 | -0.978 | 0.403 | 0.322 | 0.83 | -0.018 | 0.038 | 0.038 | 0 | 0.33 | 0.44 | 0.4 | 0.59 | 0.29 | 0.28 |
| 8 | 125238368 | rs4242365 | -1.673 | 0.58 | 0.628 | 0.99 | 0 | -0.026 | -0.026 | 1.515 | 0.36 | 0.69 | 0.42 | 0.53 | 0.32 | 0.4 |
| 8 | 125239980 | rs7815754 | -0.114 | 0.654 | 0.686 | 1.034 | -0.009 | -0.007 | -0.007 | 0.256 | 0.11 | 0.09 | 0.14 | 0.21 | 0.12 | 0.24 |
| 8 | 125240331 | rs768711 | -1.677 | 0.659 | 0.722 | 1.043 | 0.007 | 0.053 | 0.053 | 0 | 0.33 | 0.44 | 0.4 | 0.53 | 0.31 | 0.4 |
| 8 | 125241796 | rs7821423 | -1.262 | 0.736 | 0.339 | 0.456 | -0.001 | 0 | 0 | 0.48 | 0.2 | 0.2 | 0.22 | 0.22 | 0.19 | 0.27 |
| 8 | 125247147 | rs10107402 | -0.846 | 0.98 | 0.255 | 0.762 | 0.046 | 0.042 | 0.042 | 0 | 0.12 | 0.35 | 0.15 | 0.2 | 0.11 | 0.19 |
| 8 | 125251067 | rs7827309 | -0.561 | 0.749 | -0.338 | 0.593 | 0.001 | -0.005 | -0.005 | 0 | 0.41 | 0.56 | 0.4 | 0.7 | 0.4 | 0.49 |
| 8 | 125259569 | rs16899467 | -0.182 | 0.336 | 0.114 | 0.999 | -0.021 | 0.006 | 0.006 | 1.701 | 0.2 | 0.29 | 0.07 | 0.28 | 0.13 | 0.06 |
| 8 | 125263468 | rs7816704 | -0.756 | 0.558 | -0.727 | 0.615 | -0.045 | -0.004 | -0.004 | 0 | 0.76 | 0.94 | 0.62 | 0.81 | 0.73 | 0.7 |
| 8 | 125263818 | rs17360508 | 0.864 | 0.026 | -1.04 | 0.103 | -0.017 | 0.046 | 0.046 | 0 | 0.34 | 0.08 | 0.18 | 0.25 | 0.36 | 0.14 |
| 8 | 125265042 | rs7843845 | -0.55 | 0.235 | -0.372 | 0.396 | -0.002 | 0.001 | 0.001 | 0 | 0.14 | 0.09 | 0.18 | 0.13 | 0.16 | 0.27 |
| 8 | 125268122 | rs16899480 | -0.461 | 0.333 | -0.053 | 0.332 | 0.041 | -0.032 | -0.032 | 2.819 | 0.1 | 0.19 | 0.17 | 0.05 | 0.14 | 0.19 |
| 8 | 125268800 | rs17361004 | 1.469 | 0.143 | -0.516 | 0.263 | -0.059 | -0.038 | -0.038 | 0 | 0.27 | 0.01 | 0.28 | 0.24 | 0.32 | 0.1 |
| 8 | 125269095 | rs4871480 | 0.279 | 0.11 | 0.008 | 0.31 | 0.008 | -0.016 | -0.016 | 0.011 | 0.39 | 0.46 | 0.47 | 0.35 | 0.46 | 0.33 |
| 8 | 125269707 | rs4593526 | 0.272 | 0.113 | 0.01 | 0.299 | 0.008 | -0.014 | -0.014 | 0 | 0.39 | 0.46 | 0.47 | 0.37 | 0.46 | 0.33 |
| 8 | 125283389 | rs11985656 | 0.389 | 0.424 | -0.32 | 0.272 | -0.003 | 0.015 | 0.015 | 0.069 | 0.19 | 0.07 | 0.11 | 0.15 | 0.18 | 0.13 |
| 8 | 125283572 | rs13439439 | 0.665 | 0.358 | -0.179 | 0.23 | -0.001 | 0 | 0 | 0 | 0.37 | 0.36 | 0.43 | 0.36 | 0.45 | 0.32 |
| 8 | 125284359 | rs4457311 | -0.356 | 0.305 | -0.164 | 0.25 | 0.004 | 0.008 | 0.008 | 0.13 | 0.51 | 0.6 | 0.59 | 0.53 | 0.5 | 0.73 |
| 8 | 125291320 | rs4870895 | -0.593 | 0.522 | -0.058 | 0.261 | -0.009 | 0.01 | 0.01 | 0.045 | 0.82 | 0.93 | 0.89 | 0.86 | 0.82 | 0.88 |
| 8 | 125293035 | rs10755924 | 0.989 | 0.55 | 0.075 | 0.306 | 0.016 | -0.011 | -0.011 | 0.094 | 0.66 | 0.44 | 0.68 | 0.71 | 0.63 | 0.63 |
| 8 | 125293242 | rs10808533 | 0.677 | 0.642 | 0.106 | 0.356 | -0.001 | 0 | 0 | 0.011 | 0.69 | 0.66 | 0.71 | 0.71 | 0.64 | 0.64 |
| 8 | 125295256 | rs16899510 | 0.217 | 0.942 | -0.076 | 0.28 | 0.03 | 0.002 | 0.002 | 0 | 0.05 | 0.09 | 0.12 | 0.05 | 0.11 | 0.17 |
| 8 | 125296872 | rs17296667 | -0.41 | 0.853 | -0.027 | 0.311 | 0.02 | 0.001 | 0.001 | 4.209 | 0.09 | 0.1 | 0.15 | 0.09 | 0.22 | 0.24 |
| 8 | 125300254 | rs11997412 | 0.239 | 0.688 | -0.091 | 0.242 | -0.056 | 0.035 | 0.035 | 0 | 0.76 | 0.85 | 0.64 | 0.89 | 0.62 | 0.71 |
| 8 | 125309055 | rs6997326 | -1.279 | 0.603 | -0.162 | 0.198 | -0.028 | 0.068 | 0.068 | 1.109 | 0.87 | 0.97 | 0.93 | 1 | 0.87 | 0.96 |
| 8 | 125312660 | rs10956170 | 1.401 | 0.312 | -0.163 | 0.181 | 0.002 | 0.002 | 0.002 | 0 | 0.29 | 0.28 | 0.17 | 0.23 | 0.25 | 0.13 |
| 8 | 125343256 | rs9886564 | -1.159 | -0.318 | 0.092 | 0.417 | -0.002 | 0.012 | 0.012 | 0 | 0.1 | 0.16 | 0.06 | 0.17 | 0.1 | 0.14 |
| 8 | 125356918 | rs10110968 | -1.142 | -0.295 | 0.111 | 0.37 | -0.015 | 0.054 | 0.054 | 0.217 | 0.86 | 0.94 | 0.92 | 0.99 | 0.86 | 0.96 |
| 8 | 125357041 | rs10096657 | 0.133 | -0.338 | 0.13 | 0.409 | 0.005 | 0.001 | 0.001 | 0.593 | 0.06 | 0.13 | 0.04 | 0.05 | 0.08 | 0.09 |
| 8 | 125361409 | rs16899562 | -0.308 | -0.344 | 0.128 | 0.421 | 0.003 | -0.001 | -0.001 | 0 | 0.07 | 0.13 | 0.04 | 0.06 | 0.09 | 0.09 |
| 8 | 125364138 | rs4351403 | 0.451 | -0.405 | 0.246 | 0.4 | 0.004 | -0.005 | -0.005 | 1.423 | 0.4 | 0.34 | 0.43 | 0.54 | 0.32 | 0.49 |
| 8 | 125386971 | rs10091430 | -0.048 | -0.247 | 0.194 | 0.215 | -0.015 | 0.046 | 0.046 | 0 | 0.8 | 0.86 | 0.88 | 0.94 | 0.77 | 0.87 |
| 8 | 125389873 | rs7835851 | -1.061 | -0.169 | 0.311 | 0.246 | -0.001 | -0.018 | -0.018 | 0 | 0.88 | 0.87 | 0.91 | 1 | 0.85 | 0.96 |
| 8 | 125391233 | rs7386980 | 0.209 | -0.179 | 0.303 | 0.24 | -0.003 | 0.004 | 0.004 | 0 | 0.81 | 0.79 | 0.88 | 0.94 | 0.77 | 0.87 |
| 8 | 125392860 | rs4317549 | -1.337 | -0.124 | 0.241 | 0.258 | -0.013 | 0.027 | 0.027 | 0 | 0.89 | 0.94 | 0.92 | 1 | 0.85 | 0.97 |
| 8 | 125404763 | rs4612323 | 1.22 | -0.216 | 0.264 | 0.565 | -0.027 | -0.009 | -0.009 | 0 | 0.34 | 0.18 | 0.41 | 0.34 | 0.42 | 0.32 |

| | | | | | | | | | | | | | | | | |
|---|---|---|---|---|---|---|---|---|---|---|---|---|---|---|---|---|
| 8 | 125407426 | rs4617135 | 0.351 | -0.273 | 0.35 | 0.567 | 0.017 | -0.065 | -0.065 | 0 | 0.8 | 0.6 | 0.85 | 0.94 | 0.77 | 0.87 |
| 8 | 125418391 | rs4295647 | -0.51 | -0.305 | -0.036 | 0.609 | -0.011 | 0.01 | 0.01 | 0 | 0.84 | 0.92 | 0.89 | 0.88 | 0.81 | 0.89 |
| 8 | 125418855 | rs4440618 | -0.588 | -0.715 | 0.251 | 0.893 | 0.005 | -0.001 | -0.001 | 3.137 | 0.47 | 0.42 | 0.46 | 0.53 | 0.39 | 0.56 |
| 8 | 125425693 | rs4346978 | -0.388 | -0.463 | -0.057 | 1.002 | 0.006 | -0.001 | -0.001 | 0.228 | 0.43 | 0.56 | 0.44 | 0.47 | 0.31 | 0.43 |
| 8 | 125430662 | rs6994603 | -0.484 | -0.546 | -0.199 | 0.975 | -0.001 | 0 | 0 | 0.018 | 0.46 | 0.48 | 0.47 | 0.52 | 0.38 | 0.53 |
| 8 | 125437811 | rs4620270 | -1.501 | -1.261 | -0.288 | 1.111 | 0.001 | 0 | 0 | 0 | 0.3 | 0.71 | 0.23 | 0.29 | 0.17 | 0.27 |
| 8 | 125438835 | rs12675583 | -1.382 | -1.123 | -0.567 | 1.178 | 0.035 | -0.012 | -0.012 | 0.121 | 0.27 | 0.21 | 0.3 | 0.32 | 0.12 | 0.34 |
| 8 | 125450317 | rs3936120 | -0.635 | -0.423 | -0.356 | 1.174 | -0.004 | 0.002 | 0.002 | 0 | 0.08 | 0.13 | 0.15 | 0.12 | 0.04 | 0.17 |
| 8 | 125450473 | rs10956183 | -0.922 | -0.539 | -0.085 | 1.359 | -0.067 | 0.117 | 0.117 | 0.067 | 0.17 | 0.37 | 0.19 | 0.3 | 0.08 | 0.23 |
| 8 | 125455570 | rs6987358 | -0.386 | 0.362 | -1.101 | 1.127 | -0.036 | -0.03 | -0.03 | 0 | 0.17 | 0.05 | 0.3 | 0.19 | 0.23 | 0.07 |
| 8 | 125459719 | rs4512366 | -0.01 | 0.885 | -1.674 | 0.597 | -0.035 | 0.02 | 0.02 | 0 | 0.17 | 0.05 | 0.29 | 0.09 | 0.22 | 0.07 |
| 8 | 125462121 | rs4132855 | -0.518 | 0.347 | -1.564 | 0.079 | 0.002 | 0.033 | 0.033 | 0 | 0.47 | 0.67 | 0.43 | 0.3 | 0.41 | 0.51 |
| 8 | 125463083 | rs3812473 | 1.237 | -0.149 | -1.683 | 0.263 | -0.013 | 0.002 | 0.002 | 0 | 0.1 | 0.12 | 0.05 | 0.14 | 0.08 | 0.05 |
| 8 | 125463250 | rs3812475 | -0.792 | -0.171 | -1.712 | 0.252 | -0.033 | 0.004 | 0.004 | 0 | 0.57 | 0.78 | 0.48 | 0.44 | 0.5 | 0.56 |
| 8 | 125467788 | rs10956184 | 1.206 | -0.138 | -1.701 | 0.24 | -0.035 | 0.02 | 0.02 | 0 | 0.17 | 0.05 | 0.3 | 0.09 | 0.22 | 0.07 |
| 8 | 125477797 | rs10105517 | -1.599 | -0.291 | -2.283 | 0.791 | 0.006 | -0.111 | -0.111 | 0 | 0.83 | 0.99 | 0.82 | 0.7 | 0.87 | 0.79 |
| 8 | 125479475 | rs4128469 | 0.516 | -0.642 | -1.982 | 0.053 | -0.011 | -0.016 | -0.016 | 0 | 0.25 | 0.12 | 0.34 | 0.29 | 0.25 | 0.43 |
| 8 | 125479663 | rs4128468 | -0.524 | -1.194 | -1.496 | -0.235 | 0 | 0.018 | 0.018 | 0 | 0.49 | 0.36 | 0.44 | 0.44 | 0.49 | 0.56 |
| 8 | 125485201 | rs1128757 | 1.198 | -1.254 | -1.511 | -0.265 | 0.032 | 0.029 | 0.029 | 7.131 | 0.06 | 0.42 | 0.04 | 0 | 0 | 0 |
| 8 | 125485728 | rs10956185 | 1.893 | -1.27 | -1.634 | -0.275 | -0.07 | -0.001 | -0.001 | 0 | 0.16 | 0.01 | 0.29 | 0.09 | 0.22 | 0.08 |
| 8 | 125486767 | rs3812472 | -0.951 | -1.266 | -1.634 | -0.275 | -0.028 | 0.025 | 0.025 | 0.002 | 0.55 | 0.78 | 0.48 | 0.44 | 0.49 | 0.56 |
| 8 | 125493942 | rs4551335 | 1.873 | -1.266 | -1.639 | -0.273 | -0.07 | -0.007 | -0.007 | 0 | 0.17 | 0.01 | 0.29 | 0.1 | 0.22 | 0.07 |
| 8 | 125498547 | rs3812471 | -0.786 | -1.245 | -1.587 | -0.274 | -0.025 | 0.025 | 0.025 | 0.003 | 0.55 | 0.79 | 0.48 | 0.44 | 0.49 | 0.56 |
| 8 | 125531405 | rs16899649 | 1.076 | -1.152 | -1.456 | -0.126 | -0.013 | 0.02 | 0.02 | 0.139 | 0.09 | 0.13 | 0.05 | 0.19 | 0.03 | 0.35 |
| 8 | 125534041 | rs16899660 | 1.866 | -1.12 | -1.537 | -0.166 | 0.001 | -0.029 | -0.029 | 1.387 | 0.09 | 0.01 | 0.03 | 0.15 | 0.08 | 0.05 |
| 8 | 125540185 | rs11776528 | 1.355 | -0.865 | -1.46 | -0.367 | -0.026 | 0.035 | 0.035 | 0.128 | 0.14 | 0.04 | 0.1 | 0 | 0.2 | 0.06 |
| 8 | 125551567 | rs7010411 | -0.538 | -0.894 | -1.058 | 0.09 | 0.001 | 0.001 | 0.001 | 0 | 0.33 | 0.27 | 0.15 | 0.35 | 0.26 | 0.47 |
| 8 | 125552276 | rs7007115 | -0.809 | -0.79 | -1.118 | 0.073 | 0 | 0.014 | 0.014 | 0 | 0.51 | 0.35 | 0.45 | 0.45 | 0.49 | 0.56 |
| 8 | 125559807 | rs6987343 | -0.787 | -0.84 | -1.156 | 0.112 | -0.002 | 0.029 | 0.029 | 0 | 0.5 | 0.35 | 0.45 | 0.39 | 0.48 | 0.56 |
| 8 | 125565328 | rs3829037 | 0.311 | -0.495 | -1.362 | -0.038 | -0.003 | 0.021 | 0.021 | 0 | 0.23 | 0.16 | 0.11 | 0.16 | 0.24 | 0.13 |
| 8 | 125567019 | rs10956189 | 0.948 | -0.389 | -1.251 | 0.19 | -0.068 | 0.01 | 0.01 | 0 | 0.18 | 0.01 | 0.29 | 0.09 | 0.22 | 0.07 |
| 8 | 125568623 | rs3829036 | -0.032 | -0.425 | -0.71 | 0.431 | -0.002 | 0.003 | 0.003 | 0 | 0.15 | 0.16 | 0.22 | 0.14 | 0.14 | 0.04 |
| 8 | 125570144 | rs4870903 | -0.349 | -0.58 | -0.703 | 0.899 | 0.011 | -0.017 | -0.017 | 0.133 | 0.67 | 0.71 | 0.59 | 0.57 | 0.73 | 0.66 |
| 8 | 125570554 | rs6470248 | 1.236 | -0.201 | -0.939 | 0.433 | -0.046 | 0.05 | 0.05 | 0 | 0.09 | 0.01 | 0.04 | 0 | 0.11 | 0.05 |
| 8 | 125570800 | rs7846270 | 0.937 | -0.538 | -0.822 | 0.541 | -0.009 | 0.082 | 0.082 | 0.087 | 0.4 | 0.12 | 0.45 | 0.21 | 0.55 | 0.31 |
| 8 | 125571483 | rs3793398 | -0.729 | -0.62 | -0.745 | 0.796 | -0.008 | 0.034 | 0.034 | 0.129 | 0.65 | 0.75 | 0.72 | 0.73 | 0.67 | 0.64 |
| 8 | 125571586 | rs3829034 | -0.337 | -0.745 | -0.019 | 0.969 | -0.021 | 0.021 | 0.021 | 0.074 | 0.24 | 0.31 | 0.25 | 0.35 | 0.17 | 0.11 |
| 8 | 125575013 | rs4007921 | 1.5 | -0.402 | 0.073 | 0.567 | 0.016 | -0.034 | -0.034 | 0.03 | 0.1 | 0.03 | 0.03 | 0.17 | 0.09 | 0.08 |
| 8 | 125575619 | rs10110655 | 1.523 | -0.441 | 0.157 | 0.441 | -0.07 | 0.026 | 0.026 | 0.38 | 0.31 | 0.11 | 0.41 | 0.21 | 0.44 | 0.26 |
| 8 | 125578501 | rs4870904 | 0.227 | -0.488 | 0.407 | 0.399 | 0.006 | -0.009 | -0.009 | 0.594 | 0.17 | 0.19 | 0.09 | 0.16 | 0.16 | 0.31 |
| 8 | 125579788 | rs3857944 | -0.903 | -0.56 | 0.437 | 0.449 | 0.004 | -0.008 | -0.008 | 0.144 | 0.64 | 0.66 | 0.56 | 0.52 | 0.7 | 0.65 |
| 8 | 125580928 | rs2288664 | -1.299 | -0.485 | 0.544 | 0.4 | -0.015 | -0.006 | -0.006 | 0 | 0.33 | 0.56 | 0.15 | 0.34 | 0.25 | 0.4 |
| 8 | 125581871 | rs4870905 | -0.932 | -0.375 | 0.303 | 0.45 | -0.007 | -0.001 | -0.001 | 0.118 | 0.3 | 0.37 | 0.14 | 0.33 | 0.25 | 0.39 |
| 8 | 125582099 | rs4871503 | -0.146 | -0.159 | 0.121 | 0.411 | 0.006 | -0.014 | -0.014 | 0 | 0.18 | 0.23 | 0.09 | 0.16 | 0.16 | 0.31 |
| 8 | 125594916 | rs7001375 | -0.544 | -0.233 | -0.276 | 0.393 | 0.018 | -0.021 | -0.021 | 0 | 0.63 | 0.72 | 0.45 | 0.5 | 0.7 | 0.59 |
| 8 | 125600410 | rs7010757 | 1.22 | 0.391 | -0.682 | 0.275 | -0.064 | 0.027 | 0.027 | 1.014 | 0.3 | 0.07 | 0.28 | 0.2 | 0.38 | 0.21 |
| 8 | 125600818 | rs4871506 | -0.349 | -0.643 | -0.185 | 0.34 | 0.004 | 0.039 | 0.039 | 0 | 0.49 | 0.64 | 0.62 | 0.63 | 0.43 | 0.6 |
| 8 | 125600950 | rs4871507 | 0.336 | -1.34 | 0.066 | 0.376 | 0 | -0.001 | -0.001 | 0 | 0.38 | 0.37 | 0.55 | 0.55 | 0.32 | 0.51 |
| 8 | 125603298 | rs11994203 | -0.031 | -1.334 | 0.199 | 0.236 | 0 | -0.002 | -0.002 | 0 | 0.17 | 0.25 | 0.07 | 0.16 | 0.18 | 0.1 |
| 8 | 125606068 | rs4871510 | -0.464 | -0.94 | 0.424 | 0.226 | -0.018 | 0.036 | 0.036 | 0.225 | 0.57 | 0.69 | 0.65 | 0.72 | 0.5 | 0.65 |
| 8 | 125606322 | rs3857947 | 0.459 | -0.951 | 0.073 | 0.243 | 0.018 | 0.029 | 0.029 | 0 | 0.46 | 0.37 | 0.58 | 0.71 | 0.38 | 0.58 |
| 8 | 125608818 | rs7835359 | -1.821 | -1.377 | 0.906 | 0.527 | 0.002 | -0.059 | -0.059 | 0.575 | 0.18 | 0.58 | 0.09 | 0.08 | 0.14 | 0.18 |
| 8 | 125610780 | rs3857948 | -1.016 | -1.152 | 0.665 | 1.13 | 0.038 | 0.11 | 0.11 | 0.015 | 0.09 | 0.15 | 0.31 | 0.32 | 0.14 | 0.1 |
| 8 | 125610797 | rs3857949 | -1.016 | -1.152 | 0.665 | 1.108 | 0.038 | 0.105 | 0.105 | 0.015 | 0.09 | 0.15 | 0.31 | 0.32 | 0.14 | 0.1 |
| 8 | 125619203 | rs2013845 | -1.712 | -2.008 | 1.054 | 0.913 | -0.029 | 0.045 | 0.045 | 2.706 | 0.24 | 0.55 | 0.38 | 0.36 | 0.18 | 0.24 |
| 8 | 125627290 | rs4870911 | -1.287 | -1.936 | 1.509 | 1 | -0.011 | -0.016 | -0.016 | 0 | 0.2 | 0.55 | 0.25 | 0.2 | 0.15 | 0.19 |
| 8 | 125628685 | rs11994520 | -1.216 | -1.938 | 1.589 | 0.999 | -0.01 | -0.014 | -0.014 | 0 | 0.19 | 0.45 | 0.23 | 0.2 | 0.15 | 0.19 |
| 8 | 125628928 | rs11995300 | -1.304 | -1.938 | 1.474 | 0.999 | -0.012 | -0.017 | -0.017 | 0 | 0.2 | 0.52 | 0.24 | 0.2 | 0.15 | 0.19 |
| 8 | 125633124 | rs7015740 | -0.57 | -2.415 | 0.679 | 0.667 | 0.078 | -0.001 | -0.001 | 0 | 0.5 | 0.75 | 0.47 | 0.58 | 0.36 | 0.65 |
| 8 | 125635705 | rs4871518 | -0.844 | -2.365 | 0.25 | 0.306 | 0.07 | 0.077 | 0.077 | 1.633 | 0.53 | 0.89 | 0.6 | 0.73 | 0.37 | 0.71 |
| 8 | 125648122 | rs6992199 | -1.084 | -2.784 | 0.904 | 0.123 | -0.027 | -0.019 | -0.019 | 0 | 0.24 | 0.29 | 0.1 | 0.22 | 0.13 | 0.16 |
| 8 | 125648925 | rs7017138 | -1.243 | -2.848 | 1.288 | 0.091 | -0.035 | -0.025 | -0.025 | 0 | 0.27 | 0.62 | 0.13 | 0.22 | 0.13 | 0.16 |
| 8 | 125650653 | rs4871527 | 0.009 | -2.885 | 1.361 | 0.074 | -0.023 | -0.003 | -0.003 | 0 | 0.11 | 0.18 | 0.06 | 0.14 | 0.05 | 0.06 |
| 8 | 125654560 | rs4510845 | -1.377 | -2.886 | 1.102 | 0.069 | -0.049 | -0.038 | -0.038 | 0 | 0.26 | 0.6 | 0.12 | 0.22 | 0.13 | 0.16 |
| 8 | 125658852 | rs10216358 | -0.122 | -2.837 | 0.907 | 0.074 | -0.008 | -0.015 | -0.015 | 0 | 0.13 | 0.17 | 0.05 | 0.08 | 0.08 | 0.09 |
| 8 | 125660259 | rs7010266 | 0.313 | -2.822 | 0.661 | 0.083 | 0.008 | 0.011 | 0.011 | 0 | 0.11 | 0.06 | 0.04 | 0.08 | 0.08 | 0.08 |
| 8 | 125660721 | rs6470257 | -1.342 | -2.9 | 0.604 | 0.313 | -0.033 | 0.009 | 0.009 | 0.733 | 0.3 | 0.6 | 0.24 | 0.4 | 0.17 | 0.23 |
| 8 | 125661323 | rs4871529 | 0.791 | -2.693 | -0.13 | 0.423 | 0.015 | -0.048 | -0.048 | 0.149 | 0.25 | 0.16 | 0.19 | 0.46 | 0.19 | 0.43 |
| 8 | 125667226 | rs7813190 | -0.355 | -2.225 | -0.086 | 0.408 | -0.019 | -0.03 | -0.03 | 0 | 0.13 | 0.18 | 0.05 | 0.08 | 0.08 | 0.11 |

| | | | | | | | | | | | | | | | | |
|---|---|---|---|---|---|---|---|---|---|---|---|---|---|---|---|---|
| 8 | 125677005 | rs4870916 | -0.823 | -2.33 | -0.774 | 0.411 | 0.089 | 0.019 | 0.019 | 0.381 | 0.4 | 0.72 | 0.26 | 0.37 | 0.17 | 0.25 |
| 8 | 125681849 | rs7016031 | -0.616 | -1.55 | 0.023 | 0.789 | -0.087 | 0.07 | 0.07 | 1.601 | 0.16 | 0.42 | 0.3 | 0.21 | 0.07 | 0.1 |
| 8 | 125682525 | rs7845483 | -0.46 | -1.47 | -0.72 | 0.508 | 0.018 | 0.015 | 0.015 | 0.04 | 0.41 | 0.64 | 0.5 | 0.68 | 0.29 | 0.53 |
| 8 | 125683083 | rs7831518 | 0.598 | -0.547 | -0.313 | 0.852 | 0.004 | 0.027 | 0.027 | 0.324 | 0.79 | 0.59 | 0.63 | 0.73 | 0.81 | 0.75 |
| 8 | 125683183 | rs6470259 | -0.453 | -0.068 | -1.456 | 0.435 | -0.013 | 0.089 | 0.089 | 0.841 | 0.55 | 0.84 | 0.41 | 0.72 | 0.43 | 0.49 |
| 8 | 125687498 | rs3857953 | -0.259 | 0.605 | -1.055 | 0.735 | 0.003 | 0.018 | 0.018 | 0 | 0.13 | 0.17 | 0.35 | 0.25 | 0.12 | 0.2 |
| 8 | 125688407 | rs3915792 | -0.435 | 1.268 | -1.948 | 0.662 | -0.085 | -0.039 | -0.039 | 0 | 0.79 | 0.99 | 0.81 | 0.83 | 0.75 | 0.77 |
| 8 | 125691155 | rs16899960 | -0.178 | 1.689 | -1.577 | 0.854 | 0.017 | 0.087 | 0.087 | 0 | 0.11 | 0.2 | 0.18 | 0.39 | 0.15 | 0.11 |
| 8 | 125697259 | rs2382992 | 0.329 | 1.246 | -1.096 | 0.257 | 0.007 | 0.005 | 0.005 | 0 | 0.63 | 0.6 | 0.6 | 0.42 | 0.57 | 0.61 |
| 8 | 125697673 | rs2116081 | 0.211 | 1.287 | -1.137 | 0.237 | -0.003 | -0.002 | -0.002 | 1.805 | 0.64 | 0.69 | 0.61 | 0.41 | 0.54 | 0.64 |
| 8 | 125700925 | rs2382993 | -0.328 | 1.058 | -1.37 | 0.111 | 0.003 | 0.002 | 0.002 | 0 | 0.3 | 0.37 | 0.5 | 0.3 | 0.3 | 0.39 |
| 8 | 125702761 | rs3901290 | 0.406 | 0.737 | -1.339 | -0.091 | -0.001 | 0.002 | 0.002 | 0.018 | 0.13 | 0.13 | 0.24 | 0.18 | 0.13 | 0.23 |
| 8 | 125704238 | rs34410151 | 0.897 | 0.473 | -1.93 | -0.542 | -0.063 | 0.07 | 0.07 | 1.548 | 0.27 | 0.02 | 0.15 | 0.12 | 0.28 | 0.12 |
| 8 | 125706593 | rs2382995 | -0.155 | 0.579 | -1.735 | -0.435 | -0.002 | 0.056 | 0.056 | 0 | 0.44 | 0.21 | 0.4 | 0.26 | 0.43 | 0.38 |
| 8 | 125706829 | rs2163372 | -0.162 | 0.582 | -1.734 | -0.458 | -0.002 | 0.03 | 0.03 | 0 | 0.44 | 0.21 | 0.4 | 0.3 | 0.43 | 0.39 |
| 8 | 125707677 | rs12550311 | 0.366 | 0.477 | -1.909 | -0.633 | 0.026 | 0.023 | 0.023 | 0 | 0.3 | 0.15 | 0.28 | 0.18 | 0.22 | 0.26 |
| 8 | 125712420 | rs10956197 | -0.852 | 0.599 | -1.493 | -0.508 | 0.001 | -0.03 | -0.03 | 0 | 0.73 | 0.72 | 0.76 | 0.84 | 0.72 | 0.77 |
| 8 | 125713791 | rs1017342 | 0.838 | 0.596 | -1.536 | -0.466 | 0.003 | 0.085 | 0.085 | 0 | 0.29 | 0.51 | 0.35 | 0.53 | 0.29 | 0.41 |
| 8 | 125714397 | rs2116082 | -0.164 | 0.674 | -1.055 | -0.216 | 0 | -0.018 | -0.018 | 0 | 0.73 | 0.73 | 0.76 | 0.84 | 0.72 | 0.77 |
| 8 | 125724292 | rs10956198 | -0.215 | 0.354 | -1.24 | 0.212 | -0.001 | -0.001 | -0.001 | 0.887 | 0.2 | 0.19 | 0.26 | 0.18 | 0.17 | 0.23 |
| 8 | 125728351 | rs12547677 | 0.039 | 0.579 | -1.662 | 0.004 | -0.034 | 0.033 | 0.033 | 0 | 0.2 | 0.04 | 0.18 | 0.15 | 0.17 | 0.24 |
| 8 | 125728561 | rs10956200 | -0.687 | 0.619 | -0.854 | 0.028 | 0.003 | 0.005 | 0.005 | 0.074 | 0.47 | 0.55 | 0.52 | 0.36 | 0.43 | 0.54 |
| 8 | 125732857 | rs6470263 | 0.504 | -0.039 | -0.574 | 0.489 | -0.031 | 0.007 | 0.007 | 0 | 0.62 | 0.6 | 0.79 | 0.55 | 0.72 | 0.57 |
| 8 | 125736146 | rs1430849 | -0.228 | -0.358 | -0.726 | 0.544 | 0.029 | 0.019 | 0.019 | 0 | 0.8 | 0.98 | 0.92 | 0.86 | 0.84 | 0.7 |
| 8 | 125736746 | rs6993178 | 0.738 | -0.281 | -0.656 | 0.509 | -0.036 | 0.01 | 0.01 | 0.01 | 0.64 | 0.6 | 0.8 | 0.55 | 0.74 | 0.57 |
| 8 | 125742166 | rs891541 | 0.768 | -0.33 | -0.485 | 0.485 | -0.037 | 0.008 | 0.008 | 0.078 | 0.63 | 0.52 | 0.79 | 0.55 | 0.74 | 0.57 |
| 8 | 125743773 | rs733759 | 1.076 | 0.582 | -0.541 | 0.597 | 0.014 | -0.003 | -0.003 | 0.204 | 0.56 | 0.37 | 0.4 | 0.39 | 0.59 | 0.5 |
| 8 | 125747374 | rs4870923 | 0.691 | 1.012 | -1.464 | 0.405 | -0.11 | -0.067 | -0.067 | 0.048 | 0.29 | 0.02 | 0.25 | 0.29 | 0.35 | 0.37 |
| 8 | 125748913 | rs2891102 | 0.17 | 1.265 | -1.13 | 0.399 | -0.039 | -0.009 | -0.009 | 0.355 | 0.36 | 0.24 | 0.5 | 0.38 | 0.45 | 0.43 |
| 8 | 125750505 | rs6983655 | 0.837 | 1.009 | -0.618 | 0.267 | 0.007 | -0.022 | -0.022 | 0 | 0.37 | 0.44 | 0.42 | 0.23 | 0.36 | 0.28 |
| 8 | 125750897 | rs6988023 | 0.5 | 0.987 | -0.543 | 0.385 | 0.01 | 0.09 | 0.09 | 0 | 0.15 | 0.31 | 0.25 | 0.34 | 0.16 | 0.1 |
| 8 | 125751065 | rs7006084 | -0.067 | 0.906 | -0.608 | 0.338 | 0.009 | 0.013 | 0.013 | 0 | 0.53 | 0.75 | 0.67 | 0.57 | 0.52 | 0.38 |
| 8 | 125751177 | rs7006401 | 0.443 | 0.911 | -0.445 | 0.316 | 0.091 | 0.047 | 0.047 | 0 | 0.06 | 0.17 | 0.26 | 0.09 | 0.1 | 0.06 |
| 8 | 125752167 | rs10099679 | 1.298 | 0.904 | -0.604 | 0.305 | -0.018 | -0.046 | -0.046 | 0.33 | 0.33 | 0.43 | 0.19 | 0.14 | 0.26 | 0.21 |
| 8 | 125753604 | rs967943 | -1.127 | 0.855 | -0.702 | 0.295 | 0.034 | 0 | 0 | 0 | 0.83 | 0.98 | 0.95 | 0.86 | 0.86 | 0.74 |
| 8 | 125753813 | rs1367501 | 0.248 | 0.805 | -0.59 | 0.326 | 0.005 | 0.104 | 0.104 | 0 | 0.17 | 0.39 | 0.26 | 0.35 | 0.16 | 0.11 |
| 8 | 125753849 | rs1367500 | 0.096 | 0.812 | -0.611 | 0.311 | 0.039 | -0.013 | -0.013 | 0 | 0.5 | 0.82 | 0.45 | 0.49 | 0.42 | 0.32 |
| 8 | 125754151 | rs6993864 | 0.096 | 0.812 | -0.611 | 0.311 | 0.039 | -0.013 | -0.013 | 0 | 0.5 | 0.82 | 0.45 | 0.49 | 0.42 | 0.32 |
| 8 | 125755750 | rs7459793 | 0.109 | 1.135 | -0.542 | 0.205 | -0.001 | 0.003 | 0.003 | 0 | 0.42 | 0.41 | 0.48 | 0.64 | 0.43 | 0.38 |
| 8 | 125760391 | rs12675098 | -0.16 | 1.268 | -0.791 | 0.313 | -0.014 | -0.005 | -0.005 | 0 | 0.78 | 0.93 | 0.71 | 0.78 | 0.76 | 0.69 |
| 8 | 125761998 | rs4871533 | -0.186 | 1.275 | -0.827 | 0.316 | -0.018 | -0.008 | -0.008 | 0.046 | 0.78 | 0.95 | 0.7 | 0.78 | 0.76 | 0.69 |
| 8 | 125763884 | rs16900003 | 0.828 | 0.934 | -0.504 | 0.134 | 0 | -0.013 | -0.013 | 0.306 | 0.34 | 0.36 | 0.21 | 0.15 | 0.33 | 0.33 |
| 8 | 125764361 | rs6470269 | 0.149 | 1.233 | -0.681 | 0.468 | 0 | -0.013 | -0.013 | 0.16 | 0.72 | 0.77 | 0.66 | 0.49 | 0.67 | 0.62 |
| 8 | 125766271 | rs7841287 | -0.391 | 1.378 | -0.224 | 0.996 | -0.003 | 0.032 | 0.032 | 0 | 0.84 | 0.84 | 0.85 | 0.55 | 0.84 | 0.7 |
| 8 | 125766962 | rs9656810 | 0.858 | 1.585 | -0.61 | 0.972 | -0.047 | -0.07 | -0.07 | 0.039 | 0.15 | 0.01 | 0.12 | 0.25 | 0.16 | 0.23 |
| 8 | 125769891 | rs7825792 | 1.516 | 1.518 | -0.145 | 0.37 | -0.01 | -0.046 | -0.046 | 0.752 | 0.4 | 0.41 | 0.18 | 0.09 | 0.3 | 0.25 |
| 8 | 125771707 | rs13279462 | 0.744 | 1.489 | -0.309 | 0.205 | 0.007 | -0.054 | -0.054 | 0.465 | 0.16 | 0.04 | 0.12 | 0.37 | 0.16 | 0.3 |
| 8 | 125772282 | rs13270557 | 0.724 | 1.623 | -0.222 | 0.272 | 0.004 | 0 | 0 | 0 | 0.56 | 0.56 | 0.33 | 0.46 | 0.46 | 0.55 |
| 8 | 125773964 | rs7835906 | -0.029 | 1.866 | -0.341 | 0.301 | -0.099 | 0.032 | 0.032 | 0.041 | 0.19 | 0.03 | 0.35 | 0.09 | 0.27 | 0.14 |
| 8 | 125774978 | rs7011543 | -1.026 | 1.783 | 0.025 | 0.361 | -0.003 | -0.004 | -0.004 | 0 | 0.35 | 0.32 | 0.49 | 0.49 | 0.43 | 0.44 |
| 8 | 125775553 | rs6470276 | -0.825 | 1.899 | 0.165 | 0.073 | 0.052 | -0.061 | -0.061 | 0 | 0.24 | 0.4 | 0.41 | 0.09 | 0.31 | 0.15 |
| 8 | 125781412 | rs7836373 | -0.583 | 1.959 | -0.253 | 0.136 | -0.029 | -0.086 | -0.086 | 0 | 0.81 | 0.94 | 0.72 | 0.54 | 0.78 | 0.67 |
| 8 | 125787117 | rs6982303 | -0.436 | 1.734 | 0.084 | -0.006 | 0.03 | -0.04 | -0.04 | 0 | 0.23 | 0.32 | 0.37 | 0.09 | 0.29 | 0.17 |
| 8 | 125792809 | rs4870927 | -0.55 | 1.788 | -0.06 | 0.035 | -0.024 | -0.061 | -0.061 | 0 | 0.8 | 0.9 | 0.68 | 0.54 | 0.74 | 0.68 |
| 8 | 125797266 | rs11992495 | 1.722 | 1.511 | 0.034 | -0.19 | 0.032 | 0.038 | 0.038 | 0.804 | 0.39 | 0.28 | 0.17 | 0.09 | 0.26 | 0.19 |
| 8 | 125799895 | rs6985316 | 0.583 | 2.328 | -0.603 | -0.007 | -0.02 | -0.026 | -0.026 | 0 | 0.07 | 0.01 | 0.29 | 0.13 | 0.21 | 0.12 |
| 8 | 125802018 | rs3935197 | -1.327 | 2.193 | -0.512 | 0.023 | 0.023 | 0.018 | 0.018 | 0 | 0.38 | 0.51 | 0.49 | 0.48 | 0.49 | 0.5 |
| 8 | 125803141 | rs4871546 | -0.39 | 1.691 | -1.096 | 0.408 | -0.014 | -0.037 | -0.037 | 0.017 | 0.8 | 0.89 | 0.69 | 0.54 | 0.75 | 0.68 |
| 8 | 125804636 | rs10096762 | -0.095 | 1.634 | -1.047 | 0.525 | -0.002 | 0.058 | 0.058 | 0 | 0.16 | 0.29 | 0.14 | 0.36 | 0.19 | 0.33 |
| 8 | 125810701 | rs7824418 | -0.159 | 1.633 | -1.082 | 0.524 | -0.003 | 0.067 | 0.067 | 0 | 0.17 | 0.32 | 0.14 | 0.36 | 0.19 | 0.33 |
| 8 | 125815850 | rs4871550 | 0.569 | 1.97 | -1.452 | 0.021 | -0.025 | 0.055 | 0.055 | 2.558 | 0.66 | 0.78 | 0.56 | 0.18 | 0.56 | 0.37 |
| 8 | 125825915 | rs12550726 | -0.879 | 2.302 | -1.484 | 0.644 | -0.087 | -0.057 | -0.057 | 0 | 0.19 | 0.03 | 0.28 | 0.25 | 0.31 | 0.35 |
| 8 | 125831705 | rs12544599 | -0.627 | 2.61 | -1.221 | 1.054 | -0.01 | -0.008 | -0.008 | 2.551 | 0.17 | 0.16 | 0.46 | 0.39 | 0.33 | 0.26 |
| 8 | 125835989 | rs10956215 | 1.145 | 0.976 | -0.918 | -0.094 | -0.05 | -0.093 | -0.093 | 0.471 | 0.3 | 0.39 | 0.11 | 0.05 | 0.14 | 0.14 |
| 8 | 125836335 | rs10808541 | -0.057 | 1.087 | -1.011 | -0.084 | 0.036 | 0.106 | 0.106 | 0 | 0.52 | 0.72 | 0.31 | 0.28 | 0.42 | 0.46 |
| 8 | 125836386 | rs11778381 | -0.185 | 1.248 | -0.472 | 0.44 | -0.003 | -0.003 | -0.003 | 0 | 0.17 | 0.14 | 0.14 | 0.23 | 0.22 | 0.28 |
| 8 | 125839379 | rs4871555 | -0.216 | 0.879 | -0.364 | 0.488 | 0 | 0.005 | 0.005 | 0 | 0.64 | 0.73 | 0.71 | 0.68 | 0.69 | 0.69 |
| 8 | 125843240 | rs11783665 | -0.717 | 0.224 | 0.62 | 0.506 | 0.005 | 0.028 | 0.028 | 1.398 | 0.09 | 0.3 | 0.07 | 0.09 | 0.05 | 0.08 |
| 8 | 125844898 | rs7461109 | 0.753 | -0.009 | 0.172 | 0.27 | 0.023 | 0.044 | 0.044 | 0.687 | 0.18 | 0.08 | 0.11 | 0.04 | 0.11 | 0.08 |

| | | | | | | | | | | | | | | | | |
|---|---|---|---|---|---|---|---|---|---|---|---|---|---|---|---|---|
| 8 | 125845495 | rs9969440 | -0.672 | -0.338 | 0.744 | 0.539 | -0.029 | 0.072 | 0.072 | 0 | 0.17 | 0.47 | 0.12 | 0.28 | 0.09 | 0.11 |
| 8 | 125854702 | rs10956218 | -0.145 | -0.312 | 0.864 | 0.375 | -0.006 | -0.009 | -0.009 | 0 | 0.29 | 0.3 | 0.15 | 0.16 | 0.17 | 0.13 |
| 8 | 125861374 | rs7461129 | 0.131 | 0.696 | 0.573 | 0.884 | -0.04 | -0.04 | -0.04 | 0.215 | 0.15 | 0.07 | 0.34 | 0.47 | 0.34 | 0.26 |
| 8 | 125866943 | rs12547431 | -0.91 | 0.438 | 0.936 | 0.905 | 0 | 0.006 | 0.006 | 0 | 0.47 | 0.42 | 0.49 | 0.66 | 0.52 | 0.54 |
| 8 | 125872784 | rs7461652 | 0.055 | 0.53 | 0.875 | 0.888 | 0.025 | 0.028 | 0.028 | 1.536 | 0.31 | 0.23 | 0.16 | 0.19 | 0.21 | 0.27 |
| 8 | 125878316 | rs6987095 | 1.153 | 0.284 | 0.836 | 0.868 | -0.02 | 0.161 | 0.161 | 1.009 | 0.47 | 0.13 | 0.34 | 0.22 | 0.48 | 0.41 |
| 8 | 125891362 | rs6470302 | 0.763 | 0.286 | 1.214 | 0.887 | 0.031 | 0.231 | 0.231 | 0.125 | 0.93 | 0.64 | 0.77 | 0.84 | 0.97 | 0.89 |
| 8 | 125892187 | rs7016748 | -0.688 | 0.308 | 1.061 | 0.913 | -0.006 | 0.032 | 0.032 | 0 | 0.46 | 0.32 | 0.48 | 0.68 | 0.51 | 0.53 |
| 8 | 125898882 | rs7387254 | 0.674 | 0.413 | 0.752 | 1.087 | 0.028 | 0.232 | 0.232 | 4.429 | 0.93 | 0.58 | 0.83 | 0.88 | 1 | 0.94 |
| 8 | 125901416 | rs7386656 | 0.136 | 0.551 | 0.138 | 1.149 | -0.028 | -0.026 | -0.026 | 0.035 | 0.14 | 0.07 | 0.33 | 0.47 | 0.3 | 0.3 |
| 8 | 125902415 | rs6470317 | 0.872 | -0.021 | 0.51 | 0.94 | 0.023 | 0.221 | 0.221 | 4.85 | 0.93 | 0.48 | 0.82 | 0.89 | 1 | 0.94 |
| 8 | 125903495 | rs13262017 | -0.308 | 0.077 | 0.128 | 0.546 | -0.009 | -0.009 | -0.009 | 0 | 0.33 | 0.36 | 0.16 | 0.22 | 0.21 | 0.22 |
| 8 | 125908657 | rs6470323 | 0.414 | 0.291 | 0.247 | 0.588 | 0.002 | -0.002 | -0.002 | 0.134 | 0.58 | 0.55 | 0.37 | 0.38 | 0.55 | 0.57 |
| 8 | 125909352 | rs9785063 | 1.137 | -0.446 | 0.189 | 0.119 | -0.028 | 0.058 | 0.058 | 0.822 | 0.34 | 0.22 | 0.23 | 0.12 | 0.43 | 0.28 |
| 8 | 125909479 | rs10216354 | -0.477 | 0.137 | 0.34 | 0.952 | 0.001 | 0.033 | 0.033 | 2.723 | 0.74 | 0.65 | 0.62 | 0.63 | 0.76 | 0.85 |
| 8 | 125937406 | rs2555716 | -0.468 | 0.051 | 0.498 | 1.147 | 0.007 | 0.048 | 0.048 | 0 | 0.82 | 0.76 | 0.6 | 0.62 | 0.78 | 0.72 |
| 8 | 125937760 | rs11779177 | 1.009 | 0.294 | 0.212 | 0.892 | 0.02 | 0.003 | 0.003 | 0 | 0.28 | 0.22 | 0.16 | 0.27 | 0.22 | 0.3 |
| 8 | 125939953 | rs2555718 | 1.464 | 0.458 | 0.331 | 0.841 | -0.001 | -0.004 | -0.004 | 0 | 0.18 | 0.18 | 0.2 | 0.23 | 0.17 | 0.08 |
| 8 | 125940308 | rs10956224 | 0.478 | 0.398 | 0.032 | 0.668 | -0.016 | 0.02 | 0.02 | 0.004 | 0.55 | 0.28 | 0.35 | 0.38 | 0.53 | 0.6 |
| 8 | 125942505 | rs2052783 | 0.447 | 0.451 | 0.07 | 0.681 | -0.017 | 0.004 | 0.004 | 0 | 0.56 | 0.3 | 0.35 | 0.39 | 0.55 | 0.61 |
| 8 | 125944942 | rs1427083 | -0.122 | 0.456 | 0.288 | 0.672 | -0.002 | -0.003 | -0.003 | 0 | 0.63 | 0.4 | 0.39 | 0.39 | 0.62 | 0.64 |
| 8 | 125948898 | rs2077670 | 1.536 | 0.447 | -0.076 | 0.593 | -0.049 | 0.082 | 0.082 | 0.217 | 0.28 | 0.1 | 0.19 | 0.12 | 0.32 | 0.3 |
| 8 | 125949053 | rs723231 | -1.431 | 0.395 | 0.283 | 0.765 | -0.007 | 0.047 | 0.047 | 0.018 | 0.82 | 0.68 | 0.68 | 0.66 | 0.85 | 0.87 |
| 8 | 125951173 | rs16900122 | -0.16 | 0.343 | 0.4 | 0.783 | 0.015 | -0.047 | -0.047 | 0.759 | 0.09 | 0.27 | 0.05 | 0 | 0.07 | 0.03 |
| 8 | 125951673 | rs2555721 | 0.75 | 0.415 | -0.103 | 1.15 | -0.005 | -0.003 | -0.003 | 0 | 0.16 | 0.11 | 0.27 | 0.23 | 0.22 | 0.22 |
| 8 | 125951713 | rs2555722 | -0.062 | 0.609 | -0.133 | 0.841 | -0.005 | -0.006 | -0.006 | 0.102 | 0.46 | 0.34 | 0.46 | 0.5 | 0.49 | 0.55 |
| 8 | 125954124 | rs10956225 | 0.26 | 0.246 | -0.459 | 0.533 | 0.003 | 0 | 0 | 0.142 | 0.23 | 0.2 | 0.23 | 0.21 | 0.2 | 0.29 |
| 8 | 125954274 | rs2555723 | -0.061 | 0.117 | -0.327 | 0.274 | 0 | 0.003 | 0.003 | 0.019 | 0.56 | 0.59 | 0.59 | 0.72 | 0.55 | 0.61 |
| 8 | 125955401 | rs2590665 | -0.497 | -0.951 | 0.213 | -0.129 | 0.033 | -0.085 | -0.085 | 1.164 | 0.21 | 0.38 | 0.11 | 0.07 | 0.25 | 0.23 |
| 8 | 125955850 | rs2590666 | -1.716 | -1.646 | -0.275 | -0.074 | -0.006 | 0.025 | 0.025 | 0.081 | 0.38 | 0.88 | 0.49 | 0.28 | 0.34 | 0.36 |
| 8 | 125957082 | rs2590667 | -0.243 | -1.704 | 0.412 | -0.167 | 0.008 | -0.041 | -0.041 | 0 | 0.22 | 0.26 | 0.12 | 0.07 | 0.26 | 0.24 |
| 8 | 125959763 | rs1834716 | 0.154 | -1.955 | 0.144 | -0.193 | -0.01 | 0.072 | 0.072 | 0 | 0.22 | 0.32 | 0.34 | 0.42 | 0.15 | 0.31 |
| 8 | 125959812 | rs1427087 | -2.143 | -1.977 | -0.005 | -0.202 | -0.118 | -0.114 | -0.114 | 0 | 0.53 | 0.95 | 0.54 | 0.49 | 0.48 | 0.57 |
| 8 | 125964505 | rs2725879 | -0.226 | -1.702 | 0.257 | -0.143 | 0 | 0.01 | 0.01 | 0 | 0.24 | 0.24 | 0.34 | 0.42 | 0.21 | 0.34 |
| 8 | 125968156 | rs2590673 | -0.066 | -1.689 | 0.352 | -0.114 | 0.013 | 0.105 | 0.105 | 0 | 0.08 | 0.18 | 0.27 | 0.23 | 0.07 | 0.14 |
| 8 | 125968659 | rs3909714 | 0.018 | -1.718 | 0.279 | -0.152 | -0.004 | -0.097 | -0.097 | 0.01 | 0.21 | 0.57 | 0.12 | 0.07 | 0.19 | 0.2 |
| 8 | 125972165 | rs2555727 | -2.109 | -2.132 | -0.513 | -0.272 | -0.149 | -0.108 | -0.108 | 0.051 | 0.71 | 0.96 | 0.63 | 0.57 | 0.54 | 0.64 |
| 8 | 125972472 | rs12677337 | 1.463 | -2.098 | -0.102 | 0.019 | -0.046 | -0.051 | -0.051 | 0 | 0.14 | 0.01 | 0.07 | 0.2 | 0.14 | 0.2 |
| 8 | 125973689 | rs12545272 | 1.914 | -2.092 | 0.004 | -0.111 | 0.063 | 0.025 | 0.025 | 0.608 | 0.19 | 0.05 | 0.09 | 0.07 | 0.06 | 0.08 |
| 8 | 125973915 | rs2725881 | -1.19 | -1.257 | 0.722 | 0.167 | 0.001 | 0.041 | 0.041 | 0 | 0.33 | 0.47 | 0.42 | 0.42 | 0.28 | 0.36 |
| 8 | 125977492 | rs7018326 | -0.479 | -0.905 | 0.906 | -0.082 | 0.016 | -0.042 | -0.042 | 0.023 | 0.09 | 0.26 | 0.08 | 0 | 0.07 | 0.01 |
| 8 | 125989649 | rs10105106 | 0.114 | -0.941 | 1.246 | -0.042 | 0.015 | -0.015 | -0.015 | 0 | 0.06 | 0.21 | 0.05 | 0 | 0.03 | 0.01 |
| 8 | 125989698 | rs16900167 | -0.419 | -0.861 | 1.381 | 0.307 | 0.015 | 0.06 | 0.06 | 0 | 0.09 | 0.14 | 0.28 | 0.21 | 0.08 | 0.16 |
| 8 | 125990858 | rs7829785 | 0.097 | -0.818 | 1.384 | 0.273 | 0.016 | -0.015 | -0.015 | 0.721 | 0.06 | 0.22 | 0.05 | 0 | 0.03 | 0.01 |
| 8 | 125993570 | rs10481077 | 0.434 | -0.823 | 1.731 | 0.271 | -0.008 | -0.031 | -0.031 | 0 | 0.06 | 0.13 | 0.04 | 0 | 0.03 | 0.01 |
| 8 | 126013303 | rs4347009 | -0.49 | -1.847 | 1.438 | 0.029 | 0.077 | 0.001 | 0.001 | 0.945 | 0.45 | 0.7 | 0.44 | 0.36 | 0.2 | 0.47 |
| 8 | 126016176 | rs13263702 | 0.536 | -2.046 | 1.351 | -0.036 | 0.058 | -0.05 | -0.05 | 0 | 0.32 | 0.08 | 0.38 | 0.29 | 0.16 | 0.27 |
| 8 | 126023124 | rs7813917 | -0.587 | -2.071 | 1.589 | -0.168 | -0.001 | 0 | 0 | 0 | 0.45 | 0.53 | 0.44 | 0.36 | 0.26 | 0.48 |
| 8 | 126030809 | rs966946 | -1.484 | -1.857 | 1.512 | -0.309 | 0.026 | 0.002 | 0.002 | 1.211 | 0.55 | 0.73 | 0.5 | 0.36 | 0.35 | 0.5 |
| 8 | 126033450 | rs10216437 | -1.422 | -1.855 | 1.393 | -0.308 | 0.022 | -0.02 | -0.02 | 0 | 0.56 | 0.86 | 0.51 | 0.36 | 0.35 | 0.5 |
| 8 | 126034052 | rs2293985 | -0.513 | -1.861 | 1.481 | -0.308 | -0.035 | -0.011 | -0.011 | 0 | 0.1 | 0.19 | 0.02 | 0.07 | 0.03 | 0.19 |
| 8 | 126035100 | rs10112173 | -0.982 | -2.185 | 1.439 | -0.479 | 0.088 | 0.107 | 0.107 | 1.589 | 0.51 | 0.72 | 0.26 | 0.14 | 0.28 | 0.36 |
| 8 | 126042022 | rs6989556 | 0.594 | -2.194 | 1.417 | -0.333 | 0.056 | -0.042 | -0.042 | 0 | 0.32 | 0.09 | 0.38 | 0.29 | 0.17 | 0.27 |
| 8 | 126044527 | rs11542889 | -0.356 | -2.186 | 1.602 | -0.344 | -0.009 | -0.055 | -0.055 | 0 | 0.1 | 0.43 | 0.06 | 0 | 0.06 | 0.02 |
| 8 | 126045344 | rs12678891 | 1.605 | -2.024 | 1.144 | -0.143 | -0.051 | -0.044 | -0.044 | 0.114 | 0.14 | 0.01 | 0.1 | 0.19 | 0.19 | 0.19 |
| 8 | 126047864 | rs2288310 | 0.285 | -2.428 | 1.638 | -0.09 | 0.043 | -0.019 | -0.019 | 0 | 0.33 | 0.16 | 0.39 | 0.29 | 0.17 | 0.27 |
| 8 | 126063271 | rs10089574 | -0.552 | -2.499 | 1.459 | -0.136 | -0.046 | -0.012 | -0.012 | 0.337 | 0.11 | 0.36 | 0.03 | 0.07 | 0.03 | 0.2 |
| 8 | 126068873 | rs2303523 | 1.569 | -2.54 | 1.289 | -0.151 | 0.1 | 0.053 | 0.053 | 0.58 | 0.23 | 0.01 | 0.11 | 0.06 | 0.09 | 0.11 |
| 8 | 126081480 | rs10429356 | -2.142 | -2.078 | 1.676 | 0.086 | -0.005 | 0.007 | 0.007 | 2.921 | 0.22 | 0.73 | 0.32 | 0.3 | 0.12 | 0.36 |
| 8 | 126082565 | rs12546767 | 0.41 | -1.903 | 1.516 | 0.173 | 0.013 | 0.055 | 0.055 | 0.098 | 0.07 | 0.07 | 0.27 | 0.23 | 0.09 | 0.16 |
| 8 | 126089549 | rs11985518 | -1.119 | -1.988 | 1.532 | -0.07 | -0.044 | 0.004 | 0.004 | 3.972 | 0.11 | 0.47 | 0.04 | 0.07 | 0.03 | 0.2 |
| 8 | 126094499 | rs2891636 | -1.101 | -1.97 | 1.137 | -0.07 | -0.049 | -0.01 | -0.01 | 0 | 0.12 | 0.42 | 0.03 | 0.07 | 0.03 | 0.2 |
| 8 | 126096274 | rs16900369 | -1.076 | -1.966 | 1.139 | -0.068 | -0.048 | -0.009 | -0.009 | 2.121 | 0.12 | 0.42 | 0.03 | 0.07 | 0.03 | 0.2 |
| 8 | 126102820 | rs7357375 | 0.146 | -2.14 | 0.824 | 0.031 | -0.052 | 0.006 | 0.006 | 0.013 | 0.36 | 0.46 | 0.4 | 0.31 | 0.17 | 0.26 |
| 8 | 126108385 | rs6995096 | -0.351 | -2.402 | 0.862 | -0.499 | 0.018 | 0.01 | 0.01 | 0 | 0.45 | 0.61 | 0.26 | 0.29 | 0.24 | 0.46 |
| 8 | 126113303 | rs9987342 | 1.164 | -2.624 | 0.743 | -0.519 | -0.006 | 0.005 | 0.005 | 0.825 | 0.27 | 0.21 | 0.18 | 0.06 | 0.08 | 0.11 |
| 8 | 126132816 | rs7831515 | -1.256 | -2.346 | 0.394 | -0.469 | -0.003 | -0.064 | -0.064 | 0 | 0.58 | 0.96 | 0.39 | 0.37 | 0.33 | 0.54 |
| 8 | 126138197 | rs7461756 | -0.368 | -2.322 | 0.833 | -0.724 | -0.006 | 0.047 | 0.047 | 0 | 0.17 | 0.4 | 0.08 | 0.23 | 0.14 | 0.36 |
| 8 | 126156243 | rs10109490 | 1.413 | -2.497 | 0.815 | -0.789 | 0.143 | 0.101 | 0.101 | 0.504 | 0.27 | 0.04 | 0.18 | 0.06 | 0.09 | 0.13 |

| | | | | | | | | | | | | | | | | |
|---|---|---|---|---|---|---|---|---|---|---|---|---|---|---|---|---|
| 8 | 126161468 | rs6470339 | -0.415 | -2.368 | 0.877 | -0.712 | 0.165 | 0.073 | 0.073 | 0 | 0.48 | 0.79 | 0.28 | 0.29 | 0.23 | 0.49 |
| 8 | 126172340 | rs12544146 | -0.465 | -2.348 | 0.954 | -0.882 | 0.008 | -0.018 | -0.018 | 0 | 0.11 | 0.17 | 0.1 | 0.08 | 0.1 | 0.04 |
| 8 | 126172882 | rs6992545 | 0.707 | -2.601 | 1.063 | -0.756 | -0.021 | 0.003 | 0.003 | 0 | 0.1 | 0.2 | 0.05 | 0.11 | 0.08 | 0.21 |
| 8 | 126207277 | rs10089963 | 0.769 | -2.6 | 1.037 | -0.755 | -0.109 | -0.078 | -0.078 | 0.202 | 0.31 | 0.39 | 0.21 | 0.06 | 0.09 | 0.14 |
| 8 | 126216588 | rs7837996 | 0.475 | -1.897 | 0.748 | -0.443 | -0.016 | 0.001 | 0.001 | 0 | 0.1 | 0.19 | 0.05 | 0.11 | 0.08 | 0.21 |
| 8 | 126254259 | rs3955404 | -1.723 | -1.137 | -1.224 | -0.312 | -0.187 | -0.172 | -0.172 | 0.128 | 0.74 | 0.98 | 0.51 | 0.52 | 0.54 | 0.75 |
| 8 | 126264351 | rs10091027 | -1.678 | -2.086 | -0.381 | -0.419 | 0.003 | -0.012 | -0.012 | 0 | 0.3 | 0.8 | 0.17 | 0.37 | 0.23 | 0.4 |
| 8 | 126266244 | rs11994861 | -1.049 | -1.778 | -0.212 | -0.25 | -0.027 | 0.058 | 0.058 | 0 | 0.24 | 0.57 | 0.1 | 0.29 | 0.18 | 0.39 |
| 8 | 126267630 | rs7008482 | -0.872 | -1.377 | -1.251 | -0.124 | 0.016 | -0.099 | -0.099 | 0 | 0.6 | 0.94 | 0.38 | 0.43 | 0.32 | 0.61 |
| 8 | 126268020 | rs12681602 | -0.828 | 0.061 | 0.098 | 0.709 | 0.034 | 0.093 | 0.093 | 0 | 0.06 | 0.15 | 0.03 | 0.17 | 0.08 | 0.18 |
| 8 | 126272646 | rs10283134 | -1.409 | -0.384 | -0.185 | 0.506 | 0.006 | -0.005 | -0.005 | 0 | 0.28 | 0.79 | 0.13 | 0.31 | 0.19 | 0.42 |
| 8 | 126273825 | rs4350000 | 0.406 | 0.323 | -0.426 | 0.483 | -0.032 | -0.03 | -0.03 | 0.027 | 0.92 | 0.97 | 0.85 | 0.78 | 0.83 | 0.81 |
| 8 | 126275593 | rs10956234 | 0.736 | 0.292 | -0.029 | 0.415 | 0.021 | 0.051 | 0.051 | 0.282 | 0.9 | 0.85 | 0.84 | 0.76 | 0.87 | 0.81 |
| 8 | 126276327 | rs16900491 | -0.249 | 0.395 | 0.52 | 0.304 | 0.001 | -0.033 | -0.033 | 0 | 0.06 | 0.15 | 0.05 | 0 | 0.08 | 0.01 |
| 8 | 126285326 | rs10112737 | 2.173 | 0.045 | 0.713 | 0.118 | 0.007 | -0.004 | -0.004 | 2.215 | 0.68 | 0.3 | 0.73 | 0.69 | 0.73 | 0.58 |
| 8 | 126289605 | rs7816350 | 0.095 | -0.474 | 0.134 | -0.221 | 0.01 | 0.017 | 0.017 | 0 | 0.19 | 0.05 | 0.21 | 0.13 | 0.13 | 0.2 |
| 8 | 126300866 | rs7831918 | -0.577 | -0.457 | 0.246 | -0.216 | 0 | -0.045 | -0.045 | 0 | 0.08 | 0.25 | 0.07 | 0 | 0.09 | 0.01 |
| 8 | 126301986 | rs4565472 | 0.297 | -0.462 | 0.667 | -0.194 | 0.024 | 0.055 | 0.055 | 0 | 0.91 | 0.88 | 0.84 | 0.78 | 0.87 | 0.81 |
| 8 | 126303493 | rs7003102 | -0.596 | -0.662 | 0.686 | -0.292 | -0.028 | -0.038 | -0.038 | 0 | 0.27 | 0.42 | 0.25 | 0.16 | 0.16 | 0.23 |
| 8 | 126303847 | rs7007866 | -0.368 | -0.659 | 0.641 | -0.29 | 0.014 | 0.017 | 0.017 | 0 | 0.25 | 0.23 | 0.24 | 0.16 | 0.16 | 0.23 |
| 8 | 126311265 | rs10956238 | 1.598 | -0.543 | -0.064 | -0.254 | -0.002 | -0.001 | -0.001 | 0 | 0.57 | 0.23 | 0.5 | 0.62 | 0.58 | 0.55 |
| 8 | 126314626 | rs4602904 | 1.173 | 0.08 | -0.073 | -0.083 | -0.017 | -0.048 | -0.048 | 0 | 0.79 | 0.87 | 0.8 | 0.71 | 0.77 | 0.63 |
| 8 | 126315917 | rs17318327 | -0.843 | 0.344 | -0.1 | -0.231 | -0.021 | 0.021 | 0.021 | 0 | 0.07 | 0.03 | 0.07 | 0.04 | 0.13 | 0.05 |
| 8 | 126350206 | rs7820471 | 1.268 | 0.383 | -0.07 | -0.227 | -0.016 | -0.036 | -0.036 | 0.226 | 0.77 | 0.83 | 0.85 | 0.7 | 0.76 | 0.59 |
| 8 | 126351822 | rs4870938 | -0.786 | 0.158 | -0.222 | -0.358 | -0.01 | 0.033 | 0.033 | 0 | 0.22 | 0.1 | 0.26 | 0.14 | 0.23 | 0.23 |
| 8 | 126365542 | rs16900568 | 0.311 | 0.173 | -0.413 | -0.325 | 0.008 | 0.007 | 0.007 | 0 | 0.14 | 0.03 | 0.19 | 0.1 | 0.1 | 0.18 |
| 8 | 126376699 | rs13255114 | 2.331 | 0.15 | -0.368 | -0.332 | -0.037 | -0.036 | -0.036 | 0.014 | 0.48 | 0.14 | 0.53 | 0.55 | 0.52 | 0.35 |
| 8 | 126377384 | rs4871588 | -0.423 | 0.253 | -0.348 | -0.291 | -0.048 | -0.04 | -0.04 | 0 | 0.92 | 0.99 | 0.92 | 0.76 | 0.88 | 0.83 |
| 8 | 126380443 | rs4871589 | -1.273 | 0.296 | -0.229 | -0.395 | 0.011 | 0.021 | 0.021 | 0.153 | 0.29 | 0.72 | 0.32 | 0.15 | 0.24 | 0.26 |
| 8 | 126385814 | rs7000501 | 1.124 | 0.712 | -0.34 | -0.015 | 0.014 | 0.039 | 0.039 | 0 | 0.74 | 0.62 | 0.82 | 0.65 | 0.73 | 0.6 |
| 8 | 126389258 | rs10099164 | 0.911 | 1.105 | -1.353 | 0.342 | -0.003 | -0.003 | -0.003 | 0.023 | 0.72 | 0.71 | 0.79 | 0.69 | 0.66 | 0.61 |
| 8 | 126404148 | rs4332153 | -0.872 | 1.134 | -2.01 | -0.339 | -0.108 | 0.129 | 0.129 | 0 | 0.34 | 0.08 | 0.3 | 0.14 | 0.42 | 0.46 |
| 8 | 126409668 | rs4870939 | -0.925 | 1.06 | -2.063 | -0.451 | -0.08 | 0.131 | 0.131 | 0 | 0.34 | 0.12 | 0.33 | 0.12 | 0.39 | 0.46 |
| 8 | 126413669 | rs2385107 | -0.714 | -0.234 | -1.721 | 0.179 | -0.008 | 0.007 | 0.007 | 0 | 0.28 | 0.33 | 0.23 | 0.45 | 0.18 | 0.21 |
| 8 | 126416920 | rs2980872 | -0.761 | 0.136 | -1.592 | -0.45 | -0.02 | 0.042 | 0.042 | 0.148 | 0.39 | 0.65 | 0.49 | 0.83 | 0.46 | 0.4 |
| 8 | 126417614 | rs2954007 | -0.045 | -0.131 | -1.261 | 0.022 | -0.028 | -0.024 | -0.024 | 0 | 0.26 | 0.17 | 0.32 | 0.63 | 0.37 | 0.31 |
| 8 | 126419188 | rs10099227 | -0.61 | -1.385 | -1.171 | -0.431 | -0.056 | -0.074 | -0.074 | 0 | 0.22 | 0.65 | 0.15 | 0.04 | 0.06 | 0.18 |
| 8 | 126426386 | rs7822730 | -0.13 | -1.325 | -1.165 | -0.283 | 0.041 | 0.033 | 0.033 | 0 | 0.16 | 0.06 | 0.09 | 0.09 | 0.05 | 0.18 |
| 8 | 126429220 | rs7837158 | -0.181 | -1.431 | -1.036 | -0.356 | 0.024 | 0.025 | 0.025 | 0.261 | 0.17 | 0.09 | 0.21 | 0.05 | 0.04 | 0.18 |
| 8 | 126431985 | rs4559257 | -0.792 | -1.359 | -0.736 | 0.061 | -0.026 | -0.004 | -0.004 | 7.728 | 0.42 | 0.54 | 0.49 | 0.38 | 0.22 | 0.44 |
| 8 | 126437630 | rs4871593 | 0.258 | -0.687 | -1.212 | 0.191 | 0.019 | 0.04 | 0.04 | 0 | 0.47 | 0.26 | 0.34 | 0.39 | 0.39 | 0.47 |
| 8 | 126439329 | rs2385110 | 0.8 | 0.005 | -1.567 | -0.046 | -0.103 | 0.054 | 0.054 | 0 | 0.34 | 0.03 | 0.22 | 0.23 | 0.34 | 0.41 |
| 8 | 126440360 | rs10956245 | -1.074 | 0.009 | -0.087 | 0.458 | 0.025 | 0.05 | 0.05 | 0 | 0.06 | 0.32 | 0.16 | 0.07 | 0.06 | 0.07 |
| 8 | 126447308 | rs2385114 | 0.017 | 0.719 | -0.859 | 0.562 | -0.04 | -0.009 | -0.009 | 0 | 0.7 | 0.77 | 0.64 | 0.72 | 0.51 | 0.74 |
| 8 | 126448789 | rs2235108 | 0.299 | -0.297 | -1.216 | -0.324 | -0.004 | 0.05 | 0.05 | 0 | 0.3 | 0.16 | 0.17 | 0.1 | 0.32 | 0.32 |
| 8 | 126449406 | rs17405319 | 1.117 | -1.261 | -0.503 | -0.427 | 0.005 | -0.004 | -0.004 | 0 | 0.27 | 0.25 | 0.34 | 0.11 | 0.16 | 0.24 |
| 8 | 126457553 | rs13273254 | 1.044 | -1.734 | -0.937 | -0.644 | -0.021 | 0.09 | 0.09 | 1.414 | 0.23 | 0.02 | 0.31 | 0.1 | 0.18 | 0.31 |
| 8 | 126459990 | rs4871598 | -0.246 | -2.446 | -0.01 | -0.716 | -0.024 | -0.014 | -0.014 | 0 | 0.44 | 0.52 | 0.47 | 0.72 | 0.26 | 0.48 |
| 8 | 126463365 | rs10808545 | -1.018 | -2.431 | -0.266 | -1.174 | -0.014 | 0.016 | 0.016 | 0 | 0.69 | 0.75 | 0.58 | 0.84 | 0.4 | 0.75 |
| 8 | 126464388 | rs17663005 | 1.001 | -2.528 | -0.141 | -1.222 | 0.026 | 0.044 | 0.044 | 0 | 0.24 | 0.08 | 0.1 | 0.11 | 0.14 | 0.27 |
| 8 | 126465305 | rs17730649 | -0.959 | -2.288 | -0.288 | -1.143 | -0.015 | 0.02 | 0.02 | 0 | 0.69 | 0.75 | 0.58 | 0.86 | 0.4 | 0.75 |
| 8 | 126470817 | rs4518686 | -1.112 | -2.347 | -0.389 | -1.144 | -0.048 | 0.039 | 0.039 | 0 | 0.7 | 0.85 | 0.59 | 0.85 | 0.41 | 0.75 |
| 8 | 126470831 | rs12678799 | 1.776 | -2.302 | -0.367 | -0.792 | 0.005 | 0.046 | 0.046 | 0.369 | 0.22 | 0.14 | 0.33 | 0.15 | 0.18 | 0.32 |
| 8 | 126473441 | rs10956248 | -0.322 | -2.389 | 0.049 | -0.086 | 0.019 | 0.012 | 0.012 | 0 | 0.56 | 0.68 | 0.51 | 0.66 | 0.34 | 0.69 |
| 8 | 126474306 | rs2980885 | -0.741 | -2.053 | -0.493 | -0.251 | -0.003 | 0.004 | 0.004 | 0 | 0.77 | 0.87 | 0.85 | 0.81 | 0.76 | 0.91 |
| 8 | 126474356 | rs2980884 | 0.558 | -1.886 | -0.277 | -0.182 | 0.006 | 0.059 | 0.059 | 0 | 0.42 | 0.14 | 0.6 | 0.29 | 0.61 | 0.51 |
| 8 | 126479362 | rs6982502 | -0.57 | -2.113 | -0.659 | -0.357 | -0.109 | -0.014 | -0.014 | 0.525 | 0.44 | 0.92 | 0.55 | 0.58 | 0.52 | 0.46 |
| 8 | 126479522 | rs7003006 | -0.677 | -2.217 | 0.101 | -0.419 | -0.003 | -0.071 | -0.071 | 0 | 0.11 | 0.5 | 0.13 | 0.02 | 0.06 | 0.11 |
| 8 | 126481694 | rs2980876 | -0.388 | -2.214 | 0.383 | -0.453 | 0.007 | 0.012 | 0.012 | 0 | 0.6 | 0.66 | 0.78 | 0.73 | 0.67 | 0.76 |
| 8 | 126482621 | rs2954022 | 0.499 | -2.171 | -0.327 | -0.429 | -0.003 | -0.002 | -0.002 | 0 | 0.32 | 0.33 | 0.42 | 0.55 | 0.45 | 0.35 |
| 8 | 126485531 | rs7846466 | -0.737 | -2.029 | -0.826 | -0.325 | -0.014 | 0.022 | 0.022 | 0 | 0.64 | 0.93 | 0.79 | 0.73 | 0.68 | 0.77 |
| 8 | 126494317 | rs4457349 | -0.154 | -1.735 | -0.414 | 0.028 | 0.029 | 0.437 | 0.437 | 2.975 | 0.08 | 0.49 | 0.11 | 0.43 | 0.03 | 0.2 |
| 8 | 126495818 | rs10808546 | 1.028 | -1.884 | -0.733 | -0.178 | 0.026 | -0.01 | -0.01 | 0 | 0.24 | 0.31 | 0.35 | 0.12 | 0.43 | 0.22 |
| 8 | 126496775 | rs16900644 | -0.119 | -2.139 | -0.528 | -0.319 | -0.013 | 0.018 | 0.018 | 2.133 | 0.33 | 0.55 | 0.41 | 0.59 | 0.23 | 0.51 |
| 8 | 126501528 | rs7846135 | 0.291 | -1.555 | -0.556 | -0.268 | -0.001 | 0.038 | 0.038 | 1.937 | 0.29 | 0.57 | 0.43 | 0.59 | 0.25 | 0.46 |
| 8 | 126502526 | rs12677676 | 1.251 | -1.196 | -1.514 | -1.13 | -0.039 | 0.034 | 0.034 | 0.037 | 0.21 | 0.08 | 0.31 | 0.17 | 0.22 | 0.27 |
| 8 | 126514676 | rs7012891 | 0.145 | -0.983 | -2.099 | -1.618 | 0.058 | 0.127 | 0.127 | 0.167 | 0.41 | 0.79 | 0.39 | 0.81 | 0.24 | 0.44 |
| 8 | 126516988 | rs4512391 | -0.563 | -1.211 | -2.584 | -1.801 | -0.075 | 0.164 | 0.164 | 0 | 0.6 | 0.95 | 0.45 | 0.82 | 0.34 | 0.64 |
| 8 | 126523523 | rs4360309 | -1.142 | -1.077 | -2.504 | -1.876 | -0.163 | 0.135 | 0.135 | 0 | 0.78 | 0.97 | 0.76 | 0.99 | 0.49 | 0.85 |

| | | | | | | | | | | | | | | | | |
|---|---|---|---|---|---|---|---|---|---|---|---|---|---|---|---|---|
| 8 | 126525035 | rs4545126 | 0.432 | -2.079 | -1.877 | -1.808 | 0.027 | 0.046 | 0.046 | 0 | 0.33 | 0.21 | 0.28 | 0.78 | 0.2 | 0.37 |
| 8 | 126525814 | rs6470362 | -0.61 | -2.091 | -1.77 | -1.788 | 0.112 | 0.171 | 0.171 | 0.243 | 0.53 | 0.79 | 0.38 | 0.79 | 0.25 | 0.47 |
| 8 | 126526834 | rs16900667 | 0.881 | -2.414 | -1.045 | -1.792 | -0.008 | -0.026 | -0.026 | 4.453 | 0.16 | 0.23 | 0.09 | 0.1 | 0.12 | 0.07 |
| 8 | 126528731 | rs7002027 | -0.359 | -2.645 | -1.012 | -1.422 | -0.06 | 0.002 | 0.002 | 0 | 0.37 | 0.53 | 0.29 | 0.61 | 0.15 | 0.4 |
| 8 | 126534431 | rs921722 | -1.511 | -2.168 | -1.47 | -1.651 | -0.063 | 0.07 | 0.07 | 0 | 0.87 | 0.98 | 0.9 | 0.98 | 0.8 | 0.81 |
| 8 | 126534671 | rs921720 | 1.429 | -2.466 | -1.526 | -1.806 | -0.024 | -0.007 | -0.007 | 0 | 0.41 | 0.1 | 0.56 | 0.37 | 0.63 | 0.37 |
| 8 | 126539437 | rs7824301 | 0.008 | -2.498 | -1.231 | -1.9 | -0.021 | -0.036 | -0.036 | 0.275 | 0.09 | 0.39 | 0.05 | 0 | 0.02 | 0.03 |
| 8 | 126540051 | rs1551398 | 1.31 | -2.28 | -1.283 | -1.762 | -0.024 | 0.012 | 0.012 | 0.208 | 0.41 | 0.1 | 0.56 | 0.31 | 0.63 | 0.37 |
| 8 | 126541659 | rs1809167 | 0.49 | -1.726 | -0.72 | -1.331 | -0.019 | 0.018 | 0.018 | 0 | 0.25 | 0.36 | 0.24 | 0.67 | 0.11 | 0.3 |
| 8 | 126543022 | rs978533 | 0.075 | -1.002 | -0.661 | -1.479 | 0 | -0.1 | -0.1 | 0 | 0.78 | 0.7 | 0.86 | 0.98 | 0.78 | 0.78 |
| 8 | 126546389 | rs4870943 | -0.185 | -0.957 | -1.254 | -1.116 | 0 | 0.123 | 0.123 | 0 | 0.61 | 0.81 | 0.8 | 0.93 | 0.62 | 0.71 |
| 8 | 126556185 | rs969674 | -0.309 | -0.965 | -0.713 | -0.809 | 0.003 | -0.002 | -0.002 | 0 | 0.54 | 0.6 | 0.75 | 0.92 | 0.57 | 0.69 |
| 8 | 126559887 | rs6992898 | -0.87 | -1.473 | -0.002 | 0.8 | -0.082 | 0.128 | 0.128 | 0 | 0.22 | 0.45 | 0.08 | 0.59 | 0.09 | 0.26 |
| 8 | 126570279 | rs7829236 | -0.326 | -0.915 | -0.557 | 0.475 | 0.005 | 0.022 | 0.022 | 0 | 0.11 | 0.06 | 0.04 | 0.05 | 0.08 | 0.12 |
| 8 | 126576084 | rs4006555 | 0.196 | -1.249 | -0.566 | 0.468 | 0.01 | 0.016 | 0.016 | 0.385 | 0.66 | 0.73 | 0.8 | 0.79 | 0.69 | 0.74 |
| 8 | 126576712 | rs4871617 | 1.291 | -1.534 | -0.916 | 0.775 | -0.018 | 0.152 | 0.152 | 1.778 | 0.44 | 0.1 | 0.43 | 0.24 | 0.58 | 0.42 |
| 8 | 126581168 | rs897148 | -0.129 | -1.43 | -0.814 | 1.206 | -0.025 | -0.066 | -0.066 | 0 | 0.69 | 0.96 | 0.83 | 0.63 | 0.71 | 0.74 |
| 8 | 126582333 | rs17323382 | -0.436 | -1.307 | 0.299 | 1.43 | 0.006 | 0.01 | 0.01 | 0 | 0.24 | 0.36 | 0.25 | 0.26 | 0.24 | 0.31 |
| 8 | 126584451 | rs7821956 | -0.159 | -1.238 | 0.173 | 1.301 | -0.003 | 0.004 | 0.004 | 0.042 | 0.18 | 0.24 | 0.26 | 0.16 | 0.17 | 0.23 |
| 8 | 126590562 | rs1963945 | -1.371 | -1.254 | -0.053 | 1.316 | -0.011 | -0.023 | -0.023 | 0 | 0.46 | 0.83 | 0.55 | 0.42 | 0.42 | 0.55 |
| 8 | 126594738 | rs7016867 | -0.138 | -1.176 | 0.019 | 1.25 | 0.002 | -0.01 | -0.01 | 1.174 | 0.18 | 0.13 | 0.23 | 0.2 | 0.17 | 0.21 |
| 8 | 126603194 | rs4870946 | 1.441 | -1.386 | 0.537 | 1.253 | -0.003 | 0.007 | 0.007 | 0 | 0.46 | 0.55 | 0.5 | 0.36 | 0.51 | 0.43 |
| 8 | 126604637 | rs10086636 | 1.389 | -1.377 | 0.684 | 1.265 | -0.008 | 0.027 | 0.027 | 0 | 0.45 | 0.59 | 0.44 | 0.22 | 0.5 | 0.41 |
| 8 | 126611214 | rs1454616 | 1.394 | -1.391 | 0.709 | 1.069 | -0.005 | 0.016 | 0.016 | 0 | 0.46 | 0.57 | 0.5 | 0.36 | 0.51 | 0.43 |
| 8 | 126611303 | rs1454617 | 0.591 | -1.194 | 0.551 | 1.216 | 0.03 | -0.065 | -0.065 | 0.581 | 0.7 | 0.84 | 0.7 | 0.6 | 0.75 | 0.74 |
| 8 | 126614904 | rs1454621 | -0.637 | -1.264 | 0.51 | 1.254 | 0.03 | -0.033 | -0.033 | 0.038 | 0.87 | 0.94 | 0.93 | 0.8 | 0.93 | 0.96 |
| 8 | 126616615 | rs11991475 | -0.7 | -1.042 | 0.936 | 1.379 | -0.009 | 0.005 | 0.005 | 0.007 | 0.22 | 0.31 | 0.26 | 0.21 | 0.18 | 0.24 |
| 8 | 126618736 | rs4871622 | 0.448 | -1.078 | 0.654 | 1.151 | -0.002 | -0.006 | -0.006 | 0 | 0.68 | 0.36 | 0.74 | 0.78 | 0.71 | 0.84 |
| 8 | 126620703 | rs1979221 | -0.241 | -0.876 | 0.142 | 1.47 | -0.052 | -0.047 | -0.047 | 0 | 0.76 | 0.95 | 0.81 | 0.69 | 0.71 | 0.83 |
| 8 | 126621911 | rs7004305 | -1.226 | -0.959 | 0.948 | 1.642 | -0.01 | 0.023 | 0.023 | 0 | 0.26 | 0.35 | 0.33 | 0.38 | 0.2 | 0.42 |
| 8 | 126625317 | rs13282254 | -1.697 | -0.83 | 1.316 | 1.388 | -0.028 | 0.083 | 0.083 | 0.476 | 0.11 | 0.62 | 0.14 | 0.13 | 0.04 | 0.25 |
| 8 | 126629575 | rs7004902 | -1.701 | -0.727 | 1.22 | 1.469 | -0.014 | 0.037 | 0.037 | 0.137 | 0.23 | 0.62 | 0.33 | 0.41 | 0.19 | 0.31 |
| 8 | 126631628 | rs4557721 | 0.55 | -0.754 | 0.828 | 0.97 | -0.001 | -0.022 | -0.022 | 0 | 0.08 | 0.07 | 0.06 | 0 | 0.08 | 0.03 |
| 8 | 126632368 | rs1812393 | -0.717 | -0.807 | 1.343 | 1.248 | -0.004 | 0.065 | 0.065 | 0.566 | 0.06 | 0.27 | 0.16 | 0.1 | 0.03 | 0.13 |
| 8 | 126644200 | rs2124034 | -2.347 | -1.165 | 0.693 | 1.202 | 0.05 | -0.028 | -0.028 | 0 | 0.37 | 0.88 | 0.43 | 0.46 | 0.28 | 0.37 |
| 8 | 126645018 | rs11990155 | -0.829 | -1.179 | 0.862 | 1.182 | 0.003 | -0.026 | -0.026 | 0 | 0.16 | 0.12 | 0.14 | 0.34 | 0.14 | 0.19 |
| 8 | 126645347 | rs10956254 | -2.283 | -1.122 | 0.613 | 1.107 | 0.049 | -0.023 | -0.023 | 0 | 0.37 | 0.86 | 0.43 | 0.46 | 0.28 | 0.37 |
| 8 | 126648134 | rs2124036 | -2.294 | -1.179 | 0.582 | 1.198 | 0.06 | -0.052 | -0.052 | 0 | 0.34 | 0.88 | 0.39 | 0.48 | 0.23 | 0.35 |
| 8 | 126649545 | rs16900769 | -0.568 | -0.814 | 0.885 | 0.836 | -0.02 | -0.071 | -0.071 | 0.549 | 0.12 | 0.21 | 0.07 | 0 | 0.07 | 0.03 |
| 8 | 126650425 | rs11774279 | -0.235 | -0.197 | -0.527 | 0.832 | -0.002 | 0.028 | 0.028 | 0.198 | 0.51 | 0.77 | 0.45 | 0.56 | 0.4 | 0.51 |
| 8 | 126656148 | rs7819544 | -0.133 | 0.333 | -0.243 | 0.936 | 0 | -0.001 | -0.001 | 0 | 0.47 | 0.45 | 0.37 | 0.48 | 0.41 | 0.44 |
| 8 | 126656246 | rs9643207 | -0.045 | 0.393 | -0.72 | 0.926 | 0.01 | -0.022 | -0.022 | 0 | 0.44 | 0.26 | 0.35 | 0.48 | 0.36 | 0.44 |
| 8 | 126667489 | rs12678366 | -0.572 | -0.033 | -0.081 | 1.123 | -0.036 | 0.125 | 0.125 | 0.111 | 0.22 | 0.42 | 0.18 | 0.38 | 0.17 | 0.16 |
| 8 | 126671176 | rs897157 | -0.737 | 0.911 | -0.914 | 0.261 | -0.041 | 0.039 | 0.039 | 0 | 0.85 | 0.89 | 0.69 | 0.97 | 0.7 | 0.85 |
| 8 | 126672313 | rs729005 | -0.044 | 0.378 | -0.629 | 0.432 | 0.01 | 0.107 | 0.107 | 0 | 0.08 | 0.19 | 0.09 | 0.26 | 0.07 | 0.04 |
| 8 | 126672930 | rs2015243 | 0.788 | -0.203 | -1.114 | -0.12 | -0.002 | 0.058 | 0.058 | 0.853 | 0.48 | 0.33 | 0.69 | 0.74 | 0.45 | 0.57 |
| 8 | 126674443 | rs1809170 | -0.082 | 0.456 | -1.107 | 0.082 | -0.064 | 0.054 | 0.054 | 0.318 | 0.79 | 0.93 | 0.86 | 0.94 | 0.63 | 0.85 |
| 8 | 126681996 | rs2385226 | 1.181 | -0.095 | -0.936 | -0.082 | -0.027 | -0.006 | -0.006 | 0.061 | 0.59 | 0.21 | 0.62 | 0.57 | 0.49 | 0.67 |
| 8 | 126688445 | rs1159477 | 0.103 | -0.18 | -0.82 | -0.066 | -0.057 | 0.023 | 0.023 | 0 | 0.2 | 0.1 | 0.22 | 0.08 | 0.37 | 0.2 |
| 8 | 126689998 | rs11987816 | -0.343 | -0.366 | -0.144 | -0.047 | -0.036 | -0.014 | -0.014 | 0 | 0.14 | 0.44 | 0.07 | 0.12 | 0.08 | 0.08 |
| 8 | 126694158 | rs2054068 | -1.454 | -0.255 | -0.089 | -0.073 | -0.004 | -0.002 | -0.002 | 0.15 | 0.39 | 0.68 | 0.36 | 0.42 | 0.5 | 0.32 |
| 8 | 126701280 | rs7826323 | -0.422 | -0.91 | -0.383 | -0.261 | -0.051 | -0.025 | -0.025 | 0.287 | 0.15 | 0.46 | 0.07 | 0.12 | 0.08 | 0.07 |
| 8 | 126705306 | rs4871632 | 1.154 | -0.779 | -1.651 | -0.458 | 0.015 | 0.094 | 0.094 | 0.127 | 0.5 | 0.13 | 0.56 | 0.66 | 0.4 | 0.66 |
| 8 | 126707061 | rs17332697 | 0.089 | -0.61 | -1.594 | -0.539 | -0.108 | 0.086 | 0.086 | 0 | 0.28 | 0.09 | 0.27 | 0.09 | 0.46 | 0.24 |
| 8 | 126707635 | rs7846616 | -1.066 | -1.313 | -0.345 | 0.002 | -0.013 | 0 | 0 | 0.533 | 0.21 | 0.77 | 0.16 | 0.24 | 0.13 | 0.1 |
| 8 | 126708781 | rs4871633 | -0.346 | -0.693 | -0.61 | -0.138 | 0.006 | -0.027 | -0.027 | 0.201 | 0.69 | 0.89 | 0.65 | 0.64 | 0.73 | 0.65 |
| 8 | 126711943 | rs13271980 | -1.285 | -0.969 | 0.795 | -0.653 | -0.008 | -0.006 | -0.006 | 0 | 0.31 | 0.68 | 0.21 | 0.13 | 0.18 | 0.13 |
| 8 | 126717244 | rs10216366 | -1.228 | -1.013 | 1.527 | -0.88 | -0.065 | -0.147 | -0.147 | 0 | 0.3 | 0.45 | 0.13 | 0.04 | 0.19 | 0.13 |
| 8 | 126728531 | rs6998174 | -0.47 | -0.967 | 2.054 | -0.84 | 0.006 | -0.012 | -0.012 | 0 | 0.2 | 0.14 | 0.08 | 0.01 | 0.14 | 0.08 |
| 8 | 126728715 | rs7018215 | -1.001 | -0.977 | 2.349 | -0.845 | -0.063 | -0.147 | -0.147 | 0 | 0.26 | 0.44 | 0.12 | 0.04 | 0.16 | 0.12 |
| 8 | 126731757 | rs7014615 | -1.169 | -0.962 | 2.133 | -0.829 | -0.035 | -0.065 | -0.065 | 3.459 | 0.27 | 0.6 | 0.14 | 0.08 | 0.17 | 0.12 |
| 8 | 126732991 | rs10441520 | -0.429 | -0.862 | 1.836 | -0.852 | -0.019 | -0.065 | -0.065 | 0 | 0.21 | 0.23 | 0.09 | 0.01 | 0.15 | 0.08 |
| 8 | 126740960 | rs10505455 | -0.925 | -0.732 | 1.434 | -0.752 | -0.064 | -0.125 | -0.125 | 0 | 0.28 | 0.52 | 0.18 | 0.05 | 0.19 | 0.12 |
| 8 | 126749993 | rs13280027 | -1.003 | -0.711 | 0.94 | -0.851 | -0.045 | -0.111 | -0.111 | 0 | 0.29 | 0.59 | 0.14 | 0.03 | 0.19 | 0.12 |
| 8 | 126761402 | rs3087723 | 0.694 | 0.441 | -0.158 | -0.305 | -0.017 | 0.035 | 0.035 | 0 | 0.59 | 0.72 | 0.68 | 0.7 | 0.54 | 0.72 |
| 8 | 126763557 | rs4871637 | 0.618 | -0.422 | 0.24 | -0.377 | -0.018 | -0.03 | -0.03 | 0 | 0.76 | 0.71 | 0.83 | 0.9 | 0.82 | 0.91 |
| 8 | 126765322 | rs7836147 | -1.943 | -0.382 | 0.838 | -0.08 | -0.021 | 0.041 | 0.041 | 9.458 | 0.08 | 0.67 | 0.11 | 0.07 | 0.01 | 0.02 |
| 8 | 126765996 | rs11775468 | -1.272 | 0.015 | 1.032 | -0.476 | 0.024 | -0.049 | -0.049 | 0 | 0.09 | 0.21 | 0.06 | 0 | 0.11 | 0.05 |
| 8 | 126772563 | rs10956258 | -0.036 | -0.375 | -0.073 | -0.922 | -0.086 | 0.087 | 0.087 | 0.254 | 0.33 | 0.02 | 0.4 | 0.17 | 0.35 | 0.39 |

| | | | | | | | | | | | | | | | | |
|---|---|---|---|---|---|---|---|---|---|---|---|---|---|---|---|---|
| 8 | 126774170 | rs7008332 | 0.336 | -1.126 | 0.003 | -1.068 | -0.009 | -0.003 | -0.003 | 0 | 0.15 | 0.13 | 0.1 | 0.11 | 0.08 | 0.22 |
| 8 | 126774314 | rs1444003 | -0.907 | -1.259 | 0.902 | -1.096 | 0.015 | 0.077 | 0.077 | 0.065 | 0.54 | 0.74 | 0.55 | 0.32 | 0.43 | 0.62 |
| 8 | 126781758 | rs16900842 | 0.495 | -1.21 | 1.586 | -1.218 | -0.022 | -0.016 | -0.016 | 0.041 | 0.12 | 0.17 | 0.09 | 0.07 | 0.07 | 0.19 |
| 8 | 126789340 | rs6988375 | -0.35 | -1.234 | 1.731 | -1.229 | -0.064 | -0.038 | -0.038 | 0.364 | 0.16 | 0.56 | 0.11 | 0.1 | 0.07 | 0.19 |
| 8 | 126794015 | rs7003960 | 0.458 | -0.885 | 1.033 | -1.764 | 0.068 | -0.157 | -0.157 | 1.462 | 0.26 | 0.52 | 0.31 | 0.03 | 0.4 | 0.15 |
| 8 | 126795544 | rs6470379 | -1.022 | -0.934 | 0.727 | -1.568 | 0.002 | 0.001 | 0.001 | 1.38 | 0.76 | 0.77 | 0.81 | 0.25 | 0.82 | 0.74 |
| 8 | 126798409 | rs9297719 | -0.127 | -0.988 | 0.999 | -1.633 | 0.017 | -0.094 | -0.094 | 0 | 0.38 | 0.46 | 0.4 | 0.12 | 0.47 | 0.37 |
| 8 | 126799011 | rs7841870 | 0.105 | -0.944 | 1.046 | -1.612 | -0.019 | 0.026 | 0.026 | 0 | 0.35 | 0.31 | 0.35 | 0.09 | 0.46 | 0.32 |
| 8 | 126802262 | rs1443997 | 0.424 | -0.933 | 1.037 | -1.609 | -0.01 | -0.06 | -0.06 | 0 | 0.41 | 0.44 | 0.45 | 0.15 | 0.36 | 0.42 |
| 8 | 126806534 | rs1023795 | 0.405 | -0.924 | 0.797 | -1.461 | -0.007 | -0.042 | -0.042 | 0.174 | 0.41 | 0.46 | 0.45 | 0.2 | 0.36 | 0.43 |
| 8 | 126806535 | rs1023794 | 0.106 | -0.906 | 0.434 | -1.468 | 0.026 | 0.09 | 0.09 | 0.014 | 0.44 | 0.71 | 0.47 | 0.2 | 0.36 | 0.43 |
| 8 | 126812638 | rs2028709 | -0.997 | -0.967 | 0.444 | -1.416 | 0.004 | 0 | 0 | 1.547 | 0.76 | 0.78 | 0.8 | 0.28 | 0.82 | 0.74 |
| 8 | 126828417 | rs10103387 | 0.356 | -0.938 | 0.28 | -1.454 | -0.085 | 0.143 | 0.143 | 0 | 0.34 | 0.14 | 0.35 | 0.09 | 0.46 | 0.32 |
| 8 | 126830010 | rs1443993 | -0.945 | -0.917 | -0.135 | -1.236 | 0.002 | -0.002 | -0.002 | 0 | 0.76 | 0.79 | 0.81 | 0.35 | 0.82 | 0.75 |
| 8 | 126835161 | rs1373325 | 0.295 | -0.553 | -0.546 | -0.949 | -0.04 | 0.113 | 0.113 | 0.013 | 0.34 | 0.05 | 0.39 | 0.16 | 0.34 | 0.37 |
| 8 | 126843414 | rs16900891 | -0.967 | -0.337 | 0.477 | -0.255 | -0.005 | 0.002 | 0.002 | 0.994 | 0.06 | 0.26 | 0.05 | 0.03 | 0.01 | 0.1 |
| 8 | 126867367 | rs3955454 | 0.048 | -0.158 | -0.11 | 0.4 | 0 | -0.007 | -0.007 | 0 | 0.64 | 0.66 | 0.75 | 0.53 | 0.67 | 0.61 |
| 8 | 126867463 | rs3955453 | -0.003 | -0.235 | 0.005 | 0.377 | 0 | 0.03 | 0.03 | 0 | 0.68 | 0.59 | 0.78 | 0.52 | 0.7 | 0.64 |
| 8 | 126867505 | rs4006531 | 0.27 | -0.874 | -0.161 | 0.219 | 0.043 | 0.018 | 0.018 | 0.583 | 0.55 | 0.27 | 0.73 | 0.4 | 0.68 | 0.55 |
| 8 | 126868230 | rs6470391 | 0.301 | 0.134 | -0.48 | 0.361 | 0 | 0.001 | 0.001 | 0 | 0.74 | 0.74 | 0.69 | 0.39 | 0.69 | 0.74 |
| 8 | 126873484 | rs10092499 | 0.462 | 0.465 | -1.081 | -0.749 | -0.02 | 0.048 | 0.048 | 0 | 0.77 | 0.85 | 0.7 | 0.09 | 0.7 | 0.68 |
| 8 | 126888589 | rs4870957 | 0.428 | 0.338 | -0.935 | -0.812 | -0.009 | 0.03 | 0.03 | 0 | 0.75 | 0.78 | 0.7 | 0.08 | 0.7 | 0.68 |
| 8 | 126895276 | rs1491477 | 1.119 | 0.494 | -1.957 | -1.77 | -0.036 | 0.147 | 0.147 | 0.021 | 0.28 | 0.01 | 0.2 | 0.07 | 0.22 | 0.35 |
| 8 | 126898700 | rs6986812 | -0.017 | 0.955 | -1.549 | -1.519 | -0.022 | 0.054 | 0.054 | 0 | 0.76 | 0.85 | 0.7 | 0.09 | 0.7 | 0.68 |
| 8 | 126909415 | rs6999325 | 0.951 | 1.136 | -0.843 | -2.019 | 0.003 | -0.094 | -0.094 | 0.276 | 0.27 | 0.29 | 0.21 | 0.01 | 0.31 | 0.2 |
| 8 | 126912468 | rs7818391 | 0.184 | 1.216 | -0.97 | -1.934 | -0.035 | 0.073 | 0.073 | 0 | 0.24 | 0.11 | 0.29 | 0.91 | 0.3 | 0.32 |
| 8 | 126915549 | rs10505462 | 0.945 | 0.977 | -0.544 | -1.892 | 0.044 | -0.046 | -0.046 | 0 | 0.08 | 0.2 | 0.23 | 0 | 0.13 | 0.09 |
| 8 | 126927059 | rs1873549 | -0.082 | 1.068 | -0.577 | -1.379 | -0.007 | 0.02 | 0.02 | 0 | 0.23 | 0.21 | 0.3 | 0.92 | 0.27 | 0.31 |
| 8 | 126936152 | rs1388372 | 0.057 | 1.067 | -0.619 | -1.378 | -0.022 | 0.054 | 0.054 | 0 | 0.22 | 0.14 | 0.29 | 0.9 | 0.27 | 0.31 |
| 8 | 126938574 | rs10956270 | -0.192 | 1.057 | -0.631 | -1.404 | -0.008 | 0.031 | 0.031 | 0 | 0.24 | 0.21 | 0.3 | 0.92 | 0.29 | 0.32 |
| 8 | 126944294 | rs2087688 | 0.713 | 1.117 | -0.682 | -1.416 | 0.007 | -0.005 | -0.005 | 1.357 | 0.46 | 0.42 | 0.54 | 0.08 | 0.36 | 0.48 |
| 8 | 126953742 | rs729088 | -0.836 | 1.487 | -0.475 | -0.12 | 0.001 | -0.017 | -0.017 | 0 | 0.25 | 0.27 | 0.25 | 0.91 | 0.3 | 0.32 |
| 8 | 126959274 | rs1602511 | -0.682 | 1.471 | -0.658 | 0.504 | -0.003 | -0.005 | -0.005 | 0.626 | 0.18 | 0.16 | 0.1 | 0.29 | 0.2 | 0.13 |
| 8 | 126962912 | rs1491467 | -0.47 | 1.223 | -0.587 | 0.745 | -0.001 | 0.031 | 0.031 | 0.156 | 0.06 | 0.05 | 0.04 | 0.25 | 0.06 | 0.09 |
| 8 | 126964809 | rs1491466 | -0.635 | 1.602 | -0.818 | -0.007 | -0.025 | 0.038 | 0.038 | 0 | 0.27 | 0.21 | 0.21 | 0.08 | 0.36 | 0.21 |
| 8 | 126970372 | rs10481157 | -1.142 | 1.417 | -0.311 | -0.072 | 0.001 | 0.013 | 0.013 | 0 | 0.47 | 0.48 | 0.67 | 0.63 | 0.53 | 0.46 |
| 8 | 126978363 | rs4870958 | 0.302 | 0.897 | -0.033 | 0.102 | 0.012 | -0.049 | -0.049 | 0.927 | 0.2 | 0.1 | 0.44 | 0.55 | 0.17 | 0.3 |
| 8 | 126983953 | rs4576392 | -1.132 | 1.083 | 0.529 | -0.005 | -0.004 | -0.002 | -0.002 | 0 | 0.38 | 0.42 | 0.57 | 0.59 | 0.35 | 0.44 |
| 8 | 126984970 | rs967851 | -1.169 | 1.072 | 0.516 | -0.01 | -0.006 | -0.004 | -0.004 | 0 | 0.39 | 0.54 | 0.59 | 0.59 | 0.35 | 0.44 |
| 8 | 126995907 | rs13264202 | 0.546 | 1.064 | 0.195 | 0.134 | 0.009 | -0.037 | -0.037 | 0.225 | 0.18 | 0.09 | 0.33 | 0.52 | 0.14 | 0.28 |
| 8 | 126997700 | rs7017721 | -0.401 | 1.094 | 0.25 | -0.011 | 0.018 | -0.064 | -0.064 | 0.296 | 0.18 | 0.3 | 0.25 | 0.05 | 0.2 | 0.16 |
| 8 | 126997975 | rs10956274 | 0.576 | 1.128 | -0.495 | 0.25 | -0.004 | -0.104 | -0.104 | 0.069 | 0.17 | 0.03 | 0.32 | 0.51 | 0.15 | 0.29 |
| 8 | 127004405 | rs12681169 | -1.229 | 1.151 | -0.306 | -0.512 | 0.016 | 0.099 | 0.099 | 0 | 0.46 | 0.68 | 0.65 | 0.87 | 0.42 | 0.53 |
| 8 | 127006184 | rs12678127 | 0.433 | 1.078 | -0.588 | -0.042 | -0.03 | -0.08 | -0.08 | 0 | 0.16 | 0.01 | 0.32 | 0.67 | 0.15 | 0.3 |
| 8 | 127008916 | rs6470399 | -0.836 | 1.453 | -0.243 | -0.356 | -0.015 | 0.057 | 0.057 | 0 | 0.42 | 0.67 | 0.63 | 0.69 | 0.42 | 0.48 |
| 8 | 127010042 | rs16901028 | -1.179 | 1.183 | 0.343 | -0.38 | 0.038 | -0.013 | -0.013 | 0 | 0.06 | 0.23 | 0.09 | 0 | 0.07 | 0.04 |
| 8 | 127014085 | rs1155228 | -0.194 | 1.104 | -0.223 | -0.684 | 0 | 0.003 | 0.003 | 0 | 0.43 | 0.55 | 0.28 | 0.26 | 0.43 | 0.28 |
| 8 | 127015044 | rs7812787 | -1.113 | 0.658 | 0.56 | -1.024 | 0.02 | -0.015 | -0.015 | 0 | 0.06 | 0.31 | 0.06 | 0 | 0.02 | 0.02 |
| 8 | 127016559 | rs4483112 | -0.539 | 0.917 | -0.368 | -1.953 | -0.044 | 0.073 | 0.073 | 0 | 0.65 | 0.84 | 0.71 | 0.86 | 0.53 | 0.68 |
| 8 | 127018815 | rs6998016 | 0.559 | 0.342 | -0.318 | -2.661 | -0.02 | -0.116 | -0.116 | 0.078 | 0.26 | 0.42 | 0.16 | 0.01 | 0.18 | 0.21 |
| 8 | 127030253 | rs7825598 | -0.967 | 0.885 | -0.65 | -2.875 | 0.024 | -0.02 | -0.02 | 0 | 0.93 | 0.85 | 0.94 | 0.97 | 0.9 | 0.97 |
| 8 | 127031728 | rs962955 | 0.931 | 0.82 | -0.54 | -2.919 | 0.006 | -0.014 | -0.014 | 0 | 0.25 | 0.25 | 0.13 | 0.01 | 0.18 | 0.2 |
| 8 | 127032660 | rs7016046 | -0.151 | 0.796 | -0.548 | -2.901 | 0.047 | -0.033 | -0.033 | 0 | 0.91 | 0.83 | 0.91 | 0.97 | 0.84 | 0.95 |
| 8 | 127032911 | rs10113614 | 1.481 | 0.212 | -1.018 | -3.005 | -0.048 | -0.038 | -0.038 | 0 | 0.13 | 0.02 | 0.38 | 0.79 | 0.18 | 0.26 |
| 8 | 127045555 | rs13279203 | -0.138 | 0.407 | -0.668 | -3.354 | 0.003 | -0.098 | -0.098 | 5.604 | 0.66 | 0.58 | 0.78 | 0.95 | 0.66 | 0.75 |
| 8 | 127045960 | rs7460327 | 1.535 | 0.302 | -0.905 | -3.438 | -0.143 | 0.162 | 0.162 | 0.139 | 0.29 | 0.01 | 0.23 | 0.07 | 0.38 | 0.33 |
| 8 | 127050415 | rs1552706 | 0.553 | 0.225 | 0.404 | -2.752 | 0.002 | -0.131 | -0.131 | 0 | 0.65 | 0.46 | 0.75 | 0.95 | 0.66 | 0.75 |
| 8 | 127054333 | rs1491461 | 0.098 | 0.293 | 0.381 | -2.656 | 0.047 | -0.011 | -0.011 | 0 | 0.92 | 0.7 | 0.9 | 0.97 | 0.9 | 0.97 |
| 8 | 127058720 | rs12543713 | 0.961 | 0.199 | -0.199 | -2.706 | 0.014 | 0.042 | 0.042 | 0.92 | 0.21 | 0.11 | 0.09 | 0.01 | 0.16 | 0.19 |
| 8 | 127058751 | rs6984936 | -0.363 | 0.168 | -0.036 | -2.709 | -0.007 | -0.133 | -0.133 | 0.283 | 0.31 | 0.58 | 0.2 | 0.01 | 0.27 | 0.23 |
| 8 | 127071007 | rs980348 | 1.578 | -0.009 | -0.319 | -2.285 | 0.008 | 0.054 | 0.054 | 0.826 | 0.76 | 0.17 | 0.82 | 0.94 | 0.83 | 0.79 |
| 8 | 127071167 | rs980349 | 1.374 | 0.097 | 0.248 | -2.089 | -0.002 | -0.057 | -0.057 | 2.66 | 0.85 | 0.31 | 0.85 | 0.95 | 0.87 | 0.93 |
| 8 | 127071998 | rs2957795 | 1.55 | -0.119 | -0.547 | -2.282 | 0.034 | 0.109 | 0.109 | 4.226 | 0.73 | 0.16 | 0.8 | 0.96 | 0.78 | 0.9 |
| 8 | 127083695 | rs780321 | 1.532 | -0.122 | -0.49 | -2.283 | 0.032 | 0.092 | 0.092 | 3.979 | 0.73 | 0.19 | 0.81 | 0.97 | 0.78 | 0.9 |
| 8 | 127099058 | rs1872934 | -0.148 | 0.088 | -0.215 | -2.131 | 0.003 | 0.009 | 0.009 | 0 | 0.08 | 0.05 | 0.04 | 0 | 0.08 | 0.05 |
| 8 | 127103441 | rs1488024 | -1.719 | -0.166 | 0.467 | -2.287 | -0.032 | -0.148 | -0.148 | 0 | 0.23 | 0.51 | 0.13 | 0.01 | 0.18 | 0.1 |
| 8 | 127103854 | rs1542691 | -0.919 | 0.348 | -0.143 | -2.382 | 0.029 | 0.193 | 0.193 | 0.491 | 0.43 | 0.81 | 0.23 | 0.08 | 0.37 | 0.35 |
| 8 | 127107701 | rs4436090 | -0.938 | 0.634 | 0.057 | -1.655 | 0.007 | -0.053 | -0.053 | 0 | 0.11 | 0.21 | 0.06 | 0.01 | 0.1 | 0.08 |
| 8 | 127131088 | rs1386784 | -1.332 | 0.644 | -0.131 | -1.627 | 0.034 | -0.075 | -0.075 | 0 | 0.13 | 0.47 | 0.1 | 0.01 | 0.11 | 0.08 |

| | | | | | | | | | | | | | | | |
|---|---|---|---|---|---|---|---|---|---|---|---|---|---|---|---|
| 8 | 127150370 | rs7006495 | -1.233 | 0.841 | -0.939 | -1.091 | -0.063 | -0.055 | -0.055 | 0 | 0.69 | 0.99 | 0.68 | 0.33 | 0.71 | 0.65 |
| 8 | 127153541 | rs10107938 | -1.464 | 1.127 | 0.431 | -0.804 | 0.002 | -0.078 | -0.078 | 0 | 0.12 | 0.48 | 0.08 | 0 | 0.08 | 0.07 |
| 8 | 127157289 | rs1906492 | -1.743 | 1.264 | 0.095 | -0.571 | 0.032 | -0.022 | -0.022 | 2.011 | 0.17 | 0.4 | 0.3 | 0.15 | 0.21 | 0.09 |
| 8 | 127170869 | rs1545978 | -1.302 | 1.358 | 0.008 | -0.225 | 0.068 | 0.057 | 0.057 | 0 | 0.09 | 0.24 | 0.25 | 0.16 | 0.14 | 0.03 |
| 8 | 127174013 | rs4871661 | -0.864 | 1.408 | -0.254 | -0.185 | 0.034 | 0.03 | 0.03 | 0 | 0.07 | 0.13 | 0.24 | 0.16 | 0.14 | 0.03 |
| 8 | 127175463 | rs6470407 | -0.715 | 0.948 | -1.103 | -0.14 | -0.001 | -0.083 | -0.083 | 0 | 0.73 | 0.89 | 0.63 | 0.43 | 0.78 | 0.59 |
| 8 | 127176135 | rs1994469 | -0.555 | 0.945 | -0.431 | -0.501 | 0.001 | -0.017 | -0.017 | 0 | 0.29 | 0.49 | 0.21 | 0.17 | 0.27 | 0.3 |
| 8 | 127176405 | rs16901163 | -0.005 | 0.741 | -0.799 | -0.907 | -0.003 | -0.022 | -0.022 | 0.526 | 0.31 | 0.42 | 0.13 | 0.14 | 0.23 | 0.3 |
| 8 | 127177084 | rs7838729 | 0.31 | 0.612 | -0.871 | -1.006 | 0 | -0.011 | -0.011 | 0 | 0.24 | 0.24 | 0.09 | 0.13 | 0.17 | 0.27 |
| 8 | 127180144 | rs10103019 | -0.049 | 1.423 | -1.445 | -0.788 | -0.048 | -0.047 | -0.047 | 0.016 | 0.08 | 0.01 | 0.26 | 0.18 | 0.22 | 0.08 |
| 8 | 127181094 | rs2385523 | 0.255 | 0.913 | -1.374 | -1.068 | -0.042 | -0.024 | -0.024 | 0.482 | 0.17 | 0.08 | 0.18 | 0.16 | 0.22 | 0.26 |
| 8 | 127182142 | rs6999393 | -1.142 | 0.688 | -1.346 | -0.877 | -0.124 | 0.033 | 0.033 | 0.153 | 0.58 | 0.95 | 0.37 | 0.3 | 0.47 | 0.6 |
| 8 | 127188115 | rs10107324 | -0.555 | 0.511 | -0.438 | -0.627 | 0.009 | 0.034 | 0.034 | 0 | 0.25 | 0.56 | 0.4 | 0.33 | 0.28 | 0.27 |
| 8 | 127188815 | rs6470411 | -0.555 | 0.511 | -0.438 | -0.627 | 0.009 | 0.034 | 0.034 | 0 | 0.25 | 0.56 | 0.4 | 0.33 | 0.28 | 0.27 |
| 8 | 127189138 | rs7357357 | -0.708 | 0.475 | -0.407 | -0.656 | 0.002 | 0.011 | 0.011 | 0.159 | 0.27 | 0.63 | 0.41 | 0.33 | 0.28 | 0.27 |
| 8 | 127208253 | rs2891807 | -0.161 | 0.316 | -0.747 | -0.952 | -0.029 | 0.133 | 0.133 | 0 | 0.36 | 0.17 | 0.18 | 0.13 | 0.39 | 0.35 |
| 8 | 127225738 | rs1488023 | -0.479 | 0.271 | -0.089 | -0.993 | -0.001 | 0.053 | 0.053 | 0 | 0.39 | 0.25 | 0.19 | 0.13 | 0.4 | 0.36 |
| 8 | 127228137 | rs1386796 | -1.765 | 0.266 | -0.198 | -0.958 | -0.004 | -0.101 | -0.101 | 0 | 0.63 | 0.89 | 0.61 | 0.46 | 0.67 | 0.64 |
| 8 | 127239525 | rs9297731 | -0.833 | 0.214 | -0.305 | -1.126 | 0 | -0.001 | -0.001 | 0 | 0.45 | 0.55 | 0.46 | 0.45 | 0.44 | 0.52 |
| 8 | 127240653 | rs7845993 | -0.149 | 0.23 | 0.248 | -0.887 | 0.042 | 0.097 | 0.097 | 0 | 0.21 | 0.43 | 0.39 | 0.33 | 0.27 | 0.27 |
| 8 | 127247885 | rs9297732 | -1.492 | 0.223 | -0.282 | -0.715 | 0.002 | -0.111 | -0.111 | 1.253 | 0.62 | 0.89 | 0.6 | 0.47 | 0.67 | 0.63 |
| 8 | 127254484 | rs1386798 | -1.556 | 0.179 | -0.308 | -0.856 | -0.023 | -0.096 | -0.096 | 0 | 0.63 | 0.95 | 0.59 | 0.43 | 0.65 | 0.63 |
| 8 | 127263042 | rs4870964 | -0.306 | 0.185 | 0.327 | -1.039 | -0.002 | -0.004 | -0.004 | 0.412 | 0.25 | 0.26 | 0.21 | 0.29 | 0.25 | 0.29 |
| 8 | 127267251 | rs4599775 | -1.622 | 0.097 | 1.046 | -1.1 | 0 | 0 | 0 | 0 | 0.61 | 0.6 | 0.39 | 0.41 | 0.64 | 0.63 |
| 8 | 127277326 | rs10090081 | 0.018 | 0.094 | 1.31 | -1.13 | -0.012 | 0.016 | 0.016 | 0 | 0.32 | 0.22 | 0.18 | 0.12 | 0.39 | 0.34 |
| 8 | 127291414 | rs4621770 | -0.537 | 0.071 | 1.366 | -1.111 | -0.002 | -0.004 | -0.004 | 0 | 0.27 | 0.28 | 0.22 | 0.29 | 0.25 | 0.29 |
| 8 | 127292996 | rs4422742 | -1.742 | 0.093 | 1.557 | -1.129 | 0.018 | -0.002 | -0.002 | 0 | 0.62 | 0.75 | 0.42 | 0.41 | 0.64 | 0.63 |
| 8 | 127293085 | rs6990457 | -1.433 | 0.057 | 1.603 | -1.123 | -0.001 | -0.001 | -0.001 | 0 | 0.57 | 0.43 | 0.38 | 0.41 | 0.63 | 0.63 |
| 8 | 127293801 | rs4870966 | -0.264 | 0.077 | 1.382 | -1.12 | 0.003 | -0.054 | -0.054 | 0 | 0.34 | 0.33 | 0.18 | 0.12 | 0.38 | 0.34 |
| 8 | 127304374 | rs10099760 | -1.521 | 0.133 | 1.133 | -1.084 | 0.022 | 0.004 | 0.004 | 0 | 0.6 | 0.7 | 0.4 | 0.41 | 0.64 | 0.63 |
| 8 | 127304509 | rs4503066 | -0.296 | 0.134 | 1.151 | -1.1 | 0.003 | -0.07 | -0.07 | 0 | 0.36 | 0.5 | 0.2 | 0.12 | 0.38 | 0.34 |
| 8 | 127306602 | rs4557669 | 0.141 | 0.119 | 1.014 | -1.026 | -0.008 | -0.015 | -0.015 | 0 | 0.76 | 0.8 | 0.64 | 0.7 | 0.76 | 0.71 |
| 8 | 127308157 | rs7834246 | 1.574 | 0.088 | 0.899 | -1.047 | 0.011 | 0.002 | 0.002 | 0 | 0.4 | 0.36 | 0.45 | 0.58 | 0.36 | 0.38 |
| 8 | 127308560 | rs998238 | -0.229 | 0.154 | 0.956 | -1.074 | 0.012 | -0.098 | -0.098 | 0 | 0.35 | 0.42 | 0.19 | 0.11 | 0.4 | 0.34 |
| 8 | 127320627 | rs926127 | 0.211 | 0.202 | 0.272 | -0.998 | -0.019 | -0.014 | -0.014 | 0.667 | 0.72 | 0.84 | 0.61 | 0.73 | 0.69 | 0.68 |
| 8 | 127322655 | rs726477 | 0.325 | -0.117 | 0.584 | -0.985 | -0.02 | 0.032 | 0.032 | 0 | 0.27 | 0.18 | 0.14 | 0.12 | 0.33 | 0.3 |
| 8 | 127323050 | rs4130556 | -0.288 | 0.031 | 0.097 | -0.911 | -0.003 | -0.004 | -0.004 | 0.631 | 0.33 | 0.63 | 0.19 | 0.12 | 0.33 | 0.29 |
| 8 | 127335010 | rs2253162 | 1.437 | 0.141 | -0.104 | -0.849 | -0.003 | 0.03 | 0.03 | 0 | 0.42 | 0.22 | 0.43 | 0.62 | 0.41 | 0.4 |
| 8 | 127339929 | rs2467618 | 0.624 | 0.295 | -0.005 | -0.184 | -0.003 | -0.014 | -0.014 | 0 | 0.73 | 0.84 | 0.61 | 0.73 | 0.72 | 0.67 |
| 8 | 127343692 | rs10101575 | -1.286 | -0.172 | 0.177 | -0.205 | -0.012 | -0.011 | -0.011 | 1.045 | 0.31 | 0.62 | 0.21 | 0.19 | 0.24 | 0.34 |
| 8 | 127344452 | rs10090539 | -0.7 | 0.149 | -0.351 | -0.15 | -0.003 | 0.017 | 0.017 | 0 | 0.21 | 0.24 | 0.16 | 0.09 | 0.23 | 0.33 |
| 8 | 127363405 | rs10095552 | -1.307 | -0.168 | -1.332 | -0.392 | -0.122 | -0.034 | -0.034 | 0 | 0.63 | 0.97 | 0.37 | 0.29 | 0.54 | 0.61 |
| 8 | 127364695 | rs10505470 | -0.967 | -0.598 | -0.406 | -0.165 | -0.01 | 0.03 | 0.03 | 0 | 0.25 | 0.46 | 0.18 | 0.3 | 0.23 | 0.32 |
| 8 | 127366340 | rs16901318 | -0.593 | -0.431 | -0.173 | -0.145 | -0.004 | -0.007 | -0.007 | 0 | 0.23 | 0.27 | 0.16 | 0.3 | 0.26 | 0.34 |
| 8 | 127368518 | rs7004827 | -1.608 | -0.841 | -0.749 | -0.374 | 0.094 | -0.036 | -0.036 | 0 | 0.5 | 0.91 | 0.28 | 0.47 | 0.35 | 0.55 |
| 8 | 127369615 | rs7839342 | 0.999 | -1.102 | -0.461 | -0.482 | 0.04 | 0.035 | 0.035 | 0 | 0.17 | 0.07 | 0.05 | 0.06 | 0.08 | 0.19 |
| 8 | 127372598 | rs10110372 | 1.33 | -0.793 | 0.017 | -0.386 | 0.014 | 0.121 | 0.121 | 0.174 | 0.06 | 0.25 | 0.04 | 0.08 | 0 | 0.02 |
| 8 | 127380069 | rs1554962 | -1.572 | -0.517 | -0.432 | -0.459 | -0.06 | -0.013 | -0.013 | 0 | 0.55 | 0.89 | 0.34 | 0.44 | 0.51 | 0.63 |
| 8 | 127381346 | rs2293851 | 0.134 | -0.352 | -0.465 | -0.581 | -0.016 | -0.016 | -0.016 | 0 | 0.13 | 0.09 | 0.11 | 0.28 | 0.21 | 0.09 |
| 8 | 127382494 | rs7827010 | -0.686 | -0.697 | 0.025 | -0.679 | 0.004 | -0.012 | -0.012 | 0 | 0.36 | 0.51 | 0.23 | 0.33 | 0.35 | 0.37 |
| 8 | 127383089 | rs16901342 | 0.449 | -0.72 | -0.166 | -0.664 | -0.007 | -0.004 | -0.004 | 0 | 0.11 | 0.19 | 0.08 | 0.1 | 0.1 | 0.2 |
| 8 | 127386194 | rs11783521 | -1.932 | -0.637 | -0.882 | -0.618 | -0.115 | -0.055 | -0.055 | 0 | 0.56 | 0.97 | 0.35 | 0.43 | 0.51 | 0.63 |
| 8 | 127393035 | rs12549017 | -0.553 | -0.054 | -0.125 | -0.132 | 0.024 | 0.018 | 0.018 | 0.038 | 0.18 | 0.21 | 0.15 | 0.29 | 0.27 | 0.18 |
| 8 | 127393927 | rs10505471 | -1.319 | -0.029 | -0.796 | -0.149 | -0.035 | -0.013 | -0.013 | 0 | 0.38 | 0.81 | 0.3 | 0.4 | 0.41 | 0.42 |
| 8 | 127399987 | rs2736000 | 0.509 | -0.067 | -0.07 | -0.507 | -0.017 | -0.015 | -0.015 | 0 | 0.74 | 0.65 | 0.55 | 0.73 | 0.78 | 0.77 |
| 8 | 127400867 | rs4871694 | -0.081 | -0.098 | 0.036 | -0.802 | 0.002 | 0.008 | 0.008 | 0 | 0.17 | 0.11 | 0.19 | 0.1 | 0.17 | 0.16 |
| 8 | 127404270 | rs16901359 | -0.531 | -0.059 | 0.312 | -0.745 | -0.011 | -0.027 | -0.027 | 0 | 0.25 | 0.28 | 0.24 | 0.2 | 0.23 | 0.23 |
| 8 | 127406370 | rs7823015 | 0.765 | 0.128 | -0.174 | -1.106 | 0.009 | 0.029 | 0.029 | 0 | 0.29 | 0.22 | 0.15 | 0.25 | 0.29 | 0.18 |
| 8 | 127409827 | rs7841789 | -0.02 | 0.2 | -0.067 | -1.283 | 0.018 | -0.077 | -0.077 | 0.119 | 0.12 | 0.43 | 0.09 | 0 | 0.09 | 0.14 |
| 8 | 127410572 | rs10094763 | -1.082 | 0.133 | -0.339 | -1.303 | 0.021 | 0.061 | 0.061 | 0 | 0.38 | 0.67 | 0.34 | 0.21 | 0.33 | 0.38 |
| 8 | 127420218 | rs16901372 | -0.036 | 0.181 | -0.511 | -0.99 | 0.001 | 0.001 | 0.001 | 0 | 0.25 | 0.17 | 0.23 | 0.21 | 0.23 | 0.24 |
| 8 | 127422090 | rs7814955 | -1.558 | 0.289 | -0.827 | -1.026 | -0.108 | -0.089 | -0.089 | 0 | 0.7 | 0.95 | 0.51 | 0.47 | 0.65 | 0.58 |
| 8 | 127422790 | rs16901379 | -1.124 | -0.068 | 0.16 | -0.965 | -0.023 | -0.055 | -0.055 | 0.02 | 0.31 | 0.41 | 0.29 | 0.21 | 0.28 | 0.32 |
| 8 | 127425147 | rs4871704 | -0.732 | 0.015 | -0.641 | -0.725 | 0.01 | 0.02 | 0.02 | 0 | 0.25 | 0.12 | 0.23 | 0.21 | 0.23 | 0.24 |
| 8 | 127437105 | rs1076585 | -0.488 | 0.022 | -0.261 | -0.569 | -0.02 | -0.018 | -0.018 | 0.502 | 0.24 | 0.3 | 0.14 | 0.14 | 0.2 | 0.22 |
| 8 | 127437394 | rs4870971 | 0.399 | 0.177 | -0.45 | -0.954 | 0.004 | 0.012 | 0.012 | 0 | 0.28 | 0.26 | 0.13 | 0.16 | 0.25 | 0.16 |
| 8 | 127437434 | rs1078700 | 0.406 | 0.186 | -0.45 | -0.956 | 0.003 | 0.012 | 0.012 | 0 | 0.28 | 0.26 | 0.13 | 0.17 | 0.25 | 0.16 |
| 8 | 127438183 | rs2735996 | -0.708 | 0.116 | -0.175 | -0.758 | 0.001 | 0.002 | 0.002 | 0 | 0.82 | 0.84 | 0.85 | 0.89 | 0.84 | 0.76 |
| 8 | 127439415 | rs1016671 | -0.661 | 0.174 | -0.15 | -0.752 | 0 | 0.001 | 0.001 | 0 | 0.82 | 0.8 | 0.85 | 0.89 | 0.83 | 0.77 |

| | | | | | | | | | | | | | | | | |
|---|---|---|---|---|---|---|---|---|---|---|---|---|---|---|---|---|
| 8 | 127443990 | rs926128 | 1.325 | 0.528 | -0.601 | -0.898 | -0.075 | -0.095 | -0.095 | 1.399 | 0.34 | 0.05 | 0.54 | 0.52 | 0.34 | 0.45 |
| 8 | 127448900 | rs2735984 | -0.895 | 0.526 | -0.487 | -0.885 | 0.004 | 0.007 | 0.007 | 0 | 0.82 | 0.88 | 0.86 | 0.89 | 0.84 | 0.76 |
| 8 | 127457081 | rs2735974 | -0.977 | 0.708 | -0.527 | -0.891 | -0.007 | 0.011 | 0.011 | 0 | 0.82 | 0.9 | 0.85 | 0.89 | 0.81 | 0.78 |
| 8 | 127457797 | rs727373 | 0.029 | 0.691 | -0.503 | -1.027 | -0.007 | 0.059 | 0.059 | 0 | 0.54 | 0.67 | 0.71 | 0.7 | 0.53 | 0.61 |
| 8 | 127462081 | rs7008211 | 0.953 | 0.691 | -0.492 | -1.027 | -0.002 | 0.018 | 0.018 | 0 | 0.28 | 0.23 | 0.14 | 0.19 | 0.28 | 0.16 |
| 8 | 127462984 | rs2223038 | 0.044 | 0.648 | -0.5 | -1.012 | -0.009 | 0.056 | 0.056 | 0 | 0.54 | 0.66 | 0.7 | 0.7 | 0.52 | 0.61 |
| 8 | 127477665 | rs6985115 | -0.214 | 0.559 | 0.016 | -0.777 | 0.003 | -0.018 | -0.018 | 0 | 0.78 | 0.69 | 0.8 | 0.89 | 0.77 | 0.77 |
| 8 | 127478325 | rs4870976 | 0.537 | 0.606 | -0.209 | -0.652 | -0.002 | -0.014 | -0.014 | 0 | 0.61 | 0.46 | 0.69 | 0.71 | 0.6 | 0.61 |
| 8 | 127483607 | rs11784010 | -0.003 | 0.643 | -0.213 | -0.661 | -0.009 | 0.017 | 0.017 | 0 | 0.3 | 0.23 | 0.23 | 0.24 | 0.34 | 0.2 |
| 8 | 127483918 | rs3922907 | -0.092 | 0.752 | -0.448 | -0.544 | 0.017 | -0.051 | -0.051 | 0 | 0.32 | 0.48 | 0.25 | 0.24 | 0.34 | 0.18 |
| 8 | 127484015 | rs3922906 | -0.12 | 0.781 | -0.486 | -0.558 | 0.026 | -0.05 | -0.05 | 0.095 | 0.32 | 0.48 | 0.26 | 0.24 | 0.37 | 0.2 |
| 8 | 127484568 | rs4422744 | -0.387 | 0.293 | -0.795 | -0.194 | 0 | 0.088 | 0.088 | 0 | 0.49 | 0.73 | 0.38 | 0.34 | 0.55 | 0.38 |
| 8 | 127489030 | rs11989898 | -0.416 | -0.354 | -0.178 | 0.75 | -0.016 | 0.011 | 0.011 | 0 | 0.17 | 0.23 | 0.25 | 0.27 | 0.11 | 0.19 |
| 8 | 127489087 | rs16901432 | 0.413 | -0.008 | -0.447 | 0.894 | 0.024 | 0.005 | 0.005 | 0.182 | 0.1 | 0.02 | 0.04 | 0.08 | 0.05 | 0.05 |
| 8 | 127505339 | rs10505503 | -0.111 | -0.193 | -0.088 | 1.081 | 0.025 | 0.023 | 0.023 | 0.135 | 0.13 | 0.1 | 0.06 | 0.07 | 0.06 | 0.06 |
| 8 | 127513350 | rs12548156 | -0.29 | 0.124 | 0.318 | 1.616 | 0 | 0.017 | 0.017 | 0 | 0.07 | 0.05 | 0.2 | 0.27 | 0.06 | 0.15 |
| 8 | 127518368 | rs10464865 | 1.073 | -0.018 | 0.973 | 1.408 | -0.024 | 0.057 | 0.057 | 0 | 0.72 | 0.41 | 0.63 | 0.45 | 0.78 | 0.69 |
| 8 | 127524012 | rs10105985 | 1.225 | 0.607 | -0.126 | 1.01 | -0.015 | 0.155 | 0.155 | 0.232 | 0.48 | 0.08 | 0.41 | 0.17 | 0.4 | 0.47 |
| 8 | 127533573 | rs11775759 | 1.057 | 0.1 | 0.345 | 1.45 | 0.008 | -0.01 | -0.01 | 0 | 0.66 | 0.32 | 0.61 | 0.35 | 0.72 | 0.69 |
| 8 | 127538582 | rs4871716 | 1.649 | 0.161 | -0.646 | 1.132 | 0.015 | 0.06 | 0.06 | 0.174 | 0.36 | 0.04 | 0.35 | 0.14 | 0.26 | 0.32 |
| 8 | 127543114 | rs11990954 | -0.403 | 0.05 | 0.635 | 2.01 | -0.009 | -0.001 | -0.001 | 0 | 0.18 | 0.3 | 0.12 | 0.26 | 0.16 | 0.09 |
| 8 | 127546182 | rs13273870 | 1.388 | 0.585 | 0.421 | 2.278 | -0.001 | 0.021 | 0.021 | 0 | 0.85 | 0.88 | 0.74 | 0.61 | 0.87 | 0.79 |
| 8 | 127551826 | rs10156297 | 0.753 | 0.691 | 0.475 | 2.351 | -0.004 | 0.049 | 0.049 | 0 | 0.89 | 0.88 | 0.75 | 0.65 | 0.89 | 0.8 |
| 8 | 127560353 | rs7818201 | -1.852 | 0.746 | 0.598 | 1.688 | 0.013 | 0.011 | 0.011 | 0 | 0.07 | 0.2 | 0.03 | 0.05 | 0.06 | 0.02 |
| 8 | 127570511 | rs2290840 | -1.208 | 0.996 | 0.111 | 1.726 | -0.018 | -0.018 | -0.018 | 0 | 0.07 | 0.01 | 0.02 | 0.06 | 0.07 | 0.06 |
| 8 | 127575595 | rs7839958 | -0.96 | 0.985 | 0.076 | 2.685 | -0.003 | -0.043 | -0.043 | 0 | 0.11 | 0.03 | 0.19 | 0.3 | 0.12 | 0.2 |
| 8 | 127584005 | rs7832762 | -2.134 | 0.942 | 0.996 | 2.146 | 0.013 | 0.149 | 0.149 | 0.878 | 0.13 | 0.55 | 0.24 | 0.32 | 0.12 | 0.07 |
| 8 | 127585373 | rs12679253 | -1.491 | 0.935 | 1.243 | 2.141 | -0.004 | 0.064 | 0.064 | 1.449 | 0.09 | 0.14 | 0.19 | 0.32 | 0.06 | 0.07 |
| 8 | 127586371 | rs10094631 | -1.827 | 0.936 | 0.852 | 2.156 | 0.012 | 0.148 | 0.148 | 1.012 | 0.14 | 0.56 | 0.24 | 0.32 | 0.12 | 0.07 |
| 8 | 127592581 | rs6470452 | 0.602 | 1.124 | 0.066 | 1.775 | 0.012 | 0.068 | 0.068 | 0 | 0.72 | 0.47 | 0.76 | 0.5 | 0.69 | 0.62 |
| 8 | 127603549 | rs2385694 | -1.035 | 1.193 | -0.796 | 1.533 | -0.011 | -0.016 | -0.016 | 0.153 | 0.4 | 0.38 | 0.44 | 0.49 | 0.5 | 0.47 |
| 8 | 127621549 | rs13438813 | -1.372 | 0.34 | -0.604 | 1.041 | 0.006 | -0.003 | -0.003 | 0 | 0.22 | 0.22 | 0.13 | 0.27 | 0.19 | 0.29 |
| 8 | 127622258 | rs12550518 | -1.544 | 0.3 | -0.624 | 1.019 | 0.009 | -0.002 | -0.002 | 1.326 | 0.23 | 0.22 | 0.13 | 0.27 | 0.19 | 0.29 |
| 8 | 127625650 | rs11786548 | -0.208 | 0.369 | -0.669 | 0.655 | 0.014 | 0.054 | 0.054 | 0 | 0.12 | 0.01 | 0.05 | 0.02 | 0.11 | 0.22 |
| 8 | 127626359 | rs9297740 | -1.325 | 0.372 | -0.624 | 0.715 | 0.006 | 0.007 | 0.007 | 0 | 0.22 | 0.23 | 0.13 | 0.18 | 0.19 | 0.29 |
| 8 | 127642790 | rs9987338 | -1.099 | 0.655 | -0.236 | 0.938 | 0.011 | 0.035 | 0.035 | 0 | 0.09 | 0.26 | 0.08 | 0.15 | 0.08 | 0.07 |
| 8 | 127645375 | rs9987136 | -0.43 | 0.716 | -0.517 | 0.969 | -0.015 | -0.022 | -0.022 | 1.33 | 0.06 | 0.03 | 0.05 | 0.16 | 0.08 | 0.07 |
| 8 | 127647147 | rs10481080 | -0.845 | 0.239 | -0.374 | 0.753 | 0 | 0 | 0 | 0.126 | 0.18 | 0.18 | 0.07 | 0.13 | 0.14 | 0.19 |
| 8 | 127650064 | rs10110041 | -1.293 | 0.34 | 0.793 | 0.977 | 0.058 | 0.104 | 0.104 | 1.875 | 0.09 | 0.46 | 0.05 | 0.14 | 0.06 | 0.06 |
| 8 | 127651504 | rs10086465 | -0.899 | 0.339 | 1.115 | 0.978 | 0.001 | 0.076 | 0.076 | 1.651 | 0.07 | 0.24 | 0.04 | 0.14 | 0.02 | 0.05 |
| 8 | 127653484 | rs11774414 | -1.597 | 0.293 | 1.069 | 1.046 | -0.006 | 0.035 | 0.035 | 0.638 | 0.11 | 0.16 | 0.05 | 0.25 | 0.08 | 0.07 |
| 8 | 127653541 | rs11784545 | -0.226 | 0.321 | 0.873 | 0.966 | -0.017 | -0.026 | -0.026 | 0 | 0.05 | 0 | 0.03 | 0.1 | 0.06 | 0.02 |
| 8 | 127654091 | rs7461025 | 1.117 | -0.029 | 0.503 | 0.951 | 0.019 | 0.018 | 0.018 | 2.029 | 0.52 | 0.14 | 0.61 | 0.43 | 0.57 | 0.54 |
| 8 | 127656250 | rs2385685 | -1.01 | -0.017 | 0.491 | 1.021 | 0 | 0.137 | 0.137 | 0.651 | 0.24 | 0.49 | 0.22 | 0.46 | 0.25 | 0.38 |
| 8 | 127658147 | rs7011739 | -1.302 | -0.035 | 0.502 | 0.883 | 0.008 | 0.052 | 0.052 | 2.264 | 0.3 | 0.51 | 0.26 | 0.56 | 0.33 | 0.41 |
| 8 | 127659536 | rs2385686 | 0.217 | -0.049 | 0.39 | 0.343 | 0.004 | 0.038 | 0.038 | 0 | 0.47 | 0.35 | 0.47 | 0.69 | 0.46 | 0.54 |
| 8 | 127660433 | rs10101241 | -1.702 | 0.087 | 1.791 | 0.887 | -0.003 | -0.003 | -0.003 | 1.19 | 0.25 | 0.44 | 0.24 | 0.21 | 0.2 | 0.16 |
| 8 | 127661127 | rs13260097 | -1.861 | 0.325 | 2.515 | 1.301 | 0.045 | 0.115 | 0.115 | 3.394 | 0.12 | 0.54 | 0.2 | 0.14 | 0.1 | 0.03 |
| 8 | 127661447 | rs13260284 | -1.305 | 0.437 | 2.444 | 1.326 | 0.038 | 0.035 | 0.035 | 0.065 | 0.09 | 0.22 | 0.13 | 0.11 | 0.12 | 0.04 |
| 8 | 127662669 | rs10088340 | -0.8 | 0.399 | 2.066 | 1.223 | -0.003 | 0.012 | 0.012 | 0.11 | 0.09 | 0.03 | 0.06 | 0.03 | 0.11 | 0.06 |
| 8 | 127663604 | rs4871724 | -2.116 | 0.363 | 2.642 | 1.233 | 0.011 | 0.033 | 0.033 | 3.429 | 0.15 | 0.62 | 0.22 | 0.14 | 0.14 | 0.04 |
| 8 | 127665272 | rs2385688 | -2.761 | 0.35 | 2.464 | 1.217 | 0.008 | 0.004 | 0.004 | 0.298 | 0.26 | 0.78 | 0.29 | 0.17 | 0.24 | 0.1 |
| 8 | 127669242 | rs13251622 | -1.466 | 0.493 | 2.346 | 1.428 | 0.056 | 0.052 | 0.052 | 0.172 | 0.1 | 0.29 | 0.15 | 0.11 | 0.13 | 0.04 |
| 8 | 127675166 | rs10090062 | -2.51 | 0.412 | 2.394 | 2.225 | 0.039 | 0.24 | 0.24 | 1.578 | 0.13 | 0.61 | 0.24 | 0.37 | 0.14 | 0.05 |
| 8 | 127675332 | rs13259479 | -2.361 | 0.439 | 2.543 | 2.19 | 0.095 | 0.289 | 0.289 | 0 | 0.06 | 0.22 | 0.12 | 0.3 | 0.09 | 0.04 |
| 8 | 127675497 | rs13259828 | -2.644 | 0.241 | 2.488 | 2.011 | 0 | 0.054 | 0.054 | 0.705 | 0.2 | 0.58 | 0.21 | 0.28 | 0.17 | 0.11 |
| 8 | 127678326 | rs7817894 | -1.845 | 0.553 | 2.269 | 1.928 | 0.103 | 0.092 | 0.092 | 0 | 0.1 | 0.3 | 0.07 | 0.1 | 0.11 | 0.14 |
| 8 | 127680853 | rs7831824 | -2.267 | 0.599 | 2.081 | 1.946 | 0.064 | 0.095 | 0.095 | 0.429 | 0.14 | 0.47 | 0.11 | 0.17 | 0.14 | 0.18 |
| 8 | 127681578 | rs2891842 | -2.244 | 0.596 | 1.579 | 1.945 | 0.054 | 0.083 | 0.083 | 0.314 | 0.14 | 0.36 | 0.09 | 0.17 | 0.14 | 0.18 |
| 8 | 127683850 | rs7814964 | -0.88 | 0.646 | -0.075 | 1.344 | -0.002 | 0.014 | 0.014 | 0 | 0.61 | 0.74 | 0.52 | 0.65 | 0.6 | 0.59 |
| 8 | 127694107 | rs4871728 | 0.179 | 0.352 | -0.99 | 1.237 | -0.057 | -0.015 | -0.015 | 0 | 0.21 | 0.01 | 0.12 | 0.13 | 0.2 | 0.21 |
| 8 | 127696127 | rs16901647 | -0.249 | 0.298 | -0.829 | 1.251 | -0.017 | 0.004 | 0.004 | 0.026 | 0.3 | 0.08 | 0.17 | 0.2 | 0.26 | 0.26 |
| 8 | 127701327 | rs6470458 | -1.282 | 0.595 | -0.77 | 1.282 | 0 | -0.004 | -0.004 | 0.019 | 0.53 | 0.48 | 0.46 | 0.63 | 0.52 | 0.56 |
| 8 | 127701948 | rs6470459 | -0.213 | 0.718 | -1.204 | 1.517 | -0.02 | -0.079 | -0.079 | 0 | 0.19 | 0.06 | 0.26 | 0.4 | 0.25 | 0.27 |
| 8 | 127707310 | rs10956324 | -0.544 | 0.539 | -1.194 | 1.472 | 0.012 | -0.087 | -0.087 | 1.86 | 0.22 | 0.06 | 0.27 | 0.44 | 0.24 | 0.34 |
| 8 | 127707824 | rs974450 | -0.818 | 0.541 | -0.771 | 0.888 | -0.003 | 0.002 | 0.002 | 0.163 | 0.43 | 0.56 | 0.35 | 0.46 | 0.38 | 0.39 |
| 8 | 127708294 | rs974451 | -0.971 | 0.841 | -0.466 | 0.256 | 0.003 | -0.031 | -0.031 | 0 | 0.29 | 0.58 | 0.25 | 0.22 | 0.31 | 0.18 |
| 8 | 127708772 | rs4871730 | -1.65 | 0.593 | -1.162 | 0.073 | -0.026 | -0.017 | -0.017 | 0 | 0.57 | 0.81 | 0.41 | 0.47 | 0.53 | 0.52 |
| 8 | 127708875 | rs4870981 | 0.33 | 0.628 | -1.386 | -0.027 | -0.049 | -0.092 | -0.092 | 0 | 0.16 | 0.01 | 0.12 | 0.24 | 0.19 | 0.23 |

| | | | | | | | | | | | | | | | | |
|---|---|---|---|---|---|---|---|---|---|---|---|---|---|---|---|---|
| 8 | 127718806 | rs7835764 | -1.551 | 0.562 | -0.928 | -0.003 | -0.004 | 0.002 | 0.002 | 0 | 0.53 | 0.59 | 0.38 | 0.46 | 0.5 | 0.52 |
| 8 | 127719548 | rs7840169 | -1.282 | 0.59 | -0.896 | -0.058 | 0.014 | 0.005 | 0.005 | 0.256 | 0.47 | 0.75 | 0.36 | 0.44 | 0.43 | 0.41 |
| 8 | 127721159 | rs2011629 | -1.738 | 0.598 | -0.934 | -0.054 | -0.025 | -0.005 | -0.005 | 0 | 0.55 | 0.79 | 0.39 | 0.45 | 0.5 | 0.52 |
| 8 | 127724042 | rs7819715 | -1.324 | 0.507 | -0.563 | 0.333 | 0 | 0 | 0 | 0 | 0.44 | 0.54 | 0.34 | 0.44 | 0.42 | 0.4 |
| 8 | 127739184 | rs6470466 | -0.559 | 0.57 | -0.388 | 0.342 | 0.002 | -0.05 | -0.05 | 0 | 0.25 | 0.45 | 0.22 | 0.22 | 0.22 | 0.21 |
| 8 | 127740139 | rs4870982 | -1.433 | 0.533 | -0.492 | 0.492 | 0.005 | 0.02 | 0.02 | 0 | 0.52 | 0.81 | 0.38 | 0.56 | 0.47 | 0.51 |
| 8 | 127746957 | rs2029424 | -1.207 | 0.035 | -0.634 | 0.613 | 0.002 | 0.004 | 0.004 | 0 | 0.48 | 0.54 | 0.36 | 0.55 | 0.46 | 0.45 |
| 8 | 127760258 | rs1472026 | -0.418 | 0.157 | -0.985 | 0.459 | 0.001 | -0.021 | -0.021 | 0 | 0.3 | 0.16 | 0.21 | 0.27 | 0.29 | 0.35 |
| 8 | 127767127 | rs7822787 | -0.898 | 0.265 | -0.792 | 0.319 | 0.01 | 0.001 | 0.001 | 0 | 0.5 | 0.67 | 0.37 | 0.41 | 0.47 | 0.45 |
| 8 | 127768683 | rs4871739 | -0.795 | -0.866 | -0.909 | 0.082 | 0.03 | 0.082 | 0.082 | 2.098 | 0.49 | 0.83 | 0.55 | 0.64 | 0.42 | 0.63 |
| 8 | 127781440 | rs12543473 | -1.395 | -1.1 | 0.115 | 0.173 | -0.01 | 0.002 | 0.002 | 0.777 | 0.21 | 0.75 | 0.33 | 0.31 | 0.12 | 0.14 |
| 8 | 127785726 | rs7834887 | -0.329 | -1.126 | 0.6 | 0.16 | -0.007 | -0.012 | -0.012 | 0 | 0.1 | 0.13 | 0.07 | 0.06 | 0.08 | 0.05 |
| 8 | 127794263 | rs11988483 | -0.798 | -1.432 | 0.434 | -0.147 | -0.003 | -0.038 | -0.038 | 8.079 | 0.31 | 0.32 | 0.22 | 0.1 | 0.27 | 0.26 |
| 8 | 127805686 | rs7017239 | -1.223 | -1.499 | 0.817 | -0.164 | 0.001 | -0.013 | -0.013 | 0 | 0.34 | 0.63 | 0.25 | 0.1 | 0.27 | 0.25 |
| 8 | 127807071 | rs4870983 | -0.836 | -1.509 | 0.962 | -0.169 | -0.008 | -0.068 | -0.068 | 0 | 0.31 | 0.39 | 0.22 | 0.1 | 0.27 | 0.26 |
| 8 | 127808949 | rs983592 | -1.401 | -1.511 | 0.622 | -0.125 | 0.041 | 0.039 | 0.039 | 0.39 | 0.39 | 0.82 | 0.46 | 0.34 | 0.29 | 0.35 |
| 8 | 127811208 | rs7839719 | -0.459 | -0.854 | 0.101 | -0.214 | -0.001 | -0.026 | -0.026 | 4.943 | 0.26 | 0.29 | 0.19 | 0.1 | 0.24 | 0.27 |
| 8 | 127822685 | rs7842598 | -0.734 | -1.156 | -0.726 | -0.025 | 0.007 | 0.026 | 0.026 | 0.047 | 0.55 | 0.75 | 0.54 | 0.64 | 0.42 | 0.6 |
| 8 | 127830777 | rs7812779 | 0.514 | -1.247 | -0.477 | 0.072 | 0.006 | 0.003 | 0.003 | 0.048 | 0.46 | 0.31 | 0.52 | 0.6 | 0.56 | 0.35 |
| 8 | 127834830 | rs16901718 | 0.615 | -1.493 | -0.738 | 0.224 | -0.004 | -0.046 | -0.046 | 0 | 0.1 | 0.02 | 0.05 | 0.27 | 0.11 | 0.06 |
| 8 | 127839267 | rs16901732 | -0.262 | -2.055 | -0.627 | -0.103 | -0.027 | 0.023 | 0.023 | 0 | 0.14 | 0.21 | 0.08 | 0.27 | 0.08 | 0.13 |
| 8 | 127841932 | rs10956329 | -0.402 | -1.972 | -0.64 | -0.285 | -0.005 | -0.112 | -0.112 | 0 | 0.67 | 0.53 | 0.62 | 0.96 | 0.72 | 0.53 |
| 8 | 127843085 | rs11986358 | 1.633 | -1.995 | -1.275 | -0.145 | -0.18 | -0.116 | -0.116 | 0 | 0.29 | 0.03 | 0.36 | 0.37 | 0.47 | 0.19 |
| 8 | 127845389 | rs10505491 | 1.014 | -1.964 | -0.762 | -0.455 | -0.001 | -0.011 | -0.011 | 0 | 0.08 | 0.09 | 0.06 | 0.01 | 0.09 | 0.25 |
| 8 | 127845702 | rs986046 | -0.653 | -2.29 | -0.187 | -0.27 | -0.029 | 0.08 | 0.08 | 0.299 | 0.17 | 0.5 | 0.14 | 0.3 | 0.13 | 0.34 |
| 8 | 127846275 | rs16901753 | 0.079 | -2.221 | 0.038 | -0.272 | 0.003 | -0.055 | -0.055 | 0 | 0.12 | 0.4 | 0.09 | 0.01 | 0.09 | 0.25 |
| 8 | 127855936 | rs4870985 | 1.265 | -1.824 | -0.477 | -0.275 | -0.121 | -0.066 | -0.066 | 0 | 0.36 | 0.07 | 0.59 | 0.54 | 0.52 | 0.25 |
| 8 | 127856177 | rs4871743 | -1.027 | -2.004 | -0.244 | -0.52 | -0.001 | 0.032 | 0.032 | 0 | 0.75 | 0.83 | 0.83 | 0.93 | 0.78 | 0.72 |
| 8 | 127866125 | rs7833497 | -0.166 | -1.9 | -0.089 | -0.44 | 0.005 | 0.011 | 0.011 | 1.659 | 0.63 | 0.35 | 0.74 | 0.7 | 0.68 | 0.68 |
| 8 | 127868163 | rs16901797 | -0.686 | -1.784 | -0.218 | -0.302 | 0.02 | -0.024 | -0.024 | 0 | 0.34 | 0.23 | 0.19 | 0.45 | 0.27 | 0.48 |
| 8 | 127877348 | rs2893579 | -0.332 | -1.39 | -0.59 | -0.648 | 0.001 | 0.008 | 0.008 | 0.829 | 0.67 | 0.71 | 0.77 | 0.7 | 0.68 | 0.68 |
| 8 | 127881952 | rs16901814 | -0.572 | -1.069 | -0.832 | -0.542 | 0.014 | 0.017 | 0.017 | 0 | 0.24 | 0.18 | 0.14 | 0.13 | 0.17 | 0.39 |
| 8 | 127887494 | rs756100 | -0.381 | -0.089 | -1.193 | -1.238 | -0.013 | 0.021 | 0.021 | 0.049 | 0.6 | 0.67 | 0.75 | 0.99 | 0.59 | 0.65 |
| 8 | 127887560 | rs756099 | 0.356 | -0.145 | -1.24 | -0.401 | -0.002 | -0.009 | -0.009 | 0 | 0.29 | 0.24 | 0.3 | 0.39 | 0.32 | 0.44 |
| 8 | 127889918 | rs10105114 | -1.019 | 0.266 | -1.651 | -0.67 | -0.026 | 0.109 | 0.109 | 0.083 | 0.61 | 0.79 | 0.75 | 0.99 | 0.57 | 0.65 |
| 8 | 127897279 | rs17185 | 1.236 | 0.577 | -1.878 | 0.241 | -0.049 | -0.091 | -0.091 | 0 | 0.13 | 0.02 | 0.21 | 0.39 | 0.17 | 0.11 |
| 8 | 127897460 | rs4871749 | -0.078 | 0.442 | -1.453 | 0.224 | 0 | 0.004 | 0.004 | 0.192 | 0.19 | 0.17 | 0.33 | 0.23 | 0.19 | 0.16 |
| 8 | 127901649 | rs6984837 | -0.13 | 1.387 | -1.851 | 0.271 | -0.071 | -0.018 | -0.018 | 0 | 0.8 | 0.95 | 0.64 | 0.79 | 0.68 | 0.64 |
| 8 | 127902003 | rs4871750 | 1.012 | 0.792 | -1.96 | 0.09 | -0.013 | -0.008 | -0.008 | 0 | 0.52 | 0.28 | 0.39 | 0.47 | 0.48 | 0.46 |
| 8 | 127909396 | rs1565048 | 2.192 | 0.545 | -1.686 | 0.047 | -0.009 | 0.027 | 0.027 | 2.486 | 0.08 | 0.1 | 0.08 | 0.28 | 0.04 | 0.02 |
| 8 | 127916370 | rs1516937 | -0.556 | 0.985 | -1.881 | 0.103 | 0.003 | -0.002 | -0.002 | 0 | 0.33 | 0.67 | 0.34 | 0.44 | 0.31 | 0.28 |
| 8 | 127919914 | rs1516935 | -1.216 | 1.121 | -1.944 | 0.093 | -0.03 | 0.017 | 0.017 | 0 | 0.85 | 0.95 | 0.73 | 0.93 | 0.79 | 0.74 |
| 8 | 127921284 | rs13271478 | 1.637 | 0.983 | -2.451 | -0.057 | -0.043 | -0.096 | -0.096 | 0.01 | 0.32 | 0.05 | 0.27 | 0.35 | 0.32 | 0.35 |
| 8 | 127923372 | rs979201 | 0.084 | 0.612 | -2.331 | -0.13 | 0.006 | 0.001 | 0.001 | 4.636 | 0.3 | 0.71 | 0.26 | 0.33 | 0.2 | 0.19 |
| 8 | 127923720 | rs979200 | -0.711 | 0.686 | -2.402 | -0.149 | 0.05 | 0.012 | 0.012 | 0.849 | 0.47 | 0.73 | 0.35 | 0.48 | 0.36 | 0.29 |
| 8 | 127924065 | rs11995166 | -0.009 | 0.851 | -2.037 | 0.093 | -0.027 | 0.028 | 0.028 | 0 | 0.67 | 0.86 | 0.65 | 0.78 | 0.64 | 0.62 |
| 8 | 127925830 | rs4330656 | 0.204 | 0.825 | -1.546 | 0.611 | 0.017 | -0.03 | -0.03 | 3.294 | 0.42 | 0.17 | 0.5 | 0.43 | 0.37 | 0.34 |
| 8 | 127926753 | rs13270753 | 0.846 | 0.811 | -1.439 | 0.56 | 0.015 | -0.034 | -0.034 | 0.077 | 0.34 | 0.14 | 0.29 | 0.38 | 0.29 | 0.28 |
| 8 | 127928559 | rs2176339 | -0.03 | 0.717 | -1.233 | 0.651 | 0.018 | 0.046 | 0.046 | 0 | 0.44 | 0.3 | 0.57 | 0.71 | 0.38 | 0.36 |
| 8 | 127930885 | rs4871753 | 0.352 | 0.71 | -1.107 | 0.583 | -0.002 | 0.003 | 0.003 | 0 | 0.07 | 0.09 | 0.2 | 0.05 | 0.05 | 0.04 |
| 8 | 127930987 | rs4870992 | 0.352 | 0.71 | -1.076 | 0.583 | -0.002 | 0.004 | 0.004 | 0 | 0.07 | 0.09 | 0.2 | 0.05 | 0.05 | 0.04 |
| 8 | 127932940 | rs11774049 | 0.022 | 0.817 | -1.073 | 0.608 | 0.006 | -0.005 | -0.005 | 1.493 | 0.43 | 0.34 | 0.52 | 0.43 | 0.38 | 0.34 |
| 8 | 127934830 | rs4871755 | 0.999 | 0.552 | -1.122 | 0.435 | -0.001 | -0.001 | -0.001 | 0 | 0.05 | 0.02 | 0.19 | 0.04 | 0.04 | 0.02 |
| 8 | 127937839 | rs2392672 | -0.393 | 0.602 | -0.371 | 0.278 | 0.002 | 0.062 | 0.062 | 0 | 0.52 | 0.64 | 0.67 | 0.78 | 0.47 | 0.49 |
| 8 | 127938759 | rs1516952 | 0.525 | 0.384 | -0.433 | -0.114 | -0.001 | 0 | 0 | 0.324 | 0.06 | 0.05 | 0.2 | 0.05 | 0.04 | 0.04 |
| 8 | 127941276 | rs4871000 | 0.381 | 0.57 | -0.459 | -0.145 | -0.008 | 0 | 0 | 2.033 | 0.07 | 0.02 | 0.22 | 0.05 | 0.09 | 0.05 |
| 8 | 127942261 | rs763319 | 1.159 | 0.479 | -0.58 | 0.086 | 0.037 | -0.095 | -0.095 | 0 | 0.33 | 0.1 | 0.27 | 0.46 | 0.27 | 0.28 |
| 8 | 127946773 | rs3929313 | -0.191 | 0.542 | -0.326 | 0.104 | -0.001 | -0.001 | -0.001 | 0.199 | 0.49 | 0.49 | 0.57 | 0.49 | 0.43 | 0.33 |
| 8 | 127952980 | rs2839811 | 0.257 | 0.545 | -0.216 | 0.07 | 0.009 | -0.012 | -0.012 | 0.036 | 0.43 | 0.34 | 0.53 | 0.46 | 0.37 | 0.32 |
| 8 | 127968515 | rs7842079 | -0.606 | 0.638 | -0.097 | -0.012 | -0.009 | 0.048 | 0.048 | 0 | 0.54 | 0.65 | 0.67 | 0.85 | 0.51 | 0.48 |
| 8 | 127979691 | rs1400460 | 0.07 | 0.563 | -0.102 | 0.076 | 0.004 | -0.003 | -0.003 | 0.073 | 0.45 | 0.44 | 0.54 | 0.47 | 0.37 | 0.33 |
| 8 | 127980071 | rs4871768 | 0.428 | 0.611 | -0.102 | 0.013 | -0.007 | 0.007 | 0.007 | 0 | 0.07 | 0.03 | 0.22 | 0.03 | 0.09 | 0.04 |
| 8 | 127980684 | rs980426 | -0.464 | 0.828 | -0.347 | -0.331 | 0 | 0 | 0 | 0 | 0.61 | 0.45 | 0.65 | 0.57 | 0.59 | 0.52 |
| 8 | 127981249 | rs12677894 | 0.701 | 0.821 | -0.362 | -0.305 | 0.029 | -0.008 | -0.008 | 0 | 0.42 | 0.31 | 0.32 | 0.54 | 0.34 | 0.41 |
| 8 | 127986466 | rs2392683 | -1.106 | 0.891 | -0.344 | -0.459 | -0.001 | -0.026 | -0.026 | 0 | 0.68 | 0.7 | 0.78 | 0.94 | 0.66 | 0.64 |
| 8 | 127986483 | rs2392684 | 0.784 | 0.612 | -0.753 | -0.139 | 0.027 | -0.014 | -0.014 | 0 | 0.4 | 0.2 | 0.51 | 0.43 | 0.36 | 0.31 |
| 8 | 127987909 | rs1516945 | 1.197 | 0.597 | -0.761 | -0.191 | 0.025 | -0.092 | -0.092 | 0.018 | 0.35 | 0.03 | 0.29 | 0.4 | 0.28 | 0.28 |
| 8 | 127989849 | rs901592 | -0.397 | 0.581 | -0.731 | -0.173 | 0.001 | -0.006 | -0.006 | 0 | 0.62 | 0.27 | 0.66 | 0.62 | 0.61 | 0.57 |
| 8 | 127998309 | rs1516932 | 0.718 | 0.531 | -0.762 | -0.161 | -0.004 | -0.052 | -0.052 | 0 | 0.42 | 0.15 | 0.51 | 0.51 | 0.41 | 0.39 |

| | | | | | | | | | | | | | | | | |
|---|---|---|---|---|---|---|---|---|---|---|---|---|---|---|---|---|
| 8 | 128003037 | rs17762342 | 0.643 | 0.566 | -0.837 | -0.322 | -0.06 | 0.03 | 0.03 | 0 | 0.19 | 0.04 | 0.13 | 0.09 | 0.2 | 0.17 |
| 8 | 128005150 | rs961023 | -0.848 | 0.558 | -0.685 | -0.36 | -0.002 | -0.033 | -0.033 | 0 | 0.67 | 0.37 | 0.75 | 0.92 | 0.69 | 0.6 |
| 8 | 128007450 | rs7817934 | 0.61 | 0.635 | -0.675 | -0.165 | 0.006 | -0.038 | -0.038 | 0 | 0.43 | 0.21 | 0.52 | 0.5 | 0.41 | 0.39 |
| 8 | 128010274 | rs11787440 | 0.55 | 0.76 | -0.654 | -0.118 | -0.031 | 0.035 | 0.035 | 0 | 0.19 | 0.08 | 0.12 | 0.09 | 0.2 | 0.17 |
| 8 | 128010768 | rs16901896 | 0.665 | 0.7 | -0.61 | -0.072 | 0.004 | 0.128 | 0.128 | 0 | 0.06 | 0.07 | 0.1 | 0.32 | 0.08 | 0.03 |
| 8 | 128011937 | rs10086908 | -0.042 | 0.756 | -0.691 | -0.089 | -0.02 | 0.009 | 0.009 | 0 | 0.25 | 0.22 | 0.36 | 0.22 | 0.32 | 0.27 |
| 8 | 128013376 | rs4871005 | -0.364 | 0.632 | -0.758 | -0.148 | 0.002 | -0.008 | -0.008 | 0.077 | 0.63 | 0.26 | 0.66 | 0.62 | 0.62 | 0.58 |
| 8 | 128019903 | rs1516942 | -0.051 | 0.772 | -0.69 | -0.091 | -0.018 | 0.008 | 0.008 | 0 | 0.24 | 0.2 | 0.33 | 0.19 | 0.31 | 0.27 |
| 8 | 128022973 | rs9297750 | 0.492 | 0.797 | -0.906 | -0.059 | -0.037 | -0.004 | -0.004 | 0.54 | 0.21 | 0.15 | 0.32 | 0.21 | 0.25 | 0.24 |
| 8 | 128026128 | rs710885 | -0.929 | 0.588 | -0.474 | 0.202 | 0.001 | -0.053 | -0.053 | 0 | 0.71 | 0.62 | 0.78 | 0.93 | 0.69 | 0.66 |
| 8 | 128028973 | rs785005 | 0.35 | 0.327 | -0.464 | 0.417 | 0.018 | 0.199 | 0.199 | 0 | 0.06 | 0.12 | 0.1 | 0.31 | 0.07 | 0.03 |
| 8 | 128033432 | rs2015595 | 0.785 | 0.002 | -1.079 | 0.122 | -0.012 | -0.118 | -0.118 | 0 | 0.29 | 0.04 | 0.28 | 0.4 | 0.28 | 0.26 |
| 8 | 128034594 | rs931354 | 0.404 | 0.06 | -1.038 | 0.107 | -0.008 | 0.064 | 0.064 | 0 | 0.32 | 0.11 | 0.53 | 0.85 | 0.35 | 0.36 |
| 8 | 128041297 | rs16901906 | -0.426 | -0.076 | -0.22 | 0.315 | 0.001 | -0.051 | -0.051 | 0 | 0.39 | 0.57 | 0.24 | 0.08 | 0.35 | 0.26 |
| 8 | 128046529 | rs9969411 | -0.492 | -0.111 | -0.191 | 0.323 | 0.004 | -0.021 | -0.021 | 0 | 0.41 | 0.58 | 0.43 | 0.23 | 0.4 | 0.39 |
| 8 | 128049005 | rs1487236 | 0.605 | -0.194 | -0.451 | 0.237 | -0.008 | -0.005 | -0.005 | 0.121 | 0.55 | 0.39 | 0.49 | 0.47 | 0.55 | 0.57 |
| 8 | 128054522 | rs980171 | -0.669 | -0.287 | -0.412 | 0.05 | 0.008 | 0.048 | 0.048 | 0 | 0.48 | 0.62 | 0.44 | 0.23 | 0.39 | 0.4 |
| 8 | 128061786 | rs16901922 | -0.113 | 0.173 | 0.105 | 0.426 | -0.003 | 0.021 | 0.021 | 0.532 | 0.32 | 0.26 | 0.18 | 0.09 | 0.31 | 0.2 |
| 8 | 128062582 | rs7827583 | 0.267 | -0.041 | -0.173 | 0.649 | -0.02 | 0.068 | 0.068 | 0 | 0.84 | 0.71 | 0.83 | 0.65 | 0.87 | 0.86 |
| 8 | 128063756 | rs961017 | 1.243 | 0.121 | -0.969 | 0.28 | -0.021 | 0.008 | 0.008 | 1.104 | 0.47 | 0.22 | 0.42 | 0.44 | 0.52 | 0.46 |
| 8 | 128067300 | rs4288339 | 1.025 | 0.042 | -0.827 | 0.659 | 0.021 | 0.007 | 0.007 | 2.323 | 0.53 | 0.26 | 0.47 | 0.43 | 0.61 | 0.51 |
| 8 | 128067513 | rs1456308 | 0.291 | 0.515 | -1.456 | 0.539 | -0.018 | -0.009 | -0.009 | 0 | 0.68 | 0.86 | 0.79 | 0.68 | 0.7 | 0.8 |
| 8 | 128070669 | rs958653 | 0.215 | 0.804 | -1.542 | 0.701 | -0.052 | -0.084 | -0.084 | 0.132 | 0.76 | 0.93 | 0.82 | 0.64 | 0.73 | 0.78 |
| 8 | 128070847 | rs9297751 | 0.557 | 0.908 | -0.89 | 0.462 | -0.001 | 0.003 | 0.003 | 1.576 | 0.51 | 0.39 | 0.44 | 0.44 | 0.46 | 0.35 |
| 8 | 128071305 | rs9649957 | -0.489 | 1.36 | -0.591 | 0.083 | 0.012 | -0.009 | -0.009 | 0 | 0.22 | 0.29 | 0.22 | 0.11 | 0.38 | 0.2 |
| 8 | 128071922 | rs6983515 | -0.202 | 0.031 | -0.344 | 0.025 | -0.059 | -0.004 | -0.004 | 7.374 | 0.64 | 0.53 | 0.57 | 0.39 | 0.79 | 0.53 |
| 8 | 128072019 | rs7833560 | 0 | 0.191 | -0.352 | 0.009 | -0.047 | -0.006 | -0.006 | 0 | 0.37 | 0.47 | 0.44 | 0.61 | 0.23 | 0.47 |
| 8 | 128072525 | rs1031586 | -1.029 | 0.143 | -0.784 | -0.376 | -0.024 | 0.083 | 0.083 | 2.293 | 0.68 | 0.86 | 0.71 | 0.95 | 0.7 | 0.78 |
| 8 | 128075690 | rs1456314 | -0.326 | 0.076 | -0.304 | -0.37 | 0 | -0.018 | -0.018 | 0 | 0.6 | 0.47 | 0.65 | 0.66 | 0.67 | 0.65 |
| 8 | 128076545 | rs16901935 | 0.966 | -0.181 | -0.955 | 0.346 | 0.017 | 0.001 | 0.001 | 0 | 0.08 | 0.05 | 0.11 | 0.31 | 0.04 | 0.2 |
| 8 | 128079475 | rs2124598 | 1.059 | 0.396 | -1.043 | -0.698 | -0.004 | 0.018 | 0.018 | 0 | 0.26 | 0.16 | 0.15 | 0.05 | 0.24 | 0.23 |
| 8 | 128079666 | rs2392729 | 0.722 | 0.642 | -1.277 | -0.691 | -0.125 | 0.076 | 0.076 | 1.846 | 0.23 | 0.04 | 0.23 | 0.06 | 0.42 | 0.18 |
| 8 | 128081138 | rs7824923 | 0.066 | 0.604 | -1.23 | -0.732 | -0.113 | 0.124 | 0.124 | 0 | 0.31 | 0.1 | 0.28 | 0.06 | 0.48 | 0.19 |
| 8 | 128084097 | rs6993569 | 0.203 | 0.057 | -0.495 | -0.2 | 0.001 | 0.083 | 0.083 | 2.345 | 0.15 | 0.49 | 0.18 | 0.29 | 0.09 | 0.13 |
| 8 | 128084539 | rs6994316 | -0.469 | 0.383 | -1.049 | -0.534 | -0.001 | -0.015 | -0.015 | 0 | 0.73 | 0.79 | 0.73 | 0.95 | 0.75 | 0.79 |
| 8 | 128093277 | rs1031588 | 0.521 | 0.908 | -1.329 | -0.22 | -0.08 | 0.042 | 0.042 | 1.124 | 0.17 | 0.03 | 0.19 | 0.05 | 0.33 | 0.1 |
| 8 | 128093297 | rs1016343 | 0.283 | 1.12 | -1.127 | 0.382 | 0.036 | 0.065 | 0.065 | 0 | 0.14 | 0.21 | 0.12 | 0.3 | 0.22 | 0.15 |
| 8 | 128094460 | rs6981122 | -0.729 | 0.506 | -1.002 | 0.326 | -0.01 | 0.007 | 0.007 | 0 | 0.39 | 0.72 | 0.38 | 0.63 | 0.56 | 0.35 |
| 8 | 128095156 | rs13252298 | 0.487 | 0.106 | -1.918 | 0.211 | -0.036 | -0.085 | -0.085 | 0 | 0.33 | 0.03 | 0.39 | 0.36 | 0.29 | 0.51 |
| 8 | 128103937 | rs1456315 | -0.168 | 0.066 | -1.068 | 0.212 | 0 | 0.001 | 0.001 | 1.902 | 0.59 | 0.42 | 0.55 | 0.3 | 0.64 | 0.65 |
| 8 | 128106880 | rs6983561 | -0.251 | -0.16 | -0.453 | 0.634 | -0.003 | 0.222 | 0.222 | 0.433 | 0.1 | 0.53 | 0.06 | 0.21 | 0.04 | 0.1 |
| 8 | 128108993 | rs17832285 | 1.292 | -0.362 | -0.89 | 0.425 | -0.013 | -0.008 | -0.008 | 0 | 0.17 | 0.09 | 0.14 | 0.19 | 0.16 | 0.27 |
| 8 | 128109129 | rs7825340 | 0.821 | -0.087 | -0.488 | 0.858 | -0.015 | -0.015 | -0.015 | 0.02 | 0.15 | 0.09 | 0.23 | 0.27 | 0.2 | 0.19 |
| 8 | 128109287 | rs12544977 | -0.143 | -0.334 | -0.585 | 0.222 | 0.008 | -0.021 | -0.021 | 0.092 | 0.62 | 0.8 | 0.48 | 0.53 | 0.68 | 0.68 |
| 8 | 128109574 | rs7826337 | 0.4 | -0.332 | -0.645 | 0.13 | 0.002 | -0.002 | -0.002 | 0 | 0.4 | 0.62 | 0.24 | 0.44 | 0.31 | 0.57 |
| 8 | 128110252 | rs16901966 | 0.665 | -0.318 | -0.194 | 0.139 | 0.015 | 0.237 | 0.237 | 0 | 0.08 | 0.38 | 0.05 | 0.22 | 0.04 | 0.1 |
| 8 | 128114710 | rs7824785 | 0.653 | -0.31 | -0.201 | 0.143 | 0.014 | 0.234 | 0.234 | 0 | 0.08 | 0.38 | 0.05 | 0.22 | 0.04 | 0.1 |
| 8 | 128116500 | rs1456306 | 1.399 | -0.302 | -0.589 | 0.14 | -0.011 | 0.038 | 0.038 | 0 | 0.3 | 0.11 | 0.19 | 0.22 | 0.28 | 0.47 |
| 8 | 128125195 | rs10505483 | 0.131 | -0.47 | -0.275 | 0.384 | 0.01 | 0.249 | 0.249 | 0.24 | 0.09 | 0.51 | 0.05 | 0.21 | 0.04 | 0.1 |
| 8 | 128127252 | rs1456305 | -0.589 | -0.128 | -0.521 | -0.14 | 0.034 | 0.054 | 0.054 | 0.273 | 0.76 | 0.9 | 0.77 | 0.97 | 0.87 | 0.86 |
| 8 | 128128459 | rs17184796 | -0.028 | 0.727 | -0.208 | 0.681 | -0.013 | 0.013 | 0.013 | 0 | 0.21 | 0.17 | 0.23 | 0.09 | 0.36 | 0.11 |
| 8 | 128150652 | rs17446747 | 0.144 | 1.391 | -1.302 | -0.341 | -0.116 | 0.051 | 0.051 | 0.008 | 0.2 | 0.03 | 0.21 | 0.06 | 0.38 | 0.11 |
| 8 | 128162974 | rs17446916 | 0.005 | 1.147 | -1.199 | -0.286 | -0.102 | 0.07 | 0.07 | 0 | 0.22 | 0.07 | 0.25 | 0.06 | 0.44 | 0.12 |
| 8 | 128171018 | rs1378900 | -0.955 | 0.266 | -0.761 | -0.211 | -0.007 | -0.016 | -0.016 | 0.005 | 0.89 | 0.9 | 0.84 | 0.98 | 0.95 | 0.87 |
| 8 | 128172028 | rs7002343 | -0.346 | 1.019 | -0.848 | -0.331 | -0.027 | 0.036 | 0.036 | 0 | 0.24 | 0.14 | 0.24 | 0.06 | 0.35 | 0.15 |
| 8 | 128172686 | rs2124600 | -0.308 | 0.756 | -0.457 | 0.082 | 0.005 | -0.014 | -0.014 | 0.439 | 0.42 | 0.7 | 0.46 | 0.54 | 0.34 | 0.29 |
| 8 | 128178647 | rs2445606 | -0.169 | 0.802 | 0.125 | -0.009 | 0 | 0.001 | 0.001 | 0 | 0.37 | 0.37 | 0.44 | 0.56 | 0.37 | 0.32 |
| 8 | 128182451 | rs2456461 | -0.052 | 0.746 | 0.212 | -0.022 | -0.032 | -0.05 | -0.05 | 1.433 | 0.36 | 0.23 | 0.43 | 0.55 | 0.37 | 0.31 |
| 8 | 128188019 | rs2466024 | -0.409 | 0.76 | 0.051 | 0.036 | -0.002 | -0.006 | -0.006 | 0 | 0.42 | 0.65 | 0.46 | 0.56 | 0.37 | 0.33 |
| 8 | 128197080 | rs2456452 | -0.699 | 0.617 | 0.262 | 0.026 | 0.009 | -0.006 | -0.006 | 0 | 0.56 | 0.6 | 0.61 | 0.53 | 0.66 | 0.37 |
| 8 | 128197295 | rs16902008 | 1.055 | 0.438 | -0.26 | -0.599 | -0.017 | -0.047 | -0.047 | 0 | 0.13 | 0.2 | 0.05 | 0 | 0.06 | 0.02 |
| 8 | 128197819 | rs2445611 | -0.734 | 0.972 | 0.059 | 0.05 | 0.009 | -0.037 | -0.037 | 0 | 0.37 | 0.52 | 0.37 | 0.19 | 0.56 | 0.17 |
| 8 | 128209820 | rs2466032 | -0.331 | 1.125 | -0.139 | 0.481 | -0.025 | 0.015 | 0.015 | 0 | 0.17 | 0.11 | 0.16 | 0.11 | 0.28 | 0.12 |
| 8 | 128211229 | rs2466035 | -1.407 | 0.535 | -0.137 | 0.864 | -0.001 | 0.002 | 0.002 | 0.11 | 0.31 | 0.71 | 0.41 | 0.45 | 0.31 | 0.27 |
| 8 | 128213233 | rs9643217 | -0.691 | 0.229 | -0.489 | 1.392 | 0.006 | -0.005 | -0.005 | 0 | 0.08 | 0.03 | 0.27 | 0.38 | 0.07 | 0.13 |
| 8 | 128218724 | rs10094303 | -1.682 | 0.003 | 0.119 | 1.476 | -0.027 | 0.194 | 0.194 | 1.244 | 0.13 | 0.58 | 0.15 | 0.36 | 0.07 | 0.1 |
| 8 | 128226195 | rs6989575 | -0.748 | 0.526 | -1.283 | 1.423 | -0.052 | -0.107 | -0.107 | 0 | 0.28 | 0.1 | 0.28 | 0.53 | 0.31 | 0.34 |
| 8 | 128228863 | rs10087719 | -0.313 | 0.636 | -1.524 | 0.813 | -0.119 | -0.013 | -0.013 | 0 | 0.26 | 0.01 | 0.23 | 0.19 | 0.28 | 0.25 |
| 8 | 128247045 | rs2044869 | -0.803 | 0.257 | -0.971 | 1.11 | -0.001 | 0.002 | 0.002 | 0.968 | 0.32 | 0.65 | 0.4 | 0.42 | 0.29 | 0.35 |

| | | | | | | | | | | | | | | | | |
|---|---|---|---|---|---|---|---|---|---|---|---|---|---|---|---|---|
| 8 | 128251421 | rs10105730 | -0.041 | 0.312 | -0.894 | 1.044 | -0.004 | 0.008 | 0.008 | 0.737 | 0.2 | 0.2 | 0.23 | 0.12 | 0.22 | 0.21 |
| 8 | 128252993 | rs283741 | 0.409 | -0.146 | -1.373 | 1.03 | -0.014 | 0.007 | 0.007 | 0 | 0.61 | 0.73 | 0.54 | 0.57 | 0.56 | 0.56 |
| 8 | 128258492 | rs7000704 | 0.15 | -0.017 | -0.789 | 0.871 | 0.005 | -0.037 | -0.037 | 1.5 | 0.17 | 0.23 | 0.21 | 0.02 | 0.19 | 0.13 |
| 8 | 128261927 | rs6992922 | -0.421 | -0.056 | -0.733 | 1.1 | -0.002 | 0.027 | 0.027 | 0.404 | 0.26 | 0.52 | 0.36 | 0.33 | 0.22 | 0.25 |
| 8 | 128267338 | rs10099034 | -0.656 | -0.322 | -1.073 | 0.856 | -0.011 | -0.001 | -0.001 | 0 | 0.34 | 0.6 | 0.35 | 0.38 | 0.23 | 0.23 |
| 8 | 128269607 | rs283735 | 1.124 | -0.48 | -1.662 | 0.784 | 0.025 | -0.046 | -0.046 | 0 | 0.1 | 0.01 | 0.13 | 0.15 | 0.04 | 0.04 |
| 8 | 128270980 | rs4871014 | 0.146 | -0.533 | -2.218 | 0.08 | -0.01 | 0.064 | 0.064 | 2.436 | 0.73 | 0.93 | 0.49 | 0.17 | 0.75 | 0.5 |
| 8 | 128276154 | rs13281552 | 0.499 | -0.748 | -2.044 | -0.26 | -0.006 | -0.026 | -0.026 | 0 | 0.08 | 0.13 | 0.04 | 0 | 0.04 | 0.05 |
| 8 | 128278313 | rs2122835 | 0.615 | -0.879 | -2.192 | -0.318 | 0.018 | 0.135 | 0.135 | 1.709 | 0.66 | 0.88 | 0.45 | 0.13 | 0.71 | 0.48 |
| 8 | 128280808 | rs11777807 | 0.982 | -0.577 | -2.73 | -1.172 | -0.08 | 0.167 | 0.167 | 0 | 0.34 | 0.09 | 0.25 | 0.08 | 0.44 | 0.18 |
| 8 | 128281644 | rs283732 | -0.519 | -0.875 | -2.93 | -0.062 | -0.001 | -0.07 | -0.07 | 0 | 0.79 | 0.93 | 0.78 | 0.63 | 0.78 | 0.58 |
| 8 | 128281911 | rs1011387 | 0.109 | -0.908 | -2.463 | 0.418 | -0.01 | 0.123 | 0.123 | 0.085 | 0.08 | 0.14 | 0.14 | 0.35 | 0.05 | 0.11 |
| 8 | 128288920 | rs17378189 | 1.337 | -0.9 | -2.166 | 0.16 | -0.028 | 0.08 | 0.08 | 0 | 0.36 | 0.22 | 0.25 | 0.07 | 0.44 | 0.17 |
| 8 | 128289591 | rs283710 | 0.169 | -0.964 | -2.112 | 0.19 | 0.016 | 0.013 | 0.013 | 0.344 | 0.72 | 0.85 | 0.65 | 0.31 | 0.73 | 0.48 |
| 8 | 128292843 | rs924992 | 0.741 | -0.918 | -2.068 | 0.264 | -0.001 | 0.092 | 0.092 | 0.025 | 0.06 | 0.07 | 0.13 | 0.33 | 0.05 | 0.1 |
| 8 | 128295710 | rs412835 | 0.499 | -1.053 | -2.354 | -0.021 | 0.021 | 0.015 | 0.015 | 0 | 0.52 | 0.33 | 0.61 | 0.53 | 0.59 | 0.42 |
| 8 | 128301414 | rs283701 | 0.019 | -0.823 | -1.993 | 0.189 | -0.012 | -0.02 | -0.02 | 2.571 | 0.09 | 0.04 | 0.23 | 0.16 | 0.09 | 0.05 |
| 8 | 128303850 | rs10100445 | 0.052 | -1.065 | -1.814 | 0.024 | -0.024 | -0.057 | -0.057 | 0 | 0.1 | 0.59 | 0.08 | 0 | 0.02 | 0.03 |
| 8 | 128304269 | rs6984900 | 0.088 | -0.889 | -0.929 | 0.128 | 0.006 | -0.091 | -0.091 | 0 | 0.21 | 0.39 | 0.13 | 0.09 | 0.17 | 0.16 |
| 8 | 128304499 | rs924993 | 0.464 | -0.582 | -1.239 | 0.809 | 0.007 | 0.051 | 0.051 | 0 | 0.06 | 0.05 | 0.13 | 0.29 | 0.02 | 0.09 |
| 8 | 128307064 | rs11994592 | 0.504 | -0.471 | -1.208 | 1.324 | 0 | 0.076 | 0.076 | 0 | 0.06 | 0.09 | 0.04 | 0.3 | 0.02 | 0.1 |
| 8 | 128307082 | rs283718 | 1.06 | -0.355 | -1.38 | 0.95 | 0.011 | 0.085 | 0.085 | 0.144 | 0.51 | 0.23 | 0.61 | 0.32 | 0.59 | 0.49 |
| 8 | 128309965 | rs283720 | 0.571 | -0.215 | -1.057 | 1.041 | -0.003 | 0 | 0 | 0.458 | 0.64 | 0.35 | 0.66 | 0.4 | 0.71 | 0.61 |
| 8 | 128310155 | rs6996866 | 1.766 | -0.087 | -1.418 | 0.89 | -0.01 | 0.113 | 0.113 | 0 | 0.39 | 0.16 | 0.26 | 0.11 | 0.43 | 0.27 |
| 8 | 128310493 | rs283721 | -0.905 | -0.065 | -1.214 | 1.447 | -0.007 | 0.006 | 0.006 | 0.157 | 0.12 | 0.13 | 0.3 | 0.49 | 0.15 | 0.25 |
| 8 | 128323181 | rs445114 | -0.982 | -0.029 | -1.07 | 1.078 | 0.027 | -0.019 | -0.019 | 0 | 0.46 | 0.72 | 0.35 | 0.46 | 0.36 | 0.55 |
| 8 | 128339426 | rs424281 | -1.751 | 0.274 | -0.665 | 0.836 | -0.003 | -0.042 | -0.042 | 2.624 | 0.58 | 0.5 | 0.7 | 0.82 | 0.57 | 0.74 |
| 8 | 128341680 | rs587948 | -0.947 | 0.158 | -0.612 | 0.999 | -0.002 | -0.001 | -0.001 | 1.013 | 0.4 | 0.38 | 0.53 | 0.57 | 0.4 | 0.56 |
| 8 | 128342154 | rs10956359 | -0.506 | 0.235 | -0.601 | 0.659 | 0.001 | 0.007 | 0.007 | 0.141 | 0.31 | 0.26 | 0.17 | 0.13 | 0.28 | 0.31 |
| 8 | 128342866 | rs17464492 | -0.132 | 0.362 | -0.649 | 0.531 | -0.003 | 0.003 | 0.003 | 1.835 | 0.27 | 0.2 | 0.16 | 0.07 | 0.28 | 0.29 |
| 8 | 128345463 | rs672888 | -1.073 | 0.342 | -0.647 | 0.928 | -0.003 | -0.004 | -0.004 | 0 | 0.43 | 0.42 | 0.56 | 0.47 | 0.43 | 0.49 |
| 8 | 128347124 | rs391640 | 0.737 | 0.399 | -0.732 | 0.781 | -0.03 | 0.036 | 0.036 | 0.649 | 0.16 | 0.07 | 0.18 | 0.05 | 0.19 | 0.23 |
| 8 | 128353171 | rs12541832 | -0.546 | 0.326 | -0.673 | 1.01 | -0.006 | 0.004 | 0.004 | 0 | 0.27 | 0.24 | 0.46 | 0.19 | 0.33 | 0.37 |
| 8 | 128354313 | rs13271658 | 0.313 | 0.529 | -0.732 | 1.078 | 0.001 | -0.013 | -0.013 | 0 | 0.79 | 0.84 | 0.67 | 0.57 | 0.81 | 0.77 |
| 8 | 128354785 | rs10095860 | 0.753 | 0.376 | -0.649 | 1.017 | -0.002 | -0.003 | -0.003 | 0 | 0.64 | 0.59 | 0.49 | 0.28 | 0.67 | 0.55 |
| 8 | 128355019 | rs10098985 | 2.319 | 0.365 | -0.436 | 0.993 | 0 | -0.001 | -0.001 | 0.484 | 0.37 | 0.36 | 0.25 | 0.22 | 0.4 | 0.3 |
| 8 | 128356226 | rs13256275 | -1.853 | 0.374 | -0.446 | 1.085 | 0.001 | -0.002 | -0.002 | 0.427 | 0.58 | 0.49 | 0.51 | 0.64 | 0.51 | 0.6 |
| 8 | 128364973 | rs12156034 | 0.948 | 0.215 | -0.561 | 1.102 | -0.022 | -0.076 | -0.076 | 0.852 | 0.12 | 0.01 | 0.07 | 0.29 | 0.1 | 0.12 |
| 8 | 128366604 | rs11782735 | 0.92 | 0.383 | -0.476 | 1.275 | 0.001 | -0.003 | -0.003 | 0 | 0.15 | 0.14 | 0.17 | 0.25 | 0.14 | 0.19 |
| 8 | 128374117 | rs978683 | 0.504 | 0.396 | -0.576 | 1.264 | -0.014 | -0.001 | -0.001 | 0 | 0.61 | 0.7 | 0.67 | 0.64 | 0.6 | 0.63 |
| 8 | 128382357 | rs16902126 | 1.353 | 0.475 | -0.792 | 0.834 | 0.003 | 0.007 | 0.007 | 0 | 0.43 | 0.59 | 0.28 | 0.26 | 0.41 | 0.32 |
| 8 | 128382488 | rs11776260 | 0.834 | 0.551 | -0.909 | 0.554 | 0 | -0.021 | -0.021 | 0.212 | 0.15 | 0.07 | 0.16 | 0.27 | 0.14 | 0.19 |
| 8 | 128386926 | rs9643220 | -1.026 | 0.761 | -0.934 | 0.259 | -0.03 | 0.02 | 0.02 | 0.967 | 0.85 | 0.93 | 0.82 | 0.93 | 0.79 | 0.82 |
| 8 | 128387852 | rs1562430 | 1.029 | 0.288 | -0.662 | 0.363 | -0.002 | -0.056 | -0.056 | 0.457 | 0.37 | 0.46 | 0.27 | 0.18 | 0.41 | 0.23 |
| 8 | 128390660 | rs731900 | 0.426 | 0.54 | -1.219 | 0.455 | -0.052 | -0.088 | -0.088 | 0 | 0.17 | 0.01 | 0.18 | 0.32 | 0.18 | 0.2 |
| 8 | 128390957 | rs16902131 | 0.253 | 0.475 | -0.489 | -0.306 | -0.017 | -0.045 | -0.045 | 0 | 0.13 | 0.25 | 0.06 | 0.05 | 0.09 | 0.12 |
| 8 | 128396316 | rs6986543 | 0.568 | 1.032 | -1.17 | -0.262 | 0 | 0.019 | 0.019 | 0 | 0.31 | 0.31 | 0.23 | 0.19 | 0.37 | 0.18 |
| 8 | 128400176 | rs7820981 | 0.531 | 1.454 | -0.962 | 0.114 | -0.005 | -0.001 | -0.001 | 0 | 0.64 | 0.71 | 0.54 | 0.64 | 0.56 | 0.46 |
| 8 | 128401772 | rs1562871 | -0.122 | 1.565 | -1.24 | 0.313 | 0 | 0.017 | 0.017 | 0 | 0.83 | 0.81 | 0.75 | 0.66 | 0.8 | 0.62 |
| 8 | 128402907 | rs12549845 | -0.534 | 1.789 | -1.434 | -0.569 | -0.038 | 0.043 | 0.043 | 0 | 0.18 | 0.09 | 0.21 | 0.04 | 0.26 | 0.17 |
| 8 | 128403514 | rs7844673 | -0.685 | 1.002 | -0.984 | -0.311 | 0.002 | -0.018 | -0.018 | 0 | 0.08 | 0.11 | 0.03 | 0.07 | 0.07 | 0.11 |
| 8 | 128407443 | rs10505477 | 0.357 | 0.586 | -1.906 | -0.447 | -0.043 | 0.011 | 0.011 | 0.221 | 0.46 | 0.13 | 0.47 | 0.61 | 0.52 | 0.49 |
| 8 | 128408116 | rs10505476 | -0.065 | -0.028 | -1.906 | -0.858 | 0.006 | 0.017 | 0.017 | 2.704 | 0.28 | 0.73 | 0.19 | 0.19 | 0.26 | 0.16 |
| 8 | 128409511 | rs10808555 | 0.116 | -0.17 | -0.846 | -1.208 | 0.005 | 0.012 | 0.012 | 1.372 | 0.72 | 0.59 | 0.71 | 0.71 | 0.69 | 0.67 |
| 8 | 128413147 | rs10808556 | 0.937 | -0.332 | -0.724 | -1.174 | 0.005 | 0.009 | 0.009 | 3.053 | 0.61 | 0.25 | 0.66 | 0.65 | 0.61 | 0.59 |
| 8 | 128413305 | rs6983267 | 1.73 | -0.532 | -1.043 | -1.448 | -0.134 | -0.053 | -0.053 | 0 | 0.41 | 0.05 | 0.45 | 0.61 | 0.5 | 0.47 |
| 8 | 128417936 | rs10505473 | 1.022 | -0.415 | -0.005 | -1.277 | -0.004 | -0.003 | -0.003 | 0 | 0.14 | 0.21 | 0.11 | 0.13 | 0.16 | 0.08 |
| 8 | 128423127 | rs7837328 | 0.842 | -0.439 | -0.001 | -1.324 | 0.002 | 0.003 | 0.003 | 1.871 | 0.63 | 0.31 | 0.67 | 0.66 | 0.62 | 0.61 |
| 8 | 128424792 | rs7014346 | 0.389 | -0.409 | 0.565 | -1.255 | 0.002 | 0.005 | 0.005 | 0.746 | 0.7 | 0.6 | 0.71 | 0.71 | 0.67 | 0.66 |
| 8 | 128428061 | rs4871789 | 0.48 | -0.615 | -0.434 | -1.687 | 0.027 | 0.078 | 0.078 | 0 | 0.5 | 0.21 | 0.6 | 0.65 | 0.53 | 0.57 |
| 8 | 128429721 | rs6985419 | 0.082 | -0.459 | -0.28 | -1.714 | -0.011 | -0.037 | -0.037 | 0 | 0.65 | 0.51 | 0.72 | 0.79 | 0.68 | 0.66 |
| 8 | 128437686 | rs1447294 | 0.06 | -0.763 | -0.445 | -1.738 | 0.006 | 0.013 | 0.013 | 0 | 0.35 | 0.31 | 0.21 | 0.23 | 0.32 | 0.17 |
| 8 | 128440167 | rs9297756 | 0.788 | -0.772 | -0.552 | -1.753 | -0.004 | -0.002 | -0.002 | 0 | 0.17 | 0.21 | 0.12 | 0.18 | 0.18 | 0.1 |
| 8 | 128440651 | rs6995633 | -1.139 | -0.733 | -1.019 | -1.704 | 0.014 | -0.027 | -0.027 | 0.354 | 0.9 | 0.99 | 0.83 | 0.85 | 0.94 | 0.76 |
| 8 | 128440928 | rs6999921 | 1.07 | -0.929 | -1.16 | -1.78 | 0.001 | 0.002 | 0.002 | 0.003 | 0.12 | 0.1 | 0.06 | 0.04 | 0.09 | 0.09 |
| 8 | 128441170 | rs7000448 | -0.385 | -0.858 | -0.854 | -1.67 | 0.052 | 0.122 | 0.122 | 0.375 | 0.49 | 0.71 | 0.29 | 0.27 | 0.4 | 0.26 |
| 8 | 128441535 | rs4871790 | 1.065 | -0.657 | -1.177 | -1.637 | -0.097 | -0.044 | -0.044 | 0.54 | 0.34 | 0.12 | 0.48 | 0.58 | 0.47 | 0.48 |
| 8 | 128453765 | rs10090421 | 0.258 | -0.603 | -0.863 | -1.445 | -0.002 | -0.002 | -0.002 | 0 | 0.27 | 0.27 | 0.18 | 0.25 | 0.27 | 0.21 |
| 8 | 128453928 | rs12334695 | 0.245 | -0.38 | -0.919 | -1.257 | 0.015 | 0.038 | 0.038 | 1.362 | 0.46 | 0.35 | 0.26 | 0.27 | 0.41 | 0.28 |

| | | | | | | | | | | | | | | | | |
|---|---|---|---|---|---|---|---|---|---|---|---|---|---|---|---|---|
| 8 | 128455654 | rs6981397 | 0.771 | 0.695 | -1.978 | -1.182 | -0.04 | -0.049 | -0.049 | 0 | 0.25 | 0.13 | 0.4 | 0.59 | 0.31 | 0.33 |
| 8 | 128458151 | rs10109622 | 0.669 | 0.408 | -1.586 | -1.279 | -0.001 | -0.037 | -0.037 | 0.293 | 0.35 | 0.63 | 0.19 | 0.05 | 0.25 | 0.18 |
| 8 | 128458644 | rs4871791 | -0.318 | 0.649 | -1.16 | -0.637 | -0.004 | 0.058 | 0.058 | 0 | 0.44 | 0.33 | 0.53 | 0.81 | 0.46 | 0.42 |
| 8 | 128466814 | rs7841264 | -0.32 | 0.913 | -1.067 | 0.699 | 0 | -0.026 | -0.026 | 0 | 0.21 | 0.17 | 0.37 | 0.5 | 0.2 | 0.22 |
| 8 | 128467934 | rs10099905 | 0.338 | 1.365 | -0.362 | 0.972 | 0.051 | 0.031 | 0.031 | 0 | 0.88 | 0.77 | 0.74 | 0.85 | 0.82 | 0.85 |
| 8 | 128472320 | rs1447293 | 0.136 | 0.257 | -1.448 | 1.188 | 0.06 | -0.046 | -0.046 | 1.533 | 0.47 | 0.09 | 0.52 | 0.48 | 0.6 | 0.61 |
| 8 | 128475185 | rs921146 | -0.347 | -0.297 | -0.467 | 1.318 | -0.011 | 0.046 | 0.046 | 0.717 | 0.79 | 0.53 | 0.76 | 0.73 | 0.77 | 0.82 |
| 8 | 128479963 | rs4871798 | 0.331 | -0.301 | -0.348 | 1.541 | 0.079 | 0.16 | 0.16 | 1.608 | 0.9 | 0.63 | 0.81 | 0.75 | 0.87 | 0.83 |
| 8 | 128482642 | rs4871799 | 0.38 | -0.278 | -0.378 | 1.525 | 0.01 | 0.072 | 0.072 | 1.077 | 0.77 | 0.56 | 0.7 | 0.63 | 0.75 | 0.78 |
| 8 | 128483096 | rs6981424 | 1.23 | -0.019 | -0.924 | 1.403 | -0.093 | -0.002 | -0.002 | 0.274 | 0.29 | 0.08 | 0.3 | 0.26 | 0.36 | 0.43 |
| 8 | 128485038 | rs1447295 | -0.427 | 0.077 | -0.315 | 1.336 | 0.102 | 0.15 | 0.15 | 3.323 | 0.93 | 0.65 | 0.89 | 0.86 | 0.9 | 0.87 |
| 8 | 128486588 | rs9297758 | 0.016 | -0.07 | -0.201 | 1.313 | -0.015 | 0.003 | 0.003 | 0 | 0.37 | 0.31 | 0.38 | 0.34 | 0.44 | 0.26 |
| 8 | 128487522 | rs10086608 | 1.626 | -0.027 | -0.273 | 1.076 | 0.007 | -0.009 | -0.009 | 0 | 0.12 | 0.08 | 0.11 | 0.12 | 0.11 | 0.05 |
| 8 | 128491792 | rs7836840 | 0.111 | -0.198 | -0.071 | 1.033 | 0.003 | 0.002 | 0.002 | 0 | 0.58 | 0.59 | 0.54 | 0.48 | 0.47 | 0.6 |
| 8 | 128494391 | rs9643225 | -0.675 | 0.006 | -0.051 | 1.076 | 0.022 | 0.037 | 0.037 | 0 | 0.8 | 0.68 | 0.76 | 0.74 | 0.74 | 0.81 |
| 8 | 128496096 | rs9643227 | 0.367 | -0.004 | -0.293 | 1.059 | 0.003 | -0.004 | -0.004 | 0.93 | 0.56 | 0.53 | 0.46 | 0.38 | 0.49 | 0.58 |
| 8 | 128496506 | rs16902169 | 1.196 | 0.132 | -1.205 | 1.075 | -0.03 | -0.05 | -0.05 | 0 | 0.2 | 0.06 | 0.22 | 0.26 | 0.22 | 0.2 |
| 8 | 128496867 | rs13258812 | 1.353 | 0.541 | -1.171 | 0.885 | -0.037 | 0.06 | 0.06 | 0.148 | 0.25 | 0.09 | 0.23 | 0.13 | 0.28 | 0.31 |
| 8 | 128503999 | rs16902173 | 1.215 | 0.267 | -0.961 | 1.233 | -0.014 | -0.015 | -0.015 | 0 | 0.14 | 0.04 | 0.14 | 0.12 | 0.16 | 0.06 |
| 8 | 128507602 | rs1562432 | 0.769 | 0.058 | -0.776 | 1.109 | 0.01 | -0.066 | -0.066 | 0.484 | 0.19 | 0.04 | 0.15 | 0.27 | 0.13 | 0.18 |
| 8 | 128507651 | rs1562431 | 1.628 | 0.078 | -0.811 | 1.131 | 0.016 | -0.031 | -0.031 | 2.839 | 0.15 | 0.01 | 0.12 | 0.12 | 0.07 | 0.07 |
| 8 | 128513545 | rs4871808 | 0.994 | 0.104 | -0.712 | 1.055 | 0.005 | -0.078 | -0.078 | 0 | 0.19 | 0.02 | 0.15 | 0.27 | 0.13 | 0.18 |
| 8 | 128517573 | rs4242382 | 0.461 | 0.751 | 0.486 | 0.89 | 0.097 | 0.178 | 0.178 | 2.985 | 0.93 | 0.64 | 0.91 | 0.85 | 0.89 | 0.86 |
| 8 | 128519957 | rs7018386 | 0.757 | 0.816 | -0.166 | 0.67 | -0.026 | 0.068 | 0.068 | 0.158 | 0.53 | 0.19 | 0.52 | 0.65 | 0.54 | 0.43 |
| 8 | 128525985 | rs7004374 | -0.465 | 1.211 | -0.698 | 0.387 | -0.001 | -0.008 | -0.008 | 0.421 | 0.3 | 0.28 | 0.27 | 0.11 | 0.33 | 0.33 |
| 8 | 128531028 | rs13265719 | -0.452 | 1.35 | -0.772 | 0.409 | -0.004 | 0.007 | 0.007 | 0 | 0.3 | 0.26 | 0.27 | 0.21 | 0.33 | 0.3 |
| 8 | 128545383 | rs6991990 | -0.222 | 1.425 | -1.172 | 0.422 | -0.048 | 0.037 | 0.037 | 0 | 0.27 | 0.1 | 0.25 | 0.2 | 0.32 | 0.27 |
| 8 | 128545619 | rs4543510 | 0.37 | 1.451 | -0.476 | 0.925 | 0.015 | 0.08 | 0.08 | 0 | 0.77 | 0.64 | 0.65 | 0.56 | 0.73 | 0.63 |
| 8 | 128548678 | rs13258742 | -0.743 | 1.874 | -0.553 | 0.24 | 0.009 | -0.035 | -0.035 | 0 | 0.11 | 0.11 | 0.07 | 0 | 0.13 | 0.02 |
| 8 | 128550123 | rs4407842 | 0.302 | 1.733 | -0.636 | 0.392 | -0.011 | -0.007 | -0.007 | 0 | 0.58 | 0.68 | 0.6 | 0.55 | 0.55 | 0.61 |
| 8 | 128558990 | rs6470530 | 0.821 | 1.713 | -0.594 | 0.323 | 0.016 | 0.001 | 0.001 | 0 | 0.47 | 0.32 | 0.51 | 0.44 | 0.42 | 0.44 |
| 8 | 128561431 | rs7011721 | 0.277 | 1.581 | -1.121 | 0.458 | -0.039 | -0.039 | -0.039 | 0 | 0.58 | 0.79 | 0.53 | 0.53 | 0.54 | 0.6 |
| 8 | 128563299 | rs6415495 | -0.479 | 1.849 | -1.373 | 0.367 | -0.093 | 0.074 | 0.074 | 0 | 0.74 | 0.98 | 0.85 | 0.91 | 0.7 | 0.81 |
| 8 | 128563712 | rs12114450 | -0.504 | 1.42 | -0.553 | 0.889 | -0.001 | 0.017 | 0.017 | 0.074 | 0.16 | 0.18 | 0.25 | 0.35 | 0.15 | 0.21 |
| 8 | 128568712 | rs7386167 | -0.312 | 0.948 | -0.674 | -0.903 | 0.002 | -0.038 | -0.038 | 1.913 | 0.58 | 0.62 | 0.75 | 0.97 | 0.63 | 0.82 |
| 8 | 128568856 | rs4733655 | 0.493 | 1.086 | -0.817 | -0.775 | -0.004 | -0.004 | -0.004 | 0 | 0.22 | 0.18 | 0.46 | 0.24 | 0.28 | 0.29 |
| 8 | 128571059 | rs10110900 | -0.404 | 1.385 | -0.916 | -0.862 | 0.007 | -0.007 | -0.007 | 0 | 0.34 | 0.43 | 0.58 | 0.31 | 0.44 | 0.42 |
| 8 | 128571759 | rs4313118 | 0.104 | 1.474 | -0.99 | -0.534 | -0.004 | -0.017 | -0.017 | 0 | 0.24 | 0.19 | 0.17 | 0.66 | 0.19 | 0.4 |
| 8 | 128578015 | rs4733739 | -0.003 | 1.051 | -1.049 | -1.012 | 0.004 | -0.07 | -0.07 | 0.017 | 0.18 | 0.23 | 0.12 | 0 | 0.2 | 0.08 |
| 8 | 128583597 | rs7465658 | 0.066 | 1.135 | -1.54 | -0.561 | -0.042 | -0.012 | -0.012 | 0 | 0.24 | 0.1 | 0.44 | 0.26 | 0.31 | 0.34 |
| 8 | 128588822 | rs7824074 | 0.577 | 1.103 | -1.624 | -0.884 | -0.029 | 0.048 | 0.048 | 0 | 0.77 | 0.88 | 0.68 | 0.82 | 0.75 | 0.64 |
| 8 | 128592256 | rs4733807 | 0.04 | 1.015 | -1.176 | -0.831 | 0.006 | -0.006 | -0.006 | 0 | 0.13 | 0.16 | 0.28 | 0.11 | 0.17 | 0.16 |
| 8 | 128592345 | rs6470537 | -0.547 | 0.998 | -1.226 | -0.856 | -0.008 | 0.021 | 0.021 | 0 | 0.89 | 0.96 | 0.93 | 0.92 | 0.92 | 0.85 |
| 8 | 128594189 | rs6470538 | 1.042 | -0.006 | -0.891 | -1.233 | 0.006 | 0.048 | 0.048 | 0 | 0.48 | 0.35 | 0.47 | 0.24 | 0.58 | 0.4 |
| 8 | 128595878 | rs10283063 | 1.325 | 0.041 | -1.203 | -1.849 | -0.022 | 0.052 | 0.052 | 0 | 0.3 | 0.18 | 0.2 | 0.08 | 0.38 | 0.24 |
| 8 | 128597214 | rs7819582 | 0.147 | -0.316 | -1.063 | -2.015 | -0.005 | -0.01 | -0.01 | 0 | 0.37 | 0.47 | 0.45 | 0.68 | 0.34 | 0.46 |
| 8 | 128599672 | rs4551307 | 1.44 | -0.697 | -0.537 | -1.972 | -0.001 | 0 | 0 | 3.875 | 0.19 | 0.23 | 0.27 | 0.17 | 0.19 | 0.16 |
| 8 | 128600662 | rs13252781 | 0.913 | -0.58 | -0.786 | -1.978 | -0.043 | 0.132 | 0.132 | 0.304 | 0.4 | 0.18 | 0.43 | 0.19 | 0.54 | 0.39 |
| 8 | 128601372 | rs10091329 | -0.172 | -1.214 | 0.275 | -1.291 | -0.01 | -0.036 | -0.036 | 1.891 | 0.85 | 0.74 | 0.9 | 0.92 | 0.91 | 0.85 |
| 8 | 128601933 | rs13253038 | 0.727 | -1.395 | 0.325 | -1.357 | -0.023 | -0.009 | -0.009 | 0 | 0.61 | 0.5 | 0.71 | 0.35 | 0.74 | 0.56 |
| 8 | 128602212 | rs11988657 | 1.452 | -1.276 | -0.506 | -1.396 | -0.074 | 0.05 | 0.05 | 1.39 | 0.36 | 0.11 | 0.41 | 0.17 | 0.5 | 0.27 |
| 8 | 128603409 | rs4129666 | 0.811 | -1.289 | -0.454 | -0.854 | 0 | 0.004 | 0.004 | 0 | 0.13 | 0.09 | 0.07 | 0.13 | 0.12 | 0.13 |
| 8 | 128606603 | rs13259282 | -1.501 | -1.446 | -0.5 | -0.971 | -0.041 | 0.063 | 0.063 | 0 | 0.57 | 0.85 | 0.52 | 0.74 | 0.43 | 0.58 |
| 8 | 128613730 | rs4451272 | 0.311 | -1.586 | -0.106 | -1.017 | -0.006 | 0.026 | 0.026 | 0 | 0.25 | 0.19 | 0.29 | 0.18 | 0.25 | 0.3 |
| 8 | 128617544 | rs6470545 | -1.458 | -1.586 | -0.174 | -1.043 | -0.001 | -0.003 | -0.003 | 0 | 0.89 | 0.88 | 0.92 | 0.92 | 0.92 | 0.85 |
| 8 | 128617612 | rs4733597 | -0.642 | -1.65 | 0.095 | -1.049 | 0.002 | 0.026 | 0.026 | 0 | 0.41 | 0.36 | 0.36 | 0.29 | 0.37 | 0.43 |
| 8 | 128621900 | rs9643229 | -0.877 | -1.358 | 0.296 | -1.016 | 0.015 | 0.063 | 0.063 | 0.104 | 0.51 | 0.7 | 0.51 | 0.73 | 0.42 | 0.58 |
| 8 | 128623639 | rs7005795 | -1.043 | -1.151 | -0.141 | -0.869 | 0 | 0.012 | 0.012 | 0.055 | 0.53 | 0.5 | 0.49 | 0.74 | 0.43 | 0.61 |
| 8 | 128624424 | rs7388104 | 1.985 | -1.128 | -0.597 | -0.869 | -0.165 | -0.029 | -0.029 | 0.011 | 0.32 | 0.03 | 0.39 | 0.18 | 0.5 | 0.25 |
| 8 | 128626960 | rs4551310 | -1.286 | -1.182 | -0.535 | -0.838 | -0.037 | 0.02 | 0.02 | 0.51 | 0.55 | 0.91 | 0.54 | 0.7 | 0.38 | 0.62 |
| 8 | 128628601 | rs6470552 | 0.218 | -1.215 | -0.439 | -0.855 | -0.028 | -0.047 | -0.047 | 0.053 | 0.15 | 0.5 | 0.11 | 0.08 | 0.07 | 0.13 |
| 8 | 128628938 | rs6470556 | -0.57 | -0.683 | -0.343 | -0.549 | -0.006 | 0.002 | 0.002 | 0 | 0.4 | 0.39 | 0.43 | 0.62 | 0.31 | 0.48 |
| 8 | 128630589 | rs4314620 | 0.434 | -0.033 | -0.818 | -0.399 | -0.041 | 0.003 | 0.003 | 0.077 | 0.14 | 0.01 | 0.1 | 0.1 | 0.16 | 0.2 |
| 8 | 128637028 | rs9656817 | -0.506 | 0.399 | -0.259 | -0.179 | -0.048 | 0.085 | 0.085 | 4.249 | 0.61 | 0.82 | 0.41 | 0.84 | 0.44 | 0.79 |
| 8 | 128641620 | rs10956379 | -1.082 | 0.415 | -0.613 | 0.076 | -0.167 | 0.002 | 0.002 | 0.66 | 0.69 | 0.97 | 0.63 | 0.85 | 0.51 | 0.8 |
| 8 | 128643231 | rs6989963 | -0.576 | 0.371 | -0.164 | 0.466 | -0.045 | 0.094 | 0.094 | 0.402 | 0.58 | 0.85 | 0.62 | 0.84 | 0.41 | 0.69 |
| 8 | 128644955 | rs4130120 | -1.097 | 0.266 | 0.597 | 0.42 | -0.014 | -0.083 | -0.083 | 0.325 | 0.16 | 0.38 | 0.12 | 0.07 | 0.12 | 0.07 |
| 8 | 128649634 | rs10441526 | -0.925 | 0.277 | 0.535 | 0.478 | -0.082 | 0.072 | 0.072 | 1.39 | 0.63 | 0.86 | 0.67 | 0.84 | 0.49 | 0.7 |
| 8 | 128653001 | rs13252945 | 0.244 | 0.068 | 0.843 | 0.935 | 0 | 0 | 0 | 1.996 | 0.44 | 0.37 | 0.51 | 0.46 | 0.36 | 0.47 |

| | | | | | | | | | | | | | | | | |
|---|---|---|---|---|---|---|---|---|---|---|---|---|---|---|---|---|
| 8 | 128654115 | rs10956381 | -0.436 | 0.154 | 0.526 | 0.978 | 0.014 | 0.01 | 0.01 | 0 | 0.53 | 0.63 | 0.57 | 0.51 | 0.41 | 0.53 |
| 8 | 128656651 | rs7825200 | -0.982 | 0.374 | 0.027 | 0.804 | -0.002 | 0.191 | 0.191 | 0 | 0.14 | 0.32 | 0.1 | 0.45 | 0.09 | 0.23 |
| 8 | 128657277 | rs11995971 | -0.555 | 0.492 | -0.278 | -0.483 | -0.002 | -0.008 | -0.008 | 0.478 | 0.58 | 0.55 | 0.63 | 0.87 | 0.49 | 0.71 |
| 8 | 128657531 | rs7829529 | -1.049 | 0.401 | -0.257 | 0.084 | -0.013 | -0.036 | -0.036 | 0 | 0.11 | 0.2 | 0.07 | 0.07 | 0.07 | 0.06 |
| 8 | 128660096 | rs7817632 | -0.572 | 0.628 | -0.684 | 0.125 | 0 | 0.017 | 0.017 | 0 | 0.36 | 0.73 | 0.4 | 0.26 | 0.37 | 0.44 |
| 8 | 128662095 | rs4733616 | 0.155 | 0.639 | -0.118 | -0.023 | -0.001 | -0.049 | -0.049 | 0 | 0.17 | 0.18 | 0.22 | 0.03 | 0.2 | 0.18 |
| 8 | 128664590 | rs4562278 | -0.247 | 1.14 | -0.296 | 0.062 | 0.009 | -0.012 | -0.012 | 0.002 | 0.17 | 0.27 | 0.23 | 0.17 | 0.18 | 0.16 |
| 8 | 128664941 | rs10100504 | -0.193 | 1.199 | -0.162 | 0.223 | 0.016 | 0.033 | 0.033 | 0 | 0.16 | 0.21 | 0.14 | 0.29 | 0.2 | 0.27 |
| 8 | 128665346 | rs4463380 | -0.479 | 0.441 | -0.907 | 0.479 | 0.015 | -0.005 | -0.005 | 0 | 0.57 | 0.71 | 0.72 | 0.62 | 0.64 | 0.8 |
| 8 | 128675771 | rs7835137 | 0.164 | 0.164 | -0.693 | 1.005 | -0.004 | 0.029 | 0.029 | 0 | 0.89 | 0.85 | 0.85 | 0.71 | 0.88 | 0.88 |
| 8 | 128676131 | rs4133274 | -0.241 | -0.735 | -0.787 | 1.385 | 0.004 | -0.008 | -0.008 | 0 | 0.14 | 0.12 | 0.05 | 0.23 | 0.07 | 0.29 |
| 8 | 128677531 | rs7841193 | 0.494 | -0.65 | -1.794 | 0.86 | -0.121 | -0.004 | -0.004 | 0 | 0.32 | 0.06 | 0.4 | 0.21 | 0.47 | 0.31 |
| 8 | 128678986 | rs4593503 | -0.714 | -0.513 | -1.662 | 1.029 | -0.005 | 0.035 | 0.035 | 0 | 0.89 | 0.85 | 0.86 | 0.67 | 0.88 | 0.88 |
| 8 | 128680999 | rs12543549 | 0.829 | -0.682 | -1.539 | 0.399 | 0.008 | 0.071 | 0.071 | 0.433 | 0.36 | 0.14 | 0.32 | 0.26 | 0.3 | 0.26 |
| 8 | 128690629 | rs4733658 | 0.737 | -0.695 | -0.897 | 0.255 | 0.006 | 0.013 | 0.013 | 1.943 | 0.32 | 0.28 | 0.37 | 0.31 | 0.27 | 0.21 |
| 8 | 128691212 | rs13248513 | 0.053 | -1.027 | -0.78 | 0.612 | 0.003 | -0.005 | -0.005 | 0 | 0.14 | 0.14 | 0.06 | 0.18 | 0.06 | 0.27 |
| 8 | 128693521 | rs11774100 | 1.168 | -0.683 | -0.8 | 0.453 | -0.055 | 0.044 | 0.044 | 2.219 | 0.35 | 0.11 | 0.4 | 0.29 | 0.38 | 0.22 |
| 8 | 128694006 | rs4733664 | -0.154 | -0.667 | -0.318 | 0.452 | 0 | 0.001 | 0.001 | 0 | 0.52 | 0.51 | 0.49 | 0.47 | 0.44 | 0.49 |
| 8 | 128699278 | rs6990483 | -1.097 | -0.326 | -0.704 | 0.9 | -0.031 | -0.047 | -0.047 | 0 | 0.92 | 0.99 | 0.9 | 0.76 | 0.9 | 0.86 |
| 8 | 128699454 | rs10110576 | -0.076 | -0.601 | -0.109 | 0.714 | -0.001 | 0.007 | 0.007 | 0.069 | 0.54 | 0.66 | 0.54 | 0.48 | 0.45 | 0.45 |
| 8 | 128700344 | rs10505504 | 1.329 | -0.634 | -0.308 | 0.718 | -0.108 | 0.024 | 0.024 | 1.72 | 0.34 | 0.05 | 0.43 | 0.3 | 0.39 | 0.2 |
| 8 | 128702676 | rs7845292 | 0.387 | -0.644 | -0.095 | 0.711 | 0.002 | 0.018 | 0.018 | 0 | 0.49 | 0.21 | 0.49 | 0.48 | 0.45 | 0.46 |
| 8 | 128705161 | rs7840975 | -0.041 | -0.5 | 0.31 | 0.644 | -0.066 | 0.025 | 0.025 | 0.811 | 0.16 | 0.33 | 0.08 | 0.18 | 0.06 | 0.26 |
| 8 | 128706908 | rs6984323 | 0.17 | -0.211 | 0.277 | 0.15 | -0.023 | 0.028 | 0.028 | 0.018 | 0.37 | 0.28 | 0.37 | 0.27 | 0.49 | 0.41 |
| 8 | 128707095 | rs6984633 | -0.46 | 0.023 | 0.221 | 0.138 | 0.047 | -0.008 | -0.008 | 0 | 0.06 | 0.22 | 0.06 | 0 | 0.06 | 0.03 |
| 8 | 128708284 | rs10096900 | 0.448 | 0.056 | -0.198 | 0.222 | -0.001 | 0.005 | 0.005 | 0 | 0.36 | 0.33 | 0.41 | 0.3 | 0.35 | 0.2 |
| 8 | 128713173 | rs12547643 | -0.809 | -0.301 | -0.647 | -0.091 | 0.012 | 0.163 | 0.163 | 0 | 0.37 | 0.09 | 0.17 | 0.16 | 0.32 | 0.47 |
| 8 | 128713873 | rs16902328 | 0.002 | 0.143 | -0.19 | 0.392 | -0.012 | 0.011 | 0.011 | 0.336 | 0.08 | 0.02 | 0.05 | 0.07 | 0.07 | 0.17 |
| 8 | 128718068 | rs9642880 | -1.73 | 0.239 | 0.033 | 0.62 | 0.007 | 0.056 | 0.056 | 0 | 0.44 | 0.81 | 0.43 | 0.31 | 0.45 | 0.58 |
| 8 | 128718443 | rs17186926 | -0.904 | 0.624 | 0.565 | 0.722 | -0.031 | 0.002 | 0.002 | 0 | 0.13 | 0.04 | 0.11 | 0.1 | 0.21 | 0.17 |
| 8 | 128725947 | rs16902338 | -1.102 | -0.02 | 1.525 | 0.765 | -0.004 | -0.032 | -0.032 | 0 | 0.07 | 0.28 | 0.05 | 0 | 0.03 | 0.02 |
| 8 | 128739771 | rs10505505 | -1.834 | -0.031 | 1.327 | 0.751 | 0.009 | -0.047 | -0.047 | 0 | 0.1 | 0.47 | 0.08 | 0 | 0.06 | 0.02 |
| 8 | 128742851 | rs16902359 | -1.634 | 0.05 | 1.017 | 1.415 | -0.02 | 0.05 | 0.05 | 0.08 | 0.2 | 0.61 | 0.2 | 0.39 | 0.14 | 0.34 |
| 8 | 128745266 | rs16902364 | -1.649 | 0.309 | 1.053 | 1.356 | 0.057 | 0.229 | 0.229 | 0.315 | 0.07 | 0.37 | 0.1 | 0.21 | 0.05 | 0.17 |
| 8 | 128757350 | rs7462301 | -1.397 | -0.216 | 0.96 | 0.807 | -0.038 | 0.011 | 0.011 | 0.991 | 0.11 | 0.45 | 0.05 | 0.06 | 0.02 | 0.09 |
| 8 | 128758861 | rs12680047 | -1.295 | -0.251 | -0.166 | 0.944 | 0.048 | 0 | 0 | 0 | 0.44 | 0.84 | 0.34 | 0.35 | 0.37 | 0.37 |
| 8 | 128766080 | rs11992286 | -1.01 | -0.297 | 0.604 | 0.922 | 0.017 | 0.018 | 0.018 | 0 | 0.19 | 0.53 | 0.15 | 0.18 | 0.21 | 0.13 |
| 8 | 128768191 | rs7820850 | -0.38 | -0.377 | 0.338 | 1.052 | 0.006 | 0.002 | 0.002 | 0.074 | 0.18 | 0.13 | 0.14 | 0.12 | 0.13 | 0.16 |
| 8 | 128773968 | rs4483116 | -1.413 | -0.49 | 0.447 | 1.005 | 0.008 | 0.007 | 0.007 | 0 | 0.18 | 0.16 | 0.1 | 0.11 | 0.14 | 0.14 |
| 8 | 128775144 | rs10956387 | 0.631 | -0.525 | -0.746 | 0.519 | -0.092 | 0.071 | 0.071 | 0.128 | 0.34 | 0.09 | 0.37 | 0.21 | 0.38 | 0.3 |
| 8 | 128775635 | rs6981913 | -0.426 | -0.603 | 0.083 | 1.172 | 0.076 | 0.2 | 0.2 | 2.197 | 0.1 | 0.35 | 0.11 | 0.37 | 0.14 | 0.28 |
| 8 | 128779001 | rs4733560 | 0.9 | -0.469 | -0.143 | 1.119 | -0.103 | 0.057 | 0.057 | 0.071 | 0.34 | 0.09 | 0.38 | 0.22 | 0.41 | 0.29 |
| 8 | 128790616 | rs4326353 | -0.877 | -0.773 | 0.37 | 1.068 | 0.007 | 0.009 | 0.009 | 0.009 | 0.6 | 0.73 | 0.54 | 0.68 | 0.59 | 0.7 |
| 8 | 128798394 | rs4404878 | 0.733 | -0.78 | 0.173 | 0.852 | -0.086 | 0.076 | 0.076 | 0 | 0.38 | 0.12 | 0.4 | 0.22 | 0.45 | 0.3 |
| 8 | 128804840 | rs4733759 | 0.531 | -0.805 | 0.215 | 0.862 | -0.063 | 0.018 | 0.018 | 0.03 | 0.42 | 0.14 | 0.41 | 0.32 | 0.45 | 0.37 |
| 8 | 128814091 | rs6990534 | -0.93 | -0.791 | 0.605 | 1.057 | 0.014 | -0.019 | -0.019 | 0 | 0.63 | 0.69 | 0.58 | 0.51 | 0.67 | 0.71 |
| 8 | 128814765 | rs7844288 | -1.122 | -0.769 | 0.658 | 1.018 | 0.008 | -0.015 | -0.015 | 0 | 0.21 | 0.56 | 0.18 | 0.19 | 0.22 | 0.33 |
| 8 | 128818308 | rs7841347 | -0.285 | -0.976 | 0.519 | 0.827 | -0.002 | 0.01 | 0.01 | 0 | 0.53 | 0.56 | 0.44 | 0.25 | 0.46 | 0.35 |
| 8 | 128822709 | rs10108514 | 0.108 | -0.729 | 0.083 | 0.697 | -0.005 | 0.004 | 0.004 | 0.007 | 0.42 | 0.43 | 0.41 | 0.31 | 0.42 | 0.41 |
| 8 | 128834403 | rs4733789 | -0.318 | -0.6 | -0.031 | 0.943 | 0.001 | 0.002 | 0.002 | 0 | 0.56 | 0.44 | 0.7 | 0.39 | 0.54 | 0.58 |
| 8 | 128840276 | rs4733792 | -0.608 | -0.599 | -0.398 | 1.047 | -0.051 | -0.095 | -0.095 | 0.527 | 0.61 | 0.93 | 0.73 | 0.5 | 0.54 | 0.63 |
| 8 | 128844721 | rs4733798 | -0.243 | -0.484 | 0.025 | 0.974 | -0.001 | 0.005 | 0.005 | 0 | 0.56 | 0.69 | 0.66 | 0.53 | 0.51 | 0.65 |
| 8 | 128856487 | rs4484653 | -0.555 | -0.22 | -0.593 | 0.675 | -0.053 | 0.008 | 0.008 | 0 | 0.58 | 0.92 | 0.71 | 0.56 | 0.51 | 0.62 |
| 8 | 128865471 | rs4339604 | -1.165 | -0.114 | -0.433 | 0.47 | 0.006 | -0.004 | -0.004 | 2.017 | 0.1 | 0.1 | 0.09 | 0 | 0.09 | 0.08 |
| 8 | 128878635 | rs7841504 | -0.818 | -0.097 | -0.01 | 0.766 | 0.003 | 0.107 | 0.107 | 0.026 | 0.11 | 0.36 | 0.08 | 0.15 | 0.08 | 0.07 |
| 8 | 128879839 | rs4427136 | -1.427 | 0.027 | -0.892 | 0.766 | 0.011 | -0.027 | -0.027 | 0 | 0.26 | 0.8 | 0.36 | 0.4 | 0.22 | 0.27 |
| 8 | 128888319 | rs6470586 | -1.786 | 0.42 | -1.552 | 0.08 | -0.103 | 0.059 | 0.059 | 0 | 0.74 | 0.99 | 0.77 | 0.86 | 0.73 | 0.68 |
| 8 | 128889371 | rs6470588 | -1.522 | 0.978 | -1.312 | 0.319 | -0.013 | -0.017 | -0.017 | 0 | 0.39 | 0.78 | 0.5 | 0.45 | 0.43 | 0.43 |
| 8 | 128890261 | rs9656968 | 0.197 | 1.149 | -1.431 | -0.265 | -0.02 | 0.036 | 0.036 | 0 | 0.2 | 0.1 | 0.17 | 0.03 | 0.25 | 0.18 |
| 8 | 128898169 | rs4733579 | -0.953 | 0.958 | -1.596 | -0.121 | -0.034 | 0.135 | 0.135 | 0 | 0.63 | 0.9 | 0.69 | 0.83 | 0.64 | 0.69 |
| 8 | 128903273 | rs17189681 | 0.834 | 1.236 | -0.931 | 0.336 | -0.011 | 0.024 | 0.024 | 0 | 0.13 | 0.13 | 0.13 | 0.03 | 0.16 | 0.15 |
| 8 | 128906787 | rs6999585 | -0.723 | 1.342 | -1.503 | 0.538 | -0.016 | -0.065 | -0.065 | 1.303 | 0.65 | 0.87 | 0.77 | 0.47 | 0.66 | 0.69 |
| 8 | 128906881 | rs16902454 | 0.865 | 1.6 | -0.715 | 0.394 | 0.036 | -0.039 | -0.039 | 0 | 0.14 | 0.26 | 0.18 | 0.03 | 0.21 | 0.16 |
| 8 | 128909539 | rs4733806 | 1.348 | 1.378 | -0.923 | 0.368 | -0.023 | 0.139 | 0.139 | 0 | 0.34 | 0.05 | 0.22 | 0.1 | 0.31 | 0.3 |
| 8 | 128911213 | rs13260853 | 1.355 | 1.229 | -0.925 | 0.341 | -0.039 | 0.153 | 0.153 | 0 | 0.33 | 0.05 | 0.22 | 0.09 | 0.32 | 0.3 |
| 8 | 128920127 | rs959409 | -1.338 | 1.043 | 0.335 | 0.962 | 0.051 | 0.25 | 0.25 | 1.487 | 0.05 | 0.49 | 0.06 | 0.1 | 0 | 0.03 |
| 8 | 128921754 | rs13276337 | -0.779 | 1.184 | -0.197 | 0.85 | -0.063 | -0.051 | -0.051 | 0.33 | 0.66 | 0.9 | 0.75 | 0.49 | 0.63 | 0.71 |
| 8 | 128928064 | rs2555114 | -0.594 | 1.171 | -0.047 | 0.854 | -0.014 | -0.006 | -0.006 | 0.964 | 0.65 | 0.83 | 0.75 | 0.53 | 0.64 | 0.72 |
| 8 | 128928246 | rs12548864 | 1.273 | 1.132 | -0.179 | 0.599 | -0.053 | 0.151 | 0.151 | 0 | 0.35 | 0.06 | 0.24 | 0.12 | 0.35 | 0.33 |

| | | | | | | | | | | | | | | | |
|---|---|---|---|---|---|---|---|---|---|---|---|---|---|---|---|
| 8 | 128928618 | rs2721133 | -0.142 | 1.217 | 0.224 | 0.806 | 0.048 | 0.04 | 0.04 | 0 | 0.08 | 0.18 | 0.17 | 0.11 | 0.13 | 0.1 |
| 8 | 128929371 | rs7387606 | 0.879 | 1.194 | 0.156 | 0.874 | 0.016 | 0.061 | 0.061 | 0 | 0.39 | 0.23 | 0.27 | 0.2 | 0.36 | 0.37 |
| 8 | 128930415 | rs7000853 | 0.623 | 1.045 | 0.593 | 1.1 | -0.002 | 0.01 | 0.01 | 0.018 | 0.21 | 0.26 | 0.31 | 0.25 | 0.17 | 0.25 |
| 8 | 128939948 | rs10956395 | 0.718 | 0.849 | -0.107 | 0.739 | -0.045 | -0.035 | -0.035 | 0 | 0.31 | 0.11 | 0.23 | 0.47 | 0.34 | 0.3 |
| 8 | 128941536 | rs10956396 | 0.371 | 0.843 | 0.185 | 0.588 | -0.002 | 0.005 | 0.005 | 0 | 0.36 | 0.4 | 0.28 | 0.55 | 0.35 | 0.34 |
| 8 | 128944499 | rs3739379 | 0.521 | 0.918 | 0.141 | 0.611 | -0.004 | -0.025 | -0.025 | 0 | 0.33 | 0.24 | 0.27 | 0.55 | 0.35 | 0.32 |
| 8 | 128945788 | rs10441527 | 0.42 | 0.901 | 0.301 | 0.603 | 0 | -0.005 | -0.005 | 0 | 0.34 | 0.32 | 0.28 | 0.55 | 0.35 | 0.32 |
| 8 | 128946538 | rs10104535 | 0.515 | 0.902 | 0.304 | 0.603 | -0.003 | -0.022 | -0.022 | 0 | 0.33 | 0.25 | 0.27 | 0.55 | 0.35 | 0.32 |
| 8 | 128947775 | rs4733581 | -0.359 | 0.966 | 0.247 | 0.482 | 0.015 | 0.048 | 0.048 | 0.112 | 0.72 | 0.57 | 0.53 | 0.68 | 0.71 | 0.65 |
| 8 | 128947967 | rs4733582 | 0.987 | 0.998 | -0.091 | 0.469 | 0.007 | 0.121 | 0.121 | 0 | 0.37 | 0.14 | 0.24 | 0.13 | 0.35 | 0.33 |
| 8 | 128949443 | rs2392827 | 0.155 | 1.022 | 0.308 | 0.831 | 0 | 0.006 | 0.006 | 0 | 0.33 | 0.35 | 0.28 | 0.56 | 0.35 | 0.32 |
| 8 | 128951476 | rs890436 | 0.223 | 1.039 | 0.313 | 0.833 | -0.003 | -0.013 | -0.013 | 0 | 0.33 | 0.27 | 0.29 | 0.56 | 0.36 | 0.32 |
| 8 | 128965552 | rs13274181 | -0.323 | 1.263 | 0.022 | 0.751 | 0.007 | 0.019 | 0.019 | 0.05 | 0.11 | 0.17 | 0.11 | 0.16 | 0.13 | 0.06 |
| 8 | 128973665 | rs7004509 | -0.161 | 0.967 | -0.054 | 0.388 | -0.003 | -0.011 | -0.011 | 0 | 0.47 | 0.48 | 0.36 | 0.32 | 0.45 | 0.45 |
| 8 | 128978326 | rs755520 | 0.467 | 0.885 | 0.069 | 0.535 | -0.012 | -0.007 | -0.007 | 0 | 0.41 | 0.3 | 0.48 | 0.58 | 0.43 | 0.46 |
| 8 | 128978856 | rs755521 | 0.969 | 0.965 | -0.137 | 0.248 | 0.089 | 0.126 | 0.126 | 2.579 | 0.33 | 0.08 | 0.16 | 0.13 | 0.19 | 0.32 |
| 8 | 128980008 | rs752427 | 0.823 | 1.162 | 0.305 | 0.414 | 0.022 | 0.008 | 0.008 | 1.106 | 0.74 | 0.4 | 0.63 | 0.71 | 0.62 | 0.77 |
| 8 | 128980806 | rs4733584 | -0.67 | 1.372 | 0.175 | 0.401 | -0.052 | -0.018 | -0.018 | 0.077 | 0.88 | 0.95 | 0.78 | 0.89 | 0.73 | 0.89 |
| 8 | 128985644 | rs10086182 | -0.322 | 1.227 | 0.394 | 0.414 | -0.002 | -0.002 | -0.002 | 0 | 0.87 | 0.86 | 0.78 | 0.88 | 0.73 | 0.91 |
| 8 | 128988045 | rs13274301 | 0.141 | 1.157 | 0.806 | 0.438 | 0.022 | 0.087 | 0.087 | 0 | 0.1 | 0.28 | 0.12 | 0.19 | 0.1 | 0.07 |
| 8 | 128988220 | rs993725 | 1.044 | 1.159 | 0.39 | 0.438 | 0.024 | 0.01 | 0.01 | 1.073 | 0.75 | 0.37 | 0.64 | 0.7 | 0.63 | 0.83 |
| 8 | 128988967 | rs4476972 | -0.723 | 1.163 | 0.376 | 0.437 | -0.052 | -0.016 | -0.016 | 0 | 0.88 | 0.95 | 0.78 | 0.89 | 0.73 | 0.91 |
| 8 | 128989292 | rs4733809 | 0.973 | 1.157 | 0.352 | 0.439 | 0.024 | 0.01 | 0.01 | 1.073 | 0.75 | 0.36 | 0.64 | 0.7 | 0.63 | 0.83 |
| 8 | 128991152 | rs13270017 | -0.141 | 1.08 | 0.521 | 0.546 | 0.038 | 0.139 | 0.139 | 0.387 | 0.11 | 0.41 | 0.12 | 0.2 | 0.1 | 0.07 |
| 8 | 128991700 | rs17776826 | 0.59 | 0.818 | 0.008 | 0.779 | 0.061 | 0.042 | 0.042 | 2.419 | 0.34 | 0.15 | 0.19 | 0.21 | 0.24 | 0.36 |
| 8 | 128992487 | rs11993333 | 0.754 | 0.686 | -0.343 | 0.637 | -0.019 | -0.019 | -0.019 | 0 | 0.67 | 0.26 | 0.58 | 0.63 | 0.55 | 0.72 |
| 8 | 128992600 | rs12334659 | 0.465 | 0.739 | -0.143 | 0.795 | 0.015 | 0.02 | 0.02 | 0.026 | 0.45 | 0.26 | 0.61 | 0.62 | 0.55 | 0.55 |
| 8 | 128994641 | rs7013628 | -1.234 | 0.902 | 0.05 | 0.842 | 0.074 | 0.073 | 0.073 | 0 | 0.05 | 0.14 | 0.14 | 0.13 | 0.17 | 0.1 |
| 8 | 128996639 | rs4637783 | -1.127 | 0.865 | 0.049 | 0.823 | 0 | -0.001 | -0.001 | 0 | 0.85 | 0.86 | 0.89 | 0.85 | 0.84 | 0.9 |
| 8 | 129000416 | rs11786536 | 1.163 | 0.945 | -0.446 | 0.958 | -0.065 | -0.006 | -0.006 | 0.061 | 0.15 | 0.01 | 0.33 | 0.09 | 0.19 | 0.16 |
| 8 | 129002233 | rs16902508 | 1.321 | 0.609 | -0.07 | 1.13 | 0.013 | 0.011 | 0.011 | 0 | 0.08 | 0.09 | 0.14 | 0.11 | 0.16 | 0.06 |
| 8 | 129003716 | rs6995630 | 0.92 | 0.606 | -0.076 | 1.064 | -0.004 | -0.011 | -0.011 | 0 | 0.28 | 0.24 | 0.21 | 0.42 | 0.28 | 0.35 |
| 8 | 129006963 | rs7016045 | -1.356 | 0.561 | -0.091 | 0.99 | 0.003 | 0.003 | 0.003 | 0.02 | 0.86 | 0.87 | 0.9 | 0.85 | 0.84 | 0.91 |
| 8 | 129007207 | rs10956403 | 0.809 | 0.584 | -0.5 | 0.998 | 0.014 | 0.142 | 0.142 | 0 | 0.43 | 0.11 | 0.5 | 0.22 | 0.38 | 0.44 |
| 8 | 129007740 | rs16902512 | 0.509 | 1.001 | -0.189 | 1.356 | -0.009 | -0.019 | -0.019 | 0 | 0.29 | 0.24 | 0.22 | 0.45 | 0.29 | 0.4 |
| 8 | 129011513 | rs11781967 | 0.927 | 1.177 | -0.666 | 1.293 | -0.03 | -0.03 | -0.03 | 0 | 0.06 | 0.01 | 0.04 | 0.16 | 0.06 | 0.11 |
| 8 | 129012574 | rs10087240 | -0.088 | 0.895 | -0.416 | 0.73 | 0.011 | -0.005 | -0.005 | 0.104 | 0.41 | 0.5 | 0.38 | 0.59 | 0.45 | 0.48 |
| 8 | 129016070 | rs4733589 | 0.327 | 1.09 | -0.444 | 0.85 | 0.011 | 0.013 | 0.013 | 0.117 | 0.76 | 0.77 | 0.59 | 0.73 | 0.69 | 0.83 |
| 8 | 129016347 | rs1529335 | 0.509 | 1.131 | -0.693 | 0.547 | 0.006 | -0.004 | -0.004 | 0 | 0.08 | 0.12 | 0.28 | 0.07 | 0.15 | 0.07 |
| 8 | 129016936 | rs13266023 | 0.157 | 0.235 | -1.162 | 0.161 | 0.011 | 0.077 | 0.077 | 0.056 | 0.69 | 0.88 | 0.83 | 0.83 | 0.74 | 0.76 |
| 8 | 129018813 | rs4535708 | -0.652 | 0.336 | -1.017 | 0.27 | -0.004 | 0.015 | 0.015 | 0 | 0.82 | 0.91 | 0.89 | 0.88 | 0.83 | 0.89 |
| 8 | 129022078 | rs963475 | 0.175 | 0.091 | -1.261 | 0.056 | 0.031 | 0.11 | 0.11 | 0 | 0.54 | 0.8 | 0.72 | 0.7 | 0.62 | 0.54 |
| 8 | 129023025 | rs17199073 | 0.609 | -0.149 | -0.893 | 0.103 | 0.029 | 0.074 | 0.074 | 0 | 0.28 | 0.08 | 0.17 | 0.19 | 0.21 | 0.35 |
| 8 | 129025804 | rs6984736 | -1.07 | -0.151 | -0.946 | 0.102 | -0.006 | 0.015 | 0.015 | 0 | 0.82 | 0.91 | 0.89 | 0.88 | 0.84 | 0.89 |
| 8 | 129028784 | rs17199521 | 0.945 | -0.193 | -0.995 | 0.086 | 0.012 | 0.092 | 0.092 | 0 | 0.25 | 0.03 | 0.14 | 0.12 | 0.19 | 0.27 |
| 8 | 129031617 | rs1909457 | 1.746 | -0.013 | -1.396 | 0.494 | -0.097 | -0.079 | -0.079 | 0.012 | 0.2 | 0.02 | 0.26 | 0.28 | 0.37 | 0.29 |
| 8 | 129031734 | rs2033098 | -0.19 | -0.286 | -1.079 | 0.303 | 0.028 | 0.088 | 0.088 | 0 | 0.54 | 0.75 | 0.71 | 0.7 | 0.62 | 0.54 |
| 8 | 129036062 | rs7010574 | -0.279 | -0.295 | -0.743 | 0.646 | 0.007 | 0.1 | 0.1 | 2.083 | 0.09 | 0.47 | 0.07 | 0.13 | 0.04 | 0.08 |
| 8 | 129039139 | rs11783127 | -0.59 | -0.188 | -1.17 | 0.483 | 0.007 | -0.005 | -0.005 | 0 | 0.56 | 0.65 | 0.52 | 0.57 | 0.4 | 0.58 |
| 8 | 129039273 | rs16902568 | 1.066 | 0.06 | -1.804 | 0.53 | 0.001 | -0.003 | -0.003 | 0.195 | 0.1 | 0.08 | 0.23 | 0.19 | 0.06 | 0.1 |
| 8 | 129043582 | rs11985818 | 0.352 | 0.234 | -1.656 | 0.576 | 0.048 | 0.038 | 0.038 | 0 | 0.35 | 0.11 | 0.37 | 0.31 | 0.24 | 0.38 |
| 8 | 129046208 | rs2720685 | 0.179 | 0.988 | -0.865 | 0.749 | 0.046 | 0.022 | 0.022 | 0 | 0.28 | 0.44 | 0.38 | 0.31 | 0.41 | 0.28 |
| 8 | 129047325 | rs2173537 | -0.331 | 0.941 | -1.467 | 0.559 | -0.004 | 0.011 | 0.011 | 0 | 0.22 | 0.16 | 0.31 | 0.15 | 0.19 | 0.17 |
| 8 | 129050119 | rs3927165 | 0.015 | 0.903 | -1.616 | 0.823 | -0.033 | -0.032 | -0.032 | 0 | 0.38 | 0.23 | 0.45 | 0.49 | 0.49 | 0.47 |
| 8 | 129050794 | rs4733828 | -0.034 | 0.857 | -2.294 | 0.915 | -0.159 | -0.055 | -0.055 | 0 | 0.37 | 0.06 | 0.39 | 0.39 | 0.52 | 0.39 |
| 8 | 129057963 | rs11784465 | 0.186 | 0.258 | -1.416 | 1.001 | 0.011 | -0.008 | -0.008 | 0 | 0.23 | 0.19 | 0.22 | 0.27 | 0.16 | 0.25 |
| 8 | 129058356 | rs2720709 | -0.761 | 0.372 | -1.434 | 1.414 | -0.001 | 0.001 | 0.001 | 0.177 | 0.36 | 0.38 | 0.39 | 0.52 | 0.3 | 0.43 |
| 8 | 129059110 | rs11775301 | -0.372 | 0.043 | -1.506 | 1.395 | -0.004 | -0.016 | -0.016 | 0 | 0.61 | 0.35 | 0.5 | 0.52 | 0.62 | 0.56 |
| 8 | 129059568 | rs2648861 | -0.71 | 0.032 | -1.478 | 1.399 | 0.003 | -0.005 | -0.005 | 0 | 0.64 | 0.46 | 0.58 | 0.72 | 0.63 | 0.62 |
| 8 | 129063021 | rs2720662 | -0.422 | -0.163 | -1.209 | 1.453 | 0.001 | -0.002 | -0.002 | 0 | 0.35 | 0.35 | 0.39 | 0.6 | 0.24 | 0.53 |
| 8 | 129064808 | rs2648868 | -0.446 | -0.094 | -1 | 1.494 | -0.004 | 0.082 | 0.082 | 0 | 0.12 | 0.19 | 0.18 | 0.34 | 0.09 | 0.3 |
| 8 | 129066074 | rs1499374 | 0.17 | 0.02 | -1.127 | 1.401 | -0.045 | -0.019 | -0.019 | 0 | 0.4 | 0.18 | 0.35 | 0.45 | 0.48 | 0.39 |
| 8 | 129070428 | rs13250078 | 0.89 | 0.013 | -1.099 | 1.37 | -0.057 | 0.089 | 0.089 | 0 | 0.33 | 0.09 | 0.25 | 0.21 | 0.42 | 0.24 |
| 8 | 129071537 | rs2648873 | -0.924 | 0.127 | -1.002 | 1.418 | 0.009 | 0.102 | 0.102 | 0.739 | 0.07 | 0.16 | 0.11 | 0.24 | 0.06 | 0.15 |
| 8 | 129072161 | rs2648875 | -0.179 | 0.101 | -1.013 | 0.88 | -0.025 | 0.013 | 0.013 | 0.075 | 0.36 | 0.47 | 0.41 | 0.61 | 0.23 | 0.53 |
| 8 | 129072966 | rs2648876 | -1.082 | 0.32 | -0.162 | 0.851 | 0.008 | -0.017 | -0.017 | 0 | 0.74 | 0.71 | 0.68 | 0.88 | 0.68 | 0.81 |
| 8 | 129074573 | rs918179 | -0.878 | 0.218 | -0.29 | 1.083 | 0.018 | 0.074 | 0.074 | 1.17 | 0.06 | 0.2 | 0.04 | 0.07 | 0.03 | 0.11 |
| 8 | 129075832 | rs2608053 | -0.212 | 0.17 | -0.773 | 0.873 | -0.004 | 0.006 | 0.006 | 1.039 | 0.6 | 0.38 | 0.53 | 0.74 | 0.55 | 0.59 |
| 8 | 129076573 | rs13255292 | 0.894 | 0.207 | -1.466 | 0.66 | -0.055 | 0.053 | 0.053 | 0 | 0.27 | 0.04 | 0.23 | 0.19 | 0.37 | 0.23 |

| | | | | | | | | | | | | | | | | |
|---|---|---|---|---|---|---|---|---|---|---|---|---|---|---|---|---|
| 8 | 129076594 | rs2648883 | 1.318 | -0.056 | -1.318 | 1.45 | -0.015 | -0.044 | -0.044 | 1.696 | 0.15 | 0.05 | 0.05 | 0.3 | 0.13 | 0.17 |
| 8 | 129078802 | rs2608049 | -0.075 | -0.221 | -1.039 | 1.52 | -0.002 | -0.036 | -0.036 | 0 | 0.25 | 0.16 | 0.3 | 0.47 | 0.21 | 0.39 |
| 8 | 129082378 | rs2250888 | -0.207 | -0.288 | -0.647 | 1.696 | -0.001 | 0.005 | 0.005 | 0.413 | 0.26 | 0.26 | 0.31 | 0.41 | 0.21 | 0.39 |
| 8 | 129083584 | rs2720667 | -1.402 | -0.092 | -0.495 | 1.376 | -0.007 | 0.01 | 0.01 | 0 | 0.73 | 0.74 | 0.75 | 0.84 | 0.66 | 0.8 |
| 8 | 129094589 | rs1499368 | 0.332 | 0.433 | -0.599 | 1.486 | -0.003 | 0.001 | 0.001 | 0.093 | 0.06 | 0.04 | 0.24 | 0.18 | 0.08 | 0.19 |
| 8 | 129095863 | rs2648889 | -0.339 | 0.266 | -0.326 | 1.614 | -0.004 | 0.023 | 0.023 | 0.366 | 0.29 | 0.27 | 0.33 | 0.47 | 0.22 | 0.43 |
| 8 | 129099503 | rs6470602 | 0.185 | 0.241 | -0.225 | 1.464 | 0.011 | 0.032 | 0.032 | 0 | 0.49 | 0.22 | 0.64 | 0.49 | 0.52 | 0.53 |
| 8 | 129100433 | rs2648896 | 0.326 | 0.404 | 0.176 | 1.392 | -0.016 | 0.086 | 0.086 | 2.176 | 0.22 | 0.43 | 0.1 | 0.29 | 0.13 | 0.24 |
| 8 | 129102128 | rs2648898 | 0.505 | 0.419 | -0.239 | 1.4 | -0.01 | 0.033 | 0.033 | 3.818 | 0.21 | 0.26 | 0.08 | 0.29 | 0.13 | 0.26 |
| 8 | 129103032 | rs17382663 | 0.98 | 0.044 | -1.096 | 0.693 | -0.051 | 0.119 | 0.119 | 0 | 0.31 | 0.05 | 0.31 | 0.15 | 0.4 | 0.22 |
| 8 | 129105305 | rs2392938 | -0.561 | -0.545 | -0.728 | 1.254 | -0.026 | 0.072 | 0.072 | 0 | 0.11 | 0.21 | 0.2 | 0.2 | 0.06 | 0.26 |
| 8 | 129106308 | rs2648901 | 0.022 | -0.586 | -0.943 | 1.289 | 0 | 0.001 | 0.001 | 2.243 | 0.27 | 0.24 | 0.12 | 0.41 | 0.19 | 0.4 |
| 8 | 129107045 | rs2648902 | -0.797 | -0.481 | -1.62 | 0.864 | -0.002 | -0.002 | -0.002 | 0 | 0.69 | 0.63 | 0.66 | 0.71 | 0.73 | 0.76 |
| 8 | 129108697 | rs2302793 | 0.899 | 0.09 | -2.371 | 0.534 | -0.113 | 0.149 | 0.149 | 0 | 0.36 | 0.06 | 0.38 | 0.16 | 0.53 | 0.26 |
| 8 | 129111409 | rs2608032 | 0.096 | 0.037 | -2.232 | 1.184 | -0.04 | -0.094 | -0.094 | 0.244 | 0.23 | 0.01 | 0.09 | 0.3 | 0.19 | 0.22 |
| 8 | 129113456 | rs3901778 | 0.594 | -0.294 | -1.558 | 1.797 | -0.038 | 0.049 | 0.049 | 6.392 | 0.83 | 0.28 | 0.76 | 0.73 | 0.93 | 0.81 |
| 8 | 129117655 | rs4733595 | 0.977 | 0.256 | -2.795 | 0.469 | -0.106 | 0.235 | 0.235 | 1.265 | 0.43 | 0.08 | 0.34 | 0.16 | 0.51 | 0.27 |
| 8 | 129121956 | rs2608034 | -1.49 | 0.563 | -1.923 | 1.513 | -0.001 | 0.029 | 0.029 | 0.276 | 0.11 | 0.12 | 0.19 | 0.48 | 0.17 | 0.42 |
| 8 | 129124831 | rs4733833 | 1.51 | 0.418 | -2.457 | 0.756 | -0.095 | 0.179 | 0.179 | 2.952 | 0.3 | 0.03 | 0.37 | 0.04 | 0.41 | 0.22 |
| 8 | 129126175 | rs2019940 | -1.544 | 0.453 | -1.811 | 1.305 | 0.018 | 0.055 | 0.055 | 1.634 | 0.12 | 0.16 | 0.19 | 0.42 | 0.16 | 0.4 |
| 8 | 129126853 | rs5029317 | -0.162 | 0.524 | -1.834 | 1.068 | -0.011 | -0.012 | -0.012 | 0 | 0.05 | 0.03 | 0.2 | 0.09 | 0.09 | 0.12 |
| 8 | 129126887 | rs16902654 | 1.266 | 0.074 | -2.118 | 0.643 | 0.02 | 0.023 | 0.023 | 0 | 0.18 | 0.13 | 0.12 | 0.1 | 0.13 | 0.17 |
| 8 | 129127793 | rs1030105 | -0.597 | -0.044 | -1.521 | 0.659 | 0 | -0.002 | -0.002 | 2.554 | 0.36 | 0.4 | 0.49 | 0.67 | 0.31 | 0.62 |
| 8 | 129128194 | rs11984659 | -0.123 | 0.02 | -1.774 | 0.767 | -0.008 | -0.02 | -0.02 | 0.019 | 0.31 | 0.22 | 0.39 | 0.63 | 0.26 | 0.55 |
| 8 | 129138663 | rs2160766 | 1.332 | 0.063 | -1.628 | -0.345 | -0.049 | 0.156 | 0.156 | 0 | 0.35 | 0.15 | 0.23 | 0.04 | 0.42 | 0.23 |
| 8 | 129138705 | rs12544603 | 0.582 | -0.136 | -0.862 | 0.518 | 0.009 | 0.013 | 0.013 | 0 | 0.19 | 0.18 | 0.13 | 0.09 | 0.14 | 0.21 |
| 8 | 129139206 | rs17382698 | -0.592 | 0.026 | -0.899 | 0.542 | -0.015 | 0.01 | 0.01 | 0 | 0.06 | 0.01 | 0.15 | 0.04 | 0.13 | 0.07 |
| 8 | 129141856 | rs4733596 | -0.157 | -0.096 | -0.695 | 0.68 | -0.001 | -0.006 | -0.006 | 0 | 0.26 | 0.31 | 0.28 | 0.2 | 0.27 | 0.29 |
| 8 | 129145876 | rs12676630 | 0.749 | -0.534 | -0.782 | 0.712 | 0.038 | 0.029 | 0.029 | 0 | 0.21 | 0.07 | 0.12 | 0.16 | 0.14 | 0.21 |
| 8 | 129148524 | rs12680075 | 0.402 | -0.639 | -0.585 | 0.679 | 0.013 | 0.011 | 0.011 | 0.135 | 0.24 | 0.15 | 0.13 | 0.17 | 0.14 | 0.24 |
| 8 | 129150668 | rs2648832 | -0.49 | -0.553 | -0.459 | 1.142 | 0.006 | 0.001 | 0.001 | 0 | 0.32 | 0.26 | 0.31 | 0.72 | 0.25 | 0.5 |
| 8 | 129153280 | rs2542419 | -0.366 | -0.532 | -0.14 | 1.583 | 0.02 | 0.153 | 0.153 | 0 | 0.08 | 0.16 | 0.17 | 0.43 | 0.11 | 0.23 |
| 8 | 129155329 | rs4733838 | -0.834 | -0.703 | 0.029 | 0.829 | 0.002 | 0.005 | 0.005 | 0.675 | 0.59 | 0.6 | 0.54 | 0.47 | 0.44 | 0.55 |
| 8 | 129157468 | rs2254778 | -2.011 | -0.745 | -0.014 | 0.655 | -0.082 | 0.111 | 0.111 | 2.258 | 0.72 | 0.88 | 0.75 | 0.96 | 0.56 | 0.8 |
| 8 | 129158945 | rs759648 | -0.216 | -0.846 | 0.262 | 0.656 | -0.003 | -0.008 | -0.008 | 0.013 | 0.37 | 0.43 | 0.41 | 0.23 | 0.3 | 0.28 |
| 8 | 129162497 | rs10956412 | 0.786 | -0.993 | 0.293 | 0.66 | -0.001 | 0 | 0 | 0.229 | 0.27 | 0.23 | 0.18 | 0.23 | 0.17 | 0.28 |
| 8 | 129163047 | rs13279159 | 0.397 | -1.239 | 0.281 | 0.445 | 0.02 | -0.099 | -0.099 | 0 | 0.26 | 0.41 | 0.31 | 0.06 | 0.25 | 0.14 |
| 8 | 129163424 | rs2648840 | -2.337 | -1.257 | 0.211 | 0.432 | -0.084 | 0.113 | 0.113 | 2.089 | 0.72 | 0.88 | 0.75 | 0.95 | 0.56 | 0.79 |
| 8 | 129164288 | rs759651 | -0.005 | -1.274 | 0.214 | 0.529 | 0.029 | 0.002 | 0.002 | 0.02 | 0.37 | 0.23 | 0.26 | 0.35 | 0.26 | 0.35 |
| 8 | 129165056 | rs1121948 | 0.184 | -1.285 | 0.22 | 0.548 | 0.033 | -0.004 | -0.004 | 0 | 0.35 | 0.23 | 0.24 | 0.35 | 0.22 | 0.35 |
| 8 | 129165198 | rs1121946 | 1.719 | -1.243 | 0.182 | 0.504 | 0.018 | -0.044 | -0.044 | 0 | 0.22 | 0.03 | 0.15 | 0.23 | 0.16 | 0.22 |
| 8 | 129170108 | rs16893188 | 0.854 | -1.277 | 0.595 | 0.406 | 0.006 | -0.004 | -0.004 | 0 | 0.11 | 0.14 | 0.22 | 0.05 | 0.14 | 0.11 |
| 8 | 129170596 | rs4733839 | 0.277 | -1.214 | 0.383 | 0.561 | 0 | -0.054 | -0.054 | 2.179 | 0.25 | 0.53 | 0.29 | 0.12 | 0.21 | 0.14 |
| 8 | 129173046 | rs7846456 | -0.309 | -1.214 | 0.363 | 0.602 | -0.005 | 0 | 0 | 0 | 0.15 | 0.13 | 0.16 | 0.15 | 0.11 | 0.2 |
| 8 | 129174099 | rs2648833 | 0.301 | -0.987 | 0.515 | 0.763 | 0.004 | 0.203 | 0.203 | 0 | 0.08 | 0.19 | 0.15 | 0.45 | 0.08 | 0.21 |
| 8 | 129179990 | rs2648846 | -1.033 | -1.006 | 0.468 | 0.739 | -0.002 | 0.013 | 0.013 | 0.209 | 0.33 | 0.61 | 0.43 | 0.57 | 0.29 | 0.4 |
| 8 | 129184165 | rs2648850 | 0.297 | -0.917 | 0.331 | 0.827 | -0.002 | 0.01 | 0.01 | 0 | 0.08 | 0.05 | 0.14 | 0.44 | 0.08 | 0.26 |
| 8 | 129185104 | rs1902789 | 0.233 | -1.022 | 0.341 | 0.546 | 0.034 | 0.003 | 0.003 | 0.175 | 0.38 | 0.22 | 0.32 | 0.35 | 0.27 | 0.36 |
| 8 | 129192271 | rs2019960 | 0.059 | -0.912 | 0.441 | 0.549 | 0.003 | -0.1 | -0.1 | 0 | 0.25 | 0.46 | 0.28 | 0.07 | 0.21 | 0.14 |
| 8 | 129203191 | rs10492292 | 0.122 | -0.909 | 0.176 | 0.672 | 0.024 | -0.014 | -0.014 | 0.554 | 0.38 | 0.26 | 0.32 | 0.42 | 0.27 | 0.38 |
| 8 | 129205610 | rs10106296 | 0.234 | -0.81 | 0.039 | 0.691 | 0.039 | 0.006 | 0.006 | 0.07 | 0.38 | 0.22 | 0.33 | 0.35 | 0.29 | 0.36 |
| 8 | 129215846 | rs7825166 | -0.979 | -0.807 | 0.079 | 0.666 | -0.009 | -0.005 | -0.005 | 0 | 0.61 | 0.65 | 0.6 | 0.44 | 0.5 | 0.49 |
| 8 | 129218047 | rs6470612 | -1.381 | -0.702 | -0.085 | 0.866 | -0.013 | 0.047 | 0.047 | 0 | 0.95 | 0.99 | 0.87 | 0.53 | 0.93 | 0.77 |
| 8 | 129218127 | rs11997192 | 1.117 | -0.771 | -0.071 | 0.546 | 0.04 | 0.03 | 0.03 | 1.549 | 0.28 | 0.1 | 0.19 | 0.19 | 0.21 | 0.21 |
| 8 | 129218859 | rs7010390 | -0.398 | -0.345 | 0.188 | 0.713 | 0.008 | -0.065 | -0.065 | 2.348 | 0.23 | 0.4 | 0.27 | 0.1 | 0.21 | 0.14 |
| 8 | 129220073 | rs11995854 | -0.398 | -0.345 | 0.188 | 0.713 | 0.008 | -0.065 | -0.065 | 0 | 0.23 | 0.4 | 0.27 | 0.1 | 0.21 | 0.14 |
| 8 | 129224053 | rs1967314 | 0.515 | -0.702 | -0.068 | 1.348 | -0.008 | 0.114 | 0.114 | 0 | 0.91 | 0.86 | 0.85 | 0.46 | 0.93 | 0.7 |
| 8 | 129224680 | rs2909242 | 0.404 | -0.917 | 0.003 | 1.103 | 0.03 | 0.036 | 0.036 | 2.776 | 0.47 | 0.43 | 0.6 | 0.38 | 0.61 | 0.46 |
| 8 | 129225502 | rs11774172 | 0.165 | -1.489 | 0.274 | 1.041 | 0.029 | 0.037 | 0.037 | 5.066 | 0.39 | 0.23 | 0.26 | 0.21 | 0.33 | 0.22 |
| 8 | 129226886 | rs6987273 | -1.064 | -1.452 | 0.721 | 1.131 | 0.044 | 0.094 | 0.094 | 0.676 | 0.12 | 0.39 | 0.09 | 0.14 | 0.11 | 0.07 |
| 8 | 129229700 | rs2111565 | 0.176 | -1.8 | 0.301 | 1.375 | 0.009 | -0.01 | -0.01 | 0.226 | 0.8 | 0.85 | 0.8 | 0.64 | 0.86 | 0.67 |
| 8 | 129230194 | rs2909246 | 0.387 | -1.822 | 0.614 | 1.356 | -0.01 | 0.021 | 0.021 | 0 | 0.79 | 0.76 | 0.78 | 0.62 | 0.82 | 0.65 |
| 8 | 129237154 | rs1468935 | 0.216 | -1.797 | 0.712 | 1.425 | -0.006 | 0.009 | 0.009 | 0 | 0.8 | 0.78 | 0.8 | 0.61 | 0.85 | 0.66 |
| 8 | 129238669 | rs7812998 | -0.307 | -1.809 | 0.802 | 1.376 | 0.007 | 0.069 | 0.069 | 0.104 | 0.09 | 0.27 | 0.07 | 0.13 | 0.06 | 0.06 |
| 8 | 129243066 | rs1468937 | 0.352 | -1.759 | 0.629 | 1.22 | -0.015 | -0.026 | -0.026 | 0 | 0.34 | 0.47 | 0.28 | 0.2 | 0.28 | 0.25 |
| 8 | 129246415 | rs10492294 | 1.261 | -1.772 | 0.468 | 1.185 | -0.039 | 0.035 | 0.035 | 1.225 | 0.27 | 0.01 | 0.21 | 0.13 | 0.24 | 0.19 |
| 8 | 129249256 | rs2977035 | 0.741 | -1.81 | 0.551 | 1.24 | -0.047 | 0.086 | 0.086 | 0 | 0.72 | 0.57 | 0.73 | 0.5 | 0.8 | 0.6 |
| 8 | 129249391 | rs2925877 | 1.303 | -1.749 | 0.418 | 1.19 | -0.127 | 0.011 | 0.011 | 5.674 | 0.38 | 0.04 | 0.44 | 0.3 | 0.52 | 0.35 |
| 8 | 129252171 | rs17205164 | 1.265 | -1.692 | 0.504 | 1.183 | -0.088 | 0.032 | 0.032 | 5.109 | 0.38 | 0.09 | 0.45 | 0.28 | 0.52 | 0.35 |

| | | | | | | | | | | | | | | | | |
|---|---|---|---|---|---|---|---|---|---|---|---|---|---|---|---|---|
| 8 | 129257965 | rs4633023 | 1.259 | -1.662 | 0.6 | 1.283 | 0.016 | 0.243 | 0.243 | 0.022 | 0.81 | 0.53 | 0.73 | 0.49 | 0.83 | 0.61 |
| 8 | 129258525 | rs4733599 | 0.907 | -1.72 | 0.516 | 1.272 | -0.016 | 0.189 | 0.189 | 0.022 | 0.82 | 0.59 | 0.75 | 0.51 | 0.87 | 0.62 |
| 8 | 129262075 | rs7835350 | 0.405 | -1.534 | -0.421 | 0.909 | 0.014 | 0.155 | 0.155 | 6.214 | 0.36 | 0.03 | 0.24 | 0.14 | 0.3 | 0.2 |
| 8 | 129263221 | rs17205367 | 0.378 | -1.558 | -0.429 | 0.901 | 0.012 | 0.157 | 0.157 | 6.098 | 0.36 | 0.03 | 0.24 | 0.14 | 0.3 | 0.2 |
| 8 | 129264171 | rs7826937 | -0.312 | -1.531 | -0.178 | 1.076 | -0.005 | -0.062 | -0.062 | 0.629 | 0.63 | 0.85 | 0.6 | 0.49 | 0.63 | 0.54 |
| 8 | 129265059 | rs9792320 | 1.217 | -1.53 | -0.526 | 1.072 | -0.042 | 0.079 | 0.079 | 0.762 | 0.27 | 0.01 | 0.22 | 0.13 | 0.25 | 0.19 |
| 8 | 129265338 | rs7815815 | -1.08 | -1.291 | -0.374 | 1.162 | -0.005 | 0.05 | 0.05 | 0 | 0.94 | 0.98 | 0.85 | 0.62 | 0.93 | 0.74 |
| 8 | 129265938 | rs16902760 | 0.996 | -1.4 | -0.575 | 1.206 | -0.013 | -0.044 | -0.044 | 0 | 0.07 | 0 | 0.03 | 0.2 | 0.06 | 0.14 |
| 8 | 129269193 | rs4733852 | -0.644 | -1.219 | -0.353 | 1.008 | -0.012 | 0.04 | 0.04 | 0 | 0.94 | 0.97 | 0.83 | 0.62 | 0.9 | 0.73 |
| 8 | 129269466 | rs4733601 | 0.866 | -1.346 | -0.019 | 0.449 | -0.058 | 0.105 | 0.105 | 3.413 | 0.36 | 0.18 | 0.44 | 0.04 | 0.48 | 0.28 |
| 8 | 129270866 | rs13439100 | 0.053 | -1.396 | 0.786 | 1.436 | -0.04 | 0.022 | 0.022 | 0.607 | 0.11 | 0.22 | 0.05 | 0.14 | 0.06 | 0.05 |
| 8 | 129271673 | rs16902767 | -0.838 | -0.819 | 0.219 | 1.292 | 0.002 | -0.002 | -0.002 | 0 | 0.05 | 0.18 | 0.04 | 0.04 | 0.03 | 0 |
| 8 | 129271933 | rs16902768 | -0.888 | -0.819 | 0.222 | 1.292 | 0.002 | -0.002 | -0.002 | 0 | 0.05 | 0.18 | 0.04 | 0.04 | 0.03 | 0 |
| 8 | 129278573 | rs998017 | 1.708 | -0.665 | 0.156 | 1.752 | 0.042 | 0.32 | 0.32 | 0 | 0.93 | 0.77 | 0.82 | 0.65 | 0.93 | 0.8 |
| 8 | 129278791 | rs998018 | 1.72 | -0.663 | 0.153 | 1.698 | 0.042 | 0.277 | 0.277 | 0 | 0.94 | 0.78 | 0.82 | 0.65 | 0.93 | 0.82 |
| 8 | 129283324 | rs4733853 | 0.25 | -0.642 | -1.263 | 0.825 | -0.104 | 0.137 | 0.137 | 0 | 0.36 | 0.08 | 0.41 | 0.12 | 0.4 | 0.27 |
| 8 | 129284351 | rs9886419 | -1.085 | -0.805 | -1.428 | 0.903 | -0.005 | -0.056 | -0.056 | 0 | 0.69 | 0.85 | 0.6 | 0.52 | 0.66 | 0.56 |
| 8 | 129284882 | rs10094929 | 0.443 | -1.043 | -1.452 | 0.812 | 0.006 | -0.022 | -0.022 | 2.164 | 0.2 | 0.13 | 0.1 | 0.36 | 0.18 | 0.24 |
| 8 | 129284951 | rs6987802 | 0.872 | -0.986 | -1.761 | 0.75 | -0.059 | 0.069 | 0.069 | 0 | 0.3 | 0.14 | 0.4 | 0.11 | 0.35 | 0.23 |
| 8 | 129286449 | rs12056478 | 1.396 | -0.822 | -1.957 | 0.555 | -0.025 | 0.02 | 0.02 | 0.107 | 0.29 | 0.13 | 0.34 | 0.24 | 0.34 | 0.31 |
| 8 | 129286933 | rs10102883 | -1.105 | -0.549 | -2.104 | 0.807 | -0.081 | -0.049 | -0.049 | 0.696 | 0.74 | 0.95 | 0.6 | 0.68 | 0.67 | 0.73 |
| 8 | 129289112 | rs13275576 | 0.266 | -0.449 | -1.304 | 0.185 | -0.048 | -0.047 | -0.047 | 0 | 0.15 | 0.38 | 0.07 | 0.06 | 0.09 | 0.07 |
| 8 | 129289556 | rs2049824 | -0.005 | 0.31 | -2.332 | 0.667 | -0.039 | 0.059 | 0.059 | 0 | 0.53 | 0.9 | 0.49 | 0.37 | 0.48 | 0.4 |
| 8 | 129299237 | rs7016955 | -0.872 | 0.092 | -1.638 | 0.91 | 0.01 | -0.008 | -0.008 | 0 | 0.31 | 0.31 | 0.15 | 0.43 | 0.23 | 0.27 |
| 8 | 129299579 | rs6997597 | -0.3 | -0.076 | -1.412 | 0.956 | -0.022 | 0.013 | 0.013 | 0 | 0.15 | 0.21 | 0.05 | 0.16 | 0.05 | 0.14 |
| 8 | 129299649 | rs10808569 | 0.18 | 0.174 | -1.588 | 0.835 | 0.013 | -0.004 | -0.004 | 0 | 0.37 | 0.5 | 0.6 | 0.34 | 0.4 | 0.45 |
| 8 | 129303172 | rs7820941 | -0.055 | 0.098 | -1.463 | 0.947 | -0.032 | 0.022 | 0.022 | 0 | 0.16 | 0.28 | 0.07 | 0.16 | 0.06 | 0.14 |
| 8 | 129306423 | rs4733859 | -0.187 | 0.731 | -1.447 | 0.702 | 0.003 | 0.012 | 0.012 | 0 | 0.51 | 0.6 | 0.65 | 0.49 | 0.47 | 0.58 |
| 8 | 129309414 | rs7815678 | -0.01 | 0.567 | -1.828 | 0.667 | -0.039 | -0.012 | -0.012 | 0 | 0.6 | 0.82 | 0.61 | 0.57 | 0.46 | 0.67 |
| 8 | 129311824 | rs16902799 | -0.608 | -1.032 | -1.136 | 0.02 | -0.01 | -0.036 | -0.036 | 0 | 0.21 | 0.26 | 0.12 | 0.05 | 0.14 | 0.13 |
| 8 | 129312741 | rs6996426 | -0.012 | -2.118 | -1.102 | 0.236 | -0.049 | 0.055 | 0.055 | 0.807 | 0.28 | 0.4 | 0.29 | 0.62 | 0.18 | 0.36 |
| 8 | 129314766 | rs976226 | 0.499 | -2.282 | -1.258 | 0.541 | -0.04 | -0.026 | -0.026 | 0.122 | 0.09 | 0 | 0.07 | 0.15 | 0.13 | 0.15 |
| 8 | 129316050 | rs975729 | -1.55 | -2.477 | -1.023 | 0.083 | 0.03 | 0.035 | 0.035 | 0.14 | 0.54 | 0.64 | 0.41 | 0.67 | 0.36 | 0.55 |
| 8 | 129321969 | rs12114344 | 0.517 | -2.526 | -0.85 | -0.052 | -0.03 | -0.004 | -0.004 | 0 | 0.08 | 0.15 | 0.04 | 0.05 | 0.02 | 0.04 |
| 8 | 129333455 | rs4077656 | 0.3 | -2.591 | -0.521 | 0.088 | -0.03 | 0.033 | 0.033 | 0.193 | 0.09 | 0.2 | 0.05 | 0.09 | 0.02 | 0.05 |
| 8 | 129343201 | rs7014842 | -1.637 | -2.959 | 0.224 | -0.065 | 0.029 | 0.036 | 0.036 | 0 | 0.52 | 0.6 | 0.41 | 0.66 | 0.36 | 0.54 |
| 8 | 129351485 | rs4265149 | -1.932 | -2.94 | 0.237 | -0.048 | 0.035 | 0.059 | 0.059 | 0.067 | 0.55 | 0.8 | 0.43 | 0.66 | 0.37 | 0.56 |
| 8 | 129384895 | rs4265150 | -1.365 | -2.961 | -0.117 | -0.149 | 0.013 | 0.02 | 0.02 | 0 | 0.51 | 0.45 | 0.38 | 0.65 | 0.37 | 0.63 |
| 8 | 129403027 | rs1516959 | -0.387 | -2.992 | -0.23 | -0.104 | -0.006 | 0.007 | 0.007 | 0.556 | 0.29 | 0.3 | 0.3 | 0.58 | 0.22 | 0.42 |
| 8 | 129412828 | rs7841408 | -1.303 | -3.081 | 0.079 | -0.166 | -0.002 | -0.002 | -0.002 | 0.016 | 0.48 | 0.49 | 0.37 | 0.64 | 0.33 | 0.59 |
| 8 | 129426941 | rs16902898 | 0.343 | -3.395 | 0.37 | 0.265 | 0.001 | -0.004 | -0.004 | 0.026 | 0.17 | 0.14 | 0.13 | 0.28 | 0.14 | 0.21 |
| 8 | 129427518 | rs7815944 | 0.568 | -3.432 | 0.381 | 0.275 | -0.032 | 0.226 | 0.226 | 0.073 | 0.1 | 0.26 | 0.18 | 0.29 | 0.04 | 0.18 |
| 8 | 129431327 | rs7844105 | 1.162 | -3.768 | 0.038 | -0.127 | -0.001 | -0.021 | -0.021 | 0 | 0.11 | 0.12 | 0.04 | 0 | 0.07 | 0.03 |
| 8 | 129435018 | rs6998301 | 1.532 | -3.76 | 0.062 | -0.128 | 0 | -0.001 | -0.001 | 0 | 0.47 | 0.47 | 0.61 | 0.36 | 0.64 | 0.37 |
| 8 | 129439051 | rs6470627 | -0.259 | -3.739 | 0.047 | -0.105 | -0.041 | 0.106 | 0.106 | 0 | 0.84 | 0.71 | 0.81 | 0.71 | 0.92 | 0.79 |
| 8 | 129439949 | rs17803668 | 0.433 | -3.66 | -0.073 | -0.089 | 0.013 | 0.069 | 0.069 | 0 | 0.22 | 0.09 | 0.08 | 0.06 | 0.15 | 0.19 |
| 8 | 129443161 | rs1878376 | -0.65 | -3.492 | -0.214 | -0.063 | 0 | 0 | 0 | 0 | 0.4 | 0.4 | 0.22 | 0.35 | 0.29 | 0.41 |
| 8 | 129447587 | rs4733864 | 1.856 | -3.099 | -0.25 | 0.079 | -0.029 | 0 | 0 | 1.113 | 0.34 | 0.25 | 0.56 | 0.36 | 0.56 | 0.34 |
| 8 | 129456625 | rs9987116 | 1.395 | -3.173 | -0.192 | 0.328 | 0.046 | 0.045 | 0.045 | 1.954 | 0.47 | 0.35 | 0.63 | 0.36 | 0.68 | 0.42 |
| 8 | 129465024 | rs10956424 | 0.47 | -2.402 | 0.071 | 0.764 | -0.029 | 0.151 | 0.151 | 0 | 0.08 | 0.17 | 0.16 | 0.29 | 0.04 | 0.18 |
| 8 | 129473012 | rs12543535 | 0.632 | -2.48 | -0.241 | 0.449 | 0.001 | -0.001 | -0.001 | 0 | 0.75 | 0.79 | 0.88 | 0.8 | 0.8 | 0.69 |
| 8 | 129475551 | rs11775876 | -1.058 | -2.843 | 0.23 | 0.352 | -0.052 | 0.009 | 0.009 | 0.427 | 0.18 | 0.33 | 0.09 | 0.14 | 0.07 | 0.28 |
| 8 | 129475576 | rs11779444 | -1.482 | -3.347 | 0.631 | 0.462 | -0.078 | 0.045 | 0.045 | 1.002 | 0.26 | 0.54 | 0.18 | 0.32 | 0.1 | 0.37 |
| 8 | 129478011 | rs10505517 | -1.521 | -3.382 | 0.769 | 0.429 | -0.057 | 0.025 | 0.025 | 0.995 | 0.27 | 0.6 | 0.18 | 0.32 | 0.1 | 0.37 |
| 8 | 129478333 | rs11996548 | 0.346 | -3.805 | 0.372 | -0.597 | 0.023 | 0.018 | 0.018 | 0 | 0.2 | 0.16 | 0.07 | 0 | 0.11 | 0.08 |
| 8 | 129480037 | rs13282878 | -1.983 | -3.853 | 0.337 | -0.652 | 0.003 | -0.003 | -0.003 | 0 | 0.93 | 0.97 | 0.98 | 1 | 0.96 | 0.96 |
| 8 | 129481921 | rs16903004 | -1.709 | -4.095 | 0.214 | -0.502 | -0.004 | -0.003 | -0.003 | 0 | 0.49 | 0.52 | 0.37 | 0.59 | 0.33 | 0.61 |
| 8 | 129484159 | rs16903006 | 0.685 | -4.06 | 0.266 | -0.461 | -0.022 | 0.008 | 0.008 | 0 | 0.09 | 0.21 | 0.06 | 0.05 | 0.04 | 0.18 |
| 8 | 129484174 | rs1464154 | 0.911 | -3.921 | -0.229 | -0.563 | -0.025 | -0.004 | -0.004 | 0.246 | 0.63 | 0.52 | 0.69 | 0.42 | 0.75 | 0.43 |
| 8 | 129486391 | rs17805310 | 0.495 | -3.804 | -0.454 | -0.551 | 0.031 | 0.068 | 0.068 | 0 | 0.19 | 0.08 | 0.07 | 0 | 0.11 | 0.08 |
| 8 | 129487893 | rs11781913 | -0.671 | -3.488 | -0.066 | -0.286 | -0.032 | 0.022 | 0.022 | 0 | 0.3 | 0.46 | 0.3 | 0.59 | 0.22 | 0.53 |
| 8 | 129488686 | rs7831607 | 0.073 | -3.055 | -0.213 | -0.365 | -0.038 | -0.082 | -0.082 | 0 | 0.86 | 0.7 | 0.95 | 1 | 0.95 | 0.93 |
| 8 | 129492318 | rs13281279 | 1.404 | -2.003 | -1.107 | -0.709 | -0.037 | -0.041 | -0.041 | 0.02 | 0.38 | 0.12 | 0.58 | 0.46 | 0.6 | 0.42 |
| 8 | 129492753 | rs6470634 | -0.438 | -1.639 | -0.177 | -1.697 | -0.007 | -0.051 | -0.051 | 2.11 | 0.18 | 0.21 | 0.08 | 0 | 0.11 | 0.09 |
| 8 | 129503291 | rs6470636 | -0.948 | -2.28 | -0.272 | -1.99 | 0.051 | 0.068 | 0.068 | 0.447 | 0.34 | 0.2 | 0.16 | 0.17 | 0.19 | 0.35 |
| 8 | 129505512 | rs6997818 | -0.552 | -2.087 | 0.172 | -1.902 | 0.024 | 0.031 | 0.031 | 1.186 | 0.28 | 0.2 | 0.17 | 0.18 | 0.17 | 0.34 |
| 8 | 129518281 | rs1516980 | -1.377 | -2.428 | 0.651 | -2.076 | 0.001 | -0.005 | -0.005 | 0 | 0.42 | 0.61 | 0.24 | 0.22 | 0.25 | 0.45 |
| 8 | 129525114 | rs1356332 | -1.4 | -1.855 | 0.616 | -2.134 | 0.002 | -0.005 | -0.005 | 0 | 0.43 | 0.61 | 0.26 | 0.23 | 0.29 | 0.45 |
| 8 | 129530492 | rs1400485 | 0.944 | -2.206 | -0.025 | -1.991 | -0.03 | 0.058 | 0.058 | 0.221 | 0.1 | 0.01 | 0.06 | 0 | 0.1 | 0.06 |

| | | | | | | | | | | | | | | | | |
|---|---|---|---|---|---|---|---|---|---|---|---|---|---|---|---|---|
| 8 | 129532118 | rs10089519 | 1.718 | -2.306 | -0.02 | -2.143 | 0.071 | 0.224 | 0.224 | 2.388 | 0.48 | 0.08 | 0.68 | 0.77 | 0.67 | 0.47 |
| 8 | 129533646 | rs1516982 | -0.29 | -1.553 | 0.838 | -1.888 | 0.01 | -0.026 | -0.026 | 0.005 | 0.13 | 0.18 | 0.12 | 0.05 | 0.14 | 0.11 |
| 8 | 129534673 | rs16903054 | -0.285 | -1.714 | 0.808 | -1.784 | -0.016 | -0.02 | -0.02 | 0 | 0.05 | 0.11 | 0.03 | 0.01 | 0.02 | 0.07 |
| 8 | 129541377 | rs16903067 | -0.872 | -1.319 | 0.988 | -2.036 | -0.01 | -0.062 | -0.062 | 0 | 0.14 | 0.22 | 0.09 | 0 | 0.13 | 0.09 |
| 8 | 129543949 | rs10088218 | -0.538 | -1.674 | 1.734 | -2.278 | 0 | -0.01 | -0.01 | 0 | 0.14 | 0.12 | 0.09 | 0.01 | 0.1 | 0.1 |
| 8 | 129548193 | rs1516975 | -0.563 | -1.674 | 1.795 | -2.278 | -0.004 | -0.031 | -0.031 | 0.861 | 0.14 | 0.21 | 0.1 | 0.01 | 0.1 | 0.1 |
| 8 | 129550041 | rs10112057 | 0.566 | -1.676 | 1.691 | -2.281 | -0.017 | 0.047 | 0.047 | 0.079 | 0.09 | 0.01 | 0.07 | 0 | 0.09 | 0.05 |
| 8 | 129556356 | rs16903097 | -0.885 | -1.639 | 1.878 | -2.286 | -0.006 | -0.073 | -0.073 | 0 | 0.15 | 0.28 | 0.11 | 0.01 | 0.11 | 0.11 |
| 8 | 129560314 | rs10102835 | 0.244 | -1.687 | 1.897 | -2.285 | -0.001 | -0.046 | -0.046 | 0 | 0.1 | 0.16 | 0.09 | 0.01 | 0.09 | 0.05 |
| 8 | 129567181 | rs6651252 | -0.825 | -1.733 | 2.021 | -2.303 | -0.012 | -0.077 | -0.077 | 2.005 | 0.16 | 0.34 | 0.12 | 0.02 | 0.11 | 0.1 |
| 8 | 129568061 | rs1561928 | 0.277 | -1.778 | 1.894 | -2.342 | -0.002 | -0.027 | -0.027 | 0.642 | 0.86 | 0.85 | 0.92 | 1 | 0.88 | 0.92 |
| 8 | 129568078 | rs1561927 | 1.767 | -1.876 | 1.643 | -2.391 | 0.001 | -0.041 | -0.041 | 0 | 0.64 | 0.39 | 0.77 | 0.98 | 0.75 | 0.72 |
| 8 | 129577486 | rs7825794 | -1.72 | -1.898 | 0.97 | -2.262 | -0.031 | -0.063 | -0.063 | 0 | 0.32 | 0.57 | 0.17 | 0.04 | 0.15 | 0.15 |
| 8 | 129579579 | rs2395822 | 2.017 | -1.45 | 0.09 | -2.407 | 0.036 | 0.136 | 0.136 | 0.98 | 0.57 | 0.23 | 0.75 | 0.96 | 0.69 | 0.81 |
| 8 | 129586776 | rs16903111 | -0.782 | -1.047 | -0.496 | -2.192 | -0.01 | -0.056 | -0.056 | 0 | 0.21 | 0.21 | 0.1 | 0 | 0.1 | 0.09 |
| 8 | 129592535 | rs7823385 | 0.417 | -0.645 | -0.521 | -1.888 | 0 | -0.037 | -0.037 | 0 | 0.8 | 0.82 | 0.91 | 1 | 0.83 | 0.93 |
| 8 | 129596286 | rs11780763 | 0.858 | -0.863 | -1.279 | -1.635 | -0.03 | 0.118 | 0.118 | 1.94 | 0.45 | 0.05 | 0.59 | 0.75 | 0.6 | 0.54 |
| 8 | 129596877 | rs16903124 | -0.753 | -0.781 | -0.059 | -1.623 | 0.017 | 0.04 | 0.04 | 0.53 | 0.08 | 0.25 | 0.06 | 0.04 | 0.06 | 0.03 |
| 8 | 129597629 | rs10090519 | 1.068 | -0.955 | -0.728 | -1.982 | -0.029 | 0.097 | 0.097 | 1.633 | 0.4 | 0.19 | 0.61 | 0.79 | 0.56 | 0.5 |
| 8 | 129597659 | rs1372992 | 0.774 | -1.177 | 0.079 | -2.145 | -0.032 | -0.164 | -0.164 | 0.191 | 0.66 | 0.52 | 0.83 | 0.99 | 0.76 | 0.78 |
| 8 | 129604429 | rs6990201 | -0.878 | -0.894 | 0.467 | -1.807 | -0.018 | -0.073 | -0.073 | 0 | 0.2 | 0.45 | 0.16 | 0.07 | 0.13 | 0.1 |
| 8 | 129606949 | rs7006008 | -0.639 | -0.911 | 0.774 | -1.82 | 0.001 | -0.027 | -0.027 | 0 | 0.18 | 0.18 | 0.1 | 0 | 0.12 | 0.1 |
| 8 | 129607917 | rs2099898 | -1.279 | -0.832 | 0.414 | -2.56 | 0.016 | 0.058 | 0.058 | 0.124 | 0.66 | 0.82 | 0.82 | 0.93 | 0.64 | 0.72 |
| 8 | 129608359 | rs2099897 | -1.266 | -0.574 | 0.338 | -2.36 | 0.005 | 0.017 | 0.017 | 0.4 | 0.4 | 0.65 | 0.3 | 0.22 | 0.34 | 0.27 |
| 8 | 129613281 | rs1443122 | 0.077 | -0.579 | 0.366 | -2.346 | 0.008 | 0.018 | 0.018 | 0 | 0.21 | 0.45 | 0.2 | 0.22 | 0.22 | 0.17 |
| 8 | 129617924 | rs6998269 | 0.121 | -0.518 | 0.176 | -2.322 | 0.004 | 0.009 | 0.009 | 0 | 0.2 | 0.27 | 0.18 | 0.22 | 0.22 | 0.17 |
| 8 | 129620475 | rs1476165 | -1.046 | -0.512 | 0.008 | -2.319 | -0.001 | -0.003 | -0.003 | 0.176 | 0.39 | 0.6 | 0.29 | 0.22 | 0.34 | 0.27 |
| 8 | 129629940 | rs4733621 | -0.93 | -0.494 | -0.02 | -2.356 | -0.008 | -0.041 | -0.041 | 0 | 0.37 | 0.51 | 0.29 | 0.22 | 0.33 | 0.27 |
| 8 | 129630950 | rs7011171 | -1.022 | -0.434 | -0.057 | -2.401 | 0 | -0.04 | -0.04 | 0 | 0.39 | 0.51 | 0.3 | 0.22 | 0.37 | 0.28 |
| 8 | 129633809 | rs16903339 | 0.292 | -0.47 | -0.049 | -2.307 | -0.007 | -0.004 | -0.004 | 0 | 0.2 | 0.13 | 0.18 | 0.22 | 0.25 | 0.18 |
| 8 | 129659764 | rs10956433 | -1.218 | -0.605 | -0.626 | -2.425 | 0.045 | 0.079 | 0.079 | 0 | 0.63 | 0.82 | 0.83 | 0.93 | 0.68 | 0.73 |
| 8 | 129662612 | rs1432011 | 0.606 | -0.334 | -0.541 | -2.021 | 0.001 | -0.003 | -0.003 | 0 | 0.11 | 0.08 | 0.07 | 0.15 | 0.1 | 0.13 |
| 8 | 129675798 | rs10091511 | 0.358 | 0.009 | -0.666 | -1.772 | -0.018 | 0 | 0 | 0 | 0.29 | 0.23 | 0.53 | 0.7 | 0.31 | 0.46 |
| 8 | 129681741 | rs1867202 | -0.7 | -0.114 | -0.781 | -2.052 | -0.002 | 0.109 | 0.109 | 1.218 | 0.59 | 0.81 | 0.77 | 0.92 | 0.53 | 0.68 |
| 8 | 129684553 | rs1367963 | 0.179 | -0.535 | -0.433 | -1.843 | -0.011 | -0.003 | -0.003 | 0 | 0.18 | 0.2 | 0.14 | 0.19 | 0.12 | 0.27 |
| 8 | 129691246 | rs4236736 | -0.009 | -0.699 | -0.485 | -1.664 | 0.009 | 0.008 | 0.008 | 0.202 | 0.44 | 0.58 | 0.38 | 0.36 | 0.41 | 0.46 |
| 8 | 129697564 | rs1367966 | -0.134 | -0.945 | -0.26 | -1.911 | -0.001 | -0.006 | -0.006 | 0 | 0.54 | 0.51 | 0.44 | 0.25 | 0.54 | 0.47 |
| 8 | 129699660 | rs7839784 | -1.216 | -0.177 | -0.846 | -2.232 | -0.087 | 0.029 | 0.029 | 0 | 0.86 | 0.99 | 0.87 | 0.89 | 0.77 | 0.86 |
| 8 | 129703419 | rs1835756 | -0.509 | -0.614 | -0.542 | -2.34 | -0.006 | -0.014 | -0.014 | 0.415 | 0.47 | 0.58 | 0.55 | 0.72 | 0.3 | 0.5 |
| 8 | 129707097 | rs2120119 | 0.637 | -0.773 | -0.414 | -2.427 | -0.001 | 0.004 | 0.004 | 0 | 0.17 | 0.19 | 0.34 | 0.64 | 0.18 | 0.3 |
| 8 | 129714626 | rs1347001 | 1.045 | -0.764 | -0.416 | -2.222 | -0.014 | -0.008 | -0.008 | 0 | 0.13 | 0.1 | 0.33 | 0.63 | 0.19 | 0.31 |
| 8 | 129716308 | rs6470648 | -2.086 | -0.506 | -0.508 | -2.492 | -0.058 | 0.066 | 0.066 | 0.571 | 0.8 | 0.95 | 0.83 | 0.92 | 0.65 | 0.85 |
| 8 | 129725425 | rs2395851 | 0.26 | -0.646 | -0.24 | -2.556 | -0.007 | 0.001 | 0.001 | 0.086 | 0.33 | 0.41 | 0.41 | 0.65 | 0.23 | 0.41 |
| 8 | 129734161 | rs7460697 | -2.128 | -0.637 | -0.241 | -2.552 | -0.038 | 0.051 | 0.051 | 0.571 | 0.8 | 0.92 | 0.82 | 0.92 | 0.65 | 0.85 |
| 8 | 129736712 | rs9297775 | 0.862 | -0.714 | 0.028 | -2.405 | -0.008 | 0.019 | 0.019 | 0 | 0.31 | 0.22 | 0.24 | 0.19 | 0.34 | 0.32 |
| 8 | 129738118 | rs1432019 | -0.508 | -0.725 | 0.103 | -2.4 | -0.001 | -0.006 | -0.006 | 0.119 | 0.48 | 0.57 | 0.57 | 0.73 | 0.31 | 0.53 |
| 8 | 129746895 | rs6990777 | -0.812 | -0.761 | 0.124 | -2.242 | 0.021 | 0.078 | 0.078 | 0 | 0.49 | 0.68 | 0.43 | 0.27 | 0.43 | 0.43 |
| 8 | 129748718 | rs6997983 | 0.246 | -0.8 | 0.01 | -2.06 | -0.039 | -0.024 | -0.024 | 0 | 0.17 | 0.29 | 0.17 | 0.08 | 0.08 | 0.11 |
| 8 | 129795806 | rs11996973 | 0.489 | -0.622 | -0.514 | -1.992 | -0.002 | -0.001 | -0.001 | 0 | 0.16 | 0.17 | 0.17 | 0.08 | 0.09 | 0.11 |
| 8 | 129798114 | rs4733649 | -0.292 | -0.29 | -0.615 | -2.141 | 0 | -0.008 | -0.008 | 0 | 0.68 | 0.7 | 0.75 | 0.79 | 0.68 | 0.68 |
| 8 | 129800770 | rs7006281 | -1.302 | -0.134 | -1.253 | -2.332 | -0.084 | 0.116 | 0.116 | 0.745 | 0.77 | 0.97 | 0.84 | 0.93 | 0.69 | 0.8 |
| 8 | 129805974 | rs1356761 | -0.016 | -0.207 | -0.515 | -2.687 | 0.005 | -0.023 | -0.023 | 0.005 | 0.28 | 0.38 | 0.48 | 0.8 | 0.28 | 0.35 |
| 8 | 129832156 | rs6994324 | 0.218 | -0.592 | -0.345 | -2.706 | -0.017 | -0.025 | -0.025 | 0 | 0.17 | 0.19 | 0.12 | 0.06 | 0.12 | 0.23 |
| 8 | 129840739 | rs2395855 | -0.943 | -0.912 | -0.439 | -3.057 | 0.011 | 0.17 | 0.17 | 0.022 | 0.63 | 0.86 | 0.78 | 0.92 | 0.62 | 0.66 |
| 8 | 129844671 | rs4733653 | -0.412 | -0.997 | 0.033 | -3.108 | -0.004 | -0.036 | -0.036 | 0.033 | 0.57 | 0.54 | 0.67 | 0.86 | 0.58 | 0.53 |
| 8 | 129845635 | rs7844704 | -1.591 | -0.968 | -0.199 | -3.052 | -0.004 | 0.075 | 0.075 | 0.072 | 0.73 | 0.86 | 0.78 | 0.89 | 0.69 | 0.7 |
| 8 | 129857419 | rs13250924 | 1.831 | -0.937 | -0.191 | -3.019 | 0.026 | 0.035 | 0.035 | 0 | 0.12 | 0.04 | 0.04 | 0 | 0.08 | 0.1 |
| 8 | 129881299 | rs4733532 | -0.624 | -0.952 | 0.019 | -3.021 | -0.003 | -0.051 | -0.051 | 0 | 0.6 | 0.59 | 0.74 | 0.93 | 0.62 | 0.65 |
| 8 | 129888794 | rs7845615 | -0.169 | -0.911 | 0.095 | -3.003 | -0.006 | 0.006 | 0.006 | 0 | 0.28 | 0.26 | 0.49 | 0.8 | 0.32 | 0.39 |
| 8 | 129896821 | rs1519850 | -1.937 | -0.723 | 0.115 | -2.811 | -0.027 | 0.094 | 0.094 | 0.077 | 0.76 | 0.91 | 0.82 | 0.94 | 0.7 | 0.75 |
| 8 | 129901910 | rs1157136 | -0.016 | -0.925 | -0.105 | -2.573 | 0.009 | 0.081 | 0.081 | 0 | 0.46 | 0.29 | 0.31 | 0.17 | 0.42 | 0.42 |
| 8 | 129916890 | rs1519845 | -0.665 | -1.137 | 0.051 | -2.725 | -0.003 | -0.078 | -0.078 | 0 | 0.48 | 0.45 | 0.25 | 0.06 | 0.39 | 0.34 |
| 8 | 129919458 | rs1530300 | 0.216 | -1.241 | -0.159 | -2.798 | -0.013 | 0.122 | 0.122 | 0 | 0.32 | 0.09 | 0.17 | 0.04 | 0.31 | 0.25 |
| 8 | 129920280 | rs997310 | -0.378 | -1.374 | -0.081 | -2.863 | -0.021 | -0.094 | -0.094 | 0 | 0.38 | 0.53 | 0.24 | 0.08 | 0.27 | 0.24 |
| 8 | 129934111 | rs13264072 | 1.295 | -1.279 | -0.41 | -2.787 | -0.025 | 0.043 | 0.043 | 0.023 | 0.12 | 0 | 0.05 | 0.04 | 0.12 | 0.15 |
| 8 | 129945713 | rs4733666 | -1.113 | -1.293 | -0.281 | -2.804 | 0.047 | 0.176 | 0.176 | 0.095 | 0.52 | 0.71 | 0.32 | 0.12 | 0.39 | 0.4 |
| 8 | 129946154 | rs987525 | 0.163 | -1.244 | -0.103 | -2.727 | -0.013 | -0.061 | -0.061 | 0.961 | 0.27 | 0.62 | 0.22 | 0.08 | 0.19 | 0.2 |
| 8 | 129947882 | rs12548036 | -0.758 | -1.083 | -0.367 | -2.684 | 0.006 | -0.031 | -0.031 | 0.497 | 0.48 | 0.45 | 0.27 | 0.1 | 0.39 | 0.38 |
| 8 | 129949182 | rs13253056 | 0.604 | -0.928 | -0.998 | -2.472 | 0.003 | 0.031 | 0.031 | 0 | 0.15 | 0.05 | 0.07 | 0.05 | 0.14 | 0.17 |

| | | | | | | | | | | | | | | | | |
|---|---|---|---|---|---|---|---|---|---|---|---|---|---|---|---|---|
| 8 | 129950844 | rs7017252 | -0.383 | -1.017 | -0.97 | -2.531 | 0.022 | 0.096 | 0.096 | 0 | 0.43 | 0.21 | 0.24 | 0.05 | 0.38 | 0.3 |
| 8 | 129953936 | rs6995235 | -1.843 | -0.487 | -1.74 | -2.765 | -0.098 | 0.106 | 0.106 | 0.023 | 0.74 | 0.98 | 0.83 | 0.95 | 0.67 | 0.75 |
| 8 | 129954008 | rs11784358 | 0.933 | -1.358 | -1.318 | -2.795 | 0.006 | 0.024 | 0.024 | 0 | 0.13 | 0.05 | 0.08 | 0 | 0.11 | 0.13 |
| 8 | 129959316 | rs13265167 | -0.129 | -1.486 | -1.464 | -2.935 | 0.002 | 0.123 | 0.123 | 0 | 0.27 | 0.07 | 0.16 | 0.05 | 0.24 | 0.28 |
| 8 | 129964834 | rs9693430 | 0.934 | -1.632 | -1.399 | -3.078 | 0.01 | 0.014 | 0.014 | 0 | 0.11 | 0.07 | 0.04 | 0 | 0.08 | 0.06 |
| 8 | 129964873 | rs17242358 | 1.573 | -1.776 | -1.547 | -3.279 | -0.013 | 0.071 | 0.071 | 0 | 0.16 | 0.03 | 0.11 | 0.01 | 0.17 | 0.08 |
| 8 | 129970246 | rs12546244 | 0.898 | -1.751 | -1.506 | -3.23 | 0 | 0.063 | 0.063 | 0 | 0.16 | 0.04 | 0.1 | 0 | 0.15 | 0.14 |
| 8 | 129977464 | rs1470206 | -1.458 | -1.816 | -0.687 | -3.142 | -0.022 | 0.05 | 0.05 | 0 | 0.62 | 0.71 | 0.41 | 0.12 | 0.5 | 0.48 |
| 8 | 129980296 | rs993316 | -0.331 | -1.768 | -0.713 | -3.119 | 0.02 | 0.134 | 0.134 | 0.018 | 0.46 | 0.21 | 0.29 | 0.05 | 0.42 | 0.39 |
| 8 | 129987132 | rs16903937 | 0.151 | -1.789 | -0.34 | -3.039 | -0.031 | -0.073 | -0.073 | 0 | 0.14 | 0.3 | 0.07 | 0 | 0.07 | 0.1 |
| 8 | 129988219 | rs6990893 | -0.881 | -1.599 | -0.304 | -2.994 | 0.016 | 0.041 | 0.041 | 0.472 | 0.55 | 0.66 | 0.3 | 0.06 | 0.42 | 0.35 |
| 8 | 129999102 | rs7833100 | -0.621 | -2.248 | -0.512 | -2.771 | 0.047 | 0.052 | 0.052 | 0 | 0.33 | 0.21 | 0.15 | 0.04 | 0.22 | 0.26 |
| 8 | 130000524 | rs7829061 | -0.125 | -2.403 | -0.151 | -2.661 | -0.063 | -0.104 | -0.104 | 4.098 | 0.28 | 0.38 | 0.11 | 0.04 | 0.13 | 0.16 |
| 8 | 130003116 | rs748978 | 0.826 | -1.757 | -0.33 | -2.21 | -0.018 | -0.022 | -0.022 | 0 | 0.13 | 0.2 | 0.06 | 0.06 | 0.09 | 0.14 |
| 8 | 130003263 | rs894263 | -0.413 | -1.347 | -0.581 | -1.625 | 0.002 | 0.008 | 0.008 | 0 | 0.6 | 0.6 | 0.29 | 0.23 | 0.42 | 0.54 |
| 8 | 130005126 | rs11776007 | 0.413 | -1.687 | -0.347 | -0.729 | 0.057 | 0.051 | 0.051 | 0 | 0.48 | 0.4 | 0.59 | 0.69 | 0.71 | 0.5 |
| 8 | 130021961 | rs10505532 | 0.131 | -1.953 | 0.183 | -0.76 | 0.022 | 0.021 | 0.021 | 0.305 | 0.52 | 0.63 | 0.65 | 0.69 | 0.73 | 0.54 |
| 8 | 130029034 | rs1372452 | -1.077 | -2.549 | -0.15 | -1.068 | 0.026 | 0.098 | 0.098 | 0 | 0.77 | 0.96 | 0.92 | 0.9 | 0.85 | 0.8 |
| 8 | 130030130 | rs1372454 | 0.933 | -2.61 | 0.125 | -1.093 | 0.054 | 0.049 | 0.049 | 0 | 0.5 | 0.45 | 0.63 | 0.69 | 0.73 | 0.54 |
| 8 | 130042141 | rs7844734 | 0.514 | -2.632 | 0.219 | -1.11 | 0.045 | 0.048 | 0.048 | 0.582 | 0.53 | 0.69 | 0.69 | 0.76 | 0.75 | 0.56 |
| 8 | 130056077 | rs6470692 | 0.006 | -2.635 | 0.227 | -1.145 | -0.011 | -0.006 | -0.006 | 0 | 0.74 | 0.79 | 0.75 | 0.86 | 0.87 | 0.72 |
| 8 | 130056640 | rs4613963 | 0.092 | -2.635 | 0.229 | -1.148 | -0.031 | -0.016 | -0.016 | 0 | 0.74 | 0.71 | 0.71 | 0.83 | 0.86 | 0.72 |
| 8 | 130057819 | rs7843894 | 0.281 | -2.7 | 0.212 | -1.291 | -0.028 | -0.026 | -0.026 | 0 | 0.2 | 0.2 | 0.18 | 0.07 | 0.09 | 0.17 |
| 8 | 130063761 | rs4326354 | -1.056 | -2.6 | 0.252 | -1.207 | 0.03 | 0.032 | 0.032 | 0.265 | 0.5 | 0.38 | 0.32 | 0.25 | 0.29 | 0.5 |
| 8 | 130070676 | rs4286912 | -0.603 | -2.228 | 0.246 | -1.264 | -0.025 | -0.055 | -0.055 | 0 | 0.29 | 0.34 | 0.25 | 0.12 | 0.19 | 0.26 |
| 8 | 130074255 | rs10435554 | -0.114 | -2.611 | 0.58 | -0.97 | -0.022 | -0.017 | -0.017 | 0 | 0.22 | 0.22 | 0.2 | 0.12 | 0.11 | 0.21 |
| 8 | 130087172 | rs11996876 | 0.511 | -2.712 | 0.425 | -1.004 | 0.021 | 0.028 | 0.028 | 0 | 0.25 | 0.17 | 0.1 | 0.14 | 0.16 | 0.24 |
| 8 | 130095478 | rs11784932 | -0.563 | -2.518 | 0.481 | -0.901 | 0 | -0.008 | -0.008 | 1.053 | 0.45 | 0.37 | 0.3 | 0.21 | 0.25 | 0.43 |
| 8 | 130107054 | rs12548560 | -1.167 | -2.461 | 0.156 | -0.851 | 0.083 | 0.102 | 0.102 | 0 | 0.52 | 0.65 | 0.36 | 0.25 | 0.29 | 0.51 |
| 8 | 130112168 | rs4339605 | -0.779 | -2.449 | 0.312 | -0.372 | 0.039 | 0.019 | 0.019 | 0 | 0.46 | 0.57 | 0.32 | 0.37 | 0.22 | 0.49 |
| 8 | 130113940 | rs10107597 | -0.296 | -2.567 | 0.418 | -0.438 | -0.099 | 0.069 | 0.069 | 0.016 | 0.22 | 0.4 | 0.2 | 0.29 | 0.09 | 0.22 |
| 8 | 130117365 | rs13263573 | -0.291 | -2.575 | 0.506 | -0.443 | -0.1 | 0.072 | 0.072 | 0.006 | 0.22 | 0.39 | 0.2 | 0.29 | 0.08 | 0.22 |
| 8 | 130117683 | rs6984251 | -1.389 | -2.006 | 0.292 | -0.078 | 0.059 | 0.18 | 0.18 | 0 | 0.54 | 0.85 | 0.58 | 0.8 | 0.34 | 0.54 |
| 8 | 130123196 | rs6987260 | 0.012 | -2.358 | 0.242 | -0.205 | -0.064 | 0.028 | 0.028 | 0.054 | 0.2 | 0.26 | 0.2 | 0.25 | 0.07 | 0.23 |
| 8 | 130128118 | rs7013846 | -0.861 | -2.348 | -0.052 | -0.158 | 0.001 | 0.003 | 0.003 | 0 | 0.45 | 0.5 | 0.31 | 0.37 | 0.22 | 0.48 |
| 8 | 130137247 | rs6996102 | -0.544 | -2.003 | 0.255 | 0.122 | -0.093 | 0.057 | 0.057 | 0 | 0.23 | 0.48 | 0.21 | 0.29 | 0.09 | 0.22 |
| 8 | 130152109 | rs6981888 | -0.337 | -1.724 | 0.005 | 0.19 | -0.057 | 0.052 | 0.052 | 0 | 0.2 | 0.33 | 0.24 | 0.25 | 0.11 | 0.18 |
| 8 | 130173467 | rs4571700 | -0.508 | -1.726 | -0.223 | 0.1 | 0.013 | 0.006 | 0.006 | 0 | 0.43 | 0.35 | 0.3 | 0.33 | 0.27 | 0.43 |
| 8 | 130178978 | rs9643245 | -0.948 | -1.252 | -0.104 | 0.092 | 0.003 | 0.031 | 0.031 | 0 | 0.57 | 0.67 | 0.57 | 0.75 | 0.4 | 0.56 |
| 8 | 130181882 | rs7823113 | -1.074 | -1.276 | -0.148 | 0.031 | -0.018 | 0.074 | 0.074 | 0 | 0.61 | 0.83 | 0.6 | 0.8 | 0.43 | 0.59 |
| 8 | 130184121 | rs7833637 | -0.983 | -1.344 | -0.127 | 0.024 | 0.001 | 0.078 | 0.078 | 0 | 0.58 | 0.79 | 0.58 | 0.8 | 0.41 | 0.54 |
| 8 | 130185601 | rs8180979 | 0.364 | -1.647 | -0.107 | 0.333 | -0.009 | 0.019 | 0.019 | 0.02 | 0.2 | 0.26 | 0.24 | 0.33 | 0.14 | 0.16 |
| 8 | 130189014 | rs6992168 | -0.386 | -1.527 | -0.045 | 0.345 | -0.006 | 0.004 | 0.004 | 0 | 0.37 | 0.42 | 0.44 | 0.45 | 0.28 | 0.37 |
| 8 | 130208039 | rs7813493 | -0.336 | -1.482 | -0.02 | 0.37 | 0.015 | 0.154 | 0.154 | 0 | 0.17 | 0.35 | 0.35 | 0.41 | 0.15 | 0.19 |
| 8 | 130219805 | rs13280665 | -0.58 | -1.526 | -0.434 | 0.264 | 0.033 | 0.007 | 0.007 | 0 | 0.42 | 0.38 | 0.22 | 0.39 | 0.28 | 0.37 |
| 8 | 130224721 | rs7822386 | 0.282 | -1.115 | -0.724 | 0.039 | 0.043 | 0.081 | 0.081 | 0 | 0.22 | 0.11 | 0.09 | 0.07 | 0.14 | 0.24 |
| 8 | 130224742 | rs7826130 | 0.282 | -1.115 | -0.724 | 0.039 | 0.043 | 0.081 | 0.081 | 0 | 0.22 | 0.11 | 0.08 | 0.07 | 0.14 | 0.24 |
| 8 | 130232692 | rs4733683 | -0.779 | -0.91 | -0.124 | 0.584 | -0.013 | 0.011 | 0.011 | 0 | 0.34 | 0.46 | 0.45 | 0.4 | 0.3 | 0.36 |
| 8 | 130235768 | rs4733540 | -1.41 | -1.43 | -0.643 | 0.4 | -0.034 | 0.04 | 0.04 | 0 | 0.63 | 0.86 | 0.62 | 0.72 | 0.42 | 0.58 |
| 8 | 130248593 | rs10808577 | -1.58 | -0.466 | -0.73 | 0.274 | -0.057 | 0.09 | 0.09 | 0 | 0.66 | 0.88 | 0.66 | 0.82 | 0.54 | 0.6 |
| 8 | 130249805 | rs4380897 | -0.406 | -0.591 | -0.367 | 0.478 | -0.001 | 0.016 | 0.016 | 0 | 0.18 | 0.15 | 0.34 | 0.46 | 0.18 | 0.24 |
| 8 | 130250443 | rs7820834 | -0.532 | -0.661 | 0.015 | 0.442 | -0.006 | 0.003 | 0.003 | 0 | 0.43 | 0.45 | 0.48 | 0.58 | 0.36 | 0.47 |
| 8 | 130251745 | rs10093780 | 0.111 | -0.562 | 0.105 | 0.663 | -0.007 | 0.032 | 0.032 | 0 | 0.22 | 0.32 | 0.16 | 0.25 | 0.19 | 0.13 |
| 8 | 130253287 | rs13439491 | 0.113 | -0.561 | 0.107 | 0.665 | -0.007 | 0.032 | 0.032 | 0 | 0.22 | 0.32 | 0.16 | 0.25 | 0.19 | 0.13 |
| 8 | 130260946 | rs13274646 | -0.231 | -0.396 | -0.221 | 0.295 | 0.022 | 0.053 | 0.053 | 0 | 0.45 | 0.25 | 0.2 | 0.28 | 0.38 | 0.33 |
| 8 | 130261783 | rs13279101 | -1.069 | -0.693 | 0.057 | 0.573 | -0.001 | -0.009 | -0.009 | 0 | 0.54 | 0.52 | 0.55 | 0.73 | 0.44 | 0.44 |
| 8 | 130267110 | rs10086303 | 0.662 | -0.721 | -0.199 | 0.392 | 0.004 | -0.008 | -0.008 | 0 | 0.28 | 0.19 | 0.13 | 0.25 | 0.26 | 0.19 |
| 8 | 130276372 | rs4461870 | -1.801 | -0.668 | -0.044 | 0.154 | -0.004 | 0.021 | 0.021 | 0 | 0.66 | 0.74 | 0.71 | 0.81 | 0.62 | 0.67 |
| 8 | 130285470 | rs13272388 | -0.945 | -0.676 | 0.14 | 0.158 | -0.003 | -0.016 | -0.016 | 0.045 | 0.53 | 0.52 | 0.56 | 0.71 | 0.44 | 0.43 |
| 8 | 130286363 | rs10113527 | 0.893 | -0.694 | 0.121 | 0.306 | 0.005 | -0.012 | -0.012 | 0 | 0.28 | 0.2 | 0.14 | 0.25 | 0.26 | 0.19 |
| 8 | 130287026 | rs13256084 | -1.944 | -0.688 | 0.144 | 0.283 | -0.011 | 0.032 | 0.032 | 0 | 0.67 | 0.81 | 0.72 | 0.79 | 0.62 | 0.67 |
| 8 | 130291974 | rs6998433 | -2.163 | -0.765 | 0.155 | 0.245 | -0.036 | 0.05 | 0.05 | 0 | 0.69 | 0.9 | 0.73 | 0.79 | 0.62 | 0.67 |
| 8 | 130299492 | rs4480081 | 0.045 | -0.735 | 0.014 | 0.331 | 0.024 | 0.041 | 0.041 | 0 | 0.45 | 0.23 | 0.2 | 0.29 | 0.38 | 0.32 |
| 8 | 130300407 | rs10107030 | -1.85 | -0.678 | 0.057 | 0.343 | -0.007 | 0.025 | 0.025 | 0 | 0.67 | 0.79 | 0.72 | 0.79 | 0.62 | 0.67 |
| 8 | 130305930 | rs12679760 | 0.859 | -0.691 | 0.022 | 0.357 | 0.005 | -0.013 | -0.013 | 0.502 | 0.28 | 0.19 | 0.14 | 0.25 | 0.26 | 0.2 |
| 8 | 130306831 | rs13279151 | -0.245 | -0.33 | 0.341 | 0.496 | -0.001 | -0.004 | -0.004 | 0.03 | 0.53 | 0.55 | 0.55 | 0.7 | 0.44 | 0.43 |
| 8 | 130320356 | rs2568421 | 1.088 | -0.263 | 0.287 | 0.505 | -0.009 | 0.029 | 0.029 | 0 | 0.32 | 0.17 | 0.28 | 0.21 | 0.38 | 0.33 |
| 8 | 130329068 | rs919701 | 0.59 | -0.22 | 0.316 | 0.497 | 0.032 | -0.014 | -0.014 | 0.254 | 0.12 | 0.19 | 0.16 | 0.1 | 0.18 | 0.23 |
| 8 | 130344185 | rs965123 | 0.786 | -0.199 | -0.075 | 0.363 | -0.003 | 0.018 | 0.018 | 0 | 0.34 | 0.24 | 0.3 | 0.19 | 0.37 | 0.33 |

| | | | | | | | | | | | | | | | | |
|---|---|---|---|---|---|---|---|---|---|---|---|---|---|---|---|---|
| 8 | 130352075 | rs2719195 | -0.22 | -0.046 | -0.411 | 0.525 | 0.023 | 0.04 | 0.04 | 0 | 0.54 | 0.66 | 0.78 | 0.7 | 0.61 | 0.65 |
| 8 | 130353227 | rs1835851 | -0.497 | -0.199 | -0.153 | 0.718 | 0.006 | 0.032 | 0.032 | 0 | 0.2 | 0.26 | 0.43 | 0.41 | 0.23 | 0.28 |
| 8 | 130353671 | rs16904030 | -0.509 | -0.67 | 0.131 | 0.038 | -0.029 | 0.009 | 0.009 | 0 | 0.18 | 0.24 | 0.09 | 0.18 | 0.08 | 0.23 |
| 8 | 130354027 | rs2568453 | 0.198 | -0.156 | -0.391 | -0.425 | 0.005 | 0.023 | 0.023 | 0 | 0.63 | 0.79 | 0.8 | 0.81 | 0.66 | 0.66 |
| 8 | 130367280 | rs10505536 | -0.383 | -0.106 | -0.491 | -0.44 | -0.004 | -0.037 | -0.037 | 0 | 0.39 | 0.15 | 0.22 | 0.42 | 0.39 | 0.4 |
| 8 | 130367554 | rs2568440 | 0.479 | -0.299 | 0.168 | -0.722 | -0.019 | -0.059 | -0.059 | 0 | 0.87 | 0.33 | 0.87 | 0.96 | 0.91 | 0.92 |
| 8 | 130368204 | rs2579866 | 0.409 | -0.298 | -0.227 | -0.702 | -0.022 | -0.054 | -0.054 | 0 | 0.87 | 0.32 | 0.87 | 0.96 | 0.91 | 0.92 |
| 8 | 130368723 | rs2579867 | 0.247 | -0.153 | -0.422 | -0.685 | 0.006 | -0.011 | -0.011 | 1.429 | 0.72 | 0.29 | 0.81 | 0.92 | 0.75 | 0.82 |
| 8 | 130369564 | rs12544748 | 0.133 | -0.01 | -0.164 | -0.458 | -0.001 | -0.065 | -0.065 | 0 | 0.25 | 0.39 | 0.17 | 0.15 | 0.29 | 0.2 |
| 8 | 130370235 | rs7846284 | -1.14 | 0.142 | -0.532 | -0.396 | -0.014 | -0.034 | -0.034 | 0 | 0.62 | 0.76 | 0.43 | 0.46 | 0.59 | 0.56 |
| 8 | 130370414 | rs7846684 | -1.043 | 0.281 | -0.639 | -0.327 | -0.005 | -0.017 | -0.017 | 3.253 | 0.61 | 0.66 | 0.43 | 0.46 | 0.59 | 0.56 |
| 8 | 130375804 | rs11990588 | 0.767 | 0.328 | -0.89 | -0.426 | -0.007 | 0.06 | 0.06 | 0.037 | 0.27 | 0.09 | 0.2 | 0.01 | 0.25 | 0.22 |
| 8 | 130376551 | rs16904047 | 0.818 | 0.465 | -0.938 | -0.301 | -0.002 | -0.011 | -0.011 | 0 | 0.08 | 0.02 | 0.04 | 0.31 | 0.04 | 0.14 |
| 8 | 130378681 | rs3997666 | 0.121 | 0.596 | -0.63 | -0.168 | 0.004 | -0.064 | -0.064 | 0 | 0.24 | 0.39 | 0.17 | 0.14 | 0.29 | 0.2 |
| 8 | 130379330 | rs6990827 | 0.045 | 0.61 | -0.743 | 0.43 | -0.002 | -0.005 | -0.005 | 0 | 0.6 | 0.6 | 0.42 | 0.46 | 0.59 | 0.54 |
| 8 | 130382552 | rs16904052 | -0.77 | 0.417 | -0.408 | 0.22 | -0.008 | -0.04 | -0.04 | 0.071 | 0.1 | 0.13 | 0.05 | 0 | 0.06 | 0.06 |
| 8 | 130383854 | rs10090653 | -0.906 | 0.278 | -0.096 | 0.493 | -0.002 | 0 | 0 | 0 | 0.41 | 0.6 | 0.29 | 0.36 | 0.32 | 0.33 |
| 8 | 130385926 | rs10098689 | -1.079 | 0.038 | 0.323 | 0.374 | 0.024 | 0 | 0 | 0.04 | 0.44 | 0.74 | 0.3 | 0.36 | 0.32 | 0.33 |
| 8 | 130387646 | rs4601277 | -0.818 | -0.006 | 0.266 | 0.569 | -0.003 | 0.004 | 0.004 | 0 | 0.4 | 0.59 | 0.31 | 0.42 | 0.32 | 0.33 |
| 8 | 130389670 | rs6989787 | -0.05 | 0.056 | 0.768 | 0.884 | -0.003 | -0.043 | -0.043 | 0 | 0.26 | 0.15 | 0.18 | 0.37 | 0.2 | 0.24 |
| 8 | 130389988 | rs3997658 | -0.347 | 0.071 | 0.963 | 0.857 | 0.048 | 0.084 | 0.084 | 0 | 0.05 | 0.28 | 0.07 | 0.05 | 0.06 | 0.04 |
| 8 | 130395585 | rs16904061 | -0.602 | -0.048 | 1.052 | 0.816 | -0.017 | 0.037 | 0.037 | 0 | 0.3 | 0.45 | 0.2 | 0.37 | 0.2 | 0.24 |
| 8 | 130400697 | rs2250954 | -0.41 | -0.065 | 0.542 | 1.282 | -0.056 | -0.062 | -0.062 | 1.612 | 0.16 | 0.02 | 0.1 | 0.22 | 0.14 | 0.17 |
| 8 | 130402414 | rs2630520 | -1.358 | -0.095 | 1.39 | 1.557 | 0.005 | 0.052 | 0.052 | 0.248 | 0.23 | 0.34 | 0.15 | 0.35 | 0.18 | 0.21 |
| 8 | 130403930 | rs16904064 | 1.114 | -0.315 | 0.664 | 1.462 | 0.015 | 0.005 | 0.005 | 0 | 0.32 | 0.13 | 0.41 | 0.29 | 0.3 | 0.22 |
| 8 | 130405474 | rs2630519 | -0.78 | -0.098 | 0.397 | 1.267 | -0.073 | 0.001 | 0.001 | 0 | 0.63 | 0.86 | 0.62 | 0.68 | 0.53 | 0.61 |
| 8 | 130415818 | rs2579865 | -0.893 | -0.194 | 0.471 | 1.145 | -0.045 | 0.014 | 0.014 | 0 | 0.66 | 0.82 | 0.64 | 0.69 | 0.54 | 0.62 |
| 8 | 130417371 | rs2719194 | -0.656 | -0.19 | 1.086 | 1.215 | 0.004 | 0.016 | 0.016 | 0.032 | 0.26 | 0.4 | 0.18 | 0.35 | 0.22 | 0.32 |
| 8 | 130417771 | rs9297788 | 0.732 | -0.177 | 1.035 | 1.185 | -0.001 | -0.005 | -0.005 | 0 | 0.06 | 0.07 | 0.04 | 0.05 | 0.07 | 0.15 |
| 8 | 130421568 | rs13259353 | 1.088 | -0.207 | 1.062 | 1.202 | -0.011 | -0.003 | -0.003 | 3.177 | 0.34 | 0.34 | 0.43 | 0.3 | 0.29 | 0.21 |
| 8 | 130430120 | rs2630507 | -0.561 | -0.184 | 0.891 | 1.194 | -0.027 | 0.014 | 0.014 | 0 | 0.6 | 0.75 | 0.61 | 0.65 | 0.51 | 0.53 |
| 8 | 130432856 | rs7006807 | 0.002 | -0.143 | 0.926 | 1.264 | -0.001 | 0.004 | 0.004 | 0 | 0.19 | 0.2 | 0.14 | 0.39 | 0.16 | 0.17 |
| 8 | 130450346 | rs2217589 | -0.193 | 0.036 | 0.823 | 1.059 | -0.011 | 0.042 | 0.042 | 0 | 0.55 | 0.64 | 0.6 | 0.66 | 0.47 | 0.37 |
| 8 | 130454037 | rs13269801 | 1.224 | 0.047 | 0.985 | 1.239 | -0.007 | 0.006 | 0.006 | 0 | 0.33 | 0.12 | 0.41 | 0.27 | 0.32 | 0.2 |
| 8 | 130456426 | rs12681076 | -0.208 | 0.316 | 1.17 | 0.768 | 0.003 | 0.011 | 0.011 | 0 | 0.58 | 0.61 | 0.6 | 0.7 | 0.55 | 0.53 |
| 8 | 130457896 | rs7837648 | 0.395 | 0.328 | 1.172 | 0.76 | 0.001 | 0.025 | 0.025 | 0 | 0.51 | 0.53 | 0.57 | 0.65 | 0.46 | 0.39 |
| 8 | 130459030 | rs1124682 | -0.615 | 0.519 | 0.863 | 0.705 | -0.001 | 0.051 | 0.051 | 0 | 0.65 | 0.87 | 0.67 | 0.75 | 0.59 | 0.64 |
| 8 | 130460135 | rs2007003 | -0.523 | 0.514 | 1.036 | 0.742 | 0.001 | 0.004 | 0.004 | 0 | 0.64 | 0.67 | 0.65 | 0.74 | 0.6 | 0.63 |
| 8 | 130461485 | rs1897447 | 1.05 | 0.559 | 0.732 | 0.766 | 0.005 | -0.022 | -0.022 | 2.387 | 0.4 | 0.24 | 0.48 | 0.48 | 0.36 | 0.27 |
| 8 | 130465043 | rs7017767 | -0.492 | 0.805 | 0.967 | 0.907 | 0.005 | 0.072 | 0.072 | 1.792 | 0.11 | 0.53 | 0.1 | 0.12 | 0.08 | 0.11 |
| 8 | 130469412 | rs7831732 | -0.592 | 0.745 | 0.436 | 0.867 | -0.005 | -0.008 | -0.008 | 0 | 0.66 | 0.87 | 0.67 | 0.67 | 0.59 | 0.64 |
| 8 | 130471159 | rs7815330 | -0.759 | 0.894 | 0.747 | 0.879 | 0.05 | 0.018 | 0.018 | 0 | 0.16 | 0.38 | 0.12 | 0.16 | 0.19 | 0.26 |
| 8 | 130475638 | rs11783022 | 0.326 | 0.978 | 0.297 | 1.005 | 0.015 | 0.033 | 0.033 | 0 | 0.08 | 0.17 | 0.06 | 0.12 | 0.1 | 0.12 |
| 8 | 130476358 | rs17177561 | 0.1 | 0.873 | -0.641 | 0.813 | -0.012 | 0.058 | 0.058 | 0 | 0.33 | 0.14 | 0.2 | 0.22 | 0.3 | 0.48 |
| 8 | 130477823 | rs10464870 | -1.269 | 0.865 | -0.837 | 0.872 | -0.006 | -0.007 | -0.007 | 3.956 | 0.87 | 0.89 | 0.78 | 0.82 | 0.77 | 0.87 |
| 8 | 130479731 | rs11786929 | 0.493 | 0.825 | -1.007 | 0.728 | 0.013 | -0.018 | -0.018 | 0 | 0.42 | 0.21 | 0.47 | 0.48 | 0.37 | 0.27 |
| 8 | 130483338 | rs12543365 | 0.734 | 0.789 | -1.342 | 0.528 | -0.051 | -0.001 | -0.001 | 0 | 0.16 | 0.01 | 0.21 | 0.13 | 0.23 | 0.12 |
| 8 | 130483377 | rs10099383 | -0.779 | 0.22 | -1.379 | 0.237 | 0.021 | 0.022 | 0.022 | 0 | 0.49 | 0.68 | 0.44 | 0.45 | 0.45 | 0.43 |
| 8 | 130484031 | rs2395900 | 0.872 | -0.089 | -0.447 | 0.148 | -0.014 | 0.001 | 0.001 | 0.477 | 0.16 | 0.26 | 0.1 | 0.22 | 0.12 | 0.19 |
| 8 | 130486168 | rs1835847 | -1.649 | 0.209 | -0.937 | -0.31 | -0.001 | -0.028 | -0.028 | 0 | 0.62 | 0.79 | 0.54 | 0.49 | 0.62 | 0.76 |
| 8 | 130487654 | rs1108353 | -0.387 | 0.126 | -0.002 | -0.446 | 0.009 | -0.025 | -0.025 | 0.204 | 0.23 | 0.39 | 0.17 | 0.14 | 0.2 | 0.41 |
| 8 | 130491165 | rs17262815 | 1.029 | 0.22 | -0.125 | -0.456 | -0.014 | 0.019 | 0.019 | 0.038 | 0.14 | 0.11 | 0.1 | 0.05 | 0.17 | 0.3 |
| 8 | 130493148 | rs7000768 | 0.958 | 0.176 | 0.059 | -0.378 | 0.006 | -0.025 | -0.025 | 4.327 | 0.23 | 0.32 | 0.31 | 0.14 | 0.31 | 0.16 |
| 8 | 130499223 | rs16904090 | -1.114 | -0.246 | 0.735 | -0.148 | -0.002 | -0.004 | -0.004 | 0 | 0.41 | 0.58 | 0.27 | 0.34 | 0.32 | 0.59 |
| 8 | 130501930 | rs16904092 | 0.677 | -0.304 | 0.716 | -0.152 | -0.03 | 0.025 | 0.025 | 0.004 | 0.11 | 0.28 | 0.08 | 0.09 | 0.03 | 0.11 |
| 8 | 130503150 | rs16904093 | 0.13 | -0.275 | 0.694 | -0.182 | -0.021 | 0.04 | 0.04 | 0 | 0.28 | 0.13 | 0.17 | 0.25 | 0.28 | 0.48 |
| 8 | 130510068 | rs16893242 | -0.991 | -0.178 | 0.269 | -0.046 | -0.006 | -0.006 | -0.006 | 2.404 | 0.42 | 0.53 | 0.27 | 0.37 | 0.32 | 0.59 |
| 8 | 130525809 | rs7016061 | -1.106 | -0.014 | -0.046 | -0.096 | -0.005 | 0.03 | 0.03 | 0 | 0.53 | 0.67 | 0.44 | 0.6 | 0.43 | 0.66 |
| 8 | 130531990 | rs7000021 | -0.625 | -0.045 | 0.245 | -0.139 | 0 | 0.001 | 0.001 | 0 | 0.48 | 0.48 | 0.41 | 0.6 | 0.4 | 0.64 |
| 8 | 130533811 | rs7839438 | -0.347 | -0.183 | 0.203 | -0.184 | 0 | -0.029 | -0.029 | 0 | 0.23 | 0.3 | 0.13 | 0.11 | 0.2 | 0.38 |
| 8 | 130543651 | rs4733710 | 0.595 | -0.204 | -0.162 | -0.232 | -0.011 | -0.108 | -0.108 | 0 | 0.28 | 0.1 | 0.28 | 0.5 | 0.24 | 0.25 |
| 8 | 130545445 | rs4733711 | 2.163 | -0.218 | -0.408 | -0.508 | -0.053 | 0.008 | 0.008 | 0.011 | 0.31 | 0.11 | 0.46 | 0.27 | 0.43 | 0.27 |
| 8 | 130547253 | rs10505542 | 0.342 | -0.11 | -0.219 | -0.543 | -0.027 | 0.065 | 0.065 | 0 | 0.16 | 0.05 | 0.09 | 0.04 | 0.17 | 0.29 |
| 8 | 130564649 | rs4733712 | -0.689 | -0.223 | -0.275 | -0.28 | 0.004 | -0.002 | -0.002 | 0 | 0.87 | 0.97 | 0.95 | 0.87 | 0.9 | 0.89 |
| 8 | 130570669 | rs6470739 | 0.104 | -0.334 | 0.111 | -0.337 | -0.004 | 0.008 | 0.008 | 0 | 0.31 | 0.42 | 0.36 | 0.57 | 0.25 | 0.31 |
| 8 | 130572304 | rs6987588 | 1.358 | -0.248 | -0.124 | -0.5 | -0.024 | 0.056 | 0.056 | 0 | 0.4 | 0.23 | 0.49 | 0.21 | 0.5 | 0.31 |
| 8 | 130573637 | rs17179377 | 1.421 | -0.117 | 0.313 | -0.366 | -0.002 | 0.002 | 0.002 | 0 | 0.08 | 0.05 | 0.04 | 0 | 0.08 | 0.02 |
| 8 | 130585355 | rs6470742 | 0.126 | -0.214 | 0.512 | -0.3 | 0.002 | -0.016 | -0.016 | 0 | 0.26 | 0.24 | 0.33 | 0.51 | 0.23 | 0.25 |
| 8 | 130586355 | rs12677963 | 1.377 | -0.229 | 0.449 | -0.307 | -0.003 | -0.048 | -0.048 | 0 | 0.15 | 0.06 | 0.29 | 0.51 | 0.15 | 0.22 |

| | | | | | | | | | | | | | | | | |
|---|---|---|---|---|---|---|---|---|---|---|---|---|---|---|---|---|
| 8 | 130592498 | rs7011870 | -0.172 | -0.239 | 0.722 | -0.329 | -0.009 | 0.034 | 0.034 | 0 | 0.28 | 0.39 | 0.34 | 0.51 | 0.23 | 0.27 |
| 8 | 130596548 | rs2043405 | -1.013 | -0.052 | 0.724 | -0.39 | 0.013 | -0.024 | -0.024 | 0.015 | 0.87 | 0.85 | 0.92 | 0.93 | 0.85 | 0.92 |
| 8 | 130597585 | rs2163950 | 0.134 | -0.155 | 0.682 | -0.39 | -0.002 | -0.001 | -0.001 | 0 | 0.1 | 0.16 | 0.06 | 0.15 | 0.06 | 0.29 |
| 8 | 130599247 | rs7840212 | -0.895 | -0.188 | 0.769 | -0.36 | -0.005 | -0.004 | -0.004 | 0 | 0.42 | 0.5 | 0.4 | 0.64 | 0.33 | 0.56 |
| 8 | 130600611 | rs7008811 | -0.353 | 0.052 | 0.788 | -0.256 | -0.002 | 0.004 | 0.004 | 0 | 0.32 | 0.33 | 0.34 | 0.51 | 0.28 | 0.26 |
| 8 | 130600781 | rs16904121 | 0.986 | 0.062 | 0.616 | -0.263 | -0.006 | -0.081 | -0.081 | 0 | 0.2 | 0.06 | 0.29 | 0.48 | 0.18 | 0.23 |
| 8 | 130601285 | rs892353 | 1.371 | 0.074 | 0.642 | -0.264 | -0.003 | 0.006 | 0.006 | 0 | 0.08 | 0.05 | 0.05 | 0 | 0.09 | 0.03 |
| 8 | 130603247 | rs11785889 | 0.922 | 0.072 | 0.614 | -0.248 | -0.005 | -0.08 | -0.08 | 0 | 0.21 | 0.07 | 0.29 | 0.51 | 0.18 | 0.23 |
| 8 | 130604563 | rs13277237 | 0.806 | 0.172 | 0.802 | -0.184 | -0.002 | 0.004 | 0.004 | 0 | 0.55 | 0.44 | 0.56 | 0.44 | 0.57 | 0.66 |
| 8 | 130606310 | rs6470743 | 0.187 | 0.217 | 0.695 | -0.169 | -0.001 | -0.044 | -0.044 | 0 | 0.3 | 0.17 | 0.34 | 0.48 | 0.28 | 0.26 |
| 8 | 130606476 | rs1865223 | 0.65 | 0.25 | 0.382 | -0.184 | -0.003 | 0.011 | 0.011 | 0 | 0.55 | 0.39 | 0.56 | 0.43 | 0.57 | 0.66 |
| 8 | 130609320 | rs4480083 | 0.705 | 0.278 | 0.214 | -0.187 | -0.003 | 0.003 | 0.003 | 0 | 0.56 | 0.45 | 0.56 | 0.46 | 0.57 | 0.67 |
| 8 | 130620813 | rs7005206 | -0.621 | 0.622 | -0.341 | -0.317 | 0.015 | -0.037 | -0.037 | 0 | 0.87 | 0.82 | 0.92 | 0.96 | 0.85 | 0.95 |
| 8 | 130625884 | rs16893247 | 1.251 | 0.494 | -0.875 | 0.218 | -0.012 | -0.073 | -0.073 | 0.836 | 0.14 | 0.02 | 0.26 | 0.23 | 0.17 | 0.09 |
| 8 | 130639540 | rs4636162 | 0.022 | 0.779 | -0.832 | 0.215 | -0.016 | -0.07 | -0.07 | 0 | 0.27 | 0.15 | 0.38 | 0.45 | 0.31 | 0.35 |
| 8 | 130641921 | rs6470745 | 0.85 | 0.862 | -0.801 | 0.092 | 0.014 | -0.064 | -0.064 | 0 | 0.2 | 0.08 | 0.21 | 0.28 | 0.19 | 0.21 |
| 8 | 130642237 | rs6470746 | -1.544 | 0.875 | -0.896 | -0.296 | -0.003 | 0.028 | 0.028 | 0 | 0.84 | 0.9 | 0.94 | 0.93 | 0.86 | 0.96 |
| 8 | 130646164 | rs6989425 | 0.884 | 0.858 | -0.752 | -0.426 | 0.002 | -0.007 | -0.007 | 0 | 0.38 | 0.55 | 0.48 | 0.42 | 0.4 | 0.47 |
| 8 | 130656205 | rs4626563 | 1.351 | 0.812 | -0.619 | -0.452 | 0 | -0.002 | -0.002 | 0 | 0.32 | 0.29 | 0.32 | 0.19 | 0.32 | 0.33 |
| 8 | 130661901 | rs7018091 | -1.597 | 0.843 | -0.763 | -0.464 | 0 | 0.004 | 0.004 | 0 | 0.83 | 0.85 | 0.93 | 0.93 | 0.86 | 0.94 |
| 8 | 130663510 | rs12544799 | 0.924 | 0.845 | -0.752 | -0.25 | 0.002 | -0.035 | -0.035 | 1.82 | 0.34 | 0.36 | 0.35 | 0.19 | 0.37 | 0.34 |
| 8 | 130665643 | rs16904140 | 0.359 | 1.132 | -1.695 | 0.101 | 0.009 | -0.035 | -0.035 | 0 | 0.23 | 0.08 | 0.25 | 0.22 | 0.21 | 0.22 |
| 8 | 130679176 | rs6985166 | 0.858 | 1.384 | -1.486 | 0.158 | 0.008 | -0.06 | -0.06 | 0.044 | 0.29 | 0.48 | 0.34 | 0.19 | 0.3 | 0.31 |
| 8 | 130682387 | rs13267039 | 0.072 | 1.288 | -1.032 | 0.38 | 0.02 | 0.066 | 0.066 | 0.17 | 0.05 | 0.15 | 0.13 | 0.24 | 0.08 | 0.13 |
| 8 | 130685516 | rs5022680 | 0.108 | 1.043 | -1.437 | 0.176 | 0.006 | -0.024 | -0.024 | 0 | 0.17 | 0.12 | 0.23 | 0.23 | 0.18 | 0.2 |
| 8 | 130691025 | rs6470757 | 0.312 | 1.31 | -1.053 | -0.332 | 0.014 | -0.027 | -0.027 | 0 | 0.69 | 0.52 | 0.74 | 0.75 | 0.68 | 0.76 |
| 8 | 130699140 | rs10956486 | -0.276 | 1.433 | -1.849 | -0.772 | -0.061 | 0.115 | 0.115 | 0 | 0.23 | 0.03 | 0.16 | 0.01 | 0.34 | 0.23 |
| 8 | 130701417 | rs4397371 | -0.082 | 1.111 | -2.299 | -0.33 | -0.022 | -0.059 | -0.059 | 0 | 0.77 | 0.89 | 0.54 | 0.31 | 0.76 | 0.73 |
| 8 | 130702280 | rs6470759 | 0.152 | 0.54 | -2.495 | -0.989 | 0.018 | 0.17 | 0.17 | 1.407 | 0.53 | 0.74 | 0.42 | 0.15 | 0.61 | 0.43 |
| 8 | 130702811 | rs16904143 | 0.51 | 0.164 | -1.933 | -0.331 | 0.012 | -0.038 | -0.038 | 0 | 0.21 | 0.06 | 0.11 | 0.17 | 0.15 | 0.29 |
| 8 | 130704068 | rs7007679 | -0.207 | 0.259 | -0.67 | -0.363 | -0.004 | 0.016 | 0.016 | 0.498 | 0.11 | 0.52 | 0.09 | 0.14 | 0.05 | 0.1 |
| 8 | 130706301 | rs4076446 | -0.423 | 0.589 | -0.786 | -0.126 | 0.001 | 0.006 | 0.006 | 0 | 0.5 | 0.52 | 0.45 | 0.64 | 0.47 | 0.53 |
| 8 | 130708114 | rs6470761 | -0.065 | 0.215 | -0.551 | 0.164 | 0.022 | -0.013 | -0.013 | 0 | 0.28 | 0.39 | 0.36 | 0.21 | 0.36 | 0.21 |
| 8 | 130708705 | rs17274130 | -1.22 | -0.03 | -0.368 | 0.361 | 0.002 | 0.031 | 0.031 | 0 | 0.51 | 0.63 | 0.47 | 0.68 | 0.47 | 0.57 |
| 8 | 130709863 | rs7011037 | 0.183 | 0.147 | -0.104 | 0.334 | 0.003 | -0.005 | -0.005 | 0.672 | 0.25 | 0.32 | 0.32 | 0.16 | 0.33 | 0.15 |
| 8 | 130710638 | rs4733723 | -0.147 | 0.165 | -0.093 | 0.378 | 0.018 | -0.063 | -0.063 | 0 | 0.28 | 0.38 | 0.32 | 0.07 | 0.34 | 0.21 |
| 8 | 130712714 | rs11986284 | -0.313 | 0.43 | -0.158 | 0.341 | 0.012 | -0.012 | -0.012 | 0 | 0.29 | 0.38 | 0.37 | 0.19 | 0.4 | 0.21 |
| 8 | 130715510 | rs4518624 | -1.367 | 0.122 | -0.777 | 0.657 | -0.144 | 0.181 | 0.181 | 0 | 0.58 | 0.95 | 0.7 | 0.86 | 0.49 | 0.65 |
| 8 | 130718314 | rs4501553 | 0.068 | 0.057 | 0.055 | 0.759 | 0.041 | -0.009 | -0.009 | 0 | 0.25 | 0.41 | 0.36 | 0.21 | 0.3 | 0.2 |
| 8 | 130718712 | rs4130415 | -1.003 | 0.098 | -0.076 | 0.559 | 0.087 | 0.044 | 0.044 | 0.268 | 0.41 | 0.86 | 0.41 | 0.72 | 0.26 | 0.53 |
| 8 | 130724502 | rs10091174 | -0.814 | -0.188 | -0.16 | 0.434 | -0.001 | 0.008 | 0.008 | 0.04 | 0.41 | 0.6 | 0.55 | 0.71 | 0.39 | 0.54 |
| 8 | 130724556 | rs12679359 | -0.524 | -0.187 | -0.116 | 0.455 | -0.045 | 0.068 | 0.068 | 0.088 | 0.25 | 0.52 | 0.31 | 0.67 | 0.16 | 0.42 |
| 8 | 130725082 | rs10094846 | 0.216 | -0.131 | -0.176 | 0.443 | 0.043 | 0.018 | 0.018 | 0 | 0.07 | 0.32 | 0.08 | 0.05 | 0.07 | 0.08 |
| 8 | 130727016 | rs10956489 | 1.149 | -0.19 | -0.166 | 0.729 | -0.011 | 0.07 | 0.07 | 0 | 0.1 | 0.18 | 0.22 | 0.24 | 0.08 | 0.05 |
| 8 | 130732295 | rs4733729 | -2.092 | 0.131 | -0.721 | 0.509 | -0.112 | 0.074 | 0.074 | 0.051 | 0.6 | 0.96 | 0.67 | 0.79 | 0.55 | 0.65 |
| 8 | 130742744 | rs3885952 | 1.031 | 0.111 | -0.618 | 0.499 | -0.04 | 0.07 | 0.07 | 0 | 0.17 | 0.05 | 0.22 | 0.02 | 0.22 | 0.11 |
| 8 | 130754719 | rs4733737 | -0.875 | -0.002 | -0.509 | 0.968 | -0.002 | 0.008 | 0.008 | 0.625 | 0.25 | 0.72 | 0.35 | 0.45 | 0.2 | 0.26 |
| 8 | 130759038 | rs5006712 | -1.544 | 0.132 | -0.631 | 1.058 | 0.014 | 0.008 | 0.008 | 0.014 | 0.51 | 0.76 | 0.59 | 0.47 | 0.44 | 0.4 |
| 8 | 130760774 | rs4733559 | 1.078 | 0.162 | -0.908 | 1.04 | -0.076 | 0.103 | 0.103 | 0 | 0.17 | 0.01 | 0.21 | 0.02 | 0.22 | 0.11 |
| 8 | 130760850 | rs4144738 | -1.734 | 0.282 | -0.649 | 1.308 | -0.106 | 0.035 | 0.035 | 0.013 | 0.58 | 0.94 | 0.64 | 0.74 | 0.47 | 0.64 |
| 8 | 130761182 | rs4144737 | -1.275 | 0.32 | -0.614 | 1.288 | 0.005 | -0.007 | -0.007 | 0 | 0.52 | 0.81 | 0.61 | 0.48 | 0.44 | 0.4 |
| 8 | 130761644 | rs4527833 | -1.355 | 0.332 | -0.619 | 1.28 | -0.038 | -0.009 | -0.009 | 0 | 0.53 | 0.83 | 0.61 | 0.52 | 0.5 | 0.41 |
| 8 | 130764791 | rs4285452 | -1.128 | 0.448 | -0.658 | 0.857 | 0.004 | 0.004 | 0.004 | 0 | 0.49 | 0.65 | 0.42 | 0.52 | 0.45 | 0.6 |
| 8 | 130768004 | rs12680164 | -0.908 | 0.332 | -0.979 | 0.929 | 0.011 | -0.02 | -0.02 | 0 | 0.1 | 0.07 | 0.05 | 0.2 | 0.06 | 0.14 |
| 8 | 130773458 | rs12542856 | 0.12 | 0.374 | -1.116 | 1.024 | -0.002 | -0.012 | -0.012 | 0 | 0.15 | 0.09 | 0.28 | 0.32 | 0.15 | 0.12 |
| 8 | 130786737 | rs1600934 | -0.029 | 0.643 | -1.233 | 0.16 | -0.001 | -0.026 | -0.026 | 0 | 0.19 | 0.22 | 0.24 | 0.03 | 0.21 | 0.1 |
| 8 | 130788548 | rs4733741 | -0.731 | 1.001 | -2.087 | -0.007 | -0.031 | 0.033 | 0.033 | 0.838 | 0.44 | 0.91 | 0.52 | 0.7 | 0.36 | 0.56 |
| 8 | 130799184 | rs3912318 | -1.519 | 0.806 | -2.074 | 0.029 | -0.091 | -0.019 | -0.019 | 0 | 0.63 | 0.94 | 0.74 | 0.73 | 0.58 | 0.65 |
| 8 | 130809688 | rs2086639 | 0.389 | 0.607 | -1.468 | 0.08 | -0.037 | 0.086 | 0.086 | 0 | 0.15 | 0.02 | 0.19 | 0.02 | 0.17 | 0.08 |
| 8 | 130814803 | rs10098779 | -0.088 | -0.309 | -1.155 | 0.376 | -0.029 | 0.033 | 0.033 | 0.141 | 0.68 | 0.37 | 0.75 | 0.57 | 0.76 | 0.55 |
| 8 | 130821049 | rs2894493 | 1.246 | -0.429 | -1.839 | -0.117 | 0.048 | 0.072 | 0.072 | 0.575 | 0.27 | 0.12 | 0.09 | 0.1 | 0.17 | 0.14 |
| 8 | 130827576 | rs1825225 | -0.086 | -0.218 | -1.656 | 0.041 | 0.027 | -0.031 | -0.031 | 0 | 0.42 | 0.19 | 0.67 | 0.48 | 0.59 | 0.36 |
| 8 | 130828900 | rs2624802 | -0.133 | -0.236 | -1.66 | 0.033 | 0.031 | -0.03 | -0.03 | 0 | 0.42 | 0.21 | 0.68 | 0.48 | 0.59 | 0.36 |
| 8 | 130838863 | rs420331 | -1.576 | -0.228 | -1.755 | 0.207 | -0.001 | -0.044 | -0.044 | 3.281 | 0.82 | 0.95 | 0.86 | 0.79 | 0.81 | 0.7 |
| 8 | 130841905 | rs392992 | -0.112 | 0.893 | -1.774 | 0.212 | -0.081 | 0.06 | 0.06 | 0 | 0.3 | 0.1 | 0.42 | 0.16 | 0.42 | 0.17 |
| 8 | 130844400 | rs11994702 | 0.388 | 0.711 | -0.598 | 0.489 | 0.035 | 0.128 | 0.128 | 0 | 0.15 | 0.39 | 0.27 | 0.32 | 0.17 | 0.2 |
| 8 | 130849731 | rs298622 | -0.163 | 1.146 | -0.651 | -0.173 | -0.003 | -0.012 | -0.012 | 0 | 0.17 | 0.18 | 0.22 | 0.03 | 0.22 | 0.08 |
| 8 | 130853447 | rs837065 | -0.032 | 0.489 | -1.252 | -0.034 | -0.025 | -0.028 | -0.028 | 0 | 0.51 | 0.21 | 0.53 | 0.53 | 0.51 | 0.47 |
| 8 | 130856138 | rs412941 | -0.803 | 0.321 | -0.706 | -0.264 | -0.013 | -0.015 | -0.015 | 0 | 0.62 | 0.72 | 0.61 | 0.54 | 0.51 | 0.53 |

| | | | | | | | | | | | | | | | | |
|---|---|---|---|---|---|---|---|---|---|---|---|---|---|---|---|---|
| 8 | 130868363 | rs837083 | 0.666 | 0.613 | -1.323 | -0.178 | -0.01 | 0.004 | 0.004 | 0 | 0.16 | 0.15 | 0.22 | 0.02 | 0.22 | 0.06 |
| 8 | 130880015 | rs10956502 | 1.002 | 0.505 | -1.551 | 0.063 | 0.062 | -0.149 | -0.149 | 0.005 | 0.35 | 0.06 | 0.3 | 0.53 | 0.27 | 0.4 |
| 8 | 130958951 | rs16904179 | 0.576 | 1.134 | -1.062 | 0.195 | -0.008 | -0.019 | -0.019 | 0 | 0.07 | 0.03 | 0.07 | 0.26 | 0.07 | 0.23 |
| 8 | 130963259 | rs1670745 | 0.928 | 0.989 | -0.768 | -0.127 | 0.003 | -0.016 | -0.016 | 0 | 0.62 | 0.46 | 0.64 | 0.71 | 0.63 | 0.6 |
| 8 | 130971985 | rs837224 | -0.323 | 1.39 | -1.258 | -0.438 | -0.057 | 0.066 | 0.066 | 0 | 0.84 | 0.95 | 0.78 | 0.98 | 0.76 | 0.91 |
| 8 | 130980472 | rs10092658 | 1.086 | 1.434 | -1.297 | -0.5 | -0.002 | 0.003 | 0.003 | 0 | 0.06 | 0.02 | 0.04 | 0.01 | 0.06 | 0.06 |
| 8 | 130985863 | rs837230 | 1.431 | 1.464 | -2.149 | -0.377 | -0.06 | -0.067 | -0.067 | 0 | 0.23 | 0.02 | 0.25 | 0.32 | 0.28 | 0.38 |
| 8 | 130987888 | rs837231 | 0.662 | 1.257 | -2.047 | -0.277 | -0.07 | -0.091 | -0.091 | 0.451 | 0.33 | 0.09 | 0.32 | 0.45 | 0.39 | 0.46 |
| 8 | 130990662 | rs17279655 | 1.212 | 0.77 | -1.407 | -0.143 | 0 | -0.01 | -0.01 | 0.574 | 0.06 | 0.05 | 0.04 | 0.01 | 0.06 | 0.05 |
| 8 | 130992696 | rs7357390 | -0.236 | 0.499 | -1.455 | -0.322 | 0.065 | 0.025 | 0.025 | 0.008 | 0.51 | 0.88 | 0.46 | 0.53 | 0.39 | 0.48 |
| 8 | 130998316 | rs882446 | -0.236 | 0.516 | -1.363 | -0.306 | 0.067 | 0.036 | 0.036 | 0.003 | 0.51 | 0.85 | 0.46 | 0.54 | 0.39 | 0.48 |
| 8 | 131001150 | rs10100858 | 0.526 | 0.23 | -0.689 | -0.219 | -0.005 | 0.002 | 0.002 | 0 | 0.37 | 0.56 | 0.36 | 0.38 | 0.31 | 0.24 |
| 8 | 131005011 | rs921693 | 1.242 | 0.193 | -0.358 | -0.246 | -0.031 | 0.037 | 0.037 | 0 | 0.31 | 0.39 | 0.32 | 0.38 | 0.25 | 0.2 |
| 8 | 131005137 | rs17194924 | -0.106 | 0.273 | -0.948 | -0.378 | 0.001 | 0.001 | 0.001 | 0 | 0.17 | 0.17 | 0.11 | 0.16 | 0.14 | 0.28 |
| 8 | 131005345 | rs6470788 | 0.294 | 0.594 | -0.888 | -0.334 | 0.046 | 0.082 | 0.082 | 0 | 0.44 | 0.34 | 0.25 | 0.17 | 0.34 | 0.42 |
| 8 | 131010688 | rs12541347 | -0.137 | 0.893 | -1.466 | -0.508 | 0.049 | 0.025 | 0.025 | 0.049 | 0.53 | 0.84 | 0.46 | 0.56 | 0.39 | 0.52 |
| 8 | 131014233 | rs1349847 | 1.017 | 1.083 | -1.442 | -0.356 | -0.024 | 0.033 | 0.033 | 0 | 0.06 | 0 | 0.05 | 0.01 | 0.08 | 0.04 |
| 8 | 131015276 | rs6470789 | -0.289 | 1.225 | -1.501 | -0.382 | -0.104 | -0.056 | -0.056 | 0.015 | 0.6 | 0.93 | 0.51 | 0.56 | 0.49 | 0.52 |
| 8 | 131024880 | rs2060984 | -0.21 | 1.261 | -0.827 | -0.428 | -0.023 | 0.014 | 0.014 | 0 | 0.59 | 0.79 | 0.49 | 0.62 | 0.45 | 0.55 |
| 8 | 131025034 | rs2060983 | 0.105 | 1.195 | -0.019 | -0.552 | -0.001 | -0.002 | -0.002 | 0.303 | 0.54 | 0.55 | 0.43 | 0.62 | 0.4 | 0.55 |
| 8 | 131025818 | rs1466226 | -0.617 | 1.355 | -0.368 | -0.553 | -0.023 | -0.003 | -0.003 | 0.308 | 0.66 | 0.79 | 0.55 | 0.63 | 0.55 | 0.59 |
| 8 | 131027338 | rs874617 | -0.053 | 1.162 | -0.126 | -0.573 | 0.002 | 0.013 | 0.013 | 0 | 0.56 | 0.62 | 0.43 | 0.63 | 0.41 | 0.56 |
| 8 | 131030580 | rs747498 | 0.369 | 1.112 | -0.046 | -0.642 | 0.026 | 0.022 | 0.022 | 0 | 0.5 | 0.54 | 0.36 | 0.63 | 0.33 | 0.53 |
| 8 | 131033480 | rs16904190 | -0.047 | 1.109 | 0.138 | -0.409 | -0.001 | 0 | 0 | 2.034 | 0.56 | 0.49 | 0.42 | 0.55 | 0.42 | 0.56 |
| 8 | 131036253 | rs11780495 | 0.15 | 1.884 | -0.063 | -0.454 | -0.01 | 0.008 | 0.008 | 0 | 0.38 | 0.32 | 0.53 | 0.34 | 0.44 | 0.33 |
| 8 | 131039109 | rs6470791 | -0.339 | 1.881 | -0.089 | -0.472 | -0.009 | 0.021 | 0.021 | 0 | 0.45 | 0.38 | 0.59 | 0.34 | 0.54 | 0.37 |
| 8 | 131052403 | rs2395988 | -0.104 | 1.755 | 0.221 | -0.56 | -0.002 | 0.034 | 0.034 | 0 | 0.43 | 0.75 | 0.58 | 0.34 | 0.45 | 0.33 |
| 8 | 131054236 | rs4295708 | 0.79 | 1.543 | -0.026 | -0.861 | -0.012 | 0.006 | 0.006 | 0 | 0.15 | 0.05 | 0.06 | 0.14 | 0.14 | 0.31 |
| 8 | 131056738 | rs3871 | -0.204 | 1.679 | -0.137 | -0.942 | -0.015 | 0.005 | 0.005 | 0 | 0.8 | 0.88 | 0.89 | 0.83 | 0.74 | 0.58 |
| 8 | 131064299 | rs6470796 | 1.006 | 1.531 | -0.05 | -0.961 | -0.015 | 0.016 | 0.016 | 0 | 0.34 | 0.27 | 0.5 | 0.29 | 0.41 | 0.31 |
| 8 | 131065153 | rs4236749 | 1.987 | 1.331 | -0.168 | -1.05 | 0.046 | -0.12 | -0.12 | 0 | 0.28 | 0.07 | 0.23 | 0.46 | 0.17 | 0.19 |
| 8 | 131068117 | rs3758028 | 1.048 | 1.375 | -0.158 | -1.045 | -0.019 | -0.006 | -0.006 | 5.697 | 0.15 | 0.04 | 0.05 | 0.15 | 0.13 | 0.31 |
| 8 | 131074024 | rs7460645 | 0.057 | 1.352 | 0.003 | -1.051 | -0.001 | 0.06 | 0.06 | 0 | 0.45 | 0.73 | 0.62 | 0.29 | 0.51 | 0.34 |
| 8 | 131076710 | rs7387355 | -0.267 | 1.396 | 0.007 | -1.012 | -0.022 | 0.004 | 0.004 | 0 | 0.8 | 0.88 | 0.89 | 0.81 | 0.72 | 0.56 |
| 8 | 131108193 | rs7833007 | -0.798 | 1.507 | -0.061 | -1.06 | 0 | -0.01 | -0.01 | 0 | 0.63 | 0.54 | 0.47 | 0.69 | 0.59 | 0.69 |
| 8 | 131118271 | rs7837833 | 0.265 | 1.488 | -0.108 | -1.14 | 0.027 | -0.033 | -0.033 | 2.117 | 0.13 | 0.43 | 0.22 | 0.04 | 0.13 | 0.11 |
| 8 | 131123753 | rs6470801 | -1.736 | 1.492 | -0.09 | -1.141 | 0.001 | -0.005 | -0.005 | 0.234 | 0.73 | 0.73 | 0.65 | 0.74 | 0.68 | 0.8 |
| 8 | 131124559 | rs966185 | 0.132 | 1.462 | -0.232 | -1.149 | -0.008 | 0.089 | 0.089 | 0 | 0.53 | 0.22 | 0.37 | 0.72 | 0.46 | 0.65 |
| 8 | 131127831 | rs2305513 | 0.401 | 1.455 | -0.265 | -1.147 | -0.013 | 0.086 | 0.086 | 0 | 0.49 | 0.21 | 0.35 | 0.65 | 0.45 | 0.63 |
| 8 | 131129442 | rs6470802 | -0.722 | 1.488 | -0.201 | -1.144 | 0 | -0.006 | -0.006 | 0.349 | 0.63 | 0.39 | 0.46 | 0.7 | 0.59 | 0.69 |
| 8 | 131130594 | rs10956511 | 0.234 | 1.467 | -0.306 | -1.134 | 0.051 | 0.068 | 0.068 | 0 | 0.51 | 0.3 | 0.33 | 0.66 | 0.31 | 0.51 |
| 8 | 131138575 | rs2305511 | 1.718 | 1.397 | -0.375 | -1.132 | 0.032 | -0.128 | -0.128 | 0.906 | 0.31 | 0.07 | 0.26 | 0.47 | 0.21 | 0.21 |
| 8 | 131147397 | rs4733768 | -0.055 | 1.557 | -0.162 | -1.018 | 0.001 | 0.034 | 0.034 | 0 | 0.57 | 0.3 | 0.39 | 0.66 | 0.4 | 0.57 |
| 8 | 131156071 | rs6984815 | -0.085 | 1.656 | -0.218 | -0.983 | 0.072 | -0.022 | -0.022 | 0 | 0.15 | 0.36 | 0.26 | 0.07 | 0.27 | 0.23 |
| 8 | 131189480 | rs7012705 | 0.331 | 1.369 | -0.587 | -0.979 | 0.037 | 0.068 | 0.068 | 0 | 0.51 | 0.16 | 0.34 | 0.62 | 0.39 | 0.55 |
| 8 | 131190243 | rs11774633 | -0.793 | 1.387 | -0.641 | -0.985 | -0.13 | 0.017 | 0.017 | 0.427 | 0.65 | 0.92 | 0.5 | 0.7 | 0.51 | 0.61 |
| 8 | 131203755 | rs16904215 | 0.798 | 1.454 | -0.575 | -0.884 | -0.009 | -0.004 | -0.004 | 0.243 | 0.16 | 0.05 | 0.05 | 0.17 | 0.13 | 0.32 |
| 8 | 131207979 | rs11781752 | 1.137 | 1.359 | -0.605 | -0.91 | 0.049 | -0.096 | -0.096 | 0 | 0.36 | 0.11 | 0.31 | 0.46 | 0.27 | 0.24 |
| 8 | 131213570 | rs4733775 | 0.382 | 1.365 | -0.606 | -0.907 | 0.007 | 0.048 | 0.048 | 0 | 0.49 | 0.09 | 0.33 | 0.62 | 0.38 | 0.54 |
| 8 | 131217195 | rs11775292 | -0.198 | 1.391 | -0.318 | -0.9 | -0.004 | 0.019 | 0.019 | 0 | 0.57 | 0.31 | 0.4 | 0.68 | 0.43 | 0.58 |
| 8 | 131226752 | rs2303444 | 0.91 | 1.405 | -0.415 | -0.842 | -0.011 | -0.006 | -0.006 | 0.087 | 0.15 | 0.05 | 0.05 | 0.16 | 0.12 | 0.31 |
| 8 | 131243645 | rs7846662 | -0.131 | 1.634 | -0.303 | -0.888 | -0.019 | 0.004 | 0.004 | 0 | 0.79 | 0.86 | 0.89 | 0.81 | 0.73 | 0.56 |
| 8 | 131243949 | rs7834461 | -0.09 | 1.571 | -0.344 | -0.931 | -0.007 | 0.06 | 0.06 | 0 | 0.45 | 0.8 | 0.6 | 0.34 | 0.47 | 0.33 |
| 8 | 131247022 | rs10091450 | 1.809 | 1.537 | -0.028 | -0.981 | 0.001 | -0.007 | -0.007 | 0 | 0.26 | 0.28 | 0.34 | 0.26 | 0.3 | 0.19 |
| 8 | 131252758 | rs10956514 | 0.079 | 1.544 | -0.078 | -0.98 | 0.009 | 0.044 | 0.044 | 0 | 0.53 | 0.24 | 0.37 | 0.62 | 0.42 | 0.56 |
| 8 | 131256398 | rs2554374 | -0.144 | 1.55 | -0.105 | -0.979 | -0.007 | 0.058 | 0.058 | 0 | 0.45 | 0.8 | 0.6 | 0.35 | 0.47 | 0.33 |
| 8 | 131258535 | rs2554373 | 0.579 | 1.556 | 0.035 | -0.98 | 0.01 | -0.025 | -0.025 | 0 | 0.36 | 0.53 | 0.54 | 0.29 | 0.42 | 0.3 |
| 8 | 131261986 | rs1549454 | -0.303 | 1.282 | -0.065 | -0.871 | -0.017 | 0.004 | 0.004 | 0 | 0.79 | 0.85 | 0.89 | 0.81 | 0.73 | 0.56 |
| 8 | 131265218 | rs7832651 | -0.252 | 1.291 | -0.108 | -0.859 | -0.003 | 0.008 | 0.008 | 0 | 0.58 | 0.36 | 0.4 | 0.68 | 0.43 | 0.58 |
| 8 | 131269886 | rs2670876 | -0.595 | 1.317 | -0.139 | -0.847 | -0.03 | 0.024 | 0.024 | 0 | 0.89 | 0.92 | 0.81 | 0.96 | 0.8 | 0.81 |
| 8 | 131276160 | rs1008819 | 0.161 | 1.322 | -0.07 | -0.849 | 0.04 | -0.018 | -0.018 | 0 | 0.85 | 0.68 | 0.74 | 0.93 | 0.73 | 0.76 |
| 8 | 131278618 | rs10505549 | 1.957 | 1.2 | -0.241 | -0.893 | 0.024 | -0.163 | -0.163 | 0 | 0.29 | 0.03 | 0.25 | 0.46 | 0.2 | 0.2 |
| 8 | 131281826 | rs7839523 | -1.255 | 1.204 | -0.152 | -0.898 | 0.025 | -0.017 | -0.017 | 0 | 0.69 | 0.63 | 0.6 | 0.71 | 0.56 | 0.69 |
| 8 | 131290664 | rs4733571 | 0.352 | 1.145 | -0.218 | -0.904 | 0.048 | 0.074 | 0.074 | 0 | 0.5 | 0.19 | 0.35 | 0.62 | 0.38 | 0.55 |
| 8 | 131296255 | rs4733780 | -0.242 | 1.153 | -0.199 | -0.891 | -0.009 | 0.032 | 0.032 | 0 | 0.58 | 0.3 | 0.39 | 0.68 | 0.43 | 0.58 |
| 8 | 131298985 | rs16904237 | 1.07 | 1.151 | -0.321 | -0.869 | -0.013 | -0.003 | -0.003 | 0 | 0.15 | 0.05 | 0.05 | 0.15 | 0.12 | 0.31 |
| 8 | 131300036 | rs2912191 | 1.544 | 1.148 | -0.251 | -0.849 | 0 | -0.005 | -0.005 | 0 | 0.56 | 0.38 | 0.59 | 0.72 | 0.49 | 0.38 |
| 8 | 131303878 | rs16904239 | 1.921 | 1.137 | -0.68 | -0.764 | 0.017 | -0.157 | -0.157 | 0 | 0.29 | 0.03 | 0.25 | 0.46 | 0.2 | 0.2 |
| 8 | 131320963 | rs12679283 | 1.793 | 1.186 | -0.655 | -0.761 | 0.027 | -0.16 | -0.16 | 0.355 | 0.3 | 0.03 | 0.26 | 0.47 | 0.2 | 0.21 |

| | | | | | | | | | | | | | | | | |
|---|---|---|---|---|---|---|---|---|---|---|---|---|---|---|---|---|
| 8 | 131328302 | rs2670873 | 0.538 | 1.285 | -0.537 | -0.767 | -0.008 | 0.004 | 0.004 | 0 | 0.69 | 0.64 | 0.69 | 0.77 | 0.6 | 0.44 |
| 8 | 131334465 | rs1340022 | -0.4 | 1.46 | -0.394 | -0.8 | -0.008 | 0.006 | 0.006 | 0 | 0.63 | 0.42 | 0.56 | 0.71 | 0.49 | 0.67 |
| 8 | 131337836 | rs2670882 | 0.762 | 1.484 | -0.272 | -0.743 | 0.042 | -0.004 | -0.004 | 0 | 0.05 | 0.09 | 0.06 | 0.03 | 0.15 | 0.12 |
| 8 | 131349715 | rs6470816 | -0.189 | 1.309 | -0.593 | -0.795 | -0.023 | 0.049 | 0.049 | 0 | 0.44 | 0.64 | 0.5 | 0.34 | 0.54 | 0.37 |
| 8 | 131356392 | rs13258946 | -0.221 | 1.326 | -0.752 | -0.737 | -0.008 | 0.064 | 0.064 | 0 | 0.45 | 0.79 | 0.63 | 0.33 | 0.58 | 0.42 |
| 8 | 131359644 | rs16904252 | 0.829 | 1.56 | -0.235 | -0.413 | -0.024 | -0.015 | -0.015 | 0 | 0.12 | 0.03 | 0.05 | 0.1 | 0.1 | 0.3 |
| 8 | 131381098 | rs9694314 | -0.246 | 1.684 | 0.066 | -0.454 | -0.014 | 0.012 | 0.012 | 0 | 0.11 | 0.11 | 0.19 | 0.04 | 0.2 | 0.19 |
| 8 | 131387248 | rs3924416 | -0.713 | 1.656 | 0.512 | -0.386 | 0.04 | 0.044 | 0.044 | 0.049 | 0.09 | 0.29 | 0.13 | 0.1 | 0.09 | 0.07 |
| 8 | 131391874 | rs7844582 | -1.075 | 1.54 | 0.653 | -0.45 | 0.101 | -0.003 | -0.003 | 0 | 0.12 | 0.36 | 0.15 | 0.07 | 0.22 | 0.12 |
| 8 | 131404323 | rs6994854 | -1.935 | 1.473 | 0.532 | -0.549 | 0.023 | -0.038 | -0.038 | 0.046 | 0.29 | 0.63 | 0.4 | 0.12 | 0.38 | 0.26 |
| 8 | 131409885 | rs4609234 | -1.661 | 1.481 | 0.636 | -0.582 | 0.033 | -0.06 | -0.06 | 0 | 0.26 | 0.65 | 0.36 | 0.07 | 0.37 | 0.23 |
| 8 | 131411347 | rs10956520 | -0.803 | 1.565 | 0.755 | -0.496 | 0.064 | -0.004 | -0.004 | 0 | 0.11 | 0.28 | 0.14 | 0.06 | 0.22 | 0.12 |
| 8 | 131414083 | rs10090231 | -1.577 | 1.556 | 0.886 | -0.559 | 0.086 | -0.056 | -0.056 | 0 | 0.19 | 0.51 | 0.3 | 0.08 | 0.28 | 0.19 |
| 8 | 131414912 | rs11993741 | -1.563 | 1.561 | 0.881 | -0.556 | 0.089 | -0.056 | -0.056 | 0 | 0.19 | 0.51 | 0.3 | 0.08 | 0.28 | 0.19 |
| 8 | 131417595 | rs13255178 | -0.33 | 1.549 | 0.938 | -0.553 | 0.022 | 0 | 0 | 0.985 | 0.08 | 0.16 | 0.13 | 0.07 | 0.22 | 0.12 |
| 8 | 131418029 | rs16904263 | 0.475 | 1.347 | 0.868 | -0.071 | 0.001 | -0.008 | -0.008 | 0.005 | 0.15 | 0.12 | 0.04 | 0.24 | 0.12 | 0.31 |
| 8 | 131422473 | rs11784638 | -1.286 | 1.441 | 0.835 | -0.097 | 0.077 | -0.079 | -0.079 | 0.585 | 0.23 | 0.52 | 0.34 | 0.08 | 0.34 | 0.22 |
| 8 | 131430133 | rs7386870 | 1.391 | 1.095 | 0.693 | 0.243 | -0.003 | 0 | 0 | 0.361 | 0.57 | 0.34 | 0.58 | 0.57 | 0.5 | 0.46 |
| 8 | 131439767 | rs11775398 | -0.242 | 0.966 | 0.922 | 0.252 | 0.011 | 0.06 | 0.06 | 0 | 0.1 | 0.33 | 0.1 | 0.14 | 0.07 | 0.06 |
| 8 | 131444796 | rs12707941 | -0.305 | 0.576 | 0.217 | 0.016 | 0.005 | 0.006 | 0.006 | 0 | 0.57 | 0.41 | 0.56 | 0.58 | 0.59 | 0.7 |
| 8 | 131465722 | rs9643254 | 0.05 | 0.699 | 0.103 | 0.124 | 0 | 0.002 | 0.002 | 0.305 | 0.2 | 0.23 | 0.1 | 0.23 | 0.18 | 0.37 |
| 8 | 131465835 | rs4498576 | 0.8 | 1.095 | -0.411 | -0.059 | -0.039 | 0.007 | 0.007 | 0 | 0.08 | 0.03 | 0.09 | 0.04 | 0.19 | 0.08 |
| 8 | 131467319 | rs10956526 | 0.272 | 1.126 | -0.104 | 0.099 | 0.005 | -0.003 | -0.003 | 0 | 0.79 | 0.76 | 0.7 | 0.89 | 0.66 | 0.84 |
| 8 | 131469692 | rs868104 | -1.199 | 0.612 | -0.293 | 0.023 | 0.009 | 0.002 | 0.002 | 0 | 0.65 | 0.76 | 0.57 | 0.72 | 0.71 | 0.74 |
| 8 | 131473626 | rs4335138 | 1.965 | 0.551 | -0.596 | 0.024 | -0.065 | -0.075 | -0.075 | 0.028 | 0.25 | 0.01 | 0.27 | 0.31 | 0.27 | 0.26 |
| 8 | 131477907 | rs10105826 | 1.137 | 0.68 | 0.222 | -0.269 | 0.027 | 0.008 | 0.008 | 0 | 0.63 | 0.55 | 0.61 | 0.64 | 0.5 | 0.58 |
| 8 | 131480540 | rs6997195 | 2.274 | 0.65 | 0.28 | -0.303 | 0.014 | 0.001 | 0.001 | 0.053 | 0.51 | 0.38 | 0.52 | 0.53 | 0.41 | 0.55 |
| 8 | 131489046 | rs1002142 | 1.85 | 0.474 | 0.321 | -0.271 | -0.001 | 0.001 | 0.001 | 0 | 0.55 | 0.51 | 0.53 | 0.53 | 0.45 | 0.58 |
| 8 | 131489299 | rs13271059 | 0.262 | 0.523 | 0.212 | -0.193 | 0.028 | -0.01 | -0.01 | 0.902 | 0.09 | 0.12 | 0.1 | 0.05 | 0.22 | 0.08 |
| 8 | 131492228 | rs4601326 | 0.035 | 0.528 | -0.068 | -0.131 | -0.008 | -0.024 | -0.024 | 0 | 0.75 | 0.86 | 0.72 | 0.64 | 0.74 | 0.72 |
| 8 | 131493326 | rs17289507 | 1.076 | 0.86 | -0.429 | 0.02 | -0.039 | -0.059 | -0.059 | 0 | 0.1 | 0.01 | 0.04 | 0.21 | 0.11 | 0.23 |
| 8 | 131497231 | rs7386203 | 1.161 | 0.31 | -0.301 | 0.009 | 0 | 0.009 | 0.009 | 0.043 | 0.51 | 0.31 | 0.55 | 0.54 | 0.49 | 0.58 |
| 8 | 131498180 | rs1001812 | 0.757 | 0.512 | -0.23 | 0.119 | 0 | 0.003 | 0.003 | 0 | 0.5 | 0.49 | 0.57 | 0.54 | 0.48 | 0.48 |
| 8 | 131502886 | rs10956533 | 0.573 | 0.752 | -0.599 | 0.236 | -0.005 | 0.028 | 0.028 | 1.179 | 0.52 | 0.31 | 0.56 | 0.6 | 0.47 | 0.49 |
| 8 | 131506062 | rs12677494 | -0.947 | 1.241 | -0.097 | 1.118 | 0.001 | 0.032 | 0.032 | 0 | 0.24 | 0.34 | 0.29 | 0.36 | 0.25 | 0.32 |
| 8 | 131509646 | rs11781952 | -0.836 | 1.258 | 0.296 | 1.356 | -0.005 | 0.014 | 0.014 | 0.578 | 0.35 | 0.52 | 0.36 | 0.41 | 0.31 | 0.56 |
| 8 | 131511510 | rs12680421 | 0.329 | 1.603 | 0.159 | 1.342 | -0.017 | -0.019 | -0.019 | 0 | 0.16 | 0.08 | 0.28 | 0.3 | 0.29 | 0.18 |
| 8 | 131512049 | rs7007052 | -1.292 | 1.306 | 0.005 | 0.813 | 0.003 | 0.003 | 0.003 | 0 | 0.5 | 0.5 | 0.64 | 0.72 | 0.59 | 0.73 |
| 8 | 131512902 | rs9297810 | -0.468 | 1.526 | -0.534 | 0.993 | -0.053 | -0.054 | -0.054 | 0.055 | 0.27 | 0.15 | 0.48 | 0.55 | 0.39 | 0.43 |
| 8 | 131517071 | rs10087780 | -0.843 | 0.957 | -0.12 | 0.824 | 0.004 | 0.008 | 0.008 | 0 | 0.55 | 0.45 | 0.64 | 0.66 | 0.62 | 0.77 |
| 8 | 131517900 | rs6983185 | -0.979 | 0.968 | 0.347 | 1.199 | 0.007 | 0.013 | 0.013 | 0 | 0.17 | 0.25 | 0.21 | 0.23 | 0.16 | 0.31 |
| 8 | 131526147 | rs12541321 | 0.069 | 0.353 | 0.088 | 1.223 | 0.001 | -0.034 | -0.034 | 0 | 0.07 | 0.03 | 0.12 | 0.13 | 0.04 | 0.18 |
| 8 | 131528150 | rs10505550 | -0.337 | 0.335 | 0.174 | 1.25 | 0.004 | -0.006 | -0.006 | 0 | 0.08 | 0.09 | 0.12 | 0.13 | 0.04 | 0.19 |
| 8 | 131529122 | rs4354322 | 0.426 | -0.064 | -0.193 | 1.251 | 0.025 | 0.004 | 0.004 | 0 | 0.2 | 0.11 | 0.12 | 0.17 | 0.17 | 0.11 |
| 8 | 131529258 | rs4512389 | -0.535 | -0.097 | 0.243 | 1.195 | -0.013 | 0.003 | 0.003 | 0 | 0.28 | 0.36 | 0.25 | 0.31 | 0.21 | 0.3 |
| 8 | 131531571 | rs7015138 | -1.433 | -0.023 | -0.579 | 1.42 | 0.027 | 0.106 | 0.106 | 0.726 | 0.52 | 0.88 | 0.6 | 0.77 | 0.46 | 0.65 |
| 8 | 131535329 | rs4255152 | 0.006 | 0.279 | -0.211 | 1.527 | 0.002 | 0.023 | 0.023 | 0 | 0.16 | 0.26 | 0.31 | 0.37 | 0.22 | 0.21 |
| 8 | 131537955 | rs4367565 | -1.107 | 0.058 | -0.322 | 1.254 | 0.077 | -0.021 | -0.021 | 1.715 | 0.34 | 0.88 | 0.23 | 0.37 | 0.23 | 0.32 |
| 8 | 131540050 | rs10094066 | 0.466 | 0.085 | -0.313 | 1.242 | 0.005 | -0.003 | -0.003 | 0 | 0.2 | 0.04 | 0.14 | 0.16 | 0.19 | 0.14 |
| 8 | 131540586 | rs11782944 | -0.153 | 0.112 | -0.105 | 1.252 | -0.021 | -0.009 | -0.009 | 0 | 0.25 | 0.31 | 0.15 | 0.21 | 0.2 | 0.3 |
| 8 | 131540888 | rs16904287 | -0.525 | 0.433 | 0.323 | 1.354 | 0.004 | 0.095 | 0.095 | 4.346 | 0.09 | 0.57 | 0.08 | 0.16 | 0.03 | 0.02 |
| 8 | 131543633 | rs16904288 | -0.558 | 0.437 | 0.312 | 1.361 | 0 | 0.111 | 0.111 | 3.213 | 0.09 | 0.58 | 0.08 | 0.18 | 0.03 | 0.02 |
| 8 | 131544179 | rs7000049 | 0.324 | 1.326 | -0.884 | 1.232 | -0.011 | 0.006 | 0.006 | 0.12 | 0.65 | 0.72 | 0.68 | 0.71 | 0.56 | 0.76 |
| 8 | 131545671 | rs4320562 | -0.598 | 0.908 | -1.141 | 1.359 | -0.038 | -0.057 | -0.057 | 0 | 0.09 | 0.01 | 0.2 | 0.21 | 0.11 | 0.13 |
| 8 | 131550333 | rs6983862 | -0.18 | 0.724 | -0.777 | 1.399 | -0.002 | 0.009 | 0.009 | 0 | 0.19 | 0.22 | 0.13 | 0.29 | 0.16 | 0.13 |
| 8 | 131553214 | rs6981998 | -0.548 | 0.447 | -0.681 | 0.809 | -0.001 | -0.001 | -0.001 | 0.668 | 0.18 | 0.17 | 0.14 | 0.21 | 0.18 | 0.23 |
| 8 | 131554104 | rs6987139 | 0.225 | -0.416 | -0.517 | -0.364 | -0.023 | -0.083 | -0.083 | 0 | 0.37 | 0.41 | 0.15 | 0.05 | 0.23 | 0.3 |
| 8 | 131554154 | rs4615594 | -0.027 | -0.175 | -0.126 | -0.14 | -0.001 | 0.003 | 0.003 | 0 | 0.05 | 0.11 | 0.03 | 0.06 | 0.04 | 0.12 |
| 8 | 131554761 | rs10956535 | 0.001 | 0.414 | -0.033 | 0.022 | -0.001 | -0.001 | -0.001 | 0 | 0.6 | 0.38 | 0.59 | 0.47 | 0.62 | 0.59 |
| 8 | 131559901 | rs4255153 | -0.689 | 0.08 | 0.107 | -0.193 | -0.002 | -0.001 | -0.001 | 0 | 0.3 | 0.67 | 0.34 | 0.35 | 0.28 | 0.29 |
| 8 | 131576964 | rs2959319 | -0.138 | -0.104 | -0.163 | -0.118 | -0.024 | -0.015 | -0.015 | 0 | 0.31 | 0.1 | 0.29 | 0.33 | 0.3 | 0.3 |
| 8 | 131578186 | rs2975532 | -0.418 | -0.162 | 0.213 | -0.122 | -0.007 | -0.006 | -0.006 | 0 | 0.36 | 0.45 | 0.33 | 0.34 | 0.3 | 0.3 |
| 8 | 131588565 | rs2975519 | 0.076 | 0.022 | 0.631 | -0.074 | 0.005 | 0.054 | 0.054 | 0 | 0.13 | 0.21 | 0.15 | 0.28 | 0.14 | 0.17 |
| 8 | 131590662 | rs10505551 | 0.511 | -0.018 | 0.106 | -0.478 | -0.007 | 0.034 | 0.034 | 0 | 0.18 | 0.13 | 0.09 | 0.04 | 0.2 | 0.22 |
| 8 | 131592090 | rs7823789 | 0.265 | -0.06 | 0.414 | -0.496 | 0.008 | -0.109 | -0.109 | 0 | 0.2 | 0.45 | 0.12 | 0.04 | 0.2 | 0.22 |
| 8 | 131599918 | rs11988629 | -0.825 | -0.03 | 0.277 | -0.439 | 0.001 | 0 | 0 | 0 | 0.33 | 0.59 | 0.26 | 0.33 | 0.36 | 0.4 |
| 8 | 131607268 | rs7836657 | -0.129 | -0.261 | 0.286 | -0.534 | -0.003 | -0.033 | -0.033 | 0.177 | 0.1 | 0.37 | 0.06 | 0.01 | 0.05 | 0.02 |
| 8 | 131609951 | rs7014624 | 0.414 | -0.289 | 0.158 | -0.511 | -0.021 | -0.034 | -0.034 | 0 | 0.09 | 0.16 | 0.04 | 0.01 | 0.05 | 0.02 |
| 8 | 131611401 | rs12547913 | 0.556 | -0.221 | -0.03 | -0.486 | 0.001 | -0.015 | -0.015 | 0.12 | 0.12 | 0.13 | 0.13 | 0.28 | 0.12 | 0.17 |

| | | | | | | | | | | | | | | | | |
|---|---|---|---|---|---|---|---|---|---|---|---|---|---|---|---|---|
| 8 | 131620314 | rs10098680 | -0.755 | -0.291 | -0.199 | -0.465 | -0.011 | -0.004 | -0.004 | 1.129 | 0.23 | 0.37 | 0.19 | 0.22 | 0.19 | 0.27 |
| 8 | 131623388 | rs10102885 | -0.751 | 0 | -1.093 | -0.276 | 0.006 | 0.021 | 0.021 | 0 | 0.4 | 0.65 | 0.27 | 0.24 | 0.38 | 0.49 |
| 8 | 131627539 | rs4273866 | -0.798 | 0.343 | -1.122 | -0.032 | 0.004 | 0.024 | 0.024 | 0 | 0.53 | 0.73 | 0.38 | 0.45 | 0.51 | 0.67 |
| 8 | 131630914 | rs12549418 | -0.58 | 0.828 | -1.002 | 0.586 | -0.002 | -0.006 | -0.006 | 0 | 0.13 | 0.09 | 0.11 | 0.2 | 0.14 | 0.24 |
| 8 | 131636381 | rs16904323 | -0.553 | 0.925 | -1.16 | 0.468 | -0.004 | -0.006 | -0.006 | 0 | 0.13 | 0.1 | 0.12 | 0.18 | 0.16 | 0.21 |
| 8 | 131640843 | rs4295676 | 0.263 | 0.914 | -1.357 | 0.207 | -0.009 | 0.056 | 0.056 | 0.743 | 0.6 | 0.24 | 0.68 | 0.72 | 0.59 | 0.56 |
| 8 | 131641522 | rs10097323 | 0.426 | 0.812 | -1.111 | 0.144 | 0.001 | 0.044 | 0.044 | 0.649 | 0.58 | 0.28 | 0.68 | 0.72 | 0.59 | 0.56 |
| 8 | 131642423 | rs11990184 | -0.628 | 0.72 | -0.123 | -0.052 | 0.057 | -0.005 | -0.005 | 0.056 | 0.08 | 0.42 | 0.07 | 0 | 0.04 | 0.06 |
| 8 | 131642621 | rs4310223 | -0.094 | 0.727 | -0.427 | -0.077 | -0.002 | 0.009 | 0.009 | 0 | 0.71 | 0.79 | 0.8 | 0.79 | 0.68 | 0.74 |
| 8 | 131645079 | rs4460390 | 0.7 | 0.832 | -0.956 | 0.003 | -0.004 | 0.072 | 0.072 | 0.446 | 0.57 | 0.24 | 0.67 | 0.71 | 0.59 | 0.56 |
| 8 | 131649645 | rs7835450 | -1.119 | 0.621 | -0.25 | 0.722 | -0.014 | 0.022 | 0.022 | 0 | 0.15 | 0.22 | 0.18 | 0.23 | 0.09 | 0.18 |
| 8 | 131650813 | rs6986527 | 0.756 | 0.916 | -0.735 | 0.7 | 0.002 | -0.007 | -0.007 | 0 | 0.46 | 0.53 | 0.38 | 0.37 | 0.48 | 0.39 |
| 8 | 131651727 | rs13255815 | 0.453 | 1.138 | -0.837 | 0.958 | 0.001 | -0.002 | -0.002 | 0.545 | 0.57 | 0.59 | 0.62 | 0.53 | 0.65 | 0.53 |
| 8 | 131652191 | rs16904332 | -1.248 | 1.024 | -0.005 | 1.479 | 0.012 | 0.092 | 0.092 | 0.046 | 0.11 | 0.27 | 0.17 | 0.24 | 0.09 | 0.09 |
| 8 | 131654000 | rs10093085 | -0.122 | 1.06 | -0.97 | 0.496 | -0.004 | 0.033 | 0.033 | 0 | 0.44 | 0.36 | 0.42 | 0.78 | 0.41 | 0.61 |
| 8 | 131654160 | rs10106564 | 1.257 | 1.204 | -1.246 | 0.982 | 0.006 | -0.028 | -0.028 | 0 | 0.22 | 0.13 | 0.18 | 0.29 | 0.17 | 0.25 |
| 8 | 131654547 | rs4282589 | 0.409 | 1.231 | -1.226 | 0.973 | -0.009 | 0.023 | 0.023 | 0 | 0.29 | 0.33 | 0.33 | 0.54 | 0.22 | 0.33 |
| 8 | 131654755 | rs11997043 | 0.442 | 0.742 | -1.161 | 0.769 | 0.001 | 0.001 | 0.001 | 0 | 0.23 | 0.21 | 0.24 | 0.17 | 0.19 | 0.24 |
| 8 | 131657988 | rs6986586 | 0.676 | 0.712 | -1.156 | 0.743 | -0.021 | -0.018 | -0.018 | 0 | 0.2 | 0.21 | 0.07 | 0.07 | 0.11 | 0.15 |
| 8 | 131659108 | rs17312621 | -0.259 | 0.733 | -1.173 | 0.761 | 0.012 | 0.033 | 0.033 | 0 | 0.35 | 0.23 | 0.16 | 0.22 | 0.3 | 0.41 |
| 8 | 131660064 | rs4307362 | -0.209 | 0.822 | -1.181 | 0.754 | 0.017 | 0.013 | 0.013 | 0 | 0.34 | 0.23 | 0.16 | 0.32 | 0.29 | 0.42 |
| 8 | 131661241 | rs7012782 | 0.648 | 0.92 | -1.274 | 0.504 | 0 | -0.005 | -0.005 | 0 | 0.1 | 0.08 | 0.13 | 0.05 | 0.12 | 0.1 |
| 8 | 131668278 | rs7841283 | 1.041 | 0.8 | -1.259 | 0.557 | 0.007 | -0.005 | -0.005 | 0.7 | 0.16 | 0.19 | 0.1 | 0.15 | 0.18 | 0.24 |
| 8 | 131677999 | rs7813092 | -0.322 | 0.805 | -1.173 | 0.552 | -0.001 | -0.001 | -0.001 | 0 | 0.41 | 0.54 | 0.38 | 0.42 | 0.38 | 0.52 |
| 8 | 131682895 | rs6470834 | 0.412 | 0.638 | -1.129 | 0.325 | -0.007 | 0.021 | 0.021 | 0 | 0.11 | 0.08 | 0.14 | 0.05 | 0.14 | 0.1 |
| 8 | 131684705 | rs7844037 | 0.708 | 0.602 | -0.763 | 0.535 | 0.065 | 0.128 | 0.128 | 0.029 | 0.06 | 0.29 | 0.24 | 0.12 | 0.07 | 0.11 |
| 8 | 131686649 | rs10095393 | 0.001 | -0.196 | -0.695 | 0.153 | -0.058 | 0.013 | 0.013 | 0 | 0.22 | 0.44 | 0.14 | 0.22 | 0.12 | 0.25 |
| 8 | 131687912 | rs16904337 | -0.284 | -0.306 | -0.745 | 0.222 | -0.058 | 0.039 | 0.039 | 0 | 0.24 | 0.44 | 0.14 | 0.35 | 0.13 | 0.3 |
| 8 | 131691318 | rs10086514 | -1.311 | 0.025 | -1.122 | -0.086 | 0.007 | 0.013 | 0.013 | 0 | 0.63 | 0.59 | 0.47 | 0.52 | 0.6 | 0.71 |
| 8 | 131693352 | rs10101790 | -0.397 | -0.23 | -1.094 | -0.221 | -0.045 | -0.061 | -0.061 | 0 | 0.34 | 0.46 | 0.18 | 0.17 | 0.2 | 0.33 |
| 8 | 131695518 | rs11786562 | 1.97 | -0.018 | -1.28 | -0.114 | 0.003 | 0.004 | 0.004 | 0 | 0.08 | 0.03 | 0.05 | 0.06 | 0.08 | 0.06 |
| 8 | 131696746 | rs4736578 | 1.227 | 0.14 | -1.385 | -0.162 | -0.017 | 0.045 | 0.045 | 0 | 0.28 | 0.1 | 0.18 | 0.21 | 0.3 | 0.32 |
| 8 | 131697816 | rs9649963 | 0.372 | 0.186 | -1.399 | 0.136 | -0.019 | -0.04 | -0.04 | 0.085 | 0.33 | 0.19 | 0.39 | 0.43 | 0.39 | 0.43 |
| 8 | 131698706 | rs6470838 | -1.48 | 0.079 | -1.363 | 0.08 | -0.049 | -0.046 | -0.046 | 2.487 | 0.65 | 0.8 | 0.48 | 0.52 | 0.57 | 0.71 |
| 8 | 131701038 | rs10105307 | 1.522 | -0.052 | -1.231 | 0.011 | 0.005 | 0.003 | 0.003 | 0 | 0.13 | 0.12 | 0.09 | 0.14 | 0.1 | 0.17 |
| 8 | 131709637 | rs1467065 | 1.063 | 0.013 | -1.079 | 0.086 | -0.006 | -0.006 | -0.006 | 0 | 0.11 | 0.12 | 0.07 | 0.09 | 0.09 | 0.07 |
| 8 | 131713531 | rs12678126 | 2.246 | 0.09 | -1.737 | -0.085 | -0.052 | -0.083 | -0.083 | 0 | 0.14 | 0.01 | 0.2 | 0.27 | 0.15 | 0.13 |
| 8 | 131720773 | rs12056463 | 0.717 | -0.252 | -1.263 | -0.711 | 0.006 | -0.034 | -0.034 | 0.08 | 0.13 | 0.23 | 0.18 | 0.06 | 0.16 | 0.1 |
| 8 | 131721915 | rs10097031 | 1.233 | -0.372 | -1.375 | -0.798 | -0.004 | 0.022 | 0.022 | 0 | 0.27 | 0.16 | 0.17 | 0.22 | 0.29 | 0.32 |
| 8 | 131733304 | rs6470841 | 0.197 | -0.381 | -1.424 | -0.798 | -0.007 | 0.01 | 0.01 | 0 | 0.31 | 0.26 | 0.3 | 0.27 | 0.37 | 0.38 |
| 8 | 131743619 | rs925301 | -0.765 | -0.284 | -1.876 | -0.716 | 0.001 | 0.015 | 0.015 | 0.065 | 0.51 | 0.47 | 0.4 | 0.42 | 0.49 | 0.67 |
| 8 | 131745204 | rs2917062 | -1.65 | -0.196 | -2.529 | -0.518 | -0.027 | -0.03 | -0.03 | 0 | 0.68 | 0.82 | 0.68 | 0.65 | 0.64 | 0.83 |
| 8 | 131745284 | rs7813720 | -1.222 | -0.208 | -2.526 | -0.546 | -0.047 | -0.006 | -0.006 | 0 | 0.64 | 0.8 | 0.48 | 0.45 | 0.57 | 0.71 |
| 8 | 131747584 | rs12235033 | 1.802 | -0.411 | -2.473 | -0.626 | -0.004 | -0.028 | -0.028 | 0.025 | 0.06 | 0.01 | 0.05 | 0.14 | 0.04 | 0.1 |
| 8 | 131748652 | rs1588370 | -1.666 | -0.156 | -2.342 | -0.532 | -0.027 | -0.029 | -0.029 | 0 | 0.69 | 0.82 | 0.68 | 0.65 | 0.64 | 0.83 |
| 8 | 131752271 | rs10217074 | 0.156 | -0.498 | -2.588 | -1.025 | 0.015 | 0.013 | 0.013 | 0 | 0.41 | 0.38 | 0.22 | 0.17 | 0.26 | 0.34 |
| 8 | 131753029 | rs17224106 | 0.933 | -0.357 | -2.57 | -0.803 | 0.004 | 0.067 | 0.067 | 0 | 0.25 | 0.03 | 0.2 | 0.1 | 0.24 | 0.14 |
| 8 | 131756652 | rs11787478 | 1.169 | 0.117 | -2.25 | -0.502 | -0.006 | 0.02 | 0.02 | 0 | 0.16 | 0.02 | 0.16 | 0.08 | 0.17 | 0.09 |
| 8 | 131756936 | rs2889548 | 1.381 | 1.193 | -1.593 | 0.063 | -0.024 | 0.006 | 0.006 | 0 | 0.15 | 0.06 | 0.12 | 0.19 | 0.2 | 0.27 |
| 8 | 131759529 | rs1818404 | 0.632 | 1.205 | -1.792 | 0.082 | 0.064 | 0.09 | 0.09 | 0 | 0.46 | 0.32 | 0.31 | 0.21 | 0.36 | 0.4 |
| 8 | 131761999 | rs1509844 | -0.61 | 1.467 | -2.07 | 0.794 | -0.076 | -0.086 | -0.086 | 0 | 0.71 | 0.91 | 0.7 | 0.6 | 0.65 | 0.8 |
| 8 | 131763406 | rs1836626 | -0.121 | 1.362 | -2.027 | 0.653 | -0.071 | -0.012 | -0.012 | 0.001 | 0.65 | 0.82 | 0.48 | 0.43 | 0.57 | 0.67 |
| 8 | 131767789 | rs1509840 | -0.789 | 1.132 | -2.022 | 0.597 | -0.08 | -0.1 | -0.1 | 0 | 0.74 | 0.91 | 0.7 | 0.61 | 0.67 | 0.84 |
| 8 | 131769694 | rs2055831 | -0.514 | 1.133 | -1.832 | 0.441 | -0.035 | -0.041 | -0.041 | 0 | 0.71 | 0.83 | 0.7 | 0.58 | 0.68 | 0.85 |
| 8 | 131771222 | rs11786743 | 1.105 | 0.721 | -1.819 | -0.268 | -0.009 | 0.057 | 0.057 | 2.547 | 0.18 | 0.09 | 0.11 | 0 | 0.19 | 0.06 |
| 8 | 131774876 | rs16904352 | -0.233 | 0.171 | -1.341 | 0.042 | -0.023 | -0.032 | -0.032 | 0.465 | 0.43 | 0.51 | 0.33 | 0.2 | 0.34 | 0.35 |
| 8 | 131778235 | rs11774692 | 0.208 | 0.312 | -1.352 | 0.072 | 0.006 | 0.009 | 0.009 | 0 | 0.27 | 0.2 | 0.23 | 0.18 | 0.27 | 0.22 |
| 8 | 131778538 | rs1509848 | -0.372 | 0.868 | -1.614 | 0.332 | -0.008 | -0.013 | -0.013 | 0 | 0.31 | 0.28 | 0.44 | 0.39 | 0.36 | 0.57 |
| 8 | 131778855 | rs1560984 | -0.09 | 1.153 | -1.462 | 0.368 | -0.008 | -0.003 | -0.003 | 2.579 | 0.19 | 0.21 | 0.37 | 0.24 | 0.27 | 0.41 |
| 8 | 131779660 | rs6470848 | 0.289 | 0.869 | -1.305 | -0.313 | -0.004 | -0.059 | -0.059 | 0.193 | 0.23 | 0.51 | 0.18 | 0.12 | 0.22 | 0.07 |
| 8 | 131779957 | rs2572862 | 0.235 | 1.136 | -1.509 | -0.054 | 0.002 | -0.006 | -0.006 | 1.187 | 0.08 | 0.09 | 0.12 | 0.05 | 0.09 | 0.12 |
| 8 | 131780242 | rs11997892 | 0.557 | 1.177 | -2.007 | -0.013 | -0.013 | 0.03 | 0.03 | 1.916 | 0.58 | 0.73 | 0.44 | 0.29 | 0.57 | 0.28 |
| 8 | 131780258 | rs10097218 | 0.387 | 1.155 | -1.939 | 0.169 | -0.012 | 0.011 | 0.011 | 1.588 | 0.62 | 0.76 | 0.48 | 0.32 | 0.61 | 0.32 |
| 8 | 131781235 | rs17225213 | 0.449 | 1.14 | -1.915 | 0.162 | -0.007 | 0.011 | 0.011 | 1.664 | 0.62 | 0.69 | 0.47 | 0.32 | 0.61 | 0.31 |
| 8 | 131792219 | rs263247 | 0.254 | 1.164 | -2.23 | 0.263 | -0.017 | -0.014 | -0.014 | 0 | 0.65 | 0.84 | 0.57 | 0.35 | 0.66 | 0.39 |
| 8 | 131797568 | rs12546275 | 0.791 | 1.314 | -2.562 | 0.205 | -0.03 | -0.051 | -0.051 | 0 | 0.09 | 0.01 | 0.12 | 0.16 | 0.12 | 0.12 |
| 8 | 131799767 | rs12680002 | 1.014 | 1.179 | -2.35 | 0.512 | -0.008 | -0.044 | -0.044 | 0.311 | 0.12 | 0.04 | 0.23 | 0.31 | 0.14 | 0.16 |
| 8 | 131802394 | rs16904358 | 0.043 | 1.218 | -2.313 | 0.544 | -0.02 | -0.065 | -0.065 | 0.19 | 0.21 | 0.07 | 0.28 | 0.36 | 0.21 | 0.3 |
| 8 | 131812033 | rs263238 | 0.72 | 0.992 | -1.762 | 0.879 | -0.01 | -0.054 | -0.054 | 0 | 0.13 | 0.03 | 0.16 | 0.29 | 0.13 | 0.32 |

| | | | | | | | | | | | | | | | | |
|---|---|---|---|---|---|---|---|---|---|---|---|---|---|---|---|---|
| 8 | 131812287 | rs16904362 | 0.632 | 0.909 | -2.102 | 0.347 | -0.009 | 0.032 | 0.032 | 0 | 0.21 | 0.08 | 0.15 | 0.12 | 0.23 | 0.08 |
| 8 | 131815333 | rs12678910 | 0.391 | 0.974 | -2.452 | 0.392 | -0.039 | -0.049 | -0.049 | 0 | 0.14 | 0.01 | 0.22 | 0.2 | 0.13 | 0.14 |
| 8 | 131816296 | rs11986118 | 2.004 | 0.706 | -1.661 | -0.152 | -0.01 | -0.082 | -0.082 | 0 | 0.17 | 0.32 | 0.11 | 0.03 | 0.14 | 0.04 |
| 8 | 131817691 | rs17226545 | -0.564 | 0.623 | -1.21 | -0.198 | 0.005 | -0.002 | -0.002 | 0.141 | 0.6 | 0.35 | 0.58 | 0.61 | 0.63 | 0.53 |
| 8 | 131818715 | rs263234 | 1.436 | 0.583 | -1.341 | -0.217 | -0.048 | -0.114 | -0.114 | 0.012 | 0.18 | 0.01 | 0.21 | 0.3 | 0.19 | 0.37 |
| 8 | 131820892 | rs1509851 | 2.052 | 0.588 | -1.154 | -0.213 | -0.018 | -0.043 | -0.043 | 0 | 0.08 | 0.01 | 0.17 | 0.2 | 0.09 | 0.1 |
| 8 | 131821169 | rs263250 | -0.637 | 0.587 | -0.867 | -0.757 | 0.001 | 0.026 | 0.026 | 0 | 0.57 | 0.33 | 0.56 | 0.64 | 0.55 | 0.69 |
| 8 | 131822757 | rs16904364 | 0.714 | 0.741 | -1.08 | -0.726 | -0.04 | 0.05 | 0.05 | 0 | 0.29 | 0.07 | 0.17 | 0.13 | 0.27 | 0.22 |
| 8 | 131823126 | rs263252 | -1.951 | 0.716 | -0.452 | -0.557 | -0.002 | -0.034 | -0.034 | 0 | 0.81 | 0.73 | 0.82 | 0.91 | 0.82 | 0.9 |
| 8 | 131826621 | rs1435446 | 1.148 | 0.716 | -0.784 | -0.504 | -0.002 | -0.001 | -0.001 | 0 | 0.22 | 0.19 | 0.23 | 0.27 | 0.27 | 0.21 |
| 8 | 131830612 | rs166405 | -1.693 | 0.768 | -1.463 | -0.473 | -0.001 | -0.014 | -0.014 | 0 | 0.83 | 0.83 | 0.84 | 0.91 | 0.82 | 0.9 |
| 8 | 131833036 | rs3829032 | -0.129 | 0.755 | -1.851 | -0.174 | -0.023 | 0.019 | 0.019 | 0.175 | 0.26 | 0.06 | 0.14 | 0.14 | 0.21 | 0.2 |
| 8 | 131835318 | rs166403 | -0.545 | 0.794 | -1.878 | -0.172 | -0.01 | 0.008 | 0.008 | 0 | 0.84 | 0.94 | 0.82 | 0.83 | 0.81 | 0.86 |
| 8 | 131836353 | rs6997439 | 0.289 | 0.854 | -1.665 | -0.024 | 0.003 | 0.009 | 0.009 | 0 | 0.19 | 0.23 | 0.24 | 0.26 | 0.22 | 0.18 |
| 8 | 131836730 | rs263264 | 1.292 | 0.783 | -2.47 | -0.198 | -0.085 | -0.126 | -0.126 | 0.084 | 0.22 | 0.02 | 0.3 | 0.29 | 0.25 | 0.48 |
| 8 | 131843264 | rs10108440 | 0.559 | 0.75 | -2.344 | -0.133 | 0.003 | 0.012 | 0.012 | 0 | 0.2 | 0.25 | 0.24 | 0.26 | 0.22 | 0.19 |
| 8 | 131844517 | rs430659 | -0.759 | 0.2 | -1.734 | -0.365 | 0 | 0.002 | 0.002 | 4.978 | 0.59 | 0.63 | 0.67 | 0.56 | 0.6 | 0.76 |
| 8 | 131854234 | rs263257 | -0.737 | 0.197 | -1.729 | -0.361 | 0.001 | 0.003 | 0.003 | 5.893 | 0.59 | 0.63 | 0.67 | 0.55 | 0.59 | 0.75 |
| 8 | 131860807 | rs12548296 | 0.241 | 0.19 | -1.298 | -0.375 | 0 | 0.002 | 0.002 | 0.728 | 0.29 | 0.3 | 0.29 | 0.26 | 0.3 | 0.22 |
| 8 | 131865018 | rs263239 | -1.113 | 0.147 | -1.527 | -0.424 | 0.001 | 0.011 | 0.011 | 0 | 0.83 | 0.94 | 0.83 | 0.84 | 0.83 | 0.85 |
| 8 | 131865742 | rs448852 | 0.012 | -0.084 | -1.072 | -0.5 | -0.015 | -0.031 | -0.031 | 0 | 0.76 | 0.53 | 0.76 | 0.81 | 0.78 | 0.8 |
| 8 | 131866339 | rs412723 | -0.68 | 0.011 | -1.463 | -0.356 | -0.014 | -0.066 | -0.066 | 6.179 | 0.8 | 0.94 | 0.82 | 0.59 | 0.77 | 0.8 |
| 8 | 131871971 | rs12549142 | 0.094 | -0.063 | -1.402 | -0.561 | 0.002 | 0.043 | 0.043 | 3.206 | 0.48 | 0.65 | 0.4 | 0.24 | 0.49 | 0.23 |
| 8 | 131875344 | rs1588369 | 0.035 | -0.049 | -1.394 | -0.487 | 0 | -0.001 | -0.001 | 0.979 | 0.43 | 0.52 | 0.34 | 0.47 | 0.46 | 0.26 |
| 8 | 131877244 | rs11987135 | 0.18 | -0.159 | -1.137 | -0.773 | 0 | -0.033 | -0.033 | 0.003 | 0.38 | 0.48 | 0.32 | 0.24 | 0.4 | 0.21 |
| 8 | 131879068 | rs10505566 | 0.839 | 0.05 | -0.841 | -0.018 | -0.07 | -0.072 | -0.072 | 0 | 0.14 | 0.02 | 0.16 | 0.2 | 0.21 | 0.11 |
| 8 | 131883578 | rs976268 | 1.201 | -0.35 | -0.674 | -0.907 | -0.039 | -0.03 | -0.03 | 0.054 | 0.43 | 0.22 | 0.44 | 0.61 | 0.46 | 0.33 |
| 8 | 131883786 | rs976266 | 0.27 | -0.456 | -0.407 | -1.054 | -0.007 | -0.076 | -0.076 | 2.837 | 0.68 | 0.65 | 0.74 | 0.93 | 0.65 | 0.84 |
| 8 | 131884616 | rs2917048 | 0.114 | -0.411 | -0.322 | -0.939 | -0.006 | 0.06 | 0.06 | 0.025 | 0.24 | 0.35 | 0.3 | 0.32 | 0.19 | 0.51 |
| 8 | 131887274 | rs2164306 | -0.935 | -0.49 | -0.23 | -1.019 | -0.003 | 0.002 | 0.002 | 1.743 | 0.39 | 0.61 | 0.45 | 0.38 | 0.31 | 0.59 |
| 8 | 131888133 | rs7461547 | 1.297 | -0.436 | -0.373 | -1.249 | -0.007 | -0.056 | -0.056 | 0 | 0.11 | 0.21 | 0.08 | 0 | 0.09 | 0.02 |
| 8 | 131889668 | rs2860416 | 1.028 | -0.455 | -0.442 | -1.101 | -0.021 | -0.019 | -0.019 | 0 | 0.17 | 0.05 | 0.17 | 0.19 | 0.16 | 0.41 |
| 8 | 131890170 | rs7842798 | 0.734 | -0.47 | -0.387 | -1.13 | 0.005 | 0.002 | 0.002 | 0.206 | 0.56 | 0.73 | 0.56 | 0.63 | 0.56 | 0.34 |
| 8 | 131892722 | rs1369450 | 0.739 | -0.443 | -0.698 | -1.117 | 0.002 | 0.001 | 0.001 | 0.221 | 0.55 | 0.48 | 0.54 | 0.63 | 0.56 | 0.35 |
| 8 | 131893562 | rs10098645 | 1.406 | -0.542 | -0.858 | -1.114 | -0.024 | -0.006 | -0.006 | 0 | 0.42 | 0.27 | 0.43 | 0.63 | 0.44 | 0.32 |
| 8 | 131894883 | rs965814 | -0.727 | -0.633 | -0.921 | -1.262 | 0.026 | 0.083 | 0.083 | 7.184 | 0.4 | 0.8 | 0.42 | 0.22 | 0.33 | 0.51 |
| 8 | 131895923 | rs4130328 | -0.654 | -0.654 | -1.071 | -1.318 | -0.018 | -0.009 | -0.009 | 0.219 | 0.81 | 0.93 | 0.84 | 0.86 | 0.77 | 0.82 |
| 8 | 131897082 | rs4128982 | 0.226 | -0.677 | -0.999 | -1.336 | -0.001 | -0.003 | -0.003 | 0 | 0.7 | 0.67 | 0.76 | 0.86 | 0.68 | 0.8 |
| 8 | 131903134 | rs7016933 | 0.221 | -0.621 | -0.908 | -1.191 | 0 | -0.001 | -0.001 | 1.099 | 0.64 | 0.4 | 0.64 | 0.67 | 0.69 | 0.47 |
| 8 | 131905190 | rs6470860 | 1.026 | -0.616 | -1.086 | -1.226 | 0.035 | 0.015 | 0.015 | 1.618 | 0.49 | 0.19 | 0.51 | 0.62 | 0.6 | 0.37 |
| 8 | 131917676 | rs13274422 | 1.636 | -0.635 | -1.144 | -1.233 | -0.052 | -0.056 | -0.056 | 10.903 | 0.15 | 0.02 | 0.15 | 0.2 | 0.15 | 0.38 |
| 8 | 131918428 | rs13248582 | 0.536 | -0.568 | -0.709 | -1.314 | 0.041 | 0.025 | 0.025 | 1.389 | 0.5 | 0.25 | 0.52 | 0.62 | 0.58 | 0.37 |
| 8 | 131921956 | rs12547243 | -0.272 | -0.578 | -0.734 | -1.321 | 0.016 | -0.01 | -0.01 | 0 | 0.63 | 0.33 | 0.63 | 0.68 | 0.7 | 0.47 |
| 8 | 131922027 | rs12545028 | 0.79 | -0.245 | -0.421 | -1.043 | -0.013 | 0.026 | 0.026 | 0 | 0.12 | 0.07 | 0.13 | 0.06 | 0.16 | 0.13 |
| 8 | 131923069 | rs7461303 | 0.305 | -0.237 | -0.532 | -0.985 | 0.004 | 0.006 | 0.006 | 5.141 | 0.14 | 0.13 | 0.13 | 0.06 | 0.11 | 0.11 |
| 8 | 131928401 | rs7459573 | -0.709 | -0.003 | -0.276 | -0.934 | 0.002 | 0 | 0 | 0 | 0.61 | 0.65 | 0.65 | 0.64 | 0.63 | 0.44 |
| 8 | 131930260 | rs7831098 | 0.04 | 0.241 | -0.1 | -0.409 | 0.004 | 0.006 | 0.006 | 0.215 | 0.38 | 0.44 | 0.45 | 0.58 | 0.38 | 0.31 |
| 8 | 131936854 | rs6470861 | -0.867 | 0.303 | -1.325 | -0.392 | 0.013 | 0.047 | 0.047 | 0.019 | 0.52 | 0.73 | 0.59 | 0.65 | 0.54 | 0.4 |
| 8 | 131936877 | rs7459733 | 0.256 | 1.056 | -2.006 | -0.08 | -0.046 | -0.081 | -0.081 | 0.961 | 0.32 | 0.11 | 0.33 | 0.54 | 0.36 | 0.25 |
| 8 | 131945264 | rs10087086 | -0.478 | 0.97 | -1.107 | -0.221 | -0.024 | 0.033 | 0.033 | 0 | 0.1 | 0.04 | 0.06 | 0.06 | 0.16 | 0.09 |
| 8 | 131958259 | rs16929 | -0.772 | 0.942 | -1.193 | -0.194 | -0.032 | 0.045 | 0.045 | 0 | 0.17 | 0.03 | 0.24 | 0.11 | 0.23 | 0.21 |
| 8 | 131960656 | rs7815277 | -1.293 | 0.835 | -0.146 | -0.077 | 0.038 | -0.002 | -0.002 | 0 | 0.08 | 0.64 | 0.11 | 0.05 | 0.09 | 0.06 |
| 8 | 131961460 | rs4736462 | 0.984 | 0.434 | 0.268 | -0.39 | -0.02 | -0.035 | -0.035 | 0.721 | 0.84 | 0.36 | 0.8 | 0.93 | 0.89 | 0.85 |
| 8 | 131968469 | rs6988340 | 1.836 | 0.598 | 0.849 | -0.521 | 0.029 | 0.011 | 0.011 | 0.795 | 0.91 | 0.44 | 0.92 | 0.95 | 0.92 | 0.92 |
| 8 | 131970365 | rs7460718 | 1.877 | 0.544 | 0.832 | -0.474 | -0.001 | -0.004 | -0.004 | 0 | 0.89 | 0.37 | 0.88 | 0.94 | 0.91 | 0.9 |
| 8 | 131972500 | rs12156272 | 1.507 | 0.152 | 0.519 | -0.765 | 0.005 | 0 | 0 | 0.038 | 0.75 | 0.27 | 0.75 | 0.94 | 0.82 | 0.81 |
| 8 | 131987375 | rs6986982 | -1.685 | -0.186 | 0.696 | -0.877 | -0.055 | -0.08 | -0.08 | 0.795 | 0.26 | 0.52 | 0.2 | 0.08 | 0.16 | 0.23 |
| 8 | 131992819 | rs11775729 | -1.277 | -0.17 | 0.803 | -0.841 | 0.071 | 0.11 | 0.11 | 0.825 | 0.06 | 0.43 | 0.06 | 0.07 | 0.03 | 0.1 |
| 8 | 131994393 | rs16904391 | -2.144 | -0.185 | 0.702 | -0.85 | -0.017 | -0.007 | -0.007 | 0 | 0.11 | 0.56 | 0.11 | 0.07 | 0.06 | 0.14 |
| 8 | 131997225 | rs7840509 | -2.212 | -0.175 | 0.663 | -0.874 | -0.019 | -0.013 | -0.013 | 0 | 0.11 | 0.55 | 0.1 | 0.07 | 0.08 | 0.14 |
| 8 | 132000301 | rs17327852 | -0.578 | -0.214 | 0.452 | -0.906 | 0.034 | 0.08 | 0.08 | 0 | 0.16 | 0.01 | 0.11 | 0.01 | 0.09 | 0.1 |
| 8 | 132002334 | rs3750889 | -0.73 | -0.113 | -0.807 | -1.83 | -0.119 | 0.174 | 0.174 | 0.047 | 0.65 | 0.94 | 0.66 | 0.88 | 0.57 | 0.72 |
| 8 | 132005210 | rs6988204 | 0.836 | 0.025 | -0.357 | -1.698 | -0.004 | 0.01 | 0.01 | 0.04 | 0.4 | 0.41 | 0.47 | 0.8 | 0.42 | 0.49 |
| 8 | 132006323 | rs12680529 | -1.451 | -0.012 | -0.177 | -1.185 | -0.005 | 0.084 | 0.084 | 1.489 | 0.1 | 0.54 | 0.08 | 0.15 | 0.06 | 0.14 |
| 8 | 132011121 | rs1329804 | 0.446 | 0.058 | 0.705 | -0.795 | 0 | -0.006 | -0.006 | 0 | 0.43 | 0.55 | 0.49 | 0.7 | 0.46 | 0.57 |
| 8 | 132011279 | rs9297816 | 1.165 | -0.151 | 0.735 | 0.287 | -0.004 | -0.001 | -0.001 | 0 | 0.31 | 0.32 | 0.43 | 0.36 | 0.35 | 0.34 |
| 8 | 132020255 | rs12549392 | 0.479 | -0.136 | 0.707 | 0.289 | -0.005 | 0.006 | 0.006 | 0.011 | 0.4 | 0.35 | 0.42 | 0.64 | 0.43 | 0.54 |
| 8 | 132022101 | rs1028945 | -0.117 | -0.171 | 0.74 | 0.28 | -0.029 | 0.175 | 0.175 | 1.467 | 0.47 | 0.8 | 0.49 | 0.79 | 0.46 | 0.65 |
| 8 | 132022957 | rs1023095 | -0.024 | -0.166 | 0.77 | 0.281 | -0.003 | 0.069 | 0.069 | 0.701 | 0.45 | 0.71 | 0.46 | 0.74 | 0.43 | 0.61 |

| | | | | | | | | | | | | | | | | |
|---|---|---|---|---|---|---|---|---|---|---|---|---|---|---|---|---|
| 8 | 132037550 | rs10505568 | 1.161 | -0.15 | 0.428 | 0.355 | -0.001 | -0.01 | -0.01 | 0 | 0.07 | 0.02 | 0.03 | 0.3 | 0.06 | 0.16 |
| 8 | 132044385 | rs884423 | -1.103 | -0.125 | -0.015 | -0.283 | -0.096 | 0.116 | 0.116 | 0 | 0.76 | 0.96 | 0.71 | 0.91 | 0.7 | 0.78 |
| 8 | 132054152 | rs3829210 | 0.479 | -0.066 | 0.167 | -0.172 | -0.018 | 0.038 | 0.038 | 0.122 | 0.44 | 0.29 | 0.46 | 0.67 | 0.5 | 0.54 |
| 8 | 132074410 | rs7008202 | -0.556 | -0.449 | -0.313 | -0.277 | -0.138 | 0.207 | 0.207 | 0 | 0.65 | 0.95 | 0.66 | 0.9 | 0.55 | 0.74 |
| 8 | 132081007 | rs984174 | -0.601 | -0.641 | 0.414 | 0.44 | -0.007 | 0.003 | 0.003 | 0 | 0.26 | 0.7 | 0.23 | 0.44 | 0.14 | 0.33 |
| 8 | 132081604 | rs1329801 | -0.876 | -0.635 | 0.367 | 0.447 | 0.011 | -0.018 | -0.018 | 0 | 0.28 | 0.71 | 0.25 | 0.49 | 0.17 | 0.36 |
| 8 | 132092078 | rs1556161 | -0.655 | -0.465 | 0.167 | 0.178 | -0.083 | 0.162 | 0.162 | 0 | 0.65 | 0.89 | 0.65 | 0.9 | 0.56 | 0.73 |
| 8 | 132095593 | rs3897955 | 0.349 | 0.058 | 0.447 | 0.909 | 0.077 | 0.19 | 0.19 | 0.119 | 0.06 | 0.27 | 0.1 | 0.23 | 0.12 | 0.1 |
| 8 | 132106838 | rs6997019 | 1.497 | -0.076 | 0.386 | 0.894 | 0.004 | 0 | 0 | 0 | 0.07 | 0.04 | 0.01 | 0.18 | 0.04 | 0.12 |
| 8 | 132123581 | rs6988942 | 0.735 | -0.038 | 0.489 | 0.834 | 0.017 | 0.003 | 0.003 | 0.919 | 0.14 | 0.1 | 0.14 | 0.1 | 0.1 | 0.12 |
| 8 | 132131683 | rs10086390 | 0.127 | 0.244 | 0.604 | 0.618 | -0.001 | 0.139 | 0.139 | 0 | 0.11 | 0.31 | 0.09 | 0.31 | 0.07 | 0.19 |
| 8 | 132132160 | rs10096906 | -0.82 | 0.25 | 0.536 | 0.458 | -0.049 | 0.068 | 0.068 | 0 | 0.28 | 0.5 | 0.26 | 0.45 | 0.2 | 0.35 |
| 8 | 132144401 | rs13270165 | 0.72 | 0.284 | 0.124 | 0.326 | -0.012 | -0.03 | -0.03 | 2.542 | 0.4 | 0.3 | 0.43 | 0.45 | 0.41 | 0.37 |
| 8 | 132145040 | rs10099194 | -0.437 | 0.276 | -0.199 | 0.337 | -0.008 | 0.138 | 0.138 | 0 | 0.13 | 0.35 | 0.12 | 0.36 | 0.1 | 0.23 |
| 8 | 132146269 | rs4736744 | -0.908 | 0.383 | -0.238 | -0.074 | -0.019 | 0.036 | 0.036 | 0 | 0.68 | 0.83 | 0.68 | 0.91 | 0.61 | 0.72 |
| 8 | 132149744 | rs4992685 | 1.547 | -0.04 | -1.015 | 0.005 | -0.033 | 0.071 | 0.071 | 0 | 0.28 | 0.04 | 0.3 | 0.14 | 0.26 | 0.23 |
| 8 | 132150785 | rs2149890 | -0.485 | 0.161 | -0.741 | 0.084 | -0.029 | 0.043 | 0.043 | 0 | 0.27 | 0.4 | 0.24 | 0.45 | 0.2 | 0.35 |
| 8 | 132157949 | rs10101663 | 0.706 | 0.578 | -1.113 | -0.547 | 0 | -0.042 | -0.042 | 0.422 | 0.08 | 0.16 | 0.04 | 0 | 0.04 | 0.03 |
| 8 | 132160380 | rs17525431 | 0.274 | 0.769 | -1.797 | -0.209 | -0.026 | 0.012 | 0.012 | 0 | 0.34 | 0.04 | 0.33 | 0.22 | 0.3 | 0.28 |
| 8 | 132163814 | rs16904418 | -0.217 | 1.142 | -0.599 | -0.784 | -0.001 | -0.098 | -0.098 | 0.364 | 0.25 | 0.32 | 0.24 | 0 | 0.26 | 0.15 |
| 8 | 132163988 | rs4736746 | 0.416 | 1.079 | -0.65 | -0.717 | 0.005 | 0.007 | 0.007 | 0 | 0.42 | 0.33 | 0.42 | 0.62 | 0.39 | 0.52 |
| 8 | 132164319 | rs10956576 | -0.572 | 1.312 | -0.24 | -0.978 | -0.003 | -0.028 | -0.028 | 0 | 0.83 | 0.79 | 0.82 | 0.99 | 0.83 | 0.84 |
| 8 | 132165288 | rs1329796 | 0.654 | 2.01 | -0.603 | -1.477 | -0.024 | 0.04 | 0.04 | 0.007 | 0.65 | 0.74 | 0.7 | 0.83 | 0.64 | 0.62 |
| 8 | 132165751 | rs10094464 | -1.217 | 2.01 | -0.618 | -1.503 | -0.003 | -0.017 | -0.017 | 0 | 0.86 | 0.86 | 0.88 | 0.99 | 0.9 | 0.9 |
| 8 | 132167864 | rs4336594 | -0.607 | 2.193 | -0.648 | -1.531 | 0.007 | -0.046 | -0.046 | 0 | 0.83 | 0.75 | 0.84 | 0.99 | 0.84 | 0.87 |
| 8 | 132174324 | rs4587308 | 1.519 | 1.972 | -0.229 | -1.281 | 0.014 | 0.014 | 0.014 | 0 | 0.47 | 0.32 | 0.53 | 0.59 | 0.46 | 0.53 |
| 8 | 132184839 | rs7350092 | 1.292 | 1.905 | -0.108 | -1.285 | -0.007 | 0.014 | 0.014 | 0.034 | 0.15 | 0.18 | 0.12 | 0.25 | 0.13 | 0.09 |
| 8 | 132202889 | rs11991267 | 0.691 | 1.945 | 0.03 | -1.332 | -0.008 | 0.008 | 0.008 | 0 | 0.19 | 0.15 | 0.16 | 0.15 | 0.2 | 0.23 |
| 8 | 132207342 | rs4641042 | -0.667 | 1.874 | -0.026 | -1.219 | 0.012 | -0.054 | -0.054 | 0.122 | 0.82 | 0.75 | 0.81 | 0.99 | 0.8 | 0.84 |
| 8 | 132230063 | rs9643263 | -0.083 | 1.492 | -0.238 | -1.029 | -0.01 | -0.005 | -0.005 | 0 | 0.7 | 0.67 | 0.73 | 0.76 | 0.77 | 0.81 |
| 8 | 132238169 | rs4523235 | -0.226 | 1.731 | -0.058 | -1.1 | 0.038 | -0.03 | -0.03 | 0.07 | 0.24 | 0.35 | 0.21 | 0.15 | 0.31 | 0.26 |
| 8 | 132250643 | rs6470909 | 0.477 | 1.658 | -0.029 | -0.887 | 0.001 | -0.001 | -0.001 | 0 | 0.53 | 0.52 | 0.55 | 0.57 | 0.5 | 0.6 |
| 8 | 132256944 | rs7820729 | 0.924 | 2.218 | 0.101 | -1.057 | 0.005 | 0.003 | 0.003 | 0 | 0.51 | 0.49 | 0.55 | 0.57 | 0.43 | 0.55 |
| 8 | 132261701 | rs4304315 | 0.297 | 2.305 | -0.162 | -1.079 | -0.017 | 0.005 | 0.005 | 0.002 | 0.17 | 0.14 | 0.16 | 0.15 | 0.28 | 0.23 |
| 8 | 132273687 | rs6415533 | -1.32 | 2.569 | -0.263 | -1.178 | -0.018 | 0.036 | 0.036 | 0 | 0.84 | 0.93 | 0.84 | 0.99 | 0.82 | 0.83 |
| 8 | 132291603 | rs4128913 | 0.809 | 2.467 | -0.317 | -1.133 | 0.002 | 0.016 | 0.016 | 0 | 0.19 | 0.23 | 0.18 | 0.26 | 0.21 | 0.11 |
| 8 | 132292590 | rs4513159 | 1.102 | 2.019 | -0.831 | -0.981 | 0.083 | 0 | 0 | 0 | 0.45 | 0.27 | 0.46 | 0.57 | 0.33 | 0.49 |
| 8 | 132293461 | rs11779297 | 1.049 | 2.001 | -0.895 | -0.892 | -0.028 | -0.008 | -0.008 | 0.52 | 0.13 | 0.07 | 0.11 | 0.15 | 0.18 | 0.18 |
| 8 | 132301089 | rs13274156 | -0.274 | 1.531 | -0.699 | -0.915 | 0.004 | -0.009 | -0.009 | 0.045 | 0.37 | 0.53 | 0.36 | 0.28 | 0.39 | 0.28 |
| 8 | 132304508 | rs10102742 | -0.646 | 0.737 | -0.978 | -0.692 | 0.049 | 0.009 | 0.009 | 0 | 0.55 | 0.75 | 0.5 | 0.4 | 0.61 | 0.55 |
| 8 | 132306882 | rs11784039 | -0.998 | 1.048 | -0.645 | -0.708 | 0.031 | 0.067 | 0.067 | 0 | 0.47 | 0.71 | 0.38 | 0.29 | 0.57 | 0.44 |
| 8 | 132310783 | rs13277897 | -0.199 | 1.69 | -0.576 | -0.786 | -0.01 | 0.01 | 0.01 | 0 | 0.2 | 0.16 | 0.12 | 0.19 | 0.21 | 0.23 |
| 8 | 132311479 | rs4380914 | 1.677 | 1.086 | -0.989 | -0.761 | -0.02 | -0.013 | -0.013 | 0 | 0.33 | 0.34 | 0.24 | 0.18 | 0.24 | 0.21 |
| 8 | 132317922 | rs6470924 | -0.103 | 1.646 | -1.66 | -0.519 | -0.048 | -0.052 | -0.052 | 0 | 0.64 | 0.9 | 0.55 | 0.44 | 0.62 | 0.58 |
| 8 | 132328178 | rs13279605 | 0.219 | 0.9 | -1.475 | -0.73 | -0.136 | 0.127 | 0.127 | 0.778 | 0.34 | 0.06 | 0.29 | 0.14 | 0.4 | 0.22 |
| 8 | 132329300 | rs7017572 | 0.838 | 0.74 | -1.165 | -0.672 | -0.006 | -0.016 | -0.016 | 0.068 | 0.4 | 0.55 | 0.42 | 0.25 | 0.42 | 0.36 |
| 8 | 132338453 | rs10103835 | -1.087 | 0.603 | -1.056 | -0.593 | -0.001 | 0.023 | 0.023 | 0 | 0.84 | 0.83 | 0.7 | 0.5 | 0.85 | 0.72 |
| 8 | 132344697 | rs7460334 | -1.212 | 0.626 | -1.557 | -0.581 | 0.017 | -0.072 | -0.072 | 0 | 0.84 | 0.89 | 0.71 | 0.5 | 0.85 | 0.72 |
| 8 | 132346637 | rs6470938 | -1.375 | 0.649 | -1.53 | -0.557 | 0.015 | -0.072 | -0.072 | 0.856 | 0.85 | 0.97 | 0.81 | 0.53 | 0.86 | 0.73 |
| 8 | 132346996 | rs7385988 | 1.361 | 0.629 | -1.504 | -0.648 | -0.005 | -0.003 | -0.003 | 0 | 0.33 | 0.35 | 0.24 | 0.21 | 0.28 | 0.24 |
| 8 | 132350565 | rs7460903 | -1.444 | 0.587 | -1.525 | -0.601 | 0.013 | -0.082 | -0.082 | 0 | 0.86 | 0.97 | 0.9 | 0.54 | 0.86 | 0.73 |
| 8 | 132352151 | rs7845936 | 0.246 | 0.584 | -1.478 | -0.674 | 0.015 | 0.005 | 0.005 | 0 | 0.11 | 0.42 | 0.1 | 0.12 | 0.04 | 0.15 |
| 8 | 132353191 | rs6413554 | -1.483 | 0.605 | -1.505 | -0.653 | 0.013 | -0.082 | -0.082 | 0 | 0.86 | 0.96 | 0.9 | 0.54 | 0.86 | 0.73 |
| 8 | 132353735 | rs4596632 | 0.701 | 0.731 | -1.521 | -0.688 | -0.003 | 0.019 | 0.019 | 0 | 0.68 | 0.47 | 0.71 | 0.39 | 0.68 | 0.45 |
| 8 | 132355894 | rs7000465 | 0.064 | 0.958 | -1.246 | -0.502 | 0.011 | 0.001 | 0.001 | 0 | 0.12 | 0.39 | 0.1 | 0.12 | 0.04 | 0.15 |
| 8 | 132356637 | rs4574827 | 1.237 | 1.04 | -1.583 | -0.429 | 0.01 | 0.006 | 0.006 | 0 | 0.32 | 0.27 | 0.34 | 0.24 | 0.3 | 0.25 |
| 8 | 132356672 | rs4618675 | -0.602 | 1.073 | -0.924 | 0.229 | 0.045 | -0.041 | -0.041 | 0 | 0.17 | 0.49 | 0.18 | 0.15 | 0.18 | 0.28 |
| 8 | 132358549 | rs4269521 | 0.22 | 0.63 | -1.735 | 0.681 | 0.018 | -0.13 | -0.13 | 0.022 | 0.79 | 0.96 | 0.88 | 0.5 | 0.82 | 0.73 |
| 8 | 132358915 | rs4352831 | 0.407 | 0.947 | -1.369 | 0.482 | 0 | -0.015 | -0.015 | 0.287 | 0.66 | 0.69 | 0.73 | 0.56 | 0.67 | 0.73 |
| 8 | 132359898 | rs4446718 | -0.538 | 1.066 | -1.498 | -0.204 | -0.036 | 0.081 | 0.081 | 0 | 0.26 | 0.12 | 0.28 | 0.07 | 0.31 | 0.15 |
| 8 | 132367143 | rs4570136 | -0.3 | 0.177 | -1.248 | 0.571 | -0.009 | 0.012 | 0.012 | 0.643 | 0.18 | 0.2 | 0.27 | 0.39 | 0.09 | 0.41 |
| 8 | 132370660 | rs7832023 | -1.157 | 0.152 | -0.503 | 0.234 | -0.022 | 0.015 | 0.015 | 0.28 | 0.35 | 0.58 | 0.48 | 0.46 | 0.27 | 0.51 |
| 8 | 132389937 | rs6987024 | -0.417 | 0.289 | -0.364 | 0.104 | 0.015 | -0.043 | -0.043 | 0 | 0.16 | 0.28 | 0.2 | 0.07 | 0.18 | 0.1 |
| 8 | 132398697 | rs4371986 | -0.109 | 0.525 | 0.269 | -0.119 | 0.038 | -0.057 | -0.057 | 0 | 0.13 | 0.27 | 0.11 | 0.03 | 0.13 | 0.08 |
| 8 | 132401613 | rs10096114 | 1.176 | 0.263 | -0.016 | 0.214 | 0.009 | -0.013 | -0.013 | 0 | 0.53 | 0.25 | 0.43 | 0.46 | 0.6 | 0.44 |
| 8 | 132403378 | rs10808588 | 1.171 | 0.302 | -0.107 | 0.173 | 0.03 | -0.003 | -0.003 | 2.116 | 0.54 | 0.23 | 0.42 | 0.5 | 0.62 | 0.49 |
| 8 | 132403551 | rs10808589 | 1.257 | 0.939 | -0.175 | -0.061 | 0.001 | 0 | 0 | 0.424 | 0.49 | 0.19 | 0.35 | 0.49 | 0.47 | 0.43 |
| 8 | 132406960 | rs891429 | 0.96 | 0.859 | 0.187 | -0.31 | 0.01 | 0.01 | 0.01 | 1.907 | 0.51 | 0.67 | 0.45 | 0.5 | 0.47 | 0.41 |
| 8 | 132424978 | rs1449471 | 0.829 | 0.845 | 0.532 | -0.416 | -0.019 | 0.011 | 0.011 | 2.002 | 0.52 | 0.81 | 0.48 | 0.53 | 0.47 | 0.41 |

| | | | | | | | | | | | | | | | | |
|---|---|---|---|---|---|---|---|---|---|---|---|---|---|---|---|---|
| 8 | 132426760 | rs1036032 | 0.104 | 0.963 | 0.41 | -0.47 | -0.075 | 0.028 | 0.028 | 0 | 0.16 | 0.04 | 0.16 | 0.07 | 0.29 | 0.09 |
| 8 | 132428003 | rs9969634 | -0.74 | 0.686 | 0.264 | -0.579 | 0.044 | -0.076 | -0.076 | 0 | 0.84 | 0.96 | 0.71 | 0.63 | 0.89 | 0.57 |
| 8 | 132428564 | rs10111675 | -0.981 | 0.726 | 0.264 | -0.596 | -0.097 | 0.115 | 0.115 | 0 | 0.32 | 0.15 | 0.23 | 0.1 | 0.42 | 0.17 |
| 8 | 132441579 | rs2217438 | 1.592 | 0.352 | -0.125 | -1.179 | 0.004 | 0.032 | 0.032 | 0 | 0.31 | 0.41 | 0.37 | 0.51 | 0.34 | 0.32 |
| 8 | 132458949 | rs7014931 | 1.824 | 0.231 | -0.471 | -1.133 | -0.01 | 0.089 | 0.089 | 0.87 | 0.16 | 0.02 | 0.07 | 0.01 | 0.14 | 0.09 |
| 8 | 132469052 | rs273393 | 0.444 | 0.4 | 0.073 | -0.352 | 0.037 | -0.042 | -0.042 | 0.057 | 0.7 | 0.87 | 0.64 | 0.59 | 0.78 | 0.52 |
| 8 | 132471064 | rs10505573 | -1.338 | 0.638 | 0.309 | -0.498 | 0.018 | -0.034 | -0.034 | 0 | 0.37 | 0.5 | 0.38 | 0.23 | 0.42 | 0.19 |
| 8 | 132473863 | rs715969 | -0.236 | 0.609 | 0.407 | -0.124 | -0.001 | -0.001 | -0.001 | 0 | 0.17 | 0.17 | 0.19 | 0.15 | 0.12 | 0.08 |
| 8 | 132479901 | rs273429 | 2.051 | 0.563 | 0.434 | 0.009 | 0 | 0.02 | 0.02 | 1.441 | 0.52 | 0.24 | 0.43 | 0.55 | 0.53 | 0.46 |
| 8 | 132489214 | rs4736476 | 0.39 | 0.694 | 0.426 | 0.052 | 0.103 | 0.004 | 0.004 | 0 | 0.1 | 0.22 | 0.16 | 0.07 | 0.23 | 0.07 |
| 8 | 132489281 | rs1354873 | -1.179 | 0.696 | -0.023 | 0.055 | 0.035 | -0.014 | -0.014 | 0 | 0.27 | 0.39 | 0.45 | 0.23 | 0.35 | 0.16 |
| 8 | 132500886 | rs17534787 | 0.56 | 0.756 | -0.523 | 0.115 | -0.04 | -0.005 | -0.005 | 0 | 0.09 | 0.04 | 0.15 | 0.07 | 0.23 | 0.07 |
| 8 | 132507466 | rs7821997 | -1.362 | 0.734 | -0.472 | 0.104 | 0.025 | -0.024 | -0.024 | 0 | 0.3 | 0.48 | 0.46 | 0.23 | 0.37 | 0.16 |
| 8 | 132511008 | rs273403 | -0.721 | 0.513 | -0.472 | 0.239 | 0.032 | -0.051 | -0.051 | 0.995 | 0.86 | 0.98 | 0.92 | 0.78 | 0.91 | 0.64 |
| 8 | 132517335 | rs10505571 | -0.844 | 1.191 | 0.267 | 0.489 | -0.023 | -0.027 | -0.027 | 0 | 0.2 | 0.4 | 0.3 | 0.15 | 0.13 | 0.08 |
| 8 | 132521469 | rs273409 | 1.593 | 1.707 | -0.523 | 1.355 | 0.021 | -0.002 | -0.002 | 0 | 0.66 | 0.47 | 0.52 | 0.35 | 0.58 | 0.65 |
| 8 | 132527475 | rs273402 | 1.421 | 1.877 | -0.542 | 1.283 | 0.017 | -0.001 | -0.001 | 0.153 | 0.64 | 0.49 | 0.49 | 0.4 | 0.54 | 0.65 |
| 8 | 132528389 | rs2940106 | 1.291 | 1.904 | -0.527 | 1.314 | 0.006 | -0.008 | -0.008 | 0 | 0.67 | 0.58 | 0.54 | 0.73 | 0.6 | 0.77 |
| 8 | 132534844 | rs156397 | 0.821 | 0.59 | -0.131 | 0.045 | -0.031 | -0.026 | -0.026 | 0 | 0.8 | 0.58 | 0.71 | 0.81 | 0.87 | 0.85 |
| 8 | 132536085 | rs1594332 | -1.08 | 1.058 | -1.082 | -0.055 | -0.029 | 0.03 | 0.03 | 0.25 | 0.2 | 0.03 | 0.27 | 0.12 | 0.18 | 0.12 |
| 8 | 132536567 | rs1605438 | 0.022 | 0.784 | -0.643 | -0.274 | 0.004 | -0.012 | -0.012 | 0.757 | 0.45 | 0.47 | 0.35 | 0.25 | 0.44 | 0.43 |
| 8 | 132542601 | rs273420 | 0.17 | 0.431 | -0.335 | -0.799 | -0.001 | 0.015 | 0.015 | 0.365 | 0.51 | 0.43 | 0.68 | 0.78 | 0.53 | 0.68 |
| 8 | 132546350 | rs7010302 | 0.215 | 0.34 | -0.578 | -1.057 | -0.001 | -0.001 | -0.001 | 0 | 0.3 | 0.29 | 0.16 | 0.22 | 0.31 | 0.25 |
| 8 | 132555282 | rs460783 | -0.888 | 0.313 | -0.925 | -0.961 | 0.007 | 0.043 | 0.043 | 0 | 0.81 | 0.92 | 0.85 | 1 | 0.83 | 0.92 |
| 8 | 132564450 | rs457941 | 0.513 | 0.289 | -0.935 | -1.008 | -0.025 | 0.143 | 0.143 | 0.482 | 0.47 | 0.21 | 0.64 | 0.78 | 0.51 | 0.67 |
| 8 | 132564658 | rs459806 | -0.81 | 0.387 | -0.976 | -1.053 | 0.001 | -0.035 | -0.035 | 0 | 0.8 | 0.83 | 0.83 | 1 | 0.83 | 0.92 |
| 8 | 132586230 | rs4736496 | 0.578 | 0.261 | -1.19 | -1.08 | -0.015 | 0.009 | 0.009 | 0 | 0.26 | 0.18 | 0.13 | 0.22 | 0.29 | 0.25 |
| 8 | 132590744 | rs277026 | -1.045 | 0.263 | -1.776 | -1.048 | -0.044 | -0.03 | -0.03 | 0 | 0.8 | 0.99 | 0.81 | 0.79 | 0.78 | 0.72 |
| 8 | 132597684 | rs277017 | 0.664 | 0.495 | -1.146 | -1.021 | 0.002 | 0.083 | 0.083 | 0 | 0.19 | 0.05 | 0.15 | 0 | 0.17 | 0.08 |
| 8 | 132606888 | rs10095524 | -0.156 | 1 | -0.794 | -0.252 | -0.006 | 0.006 | 0.006 | 0 | 0.27 | 0.23 | 0.14 | 0.22 | 0.31 | 0.25 |
| 8 | 132620699 | rs277037 | -0.123 | 0.82 | -0.918 | 0.086 | 0.003 | 0.064 | 0.064 | 0 | 0.5 | 0.66 | 0.42 | 0.3 | 0.53 | 0.35 |
| 8 | 132620944 | rs277040 | 0.345 | 0.451 | -0.896 | -0.076 | 0.013 | -0.006 | -0.006 | 0.852 | 0.12 | 0.07 | 0.12 | 0.11 | 0.09 | 0.14 |
| 8 | 132624140 | rs6984975 | 0.785 | 0.172 | 0.122 | -0.365 | -0.018 | -0.056 | -0.056 | 0.41 | 0.1 | 0.25 | 0.08 | 0 | 0.07 | 0.01 |
| 8 | 132625999 | rs16904483 | 0.785 | 0.172 | 0.122 | -0.365 | -0.018 | -0.056 | -0.056 | 0.404 | 0.1 | 0.25 | 0.08 | 0 | 0.07 | 0.01 |
| 8 | 132627248 | rs277047 | -0.846 | -0.147 | -0.503 | -0.009 | -0.053 | -0.058 | -0.058 | 0.11 | 0.8 | 0.86 | 0.54 | 0.54 | 0.7 | 0.8 |
| 8 | 132638331 | rs950604 | 1.433 | -0.277 | -0.17 | -0.148 | 0.032 | 0.013 | 0.013 | 0 | 0.31 | 0.2 | 0.13 | 0.25 | 0.19 | 0.41 |
| 8 | 132654820 | rs12677773 | 0.788 | -0.171 | 0.045 | -0.087 | 0.004 | 0.001 | 0.001 | 0 | 0.57 | 0.48 | 0.33 | 0.44 | 0.53 | 0.58 |
| 8 | 132660486 | rs1420484 | 1.19 | -0.158 | 0.375 | -0.097 | -0.027 | -0.058 | -0.058 | 0.322 | 0.1 | 0.24 | 0.08 | 0 | 0.06 | 0.01 |
| 8 | 132668758 | rs993718 | -1.201 | -0.107 | 0.335 | -0.019 | 0.004 | -0.002 | -0.002 | 0 | 0.88 | 0.92 | 0.88 | 0.89 | 0.91 | 0.85 |
| 8 | 132679927 | rs4464950 | -1.158 | -0.124 | 0.302 | -0.028 | 0.014 | -0.005 | -0.005 | 0.502 | 0.88 | 0.93 | 0.88 | 0.89 | 0.91 | 0.85 |
| 8 | 132682208 | rs1035189 | 0.614 | 0.141 | 0.235 | -0.139 | -0.057 | -0.059 | -0.059 | 0 | 0.2 | 0.14 | 0.46 | 0.45 | 0.32 | 0.21 |
| 8 | 132685663 | rs10092594 | -0.266 | -0.097 | 0.514 | -0.15 | -0.012 | -0.019 | -0.019 | 0.018 | 0.8 | 0.82 | 0.83 | 0.92 | 0.84 | 0.84 |
| 8 | 132692795 | rs1420497 | 1.181 | 0.596 | 0.794 | -0.112 | 0.038 | 0.053 | 0.053 | 0 | 0.08 | 0.17 | 0.11 | 0.16 | 0.14 | 0.1 |
| 8 | 132696791 | rs16904501 | 1.683 | 0.341 | 1.095 | -0.218 | 0.051 | -0.01 | -0.01 | 0 | 0.31 | 0.16 | 0.13 | 0.28 | 0.18 | 0.45 |
| 8 | 132699454 | rs2058758 | 0.924 | 0.333 | 1.232 | -0.27 | 0.006 | -0.007 | -0.007 | 0 | 0.39 | 0.32 | 0.24 | 0.44 | 0.32 | 0.55 |
| 8 | 132699795 | rs6986708 | 1.965 | 0.288 | 1.485 | -0.271 | -0.009 | -0.021 | -0.021 | 0.484 | 0.08 | 0.08 | 0.04 | 0 | 0.05 | 0.01 |
| 8 | 132700563 | rs1362534 | 0.307 | 1.321 | 1.352 | -0.058 | 0 | 0.011 | 0.011 | 0 | 0.54 | 0.43 | 0.66 | 0.84 | 0.52 | 0.69 |
| 8 | 132701504 | rs1833226 | 0.273 | 1.339 | 1.656 | -0.065 | 0.009 | -0.065 | -0.065 | 0 | 0.33 | 0.43 | 0.22 | 0.09 | 0.38 | 0.16 |
| 8 | 132704717 | rs556739 | 1.309 | 1.353 | 1.602 | -0.057 | -0.037 | -0.051 | -0.051 | 0.046 | 0.15 | 0.1 | 0.41 | 0.4 | 0.19 | 0.14 |
| 8 | 132706148 | rs10105308 | 1.126 | 1.32 | 1.593 | -0.042 | 0.006 | -0.007 | -0.007 | 0 | 0.39 | 0.33 | 0.25 | 0.44 | 0.32 | 0.56 |
| 8 | 132707922 | rs16904505 | 1.326 | 1.446 | 1.58 | -0.072 | 0.015 | 0.003 | 0.003 | 0 | 0.05 | 0.08 | 0.05 | 0.07 | 0.12 | 0.08 |
| 8 | 132711614 | rs581534 | 1.322 | 1.213 | 1.593 | -0.096 | 0.007 | 0.006 | 0.006 | 0.035 | 0.12 | 0.08 | 0.11 | 0.07 | 0.09 | 0.15 |
| 8 | 132712674 | rs598630 | 1.169 | 1.216 | 1.621 | -0.102 | -0.007 | 0.005 | 0.005 | 0 | 0.39 | 0.37 | 0.25 | 0.44 | 0.32 | 0.56 |
| 8 | 132716759 | rs1345383 | 0.605 | 1.219 | 1.519 | -0.315 | -0.013 | -0.152 | -0.152 | 0.143 | 0.29 | 0.45 | 0.17 | 0.01 | 0.28 | 0.08 |
| 8 | 132718006 | rs529894 | 0.551 | 1.587 | 1.491 | 0.669 | 0.027 | 0.021 | 0.021 | 0 | 0.69 | 0.49 | 0.51 | 0.6 | 0.67 | 0.79 |
| 8 | 132718778 | rs600794 | 1.162 | 1.507 | 1.072 | 0.651 | -0.015 | 0.005 | 0.005 | 0 | 0.55 | 0.25 | 0.4 | 0.52 | 0.55 | 0.66 |
| 8 | 132719817 | rs491614 | 1.14 | 1.495 | 1.052 | 0.66 | -0.015 | 0.004 | 0.004 | 0 | 0.55 | 0.27 | 0.41 | 0.52 | 0.55 | 0.66 |
| 8 | 132721564 | rs1861515 | 0.713 | 1.341 | 1.035 | 0.802 | 0.023 | 0.024 | 0.024 | 0 | 0.08 | 0.13 | 0.14 | 0.17 | 0.15 | 0.1 |
| 8 | 132721667 | rs1514628 | -0.437 | 1.659 | 0.717 | 0.705 | -0.027 | -0.05 | -0.05 | 0.774 | 0.82 | 0.91 | 0.61 | 0.67 | 0.81 | 0.84 |
| 8 | 132723844 | rs9297830 | 0.983 | 1.612 | 1.047 | 0.605 | 0.017 | 0.028 | 0.028 | 0.176 | 0.63 | 0.51 | 0.47 | 0.46 | 0.56 | 0.61 |
| 8 | 132725731 | rs587639 | 0.705 | 1.766 | 0.685 | 0.709 | 0.011 | 0.019 | 0.019 | 0 | 0.72 | 0.58 | 0.52 | 0.52 | 0.64 | 0.73 |
| 8 | 132727074 | rs9297832 | -0.513 | 1.127 | 0.47 | 0.422 | 0.002 | -0.01 | -0.01 | 1.106 | 0.26 | 0.35 | 0.23 | 0.19 | 0.31 | 0.15 |
| 8 | 132732212 | rs925687 | -0.819 | 0.348 | 0.462 | 0.932 | -0.001 | 0.011 | 0.011 | 0 | 0.47 | 0.42 | 0.43 | 0.43 | 0.47 | 0.34 |
| 8 | 132732273 | rs925688 | -0.629 | 0.241 | 0.352 | 0.85 | -0.017 | 0.008 | 0.008 | 0 | 0.44 | 0.28 | 0.4 | 0.43 | 0.45 | 0.31 |
| 8 | 132733415 | rs13266806 | 0.357 | 0.291 | 0.52 | 0.663 | 0 | -0.042 | -0.042 | 0 | 0.21 | 0.22 | 0.17 | 0 | 0.18 | 0.09 |
| 8 | 132742230 | rs10093373 | -1.625 | 0.06 | 0.525 | 1.512 | -0.121 | -0.024 | -0.024 | 0.419 | 0.52 | 0.97 | 0.56 | 0.64 | 0.48 | 0.42 |
| 8 | 132746158 | rs6470996 | -1.424 | -0.125 | 1.127 | 1.554 | 0.087 | 0.194 | 0.194 | 0.244 | 0.1 | 0.45 | 0.14 | 0.2 | 0.09 | 0.17 |
| 8 | 132748054 | rs11992279 | -2.459 | -0.162 | 1.026 | 1.575 | -0.002 | 0.029 | 0.029 | 2.504 | 0.13 | 0.72 | 0.21 | 0.2 | 0.11 | 0.2 |
| 8 | 132776942 | rs10956608 | -0.418 | -0.239 | 1.138 | 1.639 | -0.004 | 0.007 | 0.007 | 0 | 0.32 | 0.32 | 0.34 | 0.25 | 0.34 | 0.17 |

| | | | | | | | | | | | | | | | | |
|---|---|---|---|---|---|---|---|---|---|---|---|---|---|---|---|---|
| 8 | 132782921 | rs1514633 | -0.229 | -0.228 | 1.647 | 1.768 | 0.098 | 0.247 | 0.247 | 0 | 0.06 | 0.3 | 0.12 | 0.21 | 0.06 | 0.17 |
| 8 | 132789352 | rs10956609 | -2.173 | -0.512 | 1.018 | 1.862 | -0.143 | 0.102 | 0.102 | 0 | 0.57 | 0.98 | 0.59 | 0.82 | 0.54 | 0.57 |
| 8 | 132790095 | rs2878231 | -2.122 | -0.215 | 1.329 | 1.971 | -0.021 | 0.089 | 0.089 | 0.11 | 0.16 | 0.65 | 0.19 | 0.39 | 0.09 | 0.25 |
| 8 | 132803445 | rs1514639 | -1.631 | -0.162 | 1.172 | 1.979 | -0.006 | 0.242 | 0.242 | 0 | 0.13 | 0.5 | 0.15 | 0.39 | 0.07 | 0.22 |
| 8 | 132810827 | rs10107814 | -1.55 | -0.137 | 0.424 | 1.857 | -0.053 | 0.013 | 0.013 | 0 | 0.47 | 0.92 | 0.53 | 0.66 | 0.43 | 0.43 |
| 8 | 132822037 | rs1040098 | -1.549 | -0.115 | 0.41 | 1.816 | -0.063 | -0.063 | -0.063 | 0 | 0.47 | 0.92 | 0.53 | 0.61 | 0.43 | 0.42 |
| 8 | 132827722 | rs1399359 | -0.537 | 0.04 | 0.649 | 1.669 | -0.018 | 0.056 | 0.056 | 0 | 0.32 | 0.21 | 0.23 | 0.04 | 0.36 | 0.18 |
| 8 | 132828062 | rs7815383 | -1.626 | 0.151 | 1.133 | 2.17 | 0.01 | 0.271 | 0.271 | 0 | 0.12 | 0.43 | 0.16 | 0.4 | 0.08 | 0.23 |
| 8 | 132841734 | rs7828128 | 0.135 | 0.334 | 0.647 | 1.858 | 0.049 | 0.178 | 0.178 | 0 | 0.05 | 0.22 | 0.11 | 0.21 | 0.05 | 0.17 |
| 8 | 132854416 | rs1355981 | 0.033 | 0.502 | 0.34 | 1.316 | -0.021 | 0.006 | 0.006 | 0 | 0.09 | 0.05 | 0.07 | 0.04 | 0.15 | 0.1 |
| 8 | 132855371 | rs13269138 | 0.778 | 0.407 | 0.153 | 1.364 | -0.027 | -0.022 | -0.022 | 0.689 | 0.17 | 0.03 | 0.24 | 0.19 | 0.19 | 0.08 |
| 8 | 132856646 | rs4332113 | -0.382 | 0.325 | 0.373 | 1.081 | -0.024 | 0.055 | 0.055 | 0 | 0.29 | 0.19 | 0.23 | 0.04 | 0.36 | 0.18 |
| 8 | 132862920 | rs6999964 | -1.037 | 0.042 | 0.453 | 1.564 | -0.012 | 0.046 | 0.046 | 0.792 | 0.21 | 0.26 | 0.18 | 0.6 | 0.2 | 0.4 |
| 8 | 132869766 | rs2970611 | -1.035 | 0.217 | 0.629 | 1.694 | 0.007 | 0.079 | 0.079 | 2.006 | 0.2 | 0.37 | 0.22 | 0.42 | 0.23 | 0.33 |
| 8 | 132871608 | rs11997238 | -0.215 | 0.266 | 0.558 | 1.783 | 0.044 | 0.082 | 0.082 | 0 | 0.06 | 0.2 | 0.13 | 0.18 | 0.07 | 0.11 |
| 8 | 132872192 | rs1399362 | -1.456 | 0.008 | 0.173 | 1.725 | 0.014 | 0.053 | 0.053 | 0 | 0.5 | 0.91 | 0.55 | 0.67 | 0.45 | 0.45 |
| 8 | 132874364 | rs7824023 | -1.35 | 0.3 | 0.941 | 2.05 | 0.05 | 0.078 | 0.078 | 0 | 0.08 | 0.27 | 0.14 | 0.18 | 0.08 | 0.14 |
| 8 | 132885391 | rs16904539 | 0.138 | 0.27 | 0.499 | 2.072 | 0.002 | -0.029 | -0.029 | 0 | 0.09 | 0.03 | 0.06 | 0.24 | 0.11 | 0.13 |
| 8 | 132886070 | rs7832224 | -1.71 | 0.261 | 1.144 | 1.937 | 0.002 | 0.089 | 0.089 | 0 | 0.18 | 0.4 | 0.18 | 0.3 | 0.14 | 0.2 |
| 8 | 132886443 | rs7832443 | -0.312 | 0.352 | 0.949 | 1.743 | 0.099 | 0.013 | 0.013 | 0 | 0.05 | 0.21 | 0.08 | 0.03 | 0.08 | 0.14 |
| 8 | 132886840 | rs4388438 | -2.062 | 0.305 | 0.503 | 1.966 | -0.003 | 0.049 | 0.049 | 0.116 | 0.24 | 0.33 | 0.23 | 0.56 | 0.23 | 0.31 |
| 8 | 132886989 | rs4422759 | -1.585 | 0.3 | 0.557 | 1.892 | 0.001 | 0.041 | 0.041 | 0 | 0.17 | 0.31 | 0.19 | 0.29 | 0.14 | 0.21 |
| 8 | 132887566 | rs9656990 | -0.923 | 0.717 | 0.485 | 1.768 | 0.085 | 0.072 | 0.072 | 0 | 0.07 | 0.27 | 0.16 | 0.14 | 0.1 | 0.12 |
| 8 | 132889192 | rs6471009 | 0.426 | 0.626 | -0.388 | 1.669 | -0.018 | -0.047 | -0.047 | 0.081 | 0.82 | 0.9 | 0.9 | 0.69 | 0.77 | 0.75 |
| 8 | 132897005 | rs10100214 | 0.274 | 0.136 | -0.449 | 1.389 | -0.011 | 0.021 | 0.021 | 0.733 | 0.2 | 0.22 | 0.25 | 0.35 | 0.14 | 0.37 |
| 8 | 132897432 | rs9297833 | 0.643 | 0.391 | -0.621 | 1.244 | -0.001 | -0.001 | -0.001 | 0 | 0.65 | 0.6 | 0.67 | 0.52 | 0.61 | 0.55 |
| 8 | 132897862 | rs1995317 | -0.682 | 0.43 | -0.356 | 1.316 | 0.005 | 0.018 | 0.018 | 0.066 | 0.27 | 0.38 | 0.28 | 0.43 | 0.28 | 0.33 |
| 8 | 132899760 | rs16904543 | 0.648 | 0.425 | -0.497 | 1.269 | 0.009 | -0.055 | -0.055 | 0.69 | 0.13 | 0.03 | 0.12 | 0.28 | 0.13 | 0.16 |
| 8 | 132900937 | rs2334792 | -0.698 | 0.522 | -0.406 | 1.2 | 0.007 | 0.028 | 0.028 | 0.231 | 0.21 | 0.31 | 0.27 | 0.43 | 0.25 | 0.29 |
| 8 | 132914886 | rs10101450 | -1.039 | 0.31 | -0.176 | 1.007 | -0.005 | 0.002 | 0.002 | 0 | 0.39 | 0.47 | 0.43 | 0.62 | 0.37 | 0.52 |
| 8 | 132921441 | rs1392872 | -0.194 | -0.128 | -0.351 | 0.669 | -0.001 | 0.013 | 0.013 | 0 | 0.84 | 0.87 | 0.89 | 0.93 | 0.86 | 0.75 |
| 8 | 132921874 | rs1392871 | -0.68 | -0.074 | -0.298 | 0.728 | -0.007 | 0.003 | 0.003 | 0 | 0.38 | 0.43 | 0.43 | 0.62 | 0.34 | 0.5 |
| 8 | 132922442 | rs924529 | -0.278 | -0.038 | -0.074 | 0.677 | -0.013 | 0.005 | 0.005 | 0.666 | 0.87 | 0.9 | 0.88 | 0.92 | 0.82 | 0.73 |
| 8 | 132928177 | rs12544248 | 1.273 | -0.19 | -0.295 | 0.794 | 0.002 | -0.094 | -0.094 | 0 | 0.18 | 0.01 | 0.17 | 0.33 | 0.17 | 0.15 |
| 8 | 132930694 | rs4736521 | 1.724 | -0.177 | 0.074 | 0.826 | -0.002 | 0.044 | 0.044 | 3.188 | 0.47 | 0.29 | 0.44 | 0.3 | 0.48 | 0.23 |
| 8 | 132934538 | rs7004484 | 0.387 | -0.149 | -0.04 | 0.85 | -0.021 | 0.026 | 0.026 | 0.014 | 0.25 | 0.3 | 0.25 | 0.45 | 0.21 | 0.25 |
| 8 | 132940138 | rs17560662 | 2.633 | -0.176 | -1.032 | -0.051 | -0.085 | 0.116 | 0.116 | 0.084 | 0.21 | 0.01 | 0.13 | 0.02 | 0.27 | 0.11 |
| 8 | 132947786 | rs16904553 | 1.44 | -0.27 | -0.967 | 0.107 | 0.003 | -0.072 | -0.072 | 0 | 0.21 | 0.03 | 0.17 | 0.33 | 0.21 | 0.18 |
| 8 | 132949944 | rs10956624 | 0.769 | -0.357 | -0.741 | 0.075 | -0.001 | -0.001 | -0.001 | 0 | 0.53 | 0.53 | 0.5 | 0.31 | 0.51 | 0.39 |
| 8 | 132953350 | rs10112200 | -0.176 | -0.356 | -0.663 | 0.078 | 0.004 | -0.012 | -0.012 | 0 | 0.33 | 0.28 | 0.39 | 0.62 | 0.32 | 0.5 |
| 8 | 132954930 | rs11775288 | 1.198 | -0.365 | -0.595 | 0.054 | 0.001 | 0.037 | 0.037 | 3.108 | 0.49 | 0.38 | 0.44 | 0.3 | 0.48 | 0.23 |
| 8 | 132966522 | rs16904557 | 0.437 | -0.301 | -0.934 | 0.105 | -0.007 | -0.106 | -0.106 | 0.159 | 0.29 | 0.1 | 0.31 | 0.49 | 0.3 | 0.42 |
| 8 | 132968567 | rs1501061 | -1.624 | -0.272 | -0.945 | 0.112 | -0.01 | -0.002 | -0.002 | 0 | 0.91 | 0.91 | 0.93 | 0.93 | 0.85 | 0.89 |
| 8 | 132972259 | rs7000712 | 0.828 | -0.278 | -0.953 | 0.1 | -0.001 | -0.003 | -0.003 | 0 | 0.53 | 0.53 | 0.5 | 0.31 | 0.51 | 0.39 |
| 8 | 132982824 | rs1051221 | 0.828 | -0.278 | -0.952 | 0.1 | 0 | -0.004 | -0.004 | 0 | 0.53 | 0.53 | 0.5 | 0.31 | 0.51 | 0.39 |
| 8 | 132995451 | rs1553271 | 0.209 | -0.228 | -1.251 | 0.089 | 0.002 | -0.077 | -0.077 | 0 | 0.31 | 0.14 | 0.33 | 0.49 | 0.31 | 0.47 |
| 8 | 132996059 | rs1553270 | 1.208 | 0.051 | -1.206 | -0.153 | -0.009 | -0.009 | -0.009 | 0 | 0.08 | 0.06 | 0.15 | 0.16 | 0.11 | 0.25 |
| 8 | 132996359 | rs11783016 | 2.397 | 0.97 | -1.555 | -0.297 | -0.049 | 0.102 | 0.102 | 1.101 | 0.19 | 0.01 | 0.09 | 0.02 | 0.18 | 0.09 |
| 8 | 132999936 | rs2270875 | 1.27 | 1.027 | -1.286 | -0.076 | -0.023 | -0.026 | -0.026 | 0 | 0.08 | 0.03 | 0.15 | 0.16 | 0.11 | 0.25 |
| 8 | 133023595 | rs3763568 | -1.369 | 0.933 | -1.093 | -0.432 | -0.026 | -0.004 | -0.004 | 0 | 0.91 | 0.95 | 0.94 | 0.94 | 0.85 | 0.89 |
| 8 | 133029897 | rs2133999 | -1.116 | 0.908 | -1.101 | -0.563 | -0.009 | 0.006 | 0.006 | 0 | 0.91 | 0.9 | 0.93 | 0.96 | 0.85 | 0.89 |
| 8 | 133031047 | rs1501059 | -0.016 | 0.347 | -0.883 | -0.71 | -0.001 | 0.05 | 0.05 | 5.644 | 0.54 | 0.71 | 0.51 | 0.29 | 0.51 | 0.36 |
| 8 | 133033089 | rs1604292 | -0.38 | 0.23 | -0.47 | -0.324 | 0.007 | 0.041 | 0.041 | 0 | 0.38 | 0.76 | 0.46 | 0.28 | 0.36 | 0.34 |
| 8 | 133034922 | rs7007220 | 0.245 | 0.249 | -0.504 | -0.269 | -0.001 | -0.024 | -0.024 | 0 | 0.31 | 0.52 | 0.38 | 0.28 | 0.32 | 0.29 |
| 8 | 133035196 | rs7007939 | -0.054 | 0.308 | -0.805 | -0.198 | 0 | 0.002 | 0.002 | 0.054 | 0.36 | 0.68 | 0.42 | 0.28 | 0.34 | 0.3 |
| 8 | 133037144 | rs7812433 | -0.386 | -0.425 | -0.893 | 0.035 | 0.025 | 0.011 | 0.011 | 0 | 0.46 | 0.44 | 0.51 | 0.66 | 0.35 | 0.56 |
| 8 | 133045480 | rs2202068 | -1.705 | -0.117 | -0.485 | 0.225 | 0.005 | -0.041 | -0.041 | 0 | 0.83 | 0.76 | 0.85 | 0.93 | 0.81 | 0.92 |
| 8 | 133046303 | rs7387407 | -0.424 | -0.397 | -0.74 | 0.281 | -0.004 | -0.01 | -0.01 | 1.612 | 0.71 | 0.73 | 0.77 | 0.77 | 0.72 | 0.72 |
| 8 | 133048502 | rs12682408 | 1.083 | -0.349 | -0.734 | 0.344 | 0.005 | -0.016 | -0.016 | 0.013 | 0.12 | 0.03 | 0.08 | 0.15 | 0.09 | 0.19 |
| 8 | 133048593 | rs1472571 | 0.738 | 0.03 | -0.555 | 0.397 | -0.013 | -0.068 | -0.068 | 0.003 | 0.41 | 0.16 | 0.48 | 0.51 | 0.41 | 0.39 |
| 8 | 133050900 | rs871959 | -0.504 | 0.069 | -0.438 | 0.371 | -0.008 | -0.031 | -0.031 | 0 | 0.63 | 0.51 | 0.71 | 0.75 | 0.65 | 0.68 |
| 8 | 133051551 | rs2102103 | 0.828 | 0.063 | -0.718 | 0.393 | -0.016 | -0.067 | -0.067 | 0 | 0.42 | 0.16 | 0.48 | 0.51 | 0.41 | 0.38 |
| 8 | 133053309 | rs7386782 | -1.262 | 0.073 | -0.664 | 0.397 | -0.003 | -0.009 | -0.009 | 0 | 0.69 | 0.73 | 0.77 | 0.77 | 0.71 | 0.72 |
| 8 | 133053318 | rs7386783 | -1.273 | 0.071 | -0.665 | 0.396 | -0.004 | -0.009 | -0.009 | 0 | 0.69 | 0.73 | 0.77 | 0.77 | 0.71 | 0.72 |
| 8 | 133061073 | rs12674562 | 0.698 | 0.111 | -0.904 | 0.734 | 0 | 0.011 | 0.011 | 0 | 0.07 | 0.09 | 0.15 | 0.26 | 0.05 | 0.11 |
| 8 | 133061146 | rs4304313 | 0.943 | 0.034 | -0.992 | 0.636 | -0.041 | 0.007 | 0.007 | 0 | 0.37 | 0.2 | 0.36 | 0.27 | 0.41 | 0.23 |
| 8 | 133064238 | rs7831988 | 1.594 | 0.078 | -1.268 | 0.626 | -0.078 | -0.034 | -0.034 | 0 | 0.3 | 0.05 | 0.35 | 0.23 | 0.35 | 0.19 |
| 8 | 133064496 | rs7816194 | -0.115 | -0.03 | -1.09 | 0.675 | 0.029 | 0.019 | 0.019 | 0.153 | 0.27 | 0.47 | 0.23 | 0.27 | 0.29 | 0.33 |
| 8 | 133064881 | rs7836704 | 0.552 | -0.007 | -1.054 | 0.689 | 0.027 | 0.033 | 0.033 | 0.24 | 0.22 | 0.42 | 0.19 | 0.25 | 0.23 | 0.29 |

| | | | | | | | | | | | | | | | | |
|---|---|---|---|---|---|---|---|---|---|---|---|---|---|---|---|---|
| 8 | 133065379 | rs4736537 | 0.269 | -0.856 | -0.527 | 0.64 | 0.003 | -0.008 | -0.008 | 0 | 0.07 | 0.05 | 0.15 | 0.26 | 0.05 | 0.12 |
| 8 | 133069055 | rs10956630 | 0.514 | -0.626 | -0.801 | 0.795 | 0.034 | 0.002 | 0.002 | 0 | 0.52 | 0.46 | 0.58 | 0.38 | 0.64 | 0.44 |
| 8 | 133074827 | rs3802261 | -1.079 | -0.211 | -0.19 | 1.067 | -0.017 | 0.029 | 0.029 | 0 | 0.26 | 0.68 | 0.31 | 0.42 | 0.2 | 0.36 |
| 8 | 133078261 | rs3739280 | 0.02 | -0.012 | -0.246 | 1.114 | -0.021 | 0.054 | 0.054 | 2.524 | 0.05 | 0.15 | 0.03 | 0.13 | 0.01 | 0.04 |
| 8 | 133089975 | rs2280851 | -0.016 | 0.016 | -0.42 | 0.969 | -0.01 | 0.001 | 0.001 | 0 | 0.74 | 0.75 | 0.8 | 0.67 | 0.76 | 0.76 |
| 8 | 133091905 | rs11778661 | -0.015 | 0.474 | -0.736 | 1.048 | -0.012 | 0.014 | 0.014 | 0 | 0.27 | 0.17 | 0.39 | 0.2 | 0.3 | 0.3 |
| 8 | 133092138 | rs2403730 | -0.008 | 0.472 | -0.721 | 1.046 | -0.012 | 0.013 | 0.013 | 0 | 0.27 | 0.17 | 0.39 | 0.2 | 0.3 | 0.29 |
| 8 | 133098382 | rs1966702 | 0.454 | 0.35 | 0.141 | 1.713 | 0.04 | 0.226 | 0.226 | 0 | 0.92 | 0.68 | 0.9 | 0.7 | 0.94 | 0.92 |
| 8 | 133102651 | rs891764 | 0.043 | 0.373 | 0.331 | 1.791 | 0.019 | 0.111 | 0.111 | 0 | 0.94 | 0.84 | 0.91 | 0.73 | 0.94 | 0.93 |
| 8 | 133104433 | rs2118437 | 0.054 | -0.103 | -0.116 | 1.643 | -0.001 | 0.029 | 0.029 | 0 | 0.35 | 0.13 | 0.42 | 0.24 | 0.34 | 0.37 |
| 8 | 133105833 | rs16904589 | 0.958 | 0.527 | -0.291 | 1.891 | -0.02 | -0.034 | -0.034 | 0 | 0.1 | 0.04 | 0.13 | 0.2 | 0.15 | 0.08 |
| 8 | 133106539 | rs7824804 | -0.464 | 0.444 | -0.202 | 1.721 | 0.01 | 0.078 | 0.078 | 0 | 0.95 | 0.88 | 0.91 | 0.73 | 0.94 | 0.93 |
| 8 | 133108651 | rs2469608 | -0.366 | 0.803 | 0.215 | 1.304 | 0.024 | 0.039 | 0.039 | 0 | 0.25 | 0.33 | 0.26 | 0.41 | 0.34 | 0.37 |
| 8 | 133112960 | rs2469495 | 0.689 | 0.372 | -0.063 | 0.989 | -0.002 | 0.013 | 0.013 | 0 | 0.34 | 0.24 | 0.26 | 0.3 | 0.31 | 0.25 |
| 8 | 133116251 | rs6998880 | -0.326 | 0.457 | -0.169 | 0.958 | 0 | -0.013 | -0.013 | 0.947 | 0.57 | 0.66 | 0.65 | 0.46 | 0.58 | 0.49 |
| 8 | 133118482 | rs10111409 | 0.849 | 0.315 | -0.065 | 0.982 | 0.012 | 0.012 | 0.012 | 0 | 0.34 | 0.23 | 0.39 | 0.25 | 0.3 | 0.36 |
| 8 | 133118753 | rs2469643 | -0.101 | 0.39 | -0.084 | 0.855 | 0.043 | 0.001 | 0.001 | 0.39 | 0.51 | 0.35 | 0.49 | 0.51 | 0.57 | 0.33 |
| 8 | 133120467 | rs6992214 | 0.178 | 0.715 | -0.118 | 0.965 | 0.005 | 0.065 | 0.065 | 0 | 0.06 | 0.11 | 0.12 | 0.28 | 0.09 | 0.06 |
| 8 | 133121964 | rs10105818 | -0.025 | 0.709 | -0.004 | 0.946 | 0.021 | 0.133 | 0.133 | 0 | 0.07 | 0.19 | 0.14 | 0.28 | 0.09 | 0.06 |
| 8 | 133122294 | rs2436106 | -0.198 | 0.966 | 0.006 | 0.905 | 0.006 | -0.006 | -0.006 | 0.015 | 0.59 | 0.58 | 0.52 | 0.62 | 0.5 | 0.67 |
| 8 | 133122898 | rs2403732 | 0.776 | 0.951 | 0.127 | 1.007 | 0.085 | 0.328 | 0.328 | 0 | 0.9 | 0.63 | 0.72 | 0.62 | 0.85 | 0.89 |
| 8 | 133124206 | rs7825935 | 0.201 | 1.037 | -0.644 | 0.998 | 0.01 | 0.016 | 0.016 | 0 | 0.78 | 0.72 | 0.72 | 0.57 | 0.64 | 0.84 |
| 8 | 133126844 | rs2116465 | 0.279 | 0.933 | -0.694 | 0.926 | 0.052 | 0.02 | 0.02 | 0.374 | 0.48 | 0.64 | 0.55 | 0.57 | 0.37 | 0.64 |
| 8 | 133128066 | rs2436111 | 0.09 | 0.963 | -1.221 | 0.358 | 0.001 | 0.19 | 0.19 | 0.501 | 0.46 | 0.16 | 0.32 | 0.15 | 0.53 | 0.3 |
| 8 | 133131479 | rs2436118 | 0.957 | 0.874 | -1.217 | 0.312 | -0.029 | 0.166 | 0.166 | 0.497 | 0.31 | 0.07 | 0.17 | 0 | 0.27 | 0.2 |
| 8 | 133132028 | rs11783348 | 1.163 | 0.978 | -1.205 | 0.396 | -0.025 | -0.021 | -0.021 | 0 | 0.13 | 0.08 | 0.11 | 0.15 | 0.19 | 0.09 |
| 8 | 133134877 | rs9297840 | 0.849 | 0.592 | -0.912 | 0.73 | 0.048 | 0.05 | 0.05 | 0 | 0.36 | 0.21 | 0.35 | 0.22 | 0.25 | 0.34 |
| 8 | 133136551 | rs1437824 | 0.114 | 0.47 | -1.123 | 0.838 | -0.001 | 0.115 | 0.115 | 0 | 0.87 | 0.77 | 0.82 | 0.5 | 0.85 | 0.69 |
| 8 | 133139755 | rs977939 | 0.147 | 0.557 | -1.163 | 0.65 | -0.061 | 0.168 | 0.168 | 0 | 0.44 | 0.17 | 0.3 | 0.15 | 0.48 | 0.28 |
| 8 | 133143105 | rs2436133 | 0.176 | 0.659 | -1.033 | 0.769 | -0.052 | 0.139 | 0.139 | 0.454 | 0.43 | 0.21 | 0.33 | 0.16 | 0.47 | 0.23 |
| 8 | 133143785 | rs2469510 | 0.338 | 0.554 | -0.779 | 1.283 | -0.006 | 0.095 | 0.095 | 0 | 0.88 | 0.82 | 0.86 | 0.48 | 0.88 | 0.7 |
| 8 | 133150103 | rs2469515 | 0.345 | 0.808 | -1.484 | 0.072 | -0.02 | 0.114 | 0.114 | 0.162 | 0.24 | 0.06 | 0.15 | 0.01 | 0.24 | 0.12 |
| 8 | 133150903 | rs2469615 | -0.453 | 1.5 | -1.064 | 0.633 | -0.003 | -0.002 | -0.002 | 0 | 0.36 | 0.3 | 0.49 | 0.54 | 0.53 | 0.24 |
| 8 | 133154836 | rs2436139 | -0.789 | 1.621 | -0.777 | 1.125 | -0.001 | -0.002 | -0.002 | 0 | 0.42 | 0.53 | 0.48 | 0.4 | 0.56 | 0.27 |
| 8 | 133156467 | rs2469518 | -0.793 | 1.505 | -0.657 | 1.092 | 0 | -0.001 | -0.001 | 0 | 0.42 | 0.59 | 0.48 | 0.46 | 0.55 | 0.25 |
| 8 | 133157285 | rs1864772 | 0.856 | 1.079 | -0.51 | 0.763 | -0.008 | 0.035 | 0.035 | 0.291 | 0.8 | 0.69 | 0.86 | 0.68 | 0.8 | 0.66 |
| 8 | 133157519 | rs2469520 | 0.173 | 1.058 | -0.19 | 0.769 | -0.026 | -0.027 | -0.027 | 0 | 0.34 | 0.26 | 0.42 | 0.46 | 0.46 | 0.24 |
| 8 | 133158041 | rs1864773 | -0.139 | 0.978 | -0.28 | 0.805 | -0.013 | 0.061 | 0.061 | 0.085 | 0.86 | 0.8 | 0.89 | 0.68 | 0.89 | 0.67 |
| 8 | 133162001 | rs2469522 | 1.091 | 1.638 | -0.388 | 0.497 | -0.042 | 0.007 | 0.007 | 2.794 | 0.2 | 0.05 | 0.17 | 0.17 | 0.21 | 0.09 |
| 8 | 133167640 | rs6651173 | -0.365 | 1.598 | -0.081 | 0.448 | 0 | 0 | 0 | 0.141 | 0.4 | 0.43 | 0.45 | 0.45 | 0.54 | 0.25 |
| 8 | 133168611 | rs1007215 | -0.073 | 1.621 | -0.106 | 0.445 | -0.012 | -0.011 | -0.011 | 0 | 0.34 | 0.29 | 0.41 | 0.45 | 0.47 | 0.24 |
| 8 | 133171623 | rs9693071 | 0.202 | 1.425 | 0.201 | 0.372 | 0.004 | -0.054 | -0.054 | 0 | 0.09 | 0.26 | 0.07 | 0 | 0.07 | 0.1 |
| 8 | 133174006 | rs7007544 | -0.552 | 1.443 | 0.138 | 0.36 | 0.003 | -0.103 | -0.103 | 0.986 | 0.16 | 0.32 | 0.1 | 0 | 0.15 | 0.11 |
| 8 | 133183642 | rs3779972 | -0.975 | 1.565 | -0.092 | 0.6 | 0.028 | 0.005 | 0.005 | 0 | 0.5 | 0.58 | 0.51 | 0.45 | 0.61 | 0.35 |
| 8 | 133194780 | rs10956645 | 0.506 | 1.989 | 0.137 | 0.996 | 0.081 | 0.084 | 0.084 | 0 | 0.1 | 0.21 | 0.2 | 0.19 | 0.22 | 0.16 |
| 8 | 133202844 | rs10956647 | -1.416 | 1.446 | -0.517 | 0.741 | 0.047 | -0.051 | -0.051 | 0 | 0.63 | 0.85 | 0.51 | 0.47 | 0.71 | 0.55 |
| 8 | 133208405 | rs713148 | -0.448 | 1.708 | -0.459 | 0.57 | 0.003 | 0.041 | 0.041 | 0 | 0.5 | 0.41 | 0.36 | 0.31 | 0.5 | 0.36 |
| 8 | 133208506 | rs4282550 | 0.427 | 1.7 | -0.382 | 0.387 | 0.108 | 0.032 | 0.032 | 0.471 | 0.05 | 0.33 | 0.05 | 0 | 0.07 | 0.01 |
| 8 | 133208622 | rs4388439 | 0.427 | 1.7 | -0.382 | 0.387 | 0.108 | 0.032 | 0.032 | 0.471 | 0.05 | 0.33 | 0.05 | 0 | 0.07 | 0.01 |
| 8 | 133219987 | rs17595767 | -0.236 | 1.841 | -0.677 | 0.507 | -0.121 | 0.073 | 0.073 | 0 | 0.45 | 0.04 | 0.33 | 0.31 | 0.44 | 0.31 |
| 8 | 133220509 | rs9643288 | 1.228 | 1.943 | -0.809 | 0.378 | -0.097 | -0.064 | -0.064 | 0 | 0.26 | 0.01 | 0.22 | 0.3 | 0.29 | 0.12 |
| 8 | 133223530 | rs11779500 | 0.15 | 1.87 | -0.163 | 0.933 | 0.081 | 0.064 | 0.064 | 0 | 0.12 | 0.33 | 0.15 | 0.14 | 0.19 | 0.25 |
| 8 | 133227626 | rs17595945 | -0.076 | 1.51 | -1.125 | 0.56 | -0.108 | 0.083 | 0.083 | 0 | 0.46 | 0.04 | 0.32 | 0.3 | 0.44 | 0.32 |
| 8 | 133238760 | rs6987299 | -0.41 | 1.246 | -0.856 | 0.625 | 0.003 | 0.028 | 0.028 | 0 | 0.52 | 0.35 | 0.47 | 0.42 | 0.51 | 0.4 |
| 8 | 133242547 | rs1025435 | -0.543 | 0.705 | -0.814 | 0.281 | 0.011 | 0.014 | 0.014 | 2.105 | 0.64 | 0.66 | 0.72 | 0.73 | 0.72 | 0.82 |
| 8 | 133243589 | rs10090173 | -1.278 | 1.035 | -0.439 | 0.31 | -0.011 | 0.006 | 0.006 | 0.525 | 0.35 | 0.65 | 0.5 | 0.29 | 0.46 | 0.49 |
| 8 | 133247202 | rs10092367 | -1.117 | 1.288 | -0.317 | 0.226 | 0.019 | -0.006 | -0.006 | 0 | 0.1 | 0.23 | 0.13 | 0.11 | 0.11 | 0.21 |
| 8 | 133248679 | rs930894 | -0.866 | 1.27 | -0.276 | 0.403 | 0.032 | -0.003 | -0.003 | 0 | 0.28 | 0.58 | 0.4 | 0.22 | 0.36 | 0.31 |
| 8 | 133251797 | rs1031635 | 1.714 | 1.415 | -0.131 | 0.381 | 0.052 | 0 | 0 | 1.291 | 0.76 | 0.38 | 0.64 | 0.75 | 0.62 | 0.62 |
| 8 | 133255653 | rs17596711 | -0.983 | 1.404 | -0.142 | 0.38 | 0.001 | -0.001 | -0.001 | 0.021 | 0.09 | 0.09 | 0.11 | 0.11 | 0.11 | 0.2 |
| 8 | 133262048 | rs7819270 | 0.121 | 1.39 | -0.11 | 0.391 | 0.168 | 0.047 | 0.047 | 0.005 | 0.85 | 0.46 | 0.75 | 0.86 | 0.73 | 0.82 |
| 8 | 133270890 | rs7007269 | 2.138 | 1.227 | -0.298 | 0.344 | -0.085 | 0.037 | 0.037 | 0.958 | 0.61 | 0.13 | 0.47 | 0.71 | 0.53 | 0.49 |
| 8 | 133275170 | rs1902820 | 1.995 | 1.301 | -0.296 | 0.444 | -0.111 | 0.01 | 0.01 | 1.28 | 0.63 | 0.12 | 0.48 | 0.7 | 0.53 | 0.48 |
| 8 | 133286978 | rs4736412 | -0.074 | 1.401 | -0.18 | 0.519 | 0.007 | 0.005 | 0.005 | 0 | 0.13 | 0.06 | 0.16 | 0.08 | 0.11 | 0.16 |
| 8 | 133291497 | rs13264960 | 1.923 | 1.004 | -0.408 | 0.335 | -0.085 | 0.013 | 0.013 | 1.017 | 0.59 | 0.14 | 0.46 | 0.66 | 0.53 | 0.48 |
| 8 | 133294972 | rs1457780 | 1.219 | 1.005 | -0.318 | 0.314 | 0.001 | -0.001 | -0.001 | 0 | 0.72 | 0.33 | 0.64 | 0.76 | 0.64 | 0.65 |
| 8 | 133295072 | rs1457781 | 1.895 | 1.008 | -0.341 | 0.331 | -0.095 | 0.05 | 0.05 | 0.968 | 0.59 | 0.13 | 0.57 | 0.7 | 0.53 | 0.49 |
| 8 | 133298658 | rs13249479 | 1.131 | 1.07 | -0.259 | 0.331 | 0.019 | -0.013 | -0.013 | 1.127 | 0.73 | 0.36 | 0.64 | 0.79 | 0.64 | 0.66 |
| 8 | 133306801 | rs2100644 | 1.534 | 1.066 | -0.174 | 0.345 | -0.018 | 0.03 | 0.03 | 0.252 | 0.66 | 0.27 | 0.57 | 0.81 | 0.61 | 0.66 |

| | | | | | | | | | | | | | | | | |
|---|---|---|---|---|---|---|---|---|---|---|---|---|---|---|---|---|
| 8 | 133309703 | rs7818112 | 1.483 | 1.073 | -0.194 | 0.36 | -0.005 | 0.004 | 0.004 | 0.195 | 0.66 | 0.38 | 0.58 | 0.78 | 0.61 | 0.67 |
| 8 | 133312022 | rs6471054 | -0.098 | 1.295 | 0.365 | 0.343 | 0.106 | 0.025 | 0.025 | 0 | 0.88 | 0.67 | 0.8 | 0.89 | 0.78 | 0.84 |
| 8 | 133313335 | rs1457784 | -1.109 | 1.007 | -0.205 | 0.28 | -0.002 | -0.009 | -0.009 | 0 | 0.21 | 0.24 | 0.27 | 0.18 | 0.19 | 0.3 |
| 8 | 133317111 | rs11778399 | 1.797 | 0.595 | -0.533 | 0.522 | -0.006 | 0.01 | 0.01 | 0 | 0.55 | 0.38 | 0.48 | 0.6 | 0.51 | 0.49 |
| 8 | 133319798 | rs7837201 | -1.537 | 0.773 | -0.488 | 0.538 | -0.01 | -0.006 | -0.006 | 0 | 0.29 | 0.5 | 0.36 | 0.29 | 0.21 | 0.32 |
| 8 | 133320536 | rs1457788 | -0.681 | 0.782 | -0.491 | 0.538 | -0.009 | 0.006 | 0.006 | 0 | 0.85 | 0.9 | 0.84 | 0.92 | 0.73 | 0.82 |
| 8 | 133321519 | rs4736572 | 1.906 | 0.758 | -0.496 | 0.536 | -0.006 | 0.006 | 0.006 | 0 | 0.56 | 0.39 | 0.47 | 0.59 | 0.51 | 0.5 |
| 8 | 133323802 | rs10094856 | -0.629 | 0.868 | -0.526 | 0.578 | -0.007 | 0.003 | 0.003 | 0 | 0.85 | 0.9 | 0.84 | 0.9 | 0.72 | 0.82 |
| 8 | 133325922 | rs4736413 | 1.287 | 0.802 | -0.573 | 0.562 | 0.01 | -0.003 | -0.003 | 0 | 0.63 | 0.45 | 0.49 | 0.64 | 0.52 | 0.53 |
| 8 | 133329296 | rs4736573 | 1.541 | 0.921 | -0.576 | 0.607 | -0.012 | 0.003 | 0.003 | 0 | 0.58 | 0.38 | 0.49 | 0.6 | 0.53 | 0.51 |
| 8 | 133329390 | rs12541666 | -1.251 | 0.781 | -0.728 | 0.792 | -0.005 | 0.013 | 0.013 | 0 | 0.26 | 0.5 | 0.35 | 0.29 | 0.19 | 0.31 |
| 8 | 133331038 | rs16904657 | 0.408 | 0.685 | -1.474 | 1.017 | -0.018 | -0.032 | -0.032 | 0 | 0.06 | 0 | 0.06 | 0.13 | 0.05 | 0.11 |
| 8 | 133339397 | rs13267466 | 0.492 | 0.223 | -1.753 | 1.247 | -0.059 | -0.089 | -0.089 | 3.261 | 0.19 | 0.01 | 0.12 | 0.24 | 0.17 | 0.12 |
| 8 | 133341355 | rs2896656 | -0.225 | 0.411 | -1.182 | 0.985 | 0.021 | -0.002 | -0.002 | 0.133 | 0.34 | 0.43 | 0.35 | 0.33 | 0.46 | 0.44 |
| 8 | 133344709 | rs6471060 | 0.102 | 0.603 | -1.416 | 0.811 | 0.017 | -0.007 | -0.007 | 0.175 | 0.31 | 0.38 | 0.35 | 0.24 | 0.45 | 0.44 |
| 8 | 133350326 | rs1020740 | 0.422 | -0.006 | -1.585 | 1.08 | 0.043 | -0.035 | -0.035 | 1.896 | 0.6 | 0.24 | 0.61 | 0.57 | 0.71 | 0.74 |
| 8 | 133350903 | rs6993047 | 0.121 | 0.386 | -1.615 | 1.355 | -0.037 | -0.08 | -0.08 | 0 | 0.12 | 0 | 0.08 | 0.23 | 0.12 | 0.06 |
| 8 | 133351120 | rs7824311 | -0.965 | -0.19 | -1.131 | 1.15 | -0.005 | 0.002 | 0.002 | 0 | 0.35 | 0.62 | 0.34 | 0.36 | 0.26 | 0.16 |
| 8 | 133354450 | rs13276777 | -0.921 | 0.251 | -0.334 | 1.716 | 0.029 | 0.186 | 0.186 | 0 | 0.1 | 0.23 | 0.24 | 0.41 | 0.11 | 0.14 |
| 8 | 133366787 | rs930895 | -1.376 | 0.035 | 0.011 | 1.371 | -0.002 | 0.025 | 0.025 | 0.79 | 0.25 | 0.67 | 0.33 | 0.48 | 0.24 | 0.31 |
| 8 | 133374331 | rs6984395 | -0.502 | -0.027 | 0.259 | 1.078 | 0.002 | 0.048 | 0.048 | 0 | 0.1 | 0.3 | 0.11 | 0.12 | 0.09 | 0.2 |
| 8 | 133377440 | rs2100646 | -1.548 | -0.071 | 0.548 | 1.296 | 0.022 | 0.226 | 0.226 | 0.116 | 0.19 | 0.52 | 0.34 | 0.48 | 0.19 | 0.31 |
| 8 | 133378663 | rs10095671 | 0.505 | 0.15 | 0.408 | 1.197 | 0.016 | 0.051 | 0.051 | 0 | 0.08 | 0.16 | 0.13 | 0.23 | 0.1 | 0.02 |
| 8 | 133382074 | rs1350382 | 0.107 | 0.074 | 0.333 | 1.1 | -0.019 | -0.052 | -0.052 | 0 | 0.15 | 0.33 | 0.09 | 0.05 | 0.14 | 0.19 |
| 8 | 133382498 | rs6992843 | -1.486 | 0.071 | 0.045 | 1.129 | 0.027 | 0.126 | 0.126 | 0 | 0.24 | 0.46 | 0.41 | 0.48 | 0.26 | 0.31 |
| 8 | 133383549 | rs16904676 | -0.934 | 0.107 | 0.126 | 1.087 | -0.022 | -0.01 | -0.01 | 0 | 0.23 | 0.56 | 0.19 | 0.18 | 0.22 | 0.37 |
| 8 | 133391926 | rs2198985 | 0.687 | 0.228 | 0.083 | 1.16 | -0.002 | 0.017 | 0.017 | 0 | 0.07 | 0.07 | 0.09 | 0.13 | 0.06 | 0.16 |
| 8 | 133398964 | rs2721916 | -1.638 | 0.481 | 0.578 | 1.154 | 0.015 | 0.073 | 0.073 | 0 | 0.26 | 0.56 | 0.43 | 0.49 | 0.28 | 0.3 |
| 8 | 133402654 | rs2673588 | 0.21 | 0.582 | 0.415 | 1.214 | 0.044 | 0.178 | 0.178 | 0 | 0.92 | 0.78 | 0.76 | 0.64 | 0.9 | 0.91 |
| 8 | 133404011 | rs6471065 | 1.546 | 0.552 | -0.313 | 0.455 | 0.002 | 0.01 | 0.01 | 2.306 | 0.22 | 0.12 | 0.11 | 0 | 0.19 | 0.13 |
| 8 | 133404409 | rs2673590 | 0.81 | 0.367 | -0.391 | 0.445 | -0.057 | -0.037 | -0.037 | 0 | 0.46 | 0.11 | 0.44 | 0.45 | 0.46 | 0.41 |
| 8 | 133405459 | rs2597356 | 1.092 | 0.375 | -0.425 | 0.451 | -0.093 | -0.137 | -0.137 | 0 | 0.4 | 0.02 | 0.37 | 0.45 | 0.39 | 0.4 |
| 8 | 133408442 | rs6471067 | 1.398 | 0.599 | -0.192 | 0.459 | 0 | -0.032 | -0.032 | 0.102 | 0.23 | 0.17 | 0.12 | 0 | 0.19 | 0.13 |
| 8 | 133409932 | rs2673606 | 0.411 | 0.969 | 0.194 | 1.169 | 0.031 | 0.108 | 0.108 | 0 | 0.85 | 0.67 | 0.7 | 0.63 | 0.83 | 0.9 |
| 8 | 133410500 | rs2597353 | 0.318 | 0.892 | -0.157 | 1.067 | 0.013 | 0.037 | 0.037 | 0 | 0.06 | 0.11 | 0.16 | 0.13 | 0.07 | 0.08 |
| 8 | 133416556 | rs2673562 | 0.892 | 0.829 | -0.41 | 1.072 | 0 | 0.003 | 0.003 | 0.655 | 0.69 | 0.29 | 0.58 | 0.6 | 0.65 | 0.73 |
| 8 | 133419639 | rs2721894 | 0.124 | 0.581 | -0.902 | 0.765 | -0.1 | -0.099 | -0.099 | 0.383 | 0.36 | 0.11 | 0.46 | 0.47 | 0.47 | 0.49 |
| 8 | 133423607 | rs17659499 | 0.915 | 0.525 | -1.108 | 0.687 | 0.035 | 0.077 | 0.077 | 3.339 | 0.4 | 0.2 | 0.2 | 0.13 | 0.28 | 0.28 |
| 8 | 133424083 | rs2597335 | -0.179 | 1.345 | -1.255 | 1.253 | -0.113 | -0.153 | -0.153 | 0 | 0.71 | 0.96 | 0.62 | 0.54 | 0.65 | 0.56 |
| 8 | 133427940 | rs2597331 | -0.952 | 1.542 | -0.298 | 1.015 | 0.066 | 0.087 | 0.087 | 0.657 | 0.11 | 0.36 | 0.15 | 0.26 | 0.18 | 0.08 |
| 8 | 133429049 | rs12548732 | 0.143 | 1.783 | -0.932 | 1.049 | -0.065 | -0.044 | -0.044 | 0 | 0.2 | 0.03 | 0.29 | 0.23 | 0.27 | 0.17 |
| 8 | 133434644 | rs1962131 | 0.329 | 1.307 | -0.635 | 0.727 | -0.007 | -0.002 | -0.002 | 0 | 0.28 | 0.3 | 0.18 | 0.3 | 0.25 | 0.18 |
| 8 | 133442492 | rs1515519 | 0.817 | 1.314 | -0.768 | 0.715 | -0.034 | -0.016 | -0.016 | 0 | 0.17 | 0.13 | 0.29 | 0.23 | 0.27 | 0.16 |
| 8 | 133444342 | rs2673593 | -0.026 | 0.815 | -0.375 | 0.344 | -0.008 | -0.001 | -0.001 | 0.097 | 0.3 | 0.33 | 0.2 | 0.31 | 0.29 | 0.21 |
| 8 | 133448569 | rs7834336 | 0.876 | 0.584 | -0.502 | 0.407 | -0.009 | -0.028 | -0.028 | 0 | 0.36 | 0.39 | 0.2 | 0.2 | 0.28 | 0.38 |
| 8 | 133452802 | rs16904685 | 0.704 | 0.608 | -1.191 | 0.784 | 0.02 | -0.01 | -0.01 | 0 | 0.22 | 0.13 | 0.11 | 0.26 | 0.16 | 0.1 |
| 8 | 133454051 | rs7837397 | 0.216 | 1.632 | -0.943 | 1.098 | -0.043 | -0.046 | -0.046 | 0 | 0.68 | 0.75 | 0.39 | 0.51 | 0.56 | 0.56 |
| 8 | 133456441 | rs1878016 | 0.523 | 1.58 | -0.417 | 1.051 | 0.007 | 0.018 | 0.018 | 0 | 0.84 | 0.75 | 0.78 | 0.81 | 0.83 | 0.83 |
| 8 | 133458487 | rs2721905 | 0.835 | 2.245 | -0.249 | 0.762 | 0.04 | -0.016 | -0.016 | 0.937 | 0.72 | 0.38 | 0.6 | 0.75 | 0.6 | 0.65 |
| 8 | 133464737 | rs2721908 | 0.566 | 1.881 | -0.328 | 0.89 | -0.001 | 0 | 0 | 0.094 | 0.93 | 0.92 | 0.9 | 0.95 | 0.92 | 0.88 |
| 8 | 133471941 | rs2597348 | 1.627 | 2.471 | 0.196 | 1.274 | 0.039 | 0.069 | 0.069 | 0 | 0.75 | 0.62 | 0.53 | 0.59 | 0.65 | 0.69 |
| 8 | 133472585 | rs2673586 | 0.93 | 2.5 | -0.06 | 1.161 | 0.002 | 0.005 | 0.005 | 0 | 0.87 | 0.86 | 0.87 | 0.94 | 0.87 | 0.87 |
| 8 | 133476806 | rs2673555 | 0.69 | 1.71 | -1.253 | 0.617 | -0.077 | -0.101 | -0.101 | 0.09 | 0.3 | 0.03 | 0.18 | 0.39 | 0.33 | 0.28 |
| 8 | 133479165 | rs2673559 | 0.622 | 1.752 | -0.71 | 0.82 | 0.02 | 0.065 | 0.065 | 0 | 0.85 | 0.83 | 0.61 | 0.55 | 0.74 | 0.72 |
| 8 | 133479384 | rs2673558 | -0.184 | 0.744 | -1.132 | 0.266 | 0.012 | 0.014 | 0.014 | 0.035 | 0.8 | 0.86 | 0.84 | 0.91 | 0.83 | 0.78 |
| 8 | 133481225 | rs4736586 | 0.641 | 0.976 | -1.073 | 0.416 | 0.029 | 0.089 | 0.089 | 0 | 0.08 | 0.07 | 0.29 | 0.37 | 0.19 | 0.14 |
| 8 | 133481337 | rs2673557 | 0.758 | 0.778 | -1.384 | 0.261 | -0.022 | -0.008 | -0.008 | 0.023 | 0.42 | 0.07 | 0.23 | 0.4 | 0.41 | 0.36 |
| 8 | 133482177 | rs10217015 | 0.607 | 0.725 | -0.987 | 0.228 | 0.002 | -0.004 | -0.004 | 0 | 0.37 | 0.6 | 0.33 | 0.15 | 0.29 | 0.35 |
| 8 | 133482984 | rs6471070 | 0.751 | 0.83 | -1.167 | 0.407 | 0.029 | 0.091 | 0.091 | 0.005 | 0.08 | 0.07 | 0.29 | 0.39 | 0.19 | 0.14 |
| 8 | 133487525 | rs2167171 | -0.28 | 0.648 | -1.242 | 0.22 | -0.004 | -0.004 | -0.004 | 0 | 0.92 | 0.9 | 0.89 | 0.92 | 0.91 | 0.87 |
| 8 | 133493727 | rs7002144 | 0.448 | 0.458 | -0.659 | -0.066 | -0.007 | -0.048 | -0.048 | 0 | 0.35 | 0.4 | 0.31 | 0.12 | 0.28 | 0.32 |
| 8 | 133494554 | rs3824245 | 1.024 | 0.206 | -0.894 | -0.048 | -0.005 | -0.027 | -0.027 | 0.337 | 0.27 | 0.3 | 0.26 | 0.12 | 0.2 | 0.31 |
| 8 | 133497127 | rs1901090 | 0.547 | 0.027 | -1.047 | 0.327 | -0.005 | -0.003 | -0.003 | 0 | 0.33 | 0.11 | 0.19 | 0.34 | 0.33 | 0.29 |
| 8 | 133499372 | rs10100851 | -0.731 | -0.562 | -0.371 | 0.123 | -0.084 | -0.103 | -0.103 | 0 | 0.24 | 0.49 | 0.14 | 0.05 | 0.16 | 0.12 |
| 8 | 133500118 | rs7823044 | 0.51 | -0.429 | -0.696 | 0.721 | -0.008 | 0.002 | 0.002 | 0 | 0.12 | 0.13 | 0.17 | 0.15 | 0.08 | 0.07 |
| 8 | 133505209 | rs3847171 | 0.141 | -0.558 | -1.197 | 0.616 | -0.005 | 0.028 | 0.028 | 0.098 | 0.38 | 0.25 | 0.47 | 0.31 | 0.52 | 0.46 |
| 8 | 133506298 | rs2597364 | 0.026 | -1.251 | -1.358 | 0.911 | -0.04 | 0.033 | 0.033 | 0.279 | 0.75 | 0.45 | 0.8 | 0.78 | 0.84 | 0.72 |
| 8 | 133506571 | rs10046742 | -0.148 | -1.435 | -1.275 | 1.09 | -0.006 | 0.152 | 0.152 | 0.076 | 0.11 | 0.5 | 0.09 | 0.12 | 0.05 | 0.15 |
| 8 | 133506626 | rs2673572 | 1.583 | -1.541 | -1.399 | 1.124 | 0.009 | -0.016 | -0.016 | 0.117 | 0.25 | 0.07 | 0.15 | 0.31 | 0.23 | 0.2 |

| | | | | | | | | | | | | | | | | |
|---|---|---|---|---|---|---|---|---|---|---|---|---|---|---|---|---|
| 8 | 133506801 | rs2673571 | -0.948 | -0.909 | -1.056 | 0.94 | 0.012 | 0.003 | 0.003 | 0 | 0.86 | 0.95 | 0.89 | 0.89 | 0.89 | 0.87 |
| 8 | 133507756 | rs1901088 | 0.729 | 0.077 | -1.326 | 0.853 | 0.068 | 0.011 | 0.011 | 0 | 0.11 | 0.16 | 0.25 | 0.17 | 0.2 | 0.1 |
| 8 | 133512340 | rs2673597 | -1.176 | -0.253 | -1.282 | 0.86 | 0.012 | -0.008 | -0.008 | 0.084 | 0.86 | 0.95 | 0.89 | 0.85 | 0.89 | 0.84 |
| 8 | 133512858 | rs2597362 | -0.381 | -0.33 | -1.377 | 0.777 | -0.026 | 0.042 | 0.042 | 0 | 0.63 | 0.89 | 0.7 | 0.71 | 0.64 | 0.5 |
| 8 | 133516071 | rs2673579 | -0.062 | -0.386 | -1.36 | 0.824 | 0.003 | -0.003 | -0.003 | 0 | 0.58 | 0.42 | 0.64 | 0.59 | 0.64 | 0.42 |
| 8 | 133516844 | rs13251704 | 1.388 | -0.091 | -1.297 | 0.651 | -0.071 | 0.039 | 0.039 | 0.426 | 0.23 | 0.02 | 0.19 | 0.14 | 0.25 | 0.35 |
| 8 | 133519525 | rs11779861 | -0.175 | 0.075 | -0.942 | 0.912 | 0.017 | -0.001 | -0.001 | 0 | 0.17 | 0.19 | 0.27 | 0.17 | 0.25 | 0.17 |
| 8 | 133520033 | rs16904694 | -0.275 | -0.241 | -0.53 | 1.139 | 0.076 | 0.337 | 0.337 | 0.869 | 0.05 | 0.49 | 0.06 | 0.11 | 0.01 | 0.07 |
| 8 | 133525507 | rs2597379 | 0.44 | -0.145 | -1.301 | 0.404 | 0.039 | -0.021 | -0.021 | 0.39 | 0.38 | 0.28 | 0.32 | 0.52 | 0.31 | 0.35 |
| 8 | 133526556 | rs6999947 | 0.412 | 0.189 | -1.448 | -0.05 | -0.007 | 0.051 | 0.051 | 0.227 | 0.28 | 0.1 | 0.27 | 0.14 | 0.29 | 0.41 |
| 8 | 133528075 | rs2721876 | 0.192 | -0.074 | -0.984 | 0.16 | -0.009 | -0.007 | -0.007 | 0 | 0.42 | 0.48 | 0.44 | 0.67 | 0.34 | 0.43 |
| 8 | 133528541 | rs2733162 | 0.496 | -0.472 | -2.01 | -0.801 | -0.006 | 0.177 | 0.177 | 0 | 0.32 | 0.04 | 0.21 | 0.01 | 0.28 | 0.17 |
| 8 | 133528579 | rs2673576 | 0.009 | -1.021 | -1.246 | 0.061 | -0.02 | 0.02 | 0.02 | 0 | 0.32 | 0.61 | 0.28 | 0.47 | 0.25 | 0.36 |
| 8 | 133528646 | rs10505597 | 0.42 | -0.99 | -1.028 | 0.223 | -0.006 | -0.008 | -0.008 | 0 | 0.17 | 0.24 | 0.14 | 0.18 | 0.14 | 0.15 |
| 8 | 133529038 | rs1473973 | 0.379 | -0.746 | -1.034 | 0.593 | -0.004 | 0.011 | 0.011 | 0.149 | 0.72 | 0.69 | 0.69 | 0.57 | 0.71 | 0.54 |
| 8 | 133530035 | rs2138248 | 0.264 | -0.432 | -1.188 | 0.297 | -0.002 | 0.011 | 0.011 | 0.427 | 0.72 | 0.79 | 0.62 | 0.79 | 0.66 | 0.54 |
| 8 | 133531706 | rs2738422 | 0.49 | -0.443 | -1.2 | 0.292 | 0 | 0.002 | 0.002 | 0.363 | 0.69 | 0.71 | 0.62 | 0.79 | 0.66 | 0.54 |
| 8 | 133534265 | rs2721884 | -0.229 | -0.393 | -1.575 | 0.361 | -0.061 | -0.039 | -0.039 | 0 | 0.77 | 0.98 | 0.68 | 0.79 | 0.74 | 0.55 |
| 8 | 133536338 | rs13266648 | 0.839 | -0.539 | -1.824 | 0.085 | -0.062 | 0.054 | 0.054 | 0 | 0.29 | 0.16 | 0.26 | 0.18 | 0.38 | 0.17 |
| 8 | 133540343 | rs1515509 | -0.292 | -0.496 | -1.523 | 0.302 | -0.01 | -0.001 | -0.001 | 0 | 0.77 | 0.9 | 0.61 | 0.8 | 0.74 | 0.55 |
| 8 | 133549399 | rs9297852 | 0.403 | -0.61 | -1.037 | 0.251 | -0.007 | 0.023 | 0.023 | 0.537 | 0.69 | 0.57 | 0.52 | 0.48 | 0.68 | 0.44 |
| 8 | 133562114 | rs7832599 | 0.38 | -1.326 | -1.106 | -0.639 | -0.005 | -0.044 | -0.044 | 0 | 0.36 | 0.58 | 0.2 | 0.04 | 0.19 | 0.18 |
| 8 | 133563781 | rs10956670 | 1.167 | -0.789 | -1.571 | -0.266 | -0.103 | -0.045 | -0.045 | 0 | 0.23 | 0.1 | 0.27 | 0.29 | 0.4 | 0.18 |
| 8 | 133564247 | rs10956671 | -0.937 | -0.893 | -1.69 | -0.174 | -0.076 | -0.163 | -0.163 | 2.775 | 0.73 | 0.98 | 0.55 | 0.6 | 0.71 | 0.52 |
| 8 | 133569397 | rs4621788 | -0.11 | -1.625 | -0.625 | -0.492 | -0.007 | 0.051 | 0.051 | 0.062 | 0.14 | 0.28 | 0.09 | 0.27 | 0.12 | 0.16 |
| 8 | 133573057 | rs6471080 | 0.132 | -1.401 | -0.444 | -0.693 | -0.032 | -0.091 | -0.091 | 0.474 | 0.34 | 0.52 | 0.19 | 0.04 | 0.19 | 0.18 |
| 8 | 133573931 | rs7000580 | 0.747 | -1.35 | -0.342 | -0.633 | 0.016 | -0.014 | -0.014 | 0 | 0.28 | 0.18 | 0.11 | 0.01 | 0.15 | 0.14 |
| 8 | 133577588 | rs13253244 | -0.43 | -1.075 | -1.294 | -0.096 | 0.11 | 0.126 | 0.126 | 0.692 | 0.51 | 0.88 | 0.28 | 0.32 | 0.32 | 0.37 |
| 8 | 133580193 | rs2357507 | 0.121 | -1.284 | -0.466 | -0.64 | -0.018 | -0.11 | -0.11 | 0 | 0.3 | 0.44 | 0.2 | 0.05 | 0.22 | 0.22 |
| 8 | 133583294 | rs7841268 | 0.201 | -1.351 | -0.244 | -0.675 | 0.002 | -0.027 | -0.027 | 0 | 0.11 | 0.32 | 0.09 | 0.05 | 0.09 | 0.1 |
| 8 | 133584558 | rs9297853 | 0.201 | -1.352 | -0.239 | -0.675 | 0.006 | -0.027 | -0.027 | 0 | 0.11 | 0.33 | 0.09 | 0.05 | 0.09 | 0.1 |
| 8 | 133584821 | rs16904707 | -0.097 | -1.365 | -0.308 | -0.681 | 0.008 | -0.033 | -0.033 | 0 | 0.13 | 0.42 | 0.1 | 0.05 | 0.09 | 0.1 |
| 8 | 133591952 | rs2357508 | -1.034 | -1.162 | -0.624 | -0.418 | 0.01 | 0.081 | 0.081 | 0.598 | 0.44 | 0.9 | 0.32 | 0.28 | 0.36 | 0.37 |
| 8 | 133605704 | rs7465694 | -0.436 | -0.95 | -0.194 | -0.33 | 0.049 | 0.109 | 0.109 | 0 | 0.14 | 0.44 | 0.13 | 0.24 | 0.16 | 0.12 |
| 8 | 133624912 | rs16904712 | 1.54 | -1.755 | -0.958 | 0.089 | 0.078 | -0.077 | -0.077 | 0 | 0.19 | 0.02 | 0.15 | 0.3 | 0.1 | 0.11 |
| 8 | 133629614 | rs4480107 | -1.915 | -1.738 | -1.043 | 0.027 | -0.151 | -0.073 | -0.073 | 0.861 | 0.65 | 0.95 | 0.46 | 0.59 | 0.47 | 0.49 |
| 8 | 133634814 | rs2272681 | -1.204 | -0.501 | -0.204 | 0.002 | -0.019 | 0.079 | 0.079 | 0.758 | 0.44 | 0.88 | 0.32 | 0.28 | 0.38 | 0.37 |
| 8 | 133637659 | rs2293979 | -1.212 | -0.494 | -0.231 | 0.006 | -0.021 | 0.08 | 0.08 | 0.75 | 0.45 | 0.9 | 0.32 | 0.28 | 0.38 | 0.37 |
| 8 | 133645380 | rs7834760 | 0.048 | -0.341 | 0.008 | 0.066 | 0.018 | -0.03 | -0.03 | 0.065 | 0.12 | 0.38 | 0.09 | 0.05 | 0.09 | 0.1 |
| 8 | 133649465 | rs4736598 | -0.277 | -0.246 | -0.103 | 0.382 | 0.035 | 0.128 | 0.128 | 0.669 | 0.11 | 0.29 | 0.12 | 0.35 | 0.15 | 0.13 |
| 8 | 133685650 | rs7004199 | -0.523 | -0.384 | -1.242 | 0.219 | -0.021 | 0.076 | 0.076 | 0.832 | 0.45 | 0.9 | 0.34 | 0.32 | 0.4 | 0.38 |
| 8 | 133690193 | rs6981978 | -1.044 | -0.184 | -0.177 | 0.231 | 0.037 | -0.009 | -0.009 | 0 | 0.07 | 0.59 | 0.07 | 0 | 0.02 | 0.02 |
| 8 | 133698781 | rs3909640 | -0.863 | -0.32 | -1.124 | 0.368 | -0.067 | 0.032 | 0.032 | 0 | 0.54 | 0.94 | 0.36 | 0.61 | 0.48 | 0.39 |
| 8 | 133701503 | rs7841637 | -0.577 | -0.565 | -0.635 | 0.784 | -0.025 | 0.026 | 0.026 | 0 | 0.22 | 0.6 | 0.16 | 0.23 | 0.15 | 0.16 |
| 8 | 133701531 | rs7464815 | 0.54 | -0.522 | -0.804 | 0.722 | -0.004 | -0.025 | -0.025 | 0 | 0.33 | 0.19 | 0.2 | 0.38 | 0.33 | 0.26 |
| 8 | 133704712 | rs10097529 | 0.345 | -0.505 | -0.856 | 0.821 | 0.008 | -0.034 | -0.034 | 0 | 0.15 | 0.05 | 0.09 | 0.23 | 0.14 | 0.11 |
| 8 | 133711481 | rs11780202 | -0.112 | -0.555 | -0.757 | 0.707 | 0.008 | 0.002 | 0.002 | 0 | 0.39 | 0.73 | 0.26 | 0.38 | 0.33 | 0.29 |
| 8 | 133713110 | rs4736424 | -0.033 | -0.569 | -0.842 | 0.7 | 0.01 | -0.005 | -0.005 | 2.253 | 0.4 | 0.86 | 0.26 | 0.38 | 0.33 | 0.29 |
| 8 | 133714487 | rs3843561 | 1.926 | -0.288 | -1.437 | 0.187 | -0.068 | -0.107 | -0.107 | 0 | 0.21 | 0.01 | 0.31 | 0.33 | 0.22 | 0.32 |
| 8 | 133714778 | rs3843562 | -0.154 | -0.398 | -0.79 | 0.141 | -0.007 | 0.065 | 0.065 | 0 | 0.4 | 0.86 | 0.28 | 0.3 | 0.34 | 0.26 |
| 8 | 133716772 | rs6993371 | -0.661 | -0.413 | -0.403 | 0.101 | 0.068 | 0.034 | 0.034 | 2.511 | 0.06 | 0.49 | 0.06 | 0 | 0.01 | 0.01 |
| 8 | 133717649 | rs11988034 | -1.186 | -0.454 | -0.653 | 0.071 | -0.118 | -0.034 | -0.034 | 3.532 | 0.56 | 0.95 | 0.36 | 0.6 | 0.45 | 0.36 |
| 8 | 133722281 | rs1048490 | -1.126 | -0.457 | -0.516 | 0.07 | -0.105 | -0.022 | -0.022 | 0.79 | 0.56 | 0.93 | 0.36 | 0.6 | 0.45 | 0.36 |
| 8 | 133722669 | rs2280871 | -0.929 | -0.441 | -0.216 | 0.101 | -0.007 | 0.003 | 0.003 | 1.553 | 0.23 | 0.72 | 0.16 | 0.3 | 0.12 | 0.14 |
| 8 | 133731174 | rs11996730 | -0.285 | -0.429 | -0.112 | 0.02 | 0.001 | 0.059 | 0.059 | 0 | 0.4 | 0.77 | 0.27 | 0.22 | 0.34 | 0.26 |
| 8 | 133731794 | rs2052701 | -0.037 | -0.404 | -0.213 | 0.034 | -0.001 | 0.015 | 0.015 | 0 | 0.37 | 0.65 | 0.26 | 0.22 | 0.33 | 0.25 |
| 8 | 133733145 | rs10216529 | -0.989 | -0.392 | -0.257 | 0.04 | -0.083 | -0.008 | -0.008 | 0 | 0.55 | 0.91 | 0.36 | 0.6 | 0.45 | 0.37 |
| 8 | 133734222 | rs16904724 | -0.959 | -0.193 | 0.431 | -0.066 | 0.042 | 0.005 | 0.005 | 0.143 | 0.07 | 0.4 | 0.05 | 0 | 0.02 | 0.03 |
| 8 | 133737802 | rs10283307 | -0.609 | -0.039 | 0.01 | 0.08 | -0.033 | 0.072 | 0.072 | 1.831 | 0.19 | 0.29 | 0.11 | 0.38 | 0.13 | 0.11 |
| 8 | 133745573 | rs2739015 | 0.148 | 0.318 | -0.312 | -0.573 | -0.006 | 0.004 | 0.004 | 0 | 0.94 | 0.98 | 0.95 | 1 | 0.91 | 0.95 |
| 8 | 133748265 | rs1425706 | 0.231 | 0.197 | -0.183 | -0.799 | 0.001 | -0.005 | -0.005 | 0 | 0.31 | 0.32 | 0.24 | 0.23 | 0.32 | 0.28 |
| 8 | 133757877 | rs2467970 | 0.26 | 0.195 | -0.114 | -0.795 | -0.005 | 0.006 | 0.006 | 0 | 0.31 | 0.29 | 0.24 | 0.22 | 0.32 | 0.28 |
| 8 | 133759051 | rs2739024 | 1.532 | -0.024 | -0.374 | -0.852 | -0.052 | -0.005 | -0.005 | 0 | 0.37 | 0.08 | 0.49 | 0.36 | 0.41 | 0.5 |
| 8 | 133762375 | rs2467979 | 0.399 | 0.001 | -0.318 | -0.818 | -0.006 | 0.001 | 0.001 | 0 | 0.31 | 0.28 | 0.24 | 0.31 | 0.33 | 0.29 |
| 8 | 133774068 | rs890210 | 1.777 | 0.14 | -0.466 | -0.856 | -0.021 | -0.042 | -0.042 | 0 | 0.32 | 0.06 | 0.32 | 0.36 | 0.31 | 0.45 |
| 8 | 133775851 | rs2472230 | 0.162 | 0.1 | -0.019 | -0.876 | 0.007 | -0.005 | -0.005 | 0 | 0.37 | 0.41 | 0.42 | 0.35 | 0.43 | 0.33 |
| 8 | 133777471 | rs2739030 | 0.742 | 0.194 | 0.056 | -0.827 | -0.029 | 0.015 | 0.015 | 0.057 | 0.69 | 0.47 | 0.73 | 0.68 | 0.74 | 0.78 |
| 8 | 133780141 | rs1425710 | 1.38 | -0.114 | 0.122 | -0.788 | 0.003 | -0.04 | -0.04 | 0 | 0.37 | 0.13 | 0.33 | 0.44 | 0.34 | 0.47 |
| 8 | 133782186 | rs2553600 | 1.275 | -0.119 | 0.189 | -0.79 | 0.011 | -0.013 | -0.013 | 0 | 0.38 | 0.23 | 0.33 | 0.44 | 0.34 | 0.47 |

| | | | | | | | | | | | | | | | | |
|---|---|---|---|---|---|---|---|---|---|---|---|---|---|---|---|---|
| 8 | 133786954 | rs1863349 | 0.08 | -0.135 | 0.47 | -0.88 | 0.058 | -0.036 | -0.036 | 0.066 | 0.91 | 0.68 | 0.91 | 1 | 0.9 | 0.94 |
| 8 | 133791111 | rs988068 | 0.495 | -0.178 | 0.348 | -1.077 | 0.002 | 0.008 | 0.008 | 0 | 0.64 | 0.56 | 0.69 | 0.46 | 0.64 | 0.64 |
| 8 | 133796266 | rs7839989 | 1.148 | -0.185 | 0.32 | -1.091 | -0.025 | 0.079 | 0.079 | 0 | 0.18 | 0.05 | 0.27 | 0.05 | 0.2 | 0.12 |
| 8 | 133805463 | rs1469263 | 0.743 | -0.199 | 0.32 | -1.084 | -0.001 | -0.002 | -0.002 | 2.477 | 0.62 | 0.4 | 0.68 | 0.38 | 0.63 | 0.63 |
| 8 | 133813884 | rs235426 | -0.276 | -0.196 | 0.308 | -1.088 | 0.007 | -0.099 | -0.099 | 0.131 | 0.73 | 0.63 | 0.64 | 0.95 | 0.7 | 0.82 |
| 8 | 133824377 | rs235432 | 1.012 | -0.02 | 0.384 | -1.022 | -0.001 | -0.001 | -0.001 | 7.099 | 0.45 | 0.5 | 0.42 | 0.42 | 0.43 | 0.53 |
| 8 | 133844508 | rs16904746 | 1.028 | -0.021 | 0.666 | -1.032 | -0.002 | -0.013 | -0.013 | 0 | 0.44 | 0.44 | 0.41 | 0.33 | 0.42 | 0.52 |
| 8 | 133858009 | rs756201 | 0.991 | -0.021 | 0.664 | -1.032 | -0.001 | 0 | 0 | 0 | 0.46 | 0.51 | 0.42 | 0.42 | 0.44 | 0.53 |
| 8 | 133858712 | rs4736611 | 1.803 | -0.001 | 0.635 | -1.023 | -0.069 | -0.063 | -0.063 | 0 | 0.35 | 0.03 | 0.31 | 0.33 | 0.33 | 0.46 |
| 8 | 133877427 | rs180194 | -1.122 | 0.022 | 0.706 | -1.042 | -0.002 | -0.003 | -0.003 | 0.03 | 0.55 | 0.56 | 0.58 | 0.58 | 0.55 | 0.47 |
| 8 | 133881244 | rs180204 | 0.514 | 0.032 | 0.608 | -1.049 | -0.009 | -0.003 | -0.003 | 0 | 0.17 | 0.14 | 0.14 | 0.18 | 0.25 | 0.2 |
| 8 | 133894502 | rs180217 | -0.989 | -0.365 | 0.774 | -0.869 | 0.001 | 0.001 | 0.001 | 0 | 0.54 | 0.48 | 0.58 | 0.58 | 0.55 | 0.46 |
| 8 | 133900252 | rs180223 | -0.047 | -0.403 | 0.812 | -0.83 | -0.033 | 0.074 | 0.074 | 0 | 0.58 | 0.75 | 0.58 | 0.78 | 0.55 | 0.68 |
| 8 | 133900386 | rs2069550 | -0.028 | -0.403 | 0.824 | -0.83 | -0.027 | 0.064 | 0.064 | 0 | 0.58 | 0.73 | 0.58 | 0.78 | 0.54 | 0.68 |
| 8 | 133902602 | rs909357 | 0.299 | -0.409 | 0.683 | -0.824 | 0.018 | -0.027 | -0.027 | 0.055 | 0.12 | 0.05 | 0.15 | 0.35 | 0.09 | 0.14 |
| 8 | 133904380 | rs864898 | 0.908 | -0.344 | 0.61 | -0.82 | 0.001 | 0.008 | 0.008 | 0 | 0.44 | 0.38 | 0.41 | 0.33 | 0.44 | 0.52 |
| 8 | 133909847 | rs10088696 | -0.275 | 0.076 | 0.645 | -0.842 | 0.031 | -0.043 | -0.043 | 0.003 | 0.08 | 0.26 | 0.08 | 0 | 0.09 | 0.06 |
| 8 | 133909974 | rs853326 | 0.227 | 0.083 | 0.314 | -0.81 | -0.033 | 0.076 | 0.076 | 0 | 0.58 | 0.74 | 0.58 | 0.78 | 0.55 | 0.68 |
| 8 | 133916504 | rs6984038 | -0.936 | -0.375 | 0.872 | -0.356 | 0.051 | 0.028 | 0.028 | 0 | 0.09 | 0.35 | 0.08 | 0.09 | 0.11 | 0.07 |
| 8 | 133917904 | rs853321 | 0.963 | -0.475 | -0.324 | -0.854 | 0.014 | 0.067 | 0.067 | 1.539 | 0.49 | 0.08 | 0.57 | 0.36 | 0.6 | 0.53 |
| 8 | 133918769 | rs1810396 | 0.104 | -0.059 | 0.006 | -0.593 | -0.019 | 0.041 | 0.041 | 0 | 0.68 | 0.88 | 0.79 | 0.79 | 0.69 | 0.72 |
| 8 | 133929917 | rs853308 | 1.33 | -0.046 | -0.018 | -0.538 | -0.123 | 0.026 | 0.026 | 0.014 | 0.47 | 0.04 | 0.54 | 0.35 | 0.52 | 0.47 |
| 8 | 133946089 | rs1474539 | 0.886 | -0.149 | 0.446 | -0.548 | 0.019 | 0.047 | 0.047 | 0.021 | 0.53 | 0.25 | 0.59 | 0.34 | 0.6 | 0.52 |
| 8 | 133954797 | rs12675044 | 0.577 | -0.077 | 0.525 | -0.442 | -0.005 | 0.153 | 0.153 | 0 | 0.08 | 0.16 | 0.14 | 0.44 | 0.07 | 0.17 |
| 8 | 133958037 | rs1879991 | 0.908 | -0.037 | 0.671 | -0.5 | 0.029 | 0.064 | 0.064 | 0.324 | 0.51 | 0.26 | 0.6 | 0.34 | 0.6 | 0.52 |
| 8 | 133959518 | rs2687846 | 0.908 | -0.037 | 0.671 | -0.515 | 0.029 | 0.062 | 0.062 | 0.324 | 0.51 | 0.26 | 0.6 | 0.34 | 0.6 | 0.52 |
| 8 | 133960288 | rs4736613 | -0.196 | -0.075 | 0.705 | -0.562 | -0.068 | 0.077 | 0.077 | 0.146 | 0.28 | 0.53 | 0.29 | 0.48 | 0.16 | 0.31 |
| 8 | 133962414 | rs2741228 | -0.506 | 0.025 | 0.45 | -0.528 | -0.001 | 0 | 0 | 0.58 | 0.8 | 0.83 | 0.88 | 0.82 | 0.76 | 0.83 |
| 8 | 133965346 | rs1851572 | 0.773 | 0.272 | -0.804 | -0.39 | 0.022 | 0.069 | 0.069 | 0.283 | 0.49 | 0.15 | 0.58 | 0.34 | 0.6 | 0.52 |
| 8 | 133975056 | rs7844210 | -0.08 | -0.055 | -0.703 | -0.244 | -0.052 | 0.085 | 0.085 | 0 | 0.27 | 0.48 | 0.28 | 0.5 | 0.17 | 0.3 |
| 8 | 133975283 | rs2069561 | 0.558 | 0.316 | -1.629 | -0.5 | -0.122 | -0.029 | -0.029 | 0.041 | 0.44 | 0.03 | 0.53 | 0.35 | 0.51 | 0.48 |
| 8 | 133980451 | rs6988193 | -0.579 | 0.011 | -0.444 | -0.88 | -0.045 | -0.077 | -0.077 | 0.076 | 0.23 | 0.33 | 0.15 | 0.05 | 0.12 | 0.16 |
| 8 | 133981073 | rs1879992 | 0.798 | 1.64 | -0.796 | -0.575 | 0.028 | -0.016 | -0.016 | 0 | 0.77 | 0.75 | 0.63 | 0.88 | 0.67 | 0.63 |
| 8 | 133985019 | rs1403488 | 0.568 | 1.09 | -1.667 | -1.228 | -0.031 | 0.137 | 0.137 | 0 | 0.22 | 0.04 | 0.15 | 0 | 0.22 | 0.1 |
| 8 | 133989150 | rs11777934 | 0.061 | 1.212 | -1.416 | -1.608 | 0.018 | -0.036 | -0.036 | 0 | 0.35 | 0.44 | 0.23 | 0.19 | 0.38 | 0.27 |
| 8 | 133992758 | rs4074676 | -0.952 | 1.129 | -1.105 | -1.751 | 0.02 | 0.051 | 0.051 | 0 | 0.61 | 0.77 | 0.44 | 0.2 | 0.61 | 0.38 |
| 8 | 133995459 | rs13263087 | 1.51 | 0.666 | -1.589 | -1.203 | -0.001 | -0.081 | -0.081 | 0 | 0.16 | 0 | 0.1 | 0.24 | 0.12 | 0.17 |
| 8 | 133995849 | rs10095925 | -0.72 | 0.995 | -1.279 | -1.593 | 0.005 | 0.064 | 0.064 | 0 | 0.59 | 0.77 | 0.4 | 0.2 | 0.57 | 0.38 |
| 8 | 133997374 | rs6995181 | 0.329 | 1.012 | -1.185 | -1.584 | 0.012 | -0.026 | -0.026 | 1 | 0.34 | 0.43 | 0.22 | 0.2 | 0.34 | 0.27 |
| 8 | 134000415 | rs4540390 | -0.662 | 0.931 | -1.264 | -1.588 | 0 | 0.062 | 0.062 | 0 | 0.6 | 0.77 | 0.42 | 0.19 | 0.57 | 0.37 |
| 8 | 134011224 | rs7842044 | 0.209 | 0.86 | -0.438 | -1.911 | -0.004 | -0.073 | -0.073 | 0.197 | 0.24 | 0.36 | 0.17 | 0 | 0.21 | 0.08 |
| 8 | 134016518 | rs11782027 | 0.294 | 0.882 | -0.315 | -1.929 | 0.019 | -0.093 | -0.093 | 0 | 0.28 | 0.3 | 0.22 | 0.02 | 0.34 | 0.13 |
| 8 | 134018764 | rs940075 | -0.159 | 0.878 | -0.255 | -2.187 | -0.008 | -0.097 | -0.097 | 0 | 0.23 | 0.38 | 0.2 | 0 | 0.23 | 0.06 |
| 8 | 134018991 | rs940074 | -1.102 | 0.878 | -0.314 | -2 | 0.002 | 0.119 | 0.119 | 0 | 0.48 | 0.71 | 0.38 | 0.18 | 0.5 | 0.24 |
| 8 | 134021212 | rs17693908 | 0.958 | 0.892 | -0.435 | -2.055 | -0.097 | 0.108 | 0.108 | 0 | 0.17 | 0.01 | 0.11 | 0 | 0.23 | 0.08 |
| 8 | 134027404 | rs10505603 | 0.377 | 0.823 | -0.18 | -1.953 | 0.01 | -0.01 | -0.01 | 0 | 0.24 | 0.25 | 0.17 | 0.18 | 0.27 | 0.18 |
| 8 | 134027977 | rs6471109 | -0.85 | 0.698 | -0.146 | -1.823 | 0.007 | 0.109 | 0.109 | 0 | 0.48 | 0.73 | 0.38 | 0.18 | 0.5 | 0.24 |
| 8 | 134029111 | rs6985454 | -0.486 | 0.689 | -0.064 | -1.803 | -0.001 | -0.009 | -0.009 | 0.806 | 0.44 | 0.58 | 0.35 | 0.18 | 0.46 | 0.26 |
| 8 | 134034613 | rs3758113 | -0.54 | 0.73 | -0.456 | -1.778 | -0.02 | 0.136 | 0.136 | 0 | 0.46 | 0.78 | 0.36 | 0.18 | 0.46 | 0.27 |
| 8 | 134042468 | rs6998833 | 0.611 | 0.884 | -0.388 | -1.56 | 0.001 | -0.066 | -0.066 | 0 | 0.12 | 0.25 | 0.12 | 0 | 0.11 | 0.04 |
| 8 | 134046862 | rs10108988 | 0.562 | 0.882 | -0.379 | -1.561 | 0 | -0.068 | -0.068 | 0 | 0.13 | 0.27 | 0.12 | 0 | 0.11 | 0.04 |
| 8 | 134049878 | rs10106075 | 0.655 | 0.906 | -0.401 | -1.55 | 0.002 | -0.065 | -0.065 | 0.001 | 0.12 | 0.26 | 0.12 | 0 | 0.11 | 0.04 |
| 8 | 134050678 | rs3739266 | -0.34 | 0.737 | -0.727 | -1.584 | 0.005 | 0.023 | 0.023 | 0 | 0.51 | 0.58 | 0.37 | 0.18 | 0.54 | 0.3 |
| 8 | 134056899 | rs2687837 | -0.369 | 0.696 | -0.983 | -1.237 | -0.002 | 0.021 | 0.021 | 0 | 0.65 | 0.78 | 0.76 | 0.77 | 0.64 | 0.79 |
| 8 | 134060459 | rs4736618 | 1.142 | 0.53 | -0.807 | -1.539 | -0.001 | -0.041 | -0.041 | 0 | 0.11 | 0.13 | 0.19 | 0 | 0.11 | 0.05 |
| 8 | 134060528 | rs2739172 | 0.689 | 0.152 | -1.386 | -0.861 | 0.019 | -0.07 | -0.07 | 1.512 | 0.16 | 0.07 | 0.29 | 0.53 | 0.13 | 0.37 |
| 8 | 134061712 | rs2252917 | -0.118 | 0.348 | -1.34 | -1.045 | 0.005 | 0.162 | 0.162 | 0.002 | 0.55 | 0.84 | 0.52 | 0.24 | 0.56 | 0.41 |
| 8 | 134062331 | rs2741194 | 0.3 | 0.364 | -0.8 | -0.498 | 0.005 | 0 | 0 | 0 | 0.13 | 0.11 | 0.28 | 0.49 | 0.13 | 0.4 |
| 8 | 134063532 | rs2252696 | -0.084 | 0.756 | -0.473 | -1.367 | 0 | -0.024 | -0.024 | 0 | 0.49 | 0.51 | 0.45 | 0.2 | 0.52 | 0.36 |
| 8 | 134064152 | rs1533910 | 0.413 | 0.634 | 0.092 | -1.574 | 0.002 | -0.091 | -0.091 | 0 | 0.19 | 0.27 | 0.26 | 0 | 0.2 | 0.12 |
| 8 | 134064375 | rs2702971 | 0.1 | 0.48 | 0.349 | -1.289 | -0.003 | 0.06 | 0.06 | 0.017 | 0.15 | 0.21 | 0.31 | 0.58 | 0.16 | 0.43 |
| 8 | 134064633 | rs9942754 | 0.604 | 0.299 | 0.758 | -1.376 | 0.004 | -0.019 | -0.019 | 0 | 0.05 | 0.14 | 0.01 | 0 | 0.01 | 0 |
| 8 | 134064773 | rs2252553 | 0.811 | 0.211 | 0.439 | -1.741 | -0.008 | 0.08 | 0.08 | 0 | 0.42 | 0.17 | 0.4 | 0.21 | 0.43 | 0.22 |
| 8 | 134065873 | rs10099615 | 0.844 | 0.465 | 0.399 | -1.872 | 0.002 | -0.001 | -0.001 | 0 | 0.21 | 0.24 | 0.18 | 0.21 | 0.27 | 0.24 |
| 8 | 134071833 | rs2741200 | -0.149 | 0.426 | 0.515 | -1.821 | 0.004 | -0.023 | -0.023 | 0.002 | 0.3 | 0.57 | 0.21 | 0.22 | 0.29 | 0.29 |
| 8 | 134076360 | rs2703010 | 1.848 | 0.418 | 0.318 | -1.852 | 0.008 | -0.003 | -0.003 | 0 | 0.25 | 0.19 | 0.16 | 0.21 | 0.22 | 0.14 |
| 8 | 134076725 | rs2741205 | 0.471 | 0.489 | 0.096 | -1.822 | -0.031 | -0.11 | -0.11 | 0 | 0.25 | 0.1 | 0.37 | 0.55 | 0.28 | 0.41 |
| 8 | 134084800 | rs2703004 | -0.57 | 0.47 | 0.412 | -1.961 | 0.012 | 0.055 | 0.055 | 0.214 | 0.52 | 0.72 | 0.58 | 0.79 | 0.56 | 0.77 |
| 8 | 134085354 | rs2244715 | -1.374 | 0.464 | 0.309 | -1.986 | 0.017 | 0.008 | 0.008 | 1.6 | 0.7 | 0.87 | 0.84 | 0.79 | 0.78 | 0.86 |

| | | | | | | | | | | | | | | | | |
|---|---|---|---|---|---|---|---|---|---|---|---|---|---|---|---|---|
| 8 | 134087450 | rs4301434 | 0.428 | 0.642 | 0.663 | -1.833 | 0 | -0.001 | -0.001 | 0 | 0.48 | 0.46 | 0.42 | 0.22 | 0.45 | 0.23 |
| 8 | 134091284 | rs2739166 | 0.367 | 0.66 | 0.901 | -1.808 | 0.008 | 0.041 | 0.041 | 0 | 0.51 | 0.53 | 0.44 | 0.22 | 0.46 | 0.29 |
| 8 | 134093274 | rs7009823 | 0.865 | 0.678 | 0.922 | -1.832 | -0.002 | -0.107 | -0.107 | 0.01 | 0.25 | 0.33 | 0.28 | 0 | 0.25 | 0.1 |
| 8 | 134100264 | rs870539 | 0.856 | 0.665 | 0.977 | -1.839 | 0 | -0.097 | -0.097 | 0 | 0.25 | 0.3 | 0.28 | 0 | 0.25 | 0.1 |
| 8 | 134104549 | rs17630941 | 0.591 | 0.813 | 1.048 | -1.718 | -0.015 | 0.01 | 0.01 | 0.052 | 0.23 | 0.19 | 0.17 | 0.23 | 0.28 | 0.31 |
| 8 | 134105321 | rs4443645 | -0.245 | 0.756 | 0.754 | -1.593 | -0.001 | 0 | 0 | 0 | 0.74 | 0.73 | 0.62 | 0.45 | 0.74 | 0.55 |
| 8 | 134107223 | rs2272706 | -0.243 | 1.007 | 0.636 | -1.566 | -0.016 | -0.039 | -0.039 | 0.091 | 0.77 | 0.81 | 0.65 | 0.42 | 0.74 | 0.55 |
| 8 | 134108718 | rs2739180 | 0.261 | 1.167 | 0.743 | -1.698 | 0.02 | 0.029 | 0.029 | 0 | 0.69 | 0.48 | 0.48 | 0.4 | 0.66 | 0.54 |
| 8 | 134110278 | rs1000264 | 0.389 | 0.875 | 1.027 | -1.677 | -0.001 | 0.006 | 0.006 | 0.113 | 0.51 | 0.42 | 0.33 | 0.4 | 0.49 | 0.46 |
| 8 | 134110705 | rs4388441 | 1.118 | 0.851 | 1.033 | -1.697 | 0.002 | -0.017 | -0.017 | 0 | 0.24 | 0.15 | 0.3 | 0 | 0.25 | 0.11 |
| 8 | 134111532 | rs2958668 | 0.887 | 0.839 | 1.137 | -1.766 | 0.004 | -0.004 | -0.004 | 0 | 0.24 | 0.27 | 0.17 | 0.22 | 0.27 | 0.32 |
| 8 | 134114716 | rs733735 | -0.144 | 0.79 | 1.166 | -1.796 | 0.001 | 0.025 | 0.025 | 0 | 0.5 | 0.47 | 0.48 | 0.23 | 0.53 | 0.42 |
| 8 | 134116227 | rs13256826 | 0.212 | 0.783 | 1.099 | -1.781 | 0.014 | 0.109 | 0.109 | 0.011 | 0.54 | 0.72 | 0.49 | 0.17 | 0.47 | 0.25 |
| 8 | 134118030 | rs2958674 | 0.72 | 0.791 | 1.128 | -1.771 | 0.017 | -0.019 | -0.019 | 0 | 0.25 | 0.46 | 0.18 | 0.23 | 0.27 | 0.31 |
| 8 | 134119902 | rs4736631 | 1.239 | 0.783 | 0.956 | -1.789 | 0.002 | -0.011 | -0.011 | 0.286 | 0.24 | 0.15 | 0.3 | 0 | 0.25 | 0.11 |
| 8 | 134120567 | rs2958678 | 0.758 | 0.789 | 0.866 | -1.78 | 0.005 | -0.005 | -0.005 | 0 | 0.25 | 0.34 | 0.17 | 0.23 | 0.27 | 0.33 |
| 8 | 134123385 | rs4736435 | 1.215 | 0.806 | 0.825 | -1.798 | 0.004 | 0.026 | 0.026 | 0.088 | 0.24 | 0.13 | 0.3 | 0 | 0.25 | 0.11 |
| 8 | 134125029 | rs2979040 | 1.002 | 0.817 | 0.872 | -1.748 | -0.038 | 0.009 | 0.009 | 0 | 0.23 | 0.09 | 0.15 | 0.23 | 0.27 | 0.33 |
| 8 | 134125682 | rs1133076 | 0.024 | 0.807 | 0.849 | -1.746 | -0.034 | 0.017 | 0.017 | 0.747 | 0.54 | 0.8 | 0.35 | 0.4 | 0.49 | 0.49 |
| 8 | 134130426 | rs10096431 | 1.083 | 0.848 | 0.45 | -1.741 | -0.026 | 0.008 | 0.008 | 0 | 0.23 | 0.1 | 0.15 | 0.24 | 0.27 | 0.3 |
| 8 | 134132812 | rs3779951 | 1.345 | 0.649 | 0.789 | -1.323 | 0.009 | 0.008 | 0.008 | 0.62 | 0.26 | 0.27 | 0.17 | 0.17 | 0.22 | 0.15 |
| 8 | 134138581 | rs4297024 | 0.93 | 0.64 | 0.206 | -1.093 | -0.024 | -0.018 | -0.018 | 0 | 0.05 | 0.02 | 0.17 | 0.14 | 0.08 | 0.23 |
| 8 | 134140546 | rs17705719 | 0.655 | 0.521 | 0.365 | -1.467 | 0.007 | 0.024 | 0.024 | 0.092 | 0.25 | 0.15 | 0.31 | 0 | 0.25 | 0.11 |
| 8 | 134143910 | rs17633144 | 0.698 | 0.529 | 0.37 | -1.442 | 0.009 | 0.055 | 0.055 | 0.304 | 0.26 | 0.13 | 0.31 | 0 | 0.26 | 0.11 |
| 8 | 134144113 | rs2294024 | -0.081 | 0.586 | 0.628 | -1.408 | 0 | 0.016 | 0.016 | 0 | 0.55 | 0.59 | 0.49 | 0.17 | 0.48 | 0.26 |
| 8 | 134153463 | rs10956691 | -0.102 | 0.604 | 0.585 | -1.313 | 0.005 | 0.025 | 0.025 | 0.21 | 0.52 | 0.41 | 0.48 | 0.18 | 0.46 | 0.26 |
| 8 | 134154920 | rs12549029 | -0.132 | 0.609 | 0.391 | -1.311 | -0.003 | -0.019 | -0.019 | 0 | 0.53 | 0.55 | 0.49 | 0.18 | 0.46 | 0.26 |
| 8 | 134173190 | rs11785464 | 0.087 | 0.665 | 0.215 | -1.161 | 0.005 | 0.001 | 0.001 | 0.793 | 0.58 | 0.35 | 0.38 | 0.39 | 0.6 | 0.49 |
| 8 | 134175798 | rs2737697 | -0.802 | 0.636 | 0.179 | -1.256 | -0.038 | 0.055 | 0.055 | 0 | 0.66 | 0.83 | 0.79 | 0.77 | 0.6 | 0.66 |
| 8 | 134185668 | rs4736437 | 0.064 | 0.881 | -0.086 | -1.294 | 0.001 | 0.003 | 0.003 | 0 | 0.49 | 0.52 | 0.51 | 0.17 | 0.49 | 0.26 |
| 8 | 134185894 | rs4736638 | 0.65 | 0.661 | 0.094 | -1.196 | -0.004 | -0.119 | -0.119 | 0 | 0.25 | 0.41 | 0.34 | 0 | 0.26 | 0.11 |
| 8 | 134187718 | rs4410884 | -0.626 | 0.502 | -0.204 | -1.107 | 0.013 | -0.004 | -0.004 | 2.502 | 0.43 | 0.78 | 0.67 | 0.59 | 0.42 | 0.5 |
| 8 | 134188778 | rs12156170 | 0.155 | 0.147 | 0.113 | -1.452 | -0.034 | -0.165 | -0.165 | 0 | 0.25 | 0.53 | 0.34 | 0 | 0.21 | 0.1 |
| 8 | 134189138 | rs10956695 | -0.047 | 0.61 | -0.743 | -0.92 | -0.001 | -0.01 | -0.01 | 0 | 0.49 | 0.58 | 0.49 | 0.17 | 0.49 | 0.25 |
| 8 | 134193074 | rs11782936 | -0.134 | -0.083 | -1.34 | -1.455 | 0.033 | 0.035 | 0.035 | 0.335 | 0.33 | 0.21 | 0.16 | 0 | 0.29 | 0.17 |
| 8 | 134193853 | rs13281650 | -0.258 | -0.203 | -0.976 | -1.549 | 0.009 | -0.041 | -0.041 | 0 | 0.34 | 0.3 | 0.18 | 0 | 0.29 | 0.17 |
| 8 | 134195285 | rs4588834 | 0.547 | -0.017 | -1.679 | -2.097 | -0.013 | 0.089 | 0.089 | 0.623 | 0.24 | 0.11 | 0.17 | 0.09 | 0.25 | 0.22 |
| 8 | 134196037 | rs4736640 | 0.422 | -0.113 | -1.418 | -0.866 | 0.001 | -0.004 | -0.004 | 0.021 | 0.33 | 0.34 | 0.27 | 0.51 | 0.28 | 0.3 |
| 8 | 134202992 | rs2013146 | -0.631 | -0.129 | -1.847 | -1.039 | -0.082 | -0.047 | -0.047 | 0 | 0.75 | 0.9 | 0.55 | 0.64 | 0.63 | 0.62 |
| 8 | 134205713 | rs3739262 | 0.963 | -0.189 | -1.932 | -1.034 | 0.031 | 0.089 | 0.089 | 0.398 | 0.35 | 0.14 | 0.19 | 0.15 | 0.32 | 0.25 |
| 8 | 134206279 | rs16893344 | 0.065 | 0.041 | -1.983 | -0.808 | -0.035 | 0.09 | 0.09 | 3.133 | 0.32 | 0.14 | 0.22 | 0.13 | 0.34 | 0.31 |
| 8 | 134211526 | rs2929986 | -0.075 | 0.171 | -1.536 | -0.76 | -0.013 | 0.004 | 0.004 | 0 | 0.37 | 0.25 | 0.31 | 0.61 | 0.39 | 0.45 |
| 8 | 134211965 | rs11777380 | 1.515 | -0.034 | -2.011 | -1.018 | -0.004 | 0.05 | 0.05 | 0.348 | 0.27 | 0.08 | 0.23 | 0.17 | 0.32 | 0.19 |
| 8 | 134212130 | rs11781004 | 1.244 | -0.018 | -1.871 | -0.866 | 0.003 | 0.059 | 0.059 | 0.063 | 0.31 | 0.12 | 0.24 | 0.21 | 0.35 | 0.22 |
| 8 | 134213371 | rs2930000 | -0.133 | 0.074 | -1.708 | -0.799 | 0.003 | 0.045 | 0.045 | 0 | 0.68 | 0.22 | 0.52 | 0.79 | 0.73 | 0.66 |
| 8 | 134214191 | rs4382455 | 1.203 | 0.029 | -1.745 | -0.769 | -0.039 | 0.053 | 0.053 | 0 | 0.21 | 0.09 | 0.17 | 0.06 | 0.27 | 0.2 |
| 8 | 134215112 | rs2977530 | 0.175 | -0.246 | -0.986 | -0.645 | -0.003 | 0.033 | 0.033 | 0 | 0.31 | 0.43 | 0.26 | 0.51 | 0.29 | 0.3 |
| 8 | 134215631 | rs2977531 | -0.722 | 0.311 | -0.925 | -0.854 | 0.067 | -0.028 | -0.028 | 0 | 0.92 | 0.68 | 0.66 | 1 | 0.92 | 0.94 |
| 8 | 134217705 | rs2929934 | -0.629 | 0.222 | -1.282 | -0.882 | 0.029 | -0.098 | -0.098 | 0 | 0.69 | 0.49 | 0.48 | 0.94 | 0.65 | 0.73 |
| 8 | 134218500 | rs2977533 | -0.028 | -0.16 | -1.549 | -0.466 | 0.025 | 0.1 | 0.1 | 0 | 0.47 | 0.15 | 0.36 | 0.7 | 0.46 | 0.44 |
| 8 | 134220063 | rs2977537 | -0.101 | 0.116 | -1.38 | -0.074 | 0.009 | -0.048 | -0.048 | 1.682 | 0.16 | 0.11 | 0.12 | 0.47 | 0.12 | 0.24 |
| 8 | 134221502 | rs4330674 | -0.121 | -0.096 | -0.982 | -0.758 | -0.007 | -0.035 | -0.035 | 0 | 0.54 | 0.51 | 0.4 | 0.81 | 0.58 | 0.69 |
| 8 | 134223764 | rs6992383 | -0.133 | 0.315 | -1.172 | -0.001 | -0.082 | -0.104 | -0.104 | 0 | 0.67 | 0.9 | 0.7 | 0.59 | 0.61 | 0.6 |
| 8 | 134224864 | rs13254146 | -0.71 | -0.094 | -0.161 | -0.425 | -0.028 | -0.079 | -0.079 | 0 | 0.13 | 0.39 | 0.3 | 0 | 0.07 | 0.07 |
| 8 | 134226879 | rs10956696 | 0.873 | -0.4 | -0.23 | -0.703 | -0.001 | 0.008 | 0.008 | 0.251 | 0.4 | 0.35 | 0.28 | 0.1 | 0.41 | 0.25 |
| 8 | 134228018 | rs11774069 | 0.259 | -0.366 | -0.096 | -0.633 | 0.008 | 0.096 | 0.096 | 0.012 | 0.48 | 0.33 | 0.53 | 0.1 | 0.46 | 0.28 |
| 8 | 134229535 | rs6982341 | -0.186 | -0.276 | 0.09 | -0.171 | -0.003 | -0.011 | -0.011 | 6.776 | 0.58 | 0.64 | 0.62 | 0.43 | 0.58 | 0.46 |
| 8 | 134230782 | rs10956697 | -0.445 | -0.168 | 0.039 | 0.383 | 0.004 | 0.097 | 0.097 | 0 | 0.08 | 0.14 | 0.29 | 0.32 | 0.1 | 0.13 |
| 8 | 134232303 | rs16904853 | 0.181 | -0.399 | -0.017 | 0.294 | -0.004 | 0.034 | 0.034 | 0.012 | 0.12 | 0.06 | 0.1 | 0.51 | 0.07 | 0.27 |
| 8 | 134235816 | rs2929965 | -0.47 | -0.315 | -0.261 | -0.47 | 0.003 | -0.029 | -0.029 | 0.135 | 0.58 | 0.73 | 0.5 | 0.61 | 0.58 | 0.58 |
| 8 | 134239770 | rs3739261 | 0.555 | -0.363 | 0.391 | -0.704 | 0.015 | -0.059 | -0.059 | 0 | 0.29 | 0.35 | 0.4 | 0.07 | 0.36 | 0.23 |
| 8 | 134240697 | rs2929969 | -0.388 | -0.33 | 0.312 | -0.642 | -0.007 | -0.001 | -0.001 | 0 | 0.89 | 0.91 | 0.8 | 0.7 | 0.84 | 0.82 |
| 8 | 134243550 | rs2977551 | -0.073 | -0.333 | 0.264 | -0.637 | 0 | -0.001 | -0.001 | 0 | 0.63 | 0.62 | 0.42 | 0.63 | 0.51 | 0.61 |
| 8 | 134244360 | rs2977552 | 0.206 | -0.508 | 0.336 | -0.726 | 0.014 | -0.021 | -0.021 | 0 | 0.18 | 0.11 | 0.12 | 0.51 | 0.12 | 0.27 |
| 8 | 134245409 | rs7840551 | 0.858 | -0.251 | 0.292 | -0.612 | -0.035 | 0.043 | 0.043 | 0.699 | 0.24 | 0.12 | 0.35 | 0.06 | 0.33 | 0.22 |
| 8 | 134245829 | rs7844513 | 0.801 | -0.245 | 0.461 | -0.608 | 0.019 | -0.048 | -0.048 | 0 | 0.26 | 0.3 | 0.37 | 0.06 | 0.33 | 0.23 |
| 8 | 134246718 | rs2929975 | -0.057 | 0.176 | 0.016 | -0.29 | 0.002 | 0.002 | 0.002 | 0.536 | 0.48 | 0.55 | 0.54 | 0.59 | 0.47 | 0.56 |
| 8 | 134253434 | rs17641864 | -0.268 | 0.326 | -0.518 | -0.16 | 0.005 | 0.001 | 0.001 | 0 | 0.11 | 0.09 | 0.07 | 0 | 0.09 | 0.02 |
| 8 | 134254051 | rs2272646 | 0.662 | 0.834 | -0.421 | 0.688 | 0.009 | 0.069 | 0.069 | 0 | 0.93 | 0.9 | 0.89 | 0.72 | 0.85 | 0.89 |

| | | | | | | | | | | | | | | | | |
|---|---|---|---|---|---|---|---|---|---|---|---|---|---|---|---|---|
| 8 | 134256559 | rs2233341 | 0.399 | 0.477 | -0.803 | 0.072 | 0.016 | 0.029 | 0.029 | 0 | 0.16 | 0.08 | 0.21 | 0.06 | 0.09 | 0.16 |
| 8 | 134257450 | rs2977558 | -0.021 | 0.527 | -0.359 | 0.277 | 0.044 | 0.189 | 0.189 | 0 | 0.94 | 0.84 | 0.9 | 0.74 | 0.89 | 0.9 |
| 8 | 134257697 | rs2929981 | 0.303 | 0.43 | -0.206 | 0.232 | -0.001 | 0.028 | 0.028 | 0 | 0.78 | 0.75 | 0.69 | 0.68 | 0.8 | 0.74 |
| 8 | 134260174 | rs2227262 | 0.826 | -0.094 | -1.531 | -0.391 | -0.013 | 0.013 | 0.013 | 0 | 0.2 | 0.01 | 0.08 | 0.11 | 0.15 | 0.2 |
| 8 | 134260817 | rs16904873 | 0.89 | -0.127 | -1.47 | -0.251 | 0.014 | 0.025 | 0.025 | 0 | 0.16 | 0.09 | 0.18 | 0.06 | 0.09 | 0.16 |
| 8 | 134262532 | rs2272648 | 1.265 | 0.511 | -2.319 | -0.981 | -0.048 | -0.139 | -0.139 | 0 | 0.26 | 0.08 | 0.36 | 0.58 | 0.3 | 0.42 |
| 8 | 134263453 | rs3779939 | 0.663 | 0.496 | -1.166 | -1.304 | -0.003 | -0.015 | -0.015 | 0 | 0.25 | 0.26 | 0.13 | 0.11 | 0.22 | 0.21 |
| 8 | 134264054 | rs7018287 | 0.911 | 0.454 | -1.146 | -1.333 | -0.008 | -0.066 | -0.066 | 0.631 | 0.21 | 0.34 | 0.23 | 0.06 | 0.16 | 0.16 |
| 8 | 134265495 | rs3779938 | 1.444 | 0.519 | -0.92 | -1.322 | 0.018 | 0.03 | 0.03 | 0 | 0.2 | 0.09 | 0.09 | 0.11 | 0.15 | 0.2 |
| 8 | 134266585 | rs2233334 | 0.009 | 0.443 | -0.518 | -1.331 | -0.013 | -0.048 | -0.048 | 0 | 0.41 | 0.5 | 0.34 | 0.17 | 0.31 | 0.35 |
| 8 | 134270548 | rs2272651 | -0.055 | 0.589 | -0.682 | -1.399 | -0.006 | -0.067 | -0.067 | 0.648 | 0.37 | 0.4 | 0.31 | 0.06 | 0.29 | 0.27 |
| 8 | 134270887 | rs2929984 | 0.963 | 1.125 | -0.704 | -0.729 | 0.002 | -0.01 | -0.01 | 0 | 0.23 | 0.17 | 0.31 | 0.53 | 0.22 | 0.33 |
| 8 | 134271506 | rs2233327 | 0.48 | 0.878 | -0.864 | -0.64 | 0.001 | -0.092 | -0.092 | 0.594 | 0.2 | 0.03 | 0.24 | 0.3 | 0.17 | 0.25 |
| 8 | 134271648 | rs7839335 | -0.02 | 1.178 | -0.018 | -0.898 | 0.051 | -0.063 | -0.063 | 0.666 | 0.11 | 0.25 | 0.1 | 0 | 0.17 | 0.06 |
| 8 | 134272207 | rs2929985 | -0.192 | 0.274 | -0.779 | -1.05 | 0.007 | 0.018 | 0.018 | 0.161 | 0.34 | 0.17 | 0.24 | 0.3 | 0.3 | 0.24 |
| 8 | 134272549 | rs2977560 | -1.02 | 0.66 | -0.777 | -1.129 | -0.034 | 0.033 | 0.033 | 0 | 0.91 | 0.96 | 0.87 | 1 | 0.82 | 0.98 |
| 8 | 134273868 | rs4736442 | 0.538 | 0.107 | -0.66 | -1.208 | -0.012 | -0.002 | -0.002 | 0 | 0.53 | 0.22 | 0.46 | 0.58 | 0.5 | 0.66 |
| 8 | 134275018 | rs12678229 | 1.346 | -0.045 | -0.819 | -1.21 | -0.008 | -0.056 | -0.056 | 0 | 0.37 | 0.14 | 0.37 | 0.58 | 0.38 | 0.63 |
| 8 | 134275341 | rs2976570 | 0.541 | -0.112 | -0.864 | -1.294 | -0.022 | -0.121 | -0.121 | 0 | 0.31 | 0.07 | 0.3 | 0.42 | 0.28 | 0.3 |
| 8 | 134276322 | rs2929988 | -1.994 | -0.061 | -0.848 | -1.241 | -0.061 | -0.103 | -0.103 | 0.408 | 0.67 | 0.92 | 0.61 | 0.42 | 0.6 | 0.38 |
| 8 | 134277855 | rs2977497 | -0.675 | -0.245 | 0.267 | -1.046 | 0.002 | 0.001 | 0.001 | 1.712 | 0.48 | 0.55 | 0.44 | 0.4 | 0.4 | 0.33 |
| 8 | 134281774 | rs6992807 | -0.469 | 0.285 | 1.063 | -0.973 | -0.027 | -0.127 | -0.127 | 1.272 | 0.19 | 0.4 | 0.13 | 0 | 0.12 | 0.04 |
| 8 | 134285671 | rs2977504 | 0.95 | 0.328 | 0.898 | -1.002 | 0.001 | -0.115 | -0.115 | 0 | 0.7 | 0.42 | 0.73 | 1 | 0.71 | 0.94 |
| 8 | 134294419 | rs10505606 | -1.016 | -0.017 | 0.977 | -0.914 | -0.003 | -0.157 | -0.157 | 0.1 | 0.28 | 0.41 | 0.24 | 0 | 0.27 | 0.06 |
| 8 | 134294814 | rs16904881 | -1.199 | -0.008 | 0.705 | -0.91 | -0.008 | -0.135 | -0.135 | 0 | 0.3 | 0.53 | 0.26 | 0 | 0.27 | 0.06 |
| 8 | 134295368 | rs13252851 | 0.481 | 0.414 | 0.179 | -0.353 | 0.004 | -0.032 | -0.032 | 0 | 0.1 | 0.12 | 0.13 | 0 | 0.15 | 0.02 |
| 8 | 134298492 | rs4504614 | -0.651 | 0.704 | 0.488 | -0.218 | 0.087 | -0.045 | -0.045 | 0.112 | 0.09 | 0.33 | 0.15 | 0 | 0.13 | 0.03 |
| 8 | 134298919 | rs2004066 | -0.436 | 0.362 | 0.135 | -0.027 | -0.009 | 0.022 | 0.022 | 0 | 0.37 | 0.5 | 0.27 | 0.5 | 0.34 | 0.34 |
| 8 | 134299203 | rs2977509 | -0.511 | 0.352 | 0.086 | 0.15 | -0.008 | 0.003 | 0.003 | 0 | 0.38 | 0.53 | 0.27 | 0.57 | 0.34 | 0.37 |
| 8 | 134299872 | rs2930004 | -0.886 | 0.307 | -0.348 | 0.067 | -0.024 | -0.007 | -0.007 | 0 | 0.46 | 0.81 | 0.41 | 0.51 | 0.47 | 0.35 |
| 8 | 134303881 | rs3802253 | -1.127 | 0.184 | -0.55 | -0.181 | -0.041 | -0.03 | -0.03 | 0.358 | 0.6 | 0.87 | 0.48 | 0.41 | 0.56 | 0.42 |
| 8 | 134309134 | rs10108354 | 0.077 | 0.58 | 0.238 | 0.044 | 0.038 | -0.033 | -0.033 | 0 | 0.06 | 0.19 | 0.09 | 0 | 0.1 | 0.02 |
| 8 | 134313149 | rs16904884 | -0.269 | 0.236 | -0.068 | 0.333 | -0.014 | 0.037 | 0.037 | 0 | 0.09 | 0.17 | 0.1 | 0.2 | 0.06 | 0.07 |
| 8 | 134317248 | rs6982249 | -1.343 | 0.191 | -0.685 | -0.015 | -0.044 | 0.043 | 0.043 | 0 | 0.51 | 0.87 | 0.47 | 0.42 | 0.52 | 0.35 |
| 8 | 134320263 | rs4736646 | -1.363 | 0.632 | -0.441 | 0.294 | -0.082 | -0.071 | -0.071 | 0 | 0.41 | 0.9 | 0.43 | 0.51 | 0.46 | 0.31 |
| 8 | 134323337 | rs4736648 | -1.254 | 0.828 | -0.033 | 0.162 | -0.009 | -0.021 | -0.021 | 0 | 0.31 | 0.81 | 0.33 | 0.45 | 0.34 | 0.21 |
| 8 | 134323951 | rs4521737 | -1.802 | 0.799 | -0.153 | 0.123 | -0.077 | -0.076 | -0.076 | 0 | 0.42 | 0.9 | 0.44 | 0.53 | 0.47 | 0.31 |
| 8 | 134326125 | rs16904888 | -0.024 | 1.105 | 0.879 | 0.273 | 0.022 | -0.057 | -0.057 | 0 | 0.1 | 0.23 | 0.08 | 0 | 0.11 | 0.04 |
| 8 | 134327372 | rs6989828 | -0.961 | 1.056 | 1.311 | 0.326 | 0.022 | 0.125 | 0.125 | 0 | 0.2 | 0.46 | 0.24 | 0.48 | 0.23 | 0.25 |
| 8 | 134327512 | rs13272768 | -0.212 | 1.013 | 1.109 | -0.442 | 0.02 | -0.058 | -0.058 | 0 | 0.1 | 0.24 | 0.08 | 0 | 0.11 | 0.04 |
| 8 | 134328022 | rs7012851 | -1.603 | 1.086 | 0.736 | -0.476 | 0.02 | -0.048 | -0.048 | 0 | 0.25 | 0.62 | 0.28 | 0.09 | 0.32 | 0.2 |
| 8 | 134328386 | rs7013213 | -1.711 | 1.077 | 0.689 | -0.514 | 0.021 | -0.054 | -0.054 | 1.297 | 0.24 | 0.63 | 0.28 | 0.08 | 0.31 | 0.19 |
| 8 | 134332117 | rs7005796 | -1.444 | 1.091 | 0.736 | -0.04 | 0.106 | -0.004 | -0.004 | 0 | 0.13 | 0.5 | 0.21 | 0.09 | 0.19 | 0.18 |
| 8 | 134334565 | rs13254782 | -0.845 | 0.559 | 0.037 | -0.313 | -0.008 | -0.051 | -0.051 | 0.582 | 0.12 | 0.17 | 0.08 | 0 | 0.11 | 0.13 |
| 8 | 134334904 | rs2976589 | -0.218 | 1.178 | -0.816 | -0.234 | -0.041 | -0.019 | -0.019 | 0.051 | 0.81 | 0.93 | 0.71 | 0.8 | 0.75 | 0.83 |
| 8 | 134338367 | rs11778385 | -0.552 | 0.901 | -0.695 | -0.574 | -0.009 | 0.01 | 0.01 | 0 | 0.64 | 0.76 | 0.72 | 0.35 | 0.6 | 0.46 |
| 8 | 134339152 | rs10108241 | 0.539 | 0.978 | -0.819 | -0.751 | -0.02 | 0.023 | 0.023 | 0 | 0.05 | 0.02 | 0.07 | 0 | 0.1 | 0.01 |
| 8 | 134339266 | rs10095018 | -0.307 | 0.374 | -0.106 | -1.032 | -0.039 | -0.043 | -0.043 | 0.381 | 0.1 | 0.44 | 0.03 | 0 | 0.01 | 0.08 |
| 8 | 134339994 | rs10956701 | -0.849 | 0.309 | -0.194 | -0.884 | -0.007 | 0.006 | 0.006 | 0 | 0.63 | 0.74 | 0.72 | 0.37 | 0.6 | 0.47 |
| 8 | 134346841 | rs10105270 | 0.473 | 0.427 | -0.048 | -0.849 | -0.006 | 0.001 | 0.001 | 0 | 0.06 | 0.06 | 0.08 | 0 | 0.1 | 0.01 |
| 8 | 134347512 | rs6982426 | -1.449 | 0.323 | 0.163 | -0.909 | -0.012 | -0.098 | -0.098 | 0 | 0.14 | 0.47 | 0.11 | 0 | 0.11 | 0.08 |
| 8 | 134353346 | rs11782647 | 0.501 | 0.095 | -0.191 | -1.163 | -0.059 | 0.073 | 0.073 | 0 | 0.44 | 0.08 | 0.57 | 0.37 | 0.43 | 0.39 |
| 8 | 134353632 | rs6986484 | 0.207 | 0.048 | 0.629 | -1.174 | -0.01 | -0.007 | -0.007 | 0 | 0.06 | 0.24 | 0.01 | 0 | 0.01 | 0.07 |
| 8 | 134358754 | rs4236902 | -1.58 | 0.063 | 0.49 | -1.206 | -0.02 | -0.131 | -0.131 | 0 | 0.19 | 0.56 | 0.16 | 0 | 0.17 | 0.09 |
| 8 | 134360307 | rs9649972 | -0.353 | 0.337 | 0.397 | -1.062 | 0.003 | -0.031 | -0.031 | 0 | 0.1 | 0.18 | 0.13 | 0 | 0.15 | 0.02 |
| 8 | 134360780 | rs10093875 | -0.866 | 0.417 | 0.278 | -0.738 | -0.015 | -0.005 | -0.005 | 0.125 | 0.66 | 0.78 | 0.75 | 0.4 | 0.62 | 0.49 |
| 8 | 134361379 | rs7000658 | -0.724 | 0.464 | 0.324 | -0.617 | 0 | 0.001 | 0.001 | 0.37 | 0.63 | 0.67 | 0.74 | 0.44 | 0.6 | 0.48 |
| 8 | 134364050 | rs6471119 | -0.526 | 0.335 | 0.943 | -0.996 | -0.015 | -0.052 | -0.052 | 0 | 0.11 | 0.19 | 0.08 | 0 | 0.08 | 0.06 |
| 8 | 134369383 | rs2976552 | -0.014 | 0.632 | 0.066 | -1.062 | -0.02 | 0.002 | 0.002 | 0 | 0.79 | 0.82 | 0.77 | 0.75 | 0.65 | 0.72 |
| 8 | 134371219 | rs2929959 | 1.011 | 0.573 | -0.168 | -1.121 | -0.059 | 0.129 | 0.129 | 0 | 0.58 | 0.2 | 0.6 | 0.8 | 0.48 | 0.65 |
| 8 | 134371734 | rs16904909 | 0.683 | 0.492 | -0.233 | -1.09 | -0.002 | -0.018 | -0.018 | 0 | 0.07 | 0.08 | 0.14 | 0 | 0.04 | 0.01 |
| 8 | 134380152 | rs2929929 | 0.241 | 0.661 | -0.33 | -1.014 | -0.011 | 0.012 | 0.012 | 0 | 0.72 | 0.75 | 0.71 | 0.83 | 0.58 | 0.7 |
| 8 | 134386337 | rs737438 | -0.771 | -0.275 | -0.539 | -0.837 | 0.002 | 0.002 | 0.002 | 0.008 | 0.39 | 0.57 | 0.41 | 0.32 | 0.32 | 0.26 |
| 8 | 134386699 | rs4998686 | 0.385 | -0.297 | -1.002 | -0.73 | 0.007 | -0.054 | -0.054 | 0 | 0.23 | 0.07 | 0.29 | 0.32 | 0.23 | 0.23 |
| 8 | 134394277 | rs10956705 | 0.218 | -0.311 | -1.101 | -0.506 | 0.005 | -0.102 | -0.102 | 0.197 | 0.26 | 0.1 | 0.36 | 0.53 | 0.26 | 0.31 |
| 8 | 134401509 | rs6999391 | -0.449 | -0.148 | -0.742 | -1.086 | 0.06 | -0.143 | -0.143 | 0.413 | 0.72 | 0.49 | 0.77 | 0.97 | 0.62 | 0.77 |
| 8 | 134402395 | rs13267049 | 1.395 | -0.353 | -0.935 | -1.047 | -0.023 | -0.113 | -0.113 | 0.719 | 0.36 | 0.04 | 0.35 | 0.39 | 0.31 | 0.45 |
| 8 | 134409650 | rs11779130 | 0.282 | -0.575 | -0.696 | -0.771 | 0.002 | -0.058 | -0.058 | 0 | 0.25 | 0.16 | 0.26 | 0.54 | 0.26 | 0.3 |
| 8 | 134413869 | rs1075580 | -1.256 | -0.564 | -0.537 | -0.928 | -0.032 | -0.077 | -0.077 | 0 | 0.22 | 0.37 | 0.25 | 0.05 | 0.15 | 0.07 |

| | | | | | | | | | | | | | | | | |
|---|---|---|---|---|---|---|---|---|---|---|---|---|---|---|---|---|
| 8 | 134428243 | rs2736862 | 0.587 | -0.675 | -0.685 | -1.278 | 0.018 | -0.099 | -0.099 | 0 | 0.33 | 0.38 | 0.27 | 0.03 | 0.44 | 0.26 |
| 8 | 134429018 | rs11984964 | 1.395 | -0.784 | -0.924 | -1.199 | 0 | -0.082 | -0.082 | 0 | 0.21 | 0.1 | 0.21 | 0.54 | 0.22 | 0.33 |
| 8 | 134429466 | rs2736866 | 2.312 | -0.691 | -0.951 | -1.162 | -0.061 | -0.164 | -0.164 | 0.092 | 0.23 | 0.02 | 0.27 | 0.38 | 0.19 | 0.33 |
| 8 | 134433978 | rs2737437 | -0.93 | -0.413 | -0.269 | -1.462 | 0.012 | 0.17 | 0.17 | 0 | 0.5 | 0.74 | 0.48 | 0.08 | 0.55 | 0.33 |
| 8 | 134437026 | rs7823430 | -0.314 | -0.328 | -0.283 | -1.177 | -0.011 | -0.029 | -0.029 | 0 | 0.15 | 0.23 | 0.2 | 0.05 | 0.11 | 0.07 |
| 8 | 134445332 | rs10109750 | 0.403 | -0.342 | -0.2 | -1.177 | -0.018 | -0.019 | -0.019 | 0 | 0.11 | 0.18 | 0.16 | 0.05 | 0.07 | 0.06 |
| 8 | 134446148 | rs6471124 | -0.809 | -0.281 | -0.259 | -1.142 | -0.021 | -0.06 | -0.06 | 0 | 0.21 | 0.35 | 0.24 | 0.05 | 0.15 | 0.09 |
| 8 | 134449830 | rs2736860 | -0.958 | -0.15 | -0.568 | -0.892 | 0.02 | 0.052 | 0.052 | 0 | 0.53 | 0.67 | 0.55 | 0.29 | 0.62 | 0.46 |
| 8 | 134450481 | rs2403981 | 0 | 0.497 | -0.589 | -0.656 | -0.017 | 0.012 | 0.012 | 0 | 0.69 | 0.31 | 0.64 | 0.73 | 0.54 | 0.62 |
| 8 | 134452560 | rs11781445 | 0.031 | 0.04 | -0.759 | -0.223 | -0.052 | 0.053 | 0.053 | 0 | 0.25 | 0.46 | 0.2 | 0.32 | 0.2 | 0.24 |
| 8 | 134454379 | rs13274799 | 0.847 | 0.504 | -1.956 | -0.62 | -0.04 | -0.094 | -0.094 | 0.081 | 0.46 | 0.05 | 0.46 | 0.43 | 0.36 | 0.49 |
| 8 | 134459221 | rs2037957 | -0.425 | 1.23 | -1.262 | -0.873 | -0.073 | 0.07 | 0.07 | 0 | 0.81 | 0.9 | 0.8 | 0.99 | 0.64 | 0.86 |
| 8 | 134463161 | rs7822343 | -0.424 | 0.575 | -0.456 | -0.594 | 0.038 | -0.061 | -0.061 | 0 | 0.11 | 0.44 | 0.1 | 0 | 0.06 | 0.03 |
| 8 | 134465059 | rs741900 | 0.306 | 0.544 | -1.872 | -0.32 | -0.085 | -0.133 | -0.133 | 0.038 | 0.41 | 0.03 | 0.45 | 0.52 | 0.37 | 0.46 |
| 8 | 134468864 | rs4736675 | 0.441 | 0.334 | -1.23 | -0.29 | 0.002 | -0.012 | -0.012 | 5.413 | 0.23 | 0.21 | 0.16 | 0.56 | 0.19 | 0.35 |
| 8 | 134469536 | rs11782689 | 0.782 | 0.331 | -1.262 | -0.27 | -0.014 | -0.099 | -0.099 | 0 | 0.21 | 0.08 | 0.12 | 0.55 | 0.17 | 0.31 |
| 8 | 134470631 | rs2142306 | -0.495 | 0.528 | -0.876 | -0.932 | -0.001 | 0.002 | 0.002 | 0.185 | 0.45 | 0.63 | 0.3 | 0.56 | 0.41 | 0.47 |
| 8 | 134473917 | rs2736869 | -0.072 | 0.554 | -0.71 | -1.274 | 0.008 | -0.052 | -0.052 | 0.197 | 0.75 | 0.77 | 0.82 | 0.98 | 0.68 | 0.82 |
| 8 | 134475164 | rs2736872 | -0.392 | 0.371 | -1.14 | -1.507 | -0.043 | 0.144 | 0.144 | 0 | 0.42 | 0.2 | 0.4 | 0.1 | 0.5 | 0.29 |
| 8 | 134478529 | rs6986303 | 0.067 | 0.445 | -0.735 | -1.104 | 0 | -0.001 | -0.001 | 0 | 0.31 | 0.41 | 0.33 | 0.67 | 0.26 | 0.46 |
| 8 | 134478865 | rs6990329 | 0.783 | 0.366 | -0.741 | -1.06 | 0.001 | -0.024 | -0.024 | 0 | 0.24 | 0.22 | 0.15 | 0.59 | 0.2 | 0.34 |
| 8 | 134483187 | rs16904929 | 0.412 | 0.396 | -0.352 | -0.873 | -0.001 | -0.001 | -0.001 | 0.712 | 0.07 | 0.11 | 0.15 | 0.09 | 0.07 | 0.12 |
| 8 | 134485237 | rs2737422 | -0.133 | 0.067 | -0.284 | -1.125 | 0.005 | 0.065 | 0.065 | 0 | 0.47 | 0.56 | 0.4 | 0.17 | 0.48 | 0.38 |
| 8 | 134486021 | rs2737424 | 0.14 | 0.059 | -0.178 | -1.133 | 0.002 | -0.045 | -0.045 | 0 | 0.42 | 0.44 | 0.28 | 0.08 | 0.44 | 0.27 |
| 8 | 134487723 | rs2737429 | 0.699 | 0.13 | 0.278 | -1.115 | -0.007 | -0.002 | -0.002 | 0 | 0.31 | 0.21 | 0.22 | 0.08 | 0.35 | 0.24 |
| 8 | 134487803 | rs9643299 | 0.486 | -0.072 | -0.132 | -0.782 | 0.005 | -0.006 | -0.006 | 0 | 0.18 | 0.17 | 0.18 | 0.58 | 0.17 | 0.3 |
| 8 | 134490849 | rs7838656 | 0.139 | 0.048 | 0.024 | -1.045 | -0.003 | -0.019 | -0.019 | 0 | 0.46 | 0.56 | 0.6 | 0.3 | 0.43 | 0.42 |
| 8 | 134492551 | rs2142307 | 0.944 | 0.053 | -0.268 | -1.07 | -0.017 | 0.014 | 0.014 | 0 | 0.29 | 0.23 | 0.4 | 0.25 | 0.33 | 0.25 |
| 8 | 134494524 | rs7460764 | -0.607 | 0.138 | -0.2 | -0.855 | -0.002 | 0.011 | 0.011 | 0 | 0.81 | 0.76 | 0.79 | 0.38 | 0.81 | 0.75 |
| 8 | 134497617 | rs4736678 | 1.03 | 0.545 | -0.434 | -1.104 | -0.054 | 0.074 | 0.074 | 0 | 0.27 | 0.08 | 0.17 | 0.09 | 0.33 | 0.21 |
| 8 | 134501041 | rs4736681 | 0.882 | 0.522 | -0.302 | -0.854 | -0.027 | 0.036 | 0.036 | 0 | 0.29 | 0.15 | 0.18 | 0.14 | 0.34 | 0.23 |
| 8 | 134501766 | rs11779894 | 0.912 | 0.366 | -0.38 | -0.66 | -0.022 | -0.006 | -0.006 | 0.264 | 0.28 | 0.22 | 0.4 | 0.32 | 0.34 | 0.26 |
| 8 | 134501926 | rs11996611 | 0.589 | 0.235 | -0.229 | -0.71 | 0.013 | -0.036 | -0.036 | 0.517 | 0.21 | 0.12 | 0.21 | 0.55 | 0.18 | 0.28 |
| 8 | 134502205 | rs13264936 | -0.246 | 0.227 | -0.189 | -0.706 | 0 | 0.004 | 0.004 | 0 | 0.76 | 0.75 | 0.77 | 0.45 | 0.77 | 0.69 |
| 8 | 134502322 | rs16904932 | 0.846 | 0.382 | -0.351 | -0.678 | -0.018 | 0.008 | 0.008 | 0 | 0.28 | 0.25 | 0.4 | 0.26 | 0.34 | 0.26 |
| 8 | 134505411 | rs7831227 | -0.283 | 0.062 | -0.419 | -0.127 | -0.02 | 0.053 | 0.053 | 0 | 0.23 | 0.23 | 0.22 | 0.53 | 0.2 | 0.3 |
| 8 | 134508256 | rs3739256 | 0.412 | 0.776 | -0.832 | -0.499 | 0.003 | -0.068 | -0.068 | 0.005 | 0.33 | 0.27 | 0.23 | 0 | 0.39 | 0.15 |
| 8 | 134508510 | rs3739257 | 0.294 | 0.33 | -0.949 | 0.675 | 0.012 | -0.02 | -0.02 | 0 | 0.16 | 0.09 | 0.18 | 0.52 | 0.13 | 0.24 |
| 8 | 134509215 | rs3802258 | 0.321 | 0.219 | -1.027 | 0.701 | 0.022 | -0.033 | -0.033 | 0.61 | 0.15 | 0.07 | 0.16 | 0.49 | 0.11 | 0.21 |
| 8 | 134512809 | rs11776694 | 0.049 | 0.602 | -1.043 | 0.467 | -0.002 | -0.01 | -0.01 | 0.022 | 0.2 | 0.17 | 0.19 | 0.33 | 0.24 | 0.27 |
| 8 | 134513219 | rs6995270 | -0.252 | 0.647 | -1.459 | 0.036 | -0.008 | -0.022 | -0.022 | 0 | 0.43 | 0.25 | 0.42 | 0.5 | 0.49 | 0.55 |
| 8 | 134513350 | rs10100754 | 0.925 | 0.83 | -1.27 | -0.604 | -0.007 | -0.087 | -0.087 | 0 | 0.26 | 0.25 | 0.13 | 0 | 0.2 | 0.08 |
| 8 | 134514393 | rs7000952 | -0.585 | 1.132 | -1.36 | 0.452 | -0.022 | -0.038 | -0.038 | 0 | 0.74 | 0.88 | 0.65 | 0.66 | 0.74 | 0.75 |
| 8 | 134514865 | rs4736455 | 0.024 | 0.8 | -0.932 | 0.359 | 0 | 0.003 | 0.003 | 0 | 0.22 | 0.25 | 0.21 | 0.34 | 0.26 | 0.3 |
| 8 | 134516461 | rs13281927 | -0.159 | 0.437 | -0.868 | -0.161 | 0.006 | 0.016 | 0.016 | 0 | 0.48 | 0.45 | 0.32 | 0.25 | 0.39 | 0.32 |
| 8 | 134517560 | rs11992377 | 0.373 | 0.62 | -0.879 | -0.521 | 0.003 | -0.021 | -0.021 | 1.329 | 0.19 | 0.22 | 0.14 | 0.12 | 0.19 | 0.2 |
| 8 | 134519265 | rs6989475 | 2.093 | 0.316 | -1.11 | -0.783 | 0.006 | 0.023 | 0.023 | 0 | 0.11 | 0.03 | 0.06 | 0 | 0.08 | 0.01 |
| 8 | 134519479 | rs6989914 | 0.187 | 0.258 | -0.893 | -1.293 | 0.013 | -0.148 | -0.148 | 0.373 | 0.31 | 0.45 | 0.32 | 0.04 | 0.3 | 0.18 |
| 8 | 134521802 | rs4736456 | 1.401 | 0.053 | -0.888 | -0.683 | 0.026 | -0.039 | -0.039 | 0.022 | 0.15 | 0.09 | 0.12 | 0.25 | 0.12 | 0.2 |
| 8 | 134523204 | rs11992973 | 1.303 | 0.284 | -0.672 | -0.659 | -0.001 | 0.005 | 0.005 | 0 | 0.14 | 0.13 | 0.09 | 0.12 | 0.15 | 0.18 |
| 8 | 134523882 | rs1474113 | 0.113 | 0.369 | -0.553 | 0.039 | 0.038 | 0.021 | 0.021 | 0 | 0.19 | 0.35 | 0.31 | 0.2 | 0.25 | 0.24 |
| 8 | 134527696 | rs2975739 | -0.081 | 0.448 | -0.72 | -0.272 | 0 | -0.007 | -0.007 | 0 | 0.46 | 0.51 | 0.43 | 0.32 | 0.46 | 0.48 |
| 8 | 134527871 | rs2978043 | -0.609 | 0.844 | -0.61 | -0.035 | -0.001 | 0 | 0 | 0 | 0.36 | 0.6 | 0.44 | 0.29 | 0.4 | 0.39 |
| 8 | 134531773 | rs4736690 | -0.016 | 1.014 | -0.232 | -0.65 | 0.003 | -0.02 | -0.02 | 0 | 0.17 | 0.15 | 0.24 | 0.06 | 0.21 | 0.12 |
| 8 | 134534926 | rs2945738 | 0.013 | 1.073 | -0.057 | -0.786 | 0.024 | -0.062 | -0.062 | 0 | 0.17 | 0.17 | 0.23 | 0.01 | 0.26 | 0.18 |
| 8 | 134536088 | rs4736691 | 0.721 | 0.631 | -0.037 | -0.495 | -0.006 | 0.006 | 0.006 | 0 | 0.25 | 0.28 | 0.29 | 0.28 | 0.22 | 0.33 |
| 8 | 134539196 | rs2978012 | 0.694 | 0.581 | -0.365 | -0.968 | -0.051 | 0.185 | 0.185 | 0 | 0.36 | 0.11 | 0.22 | 0.06 | 0.38 | 0.23 |
| 8 | 134539286 | rs3779925 | 0.907 | 0.358 | 0.206 | -0.794 | -0.049 | 0.006 | 0.006 | 0 | 0.13 | 0.23 | 0.07 | 0.14 | 0.04 | 0.15 |
| 8 | 134540450 | rs2945787 | -0.756 | 0.407 | 0.288 | -0.678 | 0.016 | 0.001 | 0.001 | 0 | 0.8 | 0.78 | 0.64 | 0.69 | 0.67 | 0.83 |
| 8 | 134541558 | rs16904937 | 1.506 | -0.002 | 0.81 | -0.209 | 0.032 | -0.064 | -0.064 | 0.639 | 0.21 | 0.09 | 0.14 | 0.27 | 0.17 | 0.26 |
| 8 | 134541623 | rs16904938 | -0.314 | -0.046 | 0.969 | -0.258 | 0 | 0.001 | 0.001 | 0 | 0.4 | 0.56 | 0.41 | 0.62 | 0.28 | 0.55 |
| 8 | 134544194 | rs4637803 | -0.373 | -0.155 | 0.88 | -0.252 | 0.046 | 0.032 | 0.032 | 0 | 0.42 | 0.66 | 0.41 | 0.62 | 0.27 | 0.59 |
| 8 | 134545130 | rs12545010 | 0.405 | 0.289 | 1.004 | 0.046 | 0.022 | -0.007 | -0.007 | 0 | 0.41 | 0.28 | 0.37 | 0.41 | 0.31 | 0.49 |
| 8 | 134547917 | rs1137295 | 1.04 | 0.558 | 0.876 | 0.105 | -0.031 | 0.017 | 0.017 | 0 | 0.31 | 0.41 | 0.22 | 0.45 | 0.21 | 0.34 |
| 8 | 134549109 | rs2978024 | -0.38 | 1.053 | 0.549 | -0.338 | 0.04 | 0.018 | 0.018 | 0 | 0.85 | 0.78 | 0.67 | 0.67 | 0.73 | 0.84 |
| 8 | 134551286 | rs10956709 | 0.12 | -0.167 | 0.861 | -0.069 | -0.05 | -0.02 | -0.02 | 0.165 | 0.44 | 0.53 | 0.27 | 0.59 | 0.24 | 0.5 |
| 8 | 134551406 | rs6471131 | -0.62 | 0.072 | 0.934 | 0.057 | 0.059 | -0.004 | -0.004 | 0 | 0.07 | 0.21 | 0.17 | 0.03 | 0.08 | 0.1 |
| 8 | 134556145 | rs16904939 | 0.747 | 0.515 | 0.001 | 0.214 | 0.003 | 0.028 | 0.028 | 0.061 | 0.08 | 0.09 | 0.12 | 0.32 | 0.11 | 0.09 |
| 8 | 134556356 | rs7829120 | -1.326 | 0.444 | -0.187 | 0.04 | -0.133 | 0.162 | 0.162 | 0 | 0.69 | 0.93 | 0.69 | 0.94 | 0.59 | 0.7 |

| | | | | | | | | | | | | | | | | |
|---|---|---|---|---|---|---|---|---|---|---|---|---|---|---|---|---|
| 8 | 134556752 | rs4736695 | -0.559 | 0.237 | 0.425 | 0.746 | 0.083 | 0.025 | 0.025 | 0.231 | 0.49 | 0.68 | 0.39 | 0.4 | 0.32 | 0.52 |
| 8 | 134557080 | rs11785849 | -0.511 | 1.398 | 1.297 | 1.993 | 0.032 | 0.046 | 0.046 | 0 | 0.09 | 0.13 | 0.15 | 0.32 | 0.16 | 0.1 |
| 8 | 134559333 | rs3758105 | -1.365 | 0.974 | 1.298 | 1.95 | 0.114 | 0.352 | 0.352 | 0 | 0.08 | 0.46 | 0.25 | 0.22 | 0.1 | 0.08 |
| 8 | 134562614 | rs16904945 | -0.659 | 0.51 | -0.06 | 0.817 | 0.052 | 0.003 | 0.003 | 0 | 0.12 | 0.03 | 0.04 | 0.09 | 0.03 | 0.14 |
| 8 | 134566133 | rs11775725 | -1.102 | 1.696 | 0.472 | 0.928 | 0.044 | 0.079 | 0.079 | 0 | 0.1 | 0.21 | 0.16 | 0.31 | 0.15 | 0.1 |
| 8 | 134568731 | rs12543905 | -1.257 | 1.332 | 0.41 | 0.325 | 0.027 | -0.01 | -0.01 | 0 | 0.14 | 0.42 | 0.21 | 0.08 | 0.13 | 0.07 |
| 8 | 134569528 | rs4526340 | -1.163 | 2.058 | -0.257 | 0.05 | 0.005 | -0.027 | -0.027 | 0 | 0.12 | 0.12 | 0.08 | 0 | 0.17 | 0.14 |
| 8 | 134571842 | rs7820431 | 0.353 | 1.751 | -0.752 | 0.215 | -0.001 | 0.011 | 0.011 | 0.669 | 0.55 | 0.29 | 0.61 | 0.57 | 0.54 | 0.6 |
| 8 | 134572579 | rs11776567 | 0.03 | 1.688 | -0.199 | 0.105 | -0.002 | -0.004 | -0.004 | 0 | 0.76 | 0.77 | 0.77 | 0.64 | 0.71 | 0.79 |
| 8 | 134572986 | rs7825486 | -0.51 | 1.673 | -0.267 | 0.061 | -0.019 | 0.065 | 0.065 | 0.345 | 0.16 | 0.05 | 0.09 | 0 | 0.15 | 0.15 |
| 8 | 134574971 | rs1829402 | 0.238 | 1.959 | -0.586 | 0.759 | -0.022 | 0.04 | 0.04 | 0 | 0.06 | 0.01 | 0.24 | 0.43 | 0.06 | 0.16 |
| 8 | 134579035 | rs2272675 | 0.478 | 1.65 | -0.559 | 0.374 | -0.075 | -0.024 | -0.024 | 0 | 0.68 | 0.91 | 0.76 | 0.63 | 0.61 | 0.69 |
| 8 | 134579093 | rs2272674 | 1.089 | 1.622 | 0.151 | 0.341 | -0.002 | 0.003 | 0.003 | 0.029 | 0.58 | 0.59 | 0.69 | 0.63 | 0.55 | 0.59 |
| 8 | 134586713 | rs939024 | -0.045 | 1.798 | -0.481 | 0.9 | -0.074 | -0.111 | -0.111 | 0.472 | 0.86 | 0.99 | 0.84 | 0.51 | 0.77 | 0.66 |
| 8 | 134587186 | rs1554324 | 0.646 | 1.958 | -0.601 | -0.207 | -0.005 | 0.003 | 0.003 | 0.027 | 0.59 | 0.64 | 0.54 | 0.49 | 0.54 | 0.51 |
| 8 | 134589440 | rs13272252 | 0.127 | 1.934 | -0.751 | -0.37 | -0.026 | 0.042 | 0.042 | 0 | 0.08 | 0.01 | 0.04 | 0 | 0.09 | 0.12 |
| 8 | 134598104 | rs1401346 | -0.666 | 1.913 | -0.365 | 0.212 | -0.011 | -0.038 | -0.038 | 0 | 0.88 | 0.94 | 0.86 | 0.66 | 0.86 | 0.69 |
| 8 | 134600848 | rs7825610 | 2.139 | 1.455 | -0.798 | -0.896 | 0.006 | -0.021 | -0.021 | 0 | 0.31 | 0.3 | 0.17 | 0.03 | 0.24 | 0.21 |
| 8 | 134602661 | rs9643302 | 0.001 | 1.204 | -0.34 | -0.928 | -0.001 | -0.007 | -0.007 | 0.346 | 0.21 | 0.18 | 0.13 | 0.1 | 0.19 | 0.17 |
| 8 | 134603498 | rs4288361 | 1.736 | 1.408 | -0.919 | -0.907 | 0.019 | 0.113 | 0.113 | 1.13 | 0.47 | 0.31 | 0.37 | 0.14 | 0.43 | 0.28 |
| 8 | 134604864 | rs7016568 | 2.794 | 1.413 | -0.914 | -0.92 | 0.019 | 0.079 | 0.079 | 0.609 | 0.3 | 0.2 | 0.16 | 0.03 | 0.25 | 0.22 |
| 8 | 134605669 | rs7846144 | -0.927 | 1.197 | -0.504 | -0.693 | 0 | 0 | 0 | 0 | 0.41 | 0.5 | 0.37 | 0.22 | 0.37 | 0.29 |
| 8 | 134606677 | rs2978041 | -1.383 | 1.083 | -0.544 | -0.694 | 0.016 | 0.038 | 0.038 | 0.046 | 0.34 | 0.64 | 0.2 | 0.1 | 0.26 | 0.26 |
| 8 | 134609410 | rs7830798 | -0.598 | 1.038 | -0.138 | -0.631 | -0.018 | -0.076 | -0.076 | 0 | 0.25 | 0.46 | 0.18 | 0.11 | 0.21 | 0.15 |
| 8 | 134611160 | rs6471133 | -0.53 | 1.011 | 0.091 | -0.643 | -0.018 | -0.084 | -0.084 | 0 | 0.25 | 0.44 | 0.18 | 0.11 | 0.21 | 0.15 |
| 8 | 134615750 | rs2945733 | -1.279 | 1.228 | -0.07 | -0.275 | -0.024 | -0.065 | -0.065 | 0.199 | 0.88 | 0.97 | 0.89 | 0.66 | 0.89 | 0.64 |
| 8 | 134619026 | rs6471134 | -0.088 | 1.18 | -0.23 | -0.359 | -0.03 | -0.09 | -0.09 | 0 | 0.8 | 0.96 | 0.86 | 0.66 | 0.82 | 0.58 |
| 8 | 134620862 | rs16904960 | 0.289 | 1.643 | -0.259 | -0.265 | -0.005 | 0.046 | 0.046 | 0 | 0.08 | 0.07 | 0.3 | 0.41 | 0.18 | 0.14 |
| 8 | 134621104 | rs6471136 | -0.225 | 1.417 | -0.106 | -0.379 | -0.015 | -0.08 | -0.08 | 0 | 0.24 | 0.42 | 0.18 | 0.11 | 0.21 | 0.14 |
| 8 | 134621178 | rs10956711 | 3.064 | 1.319 | -0.522 | -0.643 | 0.016 | 0.026 | 0.026 | 0.597 | 0.3 | 0.23 | 0.15 | 0.02 | 0.24 | 0.19 |
| 8 | 134621686 | rs9643306 | 0.784 | 1.481 | -0.67 | -0.302 | -0.035 | -0.023 | -0.023 | 0 | 0.16 | 0.03 | 0.2 | 0.12 | 0.19 | 0.07 |
| 8 | 134624572 | rs4736459 | -0.172 | 1.337 | -0.178 | -0.054 | -0.001 | -0.011 | -0.011 | 0 | 0.22 | 0.19 | 0.16 | 0.11 | 0.21 | 0.14 |
| 8 | 134624594 | rs2978056 | -0.764 | 1.163 | 0.121 | 0.384 | 0.004 | 0.034 | 0.034 | 0.106 | 0.16 | 0.2 | 0.29 | 0.4 | 0.21 | 0.23 |
| 8 | 134625730 | rs16904963 | 0.17 | 0.856 | 0.097 | -0.083 | -0.025 | -0.012 | -0.012 | 0 | 0.13 | 0.2 | 0.11 | 0.09 | 0.11 | 0.06 |
| 8 | 134626955 | rs2945770 | -0.387 | 1.124 | -0.129 | -0.108 | 0 | 0.005 | 0.005 | 0.254 | 0.16 | 0.14 | 0.1 | 0.09 | 0.15 | 0.1 |
| 8 | 134627403 | rs2945767 | -1.409 | 1.788 | -0.29 | 0.454 | -0.02 | -0.039 | -0.039 | 0.279 | 0.23 | 0.16 | 0.34 | 0.45 | 0.28 | 0.21 |
| 8 | 134627635 | rs2978033 | -0.858 | 1.795 | -0.31 | 0.448 | -0.063 | -0.078 | -0.078 | 0 | 0.14 | 0.04 | 0.26 | 0.37 | 0.19 | 0.16 |
| 8 | 134627756 | rs13264573 | 1.28 | 1.587 | -0.879 | 0.06 | -0.032 | 0.152 | 0.152 | 1.364 | 0.57 | 0.81 | 0.46 | 0.18 | 0.51 | 0.43 |
| 8 | 134628443 | rs10808612 | -0.075 | 1.603 | -0.903 | 0.095 | -0.103 | -0.144 | -0.144 | 1.657 | 0.74 | 0.97 | 0.78 | 0.61 | 0.73 | 0.6 |
| 8 | 134629844 | rs11989055 | -0.072 | 1.63 | -0.862 | 0.12 | -0.005 | -0.003 | -0.003 | 0.045 | 0.17 | 0.12 | 0.32 | 0.44 | 0.21 | 0.17 |
| 8 | 134632232 | rs13260215 | 0.84 | 1.661 | -0.844 | -0.242 | 0 | 0 | 0 | 0 | 0.15 | 0.15 | 0.23 | 0.14 | 0.23 | 0.1 |
| 8 | 134632509 | rs7831161 | -0.425 | 1.583 | -0.99 | -0.225 | -0.103 | -0.139 | -0.139 | 0 | 0.74 | 0.97 | 0.78 | 0.62 | 0.73 | 0.6 |
| 8 | 134635535 | rs2922462 | -1.085 | 1.351 | -0.888 | -0.13 | -0.066 | -0.135 | -0.135 | 0.433 | 0.81 | 0.99 | 0.8 | 0.62 | 0.79 | 0.65 |
| 8 | 134635688 | rs3992973 | 1.152 | 1.145 | -0.463 | -0.186 | 0.027 | 0.023 | 0.023 | 0.103 | 0.47 | 0.63 | 0.46 | 0.39 | 0.41 | 0.41 |
| 8 | 134638258 | rs3958181 | 1.717 | 0.783 | -0.187 | -0.559 | -0.017 | -0.09 | -0.09 | 1.587 | 0.39 | 0.38 | 0.2 | 0.04 | 0.27 | 0.28 |
| 8 | 134642472 | rs7008657 | 0.742 | 0.953 | 0.132 | -0.148 | -0.015 | -0.002 | -0.002 | 0 | 0.13 | 0.38 | 0.11 | 0.09 | 0.1 | 0.09 |
| 8 | 134642560 | rs2922473 | -0.837 | 1.163 | -0.005 | 0.062 | -0.001 | -0.021 | -0.021 | 0 | 0.79 | 0.82 | 0.81 | 0.6 | 0.84 | 0.67 |
| 8 | 134643435 | rs6471139 | 0.477 | 1.359 | 0.128 | -0.213 | 0.007 | -0.006 | -0.006 | 0.597 | 0.7 | 0.64 | 0.53 | 0.22 | 0.64 | 0.52 |
| 8 | 134643538 | rs2945745 | -1.125 | 1.054 | -0.284 | -0.118 | 0.002 | 0.038 | 0.038 | 0 | 0.25 | 0.35 | 0.39 | 0.56 | 0.3 | 0.26 |
| 8 | 134646398 | rs10111368 | 0.328 | 1.238 | -0.856 | -0.484 | -0.068 | 0.09 | 0.09 | 1.329 | 0.66 | 0.84 | 0.48 | 0.13 | 0.56 | 0.53 |
| 8 | 134646433 | rs16904967 | -0.337 | 1.034 | -0.902 | -0.824 | 0.008 | -0.027 | -0.027 | 0.489 | 0.27 | 0.62 | 0.3 | 0.09 | 0.29 | 0.23 |
| 8 | 134646629 | rs2945741 | -0.533 | 1.327 | -1.479 | -0.024 | -0.073 | -0.163 | -0.163 | 0 | 0.74 | 0.98 | 0.66 | 0.57 | 0.75 | 0.62 |
| 8 | 134649346 | rs2958830 | -0.812 | 1.574 | -0.829 | 0.252 | 0.056 | 0.145 | 0.145 | 0 | 0.1 | 0.22 | 0.2 | 0.45 | 0.21 | 0.19 |
| 8 | 134657960 | rs16904971 | 0.138 | 1.009 | -0.978 | -0.083 | -0.006 | -0.01 | -0.01 | 0 | 0.15 | 0.21 | 0.12 | 0.08 | 0.14 | 0.11 |
| 8 | 134660028 | rs1449070 | 0.348 | 1.322 | -1.202 | -0.372 | -0.009 | 0.005 | 0.005 | 0 | 0.14 | 0.11 | 0.12 | 0.08 | 0.19 | 0.11 |
| 8 | 134662095 | rs10505611 | 3.018 | 0.533 | -1.954 | -0.881 | 0.024 | 0.136 | 0.136 | 2.428 | 0.24 | 0.01 | 0.1 | 0.03 | 0.17 | 0.15 |
| 8 | 134663886 | rs16904973 | -1.237 | 0.795 | -1.215 | -0.651 | 0.001 | -0.022 | -0.022 | 0 | 0.36 | 0.42 | 0.26 | 0.16 | 0.38 | 0.26 |
| 8 | 134664999 | rs2922446 | -1.482 | 0.881 | -1.025 | -0.505 | -0.002 | 0.002 | 0.002 | 0.254 | 0.54 | 0.48 | 0.54 | 0.47 | 0.55 | 0.6 |
| 8 | 134667117 | rs2945744 | -0.053 | 0.554 | -1.28 | -0.28 | 0.008 | -0.016 | -0.016 | 0 | 0.32 | 0.19 | 0.36 | 0.38 | 0.27 | 0.43 |
| 8 | 134668677 | rs2860223 | -0.491 | 0.618 | -1.28 | -0.269 | 0.008 | -0.007 | -0.007 | 0.013 | 0.38 | 0.21 | 0.38 | 0.4 | 0.32 | 0.49 |
| 8 | 134673368 | rs1562676 | -0.357 | 0.83 | -0.952 | -0.212 | 0 | -0.002 | -0.002 | 0 | 0.37 | 0.35 | 0.39 | 0.39 | 0.35 | 0.48 |
| 8 | 134676027 | rs2292897 | -0.253 | 1.553 | -1.518 | -0.16 | -0.062 | -0.049 | -0.049 | 0 | 0.91 | 0.99 | 0.68 | 0.55 | 0.75 | 0.82 |
| 8 | 134676889 | rs2922500 | 0.458 | 0.833 | -1.373 | -0.296 | 0.002 | -0.031 | -0.031 | 0 | 0.3 | 0.22 | 0.36 | 0.39 | 0.29 | 0.43 |
| 8 | 134678117 | rs6981912 | 1.343 | 0.362 | -1.124 | -0.64 | -0.009 | -0.046 | -0.046 | 0 | 0.39 | 0.49 | 0.19 | 0.07 | 0.29 | 0.27 |
| 8 | 134678735 | rs2958786 | 0.638 | 0.792 | -1.043 | -0.166 | 0.011 | -0.028 | -0.028 | 0.276 | 0.28 | 0.21 | 0.33 | 0.35 | 0.25 | 0.43 |
| 8 | 134681123 | rs12681042 | 1.005 | 0.663 | -1.263 | -0.373 | 0.003 | 0.004 | 0.004 | 4.398 | 0.45 | 0.55 | 0.28 | 0.21 | 0.37 | 0.28 |
| 8 | 134684826 | rs2945757 | 1.043 | 0.879 | -1.889 | -0.052 | -0.026 | -0.057 | -0.057 | 0.009 | 0.26 | 0.02 | 0.3 | 0.26 | 0.22 | 0.41 |
| 8 | 134686065 | rs2945756 | 0.197 | 1.574 | -0.991 | 0.47 | -0.036 | -0.039 | -0.039 | 0 | 0.82 | 0.9 | 0.64 | 0.55 | 0.69 | 0.75 |
| 8 | 134689013 | rs6985794 | -0.606 | 1.139 | -0.62 | 0.3 | 0.011 | 0.02 | 0.02 | 0 | 0.11 | 0.2 | 0.19 | 0.1 | 0.13 | 0.14 |

| | | | | | | | | | | | | | | | | |
|---|---|---|---|---|---|---|---|---|---|---|---|---|---|---|---|---|
| 8 | 134689575 | rs1584448 | 0.166 | 0.863 | -0.947 | 0.078 | -0.007 | 0.043 | 0.043 | 0 | 0.47 | 0.3 | 0.5 | 0.39 | 0.47 | 0.59 |
| 8 | 134690434 | rs1975605 | 1.32 | 0.401 | -1.132 | -0.155 | 0.036 | 0.046 | 0.046 | 0.072 | 0.22 | 0.14 | 0.13 | 0.09 | 0.15 | 0.11 |
| 8 | 134691200 | rs7006459 | -0.494 | 1.107 | -0.321 | 0.613 | 0.057 | 0.079 | 0.079 | 2.918 | 0.86 | 0.81 | 0.82 | 0.63 | 0.72 | 0.8 |
| 8 | 134697809 | rs7005557 | 0.795 | 0.944 | -0.267 | 0.05 | 0.021 | 0.057 | 0.057 | 0 | 0.35 | 0.25 | 0.2 | 0.12 | 0.29 | 0.3 |
| 8 | 134700316 | rs1008744 | 1.198 | 0.759 | -0.691 | 0.153 | -0.043 | 0.006 | 0.006 | 0 | 0.31 | 0.02 | 0.48 | 0.27 | 0.29 | 0.45 |
| 8 | 134702428 | rs1610133 | -0.314 | 1.21 | 0.468 | 0.638 | 0.062 | 0.109 | 0.109 | 0.657 | 0.85 | 0.8 | 0.82 | 0.53 | 0.72 | 0.82 |
| 8 | 134702589 | rs1610131 | 0.693 | 1.131 | 0.546 | 0.561 | 0.024 | 0.008 | 0.008 | 0 | 0.7 | 0.6 | 0.75 | 0.43 | 0.59 | 0.77 |
| 8 | 134708017 | rs1868435 | -0.478 | 1.367 | 0.372 | 0.761 | 0.037 | 0.058 | 0.058 | 1.04 | 0.86 | 0.84 | 0.83 | 0.62 | 0.73 | 0.82 |
| 8 | 134708926 | rs1868431 | 0.833 | 1.101 | 0.352 | 0.671 | -0.001 | -0.001 | -0.001 | 0 | 0.14 | 0.12 | 0.07 | 0.1 | 0.13 | 0.05 |
| 8 | 134709357 | rs4612320 | 0.592 | 1.261 | 0.274 | 0.544 | 0.054 | 0.051 | 0.051 | 0 | 0.71 | 0.52 | 0.74 | 0.5 | 0.6 | 0.75 |
| 8 | 134709899 | rs10956719 | -0.019 | 0.948 | 0.096 | 0.092 | 0.013 | 0.024 | 0.024 | 0 | 0.48 | 0.59 | 0.66 | 0.38 | 0.45 | 0.53 |
| 8 | 134716682 | rs2958810 | 0.199 | 1.363 | 0.52 | -0.065 | 0 | -0.007 | -0.007 | 0 | 0.58 | 0.36 | 0.38 | 0.2 | 0.54 | 0.44 |
| 8 | 134722557 | rs11780515 | -0.069 | 1.34 | 0.488 | -0.157 | 0.023 | 0.03 | 0.03 | 0 | 0.12 | 0.3 | 0.1 | 0.12 | 0.11 | 0.13 |
| 8 | 134730076 | rs7819398 | -0.946 | 1.452 | -0.178 | -0.358 | -0.04 | -0.012 | -0.012 | 0 | 0.73 | 0.9 | 0.59 | 0.72 | 0.68 | 0.57 |
| 8 | 134730103 | rs7819295 | 0.729 | 1.5 | -0.048 | -1.156 | 0.012 | 0.109 | 0.109 | 0 | 0.5 | 0.36 | 0.29 | 0.08 | 0.45 | 0.29 |
| 8 | 134733093 | rs4736713 | 0.375 | 1.48 | -0.249 | -0.589 | -0.044 | 0.106 | 0.106 | 0 | 0.27 | 0.09 | 0.4 | 0.18 | 0.32 | 0.44 |
| 8 | 134736033 | rs2922494 | -0.534 | 1.251 | -0.032 | -0.293 | 0.021 | -0.066 | -0.066 | 0 | 0.28 | 0.55 | 0.4 | 0.17 | 0.32 | 0.39 |
| 8 | 134756506 | rs2922464 | -0.485 | 1.037 | -0.19 | 0.347 | -0.01 | 0.045 | 0.045 | 0 | 0.85 | 0.8 | 0.76 | 0.64 | 0.9 | 0.8 |
| 8 | 134773993 | rs16904985 | 0.064 | 1.814 | -0.322 | 0.781 | -0.039 | -0.052 | -0.052 | 0.444 | 0.1 | 0.01 | 0.07 | 0.39 | 0.14 | 0.09 |
| 8 | 134783002 | rs17730982 | 0.02 | 1.589 | 0.167 | -0.356 | -0.006 | 0.003 | 0.003 | 0 | 0.06 | 0.04 | 0.02 | 0 | 0.06 | 0.02 |
| 8 | 134783185 | rs4301435 | -0.035 | 1.742 | -0.436 | -0.275 | -0.068 | -0.038 | -0.038 | 0 | 0.13 | 0.01 | 0.23 | 0.17 | 0.23 | 0.37 |
| 8 | 134784746 | rs6981181 | -0.944 | 1.124 | 0.569 | -0.437 | 0.049 | -0.057 | -0.057 | 0 | 0.11 | 0.42 | 0.05 | 0 | 0.06 | 0.02 |
| 8 | 134785575 | rs7830468 | -0.565 | 1.252 | 0.538 | -0.932 | 0.021 | -0.104 | -0.104 | 0 | 0.23 | 0.4 | 0.17 | 0.02 | 0.28 | 0.2 |
| 8 | 134785709 | rs7814358 | -0.823 | 1.052 | 0.761 | -0.91 | 0.015 | -0.112 | -0.112 | 3.333 | 0.19 | 0.39 | 0.12 | 0.01 | 0.18 | 0.14 |
| 8 | 134786518 | rs4452783 | -0.585 | 0.536 | 0.317 | -0.833 | 0.003 | 0.017 | 0.017 | 0 | 0.53 | 0.43 | 0.53 | 0.67 | 0.59 | 0.65 |
| 8 | 134797424 | rs4563869 | -0.898 | 0.339 | 0.242 | -0.943 | 0.02 | 0.051 | 0.051 | 0 | 0.58 | 0.74 | 0.66 | 0.71 | 0.65 | 0.68 |
| 8 | 134801669 | rs4512362 | 0.181 | 0.752 | -0.077 | -0.346 | -0.032 | -0.05 | -0.05 | 0 | 0.27 | 0.19 | 0.4 | 0.49 | 0.34 | 0.27 |
| 8 | 134801874 | rs17731209 | 0.694 | 0.49 | -0.021 | -0.122 | 0.003 | -0.018 | -0.018 | 0.042 | 0.05 | 0.07 | 0.04 | 0 | 0.06 | 0.01 |
| 8 | 134804170 | rs6988999 | 0.238 | 0.435 | -0.091 | -0.108 | -0.001 | 0.002 | 0.002 | 0 | 0.15 | 0.15 | 0.15 | 0.23 | 0.19 | 0.35 |
| 8 | 134815703 | rs10956725 | -0.179 | 0.401 | -0.087 | -0.188 | -0.001 | 0.003 | 0.003 | 0.057 | 0.33 | 0.4 | 0.38 | 0.44 | 0.28 | 0.26 |
| 8 | 134817433 | rs7010463 | 0.322 | 0.649 | -0.32 | -0.317 | 0.004 | -0.018 | -0.018 | 0.327 | 0.05 | 0.07 | 0.04 | 0 | 0.06 | 0.01 |
| 8 | 134822433 | rs11786842 | 0.712 | 0.511 | -0.624 | -0.348 | 0.003 | 0.045 | 0.045 | 0 | 0.51 | 0.4 | 0.4 | 0.28 | 0.47 | 0.36 |
| 8 | 134824398 | rs4399592 | 0.19 | 0.48 | -0.692 | -0.307 | -0.018 | -0.04 | -0.04 | 0 | 0.31 | 0.22 | 0.49 | 0.5 | 0.39 | 0.26 |
| 8 | 134829154 | rs2404038 | 0.647 | 0.257 | -0.823 | -0.409 | 0 | 0 | 0 | 0 | 0.46 | 0.43 | 0.31 | 0.28 | 0.44 | 0.35 |
| 8 | 134832652 | rs2134672 | 0.188 | 0.068 | -0.31 | -0.21 | -0.028 | 0.08 | 0.08 | 0 | 0.85 | 0.63 | 0.93 | 0.78 | 0.92 | 0.63 |
| 8 | 134835998 | rs13255038 | -0.182 | 0.483 | -1.581 | -1.024 | -0.047 | 0.151 | 0.151 | 0.012 | 0.39 | 0.13 | 0.48 | 0.13 | 0.54 | 0.26 |
| 8 | 134838488 | rs13275988 | -0.178 | 0.461 | -1.224 | -1.055 | -0.017 | 0.084 | 0.084 | 0.045 | 0.38 | 0.24 | 0.49 | 0.13 | 0.54 | 0.26 |
| 8 | 134842573 | rs4897689 | -0.239 | 0.538 | -0.892 | -0.681 | -0.004 | 0.007 | 0.007 | 0.258 | 0.09 | 0.17 | 0.05 | 0.1 | 0.07 | 0.06 |
| 8 | 134846363 | rs4897731 | -0.053 | 0.678 | -1.255 | -0.668 | -0.002 | -0.002 | -0.002 | 0 | 0.08 | 0.06 | 0.05 | 0.1 | 0.08 | 0.06 |
| 8 | 134846732 | rs4897732 | -0.087 | 0.095 | -1.764 | -0.74 | 0.018 | -0.012 | -0.012 | 0 | 0.24 | 0.03 | 0.16 | 0.23 | 0.21 | 0.42 |
| 8 | 134851366 | rs924768 | 0.168 | 0.531 | -1.349 | -1.69 | -0.021 | 0.057 | 0.057 | 0 | 0.41 | 0.34 | 0.55 | 0.12 | 0.45 | 0.29 |
| 8 | 134854031 | rs7812695 | 0.274 | 0.338 | -0.733 | -1.66 | 0.04 | 0.004 | 0.004 | 0 | 0.05 | 0.2 | 0.03 | 0 | 0.04 | 0.02 |
| 8 | 134856613 | rs10113320 | -0.649 | 0.029 | -1.648 | -1.899 | -0.023 | 0.066 | 0.066 | 0 | 0.62 | 0.98 | 0.73 | 0.19 | 0.66 | 0.39 |
| 8 | 134861390 | rs10090091 | 0.725 | -0.233 | -1.322 | -2.557 | 0.004 | 0.074 | 0.074 | 0.119 | 0.3 | 0.13 | 0.15 | 0.01 | 0.28 | 0.17 |
| 8 | 134863691 | rs11996535 | 0.289 | -0.236 | -0.601 | -2.599 | 0.101 | 0.046 | 0.046 | 4.003 | 0.05 | 0.31 | 0.04 | 0 | 0.04 | 0 |
| 8 | 134865952 | rs7845379 | 0.081 | -0.537 | -0.598 | -2.865 | -0.012 | -0.128 | -0.128 | 0.08 | 0.41 | 0.5 | 0.21 | 0.01 | 0.33 | 0.21 |
| 8 | 134868049 | rs17670773 | 0.385 | -0.707 | -0.645 | -2.702 | -0.001 | 0.008 | 0.008 | 0 | 0.1 | 0.06 | 0.05 | 0.04 | 0.08 | 0.07 |
| 8 | 134870846 | rs16905007 | -1.023 | -0.77 | -0.264 | -2.74 | -0.023 | -0.101 | -0.101 | 0 | 0.15 | 0.55 | 0.1 | 0 | 0.09 | 0.04 |
| 8 | 134871788 | rs10109162 | -1.12 | -0.767 | -0.694 | -2.944 | -0.142 | 0.245 | 0.245 | 0.151 | 0.6 | 0.96 | 0.33 | 0.05 | 0.46 | 0.3 |
| 8 | 134871987 | rs6983078 | 0.58 | -0.763 | -0.65 | -2.948 | 0.028 | 0.047 | 0.047 | 0 | 0.37 | 0.28 | 0.19 | 0.01 | 0.31 | 0.19 |
| 8 | 134873886 | rs6993205 | -1.296 | -0.764 | -0.667 | -2.947 | -0.162 | 0.212 | 0.212 | 0.574 | 0.63 | 0.96 | 0.34 | 0.05 | 0.47 | 0.31 |
| 8 | 134876452 | rs4897697 | -1.325 | -0.452 | -0.278 | -2.624 | 0.002 | -0.01 | -0.01 | 0 | 0.26 | 0.67 | 0.15 | 0.04 | 0.17 | 0.12 |
| 8 | 134885463 | rs6998502 | 0.462 | -0.639 | -0.46 | -2.617 | 0.037 | 0.028 | 0.028 | 0.46 | 0.39 | 0.3 | 0.16 | 0.01 | 0.26 | 0.2 |
| 8 | 134886783 | rs924766 | 0.308 | -0.305 | -0.595 | -2.394 | -0.001 | -0.004 | -0.004 | 0 | 0.1 | 0.11 | 0.06 | 0.07 | 0.07 | 0.06 |
| 8 | 134901186 | rs6471165 | -0.935 | -0.222 | -0.81 | -2.384 | -0.132 | 0.187 | 0.187 | 5.265 | 0.63 | 0.96 | 0.32 | 0.08 | 0.42 | 0.31 |
| 8 | 134906698 | rs4897729 | -0.35 | 0.026 | -0.363 | -2.16 | -0.001 | 0 | 0 | 0 | 0.14 | 0.12 | 0.08 | 0.05 | 0.14 | 0.07 |
| 8 | 134911143 | rs1503759 | -1.075 | -0.091 | 0.057 | -2.28 | 0.019 | -0.046 | -0.046 | 0.194 | 0.1 | 0.49 | 0.06 | 0 | 0.04 | 0.02 |
| 8 | 134911488 | rs4897730 | -1.713 | -0.103 | -0.046 | -2.329 | -0.001 | -0.015 | -0.015 | 0 | 0.27 | 0.7 | 0.19 | 0.05 | 0.24 | 0.13 |
| 8 | 134914331 | rs1503758 | -1.581 | -0.156 | 0.094 | -2.27 | 0 | -0.098 | -0.098 | 0 | 0.15 | 0.57 | 0.11 | 0 | 0.12 | 0.06 |
| 8 | 134915089 | rs7846118 | -1.446 | -0.16 | 0.07 | -2.272 | -0.002 | -0.101 | -0.101 | 0 | 0.15 | 0.56 | 0.11 | 0 | 0.12 | 0.06 |
| 8 | 134918258 | rs7014384 | -0.462 | -0.149 | -0.562 | -2.403 | 0.089 | 0.265 | 0.265 | 7.045 | 0.53 | 0.86 | 0.27 | 0.01 | 0.34 | 0.24 |
| 8 | 134927281 | rs2896730 | -0.772 | -0.07 | -0.201 | -2.1 | -0.051 | 0.09 | 0.09 | 3.257 | 0.62 | 0.83 | 0.36 | 0.19 | 0.5 | 0.31 |
| 8 | 134929444 | rs10109095 | -1.114 | 0.023 | -0.3 | -1.988 | 0.002 | -0.016 | -0.016 | 0 | 0.25 | 0.64 | 0.19 | 0.17 | 0.24 | 0.13 |
| 8 | 134932987 | rs6999388 | -0.401 | 0.159 | -0.482 | -1.147 | 0.004 | 0.029 | 0.029 | 2.369 | 0.55 | 0.65 | 0.3 | 0.35 | 0.43 | 0.31 |
| 8 | 134938386 | rs11994154 | -0.418 | 0.14 | -0.74 | -1.145 | -0.028 | 0.04 | 0.04 | 1.32 | 0.57 | 0.74 | 0.32 | 0.36 | 0.46 | 0.31 |
| 8 | 134940844 | rs963164 | 0.1 | 0.23 | -0.587 | -1.514 | -0.005 | -0.036 | -0.036 | 0 | 0.07 | 0.16 | 0.03 | 0 | 0.03 | 0.03 |
| 8 | 134942372 | rs4897645 | 0.162 | 0.375 | -1.125 | -1.486 | -0.055 | 0.067 | 0.067 | 2.256 | 0.45 | 0.08 | 0.33 | 0.3 | 0.42 | 0.31 |
| 8 | 134943118 | rs10109080 | -0.067 | 0.351 | -0.979 | -1.389 | -0.002 | 0.052 | 0.052 | 0.907 | 0.51 | 0.6 | 0.4 | 0.3 | 0.46 | 0.31 |
| 8 | 134944520 | rs1587768 | -0.557 | 0.084 | -0.738 | -1.835 | -0.01 | -0.053 | -0.053 | 1.186 | 0.14 | 0.18 | 0.06 | 0 | 0.08 | 0.04 |

| | | | | | | | | | | | | | | | | |
|---|---|---|---|---|---|---|---|---|---|---|---|---|---|---|---|---|
| 8 | 134945681 | rs1961455 | -0.636 | 0.068 | -1.227 | -2.512 | 0.054 | 0.121 | 0.121 | 1.526 | 0.5 | 0.66 | 0.25 | 0.01 | 0.32 | 0.22 |
| 8 | 134947433 | rs7464119 | -0.799 | -0.021 | -1.561 | -2.686 | -0.047 | 0.102 | 0.102 | 0 | 0.78 | 0.95 | 0.82 | 1 | 0.76 | 0.83 |
| 8 | 134948065 | rs950757 | 1.031 | -0.389 | -1.977 | -2.652 | -0.087 | 0.073 | 0.073 | 1.384 | 0.42 | 0.15 | 0.54 | 0.72 | 0.51 | 0.69 |
| 8 | 134953726 | rs10113469 | 0.45 | -0.618 | -1.549 | -2.73 | -0.013 | -0.028 | -0.028 | 6.882 | 0.08 | 0.12 | 0.05 | 0 | 0.04 | 0.05 |
| 8 | 134954618 | rs1503753 | -0.028 | -0.859 | -0.638 | -2.859 | -0.036 | -0.061 | -0.061 | 0.407 | 0.18 | 0.36 | 0.09 | 0.05 | 0.1 | 0.03 |
| 8 | 134955476 | rs12677718 | -1.442 | -0.809 | -0.363 | -2.879 | -0.013 | 0.027 | 0.027 | 0 | 0.77 | 0.87 | 0.81 | 1 | 0.77 | 0.83 |
| 8 | 134957089 | rs1317081 | 1.785 | -0.695 | -0.543 | -3.003 | -0.07 | 0.021 | 0.021 | 0.516 | 0.35 | 0.15 | 0.51 | 0.72 | 0.46 | 0.67 |
| 8 | 134957296 | rs1014935 | 0.96 | -0.81 | -0.214 | -3.045 | 0.006 | 0.054 | 0.054 | 2.453 | 0.05 | 0.25 | 0.04 | 0.05 | 0.01 | 0.01 |
| 8 | 134957528 | rs10956733 | 0.256 | -0.827 | -0.098 | -3.327 | 0 | 0.001 | 0.001 | 0 | 0.55 | 0.59 | 0.63 | 0.76 | 0.58 | 0.71 |
| 8 | 134958011 | rs6982919 | 1.177 | -0.719 | -0.093 | -3.227 | 0 | -0.022 | -0.022 | 0.24 | 0.12 | 0.11 | 0.05 | 0 | 0.09 | 0.02 |
| 8 | 134965071 | rs4897646 | 0.152 | -0.783 | -0.121 | -3.341 | -0.01 | 0.04 | 0.04 | 0 | 0.34 | 0.21 | 0.21 | 0 | 0.33 | 0.21 |
| 8 | 134967483 | rs7815319 | -0.315 | -0.695 | -0.077 | -3.172 | -0.034 | -0.136 | -0.136 | 5.795 | 0.2 | 0.45 | 0.11 | 0.01 | 0.12 | 0.03 |
| 8 | 134968245 | rs7008637 | -0.895 | -0.615 | -0.099 | -3.203 | -0.048 | -0.174 | -0.174 | 0 | 0.26 | 0.54 | 0.13 | 0.01 | 0.14 | 0.08 |
| 8 | 134969520 | rs1354365 | -1.414 | -0.515 | -0.315 | -3.166 | 0.001 | -0.006 | -0.006 | 1.163 | 0.63 | 0.6 | 0.42 | 0.25 | 0.57 | 0.35 |
| 8 | 134969926 | rs7837524 | 0.467 | -0.346 | -0.375 | -3.049 | -0.007 | -0.061 | -0.061 | 0 | 0.15 | 0.23 | 0.09 | 0 | 0.11 | 0.03 |
| 8 | 134974025 | rs1350337 | 0.089 | 0.061 | -0.849 | -2.799 | -0.001 | -0.002 | -0.002 | 0 | 0.36 | 0.28 | 0.28 | 0.24 | 0.38 | 0.25 |
| 8 | 134977914 | rs12676561 | -1.131 | 0.178 | -1.231 | -2.377 | 0.052 | 0.14 | 0.14 | 1.854 | 0.45 | 0.85 | 0.31 | 0.26 | 0.32 | 0.16 |
| 8 | 134993530 | rs1125044 | -0.746 | -0.325 | -0.855 | -2.631 | -0.022 | 0.071 | 0.071 | 1.999 | 0.59 | 0.77 | 0.31 | 0.01 | 0.43 | 0.31 |
| 8 | 134999918 | rs7813257 | 0.302 | -0.869 | -0.55 | -2.917 | -0.001 | -0.028 | -0.028 | 0 | 0.19 | 0.2 | 0.05 | 0.01 | 0.07 | 0.08 |
| 8 | 135003093 | rs11992450 | -0.673 | -0.934 | -0.019 | -2.978 | 0.053 | 0.006 | 0.006 | 0.999 | 0.07 | 0.36 | 0.06 | 0 | 0.04 | 0.03 |
| 8 | 135005295 | rs1125718 | -1.06 | -1.011 | -0.093 | -2.804 | 0.018 | 0.015 | 0.015 | 1.037 | 0.41 | 0.65 | 0.25 | 0.29 | 0.28 | 0.15 |
| 8 | 135005442 | rs1125719 | -1.083 | -0.929 | -0.117 | -2.795 | -0.012 | 0.008 | 0.008 | 0.729 | 0.63 | 0.7 | 0.41 | 0.29 | 0.53 | 0.32 |
| 8 | 135007667 | rs6982411 | -1.301 | -0.94 | -0.227 | -2.763 | -0.022 | 0.008 | 0.008 | 1.602 | 0.65 | 0.74 | 0.41 | 0.3 | 0.52 | 0.32 |
| 8 | 135012293 | rs17746690 | 0.444 | -0.496 | 0.36 | -2.375 | 0.001 | -0.074 | -0.074 | 0.601 | 0.29 | 0.29 | 0.22 | 0 | 0.28 | 0.2 |
| 8 | 135014817 | rs7844164 | -1.128 | -0.548 | 0.163 | -2.305 | -0.007 | 0.073 | 0.073 | 0.044 | 0.52 | 0.69 | 0.41 | 0.28 | 0.49 | 0.27 |
| 8 | 135018790 | rs1379438 | 1.192 | -0.372 | 0.155 | -2.483 | -0.05 | 0.139 | 0.139 | 0 | 0.22 | 0.04 | 0.13 | 0 | 0.23 | 0.16 |
| 8 | 135021967 | rs13267601 | -1.088 | -0.233 | 0.856 | -2.252 | 0.045 | 0.067 | 0.067 | 0 | 0.16 | 0.43 | 0.19 | 0.18 | 0.19 | 0.1 |
| 8 | 135025593 | rs16905048 | -1.591 | -0.342 | 1.097 | -2.275 | -0.016 | -0.029 | -0.029 | 0.002 | 0.24 | 0.47 | 0.15 | 0.15 | 0.2 | 0.08 |
| 8 | 135026160 | rs7819815 | -0.964 | -0.364 | 1.173 | -2.248 | -0.007 | -0.001 | -0.001 | 0 | 0.16 | 0.29 | 0.08 | 0.13 | 0.12 | 0.06 |
| 8 | 135026266 | rs7820063 | -1.099 | -0.421 | 1.227 | -2.238 | -0.005 | 0.001 | 0.001 | 0.073 | 0.17 | 0.26 | 0.11 | 0.16 | 0.14 | 0.06 |
| 8 | 135029384 | rs16905051 | 0.495 | -0.422 | 1.297 | -2.512 | 0.001 | -0.066 | -0.066 | 0.508 | 0.3 | 0.28 | 0.21 | 0 | 0.31 | 0.2 |
| 8 | 135031503 | rs4897625 | -0.975 | -0.474 | 1.491 | -2.498 | -0.001 | -0.025 | -0.025 | 0 | 0.48 | 0.49 | 0.35 | 0.15 | 0.46 | 0.28 |
| 8 | 135035923 | rs10505616 | 0.51 | -0.437 | 1.35 | -2.479 | -0.004 | -0.036 | -0.036 | 0.015 | 0.3 | 0.23 | 0.21 | 0 | 0.31 | 0.2 |
| 8 | 135038599 | rs4897627 | 1.092 | -0.436 | 1.15 | -2.495 | -0.072 | 0.161 | 0.161 | 0 | 0.25 | 0.02 | 0.14 | 0 | 0.26 | 0.17 |
| 8 | 135039549 | rs16905052 | -0.742 | -0.436 | 1.506 | -2.456 | 0.036 | -0.013 | -0.013 | 1.152 | 0.05 | 0.22 | 0.07 | 0 | 0.05 | 0.03 |
| 8 | 135040165 | rs16905054 | -0.888 | -0.438 | 1.576 | -2.456 | -0.018 | 0.012 | 0.012 | 1.328 | 0.16 | 0.29 | 0.13 | 0.14 | 0.12 | 0.09 |
| 8 | 135044823 | rs17675353 | 0.584 | -0.382 | 1.345 | -2.48 | -0.017 | 0.035 | 0.035 | 0 | 0.29 | 0.17 | 0.2 | 0 | 0.31 | 0.2 |
| 8 | 135045018 | rs17675396 | 0.584 | -0.382 | 1.345 | -2.48 | -0.017 | 0.035 | 0.035 | 0 | 0.29 | 0.17 | 0.2 | 0 | 0.31 | 0.2 |
| 8 | 135047450 | rs17748159 | 0.865 | -0.415 | 1.152 | -2.463 | -0.027 | 0.116 | 0.116 | 0 | 0.27 | 0.07 | 0.15 | 0 | 0.27 | 0.17 |
| 8 | 135048602 | rs16905057 | -1.663 | -0.61 | 1.69 | -1.939 | -0.033 | 0.018 | 0.018 | 0.047 | 0.2 | 0.44 | 0.15 | 0.19 | 0.15 | 0.09 |
| 8 | 135053994 | rs1457480 | -1.801 | -0.571 | 1.585 | -2.283 | -0.034 | -0.134 | -0.134 | 0.02 | 0.19 | 0.43 | 0.11 | 0 | 0.1 | 0.06 |
| 8 | 135062170 | rs16905062 | -1.183 | -0.58 | 1.712 | -2.286 | -0.018 | -0.058 | -0.058 | 0 | 0.14 | 0.18 | 0.04 | 0 | 0.06 | 0.06 |
| 8 | 135064801 | rs16905064 | 0.017 | -0.607 | 1.859 | -2.299 | 0.003 | -0.007 | -0.007 | 0.258 | 0.08 | 0.05 | 0.02 | 0 | 0.03 | 0.03 |
| 8 | 135065947 | rs1117497 | -1.161 | -0.599 | 1.839 | -2.295 | -0.018 | -0.058 | -0.058 | 0 | 0.14 | 0.18 | 0.04 | 0.01 | 0.06 | 0.05 |
| 8 | 135068340 | rs6471186 | -2.239 | -0.641 | 1.729 | -2.323 | -0.04 | -0.128 | -0.128 | 0 | 0.22 | 0.57 | 0.13 | 0 | 0.13 | 0.07 |
| 8 | 135077117 | rs7836259 | -1.756 | -0.32 | 1.636 | -1.965 | 0.028 | -0.033 | -0.033 | 0.631 | 0.09 | 0.4 | 0.08 | 0 | 0.07 | 0.02 |
| 8 | 135080135 | rs7835234 | -1.504 | -0.201 | 0.694 | -2.395 | -0.126 | -0.02 | -0.02 | 0.207 | 0.72 | 0.95 | 0.45 | 0.18 | 0.58 | 0.37 |
| 8 | 135082117 | rs10087922 | -2.199 | -0.287 | 1.303 | -2.411 | 0.009 | 0.041 | 0.041 | 0.269 | 0.2 | 0.54 | 0.19 | 0.18 | 0.19 | 0.1 |
| 8 | 135092183 | rs10093555 | -1.698 | -0.522 | 1.48 | -2.291 | 0.057 | 0.202 | 0.202 | 0.592 | 0.1 | 0.58 | 0.13 | 0.22 | 0.11 | 0.07 |
| 8 | 135093701 | rs1870126 | -0.654 | -0.744 | 1.205 | -2.56 | -0.033 | 0.047 | 0.047 | 0 | 0.59 | 0.76 | 0.38 | 0.22 | 0.5 | 0.32 |
| 8 | 135094049 | rs16905068 | -1.018 | -0.745 | 1.401 | -2.56 | 0.026 | -0.008 | -0.008 | 0 | 0.07 | 0.34 | 0.04 | 0 | 0.02 | 0.02 |
| 8 | 135094936 | rs6981423 | -1.707 | -0.705 | 1.019 | -2.548 | -0.142 | -0.034 | -0.034 | 0.002 | 0.72 | 0.95 | 0.42 | 0.22 | 0.55 | 0.36 |
| 8 | 135098256 | rs1457461 | -2.297 | -0.721 | 1.229 | -2.406 | -0.006 | 0.011 | 0.011 | 0.503 | 0.2 | 0.71 | 0.17 | 0.21 | 0.18 | 0.11 |
| 8 | 135099778 | rs1457463 | -0.898 | -0.492 | 0.823 | -2.356 | -0.089 | 0.04 | 0.04 | 0 | 0.64 | 0.91 | 0.41 | 0.22 | 0.54 | 0.36 |
| 8 | 135107424 | rs16905072 | 0.154 | -0.438 | 1.102 | -2.131 | -0.01 | -0.03 | -0.03 | 0.393 | 0.06 | 0.11 | 0.02 | 0 | 0.03 | 0.02 |
| 8 | 135122155 | rs10106881 | -1.191 | -0.241 | 1.705 | -2.036 | 0.095 | 0.15 | 0.15 | 0 | 0.07 | 0.28 | 0.11 | 0.14 | 0.1 | 0.06 |
| 8 | 135122702 | rs1123762 | -2.056 | -0.27 | 1.811 | -2.097 | 0.064 | 0.096 | 0.096 | 0.017 | 0.11 | 0.39 | 0.14 | 0.14 | 0.14 | 0.07 |
| 8 | 135122889 | rs1124525 | -0.759 | -0.269 | 1.639 | -2.371 | -0.071 | 0.056 | 0.056 | 0 | 0.62 | 0.88 | 0.39 | 0.22 | 0.51 | 0.37 |
| 8 | 135129799 | rs2404087 | -1.042 | -0.219 | 1.638 | -2.286 | 0.095 | 0.15 | 0.15 | 0 | 0.07 | 0.27 | 0.11 | 0.14 | 0.1 | 0.06 |
| 8 | 135137056 | rs16905081 | 0.039 | -0.584 | 1.283 | -2.523 | 0.007 | -0.011 | -0.011 | 0 | 0.13 | 0.13 | 0.05 | 0 | 0.05 | 0.04 |
| 8 | 135137585 | rs16905082 | 0.389 | -0.588 | 1.225 | -2.522 | 0.027 | 0.026 | 0.026 | 0 | 0.12 | 0.07 | 0.04 | 0 | 0.04 | 0.03 |
| 8 | 135143521 | rs4897681 | -0.866 | -0.581 | 1.133 | -2.574 | -0.054 | 0.054 | 0.054 | 0 | 0.62 | 0.84 | 0.39 | 0.22 | 0.51 | 0.37 |
| 8 | 135143604 | rs6471191 | -1.753 | -0.513 | 1.391 | -2.459 | -0.011 | -0.016 | -0.016 | 0 | 0.26 | 0.68 | 0.19 | 0.14 | 0.2 | 0.11 |
| 8 | 135147391 | rs2404088 | -1.102 | -0.474 | 1.21 | -2.406 | -0.068 | -0.089 | -0.089 | 0 | 0.2 | 0.54 | 0.09 | 0.07 | 0.1 | 0.07 |
| 8 | 135150030 | rs6992223 | -1.511 | -0.445 | 0.766 | -2.404 | -0.001 | -0.001 | -0.001 | 0 | 0.25 | 0.73 | 0.19 | 0.21 | 0.19 | 0.12 |
| 8 | 135151600 | rs1840936 | -1.627 | -0.264 | 0.384 | -2.482 | -0.145 | -0.024 | -0.024 | 0.058 | 0.72 | 0.95 | 0.4 | 0.22 | 0.55 | 0.41 |
| 8 | 135154825 | rs1579879 | -0.917 | -0.188 | 0.802 | -2.373 | -0.017 | 0.037 | 0.037 | 0.015 | 0.2 | 0.44 | 0.16 | 0.22 | 0.15 | 0.11 |
| 8 | 135155736 | rs1596948 | -0.502 | 0.037 | 0.508 | -2.17 | -0.048 | 0.049 | 0.049 | 0 | 0.63 | 0.8 | 0.36 | 0.22 | 0.49 | 0.37 |
| 8 | 135157479 | rs11987768 | -0.178 | -0.079 | 0.809 | -2.279 | 0 | -0.01 | -0.01 | 0.693 | 0.55 | 0.59 | 0.33 | 0.18 | 0.41 | 0.29 |

| | | | | | | | | | | | | | | | | |
|---|---|---|---|---|---|---|---|---|---|---|---|---|---|---|---|---|
| 8 | 135162202 | rs895598 | 0.358 | -0.314 | 0.731 | -2.185 | 0.014 | 0.048 | 0.048 | 0 | 0.44 | 0.67 | 0.31 | 0.16 | 0.38 | 0.23 |
| 8 | 135164948 | rs10095649 | -0.622 | -0.267 | 0.676 | -2.187 | -0.013 | 0.002 | 0.002 | 0.192 | 0.2 | 0.48 | 0.14 | 0.18 | 0.11 | 0.13 |
| 8 | 135165772 | rs11783369 | -0.427 | -0.257 | 0.456 | -2.169 | -0.008 | 0.009 | 0.009 | 0.054 | 0.18 | 0.38 | 0.14 | 0.17 | 0.11 | 0.13 |
| 8 | 135171724 | rs17757570 | 0.599 | -0.187 | -0.71 | -2.103 | 0.016 | 0.007 | 0.007 | 0 | 0.39 | 0.34 | 0.23 | 0.03 | 0.35 | 0.2 |
| 8 | 135172796 | rs7002928 | -0.376 | 0.617 | -1.237 | -1.462 | -0.131 | 0.062 | 0.062 | 0 | 0.66 | 0.91 | 0.4 | 0.2 | 0.48 | 0.35 |
| 8 | 135174385 | rs1584260 | -0.812 | 0.626 | -0.465 | -1.249 | 0.014 | -0.024 | -0.024 | 0 | 0.26 | 0.5 | 0.28 | 0.27 | 0.23 | 0.25 |
| 8 | 135174629 | rs1584262 | -0.869 | 0.595 | -1.299 | -2.155 | -0.082 | 0.067 | 0.067 | 0 | 0.66 | 0.89 | 0.47 | 0.27 | 0.5 | 0.46 |
| 8 | 135192044 | rs4406434 | -0.381 | 0.022 | -0.904 | -2.515 | 0 | 0 | 0 | 0 | 0.25 | 0.3 | 0.14 | 0.26 | 0.16 | 0.19 |
| 8 | 135201068 | rs16905106 | -0.506 | -0.776 | -0.651 | -2.152 | -0.005 | -0.009 | -0.009 | 0 | 0.57 | 0.55 | 0.33 | 0.36 | 0.39 | 0.46 |
| 8 | 135205136 | rs10094837 | 0.801 | -0.894 | -0.737 | -1.546 | 0 | 0.001 | 0.001 | 0 | 0.17 | 0.24 | 0.21 | 0.32 | 0.12 | 0.2 |
| 8 | 135207115 | rs16905109 | 0.588 | -0.847 | -0.345 | -1.623 | -0.004 | -0.04 | -0.04 | 0.008 | 0.09 | 0.14 | 0.03 | 0 | 0.05 | 0.06 |
| 8 | 135208887 | rs16905111 | 0.099 | -0.853 | -0.761 | -1.606 | 0.006 | -0.018 | -0.018 | 0.349 | 0.23 | 0.24 | 0.23 | 0.32 | 0.17 | 0.23 |
| 8 | 135214632 | rs16905118 | 1.956 | -0.703 | -0.886 | -1.749 | 0.007 | -0.009 | -0.009 | 0 | 0.08 | 0.04 | 0.04 | 0.1 | 0.02 | 0.11 |
| 8 | 135215172 | rs17686445 | 1.869 | -0.533 | -0.792 | -1.907 | 0.001 | 0.04 | 0.04 | 0 | 0.24 | 0.13 | 0.14 | 0 | 0.21 | 0.14 |
| 8 | 135217316 | rs13266721 | 0.792 | -0.214 | -0.79 | -1.623 | 0.007 | -0.026 | -0.026 | 0 | 0.07 | 0.07 | 0.14 | 0 | 0.09 | 0.07 |
| 8 | 135218576 | rs4400414 | -1.815 | -0.038 | -1.093 | -0.738 | -0.136 | -0.048 | -0.048 | 0.593 | 0.79 | 0.98 | 0.69 | 0.79 | 0.58 | 0.69 |
| 8 | 135226098 | rs6471199 | 0.502 | -1.02 | -0.771 | -1.017 | 0.009 | 0.021 | 0.021 | 0 | 0.14 | 0.09 | 0.05 | 0 | 0.1 | 0.06 |
| 8 | 135229994 | rs16905126 | -0.653 | -1.17 | -0.333 | -0.819 | 0.028 | 0 | 0 | 0 | 0.42 | 0.63 | 0.3 | 0.41 | 0.22 | 0.34 |
| 8 | 135236877 | rs6471201 | -1.445 | -0.699 | -0.888 | -0.786 | -0.193 | -0.173 | -0.173 | 0 | 0.73 | 0.98 | 0.67 | 0.48 | 0.49 | 0.56 |
| 8 | 135241108 | rs7008781 | 1.064 | -0.8 | -0.072 | -1.017 | -0.007 | -0.093 | -0.093 | 0 | 0.28 | 0.32 | 0.17 | 0.01 | 0.22 | 0.15 |
| 8 | 135245260 | rs7012883 | 1.133 | -0.45 | -0.634 | -0.878 | -0.008 | 0.03 | 0.03 | 0 | 0.08 | 0.02 | 0.04 | 0 | 0.06 | 0.07 |
| 8 | 135249414 | rs9650120 | -0.684 | -0.469 | -0.432 | -0.621 | -0.009 | -0.009 | -0.009 | 0.325 | 0.67 | 0.67 | 0.51 | 0.53 | 0.48 | 0.47 |
| 8 | 135249548 | rs6996817 | -0.941 | -0.431 | -0.887 | -0.556 | -0.173 | -0.157 | -0.157 | 0.823 | 0.69 | 0.98 | 0.53 | 0.57 | 0.45 | 0.46 |
| 8 | 135250072 | rs4628283 | 0.303 | -0.82 | -0.885 | -0.546 | 0.052 | -0.032 | -0.032 | 0 | 0.3 | 0.15 | 0.22 | 0.34 | 0.19 | 0.22 |
| 8 | 135252366 | rs2028939 | -0.922 | -0.493 | -0.902 | -0.633 | -0.142 | -0.112 | -0.112 | 0 | 0.69 | 0.91 | 0.54 | 0.57 | 0.48 | 0.47 |
| 8 | 135262393 | rs11990826 | 0.066 | -0.726 | -0.638 | -0.952 | 0.011 | -0.023 | -0.023 | 0 | 0.11 | 0.36 | 0.05 | 0.04 | 0.05 | 0.08 |
| 8 | 135262880 | rs4897704 | -0.834 | -0.404 | -1.599 | -0.663 | -0.141 | -0.139 | -0.139 | 0 | 0.69 | 0.91 | 0.5 | 0.53 | 0.47 | 0.47 |
| 8 | 135269674 | rs4581088 | -0.166 | -0.837 | -0.974 | -0.297 | 0.017 | -0.015 | -0.015 | 0 | 0.28 | 0.26 | 0.23 | 0.38 | 0.18 | 0.23 |
| 8 | 135271200 | rs1596953 | -0.323 | 0.112 | -0.916 | -0.795 | -0.001 | -0.009 | -0.009 | 0 | 0.27 | 0.63 | 0.28 | 0.17 | 0.29 | 0.16 |
| 8 | 135278536 | rs4500107 | 0.216 | -0.045 | -1.094 | -1.249 | 0.003 | 0.042 | 0.042 | 0 | 0.46 | 0.37 | 0.62 | 0.84 | 0.53 | 0.66 |
| 8 | 135283869 | rs4298516 | -0.771 | -0.187 | -1.122 | -1.409 | -0.003 | -0.058 | -0.058 | 0 | 0.71 | 0.7 | 0.88 | 0.99 | 0.85 | 0.86 |
| 8 | 135288110 | rs10109387 | 1.25 | 0.327 | -1.479 | -1.205 | 0.003 | 0.012 | 0.012 | 0 | 0.26 | 0.16 | 0.26 | 0.22 | 0.22 | 0.18 |
| 8 | 135288668 | rs895594 | 1.549 | 0.577 | -1.943 | -1.136 | -0.09 | -0.101 | -0.101 | 0 | 0.13 | 0.01 | 0.34 | 0.62 | 0.29 | 0.46 |
| 8 | 135290290 | rs13256457 | -0.941 | -0.348 | -1.575 | -1.093 | -0.01 | -0.064 | -0.064 | 0 | 0.74 | 0.7 | 0.89 | 0.99 | 0.87 | 0.86 |
| 8 | 135294755 | rs4401895 | 0.521 | 0.198 | -1.493 | -0.265 | 0.005 | -0.007 | -0.007 | 0.706 | 0.12 | 0.09 | 0.08 | 0.14 | 0.12 | 0.05 |
| 8 | 135303085 | rs10098827 | -0.281 | 0.486 | -1.437 | -0.261 | -0.017 | -0.007 | -0.007 | 0.053 | 0.4 | 0.48 | 0.36 | 0.31 | 0.34 | 0.23 |
| 8 | 135307548 | rs1596959 | -0.239 | 1.359 | -2.256 | -0.043 | -0.144 | -0.075 | -0.075 | 0.208 | 0.27 | 0.03 | 0.44 | 0.28 | 0.42 | 0.53 |
| 8 | 135308849 | rs4308771 | -0.23 | 0.924 | -2.485 | -0.855 | -0.009 | 0.065 | 0.065 | 0 | 0.76 | 0.9 | 0.85 | 1 | 0.75 | 0.86 |
| 8 | 135313764 | rs1867949 | -0.536 | 1.385 | -2.345 | 0 | -0.078 | -0.067 | -0.067 | 1.489 | 0.38 | 0.11 | 0.54 | 0.44 | 0.51 | 0.63 |
| 8 | 135319528 | rs11787445 | 0.277 | 1.744 | -2.37 | -0.123 | -0.056 | 0.013 | 0.013 | 0 | 0.16 | 0.03 | 0.16 | 0.14 | 0.23 | 0.1 |
| 8 | 135326486 | rs12679451 | -0.911 | 1.704 | -1.476 | -0.075 | -0.001 | 0 | 0 | 0 | 0.44 | 0.4 | 0.58 | 0.43 | 0.6 | 0.65 |
| 8 | 135328582 | rs10100237 | -0.118 | 1.929 | -1.346 | 0.504 | -0.063 | -0.05 | -0.05 | 0 | 0.23 | 0.1 | 0.38 | 0.29 | 0.36 | 0.53 |
| 8 | 135336557 | rs4584173 | -0.211 | 1.638 | -1.105 | 0.339 | 0.004 | 0 | 0 | 0 | 0.28 | 0.3 | 0.55 | 0.65 | 0.4 | 0.57 |
| 8 | 135337294 | rs12541052 | -0.202 | 1.215 | -0.899 | 0.13 | -0.001 | 0.011 | 0.011 | 3.225 | 0.78 | 0.83 | 0.77 | 0.85 | 0.79 | 0.88 |
| 8 | 135339383 | rs6471216 | -0.235 | 1.009 | -0.156 | 0.125 | -0.023 | -0.019 | -0.019 | 0 | 0.29 | 0.61 | 0.16 | 0.15 | 0.18 | 0.18 |
| 8 | 135343419 | rs4412399 | -0.427 | 1.075 | -0.201 | 0.264 | 0.042 | -0.019 | -0.019 | 0 | 0.31 | 0.86 | 0.24 | 0.23 | 0.2 | 0.19 |
| 8 | 135346972 | rs7831261 | -0.572 | 1.072 | -0.26 | 0.212 | 0.038 | -0.023 | -0.023 | 0.268 | 0.33 | 0.89 | 0.25 | 0.23 | 0.21 | 0.19 |
| 8 | 135348578 | rs7463966 | -0.379 | 1.057 | -0.077 | 0.207 | 0.01 | 0 | 0 | 0 | 0.31 | 0.71 | 0.22 | 0.23 | 0.19 | 0.2 |
| 8 | 135353633 | rs11997394 | -1.973 | 0.997 | 0.601 | 0.171 | 0.057 | 0.037 | 0.037 | 2.994 | 0.07 | 0.56 | 0.07 | 0.02 | 0.03 | 0.07 |
| 8 | 135357247 | rs13273294 | 1.006 | 1.495 | 0.059 | 0.306 | -0.072 | -0.079 | -0.079 | 0 | 0.51 | 0.06 | 0.53 | 0.6 | 0.52 | 0.61 |
| 8 | 135367334 | rs4554499 | -0.463 | 1.533 | 0.21 | 0.324 | -0.027 | -0.012 | -0.012 | 0 | 0.85 | 0.91 | 0.76 | 0.81 | 0.74 | 0.82 |
| 8 | 135372035 | rs6471220 | 0.393 | 1.497 | 0.33 | 0.313 | -0.022 | 0.002 | 0.002 | 0 | 0.27 | 0.34 | 0.17 | 0.2 | 0.18 | 0.14 |
| 8 | 135372184 | rs6471221 | -0.143 | 1.495 | 0.356 | 0.309 | 0.006 | 0 | 0 | 0 | 0.83 | 0.77 | 0.75 | 0.81 | 0.74 | 0.82 |
| 8 | 135372622 | rs4486240 | 1.069 | 1.494 | 0.275 | 0.307 | -0.068 | -0.076 | -0.076 | 0 | 0.52 | 0.07 | 0.53 | 0.6 | 0.52 | 0.62 |
| 8 | 135373474 | rs7812511 | 0.547 | 1.615 | 0.452 | 0.338 | 0.042 | 0.003 | 0.003 | 0.221 | 0.79 | 0.54 | 0.71 | 0.79 | 0.7 | 0.76 |
| 8 | 135385238 | rs10956757 | 0.413 | 1.735 | 0.289 | 0.161 | 0 | -0.001 | -0.001 | 0 | 0.57 | 0.54 | 0.59 | 0.68 | 0.54 | 0.66 |
| 8 | 135389370 | rs7011517 | 0.67 | 1.445 | -0.119 | 0.051 | -0.001 | 0.008 | 0.008 | 0 | 0.21 | 0.26 | 0.16 | 0.2 | 0.18 | 0.16 |
| 8 | 135389808 | rs4897638 | -0.906 | 1.442 | -1.095 | 0.093 | 0.002 | 0.021 | 0.021 | 0 | 0.46 | 0.66 | 0.43 | 0.32 | 0.47 | 0.34 |
| 8 | 135394067 | rs6415556 | -1.172 | 1.474 | -1.109 | 0.073 | 0.003 | 0.01 | 0.01 | 0.055 | 0.51 | 0.92 | 0.54 | 0.35 | 0.59 | 0.43 |
| 8 | 135394086 | rs6471223 | -1.172 | 1.474 | -1.109 | 0.073 | 0.003 | 0.01 | 0.01 | 0.055 | 0.51 | 0.92 | 0.54 | 0.35 | 0.59 | 0.43 |
| 8 | 135397468 | rs4897719 | -1.06 | 1.561 | -1.102 | 0.144 | 0.019 | 0.031 | 0.031 | 0 | 0.48 | 0.67 | 0.51 | 0.32 | 0.59 | 0.38 |
| 8 | 135398029 | rs3923951 | -0.567 | 1.02 | -0.99 | 0.051 | -0.03 | 0.046 | 0.046 | 0 | 0.21 | 0.06 | 0.23 | 0.12 | 0.25 | 0.17 |
| 8 | 135409616 | rs10808622 | -0.016 | 1.15 | -0.926 | 0.31 | -0.024 | 0.006 | 0.006 | 0 | 0.77 | 0.89 | 0.83 | 0.78 | 0.76 | 0.77 |
| 8 | 135412090 | rs3934606 | 0.158 | 1.261 | -1.277 | 0.676 | -0.009 | -0.021 | -0.021 | 0 | 0.14 | 0.11 | 0.24 | 0.19 | 0.18 | 0.07 |
| 8 | 135413853 | rs6471235 | 0.054 | 0.814 | -1.277 | 0.384 | -0.004 | -0.005 | -0.005 | 0.949 | 0.37 | 0.31 | 0.43 | 0.57 | 0.42 | 0.44 |
| 8 | 135416927 | rs11785168 | 1.076 | 0.457 | -1.335 | -0.172 | 0.005 | 0.025 | 0.025 | 0 | 0.18 | 0.11 | 0.12 | 0.04 | 0.14 | 0.17 |
| 8 | 135419683 | rs7011450 | 1.35 | 0.351 | -1.361 | -0.192 | -0.001 | 0.004 | 0.004 | 0 | 0.25 | 0.26 | 0.2 | 0.38 | 0.24 | 0.37 |
| 8 | 135420488 | rs4897723 | -0.68 | -0.016 | -1.206 | -0.42 | 0.025 | 0.005 | 0.005 | 0 | 0.65 | 0.5 | 0.58 | 0.62 | 0.57 | 0.64 |
| 8 | 135422805 | rs9785079 | 1.591 | 0.05 | -1.217 | -0.349 | 0.001 | -0.012 | -0.012 | 0.023 | 0.23 | 0.2 | 0.2 | 0.38 | 0.24 | 0.37 |

| | | | | | | | | | | | | | | | | |
|---|---|---|---|---|---|---|---|---|---|---|---|---|---|---|---|---|
| 8 | 135422909 | rs7003297 | 0.324 | 0.074 | -1.208 | -0.342 | 0.011 | 0.006 | 0.006 | 0 | 0.43 | 0.35 | 0.32 | 0.43 | 0.38 | 0.54 |
| 8 | 135427284 | rs10448002 | 0.781 | 0.321 | -1.248 | -0.386 | 0.004 | 0.057 | 0.057 | 0 | 0.18 | 0.09 | 0.12 | 0.05 | 0.14 | 0.17 |
| 8 | 135432541 | rs11775881 | 0.479 | 0.37 | -0.473 | -0.272 | 0.007 | -0.024 | -0.024 | 0 | 0.24 | 0.26 | 0.18 | 0.15 | 0.25 | 0.25 |
| 8 | 135433894 | rs7844624 | -0.442 | 0.247 | -0.402 | -0.25 | 0.008 | -0.011 | -0.011 | 0 | 0.39 | 0.48 | 0.37 | 0.31 | 0.43 | 0.32 |
| 8 | 135437151 | rs7012497 | 0.686 | 0.371 | -0.482 | -0.196 | -0.004 | -0.011 | -0.011 | 0 | 0.16 | 0.15 | 0.17 | 0.16 | 0.18 | 0.07 |
| 8 | 135441161 | rs10956760 | -1.41 | 0.178 | -0.974 | -0.254 | -0.038 | 0.006 | 0.006 | 0 | 0.79 | 0.91 | 0.69 | 0.78 | 0.74 | 0.79 |
| 8 | 135442374 | rs7840306 | 1.753 | -0.133 | -0.898 | -0.346 | 0.004 | -0.054 | -0.054 | 0 | 0.24 | 0.12 | 0.28 | 0.41 | 0.25 | 0.39 |
| 8 | 135444101 | rs11775304 | -0.118 | -0.121 | -0.79 | -0.39 | -0.017 | -0.017 | -0.017 | 0 | 0.61 | 0.71 | 0.5 | 0.56 | 0.53 | 0.7 |
| 8 | 135445253 | rs13273819 | 1.396 | 0.07 | -0.669 | -0.114 | 0.006 | -0.024 | -0.024 | 0 | 0.25 | 0.18 | 0.29 | 0.41 | 0.25 | 0.4 |
| 8 | 135456499 | rs12155608 | 0.432 | 0.501 | -0.465 | -0.075 | -0.008 | 0.015 | 0.015 | 0 | 0.66 | 0.77 | 0.64 | 0.72 | 0.67 | 0.72 |
| 8 | 135460497 | rs6415559 | -0.359 | 0.556 | -0.43 | -0.014 | 0.004 | 0 | 0 | 0 | 0.74 | 0.76 | 0.66 | 0.72 | 0.7 | 0.76 |
| 8 | 135468912 | rs7816086 | -0.22 | 0.673 | -0.761 | -0.12 | -0.071 | 0.046 | 0.046 | 0 | 0.73 | 0.97 | 0.68 | 0.81 | 0.71 | 0.72 |
| 8 | 135483883 | rs6994619 | -0.905 | 0.838 | 0.07 | 0.287 | 0.053 | 0.081 | 0.081 | 0 | 0.1 | 0.3 | 0.09 | 0.16 | 0.12 | 0.06 |
| 8 | 135487835 | rs11774192 | 0.697 | 1.371 | -0.497 | 0.104 | -0.024 | 0.007 | 0.007 | 0 | 0.13 | 0.05 | 0.25 | 0.1 | 0.22 | 0.16 |
| 8 | 135489339 | rs4243853 | 0.545 | 1.332 | -0.772 | 0.748 | -0.019 | -0.003 | -0.003 | 0.376 | 0.07 | 0.02 | 0.11 | 0.29 | 0.1 | 0.15 |
| 8 | 135490539 | rs11538239 | -1.473 | 0.727 | -0.985 | 0.16 | 0.012 | -0.022 | -0.022 | 0.587 | 0.74 | 0.83 | 0.7 | 0.57 | 0.76 | 0.61 |
| 8 | 135491990 | rs12544587 | 0.804 | 0.912 | -0.639 | 0.148 | 0.001 | -0.001 | -0.001 | 0 | 0.16 | 0.13 | 0.31 | 0.24 | 0.28 | 0.19 |
| 8 | 135494243 | rs5025124 | 0.456 | 0.572 | -0.826 | -0.428 | -0.001 | -0.011 | -0.011 | 4.254 | 0.28 | 0.23 | 0.22 | 0.29 | 0.3 | 0.27 |
| 8 | 135496060 | rs10103826 | 0.852 | 0.17 | -0.914 | -0.579 | -0.008 | 0.059 | 0.059 | 0 | 0.2 | 0.08 | 0.12 | 0 | 0.15 | 0.12 |
| 8 | 135497576 | rs7828867 | -0.06 | 0.104 | -0.497 | -0.596 | 0.001 | -0.013 | -0.013 | 0 | 0.44 | 0.51 | 0.45 | 0.28 | 0.52 | 0.33 |
| 8 | 135497615 | rs11786204 | -0.063 | 0.195 | -0.517 | -0.408 | -0.005 | -0.023 | -0.023 | 0 | 0.28 | 0.18 | 0.23 | 0.29 | 0.24 | 0.27 |
| 8 | 135499383 | rs4391381 | -0.264 | -0.006 | -0.045 | -0.704 | -0.013 | -0.043 | -0.043 | 0.11 | 0.16 | 0.33 | 0.11 | 0.08 | 0.14 | 0.18 |
| 8 | 135499696 | rs10113404 | 0.536 | -0.039 | -0.219 | -1.019 | -0.003 | 0.032 | 0.032 | 0 | 0.24 | 0.12 | 0.27 | 0 | 0.19 | 0.16 |
| 8 | 135500509 | rs4909615 | 0.6 | -0.04 | -0.194 | -1.02 | -0.003 | 0.018 | 0.018 | 0 | 0.23 | 0.14 | 0.27 | 0 | 0.19 | 0.16 |
| 8 | 135500567 | rs4909616 | 0.685 | -0.044 | -0.239 | -1.226 | -0.003 | 0.021 | 0.021 | 0 | 0.24 | 0.19 | 0.16 | 0.05 | 0.26 | 0.25 |
| 8 | 135500832 | rs7812708 | -2.453 | -0.017 | -0.423 | -1.069 | -0.001 | -0.134 | -0.134 | 1.402 | 0.75 | 0.97 | 0.71 | 0.5 | 0.75 | 0.59 |
| 8 | 135503898 | rs7010252 | 0.848 | 0.049 | -0.049 | -0.977 | -0.02 | -0.018 | -0.018 | 0 | 0.08 | 0.25 | 0.04 | 0.03 | 0.05 | 0.02 |
| 8 | 135506043 | rs4424215 | -0.396 | 0.146 | -0.061 | -0.933 | -0.001 | 0 | 0 | 0 | 0.37 | 0.62 | 0.4 | 0.26 | 0.3 | 0.22 |
| 8 | 135506073 | rs4076195 | -0.716 | 0.039 | 0.007 | -0.911 | 0.001 | -0.004 | -0.004 | 0 | 0.47 | 0.59 | 0.52 | 0.45 | 0.48 | 0.34 |
| 8 | 135506830 | rs11776162 | -0.016 | -0.456 | -0.176 | -1.434 | -0.01 | -0.071 | -0.071 | 0 | 0.59 | 0.52 | 0.63 | 0.92 | 0.68 | 0.65 |
| 8 | 135508883 | rs894351 | 1.869 | -0.215 | -0.325 | -1.43 | -0.042 | 0.07 | 0.07 | 0.363 | 0.15 | 0.03 | 0.13 | 0.04 | 0.19 | 0.15 |
| 8 | 135512875 | rs754234 | 1.883 | -0.22 | -0.39 | -1.081 | -0.029 | -0.074 | -0.074 | 0.197 | 0.19 | 0.05 | 0.27 | 0.42 | 0.28 | 0.38 |
| 8 | 135516280 | rs4366044 | 0.322 | -0.514 | -0.071 | -1.377 | -0.033 | 0.23 | 0.23 | 0 | 0.13 | 0.39 | 0.1 | 0.26 | 0.06 | 0.06 |
| 8 | 135517527 | rs2083683 | -1.352 | -0.574 | -0.134 | -1.51 | 0.004 | 0.044 | 0.044 | 2.322 | 0.74 | 0.84 | 0.83 | 0.96 | 0.76 | 0.79 |
| 8 | 135517955 | rs4909834 | 0.025 | -0.572 | -0.117 | -1.505 | -0.009 | -0.001 | -0.001 | 0 | 0.6 | 0.45 | 0.73 | 0.7 | 0.7 | 0.72 |
| 8 | 135519914 | rs7001459 | 0.26 | -0.416 | 0.032 | -1.393 | -0.034 | 0.218 | 0.218 | 0 | 0.13 | 0.39 | 0.1 | 0.26 | 0.06 | 0.06 |
| 8 | 135522921 | rs6578007 | 1.936 | -0.407 | -0.062 | -1.452 | -0.035 | 0.07 | 0.07 | 0 | 0.13 | 0.02 | 0.23 | 0 | 0.15 | 0.12 |
| 8 | 135522989 | rs6578009 | -1.4 | -0.442 | -0.088 | -1.493 | -0.028 | -0.062 | -0.062 | 0 | 0.67 | 0.93 | 0.63 | 0.3 | 0.64 | 0.47 |
| 8 | 135523766 | rs17688465 | 1.735 | -0.464 | 0.041 | -1.512 | -0.012 | 0.026 | 0.026 | 0 | 0.22 | 0.16 | 0.13 | 0 | 0.23 | 0.13 |
| 8 | 135523829 | rs6988595 | 1.127 | -0.422 | 0.342 | -1.331 | -0.027 | -0.019 | -0.019 | 0.193 | 0.1 | 0.19 | 0.03 | 0.04 | 0.05 | 0.03 |
| 8 | 135524797 | rs3739429 | 0.346 | -0.405 | 0.2 | -1.313 | -0.034 | 0.214 | 0.214 | 0 | 0.13 | 0.39 | 0.1 | 0.26 | 0.06 | 0.06 |
| 8 | 135526317 | rs4909858 | -0.931 | -0.39 | 0.078 | -1.349 | 0.01 | 0.021 | 0.021 | 0 | 0.45 | 0.76 | 0.5 | 0.3 | 0.41 | 0.34 |
| 8 | 135528095 | rs6578041 | -0.783 | -0.367 | 0.045 | -1.277 | 0.005 | 0.005 | 0.005 | 0 | 0.44 | 0.64 | 0.5 | 0.34 | 0.41 | 0.34 |
| 8 | 135534990 | rs4909874 | 0.725 | -0.334 | 0.026 | -1.287 | -0.022 | 0.052 | 0.052 | 0 | 0.29 | 0.17 | 0.38 | 0.04 | 0.34 | 0.28 |
| 8 | 135536234 | rs6992913 | 0.167 | -0.237 | 0.175 | -1.15 | -0.037 | 0.244 | 0.244 | 0 | 0.14 | 0.43 | 0.11 | 0.26 | 0.06 | 0.06 |
| 8 | 135537773 | rs11994231 | 0.156 | -0.15 | 0.013 | -1.118 | -0.01 | 0.017 | 0.017 | 0.039 | 0.3 | 0.59 | 0.26 | 0.3 | 0.26 | 0.22 |
| 8 | 135540632 | rs16905165 | 0.999 | -0.171 | -0.05 | -1.137 | 0 | -0.015 | -0.015 | 0 | 0.18 | 0.11 | 0.31 | 0.26 | 0.17 | 0.17 |
| 8 | 135540786 | rs6578149 | 0.233 | -0.346 | -0.053 | -1.118 | -0.009 | 0.013 | 0.013 | 0 | 0.43 | 0.33 | 0.43 | 0.43 | 0.52 | 0.56 |
| 8 | 135541831 | rs7011356 | 0.217 | -0.423 | -0.091 | -1.291 | 0.019 | 0.028 | 0.028 | 2.759 | 0.55 | 0.36 | 0.64 | 0.36 | 0.66 | 0.61 |
| 8 | 135542084 | rs1550582 | 1.664 | -0.328 | -0.294 | -1.122 | -0.009 | -0.019 | -0.019 | 0.712 | 0.19 | 0.11 | 0.28 | 0.36 | 0.28 | 0.36 |
| 8 | 135543194 | rs1020236 | 0.683 | -0.346 | -0.354 | -1.177 | -0.003 | -0.033 | -0.033 | 0.856 | 0.25 | 0.27 | 0.17 | 0.07 | 0.25 | 0.2 |
| 8 | 135549042 | rs2119879 | 0.476 | -0.136 | -0.289 | -0.887 | -0.011 | 0.012 | 0.012 | 0 | 0.35 | 0.55 | 0.42 | 0.67 | 0.34 | 0.46 |
| 8 | 135550073 | rs16905167 | 0.255 | -0.072 | -0.622 | -1.029 | -0.007 | 0.015 | 0.015 | 0.18 | 0.35 | 0.24 | 0.56 | 0.35 | 0.42 | 0.44 |
| 8 | 135552427 | rs2315833 | -2.268 | 0.075 | -0.451 | -1.343 | -0.043 | 0.075 | 0.075 | 0.154 | 0.86 | 0.98 | 0.88 | 0.97 | 0.81 | 0.84 |
| 8 | 135553329 | rs9324581 | -0.164 | 0.098 | 0.065 | -0.876 | 0.006 | 0.033 | 0.033 | 0 | 0.49 | 0.25 | 0.65 | 0.67 | 0.51 | 0.6 |
| 8 | 135558301 | rs1530318 | 0.558 | 0.295 | -0.258 | -0.671 | 0.001 | -0.041 | -0.041 | 0 | 0.2 | 0.1 | 0.33 | 0.23 | 0.18 | 0.17 |
| 8 | 135564627 | rs4909909 | 0.447 | 0.402 | -0.133 | -0.653 | -0.006 | 0.011 | 0.011 | 0 | 0.34 | 0.37 | 0.4 | 0.67 | 0.34 | 0.46 |
| 8 | 135565255 | rs4909911 | 0.447 | 0.402 | -0.133 | -0.653 | -0.006 | 0.011 | 0.011 | 0 | 0.34 | 0.37 | 0.4 | 0.67 | 0.34 | 0.46 |
| 8 | 135565420 | rs6578231 | -0.555 | 0.405 | -0.12 | -0.769 | -0.002 | 0.001 | 0.001 | 0 | 0.54 | 0.55 | 0.69 | 0.67 | 0.52 | 0.6 |
| 8 | 135565904 | rs7008121 | 0.254 | 0.513 | -0.23 | -0.307 | -0.003 | 0.005 | 0.005 | 0 | 0.33 | 0.34 | 0.39 | 0.67 | 0.34 | 0.46 |
| 8 | 135571916 | rs6986489 | -0.357 | 0.628 | -0.477 | -1.117 | 0.003 | 0.096 | 0.096 | 0 | 0.55 | 0.66 | 0.55 | 0.91 | 0.55 | 0.63 |
| 8 | 135572715 | rs4909483 | -0.627 | 0.477 | -0.425 | -1.215 | -0.002 | -0.001 | -0.001 | 0 | 0.58 | 0.56 | 0.73 | 0.59 | 0.65 | 0.62 |
| 8 | 135573657 | rs6993032 | 0.239 | 0.513 | -0.373 | -1.153 | 0.006 | 0.002 | 0.002 | 0 | 0.39 | 0.52 | 0.44 | 0.59 | 0.46 | 0.48 |
| 8 | 135574196 | rs721000 | -1.294 | 0.496 | -0.305 | -1.094 | 0 | -0.005 | -0.005 | 2.392 | 0.74 | 0.74 | 0.84 | 0.91 | 0.75 | 0.77 |
| 8 | 135578218 | rs9693237 | 1.23 | 0.59 | 0.206 | -1 | 0 | 0.019 | 0.019 | 1.379 | 0.17 | 0.21 | 0.33 | 0.54 | 0.25 | 0.36 |
| 8 | 135584316 | rs6998749 | 1.339 | 0.593 | 0.145 | -0.998 | -0.009 | -0.036 | -0.036 | 0.051 | 0.17 | 0.12 | 0.31 | 0.54 | 0.25 | 0.36 |
| 8 | 135586279 | rs7003440 | 1.009 | 0.534 | 0.202 | -1.21 | -0.006 | -0.004 | -0.004 | 0 | 0.21 | 0.19 | 0.11 | 0.02 | 0.21 | 0.11 |
| 8 | 135589494 | rs16905185 | 2.031 | 0.528 | 0.024 | -1.047 | -0.006 | -0.072 | -0.072 | 0 | 0.14 | 0.02 | 0.1 | 0.34 | 0.09 | 0.15 |
| 8 | 135593713 | rs7841159 | -0.205 | 0.496 | 0.461 | -1.097 | 0.002 | 0.004 | 0.004 | 7.072 | 0.38 | 0.53 | 0.62 | 0.54 | 0.44 | 0.5 |

| | | | | | | | | | | | | | | | | |
|---|---|---|---|---|---|---|---|---|---|---|---|---|---|---|---|---|
| 8 | 135598675 | rs11988811 | 0.998 | 0.62 | 0.519 | -0.9 | 0.005 | 0.104 | 0.104 | 1.963 | 0.16 | 0.3 | 0.32 | 0.54 | 0.2 | 0.35 |
| 8 | 135607485 | rs7829476 | 1.266 | 0.651 | 0.379 | -1.236 | -0.03 | 0.043 | 0.043 | 0 | 0.16 | 0.11 | 0.09 | 0.02 | 0.2 | 0.11 |
| 8 | 135607823 | rs2119880 | 1.393 | 0.771 | 0.386 | -1.174 | 0.002 | 0.044 | 0.044 | 2.682 | 0.15 | 0.19 | 0.32 | 0.54 | 0.22 | 0.36 |
| 8 | 135610120 | rs16905193 | 1.107 | 0.76 | 0.462 | -1.19 | 0.006 | 0.106 | 0.106 | 2.509 | 0.16 | 0.31 | 0.33 | 0.54 | 0.22 | 0.36 |
| 8 | 135611117 | rs6982146 | 1.415 | 0.77 | 0.365 | -1.185 | 0.002 | 0.044 | 0.044 | 2.557 | 0.14 | 0.19 | 0.32 | 0.54 | 0.22 | 0.36 |
| 8 | 135611154 | rs12549256 | 1.717 | 0.592 | 0.374 | -1.213 | 0.012 | -0.047 | -0.047 | 0.054 | 0.17 | 0.06 | 0.1 | 0.34 | 0.1 | 0.15 |
| 8 | 135612595 | rs894343 | -1.338 | 0.709 | 0.451 | -1.426 | 0 | 0.008 | 0.008 | 0 | 0.7 | 0.76 | 0.81 | 0.91 | 0.69 | 0.76 |
| 8 | 135612745 | rs894344 | 0.165 | 0.782 | 0.474 | -1.392 | 0.009 | 0.01 | 0.01 | 2.743 | 0.35 | 0.42 | 0.62 | 0.55 | 0.43 | 0.5 |
| 8 | 135614972 | rs3739422 | 0.951 | 0.813 | 0.323 | -1.482 | -0.001 | 0.005 | 0.005 | 2.578 | 0.26 | 0.29 | 0.39 | 0.55 | 0.28 | 0.46 |
| 8 | 135617960 | rs7821828 | 2.053 | 0.853 | 0.202 | -1.466 | 0.021 | -0.078 | -0.078 | 0 | 0.16 | 0.02 | 0.09 | 0.33 | 0.1 | 0.16 |
| 8 | 135621369 | rs11992096 | 0.531 | 0.91 | 0.278 | -1.496 | -0.051 | 0.065 | 0.065 | 0 | 0.25 | 0.17 | 0.32 | 0.02 | 0.34 | 0.17 |
| 8 | 135624103 | rs17770304 | 1.459 | 0.914 | 0.258 | -1.472 | -0.051 | 0.07 | 0.07 | 0 | 0.16 | 0.08 | 0.09 | 0.02 | 0.2 | 0.11 |
| 8 | 135631181 | rs17770844 | 1.971 | 1.098 | 0.179 | -1.516 | -0.006 | 0 | 0 | 0 | 0.06 | 0.07 | 0.03 | 0 | 0.08 | 0.05 |
| 8 | 135635174 | rs11990335 | 1.273 | 0.93 | 0.46 | -1.495 | 0.016 | -0.048 | -0.048 | 0 | 0.09 | 0.16 | 0.24 | 0 | 0.13 | 0.06 |
| 8 | 135635481 | rs1077224 | 0.427 | 0.889 | 0.253 | -1.49 | -0.031 | -0.013 | -0.013 | 0 | 0.34 | 0.14 | 0.18 | 0.35 | 0.3 | 0.27 |
| 8 | 135639045 | rs966659 | -1.436 | 1.261 | -0.056 | -1.717 | 0.001 | 0.007 | 0.007 | 0 | 0.71 | 0.77 | 0.82 | 0.9 | 0.72 | 0.77 |
| 8 | 135644369 | rs4909330 | -1.327 | 1.294 | -0.266 | -1.701 | -0.008 | -0.07 | -0.07 | 1.178 | 0.7 | 0.64 | 0.81 | 0.9 | 0.72 | 0.77 |
| 8 | 135648040 | rs2028556 | -0.314 | 1.251 | -1.012 | -1.304 | -0.002 | 0.194 | 0.194 | 0 | 0.49 | 0.25 | 0.7 | 0.88 | 0.5 | 0.66 |
| 8 | 135648397 | rs2028557 | 0.068 | 0.94 | -1.005 | -1.132 | 0.022 | 0.153 | 0.153 | 0 | 0.42 | 0.21 | 0.47 | 0.88 | 0.38 | 0.6 |
| 8 | 135649848 | rs12541381 | 0.037 | 1.225 | -1.102 | 0.009 | -0.038 | -0.065 | -0.065 | 0.006 | 0.19 | 0.06 | 0.22 | 0.31 | 0.26 | 0.28 |
| 8 | 135650071 | rs7831658 | 0.113 | 0.886 | -0.172 | -0.848 | 0.015 | -0.066 | -0.066 | 0 | 0.11 | 0.32 | 0.18 | 0 | 0.11 | 0.04 |
| 8 | 135654340 | rs11166643 | -0.312 | 0.613 | -0.483 | -1.012 | 0.021 | 0.066 | 0.066 | 0 | 0.5 | 0.4 | 0.59 | 0.83 | 0.43 | 0.65 |
| 8 | 135655107 | rs17699449 | 0.988 | 0.9 | -0.885 | -1.138 | -0.039 | 0.046 | 0.046 | 0 | 0.08 | 0 | 0.06 | 0 | 0.11 | 0.06 |
| 8 | 135661484 | rs1583969 | -1.463 | 1.023 | -0.935 | -1.429 | -0.019 | 0.029 | 0.029 | 0 | 0.74 | 0.87 | 0.75 | 0.83 | 0.67 | 0.77 |
| 8 | 135664673 | rs7000796 | -0.518 | 0.585 | -0.619 | -1.21 | 0.003 | 0.004 | 0.004 | 0 | 0.42 | 0.47 | 0.51 | 0.51 | 0.43 | 0.59 |
| 8 | 135669222 | rs16905212 | 0.344 | 0.601 | -0.264 | -1.16 | -0.01 | -0.055 | -0.055 | 0 | 0.3 | 0.16 | 0.43 | 0.51 | 0.31 | 0.53 |
| 8 | 135669810 | rs17778003 | 1.129 | 0.661 | -0.096 | -1.053 | 0.017 | -0.043 | -0.043 | 0.952 | 0.08 | 0.15 | 0.06 | 0 | 0.09 | 0.06 |
| 8 | 135670229 | rs2467026 | -0.788 | 0.591 | -0.342 | -0.746 | -0.028 | 0.023 | 0.023 | 0 | 0.66 | 0.84 | 0.67 | 0.73 | 0.6 | 0.56 |
| 8 | 135676891 | rs16905215 | 0.671 | 0.527 | -0.284 | -0.714 | -0.017 | 0.007 | 0.007 | 0 | 0.33 | 0.19 | 0.33 | 0.29 | 0.4 | 0.44 |
| 8 | 135678804 | rs9785140 | 0.627 | 0.528 | -0.179 | -0.718 | -0.005 | 0.004 | 0.004 | 0 | 0.34 | 0.25 | 0.34 | 0.29 | 0.4 | 0.44 |
| 8 | 135693723 | rs10505627 | 0.627 | 0.528 | -0.179 | -0.718 | -0.005 | 0.004 | 0.004 | 0 | 0.34 | 0.25 | 0.34 | 0.29 | 0.41 | 0.44 |
| 8 | 135697951 | rs7829427 | -0.891 | 0.521 | -0.52 | -0.778 | 0.008 | 0.006 | 0.006 | 0.436 | 0.71 | 0.82 | 0.8 | 0.67 | 0.77 | 0.83 |
| 8 | 135703504 | rs4909347 | 0.776 | 0.478 | -0.247 | -0.672 | -0.016 | -0.058 | -0.058 | 0 | 0.3 | 0.16 | 0.38 | 0.41 | 0.3 | 0.34 |
| 8 | 135712067 | rs4909609 | 0.265 | 0.485 | -0.112 | -0.66 | -0.001 | 0.001 | 0.001 | 0 | 0.35 | 0.36 | 0.44 | 0.41 | 0.36 | 0.4 |
| 8 | 135731578 | rs9969668 | 1.468 | 0.52 | -0.069 | -0.694 | 0.029 | -0.053 | -0.053 | 0 | 0.1 | 0.16 | 0.06 | 0 | 0.13 | 0.06 |
| 8 | 135733982 | rs1806812 | -0.664 | -0.067 | 0.076 | -0.646 | -0.072 | 0.022 | 0.022 | 0.183 | 0.74 | 0.56 | 0.79 | 0.66 | 0.84 | 0.84 |
| 8 | 135734604 | rs1806765 | -0.669 | -0.069 | 0.152 | -0.647 | -0.054 | 0.015 | 0.015 | 0.252 | 0.74 | 0.49 | 0.78 | 0.66 | 0.84 | 0.84 |
| 8 | 135736517 | rs16905222 | 0.902 | -0.037 | 0.07 | -0.64 | 0.011 | -0.02 | -0.02 | 0 | 0.05 | 0.18 | 0.05 | 0 | 0.06 | 0.06 |
| 8 | 135738044 | rs7817804 | -1.917 | -0.073 | -0.641 | -0.645 | 0.028 | -0.09 | -0.09 | 0 | 0.83 | 0.99 | 0.87 | 0.66 | 0.9 | 0.9 |
| 8 | 135739157 | rs13278719 | 0.657 | -0.038 | -0.704 | -0.719 | -0.011 | 0.015 | 0.015 | 0.051 | 0.34 | 0.24 | 0.34 | 0.27 | 0.4 | 0.45 |
| 8 | 135744460 | rs4909630 | 0.057 | 0.06 | -0.87 | -0.678 | -0.041 | -0.029 | -0.029 | 0.285 | 0.41 | 0.28 | 0.44 | 0.41 | 0.43 | 0.39 |
| 8 | 135745432 | rs4243863 | -0.176 | 0.288 | -0.851 | -0.381 | -0.004 | -0.015 | -0.015 | 0 | 0.38 | 0.64 | 0.47 | 0.41 | 0.38 | 0.39 |
| 8 | 135745502 | rs4388416 | 0.118 | 0.333 | -1.436 | -0.262 | -0.016 | -0.024 | -0.024 | 0.025 | 0.13 | 0 | 0.08 | 0.12 | 0.11 | 0.21 |
| 8 | 135748474 | rs11775800 | 0.306 | 0.346 | -1.098 | -0.173 | -0.018 | -0.032 | -0.032 | 0.181 | 0.3 | 0.22 | 0.45 | 0.39 | 0.34 | 0.35 |
| 8 | 135750322 | rs13265479 | 0.576 | 0.776 | -0.845 | -0.154 | -0.028 | -0.005 | -0.005 | 0 | 0.24 | 0.18 | 0.24 | 0.25 | 0.4 | 0.39 |
| 8 | 135751142 | rs4395854 | -0.714 | 0.017 | -0.991 | -0.172 | 0.017 | -0.024 | -0.024 | 0.005 | 0.83 | 0.88 | 0.84 | 0.64 | 0.89 | 0.86 |
| 8 | 135751481 | rs7821456 | -0.489 | -0.038 | -0.791 | -0.206 | 0.001 | -0.002 | -0.002 | 0 | 0.81 | 0.81 | 0.83 | 0.64 | 0.88 | 0.84 |
| 8 | 135752635 | rs13272667 | 0.816 | 0.213 | -0.953 | -0.287 | -0.034 | 0.041 | 0.041 | 0 | 0.08 | 0.02 | 0.04 | 0 | 0.09 | 0.07 |
| 8 | 135755248 | rs16905236 | 0.366 | 0.158 | -0.39 | -0.304 | 0.021 | -0.008 | -0.008 | 0 | 0.29 | 0.54 | 0.28 | 0.24 | 0.4 | 0.41 |
| 8 | 135760460 | rs13277671 | 0.188 | 0.087 | -0.211 | -0.351 | 0.04 | -0.013 | -0.013 | 0 | 0.31 | 0.46 | 0.29 | 0.25 | 0.4 | 0.42 |
| 8 | 135762452 | rs7013711 | -0.342 | -0.121 | -0.371 | -0.374 | 0.005 | -0.024 | -0.024 | 4.157 | 0.74 | 0.83 | 0.79 | 0.65 | 0.78 | 0.75 |
| 8 | 135764140 | rs12547792 | 1.118 | -0.141 | -0.352 | -0.387 | 0.019 | -0.004 | -0.004 | 0 | 0.41 | 0.27 | 0.5 | 0.4 | 0.37 | 0.32 |
| 8 | 135766277 | rs17707545 | 0.781 | -0.217 | -0.241 | -0.346 | 0.004 | -0.001 | -0.001 | 0.531 | 0.05 | 0.04 | 0.04 | 0 | 0.03 | 0.06 |
| 8 | 135768759 | rs6995897 | 1.12 | -0.051 | -0.346 | -0.447 | 0.037 | -0.015 | -0.015 | 0 | 0.39 | 0.24 | 0.48 | 0.4 | 0.33 | 0.31 |
| 8 | 135769607 | rs12681690 | -0.079 | 0.04 | -0.063 | -0.429 | 0 | 0.005 | 0.005 | 0 | 0.7 | 0.63 | 0.76 | 0.64 | 0.72 | 0.75 |
| 8 | 135771473 | rs1986890 | 0.451 | 0.002 | 0.013 | -0.446 | 0.011 | 0.005 | 0.005 | 0 | 0.48 | 0.5 | 0.54 | 0.4 | 0.38 | 0.41 |
| 8 | 135771497 | rs1986891 | 0.352 | -0.007 | -0.002 | -0.45 | 0.068 | 0.03 | 0.03 | 0.066 | 0.5 | 0.66 | 0.57 | 0.4 | 0.38 | 0.41 |
| 8 | 135771896 | rs7830265 | 0.662 | 0.044 | -0.269 | -0.412 | 0.036 | 0.011 | 0.011 | 0.028 | 0.46 | 0.38 | 0.52 | 0.4 | 0.36 | 0.41 |
| 8 | 135773607 | rs7842889 | -0.248 | 0.247 | -0.521 | -0.266 | -0.043 | -0.051 | -0.051 | 0.295 | 0.74 | 0.92 | 0.79 | 0.65 | 0.74 | 0.76 |
| 8 | 135787891 | rs7832498 | -0.895 | 0.429 | 0.113 | -0.366 | 0.017 | -0.012 | -0.012 | 0 | 0.07 | 0.3 | 0.05 | 0 | 0.03 | 0.06 |
| 8 | 135792825 | rs16905249 | 0.284 | 0.766 | -0.256 | -0.459 | 0.054 | -0.033 | -0.033 | 0 | 0.27 | 0.5 | 0.23 | 0.17 | 0.37 | 0.39 |
| 8 | 135793642 | rs12679681 | -0.672 | 0.272 | -0.599 | -0.264 | 0.007 | 0.012 | 0.012 | 0 | 0.45 | 0.67 | 0.4 | 0.59 | 0.56 | 0.64 |
| 8 | 135793846 | rs12679206 | 0.384 | 0.394 | -0.597 | -0.249 | 0.008 | 0.005 | 0.005 | 0 | 0.27 | 0.26 | 0.29 | 0.58 | 0.41 | 0.51 |
| 8 | 135793900 | rs11777778 | 0.979 | -0.1 | -1.591 | -0.37 | -0.035 | -0.096 | -0.096 | 0.074 | 0.39 | 0.02 | 0.52 | 0.41 | 0.34 | 0.3 |
| 8 | 135794496 | rs7828209 | 0.18 | -0.223 | -1.114 | -0.429 | 0.014 | 0.014 | 0.014 | 0 | 0.53 | 0.57 | 0.5 | 0.39 | 0.61 | 0.6 |
| 8 | 135796966 | rs10505630 | -1.405 | 1.382 | -0.765 | -0.525 | -0.004 | 0.003 | 0.003 | 0.223 | 0.39 | 0.61 | 0.37 | 0.31 | 0.49 | 0.4 |
| 8 | 135802892 | rs7831733 | -0.675 | 2.039 | -0.4 | -0.482 | 0.042 | -0.028 | -0.028 | 0 | 0.13 | 0.25 | 0.13 | 0.04 | 0.2 | 0.11 |
| 8 | 135805476 | rs10099070 | -0.412 | 2.028 | -0.446 | -0.509 | -0.013 | 0.013 | 0.013 | 0 | 0.15 | 0.11 | 0.15 | 0.07 | 0.22 | 0.13 |
| 8 | 135809634 | rs10106201 | -1.704 | 2.151 | -0.294 | -0.598 | 0.029 | -0.089 | -0.089 | 1.039 | 0.24 | 0.42 | 0.2 | 0.05 | 0.27 | 0.19 |

| | | | | | | | | | | | | | | | |
|---|---|---|---|---|---|---|---|---|---|---|---|---|---|---|---|
| 8 | 135812548 | rs10095483 | -0.405 | 2.185 | -0.35 | -0.58 | -0.009 | 0.011 | 0.011 | 0 | 0.16 | 0.13 | 0.15 | 0.04 | 0.22 | 0.12 |
| 8 | 135813693 | rs16905252 | 0.036 | 2.154 | -0.651 | -0.55 | 0.017 | -0.032 | -0.032 | 0 | 0.08 | 0.14 | 0.09 | 0.01 | 0.13 | 0.08 |
| 8 | 135837130 | rs7017546 | -0.764 | 1.873 | -0.901 | -0.454 | 0.031 | -0.103 | -0.103 | 0.145 | 0.25 | 0.47 | 0.2 | 0.05 | 0.27 | 0.19 |
| 8 | 135840879 | rs3735773 | 0.331 | 1.868 | -1.087 | -0.308 | -0.032 | -0.035 | -0.035 | 0 | 0.1 | 0.03 | 0.11 | 0.26 | 0.16 | 0.08 |
| 8 | 135847101 | rs733122 | -1.416 | 1.872 | -1.064 | -0.425 | -0.001 | 0.014 | 0.014 | 0.03 | 0.42 | 0.62 | 0.39 | 0.31 | 0.53 | 0.41 |
| 8 | 135847442 | rs737196 | -1.099 | 1.829 | -1.023 | -0.897 | 0.019 | -0.087 | -0.087 | 0 | 0.31 | 0.52 | 0.28 | 0.06 | 0.37 | 0.33 |
| 8 | 135856404 | rs6987233 | -1.037 | 1.667 | -1.913 | -0.167 | -0.033 | -0.067 | -0.067 | 0.294 | 0.54 | 0.97 | 0.55 | 0.58 | 0.56 | 0.5 |
| 8 | 135864228 | rs2205362 | -1.208 | 2.214 | -0.564 | 0.047 | 0.069 | -0.08 | -0.08 | 0.344 | 0.25 | 0.45 | 0.32 | 0.13 | 0.35 | 0.34 |
| 8 | 135866471 | rs17519106 | -0.742 | 1.355 | -1.185 | 0.283 | 0.037 | -0.052 | -0.052 | 0 | 0.1 | 0.17 | 0.1 | 0 | 0.13 | 0.15 |
| 8 | 135870334 | rs1000394 | -0.394 | 0.121 | -1.685 | 0.706 | 0.031 | -0.026 | -0.026 | 0 | 0.69 | 0.82 | 0.79 | 0.64 | 0.8 | 0.67 |
| 8 | 135871032 | rs12542413 | -0.699 | 1.585 | -1.605 | 0.508 | -0.054 | 0.043 | 0.043 | 2.952 | 0.3 | 0.17 | 0.46 | 0.23 | 0.52 | 0.48 |
| 8 | 135878115 | rs4909799 | -0.82 | 1.102 | -1.421 | 1.371 | -0.009 | -0.026 | -0.026 | 0.047 | 0.07 | 0 | 0.21 | 0.3 | 0.08 | 0.17 |
| 8 | 135878725 | rs16905258 | -0.279 | 1.098 | -1.642 | 1.534 | -0.012 | -0.031 | -0.031 | 0 | 0.06 | 0.01 | 0.13 | 0.12 | 0.09 | 0.13 |
| 8 | 135879159 | rs4909801 | -1.206 | 1.653 | -0.9 | 1.1 | 0.017 | 0.04 | 0.04 | 0.12 | 0.21 | 0.26 | 0.41 | 0.41 | 0.27 | 0.33 |
| 8 | 135882007 | rs7834489 | -0.899 | 1.289 | -1.029 | 0.421 | 0.028 | 0.038 | 0.038 | 0 | 0.56 | 0.76 | 0.67 | 0.69 | 0.61 | 0.65 |
| 8 | 135883928 | rs6992901 | 0.159 | 1.385 | -0.63 | 0.753 | 0.009 | 0.017 | 0.017 | 0 | 0.12 | 0.2 | 0.16 | 0.17 | 0.15 | 0.11 |
| 8 | 135884188 | rs4243894 | -1.054 | 1.361 | -0.342 | 0.537 | 0.021 | 0.01 | 0.01 | 0 | 0.37 | 0.56 | 0.59 | 0.59 | 0.45 | 0.39 |
| 8 | 135888570 | rs4434578 | -0.566 | 0.918 | -0.371 | 0.438 | -0.011 | -0.018 | -0.018 | 0 | 0.23 | 0.39 | 0.21 | 0.18 | 0.2 | 0.14 |
| 8 | 135889561 | rs4624986 | 0.475 | 1.134 | -0.994 | 0.573 | -0.024 | -0.033 | -0.033 | 0 | 0.1 | 0.06 | 0.15 | 0.17 | 0.15 | 0.11 |
| 8 | 135894058 | rs4909820 | 0.033 | 1.195 | -0.967 | 0.625 | -0.001 | 0.017 | 0.017 | 0.037 | 0.14 | 0.15 | 0.37 | 0.41 | 0.23 | 0.26 |
| 8 | 135894974 | rs17519875 | 0.463 | 1.194 | -1.013 | 0.706 | -0.022 | -0.032 | -0.032 | 0.001 | 0.1 | 0.06 | 0.14 | 0.18 | 0.14 | 0.11 |
| 8 | 135895391 | rs1985221 | -0.377 | 0.929 | -0.776 | 0.227 | 0 | 0 | 0 | 0 | 0.35 | 0.39 | 0.57 | 0.59 | 0.43 | 0.39 |
| 8 | 135899630 | rs4601273 | -0.244 | 0.637 | -0.873 | 0.114 | 0.021 | 0.008 | 0.008 | 0 | 0.11 | 0.01 | 0.03 | 0.09 | 0.04 | 0.17 |
| 8 | 135907056 | rs7844371 | 0.139 | 0.668 | -0.204 | -0.186 | -0.004 | -0.073 | -0.073 | 0.049 | 0.82 | 0.62 | 0.88 | 0.94 | 0.86 | 0.83 |
| 8 | 135908515 | rs7462577 | 0.418 | 0.457 | -0.761 | -0.681 | -0.006 | -0.01 | -0.01 | 2.422 | 0.38 | 0.33 | 0.31 | 0.19 | 0.28 | 0.17 |
| 8 | 135910193 | rs10097238 | 0.901 | 1.064 | -1.531 | -0.23 | -0.064 | -0.12 | -0.12 | 0 | 0.25 | 0.08 | 0.28 | 0.46 | 0.35 | 0.41 |
| 8 | 135918442 | rs4909841 | 0.125 | 0.807 | -1.553 | -0.574 | -0.027 | -0.029 | -0.029 | 0.016 | 0.44 | 0.33 | 0.47 | 0.47 | 0.46 | 0.38 |
| 8 | 135918992 | rs7464953 | 0.683 | 0.604 | -1.329 | -0.929 | -0.004 | 0.111 | 0.111 | 0 | 0.5 | 0.23 | 0.46 | 0.65 | 0.55 | 0.52 |
| 8 | 135923654 | rs10111298 | 0.376 | 1.113 | -0.824 | -0.196 | 0.013 | 0.019 | 0.019 | 0 | 0.16 | 0.22 | 0.28 | 0.28 | 0.23 | 0.25 |
| 8 | 135945925 | rs11166887 | 0.423 | 1.111 | -0.827 | -0.126 | -0.009 | 0.03 | 0.03 | 2.678 | 0.36 | 0.25 | 0.37 | 0.29 | 0.37 | 0.35 |
| 8 | 135953131 | rs10110613 | -0.826 | 1.144 | -0.431 | 0.12 | -0.001 | -0.001 | -0.001 | 0.135 | 0.67 | 0.66 | 0.62 | 0.7 | 0.72 | 0.66 |
| 8 | 135954467 | rs3925094 | -0.824 | 1.143 | -0.576 | 0.12 | -0.019 | -0.025 | -0.025 | 0 | 0.67 | 0.55 | 0.61 | 0.7 | 0.72 | 0.66 |
| 8 | 135958538 | rs7011963 | 0.142 | 1.412 | -0.594 | 0.422 | 0.034 | 0.093 | 0.093 | 0.056 | 0.18 | 0.26 | 0.21 | 0.4 | 0.28 | 0.18 |
| 8 | 135960258 | rs11166907 | 0.036 | 1.192 | -0.656 | 0.301 | 0 | 0.002 | 0.002 | 0 | 0.12 | 0.08 | 0.06 | 0.06 | 0.08 | 0.11 |
| 8 | 135961826 | rs7016799 | 0.099 | 1.271 | -0.736 | 0.425 | 0.004 | 0.045 | 0.045 | 0 | 0.19 | 0.26 | 0.19 | 0.41 | 0.26 | 0.18 |
| 8 | 135962461 | rs4909849 | -0.161 | 1.29 | -0.711 | 0.292 | -0.002 | -0.004 | -0.004 | 0.309 | 0.29 | 0.3 | 0.33 | 0.48 | 0.36 | 0.32 |
| 8 | 135962651 | rs11785063 | 0.08 | 1.04 | -0.964 | 0.043 | 0.005 | 0.026 | 0.026 | 5.184 | 0.39 | 0.3 | 0.27 | 0.24 | 0.33 | 0.36 |
| 8 | 135964558 | rs12679813 | 0.034 | 0.877 | -1.177 | 0.23 | -0.008 | -0.03 | -0.03 | 0 | 0.25 | 0.17 | 0.31 | 0.28 | 0.22 | 0.21 |
| 8 | 135965941 | rs4909851 | 0.548 | 0.538 | -1.14 | -0.233 | -0.013 | -0.021 | -0.021 | 0 | 0.4 | 0.29 | 0.38 | 0.56 | 0.47 | 0.48 |
| 8 | 135967706 | rs11994368 | 0.564 | 0.43 | -1.203 | -0.297 | 0 | -0.027 | -0.027 | 2.424 | 0.26 | 0.18 | 0.31 | 0.27 | 0.21 | 0.21 |
| 8 | 135968839 | rs7461441 | 0.375 | 0.388 | -0.875 | -0.327 | -0.006 | 0.025 | 0.025 | 0 | 0.28 | 0.47 | 0.33 | 0.27 | 0.21 | 0.21 |
| 8 | 135969632 | rs7820325 | -0.133 | 0.572 | -1.223 | -0.274 | 0.008 | 0.021 | 0.021 | 0.704 | 0.68 | 0.77 | 0.73 | 0.81 | 0.69 | 0.7 |
| 8 | 135971872 | rs11784641 | 0.471 | 0.989 | -1.839 | -0.254 | -0.034 | 0.007 | 0.007 | 0 | 0.15 | 0.04 | 0.13 | 0.1 | 0.26 | 0.11 |
| 8 | 135972481 | rs11987389 | -0.304 | 0.662 | -1.658 | -0.506 | 0 | 0.015 | 0.015 | 0.069 | 0.75 | 0.9 | 0.76 | 0.81 | 0.74 | 0.72 |
| 8 | 135977336 | rs7813240 | -0.352 | 0.664 | -1.906 | -0.809 | -0.056 | -0.041 | -0.041 | 0.055 | 0.71 | 0.92 | 0.64 | 0.4 | 0.57 | 0.6 |
| 8 | 135980247 | rs6992707 | 1.244 | 0.404 | -2.359 | -1.401 | -0.062 | 0.115 | 0.115 | 0 | 0.22 | 0.06 | 0.22 | 0.02 | 0.24 | 0.11 |
| 8 | 135986892 | rs7465467 | 0.394 | -0.06 | -1.698 | -1.635 | -0.044 | -0.063 | -0.063 | 0 | 0.18 | 0.34 | 0.13 | 0.06 | 0.09 | 0.13 |
| 8 | 136005077 | rs6998599 | -1.019 | 0.032 | -1.893 | -1.682 | -0.059 | -0.041 | -0.041 | 0 | 0.71 | 0.92 | 0.63 | 0.4 | 0.56 | 0.61 |
| 8 | 136005606 | rs11778431 | 1.536 | -0.041 | -1.847 | -1.749 | -0.032 | 0.069 | 0.069 | 0 | 0.22 | 0.11 | 0.22 | 0.02 | 0.23 | 0.11 |
| 8 | 136006630 | rs7014470 | 0.271 | -0.192 | -1.49 | -1.75 | 0.004 | 0.023 | 0.023 | 0 | 0.51 | 0.57 | 0.5 | 0.34 | 0.47 | 0.49 |
| 8 | 136009494 | rs6578047 | -0.121 | -0.398 | -1.398 | -1.741 | -0.014 | -0.065 | -0.065 | 0 | 0.25 | 0.35 | 0.22 | 0.06 | 0.21 | 0.19 |
| 8 | 136017359 | rs4909854 | 0.79 | 0.111 | -1.681 | -1.734 | -0.012 | 0.006 | 0.006 | 0.508 | 0.32 | 0.1 | 0.42 | 0.66 | 0.4 | 0.49 |
| 8 | 136017930 | rs7462529 | -0.863 | 0.702 | -1.347 | -1.206 | -0.053 | 0.076 | 0.076 | 0 | 0.76 | 0.88 | 0.75 | 0.98 | 0.69 | 0.89 |
| 8 | 136024190 | rs12679789 | 0.49 | 0.27 | -1.083 | -0.951 | -0.017 | 0.012 | 0.012 | 0 | 0.22 | 0.32 | 0.19 | 0.32 | 0.11 | 0.31 |
| 8 | 136032287 | rs10113126 | -0.088 | 0.978 | -1.445 | -1.029 | -0.043 | 0.045 | 0.045 | 0 | 0.69 | 0.79 | 0.66 | 0.98 | 0.56 | 0.83 |
| 8 | 136038123 | rs6578068 | -0.187 | 0.788 | -1.247 | 0.207 | -0.058 | -0.042 | -0.042 | 0 | 0.66 | 0.81 | 0.57 | 0.56 | 0.57 | 0.59 |
| 8 | 136039314 | rs6987207 | -0.144 | -0.017 | -0.794 | 0.063 | -0.084 | 0.144 | 0.144 | 0 | 0.22 | 0.49 | 0.2 | 0.43 | 0.08 | 0.32 |
| 8 | 136039623 | rs6578070 | 0.55 | -0.01 | -0.855 | -0.257 | 0.004 | 0.114 | 0.114 | 0 | 0.45 | 0.28 | 0.37 | 0.14 | 0.51 | 0.27 |
| 8 | 136042131 | rs6578071 | -0.369 | 0.049 | -0.671 | 0.235 | -0.048 | -0.008 | -0.008 | 0 | 0.66 | 0.85 | 0.56 | 0.67 | 0.59 | 0.59 |
| 8 | 136046135 | rs1480806 | -0.245 | -0.011 | -0.481 | 0.526 | -0.118 | -0.052 | -0.052 | 0 | 0.63 | 0.94 | 0.55 | 0.64 | 0.57 | 0.57 |
| 8 | 136050386 | rs4909459 | -0.02 | 0.375 | -0.39 | 0.465 | 0.002 | -0.003 | -0.003 | 0 | 0.39 | 0.37 | 0.35 | 0.52 | 0.32 | 0.43 |
| 8 | 136050413 | rs10875449 | -0.41 | 0.734 | -0.28 | 0.896 | 0.021 | 0.117 | 0.117 | 0 | 0.17 | 0.35 | 0.32 | 0.36 | 0.17 | 0.27 |
| 8 | 136053640 | rs2086438 | 0.235 | 0.275 | -1.163 | 0.52 | 0.006 | -0.002 | -0.002 | 0.249 | 0.35 | 0.13 | 0.34 | 0.38 | 0.33 | 0.37 |
| 8 | 136055420 | rs6999421 | -0.027 | -0.036 | -0.706 | -0.085 | -0.001 | -0.036 | -0.036 | 0 | 0.42 | 0.44 | 0.5 | 0.16 | 0.59 | 0.36 |
| 8 | 136058795 | rs4909871 | 0.696 | -0.477 | -1.011 | 0.485 | -0.003 | -0.065 | -0.065 | 0 | 0.42 | 0.11 | 0.32 | 0.42 | 0.32 | 0.43 |
| 8 | 136063977 | rs4909872 | 0.602 | -0.707 | -0.157 | 0.339 | -0.041 | -0.061 | -0.061 | 0.221 | 0.87 | 0.44 | 0.91 | 0.97 | 0.94 | 0.96 |
| 8 | 136068182 | rs6981389 | -1.774 | -0.782 | -0.147 | 0.363 | -0.049 | 0.084 | 0.084 | 0 | 0.19 | 0.64 | 0.22 | 0.44 | 0.1 | 0.22 |
| 8 | 136073944 | rs1383453 | 0.501 | -0.654 | -0.249 | 0.305 | -0.08 | 0.137 | 0.137 | 0.074 | 0.38 | 0.11 | 0.45 | 0.13 | 0.58 | 0.35 |
| 8 | 136077967 | rs7846098 | 0.144 | -0.673 | -0.061 | 0.398 | -0.02 | 0.179 | 0.179 | 0 | 0.07 | 0.2 | 0.15 | 0.41 | 0.04 | 0.17 |

| | | | | | | | | | | | | | | | | |
|---|---|---|---|---|---|---|---|---|---|---|---|---|---|---|---|---|
| 8 | 136085856 | rs12682229 | 0.286 | -0.747 | -0.144 | 0.279 | 0.007 | 0.001 | 0.001 | 0 | 0.49 | 0.39 | 0.35 | 0.46 | 0.35 | 0.47 |
| 8 | 136086741 | rs6578122 | 0.587 | -0.68 | -0.009 | 0.384 | -0.018 | 0.304 | 0.304 | 0 | 0.91 | 0.61 | 0.83 | 0.59 | 0.96 | 0.83 |
| 8 | 136089776 | rs1383480 | 1.413 | -0.79 | -0.115 | 0.307 | -0.093 | 0.158 | 0.158 | 0.023 | 0.81 | 0.48 | 0.8 | 0.58 | 0.93 | 0.79 |
| 8 | 136097222 | rs4909463 | 0.689 | -0.723 | -0.214 | 0.273 | 0.004 | -0.002 | -0.002 | 0 | 0.44 | 0.33 | 0.35 | 0.46 | 0.35 | 0.44 |
| 8 | 136124463 | rs1480804 | 0.17 | -0.631 | -0.063 | 0.298 | -0.039 | -0.041 | -0.041 | 0 | 0.93 | 0.84 | 0.98 | 1 | 0.99 | 0.97 |
| 8 | 136132998 | rs1480803 | 0.972 | -0.659 | -0.264 | 0.578 | -0.09 | 0.208 | 0.208 | 0.068 | 0.83 | 0.52 | 0.81 | 0.57 | 0.94 | 0.78 |
| 8 | 136133575 | rs10104990 | 0.304 | -0.455 | -0.302 | 0.414 | -0.059 | 0.08 | 0.08 | 0 | 0.37 | 0.18 | 0.47 | 0.16 | 0.54 | 0.29 |
| 8 | 136140833 | rs4909467 | 0.292 | -0.085 | -0.333 | 0.313 | 0.011 | 0.024 | 0.024 | 0 | 0.5 | 0.35 | 0.35 | 0.4 | 0.41 | 0.48 |
| 8 | 136147715 | rs4909890 | -0.455 | 0.362 | -0.018 | 0.165 | 0.019 | 0.095 | 0.095 | 1.823 | 0.49 | 0.32 | 0.63 | 0.73 | 0.61 | 0.69 |
| 8 | 136150965 | rs971448 | 0.265 | 0.149 | 0.502 | 0.379 | 0.07 | 0.399 | 0.399 | 0.771 | 0.95 | 0.61 | 0.89 | 0.76 | 0.98 | 0.97 |
| 8 | 136153151 | rs16905290 | 0.496 | 0.233 | 0.364 | 0.353 | -0.001 | -0.024 | -0.024 | 0 | 0.15 | 0.1 | 0.3 | 0.53 | 0.16 | 0.39 |
| 8 | 136179139 | rs17588701 | -0.557 | 0.511 | -0.226 | -0.17 | -0.023 | 0.072 | 0.072 | 0.839 | 0.43 | 0.21 | 0.58 | 0.72 | 0.5 | 0.63 |
| 8 | 136187340 | rs11167077 | -1.405 | 0.49 | -0.864 | -0.452 | -0.009 | 0.136 | 0.136 | 0.569 | 0.61 | 0.83 | 0.79 | 0.97 | 0.67 | 0.77 |
| 8 | 136193650 | rs10156288 | -0.039 | 0.464 | -0.388 | 0.221 | 0.029 | 0.271 | 0.271 | 4.241 | 0.05 | 0.3 | 0.11 | 0.22 | 0.01 | 0.03 |
| 8 | 136194794 | rs17589261 | 0.071 | 1.016 | -1.47 | 0.402 | -0.048 | -0.092 | -0.092 | 0 | 0.17 | 0.07 | 0.37 | 0.51 | 0.24 | 0.47 |
| 8 | 136200559 | rs7002465 | -0.625 | 0.622 | -1.464 | 0.14 | -0.043 | -0.043 | -0.043 | 0 | 0.39 | 0.24 | 0.56 | 0.52 | 0.56 | 0.34 |
| 8 | 136202918 | rs13269684 | 0.991 | 0.795 | -1.601 | -0.107 | -0.012 | -0.079 | -0.079 | 0 | 0.15 | 0.07 | 0.26 | 0.44 | 0.16 | 0.36 |
| 8 | 136203731 | rs6578220 | -1.293 | 0.515 | -1.025 | -0.193 | 0 | -0.045 | -0.045 | 0 | 0.67 | 0.72 | 0.83 | 0.96 | 0.73 | 0.73 |
| 8 | 136203934 | rs7820620 | -0.15 | 0.72 | -2.079 | -0.269 | 0.002 | -0.073 | -0.073 | 0.08 | 0.27 | 0.09 | 0.32 | 0.42 | 0.29 | 0.23 |
| 8 | 136204327 | rs4909899 | -0.945 | 0.485 | -1.585 | -0.024 | -0.002 | -0.088 | -0.088 | 0.587 | 0.65 | 0.47 | 0.8 | 0.96 | 0.73 | 0.73 |
| 8 | 136206652 | rs6998848 | -0.523 | 0.537 | -1.551 | 1.335 | 0.013 | -0.053 | -0.053 | 0.017 | 0.18 | 0.09 | 0.26 | 0.4 | 0.19 | 0.2 |
| 8 | 136212232 | rs1157331 | 0.559 | 0.39 | -1.787 | 0.89 | 0.137 | 0.004 | 0.004 | 0.01 | 0.49 | 0.13 | 0.34 | 0.4 | 0.33 | 0.43 |
| 8 | 136225420 | rs4909472 | 0.402 | 0.853 | -0.87 | 0.85 | -0.004 | 0 | 0 | 0.026 | 0.57 | 0.5 | 0.44 | 0.5 | 0.39 | 0.5 |
| 8 | 136225944 | rs964043 | 0.321 | 1.42 | -1.472 | 0.596 | -0.056 | -0.012 | -0.012 | 0 | 0.11 | 0.01 | 0.11 | 0.09 | 0.19 | 0.16 |
| 8 | 136238597 | rs7016887 | 0.323 | 0.544 | -1.145 | 0.579 | 0.051 | 0.074 | 0.074 | 0 | 0.55 | 0.71 | 0.43 | 0.65 | 0.34 | 0.52 |
| 8 | 136238694 | rs6578225 | 0.392 | 0.341 | -1.164 | 0.633 | 0.085 | 0.084 | 0.084 | 0 | 0.54 | 0.76 | 0.38 | 0.64 | 0.31 | 0.51 |
| 8 | 136266912 | rs11782903 | -0.187 | 0.757 | -0.616 | 0.405 | -0.001 | 0.016 | 0.016 | 0 | 0.42 | 0.19 | 0.65 | 0.6 | 0.58 | 0.56 |
| 8 | 136267693 | rs6578226 | 0.052 | 0.315 | 0.072 | 0.093 | -0.001 | 0.034 | 0.034 | 0.698 | 0.83 | 0.77 | 0.84 | 0.69 | 0.81 | 0.83 |
| 8 | 136270473 | rs7004726 | 0.932 | -0.108 | 0.226 | -0.031 | -0.01 | 0.032 | 0.032 | 0.346 | 0.76 | 0.28 | 0.79 | 0.57 | 0.79 | 0.78 |
| 8 | 136280698 | rs6984612 | -0.7 | 0.01 | 0.372 | -0.206 | 0.016 | 0.104 | 0.104 | 0.031 | 0.1 | 0.36 | 0.09 | 0.23 | 0.07 | 0.15 |
| 8 | 136293806 | rs10095580 | -0.685 | 0.025 | 0.685 | -0.128 | 0.025 | 0.123 | 0.123 | 0 | 0.1 | 0.38 | 0.08 | 0.23 | 0.08 | 0.13 |
| 8 | 136294570 | rs7831546 | 0.124 | 0.015 | 0.689 | -0.15 | 0 | 0.024 | 0.024 | 0 | 0.06 | 0.07 | 0.06 | 0.23 | 0.05 | 0.13 |
| 8 | 136294660 | rs10464988 | 0.529 | -0.259 | 0.802 | -0.346 | -0.042 | 0.002 | 0.002 | 0 | 0.39 | 0.09 | 0.63 | 0.6 | 0.58 | 0.53 |
| 8 | 136294856 | rs10099533 | 1.382 | -0.751 | 0.464 | -0.606 | -0.001 | 0.221 | 0.221 | 0 | 0.3 | 0.01 | 0.13 | 0.01 | 0.21 | 0.21 |
| 8 | 136298197 | rs4354348 | 0.1 | -0.688 | 1.237 | -0.207 | -0.004 | 0.01 | 0.01 | 0 | 0.89 | 0.77 | 0.9 | 0.86 | 0.88 | 0.93 |
| 8 | 136317085 | rs7840823 | 1.571 | -0.695 | 1.271 | -0.209 | -0.01 | 0.082 | 0.082 | 0.25 | 0.79 | 0.4 | 0.8 | 0.62 | 0.81 | 0.79 |
| 8 | 136320986 | rs12114371 | 1.496 | -0.687 | 1.249 | -0.222 | -0.004 | 0.056 | 0.056 | 0.13 | 0.8 | 0.51 | 0.83 | 0.64 | 0.81 | 0.79 |
| 8 | 136324034 | rs1455476 | 0.065 | -0.789 | 1.231 | -0.239 | -0.004 | 0 | 0 | 0 | 0.89 | 0.82 | 0.9 | 0.87 | 0.88 | 0.93 |
| 8 | 136330821 | rs2925046 | 1.627 | -0.818 | 1.34 | -0.253 | -0.026 | 0.068 | 0.068 | 0.395 | 0.79 | 0.45 | 0.82 | 0.62 | 0.83 | 0.79 |
| 8 | 136337966 | rs1350031 | -0.21 | -0.789 | 1.142 | -0.251 | 0 | -0.009 | -0.009 | 0.219 | 0.91 | 0.92 | 0.91 | 0.98 | 0.88 | 0.96 |
| 8 | 136355603 | rs7825136 | 1.252 | -0.793 | 1.226 | -0.176 | -0.01 | 0.05 | 0.05 | 0.003 | 0.81 | 0.68 | 0.82 | 0.62 | 0.83 | 0.8 |
| 8 | 136358469 | rs882615 | 1.602 | -0.759 | 1.237 | -0.243 | -0.023 | 0.013 | 0.013 | 0.293 | 0.74 | 0.41 | 0.76 | 0.62 | 0.8 | 0.79 |
| 8 | 136361527 | rs2124279 | 1.778 | -0.758 | 1.053 | -0.17 | 0 | -0.001 | -0.001 | 0 | 0.73 | 0.24 | 0.75 | 0.61 | 0.8 | 0.78 |
| 8 | 136366399 | rs1110710 | 0.19 | -0.536 | 0.84 | -0.072 | -0.003 | 0.006 | 0.006 | 0 | 0.9 | 0.83 | 0.9 | 0.85 | 0.88 | 0.93 |
| 8 | 136368501 | rs7843508 | -0.461 | -0.3 | 0.243 | -0.196 | -0.035 | 0.103 | 0.103 | 0.158 | 0.6 | 0.85 | 0.35 | 0.21 | 0.37 | 0.32 |
| 8 | 136374029 | rs1455469 | 0.275 | -0.581 | 0.894 | -0.499 | 0.018 | -0.014 | -0.014 | 0.121 | 0.45 | 0.59 | 0.22 | 0.03 | 0.24 | 0.21 |
| 8 | 136377986 | rs1455474 | -0.848 | -0.878 | 0.736 | -0.184 | -0.067 | -0.075 | -0.075 | 0 | 0.61 | 0.86 | 0.45 | 0.48 | 0.4 | 0.39 |
| 8 | 136378593 | rs6578233 | 0.62 | -0.948 | 1.148 | -0.226 | 0.007 | -0.031 | -0.031 | 0 | 0.41 | 0.37 | 0.2 | 0.02 | 0.24 | 0.19 |
| 8 | 136390657 | rs10505636 | -0.877 | -0.769 | 0.458 | -0.107 | -0.079 | -0.086 | -0.086 | 1.943 | 0.62 | 0.9 | 0.45 | 0.48 | 0.4 | 0.39 |
| 8 | 136404884 | rs1378471 | -0.684 | -0.7 | 0.448 | -0.039 | -0.019 | 0.023 | 0.023 | 0.692 | 0.58 | 0.83 | 0.36 | 0.36 | 0.35 | 0.3 |
| 8 | 136417980 | rs930836 | -0.719 | -0.405 | 0.385 | -0.286 | -0.05 | 0.053 | 0.053 | 0.294 | 0.58 | 0.86 | 0.37 | 0.34 | 0.39 | 0.31 |
| 8 | 136418675 | rs16905364 | -0.177 | -0.418 | 0.622 | -0.794 | 0.023 | 0.02 | 0.02 | 0.064 | 0.47 | 0.64 | 0.25 | 0.03 | 0.3 | 0.23 |
| 8 | 136428948 | rs897407 | -0.142 | -0.262 | 0.653 | -0.716 | -0.006 | -0.082 | -0.082 | 0.054 | 0.45 | 0.58 | 0.25 | 0.03 | 0.3 | 0.23 |
| 8 | 136444089 | rs7829575 | -1.862 | -0.495 | 0.979 | -0.142 | -0.046 | 0.009 | 0.009 | 0.344 | 0.16 | 0.5 | 0.17 | 0.15 | 0.11 | 0.12 |
| 8 | 136449715 | rs10106123 | -1.737 | -0.552 | 1.459 | -0.165 | -0.05 | -0.048 | -0.048 | 0 | 0.1 | 0.4 | 0.07 | 0.01 | 0.03 | 0.04 |
| 8 | 136450246 | rs7003412 | -2.19 | -0.485 | 1.425 | -0.152 | -0.043 | 0 | 0 | 0.298 | 0.16 | 0.58 | 0.19 | 0.15 | 0.11 | 0.11 |
| 8 | 136455412 | rs736647 | -0.56 | -0.377 | 0.513 | -0.771 | -0.024 | 0.099 | 0.099 | 0 | 0.69 | 0.86 | 0.82 | 0.96 | 0.73 | 0.82 |
| 8 | 136475038 | rs12679393 | -0.401 | -0.403 | 0.973 | -0.298 | 0.006 | 0.085 | 0.085 | 0.114 | 0.07 | 0.18 | 0.12 | 0.19 | 0.06 | 0.08 |
| 8 | 136500242 | rs12678148 | 0.644 | -0.714 | 1.433 | -0.668 | 0.026 | 0.116 | 0.116 | 0.059 | 0.52 | 0.3 | 0.62 | 0.78 | 0.61 | 0.68 |
| 8 | 136500554 | rs2873420 | 0.636 | -0.715 | 1.433 | -0.668 | 0.025 | 0.115 | 0.115 | 0.045 | 0.52 | 0.31 | 0.62 | 0.78 | 0.61 | 0.68 |
| 8 | 136522558 | rs11779943 | 1.604 | -0.676 | 0.905 | -0.003 | -0.082 | -0.066 | -0.066 | 0 | 0.35 | 0.1 | 0.45 | 0.38 | 0.44 | 0.55 |
| 8 | 136523497 | rs10104847 | -1.215 | -0.697 | 1.11 | -0.018 | 0.075 | 0.048 | 0.048 | 0.086 | 0.05 | 0.4 | 0.08 | 0 | 0.04 | 0.03 |
| 8 | 136526248 | rs10092358 | 1.6 | -0.68 | 0.781 | 0.006 | -0.085 | -0.071 | -0.071 | 0.436 | 0.36 | 0.09 | 0.45 | 0.39 | 0.44 | 0.56 |
| 8 | 136539132 | rs16905387 | 1.612 | -0.699 | 0.742 | 0.038 | -0.122 | -0.155 | -0.155 | 0 | 0.35 | 0.03 | 0.44 | 0.45 | 0.44 | 0.56 |
| 8 | 136563532 | rs10095429 | 0.232 | -0.667 | 1.081 | 0.04 | 0.011 | 0.059 | 0.059 | 0.051 | 0.55 | 0.35 | 0.63 | 0.78 | 0.63 | 0.68 |
| 8 | 136569244 | rs12681761 | -1.23 | -0.791 | 1.1 | -0.037 | -0.033 | 0.05 | 0.05 | 0 | 0.16 | 0.45 | 0.19 | 0.18 | 0.11 | 0.14 |
| 8 | 136570502 | rs2317280 | -0.795 | -0.846 | 0.9 | -0.11 | -0.005 | 0.044 | 0.044 | 0 | 0.69 | 0.81 | 0.82 | 0.96 | 0.73 | 0.82 |
| 8 | 136580911 | rs16905413 | 0.166 | -0.845 | 0.96 | -0.041 | 0 | 0.013 | 0.013 | 0 | 0.15 | 0.18 | 0.17 | 0.33 | 0.16 | 0.12 |
| 8 | 136581105 | rs11781018 | 1.705 | -0.863 | 0.924 | -0.086 | -0.045 | -0.072 | -0.072 | 0 | 0.37 | 0.13 | 0.46 | 0.45 | 0.46 | 0.55 |

| | | | | | | | | | | | | | | | | |
|---|---|---|---|---|---|---|---|---|---|---|---|---|---|---|---|---|
| 8 | 136614336 | rs10088630 | -0.265 | -0.88 | 1.211 | -0.131 | -0.005 | -0.135 | -0.135 | 0.015 | 0.63 | 0.5 | 0.75 | 0.96 | 0.68 | 0.76 |
| 8 | 136617235 | rs6983445 | -0.645 | -0.857 | 1.418 | -0.105 | 0.048 | 0.024 | 0.024 | 0 | 0.05 | 0.3 | 0.07 | 0 | 0.04 | 0.04 |
| 8 | 136623771 | rs4573318 | 0.649 | -0.927 | 0.947 | -0.104 | 0.017 | 0.004 | 0.004 | 0.01 | 0.81 | 0.68 | 0.77 | 0.81 | 0.79 | 0.87 |
| 8 | 136624184 | rs11166605 | 1.792 | -0.945 | 0.906 | -0.102 | -0.072 | -0.102 | -0.102 | 3.661 | 0.37 | 0.1 | 0.36 | 0.45 | 0.46 | 0.55 |
| 8 | 136636759 | rs4286981 | -1.011 | -0.901 | 0.925 | -0.087 | 0.031 | -0.04 | -0.04 | 0 | 0.91 | 0.83 | 0.88 | 0.99 | 0.85 | 0.97 |
| 8 | 136639174 | rs9657441 | 0.018 | -0.914 | 0.914 | -0.082 | -0.03 | 0.158 | 0.158 | 0 | 0.09 | 0.21 | 0.09 | 0.32 | 0.02 | 0.08 |
| 8 | 136646548 | rs4909494 | -0.07 | -0.884 | 0.548 | -0.259 | 0.007 | -0.097 | -0.097 | 0 | 0.58 | 0.52 | 0.66 | 0.96 | 0.54 | 0.74 |
| 8 | 136650653 | rs10081435 | -1.451 | -0.831 | 0.885 | 0.175 | -0.051 | 0.165 | 0.165 | 0.098 | 0.19 | 0.36 | 0.19 | 0.51 | 0.08 | 0.18 |
| 8 | 136659426 | rs3184618 | -0.697 | -0.738 | 0.562 | 0.136 | 0.03 | -0.039 | -0.039 | 0 | 0.91 | 0.85 | 0.89 | 0.99 | 0.85 | 0.96 |
| 8 | 136663746 | rs10104139 | -1.502 | -0.962 | 0.401 | 0.253 | -0.058 | 0.173 | 0.173 | 0.011 | 0.2 | 0.39 | 0.2 | 0.51 | 0.08 | 0.19 |
| 8 | 136666924 | rs10110397 | -0.728 | -0.572 | -0.181 | -0.062 | 0.01 | 0.07 | 0.07 | 0.039 | 0.74 | 0.9 | 0.85 | 0.98 | 0.81 | 0.77 |
| 8 | 136667194 | rs13254363 | 1.067 | -0.062 | 0.108 | 0.383 | -0.006 | -0.001 | -0.001 | 0 | 0.44 | 0.34 | 0.51 | 0.48 | 0.45 | 0.54 |
| 8 | 136671060 | rs4243847 | 0.666 | -0.042 | -0.13 | 0.692 | -0.035 | 0.24 | 0.24 | 0.056 | 0.87 | 0.66 | 0.85 | 0.48 | 0.93 | 0.85 |
| 8 | 136683843 | rs3935903 | -0.85 | 0.185 | -0.613 | 0.516 | 0.005 | -0.004 | -0.004 | 0 | 0.58 | 0.53 | 0.48 | 0.56 | 0.56 | 0.52 |
| 8 | 136686466 | rs7839810 | -0.761 | 0.695 | -0.124 | 1.683 | -0.002 | -0.002 | -0.002 | 0 | 0.17 | 0.18 | 0.18 | 0.19 | 0.25 | 0.12 |
| 8 | 136690813 | rs7002795 | 0.126 | 0.297 | -1.018 | 1.302 | -0.013 | -0.125 | -0.125 | 0 | 0.85 | 1 | 0.8 | 0.6 | 0.89 | 0.6 |
| 8 | 136695516 | rs7462758 | 0.128 | 0.31 | -0.49 | 0.501 | 0.002 | -0.044 | -0.044 | 0 | 0.16 | 0.29 | 0.08 | 0.07 | 0.13 | 0.11 |
| 8 | 136698907 | rs4075284 | -0.408 | 0.359 | -0.581 | 0.199 | 0 | -0.017 | -0.017 | 0 | 0.23 | 0.22 | 0.14 | 0.08 | 0.19 | 0.21 |
| 8 | 136705975 | rs6984541 | -0.722 | 0.402 | -0.403 | 0.103 | -0.001 | 0 | 0 | 0 | 0.32 | 0.31 | 0.37 | 0.5 | 0.26 | 0.54 |
| 8 | 136715743 | rs4909303 | -0.073 | 0.569 | -0.734 | -0.544 | -0.01 | 0.046 | 0.046 | 0 | 0.34 | 0.21 | 0.43 | 0.72 | 0.38 | 0.59 |
| 8 | 136716059 | rs7821328 | -0.053 | 0.543 | -0.709 | -0.556 | -0.032 | 0.072 | 0.072 | 0 | 0.34 | 0.1 | 0.42 | 0.72 | 0.38 | 0.6 |
| 8 | 136719754 | rs7823639 | -1.353 | 0.283 | -0.125 | -0.93 | -0.011 | 0.041 | 0.041 | 0 | 0.66 | 0.74 | 0.62 | 0.79 | 0.59 | 0.82 |
| 8 | 136721432 | rs7846197 | 0.09 | 0.267 | -0.004 | -0.855 | -0.014 | -0.059 | -0.059 | 0 | 0.24 | 0.59 | 0.15 | 0.1 | 0.16 | 0.14 |
| 8 | 136722239 | rs7004576 | -0.226 | 0.272 | -0.006 | -0.917 | 0.001 | 0.101 | 0.101 | 0 | 0.43 | 0.21 | 0.48 | 0.72 | 0.43 | 0.68 |
| 8 | 136725653 | rs11984564 | 0.062 | 0.445 | -0.205 | -1.235 | -0.002 | -0.06 | -0.06 | 0 | 0.29 | 0.61 | 0.18 | 0.07 | 0.22 | 0.15 |
| 8 | 136726750 | rs7814248 | 0.026 | 0.431 | -0.259 | -1.246 | 0.001 | 0.107 | 0.107 | 0 | 0.44 | 0.22 | 0.48 | 0.73 | 0.43 | 0.68 |
| 8 | 136730241 | rs12675240 | 1.389 | 0.415 | -0.239 | -1.235 | -0.005 | 0.023 | 0.023 | 0 | 0.16 | 0.08 | 0.07 | 0.07 | 0.13 | 0.13 |
| 8 | 136733007 | rs7830941 | -1.586 | 0.436 | -0.291 | -1.191 | -0.016 | 0.048 | 0.048 | 0.424 | 0.73 | 0.85 | 0.67 | 0.8 | 0.66 | 0.84 |
| 8 | 136735506 | rs7460819 | -1.454 | 0.441 | -0.244 | -1.203 | -0.009 | 0.046 | 0.046 | 0.892 | 0.72 | 0.84 | 0.67 | 0.8 | 0.66 | 0.83 |
| 8 | 136738372 | rs11993063 | 0.107 | 0.38 | 0.015 | -1.269 | -0.002 | -0.11 | -0.11 | 0 | 0.29 | 0.42 | 0.17 | 0.07 | 0.23 | 0.16 |
| 8 | 136738670 | rs10088869 | 0.612 | 0.439 | 0.127 | -1.163 | 0.015 | -0.085 | -0.085 | 0 | 0.13 | 0.37 | 0.1 | 0 | 0.09 | 0.02 |
| 8 | 136739516 | rs6981947 | -1.221 | 0.445 | -0.223 | -0.976 | 0.001 | -0.022 | -0.022 | 0 | 0.72 | 0.67 | 0.65 | 0.79 | 0.66 | 0.83 |
| 8 | 136743544 | rs7832101 | -1.311 | 0.486 | -0.909 | -0.926 | -0.008 | 0.032 | 0.032 | 0.177 | 0.73 | 0.84 | 0.71 | 0.81 | 0.66 | 0.87 |
| 8 | 136746031 | rs4268160 | 0.117 | 0.592 | -1.282 | -0.775 | -0.035 | 0.058 | 0.058 | 0 | 0.34 | 0.12 | 0.44 | 0.69 | 0.38 | 0.58 |
| 8 | 136748778 | rs9657442 | 0.718 | 0.348 | -1.29 | -0.295 | -0.028 | -0.069 | -0.069 | 0.175 | 0.19 | 0.02 | 0.16 | 0.2 | 0.2 | 0.18 |
| 8 | 136752206 | rs4354346 | -1.077 | 0.391 | -0.84 | -0.695 | 0.007 | 0.013 | 0.013 | 0 | 0.81 | 0.85 | 0.8 | 0.84 | 0.81 | 0.93 |
| 8 | 136755003 | rs6982188 | -0.91 | 0.342 | -0.864 | -0.719 | 0.011 | 0.018 | 0.018 | 0 | 0.8 | 0.85 | 0.8 | 0.84 | 0.81 | 0.93 |
| 8 | 136755673 | rs6577666 | 0.378 | 0.505 | -0.232 | -0.696 | 0.041 | -0.073 | -0.073 | 0 | 0.11 | 0.38 | 0.11 | 0 | 0.14 | 0.05 |
| 8 | 136756510 | rs4909515 | -0.534 | 0.692 | -0.547 | -0.835 | -0.01 | 0.091 | 0.091 | 0 | 0.62 | 0.88 | 0.69 | 0.72 | 0.65 | 0.79 |
| 8 | 136758163 | rs7008053 | 1.142 | 0.694 | -0.405 | -0.832 | 0.011 | -0.04 | -0.04 | 0 | 0.09 | 0.16 | 0.09 | 0 | 0.14 | 0.05 |
| 8 | 136758835 | rs4909306 | -0.487 | 0.65 | -0.504 | -0.838 | 0.011 | 0.058 | 0.058 | 0 | 0.6 | 0.74 | 0.68 | 0.73 | 0.64 | 0.79 |
| 8 | 136759839 | rs7828976 | -0.579 | 0.665 | -0.775 | -0.385 | -0.021 | -0.02 | -0.02 | 0 | 0.62 | 0.89 | 0.7 | 0.64 | 0.64 | 0.79 |
| 8 | 136767239 | rs7837809 | -0.621 | 0.664 | 0.01 | -0.384 | 0.036 | -0.118 | -0.118 | 0 | 0.17 | 0.44 | 0.17 | 0.02 | 0.17 | 0.12 |
| 8 | 136769999 | rs6577669 | 0.285 | 0.397 | -0.32 | -0.154 | -0.002 | 0.01 | 0.01 | 0 | 0.41 | 0.34 | 0.59 | 0.63 | 0.43 | 0.66 |
| 8 | 136770319 | rs4506254 | 0.697 | 0.134 | -0.352 | -0.4 | 0.009 | -0.002 | -0.002 | 0 | 0.19 | 0.16 | 0.06 | 0.16 | 0.09 | 0.07 |
| 8 | 136770869 | rs7822171 | 1.03 | 0.07 | -0.52 | 0.043 | -0.001 | -0.001 | -0.001 | 0.114 | 0.13 | 0.12 | 0.1 | 0.14 | 0.07 | 0.07 |
| 8 | 136770912 | rs4335161 | 0.006 | 0.182 | -0.343 | -0.058 | -0.004 | 0.015 | 0.015 | 0 | 0.43 | 0.37 | 0.6 | 0.66 | 0.46 | 0.71 |
| 8 | 136785992 | rs4406435 | 0.066 | 0.308 | -0.159 | 0.366 | 0.001 | -0.003 | -0.003 | 0.03 | 0.39 | 0.38 | 0.54 | 0.46 | 0.34 | 0.56 |
| 8 | 136788138 | rs4909520 | -0.105 | 0.039 | 0.348 | -0.189 | -0.001 | 0.011 | 0.011 | 0 | 0.51 | 0.54 | 0.58 | 0.69 | 0.48 | 0.72 |
| 8 | 136789772 | rs6991363 | 0.866 | 0.031 | 0.426 | -0.026 | 0 | -0.01 | -0.01 | 0 | 0.29 | 0.29 | 0.45 | 0.43 | 0.3 | 0.55 |
| 8 | 136792211 | rs10096833 | -0.907 | 0.333 | 0.479 | -0.338 | 0.005 | -0.003 | -0.003 | 0 | 0.73 | 0.8 | 0.73 | 0.69 | 0.73 | 0.84 |
| 8 | 136794579 | rs10464971 | 0.256 | 0.39 | 0.665 | -0.339 | 0.113 | 0.146 | 0.146 | 0 | 0.06 | 0.3 | 0.08 | 0.17 | 0.09 | 0.1 |
| 8 | 136807014 | rs12676004 | -0.463 | 0.285 | 0.291 | -0.398 | -0.006 | 0.064 | 0.064 | 0 | 0.47 | 0.64 | 0.31 | 0.29 | 0.47 | 0.35 |
| 8 | 136809038 | rs10090960 | -0.321 | 0.284 | 0.308 | -0.443 | 0.003 | -0.011 | -0.011 | 0 | 0.31 | 0.64 | 0.24 | 0.2 | 0.37 | 0.29 |
| 8 | 136811545 | rs11784836 | -0.347 | 0.249 | 0.718 | -0.323 | 0.029 | -0.009 | -0.009 | 0 | 0.31 | 0.47 | 0.29 | 0.31 | 0.38 | 0.22 |
| 8 | 136812571 | rs11785754 | 1.097 | 0.223 | 0.672 | -0.317 | 0.002 | -0.032 | -0.032 | 0 | 0.34 | 0.25 | 0.47 | 0.42 | 0.33 | 0.62 |
| 8 | 136816446 | rs11166618 | -1.889 | 0.139 | 0.562 | -0.277 | -0.017 | 0.005 | 0.005 | 0 | 0.85 | 0.93 | 0.83 | 0.83 | 0.8 | 0.91 |
| 8 | 136817274 | rs9644463 | -1.888 | 0.145 | 0.55 | -0.286 | -0.017 | 0.004 | 0.004 | 0 | 0.85 | 0.93 | 0.83 | 0.83 | 0.8 | 0.91 |
| 8 | 136822043 | rs4637882 | 0.227 | -0.021 | 0.808 | -0.337 | -0.001 | 0.005 | 0.005 | 0 | 0.55 | 0.62 | 0.56 | 0.53 | 0.44 | 0.69 |
| 8 | 136829636 | rs10086507 | 1.386 | 0.133 | 0.664 | -0.437 | 0.016 | -0.045 | -0.045 | 3.341 | 0.13 | 0.16 | 0.12 | 0 | 0.22 | 0.05 |
| 8 | 136832856 | rs12678843 | -0.408 | 0.019 | 0.613 | -0.428 | 0.003 | 0.005 | 0.005 | 0.051 | 0.45 | 0.44 | 0.28 | 0.24 | 0.36 | 0.2 |
| 8 | 136835925 | rs7830100 | 0.153 | -0.249 | 0.589 | -0.256 | -0.045 | -0.018 | -0.018 | 0 | 0.31 | 0.39 | 0.19 | 0.21 | 0.14 | 0.15 |
| 8 | 136838800 | rs4345596 | 0.634 | -0.075 | 0.428 | -0.006 | -0.001 | 0.002 | 0.002 | 0 | 0.46 | 0.41 | 0.52 | 0.36 | 0.53 | 0.57 |
| 8 | 136839229 | rs4288409 | -0.65 | 0.142 | 0.256 | 0.125 | -0.015 | -0.017 | -0.017 | 0 | 0.78 | 0.83 | 0.7 | 0.57 | 0.67 | 0.71 |
| 8 | 136843565 | rs4562356 | 1.158 | -0.291 | 0.288 | -0.033 | -0.003 | -0.002 | -0.002 | 0.142 | 0.12 | 0.11 | 0.09 | 0.14 | 0.05 | 0.06 |
| 8 | 136850133 | rs4351452 | 0.038 | -0.495 | 0.315 | -0.179 | 0.027 | 0.013 | 0.013 | 0 | 0.3 | 0.21 | 0.16 | 0.21 | 0.14 | 0.14 |
| 8 | 136855695 | rs7841070 | 1.447 | -0.259 | 0.098 | -0.15 | -0.006 | 0.031 | 0.031 | 0 | 0.16 | 0.01 | 0.06 | 0.08 | 0.09 | 0.08 |
| 8 | 136868367 | rs4339682 | -0.74 | 0.197 | 0.642 | -0.106 | 0.005 | -0.002 | -0.002 | 0 | 0.35 | 0.6 | 0.31 | 0.37 | 0.39 | 0.29 |
| 8 | 136868522 | rs4909528 | 0.786 | -0.223 | 0.691 | 0.048 | -0.045 | 0.014 | 0.014 | 0 | 0.13 | 0.23 | 0.1 | 0.13 | 0.04 | 0.06 |

| | | | | | | | | | | | | | | | | |
|---|---|---|---|---|---|---|---|---|---|---|---|---|---|---|---|---|
| 8 | 136871355 | rs7460887 | 1.051 | -0.047 | 0.59 | 0.429 | 0.002 | -0.01 | -0.01 | 0.016 | 0.3 | 0.19 | 0.4 | 0.34 | 0.27 | 0.47 |
| 8 | 136871739 | rs12680546 | 0.957 | 0.129 | -0.178 | 0.137 | -0.061 | 0.103 | 0.103 | 0 | 0.3 | 0.07 | 0.18 | 0.08 | 0.28 | 0.14 |
| 8 | 136874323 | rs4552942 | 0.79 | 0.108 | -0.049 | 0.127 | -0.022 | 0.093 | 0.093 | 0.425 | 0.31 | 0.11 | 0.19 | 0.08 | 0.28 | 0.14 |
| 8 | 136878098 | rs7826824 | 0.126 | 0.158 | -0.015 | 0.201 | 0.013 | 0.001 | 0.001 | 0 | 0.44 | 0.68 | 0.52 | 0.55 | 0.39 | 0.64 |
| 8 | 136880836 | rs7818542 | 0.089 | 0.179 | 0.175 | 0.245 | 0.01 | 0.034 | 0.034 | 0 | 0.07 | 0.14 | 0.07 | 0.19 | 0.08 | 0.12 |
| 8 | 136885072 | rs4433188 | -1.12 | 0.124 | 0.096 | 0.178 | -0.04 | 0.017 | 0.017 | 0 | 0.78 | 0.93 | 0.77 | 0.79 | 0.69 | 0.83 |
| 8 | 136887474 | rs11785202 | 0.554 | 0.237 | 0.228 | 0.249 | 0.004 | -0.003 | -0.003 | 0 | 0.38 | 0.59 | 0.46 | 0.36 | 0.31 | 0.53 |
| 8 | 136901085 | rs2317571 | 0.204 | 0.173 | 0.102 | 0.206 | -0.006 | -0.004 | -0.004 | 0 | 0.41 | 0.33 | 0.31 | 0.44 | 0.39 | 0.3 |
| 8 | 136902755 | rs13276788 | 0.669 | 0.116 | 0.095 | 0.197 | 0.002 | -0.002 | -0.002 | 0 | 0.38 | 0.58 | 0.46 | 0.37 | 0.3 | 0.52 |
| 8 | 136918435 | rs4463457 | -0.126 | 0.058 | 0.219 | 0.091 | 0.001 | 0.001 | 0.001 | 0 | 0.43 | 0.4 | 0.32 | 0.43 | 0.4 | 0.3 |
| 8 | 136925074 | rs6997347 | 0.619 | 0.131 | -0.274 | -0.303 | -0.013 | 0.096 | 0.096 | 0 | 0.31 | 0.11 | 0.2 | 0.08 | 0.29 | 0.13 |
| 8 | 136932517 | rs13275610 | 0.822 | 0.581 | 0.259 | 0.191 | -0.006 | -0.018 | -0.018 | 0 | 0.29 | 0.19 | 0.46 | 0.59 | 0.3 | 0.51 |
| 8 | 136932865 | rs1108711 | -0.06 | 0.531 | 0.366 | 0.063 | 0.018 | 0.02 | 0.02 | 0 | 0.72 | 0.55 | 0.74 | 0.74 | 0.64 | 0.77 |
| 8 | 136933622 | rs2317547 | -0.51 | 0.618 | 0.125 | 0.013 | -0.007 | 0.003 | 0.003 | 0 | 0.77 | 0.83 | 0.78 | 0.83 | 0.68 | 0.8 |
| 8 | 136938641 | rs2978220 | -0.448 | 0.657 | 0.128 | 0.027 | -0.002 | 0.003 | 0.003 | 0 | 0.76 | 0.75 | 0.78 | 0.83 | 0.67 | 0.81 |
| 8 | 136941348 | rs962157 | 0.536 | 0.537 | 0.014 | -0.053 | 0.029 | 0.151 | 0.151 | 0 | 0.41 | 0.15 | 0.26 | 0.14 | 0.34 | 0.26 |
| 8 | 136944696 | rs2317534 | -1.148 | 0.61 | 0.73 | 0.618 | 0.103 | 0.163 | 0.163 | 0.535 | 0.05 | 0.35 | 0.05 | 0.08 | 0.04 | 0.04 |
| 8 | 136957864 | rs1947181 | 0.636 | 0.55 | 0.142 | 0.411 | -0.001 | -0.001 | -0.001 | 0.5 | 0.69 | 0.28 | 0.69 | 0.76 | 0.64 | 0.77 |
| 8 | 136964642 | rs7016929 | 0.009 | 0.035 | -0.168 | -0.139 | 0.037 | 0.071 | 0.071 | 0 | 0.4 | 0.19 | 0.23 | 0.31 | 0.3 | 0.27 |
| 8 | 136974985 | rs13257256 | 0.117 | -0.083 | 0.249 | -0.245 | 0.031 | 0.101 | 0.101 | 0 | 0.38 | 0.08 | 0.21 | 0.3 | 0.3 | 0.28 |
| 8 | 136978770 | rs7012670 | 0.122 | -0.093 | 0.324 | -0.254 | 0.043 | 0.092 | 0.092 | 0 | 0.4 | 0.2 | 0.23 | 0.3 | 0.3 | 0.27 |
| 8 | 136985475 | rs2952054 | -0.141 | -0.197 | 0.747 | -0.321 | -0.018 | -0.053 | -0.053 | 0.298 | 0.43 | 0.51 | 0.27 | 0.22 | 0.29 | 0.27 |
| 8 | 136986585 | rs2122055 | -0.037 | -0.112 | 0.308 | -0.27 | 0.036 | 0.063 | 0.063 | 0 | 0.42 | 0.3 | 0.25 | 0.24 | 0.28 | 0.27 |
| 8 | 136991746 | rs1470615 | 0.269 | -0.03 | 0.109 | -0.273 | 0.055 | 0.157 | 0.157 | 0 | 0.38 | 0.12 | 0.22 | 0.1 | 0.28 | 0.23 |
| 8 | 136994952 | rs12335000 | -0.88 | 0.391 | 0.039 | -0.114 | 0.011 | 0.001 | 0.001 | 0 | 0.61 | 0.57 | 0.53 | 0.51 | 0.52 | 0.5 |
| 8 | 136996193 | rs2873443 | -0.552 | 0.531 | 0.143 | -0.022 | 0.003 | 0.021 | 0.021 | 0 | 0.22 | 0.34 | 0.31 | 0.41 | 0.25 | 0.28 |
| 8 | 136999297 | rs748843 | -0.977 | 0.446 | 0.012 | -0.065 | 0.016 | 0.004 | 0.004 | 0 | 0.63 | 0.57 | 0.54 | 0.51 | 0.53 | 0.51 |
| 8 | 136999912 | rs2084477 | -1.388 | 0.742 | -0.016 | 0.612 | 0.034 | 0.076 | 0.076 | 0.579 | 0.06 | 0.31 | 0.06 | 0.13 | 0.02 | 0.04 |
| 8 | 137007398 | rs6577698 | -0.726 | 0.741 | -0.167 | 0.482 | 0.001 | 0.009 | 0.009 | 0 | 0.21 | 0.27 | 0.29 | 0.27 | 0.23 | 0.23 |
| 8 | 137021343 | rs13259113 | -0.709 | 0.641 | -0.82 | 0.831 | -0.011 | -0.069 | -0.069 | 2.36 | 0.13 | 0.02 | 0.22 | 0.25 | 0.14 | 0.23 |
| 8 | 137023806 | rs10505642 | -0.406 | 0.693 | -0.663 | 0.324 | 0.005 | 0.054 | 0.054 | 0 | 0.48 | 0.35 | 0.64 | 0.66 | 0.53 | 0.64 |
| 8 | 137041406 | rs1159345 | 0.633 | 0.657 | -0.14 | 0.167 | -0.002 | -0.037 | -0.037 | 0.119 | 0.47 | 0.46 | 0.31 | 0.12 | 0.4 | 0.3 |
| 8 | 137042133 | rs13248990 | -0.738 | 0.589 | -0.041 | 0.077 | 0.037 | -0.025 | -0.025 | 0.248 | 0.68 | 0.54 | 0.6 | 0.29 | 0.62 | 0.53 |
| 8 | 137043988 | rs7001747 | 0.691 | 0.576 | 0.232 | 0.054 | -0.007 | 0.003 | 0.003 | 0 | 0.31 | 0.29 | 0.38 | 0.71 | 0.38 | 0.47 |
| 8 | 137044987 | rs11989223 | 0.998 | 0.555 | 0.217 | 0.053 | 0.008 | -0.001 | -0.001 | 0 | 0.48 | 0.39 | 0.3 | 0.12 | 0.4 | 0.31 |
| 8 | 137047614 | rs1900811 | 0.462 | 0.33 | -0.02 | 0.054 | 0.005 | 0.032 | 0.032 | 0 | 0.07 | 0.09 | 0.13 | 0.22 | 0.07 | 0.06 |
| 8 | 137050790 | rs2044661 | 1.115 | 0.146 | 0.403 | -0.035 | -0.007 | 0.01 | 0.01 | 0 | 0.7 | 0.45 | 0.54 | 0.61 | 0.71 | 0.72 |
| 8 | 137056293 | rs10505647 | 1.282 | 0.035 | 0.235 | -0.044 | 0 | 0.002 | 0.002 | 0 | 0.25 | 0.24 | 0.26 | 0.49 | 0.32 | 0.41 |
| 8 | 137062557 | rs4909552 | 0.05 | -0.001 | 0.018 | -0.049 | -0.049 | -0.007 | -0.007 | 0 | 0.17 | 0.01 | 0.27 | 0.17 | 0.2 | 0.18 |
| 8 | 137079029 | rs4909560 | -0.082 | 0.107 | 0.107 | -0.032 | -0.004 | 0.009 | 0.009 | 0 | 0.17 | 0.17 | 0.29 | 0.17 | 0.18 | 0.17 |
| 8 | 137091947 | rs2166944 | 1.487 | -0.282 | -0.261 | 0.34 | 0.009 | 0.068 | 0.068 | 0 | 0.35 | 0.17 | 0.26 | 0.11 | 0.35 | 0.25 |
| 8 | 137095292 | rs4909564 | -1.441 | -0.012 | -0.208 | 0.369 | 0.001 | 0.006 | 0.006 | 0 | 0.62 | 0.74 | 0.73 | 0.89 | 0.64 | 0.75 |
| 8 | 137096515 | rs1562635 | -0.174 | 0.302 | -0.526 | 0.648 | -0.005 | 0.042 | 0.042 | 0 | 0.42 | 0.31 | 0.43 | 0.72 | 0.44 | 0.58 |
| 8 | 137112133 | rs12548244 | 0.638 | 0.319 | -0.775 | 0.752 | -0.013 | -0.033 | -0.033 | 0 | 0.33 | 0.21 | 0.29 | 0.49 | 0.37 | 0.49 |
| 8 | 137112282 | rs1448697 | -0.687 | -0.02 | -0.735 | 1.168 | -0.012 | -0.058 | -0.058 | 0 | 0.1 | 0.01 | 0.13 | 0.24 | 0.07 | 0.09 |
| 8 | 137118961 | rs4909568 | 0.559 | 0.123 | -0.427 | 1.469 | 0.005 | 0.04 | 0.04 | 0 | 0.8 | 0.75 | 0.65 | 0.59 | 0.79 | 0.74 |
| 8 | 137123266 | rs12375319 | 0.292 | -0.131 | -0.93 | 1.33 | 0.003 | -0.035 | -0.035 | 0 | 0.22 | 0.06 | 0.16 | 0.21 | 0.25 | 0.39 |
| 8 | 137125320 | rs4551400 | 1.067 | -0.4 | -0.34 | 1.47 | 0.002 | 0.003 | 0.003 | 0 | 0.54 | 0.43 | 0.49 | 0.28 | 0.55 | 0.31 |
| 8 | 137128194 | rs4562359 | 0.574 | -0.523 | -0.474 | 1.473 | 0.005 | -0.033 | -0.033 | 0 | 0.22 | 0.06 | 0.16 | 0.21 | 0.25 | 0.4 |
| 8 | 137128340 | rs11775576 | 1.387 | -0.529 | -0.251 | 1.479 | 0.017 | 0.053 | 0.053 | 0 | 0.49 | 0.35 | 0.39 | 0.09 | 0.51 | 0.28 |
| 8 | 137129374 | rs6987454 | -0.644 | -0.494 | 0.291 | 1.847 | -0.008 | 0.072 | 0.072 | 0 | 0.22 | 0.35 | 0.33 | 0.49 | 0.2 | 0.29 |
| 8 | 137141233 | rs6577710 | 1.03 | -0.474 | 0.319 | 1.641 | -0.018 | 0.041 | 0.041 | 0 | 0.73 | 0.51 | 0.57 | 0.3 | 0.77 | 0.67 |
| 8 | 137142227 | rs16905510 | -0.184 | -0.253 | 0.422 | 1.491 | 0.002 | -0.003 | -0.003 | 0 | 0.16 | 0.22 | 0.29 | 0.16 | 0.17 | 0.18 |
| 8 | 137155913 | rs4909329 | -0.35 | -0.486 | 0.167 | 1.289 | 0 | 0 | 0 | 0 | 0.37 | 0.4 | 0.4 | 0.34 | 0.35 | 0.36 |
| 8 | 137156450 | rs10505648 | 1.114 | -0.665 | 0.328 | 1.128 | -0.006 | 0.077 | 0.077 | 0 | 0.44 | 0.28 | 0.36 | 0.09 | 0.52 | 0.27 |
| 8 | 137164996 | rs2317727 | 0.22 | -0.752 | 0.462 | 1.411 | -0.03 | -0.081 | -0.081 | 2.891 | 0.32 | 0.13 | 0.43 | 0.52 | 0.36 | 0.35 |
| 8 | 137165362 | rs2317726 | -0.889 | -0.771 | 0.515 | 1.441 | -0.001 | -0.008 | -0.008 | 0.152 | 0.51 | 0.52 | 0.58 | 0.79 | 0.47 | 0.66 |
| 8 | 137171813 | rs4415351 | 1.078 | -0.856 | 0.279 | 1.559 | 0.014 | -0.035 | -0.035 | 0 | 0.13 | 0.04 | 0.13 | 0.21 | 0.11 | 0.25 |
| 8 | 137185136 | rs12676891 | 1.288 | -0.854 | 0.256 | 1.56 | 0.007 | -0.053 | -0.053 | 1.024 | 0.12 | 0.02 | 0.13 | 0.22 | 0.11 | 0.25 |
| 8 | 137196003 | rs7835733 | 1.528 | -0.718 | 0.218 | 1.171 | -0.013 | 0.015 | 0.015 | 0 | 0.33 | 0.24 | 0.2 | 0.03 | 0.36 | 0.21 |
| 8 | 137201005 | rs6981818 | -0.392 | -0.724 | 0.162 | 1.251 | 0.003 | 0.078 | 0.078 | 0 | 0.5 | 0.38 | 0.63 | 0.79 | 0.5 | 0.67 |
| 8 | 137210949 | rs2873494 | 0.547 | -0.71 | 0.046 | 1.384 | -0.032 | -0.036 | -0.036 | 0 | 0.32 | 0.13 | 0.44 | 0.34 | 0.36 | 0.37 |
| 8 | 137211259 | rs7007471 | -1.42 | -0.707 | 0.091 | 1.321 | 0 | 0.001 | 0.001 | 0 | 0.89 | 0.89 | 0.91 | 0.95 | 0.87 | 0.96 |
| 8 | 137218971 | rs4909582 | -0.91 | -0.554 | 0.309 | 1.593 | -0.001 | 0.003 | 0.003 | 0 | 0.53 | 0.57 | 0.65 | 0.79 | 0.5 | 0.68 |
| 8 | 137240873 | rs16905528 | 0.506 | -0.439 | -0.312 | 1.865 | -0.001 | -0.039 | -0.039 | 0 | 0.08 | 0 | 0.12 | 0.25 | 0.08 | 0.21 |
| 8 | 137267394 | rs11781934 | -0.144 | -0.429 | -0.219 | 1.626 | -0.011 | -0.008 | -0.008 | 0 | 0.65 | 0.76 | 0.53 | 0.66 | 0.62 | 0.61 |
| 8 | 137275863 | rs11166644 | 0.448 | -0.712 | -0.459 | 1.962 | -0.012 | 0.015 | 0.015 | 0 | 0.15 | 0.21 | 0.15 | 0.22 | 0.13 | 0.26 |
| 8 | 137278841 | rs13272742 | 0.43 | -0.701 | -0.455 | 1.789 | -0.012 | 0.014 | 0.014 | 0 | 0.15 | 0.21 | 0.15 | 0.22 | 0.13 | 0.26 |
| 8 | 137281470 | rs11166646 | -0.779 | -0.816 | -0.371 | 1.643 | -0.037 | 0 | 0 | 0 | 0.28 | 0.43 | 0.34 | 0.71 | 0.23 | 0.5 |

| | | | | | | | | | | | | | | | | |
|---|---|---|---|---|---|---|---|---|---|---|---|---|---|---|---|---|
| 8 | 137282949 | rs7844595 | -0.113 | -0.59 | -0.539 | 1.763 | -0.023 | 0.075 | 0.075 | 0 | 0.19 | 0.32 | 0.21 | 0.55 | 0.15 | 0.27 |
| 8 | 137286723 | rs13260793 | 0.71 | -0.762 | -1.336 | 1.33 | 0.006 | 0.001 | 0.001 | 0 | 0.12 | 0.07 | 0.11 | 0.12 | 0.09 | 0.13 |
| 8 | 137286985 | rs1492985 | -1.164 | -0.58 | -1.453 | 1.004 | 0.033 | 0.029 | 0.029 | 0 | 0.69 | 0.53 | 0.61 | 0.55 | 0.65 | 0.73 |
| 8 | 137290624 | rs17620825 | 0.58 | -0.351 | -2.571 | 0.939 | -0.063 | 0.004 | 0.004 | 0 | 0.35 | 0.09 | 0.35 | 0.22 | 0.41 | 0.37 |
| 8 | 137292810 | rs7832973 | 0.446 | 0.108 | -1.941 | 1.487 | 0.006 | 0.001 | 0.001 | 0 | 0.12 | 0.07 | 0.11 | 0.12 | 0.09 | 0.13 |
| 8 | 137297600 | rs13250040 | -0.32 | 0.103 | -1.944 | 1.13 | -0.015 | -0.006 | -0.006 | 0 | 0.62 | 0.24 | 0.57 | 0.55 | 0.62 | 0.73 |
| 8 | 137302684 | rs13272404 | -0.003 | 0.285 | -2.124 | 0.26 | -0.049 | 0.167 | 0.167 | 0 | 0.47 | 0.1 | 0.3 | 0.22 | 0.48 | 0.43 |
| 8 | 137309449 | rs2201229 | -0.574 | 0.11 | -1.547 | 0.907 | 0.014 | -0.043 | -0.043 | 0 | 0.18 | 0.11 | 0.31 | 0.5 | 0.16 | 0.37 |
| 8 | 137311996 | rs1389081 | 0.044 | -0.315 | -0.634 | 0.508 | -0.007 | -0.035 | -0.035 | 0 | 0.93 | 0.74 | 0.93 | 0.99 | 0.96 | 0.88 |
| 8 | 137313825 | rs1873817 | -0.684 | -0.997 | -0.656 | -0.696 | 0.004 | 0.001 | 0.001 | 0 | 0.48 | 0.42 | 0.45 | 0.52 | 0.36 | 0.52 |
| 8 | 137316285 | rs993325 | 1.084 | -1.011 | -0.53 | -0.715 | 0.019 | -0.009 | -0.009 | 0.057 | 0.43 | 0.16 | 0.43 | 0.46 | 0.58 | 0.34 |
| 8 | 137317540 | rs9644476 | 0.963 | 0.107 | -0.955 | -0.835 | -0.054 | -0.032 | -0.032 | 0.673 | 0.11 | 0.03 | 0.15 | 0.22 | 0.15 | 0.08 |
| 8 | 137318613 | rs902957 | -0.948 | -0.066 | -0.664 | -0.851 | -0.019 | 0.052 | 0.052 | 0 | 0.56 | 0.66 | 0.64 | 0.75 | 0.5 | 0.62 |
| 8 | 137319350 | rs923791 | 0.364 | 0.053 | -0.73 | -0.858 | 0.004 | -0.028 | -0.028 | 0 | 0.19 | 0.14 | 0.32 | 0.53 | 0.19 | 0.38 |
| 8 | 137321806 | rs16905602 | 0.686 | 0.166 | -0.372 | -0.93 | 0.002 | 0.001 | 0.001 | 0 | 0.08 | 0.07 | 0.13 | 0.21 | 0.11 | 0.06 |
| 8 | 137340282 | rs7826564 | 0.246 | -0.721 | 0.506 | -1.194 | 0.022 | 0.002 | 0.002 | 0 | 0.27 | 0.19 | 0.12 | 0.02 | 0.17 | 0.16 |
| 8 | 137343147 | rs7822138 | 0.397 | -0.863 | 0.579 | -1.281 | -0.009 | -0.071 | -0.071 | 0 | 0.65 | 0.54 | 0.78 | 0.96 | 0.76 | 0.7 |
| 8 | 137346996 | rs16905668 | 0.285 | -0.841 | 0.261 | -1.258 | 0.037 | 0.117 | 0.117 | 0 | 0.25 | 0.09 | 0.1 | 0.02 | 0.17 | 0.15 |
| 8 | 137350456 | rs4909593 | 0.686 | -0.84 | 0.403 | -1.23 | -0.039 | -0.13 | -0.13 | 0 | 0.63 | 0.47 | 0.8 | 0.98 | 0.78 | 0.72 |
| 8 | 137351689 | rs16905672 | 0.082 | -0.568 | 0.055 | -0.939 | -0.004 | -0.052 | -0.052 | 2.099 | 0.26 | 0.23 | 0.13 | 0.02 | 0.17 | 0.17 |
| 8 | 137353807 | rs6984599 | 0.056 | -0.571 | -0.063 | -0.943 | -0.006 | -0.057 | -0.057 | 0.003 | 0.26 | 0.26 | 0.12 | 0.02 | 0.17 | 0.16 |
| 8 | 137354700 | rs7842238 | 0.839 | -0.336 | 0.524 | -0.48 | 0 | -0.067 | -0.067 | 0.957 | 0.81 | 0.44 | 0.71 | 0.88 | 0.85 | 0.8 |
| 8 | 137358265 | rs13265887 | -0.245 | -0.239 | 0.501 | 0.317 | 0.002 | 0.021 | 0.021 | 0 | 0.23 | 0.22 | 0.25 | 0.49 | 0.26 | 0.28 |
| 8 | 137383553 | rs215153 | 1.007 | -0.595 | 0.992 | 0.979 | -0.016 | 0.124 | 0.124 | 0 | 0.54 | 0.16 | 0.42 | 0.24 | 0.54 | 0.43 |
| 8 | 137390958 | rs302298 | -0.002 | -0.506 | 1.302 | 1.015 | 0.01 | 0.081 | 0.081 | 0 | 0.76 | 0.66 | 0.73 | 0.52 | 0.74 | 0.71 |
| 8 | 137408734 | rs7835274 | 1.859 | -0.494 | -0.038 | 1.079 | -0.116 | -0.084 | -0.084 | 0 | 0.25 | 0.05 | 0.31 | 0.34 | 0.35 | 0.28 |
| 8 | 137419150 | rs6577727 | -1.201 | -0.837 | -0.18 | 0.833 | -0.118 | -0.04 | -0.04 | 0.003 | 0.52 | 0.95 | 0.6 | 0.6 | 0.47 | 0.56 |
| 8 | 137424202 | rs17566877 | 0.2 | -0.858 | -0.013 | 0.863 | 0.001 | -0.001 | -0.001 | 0.249 | 0.2 | 0.17 | 0.2 | 0.17 | 0.2 | 0.18 |
| 8 | 137426590 | rs7833279 | 0.414 | -0.654 | 0.155 | 0.232 | 0.001 | 0.036 | 0.036 | 0.071 | 0.48 | 0.63 | 0.67 | 0.65 | 0.53 | 0.59 |
| 8 | 137428405 | rs2119235 | 2.066 | -0.642 | -0.227 | 0.272 | -0.078 | -0.102 | -0.102 | 0 | 0.26 | 0.05 | 0.32 | 0.36 | 0.32 | 0.35 |
| 8 | 137430757 | rs1545201 | -0.419 | -0.78 | -0.016 | 0.179 | 0.012 | 0.028 | 0.028 | 0 | 0.72 | 0.95 | 0.88 | 0.8 | 0.78 | 0.79 |
| 8 | 137445034 | rs28897205 | 0.027 | -0.834 | 0.185 | -0.081 | 0 | -0.002 | -0.002 | 0 | 0.21 | 0.31 | 0.23 | 0.17 | 0.21 | 0.17 |
| 8 | 137464965 | rs7012640 | 0.003 | -0.788 | -0.259 | -0.063 | 0.001 | -0.002 | -0.002 | 0 | 0.21 | 0.31 | 0.24 | 0.17 | 0.22 | 0.17 |
| 8 | 137465022 | rs1371749 | -0.57 | -0.987 | -0.806 | -0.355 | 0.061 | 0.123 | 0.123 | 0 | 0.75 | 0.97 | 0.91 | 0.92 | 0.8 | 0.84 |
| 8 | 137485507 | rs4909596 | 0.153 | -0.521 | -0.967 | -0.028 | -0.006 | 0.013 | 0.013 | 0 | 0.63 | 0.83 | 0.7 | 0.71 | 0.65 | 0.77 |
| 8 | 137488037 | rs13269920 | 0.183 | -0.442 | -0.424 | -0.277 | -0.005 | -0.031 | -0.031 | 0.1 | 0.15 | 0.2 | 0.14 | 0.12 | 0.12 | 0.14 |
| 8 | 137492260 | rs875367 | 0.233 | -0.377 | -0.723 | -0.408 | -0.021 | 0.063 | 0.063 | 0 | 0.18 | 0.03 | 0.15 | 0.03 | 0.18 | 0.14 |
| 8 | 137493215 | rs875366 | 0.084 | -0.51 | -0.51 | -0.295 | 0.047 | 0.022 | 0.022 | 0 | 0.53 | 0.26 | 0.64 | 0.55 | 0.62 | 0.61 |
| 8 | 137493361 | rs1812882 | 1.378 | -0.432 | -0.832 | -0.255 | -0.134 | -0.135 | -0.135 | 1.426 | 0.32 | 0.08 | 0.48 | 0.52 | 0.44 | 0.43 |
| 8 | 137509472 | rs2932212 | -0.249 | -0.701 | 0.079 | -0.157 | -0.03 | 0.006 | 0.006 | 0 | 0.66 | 0.54 | 0.78 | 0.67 | 0.74 | 0.72 |
| 8 | 137511102 | rs1371746 | -0.466 | -0.639 | -0.38 | -0.126 | 0.002 | -0.001 | -0.001 | 1.12 | 0.68 | 0.71 | 0.79 | 0.67 | 0.74 | 0.76 |
| 8 | 137518037 | rs1899065 | -0.687 | -0.215 | 0.337 | -0.332 | -0.014 | -0.052 | -0.052 | 0 | 0.17 | 0.35 | 0.14 | 0.14 | 0.12 | 0.18 |
| 8 | 137522988 | rs1866698 | -0.314 | -0.217 | -0.909 | -0.177 | 0 | 0.004 | 0.004 | 0 | 0.09 | 0.03 | 0.09 | 0.04 | 0.09 | 0.14 |
| 8 | 137523046 | rs1971833 | -0.817 | -0.18 | -0.877 | -0.162 | 0.015 | 0.02 | 0.02 | 0 | 0.27 | 0.74 | 0.26 | 0.17 | 0.19 | 0.34 |
| 8 | 137526799 | rs2660665 | -0.031 | 0.441 | -0.671 | 0.111 | -0.001 | 0 | 0 | 0 | 0.42 | 0.54 | 0.51 | 0.53 | 0.43 | 0.66 |
| 8 | 137529123 | rs4909345 | -0.26 | 0.171 | -0.486 | -0.409 | 0.002 | -0.023 | -0.023 | 0 | 0.58 | 0.54 | 0.68 | 0.78 | 0.59 | 0.76 |
| 8 | 137538931 | rs1978894 | -0.469 | 0.188 | -0.175 | -0.467 | -0.043 | 0.052 | 0.052 | 0 | 0.63 | 0.84 | 0.72 | 0.78 | 0.61 | 0.77 |
| 8 | 137543379 | rs7837030 | 0.197 | 0.02 | -0.023 | -0.434 | 0.04 | 0.053 | 0.053 | 0.118 | 0.49 | 0.75 | 0.56 | 0.66 | 0.45 | 0.68 |
| 8 | 137545501 | rs1444027 | 1.046 | 0.2 | -0.259 | -0.433 | -0.05 | -0.185 | -0.185 | 0.001 | 0.36 | 0.02 | 0.39 | 0.53 | 0.32 | 0.63 |
| 8 | 137553132 | rs2587851 | 0.533 | 0.188 | -0.248 | -0.308 | 0.001 | -0.007 | -0.007 | 0 | 0.06 | 0.05 | 0.13 | 0.11 | 0.07 | 0.04 |
| 8 | 137554206 | rs2660666 | -1.116 | 0.15 | -0.1 | -0.452 | -0.003 | 0.003 | 0.003 | 0 | 0.27 | 0.7 | 0.33 | 0.31 | 0.24 | 0.15 |
| 8 | 137561504 | rs305262 | 0.862 | 0.143 | -0.09 | -0.455 | -0.003 | 0.003 | 0.003 | 0 | 0.73 | 0.3 | 0.67 | 0.69 | 0.76 | 0.84 |
| 8 | 137564311 | rs1837696 | -0.509 | 0.145 | 0.352 | -0.467 | 0.032 | 0.08 | 0.08 | 0.573 | 0.07 | 0.41 | 0.09 | 0.09 | 0.02 | 0.02 |
| 8 | 137586334 | rs1372069 | -0.054 | 0.22 | 0.399 | -0.468 | 0.003 | -0.046 | -0.046 | 1.333 | 0.16 | 0.31 | 0.15 | 0.09 | 0.16 | 0.09 |
| 8 | 137595285 | rs7815091 | -0.304 | 0.217 | 0.53 | -0.456 | 0.045 | 0.096 | 0.096 | 0.085 | 0.07 | 0.46 | 0.09 | 0.09 | 0.02 | 0.02 |
| 8 | 137600653 | rs305234 | 0.635 | 0.3 | 0.443 | -0.493 | -0.012 | 0.026 | 0.026 | 0 | 0.38 | 0.27 | 0.28 | 0.16 | 0.44 | 0.21 |
| 8 | 137604801 | rs305230 | 0.269 | 0.431 | 1.188 | -0.068 | 0 | 0.004 | 0.004 | 0 | 0.92 | 0.91 | 0.83 | 0.78 | 0.92 | 0.94 |
| 8 | 137613619 | rs305241 | 0.46 | 0.389 | 1.1 | -0.028 | 0 | 0.002 | 0.002 | 0 | 0.79 | 0.75 | 0.76 | 0.77 | 0.78 | 0.87 |
| 8 | 137617180 | rs305246 | 0.316 | 0.391 | 0.813 | -0.003 | -0.008 | -0.019 | -0.019 | 0 | 0.8 | 0.84 | 0.76 | 0.77 | 0.78 | 0.87 |
| 8 | 137653054 | rs305348 | 0.977 | 0.402 | 0.946 | 0.055 | -0.007 | 0.016 | 0.016 | 0 | 0.74 | 0.34 | 0.67 | 0.68 | 0.76 | 0.84 |
| 8 | 137664271 | rs305366 | 0.988 | 0.417 | 0.3 | 0.064 | -0.004 | 0.008 | 0.008 | 0 | 0.74 | 0.3 | 0.69 | 0.68 | 0.76 | 0.84 |
| 8 | 137670528 | rs305297 | 0.442 | 0.87 | -0.7 | 0.181 | -0.008 | -0.018 | -0.018 | 0 | 0.81 | 0.83 | 0.76 | 0.77 | 0.78 | 0.87 |
| 8 | 137672952 | rs305301 | 0.437 | 0.876 | -0.699 | 0.184 | -0.007 | -0.016 | -0.016 | 0 | 0.81 | 0.82 | 0.76 | 0.77 | 0.78 | 0.87 |
| 8 | 137677083 | rs305306 | 0.216 | 0.898 | -0.926 | -0.125 | -0.002 | 0.003 | 0.003 | 0 | 0.69 | 0.73 | 0.63 | 0.73 | 0.69 | 0.74 |
| 8 | 137677221 | rs10102048 | -0.517 | 0.655 | -1.433 | -0.192 | -0.014 | -0.016 | -0.016 | 0.364 | 0.12 | 0.03 | 0.14 | 0.13 | 0.11 | 0.11 |
| 8 | 137677675 | rs894091 | 0.292 | 0.88 | -1.611 | -0.297 | -0.005 | -0.002 | -0.002 | 0.088 | 0.21 | 0.22 | 0.31 | 0.22 | 0.33 | 0.25 |
| 8 | 137677896 | rs894089 | -0.339 | 0.579 | -0.923 | -0.852 | 0.012 | -0.021 | -0.021 | 0 | 0.31 | 0.49 | 0.37 | 0.22 | 0.36 | 0.27 |
| 8 | 137679805 | rs6991548 | 0.356 | 0.049 | -1.225 | -0.654 | -0.011 | -0.019 | -0.019 | 0 | 0.21 | 0.27 | 0.17 | 0.18 | 0.17 | 0.17 |
| 8 | 137682484 | rs305307 | -0.776 | -0.279 | -1.802 | -0.378 | -0.006 | 0.074 | 0.074 | 0.715 | 0.78 | 0.61 | 0.7 | 0.55 | 0.82 | 0.68 |

| | | | | | | | | | | | | | | | | |
|---|---|---|---|---|---|---|---|---|---|---|---|---|---|---|---|---|
| 8 | 137687538 | rs894087 | 1.586 | -0.314 | -1.651 | -0.441 | 0.003 | 0.001 | 0.001 | 0 | 0.2 | 0.24 | 0.3 | 0.22 | 0.29 | 0.25 |
| 8 | 137688230 | rs732943 | 1.732 | -0.322 | -1.652 | -0.445 | -0.012 | -0.008 | -0.008 | 0 | 0.19 | 0.18 | 0.29 | 0.21 | 0.29 | 0.28 |
| 8 | 137697835 | rs305276 | -0.309 | -0.37 | -1.386 | -0.224 | -0.007 | 0.003 | 0.003 | 0 | 0.76 | 0.69 | 0.74 | 0.73 | 0.8 | 0.81 |
| 8 | 137703525 | rs2318172 | 1.709 | -0.4 | -1.377 | -0.276 | -0.011 | -0.071 | -0.071 | 0.7 | 0.11 | 0.26 | 0.07 | 0 | 0.07 | 0.02 |
| 8 | 137720395 | rs16906244 | 1.366 | -0.453 | -1.417 | -0.302 | -0.02 | -0.083 | -0.083 | 0 | 0.13 | 0.33 | 0.07 | 0 | 0.07 | 0.02 |
| 8 | 137727717 | rs2577049 | 1.067 | -0.587 | -1.602 | -0.417 | -0.107 | -0.13 | -0.13 | 0.008 | 0.43 | 0.04 | 0.36 | 0.51 | 0.43 | 0.54 |
| 8 | 137730280 | rs2577047 | -0.487 | -0.65 | -1.556 | -0.449 | -0.014 | 0.128 | 0.128 | 0 | 0.88 | 0.64 | 0.81 | 0.77 | 0.93 | 0.88 |
| 8 | 137734805 | rs17587101 | 1.79 | -0.591 | -1.703 | -0.436 | -0.02 | 0.062 | 0.062 | 0.127 | 0.15 | 0.03 | 0.07 | 0.05 | 0.14 | 0.08 |
| 8 | 137742872 | rs2262343 | 0.48 | -0.583 | -1.7 | -0.458 | -0.001 | -0.002 | -0.002 | 0.55 | 0.55 | 0.37 | 0.43 | 0.52 | 0.5 | 0.56 |
| 8 | 137745177 | rs2245160 | 0.485 | -0.579 | -1.772 | -0.459 | -0.051 | -0.112 | -0.112 | 0.796 | 0.58 | 0.07 | 0.43 | 0.56 | 0.57 | 0.62 |
| 8 | 137747692 | rs4909606 | 2.566 | -0.515 | -1.749 | -0.463 | -0.05 | -0.055 | -0.055 | 0 | 0.12 | 0.01 | 0.11 | 0.22 | 0.12 | 0.21 |
| 8 | 137754784 | rs3850497 | 1.437 | -0.613 | -1.524 | -0.509 | 0.001 | 0.035 | 0.035 | 0.03 | 0.15 | 0.11 | 0.08 | 0.05 | 0.14 | 0.08 |
| 8 | 137769663 | rs10505664 | 1.489 | -0.637 | -1.485 | -0.507 | -0.02 | -0.077 | -0.077 | 4.731 | 0.12 | 0.26 | 0.08 | 0 | 0.07 | 0.02 |
| 8 | 137771602 | rs2610082 | 1.086 | -0.598 | -1.562 | -0.459 | -0.052 | -0.066 | -0.066 | 0 | 0.44 | 0.12 | 0.36 | 0.51 | 0.43 | 0.54 |
| 8 | 137772759 | rs2610084 | 0.135 | -0.633 | -1.429 | -0.451 | -0.04 | 0.005 | 0.005 | 0.153 | 0.77 | 0.44 | 0.73 | 0.77 | 0.86 | 0.85 |
| 8 | 137776239 | rs17594630 | 1.515 | -0.657 | -1.439 | -0.48 | -0.008 | 0.064 | 0.064 | 0 | 0.16 | 0.06 | 0.08 | 0.05 | 0.14 | 0.08 |
| 8 | 137776731 | rs2582478 | 0.131 | -0.662 | -1.434 | -0.477 | -0.04 | 0.004 | 0.004 | 0.203 | 0.77 | 0.44 | 0.73 | 0.77 | 0.86 | 0.85 |
| 8 | 137779385 | rs2582475 | 1.146 | -0.655 | -1.45 | -0.475 | -0.055 | -0.068 | -0.068 | 0 | 0.44 | 0.11 | 0.36 | 0.51 | 0.43 | 0.54 |
| 8 | 137784415 | rs2253765 | -0.413 | -0.552 | -1.3 | -0.398 | -0.021 | 0.098 | 0.098 | 0 | 0.89 | 0.69 | 0.8 | 0.78 | 0.93 | 0.88 |
| 8 | 137787569 | rs2610093 | -0.09 | -0.459 | -1.648 | -0.344 | -0.04 | 0.004 | 0.004 | 0.115 | 0.77 | 0.44 | 0.73 | 0.77 | 0.86 | 0.85 |
| 8 | 137790603 | rs17595046 | 1.638 | -0.429 | -1.68 | -0.347 | -0.02 | 0.061 | 0.061 | 0.055 | 0.14 | 0.02 | 0.07 | 0.05 | 0.14 | 0.08 |
| 8 | 137791478 | rs17595144 | 1.415 | -0.424 | -1.677 | -0.345 | -0.008 | 0.064 | 0.064 | 0 | 0.16 | 0.06 | 0.08 | 0.05 | 0.14 | 0.08 |
| 8 | 137798133 | rs2582465 | 1.142 | -0.447 | -1.73 | -0.358 | -0.119 | -0.139 | -0.139 | 0 | 0.4 | 0.04 | 0.35 | 0.51 | 0.43 | 0.54 |
| 8 | 137801130 | rs2249732 | 1.22 | -0.445 | -1.692 | -0.336 | -0.063 | -0.075 | -0.075 | 0.402 | 0.41 | 0.11 | 0.36 | 0.51 | 0.43 | 0.54 |
| 8 | 137802814 | rs3850500 | 1.645 | -0.423 | -1.624 | -0.294 | 0.007 | 0.002 | 0.002 | 0 | 0.21 | 0.27 | 0.3 | 0.21 | 0.29 | 0.23 |
| 8 | 137806774 | rs2613837 | 0.813 | -0.464 | -1.66 | -0.341 | 0.032 | 0.032 | 0.032 | 1.107 | 0.61 | 0.29 | 0.61 | 0.72 | 0.7 | 0.78 |
| 8 | 137806865 | rs2582458 | 1.253 | -0.42 | -1.739 | -0.351 | -0.069 | -0.091 | -0.091 | 0 | 0.4 | 0.1 | 0.35 | 0.51 | 0.42 | 0.55 |
| 8 | 137808181 | rs2610104 | 0.688 | -0.395 | -1.53 | -0.188 | -0.001 | -0.056 | -0.056 | 2.742 | 0.56 | 0.16 | 0.44 | 0.56 | 0.57 | 0.64 |
| 8 | 137811273 | rs2245810 | 0.623 | -0.38 | -1.54 | -0.18 | -0.001 | -0.061 | -0.061 | 1.034 | 0.55 | 0.13 | 0.43 | 0.56 | 0.57 | 0.64 |
| 8 | 137811319 | rs6995884 | 1.321 | -0.393 | -1.479 | -0.193 | 0.008 | 0.003 | 0.003 | 0 | 0.22 | 0.3 | 0.28 | 0.21 | 0.29 | 0.22 |
| 8 | 137822965 | rs10505665 | -0.198 | -0.524 | -0.985 | -0.113 | -0.056 | -0.06 | -0.06 | 0 | 0.19 | 0.49 | 0.18 | 0.13 | 0.12 | 0.12 |
| 8 | 137838235 | rs7842212 | -0.386 | 0.02 | -0.875 | -0.143 | 0 | 0.002 | 0.002 | 0 | 0.38 | 0.37 | 0.28 | 0.33 | 0.34 | 0.41 |
| 8 | 137838984 | rs12114965 | 1.211 | -0.033 | -0.945 | -0.254 | -0.002 | 0.027 | 0.027 | 0.046 | 0.15 | 0.12 | 0.09 | 0.05 | 0.15 | 0.08 |
| 8 | 137840400 | rs2681672 | 0.597 | -0.112 | -0.671 | -0.167 | -0.005 | -0.005 | -0.005 | 0.514 | 0.58 | 0.48 | 0.65 | 0.6 | 0.63 | 0.58 |
| 8 | 137843383 | rs10093517 | -0.509 | -0.09 | -0.835 | -0.154 | 0.001 | 0.008 | 0.008 | 0 | 0.38 | 0.34 | 0.28 | 0.33 | 0.34 | 0.41 |
| 8 | 137846185 | rs17650786 | 1.77 | -0.027 | -0.727 | -0.236 | -0.001 | -0.016 | -0.016 | 0.005 | 0.08 | 0.05 | 0.05 | 0 | 0.06 | 0.02 |
| 8 | 137847061 | rs7825584 | 1.77 | -0.027 | -0.714 | -0.236 | -0.002 | -0.021 | -0.021 | 0 | 0.08 | 0.06 | 0.05 | 0 | 0.06 | 0.02 |
| 8 | 137848316 | rs17650875 | 0.008 | -0.086 | -0.74 | -0.08 | 0 | 0.003 | 0.003 | 0 | 0.31 | 0.31 | 0.24 | 0.34 | 0.28 | 0.4 |
| 8 | 137850503 | rs7016048 | -0.979 | -0.061 | -0.721 | -0.062 | -0.01 | 0.005 | 0.005 | 0 | 0.89 | 0.83 | 0.89 | 0.93 | 0.91 | 0.98 |
| 8 | 137851655 | rs17597626 | 1.768 | 0.014 | -0.809 | -0.079 | -0.001 | -0.014 | -0.014 | 0 | 0.08 | 0.06 | 0.05 | 0 | 0.06 | 0.02 |
| 8 | 137852251 | rs10092767 | -0.008 | 0.045 | -0.837 | -0.029 | 0 | 0.001 | 0.001 | 0 | 0.3 | 0.31 | 0.24 | 0.34 | 0.28 | 0.4 |
| 8 | 137857327 | rs10107003 | 1.288 | 0.068 | -0.797 | 0.033 | -0.001 | 0 | 0 | 0 | 0.15 | 0.16 | 0.15 | 0.29 | 0.13 | 0.3 |
| 8 | 137876402 | rs17598411 | 1.212 | 0.065 | -0.826 | -0.063 | -0.001 | 0.017 | 0.017 | 0.008 | 0.15 | 0.11 | 0.08 | 0.05 | 0.15 | 0.08 |
| 8 | 137886896 | rs3850504 | 1.393 | -0.033 | -1.201 | -0.166 | -0.024 | 0.01 | 0.01 | 0 | 0.31 | 0.08 | 0.25 | 0.27 | 0.31 | 0.34 |
| 8 | 137894507 | rs17045 | 0.713 | -0.012 | -0.714 | -0.222 | -0.004 | 0.058 | 0.058 | 0.475 | 0.21 | 0.07 | 0.13 | 0.05 | 0.21 | 0.09 |
| 8 | 137899228 | rs3850507 | -0.832 | 0.007 | -0.702 | -0.161 | -0.003 | 0.051 | 0.051 | 0 | 0.8 | 0.69 | 0.78 | 0.64 | 0.83 | 0.65 |
| 8 | 137905065 | rs12676794 | 1.009 | -0.085 | -0.728 | -0.21 | 0.001 | -0.007 | -0.007 | 0 | 0.17 | 0.16 | 0.16 | 0.28 | 0.14 | 0.33 |
| 8 | 137907651 | rs16906528 | 1.889 | -0.1 | -0.989 | -0.459 | 0.006 | 0.004 | 0.004 | 0.024 | 0.14 | 0.06 | 0.08 | 0 | 0.12 | 0.06 |
| 8 | 137913324 | rs6981340 | -0.71 | 0.524 | -0.843 | -0.017 | -0.003 | 0.049 | 0.049 | 0 | 0.81 | 0.7 | 0.78 | 0.63 | 0.83 | 0.65 |
| 8 | 137938502 | rs4909614 | -0.108 | 0.131 | -0.848 | -0.243 | -0.006 | -0.023 | -0.023 | 0.023 | 0.74 | 0.7 | 0.81 | 0.87 | 0.77 | 0.89 |
| 8 | 137943303 | rs1846525 | -1.368 | 0.349 | -0.474 | 0.284 | -0.005 | 0.021 | 0.021 | 0 | 0.86 | 0.76 | 0.87 | 0.73 | 0.89 | 0.75 |
| 8 | 137943783 | rs2019131 | 0.647 | 0.645 | -0.455 | 0.375 | -0.036 | -0.037 | -0.037 | 0 | 0.26 | 0.15 | 0.34 | 0.3 | 0.39 | 0.28 |
| 8 | 137949781 | rs1499446 | -0.05 | 0.658 | -0.273 | 0.008 | 0.007 | 0.032 | 0.032 | 0 | 0.5 | 0.39 | 0.49 | 0.43 | 0.47 | 0.44 |
| 8 | 137977335 | rs1499451 | -0.486 | 0.844 | -0.677 | 0.082 | -0.002 | -0.009 | -0.009 | 0 | 0.49 | 0.49 | 0.42 | 0.24 | 0.59 | 0.33 |
| 8 | 137979848 | rs6577751 | -1.341 | 0.612 | -0.856 | 0.491 | 0.025 | -0.055 | -0.055 | 0 | 0.69 | 0.81 | 0.71 | 0.61 | 0.77 | 0.6 |
| 8 | 137991169 | rs7845945 | -0.331 | 0.962 | -0.552 | 0.473 | -0.017 | 0.054 | 0.054 | 0 | 0.42 | 0.71 | 0.35 | 0.23 | 0.46 | 0.3 |
| 8 | 137995401 | rs1499449 | 1.135 | 0.858 | -0.624 | 0.462 | -0.04 | -0.116 | -0.116 | 0.159 | 0.39 | 0.07 | 0.35 | 0.39 | 0.36 | 0.43 |
| 8 | 138005370 | rs10090384 | 0.784 | 1.14 | -0.404 | 0.608 | -0.001 | 0.001 | 0.001 | 0 | 0.18 | 0.16 | 0.28 | 0.38 | 0.18 | 0.26 |
| 8 | 138006031 | rs10103797 | 0.942 | 1.141 | -0.646 | 0.618 | -0.008 | -0.029 | -0.029 | 0 | 0.17 | 0.09 | 0.27 | 0.38 | 0.18 | 0.26 |
| 8 | 138018770 | rs4366108 | -0.455 | 1.241 | -0.911 | 0.599 | 0.012 | -0.044 | -0.044 | 0 | 0.69 | 0.66 | 0.78 | 0.88 | 0.63 | 0.79 |
| 8 | 138020661 | rs6988114 | 1.064 | 1.107 | -1.195 | 0.801 | -0.019 | -0.052 | -0.052 | 0 | 0.42 | 0.07 | 0.32 | 0.35 | 0.36 | 0.42 |
| 8 | 138040724 | rs16906685 | 0.631 | 1.653 | -1.167 | 0.585 | 0.003 | 0.032 | 0.032 | 0.13 | 0.49 | 0.15 | 0.58 | 0.53 | 0.56 | 0.55 |
| 8 | 138040743 | rs9324441 | 0.628 | 1.734 | -0.922 | 0.627 | 0.001 | 0.047 | 0.047 | 0 | 0.19 | 0.35 | 0.23 | 0.36 | 0.18 | 0.24 |
| 8 | 138041153 | rs10096434 | 0.068 | 1.825 | -0.666 | 0.369 | -0.015 | -0.062 | -0.062 | 0 | 0.7 | 0.62 | 0.83 | 0.89 | 0.74 | 0.81 |
| 8 | 138046288 | rs6986650 | -0.788 | 2.136 | -1.096 | 0.891 | 0.011 | -0.001 | -0.001 | 0.345 | 0.06 | 0.06 | 0.17 | 0.12 | 0.19 | 0.12 |
| 8 | 138055195 | rs10096383 | -0.753 | 1.426 | -1.646 | 0.578 | 0.01 | 0.083 | 0.083 | 0.899 | 0.51 | 0.85 | 0.54 | 0.71 | 0.55 | 0.52 |
| 8 | 138056227 | rs9650553 | -0.549 | 1.419 | -0.56 | 0.567 | 0.014 | 0.025 | 0.025 | 0.01 | 0.38 | 0.47 | 0.38 | 0.58 | 0.43 | 0.39 |
| 8 | 138058748 | rs12542475 | -0.305 | 1.238 | -0.285 | 0.546 | 0.001 | 0.006 | 0.006 | 0 | 0.3 | 0.27 | 0.31 | 0.65 | 0.32 | 0.36 |
| 8 | 138062943 | rs11784156 | -0.321 | 1.112 | -0.171 | 0.439 | -0.001 | 0.016 | 0.016 | 0 | 0.14 | 0.07 | 0.05 | 0.08 | 0.1 | 0.07 |

| | | | | | | | | | | | | | | | | |
|---|---|---|---|---|---|---|---|---|---|---|---|---|---|---|---|---|
| 8 | 138075823 | rs7014510 | -0.768 | 0.987 | 0.988 | 0.699 | -0.005 | 0.082 | 0.082 | 0 | 0.16 | 0.24 | 0.18 | 0.55 | 0.15 | 0.29 |
| 8 | 138081290 | rs4295697 | -0.974 | 1.214 | 1.393 | 0.608 | 0.13 | 0.094 | 0.094 | 0.141 | 0.07 | 0.39 | 0.11 | 0.04 | 0.12 | 0.02 |
| 8 | 138103060 | rs6988964 | -2.286 | 0.782 | 0.913 | 0.425 | -0.008 | -0.003 | -0.003 | 0 | 0.42 | 0.9 | 0.39 | 0.64 | 0.38 | 0.39 |
| 8 | 138105524 | rs16906754 | -1.429 | 0.681 | 1.372 | 0.526 | -0.011 | -0.044 | -0.044 | 0 | 0.18 | 0.24 | 0.12 | 0.08 | 0.11 | 0.09 |
| 8 | 138110573 | rs12541799 | -0.419 | 0.678 | 1.479 | 0.528 | -0.022 | -0.043 | -0.043 | 0.582 | 0.08 | 0.15 | 0.06 | 0 | 0.03 | 0.02 |
| 8 | 138111042 | rs16906756 | -0.57 | 0.681 | 1.622 | 0.607 | -0.011 | 0.112 | 0.112 | 0 | 0.15 | 0.23 | 0.11 | 0.52 | 0.12 | 0.28 |
| 8 | 138114639 | rs16906760 | -1.542 | 0.743 | 1.631 | 0.449 | 0.083 | 0.062 | 0.062 | 0.41 | 0.08 | 0.44 | 0.12 | 0.04 | 0.13 | 0.03 |
| 8 | 138121678 | rs11780634 | -2.047 | 0.824 | 1.58 | 0.511 | 0 | -0.037 | -0.037 | 0.51 | 0.25 | 0.41 | 0.23 | 0.15 | 0.24 | 0.11 |
| 8 | 138135202 | rs4527908 | -0.662 | 0.459 | 1.127 | 0.16 | -0.016 | 0.196 | 0.196 | 0 | 0.2 | 0.5 | 0.15 | 0.54 | 0.18 | 0.3 |
| 8 | 138135936 | rs7827430 | -2.099 | 0.444 | 1.062 | 0.099 | 0.037 | -0.004 | -0.004 | 0.108 | 0.39 | 0.83 | 0.29 | 0.66 | 0.3 | 0.39 |
| 8 | 138145239 | rs16906787 | 0.077 | 0.587 | 1.131 | 0.153 | 0.017 | 0.022 | 0.022 | 0 | 0.09 | 0.04 | 0.05 | 0 | 0.04 | 0.03 |
| 8 | 138146684 | rs16906788 | 0.43 | 0.604 | 1.154 | 0.178 | 0.004 | 0.001 | 0.001 | 0 | 0.07 | 0.07 | 0.07 | 0.11 | 0.08 | 0.04 |
| 8 | 138148917 | rs16906793 | -0.219 | 0.54 | 1.166 | 0.179 | -0.014 | 0.277 | 0.277 | 0 | 0.16 | 0.47 | 0.1 | 0.51 | 0.12 | 0.29 |
| 8 | 138153441 | rs10282908 | -0.147 | 0.626 | 0.855 | 0.049 | -0.025 | -0.053 | -0.053 | 0.263 | 0.11 | 0.17 | 0.07 | 0 | 0.04 | 0.03 |
| 8 | 138159231 | rs13248077 | -0.322 | 0.828 | 0.733 | 0.145 | 0.067 | 0.073 | 0.073 | 0 | 0.08 | 0.23 | 0.11 | 0.15 | 0.13 | 0.04 |
| 8 | 138170251 | rs3112824 | 1.865 | 0.697 | 0.521 | 0.251 | 0.001 | -0.008 | -0.008 | 0 | 0.6 | 0.12 | 0.66 | 0.36 | 0.63 | 0.62 |
| 8 | 138179102 | rs11166687 | 0.306 | 0.792 | 0.225 | 0.008 | -0.001 | 0 | 0 | 0 | 0.09 | 0.06 | 0.07 | 0.12 | 0.07 | 0.03 |
| 8 | 138179125 | rs1012332 | -1.632 | 0.905 | 0.345 | -0.108 | 0.005 | -0.031 | -0.031 | 0 | 0.24 | 0.35 | 0.22 | 0.16 | 0.24 | 0.08 |
| 8 | 138181704 | rs2978607 | 1.875 | 1.03 | -0.382 | -0.239 | 0.011 | 0.02 | 0.02 | 0 | 0.33 | 0.04 | 0.36 | 0.26 | 0.28 | 0.48 |
| 8 | 138182862 | rs10088280 | 0.21 | 0.926 | 0.359 | -0.152 | -0.029 | -0.052 | -0.052 | 0.101 | 0.1 | 0.17 | 0.06 | 0 | 0.03 | 0.03 |
| 8 | 138189660 | rs11785515 | -1.384 | 1.133 | 0.284 | -0.151 | 0.022 | -0.025 | -0.025 | 6.212 | 0.21 | 0.38 | 0.24 | 0.16 | 0.2 | 0.08 |
| 8 | 138197117 | rs4626644 | -1.79 | 1.186 | 0.289 | -0.127 | 0.031 | -0.043 | -0.043 | 0 | 0.25 | 0.43 | 0.3 | 0.16 | 0.28 | 0.09 |
| 8 | 138199228 | rs11780467 | -1.276 | 1.081 | 0.278 | -0.128 | -0.011 | -0.046 | -0.046 | 9.449 | 0.22 | 0.34 | 0.19 | 0.12 | 0.18 | 0.07 |
| 8 | 138203140 | rs4909633 | -0.149 | 1.039 | 0.301 | 0.305 | 0.003 | 0.04 | 0.04 | 0 | 0.83 | 0.73 | 0.81 | 0.67 | 0.82 | 0.72 |
| 8 | 138209669 | rs7012808 | -0.638 | 1.106 | 0.106 | 0.073 | 0.004 | -0.048 | -0.048 | 0 | 0.15 | 0.25 | 0.14 | 0.04 | 0.17 | 0.05 |
| 8 | 138213836 | rs16906838 | 0.32 | 1.005 | -0.15 | 0.037 | -0.007 | -0.033 | -0.033 | 1.889 | 0.11 | 0.11 | 0.06 | 0 | 0.05 | 0.06 |
| 8 | 138213978 | rs9644431 | 1.348 | 0.983 | -0.336 | 0.103 | 0.001 | -0.022 | -0.022 | 0.523 | 0.57 | 0.3 | 0.58 | 0.47 | 0.57 | 0.65 |
| 8 | 138219589 | rs7830180 | 0.387 | 0.722 | -0.405 | 0.449 | 0.001 | 0.002 | 0.002 | 0 | 0.87 | 0.82 | 0.89 | 0.86 | 0.87 | 0.88 |
| 8 | 138220815 | rs4909356 | 0.71 | 0.879 | -0.713 | 0.513 | -0.007 | -0.032 | -0.032 | 0 | 0.75 | 0.65 | 0.83 | 0.86 | 0.8 | 0.82 |
| 8 | 138222987 | rs4481636 | 0.278 | 1.135 | -1.166 | 0.992 | 0.005 | -0.01 | -0.01 | 0.636 | 0.43 | 0.36 | 0.49 | 0.47 | 0.38 | 0.59 |
| 8 | 138223632 | rs4243865 | 0.42 | 1.184 | -1.19 | 0.967 | -0.013 | -0.015 | -0.015 | 0 | 0.94 | 0.97 | 0.95 | 0.92 | 0.93 | 0.96 |
| 8 | 138228075 | rs4909357 | -0.099 | 0.94 | -1.566 | 1.164 | -0.012 | -0.089 | -0.089 | 2.074 | 0.22 | 0.09 | 0.31 | 0.41 | 0.26 | 0.38 |
| 8 | 138231791 | rs17668646 | 0.313 | 0.023 | -1.993 | 0.294 | 0.026 | 0.018 | 0.018 | 0 | 0.18 | 0.1 | 0.06 | 0 | 0.07 | 0.06 |
| 8 | 138235253 | rs10112576 | 0.931 | 0.176 | -1.242 | -0.27 | 0.011 | -0.054 | -0.054 | 0.25 | 0.15 | 0.34 | 0.1 | 0.05 | 0.15 | 0.14 |
| 8 | 138239862 | rs13282991 | 0.271 | 0.951 | -0.99 | -0.093 | 0.004 | 0.019 | 0.019 | 0 | 0.25 | 0.18 | 0.14 | 0.17 | 0.21 | 0.13 |
| 8 | 138241127 | rs4313191 | 0.184 | 0.881 | -1.06 | -0.067 | -0.001 | 0.007 | 0.007 | 0 | 0.38 | 0.44 | 0.41 | 0.46 | 0.41 | 0.52 |
| 8 | 138241598 | rs9644432 | 1.644 | 1.16 | -0.865 | -0.284 | 0 | -0.008 | -0.008 | 0.073 | 0.24 | 0.23 | 0.38 | 0.3 | 0.22 | 0.26 |
| 8 | 138243039 | rs9324442 | -0.532 | 1.169 | -0.856 | -0.275 | 0.003 | 0.003 | 0.003 | 0 | 0.63 | 0.6 | 0.55 | 0.63 | 0.62 | 0.65 |
| 8 | 138243207 | rs9324443 | -0.185 | 1.221 | -0.741 | -0.586 | 0.002 | -0.007 | -0.007 | 0 | 0.7 | 0.75 | 0.87 | 0.93 | 0.77 | 0.86 |
| 8 | 138243618 | rs16906888 | 0.436 | 1.328 | -0.278 | -0.481 | 0.027 | -0.007 | -0.007 | 0 | 0.22 | 0.31 | 0.17 | 0.22 | 0.29 | 0.22 |
| 8 | 138246221 | rs6987191 | 1.285 | 1.376 | -0.26 | -0.432 | 0.029 | 0.034 | 0.034 | 0 | 0.07 | 0.12 | 0.08 | 0.17 | 0.12 | 0.08 |
| 8 | 138253889 | rs6577763 | 0.317 | 1.09 | 0.19 | -0.693 | 0.007 | 0.032 | 0.032 | 3.712 | 0.55 | 0.4 | 0.73 | 0.88 | 0.6 | 0.72 |
| 8 | 138254326 | rs4909638 | -0.563 | 1.072 | 0.488 | -0.732 | 0.033 | -0.024 | -0.024 | 0 | 0.86 | 0.71 | 0.88 | 0.93 | 0.83 | 0.91 |
| 8 | 138260171 | rs11780629 | 0.38 | 1.067 | 0.475 | -0.741 | 0.007 | 0.032 | 0.032 | 3.712 | 0.55 | 0.4 | 0.73 | 0.88 | 0.61 | 0.72 |
| 8 | 138262220 | rs7006748 | -0.541 | 1.066 | 0.465 | -0.731 | 0.033 | -0.024 | -0.024 | 0 | 0.86 | 0.71 | 0.88 | 0.93 | 0.83 | 0.91 |
| 8 | 138263180 | rs7818464 | 1.525 | 1.086 | 0.39 | -0.691 | 0.019 | 0.022 | 0.022 | 0 | 0.07 | 0.09 | 0.08 | 0.17 | 0.12 | 0.08 |
| 8 | 138264020 | rs7824123 | 1.143 | 1.13 | 0.37 | -0.687 | 0.019 | 0.044 | 0.044 | 3.103 | 0.43 | 0.17 | 0.59 | 0.64 | 0.37 | 0.6 |
| 8 | 138264848 | rs7018106 | 0.391 | 1.036 | 0.539 | -0.787 | -0.006 | 0.007 | 0.007 | 0.287 | 0.67 | 0.37 | 0.72 | 0.81 | 0.56 | 0.74 |
| 8 | 138266983 | rs4634676 | 0.349 | 0.816 | 0.508 | -0.779 | -0.002 | -0.061 | -0.061 | 0 | 0.56 | 0.51 | 0.75 | 0.88 | 0.6 | 0.72 |
| 8 | 138271518 | rs7002431 | 1.43 | 0.699 | 0.119 | -0.656 | 0 | -0.011 | -0.011 | 0 | 0.14 | 0.16 | 0.09 | 0.05 | 0.15 | 0.14 |
| 8 | 138273650 | rs4446770 | -0.269 | 0.743 | 0.291 | -0.701 | 0.015 | -0.041 | -0.041 | 0.181 | 0.77 | 0.59 | 0.85 | 0.88 | 0.78 | 0.79 |
| 8 | 138273952 | rs16906917 | 1.846 | 0.548 | 0.009 | -0.244 | -0.006 | -0.056 | -0.056 | 3.034 | 0.23 | 0.17 | 0.35 | 0.39 | 0.22 | 0.26 |
| 8 | 138280429 | rs4330721 | -0.83 | 0.765 | 0.475 | -0.485 | -0.014 | -0.011 | -0.011 | 0 | 0.92 | 0.96 | 0.96 | 0.93 | 0.93 | 0.94 |
| 8 | 138281807 | rs6987993 | -0.981 | 0.79 | 0.5 | -0.473 | -0.009 | -0.007 | -0.007 | 0 | 0.93 | 0.95 | 0.96 | 0.93 | 0.93 | 0.94 |
| 8 | 138282361 | rs11774039 | 1.343 | 0.722 | 0.397 | -0.495 | 0.017 | 0.045 | 0.045 | 3.374 | 0.43 | 0.19 | 0.59 | 0.64 | 0.37 | 0.6 |
| 8 | 138286806 | rs11997699 | -0.364 | 0.721 | 0.999 | -0.203 | 0.086 | 0.017 | 0.017 | 0 | 0.18 | 0.35 | 0.21 | 0.24 | 0.33 | 0.13 |
| 8 | 138287866 | rs4410940 | 1.111 | 0.55 | 1.049 | -0.268 | -0.007 | 0.014 | 0.014 | 0.746 | 0.58 | 0.44 | 0.69 | 0.69 | 0.53 | 0.75 |
| 8 | 138295559 | rs11166700 | 0.314 | 1.063 | 1.307 | 0.044 | 0.117 | 0.068 | 0.068 | 0 | 0.81 | 0.65 | 0.75 | 0.69 | 0.62 | 0.82 |
| 8 | 138300484 | rs16906944 | -0.467 | 0.943 | 1.383 | 0.046 | 0.004 | -0.051 | -0.051 | 0.01 | 0.09 | 0.25 | 0.05 | 0 | 0.05 | 0.02 |
| 8 | 138304288 | rs10505671 | -0.075 | 0.974 | 1.238 | 0.061 | 0 | -0.046 | -0.046 | 2.824 | 0.08 | 0.18 | 0.04 | 0 | 0.05 | 0.02 |
| 8 | 138305248 | rs4909642 | 0.355 | 1.037 | 0.987 | 0.141 | 0.087 | 0.044 | 0.044 | 0 | 0.81 | 0.66 | 0.76 | 0.7 | 0.64 | 0.81 |
| 8 | 138309026 | rs4909643 | 1.041 | 0.818 | 0.54 | 0.183 | 0.002 | 0.001 | 0.001 | 0 | 0.71 | 0.33 | 0.59 | 0.64 | 0.62 | 0.74 |
| 8 | 138309596 | rs4909644 | 0.782 | 0.867 | 0.833 | 0.278 | 0.059 | 0.055 | 0.055 | 0 | 0.76 | 0.43 | 0.62 | 0.62 | 0.64 | 0.75 |
| 8 | 138309956 | rs17669995 | 0.133 | 0.463 | 0.302 | -0.101 | 0.045 | 0.111 | 0.111 | 1.352 | 0.25 | 0.07 | 0.09 | 0 | 0.12 | 0.09 |
| 8 | 138312015 | rs4413808 | 0.72 | 0.601 | 0.344 | 0.004 | 0.036 | 0.035 | 0.035 | 0.003 | 0.75 | 0.39 | 0.61 | 0.62 | 0.64 | 0.75 |
| 8 | 138315178 | rs4288411 | 1.073 | 0.652 | -0.2 | 0.164 | -0.007 | 0.074 | 0.074 | 0 | 0.5 | 0.25 | 0.52 | 0.62 | 0.53 | 0.67 |
| 8 | 138315997 | rs7829832 | 0.833 | 0.658 | -0.174 | 0.066 | 0.004 | 0.081 | 0.081 | 0 | 0.54 | 0.29 | 0.67 | 0.68 | 0.62 | 0.69 |
| 8 | 138325276 | rs13261938 | -0.821 | 0.314 | 0.228 | 0.099 | -0.01 | -0.017 | -0.017 | 0 | 0.23 | 0.7 | 0.18 | 0.13 | 0.11 | 0.1 |
| 8 | 138333969 | rs3887806 | -0.276 | -0.084 | 0.001 | -0.656 | 0.005 | -0.007 | -0.007 | 1.353 | 0.41 | 0.39 | 0.26 | 0.15 | 0.28 | 0.26 |

| | | | | | | | | | | | | | | | |
|---|---|---|---|---|---|---|---|---|---|---|---|---|---|---|---|
| 8 | 138337729 | rs7837472 | 1.141 | -0.117 | -0.611 | -0.69 | 0.001 | 0.07 | 0.07 | 0.045 | 0.22 | 0.13 | 0.15 | 0.05 | 0.19 | 0.15 |
| 8 | 138338162 | rs6577777 | -0.458 | 0.01 | -0.65 | -0.527 | 0.068 | 0.216 | 0.216 | 1.643 | 0.45 | 0.85 | 0.31 | 0.17 | 0.28 | 0.26 |
| 8 | 138353376 | rs12114107 | -0.449 | 0.324 | 0.022 | 0.19 | -0.011 | -0.008 | -0.008 | 0 | 0.26 | 0.3 | 0.19 | 0.28 | 0.19 | 0.24 |
| 8 | 138354352 | rs13279979 | 0.55 | 0.472 | -0.222 | 0.638 | 0.009 | 0.009 | 0.009 | 0 | 0.51 | 0.6 | 0.6 | 0.51 | 0.57 | 0.58 |
| 8 | 138364362 | rs13249103 | 0.673 | 0.407 | 0.169 | 0.598 | 0.008 | 0.013 | 0.013 | 0 | 0.49 | 0.28 | 0.55 | 0.51 | 0.56 | 0.56 |
| 8 | 138368191 | rs9324446 | -1.369 | 0.443 | -0.124 | 0.634 | -0.029 | -0.01 | -0.01 | 1.855 | 0.77 | 0.89 | 0.78 | 0.8 | 0.75 | 0.78 |
| 8 | 138372696 | rs10097104 | -0.505 | 0.408 | 0.213 | 0.529 | -0.028 | -0.015 | -0.015 | 0.298 | 0.27 | 0.53 | 0.22 | 0.3 | 0.2 | 0.22 |
| 8 | 138376355 | rs16907008 | 0.576 | 0.685 | -0.037 | 0.382 | -0.005 | -0.045 | -0.045 | 1.11 | 0.1 | 0.19 | 0.06 | 0.02 | 0.08 | 0.03 |
| 8 | 138376514 | rs6421006 | -0.839 | 0.844 | -0.504 | 0.851 | -0.035 | -0.053 | -0.053 | 0 | 0.77 | 0.9 | 0.79 | 0.75 | 0.75 | 0.79 |
| 8 | 138379427 | rs4909649 | -0.752 | 0.746 | -0.193 | 0.918 | -0.001 | -0.001 | -0.001 | 0.701 | 0.82 | 0.8 | 0.82 | 0.74 | 0.81 | 0.81 |
| 8 | 138379666 | rs4909364 | 0.329 | 0.872 | -0.306 | 0.839 | -0.019 | -0.003 | -0.003 | 0 | 0.14 | 0.06 | 0.14 | 0.12 | 0.18 | 0.07 |
| 8 | 138379927 | rs7017772 | 0.564 | 0.241 | -0.417 | 0.465 | -0.007 | -0.003 | -0.003 | 0 | 0.35 | 0.26 | 0.41 | 0.43 | 0.4 | 0.46 |
| 8 | 138382230 | rs4579564 | -0.2 | 0.084 | -0.06 | 0.437 | 0.003 | 0.005 | 0.005 | 0 | 0.66 | 0.61 | 0.67 | 0.64 | 0.63 | 0.74 |
| 8 | 138386546 | rs12541665 | -1.304 | 0.087 | -0.006 | 0.446 | 0.001 | 0.014 | 0.014 | 0 | 0.79 | 0.67 | 0.81 | 0.78 | 0.81 | 0.81 |
| 8 | 138387871 | rs11787252 | 0.781 | 0.026 | 0.025 | 0.382 | -0.045 | -0.033 | -0.033 | 0.059 | 0.33 | 0.12 | 0.39 | 0.43 | 0.39 | 0.46 |
| 8 | 138391587 | rs17626729 | 1.307 | 0.247 | -0.337 | 0.233 | -0.033 | 0.023 | 0.023 | 0 | 0.16 | 0.07 | 0.28 | 0.08 | 0.29 | 0.13 |
| 8 | 138404925 | rs4909666 | 0.19 | 0.162 | -0.231 | 0.22 | 0.008 | 0.013 | 0.013 | 0 | 0.46 | 0.41 | 0.39 | 0.33 | 0.42 | 0.35 |
| 8 | 138414687 | rs2960117 | 0.123 | 0.193 | -0.123 | 0.319 | 0.006 | 0.009 | 0.009 | 0 | 0.37 | 0.39 | 0.43 | 0.45 | 0.4 | 0.47 |
| 8 | 138420975 | rs10109851 | 1.02 | 0.133 | -0.218 | 0.288 | -0.004 | -0.003 | -0.003 | 0 | 0.31 | 0.33 | 0.25 | 0.23 | 0.24 | 0.28 |
| 8 | 138421223 | rs1514212 | -0.853 | 0.174 | -0.459 | 0.307 | -0.012 | -0.006 | -0.006 | 0 | 0.69 | 0.73 | 0.68 | 0.68 | 0.64 | 0.75 |
| 8 | 138426274 | rs2943191 | 0.148 | 0.123 | -0.368 | 0.315 | -0.017 | -0.022 | -0.022 | 0.256 | 0.35 | 0.23 | 0.41 | 0.45 | 0.4 | 0.47 |
| 8 | 138429552 | rs2960100 | -1.627 | 0.223 | -0.279 | 0.32 | 0 | -0.001 | -0.001 | 0 | 0.8 | 0.83 | 0.83 | 0.8 | 0.82 | 0.81 |
| 8 | 138437870 | rs2960106 | -0.652 | 0.325 | -0.312 | 0.347 | -0.006 | -0.006 | -0.006 | 0 | 0.68 | 0.75 | 0.68 | 0.66 | 0.66 | 0.77 |
| 8 | 138444604 | rs2960109 | 0.728 | 0.493 | -0.182 | 0.221 | -0.002 | -0.006 | -0.006 | 0 | 0.35 | 0.37 | 0.29 | 0.21 | 0.31 | 0.3 |
| 8 | 138449358 | rs1514194 | 0.314 | 0.633 | -0.635 | 0.326 | -0.076 | -0.03 | -0.03 | 0 | 0.25 | 0.1 | 0.38 | 0.29 | 0.4 | 0.2 |
| 8 | 138454656 | rs2960110 | 1.558 | 0.187 | -0.839 | -0.108 | -0.009 | 0.109 | 0.109 | 0.278 | 0.2 | 0.03 | 0.18 | 0.01 | 0.2 | 0.15 |
| 8 | 138455004 | rs9324449 | 1.452 | 0.24 | -0.649 | 0.184 | 0.018 | -0.017 | -0.017 | 0 | 0.18 | 0.15 | 0.12 | 0.27 | 0.11 | 0.32 |
| 8 | 138456460 | rs11166722 | 2.013 | 0.789 | -0.809 | 0.145 | -0.024 | 0.021 | 0.021 | 0 | 0.1 | 0.04 | 0.1 | 0.03 | 0.19 | 0.04 |
| 8 | 138457891 | rs17679798 | 1.608 | 0.616 | -0.815 | 0.169 | 0.031 | -0.067 | -0.067 | 0.747 | 0.18 | 0.03 | 0.11 | 0.26 | 0.11 | 0.34 |
| 8 | 138459740 | rs13438943 | 0.859 | 0.674 | -0.491 | 0.218 | -0.004 | -0.004 | -0.004 | 0.097 | 0.31 | 0.28 | 0.3 | 0.33 | 0.33 | 0.21 |
| 8 | 138461211 | rs10505679 | 1.088 | 0.785 | -0.443 | 0.231 | -0.005 | -0.001 | -0.001 | 0.621 | 0.15 | 0.12 | 0.27 | 0.18 | 0.24 | 0.15 |
| 8 | 138464355 | rs964944 | -0.006 | 0.65 | -0.302 | 0.199 | 0 | 0.001 | 0.001 | 0.913 | 0.51 | 0.51 | 0.44 | 0.6 | 0.45 | 0.55 |
| 8 | 138466049 | rs11780660 | 0.554 | 0.802 | -0.618 | 0.253 | -0.017 | -0.006 | -0.006 | 0.17 | 0.34 | 0.25 | 0.44 | 0.4 | 0.41 | 0.31 |
| 8 | 138467144 | rs9693212 | -0.187 | 0.646 | -0.588 | 0.268 | 0.005 | -0.002 | -0.002 | 0 | 0.46 | 0.43 | 0.57 | 0.51 | 0.58 | 0.36 |
| 8 | 138468688 | rs4909369 | 0.694 | 0.632 | -0.604 | 0.272 | 0.001 | 0.001 | 0.001 | 0 | 0.32 | 0.32 | 0.31 | 0.33 | 0.33 | 0.21 |
| 8 | 138469175 | rs16907232 | 0.763 | 0.548 | -0.799 | 0.275 | 0.03 | -0.054 | -0.054 | 0 | 0.37 | 0.17 | 0.4 | 0.51 | 0.29 | 0.51 |
| 8 | 138473321 | rs1399173 | -0.049 | 0.583 | -0.715 | 0.253 | 0.028 | -0.01 | -0.01 | 0 | 0.45 | 0.29 | 0.57 | 0.51 | 0.58 | 0.36 |
| 8 | 138475068 | rs6981649 | 1.913 | 0.49 | -0.789 | 0.348 | -0.03 | -0.016 | -0.016 | 0 | 0.12 | 0.03 | 0.12 | 0.09 | 0.16 | 0.05 |
| 8 | 138476014 | rs11166725 | 1.791 | 0.503 | -0.824 | 0.332 | -0.007 | -0.021 | -0.021 | 0 | 0.19 | 0.09 | 0.17 | 0.23 | 0.18 | 0.16 |
| 8 | 138477671 | rs1514208 | 0.425 | 0.255 | -0.678 | 0.374 | 0.011 | -0.035 | -0.035 | 0 | 0.3 | 0.19 | 0.23 | 0.38 | 0.26 | 0.4 |
| 8 | 138487768 | rs17681845 | 1.966 | 0.295 | -0.782 | 0.379 | -0.052 | -0.034 | -0.034 | 0 | 0.11 | 0.01 | 0.12 | 0.09 | 0.16 | 0.05 |
| 8 | 138488244 | rs11166726 | 0.613 | 0.256 | -0.862 | 0.217 | -0.052 | -0.013 | -0.013 | 0 | 0.37 | 0.2 | 0.51 | 0.41 | 0.47 | 0.35 |
| 8 | 138495720 | rs6577800 | -0.521 | 0.091 | -0.656 | 0.176 | 0.005 | 0.006 | 0.006 | 1.335 | 0.67 | 0.34 | 0.74 | 0.78 | 0.73 | 0.75 |
| 8 | 138504602 | rs925648 | 1.609 | 0.15 | -0.81 | 0.342 | 0.033 | -0.042 | -0.042 | 0 | 0.18 | 0.08 | 0.1 | 0.28 | 0.1 | 0.34 |
| 8 | 138505969 | rs4053409 | -0.027 | 0.281 | -0.37 | 0.245 | 0 | 0.005 | 0.005 | 1.206 | 0.5 | 0.52 | 0.43 | 0.61 | 0.44 | 0.54 |
| 8 | 138506934 | rs1849692 | 1.587 | 0.187 | -0.267 | 0.271 | -0.017 | 0.013 | 0.013 | 0 | 0.19 | 0.23 | 0.11 | 0.29 | 0.1 | 0.35 |
| 8 | 138511344 | rs1399175 | 1.872 | -0.025 | -0.579 | 0.003 | 0.009 | 0.085 | 0.085 | 0.273 | 0.17 | 0.03 | 0.11 | 0.01 | 0.16 | 0.1 |
| 8 | 138516786 | rs7816962 | 1.122 | -0.012 | -0.628 | 0.044 | 0.005 | -0.113 | -0.113 | 0.483 | 0.36 | 0.12 | 0.27 | 0.51 | 0.28 | 0.5 |
| 8 | 138522685 | rs16907427 | 1.989 | -0.06 | -0.617 | 0.059 | 0.03 | -0.067 | -0.067 | 0 | 0.18 | 0.03 | 0.1 | 0.28 | 0.1 | 0.35 |
| 8 | 138526638 | rs2943195 | -1.36 | -0.054 | -0.253 | 0.097 | -0.026 | 0.032 | 0.032 | 0.556 | 0.84 | 0.69 | 0.87 | 0.79 | 0.87 | 0.83 |
| 8 | 138534526 | rs11776612 | 1.981 | 0.167 | -0.297 | 0.134 | -0.03 | -0.03 | -0.03 | 0 | 0.18 | 0.09 | 0.17 | 0.23 | 0.19 | 0.14 |
| 8 | 138538597 | rs6577808 | 0.816 | 0.232 | -0.174 | 0.145 | -0.001 | -0.001 | -0.001 | 0 | 0.31 | 0.29 | 0.31 | 0.32 | 0.34 | 0.19 |
| 8 | 138542450 | rs7815475 | 1.276 | 0.294 | -0.341 | 0.149 | 0.025 | -0.001 | -0.001 | 0.513 | 0.18 | 0.24 | 0.34 | 0.18 | 0.27 | 0.19 |
| 8 | 138547199 | rs17685635 | 1.793 | 0.339 | -0.395 | 0.546 | -0.027 | -0.012 | -0.012 | 0 | 0.11 | 0.03 | 0.12 | 0.09 | 0.15 | 0.05 |
| 8 | 138547529 | rs1514203 | 1.648 | 0.388 | -0.365 | 0.457 | -0.021 | -0.02 | -0.02 | 0 | 0.18 | 0.11 | 0.17 | 0.23 | 0.19 | 0.14 |
| 8 | 138551517 | rs12156116 | -0.352 | 0.342 | -0.486 | 0.617 | 0 | 0.014 | 0.014 | 1.458 | 0.5 | 0.47 | 0.45 | 0.68 | 0.48 | 0.58 |
| 8 | 138553664 | rs7824808 | 0.55 | 0.446 | -0.301 | 0.532 | 0.009 | -0.005 | -0.005 | 0 | 0.33 | 0.34 | 0.35 | 0.42 | 0.25 | 0.51 |
| 8 | 138558314 | rs12550796 | 0.479 | 0.942 | -0.065 | 1.093 | 0 | 0.001 | 0.001 | 0.073 | 0.14 | 0.1 | 0.15 | 0.18 | 0.16 | 0.05 |
| 8 | 138562411 | rs7843558 | -0.142 | 0.753 | -0.196 | 0.875 | 0.015 | 0.027 | 0.027 | 1.066 | 0.33 | 0.42 | 0.34 | 0.41 | 0.35 | 0.17 |
| 8 | 138567221 | rs9644434 | -0.197 | 0.801 | -0.203 | 1.007 | 0.03 | 0.054 | 0.054 | 0.044 | 0.29 | 0.45 | 0.34 | 0.52 | 0.34 | 0.15 |
| 8 | 138577014 | rs6992974 | -0.365 | -0.155 | 0.261 | 0.74 | -0.045 | 0.007 | 0.007 | 0 | 0.16 | 0.28 | 0.21 | 0.15 | 0.1 | 0.14 |
| 8 | 138585238 | rs7844139 | -1.021 | -0.307 | 0.009 | 0.816 | -0.02 | 0.005 | 0.005 | 0 | 0.25 | 0.26 | 0.31 | 0.29 | 0.16 | 0.21 |
| 8 | 138585743 | rs7822502 | -1.021 | -0.307 | 0.009 | 0.821 | -0.02 | 0.004 | 0.004 | 0 | 0.25 | 0.26 | 0.31 | 0.28 | 0.16 | 0.21 |
| 8 | 138588027 | rs13269649 | -1.324 | -0.346 | -0.181 | 0.698 | 0 | 0 | 0 | 2.237 | 0.74 | 0.73 | 0.65 | 0.64 | 0.63 | 0.59 |
| 8 | 138589339 | rs13254131 | 0.77 | -0.484 | 0.074 | 0.634 | 0 | 0.012 | 0.012 | 0 | 0.46 | 0.33 | 0.33 | 0.38 | 0.46 | 0.38 |
| 8 | 138589542 | rs10156236 | -0.1 | -0.62 | 0.243 | 0.443 | -0.007 | 0.002 | 0.002 | 0 | 0.13 | 0.21 | 0.13 | 0.14 | 0.09 | 0.02 |
| 8 | 138590324 | rs6996902 | 0.349 | -0.508 | 0.076 | 0.395 | -0.027 | -0.002 | -0.002 | 0 | 0.14 | 0.19 | 0.19 | 0.13 | 0.07 | 0.16 |
| 8 | 138593875 | rs13260366 | 0.541 | -0.278 | -0.234 | 0.061 | -0.002 | -0.005 | -0.005 | 0.016 | 0.11 | 0.07 | 0.09 | 0 | 0.14 | 0.04 |
| 8 | 138596758 | rs4599850 | 1.123 | -0.362 | -0.252 | 0.304 | 0 | 0.007 | 0.007 | 0 | 0.18 | 0.21 | 0.28 | 0.3 | 0.17 | 0.18 |

| | | | | | | | | | | | | | | | | |
|---|---|---|---|---|---|---|---|---|---|---|---|---|---|---|---|---|
| 8 | 138598197 | rs9650558 | 0.041 | -0.562 | -0.288 | 0.236 | -0.004 | -0.001 | -0.001 | 0.487 | 0.25 | 0.25 | 0.2 | 0.25 | 0.19 | 0.18 |
| 8 | 138599067 | rs868421 | 0.516 | -0.462 | -0.833 | -0.318 | 0.003 | 0.006 | 0.006 | 0.177 | 0.3 | 0.28 | 0.18 | 0.08 | 0.28 | 0.21 |
| 8 | 138599275 | rs907991 | -1.183 | -0.108 | -1.074 | -0.072 | -0.039 | -0.029 | -0.029 | 0.066 | 0.73 | 0.84 | 0.67 | 0.63 | 0.65 | 0.58 |
| 8 | 138601192 | rs949449 | 0.431 | -0.672 | -1.05 | -0.545 | 0.034 | 0.032 | 0.032 | 0 | 0.24 | 0.12 | 0.1 | 0 | 0.16 | 0.15 |
| 8 | 138612206 | rs12334660 | 0.03 | -0.57 | -1.187 | -0.349 | 0.015 | 0.048 | 0.048 | 0 | 0.26 | 0.1 | 0.2 | 0.14 | 0.25 | 0.12 |
| 8 | 138612383 | rs11992976 | 0.024 | -0.813 | -0.945 | -0.257 | 0.003 | 0.018 | 0.018 | 0 | 0.3 | 0.35 | 0.35 | 0.65 | 0.31 | 0.37 |
| 8 | 138612866 | rs17634990 | 1.058 | -0.908 | -1.131 | -0.498 | 0.011 | 0.036 | 0.036 | 0 | 0.2 | 0.03 | 0.17 | 0.08 | 0.16 | 0.13 |
| 8 | 138616654 | rs10086487 | 0.494 | -0.953 | -0.765 | -0.5 | -0.02 | -0.045 | -0.045 | 0 | 0.28 | 0.39 | 0.23 | 0.14 | 0.25 | 0.12 |
| 8 | 138617100 | rs11992583 | 0.926 | -0.949 | -0.765 | -0.492 | -0.004 | 0.093 | 0.093 | 0 | 0.23 | 0.35 | 0.23 | 0.52 | 0.21 | 0.36 |
| 8 | 138619522 | rs6991627 | -0.607 | -0.894 | -1.018 | -0.473 | 0.024 | 0.051 | 0.051 | 0 | 0.51 | 0.75 | 0.47 | 0.66 | 0.46 | 0.48 |
| 8 | 138620757 | rs7831168 | -1.813 | -0.919 | -1.044 | -0.501 | -0.109 | -0.025 | -0.025 | 0 | 0.73 | 0.92 | 0.65 | 0.74 | 0.63 | 0.61 |
| 8 | 138621005 | rs10875408 | -0.607 | -0.853 | -0.753 | -0.412 | 0.009 | 0.014 | 0.014 | 0 | 0.5 | 0.51 | 0.44 | 0.66 | 0.46 | 0.48 |
| 8 | 138624129 | rs11985328 | 0.367 | -0.513 | -0.818 | 0.062 | 0.004 | 0.132 | 0.132 | 0 | 0.24 | 0.42 | 0.24 | 0.52 | 0.22 | 0.36 |
| 8 | 138626268 | rs7000472 | -1.052 | -0.089 | -1.452 | 0.012 | -0.071 | -0.018 | -0.018 | 0 | 0.73 | 0.86 | 0.64 | 0.74 | 0.64 | 0.62 |
| 8 | 138627139 | rs3890693 | -0.016 | -1.073 | -0.964 | -0.465 | -0.029 | -0.029 | -0.029 | 0 | 0.28 | 0.69 | 0.21 | 0.13 | 0.14 | 0.16 |
| 8 | 138627984 | rs7014826 | 0.27 | -0.675 | -1.299 | -0.414 | 0.017 | 0.046 | 0.046 | 0 | 0.3 | 0.13 | 0.24 | 0.17 | 0.27 | 0.28 |
| 8 | 138629624 | rs11166739 | -0.581 | 0.023 | -0.906 | -0.634 | -0.001 | -0.047 | -0.047 | 0 | 0.78 | 0.76 | 0.87 | 1 | 0.78 | 0.91 |
| 8 | 138629984 | rs4620319 | 0.357 | 0.005 | -0.838 | -0.119 | -0.006 | -0.001 | -0.001 | 0.154 | 0.27 | 0.27 | 0.4 | 0.26 | 0.42 | 0.47 |
| 8 | 138633720 | rs4909680 | 0.7 | -0.777 | -1.286 | 0.3 | 0.069 | -0.02 | -0.02 | 0 | 0.36 | 0.09 | 0.22 | 0.32 | 0.25 | 0.31 |
| 8 | 138634023 | rs4909681 | -0.686 | -0.313 | -1.344 | -0.329 | -0.034 | 0.058 | 0.058 | 0 | 0.76 | 0.88 | 0.85 | 0.99 | 0.7 | 0.89 |
| 8 | 138638199 | rs10090365 | 0.282 | -0.464 | -0.404 | 0.124 | 0 | -0.001 | -0.001 | 0 | 0.36 | 0.35 | 0.5 | 0.31 | 0.44 | 0.46 |
| 8 | 138640055 | rs10110872 | -0.608 | -0.437 | -0.567 | -0.028 | -0.027 | 0.042 | 0.042 | 0 | 0.74 | 0.85 | 0.84 | 0.99 | 0.69 | 0.89 |
| 8 | 138655344 | rs4623466 | 0.304 | 0.001 | -0.126 | 0.362 | -0.01 | -0.004 | -0.004 | 0 | 0.32 | 0.3 | 0.43 | 0.32 | 0.36 | 0.43 |
| 8 | 138657388 | rs6577815 | -0.39 | -0.291 | 0.32 | 0.414 | -0.001 | -0.004 | -0.004 | 0.339 | 0.41 | 0.71 | 0.51 | 0.4 | 0.41 | 0.56 |
| 8 | 138659257 | rs4463463 | -0.299 | -0.295 | 0.464 | 0.412 | -0.004 | -0.005 | -0.005 | 0.283 | 0.38 | 0.35 | 0.47 | 0.41 | 0.4 | 0.56 |
| 8 | 138664030 | rs4487812 | -0.063 | -0.37 | 0.521 | 0.746 | -0.004 | -0.017 | -0.017 | 0 | 0.38 | 0.31 | 0.47 | 0.48 | 0.38 | 0.54 |
| 8 | 138669620 | rs12676177 | -0.755 | -0.563 | 0.321 | 0.657 | -0.015 | 0.02 | 0.02 | 0 | 0.73 | 0.74 | 0.73 | 0.84 | 0.64 | 0.85 |
| 8 | 138675098 | rs12549338 | 1.472 | -0.605 | 0.326 | 0.654 | 0.018 | -0.026 | -0.026 | 0.106 | 0.3 | 0.06 | 0.21 | 0.28 | 0.25 | 0.2 |
| 8 | 138688770 | rs12545584 | 0.438 | -0.767 | 0.595 | 0.617 | 0 | -0.003 | -0.003 | 1.129 | 0.6 | 0.31 | 0.57 | 0.63 | 0.58 | 0.71 |
| 8 | 138692245 | rs4469495 | 0.439 | -0.789 | 0.512 | 0.607 | 0 | -0.007 | -0.007 | 0.053 | 0.6 | 0.44 | 0.58 | 0.63 | 0.58 | 0.71 |
| 8 | 138692559 | rs7835880 | -0.966 | -0.906 | 0.481 | 0.538 | -0.003 | 0.001 | 0.001 | 0 | 0.77 | 0.72 | 0.74 | 0.84 | 0.63 | 0.9 |
| 8 | 138694431 | rs4416856 | 0.377 | -0.749 | 0.392 | 0.68 | -0.038 | 0.042 | 0.042 | 0 | 0.13 | 0.3 | 0.15 | 0.21 | 0.06 | 0.14 |
| 8 | 138702381 | rs4389979 | 0.971 | -0.679 | 0.181 | 0.562 | -0.083 | -0.058 | -0.058 | 0 | 0.28 | 0.08 | 0.35 | 0.34 | 0.35 | 0.49 |
| 8 | 138703757 | rs11166747 | 1.548 | -0.776 | 0.367 | 0.554 | -0.007 | 0.003 | 0.003 | 0 | 0.32 | 0.22 | 0.31 | 0.35 | 0.25 | 0.2 |
| 8 | 138705476 | rs4495466 | -0.002 | -0.439 | 0.685 | 0.962 | -0.036 | 0.016 | 0.016 | 0 | 0.13 | 0.29 | 0.08 | 0.16 | 0.05 | 0.16 |
| 8 | 138709985 | rs11786747 | 0.345 | -0.25 | 0.699 | 1.004 | -0.005 | -0.016 | -0.016 | 0 | 0.59 | 0.63 | 0.57 | 0.5 | 0.6 | 0.34 |
| 8 | 138723448 | rs4909385 | 0.516 | -0.062 | 0.71 | 0.931 | -0.009 | -0.003 | -0.003 | 0 | 0.58 | 0.3 | 0.51 | 0.55 | 0.54 | 0.41 |
| 8 | 138737357 | rs6577825 | 0.441 | -0.102 | 0.13 | 0.894 | -0.058 | -0.023 | -0.023 | 0 | 0.28 | 0.1 | 0.34 | 0.29 | 0.33 | 0.47 |
| 8 | 138739706 | rs7006635 | 0.518 | -0.129 | 0.39 | 0.912 | 0.006 | 0.002 | 0.002 | 0 | 0.59 | 0.48 | 0.53 | 0.44 | 0.54 | 0.4 |
| 8 | 138744981 | rs4909692 | 0.406 | 0.006 | 0.308 | 0.805 | -0.053 | -0.051 | -0.051 | 0 | 0.28 | 0.11 | 0.35 | 0.38 | 0.34 | 0.51 |
| 8 | 138749970 | rs12678736 | 0.813 | 0.232 | 0.533 | 0.874 | -0.025 | -0.032 | -0.032 | 0.123 | 0.6 | 0.29 | 0.5 | 0.47 | 0.54 | 0.39 |
| 8 | 138750268 | rs4631498 | -0.175 | 0.431 | -0.352 | 0.98 | -0.001 | 0.029 | 0.029 | 0.084 | 0.89 | 0.71 | 0.9 | 0.84 | 0.87 | 0.88 |
| 8 | 138759742 | rs10106108 | 0.292 | 0.354 | -0.697 | 0.737 | 0.003 | 0.004 | 0.004 | 0 | 0.63 | 0.58 | 0.53 | 0.47 | 0.54 | 0.39 |
| 8 | 138769527 | rs6981722 | 0.988 | 0.178 | -0.978 | 0.542 | 0 | -0.044 | -0.044 | 1.57 | 0.61 | 0.15 | 0.63 | 0.48 | 0.61 | 0.65 |
| 8 | 138769669 | rs6981916 | -0.364 | 0.12 | -0.643 | 0.55 | -0.01 | -0.018 | -0.018 | 0 | 0.87 | 0.81 | 0.87 | 0.91 | 0.87 | 0.83 |
| 8 | 138771147 | rs4129321 | -0.605 | 0.089 | -0.61 | 0.553 | 0.001 | 0.035 | 0.035 | 0 | 0.27 | 0.66 | 0.24 | 0.43 | 0.26 | 0.19 |
| 8 | 138775160 | rs11781661 | 0.588 | -0.305 | -0.797 | 0.322 | -0.007 | 0.006 | 0.006 | 0 | 0.8 | 0.78 | 0.83 | 0.74 | 0.82 | 0.7 |
| 8 | 138775796 | rs9324461 | -0.535 | -0.066 | -0.617 | 0.026 | 0.008 | 0.02 | 0.02 | 0 | 0.31 | 0.45 | 0.35 | 0.35 | 0.31 | 0.21 |
| 8 | 138776768 | rs10108925 | -0.318 | -0.084 | -0.651 | 0.019 | -0.011 | -0.024 | -0.024 | 0.021 | 0.86 | 0.79 | 0.88 | 0.91 | 0.87 | 0.83 |
| 8 | 138777847 | rs10112884 | 0.84 | 0.102 | -1.504 | 0.102 | 0.004 | 0 | 0 | 0 | 0.56 | 0.29 | 0.53 | 0.56 | 0.56 | 0.63 |
| 8 | 138777955 | rs12543628 | 0.88 | 0.128 | -1.836 | 0.115 | 0.007 | -0.002 | -0.002 | 0 | 0.55 | 0.21 | 0.52 | 0.56 | 0.56 | 0.62 |
| 8 | 138788813 | rs7005722 | -0.25 | 0.705 | -2.148 | 0.071 | -0.088 | -0.011 | -0.011 | 0 | 0.21 | 0.03 | 0.26 | 0.17 | 0.26 | 0.17 |
| 8 | 138791415 | rs4548214 | 0.457 | 0.675 | -1.924 | -0.082 | -0.115 | -0.107 | -0.107 | 0 | 0.45 | 0.11 | 0.36 | 0.45 | 0.5 | 0.28 |
| 8 | 138791736 | rs4548215 | 0.714 | 0.233 | -0.708 | -0.626 | 0.03 | 0.038 | 0.038 | 2.301 | 0.7 | 0.29 | 0.81 | 0.85 | 0.82 | 0.66 |
| 8 | 138796025 | rs7830879 | -1.252 | 0.853 | 0.283 | -0.082 | 0.062 | 0.073 | 0.073 | 0 | 0.07 | 0.29 | 0.07 | 0.07 | 0.08 | 0.07 |
| 8 | 138796799 | rs7462989 | 0.591 | 0.951 | 0.188 | -0.48 | 0.004 | -0.085 | -0.085 | 0 | 0.79 | 0.49 | 0.85 | 0.87 | 0.81 | 0.8 |
| 8 | 138801264 | rs11787188 | -1.168 | 0.841 | -0.206 | -0.548 | 0 | -0.001 | -0.001 | 1.793 | 0.11 | 0.1 | 0.09 | 0.08 | 0.13 | 0.09 |
| 8 | 138806916 | rs13267220 | -0.533 | 1.265 | -0.646 | -1.244 | -0.023 | 0.067 | 0.067 | 0 | 0.21 | 0.1 | 0.1 | 0 | 0.19 | 0.08 |
| 8 | 138810922 | rs7822022 | 0.468 | 1.098 | -0.206 | -1.047 | 0.003 | 0.012 | 0.012 | 0.036 | 0.89 | 0.86 | 0.75 | 0.84 | 0.86 | 0.72 |
| 8 | 138813360 | rs6577842 | 0.614 | 1.043 | 0.202 | -1.086 | 0.005 | 0.014 | 0.014 | 0.029 | 0.89 | 0.81 | 0.73 | 0.84 | 0.86 | 0.73 |
| 8 | 138816115 | rs949983 | 0.617 | 1.392 | -0.1 | -1.195 | -0.074 | 0.077 | 0.077 | 0 | 0.72 | 0.95 | 0.68 | 0.85 | 0.69 | 0.65 |
| 8 | 138821460 | rs7845645 | -1.329 | 1.223 | 0.492 | -2.05 | 0.024 | 0.06 | 0.06 | 0 | 0.49 | 0.62 | 0.58 | 0.17 | 0.56 | 0.45 |
| 8 | 138833669 | rs11988641 | -0.766 | 1.317 | 0.884 | -2.069 | 0.021 | -0.064 | -0.064 | 6.405 | 0.35 | 0.52 | 0.48 | 0.14 | 0.39 | 0.4 |
| 8 | 138843225 | rs1511852 | -0.219 | 0.575 | 0.359 | -2.531 | -0.043 | 0.143 | 0.143 | 0 | 0.4 | 0.21 | 0.45 | 0.14 | 0.42 | 0.43 |
| 8 | 138844671 | rs12545671 | -0.226 | 0.555 | 0.302 | -2.53 | -0.091 | 0.154 | 0.154 | 0 | 0.39 | 0.1 | 0.43 | 0.16 | 0.42 | 0.43 |
| 8 | 138844863 | rs979289 | 0.209 | 0.877 | 0.789 | -2.666 | 0.069 | -0.113 | -0.113 | 0 | 0.16 | 0.36 | 0.15 | 0.01 | 0.21 | 0.07 |
| 8 | 138846091 | rs7842171 | -1.175 | 0.884 | 0.785 | -2.423 | 0 | -0.006 | -0.006 | 0 | 0.57 | 0.55 | 0.63 | 0.22 | 0.63 | 0.5 |
| 8 | 138846311 | rs10098705 | -0.052 | 0.898 | 0.793 | -2.515 | 0.069 | -0.13 | -0.13 | 1.694 | 0.17 | 0.44 | 0.17 | 0.01 | 0.21 | 0.07 |
| 8 | 138849473 | rs13282624 | -1 | 0.703 | 0.779 | -2.399 | 0.002 | -0.01 | -0.01 | 0 | 0.56 | 0.56 | 0.6 | 0.18 | 0.63 | 0.49 |

| | | | | | | | | | | | | | | | | |
|---|---|---|---|---|---|---|---|---|---|---|---|---|---|---|---|---|
| 8 | 138864096 | rs7386095 | 0.709 | 0.177 | -0.163 | -2.394 | -0.03 | 0.213 | 0.213 | 0.532 | 0.49 | 0.13 | 0.5 | 0.8 | 0.48 | 0.64 |
| 8 | 138866091 | rs4909709 | 0.646 | 0.167 | 0.231 | -2.399 | 0.004 | 0.109 | 0.109 | 0.075 | 0.5 | 0.27 | 0.51 | 0.8 | 0.48 | 0.64 |
| 8 | 138866267 | rs2326132 | 0.348 | 0.2 | 0.12 | -2.543 | -0.021 | 0.024 | 0.024 | 0 | 0.16 | 0.1 | 0.14 | 0.05 | 0.23 | 0.13 |
| 8 | 138866496 | rs2326130 | -0.507 | 0.103 | 0.611 | -2.58 | 0.007 | 0.009 | 0.009 | 0 | 0.7 | 0.72 | 0.7 | 0.85 | 0.73 | 0.77 |
| 8 | 138871092 | rs16907783 | 0.53 | 0.199 | 0.451 | -2.277 | -0.015 | -0.043 | -0.043 | 0 | 0.11 | 0.02 | 0.21 | 0.17 | 0.12 | 0.28 |
| 8 | 138875499 | rs907461 | 0.961 | 0.268 | 0.769 | -2.636 | 0.003 | 0.072 | 0.072 | 0.007 | 0.5 | 0.32 | 0.52 | 0.8 | 0.48 | 0.65 |
| 8 | 138876628 | rs1995983 | 0.428 | 0.186 | 0.491 | -2.777 | -0.009 | 0.006 | 0.006 | 0 | 0.66 | 0.63 | 0.61 | 0.89 | 0.51 | 0.7 |
| 8 | 138877155 | rs10095685 | -0.125 | 0.142 | 0.538 | -2.775 | -0.034 | -0.04 | -0.04 | 0.077 | 0.15 | 0.26 | 0.1 | 0.09 | 0.07 | 0.07 |
| 8 | 138877271 | rs10095814 | -0.686 | 0.19 | 0.38 | -2.765 | 0.003 | -0.026 | -0.026 | 0 | 0.8 | 0.66 | 0.74 | 0.91 | 0.73 | 0.82 |
| 8 | 138881439 | rs1949141 | -0.861 | -0.14 | 0.489 | -2.67 | -0.003 | -0.025 | -0.025 | 0 | 0.81 | 0.67 | 0.75 | 0.91 | 0.78 | 0.86 |
| 8 | 138884435 | rs1511851 | 0.528 | -0.088 | 1.093 | -2.435 | 0.021 | -0.037 | -0.037 | 0 | 0.77 | 0.65 | 0.79 | 0.93 | 0.71 | 0.81 |
| 8 | 138885943 | rs4909712 | -1.4 | -0.108 | 1.117 | -2.442 | 0.003 | -0.085 | -0.085 | 0 | 0.3 | 0.5 | 0.25 | 0.1 | 0.29 | 0.19 |
| 8 | 138887431 | rs972679 | 1.62 | -0.089 | 0.813 | -2.429 | -0.016 | 0.129 | 0.129 | 1.445 | 0.61 | 0.16 | 0.66 | 0.84 | 0.64 | 0.75 |
| 8 | 138890763 | rs17689136 | 1.632 | -0.064 | 0.519 | -2.299 | -0.04 | 0.147 | 0.147 | 1.542 | 0.61 | 0.13 | 0.66 | 0.84 | 0.65 | 0.75 |
| 8 | 138902764 | rs907458 | 0.258 | 0.144 | 0.688 | -2.275 | 0.002 | -0.016 | -0.016 | 0.045 | 0.14 | 0.13 | 0.12 | 0.01 | 0.21 | 0.12 |
| 8 | 138911999 | rs2013354 | 0.464 | -0.043 | 0.61 | -2.213 | 0.016 | -0.029 | -0.029 | 0 | 0.77 | 0.68 | 0.8 | 0.93 | 0.71 | 0.82 |
| 8 | 138912997 | rs2668096 | -0.689 | -0.159 | 0.661 | -2.248 | -0.045 | -0.045 | -0.045 | 0 | 0.15 | 0.52 | 0.14 | 0.09 | 0.07 | 0.07 |
| 8 | 138915097 | rs9644435 | 1.61 | -0.161 | 0.579 | -2.265 | -0.061 | 0.136 | 0.136 | 1.754 | 0.61 | 0.07 | 0.65 | 0.84 | 0.64 | 0.75 |
| 8 | 138915665 | rs2668100 | -0.642 | -0.147 | 0.735 | -2.193 | -0.061 | -0.043 | -0.043 | 0 | 0.15 | 0.41 | 0.11 | 0.09 | 0.06 | 0.07 |
| 8 | 138915718 | rs2705302 | -0.666 | -0.097 | 0.634 | -2.183 | 0.016 | 0.021 | 0.021 | 0 | 0.91 | 0.7 | 0.92 | 0.94 | 0.93 | 0.94 |
| 8 | 138923107 | rs16907984 | 0.152 | -0.15 | 0.635 | -2.173 | -0.001 | 0.001 | 0.001 | 0.434 | 0.12 | 0.12 | 0.09 | 0.1 | 0.05 | 0.07 |
| 8 | 138927680 | rs2221844 | 0.998 | -0.062 | 0.37 | -2.135 | -0.021 | -0.016 | -0.016 | 0.015 | 0.76 | 0.33 | 0.8 | 0.85 | 0.87 | 0.88 |
| 8 | 138928954 | rs12545857 | 1.217 | -0.006 | -0.303 | -2.087 | -0.025 | 0.124 | 0.124 | 1.538 | 0.62 | 0.15 | 0.66 | 0.83 | 0.64 | 0.76 |
| 8 | 138930779 | rs1858367 | 0.725 | -0.137 | 0.402 | -1.782 | -0.016 | -0.027 | -0.027 | 0.022 | 0.86 | 0.65 | 0.87 | 0.9 | 0.92 | 0.94 |
| 8 | 138931602 | rs7003232 | -0.803 | -0.254 | -0.525 | -1.676 | 0.032 | 0.019 | 0.019 | 0 | 0.17 | 0.11 | 0.22 | 0.12 | 0.11 | 0.29 |
| 8 | 138932113 | rs2203112 | -0.755 | 0.28 | 0.28 | -2.524 | 0.018 | -0.067 | -0.067 | 0 | 0.22 | 0.34 | 0.21 | 0.07 | 0.28 | 0.21 |
| 8 | 138932176 | rs1398034 | 0.214 | -0.292 | -0.195 | -3.101 | 0.034 | 0.062 | 0.062 | 0 | 0.31 | 0.16 | 0.41 | 0.83 | 0.23 | 0.46 |
| 8 | 138935483 | rs2668109 | 0.212 | -0.116 | 0.62 | -3.13 | 0.023 | -0.054 | -0.054 | 0 | 0.77 | 0.64 | 0.71 | 0.85 | 0.72 | 0.8 |
| 8 | 138967803 | rs2705325 | 0.597 | -0.22 | 0.555 | -3.209 | 0 | -0.094 | -0.094 | 0.091 | 0.49 | 0.53 | 0.3 | 0.02 | 0.51 | 0.35 |
| 8 | 138973526 | rs6999779 | 0.1 | 0.02 | 0.2 | -2.897 | 0.016 | -0.005 | -0.005 | 0 | 0.13 | 0.15 | 0.25 | 0.15 | 0.2 | 0.14 |
| 8 | 138985519 | rs4392939 | -0.285 | -0.129 | 0.288 | -3.266 | -0.007 | 0.101 | 0.101 | 0 | 0.48 | 0.24 | 0.57 | 0.99 | 0.51 | 0.69 |
| 8 | 138993531 | rs3861780 | -0.212 | -0.043 | 0.253 | -3.247 | -0.007 | 0.019 | 0.019 | 0 | 0.47 | 0.34 | 0.56 | 0.99 | 0.51 | 0.71 |
| 8 | 139009683 | rs10103652 | -0.043 | 0.162 | 0.13 | -3.069 | -0.007 | -0.013 | -0.013 | 0.19 | 0.63 | 0.31 | 0.62 | 0.99 | 0.62 | 0.79 |
| 8 | 139023474 | rs13270582 | -0.218 | 0.346 | 0.79 | -3.042 | 0 | -0.051 | -0.051 | 0 | 0.37 | 0.33 | 0.26 | 0.03 | 0.36 | 0.29 |
| 8 | 139024326 | rs7840352 | 0.257 | 0.264 | 1.188 | -3.081 | 0.001 | 0.061 | 0.061 | 0 | 0.6 | 0.3 | 0.71 | 0.96 | 0.64 | 0.68 |
| 8 | 139025160 | rs13282681 | -0.104 | 0.315 | 1.259 | -3.096 | -0.013 | 0.062 | 0.062 | 0 | 0.34 | 0.17 | 0.19 | 0.03 | 0.35 | 0.24 |
| 8 | 139028105 | rs6577862 | -0.328 | 0.427 | 0.793 | -3.035 | 0.003 | -0.123 | -0.123 | 0 | 0.38 | 0.45 | 0.26 | 0.03 | 0.36 | 0.29 |
| 8 | 139048470 | rs11166774 | -0.03 | 0.634 | 0.088 | -2.84 | -0.005 | 0.007 | 0.007 | 0 | 0.63 | 0.74 | 0.8 | 0.94 | 0.64 | 0.71 |
| 8 | 139051681 | rs9657452 | 0.167 | 0.643 | 0.071 | -2.891 | -0.001 | -0.063 | -0.063 | 0.099 | 0.59 | 0.54 | 0.75 | 0.94 | 0.67 | 0.67 |
| 8 | 139051846 | rs9324470 | -0.257 | 0.58 | -0.006 | -2.872 | -0.001 | -0.053 | -0.053 | 0 | 0.4 | 0.43 | 0.21 | 0.06 | 0.33 | 0.27 |
| 8 | 139062339 | rs7005095 | -0.07 | 0.493 | -0.043 | -3.051 | -0.003 | -0.076 | -0.076 | 0.379 | 0.42 | 0.48 | 0.2 | 0.05 | 0.31 | 0.31 |
| 8 | 139069173 | rs982842 | 0.016 | 0.486 | -0.575 | -3.139 | -0.059 | 0.118 | 0.118 | 0 | 0.68 | 0.88 | 0.74 | 1 | 0.55 | 0.8 |
| 8 | 139074250 | rs4332169 | 0.273 | 0.659 | 0.158 | -3.149 | 0.014 | -0.118 | -0.118 | 0 | 0.81 | 0.61 | 0.86 | 1 | 0.79 | 0.82 |
| 8 | 139075196 | rs6990911 | 0.234 | 0.699 | -0.016 | -3.537 | -0.005 | -0.018 | -0.018 | 0 | 0.19 | 0.16 | 0.19 | 0.01 | 0.24 | 0.11 |
| 8 | 139075826 | rs1325058 | 1.117 | 0.349 | 0.291 | -4.131 | 0.003 | 0.03 | 0.03 | 0.014 | 0.49 | 0.37 | 0.61 | 0.99 | 0.49 | 0.65 |
| 8 | 139077206 | rs1028777 | -0.552 | 0.347 | 0.3 | -4.113 | -0.01 | 0.02 | 0.02 | 0 | 0.32 | 0.25 | 0.27 | 0.01 | 0.32 | 0.19 |
| 8 | 139084283 | rs10111139 | 0.451 | 0.34 | 0.819 | -4.125 | 0.01 | -0.126 | -0.126 | 0 | 0.66 | 0.44 | 0.7 | 0.99 | 0.58 | 0.76 |
| 8 | 139084636 | rs7816490 | 0.649 | -0.621 | 0.985 | -3.151 | -0.013 | -0.013 | -0.013 | 1.21 | 0.72 | 0.42 | 0.59 | 0.81 | 0.83 | 0.69 |
| 8 | 139087040 | rs1325054 | 0.181 | -0.174 | 0.628 | -3.087 | 0.002 | -0.027 | -0.027 | 0 | 0.14 | 0.13 | 0.11 | 0 | 0.22 | 0.15 |
| 8 | 139089126 | rs10109789 | -0.641 | -0.785 | 0.208 | -3.224 | 0.015 | 0.057 | 0.057 | 0 | 0.46 | 0.33 | 0.59 | 0.21 | 0.56 | 0.46 |
| 8 | 139089532 | rs11166780 | 0.513 | -0.941 | 0.871 | -2.492 | -0.004 | 0.01 | 0.01 | 0 | 0.11 | 0.11 | 0.32 | 0.19 | 0.08 | 0.17 |
| 8 | 139090718 | rs10096264 | -0.663 | -1.09 | 0.612 | -2.612 | -0.042 | -0.007 | -0.007 | 0 | 0.23 | 0.4 | 0.39 | 0.2 | 0.13 | 0.22 |
| 8 | 139090974 | rs1570770 | -0.752 | -1.081 | 0.488 | -2.817 | 0.004 | 0.01 | 0.01 | 0 | 0.48 | 0.43 | 0.56 | 0.21 | 0.4 | 0.38 |
| 8 | 139092646 | rs1022900 | 0.28 | -1.457 | 1.117 | -2.055 | 0 | -0.037 | -0.037 | 0.293 | 0.59 | 0.43 | 0.61 | 0.85 | 0.57 | 0.77 |
| 8 | 139101291 | rs1033917 | -0.18 | -1.475 | 1.258 | -1.953 | -0.012 | 0.026 | 0.026 | 0 | 0.66 | 0.69 | 0.67 | 0.86 | 0.59 | 0.81 |
| 8 | 139102267 | rs11786500 | 0.47 | -1.376 | 0.71 | -1.988 | -0.008 | 0.051 | 0.051 | 0 | 0.09 | 0.02 | 0.04 | 0 | 0.07 | 0.04 |
| 8 | 139102790 | rs1408762 | -0.029 | -1.311 | 0.936 | -1.944 | 0.011 | -0.07 | -0.07 | 0 | 0.63 | 0.47 | 0.63 | 0.86 | 0.59 | 0.81 |
| 8 | 139109143 | rs1830877 | 0.708 | -1.12 | 0.776 | -1.6 | -0.001 | 0.009 | 0.009 | 0 | 0.46 | 0.37 | 0.53 | 0.85 | 0.44 | 0.7 |
| 8 | 139113320 | rs10429384 | 0.72 | -1.077 | 0.527 | -1.432 | -0.002 | 0.039 | 0.039 | 0 | 0.46 | 0.35 | 0.53 | 0.8 | 0.44 | 0.69 |
| 8 | 139116305 | rs7815520 | -0.018 | -0.195 | 0.426 | -0.936 | -0.019 | 0.036 | 0.036 | 0 | 0.15 | 0.29 | 0.14 | 0.18 | 0.12 | 0.12 |
| 8 | 139117698 | rs1886578 | 0.745 | 0.476 | 0.45 | -1.017 | 0.022 | -0.053 | -0.053 | 0 | 0.61 | 0.46 | 0.59 | 0.79 | 0.5 | 0.71 |
| 8 | 139123897 | rs10109073 | 0.224 | 0.879 | -0.203 | -1.066 | -0.043 | 0.026 | 0.026 | 0 | 0.71 | 0.82 | 0.72 | 1 | 0.55 | 0.88 |
| 8 | 139140208 | rs7462761 | 0.48 | 1.176 | 0.025 | -0.046 | 0.036 | -0.021 | -0.021 | 0.809 | 0.75 | 0.58 | 0.68 | 0.78 | 0.62 | 0.74 |
| 8 | 139140364 | rs16908252 | 0.791 | 1.191 | -0.486 | 0.27 | -0.002 | 0.002 | 0.002 | 0.593 | 0.61 | 0.59 | 0.39 | 0.23 | 0.45 | 0.44 |
| 8 | 139140561 | rs16908261 | 0.042 | 0.841 | -0.106 | 0.312 | -0.048 | 0.055 | 0.055 | 0.711 | 0.11 | 0.27 | 0.09 | 0.21 | 0.05 | 0.17 |
| 8 | 139141106 | rs7463648 | 0.141 | 0.655 | -0.038 | -0.978 | 0.004 | -0.116 | -0.116 | 0 | 0.57 | 0.54 | 0.66 | 0.98 | 0.55 | 0.78 |
| 8 | 139141229 | rs7816669 | 0.236 | 0.374 | 0.167 | -0.836 | -0.003 | -0.025 | -0.025 | 0 | 0.36 | 0.44 | 0.29 | 0.22 | 0.3 | 0.3 |
| 8 | 139145480 | rs7461621 | -1.135 | 0.252 | 0.066 | -1 | -0.002 | -0.012 | -0.012 | 0 | 0.87 | 0.87 | 0.87 | 1 | 0.78 | 0.94 |
| 8 | 139149406 | rs6577876 | 0.939 | 0.21 | 0.352 | -1.039 | 0.001 | -0.064 | -0.064 | 1.113 | 0.3 | 0.31 | 0.2 | 0.01 | 0.26 | 0.16 |

| | | | | | | | | | | | | | | | | |
|---|---|---|---|---|---|---|---|---|---|---|---|---|---|---|---|---|
| 8 | 139152091 | rs7828715 | 0.85 | -0.361 | 0.682 | -1.071 | -0.007 | 0.006 | 0.006 | 1.204 | 0.64 | 0.39 | 0.62 | 0.56 | 0.65 | 0.6 |
| 8 | 139152515 | rs11996097 | 0.148 | -0.46 | 0.746 | -1.029 | -0.018 | 0.182 | 0.182 | 0 | 0.08 | 0.29 | 0.12 | 0.21 | 0.04 | 0.14 |
| 8 | 139158169 | rs2280849 | 0.672 | -0.425 | 0.518 | -0.886 | -0.009 | -0.002 | -0.002 | 0 | 0.16 | 0.16 | 0.12 | 0.24 | 0.1 | 0.2 |
| 8 | 139159247 | rs4501627 | 1.219 | -0.507 | 0.442 | -0.987 | -0.018 | -0.039 | -0.039 | 0 | 0.34 | 0.18 | 0.42 | 0.53 | 0.38 | 0.43 |
| 8 | 139160199 | rs16908366 | 1.028 | -0.569 | 0.304 | -1.477 | -0.001 | -0.001 | -0.001 | 0 | 0.2 | 0.2 | 0.19 | 0.23 | 0.17 | 0.25 |
| 8 | 139160613 | rs6577879 | 0.064 | -0.563 | 0.274 | -1.497 | 0.001 | 0.002 | 0.002 | 0.871 | 0.71 | 0.68 | 0.74 | 0.76 | 0.69 | 0.73 |
| 8 | 139161806 | rs11985645 | -1.663 | -0.488 | 0.247 | -1.483 | -0.003 | -0.01 | -0.01 | 0 | 0.87 | 0.88 | 0.87 | 1 | 0.78 | 0.95 |
| 8 | 139164192 | rs3750306 | 1.578 | -0.474 | 0.245 | -1.317 | -0.017 | -0.043 | -0.043 | 1.141 | 0.34 | 0.18 | 0.42 | 0.53 | 0.38 | 0.43 |
| 8 | 139164570 | rs3750307 | 0.727 | -0.647 | 0.156 | -1.413 | 0 | -0.004 | -0.004 | 0 | 0.22 | 0.21 | 0.19 | 0.23 | 0.17 | 0.26 |
| 8 | 139165068 | rs3763590 | 1.496 | -0.55 | -0.07 | -1.299 | -0.022 | -0.052 | -0.052 | 1.05 | 0.33 | 0.16 | 0.41 | 0.53 | 0.38 | 0.43 |
| 8 | 139165289 | rs7835830 | 0.297 | -0.396 | -0.173 | -1.6 | 0.001 | 0.003 | 0.003 | 0.913 | 0.7 | 0.68 | 0.74 | 0.77 | 0.69 | 0.73 |
| 8 | 139173342 | rs11166794 | 0.332 | -0.321 | -0.223 | -1.453 | -0.008 | -0.011 | -0.011 | 0 | 0.7 | 0.62 | 0.74 | 0.73 | 0.68 | 0.76 |
| 8 | 139175182 | rs17655198 | -0.232 | -0.025 | -0.761 | -1.675 | -0.032 | 0.027 | 0.027 | 0 | 0.28 | 0.21 | 0.25 | 0.01 | 0.36 | 0.18 |
| 8 | 139175349 | rs3898456 | -0.495 | -0.104 | -0.711 | -1.568 | -0.007 | 0.037 | 0.037 | 0.168 | 0.29 | 0.22 | 0.2 | 0.13 | 0.31 | 0.17 |
| 8 | 139175625 | rs3897732 | -0.436 | -0.606 | -0.837 | -0.875 | 0.002 | -0.002 | -0.002 | 0 | 0.41 | 0.41 | 0.46 | 0.57 | 0.31 | 0.45 |
| 8 | 139177269 | rs7814738 | -0.119 | -0.317 | -0.78 | -1.505 | 0.006 | 0.119 | 0.119 | 0 | 0.66 | 0.88 | 0.78 | 0.98 | 0.71 | 0.79 |
| 8 | 139178416 | rs7835663 | 0.223 | -0.08 | -0.32 | -1.127 | -0.004 | -0.024 | -0.024 | 0 | 0.56 | 0.51 | 0.65 | 0.79 | 0.65 | 0.62 |
| 8 | 139186240 | rs4493955 | 0.384 | 0.387 | -0.668 | -0.83 | 0.007 | 0.037 | 0.037 | 0 | 0.47 | 0.41 | 0.59 | 0.78 | 0.58 | 0.61 |
| 8 | 139188250 | rs7824771 | 0.053 | 0.386 | -0.675 | -0.853 | 0.009 | -0.05 | -0.05 | 0.004 | 0.49 | 0.56 | 0.65 | 0.98 | 0.59 | 0.71 |
| 8 | 139191312 | rs4455890 | -0.163 | 0.318 | -1.129 | -0.513 | 0.014 | 0.024 | 0.024 | 0 | 0.47 | 0.56 | 0.36 | 0.21 | 0.33 | 0.38 |
| 8 | 139195589 | rs11780230 | -0.12 | 0.586 | -1.447 | -0.092 | 0.002 | 0.054 | 0.054 | 0 | 0.41 | 0.27 | 0.5 | 0.23 | 0.44 | 0.44 |
| 8 | 139196987 | rs16908495 | -0.399 | 0.167 | -0.847 | -0.241 | -0.022 | -0.032 | -0.032 | 0 | 0.06 | 0.23 | 0.03 | 0 | 0.02 | 0 |
| 8 | 139197874 | rs4520207 | -0.224 | -0.308 | -1.139 | -0.169 | 0.017 | 0.003 | 0.003 | 0 | 0.44 | 0.43 | 0.28 | 0.43 | 0.4 | 0.35 |
| 8 | 139198651 | rs11783192 | -0.379 | -0.145 | -0.357 | 0.024 | 0.004 | -0.015 | -0.015 | 0 | 0.57 | 0.48 | 0.66 | 0.76 | 0.58 | 0.7 |
| 8 | 139199407 | rs12550019 | 0.729 | -0.15 | -0.407 | 0.022 | 0.028 | -0.016 | -0.016 | 0 | 0.39 | 0.21 | 0.26 | 0.43 | 0.39 | 0.35 |
| 8 | 139199667 | rs7011071 | -0.637 | 0.122 | -0.199 | -0.297 | -0.004 | 0 | 0 | 0 | 0.39 | 0.6 | 0.58 | 0.44 | 0.33 | 0.42 |
| 8 | 139200513 | rs733518 | 0.22 | 0.058 | -0.071 | -0.287 | 0.001 | -0.011 | -0.011 | 1.269 | 0.53 | 0.27 | 0.42 | 0.55 | 0.51 | 0.54 |
| 8 | 139201036 | rs744881 | 1.603 | -0.5 | -0.5 | -0.65 | -0.046 | 0.069 | 0.069 | 0 | 0.16 | 0.03 | 0.15 | 0.02 | 0.27 | 0.06 |
| 8 | 139202095 | rs4579575 | 1.507 | -0.437 | -0.048 | -0.586 | 0.003 | 0.017 | 0.017 | 0 | 0.17 | 0.25 | 0.38 | 0.28 | 0.19 | 0.28 |
| 8 | 139202536 | rs6984968 | 0.445 | -0.465 | 0.057 | -0.631 | -0.013 | -0.036 | -0.036 | 0.003 | 0.22 | 0.33 | 0.2 | 0.14 | 0.15 | 0.16 |
| 8 | 139206628 | rs876369 | -0.303 | -0.455 | -0.007 | -0.585 | 0.001 | 0.009 | 0.009 | 0.01 | 0.57 | 0.64 | 0.66 | 0.82 | 0.44 | 0.69 |
| 8 | 139207792 | rs4634696 | 0.614 | -0.525 | -0.109 | -0.519 | -0.021 | 0.039 | 0.039 | 0.449 | 0.23 | 0.51 | 0.39 | 0.27 | 0.19 | 0.28 |
| 8 | 139210666 | rs16908562 | 2.199 | -0.496 | -0.52 | -0.53 | 0.053 | -0.086 | -0.086 | 0.921 | 0.21 | 0.06 | 0.1 | 0.38 | 0.14 | 0.34 |
| 8 | 139210676 | rs7845372 | 1.421 | -0.454 | -0.384 | -0.49 | 0 | -0.001 | -0.001 | 0 | 0.15 | 0.11 | 0.16 | 0.11 | 0.11 | 0.12 |
| 8 | 139211781 | rs6989344 | 0.111 | -0.421 | -0.471 | -0.286 | 0 | -0.001 | -0.001 | 0 | 0.54 | 0.51 | 0.44 | 0.51 | 0.51 | 0.53 |
| 8 | 139215025 | rs11995943 | -0.938 | -0.471 | -0.246 | 0.121 | -0.005 | 0.001 | 0.001 | 0 | 0.77 | 0.82 | 0.84 | 0.84 | 0.74 | 0.83 |
| 8 | 139221664 | rs11166797 | 1.294 | -0.531 | -0.201 | 0.31 | 0.065 | -0.089 | -0.089 | 0 | 0.21 | 0.06 | 0.1 | 0.38 | 0.13 | 0.29 |
| 8 | 139226808 | rs7462490 | -0.531 | 0.57 | 0.454 | 0.311 | 0.015 | 0.001 | 0.001 | 0.006 | 0.81 | 0.78 | 0.85 | 0.83 | 0.77 | 0.84 |
| 8 | 139228642 | rs7001598 | 0.305 | 0.506 | 0.353 | 0.42 | 0.008 | 0.028 | 0.028 | 0.196 | 0.72 | 0.52 | 0.76 | 0.7 | 0.73 | 0.76 |
| 8 | 139232371 | rs10093007 | -0.528 | 1.117 | 0.047 | 0.172 | -0.006 | 0 | 0 | 0 | 0.82 | 0.84 | 0.85 | 0.85 | 0.77 | 0.83 |
| 8 | 139233180 | rs6989502 | 0.771 | 0.88 | 0.513 | 0.585 | -0.013 | -0.005 | -0.005 | 2.227 | 0.36 | 0.35 | 0.21 | 0.3 | 0.26 | 0.44 |
| 8 | 139236845 | rs6577893 | -1.327 | 1.492 | 1.163 | 0.5 | 0.002 | 0.12 | 0.12 | 0 | 0.16 | 0.44 | 0.22 | 0.28 | 0.15 | 0.11 |
| 8 | 139237014 | rs6577894 | -1.299 | 1.794 | 1.204 | 0.355 | 0.014 | -0.014 | -0.014 | 0.462 | 0.31 | 0.45 | 0.41 | 0.29 | 0.37 | 0.18 |
| 8 | 139240445 | rs7017613 | -1.051 | 1.764 | 0.718 | 0.363 | 0.034 | -0.05 | -0.05 | 0 | 0.1 | 0.24 | 0.14 | 0.01 | 0.14 | 0.04 |
| 8 | 139240592 | rs7845202 | 0.063 | 1.783 | 0.55 | 0.374 | 0.008 | -0.025 | -0.025 | 0 | 0.07 | 0.09 | 0.1 | 0.01 | 0.11 | 0.04 |
| 8 | 139241611 | rs11166799 | -1.513 | 1.834 | 0.383 | 0.427 | 0.01 | -0.006 | -0.006 | 0 | 0.33 | 0.46 | 0.45 | 0.32 | 0.41 | 0.18 |
| 8 | 139241826 | rs6988219 | -1.324 | 1.595 | -0.075 | 0.532 | 0.007 | -0.009 | -0.009 | 0 | 0.31 | 0.45 | 0.43 | 0.3 | 0.38 | 0.17 |
| 8 | 139244132 | rs9969442 | -0.263 | 1.33 | -0.112 | 0.242 | 0.017 | -0.115 | -0.115 | 0.526 | 0.2 | 0.37 | 0.22 | 0.01 | 0.22 | 0.07 |
| 8 | 139244798 | rs16908658 | 0.002 | 1.549 | -0.757 | 0.461 | -0.05 | -0.053 | -0.053 | 0.575 | 0.18 | 0.05 | 0.13 | 0.23 | 0.18 | 0.19 |
| 8 | 139247027 | rs6577898 | -0.919 | 0.932 | 0.053 | 0.192 | 0.004 | -0.019 | -0.019 | 0 | 0.24 | 0.39 | 0.27 | 0.23 | 0.26 | 0.12 |
| 8 | 139253064 | rs872501 | -1.02 | 0.716 | 0.071 | 0.186 | 0.008 | -0.041 | -0.041 | 0.054 | 0.17 | 0.43 | 0.23 | 0.08 | 0.13 | 0.25 |
| 8 | 139253805 | rs4243888 | 0.173 | 0.254 | -0.533 | 0.391 | 0.011 | -0.004 | -0.004 | 0 | 0.54 | 0.42 | 0.43 | 0.49 | 0.37 | 0.48 |
| 8 | 139255733 | rs6577900 | -0.719 | 0.638 | -0.644 | 0.026 | 0 | -0.007 | -0.007 | 0.299 | 0.3 | 0.34 | 0.37 | 0.79 | 0.35 | 0.43 |
| 8 | 139261705 | rs7460570 | -0.102 | 0.865 | -0.768 | -1.308 | 0.013 | -0.081 | -0.081 | 0.989 | 0.38 | 0.55 | 0.23 | 0.07 | 0.35 | 0.37 |
| 8 | 139262090 | rs1996030 | 0.091 | 0.365 | -1.27 | -1.085 | 0.027 | -0.016 | -0.016 | 0 | 0.31 | 0.2 | 0.43 | 0.63 | 0.32 | 0.35 |
| 8 | 139263611 | rs7461019 | -0.86 | -0.307 | -1.722 | -1.465 | -0.063 | -0.034 | -0.034 | 0 | 0.71 | 0.98 | 0.71 | 0.73 | 0.71 | 0.73 |
| 8 | 139263840 | rs6577902 | 1.141 | 0.352 | -1.869 | -1.522 | 0.022 | -0.078 | -0.078 | 0 | 0.23 | 0.57 | 0.15 | 0.08 | 0.18 | 0.23 |
| 8 | 139269984 | rs1033129 | -0.163 | 0.277 | -1.805 | -0.908 | 0.015 | -0.059 | -0.059 | 0.692 | 0.39 | 0.5 | 0.36 | 0.16 | 0.4 | 0.39 |
| 8 | 139271002 | rs16908717 | -0.162 | 0.222 | -1.239 | -0.834 | 0.034 | -0.016 | -0.016 | 0 | 0.14 | 0.31 | 0.13 | 0.08 | 0.17 | 0.19 |
| 8 | 139271579 | rs7816485 | 0.996 | 0.287 | -1.096 | -0.833 | 0.011 | -0.035 | -0.035 | 0 | 0.24 | 0.44 | 0.25 | 0.15 | 0.18 | 0.25 |
| 8 | 139274137 | rs4909414 | 0.262 | 0.495 | -1.375 | -0.711 | 0.006 | -0.051 | -0.051 | 0 | 0.33 | 0.21 | 0.34 | 0.51 | 0.35 | 0.29 |
| 8 | 139278747 | rs4909768 | -0.338 | 0.17 | -1.372 | -0.49 | -0.006 | 0.008 | 0.008 | 0 | 0.29 | 0.14 | 0.3 | 0.27 | 0.35 | 0.33 |
| 8 | 139282109 | rs1398295 | 0.949 | -0.044 | -1.838 | -0.376 | -0.066 | 0.046 | 0.046 | 2.238 | 0.37 | 0.02 | 0.32 | 0.24 | 0.38 | 0.22 |
| 8 | 139286075 | rs16908741 | -1.421 | 0.02 | -1.314 | 0.15 | 0.006 | -0.013 | -0.013 | 0.713 | 0.66 | 0.79 | 0.6 | 0.66 | 0.67 | 0.73 |
| 8 | 139287998 | rs12545802 | -0.84 | 0.343 | -0.894 | 0.028 | 0.016 | 0.056 | 0.056 | 0.386 | 0.49 | 0.67 | 0.45 | 0.31 | 0.53 | 0.59 |
| 8 | 139289506 | rs16908758 | 0.452 | 0.221 | -0.453 | 0.197 | -0.002 | -0.025 | -0.025 | 0 | 0.1 | 0.1 | 0.08 | 0.01 | 0.11 | 0.12 |
| 8 | 139291730 | rs1398296 | 0.178 | 0.686 | -0.25 | 0.467 | -0.003 | -0.027 | -0.027 | 0.159 | 0.32 | 0.46 | 0.44 | 0.21 | 0.3 | 0.42 |
| 8 | 139293824 | rs1398297 | -0.465 | 0.809 | 0.185 | 0.981 | 0.044 | 0.043 | 0.043 | 0 | 0.89 | 0.67 | 0.92 | 0.87 | 0.88 | 0.88 |
| 8 | 139298819 | rs10505698 | -0.556 | 0.799 | -0.221 | 1.035 | -0.04 | -0.006 | -0.006 | 0 | 0.22 | 0.06 | 0.1 | 0.2 | 0.25 | 0.25 |

| | | | | | | | | | | | | | | | | |
|---|---|---|---|---|---|---|---|---|---|---|---|---|---|---|---|---|
| 8 | 139299165 | rs1512406 | 0.949 | 0.78 | -0.261 | 1.342 | 0.005 | 0.014 | 0.014 | 0 | 0.66 | 0.62 | 0.76 | 0.44 | 0.64 | 0.56 |
| 8 | 139302217 | rs7831210 | 1.053 | 0.175 | 0.303 | 1.216 | 0.008 | 0.067 | 0.067 | 0 | 0.49 | 0.38 | 0.42 | 0.27 | 0.44 | 0.35 |
| 8 | 139307042 | rs1512384 | 0.889 | -0.395 | 1.518 | 1.712 | 0.066 | 0.529 | 0.529 | 0 | 0.95 | 0.72 | 0.63 | 0.56 | 0.94 | 0.91 |
| 8 | 139309169 | rs1398293 | -0.611 | -0.394 | 1.724 | 1.381 | -0.002 | -0.011 | -0.011 | 1.793 | 0.32 | 0.36 | 0.18 | 0.22 | 0.3 | 0.44 |
| 8 | 139310880 | rs12542289 | -0.751 | -0.405 | 1.383 | 1.02 | 0.002 | -0.05 | -0.05 | 0 | 0.36 | 0.48 | 0.22 | 0.22 | 0.35 | 0.44 |
| 8 | 139316223 | rs1567689 | 0.03 | -0.21 | 0.861 | 1.116 | -0.018 | 0.056 | 0.056 | 0.006 | 0.94 | 0.88 | 0.97 | 0.83 | 0.97 | 0.97 |
| 8 | 139320385 | rs1022203 | -0.012 | -0.204 | 0.863 | 1.022 | -0.018 | 0.03 | 0.03 | 0.023 | 0.94 | 0.87 | 0.97 | 0.88 | 0.97 | 0.98 |
| 8 | 139321179 | rs6577909 | 0.782 | -0.447 | 1.311 | 0.799 | 0.002 | -0.009 | -0.009 | 0 | 0.6 | 0.5 | 0.44 | 0.32 | 0.61 | 0.49 |
| 8 | 139323883 | rs10875424 | -0.196 | -0.449 | 1.227 | 0.771 | -0.015 | 0.063 | 0.063 | 0 | 0.95 | 0.81 | 0.97 | 0.89 | 0.99 | 0.98 |
| 8 | 139327776 | rs6577913 | 0.316 | -0.691 | 1.479 | 0.761 | -0.022 | 0.015 | 0.015 | 0 | 0.72 | 0.54 | 0.51 | 0.38 | 0.76 | 0.69 |
| 8 | 139328196 | rs11166805 | 1.659 | -0.758 | 1.178 | 0.575 | 0.023 | 0.183 | 0.183 | 0 | 0.51 | 0.2 | 0.36 | 0.19 | 0.56 | 0.4 |
| 8 | 139337546 | rs1512394 | 1.068 | -0.83 | 1.382 | 0.924 | -0.013 | 0.24 | 0.24 | 0 | 0.9 | 0.64 | 0.86 | 0.71 | 0.96 | 0.91 |
| 8 | 139337902 | rs11782223 | -0.309 | -1.111 | 1.231 | 0.751 | 0.004 | 0.007 | 0.007 | 0.012 | 0.23 | 0.21 | 0.14 | 0.18 | 0.19 | 0.35 |
| 8 | 139339358 | rs1512396 | -1.036 | -1.096 | 1.222 | 0.724 | 0.005 | -0.009 | -0.009 | 0 | 0.4 | 0.33 | 0.48 | 0.55 | 0.39 | 0.53 |
| 8 | 139353715 | rs10105423 | 1.103 | -1.036 | 1.239 | 0.886 | 0.031 | 0.026 | 0.026 | 0 | 0.57 | 0.29 | 0.47 | 0.32 | 0.63 | 0.44 |
| 8 | 139357251 | rs12546144 | 1.085 | -1.03 | 1.279 | 0.845 | 0.013 | 0.009 | 0.009 | 0.126 | 0.58 | 0.37 | 0.48 | 0.34 | 0.63 | 0.42 |
| 8 | 139374614 | rs4909779 | 0.591 | -0.944 | 1.135 | 0.983 | 0.001 | -0.021 | -0.021 | 1.149 | 0.18 | 0.13 | 0.34 | 0.36 | 0.18 | 0.21 |
| 8 | 139381546 | rs340708 | 0.837 | -0.724 | 0.98 | 0.927 | 0.066 | 0.16 | 0.16 | 0 | 0.52 | 0.23 | 0.39 | 0.16 | 0.61 | 0.36 |
| 8 | 139381863 | rs908843 | -0.459 | -0.713 | 0.84 | 0.891 | -0.014 | -0.007 | -0.007 | 0 | 0.41 | 0.51 | 0.51 | 0.65 | 0.37 | 0.58 |
| 8 | 139387080 | rs7815456 | 0.544 | -0.607 | -0.159 | 0.873 | 0.003 | 0.047 | 0.047 | 0.636 | 0.3 | 0.16 | 0.17 | 0.26 | 0.3 | 0.43 |
| 8 | 139390139 | rs4909781 | -0.627 | -0.357 | -0.11 | 0.18 | 0.051 | 0.114 | 0.114 | 0 | 0.47 | 0.73 | 0.61 | 0.84 | 0.38 | 0.65 |
| 8 | 139394114 | rs1519370 | -0.042 | 0.021 | -0.812 | 0.154 | 0.016 | -0.003 | -0.003 | 0.189 | 0.37 | 0.29 | 0.46 | 0.64 | 0.34 | 0.54 |
| 8 | 139396960 | rs17667285 | 0.042 | 0.561 | -0.985 | -0.588 | -0.012 | 0.039 | 0.039 | 0 | 0.21 | 0.05 | 0.13 | 0.1 | 0.2 | 0.19 |
| 8 | 139398544 | rs12679167 | -0.296 | 0.608 | -0.656 | -0.53 | -0.002 | -0.008 | -0.008 | 0 | 0.23 | 0.29 | 0.18 | 0.12 | 0.24 | 0.18 |
| 8 | 139399116 | rs2056277 | 0.111 | 0.844 | -1.401 | -0.411 | -0.013 | 0.019 | 0.019 | 0 | 0.21 | 0.11 | 0.16 | 0.12 | 0.24 | 0.17 |
| 8 | 139399624 | rs7837068 | -0.419 | 1.446 | -0.743 | -0.286 | 0.01 | -0.064 | -0.064 | 0 | 0.2 | 0.24 | 0.18 | 0.03 | 0.24 | 0.17 |
| 8 | 139403283 | rs908841 | -0.911 | 1.463 | -1.515 | 0.369 | -0.057 | -0.029 | -0.029 | 0 | 0.5 | 0.93 | 0.79 | 0.55 | 0.54 | 0.47 |
| 8 | 139405963 | rs16909084 | -0.435 | 1.991 | -0.642 | 0.616 | -0.002 | -0.023 | -0.023 | 5.976 | 0.08 | 0.12 | 0.04 | 0 | 0.06 | 0.02 |
| 8 | 139413333 | rs2056412 | -0.991 | 2.241 | 0.002 | 0.928 | 0.016 | 0.158 | 0.158 | 0 | 0.09 | 0.15 | 0.36 | 0.45 | 0.11 | 0.23 |
| 8 | 139422252 | rs6577921 | -1.338 | 2.254 | 0.931 | 1.076 | 0.02 | -0.008 | -0.008 | 0 | 0.24 | 0.54 | 0.37 | 0.21 | 0.33 | 0.28 |
| 8 | 139424537 | rs4909424 | -1.838 | 2.297 | 0.822 | 1.103 | -0.004 | -0.002 | -0.002 | 0 | 0.3 | 0.71 | 0.6 | 0.36 | 0.41 | 0.24 |
| 8 | 139440366 | rs10101262 | -1.428 | 2.01 | 0.97 | 0.814 | -0.019 | 0.05 | 0.05 | 1.266 | 0.43 | 0.77 | 0.51 | 0.28 | 0.56 | 0.3 |
| 8 | 139446282 | rs925938 | -0.174 | 1.991 | 1.434 | 0.699 | -0.019 | 0.023 | 0.023 | 0 | 0.28 | 0.19 | 0.35 | 0.02 | 0.44 | 0.15 |
| 8 | 139448798 | rs1160458 | -1.641 | 1.933 | 1.193 | 0.888 | -0.052 | -0.036 | -0.036 | 9.492 | 0.48 | 0.91 | 0.79 | 0.54 | 0.6 | 0.41 |
| 8 | 139453936 | rs985679 | -1.313 | 1.918 | 2.157 | 1.257 | -0.009 | 0.02 | 0.02 | 2.661 | 0.39 | 0.71 | 0.68 | 0.28 | 0.5 | 0.3 |
| 8 | 139455120 | rs1519386 | -1.535 | 1.995 | 2.125 | 1.154 | -0.048 | 0.063 | 0.063 | 2.697 | 0.43 | 0.85 | 0.52 | 0.33 | 0.55 | 0.38 |
| 8 | 139462616 | rs1589228 | -1.478 | 1.976 | 2.17 | 0.867 | -0.038 | 0.139 | 0.139 | 2.68 | 0.43 | 0.84 | 0.45 | 0.25 | 0.53 | 0.37 |
| 8 | 139467139 | rs4397407 | -0.985 | 2.031 | 2.99 | 0.847 | 0.009 | -0.085 | -0.085 | 0 | 0.36 | 0.57 | 0.39 | 0.09 | 0.47 | 0.25 |
| 8 | 139472163 | rs1519372 | 0.329 | 2.088 | 3.248 | 0.845 | -0.038 | 0.082 | 0.082 | 0 | 0.69 | 0.83 | 0.71 | 0.99 | 0.56 | 0.77 |
| 8 | 139478382 | rs4909427 | 0.765 | 2.064 | 3.348 | 0.868 | 0.007 | -0.064 | -0.064 | 2.887 | 0.64 | 0.4 | 0.65 | 0.9 | 0.54 | 0.76 |
| 8 | 139478670 | rs16909189 | -1.53 | 2.062 | 3.28 | 0.86 | 0.075 | 0.087 | 0.087 | 0 | 0.05 | 0.22 | 0.06 | 0.16 | 0.06 | 0.12 |
| 8 | 139480820 | rs13249380 | 1.023 | 1.955 | 2.884 | 0.951 | -0.028 | 0.077 | 0.077 | 0.982 | 0.6 | 0.28 | 0.61 | 0.83 | 0.47 | 0.65 |
| 8 | 139481671 | rs11166825 | 0.947 | 1.956 | 2.753 | 0.952 | -0.037 | 0.114 | 0.114 | 1.319 | 0.6 | 0.24 | 0.6 | 0.83 | 0.47 | 0.64 |
| 8 | 139483441 | rs13251646 | 1.264 | 1.932 | 2.318 | 1.01 | -0.119 | -0.048 | -0.048 | 0 | 0.56 | 0.06 | 0.3 | 0.5 | 0.45 | 0.57 |
| 8 | 139499127 | rs6577927 | 2.505 | 1.958 | 3.285 | 1.321 | 0.008 | -0.002 | -0.002 | 1.241 | 0.84 | 0.25 | 0.8 | 0.83 | 0.83 | 0.81 |
| 8 | 139513971 | rs6992821 | -3.22 | 1.354 | 3.799 | 1.263 | 0.07 | -0.067 | -0.067 | 0.089 | 0.14 | 0.44 | 0.14 | 0.03 | 0.15 | 0.2 |
| 8 | 139518806 | rs4909430 | -3.094 | 1.19 | 3.821 | 1.275 | 0.05 | -0.061 | -0.061 | 0.288 | 0.14 | 0.44 | 0.13 | 0.03 | 0.14 | 0.2 |
| 8 | 139518938 | rs1519379 | -4.182 | 1.153 | 3.79 | 1.396 | 0.008 | 0.007 | 0.007 | 2.304 | 0.2 | 0.89 | 0.23 | 0.19 | 0.15 | 0.25 |
| 8 | 139537896 | rs7816642 | -2.465 | 1.052 | 4.03 | 1.814 | 0.032 | 0.178 | 0.178 | 0 | 0.09 | 0.35 | 0.14 | 0.19 | 0.06 | 0.24 |
| 8 | 139551812 | rs4909806 | -2.781 | 1.061 | 3.572 | 1.657 | -0.005 | 0.014 | 0.014 | 0 | 0.07 | 0.05 | 0.06 | 0.01 | 0.08 | 0.04 |
| 8 | 139554865 | rs7005917 | -3.246 | 1.037 | 3.869 | 1.683 | 0.011 | 0.044 | 0.044 | 0.071 | 0.11 | 0.36 | 0.14 | 0.17 | 0.09 | 0.12 |
| 8 | 139561394 | rs7014706 | -2.639 | 1.151 | 3.752 | 1.116 | 0.021 | -0.042 | -0.042 | 0 | 0.1 | 0.47 | 0.09 | 0.01 | 0.06 | 0.02 |
| 8 | 139566915 | rs4588859 | -3.152 | 0.983 | 3.76 | 1.333 | 0.033 | 0.145 | 0.145 | 0 | 0.12 | 0.39 | 0.17 | 0.21 | 0.1 | 0.26 |
| 8 | 139589488 | rs4909431 | -3.025 | 1.02 | 3.359 | 0.954 | 0.006 | -0.081 | -0.081 | 0 | 0.15 | 0.54 | 0.14 | 0.03 | 0.12 | 0.19 |
| 8 | 139593666 | rs6577934 | -2.907 | 0.86 | 3.422 | 0.926 | -0.011 | 0.013 | 0.013 | 3.763 | 0.17 | 0.56 | 0.24 | 0.15 | 0.13 | 0.12 |
| 8 | 139593796 | rs7017524 | -2.579 | 0.864 | 3.581 | 0.949 | 0.023 | 0.064 | 0.064 | 2.245 | 0.13 | 0.39 | 0.21 | 0.15 | 0.11 | 0.11 |
| 8 | 139598820 | rs7465307 | -2.144 | 0.857 | 3.153 | 0.936 | -0.008 | -0.038 | -0.038 | 0 | 0.12 | 0.23 | 0.12 | 0.04 | 0.08 | 0.17 |
| 8 | 139603111 | rs10091528 | -2.664 | 0.868 | 2.607 | 0.987 | 0.042 | 0.134 | 0.134 | 2.19 | 0.08 | 0.35 | 0.18 | 0.16 | 0.08 | 0.09 |
| 8 | 139604673 | rs10096258 | -2.823 | 0.841 | 1.468 | 0.951 | 0.009 | 0.005 | 0.005 | 1.673 | 0.29 | 0.95 | 0.37 | 0.21 | 0.2 | 0.27 |
| 8 | 139609087 | rs7832390 | -2.576 | 1.314 | 2.287 | 0.892 | 0.015 | -0.03 | -0.03 | 0 | 0.2 | 0.39 | 0.27 | 0.16 | 0.2 | 0.27 |
| 8 | 139620049 | rs12681582 | 2.816 | 1.227 | 3.236 | 1.168 | 0.001 | -0.003 | -0.003 | 0 | 0.75 | 0.25 | 0.56 | 0.7 | 0.76 | 0.63 |
| 8 | 139622885 | rs4341165 | 2.762 | 1.233 | 3.992 | 1.171 | -0.003 | 0.01 | 0.01 | 0.299 | 0.76 | 0.42 | 0.59 | 0.7 | 0.76 | 0.63 |
| 8 | 139623525 | rs7012042 | -2.87 | 1.237 | 3.989 | 1.144 | -0.006 | -0.077 | -0.077 | 0 | 0.2 | 0.39 | 0.23 | 0.1 | 0.19 | 0.24 |
| 8 | 139628889 | rs4076991 | 3.181 | 1.091 | 4.328 | 1.093 | 0.03 | 0.128 | 0.128 | 0.285 | 0.87 | 0.51 | 0.76 | 0.79 | 0.89 | 0.86 |
| 8 | 139634170 | rs4623432 | 3.502 | 1.064 | 4.301 | 1.157 | -0.018 | 0.087 | 0.087 | 0 | 0.85 | 0.49 | 0.75 | 0.8 | 0.92 | 0.85 |
| 8 | 139634852 | rs4576426 | 3.59 | 1.055 | 4.398 | 1.172 | -0.017 | 0.025 | 0.025 | 0.049 | 0.82 | 0.36 | 0.72 | 0.78 | 0.86 | 0.83 |
| 8 | 139637872 | rs6983378 | 3.671 | 1.147 | 4.417 | 1.181 | -0.012 | 0.013 | 0.013 | 0.167 | 0.82 | 0.29 | 0.7 | 0.78 | 0.87 | 0.83 |
| 8 | 139639729 | rs10094167 | 3.547 | 1.148 | 4.41 | 1.181 | -0.014 | 0.026 | 0.026 | 0.023 | 0.83 | 0.35 | 0.72 | 0.78 | 0.87 | 0.84 |
| 8 | 139642327 | rs4301456 | 3.328 | 1.184 | 4.417 | 1.224 | -0.049 | 0.062 | 0.062 | 1.988 | 0.86 | 0.46 | 0.79 | 0.83 | 0.96 | 0.96 |

| | | | | | | | | | | | | | | | | |
|---|---|---|---|---|---|---|---|---|---|---|---|---|---|---|---|---|
| 8 | 139642974 | rs4131277 | -3.703 | 1.148 | 4.204 | 1.181 | -0.018 | 0.059 | 0.059 | 0.013 | 0.16 | 0.54 | 0.25 | 0.2 | 0.08 | 0.14 |
| 8 | 139651182 | rs4909818 | 3.254 | 1.164 | 4.276 | 1.07 | 0.006 | -0.05 | -0.05 | 0 | 0.86 | 0.54 | 0.87 | 0.97 | 0.93 | 0.9 |
| 8 | 139652806 | rs10097597 | 2.831 | 1.18 | 4.302 | 1.08 | 0.027 | -0.017 | -0.017 | 0.16 | 0.89 | 0.55 | 0.89 | 0.97 | 0.94 | 0.89 |
| 8 | 139658057 | rs9324491 | -3.068 | 1.152 | 4.093 | 1.07 | -0.001 | -0.013 | -0.013 | 0 | 0.08 | 0.15 | 0.08 | 0.03 | 0.05 | 0.1 |
| 8 | 139658747 | rs4909439 | 2.839 | 1.123 | 4.164 | 1.1 | -0.006 | -0.019 | -0.019 | 0 | 0.83 | 0.7 | 0.79 | 0.9 | 0.87 | 0.7 |
| 8 | 139659737 | rs4909440 | 2.21 | 1.121 | 4.164 | 1.096 | -0.01 | -0.026 | -0.026 | 0.062 | 0.87 | 0.73 | 0.84 | 0.93 | 0.91 | 0.8 |
| 8 | 139662602 | rs4909825 | -2.188 | 1.151 | 3.753 | 1.239 | 0 | 0.026 | 0.026 | 0 | 0.06 | 0.08 | 0.17 | 0.2 | 0.05 | 0.18 |
| 8 | 139664209 | rs7460457 | 3.872 | 1.068 | 3.858 | 1.125 | 0.018 | -0.021 | -0.021 | 0.73 | 0.72 | 0.11 | 0.56 | 0.71 | 0.79 | 0.6 |
| 8 | 139665874 | rs4073446 | 2.966 | 1.051 | 4.249 | 1.11 | -0.033 | -0.056 | -0.056 | 0.118 | 0.81 | 0.37 | 0.79 | 0.93 | 0.9 | 0.79 |
| 8 | 139668032 | rs11166832 | 3.819 | 0.961 | 3.681 | 1.05 | -0.016 | -0.05 | -0.05 | 0.717 | 0.7 | 0.05 | 0.55 | 0.73 | 0.76 | 0.59 |
| 8 | 139669063 | rs7002247 | 3.047 | 0.894 | 4.512 | 1.056 | -0.019 | 0.036 | 0.036 | 0.321 | 0.8 | 0.43 | 0.67 | 0.76 | 0.88 | 0.75 |
| 8 | 139669119 | rs6989017 | 2.454 | 0.91 | 4.634 | 1.065 | 0.004 | 0.103 | 0.103 | 0.645 | 0.84 | 0.48 | 0.69 | 0.76 | 0.9 | 0.77 |
| 8 | 139669569 | rs7460303 | 1.899 | 0.924 | 4.589 | 0.977 | 0.027 | 0.101 | 0.101 | 0 | 0.88 | 0.66 | 0.71 | 0.77 | 0.9 | 0.77 |
| 8 | 139669711 | rs12547337 | -3.161 | 0.864 | 4.578 | 1.09 | 0.066 | 0.078 | 0.078 | 0 | 0.06 | 0.32 | 0.07 | 0.04 | 0.06 | 0.15 |
| 8 | 139674972 | rs4549820 | 2.601 | 0.902 | 4.237 | 0.886 | 0.022 | 0.06 | 0.06 | 0 | 0.78 | 0.43 | 0.58 | 0.73 | 0.77 | 0.61 |
| 8 | 139675763 | rs13272198 | 3.788 | 0.91 | 3.818 | 1.043 | -0.016 | -0.05 | -0.05 | 1.059 | 0.66 | 0.07 | 0.49 | 0.66 | 0.67 | 0.56 |
| 8 | 139675869 | rs11166835 | 2.796 | 0.932 | 3.852 | 1.061 | -0.001 | 0.014 | 0.014 | 0 | 0.74 | 0.33 | 0.54 | 0.69 | 0.75 | 0.57 |
| 8 | 139681125 | rs9657455 | 2.632 | 0.933 | 3.171 | 0.985 | -0.002 | -0.001 | -0.001 | 0 | 0.73 | 0.29 | 0.47 | 0.73 | 0.72 | 0.71 |
| 8 | 139686706 | rs6997633 | 2.793 | 1.158 | 2.426 | 0.717 | -0.029 | 0.031 | 0.031 | 0.02 | 0.64 | 0.11 | 0.45 | 0.73 | 0.59 | 0.57 |
| 8 | 139686742 | rs7844565 | -1.976 | 1.225 | 2.645 | 0.629 | -0.008 | 0.008 | 0.008 | 0 | 0.12 | 0.09 | 0.12 | 0.05 | 0.19 | 0.19 |
| 8 | 139689876 | rs7842696 | 1.85 | 1.19 | 3.706 | 0.608 | 0.024 | 0.011 | 0.011 | 0 | 0.83 | 0.57 | 0.76 | 0.83 | 0.81 | 0.77 |
| 8 | 139693463 | rs11777150 | -2.69 | 1.29 | 3.508 | 0.525 | -0.011 | 0.023 | 0.023 | 0 | 0.17 | 0.1 | 0.2 | 0.06 | 0.2 | 0.22 |
| 8 | 139693681 | rs7839680 | -1.92 | 1.275 | 4.038 | 0.536 | 0.006 | 0.006 | 0.006 | 0 | 0.14 | 0.2 | 0.21 | 0.16 | 0.19 | 0.23 |
| 8 | 139694420 | rs7844188 | 2.988 | 1.315 | 4.19 | 0.463 | -0.003 | -0.029 | -0.029 | 0 | 0.77 | 0.63 | 0.74 | 0.86 | 0.77 | 0.75 |
| 8 | 139696984 | rs11787079 | 3.133 | 1.019 | 4.39 | 0.236 | 0.005 | -0.025 | -0.025 | 0.665 | 0.63 | 0.42 | 0.62 | 0.79 | 0.57 | 0.53 |
| 8 | 139700911 | rs4909443 | -3.743 | 1.072 | 4.288 | 0.2 | -0.001 | 0.014 | 0.014 | 0 | 0.43 | 0.64 | 0.4 | 0.21 | 0.46 | 0.5 |
| 8 | 139701209 | rs4909444 | -2.651 | 1.099 | 4.189 | 0.22 | 0.005 | -0.001 | -0.001 | 0 | 0.24 | 0.25 | 0.3 | 0.18 | 0.34 | 0.4 |
| 8 | 139706888 | rs10109299 | -2.742 | 0.96 | 3.714 | 0.217 | -0.042 | -0.085 | -0.085 | 0.151 | 0.16 | 0.35 | 0.08 | 0.02 | 0.07 | 0.08 |
| 8 | 139708781 | rs10107433 | -3.265 | 1.018 | 3.792 | 0.188 | -0.034 | -0.056 | -0.056 | 0 | 0.21 | 0.52 | 0.16 | 0.13 | 0.15 | 0.14 |
| 8 | 139709805 | rs7835385 | 1.617 | 1.146 | 3.571 | 0.368 | 0.008 | 0.025 | 0.025 | 0.051 | 0.89 | 0.64 | 0.9 | 0.92 | 0.9 | 0.88 |
| 8 | 139711869 | rs4574887 | 2.628 | 1.127 | 3.375 | 0.334 | 0 | -0.006 | -0.006 | 0.046 | 0.69 | 0.64 | 0.71 | 0.78 | 0.67 | 0.58 |
| 8 | 139712346 | rs4909836 | -1.698 | 1.034 | 3.013 | 0.343 | -0.014 | 0.013 | 0.013 | 0 | 0.08 | 0.16 | 0.09 | 0.12 | 0.03 | 0.1 |
| 8 | 139725718 | rs12681358 | 1.943 | 1.114 | 3.134 | 0.386 | -0.014 | 0.04 | 0.04 | 0 | 0.82 | 0.58 | 0.81 | 0.79 | 0.87 | 0.8 |
| 8 | 139727981 | rs9324493 | -1.535 | 1.401 | 2.85 | 0.315 | 0.027 | -0.005 | -0.005 | 0 | 0.1 | 0.25 | 0.11 | 0.05 | 0.14 | 0.08 |
| 8 | 139730104 | rs11778635 | -1.252 | 1.565 | 2.369 | 0.236 | -0.037 | 0.058 | 0.058 | 0.003 | 0.16 | 0.05 | 0.16 | 0.02 | 0.22 | 0.21 |
| 8 | 139731297 | rs11166838 | -2.084 | 1.293 | 2.482 | 0.348 | 0.001 | -0.008 | -0.008 | 0 | 0.43 | 0.59 | 0.45 | 0.24 | 0.49 | 0.4 |
| 8 | 139737297 | rs12550730 | 1.739 | 2.028 | 1.564 | 0.955 | -0.021 | 0.003 | 0.003 | 0.089 | 0.6 | 0.31 | 0.53 | 0.62 | 0.51 | 0.6 |
| 8 | 139738961 | rs4075073 | -2.263 | 2.045 | 2.258 | 1.065 | 0.001 | 0.003 | 0.003 | 0 | 0.35 | 0.55 | 0.39 | 0.38 | 0.38 | 0.37 |
| 8 | 139745582 | rs10096388 | -2.378 | 1.998 | 2.9 | 1.255 | 0 | 0.002 | 0.002 | 0 | 0.35 | 0.66 | 0.42 | 0.33 | 0.35 | 0.36 |
| 8 | 139749638 | rs4409435 | -2.454 | 1.804 | 3.272 | 1.594 | 0.005 | 0.03 | 0.03 | 1.232 | 0.33 | 0.47 | 0.45 | 0.49 | 0.31 | 0.41 |
| 8 | 139749799 | rs10111520 | -0.738 | 1.869 | 2.833 | 1.387 | -0.004 | -0.012 | -0.012 | 4.373 | 0.06 | 0.01 | 0.11 | 0.05 | 0.05 | 0.05 |
| 8 | 139751900 | rs10109866 | 2.167 | 3.109 | 3.397 | 1.282 | 0.016 | 0.005 | 0.005 | 0.823 | 0.81 | 0.27 | 0.8 | 0.76 | 0.79 | 0.85 |
| 8 | 139752536 | rs7835533 | -1.306 | 3.037 | 3.096 | 1.229 | -0.029 | 0.029 | 0.029 | 0 | 0.11 | 0.04 | 0.15 | 0.04 | 0.16 | 0.11 |
| 8 | 139752908 | rs7839333 | -1.356 | 3.035 | 3.116 | 1.23 | -0.016 | 0.018 | 0.018 | 0 | 0.11 | 0.07 | 0.16 | 0.05 | 0.16 | 0.11 |
| 8 | 139764396 | rs9324496 | 2.781 | 3.304 | 3.335 | 1.517 | -0.033 | 0.001 | 0.001 | 0.018 | 0.69 | 0.23 | 0.63 | 0.64 | 0.62 | 0.7 |
| 8 | 139764832 | rs2318340 | 1.75 | 3.382 | 3.607 | 1.514 | 0.047 | -0.051 | -0.051 | 1.343 | 0.87 | 0.78 | 0.88 | 0.98 | 0.79 | 0.94 |
| 8 | 139768085 | rs7825723 | 2.466 | 3.39 | 3.64 | 1.107 | 0.026 | -0.021 | -0.021 | 0 | 0.77 | 0.73 | 0.71 | 0.87 | 0.63 | 0.77 |
| 8 | 139768859 | rs7831232 | 1.204 | 3.791 | 3.387 | 0.994 | 0.047 | -0.029 | -0.029 | 0 | 0.93 | 0.87 | 0.91 | 1 | 0.82 | 0.94 |
| 8 | 139769993 | rs4736185 | 2.448 | 3.708 | 3.601 | 0.996 | 0.069 | -0.092 | -0.092 | 0 | 0.78 | 0.54 | 0.79 | 0.94 | 0.73 | 0.79 |
| 8 | 139770716 | rs7824025 | 1.173 | 3.563 | 2.854 | 0.975 | 0.006 | -0.019 | -0.019 | 0 | 0.93 | 0.87 | 0.94 | 0.99 | 0.91 | 0.95 |
| 8 | 139774092 | rs13279646 | -2.024 | 3.527 | 2.959 | 1.125 | 0.096 | 0.02 | 0.02 | 0.159 | 0.17 | 0.49 | 0.21 | 0.18 | 0.23 | 0.25 |
| 8 | 139774409 | rs6577946 | 1.817 | 3.194 | 1.917 | 1.156 | -0.016 | 0.018 | 0.018 | 0 | 0.69 | 0.28 | 0.67 | 0.81 | 0.61 | 0.68 |
| 8 | 139781866 | rs7844608 | -1.723 | 2.751 | 1.897 | 1.547 | 0.022 | 0.053 | 0.053 | 0 | 0.13 | 0.34 | 0.13 | 0.21 | 0.17 | 0.22 |
| 8 | 139782538 | rs4736183 | -1.493 | 2.391 | 1.099 | 1.363 | 0.011 | 0.029 | 0.029 | 0 | 0.33 | 0.6 | 0.43 | 0.46 | 0.4 | 0.51 |
| 8 | 139785792 | rs5019355 | -2.018 | 1.83 | 1.448 | 0.795 | 0.043 | -0.014 | -0.014 | 0 | 0.07 | 0.36 | 0.11 | 0 | 0.04 | 0.1 |
| 8 | 139818903 | rs11779129 | -1.704 | 1.826 | 0.962 | 0.362 | -0.022 | 0.08 | 0.08 | 0.998 | 0.3 | 0.87 | 0.32 | 0.09 | 0.27 | 0.43 |
| 8 | 139821059 | rs10090299 | 1.791 | 1.701 | 1.502 | 0.897 | -0.004 | -0.004 | -0.004 | 0.054 | 0.6 | 0.37 | 0.43 | 0.57 | 0.53 | 0.39 |
| 8 | 139822483 | rs1879051 | 1.929 | 1.704 | 1.727 | 0.895 | -0.028 | -0.021 | -0.021 | 1.089 | 0.58 | 0.26 | 0.41 | 0.57 | 0.53 | 0.39 |
| 8 | 139824057 | rs10101430 | 1.759 | 1.793 | 1.899 | 0.968 | 0.01 | 0.009 | 0.009 | 0.008 | 0.64 | 0.41 | 0.44 | 0.57 | 0.55 | 0.44 |
| 8 | 139825393 | rs6577951 | 0.68 | 1.578 | 1.726 | 0.878 | 0.002 | 0.039 | 0.039 | 0 | 0.85 | 0.77 | 0.72 | 0.66 | 0.82 | 0.79 |
| 8 | 139826029 | rs10481409 | 0.778 | 1.581 | 1.67 | 0.873 | 0.002 | 0.075 | 0.075 | 0 | 0.84 | 0.67 | 0.7 | 0.66 | 0.81 | 0.78 |
| 8 | 139830796 | rs1320279 | 0.056 | 1.7 | 0.98 | 0.968 | -0.032 | -0.022 | -0.022 | 0 | 0.94 | 0.99 | 0.89 | 0.75 | 0.88 | 0.93 |
| 8 | 139831266 | rs1316523 | -1.077 | 1.541 | 1.076 | 0.614 | -0.018 | 0.025 | 0.025 | 0 | 0.19 | 0.11 | 0.21 | 0.05 | 0.23 | 0.3 |
| 8 | 139838912 | rs2292927 | 0.509 | 1.189 | 0.587 | 0.358 | 0.004 | -0.013 | -0.013 | 1.522 | 0.8 | 0.87 | 0.82 | 0.76 | 0.81 | 0.88 |
| 8 | 139839361 | rs3750282 | 1.128 | 0.929 | 0.324 | 0.176 | -0.006 | 0.015 | 0.015 | 0 | 0.57 | 0.42 | 0.58 | 0.69 | 0.58 | 0.52 |
| 8 | 139842957 | rs4736009 | 0.921 | 0.84 | 0.098 | 0.249 | 0.001 | -0.005 | -0.005 | 0 | 0.6 | 0.46 | 0.63 | 0.7 | 0.6 | 0.56 |
| 8 | 139846822 | rs6995718 | 0.888 | 0.851 | -0.261 | 0.263 | -0.002 | 0.003 | 0.003 | 0 | 0.61 | 0.42 | 0.6 | 0.7 | 0.61 | 0.56 |
| 8 | 139859313 | rs2318331 | 0.354 | 0.8 | -1.118 | -0.327 | 0.01 | 0.129 | 0.129 | 1.257 | 0.52 | 0.2 | 0.42 | 0.7 | 0.57 | 0.63 |
| 8 | 139862185 | rs9644504 | -0.298 | -0.226 | -1.175 | -0.656 | 0.047 | 0.008 | 0.008 | 0.386 | 0.28 | 0.12 | 0.25 | 0.25 | 0.15 | 0.18 |

| | | | | | | | | | | | | | | | | |
|---|---|---|---|---|---|---|---|---|---|---|---|---|---|---|---|---|
| 8 | 139863048 | rs6577956 | 1.01 | -0.05 | -0.483 | -1.146 | 0.028 | 0.04 | 0.04 | 0.574 | 0.49 | 0.36 | 0.47 | 0.7 | 0.64 | 0.7 |
| 8 | 139866268 | rs7833411 | 1.097 | 0.726 | 0.045 | -1.626 | -0.005 | 0.029 | 0.029 | 0.034 | 0.45 | 0.37 | 0.49 | 0.75 | 0.55 | 0.64 |
| 8 | 139867676 | rs11777748 | -0.03 | 0.662 | -0.135 | -1.733 | 0.043 | 0.042 | 0.042 | 0.024 | 0.32 | 0.12 | 0.25 | 0.23 | 0.2 | 0.25 |
| 8 | 139869685 | rs12675384 | -0.819 | 0.903 | 0.279 | -1.598 | -0.004 | -0.026 | -0.026 | 0 | 0.36 | 0.61 | 0.44 | 0.24 | 0.3 | 0.27 |
| 8 | 139872620 | rs1574172 | 0.481 | 1.054 | 0.768 | -1.404 | -0.009 | -0.07 | -0.07 | 0 | 0.81 | 0.72 | 0.81 | 0.98 | 0.86 | 0.92 |
| 8 | 139875116 | rs4736078 | 1.163 | 1.002 | 0.051 | -1.42 | 0.001 | -0.09 | -0.09 | 1.34 | 0.75 | 0.34 | 0.71 | 0.98 | 0.76 | 0.88 |
| 8 | 139876185 | rs11166852 | -0.128 | 0.77 | -0.104 | -1.283 | 0 | -0.001 | -0.001 | 0.763 | 0.89 | 0.92 | 0.92 | 0.98 | 0.91 | 0.95 |
| 8 | 139876426 | rs11775756 | 0.729 | 1.112 | -0.538 | -0.541 | -0.001 | -0.001 | -0.001 | 0 | 0.68 | 0.32 | 0.6 | 0.79 | 0.63 | 0.79 |
| 8 | 139879572 | rs16909722 | 0.049 | 0.718 | -0.437 | -0.548 | -0.002 | 0.001 | 0.001 | 0 | 0.19 | 0.2 | 0.14 | 0.25 | 0.12 | 0.18 |
| 8 | 139880901 | rs11991356 | -0.762 | 0.758 | -0.932 | -1.865 | 0.015 | 0.124 | 0.124 | 0.634 | 0.4 | 0.77 | 0.27 | 0.02 | 0.29 | 0.14 |
| 8 | 139883488 | rs13265526 | -0.93 | 0.831 | -0.287 | -1.939 | 0.001 | -0.043 | -0.043 | 0 | 0.39 | 0.59 | 0.28 | 0.02 | 0.3 | 0.16 |
| 8 | 139884509 | rs6988229 | -0.473 | 0.927 | 0.064 | -1.995 | 0.025 | -0.122 | -0.122 | 0 | 0.21 | 0.6 | 0.18 | 0.01 | 0.19 | 0.08 |
| 8 | 139886780 | rs6998323 | 0.154 | 1.17 | 0.78 | -1.705 | 0.036 | -0.027 | -0.027 | 0.057 | 0.8 | 0.66 | 0.87 | 0.88 | 0.77 | 0.83 |
| 8 | 139886837 | rs7011683 | 1.366 | 0.898 | -0.036 | -2.163 | 0.025 | 0.122 | 0.122 | 0 | 0.48 | 0.3 | 0.6 | 0.85 | 0.46 | 0.7 |
| 8 | 139888196 | rs12549200 | -1.537 | 0.917 | 0.841 | -1.772 | 0.014 | -0.03 | -0.03 | 0 | 0.18 | 0.41 | 0.17 | 0.12 | 0.18 | 0.21 |
| 8 | 139894529 | rs12541806 | 0.674 | 0.868 | 1.132 | -1.784 | 0.021 | -0.05 | -0.05 | 0.391 | 0.77 | 0.4 | 0.77 | 0.9 | 0.72 | 0.8 |
| 8 | 139901178 | rs2318345 | -0.074 | 0.918 | 1.468 | -1.712 | 0.128 | 0.025 | 0.025 | 0.019 | 0.05 | 0.34 | 0.06 | 0.01 | 0.1 | 0.03 |
| 8 | 139902305 | rs7013414 | 0.705 | 0.948 | 1.584 | -1.73 | 0.121 | 0.02 | 0.02 | 0.128 | 0.93 | 0.6 | 0.89 | 1 | 0.91 | 0.94 |
| 8 | 139904247 | rs7000425 | 1.202 | 0.658 | 1.554 | -1.793 | 0.004 | -0.039 | -0.039 | 0.001 | 0.69 | 0.42 | 0.75 | 0.86 | 0.68 | 0.71 |
| 8 | 139906088 | rs7006103 | -0.866 | 0.656 | 1.749 | -1.79 | -0.014 | -0.022 | -0.022 | 0 | 0.25 | 0.35 | 0.14 | 0.14 | 0.17 | 0.23 |
| 8 | 139906733 | rs1879046 | -0.504 | 0.657 | 1.708 | -1.789 | 0 | -0.001 | -0.001 | 0 | 0.23 | 0.22 | 0.13 | 0.14 | 0.17 | 0.23 |
| 8 | 139910597 | rs2318346 | 1.261 | 0.718 | 1.776 | -1.759 | 0 | -0.005 | -0.005 | 0.578 | 0.69 | 0.31 | 0.74 | 0.86 | 0.68 | 0.71 |
| 8 | 139911539 | rs2318348 | 0.725 | 0.704 | 1.564 | -1.738 | -0.003 | -0.038 | -0.038 | 0 | 0.75 | 0.57 | 0.8 | 0.85 | 0.78 | 0.74 |
| 8 | 139914637 | rs7007907 | 1.461 | 0.543 | 0.185 | -1.591 | -0.088 | 0.027 | 0.027 | 0 | 0.4 | 0.03 | 0.4 | 0.35 | 0.48 | 0.28 |
| 8 | 139916909 | rs11780422 | 0.296 | 0.459 | 0.452 | -1.6 | -0.001 | -0.003 | -0.003 | 0 | 0.22 | 0.21 | 0.13 | 0.15 | 0.18 | 0.21 |
| 8 | 139919267 | rs6986041 | 0.391 | 0.574 | 0.95 | -1.273 | 0 | 0.003 | 0.003 | 0 | 0.62 | 0.45 | 0.56 | 0.47 | 0.65 | 0.49 |
| 8 | 139920060 | rs12681025 | 1.341 | 0.842 | 0.419 | -1.183 | -0.025 | -0.075 | -0.075 | 0.12 | 0.22 | 0.08 | 0.32 | 0.5 | 0.19 | 0.42 |
| 8 | 139921285 | rs4736204 | 0.126 | 0.651 | 1.142 | -1.52 | 0.041 | 0.107 | 0.107 | 0 | 0.35 | 0.14 | 0.19 | 0.11 | 0.25 | 0.25 |
| 8 | 139924336 | rs1316448 | 0.44 | 0.579 | 0.841 | -1.417 | 0.014 | 0.048 | 0.048 | 0 | 0.55 | 0.33 | 0.68 | 0.85 | 0.6 | 0.7 |
| 8 | 139926589 | rs2318349 | 0.3 | 0.969 | 1.044 | -1.3 | -0.004 | 0.026 | 0.026 | 0 | 0.58 | 0.33 | 0.68 | 0.85 | 0.57 | 0.72 |
| 8 | 139944993 | rs11166859 | 2.814 | 1.316 | 2.82 | 0.246 | 0.089 | 0.03 | 0.03 | 0.306 | 0.91 | 0.61 | 0.86 | 0.95 | 0.9 | 0.81 |
| 8 | 139951715 | rs11166863 | 1.074 | 1.541 | 2.91 | 0.207 | 0.115 | -0.061 | -0.061 | 2.271 | 0.9 | 0.72 | 0.85 | 1 | 0.81 | 0.87 |
| 8 | 139954311 | rs13259487 | 0.57 | 1.634 | 2.953 | 0.312 | 0.105 | -0.021 | -0.021 | 0.154 | 0.85 | 0.54 | 0.82 | 0.88 | 0.82 | 0.83 |
| 8 | 139962091 | rs2135461 | -1.211 | 1.591 | 2.951 | 0.409 | 0.073 | -0.039 | -0.039 | 0.58 | 0.09 | 0.42 | 0.07 | 0 | 0.09 | 0.1 |
| 8 | 139962250 | rs2318822 | -0.844 | 1.593 | 2.764 | 0.415 | 0.037 | -0.044 | -0.044 | 0.004 | 0.08 | 0.27 | 0.05 | 0 | 0.09 | 0.1 |
| 8 | 139964213 | rs10091597 | -0.893 | 1.602 | 2.711 | 0.419 | 0.047 | -0.041 | -0.041 | 0.248 | 0.09 | 0.33 | 0.06 | 0 | 0.09 | 0.1 |
| 8 | 139975998 | rs6577977 | 0.921 | 1.723 | 2.398 | 1.256 | 0.033 | 0.086 | 0.086 | 0 | 0.81 | 0.68 | 0.64 | 0.64 | 0.76 | 0.68 |
| 8 | 139976182 | rs7834262 | -0.031 | 1.425 | 2.265 | 0.821 | -0.006 | 0.017 | 0.017 | 0 | 0.2 | 0.17 | 0.12 | 0.17 | 0.2 | 0.24 |
| 8 | 139976312 | rs6577978 | 1.709 | 1.38 | 3.168 | 1.284 | 0.024 | -0.101 | -0.101 | 3.668 | 0.83 | 0.69 | 0.8 | 1 | 0.82 | 0.86 |
| 8 | 139977470 | rs7817046 | 1.966 | 0.592 | 2.984 | 2.022 | -0.002 | 0.05 | 0.05 | 0 | 0.85 | 0.78 | 0.76 | 0.65 | 0.87 | 0.79 |
| 8 | 139989927 | rs2135455 | 2.332 | 0.636 | 2.586 | 1.979 | -0.003 | 0.018 | 0.018 | 0 | 0.82 | 0.77 | 0.74 | 0.65 | 0.84 | 0.66 |
| 8 | 139991388 | rs13262435 | -2.078 | 0.579 | 2.523 | 1.972 | 0.022 | 0.184 | 0.184 | 0 | 0.06 | 0.27 | 0.1 | 0.22 | 0.04 | 0.05 |
| 8 | 139996693 | rs4736198 | 1.276 | 0.708 | 1.748 | 1.337 | -0.01 | -0.047 | -0.047 | 0.157 | 0.89 | 0.85 | 0.95 | 1 | 0.94 | 0.95 |
| 8 | 140006609 | rs2318832 | 2.019 | 0.575 | 1.431 | 1.345 | 0.029 | 0.036 | 0.036 | 0.655 | 0.74 | 0.2 | 0.69 | 0.81 | 0.87 | 0.76 |
| 8 | 140007284 | rs4297062 | -2.599 | 0.922 | 2.656 | 1.422 | 0.11 | 0.097 | 0.097 | 0 | 0.09 | 0.47 | 0.22 | 0.13 | 0.1 | 0.13 |
| 8 | 140011514 | rs6577982 | -2.815 | 0.978 | 3.221 | 1.382 | 0.066 | 0.035 | 0.035 | 0 | 0.11 | 0.38 | 0.25 | 0.13 | 0.13 | 0.15 |
| 8 | 140013519 | rs1027787 | 1.685 | 0.904 | 3.347 | 1.223 | -0.004 | -0.038 | -0.038 | 0 | 0.91 | 0.84 | 0.95 | 1 | 0.95 | 0.96 |
| 8 | 140015639 | rs16909936 | 2.989 | 0.915 | 3.988 | 1.534 | 0.004 | -0.023 | -0.023 | 0 | 0.74 | 0.56 | 0.66 | 0.7 | 0.78 | 0.69 |
| 8 | 140018998 | rs2873810 | -2.96 | 0.738 | 3.881 | 1.561 | 0.006 | 0.007 | 0.007 | 0 | 0.18 | 0.3 | 0.28 | 0.28 | 0.17 | 0.29 |
| 8 | 140021020 | rs12680976 | -1.947 | 0.838 | 3.795 | 2.105 | -0.004 | 0.098 | 0.098 | 0.299 | 0.1 | 0.13 | 0.14 | 0.51 | 0.1 | 0.24 |
| 8 | 140024361 | rs1588022 | -2.494 | 0.935 | 3.873 | 2.02 | 0.037 | 0.058 | 0.058 | 0 | 0.14 | 0.27 | 0.28 | 0.28 | 0.16 | 0.23 |
| 8 | 140024703 | rs13257708 | -2.263 | 1.062 | 3.909 | 2.578 | 0 | 0.113 | 0.113 | 0.285 | 0.11 | 0.18 | 0.16 | 0.44 | 0.12 | 0.28 |
| 8 | 140025425 | rs12680224 | -3.077 | 1.057 | 3.854 | 2.573 | -0.006 | 0.102 | 0.102 | 0 | 0.15 | 0.25 | 0.16 | 0.44 | 0.13 | 0.3 |
| 8 | 140028933 | rs2318830 | 2.003 | 1.063 | 3.294 | 2.394 | 0.003 | -0.017 | -0.017 | 0 | 0.86 | 0.87 | 0.9 | 1 | 0.9 | 0.91 |
| 8 | 140029711 | rs1032928 | -2.928 | 1.078 | 3.395 | 2.506 | 0.026 | 0.077 | 0.077 | 0 | 0.12 | 0.27 | 0.23 | 0.29 | 0.15 | 0.26 |
| 8 | 140031004 | rs10095842 | 2.314 | 1.051 | 3.303 | 2.403 | 0.001 | 0.01 | 0.01 | 0.363 | 0.8 | 0.81 | 0.71 | 0.55 | 0.83 | 0.71 |
| 8 | 140056945 | rs13267587 | -2.888 | 1.058 | 1.79 | 2.47 | -0.003 | 0.002 | 0.002 | 1.81 | 0.3 | 0.67 | 0.43 | 0.63 | 0.23 | 0.41 |
| 8 | 140061723 | rs1827915 | 1.176 | 1.522 | 0.759 | 2.678 | -0.016 | -0.007 | -0.007 | 0 | 0.93 | 0.96 | 0.8 | 0.91 | 0.87 | 0.86 |
| 8 | 140062905 | rs1827916 | -1.533 | 1.549 | 0.708 | 2.799 | -0.033 | -0.019 | -0.019 | 0 | 0.08 | 0 | 0.2 | 0.45 | 0.08 | 0.22 |
| 8 | 140068246 | rs10099744 | -1.01 | 1.427 | 1.03 | 2.171 | -0.007 | -0.035 | -0.035 | 0 | 0.08 | 0.16 | 0.09 | 0.01 | 0.05 | 0.08 |
| 8 | 140073693 | rs1395316 | 0.958 | 1.406 | 0.835 | 2.197 | 0.019 | -0.018 | -0.018 | 0 | 0.85 | 0.72 | 0.68 | 0.9 | 0.83 | 0.78 |
| 8 | 140081806 | rs16910069 | -1.72 | 1.288 | 0.068 | 2.384 | -0.019 | 0.017 | 0.017 | 0 | 0.1 | 0.22 | 0.05 | 0.19 | 0.07 | 0.09 |
| 8 | 140093896 | rs16910093 | -1.791 | 1.813 | -0.627 | 2.789 | -0.02 | 0.001 | 0.001 | 0 | 0.06 | 0.01 | 0.14 | 0.33 | 0.09 | 0.15 |
| 8 | 140096660 | rs13257220 | -1.665 | 2.204 | -0.597 | 2.49 | -0.007 | 0.002 | 0.002 | 0 | 0.14 | 0.12 | 0.35 | 0.52 | 0.27 | 0.31 |
| 8 | 140105131 | rs10875434 | -1.65 | 2.594 | -0.876 | 2.458 | -0.045 | -0.06 | -0.06 | 0 | 0.2 | 0.11 | 0.4 | 0.48 | 0.33 | 0.48 |
| 8 | 140109524 | rs2202559 | -0.564 | 1.585 | -0.317 | 2.179 | -0.03 | 0.01 | 0.01 | 0.005 | 0.26 | 0.54 | 0.26 | 0.28 | 0.19 | 0.25 |
| 8 | 140112894 | rs6577989 | -1.279 | 1.909 | 0.659 | 1.601 | -0.003 | -0.019 | -0.019 | 0.007 | 0.37 | 0.57 | 0.33 | 0.26 | 0.32 | 0.31 |
| 8 | 140112914 | rs13248142 | -0.921 | 1.805 | 1.013 | 1.516 | 0 | -0.001 | -0.001 | 0 | 0.11 | 0.11 | 0.1 | 0.04 | 0.12 | 0.19 |
| 8 | 140116013 | rs17750756 | 0.984 | 1.613 | -0.092 | 0.595 | -0.018 | 0.057 | 0.057 | 0.575 | 0.13 | 0.02 | 0.09 | 0 | 0.12 | 0.03 |

| | | | | | | | | | | | | | | | | |
|---|---|---|---|---|---|---|---|---|---|---|---|---|---|---|---|---|
| 8 | 140118795 | rs13265485 | -1.445 | 1.573 | 0.066 | 0.633 | 0.004 | 0.046 | 0.046 | 0.253 | 0.38 | 0.79 | 0.35 | 0.29 | 0.32 | 0.32 |
| 8 | 140121236 | rs12548622 | -0.928 | 1.489 | 0.468 | 0.611 | -0.002 | -0.02 | -0.02 | 0.235 | 0.13 | 0.18 | 0.1 | 0.04 | 0.13 | 0.2 |
| 8 | 140127523 | rs6577993 | -1.475 | 1.639 | -0.493 | -0.276 | -0.029 | 0.084 | 0.084 | 0.197 | 0.6 | 0.9 | 0.51 | 0.26 | 0.55 | 0.39 |
| 8 | 140129789 | rs12543015 | -0.254 | 1.477 | -0.296 | -0.358 | -0.007 | -0.041 | -0.041 | 0.021 | 0.26 | 0.27 | 0.15 | 0.04 | 0.2 | 0.22 |
| 8 | 140132539 | rs11987078 | -0.453 | 1.412 | -0.333 | -0.121 | -0.005 | -0.023 | -0.023 | 1.276 | 0.27 | 0.27 | 0.15 | 0.07 | 0.2 | 0.24 |
| 8 | 140132614 | rs11997682 | 1.012 | 1.404 | -0.29 | -0.216 | -0.009 | -0.046 | -0.046 | 0 | 0.15 | 0.15 | 0.06 | 0 | 0.07 | 0.03 |
| 8 | 140132848 | rs11166878 | -0.333 | 1.405 | -0.29 | -0.214 | -0.005 | -0.036 | -0.036 | 0 | 0.26 | 0.26 | 0.14 | 0.04 | 0.2 | 0.22 |
| 8 | 140137118 | rs7815794 | -0.617 | 1.387 | -0.026 | -0.179 | -0.017 | -0.012 | -0.012 | 0 | 0.28 | 0.53 | 0.26 | 0.23 | 0.2 | 0.23 |
| 8 | 140138934 | rs4736209 | -1.707 | 1.602 | -0.044 | -0.271 | 0.033 | 0.03 | 0.03 | 0 | 0.22 | 0.56 | 0.43 | 0.28 | 0.25 | 0.29 |
| 8 | 140141910 | rs7462091 | -1.586 | 0.748 | -0.908 | -0.008 | 0.015 | -0.086 | -0.086 | 0.774 | 0.88 | 0.99 | 0.94 | 0.57 | 0.92 | 0.76 |
| 8 | 140149457 | rs6981713 | 0.084 | 0.744 | -0.575 | -0.183 | -0.023 | -0.089 | -0.089 | 0 | 0.26 | 0.35 | 0.15 | 0.04 | 0.2 | 0.22 |
| 8 | 140149784 | rs13264432 | 0.613 | 0.795 | -0.621 | -0.127 | -0.001 | 0 | 0 | 7.972 | 0.11 | 0.11 | 0.09 | 0.04 | 0.12 | 0.19 |
| 8 | 140153172 | rs11775767 | -0.549 | 0.891 | -0.286 | -0.28 | -0.006 | 0.034 | 0.034 | 0 | 0.4 | 0.67 | 0.35 | 0.08 | 0.4 | 0.3 |
| 8 | 140153347 | rs11775824 | 2.481 | 0.92 | -0.358 | -0.304 | -0.03 | 0.038 | 0.038 | 0 | 0.1 | 0.05 | 0.1 | 0 | 0.14 | 0.05 |
| 8 | 140154013 | rs7464161 | 0.214 | 0.825 | -0.181 | -0.271 | -0.012 | -0.041 | -0.041 | 0.615 | 0.24 | 0.52 | 0.22 | 0.21 | 0.17 | 0.12 |
| 8 | 140158294 | rs6577995 | -0.559 | 0.818 | -0.273 | -0.288 | 0.013 | 0.107 | 0.107 | 0.202 | 0.4 | 0.75 | 0.34 | 0.21 | 0.3 | 0.15 |
| 8 | 140159775 | rs13278167 | -0.558 | 0.842 | -0.268 | -0.29 | 0.009 | 0.118 | 0.118 | 0.187 | 0.4 | 0.78 | 0.34 | 0.21 | 0.3 | 0.15 |
| 8 | 140162809 | rs7010100 | -0.935 | 0.836 | 0.004 | -0.294 | -0.006 | 0.021 | 0.021 | 0.328 | 0.45 | 0.59 | 0.55 | 0.27 | 0.45 | 0.37 |
| 8 | 140165114 | rs3935646 | -0.96 | 0.817 | 0.171 | -0.285 | -0.001 | -0.001 | -0.001 | 0 | 0.45 | 0.52 | 0.55 | 0.27 | 0.45 | 0.36 |
| 8 | 140165476 | rs7460675 | 1.544 | 0.788 | 0.296 | -0.313 | -0.014 | -0.052 | -0.052 | 2.895 | 0.16 | 0.21 | 0.05 | 0.01 | 0.08 | 0.06 |
| 8 | 140169439 | rs11781079 | 0.704 | 0.815 | 0.319 | -0.295 | -0.025 | 0.032 | 0.032 | 0 | 0.2 | 0.14 | 0.19 | 0.07 | 0.27 | 0.24 |
| 8 | 140174644 | rs4736212 | -0.987 | 0.81 | 0.334 | -0.297 | -0.001 | -0.001 | -0.001 | 0 | 0.44 | 0.53 | 0.55 | 0.27 | 0.45 | 0.37 |
| 8 | 140180024 | rs6997709 | 0.7 | 0.85 | 0.316 | -0.291 | -0.022 | 0.029 | 0.029 | 0 | 0.2 | 0.14 | 0.19 | 0.07 | 0.27 | 0.24 |
| 8 | 140188319 | rs12548575 | -0.223 | 0.838 | 0.324 | -0.316 | 0.01 | -0.067 | -0.067 | 0 | 0.37 | 0.48 | 0.32 | 0.07 | 0.39 | 0.27 |
| 8 | 140188963 | rs7465545 | 1.711 | 0.842 | 0.307 | -0.315 | 0.002 | -0.07 | -0.07 | 0.176 | 0.15 | 0.18 | 0.11 | 0 | 0.12 | 0.03 |
| 8 | 140190193 | rs11786307 | -1.314 | 0.829 | 0.282 | -0.291 | -0.002 | -0.005 | -0.005 | 0 | 0.58 | 0.56 | 0.65 | 0.27 | 0.57 | 0.4 |
| 8 | 140200107 | rs11166892 | 0.608 | 0.807 | 0.072 | -0.293 | -0.022 | 0.03 | 0.03 | 0 | 0.21 | 0.14 | 0.19 | 0.07 | 0.27 | 0.24 |
| 8 | 140213324 | rs7462825 | -1.458 | 0.803 | 0.049 | -0.183 | -0.012 | 0.017 | 0.017 | 0.008 | 0.62 | 0.88 | 0.7 | 0.29 | 0.56 | 0.4 |
| 8 | 140214015 | rs6578010 | -1.411 | 0.733 | 0.051 | -0.217 | 0.013 | 0.034 | 0.034 | 0.005 | 0.6 | 0.74 | 0.68 | 0.29 | 0.56 | 0.4 |
| 8 | 140217176 | rs6989522 | 2.151 | 0.759 | -0.023 | -0.246 | -0.033 | 0.049 | 0.049 | 0 | 0.12 | 0.05 | 0.11 | 0 | 0.15 | 0.05 |
| 8 | 140218012 | rs4133282 | 1.237 | 0.672 | 0.039 | -0.218 | 0.006 | 0.004 | 0.004 | 0 | 0.08 | 0.1 | 0.07 | 0.07 | 0.11 | 0.19 |
| 8 | 140218260 | rs4736218 | -2.006 | 0.564 | -0.113 | -0.024 | 0.004 | -0.061 | -0.061 | 3.187 | 0.9 | 0.99 | 0.95 | 0.55 | 0.92 | 0.77 |
| 8 | 140223383 | rs6988320 | 2.145 | 0.568 | -0.13 | -0.049 | -0.054 | -0.064 | -0.064 | 0 | 0.28 | 0.04 | 0.25 | 0.26 | 0.35 | 0.38 |
| 8 | 140231770 | rs11775898 | -0.737 | 0.622 | 0.55 | -0.664 | 0.002 | 0.043 | 0.043 | 1.422 | 0.52 | 0.6 | 0.6 | 0.23 | 0.46 | 0.22 |
| 8 | 140232935 | rs4556079 | 0.672 | 0.632 | 0.455 | -0.705 | -0.058 | -0.016 | -0.016 | 0 | 0.19 | 0.11 | 0.42 | 0.21 | 0.25 | 0.11 |
| 8 | 140234888 | rs1986626 | -0.557 | 0.596 | 0.606 | -0.676 | 0.004 | 0.069 | 0.069 | 1.984 | 0.49 | 0.36 | 0.57 | 0.21 | 0.45 | 0.19 |
| 8 | 140236246 | rs12548929 | 0.671 | 0.667 | 0.545 | -0.618 | 0.017 | -0.014 | -0.014 | 0 | 0.1 | 0.18 | 0.09 | 0.04 | 0.11 | 0.19 |
| 8 | 140238440 | rs17694995 | 1.76 | 0.635 | 0.513 | -0.627 | 0 | -0.023 | -0.023 | 0 | 0.14 | 0.11 | 0.1 | 0 | 0.12 | 0.03 |
| 8 | 140239452 | rs7017449 | -0.165 | 0.641 | 0.526 | -0.628 | -0.031 | 0.041 | 0.041 | 0 | 0.34 | 0.24 | 0.53 | 0.21 | 0.37 | 0.14 |
| 8 | 140247495 | rs11991162 | 0.544 | 0.216 | 0.803 | -0.528 | -0.027 | -0.16 | -0.16 | 0 | 0.3 | 0.43 | 0.17 | 0.02 | 0.2 | 0.09 |
| 8 | 140247662 | rs11781101 | -0.451 | 0.59 | 0.88 | -0.152 | 0.002 | -0.001 | -0.001 | 0 | 0.29 | 0.3 | 0.51 | 0.25 | 0.39 | 0.32 |
| 8 | 140258348 | rs4736224 | -0.119 | 0.696 | 0.469 | -0.102 | -0.032 | -0.001 | -0.001 | 0.449 | 0.27 | 0.22 | 0.5 | 0.25 | 0.38 | 0.32 |
| 8 | 140259225 | rs7004938 | -0.085 | 0.766 | -0.381 | -0.109 | -0.055 | 0.01 | 0.01 | 0 | 0.27 | 0.16 | 0.49 | 0.22 | 0.38 | 0.32 |
| 8 | 140259539 | rs11783351 | -0.114 | 0.791 | -0.49 | -0.108 | -0.069 | 0.01 | 0.01 | 0 | 0.27 | 0.14 | 0.49 | 0.22 | 0.38 | 0.32 |
| 8 | 140264369 | rs4736226 | 0.216 | 0.024 | -0.744 | 0.01 | -0.002 | 0.003 | 0.003 | 3.967 | 0.48 | 0.42 | 0.41 | 0.47 | 0.52 | 0.5 |
| 8 | 140264728 | rs4440614 | -0.671 | -0.004 | -0.731 | 0.091 | -0.015 | -0.006 | -0.006 | 0 | 0.8 | 0.57 | 0.75 | 0.85 | 0.88 | 0.89 |
| 8 | 140265594 | rs7814156 | -1.002 | 0.109 | -0.784 | 0.183 | 0 | 0.012 | 0.012 | 0.655 | 0.82 | 0.62 | 0.76 | 0.85 | 0.89 | 0.89 |
| 8 | 140267889 | rs12675641 | 0.366 | 0.173 | -1.737 | 0.139 | -0.048 | -0.016 | -0.016 | 0 | 0.31 | 0.08 | 0.34 | 0.38 | 0.36 | 0.39 |
| 8 | 140270356 | rs2873902 | -0.02 | -0.284 | -0.868 | 0.579 | -0.006 | 0.005 | 0.005 | 0 | 0.74 | 0.68 | 0.6 | 0.42 | 0.77 | 0.58 |
| 8 | 140272646 | rs10113442 | -1.16 | 0.275 | -0.473 | -0.007 | 0.004 | -0.041 | -0.041 | 0.598 | 0.35 | 0.54 | 0.53 | 0.19 | 0.33 | 0.21 |
| 8 | 140274384 | rs10216876 | -0.685 | 0.24 | -0.314 | -0.047 | 0 | -0.003 | -0.003 | 0.765 | 0.32 | 0.34 | 0.51 | 0.18 | 0.33 | 0.21 |
| 8 | 140277409 | rs7834945 | -0.814 | 0.057 | -0.595 | -0.062 | -0.071 | -0.048 | -0.048 | 0 | 0.18 | 0.43 | 0.25 | 0.1 | 0.09 | 0.1 |
| 8 | 140278162 | rs17701262 | -0.075 | 1.058 | -1.133 | 0.204 | -0.101 | 0.026 | 0.026 | 0 | 0.17 | 0.02 | 0.29 | 0.09 | 0.26 | 0.11 |
| 8 | 140280887 | rs4736230 | -0.368 | 1.177 | -1.015 | 0.238 | -0.007 | 0.003 | 0.003 | 0 | 0.21 | 0.2 | 0.32 | 0.11 | 0.31 | 0.15 |
| 8 | 140281666 | rs7820230 | -0.297 | -0.051 | -1.33 | 0.215 | 0.03 | 0.031 | 0.031 | 0.128 | 0.37 | 0.28 | 0.17 | 0.25 | 0.29 | 0.41 |
| 8 | 140282987 | rs11166903 | 0.065 | 0.142 | -1.678 | 0.505 | -0.028 | 0.002 | 0.002 | 0 | 0.24 | 0.04 | 0.27 | 0.18 | 0.27 | 0.08 |
| 8 | 140293281 | rs2319075 | 0.551 | -0.345 | 0.267 | 1.278 | 0.017 | 0.293 | 0.293 | 0.255 | 0.91 | 0.53 | 0.92 | 0.73 | 0.98 | 0.9 |
| 8 | 140293780 | rs7004320 | 0.231 | -0.34 | 0.275 | 1.283 | 0.01 | 0.012 | 0.012 | 0 | 0.94 | 0.74 | 0.97 | 0.96 | 0.98 | 0.93 |
| 8 | 140299457 | rs2873896 | 1.316 | -0.251 | 0.318 | 1.177 | 0.012 | 0.201 | 0.201 | 0 | 0.89 | 0.54 | 0.9 | 0.73 | 0.94 | 0.89 |
| 8 | 140300344 | rs4636184 | 0.718 | -0.03 | -0.128 | 0.422 | 0.001 | 0.029 | 0.029 | 0.119 | 0.43 | 0.33 | 0.39 | 0.69 | 0.39 | 0.69 |
| 8 | 140301753 | rs4736239 | 0.147 | -0.704 | -0.203 | 0.64 | -0.009 | -0.005 | -0.005 | 0.431 | 0.64 | 0.42 | 0.62 | 0.7 | 0.68 | 0.8 |
| 8 | 140304087 | rs2319093 | 0.566 | -0.904 | -0.331 | 0.734 | -0.051 | 0.138 | 0.138 | 0 | 0.85 | 0.48 | 0.83 | 0.73 | 0.93 | 0.74 |
| 8 | 140308182 | rs13270196 | 0.601 | -0.948 | -0.296 | 0.683 | 0.02 | -0.019 | -0.019 | 0.035 | 0.91 | 0.71 | 0.94 | 0.97 | 0.94 | 0.93 |
| 8 | 140309909 | rs4445205 | -0.116 | -0.972 | -0.518 | 0.768 | -0.006 | -0.04 | -0.04 | 0 | 0.64 | 0.58 | 0.64 | 0.8 | 0.64 | 0.84 |
| 8 | 140318368 | rs6993541 | 0.379 | -0.791 | -1.014 | 0.959 | 0.016 | -0.01 | -0.01 | 0 | 0.56 | 0.36 | 0.43 | 0.51 | 0.6 | 0.65 |
| 8 | 140330596 | rs766625 | 0.654 | -0.526 | -1.931 | 0.658 | -0.036 | 0.044 | 0.044 | 0.058 | 0.27 | 0.06 | 0.31 | 0.17 | 0.33 | 0.08 |
| 8 | 140333076 | rs994769 | 0.589 | 0.136 | -1.948 | 0.606 | -0.006 | -0.107 | -0.107 | 0.211 | 0.25 | 0.01 | 0.26 | 0.34 | 0.17 | 0.44 |
| 8 | 140333323 | rs994770 | 0.08 | 0.119 | -1.023 | 0.924 | 0 | -0.001 | -0.001 | 0.236 | 0.56 | 0.55 | 0.52 | 0.49 | 0.59 | 0.6 |
| 8 | 140337281 | rs6981027 | -0.92 | 0.096 | -1.323 | 1.018 | 0.018 | -0.043 | -0.043 | 0.194 | 0.25 | 0.14 | 0.21 | 0.38 | 0.21 | 0.28 |

| | | | | | | | | | | | | | | | | |
|---|---|---|---|---|---|---|---|---|---|---|---|---|---|---|---|---|
| 8 | 140340558 | rs6997652 | -0.983 | 0.07 | -0.535 | 0.365 | -0.009 | -0.021 | -0.021 | 0 | 0.26 | 0.43 | 0.29 | 0.22 | 0.25 | 0.22 |
| 8 | 140343359 | rs11781232 | 0.74 | -0.03 | -0.363 | 0.008 | 0.001 | -0.004 | -0.004 | 0 | 0.66 | 0.62 | 0.56 | 0.67 | 0.63 | 0.68 |
| 8 | 140345180 | rs1031824 | 1.508 | -0.039 | -0.216 | -0.012 | 0.002 | 0.039 | 0.039 | 0 | 0.51 | 0.26 | 0.4 | 0.56 | 0.54 | 0.61 |
| 8 | 140348321 | rs3843560 | 1.028 | 0.257 | -0.04 | 0.1 | 0.004 | 0.006 | 0.006 | 2.109 | 0.07 | 0.08 | 0.13 | 0.1 | 0.09 | 0.03 |
| 8 | 140348367 | rs3847167 | 1.085 | 0.177 | 0.439 | 0.158 | 0.009 | 0.003 | 0.003 | 0.064 | 0.6 | 0.48 | 0.42 | 0.55 | 0.58 | 0.65 |
| 8 | 140349289 | rs7845540 | 1.254 | 0.215 | 0.535 | 0.177 | -0.001 | -0.001 | -0.001 | 0 | 0.58 | 0.38 | 0.42 | 0.55 | 0.57 | 0.64 |
| 8 | 140349339 | rs7832666 | 0.759 | 0.2 | 0.444 | 0.15 | 0.005 | -0.003 | -0.003 | 0 | 0.67 | 0.58 | 0.57 | 0.65 | 0.67 | 0.67 |
| 8 | 140349351 | rs7844572 | 1.117 | 0.212 | -0.156 | 0.155 | -0.017 | 0.001 | 0.001 | 0 | 0.57 | 0.3 | 0.42 | 0.55 | 0.57 | 0.64 |
| 8 | 140351500 | rs16910485 | -1.111 | 0.293 | -0.31 | 0.015 | 0 | 0.004 | 0.004 | 0 | 0.15 | 0.13 | 0.15 | 0.11 | 0.16 | 0.16 |
| 8 | 140353842 | rs7833826 | -0.931 | 0.198 | -0.634 | -0.222 | -0.002 | 0.027 | 0.027 | 1.103 | 0.18 | 0.11 | 0.14 | 0.11 | 0.2 | 0.16 |
| 8 | 140354650 | rs4736251 | -0.414 | 0.344 | -0.335 | -0.426 | 0.002 | -0.009 | -0.009 | 0 | 0.36 | 0.59 | 0.46 | 0.31 | 0.43 | 0.42 |
| 8 | 140358700 | rs6578026 | -0.429 | 0.241 | -0.821 | -0.525 | -0.029 | -0.004 | -0.004 | 0 | 0.78 | 0.9 | 0.72 | 0.74 | 0.81 | 0.78 |
| 8 | 140360853 | rs3857922 | 0.54 | -0.149 | -0.085 | -1.057 | 0 | -0.001 | -0.001 | 1.437 | 0.33 | 0.3 | 0.26 | 0.49 | 0.27 | 0.29 |
| 8 | 140369906 | rs16910519 | 0.719 | -0.177 | -0.162 | -1.071 | 0.014 | -0.044 | -0.044 | 0 | 0.31 | 0.16 | 0.24 | 0.49 | 0.27 | 0.29 |
| 8 | 140375282 | rs10505712 | 0.325 | -0.177 | -0.232 | -1.071 | -0.024 | 0.02 | 0.02 | 0 | 0.34 | 0.54 | 0.27 | 0.49 | 0.27 | 0.29 |
| 8 | 140375578 | rs4577935 | -1.269 | 0.111 | -0.439 | -1.094 | -0.029 | 0.04 | 0.04 | 0 | 0.82 | 0.99 | 0.83 | 0.92 | 0.89 | 0.88 |
| 8 | 140383483 | rs12678674 | 0.366 | -0.038 | -0.211 | -1.06 | -0.025 | 0.022 | 0.022 | 0 | 0.34 | 0.5 | 0.27 | 0.49 | 0.27 | 0.29 |
| 8 | 140388179 | rs2467180 | 0.525 | -0.064 | -0.662 | -1.008 | -0.061 | 0.1 | 0.1 | 0.221 | 0.4 | 0.13 | 0.42 | 0.25 | 0.52 | 0.45 |
| 8 | 140393623 | rs1382023 | 0.251 | -0.081 | -0.337 | -0.805 | 0.01 | -0.007 | -0.007 | 0.373 | 0.36 | 0.65 | 0.28 | 0.49 | 0.3 | 0.3 |
| 8 | 140398392 | rs7845764 | 0.174 | -0.073 | -0.581 | -0.801 | 0.024 | -0.014 | -0.014 | 0 | 0.37 | 0.73 | 0.29 | 0.5 | 0.31 | 0.3 |
| 8 | 140408107 | rs922382 | 0.863 | 0.179 | -1.281 | -0.687 | -0.02 | -0.104 | -0.104 | 0 | 0.3 | 0.04 | 0.23 | 0.49 | 0.3 | 0.3 |
| 8 | 140415986 | rs2468719 | -0.742 | -0.009 | -1.392 | -0.732 | -0.035 | -0.016 | -0.016 | 0 | 0.79 | 0.9 | 0.72 | 0.74 | 0.8 | 0.77 |
| 8 | 140417049 | rs899073 | 0.048 | 0.088 | -1.352 | -0.893 | -0.005 | 0.197 | 0.197 | 0 | 0.49 | 0.83 | 0.53 | 0.25 | 0.55 | 0.47 |
| 8 | 140421115 | rs10435688 | 0.254 | -0.149 | -0.85 | -0.431 | 0.03 | -0.018 | -0.018 | 2.619 | 0.33 | 0.22 | 0.2 | 0.48 | 0.22 | 0.33 |
| 8 | 140444375 | rs17786072 | 0.314 | -0.455 | -0.733 | -0.6 | 0.031 | 0.031 | 0.031 | 2.36 | 0.35 | 0.24 | 0.23 | 0.2 | 0.25 | 0.23 |
| 8 | 140446725 | rs11990329 | 0.914 | 0.198 | -0.921 | -0.536 | -0.036 | -0.023 | -0.023 | 0 | 0.28 | 0.15 | 0.41 | 0.6 | 0.41 | 0.43 |
| 8 | 140452253 | rs2467184 | -0.649 | 0.01 | -0.331 | -0.859 | -0.008 | 0.017 | 0.017 | 0 | 0.72 | 0.74 | 0.69 | 0.8 | 0.7 | 0.7 |
| 8 | 140459422 | rs1600047 | -0.694 | -0.078 | -0.266 | -0.384 | -0.003 | -0.011 | -0.011 | 0 | 0.8 | 0.82 | 0.74 | 0.76 | 0.79 | 0.71 |
| 8 | 140462221 | rs2467182 | -0.023 | -0.58 | -0.068 | -0.807 | -0.001 | 0.005 | 0.005 | 0 | 0.64 | 0.66 | 0.65 | 0.76 | 0.63 | 0.65 |
| 8 | 140465267 | rs17786392 | 0.891 | -0.51 | -0.135 | -0.894 | 0.011 | -0.072 | -0.072 | 0 | 0.22 | 0.24 | 0.25 | 0.01 | 0.26 | 0.24 |
| 8 | 140465847 | rs1827530 | 0.791 | -1.014 | -0.177 | -1.007 | -0.069 | -0.064 | -0.064 | 0 | 0.24 | 0.29 | 0.16 | 0.09 | 0.1 | 0.15 |
| 8 | 140467417 | rs2447560 | 0.504 | -0.772 | -0.517 | -0.817 | -0.052 | 0.002 | 0.002 | 0 | 0.37 | 0.16 | 0.47 | 0.66 | 0.52 | 0.48 |
| 8 | 140469250 | rs2455790 | 0.473 | -0.761 | -0.517 | -0.812 | -0.045 | 0.005 | 0.005 | 0 | 0.37 | 0.18 | 0.48 | 0.66 | 0.52 | 0.48 |
| 8 | 140478970 | rs2045855 | 1.267 | -1.073 | -0.559 | -0.76 | -0.036 | -0.026 | -0.026 | 0 | 0.19 | 0.2 | 0.15 | 0.08 | 0.09 | 0.12 |
| 8 | 140482534 | rs2468731 | 0.349 | -0.434 | -1.183 | -0.51 | -0.121 | -0.044 | -0.044 | 7.235 | 0.34 | 0.07 | 0.43 | 0.65 | 0.46 | 0.45 |
| 8 | 140489319 | rs2471095 | 0.112 | -1.032 | -0.758 | -0.38 | -0.012 | 0.004 | 0.004 | 0.23 | 0.4 | 0.25 | 0.68 | 0.62 | 0.53 | 0.58 |
| 8 | 140497775 | rs1382022 | -1.5 | -1.293 | -0.813 | -0.278 | 0.016 | -0.02 | -0.02 | 0 | 0.86 | 0.92 | 0.93 | 0.74 | 0.9 | 0.93 |
| 8 | 140497806 | rs10101763 | 0.954 | -1.326 | -1.28 | -0.544 | 0.005 | 0.072 | 0.072 | 0 | 0.22 | 0.03 | 0.11 | 0.08 | 0.19 | 0.17 |
| 8 | 140501230 | rs1870977 | 0.363 | -0.455 | -1.413 | -0.157 | -0.061 | 0.079 | 0.079 | 0 | 0.38 | 0.14 | 0.44 | 0.25 | 0.46 | 0.45 |
| 8 | 140503539 | rs2471096 | 0.405 | 0.063 | -0.279 | 0.263 | 0.036 | 0.129 | 0.129 | 0 | 0.16 | 0.33 | 0.35 | 0.41 | 0.2 | 0.26 |
| 8 | 140505503 | rs17725186 | 0.795 | 0.097 | -0.889 | -0.208 | -0.039 | 0.081 | 0.081 | 0 | 0.31 | 0.09 | 0.36 | 0.16 | 0.39 | 0.47 |
| 8 | 140508079 | rs3934864 | 1.294 | -0.696 | -0.792 | 0.31 | -0.027 | -0.046 | -0.046 | 0 | 0.13 | 0.05 | 0.21 | 0.27 | 0.17 | 0.24 |
| 8 | 140508781 | rs2468670 | 1.145 | -1.079 | -0.976 | 0.231 | 0.042 | -0.096 | -0.096 | 0 | 0.19 | 0.05 | 0.14 | 0.39 | 0.11 | 0.12 |
| 8 | 140510093 | rs12548447 | 0.989 | -1.054 | -0.946 | 0.137 | -0.073 | 0.069 | 0.069 | 0 | 0.35 | 0.12 | 0.46 | 0.22 | 0.48 | 0.44 |
| 8 | 140510694 | rs17787479 | 1.595 | -1.323 | -1.005 | 0.02 | -0.006 | 0.065 | 0.065 | 0 | 0.19 | 0.02 | 0.1 | 0.07 | 0.18 | 0.14 |
| 8 | 140511074 | rs4236885 | 0.341 | -1.363 | -0.7 | -0.007 | -0.011 | -0.035 | -0.035 | 0 | 0.31 | 0.6 | 0.16 | 0.11 | 0.23 | 0.21 |
| 8 | 140511083 | rs4736261 | 1.488 | -1.323 | -0.828 | 0.03 | 0.008 | 0.05 | 0.05 | 0 | 0.2 | 0.06 | 0.11 | 0.11 | 0.18 | 0.17 |
| 8 | 140511526 | rs2447406 | 1.211 | -1.255 | -0.807 | 0.141 | 0.042 | -0.099 | -0.099 | 0 | 0.19 | 0.05 | 0.14 | 0.4 | 0.11 | 0.12 |
| 8 | 140514193 | rs13252879 | 1.703 | -1.104 | -0.89 | -0.148 | -0.045 | 0.016 | 0.016 | 0 | 0.23 | 0.09 | 0.32 | 0.21 | 0.33 | 0.4 |
| 8 | 140516541 | rs6578030 | 1.81 | -1.381 | -0.128 | 0.048 | -0.013 | -0.042 | -0.042 | 0.749 | 0.08 | 0.18 | 0.01 | 0 | 0.05 | 0.04 |
| 8 | 140519341 | rs2471083 | -0.077 | -1.381 | -0.356 | 0.097 | 0.02 | 0.014 | 0.014 | 0 | 0.33 | 0.24 | 0.43 | 0.66 | 0.28 | 0.36 |
| 8 | 140521806 | rs10109373 | -0.752 | -0.608 | 0.189 | 0.183 | -0.005 | -0.003 | -0.003 | 0 | 0.45 | 0.53 | 0.37 | 0.38 | 0.4 | 0.45 |
| 8 | 140525121 | rs4736077 | 0.999 | -0.24 | 0.037 | 0.362 | 0.018 | 0.041 | 0.041 | 0 | 0.12 | 0.22 | 0.23 | 0.23 | 0.18 | 0.24 |
| 8 | 140526489 | rs2447356 | 0.044 | -0.521 | 0.4 | 0.415 | -0.001 | -0.001 | -0.001 | 0 | 0.6 | 0.59 | 0.71 | 0.69 | 0.71 | 0.75 |
| 8 | 140529378 | rs16893589 | 0.919 | -0.648 | 0.408 | 0.398 | 0.011 | -0.008 | -0.008 | 0.045 | 0.19 | 0.17 | 0.15 | 0.28 | 0.1 | 0.12 |
| 8 | 140533774 | rs2447372 | 0.228 | -0.682 | 0.476 | 0.361 | -0.023 | -0.015 | -0.015 | 0.092 | 0.61 | 0.53 | 0.71 | 0.65 | 0.72 | 0.73 |
| 8 | 140534479 | rs2447376 | 0.873 | -0.644 | 0.571 | 0.373 | -0.018 | -0.001 | -0.001 | 0 | 0.4 | 0.31 | 0.48 | 0.42 | 0.49 | 0.46 |
| 8 | 140542270 | rs2447395 | -0.01 | -0.657 | 0.775 | 0.37 | -0.003 | -0.006 | -0.006 | 0 | 0.59 | 0.47 | 0.61 | 0.7 | 0.59 | 0.58 |
| 8 | 140545742 | rs4380912 | 1.268 | -0.643 | 0.613 | 0.377 | -0.012 | -0.022 | -0.022 | 2.458 | 0.19 | 0.12 | 0.23 | 0.23 | 0.23 | 0.27 |
| 8 | 140549752 | rs4736270 | 0.978 | -0.663 | 0.28 | 0.326 | 0.011 | 0.032 | 0.032 | 0 | 0.19 | 0.09 | 0.11 | 0.07 | 0.17 | 0.14 |
| 8 | 140551003 | rs2468708 | 0.872 | -0.682 | 0.297 | 0.339 | -0.025 | -0.001 | -0.001 | 0 | 0.4 | 0.24 | 0.47 | 0.42 | 0.49 | 0.45 |
| 8 | 140555844 | rs2542474 | -1.652 | -0.676 | 0.202 | 0.198 | -0.001 | -0.053 | -0.053 | 0 | 0.83 | 0.99 | 0.85 | 0.76 | 0.88 | 0.86 |
| 8 | 140556571 | rs2615387 | 0.032 | -0.531 | 0.398 | 0.373 | -0.005 | 0 | 0 | 0 | 0.59 | 0.41 | 0.68 | 0.6 | 0.71 | 0.71 |
| 8 | 140559736 | rs2615391 | 0.05 | -0.555 | 0.413 | 0.361 | -0.024 | -0.002 | -0.002 | 0.038 | 0.59 | 0.5 | 0.69 | 0.6 | 0.7 | 0.71 |
| 8 | 140564192 | rs969124 | -0.086 | -0.542 | 0.398 | 0.368 | 0.043 | -0.01 | -0.01 | 0 | 0.61 | 0.77 | 0.72 | 0.6 | 0.7 | 0.72 |
| 8 | 140567626 | rs2261712 | 0.777 | -0.505 | 0.373 | 0.379 | 0 | -0.001 | -0.001 | 0.154 | 0.4 | 0.39 | 0.47 | 0.37 | 0.46 | 0.43 |
| 8 | 140569221 | rs2615401 | 0.802 | -0.461 | 0.326 | 0.411 | -0.027 | 0.023 | 0.023 | 0 | 0.4 | 0.24 | 0.47 | 0.37 | 0.48 | 0.45 |
| 8 | 140571010 | rs13265550 | 1.164 | -0.426 | -0.176 | 0.443 | 0 | 0.001 | 0.001 | 0.009 | 0.19 | 0.2 | 0.24 | 0.23 | 0.23 | 0.28 |
| 8 | 140576819 | rs2542454 | 0.416 | -0.451 | -0.268 | 0.414 | -0.025 | 0.045 | 0.045 | 0 | 0.44 | 0.71 | 0.52 | 0.37 | 0.49 | 0.46 |

| | | | | | | | | | | | | | | | | |
|---|---|---|---|---|---|---|---|---|---|---|---|---|---|---|---|---|
| 8 | 140577572 | rs1833582 | -1.177 | -0.547 | -0.388 | 0.415 | 0.03 | -0.009 | -0.009 | 1.795 | 0.78 | 0.86 | 0.84 | 0.73 | 0.89 | 0.86 |
| 8 | 140577858 | rs1862222 | -0.767 | -0.53 | -0.473 | 0.384 | 0.023 | -0.002 | -0.002 | 1.115 | 0.75 | 0.87 | 0.82 | 0.73 | 0.84 | 0.84 |
| 8 | 140578820 | rs16910949 | 1.588 | -0.348 | -0.901 | 0.453 | 0.01 | -0.03 | -0.03 | 0 | 0.12 | 0.03 | 0.08 | 0.12 | 0.12 | 0.1 |
| 8 | 140584877 | rs10101646 | 0.898 | -0.334 | -0.838 | 0.418 | 0.027 | 0.021 | 0.021 | 0 | 0.21 | 0.47 | 0.26 | 0.18 | 0.23 | 0.26 |
| 8 | 140585284 | rs2615405 | -1.531 | -0.222 | -1.738 | 0.351 | 0.01 | -0.019 | -0.019 | 0 | 0.81 | 0.99 | 0.87 | 0.83 | 0.89 | 0.88 |
| 8 | 140585403 | rs2542450 | 0.975 | 0.68 | -1.281 | 0.712 | -0.047 | -0.038 | -0.038 | 0 | 0.24 | 0.07 | 0.33 | 0.37 | 0.32 | 0.41 |
| 8 | 140585526 | rs11166916 | -1.062 | 0.704 | -0.482 | 0.978 | -0.001 | 0.013 | 0.013 | 0 | 0.33 | 0.38 | 0.34 | 0.42 | 0.35 | 0.39 |
| 8 | 140587030 | rs11775044 | -0.754 | 0.616 | -0.068 | 0.769 | -0.005 | 0.011 | 0.011 | 0 | 0.17 | 0.26 | 0.14 | 0.19 | 0.18 | 0.15 |
| 8 | 140590599 | rs2615412 | -0.837 | 0.926 | 0.134 | 0.701 | 0.019 | 0.033 | 0.033 | 0.296 | 0.23 | 0.38 | 0.21 | 0.32 | 0.25 | 0.27 |
| 8 | 140590707 | rs6578037 | 0.792 | 0.832 | -0.654 | -0.059 | 0.032 | -0.035 | -0.035 | 0 | 0.38 | 0.14 | 0.32 | 0.6 | 0.33 | 0.46 |
| 8 | 140590799 | rs2615413 | -0.922 | 0.826 | -0.547 | -0.459 | -0.051 | 0.1 | 0.1 | 0 | 0.73 | 0.93 | 0.63 | 0.95 | 0.68 | 0.75 |
| 8 | 140591581 | rs2542443 | 0.021 | 0.903 | -0.101 | -0.253 | 0 | -0.011 | -0.011 | 3.469 | 0.1 | 0.14 | 0.07 | 0 | 0.12 | 0.02 |
| 8 | 140592229 | rs10097949 | 0.114 | 0.888 | 0.21 | -0.254 | 0.001 | -0.056 | -0.056 | 0 | 0.1 | 0.29 | 0.08 | 0.01 | 0.1 | 0.02 |
| 8 | 140595280 | rs2542440 | 0.029 | 0.918 | -0.059 | -0.246 | 0 | -0.047 | -0.047 | 0 | 0.1 | 0.22 | 0.09 | 0 | 0.11 | 0.02 |
| 8 | 140595992 | rs6981756 | 0.813 | 0.955 | -0.34 | -0.213 | 0.032 | 0.008 | 0.008 | 0 | 0.4 | 0.14 | 0.33 | 0.62 | 0.34 | 0.47 |
| 8 | 140598664 | rs16911026 | 0.144 | 0.838 | -0.248 | -0.329 | -0.002 | -0.028 | -0.028 | 0 | 0.09 | 0.14 | 0.08 | 0 | 0.1 | 0.02 |
| 8 | 140599226 | rs2111573 | 0.86 | 0.923 | -0.007 | 0.081 | 0.028 | 0.097 | 0.097 | 0.87 | 0.41 | 0.2 | 0.15 | 0.26 | 0.35 | 0.44 |
| 8 | 140602616 | rs2542436 | -0.786 | 1.031 | 0.138 | 0.433 | 0.011 | 0.144 | 0.144 | 0 | 0.14 | 0.27 | 0.28 | 0.45 | 0.19 | 0.08 |
| 8 | 140604643 | rs16911045 | 0.286 | 1.093 | 0.583 | 0.244 | 0.022 | 0.083 | 0.083 | 0 | 0.16 | 0.3 | 0.16 | 0.32 | 0.19 | 0.25 |
| 8 | 140607542 | rs2261824 | 0.113 | 1.074 | 0.343 | 0.102 | -0.002 | 0.016 | 0.016 | 0 | 0.54 | 0.6 | 0.44 | 0.66 | 0.5 | 0.52 |
| 8 | 140608316 | rs7003899 | 0.957 | 0.895 | 0.201 | 0.225 | 0.011 | 0.017 | 0.017 | 0 | 0.41 | 0.34 | 0.16 | 0.24 | 0.32 | 0.46 |
| 8 | 140610241 | rs2430822 | -0.399 | 0.956 | 0.151 | 0.267 | 0.001 | 0.001 | 0.001 | 0 | 0.2 | 0.22 | 0.32 | 0.73 | 0.25 | 0.33 |
| 8 | 140611754 | rs2431300 | 0.68 | 0.886 | -0.217 | -0.16 | -0.003 | -0.004 | -0.004 | 4.171 | 0.46 | 0.52 | 0.35 | 0.69 | 0.38 | 0.53 |
| 8 | 140611843 | rs2447342 | 0.138 | 1.011 | -0.091 | -0.644 | 0.003 | -0.072 | -0.072 | 0 | 0.61 | 0.58 | 0.48 | 0.95 | 0.58 | 0.77 |
| 8 | 140618792 | rs2545432 | -0.076 | 1.157 | -0.734 | -0.689 | -0.03 | 0.149 | 0.149 | 0 | 0.37 | 0.16 | 0.39 | 0.06 | 0.35 | 0.22 |
| 8 | 140626391 | rs2471093 | -0.447 | 0.983 | -0.035 | 0.444 | -0.005 | 0.124 | 0.124 | 0 | 0.12 | 0.17 | 0.19 | 0.6 | 0.11 | 0.38 |
| 8 | 140630990 | rs2615374 | -0.854 | 0.92 | -0.78 | 0.039 | -0.007 | 0.055 | 0.055 | 3.36 | 0.61 | 0.88 | 0.66 | 0.7 | 0.64 | 0.66 |
| 8 | 140632384 | rs2542425 | 0.383 | 0.966 | -0.606 | 0.535 | -0.025 | 0.077 | 0.077 | 0.001 | 0.38 | 0.27 | 0.49 | 0.08 | 0.42 | 0.24 |
| 8 | 140635346 | rs2471094 | -0.554 | 1.04 | -0.625 | 0.701 | -0.004 | 0.116 | 0.116 | 0.693 | 0.22 | 0.49 | 0.24 | 0.55 | 0.23 | 0.44 |
| 8 | 140637124 | rs2542422 | -0.74 | 1.061 | -1.673 | 0.629 | -0.049 | 0.008 | 0.008 | 2.206 | 0.61 | 0.97 | 0.74 | 0.69 | 0.65 | 0.67 |
| 8 | 140640634 | rs2014712 | -0.782 | 1.353 | -1.17 | 0.83 | -0.02 | -0.029 | -0.029 | 0 | 0.18 | 0.1 | 0.13 | 0.29 | 0.27 | 0.34 |
| 8 | 140645427 | rs2615377 | 0.044 | 1.055 | -1.043 | 0.279 | -0.011 | 0.048 | 0.048 | 0.342 | 0.45 | 0.38 | 0.47 | 0.12 | 0.47 | 0.28 |
| 8 | 140646990 | rs888345 | 0.251 | 1.435 | -0.95 | 1.056 | 0.026 | 0.071 | 0.071 | 0 | 0.89 | 0.84 | 0.92 | 0.7 | 0.81 | 0.69 |
| 8 | 140658801 | rs3780051 | 0.204 | 0.709 | -2.146 | 0.846 | -0.131 | -0.089 | -0.089 | 1.074 | 0.48 | 0.03 | 0.32 | 0.42 | 0.53 | 0.58 |
| 8 | 140661285 | rs2545457 | 0.364 | -0.286 | -2.134 | 0.12 | 0.008 | 0.035 | 0.035 | 2.581 | 0.53 | 0.8 | 0.69 | 0.64 | 0.61 | 0.42 |
| 8 | 140662310 | rs882555 | 1.028 | -2.059 | -2.38 | -0.718 | 0.081 | 0.225 | 0.225 | 2.866 | 0.5 | 0.21 | 0.49 | 0.13 | 0.64 | 0.46 |
| 8 | 140662811 | rs3780047 | -0.811 | -2.32 | -1.553 | -0.195 | -0.056 | 0.091 | 0.091 | 0.245 | 0.17 | 0.63 | 0.21 | 0.57 | 0.05 | 0.21 |
| 8 | 140663612 | rs12544666 | 0.869 | -1.937 | -1.59 | -0.315 | 0 | -0.019 | -0.019 | 0 | 0.12 | 0.12 | 0.04 | 0.01 | 0.08 | 0.15 |
| 8 | 140668477 | rs3780045 | -0.197 | -1.967 | -1.047 | -0.062 | -0.016 | 0.152 | 0.152 | 0 | 0.05 | 0.16 | 0.1 | 0.27 | 0.01 | 0.13 |
| 8 | 140669745 | rs888349 | -0.218 | -2.178 | -1.022 | -0.994 | 0.002 | -0.017 | -0.017 | 0.159 | 0.75 | 0.75 | 0.86 | 0.97 | 0.8 | 0.85 |
| 8 | 140669967 | rs759656 | -1.127 | -2.724 | -0.562 | -1.507 | 0.005 | -0.001 | -0.001 | 0 | 0.36 | 0.64 | 0.35 | 0.87 | 0.23 | 0.52 |
| 8 | 140670248 | rs2258791 | -0.592 | -2.781 | 0.415 | -1.325 | -0.048 | 0.008 | 0.008 | 0 | 0.25 | 0.3 | 0.22 | 0.73 | 0.11 | 0.35 |
| 8 | 140670853 | rs885725 | -1.657 | -3.094 | -0.203 | -1.717 | 0.092 | 0.313 | 0.313 | 0.334 | 0.5 | 0.95 | 0.44 | 0.88 | 0.3 | 0.63 |
| 8 | 140670930 | rs885724 | -1.478 | -3.089 | 0.148 | -1.622 | 0.089 | 0.165 | 0.165 | 0.154 | 0.48 | 0.77 | 0.44 | 0.86 | 0.3 | 0.62 |
| 8 | 140673111 | rs7822827 | -0.641 | -3.313 | 0.41 | -1.617 | -0.012 | -0.018 | -0.018 | 1.825 | 0.31 | 0.37 | 0.22 | 0.2 | 0.21 | 0.41 |
| 8 | 140676100 | rs3824281 | -0.333 | -3.323 | 0.529 | -1.615 | -0.06 | 0.183 | 0.183 | 0 | 0.16 | 0.39 | 0.21 | 0.65 | 0.08 | 0.26 |
| 8 | 140676452 | rs3780040 | -1.699 | -3.307 | 0.418 | -1.626 | 0.113 | 0.213 | 0.213 | 0.033 | 0.48 | 0.79 | 0.43 | 0.86 | 0.29 | 0.63 |
| 8 | 140678419 | rs1159873 | -0.693 | -2.859 | 0.589 | -1.383 | -0.002 | -0.004 | -0.004 | 0.204 | 0.3 | 0.3 | 0.21 | 0.2 | 0.21 | 0.36 |
| 8 | 140685621 | rs10110946 | -0.208 | -2.625 | 0.328 | -1.568 | 0.015 | 0.002 | 0.002 | 0 | 0.64 | 0.77 | 0.69 | 0.34 | 0.73 | 0.68 |
| 8 | 140686841 | rs7828107 | 1.17 | -2.26 | 0.315 | -1.409 | -0.075 | 0.101 | 0.101 | 0 | 0.33 | 0.19 | 0.44 | 0.1 | 0.5 | 0.3 |
| 8 | 140687274 | rs7846299 | 0.114 | -2.322 | 0.309 | -1.333 | 0.036 | 0.073 | 0.073 | 2.231 | 0.5 | 0.29 | 0.61 | 0.35 | 0.58 | 0.57 |
| 8 | 140694111 | rs2111578 | 0.135 | -2.319 | 0.475 | -1.327 | 0.01 | 0.018 | 0.018 | 3.836 | 0.49 | 0.44 | 0.58 | 0.36 | 0.58 | 0.6 |
| 8 | 140695375 | rs3780032 | 0.478 | -2.095 | 0.082 | -1.413 | -0.015 | 0.166 | 0.166 | 0.982 | 0.44 | 0.2 | 0.46 | 0.15 | 0.54 | 0.47 |
| 8 | 140703184 | rs16911238 | 0.772 | -2.063 | 0.079 | -1.356 | 0.001 | -0.004 | -0.004 | 0.014 | 0.13 | 0.13 | 0.03 | 0.05 | 0.04 | 0.17 |
| 8 | 140703236 | rs11166920 | 1.419 | -2.169 | 0.204 | -1.332 | -0.08 | 0.067 | 0.067 | 0.216 | 0.31 | 0.12 | 0.45 | 0.15 | 0.49 | 0.29 |
| 8 | 140707755 | rs11166921 | -0.39 | -2.364 | 0.483 | -1.338 | 0.042 | -0.002 | -0.002 | 0 | 0.57 | 0.81 | 0.68 | 0.42 | 0.65 | 0.61 |
| 8 | 140708717 | rs1619829 | 0.041 | -2.225 | 0.287 | -1.259 | 0.001 | 0.001 | 0.001 | 1.666 | 0.5 | 0.53 | 0.6 | 0.41 | 0.58 | 0.59 |
| 8 | 140710362 | rs13278664 | 0.744 | -2.005 | -0.102 | -1.127 | -0.009 | 0.043 | 0.043 | 0 | 0.38 | 0.31 | 0.5 | 0.15 | 0.55 | 0.3 |
| 8 | 140712201 | rs199212 | -0.155 | -1.39 | -0.008 | -0.38 | 0.027 | 0.261 | 0.261 | 0 | 0.06 | 0.32 | 0.13 | 0.23 | 0.05 | 0.13 |
| 8 | 140720163 | rs7826309 | 0.911 | -1.374 | -1.355 | -0.824 | -0.003 | 0.161 | 0.161 | 0.747 | 0.45 | 0.19 | 0.33 | 0.2 | 0.53 | 0.59 |
| 8 | 140720412 | rs83520 | 0.181 | -1.766 | -1.199 | -0.616 | 0.007 | -0.001 | -0.001 | 0 | 0.6 | 0.66 | 0.67 | 0.45 | 0.66 | 0.69 |
| 8 | 140722178 | rs124672 | -0.474 | -1.227 | -1.914 | 0.208 | 0.084 | -0.029 | -0.029 | 2.654 | 0.43 | 0.89 | 0.42 | 0.57 | 0.3 | 0.33 |
| 8 | 140726189 | rs199219 | -0.713 | -1.508 | -0.896 | 0.107 | -0.048 | -0.013 | -0.013 | 1.366 | 0.26 | 0.59 | 0.13 | 0.23 | 0.11 | 0.29 |
| 8 | 140726933 | rs1946213 | 0.605 | -1.033 | -2.053 | 0.166 | -0.142 | -0.035 | -0.035 | 0.336 | 0.35 | 0.06 | 0.42 | 0.39 | 0.56 | 0.42 |
| 8 | 140733440 | rs4736295 | 0.034 | -1.225 | -0.929 | -0.181 | -0.001 | 0.021 | 0.021 | 0.042 | 0.06 | 0.19 | 0.17 | 0.08 | 0.03 | 0.06 |
| 8 | 140733497 | rs4736088 | -1.066 | -0.937 | -0.959 | -0.27 | -0.001 | -0.001 | -0.001 | 0.525 | 0.09 | 0.12 | 0.19 | 0.08 | 0.08 | 0.08 |
| 8 | 140734056 | rs199214 | -0.398 | -1.383 | -0.555 | -0.481 | 0.047 | 0.063 | 0.063 | 0 | 0.25 | 0.08 | 0.11 | 0.14 | 0.16 | 0.25 |
| 8 | 140741551 | rs3739373 | 0.37 | -1.232 | -0.482 | -0.518 | -0.003 | 0.004 | 0.004 | 0.029 | 0.06 | 0.1 | 0.16 | 0.1 | 0.03 | 0.06 |
| 8 | 140742121 | rs199228 | -1.44 | -1.366 | -0.294 | -0.571 | -0.005 | -0.006 | -0.006 | 0 | 0.33 | 0.6 | 0.31 | 0.21 | 0.19 | 0.29 |

| | | | | | | | | | | | | | | | | |
|---|---|---|---|---|---|---|---|---|---|---|---|---|---|---|---|---|
| 8 | 140743200 | rs199226 | -0.592 | -1.318 | 0.395 | -0.585 | -0.035 | -0.059 | -0.059 | 0 | 0.25 | 0.32 | 0.13 | 0.11 | 0.16 | 0.22 |
| 8 | 140744726 | rs199225 | -0.812 | -1.316 | 0.381 | -0.585 | -0.063 | -0.098 | -0.098 | 0 | 0.27 | 0.43 | 0.15 | 0.12 | 0.16 | 0.23 |
| 8 | 140748094 | rs1675436 | -0.691 | -1.315 | 0.442 | -0.584 | -0.022 | -0.038 | -0.038 | 1.804 | 0.26 | 0.3 | 0.14 | 0.12 | 0.16 | 0.23 |
| 8 | 140750045 | rs898662 | 0.731 | -1.452 | 0.468 | -0.58 | -0.029 | -0.037 | -0.037 | 0.056 | 0.73 | 0.41 | 0.85 | 0.88 | 0.84 | 0.78 |
| 8 | 140751896 | rs10093271 | 0.653 | -1.423 | 0.476 | -0.568 | 0.015 | 0.015 | 0.015 | 0 | 0.75 | 0.82 | 0.88 | 0.89 | 0.84 | 0.79 |
| 8 | 140752288 | rs7839776 | 0.569 | -1.413 | 0.393 | -0.563 | 0.037 | 0.045 | 0.045 | 0 | 0.76 | 0.86 | 0.89 | 0.89 | 0.85 | 0.79 |
| 8 | 140758956 | rs7843846 | 0.192 | -1.459 | 0.245 | -0.582 | 0.013 | 0.021 | 0.021 | 0.265 | 0.13 | 0.09 | 0.07 | 0.09 | 0.09 | 0.09 |
| 8 | 140762483 | rs11992718 | -1.991 | -1.664 | 0.327 | -0.74 | -0.002 | 0 | 0 | 1.113 | 0.35 | 0.64 | 0.18 | 0.2 | 0.2 | 0.27 |
| 8 | 140771011 | rs11773968 | -0.332 | -1.735 | 0.237 | -0.797 | 0.026 | 0.029 | 0.029 | 0 | 0.12 | 0.02 | 0.03 | 0.03 | 0.06 | 0.1 |
| 8 | 140771161 | rs11781035 | -1.278 | -1.749 | 0.411 | -0.782 | -0.055 | -0.068 | -0.068 | 0.138 | 0.26 | 0.55 | 0.14 | 0.09 | 0.15 | 0.19 |
| 8 | 140777251 | rs898670 | -1.965 | -1.659 | 0.369 | -0.67 | -0.02 | -0.017 | -0.017 | 0.155 | 0.34 | 0.56 | 0.32 | 0.21 | 0.21 | 0.26 |
| 8 | 140785732 | rs7820434 | 0.315 | -1.556 | 0.384 | -0.624 | -0.001 | -0.005 | -0.005 | 0 | 0.11 | 0.12 | 0.07 | 0.05 | 0.08 | 0.08 |
| 8 | 140786172 | rs898668 | -1.796 | -1.625 | 0.493 | -0.648 | -0.018 | -0.013 | -0.013 | 0.056 | 0.3 | 0.61 | 0.33 | 0.21 | 0.19 | 0.26 |
| 8 | 140788299 | rs2126128 | 0.2 | -1.968 | 0.226 | -0.785 | -0.001 | 0.001 | 0.001 | 0 | 0.44 | 0.55 | 0.54 | 0.35 | 0.56 | 0.55 |
| 8 | 140789847 | rs12676068 | -0.091 | -1.886 | 0.114 | -0.438 | 0.028 | -0.011 | -0.011 | 0 | 0.16 | 0.04 | 0.15 | 0.18 | 0.1 | 0.18 |
| 8 | 140792787 | rs7009797 | -1.379 | -2.11 | 0.278 | -0.652 | -0.014 | -0.034 | -0.034 | 0 | 0.72 | 0.82 | 0.74 | 0.55 | 0.72 | 0.77 |
| 8 | 140795752 | rs11166926 | 1.06 | -1.947 | 0.181 | -0.573 | -0.1 | 0.106 | 0.106 | 0.011 | 0.36 | 0.07 | 0.32 | 0.22 | 0.51 | 0.5 |
| 8 | 140796420 | rs11166927 | 0.94 | -1.945 | 0.283 | -0.577 | -0.074 | 0.108 | 0.108 | 0 | 0.37 | 0.11 | 0.32 | 0.22 | 0.51 | 0.5 |
| 8 | 140805932 | rs7820154 | -1.616 | -1.953 | 0.91 | -0.49 | -0.038 | 0.069 | 0.069 | 0 | 0.19 | 0.47 | 0.35 | 0.29 | 0.12 | 0.2 |
| 8 | 140806026 | rs7816574 | -1.189 | -1.974 | 0.924 | -0.495 | -0.065 | 0.019 | 0.019 | 0.015 | 0.16 | 0.48 | 0.17 | 0.15 | 0.06 | 0.12 |
| 8 | 140812347 | rs12679196 | -0.994 | -1.913 | 0.94 | -0.491 | 0.008 | -0.02 | -0.02 | 0 | 0.15 | 0.08 | 0.3 | 0.28 | 0.11 | 0.18 |
| 8 | 140817192 | rs4736007 | -0.059 | -2.032 | 1.058 | -0.454 | 0.006 | 0.016 | 0.016 | 0 | 0.5 | 0.64 | 0.64 | 0.5 | 0.58 | 0.68 |
| 8 | 140819035 | rs7836476 | 0.868 | -1.9 | 0.938 | -0.476 | 0.017 | -0.052 | -0.052 | 0 | 0.38 | 0.47 | 0.35 | 0.21 | 0.52 | 0.5 |
| 8 | 140819819 | rs2242181 | 0.923 | -1.906 | 0.997 | -0.479 | -0.007 | 0.01 | 0.01 | 0.255 | 0.37 | 0.32 | 0.34 | 0.21 | 0.51 | 0.51 |
| 8 | 140821181 | rs1109796 | 0.388 | -1.653 | 0.888 | -0.35 | -0.018 | 0.036 | 0.036 | 0 | 0.06 | 0.11 | 0.1 | 0.14 | 0.01 | 0.11 |
| 8 | 140821491 | rs1109797 | 0.388 | -1.672 | 0.861 | -0.35 | -0.021 | 0.034 | 0.034 | 0 | 0.06 | 0.1 | 0.1 | 0.14 | 0.01 | 0.11 |
| 8 | 140822965 | rs4736008 | 0.308 | -1.959 | 0.473 | -0.566 | -0.001 | -0.001 | -0.001 | 0 | 0.46 | 0.5 | 0.62 | 0.5 | 0.58 | 0.66 |
| 8 | 140830241 | rs9324514 | -1.019 | -1.844 | 0.245 | -0.558 | -0.001 | 0.088 | 0.088 | 0 | 0.56 | 0.73 | 0.64 | 0.74 | 0.43 | 0.49 |
| 8 | 140831320 | rs2035087 | -0.942 | -1.935 | 0.03 | -0.609 | -0.003 | 0.176 | 0.176 | 0 | 0.53 | 0.94 | 0.62 | 0.74 | 0.39 | 0.49 |
| 8 | 140841314 | rs7844870 | -1.027 | -2.325 | 0.139 | -0.795 | -0.073 | -0.017 | -0.017 | 0.017 | 0.25 | 0.55 | 0.36 | 0.22 | 0.11 | 0.17 |
| 8 | 140842723 | rs11985404 | -0.481 | -2.337 | -0.294 | -0.83 | -0.077 | 0.031 | 0.031 | 0.135 | 0.19 | 0.47 | 0.33 | 0.22 | 0.06 | 0.17 |
| 8 | 140843167 | rs10111401 | -1.703 | -2.214 | -0.643 | -0.892 | -0.111 | 0.117 | 0.117 | 0 | 0.59 | 0.96 | 0.65 | 0.74 | 0.45 | 0.5 |
| 8 | 140847892 | rs876745 | -1.479 | -2.211 | -0.603 | -0.848 | 0.099 | 0.075 | 0.075 | 0.211 | 0.41 | 0.86 | 0.45 | 0.28 | 0.25 | 0.31 |
| 8 | 140848850 | rs11782640 | -0.674 | -2.209 | -0.531 | -0.843 | -0.074 | -0.014 | -0.014 | 0 | 0.25 | 0.47 | 0.34 | 0.22 | 0.12 | 0.18 |
| 8 | 140850709 | rs10100605 | 1.382 | -2.032 | -0.721 | -0.775 | -0.047 | -0.099 | -0.099 | 0 | 0.16 | 0.05 | 0.18 | 0.46 | 0.18 | 0.17 |
| 8 | 140851618 | rs7812357 | 0.393 | -2.012 | -0.642 | -0.826 | 0 | -0.003 | -0.003 | 0 | 0.13 | 0.13 | 0.09 | 0.07 | 0.13 | 0.13 |
| 8 | 140852638 | rs898806 | -0.334 | -1.937 | -0.611 | -0.772 | -0.077 | 0.017 | 0.017 | 0.022 | 0.2 | 0.48 | 0.32 | 0.21 | 0.07 | 0.17 |
| 8 | 140855827 | rs10100424 | 0.557 | -1.478 | -1.656 | -0.688 | -0.004 | -0.005 | -0.005 | 0 | 0.24 | 0.15 | 0.29 | 0.67 | 0.21 | 0.31 |
| 8 | 140857481 | rs898807 | -0.798 | -0.689 | -1.868 | -0.225 | -0.089 | 0.142 | 0.142 | 0 | 0.57 | 0.94 | 0.63 | 0.75 | 0.44 | 0.52 |
| 8 | 140862119 | rs2054313 | -0.637 | 0.446 | -0.63 | 0.421 | -0.007 | 0.009 | 0.009 | 0 | 0.12 | 0.11 | 0.09 | 0.07 | 0.15 | 0.13 |
| 8 | 140863075 | rs1473416 | -0.782 | 0.435 | -0.576 | 0.513 | 0.015 | 0.025 | 0.025 | 0 | 0.13 | 0.19 | 0.23 | 0.28 | 0.22 | 0.26 |
| 8 | 140869942 | rs11786354 | -0.236 | 0.699 | -0.803 | 0.395 | 0 | 0.001 | 0.001 | 0 | 0.46 | 0.55 | 0.33 | 0.75 | 0.35 | 0.36 |
| 8 | 140874687 | rs729855 | 0.186 | 0.472 | -1.228 | 0.738 | 0.016 | -0.024 | -0.024 | 0 | 0.17 | 0.06 | 0.06 | 0.27 | 0.12 | 0.13 |
| 8 | 140875002 | rs729854 | 0.368 | 0.326 | -0.54 | 0.653 | 0.001 | 0.025 | 0.025 | 0 | 0.25 | 0.27 | 0.23 | 0.47 | 0.22 | 0.23 |
| 8 | 140876251 | rs6987973 | -1.151 | 0.21 | -1.454 | -0.011 | -0.132 | 0.021 | 0.021 | 0 | 0.68 | 0.92 | 0.56 | 0.83 | 0.51 | 0.67 |
| 8 | 140882517 | rs922280 | 0.897 | 0.709 | -1.655 | -0.039 | -0.036 | -0.022 | -0.022 | 0.625 | 0.28 | 0.07 | 0.39 | 0.22 | 0.32 | 0.35 |
| 8 | 140882563 | rs4236865 | 0.926 | 0.691 | -1.668 | -0.048 | -0.04 | -0.026 | -0.026 | 0 | 0.28 | 0.06 | 0.39 | 0.22 | 0.32 | 0.35 |
| 8 | 140886603 | rs2010771 | 0.3 | 0.257 | -1.127 | -0.092 | 0.009 | -0.005 | -0.005 | 0.227 | 0.35 | 0.28 | 0.4 | 0.32 | 0.32 | 0.34 |
| 8 | 140887602 | rs13254009 | 1.224 | 1.073 | -1.437 | 0.354 | -0.067 | -0.062 | -0.062 | 0 | 0.17 | 0.06 | 0.14 | 0.33 | 0.25 | 0.15 |
| 8 | 140887640 | rs1870810 | 0.189 | 0.297 | -0.841 | -0.013 | -0.001 | -0.001 | -0.001 | 0 | 0.62 | 0.63 | 0.62 | 0.66 | 0.65 | 0.57 |
| 8 | 140891088 | rs4736110 | 0.366 | 0.154 | -0.722 | -0.578 | -0.006 | -0.052 | -0.052 | 0.456 | 0.7 | 0.58 | 0.7 | 0.89 | 0.73 | 0.67 |
| 8 | 140892871 | rs7819785 | -0.38 | -0.301 | -0.543 | -0.099 | -0.069 | 0.022 | 0.022 | 0 | 0.28 | 0.4 | 0.11 | 0.4 | 0.13 | 0.21 |
| 8 | 140898666 | rs922279 | 0.587 | 0.244 | -1.434 | -0.165 | 0.003 | 0.002 | 0.002 | 0 | 0.48 | 0.1 | 0.41 | 0.51 | 0.4 | 0.55 |
| 8 | 140901915 | rs12679771 | -0.322 | 0.435 | -0.36 | -0.356 | 0.038 | -0.058 | -0.058 | 3.814 | 0.2 | 0.51 | 0.36 | 0.08 | 0.21 | 0.26 |
| 8 | 140910308 | rs4736112 | -0.399 | 0.153 | -0.162 | 0.047 | -0.004 | 0.003 | 0.003 | 0 | 0.43 | 0.43 | 0.36 | 0.63 | 0.42 | 0.41 |
| 8 | 140917457 | rs1075493 | 0.35 | 0.032 | 0.064 | 0.006 | -0.008 | 0.014 | 0.014 | 0 | 0.26 | 0.33 | 0.22 | 0.38 | 0.29 | 0.16 |
| 8 | 140921339 | rs11986292 | 0.03 | -0.176 | 0.308 | -0.052 | -0.029 | 0.046 | 0.046 | 0 | 0.22 | 0.48 | 0.17 | 0.3 | 0.15 | 0.27 |
| 8 | 140922699 | rs1551806 | 0.428 | -0.169 | 0.312 | -0.046 | -0.006 | 0.027 | 0.027 | 0 | 0.25 | 0.34 | 0.22 | 0.38 | 0.28 | 0.16 |
| 8 | 140924076 | rs2035131 | -1.594 | 0.151 | -0.122 | -0.133 | -0.136 | -0.061 | -0.061 | 0 | 0.56 | 0.97 | 0.66 | 0.68 | 0.53 | 0.54 |
| 8 | 140926772 | rs2306078 | -0.356 | 0.227 | 0.19 | -0.116 | -0.003 | 0.011 | 0.011 | 0 | 0.78 | 0.72 | 0.84 | 0.7 | 0.81 | 0.71 |
| 8 | 140928764 | rs2290928 | -0.735 | 0.192 | 0.161 | -0.08 | 0.005 | -0.005 | -0.005 | 7.694 | 0.35 | 0.46 | 0.48 | 0.39 | 0.37 | 0.26 |
| 8 | 140932105 | rs10097265 | 0.442 | 0.155 | -0.268 | -0.236 | -0.006 | 0.016 | 0.016 | 0.227 | 0.26 | 0.21 | 0.34 | 0.17 | 0.25 | 0.38 |
| 8 | 140934326 | rs13261649 | 0.938 | 0.011 | -0.633 | -0.248 | 0.008 | 0.036 | 0.036 | 0.385 | 0.22 | 0.06 | 0.1 | 0.17 | 0.16 | 0.27 |
| 8 | 140934805 | rs2085925 | 0.12 | 0.01 | -0.732 | -0.617 | 0 | -0.019 | -0.019 | 0 | 0.27 | 0.22 | 0.18 | 0.05 | 0.3 | 0.18 |
| 8 | 140937473 | rs6578053 | -1.425 | 0.021 | -1.434 | -0.259 | -0.13 | -0.115 | -0.115 | 0.192 | 0.66 | 0.97 | 0.62 | 0.71 | 0.59 | 0.58 |
| 8 | 140943503 | rs2233230 | -1.203 | -0.296 | -1.255 | -0.108 | -0.179 | -0.093 | -0.093 | 0 | 0.61 | 0.97 | 0.59 | 0.7 | 0.51 | 0.56 |
| 8 | 140950509 | rs7011240 | 1.429 | -0.25 | -0.986 | -0.047 | -0.012 | -0.013 | -0.013 | 0 | 0.21 | 0.06 | 0.09 | 0.23 | 0.17 | 0.25 |
| 8 | 140953959 | rs2665946 | 0.782 | -0.218 | -0.508 | -0.083 | 0.001 | -0.002 | -0.002 | 0 | 0.25 | 0.25 | 0.35 | 0.25 | 0.26 | 0.36 |
| 8 | 140954213 | rs2665945 | -0.541 | -0.429 | -0.159 | -0.073 | -0.001 | 0 | 0 | 1.212 | 0.3 | 0.65 | 0.43 | 0.39 | 0.3 | 0.26 |

| | | | | | | | | | | | | | | | | |
|---|---|---|---|---|---|---|---|---|---|---|---|---|---|---|---|---|
| 8 | 140956133 | rs6995955 | -0.444 | -0.507 | -0.254 | -0.135 | -0.007 | 0.001 | 0.001 | 0.799 | 0.46 | 0.48 | 0.26 | 0.62 | 0.39 | 0.42 |
| 8 | 140962615 | rs1542502 | 0.755 | -0.346 | -0.421 | -0.198 | 0.003 | 0.003 | 0.003 | 0 | 0.27 | 0.27 | 0.14 | 0.25 | 0.19 | 0.25 |
| 8 | 140963102 | rs6999707 | 0.084 | -0.623 | -0.109 | -0.333 | 0.039 | -0.009 | -0.009 | 0 | 0.4 | 0.24 | 0.21 | 0.64 | 0.28 | 0.39 |
| 8 | 140963276 | rs2665917 | -1.085 | -0.389 | -0.626 | -0.444 | -0.144 | -0.006 | -0.006 | 0.023 | 0.59 | 0.95 | 0.6 | 0.71 | 0.51 | 0.6 |
| 8 | 140966715 | rs7834946 | -0.161 | -0.104 | 0.681 | -0.343 | 0.013 | -0.074 | -0.074 | 0.354 | 0.12 | 0.33 | 0.1 | 0 | 0.13 | 0.07 |
| 8 | 140969818 | rs732803 | -0.412 | -0.201 | 0.486 | -0.369 | 0.007 | 0.006 | 0.006 | 0 | 0.55 | 0.61 | 0.31 | 0.66 | 0.46 | 0.52 |
| 8 | 140970457 | rs2614733 | -0.687 | -0.213 | 0.152 | -0.402 | -0.052 | -0.004 | -0.004 | 0 | 0.59 | 0.76 | 0.59 | 0.71 | 0.54 | 0.62 |
| 8 | 140973610 | rs2665926 | -0.978 | -0.17 | -0.196 | -0.425 | -0.176 | -0.021 | -0.021 | 0.039 | 0.64 | 0.95 | 0.61 | 0.77 | 0.55 | 0.63 |
| 8 | 140977205 | rs2665931 | -0.918 | -0.051 | 0.532 | -0.432 | 0.044 | -0.122 | -0.122 | 0 | 0.18 | 0.53 | 0.38 | 0.02 | 0.22 | 0.17 |
| 8 | 140980456 | rs10875447 | 0.827 | -0.405 | 0.17 | -0.275 | 0.007 | 0.027 | 0.027 | 0 | 0.35 | 0.25 | 0.17 | 0.28 | 0.34 | 0.35 |
| 8 | 140982636 | rs6981165 | -0.043 | -0.649 | 1.042 | -0.209 | -0.022 | 0.059 | 0.059 | 0 | 0.16 | 0.22 | 0.12 | 0.41 | 0.08 | 0.1 |
| 8 | 140983577 | rs2927052 | -1.118 | -0.641 | 1.156 | -0.201 | 0.022 | 0.022 | 0.022 | 0 | 0.13 | 0.43 | 0.31 | 0.08 | 0.12 | 0.17 |
| 8 | 140984964 | rs2665951 | -0.98 | -0.613 | 1.155 | -0.187 | 0.03 | 0.03 | 0.03 | 0 | 0.12 | 0.42 | 0.31 | 0.08 | 0.12 | 0.17 |
| 8 | 140986842 | rs4736121 | 1.165 | -0.171 | 0.207 | -0.409 | 0.001 | -0.023 | -0.023 | 0 | 0.58 | 0.3 | 0.47 | 0.49 | 0.61 | 0.48 |
| 8 | 140989919 | rs7821395 | 0.215 | 0.296 | 0.45 | -0.256 | -0.008 | -0.012 | -0.012 | 0 | 0.72 | 0.77 | 0.61 | 0.59 | 0.64 | 0.59 |
| 8 | 140990501 | rs2085926 | -0.053 | 0.297 | 0.166 | -0.3 | -0.078 | -0.113 | -0.113 | 0 | 0.75 | 0.97 | 0.78 | 0.59 | 0.68 | 0.64 |
| 8 | 140990677 | rs898809 | 1.669 | 0.026 | 0.16 | -0.444 | -0.023 | -0.023 | -0.023 | 0 | 0.48 | 0.19 | 0.41 | 0.47 | 0.51 | 0.49 |
| 8 | 140990959 | rs13251519 | -0.57 | 0 | 0.667 | -0.468 | -0.044 | -0.032 | -0.032 | 0 | 0.2 | 0.43 | 0.17 | 0.11 | 0.12 | 0.1 |
| 8 | 140991284 | rs4736126 | 1.502 | -0.115 | 0.242 | -0.379 | -0.023 | -0.029 | -0.029 | 0.031 | 0.48 | 0.19 | 0.4 | 0.47 | 0.5 | 0.5 |
| 8 | 140994914 | rs1870807 | -0.976 | -0.028 | 0.746 | -0.29 | 0.041 | 0.068 | 0.068 | 0 | 0.08 | 0.3 | 0.22 | 0.07 | 0.08 | 0.1 |
| 8 | 140995243 | rs1870805 | 0.77 | 0.009 | 0.252 | -0.34 | -0.001 | -0.002 | -0.002 | 1.525 | 0.61 | 0.57 | 0.64 | 0.59 | 0.59 | 0.65 |
| 8 | 140997024 | rs1542501 | 1.711 | 0.344 | -0.462 | -0.242 | -0.065 | -0.109 | -0.109 | 0 | 0.47 | 0.04 | 0.32 | 0.51 | 0.4 | 0.52 |
| 8 | 140997950 | rs2927051 | -0.111 | 0.384 | -0.429 | -0.229 | -0.071 | -0.075 | -0.075 | 6.267 | 0.73 | 0.93 | 0.71 | 0.62 | 0.62 | 0.66 |
| 8 | 141004455 | rs1870809 | -1.463 | 0.527 | 0.204 | -0.161 | 0.004 | 0.052 | 0.052 | 3.114 | 0.26 | 0.84 | 0.38 | 0.11 | 0.23 | 0.14 |
| 8 | 141005450 | rs2665941 | -0.835 | 1.149 | 0.574 | 0.205 | 0.112 | 0.044 | 0.044 | 2.259 | 0.1 | 0.36 | 0.29 | 0.06 | 0.14 | 0.08 |
| 8 | 141015327 | rs10093149 | 0.184 | 1.249 | -0.004 | 0.3 | -0.01 | -0.009 | -0.009 | 0 | 0.78 | 0.81 | 0.56 | 0.66 | 0.62 | 0.64 |
| 8 | 141015586 | rs2011620 | -0.423 | 0.7 | 0.515 | -0.082 | -0.002 | -0.051 | -0.051 | 0 | 0.17 | 0.27 | 0.13 | 0.05 | 0.17 | 0.11 |
| 8 | 141018696 | rs6988665 | -1.528 | 0.342 | 0.645 | -0.398 | 0.012 | 0.019 | 0.019 | 1.352 | 0.32 | 0.74 | 0.23 | 0.09 | 0.21 | 0.18 |
| 8 | 141018901 | rs12549400 | 0.974 | 0.733 | 0.725 | -0.21 | 0.017 | 0.017 | 0.017 | 0.001 | 0.66 | 0.47 | 0.48 | 0.62 | 0.58 | 0.59 |
| 8 | 141020809 | rs6578056 | -0.747 | 0.372 | 1.27 | -0.022 | -0.013 | -0.027 | -0.027 | 0.619 | 0.13 | 0.47 | 0.1 | 0.04 | 0.04 | 0.06 |
| 8 | 141040170 | rs6982545 | -0.175 | 0.404 | 1.363 | -0.006 | 0.036 | -0.035 | -0.035 | 0 | 0.12 | 0.28 | 0.1 | 0.07 | 0.13 | 0.04 |
| 8 | 141040629 | rs11783626 | -0.616 | 0.34 | 1.477 | 0.014 | 0.002 | -0.007 | -0.007 | 0 | 0.11 | 0.41 | 0.1 | 0.05 | 0.05 | 0.06 |
| 8 | 141042931 | rs6998315 | 0.185 | 0.679 | 0.491 | 0.327 | -0.024 | -0.061 | -0.061 | 0 | 0.82 | 0.93 | 0.58 | 0.61 | 0.76 | 0.64 |
| 8 | 141045732 | rs6578061 | 1.612 | 0.356 | 0.27 | 0.132 | -0.043 | -0.069 | -0.069 | 0 | 0.54 | 0.1 | 0.36 | 0.5 | 0.57 | 0.56 |
| 8 | 141056362 | rs10100512 | 0.091 | 0.279 | 1.042 | 0.02 | 0.034 | -0.053 | -0.053 | 0.035 | 0.11 | 0.26 | 0.1 | 0.01 | 0.13 | 0.04 |
| 8 | 141059415 | rs7843613 | -0.071 | -0.025 | 0.474 | 0.322 | -0.031 | -0.08 | -0.08 | 0 | 0.83 | 0.95 | 0.59 | 0.53 | 0.77 | 0.61 |
| 8 | 141061798 | rs4374967 | 1.632 | -0.337 | 0.887 | 0.201 | 0.02 | -0.003 | -0.003 | 0 | 0.54 | 0.3 | 0.4 | 0.45 | 0.61 | 0.52 |
| 8 | 141068140 | rs4629846 | 0.449 | -0.373 | 0.491 | 0.152 | -0.064 | -0.093 | -0.093 | 0 | 0.78 | 0.93 | 0.61 | 0.57 | 0.69 | 0.62 |
| 8 | 141070923 | rs9774091 | -0.471 | -0.497 | 1.394 | 0.177 | -0.081 | -0.044 | -0.044 | 0 | 0.18 | 0.32 | 0.05 | 0.09 | 0.03 | 0.07 |
| 8 | 141071527 | rs11991065 | -0.737 | -0.508 | 1.582 | 0.174 | -0.114 | -0.033 | -0.033 | 0 | 0.19 | 0.46 | 0.07 | 0.11 | 0.03 | 0.07 |
| 8 | 141073802 | rs6578065 | 0.611 | -0.413 | 1.305 | 0.221 | -0.073 | -0.004 | -0.004 | 0.879 | 0.79 | 0.6 | 0.77 | 0.83 | 0.93 | 0.88 |
| 8 | 141074664 | rs7005840 | 0.788 | -0.288 | 1.378 | 0.186 | -0.071 | -0.008 | -0.008 | 0 | 0.78 | 0.6 | 0.77 | 0.83 | 0.91 | 0.86 |
| 8 | 141076197 | rs883613 | -0.859 | 0.087 | 0.934 | -0.105 | -0.014 | -0.012 | -0.012 | 0 | 0.27 | 0.23 | 0.33 | 0.38 | 0.37 | 0.36 |
| 8 | 141083615 | rs10875448 | 1.062 | 0.117 | 1.209 | -0.035 | 0.006 | 0.004 | 0.004 | 0 | 0.58 | 0.52 | 0.5 | 0.54 | 0.54 | 0.54 |
| 8 | 141085094 | rs13261464 | 1.662 | 0.13 | 0.567 | -0.062 | -0.111 | -0.069 | -0.069 | 0 | 0.48 | 0.04 | 0.38 | 0.43 | 0.46 | 0.41 |
| 8 | 141086326 | rs7005952 | -0.337 | 0.102 | 0.855 | -0.341 | 0 | 0 | 0 | 0 | 0.14 | 0.14 | 0.06 | 0.1 | 0.07 | 0.07 |
| 8 | 141086363 | rs885368 | -0.039 | 0.252 | 0.899 | -0.296 | 0.017 | 0.014 | 0.014 | 0 | 0.77 | 0.73 | 0.6 | 0.65 | 0.66 | 0.63 |
| 8 | 141090700 | rs10046672 | 1.639 | 0.241 | 0.894 | -0.358 | -0.028 | -0.051 | -0.051 | 0 | 0.51 | 0.1 | 0.42 | 0.5 | 0.51 | 0.49 |
| 8 | 141092683 | rs12542612 | 0.071 | 0.615 | 1.348 | -0.146 | 0.045 | 0.057 | 0.057 | 0 | 0.78 | 0.64 | 0.59 | 0.63 | 0.66 | 0.67 |
| 8 | 141093505 | rs4551331 | -0.871 | 0.537 | 0.922 | -0.385 | -0.033 | -0.071 | -0.071 | 0.339 | 0.22 | 0.53 | 0.22 | 0.08 | 0.15 | 0.11 |
| 8 | 141094804 | rs13282606 | 1.664 | 0.675 | -0.076 | -0.436 | -0.026 | -0.029 | -0.029 | 0.013 | 0.51 | 0.12 | 0.38 | 0.54 | 0.48 | 0.49 |
| 8 | 141095080 | rs10096038 | -0.313 | 1.029 | -0.238 | -0.168 | -0.055 | -0.104 | -0.104 | 5.313 | 0.8 | 0.95 | 0.71 | 0.64 | 0.75 | 0.76 |
| 8 | 141096207 | rs6578067 | 0.067 | 0.851 | -0.194 | -0.169 | -0.05 | -0.053 | -0.053 | 0.362 | 0.77 | 0.88 | 0.64 | 0.61 | 0.66 | 0.64 |
| 8 | 141099377 | rs6988625 | -0.616 | 0.37 | 0.042 | -0.425 | -0.024 | -0.04 | -0.04 | 0.706 | 0.18 | 0.29 | 0.11 | 0.07 | 0.13 | 0.14 |
| 8 | 141100742 | rs6985153 | -1.009 | 0.332 | -0.434 | -0.726 | -0.004 | -0.035 | -0.035 | 0 | 0.28 | 0.61 | 0.25 | 0.04 | 0.19 | 0.19 |
| 8 | 141100784 | rs6985285 | 0.166 | 0.336 | -1.107 | -0.607 | -0.011 | -0.019 | -0.019 | 0 | 0.72 | 0.79 | 0.56 | 0.65 | 0.67 | 0.61 |
| 8 | 141100950 | rs4736135 | -0.346 | 0.4 | -1.429 | -0.525 | -0.04 | -0.085 | -0.085 | 1.479 | 0.79 | 0.92 | 0.65 | 0.66 | 0.75 | 0.76 |
| 8 | 141104817 | rs12547311 | 1.279 | 0.361 | -1.579 | -0.731 | -0.116 | -0.065 | -0.065 | 1.208 | 0.43 | 0.02 | 0.3 | 0.6 | 0.44 | 0.42 |
| 8 | 141105774 | rs2270409 | 0.727 | 0.239 | -1.671 | -2.049 | 0.041 | 0.068 | 0.068 | 0.273 | 0.35 | 0.22 | 0.3 | 0.84 | 0.24 | 0.43 |
| 8 | 141106204 | rs4289794 | 0.396 | 0.451 | -1.003 | -2.196 | -0.013 | 0.042 | 0.042 | 0 | 0.67 | 0.74 | 0.56 | 0.89 | 0.61 | 0.67 |
| 8 | 141107977 | rs3802217 | -0.431 | 0.6 | -0.95 | -2.249 | -0.024 | 0.04 | 0.04 | 1.378 | 0.74 | 0.82 | 0.62 | 0.88 | 0.66 | 0.67 |
| 8 | 141113716 | rs4460346 | 1.386 | 0.487 | -0.968 | -2.449 | 0.004 | 0.044 | 0.044 | 0.1 | 0.39 | 0.32 | 0.4 | 0.86 | 0.38 | 0.4 |
| 8 | 141113741 | rs4297022 | 0.513 | 0.71 | -0.968 | -1.951 | -0.069 | 0.14 | 0.14 | 0 | 0.64 | 0.86 | 0.57 | 0.87 | 0.57 | 0.62 |
| 8 | 141121745 | rs10101006 | -0.175 | 0.906 | -0.298 | -1.709 | 0.005 | -0.038 | -0.038 | 1.645 | 0.73 | 0.66 | 0.62 | 0.85 | 0.71 | 0.75 |
| 8 | 141128357 | rs4736138 | 0.017 | 0.911 | -0.349 | -1.705 | 0.007 | -0.042 | -0.042 | 0 | 0.27 | 0.36 | 0.38 | 0.15 | 0.29 | 0.25 |
| 8 | 141128606 | rs4736139 | -1.249 | 0.815 | -0.209 | -1.932 | -0.005 | 0.121 | 0.121 | 0 | 0.46 | 0.76 | 0.53 | 0.16 | 0.46 | 0.39 |
| 8 | 141132306 | rs6987302 | 0.103 | 0.904 | 0.164 | -1.495 | 0.015 | -0.043 | -0.043 | 0 | 0.09 | 0.24 | 0.08 | 0.01 | 0.08 | 0.1 |
| 8 | 141133631 | rs6993130 | -0.547 | 0.952 | -0.536 | -1.502 | -0.005 | 0.033 | 0.033 | 0.647 | 0.37 | 0.69 | 0.48 | 0.16 | 0.38 | 0.35 |
| 8 | 141142549 | rs10216841 | -0.635 | 0.738 | -0.688 | -0.879 | 0.016 | 0.008 | 0.008 | 0.471 | 0.57 | 0.59 | 0.75 | 0.37 | 0.67 | 0.58 |

| | | | | | | | | | | | | | | | | |
|---|---|---|---|---|---|---|---|---|---|---|---|---|---|---|---|---|
| 8 | 141143227 | rs11985315 | -0.189 | 1.494 | -0.777 | -1.197 | 0.001 | 0.01 | 0.01 | 0.188 | 0.44 | 0.59 | 0.49 | 0.36 | 0.45 | 0.48 |
| 8 | 141149918 | rs7006969 | -0.272 | 1.6 | -0.792 | -1.262 | 0 | 0.003 | 0.003 | 0 | 0.45 | 0.55 | 0.48 | 0.36 | 0.46 | 0.45 |
| 8 | 141150169 | rs6992848 | -0.393 | 1.654 | -0.722 | -1.558 | 0.096 | -0.103 | -0.103 | 0 | 0.18 | 0.41 | 0.32 | 0.01 | 0.27 | 0.14 |
| 8 | 141160226 | rs11774304 | 0.493 | 1.694 | -1.055 | -1.512 | -0.04 | 0.038 | 0.038 | 3.792 | 0.14 | 0.05 | 0.26 | 0.01 | 0.21 | 0.09 |
| 8 | 141163406 | rs6996708 | -1.206 | 1.422 | -1.275 | -0.739 | 0.018 | -0.152 | -0.152 | 0 | 0.62 | 0.98 | 0.82 | 0.55 | 0.71 | 0.59 |
| 8 | 141166373 | rs4736143 | -0.918 | 1.63 | -0.467 | -1.051 | 0.053 | -0.114 | -0.114 | 0.161 | 0.26 | 0.55 | 0.4 | 0.01 | 0.35 | 0.18 |
| 8 | 141168773 | rs6578074 | -0.623 | 1.424 | -0.654 | -1.32 | -0.003 | -0.014 | -0.014 | 0 | 0.87 | 0.88 | 0.86 | 1 | 0.82 | 0.94 |
| 8 | 141171050 | rs4567012 | -0.436 | 1.299 | -0.693 | -1.352 | -0.003 | -0.008 | -0.008 | 0 | 0.86 | 0.88 | 0.86 | 1 | 0.82 | 0.94 |
| 8 | 141177398 | rs6988524 | 0.908 | 1.352 | -0.696 | -1.461 | 0.04 | -0.13 | -0.13 | 0.792 | 0.68 | 0.45 | 0.54 | 0.99 | 0.58 | 0.8 |
| 8 | 141177627 | rs6578076 | -0.108 | 1.408 | -0.398 | -1.46 | 0.024 | -0.106 | -0.106 | 4.98 | 0.84 | 0.63 | 0.81 | 1 | 0.79 | 0.9 |
| 8 | 141178125 | rs4262308 | 0.725 | 1.336 | -0.43 | -1.437 | 0.059 | -0.153 | -0.153 | 2.508 | 0.69 | 0.52 | 0.55 | 0.99 | 0.58 | 0.8 |
| 8 | 141180368 | rs7017708 | 0.344 | 0.957 | -0.72 | -1.069 | -0.002 | 0.013 | 0.013 | 0.245 | 0.37 | 0.42 | 0.43 | 0.52 | 0.38 | 0.43 |
| 8 | 141184000 | rs12682134 | 0.479 | 0.891 | -1.507 | -0.969 | -0.027 | 0.002 | 0.002 | 2.075 | 0.46 | 0.09 | 0.38 | 0.39 | 0.41 | 0.47 |
| 8 | 141188691 | rs7816828 | 0.059 | 0.82 | -1.47 | -1.545 | -0.019 | 0.017 | 0.017 | 0.076 | 0.65 | 0.26 | 0.65 | 0.99 | 0.55 | 0.77 |
| 8 | 141198629 | rs11776464 | 0.484 | 1.199 | -1.748 | -1.222 | -0.067 | -0.005 | -0.005 | 0.542 | 0.45 | 0.09 | 0.38 | 0.36 | 0.46 | 0.47 |
| 8 | 141203709 | rs3935989 | -0.099 | 1.131 | -1.973 | -2.14 | -0.106 | 0.152 | 0.152 | 5.407 | 0.31 | 0.09 | 0.42 | 0.01 | 0.41 | 0.18 |
| 8 | 141204995 | rs28575441 | -1.562 | 0.571 | -1.68 | -1.553 | 0.027 | -0.051 | -0.051 | 0 | 0.74 | 0.91 | 0.88 | 0.59 | 0.8 | 0.68 |
| 8 | 141210214 | rs13259162 | 0.12 | 0.35 | -1.31 | -1.831 | 0.023 | -0.003 | -0.003 | 0 | 0.39 | 0.32 | 0.44 | 0.58 | 0.38 | 0.42 |
| 8 | 141213288 | rs10088351 | -1.351 | 0.483 | -1.281 | -1.356 | 0.027 | -0.053 | -0.053 | 0 | 0.74 | 0.91 | 0.88 | 0.59 | 0.8 | 0.68 |
| 8 | 141219996 | rs4463400 | 0.153 | 1.139 | -0.751 | -1.445 | 0.031 | -0.045 | -0.045 | 0.004 | 0.14 | 0.19 | 0.28 | 0 | 0.24 | 0.09 |
| 8 | 141220019 | rs4407863 | -1.001 | -0.093 | -0.948 | -1.132 | 0.02 | -0.05 | -0.05 | 0 | 0.75 | 0.9 | 0.87 | 0.6 | 0.81 | 0.66 |
| 8 | 141222648 | rs6578080 | 0.185 | 0.31 | -0.862 | -1.504 | -0.03 | 0.227 | 0.227 | 0 | 0.41 | 0.11 | 0.47 | 0.01 | 0.53 | 0.22 |
| 8 | 141224852 | rs4397386 | 0.246 | 0.316 | -0.917 | -1.509 | -0.06 | 0.285 | 0.285 | 0 | 0.41 | 0.06 | 0.46 | 0.01 | 0.53 | 0.22 |
| 8 | 141227581 | rs4286926 | -0.419 | -0.202 | -0.171 | -1.213 | 0.023 | 0.005 | 0.005 | 0 | 0.58 | 0.65 | 0.79 | 0.55 | 0.73 | 0.62 |
| 8 | 141234162 | rs10448121 | -0.225 | -0.185 | 0.043 | -1.207 | -0.014 | -0.004 | -0.004 | 2.046 | 0.55 | 0.51 | 0.76 | 0.55 | 0.7 | 0.64 |
| 8 | 141242366 | rs7822130 | -0.155 | -0.069 | -0.241 | -1.134 | -0.005 | -0.001 | -0.001 | 0 | 0.56 | 0.56 | 0.76 | 0.51 | 0.7 | 0.63 |
| 8 | 141242865 | rs9324519 | -0.422 | 0.098 | -0.823 | -1.026 | 0.041 | -0.007 | -0.007 | 0 | 0.59 | 0.71 | 0.78 | 0.51 | 0.72 | 0.64 |
| 8 | 141259076 | rs6578086 | -0.219 | 0.937 | -1.14 | -1.851 | 0.002 | 0.023 | 0.023 | 0 | 0.47 | 0.36 | 0.47 | 0.02 | 0.45 | 0.28 |
| 8 | 141259245 | rs4736023 | -0.729 | 1.329 | -0.994 | -1.754 | 0.006 | -0.047 | -0.047 | 0.993 | 0.16 | 0.27 | 0.1 | 0.05 | 0.15 | 0.18 |
| 8 | 141260967 | rs7017431 | -0.283 | 2.288 | -1.44 | -1.35 | -0.074 | -0.051 | -0.051 | 0 | 0.86 | 0.97 | 0.85 | 0.75 | 0.71 | 0.78 |
| 8 | 141262869 | rs11166964 | 0.378 | 1.733 | -1.521 | -1.557 | -0.026 | 0.048 | 0.048 | 0 | 0.09 | 0.02 | 0.2 | 0 | 0.12 | 0.07 |
| 8 | 141263015 | rs11166965 | 0.378 | 1.734 | -1.521 | -1.557 | -0.024 | 0.048 | 0.048 | 0 | 0.09 | 0.02 | 0.19 | 0 | 0.12 | 0.07 |
| 8 | 141265687 | rs4078541 | -1.082 | 1.507 | -0.928 | -1.681 | -0.011 | -0.1 | -0.1 | 0.817 | 0.18 | 0.34 | 0.25 | 0.02 | 0.17 | 0.17 |
| 8 | 141271075 | rs11166967 | -0.125 | 1.591 | -0.958 | -1.678 | 0.001 | -0.025 | -0.025 | 0 | 0.13 | 0.14 | 0.24 | 0.02 | 0.17 | 0.17 |
| 8 | 141274104 | rs4076117 | 1.205 | 0.852 | -1.549 | -2.593 | 0.041 | 0.099 | 0.099 | 0 | 0.33 | 0.16 | 0.21 | 0 | 0.25 | 0.09 |
| 8 | 141275157 | rs6578089 | -0.537 | 0.946 | -1.31 | -2.447 | -0.006 | -0.108 | -0.108 | 0.051 | 0.19 | 0.38 | 0.27 | 0.02 | 0.19 | 0.17 |
| 8 | 141280413 | rs10464982 | 0.676 | 0.572 | -1.043 | -2.44 | 0.015 | 0.019 | 0.019 | 0 | 0.44 | 0.38 | 0.43 | 0.73 | 0.37 | 0.42 |
| 8 | 141281761 | rs4236871 | -0.025 | 1.311 | -1.102 | -2.275 | -0.014 | 0.024 | 0.024 | 0 | 0.67 | 0.78 | 0.62 | 0.23 | 0.65 | 0.48 |
| 8 | 141281786 | rs4736166 | 0.848 | 1.189 | -1.469 | -2.928 | -0.029 | 0.05 | 0.05 | 0 | 0.22 | 0.14 | 0.23 | 0.02 | 0.29 | 0.2 |
| 8 | 141281899 | rs4469435 | 0.45 | 1.042 | -0.938 | -2.94 | -0.007 | -0.084 | -0.084 | 0 | 0.25 | 0.31 | 0.18 | 0 | 0.2 | 0.16 |
| 8 | 141284382 | rs4736167 | 0.395 | -0.044 | -0.997 | -2.55 | -0.005 | -0.006 | -0.006 | 0 | 0.66 | 0.56 | 0.5 | 0.25 | 0.67 | 0.51 |
| 8 | 141285098 | rs4317547 | -0.492 | 0.158 | -1.08 | -2.369 | -0.014 | -0.038 | -0.038 | 0 | 0.75 | 0.9 | 0.56 | 0.27 | 0.72 | 0.59 |
| 8 | 141288769 | rs4075647 | 0.263 | -0.299 | -0.981 | -2.316 | 0.011 | -0.028 | -0.028 | 0 | 0.24 | 0.15 | 0.43 | 0.73 | 0.23 | 0.41 |
| 8 | 141290106 | rs4560764 | -0.024 | -0.139 | -0.785 | -2.495 | -0.008 | -0.147 | -0.147 | 0 | 0.34 | 0.44 | 0.23 | 0.01 | 0.33 | 0.2 |
| 8 | 141297613 | rs6999767 | -0.739 | 0.191 | -1.251 | -1.531 | -0.03 | -0.055 | -0.055 | 0.548 | 0.7 | 0.9 | 0.78 | 0.65 | 0.67 | 0.79 |
| 8 | 141312541 | rs4311638 | -0.617 | 0.491 | -0.518 | -1.771 | 0.008 | 0.026 | 0.026 | 0 | 0.54 | 0.75 | 0.67 | 0.65 | 0.52 | 0.68 |
| 8 | 141327474 | rs10091104 | 0.762 | 0.446 | -0.319 | -1.808 | -0.006 | 0.012 | 0.012 | 0.024 | 0.33 | 0.25 | 0.26 | 0.01 | 0.37 | 0.19 |
| 8 | 141330721 | rs2049236 | -0.923 | 0.443 | -0.298 | -1.792 | 0.008 | -0.006 | -0.006 | 0 | 0.61 | 0.71 | 0.71 | 0.64 | 0.6 | 0.69 |
| 8 | 141332262 | rs1568605 | -0.367 | 0.413 | -0.466 | -1.795 | 0.001 | 0.004 | 0.004 | 0 | 0.54 | 0.56 | 0.65 | 0.64 | 0.52 | 0.57 |
| 8 | 141339278 | rs7010142 | 0.161 | 0.524 | -0.63 | -1.662 | -0.002 | 0.037 | 0.037 | 0 | 0.22 | 0.24 | 0.4 | 0.63 | 0.17 | 0.38 |
| 8 | 141344108 | rs11166972 | 0.442 | 0.627 | -1.269 | -1.606 | 0.001 | -0.021 | -0.021 | 0 | 0.18 | 0.11 | 0.37 | 0.63 | 0.15 | 0.36 |
| 8 | 141345638 | rs7835808 | -0.276 | 0.514 | -1.41 | -1.144 | -0.003 | 0 | 0 | 0.229 | 0.59 | 0.77 | 0.72 | 0.65 | 0.57 | 0.67 |
| 8 | 141355502 | rs6993246 | -0.576 | 0.626 | -0.542 | -1.205 | -0.017 | -0.161 | -0.161 | 0 | 0.26 | 0.51 | 0.21 | 0.02 | 0.2 | 0.28 |
| 8 | 141360167 | rs7839705 | -0.518 | 0.846 | -0.933 | -1.506 | -0.001 | -0.032 | -0.032 | 0 | 0.39 | 0.34 | 0.22 | 0.01 | 0.35 | 0.36 |
| 8 | 141360600 | rs6578099 | -0.548 | 0.913 | -1.866 | -1.286 | -0.046 | -0.08 | -0.08 | 0 | 0.79 | 0.99 | 0.88 | 0.77 | 0.78 | 0.77 |
| 8 | 141362279 | rs7815187 | -0.245 | 0.948 | -1.766 | -1.176 | 0 | -0.035 | -0.035 | 0 | 0.75 | 0.84 | 0.83 | 0.71 | 0.78 | 0.72 |
| 8 | 141371593 | rs909055 | -0.346 | 0.84 | -1.485 | -1.167 | 0.01 | -0.078 | -0.078 | 0 | 0.66 | 0.88 | 0.78 | 0.6 | 0.69 | 0.57 |
| 8 | 141371655 | rs884900 | -0.16 | 1.518 | -0.807 | -1.474 | 0.136 | -0.075 | -0.075 | 0 | 0.13 | 0.31 | 0.21 | 0.01 | 0.3 | 0.1 |
| 8 | 141372344 | rs10110999 | 1.406 | 1.173 | -1.055 | -1.782 | 0.004 | 0.158 | 0.158 | 0.555 | 0.29 | 0.06 | 0.15 | 0.01 | 0.21 | 0.11 |
| 8 | 141386311 | rs7834518 | -0.778 | 1.103 | -0.972 | -1.77 | 0.014 | -0.083 | -0.083 | 0.887 | 0.64 | 0.86 | 0.78 | 0.6 | 0.68 | 0.56 |
| 8 | 141407698 | rs3779658 | 0.672 | 1.156 | -0.835 | -1.667 | 0.009 | -0.093 | -0.093 | 0 | 0.35 | 0.11 | 0.22 | 0.42 | 0.31 | 0.44 |
| 8 | 141410497 | rs11778051 | 0.462 | 1.139 | -0.801 | -1.706 | 0.028 | 0.042 | 0.042 | 0.082 | 0.71 | 0.65 | 0.42 | 0.44 | 0.55 | 0.58 |
| 8 | 141411588 | rs13265944 | 0.29 | 1.301 | -0.915 | -1.194 | 0.001 | -0.106 | -0.106 | 0.27 | 0.35 | 0.09 | 0.22 | 0.4 | 0.31 | 0.44 |
| 8 | 141441981 | rs4515527 | 0.48 | 1.734 | -0.52 | -1.346 | 0.13 | -0.086 | -0.086 | 0 | 0.86 | 0.62 | 0.79 | 1 | 0.72 | 0.9 |
| 8 | 141458494 | rs12681305 | -0.657 | 0.98 | -1.106 | -0.837 | 0.001 | -0.002 | -0.002 | 0 | 0.56 | 0.56 | 0.41 | 0.53 | 0.56 | 0.58 |
| 8 | 141459861 | rs7841423 | -0.215 | 1.064 | -1.086 | -0.87 | 0.01 | -0.019 | -0.019 | 0.003 | 0.07 | 0.1 | 0.07 | 0 | 0.08 | 0.05 |
| 8 | 141461062 | rs3735803 | -0.181 | 1.061 | -1.102 | -0.868 | 0.011 | 0.01 | 0.01 | 0.31 | 0.49 | 0.44 | 0.34 | 0.52 | 0.48 | 0.53 |
| 8 | 141461266 | rs3735801 | -0.668 | 0.903 | -1.497 | -0.852 | 0 | -0.001 | -0.001 | 0 | 0.56 | 0.56 | 0.41 | 0.53 | 0.56 | 0.57 |
| 8 | 141467960 | rs3735799 | 0.299 | 0.345 | -1.274 | -0.552 | 0.001 | 0.003 | 0.003 | 0.29 | 0.49 | 0.51 | 0.64 | 0.49 | 0.47 | 0.46 |

| | | | | | | | | | | | | | | | | |
|---|---|---|---|---|---|---|---|---|---|---|---|---|---|---|---|---|
| 8 | 141483528 | rs4961282 | 0.545 | -0.576 | -0.763 | -0.613 | -0.075 | -0.121 | -0.121 | 3.752 | 0.76 | 0.5 | 0.88 | 0.96 | 0.86 | 0.81 |
| 8 | 141506903 | rs11780595 | 0.124 | -0.112 | -2.012 | -0.333 | 0.009 | 0.053 | 0.053 | 1.441 | 0.44 | 0.09 | 0.59 | 0.68 | 0.63 | 0.48 |
| 8 | 141514184 | rs4961309 | -0.013 | -0.519 | -1.089 | -0.464 | 0.004 | 0.035 | 0.035 | 0 | 0.83 | 0.71 | 0.82 | 0.76 | 0.83 | 0.75 |
| 8 | 141518037 | rs1057083 | -0.147 | -0.545 | -0.985 | -0.477 | -0.008 | -0.028 | -0.028 | 0.923 | 0.84 | 0.91 | 0.84 | 0.76 | 0.84 | 0.75 |
| 8 | 141520581 | rs6578111 | -0.147 | -0.545 | -0.985 | -0.477 | -0.008 | -0.028 | -0.028 | 0.171 | 0.84 | 0.91 | 0.84 | 0.76 | 0.84 | 0.75 |
| 8 | 141526038 | rs4246131 | 0.052 | -0.455 | -1.026 | -0.485 | 0 | 0.037 | 0.037 | 0.001 | 0.83 | 0.72 | 0.83 | 0.76 | 0.85 | 0.75 |
| 8 | 141526231 | rs4961323 | 0.135 | -0.467 | -1.026 | -0.486 | 0.005 | 0.033 | 0.033 | 0 | 0.83 | 0.72 | 0.82 | 0.76 | 0.83 | 0.75 |
| 8 | 141530057 | rs10109589 | -0.417 | -0.637 | -0.701 | -1.059 | -0.01 | -0.031 | -0.031 | 0 | 0.18 | 0.2 | 0.11 | 0.07 | 0.13 | 0.09 |
| 8 | 141533818 | rs11595 | 0.65 | -0.375 | -0.731 | -1.077 | 0.006 | 0.024 | 0.024 | 0.367 | 0.49 | 0.37 | 0.61 | 0.64 | 0.55 | 0.53 |
| 8 | 141534198 | rs2977451 | -0.329 | -0.309 | -0.238 | -1.273 | -0.001 | -0.066 | -0.066 | 0 | 0.83 | 0.74 | 0.9 | 0.99 | 0.82 | 0.89 |
| 8 | 141537241 | rs6578113 | -0.033 | -0.333 | 0.152 | -1.163 | 0 | -0.027 | -0.027 | 0 | 0.08 | 0.15 | 0.04 | 0.04 | 0.07 | 0.06 |
| 8 | 141537406 | rs2235117 | -1.452 | -0.443 | 0.46 | -1.219 | -0.001 | 0.057 | 0.057 | 0 | 0.2 | 0.55 | 0.28 | 0.3 | 0.19 | 0.29 |
| 8 | 141538846 | rs2977453 | -1.503 | -0.463 | 0.463 | -1.237 | -0.002 | 0.052 | 0.052 | 0 | 0.2 | 0.57 | 0.28 | 0.3 | 0.19 | 0.3 |
| 8 | 141539854 | rs2944760 | -1.51 | -0.452 | 0.441 | -1.255 | -0.004 | 0.04 | 0.04 | 0 | 0.21 | 0.62 | 0.29 | 0.3 | 0.19 | 0.29 |
| 8 | 141541288 | rs2977461 | -1.277 | -0.464 | 0.477 | -1.362 | -0.013 | -0.072 | -0.072 | 0 | 0.17 | 0.62 | 0.12 | 0.04 | 0.13 | 0.1 |
| 8 | 141541888 | rs1060832 | -0.658 | -0.394 | 0.404 | -1.321 | 0.024 | -0.059 | -0.059 | 0.001 | 0.13 | 0.34 | 0.09 | 0.04 | 0.12 | 0.1 |
| 8 | 141547226 | rs10096598 | 0.235 | 0.392 | -0.766 | -1.117 | -0.063 | -0.026 | -0.026 | 0 | 0.27 | 0.08 | 0.29 | 0.33 | 0.27 | 0.27 |
| 8 | 141549291 | rs2977469 | -1.259 | 0.159 | -0.334 | -1.202 | 0 | 0.002 | 0.002 | 0 | 0.5 | 0.54 | 0.57 | 0.64 | 0.48 | 0.55 |
| 8 | 141551407 | rs2293939 | 0.555 | 0.036 | -0.648 | -1.488 | -0.078 | -0.054 | -0.054 | 0 | 0.19 | 0.06 | 0.28 | 0.36 | 0.27 | 0.24 |
| 8 | 141554959 | rs2977477 | -1.678 | -0.001 | -0.86 | -1.723 | -0.057 | 0.083 | 0.083 | 0 | 0.86 | 0.97 | 0.9 | 0.99 | 0.79 | 0.91 |
| 8 | 141555862 | rs3864659 | 0.01 | -0.11 | -0.527 | -1.756 | 0.001 | -0.067 | -0.067 | 0 | 0.14 | 0.31 | 0.1 | 0.04 | 0.12 | 0.09 |
| 8 | 141560634 | rs2944765 | -0.14 | 0.429 | -1.117 | -1.645 | -0.11 | -0.012 | -0.012 | 0 | 0.65 | 0.92 | 0.61 | 0.63 | 0.49 | 0.63 |
| 8 | 141561357 | rs2292780 | 0.957 | 0.411 | -1.018 | -1.654 | -0.003 | -0.025 | -0.025 | 0 | 0.08 | 0.15 | 0.04 | 0.04 | 0.07 | 0.04 |
| 8 | 141564525 | rs2977482 | -0.117 | 0.553 | -1.443 | -1.584 | -0.114 | -0.023 | -0.023 | 0.028 | 0.66 | 0.92 | 0.62 | 0.6 | 0.5 | 0.62 |
| 8 | 141566311 | rs2271738 | 1.724 | 0.185 | -1.302 | -1.32 | 0.003 | 0.005 | 0.005 | 0.048 | 0.32 | 0.29 | 0.58 | 0.64 | 0.33 | 0.5 |
| 8 | 141569310 | rs2271735 | 0.458 | 0.835 | -1.48 | -0.644 | -0.097 | -0.045 | -0.045 | 0.71 | 0.19 | 0.03 | 0.16 | 0.28 | 0.28 | 0.23 |
| 8 | 141572059 | rs2977489 | -1.606 | 0.979 | -1.301 | -0.638 | -0.038 | 0.035 | 0.035 | 0 | 0.86 | 0.94 | 0.79 | 0.93 | 0.83 | 0.93 |
| 8 | 141572167 | rs3928672 | 0.13 | 0.676 | -1.153 | -0.572 | -0.008 | 0.015 | 0.015 | 0.393 | 0.28 | 0.1 | 0.17 | 0.29 | 0.25 | 0.23 |
| 8 | 141572309 | rs2292775 | 0.084 | 0.187 | -0.309 | -0.696 | 0.032 | -0.007 | -0.007 | 0 | 0.09 | 0.38 | 0.05 | 0.03 | 0.06 | 0.04 |
| 8 | 141573358 | rs2977490 | -0.507 | 0.35 | -1.251 | -0.446 | -0.001 | 0.03 | 0.03 | 1.133 | 0.51 | 0.61 | 0.37 | 0.59 | 0.46 | 0.45 |
| 8 | 141582896 | rs13276958 | -0.615 | 0.837 | -1.588 | -0.937 | -0.079 | 0.121 | 0.121 | 0 | 0.29 | 0.14 | 0.24 | 0.05 | 0.45 | 0.24 |
| 8 | 141584017 | rs7824304 | -0.358 | 0.421 | -0.974 | -1.016 | -0.003 | -0.023 | -0.023 | 0.775 | 0.26 | 0.24 | 0.22 | 0.04 | 0.34 | 0.28 |
| 8 | 141587404 | rs9694342 | -0.522 | 0.299 | -0.663 | -1.079 | 0.006 | -0.093 | -0.093 | 0 | 0.28 | 0.35 | 0.23 | 0.04 | 0.34 | 0.28 |
| 8 | 141592657 | rs7825416 | -0.371 | -0.284 | -0.658 | 0.022 | 0.003 | 0.035 | 0.035 | 0 | 0.86 | 0.83 | 0.84 | 0.68 | 0.84 | 0.79 |
| 8 | 141594881 | rs7009635 | 0.223 | -1.271 | -1.586 | -0.305 | -0.014 | -0.042 | -0.042 | 0.023 | 0.41 | 0.07 | 0.34 | 0.39 | 0.62 | 0.38 |
| 8 | 141595023 | rs6985156 | 0.527 | -0.59 | -0.948 | -0.589 | 0 | 0.002 | 0.002 | 0.496 | 0.12 | 0.06 | 0.06 | 0 | 0.12 | 0.05 |
| 8 | 141601542 | rs7843258 | -0.452 | -0.642 | -0.734 | -0.044 | 0.006 | 0.076 | 0.076 | 0 | 0.84 | 0.78 | 0.83 | 0.58 | 0.83 | 0.77 |
| 8 | 141606989 | rs4961271 | -0.247 | -0.656 | -0.553 | -0.051 | 0.008 | 0.098 | 0.098 | 0 | 0.83 | 0.75 | 0.81 | 0.58 | 0.82 | 0.76 |
| 8 | 141621038 | rs7001653 | 0.78 | -1.115 | -0.472 | -0.398 | -0.018 | 0.017 | 0.017 | 0.238 | 0.46 | 0.26 | 0.45 | 0.45 | 0.5 | 0.4 |
| 8 | 141622537 | rs2176397 | 0.579 | -1.064 | 0.255 | -0.411 | -0.004 | -0.029 | -0.029 | 0 | 0.35 | 0.37 | 0.36 | 0.13 | 0.28 | 0.34 |
| 8 | 141625045 | rs10087629 | 0.574 | -1.045 | 0.26 | -0.373 | 0.001 | 0.026 | 0.026 | 0.172 | 0.49 | 0.32 | 0.46 | 0.45 | 0.52 | 0.45 |
| 8 | 141630172 | rs11786030 | -1.044 | -0.761 | 0.257 | -0.066 | 0.009 | 0.078 | 0.078 | 0 | 0.87 | 0.86 | 0.86 | 0.58 | 0.83 | 0.8 |
| 8 | 141634540 | rs1878478 | 1.248 | -1.168 | 0.298 | -0.457 | -0.054 | -0.098 | -0.098 | 0 | 0.26 | 0.45 | 0.33 | 0.1 | 0.17 | 0.25 |
| 8 | 141642392 | rs4961278 | 0.239 | -0.734 | 0.436 | 0.057 | 0.016 | 0.06 | 0.06 | 0 | 0.74 | 0.63 | 0.77 | 0.5 | 0.71 | 0.7 |
| 8 | 141652465 | rs6578128 | -0.42 | -0.419 | 0.111 | 0.29 | -0.004 | 0 | 0 | 0 | 0.83 | 0.86 | 0.84 | 0.55 | 0.82 | 0.76 |
| 8 | 141660770 | rs10108278 | 0.838 | -0.845 | 0.279 | 0.19 | -0.001 | -0.007 | -0.007 | 0.84 | 0.67 | 0.31 | 0.72 | 0.53 | 0.7 | 0.67 |
| 8 | 141675931 | rs13273096 | 0.62 | -1.393 | 0.48 | -0.275 | -0.005 | 0.021 | 0.021 | 0.107 | 0.45 | 0.34 | 0.46 | 0.39 | 0.52 | 0.44 |
| 8 | 141677324 | rs7016497 | -0.476 | -1.487 | 0.519 | -0.326 | -0.001 | 0.004 | 0.004 | 0 | 0.81 | 0.85 | 0.85 | 0.57 | 0.82 | 0.74 |
| 8 | 141677470 | rs11166992 | 0.638 | -1.363 | 0.631 | -0.225 | -0.033 | 0.037 | 0.037 | 0.278 | 0.44 | 0.2 | 0.44 | 0.41 | 0.52 | 0.44 |
| 8 | 141701476 | rs7820179 | 0.255 | -1.464 | 0.773 | -0.285 | -0.001 | 0.03 | 0.03 | 0.122 | 0.48 | 0.67 | 0.49 | 0.48 | 0.52 | 0.46 |
| 8 | 141704996 | rs11785444 | 0.966 | -0.981 | 0.242 | -0.834 | 0.002 | 0.008 | 0.008 | 0.063 | 0.17 | 0.1 | 0.2 | 0.18 | 0.12 | 0.22 |
| 8 | 141706320 | rs4961287 | 0.642 | -2.151 | 0.302 | -1.773 | 0.002 | 0.18 | 0.18 | 1.146 | 0.36 | 0.12 | 0.35 | 0.08 | 0.34 | 0.28 |
| 8 | 141708394 | rs4596627 | -0.925 | -2.155 | 0.532 | -1.566 | 0 | 0.034 | 0.034 | 0 | 0.85 | 0.84 | 0.73 | 0.51 | 0.87 | 0.73 |
| 8 | 141713774 | rs10109684 | -0.237 | -2.178 | 0.933 | -1.572 | 0.001 | 0.133 | 0.133 | 0 | 0.81 | 0.56 | 0.67 | 0.43 | 0.85 | 0.71 |
| 8 | 141762659 | rs6993266 | -1.197 | -2.497 | -0.391 | -1.798 | -0.174 | -0.048 | -0.048 | 0 | 0.71 | 0.97 | 0.71 | 0.66 | 0.49 | 0.62 |
| 8 | 141774282 | rs7840381 | -0.602 | -1.934 | 0.227 | -1.365 | 0.02 | 0.079 | 0.079 | 0.003 | 0.48 | 0.26 | 0.6 | 0.58 | 0.43 | 0.53 |
| 8 | 141826739 | rs10090774 | -1.06 | -1.832 | 0.003 | -1.532 | -0.021 | 0.025 | 0.025 | 0.715 | 0.55 | 0.83 | 0.67 | 0.61 | 0.46 | 0.58 |
| 8 | 141831989 | rs9650572 | -1.102 | -1.832 | 0.032 | -1.533 | -0.023 | 0.023 | 0.023 | 0.297 | 0.55 | 0.85 | 0.67 | 0.61 | 0.46 | 0.58 |
| 8 | 141847249 | rs7831543 | 0.208 | -1.886 | -0.018 | -1.575 | 0.026 | 0.088 | 0.088 | 2.634 | 0.37 | 0.21 | 0.34 | 0.08 | 0.32 | 0.28 |
| 8 | 141857322 | rs4327896 | -0.781 | -1.877 | 0 | -1.562 | 0.012 | 0.017 | 0.017 | 0.399 | 0.51 | 0.36 | 0.61 | 0.53 | 0.45 | 0.56 |
| 8 | 141858620 | rs10087782 | 0.871 | -1.818 | -0.15 | -1.537 | -0.013 | 0.039 | 0.039 | 0.336 | 0.47 | 0.19 | 0.34 | 0.39 | 0.55 | 0.43 |
| 8 | 141902493 | rs4961237 | 0.239 | -1.786 | -0.24 | -1.584 | 0.021 | 0.082 | 0.082 | 2.022 | 0.36 | 0.22 | 0.33 | 0.06 | 0.32 | 0.28 |
| 8 | 141928350 | rs10111363 | 0.965 | -1.622 | -0.281 | -1.439 | 0.005 | 0.04 | 0.04 | 0.275 | 0.46 | 0.21 | 0.35 | 0.4 | 0.55 | 0.42 |
| 8 | 141932083 | rs12678285 | 0.65 | -1.665 | -0.534 | -1.454 | 0.002 | 0.071 | 0.071 | 0.004 | 0.13 | 0.15 | 0.26 | 0.43 | 0.13 | 0.26 |
| 8 | 141956904 | rs12677559 | -0.237 | -1.379 | -0.37 | -1.322 | 0.017 | 0.02 | 0.02 | 0.147 | 0.48 | 0.27 | 0.57 | 0.5 | 0.44 | 0.56 |
| 8 | 141965154 | rs10092810 | 0.444 | -1.39 | -1.371 | -1.271 | -0.031 | 0.025 | 0.025 | 0.329 | 0.46 | 0.12 | 0.34 | 0.4 | 0.55 | 0.42 |
| 8 | 141966167 | rs7007003 | 1.153 | -1.44 | -1.432 | -1.336 | -0.037 | 0.049 | 0.049 | 0 | 0.09 | 0.01 | 0.05 | 0 | 0.13 | 0.06 |
| 8 | 142003311 | rs11167018 | 0.127 | -1.042 | -2.173 | -1.194 | -0.032 | 0.024 | 0.024 | 0.427 | 0.46 | 0.12 | 0.34 | 0.4 | 0.55 | 0.42 |
| 8 | 142004114 | rs7816620 | 0.094 | -1.007 | -2.158 | -1.176 | -0.028 | 0.028 | 0.028 | 0.188 | 0.46 | 0.14 | 0.34 | 0.4 | 0.55 | 0.42 |

| | | | | | | | | | | | | | | | | |
|---|---|---|---|---|---|---|---|---|---|---|---|---|---|---|---|---|
| 8 | 142004193 | rs7813058 | 0.077 | -0.981 | -2.317 | -1.163 | -0.047 | 0.015 | 0.015 | 0.244 | 0.47 | 0.12 | 0.34 | 0.4 | 0.55 | 0.42 |
| 8 | 142022686 | rs12545416 | 0.276 | -0.437 | -1.739 | -1.35 | 0.004 | 0.047 | 0.047 | 0 | 0.27 | 0.17 | 0.34 | 0.05 | 0.28 | 0.33 |
| 8 | 142024236 | rs7840147 | 0.867 | -0.973 | -1.548 | -1.327 | -0.035 | -0.037 | -0.037 | 0.129 | 0.28 | 0.28 | 0.11 | 0.13 | 0.07 | 0.18 |
| 8 | 142025150 | rs11167020 | -0.439 | 0.134 | -1.042 | -0.502 | 0.019 | 0.064 | 0.064 | 0 | 0.73 | 0.57 | 0.66 | 0.46 | 0.71 | 0.73 |
| 8 | 142025644 | rs11167021 | 0.045 | 0.238 | -1.085 | -0.084 | -0.001 | 0.022 | 0.022 | 0.066 | 0.5 | 0.42 | 0.33 | 0.4 | 0.5 | 0.42 |
| 8 | 142038599 | rs1550955 | -0.531 | -0.097 | -1.18 | -0.406 | 0.034 | 0.072 | 0.072 | 0 | 0.52 | 0.68 | 0.42 | 0.3 | 0.42 | 0.58 |
| 8 | 142039892 | rs7001816 | 0.042 | -0.3 | -1.206 | -0.463 | -0.027 | -0.036 | -0.036 | 0 | 0.22 | 0.11 | 0.28 | 0.29 | 0.31 | 0.37 |
| 8 | 142041794 | rs3887342 | -0.57 | -0.392 | -0.645 | -0.491 | -0.007 | -0.016 | -0.016 | 0 | 0.36 | 0.5 | 0.35 | 0.3 | 0.37 | 0.46 |
| 8 | 142046065 | rs306983 | -0.446 | 0.072 | -0.845 | -0.488 | -0.004 | 0.001 | 0.001 | 0.357 | 0.95 | 0.96 | 0.96 | 1 | 0.93 | 0.99 |
| 8 | 142054224 | rs306974 | -0.124 | 0.072 | -0.79 | -0.497 | 0 | -0.008 | -0.008 | 0.148 | 0.94 | 0.92 | 0.95 | 0.99 | 0.93 | 0.98 |
| 8 | 142055976 | rs306991 | -0.173 | 0.076 | -0.762 | -0.495 | 0.002 | -0.012 | -0.012 | 0 | 0.95 | 0.92 | 0.95 | 0.99 | 0.93 | 0.98 |
| 8 | 142072158 | rs306966 | 0.011 | -0.992 | -0.712 | -0.484 | 0.018 | 0.008 | 0.008 | 0.001 | 0.58 | 0.44 | 0.77 | 0.35 | 0.72 | 0.59 |
| 8 | 142072614 | rs306967 | 0.968 | -1.028 | -1.301 | -0.779 | -0.077 | 0.122 | 0.122 | 0 | 0.33 | 0.11 | 0.37 | 0.11 | 0.58 | 0.26 |
| 8 | 142073846 | rs1588459 | 0.033 | -0.953 | -1.115 | -0.57 | -0.017 | -0.055 | -0.055 | 1.944 | 0.11 | 0.03 | 0.29 | 0.23 | 0.08 | 0.24 |
| 8 | 142081677 | rs12115135 | -0.29 | -1.175 | 0.098 | -0.553 | -0.033 | 0.092 | 0.092 | 1.409 | 0.12 | 0.51 | 0.05 | 0.14 | 0.03 | 0.08 |
| 8 | 142084221 | rs1567467 | -1.231 | -1.177 | -0.327 | -0.886 | -0.081 | 0.124 | 0.124 | 0 | 0.67 | 0.87 | 0.66 | 0.9 | 0.46 | 0.75 |
| 8 | 142091853 | rs13252126 | 0.355 | -1.984 | 0.117 | -0.947 | 0.001 | 0.013 | 0.013 | 0.259 | 0.25 | 0.21 | 0.32 | 0.67 | 0.26 | 0.39 |
| 8 | 142094777 | rs12334878 | -0.859 | -2.011 | 0.211 | -1.236 | 0.012 | 0.102 | 0.102 | 0 | 0.43 | 0.68 | 0.55 | 0.9 | 0.37 | 0.53 |
| 8 | 142097004 | rs10089264 | -0.307 | -1.968 | 0.852 | -1.017 | -0.021 | 0.09 | 0.09 | 0.603 | 0.14 | 0.35 | 0.18 | 0.22 | 0.07 | 0.12 |
| 8 | 142104071 | rs1107258 | 0.466 | -1.934 | 0.62 | -0.745 | -0.003 | -0.02 | -0.02 | 1.335 | 0.24 | 0.15 | 0.31 | 0.63 | 0.25 | 0.43 |
| 8 | 142104108 | rs2005358 | -0.661 | -1.9 | 0.57 | -0.871 | 0.005 | 0.017 | 0.017 | 0 | 0.38 | 0.59 | 0.5 | 0.84 | 0.33 | 0.55 |
| 8 | 142104814 | rs4961329 | -0.489 | -1.803 | 0.67 | -0.581 | -0.011 | 0.008 | 0.008 | 0 | 0.32 | 0.44 | 0.4 | 0.66 | 0.28 | 0.53 |
| 8 | 142104944 | rs4961252 | -0.686 | -1.2 | -0.583 | -0.584 | 0.031 | 0.057 | 0.057 | 0 | 0.46 | 0.74 | 0.5 | 0.68 | 0.4 | 0.57 |
| 8 | 142106228 | rs11167023 | -0.328 | -0.511 | -0.097 | 0.437 | 0.033 | 0.166 | 0.166 | 0.144 | 0.17 | 0.34 | 0.32 | 0.38 | 0.18 | 0.34 |
| 8 | 142108239 | rs11167024 | -0.552 | -0.213 | -1.347 | 0.289 | 0.026 | -0.036 | -0.036 | 1.199 | 0.42 | 0.76 | 0.48 | 0.48 | 0.34 | 0.52 |
| 8 | 142109356 | rs4961331 | -0.533 | -0.188 | -1.378 | 0.275 | 0.023 | -0.026 | -0.026 | 0 | 0.42 | 0.74 | 0.48 | 0.48 | 0.34 | 0.52 |
| 8 | 142122693 | rs11989678 | -0.043 | 0.416 | -1.46 | 0.156 | -0.01 | 0.049 | 0.049 | 0 | 0.55 | 0.75 | 0.45 | 0.33 | 0.49 | 0.53 |
| 8 | 142125305 | rs2221770 | 0.334 | -0.271 | -1.273 | -0.223 | -0.002 | 0.061 | 0.061 | 0 | 0.46 | 0.68 | 0.42 | 0.26 | 0.47 | 0.47 |
| 8 | 142131002 | rs11785437 | 0.398 | -0.687 | -0.603 | -0.456 | -0.006 | 0.022 | 0.022 | 0 | 0.4 | 0.34 | 0.36 | 0.17 | 0.44 | 0.36 |
| 8 | 142132965 | rs1467073 | -0.988 | -1.184 | -1.07 | -0.203 | 0.063 | -0.096 | -0.096 | 0.728 | 0.78 | 0.95 | 0.76 | 0.5 | 0.87 | 0.69 |
| 8 | 142134478 | rs7462379 | 0.807 | -0.546 | -1.135 | -1.151 | 0 | -0.048 | -0.048 | 0 | 0.33 | 0.3 | 0.27 | 0.03 | 0.33 | 0.19 |
| 8 | 142137634 | rs7009780 | -0.138 | -0.431 | -0.848 | -0.964 | 0.006 | 0.051 | 0.051 | 0 | 0.49 | 0.65 | 0.43 | 0.26 | 0.46 | 0.45 |
| 8 | 142137704 | rs6988905 | -0.125 | -0.448 | -0.869 | -0.962 | 0.006 | 0.042 | 0.042 | 0 | 0.49 | 0.64 | 0.43 | 0.26 | 0.46 | 0.45 |
| 8 | 142146708 | rs307772 | -0.662 | 0.061 | -1.156 | -1.251 | -0.018 | 0.085 | 0.085 | 0 | 0.79 | 0.93 | 0.85 | 0.99 | 0.76 | 0.92 |
| 8 | 142160876 | rs170088 | 0.526 | -0.272 | 0.241 | -0.682 | -0.022 | 0.229 | 0.229 | 0 | 0.17 | 0.58 | 0.29 | 0.46 | 0.11 | 0.2 |
| 8 | 142161064 | rs3816063 | 0.748 | -0.183 | -0.4 | -1.298 | -0.002 | 0.001 | 0.001 | 0 | 0.28 | 0.25 | 0.24 | 0.2 | 0.38 | 0.27 |
| 8 | 142161873 | rs2304284 | 1.523 | -0.236 | -0.184 | -1.222 | -0.01 | 0.129 | 0.129 | 0 | 0.14 | 0.17 | 0.24 | 0.46 | 0.11 | 0.2 |
| 8 | 142161888 | rs2304283 | 1.523 | -0.236 | -0.184 | -1.222 | -0.01 | 0.129 | 0.129 | 0 | 0.14 | 0.17 | 0.24 | 0.46 | 0.11 | 0.2 |
| 8 | 142170884 | rs2289001 | 0.747 | -0.465 | -0.252 | -1.323 | 0.029 | 0.001 | 0.001 | 0 | 0.35 | 0.13 | 0.34 | 0.34 | 0.27 | 0.45 |
| 8 | 142173356 | rs428300 | -0.688 | -0.184 | 0.009 | -1.177 | -0.003 | -0.022 | -0.022 | 0 | 0.67 | 0.73 | 0.6 | 0.53 | 0.65 | 0.74 |
| 8 | 142177251 | rs372758 | -0.297 | 0.257 | 0.64 | -0.381 | 0.013 | 0.071 | 0.071 | 0 | 0.07 | 0.22 | 0.05 | 0.14 | 0.04 | 0.1 |
| 8 | 142182417 | rs3739230 | 0.184 | 0.704 | 0.552 | -0.526 | -0.015 | -0.043 | -0.043 | 0 | 0.14 | 0.21 | 0.21 | 0.03 | 0.12 | 0.12 |
| 8 | 142182644 | rs4961333 | -0.76 | 0.873 | 0.252 | -0.6 | 0.007 | -0.038 | -0.038 | 0 | 0.35 | 0.53 | 0.42 | 0.22 | 0.39 | 0.34 |
| 8 | 142188039 | rs307731 | -0.5 | 0.745 | 0.257 | -0.375 | 0.009 | 0.002 | 0.002 | 0.317 | 0.32 | 0.57 | 0.28 | 0.33 | 0.37 | 0.34 |
| 8 | 142191032 | rs2278448 | 0.193 | 1.014 | -0.907 | -0.233 | -0.058 | -0.033 | -0.033 | 0.506 | 0.26 | 0.1 | 0.29 | 0.34 | 0.39 | 0.34 |
| 8 | 142192569 | rs307739 | -0.536 | 0.551 | -0.984 | -0.861 | 0.019 | 0.161 | 0.161 | 0 | 0.49 | 0.76 | 0.54 | 0.8 | 0.52 | 0.56 |
| 8 | 142197681 | rs3922678 | -0.509 | 0.49 | -0.595 | -0.863 | 0.002 | 0.001 | 0.001 | 0 | 0.54 | 0.57 | 0.71 | 0.79 | 0.63 | 0.64 |
| 8 | 142198596 | rs7001673 | -0.858 | 0.755 | -0.005 | -0.079 | 0.027 | 0.015 | 0.015 | 0 | 0.48 | 0.62 | 0.44 | 0.6 | 0.39 | 0.53 |
| 8 | 142200731 | rs2288997 | -0.267 | 0.403 | -0.053 | 0.407 | 0.018 | -0.081 | -0.081 | 0 | 0.3 | 0.11 | 0.29 | 0.46 | 0.23 | 0.4 |
| 8 | 142201477 | rs13270024 | -0.208 | 1.047 | 0.594 | 0.258 | 0.128 | 0.086 | 0.086 | 0 | 0.07 | 0.25 | 0.11 | 0.15 | 0.19 | 0.03 |
| 8 | 142205964 | rs10102728 | -1.4 | 0.576 | 0.756 | -0.006 | 0.005 | -0.035 | -0.035 | 0 | 0.59 | 0.5 | 0.55 | 0.8 | 0.6 | 0.58 |
| 8 | 142209543 | rs6578153 | -0.875 | 0.529 | 0.676 | -0.022 | 0.043 | 0.063 | 0.063 | 0 | 0.52 | 0.27 | 0.45 | 0.67 | 0.42 | 0.6 |
| 8 | 142219810 | rs3739242 | 0.622 | 1.088 | 0.749 | 0.195 | -0.015 | -0.018 | -0.018 | 0 | 0.06 | 0.01 | 0.09 | 0.17 | 0.18 | 0.04 |
| 8 | 142221414 | rs3739240 | -1.458 | 0.906 | 0.967 | 0.077 | 0.001 | 0.023 | 0.023 | 0 | 0.57 | 0.64 | 0.56 | 0.81 | 0.6 | 0.64 |
| 8 | 142221874 | rs3739239 | 0.596 | 0.786 | 0.621 | 0.294 | -0.013 | -0.089 | -0.089 | 0 | 0.14 | 0.02 | 0.12 | 0.36 | 0.12 | 0.23 |
| 8 | 142222445 | rs3739238 | -1.431 | 1.28 | 1.015 | 0.551 | -0.022 | 0.11 | 0.11 | 0 | 0.59 | 0.84 | 0.58 | 0.81 | 0.59 | 0.65 |
| 8 | 142225990 | rs2288998 | -1.017 | 1.263 | 1.294 | 0.616 | 0.014 | 0.106 | 0.106 | 0 | 0.21 | 0.35 | 0.18 | 0.44 | 0.26 | 0.21 |
| 8 | 142227083 | rs7813723 | 0.173 | 1.407 | 0.635 | 0.553 | 0.023 | -0.04 | -0.04 | 3.47 | 0.15 | 0.08 | 0.1 | 0.24 | 0.13 | 0.17 |
| 8 | 142228909 | rs753778 | 0.139 | 1.432 | 0.685 | 0.548 | -0.005 | 0.003 | 0.003 | 0 | 0.34 | 0.37 | 0.37 | 0.4 | 0.29 | 0.44 |
| 8 | 142229543 | rs2304282 | -1.112 | 1.633 | 0.276 | 0.58 | -0.003 | 0.091 | 0.091 | 0 | 0.58 | 0.76 | 0.58 | 0.82 | 0.6 | 0.64 |
| 8 | 142233819 | rs13281845 | 0.658 | 1.696 | 0.101 | 0.763 | -0.014 | -0.016 | -0.016 | 0 | 0.06 | 0.01 | 0.08 | 0.18 | 0.18 | 0.04 |
| 8 | 142237398 | rs12547917 | 0.494 | 1.717 | 0.175 | 0.701 | 0.027 | 0.03 | 0.03 | 0 | 0.48 | 0.26 | 0.49 | 0.38 | 0.58 | 0.41 |
| 8 | 142239210 | rs7018178 | -0.807 | 1.432 | -0.26 | 0.334 | -0.003 | 0.028 | 0.028 | 0 | 0.55 | 0.66 | 0.55 | 0.85 | 0.53 | 0.61 |
| 8 | 142239377 | rs6994692 | 0.558 | 0.912 | -0.467 | 0.566 | 0.015 | -0.055 | -0.055 | 0.037 | 0.13 | 0.06 | 0.08 | 0.25 | 0.09 | 0.16 |
| 8 | 142240768 | rs11782278 | 0.707 | 0.928 | -0.477 | 0.535 | 0.006 | -0.073 | -0.073 | 0 | 0.13 | 0.05 | 0.08 | 0.24 | 0.09 | 0.15 |
| 8 | 142241238 | rs3824233 | 1.442 | 1.077 | -0.508 | 0.364 | -0.051 | 0.095 | 0.095 | 0 | 0.35 | 0.07 | 0.23 | 0.15 | 0.37 | 0.29 |
| 8 | 142245464 | rs10875468 | -0.326 | 1.174 | -0.016 | 0.659 | 0.091 | -0.002 | -0.002 | 0 | 0.94 | 0.7 | 0.79 | 1 | 0.92 | 0.97 |
| 8 | 142246550 | rs11782904 | 0.455 | 1.16 | -0.984 | 0.74 | 0.014 | -0.05 | -0.05 | 1.707 | 0.13 | 0.05 | 0.08 | 0.23 | 0.09 | 0.15 |
| 8 | 142246961 | rs13276184 | -0.433 | 1.026 | -0.964 | 0.619 | -0.011 | -0.032 | -0.032 | 0 | 0.19 | 0.15 | 0.16 | 0.39 | 0.24 | 0.19 |

| | | | | | | | | | | | | | | | | |
|---|---|---|---|---|---|---|---|---|---|---|---|---|---|---|---|---|
| 8 | 142247390 | rs12550677 | 1.06 | 1.338 | -1.205 | 0.734 | -0.037 | 0.123 | 0.123 | 0 | 0.39 | 0.12 | 0.26 | 0.15 | 0.41 | 0.35 |
| 8 | 142247539 | rs4437700 | 1.149 | 2.077 | -0.536 | 1.035 | 0.011 | 0.002 | 0.002 | 0.12 | 0.64 | 0.53 | 0.6 | 0.6 | 0.57 | 0.62 |
| 8 | 142248021 | rs7014142 | -0.6 | 1.596 | -1.356 | 1.242 | 0.017 | 0.017 | 0.017 | 0 | 0.24 | 0.2 | 0.12 | 0.17 | 0.17 | 0.25 |
| 8 | 142251553 | rs2369481 | -0.911 | 2.282 | -0.865 | 1.58 | -0.01 | 0.015 | 0.015 | 0 | 0.06 | 0.02 | 0.09 | 0.26 | 0.18 | 0.08 |
| 8 | 142256716 | rs11778032 | 0.649 | 1.379 | -0.567 | 0.729 | -0.003 | -0.017 | -0.017 | 0 | 0.8 | 0.78 | 0.67 | 0.8 | 0.82 | 0.78 |
| 8 | 142257506 | rs7016433 | 0.276 | 1.547 | -0.738 | 0.296 | -0.002 | 0.014 | 0.014 | 0 | 0.72 | 0.78 | 0.77 | 0.87 | 0.7 | 0.72 |
| 8 | 142259829 | rs2060357 | 0.53 | 1.047 | -1.003 | 0.429 | 0.023 | -0.022 | -0.022 | 0 | 0.34 | 0.23 | 0.31 | 0.38 | 0.3 | 0.28 |
| 8 | 142260995 | rs10093988 | 0.552 | 0.982 | -0.973 | 0.396 | 0.024 | -0.028 | -0.028 | 0 | 0.34 | 0.18 | 0.31 | 0.38 | 0.3 | 0.28 |
| 8 | 142269874 | rs11167042 | -0.195 | 1.149 | -0.517 | -0.124 | -0.001 | 0.015 | 0.015 | 0.089 | 0.58 | 0.4 | 0.62 | 0.86 | 0.59 | 0.63 |
| 8 | 142274632 | rs1865246 | 0.083 | 0.987 | 0.005 | -0.252 | 0.032 | 0.087 | 0.087 | 0.058 | 0.26 | 0.46 | 0.35 | 0.48 | 0.3 | 0.35 |
| 8 | 142279412 | rs6992286 | 0.504 | 0.845 | -0.003 | -0.308 | 0.013 | -0.004 | -0.004 | 0 | 0.4 | 0.65 | 0.38 | 0.39 | 0.32 | 0.29 |
| 8 | 142295965 | rs7819263 | 0.047 | 1.027 | 0.176 | -0.212 | -0.001 | 0 | 0 | 0 | 0.56 | 0.61 | 0.64 | 0.45 | 0.58 | 0.45 |
| 8 | 142300315 | rs7827290 | 0.37 | 0.878 | 0.202 | -0.29 | -0.015 | 0.071 | 0.071 | 0.183 | 0.34 | 0.16 | 0.28 | 0.14 | 0.35 | 0.34 |
| 8 | 142302024 | rs12677618 | 1.389 | 0.007 | -0.468 | -0.575 | -0.022 | -0.051 | -0.051 | 0 | 0.18 | 0.07 | 0.12 | 0.34 | 0.27 | 0.25 |
| 8 | 142302260 | rs919873 | 1.728 | -0.174 | -0.408 | -0.628 | 0.001 | 0.002 | 0.002 | 0.18 | 0.16 | 0.14 | 0.22 | 0.06 | 0.06 | 0.05 |
| 8 | 142304722 | rs7828326 | 0.799 | -0.116 | -0.212 | -0.578 | 0.011 | 0.233 | 0.233 | 0.084 | 0.08 | 0.22 | 0.06 | 0.42 | 0.04 | 0.2 |
| 8 | 142305792 | rs6999890 | 0.355 | -0.025 | -0.482 | -0.665 | 0.013 | 0.005 | 0.005 | 0 | 0.53 | 0.29 | 0.53 | 0.48 | 0.63 | 0.61 |
| 8 | 142310657 | rs11786212 | 0.336 | 0.177 | -0.783 | -0.897 | -0.054 | 0.165 | 0.165 | 0.057 | 0.42 | 0.11 | 0.5 | 0.18 | 0.54 | 0.42 |
| 8 | 142315838 | rs7831477 | 0.305 | 0.173 | -0.915 | -1.006 | -0.099 | 0.131 | 0.131 | 0 | 0.39 | 0.08 | 0.48 | 0.18 | 0.52 | 0.41 |
| 8 | 142323189 | rs7818687 | 0.147 | -0.291 | -0.366 | -0.948 | 0.034 | -0.014 | -0.014 | 0.081 | 0.06 | 0.25 | 0.05 | 0 | 0.04 | 0.04 |
| 8 | 142324925 | rs2117149 | 0.276 | -0.681 | -0.887 | -1.505 | 0.006 | 0.077 | 0.077 | 0 | 0.55 | 0.33 | 0.51 | 0.14 | 0.53 | 0.52 |
| 8 | 142327998 | rs878422 | 0.888 | -0.68 | -1.121 | -1.132 | -0.066 | 0.256 | 0.256 | 0 | 0.47 | 0.07 | 0.46 | 0.14 | 0.49 | 0.48 |
| 8 | 142328020 | rs878421 | -0.105 | -0.816 | -1.09 | -0.788 | 0.047 | -0.018 | -0.018 | 0 | 0.66 | 0.29 | 0.78 | 0.6 | 0.85 | 0.72 |
| 8 | 142333425 | rs10875469 | 2.454 | -0.31 | -1.353 | -0.82 | -0.081 | 0.025 | 0.025 | 0 | 0.22 | 0.02 | 0.34 | 0.13 | 0.28 | 0.3 |
| 8 | 142334326 | rs10086688 | 1.07 | -0.601 | -0.434 | 0.077 | -0.021 | -0.006 | -0.006 | 0.228 | 0.1 | 0.05 | 0.24 | 0.43 | 0.19 | 0.17 |
| 8 | 142337734 | rs6985508 | -0.393 | -0.62 | -0.226 | 0.034 | 0.042 | 0.075 | 0.075 | 1.093 | 0.19 | 0.29 | 0.33 | 0.47 | 0.35 | 0.24 |
| 8 | 142341758 | rs13279009 | -0.743 | -0.65 | -0.179 | -0.405 | 0.002 | 0.013 | 0.013 | 0 | 0.45 | 0.4 | 0.46 | 0.86 | 0.47 | 0.48 |
| 8 | 142355742 | rs4961364 | -0.722 | -0.147 | -1.226 | 0.199 | -0.012 | 0.002 | 0.002 | 0 | 0.89 | 0.95 | 0.77 | 0.56 | 0.81 | 0.88 |
| 8 | 142359368 | rs7386745 | 0.735 | -0.334 | -1.373 | -0.164 | -0.007 | 0.237 | 0.237 | 0 | 0.5 | 0.08 | 0.46 | 0.17 | 0.51 | 0.54 |
| 8 | 142364163 | rs4961374 | 0.349 | -0.769 | -1.253 | -0.087 | 0.005 | -0.021 | -0.021 | 0 | 0.75 | 0.81 | 0.82 | 0.51 | 0.78 | 0.66 |
| 8 | 142366200 | rs7843131 | 0.21 | -0.76 | -1.28 | 0.007 | 0.044 | -0.021 | -0.021 | 0.029 | 0.62 | 0.23 | 0.73 | 0.51 | 0.73 | 0.71 |
| 8 | 142373509 | rs7830293 | 0.638 | -0.522 | -1.108 | 0.575 | 0.008 | 0.015 | 0.015 | 0 | 0.1 | 0.08 | 0.24 | 0.43 | 0.07 | 0.2 |
| 8 | 142373969 | rs13260421 | 0.871 | -0.348 | -1.605 | -0.58 | -0.141 | 0.134 | 0.134 | 0 | 0.39 | 0.04 | 0.38 | 0.17 | 0.44 | 0.44 |
| 8 | 142382757 | rs10112201 | -1.158 | -0.849 | 0.145 | -0.034 | 0.077 | 0.017 | 0.017 | 0.643 | 0.07 | 0.4 | 0.05 | 0 | 0.04 | 0.02 |
| 8 | 142414326 | rs6578177 | -0.254 | -1.307 | -0.677 | 0.091 | 0.032 | -0.001 | -0.001 | 0.971 | 0.4 | 0.65 | 0.41 | 0.6 | 0.25 | 0.39 |
| 8 | 142418127 | rs7465330 | -0.176 | -1.374 | -1.333 | 0.261 | 0.105 | -0.064 | -0.064 | 0.012 | 0.35 | 0.11 | 0.34 | 0.6 | 0.21 | 0.36 |
| 8 | 142429135 | rs7828919 | -0.641 | -0.564 | -0.137 | 0.283 | 0.006 | -0.006 | -0.006 | 0 | 0.2 | 0.47 | 0.14 | 0.16 | 0.18 | 0.14 |
| 8 | 142437352 | rs9987318 | 0.808 | -0.517 | -1.207 | 0.07 | -0.078 | 0.048 | 0.048 | 0.403 | 0.36 | 0.13 | 0.38 | 0.28 | 0.46 | 0.47 |
| 8 | 142442538 | rs6987971 | -0.325 | -0.869 | -0.253 | 0.066 | -0.006 | -0.001 | -0.001 | 0.151 | 0.57 | 0.52 | 0.52 | 0.43 | 0.65 | 0.64 |
| 8 | 142443189 | rs7836875 | -0.335 | -0.887 | -0.258 | 0.076 | -0.006 | -0.001 | -0.001 | 1.953 | 0.57 | 0.52 | 0.52 | 0.43 | 0.65 | 0.62 |
| 8 | 142446923 | rs11782383 | 0.961 | -0.8 | -0.415 | -0.072 | -0.04 | 0.039 | 0.039 | 0 | 0.35 | 0.21 | 0.28 | 0.27 | 0.43 | 0.43 |
| 8 | 142447850 | rs1137665 | -0.482 | -1.281 | -0.207 | 0.317 | 0 | -0.001 | -0.001 | 0 | 0.35 | 0.61 | 0.2 | 0.33 | 0.29 | 0.21 |
| 8 | 142452509 | rs4075035 | 1.62 | -1.128 | -0.281 | 0.48 | 0.014 | -0.025 | -0.025 | 0 | 0.14 | 0.06 | 0.05 | 0.16 | 0.11 | 0.05 |
| 8 | 142453585 | rs7816220 | 0.754 | -1.104 | 0.315 | 0.331 | 0.003 | -0.012 | -0.012 | 8.457 | 0.39 | 0.45 | 0.31 | 0.27 | 0.43 | 0.43 |
| 8 | 142458053 | rs6578185 | 0.817 | -0.931 | 0.006 | 0.318 | -0.046 | 0.041 | 0.041 | 0 | 0.38 | 0.24 | 0.3 | 0.3 | 0.47 | 0.44 |
| 8 | 142459032 | rs11167062 | -0.514 | -1.197 | 0.202 | 0.334 | 0.007 | -0.001 | -0.001 | 2.521 | 0.38 | 0.3 | 0.22 | 0.32 | 0.35 | 0.2 |
| 8 | 142460269 | rs11776933 | 0.278 | -1.29 | -0.023 | 0.309 | -0.045 | -0.005 | -0.005 | 0 | 0.21 | 0.03 | 0.14 | 0.15 | 0.25 | 0.15 |
| 8 | 142460495 | rs11781250 | -0.708 | -1.363 | 0.008 | 0.285 | 0.01 | -0.001 | -0.001 | 0 | 0.38 | 0.28 | 0.22 | 0.32 | 0.35 | 0.2 |
| 8 | 142460808 | rs7836905 | 0.732 | -1.034 | -0.147 | 0.638 | 0.078 | -0.015 | -0.015 | 0.248 | 0.53 | 0.18 | 0.4 | 0.45 | 0.67 | 0.57 |
| 8 | 142461694 | rs7842575 | -1.195 | -0.806 | 0.247 | 0.543 | 0.001 | -0.01 | -0.01 | 0 | 0.8 | 0.85 | 0.55 | 0.64 | 0.83 | 0.62 |
| 8 | 142464967 | rs13248753 | 1.917 | -0.666 | -0.16 | 0.383 | -0.082 | 0.09 | 0.09 | 0.247 | 0.33 | 0.07 | 0.24 | 0.18 | 0.4 | 0.35 |
| 8 | 142466821 | rs11777747 | 0.418 | -0.541 | -0.096 | 0.517 | -0.056 | -0.028 | -0.028 | 0.053 | 0.15 | 0.01 | 0.11 | 0.13 | 0.2 | 0.13 |
| 8 | 142467575 | rs11782199 | -0.127 | -0.68 | 0.409 | 0.589 | -0.023 | 0.038 | 0.038 | 0.607 | 0.71 | 0.49 | 0.47 | 0.58 | 0.77 | 0.6 |
| 8 | 142468255 | rs7845743 | 0.839 | -0.575 | 0.517 | 0.648 | -0.002 | 0.005 | 0.005 | 0.548 | 0.56 | 0.46 | 0.37 | 0.45 | 0.57 | 0.48 |
| 8 | 142476529 | rs7816228 | 0.967 | -0.577 | 0.384 | 0.663 | 0 | -0.029 | -0.029 | 0 | 0.19 | 0.12 | 0.39 | 0.42 | 0.17 | 0.35 |
| 8 | 142480610 | rs4907398 | 1.274 | -0.574 | 1.02 | 0.453 | 0.042 | -0.016 | -0.016 | 0 | 0.91 | 0.52 | 0.91 | 0.99 | 0.95 | 0.98 |
| 8 | 142484621 | rs10102492 | 1.902 | -0.57 | 0.287 | 0.39 | -0.107 | 0.093 | 0.093 | 0.323 | 0.34 | 0.05 | 0.25 | 0.18 | 0.39 | 0.33 |
| 8 | 142486625 | rs2613646 | 0.971 | -0.659 | 0.422 | 0.516 | -0.02 | 0.02 | 0.02 | 2.262 | 0.5 | 0.16 | 0.31 | 0.39 | 0.48 | 0.41 |
| 8 | 142488994 | rs7816304 | -0.23 | -0.751 | 0.377 | 0.523 | -0.046 | -0.033 | -0.033 | 0.007 | 0.4 | 0.16 | 0.58 | 0.6 | 0.47 | 0.58 |
| 8 | 142489410 | rs2748418 | 1.887 | -0.7 | 0.295 | 0.538 | -0.104 | 0.086 | 0.086 | 0.289 | 0.33 | 0.06 | 0.26 | 0.19 | 0.39 | 0.32 |
| 8 | 142490120 | rs2748416 | 1.441 | -0.578 | 1.039 | 0.738 | 0.056 | 0.294 | 0.294 | 2.778 | 0.9 | 0.53 | 0.91 | 0.78 | 0.92 | 0.9 |
| 8 | 142490166 | rs7838192 | -0.235 | -0.305 | 0.874 | 0.739 | -0.012 | 0.007 | 0.007 | 0 | 0.18 | 0.11 | 0.15 | 0.14 | 0.27 | 0.12 |
| 8 | 142491238 | rs7014345 | -0.627 | -0.366 | 0.809 | 0.733 | 0.005 | 0 | 0 | 0 | 0.43 | 0.44 | 0.61 | 0.6 | 0.5 | 0.56 |
| 8 | 142496058 | rs12056882 | -0.372 | -0.486 | 0.366 | 0.928 | 0.036 | -0.007 | -0.007 | 0 | 0.39 | 0.25 | 0.49 | 0.44 | 0.28 | 0.46 |
| 8 | 142496142 | rs12056884 | -0.554 | -0.48 | 0.386 | 0.941 | 0.039 | 0.006 | 0.006 | 0 | 0.43 | 0.26 | 0.52 | 0.44 | 0.33 | 0.47 |
| 8 | 142501186 | rs10099308 | -0.491 | 0.073 | 0.296 | 0.937 | -0.015 | -0.025 | -0.025 | 0 | 0.39 | 0.17 | 0.54 | 0.6 | 0.41 | 0.55 |
| 8 | 142503997 | rs11167065 | -0.396 | 0.107 | 0.348 | 1.094 | 0.007 | 0.034 | 0.034 | 0 | 0.84 | 0.78 | 0.84 | 0.67 | 0.79 | 0.89 |
| 8 | 142504796 | rs2748406 | 1.752 | -0.169 | 0.057 | 0.878 | -0.102 | 0.138 | 0.138 | 0.803 | 0.38 | 0.09 | 0.28 | 0.19 | 0.43 | 0.3 |
| 8 | 142505578 | rs2748404 | 0.298 | -0.117 | 0.549 | 1.182 | 0.037 | 0.116 | 0.116 | 0 | 0.79 | 0.64 | 0.8 | 0.47 | 0.71 | 0.76 |

| | | | | | | | | | | | | | | | | |
|---|---|---|---|---|---|---|---|---|---|---|---|---|---|---|---|---|
| 8 | 142506121 | rs2748402 | 0.598 | -0.209 | 0.482 | 1.195 | 0.03 | 0.135 | 0.135 | 0 | 0.06 | 0.14 | 0.04 | 0.19 | 0.08 | 0.12 |
| 8 | 142507908 | rs2748423 | 0.358 | -0.223 | 0.553 | 1.186 | 0.034 | 0.161 | 0.161 | 0 | 0.08 | 0.26 | 0.05 | 0.19 | 0.08 | 0.12 |
| 8 | 142509050 | rs7015884 | -0.004 | -0.363 | 0.591 | 1.111 | -0.002 | -0.001 | -0.001 | 0 | 0.35 | 0.31 | 0.48 | 0.27 | 0.27 | 0.45 |
| 8 | 142509280 | rs2613636 | -0.96 | -0.268 | 0.263 | 1.148 | 0.044 | -0.011 | -0.011 | 0 | 0.47 | 0.69 | 0.56 | 0.47 | 0.36 | 0.59 |
| 8 | 142515399 | rs2748414 | 1.271 | -0.07 | 0.245 | 1.208 | -0.031 | 0.021 | 0.021 | 0.587 | 0.45 | 0.26 | 0.31 | 0.38 | 0.48 | 0.42 |
| 8 | 142517222 | rs2748421 | 1.156 | -0.07 | 0.206 | 1.208 | -0.029 | 0.02 | 0.02 | 0.602 | 0.45 | 0.26 | 0.31 | 0.39 | 0.48 | 0.42 |
| 8 | 142518112 | rs6578196 | -0.395 | 0.163 | 0.263 | 1.076 | -0.001 | 0.009 | 0.009 | 0.024 | 0.25 | 0.32 | 0.45 | 0.27 | 0.22 | 0.44 |
| 8 | 142527473 | rs7386137 | -0.224 | 0.273 | -0.199 | 1.147 | 0.002 | -0.013 | -0.013 | 0 | 0.23 | 0.15 | 0.43 | 0.28 | 0.21 | 0.43 |
| 8 | 142529211 | rs12235081 | 0.203 | 0.695 | -0.264 | 1.254 | -0.01 | -0.028 | -0.028 | 0.996 | 0.19 | 0.12 | 0.37 | 0.28 | 0.18 | 0.39 |
| 8 | 142529703 | rs5019176 | 0.44 | 0.279 | -0.25 | 1.223 | 0.006 | -0.019 | -0.019 | 0.044 | 0.17 | 0.1 | 0.24 | 0.17 | 0.15 | 0.35 |
| 8 | 142530251 | rs6994715 | 1.478 | -0.005 | -0.482 | 0.673 | -0.004 | 0.089 | 0.089 | 1.589 | 0.47 | 0.21 | 0.43 | 0.33 | 0.5 | 0.33 |
| 8 | 142530711 | rs873225 | -0.476 | -0.032 | -0.024 | 1.257 | 0.003 | -0.01 | -0.01 | 0 | 0.27 | 0.24 | 0.35 | 0.5 | 0.23 | 0.52 |
| 8 | 142530964 | rs873224 | 0.098 | -0.001 | -0.133 | 1.257 | -0.002 | 0.002 | 0.002 | 0 | 0.06 | 0.04 | 0.09 | 0.02 | 0.08 | 0.13 |
| 8 | 142532935 | rs4907333 | 0.15 | -0.019 | -0.034 | 1.422 | 0.012 | -0.031 | -0.031 | 0 | 0.21 | 0.15 | 0.26 | 0.49 | 0.15 | 0.39 |
| 8 | 142537715 | rs4907374 | 0.396 | 0.362 | 0.823 | 0.937 | 0.033 | 0.007 | 0.007 | 2.146 | 0.84 | 0.65 | 0.83 | 0.84 | 0.77 | 0.86 |
| 8 | 142544410 | rs2085841 | 0.459 | -0.204 | 0.227 | 0.764 | 0.01 | 0.014 | 0.014 | 0 | 0.35 | 0.2 | 0.33 | 0.21 | 0.35 | 0.4 |
| 8 | 142549778 | rs11167068 | 0.39 | -0.379 | 0.781 | 0.405 | -0.015 | -0.028 | -0.028 | 0 | 0.83 | 0.61 | 0.72 | 0.87 | 0.85 | 0.77 |
| 8 | 142556805 | rs1566080 | -0.752 | -0.743 | 0.812 | 0.357 | -0.021 | 0.032 | 0.032 | 0 | 0.62 | 0.67 | 0.74 | 0.78 | 0.57 | 0.72 |
| 8 | 142564072 | rs12334747 | 0.752 | -0.666 | 1.186 | 0.29 | 0.025 | 0.012 | 0.012 | 3.605 | 0.95 | 0.7 | 0.94 | 1 | 0.97 | 1 |
| 8 | 142565478 | rs13271156 | 0.139 | -0.891 | 1.322 | 0.423 | -0.008 | -0.01 | -0.01 | 0 | 0.44 | 0.3 | 0.26 | 0.52 | 0.49 | 0.35 |
| 8 | 142566820 | rs10099507 | 0.692 | -1.021 | 0.958 | 0.367 | 0.013 | -0.009 | -0.009 | 0 | 0.39 | 0.26 | 0.46 | 0.35 | 0.36 | 0.42 |
| 8 | 142568559 | rs13266025 | 0.149 | -0.947 | 1.061 | 0.392 | 0.019 | 0.003 | 0.003 | 0 | 0.45 | 0.33 | 0.46 | 0.63 | 0.54 | 0.54 |
| 8 | 142569536 | rs871952 | 0.688 | -0.652 | 1.213 | 0.175 | -0.061 | -0.119 | -0.119 | 0 | 0.84 | 0.54 | 0.92 | 0.98 | 0.9 | 0.97 |
| 8 | 142571042 | rs13267815 | 0.688 | -0.649 | 1.213 | 0.177 | -0.061 | -0.12 | -0.12 | 0.037 | 0.84 | 0.53 | 0.92 | 0.98 | 0.9 | 0.97 |
| 8 | 142572865 | rs10112716 | 0.523 | -0.593 | 0.723 | 0.205 | -0.044 | -0.089 | -0.089 | 0 | 0.85 | 0.66 | 0.93 | 0.98 | 0.91 | 0.97 |
| 8 | 142576687 | rs13277303 | 0.522 | 0.019 | -0.137 | 0.414 | -0.058 | 0.025 | 0.025 | 0 | 0.27 | 0.1 | 0.16 | 0.25 | 0.32 | 0.27 |
| 8 | 142577813 | rs307393 | -0.44 | 0.021 | -0.193 | 0.266 | 0.002 | 0.011 | 0.011 | 0.087 | 0.91 | 0.94 | 0.91 | 0.95 | 0.92 | 0.9 |
| 8 | 142578923 | rs11785794 | 0.947 | -0.364 | -0.436 | 0.503 | -0.053 | 0.013 | 0.013 | 1.775 | 0.48 | 0.09 | 0.42 | 0.33 | 0.5 | 0.48 |
| 8 | 142579169 | rs898994 | 1.464 | -0.454 | 1.35 | 0.714 | -0.031 | -0.023 | -0.023 | 0 | 0.8 | 0.36 | 0.7 | 0.85 | 0.87 | 0.88 |
| 8 | 142580648 | rs876096 | -2.236 | -0.52 | 1.455 | 0.841 | -0.025 | 0.01 | 0.01 | 5.094 | 0.22 | 0.62 | 0.35 | 0.26 | 0.16 | 0.25 |
| 8 | 142580844 | rs876097 | 2.228 | -0.425 | 1.382 | 0.795 | 0.017 | -0.019 | -0.019 | 0 | 0.66 | 0.23 | 0.52 | 0.59 | 0.71 | 0.65 |
| 8 | 142581653 | rs4425787 | 1.947 | -0.418 | 1.398 | 0.498 | -0.014 | -0.043 | -0.043 | 0 | 0.69 | 0.52 | 0.55 | 0.83 | 0.74 | 0.75 |
| 8 | 142582850 | rs875056 | -0.485 | -0.319 | 1.305 | 0.561 | -0.025 | -0.025 | -0.025 | 0 | 0.09 | 0.01 | 0.26 | 0.11 | 0.1 | 0.14 |
| 8 | 142584729 | rs4907356 | 1.091 | -0.251 | 1.529 | 0.652 | -0.012 | 0.006 | 0.006 | 2.458 | 0.74 | 0.33 | 0.6 | 0.69 | 0.82 | 0.79 |
| 8 | 142593586 | rs13252791 | 1.544 | -0.147 | 0.018 | 0.473 | 0.022 | -0.001 | -0.001 | 0 | 0.63 | 0.26 | 0.48 | 0.62 | 0.72 | 0.68 |
| 8 | 142593773 | rs670677 | 0.622 | 0.721 | -0.288 | 0.029 | 0 | -0.001 | -0.001 | 0.044 | 0.87 | 0.86 | 0.89 | 0.94 | 0.9 | 0.91 |
| 8 | 142598202 | rs7819283 | 0.73 | 0.929 | -0.591 | 0.075 | 0 | -0.008 | -0.008 | 0 | 0.79 | 0.76 | 0.68 | 0.91 | 0.76 | 0.78 |
| 8 | 142604299 | rs11779803 | -0.371 | 0.57 | -0.895 | 0.277 | 0 | -0.007 | -0.007 | 0 | 0.56 | 0.44 | 0.33 | 0.67 | 0.45 | 0.53 |
| 8 | 142604964 | rs10092704 | -0.502 | 0.453 | -1.216 | 0.778 | 0.008 | -0.05 | -0.05 | 2.375 | 0.29 | 0.15 | 0.18 | 0.39 | 0.28 | 0.15 |
| 8 | 142607603 | rs898987 | -0.337 | 0.332 | -1.247 | 0.872 | 0.009 | -0.048 | -0.048 | 1.422 | 0.17 | 0.09 | 0.09 | 0.39 | 0.14 | 0.11 |
| 8 | 142609665 | rs307401 | 0.961 | 0.181 | -0.855 | 0.762 | 0.004 | -0.001 | -0.001 | 0 | 0.75 | 0.77 | 0.86 | 0.75 | 0.8 | 0.89 |
| 8 | 142613054 | rs307415 | -1.278 | -0.206 | 0.334 | 1.233 | -0.04 | 0.192 | 0.192 | 0.281 | 0.15 | 0.34 | 0.22 | 0.34 | 0.08 | 0.23 |
| 8 | 142613212 | rs307416 | -1.533 | 0.838 | 0.257 | 0.559 | 0.018 | 0.023 | 0.023 | 1.629 | 0.33 | 0.53 | 0.39 | 0.37 | 0.39 | 0.4 |
| 8 | 142613531 | rs9644542 | -1.579 | 0.808 | 0.638 | 0.551 | 0.027 | 0.034 | 0.034 | 1.901 | 0.33 | 0.5 | 0.39 | 0.37 | 0.4 | 0.4 |
| 8 | 142620990 | rs307426 | -0.23 | 0.88 | 1.035 | 0.536 | 0.073 | -0.095 | -0.095 | 0.009 | 0.18 | 0.32 | 0.18 | 0.01 | 0.29 | 0.16 |
| 8 | 142621485 | rs1004380 | -1.631 | 0.153 | 0.564 | 0.135 | -0.023 | 0.002 | 0.002 | 0 | 0.59 | 0.45 | 0.66 | 0.53 | 0.74 | 0.59 |
| 8 | 142629143 | rs1496863 | -1.711 | -0.014 | 0.372 | 0.374 | 0.103 | 0.023 | 0.023 | 0 | 0.61 | 0.81 | 0.72 | 0.66 | 0.75 | 0.6 |
| 8 | 142633778 | rs4907370 | 0.276 | -0.413 | 0.457 | 0.338 | -0.019 | 0.025 | 0.025 | 0 | 0.8 | 0.76 | 0.91 | 0.52 | 0.93 | 0.8 |
| 8 | 142639657 | rs7846428 | 0.304 | 0.159 | 0.209 | 0.232 | -0.078 | 0.024 | 0.024 | 0.072 | 0.2 | 0.08 | 0.27 | 0.16 | 0.36 | 0.22 |
| 8 | 142647889 | rs10107586 | -1.004 | -0.062 | 0.564 | 0.217 | -0.014 | -0.011 | -0.011 | 1.611 | 0.43 | 0.47 | 0.64 | 0.32 | 0.68 | 0.41 |
| 8 | 142649191 | rs7835431 | -0.682 | -0.005 | 0.831 | 0.552 | -0.085 | 0.126 | 0.126 | 0 | 0.21 | 0.37 | 0.07 | 0.37 | 0.07 | 0.22 |
| 8 | 142654591 | rs6994100 | -0.301 | 0.566 | 0.658 | 0.544 | -0.014 | 0.014 | 0.014 | 0.506 | 0.29 | 0.21 | 0.28 | 0.22 | 0.34 | 0.25 |
| 8 | 142655880 | rs11778217 | 1.941 | 0.614 | 0.396 | 0.519 | 0.075 | 0.034 | 0.034 | 0 | 0.42 | 0.06 | 0.31 | 0.32 | 0.32 | 0.45 |
| 8 | 142656636 | rs7006068 | 0.744 | 0.595 | 0.847 | 0.713 | -0.014 | -0.022 | -0.022 | 0.126 | 0.65 | 0.31 | 0.67 | 0.46 | 0.64 | 0.67 |
| 8 | 142665525 | rs10111326 | -0.972 | -0.095 | 1.062 | 1.141 | 0.087 | 0.578 | 0.578 | 0.393 | 0.05 | 0.45 | 0.06 | 0.3 | 0.02 | 0.07 |
| 8 | 142667068 | rs2379250 | 0.968 | -0.097 | 0.741 | 1.147 | 0.04 | 0.457 | 0.457 | 3.024 | 0.94 | 0.45 | 0.94 | 0.7 | 0.98 | 0.93 |
| 8 | 142671483 | rs10089652 | 1.308 | 0.619 | -0.427 | 0.878 | -0.056 | 0.107 | 0.107 | 0 | 0.49 | 0.08 | 0.41 | 0.34 | 0.5 | 0.48 |
| 8 | 142674928 | rs7837686 | 1.017 | 0.944 | -0.121 | 0.81 | -0.021 | -0.005 | -0.005 | 0 | 0.56 | 0.26 | 0.49 | 0.46 | 0.55 | 0.57 |
| 8 | 142677833 | rs10100606 | -0.698 | 0.785 | 0.212 | 0.453 | 0.007 | -0.014 | -0.014 | 0 | 0.24 | 0.34 | 0.21 | 0.22 | 0.2 | 0.24 |
| 8 | 142683197 | rs12547611 | -0.495 | 0.391 | -0.568 | 0.341 | 0.012 | 0.006 | 0.006 | 0.101 | 0.28 | 0.11 | 0.21 | 0.27 | 0.2 | 0.32 |
| 8 | 142684445 | rs4907391 | 0.518 | 0.234 | 0.099 | 0.692 | 0.007 | -0.005 | -0.005 | 3.274 | 0.7 | 0.7 | 0.77 | 0.64 | 0.8 | 0.67 |
| 8 | 142684783 | rs13248484 | 1.097 | -0.389 | 0.466 | 0.999 | 0.033 | 0.039 | 0.039 | 0 | 0.62 | 0.29 | 0.67 | 0.21 | 0.77 | 0.59 |
| 8 | 142689315 | rs868480 | -0.47 | -0.315 | 0.244 | 0.968 | 0.018 | -0.013 | -0.013 | 0 | 0.3 | 0.21 | 0.22 | 0.37 | 0.2 | 0.33 |
| 8 | 142696491 | rs7004533 | 0.196 | 0.201 | -0.384 | 0.471 | -0.028 | 0.043 | 0.043 | 0.311 | 0.15 | 0.05 | 0.11 | 0.01 | 0.21 | 0.1 |
| 8 | 142696638 | rs10106557 | 1.061 | -0.043 | -0.28 | 0.548 | -0.008 | 0.206 | 0.206 | 0 | 0.48 | 0.13 | 0.55 | 0.19 | 0.56 | 0.5 |
| 8 | 142698496 | rs903965 | 1.019 | -0.104 | 0.763 | 0.989 | 0.028 | 0.587 | 0.587 | 2.18 | 0.93 | 0.46 | 0.89 | 0.57 | 0.97 | 0.94 |
| 8 | 142705898 | rs903956 | -0.585 | 0.447 | 0.179 | 0.324 | 0.005 | -0.025 | -0.025 | 0.251 | 0.42 | 0.5 | 0.39 | 0.29 | 0.49 | 0.36 |
| 8 | 142706843 | rs7844529 | 0.609 | 0.064 | -0.097 | 0.304 | -0.003 | -0.001 | -0.001 | 0 | 0.61 | 0.57 | 0.62 | 0.35 | 0.69 | 0.56 |
| 8 | 142710386 | rs903957 | 0.328 | 0.39 | 0.013 | 0.524 | 0.009 | -0.002 | -0.002 | 0 | 0.15 | 0.18 | 0.19 | 0.15 | 0.17 | 0.16 |

| | | | | | | | | | | | | | | | | |
|---|---|---|---|---|---|---|---|---|---|---|---|---|---|---|---|---|
| 8 | 142710708 | rs754396 | -0.471 | 0.285 | -0.134 | 0.57 | 0.007 | -0.002 | -0.002 | 0 | 0.53 | 0.47 | 0.63 | 0.54 | 0.62 | 0.47 |
| 8 | 142712678 | rs11167086 | -1.167 | 0.186 | -0.878 | 0.821 | 0.025 | 0.008 | 0.008 | 0 | 0.7 | 0.85 | 0.76 | 0.79 | 0.76 | 0.6 |
| 8 | 142713568 | rs6982668 | -0.248 | 0.678 | -0.382 | 1.254 | 0.004 | 0.004 | 0.004 | 0 | 0.42 | 0.46 | 0.51 | 0.53 | 0.51 | 0.27 |
| 8 | 142715006 | rs4907386 | -0.606 | 0.732 | -0.393 | 1.107 | 0.003 | -0.003 | -0.003 | 0 | 0.45 | 0.47 | 0.54 | 0.63 | 0.53 | 0.35 |
| 8 | 142715130 | rs4907385 | -0.037 | 0.635 | -0.41 | 1.113 | 0.014 | -0.005 | -0.005 | 0 | 0.39 | 0.46 | 0.5 | 0.39 | 0.49 | 0.24 |
| 8 | 142715381 | rs4907384 | -0.209 | 0.854 | -0.757 | 0.526 | -0.002 | 0.001 | 0.001 | 0 | 0.45 | 0.6 | 0.54 | 0.63 | 0.53 | 0.35 |
| 8 | 142726007 | rs7460203 | -0.711 | 1.113 | -0.706 | 0.528 | 0.015 | 0.016 | 0.016 | 0 | 0.33 | 0.54 | 0.5 | 0.59 | 0.4 | 0.29 |
| 8 | 142728247 | rs11785008 | 0.199 | 1.324 | -1.056 | -0.38 | -0.068 | 0.121 | 0.121 | 0.722 | 0.26 | 0.07 | 0.23 | 0.03 | 0.29 | 0.16 |
| 8 | 142729853 | rs12549250 | -1.035 | 1.217 | -0.479 | -0.249 | 0.032 | 0.011 | 0.011 | 0.971 | 0.51 | 0.44 | 0.68 | 0.61 | 0.64 | 0.33 |
| 8 | 142730346 | rs10092983 | -0.566 | 1.342 | -0.561 | 0.165 | -0.001 | 0 | 0 | 0 | 0.39 | 0.42 | 0.31 | 0.32 | 0.39 | 0.24 |
| 8 | 142730519 | rs10111342 | -1.389 | 1.085 | -0.563 | -0.057 | 0.027 | -0.001 | -0.001 | 2.959 | 0.57 | 0.67 | 0.74 | 0.61 | 0.68 | 0.35 |
| 8 | 142731374 | rs10097833 | 0.913 | 1.248 | -0.955 | -0.092 | -0.098 | 0.115 | 0.115 | 0 | 0.22 | 0.02 | 0.2 | 0.05 | 0.24 | 0.16 |
| 8 | 142731821 | rs7820007 | -0.994 | 1.042 | -0.499 | -0.026 | 0.006 | -0.001 | -0.001 | 0.838 | 0.48 | 0.55 | 0.69 | 0.49 | 0.57 | 0.31 |
| 8 | 142733541 | rs4976944 | -0.193 | 1.362 | -0.514 | 0.466 | -0.074 | -0.073 | -0.073 | 0 | 0.16 | 0.04 | 0.44 | 0.27 | 0.27 | 0.11 |
| 8 | 142736531 | rs4072286 | -1.296 | 1.323 | -0.361 | 0.342 | 0.01 | 0.031 | 0.031 | 0 | 0.29 | 0.61 | 0.54 | 0.55 | 0.37 | 0.21 |
| 8 | 142740731 | rs4074112 | 1.042 | 0.975 | -0.954 | 0.245 | 0.025 | -0.061 | -0.061 | 0 | 0.34 | 0.13 | 0.23 | 0.46 | 0.31 | 0.65 |
| 8 | 142745213 | rs12545322 | -0.647 | 1.42 | -0.01 | 0.69 | -0.051 | -0.066 | -0.066 | 0 | 0.16 | 0.08 | 0.47 | 0.39 | 0.29 | 0.16 |
| 8 | 142745374 | rs12676599 | 1.346 | 1.298 | 0.247 | 0.513 | -0.015 | -0.03 | -0.03 | 0.001 | 0.69 | 0.27 | 0.43 | 0.5 | 0.62 | 0.78 |
| 8 | 142827997 | rs11167091 | 1.12 | 1.301 | 0.485 | -0.278 | 0.018 | -0.002 | -0.002 | 1.254 | 0.71 | 0.45 | 0.53 | 0.75 | 0.67 | 0.82 |
| 8 | 142832285 | rs12156174 | 0.516 | 1.276 | -0.394 | -0.512 | -0.089 | 0.088 | 0.088 | 0.001 | 0.21 | 0.01 | 0.14 | 0.06 | 0.26 | 0.15 |
| 8 | 142834855 | rs11785612 | 1.268 | 1.478 | 0.082 | -0.217 | -0.007 | 0.004 | 0.004 | 0 | 0.7 | 0.29 | 0.52 | 0.75 | 0.68 | 0.82 |
| 8 | 142837649 | rs13274472 | 0.497 | 1.506 | 0.134 | 0.087 | 0.019 | 0.024 | 0.024 | 0 | 0.85 | 0.81 | 0.6 | 0.76 | 0.78 | 0.88 |
| 8 | 142839492 | rs9694100 | -0.128 | 0.816 | -0.11 | -0.342 | 0.021 | -0.109 | -0.109 | 0 | 0.36 | 0.52 | 0.27 | 0.06 | 0.41 | 0.22 |
| 8 | 142841497 | rs12682375 | 0.86 | 0.835 | -1.021 | -0.035 | 0.002 | 0.068 | 0.068 | 0 | 0.49 | 0.21 | 0.49 | 0.63 | 0.49 | 0.67 |
| 8 | 142847158 | rs9774266 | 1.029 | 0.52 | -0.65 | 0.991 | 0.007 | -0.043 | -0.043 | 0 | 0.9 | 0.97 | 0.82 | 0.7 | 0.92 | 0.89 |
| 8 | 142857919 | rs9694951 | -1.3 | 0.51 | -0.062 | 0.234 | -0.015 | -0.067 | -0.067 | 0 | 0.11 | 0.27 | 0.09 | 0 | 0.07 | 0.05 |
| 8 | 142864061 | rs9720781 | -0.488 | 0.725 | -0.298 | 0.208 | 0 | -0.013 | -0.013 | 0 | 0.4 | 0.53 | 0.33 | 0.29 | 0.54 | 0.4 |
| 8 | 142866345 | rs9324566 | 0.498 | 0.458 | -1.01 | 0.702 | 0.003 | 0.001 | 0.001 | 1.102 | 0.51 | 0.12 | 0.51 | 0.43 | 0.48 | 0.4 |
| 8 | 142870593 | rs9324568 | 0.616 | 0.632 | -0.866 | -0.048 | 0.003 | 0.079 | 0.079 | 0.016 | 0.49 | 0.3 | 0.45 | 0.67 | 0.5 | 0.49 |
| 8 | 142877020 | rs9324570 | 0.616 | 0.46 | -0.287 | 0.212 | -0.011 | 0.016 | 0.016 | 0 | 0.78 | 0.73 | 0.67 | 0.69 | 0.8 | 0.71 |
| 8 | 142878259 | rs9772881 | 0.397 | 1.095 | -0.552 | 0.109 | -0.025 | -0.009 | -0.009 | 0.607 | 0.94 | 0.99 | 0.78 | 0.92 | 0.9 | 0.92 |
| 8 | 142883316 | rs7836736 | -0.858 | 1.254 | -0.035 | -0.47 | 0.027 | -0.111 | -0.111 | 0 | 0.31 | 0.47 | 0.53 | 0.06 | 0.42 | 0.43 |
| 8 | 142884265 | rs6583654 | -0.911 | 1.247 | -0.027 | -0.523 | 0.021 | -0.096 | -0.096 | 0.066 | 0.31 | 0.44 | 0.52 | 0.05 | 0.42 | 0.43 |
| 8 | 142885204 | rs10110226 | -0.663 | 1.298 | -0.157 | -0.604 | -0.077 | 0.14 | 0.14 | 0 | 0.28 | 0.09 | 0.47 | 0.02 | 0.37 | 0.37 |
| 8 | 142895919 | rs13256308 | 0.603 | 1.352 | 0.248 | 0.628 | 0.007 | 0.264 | 0.264 | 0 | 0.95 | 0.86 | 0.91 | 0.57 | 0.94 | 0.93 |
| 8 | 142897314 | rs4917230 | -0.569 | 1.407 | 0.158 | 0.563 | -0.074 | 0.131 | 0.131 | 0 | 0.26 | 0.09 | 0.48 | 0.04 | 0.37 | 0.36 |
| 8 | 142897992 | rs6985874 | 0.989 | 1.384 | 0.428 | 0.633 | 0.044 | 0.452 | 0.452 | 0 | 0.94 | 0.76 | 0.9 | 0.55 | 0.94 | 0.92 |
| 8 | 142921869 | rs6583614 | -0.893 | 1.337 | 0.506 | 0.583 | 0.011 | 0.298 | 0.298 | 0 | 0.06 | 0.15 | 0.09 | 0.43 | 0.06 | 0.07 |
| 8 | 142923384 | rs7012875 | -1.284 | 1.323 | 0.556 | 0.419 | -0.009 | -0.015 | -0.015 | 2.266 | 0.32 | 0.31 | 0.55 | 0.44 | 0.43 | 0.42 |
| 8 | 142926997 | rs9785081 | -1.385 | 1.282 | 0.866 | 0.399 | 0.015 | 0.032 | 0.032 | 0 | 0.33 | 0.42 | 0.56 | 0.46 | 0.43 | 0.43 |
| 8 | 142930514 | rs1446939 | -1.579 | 1.262 | 0.985 | 0.407 | 0.022 | 0.025 | 0.025 | 0 | 0.37 | 0.56 | 0.6 | 0.47 | 0.49 | 0.49 |
| 8 | 142932284 | rs6985498 | -1.063 | 1.331 | 0.816 | 0.297 | 0.004 | -0.056 | -0.056 | 4.988 | 0.3 | 0.36 | 0.5 | 0.04 | 0.42 | 0.42 |
| 8 | 142934184 | rs11988083 | -0.601 | 1.298 | 0.595 | 0.315 | -0.066 | 0.114 | 0.114 | 0 | 0.25 | 0.1 | 0.45 | 0.02 | 0.37 | 0.34 |
| 8 | 142940178 | rs4917320 | -0.895 | 0.798 | 0.178 | -0.309 | -0.003 | 0.023 | 0.023 | 0.014 | 0.49 | 0.41 | 0.58 | 0.02 | 0.53 | 0.4 |
| 8 | 142942631 | rs4917239 | -0.574 | 1.018 | 0.037 | -0.411 | -0.002 | -0.056 | -0.056 | 0 | 0.14 | 0.21 | 0.27 | 0.02 | 0.14 | 0.24 |
| 8 | 142946225 | rs4917238 | -1.722 | 1.028 | -0.008 | -0.269 | -0.002 | 0 | 0 | 0.061 | 0.55 | 0.57 | 0.68 | 0.47 | 0.6 | 0.48 |
| 8 | 142949112 | rs6583640 | -0.142 | 0.788 | -0.286 | -0.326 | -0.034 | 0.154 | 0.154 | 2.103 | 0.34 | 0.14 | 0.3 | 0 | 0.39 | 0.15 |
| 8 | 142949616 | rs4917236 | -1.139 | 0.632 | -0.239 | -0.169 | 0 | -0.036 | -0.036 | 0 | 0.49 | 0.53 | 0.6 | 0.04 | 0.54 | 0.4 |
| 8 | 142950775 | rs1992805 | -0.574 | 0.847 | -0.948 | 0.223 | -0.054 | -0.111 | -0.111 | 0.118 | 0.27 | 0.13 | 0.47 | 0.44 | 0.37 | 0.36 |
| 8 | 142972130 | rs11167103 | -0.921 | 0.981 | -0.377 | 0.249 | -0.001 | 0.004 | 0.004 | 0.107 | 0.5 | 0.47 | 0.61 | 0.54 | 0.53 | 0.43 |
| 8 | 142972383 | rs12547538 | 1.015 | 0.896 | -0.745 | 0.153 | 0.019 | 0.067 | 0.067 | 2.303 | 0.21 | 0.1 | 0.16 | 0.01 | 0.17 | 0.09 |
| 8 | 142980109 | rs2028930 | -0.815 | 1.143 | -0.536 | 0.364 | 0 | 0 | 0 | 0 | 0.5 | 0.45 | 0.61 | 0.54 | 0.53 | 0.42 |
| 8 | 142986040 | rs11167105 | 0.834 | 2.022 | -0.254 | 0.902 | 0.015 | 0.019 | 0.019 | 0 | 0.68 | 0.44 | 0.56 | 0.47 | 0.61 | 0.62 |
| 8 | 142986188 | rs11167106 | -0.467 | 2.164 | -0.273 | 1.042 | -0.156 | -0.184 | -0.184 | 0 | 0.26 | 0.03 | 0.39 | 0.46 | 0.39 | 0.3 |
| 8 | 142986509 | rs6583548 | 0.828 | 2.035 | -0.252 | 1.02 | 0.019 | 0.019 | 0.019 | 0 | 0.68 | 0.45 | 0.56 | 0.47 | 0.6 | 0.62 |
| 8 | 142989330 | rs9644548 | 0.802 | 2.006 | -0.249 | 0.995 | 0.018 | 0.007 | 0.007 | 0 | 0.69 | 0.45 | 0.56 | 0.64 | 0.6 | 0.62 |
| 8 | 142990575 | rs4917328 | 0.334 | 2.156 | -0.172 | 1.013 | 0.026 | -0.072 | -0.072 | 0.12 | 0.74 | 0.57 | 0.63 | 0.89 | 0.71 | 0.67 |
| 8 | 142990646 | rs4917329 | 1.725 | 2.067 | 0.749 | 1.686 | 0.142 | 0.207 | 0.207 | 0 | 0.92 | 0.66 | 0.88 | 0.75 | 0.86 | 0.94 |
| 8 | 142997063 | rs7463711 | 1.598 | 1.903 | 0.554 | 1.645 | 0.081 | 0.204 | 0.204 | 0.117 | 0.89 | 0.55 | 0.87 | 0.74 | 0.91 | 0.75 |
| 8 | 143010636 | rs3937642 | -0.687 | 1.37 | -0.054 | 1.741 | -0.001 | 0.012 | 0.012 | 0 | 0.14 | 0.32 | 0.13 | 0.18 | 0.12 | 0.19 |
| 8 | 143019293 | rs935459 | -0.163 | 1.394 | -0.28 | 1.82 | 0.023 | 0.088 | 0.088 | 0 | 0.05 | 0.07 | 0.05 | 0.22 | 0.1 | 0.07 |
| 8 | 143020731 | rs7845056 | 1.076 | 1.2 | -0.712 | 1.909 | -0.03 | 0.031 | 0.031 | 1.098 | 0.54 | 0.11 | 0.66 | 0.4 | 0.6 | 0.62 |
| 8 | 143034048 | rs4917217 | 1.499 | 1.997 | 0.22 | 1.959 | 0.028 | 0.097 | 0.097 | 0 | 0.8 | 0.62 | 0.82 | 0.6 | 0.77 | 0.75 |
| 8 | 143038945 | rs10087678 | 0.441 | 1.853 | 0.018 | 1.461 | -0.001 | -0.001 | -0.001 | 2.649 | 0.87 | 0.85 | 0.9 | 0.82 | 0.88 | 0.82 |
| 8 | 143042604 | rs4130074 | 0.898 | 1.899 | 0.043 | 1.434 | 0.054 | 0.057 | 0.057 | 0 | 0.79 | 0.58 | 0.82 | 0.7 | 0.72 | 0.75 |
| 8 | 143046546 | rs10106516 | -1.399 | 1.719 | 0.537 | 1.673 | 0.087 | 0.134 | 0.134 | 0.326 | 0.2 | 0.56 | 0.48 | 0.38 | 0.23 | 0.22 |
| 8 | 143048683 | rs1542758 | -1.2 | 1.55 | 1.625 | 1.679 | 0.004 | 0.213 | 0.213 | 0.235 | 0.13 | 0.54 | 0.4 | 0.36 | 0.09 | 0.13 |
| 8 | 143057986 | rs12676273 | -0.996 | 1.221 | 1.906 | 2.012 | 0.041 | 0.388 | 0.388 | 0 | 0.11 | 0.43 | 0.34 | 0.41 | 0.09 | 0.13 |
| 8 | 143071927 | rs6981779 | -0.188 | 0.701 | 0.657 | 1.283 | -0.001 | 0.015 | 0.015 | 0.069 | 0.51 | 0.53 | 0.6 | 0.59 | 0.55 | 0.66 |

| | | | | | | | | | | | | | | | | |
|---|---|---|---|---|---|---|---|---|---|---|---|---|---|---|---|---|
| 8 | 143072068 | rs7004581 | -0.938 | 0.444 | 0.191 | 0.942 | -0.018 | -0.034 | -0.034 | 0.235 | 0.72 | 0.88 | 0.75 | 0.69 | 0.76 | 0.78 |
| 8 | 143074699 | rs2101663 | 0.321 | 0.59 | 0.366 | 1.006 | 0.007 | -0.006 | -0.006 | 0 | 0.05 | 0.09 | 0.09 | 0.01 | 0.12 | 0.08 |
| 8 | 143075220 | rs1565623 | -0.686 | 0.5 | 0.921 | 1.079 | 0.051 | 0.15 | 0.15 | 0 | 0.16 | 0.37 | 0.41 | 0.42 | 0.21 | 0.2 |
| 8 | 143078930 | rs9772780 | 0.86 | 0.281 | 0.502 | 0.664 | 0.001 | 0.041 | 0.041 | 0 | 0.41 | 0.26 | 0.28 | 0.19 | 0.47 | 0.52 |
| 8 | 143086093 | rs750195 | 1.019 | -0.424 | -0.169 | 0.489 | -0.053 | -0.029 | -0.029 | 0.061 | 0.29 | 0.15 | 0.28 | 0.31 | 0.41 | 0.42 |
| 8 | 143087187 | rs4917274 | -0.823 | -1.086 | -0.146 | -0.106 | -0.019 | 0.012 | 0.012 | 0 | 0.71 | 0.81 | 0.77 | 0.98 | 0.66 | 0.8 |
| 8 | 143092708 | rs731794 | 0.441 | -1.321 | 0.28 | 0.297 | -0.033 | 0.013 | 0.013 | 0 | 0.17 | 0.16 | 0.27 | 0.25 | 0.08 | 0.22 |
| 8 | 143101061 | rs7003047 | -0.358 | -1.726 | 0.22 | 0.108 | -0.039 | 0.026 | 0.026 | 0.194 | 0.33 | 0.57 | 0.3 | 0.53 | 0.16 | 0.33 |
| 8 | 143103398 | rs1492366 | 1.052 | -1.751 | 0.265 | 0.046 | 0 | 0 | 0 | 0 | 0.41 | 0.49 | 0.36 | 0.42 | 0.51 | 0.42 |
| 8 | 143107657 | rs7816988 | -1.134 | -1.6 | 0.182 | -0.036 | -0.023 | 0.064 | 0.064 | 0 | 0.79 | 0.91 | 0.71 | 0.95 | 0.79 | 0.85 |
| 8 | 143108079 | rs11167118 | 0.431 | -1.668 | 0.211 | 0.039 | -0.003 | 0.002 | 0.002 | 0 | 0.62 | 0.65 | 0.48 | 0.64 | 0.58 | 0.62 |
| 8 | 143110435 | rs12677032 | 0.838 | -1.579 | 0.141 | 0.101 | 0 | 0.001 | 0.001 | 0 | 0.1 | 0.11 | 0.14 | 0.31 | 0.08 | 0.13 |
| 8 | 143114379 | rs6583565 | -0.576 | -1.671 | 0.107 | 0.011 | -0.026 | 0.042 | 0.042 | 0.501 | 0.73 | 0.85 | 0.63 | 0.95 | 0.67 | 0.75 |
| 8 | 143116334 | rs4917227 | -0.161 | -1.396 | 0.442 | 0.387 | -0.008 | 0.052 | 0.052 | 0.058 | 0.12 | 0.24 | 0.15 | 0.31 | 0.09 | 0.13 |
| 8 | 143116776 | rs11987321 | 0.547 | -1.405 | 0.283 | 0.258 | -0.012 | -0.001 | -0.001 | 0 | 0.21 | 0.17 | 0.13 | 0.22 | 0.08 | 0.21 |
| 8 | 143120934 | rs13272465 | 0.328 | -0.297 | 0.346 | 0.714 | 0.001 | 0.005 | 0.005 | 0.018 | 0.35 | 0.38 | 0.46 | 0.45 | 0.43 | 0.43 |
| 8 | 143121176 | rs4917298 | -0.29 | -0.262 | 0.413 | 0.675 | 0 | -0.007 | -0.007 | 0 | 0.46 | 0.57 | 0.6 | 0.75 | 0.48 | 0.55 |
| 8 | 143121557 | rs4917299 | 0.295 | -0.29 | 0.131 | 0.679 | 0.001 | -0.016 | -0.016 | 0 | 0.09 | 0.07 | 0.14 | 0.25 | 0.06 | 0.12 |
| 8 | 143121671 | rs4917300 | -0.439 | -0.287 | 0.132 | 0.682 | 0.001 | 0.023 | 0.023 | 0.02 | 0.47 | 0.65 | 0.63 | 0.69 | 0.51 | 0.55 |
| 8 | 143123669 | rs7844494 | -0.202 | -0.486 | -0.519 | 0.935 | -0.005 | 0.031 | 0.031 | 0.165 | 0.11 | 0.19 | 0.15 | 0.33 | 0.07 | 0.13 |
| 8 | 143126970 | rs6993344 | 0.068 | -0.275 | -0.969 | 0.843 | -0.04 | -0.042 | -0.042 | 0 | 0.38 | 0.25 | 0.54 | 0.46 | 0.58 | 0.49 |
| 8 | 143128774 | rs7843263 | 0.167 | -0.038 | -1.764 | 0.896 | -0.115 | -0.11 | -0.11 | 0.107 | 0.35 | 0.13 | 0.47 | 0.4 | 0.47 | 0.48 |
| 8 | 143138194 | rs6986353 | 0.325 | -0.194 | -0.777 | 0.952 | 0.003 | -0.007 | -0.007 | 0 | 0.89 | 0.92 | 0.83 | 0.8 | 0.9 | 0.89 |
| 8 | 143145465 | rs7010485 | 0.769 | -0.472 | -0.559 | 0.797 | -0.021 | 0.021 | 0.021 | 0 | 0.89 | 0.82 | 0.77 | 0.82 | 0.94 | 0.81 |
| 8 | 143147535 | rs7814958 | 0.212 | -0.329 | -0.267 | 0.153 | 0.01 | 0.033 | 0.033 | 0 | 0.44 | 0.62 | 0.47 | 0.16 | 0.38 | 0.41 |
| 8 | 143147708 | rs6583572 | 0.532 | -0.278 | -0.26 | 0.149 | -0.003 | -0.051 | -0.051 | 0 | 0.41 | 0.43 | 0.44 | 0.13 | 0.37 | 0.33 |
| 8 | 143156531 | rs4244246 | -0.09 | -0.244 | -0.375 | 0.145 | 0.031 | 0.045 | 0.045 | 0 | 0.49 | 0.31 | 0.64 | 0.7 | 0.57 | 0.47 |
| 8 | 143158487 | rs4917312 | -0.181 | -0.23 | -0.44 | 0.062 | 0.043 | 0.084 | 0.084 | 0 | 0.5 | 0.27 | 0.48 | 0.73 | 0.58 | 0.57 |
| 8 | 143160140 | rs10090087 | -0.186 | -0.217 | -0.487 | 0.086 | 0.045 | 0.086 | 0.086 | 0 | 0.5 | 0.23 | 0.48 | 0.73 | 0.58 | 0.56 |
| 8 | 143162308 | rs7824033 | 0.28 | 0.004 | -1.132 | -0.043 | -0.146 | -0.087 | -0.087 | 0 | 0.36 | 0.04 | 0.54 | 0.68 | 0.49 | 0.41 |
| 8 | 143162402 | rs11777775 | -0.156 | 0.495 | -0.159 | -0.561 | 0.001 | -0.026 | -0.026 | 0.374 | 0.33 | 0.41 | 0.28 | 0.16 | 0.28 | 0.26 |
| 8 | 143182515 | rs7386204 | 0.586 | 0.859 | 0.129 | -0.525 | -0.014 | 0.004 | 0.004 | 0.261 | 0.65 | 0.28 | 0.58 | 0.69 | 0.6 | 0.58 |
| 8 | 143184571 | rs7819659 | 0.34 | 0.81 | 0.766 | -0.563 | 0.024 | 0.028 | 0.028 | 0.051 | 0.78 | 0.67 | 0.72 | 0.7 | 0.73 | 0.71 |
| 8 | 143187128 | rs744769 | 0.716 | 0.496 | -0.091 | -1.313 | 0.022 | 0.038 | 0.038 | 0.558 | 0.33 | 0.27 | 0.18 | 0.05 | 0.27 | 0.25 |
| 8 | 143187847 | rs1873701 | 0.346 | 0.574 | -0.215 | -1.573 | -0.004 | -0.001 | -0.001 | 0 | 0.33 | 0.32 | 0.48 | 0.65 | 0.34 | 0.38 |
| 8 | 143190003 | rs902829 | -0.247 | 0.638 | -0.46 | -1.544 | 0.008 | -0.006 | -0.006 | 0 | 0.66 | 0.66 | 0.67 | 0.7 | 0.62 | 0.63 |
| 8 | 143190271 | rs902832 | 0.079 | 1.197 | -1.096 | -0.957 | -0.024 | -0.011 | -0.011 | 0 | 0.31 | 0.17 | 0.47 | 0.64 | 0.34 | 0.38 |
| 8 | 143195007 | rs4917314 | 0.259 | 1.969 | -1.01 | -0.415 | -0.037 | -0.038 | -0.038 | 0 | 0.07 | 0.01 | 0.12 | 0.19 | 0.17 | 0.18 |
| 8 | 143199039 | rs11778623 | 0.207 | 1.539 | -1.232 | -0.394 | -0.061 | -0.083 | -0.083 | 0 | 0.13 | 0.01 | 0.12 | 0.27 | 0.18 | 0.18 |
| 8 | 143199529 | rs7017932 | -0.037 | 1.328 | -0.237 | -0.466 | 0.036 | 0.106 | 0.106 | 0 | 0.83 | 0.59 | 0.85 | 0.72 | 0.82 | 0.81 |
| 8 | 143200557 | rs13253406 | -0.035 | 0.828 | -0.664 | -1.956 | -0.001 | -0.005 | -0.005 | 0 | 0.57 | 0.43 | 0.41 | 0.15 | 0.52 | 0.43 |
| 8 | 143202173 | rs6981017 | 0.013 | 0.902 | -0.616 | -1.917 | 0.001 | -0.038 | -0.038 | 0 | 0.57 | 0.51 | 0.41 | 0.15 | 0.52 | 0.43 |
| 8 | 143205583 | rs7011064 | 0.545 | 0.386 | -0.823 | -1.602 | -0.001 | -0.006 | -0.006 | 0 | 0.16 | 0.15 | 0.16 | 0.1 | 0.14 | 0.11 |
| 8 | 143206611 | rs10087556 | 0.194 | 0.457 | -0.99 | -1.733 | -0.002 | -0.026 | -0.026 | 0 | 0.34 | 0.33 | 0.31 | 0.1 | 0.34 | 0.31 |
| 8 | 143207378 | rs7835207 | -0.679 | 0.737 | -1.557 | -1.316 | -0.025 | -0.026 | -0.026 | 0 | 0.83 | 0.97 | 0.84 | 0.85 | 0.85 | 0.89 |
| 8 | 143207696 | rs747769 | -0.137 | 1.067 | -1.745 | -1.778 | -0.038 | 0.053 | 0.053 | 3.277 | 0.31 | 0.13 | 0.53 | 0.82 | 0.39 | 0.54 |
| 8 | 143208340 | rs747898 | -0.602 | 0.907 | -1.39 | -2.078 | 0.037 | 0.12 | 0.12 | 0.052 | 0.57 | 0.26 | 0.77 | 0.97 | 0.7 | 0.73 |
| 8 | 143211177 | rs747551 | -0.841 | 1.479 | -0.409 | -0.998 | -0.01 | 0.008 | 0.008 | 0.293 | 0.32 | 0.28 | 0.52 | 0.76 | 0.4 | 0.53 |
| 8 | 143233571 | rs7817259 | 0.068 | 1.674 | -0.106 | -0.425 | 0.001 | 0.004 | 0.004 | 0 | 0.8 | 0.79 | 0.68 | 0.35 | 0.71 | 0.7 |
| 8 | 143233677 | rs10085978 | 0.913 | 1.529 | 0.033 | -0.616 | -0.007 | 0.038 | 0.038 | 0.017 | 0.6 | 0.3 | 0.48 | 0.21 | 0.55 | 0.57 |
| 8 | 143234609 | rs7465378 | 0.591 | 1.365 | -0.233 | -0.429 | 0.001 | 0.007 | 0.007 | 0.335 | 0.16 | 0.13 | 0.17 | 0.14 | 0.16 | 0.13 |
| 8 | 143240263 | rs7463731 | 0.653 | 1.837 | 0.438 | 0.064 | 0.094 | 0.088 | 0.088 | 0 | 0.77 | 0.52 | 0.68 | 0.35 | 0.68 | 0.7 |
| 8 | 143247678 | rs11167132 | 0.636 | 0.994 | 0.249 | -0.02 | -0.013 | 0.03 | 0.03 | 0 | 0.77 | 0.65 | 0.75 | 0.35 | 0.79 | 0.74 |
| 8 | 143247742 | rs11781880 | 0.644 | 0.859 | -0.278 | -0.991 | -0.017 | 0.081 | 0.081 | 0 | 0.4 | 0.22 | 0.3 | 0.06 | 0.42 | 0.23 |
| 8 | 143249252 | rs6583595 | 0.759 | 0.705 | -0.038 | 0.689 | 0.005 | -0.043 | -0.043 | 0.908 | 0.77 | 0.87 | 0.78 | 0.52 | 0.8 | 0.71 |
| 8 | 143249727 | rs7823353 | -1.262 | 1.031 | 1.1 | 0.886 | 0.092 | 0.344 | 0.344 | 0.079 | 0.08 | 0.35 | 0.14 | 0.37 | 0.07 | 0.15 |
| 8 | 143257317 | rs7846108 | -1.592 | 0.939 | 1.085 | 0.683 | 0.033 | 0.037 | 0.037 | 0.134 | 0.1 | 0.51 | 0.26 | 0.13 | 0.09 | 0.16 |
| 8 | 143258503 | rs11167133 | 0.616 | 1.078 | 0.687 | 1.198 | -0.024 | -0.108 | -0.108 | 3.074 | 0.79 | 0.92 | 0.79 | 0.56 | 0.79 | 0.67 |
| 8 | 143259455 | rs7828532 | -1.827 | 0.895 | 1.531 | 1.05 | 0.082 | 0.143 | 0.143 | 2.816 | 0.06 | 0.5 | 0.06 | 0.11 | 0.04 | 0.06 |
| 8 | 143265173 | rs3935507 | 0.773 | 0.694 | 0.181 | 1.469 | 0.009 | -0.049 | -0.049 | 0.014 | 0.79 | 0.86 | 0.82 | 0.63 | 0.81 | 0.69 |
| 8 | 143270729 | rs3923878 | -1.466 | 0.437 | 0.405 | 1.686 | -0.019 | 0.093 | 0.093 | 0.097 | 0.15 | 0.31 | 0.32 | 0.41 | 0.09 | 0.17 |
| 8 | 143274934 | rs7465207 | -1.561 | 0.428 | 0.453 | 1.746 | -0.014 | 0.216 | 0.216 | 0.687 | 0.09 | 0.32 | 0.21 | 0.41 | 0.04 | 0.11 |
| 8 | 143291396 | rs4345692 | 1.595 | 0.525 | 0.804 | 1.636 | -0.007 | 0.084 | 0.084 | 1.055 | 0.87 | 0.42 | 0.81 | 0.83 | 0.93 | 0.87 |
| 8 | 143293307 | rs4976967 | -0.41 | 1.218 | -0.209 | 1.034 | -0.075 | 0.151 | 0.151 | 0 | 0.27 | 0.04 | 0.26 | 0.03 | 0.32 | 0.31 |
| 8 | 143293558 | rs11992923 | 0.555 | 0.947 | 0.164 | 1.387 | -0.027 | -0.038 | -0.038 | 0.232 | 0.93 | 0.98 | 0.87 | 0.83 | 0.93 | 0.87 |
| 8 | 143297620 | rs6583602 | 0.248 | 0.934 | 0.272 | 1.402 | -0.001 | 0.023 | 0.023 | 0 | 0.94 | 0.93 | 0.87 | 0.83 | 0.93 | 0.87 |
| 8 | 143300157 | rs4327969 | 0.284 | 0.934 | 0.394 | 1.436 | 0.011 | 0.034 | 0.034 | 0 | 0.94 | 0.86 | 0.86 | 0.86 | 0.93 | 0.87 |
| 8 | 143301529 | rs11167134 | 1.426 | 0.732 | -0.108 | 0.969 | 0.014 | 0.092 | 0.092 | 0.58 | 0.41 | 0.15 | 0.27 | 0.21 | 0.42 | 0.24 |
| 8 | 143302166 | rs7463863 | -0.823 | 0.745 | 0.289 | 1.11 | 0.023 | 0.033 | 0.033 | 0.084 | 0.5 | 0.29 | 0.53 | 0.59 | 0.51 | 0.59 |

| | | | | | | | | | | | | | | | | |
|---|---|---|---|---|---|---|---|---|---|---|---|---|---|---|---|---|
| 8 | 143302580 | rs6583604 | -0.975 | 0.722 | 0.322 | 1.117 | 0.009 | 0.036 | 0.036 | 0.02 | 0.53 | 0.68 | 0.59 | 0.65 | 0.52 | 0.63 |
| 8 | 143302889 | rs11786405 | 1.393 | 0.73 | 0.319 | 1.119 | 0.01 | 0.092 | 0.092 | 0.609 | 0.4 | 0.13 | 0.27 | 0.21 | 0.42 | 0.24 |
| 8 | 143305232 | rs7016042 | -0.118 | 0.766 | 0.259 | 1.162 | 0.018 | 0.048 | 0.048 | 0 | 0.93 | 0.82 | 0.86 | 0.86 | 0.93 | 0.87 |
| 8 | 143305508 | rs11167135 | 1.202 | 0.677 | 0.076 | 0.952 | -0.009 | 0.087 | 0.087 | 0.605 | 0.4 | 0.08 | 0.26 | 0.22 | 0.42 | 0.24 |
| 8 | 143310815 | rs11167136 | -0.731 | 0.552 | 0.427 | 1.061 | 0.004 | 0.002 | 0.002 | 0 | 0.5 | 0.57 | 0.57 | 0.5 | 0.43 | 0.61 |
| 8 | 143312933 | rs4129585 | -1.67 | 0.625 | 0.104 | 1.077 | -0.068 | 0.008 | 0.008 | 0 | 0.65 | 0.93 | 0.73 | 0.74 | 0.55 | 0.79 |
| 8 | 143319470 | rs7830520 | -0.053 | 0.27 | 0.647 | 1.004 | 0.002 | -0.035 | -0.035 | 0 | 0.17 | 0.46 | 0.23 | 0.08 | 0.15 | 0.14 |
| 8 | 143323109 | rs7822538 | -1.23 | 0.303 | 0.406 | 1.033 | -0.002 | -0.018 | -0.018 | 0 | 0.54 | 0.81 | 0.64 | 0.51 | 0.43 | 0.62 |
| 8 | 143326402 | rs8180914 | -1.027 | 0.463 | 0.447 | 1.113 | 0.015 | 0.004 | 0.004 | 0.306 | 0.51 | 0.7 | 0.63 | 0.5 | 0.43 | 0.61 |
| 8 | 143345885 | rs12547214 | -0.84 | 0.3 | 0.469 | 0.736 | 0 | -0.024 | -0.024 | 0 | 0.38 | 0.53 | 0.37 | 0.2 | 0.33 | 0.34 |
| 8 | 143347243 | rs6583612 | -0.478 | 0.048 | 0.619 | 0.716 | -0.02 | -0.046 | -0.046 | 0 | 0.21 | 0.6 | 0.24 | 0.08 | 0.15 | 0.16 |
| 8 | 143350750 | rs4976955 | -0.468 | 0.043 | 0.645 | 0.697 | -0.02 | -0.049 | -0.049 | 0 | 0.21 | 0.59 | 0.25 | 0.08 | 0.15 | 0.16 |
| 8 | 143374261 | rs4325021 | -0.604 | -0.082 | 1.155 | 1.632 | -0.046 | 0.044 | 0.044 | 0 | 0.25 | 0.42 | 0.25 | 0.31 | 0.17 | 0.28 |
| 8 | 143376802 | rs7832163 | -1.231 | 0.516 | 0.952 | 1.517 | 0.002 | 0.004 | 0.004 | 1.097 | 0.43 | 0.47 | 0.41 | 0.46 | 0.42 | 0.54 |
| 8 | 143384314 | rs10102944 | -1.562 | 0.477 | 1.068 | 1.561 | -0.017 | 0.035 | 0.035 | 1.625 | 0.47 | 0.78 | 0.54 | 0.57 | 0.45 | 0.6 |
| 8 | 143388250 | rs4581170 | 0.721 | 0.589 | 0.807 | 1.587 | -0.025 | 0.05 | 0.05 | 0 | 0.69 | 0.56 | 0.7 | 0.53 | 0.77 | 0.69 |
| 8 | 143394208 | rs11167137 | -1.706 | 0.357 | 0.939 | 1.338 | -0.004 | 0.043 | 0.043 | 0.686 | 0.51 | 0.84 | 0.55 | 0.55 | 0.44 | 0.59 |
| 8 | 143394496 | rs4325020 | -1.581 | 0.356 | 1.008 | 1.342 | -0.003 | 0.016 | 0.016 | 0.203 | 0.49 | 0.76 | 0.53 | 0.51 | 0.44 | 0.56 |
| 8 | 143398255 | rs13261479 | -0.373 | 0.034 | 1.008 | 1.379 | -0.037 | 0.043 | 0.043 | 0 | 0.25 | 0.4 | 0.26 | 0.33 | 0.16 | 0.28 |
| 8 | 143403693 | rs13266463 | -2.33 | 0.357 | 0.365 | 1.29 | 0.004 | 0.042 | 0.042 | 0.534 | 0.63 | 0.85 | 0.75 | 0.75 | 0.61 | 0.8 |
| 8 | 143405289 | rs7814338 | -1.005 | -0.402 | 1.441 | 1.643 | -0.061 | 0.043 | 0.043 | 0 | 0.27 | 0.51 | 0.26 | 0.3 | 0.17 | 0.27 |
| 8 | 143408785 | rs7000748 | -1.897 | -0.286 | 1.432 | 1.656 | -0.013 | 0.008 | 0.008 | 0 | 0.42 | 0.55 | 0.45 | 0.54 | 0.33 | 0.44 |
| 8 | 143413136 | rs10435683 | -1.831 | -0.291 | 1.465 | 1.678 | -0.015 | 0.007 | 0.007 | 0 | 0.42 | 0.54 | 0.44 | 0.47 | 0.32 | 0.34 |
| 8 | 143416412 | rs13265349 | -1.059 | -0.344 | 1.564 | 1.776 | 0.002 | 0.004 | 0.004 | 1.267 | 0.16 | 0.19 | 0.19 | 0.28 | 0.14 | 0.29 |
| 8 | 143425720 | rs10100935 | -2.261 | -0.389 | 1.561 | 1.806 | 0.042 | 0.002 | 0.002 | 0.099 | 0.45 | 0.86 | 0.48 | 0.63 | 0.34 | 0.5 |
| 8 | 143436034 | rs7814359 | -1.343 | -0.362 | 1.688 | 1.804 | -0.029 | 0.029 | 0.029 | 0 | 0.29 | 0.63 | 0.29 | 0.35 | 0.2 | 0.21 |
| 8 | 143443810 | rs12682505 | -1.164 | -0.363 | 1.755 | 1.826 | -0.045 | 0.026 | 0.026 | 0 | 0.28 | 0.57 | 0.27 | 0.26 | 0.18 | 0.17 |
| 8 | 143458833 | rs7013121 | -1.295 | -0.349 | 1.776 | 1.816 | -0.031 | 0.035 | 0.035 | 0 | 0.28 | 0.61 | 0.29 | 0.35 | 0.2 | 0.21 |
| 8 | 143475927 | rs7387891 | -2.275 | -0.28 | 1.623 | 1.656 | 0.024 | -0.021 | -0.021 | 0.048 | 0.44 | 0.85 | 0.49 | 0.6 | 0.35 | 0.51 |
| 8 | 143480669 | rs6583623 | -1.16 | -0.135 | 1.581 | 1.629 | 0.012 | 0.014 | 0.014 | 0.558 | 0.16 | 0.25 | 0.22 | 0.29 | 0.17 | 0.24 |
| 8 | 143488721 | rs10109771 | 0.627 | -0.204 | 1.572 | 1.689 | 0.004 | 0.005 | 0.005 | 0.978 | 0.86 | 0.83 | 0.83 | 0.77 | 0.86 | 0.82 |
| 8 | 143490718 | rs7387733 | 0.708 | -0.203 | 1.585 | 1.692 | 0.009 | 0.016 | 0.016 | 0.743 | 0.85 | 0.79 | 0.81 | 0.71 | 0.84 | 0.78 |
| 8 | 143491473 | rs9324588 | -0.399 | -0.283 | 1.964 | 1.587 | -0.001 | -0.078 | -0.078 | 0 | 0.13 | 0.35 | 0.11 | 0 | 0.1 | 0.1 |
| 8 | 143497052 | rs6583630 | 0.869 | -0.377 | 2.089 | 1.755 | 0.008 | 0.016 | 0.016 | 0.706 | 0.85 | 0.79 | 0.81 | 0.71 | 0.84 | 0.78 |
| 8 | 143497202 | rs4073866 | 2.24 | -0.476 | 2.092 | 1.709 | 0.026 | -0.019 | -0.019 | 0 | 0.59 | 0.19 | 0.54 | 0.4 | 0.68 | 0.52 |
| 8 | 143500243 | rs6991885 | 2.023 | -0.482 | 2.155 | 1.744 | -0.003 | 0.012 | 0.012 | 1.042 | 0.65 | 0.37 | 0.57 | 0.42 | 0.7 | 0.63 |
| 8 | 143504469 | rs9324591 | 1.786 | -0.482 | 2.155 | 1.744 | -0.003 | 0.011 | 0.011 | 1.085 | 0.65 | 0.37 | 0.57 | 0.42 | 0.7 | 0.63 |
| 8 | 143511806 | rs9801760 | 0.545 | -0.428 | 1.995 | 1.647 | 0.003 | -0.001 | -0.001 | 0 | 0.87 | 0.84 | 0.83 | 0.8 | 0.86 | 0.82 |
| 8 | 143514434 | rs7827372 | 1.245 | -0.402 | 1.709 | 1.666 | 0.004 | -0.05 | -0.05 | 0 | 0.76 | 0.5 | 0.74 | 0.79 | 0.79 | 0.73 |
| 8 | 143516063 | rs9801746 | 1.635 | -0.294 | 1.048 | 1.706 | 0.024 | -0.032 | -0.032 | 0 | 0.6 | 0.19 | 0.63 | 0.41 | 0.69 | 0.52 |
| 8 | 143522536 | rs11779434 | 0.735 | 0.43 | 1.312 | 2.021 | 0.06 | 0.182 | 0.182 | 0 | 0.94 | 0.8 | 0.77 | 0.7 | 0.92 | 0.86 |
| 8 | 143528511 | rs13275839 | -0.476 | -0.04 | 1.5 | 1.303 | -0.005 | -0.056 | -0.056 | 0 | 0.1 | 0.27 | 0.07 | 0 | 0.06 | 0.08 |
| 8 | 143535189 | rs12682623 | -1.137 | -0.25 | 1.352 | 1.744 | -0.031 | 0.071 | 0.071 | 0.084 | 0.16 | 0.32 | 0.12 | 0.33 | 0.11 | 0.24 |
| 8 | 143549588 | rs6583634 | -1.065 | 0.06 | 1.193 | 1.553 | 0.004 | 0.058 | 0.058 | 0 | 0.18 | 0.3 | 0.29 | 0.28 | 0.18 | 0.14 |
| 8 | 143553868 | rs7822554 | 0.777 | -0.131 | 0.771 | 1.179 | -0.028 | -0.022 | -0.022 | 0 | 0.75 | 0.54 | 0.69 | 0.78 | 0.81 | 0.61 |
| 8 | 143565607 | rs10875483 | 0.303 | 0.291 | 0.221 | 1.278 | -0.054 | -0.083 | -0.083 | 0 | 0.82 | 0.99 | 0.79 | 0.78 | 0.8 | 0.84 |
| 8 | 143566729 | rs4074758 | -1.035 | 0.318 | 0.956 | 1.318 | 0.015 | 0.028 | 0.028 | 1.349 | 0.18 | 0.56 | 0.3 | 0.26 | 0.17 | 0.25 |
| 8 | 143567370 | rs7386480 | -1.503 | 0.078 | 0.925 | 1.199 | -0.008 | -0.003 | -0.003 | 0 | 0.22 | 0.62 | 0.32 | 0.26 | 0.18 | 0.25 |
| 8 | 143569216 | rs7462966 | 1.518 | 0.374 | 0.63 | 1.267 | 0.008 | 0.025 | 0.025 | 0.686 | 0.64 | 0.48 | 0.49 | 0.52 | 0.63 | 0.58 |
| 8 | 143585398 | rs7386358 | 1.145 | 0.132 | 0.251 | 0.823 | -0.005 | 0 | 0 | 0 | 0.75 | 0.62 | 0.74 | 0.78 | 0.77 | 0.81 |
| 8 | 143587121 | rs12546526 | 0.378 | -0.208 | -0.753 | 0.949 | -0.003 | -0.07 | -0.07 | 0 | 0.83 | 0.99 | 0.83 | 0.8 | 0.88 | 0.84 |
| 8 | 143596497 | rs9657474 | -0.593 | -0.136 | -0.289 | 0.386 | -0.014 | -0.021 | -0.021 | 0 | 0.31 | 0.41 | 0.28 | 0.12 | 0.24 | 0.23 |
| 8 | 143608639 | rs10283377 | -0.981 | 0.226 | 0.104 | 1.3 | 0.006 | 0.11 | 0.11 | 0 | 0.15 | 0.36 | 0.28 | 0.35 | 0.15 | 0.19 |
| 8 | 143609913 | rs7838992 | 0.416 | 0.107 | -0.428 | 0.737 | 0.001 | 0.014 | 0.014 | 0.201 | 0.6 | 0.35 | 0.77 | 0.65 | 0.63 | 0.58 |
| 8 | 143612149 | rs7846567 | -0.884 | -0.244 | -1.245 | -0.322 | -0.143 | 0.26 | 0.26 | 0.6 | 0.64 | 0.94 | 0.58 | 0.99 | 0.53 | 0.79 |
| 8 | 143615524 | rs4394435 | -0.653 | -0.321 | -0.337 | 0.415 | -0.003 | -0.005 | -0.005 | 0 | 0.31 | 0.35 | 0.36 | 0.7 | 0.29 | 0.5 |
| 8 | 143620393 | rs7462518 | -0.845 | -0.691 | 0.343 | 0.649 | -0.024 | 0.142 | 0.142 | 0 | 0.15 | 0.45 | 0.13 | 0.22 | 0.12 | 0.28 |
| 8 | 143620400 | rs7460047 | -0.153 | -0.569 | -0.261 | 0.696 | -0.009 | -0.073 | -0.073 | 0 | 0.11 | 0.01 | 0.07 | 0.22 | 0.11 | 0.28 |
| 8 | 143628958 | rs7387708 | -0.468 | -0.508 | 0.313 | -0.208 | -0.005 | 0.011 | 0.011 | 0.21 | 0.57 | 0.43 | 0.51 | 0.85 | 0.54 | 0.68 |
| 8 | 143633066 | rs7387754 | -0.731 | -0.613 | 0.841 | -0.228 | -0.008 | 0.012 | 0.012 | 0.78 | 0.59 | 0.67 | 0.51 | 0.84 | 0.52 | 0.67 |
| 8 | 143642776 | rs12680736 | 0.531 | -0.864 | 0.656 | -0.173 | -0.015 | -0.053 | -0.053 | 0.37 | 0.35 | 0.08 | 0.23 | 0.59 | 0.33 | 0.44 |
| 8 | 143643575 | rs13269784 | -1.297 | -0.858 | 0.865 | -0.274 | -0.059 | 0.109 | 0.109 | 2.054 | 0.65 | 0.81 | 0.54 | 0.99 | 0.54 | 0.78 |
| 8 | 143647801 | rs7460370 | -0.808 | -0.748 | 0.992 | -0.166 | -0.023 | 0.042 | 0.042 | 0.853 | 0.6 | 0.72 | 0.52 | 0.84 | 0.51 | 0.67 |
| 8 | 143653364 | rs4977106 | 0.229 | -0.778 | 0.928 | -0.152 | 0.002 | 0 | 0 | 0 | 0.4 | 0.42 | 0.26 | 0.58 | 0.34 | 0.43 |
| 8 | 143656021 | rs6996219 | -0.687 | -0.229 | 1.665 | 0.15 | -0.028 | -0.023 | -0.023 | 0.055 | 0.23 | 0.53 | 0.28 | 0.26 | 0.17 | 0.23 |
| 8 | 143660032 | rs4073782 | -0.427 | -0.096 | 1.505 | 0.198 | -0.004 | -0.012 | -0.012 | 0 | 0.21 | 0.43 | 0.28 | 0.26 | 0.18 | 0.23 |
| 8 | 143664312 | rs3934486 | -0.631 | -0.068 | 1.379 | 0.193 | -0.009 | -0.016 | -0.016 | 0 | 0.22 | 0.53 | 0.29 | 0.27 | 0.18 | 0.24 |
| 8 | 143677819 | rs4977101 | -0.333 | -0.047 | 1.06 | 0.185 | -0.002 | -0.014 | -0.014 | 0 | 0.2 | 0.34 | 0.28 | 0.23 | 0.19 | 0.23 |
| 8 | 143682428 | rs7835613 | 0.975 | -0.287 | 0.075 | -0.179 | 0.071 | 0.115 | 0.115 | 0 | 0.41 | 0.21 | 0.2 | 0.19 | 0.28 | 0.29 |

| | | | | | | | | | | | | | | | | |
|---|---|---|---|---|---|---|---|---|---|---|---|---|---|---|---|---|
| 8 | 143682867 | rs12546658 | -0.839 | 0.149 | -0.078 | -0.173 | -0.081 | 0.092 | 0.092 | 0 | 0.75 | 0.9 | 0.6 | 0.99 | 0.6 | 0.84 |
| 8 | 143686995 | rs9324593 | -0.132 | 0.077 | 0.194 | -0.053 | -0.002 | 0.028 | 0.028 | 0.067 | 0.15 | 0.23 | 0.12 | 0.59 | 0.13 | 0.34 |
| 8 | 143688896 | rs10103454 | -0.881 | 0.01 | -0.058 | -0.134 | -0.079 | 0.094 | 0.094 | 0 | 0.74 | 0.86 | 0.6 | 0.99 | 0.6 | 0.85 |
| 8 | 143690833 | rs10110456 | 0.285 | 0.059 | -0.096 | -0.014 | 0.006 | -0.006 | -0.006 | 0 | 0.12 | 0.11 | 0.08 | 0.6 | 0.09 | 0.33 |
| 8 | 143694184 | rs10097505 | -0.356 | 0.225 | 0.157 | 0.403 | -0.013 | -0.006 | -0.006 | 4.468 | 0.38 | 0.42 | 0.5 | 0.4 | 0.48 | 0.38 |
| 8 | 143707473 | rs28473387 | -0.852 | 0.319 | 0.03 | -0.084 | 0.014 | 0.009 | 0.009 | 0 | 0.49 | 0.54 | 0.72 | 0.22 | 0.66 | 0.35 |
| 8 | 143716030 | rs4545075 | 0.744 | 0.46 | -0.046 | -0.053 | 0.003 | 0.001 | 0.001 | 0 | 0.5 | 0.49 | 0.29 | 0.49 | 0.34 | 0.53 |
| 8 | 143728375 | rs9297973 | -0.058 | 0.676 | -0.192 | -0.293 | -0.002 | 0.025 | 0.025 | 0 | 0.76 | 0.74 | 0.71 | 0.53 | 0.78 | 0.7 |
| 8 | 143733540 | rs4327844 | 0.042 | -0.159 | 0.153 | -0.1 | 0.038 | 0.007 | 0.007 | 0 | 0.54 | 0.72 | 0.32 | 0.51 | 0.39 | 0.55 |
| 8 | 143738529 | rs4587307 | 0.192 | -0.152 | 0.196 | -0.091 | 0.043 | 0.005 | 0.005 | 0 | 0.53 | 0.67 | 0.3 | 0.49 | 0.35 | 0.55 |
| 8 | 143740596 | rs6988393 | -0.05 | -0.151 | 0.196 | -0.091 | 0.033 | 0.004 | 0.004 | 0 | 0.55 | 0.72 | 0.32 | 0.52 | 0.39 | 0.56 |
| 8 | 143741170 | rs3735993 | 0.394 | -0.162 | 0.338 | -0.096 | 0.006 | 0.001 | 0.001 | 0 | 0.51 | 0.46 | 0.29 | 0.48 | 0.35 | 0.54 |
| 8 | 143742683 | rs3735994 | -0.054 | -0.161 | 0.334 | -0.096 | 0.033 | 0.004 | 0.004 | 0 | 0.55 | 0.72 | 0.32 | 0.52 | 0.39 | 0.56 |
| 8 | 143743611 | rs2976400 | 0.188 | -0.162 | 0.346 | -0.096 | 0.038 | 0.004 | 0.004 | 0 | 0.53 | 0.66 | 0.3 | 0.49 | 0.35 | 0.55 |
| 8 | 143744931 | rs750530 | 0.188 | -0.14 | 0.321 | -0.096 | 0.041 | 0.005 | 0.005 | 0 | 0.53 | 0.67 | 0.3 | 0.49 | 0.35 | 0.55 |
| 8 | 143746050 | rs2976399 | 0.39 | -0.175 | 0.205 | -0.093 | 0.006 | 0.001 | 0.001 | 0 | 0.51 | 0.47 | 0.29 | 0.48 | 0.35 | 0.56 |
| 8 | 143746069 | rs3735999 | 0.061 | 0.003 | -0.624 | -0.019 | 0.032 | 0.004 | 0.004 | 0 | 0.55 | 0.72 | 0.32 | 0.52 | 0.39 | 0.59 |
| 8 | 143746701 | rs3802232 | 0.061 | 0.003 | -0.651 | -0.019 | 0.033 | 0.004 | 0.004 | 0 | 0.55 | 0.72 | 0.32 | 0.52 | 0.39 | 0.56 |
| 8 | 143751864 | rs2978974 | -0.249 | 0.402 | -0.512 | -0.08 | 0.01 | -0.005 | -0.005 | 0 | 0.33 | 0.54 | 0.25 | 0.29 | 0.35 | 0.3 |
| 8 | 143757037 | rs2920283 | -0.627 | 0.13 | -0.953 | -0.076 | 0.022 | 0.121 | 0.121 | 0 | 0.48 | 0.35 | 0.5 | 0.34 | 0.45 | 0.41 |
| 8 | 143761931 | rs2294008 | -0.643 | 0.133 | -0.922 | -0.075 | 0.023 | 0.119 | 0.119 | 0 | 0.48 | 0.37 | 0.5 | 0.34 | 0.45 | 0.41 |
| 8 | 143762724 | rs2976391 | 0.915 | 0.099 | -0.531 | 0.022 | 0.003 | -0.005 | -0.005 | 0 | 0.43 | 0.42 | 0.43 | 0.33 | 0.47 | 0.36 |
| 8 | 143762932 | rs2976392 | -0.506 | 0.068 | -0.569 | 0.08 | 0.023 | 0.121 | 0.121 | 0 | 0.48 | 0.37 | 0.51 | 0.34 | 0.45 | 0.41 |
| 8 | 143763043 | rs2920298 | -0.482 | 0.061 | -0.562 | 0.077 | 0.022 | 0.125 | 0.125 | 0 | 0.48 | 0.35 | 0.51 | 0.34 | 0.45 | 0.41 |
| 8 | 143763690 | rs10216533 | -0.474 | 0.059 | -0.562 | 0.076 | 0.021 | 0.124 | 0.124 | 0 | 0.48 | 0.36 | 0.51 | 0.34 | 0.45 | 0.41 |
| 8 | 143763750 | rs2976395 | -0.474 | 0.059 | -0.562 | 0.076 | 0.021 | 0.124 | 0.124 | 0 | 0.48 | 0.35 | 0.51 | 0.34 | 0.45 | 0.41 |
| 8 | 143763757 | rs1045547 | -0.474 | 0.059 | -0.562 | 0.076 | 0.021 | 0.124 | 0.124 | 0 | 0.48 | 0.35 | 0.51 | 0.34 | 0.45 | 0.41 |
| 8 | 143763958 | rs1045574 | -0.474 | 0.059 | -0.562 | 0.076 | 0.021 | 0.124 | 0.124 | 0 | 0.48 | 0.35 | 0.51 | 0.34 | 0.45 | 0.41 |
| 8 | 143764001 | rs2976396 | -0.474 | 0.059 | -0.558 | 0.076 | 0.021 | 0.126 | 0.126 | 0 | 0.48 | 0.35 | 0.51 | 0.34 | 0.45 | 0.41 |
| 8 | 143764613 | rs2976397 | -0.322 | -0.06 | -0.587 | 0.042 | -0.035 | 0.218 | 0.218 | 0 | 0.45 | 0.15 | 0.48 | 0.23 | 0.43 | 0.37 |
| 8 | 143765885 | rs12155758 | 0.102 | -0.091 | -0.655 | -0.25 | -0.058 | 0.065 | 0.065 | 0 | 0.2 | 0.08 | 0.33 | 0.13 | 0.25 | 0.22 |
| 8 | 143780775 | rs1435453 | 0.996 | -0.355 | -0.463 | -0.1 | 0.003 | -0.005 | -0.005 | 0 | 0.42 | 0.42 | 0.43 | 0.33 | 0.47 | 0.36 |
| 8 | 143784885 | rs2244152 | 0.806 | -0.39 | -0.481 | -0.057 | -0.008 | -0.005 | -0.005 | 0.043 | 0.44 | 0.59 | 0.45 | 0.42 | 0.47 | 0.4 |
| 8 | 143785345 | rs2585153 | 0.818 | -0.387 | -0.472 | -0.057 | -0.008 | -0.006 | -0.006 | 0.021 | 0.43 | 0.59 | 0.45 | 0.42 | 0.47 | 0.4 |
| 8 | 143785659 | rs2164308 | 0.802 | -0.361 | -0.459 | -0.044 | -0.008 | -0.006 | -0.006 | 0 | 0.44 | 0.59 | 0.45 | 0.42 | 0.47 | 0.4 |
| 8 | 143786835 | rs6471588 | -0.678 | -0.477 | -0.603 | -0.119 | 0.01 | -0.018 | -0.018 | 0 | 0.72 | 0.75 | 0.84 | 0.58 | 0.8 | 0.68 |
| 8 | 143806896 | rs2585140 | 0.87 | -0.375 | -0.694 | -0.08 | -0.013 | -0.01 | -0.01 | 0 | 0.44 | 0.65 | 0.46 | 0.42 | 0.47 | 0.4 |
| 8 | 143807651 | rs3758079 | 0.831 | -0.406 | -0.789 | -0.146 | -0.094 | 0.026 | 0.026 | 0 | 0.16 | 0.02 | 0.32 | 0.16 | 0.27 | 0.23 |
| 8 | 143808951 | rs2585138 | 0.679 | -0.141 | -0.509 | 0.075 | -0.01 | -0.009 | -0.009 | 0 | 0.44 | 0.64 | 0.46 | 0.42 | 0.47 | 0.4 |
| 8 | 143819383 | rs7828042 | 0.026 | -0.118 | -0.59 | 0.136 | -0.046 | 0.045 | 0.045 | 0 | 0.3 | 0.2 | 0.4 | 0.28 | 0.35 | 0.36 |
| 8 | 143822028 | rs34956412 | 0.486 | -0.121 | -0.597 | 0.132 | -0.068 | 0.072 | 0.072 | 0 | 0.26 | 0.12 | 0.35 | 0.19 | 0.31 | 0.32 |
| 8 | 143824373 | rs3758081 | 0.625 | -0.095 | -0.58 | 0.145 | -0.092 | 0.06 | 0.06 | 0 | 0.25 | 0.05 | 0.35 | 0.19 | 0.31 | 0.32 |
| 8 | 143824483 | rs3758082 | 0.08 | -0.052 | -0.566 | 0.169 | -0.046 | 0.044 | 0.044 | 0 | 0.29 | 0.2 | 0.4 | 0.28 | 0.35 | 0.36 |
| 8 | 143828773 | rs2585184 | 0.298 | -0.114 | -0.535 | 0.228 | 0.004 | -0.006 | -0.006 | 1.557 | 0.29 | 0.28 | 0.16 | 0.42 | 0.2 | 0.28 |
| 8 | 143830359 | rs4736323 | 0.36 | 0.13 | -0.679 | 0.179 | -0.059 | 0.072 | 0.072 | 0 | 0.26 | 0.12 | 0.35 | 0.19 | 0.31 | 0.32 |
| 8 | 143842901 | rs1036385 | 0.542 | 0.401 | -0.265 | 0.221 | 0.006 | 0.003 | 0.003 | 0 | 0.35 | 0.42 | 0.38 | 0.29 | 0.38 | 0.32 |
| 8 | 143845269 | rs7822193 | 0.578 | 0.454 | -0.431 | 0.265 | 0.008 | -0.007 | -0.007 | 0 | 0.35 | 0.44 | 0.39 | 0.2 | 0.4 | 0.33 |
| 8 | 143848210 | rs1369453 | -0.025 | 0.535 | -0.76 | 0.163 | -0.008 | 0.021 | 0.021 | 0 | 0.29 | 0.21 | 0.2 | 0.2 | 0.32 | 0.33 |
| 8 | 143854600 | rs3808493 | -0.036 | -0.33 | -0.691 | 0.979 | 0.01 | -0.02 | -0.02 | 0 | 0.74 | 0.78 | 0.84 | 0.53 | 0.83 | 0.72 |
| 8 | 143859181 | rs2585126 | 0.601 | -0.48 | -0.394 | 1.291 | -0.002 | 0.048 | 0.048 | 0.527 | 0.86 | 0.86 | 0.74 | 0.53 | 0.89 | 0.77 |
| 8 | 143871387 | rs11988843 | 0.04 | -0.817 | 0.329 | 1.136 | -0.023 | 0.037 | 0.037 | 0 | 0.11 | 0.22 | 0.09 | 0.22 | 0.05 | 0.1 |
| 8 | 143879927 | rs2717611 | -0.148 | -0.677 | 0.392 | 0.972 | -0.002 | 0.012 | 0.012 | 0 | 0.45 | 0.4 | 0.48 | 0.28 | 0.47 | 0.38 |
| 8 | 143882962 | rs11783855 | -0.031 | -0.901 | 0.249 | 0.917 | -0.028 | -0.012 | -0.012 | 0 | 0.4 | 0.2 | 0.48 | 0.6 | 0.5 | 0.56 |
| 8 | 143886601 | rs10956986 | -1.2 | -0.728 | 0.705 | 1.121 | 0.001 | -0.034 | -0.034 | 0.354 | 0.63 | 0.45 | 0.59 | 0.84 | 0.6 | 0.65 |
| 8 | 143897882 | rs2717586 | -1.272 | -0.807 | 0.341 | 1.091 | -0.009 | 0.041 | 0.041 | 0 | 0.64 | 0.75 | 0.62 | 0.84 | 0.6 | 0.67 |
| 8 | 143900017 | rs2516380 | -0.004 | -0.742 | 0.291 | 1.357 | -0.011 | 0.017 | 0.017 | 0 | 0.1 | 0.15 | 0.09 | 0.24 | 0.05 | 0.08 |
| 8 | 143901987 | rs7846202 | 0.28 | -0.725 | 0.136 | 1.367 | 0.003 | -0.005 | -0.005 | 0 | 0.09 | 0.09 | 0.07 | 0.24 | 0.05 | 0.08 |
| 8 | 143902692 | rs378433 | -0.806 | -0.711 | 0.197 | 1.205 | -0.001 | 0.033 | 0.033 | 0 | 0.16 | 0.21 | 0.13 | 0.56 | 0.14 | 0.3 |
| 8 | 143903291 | rs369895 | 0.078 | -0.535 | 0.336 | 0.381 | -0.005 | 0.023 | 0.023 | 0.001 | 0.54 | 0.61 | 0.7 | 0.88 | 0.54 | 0.63 |
| 8 | 143903463 | rs2717591 | 0.278 | -0.634 | -0.083 | 0.067 | 0.008 | 0.012 | 0.012 | 0.018 | 0.62 | 0.72 | 0.64 | 0.23 | 0.69 | 0.39 |
| 8 | 143903612 | rs439747 | -0.773 | -0.297 | 0.053 | -0.324 | 0 | -0.013 | -0.013 | 0.148 | 0.6 | 0.59 | 0.62 | 0.91 | 0.6 | 0.68 |
| 8 | 143906396 | rs2003045 | 0.253 | -0.43 | -0.161 | -0.009 | 0.007 | 0.012 | 0.012 | 0 | 0.52 | 0.54 | 0.5 | 0.36 | 0.48 | 0.44 |
| 8 | 143912840 | rs7816010 | 0.633 | -0.549 | 0.183 | -0.022 | -0.019 | 0.043 | 0.043 | 0.521 | 0.1 | 0.27 | 0.04 | 0.09 | 0.04 | 0.07 |
| 8 | 143920297 | rs4736337 | -0.136 | -0.339 | -0.002 | 0.252 | 0.005 | 0.107 | 0.107 | 0 | 0.12 | 0.2 | 0.16 | 0.48 | 0.11 | 0.22 |
| 8 | 143920340 | rs4736338 | -0.4 | -0.369 | -0.094 | -0.032 | -0.005 | 0.004 | 0.004 | 0 | 0.61 | 0.66 | 0.65 | 0.86 | 0.59 | 0.66 |
| 8 | 143921804 | rs3750246 | 1.596 | -0.964 | -0.553 | -0.047 | -0.05 | 0.062 | 0.062 | 1.679 | 0.37 | 0.03 | 0.44 | 0.29 | 0.43 | 0.37 |
| 8 | 143922620 | rs3764795 | 0.853 | -0.94 | -0.314 | 0.142 | -0.012 | 0.144 | 0.144 | 0 | 0.09 | 0.17 | 0.13 | 0.48 | 0.05 | 0.21 |
| 8 | 143923250 | rs3819495 | -1.071 | -1.001 | -0.088 | -0.045 | -0.082 | 0.093 | 0.093 | 0.519 | 0.23 | 0.58 | 0.18 | 0.56 | 0.09 | 0.29 |
| 8 | 143923891 | rs3819496 | 1.601 | -0.908 | -0.535 | -0.01 | -0.051 | 0.017 | 0.017 | 2.113 | 0.37 | 0.03 | 0.44 | 0.32 | 0.42 | 0.38 |

| | | | | | | | | | | | | | | | | |
|---|---|---|---|---|---|---|---|---|---|---|---|---|---|---|---|---|
| 8 | 143928233 | rs2078674 | 0.025 | -0.853 | -0.078 | -0.1 | 0.002 | 0.064 | 0.064 | 0.329 | 0.55 | 0.34 | 0.59 | 0.84 | 0.52 | 0.65 |
| 8 | 143949604 | rs10098023 | -0.948 | -1.119 | -0.626 | -0.558 | -0.004 | 0.002 | 0.002 | 0 | 0.73 | 0.69 | 0.66 | 0.85 | 0.6 | 0.71 |
| 8 | 143950026 | rs7016088 | -0.993 | -1.114 | -0.609 | -0.556 | -0.007 | 0.004 | 0.004 | 0 | 0.74 | 0.7 | 0.67 | 0.85 | 0.6 | 0.72 |
| 8 | 143950214 | rs7005638 | -0.959 | -1.113 | -0.611 | -0.556 | -0.003 | 0.002 | 0.002 | 0 | 0.74 | 0.69 | 0.67 | 0.85 | 0.6 | 0.72 |
| 8 | 143950352 | rs7016594 | -1.131 | -1.121 | -0.57 | -0.56 | -0.075 | 0.035 | 0.035 | 0 | 0.75 | 0.84 | 0.68 | 0.85 | 0.6 | 0.72 |
| 8 | 143950501 | rs4548164 | -0.956 | -1.117 | -0.583 | -0.557 | -0.003 | 0.002 | 0.002 | 0 | 0.74 | 0.69 | 0.67 | 0.85 | 0.6 | 0.72 |
| 8 | 143950796 | rs4464947 | 0.653 | -1.164 | -0.525 | -0.554 | -0.043 | -0.106 | -0.106 | 0 | 0.17 | 0.26 | 0.06 | 0 | 0.08 | 0.07 |
| 8 | 143951849 | rs4736346 | 0.608 | -1.144 | -0.603 | -0.558 | 0.002 | 0.057 | 0.057 | 0.311 | 0.55 | 0.35 | 0.59 | 0.85 | 0.52 | 0.65 |
| 8 | 143952700 | rs4736311 | -1.088 | -1.141 | -0.59 | -0.563 | -0.062 | 0.032 | 0.032 | 0 | 0.74 | 0.81 | 0.67 | 0.85 | 0.59 | 0.72 |
| 8 | 143953937 | rs4736312 | -0.912 | -1.143 | -0.559 | -0.564 | 0.004 | -0.003 | -0.003 | 0 | 0.73 | 0.66 | 0.66 | 0.85 | 0.59 | 0.72 |
| 8 | 143954223 | rs1134096 | -0.912 | -1.143 | -0.559 | -0.564 | 0.004 | -0.003 | -0.003 | 0.014 | 0.73 | 0.66 | 0.66 | 0.85 | 0.59 | 0.71 |
| 8 | 143954290 | rs1134095 | -0.912 | -1.143 | -0.559 | -0.564 | 0.004 | -0.003 | -0.003 | 0 | 0.73 | 0.66 | 0.66 | 0.85 | 0.59 | 0.72 |
| 8 | 143954747 | rs7003319 | -0.912 | -1.143 | -0.559 | -0.564 | 0.004 | -0.003 | -0.003 | 0 | 0.73 | 0.66 | 0.66 | 0.85 | 0.59 | 0.72 |
| 8 | 143954769 | rs5017238 | -0.912 | -1.143 | -0.559 | -0.564 | 0.004 | -0.003 | -0.003 | 3.229 | 0.73 | 0.66 | 0.66 | 0.85 | 0.59 | 0.72 |
| 8 | 143955095 | rs5303 | -1.133 | -1.135 | -0.721 | -0.56 | -0.077 | 0.035 | 0.035 | 0 | 0.75 | 0.84 | 0.68 | 0.85 | 0.59 | 0.72 |
| 8 | 143955657 | rs5297 | 0.243 | -1.175 | -0.639 | -0.541 | -0.055 | -0.121 | -0.121 | 0 | 0.18 | 0.32 | 0.07 | 0 | 0.08 | 0.07 |
| 8 | 143959272 | rs9657021 | 0.855 | -1.271 | -0.68 | -0.358 | -0.025 | 0.07 | 0.07 | 2.181 | 0.11 | 0.44 | 0.05 | 0.09 | 0.04 | 0.08 |
| 8 | 143961005 | rs6410 | 0.197 | -1.367 | -0.839 | -0.39 | 0.015 | -0.012 | -0.012 | 0.367 | 0.62 | 0.42 | 0.63 | 0.75 | 0.56 | 0.64 |
| 8 | 143961293 | rs5280 | 2.189 | -1.415 | -0.901 | -0.16 | 0.018 | 0.064 | 0.064 | 0 | 0.06 | 0.02 | 0.1 | 0.44 | 0.02 | 0.19 |
| 8 | 143967432 | rs4736349 | 1.596 | -1.462 | -0.941 | -0.253 | -0.026 | 0.073 | 0.073 | 1.986 | 0.46 | 0.09 | 0.55 | 0.75 | 0.48 | 0.57 |
| 8 | 143969032 | rs4736350 | 0.058 | -1.444 | -0.878 | -0.242 | 0.012 | -0.009 | -0.009 | 0 | 0.63 | 0.38 | 0.62 | 0.75 | 0.55 | 0.63 |
| 8 | 143977819 | rs4641039 | -1.249 | -1.431 | -0.915 | -0.257 | -0.104 | 0.04 | 0.04 | 0 | 0.75 | 0.89 | 0.68 | 0.84 | 0.59 | 0.72 |
| 8 | 143982392 | rs7011830 | -1.223 | -1.427 | -0.9 | -0.256 | -0.073 | 0.034 | 0.034 | 0 | 0.75 | 0.84 | 0.68 | 0.84 | 0.59 | 0.72 |
| 8 | 143982753 | rs4736317 | -1.252 | -1.427 | -0.917 | -0.255 | -0.093 | 0.038 | 0.038 | 0 | 0.75 | 0.86 | 0.68 | 0.84 | 0.59 | 0.72 |
| 8 | 143982831 | rs4736318 | 1.62 | -1.451 | -1.067 | -0.267 | -0.031 | 0.072 | 0.072 | 1.79 | 0.45 | 0.08 | 0.55 | 0.75 | 0.48 | 0.57 |
| 8 | 143984880 | rs4736319 | -0.652 | -1.389 | -0.792 | 0.113 | -0.06 | 0.07 | 0.07 | 0 | 0.24 | 0.3 | 0.17 | 0.43 | 0.1 | 0.27 |
| 8 | 143986701 | rs6471580 | 0.893 | -1.365 | -0.79 | -0.174 | 0 | -0.002 | -0.002 | 0 | 0.56 | 0.66 | 0.62 | 0.84 | 0.52 | 0.66 |
| 8 | 143987191 | rs6987382 | 0.893 | -1.365 | -0.772 | -0.174 | 0 | 0.001 | 0.001 | 0.285 | 0.56 | 0.66 | 0.62 | 0.84 | 0.52 | 0.66 |
| 8 | 143987310 | rs4736358 | 0.298 | -1.153 | -0.738 | 0.099 | -0.012 | 0.345 | 0.345 | 0 | 0.06 | 0.23 | 0.11 | 0.5 | 0.03 | 0.21 |
| 8 | 143987427 | rs4736359 | -1.124 | -1.134 | -1.005 | 0.069 | -0.154 | 0.035 | 0.035 | 0 | 0.75 | 0.94 | 0.68 | 0.84 | 0.59 | 0.72 |
| 8 | 143992218 | rs3802228 | -0.37 | -1.154 | -0.914 | 0.086 | -0.112 | -0.029 | -0.029 | 3.021 | 0.69 | 0.89 | 0.58 | 0.34 | 0.57 | 0.51 |
| 8 | 143993640 | rs6433 | -1.154 | -1.112 | -0.929 | -0.026 | -0.148 | 0.033 | 0.033 | 0 | 0.75 | 0.94 | 0.69 | 0.84 | 0.59 | 0.72 |
| 8 | 143993765 | rs6431 | -1.154 | -1.112 | -0.929 | -0.026 | -0.148 | 0.033 | 0.033 | 0 | 0.75 | 0.94 | 0.69 | 0.84 | 0.59 | 0.72 |
| 8 | 143995743 | rs4543 | 0.141 | -1.343 | -0.648 | -0.147 | -0.047 | -0.106 | -0.106 | 0.401 | 0.18 | 0.25 | 0.06 | 0 | 0.08 | 0.07 |
| 8 | 143999600 | rs1799998 | 1.564 | -1.371 | -0.782 | -0.183 | 0.003 | 0.118 | 0.118 | 0 | 0.46 | 0.19 | 0.47 | 0.29 | 0.49 | 0.39 |
| 8 | 143999726 | rs10087214 | 1.609 | -1.374 | -0.784 | -0.185 | 0 | 0.114 | 0.114 | 0 | 0.46 | 0.19 | 0.47 | 0.29 | 0.48 | 0.37 |
| 8 | 143999901 | rs11781082 | 1.552 | -0.89 | -1.038 | -0.234 | -0.102 | 0 | 0 | 2.227 | 0.15 | 0.02 | 0.23 | 0.11 | 0.26 | 0.07 |
| 8 | 144002184 | rs7844961 | 0.812 | -1.613 | -0.553 | -0.086 | -0.035 | -0.094 | -0.094 | 0 | 0.15 | 0.21 | 0.06 | 0 | 0.08 | 0.07 |
| 8 | 144002283 | rs7831617 | 1.674 | -1.545 | -0.773 | -0.017 | 0.006 | 0.118 | 0.118 | 0 | 0.46 | 0.19 | 0.47 | 0.29 | 0.48 | 0.39 |
| 8 | 144005285 | rs7011901 | -1.215 | -1.48 | -0.794 | 0.161 | -0.003 | 0.137 | 0.137 | 0 | 0.95 | 0.94 | 0.9 | 0.48 | 0.98 | 0.81 |
| 8 | 144006073 | rs7016924 | -1.551 | -1.602 | -0.944 | 0.2 | -0.023 | 0.155 | 0.155 | 0 | 0.49 | 0.91 | 0.44 | 0.21 | 0.5 | 0.46 |
| 8 | 144007104 | rs6988985 | -0.348 | -1.966 | -1.291 | -0.349 | -0.099 | -0.021 | -0.021 | 0 | 0.63 | 0.93 | 0.54 | 0.32 | 0.51 | 0.52 |
| 8 | 144007939 | rs6981918 | 0.028 | -1.606 | -0.591 | 0.037 | 0.041 | 0.111 | 0.111 | 2.588 | 0.05 | 0.39 | 0.05 | 0.03 | 0.01 | 0.09 |
| 8 | 144008181 | rs11775009 | 0.992 | -1.595 | -0.733 | 0.02 | -0.102 | 0.046 | 0.046 | 0 | 0.27 | 0.13 | 0.31 | 0.16 | 0.41 | 0.27 |
| 8 | 144035121 | rs4074074 | -0.72 | -1.507 | -0.511 | 0.024 | -0.001 | -0.021 | -0.021 | 0 | 0.43 | 0.51 | 0.41 | 0.19 | 0.5 | 0.47 |
| 8 | 144043126 | rs9692931 | -0.78 | -1.335 | -0.45 | 0.187 | -0.02 | 0.191 | 0.191 | 0 | 0.94 | 0.86 | 0.89 | 0.48 | 0.97 | 0.84 |
| 8 | 144052791 | rs10090696 | 1.678 | -1.34 | -0.516 | 0.136 | -0.053 | 0.092 | 0.092 | 0 | 0.47 | 0.02 | 0.45 | 0.29 | 0.47 | 0.37 |
| 8 | 144055179 | rs9330408 | 1.131 | -1.325 | -0.46 | 0.153 | -0.033 | 0.024 | 0.024 | 0 | 0.75 | 0.33 | 0.79 | 0.41 | 0.89 | 0.67 |
| 8 | 144056123 | rs9694474 | -0.254 | -0.765 | -0.941 | -0.247 | 0.001 | 0.139 | 0.139 | 0 | 0.44 | 0.71 | 0.42 | 0.13 | 0.5 | 0.38 |
| 8 | 144067080 | rs13248022 | -0.184 | -1.122 | -0.825 | 0.379 | -0.014 | 0.059 | 0.059 | 0.036 | 0.43 | 0.63 | 0.28 | 0.18 | 0.4 | 0.44 |
| 8 | 144075281 | rs4424237 | 0.696 | -0.713 | -1.614 | 0.767 | -0.043 | 0.005 | 0.005 | 0 | 0.43 | 0.14 | 0.64 | 0.61 | 0.51 | 0.38 |
| 8 | 144079588 | rs12675512 | 0.696 | -0.713 | -1.614 | 0.756 | -0.043 | 0.009 | 0.009 | 0 | 0.43 | 0.15 | 0.64 | 0.63 | 0.51 | 0.39 |
| 8 | 144093864 | rs4270938 | 0.695 | -1.001 | -0.84 | 0.366 | 0.003 | 0.148 | 0.148 | 0.001 | 0.51 | 0.13 | 0.51 | 0.25 | 0.58 | 0.42 |
| 8 | 144103435 | rs4464948 | -0.01 | -0.593 | 0.413 | 0.626 | 0.013 | 0 | 0 | 0 | 0.94 | 0.71 | 0.97 | 1 | 0.98 | 0.98 |
| 8 | 144107977 | rs10283236 | 0.5 | -0.53 | 0.088 | 0.838 | -0.027 | 0.029 | 0.029 | 0 | 0.84 | 0.53 | 0.89 | 0.8 | 0.9 | 0.69 |
| 8 | 144110454 | rs4977145 | -0.373 | -0.213 | -0.243 | 1.083 | -0.015 | -0.031 | -0.031 | 0 | 0.29 | 0.21 | 0.35 | 0.54 | 0.33 | 0.21 |
| 8 | 144124512 | rs11136299 | -0.712 | -1.203 | 0.847 | 1.166 | -0.004 | -0.006 | -0.006 | 0 | 0.44 | 0.48 | 0.33 | 0.42 | 0.36 | 0.3 |
| 8 | 144124609 | rs11136300 | 0.397 | -1.296 | 0.935 | 1.211 | 0.004 | 0.064 | 0.064 | 0 | 0.59 | 0.74 | 0.69 | 0.73 | 0.64 | 0.71 |
| 8 | 144134961 | rs13248517 | 0.272 | -1.339 | 0.084 | 1.013 | -0.088 | 0.052 | 0.052 | 0 | 0.13 | 0.01 | 0.12 | 0.09 | 0.24 | 0.11 |
| 8 | 144134969 | rs7823214 | -0.16 | -1.315 | 0.447 | 1.025 | 0.068 | -0.092 | -0.092 | 0 | 0.82 | 0.69 | 0.85 | 0.91 | 0.76 | 0.86 |
| 8 | 144135387 | rs4977119 | -0.112 | -1.316 | 0.458 | 1.025 | 0.068 | -0.095 | -0.095 | 0 | 0.81 | 0.66 | 0.85 | 0.91 | 0.76 | 0.86 |
| 8 | 144137403 | rs10103088 | -0.13 | -1.319 | 0.467 | 1.023 | 0.072 | -0.097 | -0.097 | 0 | 0.81 | 0.66 | 0.85 | 0.91 | 0.76 | 0.86 |
| 8 | 144137783 | rs4977120 | -0.143 | -1.306 | 0.473 | 1.03 | 0.073 | -0.096 | -0.096 | 2.198 | 0.81 | 0.66 | 0.85 | 0.91 | 0.76 | 0.85 |
| 8 | 144146404 | rs12675829 | 1.708 | -1.644 | 0.136 | 1.256 | -0.105 | 0.119 | 0.119 | 0.105 | 0.41 | 0.05 | 0.42 | 0.18 | 0.57 | 0.4 |
| 8 | 144157565 | rs7462934 | 0.169 | -1.743 | 0.367 | 1.364 | -0.054 | 0.022 | 0.022 | 1.071 | 0.27 | 0.36 | 0.19 | 0.33 | 0.12 | 0.15 |
| 8 | 144158342 | rs7824044 | -0.395 | -1.708 | 0.184 | 1.35 | 0.069 | -0.086 | -0.086 | 0.342 | 0.82 | 0.7 | 0.85 | 0.91 | 0.76 | 0.87 |
| 8 | 144169091 | rs4246828 | 0.387 | -1.719 | -0.181 | 1.446 | -0.001 | -0.011 | -0.011 | 1.709 | 0.74 | 0.31 | 0.81 | 0.84 | 0.75 | 0.78 |
| 8 | 144170824 | rs6988480 | 0.943 | -1.555 | -0.204 | 1.527 | 0.042 | 0.014 | 0.014 | 0.21 | 0.49 | 0.14 | 0.64 | 0.5 | 0.64 | 0.62 |
| 8 | 144179460 | rs4977114 | 0.156 | -0.871 | 0.035 | 1.753 | -0.007 | 0.102 | 0.102 | 0 | 0.07 | 0.09 | 0.22 | 0.32 | 0.06 | 0.22 |

| | | | | | | | | | | | | | | | | |
|---|---|---|---|---|---|---|---|---|---|---|---|---|---|---|---|---|
| 8 | 144183966 | rs7846671 | 0.036 | -0.61 | 0.072 | 1.009 | 0.081 | -0.093 | -0.093 | 0 | 0.82 | 0.67 | 0.86 | 0.91 | 0.76 | 0.85 |
| 8 | 144201854 | rs12680363 | -0.568 | -1.252 | -0.798 | 0.781 | 0.001 | 0.003 | 0.003 | 0.036 | 0.27 | 0.24 | 0.18 | 0.21 | 0.11 | 0.16 |
| 8 | 144216966 | rs13281215 | -0.324 | -0.493 | -1.044 | 1.031 | 0.007 | -0.002 | -0.002 | 0 | 0.28 | 0.16 | 0.21 | 0.26 | 0.22 | 0.18 |
| 8 | 144218678 | rs7845556 | -0.235 | -1.059 | -0.277 | 0.858 | -0.017 | -0.003 | -0.003 | 0 | 0.9 | 0.64 | 0.95 | 0.94 | 0.96 | 0.88 |
| 8 | 144223556 | rs11990514 | 1.001 | -1.436 | -0.334 | 1.474 | 0.082 | -0.016 | -0.016 | 0 | 0.53 | 0.13 | 0.59 | 0.48 | 0.68 | 0.48 |
| 8 | 144224593 | rs11782029 | 0.596 | -1.485 | -0.002 | 1.58 | -0.054 | 0.056 | 0.056 | 0 | 0.82 | 0.41 | 0.81 | 0.74 | 0.9 | 0.66 |
| 8 | 144225931 | rs4977136 | 0.676 | -1.483 | 0.004 | 1.577 | -0.052 | 0.05 | 0.05 | 0 | 0.81 | 0.36 | 0.81 | 0.72 | 0.9 | 0.66 |
| 8 | 144226526 | rs6994704 | 0.501 | -0.764 | -0.711 | 1.467 | -0.089 | -0.078 | -0.078 | 0.247 | 0.17 | 0.01 | 0.31 | 0.27 | 0.24 | 0.28 |
| 8 | 144227728 | rs7461733 | -0.763 | -1.401 | -0.228 | 1.506 | -0.019 | 0.008 | 0.008 | 0 | 0.9 | 0.64 | 0.95 | 0.92 | 0.96 | 0.88 |
| 8 | 144229103 | rs4075410 | 0.792 | -1.262 | -0.859 | 1.517 | 0.09 | 0.029 | 0.029 | 0 | 0.52 | 0.13 | 0.58 | 0.45 | 0.68 | 0.48 |
| 8 | 144229984 | rs11782545 | -0.159 | -0.456 | -0.53 | 1.212 | 0.007 | 0.025 | 0.025 | 0 | 0.27 | 0.19 | 0.13 | 0.07 | 0.21 | 0.17 |
| 8 | 144231514 | rs4977138 | 0.911 | -0.133 | 0.143 | 2.234 | -0.009 | 0.172 | 0.172 | 0.531 | 0.92 | 0.77 | 0.86 | 0.74 | 0.94 | 0.79 |
| 8 | 144234489 | rs6420183 | 1.164 | -0.124 | 0.078 | 2.138 | -0.015 | 0.178 | 0.178 | 0.292 | 0.9 | 0.77 | 0.86 | 0.71 | 0.93 | 0.78 |
| 8 | 144236881 | rs7834727 | -1.73 | -0.263 | 0.372 | 2.476 | -0.024 | 0.216 | 0.216 | 1.757 | 0.12 | 0.25 | 0.14 | 0.38 | 0.07 | 0.28 |
| 8 | 144237145 | rs4977116 | 1.376 | -0.089 | 0.185 | 2.943 | -0.002 | -0.006 | -0.006 | 0 | 0.86 | 0.9 | 0.81 | 0.66 | 0.84 | 0.73 |
| 8 | 144239099 | rs11787024 | -0.117 | 0.176 | -0.574 | 2.034 | -0.036 | 0.02 | 0.02 | 0.077 | 0.34 | 0.18 | 0.2 | 0.26 | 0.3 | 0.15 |
| 8 | 144239660 | rs7007949 | -0.57 | -0.473 | -0.069 | 1.156 | -0.007 | 0.051 | 0.051 | 0 | 0.53 | 0.68 | 0.33 | 0.28 | 0.4 | 0.33 |
| 8 | 144239859 | rs10109061 | -0.579 | -0.433 | -0.1 | 1.144 | -0.012 | 0.057 | 0.057 | 0 | 0.53 | 0.72 | 0.33 | 0.28 | 0.4 | 0.33 |
| 8 | 144258705 | rs7829800 | -0.78 | -0.524 | -0.392 | 1.072 | -0.035 | 0.014 | 0.014 | 0.036 | 0.68 | 0.79 | 0.79 | 0.74 | 0.7 | 0.87 |
| 8 | 144258826 | rs7843980 | 1.006 | 0.007 | -1.159 | 0.579 | -0.064 | 0.063 | 0.063 | 0 | 0.18 | 0.03 | 0.16 | 0.04 | 0.28 | 0.2 |
| 8 | 144261356 | rs7386087 | 0.151 | -0.342 | -0.84 | 0.711 | 0.006 | -0.045 | -0.045 | 0 | 0.32 | 0.11 | 0.41 | 0.66 | 0.33 | 0.51 |
| 8 | 144263147 | rs11781387 | 0.204 | 0.055 | -0.157 | 0.393 | -0.014 | -0.015 | -0.015 | 0.296 | 0.4 | 0.57 | 0.34 | 0.3 | 0.3 | 0.25 |
| 8 | 144266114 | rs13255714 | 0.867 | -0.147 | 0.087 | 0.361 | -0.022 | -0.033 | -0.033 | 5.969 | 0.13 | 0.17 | 0.08 | 0.04 | 0.09 | 0.16 |
| 8 | 144280266 | rs9650460 | 0.603 | -0.426 | -0.405 | 0.587 | -0.001 | 0.041 | 0.041 | 0.027 | 0.46 | 0.34 | 0.46 | 0.4 | 0.47 | 0.3 |
| 8 | 144282591 | rs12548918 | -0.42 | -0.383 | -0.617 | 1.021 | -0.001 | 0.036 | 0.036 | 0.028 | 0.18 | 0.16 | 0.23 | 0.39 | 0.2 | 0.35 |
| 8 | 144283651 | rs7388577 | 0.645 | -0.351 | -0.804 | 1.184 | -0.02 | -0.021 | -0.021 | 0 | 0.08 | 0.01 | 0.13 | 0.32 | 0.1 | 0.19 |
| 8 | 144283990 | rs7464213 | 1.269 | -0.43 | -1.14 | 0.843 | -0.047 | -0.023 | -0.023 | 0 | 0.17 | 0.02 | 0.19 | 0.18 | 0.22 | 0.19 |
| 8 | 144284479 | rs7815797 | 1.257 | -0.437 | -1.141 | 0.84 | -0.047 | -0.023 | -0.023 | 0 | 0.17 | 0.02 | 0.2 | 0.18 | 0.22 | 0.19 |
| 8 | 144288864 | rs7386576 | 0.272 | -0.512 | -0.897 | 1.03 | 0.004 | 0.092 | 0.092 | 0.091 | 0.08 | 0.1 | 0.14 | 0.29 | 0.1 | 0.17 |
| 8 | 144294773 | rs11136268 | 1.127 | -0.495 | -1.166 | 0.987 | -0.012 | -0.013 | -0.013 | 0 | 0.08 | 0.01 | 0.09 | 0.06 | 0.1 | 0.14 |
| 8 | 144297667 | rs12545739 | 0.545 | -0.819 | -0.607 | 0.485 | -0.038 | 0.022 | 0.022 | 0.341 | 0.67 | 0.45 | 0.74 | 0.65 | 0.74 | 0.65 |
| 8 | 144303418 | rs10087900 | 0.938 | -0.494 | -1.205 | -0.008 | -0.018 | 0.012 | 0.012 | 0 | 0.43 | 0.23 | 0.43 | 0.44 | 0.45 | 0.34 |
| 8 | 144304941 | rs7460860 | -0.366 | -0.118 | -0.751 | 0.344 | 0.001 | 0.016 | 0.016 | 0 | 0.34 | 0.53 | 0.46 | 0.51 | 0.4 | 0.48 |
| 8 | 144309595 | rs6990605 | 0.141 | 0.51 | -1.282 | -0.275 | -0.014 | 0.044 | 0.044 | 0.279 | 0.33 | 0.23 | 0.2 | 0.23 | 0.3 | 0.17 |
| 8 | 144310057 | rs6987719 | 0.58 | 0.402 | -1.371 | -0.543 | 0.003 | 0.003 | 0.003 | 0.224 | 0.22 | 0.18 | 0.12 | 0.18 | 0.16 | 0.11 |
| 8 | 144312990 | rs7011172 | -0.077 | 0.511 | -0.738 | -0.573 | -0.003 | -0.012 | -0.012 | 0 | 0.31 | 0.4 | 0.17 | 0.24 | 0.24 | 0.13 |
| 8 | 144315939 | rs7386971 | 0.581 | 0.137 | -1.938 | -1.092 | -0.048 | 0.044 | 0.044 | 0.551 | 0.38 | 0.14 | 0.64 | 0.68 | 0.46 | 0.64 |
| 8 | 144316984 | rs7387173 | 0.619 | 0.141 | -2.025 | -1.089 | -0.087 | 0.03 | 0.03 | 0.706 | 0.38 | 0.06 | 0.64 | 0.68 | 0.46 | 0.64 |
| 8 | 144323692 | rs13277059 | 2.248 | 0.148 | -2.057 | -1.125 | -0.006 | -0.008 | -0.008 | 0 | 0.05 | 0.02 | 0.04 | 0.09 | 0.06 | 0.05 |
| 8 | 144327634 | rs34029908 | 2.248 | 0.148 | -2.057 | -1.125 | -0.006 | -0.008 | -0.008 | 0 | 0.05 | 0.02 | 0.04 | 0.09 | 0.06 | 0.05 |
| 8 | 144330452 | rs7464066 | 0.679 | 0.116 | -2.107 | -1.153 | -0.08 | 0.034 | 0.034 | 0.7 | 0.38 | 0.07 | 0.64 | 0.68 | 0.46 | 0.65 |
| 8 | 144333315 | rs4570175 | 0.639 | 0.035 | -1.697 | -1.199 | -0.032 | 0.042 | 0.042 | 0.335 | 0.38 | 0.16 | 0.65 | 0.68 | 0.46 | 0.65 |
| 8 | 144335295 | rs2272631 | -1.332 | 0.051 | -1.857 | -1.287 | -0.02 | 0.063 | 0.063 | 0.834 | 0.77 | 0.94 | 0.9 | 0.85 | 0.81 | 0.92 |
| 8 | 144337633 | rs7387080 | 0.7 | 0.078 | -1.773 | -1.162 | -0.08 | 0.037 | 0.037 | 0.826 | 0.38 | 0.07 | 0.64 | 0.68 | 0.46 | 0.65 |
| 8 | 144339053 | rs10090179 | 0.7 | 0.078 | -1.773 | -1.164 | -0.08 | 0.034 | 0.034 | 0.826 | 0.38 | 0.07 | 0.64 | 0.68 | 0.45 | 0.65 |
| 8 | 144344387 | rs3750213 | -1.404 | 0.068 | -1.589 | -1.006 | -0.009 | 0.028 | 0.028 | 0 | 0.82 | 0.9 | 0.91 | 0.85 | 0.86 | 0.92 |
| 8 | 144351394 | rs7819483 | 1.221 | 0.02 | -1.393 | -1.071 | -0.002 | -0.072 | -0.072 | 0.008 | 0.31 | 0.53 | 0.19 | 0.09 | 0.3 | 0.22 |
| 8 | 144357772 | rs7387207 | 0.432 | 0.047 | -1.721 | -0.971 | -0.006 | 0.075 | 0.075 | 0.115 | 0.45 | 0.26 | 0.67 | 0.68 | 0.49 | 0.66 |
| 8 | 144358381 | rs1056148 | 0.578 | 0.045 | -1.753 | -0.972 | -0.067 | 0.08 | 0.08 | 0.342 | 0.43 | 0.07 | 0.66 | 0.66 | 0.48 | 0.66 |
| 8 | 144364579 | rs11782193 | 0.528 | 0.064 | -1.748 | -0.962 | -0.027 | 0.086 | 0.086 | 0.115 | 0.44 | 0.19 | 0.66 | 0.67 | 0.48 | 0.66 |
| 8 | 144376881 | rs6651422 | 0.236 | 0.082 | -1.74 | -0.956 | -0.018 | 0.088 | 0.088 | 0.071 | 0.45 | 0.24 | 0.66 | 0.68 | 0.48 | 0.66 |
| 8 | 144379252 | rs3750217 | 0.626 | 0.085 | -1.826 | -0.95 | -0.133 | 0.045 | 0.045 | 0.328 | 0.43 | 0.04 | 0.66 | 0.67 | 0.49 | 0.66 |
| 8 | 144379934 | rs7386378 | 0.595 | 0.063 | -1.824 | -0.949 | -0.121 | 0.052 | 0.052 | 0.349 | 0.43 | 0.05 | 0.65 | 0.67 | 0.48 | 0.66 |
| 8 | 144387485 | rs1122242 | 0.757 | -0.061 | -1.894 | -0.982 | -0.133 | 0.032 | 0.032 | 0.007 | 0.41 | 0.05 | 0.63 | 0.67 | 0.46 | 0.66 |
| 8 | 144387833 | rs9644251 | 0.737 | 0.162 | -1.8 | -0.245 | -0.005 | 0 | 0 | 0 | 0.33 | 0.26 | 0.41 | 0.31 | 0.35 | 0.57 |
| 8 | 144389516 | rs11136284 | -1.172 | -0.002 | -1.015 | -0.436 | -0.026 | 0.018 | 0.018 | 0 | 0.83 | 0.94 | 0.92 | 0.84 | 0.86 | 0.93 |
| 8 | 144406627 | rs3829018 | 0.135 | 0.129 | -0.656 | -0.344 | -0.005 | -0.001 | -0.001 | 0 | 0.09 | 0.24 | 0.08 | 0.09 | 0.09 | 0.05 |
| 8 | 144411759 | rs3812436 | 0.112 | 0.237 | -0.534 | -0.632 | 0.001 | 0.01 | 0.01 | 0 | 0.51 | 0.53 | 0.64 | 0.73 | 0.57 | 0.66 |
| 8 | 144411767 | rs2467894 | 0.221 | -0.265 | -0.625 | -0.317 | -0.034 | -0.027 | -0.027 | 0 | 0.45 | 0.83 | 0.55 | 0.52 | 0.51 | 0.64 |
| 8 | 144413416 | rs2450772 | -0.248 | -0.356 | 0.999 | -0.58 | 0.022 | 0.022 | 0.022 | 0.003 | 0.11 | 0.46 | 0.09 | 0.08 | 0.11 | 0.07 |
| 8 | 144420930 | rs10866910 | 1.52 | -0.548 | 0.893 | -0.657 | 0.023 | 0.023 | 0.023 | 5.415 | 0.67 | 0.19 | 0.78 | 0.7 | 0.76 | 0.86 |
| 8 | 144425247 | rs7837655 | 0.564 | -0.486 | 0.87 | -0.665 | -0.001 | 0.006 | 0.006 | 0.398 | 0.76 | 0.79 | 0.85 | 0.78 | 0.82 | 0.91 |
| 8 | 144434556 | rs2450749 | 0.038 | -0.439 | 0.72 | -0.623 | -0.001 | 0.029 | 0.029 | 0.307 | 0.8 | 0.87 | 0.89 | 0.86 | 0.85 | 0.93 |
| 8 | 144438516 | rs1017943 | -0.483 | -0.513 | 0.805 | -0.601 | -0.018 | -0.014 | -0.014 | 0 | 0.12 | 0.64 | 0.1 | 0.08 | 0.08 | 0.06 |
| 8 | 144446285 | rs2467938 | -0.282 | -0.273 | 0.43 | -0.485 | -0.026 | -0.026 | -0.026 | 0 | 0.82 | 0.96 | 0.91 | 0.78 | 0.87 | 0.93 |
| 8 | 144449191 | rs2467950 | 0.094 | -0.036 | 0.568 | -0.399 | -0.001 | 0.026 | 0.026 | 0 | 0.81 | 0.88 | 0.88 | 0.86 | 0.85 | 0.93 |
| 8 | 144449192 | rs2450764 | 0.094 | -0.036 | 0.568 | -0.399 | -0.001 | 0.026 | 0.026 | 0 | 0.81 | 0.88 | 0.88 | 0.86 | 0.85 | 0.93 |
| 8 | 144462169 | rs3812440 | 0.339 | -0.126 | 0.332 | -0.058 | -0.02 | -0.03 | -0.03 | 0 | 0.74 | 0.58 | 0.82 | 0.78 | 0.82 | 0.91 |
| 8 | 144465285 | rs3814772 | -0.481 | 0.402 | 0.27 | 0.606 | -0.011 | -0.008 | -0.008 | 0 | 0.1 | 0.18 | 0.07 | 0.08 | 0.09 | 0.07 |

| | | | | | | | | | | | | | | | | |
|---|---|---|---|---|---|---|---|---|---|---|---|---|---|---|---|---|
| 8 | 144475424 | rs901764 | 0.248 | 0.323 | -0.028 | 0.657 | -0.02 | 0.01 | 0.01 | 0 | 0.82 | 0.7 | 0.82 | 0.76 | 0.9 | 0.86 |
| 8 | 144477853 | rs901770 | 0.408 | 0.327 | 0.014 | 0.498 | -0.024 | -0.026 | -0.026 | 0.02 | 0.89 | 0.75 | 0.92 | 0.96 | 0.92 | 0.94 |
| 8 | 144482856 | rs4873792 | -0.672 | 1.047 | -0.36 | 0.361 | -0.041 | 0.021 | 0.021 | 0.912 | 0.34 | 0.23 | 0.52 | 0.28 | 0.44 | 0.47 |
| 8 | 144483703 | rs4336628 | -0.683 | 1.2 | -0.053 | 0.471 | 0.012 | 0.002 | 0.002 | 1.61 | 0.25 | 0.25 | 0.4 | 0.25 | 0.37 | 0.39 |
| 8 | 144484673 | rs879740 | -0.31 | 1.05 | -0.174 | 0.444 | 0.001 | -0.006 | -0.006 | 0 | 0.11 | 0.25 | 0.12 | 0.1 | 0.09 | 0.16 |
| 8 | 144494963 | rs875859 | 0.421 | 1.09 | -0.501 | 0.65 | 0 | -0.004 | -0.004 | 0 | 0.54 | 0.45 | 0.37 | 0.55 | 0.5 | 0.33 |
| 8 | 144508768 | rs11783095 | -0.414 | 1.396 | -0.98 | 0.659 | -0.011 | -0.021 | -0.021 | 0 | 0.36 | 0.31 | 0.43 | 0.48 | 0.4 | 0.54 |
| 8 | 144510023 | rs2280930 | 0.553 | 1.515 | -1.164 | 0.727 | -0.058 | -0.145 | -0.145 | 0.034 | 0.22 | 0.04 | 0.16 | 0.41 | 0.24 | 0.25 |
| 8 | 144511199 | rs4874114 | 0.576 | 1.522 | -1.166 | 0.731 | -0.068 | -0.156 | -0.156 | 0.15 | 0.22 | 0.02 | 0.16 | 0.41 | 0.24 | 0.25 |
| 8 | 144514383 | rs1994448 | -0.473 | 1.362 | -1.027 | 0.72 | -0.019 | -0.038 | -0.038 | 0.167 | 0.35 | 0.28 | 0.43 | 0.45 | 0.4 | 0.57 |
| 8 | 144518049 | rs11136295 | 0.176 | 1.818 | -1.253 | 0.546 | -0.047 | -0.046 | -0.046 | 0.043 | 0.15 | 0.04 | 0.16 | 0.28 | 0.22 | 0.05 |
| 8 | 144526905 | rs10464952 | -0.319 | 1.798 | -1.235 | 0.492 | -0.011 | -0.027 | -0.027 | 0.204 | 0.37 | 0.28 | 0.45 | 0.51 | 0.37 | 0.58 |
| 8 | 144530651 | rs7835012 | 0.204 | 1.782 | -1.165 | 0.484 | 0 | 0 | 0 | 0 | 0.62 | 0.63 | 0.55 | 0.49 | 0.63 | 0.42 |
| 8 | 144543160 | rs4874137 | 0.725 | 1.41 | -0.659 | -0.134 | -0.006 | -0.019 | -0.019 | 1.314 | 0.44 | 0.47 | 0.37 | 0.18 | 0.38 | 0.36 |
| 8 | 144548806 | rs369051 | 0.388 | 1.883 | -0.473 | 0.105 | 0.02 | -0.017 | -0.017 | 0 | 0.83 | 0.75 | 0.7 | 0.9 | 0.78 | 0.65 |
| 8 | 144557036 | rs11136297 | -0.413 | 1.795 | -0.542 | -0.402 | 0.014 | -0.04 | -0.04 | 0 | 0.23 | 0.28 | 0.25 | 0.07 | 0.28 | 0.19 |
| 8 | 144558158 | rs11986221 | 0.667 | 1.034 | -0.926 | -0.916 | 0.015 | 0.02 | 0.02 | 1.057 | 0.3 | 0.24 | 0.29 | 0.18 | 0.28 | 0.39 |
| 8 | 144559165 | rs11780582 | 0.042 | 1.265 | -0.893 | -1.042 | -0.03 | 0.048 | 0.048 | 0.13 | 0.21 | 0.12 | 0.22 | 0.07 | 0.24 | 0.18 |
| 8 | 144560049 | rs13257849 | -0.662 | 1.071 | -0.511 | -0.998 | -0.015 | -0.04 | -0.04 | 0 | 0.59 | 0.49 | 0.61 | 0.78 | 0.69 | 0.78 |
| 8 | 144565905 | rs380904 | -0.118 | 1.263 | -0.196 | -1.071 | 0.007 | -0.003 | -0.003 | 0 | 0.75 | 0.71 | 0.77 | 0.75 | 0.73 | 0.82 |
| 8 | 144566969 | rs2294119 | 1.54 | 1.146 | -0.919 | -0.744 | -0.038 | -0.088 | -0.088 | 0 | 0.13 | 0.03 | 0.1 | 0.46 | 0.15 | 0.17 |
| 8 | 144568375 | rs12542047 | 1.153 | 1.252 | -1.044 | -0.982 | -0.089 | 0.059 | 0.059 | 0.638 | 0.18 | 0.01 | 0.2 | 0.07 | 0.23 | 0.2 |
| 8 | 144571431 | rs7013633 | 2.695 | 0.926 | -1.058 | -0.973 | 0.079 | 0.071 | 0.071 | 5.393 | 0.26 | 0.03 | 0.26 | 0.1 | 0.19 | 0.27 |
| 8 | 144573123 | rs11779488 | 2.345 | 1.002 | -1.082 | -0.997 | 0.03 | 0.086 | 0.086 | 2.856 | 0.28 | 0.04 | 0.27 | 0.09 | 0.26 | 0.29 |
| 8 | 144573675 | rs448340 | -0.561 | 1.363 | -0.498 | -0.407 | -0.015 | -0.029 | -0.029 | 0.929 | 0.38 | 0.3 | 0.42 | 0.5 | 0.4 | 0.52 |
| 8 | 144573910 | rs388357 | 0.166 | 1.311 | 0.09 | -0.778 | 0.017 | -0.128 | -0.128 | 0 | 0.69 | 0.44 | 0.69 | 0.93 | 0.69 | 0.68 |
| 8 | 144584780 | rs896945 | 0.395 | 1.152 | -0.196 | -0.485 | 0 | 0.003 | 0.003 | 0.174 | 0.55 | 0.57 | 0.56 | 0.67 | 0.5 | 0.62 |
| 8 | 144611729 | rs4242488 | 0.531 | 1.171 | -0.192 | -0.463 | 0 | -0.003 | -0.003 | 0 | 0.55 | 0.53 | 0.56 | 0.67 | 0.5 | 0.62 |
| 8 | 144613084 | rs2382954 | 0.698 | 1.139 | -0.138 | -0.405 | 0.031 | 0.03 | 0.03 | 0.826 | 0.13 | 0.27 | 0.19 | 0.12 | 0.15 | 0.19 |
| 8 | 144613680 | rs3889129 | 1.263 | 1.1 | -0.242 | -0.411 | 0.053 | -0.021 | -0.021 | 0.201 | 0.42 | 0.26 | 0.38 | 0.55 | 0.35 | 0.42 |
| 8 | 144620183 | rs4874147 | -0.618 | 1.287 | -0.418 | -0.592 | 0.008 | -0.061 | -0.061 | 0.112 | 0.74 | 0.69 | 0.76 | 0.93 | 0.74 | 0.79 |
| 8 | 144620342 | rs1318196 | 0.664 | 0.447 | -0.379 | -0.333 | 0.003 | 0.003 | 0.003 | 0.004 | 0.42 | 0.58 | 0.47 | 0.21 | 0.35 | 0.45 |
| 8 | 144620855 | rs4873802 | 2.256 | 0.493 | -1.021 | -0.289 | 0.08 | 0.084 | 0.084 | 5.262 | 0.26 | 0.03 | 0.25 | 0.08 | 0.2 | 0.25 |
| 8 | 144621035 | rs3750208 | 0.815 | 0.852 | -1.617 | -0.112 | -0.032 | -0.033 | -0.033 | 0 | 0.12 | 0.04 | 0.17 | 0.13 | 0.15 | 0.19 |
| 8 | 144621839 | rs964196 | 0.689 | 0.862 | -1.586 | -0.107 | 0.002 | 0.002 | 0.002 | 0.938 | 0.13 | 0.16 | 0.17 | 0.13 | 0.15 | 0.19 |
| 8 | 144623574 | rs2242093 | 1.251 | 0.357 | -1.882 | -0.029 | -0.047 | -0.108 | -0.108 | 0 | 0.14 | 0.01 | 0.1 | 0.46 | 0.16 | 0.17 |
| 8 | 144624438 | rs6991958 | 1.198 | -0.552 | -1.49 | -0.36 | 0.057 | 0.058 | 0.058 | 5.363 | 0.29 | 0.15 | 0.27 | 0.08 | 0.2 | 0.25 |
| 8 | 144641370 | rs2305492 | 0.674 | 0.731 | -1.664 | 0.419 | -0.127 | -0.121 | -0.121 | 0 | 0.2 | 0.03 | 0.28 | 0.43 | 0.34 | 0.25 |
| 8 | 144645460 | rs10099003 | 0.115 | 0.48 | -1.483 | -0.333 | -0.016 | 0.015 | 0.015 | 0 | 0.23 | 0.2 | 0.17 | 0.04 | 0.27 | 0.17 |
| 8 | 144645513 | rs13266821 | 0.81 | 0.054 | -1.84 | -0.093 | -0.021 | -0.095 | -0.095 | 0 | 0.14 | 0.01 | 0.18 | 0.31 | 0.12 | 0.22 |
| 8 | 144647928 | rs12541647 | 0.416 | -0.057 | -1.248 | -0.148 | 0 | 0.001 | 0.001 | 0 | 0.18 | 0.12 | 0.21 | 0.31 | 0.12 | 0.21 |
| 8 | 144648509 | rs12168 | 0.488 | -0.385 | -1.177 | -0.669 | -0.039 | 0.068 | 0.068 | 0 | 0.17 | 0.11 | 0.13 | 0.04 | 0.22 | 0.17 |
| 8 | 144656187 | rs12546272 | -0.489 | -1.599 | 0.244 | -0.109 | -0.027 | -0.01 | -0.01 | 0 | 0.64 | 0.56 | 0.67 | 0.76 | 0.76 | 0.6 |
| 8 | 144656754 | rs4606077 | -0.489 | -1.599 | 0.257 | -0.109 | -0.022 | -0.007 | -0.007 | 0 | 0.64 | 0.57 | 0.68 | 0.76 | 0.76 | 0.6 |
| 8 | 144657600 | rs2290416 | -0.183 | -1.898 | -0.249 | -0.249 | 0.004 | -0.035 | -0.035 | 0 | 0.14 | 0.03 | 0.07 | 0.15 | 0.09 | 0.17 |
| 8 | 144660046 | rs896954 | 1.273 | -1.938 | -0.101 | -0.284 | 0.074 | 0.064 | 0.064 | 0.485 | 0.24 | 0.07 | 0.26 | 0.09 | 0.17 | 0.24 |
| 8 | 144661564 | rs4874156 | 0.694 | -1.974 | 0.231 | 0.082 | -0.001 | 0.172 | 0.172 | 0 | 0.12 | 0.23 | 0.17 | 0.35 | 0.09 | 0.16 |
| 8 | 144662353 | rs1062391 | 1.074 | -2.016 | 0.175 | -0.009 | 0.064 | 0.019 | 0.019 | 0 | 0.46 | 0.1 | 0.45 | 0.4 | 0.62 | 0.41 |
| 8 | 144663075 | rs1809148 | -0.292 | -1.961 | 0.389 | 0.04 | -0.008 | 0.098 | 0.098 | 0 | 0.17 | 0.32 | 0.2 | 0.35 | 0.13 | 0.17 |
| 8 | 144664299 | rs3793371 | 1.291 | -1.928 | 0.24 | 0.061 | -0.01 | -0.046 | -0.046 | 0 | 0.09 | 0.03 | 0.05 | 0.15 | 0.07 | 0.16 |
| 8 | 144664594 | rs3793368 | -0.975 | -1.975 | 0.491 | 0.003 | -0.076 | -0.071 | -0.071 | 0 | 0.72 | 0.45 | 0.7 | 0.91 | 0.82 | 0.76 |
| 8 | 144669392 | rs2382962 | -0.294 | -0.933 | 1.013 | 0.552 | 0.017 | 0.25 | 0.25 | 0 | 0.88 | 0.54 | 0.81 | 0.66 | 0.92 | 0.85 |
| 8 | 144670950 | rs4874159 | 0.567 | -0.928 | 0.994 | 0.508 | -0.015 | 0.14 | 0.14 | 0.088 | 0.79 | 0.49 | 0.75 | 0.51 | 0.85 | 0.69 |
| 8 | 144671922 | rs3812448 | 0.538 | -0.927 | 1.021 | 0.508 | -0.014 | 0.137 | 0.137 | 0.061 | 0.79 | 0.48 | 0.75 | 0.51 | 0.85 | 0.69 |
| 8 | 144681777 | rs10282929 | 1.179 | -0.891 | 0.821 | 0.519 | -0.002 | 0.006 | 0.006 | 0.36 | 0.72 | 0.4 | 0.72 | 0.51 | 0.8 | 0.67 |
| 8 | 144686512 | rs755217 | -0.171 | -0.37 | 0.407 | 0.481 | 0.03 | 0.1 | 0.1 | 0 | 0.14 | 0.21 | 0.22 | 0.37 | 0.17 | 0.25 |
| 8 | 144687092 | rs896962 | -0.171 | -0.37 | 0.407 | 0.493 | 0.03 | 0.105 | 0.105 | 0 | 0.14 | 0.21 | 0.22 | 0.38 | 0.17 | 0.25 |
| 8 | 144689146 | rs2242089 | -0.382 | -0.487 | 0.274 | 0.406 | 0.017 | 0.056 | 0.056 | 0 | 0.16 | 0.18 | 0.22 | 0.37 | 0.2 | 0.27 |
| 8 | 144689289 | rs11549789 | -0.192 | -0.508 | -0.067 | 0.18 | -0.002 | -0.023 | -0.023 | 0.891 | 0.07 | 0.09 | 0.03 | 0 | 0.05 | 0.02 |
| 8 | 144690886 | rs10097337 | 1.214 | -0.641 | -0.057 | 0.249 | -0.003 | -0.021 | -0.021 | 0 | 0.66 | 0.31 | 0.67 | 0.49 | 0.68 | 0.56 |
| 8 | 144692333 | rs4403422 | -0.346 | -0.925 | -0.613 | -0.482 | 0.009 | 0.012 | 0.012 | 0 | 0.52 | 0.54 | 0.44 | 0.2 | 0.55 | 0.48 |
| 8 | 144694067 | rs4873809 | -1.109 | -1.102 | -0.642 | 0.48 | 0 | -0.006 | -0.006 | 0.001 | 0.31 | 0.69 | 0.31 | 0.67 | 0.28 | 0.46 |
| 8 | 144695499 | rs2242086 | -0.34 | -1.488 | -0.45 | 0.613 | -0.001 | 0.016 | 0.016 | 0 | 0.19 | 0.16 | 0.22 | 0.54 | 0.21 | 0.33 |
| 8 | 144697041 | rs1049832 | 1.17 | -1.598 | -1.395 | 0.096 | 0.063 | 0.185 | 0.185 | 0 | 0.49 | 0.09 | 0.46 | 0.23 | 0.62 | 0.38 |
| 8 | 144697653 | rs7817301 | 0.015 | -1.259 | -0.41 | 0.359 | -0.01 | -0.002 | -0.002 | 0.007 | 0.8 | 0.77 | 0.76 | 0.9 | 0.89 | 0.83 |
| 8 | 144722192 | rs4873814 | -0.719 | -1.031 | 0.104 | 1.322 | -0.002 | 0.125 | 0.125 | 0 | 0.14 | 0.23 | 0.2 | 0.46 | 0.15 | 0.3 |
| 8 | 144725063 | rs4873815 | -1.49 | -1.554 | 0.319 | 0.829 | 0.003 | 0.001 | 0.001 | 0 | 0.44 | 0.56 | 0.51 | 0.8 | 0.35 | 0.59 |
| 8 | 144732450 | rs3750204 | -0.791 | -1.371 | 0.407 | 0.838 | -0.014 | 0.01 | 0.01 | 0 | 0.31 | 0.36 | 0.31 | 0.37 | 0.23 | 0.27 |
| 8 | 144735347 | rs921669 | -0.056 | -1.297 | -0.133 | 0.533 | 0.009 | 0.019 | 0.019 | 0 | 0.17 | 0.04 | 0.21 | 0.11 | 0.12 | 0.1 |

| | | | | | | | | | | | | | | | | |
|---|---|---|---|---|---|---|---|---|---|---|---|---|---|---|---|---|
| 8 | 144756386 | rs12677473 | -0.869 | -1.986 | 0.105 | 0.224 | -0.003 | -0.003 | -0.003 | 0 | 0.36 | 0.41 | 0.28 | 0.76 | 0.25 | 0.54 |
| 8 | 144764119 | rs4875034 | 0.988 | -1.713 | 0.138 | 0.263 | 0.004 | 0 | 0 | 2.446 | 0.57 | 0.44 | 0.59 | 0.6 | 0.68 | 0.62 |
| 8 | 144766758 | rs28548729 | -0.724 | -2.245 | 0.048 | 0.446 | 0.019 | -0.014 | -0.014 | 2.211 | 0.31 | 0.19 | 0.32 | 0.34 | 0.24 | 0.33 |
| 8 | 144784656 | rs11136317 | -0.628 | -2.076 | -1.107 | 0.123 | 0.002 | -0.02 | -0.02 | 0 | 0.23 | 0.21 | 0.23 | 0.29 | 0.22 | 0.28 |
| 8 | 144788265 | rs7829491 | 0.606 | -2.677 | -1.515 | -0.039 | 0.079 | -0.143 | -0.143 | 0.215 | 0.22 | 0.04 | 0.08 | 0.48 | 0.1 | 0.2 |
| 8 | 144790997 | rs6980774 | 0.38 | -2.612 | -1.545 | -0.246 | -0.004 | -0.036 | -0.036 | 0 | 0.75 | 0.81 | 0.77 | 0.68 | 0.74 | 0.56 |
| 8 | 144791126 | rs10100154 | 0.389 | -2.621 | -1.551 | -0.246 | -0.003 | -0.036 | -0.036 | 0 | 0.75 | 0.8 | 0.77 | 0.68 | 0.74 | 0.56 |
| 8 | 144799465 | rs11776672 | 0.478 | -3.427 | -1.635 | -0.602 | 0.16 | -0.111 | -0.111 | 0 | 0.61 | 0.13 | 0.63 | 0.48 | 0.81 | 0.68 |
| 8 | 144801681 | rs11778657 | 0.546 | -3.281 | -1.86 | -0.661 | 0.149 | -0.135 | -0.135 | 0 | 0.57 | 0.09 | 0.62 | 0.44 | 0.78 | 0.65 |
| 8 | 144807456 | rs4072459 | 0.238 | -3.064 | -1.173 | -0.565 | 0.003 | -0.006 | -0.006 | 0.991 | 0.65 | 0.38 | 0.68 | 0.48 | 0.8 | 0.72 |
| 8 | 144822037 | rs7831467 | -0.047 | -2.785 | -1.99 | -0.443 | 0.064 | -0.078 | -0.078 | 0 | 0.3 | 0.12 | 0.27 | 0.52 | 0.18 | 0.28 |
| 8 | 144870701 | rs11136325 | 0.525 | -1.997 | -2.33 | -1.377 | -0.064 | 0.295 | 0.295 | 0 | 0.38 | 0.02 | 0.44 | 0.01 | 0.58 | 0.24 |
| 8 | 144880924 | rs7822430 | 0.345 | -2.015 | -1.891 | -1.427 | 0.039 | 0.305 | 0.305 | 0 | 0.45 | 0.11 | 0.47 | 0.02 | 0.6 | 0.27 |
| 8 | 144882423 | rs4874100 | 0.271 | -1.604 | -1.612 | -1.212 | 0.006 | -0.009 | -0.009 | 0 | 0.91 | 0.87 | 0.94 | 0.98 | 0.9 | 0.96 |
| 8 | 144882710 | rs11786637 | 0.514 | -1.625 | -1.483 | -1.179 | 0.027 | -0.015 | -0.015 | 0.042 | 0.91 | 0.77 | 0.92 | 0.95 | 0.89 | 0.96 |
| 8 | 144893084 | rs6558394 | 0.108 | -2.081 | -0.945 | -0.914 | -0.001 | 0 | 0 | 0 | 0.53 | 0.47 | 0.67 | 0.35 | 0.63 | 0.62 |
| 8 | 144894496 | rs4875060 | 0.085 | -1.906 | -1.224 | -0.884 | 0.029 | -0.018 | -0.018 | 0 | 0.92 | 0.8 | 0.93 | 0.98 | 0.9 | 0.97 |
| 8 | 144905947 | rs6985555 | 0.5 | -2.012 | -0.93 | -0.843 | 0.004 | -0.004 | -0.004 | 0 | 0.34 | 0.34 | 0.3 | 0.62 | 0.28 | 0.28 |
| 8 | 144906112 | rs4875065 | -0.183 | -1.676 | -2.142 | -0.853 | 0.021 | -0.009 | -0.009 | 0 | 0.93 | 0.86 | 0.94 | 0.98 | 0.9 | 0.96 |
| 8 | 144912303 | rs13252406 | 0.76 | -1.756 | -2.33 | -0.942 | -0.083 | 0.289 | 0.289 | 0 | 0.37 | 0.02 | 0.44 | 0.01 | 0.56 | 0.26 |
| 8 | 144927272 | rs11778556 | -0.31 | -0.835 | -1.68 | -0.264 | 0.04 | -0.012 | -0.012 | 0 | 0.63 | 0.27 | 0.67 | 0.37 | 0.72 | 0.72 |
| 8 | 144928660 | rs6985307 | 0.017 | -0.254 | -1.996 | 0.004 | 0.061 | 0.048 | 0.048 | 0 | 0.51 | 0.11 | 0.58 | 0.34 | 0.57 | 0.67 |
| 8 | 144929811 | rs11779128 | 0.054 | -0.251 | -1.983 | 0.019 | 0.072 | 0.055 | 0.055 | 0 | 0.5 | 0.11 | 0.58 | 0.34 | 0.57 | 0.66 |
| 8 | 144941181 | rs7839934 | -0.042 | 0.561 | -1.563 | 0.242 | 0.021 | 0.019 | 0.019 | 0.003 | 0.34 | 0.23 | 0.3 | 0.27 | 0.26 | 0.41 |
| 8 | 144948861 | rs9314419 | 0.1 | 1.292 | -1.227 | 1.054 | -0.028 | 0.072 | 0.072 | 0 | 0.8 | 0.74 | 0.87 | 0.57 | 0.87 | 0.74 |
| 8 | 144950720 | rs7008775 | -0.805 | 2.054 | -1.571 | 0.657 | 0.041 | 0 | 0 | 0 | 0.05 | 0.07 | 0.19 | 0.04 | 0.19 | 0.16 |
| 8 | 144956400 | rs4977189 | -0.948 | 1.494 | -2.004 | 0.602 | -0.036 | -0.009 | -0.009 | 0 | 0.31 | 0.21 | 0.38 | 0.34 | 0.36 | 0.42 |
| 8 | 144956464 | rs4977188 | -0.39 | -0.204 | -2.036 | 0.72 | 0.052 | -0.027 | -0.027 | 0 | 0.21 | 0.11 | 0.16 | 0.29 | 0.07 | 0.22 |
| 8 | 145677011 | rs2721195 | 0.745 | 1.755 | 0.778 | 1.849 | 0.012 | -0.004 | -0.004 | 0 | 0.63 | 0.59 | 0.4 | 0.25 | 0.52 | 0.42 |
| 8 | 145682115 | rs748193 | -0.863 | 1.767 | 0.655 | 1.924 | -0.116 | -0.104 | -0.104 | 2.053 | 0.34 | 0.09 | 0.51 | 0.56 | 0.48 | 0.57 |
| 8 | 145687475 | rs10108150 | -0.865 | 1.771 | 0.614 | 1.939 | -0.097 | -0.085 | -0.085 | 0.02 | 0.34 | 0.13 | 0.51 | 0.55 | 0.48 | 0.57 |
| 8 | 145701453 | rs750472 | 0.881 | 1.676 | 1.003 | 1.859 | 0.009 | -0.004 | -0.004 | 0.004 | 0.62 | 0.59 | 0.38 | 0.25 | 0.5 | 0.42 |
| 8 | 145705213 | rs4925810 | 1.288 | 1.613 | 1.382 | 1.77 | 0.04 | 0.164 | 0.164 | 0.758 | 0.88 | 0.55 | 0.81 | 0.74 | 0.92 | 0.98 |
| 8 | 145724275 | rs3816732 | -0.899 | 1.634 | 1.031 | 1.603 | -0.106 | -0.132 | -0.132 | 1.731 | 0.35 | 0.09 | 0.5 | 0.54 | 0.46 | 0.57 |
| 8 | 145728735 | rs13254911 | -0.937 | 1.572 | 0.936 | 1.571 | -0.135 | -0.162 | -0.162 | 0 | 0.36 | 0.06 | 0.49 | 0.53 | 0.46 | 0.57 |
| 8 | 145785009 | rs11997608 | -1.079 | 1.132 | 1.743 | 1.139 | -0.024 | -0.037 | -0.037 | 0.086 | 0.09 | 0.31 | 0.14 | 0.01 | 0.02 | 0.04 |
| 8 | 145818782 | rs11988651 | -1.062 | 1.507 | 1.482 | 1.179 | -0.003 | -0.033 | -0.033 | 0.017 | 0.09 | 0.36 | 0.14 | 0.01 | 0.02 | 0.04 |
| 8 | 145846570 | rs7010259 | 0.646 | 2.146 | 1.304 | 1.953 | 0.002 | 0.013 | 0.013 | 2.625 | 0.74 | 0.72 | 0.52 | 0.47 | 0.68 | 0.59 |
| 9 | 279753 | rs10967642 | -1.84 | 0.319 | 0.976 | -1.828 | 0.003 | -0.055 | -0.055 | 0 | 0.11 | 0.35 | 0.14 | 0.01 | 0.06 | 0.09 |
| 9 | 286593 | rs529208 | 1.93 | 0.927 | 1.195 | -2.057 | 0.004 | 0.028 | 0.028 | 0 | 0.54 | 0.39 | 0.52 | 0.72 | 0.48 | 0.57 |
| 9 | 289061 | rs6476030 | -0.97 | 1.326 | 1.365 | -1.863 | 0.014 | -0.012 | -0.012 | 3.086 | 0.05 | 0.18 | 0.1 | 0.02 | 0.03 | 0.05 |
| 9 | 290670 | rs7030730 | -1.806 | 1.383 | 1.408 | -1.944 | -0.005 | 0.004 | 0.004 | 0 | 0.38 | 0.61 | 0.44 | 0.25 | 0.44 | 0.34 |
| 9 | 293360 | rs943628 | -1.145 | 1.577 | 1.384 | -1.959 | 0.026 | -0.063 | -0.063 | 0 | 0.19 | 0.31 | 0.31 | 0.05 | 0.24 | 0.16 |
| 9 | 299315 | rs785857 | 1.452 | 1.413 | 1.058 | -1.932 | 0.026 | -0.011 | -0.011 | 0 | 0.7 | 0.6 | 0.61 | 0.76 | 0.61 | 0.72 |
| 9 | 303727 | rs10757847 | 2.011 | 1.397 | 0.87 | -1.974 | 0.02 | -0.027 | -0.027 | 0 | 0.62 | 0.53 | 0.57 | 0.75 | 0.53 | 0.65 |
| 9 | 316515 | rs7024448 | -0.993 | 0.311 | 0.579 | -1.973 | -0.008 | -0.067 | -0.067 | 0 | 0.27 | 0.51 | 0.37 | 0.13 | 0.26 | 0.17 |
| 9 | 318153 | rs1980876 | -1.665 | -0.183 | 0.629 | -2.614 | -0.003 | 0.042 | 0.042 | 0 | 0.53 | 0.66 | 0.5 | 0.14 | 0.46 | 0.35 |
| 9 | 318207 | rs4294238 | 0.524 | -0.029 | 0.674 | -2.523 | -0.001 | -0.044 | -0.044 | 0.451 | 0.24 | 0.19 | 0.13 | 0.01 | 0.19 | 0.18 |
| 9 | 326767 | rs1556022 | 1.739 | -0.145 | 0.033 | -2.683 | -0.002 | 0.003 | 0.003 | 0 | 0.34 | 0.26 | 0.39 | 0.7 | 0.41 | 0.51 |
| 9 | 329462 | rs10970611 | 0.073 | -0.181 | 0.148 | -2.723 | 0.012 | -0.025 | -0.025 | 0 | 0.18 | 0.33 | 0.23 | 0.12 | 0.21 | 0.09 |
| 9 | 329830 | rs10970638 | 0.493 | -0.148 | 0.23 | -2.692 | 0.001 | -0.122 | -0.122 | 0.338 | 0.29 | 0.36 | 0.25 | 0.01 | 0.24 | 0.21 |
| 9 | 331490 | rs10813766 | 0.744 | -0.17 | -0.155 | -2.484 | 0.004 | 0.009 | 0.009 | 0 | 0.62 | 0.61 | 0.64 | 0.71 | 0.65 | 0.77 |
| 9 | 334337 | rs10970979 | 0.187 | -0.025 | -0.078 | -2.73 | 0.004 | 0.05 | 0.05 | 0 | 0.34 | 0.21 | 0.21 | 0.04 | 0.27 | 0.28 |
| 9 | 334708 | rs12335694 | -0.444 | -0.056 | 0.566 | -2.587 | 0.007 | -0.015 | -0.015 | 0 | 0.08 | 0.34 | 0.13 | 0 | 0.05 | 0.04 |
| 9 | 335387 | rs17722518 | -0.405 | 0.064 | 0.18 | -2.539 | 0.005 | 0.005 | 0.005 | 0.768 | 0.07 | 0.15 | 0.06 | 0.08 | 0.08 | 0.01 |
| 9 | 335545 | rs7854418 | 1.463 | 0.173 | -0.059 | -2.644 | -0.001 | 0.002 | 0.002 | 0.01 | 0.35 | 0.31 | 0.5 | 0.71 | 0.45 | 0.53 |
| 9 | 336026 | rs12377330 | 0.065 | -0.268 | -0.103 | -2.81 | -0.001 | 0.121 | 0.121 | 0.127 | 0.38 | 0.16 | 0.23 | 0.04 | 0.29 | 0.32 |
| 9 | 336186 | rs4741814 | 0.551 | -0.05 | -0.212 | -2.294 | -0.001 | -0.001 | -0.001 | 0 | 0.4 | 0.4 | 0.56 | 0.79 | 0.53 | 0.55 |
| 9 | 344508 | rs12347078 | -1.985 | -0.485 | -0.123 | -2.242 | 0.01 | -0.005 | -0.005 | 0 | 0.14 | 0.84 | 0.1 | 0 | 0.03 | 0.06 |
| 9 | 345900 | rs13292760 | 0.462 | -0.508 | -0.373 | -2.699 | -0.014 | 0.093 | 0.093 | 0 | 0.24 | 0.09 | 0.1 | 0.01 | 0.18 | 0.18 |
| 9 | 347559 | rs1410216 | 0.696 | -0.85 | 0.521 | -2.891 | -0.022 | -0.048 | -0.048 | 0.989 | 0.79 | 0.62 | 0.83 | 0.85 | 0.85 | 0.87 |
| 9 | 348772 | rs13289414 | 0.572 | -0.804 | 0.819 | -2.873 | 0 | 0.024 | 0.024 | 0 | 0.24 | 0.16 | 0.11 | 0.01 | 0.18 | 0.18 |
| 9 | 349518 | rs7855377 | -0.424 | -0.933 | 1.326 | -2.989 | -0.003 | -0.085 | -0.085 | 0 | 0.34 | 0.62 | 0.26 | 0.01 | 0.25 | 0.25 |
| 9 | 349557 | rs7855392 | -0.494 | -0.928 | 1.384 | -2.987 | 0.002 | -0.041 | -0.041 | 0 | 0.35 | 0.66 | 0.27 | 0.01 | 0.24 | 0.25 |
| 9 | 354334 | rs943625 | 0.565 | -0.918 | 1.683 | -2.975 | 0 | -0.035 | -0.035 | 0 | 0.24 | 0.24 | 0.11 | 0.01 | 0.18 | 0.18 |
| 9 | 354680 | rs1887528 | 1.2 | -0.972 | 1.539 | -2.936 | 0.021 | 0.124 | 0.124 | 1.447 | 0.49 | 0.3 | 0.63 | 0.82 | 0.62 | 0.66 |
| 9 | 355718 | rs4741827 | 0.773 | -0.689 | 1.684 | -2.767 | -0.017 | -0.018 | -0.018 | 0 | 0.79 | 0.67 | 0.84 | 0.82 | 0.86 | 0.86 |
| 9 | 355793 | rs4741828 | 0.945 | -0.65 | 1.283 | -2.712 | 0.013 | 0.074 | 0.074 | 0 | 0.54 | 0.35 | 0.64 | 0.81 | 0.64 | 0.67 |
| 9 | 356655 | rs10972190 | 1.047 | -0.624 | 1.202 | -2.718 | 0.022 | 0.122 | 0.122 | 0 | 0.53 | 0.29 | 0.64 | 0.82 | 0.64 | 0.67 |

| | | | | | | | | | | | | | | | | |
|---|---|---|---|---|---|---|---|---|---|---|---|---|---|---|---|---|
| 9 | 357653 | rs12551785 | 0.598 | -0.536 | 0.687 | -2.771 | -0.024 | 0.169 | 0.169 | 0.063 | 0.26 | 0.02 | 0.17 | 0.01 | 0.18 | 0.2 |
| 9 | 359338 | rs10972266 | 0.211 | -0.609 | 1.126 | -2.748 | -0.001 | -0.024 | -0.024 | 1.883 | 0.3 | 0.27 | 0.27 | 0.01 | 0.26 | 0.25 |
| 9 | 362209 | rs4741833 | 0.942 | -0.552 | 0.992 | -2.71 | 0.006 | 0.263 | 0.263 | 0 | 0.53 | 0.15 | 0.63 | 0.85 | 0.65 | 0.67 |
| 9 | 362917 | rs750587 | 0.718 | -0.491 | 1.965 | -2.595 | 0.013 | -0.014 | -0.014 | 0 | 0.84 | 0.54 | 0.86 | 0.85 | 0.87 | 0.87 |
| 9 | 366281 | rs11791465 | 0.535 | -0.631 | 2.493 | -2.715 | 0.01 | 0.022 | 0.022 | 0 | 0.27 | 0.16 | 0.2 | 0.01 | 0.21 | 0.19 |
| 9 | 366911 | rs10814269 | -0.337 | -0.643 | 2.632 | -2.735 | 0.001 | -0.046 | -0.046 | 0 | 0.34 | 0.65 | 0.3 | 0.01 | 0.25 | 0.22 |
| 9 | 367030 | rs10972564 | -1.399 | -0.538 | 2.591 | -2.805 | -0.011 | -0.074 | -0.074 | 0.577 | 0.16 | 0.69 | 0.15 | 0.03 | 0.09 | 0.2 |
| 9 | 370244 | rs913703 | -0.154 | -0.356 | 2.959 | -2.476 | 0.013 | 0.017 | 0.017 | 0 | 0.32 | 0.29 | 0.21 | 0.08 | 0.23 | 0.25 |
| 9 | 370439 | rs751352 | -0.397 | -0.344 | 2.834 | -2.475 | -0.002 | -0.05 | -0.05 | 0 | 0.36 | 0.61 | 0.27 | 0.08 | 0.26 | 0.27 |
| 9 | 372341 | rs4436173 | -2.127 | -0.291 | 2.841 | -2.419 | 0.079 | 0.022 | 0.022 | 1.655 | 0.06 | 0.39 | 0.06 | 0 | 0.03 | 0.03 |
| 9 | 376538 | rs10973014 | -0.613 | -0.315 | 2.257 | -2.448 | 0.022 | 0.128 | 0.128 | 0.318 | 0.38 | 0.77 | 0.3 | 0.08 | 0.28 | 0.28 |
| 9 | 384134 | rs954872 | 0.374 | -0.144 | 2.063 | -2.1 | -0.018 | 0 | 0 | 0 | 0.89 | 0.8 | 0.93 | 0.91 | 0.94 | 0.96 |
| 9 | 384150 | rs954871 | 0.152 | -0.126 | 2.017 | -2.091 | -0.015 | 0.002 | 0.002 | 0 | 0.9 | 0.82 | 0.93 | 0.91 | 0.94 | 0.96 |
| 9 | 388167 | rs10814626 | -0.596 | 0.124 | 2.338 | -2.122 | -0.001 | -0.046 | -0.046 | 2.08 | 0.34 | 0.65 | 0.29 | 0.08 | 0.28 | 0.26 |
| 9 | 389285 | rs10814632 | -0.599 | 0.111 | 1.983 | -2.144 | 0.042 | 0.111 | 0.111 | 0.591 | 0.35 | 0.79 | 0.28 | 0.08 | 0.25 | 0.24 |
| 9 | 390228 | rs7028802 | -0.537 | 0.239 | 1.862 | -2.084 | 0.011 | -0.01 | -0.01 | 0 | 0.07 | 0.15 | 0.13 | 0.07 | 0.08 | 0.11 |
| 9 | 390512 | rs2297075 | -0.033 | 0.276 | 2.127 | -2.016 | -0.011 | -0.073 | -0.073 | 0.403 | 0.31 | 0.4 | 0.25 | 0.1 | 0.25 | 0.24 |
| 9 | 395250 | rs10739001 | -0.475 | 0.593 | 0.949 | -1.642 | -0.008 | 0.105 | 0.105 | 0 | 0.47 | 0.71 | 0.54 | 0.76 | 0.48 | 0.44 |
| 9 | 396232 | rs10973946 | -0.859 | 0.563 | 0.435 | -1.723 | 0.001 | 0.175 | 0.175 | 0 | 0.52 | 0.82 | 0.61 | 0.78 | 0.55 | 0.55 |
| 9 | 396293 | rs10973952 | 0.002 | 0.516 | 0.419 | -1.75 | 0.033 | 0.097 | 0.097 | 0.241 | 0.34 | 0.76 | 0.27 | 0.1 | 0.24 | 0.23 |
| 9 | 396380 | rs10814731 | -0.265 | 0.661 | 0.004 | -1.575 | -0.031 | 0.11 | 0.11 | 0 | 0.45 | 0.2 | 0.52 | 0.78 | 0.5 | 0.52 |
| 9 | 398673 | rs12346176 | -0.322 | 0.021 | 0.277 | -1.384 | -0.007 | -0.12 | -0.12 | 0.089 | 0.34 | 0.46 | 0.29 | 0.1 | 0.31 | 0.28 |
| 9 | 407227 | rs4741867 | -1.054 | 0.153 | 0.473 | 0.06 | 0.007 | 0 | 0 | 0.987 | 0.4 | 0.55 | 0.52 | 0.51 | 0.47 | 0.44 |
| 9 | 413241 | rs12235927 | -0.916 | 0.211 | 0.639 | 0.168 | 0.001 | 0.033 | 0.033 | 0 | 0.31 | 0.46 | 0.43 | 0.49 | 0.35 | 0.4 |
| 9 | 418003 | rs10814836 | -0.864 | -0.337 | 1.133 | 0.396 | -0.037 | 0.315 | 0.315 | 0 | 0.14 | 0.44 | 0.19 | 0.47 | 0.11 | 0.23 |
| 9 | 419153 | rs10758556 | 1.105 | -0.603 | 0.745 | 0.345 | -0.004 | -0.003 | -0.003 | 0 | 0.55 | 0.51 | 0.49 | 0.42 | 0.57 | 0.52 |
| 9 | 422157 | rs10217719 | 0.765 | -1.039 | 0.471 | -0.493 | 0.012 | 0.038 | 0.038 | 0 | 0.6 | 0.32 | 0.65 | 0.95 | 0.66 | 0.65 |
| 9 | 423052 | rs7853977 | 1.639 | -1.246 | 0.327 | -0.049 | 0 | 0.004 | 0.004 | 0 | 0.46 | 0.26 | 0.46 | 0.48 | 0.54 | 0.48 |
| 9 | 424327 | rs6476810 | -0.408 | -1.238 | 1.116 | 0.044 | -0.015 | 0.124 | 0.124 | 0 | 0.13 | 0.23 | 0.2 | 0.47 | 0.11 | 0.17 |
| 9 | 424875 | rs10814854 | -1.27 | -1.464 | 0.905 | -0.136 | -0.013 | 0.118 | 0.118 | 0 | 0.23 | 0.41 | 0.32 | 0.45 | 0.18 | 0.27 |
| 9 | 425228 | rs10733505 | 0.402 | -1.629 | 1.094 | -0.41 | -0.023 | -0.173 | -0.173 | 0 | 0.69 | 0.52 | 0.72 | 0.99 | 0.74 | 0.67 |
| 9 | 426215 | rs12353411 | -0.15 | -1.59 | 0.797 | -0.39 | 0 | -0.04 | -0.04 | 0 | 0.07 | 0.15 | 0.06 | 0 | 0.07 | 0.02 |
| 9 | 428706 | rs2297082 | 2.264 | -1.591 | 0.525 | -0.318 | -0.083 | -0.102 | -0.102 | 0.476 | 0.44 | 0.06 | 0.43 | 0.53 | 0.55 | 0.38 |
| 9 | 431860 | rs1329376 | 1.132 | -1.583 | 0.955 | -0.757 | -0.009 | -0.096 | -0.096 | 0 | 0.67 | 0.45 | 0.81 | 0.98 | 0.77 | 0.68 |
| 9 | 432918 | rs7025078 | 0.296 | -1.592 | 1.094 | -0.772 | -0.034 | -0.153 | -0.153 | 3.022 | 0.74 | 0.52 | 0.84 | 0.98 | 0.84 | 0.71 |
| 9 | 433048 | rs10758582 | 1.704 | -1.588 | 1.123 | -0.768 | 0.042 | 0.222 | 0.222 | 0.243 | 0.57 | 0.18 | 0.73 | 0.98 | 0.7 | 0.65 |
| 9 | 433347 | rs7040842 | 0.998 | -1.386 | 1.027 | -0.7 | -0.009 | -0.127 | -0.127 | 0.051 | 0.68 | 0.51 | 0.81 | 0.98 | 0.76 | 0.67 |
| 9 | 438295 | rs4740757 | 1.408 | -1.406 | 0.927 | -0.664 | 0.016 | 0.124 | 0.124 | 0.014 | 0.62 | 0.21 | 0.73 | 0.96 | 0.7 | 0.66 |
| 9 | 438779 | rs10974442 | -1.178 | -1.456 | 1.325 | -0.669 | 0.012 | -0.047 | -0.047 | 0 | 0.09 | 0.29 | 0.07 | 0 | 0.07 | 0.01 |
| 9 | 440683 | rs10758602 | 1.054 | -1.487 | 1.247 | -0.728 | 0.012 | 0 | 0 | 0 | 0.69 | 0.3 | 0.78 | 0.96 | 0.78 | 0.71 |
| 9 | 442030 | rs1887958 | -1.72 | -1.689 | 1.188 | -0.407 | -0.008 | 0.024 | 0.024 | 0 | 0.32 | 0.4 | 0.43 | 0.41 | 0.29 | 0.44 |
| 9 | 444116 | rs10974488 | -0.117 | -1.526 | 1.36 | -0.455 | 0.026 | -0.027 | -0.027 | 0 | 0.07 | 0.2 | 0.05 | 0 | 0.07 | 0.01 |
| 9 | 447887 | rs4741955 | 1.683 | -1.547 | 1.274 | -0.519 | 0.027 | 0.188 | 0.188 | 0.05 | 0.61 | 0.2 | 0.76 | 0.98 | 0.69 | 0.61 |
| 9 | 447960 | rs16908874 | -1.025 | -1.573 | 1.39 | -0.529 | 0.01 | -0.031 | -0.031 | 0.394 | 0.08 | 0.21 | 0.06 | 0.01 | 0.07 | 0.2 |
| 9 | 448429 | rs4741958 | 1.057 | -1.563 | 1.371 | -0.526 | -0.005 | -0.081 | -0.081 | 7.441 | 0.69 | 0.41 | 0.82 | 1 | 0.76 | 0.81 |
| 9 | 449373 | rs10974514 | -1.838 | -1.612 | 1.47 | -0.512 | -0.004 | -0.077 | -0.077 | 0 | 0.2 | 0.32 | 0.12 | 0.02 | 0.18 | 0.26 |
| 9 | 449798 | rs10491684 | -0.248 | -1.517 | 1.472 | -0.482 | 0.006 | -0.041 | -0.041 | 0 | 0.07 | 0.17 | 0.05 | 0 | 0.08 | 0.01 |
| 9 | 450420 | rs10814967 | 1.476 | -1.522 | 0.85 | -0.49 | 0.018 | 0.089 | 0.089 | 0 | 0.62 | 0.28 | 0.77 | 0.99 | 0.7 | 0.61 |
| 9 | 452926 | rs16909299 | -1.439 | -1.388 | 0.88 | -0.396 | 0.003 | -0.042 | -0.042 | 0 | 0.14 | 0.21 | 0.1 | 0.01 | 0.14 | 0.21 |
| 9 | 457305 | rs12682895 | -0.614 | -1.463 | 0.937 | 0.02 | 0.002 | 0.011 | 0.011 | 0 | 0.17 | 0.19 | 0.32 | 0.41 | 0.13 | 0.2 |
| 9 | 459024 | rs7853027 | 1.335 | -1.428 | 0.822 | -0.147 | 0.012 | 0.06 | 0.06 | 0.155 | 0.66 | 0.28 | 0.77 | 0.96 | 0.75 | 0.58 |
| 9 | 460072 | rs10758625 | 1.794 | -1.469 | 1.169 | 0.011 | 0.024 | -0.008 | -0.008 | 0 | 0.57 | 0.38 | 0.5 | 0.59 | 0.69 | 0.61 |
| 9 | 460877 | rs10814999 | 0.492 | -1.429 | 1.305 | -0.375 | -0.021 | -0.141 | -0.141 | 0 | 0.74 | 0.52 | 0.81 | 0.98 | 0.82 | 0.8 |
| 9 | 464920 | rs3780338 | 0.223 | -1.499 | 1.196 | -0.37 | -0.001 | -0.041 | -0.041 | 1.14 | 0.78 | 0.84 | 0.84 | 0.97 | 0.83 | 0.8 |
| 9 | 466816 | rs4742002 | 0.172 | -1.557 | 1.178 | -0.369 | 0 | -0.037 | -0.037 | 0.575 | 0.78 | 0.85 | 0.88 | 1 | 0.83 | 0.81 |
| 9 | 468705 | rs10974637 | -1.603 | -1.506 | 1.001 | -0.344 | -0.005 | -0.018 | -0.018 | 0 | 0.08 | 0.16 | 0.05 | 0.02 | 0.06 | 0.2 |
| 9 | 469245 | rs1329381 | 1.085 | -1.6 | 1.041 | -0.414 | 0.017 | -0.01 | -0.01 | 0 | 0.7 | 0.31 | 0.79 | 0.97 | 0.83 | 0.75 |
| 9 | 469265 | rs10974644 | -0.574 | -1.462 | 1.019 | -0.33 | 0.007 | -0.008 | -0.008 | 0 | 0.05 | 0.13 | 0.04 | 0.02 | 0.05 | 0.19 |
| 9 | 469767 | rs10815034 | 1.702 | -1.581 | 0.873 | -0.422 | 0.064 | 0.165 | 0.165 | 0.003 | 0.6 | 0.19 | 0.73 | 0.95 | 0.77 | 0.51 |
| 9 | 473379 | rs7867447 | -2.4 | -1.191 | 1.69 | 0.319 | -0.033 | -0.053 | -0.053 | 0.192 | 0.17 | 0.73 | 0.15 | 0.03 | 0.06 | 0.17 |
| 9 | 478153 | rs2282039 | -0.705 | -0.71 | 0.978 | 1.112 | -0.019 | -0.1 | -0.1 | 0.003 | 0.13 | 0.01 | 0.32 | 0.46 | 0.14 | 0.19 |
| 9 | 478530 | rs2282038 | -1.588 | -0.745 | 1.428 | 1.101 | -0.027 | 0.106 | 0.106 | 0.091 | 0.23 | 0.5 | 0.39 | 0.48 | 0.19 | 0.35 |
| 9 | 483018 | rs4742040 | -0.808 | -0.364 | 0.978 | 2.043 | 0.015 | -0.042 | -0.042 | 0 | 0.11 | 0.06 | 0.2 | 0.42 | 0.08 | 0.16 |
| 9 | 483798 | rs2360717 | -1.696 | -0.363 | 2.217 | 2.229 | -0.013 | 0.215 | 0.215 | 1 | 0.16 | 0.35 | 0.25 | 0.44 | 0.15 | 0.31 |
| 9 | 484745 | rs4742043 | -1.996 | -0.366 | 3.218 | 2.435 | -0.037 | 0.155 | 0.155 | 0.607 | 0.18 | 0.4 | 0.24 | 0.4 | 0.13 | 0.3 |
| 9 | 484786 | rs4742044 | -1.407 | -0.444 | 2.73 | 1.962 | -0.01 | 0.09 | 0.09 | 0.717 | 0.15 | 0.24 | 0.23 | 0.42 | 0.15 | 0.31 |
| 9 | 489338 | rs10758653 | -1.55 | -0.505 | 2.573 | 1.792 | -0.022 | 0.096 | 0.096 | 0.012 | 0.22 | 0.32 | 0.27 | 0.44 | 0.18 | 0.36 |
| 9 | 500979 | rs10118798 | -3.192 | -0.391 | 3.115 | 1.062 | -0.03 | -0.051 | -0.051 | 0 | 0.11 | 0.77 | 0.1 | 0 | 0.06 | 0.13 |
| 9 | 512589 | rs1410968 | -2.556 | -0.293 | 3.238 | 1.098 | 0.119 | 0.076 | 0.076 | 0.479 | 0.05 | 0.46 | 0.06 | 0 | 0.07 | 0.12 |

| | | | | | | | | | | | | | | | | |
|---|---|---|---|---|---|---|---|---|---|---|---|---|---|---|---|---|
| 9 | 516413 | rs12343458 | -2.166 | -0.374 | 2.369 | 0.743 | -0.011 | -0.05 | -0.05 | 0.091 | 0.16 | 0.74 | 0.13 | 0 | 0.14 | 0.23 |
| 9 | 516638 | rs12343559 | -1.329 | -0.336 | 2.551 | 0.759 | 0.04 | -0.064 | -0.064 | 0 | 0.11 | 0.44 | 0.1 | 0 | 0.14 | 0.11 |
| 9 | 517715 | rs1928415 | -2.172 | -0.362 | 1.79 | 0.733 | -0.007 | -0.045 | -0.045 | 0 | 0.16 | 0.75 | 0.14 | 0 | 0.16 | 0.23 |
| 9 | 518675 | rs7026716 | -0.459 | 0.8 | 0.649 | -0.218 | -0.083 | 0.1 | 0.1 | 0 | 0.27 | 0.09 | 0.32 | 0 | 0.43 | 0.2 |
| 9 | 518755 | rs10815156 | 0.682 | 0.683 | 0.654 | 0.695 | -0.007 | -0.014 | -0.014 | 0 | 0.58 | 0.67 | 0.32 | 0.57 | 0.53 | 0.64 |
| 9 | 520886 | rs2026075 | 0.93 | 0.577 | 1.461 | 1.593 | 0 | -0.001 | -0.001 | 0 | 0.73 | 0.66 | 0.56 | 0.61 | 0.71 | 0.74 |
| 9 | 523495 | rs7853150 | -0.258 | 0.668 | 2.267 | 1.675 | 0.028 | -0.017 | -0.017 | 0 | 0.07 | 0.28 | 0.06 | 0 | 0.07 | 0.09 |
| 9 | 527180 | rs10974992 | -1.929 | 0.66 | 2.048 | 1.873 | -0.008 | 0.047 | 0.047 | 0 | 0.24 | 0.66 | 0.45 | 0.39 | 0.28 | 0.35 |
| 9 | 537358 | rs10120392 | -1.429 | 0.901 | 1.912 | 1.96 | 0.026 | 0.127 | 0.127 | 0 | 0.14 | 0.33 | 0.17 | 0.32 | 0.15 | 0.12 |
| 9 | 541031 | rs9299011 | -1.619 | 0.684 | 2.294 | 1.891 | -0.022 | 0.071 | 0.071 | 0.437 | 0.15 | 0.27 | 0.15 | 0.31 | 0.15 | 0.19 |
| 9 | 542531 | rs4742073 | -1.339 | 0.641 | 2.373 | 2.246 | -0.007 | 0.049 | 0.049 | 0.039 | 0.16 | 0.38 | 0.44 | 0.25 | 0.18 | 0.2 |
| 9 | 544405 | rs16919380 | -0.515 | 0.666 | 2.457 | 2.026 | -0.002 | -0.022 | -0.022 | 1.202 | 0.08 | 0.14 | 0.15 | 0 | 0.08 | 0.09 |
| 9 | 544519 | rs9408628 | 0.597 | 0.654 | 2.418 | 2.031 | -0.001 | -0.014 | -0.014 | 0 | 0.06 | 0.05 | 0.03 | 0 | 0.06 | 0.09 |
| 9 | 546198 | rs10758683 | -1.883 | 0.566 | 2.488 | 2.545 | -0.019 | 0.127 | 0.127 | 0 | 0.19 | 0.42 | 0.47 | 0.47 | 0.18 | 0.21 |
| 9 | 549518 | rs2279619 | -0.342 | 0.668 | 2.239 | 2.515 | -0.009 | -0.04 | -0.04 | 0.014 | 0.18 | 0.09 | 0.45 | 0.31 | 0.22 | 0.19 |
| 9 | 553248 | rs2641987 | -0.764 | 0.734 | 2.224 | 2.33 | -0.002 | -0.023 | -0.023 | 0 | 0.94 | 0.94 | 0.96 | 1 | 0.93 | 0.91 |
| 9 | 555653 | rs2641984 | 2.317 | 0.717 | 2.423 | 2.416 | 0.009 | -0.025 | -0.025 | 0 | 0.56 | 0.28 | 0.41 | 0.46 | 0.52 | 0.62 |
| 9 | 555772 | rs2641983 | 2.576 | 0.634 | 2.233 | 2.418 | -0.014 | 0.13 | 0.13 | 0.108 | 0.53 | 0.06 | 0.38 | 0.25 | 0.53 | 0.6 |
| 9 | 557617 | rs3739592 | -0.838 | 0.682 | 2.123 | 2.342 | -0.006 | -0.014 | -0.014 | 0 | 0.12 | 0.08 | 0.12 | 0.28 | 0.16 | 0.12 |
| 9 | 558343 | rs2804261 | 1.793 | 0.669 | 2.308 | 2.323 | 0 | -0.004 | -0.004 | 0 | 0.62 | 0.62 | 0.47 | 0.46 | 0.57 | 0.63 |
| 9 | 559714 | rs2804262 | 1.139 | 0.688 | 2.273 | 2.3 | 0 | 0.002 | 0.002 | 0 | 0.83 | 0.86 | 0.85 | 0.72 | 0.79 | 0.8 |
| 9 | 564836 | rs16919733 | -0.609 | 0.663 | 2.203 | 2.083 | 0 | 0 | 0 | 0 | 0.22 | 0.25 | 0.39 | 0.26 | 0.23 | 0.17 |
| 9 | 569992 | rs881985 | -1.302 | 0.592 | 2.096 | 1.877 | -0.004 | -0.014 | -0.014 | 0.41 | 0.14 | 0.11 | 0.26 | 0.34 | 0.18 | 0.13 |
| 9 | 582371 | rs9299021 | -0.794 | 0.842 | 2.541 | 1.908 | -0.001 | 0.061 | 0.061 | 0 | 0.23 | 0.39 | 0.38 | 0.36 | 0.25 | 0.19 |
| 9 | 585409 | rs7033716 | -0.89 | 0.906 | 2.368 | 1.898 | 0 | 0.011 | 0.011 | 0 | 0.23 | 0.28 | 0.4 | 0.36 | 0.3 | 0.21 |
| 9 | 589511 | rs2804280 | 1.075 | 0.821 | 2.289 | 1.926 | -0.001 | -0.026 | -0.026 | 2.163 | 0.8 | 0.89 | 0.72 | 0.66 | 0.78 | 0.79 |
| 9 | 589875 | rs11795359 | -0.479 | 0.957 | 2.576 | 2.114 | -0.002 | 0.019 | 0.019 | 0 | 0.2 | 0.24 | 0.35 | 0.34 | 0.23 | 0.17 |
| 9 | 592889 | rs10491598 | -1.131 | 0.873 | 2.725 | 2.067 | -0.012 | 0.053 | 0.053 | 0.165 | 0.25 | 0.51 | 0.39 | 0.36 | 0.27 | 0.19 |
| 9 | 607738 | rs7850218 | 1.097 | 0.74 | 2.59 | 1.97 | 0.002 | 0.062 | 0.062 | 0 | 0.81 | 0.67 | 0.69 | 0.44 | 0.77 | 0.77 |
| 9 | 608622 | rs2804309 | 1.24 | 0.734 | 2.752 | 1.966 | 0.008 | 0.083 | 0.083 | 0 | 0.78 | 0.55 | 0.66 | 0.44 | 0.73 | 0.76 |
| 9 | 613908 | rs7028675 | 1.381 | 0.679 | 2.79 | 1.941 | -0.01 | 0.045 | 0.045 | 0 | 0.75 | 0.35 | 0.65 | 0.44 | 0.75 | 0.77 |
| 9 | 623848 | rs10975462 | -0.524 | 0.767 | 2.7 | 1.934 | -0.006 | 0.026 | 0.026 | 0 | 0.2 | 0.26 | 0.34 | 0.34 | 0.22 | 0.19 |
| 9 | 626096 | rs16921323 | -2.053 | 0.816 | 3.022 | 2.056 | 0.058 | 0.421 | 0.421 | 1.187 | 0.06 | 0.45 | 0.05 | 0.24 | 0.01 | 0.02 |
| 9 | 628788 | rs12338965 | -0.104 | 0.829 | 2.692 | 1.938 | -0.001 | -0.007 | -0.007 | 0.067 | 0.15 | 0.13 | 0.2 | 0.01 | 0.16 | 0.07 |
| 9 | 634160 | rs6477056 | -1.87 | 1.597 | 3.245 | 2.447 | 0.006 | -0.006 | -0.006 | 0 | 0.37 | 0.42 | 0.46 | 0.33 | 0.44 | 0.29 |
| 9 | 635677 | rs7047127 | -1.052 | 1.219 | 2.725 | 2.032 | 0.005 | 0.022 | 0.022 | 0 | 0.2 | 0.22 | 0.23 | 0.33 | 0.25 | 0.21 |
| 9 | 649553 | rs16921797 | -0.023 | 1.217 | 2.107 | 1.507 | -0.001 | -0.032 | -0.032 | 0.317 | 0.15 | 0.15 | 0.21 | 0 | 0.15 | 0.06 |
| 9 | 651461 | rs9407334 | -0.459 | 1.371 | 1.818 | 1.733 | -0.03 | -0.05 | -0.05 | 0.195 | 0.15 | 0.07 | 0.19 | 0.31 | 0.22 | 0.2 |
| 9 | 653505 | rs7851766 | -0.489 | 1.36 | 1.871 | 1.728 | -0.021 | -0.036 | -0.036 | 0.055 | 0.15 | 0.09 | 0.19 | 0.31 | 0.22 | 0.2 |
| 9 | 654101 | rs7021495 | -1.193 | 1.12 | 2.468 | 1.822 | -0.036 | 0.079 | 0.079 | 0 | 0.26 | 0.53 | 0.46 | 0.57 | 0.22 | 0.21 |
| 9 | 658446 | rs4742197 | -0.849 | 1.283 | 2.069 | 1.954 | 0.001 | 0.025 | 0.025 | 0.111 | 0.17 | 0.25 | 0.24 | 0.53 | 0.24 | 0.24 |
| 9 | 659722 | rs9408655 | 0 | 1.772 | 1.771 | 1.3 | 0.016 | -0.035 | -0.035 | 0 | 0.88 | 0.85 | 0.89 | 0.97 | 0.83 | 0.93 |
| 9 | 661057 | rs10815440 | 0 | 1.772 | 1.771 | 1.3 | 0.016 | -0.035 | -0.035 | 0 | 0.88 | 0.85 | 0.89 | 0.97 | 0.83 | 0.93 |
| 9 | 666389 | rs10975653 | 2.427 | 1.31 | 1.225 | 0.985 | -0.01 | 0.059 | 0.059 | 0 | 0.41 | 0.17 | 0.2 | 0.3 | 0.34 | 0.48 |
| 9 | 666970 | rs10975657 | 2.44 | 1.309 | 1.225 | 0.985 | -0.011 | 0.059 | 0.059 | 0 | 0.41 | 0.17 | 0.2 | 0.3 | 0.34 | 0.48 |
| 9 | 674275 | rs11791986 | -1.359 | 1.207 | 2.014 | 1.226 | -0.041 | 0.163 | 0.163 | 0 | 0.15 | 0.32 | 0.27 | 0.35 | 0.12 | 0.19 |
| 9 | 675419 | rs4742220 | -1.872 | 1.252 | 2.185 | 1.111 | -0.01 | 0.003 | 0.003 | 0 | 0.33 | 0.56 | 0.48 | 0.64 | 0.33 | 0.39 |
| 9 | 678088 | rs4742226 | -1.855 | 1.358 | 2.082 | 1.097 | 0.001 | -0.016 | -0.016 | 0.251 | 0.38 | 0.79 | 0.57 | 0.6 | 0.39 | 0.34 |
| 9 | 678837 | rs12352279 | -0.81 | 1.273 | 2.242 | 1.148 | -0.037 | 0.211 | 0.211 | 0 | 0.13 | 0.33 | 0.25 | 0.33 | 0.09 | 0.2 |
| 9 | 679901 | rs1885924 | -2.497 | 1.328 | 2.054 | 0.932 | 0.021 | 0.149 | 0.149 | 0 | 0.5 | 0.72 | 0.75 | 0.9 | 0.58 | 0.56 |
| 9 | 680323 | rs10975792 | 0.463 | 1.384 | 1.959 | 0.966 | 0.004 | -0.002 | -0.002 | 0 | 0.06 | 0.05 | 0.05 | 0.03 | 0.09 | 0.06 |
| 9 | 680585 | rs10975803 | 0.432 | 1.384 | 1.959 | 0.966 | 0.004 | -0.002 | -0.002 | 1.075 | 0.06 | 0.05 | 0.06 | 0.03 | 0.09 | 0.06 |
| 9 | 680667 | rs7035584 | -1.008 | 1.365 | 2.241 | 1.009 | 0.025 | 0.01 | 0.01 | 0 | 0.24 | 0.57 | 0.31 | 0.26 | 0.26 | 0.15 |
| 9 | 680714 | rs10815466 | -1.439 | 1.369 | 1.905 | 1.084 | 0.004 | 0.162 | 0.162 | 0 | 0.19 | 0.43 | 0.31 | 0.36 | 0.18 | 0.26 |
| 9 | 680763 | rs9408671 | -0.473 | 1.298 | 1.269 | 0.594 | -0.044 | -0.066 | -0.066 | 0 | 0.17 | 0.06 | 0.2 | 0.31 | 0.23 | 0.2 |
| 9 | 680791 | rs17431629 | 0.802 | 1.323 | 1.228 | 0.593 | -0.039 | 0.076 | 0.076 | 0 | 0.13 | 0.02 | 0.19 | 0 | 0.15 | 0.06 |
| 9 | 680905 | rs1323263 | -0.958 | 1.249 | 1.314 | 0.636 | -0.036 | -0.006 | -0.006 | 0 | 0.31 | 0.17 | 0.41 | 0.32 | 0.38 | 0.26 |
| 9 | 686170 | rs9408678 | -1.089 | 0.959 | 1.741 | 0.868 | -0.031 | 0.109 | 0.109 | 0 | 0.17 | 0.32 | 0.29 | 0.55 | 0.14 | 0.31 |
| 9 | 687205 | rs7034698 | -1.259 | 0.981 | 1.705 | 0.619 | -0.032 | 0.073 | 0.073 | 0.057 | 0.21 | 0.36 | 0.33 | 0.57 | 0.17 | 0.35 |
| 9 | 687265 | rs7044480 | -1.407 | 0.958 | 1.818 | 0.612 | -0.042 | 0.082 | 0.082 | 0.17 | 0.22 | 0.4 | 0.34 | 0.57 | 0.17 | 0.35 |
| 9 | 689340 | rs9299038 | -1.192 | 1.064 | 1.793 | 0.657 | -0.022 | 0.116 | 0.116 | 0.3 | 0.18 | 0.38 | 0.32 | 0.57 | 0.14 | 0.33 |
| 9 | 693182 | rs17432514 | -0.19 | 1.475 | 1.7 | 0.576 | -0.006 | 0.023 | 0.023 | 0 | 0.17 | 0.09 | 0.24 | 0.01 | 0.21 | 0.07 |
| 9 | 695607 | rs732119 | 0.453 | 1.917 | 1.742 | 0.519 | 0.027 | 0.036 | 0.036 | 0 | 0.82 | 0.71 | 0.8 | 0.67 | 0.76 | 0.8 |
| 9 | 696683 | rs7854502 | 1.323 | 1.677 | 1.257 | 0.488 | -0.014 | 0.018 | 0.018 | 0 | 0.6 | 0.3 | 0.54 | 0.66 | 0.57 | 0.73 |
| 9 | 706479 | rs7019865 | -0.714 | 2.503 | 1.416 | 0.14 | 0.03 | -0.088 | -0.088 | 0 | 0.2 | 0.25 | 0.24 | 0.01 | 0.22 | 0.07 |
| 9 | 712156 | rs912174 | -0.353 | 2.533 | 1.117 | 0.623 | 0.003 | 0 | 0 | 0 | 0.16 | 0.25 | 0.31 | 0.19 | 0.21 | 0.07 |
| 9 | 712599 | rs11789987 | 1.253 | 1.957 | 0.089 | 0.131 | -0.004 | 0.193 | 0.193 | 0.009 | 0.43 | 0.09 | 0.23 | 0.13 | 0.41 | 0.46 |
| 9 | 716570 | rs4740866 | -1.082 | 1.77 | 0.996 | 0.952 | 0.017 | 0.175 | 0.175 | 0.009 | 0.11 | 0.33 | 0.24 | 0.63 | 0.1 | 0.18 |
| 9 | 720179 | rs7043585 | -0.548 | 1.737 | 1.094 | 0.469 | 0.001 | 0.015 | 0.015 | 0 | 0.73 | 0.76 | 0.73 | 0.81 | 0.75 | 0.71 |

| | | | | | | | | | | | | | | | | |
|---|---|---|---|---|---|---|---|---|---|---|---|---|---|---|---|---|
| 9 | 720832 | rs4742292 | -0.062 | 1.024 | 0.86 | -0.049 | 0 | 0.006 | 0.006 | 0 | 0.64 | 0.64 | 0.6 | 0.33 | 0.68 | 0.5 |
| 9 | 721717 | rs10815523 | 0.922 | 1.091 | 0.586 | -0.262 | -0.044 | 0.218 | 0.218 | 0 | 0.46 | 0.1 | 0.36 | 0.15 | 0.51 | 0.46 |
| 9 | 724217 | rs10815530 | 0.95 | 1.379 | 0.615 | -0.267 | -0.078 | 0.216 | 0.216 | 0 | 0.46 | 0.1 | 0.25 | 0.18 | 0.51 | 0.44 |
| 9 | 724287 | rs10758839 | 0.882 | 0.975 | 0.659 | -0.258 | -0.027 | 0.213 | 0.213 | 0 | 0.46 | 0.16 | 0.25 | 0.18 | 0.45 | 0.44 |
| 9 | 725147 | rs12685329 | -0.12 | 0.983 | 1.18 | -0.082 | 0.011 | -0.002 | -0.002 | 0 | 0.17 | 0.35 | 0.31 | 0.18 | 0.17 | 0.06 |
| 9 | 733049 | rs10815567 | 0.839 | 0.682 | 0.645 | -0.263 | 0.007 | 0.016 | 0.016 | 0 | 0.41 | 0.31 | 0.33 | 0.36 | 0.38 | 0.36 |
| 9 | 734679 | rs10429625 | 0.359 | 0.792 | 0.703 | -0.431 | 0.014 | 0.151 | 0.151 | 0 | 0.51 | 0.22 | 0.39 | 0.17 | 0.55 | 0.4 |
| 9 | 737436 | rs10739127 | 0.988 | 0.741 | 0.635 | -0.462 | 0 | 0.091 | 0.091 | 0 | 0.38 | 0.14 | 0.19 | 0.16 | 0.37 | 0.35 |
| 9 | 738238 | rs2296052 | -0.76 | 0.579 | 0.75 | -0.532 | -0.045 | 0.011 | 0.011 | 0 | 0.64 | 0.9 | 0.58 | 0.33 | 0.6 | 0.54 |
| 9 | 738940 | rs10491590 | -0.362 | 0.406 | 0.94 | -0.547 | -0.001 | -0.058 | -0.058 | 0.108 | 0.11 | 0.46 | 0.09 | 0 | 0.06 | 0.12 |
| 9 | 741001 | rs7027152 | 0.605 | 0.627 | 0.418 | -0.398 | 0 | -0.004 | -0.004 | 0 | 0.39 | 0.39 | 0.23 | 0.17 | 0.36 | 0.38 |
| 9 | 742884 | rs12377726 | 0.229 | 0.438 | -0.031 | -0.787 | 0.006 | -0.012 | -0.012 | 0 | 0.33 | 0.31 | 0.14 | 0.06 | 0.2 | 0.26 |
| 9 | 743046 | rs10815596 | -0.071 | 0.73 | 0.172 | 0.017 | -0.002 | 0.025 | 0.025 | 0 | 0.49 | 0.38 | 0.41 | 0.41 | 0.47 | 0.52 |
| 9 | 745289 | rs17369029 | 0.155 | 0.844 | -0.028 | -0.367 | -0.006 | 0.002 | 0.002 | 0 | 0.18 | 0.13 | 0.2 | 0 | 0.19 | 0.07 |
| 9 | 745413 | rs3739586 | -1.119 | 0.741 | -0.244 | 0.218 | -0.032 | -0.101 | -0.101 | 0 | 0.69 | 0.92 | 0.63 | 0.57 | 0.67 | 0.61 |
| 9 | 750247 | rs10815610 | 0.227 | 0.693 | -0.005 | 0.137 | 0.024 | -0.026 | -0.026 | 0 | 0.53 | 0.24 | 0.49 | 0.54 | 0.59 | 0.46 |
| 9 | 750366 | rs10758866 | -0.029 | 0.689 | -0.123 | 0.136 | 0.006 | -0.014 | -0.014 | 0.016 | 0.56 | 0.37 | 0.5 | 0.54 | 0.59 | 0.46 |
| 9 | 750892 | rs962817 | -1.273 | 0.744 | -0.491 | -0.011 | 0.018 | 0.124 | 0.124 | 0.15 | 0.34 | 0.82 | 0.33 | 0.02 | 0.25 | 0.21 |
| 9 | 751359 | rs7865413 | -1.29 | 0.745 | -0.628 | -0.011 | 0.014 | 0.168 | 0.168 | 0.187 | 0.34 | 0.85 | 0.33 | 0.02 | 0.25 | 0.21 |
| 9 | 753218 | rs4742339 | -1.29 | 0.774 | -0.698 | 0.12 | -0.041 | -0.11 | -0.11 | 0.342 | 0.7 | 0.91 | 0.63 | 0.56 | 0.67 | 0.6 |
| 9 | 754527 | rs7858284 | 1.149 | 0.834 | -0.659 | 0.268 | -0.089 | -0.159 | -0.159 | 1.36 | 0.36 | 0.06 | 0.29 | 0.55 | 0.42 | 0.39 |
| 9 | 755034 | rs12005362 | -0.603 | 0.453 | 0.014 | 0.092 | 0 | 0.001 | 0.001 | 0 | 0.09 | 0.2 | 0.05 | 0.05 | 0.06 | 0.1 |
| 9 | 756415 | rs1885927 | -1.359 | 0.432 | -0.096 | 0.093 | -0.012 | -0.054 | -0.054 | 0.308 | 0.28 | 0.63 | 0.28 | 0.06 | 0.2 | 0.13 |
| 9 | 761569 | rs1917 | 0.08 | 0.595 | -0.157 | 0.542 | 0 | 0.006 | 0.006 | 0.002 | 0.15 | 0.17 | 0.26 | 0.19 | 0.15 | 0.18 |
| 9 | 761813 | rs6477206 | -0.684 | 1.388 | -0.78 | 0.993 | -0.05 | -0.012 | -0.012 | 0 | 0.62 | 0.84 | 0.59 | 0.62 | 0.59 | 0.53 |
| 9 | 762970 | rs7469926 | 0.012 | 0.674 | -0.126 | 0.612 | 0 | -0.001 | -0.001 | 0 | 0.14 | 0.14 | 0.21 | 0.12 | 0.14 | 0.19 |
| 9 | 764128 | rs17436244 | 0.735 | 0.67 | -0.248 | 0.6 | -0.039 | -0.014 | -0.014 | 0 | 0.1 | 0.03 | 0.2 | 0.12 | 0.15 | 0.19 |
| 9 | 770667 | rs11534212 | 0.22 | 0.212 | -0.295 | 0.541 | 0.001 | 0.001 | 0.001 | 0.273 | 0.56 | 0.32 | 0.51 | 0.45 | 0.58 | 0.45 |
| 9 | 776349 | rs2370197 | -0.6 | 0.1 | -0.5 | 0.565 | -0.036 | -0.073 | -0.073 | 0.461 | 0.67 | 0.82 | 0.62 | 0.52 | 0.62 | 0.55 |
| 9 | 781115 | rs10976526 | 1.063 | 0.021 | -0.66 | 0.38 | -0.027 | -0.005 | -0.005 | 0.72 | 0.39 | 0.13 | 0.28 | 0.33 | 0.39 | 0.25 |
| 9 | 783574 | rs10815670 | 0.37 | 0.009 | -0.22 | 0.375 | 0.016 | 0.005 | 0.005 | 0 | 0.51 | 0.29 | 0.5 | 0.46 | 0.56 | 0.46 |
| 9 | 783717 | rs11794001 | 0.284 | 0.119 | -0.254 | 0.413 | -0.029 | -0.02 | -0.02 | 0 | 0.14 | 0.05 | 0.21 | 0.15 | 0.17 | 0.2 |
| 9 | 784064 | rs1853426 | -1.169 | 0.239 | -0.109 | 0.364 | 0.005 | 0.017 | 0.017 | 0 | 0.24 | 0.76 | 0.32 | 0.18 | 0.21 | 0.3 |
| 9 | 787050 | rs6477235 | 0.073 | 0.137 | -0.475 | 0.597 | 0 | -0.03 | -0.03 | 0 | 0.54 | 0.75 | 0.59 | 0.5 | 0.58 | 0.5 |
| 9 | 788730 | rs4742390 | 0.664 | -0.216 | 0.049 | 0.8 | -0.001 | 0 | 0 | 0 | 0.58 | 0.49 | 0.51 | 0.46 | 0.68 | 0.46 |
| 9 | 791067 | rs10758904 | 1.413 | 0.031 | -0.193 | 0.894 | 0.031 | 0.105 | 0.105 | 0.51 | 0.5 | 0.18 | 0.3 | 0.33 | 0.55 | 0.33 |
| 9 | 791133 | rs10758905 | 0.693 | 0.244 | -0.04 | 0.989 | 0.003 | -0.003 | -0.003 | 0.025 | 0.61 | 0.39 | 0.52 | 0.51 | 0.7 | 0.58 |
| 9 | 791858 | rs10481615 | 0.75 | 0.836 | -0.58 | 1.523 | -0.005 | 0.056 | 0.056 | 0.388 | 0.76 | 0.69 | 0.63 | 0.51 | 0.79 | 0.71 |
| 9 | 792777 | rs17371714 | -0.536 | 1.18 | -1.036 | 1.428 | -0.012 | -0.01 | -0.01 | 0 | 0.07 | 0.04 | 0.16 | 0.15 | 0.15 | 0.2 |
| 9 | 794761 | rs16924003 | -0.228 | 0.774 | -0.894 | 1.762 | 0.006 | 0.056 | 0.056 | 2.265 | 0.07 | 0.11 | 0.16 | 0.14 | 0.08 | 0.03 |
| 9 | 798773 | rs10815704 | -0.781 | 0.766 | -1.259 | 1.48 | -0.037 | -0.082 | -0.082 | 0.052 | 0.13 | 0.02 | 0.16 | 0.4 | 0.15 | 0.27 |
| 9 | 800919 | rs7019952 | -0.855 | 0.097 | -0.853 | 0.364 | 0.001 | -0.003 | -0.003 | 0.322 | 0.34 | 0.67 | 0.34 | 0.19 | 0.27 | 0.31 |
| 9 | 801602 | rs1323267 | -1.296 | 0.36 | -0.9 | 0.393 | 0.008 | 0.101 | 0.101 | 0.014 | 0.48 | 0.81 | 0.64 | 0.72 | 0.46 | 0.53 |
| 9 | 802801 | rs2370200 | -0.402 | 0.684 | -0.448 | 0.607 | 0.012 | -0.003 | -0.003 | 0 | 0.16 | 0.16 | 0.21 | 0.13 | 0.27 | 0.15 |
| 9 | 804436 | rs4742419 | -1.703 | 0.335 | -0.946 | 0.127 | -0.077 | 0.095 | 0.095 | 0 | 0.55 | 0.89 | 0.68 | 0.72 | 0.53 | 0.5 |
| 9 | 811618 | rs7035551 | -0.38 | 0.462 | -0.745 | 0.237 | -0.008 | -0.001 | -0.001 | 0 | 0.27 | 0.21 | 0.52 | 0.71 | 0.36 | 0.39 |
| 9 | 816742 | rs4742429 | 1.453 | 0.504 | -0.713 | 0.446 | 0 | -0.016 | -0.016 | 0 | 0.12 | 0.07 | 0.32 | 0.63 | 0.12 | 0.24 |
| 9 | 818693 | rs10815756 | -1.754 | 0.619 | -0.726 | 0.293 | -0.063 | 0.135 | 0.135 | 0 | 0.54 | 0.89 | 0.67 | 0.74 | 0.51 | 0.52 |
| 9 | 822594 | rs4141756 | -0.39 | 0.438 | 0.431 | 0.206 | 0.008 | -0.057 | -0.057 | 0.485 | 0.3 | 0.38 | 0.3 | 0.1 | 0.36 | 0.18 |
| 9 | 824661 | rs10976808 | 0.229 | 0.552 | 0.939 | 1.489 | -0.008 | 0.053 | 0.053 | 0.069 | 0.17 | 0.21 | 0.15 | 0.48 | 0.1 | 0.24 |
| 9 | 833828 | rs16924648 | 0.782 | 0.598 | 1.229 | 1.759 | 0.007 | -0.012 | -0.012 | 0 | 0.16 | 0.09 | 0.18 | 0.49 | 0.09 | 0.21 |
| 9 | 839141 | rs17372837 | -0.15 | 0.877 | 1.137 | 1.333 | -0.011 | 0.061 | 0.061 | 0 | 0.27 | 0.16 | 0.24 | 0.11 | 0.31 | 0.21 |
| 9 | 841152 | rs912062 | 0.798 | 0.864 | 1.751 | 1.502 | 0.013 | 0.173 | 0.173 | 0.094 | 0.92 | 0.79 | 0.81 | 0.79 | 0.93 | 0.87 |
| 9 | 843382 | rs13295838 | 1.35 | 0.701 | 1.273 | 1.214 | -0.081 | 0.087 | 0.087 | 0 | 0.43 | 0.11 | 0.32 | 0.23 | 0.51 | 0.4 |
| 9 | 843763 | rs4338164 | 0.974 | 0.541 | 1.383 | 1.254 | 0.017 | -0.031 | -0.031 | 0 | 0.2 | 0.09 | 0.2 | 0.43 | 0.12 | 0.26 |
| 9 | 844001 | rs7862426 | 0.027 | 0.663 | 1.64 | 1.179 | 0 | 0 | 0 | 0 | 0.24 | 0.25 | 0.27 | 0.17 | 0.26 | 0.22 |
| 9 | 845516 | rs7040024 | -0.025 | 0.669 | 1.414 | 1.182 | 0 | 0.015 | 0.015 | 0 | 0.24 | 0.26 | 0.26 | 0.17 | 0.26 | 0.22 |
| 9 | 846480 | rs16924786 | 0.886 | 0.509 | 1.462 | 1.458 | 0.014 | -0.015 | -0.015 | 0.015 | 0.2 | 0.15 | 0.2 | 0.4 | 0.11 | 0.25 |
| 9 | 847547 | rs2146532 | -0.254 | 0.856 | 2.15 | 1.412 | -0.022 | 0.087 | 0.087 | 0 | 0.72 | 0.5 | 0.61 | 0.4 | 0.8 | 0.64 |
| 9 | 853004 | rs4740943 | -0.469 | 2.156 | 2.296 | 1.675 | 0.003 | -0.002 | -0.002 | 0.275 | 0.45 | 0.46 | 0.49 | 0.6 | 0.39 | 0.48 |
| 9 | 855091 | rs7029148 | 0.864 | 0.987 | 1.436 | 0.782 | -0.001 | 0.016 | 0.016 | 0 | 0.47 | 0.33 | 0.35 | 0.23 | 0.54 | 0.43 |
| 9 | 857511 | rs10739166 | 1.169 | 1.078 | 0.566 | 0.842 | -0.031 | 0.069 | 0.069 | 0 | 0.42 | 0.25 | 0.31 | 0.22 | 0.48 | 0.38 |
| 9 | 863635 | rs755383 | 1.057 | 0.799 | 0.766 | 0.79 | 0 | -0.003 | -0.003 | 0.155 | 0.61 | 0.35 | 0.55 | 0.65 | 0.62 | 0.59 |
| 9 | 864122 | rs749427 | -0.692 | 0.614 | 0.954 | 0.503 | 0.038 | -0.041 | -0.041 | 0 | 0.09 | 0.25 | 0.07 | 0.01 | 0.09 | 0.08 |
| 9 | 866617 | rs4742515 | 0.428 | 0.677 | 0.806 | 0.346 | 0.048 | 0.011 | 0.011 | 0 | 0.86 | 0.67 | 0.86 | 0.87 | 0.82 | 0.86 |
| 9 | 866725 | rs4742516 | -0.575 | 0.812 | 0.467 | 0.357 | -0.004 | 0.001 | 0.001 | 0 | 0.56 | 0.62 | 0.55 | 0.58 | 0.46 | 0.57 |
| 9 | 872418 | rs6477322 | 0.289 | 0.277 | 0.049 | 0.071 | 0.013 | -0.005 | -0.005 | 0 | 0.19 | 0.17 | 0.11 | 0 | 0.13 | 0.13 |
| 9 | 874129 | rs7047524 | -0.466 | 0.352 | -0.033 | 0.126 | 0.016 | 0.01 | 0.01 | 0 | 0.9 | 0.86 | 0.92 | 0.86 | 0.86 | 0.88 |
| 9 | 875364 | rs2370214 | -0.26 | 0.408 | -0.774 | 0.278 | 0.006 | 0.07 | 0.07 | 0 | 0.72 | 0.83 | 0.82 | 0.85 | 0.73 | 0.75 |

| | | | | | | | | | | | | | | | | |
|---|---|---|---|---|---|---|---|---|---|---|---|---|---|---|---|---|
| 9 | 875947 | rs912058 | 1.07 | 0.559 | -1.192 | 0.516 | -0.033 | -0.054 | -0.054 | 0.287 | 0.26 | 0.11 | 0.26 | 0.36 | 0.26 | 0.33 |
| 9 | 876308 | rs10977244 | 1.204 | 0.792 | -1.174 | 0.78 | -0.069 | -0.027 | -0.027 | 0.412 | 0.3 | 0.13 | 0.32 | 0.31 | 0.38 | 0.3 |
| 9 | 877032 | rs1998437 | -0.8 | 0.389 | -0.196 | 0.752 | -0.008 | 0.029 | 0.029 | 0 | 0.08 | 0.12 | 0.19 | 0.18 | 0.08 | 0.11 |
| 9 | 877871 | rs7875857 | -0.761 | 0.399 | -0.314 | 0.724 | -0.003 | 0.015 | 0.015 | 0 | 0.08 | 0.09 | 0.19 | 0.18 | 0.08 | 0.11 |
| 9 | 880777 | rs10733547 | 0.543 | 0.417 | -0.366 | 0.567 | 0.01 | 0.029 | 0.029 | 0 | 0.72 | 0.81 | 0.84 | 0.82 | 0.73 | 0.76 |
| 9 | 883226 | rs10977270 | 1.415 | 0.437 | -0.552 | 0.549 | -0.001 | -0.008 | -0.008 | 0 | 0.61 | 0.11 | 0.56 | 0.65 | 0.61 | 0.69 |
| 9 | 897949 | rs7859987 | 0.565 | 0.644 | -0.115 | 0.547 | -0.003 | 0.001 | 0.001 | 0.002 | 0.78 | 0.82 | 0.83 | 0.78 | 0.74 | 0.8 |
| 9 | 900015 | rs7875002 | 0.714 | 0.697 | 0.121 | 0.506 | -0.001 | -0.002 | -0.002 | 0.009 | 0.73 | 0.74 | 0.8 | 0.78 | 0.69 | 0.78 |
| 9 | 902302 | rs2057429 | 0.988 | 1.147 | 0.432 | 0.481 | 0.004 | -0.018 | -0.018 | 0 | 0.75 | 0.32 | 0.76 | 0.88 | 0.76 | 0.8 |
| 9 | 902888 | rs10816007 | 2.344 | 1.614 | 0.678 | 1.093 | -0.014 | -0.016 | -0.016 | 1.819 | 0.76 | 0.21 | 0.67 | 0.76 | 0.73 | 0.77 |
| 9 | 905068 | rs10977402 | -2.43 | 0.934 | 0.853 | 1.057 | 0.04 | -0.011 | -0.011 | 0 | 0.25 | 0.78 | 0.32 | 0.22 | 0.13 | 0.22 |
| 9 | 907561 | rs4742545 | 2.144 | 1.255 | 2.052 | 1.162 | 0.175 | 0.109 | 0.109 | 0.696 | 0.93 | 0.68 | 0.88 | 0.9 | 0.87 | 0.91 |
| 9 | 907988 | rs2370218 | 2.641 | 1.085 | 1.862 | 1.116 | 0.001 | -0.014 | -0.014 | 0 | 0.73 | 0.22 | 0.66 | 0.73 | 0.75 | 0.7 |
| 9 | 910700 | rs16925796 | -1.415 | 1.115 | 2.284 | 1.205 | -0.014 | 0.107 | 0.107 | 0 | 0.05 | 0.15 | 0.19 | 0.16 | 0.02 | 0.01 |
| 9 | 916220 | rs1322984 | -2.838 | 1.004 | 2.641 | 1.086 | -0.045 | 0.016 | 0.016 | 0.601 | 0.19 | 0.41 | 0.27 | 0.17 | 0.09 | 0.2 |
| 9 | 917214 | rs9299086 | -2.328 | 1.014 | 2.825 | 1.101 | -0.01 | 0.127 | 0.127 | 0.217 | 0.08 | 0.26 | 0.19 | 0.16 | 0.03 | 0.01 |
| 9 | 917667 | rs2370220 | 0.809 | 1.232 | 2.599 | 0.943 | 0.143 | 0.024 | 0.024 | 0.376 | 0.88 | 0.67 | 0.79 | 0.89 | 0.77 | 0.9 |
| 9 | 923342 | rs11791208 | -2.462 | 1.115 | 2.25 | 0.853 | -0.034 | -0.092 | -0.092 | 0 | 0.15 | 0.38 | 0.11 | 0.01 | 0.08 | 0.19 |
| 9 | 923402 | rs10739185 | 1.594 | 1.315 | 2.273 | 1.012 | 0.118 | 0.124 | 0.124 | 0.917 | 0.83 | 0.54 | 0.61 | 0.73 | 0.75 | 0.9 |
| 9 | 924525 | rs10977534 | -2.009 | 1.046 | 1.673 | 0.879 | -0.018 | -0.02 | -0.02 | 0 | 0.12 | 0.25 | 0.1 | 0.05 | 0.08 | 0.19 |
| 9 | 930033 | rs10977581 | 2.483 | 1.392 | 1.001 | 1.114 | -0.048 | -0.044 | -0.044 | 0.203 | 0.65 | 0.17 | 0.61 | 0.59 | 0.61 | 0.71 |
| 9 | 930145 | rs10816063 | 2.315 | 1.371 | 0.706 | 0.758 | -0.068 | 0.086 | 0.086 | 0.643 | 0.61 | 0.17 | 0.62 | 0.74 | 0.57 | 0.65 |
| 9 | 932251 | rs10816069 | -1.182 | 1.21 | 2.176 | 0.805 | 0.003 | -0.082 | -0.082 | 0 | 0.13 | 0.33 | 0.13 | 0 | 0.12 | 0.06 |
| 9 | 932504 | rs10816070 | -2.254 | 1.186 | 2.252 | 0.935 | -0.027 | 0.042 | 0.042 | 1.631 | 0.19 | 0.59 | 0.33 | 0.21 | 0.16 | 0.09 |
| 9 | 932875 | rs7863679 | -2.267 | 1.185 | 2.272 | 0.935 | -0.027 | 0.038 | 0.038 | 1.814 | 0.19 | 0.61 | 0.33 | 0.21 | 0.16 | 0.09 |
| 9 | 934261 | rs10977617 | -1.713 | 1.286 | 2.369 | 0.72 | -0.015 | -0.009 | -0.009 | 0.123 | 0.09 | 0.16 | 0.05 | 0.05 | 0.04 | 0.17 |
| 9 | 939156 | rs7856742 | -2.352 | 1.263 | 2.388 | 0.696 | -0.008 | -0.026 | -0.026 | 0 | 0.22 | 0.7 | 0.19 | 0.05 | 0.16 | 0.23 |
| 9 | 939653 | rs10491584 | -1.341 | 1.446 | 2.311 | 0.714 | 0.015 | -0.091 | -0.091 | 0.176 | 0.13 | 0.47 | 0.13 | 0 | 0.12 | 0.06 |
| 9 | 953465 | rs10759073 | 1.866 | 1.387 | 2.565 | 0.682 | 0.014 | -0.021 | -0.021 | 2.782 | 0.72 | 0.48 | 0.62 | 0.81 | 0.71 | 0.71 |
| 9 | 955794 | rs364477 | 1.625 | 1.827 | 3.155 | 1.022 | 0.062 | 0.066 | 0.066 | 0 | 0.86 | 0.68 | 0.7 | 0.82 | 0.81 | 0.94 |
| 9 | 958073 | rs10123824 | 2.467 | 1.825 | 3.235 | 1.011 | 0.041 | 0.029 | 0.029 | 0 | 0.82 | 0.65 | 0.67 | 0.81 | 0.78 | 0.8 |
| 9 | 960479 | rs279904 | 0.885 | 1.832 | 2.953 | 0.929 | 0.009 | -0.03 | -0.03 | 2.256 | 0.89 | 0.85 | 0.88 | 1 | 0.83 | 0.95 |
| 9 | 962184 | rs279903 | 3.108 | 1.928 | 3.379 | 1.256 | 0.049 | 0.09 | 0.09 | 0 | 0.81 | 0.45 | 0.62 | 0.74 | 0.78 | 0.77 |
| 9 | 967606 | rs12336949 | -2.541 | 2.165 | 3.094 | 1.339 | 0.035 | 0.145 | 0.145 | 0 | 0.13 | 0.27 | 0.29 | 0.29 | 0.13 | 0.22 |
| 9 | 969044 | rs4742608 | -2.82 | 2.163 | 3.09 | 1.338 | 0.039 | 0.192 | 0.192 | 0.02 | 0.15 | 0.47 | 0.31 | 0.29 | 0.14 | 0.22 |
| 9 | 972476 | rs408200 | 1.843 | 3.044 | 2.722 | 1.718 | 0.096 | 0.166 | 0.166 | 0 | 0.79 | 0.55 | 0.52 | 0.58 | 0.68 | 0.86 |
| 9 | 975112 | rs279910 | 2.357 | 3.039 | 1.953 | 1.549 | 0.024 | 0.009 | 0.009 | 0 | 0.71 | 0.55 | 0.58 | 0.72 | 0.69 | 0.71 |
| 9 | 978833 | rs279888 | 2.421 | 2.82 | 1.785 | 1.901 | 0.069 | 0.052 | 0.052 | 0 | 0.69 | 0.49 | 0.49 | 0.48 | 0.57 | 0.69 |
| 9 | 985569 | rs12005831 | -1.569 | 2.976 | 1.255 | 2.016 | 0.124 | 0.107 | 0.107 | 0.002 | 0.15 | 0.49 | 0.18 | 0.25 | 0.23 | 0.2 |
| 9 | 986085 | rs279880 | 1.821 | 2.749 | 1.225 | 1.95 | 0.061 | 0.043 | 0.043 | 0 | 0.71 | 0.42 | 0.51 | 0.57 | 0.59 | 0.71 |
| 9 | 990840 | rs7021244 | 0.012 | 1.779 | -0.218 | 1.623 | -0.018 | -0.002 | -0.002 | 0 | 0.24 | 0.01 | 0.18 | 0.23 | 0.22 | 0.35 |
| 9 | 1000520 | rs7047346 | -1.439 | 2.221 | 1.027 | 1.978 | 0.117 | 0.257 | 0.257 | 0.806 | 0.08 | 0.37 | 0.28 | 0.22 | 0.1 | 0.13 |
| 9 | 1001309 | rs12238866 | -0.636 | 2.418 | 1.057 | 1.537 | -0.008 | -0.005 | -0.005 | 0 | 0.11 | 0.1 | 0.14 | 0.09 | 0.18 | 0.09 |
| 9 | 1001397 | rs7033404 | -1.982 | 2.418 | 1.025 | 1.599 | 0.08 | 0.059 | 0.059 | 0 | 0.19 | 0.55 | 0.26 | 0.22 | 0.28 | 0.21 |
| 9 | 1002280 | rs12343006 | -1.451 | 1.618 | 0.767 | 1.387 | 0.131 | 0.142 | 0.142 | 0.854 | 0.07 | 0.37 | 0.11 | 0.13 | 0.1 | 0.12 |
| 9 | 1012154 | rs10738147 | -0.261 | 1.073 | -0.805 | 1.155 | 0.097 | 0.049 | 0.049 | 0 | 0.5 | 0.83 | 0.5 | 0.6 | 0.61 | 0.39 |
| 9 | 1016842 | rs1588303 | 0.654 | 1.652 | -0.389 | 1.335 | 0.003 | 0.003 | 0.003 | 0.111 | 0.68 | 0.35 | 0.52 | 0.65 | 0.62 | 0.72 |
| 9 | 1019132 | rs1509198 | 0.814 | 1.551 | -0.501 | 1.363 | 0.056 | 0.058 | 0.058 | 0 | 0.43 | 0.28 | 0.29 | 0.33 | 0.34 | 0.57 |
| 9 | 1019207 | rs10958885 | 0.749 | 1.437 | 0.345 | 1.457 | -0.004 | -0.002 | -0.002 | 1.903 | 0.28 | 0.23 | 0.25 | 0.3 | 0.29 | 0.15 |
| 9 | 1021801 | rs10756003 | 0.984 | 1.474 | 0.325 | 1.475 | -0.042 | -0.035 | -0.035 | 0 | 0.25 | 0.1 | 0.23 | 0.3 | 0.28 | 0.15 |
| 9 | 1022010 | rs10738153 | 0.805 | 1.978 | 0.843 | 1.679 | 0.076 | 0.078 | 0.078 | 0 | 0.91 | 0.78 | 0.87 | 0.85 | 0.87 | 0.89 |
| 9 | 1024471 | rs10756006 | 0.98 | 1.522 | 0.282 | 1.514 | 0.041 | 0.043 | 0.043 | 0 | 0.4 | 0.28 | 0.29 | 0.32 | 0.34 | 0.56 |
| 9 | 1032567 | rs1016565 | -1.27 | 1.807 | 0.291 | 1.715 | -0.013 | -0.046 | -0.046 | 0 | 0.36 | 0.8 | 0.5 | 0.43 | 0.33 | 0.29 |
| 9 | 1041524 | rs2279989 | -0.691 | 1.932 | 0.303 | 1.758 | -0.01 | -0.006 | -0.006 | 0 | 0.07 | 0.03 | 0.03 | 0.06 | 0.08 | 0.1 |
| 9 | 1042166 | rs2279988 | 1.311 | 2.303 | 0.839 | 1.916 | 0.046 | 0.072 | 0.072 | 0.017 | 0.71 | 0.43 | 0.5 | 0.52 | 0.65 | 0.72 |
| 9 | 1043752 | rs6474536 | 0.629 | 1.662 | 0.823 | 1.694 | -0.01 | 0.029 | 0.029 | 0 | 0.82 | 0.69 | 0.79 | 0.76 | 0.81 | 0.9 |
| 9 | 1047343 | rs4740448 | -0.743 | 1.442 | 0.768 | 1.768 | 0.023 | 0.117 | 0.117 | 0 | 0.23 | 0.41 | 0.44 | 0.56 | 0.23 | 0.36 |
| 9 | 1050981 | rs7869989 | -0.326 | 0.974 | -0.248 | 0.58 | 0.007 | 0.087 | 0.087 | 1.473 | 0.53 | 0.66 | 0.35 | 0.23 | 0.48 | 0.5 |
| 9 | 1052722 | rs6474550 | -0.183 | 0.327 | -0.62 | -0.103 | 0.053 | 0.12 | 0.12 | 0.29 | 0.47 | 0.72 | 0.28 | 0.19 | 0.39 | 0.43 |
| 9 | 1053465 | rs869939 | -0.546 | -0.181 | 0.315 | -0.198 | -0.003 | 0.019 | 0.019 | 0 | 0.11 | 0.39 | 0.12 | 0.08 | 0.05 | 0.08 |
| 9 | 1053910 | rs2275516 | 0.472 | -0.376 | -0.066 | -0.552 | 0.013 | 0.023 | 0.023 | 0 | 0.2 | 0.19 | 0.1 | 0.04 | 0.18 | 0.14 |
| 9 | 1054991 | rs10959107 | 0.525 | -0.414 | -0.132 | -0.553 | 0.02 | 0.038 | 0.038 | 0 | 0.19 | 0.13 | 0.08 | 0.04 | 0.18 | 0.14 |
| 9 | 1057318 | rs9140 | -1.748 | -0.03 | -0.315 | -0.246 | -0.073 | 0.058 | 0.058 | 0.291 | 0.7 | 0.89 | 0.71 | 0.84 | 0.62 | 0.79 |
| 9 | 1057351 | rs4741030 | -0.216 | -0.651 | 0.646 | 0.131 | -0.023 | 0.01 | 0.01 | 0 | 0.11 | 0.2 | 0.13 | 0.1 | 0.05 | 0.1 |
| 9 | 1057797 | rs10809097 | -1.099 | -0.721 | 0.542 | 0.221 | -0.056 | 0.004 | 0.004 | 0.004 | 0.21 | 0.57 | 0.18 | 0.22 | 0.14 | 0.17 |
| 9 | 1062999 | rs959481 | -0.177 | -0.398 | 0.251 | 0.476 | 0.01 | -0.001 | -0.001 | 0.227 | 0.48 | 0.51 | 0.51 | 0.46 | 0.56 | 0.62 |
| 9 | 1064985 | rs7039375 | -0.224 | -0.412 | 0.125 | 0.42 | 0.008 | 0.004 | 0.004 | 0 | 0.51 | 0.63 | 0.53 | 0.6 | 0.59 | 0.63 |
| 9 | 1069663 | rs7857644 | -1.594 | -0.471 | -0.921 | 0.45 | 0.005 | 0.011 | 0.011 | 0 | 0.72 | 0.9 | 0.64 | 0.78 | 0.73 | 0.75 |
| 9 | 1070591 | rs10738171 | -0.611 | 0.006 | -0.826 | 0.3 | 0.036 | 0.016 | 0.016 | 0 | 0.53 | 0.77 | 0.54 | 0.61 | 0.59 | 0.63 |

| | | | | | | | | | | | | | | | | |
|---|---|---|---|---|---|---|---|---|---|---|---|---|---|---|---|---|
| 9 | 1071814 | rs7043337 | -0.419 | 0.615 | -0.494 | -0.062 | 0.007 | -0.009 | -0.009 | 0 | 0.33 | 0.6 | 0.27 | 0.28 | 0.35 | 0.33 |
| 9 | 1076795 | rs10959226 | -0.093 | 0.912 | -2.419 | 0.104 | -0.064 | 0.004 | 0.004 | 0 | 0.29 | 0.09 | 0.23 | 0.27 | 0.35 | 0.3 |
| 9 | 1077862 | rs17641200 | -0.235 | 0.41 | -1.687 | 0.035 | -0.012 | 0.02 | 0.02 | 0 | 0.14 | 0.01 | 0.06 | 0.08 | 0.13 | 0.07 |
| 9 | 1080866 | rs4497022 | 0.03 | 0.505 | -2.057 | -0.488 | -0.018 | 0.16 | 0.16 | 2.262 | 0.69 | 0.93 | 0.82 | 0.9 | 0.68 | 0.78 |
| 9 | 1081038 | rs7859614 | 0.304 | 0.385 | -2.006 | 0.152 | 0.065 | -0.044 | -0.044 | 0 | 0.42 | 0.81 | 0.27 | 0.46 | 0.28 | 0.47 |
| 9 | 1081794 | rs10809172 | 0.639 | 0.33 | -0.73 | 0.201 | -0.025 | 0.009 | 0.009 | 0 | 0.35 | 0.52 | 0.22 | 0.35 | 0.27 | 0.45 |
| 9 | 1082544 | rs10156555 | 0.131 | 0.257 | -0.193 | 0.229 | 0.042 | -0.029 | -0.029 | 0 | 0.42 | 0.76 | 0.28 | 0.45 | 0.31 | 0.49 |
| 9 | 1085999 | rs4741056 | 0.965 | 0.108 | 0.488 | 0.85 | 0.016 | -0.009 | -0.009 | 0 | 0.3 | 0.25 | 0.16 | 0.32 | 0.21 | 0.41 |
| 9 | 1087874 | rs12353456 | -0.986 | -0.107 | 0.83 | 0.239 | 0.06 | 0.003 | 0.003 | 0.053 | 0.07 | 0.36 | 0.04 | 0 | 0.02 | 0.02 |
| 9 | 1089920 | rs7020419 | -0.022 | 0.007 | 0.513 | 0.033 | 0.007 | 0.013 | 0.013 | 0 | 0.41 | 0.46 | 0.66 | 0.56 | 0.5 | 0.38 |
| 9 | 1090529 | rs10738191 | -0.961 | 0.099 | 0.305 | -0.01 | -0.032 | 0.053 | 0.053 | 0.084 | 0.8 | 0.98 | 0.89 | 0.9 | 0.8 | 0.83 |
| 9 | 1090566 | rs10809197 | 0.954 | 0.085 | 0.424 | 0.007 | 0.001 | -0.001 | -0.001 | 0 | 0.35 | 0.35 | 0.2 | 0.32 | 0.28 | 0.42 |
| 9 | 1090919 | rs10738192 | 0.397 | 0.394 | 1.353 | 0.23 | -0.007 | -0.034 | -0.034 | 0 | 0.74 | 0.64 | 0.84 | 0.81 | 0.78 | 0.8 |
| 9 | 1094644 | rs1396633 | -0.439 | 0.424 | -0.006 | 0.342 | -0.044 | 0.029 | 0.029 | 0 | 0.82 | 0.98 | 0.88 | 0.89 | 0.79 | 0.84 |
| 9 | 1094973 | rs1396634 | 0.269 | 0.507 | 0.126 | 0.4 | 0.005 | -0.035 | -0.035 | 0 | 0.74 | 0.52 | 0.75 | 0.76 | 0.76 | 0.81 |
| 9 | 1095101 | rs10959331 | -1.268 | 0.593 | 0.344 | 0.787 | 0.079 | 0.114 | 0.114 | 1.161 | 0.05 | 0.33 | 0.11 | 0.13 | 0.02 | 0.03 |
| 9 | 1095922 | rs1509186 | -0.225 | 0.826 | -0.958 | 0.782 | -0.07 | -0.096 | -0.096 | 0 | 0.17 | 0.01 | 0.3 | 0.33 | 0.18 | 0.16 |
| 9 | 1100738 | rs1355142 | -0.984 | -0.03 | 0.14 | 0.299 | -0.067 | 0.016 | 0.016 | 0.149 | 0.17 | 0.41 | 0.14 | 0.16 | 0.04 | 0.04 |
| 9 | 1102058 | rs10809246 | -0.865 | 0.473 | -0.587 | -0.611 | 0.002 | 0.043 | 0.043 | 0.019 | 0.28 | 0.75 | 0.25 | 0.09 | 0.27 | 0.25 |
| 9 | 1102943 | rs10756149 | 0.821 | 0.33 | -0.594 | -1.517 | 0 | 0.025 | 0.025 | 0 | 0.57 | 0.4 | 0.67 | 0.91 | 0.6 | 0.58 |
| 9 | 1103477 | rs10809249 | 1.265 | 0.627 | -1.162 | -1.256 | 0.017 | 0.012 | 0.012 | 0 | 0.32 | 0.13 | 0.36 | 0.26 | 0.31 | 0.17 |
| 9 | 1107806 | rs10118075 | -0.812 | 0.581 | -0.95 | -1.289 | 0.019 | 0.15 | 0.15 | 0.078 | 0.5 | 0.81 | 0.63 | 0.75 | 0.56 | 0.66 |
| 9 | 1116335 | rs4146595 | -0.406 | 0.551 | 0.775 | -1.525 | -0.009 | -0.134 | -0.134 | 0.03 | 0.74 | 0.66 | 0.85 | 0.97 | 0.82 | 0.79 |
| 9 | 1117945 | rs4741086 | 0.377 | 0.762 | 0.788 | -1.692 | -0.027 | -0.064 | -0.064 | 2.229 | 0.16 | 0.33 | 0.13 | 0.06 | 0.12 | 0.21 |
| 9 | 1118896 | rs7025111 | 0.64 | 2.034 | 1.309 | -1.69 | 0.05 | 0.117 | 0.117 | 0.021 | 0.57 | 0.3 | 0.72 | 0.92 | 0.71 | 0.59 |
| 9 | 1121085 | rs10959548 | 1.32 | 1.174 | 0.889 | -1.755 | 0.031 | 0.052 | 0.052 | 0 | 0.36 | 0.18 | 0.32 | 0.22 | 0.32 | 0.19 |
| 9 | 1121353 | rs7861975 | 0.341 | 1.175 | 0.863 | -1.775 | -0.045 | 0.001 | 0.001 | 0 | 0.26 | 0.19 | 0.43 | 0.7 | 0.4 | 0.42 |
| 9 | 1123530 | rs10809358 | 1.594 | 1.114 | 0.879 | -1.751 | 0.01 | 0.022 | 0.022 | 0 | 0.33 | 0.24 | 0.3 | 0.21 | 0.3 | 0.15 |
| 9 | 1124087 | rs4740471 | -1.178 | 1.204 | 0.744 | -1.725 | -0.002 | 0 | 0 | 0 | 0.31 | 0.66 | 0.4 | 0.69 | 0.26 | 0.34 |
| 9 | 1125972 | rs10809367 | 1.087 | 1.26 | 0.626 | -1.731 | -0.065 | 0.082 | 0.082 | 0.106 | 0.48 | 0.09 | 0.47 | 0.29 | 0.58 | 0.46 |
| 9 | 1128641 | rs7859005 | -0.678 | 1.353 | 0.8 | -1.765 | -0.001 | 0 | 0 | 0.051 | 0.33 | 0.33 | 0.47 | 0.7 | 0.44 | 0.46 |
| 9 | 1129674 | rs4237127 | -1.101 | 1.331 | 0.669 | -1.706 | -0.014 | 0.047 | 0.047 | 0 | 0.8 | 0.92 | 0.88 | 0.97 | 0.82 | 0.79 |
| 9 | 1130580 | rs4741117 | -1.333 | 1.325 | 0.608 | -1.698 | -0.03 | 0.073 | 0.073 | 0 | 0.82 | 0.96 | 0.89 | 0.97 | 0.82 | 0.8 |
| 9 | 1142276 | rs953166 | -0.15 | 1.301 | 0.74 | -1.712 | 0 | -0.12 | -0.12 | 0 | 0.67 | 0.55 | 0.76 | 0.92 | 0.69 | 0.6 |
| 9 | 1142653 | rs2375254 | -0.464 | 1.4 | 0.23 | -1.478 | -0.058 | 0.007 | 0.007 | 0.005 | 0.31 | 0.2 | 0.42 | 0.68 | 0.41 | 0.41 |
| 9 | 1152096 | rs12553921 | 0.527 | 1.323 | 0.073 | -1.392 | -0.089 | 0.046 | 0.046 | 0.262 | 0.2 | 0.03 | 0.19 | 0.07 | 0.28 | 0.31 |
| 9 | 1152396 | rs1337619 | 1.566 | 0.767 | 0.277 | -1.623 | 0.027 | 0.035 | 0.035 | 0 | 0.31 | 0.1 | 0.27 | 0.22 | 0.27 | 0.16 |
| 9 | 1153978 | rs10809481 | 1.518 | 0.73 | 0.201 | -1.64 | 0.01 | 0.023 | 0.023 | 0 | 0.32 | 0.05 | 0.27 | 0.23 | 0.27 | 0.16 |
| 9 | 1158171 | rs10738237 | 0.497 | 0.795 | 0.552 | -1.646 | 0.023 | 0.028 | 0.028 | 0 | 0.48 | 0.58 | 0.31 | 0.29 | 0.36 | 0.35 |
| 9 | 1158199 | rs4741151 | -0.892 | 0.909 | 0.292 | -1.56 | -0.038 | 0.064 | 0.064 | 0 | 0.79 | 0.9 | 0.77 | 0.97 | 0.76 | 0.8 |
| 9 | 1163465 | rs10809512 | -0.593 | 0.833 | 0.533 | -1.555 | -0.001 | 0.003 | 0.003 | 0 | 0.45 | 0.59 | 0.57 | 0.74 | 0.48 | 0.64 |
| 9 | 1169485 | rs2375256 | -0.647 | 0.845 | 0.496 | -1.572 | -0.001 | 0.012 | 0.012 | 0 | 0.46 | 0.6 | 0.58 | 0.75 | 0.48 | 0.63 |
| 9 | 1170012 | rs2152850 | -0.744 | 1.025 | -0.016 | -1.436 | -0.035 | 0.064 | 0.064 | 0 | 0.78 | 0.89 | 0.86 | 0.97 | 0.76 | 0.79 |
| 9 | 1170227 | rs7848270 | -0.297 | 1.035 | 0.013 | -1.374 | -0.009 | 0.01 | 0.01 | 0.15 | 0.35 | 0.33 | 0.47 | 0.69 | 0.42 | 0.43 |
| 9 | 1174357 | rs10960072 | 0.394 | 0.273 | -0.237 | -1.058 | 0.022 | 0.057 | 0.057 | 0 | 0.38 | 0.24 | 0.28 | 0.12 | 0.35 | 0.3 |
| 9 | 1183695 | rs13297826 | -0.196 | 0.518 | -0.799 | -0.715 | 0.007 | 0.093 | 0.093 | 0 | 0.55 | 0.84 | 0.35 | 0.25 | 0.46 | 0.46 |
| 9 | 1183870 | rs12342237 | -1.184 | 0.194 | -0.371 | -0.926 | -0.005 | 0.015 | 0.015 | 1.742 | 0.14 | 0.63 | 0.13 | 0.14 | 0.11 | 0.05 |
| 9 | 1184213 | rs10125101 | -0.426 | 0.124 | 0.31 | -0.885 | 0.033 | 0.101 | 0.101 | 0 | 0.07 | 0.29 | 0.07 | 0.12 | 0.06 | 0.03 |
| 9 | 1189365 | rs17641744 | 0.248 | 0.134 | 0.2 | -0.904 | 0.038 | 0.118 | 0.118 | 0 | 0.45 | 0.23 | 0.23 | 0.14 | 0.4 | 0.4 |
| 9 | 1196570 | rs1933040 | 0.343 | 0.311 | -0.024 | -1.08 | 0.017 | 0.024 | 0.024 | 0 | 0.45 | 0.38 | 0.26 | 0.12 | 0.36 | 0.4 |
| 9 | 1197071 | rs7029687 | -0.031 | 0.977 | -0.832 | -1.357 | -0.055 | 0.049 | 0.049 | 0 | 0.26 | 0.18 | 0.24 | 0.03 | 0.41 | 0.23 |
| 9 | 1197632 | rs10809638 | -0.092 | 0.563 | -0.574 | -1.312 | 0.001 | -0.007 | -0.007 | 0 | 0.61 | 0.57 | 0.37 | 0.21 | 0.53 | 0.65 |
| 9 | 1197716 | rs10960353 | 0.699 | 0.147 | -1.23 | -1.71 | -0.047 | 0.115 | 0.115 | 0 | 0.3 | 0.08 | 0.24 | 0.08 | 0.35 | 0.17 |
| 9 | 1199448 | rs10809642 | -0.695 | -0.231 | -0.156 | -1.261 | -0.029 | -0.011 | -0.011 | 1.384 | 0.23 | 0.36 | 0.16 | 0.22 | 0.19 | 0.35 |
| 9 | 1200011 | rs7863802 | -0.776 | -0.269 | -0.175 | -1.285 | -0.03 | -0.031 | -0.031 | 0.258 | 0.23 | 0.36 | 0.14 | 0.18 | 0.19 | 0.35 |
| 9 | 1200108 | rs10960360 | -0.147 | -0.264 | -0.081 | -1.27 | 0.001 | -0.029 | -0.029 | 0 | 0.07 | 0.16 | 0.06 | 0.01 | 0.07 | 0.05 |
| 9 | 1200946 | rs12341136 | 0.014 | -0.267 | -0.07 | -1.27 | 0.01 | -0.029 | -0.029 | 0.781 | 0.06 | 0.15 | 0.06 | 0.01 | 0.08 | 0.06 |
| 9 | 1201091 | rs867579 | -0.973 | -0.273 | 0.421 | -1.357 | -0.008 | -0.022 | -0.022 | 0 | 0.21 | 0.48 | 0.16 | 0.2 | 0.2 | 0.28 |
| 9 | 1201156 | rs12342106 | -1.138 | -0.331 | 0.764 | -1.372 | -0.013 | -0.026 | -0.026 | 7.806 | 0.24 | 0.57 | 0.19 | 0.2 | 0.22 | 0.29 |
| 9 | 1201248 | rs10960372 | 0.136 | -0.397 | 0.701 | -1.375 | 0.017 | -0.002 | -0.002 | 0.077 | 0.14 | 0.1 | 0.07 | 0.18 | 0.1 | 0.24 |
| 9 | 1202371 | rs10809650 | 1.406 | -0.338 | 0.31 | -1.252 | -0.047 | -0.07 | -0.07 | 0 | 0.2 | 0.11 | 0.46 | 0.53 | 0.29 | 0.21 |
| 9 | 1202824 | rs12353067 | -0.609 | -0.525 | 0.546 | -1.183 | -0.029 | -0.023 | -0.023 | 0 | 0.22 | 0.37 | 0.16 | 0.2 | 0.19 | 0.29 |
| 9 | 1204386 | rs16925544 | 0.975 | -0.715 | 0.344 | -1.334 | 0.006 | -0.002 | -0.002 | 0 | 0.2 | 0.19 | 0.14 | 0.21 | 0.14 | 0.17 |
| 9 | 1207469 | rs10960412 | 0.706 | -0.855 | 0.334 | -1.209 | 0.008 | 0.001 | 0.001 | 3.876 | 0.12 | 0.1 | 0.05 | 0.15 | 0.08 | 0.23 |
| 9 | 1209190 | rs12343642 | -0.257 | -1.283 | 0.436 | -1.233 | 0.003 | -0.044 | -0.044 | 0 | 0.1 | 0.2 | 0.09 | 0.01 | 0.12 | 0.06 |
| 9 | 1209265 | rs10809665 | -0.78 | -1.269 | 0.377 | -1.251 | -0.03 | -0.039 | -0.039 | 0.221 | 0.23 | 0.57 | 0.17 | 0.16 | 0.18 | 0.29 |
| 9 | 1213024 | rs10809682 | -0.057 | -0.829 | -0.113 | -1.161 | 0.001 | 0.032 | 0.032 | 0.706 | 0.47 | 0.54 | 0.35 | 0.13 | 0.51 | 0.39 |
| 9 | 1214563 | rs1890126 | 1.433 | -0.716 | -0.583 | -1.117 | -0.085 | -0.116 | -0.116 | 0.011 | 0.2 | 0.06 | 0.45 | 0.51 | 0.3 | 0.19 |
| 9 | 1216007 | rs12236917 | 1.404 | -0.708 | -0.581 | -1.174 | -0.081 | -0.108 | -0.108 | 0.003 | 0.2 | 0.07 | 0.46 | 0.52 | 0.3 | 0.21 |

| | | | | | | | | | | | | | | | | |
|---|---|---|---|---|---|---|---|---|---|---|---|---|---|---|---|---|
| 9 | 1216635 | rs3920797 | -0.596 | -0.717 | -0.551 | -1.159 | 0.018 | -0.004 | -0.004 | 0.212 | 0.68 | 0.72 | 0.81 | 0.63 | 0.8 | 0.63 |
| 9 | 1218440 | rs10809695 | 0.723 | -0.37 | -1.348 | -1.26 | 0.008 | 0.077 | 0.077 | 0 | 0.35 | 0.19 | 0.22 | 0.08 | 0.34 | 0.33 |
| 9 | 1218605 | rs10960472 | 0.673 | -0.33 | -1.325 | -1.253 | 0.015 | -0.019 | -0.019 | 0.013 | 0.06 | 0.11 | 0.05 | 0.01 | 0.11 | 0.06 |
| 9 | 1218816 | rs10960475 | 0.547 | -0.323 | -1.735 | -1.066 | 0.009 | 0.072 | 0.072 | 0 | 0.35 | 0.2 | 0.22 | 0.08 | 0.34 | 0.34 |
| 9 | 1219329 | rs10960480 | 0.325 | -0.219 | -1.423 | -0.041 | -0.134 | -0.12 | -0.12 | 0 | 0.33 | 0.08 | 0.53 | 0.52 | 0.43 | 0.23 |
| 9 | 1221729 | rs12339290 | 0.009 | -0.453 | -0.458 | -0.142 | -0.017 | -0.007 | -0.007 | 0 | 0.11 | 0.12 | 0.05 | 0.08 | 0.05 | 0.13 |
| 9 | 1221840 | rs6474685 | -0.406 | -0.842 | -0.569 | -0.148 | 0.005 | -0.004 | -0.004 | 2.759 | 0.87 | 0.9 | 0.9 | 0.86 | 0.88 | 0.83 |
| 9 | 1222087 | rs12345035 | 0.192 | -0.93 | -0.603 | -0.189 | -0.009 | -0.003 | -0.003 | 0 | 0.12 | 0.12 | 0.06 | 0.09 | 0.05 | 0.14 |
| 9 | 1222118 | rs7874807 | 0.879 | -1.099 | 0.504 | -0.315 | -0.053 | -0.023 | -0.023 | 2.112 | 0.72 | 0.57 | 0.82 | 0.77 | 0.83 | 0.68 |
| 9 | 1223322 | rs10738265 | 1.122 | -1.277 | 1.279 | -0.402 | -0.03 | -0.015 | -0.015 | 0.432 | 0.7 | 0.42 | 0.82 | 0.76 | 0.82 | 0.68 |
| 9 | 1230377 | rs17709273 | 0.544 | -1.256 | 1.026 | -0.247 | -0.017 | -0.026 | -0.026 | 4.543 | 0.06 | 0.01 | 0.09 | 0.09 | 0.09 | 0.1 |
| 9 | 1230994 | rs10756345 | 0.135 | -1.369 | 1.444 | -0.482 | -0.019 | -0.027 | -0.027 | 0 | 0.83 | 0.67 | 0.88 | 0.89 | 0.87 | 0.82 |
| 9 | 1231408 | rs10756347 | 0.412 | -1.376 | 1.338 | -0.485 | -0.029 | -0.038 | -0.038 | 0 | 0.82 | 0.55 | 0.87 | 0.89 | 0.87 | 0.82 |
| 9 | 1235507 | rs13293948 | 0.955 | -1.443 | 0.889 | -0.854 | 0 | 0.051 | 0.051 | 0 | 0.27 | 0.18 | 0.19 | 0.09 | 0.24 | 0.28 |
| 9 | 1235703 | rs10429609 | -0.403 | -1.25 | 1.33 | -0.809 | -0.003 | -0.018 | -0.018 | 0 | 0.84 | 0.73 | 0.87 | 0.89 | 0.86 | 0.82 |
| 9 | 1235993 | rs945862 | -0.539 | -1.236 | 1.037 | -0.805 | -0.002 | -0.006 | -0.006 | 0 | 0.85 | 0.81 | 0.88 | 0.89 | 0.86 | 0.82 |
| 9 | 1239644 | rs12346252 | 0.342 | -1.341 | 0.949 | -0.741 | -0.022 | -0.006 | -0.006 | 0 | 0.14 | 0.22 | 0.08 | 0.13 | 0.08 | 0.14 |
| 9 | 1239776 | rs4741213 | 1.024 | -1.363 | 0.16 | -0.689 | -0.039 | 0.038 | 0.038 | 0 | 0.41 | 0.15 | 0.61 | 0.67 | 0.56 | 0.4 |
| 9 | 1241530 | rs1556786 | -0.149 | -1.228 | -0.045 | -0.814 | 0.006 | 0.031 | 0.031 | 0.394 | 0.6 | 0.63 | 0.7 | 0.82 | 0.66 | 0.54 |
| 9 | 1243842 | rs4741218 | -0.268 | -0.738 | -0.793 | -0.304 | 0.033 | 0.097 | 0.097 | 0.095 | 0.48 | 0.88 | 0.33 | 0.31 | 0.36 | 0.48 |
| 9 | 1252572 | rs9298675 | -0.421 | -0.641 | -0.652 | -0.284 | 0.002 | -0.012 | -0.012 | 0.057 | 0.87 | 0.99 | 0.93 | 0.91 | 0.9 | 0.82 |
| 9 | 1258510 | rs10491875 | 0.169 | -0.71 | 0.061 | -0.482 | 0.007 | 0.072 | 0.072 | 0 | 0.36 | 0.21 | 0.26 | 0.17 | 0.32 | 0.4 |
| 9 | 1265994 | rs7037665 | -0.398 | -0.592 | -0.407 | -0.445 | 0.011 | 0.099 | 0.099 | 0 | 0.44 | 0.85 | 0.34 | 0.29 | 0.38 | 0.46 |
| 9 | 1267493 | rs10809798 | -0.318 | -0.607 | -0.383 | -0.409 | 0.039 | 0.087 | 0.087 | 0 | 0.43 | 0.84 | 0.32 | 0.28 | 0.33 | 0.46 |
| 9 | 1270676 | rs882092 | -0.153 | -0.776 | 0.076 | -0.338 | -0.005 | -0.03 | -0.03 | 0 | 0.44 | 0.58 | 0.3 | 0.19 | 0.33 | 0.41 |
| 9 | 1270931 | rs885043 | -0.231 | -0.592 | -0.778 | -0.254 | 0.071 | 0.102 | 0.102 | 0 | 0.47 | 0.78 | 0.33 | 0.31 | 0.36 | 0.45 |
| 9 | 1275707 | rs1582167 | -0.242 | -0.558 | -1.061 | -0.136 | 0.097 | 0.109 | 0.109 | 0 | 0.47 | 0.83 | 0.31 | 0.31 | 0.32 | 0.46 |
| 9 | 1279854 | rs7871622 | -0.233 | 0.297 | -0.828 | -0.043 | -0.013 | 0 | 0 | 0.042 | 0.31 | 0.22 | 0.53 | 0.65 | 0.4 | 0.4 |
| 9 | 1281453 | rs1415612 | 0.492 | 0.618 | -0.996 | -0.123 | -0.016 | 0.004 | 0.004 | 0 | 0.61 | 0.71 | 0.39 | 0.37 | 0.47 | 0.56 |
| 9 | 1282293 | rs10960773 | 0.545 | -0.428 | -0.59 | -0.763 | -0.057 | -0.011 | -0.011 | 0 | 0.22 | 0.34 | 0.14 | 0.23 | 0.09 | 0.18 |
| 9 | 1283014 | rs10809833 | 1.206 | -0.307 | -0.933 | -0.44 | -0.1 | -0.133 | -0.133 | 0.497 | 0.18 | 0.04 | 0.41 | 0.54 | 0.3 | 0.25 |
| 9 | 1285715 | rs7848287 | -0.838 | -0.453 | -0.182 | -0.638 | -0.012 | 0.042 | 0.042 | 0.008 | 0.57 | 0.63 | 0.71 | 0.87 | 0.53 | 0.61 |
| 9 | 1288255 | rs10960798 | 1.164 | -0.529 | -0.095 | -0.643 | -0.015 | 0.08 | 0.08 | 0 | 0.38 | 0.25 | 0.23 | 0.13 | 0.37 | 0.38 |
| 9 | 1288574 | rs7870599 | 0.879 | -0.777 | 0.189 | -0.559 | -0.052 | -0.001 | -0.001 | 1.23 | 0.2 | 0.35 | 0.15 | 0.23 | 0.09 | 0.2 |
| 9 | 1290551 | rs4314695 | -0.886 | -0.775 | 0.194 | -0.577 | -0.008 | 0.03 | 0.03 | 0.019 | 0.56 | 0.61 | 0.7 | 0.87 | 0.52 | 0.61 |
| 9 | 1291196 | rs4373571 | 0.1 | -0.774 | 0.267 | -0.559 | -0.005 | 0.001 | 0.001 | 0.07 | 0.36 | 0.28 | 0.56 | 0.64 | 0.42 | 0.41 |
| 9 | 1293445 | rs9298685 | 0.793 | -0.779 | 0.291 | -0.528 | -0.063 | -0.008 | -0.008 | 0 | 0.21 | 0.39 | 0.16 | 0.23 | 0.09 | 0.2 |
| 9 | 1294000 | rs10122927 | 1.089 | -0.905 | 0.401 | -0.59 | 0.042 | -0.012 | -0.012 | 0 | 0.17 | 0.06 | 0.12 | 0.23 | 0.09 | 0.2 |
| 9 | 1294441 | rs4618752 | 0.198 | -0.832 | 0.379 | -0.606 | -0.015 | -0.006 | -0.006 | 0 | 0.34 | 0.22 | 0.55 | 0.63 | 0.41 | 0.41 |
| 9 | 1294789 | rs9969818 | 0.732 | -0.767 | 0.578 | -0.489 | -0.004 | 0 | 0 | 0 | 0.18 | 0.17 | 0.13 | 0.23 | 0.09 | 0.2 |
| 9 | 1304815 | rs7864098 | 0.832 | -0.594 | -0.267 | -0.409 | 0.029 | -0.004 | -0.004 | 0 | 0.18 | 0.1 | 0.12 | 0.23 | 0.09 | 0.2 |
| 9 | 1308288 | rs1340010 | 0.049 | -0.359 | -0.267 | -0.179 | -0.01 | 0.002 | 0.002 | 0.347 | 0.33 | 0.23 | 0.54 | 0.67 | 0.4 | 0.38 |
| 9 | 1309997 | rs7028809 | 0.327 | -0.829 | 0.014 | -0.077 | -0.011 | 0.001 | 0.001 | 0 | 0.22 | 0.28 | 0.22 | 0.26 | 0.14 | 0.23 |
| 9 | 1310080 | rs4741252 | 0.06 | -0.982 | -0.445 | -0.004 | 0.018 | 0.009 | 0.009 | 1.239 | 0.58 | 0.27 | 0.56 | 0.4 | 0.64 | 0.67 |
| 9 | 1310475 | rs7048584 | -0.86 | -1.108 | -0.497 | -0.35 | -0.051 | 0.011 | 0.011 | 0 | 0.69 | 0.84 | 0.49 | 0.31 | 0.58 | 0.66 |
| 9 | 1311050 | rs7041725 | -0.762 | -0.977 | -0.007 | -1.072 | 0.002 | -0.053 | -0.053 | 5.11 | 0.67 | 0.58 | 0.78 | 0.91 | 0.68 | 0.85 |
| 9 | 1311310 | rs16927344 | -0.19 | -1.305 | -0.013 | -0.927 | -0.05 | -0.023 | -0.023 | 0 | 0.14 | 0.23 | 0.04 | 0.08 | 0.04 | 0.17 |
| 9 | 1315843 | rs10809892 | -0.676 | -0.556 | -0.455 | -1.206 | -0.002 | 0.008 | 0.008 | 0 | 0.45 | 0.46 | 0.61 | 0.82 | 0.51 | 0.55 |
| 9 | 1315952 | rs10809893 | -0.4 | -0.676 | -0.349 | -1.147 | -0.034 | 0.052 | 0.052 | 0.813 | 0.41 | 0.27 | 0.6 | 0.78 | 0.51 | 0.53 |
| 9 | 1316816 | rs7044362 | -0.66 | -0.632 | -0.449 | -1.075 | -0.003 | 0.005 | 0.005 | 0.563 | 0.43 | 0.44 | 0.61 | 0.75 | 0.5 | 0.55 |
| 9 | 1317163 | rs10960938 | 0.158 | -0.535 | -1.2 | -1.173 | 0.023 | 0.054 | 0.054 | 0.043 | 0.24 | 0.11 | 0.13 | 0.17 | 0.19 | 0.29 |
| 9 | 1319316 | rs7853956 | 0.376 | -0.604 | -1.342 | -1.3 | 0.024 | 0.083 | 0.083 | 2.653 | 0.25 | 0.09 | 0.09 | 0.09 | 0.17 | 0.31 |
| 9 | 1320351 | rs17740968 | 1.604 | -0.044 | -1.318 | -1.196 | -0.038 | 0.032 | 0.032 | 0 | 0.06 | 0.01 | 0.15 | 0 | 0.14 | 0.03 |
| 9 | 1321518 | rs7869745 | -1.141 | -0.56 | -0.948 | -1.307 | -0.01 | 0.071 | 0.071 | 2.258 | 0.47 | 0.7 | 0.6 | 0.72 | 0.47 | 0.53 |
| 9 | 1321793 | rs10809905 | 0.742 | -0.41 | -0.646 | -1.341 | -0.031 | -0.039 | -0.039 | 0 | 0.18 | 0.13 | 0.47 | 0.61 | 0.3 | 0.21 |
| 9 | 1322136 | rs10491865 | -2.473 | -0.992 | -0.721 | -1.855 | -0.001 | -0.008 | -0.008 | 0.307 | 0.72 | 0.74 | 0.72 | 0.92 | 0.67 | 0.84 |
| 9 | 1322730 | rs9792626 | -0.695 | -1.431 | -0.353 | -2.082 | 0 | -0.049 | -0.049 | 1.404 | 0.32 | 0.5 | 0.27 | 0.2 | 0.29 | 0.4 |
| 9 | 1323164 | rs4740541 | -2.632 | -1.116 | -0.305 | -1.885 | -0.004 | 0.002 | 0.002 | 0 | 0.7 | 0.76 | 0.73 | 0.94 | 0.64 | 0.82 |
| 9 | 1324127 | rs1340007 | -0.402 | -0.679 | -0.159 | -1.702 | -0.057 | 0.105 | 0.105 | 0 | 0.31 | 0.15 | 0.27 | 0.04 | 0.4 | 0.47 |
| 9 | 1324647 | rs1538714 | 0.618 | -0.635 | 0.309 | -1.004 | -0.01 | -0.042 | -0.042 | 0 | 0.13 | 0.09 | 0.36 | 0.68 | 0.16 | 0.16 |
| 9 | 1328424 | rs16927623 | 0.827 | -0.575 | 0.191 | -0.876 | -0.015 | -0.048 | -0.048 | 0 | 0.13 | 0.06 | 0.35 | 0.62 | 0.16 | 0.16 |
| 9 | 1328742 | rs10491864 | 0.342 | -1.139 | 0.359 | -1.173 | -0.004 | 0.018 | 0.018 | 2.837 | 0.16 | 0.22 | 0.36 | 0.64 | 0.15 | 0.16 |
| 9 | 1330623 | rs10809918 | -0.038 | -1.406 | 0.088 | -1.356 | 0.037 | 0.021 | 0.021 | 0 | 0.26 | 0.13 | 0.39 | 0.78 | 0.2 | 0.26 |
| 9 | 1331350 | rs7875816 | 0.12 | -1.426 | 0.319 | -1.855 | 0.005 | -0.006 | -0.006 | 1.024 | 0.19 | 0.14 | 0.1 | 0.03 | 0.17 | 0.29 |
| 9 | 1332257 | rs999924 | -0.552 | -1.439 | 0.515 | -1.765 | -0.012 | 0.015 | 0.015 | 0 | 0.31 | 0.4 | 0.45 | 0.63 | 0.3 | 0.43 |
| 9 | 1334190 | rs10809921 | 0.734 | -1.268 | -0.242 | -1.709 | 0.006 | -0.012 | -0.012 | 0 | 0.05 | 0.04 | 0.02 | 0.13 | 0.03 | 0.09 |
| 9 | 1336460 | rs4741290 | 1.467 | -1.149 | -0.386 | -1.618 | -0.003 | 0.009 | 0.009 | 7.442 | 0.1 | 0.07 | 0.29 | 0.59 | 0.15 | 0.16 |
| 9 | 1337835 | rs7855990 | 0.671 | -1.184 | -0.528 | -1.94 | -0.008 | 0.074 | 0.074 | 0 | 0.14 | 0.04 | 0.07 | 0.01 | 0.13 | 0.22 |
| 9 | 1339942 | rs12345947 | -1.038 | -1.526 | -0.245 | -2.145 | 0.04 | 0.034 | 0.034 | 1.896 | 0.42 | 0.62 | 0.2 | 0.31 | 0.25 | 0.5 |

| | | | | | | | | | | | | | | | | |
|---|---|---|---|---|---|---|---|---|---|---|---|---|---|---|---|---|
| 9 | 1340735 | rs10756470 | -1.387 | -1.426 | -0.276 | -2.089 | 0.048 | 0.129 | 0.129 | 0 | 0.53 | 0.68 | 0.49 | 0.92 | 0.39 | 0.66 |
| 9 | 1342328 | rs10961025 | 0.094 | -1.563 | 0.034 | -2.028 | -0.002 | 0.053 | 0.053 | 0 | 0.08 | 0.23 | 0.05 | 0.15 | 0.05 | 0.15 |
| 9 | 1342663 | rs16927838 | 0.997 | -1.449 | -0.798 | -1.999 | 0.042 | -0.029 | -0.029 | 0 | 0.14 | 0.03 | 0.05 | 0.18 | 0.06 | 0.12 |
| 9 | 1342763 | rs2226014 | -2.017 | -0.891 | -0.831 | -2.169 | -0.028 | 0.109 | 0.109 | 0 | 0.68 | 0.83 | 0.75 | 0.95 | 0.64 | 0.82 |
| 9 | 1346388 | rs10809943 | -0.976 | -0.953 | -0.701 | -2.018 | 0.063 | 0.179 | 0.179 | 0 | 0.52 | 0.71 | 0.49 | 0.93 | 0.39 | 0.64 |
| 9 | 1347094 | rs12684796 | 1.091 | -0.803 | -0.515 | -1.468 | -0.001 | 0.027 | 0.027 | 7.146 | 0.11 | 0.11 | 0.3 | 0.6 | 0.16 | 0.17 |
| 9 | 1347581 | rs7037591 | 0.673 | -0.882 | -0.679 | -1.806 | 0.035 | -0.02 | -0.02 | 5.486 | 0.15 | 0.03 | 0.05 | 0.17 | 0.07 | 0.12 |
| 9 | 1350453 | rs7866259 | 1.572 | -0.834 | -0.035 | -2.099 | 0.013 | -0.074 | -0.074 | 0 | 0.3 | 0.39 | 0.27 | 0.08 | 0.36 | 0.22 |
| 9 | 1351141 | rs7031390 | -0.823 | -1.061 | 0.05 | -2.327 | -0.009 | 0.064 | 0.064 | 0 | 0.56 | 0.69 | 0.67 | 0.92 | 0.51 | 0.57 |
| 9 | 1353353 | rs7021191 | 1.131 | -0.96 | -0.026 | -2.162 | -0.005 | -0.016 | -0.016 | 0.319 | 0.12 | 0.08 | 0.3 | 0.73 | 0.16 | 0.23 |
| 9 | 1354036 | rs7871293 | 0.578 | -1.142 | 0.297 | -2.301 | -0.014 | -0.017 | -0.017 | 0.379 | 0.66 | 0.45 | 0.59 | 0.27 | 0.7 | 0.54 |
| 9 | 1355029 | rs1412252 | 0.862 | -0.984 | 0.272 | -2.294 | -0.013 | 0.064 | 0.064 | 10.921 | 0.45 | 0.7 | 0.59 | 0.82 | 0.53 | 0.46 |
| 9 | 1355061 | rs1538721 | -0.315 | -0.808 | 0.022 | -2.157 | -0.012 | 0.079 | 0.079 | 4.647 | 0.35 | 0.16 | 0.31 | 0.17 | 0.32 | 0.32 |
| 9 | 1356482 | rs9406460 | 1.003 | -0.817 | 0.77 | -1.729 | 0.043 | 0.092 | 0.092 | 0 | 0.53 | 0.29 | 0.52 | 0.21 | 0.63 | 0.43 |
| 9 | 1358244 | rs682927 | 1.307 | -0.762 | 0.341 | -1.672 | 0.026 | 0.109 | 0.109 | 0 | 0.5 | 0.31 | 0.36 | 0.19 | 0.56 | 0.43 |
| 9 | 1358574 | rs650902 | 0.644 | -1.146 | 0.301 | -1.472 | -0.051 | -0.06 | -0.06 | 0 | 0.65 | 0.54 | 0.82 | 0.82 | 0.8 | 0.63 |
| 9 | 1360104 | rs16928128 | 0.131 | -0.166 | -0.09 | -0.735 | -0.012 | -0.015 | -0.015 | 1.871 | 0.11 | 0.09 | 0.32 | 0.65 | 0.16 | 0.17 |
| 9 | 1362503 | rs10961113 | -0.94 | -0.922 | 0.227 | -1.702 | 0.001 | -0.023 | -0.023 | 0 | 0.14 | 0.15 | 0.05 | 0.01 | 0.1 | 0.12 |
| 9 | 1367182 | rs2375528 | -1.323 | -1.152 | 0.575 | -2.013 | -0.024 | -0.127 | -0.127 | 0 | 0.26 | 0.58 | 0.16 | 0.01 | 0.18 | 0.2 |
| 9 | 1373944 | rs646785 | 1.432 | -1.01 | 0.364 | -2.082 | -0.001 | -0.082 | -0.082 | 0 | 0.62 | 0.34 | 0.67 | 0.99 | 0.66 | 0.73 |
| 9 | 1383446 | rs17743398 | -0.551 | -0.481 | -0.73 | -1.764 | 0.004 | 0.013 | 0.013 | 0 | 0.14 | 0.08 | 0.16 | 0.01 | 0.12 | 0.11 |
| 9 | 1383919 | rs998573 | -0.525 | -0.524 | -1.044 | -1.878 | -0.004 | 0.054 | 0.054 | 0 | 0.23 | 0.13 | 0.15 | 0 | 0.24 | 0.19 |
| 9 | 1385087 | rs10961225 | -0.802 | -0.209 | -0.809 | -1.602 | -0.01 | -0.164 | -0.164 | 0.319 | 0.39 | 0.55 | 0.34 | 0.01 | 0.35 | 0.32 |
| 9 | 1387193 | rs627024 | 0.517 | 0.038 | -1.482 | -0.294 | -0.016 | -0.088 | -0.088 | 0 | 0.82 | 0.99 | 0.81 | 0.72 | 0.85 | 0.84 |
| 9 | 1387954 | rs677079 | -0.439 | -0.056 | -1.348 | -0.89 | 0.013 | 0.026 | 0.026 | 0 | 0.14 | 0.07 | 0.07 | 0.07 | 0.1 | 0.12 |
| 9 | 1391887 | rs590938 | -0.089 | -0.081 | -1.374 | -0.915 | 0.014 | 0.032 | 0.032 | 0 | 0.11 | 0.02 | 0.05 | 0.07 | 0.08 | 0.07 |
| 9 | 1393733 | rs10961278 | -0.13 | 0.65 | -1.377 | -0.927 | -0.006 | 0.019 | 0.019 | 0 | 0.11 | 0.07 | 0.07 | 0.06 | 0.09 | 0.1 |
| 9 | 1403492 | rs659105 | 0.835 | 1.56 | -0.708 | -1.347 | -0.014 | 0.066 | 0.066 | 0.812 | 0.57 | 0.23 | 0.42 | 0.27 | 0.53 | 0.48 |
| 9 | 1403749 | rs6474798 | -0.064 | 1.161 | 0.256 | -1.657 | 0.034 | 0.012 | 0.012 | 0.337 | 0.74 | 0.64 | 0.46 | 0.34 | 0.66 | 0.6 |
| 9 | 1404008 | rs6474799 | -0.884 | 1.047 | 0.696 | -1.561 | 0.011 | -0.069 | -0.069 | 0 | 0.25 | 0.65 | 0.18 | 0.09 | 0.23 | 0.18 |
| 9 | 1405732 | rs10810061 | -0.867 | 1.054 | 1.148 | -1.504 | 0.011 | -0.05 | -0.05 | 0 | 0.25 | 0.65 | 0.18 | 0.11 | 0.24 | 0.19 |
| 9 | 1411571 | rs1412248 | -0.498 | 1.004 | 1.408 | -1.516 | 0.026 | -0.073 | -0.073 | 2.639 | 0.21 | 0.37 | 0.15 | 0.08 | 0.21 | 0.14 |
| 9 | 1413376 | rs7873128 | -0.008 | 0.468 | 1.1 | -1.764 | -0.001 | -0.003 | -0.003 | 0 | 0.73 | 0.79 | 0.57 | 0.33 | 0.71 | 0.58 |
| 9 | 1413700 | rs10119492 | 0.013 | 0.881 | 1.425 | -1.322 | -0.001 | 0.113 | 0.113 | 0 | 0.16 | 0.37 | 0.36 | 0.58 | 0.14 | 0.28 |
| 9 | 1416909 | rs16928755 | 0.637 | 0.936 | 1.164 | -1.301 | 0 | 0.027 | 0.027 | 0 | 0.13 | 0.11 | 0.34 | 0.55 | 0.14 | 0.25 |
| 9 | 1417953 | rs10120239 | -0.823 | 0.704 | 0.629 | -1.597 | -0.001 | -0.023 | -0.023 | 0 | 0.25 | 0.33 | 0.12 | 0.08 | 0.18 | 0.14 |
| 9 | 1428059 | rs10961410 | 0.668 | 0.85 | 0.234 | -1.475 | -0.021 | 0.041 | 0.041 | 0 | 0.13 | 0.03 | 0.06 | 0.07 | 0.12 | 0.11 |
| 9 | 1429660 | rs10961423 | -0.189 | 0.916 | 0.314 | -1.511 | -0.003 | 0.037 | 0.037 | 0 | 0.18 | 0.08 | 0.09 | 0.08 | 0.16 | 0.12 |
| 9 | 1435496 | rs880013 | -1.524 | 1.008 | 0.493 | -1.611 | -0.008 | 0.033 | 0.033 | 0.863 | 0.44 | 0.7 | 0.63 | 0.71 | 0.45 | 0.44 |
| 9 | 1436134 | rs876977 | 0.122 | 1.038 | 0.235 | -1.606 | -0.004 | 0.005 | 0.005 | 0.019 | 0.2 | 0.22 | 0.49 | 0.66 | 0.21 | 0.3 |
| 9 | 1436383 | rs876976 | -1.384 | 1.851 | -0.218 | -1.822 | 0.017 | -0.096 | -0.096 | 0 | 0.32 | 0.39 | 0.22 | 0.03 | 0.36 | 0.26 |
| 9 | 1438677 | rs1412260 | -0.379 | 1.79 | -0.438 | -1.652 | 0.018 | -0.044 | -0.044 | 0 | 0.17 | 0.25 | 0.12 | 0.03 | 0.22 | 0.12 |
| 9 | 1444746 | rs9406475 | -0.85 | 1.38 | -1.212 | -1.773 | -0.035 | 0.142 | 0.142 | 0 | 0.31 | 0.07 | 0.17 | 0.09 | 0.3 | 0.22 |
| 9 | 1446149 | rs604829 | 0.381 | 1.341 | -1.087 | -1.576 | -0.01 | 0.048 | 0.048 | 0 | 0.13 | 0.02 | 0.07 | 0.04 | 0.12 | 0.08 |
| 9 | 1446459 | rs4741361 | -0.476 | 1.452 | -1.241 | -1.817 | -0.039 | -0.014 | -0.014 | 0 | 0.24 | 0.19 | 0.52 | 0.68 | 0.36 | 0.42 |
| 9 | 1447011 | rs10122783 | 1.373 | 0.816 | -0.685 | -2.299 | 0.007 | 0.047 | 0.047 | 0.028 | 0.45 | 0.31 | 0.33 | 0.2 | 0.43 | 0.34 |
| 9 | 1448247 | rs9407572 | 0.399 | 0.983 | -0.721 | -2.212 | -0.003 | 0.024 | 0.024 | 0 | 0.18 | 0.13 | 0.11 | 0.04 | 0.18 | 0.11 |
| 9 | 1449084 | rs13294519 | 0.647 | 1.164 | -0.734 | -2.228 | -0.007 | -0.028 | -0.028 | 0 | 0.15 | 0.11 | 0.45 | 0.68 | 0.21 | 0.39 |
| 9 | 1449326 | rs10756548 | 0.318 | 1.195 | -0.807 | -2.227 | -0.002 | 0.026 | 0.026 | 0 | 0.19 | 0.13 | 0.11 | 0.04 | 0.18 | 0.11 |
| 9 | 1450274 | rs7037453 | 0.569 | 0.958 | -0.789 | -2.183 | -0.004 | 0.01 | 0.01 | 3.104 | 0.81 | 0.86 | 0.82 | 0.93 | 0.8 | 0.86 |
| 9 | 1450428 | rs7036822 | 0.045 | 0.974 | -0.755 | -2.144 | -0.002 | -0.001 | -0.001 | 0.021 | 0.85 | 0.87 | 0.84 | 0.93 | 0.83 | 0.86 |
| 9 | 1453731 | rs1332004 | 1.368 | 0.878 | -0.677 | -2.224 | 0.015 | 0.048 | 0.048 | 1.652 | 0.46 | 0.31 | 0.32 | 0.19 | 0.42 | 0.35 |
| 9 | 1454200 | rs4142436 | -0.552 | 0.956 | -0.492 | -2.195 | -0.001 | -0.04 | -0.04 | 1.046 | 0.87 | 0.8 | 0.94 | 0.91 | 0.91 | 0.87 |
| 9 | 1455409 | rs10810165 | 0.198 | 1.067 | -0.862 | -2.075 | -0.006 | -0.011 | -0.011 | 0 | 0.2 | 0.21 | 0.15 | 0.04 | 0.26 | 0.15 |
| 9 | 1457305 | rs10961539 | 0.38 | 0.582 | -1.272 | -1.644 | 0.02 | 0.052 | 0.052 | 2.054 | 0.47 | 0.34 | 0.32 | 0.21 | 0.42 | 0.36 |
| 9 | 1460365 | rs2210421 | 0.036 | 0.524 | -2.089 | -1.166 | 0.026 | 0.034 | 0.034 | 0.334 | 0.42 | 0.25 | 0.43 | 0.27 | 0.45 | 0.2 |
| 9 | 1460427 | rs10961548 | -0.476 | 0.736 | -2.812 | -1.839 | -0.107 | 0.224 | 0.224 | 0.191 | 0.34 | 0.05 | 0.2 | 0.05 | 0.39 | 0.29 |
| 9 | 1463618 | rs10124818 | -0.746 | -0.32 | -2.711 | -1.618 | 0.062 | 0.101 | 0.101 | 0.617 | 0.33 | 0.83 | 0.26 | 0.14 | 0.19 | 0.17 |
| 9 | 1463904 | rs1339488 | -0.857 | -0.319 | -3.769 | -2.359 | -0.027 | 0.131 | 0.131 | 2.336 | 0.69 | 0.94 | 0.76 | 0.92 | 0.74 | 0.75 |
| 9 | 1464762 | rs7857963 | -0.1 | -0.582 | -3.246 | -2.458 | 0.001 | -0.015 | -0.015 | 0.293 | 0.27 | 0.69 | 0.22 | 0.08 | 0.15 | 0.14 |
| 9 | 1466336 | rs10961586 | 0.955 | 0.264 | -3.716 | -2.767 | -0.116 | 0.139 | 0.139 | 0.475 | 0.23 | 0.04 | 0.28 | 0.02 | 0.38 | 0.17 |
| 9 | 1468035 | rs16929726 | -0.303 | -0.19 | -2.541 | -2.789 | -0.008 | -0.032 | -0.032 | 1.796 | 0.2 | 0.24 | 0.15 | 0.09 | 0.15 | 0.07 |
| 9 | 1468923 | rs16929773 | 0.677 | -0.267 | -2.493 | -2.785 | 0.008 | 0.007 | 0.007 | 1.379 | 0.12 | 0.09 | 0.11 | 0.03 | 0.08 | 0.06 |
| 9 | 1472542 | rs7041813 | -1.509 | -0.263 | -2.373 | -2.943 | -0.024 | 0.116 | 0.116 | 1.931 | 0.71 | 0.95 | 0.76 | 0.91 | 0.76 | 0.8 |
| 9 | 1474523 | rs10961637 | 0.475 | -0.398 | -2.429 | -2.907 | -0.072 | 0.13 | 0.13 | 0.117 | 0.39 | 0.11 | 0.53 | 0.79 | 0.58 | 0.6 |
| 9 | 1475567 | rs10119014 | -0.563 | -0.533 | -1.537 | -2.771 | -0.029 | -0.105 | -0.105 | 0 | 0.25 | 0.53 | 0.18 | 0.11 | 0.17 | 0.17 |
| 9 | 1481609 | rs12343051 | -0.974 | -0.68 | -2.093 | -2.788 | 0.032 | 0.045 | 0.045 | 0.098 | 0.31 | 0.81 | 0.2 | 0.11 | 0.17 | 0.17 |
| 9 | 1484527 | rs4741416 | -0.489 | -0.492 | -2.295 | -2.605 | 0.008 | 0.156 | 0.156 | 5.513 | 0.56 | 0.2 | 0.64 | 0.87 | 0.64 | 0.72 |
| 9 | 1486723 | rs7859959 | 0.466 | -0.398 | -2.311 | -2.01 | -0.085 | 0.062 | 0.062 | 0.555 | 0.38 | 0.12 | 0.51 | 0.73 | 0.53 | 0.56 |

| | | | | | | | | | | | | | | | | |
|---|---|---|---|---|---|---|---|---|---|---|---|---|---|---|---|---|
| 9 | 1487108 | rs7864191 | -1.486 | -0.406 | -2.347 | -2.178 | -0.027 | 0.119 | 0.119 | 3.137 | 0.69 | 0.93 | 0.72 | 0.91 | 0.7 | 0.74 |
| 9 | 1488208 | rs2375535 | -0.651 | -0.501 | -2.188 | -2.085 | -0.024 | -0.049 | -0.049 | 0 | 0.19 | 0.68 | 0.09 | 0.1 | 0.09 | 0.12 |
| 9 | 1488355 | rs10124962 | -1.516 | -0.514 | -2.341 | -2.042 | -0.006 | 0.082 | 0.082 | 2.385 | 0.68 | 0.9 | 0.7 | 0.89 | 0.68 | 0.75 |
| 9 | 1488655 | rs10961710 | 0.688 | 0.226 | -2.858 | -2.066 | -0.112 | -0.057 | -0.057 | 5.591 | 0.25 | 0.04 | 0.19 | 0.7 | 0.34 | 0.48 |
| 9 | 1489454 | rs10961719 | 0.367 | 0.215 | -2.811 | -2.131 | -0.016 | 0.081 | 0.081 | 4.028 | 0.27 | 0.12 | 0.41 | 0.09 | 0.28 | 0.17 |
| 9 | 1490577 | rs7854400 | -1.875 | 0.167 | -2.709 | -1.96 | -0.007 | 0.072 | 0.072 | 1.955 | 0.68 | 0.86 | 0.71 | 0.92 | 0.68 | 0.76 |
| 9 | 1490830 | rs10810253 | 0.869 | 0.557 | -2.988 | -1.627 | -0.107 | 0.049 | 0.049 | 2.68 | 0.26 | 0.04 | 0.28 | 0.09 | 0.34 | 0.3 |
| 9 | 1491213 | rs12379509 | 0.105 | -0.017 | -2.709 | -1.793 | -0.134 | 0.086 | 0.086 | 0.672 | 0.39 | 0.07 | 0.39 | 0.75 | 0.52 | 0.57 |
| 9 | 1492266 | rs10961743 | -0.815 | -0.187 | -1.202 | -1.214 | -0.038 | -0.037 | -0.037 | 0 | 0.27 | 0.67 | 0.32 | 0.2 | 0.16 | 0.2 |
| 9 | 1495780 | rs7846887 | -0.003 | 0.399 | -1.37 | -0.888 | -0.003 | 0.003 | 0.003 | 0 | 0.08 | 0.05 | 0.18 | 0.05 | 0.1 | 0.06 |
| 9 | 1496230 | rs4740589 | -0.398 | 0.817 | -1.335 | -0.96 | -0.006 | 0.004 | 0.004 | 0 | 0.28 | 0.3 | 0.5 | 0.19 | 0.35 | 0.3 |
| 9 | 1499001 | rs10756623 | -0.055 | 0.512 | -1.965 | -1.162 | -0.187 | 0.014 | 0.014 | 0 | 0.34 | 0.05 | 0.35 | 0.76 | 0.45 | 0.53 |
| 9 | 1499641 | rs1331999 | -1.198 | 0.041 | -1.287 | -1.441 | 0.018 | 0.062 | 0.062 | 0 | 0.55 | 0.31 | 0.73 | 0.86 | 0.61 | 0.74 |
| 9 | 1501815 | rs2095023 | -1.091 | -0.223 | -0.981 | -1.442 | -0.005 | -0.031 | -0.031 | 0 | 0.57 | 0.48 | 0.72 | 0.83 | 0.6 | 0.71 |
| 9 | 1501977 | rs10961794 | 0.188 | -0.264 | -1.039 | -1.446 | -0.04 | -0.037 | -0.037 | 0 | 0.17 | 0.25 | 0.34 | 0.1 | 0.1 | 0.2 |
| 9 | 1502113 | rs7025212 | -1.932 | -0.626 | -1.329 | -1.61 | -0.051 | 0.046 | 0.046 | 0.977 | 0.72 | 0.96 | 0.83 | 0.87 | 0.74 | 0.75 |
| 9 | 1503356 | rs1576748 | -1.723 | -0.7 | -0.645 | -1.644 | 0.001 | 0.006 | 0.006 | 1.106 | 0.71 | 0.78 | 0.82 | 0.87 | 0.74 | 0.74 |
| 9 | 1505268 | rs955956 | -0.893 | -0.179 | -0.726 | -1.74 | -0.009 | 0.137 | 0.137 | 0.205 | 0.48 | 0.78 | 0.64 | 0.18 | 0.52 | 0.34 |
| 9 | 1505457 | rs1331998 | -0.732 | -0.175 | -0.693 | -1.771 | 0 | 0.115 | 0.115 | 0.714 | 0.44 | 0.76 | 0.61 | 0.17 | 0.46 | 0.31 |
| 9 | 1506101 | rs10756631 | -0.741 | -0.153 | -0.249 | -1.805 | 0.001 | -0.027 | -0.027 | 0 | 0.44 | 0.53 | 0.62 | 0.18 | 0.52 | 0.34 |
| 9 | 1507712 | rs16930214 | 0.159 | -0.137 | -0.09 | -1.741 | -0.001 | -0.027 | -0.027 | 0 | 0.06 | 0.14 | 0.04 | 0.02 | 0.05 | 0.03 |
| 9 | 1510230 | rs1412254 | -2.328 | -0.046 | -0.257 | -1.796 | 0.004 | 0.076 | 0.076 | 0 | 0.76 | 0.96 | 0.91 | 0.91 | 0.83 | 0.75 |
| 9 | 1510495 | rs10491858 | 1.253 | -0.164 | -0.043 | -1.79 | -0.001 | 0 | 0 | 0 | 0.11 | 0.14 | 0.07 | 0.07 | 0.14 | 0.03 |
| 9 | 1511184 | rs7019932 | 1.121 | -0.175 | 0.052 | -1.795 | 0.001 | -0.013 | -0.013 | 0 | 0.11 | 0.2 | 0.08 | 0.07 | 0.14 | 0.03 |
| 9 | 1511341 | rs9918936 | 0.258 | -0.208 | 0.087 | -1.894 | -0.013 | 0.019 | 0.019 | 0 | 0.32 | 0.24 | 0.53 | 0.14 | 0.4 | 0.3 |
| 9 | 1512169 | rs10961842 | 1.832 | -0.354 | 0.196 | -1.954 | -0.015 | -0.003 | -0.003 | 1.261 | 0.08 | 0.08 | 0.21 | 0.07 | 0.06 | 0.12 |
| 9 | 1512232 | rs10491857 | 1.173 | -0.131 | 0.095 | -1.66 | 0.002 | -0.003 | -0.003 | 0 | 0.11 | 0.18 | 0.07 | 0.09 | 0.14 | 0.03 |
| 9 | 1512369 | rs10810330 | -0.665 | -0.14 | 0.094 | -1.721 | 0.001 | -0.037 | -0.037 | 0 | 0.45 | 0.53 | 0.6 | 0.21 | 0.5 | 0.34 |
| 9 | 1513074 | rs6474880 | -2.199 | -0.166 | -0.114 | -1.722 | -0.003 | 0.081 | 0.081 | 0.147 | 0.75 | 0.95 | 0.88 | 0.9 | 0.81 | 0.74 |
| 9 | 1513849 | rs1999435 | 0.226 | -0.103 | -0.022 | -1.641 | -0.004 | 0.013 | 0.013 | 0 | 0.32 | 0.25 | 0.51 | 0.14 | 0.36 | 0.31 |
| 9 | 1513917 | rs2050757 | 0.443 | -0.111 | 0.009 | -1.625 | -0.003 | 0.01 | 0.01 | 0 | 0.31 | 0.24 | 0.51 | 0.14 | 0.35 | 0.3 |
| 9 | 1519043 | rs7022816 | -0.348 | -0.564 | 0.089 | -1.08 | -0.016 | -0.007 | -0.007 | 0 | 0.38 | 0.54 | 0.28 | 0.68 | 0.33 | 0.42 |
| 9 | 1520317 | rs10756653 | -0.264 | -0.553 | -0.031 | -1.12 | 0.005 | 0.05 | 0.05 | 0.087 | 0.38 | 0.24 | 0.3 | 0.76 | 0.39 | 0.42 |
| 9 | 1521178 | rs7856537 | -2.22 | -0.786 | -0.265 | -1.237 | 0.003 | 0.004 | 0.004 | 0 | 0.83 | 0.87 | 0.9 | 0.9 | 0.88 | 0.77 |
| 9 | 1521204 | rs9886784 | -0.853 | -0.86 | -0.141 | -1.244 | -0.018 | -0.013 | -0.013 | 0 | 0.75 | 0.54 | 0.84 | 0.81 | 0.81 | 0.74 |
| 9 | 1521294 | rs10961888 | 0.627 | -0.843 | -0.153 | -1.085 | -0.003 | 0.002 | 0.002 | 0 | 0.32 | 0.27 | 0.53 | 0.23 | 0.39 | 0.41 |
| 9 | 1523296 | rs7048299 | 1.138 | -0.803 | -0.137 | -0.943 | 0.004 | 0.019 | 0.019 | 0 | 0.07 | 0.21 | 0.04 | 0.09 | 0.06 | 0.02 |
| 9 | 1526250 | rs10810355 | 0.977 | -0.908 | -0.327 | -0.978 | -0.005 | -0.096 | -0.096 | 0.011 | 0.28 | 0.1 | 0.22 | 0.58 | 0.27 | 0.38 |
| 9 | 1526834 | rs1888838 | 1.003 | -0.912 | -0.34 | -0.976 | -0.015 | -0.056 | -0.056 | 0 | 0.28 | 0.07 | 0.21 | 0.67 | 0.26 | 0.38 |
| 9 | 1527191 | rs10961907 | 0.289 | -0.899 | -0.231 | -1.098 | 0.009 | -0.001 | -0.001 | 0 | 0.35 | 0.29 | 0.26 | 0.68 | 0.33 | 0.4 |
| 9 | 1527227 | rs10961908 | 0.29 | -0.901 | -0.232 | -1.099 | 0.009 | -0.001 | -0.001 | 0 | 0.35 | 0.29 | 0.26 | 0.68 | 0.33 | 0.4 |
| 9 | 1527703 | rs10961911 | -0.234 | -0.951 | 0.03 | -1.126 | -0.011 | -0.009 | -0.009 | 0.084 | 0.39 | 0.58 | 0.29 | 0.7 | 0.33 | 0.41 |
| 9 | 1528388 | rs944608 | -0.245 | -0.819 | -0.484 | -1.226 | 0.005 | 0.093 | 0.093 | 0 | 0.48 | 0.66 | 0.66 | 0.22 | 0.54 | 0.34 |
| 9 | 1533064 | rs1114132 | -0.874 | -0.981 | 0.069 | -1.031 | -0.003 | -0.017 | -0.017 | 0.085 | 0.61 | 0.61 | 0.76 | 0.85 | 0.68 | 0.68 |
| 9 | 1533757 | rs1745886 | -0.347 | -1.152 | 0.053 | -0.953 | 0.007 | 0.014 | 0.014 | 0 | 0.49 | 0.44 | 0.66 | 0.75 | 0.59 | 0.53 |
| 9 | 1534205 | rs1796902 | -0.348 | -1.16 | 0.007 | -0.99 | -0.002 | -0.009 | -0.009 | 0.069 | 0.49 | 0.51 | 0.67 | 0.74 | 0.6 | 0.52 |
| 9 | 1540499 | rs7045593 | 0.154 | -1.128 | 0.063 | -0.95 | -0.047 | -0.055 | -0.055 | 0 | 0.21 | 0.47 | 0.15 | 0.14 | 0.14 | 0.18 |
| 9 | 1550403 | rs17723131 | 1.264 | -0.996 | -0.284 | -0.708 | 0.004 | -0.003 | -0.003 | 0 | 0.08 | 0.05 | 0.19 | 0.15 | 0.06 | 0.11 |
| 9 | 1553462 | rs2486442 | -1.163 | -0.99 | -0.352 | -1.415 | -0.002 | -0.029 | -0.029 | 0 | 0.66 | 0.66 | 0.79 | 0.91 | 0.74 | 0.7 |
| 9 | 1554468 | rs771918 | -1.245 | -0.98 | -0.429 | -1.41 | 0.005 | 0.007 | 0.007 | 0.29 | 0.67 | 0.73 | 0.8 | 0.91 | 0.74 | 0.7 |
| 9 | 1555134 | rs771922 | -1.219 | -0.981 | -0.434 | -1.373 | -0.001 | -0.01 | -0.01 | 0 | 0.67 | 0.71 | 0.8 | 0.89 | 0.74 | 0.7 |
| 9 | 1556054 | rs7873915 | 0.07 | -0.897 | -0.636 | -1.26 | -0.011 | -0.007 | -0.007 | 0 | 0.09 | 0.25 | 0.05 | 0.07 | 0.06 | 0.02 |
| 9 | 1556594 | rs771927 | -1.407 | -0.471 | -1.389 | -1.469 | -0.081 | 0.096 | 0.096 | 2.037 | 0.66 | 0.97 | 0.79 | 0.85 | 0.7 | 0.62 |
| 9 | 1565742 | rs771667 | 0.234 | -0.56 | -1.423 | -1.694 | -0.149 | 0.001 | 0.001 | 0 | 0.34 | 0.11 | 0.43 | 0.68 | 0.5 | 0.35 |
| 9 | 1567276 | rs12551843 | 1.498 | 0.193 | -1.65 | -1.975 | -0.114 | 0.034 | 0.034 | 0 | 0.16 | 0.03 | 0.26 | 0.08 | 0.28 | 0.15 |
| 9 | 1568904 | rs771663 | -1.375 | -0.685 | -1.37 | -2.033 | 0.04 | 0.182 | 0.182 | 0.32 | 0.62 | 0.88 | 0.81 | 0.89 | 0.73 | 0.64 |
| 9 | 1568941 | rs771662 | -0.678 | -0.707 | -0.704 | -2.052 | 0.029 | 0.05 | 0.05 | 0.035 | 0.53 | 0.37 | 0.73 | 0.87 | 0.68 | 0.61 |
| 9 | 1577505 | rs2785727 | -1.127 | -0.823 | -0.697 | -1.99 | 0.024 | 0.146 | 0.146 | 0.068 | 0.58 | 0.76 | 0.75 | 0.89 | 0.69 | 0.6 |
| 9 | 1583103 | rs7874725 | 1.137 | -0.613 | -0.234 | -1.789 | 0.007 | -0.011 | -0.011 | 0 | 0.07 | 0.2 | 0.04 | 0.04 | 0.06 | 0.02 |
| 9 | 1591811 | rs771912 | -0.253 | -0.892 | -0.565 | -2.049 | -0.008 | 0.041 | 0.041 | 0 | 0.43 | 0.59 | 0.64 | 0.77 | 0.54 | 0.46 |
| 9 | 1591888 | rs7873584 | 0.079 | -0.941 | -0.286 | -1.811 | -0.016 | -0.021 | -0.021 | 0.011 | 0.19 | 0.32 | 0.13 | 0.14 | 0.14 | 0.16 |
| 9 | 1596030 | rs10810459 | 1.033 | -0.762 | -0.125 | -1.664 | -0.023 | -0.022 | -0.022 | 0 | 0.13 | 0.28 | 0.1 | 0.09 | 0.09 | 0.15 |
| 9 | 1598874 | rs1853418 | 0.243 | -0.814 | -0.297 | -2 | -0.02 | 0.029 | 0.029 | 0 | 0.37 | 0.28 | 0.6 | 0.75 | 0.52 | 0.44 |
| 9 | 1599073 | rs1853419 | 1.453 | -0.927 | -0.211 | -2.076 | -0.012 | -0.029 | -0.029 | 0 | 0.06 | 0.16 | 0.03 | 0 | 0.03 | 0.01 |
| 9 | 1599200 | rs9298744 | -0.344 | -0.924 | -0.124 | -2.066 | 0.001 | -0.037 | -0.037 | 0 | 0.43 | 0.53 | 0.64 | 0.75 | 0.55 | 0.46 |
| 9 | 1600210 | rs10283925 | -0.97 | -0.972 | -0.206 | -2.121 | 0.019 | 0.099 | 0.099 | 0 | 0.56 | 0.71 | 0.73 | 0.84 | 0.64 | 0.61 |
| 9 | 1603814 | rs702160 | -0.493 | -1.005 | -0.198 | -2.151 | 0.001 | 0.102 | 0.102 | 0 | 0.46 | 0.73 | 0.66 | 0.76 | 0.55 | 0.46 |
| 9 | 1605103 | rs1969771 | 0.099 | -1.185 | -0.537 | -2.52 | 0.014 | 0.089 | 0.089 | 0 | 0.46 | 0.29 | 0.65 | 0.75 | 0.6 | 0.45 |
| 9 | 1605827 | rs702167 | -1.221 | -0.968 | -0.387 | -2.212 | 0.025 | 0.17 | 0.17 | 0.047 | 0.55 | 0.8 | 0.74 | 0.85 | 0.63 | 0.6 |

| | | | | | | | | | | | | | | | | |
|---|---|---|---|---|---|---|---|---|---|---|---|---|---|---|---|---|
| 9 | 1607013 | rs702172 | 1.286 | -1.055 | 0.055 | -1.907 | -0.009 | -0.029 | -0.029 | 0.206 | 0.05 | 0.17 | 0.03 | 0 | 0.03 | 0.01 |
| 9 | 1607940 | rs697047 | -0.76 | -0.973 | -0.014 | -1.979 | -0.015 | 0.076 | 0.076 | 0 | 0.44 | 0.71 | 0.66 | 0.76 | 0.53 | 0.45 |
| 9 | 1610367 | rs702176 | -2.433 | -1.047 | -0.498 | -2.017 | -0.077 | 0.123 | 0.123 | 1.828 | 0.69 | 0.98 | 0.79 | 0.88 | 0.72 | 0.63 |
| 9 | 1610861 | rs7030942 | -0.405 | -1.045 | -0.284 | -2.041 | -0.016 | 0.056 | 0.056 | 0 | 0.43 | 0.33 | 0.6 | 0.76 | 0.53 | 0.45 |
| 9 | 1614282 | rs702177 | -0.894 | -0.981 | -0.238 | -1.857 | -0.054 | -0.098 | -0.098 | 0 | 0.27 | 0.54 | 0.17 | 0.12 | 0.19 | 0.19 |
| 9 | 1616251 | rs702179 | 1.107 | -0.715 | -0.897 | -1.672 | -0.011 | -0.029 | -0.029 | 0 | 0.06 | 0.16 | 0.03 | 0 | 0.03 | 0.01 |
| 9 | 1616918 | rs945657 | 1.216 | -0.034 | -1.275 | -1.683 | -0.078 | 0.038 | 0.038 | 0 | 0.13 | 0.03 | 0.26 | 0.07 | 0.24 | 0.13 |
| 9 | 1617716 | rs10962274 | 1.277 | -0.255 | -1.036 | -1.513 | 0.014 | 0.009 | 0.009 | 0.313 | 0.11 | 0.06 | 0.09 | 0.1 | 0.09 | 0.15 |
| 9 | 1625303 | rs1537966 | -1.499 | -0.289 | -1.183 | -1.743 | 0.007 | 0.172 | 0.172 | 0 | 0.59 | 0.78 | 0.74 | 0.88 | 0.66 | 0.62 |
| 9 | 1626977 | rs12685706 | 0.849 | 0.112 | -1.226 | -1.118 | -0.042 | -0.068 | -0.068 | 0 | 0.11 | 0.02 | 0.11 | 0.53 | 0.14 | 0.17 |
| 9 | 1628468 | rs702188 | -1.067 | -0.205 | -0.676 | -0.872 | -0.001 | 0.026 | 0.026 | 0 | 0.27 | 0.44 | 0.19 | 0.47 | 0.28 | 0.22 |
| 9 | 1629344 | rs771668 | 1.257 | -0.308 | -0.693 | -0.89 | -0.01 | -0.03 | -0.03 | 0 | 0.06 | 0.16 | 0.03 | 0 | 0.03 | 0.01 |
| 9 | 1630823 | rs702189 | -2.364 | -0.423 | -0.912 | -1.041 | -0.053 | 0.174 | 0.174 | 0.195 | 0.63 | 0.96 | 0.77 | 0.86 | 0.69 | 0.62 |
| 9 | 1632203 | rs9407731 | -0.209 | -0.247 | -0.767 | -0.773 | 0.003 | 0.04 | 0.04 | 0 | 0.19 | 0.29 | 0.15 | 0.44 | 0.22 | 0.19 |
| 9 | 1634618 | rs7869423 | -0.547 | -0.31 | -0.48 | -1.017 | -0.003 | -0.044 | -0.044 | 0 | 0.35 | 0.5 | 0.43 | 0.2 | 0.37 | 0.27 |
| 9 | 1635485 | rs2066118 | -0.641 | -0.349 | -0.538 | -1.052 | -0.003 | -0.015 | -0.015 | 0 | 0.37 | 0.51 | 0.45 | 0.27 | 0.38 | 0.26 |
| 9 | 1639619 | rs771889 | -0.662 | -0.31 | 0.026 | -1.036 | 0.012 | 0.033 | 0.033 | 0 | 0.28 | 0.47 | 0.34 | 0.59 | 0.32 | 0.35 |
| 9 | 1639935 | rs796424 | -0.807 | -0.303 | -0.174 | -1.033 | 0.006 | 0.021 | 0.021 | 0 | 0.29 | 0.56 | 0.35 | 0.59 | 0.32 | 0.35 |
| 9 | 1641933 | rs7847664 | -0.366 | -0.111 | -0.075 | -1.055 | 0 | -0.006 | -0.006 | 0 | 0.33 | 0.33 | 0.41 | 0.2 | 0.35 | 0.27 |
| 9 | 1644465 | rs10126007 | 0.091 | -0.116 | -0.131 | -1.135 | -0.029 | 0.073 | 0.073 | 0.039 | 0.23 | 0.11 | 0.32 | 0.08 | 0.27 | 0.18 |
| 9 | 1654030 | rs13289158 | 0.087 | -0.132 | -0.153 | -1.089 | -0.06 | 0.054 | 0.054 | 0 | 0.23 | 0.07 | 0.33 | 0.14 | 0.31 | 0.2 |
| 9 | 1654460 | rs771901 | -0.51 | -0.316 | 0.362 | -0.907 | 0.012 | 0.035 | 0.035 | 0 | 0.25 | 0.42 | 0.32 | 0.6 | 0.31 | 0.35 |
| 9 | 1659913 | rs2181515 | -0.574 | -0.657 | 0.344 | -0.968 | -0.073 | -0.111 | -0.111 | 0 | 0.19 | 0.38 | 0.14 | 0.02 | 0.09 | 0.08 |
| 9 | 1661432 | rs2476245 | -1.452 | -0.201 | 0.65 | -0.576 | 0.014 | 0.044 | 0.044 | 0 | 0.4 | 0.77 | 0.43 | 0.69 | 0.4 | 0.44 |
| 9 | 1661825 | rs2476244 | -0.524 | -0.104 | 0.387 | -0.478 | 0.008 | 0.033 | 0.033 | 0 | 0.25 | 0.41 | 0.33 | 0.6 | 0.3 | 0.35 |
| 9 | 1663954 | rs1885917 | 0.711 | -0.126 | 0.351 | -0.495 | -0.017 | -0.001 | -0.001 | 0.039 | 0.13 | 0.23 | 0.1 | 0.09 | 0.1 | 0.09 |
| 9 | 1666693 | rs2485175 | -0.657 | -0.202 | -0.087 | -0.328 | -0.002 | 0.007 | 0.007 | 0 | 0.24 | 0.39 | 0.3 | 0.67 | 0.26 | 0.4 |
| 9 | 1667544 | rs10962498 | 0.243 | -0.014 | -0.474 | -0.37 | -0.13 | 0.044 | 0.044 | 1.035 | 0.26 | 0.05 | 0.28 | 0.13 | 0.37 | 0.18 |
| 9 | 1667575 | rs2485178 | -0.552 | -0.034 | -0.294 | -0.326 | -0.001 | 0.024 | 0.024 | 0 | 0.24 | 0.42 | 0.3 | 0.67 | 0.26 | 0.4 |
| 9 | 1668272 | rs10962501 | 0.088 | 0.221 | -0.593 | -0.17 | -0.122 | 0.034 | 0.034 | 0.842 | 0.26 | 0.05 | 0.28 | 0.14 | 0.37 | 0.18 |
| 9 | 1668830 | rs12683735 | -0.384 | 0.26 | -0.011 | 0.206 | 0.021 | 0.098 | 0.098 | 0 | 0.18 | 0.33 | 0.17 | 0.51 | 0.23 | 0.27 |
| 9 | 1674726 | rs10962521 | -0.326 | 0.251 | -0.46 | 0.046 | -0.076 | 0.027 | 0.027 | 0.809 | 0.29 | 0.13 | 0.3 | 0.19 | 0.4 | 0.19 |
| 9 | 1676766 | rs4740636 | 0.498 | 0.312 | -0.544 | -0.259 | -0.127 | 0.058 | 0.058 | 0 | 0.22 | 0.04 | 0.26 | 0.11 | 0.33 | 0.15 |
| 9 | 1678482 | rs945663 | -0.425 | -0.47 | 0.36 | -0.653 | -0.049 | -0.066 | -0.066 | 0 | 0.14 | 0.32 | 0.18 | 0.04 | 0.09 | 0.13 |
| 9 | 1679714 | rs10124068 | -0.605 | -0.566 | 0.716 | -0.63 | 0.002 | -0.008 | -0.008 | 0 | 0.4 | 0.47 | 0.43 | 0.62 | 0.54 | 0.43 |
| 9 | 1683466 | rs523006 | 0.005 | -0.534 | 0.916 | -0.494 | -0.008 | -0.01 | -0.01 | 0 | 0.13 | 0.16 | 0.16 | 0.05 | 0.08 | 0.12 |
| 9 | 1685649 | rs2760397 | -2.595 | -0.651 | 0.389 | -0.671 | 0.003 | 0.109 | 0.109 | 0 | 0.65 | 0.9 | 0.78 | 0.82 | 0.71 | 0.71 |
| 9 | 1686322 | rs10810620 | -0.335 | -0.698 | 0.666 | -0.673 | -0.014 | 0.02 | 0.02 | 0 | 0.4 | 0.3 | 0.4 | 0.62 | 0.53 | 0.42 |
| 9 | 1688398 | rs4740637 | -0.59 | -0.722 | 0.958 | -0.635 | 0.01 | 0.03 | 0.03 | 0 | 0.45 | 0.34 | 0.44 | 0.63 | 0.57 | 0.45 |
| 9 | 1689357 | rs1323243 | -0.56 | -0.638 | 1.157 | -0.509 | -0.044 | 0.009 | 0.009 | 0 | 0.17 | 0.48 | 0.34 | 0.2 | 0.14 | 0.26 |
| 9 | 1697268 | rs7039730 | -0.574 | -0.625 | 0.647 | -0.527 | 0.002 | 0.028 | 0.028 | 0 | 0.43 | 0.23 | 0.42 | 0.6 | 0.57 | 0.38 |
| 9 | 1699307 | rs2485153 | -0.236 | -0.561 | 0.727 | -0.478 | -0.042 | -0.055 | -0.055 | 0 | 0.14 | 0.32 | 0.19 | 0.05 | 0.09 | 0.11 |
| 9 | 1704023 | rs771386 | -2.2 | -0.634 | -0.014 | -0.565 | -0.005 | 0.128 | 0.128 | 0 | 0.62 | 0.9 | 0.77 | 0.82 | 0.69 | 0.64 |
| 9 | 1705519 | rs7852382 | 0.47 | -0.019 | -0.111 | -0.36 | -0.057 | -0.092 | -0.092 | 0.015 | 0.15 | 0.03 | 0.18 | 0.47 | 0.24 | 0.19 |
| 9 | 1707496 | rs7029758 | -0.718 | -0.215 | -0.048 | -0.55 | -0.055 | 0.06 | 0.06 | 0 | 0.39 | 0.16 | 0.58 | 0.71 | 0.54 | 0.48 |
| 9 | 1707684 | rs12377978 | -0.838 | -0.209 | -0.042 | -0.555 | -0.034 | 0.067 | 0.067 | 0 | 0.4 | 0.17 | 0.59 | 0.71 | 0.56 | 0.48 |
| 9 | 1708902 | rs10962807 | -0.541 | -0.186 | -0.073 | -0.548 | -0.059 | 0.05 | 0.05 | 0 | 0.37 | 0.16 | 0.58 | 0.71 | 0.53 | 0.48 |
| 9 | 1709013 | rs510169 | -2.103 | -0.286 | 0.159 | -0.571 | -0.001 | -0.007 | -0.007 | 0 | 0.55 | 0.48 | 0.71 | 0.76 | 0.64 | 0.59 |
| 9 | 1709308 | rs512774 | -0.282 | -0.29 | -0.04 | -0.531 | -0.03 | -0.042 | -0.042 | 0.507 | 0.15 | 0.23 | 0.11 | 0.05 | 0.08 | 0.12 |
| 9 | 1709649 | rs7038689 | -0.502 | 0.17 | -0.507 | -0.291 | -0.155 | -0.081 | -0.081 | 0 | 0.35 | 0.07 | 0.44 | 0.55 | 0.48 | 0.34 |
| 9 | 1712303 | rs10962829 | -0.004 | 0.325 | -0.356 | 0.536 | -0.028 | -0.041 | -0.041 | 0.46 | 0.15 | 0.09 | 0.16 | 0.22 | 0.22 | 0.14 |
| 9 | 1712474 | rs496867 | -0.833 | -0.437 | -0.503 | -0.608 | 0.052 | 0.063 | 0.063 | 0 | 0.35 | 0.73 | 0.42 | 0.22 | 0.26 | 0.33 |
| 9 | 1712562 | rs574865 | -0.11 | -0.414 | -0.148 | -0.759 | -0.024 | -0.095 | -0.095 | 0.025 | 0.24 | 0.37 | 0.11 | 0.04 | 0.17 | 0.16 |
| 9 | 1714233 | rs1038050 | -1.134 | -0.613 | -0.581 | -0.493 | 0.038 | 0.045 | 0.045 | 0 | 0.6 | 0.81 | 0.64 | 0.63 | 0.71 | 0.6 |
| 9 | 1715155 | rs508876 | -0.332 | -0.771 | 0.042 | -0.71 | -0.08 | -0.115 | -0.115 | 0 | 0.27 | 0.52 | 0.11 | 0.09 | 0.15 | 0.15 |
| 9 | 1715256 | rs507970 | 0.117 | -0.574 | 0.139 | -0.553 | -0.016 | -0.018 | -0.018 | 0.068 | 0.1 | 0.17 | 0.04 | 0.06 | 0.05 | 0.05 |
| 9 | 1715820 | rs7020507 | -0.225 | 0.058 | 0.363 | -0.352 | 0.141 | 0.13 | 0.13 | 0.59 | 0.07 | 0.32 | 0.21 | 0.13 | 0.14 | 0.15 |
| 9 | 1717960 | rs2095614 | -2.108 | -0.118 | -0.15 | -0.443 | -0.039 | -0.03 | -0.03 | 1.621 | 0.71 | 0.98 | 0.69 | 0.68 | 0.76 | 0.65 |
| 9 | 1718076 | rs1809342 | 0.087 | -0.347 | 0.111 | -0.598 | -0.038 | -0.063 | -0.063 | 0 | 0.27 | 0.39 | 0.1 | 0.1 | 0.18 | 0.17 |
| 9 | 1718324 | rs1382183 | -1.347 | -0.364 | 0.423 | -0.632 | 0.005 | 0.003 | 0.003 | 0 | 0.62 | 0.57 | 0.48 | 0.55 | 0.6 | 0.47 |
| 9 | 1720593 | rs2616833 | -1.137 | -0.372 | 0.448 | -0.648 | 0.006 | 0.001 | 0.001 | 0 | 0.6 | 0.55 | 0.46 | 0.54 | 0.59 | 0.47 |
| 9 | 1720702 | rs2616832 | -1.11 | -0.379 | 0.441 | -0.651 | 0.006 | 0 | 0 | 0 | 0.6 | 0.54 | 0.46 | 0.54 | 0.6 | 0.48 |
| 9 | 1721778 | rs2616829 | 0.417 | -0.272 | 0.524 | -0.655 | -0.028 | -0.051 | -0.051 | 0.008 | 0.26 | 0.34 | 0.09 | 0.1 | 0.17 | 0.17 |
| 9 | 1722019 | rs10810734 | 0.156 | -0.226 | 0.578 | -0.627 | -0.016 | -0.024 | -0.024 | 6.448 | 0.35 | 0.28 | 0.38 | 0.44 | 0.42 | 0.29 |
| 9 | 1725777 | rs521124 | -0.014 | -0.34 | 0.707 | -0.574 | -0.031 | -0.065 | -0.065 | 0 | 0.27 | 0.4 | 0.11 | 0.1 | 0.18 | 0.18 |
| 9 | 1731076 | rs10962915 | 0.715 | -0.178 | 0.278 | -0.637 | -0.026 | -0.054 | -0.054 | 0 | 0.28 | 0.18 | 0.44 | 0.46 | 0.35 | 0.28 |
| 9 | 1732880 | rs6475120 | -0.99 | -0.081 | 0.331 | -0.579 | -0.015 | 0.014 | 0.014 | 0 | 0.57 | 0.71 | 0.56 | 0.57 | 0.53 | 0.46 |
| 9 | 1734768 | rs519864 | -1.257 | -0.06 | 0.244 | -0.55 | -0.038 | 0.001 | 0.001 | 0 | 0.59 | 0.81 | 0.58 | 0.55 | 0.55 | 0.47 |
| 9 | 1739568 | rs9298771 | -0.412 | -0.213 | 0.16 | -0.542 | 0 | -0.001 | -0.001 | 0 | 0.44 | 0.57 | 0.41 | 0.49 | 0.48 | 0.38 |

| | | | | | | | | | | | | | | | | |
|---|---|---|---|---|---|---|---|---|---|---|---|---|---|---|---|---|
| 9 | 1740278 | rs4741586 | 0.842 | -0.208 | 0.107 | -0.534 | -0.044 | -0.084 | -0.084 | 0.122 | 0.24 | 0.12 | 0.43 | 0.45 | 0.34 | 0.25 |
| 9 | 1742374 | rs2077415 | 0.601 | -0.222 | 0.291 | -0.541 | -0.003 | -0.003 | -0.003 | 0.278 | 0.26 | 0.28 | 0.44 | 0.45 | 0.34 | 0.26 |
| 9 | 1745174 | rs12344952 | 0.639 | -0.22 | 0.294 | -0.541 | -0.005 | -0.006 | -0.006 | 2.143 | 0.25 | 0.26 | 0.44 | 0.45 | 0.34 | 0.26 |
| 9 | 1746224 | rs4571782 | 1.632 | -0.327 | 0.426 | -0.662 | -0.022 | -0.05 | -0.05 | 0.97 | 0.16 | 0.23 | 0.06 | 0.06 | 0.13 | 0.12 |
| 9 | 1746287 | rs4297077 | 1.632 | -0.326 | 0.426 | -0.662 | -0.02 | -0.05 | -0.05 | 0.473 | 0.16 | 0.23 | 0.06 | 0.06 | 0.13 | 0.12 |
| 9 | 1757274 | rs7849487 | 0.068 | -0.078 | 0.311 | -0.492 | 0.01 | 0.04 | 0.04 | 0 | 0.28 | 0.41 | 0.35 | 0.43 | 0.36 | 0.25 |
| 9 | 1759486 | rs4741603 | 0.834 | -0.055 | 0.173 | -0.476 | -0.064 | -0.116 | -0.116 | 0 | 0.23 | 0.06 | 0.31 | 0.43 | 0.34 | 0.24 |
| 9 | 1759811 | rs486815 | 1.598 | -0.03 | -0.099 | -0.611 | -0.098 | 0.02 | 0.02 | 0 | 0.36 | 0.02 | 0.23 | 0.29 | 0.32 | 0.42 |
| 9 | 1761505 | rs10810799 | 0.503 | 0.11 | 0.424 | -0.319 | -0.036 | -0.072 | -0.072 | 0 | 0.24 | 0.13 | 0.31 | 0.44 | 0.35 | 0.24 |
| 9 | 1762597 | rs4741604 | -0.041 | 0.133 | 0.431 | -0.306 | 0.014 | 0.053 | 0.053 | 0 | 0.28 | 0.38 | 0.35 | 0.44 | 0.36 | 0.24 |
| 9 | 1773538 | rs10963188 | 0.316 | 0.204 | 0.33 | -0.297 | -0.002 | 0.002 | 0.002 | 0 | 0.25 | 0.21 | 0.32 | 0.44 | 0.34 | 0.25 |
| 9 | 1784090 | rs7035491 | -0.315 | 0.091 | 0.181 | -0.522 | 0.007 | -0.015 | -0.015 | 0.199 | 0.36 | 0.3 | 0.33 | 0.43 | 0.34 | 0.28 |
| 9 | 1784948 | rs13288107 | 1.921 | 0.277 | 0.091 | -0.418 | 0.002 | -0.001 | -0.001 | 0.884 | 0.15 | 0.12 | 0.16 | 0.09 | 0.22 | 0.11 |
| 9 | 1785717 | rs494566 | 1.765 | 0.138 | -0.129 | -0.445 | -0.063 | -0.066 | -0.066 | 3.03 | 0.33 | 0.03 | 0.37 | 0.36 | 0.29 | 0.46 |
| 9 | 1786335 | rs7028588 | -0.703 | 0.091 | 0.36 | -0.327 | 0.025 | 0.047 | 0.047 | 0 | 0.34 | 0.48 | 0.4 | 0.46 | 0.43 | 0.26 |
| 9 | 1786681 | rs7048135 | -0.507 | 0.046 | 0.156 | -0.285 | -0.005 | 0.116 | 0.116 | 0 | 0.19 | 0.38 | 0.24 | 0.37 | 0.19 | 0.17 |
| 9 | 1789663 | rs3843567 | 1.82 | 0.303 | -0.56 | -0.407 | -0.008 | 0.008 | 0.008 | 0 | 0.15 | 0.09 | 0.17 | 0.09 | 0.23 | 0.1 |
| 9 | 1790488 | rs3847223 | -1.334 | 0.102 | -0.395 | -0.473 | 0 | -0.001 | -0.001 | 0.909 | 0.54 | 0.54 | 0.51 | 0.51 | 0.57 | 0.38 |
| 9 | 1790489 | rs3847224 | 0.416 | 0.262 | -0.799 | -0.363 | -0.003 | -0.006 | -0.006 | 0.003 | 0.15 | 0.11 | 0.19 | 0.38 | 0.18 | 0.16 |
| 9 | 1791880 | rs12553036 | -0.307 | 0.219 | -0.834 | -0.757 | -0.005 | 0.02 | 0.02 | 0 | 0.36 | 0.29 | 0.29 | 0.11 | 0.38 | 0.23 |
| 9 | 1792147 | rs2068428 | 0.976 | 0.636 | -0.697 | -0.177 | -0.008 | 0.008 | 0.008 | 0 | 0.15 | 0.08 | 0.17 | 0.07 | 0.23 | 0.1 |
| 9 | 1793162 | rs7852096 | 0.078 | -0.215 | -0.394 | -0.001 | -0.031 | -0.021 | -0.021 | 0 | 0.35 | 0.45 | 0.31 | 0.36 | 0.24 | 0.37 |
| 9 | 1793408 | rs3912096 | 0.263 | -0.029 | -1.042 | -0.007 | 0.037 | 0.126 | 0.126 | 0 | 0.57 | 0.18 | 0.62 | 0.78 | 0.64 | 0.48 |
| 9 | 1794330 | rs10756919 | -1.887 | -0.111 | -0.414 | 0.952 | 0.021 | -0.044 | -0.044 | 0 | 0.27 | 0.78 | 0.2 | 0.46 | 0.16 | 0.29 |
| 9 | 1795468 | rs3847226 | -1.232 | 0.024 | -0.019 | 0.932 | -0.003 | 0.001 | 0.001 | 0 | 0.15 | 0.15 | 0.16 | 0.35 | 0.17 | 0.2 |
| 9 | 1795548 | rs4741606 | -1.712 | 0.042 | 0.246 | 0.872 | 0 | 0.032 | 0.032 | 0 | 0.18 | 0.24 | 0.19 | 0.41 | 0.2 | 0.28 |
| 9 | 1798170 | rs3847227 | -2.879 | -0.096 | 1.004 | 0.801 | -0.003 | 0.01 | 0.01 | 0.97 | 0.27 | 0.69 | 0.23 | 0.5 | 0.23 | 0.36 |
| 9 | 1798324 | rs16934185 | 0.032 | -0.088 | 0.935 | 0.741 | -0.002 | -0.004 | -0.004 | 5.466 | 0.06 | 0.07 | 0.04 | 0 | 0.1 | 0.06 |
| 9 | 1798805 | rs3847228 | 1.373 | 0.19 | 0.372 | 0.373 | -0.149 | 0.065 | 0.065 | 0.541 | 0.35 | 0.03 | 0.25 | 0.17 | 0.41 | 0.26 |
| 9 | 1799943 | rs16934200 | -2.106 | 0.032 | 1.617 | 0.998 | 0 | 0.001 | 0.001 | 0 | 0.11 | 0.12 | 0.05 | 0.09 | 0.12 | 0.13 |
| 9 | 1809976 | rs10810883 | -0.542 | 0.049 | 1.593 | 0.981 | 0.001 | 0.08 | 0.08 | 0 | 0.08 | 0.08 | 0.12 | 0.35 | 0.07 | 0.16 |
| 9 | 1815727 | rs10117983 | -3 | 0.02 | 1.715 | 0.974 | -0.024 | 0.076 | 0.076 | 1.439 | 0.22 | 0.64 | 0.2 | 0.48 | 0.11 | 0.3 |
| 9 | 1816322 | rs10756927 | -3.703 | 0.041 | 2.305 | 0.976 | 0.013 | -0.033 | -0.033 | 0 | 0.28 | 0.76 | 0.24 | 0.48 | 0.22 | 0.36 |
| 9 | 1816832 | rs12346108 | -3.049 | 0.246 | 2.337 | 1.173 | -0.053 | 0.01 | 0.01 | 3.1 | 0.14 | 0.56 | 0.07 | 0.13 | 0.04 | 0.15 |
| 9 | 1818518 | rs1382189 | -2.624 | 0.259 | 2.345 | 1.166 | -0.007 | 0.015 | 0.015 | 2.036 | 0.09 | 0.53 | 0.06 | 0.05 | 0.02 | 0.08 |
| 9 | 1826357 | rs12686439 | -1.648 | 0.589 | 2.326 | 1.186 | 0.063 | 0.15 | 0.15 | 0 | 0.11 | 0.21 | 0.16 | 0.34 | 0.19 | 0.17 |
| 9 | 1830286 | rs10283612 | -3.205 | -0.244 | 2.368 | 1.213 | -0.021 | 0.084 | 0.084 | 1.506 | 0.2 | 0.67 | 0.19 | 0.48 | 0.11 | 0.3 |
| 9 | 1835334 | rs10963540 | -1.596 | -0.068 | 2.093 | 1.339 | 0.034 | 0.313 | 0.313 | 0 | 0.09 | 0.28 | 0.14 | 0.35 | 0.07 | 0.15 |
| 9 | 1836746 | rs10116883 | -0.948 | -0.458 | 1.515 | 0.293 | -0.018 | 0.018 | 0.018 | 0.012 | 0.05 | 0.13 | 0.01 | 0.08 | 0.02 | 0.07 |
| 9 | 1839259 | rs10963567 | -1.931 | -0.368 | 1.727 | 0.094 | 0.066 | 0.004 | 0.004 | 1.079 | 0.08 | 0.36 | 0.07 | 0.06 | 0.11 | 0.12 |
| 9 | 1840337 | rs12005947 | -1.522 | -0.345 | 1.797 | 0.105 | 0.066 | 0.01 | 0.01 | 1.069 | 0.08 | 0.31 | 0.06 | 0.05 | 0.1 | 0.12 |
| 9 | 1842265 | rs7019171 | -2.04 | -0.313 | 1.833 | 0.054 | 0.039 | -0.004 | -0.004 | 0 | 0.09 | 0.34 | 0.05 | 0.06 | 0.1 | 0.1 |
| 9 | 1845946 | rs7847373 | -2.589 | -1.072 | 1.376 | -0.107 | -0.026 | -0.032 | -0.032 | 0 | 0.23 | 0.7 | 0.13 | 0.12 | 0.14 | 0.2 |
| 9 | 1849664 | rs7019385 | -2.174 | -1.076 | 1.613 | -0.123 | -0.029 | -0.08 | -0.08 | 0 | 0.13 | 0.66 | 0.11 | 0.01 | 0.08 | 0.06 |
| 9 | 1866537 | rs7860714 | -1.247 | -1.233 | 0.58 | 0.024 | -0.088 | -0.146 | -0.146 | 0 | 0.67 | 0.97 | 0.61 | 0.45 | 0.67 | 0.6 |
| 9 | 1867013 | rs10963648 | -1.888 | -1.091 | 2.281 | 0.486 | 0.032 | 0 | 0 | 0 | 0.1 | 0.47 | 0.1 | 0.05 | 0.08 | 0.1 |
| 9 | 1868585 | rs10963660 | -2.812 | -1.151 | 2.184 | 0.507 | -0.045 | -0.028 | -0.028 | 0.847 | 0.22 | 0.61 | 0.11 | 0.16 | 0.12 | 0.19 |
| 9 | 1872957 | rs17724855 | -1.359 | -1.206 | 2.109 | 0.747 | -0.034 | 0.008 | 0.008 | 0.257 | 0.07 | 0.17 | 0.03 | 0.1 | 0.02 | 0.09 |
| 9 | 1875165 | rs7048797 | -1.595 | -1.209 | 2.014 | 0.731 | -0.084 | -0.036 | -0.036 | 0 | 0.15 | 0.35 | 0.05 | 0.1 | 0.04 | 0.09 |
| 9 | 1876048 | rs7022128 | -1.801 | -1.179 | 1.429 | 0.758 | -0.089 | -0.027 | -0.027 | 0.249 | 0.19 | 0.58 | 0.07 | 0.15 | 0.05 | 0.15 |
| 9 | 1876496 | rs10963717 | -0.197 | -0.957 | 0.663 | 0.742 | -0.068 | 0.002 | 0.002 | 0.487 | 0.22 | 0.09 | 0.27 | 0.19 | 0.36 | 0.19 |
| 9 | 1876592 | rs10811009 | -1.688 | -1.689 | 1.559 | 1.049 | -0.095 | -0.006 | -0.006 | 0.214 | 0.16 | 0.46 | 0.06 | 0.15 | 0.05 | 0.14 |
| 9 | 1878648 | rs7862396 | -0.461 | -1.116 | 0.857 | 1.137 | -0.005 | -0.002 | -0.002 | 0 | 0.14 | 0.13 | 0.06 | 0.09 | 0.1 | 0.22 |
| 9 | 1883627 | rs10963744 | -0.258 | -1.233 | 0.814 | 1.767 | -0.024 | -0.075 | -0.075 | 0 | 0.11 | 0.02 | 0.18 | 0.35 | 0.12 | 0.09 |
| 9 | 1884163 | rs2039317 | -1.571 | -1.367 | 1.01 | 1.736 | -0.007 | 0.046 | 0.046 | 0 | 0.25 | 0.33 | 0.29 | 0.46 | 0.25 | 0.19 |
| 9 | 1884934 | rs10811026 | -1.902 | -1.42 | 0.919 | 1.292 | -0.003 | -0.004 | -0.004 | 0 | 0.32 | 0.33 | 0.47 | 0.73 | 0.29 | 0.42 |
| 9 | 1887124 | rs4741619 | -2.041 | -1.325 | 0.063 | 1.297 | 0.066 | 0.156 | 0.156 | 0 | 0.47 | 0.79 | 0.55 | 0.78 | 0.41 | 0.56 |
| 9 | 1887281 | rs4741620 | -1.612 | -0.043 | 0.447 | 2.238 | -0.073 | -0.076 | -0.076 | 1.811 | 0.23 | 0.04 | 0.44 | 0.71 | 0.29 | 0.27 |
| 9 | 1900689 | rs10122055 | -2.084 | -0.247 | 0.601 | 1.712 | -0.012 | -0.023 | -0.023 | 0 | 0.3 | 0.58 | 0.47 | 0.78 | 0.28 | 0.27 |
| 9 | 1908927 | rs4741629 | -1.68 | 0.25 | -0.732 | 1.868 | -0.004 | 0.021 | 0.021 | 0 | 0.27 | 0.18 | 0.47 | 0.8 | 0.3 | 0.27 |
| 9 | 1909586 | rs10811093 | -1.037 | -0.306 | -0.164 | 1.788 | -0.019 | 0.037 | 0.037 | 0 | 0.16 | 0.22 | 0.25 | 0.3 | 0.13 | 0.2 |
| 9 | 1915483 | rs4551422 | 1.052 | 0.248 | -0.484 | 1.463 | 0 | 0.002 | 0.002 | 0 | 0.94 | 0.97 | 0.95 | 0.87 | 0.94 | 0.85 |
| 9 | 1919833 | rs10811114 | -0.533 | 0.46 | -0.324 | 1.183 | 0.004 | -0.002 | -0.002 | 0 | 0.34 | 0.41 | 0.52 | 0.7 | 0.39 | 0.35 |
| 9 | 1923359 | rs10512035 | -0.955 | -0.199 | 0.983 | 1.244 | 0.03 | 0.088 | 0.088 | 0 | 0.1 | 0.46 | 0.15 | 0.08 | 0.06 | 0.03 |
| 9 | 1945517 | rs7047972 | -1.656 | -0.842 | 1.477 | 1.349 | -0.05 | 0.101 | 0.101 | 0 | 0.13 | 0.28 | 0.24 | 0.16 | 0.06 | 0.12 |
| 9 | 1950813 | rs7030343 | -1.831 | -1.393 | 1.428 | 1.901 | -0.022 | 0.016 | 0.016 | 1.416 | 0.21 | 0.71 | 0.38 | 0.31 | 0.09 | 0.24 |
| 9 | 1950944 | rs10964130 | -1.405 | -2.337 | 1.611 | 2.231 | -0.079 | 0.072 | 0.072 | 0.088 | 0.26 | 0.49 | 0.39 | 0.57 | 0.14 | 0.34 |
| 9 | 1953536 | rs3935922 | -2.17 | -2.197 | 2.094 | 3.311 | -0.003 | 0.399 | 0.399 | 0 | 0.08 | 0.23 | 0.24 | 0.45 | 0.02 | 0.17 |
| 9 | 1957974 | rs7857865 | -1.02 | -1.743 | 1.165 | 2.62 | -0.018 | 0.105 | 0.105 | 0 | 0.51 | 0.79 | 0.66 | 0.67 | 0.44 | 0.52 |

| | | | | | | | | | | | | | | | | |
|---|---|---|---|---|---|---|---|---|---|---|---|---|---|---|---|---|
| 9 | 1958885 | rs7020398 | -1.707 | -1.456 | 1.11 | 2.868 | 0.001 | 0.373 | 0.373 | 0.061 | 0.08 | 0.24 | 0.25 | 0.43 | 0.03 | 0.17 |
| 9 | 1960788 | rs1538533 | -1.412 | -1.444 | 1.123 | 2.839 | 0.007 | 0.373 | 0.373 | 0 | 0.07 | 0.22 | 0.23 | 0.38 | 0.02 | 0.13 |
| 9 | 1962977 | rs10964196 | -1.543 | -1.429 | 1.134 | 2.824 | 0.011 | 0.415 | 0.415 | 0 | 0.07 | 0.23 | 0.23 | 0.42 | 0.03 | 0.13 |
| 9 | 1965945 | rs4562384 | -0.429 | -1.44 | 0.049 | 1.845 | 0.025 | 0.048 | 0.048 | 0.197 | 0.41 | 0.79 | 0.41 | 0.22 | 0.32 | 0.35 |
| 9 | 1966130 | rs4014085 | -0.745 | -1.121 | -0.438 | 2.129 | -0.05 | 0.059 | 0.059 | 0.895 | 0.47 | 0.92 | 0.65 | 0.6 | 0.41 | 0.47 |
| 9 | 1979053 | rs16936187 | -0.659 | -1.038 | 0.113 | 1.843 | -0.008 | 0.044 | 0.044 | 0 | 0.09 | 0.11 | 0.08 | 0.22 | 0.07 | 0.11 |
| 9 | 1979764 | rs1416621 | -0.935 | -0.956 | 0.09 | 0.721 | 0.003 | 0.007 | 0.007 | 0.045 | 0.49 | 0.49 | 0.46 | 0.23 | 0.4 | 0.43 |
| 9 | 1980819 | rs7858803 | -1.306 | -0.614 | 1.012 | 1.179 | -0.002 | 0.044 | 0.044 | 0.046 | 0.19 | 0.33 | 0.23 | 0.28 | 0.16 | 0.2 |
| 9 | 1980918 | rs7847882 | -1.198 | -0.614 | 1.016 | 1.182 | -0.015 | 0.044 | 0.044 | 0.107 | 0.18 | 0.33 | 0.23 | 0.28 | 0.13 | 0.2 |
| 9 | 1981634 | rs10120400 | -0.995 | -0.637 | 1.166 | 1.198 | 0 | 0.138 | 0.138 | 1.934 | 0.08 | 0.19 | 0.09 | 0.26 | 0.04 | 0.09 |
| 9 | 1981979 | rs16936230 | -0.91 | -0.813 | 1.508 | 0.927 | 0.013 | -0.015 | -0.015 | 0 | 0.05 | 0.18 | 0.04 | 0 | 0.03 | 0.01 |
| 9 | 1982607 | rs7027936 | -2.412 | -0.12 | 1.545 | 0.817 | 0.013 | 0 | 0 | 1.157 | 0.27 | 0.52 | 0.24 | 0.26 | 0.23 | 0.25 |
| 9 | 1989033 | rs7864828 | -2.636 | -0.153 | 1.359 | 0.865 | -0.011 | 0.021 | 0.021 | 2.809 | 0.21 | 0.65 | 0.19 | 0.3 | 0.16 | 0.17 |
| 9 | 1992516 | rs7872017 | -1.446 | -0.183 | 1.389 | 0.821 | -0.015 | -0.018 | -0.018 | 0 | 0.1 | 0.32 | 0.11 | 0.07 | 0.06 | 0.03 |
| 9 | 1994144 | rs10964314 | -0.897 | -0.02 | 1.342 | 0.944 | 0.007 | 0.153 | 0.153 | 2.249 | 0.07 | 0.21 | 0.09 | 0.23 | 0.04 | 0.1 |
| 9 | 1994871 | rs10964319 | -1.105 | -0.033 | 1.347 | 0.949 | 0.02 | 0.081 | 0.081 | 2.582 | 0.11 | 0.23 | 0.11 | 0.26 | 0.1 | 0.13 |
| 9 | 1997332 | rs17529312 | -1.45 | 0.015 | 1.314 | 0.904 | 0.01 | 0.014 | 0.014 | 0.081 | 0.2 | 0.27 | 0.19 | 0.28 | 0.2 | 0.24 |
| 9 | 1998493 | rs7852330 | -0.098 | 0.107 | 0.948 | 0.75 | 0 | -0.01 | -0.01 | 0.022 | 0.24 | 0.3 | 0.26 | 0.18 | 0.22 | 0.23 |
| 9 | 1999277 | rs10119055 | -1.463 | 0.12 | 0.793 | 0.835 | 0.008 | 0.001 | 0.001 | 0 | 0.42 | 0.53 | 0.45 | 0.43 | 0.42 | 0.41 |
| 9 | 2000062 | rs10964337 | -0.117 | 0.19 | 0.911 | 0.529 | -0.039 | -0.012 | -0.012 | 0 | 0.15 | 0.01 | 0.26 | 0.12 | 0.15 | 0.19 |
| 9 | 2007485 | rs7853498 | -1.519 | 0.406 | 1.589 | 0.858 | -0.011 | -0.019 | -0.019 | 0 | 0.09 | 0.33 | 0.06 | 0.03 | 0.05 | 0.02 |
| 9 | 2013580 | rs4741636 | -0.41 | 0.742 | 1.076 | 0.985 | 0 | 0.004 | 0.004 | 0.063 | 0.41 | 0.45 | 0.43 | 0.24 | 0.37 | 0.37 |
| 9 | 2013962 | rs7040790 | -0.396 | 0.862 | 1.148 | 1.079 | 0 | 0.005 | 0.005 | 0.068 | 0.41 | 0.43 | 0.43 | 0.25 | 0.36 | 0.35 |
| 9 | 2024076 | rs6475412 | -2.883 | 1.529 | 2.319 | 2.172 | 0.09 | 0.16 | 0.16 | 0.146 | 0.06 | 0.26 | 0.16 | 0.17 | 0.07 | 0.05 |
| 9 | 2028237 | rs16936751 | -2.903 | 1.424 | 1.534 | 2.373 | 0.054 | 0.134 | 0.134 | 0.019 | 0.07 | 0.26 | 0.26 | 0.22 | 0.08 | 0.08 |
| 9 | 2029454 | rs7864163 | -2.426 | 1.964 | 1.265 | 2.409 | 0.088 | 0.103 | 0.103 | 0 | 0.05 | 0.24 | 0.25 | 0.13 | 0.07 | 0.06 |
| 9 | 2034440 | rs10964500 | -2.885 | 1.898 | 1.055 | 2.44 | 0.064 | 0.061 | 0.061 | 0.452 | 0.13 | 0.26 | 0.34 | 0.34 | 0.21 | 0.22 |
| 9 | 2034908 | rs7862617 | -2.849 | 1.702 | 1.025 | 2.358 | 0.052 | 0.051 | 0.051 | 0.052 | 0.13 | 0.24 | 0.31 | 0.3 | 0.19 | 0.21 |
| 9 | 2036252 | rs12554293 | -2.144 | 1.554 | 1 | 1.821 | 0.021 | 0.026 | 0.026 | 0.017 | 0.09 | 0.2 | 0.24 | 0.14 | 0.11 | 0.05 |
| 9 | 2039973 | rs10964527 | -1.658 | 1.406 | 0.604 | 1.918 | 0.001 | 0.064 | 0.064 | 0.01 | 0.05 | 0.07 | 0.12 | 0.44 | 0.09 | 0.16 |
| 9 | 2039983 | rs10964528 | -1.743 | 1.001 | 0.721 | 1.426 | 0 | 0.026 | 0.026 | 0.381 | 0.3 | 0.6 | 0.3 | 0.52 | 0.23 | 0.3 |
| 9 | 2040973 | rs7865562 | -1.43 | 0.513 | 0.613 | 0.738 | 0.039 | 0.087 | 0.087 | 0 | 0.05 | 0.22 | 0.13 | 0.09 | 0.04 | 0.02 |
| 9 | 2042083 | rs3793482 | -1.31 | 0.117 | -0.298 | 0.204 | -0.018 | 0.116 | 0.116 | 0.019 | 0.23 | 0.02 | 0.16 | 0.04 | 0.19 | 0.12 |
| 9 | 2048361 | rs10122060 | -1.812 | 0.196 | 0.968 | 0.754 | 0.02 | 0.026 | 0.026 | 0 | 0.19 | 0.35 | 0.2 | 0.26 | 0.18 | 0.15 |
| 9 | 2048876 | rs7028212 | -2.72 | 0.014 | 1.485 | 0.745 | 0 | -0.004 | -0.004 | 0.148 | 0.3 | 0.72 | 0.4 | 0.36 | 0.25 | 0.21 |
| 9 | 2052014 | rs7036529 | -2.244 | -0.053 | 1.857 | 0.726 | 0.001 | 0.007 | 0.007 | 0 | 0.23 | 0.65 | 0.32 | 0.3 | 0.19 | 0.19 |
| 9 | 2054409 | rs3750417 | -1.964 | -0.079 | 2.107 | 0.719 | 0.014 | 0.004 | 0.004 | 0 | 0.21 | 0.29 | 0.24 | 0.25 | 0.2 | 0.16 |
| 9 | 2056563 | rs3829069 | -3.243 | -0.378 | 1.938 | 0.575 | 0.013 | 0.012 | 0.012 | 0.022 | 0.34 | 0.78 | 0.39 | 0.32 | 0.24 | 0.22 |
| 9 | 2058133 | rs6475447 | -3.221 | -0.481 | 2.014 | 0.671 | 0.014 | 0.002 | 0.002 | 0 | 0.33 | 0.74 | 0.44 | 0.35 | 0.24 | 0.22 |
| 9 | 2062213 | rs7853411 | -1.706 | -0.311 | 2.87 | 0.677 | -0.013 | -0.102 | -0.102 | 0.047 | 0.16 | 0.37 | 0.08 | 0.02 | 0.09 | 0.08 |
| 9 | 2064795 | rs7035898 | -2.56 | -0.313 | 3.055 | 0.7 | 0.003 | -0.001 | -0.001 | 1.141 | 0.22 | 0.66 | 0.28 | 0.25 | 0.18 | 0.17 |
| 9 | 2066976 | rs13300103 | -1.979 | -0.667 | 3.519 | 0.466 | 0.037 | 0.142 | 0.142 | 0.481 | 0.16 | 0.46 | 0.36 | 0.34 | 0.18 | 0.23 |
| 9 | 2067703 | rs12002861 | -1.994 | -0.675 | 3.54 | 0.447 | 0.031 | 0.125 | 0.125 | 0.581 | 0.16 | 0.47 | 0.37 | 0.34 | 0.18 | 0.23 |
| 9 | 2068747 | rs10121854 | -2.009 | -0.695 | 3.629 | 0.355 | -0.006 | -0.052 | -0.052 | 0 | 0.17 | 0.24 | 0.09 | 0.02 | 0.11 | 0.07 |
| 9 | 2072356 | rs4741641 | -1.42 | -0.613 | 3.647 | 0.461 | 0.029 | 0.119 | 0.119 | 0 | 0.05 | 0.09 | 0.19 | 0.29 | 0.09 | 0.11 |
| 9 | 2073292 | rs13288443 | -0.532 | -0.638 | 3.833 | 0.379 | 0 | -0.008 | -0.008 | 0 | 0.09 | 0.13 | 0.15 | 0.04 | 0.1 | 0.12 |
| 9 | 2073839 | rs2066111 | -3.014 | -0.668 | 3.949 | 0.36 | -0.002 | -0.001 | -0.001 | 0.115 | 0.29 | 0.71 | 0.43 | 0.31 | 0.27 | 0.29 |
| 9 | 2074070 | rs2066110 | -2.301 | -0.649 | 3.905 | 0.264 | -0.022 | -0.097 | -0.097 | 0.431 | 0.17 | 0.38 | 0.1 | 0.02 | 0.1 | 0.05 |
| 9 | 2079250 | rs7035071 | -1.761 | -0.716 | 3.473 | 0.268 | -0.001 | -0.006 | -0.006 | 0 | 0.16 | 0.13 | 0.2 | 0.05 | 0.17 | 0.14 |
| 9 | 2081292 | rs10123771 | -2.854 | -0.756 | 3.459 | 0.247 | -0.015 | -0.076 | -0.076 | 0 | 0.27 | 0.6 | 0.27 | 0.06 | 0.22 | 0.18 |
| 9 | 2089841 | rs10811409 | -1.849 | -0.808 | 3.21 | 0.25 | 0.011 | 0.066 | 0.066 | 0 | 0.16 | 0.03 | 0.06 | 0.02 | 0.12 | 0.06 |
| 9 | 2091914 | rs17725419 | -1.644 | -0.791 | 3.622 | 0.438 | 0.042 | 0.123 | 0.123 | 0 | 0.06 | 0.2 | 0.19 | 0.21 | 0.07 | 0.1 |
| 9 | 2095771 | rs3793489 | -1.565 | -1.043 | 2.424 | 0.173 | 0.008 | 0.021 | 0.021 | 0 | 0.11 | 0.08 | 0.15 | 0.03 | 0.11 | 0.1 |
| 9 | 2097585 | rs2275419 | -2.632 | -1.147 | 2.645 | 0.665 | -0.004 | 0.005 | 0.005 | 0.114 | 0.23 | 0.66 | 0.3 | 0.36 | 0.15 | 0.18 |
| 9 | 2100396 | rs10811426 | -3.048 | -1.135 | 2.517 | 0.675 | 0.039 | -0.018 | -0.018 | 0.022 | 0.36 | 0.82 | 0.39 | 0.4 | 0.25 | 0.27 |
| 9 | 2106705 | rs10757167 | -2.486 | -1.28 | 3.15 | 0.675 | -0.052 | -0.138 | -0.138 | 1.095 | 0.24 | 0.39 | 0.12 | 0.02 | 0.15 | 0.1 |
| 9 | 2109792 | rs12336319 | -2.238 | -1.208 | 3.375 | 0.893 | 0.072 | 0.379 | 0.379 | 0 | 0.08 | 0.41 | 0.16 | 0.38 | 0.07 | 0.1 |
| 9 | 2110581 | rs1962292 | -3.178 | -1.253 | 3.236 | 0.871 | -0.023 | 0.014 | 0.014 | 0 | 0.29 | 0.39 | 0.24 | 0.41 | 0.21 | 0.21 |
| 9 | 2111748 | rs3793495 | -1.798 | -0.943 | 2.935 | 0.999 | 0.045 | 0.261 | 0.261 | 0 | 0.07 | 0.29 | 0.15 | 0.34 | 0.06 | 0.1 |
| 9 | 2114024 | rs10511434 | -2.824 | -0.912 | 2.854 | 1.024 | 0.003 | 0.201 | 0.201 | 0 | 0.11 | 0.31 | 0.17 | 0.41 | 0.08 | 0.14 |
| 9 | 2115271 | rs1572744 | -2.6 | -0.797 | 1.905 | 1.065 | -0.008 | 0.009 | 0.009 | 0 | 0.26 | 0.64 | 0.25 | 0.41 | 0.17 | 0.17 |
| 9 | 2116414 | rs7020980 | -2.006 | -0.331 | 1.306 | 1.187 | 0.009 | -0.043 | -0.043 | 0 | 0.25 | 0.82 | 0.26 | 0.41 | 0.18 | 0.18 |
| 9 | 2120825 | rs2001776 | -1.765 | -0.417 | 2.127 | 1.777 | -0.019 | 0.229 | 0.229 | 0.007 | 0.13 | 0.47 | 0.22 | 0.37 | 0.07 | 0.17 |
| 9 | 2121371 | rs10757185 | -2.29 | -0.316 | 2.547 | 1.566 | -0.022 | 0.053 | 0.053 | 0.235 | 0.23 | 0.7 | 0.27 | 0.46 | 0.14 | 0.21 |
| 9 | 2122780 | rs10757188 | -1.79 | -0.161 | 2.78 | 2.088 | 0.005 | 0.143 | 0.143 | 0 | 0.17 | 0.39 | 0.22 | 0.56 | 0.16 | 0.22 |
| 9 | 2123435 | rs7866316 | -2.155 | 0.015 | 2.598 | 2.051 | -0.008 | 0.182 | 0.182 | 0 | 0.21 | 0.5 | 0.26 | 0.48 | 0.19 | 0.22 |
| 9 | 2123639 | rs3829070 | -1.811 | -0.378 | 2.524 | 1.807 | -0.044 | 0.158 | 0.158 | 0 | 0.2 | 0.53 | 0.23 | 0.56 | 0.12 | 0.24 |
| 9 | 2131534 | rs6475524 | -2.618 | 0.014 | 2.351 | 1.083 | 0.033 | -0.004 | -0.004 | 0.252 | 0.1 | 0.48 | 0.08 | 0.02 | 0.07 | 0.02 |
| 9 | 2137725 | rs13283799 | -2.497 | 0.015 | 2.092 | 1.067 | 0.017 | -0.021 | -0.021 | 0.164 | 0.11 | 0.45 | 0.06 | 0.02 | 0.06 | 0.03 |

| | | | | | | | | | | | | | | | | |
|---|---|---|---|---|---|---|---|---|---|---|---|---|---|---|---|---|
| 9 | 2141357 | rs7868484 | -1.225 | 0.232 | 1.133 | 1.32 | 0.003 | 0.01 | 0.01 | 0 | 0.26 | 0.71 | 0.41 | 0.4 | 0.26 | 0.39 |
| 9 | 2142191 | rs10811530 | 0.953 | 0.398 | 0.448 | 0.659 | -0.002 | -0.001 | -0.001 | 0.21 | 0.56 | 0.42 | 0.44 | 0.51 | 0.5 | 0.47 |
| 9 | 2143221 | rs2376308 | -0.112 | 0.166 | -0.33 | 0.485 | -0.014 | 0.042 | 0.042 | 0 | 0.72 | 0.95 | 0.76 | 0.83 | 0.73 | 0.78 |
| 9 | 2143543 | rs2376305 | 0.643 | 0.132 | -0.098 | 0.451 | 0.005 | 0.01 | 0.01 | 0 | 0.59 | 0.65 | 0.66 | 0.8 | 0.64 | 0.71 |
| 9 | 2148213 | rs10964978 | 1.219 | 0.13 | -1.013 | 0.407 | -0.015 | -0.061 | -0.061 | 0 | 0.47 | 0.14 | 0.42 | 0.47 | 0.53 | 0.37 |
| 9 | 2151194 | rs10811540 | 1.123 | 0.159 | -1.24 | 0.495 | -0.031 | 0.013 | 0.013 | 0 | 0.49 | 0.17 | 0.4 | 0.48 | 0.54 | 0.43 |
| 9 | 2154771 | rs17386332 | -0.099 | -0.222 | -1.322 | 0.58 | 0.006 | 0.017 | 0.017 | 0.27 | 0.18 | 0.01 | 0.11 | 0.12 | 0.15 | 0.15 |
| 9 | 2155164 | rs3793503 | 0.491 | -0.033 | -0.562 | 0.754 | -0.031 | -0.061 | -0.061 | 0.965 | 0.83 | 0.92 | 0.75 | 0.72 | 0.79 | 0.68 |
| 9 | 2157572 | rs7025777 | -1.221 | -0.175 | 0.265 | 0.293 | 0.008 | -0.032 | -0.032 | 0.095 | 0.09 | 0.4 | 0.1 | 0 | 0.03 | 0.02 |
| 9 | 2157934 | rs10965030 | 0.619 | -0.366 | -0.403 | 0.38 | 0.011 | 0.009 | 0.009 | 0.121 | 0.6 | 0.33 | 0.58 | 0.71 | 0.66 | 0.61 |
| 9 | 2160105 | rs3793504 | -0.635 | -0.34 | -0.12 | 0.8 | -0.019 | 0.08 | 0.08 | 0.048 | 0.26 | 0.61 | 0.45 | 0.45 | 0.19 | 0.22 |
| 9 | 2161949 | rs2274095 | -0.351 | -0.375 | -0.474 | 0.62 | -0.008 | -0.002 | -0.002 | 0 | 0.31 | 0.64 | 0.2 | 0.24 | 0.24 | 0.34 |
| 9 | 2165758 | rs10491695 | 0.679 | 0.013 | -0.918 | 0.518 | 0.002 | -0.005 | -0.005 | 1.432 | 0.37 | 0.38 | 0.42 | 0.28 | 0.4 | 0.29 |
| 9 | 2167498 | rs3793508 | 0.389 | -0.181 | -1.451 | 0.665 | -0.011 | 0.013 | 0.013 | 0.263 | 0.25 | 0.31 | 0.14 | 0.26 | 0.2 | 0.33 |
| 9 | 2167664 | rs3829074 | 0.213 | 0.249 | -1.929 | 0.781 | -0.06 | -0.098 | -0.098 | 0 | 0.66 | 0.95 | 0.63 | 0.55 | 0.65 | 0.62 |
| 9 | 2170884 | rs3793514 | -0.638 | -0.221 | -1.193 | 1.013 | -0.003 | 0.005 | 0.005 | 0 | 0.08 | 0.09 | 0.15 | 0.18 | 0.04 | 0.16 |
| 9 | 2172011 | rs16937826 | -0.465 | -0.837 | -1.095 | 1.242 | -0.006 | 0.006 | 0.006 | 0 | 0.1 | 0.08 | 0.05 | 0.14 | 0.06 | 0.13 |
| 9 | 2172103 | rs10965085 | 0.665 | -0.845 | -1.091 | 0.579 | -0.005 | -0.013 | -0.013 | 0 | 0.42 | 0.33 | 0.56 | 0.49 | 0.47 | 0.37 |
| 9 | 2172726 | rs7858891 | 0.283 | -0.596 | -1.096 | 0.609 | 0.002 | -0.001 | -0.001 | 0 | 0.09 | 0.1 | 0.16 | 0.22 | 0.06 | 0.19 |
| 9 | 2173296 | rs7048976 | 0.247 | -1.079 | -1.413 | 0.404 | 0.002 | 0.026 | 0.026 | 0 | 0.47 | 0.41 | 0.7 | 0.67 | 0.5 | 0.54 |
| 9 | 2173478 | rs12345422 | -0.35 | -1.422 | -1.342 | 0.038 | -0.01 | -0.001 | -0.001 | 0.203 | 0.2 | 0.24 | 0.21 | 0.23 | 0.1 | 0.23 |
| 9 | 2173590 | rs1886261 | -0.354 | -2.047 | -1.565 | -0.57 | -0.029 | -0.031 | -0.031 | 0 | 0.24 | 0.49 | 0.15 | 0.15 | 0.15 | 0.21 |
| 9 | 2173752 | rs2376230 | 0.999 | -2.109 | -0.876 | -0.61 | -0.002 | -0.02 | -0.02 | 0.75 | 0.09 | 0.1 | 0.04 | 0.01 | 0.07 | 0.08 |
| 9 | 2175247 | rs10757230 | 0.352 | -2.149 | -0.504 | -0.64 | -0.056 | -0.022 | -0.022 | 0 | 0.76 | 0.57 | 0.75 | 0.77 | 0.86 | 0.75 |
| 9 | 2177391 | rs9298820 | -0.511 | -2.572 | -0.157 | -0.857 | 0.014 | -0.018 | -0.018 | 0 | 0.65 | 0.6 | 0.76 | 0.71 | 0.59 | 0.66 |
| 9 | 2177492 | rs10491697 | -0.576 | -2.68 | 0 | -0.799 | -0.063 | 0.012 | 0.012 | 0.026 | 0.31 | 0.56 | 0.35 | 0.36 | 0.16 | 0.35 |
| 9 | 2180149 | rs1547635 | -0.633 | -2.701 | 0.196 | -0.63 | -0.064 | 0 | 0 | 0.821 | 0.26 | 0.62 | 0.24 | 0.24 | 0.11 | 0.31 |
| 9 | 2182062 | rs2281786 | -0.839 | -2.515 | -0.005 | -0.803 | -0.004 | 0.002 | 0.002 | 0 | 0.71 | 0.73 | 0.8 | 0.74 | 0.62 | 0.72 |
| 9 | 2184301 | rs11787578 | 0.097 | -2.729 | 0.259 | -0.836 | -0.049 | 0.003 | 0.003 | 0 | 0.22 | 0.32 | 0.22 | 0.24 | 0.09 | 0.27 |
| 9 | 2184771 | rs7871591 | -0.902 | -2.659 | 0.293 | -0.845 | -0.012 | 0.008 | 0.008 | 0.032 | 0.28 | 0.62 | 0.27 | 0.36 | 0.18 | 0.42 |
| 9 | 2184925 | rs7860758 | -1.363 | -2.681 | -0.017 | -0.789 | -0.098 | 0.041 | 0.041 | 0 | 0.74 | 0.98 | 0.87 | 0.85 | 0.69 | 0.82 |
| 9 | 2187750 | rs2147263 | -1.414 | -2.615 | 0.131 | -0.731 | -0.025 | -0.024 | -0.024 | 0.057 | 0.58 | 0.75 | 0.45 | 0.53 | 0.5 | 0.54 |
| 9 | 2188383 | rs10116703 | 0.938 | -2.503 | 0.17 | -0.662 | -0.057 | -0.059 | -0.059 | 0 | 0.45 | 0.14 | 0.57 | 0.49 | 0.52 | 0.41 |
| 9 | 2190296 | rs7859695 | -0.721 | -2.673 | 0.494 | -0.334 | -0.036 | 0.108 | 0.108 | 0.003 | 0.2 | 0.56 | 0.24 | 0.37 | 0.12 | 0.33 |
| 9 | 2191189 | rs10965149 | 0.662 | -2.487 | 0.754 | -0.306 | -0.047 | 0.001 | 0.001 | 0.013 | 0.2 | 0.3 | 0.2 | 0.21 | 0.08 | 0.26 |
| 9 | 2191246 | rs3818384 | -1.101 | -2.336 | 0.632 | -0.285 | 0.07 | 0.114 | 0.114 | 0 | 0.08 | 0.35 | 0.08 | 0.14 | 0.08 | 0.09 |
| 9 | 2193975 | rs11789313 | 0.357 | -2.432 | 0.538 | -0.344 | -0.041 | -0.001 | -0.001 | 0 | 0.22 | 0.3 | 0.22 | 0.24 | 0.1 | 0.27 |
| 9 | 2194227 | rs4741652 | -0.574 | -2.108 | -0.171 | -0.399 | -0.005 | -0.01 | -0.01 | 0 | 0.75 | 0.85 | 0.87 | 0.7 | 0.74 | 0.73 |
| 9 | 2194805 | rs1535844 | -0.232 | -2.182 | -0.218 | -0.484 | -0.033 | -0.02 | -0.02 | 0 | 0.36 | 0.55 | 0.28 | 0.23 | 0.18 | 0.35 |
| 9 | 2194911 | rs7022827 | -0.302 | -2.182 | -0.192 | -0.478 | -0.011 | -0.007 | -0.007 | 0 | 0.36 | 0.61 | 0.28 | 0.24 | 0.19 | 0.35 |
| 9 | 2195420 | rs10965167 | 1.114 | -1.825 | -0.698 | -0.27 | -0.089 | -0.094 | -0.094 | 0 | 0.36 | 0.1 | 0.56 | 0.45 | 0.56 | 0.4 |
| 9 | 2195846 | rs7865072 | -0.229 | -1.271 | -0.222 | -0.159 | -0.003 | -0.006 | -0.006 | 0 | 0.75 | 0.89 | 0.82 | 0.74 | 0.75 | 0.66 |
| 9 | 2196183 | rs10757258 | 0.016 | -1.409 | 0.173 | -0.622 | -0.008 | -0.035 | -0.035 | 0 | 0.2 | 0.3 | 0.1 | 0.1 | 0.16 | 0.23 |
| 9 | 2196566 | rs1886263 | 0.496 | -1.003 | -0.585 | -0.499 | -0.007 | -0.009 | -0.009 | 0.021 | 0.26 | 0.12 | 0.23 | 0.28 | 0.24 | 0.27 |
| 9 | 2196734 | rs1886264 | -0.589 | -0.757 | 0.072 | -0.398 | 0.006 | 0.005 | 0.005 | 0 | 0.52 | 0.54 | 0.35 | 0.44 | 0.53 | 0.4 |
| 9 | 2196932 | rs7031622 | -0.615 | 0.277 | 0.187 | -0.356 | 0.059 | -0.086 | -0.086 | 0 | 0.16 | 0.44 | 0.11 | 0.01 | 0.14 | 0.08 |
| 9 | 2197557 | rs7874004 | 0.368 | 0.156 | -0.223 | 0.045 | -0.001 | -0.001 | -0.001 | 0 | 0.2 | 0.22 | 0.09 | 0.23 | 0.19 | 0.14 |
| 9 | 2198627 | rs10965177 | -0.35 | 0.252 | 0.135 | -0.136 | 0.013 | -0.019 | -0.019 | 0 | 0.34 | 0.44 | 0.27 | 0.25 | 0.35 | 0.28 |
| 9 | 2203544 | rs7049212 | -0.214 | -0.102 | -0.476 | -0.129 | -0.001 | -0.001 | -0.001 | 0 | 0.52 | 0.57 | 0.3 | 0.4 | 0.41 | 0.46 |
| 9 | 2203938 | rs6475600 | -0.233 | -0.975 | -1.003 | -0.898 | 0.019 | 0.023 | 0.023 | 0 | 0.42 | 0.35 | 0.31 | 0.17 | 0.3 | 0.4 |
| 9 | 2205924 | rs7044370 | 0.432 | -0.486 | -1.03 | -0.863 | 0.016 | 0.082 | 0.082 | 0 | 0.3 | 0.15 | 0.21 | 0.09 | 0.25 | 0.31 |
| 9 | 2208793 | rs10965207 | 0.485 | -0.415 | -0.884 | -0.511 | -0.002 | -0.007 | -0.007 | 0.272 | 0.14 | 0.11 | 0.09 | 0.27 | 0.14 | 0.11 |
| 9 | 2208810 | rs7026521 | -0.264 | -0.535 | -0.711 | -0.987 | -0.012 | -0.034 | -0.034 | 0.286 | 0.16 | 0.21 | 0.16 | 0.04 | 0.09 | 0.26 |
| 9 | 2210547 | rs942401 | 0.209 | -0.247 | -0.809 | -0.942 | -0.015 | -0.021 | -0.021 | 0 | 0.18 | 0.12 | 0.15 | 0.34 | 0.23 | 0.13 |
| 9 | 2210583 | rs942402 | 0.252 | -0.241 | -0.806 | -0.939 | -0.015 | -0.021 | -0.021 | 0.041 | 0.18 | 0.12 | 0.14 | 0.34 | 0.23 | 0.13 |
| 9 | 2210993 | rs7863990 | 0.734 | -0.752 | -0.313 | -1.432 | -0.024 | -0.084 | -0.084 | 0 | 0.35 | 0.45 | 0.23 | 0.09 | 0.25 | 0.25 |
| 9 | 2212546 | rs1535842 | -0.833 | -0.876 | -0.356 | -1.404 | -0.068 | -0.056 | -0.056 | 1.748 | 0.62 | 0.85 | 0.46 | 0.47 | 0.49 | 0.48 |
| 9 | 2212574 | rs1535841 | -0.32 | -0.697 | -0.198 | -1.243 | 0 | -0.002 | -0.002 | 0.726 | 0.28 | 0.26 | 0.22 | 0.38 | 0.27 | 0.31 |
| 9 | 2213818 | rs7037884 | 0.487 | -0.702 | -0.097 | -1.149 | -0.017 | -0.016 | -0.016 | 0 | 0.11 | 0.14 | 0.08 | 0.04 | 0.05 | 0.17 |
| 9 | 2214693 | rs1075039 | 0.444 | -1.024 | -0.355 | -1.374 | -0.012 | 0.009 | 0.009 | 0 | 0.45 | 0.25 | 0.31 | 0.43 | 0.43 | 0.3 |
| 9 | 2217178 | rs12350147 | 0.615 | -0.962 | -0.209 | -1.465 | -0.005 | -0.026 | -0.026 | 0 | 0.12 | 0.13 | 0.08 | 0.07 | 0.12 | 0.02 |
| 9 | 2220211 | rs7025331 | -0.318 | -0.643 | -0.346 | -1.344 | -0.073 | -0.091 | -0.091 | 0.116 | 0.63 | 0.88 | 0.49 | 0.51 | 0.51 | 0.53 |
| 9 | 2220796 | rs2093833 | 1.833 | -0.818 | -0.423 | -1.483 | -0.028 | 0.063 | 0.063 | 0.234 | 0.27 | 0.06 | 0.21 | 0.11 | 0.21 | 0.17 |
| 9 | 2220915 | rs10757276 | 0.797 | -0.965 | 0.262 | -1.504 | 0 | -0.002 | -0.002 | 0 | 0.21 | 0.23 | 0.1 | 0.27 | 0.17 | 0.15 |
| 9 | 2227217 | rs7024071 | 0.432 | -0.921 | 0.433 | -1.499 | -0.004 | -0.028 | -0.028 | 0 | 0.39 | 0.5 | 0.38 | 0.24 | 0.34 | 0.37 |
| 9 | 2227896 | rs10491703 | 0.295 | -0.719 | -0.114 | -1.512 | -0.067 | -0.085 | -0.085 | 0.113 | 0.34 | 0.09 | 0.52 | 0.49 | 0.49 | 0.46 |
| 9 | 2228689 | rs7045699 | 0.334 | -0.783 | 0.037 | -1.435 | 0.018 | -0.022 | -0.022 | 0.6 | 0.87 | 0.67 | 0.81 | 0.88 | 0.84 | 0.77 |
| 9 | 2229436 | rs2093836 | -0.516 | -0.804 | -0.358 | -1.428 | -0.011 | -0.022 | -0.022 | 0 | 0.9 | 0.75 | 0.85 | 0.95 | 0.91 | 0.77 |
| 9 | 2229694 | rs4741655 | 0.437 | -0.873 | -0.463 | -1.447 | -0.001 | 0 | 0 | 0.465 | 0.65 | 0.34 | 0.7 | 0.61 | 0.7 | 0.62 |

| | | | | | | | | | | | | | | | | |
|---|---|---|---|---|---|---|---|---|---|---|---|---|---|---|---|---|
| 9 | 2231490 | rs4741656 | -0.094 | -0.253 | -1.153 | -1.776 | -0.015 | 0.098 | 0.098 | 0 | 0.56 | 0.69 | 0.55 | 0.82 | 0.58 | 0.53 |
| 9 | 2234367 | rs942404 | 0.221 | -0.448 | -0.786 | -1.291 | 0.006 | 0.015 | 0.015 | 0 | 0.91 | 0.96 | 0.87 | 0.94 | 0.92 | 0.78 |
| 9 | 2239429 | rs4741657 | -0.176 | -0.564 | -0.676 | -1.659 | 0.028 | 0.031 | 0.031 | 1.758 | 0.59 | 0.67 | 0.66 | 0.81 | 0.74 | 0.6 |
| 9 | 2241091 | rs10965321 | -0.078 | 0.568 | -0.934 | -1.158 | -0.034 | -0.007 | -0.007 | 0.207 | 0.21 | 0.16 | 0.45 | 0.74 | 0.31 | 0.36 |
| 9 | 2241994 | rs4741658 | -0.03 | 0.299 | -1.127 | -1.099 | -0.078 | -0.05 | -0.05 | 0 | 0.19 | 0.02 | 0.41 | 0.78 | 0.27 | 0.36 |
| 9 | 2245401 | rs2181829 | -1.365 | 0.026 | -1.257 | -1.51 | -0.038 | 0.075 | 0.075 | 0 | 0.81 | 0.98 | 0.91 | 0.94 | 0.82 | 0.88 |
| 9 | 2247072 | rs6475635 | 0.143 | -0.146 | -0.624 | -1.702 | 0 | 0.088 | 0.088 | 0 | 0.58 | 0.77 | 0.48 | 0.14 | 0.59 | 0.42 |
| 9 | 2247904 | rs10965339 | 0.322 | -0.239 | -0.131 | -1.743 | -0.001 | -0.017 | -0.017 | 0.522 | 0.56 | 0.58 | 0.47 | 0.14 | 0.59 | 0.42 |
| 9 | 2250385 | rs7044488 | 1.31 | -0.409 | -0.655 | -2.15 | -0.041 | 0.107 | 0.107 | 0.102 | 0.32 | 0.12 | 0.27 | 0.09 | 0.35 | 0.2 |
| 9 | 2250413 | rs7041362 | 0.61 | -0.214 | -0.428 | -1.836 | -0.011 | -0.027 | -0.027 | 0 | 0.19 | 0.07 | 0.4 | 0.77 | 0.2 | 0.4 |
| 9 | 2252454 | rs10965349 | 0.579 | -0.215 | -0.385 | -1.84 | -0.003 | -0.013 | -0.013 | 3.099 | 0.19 | 0.11 | 0.4 | 0.79 | 0.21 | 0.4 |
| 9 | 2252617 | rs10811711 | 0.606 | -0.216 | -0.386 | -1.841 | -0.003 | -0.013 | -0.013 | 0 | 0.19 | 0.1 | 0.4 | 0.79 | 0.2 | 0.38 |
| 9 | 2253087 | rs942397 | -1.817 | -0.174 | -0.464 | -1.865 | -0.022 | 0.067 | 0.067 | 0 | 0.81 | 0.99 | 0.92 | 0.94 | 0.84 | 0.91 |
| 9 | 2253678 | rs942398 | -1.716 | -0.14 | -0.45 | -1.77 | -0.019 | 0.053 | 0.053 | 0 | 0.8 | 0.99 | 0.92 | 0.91 | 0.84 | 0.9 |
| 9 | 2254498 | rs7042935 | -1.455 | -0.154 | -0.234 | -1.777 | 0.001 | -0.001 | -0.001 | 0 | 0.79 | 0.8 | 0.91 | 0.91 | 0.84 | 0.9 |
| 9 | 2255397 | rs1555646 | -0.196 | -0.269 | -0.03 | -1.814 | -0.001 | -0.015 | -0.015 | 0.174 | 0.58 | 0.56 | 0.5 | 0.14 | 0.63 | 0.52 |
| 9 | 2256092 | rs16938145 | 0.226 | -0.484 | 0.168 | -1.952 | 0.003 | -0.081 | -0.081 | 0 | 0.21 | 0.32 | 0.23 | 0.06 | 0.2 | 0.3 |
| 9 | 2261052 | rs7847256 | 0.405 | -0.519 | 0.216 | -1.3 | -0.013 | -0.019 | -0.019 | 0.096 | 0.18 | 0.08 | 0.39 | 0.78 | 0.22 | 0.38 |
| 9 | 2263591 | rs10125336 | -1.187 | -0.618 | 0.338 | -0.758 | 0.02 | -0.018 | -0.018 | 0 | 0.79 | 0.91 | 0.86 | 0.7 | 0.84 | 0.84 |
| 9 | 2264009 | rs12552398 | 1.149 | -0.616 | 0.299 | -0.753 | 0.019 | -0.008 | -0.008 | 0 | 0.2 | 0.07 | 0.14 | 0.24 | 0.16 | 0.16 |
| 9 | 2267871 | rs7872273 | 0.881 | -0.618 | 0.219 | -1.282 | -0.052 | -0.081 | -0.081 | 0 | 0.15 | 0.02 | 0.37 | 0.74 | 0.2 | 0.36 |
| 9 | 2270046 | rs7023476 | -1.243 | -0.68 | 0.762 | -1.546 | -0.001 | -0.014 | -0.014 | 0.664 | 0.54 | 0.59 | 0.58 | 0.88 | 0.47 | 0.58 |
| 9 | 2270173 | rs17471115 | 0.159 | -0.383 | 0.558 | -1.377 | -0.006 | 0.026 | 0.026 | 0 | 0.1 | 0.03 | 0.05 | 0 | 0.09 | 0.02 |
| 9 | 2271325 | rs17405547 | 1.197 | -0.358 | 0.561 | -1.309 | -0.036 | 0.078 | 0.078 | 0 | 0.2 | 0.01 | 0.1 | 0.06 | 0.18 | 0.13 |
| 9 | 2272512 | rs10811756 | -0.271 | -0.216 | 0.724 | -1.252 | 0.001 | 0.005 | 0.005 | 0 | 0.41 | 0.42 | 0.52 | 0.83 | 0.4 | 0.57 |
| 9 | 2275029 | rs7869041 | -0.234 | 0.019 | 0.028 | -1.071 | -0.018 | -0.091 | -0.091 | 0.906 | 0.36 | 0.54 | 0.2 | 0.11 | 0.26 | 0.23 |
| 9 | 2282951 | rs1324208 | -0.311 | 0.003 | -0.507 | -0.379 | -0.004 | -0.018 | -0.018 | 0 | 0.25 | 0.12 | 0.36 | 0.69 | 0.27 | 0.49 |
| 9 | 2285443 | rs10811770 | 0.252 | 0.095 | -0.559 | -0.659 | 0.035 | 0.119 | 0.119 | 0 | 0.49 | 0.32 | 0.48 | 0.23 | 0.56 | 0.33 |
| 9 | 2290590 | rs9644866 | -0.546 | 0.391 | -0.233 | -0.552 | 0.028 | -0.047 | -0.047 | 0 | 0.23 | 0.51 | 0.28 | 0.14 | 0.26 | 0.19 |
| 9 | 2291635 | rs744216 | -0.867 | -0.32 | -0.631 | -0.37 | 0.017 | 0.031 | 0.031 | 0.004 | 0.43 | 0.68 | 0.43 | 0.31 | 0.37 | 0.41 |
| 9 | 2293921 | rs6475688 | 0.024 | -0.419 | -1.402 | -0.444 | 0.044 | 0.045 | 0.045 | 0.014 | 0.59 | 0.74 | 0.54 | 0.28 | 0.69 | 0.5 |
| 9 | 2295503 | rs1555644 | 1.285 | -0.395 | -1.046 | -0.592 | -0.036 | 0.093 | 0.093 | 0 | 0.34 | 0.15 | 0.24 | 0.1 | 0.41 | 0.22 |
| 9 | 2299042 | rs12554219 | 1.247 | -0.509 | -1.151 | -0.55 | 0.041 | -0.016 | -0.016 | 0 | 0.19 | 0.06 | 0.15 | 0.16 | 0.12 | 0.22 |
| 9 | 2299220 | rs10965506 | 1.084 | -0.397 | -1.087 | -0.565 | -0.042 | 0.11 | 0.11 | 0 | 0.36 | 0.17 | 0.25 | 0.1 | 0.43 | 0.3 |
| 9 | 2300565 | rs13292475 | 1.265 | -0.277 | -1.095 | -0.531 | -0.041 | 0.101 | 0.101 | 0 | 0.33 | 0.1 | 0.22 | 0.1 | 0.39 | 0.21 |
| 9 | 2301610 | rs12684262 | 0.702 | -0.545 | -0.713 | -0.384 | -0.018 | -0.004 | -0.004 | 0 | 0.22 | 0.25 | 0.16 | 0.16 | 0.13 | 0.27 |
| 9 | 2301692 | rs10965522 | 0.039 | 0.013 | -0.352 | -0.293 | 0.039 | -0.036 | -0.036 | 0 | 0.22 | 0.45 | 0.27 | 0.14 | 0.25 | 0.18 |
| 9 | 2302693 | rs7032999 | 0.224 | 0.245 | -0.41 | -0.142 | -0.005 | 0.028 | 0.028 | 0 | 0.43 | 0.66 | 0.38 | 0.24 | 0.45 | 0.47 |
| 9 | 2303095 | rs7036558 | 1.323 | 0.133 | 0.036 | -0.382 | 0.013 | -0.05 | -0.05 | 0 | 0.86 | 0.56 | 0.86 | 0.94 | 0.86 | 0.96 |
| 9 | 2305679 | rs10965543 | 0.172 | 0.689 | -0.676 | 0.141 | -0.001 | 0.017 | 0.017 | 0 | 0.17 | 0.19 | 0.32 | 0.54 | 0.28 | 0.23 |
| 9 | 2312205 | rs10965569 | -0.096 | 0.288 | -1.316 | -0.332 | -0.074 | 0.024 | 0.024 | 0 | 0.34 | 0.17 | 0.55 | 0.73 | 0.47 | 0.4 |
| 9 | 2320311 | rs10448215 | 0.946 | 0.164 | -1.939 | -0.913 | -0.076 | 0.156 | 0.156 | 0.396 | 0.43 | 0.07 | 0.16 | 0.21 | 0.34 | 0.43 |
| 9 | 2321242 | rs12553062 | 1.29 | 0.02 | -1.744 | -0.968 | -0.008 | 0.001 | 0.001 | 0.025 | 0.11 | 0.09 | 0.18 | 0.07 | 0.16 | 0.2 |
| 9 | 2322572 | rs7021419 | -0.644 | -0.133 | -1.371 | -0.952 | 0.042 | 0.048 | 0.048 | 0 | 0.45 | 0.67 | 0.53 | 0.24 | 0.38 | 0.38 |
| 9 | 2323909 | rs12380394 | 0.977 | 0.162 | -1.155 | -0.783 | -0.007 | 0.003 | 0.003 | 0 | 0.17 | 0.2 | 0.33 | 0.57 | 0.3 | 0.23 |
| 9 | 2324558 | rs16938404 | 1.809 | 0.067 | -1.318 | -0.832 | 0.003 | -0.02 | -0.02 | 2.295 | 0.19 | 0.06 | 0.23 | 0.17 | 0.17 | 0.16 |
| 9 | 2324646 | rs17473202 | 0.863 | 0.209 | -0.545 | -0.787 | 0.008 | 0.104 | 0.104 | 0 | 0.42 | 0.19 | 0.16 | 0.2 | 0.32 | 0.39 |
| 9 | 2325739 | rs1002780 | -0.136 | 0.221 | -0.102 | -0.684 | -0.007 | -0.007 | -0.007 | 0.037 | 0.6 | 0.4 | 0.52 | 0.76 | 0.65 | 0.63 |
| 9 | 2329724 | rs1571837 | 0.529 | 0.508 | -0.115 | -0.267 | -0.005 | -0.006 | -0.006 | 0 | 0.19 | 0.17 | 0.34 | 0.49 | 0.29 | 0.21 |
| 9 | 2330643 | rs10125660 | -0.235 | -0.137 | 0.159 | -1.139 | -0.023 | -0.083 | -0.083 | 0.082 | 0.28 | 0.5 | 0.29 | 0.11 | 0.26 | 0.3 |
| 9 | 2334748 | rs1109816 | 0.517 | -0.077 | 0.14 | -1.176 | 0.003 | -0.04 | -0.04 | 0 | 0.39 | 0.47 | 0.36 | 0.18 | 0.4 | 0.44 |
| 9 | 2338786 | rs6475728 | -1.767 | 0.275 | -0.762 | -0.986 | -0.03 | -0.045 | -0.045 | 0.027 | 0.84 | 0.95 | 0.83 | 0.71 | 0.82 | 0.76 |
| 9 | 2340310 | rs4741676 | -0.308 | 0.246 | -0.487 | -1.009 | 0.007 | 0.015 | 0.015 | 0 | 0.66 | 0.76 | 0.75 | 0.72 | 0.71 | 0.7 |
| 9 | 2342136 | rs1331817 | 0.409 | 0.493 | -0.686 | -0.491 | -0.012 | -0.059 | -0.059 | 0 | 0.26 | 0.16 | 0.38 | 0.54 | 0.3 | 0.25 |
| 9 | 2361503 | rs4475544 | 0.41 | 0.067 | 0.288 | -0.746 | -0.016 | -0.05 | -0.05 | 0 | 0.14 | 0.2 | 0.07 | 0 | 0.08 | 0.07 |
| 9 | 2361656 | rs10123371 | 0.186 | 0.266 | 0.415 | -0.86 | 0.051 | 0.026 | 0.026 | 0 | 0.12 | 0.34 | 0.11 | 0.1 | 0.14 | 0.24 |
| 9 | 2363667 | rs2225500 | 0.734 | 0.393 | -0.463 | -0.918 | -0.008 | 0.066 | 0.066 | 0 | 0.49 | 0.16 | 0.59 | 0.63 | 0.54 | 0.43 |
| 9 | 2363874 | rs2183810 | 0.734 | 0.393 | -0.463 | -0.919 | -0.008 | 0.063 | 0.063 | 0 | 0.49 | 0.17 | 0.59 | 0.63 | 0.54 | 0.42 |
| 9 | 2379343 | rs1113842 | -0.424 | 0.448 | -0.145 | -0.991 | -0.01 | -0.088 | -0.088 | 0 | 0.28 | 0.46 | 0.2 | 0.11 | 0.26 | 0.32 |
| 9 | 2379389 | rs1113841 | 0.9 | 0.428 | -0.496 | -0.979 | -0.031 | 0.045 | 0.045 | 0 | 0.47 | 0.23 | 0.49 | 0.61 | 0.51 | 0.42 |
| 9 | 2380408 | rs1331824 | 0.063 | 0.272 | 0.031 | -1.119 | -0.001 | -0.002 | -0.002 | 1.608 | 0.59 | 0.46 | 0.67 | 0.71 | 0.63 | 0.66 |
| 9 | 2380595 | rs1331825 | 0.957 | 0.539 | -0.347 | -1.106 | -0.004 | 0.043 | 0.043 | 0 | 0.46 | 0.19 | 0.43 | 0.61 | 0.44 | 0.41 |
| 9 | 2381672 | rs1023197 | -0.798 | 0.8 | -0.099 | -0.907 | -0.015 | -0.008 | -0.008 | 0 | 0.77 | 0.85 | 0.76 | 0.72 | 0.72 | 0.76 |
| 9 | 2382247 | rs4741690 | 0.162 | 0.81 | -0.034 | -0.886 | -0.014 | 0.002 | 0.002 | 0 | 0.61 | 0.81 | 0.5 | 0.6 | 0.45 | 0.48 |
| 9 | 2384754 | rs11794776 | 0.697 | 0.412 | -0.427 | -1.062 | 0.003 | -0.054 | -0.054 | 0.011 | 0.18 | 0.09 | 0.25 | 0.51 | 0.19 | 0.25 |
| 9 | 2385314 | rs10491708 | 0.943 | 0.61 | -0.559 | -1.093 | -0.044 | -0.024 | -0.024 | 1.434 | 0.08 | 0.02 | 0.19 | 0.1 | 0.16 | 0.25 |
| 9 | 2385584 | rs10966074 | 0.247 | 0.484 | 0.171 | -1.197 | 0.007 | 0.019 | 0.019 | 0 | 0.51 | 0.6 | 0.42 | 0.19 | 0.43 | 0.49 |
| 9 | 2386539 | rs4740678 | 0.562 | 0.33 | 0.153 | -1.126 | -0.005 | 0.042 | 0.042 | 0 | 0.2 | 0.2 | 0.28 | 0.51 | 0.19 | 0.23 |
| 9 | 2387137 | rs999644 | 0.752 | 0.253 | 0.391 | -1.236 | 0.011 | -0.001 | -0.001 | 0 | 0.4 | 0.38 | 0.21 | 0.09 | 0.27 | 0.24 |

| | | | | | | | | | | | | | | | | |
|---|---|---|---|---|---|---|---|---|---|---|---|---|---|---|---|---|
| 9 | 2388044 | rs1571838 | -0.718 | 0.559 | 0.468 | -1.11 | 0.014 | -0.003 | -0.003 | 0 | 0.71 | 0.69 | 0.68 | 0.72 | 0.62 | 0.74 |
| 9 | 2388302 | rs12342955 | 1.639 | -0.164 | 0.196 | -1.406 | -0.008 | -0.032 | -0.032 | 0 | 0.26 | 0.34 | 0.17 | 0.09 | 0.2 | 0.17 |
| 9 | 2391261 | rs10811970 | 1.254 | -0.299 | 0.406 | -1.254 | -0.075 | 0.072 | 0.072 | 0 | 0.32 | 0.05 | 0.35 | 0.2 | 0.38 | 0.41 |
| 9 | 2394917 | rs959379 | -0.458 | -0.063 | 0.763 | -0.994 | 0.015 | -0.006 | -0.006 | 0.388 | 0.69 | 0.48 | 0.66 | 0.68 | 0.68 | 0.71 |
| 9 | 2396673 | rs7025116 | -0.725 | -0.059 | 0.406 | -0.992 | 0.004 | -0.001 | -0.001 | 0.018 | 0.71 | 0.64 | 0.68 | 0.68 | 0.68 | 0.71 |
| 9 | 2397564 | rs7870386 | 0.428 | -0.082 | 0.325 | -1.025 | 0.014 | 0.06 | 0.06 | 0 | 0.52 | 0.27 | 0.6 | 0.67 | 0.59 | 0.65 |
| 9 | 2407101 | rs11793711 | 0.373 | -0.051 | 0.427 | -0.215 | 0.01 | 0.144 | 0.144 | 0 | 0.17 | 0.35 | 0.23 | 0.39 | 0.16 | 0.21 |
| 9 | 2408770 | rs7038629 | 0.188 | -0.306 | 0.364 | -0.358 | -0.025 | -0.075 | -0.075 | 0 | 0.19 | 0.29 | 0.1 | 0.04 | 0.16 | 0.1 |
| 9 | 2409096 | rs7030433 | -1.085 | -0.289 | 0.385 | -0.358 | 0.017 | -0.01 | -0.01 | 0 | 0.38 | 0.7 | 0.34 | 0.44 | 0.34 | 0.31 |
| 9 | 2411572 | rs7025994 | 0.309 | -0.172 | 0.394 | -0.244 | 0.019 | 0.163 | 0.163 | 0 | 0.17 | 0.35 | 0.27 | 0.43 | 0.18 | 0.23 |
| 9 | 2412296 | rs1571840 | 0.123 | -0.168 | 0.39 | -0.244 | 0.012 | 0.146 | 0.146 | 0 | 0.19 | 0.36 | 0.27 | 0.43 | 0.18 | 0.22 |
| 9 | 2414600 | rs2376120 | 0.04 | 0.275 | -0.091 | 0.083 | -0.001 | -0.001 | -0.001 | 0 | 0.55 | 0.41 | 0.56 | 0.48 | 0.54 | 0.59 |
| 9 | 2419538 | rs872039 | -0.89 | 0.394 | -0.115 | -0.341 | 0.022 | 0.016 | 0.016 | 0 | 0.73 | 0.6 | 0.7 | 0.66 | 0.69 | 0.76 |
| 9 | 2419927 | rs914491 | 0.694 | 0.118 | -0.445 | -0.836 | -0.046 | 0.053 | 0.053 | 0 | 0.36 | 0.16 | 0.39 | 0.21 | 0.39 | 0.46 |
| 9 | 2421084 | rs16906059 | -0.282 | -0.026 | 0.121 | -0.573 | -0.018 | -0.06 | -0.06 | 0.058 | 0.16 | 0.24 | 0.08 | 0.01 | 0.11 | 0.07 |
| 9 | 2421690 | rs9696013 | -0.174 | 0.104 | -0.311 | -0.518 | -0.02 | -0.065 | -0.065 | 0 | 0.14 | 0.22 | 0.06 | 0 | 0.07 | 0.05 |
| 9 | 2425428 | rs10812032 | -0.209 | 0.225 | -0.803 | -0.655 | 0.003 | -0.005 | -0.005 | 0.858 | 0.56 | 0.41 | 0.69 | 0.83 | 0.57 | 0.7 |
| 9 | 2425663 | rs10812034 | -0.078 | 0.587 | -1.18 | -0.134 | -0.018 | -0.037 | -0.037 | 0 | 0.37 | 0.21 | 0.39 | 0.43 | 0.35 | 0.49 |
| 9 | 2426100 | rs10966251 | 0.778 | 0.67 | -1.049 | -0.037 | 0 | 0.008 | 0.008 | 0.664 | 0.19 | 0.2 | 0.3 | 0.4 | 0.22 | 0.19 |
| 9 | 2428377 | rs10812036 | 0.502 | 0.7 | -0.969 | -0.32 | 0.005 | -0.002 | -0.002 | 1.959 | 0.48 | 0.52 | 0.32 | 0.17 | 0.44 | 0.34 |
| 9 | 2429859 | rs2210396 | -0.009 | 0.692 | -1.262 | 0.001 | -0.012 | -0.03 | -0.03 | 0 | 0.32 | 0.21 | 0.45 | 0.53 | 0.31 | 0.46 |
| 9 | 2430911 | rs2376117 | 0.989 | 0.696 | -1.244 | -0.319 | -0.011 | -0.035 | -0.035 | 0.036 | 0.18 | 0.13 | 0.28 | 0.39 | 0.22 | 0.19 |
| 9 | 2431272 | rs1155008 | -0.004 | 0.744 | -1.205 | -0.332 | -0.002 | -0.006 | -0.006 | 0 | 0.31 | 0.3 | 0.35 | 0.44 | 0.3 | 0.46 |
| 9 | 2432562 | rs10966259 | 0.717 | 0.643 | -1.429 | -0.4 | 0.002 | 0.131 | 0.131 | 0 | 0.45 | 0.27 | 0.32 | 0.13 | 0.45 | 0.47 |
| 9 | 2435209 | rs2889293 | 0.61 | 0.568 | -0.927 | 0.008 | 0 | -0.012 | -0.012 | 0.239 | 0.59 | 0.46 | 0.43 | 0.24 | 0.6 | 0.55 |
| 9 | 2437565 | rs12338622 | -0.043 | 1.053 | -1.408 | 0.254 | -0.032 | -0.055 | -0.055 | 0 | 0.22 | 0.13 | 0.32 | 0.4 | 0.26 | 0.21 |
| 9 | 2442052 | rs962606 | -0.27 | 0.726 | -1.246 | -0.98 | 0.023 | 0.082 | 0.082 | 0.813 | 0.52 | 0.68 | 0.67 | 0.92 | 0.59 | 0.6 |
| 9 | 2447910 | rs10757471 | -0.254 | 0.752 | -1.306 | -0.488 | -0.01 | -0.03 | -0.03 | 0 | 0.36 | 0.33 | 0.45 | 0.52 | 0.36 | 0.38 |
| 9 | 2451856 | rs10757474 | 0.035 | 0.721 | -1.037 | -0.935 | -0.001 | 0 | 0 | 0 | 0.39 | 0.37 | 0.27 | 0.39 | 0.33 | 0.23 |
| 9 | 2456676 | rs10812051 | 0.123 | 0.821 | -0.942 | -1.092 | 0.001 | 0.017 | 0.017 | 0 | 0.2 | 0.24 | 0.4 | 0.51 | 0.24 | 0.49 |
| 9 | 2459387 | rs1158989 | 1.276 | 0.503 | -0.772 | -1.412 | 0.004 | -0.016 | -0.016 | 0.064 | 0.28 | 0.19 | 0.21 | 0.39 | 0.27 | 0.19 |
| 9 | 2459760 | rs954624 | 1.19 | 0.905 | -0.901 | -1.312 | -0.047 | -0.024 | -0.024 | 0 | 0.12 | 0.02 | 0.18 | 0.1 | 0.19 | 0.27 |
| 9 | 2460468 | rs10491712 | 1.259 | 0.778 | -0.821 | -1.4 | -0.031 | 0.067 | 0.067 | 0 | 0.21 | 0.07 | 0.19 | 0.1 | 0.25 | 0.17 |
| 9 | 2466509 | rs10115153 | -1.245 | 0.819 | 0.042 | -1.148 | 0.009 | -0.054 | -0.054 | 0.295 | 0.88 | 0.84 | 0.91 | 1 | 0.86 | 0.95 |
| 9 | 2468858 | rs914492 | 0.638 | 0.81 | -0.1 | -0.918 | 0 | 0.099 | 0.099 | 0 | 0.06 | 0.08 | 0.21 | 0.41 | 0.06 | 0.23 |
| 9 | 2469509 | rs1331458 | 0.884 | 0.615 | -0.208 | -1.285 | -0.021 | -0.014 | -0.014 | 0 | 0.57 | 0.28 | 0.46 | 0.48 | 0.6 | 0.46 |
| 9 | 2469742 | rs2150720 | 0.813 | 0.762 | -0.202 | -1.39 | 0 | 0.002 | 0.002 | 0 | 0.68 | 0.33 | 0.6 | 0.55 | 0.69 | 0.67 |
| 9 | 2470058 | rs4741713 | 1.047 | 0.958 | -0.731 | -1.354 | -0.022 | -0.045 | -0.045 | 0 | 0.16 | 0.06 | 0.18 | 0.32 | 0.21 | 0.17 |
| 9 | 2472026 | rs7041090 | -0.48 | 0.906 | -0.304 | -1.517 | 0.001 | 0.018 | 0.018 | 0 | 0.35 | 0.41 | 0.48 | 0.53 | 0.28 | 0.5 |
| 9 | 2472410 | rs6475804 | -0.403 | 0.969 | -0.116 | -1.58 | 0 | 0 | 0 | 1.327 | 0.7 | 0.68 | 0.74 | 0.66 | 0.64 | 0.78 |
| 9 | 2475688 | rs2150721 | 0.909 | 0.6 | -0.347 | -1.53 | 0.041 | 0.114 | 0.114 | 0 | 0.39 | 0.16 | 0.27 | 0.13 | 0.35 | 0.31 |
| 9 | 2476620 | rs12352307 | -0.027 | 0.614 | -0.052 | -1.326 | -0.004 | 0.031 | 0.031 | 0 | 0.27 | 0.33 | 0.36 | 0.5 | 0.2 | 0.39 |
| 9 | 2482283 | rs2376112 | 1.599 | 0.771 | -0.698 | -1.61 | -0.073 | 0.144 | 0.144 | 0 | 0.34 | 0.05 | 0.32 | 0.12 | 0.37 | 0.33 |
| 9 | 2482545 | rs2889291 | -1.478 | 0.717 | -0.178 | -1.478 | -0.002 | -0.013 | -0.013 | 0 | 0.87 | 0.87 | 0.92 | 1 | 0.84 | 0.94 |
| 9 | 2485202 | rs2026344 | 0.21 | 0.579 | 0.094 | -1.375 | -0.004 | 0.007 | 0.007 | 0 | 0.63 | 0.53 | 0.71 | 0.61 | 0.63 | 0.72 |
| 9 | 2487356 | rs10966459 | -0.177 | 0.656 | 0.19 | -1.348 | -0.004 | -0.017 | -0.017 | 0.171 | 0.07 | 0.08 | 0.03 | 0 | 0.04 | 0.02 |
| 9 | 2490339 | rs10966485 | 0.2 | 0.974 | 0.452 | -1.361 | -0.001 | -0.053 | -0.053 | 0 | 0.1 | 0.2 | 0.05 | 0 | 0.05 | 0.05 |
| 9 | 2490690 | rs1571808 | -0.082 | 1.096 | 0.467 | -1.359 | 0.011 | 0.024 | 0.024 | 0 | 0.71 | 0.6 | 0.73 | 0.61 | 0.66 | 0.76 |
| 9 | 2493123 | rs4237145 | -0.823 | 1.206 | 0.65 | -1.302 | 0.044 | -0.074 | -0.074 | 0.594 | 0.87 | 0.72 | 0.88 | 1 | 0.82 | 0.94 |
| 9 | 2493381 | rs1002393 | 0.078 | 1.151 | 0.17 | -1.389 | -0.007 | 0.075 | 0.075 | 0 | 0.19 | 0.27 | 0.32 | 0.49 | 0.16 | 0.36 |
| 9 | 2493528 | rs1002394 | 1.105 | 1.202 | -0.316 | -1.394 | -0.055 | 0.194 | 0.194 | 0 | 0.4 | 0.13 | 0.35 | 0.11 | 0.45 | 0.38 |
| 9 | 2495675 | rs4559295 | -0.286 | 0.877 | -0.139 | -1.011 | 0.035 | -0.04 | -0.04 | 0 | 0.36 | 0.21 | 0.25 | 0.47 | 0.27 | 0.42 |
| 9 | 2495694 | rs875587 | -0.846 | 0.929 | 0.224 | -1.057 | 0.015 | 0.032 | 0.032 | 0 | 0.43 | 0.34 | 0.5 | 0.88 | 0.35 | 0.57 |
| 9 | 2495870 | rs1571812 | -0.179 | 0.907 | 0.009 | -1.039 | 0.035 | -0.063 | -0.063 | 0 | 0.36 | 0.14 | 0.25 | 0.47 | 0.27 | 0.42 |
| 9 | 2496567 | rs872257 | 0.207 | 1 | 0.115 | -1.119 | 0 | 0.005 | 0.005 | 0 | 0.75 | 0.31 | 0.61 | 0.59 | 0.73 | 0.75 |
| 9 | 2499251 | rs10812126 | 1.373 | 1.084 | -0.055 | -1.241 | -0.11 | 0.206 | 0.206 | 0 | 0.4 | 0.11 | 0.4 | 0.11 | 0.49 | 0.35 |
| 9 | 2499273 | rs7848645 | 0.696 | 0.762 | 0.342 | -1.292 | -0.004 | -0.003 | -0.003 | 0.441 | 0.13 | 0.17 | 0.06 | 0.08 | 0.07 | 0.19 |
| 9 | 2499345 | rs6475827 | 0.226 | 0.754 | 0.818 | -1.33 | -0.004 | -0.102 | -0.102 | 0.041 | 0.71 | 0.5 | 0.83 | 0.92 | 0.73 | 0.73 |
| 9 | 2500687 | rs10812133 | 1.101 | 0.772 | 0.763 | -1.322 | -0.015 | -0.059 | -0.059 | 0 | 0.18 | 0.06 | 0.17 | 0.4 | 0.19 | 0.19 |
| 9 | 2500856 | rs1105016 | 0.733 | 0.745 | 0.747 | -1.335 | -0.004 | -0.003 | -0.003 | 0 | 0.13 | 0.16 | 0.06 | 0.08 | 0.07 | 0.18 |
| 9 | 2501314 | rs17479130 | 1.895 | 0.746 | 0.674 | -1.343 | -0.076 | 0.192 | 0.192 | 0 | 0.39 | 0.1 | 0.36 | 0.12 | 0.44 | 0.33 |
| 9 | 2503034 | rs10812141 | 0.254 | 0.874 | 0.714 | -1.313 | -0.006 | -0.042 | -0.042 | 2.974 | 0.24 | 0.13 | 0.19 | 0.39 | 0.23 | 0.25 |
| 9 | 2503549 | rs10738699 | -0.963 | 0.776 | 0.753 | -1.445 | 0.02 | -0.078 | -0.078 | 1.351 | 0.85 | 0.73 | 0.89 | 1 | 0.82 | 0.91 |
| 9 | 2505393 | rs944546 | -1.247 | 0.875 | 0.601 | -1.388 | -0.007 | -0.057 | -0.057 | 0 | 0.45 | 0.47 | 0.52 | 0.88 | 0.35 | 0.58 |
| 9 | 2506794 | rs4741719 | -0.687 | 0.912 | 0.792 | -1.187 | -0.008 | -0.04 | -0.04 | 0 | 0.33 | 0.4 | 0.47 | 0.8 | 0.28 | 0.4 |
| 9 | 2508256 | rs4741720 | 0.557 | 1.019 | 1.08 | -1.226 | 0.015 | 0.097 | 0.097 | 0 | 0.74 | 0.53 | 0.64 | 0.59 | 0.73 | 0.71 |
| 9 | 2508296 | rs4741721 | 0.728 | 0.871 | 0.835 | -0.951 | -0.008 | -0.037 | -0.037 | 0.387 | 0.19 | 0.11 | 0.18 | 0.4 | 0.2 | 0.19 |
| 9 | 2508727 | rs11999258 | 0.176 | 0.881 | 0.712 | -0.94 | 0 | 0.193 | 0.193 | 0 | 0.06 | 0.11 | 0.23 | 0.41 | 0.05 | 0.15 |
| 9 | 2509947 | rs1970074 | 0.606 | 0.847 | 0.489 | -1.015 | 0.001 | 0.026 | 0.026 | 0 | 0.2 | 0.25 | 0.19 | 0.4 | 0.2 | 0.18 |

| | | | | | | | | | | | | | | | | |
|---|---|---|---|---|---|---|---|---|---|---|---|---|---|---|---|---|
| 9 | 2515607 | rs7042212 | -0.833 | 0.791 | 0.665 | -1.283 | 0.001 | -0.045 | -0.045 | 0.176 | 0.86 | 0.81 | 0.93 | 1 | 0.85 | 0.93 |
| 9 | 2522475 | rs10966757 | -0.79 | 0.712 | 0.728 | -1.317 | -0.015 | 0.109 | 0.109 | 0.027 | 0.27 | 0.5 | 0.41 | 0.49 | 0.22 | 0.41 |
| 9 | 2522863 | rs4740687 | -0.996 | 0.678 | 0.328 | -1.704 | 0 | 0.13 | 0.13 | 1.115 | 0.49 | 0.67 | 0.63 | 0.93 | 0.46 | 0.63 |
| 9 | 2524146 | rs7864837 | 0.238 | 0.905 | 0.758 | -1.537 | 0.03 | -0.055 | -0.055 | 0 | 0.28 | 0.37 | 0.5 | 0.85 | 0.31 | 0.34 |
| 9 | 2527980 | rs7864942 | 0.165 | 0.853 | 0.741 | -1.448 | -0.001 | 0.019 | 0.019 | 0 | 0.17 | 0.16 | 0.34 | 0.5 | 0.19 | 0.36 |
| 9 | 2530047 | rs7857522 | -0.284 | 0.857 | 0.719 | -1.931 | 0.019 | -0.072 | -0.072 | 0 | 0.76 | 0.66 | 0.76 | 0.98 | 0.67 | 0.87 |
| 9 | 2531098 | rs693007 | -0.477 | 0.723 | 0.799 | -1.991 | 0.001 | -0.061 | -0.061 | 0 | 0.43 | 0.49 | 0.58 | 0.92 | 0.41 | 0.62 |
| 9 | 2533445 | rs963197 | 1.802 | 0.559 | 0.716 | -2.064 | 0.042 | 0.086 | 0.086 | 0 | 0.29 | 0.15 | 0.17 | 0.05 | 0.24 | 0.24 |
| 9 | 2533850 | rs10491717 | 1.305 | 0.544 | 0.722 | -2.082 | 0.052 | 0.104 | 0.104 | 0 | 0.33 | 0.17 | 0.19 | 0.05 | 0.27 | 0.29 |
| 9 | 2535524 | rs655568 | -0.785 | 0.778 | 0.845 | -2.154 | 0.026 | 0.098 | 0.098 | 0 | 0.52 | 0.66 | 0.72 | 0.93 | 0.53 | 0.63 |
| 9 | 2535695 | rs656414 | -1.607 | 0.79 | 0.805 | -2.145 | 0.003 | -0.028 | -0.028 | 0.63 | 0.85 | 0.83 | 0.91 | 0.98 | 0.8 | 0.92 |
| 9 | 2535913 | rs475384 | 0.656 | 0.827 | 0.852 | -2.113 | -0.005 | -0.01 | -0.01 | 0 | 0.17 | 0.13 | 0.35 | 0.51 | 0.18 | 0.37 |
| 9 | 2535947 | rs2297352 | 1.686 | 0.838 | 1.058 | -2.117 | -0.003 | -0.014 | -0.014 | 0 | 0.22 | 0.16 | 0.19 | 0.42 | 0.21 | 0.21 |
| 9 | 2536241 | rs499532 | -0.228 | 0.829 | 0.875 | -2.059 | 0.002 | -0.002 | -0.002 | 0 | 0.58 | 0.57 | 0.68 | 0.56 | 0.56 | 0.67 |
| 9 | 2536666 | rs7864004 | -0.099 | 0.871 | 0.844 | -2.021 | 0.01 | -0.052 | -0.052 | 0 | 0.14 | 0.25 | 0.17 | 0.02 | 0.16 | 0.09 |
| 9 | 2536828 | rs10812210 | 1.848 | 0.875 | 0.665 | -1.949 | -0.031 | 0.061 | 0.061 | 0 | 0.05 | 0.01 | 0.26 | 0.42 | 0.11 | 0.13 |
| 9 | 2538201 | rs559776 | 0.335 | 0.791 | 0.787 | -2.005 | 0.014 | 0.01 | 0.01 | 0 | 0.48 | 0.38 | 0.52 | 0.56 | 0.43 | 0.62 |
| 9 | 2538431 | rs591642 | -0.669 | 0.825 | 0.577 | -2.034 | -0.002 | -0.001 | -0.001 | 1.304 | 0.28 | 0.29 | 0.5 | 0.53 | 0.32 | 0.43 |
| 9 | 2538476 | rs13299915 | 0.642 | 0.986 | 0.739 | -1.884 | 0 | -0.013 | -0.013 | 0.179 | 0.1 | 0.12 | 0.15 | 0.02 | 0.13 | 0.05 |
| 9 | 2539222 | rs2297350 | 0.673 | 0.973 | 0.738 | -1.913 | 0.013 | 0.133 | 0.133 | 0.189 | 0.38 | 0.19 | 0.31 | 0.07 | 0.37 | 0.29 |
| 9 | 2539495 | rs482376 | 0.094 | 1.197 | 0.96 | -1.8 | 0.001 | -0.032 | -0.032 | 0 | 0.75 | 0.72 | 0.76 | 0.97 | 0.67 | 0.86 |
| 9 | 2540365 | rs620949 | -0.107 | 1.228 | 0.959 | -1.7 | 0.026 | 0.075 | 0.075 | 0 | 0.75 | 0.58 | 0.6 | 0.55 | 0.68 | 0.75 |
| 9 | 2541567 | rs10966854 | 0.413 | 1.02 | 0.617 | -1.674 | 0.007 | -0.013 | -0.013 | 0 | 0.38 | 0.34 | 0.47 | 0.48 | 0.36 | 0.4 |
| 9 | 2544610 | rs1512712 | -0.267 | 1.02 | 0.627 | -1.688 | -0.001 | 0 | 0 | 3.03 | 0.48 | 0.55 | 0.64 | 0.49 | 0.51 | 0.46 |
| 9 | 2547156 | rs4741729 | -0.034 | 1.025 | 0.606 | -1.651 | -0.003 | -0.003 | -0.003 | 0 | 0.45 | 0.42 | 0.61 | 0.49 | 0.49 | 0.44 |
| 9 | 2547785 | rs4741730 | -0.17 | 1.249 | 0.584 | -1.618 | 0.038 | 0.114 | 0.114 | 0 | 0.16 | 0.28 | 0.43 | 0.44 | 0.21 | 0.2 |
| 9 | 2548026 | rs4741732 | 0.857 | 1.225 | 0.484 | -1.595 | 0.021 | 0.146 | 0.146 | 0 | 0.08 | 0.14 | 0.29 | 0.42 | 0.12 | 0.15 |
| 9 | 2549364 | rs1398433 | 0.505 | 1.209 | 0.509 | -1.572 | 0.019 | -0.029 | -0.029 | 0.086 | 0.38 | 0.28 | 0.46 | 0.48 | 0.34 | 0.41 |
| 9 | 2550811 | rs520752 | 0.762 | 1.058 | 0.432 | -1.652 | 0.006 | -0.03 | -0.03 | 0 | 0.34 | 0.2 | 0.25 | 0.49 | 0.28 | 0.49 |
| 9 | 2551526 | rs12348301 | 0.691 | 1.044 | 0.263 | -1.327 | 0.029 | -0.036 | -0.036 | 0.01 | 0.08 | 0.14 | 0.14 | 0.02 | 0.12 | 0.04 |
| 9 | 2551594 | rs10966901 | 0.197 | 0.727 | -0.131 | -1.183 | 0.01 | -0.02 | -0.02 | 0.715 | 0.39 | 0.33 | 0.47 | 0.48 | 0.36 | 0.37 |
| 9 | 2558513 | rs542859 | -0.296 | 0.102 | 0.132 | -1.867 | -0.001 | -0.023 | -0.023 | 0 | 0.65 | 0.35 | 0.69 | 0.93 | 0.71 | 0.71 |
| 9 | 2559492 | rs1590979 | 0.275 | 0.323 | 0.387 | -2.17 | 0 | -0.041 | -0.041 | 0.344 | 0.48 | 0.49 | 0.59 | 0.92 | 0.45 | 0.61 |
| 9 | 2563876 | rs1707299 | 0.388 | 0.278 | 0.427 | -2.192 | -0.004 | 0.006 | 0.006 | 0 | 0.46 | 0.39 | 0.59 | 0.92 | 0.44 | 0.6 |
| 9 | 2565685 | rs1512715 | 0.577 | 0.493 | 0.539 | -2.107 | 0.026 | -0.074 | -0.074 | 0 | 0.17 | 0.3 | 0.13 | 0.02 | 0.24 | 0.09 |
| 9 | 2568043 | rs7858086 | 0.509 | 0.291 | 0.552 | -2.113 | 0.016 | -0.033 | -0.033 | 0 | 0.1 | 0.15 | 0.16 | 0.01 | 0.14 | 0.05 |
| 9 | 2569655 | rs10738726 | 0.588 | -0.445 | -0.491 | -2.34 | -0.086 | 0.156 | 0.156 | 0 | 0.41 | 0.1 | 0.52 | 0.84 | 0.45 | 0.4 |
| 9 | 2573499 | rs7860538 | 0.291 | -0.391 | -0.176 | -2.278 | -0.003 | -0.031 | -0.031 | 0 | 0.35 | 0.32 | 0.29 | 0.07 | 0.32 | 0.3 |
| 9 | 2574919 | rs7847026 | 0.352 | 0.043 | -0.463 | -1.781 | 0.015 | -0.057 | -0.057 | 0 | 0.16 | 0.26 | 0.12 | 0.01 | 0.21 | 0.09 |
| 9 | 2580421 | rs10757565 | 0.367 | -0.11 | -0.945 | -1.397 | -0.007 | -0.009 | -0.009 | 0 | 0.55 | 0.34 | 0.41 | 0.47 | 0.59 | 0.5 |
| 9 | 2582380 | rs10967052 | 0.779 | 0.042 | -1.13 | -1.326 | 0.005 | -0.035 | -0.035 | 0 | 0.2 | 0.15 | 0.2 | 0.4 | 0.2 | 0.22 |
| 9 | 2583439 | rs4741738 | 0.562 | -0.034 | -0.786 | -1.372 | -0.008 | 0.03 | 0.03 | 0.542 | 0.21 | 0.32 | 0.21 | 0.4 | 0.2 | 0.22 |
| 9 | 2593528 | rs986567 | 0.226 | 0.219 | -0.635 | -1.528 | 0.017 | -0.045 | -0.045 | 0 | 0.12 | 0.2 | 0.16 | 0.01 | 0.15 | 0.06 |
| 9 | 2596569 | rs1454637 | 0.454 | 0.138 | -0.806 | -1.657 | -0.041 | -0.03 | -0.03 | 0.084 | 0.59 | 0.2 | 0.42 | 0.46 | 0.59 | 0.52 |
| 9 | 2596783 | rs1156847 | -0.366 | 0.114 | -0.642 | -1.599 | 0.004 | 0.051 | 0.051 | 0 | 0.71 | 0.42 | 0.59 | 0.47 | 0.73 | 0.58 |
| 9 | 2598844 | rs10812321 | 0.121 | 0.072 | -0.973 | -1.451 | -0.02 | -0.053 | -0.053 | 0.133 | 0.29 | 0.13 | 0.34 | 0.46 | 0.31 | 0.27 |
| 9 | 2599564 | rs882259 | 0.962 | 0.029 | -1.021 | -1.564 | 0.008 | 0.17 | 0.17 | 0 | 0.4 | 0.13 | 0.24 | 0.06 | 0.41 | 0.32 |
| 9 | 2600052 | rs728841 | 1.278 | 0.038 | -1.015 | -1.495 | 0 | -0.076 | -0.076 | 5.18 | 0.19 | 0.07 | 0.2 | 0.45 | 0.19 | 0.21 |
| 9 | 2600253 | rs729367 | -0.477 | 0.112 | -0.82 | -1.366 | 0.015 | 0.083 | 0.083 | 0 | 0.72 | 0.49 | 0.61 | 0.53 | 0.75 | 0.59 |
| 9 | 2602831 | rs10812325 | -0.547 | 0.069 | -1.061 | -1.448 | 0.029 | 0.131 | 0.131 | 0 | 0.76 | 0.56 | 0.57 | 0.52 | 0.77 | 0.61 |
| 9 | 2603361 | rs10812326 | 0.126 | 0.078 | -1.082 | -1.538 | -0.001 | 0.012 | 0.012 | 0 | 0.54 | 0.39 | 0.42 | 0.08 | 0.56 | 0.38 |
| 9 | 2605023 | rs12350335 | 0.849 | -0.057 | -1.656 | -1.311 | 0.013 | 0.152 | 0.152 | 0 | 0.4 | 0.16 | 0.24 | 0.06 | 0.41 | 0.32 |
| 9 | 2607430 | rs10812332 | 0.522 | -0.002 | -1.241 | -0.676 | 0.003 | -0.019 | -0.019 | 0 | 0.2 | 0.16 | 0.21 | 0.45 | 0.2 | 0.21 |
| 9 | 2614294 | rs9969780 | -0.496 | 0.048 | -1.127 | -0.686 | -0.004 | 0.01 | 0.01 | 0.274 | 0.64 | 0.61 | 0.48 | 0.47 | 0.65 | 0.58 |
| 9 | 2616042 | rs11790563 | 0.778 | -0.307 | -1.235 | -0.765 | -0.015 | 0.012 | 0.012 | 0 | 0.34 | 0.31 | 0.25 | 0.08 | 0.43 | 0.32 |
| 9 | 2617099 | rs10967186 | 1.031 | -0.299 | -1.305 | -0.78 | -0.068 | 0.085 | 0.085 | 0 | 0.32 | 0.15 | 0.22 | 0.08 | 0.43 | 0.32 |
| 9 | 2617165 | rs10967188 | -0.098 | -0.401 | -1.357 | -0.774 | -0.024 | 0.009 | 0.009 | 0.959 | 0.29 | 0.36 | 0.23 | 0.39 | 0.19 | 0.24 |
| 9 | 2618721 | rs4741745 | -0.505 | -0.016 | -1.527 | -0.501 | -0.002 | 0 | 0 | 0 | 0.59 | 0.59 | 0.46 | 0.44 | 0.6 | 0.56 |
| 9 | 2618969 | rs4741746 | -0.105 | -0.084 | -1.131 | 0.325 | 0.007 | -0.012 | -0.012 | 0 | 0.19 | 0.17 | 0.18 | 0.41 | 0.16 | 0.2 |
| 9 | 2621030 | rs1454626 | -0.617 | 0.597 | -1.141 | 0.182 | -0.005 | -0.013 | -0.013 | 0 | 0.54 | 0.52 | 0.76 | 0.78 | 0.69 | 0.6 |
| 9 | 2622134 | rs34881325 | -0.723 | 2.112 | -1.333 | 1.239 | -0.054 | -0.058 | -0.058 | 0.012 | 0.23 | 0.12 | 0.43 | 0.42 | 0.43 | 0.25 |
| 9 | 2622278 | rs2219143 | -0.632 | 2.248 | -1.402 | 1.279 | -0.073 | -0.076 | -0.076 | 0 | 0.22 | 0.1 | 0.43 | 0.4 | 0.42 | 0.23 |
| 9 | 2623615 | rs7032549 | -0.957 | 1.687 | -1.131 | 1.027 | -0.004 | -0.013 | -0.013 | 0 | 0.36 | 0.28 | 0.68 | 0.58 | 0.56 | 0.31 |
| 9 | 2624433 | rs13290598 | -1.138 | 1.957 | -0.336 | 0.765 | 0.028 | 0.108 | 0.108 | 0 | 0.14 | 0.22 | 0.36 | 0.49 | 0.21 | 0.2 |
| 9 | 2624874 | rs1545565 | -0.301 | 0.713 | -0.052 | 0.115 | -0.017 | 0.02 | 0.02 | 0 | 0.86 | 0.83 | 0.9 | 0.78 | 0.91 | 0.9 |
| 9 | 2625172 | rs10967213 | -0.443 | 0.762 | -0.368 | -0.477 | 0.004 | -0.043 | -0.043 | 2.738 | 0.33 | 0.4 | 0.28 | 0.13 | 0.38 | 0.14 |
| 9 | 2626313 | rs1545568 | -0.866 | 0.363 | -0.317 | -0.524 | 0.038 | 0.025 | 0.025 | 0.214 | 0.56 | 0.79 | 0.74 | 0.59 | 0.64 | 0.39 |
| 9 | 2626851 | rs4741748 | -0.36 | 0.417 | -0.236 | -0.468 | 0.002 | -0.002 | -0.002 | 0 | 0.45 | 0.39 | 0.65 | 0.49 | 0.57 | 0.35 |
| 9 | 2629324 | rs10967228 | 0.511 | 0.077 | 0.328 | -0.193 | -0.009 | 0.022 | 0.022 | 0 | 0.12 | 0.17 | 0.07 | 0.19 | 0.08 | 0.08 |

| | | | | | | | | | | | | | | | | |
|---|---|---|---|---|---|---|---|---|---|---|---|---|---|---|---|---|
| 9 | 2631092 | rs2375994 | -0.982 | 0.058 | 0.224 | -0.592 | 0.011 | 0.066 | 0.066 | 0.071 | 0.44 | 0.73 | 0.62 | 0.74 | 0.39 | 0.41 |
| 9 | 2631680 | rs7030282 | 0.099 | 0.015 | 0.584 | -0.446 | -0.02 | 0.046 | 0.046 | 0 | 0.14 | 0.27 | 0.08 | 0.21 | 0.09 | 0.06 |
| 9 | 2633460 | rs10812378 | 0.268 | 0.216 | 0.823 | -0.818 | -0.02 | 0.011 | 0.011 | 1.025 | 0.06 | 0.24 | 0.03 | 0.06 | 0.01 | 0.03 |
| 9 | 2633548 | rs10812379 | -0.335 | 0.449 | 0.699 | -1.234 | 0.013 | -0.12 | -0.12 | 0 | 0.31 | 0.47 | 0.23 | 0.06 | 0.35 | 0.12 |
| 9 | 2633562 | rs10812380 | 0.805 | 0.523 | 0.408 | -1.226 | -0.05 | -0.074 | -0.074 | 0.496 | 0.18 | 0.06 | 0.47 | 0.63 | 0.25 | 0.24 |
| 9 | 2636161 | rs7035616 | -0.609 | 0.526 | 0.645 | -1.296 | 0.009 | -0.126 | -0.126 | 0 | 0.35 | 0.5 | 0.31 | 0.06 | 0.38 | 0.15 |
| 9 | 2636492 | rs10812382 | -0.279 | 0.398 | 1.037 | -0.78 | 0.009 | 0.019 | 0.019 | 0.097 | 0.5 | 0.57 | 0.71 | 0.7 | 0.61 | 0.37 |
| 9 | 2637551 | rs4740698 | -0.562 | 0.411 | 0.969 | -0.72 | 0.002 | 0.002 | 0.002 | 0.048 | 0.53 | 0.57 | 0.72 | 0.69 | 0.63 | 0.39 |
| 9 | 2640759 | rs3780181 | -1.012 | 0.498 | 1.625 | -0.078 | -0.007 | 0 | 0 | 0 | 0.1 | 0.25 | 0.15 | 0.11 | 0.05 | 0.07 |
| 9 | 2640805 | rs10967306 | -0.513 | 0.508 | 0.976 | -0.112 | 0.006 | -0.007 | -0.007 | 0 | 0.11 | 0.08 | 0.06 | 0.19 | 0.08 | 0.23 |
| 9 | 2644954 | rs11789583 | -0.869 | 1.418 | 0.886 | -0.635 | -0.037 | 0.031 | 0.031 | 0 | 0.13 | 0.07 | 0.08 | 0 | 0.2 | 0.07 |
| 9 | 2647760 | rs6475888 | -1.664 | 1.4 | 1.299 | -0.705 | 0.059 | -0.09 | -0.09 | 0 | 0.18 | 0.26 | 0.13 | 0.01 | 0.24 | 0.09 |
| 9 | 2648773 | rs6148 | -0.877 | 1.416 | 1.234 | -0.71 | -0.069 | 0.067 | 0.067 | 0 | 0.13 | 0.03 | 0.08 | 0 | 0.21 | 0.07 |
| 9 | 2649217 | rs7865617 | -1.787 | 1.371 | 1.468 | -0.731 | 0.061 | -0.116 | -0.116 | 0 | 0.2 | 0.35 | 0.14 | 0.01 | 0.23 | 0.09 |
| 9 | 2651962 | rs6145 | -1.555 | 1.43 | 1.337 | -0.723 | 0.096 | -0.108 | -0.108 | 0.19 | 0.17 | 0.35 | 0.13 | 0.01 | 0.23 | 0.07 |
| 9 | 2652301 | rs17712698 | -1.219 | 1.452 | 0.987 | -0.711 | -0.042 | 0.037 | 0.037 | 0 | 0.14 | 0.06 | 0.08 | 0 | 0.21 | 0.07 |
| 9 | 2652589 | rs35075133 | -0.818 | 1.449 | 0.939 | -0.71 | 0.013 | -0.031 | -0.031 | 0 | 0.13 | 0.16 | 0.1 | 0 | 0.2 | 0.05 |
| 9 | 2654419 | rs8210 | -1.213 | 1.475 | 0.736 | -0.67 | -0.019 | 0.006 | 0.006 | 0 | 0.14 | 0.1 | 0.09 | 0 | 0.21 | 0.07 |
| 9 | 2655520 | rs7040916 | -1.699 | 1.44 | 0.526 | -0.664 | 0.089 | -0.118 | -0.118 | 1.299 | 0.18 | 0.4 | 0.13 | 0.01 | 0.23 | 0.07 |
| 9 | 2658587 | rs11790283 | -0.367 | 1.398 | 0.306 | -0.641 | -0.073 | 0.065 | 0.065 | 0 | 0.11 | 0.01 | 0.08 | 0 | 0.2 | 0.05 |
| 9 | 2661766 | rs7045124 | -1.322 | 1.323 | 0.435 | -0.612 | 0.075 | -0.124 | -0.124 | 0 | 0.2 | 0.4 | 0.14 | 0.01 | 0.23 | 0.08 |
| 9 | 2668085 | rs1470943 | -0.99 | 0.639 | -0.352 | -0.768 | 0.018 | 0.088 | 0.088 | 0.368 | 0.38 | 0.72 | 0.37 | 0.13 | 0.32 | 0.17 |
| 9 | 2668187 | rs4741756 | -0.221 | 0.815 | -0.186 | -0.054 | -0.001 | -0.003 | -0.003 | 4.49 | 0.27 | 0.29 | 0.22 | 0.31 | 0.3 | 0.34 |
| 9 | 2674110 | rs7859818 | 0.201 | 0.692 | -0.114 | 0.001 | -0.002 | 0.004 | 0.004 | 0 | 0.09 | 0.12 | 0.09 | 0.13 | 0.06 | 0.06 |
| 9 | 2679427 | rs11794349 | -0.812 | 0.597 | 0.704 | -0.119 | 0.034 | -0.028 | -0.028 | 0 | 0.1 | 0.4 | 0.16 | 0.04 | 0.1 | 0.06 |
| 9 | 2679579 | rs10125071 | -1.519 | 0.646 | 0.715 | 0.122 | 0.025 | 0.016 | 0.016 | 0 | 0.46 | 0.74 | 0.42 | 0.45 | 0.41 | 0.47 |
| 9 | 2681430 | rs7870360 | -0.968 | 0.647 | 0.629 | 0.367 | 0.003 | 0.003 | 0.003 | 0 | 0.16 | 0.25 | 0.19 | 0.2 | 0.14 | 0.12 |
| 9 | 2682322 | rs11794855 | -0.078 | 0.624 | 0.651 | 0.348 | -0.004 | -0.002 | -0.002 | 0.215 | 0.11 | 0.09 | 0.12 | 0.13 | 0.11 | 0.09 |
| 9 | 2684117 | rs10967515 | -0.928 | 0.642 | 0.496 | 0.545 | 0.01 | 0.006 | 0.006 | 0 | 0.17 | 0.25 | 0.18 | 0.19 | 0.17 | 0.16 |
| 9 | 2691186 | rs10738760 | -1.517 | 1.037 | 0.341 | 0.479 | -0.007 | 0.002 | 0.002 | 0 | 0.48 | 0.76 | 0.52 | 0.59 | 0.5 | 0.52 |
| 9 | 2691446 | rs10812474 | 0.166 | 0.585 | 0.015 | 0.417 | 0.005 | -0.019 | -0.019 | 0 | 0.22 | 0.11 | 0.15 | 0.27 | 0.22 | 0.31 |
| 9 | 2691951 | rs10122587 | -0.414 | 0.659 | -0.007 | 0.384 | -0.016 | 0.01 | 0.01 | 0 | 0.33 | 0.44 | 0.25 | 0.4 | 0.29 | 0.38 |
| 9 | 2694436 | rs748786 | -0.232 | 0.741 | -0.442 | 0.098 | -0.019 | 0.035 | 0.035 | 0 | 0.12 | 0.07 | 0.06 | 0.03 | 0.14 | 0.06 |
| 9 | 2696555 | rs930811 | -0.266 | 0.622 | -0.148 | -0.116 | 0.005 | 0.022 | 0.022 | 2.997 | 0.54 | 0.52 | 0.6 | 0.69 | 0.43 | 0.46 |
| 9 | 2699004 | rs1884441 | 0.231 | 0.151 | 0.394 | 0.028 | -0.014 | 0.166 | 0.166 | 0 | 0.17 | 0.35 | 0.43 | 0.52 | 0.16 | 0.17 |
| 9 | 2699606 | rs7034098 | -0.226 | 0.665 | 0.586 | -0.129 | 0.019 | 0.13 | 0.13 | 0 | 0.21 | 0.37 | 0.52 | 0.54 | 0.21 | 0.2 |
| 9 | 2700620 | rs1454629 | 0.254 | 0.351 | 0.374 | -0.365 | -0.006 | 0.116 | 0.116 | 0.475 | 0.38 | 0.08 | 0.2 | 0.19 | 0.33 | 0.35 |
| 9 | 2702703 | rs7855332 | -0.296 | 0.853 | 0.767 | -0.338 | -0.003 | 0.013 | 0.013 | 1.647 | 0.56 | 0.34 | 0.63 | 0.71 | 0.51 | 0.53 |
| 9 | 2704399 | rs7032456 | -0.911 | 0.679 | 0.832 | -0.274 | -0.005 | 0.008 | 0.008 | 2.175 | 0.63 | 0.62 | 0.74 | 0.76 | 0.56 | 0.56 |
| 9 | 2705874 | rs1412693 | -0.942 | 0.717 | 0.806 | -0.276 | -0.006 | 0.01 | 0.01 | 2.076 | 0.63 | 0.63 | 0.74 | 0.76 | 0.57 | 0.58 |
| 9 | 2710537 | rs897165 | -1.056 | 0.56 | 0.672 | -0.266 | -0.027 | 0.028 | 0.028 | 3.665 | 0.64 | 0.78 | 0.75 | 0.73 | 0.53 | 0.56 |
| 9 | 2716921 | rs9969833 | -0.906 | 0.752 | 1.348 | 0.125 | 0.028 | 0.061 | 0.061 | 0.03 | 0.05 | 0.32 | 0.04 | 0.02 | 0 | 0.01 |
| 9 | 2721249 | rs10812526 | -0.54 | 0.909 | 0.967 | -0.276 | 0.005 | 0.028 | 0.028 | 0.516 | 0.45 | 0.37 | 0.56 | 0.73 | 0.45 | 0.41 |
| 9 | 2725283 | rs1006698 | -1.386 | 0.814 | 1.447 | -0.302 | -0.007 | 0.102 | 0.102 | 0.326 | 0.53 | 0.76 | 0.69 | 0.76 | 0.53 | 0.43 |
| 9 | 2730988 | rs10967775 | -0.05 | 0.627 | 1.327 | -0.514 | -0.02 | 0.034 | 0.034 | 0 | 0.11 | 0.05 | 0.15 | 0.02 | 0.11 | 0.08 |
| 9 | 2731519 | rs12352612 | -1.164 | 0.752 | 1.589 | -0.401 | 0.097 | 0.009 | 0.009 | 0.122 | 0.08 | 0.39 | 0.14 | 0.02 | 0.07 | 0.04 |
| 9 | 2734038 | rs13294756 | 0.083 | 0.636 | 1.35 | -0.357 | -0.003 | 0.002 | 0.002 | 0 | 0.17 | 0.14 | 0.11 | 0.18 | 0.18 | 0.11 |
| 9 | 2735053 | rs7048820 | -1.8 | 0.579 | 1.156 | -0.306 | -0.009 | 0.125 | 0.125 | 0.174 | 0.51 | 0.76 | 0.57 | 0.72 | 0.51 | 0.39 |
| 9 | 2735415 | rs7029249 | -0.543 | 0.4 | 1.337 | -0.162 | -0.001 | 0.017 | 0.017 | 0 | 0.33 | 0.58 | 0.46 | 0.54 | 0.34 | 0.28 |
| 9 | 2737697 | rs1470945 | -0.325 | 0.349 | 1.31 | -0.123 | -0.01 | 0.017 | 0.017 | 0 | 0.31 | 0.64 | 0.38 | 0.52 | 0.27 | 0.24 |
| 9 | 2741196 | rs10738768 | 0.536 | 0.385 | 1.062 | 0.004 | -0.009 | 0.153 | 0.153 | 0 | 0.2 | 0.4 | 0.33 | 0.49 | 0.2 | 0.22 |
| 9 | 2742771 | rs2034764 | -1.618 | 0.403 | 0.741 | -0.225 | 0.005 | 0.107 | 0.107 | 0.648 | 0.49 | 0.74 | 0.63 | 0.71 | 0.49 | 0.36 |
| 9 | 2744472 | rs10967908 | -0.557 | 0.408 | 1.015 | -0.147 | -0.026 | -0.022 | -0.022 | 0 | 0.21 | 0.39 | 0.1 | 0.11 | 0.13 | 0.09 |
| 9 | 2746270 | rs1547501 | 0.516 | 0.413 | 0.56 | -0.056 | -0.002 | 0.005 | 0.005 | 0.48 | 0.8 | 0.82 | 0.8 | 0.86 | 0.82 | 0.88 |
| 9 | 2746548 | rs10812577 | 0.021 | 0.319 | 0.468 | -0.003 | 0.001 | -0.025 | -0.025 | 0.6 | 0.32 | 0.27 | 0.43 | 0.5 | 0.37 | 0.29 |
| 9 | 2746594 | rs7852307 | 0.226 | 0.243 | 0.451 | -0.066 | -0.001 | 0.008 | 0.008 | 0.414 | 0.8 | 0.81 | 0.8 | 0.91 | 0.82 | 0.88 |
| 9 | 2751726 | rs1378062 | 0.849 | 0.09 | 0.204 | -0.21 | -0.001 | 0.014 | 0.014 | 0 | 0.49 | 0.58 | 0.43 | 0.41 | 0.45 | 0.6 |
| 9 | 2753505 | rs2375957 | 0.992 | 0.341 | -0.907 | -0.271 | -0.039 | 0.061 | 0.061 | 0.017 | 0.46 | 0.08 | 0.38 | 0.36 | 0.47 | 0.59 |
| 9 | 2756756 | rs7857841 | 0.541 | -0.109 | -0.374 | -0.335 | -0.01 | -0.035 | -0.035 | 0 | 0.84 | 0.79 | 0.92 | 0.97 | 0.89 | 0.92 |
| 9 | 2759058 | rs7854358 | 0.339 | -0.241 | -1.018 | -0.182 | 0.056 | 0.082 | 0.082 | 0 | 0.55 | 0.28 | 0.37 | 0.25 | 0.66 | 0.68 |
| 9 | 2762721 | rs958505 | 0.167 | -0.146 | -0.73 | 0.359 | -0.005 | -0.117 | -0.117 | 1.815 | 0.8 | 0.97 | 0.73 | 0.57 | 0.81 | 0.79 |
| 9 | 2765753 | rs2168134 | 0.633 | -0.16 | -0.167 | -0.031 | 0.007 | -0.01 | -0.01 | 0.062 | 0.64 | 0.64 | 0.63 | 0.53 | 0.65 | 0.7 |
| 9 | 2783788 | rs1060812 | 0.574 | -0.131 | -0.188 | -0.038 | 0.009 | -0.023 | -0.023 | 0.034 | 0.64 | 0.76 | 0.63 | 0.54 | 0.61 | 0.7 |
| 9 | 2783928 | rs3739459 | -0.15 | 0.228 | 0.135 | -0.48 | 0.007 | -0.056 | -0.056 | 0 | 0.14 | 0.23 | 0.1 | 0.03 | 0.13 | 0.19 |
| 9 | 2787856 | rs10757686 | 1.466 | 0.059 | 0.002 | -0.518 | -0.029 | -0.026 | -0.026 | 0 | 0.44 | 0.21 | 0.44 | 0.45 | 0.48 | 0.47 |
| 9 | 2790307 | rs1454635 | 1.06 | 0.298 | 0.563 | 0.069 | -0.005 | 0.003 | 0.003 | 0 | 0.58 | 0.41 | 0.53 | 0.48 | 0.59 | 0.64 |
| 9 | 2795953 | rs6476022 | 0.647 | 0.453 | -0.18 | 0.135 | 0.013 | -0.037 | -0.037 | 0.707 | 0.64 | 0.75 | 0.57 | 0.51 | 0.62 | 0.71 |
| 9 | 2800854 | rs870652 | 0.371 | 0.616 | -0.323 | 0.312 | -0.015 | -0.095 | -0.095 | 3.755 | 0.73 | 0.92 | 0.63 | 0.55 | 0.7 | 0.71 |
| 9 | 2801858 | rs7032686 | 0.448 | 0.886 | 0.415 | 0.552 | -0.05 | -0.041 | -0.041 | 0 | 0.67 | 0.77 | 0.4 | 0.51 | 0.52 | 0.64 |

| | | | | | | | | | | | | | | | | |
|---|---|---|---|---|---|---|---|---|---|---|---|---|---|---|---|---|
| 9 | 2804393 | rs12171 | 0.162 | 0.966 | 1.115 | 0.449 | 0.023 | 0.008 | 0.008 | 0.109 | 0.06 | 0.21 | 0.09 | 0.05 | 0.04 | 0.1 |
| 9 | 2809548 | rs7850317 | -0.261 | 1.057 | 0.759 | 0.483 | -0.043 | -0.027 | -0.027 | 0 | 0.73 | 0.83 | 0.59 | 0.61 | 0.57 | 0.71 |
| 9 | 2817887 | rs10812738 | -1.305 | 0.814 | 1.093 | 0.322 | -0.006 | -0.023 | -0.023 | 0 | 0.13 | 0.51 | 0.13 | 0.08 | 0.09 | 0.12 |
| 9 | 2825965 | rs10968361 | 0.566 | 0.831 | 0.764 | 0.46 | -0.014 | -0.012 | -0.012 | 8.375 | 0.6 | 0.37 | 0.41 | 0.52 | 0.49 | 0.59 |
| 9 | 2827752 | rs10757715 | 0.984 | 0.676 | 0.64 | 0.435 | -0.053 | -0.038 | -0.038 | 0 | 0.53 | 0.15 | 0.38 | 0.45 | 0.49 | 0.55 |
| 9 | 2831891 | rs10125846 | 0.341 | 1.322 | 1.015 | 0.562 | 0.04 | 0.022 | 0.022 | 0.001 | 0.66 | 0.56 | 0.48 | 0.56 | 0.53 | 0.63 |
| 9 | 2833267 | rs10217194 | 0.239 | 1.15 | 0.953 | 0.616 | 0.025 | 0.012 | 0.012 | 0 | 0.64 | 0.58 | 0.47 | 0.55 | 0.51 | 0.62 |
| 9 | 2840829 | rs10116824 | 0.678 | 1.176 | 0.693 | 0.55 | -0.027 | -0.025 | -0.025 | 0.006 | 0.59 | 0.3 | 0.39 | 0.5 | 0.47 | 0.59 |
| 9 | 2847195 | rs4741769 | 0.754 | 1.038 | -0.119 | 0.59 | -0.013 | -0.024 | -0.024 | 0.271 | 0.54 | 0.26 | 0.3 | 0.44 | 0.48 | 0.55 |
| 9 | 2851026 | rs12001239 | -1.177 | 0.709 | 0.139 | 0.677 | 0.031 | 0.027 | 0.027 | 0.476 | 0.08 | 0.31 | 0.1 | 0.05 | 0.07 | 0.03 |
| 9 | 2852747 | rs7026256 | 0.388 | 1.229 | -1.176 | 0.803 | -0.082 | -0.079 | -0.079 | 20.692 | 0.66 | 0.85 | 0.45 | 0.53 | 0.56 | 0.65 |
| 9 | 2864428 | rs10968651 | -0.571 | 1.042 | -0.536 | 0.76 | 0.004 | 0.01 | 0.01 | 0 | 0.38 | 0.51 | 0.29 | 0.4 | 0.35 | 0.42 |
| 9 | 2864819 | rs10968653 | -0.776 | 0.857 | -0.327 | 0.607 | -0.001 | -0.001 | -0.001 | 0.15 | 0.42 | 0.57 | 0.31 | 0.39 | 0.38 | 0.42 |
| 9 | 2871439 | rs7848307 | -0.724 | 0.752 | -0.105 | 0.684 | 0.008 | -0.002 | -0.002 | 0.512 | 0.4 | 0.45 | 0.29 | 0.37 | 0.38 | 0.42 |
| 9 | 2890451 | rs7037565 | -0.515 | 0.648 | -0.154 | 0.583 | -0.064 | -0.052 | -0.052 | 0 | 0.33 | 0.11 | 0.25 | 0.37 | 0.35 | 0.35 |
| 9 | 2900843 | rs13297517 | -0.517 | 0.502 | -0.009 | 0.671 | -0.124 | -0.147 | -0.147 | 0 | 0.33 | 0.04 | 0.31 | 0.44 | 0.34 | 0.33 |
| 9 | 2906368 | rs12346338 | -1.769 | -0.202 | 0.848 | 0.285 | -0.06 | -0.062 | -0.062 | 0.407 | 0.11 | 0.69 | 0.06 | 0.01 | 0.01 | 0.05 |
| 9 | 2909883 | rs7021223 | -2.022 | -0.145 | 1.204 | 0.282 | -0.043 | -0.038 | -0.038 | 0 | 0.13 | 0.62 | 0.07 | 0.05 | 0.05 | 0.14 |
| 9 | 2913620 | rs7865808 | -2.169 | -0.175 | 0.852 | 0.281 | 0.012 | 0.012 | 0.012 | 4.983 | 0.15 | 0.84 | 0.1 | 0.05 | 0.05 | 0.09 |
| 9 | 2916741 | rs10117905 | -0.846 | 0.388 | 0.423 | 0.078 | 0.009 | -0.007 | -0.007 | 4.08 | 0.4 | 0.55 | 0.46 | 0.33 | 0.44 | 0.46 |
| 9 | 2918656 | rs1501576 | -1.252 | 0.274 | 0.412 | -0.576 | -0.018 | -0.072 | -0.072 | 0 | 0.31 | 0.54 | 0.18 | 0.1 | 0.27 | 0.29 |
| 9 | 2929424 | rs10969013 | -1.817 | -0.079 | 0.165 | -0.965 | 0.037 | 0.037 | 0.037 | 0 | 0.42 | 0.71 | 0.22 | 0.04 | 0.28 | 0.37 |
| 9 | 2931315 | rs10121692 | -1.37 | 0.165 | 0.705 | -0.906 | -0.028 | -0.135 | -0.135 | 0 | 0.29 | 0.48 | 0.17 | 0.04 | 0.23 | 0.29 |
| 9 | 2931341 | rs13292810 | -1.055 | 0.137 | 0.702 | -0.921 | -0.02 | -0.134 | -0.134 | 0 | 0.27 | 0.46 | 0.16 | 0.03 | 0.22 | 0.24 |
| 9 | 2932710 | rs13285246 | -1.265 | 0.936 | 0.396 | -1.006 | 0 | -0.027 | -0.027 | 0 | 0.29 | 0.33 | 0.23 | 0.04 | 0.34 | 0.34 |
| 9 | 2933001 | rs7875771 | -1.464 | 1.343 | 0.598 | -1.155 | -0.003 | -0.095 | -0.095 | 0.105 | 0.29 | 0.41 | 0.16 | 0.03 | 0.27 | 0.29 |
| 9 | 2933171 | rs10813032 | -0.953 | 0.695 | 0.427 | -1.279 | -0.013 | -0.084 | -0.084 | 2.622 | 0.24 | 0.35 | 0.14 | 0.04 | 0.19 | 0.16 |
| 9 | 2933816 | rs10114469 | -0.656 | -0.012 | -0.044 | -0.648 | 0.002 | -0.066 | -0.066 | 0.038 | 0.75 | 0.89 | 0.78 | 0.63 | 0.72 | 0.73 |
| 9 | 2933896 | rs10123521 | 0.598 | 0.424 | 0.292 | -0.988 | -0.001 | 0.003 | 0.003 | 0 | 0.5 | 0.52 | 0.48 | 0.49 | 0.53 | 0.54 |
| 9 | 2934146 | rs10969031 | -0.296 | -0.004 | 0.468 | -0.57 | 0.003 | 0.015 | 0.015 | 0 | 0.69 | 0.68 | 0.57 | 0.57 | 0.64 | 0.6 |
| 9 | 2936864 | rs10156534 | 0.128 | 0.564 | 0.839 | -0.608 | 0.021 | 0.014 | 0.014 | 0 | 0.24 | 0.3 | 0.34 | 0.47 | 0.28 | 0.24 |
| 9 | 2937654 | rs3847230 | -0.023 | 0.451 | 0.784 | -0.65 | 0.009 | 0.007 | 0.007 | 0 | 0.27 | 0.3 | 0.36 | 0.49 | 0.29 | 0.25 |
| 9 | 2942047 | rs10757834 | 0.347 | 0.185 | 0.599 | -0.687 | -0.004 | -0.035 | -0.035 | 0 | 0.3 | 0.14 | 0.39 | 0.46 | 0.32 | 0.39 |
| 9 | 2949336 | rs10813072 | 0.153 | 0.214 | 0.851 | -0.769 | 0.002 | -0.005 | -0.005 | 0 | 0.32 | 0.28 | 0.4 | 0.49 | 0.34 | 0.41 |
| 9 | 2949580 | rs17799828 | 0.116 | 0.148 | 1.288 | -0.916 | 0.003 | 0 | 0 | 0 | 0.3 | 0.3 | 0.36 | 0.48 | 0.31 | 0.38 |
| 9 | 2951475 | rs3913466 | -0.099 | -0.122 | 1.619 | -1.058 | -0.011 | -0.008 | -0.008 | 0 | 0.36 | 0.57 | 0.42 | 0.49 | 0.34 | 0.41 |
| 9 | 2951623 | rs10813084 | 0.905 | -0.123 | 1.527 | -1.061 | -0.006 | -0.013 | -0.013 | 0 | 0.27 | 0.23 | 0.4 | 0.46 | 0.33 | 0.38 |
| 9 | 2954622 | rs3913468 | 0.104 | -0.102 | 1.572 | -1.239 | 0.004 | 0.006 | 0.006 | 0 | 0.33 | 0.3 | 0.41 | 0.51 | 0.34 | 0.41 |
| 9 | 2958182 | rs7846783 | -1.322 | -0.03 | 0.862 | -1.61 | 0.041 | 0.18 | 0.18 | 0 | 0.65 | 0.92 | 0.8 | 0.97 | 0.7 | 0.79 |
| 9 | 2963074 | rs10969226 | -1.899 | -0.256 | 1.27 | -1.529 | 0.017 | 0.006 | 0.006 | 1.572 | 0.08 | 0.48 | 0.04 | 0 | 0.03 | 0.01 |
| 9 | 2963183 | rs10969227 | -1.134 | -0.243 | 1.261 | -1.522 | 0.042 | 0.034 | 0.034 | 0 | 0.05 | 0.34 | 0.03 | 0 | 0.03 | 0.01 |
| 9 | 2964750 | rs3858034 | -1.105 | -0.182 | 1.066 | -1.621 | -0.019 | -0.121 | -0.121 | 0.167 | 0.62 | 0.53 | 0.76 | 0.97 | 0.66 | 0.79 |
| 9 | 2964981 | rs10969237 | 0.447 | 0.015 | 0.942 | -1.361 | 0.007 | -0.03 | -0.03 | 0.12 | 0.31 | 0.2 | 0.42 | 0.43 | 0.33 | 0.42 |
| 9 | 2968044 | rs3858035 | -1.104 | 0.028 | 0.736 | -1.869 | 0.027 | 0.182 | 0.182 | 0 | 0.65 | 0.89 | 0.79 | 0.97 | 0.68 | 0.8 |
| 9 | 2968451 | rs1393037 | -0.908 | 0.03 | 0.882 | -1.868 | 0.025 | 0.052 | 0.052 | 0 | 0.63 | 0.75 | 0.78 | 0.97 | 0.66 | 0.79 |
| 9 | 2982931 | rs7044150 | -0.845 | 0.722 | -0.573 | -1.624 | -0.056 | 0.233 | 0.233 | 0 | 0.65 | 0.95 | 0.78 | 0.97 | 0.63 | 0.81 |
| 9 | 2985743 | rs1393040 | -1.369 | 0.722 | -0.627 | -1.64 | -0.046 | 0.199 | 0.199 | 0 | 0.7 | 0.98 | 0.82 | 0.97 | 0.69 | 0.83 |
| 9 | 2987835 | rs3847233 | -1.018 | 0.772 | -0.36 | -1.573 | 0.012 | 0.141 | 0.141 | 0 | 0.63 | 0.83 | 0.78 | 0.93 | 0.64 | 0.79 |
| 9 | 2989907 | rs4741798 | -0.642 | 0.839 | -0.343 | -1.579 | 0 | 0.15 | 0.15 | 0 | 0.58 | 0.76 | 0.75 | 0.93 | 0.58 | 0.77 |
| 9 | 2994084 | rs7041442 | -0.465 | 1.474 | 0.485 | -0.717 | 0.005 | 0.002 | 0.002 | 0.398 | 0.22 | 0.19 | 0.31 | 0.43 | 0.33 | 0.31 |
| 9 | 3010878 | rs7040143 | 0.515 | 0.893 | 0.002 | -1.902 | 0.014 | 0.088 | 0.088 | 0 | 0.49 | 0.32 | 0.51 | 0 | 0.54 | 0.25 |
| 9 | 3011890 | rs7855587 | 0.258 | 0.752 | 0.428 | -1.932 | -0.026 | 0.051 | 0.051 | 0.534 | 0.62 | 0.7 | 0.57 | 0.1 | 0.52 | 0.32 |
| 9 | 3012372 | rs10123800 | -0.75 | 0.789 | 0.683 | -2.006 | 0.08 | 0.033 | 0.033 | 0 | 0.09 | 0.38 | 0.12 | 0.09 | 0.11 | 0.11 |
| 9 | 3015465 | rs10813222 | 0.645 | 0.627 | 0.609 | -1.967 | 0.006 | -0.079 | -0.079 | 0.26 | 0.57 | 0.45 | 0.55 | 0.1 | 0.52 | 0.33 |
| 9 | 3022592 | rs4741801 | 0.3 | 0.788 | 0.343 | -1.719 | -0.017 | 0.03 | 0.03 | 0.254 | 0.63 | 0.66 | 0.57 | 0.12 | 0.54 | 0.37 |
| 9 | 3030814 | rs7033621 | 1.098 | 0.515 | 0.693 | -1.937 | -0.004 | -0.118 | -0.118 | 1.094 | 0.42 | 0.49 | 0.41 | 0.01 | 0.38 | 0.25 |
| 9 | 3034077 | rs2376770 | 0.863 | 0.251 | 1.065 | -1.284 | -0.021 | 0.006 | 0.006 | 0 | 0.76 | 0.39 | 0.69 | 0.67 | 0.82 | 0.81 |
| 9 | 3037903 | rs7860862 | 0.27 | 0.233 | 1.319 | -1.234 | 0.001 | 0.013 | 0.013 | 0 | 0.81 | 0.87 | 0.82 | 0.84 | 0.82 | 0.85 |
| 9 | 3044447 | rs6476195 | -2.096 | 0.198 | 1.407 | -1.187 | 0.048 | 0.247 | 0.247 | 4.386 | 0.06 | 0.54 | 0.14 | 0.15 | 0.01 | 0.01 |
| 9 | 3044503 | rs6476196 | 0.559 | 0.253 | 1.032 | -1.289 | -0.019 | 0.052 | 0.052 | 0 | 0.77 | 0.96 | 0.77 | 0.81 | 0.78 | 0.76 |
| 9 | 3049080 | rs3904463 | -2.2 | 0.309 | 0.978 | -1.423 | -0.009 | 0.12 | 0.12 | 0 | 0.62 | 0.85 | 0.65 | 0.98 | 0.62 | 0.76 |
| 9 | 3050066 | rs13298031 | -0.572 | 0.373 | 1.115 | -1.315 | -0.015 | 0.043 | 0.043 | 0 | 0.34 | 0.28 | 0.37 | 0.75 | 0.4 | 0.52 |
| 9 | 3050940 | rs2327695 | -2.254 | 0.34 | 0.808 | -1.368 | -0.027 | 0.179 | 0.179 | 0 | 0.63 | 0.89 | 0.64 | 0.95 | 0.6 | 0.76 |
| 9 | 3061937 | rs4404971 | 0.849 | 0.379 | 1.576 | -0.969 | -0.019 | -0.06 | -0.06 | 0.127 | 0.73 | 0.55 | 0.73 | 0.81 | 0.78 | 0.73 |
| 9 | 3064335 | rs10757975 | 1.21 | 0.357 | 1.519 | -0.856 | -0.012 | -0.003 | -0.003 | 0 | 0.7 | 0.4 | 0.62 | 0.66 | 0.75 | 0.72 |
| 9 | 3066302 | rs2327719 | 0.798 | 0.508 | 0.857 | -0.832 | 0 | -0.004 | -0.004 | 0 | 0.72 | 0.79 | 0.66 | 0.66 | 0.75 | 0.75 |
| 9 | 3070489 | rs3904474 | 0.499 | 0.93 | 0.49 | -0.278 | -0.053 | -0.058 | -0.058 | 0 | 0.11 | 0.02 | 0.17 | 0.52 | 0.22 | 0.31 |
| 9 | 3073208 | rs3904473 | -2.189 | 0.319 | 1.711 | -0.465 | 0.059 | 0.259 | 0.259 | 4.875 | 0.06 | 0.5 | 0.11 | 0.15 | 0 | 0.01 |
| 9 | 3077843 | rs10511442 | -0.132 | 0.127 | 1.54 | -1.045 | 0.005 | 0.01 | 0.01 | 0 | 0.18 | 0.13 | 0.1 | 0.09 | 0.13 | 0.17 |

| | | | | | | | | | | | | | | | | |
|---|---|---|---|---|---|---|---|---|---|---|---|---|---|---|---|---|
| 9 | 3080838 | rs3899009 | 1.298 | 0.204 | 2.07 | -1.075 | 0.065 | -0.029 | -0.029 | 0 | 0.92 | 0.66 | 0.93 | 1 | 0.93 | 0.93 |
| 9 | 3080970 | rs11789022 | 1.994 | 0.38 | 1.193 | -0.961 | 0.018 | 0.159 | 0.159 | 0 | 0.38 | 0.11 | 0.38 | 0.12 | 0.36 | 0.22 |
| 9 | 3081886 | rs3847257 | 1.922 | 0.155 | 2.005 | -0.38 | 0.036 | -0.062 | -0.062 | 0 | 0.89 | 0.45 | 0.91 | 1 | 0.92 | 0.89 |
| 9 | 3088774 | rs10813363 | 1.481 | 0.073 | 1.426 | -0.18 | -0.006 | 0.018 | 0.018 | 0 | 0.7 | 0.37 | 0.68 | 0.6 | 0.76 | 0.68 |
| 9 | 3096420 | rs17727514 | -0.168 | 0.345 | 1.36 | -0.015 | 0.015 | 0.041 | 0.041 | 0 | 0.19 | 0.2 | 0.31 | 0.54 | 0.3 | 0.39 |
| 9 | 3096561 | rs12377888 | -0.099 | 0.092 | 1.944 | -0.237 | -0.018 | -0.047 | -0.047 | 0 | 0.11 | 0.2 | 0.05 | 0.02 | 0.08 | 0.08 |
| 9 | 3097453 | rs7021672 | 1.161 | -0.081 | 2.388 | -0.288 | -0.001 | -0.001 | -0.001 | 2.145 | 0.75 | 0.71 | 0.75 | 0.73 | 0.77 | 0.72 |
| 9 | 3099009 | rs4741806 | -0.449 | -0.006 | 2.36 | -0.205 | 0.002 | 0.047 | 0.047 | 0 | 0.09 | 0.12 | 0.11 | 0.17 | 0.09 | 0.14 |
| 9 | 3101606 | rs17800615 | 0.153 | 0.01 | 2.362 | -0.234 | 0.002 | 0.015 | 0.015 | 0 | 0.19 | 0.18 | 0.31 | 0.54 | 0.3 | 0.38 |
| 9 | 3101626 | rs9696699 | 0.592 | -0.315 | 2.101 | -0.346 | -0.003 | 0.006 | 0.006 | 0 | 0.85 | 0.84 | 0.86 | 0.9 | 0.85 | 0.85 |
| 9 | 3121992 | rs2078432 | -0.078 | -0.197 | 2.309 | -0.193 | 0.018 | 0.019 | 0.019 | 0.099 | 0.26 | 0.33 | 0.34 | 0.68 | 0.34 | 0.43 |
| 9 | 3126263 | rs626533 | 1.104 | -0.206 | 2.366 | -0.199 | -0.024 | 0.036 | 0.036 | 0.464 | 0.82 | 0.71 | 0.85 | 0.81 | 0.87 | 0.79 |
| 9 | 3128374 | rs506826 | -0.329 | -0.188 | 1.849 | -0.22 | -0.02 | 0.007 | 0.007 | 0 | 0.1 | 0.05 | 0.13 | 0.1 | 0.14 | 0.12 |
| 9 | 3130428 | rs657877 | 0.82 | -0.339 | 1.995 | -0.197 | -0.01 | 0.028 | 0.028 | 0.451 | 0.89 | 0.83 | 0.88 | 0.87 | 0.94 | 0.84 |
| 9 | 3131352 | rs1571974 | 1.581 | -0.287 | 2.042 | -0.176 | -0.001 | -0.001 | -0.001 | 0 | 0.71 | 0.69 | 0.72 | 0.75 | 0.74 | 0.69 |
| 9 | 3136949 | rs928449 | 1.976 | -0.533 | 2.111 | -0.159 | 0.032 | 0.094 | 0.094 | 0 | 0.46 | 0.35 | 0.38 | 0.07 | 0.4 | 0.25 |
| 9 | 3137117 | rs662562 | -0.015 | -0.415 | 2.1 | -0.081 | 0.047 | 0.032 | 0.032 | 0.105 | 0.26 | 0.34 | 0.34 | 0.68 | 0.37 | 0.43 |
| 9 | 3140096 | rs494408 | 1.09 | -0.466 | 2 | 0.001 | -0.009 | 0.009 | 0.009 | 0.243 | 0.81 | 0.78 | 0.84 | 0.79 | 0.87 | 0.79 |
| 9 | 3151278 | rs1983765 | -0.036 | -0.188 | 2.028 | 0.056 | 0.034 | 0.027 | 0.027 | 0.318 | 0.25 | 0.31 | 0.31 | 0.67 | 0.36 | 0.42 |
| 9 | 3153178 | rs10813578 | -1.415 | -0.323 | 1.877 | -0.384 | -0.013 | -0.024 | -0.024 | 0.433 | 0.2 | 0.41 | 0.18 | 0.17 | 0.17 | 0.2 |
| 9 | 3154726 | rs12380711 | -0.463 | -0.244 | 1.415 | -0.262 | -0.002 | 0.003 | 0.003 | 0 | 0.09 | 0.09 | 0.12 | 0.09 | 0.1 | 0.11 |
| 9 | 3155802 | rs10970366 | 0.835 | -0.246 | 1.331 | -0.259 | 0.013 | -0.003 | -0.003 | 0 | 0.08 | 0.02 | 0.03 | 0.07 | 0.07 | 0.09 |
| 9 | 3160003 | rs10970390 | -0.435 | 0.017 | 1.554 | -0.161 | -0.009 | 0.007 | 0.007 | 0 | 0.09 | 0.05 | 0.11 | 0.09 | 0.1 | 0.11 |
| 9 | 3168100 | rs6476300 | -1.51 | -0.111 | 1.739 | -0.282 | 0.012 | 0.02 | 0.02 | 0 | 0.51 | 0.54 | 0.59 | 0.84 | 0.6 | 0.74 |
| 9 | 3169779 | rs10813625 | -0.891 | 0.159 | 1.476 | -0.216 | -0.023 | 0.019 | 0.019 | 0 | 0.13 | 0.05 | 0.13 | 0.11 | 0.15 | 0.16 |
| 9 | 3169820 | rs1322228 | -0.437 | 0.216 | 1.685 | -0.054 | 0.008 | -0.005 | -0.005 | 0 | 0.35 | 0.32 | 0.42 | 0.7 | 0.44 | 0.56 |
| 9 | 3176796 | rs10813628 | 0.631 | -0.153 | 0.672 | -0.372 | -0.001 | 0.023 | 0.023 | 0 | 0.54 | 0.62 | 0.55 | 0.15 | 0.51 | 0.4 |
| 9 | 3181278 | rs7862467 | 0.655 | -0.147 | 0.541 | -0.319 | -0.001 | -0.019 | -0.019 | 0.646 | 0.55 | 0.57 | 0.55 | 0.18 | 0.51 | 0.41 |
| 9 | 3182850 | rs7036970 | -0.888 | -0.041 | 0.256 | -0.289 | -0.001 | 0.009 | 0.009 | 0 | 0.1 | 0.07 | 0.11 | 0.09 | 0.11 | 0.15 |
| 9 | 3183137 | rs4740717 | -0.145 | -0.012 | 0.104 | -0.099 | 0.01 | -0.01 | -0.01 | 0.47 | 0.67 | 0.64 | 0.65 | 0.26 | 0.59 | 0.55 |
| 9 | 3184864 | rs10970516 | -1.205 | -0.356 | -0.084 | 0.565 | -0.003 | 0.022 | 0.022 | 0 | 0.1 | 0.02 | 0.08 | 0.08 | 0.08 | 0.13 |
| 9 | 3184967 | rs7875874 | 1.217 | -0.373 | 0.174 | 0.685 | -0.046 | 0.001 | 0.001 | 0.506 | 0.73 | 0.55 | 0.85 | 0.72 | 0.85 | 0.72 |
| 9 | 3189170 | rs10970529 | -1.212 | 0.293 | -0.646 | 0.726 | -0.032 | 0.006 | 0.006 | 0 | 0.12 | 0.02 | 0.14 | 0.09 | 0.17 | 0.07 |
| 9 | 3192508 | rs10970548 | -1.219 | -0.017 | -0.62 | 1.351 | -0.003 | -0.056 | -0.056 | 0 | 0.13 | 0.02 | 0.17 | 0.28 | 0.13 | 0.06 |
| 9 | 3192845 | rs12342853 | -1.431 | -0.072 | 0.595 | 1.307 | 0 | 0.101 | 0.101 | 0 | 0.14 | 0.26 | 0.19 | 0.29 | 0.13 | 0.06 |
| 9 | 3192985 | rs17801047 | -2.074 | -0.395 | 0.943 | 1.606 | -0.02 | 0.107 | 0.107 | 0 | 0.23 | 0.39 | 0.32 | 0.49 | 0.22 | 0.16 |
| 9 | 3193321 | rs1322231 | 1.731 | -1.099 | 1.235 | 1.253 | -0.035 | 0.002 | 0.002 | 2.217 | 0.78 | 0.6 | 0.79 | 0.73 | 0.87 | 0.71 |
| 9 | 3193862 | rs2093451 | -1.474 | -0.945 | 1.409 | 1.233 | 0.008 | 0.043 | 0.043 | 0.06 | 0.09 | 0.28 | 0.14 | 0.17 | 0.09 | 0.1 |
| 9 | 3194613 | rs12236127 | -1.729 | -1.368 | 1.596 | 1.513 | -0.014 | 0.06 | 0.06 | 0 | 0.19 | 0.22 | 0.23 | 0.43 | 0.15 | 0.17 |
| 9 | 3197716 | rs4741815 | -2.269 | -1.896 | 2.06 | 1.38 | -0.013 | 0.004 | 0.004 | 0 | 0.37 | 0.6 | 0.34 | 0.53 | 0.22 | 0.33 |
| 9 | 3202370 | rs7873419 | -2.115 | -1.963 | 2.28 | 1.369 | -0.037 | 0.012 | 0.012 | 0 | 0.36 | 0.48 | 0.32 | 0.55 | 0.22 | 0.33 |
| 9 | 3204080 | rs1541125 | -2.316 | -2.03 | 2.384 | 1.343 | 0.022 | -0.007 | -0.007 | 0 | 0.37 | 0.71 | 0.34 | 0.55 | 0.22 | 0.33 |
| 9 | 3204115 | rs10813672 | -1.6 | -2.055 | 2.356 | 1.37 | -0.02 | 0.038 | 0.038 | 0 | 0.26 | 0.31 | 0.26 | 0.48 | 0.17 | 0.24 |
| 9 | 3207669 | rs7868169 | -1.653 | -2.143 | 2.297 | 1.501 | -0.051 | 0.088 | 0.088 | 0 | 0.26 | 0.4 | 0.27 | 0.44 | 0.16 | 0.25 |
| 9 | 3208361 | rs10116401 | -2.199 | -2.255 | 2.182 | 1.406 | -0.052 | 0.05 | 0.05 | 0 | 0.32 | 0.54 | 0.22 | 0.48 | 0.17 | 0.3 |
| 9 | 3208472 | rs580157 | 1.027 | -2.006 | 1.195 | 0.513 | 0.015 | 0.01 | 0.01 | 0 | 0.48 | 0.26 | 0.54 | 0.43 | 0.6 | 0.44 |
| 9 | 3208516 | rs580944 | 1.024 | -1.917 | 1.217 | 0.335 | -0.041 | -0.045 | -0.045 | 0 | 0.85 | 0.47 | 0.85 | 0.94 | 0.91 | 0.89 |
| 9 | 3212494 | rs2104168 | -1.392 | -1.64 | 0.659 | 0.37 | -0.025 | -0.02 | -0.02 | 0 | 0.22 | 0.66 | 0.21 | 0.15 | 0.16 | 0.14 |
| 9 | 3220602 | rs17658489 | 0.554 | -1.335 | -0.479 | 0.055 | -0.04 | 0.1 | 0.1 | 0 | 0.24 | 0.08 | 0.14 | 0.08 | 0.24 | 0.32 |
| 9 | 3221674 | rs1281813 | 1.618 | -1.634 | -0.226 | 0.539 | 0.064 | -0.06 | -0.06 | 0 | 0.6 | 0.15 | 0.69 | 0.47 | 0.74 | 0.7 |
| 9 | 3223166 | rs10813760 | -0.444 | -1.678 | -0.076 | 0.883 | 0.005 | -0.054 | -0.054 | 0 | 0.17 | 0.1 | 0.11 | 0.43 | 0.12 | 0.17 |
| 9 | 3223441 | rs16915985 | -0.426 | -1.872 | 0.23 | 0.536 | 0.003 | 0.001 | 0.001 | 0 | 0.11 | 0.09 | 0.08 | 0.02 | 0.08 | 0.1 |
| 9 | 3225765 | rs687511 | 1.597 | -2.054 | 1.077 | 0.448 | -0.054 | -0.078 | -0.078 | 0 | 0.83 | 0.41 | 0.86 | 0.98 | 0.92 | 0.92 |
| 9 | 3226462 | rs684434 | 1.252 | -2.057 | 1.676 | 0.447 | -0.046 | -0.077 | -0.077 | 0.006 | 0.85 | 0.62 | 0.88 | 0.98 | 0.92 | 0.91 |
| 9 | 3228700 | rs643429 | 2.218 | -2.229 | 1.236 | 0.641 | 0.085 | -0.123 | -0.123 | 0 | 0.61 | 0.11 | 0.71 | 0.54 | 0.75 | 0.75 |
| 9 | 3236394 | rs620745 | 1.735 | -1.968 | 1.771 | 0.392 | -0.065 | -0.091 | -0.091 | 0 | 0.82 | 0.41 | 0.85 | 0.98 | 0.9 | 0.91 |
| 9 | 3237890 | rs832307 | 1.841 | -1.967 | 1.778 | 0.39 | -0.034 | -0.089 | -0.089 | 0 | 0.81 | 0.4 | 0.84 | 0.98 | 0.87 | 0.91 |
| 9 | 3239762 | rs7856125 | -1.216 | -1.911 | 1.618 | 0.424 | -0.01 | -0.037 | -0.037 | 0.526 | 0.14 | 0.2 | 0.12 | 0.02 | 0.12 | 0.09 |
| 9 | 3242834 | rs515065 | 1.366 | -2.041 | 0.623 | 0.102 | 0.053 | 0.194 | 0.194 | 0.183 | 0.5 | 0.19 | 0.56 | 0.77 | 0.6 | 0.58 |
| 9 | 3246491 | rs535614 | 1.05 | -1.869 | 1.444 | 0.334 | -0.039 | -0.022 | -0.022 | 0 | 0.8 | 0.45 | 0.82 | 0.87 | 0.86 | 0.88 |
| 9 | 3248543 | rs636548 | 1.328 | -1.9 | 1.417 | 0.306 | -0.037 | -0.024 | -0.024 | 0 | 0.79 | 0.41 | 0.82 | 0.86 | 0.87 | 0.85 |
| 9 | 3255945 | rs594370 | 1.212 | -1.928 | 1.555 | 0.16 | -0.04 | -0.059 | -0.059 | 0.36 | 0.8 | 0.51 | 0.83 | 0.9 | 0.87 | 0.9 |
| 9 | 3277260 | rs2304929 | 0.178 | -2.038 | 0.682 | 0.526 | 0.001 | -0.09 | -0.09 | 3.363 | 0.12 | 0 | 0.07 | 0.36 | 0.08 | 0.15 |
| 9 | 3284500 | rs2991328 | -2.528 | -2.105 | 0.429 | 0.301 | -0.168 | -0.079 | -0.079 | 0 | 0.64 | 0.97 | 0.53 | 0.64 | 0.46 | 0.56 |
| 9 | 3301887 | rs2986687 | -1.201 | -2.181 | 1.058 | 0.196 | -0.056 | -0.042 | -0.042 | 0.177 | 0.2 | 0.44 | 0.16 | 0.13 | 0.12 | 0.15 |
| 9 | 3307745 | rs2986697 | -0.549 | -2.161 | 0.939 | 0.265 | -0.014 | -0.009 | -0.009 | 0 | 0.16 | 0.19 | 0.13 | 0.12 | 0.11 | 0.15 |
| 9 | 3322283 | rs3012708 | -1.424 | -2.174 | 1.069 | 0.087 | 0.007 | 0.014 | 0.014 | 1.494 | 0.25 | 0.74 | 0.22 | 0.12 | 0.19 | 0.2 |
| 9 | 3327260 | rs2986676 | -1.953 | -2.238 | 0.899 | 0.224 | 0.028 | -0.106 | -0.106 | 0.512 | 0.4 | 0.88 | 0.27 | 0.45 | 0.28 | 0.3 |
| 9 | 3332974 | rs2991317 | -0.948 | -2.181 | 0.871 | -0.038 | -0.002 | -0.012 | -0.012 | 0 | 0.21 | 0.22 | 0.14 | 0.12 | 0.19 | 0.19 |

| | | | | | | | | | | | | | | | | |
|---|---|---|---|---|---|---|---|---|---|---|---|---|---|---|---|---|
| 9 | 3334185 | rs10971452 | 0.629 | -2.078 | -0.024 | -0.128 | 0.018 | 0.094 | 0.094 | 0 | 0.21 | 0.07 | 0.2 | 0.11 | 0.16 | 0.26 |
| 9 | 3335679 | rs10121595 | -0.329 | -1.903 | 0.717 | 0.582 | 0.015 | -0.049 | -0.049 | 3.271 | 0.17 | 0.1 | 0.09 | 0.43 | 0.1 | 0.17 |
| 9 | 3345623 | rs2991321 | -1.627 | -1.138 | 0.195 | -0.25 | -0.002 | 0.009 | 0.009 | 4.405 | 0.23 | 0.74 | 0.21 | 0.12 | 0.21 | 0.18 |
| 9 | 3349487 | rs3012675 | -2.104 | -1.171 | 0.597 | -0.234 | 0.015 | -0.093 | -0.093 | 0.047 | 0.14 | 0.55 | 0.09 | 0 | 0.11 | 0.04 |
| 9 | 3353579 | rs3012677 | -1.886 | -1.407 | 0.605 | -0.178 | -0.005 | -0.005 | -0.005 | 0 | 0.19 | 0.78 | 0.2 | 0.14 | 0.11 | 0.19 |
| 9 | 3363069 | rs13296217 | -0.538 | -1.442 | 0.428 | -0.201 | -0.022 | 0.015 | 0.015 | 0 | 0.06 | 0.01 | 0.03 | 0.03 | 0.05 | 0.09 |
| 9 | 3364535 | rs10511447 | -1.039 | -1.315 | 1.022 | -0.207 | -0.008 | 0.01 | 0.01 | 0.212 | 0.08 | 0.14 | 0.1 | 0.11 | 0.04 | 0.08 |
| 9 | 3370160 | rs2920359 | -1.773 | -2.205 | -0.169 | -0.273 | 0.042 | 0.016 | 0.016 | 0.322 | 0.54 | 0.94 | 0.43 | 0.62 | 0.3 | 0.58 |
| 9 | 3380518 | rs1373051 | -1.757 | -1.655 | -0.187 | -0.944 | -0.033 | 0.048 | 0.048 | 0 | 0.91 | 1 | 0.92 | 1 | 0.89 | 0.98 |
| 9 | 3384794 | rs10971685 | -0.189 | -2.067 | 0.568 | -0.907 | -0.011 | -0.019 | -0.019 | 0 | 0.28 | 0.35 | 0.21 | 0.18 | 0.17 | 0.37 |
| 9 | 3394094 | rs16917918 | -0.867 | -2.699 | 0.956 | -0.548 | -0.016 | 0.005 | 0.005 | 0 | 0.39 | 0.5 | 0.28 | 0.57 | 0.22 | 0.48 |
| 9 | 3395848 | rs3739461 | -0.9 | -2.86 | 1.371 | -0.458 | 0.018 | -0.005 | -0.005 | 0 | 0.38 | 0.63 | 0.3 | 0.57 | 0.2 | 0.39 |
| 9 | 3408858 | rs12341737 | -1.308 | -2.306 | 2.07 | 0.184 | 0.001 | 0.051 | 0.051 | 4.83 | 0.14 | 0.5 | 0.16 | 0.13 | 0.08 | 0.08 |
| 9 | 3416639 | rs11795111 | -0.01 | -2.327 | 1.774 | 0.185 | -0.014 | 0.009 | 0.009 | 0 | 0.07 | 0 | 0.03 | 0.04 | 0.05 | 0.04 |
| 9 | 3427166 | rs7854273 | -1.919 | -2.242 | 2.044 | 0.201 | -0.035 | 0.001 | 0.001 | 3.467 | 0.18 | 0.61 | 0.22 | 0.16 | 0.11 | 0.17 |
| 9 | 3432447 | rs7864440 | -1.702 | -2.173 | 2.083 | 0.292 | -0.03 | 0.008 | 0.008 | 3.737 | 0.17 | 0.61 | 0.21 | 0.16 | 0.09 | 0.16 |
| 9 | 3440355 | rs7864459 | -1.031 | -2.13 | 2.098 | 0.304 | 0.007 | 0.039 | 0.039 | 3.835 | 0.13 | 0.37 | 0.17 | 0.14 | 0.09 | 0.08 |
| 9 | 3446277 | rs10758256 | -2.078 | -2.22 | 1.003 | 0.231 | 0.092 | 0.07 | 0.07 | 0.792 | 0.4 | 0.89 | 0.42 | 0.75 | 0.2 | 0.35 |
| 9 | 3454967 | rs7852899 | -1.368 | -2.158 | 1.697 | 0.723 | -0.029 | 0.088 | 0.088 | 0 | 0.19 | 0.54 | 0.29 | 0.65 | 0.11 | 0.19 |
| 9 | 3468075 | rs12236983 | -1.251 | -2.053 | 1.437 | 0.552 | -0.033 | 0.016 | 0.016 | 0.462 | 0.27 | 0.61 | 0.28 | 0.65 | 0.13 | 0.22 |
| 9 | 3479911 | rs7026025 | -1.189 | -1.853 | 1.411 | 0.683 | -0.09 | 0.133 | 0.133 | 0 | 0.22 | 0.58 | 0.2 | 0.55 | 0.07 | 0.25 |
| 9 | 3496660 | rs881167 | -0.145 | -1.791 | 1.454 | 0.716 | -0.027 | 0.091 | 0.091 | 0.027 | 0.14 | 0.19 | 0.16 | 0.54 | 0.06 | 0.15 |
| 9 | 3500471 | rs12340274 | -1.149 | -1.793 | 1.411 | 0.693 | -0.095 | 0.159 | 0.159 | 0 | 0.21 | 0.57 | 0.2 | 0.54 | 0.07 | 0.24 |
| 9 | 3508131 | rs10814189 | -0.403 | -1.718 | 0.796 | 0.73 | -0.054 | 0.174 | 0.174 | 0 | 0.16 | 0.34 | 0.18 | 0.54 | 0.07 | 0.15 |
| 9 | 3517672 | rs7046581 | -0.228 | -1.634 | -0.09 | 0.102 | -0.025 | 0.082 | 0.082 | 0.23 | 0.85 | 0.98 | 0.88 | 1 | 0.84 | 0.97 |
| 9 | 3536260 | rs10814219 | -1.56 | -1.514 | 0.693 | 0.979 | -0.051 | 0.05 | 0.05 | 0 | 0.24 | 0.68 | 0.2 | 0.51 | 0.08 | 0.25 |
| 9 | 3547259 | rs10814233 | -0.766 | -0.465 | -0.236 | 1.496 | -0.03 | 0.108 | 0.108 | 0 | 0.15 | 0.24 | 0.16 | 0.51 | 0.07 | 0.17 |
| 9 | 3573936 | rs951777 | -0.2 | 0.437 | -0.571 | 0.956 | 0.007 | -0.003 | -0.003 | 0 | 0.4 | 0.48 | 0.47 | 0.34 | 0.47 | 0.37 |
| 9 | 3574369 | rs2488292 | 0.061 | 0.327 | -1.176 | 0.667 | -0.02 | 0.04 | 0.04 | 0 | 0.52 | 0.24 | 0.64 | 0.65 | 0.56 | 0.65 |
| 9 | 3575331 | rs7026527 | 0.644 | -0.361 | -0.222 | 0.759 | -0.043 | 0.025 | 0.025 | 0.154 | 0.81 | 0.46 | 0.81 | 0.78 | 0.91 | 0.81 |
| 9 | 3576434 | rs2184239 | 0.481 | -0.696 | -0.805 | 1.092 | 0.016 | -0.046 | -0.046 | 0 | 0.16 | 0.07 | 0.1 | 0.37 | 0.11 | 0.18 |
| 9 | 3578094 | rs10758318 | -0.476 | -0.637 | -0.07 | 0.908 | 0.023 | 0.104 | 0.104 | 0 | 0.89 | 0.71 | 0.91 | 0.83 | 0.9 | 0.92 |
| 9 | 3579891 | rs10814271 | 0.817 | -0.175 | -0.724 | 0.009 | -0.048 | -0.018 | -0.018 | 0 | 0.3 | 0.16 | 0.51 | 0.35 | 0.41 | 0.43 |
| 9 | 3579989 | rs10758319 | 0.91 | -0.221 | -0.851 | -0.021 | -0.093 | -0.048 | -0.048 | 1.632 | 0.28 | 0.08 | 0.5 | 0.35 | 0.41 | 0.43 |
| 9 | 3584207 | rs1001388 | 0.966 | -0.093 | -0.404 | 0.112 | 0.019 | 0.081 | 0.081 | 0 | 0.1 | 0.25 | 0.09 | 0.17 | 0.11 | 0.09 |
| 9 | 3584543 | rs1001389 | 0.438 | -0.498 | -1.069 | -0.035 | -0.042 | -0.002 | -0.002 | 0 | 0.44 | 0.21 | 0.6 | 0.45 | 0.49 | 0.63 |
| 9 | 3585271 | rs10972572 | 0.906 | -0.318 | -0.776 | 0.665 | 0.016 | -0.042 | -0.042 | 0 | 0.17 | 0.08 | 0.11 | 0.39 | 0.12 | 0.18 |
| 9 | 3589867 | rs10814278 | -0.492 | -0.016 | -0.288 | 0.525 | 0.005 | 0.078 | 0.078 | 0 | 0.89 | 0.73 | 0.91 | 0.82 | 0.91 | 0.9 |
| 9 | 3590468 | rs10814280 | 0.204 | 0.085 | -0.974 | -0.256 | -0.025 | 0.06 | 0.06 | 0.031 | 0.32 | 0.23 | 0.27 | 0.13 | 0.38 | 0.21 |
| 9 | 3593961 | rs10814284 | -0.056 | 0.166 | -0.841 | 0.004 | 0 | 0.01 | 0.01 | 0 | 0.36 | 0.4 | 0.47 | 0.46 | 0.35 | 0.49 |
| 9 | 3594049 | rs10972598 | 0.305 | 0.126 | -0.469 | 0.169 | 0.004 | 0.003 | 0.003 | 0 | 0.28 | 0.38 | 0.43 | 0.33 | 0.3 | 0.39 |
| 9 | 3595084 | rs7860641 | 0.962 | -0.13 | -0.672 | 0.241 | 0.007 | -0.005 | -0.005 | 0 | 0.26 | 0.17 | 0.11 | 0.34 | 0.22 | 0.24 |
| 9 | 3599498 | rs11787813 | 1.583 | -0.042 | -0.867 | 0.548 | -0.02 | -0.044 | -0.044 | 0 | 0.08 | 0.02 | 0.09 | 0.24 | 0.05 | 0.11 |
| 9 | 3601355 | rs10758333 | -0.313 | -0.165 | -0.588 | 0.147 | 0.013 | 0.015 | 0.015 | 0 | 0.48 | 0.54 | 0.48 | 0.59 | 0.48 | 0.53 |
| 9 | 3602018 | rs1334069 | 0.959 | 0.08 | -0.791 | -0.146 | -0.039 | -0.051 | -0.051 | 0 | 0.08 | 0.01 | 0.18 | 0.18 | 0.11 | 0.17 |
| 9 | 3603238 | rs7856970 | 1.529 | -0.124 | -0.511 | -0.275 | 0.015 | 0.002 | 0.002 | 0 | 0.16 | 0.22 | 0.34 | 0.17 | 0.24 | 0.28 |
| 9 | 3607242 | rs10972661 | 1.277 | -0.101 | -0.505 | -0.246 | 0.006 | -0.039 | -0.039 | 0.049 | 0.21 | 0.07 | 0.1 | 0.34 | 0.2 | 0.2 |
| 9 | 3631008 | rs17728981 | 1.574 | -0.142 | -0.728 | -0.207 | 0.013 | -0.035 | -0.035 | 0.451 | 0.18 | 0.07 | 0.09 | 0.33 | 0.15 | 0.16 |
| 9 | 3639538 | rs10814372 | -0.034 | -0.204 | -0.702 | -0.425 | 0.015 | -0.006 | -0.006 | 2.337 | 0.45 | 0.54 | 0.45 | 0.53 | 0.43 | 0.48 |
| 9 | 3646026 | rs10972874 | 0.871 | -0.699 | -0.758 | -0.958 | -0.043 | 0.146 | 0.146 | 0 | 0.34 | 0.15 | 0.21 | 0.01 | 0.39 | 0.21 |
| 9 | 3649190 | rs1537524 | -0.04 | -1.658 | -0.46 | -0.642 | -0.044 | 0.027 | 0.027 | 0 | 0.81 | 0.66 | 0.87 | 0.82 | 0.94 | 0.84 |
| 9 | 3650174 | rs1548299 | -0.356 | -1.206 | -0.22 | -0.61 | 0.007 | 0.009 | 0.009 | 0 | 0.53 | 0.52 | 0.71 | 0.8 | 0.65 | 0.67 |
| 9 | 3650939 | rs12379966 | 1.007 | -1.219 | 0.202 | -0.249 | 0.018 | 0.068 | 0.068 | 0 | 0.11 | 0.12 | 0.12 | 0.36 | 0.18 | 0.19 |
| 9 | 3651396 | rs10814387 | -0.915 | -1.215 | -0.263 | -0.354 | 0 | -0.001 | -0.001 | 0 | 0.4 | 0.55 | 0.34 | 0.58 | 0.3 | 0.41 |
| 9 | 3670015 | rs17729056 | 0.532 | -1.364 | -0.508 | -0.26 | 0.011 | -0.046 | -0.046 | 0.267 | 0.22 | 0.11 | 0.12 | 0.36 | 0.18 | 0.17 |
| 9 | 3670864 | rs2380874 | -0.708 | -1.191 | -0.33 | -0.176 | 0.048 | -0.018 | -0.018 | 2.353 | 0.41 | 0.73 | 0.35 | 0.57 | 0.3 | 0.38 |
| 9 | 3673041 | rs10511452 | 0.309 | -1.401 | -0.037 | -0.28 | 0.013 | -0.029 | -0.029 | 0 | 0.35 | 0.34 | 0.42 | 0.42 | 0.22 | 0.4 |
| 9 | 3676408 | rs7041297 | -0.969 | -0.862 | 0.897 | 0.391 | 0.011 | -0.003 | -0.003 | 0 | 0.39 | 0.61 | 0.42 | 0.59 | 0.26 | 0.36 |
| 9 | 3676560 | rs7040514 | 0.73 | -0.773 | 1.073 | 0.438 | -0.016 | 0.008 | 0.008 | 0.039 | 0.2 | 0.29 | 0.29 | 0.23 | 0.11 | 0.18 |
| 9 | 3677809 | rs10973024 | 1.118 | -0.728 | 0.865 | 0.466 | 0.003 | -0.005 | -0.005 | 1.942 | 0.18 | 0.19 | 0.28 | 0.23 | 0.11 | 0.17 |
| 9 | 3678039 | rs10814428 | -0.297 | -0.725 | 0.941 | 0.471 | -0.001 | 0.085 | 0.085 | 0 | 0.15 | 0.23 | 0.12 | 0.36 | 0.15 | 0.17 |
| 9 | 3681072 | rs10738972 | -0.314 | -0.697 | 0.883 | 0.458 | -0.012 | 0.036 | 0.036 | 0 | 0.31 | 0.44 | 0.4 | 0.59 | 0.25 | 0.35 |
| 9 | 3684027 | rs7040907 | -0.023 | -0.685 | 0.438 | 0.486 | 0.014 | 0.15 | 0.15 | 0 | 0.12 | 0.25 | 0.18 | 0.36 | 0.14 | 0.16 |
| 9 | 3684635 | rs2039470 | 0.853 | -0.716 | 0.506 | 0.471 | -0.013 | 0.008 | 0.008 | 0 | 0.2 | 0.29 | 0.22 | 0.23 | 0.11 | 0.19 |
| 9 | 3684798 | rs11794229 | 1.066 | -0.636 | 0.31 | 0.311 | -0.117 | 0.021 | 0.021 | 0 | 0.21 | 0.03 | 0.28 | 0.2 | 0.36 | 0.33 |
| 9 | 3686568 | rs10758400 | -0.29 | -0.638 | 0.322 | 0.319 | -0.01 | 0.019 | 0.019 | 0 | 0.32 | 0.54 | 0.4 | 0.59 | 0.25 | 0.35 |
| 9 | 3687942 | rs10973079 | 0.027 | -0.619 | 0.257 | 0.327 | 0.036 | 0.24 | 0.24 | 0 | 0.13 | 0.34 | 0.13 | 0.36 | 0.14 | 0.16 |
| 9 | 3688348 | rs7466772 | 1.06 | -0.502 | -0.515 | 0.357 | 0.029 | 0.147 | 0.147 | 0 | 0.08 | 0.12 | 0.11 | 0.36 | 0.14 | 0.16 |
| 9 | 3689790 | rs11789122 | 0.726 | -0.611 | -1.234 | -0.387 | -0.117 | 0.021 | 0.021 | 0 | 0.21 | 0.03 | 0.28 | 0.21 | 0.36 | 0.33 |

| | | | | | | | | | | | | | | | | |
|---|---|---|---|---|---|---|---|---|---|---|---|---|---|---|---|---|
| 9 | 3692923 | rs6476583 | -0.052 | -1.22 | -0.229 | 0.333 | 0.004 | -0.009 | -0.009 | 0 | 0.33 | 0.62 | 0.42 | 0.58 | 0.25 | 0.35 |
| 9 | 3698029 | rs13283774 | -0.444 | -0.551 | -0.436 | 0.478 | -0.003 | -0.003 | -0.003 | 0 | 0.06 | 0.06 | 0.02 | 0 | 0.01 | 0.02 |
| 9 | 3701629 | rs7035633 | 0.469 | -0.742 | 0.049 | 0.994 | -0.056 | 0.105 | 0.105 | 0 | 0.79 | 0.63 | 0.86 | 0.68 | 0.9 | 0.81 |
| 9 | 3701667 | rs7038842 | 0.748 | -0.314 | -0.236 | 0.785 | -0.001 | 0.012 | 0.012 | 0 | 0.53 | 0.45 | 0.45 | 0.37 | 0.51 | 0.52 |
| 9 | 3701912 | rs10511453 | -0.15 | -0.48 | -0.345 | 1.023 | 0.026 | -0.063 | -0.063 | 0.493 | 0.18 | 0.05 | 0.11 | 0.25 | 0.11 | 0.15 |
| 9 | 3705373 | rs933036 | 1.274 | -0.402 | -0.107 | 0.983 | 0.009 | 0.023 | 0.023 | 0 | 0.54 | 0.45 | 0.55 | 0.35 | 0.58 | 0.56 |
| 9 | 3707775 | rs7872019 | 0.208 | -0.207 | 0.074 | 0.944 | 0.031 | 0.01 | 0.01 | 0 | 0.74 | 0.59 | 0.7 | 0.71 | 0.69 | 0.75 |
| 9 | 3710068 | rs7871296 | -0.436 | -0.096 | 0.228 | 0.894 | 0.074 | 0.019 | 0.019 | 0 | 0.95 | 0.77 | 0.97 | 1 | 0.96 | 0.98 |
| 9 | 3714403 | rs7031446 | 1.56 | -0.211 | -0.007 | 0.925 | 0.013 | 0.101 | 0.101 | 0 | 0.51 | 0.4 | 0.53 | 0.25 | 0.51 | 0.46 |
| 9 | 3716878 | rs3954950 | 0.598 | -0.244 | -0.079 | 0.966 | -0.004 | -0.007 | -0.007 | 0 | 0.18 | 0.2 | 0.24 | 0.29 | 0.24 | 0.24 |
| 9 | 3723221 | rs13283466 | 0.708 | -0.397 | -0.447 | 1.003 | -0.007 | -0.08 | -0.08 | 0.002 | 0.12 | 0.01 | 0.11 | 0.22 | 0.09 | 0.12 |
| 9 | 3726443 | rs4741847 | 0.524 | -0.204 | -0.137 | 0.776 | 0 | 0.01 | 0.01 | 0 | 0.54 | 0.51 | 0.57 | 0.39 | 0.53 | 0.52 |
| 9 | 3735490 | rs10973279 | 0.046 | -0.17 | -0.316 | 0.973 | -0.009 | 0.029 | 0.029 | 0 | 0.05 | 0.09 | 0.06 | 0.15 | 0.03 | 0.06 |
| 9 | 3741507 | rs7875110 | 0.547 | -0.009 | -0.603 | 0.623 | -0.002 | 0.005 | 0.005 | 0 | 0.61 | 0.51 | 0.68 | 0.52 | 0.6 | 0.68 |
| 9 | 3743938 | rs10511454 | -0.41 | 0.302 | -0.993 | 0.82 | -0.036 | -0.049 | -0.049 | 0.196 | 0.12 | 0.06 | 0.15 | 0.23 | 0.15 | 0.27 |
| 9 | 3744144 | rs7046742 | 0.268 | -0.253 | -0.822 | 1.086 | -0.007 | 0.033 | 0.033 | 0 | 0.86 | 0.82 | 0.8 | 0.74 | 0.9 | 0.91 |
| 9 | 3746413 | rs4147122 | -0.131 | -0.269 | -0.825 | 1.082 | -0.013 | 0.043 | 0.043 | 0 | 0.89 | 0.85 | 0.83 | 0.75 | 0.92 | 0.93 |
| 9 | 3747376 | rs4740731 | 0.944 | -0.466 | -0.878 | 0.94 | -0.003 | -0.013 | -0.013 | 0 | 0.74 | 0.73 | 0.64 | 0.51 | 0.74 | 0.63 |
| 9 | 3755135 | rs4626661 | -0.719 | -0.836 | -1.37 | 0.672 | 0.013 | -0.023 | -0.023 | 2.432 | 0.45 | 0.27 | 0.37 | 0.48 | 0.39 | 0.51 |
| 9 | 3755938 | rs7868939 | -0.671 | -0.895 | -1.363 | 0.649 | 0.009 | -0.02 | -0.02 | 0.043 | 0.44 | 0.29 | 0.37 | 0.47 | 0.39 | 0.48 |
| 9 | 3756613 | rs11791065 | 0.249 | -0.656 | -1.426 | 0.592 | 0.003 | 0.002 | 0.002 | 0 | 0.27 | 0.21 | 0.18 | 0.26 | 0.24 | 0.24 |
| 9 | 3756990 | rs4740732 | 0.229 | -0.326 | -1.507 | 0.597 | -0.004 | -0.002 | -0.002 | 0.056 | 0.27 | 0.27 | 0.18 | 0.26 | 0.23 | 0.25 |
| 9 | 3757758 | rs13286706 | 0.607 | 0.44 | -2.594 | 0.455 | -0.026 | -0.023 | -0.023 | 0 | 0.2 | 0.14 | 0.31 | 0.26 | 0.29 | 0.17 |
| 9 | 3758867 | rs2183689 | 0.269 | -0.261 | -2.382 | 0.042 | 0.023 | 0.055 | 0.055 | 0 | 0.33 | 0.2 | 0.29 | 0.2 | 0.29 | 0.28 |
| 9 | 3761548 | rs4740734 | -0.355 | -0.379 | -1.911 | 0.793 | 0.013 | -0.006 | -0.006 | 0 | 0.17 | 0.11 | 0.13 | 0.21 | 0.13 | 0.25 |
| 9 | 3761909 | rs954980 | -0.635 | -0.174 | -1.865 | 0.728 | 0.028 | -0.022 | -0.022 | 3.446 | 0.35 | 0.19 | 0.33 | 0.49 | 0.32 | 0.43 |
| 9 | 3765198 | rs927584 | -0.383 | 0.427 | -2.316 | 0.885 | -0.065 | -0.076 | -0.076 | 0 | 0.38 | 0.12 | 0.36 | 0.42 | 0.49 | 0.53 |
| 9 | 3765395 | rs6476642 | 0.509 | -0.653 | -2.065 | 0.785 | -0.001 | -0.001 | -0.001 | 0 | 0.28 | 0.31 | 0.45 | 0.3 | 0.28 | 0.2 |
| 9 | 3772674 | rs10814589 | -0.233 | -0.987 | -2.064 | 0.509 | 0.003 | -0.017 | -0.017 | 1.596 | 0.33 | 0.34 | 0.44 | 0.44 | 0.31 | 0.43 |
| 9 | 3773699 | rs12351927 | 1.592 | -0.919 | -2.066 | 0.491 | 0.001 | -0.013 | -0.013 | 0 | 0.1 | 0.11 | 0.08 | 0 | 0.09 | 0.01 |
| 9 | 3774843 | rs1455181 | 1.259 | -0.795 | -2.143 | 0.54 | -0.073 | -0.019 | -0.019 | 0 | 0.27 | 0.11 | 0.3 | 0.25 | 0.39 | 0.28 |
| 9 | 3775583 | rs1455180 | -1.1 | -0.911 | -1.41 | 0.539 | 0.003 | 0.012 | 0.012 | 0 | 0.9 | 0.89 | 0.9 | 0.87 | 0.9 | 0.81 |
| 9 | 3777523 | rs10814596 | 0.725 | -1.045 | -1.011 | 0.473 | -0.002 | 0.004 | 0.004 | 0 | 0.21 | 0.24 | 0.32 | 0.29 | 0.19 | 0.21 |
| 9 | 3779330 | rs10758451 | 0.752 | -1.323 | -0.926 | 0.42 | -0.019 | 0.005 | 0.005 | 0.055 | 0.24 | 0.29 | 0.16 | 0.32 | 0.2 | 0.25 |
| 9 | 3781983 | rs7039314 | 0.393 | -1.366 | -1.71 | 0.781 | -0.021 | -0.028 | -0.028 | 0 | 0.43 | 0.16 | 0.45 | 0.43 | 0.54 | 0.42 |
| 9 | 3786161 | rs10973533 | 0.019 | -1.347 | -1.203 | 0.796 | 0.026 | -0.014 | -0.014 | 0 | 0.15 | 0.08 | 0.08 | 0.26 | 0.08 | 0.18 |
| 9 | 3791433 | rs7047408 | -0.011 | -0.81 | -0.961 | 0.491 | 0.014 | 0.047 | 0.047 | 0.033 | 0.51 | 0.36 | 0.56 | 0.67 | 0.52 | 0.46 |
| 9 | 3801316 | rs2124210 | 0.438 | 0.744 | 0.659 | 1.226 | 0.081 | 0.023 | 0.023 | 0 | 0.83 | 0.49 | 0.82 | 0.86 | 0.82 | 0.88 |
| 9 | 3808284 | rs7033413 | 0.061 | 0.261 | 0.688 | 1.246 | -0.002 | -0.003 | -0.003 | 0 | 0.86 | 0.88 | 0.86 | 0.84 | 0.86 | 0.89 |
| 9 | 3809726 | rs1537162 | 0.643 | 0.183 | 0.599 | 1.198 | -0.005 | -0.012 | -0.012 | 0 | 0.83 | 0.89 | 0.85 | 0.81 | 0.84 | 0.85 |
| 9 | 3810196 | rs10973640 | -0.152 | -0.072 | 0.704 | 1.276 | -0.01 | 0.005 | 0.005 | 2.711 | 0.18 | 0.21 | 0.27 | 0.26 | 0.1 | 0.28 |
| 9 | 3810281 | rs7870234 | 1.036 | -0.035 | 0.759 | 1.247 | -0.024 | 0.036 | 0.036 | 1.237 | 0.64 | 0.46 | 0.58 | 0.55 | 0.74 | 0.59 |
| 9 | 3817537 | rs2380902 | 1.008 | -0.259 | 0.524 | 1.132 | 0.008 | -0.004 | -0.004 | 0 | 0.69 | 0.23 | 0.72 | 0.66 | 0.82 | 0.73 |
| 9 | 3823277 | rs1411880 | -0.057 | -0.248 | 0.323 | 1.11 | -0.053 | -0.003 | -0.003 | 0.022 | 0.83 | 0.71 | 0.84 | 0.83 | 0.93 | 0.83 |
| 9 | 3824004 | rs7049103 | 0.319 | 0.556 | -0.036 | 1.322 | 0 | 0.016 | 0.016 | 0 | 0.11 | 0.14 | 0.11 | 0.18 | 0.11 | 0.12 |
| 9 | 3826081 | rs3739618 | -0.952 | 1.4 | 0.395 | 1.583 | 0.083 | 0.143 | 0.143 | 0 | 0.09 | 0.29 | 0.3 | 0.27 | 0.21 | 0.09 |
| 9 | 3826710 | rs12376878 | 0.45 | 0.961 | -0.512 | 0.602 | -0.023 | 0.067 | 0.067 | 0 | 0.47 | 0.16 | 0.38 | 0.32 | 0.45 | 0.48 |
| 9 | 3827075 | rs10973690 | 0.034 | 0.846 | -0.352 | 0.695 | -0.033 | -0.056 | -0.056 | 0 | 0.34 | 0.19 | 0.47 | 0.53 | 0.47 | 0.36 |
| 9 | 3827120 | rs1331260 | -0.016 | 0.896 | -0.239 | 0.276 | 0.015 | -0.003 | -0.003 | 0 | 0.37 | 0.44 | 0.32 | 0.32 | 0.37 | 0.48 |
| 9 | 3827824 | rs734752 | 0.34 | 1.075 | -1.036 | 0.292 | -0.048 | -0.057 | -0.057 | 0.674 | 0.3 | 0.08 | 0.26 | 0.34 | 0.3 | 0.47 |
| 9 | 3828696 | rs12001442 | 0.57 | 1.048 | -0.688 | -0.272 | 0 | -0.029 | -0.029 | 0 | 0.13 | 0.14 | 0.09 | 0 | 0.18 | 0.05 |
| 9 | 3829212 | rs7029652 | -0.18 | 1.091 | -0.664 | -0.507 | -0.003 | -0.014 | -0.014 | 1.414 | 0.2 | 0.23 | 0.12 | 0 | 0.22 | 0.09 |
| 9 | 3832812 | rs9644834 | 0.167 | 1.502 | -0.702 | 0.494 | -0.012 | 0.013 | 0.013 | 0 | 0.1 | 0.07 | 0.23 | 0.45 | 0.15 | 0.17 |
| 9 | 3833480 | rs7024250 | 0.462 | 0.358 | -0.903 | 0.004 | 0.01 | -0.003 | -0.003 | 0 | 0.18 | 0.21 | 0.13 | 0.18 | 0.08 | 0.19 |
| 9 | 3838888 | rs10973736 | 1.423 | 0.713 | -1.406 | -0.366 | -0.01 | -0.003 | -0.003 | 0.005 | 0.2 | 0.06 | 0.17 | 0.16 | 0.22 | 0.13 |
| 9 | 3838929 | rs10973737 | 1.2 | 0.464 | -1.587 | -0.603 | 0.013 | 0.018 | 0.018 | 0.008 | 0.18 | 0.14 | 0.08 | 0.09 | 0.12 | 0.27 |
| 9 | 3840882 | rs4612398 | -0.356 | 0.693 | -1.129 | -0.792 | 0 | 0.001 | 0.001 | 0 | 0.48 | 0.58 | 0.65 | 0.73 | 0.49 | 0.54 |
| 9 | 3842939 | rs1378325 | 1.406 | 0.987 | -1.605 | -0.725 | -0.087 | -0.062 | -0.062 | 0 | 0.24 | 0.01 | 0.18 | 0.21 | 0.27 | 0.15 |
| 9 | 3843017 | rs10973755 | 0.662 | 0.966 | -1.178 | -0.592 | 0.003 | 0.062 | 0.062 | 0 | 0.15 | 0.15 | 0.28 | 0.52 | 0.23 | 0.29 |
| 9 | 3844565 | rs12685627 | -0.135 | 1.035 | -1.415 | -0.786 | -0.014 | -0.035 | -0.035 | 0.497 | 0.25 | 0.15 | 0.43 | 0.55 | 0.34 | 0.35 |
| 9 | 3845784 | rs10973760 | 0.766 | 0.736 | -1.203 | -0.58 | -0.005 | 0.004 | 0.004 | 0 | 0.14 | 0.09 | 0.29 | 0.49 | 0.21 | 0.26 |
| 9 | 3848687 | rs7046660 | -0.553 | 0.025 | -0.908 | -0.915 | -0.018 | -0.012 | -0.012 | 0 | 0.67 | 0.33 | 0.69 | 0.73 | 0.81 | 0.61 |
| 9 | 3851002 | rs11788034 | 1.285 | 0.048 | -0.917 | -0.691 | -0.003 | -0.052 | -0.052 | 0.19 | 0.12 | 0.04 | 0.23 | 0.5 | 0.13 | 0.24 |
| 9 | 3851138 | rs10758497 | -0.775 | -0.286 | -0.365 | -1.291 | -0.038 | -0.015 | -0.015 | 0 | 0.72 | 0.54 | 0.71 | 0.75 | 0.83 | 0.63 |
| 9 | 3853657 | rs4741860 | 0.285 | -0.262 | -0.423 | -1.192 | -0.008 | -0.005 | -0.005 | 0 | 0.43 | 0.35 | 0.43 | 0.09 | 0.48 | 0.49 |
| 9 | 3854070 | rs10758498 | 0.323 | -0.096 | -0.445 | -1.04 | -0.02 | 0.06 | 0.06 | 0 | 0.42 | 0.27 | 0.47 | 0.7 | 0.48 | 0.41 |
| 9 | 3855616 | rs2281731 | -1.195 | 0.208 | -0.181 | -0.352 | 0.001 | 0.001 | 0.001 | 1.316 | 0.65 | 0.64 | 0.78 | 0.79 | 0.74 | 0.82 |
| 9 | 3862693 | rs10973877 | 0.578 | 0.497 | -0.104 | -0.367 | 0.001 | 0.018 | 0.018 | 0.415 | 0.51 | 0.44 | 0.61 | 0.9 | 0.53 | 0.5 |
| 9 | 3864491 | rs7867243 | 0.344 | 0.716 | -0.128 | 0.977 | -0.026 | 0.008 | 0.008 | 1.432 | 0.05 | 0.01 | 0.23 | 0.23 | 0.09 | 0.07 |

| | | | | | | | | | | | | | | | | |
|---|---|---|---|---|---|---|---|---|---|---|---|---|---|---|---|---|
| 9 | 3865097 | rs10739002 | 1.051 | 0.646 | 0.189 | 0.673 | -0.007 | -0.002 | -0.002 | 0 | 0.66 | 0.73 | 0.57 | 0.67 | 0.63 | 0.71 |
| 9 | 3865283 | rs16919809 | 0.294 | 0.582 | 0.646 | 0.895 | -0.002 | 0.08 | 0.08 | 0 | 0.14 | 0.22 | 0.27 | 0.47 | 0.16 | 0.26 |
| 9 | 3866849 | rs6476722 | 0.393 | 0.685 | 0.894 | 0.314 | 0.032 | -0.025 | -0.025 | 0 | 0.93 | 0.77 | 0.96 | 1 | 0.92 | 0.94 |
| 9 | 3869844 | rs1475223 | 0.69 | 0.735 | 1.021 | 0.332 | 0.011 | -0.103 | -0.103 | 0.041 | 0.81 | 0.67 | 0.85 | 0.98 | 0.81 | 0.85 |
| 9 | 3879097 | rs10814737 | 1.232 | 0.703 | 1.145 | 0.453 | 0.022 | -0.037 | -0.037 | 0 | 0.91 | 0.65 | 0.95 | 1 | 0.93 | 0.95 |
| 9 | 3880627 | rs10973995 | -0.081 | 0.787 | 0.708 | 0.75 | -0.01 | 0.004 | 0.004 | 0 | 0.16 | 0.1 | 0.13 | 0.17 | 0.18 | 0.17 |
| 9 | 3880643 | rs10973996 | -0.079 | 0.795 | 0.712 | 0.754 | -0.01 | 0.004 | 0.004 | 0 | 0.16 | 0.1 | 0.13 | 0.17 | 0.18 | 0.17 |
| 9 | 3883042 | rs895229 | -0.413 | 0.792 | 0.948 | 0.8 | 0.012 | -0.001 | -0.001 | 0 | 0.17 | 0.28 | 0.15 | 0.19 | 0.19 | 0.17 |
| 9 | 3891863 | rs1869690 | 1.557 | 0.767 | 0.426 | 0.738 | 0.003 | 0.026 | 0.026 | 0.391 | 0.62 | 0.35 | 0.75 | 0.72 | 0.65 | 0.56 |
| 9 | 3895172 | rs1547127 | 1.243 | 1.016 | -0.065 | 0.899 | 0.007 | -0.025 | -0.025 | 0 | 0.76 | 0.71 | 0.83 | 0.82 | 0.74 | 0.78 |
| 9 | 3897225 | rs10974064 | 0.282 | 1.1 | -0.363 | 1.333 | -0.045 | -0.059 | -0.059 | 0 | 0.09 | 0 | 0.1 | 0.17 | 0.1 | 0.09 |
| 9 | 3898987 | rs10122183 | 1.213 | 0.722 | 0.115 | 0.95 | 0 | -0.026 | -0.026 | 0.373 | 0.77 | 0.74 | 0.85 | 0.84 | 0.76 | 0.8 |
| 9 | 3900982 | rs10974067 | -0.307 | 0.799 | -0.409 | 1.012 | -0.04 | -0.003 | -0.003 | 1.313 | 0.16 | 0.04 | 0.13 | 0.17 | 0.19 | 0.17 |
| 9 | 3901415 | rs10974071 | -0.223 | 0.761 | -0.405 | 1.022 | -0.039 | -0.007 | -0.007 | 1.666 | 0.15 | 0.03 | 0.12 | 0.17 | 0.18 | 0.16 |
| 9 | 3908403 | rs702268 | 1.287 | 0.587 | 0.029 | 1.541 | -0.003 | 0.014 | 0.014 | 4.263 | 0.85 | 0.85 | 0.89 | 0.78 | 0.87 | 0.69 |
| 9 | 3908583 | rs16919945 | -0.236 | 0.602 | -0.05 | 1.529 | -0.021 | -0.018 | -0.018 | 0.047 | 0.09 | 0.02 | 0.07 | 0.15 | 0.1 | 0.14 |
| 9 | 3909453 | rs453166 | 1.947 | 0.707 | 0.151 | 1.592 | 0.006 | 0.032 | 0.032 | 2.831 | 0.73 | 0.67 | 0.79 | 0.59 | 0.72 | 0.49 |
| 9 | 3910136 | rs519644 | 1.153 | 0.706 | 0.166 | 1.588 | 0.023 | 0.064 | 0.064 | 2.168 | 0.86 | 0.78 | 0.86 | 0.76 | 0.83 | 0.65 |
| 9 | 3911976 | rs570649 | 1.828 | 0.627 | -0.018 | 1.565 | 0 | 0.005 | 0.005 | 4.641 | 0.7 | 0.72 | 0.72 | 0.53 | 0.71 | 0.49 |
| 9 | 3915436 | rs10974127 | -0.454 | 0.531 | -0.099 | 1.188 | -0.019 | -0.028 | -0.028 | 0 | 0.09 | 0.02 | 0.07 | 0.17 | 0.1 | 0.14 |
| 9 | 3920688 | rs487735 | 1.16 | 0.415 | 0.089 | 1.079 | -0.014 | -0.008 | -0.008 | 0 | 0.85 | 0.83 | 0.92 | 0.84 | 0.91 | 0.83 |
| 9 | 3928115 | rs624290 | 1.379 | 0.824 | 0.696 | 1.46 | 0.012 | 0.127 | 0.127 | 0 | 0.88 | 0.63 | 0.87 | 0.75 | 0.89 | 0.83 |
| 9 | 3931626 | rs681080 | 0.625 | 0.977 | 0.332 | 0.669 | 0.001 | -0.108 | -0.108 | 1.613 | 0.68 | 0.5 | 0.81 | 0.91 | 0.72 | 0.75 |
| 9 | 3934738 | rs581222 | 0.282 | 1.055 | 0.571 | 0.618 | 0.016 | -0.085 | -0.085 | 0 | 0.73 | 0.56 | 0.83 | 0.93 | 0.73 | 0.77 |
| 9 | 3935560 | rs7870379 | -0.185 | 1.158 | 0.522 | 0.608 | 0.003 | 0.011 | 0.011 | 0 | 0.23 | 0.21 | 0.12 | 0.07 | 0.2 | 0.21 |
| 9 | 3936301 | rs662234 | 0.089 | 1.126 | 0.596 | 0.613 | -0.002 | -0.027 | -0.027 | 0.799 | 0.21 | 0.26 | 0.13 | 0.08 | 0.2 | 0.21 |
| 9 | 3937288 | rs10974212 | -1.027 | 1.795 | 0.666 | 1.296 | 0.116 | 0.269 | 0.269 | 0 | 0.15 | 0.43 | 0.25 | 0.4 | 0.25 | 0.22 |
| 9 | 3940862 | rs12344176 | -1.314 | 1.682 | 0.41 | 1.345 | 0.067 | 0.059 | 0.059 | 0.238 | 0.15 | 0.59 | 0.27 | 0.22 | 0.22 | 0.16 |
| 9 | 3940874 | rs564618 | -1.421 | 1.679 | 0.287 | 1.283 | -0.075 | 0 | 0 | 0.125 | 0.37 | 0.92 | 0.38 | 0.27 | 0.36 | 0.33 |
| 9 | 3941263 | rs628338 | -0.251 | 1.666 | 0.285 | 1.281 | -0.001 | -0.027 | -0.027 | 0 | 0.25 | 0.28 | 0.14 | 0.05 | 0.21 | 0.2 |
| 9 | 3941973 | rs533520 | -1.283 | 1.684 | 0.051 | 1.296 | -0.033 | 0.021 | 0.021 | 0.117 | 0.35 | 0.84 | 0.37 | 0.26 | 0.34 | 0.32 |
| 9 | 3942136 | rs12554821 | 0.348 | 1.526 | 0.033 | 1.314 | 0.02 | 0.037 | 0.037 | 0 | 0.2 | 0.14 | 0.1 | 0.06 | 0.15 | 0.18 |
| 9 | 3943449 | rs1445339 | -0.779 | 1.845 | 0.188 | 1.495 | 0.057 | 0.135 | 0.135 | 0.16 | 0.1 | 0.59 | 0.23 | 0.19 | 0.13 | 0.12 |
| 9 | 3946361 | rs1445338 | -1.352 | 1.542 | -0.773 | 1.164 | 0.027 | -0.01 | -0.01 | 0.155 | 0.52 | 0.76 | 0.48 | 0.45 | 0.57 | 0.42 |
| 9 | 3948365 | rs1017568 | -0.661 | 1.704 | -1.085 | 1.326 | -0.051 | -0.055 | -0.055 | 0.012 | 0.1 | 0.01 | 0.11 | 0.25 | 0.18 | 0.21 |
| 9 | 3949244 | rs17701244 | -1.339 | 1.858 | -1.072 | 1.244 | -0.051 | -0.018 | -0.018 | 0 | 0.17 | 0.05 | 0.17 | 0.26 | 0.26 | 0.25 |
| 9 | 3952421 | rs592310 | -0.736 | 1.671 | -0.362 | 1.579 | 0.078 | 0.13 | 0.13 | 0 | 0.17 | 0.53 | 0.23 | 0.25 | 0.21 | 0.19 |
| 9 | 3952892 | rs10974256 | -0.421 | 1.538 | -0.821 | 1.1 | 0 | -0.001 | -0.001 | 0 | 0.3 | 0.25 | 0.29 | 0.31 | 0.34 | 0.2 |
| 9 | 3957866 | rs12238888 | -1.33 | 2.244 | -1.023 | 1.513 | 0.01 | 0.001 | 0.001 | 0.464 | 0.09 | 0.04 | 0.14 | 0.25 | 0.24 | 0.1 |
| 9 | 3958355 | rs11791772 | -0.739 | 0.53 | -1.56 | 0.783 | 0.034 | 0.051 | 0.051 | 1.478 | 0.17 | 0.04 | 0.1 | 0.06 | 0.09 | 0.13 |
| 9 | 3958518 | rs528938 | -0.491 | -0.048 | -2.404 | 0.326 | 0.052 | 0.205 | 0.205 | 0 | 0.46 | 0.83 | 0.6 | 0.87 | 0.37 | 0.37 |
| 9 | 3958553 | rs10814808 | -0.486 | 0.429 | -1.945 | 1.635 | -0.04 | 0.013 | 0.013 | 0 | 0.07 | 0.01 | 0.13 | 0.4 | 0.13 | 0.11 |
| 9 | 3959104 | rs2839895 | -1.205 | -0.249 | -1.642 | 0.88 | 0.003 | 0.005 | 0.005 | 0.055 | 0.12 | 0.09 | 0.19 | 0.48 | 0.1 | 0.13 |
| 9 | 3961492 | rs635171 | -0.71 | -0.513 | -1.859 | -0.428 | 0.034 | 0.142 | 0.142 | 0 | 0.38 | 0.8 | 0.29 | 0.03 | 0.34 | 0.1 |
| 9 | 3962727 | rs2182633 | -1.207 | -0.214 | -1.67 | -0.085 | -0.007 | -0.031 | -0.031 | 0 | 0.16 | 0.12 | 0.23 | 0.53 | 0.17 | 0.18 |
| 9 | 3962983 | rs789263 | -0.674 | -0.587 | -0.987 | -0.773 | -0.002 | -0.083 | -0.083 | 0 | 0.4 | 0.43 | 0.32 | 0.03 | 0.4 | 0.15 |
| 9 | 3965689 | rs510807 | -1.137 | -0.613 | -0.378 | -0.408 | 0 | 0.001 | 0.001 | 0 | 0.46 | 0.48 | 0.45 | 0.48 | 0.48 | 0.27 |
| 9 | 3965805 | rs509879 | -1.887 | -0.858 | -0.835 | -0.744 | -0.024 | -0.016 | -0.016 | 0.967 | 0.61 | 0.77 | 0.5 | 0.48 | 0.56 | 0.31 |
| 9 | 3967645 | rs12552476 | 0.721 | -0.787 | -0.729 | -0.702 | 0.004 | -0.011 | -0.011 | 2.697 | 0.12 | 0.08 | 0.04 | 0 | 0.08 | 0.04 |
| 9 | 3967758 | rs10974265 | 0.196 | -0.842 | -0.944 | -0.913 | -0.023 | -0.085 | -0.085 | 0 | 0.15 | 0.03 | 0.2 | 0.47 | 0.18 | 0.15 |
| 9 | 3969423 | rs661726 | -2.045 | -0.855 | -0.848 | -1.043 | -0.032 | -0.003 | -0.003 | 0 | 0.61 | 0.77 | 0.5 | 0.39 | 0.55 | 0.3 |
| 9 | 3969842 | rs663542 | -2.008 | -0.886 | -0.78 | -0.991 | -0.024 | -0.005 | -0.005 | 0 | 0.61 | 0.78 | 0.5 | 0.4 | 0.57 | 0.3 |
| 9 | 3970103 | rs675148 | -2.005 | -0.905 | -0.789 | -1.008 | -0.029 | 0.001 | 0.001 | 0 | 0.59 | 0.77 | 0.5 | 0.4 | 0.56 | 0.29 |
| 9 | 3971303 | rs7045016 | 0.46 | -0.614 | -1.081 | -0.955 | 0.003 | -0.013 | -0.013 | 0 | 0.12 | 0.09 | 0.04 | 0 | 0.08 | 0.04 |
| 9 | 3971746 | rs585217 | 0.251 | -0.502 | -1.352 | -0.789 | 0.001 | -0.049 | -0.049 | 0 | 0.22 | 0.29 | 0.18 | 0.01 | 0.24 | 0.08 |
| 9 | 3971787 | rs10758547 | -0.35 | 0.281 | -1.859 | 0.059 | -0.008 | -0.02 | -0.02 | 0.353 | 0.17 | 0.13 | 0.23 | 0.46 | 0.22 | 0.18 |
| 9 | 3975129 | rs657821 | -0.515 | -0.104 | -1.688 | -0.689 | 0.037 | 0.073 | 0.073 | 1.26 | 0.36 | 0.8 | 0.25 | 0.02 | 0.27 | 0.12 |
| 9 | 3981076 | rs627947 | -0.744 | 0.389 | -1.991 | 0.041 | -0.017 | -0.04 | -0.04 | 0 | 0.44 | 0.94 | 0.46 | 0.54 | 0.39 | 0.35 |
| 9 | 3981646 | rs12351382 | -1.165 | 0.604 | -0.498 | -0.497 | 0.06 | -0.077 | -0.077 | 0 | 0.13 | 0.43 | 0.11 | 0.01 | 0.13 | 0.07 |
| 9 | 3983003 | rs657076 | -0.565 | 0.758 | -1.434 | -0.454 | -0.038 | -0.046 | -0.046 | 0 | 0.59 | 0.88 | 0.51 | 0.48 | 0.59 | 0.47 |
| 9 | 3985733 | rs10814812 | 0.116 | 0.841 | -0.573 | -0.674 | 0.02 | 0.035 | 0.035 | 1.778 | 0.37 | 0.3 | 0.23 | 0.24 | 0.31 | 0.23 |
| 9 | 3986123 | rs494215 | -0.427 | 1.142 | 0.454 | -0.379 | 0.011 | -0.003 | -0.003 | 0 | 0.26 | 0.3 | 0.24 | 0.27 | 0.32 | 0.3 |
| 9 | 3986952 | rs635884 | -0.347 | 0.694 | -0.09 | -0.276 | -0.02 | -0.004 | -0.004 | 0 | 0.23 | 0.14 | 0.21 | 0.26 | 0.25 | 0.26 |
| 9 | 4002823 | rs500044 | 1.215 | 0.239 | 0.388 | -0.318 | 0.002 | 0.001 | 0.001 | 0.012 | 0.84 | 0.86 | 0.85 | 0.97 | 0.88 | 0.94 |
| 9 | 4005108 | rs16920327 | -0.834 | 0.563 | 0.347 | -0.272 | -0.003 | -0.002 | -0.002 | 0 | 0.09 | 0.11 | 0.11 | 0.13 | 0.09 | 0.23 |
| 9 | 4005592 | rs702263 | -1.174 | 0.459 | 1.03 | -0.258 | 0.018 | 0.051 | 0.051 | 0.037 | 0.11 | 0.33 | 0.15 | 0.17 | 0.09 | 0.23 |
| 9 | 4008954 | rs660812 | 1.055 | 0.309 | 0.732 | -0.445 | -0.003 | 0.029 | 0.029 | 0 | 0.86 | 0.92 | 0.88 | 0.98 | 0.88 | 0.93 |
| 9 | 4020748 | rs627938 | -1.144 | -0.041 | 1.386 | -0.528 | -0.05 | -0.094 | -0.094 | 0 | 0.18 | 0.43 | 0.15 | 0.04 | 0.13 | 0.09 |
| 9 | 4023999 | rs12344339 | -0.221 | 0.678 | -0.044 | -0.818 | -0.055 | 0.077 | 0.077 | 0 | 0.3 | 0.12 | 0.21 | 0.14 | 0.4 | 0.3 |

| | | | | | | | | | | | | | | | | |
|---|---|---|---|---|---|---|---|---|---|---|---|---|---|---|---|---|
| 9 | 4028842 | rs17703489 | -0.113 | 0.785 | -0.147 | -0.867 | -0.074 | 0.086 | 0.086 | 0 | 0.25 | 0.08 | 0.16 | 0.12 | 0.31 | 0.29 |
| 9 | 4030806 | rs16920428 | -0.054 | 0.457 | 0.562 | -0.944 | -0.019 | -0.037 | -0.037 | 0 | 0.07 | 0.18 | 0.05 | 0 | 0.03 | 0.02 |
| 9 | 4032693 | rs16920433 | -0.395 | 0.605 | 0.559 | -1.061 | 0.003 | -0.046 | -0.046 | 0 | 0.1 | 0.21 | 0.1 | 0 | 0.1 | 0.06 |
| 9 | 4032747 | rs606215 | -0.526 | 0.245 | 0.411 | -0.603 | 0.009 | -0.01 | -0.01 | 0.284 | 0.35 | 0.47 | 0.28 | 0.37 | 0.4 | 0.42 |
| 9 | 4034898 | rs7853251 | -0.923 | -0.498 | 0.206 | -0.011 | -0.035 | -0.003 | -0.003 | 0 | 0.32 | 0.56 | 0.4 | 0.33 | 0.23 | 0.46 |
| 9 | 4035710 | rs10814817 | -0.513 | -0.059 | -0.789 | 0.137 | 0.006 | -0.017 | -0.017 | 0.064 | 0.33 | 0.11 | 0.45 | 0.65 | 0.31 | 0.34 |
| 9 | 4036449 | rs515380 | -0.994 | -0.526 | -0.651 | 0.276 | 0.006 | 0.002 | 0.002 | 0 | 0.44 | 0.59 | 0.6 | 0.55 | 0.38 | 0.44 |
| 9 | 4036616 | rs16920450 | 0.283 | -0.527 | -0.813 | 0.611 | -0.015 | -0.02 | -0.02 | 0 | 0.08 | 0 | 0.21 | 0.38 | 0.07 | 0.19 |
| 9 | 4037925 | rs571317 | -0.487 | -0.602 | -0.383 | 0.446 | -0.018 | -0.063 | -0.063 | 1.252 | 0.34 | 0.49 | 0.33 | 0.14 | 0.27 | 0.23 |
| 9 | 4038228 | rs895236 | 0.288 | -0.448 | -0.186 | 0.589 | -0.004 | 0.047 | 0.047 | 0 | 0.08 | 0.07 | 0.27 | 0.38 | 0.12 | 0.2 |
| 9 | 4038257 | rs601370 | -0.873 | -0.651 | -0.405 | 0.448 | -0.033 | 0.088 | 0.088 | 0 | 0.29 | 0.53 | 0.38 | 0.51 | 0.22 | 0.32 |
| 9 | 4038998 | rs2380931 | 0.296 | -0.6 | -0.42 | 0.431 | 0.003 | 0.103 | 0.103 | 0 | 0.08 | 0.09 | 0.27 | 0.41 | 0.12 | 0.2 |
| 9 | 4039027 | rs614782 | -1.279 | -0.493 | -1.019 | 0.009 | -0.017 | -0.013 | -0.013 | 1.167 | 0.81 | 0.93 | 0.83 | 0.81 | 0.78 | 0.8 |
| 9 | 4039637 | rs4741875 | 1.71 | -0.954 | -1.04 | -0.107 | -0.082 | 0.065 | 0.065 | 0 | 0.33 | 0.07 | 0.2 | 0.26 | 0.38 | 0.4 |
| 9 | 4039727 | rs501461 | -0.307 | -0.898 | -0.489 | -0.09 | -0.007 | 0.016 | 0.016 | 0 | 0.66 | 0.78 | 0.6 | 0.8 | 0.61 | 0.68 |
| 9 | 4040641 | rs632562 | -0.375 | -0.929 | -0.371 | 0.248 | -0.03 | 0.066 | 0.066 | 0 | 0.28 | 0.5 | 0.37 | 0.55 | 0.22 | 0.31 |
| 9 | 4043590 | rs676472 | -0.028 | -1.022 | -0.366 | -0.029 | -0.083 | -0.08 | -0.08 | 0 | 0.24 | 0.49 | 0.13 | 0.14 | 0.11 | 0.1 |
| 9 | 4046569 | rs1536468 | -0.94 | -0.738 | -1.391 | 0.229 | 0.057 | -0.067 | -0.067 | 0.063 | 0.48 | 0.9 | 0.57 | 0.46 | 0.37 | 0.4 |
| 9 | 4046886 | rs4741877 | 1.436 | -0.549 | -1.75 | 0.18 | -0.132 | 0.033 | 0.033 | 0 | 0.33 | 0.04 | 0.25 | 0.26 | 0.41 | 0.41 |
| 9 | 4047947 | rs10733504 | -0.836 | -0.28 | -1.28 | 0.408 | -0.039 | -0.065 | -0.065 | 0 | 0.81 | 0.97 | 0.84 | 0.74 | 0.78 | 0.81 |
| 9 | 4048636 | rs6476798 | -0.325 | -0.283 | -1.001 | 0.39 | -0.01 | -0.019 | -0.019 | 0 | 0.76 | 0.81 | 0.79 | 0.7 | 0.73 | 0.77 |
| 9 | 4050630 | rs10814822 | -0.346 | -0.866 | -1.584 | 0.309 | -0.027 | -0.134 | -0.134 | 0.159 | 0.25 | 0.01 | 0.18 | 0.35 | 0.21 | 0.27 |
| 9 | 4051387 | rs11791472 | -0.271 | -0.895 | -1.226 | 0.097 | 0.008 | -0.039 | -0.039 | 0 | 0.32 | 0.18 | 0.28 | 0.37 | 0.33 | 0.25 |
| 9 | 4051921 | rs6476800 | -0.224 | -0.441 | -0.645 | -0.2 | 0.026 | -0.072 | -0.072 | 0.589 | 0.72 | 0.51 | 0.66 | 0.92 | 0.66 | 0.78 |
| 9 | 4055471 | rs7863393 | -0.449 | -0.397 | -0.531 | -0.271 | 0.027 | -0.098 | -0.098 | 0 | 0.74 | 0.65 | 0.81 | 0.96 | 0.69 | 0.77 |
| 9 | 4058403 | rs10974307 | 1.72 | -0.644 | -1.132 | 0.103 | -0.089 | 0.005 | 0.005 | 0 | 0.32 | 0.03 | 0.16 | 0.31 | 0.32 | 0.46 |
| 9 | 4059114 | rs7864487 | 0.051 | -0.691 | -0.733 | -0.013 | 0.011 | -0.093 | -0.093 | 0 | 0.68 | 0.52 | 0.75 | 0.94 | 0.66 | 0.71 |
| 9 | 4064137 | rs7035962 | 0.601 | -0.733 | -1.005 | 0.125 | -0.016 | -0.002 | -0.002 | 0 | 0.61 | 0.11 | 0.52 | 0.66 | 0.65 | 0.63 |
| 9 | 4065171 | rs10758550 | 0.476 | -0.705 | -1.007 | 0.101 | -0.015 | 0.016 | 0.016 | 0.06 | 0.62 | 0.12 | 0.52 | 0.69 | 0.66 | 0.69 |
| 9 | 4068002 | rs10758551 | 0.935 | -0.841 | -0.551 | -0.015 | -0.002 | 0.003 | 0.003 | 0 | 0.39 | 0.61 | 0.41 | 0.58 | 0.33 | 0.48 |
| 9 | 4069657 | rs2027394 | -0.213 | -0.972 | -0.396 | 0.233 | 0.022 | -0.018 | -0.018 | 1.509 | 0.36 | 0.23 | 0.39 | 0.38 | 0.35 | 0.24 |
| 9 | 4071952 | rs7870442 | -0.773 | -0.953 | -0.606 | 0.174 | 0.064 | 0.066 | 0.066 | 0 | 0.45 | 0.79 | 0.67 | 0.64 | 0.39 | 0.34 |
| 9 | 4078864 | rs10814830 | 0.547 | -0.83 | -1.122 | 0.104 | -0.01 | -0.11 | -0.11 | 0.081 | 0.23 | 0.05 | 0.3 | 0.38 | 0.24 | 0.21 |
| 9 | 4079470 | rs4741887 | -0.964 | -0.794 | -0.987 | -0.433 | 0.006 | 0.03 | 0.03 | 0 | 0.52 | 0.66 | 0.7 | 0.66 | 0.47 | 0.43 |
| 9 | 4080699 | rs3925024 | 0.289 | -0.816 | -0.757 | -0.39 | -0.011 | 0.026 | 0.026 | 0 | 0.27 | 0.45 | 0.38 | 0.3 | 0.22 | 0.21 |
| 9 | 4080755 | rs7028492 | 0.908 | -0.362 | -1.06 | -0.478 | 0.009 | -0.037 | -0.037 | 0 | 0.1 | 0.12 | 0.11 | 0 | 0.13 | 0.08 |
| 9 | 4080780 | rs7875452 | 2.149 | -0.45 | -1.355 | -0.542 | -0.101 | -0.053 | -0.053 | 0 | 0.28 | 0.02 | 0.13 | 0.31 | 0.29 | 0.43 |
| 9 | 4083297 | rs7030223 | 0.552 | -0.447 | -0.952 | -0.422 | 0.018 | -0.072 | -0.072 | 0 | 0.29 | 0.13 | 0.36 | 0.37 | 0.27 | 0.24 |
| 9 | 4083587 | rs7027277 | 0.605 | -0.449 | -0.957 | -0.405 | 0.013 | -0.078 | -0.078 | 0 | 0.29 | 0.11 | 0.36 | 0.37 | 0.27 | 0.24 |
| 9 | 4083665 | rs6476804 | -0.354 | -0.287 | -0.602 | -0.553 | 0.001 | 0 | 0 | 0 | 0.4 | 0.37 | 0.47 | 0.37 | 0.4 | 0.32 |
| 9 | 4084390 | rs10814832 | -0.336 | -0.189 | -0.775 | -0.536 | 0.001 | 0 | 0 | 0 | 0.39 | 0.3 | 0.47 | 0.35 | 0.39 | 0.32 |
| 9 | 4086370 | rs10814834 | -0.388 | -0.061 | -0.928 | -0.61 | -0.009 | 0.016 | 0.016 | 0 | 0.42 | 0.24 | 0.48 | 0.33 | 0.42 | 0.33 |
| 9 | 4086553 | rs1556987 | 1.033 | -0.241 | -1.252 | -0.505 | -0.001 | -0.092 | -0.092 | 0.05 | 0.25 | 0.04 | 0.32 | 0.32 | 0.24 | 0.23 |
| 9 | 4088131 | rs2380935 | -1.614 | 0.279 | -1.012 | -0.543 | 0.017 | 0.029 | 0.029 | 0 | 0.65 | 0.84 | 0.82 | 0.69 | 0.66 | 0.51 |
| 9 | 4088648 | rs7849831 | -0.614 | 0.293 | -0.974 | -0.473 | -0.004 | -0.007 | -0.007 | 0.009 | 0.31 | 0.74 | 0.44 | 0.34 | 0.3 | 0.21 |
| 9 | 4088934 | rs10814835 | -0.746 | 0.411 | -1.055 | -0.834 | 0.004 | 0.008 | 0.008 | 0 | 0.5 | 0.47 | 0.52 | 0.35 | 0.52 | 0.33 |
| 9 | 4089046 | rs10974326 | -0.191 | 0.382 | -1.598 | -0.557 | -0.011 | 0.003 | 0.003 | 0 | 0.39 | 0.2 | 0.41 | 0.33 | 0.39 | 0.3 |
| 9 | 4090601 | rs10814837 | 0.676 | 0.124 | -1.581 | -0.718 | 0.064 | -0.059 | -0.059 | 0 | 0.37 | 0.14 | 0.36 | 0.47 | 0.29 | 0.31 |
| 9 | 4090724 | rs17273930 | 0.099 | 0.562 | -1.693 | -1.032 | -0.027 | 0.056 | 0.056 | 0 | 0.23 | 0.14 | 0.18 | 0.06 | 0.28 | 0.14 |
| 9 | 4091519 | rs2755087 | -1.045 | 0.206 | -1.305 | -0.824 | 0.041 | -0.035 | -0.035 | 0 | 0.61 | 0.75 | 0.56 | 0.51 | 0.69 | 0.67 |
| 9 | 4093295 | rs484781 | -0.014 | 0.389 | -0.994 | -1.161 | -0.002 | -0.086 | -0.086 | 0 | 0.25 | 0.49 | 0.18 | 0.07 | 0.27 | 0.15 |
| 9 | 4094202 | rs646827 | 0.123 | 0.369 | -1.035 | -1.171 | -0.003 | -0.101 | -0.101 | 0.176 | 0.25 | 0.48 | 0.18 | 0.07 | 0.27 | 0.16 |
| 9 | 4094225 | rs565114 | -1.019 | 0.486 | -1.093 | -1.226 | -0.03 | -0.035 | -0.035 | 0 | 0.72 | 0.84 | 0.72 | 0.56 | 0.68 | 0.46 |
| 9 | 4094321 | rs564275 | -0.068 | 0.429 | -1.134 | -1.23 | -0.005 | -0.003 | -0.003 | 1.804 | 0.62 | 0.6 | 0.61 | 0.55 | 0.57 | 0.44 |
| 9 | 4096451 | rs806034 | 0.115 | 0.308 | -0.75 | -1.172 | 0.043 | -0.049 | -0.049 | 0 | 0.1 | 0.39 | 0.09 | 0 | 0.11 | 0.05 |
| 9 | 4097131 | rs4740750 | 1.323 | 0.348 | -0.988 | -1.097 | -0.003 | 0.015 | 0.015 | 0.276 | 0.14 | 0.06 | 0.09 | 0.08 | 0.16 | 0.11 |
| 9 | 4097344 | rs806032 | -0.721 | 0.38 | -0.99 | -1.067 | -0.002 | -0.002 | -0.002 | 0 | 0.7 | 0.72 | 0.68 | 0.55 | 0.65 | 0.46 |
| 9 | 4099510 | rs487732 | -0.742 | 0.466 | -1.023 | -1.134 | -0.028 | -0.032 | -0.032 | 0 | 0.7 | 0.82 | 0.7 | 0.56 | 0.66 | 0.46 |
| 9 | 4099988 | rs483774 | -0.68 | 0.245 | -0.616 | -1.095 | 0.044 | -0.13 | -0.13 | 0 | 0.17 | 0.46 | 0.18 | 0 | 0.21 | 0.06 |
| 9 | 4100236 | rs806031 | -0.164 | 0.355 | -1.029 | -1.058 | 0.027 | -0.001 | -0.001 | 0.155 | 0.49 | 0.66 | 0.39 | 0.49 | 0.39 | 0.31 |
| 9 | 4102339 | rs10974334 | 1.205 | 0.416 | -0.7 | -1.022 | -0.001 | -0.013 | -0.013 | 0 | 0.81 | 0.22 | 0.73 | 0.95 | 0.77 | 0.94 |
| 9 | 4103200 | rs10814843 | 0.323 | 0.448 | -0.513 | -1.035 | 0.015 | -0.05 | -0.05 | 0 | 0.87 | 0.42 | 0.82 | 0.96 | 0.88 | 0.95 |
| 9 | 4103703 | rs10739033 | 1.138 | 0.405 | -0.548 | -1.046 | 0.019 | -0.068 | -0.068 | 0 | 0.83 | 0.36 | 0.75 | 0.96 | 0.8 | 0.94 |
| 9 | 4103815 | rs10974337 | 0.537 | 0.617 | -0.853 | -0.875 | -0.003 | -0.009 | -0.009 | 0.075 | 0.24 | 0.18 | 0.28 | 0.28 | 0.26 | 0.22 |
| 9 | 4104139 | rs4741891 | 0.753 | 0.601 | -0.92 | -0.871 | -0.018 | -0.044 | -0.044 | 0 | 0.23 | 0.07 | 0.27 | 0.27 | 0.25 | 0.22 |
| 9 | 4106866 | rs2064986 | -0.561 | -0.007 | -1.105 | -1.164 | -0.038 | -0.084 | -0.084 | 0 | 0.77 | 0.41 | 0.8 | 0.96 | 0.86 | 0.9 |
| 9 | 4109656 | rs12345723 | 0.752 | 0.182 | -1.352 | -1.051 | -0.003 | -0.011 | -0.011 | 0.763 | 0.23 | 0.18 | 0.27 | 0.27 | 0.25 | 0.22 |
| 9 | 4109906 | rs10814846 | 0.597 | 0.201 | -1.633 | -0.932 | -0.112 | 0.016 | 0.016 | 0 | 0.47 | 0.06 | 0.44 | 0.67 | 0.49 | 0.65 |
| 9 | 4117602 | rs537484 | -0.627 | 0.174 | -1.293 | -1.089 | 0.026 | 0.128 | 0.128 | 0 | 0.49 | 0.78 | 0.5 | 0.25 | 0.46 | 0.34 |

| | | | | | | | | | | | | | | | | |
|---|---|---|---|---|---|---|---|---|---|---|---|---|---|---|---|---|
| 9 | 4118111 | rs6415788 | -0.196 | 0.377 | -0.822 | -1.175 | 0.002 | -0.042 | -0.042 | 0 | 0.6 | 0.6 | 0.76 | 0.87 | 0.63 | 0.56 |
| 9 | 4119049 | rs13301186 | 0.102 | 0.195 | -0.692 | -1.247 | -0.001 | 0.012 | 0.012 | 0 | 0.31 | 0.27 | 0.31 | 0.21 | 0.33 | 0.29 |
| 9 | 4121399 | rs806050 | 0.047 | -0.046 | -0.409 | -1.206 | -0.002 | 0.017 | 0.017 | 0 | 0.53 | 0.42 | 0.7 | 0.83 | 0.57 | 0.51 |
| 9 | 4122302 | rs499357 | 1.088 | 0.143 | -0.292 | -1.033 | 0 | -0.014 | -0.014 | 0 | 0.3 | 0.27 | 0.47 | 0.62 | 0.33 | 0.28 |
| 9 | 4124377 | rs7872188 | 0.465 | 0.162 | -0.418 | -1.167 | 0.039 | -0.042 | -0.042 | 1.262 | 0.29 | 0.39 | 0.24 | 0.16 | 0.37 | 0.37 |
| 9 | 4131657 | rs1340252 | 0.838 | -0.041 | -0.391 | -1.051 | -0.006 | 0.01 | 0.01 | 0 | 0.32 | 0.39 | 0.41 | 0.6 | 0.32 | 0.29 |
| 9 | 4136715 | rs611698 | -0.537 | -0.016 | -0.66 | -1.096 | 0.045 | 0.062 | 0.062 | 1.111 | 0.63 | 0.86 | 0.74 | 0.79 | 0.71 | 0.68 |
| 9 | 4143590 | rs2791757 | 0.365 | -0.103 | -0.584 | -1.084 | 0.036 | -0.035 | -0.035 | 0 | 0.31 | 0.41 | 0.32 | 0.19 | 0.4 | 0.41 |
| 9 | 4143674 | rs2755084 | -1.241 | -0.143 | -0.584 | -0.989 | -0.01 | 0.031 | 0.031 | 0 | 0.7 | 0.56 | 0.59 | 0.4 | 0.69 | 0.73 |
| 9 | 4145545 | rs13299873 | -0.12 | -0.668 | -0.565 | -0.797 | 0.001 | 0.013 | 0.013 | 0.057 | 0.3 | 0.18 | 0.33 | 0.24 | 0.3 | 0.29 |
| 9 | 4145587 | rs13299289 | 0.121 | -0.722 | -0.607 | -0.791 | 0.005 | 0.02 | 0.02 | 0.115 | 0.28 | 0.14 | 0.25 | 0.21 | 0.27 | 0.27 |
| 9 | 4146903 | rs6476808 | -0.197 | -0.562 | -0.45 | -0.699 | 0.011 | 0.016 | 0.016 | 0 | 0.38 | 0.31 | 0.35 | 0.24 | 0.31 | 0.29 |
| 9 | 4149475 | rs982820 | -0.486 | -0.565 | 0.142 | -0.404 | -0.035 | -0.038 | -0.038 | 0 | 0.27 | 0.39 | 0.24 | 0.11 | 0.18 | 0.23 |
| 9 | 4150202 | rs10125159 | -0.372 | -0.607 | 0.545 | -0.363 | -0.052 | -0.043 | -0.043 | 0.516 | 0.26 | 0.56 | 0.26 | 0.11 | 0.16 | 0.2 |
| 9 | 4156206 | rs927320 | 0.042 | -0.366 | 0.631 | 0.248 | -0.021 | 0.054 | 0.054 | 0 | 0.21 | 0.39 | 0.35 | 0.31 | 0.18 | 0.18 |
| 9 | 4157226 | rs7026219 | -0.712 | -0.152 | 0.157 | -0.245 | 0.001 | -0.007 | -0.007 | 0 | 0.46 | 0.54 | 0.48 | 0.28 | 0.47 | 0.51 |
| 9 | 4159313 | rs10758559 | 0.453 | -0.183 | 0.066 | -0.47 | -0.026 | 0.121 | 0.121 | 0 | 0.55 | 0.23 | 0.63 | 0.82 | 0.56 | 0.57 |
| 9 | 4160364 | rs1983753 | -0.378 | -0.193 | 0.242 | -0.489 | -0.035 | -0.14 | -0.14 | 0 | 0.8 | 0.52 | 0.88 | 0.99 | 0.86 | 0.93 |
| 9 | 4162147 | rs4741897 | 0.174 | -0.142 | 0.035 | -0.395 | -0.01 | 0.041 | 0.041 | 0 | 0.62 | 0.3 | 0.7 | 0.82 | 0.65 | 0.58 |
| 9 | 4163904 | rs7847753 | 0.27 | -0.057 | -0.157 | -0.317 | -0.005 | 0.003 | 0.003 | 0.051 | 0.25 | 0.25 | 0.26 | 0.16 | 0.33 | 0.4 |
| 9 | 4165934 | rs10758561 | 0.782 | -0.399 | -0.559 | -0.31 | -0.01 | 0.095 | 0.095 | 0.234 | 0.49 | 0.1 | 0.51 | 0.69 | 0.46 | 0.45 |
| 9 | 4169816 | rs4740753 | 0.879 | -0.301 | -0.468 | -0.031 | -0.05 | 0.006 | 0.006 | 2.013 | 0.57 | 0.09 | 0.52 | 0.64 | 0.47 | 0.46 |
| 9 | 4170113 | rs7853833 | -0.386 | -0.26 | 0.761 | -0.061 | 0.039 | 0.009 | 0.009 | 0.341 | 0.33 | 0.48 | 0.38 | 0.34 | 0.42 | 0.53 |
| 9 | 4170979 | rs7865812 | 0.955 | -0.452 | 0.878 | -0.087 | -0.024 | 0.019 | 0.019 | 0 | 0.55 | 0.23 | 0.56 | 0.65 | 0.49 | 0.43 |
| 9 | 4171953 | rs10739036 | 0.351 | -0.346 | 1.058 | -0.077 | 0.021 | -0.03 | -0.03 | 0 | 0.62 | 0.51 | 0.65 | 0.78 | 0.53 | 0.51 |
| 9 | 4172087 | rs10974362 | -0.042 | -0.472 | 1.094 | -0.273 | 0.009 | 0.046 | 0.046 | 0.009 | 0.07 | 0.15 | 0.11 | 0.15 | 0.07 | 0.11 |
| 9 | 4172899 | rs927316 | -0.645 | -0.846 | 0.641 | -0.649 | 0.004 | 0.005 | 0.005 | 0 | 0.42 | 0.59 | 0.29 | 0.24 | 0.33 | 0.27 |
| 9 | 4175223 | rs7044458 | -0.84 | -0.833 | 0.82 | -0.704 | 0 | -0.036 | -0.036 | 0.176 | 0.15 | 0.41 | 0.14 | 0.09 | 0.15 | 0.08 |
| 9 | 4175324 | rs10814856 | 1.161 | -0.529 | 0.566 | -0.081 | 0.004 | -0.017 | -0.017 | 1.136 | 0.33 | 0.23 | 0.4 | 0.4 | 0.3 | 0.45 |
| 9 | 4176178 | rs4741904 | -1.227 | -0.495 | 0.292 | 0.096 | -0.03 | -0.02 | -0.02 | 0 | 0.78 | 0.87 | 0.71 | 0.75 | 0.71 | 0.72 |
| 9 | 4177109 | rs11790620 | 0.169 | -0.678 | 0.315 | -0.09 | -0.006 | -0.024 | -0.024 | 0 | 0.29 | 0.31 | 0.17 | 0.13 | 0.27 | 0.1 |
| 9 | 4177749 | rs714867 | -1.256 | -0.56 | -0.005 | 0.133 | -0.058 | -0.131 | -0.131 | 0.66 | 0.64 | 0.93 | 0.57 | 0.59 | 0.65 | 0.47 |
| 9 | 4178016 | rs2380941 | -0.935 | -0.55 | 0.607 | 0.178 | 0.011 | 0.069 | 0.069 | 0 | 0.14 | 0.32 | 0.15 | 0.29 | 0.17 | 0.1 |
| 9 | 4179609 | rs7045544 | 1.25 | -0.601 | 0.059 | 0.109 | -0.013 | -0.067 | -0.067 | 0 | 0.3 | 0.11 | 0.42 | 0.37 | 0.28 | 0.47 |
| 9 | 4180887 | rs10814859 | 0.714 | -0.795 | -0.012 | -0.011 | -0.013 | -0.091 | -0.091 | 0 | 0.36 | 0.06 | 0.43 | 0.37 | 0.31 | 0.56 |
| 9 | 4183103 | rs7040304 | -0.207 | -0.752 | 0.301 | -0.169 | 0.009 | -0.004 | -0.004 | 0 | 0.64 | 0.55 | 0.64 | 0.73 | 0.58 | 0.67 |
| 9 | 4183453 | rs10119355 | 1.138 | -0.926 | 0.382 | -0.263 | -0.036 | -0.034 | -0.034 | 0.473 | 0.23 | 0.19 | 0.28 | 0.35 | 0.34 | 0.28 |
| 9 | 4184508 | rs10814865 | 1.074 | -0.98 | 0.614 | -0.216 | 0.008 | 0.106 | 0.106 | 0 | 0.07 | 0.17 | 0.1 | 0.24 | 0.08 | 0.1 |
| 9 | 4185934 | rs10123353 | -0.312 | -1.025 | 0.529 | -0.322 | 0.007 | 0.006 | 0.006 | 0 | 0.55 | 0.64 | 0.49 | 0.64 | 0.6 | 0.43 |
| 9 | 4193350 | rs10814871 | 1.063 | -1.026 | 0.376 | -0.322 | -0.003 | -0.046 | -0.046 | 0 | 0.32 | 0.06 | 0.44 | 0.36 | 0.3 | 0.52 |
| 9 | 4193531 | rs4741910 | 0.964 | -1.033 | 0.471 | -0.32 | 0.01 | -0.017 | -0.017 | 0.09 | 0.33 | 0.14 | 0.44 | 0.35 | 0.3 | 0.51 |
| 9 | 4194762 | rs10814874 | 0.076 | -0.827 | 0.71 | -0.247 | -0.003 | -0.003 | -0.003 | 0 | 0.46 | 0.46 | 0.52 | 0.35 | 0.4 | 0.57 |
| 9 | 4198077 | rs7341838 | -0.125 | -0.73 | 0.712 | -0.271 | 0.009 | -0.055 | -0.055 | 0 | 0.1 | 0.29 | 0.07 | 0 | 0.1 | 0.02 |
| 9 | 4200150 | rs4741914 | 0.166 | -0.737 | 0.258 | -0.022 | -0.001 | -0.003 | -0.003 | 0 | 0.44 | 0.56 | 0.5 | 0.3 | 0.39 | 0.52 |
| 9 | 4203214 | rs10974390 | 0.977 | -0.358 | 0.554 | 0.197 | -0.003 | 0.065 | 0.065 | 0 | 0.06 | 0.08 | 0.09 | 0.26 | 0.08 | 0.1 |
| 9 | 4203442 | rs2093322 | 0.724 | -0.371 | 0.609 | -0.01 | -0.009 | -0.08 | -0.08 | 0.339 | 0.88 | 0.49 | 0.92 | 1 | 0.91 | 0.99 |
| 9 | 4205498 | rs2068911 | 0.628 | -0.136 | 0.151 | 0.068 | -0.007 | -0.007 | -0.007 | 0 | 0.26 | 0.28 | 0.29 | 0.37 | 0.34 | 0.28 |
| 9 | 4206388 | rs7048915 | -0.107 | -0.647 | 0.447 | -0.371 | -0.029 | -0.168 | -0.168 | 0.059 | 0.69 | 0.51 | 0.83 | 0.97 | 0.75 | 0.94 |
| 9 | 4206671 | rs6476823 | 0.066 | -1.11 | -0.127 | -0.473 | 0.016 | 0.015 | 0.015 | 0 | 0.32 | 0.18 | 0.21 | 0.29 | 0.26 | 0.16 |
| 9 | 4206769 | rs6476824 | 0.122 | -0.714 | -0.16 | -0.355 | -0.021 | 0.001 | 0.001 | 0 | 0.61 | 0.22 | 0.64 | 0.65 | 0.56 | 0.7 |
| 9 | 4207526 | rs7864894 | -0.711 | -0.83 | -0.297 | -0.352 | -0.021 | -0.015 | -0.015 | 0 | 0.57 | 0.86 | 0.47 | 0.56 | 0.6 | 0.45 |
| 9 | 4207680 | rs7850913 | -0.618 | -0.872 | 0.261 | -0.373 | 0.001 | 0.002 | 0.002 | 0 | 0.54 | 0.52 | 0.44 | 0.56 | 0.6 | 0.45 |
| 9 | 4208429 | rs10974395 | 0.388 | -0.823 | 0.242 | -0.376 | -0.052 | -0.005 | -0.005 | 0 | 0.31 | 0.2 | 0.28 | 0.25 | 0.43 | 0.29 |
| 9 | 4209053 | rs7030766 | 0.42 | -0.891 | 0.575 | -0.319 | -0.051 | 0.006 | 0.006 | 0 | 0.27 | 0.47 | 0.2 | 0.33 | 0.21 | 0.16 |
| 9 | 4211589 | rs12337783 | 0.078 | -0.853 | 0.321 | -0.415 | -0.003 | -0.078 | -0.078 | 0 | 0.12 | 0.31 | 0.08 | 0 | 0.12 | 0.01 |
| 9 | 4212601 | rs12554508 | 2.159 | -0.873 | -0.131 | -0.415 | -0.059 | -0.102 | -0.102 | 0 | 0.27 | 0.02 | 0.44 | 0.33 | 0.24 | 0.49 |
| 9 | 4214311 | rs13292114 | 0.959 | -0.736 | 0.111 | -0.367 | 0 | 0.007 | 0.007 | 0 | 0.1 | 0.09 | 0.05 | 0.09 | 0.1 | 0.03 |
| 9 | 4215542 | rs12006019 | 1.098 | -0.858 | 0.085 | -0.414 | -0.008 | -0.036 | -0.036 | 0 | 0.2 | 0.26 | 0.1 | 0.03 | 0.13 | 0.06 |
| 9 | 4217028 | rs736893 | 1.384 | -0.701 | -0.343 | -0.252 | -0.101 | -0.084 | -0.084 | 0.763 | 0.21 | 0.04 | 0.21 | 0.26 | 0.33 | 0.28 |
| 9 | 4218178 | rs10758578 | 0.8 | -0.802 | -0.075 | -0.348 | -0.004 | 0.013 | 0.013 | 0 | 0.42 | 0.31 | 0.53 | 0.34 | 0.42 | 0.52 |
| 9 | 4218631 | rs10121596 | 0.201 | -0.991 | 0.011 | -0.345 | -0.044 | -0.057 | -0.057 | 0 | 0.28 | 0.47 | 0.16 | 0.13 | 0.18 | 0.08 |
| 9 | 4219259 | rs732957 | 1.671 | -1.035 | -0.293 | -0.244 | 0.005 | -0.017 | -0.017 | 0 | 0.07 | 0.04 | 0.08 | 0.25 | 0.07 | 0.09 |
| 9 | 4221298 | rs7873463 | 0.209 | -0.988 | -0.186 | -0.329 | 0.02 | 0.006 | 0.006 | 0 | 0.64 | 0.28 | 0.73 | 0.62 | 0.72 | 0.83 |
| 9 | 4226630 | rs10814891 | 0.083 | -0.63 | -0.586 | -0.316 | -0.025 | 0.008 | 0.008 | 0 | 0.34 | 0.22 | 0.27 | 0.25 | 0.45 | 0.26 |
| 9 | 4228550 | rs2890551 | 0.505 | -0.365 | -0.818 | -0.076 | -0.001 | 0 | 0 | 1.587 | 0.4 | 0.5 | 0.53 | 0.43 | 0.32 | 0.59 |
| 9 | 4232871 | rs10491551 | 0.466 | -0.094 | -1.328 | -0.396 | 0.002 | 0.001 | 0.001 | 0.018 | 0.11 | 0.09 | 0.06 | 0.07 | 0.08 | 0.02 |
| 9 | 4234586 | rs10814895 | 0.748 | -0.044 | -1.932 | -0.398 | -0.069 | -0.017 | -0.017 | 0 | 0.29 | 0.06 | 0.2 | 0.23 | 0.33 | 0.27 |
| 9 | 4237041 | rs10974406 | 0.712 | -0.136 | -1.649 | -0.071 | -0.032 | -0.068 | -0.068 | 1.417 | 0.36 | 0.06 | 0.54 | 0.65 | 0.32 | 0.62 |
| 9 | 4239545 | rs11999791 | -0.518 | 0.378 | -0.747 | 0.306 | 0.006 | 0.017 | 0.017 | 0 | 0.44 | 0.7 | 0.46 | 0.32 | 0.53 | 0.37 |

| | | | | | | | | | | | | | | | | |
|---|---|---|---|---|---|---|---|---|---|---|---|---|---|---|---|---|
| 9 | 4240743 | rs10120959 | -0.75 | 0.363 | -0.359 | 0.389 | 0.031 | -0.002 | -0.002 | 0.228 | 0.48 | 0.66 | 0.54 | 0.41 | 0.58 | 0.4 |
| 9 | 4243162 | rs911490 | 1.803 | 1.026 | 0.893 | 0.38 | 0.076 | -0.04 | -0.04 | 0 | 0.88 | 0.42 | 0.85 | 0.95 | 0.82 | 0.94 |
| 9 | 4257339 | rs6476830 | 1.214 | 0.914 | 1.164 | 0.291 | 0.079 | -0.048 | -0.048 | 0.109 | 0.89 | 0.62 | 0.89 | 0.98 | 0.84 | 0.94 |
| 9 | 4258586 | rs10122830 | -1.127 | 0.922 | 1.13 | 0.29 | 0.025 | -0.035 | -0.035 | 0.451 | 0.11 | 0.18 | 0.1 | 0.02 | 0.16 | 0.06 |
| 9 | 4261496 | rs12378556 | 1.035 | 0.967 | 0.271 | 0.468 | 0.069 | 0.008 | 0.008 | 0 | 0.39 | 0.09 | 0.22 | 0.41 | 0.26 | 0.49 |
| 9 | 4262516 | rs12380197 | 0.251 | 1.404 | 0.332 | 0.635 | -0.001 | 0 | 0 | 0 | 0.17 | 0.16 | 0.24 | 0.25 | 0.2 | 0.2 |
| 9 | 4265363 | rs7037716 | -1.201 | 1.371 | 0.673 | 0.578 | 0.009 | 0.029 | 0.029 | 0 | 0.53 | 0.68 | 0.7 | 0.57 | 0.59 | 0.47 |
| 9 | 4265865 | rs10974423 | 1.708 | 1.325 | 0.317 | 0.571 | 0.053 | 0.001 | 0.001 | 0 | 0.37 | 0.09 | 0.21 | 0.41 | 0.25 | 0.47 |
| 9 | 4267787 | rs7045959 | 0.009 | 1.378 | 0.556 | 0.642 | 0.002 | 0.011 | 0.011 | 0.026 | 0.34 | 0.38 | 0.45 | 0.32 | 0.39 | 0.26 |
| 9 | 4269407 | rs10511458 | 0.698 | 1.564 | 0.602 | 0.348 | 0.035 | -0.019 | -0.019 | 0 | 0.73 | 0.62 | 0.67 | 0.73 | 0.64 | 0.73 |
| 9 | 4270669 | rs10814906 | 0.066 | 1.608 | 0.453 | 0.349 | 0.001 | 0 | 0 | 0 | 0.18 | 0.17 | 0.23 | 0.25 | 0.2 | 0.21 |
| 9 | 4270682 | rs10758589 | 0.571 | 1.489 | 0.459 | 0.367 | 0.026 | -0.014 | -0.014 | 0 | 0.73 | 0.67 | 0.68 | 0.73 | 0.64 | 0.73 |
| 9 | 4272041 | rs4740762 | 0.597 | 1.377 | 0.407 | 0.394 | 0.035 | -0.02 | -0.02 | 0 | 0.73 | 0.58 | 0.68 | 0.73 | 0.65 | 0.72 |
| 9 | 4277721 | rs10758590 | 0.375 | 0.725 | -0.019 | 0.722 | 0.003 | 0.005 | 0.005 | 0.041 | 0.62 | 0.25 | 0.5 | 0.74 | 0.62 | 0.7 |
| 9 | 4283597 | rs3892341 | 0.068 | 0.559 | 0.349 | 0.825 | 0.035 | -0.017 | -0.017 | 0.189 | 0.07 | 0.2 | 0.05 | 0.01 | 0.08 | 0.05 |
| 9 | 4284369 | rs10974436 | -1.332 | 0.462 | 0.657 | 0.693 | 0.02 | -0.081 | -0.081 | 0.818 | 0.14 | 0.44 | 0.11 | 0.02 | 0.12 | 0.06 |
| 9 | 4289050 | rs7034200 | 0.536 | 0.622 | 0.276 | 0.939 | 0 | 0 | 0 | 0.43 | 0.59 | 0.65 | 0.53 | 0.47 | 0.52 | 0.57 |
| 9 | 4289747 | rs2380950 | 0.969 | 0.55 | 0.107 | 0.864 | 0.006 | -0.048 | -0.048 | 0 | 0.88 | 0.59 | 0.9 | 0.95 | 0.91 | 0.92 |
| 9 | 4290541 | rs10116772 | 1.297 | 0.599 | 0.073 | 0.914 | 0 | 0.004 | 0.004 | 0 | 0.51 | 0.52 | 0.49 | 0.44 | 0.45 | 0.52 |
| 9 | 4291268 | rs6476842 | -0.387 | 0.749 | -0.767 | 1.045 | -0.048 | -0.061 | -0.061 | 0 | 0.36 | 0.09 | 0.35 | 0.41 | 0.45 | 0.39 |
| 9 | 4291928 | rs10974438 | 1.746 | 0.752 | -1.043 | 1.138 | -0.056 | 0.018 | 0.018 | 0 | 0.42 | 0.09 | 0.26 | 0.4 | 0.38 | 0.47 |
| 9 | 4292012 | rs7043265 | 0.251 | 0.992 | -0.751 | 1.03 | -0.067 | -0.072 | -0.072 | 0 | 0.14 | 0.02 | 0.21 | 0.28 | 0.21 | 0.18 |
| 9 | 4295583 | rs10758594 | -0.747 | 0.632 | -0.491 | 0.839 | -0.043 | -0.022 | -0.022 | 0 | 0.43 | 0.2 | 0.39 | 0.42 | 0.55 | 0.44 |
| 9 | 4300862 | rs4741942 | 1.31 | 0.159 | 0.495 | 0.608 | -0.014 | -0.018 | -0.018 | 0.268 | 0.88 | 0.37 | 0.79 | 0.95 | 0.91 | 0.95 |
| 9 | 4302988 | rs7861665 | 1.211 | 0.147 | 0.484 | 0.618 | -0.021 | -0.003 | -0.003 | 0.001 | 0.89 | 0.38 | 0.79 | 0.95 | 0.92 | 0.95 |
| 9 | 4303682 | rs7018983 | 1.026 | -0.082 | -0.105 | 0.58 | -0.003 | -0.002 | -0.002 | 0 | 0.65 | 0.36 | 0.72 | 0.77 | 0.73 | 0.79 |
| 9 | 4304915 | rs4740766 | -1.48 | 0.183 | 0.324 | 0.711 | -0.002 | -0.002 | -0.002 | 0.213 | 0.44 | 0.44 | 0.6 | 0.39 | 0.41 | 0.44 |
| 9 | 4305320 | rs16921094 | -0.526 | 0.206 | 0.632 | 0.714 | 0.022 | -0.033 | -0.033 | 0 | 0.08 | 0.39 | 0.07 | 0.01 | 0.06 | 0.01 |
| 9 | 4305959 | rs10814919 | 0.066 | 0.212 | 0.277 | 0.715 | -0.013 | -0.006 | -0.006 | 0.046 | 0.85 | 0.92 | 0.63 | 0.79 | 0.8 | 0.76 |
| 9 | 4307572 | rs10814921 | 1.851 | 0.189 | 0.181 | 0.668 | -0.104 | -0.066 | -0.066 | 0.196 | 0.47 | 0.11 | 0.31 | 0.56 | 0.5 | 0.54 |
| 9 | 4307589 | rs12377521 | 0.371 | 0.303 | 0.193 | 0.535 | 0.003 | -0.031 | -0.031 | 0 | 0.1 | 0.01 | 0.15 | 0.17 | 0.06 | 0.15 |
| 9 | 4307733 | rs10739044 | -0.349 | 0.303 | 0.229 | 0.539 | 0.004 | -0.002 | -0.002 | 0.141 | 0.9 | 0.94 | 0.87 | 0.83 | 0.94 | 0.86 |
| 9 | 4308010 | rs10974448 | -0.044 | 0.293 | 0.539 | 0.532 | 0.003 | 0.004 | 0.004 | 0 | 0.25 | 0.2 | 0.1 | 0.13 | 0.2 | 0.17 |
| 9 | 4309006 | rs4567095 | -1.541 | 0.415 | 1.163 | 0.619 | 0.025 | -0.036 | -0.036 | 0.988 | 0.23 | 0.48 | 0.43 | 0.2 | 0.27 | 0.25 |
| 9 | 4310558 | rs12336107 | -1.912 | 0.385 | 1.186 | 0.572 | 0.03 | -0.017 | -0.017 | 0 | 0.18 | 0.67 | 0.46 | 0.1 | 0.27 | 0.15 |
| 9 | 4311777 | rs6476846 | -0.345 | 0.311 | 1.273 | 0.435 | 0.002 | -0.008 | -0.008 | 1.025 | 0.88 | 0.86 | 0.75 | 0.97 | 0.82 | 0.89 |
| 9 | 4311864 | rs10974450 | 1.824 | 0.214 | 1.326 | 0.525 | 0.001 | 0.016 | 0.016 | 0 | 0.68 | 0.3 | 0.5 | 0.81 | 0.68 | 0.71 |
| 9 | 4311892 | rs6476847 | 1.047 | 0.277 | 1.575 | 0.583 | 0.034 | -0.099 | -0.099 | 0 | 0.8 | 0.57 | 0.68 | 0.97 | 0.74 | 0.86 |
| 9 | 4314752 | rs7038304 | -1.07 | 0.09 | 0.953 | 0.926 | 0.013 | -0.007 | -0.007 | 0.223 | 0.43 | 0.53 | 0.42 | 0.44 | 0.34 | 0.35 |
| 9 | 4316336 | rs10739046 | -1.194 | 0.004 | 0.698 | 1.2 | 0.087 | 0.007 | 0.007 | 0 | 0.46 | 0.79 | 0.47 | 0.54 | 0.34 | 0.38 |
| 9 | 4316407 | rs10733510 | -0.697 | 0.059 | 0.677 | 1.098 | -0.003 | -0.001 | -0.001 | 0 | 0.88 | 0.88 | 0.78 | 0.97 | 0.82 | 0.89 |
| 9 | 4316859 | rs10814929 | -1.135 | 0.048 | 0.711 | 1.156 | 0.035 | -0.016 | -0.016 | 0 | 0.44 | 0.6 | 0.43 | 0.44 | 0.34 | 0.35 |
| 9 | 4317128 | rs10974453 | -0.721 | -0.003 | 0.492 | 1.279 | 0.009 | 0 | 0 | 0.01 | 0.38 | 0.32 | 0.26 | 0.35 | 0.28 | 0.32 |
| 9 | 4317825 | rs960485 | 1.041 | 0.143 | -0.962 | 1.15 | -0.102 | -0.083 | -0.083 | 0 | 0.45 | 0.08 | 0.33 | 0.57 | 0.49 | 0.46 |
| 9 | 4319915 | rs10814931 | -0.697 | -0.078 | 0.262 | 1.502 | -0.023 | 0.039 | 0.039 | 0 | 0.24 | 0.33 | 0.21 | 0.31 | 0.18 | 0.26 |
| 9 | 4321558 | rs11789633 | -0.842 | 0.476 | 0.097 | 1.676 | -0.007 | -0.009 | -0.009 | 0 | 0.09 | 0.07 | 0.14 | 0.23 | 0.16 | 0.17 |
| 9 | 4323880 | rs10814934 | -1.778 | 0.152 | 0.811 | 2.001 | -0.003 | 0.125 | 0.125 | 0.257 | 0.16 | 0.28 | 0.25 | 0.36 | 0.18 | 0.19 |
| 9 | 4324137 | rs10814935 | -1.929 | -0.098 | 1.699 | 1.83 | -0.037 | 0.088 | 0.088 | 3.195 | 0.1 | 0.45 | 0.12 | 0.12 | 0.02 | 0.03 |
| 9 | 4326425 | rs10758606 | -2.334 | -0.011 | 1.641 | 1.841 | -0.012 | 0.049 | 0.049 | 2.27 | 0.19 | 0.57 | 0.27 | 0.35 | 0.19 | 0.19 |
| 9 | 4327048 | rs10122246 | -0.808 | 0.242 | 1.757 | 1.857 | 0.027 | 0.035 | 0.035 | 0 | 0.1 | 0.16 | 0.16 | 0.24 | 0.19 | 0.16 |
| 9 | 4327105 | rs10435804 | -0.596 | 0.444 | 1.337 | 2.548 | -0.012 | -0.01 | -0.01 | 0.207 | 0.05 | 0.02 | 0.06 | 0.18 | 0.05 | 0.08 |
| 9 | 4327938 | rs12351393 | -2.062 | 0.452 | 1.703 | 2.458 | 0.006 | 0.094 | 0.094 | 1.877 | 0.18 | 0.53 | 0.28 | 0.36 | 0.2 | 0.2 |
| 9 | 4328940 | rs10974470 | -1.921 | 0.141 | 1.348 | 2.084 | -0.022 | 0.037 | 0.037 | 0.308 | 0.24 | 0.55 | 0.26 | 0.29 | 0.2 | 0.13 |
| 9 | 4329033 | rs10974471 | -1.366 | 0.148 | 1.492 | 2.052 | -0.022 | -0.01 | -0.01 | 0 | 0.17 | 0.54 | 0.2 | 0.15 | 0.15 | 0.05 |
| 9 | 4329209 | rs3895473 | 0.214 | 0.087 | 0.142 | 1.955 | 0 | 0.001 | 0.001 | 0.138 | 0.23 | 0.19 | 0.31 | 0.36 | 0.22 | 0.39 |
| 9 | 4329354 | rs10814941 | 1.682 | 0.046 | 0.383 | 2.047 | -0.011 | 0.036 | 0.036 | 0.168 | 0.78 | 0.36 | 0.73 | 0.69 | 0.81 | 0.88 |
| 9 | 4330843 | rs10814942 | -0.215 | 0.052 | -0.423 | 2.039 | -0.008 | -0.053 | -0.053 | 0 | 0.17 | 0.04 | 0.32 | 0.26 | 0.16 | 0.15 |
| 9 | 4331717 | rs12378782 | 1.408 | -0.149 | -0.344 | 2.19 | 0.03 | -0.073 | -0.073 | 0 | 0.72 | 0.23 | 0.55 | 0.54 | 0.81 | 0.75 |
| 9 | 4331760 | rs13286735 | -0.334 | 0.048 | -0.483 | 1.827 | -0.001 | -0.055 | -0.055 | 0 | 0.17 | 0.04 | 0.32 | 0.26 | 0.16 | 0.15 |
| 9 | 4332667 | rs4305953 | -0.686 | -0.003 | -0.639 | 1.728 | -0.003 | -0.065 | -0.065 | 0.051 | 0.23 | 0.04 | 0.36 | 0.29 | 0.21 | 0.17 |
| 9 | 4334606 | rs4741961 | 1.305 | -0.365 | 0.217 | 1.977 | -0.026 | 0.058 | 0.058 | 0 | 0.82 | 0.38 | 0.86 | 0.66 | 0.83 | 0.77 |
| 9 | 4335668 | rs4131424 | -0.865 | -0.2 | 0.636 | 1.995 | 0.001 | -0.003 | -0.003 | 0.014 | 0.13 | 0.21 | 0.09 | 0.13 | 0.15 | 0.13 |
| 9 | 4338987 | rs10974480 | -0.838 | -0.047 | -0.155 | 1.936 | -0.029 | -0.011 | -0.011 | 0 | 0.34 | 0.08 | 0.34 | 0.23 | 0.35 | 0.26 |
| 9 | 4340081 | rs10814943 | -1.393 | -0.091 | 0.72 | 1.88 | 0.001 | 0.003 | 0.003 | 0 | 0.45 | 0.6 | 0.45 | 0.56 | 0.47 | 0.49 |
| 9 | 4345431 | rs12376666 | -1.453 | -0.29 | 1.157 | 1.928 | -0.025 | 0.024 | 0.024 | 0 | 0.23 | 0.38 | 0.16 | 0.35 | 0.2 | 0.29 |
| 9 | 4345557 | rs13295703 | 0.309 | -0.052 | 0.616 | 2.086 | -0.031 | -0.038 | -0.038 | 0 | 0.06 | 0 | 0.09 | 0.17 | 0.13 | 0.06 |
| 9 | 4347606 | rs10974484 | 1.656 | -0.212 | 0.727 | 1.888 | 0.024 | 0.017 | 0.017 | 0 | 0.47 | 0.24 | 0.51 | 0.44 | 0.54 | 0.44 |
| 9 | 4350760 | rs10974487 | -0.137 | 0.125 | 0.457 | 2.039 | -0.037 | -0.086 | -0.086 | 0 | 0.26 | 0.06 | 0.34 | 0.29 | 0.26 | 0.31 |
| 9 | 4352222 | rs10974489 | -0.14 | 0.131 | 0.46 | 1.911 | -0.037 | -0.058 | -0.058 | 0.108 | 0.26 | 0.06 | 0.34 | 0.26 | 0.26 | 0.31 |

| | | | | | | | | | | | | | | | | |
|---|---|---|---|---|---|---|---|---|---|---|---|---|---|---|---|---|
| 9 | 4356048 | rs10814947 | 1.244 | 0.28 | 0.894 | 1.645 | 0.029 | 0.011 | 0.011 | 0 | 0.5 | 0.34 | 0.51 | 0.5 | 0.56 | 0.47 |
| 9 | 4357693 | rs12005536 | -0.671 | -0.231 | 1.088 | 1.538 | 0.028 | 0.049 | 0.049 | 0.131 | 0.25 | 0.19 | 0.1 | 0.13 | 0.18 | 0.16 |
| 9 | 4358255 | rs4237154 | 0.511 | -0.166 | 1.396 | 1.631 | 0.082 | 0.096 | 0.096 | 0 | 0.95 | 0.7 | 0.96 | 0.97 | 0.96 | 0.9 |
| 9 | 4359689 | rs17741020 | 0.445 | 0.074 | 0.94 | 1.745 | -0.063 | -0.102 | -0.102 | 0 | 0.2 | 0.07 | 0.37 | 0.37 | 0.31 | 0.27 |
| 9 | 4360058 | rs6476853 | 1.575 | -0.083 | 1.123 | 1.647 | -0.003 | 0.003 | 0.003 | 0 | 0.47 | 0.38 | 0.48 | 0.47 | 0.47 | 0.47 |
| 9 | 4362167 | rs4284081 | 0.332 | 0.589 | 0.885 | 1.507 | -0.022 | -0.016 | -0.016 | 0 | 0.72 | 0.74 | 0.43 | 0.49 | 0.53 | 0.6 |
| 9 | 4362624 | rs10758611 | 1.091 | 0.653 | 1.119 | 1.211 | 0.102 | 0.014 | 0.014 | 0 | 0.93 | 0.68 | 0.91 | 0.98 | 0.87 | 0.88 |
| 9 | 4364108 | rs10733512 | 1.569 | 0.589 | 1.179 | 1.322 | 0.006 | 0.01 | 0.01 | 0 | 0.47 | 0.37 | 0.32 | 0.42 | 0.36 | 0.46 |
| 9 | 4364438 | rs4304371 | -0.262 | 0.705 | 1.038 | 1.382 | -0.048 | -0.064 | -0.064 | 0 | 0.78 | 0.86 | 0.46 | 0.57 | 0.57 | 0.69 |
| 9 | 4368838 | rs10814948 | 1.335 | 0.012 | 0.391 | 1.179 | 0.008 | 0.012 | 0.012 | 0 | 0.46 | 0.37 | 0.28 | 0.36 | 0.31 | 0.45 |
| 9 | 4371339 | rs17717668 | -0.21 | -0.139 | 0.204 | 0.988 | -0.004 | 0.03 | 0.03 | 0.235 | 0.21 | 0.01 | 0.05 | 0.14 | 0.17 | 0.16 |
| 9 | 4372189 | rs6476856 | 0.755 | 0.118 | 0.358 | 1.231 | -0.002 | -0.005 | -0.005 | 0 | 0.44 | 0.41 | 0.29 | 0.35 | 0.32 | 0.43 |
| 9 | 4374278 | rs4741967 | 0.106 | 0.367 | -0.527 | 1.496 | -0.137 | -0.128 | -0.128 | 0 | 0.37 | 0.05 | 0.58 | 0.46 | 0.51 | 0.35 |
| 9 | 4378727 | rs6476858 | -0.573 | 0.059 | 0.224 | 1.086 | 0.01 | 0.028 | 0.028 | 0 | 0.6 | 0.73 | 0.74 | 0.67 | 0.7 | 0.58 |
| 9 | 4380096 | rs3931525 | -1.923 | 0.026 | 1.273 | 1.852 | 0.027 | 0.223 | 0.223 | 0.044 | 0.1 | 0.32 | 0.18 | 0.24 | 0.07 | 0.13 |
| 9 | 4383219 | rs4474078 | 0.614 | -0.089 | 1.916 | 1.686 | -0.003 | 0.132 | 0.132 | 0.331 | 0.88 | 0.78 | 0.84 | 0.66 | 0.9 | 0.82 |
| 9 | 4383842 | rs10739055 | 1.671 | -0.24 | 1.789 | 1.511 | -0.016 | 0.103 | 0.103 | 0.682 | 0.77 | 0.45 | 0.78 | 0.64 | 0.83 | 0.71 |
| 9 | 4384289 | rs10739056 | 1.575 | -0.259 | 1.631 | 1.465 | -0.014 | 0.054 | 0.054 | 0 | 0.74 | 0.41 | 0.77 | 0.64 | 0.82 | 0.7 |
| 9 | 4392649 | rs4317631 | 1.316 | -0.071 | 1.291 | 1.628 | -0.008 | 0.167 | 0.167 | 1.028 | 0.79 | 0.63 | 0.8 | 0.64 | 0.85 | 0.74 |
| 9 | 4394771 | rs10511457 | -1.19 | -0.057 | 0.897 | 1.607 | -0.008 | 0.144 | 0.144 | 0.747 | 0.2 | 0.34 | 0.21 | 0.36 | 0.15 | 0.26 |
| 9 | 4400991 | rs4401921 | -1.228 | -0.364 | 0.636 | 1.24 | -0.039 | 0.055 | 0.055 | 5.322 | 0.28 | 0.46 | 0.25 | 0.34 | 0.2 | 0.23 |
| 9 | 4410854 | rs4237157 | 0.02 | -0.591 | 0.649 | 1.281 | 0.022 | 0.049 | 0.049 | 0 | 0.57 | 0.63 | 0.75 | 0.77 | 0.65 | 0.8 |
| 9 | 4411383 | rs7851353 | 0.042 | -0.356 | -0.068 | 0.915 | 0.003 | 0.126 | 0.126 | 0 | 0.49 | 0.73 | 0.68 | 0.68 | 0.52 | 0.58 |
| 9 | 4412949 | rs10122237 | 0.871 | 0.215 | 0.44 | 0.733 | -0.002 | 0 | 0 | 0 | 0.34 | 0.52 | 0.52 | 0.6 | 0.33 | 0.42 |
| 9 | 4413449 | rs10758621 | -0.029 | 0.083 | 0.336 | 0.716 | -0.008 | 0.107 | 0.107 | 0 | 0.48 | 0.71 | 0.67 | 0.66 | 0.51 | 0.58 |
| 9 | 4415824 | rs13295153 | 0.054 | -0.303 | 0.49 | 0.795 | 0.048 | 0.001 | 0.001 | 0 | 0.51 | 0.33 | 0.62 | 0.49 | 0.64 | 0.73 |
| 9 | 4417219 | rs4740779 | -1.139 | -0.623 | 0.14 | 0.65 | 0.016 | 0.062 | 0.062 | 0 | 0.65 | 0.79 | 0.77 | 0.78 | 0.69 | 0.82 |
| 9 | 4417406 | rs10974525 | 0.081 | -0.551 | -0.186 | 0.733 | 0.105 | -0.033 | -0.033 | 0.087 | 0.51 | 0.15 | 0.61 | 0.5 | 0.65 | 0.72 |
| 9 | 4419154 | rs10814969 | -0.253 | -0.12 | -0.239 | -0.029 | 0 | -0.008 | -0.008 | 0.29 | 0.51 | 0.44 | 0.38 | 0.23 | 0.54 | 0.53 |
| 9 | 4420767 | rs7039988 | 0.838 | -0.184 | -0.22 | -0.451 | -0.02 | -0.101 | -0.101 | 0.206 | 0.25 | 0.29 | 0.14 | 0.01 | 0.19 | 0.08 |
| 9 | 4420971 | rs13293594 | 0.856 | 0.18 | -0.837 | -0.037 | -0.035 | -0.021 | -0.021 | 0 | 0.14 | 0.04 | 0.18 | 0.13 | 0.15 | 0.29 |
| 9 | 4423789 | rs10974528 | 0.892 | 0.101 | -1.245 | -0.078 | -0.001 | 0.074 | 0.074 | 0 | 0.23 | 0.1 | 0.21 | 0.08 | 0.24 | 0.09 |
| 9 | 4425033 | rs10974530 | 0.522 | 0.469 | -0.453 | 0.137 | 0.04 | -0.007 | -0.007 | 0 | 0.15 | 0.25 | 0.22 | 0.12 | 0.18 | 0.25 |
| 9 | 4426991 | rs16921295 | -0.024 | 0.231 | -0.458 | 0.117 | -0.026 | -0.015 | -0.015 | 0.912 | 0.68 | 0.49 | 0.6 | 0.26 | 0.74 | 0.66 |
| 9 | 4428615 | rs10974533 | -0.045 | 0.109 | -0.244 | 0.127 | 0.011 | -0.044 | -0.044 | 0 | 0.18 | 0.34 | 0.19 | 0.11 | 0.13 | 0.24 |
| 9 | 4429166 | rs4562386 | -0.381 | 0.097 | -0.121 | 0.106 | 0.014 | -0.049 | -0.049 | 0 | 0.2 | 0.42 | 0.21 | 0.12 | 0.15 | 0.24 |
| 9 | 4429762 | rs10814970 | -0.227 | 0.196 | -0.645 | 0.24 | 0.008 | -0.01 | -0.01 | 0 | 0.22 | 0.3 | 0.29 | 0.19 | 0.18 | 0.25 |
| 9 | 4430975 | rs17753881 | 1.53 | -0.537 | -1.136 | 0.34 | 0.013 | 0.077 | 0.077 | 0.123 | 0.28 | 0.08 | 0.11 | 0.14 | 0.26 | 0.15 |
| 9 | 4434601 | rs7847689 | 0.902 | -0.583 | -1.087 | 0.712 | 0.03 | -0.048 | -0.048 | 0 | 0.33 | 0.18 | 0.25 | 0.43 | 0.3 | 0.23 |
| 9 | 4438240 | rs2225360 | -1.04 | -0.786 | -0.868 | 0.621 | 0.006 | 0.025 | 0.025 | 0.018 | 0.79 | 0.65 | 0.79 | 0.7 | 0.76 | 0.81 |
| 9 | 4438264 | rs10814971 | 1.737 | -0.727 | -1.356 | 0.5 | -0.001 | 0.055 | 0.055 | 0 | 0.27 | 0.13 | 0.1 | 0.13 | 0.24 | 0.15 |
| 9 | 4438435 | rs7874211 | 0.92 | -0.701 | -1.622 | 0.36 | -0.014 | 0.14 | 0.14 | 0.348 | 0.38 | 0.12 | 0.17 | 0.14 | 0.35 | 0.21 |
| 9 | 4441054 | rs1590929 | 0.329 | -0.824 | -1.563 | 0.432 | 0.005 | -0.008 | -0.008 | 0 | 0.46 | 0.3 | 0.4 | 0.51 | 0.38 | 0.48 |
| 9 | 4441460 | rs4741989 | 1.174 | -0.793 | -1.584 | 0.46 | -0.051 | -0.028 | -0.028 | 0.432 | 0.14 | 0.02 | 0.17 | 0.14 | 0.12 | 0.19 |
| 9 | 4442884 | rs10974540 | 2.011 | -0.797 | -1.573 | 0.451 | -0.011 | 0.07 | 0.07 | 0 | 0.26 | 0.09 | 0.1 | 0.13 | 0.24 | 0.15 |
| 9 | 4443086 | rs4625083 | 0.816 | -0.785 | -1.557 | 0.456 | -0.017 | 0.068 | 0.068 | 0 | 0.41 | 0.26 | 0.28 | 0.27 | 0.36 | 0.34 |
| 9 | 4446307 | rs7855993 | 1.26 | -0.728 | -1.528 | 0.492 | -0.017 | -0.002 | -0.002 | 1.903 | 0.14 | 0.09 | 0.17 | 0.13 | 0.12 | 0.18 |
| 9 | 4447405 | rs10814973 | 1.742 | -0.713 | -1.473 | 0.52 | 0.002 | 0.041 | 0.041 | 0 | 0.27 | 0.15 | 0.1 | 0.17 | 0.24 | 0.17 |
| 9 | 4455794 | rs10974547 | -0.163 | -0.918 | -0.026 | 0.702 | 0.023 | 0.021 | 0.021 | 0 | 0.64 | 0.56 | 0.6 | 0.59 | 0.56 | 0.57 |
| 9 | 4456373 | rs10814978 | 0.004 | -0.969 | -0.125 | 0.681 | 0.019 | 0.016 | 0.016 | 0.012 | 0.6 | 0.36 | 0.6 | 0.63 | 0.59 | 0.72 |
| 9 | 4457091 | rs4333659 | -0.665 | -0.221 | 0.337 | 0.849 | 0.006 | 0.054 | 0.054 | 0 | 0.79 | 0.59 | 0.79 | 0.71 | 0.78 | 0.82 |
| 9 | 4458297 | rs7046512 | -0.219 | 0.208 | -0.389 | 0.054 | -0.011 | 0.002 | 0.002 | 0 | 0.18 | 0.17 | 0.21 | 0.13 | 0.22 | 0.29 |
| 9 | 4458374 | rs13297783 | 0.333 | 0.385 | -0.526 | 0.041 | -0.06 | -0.005 | -0.005 | 0 | 0.15 | 0.04 | 0.19 | 0.1 | 0.22 | 0.24 |
| 9 | 4460786 | rs6476867 | 1.134 | -0.644 | -0.331 | 0.214 | -0.041 | 0.066 | 0.066 | 0 | 0.46 | 0.22 | 0.39 | 0.36 | 0.44 | 0.47 |
| 9 | 4461536 | rs7048574 | 0.034 | -0.683 | -0.173 | 0.202 | -0.003 | 0.013 | 0.013 | 0.192 | 0.63 | 0.52 | 0.58 | 0.48 | 0.57 | 0.71 |
| 9 | 4462265 | rs4741996 | 1.225 | -0.635 | -0.269 | -0.286 | -0.069 | 0.125 | 0.125 | 0 | 0.45 | 0.15 | 0.4 | 0.28 | 0.44 | 0.39 |
| 9 | 4463421 | rs10974552 | 0.197 | -0.517 | -0.113 | -0.153 | -0.006 | -0.004 | -0.004 | 0 | 0.62 | 0.54 | 0.59 | 0.4 | 0.57 | 0.63 |
| 9 | 4476238 | rs954864 | 0.545 | -0.19 | -0.788 | -0.203 | -0.003 | -0.002 | -0.002 | 0 | 0.15 | 0.1 | 0.19 | 0.15 | 0.2 | 0.17 |
| 9 | 4479464 | rs10814983 | 0.366 | -0.313 | -0.661 | -0.144 | 0.017 | -0.01 | -0.01 | 0 | 0.62 | 0.26 | 0.54 | 0.43 | 0.65 | 0.57 |
| 9 | 4481668 | rs4740788 | 0.099 | 0.028 | -0.077 | -0.186 | 0.017 | -0.003 | -0.003 | 0 | 0.13 | 0.26 | 0.2 | 0.1 | 0.12 | 0.24 |
| 9 | 4482706 | rs10491734 | -0.759 | -0.071 | -1.003 | -0.734 | -0.111 | 0.125 | 0.125 | 0 | 0.58 | 0.9 | 0.68 | 0.75 | 0.56 | 0.66 |
| 9 | 4483189 | rs10491733 | -0.3 | -0.181 | -0.49 | -0.73 | 0.003 | 0.027 | 0.027 | 0.007 | 0.44 | 0.57 | 0.58 | 0.73 | 0.49 | 0.55 |
| 9 | 4487024 | rs2150192 | 0.885 | 0.732 | -0.413 | -0.146 | 0.019 | 0.027 | 0.027 | 0 | 0.51 | 0.43 | 0.3 | 0.32 | 0.44 | 0.34 |
| 9 | 4487374 | rs2183472 | 0.709 | 1.269 | -0.432 | -0.112 | -0.088 | 0.058 | 0.058 | 0 | 0.84 | 0.97 | 0.77 | 0.89 | 0.78 | 0.73 |
| 9 | 4489494 | rs1360329 | 0.594 | 1.392 | -0.251 | -0.035 | 0.043 | -0.039 | -0.039 | 0 | 0.84 | 0.8 | 0.77 | 0.9 | 0.78 | 0.8 |
| 9 | 4499361 | rs7021409 | -0.576 | -0.309 | -0.431 | 0.164 | -0.021 | -0.005 | -0.005 | 0 | 0.33 | 0.43 | 0.15 | 0.37 | 0.24 | 0.45 |
| 9 | 4501492 | rs7023050 | -0.436 | 0.077 | -0.021 | -0.125 | -0.005 | 0.013 | 0.013 | 0 | 0.72 | 0.79 | 0.56 | 0.82 | 0.69 | 0.76 |
| 9 | 4505544 | rs10814995 | 0.01 | 0.505 | -0.477 | 0.214 | 0.025 | 0.009 | 0.009 | 0 | 0.28 | 0.3 | 0.43 | 0.27 | 0.32 | 0.16 |
| 9 | 4507513 | rs16921385 | 0.121 | 0.14 | -1.539 | -0.476 | -0.056 | 0.091 | 0.091 | 0 | 0.33 | 0.06 | 0.16 | 0.21 | 0.36 | 0.43 |

| | | | | | | | | | | | | | | | | |
|---|---|---|---|---|---|---|---|---|---|---|---|---|---|---|---|---|
| 9 | 4509590 | rs10974584 | 0.646 | 0.095 | -1.253 | -0.339 | -0.014 | 0.002 | 0.002 | 0 | 0.27 | 0.15 | 0.38 | 0.2 | 0.32 | 0.16 |
| 9 | 4511788 | rs2150195 | -0.755 | -0.876 | -1.717 | -0.766 | 0.001 | -0.057 | -0.057 | 0.237 | 0.68 | 0.79 | 0.62 | 0.47 | 0.66 | 0.61 |
| 9 | 4512200 | rs7041093 | -0.684 | -0.655 | -1.869 | -0.756 | 0.022 | -0.047 | -0.047 | 0 | 0.7 | 0.8 | 0.65 | 0.55 | 0.73 | 0.64 |
| 9 | 4515470 | rs10974587 | 1.024 | -0.651 | -1.723 | -0.828 | -0.013 | 0.007 | 0.007 | 0 | 0.29 | 0.07 | 0.16 | 0.28 | 0.29 | 0.41 |
| 9 | 4515982 | rs12351328 | -0.748 | -0.633 | -0.893 | -0.783 | 0.037 | 0.034 | 0.034 | 5.767 | 0.05 | 0.33 | 0.04 | 0 | 0 | 0 |
| 9 | 4516768 | rs17755777 | 1.484 | -0.519 | -1.068 | -0.761 | -0.063 | -0.031 | -0.031 | 0 | 0.2 | 0.04 | 0.34 | 0.2 | 0.26 | 0.15 |
| 9 | 4526525 | rs7864309 | 0.515 | -0.307 | -1.134 | -1.064 | -0.049 | 0.029 | 0.029 | 0 | 0.32 | 0.14 | 0.42 | 0.19 | 0.37 | 0.18 |
| 9 | 4526642 | rs4742004 | 0.726 | -0.672 | -1.476 | -1.179 | -0.081 | -0.008 | -0.008 | 0.364 | 0.33 | 0.04 | 0.21 | 0.29 | 0.36 | 0.41 |
| 9 | 4526796 | rs10815004 | 0.491 | -0.717 | -1.19 | -1.138 | -0.005 | 0.004 | 0.004 | 0 | 0.36 | 0.34 | 0.47 | 0.2 | 0.44 | 0.21 |
| 9 | 4527201 | rs6476874 | -1.068 | -1.026 | -1.205 | -0.962 | -0.038 | 0.091 | 0.091 | 0 | 0.79 | 0.67 | 0.72 | 0.5 | 0.84 | 0.67 |
| 9 | 4527624 | rs17756150 | 1.995 | -0.575 | -1.382 | -0.775 | 0.017 | 0.028 | 0.028 | 0 | 0.06 | 0 | 0.02 | 0 | 0.05 | 0.05 |
| 9 | 4529671 | rs6476875 | 1.132 | -0.143 | -1.439 | -0.804 | -0.099 | -0.012 | -0.012 | 0 | 0.26 | 0.05 | 0.4 | 0.2 | 0.34 | 0.16 |
| 9 | 4539377 | rs7848533 | -0.437 | -0.306 | -1.265 | -1.049 | 0.002 | -0.001 | -0.001 | 0.079 | 0.53 | 0.41 | 0.54 | 0.43 | 0.49 | 0.53 |
| 9 | 4541368 | rs10974611 | 1.015 | 0.192 | -1.627 | -0.661 | -0.015 | -0.075 | -0.075 | 0 | 0.15 | 0 | 0.1 | 0.25 | 0.11 | 0.32 |
| 9 | 4545973 | rs16921457 | 0.938 | 0.842 | -1.616 | -0.38 | -0.04 | -0.064 | -0.064 | 0 | 0.12 | 0.02 | 0.11 | 0.23 | 0.13 | 0.28 |
| 9 | 4545998 | rs10815016 | 0.541 | 0.642 | -0.923 | -0.552 | 0.003 | -0.062 | -0.062 | 0 | 0.12 | 0.22 | 0.05 | 0 | 0.1 | 0.09 |
| 9 | 4546594 | rs10815017 | 0.721 | 0.697 | -1.25 | -0.012 | -0.029 | 0.002 | 0.002 | 0 | 0.23 | 0.12 | 0.37 | 0.18 | 0.28 | 0.13 |
| 9 | 4550752 | rs10739065 | -0.602 | -0.439 | -0.952 | 0.082 | 0.014 | -0.018 | -0.018 | 0 | 0.39 | 0.37 | 0.59 | 0.59 | 0.32 | 0.41 |
| 9 | 4555305 | rs7856675 | 0.012 | -0.007 | -0.787 | -0.706 | 0.006 | -0.035 | -0.035 | 0 | 0.08 | 0.25 | 0.06 | 0 | 0.07 | 0.05 |
| 9 | 4555573 | rs3780415 | 0.375 | -0.102 | -1.873 | -0.761 | -0.02 | 0.229 | 0.229 | 0.016 | 0.48 | 0.09 | 0.5 | 0.84 | 0.6 | 0.68 |
| 9 | 4556448 | rs10974619 | -0.212 | -0.01 | -1.028 | -0.546 | 0.001 | 0.071 | 0.071 | 0 | 0.36 | 0.23 | 0.38 | 0.07 | 0.33 | 0.33 |
| 9 | 4559775 | rs4742007 | -0.691 | 0.814 | -0.005 | 1.051 | 0.013 | 0.059 | 0.059 | 0 | 0.14 | 0.19 | 0.3 | 0.46 | 0.28 | 0.19 |
| 9 | 4561374 | rs10974624 | -0.616 | 0.814 | -0.26 | 1.051 | -0.025 | -0.025 | -0.025 | 0 | 0.14 | 0.11 | 0.3 | 0.47 | 0.28 | 0.19 |
| 9 | 4564432 | rs2228622 | 0.733 | 0.568 | -0.118 | 0.873 | 0.039 | 0.149 | 0.149 | 0 | 0.49 | 0.19 | 0.46 | 0.24 | 0.41 | 0.43 |
| 9 | 4566210 | rs7022772 | -0.945 | 0.482 | -0.319 | 0.884 | -0.076 | -0.055 | -0.055 | 0.358 | 0.73 | 0.98 | 0.84 | 0.73 | 0.74 | 0.66 |
| 9 | 4566863 | rs11792309 | 0.77 | 0.378 | -0.184 | 0.751 | 0.035 | 0.159 | 0.159 | 0 | 0.5 | 0.17 | 0.46 | 0.24 | 0.41 | 0.46 |
| 9 | 4571728 | rs7864496 | 0.706 | 0.358 | -0.123 | 0.867 | 0.013 | 0.109 | 0.109 | 0 | 0.52 | 0.22 | 0.46 | 0.26 | 0.42 | 0.46 |
| 9 | 4572480 | rs3780412 | 0.536 | 0.525 | -0.15 | 0.933 | 0.001 | 0.03 | 0.03 | 0 | 0.55 | 0.58 | 0.5 | 0.26 | 0.43 | 0.46 |
| 9 | 4577346 | rs6476879 | 0.688 | 0.14 | -0.108 | 0.373 | 0.006 | 0.033 | 0.033 | 0 | 0.51 | 0.43 | 0.64 | 0.64 | 0.57 | 0.35 |
| 9 | 4577932 | rs188537 | 0.105 | 0.158 | -0.854 | 0.157 | -0.003 | 0.037 | 0.037 | 0 | 0.68 | 0.75 | 0.83 | 0.85 | 0.72 | 0.61 |
| 9 | 4598379 | rs2771 | 0.79 | -0.57 | -0.313 | 0.987 | -0.004 | 0 | 0 | 0 | 0.68 | 0.36 | 0.58 | 0.28 | 0.74 | 0.54 |
| 9 | 4600804 | rs3739642 | -0.763 | -0.599 | 0.341 | 1.977 | -0.007 | 0.029 | 0.029 | 0 | 0.13 | 0.12 | 0.2 | 0.34 | 0.07 | 0.2 |
| 9 | 4602356 | rs3780407 | -0.345 | -0.597 | 0.297 | 2.061 | 0.004 | -0.006 | -0.006 | 0.179 | 0.07 | 0.03 | 0.1 | 0.32 | 0.04 | 0.14 |
| 9 | 4604807 | rs301453 | 0.151 | -0.774 | 1.559 | 2.332 | 0.007 | 0.062 | 0.062 | 0 | 0.9 | 0.81 | 0.86 | 0.74 | 0.86 | 0.84 |
| 9 | 4611901 | rs4740791 | -2.078 | -0.853 | 2.145 | 2.258 | -0.04 | 0.02 | 0.02 | 0 | 0.17 | 0.44 | 0.21 | 0.19 | 0.07 | 0.2 |
| 9 | 4615020 | rs3824441 | -0.677 | -0.785 | 1.894 | 2.326 | -0.004 | 0.052 | 0.052 | 0 | 0.07 | 0.13 | 0.11 | 0.26 | 0.05 | 0.14 |
| 9 | 4622453 | rs10974657 | -0.598 | -0.807 | 1.839 | 2.308 | -0.001 | 0.023 | 0.023 | 0 | 0.07 | 0.08 | 0.1 | 0.26 | 0.05 | 0.14 |
| 9 | 4629595 | rs4742027 | -1.347 | -1.025 | 2.027 | 2.385 | -0.017 | 0.002 | 0.002 | 0 | 0.1 | 0.21 | 0.19 | 0.06 | 0.05 | 0.1 |
| 9 | 4630675 | rs10815039 | -2.547 | -1.089 | 2.487 | 2.737 | -0.038 | 0.159 | 0.159 | 0 | 0.2 | 0.43 | 0.33 | 0.41 | 0.11 | 0.23 |
| 9 | 4630762 | rs10974672 | -1.012 | -1.076 | 2.266 | 2.717 | -0.008 | 0 | 0 | 0 | 0.09 | 0.1 | 0.18 | 0.06 | 0.05 | 0.1 |
| 9 | 4633229 | rs10815041 | -2.461 | -1.095 | 2.395 | 2.735 | -0.046 | 0.147 | 0.147 | 0.107 | 0.21 | 0.52 | 0.34 | 0.41 | 0.11 | 0.23 |
| 9 | 4633488 | rs10974682 | -1.281 | -1.021 | 2.179 | 2.738 | -0.003 | 0.165 | 0.165 | 0 | 0.09 | 0.22 | 0.14 | 0.34 | 0.06 | 0.13 |
| 9 | 4635232 | rs16921669 | -1.454 | -1.02 | 2.099 | 2.738 | 0.002 | 0.217 | 0.217 | 0 | 0.1 | 0.31 | 0.14 | 0.34 | 0.06 | 0.13 |
| 9 | 4640212 | rs4742035 | -2.104 | -1.298 | 1.135 | 2.499 | -0.012 | 0.021 | 0.021 | 0.092 | 0.24 | 0.25 | 0.26 | 0.41 | 0.14 | 0.24 |
| 9 | 4640712 | rs10815044 | -1.756 | -1.316 | 1.215 | 2.007 | -0.027 | -0.005 | -0.005 | 0 | 0.18 | 0.24 | 0.15 | 0.14 | 0.08 | 0.11 |
| 9 | 4644477 | rs10815051 | 0.506 | -1.265 | -0.084 | 1.592 | 0.006 | 0.007 | 0.007 | 0 | 0.58 | 0.17 | 0.54 | 0.37 | 0.58 | 0.52 |
| 9 | 4645935 | rs301468 | -1.078 | -1.667 | -0.849 | 1.056 | -0.128 | 0.017 | 0.017 | 0 | 0.61 | 0.94 | 0.57 | 0.71 | 0.52 | 0.54 |
| 9 | 4651478 | rs1385456 | -0.162 | -2.006 | 0.01 | 1.113 | -0.002 | 0.001 | 0.001 | 0 | 0.39 | 0.43 | 0.37 | 0.58 | 0.42 | 0.48 |
| 9 | 4654195 | rs392979 | 0.702 | -1.999 | 0.28 | 1.021 | -0.013 | -0.02 | -0.02 | 0.382 | 0.15 | 0.18 | 0.1 | 0.06 | 0.09 | 0.03 |
| 9 | 4657040 | rs2146423 | -0.135 | -1.877 | -0.01 | 1.139 | -0.005 | -0.005 | -0.005 | 0 | 0.4 | 0.45 | 0.36 | 0.64 | 0.42 | 0.48 |
| 9 | 4658449 | rs1385452 | 0.367 | -1.931 | -0.443 | 1.383 | 0.004 | 0.011 | 0.011 | 0 | 0.27 | 0.3 | 0.25 | 0.36 | 0.36 | 0.35 |
| 9 | 4659111 | rs7047933 | -0.851 | -1.288 | -1.139 | 0.854 | -0.07 | 0.096 | 0.096 | 0 | 0.59 | 0.87 | 0.55 | 0.78 | 0.51 | 0.55 |
| 9 | 4661853 | rs301489 | -0.617 | -1.34 | -1.655 | 0.747 | -0.024 | 0.051 | 0.051 | 0.222 | 0.55 | 0.88 | 0.54 | 0.66 | 0.46 | 0.56 |
| 9 | 4662065 | rs301488 | -0.634 | -1.556 | -0.716 | 1.062 | -0.008 | 0.13 | 0.13 | 0 | 0.19 | 0.36 | 0.14 | 0.45 | 0.18 | 0.3 |
| 9 | 4662369 | rs301487 | -0.85 | -0.285 | -1.777 | 0.23 | -0.08 | 0.055 | 0.055 | 2.684 | 0.61 | 0.9 | 0.52 | 0.71 | 0.42 | 0.77 |
| 9 | 4664749 | rs1485202 | -0.122 | -0.419 | -0.788 | 0.907 | 0.006 | 0.137 | 0.137 | 0 | 0.1 | 0.2 | 0.15 | 0.38 | 0.11 | 0.35 |
| 9 | 4664852 | rs2295963 | 0.611 | -0.721 | -1.111 | 0.195 | 0.002 | 0.038 | 0.038 | 0 | 0.26 | 0.2 | 0.17 | 0.06 | 0.18 | 0.1 |
| 9 | 4671022 | rs1385453 | -1.012 | -0.97 | -0.452 | 0.33 | 0.011 | 0.003 | 0.003 | 0.77 | 0.44 | 0.64 | 0.36 | 0.59 | 0.31 | 0.57 |
| 9 | 4671409 | rs6476893 | -1.001 | -1.058 | -0.417 | 0.334 | 0.038 | 0.021 | 0.021 | 1.233 | 0.46 | 0.77 | 0.38 | 0.6 | 0.31 | 0.57 |
| 9 | 4677454 | rs10974724 | 0.751 | -1.031 | -0.489 | 0.495 | -0.035 | -0.033 | -0.033 | 0.035 | 0.08 | 0.01 | 0.13 | 0.37 | 0.12 | 0.35 |
| 9 | 4678754 | rs7848725 | -0.913 | -0.924 | -0.376 | 0.408 | 0.016 | 0.004 | 0.004 | 1.324 | 0.42 | 0.67 | 0.35 | 0.6 | 0.28 | 0.54 |
| 9 | 4679906 | rs2295964 | -1.203 | -1.012 | -0.583 | 0.251 | 0.042 | 0.017 | 0.017 | 2.908 | 0.46 | 0.84 | 0.4 | 0.6 | 0.32 | 0.56 |
| 9 | 4680653 | rs301486 | 0.721 | -0.772 | -0.998 | -0.137 | -0.008 | 0.042 | 0.042 | 0 | 0.18 | 0.08 | 0.15 | 0.04 | 0.17 | 0.14 |
| 9 | 4691570 | rs3824440 | 1.169 | -0.879 | -0.977 | -0.329 | -0.014 | 0.015 | 0.015 | 0 | 0.17 | 0.14 | 0.16 | 0.06 | 0.17 | 0.08 |
| 9 | 4691765 | rs3780397 | 1.017 | -0.792 | -1.106 | -0.218 | 0.002 | 0.02 | 0.02 | 0 | 0.08 | 0.03 | 0.11 | 0.37 | 0.09 | 0.32 |
| 9 | 4695004 | rs12685759 | 1.448 | -0.922 | -1.186 | -0.285 | 0.009 | -0.039 | -0.039 | 0 | 0.12 | 0 | 0.06 | 0.18 | 0.06 | 0.16 |
| 9 | 4695068 | rs7874378 | -0.282 | -1.126 | -0.634 | -0.196 | -0.005 | -0.032 | -0.032 | 0.048 | 0.34 | 0.54 | 0.25 | 0.24 | 0.23 | 0.24 |
| 9 | 4696919 | rs2295970 | -1.071 | -1.126 | -0.573 | -0.318 | 0.018 | 0.012 | 0.012 | 2.203 | 0.44 | 0.76 | 0.38 | 0.61 | 0.32 | 0.55 |
| 9 | 4698594 | rs10122598 | -0.29 | -1.092 | -0.57 | -0.411 | -0.005 | -0.047 | -0.047 | 0 | 0.34 | 0.54 | 0.25 | 0.15 | 0.23 | 0.23 |

| | | | | | | | | | | | | | | | | |
|---|---|---|---|---|---|---|---|---|---|---|---|---|---|---|---|---|
| 9 | 4704420 | rs1331191 | -1.859 | -0.955 | -0.672 | -0.661 | -0.056 | 0.005 | 0.005 | 1.926 | 0.63 | 0.87 | 0.54 | 0.65 | 0.49 | 0.69 |
| 9 | 4706214 | rs2181341 | -0.776 | -0.887 | -0.26 | -0.278 | -0.001 | -0.001 | -0.001 | 0 | 0.41 | 0.55 | 0.39 | 0.6 | 0.31 | 0.55 |
| 9 | 4706366 | rs3174298 | -0.953 | -0.82 | -0.707 | -0.264 | 0.016 | 0.021 | 0.021 | 2.088 | 0.43 | 0.72 | 0.41 | 0.61 | 0.31 | 0.55 |
| 9 | 4709784 | rs1048020 | 0.175 | -0.405 | -0.481 | -0.578 | -0.014 | 0.015 | 0.015 | 0 | 0.16 | 0.09 | 0.1 | 0 | 0.18 | 0.06 |
| 9 | 4713213 | rs296827 | 0.226 | -0.49 | 0.186 | -0.673 | 0.016 | -0.049 | -0.049 | 0 | 0.09 | 0.22 | 0.09 | 0.01 | 0.09 | 0.11 |
| 9 | 4715847 | rs403860 | -1.227 | -0.992 | 0.399 | -0.229 | 0.008 | 0.005 | 0.005 | 3.755 | 0.42 | 0.6 | 0.31 | 0.28 | 0.32 | 0.37 |
| 9 | 4716078 | rs12684167 | -0.486 | -1.1 | 0.413 | -0.077 | -0.002 | 0.002 | 0.002 | 1.598 | 0.3 | 0.28 | 0.21 | 0.27 | 0.23 | 0.25 |
| 9 | 4717365 | rs463684 | 1.393 | -0.91 | -0.159 | -0.228 | -0.004 | 0.001 | 0.001 | 0 | 0.06 | 0.05 | 0.07 | 0.07 | 0.05 | 0.01 |
| 9 | 4717384 | rs7856334 | 0.572 | -0.931 | -0.189 | -0.276 | -0.046 | 0.07 | 0.07 | 0.051 | 0.14 | 0.01 | 0.09 | 0 | 0.17 | 0.06 |
| 9 | 4718907 | rs456374 | -0.374 | -0.852 | 0.409 | 0.054 | -0.009 | -0.003 | -0.003 | 0.089 | 0.37 | 0.47 | 0.29 | 0.29 | 0.28 | 0.26 |
| 9 | 4718921 | rs465793 | -0.91 | -0.906 | 0.059 | 0.055 | 0.01 | 0.009 | 0.009 | 1.261 | 0.47 | 0.62 | 0.37 | 0.35 | 0.39 | 0.38 |
| 9 | 4719031 | rs16921966 | -0.082 | -0.832 | 0.245 | 0.17 | -0.022 | -0.005 | -0.005 | 0.491 | 0.33 | 0.52 | 0.23 | 0.22 | 0.23 | 0.25 |
| 9 | 4720263 | rs458853 | -0.332 | -0.195 | -0.249 | 0.147 | -0.018 | 0 | 0 | 0.129 | 0.61 | 0.69 | 0.48 | 0.6 | 0.55 | 0.58 |
| 9 | 4722462 | rs2274821 | 0.709 | 0.18 | -0.785 | 0.462 | -0.04 | -0.075 | -0.075 | 0.605 | 0.14 | 0.02 | 0.14 | 0.38 | 0.23 | 0.27 |
| 9 | 4725823 | rs2039300 | 0.097 | -0.245 | -0.85 | -0.154 | -0.056 | 0.044 | 0.044 | 0 | 0.39 | 0.11 | 0.57 | 0.73 | 0.58 | 0.57 |
| 9 | 4730743 | rs429491 | -0.307 | 0.323 | -0.114 | 0.097 | -0.008 | -0.017 | -0.017 | 0 | 0.87 | 0.82 | 0.86 | 0.94 | 0.9 | 0.87 |
| 9 | 4731040 | rs439486 | -0.206 | 0.674 | -0.402 | 0.347 | 0 | -0.004 | -0.004 | 0 | 0.08 | 0.14 | 0.07 | 0.07 | 0.07 | 0.03 |
| 9 | 4736508 | rs412899 | -0.073 | 0.864 | -1.121 | 0.422 | 0.003 | -0.002 | -0.002 | 1.846 | 0.08 | 0.11 | 0.08 | 0.07 | 0.1 | 0.04 |
| 9 | 4737890 | rs435100 | -0.371 | 0.294 | -1.157 | 0.563 | -0.025 | 0 | 0 | 0.017 | 0.34 | 0.3 | 0.51 | 0.36 | 0.48 | 0.3 |
| 9 | 4740648 | rs1756867 | 0.125 | -0.509 | -1.59 | 0.752 | 0.05 | -0.052 | -0.052 | 0 | 0.59 | 0.2 | 0.73 | 0.5 | 0.69 | 0.51 |
| 9 | 4741883 | rs3808857 | -0.476 | -1.434 | -1.489 | 0.682 | 0.039 | -0.005 | -0.005 | 0 | 0.17 | 0.07 | 0.14 | 0.13 | 0.06 | 0.22 |
| 9 | 4742323 | rs378117 | 0.944 | -1.99 | -0.444 | 0.756 | -0.066 | 0.013 | 0.013 | 2.324 | 0.74 | 0.43 | 0.76 | 0.71 | 0.91 | 0.67 |
| 9 | 4743341 | rs12352353 | 0.374 | -2.512 | -0.426 | -0.193 | 0.013 | 0.02 | 0.02 | 0 | 0.21 | 0.12 | 0.19 | 0.05 | 0.15 | 0.16 |
| 9 | 4743477 | rs12352385 | 0.374 | -2.512 | -0.426 | -0.193 | 0.013 | 0.02 | 0.02 | 0 | 0.21 | 0.12 | 0.19 | 0.05 | 0.15 | 0.16 |
| 9 | 4743626 | rs396861 | -0.088 | -2.606 | 0.272 | -0.239 | -0.049 | -0.106 | -0.106 | 0.32 | 0.74 | 0.46 | 0.77 | 0.95 | 0.84 | 0.84 |
| 9 | 4746363 | rs4742045 | -0.341 | -2.13 | -0.291 | -0.529 | 0 | 0.003 | 0.003 | 0 | 0.44 | 0.41 | 0.49 | 0.2 | 0.45 | 0.38 |
| 9 | 4749459 | rs11790285 | 0.703 | -1.797 | -0.044 | 0.058 | -0.045 | -0.002 | -0.002 | 0 | 0.2 | 0.09 | 0.26 | 0.21 | 0.29 | 0.23 |
| 9 | 4750996 | rs10974763 | 0.317 | -2.04 | -0.073 | 0.265 | 0.05 | -0.047 | -0.047 | 0.121 | 0.18 | 0.08 | 0.14 | 0.22 | 0.06 | 0.21 |
| 9 | 4751056 | rs4742046 | 0.506 | -1.091 | 0.419 | -0.01 | -0.052 | 0.033 | 0.033 | 0 | 0.2 | 0.08 | 0.26 | 0.15 | 0.28 | 0.22 |
| 9 | 4757649 | rs10123382 | 0.234 | -0.762 | 0.315 | -0.006 | -0.004 | 0.002 | 0.002 | 0 | 0.22 | 0.22 | 0.29 | 0.15 | 0.33 | 0.24 |
| 9 | 4758022 | rs1756871 | -0.054 | -0.743 | 0.302 | 0.064 | 0.012 | -0.014 | -0.014 | 0 | 0.73 | 0.4 | 0.56 | 0.76 | 0.65 | 0.7 |
| 9 | 4763176 | rs385893 | 0.194 | -1.008 | -0.49 | -0.255 | -0.009 | 0.079 | 0.079 | 0 | 0.41 | 0.25 | 0.43 | 0.78 | 0.49 | 0.53 |
| 9 | 4763368 | rs409950 | 0.344 | -0.795 | -1.061 | -0.186 | 0.018 | -0.049 | -0.049 | 0 | 0.19 | 0.08 | 0.11 | 0.19 | 0.17 | 0.17 |
| 9 | 4766022 | rs10815074 | -0.643 | -1.229 | -1.079 | -0.776 | 0.027 | 0.041 | 0.041 | 0.048 | 0.76 | 0.8 | 0.84 | 0.93 | 0.83 | 0.87 |
| 9 | 4767341 | rs420470 | -0.539 | -0.727 | -1.107 | -0.903 | 0.016 | 0.042 | 0.042 | 0 | 0.73 | 0.78 | 0.8 | 0.93 | 0.77 | 0.85 |
| 9 | 4767677 | rs10974772 | 1.401 | 0.246 | -0.666 | -0.499 | 0.021 | 0.069 | 0.069 | 0.403 | 0.15 | 0.17 | 0.26 | 0.58 | 0.22 | 0.32 |
| 9 | 4769647 | rs407504 | -0.776 | -0.122 | -0.918 | -0.756 | 0.019 | 0.028 | 0.028 | 0 | 0.78 | 0.81 | 0.84 | 0.93 | 0.84 | 0.88 |
| 9 | 4770942 | rs1756875 | -0.068 | -0.195 | -1.064 | -0.89 | 0.011 | 0.097 | 0.097 | 0 | 0.54 | 0.73 | 0.49 | 0.76 | 0.57 | 0.67 |
| 9 | 4773074 | rs296862 | -0.504 | -0.706 | -0.509 | -0.715 | 0.013 | 0.011 | 0.011 | 0.099 | 0.78 | 0.84 | 0.85 | 0.93 | 0.84 | 0.88 |
| 9 | 4775634 | rs172459 | -0.528 | -0.908 | -0.534 | -0.707 | 0.018 | 0.016 | 0.016 | 0.031 | 0.78 | 0.85 | 0.84 | 0.93 | 0.85 | 0.88 |
| 9 | 4784933 | rs7025754 | 0.06 | -1.022 | -0.215 | -0.638 | -0.032 | 0.034 | 0.034 | 0 | 0.18 | 0.31 | 0.1 | 0.15 | 0.13 | 0.26 |
| 9 | 4785307 | rs420565 | -0.283 | -0.59 | -0.769 | -0.578 | 0.036 | 0.059 | 0.059 | 0 | 0.78 | 0.89 | 0.84 | 0.92 | 0.85 | 0.86 |
| 9 | 4786985 | rs296854 | -0.107 | -0.604 | -0.702 | -0.584 | 0.017 | 0.015 | 0.015 | 0.029 | 0.76 | 0.83 | 0.83 | 0.92 | 0.85 | 0.85 |
| 9 | 4790165 | rs4740801 | -0.067 | -0.01 | -0.654 | -0.463 | 0 | 0.005 | 0.005 | 0 | 0.61 | 0.58 | 0.56 | 0.72 | 0.57 | 0.56 |
| 9 | 4792339 | rs423955 | 0.31 | -0.514 | -0.661 | -0.781 | 0.014 | 0.021 | 0.021 | 0 | 0.61 | 0.3 | 0.6 | 0.78 | 0.68 | 0.55 |
| 9 | 4795595 | rs10974788 | 0.729 | -0.299 | -0.641 | -0.527 | -0.025 | -0.029 | -0.029 | 0.958 | 0.16 | 0.05 | 0.23 | 0.17 | 0.18 | 0.35 |
| 9 | 4796225 | rs7851250 | 0.056 | -0.41 | 0.319 | -0.601 | -0.001 | 0.003 | 0.003 | 0 | 0.19 | 0.41 | 0.26 | 0.16 | 0.17 | 0.34 |
| 9 | 4806233 | rs10974795 | 0.354 | 0.385 | 0.165 | -0.46 | -0.006 | -0.003 | -0.003 | 0 | 0.18 | 0.19 | 0.33 | 0.19 | 0.23 | 0.37 |
| 9 | 4808069 | rs10758654 | -0.202 | 0.196 | 0.476 | -0.46 | 0.023 | 0.047 | 0.047 | 0 | 0.23 | 0.42 | 0.38 | 0.28 | 0.25 | 0.43 |
| 9 | 4819421 | rs377186 | -0.08 | 0.128 | -0.105 | -0.841 | -0.035 | 0.016 | 0.016 | 0.648 | 0.39 | 0.18 | 0.47 | 0.26 | 0.44 | 0.49 |
| 9 | 4821348 | rs7038267 | 0.553 | 0.437 | 0.496 | -0.604 | 0.008 | 0 | 0 | 0 | 0.18 | 0.21 | 0.32 | 0.17 | 0.22 | 0.36 |
| 9 | 4824003 | rs404464 | 1.24 | 0.369 | 0.049 | -0.53 | 0.024 | 0.123 | 0.123 | 0 | 0.4 | 0.08 | 0.31 | 0.15 | 0.39 | 0.19 |
| 9 | 4826618 | rs7871570 | 0.756 | 0.242 | 0.289 | -0.547 | 0.012 | -0.002 | -0.002 | 0 | 0.18 | 0.24 | 0.31 | 0.16 | 0.22 | 0.36 |
| 9 | 4830849 | rs531122 | -0.13 | -0.004 | 0.19 | -0.717 | -0.053 | -0.133 | -0.133 | 0 | 0.76 | 0.38 | 0.86 | 1 | 0.86 | 0.87 |
| 9 | 4832888 | rs439975 | 0.015 | 0.049 | 0.059 | -0.701 | -0.037 | -0.043 | -0.043 | 0.254 | 0.72 | 0.39 | 0.82 | 0.92 | 0.84 | 0.8 |
| 9 | 4833437 | rs458552 | 0.755 | 0.193 | -0.5 | -0.695 | 0.02 | 0.104 | 0.104 | 0.071 | 0.53 | 0.1 | 0.5 | 0.75 | 0.61 | 0.44 |
| 9 | 4834034 | rs466547 | -0.749 | 0.232 | 0.103 | -0.45 | 0 | -0.027 | -0.027 | 0 | 0.82 | 0.83 | 0.91 | 1 | 0.87 | 0.87 |
| 9 | 4834394 | rs457287 | -0.514 | 0.235 | -0.087 | -0.44 | -0.048 | -0.128 | -0.128 | 0 | 0.8 | 0.62 | 0.88 | 1 | 0.87 | 0.87 |
| 9 | 4835574 | rs3780359 | 1.202 | 1.105 | -0.976 | 0.193 | -0.062 | -0.072 | -0.072 | 0 | 0.12 | 0.03 | 0.18 | 0.6 | 0.19 | 0.24 |
| 9 | 4836566 | rs7860776 | 0.066 | 1.068 | -1.158 | -0.346 | -0.026 | 0.025 | 0.025 | 0.226 | 0.25 | 0.13 | 0.36 | 0.15 | 0.27 | 0.41 |
| 9 | 4836621 | rs10974807 | -0.522 | 0.267 | -1.65 | -0.092 | 0.063 | -0.019 | -0.019 | 0.501 | 0.35 | 0.11 | 0.33 | 0.33 | 0.23 | 0.45 |
| 9 | 4836811 | rs7861943 | 0.008 | 0.448 | -1.17 | 0.004 | -0.007 | -0.053 | -0.053 | 1.285 | 0.29 | 0.12 | 0.46 | 0.71 | 0.28 | 0.55 |
| 9 | 4840063 | rs295263 | 0.069 | -0.139 | -0.967 | -0.497 | 0.019 | 0.015 | 0.015 | 0 | 0.8 | 0.86 | 0.9 | 1 | 0.87 | 0.87 |
| 9 | 4842061 | rs10815093 | 0.272 | -0.024 | -0.129 | -0.446 | 0.025 | -0.01 | -0.01 | 0 | 0.29 | 0.46 | 0.42 | 0.28 | 0.3 | 0.49 |
| 9 | 4844459 | rs159432 | 0.383 | -0.222 | -0.004 | -0.453 | 0.02 | -0.017 | -0.017 | 0 | 0.29 | 0.45 | 0.4 | 0.24 | 0.28 | 0.47 |
| 9 | 4844507 | rs295259 | -0.077 | -0.327 | 0.093 | -0.541 | 0.019 | 0.018 | 0.018 | 0 | 0.81 | 0.89 | 0.92 | 1 | 0.88 | 0.87 |
| 9 | 4844704 | rs2236497 | 0.958 | -0.136 | -0.01 | -0.538 | -0.015 | -0.01 | -0.01 | 0 | 0.14 | 0.11 | 0.18 | 0.56 | 0.19 | 0.22 |
| 9 | 4845242 | rs295258 | 0.805 | -0.144 | 0.029 | -0.54 | -0.001 | 0.006 | 0.006 | 0 | 0.65 | 0.61 | 0.7 | 0.44 | 0.68 | 0.65 |
| 9 | 4848630 | rs295254 | 1.597 | -0.279 | -0.116 | -0.579 | -0.05 | 0.018 | 0.018 | 0 | 0.35 | 0.04 | 0.3 | 0.2 | 0.4 | 0.19 |

| | | | | | | | | | | | | | | | | |
|---|---|---|---|---|---|---|---|---|---|---|---|---|---|---|---|---|
| 9 | 4848875 | rs159433 | 0.134 | -0.373 | 0.172 | -0.625 | 0.024 | 0.021 | 0.021 | 0 | 0.8 | 0.87 | 0.91 | 1 | 0.88 | 0.87 |
| 9 | 4851515 | rs10115106 | 0.48 | -0.351 | 0.3 | -0.576 | 0.016 | -0.004 | -0.004 | 0.176 | 0.28 | 0.46 | 0.41 | 0.25 | 0.28 | 0.47 |
| 9 | 4855514 | rs457417 | 0.309 | -0.31 | 0.173 | -0.633 | -0.028 | -0.094 | -0.094 | 0 | 0.78 | 0.63 | 0.89 | 1 | 0.87 | 0.88 |
| 9 | 4856234 | rs1887430 | 0.217 | -0.041 | -0.243 | -0.255 | -0.001 | -0.005 | -0.005 | 0 | 0.54 | 0.51 | 0.51 | 0.74 | 0.59 | 0.4 |
| 9 | 4860300 | rs15583 | 0.72 | 0.565 | -0.228 | 0.217 | 0.009 | 0.067 | 0.067 | 0 | 0.7 | 0.47 | 0.68 | 0.43 | 0.7 | 0.69 |
| 9 | 4860569 | rs1260073 | -0.085 | 0.273 | -0.375 | -0.282 | -0.018 | 0.027 | 0.027 | 0.667 | 0.66 | 0.77 | 0.59 | 0.77 | 0.62 | 0.46 |
| 9 | 4860943 | rs295918 | 0.448 | 0.212 | -0.268 | -0.277 | 0.005 | -0.008 | -0.008 | 0 | 0.29 | 0.32 | 0.47 | 0.24 | 0.33 | 0.48 |
| 9 | 4861479 | rs4740804 | -0.52 | 0.479 | 0.03 | 0.272 | 0.023 | 0.101 | 0.101 | 0 | 0.23 | 0.44 | 0.27 | 0.62 | 0.25 | 0.24 |
| 9 | 4863952 | rs10758661 | 0.081 | 0.401 | -0.449 | 0.322 | -0.071 | -0.139 | -0.139 | 0.298 | 0.8 | 0.96 | 0.78 | 0.46 | 0.76 | 0.75 |
| 9 | 4864891 | rs7041623 | 0.826 | 0.466 | -0.115 | 0.401 | 0.014 | 0.041 | 0.041 | 0 | 0.7 | 0.58 | 0.69 | 0.36 | 0.7 | 0.64 |
| 9 | 4866010 | rs7032487 | 0.996 | 0.643 | -0.995 | 0.234 | -0.012 | 0.005 | 0.005 | 0 | 0.62 | 0.34 | 0.6 | 0.33 | 0.56 | 0.62 |
| 9 | 4866840 | rs7850837 | -0.37 | 0.512 | -1.447 | 0.417 | 0.006 | 0.014 | 0.014 | 0.326 | 0.42 | 0.23 | 0.47 | 0.39 | 0.4 | 0.64 |
| 9 | 4869801 | rs10974824 | -0.789 | 0.399 | -0.982 | 0.512 | -0.001 | 0.006 | 0.006 | 0 | 0.08 | 0.08 | 0.09 | 0.13 | 0.08 | 0.11 |
| 9 | 4876202 | rs10815099 | 0.18 | 0.475 | -1.305 | 0.14 | -0.029 | 0.149 | 0.149 | 0 | 0.46 | 0.19 | 0.35 | 0.17 | 0.52 | 0.23 |
| 9 | 4876997 | rs913259 | -0.743 | 0.047 | -0.32 | 0.972 | -0.012 | 0.097 | 0.097 | 3.173 | 0.24 | 0.62 | 0.3 | 0.52 | 0.23 | 0.22 |
| 9 | 4883257 | rs10117338 | -1.655 | -0.48 | 0.573 | 0.72 | -0.037 | 0.004 | 0.004 | 1.975 | 0.11 | 0.59 | 0.08 | 0.06 | 0.04 | 0.05 |
| 9 | 4883937 | rs10758662 | -0.443 | -0.863 | 0.056 | 1.05 | 0.013 | -0.016 | -0.016 | 0 | 0.85 | 0.9 | 0.93 | 0.77 | 0.9 | 0.81 |
| 9 | 4888355 | rs10117872 | -1.209 | -0.784 | 1.133 | 0.938 | 0.008 | 0.07 | 0.07 | 0 | 0.07 | 0.36 | 0.04 | 0.05 | 0.01 | 0.05 |
| 9 | 4888441 | rs9969783 | 0.637 | -0.426 | 0.061 | 0.609 | -0.044 | -0.047 | -0.047 | 0 | 0.46 | 0.1 | 0.5 | 0.48 | 0.53 | 0.31 |
| 9 | 4895391 | rs10974837 | 0.279 | -0.451 | 0.375 | 0.53 | 0 | 0 | 0 | 0 | 0.49 | 0.49 | 0.5 | 0.49 | 0.49 | 0.28 |
| 9 | 4895668 | rs6476919 | 0.427 | -0.568 | 0.498 | 0.553 | -0.016 | 0.017 | 0.017 | 0.051 | 0.32 | 0.27 | 0.39 | 0.21 | 0.4 | 0.44 |
| 9 | 4902707 | rs7022229 | -0.778 | -0.728 | 0.445 | 0.626 | -0.004 | 0.002 | 0.002 | 0.997 | 0.84 | 0.84 | 0.92 | 0.75 | 0.9 | 0.79 |
| 9 | 4903289 | rs6476924 | -0.728 | -0.755 | 0.465 | 0.616 | -0.009 | 0.005 | 0.005 | 0.89 | 0.83 | 0.83 | 0.92 | 0.76 | 0.9 | 0.8 |
| 9 | 4903818 | rs12006302 | -0.343 | -0.834 | 0.529 | 0.628 | 0.029 | 0.149 | 0.149 | 2.4 | 0.05 | 0.33 | 0.03 | 0.08 | 0.01 | 0.15 |
| 9 | 4904969 | rs10815105 | -0.13 | -0.853 | 0.402 | 0.641 | -0.023 | 0.017 | 0.017 | 0.659 | 0.8 | 0.77 | 0.9 | 0.72 | 0.89 | 0.73 |
| 9 | 4905038 | rs17803986 | 0.647 | -0.325 | -0.188 | 0.475 | 0.042 | -0.025 | -0.025 | 0 | 0.49 | 0.13 | 0.61 | 0.53 | 0.57 | 0.33 |
| 9 | 4907617 | rs7870918 | 0.045 | -0.662 | 0.118 | 0.622 | -0.061 | 0.083 | 0.083 | 0.341 | 0.77 | 0.45 | 0.89 | 0.63 | 0.89 | 0.58 |
| 9 | 4908182 | rs2381180 | 0.855 | -0.412 | -0.15 | 0.635 | -0.031 | -0.019 | -0.019 | 0 | 0.45 | 0.17 | 0.57 | 0.47 | 0.52 | 0.28 |
| 9 | 4909719 | rs7864244 | 0.203 | -0.194 | 0.053 | 0.729 | -0.009 | 0.018 | 0.018 | 0 | 0.32 | 0.29 | 0.33 | 0.15 | 0.37 | 0.31 |
| 9 | 4921639 | rs715769 | 0.515 | -0.207 | -0.158 | 0.695 | -0.037 | 0.066 | 0.066 | 0.087 | 0.29 | 0.16 | 0.3 | 0.11 | 0.37 | 0.29 |
| 9 | 4921824 | rs1410004 | 0.861 | -0.159 | -0.504 | 0.409 | -0.036 | -0.066 | -0.066 | 0.305 | 0.34 | 0.19 | 0.51 | 0.52 | 0.41 | 0.26 |
| 9 | 4926113 | rs10815121 | 0.658 | 0.05 | -0.678 | 0.411 | -0.032 | 0.047 | 0.047 | 0.012 | 0.3 | 0.17 | 0.35 | 0.16 | 0.37 | 0.3 |
| 9 | 4930487 | rs10974859 | 0.818 | -0.01 | -0.488 | 0.742 | -0.012 | -0.013 | -0.013 | 0 | 0.33 | 0.29 | 0.39 | 0.44 | 0.38 | 0.28 |
| 9 | 4934896 | rs7023498 | 0.799 | -0.116 | -0.486 | 0.698 | -0.026 | 0.03 | 0.03 | 0.008 | 0.3 | 0.17 | 0.34 | 0.17 | 0.36 | 0.3 |
| 9 | 4934926 | rs10815126 | 1.465 | -0.247 | -0.786 | 0.636 | -0.117 | -0.102 | -0.102 | 0.203 | 0.27 | 0.04 | 0.35 | 0.4 | 0.34 | 0.23 |
| 9 | 4935926 | rs1571221 | 0.361 | -0.125 | -0.629 | 0.541 | 0.042 | -0.005 | -0.005 | 1.992 | 0.58 | 0.33 | 0.7 | 0.6 | 0.71 | 0.53 |
| 9 | 4937331 | rs16922344 | 0.725 | -0.041 | -0.604 | 0.421 | 0 | -0.001 | -0.001 | 0 | 0.31 | 0.32 | 0.36 | 0.19 | 0.37 | 0.32 |
| 9 | 4937872 | rs7025867 | 0.868 | -0.031 | -0.853 | 0.291 | -0.025 | 0.032 | 0.032 | 0 | 0.3 | 0.18 | 0.34 | 0.18 | 0.36 | 0.3 |
| 9 | 4938232 | rs2381181 | 0.166 | -0.576 | -0.593 | 0.288 | 0.046 | 0.042 | 0.042 | 1.466 | 0.68 | 0.91 | 0.85 | 0.77 | 0.76 | 0.71 |
| 9 | 4939569 | rs1327489 | 0.715 | -0.413 | -0.005 | 0.7 | -0.052 | 0.074 | 0.074 | 1.134 | 0.78 | 0.54 | 0.79 | 0.64 | 0.85 | 0.61 |
| 9 | 4943454 | rs10974873 | 0.678 | -0.237 | -0.168 | 0.62 | -0.036 | 0.045 | 0.045 | 0 | 0.73 | 0.57 | 0.78 | 0.64 | 0.78 | 0.55 |
| 9 | 4946989 | rs7021993 | -0.759 | 0.086 | -0.798 | 0.07 | 0.003 | 0.034 | 0.034 | 0.084 | 0.35 | 0.82 | 0.46 | 0.23 | 0.35 | 0.4 |
| 9 | 4955050 | rs7037047 | -0.49 | 0.151 | -0.071 | 0.046 | 0.019 | -0.055 | -0.055 | 0.249 | 0.32 | 0.45 | 0.43 | 0.15 | 0.37 | 0.29 |
| 9 | 4959024 | rs6476931 | 0.575 | -0.049 | -0.393 | -0.218 | 0.002 | -0.046 | -0.046 | 0 | 0.6 | 0.5 | 0.53 | 0.85 | 0.57 | 0.65 |
| 9 | 4959331 | rs2381194 | 0.027 | 0.253 | 0.062 | -0.073 | 0.081 | -0.029 | -0.029 | 0 | 0.1 | 0.33 | 0.25 | 0.01 | 0.13 | 0.07 |
| 9 | 4964774 | rs10974892 | 0.473 | 0.012 | -1.036 | 0.115 | -0.098 | -0.108 | -0.108 | 0 | 0.37 | 0.09 | 0.35 | 0.44 | 0.45 | 0.33 |
| 9 | 4970909 | rs7870381 | -0.144 | 0.257 | -0.605 | -0.168 | 0.021 | -0.051 | -0.051 | 1.376 | 0.35 | 0.48 | 0.45 | 0.16 | 0.42 | 0.23 |
| 9 | 4972398 | rs7862852 | -1.078 | -0.068 | -0.505 | 0.16 | -0.002 | -0.004 | -0.004 | 0 | 0.65 | 0.71 | 0.69 | 0.66 | 0.65 | 0.65 |
| 9 | 4974441 | rs1327495 | 0.055 | 0.09 | -0.841 | -0.176 | -0.004 | 0.004 | 0.004 | 0 | 0.32 | 0.29 | 0.44 | 0.17 | 0.41 | 0.23 |
| 9 | 4981602 | rs10758669 | -0.454 | -0.448 | -1.363 | -0.226 | -0.01 | 0.045 | 0.045 | 0 | 0.62 | 0.86 | 0.57 | 0.63 | 0.65 | 0.61 |
| 9 | 4984549 | rs1887429 | -0.033 | -0.545 | -0.905 | -1.17 | 0.013 | 0.021 | 0.021 | 0.243 | 0.3 | 0.65 | 0.24 | 0.15 | 0.28 | 0.24 |
| 9 | 4985879 | rs2274471 | 0.507 | -1.277 | -1.026 | -1.107 | 0.018 | 0.033 | 0.033 | 0 | 0.25 | 0.12 | 0.2 | 0.17 | 0.26 | 0.15 |
| 9 | 5003338 | rs4372063 | -1.032 | -1.451 | -1.541 | -1.082 | -0.128 | -0.126 | -0.126 | 0.438 | 0.67 | 0.93 | 0.63 | 0.54 | 0.61 | 0.59 |
| 9 | 5008070 | rs7030260 | -0.095 | -1.562 | -0.659 | -1.313 | -0.004 | -0.015 | -0.015 | 0 | 0.33 | 0.66 | 0.39 | 0.26 | 0.27 | 0.31 |
| 9 | 5021514 | rs7034753 | -1.278 | -1.626 | -0.583 | -1.319 | -0.147 | -0.059 | -0.059 | 0.089 | 0.63 | 0.94 | 0.62 | 0.46 | 0.58 | 0.51 |
| 9 | 5022807 | rs7851556 | 0.004 | -1.694 | -0.251 | -1.274 | -0.005 | -0.036 | -0.036 | 0 | 0.31 | 0.5 | 0.38 | 0.27 | 0.27 | 0.31 |
| 9 | 5055434 | rs1536800 | 0.14 | -1.89 | -0.516 | -1.437 | -0.007 | 0.034 | 0.034 | 0 | 0.23 | 0.11 | 0.3 | 0.18 | 0.22 | 0.22 |
| 9 | 5063701 | rs10815149 | -0.299 | -1.651 | -0.216 | -1.318 | -0.012 | -0.041 | -0.041 | 0.274 | 0.34 | 0.51 | 0.38 | 0.28 | 0.28 | 0.3 |
| 9 | 5072846 | rs10974947 | 0.531 | -1.229 | 0.038 | -1.192 | -0.002 | -0.007 | -0.007 | 1.657 | 0.24 | 0.22 | 0.19 | 0.17 | 0.26 | 0.19 |
| 9 | 5081585 | rs7034539 | 0.408 | -1.697 | 0.129 | -1.253 | -0.03 | -0.057 | -0.057 | 0 | 0.26 | 0.41 | 0.21 | 0.16 | 0.25 | 0.19 |
| 9 | 5081780 | rs2230724 | -1.236 | -1.757 | -0.123 | -1.29 | -0.126 | -0.007 | -0.007 | 0 | 0.58 | 0.93 | 0.59 | 0.44 | 0.52 | 0.49 |
| 9 | 5083173 | rs1410779 | 1.174 | -1.763 | -0.049 | -1.272 | 0.014 | 0.021 | 0.021 | 0 | 0.2 | 0.04 | 0.16 | 0.16 | 0.22 | 0.15 |
| 9 | 5097544 | rs3780372 | -0.067 | -1.881 | 0.042 | -1.356 | 0.002 | -0.037 | -0.037 | 0 | 0.31 | 0.47 | 0.39 | 0.27 | 0.29 | 0.3 |
| 9 | 5100628 | rs7870694 | -0.395 | -1.879 | 0.049 | -1.355 | 0.001 | 0.004 | 0.004 | 0 | 0.34 | 0.69 | 0.41 | 0.27 | 0.29 | 0.3 |
| 9 | 5112519 | rs3780379 | 1.185 | -1.87 | -0.06 | -1.278 | 0.005 | -0.017 | -0.017 | 0 | 0.2 | 0.03 | 0.16 | 0.21 | 0.22 | 0.15 |
| 9 | 5112844 | rs10815158 | -1.325 | -1.894 | -0.092 | -1.241 | -0.139 | -0.1 | -0.1 | 2.64 | 0.58 | 0.93 | 0.6 | 0.55 | 0.55 | 0.56 |
| 9 | 5113577 | rs3824433 | -0.24 | -1.716 | 0.173 | -1.266 | -0.001 | -0.021 | -0.021 | 0.957 | 0.32 | 0.58 | 0.39 | 0.26 | 0.29 | 0.3 |
| 9 | 5114522 | rs884132 | 0.231 | -1.714 | 0.116 | -1.265 | -0.019 | 0.029 | 0.029 | 0 | 0.28 | 0.15 | 0.36 | 0.26 | 0.29 | 0.26 |
| 9 | 5150058 | rs10815167 | -0.516 | -1.52 | 0.605 | -0.912 | -0.002 | -0.016 | -0.016 | 2.112 | 0.33 | 0.58 | 0.39 | 0.26 | 0.29 | 0.3 |

| | | | | | | | | | | | | | | | | |
|---|---|---|---|---|---|---|---|---|---|---|---|---|---|---|---|---|
| 9 | 5156061 | rs10974985 | 0.04 | -1.413 | 0.73 | -0.833 | -0.002 | -0.001 | -0.001 | 0 | 0.27 | 0.23 | 0.28 | 0.23 | 0.26 | 0.2 |
| 9 | 5174638 | rs7029084 | -0.585 | -1.073 | 0.932 | -0.857 | 0.001 | -0.016 | -0.016 | 0 | 0.31 | 0.59 | 0.39 | 0.26 | 0.29 | 0.27 |
| 9 | 5181467 | rs7047795 | -0.807 | -1.084 | 0.9 | -0.863 | 0 | 0.042 | 0.042 | 0 | 0.33 | 0.78 | 0.41 | 0.26 | 0.29 | 0.27 |
| 9 | 5181610 | rs7036536 | -0.636 | -1.079 | 0.858 | -0.86 | -0.001 | -0.003 | -0.003 | 0 | 0.32 | 0.65 | 0.4 | 0.26 | 0.29 | 0.27 |
| 9 | 5183892 | rs7045342 | -0.584 | -1.075 | 0.822 | -0.887 | 0 | 0.001 | 0.001 | 0 | 0.29 | 0.69 | 0.36 | 0.26 | 0.25 | 0.29 |
| 9 | 5188078 | rs10758677 | -1.802 | -1.214 | 0.65 | -0.926 | -0.036 | 0.049 | 0.049 | 0 | 0.54 | 0.86 | 0.6 | 0.38 | 0.51 | 0.48 |
| 9 | 5189773 | rs11999802 | -0.345 | -1.03 | 0.88 | -0.835 | -0.003 | -0.02 | -0.02 | 4.746 | 0.27 | 0.55 | 0.35 | 0.26 | 0.26 | 0.31 |
| 9 | 5191297 | rs2031903 | 0.537 | -0.961 | 0.927 | -0.766 | 0.007 | 0.005 | 0.005 | 0 | 0.21 | 0.03 | 0.14 | 0.13 | 0.23 | 0.13 |
| 9 | 5196253 | rs2224571 | -0.654 | -0.538 | 1.645 | -0.829 | 0 | -0.014 | -0.014 | 0 | 0.28 | 0.38 | 0.37 | 0.25 | 0.29 | 0.31 |
| 9 | 5200127 | rs16922779 | -0.666 | -0.538 | 1.651 | -0.863 | 0.001 | -0.027 | -0.027 | 0.424 | 0.29 | 0.42 | 0.37 | 0.25 | 0.29 | 0.32 |
| 9 | 5213687 | rs10975003 | -1.42 | -0.466 | 1.388 | -0.912 | 0.007 | 0.022 | 0.022 | 0 | 0.4 | 0.69 | 0.44 | 0.23 | 0.35 | 0.38 |
| 9 | 5229116 | rs10114531 | -1.285 | -0.315 | 1.006 | -0.746 | 0.012 | 0.075 | 0.075 | 0 | 0.37 | 0.8 | 0.43 | 0.23 | 0.32 | 0.33 |
| 9 | 5230447 | rs2104166 | -1.21 | -0.31 | 0.995 | -0.786 | -0.004 | -0.042 | -0.042 | 0.439 | 0.35 | 0.55 | 0.4 | 0.22 | 0.31 | 0.32 |
| 9 | 5238460 | rs10118434 | -0.714 | -0.175 | 1.275 | -0.455 | 0.002 | -0.013 | -0.013 | 0.012 | 0.27 | 0.29 | 0.34 | 0.25 | 0.29 | 0.26 |
| 9 | 5248742 | rs6476953 | -1.191 | -0.156 | 0.414 | -0.466 | 0.007 | 0.031 | 0.031 | 0 | 0.31 | 0.78 | 0.39 | 0.25 | 0.29 | 0.27 |
| 9 | 5267043 | rs1575284 | -0.612 | -0.116 | 0.13 | -0.377 | -0.04 | -0.021 | -0.021 | 0 | 0.23 | 0.48 | 0.15 | 0.24 | 0.2 | 0.19 |
| 9 | 5270239 | rs16922908 | 0.183 | 0.448 | -0.622 | -0.157 | -0.006 | -0.012 | -0.012 | 0 | 0.14 | 0.02 | 0.06 | 0.14 | 0.14 | 0.14 |
| 9 | 5270603 | rs2381216 | -0.582 | 0.614 | -0.644 | -0.084 | -0.031 | -0.017 | -0.017 | 0 | 0.05 | 0.01 | 0.2 | 0.09 | 0.08 | 0.16 |
| 9 | 5272381 | rs12683801 | 0.134 | 0.579 | -0.662 | -0.297 | -0.02 | 0.052 | 0.052 | 0 | 0.31 | 0.18 | 0.42 | 0.17 | 0.39 | 0.22 |
| 9 | 5273180 | rs7861755 | -0.498 | 0.643 | -0.545 | -0.112 | 0.002 | 0.025 | 0.025 | 0 | 0.48 | 0.58 | 0.52 | 0.31 | 0.53 | 0.37 |
| 9 | 5274464 | rs10975037 | -0.326 | 0.652 | -0.749 | 0.085 | 0.005 | 0.037 | 0.037 | 0.523 | 0.37 | 0.25 | 0.46 | 0.26 | 0.44 | 0.26 |
| 9 | 5274648 | rs12684204 | -0.364 | 0.308 | -0.443 | 0.426 | 0.016 | 0.001 | 0.001 | 0.171 | 0.36 | 0.29 | 0.45 | 0.4 | 0.33 | 0.26 |
| 9 | 5277261 | rs2146043 | 0.4 | 0.188 | -0.683 | 0.59 | 0.061 | -0.026 | -0.026 | 0.529 | 0.23 | 0.06 | 0.26 | 0.32 | 0.18 | 0.17 |
| 9 | 5277752 | rs13288217 | -0.089 | 0.902 | -0.475 | 0.312 | 0.002 | -0.008 | -0.008 | 0.293 | 0.22 | 0.29 | 0.16 | 0.15 | 0.2 | 0.2 |
| 9 | 5289943 | rs2381217 | 0.081 | 0.912 | -0.332 | 0.353 | -0.002 | 0.002 | 0.002 | 0.334 | 0.21 | 0.21 | 0.16 | 0.15 | 0.19 | 0.2 |
| 9 | 5291710 | rs10975041 | 0.096 | 0.883 | -0.949 | 0.567 | -0.046 | -0.047 | -0.047 | 0 | 0.13 | 0.02 | 0.15 | 0.17 | 0.12 | 0.28 |
| 9 | 5292257 | rs1407374 | -0.07 | 0.596 | -0.049 | 0.598 | 0.013 | -0.036 | -0.036 | 0 | 0.23 | 0.44 | 0.18 | 0.15 | 0.19 | 0.2 |
| 9 | 5297563 | rs7047392 | 0.522 | 0.581 | -0.257 | 0.674 | 0.009 | -0.008 | -0.008 | 0 | 0.36 | 0.29 | 0.33 | 0.51 | 0.31 | 0.47 |
| 9 | 5299385 | rs10975043 | -0.706 | 0.588 | -0.42 | 0.506 | -0.064 | 0.011 | 0.011 | 0 | 0.6 | 0.85 | 0.58 | 0.71 | 0.5 | 0.67 |
| 9 | 5306824 | rs3758239 | 0.006 | 0.767 | -0.296 | 0.501 | 0.002 | -0.004 | -0.004 | 0.541 | 0.22 | 0.3 | 0.15 | 0.19 | 0.19 | 0.2 |
| 9 | 5315460 | rs4742078 | 0.107 | 0.393 | -0.537 | 0.691 | 0.012 | -0.008 | -0.008 | 0 | 0.38 | 0.31 | 0.42 | 0.51 | 0.32 | 0.48 |
| 9 | 5317534 | rs7848509 | -0.056 | 0.998 | -0.719 | 0.418 | 0.002 | 0 | 0 | 0 | 0.35 | 0.65 | 0.28 | 0.3 | 0.38 | 0.23 |
| 9 | 5318770 | rs10491649 | -0.48 | 1.025 | -0.724 | 0.148 | -0.021 | 0.046 | 0.046 | 1.527 | 0.34 | 0.27 | 0.4 | 0.23 | 0.42 | 0.39 |
| 9 | 5322687 | rs4740810 | 0.249 | 0.36 | -0.566 | 0.228 | -0.003 | 0 | 0 | 0 | 0.34 | 0.32 | 0.34 | 0.43 | 0.26 | 0.43 |
| 9 | 5324922 | rs6476962 | 0.498 | 0.388 | -0.707 | 0.262 | 0.02 | -0.017 | -0.017 | 0 | 0.31 | 0.21 | 0.32 | 0.43 | 0.25 | 0.41 |
| 9 | 5335470 | rs2273782 | 0.359 | 0.381 | -0.222 | 0.32 | -0.016 | 0.009 | 0.009 | 0 | 0.33 | 0.3 | 0.33 | 0.42 | 0.26 | 0.45 |
| 9 | 5340596 | rs1322220 | -0.123 | 0.645 | -0.291 | 0.198 | -0.001 | 0 | 0 | 0 | 0.46 | 0.55 | 0.45 | 0.48 | 0.42 | 0.57 |
| 9 | 5354681 | rs10815188 | -0.088 | 0.309 | -0.597 | 0.204 | 0.046 | 0.004 | 0.004 | 0.059 | 0.46 | 0.62 | 0.52 | 0.52 | 0.4 | 0.47 |
| 9 | 5356333 | rs7862502 | -1.83 | 0.436 | -0.686 | 0.195 | -0.084 | -0.082 | -0.082 | 0 | 0.69 | 0.95 | 0.76 | 0.67 | 0.68 | 0.81 |
| 9 | 5356674 | rs10123951 | -0.158 | 0.44 | -0.596 | 0.219 | 0.001 | -0.003 | -0.003 | 0.413 | 0.24 | 0.33 | 0.23 | 0.16 | 0.27 | 0.35 |
| 9 | 5360293 | rs2181146 | 1.676 | 0.269 | -0.641 | 0.17 | -0.018 | 0.01 | 0.01 | 0.665 | 0.21 | 0.23 | 0.2 | 0.32 | 0.12 | 0.2 |
| 9 | 5362028 | rs2236320 | 0.231 | 0.654 | -0.793 | 0.177 | -0.022 | 0.005 | 0.005 | 1.497 | 0.2 | 0.15 | 0.21 | 0.15 | 0.24 | 0.3 |
| 9 | 5364592 | rs7874937 | 0.062 | 0.451 | -0.664 | 0.161 | 0 | -0.002 | -0.002 | 0.053 | 0.21 | 0.28 | 0.22 | 0.15 | 0.23 | 0.3 |
| 9 | 5373441 | rs10975080 | 0.476 | 0.44 | -1.053 | 0.196 | -0.055 | -0.006 | -0.006 | 0.089 | 0.2 | 0.03 | 0.28 | 0.2 | 0.26 | 0.25 |
| 9 | 5377177 | rs10975083 | -0.006 | 0.449 | -0.694 | 0.228 | 0.001 | -0.01 | -0.01 | 0.031 | 0.24 | 0.32 | 0.31 | 0.24 | 0.26 | 0.24 |
| 9 | 5381870 | rs2381277 | -1.646 | 0.26 | -1.199 | 0.29 | -0.127 | -0.077 | -0.077 | 0 | 0.69 | 0.96 | 0.75 | 0.7 | 0.65 | 0.82 |
| 9 | 5382192 | rs16923101 | 1.722 | 0.059 | -0.962 | 0.243 | 0.023 | -0.024 | -0.024 | 0 | 0.2 | 0.09 | 0.21 | 0.32 | 0.15 | 0.2 |
| 9 | 5384832 | rs7027829 | -1.121 | 0.102 | -0.727 | -0.063 | -0.007 | 0.046 | 0.046 | 4.178 | 0.45 | 0.88 | 0.52 | 0.37 | 0.45 | 0.59 |
| 9 | 5385642 | rs10815201 | 0.15 | 0.309 | -0.203 | 0.041 | -0.002 | -0.008 | -0.008 | 0 | 0.22 | 0.24 | 0.27 | 0.19 | 0.23 | 0.25 |
| 9 | 5385783 | rs10975086 | -0.149 | -0.152 | -0.152 | -0.015 | 0.003 | -0.012 | -0.012 | 0 | 0.23 | 0.36 | 0.23 | 0.16 | 0.23 | 0.31 |
| 9 | 5386384 | rs10975087 | 0.16 | -0.352 | -0.46 | 0.378 | 0.039 | -0.013 | -0.013 | 0 | 0.35 | 0.15 | 0.19 | 0.37 | 0.29 | 0.34 |
| 9 | 5389075 | rs10815203 | -0.251 | -0.618 | 0.227 | 0.155 | -0.034 | -0.031 | -0.031 | 0.886 | 0.27 | 0.44 | 0.25 | 0.18 | 0.22 | 0.36 |
| 9 | 5389652 | rs10975090 | -0.379 | -0.709 | 0.256 | 0.017 | 0.003 | -0.017 | -0.017 | 0 | 0.24 | 0.3 | 0.28 | 0.16 | 0.27 | 0.25 |
| 9 | 5392199 | rs3739636 | -0.962 | -0.627 | 0.297 | 0.086 | -0.008 | -0.015 | -0.015 | 11.04 | 0.41 | 0.52 | 0.47 | 0.36 | 0.42 | 0.57 |
| 9 | 5392243 | rs3739637 | 0.152 | -0.782 | 0.219 | 0.08 | -0.043 | -0.035 | -0.035 | 0 | 0.26 | 0.45 | 0.24 | 0.17 | 0.2 | 0.33 |
| 9 | 5407628 | rs10283595 | -2.202 | -0.982 | -0.405 | 0.192 | -0.137 | -0.162 | -0.162 | 0 | 0.6 | 0.97 | 0.58 | 0.53 | 0.51 | 0.79 |
| 9 | 5407803 | rs10283755 | -0.162 | -0.523 | 0.301 | 0.44 | -0.053 | -0.034 | -0.034 | 0 | 0.28 | 0.52 | 0.26 | 0.21 | 0.2 | 0.34 |
| 9 | 5410624 | rs1535453 | 0.892 | -0.524 | 0.141 | 0.354 | -0.009 | 0.009 | 0.009 | 1.793 | 0.1 | 0.13 | 0.06 | 0.16 | 0.06 | 0.2 |
| 9 | 5410723 | rs1535454 | 0.932 | -0.473 | 0.076 | 0.379 | -0.005 | 0.006 | 0.006 | 1.581 | 0.1 | 0.09 | 0.05 | 0.16 | 0.05 | 0.2 |
| 9 | 5412546 | rs10815211 | 0.091 | -0.233 | 0.098 | 0.01 | -0.029 | -0.004 | -0.004 | 0 | 0.25 | 0.38 | 0.24 | 0.25 | 0.2 | 0.34 |
| 9 | 5415874 | rs10975105 | 0.427 | -0.173 | 0.095 | 0.037 | -0.021 | 0.019 | 0.019 | 0 | 0.17 | 0.09 | 0.25 | 0.12 | 0.26 | 0.25 |
| 9 | 5417680 | rs11789768 | 0.406 | -0.155 | 0.103 | -0.005 | -0.02 | 0.025 | 0.025 | 0 | 0.17 | 0.09 | 0.25 | 0.09 | 0.26 | 0.25 |
| 9 | 5418191 | rs10815213 | -0.547 | -0.601 | 0.466 | 0.452 | -0.019 | -0.004 | -0.004 | 0 | 0.27 | 0.37 | 0.26 | 0.28 | 0.22 | 0.36 |
| 9 | 5418334 | rs10815214 | -0.806 | -0.621 | 0.025 | 0.467 | -0.037 | -0.012 | -0.012 | 0 | 0.3 | 0.55 | 0.28 | 0.28 | 0.22 | 0.36 |
| 9 | 5423103 | rs10491647 | -0.348 | -0.065 | -0.834 | 0.55 | -0.016 | -0.055 | -0.055 | 0.018 | 0.91 | 0.99 | 0.87 | 0.77 | 0.93 | 0.76 |
| 9 | 5435709 | rs1322250 | 0.172 | 0.016 | -0.438 | 0.429 | -0.008 | -0.003 | -0.003 | 0.057 | 0.63 | 0.6 | 0.69 | 0.67 | 0.69 | 0.49 |
| 9 | 5436461 | rs2295870 | 0.298 | 0.101 | -0.489 | 0.469 | -0.018 | -0.01 | -0.01 | 0 | 0.61 | 0.53 | 0.68 | 0.66 | 0.67 | 0.46 |
| 9 | 5438435 | rs911760 | -0.452 | 0.655 | -0.248 | 0.446 | 0.011 | 0.006 | 0.006 | 0 | 0.2 | 0.33 | 0.22 | 0.23 | 0.19 | 0.28 |
| 9 | 5442487 | rs822349 | 0.679 | 0.203 | -0.367 | -0.035 | -0.004 | 0.033 | 0.033 | 0.057 | 0.48 | 0.31 | 0.45 | 0.24 | 0.5 | 0.46 |

| | | | | | | | | | | | | | | | | |
|---|---|---|---|---|---|---|---|---|---|---|---|---|---|---|---|---|
| 9 | 5445990 | rs4742095 | 0.756 | -0.027 | -0.419 | -0.266 | -0.005 | 0.006 | 0.006 | 0.429 | 0.13 | 0.17 | 0.14 | 0.17 | 0.12 | 0.13 |
| 9 | 5446078 | rs10975121 | -0.098 | -0.17 | -0.048 | -0.441 | -0.008 | -0.027 | -0.027 | 0 | 0.15 | 0.25 | 0.09 | 0.08 | 0.12 | 0.17 |
| 9 | 5448218 | rs822335 | -0.176 | -0.105 | -0.024 | -0.239 | -0.001 | -0.001 | -0.001 | 0.096 | 0.65 | 0.61 | 0.55 | 0.34 | 0.63 | 0.65 |
| 9 | 5453172 | rs822339 | -0.334 | -0.078 | -0.4 | -0.053 | -0.047 | -0.097 | -0.097 | 1.465 | 0.81 | 0.94 | 0.67 | 0.45 | 0.72 | 0.77 |
| 9 | 5455632 | rs4742097 | -1.974 | -0.366 | 0.041 | -0.369 | 0.006 | -0.052 | -0.052 | 0.021 | 0.58 | 0.47 | 0.64 | 0.92 | 0.56 | 0.62 |
| 9 | 5457733 | rs17804441 | -0.171 | -0.2 | 0.058 | -0.045 | 0.017 | -0.034 | -0.034 | 0 | 0.26 | 0.12 | 0.25 | 0.32 | 0.24 | 0.33 |
| 9 | 5458095 | rs10481593 | 0.22 | -0.039 | 0.158 | 0.059 | 0.011 | 0.111 | 0.111 | 0 | 0.14 | 0.27 | 0.25 | 0.49 | 0.22 | 0.13 |
| 9 | 5459419 | rs1411262 | -0.879 | -0.068 | 0.092 | 0.029 | 0.016 | 0.097 | 0.097 | 0.015 | 0.24 | 0.46 | 0.37 | 0.55 | 0.29 | 0.23 |
| 9 | 5460910 | rs7028894 | -1.695 | -0.018 | 0.042 | -0.51 | -0.001 | -0.012 | -0.012 | 0 | 0.52 | 0.61 | 0.62 | 0.87 | 0.53 | 0.56 |
| 9 | 5464065 | rs10125854 | -0.128 | -0.042 | -0.021 | -0.046 | -0.003 | 0.005 | 0.005 | 0 | 0.08 | 0.06 | 0.04 | 0.06 | 0.07 | 0.1 |
| 9 | 5465130 | rs2890658 | 0.204 | -0.62 | 0.12 | 0.344 | -0.022 | -0.001 | -0.001 | 0 | 0.14 | 0.19 | 0.17 | 0.15 | 0.1 | 0.18 |
| 9 | 5465262 | rs16923173 | -0.836 | -0.237 | 0.461 | 0.303 | 0.011 | 0.008 | 0.008 | 0 | 0.08 | 0.27 | 0.13 | 0.08 | 0.04 | 0.07 |
| 9 | 5467955 | rs2297136 | -1.097 | -0.266 | -0.242 | 0.206 | 0.001 | -0.014 | -0.014 | 0 | 0.6 | 0.66 | 0.66 | 0.81 | 0.55 | 0.67 |
| 9 | 5468257 | rs4143815 | -0.988 | -0.376 | -0.615 | 1.369 | -0.079 | -0.155 | -0.155 | 0.02 | 0.22 | 0.01 | 0.43 | 0.57 | 0.33 | 0.2 |
| 9 | 5470954 | rs4742100 | -0.955 | -0.458 | -0.282 | 1.459 | -0.005 | -0.021 | -0.021 | 0 | 0.21 | 0.16 | 0.39 | 0.53 | 0.27 | 0.2 |
| 9 | 5483659 | rs10815236 | -1.118 | -0.789 | -0.411 | 1.126 | 0.006 | 0.002 | 0.002 | 0 | 0.31 | 0.2 | 0.4 | 0.28 | 0.34 | 0.47 |
| 9 | 5484418 | rs11790653 | -0.33 | -0.886 | -0.449 | 1.072 | -0.002 | 0 | 0 | 0 | 0.13 | 0.13 | 0.08 | 0.15 | 0.09 | 0.17 |
| 9 | 5485166 | rs10815237 | -1.645 | -1.117 | -0.03 | 1.435 | 0.014 | 0.002 | 0.002 | 0 | 0.4 | 0.38 | 0.55 | 0.62 | 0.36 | 0.5 |
| 9 | 5486106 | rs1556188 | -2.27 | -0.912 | 0.117 | 1.156 | 0.012 | 0.029 | 0.029 | 0 | 0.51 | 0.47 | 0.59 | 0.74 | 0.47 | 0.67 |
| 9 | 5486856 | rs10435744 | -1.079 | -0.697 | 0.235 | 1.424 | -0.006 | 0.017 | 0.017 | 0 | 0.19 | 0.2 | 0.19 | 0.43 | 0.11 | 0.2 |
| 9 | 5493994 | rs12338205 | -0.96 | -0.622 | 0.563 | 0.918 | 0.013 | -0.002 | -0.002 | 0 | 0.2 | 0.45 | 0.42 | 0.18 | 0.23 | 0.3 |
| 9 | 5494438 | rs7034893 | -1.821 | -0.7 | 0.531 | 0.944 | -0.001 | -0.021 | -0.021 | 1.144 | 0.3 | 0.57 | 0.5 | 0.29 | 0.31 | 0.39 |
| 9 | 5497531 | rs12376596 | -1.047 | -0.728 | 0.519 | 0.929 | -0.001 | -0.007 | -0.007 | 0 | 0.22 | 0.35 | 0.4 | 0.19 | 0.24 | 0.3 |
| 9 | 5500300 | rs10975148 | -1.785 | -0.681 | 0.289 | 0.959 | -0.009 | -0.027 | -0.027 | 0 | 0.33 | 0.56 | 0.51 | 0.3 | 0.31 | 0.39 |
| 9 | 5502492 | rs10975152 | -1.989 | -0.361 | 0.409 | 1.144 | -0.011 | -0.002 | -0.002 | 0.285 | 0.39 | 0.57 | 0.59 | 0.57 | 0.35 | 0.47 |
| 9 | 5509091 | rs3808847 | -0.375 | -0.239 | -0.065 | 1.161 | 0 | 0.007 | 0.007 | 0 | 0.09 | 0.03 | 0.07 | 0.11 | 0.07 | 0.08 |
| 9 | 5510644 | rs16923189 | -1.011 | -0.266 | -0.148 | 0.995 | -0.001 | 0.023 | 0.023 | 0.014 | 0.27 | 0.23 | 0.44 | 0.18 | 0.3 | 0.36 |
| 9 | 5512459 | rs16923193 | -0.918 | -0.203 | -0.377 | 1.089 | -0.005 | 0.017 | 0.017 | 0 | 0.12 | 0.08 | 0.13 | 0.34 | 0.14 | 0.14 |
| 9 | 5512522 | rs16923195 | -1.65 | 0.009 | -0.245 | 0.123 | -0.002 | -0.002 | -0.002 | 0 | 0.31 | 0.34 | 0.48 | 0.3 | 0.27 | 0.38 |
| 9 | 5512786 | rs16923198 | -0.656 | 0.115 | -0.175 | 0.298 | -0.008 | -0.002 | -0.002 | 0 | 0.19 | 0.25 | 0.37 | 0.19 | 0.15 | 0.26 |
| 9 | 5514372 | rs1360238 | -0.992 | 0.215 | -0.095 | 0.395 | -0.001 | 0.001 | 0.001 | 0.005 | 0.29 | 0.3 | 0.47 | 0.2 | 0.32 | 0.37 |
| 9 | 5518260 | rs12001295 | -1.49 | 0.148 | 0.284 | 0.603 | -0.007 | 0.02 | 0.02 | 0 | 0.09 | 0.31 | 0.08 | 0.14 | 0.06 | 0.08 |
| 9 | 5523773 | rs16923201 | -1.608 | 0.186 | -0.001 | 0.422 | -0.009 | -0.027 | -0.027 | 0.249 | 0.1 | 0.19 | 0.06 | 0.03 | 0.09 | 0.06 |
| 9 | 5526124 | rs4237162 | -0.913 | 0.358 | -0.555 | 0.149 | -0.014 | 0.029 | 0.029 | 0 | 0.64 | 0.82 | 0.52 | 0.78 | 0.54 | 0.8 |
| 9 | 5532871 | rs4742104 | -0.059 | 0.215 | -0.513 | 0.783 | 0.002 | -0.031 | -0.031 | 0 | 0.21 | 0.13 | 0.12 | 0.43 | 0.16 | 0.44 |
| 9 | 5533831 | rs10435814 | -0.825 | 0.484 | -0.588 | 0.326 | -0.061 | 0.098 | 0.098 | 0 | 0.27 | 0.01 | 0.2 | 0.11 | 0.26 | 0.27 |
| 9 | 5534087 | rs10435815 | -1.413 | 0.428 | -0.65 | 0.265 | -0.097 | 0.157 | 0.157 | 0.01 | 0.33 | 0.01 | 0.23 | 0.11 | 0.33 | 0.29 |
| 9 | 5534243 | rs4143813 | -1.091 | 0.19 | -0.066 | 0.609 | 0.007 | 0.103 | 0.103 | 2.074 | 0.13 | 0.67 | 0.21 | 0.24 | 0.11 | 0.06 |
| 9 | 5544405 | rs10975175 | -1.686 | 0.177 | 0.371 | 0.561 | 0.004 | 0.016 | 0.016 | 0 | 0.34 | 0.46 | 0.39 | 0.39 | 0.35 | 0.3 |
| 9 | 5546496 | rs7028800 | -0.251 | 0.247 | 0.876 | 0.754 | 0.115 | 0.328 | 0.328 | 0 | 0.07 | 0.3 | 0.18 | 0.29 | 0.1 | 0.06 |
| 9 | 5551261 | rs7869201 | -1.025 | 0.182 | 0.444 | 0.541 | -0.015 | 0.034 | 0.034 | 0 | 0.26 | 0.63 | 0.34 | 0.38 | 0.22 | 0.17 |
| 9 | 5555023 | rs7871338 | -0.941 | 0.185 | 0.364 | 0.635 | 0.001 | 0.032 | 0.032 | 0 | 0.3 | 0.41 | 0.37 | 0.39 | 0.29 | 0.26 |
| 9 | 5556786 | rs1009759 | -1.225 | 0.173 | 0.044 | 0.656 | -0.001 | -0.006 | -0.006 | 0 | 0.37 | 0.69 | 0.39 | 0.49 | 0.34 | 0.31 |
| 9 | 5557708 | rs7854413 | -1.43 | -0.581 | 0.708 | 0.309 | -0.038 | -0.031 | -0.031 | 0 | 0.24 | 0.64 | 0.13 | 0.12 | 0.1 | 0.25 |
| 9 | 5559893 | rs10975179 | -1.103 | -0.375 | 0.945 | 0.419 | -0.042 | -0.028 | -0.028 | 0 | 0.21 | 0.44 | 0.11 | 0.12 | 0.11 | 0.22 |
| 9 | 5564748 | rs7858319 | -1.186 | -0.41 | 0.904 | 0.464 | -0.013 | -0.004 | -0.004 | 0.369 | 0.19 | 0.19 | 0.09 | 0.14 | 0.11 | 0.24 |
| 9 | 5568222 | rs7031529 | -1.183 | -0.275 | 0.532 | 0.445 | -0.043 | -0.03 | -0.03 | 0 | 0.2 | 0.41 | 0.11 | 0.09 | 0.11 | 0.22 |
| 9 | 5579041 | rs7025291 | -0.477 | 0.089 | -0.637 | 0.385 | 0.024 | 0.092 | 0.092 | 0 | 0.51 | 0.73 | 0.61 | 0.69 | 0.54 | 0.56 |
| 9 | 5579213 | rs7024586 | -0.665 | -0.098 | -0.24 | 0.787 | -0.003 | -0.008 | -0.008 | 0 | 0.24 | 0.61 | 0.32 | 0.21 | 0.2 | 0.13 |
| 9 | 5581140 | rs1536927 | -1.072 | 0.014 | -0.61 | 1.419 | 0.025 | -0.038 | -0.038 | 0 | 0.19 | 0.06 | 0.12 | 0.3 | 0.12 | 0.38 |
| 9 | 5593419 | rs10975190 | -0.394 | -0.022 | -0.378 | 1.536 | 0 | 0.001 | 0.001 | 0 | 0.14 | 0.14 | 0.12 | 0.27 | 0.07 | 0.31 |
| 9 | 5607230 | rs10815252 | -0.907 | -0.181 | -0.35 | 1.298 | 0.036 | 0.01 | 0.01 | 0 | 0.18 | 0.06 | 0.11 | 0.19 | 0.1 | 0.32 |
| 9 | 5607774 | rs10975196 | -1.149 | -0.325 | 0.622 | 1.21 | -0.043 | -0.024 | -0.024 | 0 | 0.22 | 0.31 | 0.16 | 0.2 | 0.12 | 0.33 |
| 9 | 5608026 | rs7036748 | -1.272 | -0.324 | 0.866 | 1.196 | -0.058 | -0.039 | -0.039 | 0.216 | 0.23 | 0.42 | 0.18 | 0.2 | 0.13 | 0.33 |
| 9 | 5610320 | rs4740819 | -0.693 | -0.037 | 0.557 | 0.831 | 0.004 | -0.005 | -0.005 | 0 | 0.35 | 0.32 | 0.4 | 0.6 | 0.26 | 0.41 |
| 9 | 5610381 | rs4740820 | -0.817 | -0.042 | 0.551 | 0.822 | 0.006 | -0.006 | -0.006 | 0.867 | 0.35 | 0.3 | 0.4 | 0.6 | 0.26 | 0.41 |
| 9 | 5610501 | rs4740821 | -1.64 | 0.038 | 0.462 | 0.42 | 0.029 | 0.017 | 0.017 | 0 | 0.52 | 0.67 | 0.53 | 0.6 | 0.42 | 0.51 |
| 9 | 5611912 | rs7037208 | -1.058 | 0.247 | 0.302 | 0.7 | -0.01 | 0.001 | 0.001 | 2.227 | 0.4 | 0.4 | 0.43 | 0.6 | 0.3 | 0.47 |
| 9 | 5621588 | rs10975202 | -0.722 | 0.217 | 0.139 | 0.823 | -0.017 | 0.017 | 0.017 | 1.918 | 0.34 | 0.41 | 0.43 | 0.58 | 0.26 | 0.4 |
| 9 | 5625118 | rs10122199 | -0.861 | 0.043 | 0.048 | 0.853 | -0.024 | -0.02 | -0.02 | 0 | 0.38 | 0.44 | 0.45 | 0.72 | 0.26 | 0.44 |
| 9 | 5632721 | rs4740823 | -0.697 | 0.595 | -0.329 | 0.907 | -0.007 | -0.033 | -0.033 | 0 | 0.35 | 0.13 | 0.38 | 0.61 | 0.3 | 0.44 |
| 9 | 5639227 | rs7019690 | -0.59 | 0.077 | 0.097 | 0.332 | -0.025 | -0.104 | -0.104 | 0 | 0.15 | 0.39 | 0.1 | 0 | 0.08 | 0.07 |
| 9 | 5644555 | rs7860397 | -1.266 | 0.1 | 0.096 | 0.426 | 0.013 | 0.025 | 0.025 | 0 | 0.49 | 0.59 | 0.46 | 0.56 | 0.39 | 0.49 |
| 9 | 5692233 | rs10975244 | -0.376 | -0.08 | 0.011 | 0.906 | -0.003 | -0.006 | -0.006 | 0 | 0.34 | 0.17 | 0.32 | 0.59 | 0.31 | 0.43 |
| 9 | 5696021 | rs2760415 | -0.032 | -0.333 | -0.067 | 0.663 | -0.003 | 0.019 | 0.019 | 6.112 | 0.19 | 0.37 | 0.23 | 0.27 | 0.19 | 0.19 |
| 9 | 5700676 | rs16923398 | 1.593 | -0.227 | -0.326 | 0.605 | -0.04 | -0.051 | -0.051 | 0 | 0.08 | 0.01 | 0.14 | 0.25 | 0.1 | 0.13 |
| 9 | 5703394 | rs7047997 | -0.36 | -0.409 | -0.116 | 0.51 | 0 | 0.004 | 0.004 | 5.724 | 0.2 | 0.26 | 0.23 | 0.26 | 0.2 | 0.21 |
| 9 | 5703443 | rs7028052 | -1.262 | -0.461 | -0.342 | 0.457 | 0.01 | 0.012 | 0.012 | 4.57 | 0.27 | 0.84 | 0.3 | 0.26 | 0.24 | 0.21 |
| 9 | 5715597 | rs17292138 | -0.634 | -0.425 | -0.034 | 0.761 | 0.005 | 0.011 | 0.011 | 0 | 0.39 | 0.23 | 0.34 | 0.6 | 0.32 | 0.48 |

| | | | | | | | | | | | | | | | | |
|---|---|---|---|---|---|---|---|---|---|---|---|---|---|---|---|---|
| 9 | 5719116 | rs10975258 | -0.746 | -0.614 | -0.261 | 0.825 | -0.08 | 0.049 | 0.049 | 0 | 0.35 | 0.12 | 0.44 | 0.75 | 0.39 | 0.4 |
| 9 | 5722220 | rs10975262 | -1.545 | -0.88 | -0.418 | 0.688 | -0.046 | 0.271 | 0.271 | 0.886 | 0.51 | 0.11 | 0.48 | 0.83 | 0.43 | 0.62 |
| 9 | 5728968 | rs10815274 | -1.869 | -0.888 | -0.202 | 0.634 | 0.004 | 0.087 | 0.087 | 0.252 | 0.53 | 0.3 | 0.52 | 0.84 | 0.47 | 0.62 |
| 9 | 5731062 | rs4742118 | -0.944 | -1.271 | 0.109 | 1.012 | -0.02 | -0.037 | -0.037 | 0 | 0.38 | 0.1 | 0.33 | 0.58 | 0.32 | 0.48 |
| 9 | 5742085 | rs4742120 | -0.743 | -1.346 | 0.099 | 1.097 | -0.022 | -0.103 | -0.103 | 0.102 | 0.31 | 0.11 | 0.29 | 0.5 | 0.28 | 0.26 |
| 9 | 5742450 | rs7869667 | 0.965 | -1.374 | 0.177 | 1.119 | -0.009 | -0.004 | -0.004 | 0 | 0.12 | 0.14 | 0.04 | 0.08 | 0.06 | 0.25 |
| 9 | 5747187 | rs10815278 | -1.255 | -1.674 | 0.366 | 1.267 | -0.001 | 0.016 | 0.016 | 0.062 | 0.33 | 0.6 | 0.33 | 0.41 | 0.29 | 0.21 |
| 9 | 5748218 | rs10975270 | -0.726 | -1.583 | 0.415 | 1.13 | -0.033 | -0.119 | -0.119 | 0.368 | 0.3 | 0.07 | 0.29 | 0.49 | 0.27 | 0.26 |
| 9 | 5750198 | rs2031179 | -1.149 | -1.568 | 0.427 | 1.115 | -0.002 | 0.02 | 0.02 | 0 | 0.34 | 0.61 | 0.35 | 0.49 | 0.29 | 0.25 |
| 9 | 5756067 | rs7030124 | -1.563 | -1.593 | 0.806 | 1.092 | -0.004 | -0.003 | -0.003 | 0 | 0.47 | 0.49 | 0.38 | 0.58 | 0.36 | 0.5 |
| 9 | 5762600 | rs3739646 | 1.115 | -1.599 | 0.724 | 1.102 | 0.034 | -0.008 | -0.008 | 0 | 0.11 | 0.01 | 0.03 | 0.07 | 0.04 | 0.23 |
| 9 | 5762894 | rs1543527 | -1.547 | -1.561 | 0.912 | 1.041 | -0.004 | -0.004 | -0.004 | 0 | 0.48 | 0.5 | 0.38 | 0.58 | 0.36 | 0.5 |
| 9 | 5765984 | rs754241 | -1.495 | -1.538 | 0.606 | 1.017 | 0.001 | 0.001 | 0.001 | 0 | 0.47 | 0.51 | 0.39 | 0.58 | 0.35 | 0.49 |
| 9 | 5774225 | rs3739648 | -0.406 | -1.485 | -0.266 | 1.096 | -0.018 | -0.058 | -0.058 | 1.339 | 0.31 | 0.12 | 0.3 | 0.59 | 0.29 | 0.43 |
| 9 | 5777553 | rs16923507 | -0.571 | -1.469 | -0.234 | 1.026 | -0.005 | -0.032 | -0.032 | 0.245 | 0.32 | 0.16 | 0.32 | 0.58 | 0.29 | 0.44 |
| 9 | 5786850 | rs1028702 | -0.589 | -1.44 | -0.179 | 1.157 | 0 | -0.073 | -0.073 | 0.129 | 0.33 | 0.14 | 0.31 | 0.53 | 0.28 | 0.43 |
| 9 | 5791151 | rs1342854 | -0.248 | -1.496 | -0.249 | 1.166 | -0.016 | -0.03 | -0.03 | 0.967 | 0.24 | 0.19 | 0.4 | 0.59 | 0.28 | 0.21 |
| 9 | 5792106 | rs10975282 | -0.61 | -1.292 | -0.044 | 0.966 | 0.007 | 0.044 | 0.044 | 1.208 | 0.36 | 0.23 | 0.46 | 0.74 | 0.34 | 0.45 |
| 9 | 5792156 | rs10975283 | -0.405 | -1.282 | -0.133 | 0.969 | 0.002 | 0.023 | 0.023 | 1.343 | 0.36 | 0.19 | 0.43 | 0.68 | 0.32 | 0.44 |
| 9 | 5801121 | rs3824444 | -0.309 | -1.3 | -0.254 | 0.838 | -0.008 | 0.041 | 0.041 | 0.668 | 0.34 | 0.15 | 0.42 | 0.73 | 0.31 | 0.44 |
| 9 | 5820306 | rs10975302 | -0.609 | -1.054 | -0.193 | 0.941 | 0.02 | 0.13 | 0.13 | 0.023 | 0.45 | 0.17 | 0.44 | 0.73 | 0.35 | 0.52 |
| 9 | 5820878 | rs7861988 | 0.714 | -1.046 | -0.215 | 0.932 | 0.017 | 0.039 | 0.039 | 0 | 0.52 | 0.56 | 0.54 | 0.27 | 0.65 | 0.48 |
| 9 | 5825768 | rs10975303 | 1.154 | -1.057 | -1.328 | 0.732 | -0.019 | -0.016 | -0.016 | 0 | 0.45 | 0.09 | 0.55 | 0.43 | 0.62 | 0.49 |
| 9 | 5833337 | rs10975309 | -0.737 | -0.676 | -1.261 | 0.911 | 0.011 | -0.055 | -0.055 | 0.564 | 0.44 | 0.13 | 0.33 | 0.52 | 0.33 | 0.5 |
| 9 | 5845105 | rs10758710 | 0.792 | -0.345 | -0.52 | 0.605 | 0.005 | 0.037 | 0.037 | 0 | 0.58 | 0.36 | 0.53 | 0.33 | 0.62 | 0.66 |
| 9 | 5850834 | rs4289865 | 0.74 | -0.401 | -0.374 | 0.628 | 0.001 | 0.007 | 0.007 | 0 | 0.61 | 0.4 | 0.53 | 0.33 | 0.65 | 0.69 |
| 9 | 5854501 | rs1418746 | 0.163 | -0.641 | 0.169 | 0.894 | -0.01 | -0.006 | -0.006 | 0.408 | 0.8 | 0.76 | 0.93 | 0.85 | 0.91 | 0.87 |
| 9 | 5856209 | rs6477005 | 0.498 | -0.509 | 0.906 | 0.997 | 0.058 | 0.171 | 0.171 | 0 | 0.81 | 0.53 | 0.61 | 0.5 | 0.74 | 0.82 |
| 9 | 5859800 | rs7041923 | -0.249 | -0.445 | 0.912 | 1.06 | 0.066 | 0.205 | 0.205 | 0 | 0.17 | 0.42 | 0.38 | 0.5 | 0.24 | 0.18 |
| 9 | 5861604 | rs7044017 | -0.582 | -0.491 | 0.936 | 1.05 | 0.064 | 0.175 | 0.175 | 0 | 0.19 | 0.43 | 0.39 | 0.5 | 0.26 | 0.18 |
| 9 | 5864988 | rs1418742 | 0.701 | -0.943 | 0.54 | 1.178 | -0.017 | 0.004 | 0.004 | 0 | 0.79 | 0.72 | 0.82 | 0.75 | 0.9 | 0.85 |
| 9 | 5870703 | rs4740826 | 0.503 | -1.377 | 0.528 | 1.082 | -0.061 | 0.061 | 0.061 | 0.145 | 0.74 | 0.47 | 0.74 | 0.6 | 0.86 | 0.77 |
| 9 | 5872949 | rs10758713 | -0.285 | -1.47 | 0.564 | 1.039 | 0.045 | -0.076 | -0.076 | 0 | 0.82 | 0.97 | 0.9 | 0.71 | 0.9 | 0.79 |
| 9 | 5875294 | rs10739078 | 1.411 | -0.692 | 0.573 | 0.985 | 0.002 | 0.107 | 0.107 | 0.661 | 0.6 | 0.24 | 0.47 | 0.2 | 0.64 | 0.66 |
| 9 | 5875690 | rs6477016 | -0.21 | -0.887 | 0.359 | 1.053 | 0.045 | -0.078 | -0.078 | 1.522 | 0.81 | 0.98 | 0.9 | 0.7 | 0.89 | 0.78 |
| 9 | 5877948 | rs11791517 | 0.917 | -1.047 | 0.437 | 0.851 | 0.004 | -0.003 | -0.003 | 0 | 0.62 | 0.37 | 0.65 | 0.53 | 0.75 | 0.58 |
| 9 | 5884281 | rs10758717 | -0.936 | -1.083 | 0.736 | 0.846 | 0.01 | 0.01 | 0.01 | 0 | 0.34 | 0.3 | 0.24 | 0.12 | 0.25 | 0.33 |
| 9 | 5890270 | rs2233173 | -0.306 | -1.101 | 0.833 | 1.003 | -0.016 | 0.031 | 0.031 | 0 | 0.9 | 0.84 | 0.94 | 0.75 | 0.94 | 0.76 |
| 9 | 5894344 | rs1052994 | 0.843 | -0.356 | 0.791 | 1.424 | 0.029 | 0.07 | 0.07 | 0 | 0.11 | 0.13 | 0.36 | 0.45 | 0.24 | 0.12 |
| 9 | 5895586 | rs10975339 | 1.153 | -0.953 | 0.38 | 1.019 | 0 | 0.036 | 0.036 | 0.527 | 0.61 | 0.16 | 0.51 | 0.31 | 0.65 | 0.6 |
| 9 | 5898829 | rs16923640 | -0.056 | -0.874 | 0.511 | 1.05 | 0.01 | 0.034 | 0.034 | 0 | 0.26 | 0.35 | 0.41 | 0.43 | 0.27 | 0.15 |
| 9 | 5899749 | rs4554522 | 0.019 | -1.465 | 0.379 | 0.751 | 0.019 | 0.036 | 0.036 | 1.687 | 0.25 | 0.11 | 0.18 | 0.1 | 0.19 | 0.19 |
| 9 | 5901100 | rs10815303 | 1.042 | -1.549 | 0.173 | 0.764 | 0.007 | 0 | 0 | 2.421 | 0.17 | 0 | 0.12 | 0.1 | 0.12 | 0.14 |
| 9 | 5909152 | rs1056796 | 1.214 | -1.46 | 0.347 | 0.72 | 0.048 | 0.009 | 0.009 | 0 | 0.49 | 0.38 | 0.56 | 0.53 | 0.69 | 0.5 |
| 9 | 5909259 | rs1056798 | 1.223 | -1.461 | 0.397 | 0.719 | 0.065 | 0.012 | 0.012 | 0 | 0.49 | 0.35 | 0.56 | 0.53 | 0.69 | 0.5 |
| 9 | 5913668 | rs10758719 | -0.873 | -1.608 | 0.625 | 0.679 | 0.029 | 0.03 | 0.03 | 0.874 | 0.4 | 0.3 | 0.29 | 0.13 | 0.21 | 0.38 |
| 9 | 5915835 | rs10975346 | -0.58 | -1.659 | 0.677 | 0.655 | 0.072 | 0.096 | 0.096 | 1.556 | 0.37 | 0.13 | 0.27 | 0.1 | 0.2 | 0.23 |
| 9 | 5931216 | rs1478938 | -0.116 | -1.348 | 1.008 | 0.861 | -0.003 | 0.154 | 0.154 | 0 | 0.88 | 0.64 | 0.84 | 0.66 | 0.9 | 0.89 |
| 9 | 5932616 | rs10815311 | -0.594 | -1.478 | 0.825 | 0.706 | 0.029 | 0.03 | 0.03 | 0.503 | 0.4 | 0.3 | 0.29 | 0.14 | 0.22 | 0.38 |
| 9 | 5939792 | rs10975360 | -0.584 | -1.481 | 0.979 | 0.699 | 0.011 | 0.009 | 0.009 | 0.54 | 0.4 | 0.35 | 0.29 | 0.14 | 0.22 | 0.37 |
| 9 | 5944012 | rs10815315 | -0.335 | -1.449 | 0.811 | 0.703 | 0.086 | 0.115 | 0.115 | 1.761 | 0.37 | 0.11 | 0.27 | 0.11 | 0.22 | 0.24 |
| 9 | 5956318 | rs1984272 | 0.763 | -1.422 | 0.901 | 0.886 | 0.025 | -0.006 | -0.006 | 0 | 0.56 | 0.39 | 0.6 | 0.53 | 0.76 | 0.54 |
| 9 | 5966898 | rs13289592 | -0.099 | -1.354 | 0.822 | 0.709 | 0.002 | 0.022 | 0.022 | 0 | 0.14 | 0.05 | 0.09 | 0 | 0.14 | 0.06 |
| 9 | 5971977 | rs10815327 | 0.134 | -1.393 | 0.881 | 0.838 | 0.086 | 0.087 | 0.087 | 3.564 | 0.32 | 0.09 | 0.22 | 0.12 | 0.16 | 0.21 |
| 9 | 6009761 | rs343477 | -1.481 | -1.225 | 1.114 | 1.077 | 0.048 | 0.028 | 0.028 | 0 | 0.54 | 0.66 | 0.36 | 0.32 | 0.32 | 0.51 |
| 9 | 6011335 | rs7425 | -0.179 | -1.226 | 1.117 | 1.203 | 0.094 | 0.086 | 0.086 | 1.877 | 0.38 | 0.09 | 0.27 | 0.16 | 0.23 | 0.25 |
| 9 | 6011757 | rs10758736 | -0.411 | -1.192 | 1.237 | 1.186 | 0.048 | 0.045 | 0.045 | 0.664 | 0.41 | 0.27 | 0.29 | 0.17 | 0.23 | 0.39 |
| 9 | 6033278 | rs12378118 | 0.538 | 0.55 | 1.111 | 1.773 | 0.064 | 0.076 | 0.076 | 0 | 0.07 | 0.12 | 0.15 | 0.29 | 0.16 | 0.08 |
| 9 | 6034076 | rs899381 | -1.38 | -0.016 | 1.345 | 1.544 | 0.024 | 0.318 | 0.318 | 0.948 | 0.07 | 0.44 | 0.05 | 0.22 | 0.02 | 0.27 |
| 9 | 6042708 | rs10815345 | -0.746 | 0.1 | 0.921 | 1.135 | 0.051 | 0.031 | 0.031 | 0 | 0.49 | 0.59 | 0.35 | 0.31 | 0.3 | 0.54 |
| 9 | 6049547 | rs10975412 | 1.339 | 0.243 | 0.351 | 0.407 | 0.003 | -0.002 | -0.002 | 0.289 | 0.2 | 0.18 | 0.14 | 0.02 | 0.19 | 0.24 |
| 9 | 6051924 | rs10975416 | 1.495 | 0.257 | 0.274 | 0.414 | 0.007 | 0.023 | 0.023 | 0.349 | 0.19 | 0.14 | 0.15 | 0.02 | 0.2 | 0.25 |
| 9 | 6067122 | rs10739083 | 0.445 | 0.186 | 0.041 | 0.387 | 0.054 | 0.139 | 0.139 | 0.046 | 0.4 | 0.09 | 0.32 | 0.14 | 0.3 | 0.29 |
| 9 | 6070342 | rs343494 | -0.211 | 0.346 | 0.796 | 0.486 | 0.01 | -0.004 | -0.004 | 1.497 | 0.65 | 0.59 | 0.5 | 0.27 | 0.51 | 0.57 |
| 9 | 6071843 | rs7872100 | -0.107 | 0.287 | 0.761 | 0.466 | 0.01 | 0.103 | 0.103 | 0 | 0.14 | 0.34 | 0.23 | 0.25 | 0.14 | 0.21 |
| 9 | 6072597 | rs343476 | -1.906 | 0.37 | 0.36 | 0.238 | 0.006 | 0.012 | 0.012 | 0.436 | 0.81 | 0.85 | 0.85 | 0.98 | 0.81 | 0.76 |
| 9 | 6085078 | rs340933 | -1.731 | 0.375 | 0.311 | 0.234 | -0.009 | -0.041 | -0.041 | 0 | 0.8 | 0.76 | 0.85 | 0.98 | 0.82 | 0.77 |
| 9 | 6088457 | rs10120134 | 0.878 | 1.085 | 0.351 | 0.738 | 0.046 | 0.052 | 0.052 | 0 | 0.12 | 0.17 | 0.2 | 0.32 | 0.21 | 0.09 |
| 9 | 6089527 | rs4740837 | 0.324 | 1.014 | -0.047 | 0.522 | -0.027 | -0.017 | -0.017 | 0 | 0.13 | 0.01 | 0.19 | 0.12 | 0.13 | 0.18 |

| | | | | | | | | | | | | | | | | |
|---|---|---|---|---|---|---|---|---|---|---|---|---|---|---|---|---|
| 9 | 6098662 | rs340892 | 0.278 | 0.878 | 0.147 | 0.571 | 0.021 | 0.009 | 0.009 | 5.768 | 0.31 | 0.35 | 0.49 | 0.67 | 0.47 | 0.43 |
| 9 | 6100169 | rs340890 | 1.919 | 0.765 | -0.028 | 0.778 | -0.024 | -0.021 | -0.021 | 0.766 | 0.17 | 0.06 | 0.27 | 0.26 | 0.24 | 0.31 |
| 9 | 6118174 | rs6477037 | 0.177 | 0.329 | -0.032 | 0.473 | 0.005 | 0.016 | 0.016 | 0 | 0.11 | 0.24 | 0.19 | 0.12 | 0.12 | 0.1 |
| 9 | 6119329 | rs10975444 | 0.795 | 0.164 | -0.244 | 0.366 | 0.08 | 0.049 | 0.049 | 1.421 | 0.34 | 0.07 | 0.13 | 0.19 | 0.21 | 0.18 |
| 9 | 6122556 | rs693838 | -0.151 | 0.073 | 0.103 | 0.032 | -0.021 | -0.051 | -0.051 | 0 | 0.69 | 0.52 | 0.65 | 0.87 | 0.69 | 0.66 |
| 9 | 6133074 | rs6477040 | 0.007 | 0.206 | -0.2 | 0.103 | 0.014 | 0.019 | 0.019 | 0 | 0.11 | 0.24 | 0.19 | 0.11 | 0.14 | 0.1 |
| 9 | 6134999 | rs376382 | -0.204 | 0.199 | -0.282 | 0.062 | -0.018 | -0.06 | -0.06 | 0 | 0.69 | 0.49 | 0.64 | 0.87 | 0.68 | 0.66 |
| 9 | 6142157 | rs544253 | 0.855 | 0.639 | -1.005 | 0.042 | -0.064 | -0.064 | -0.064 | 0 | 0.21 | 0.03 | 0.31 | 0.35 | 0.29 | 0.41 |
| 9 | 6142948 | rs413382 | -0.757 | -0.009 | -0.183 | -0.334 | -0.007 | -0.055 | -0.055 | 0 | 0.8 | 0.76 | 0.84 | 0.99 | 0.8 | 0.76 |
| 9 | 6145022 | rs387149 | 0.382 | 0.08 | 0.042 | 0.239 | -0.01 | 0.036 | 0.036 | 1.758 | 0.64 | 0.56 | 0.67 | 0.49 | 0.63 | 0.75 |
| 9 | 6145491 | rs12349559 | 1.063 | -0.402 | 0.302 | -0.093 | -0.001 | 0 | 0 | 0 | 0.11 | 0.14 | 0.07 | 0.1 | 0.08 | 0.07 |
| 9 | 6149006 | rs10975463 | 0.017 | -0.356 | 0.337 | -0.356 | -0.002 | -0.005 | -0.005 | 0 | 0.09 | 0.08 | 0.07 | 0 | 0.09 | 0.04 |
| 9 | 6158778 | rs10117792 | 1.46 | -0.386 | 0.536 | -0.173 | -0.009 | 0.018 | 0.018 | 0.828 | 0.1 | 0.18 | 0.06 | 0.15 | 0.06 | 0.07 |
| 9 | 6161686 | rs10124250 | -1.466 | -0.506 | 0.915 | -0.484 | 0.011 | 0.039 | 0.039 | 0 | 0.49 | 0.63 | 0.39 | 0.22 | 0.43 | 0.38 |
| 9 | 6163823 | rs10119713 | -1.513 | -0.52 | 0.929 | -0.488 | 0.012 | 0.034 | 0.034 | 0 | 0.49 | 0.64 | 0.39 | 0.3 | 0.43 | 0.38 |
| 9 | 6164771 | rs7863536 | 1.769 | -0.38 | 0.412 | -0.459 | -0.003 | -0.001 | -0.001 | 0.223 | 0.1 | 0.12 | 0.06 | 0.08 | 0.06 | 0.07 |
| 9 | 6166653 | rs2079 | -0.79 | -0.402 | 0.465 | -0.482 | -0.007 | 0.076 | 0.076 | 0.337 | 0.37 | 0.2 | 0.32 | 0.15 | 0.37 | 0.27 |
| 9 | 6170162 | rs10118918 | 1.417 | -0.325 | 0.758 | -0.503 | -0.003 | 0.006 | 0.006 | 0.621 | 0.1 | 0.13 | 0.06 | 0.15 | 0.06 | 0.08 |
| 9 | 6176770 | rs4742165 | -0.76 | -0.563 | 1 | -0.865 | -0.029 | -0.017 | -0.017 | 0 | 0.87 | 0.79 | 0.93 | 0.91 | 0.94 | 0.93 |
| 9 | 6184165 | rs2890704 | -0.434 | -0.546 | 1.055 | -1.054 | -0.02 | -0.058 | -0.058 | 0 | 0.84 | 0.73 | 0.94 | 0.95 | 0.91 | 0.91 |
| 9 | 6185360 | rs13291323 | 0.732 | -0.563 | 0.783 | -1.106 | -0.004 | 0.033 | 0.033 | 0 | 0.06 | 0.01 | 0.04 | 0 | 0.05 | 0.01 |
| 9 | 6188652 | rs1412426 | 1.311 | -0.773 | 0.972 | -1.397 | 0.039 | 0.101 | 0.101 | 0 | 0.58 | 0.25 | 0.74 | 0.92 | 0.69 | 0.71 |
| 9 | 6201364 | rs2150970 | -0.42 | -0.652 | 1.352 | -1.149 | -0.024 | -0.053 | -0.053 | 0 | 0.84 | 0.7 | 0.93 | 0.93 | 0.91 | 0.91 |
| 9 | 6209697 | rs992969 | 0.388 | -0.671 | 1.265 | -1.194 | -0.006 | -0.066 | -0.066 | 0.463 | 0.69 | 0.67 | 0.83 | 0.95 | 0.75 | 0.78 |
| 9 | 6210099 | rs3939286 | 0.71 | -0.646 | 1.201 | -1.182 | -0.022 | -0.125 | -0.125 | 0 | 0.66 | 0.49 | 0.8 | 0.95 | 0.75 | 0.77 |
| 9 | 6213468 | rs7848215 | -0.776 | -0.474 | 0.606 | -1.103 | -0.006 | -0.055 | -0.055 | 0 | 0.35 | 0.64 | 0.2 | 0.08 | 0.26 | 0.27 |
| 9 | 6219176 | rs2066362 | -0.36 | -0.563 | 0.668 | -1.049 | -0.025 | -0.124 | -0.124 | 0 | 0.22 | 0.44 | 0.13 | 0 | 0.17 | 0.21 |
| 9 | 6219845 | rs1891385 | 0.325 | -0.617 | -0.091 | -0.593 | -0.022 | -0.065 | -0.065 | 0.118 | 0.13 | 0.01 | 0.15 | 0.22 | 0.13 | 0.15 |
| 9 | 6221246 | rs16924144 | 1.187 | -0.191 | -0.555 | -0.475 | 0.008 | 0.007 | 0.007 | 0 | 0.36 | 0.13 | 0.39 | 0.31 | 0.33 | 0.39 |
| 9 | 6222110 | rs11794419 | 0.512 | -0.202 | 0.134 | -0.585 | -0.019 | -0.049 | -0.049 | 0 | 0.09 | 0.24 | 0.04 | 0 | 0.03 | 0.08 |
| 9 | 6225535 | rs10435816 | -0.212 | 0.311 | 0.195 | -0.687 | 0.005 | 0.049 | 0.049 | 1.874 | 0.27 | 0.61 | 0.27 | 0.49 | 0.25 | 0.31 |
| 9 | 6227752 | rs2210463 | -0.083 | 0.319 | 0.341 | -0.683 | 0.014 | 0.083 | 0.083 | 1.75 | 0.26 | 0.56 | 0.25 | 0.49 | 0.25 | 0.3 |
| 9 | 6229417 | rs16924159 | 1.691 | 0.236 | -0.276 | -0.599 | 0.004 | 0.003 | 0.003 | 0 | 0.32 | 0.13 | 0.38 | 0.27 | 0.32 | 0.38 |
| 9 | 6230912 | rs16924161 | 1.108 | 0.573 | 0.023 | -0.573 | -0.001 | 0.076 | 0.076 | 0 | 0.06 | 0.03 | 0.17 | 0.46 | 0.17 | 0.16 |
| 9 | 6231239 | rs12551256 | 0.713 | 0.478 | -0.03 | -0.777 | -0.005 | -0.023 | -0.023 | 0 | 0.48 | 0.16 | 0.55 | 0.47 | 0.47 | 0.52 |
| 9 | 6232242 | rs10815388 | 0.666 | 0.251 | 0.793 | -0.609 | -0.001 | 0.023 | 0.023 | 2.658 | 0.67 | 0.41 | 0.69 | 0.48 | 0.7 | 0.66 |
| 9 | 6233221 | rs10975501 | -1.119 | 0.331 | 0.691 | -0.559 | -0.002 | -0.036 | -0.036 | 0 | 0.93 | 0.82 | 0.93 | 1 | 0.95 | 0.99 |
| 9 | 6242950 | rs4742170 | 0.601 | 0.18 | 0.46 | -0.643 | 0 | -0.014 | -0.014 | 0 | 0.66 | 0.3 | 0.68 | 0.48 | 0.69 | 0.66 |
| 9 | 6247430 | rs7037276 | -1.207 | 0.123 | 0.582 | -0.397 | 0 | -0.018 | -0.018 | 0 | 0.94 | 0.94 | 0.95 | 1 | 0.94 | 0.98 |
| 9 | 6247693 | rs10975516 | 0.205 | -0.232 | 0.471 | -0.88 | 0.001 | -0.003 | -0.003 | 0 | 0.39 | 0.6 | 0.47 | 0.46 | 0.32 | 0.41 |
| 9 | 6251507 | rs1330383 | 0.474 | -0.271 | 0.49 | -0.879 | -0.008 | 0.011 | 0.011 | 0 | 0.37 | 0.41 | 0.45 | 0.46 | 0.32 | 0.41 |
| 9 | 6253571 | rs10975519 | 0.423 | -0.26 | 0.552 | -0.863 | -0.008 | 0.008 | 0.008 | 0 | 0.38 | 0.42 | 0.45 | 0.44 | 0.32 | 0.41 |
| 9 | 6255881 | rs1332290 | -0.009 | -0.31 | 0.692 | -0.931 | 0.059 | -0.005 | -0.005 | 0 | 0.53 | 0.25 | 0.48 | 0.57 | 0.64 | 0.58 |
| 9 | 6256292 | rs1048274 | 0.367 | -0.336 | 0.733 | -0.916 | -0.007 | 0.006 | 0.006 | 0 | 0.39 | 0.42 | 0.45 | 0.43 | 0.32 | 0.4 |
| 9 | 6266440 | rs2026991 | -0.011 | -0.287 | 0.214 | -0.936 | -0.012 | -0.073 | -0.073 | 0 | 0.88 | 0.4 | 0.9 | 1 | 0.93 | 0.97 |
| 9 | 6272766 | rs10815398 | 0.057 | -0.314 | -0.053 | -0.915 | 0.067 | -0.03 | -0.03 | 0.027 | 0.52 | 0.15 | 0.46 | 0.55 | 0.64 | 0.59 |
| 9 | 6280786 | rs2169285 | 0.087 | -0.243 | 0.274 | -0.921 | -0.02 | -0.076 | -0.076 | 0.601 | 0.88 | 0.35 | 0.9 | 1 | 0.93 | 0.96 |
| 9 | 6293715 | rs10815402 | 0.751 | -0.353 | 0.067 | -0.901 | 0.019 | -0.07 | -0.07 | 0.173 | 0.33 | 0.15 | 0.43 | 0.44 | 0.29 | 0.38 |
| 9 | 6306896 | rs12237914 | 0.159 | -0.047 | -0.459 | -1.158 | -0.024 | -0.052 | -0.052 | 0 | 0.37 | 0.18 | 0.34 | 0.57 | 0.39 | 0.43 |
| 9 | 6324744 | rs16924328 | 0.86 | -0.13 | -0.8 | -0.894 | -0.019 | -0.084 | -0.084 | 0.601 | 0.14 | 0 | 0.17 | 0.27 | 0.12 | 0.17 |
| 9 | 6330380 | rs1052335 | 1.276 | -0.518 | -0.945 | -1.044 | 0.021 | -0.021 | -0.021 | 0 | 0.25 | 0.07 | 0.27 | 0.23 | 0.19 | 0.26 |
| 9 | 6331324 | rs2890707 | -1.57 | -0.387 | 0.268 | -1.081 | -0.005 | -0.031 | -0.031 | 0.145 | 0.94 | 0.79 | 0.94 | 1 | 0.97 | 0.98 |
| 9 | 6331610 | rs16924356 | 1.018 | -0.316 | 0.604 | -1.198 | 0.006 | -0.063 | -0.063 | 0 | 0.21 | 0.23 | 0.17 | 0 | 0.28 | 0.15 |
| 9 | 6342328 | rs731585 | -0.189 | -0.353 | 0.771 | -1.224 | 0 | -0.042 | -0.042 | 0 | 0.69 | 0.28 | 0.76 | 1 | 0.68 | 0.81 |
| 9 | 6342855 | rs2381438 | -1.36 | -0.327 | 0.769 | -1.162 | -0.007 | -0.034 | -0.034 | 0 | 0.93 | 0.77 | 0.94 | 1 | 0.96 | 0.98 |
| 9 | 6351111 | rs16924428 | 0.949 | -0.235 | 0.56 | -1.159 | 0.005 | -0.067 | -0.067 | 1.344 | 0.22 | 0.23 | 0.17 | 0 | 0.28 | 0.16 |
| 9 | 6395247 | rs7025295 | 0.062 | -0.121 | 0.453 | -1.032 | -0.015 | 0.026 | 0.026 | 3.584 | 0.34 | 0.21 | 0.34 | 0.26 | 0.4 | 0.32 |
| 9 | 6401077 | rs4742194 | 0.81 | -0.098 | -0.241 | -1.164 | -0.017 | -0.079 | -0.079 | 0 | 0.24 | 0.1 | 0.26 | 0.52 | 0.23 | 0.31 |
| 9 | 6407843 | rs10739097 | 0.501 | -0.1 | 0.047 | -0.987 | 0.001 | 0.036 | 0.036 | 1.733 | 0.57 | 0.31 | 0.59 | 0.76 | 0.56 | 0.67 |
| 9 | 6408429 | rs10125313 | -0.82 | 0.136 | 0.416 | -0.566 | -0.013 | -0.039 | -0.039 | 0 | 0.93 | 0.75 | 0.94 | 1 | 0.96 | 0.98 |
| 9 | 6416844 | rs10975583 | 0.348 | -0.177 | 0.145 | -0.786 | -0.016 | -0.005 | -0.005 | 0.044 | 0.71 | 0.33 | 0.76 | 0.74 | 0.82 | 0.83 |
| 9 | 6423233 | rs1156747 | -0.155 | -0.156 | 0.255 | -0.792 | -0.03 | 0.014 | 0.014 | 0.326 | 0.23 | 0.59 | 0.25 | 0.25 | 0.18 | 0.15 |
| 9 | 6427721 | rs7043824 | -1.342 | -0.188 | 0.043 | -0.781 | 0.006 | 0.007 | 0.007 | 0 | 0.94 | 0.96 | 0.96 | 0.99 | 0.96 | 0.98 |
| 9 | 6430598 | rs4742202 | 0.346 | -0.168 | 0.284 | -0.791 | -0.038 | -0.011 | -0.011 | 0.155 | 0.72 | 0.43 | 0.77 | 0.74 | 0.82 | 0.84 |
| 9 | 6432353 | rs10739104 | -1.511 | -0.099 | -0.201 | -0.759 | 0.005 | 0.005 | 0.005 | 0 | 0.94 | 0.96 | 0.96 | 0.99 | 0.96 | 0.98 |
| 9 | 6440123 | rs7033570 | 0.882 | -0.119 | -0.28 | -0.709 | -0.004 | -0.066 | -0.066 | 0.999 | 0.17 | 0 | 0.12 | 0.21 | 0.11 | 0.14 |
| 9 | 6450183 | rs12554047 | 1.515 | -0.171 | -0.408 | -0.961 | 0.002 | -0.006 | -0.006 | 0 | 0.19 | 0.11 | 0.19 | 0.18 | 0.18 | 0.25 |
| 9 | 6455048 | rs401733 | 0.213 | -0.229 | -0.019 | -1.142 | 0.014 | 0.01 | 0.01 | 0 | 0.53 | 0.37 | 0.53 | 0.63 | 0.62 | 0.58 |
| 9 | 6459274 | rs524888 | 0.485 | -0.213 | -0.3 | -1.163 | 0.002 | -0.001 | -0.001 | 0.36 | 0.46 | 0.26 | 0.42 | 0.61 | 0.54 | 0.57 |

| | | | | | | | | | | | | | | | | |
|---|---|---|---|---|---|---|---|---|---|---|---|---|---|---|---|---|
| 9 | 6462585 | rs16924591 | 1.082 | -0.182 | -0.485 | -1.159 | -0.016 | -0.08 | -0.08 | 0.148 | 0.17 | 0 | 0.19 | 0.25 | 0.13 | 0.14 |
| 9 | 6469651 | rs12346455 | -0.406 | -0.164 | -0.402 | -1.154 | -0.018 | -0.008 | -0.008 | 0 | 0.3 | 0.65 | 0.29 | 0.25 | 0.21 | 0.18 |
| 9 | 6472714 | rs6477087 | -0.375 | -0.241 | -0.429 | -1.146 | -0.029 | -0.012 | -0.012 | 0.03 | 0.29 | 0.62 | 0.28 | 0.24 | 0.2 | 0.18 |
| 9 | 6473630 | rs7860627 | 1.297 | -0.167 | -0.588 | -1.196 | -0.036 | 0.012 | 0.012 | 0 | 0.23 | 0.05 | 0.3 | 0.15 | 0.27 | 0.24 |
| 9 | 6479626 | rs10115883 | -0.17 | -0.821 | -0.088 | -1.214 | -0.032 | 0.011 | 0.011 | 0.206 | 0.24 | 0.59 | 0.25 | 0.24 | 0.17 | 0.16 |
| 9 | 6491371 | rs13285035 | 1.176 | -0.895 | -0.205 | -0.88 | -0.013 | -0.032 | -0.032 | 1.667 | 0.06 | 0.13 | 0.04 | 0 | 0.03 | 0.01 |
| 9 | 6495789 | rs16924634 | 0.458 | -0.79 | -0.519 | -0.795 | 0.001 | -0.002 | -0.002 | 0.033 | 0.18 | 0.16 | 0.2 | 0.24 | 0.13 | 0.15 |
| 9 | 6506038 | rs33962342 | 0.692 | -0.715 | -1.25 | -0.752 | -0.017 | -0.079 | -0.079 | 3.739 | 0.16 | 0.01 | 0.18 | 0.24 | 0.13 | 0.15 |
| 9 | 6515433 | rs10118449 | 0.008 | -0.387 | -0.541 | -0.556 | -0.026 | 0.005 | 0.005 | 0.018 | 0.25 | 0.41 | 0.25 | 0.26 | 0.18 | 0.17 |
| 9 | 6532344 | rs10975629 | 0.364 | 0.048 | -0.695 | -0.769 | 0 | -0.007 | -0.007 | 0 | 0.46 | 0.44 | 0.54 | 0.7 | 0.5 | 0.57 |
| 9 | 6532477 | rs7848919 | -0.471 | -0.043 | -0.813 | -0.973 | -0.001 | -0.021 | -0.021 | 0 | 0.7 | 0.72 | 0.78 | 0.97 | 0.68 | 0.72 |
| 9 | 6534845 | rs2282160 | 0.029 | 0.147 | -0.991 | -0.554 | -0.011 | -0.008 | -0.008 | 0 | 0.44 | 0.34 | 0.5 | 0.46 | 0.45 | 0.43 |
| 9 | 6535057 | rs2282158 | 0.916 | 0.219 | -1.182 | -0.587 | 0.008 | -0.047 | -0.047 | 0 | 0.22 | 0.05 | 0.21 | 0.26 | 0.19 | 0.14 |
| 9 | 6535975 | rs3765555 | 1.504 | 0.36 | -1.095 | -0.552 | -0.067 | 0.001 | 0.001 | 0 | 0.19 | 0.05 | 0.28 | 0.17 | 0.25 | 0.25 |
| 9 | 6538417 | rs6477088 | 0.53 | 0.34 | -0.957 | -0.719 | -0.004 | 0.121 | 0.121 | 0 | 0.25 | 0.42 | 0.25 | 0.53 | 0.21 | 0.33 |
| 9 | 6539087 | rs4742213 | -0.012 | 0.276 | -1.544 | -0.776 | -0.003 | 0.001 | 0.001 | 0 | 0.47 | 0.46 | 0.52 | 0.44 | 0.47 | 0.41 |
| 9 | 6540540 | rs2297442 | 1.307 | 0.464 | -2.195 | -0.782 | -0.031 | 0.003 | 0.003 | 0 | 0.21 | 0.15 | 0.29 | 0.18 | 0.27 | 0.26 |
| 9 | 6544766 | rs4419859 | -0.747 | 0.478 | -2.259 | -0.941 | -0.033 | 0.14 | 0.14 | 0 | 0.69 | 0.89 | 0.71 | 0.98 | 0.6 | 0.72 |
| 9 | 6549727 | rs12347456 | -0.437 | 0.525 | -1.859 | -0.246 | 0.018 | 0.027 | 0.027 | 0 | 0.5 | 0.3 | 0.52 | 0.45 | 0.52 | 0.47 |
| 9 | 6552278 | rs16924682 | 0.102 | 0.326 | -1.906 | 0.044 | 0.008 | -0.027 | -0.027 | 0.397 | 0.24 | 0.05 | 0.21 | 0.26 | 0.2 | 0.17 |
| 9 | 6554558 | rs4740848 | -0.22 | 0.443 | -1.51 | 0.173 | 0.001 | 0.006 | 0.006 | 0 | 0.48 | 0.44 | 0.54 | 0.62 | 0.52 | 0.58 |
| 9 | 6555079 | rs1929933 | 0.608 | 0.796 | -2.358 | -0.489 | -0.029 | 0.133 | 0.133 | 0 | 0.3 | 0.07 | 0.17 | 0.08 | 0.29 | 0.17 |
| 9 | 6555183 | rs10815443 | -0.052 | 0.967 | -2.442 | -0.031 | -0.031 | 0.005 | 0.005 | 0 | 0.31 | 0.21 | 0.38 | 0.29 | 0.36 | 0.41 |
| 9 | 6555311 | rs10125618 | -0.025 | 0.325 | -1.857 | -0.194 | -0.011 | -0.001 | -0.001 | 1.734 | 0.72 | 0.8 | 0.53 | 0.3 | 0.66 | 0.5 |
| 9 | 6555417 | rs10975640 | -0.018 | 0.294 | -1.284 | -0.563 | -0.017 | 0.013 | 0.013 | 0.126 | 0.28 | 0.22 | 0.34 | 0.25 | 0.32 | 0.43 |
| 9 | 6556948 | rs4333664 | -0.577 | 0.015 | -1.154 | -0.904 | 0.003 | 0.035 | 0.035 | 0 | 0.46 | 0.71 | 0.41 | 0.42 | 0.43 | 0.53 |
| 9 | 6567841 | rs7870809 | -0.261 | 0.084 | -0.941 | -1.412 | -0.008 | 0.031 | 0.031 | 0.498 | 0.67 | 0.85 | 0.63 | 0.23 | 0.67 | 0.59 |
| 9 | 6571569 | rs10815447 | -0.605 | -0.354 | -0.137 | -1.523 | -0.001 | -0.007 | -0.007 | 0 | 0.58 | 0.6 | 0.52 | 0.17 | 0.57 | 0.45 |
| 9 | 6581461 | rs10815450 | 0.814 | -0.48 | -0.712 | -1.788 | -0.005 | 0.129 | 0.129 | 1.256 | 0.26 | 0.07 | 0.16 | 0 | 0.28 | 0.1 |
| 9 | 6587454 | rs7021684 | -0.598 | -0.402 | -0.603 | -1.695 | 0 | -0.024 | -0.024 | 0 | 0.56 | 0.5 | 0.52 | 0.17 | 0.57 | 0.45 |
| 9 | 6594978 | rs11789777 | 0.334 | 0.04 | -1.352 | -1.14 | -0.04 | 0.033 | 0.033 | 0 | 0.23 | 0.07 | 0.3 | 0.16 | 0.26 | 0.28 |
| 9 | 6602821 | rs7030760 | 0.548 | 0.133 | -1.066 | -1.275 | -0.018 | 0.104 | 0.104 | 0 | 0.23 | 0.05 | 0.15 | 0 | 0.27 | 0.09 |
| 9 | 6609578 | rs10125015 | -0.213 | -0.021 | -0.558 | -0.485 | 0 | 0.013 | 0.013 | 0 | 0.28 | 0.26 | 0.19 | 0.09 | 0.27 | 0.34 |
| 9 | 6618004 | rs10120677 | -0.453 | -0.439 | 0.035 | 0.558 | -0.044 | 0.141 | 0.141 | 0 | 0.2 | 0.42 | 0.3 | 0.46 | 0.14 | 0.25 |
| 9 | 6618018 | rs10124449 | -0.53 | -0.457 | 0.18 | 0.549 | -0.031 | 0.104 | 0.104 | 0 | 0.19 | 0.32 | 0.29 | 0.46 | 0.14 | 0.25 |
| 9 | 6620059 | rs2118653 | -1.011 | -0.4 | -1.244 | 0.166 | -0.15 | 0.081 | 0.081 | 0 | 0.61 | 0.95 | 0.71 | 0.8 | 0.53 | 0.75 |
| 9 | 6624484 | rs2578268 | -0.738 | -0.461 | -1.246 | 0.297 | -0.103 | 0.105 | 0.105 | 1.434 | 0.55 | 0.96 | 0.69 | 0.74 | 0.51 | 0.66 |
| 9 | 6628916 | rs2773507 | -0.02 | -0.713 | -0.487 | 0.368 | -0.015 | -0.009 | -0.009 | 0.645 | 0.18 | 0.26 | 0.14 | 0.16 | 0.13 | 0.11 |
| 9 | 6631611 | rs934854 | -0.548 | -0.732 | 0.062 | 0.513 | -0.031 | 0.062 | 0.062 | 0 | 0.32 | 0.5 | 0.41 | 0.54 | 0.26 | 0.29 |
| 9 | 6631702 | rs934855 | -0.381 | -0.777 | -0.271 | 0.511 | -0.002 | 0.004 | 0.004 | 0 | 0.3 | 0.32 | 0.39 | 0.52 | 0.25 | 0.29 |
| 9 | 6635045 | rs7031908 | 0.537 | -0.623 | -0.242 | 0.444 | 0.001 | 0 | 0 | 0 | 0.23 | 0.17 | 0.17 | 0.18 | 0.25 | 0.31 |
| 9 | 6641802 | rs1755606 | -1.194 | -0.532 | -0.778 | 0.373 | -0.044 | 0.168 | 0.168 | 0.229 | 0.54 | 0.89 | 0.69 | 0.76 | 0.5 | 0.64 |
| 9 | 6644932 | rs1755617 | -0.635 | -0.226 | 0.005 | 1.093 | -0.003 | 0.019 | 0.019 | 0 | 0.27 | 0.31 | 0.39 | 0.54 | 0.25 | 0.28 |
| 9 | 6654000 | rs10975714 | -0.502 | -0.488 | 0.311 | 1.22 | -0.037 | 0.135 | 0.135 | 0 | 0.21 | 0.42 | 0.3 | 0.47 | 0.15 | 0.35 |
| 9 | 6654050 | rs11999442 | 0.426 | 0.298 | 0.052 | 0.597 | -0.009 | -0.003 | -0.003 | 0 | 0.11 | 0.12 | 0.1 | 0 | 0.18 | 0.04 |
| 9 | 6654203 | rs10815461 | -1.254 | 1.009 | -0.242 | 0.47 | -0.008 | 0.01 | 0.01 | 0 | 0.42 | 0.62 | 0.54 | 0.71 | 0.47 | 0.47 |
| 9 | 6655056 | rs7031325 | -1.175 | 0.124 | -0.254 | 0.422 | 0.03 | 0.031 | 0.031 | 0.016 | 0.4 | 0.77 | 0.49 | 0.71 | 0.34 | 0.46 |
| 9 | 6657192 | rs10758801 | -0.74 | 0.127 | -0.051 | 0.68 | 0.007 | -0.028 | -0.028 | 0 | 0.32 | 0.27 | 0.38 | 0.48 | 0.3 | 0.39 |
| 9 | 6658080 | rs7868175 | -0.381 | 0.301 | 0.178 | 0.379 | 0.002 | -0.002 | -0.002 | 0.793 | 0.18 | 0.23 | 0.16 | 0.15 | 0.2 | 0.12 |
| 9 | 6664282 | rs2918182 | -0.281 | 0.159 | -0.245 | -0.11 | 0.009 | 0.015 | 0.015 | 0 | 0.55 | 0.61 | 0.5 | 0.27 | 0.62 | 0.45 |
| 9 | 6664902 | rs1624139 | -0.441 | -0.089 | -0.318 | 0.336 | 0.008 | -0.021 | -0.021 | 0 | 0.65 | 0.69 | 0.76 | 0.51 | 0.68 | 0.65 |
| 9 | 6665010 | rs1658972 | 1.588 | 0.242 | -0.256 | 0.124 | -0.003 | -0.004 | -0.004 | 0.687 | 0.18 | 0.2 | 0.14 | 0.12 | 0.16 | 0.28 |
| 9 | 6669474 | rs820505 | -0.609 | 0.133 | -0.194 | 0.216 | -0.001 | 0.002 | 0.002 | 0 | 0.66 | 0.65 | 0.76 | 0.51 | 0.7 | 0.66 |
| 9 | 6673285 | rs12347930 | 0.77 | 0.146 | -0.138 | 0.255 | 0.02 | 0.073 | 0.073 | 0 | 0.08 | 0.27 | 0.09 | 0.15 | 0.08 | 0.09 |
| 9 | 6673465 | rs7871969 | 0.841 | 0.149 | -0.248 | 0.256 | 0.007 | 0.048 | 0.048 | 0 | 0.08 | 0.2 | 0.08 | 0.15 | 0.08 | 0.09 |
| 9 | 6673830 | rs1759414 | -1.599 | 0.182 | -0.817 | 0.252 | 0.004 | -0.055 | -0.055 | 0 | 0.74 | 0.93 | 0.87 | 0.66 | 0.78 | 0.75 |
| 9 | 6673957 | rs2578275 | 0.028 | 0.346 | -0.547 | -0.105 | -0.007 | 0.035 | 0.035 | 0 | 0.46 | 0.65 | 0.5 | 0.27 | 0.5 | 0.4 |
| 9 | 6679992 | rs12003721 | 0.521 | 0.452 | -0.536 | -0.047 | -0.006 | 0.11 | 0.11 | 0 | 0.18 | 0.26 | 0.29 | 0.49 | 0.15 | 0.31 |
| 9 | 6680477 | rs1759402 | -1.135 | 0.437 | -0.857 | -0.073 | -0.017 | 0.091 | 0.091 | 0 | 0.65 | 0.91 | 0.79 | 0.81 | 0.66 | 0.71 |
| 9 | 6682871 | rs7046643 | 0.408 | 0.582 | -0.727 | -0.353 | -0.008 | 0.015 | 0.015 | 0 | 0.28 | 0.49 | 0.35 | 0.31 | 0.25 | 0.35 |
| 9 | 6684955 | rs17501756 | 1.107 | 0.623 | -1.011 | -0.502 | 0.005 | -0.062 | -0.062 | 0 | 0.2 | 0.22 | 0.17 | 0 | 0.25 | 0.05 |
| 9 | 6695397 | rs1094038 | -0.948 | 0.62 | -1.461 | -0.438 | -0.026 | 0.109 | 0.109 | 0 | 0.66 | 0.92 | 0.77 | 0.82 | 0.65 | 0.7 |
| 9 | 6695472 | rs1094039 | 0.127 | 1.154 | -1.179 | -0.21 | -0.026 | 0.075 | 0.075 | 0 | 0.38 | 0.25 | 0.42 | 0.13 | 0.44 | 0.31 |
| 9 | 6695502 | rs1094040 | 1.076 | 1.012 | -1.413 | 0.025 | -0.019 | 0.013 | 0.013 | 0 | 0.18 | 0.06 | 0.25 | 0.13 | 0.19 | 0.26 |
| 9 | 6697128 | rs10739110 | -0.025 | 1.156 | -0.819 | 0.579 | 0.019 | 0.039 | 0.039 | 0 | 0.33 | 0.41 | 0.31 | 0.46 | 0.38 | 0.34 |
| 9 | 6704475 | rs820493 | -0.431 | 1.163 | -1.516 | 0.354 | -0.078 | -0.064 | -0.064 | 0.057 | 0.82 | 0.99 | 0.73 | 0.74 | 0.75 | 0.8 |
| 9 | 6722102 | rs2996060 | 1.029 | 1.135 | -1.051 | -0.273 | 0.003 | -0.042 | -0.042 | 0 | 0.17 | 0.16 | 0.15 | 0 | 0.24 | 0.04 |
| 9 | 6722212 | rs2990659 | -0.441 | 0.758 | -1.004 | -0.285 | -0.025 | 0.019 | 0.019 | 0 | 0.57 | 0.65 | 0.54 | 0.36 | 0.51 | 0.48 |
| 9 | 6726613 | rs4742247 | 0.482 | 0.525 | -0.756 | -0.519 | 0.001 | 0.002 | 0.002 | 0 | 0.25 | 0.22 | 0.34 | 0.2 | 0.21 | 0.3 |

| | | | | | | | | | | | | | | | | |
|---|---|---|---|---|---|---|---|---|---|---|---|---|---|---|---|---|
| 9 | 6739489 | rs2988413 | -0.22 | 0.428 | -0.583 | -0.555 | 0 | -0.011 | -0.011 | 0 | 0.45 | 0.52 | 0.5 | 0.2 | 0.45 | 0.3 |
| 9 | 6743440 | rs4008332 | 0.417 | -0.099 | -0.533 | -0.37 | -0.009 | 0.001 | 0.001 | 0 | 0.16 | 0.15 | 0.25 | 0.2 | 0.06 | 0.18 |
| 9 | 6744721 | rs7036872 | -0.828 | 0.04 | -0.457 | -0.246 | 0.015 | 0.001 | 0.001 | 0 | 0.42 | 0.61 | 0.46 | 0.43 | 0.33 | 0.46 |
| 9 | 6759209 | rs16924852 | 0.744 | -0.287 | -0.366 | -0.39 | 0.006 | -0.004 | -0.004 | 0 | 0.13 | 0.08 | 0.22 | 0.19 | 0.05 | 0.23 |
| 9 | 6760919 | rs6477107 | -0.978 | -0.143 | -0.338 | -0.343 | 0.021 | 0.007 | 0.007 | 0 | 0.43 | 0.61 | 0.45 | 0.39 | 0.31 | 0.45 |
| 9 | 6763072 | rs12001316 | -0.241 | -0.356 | -0.463 | -0.435 | 0.029 | -0.046 | -0.046 | 0 | 0.21 | 0.12 | 0.28 | 0.38 | 0.12 | 0.32 |
| 9 | 6764157 | rs10975823 | 0.95 | -0.297 | -0.179 | -0.53 | -0.019 | -0.069 | -0.069 | 0 | 0.15 | 0.2 | 0.08 | 0 | 0.09 | 0.1 |
| 9 | 6769653 | rs2990658 | 1.552 | -0.09 | -0.479 | -1.016 | 0.001 | -0.055 | -0.055 | 0 | 0.26 | 0.24 | 0.24 | 0 | 0.38 | 0.1 |
| 9 | 6769692 | rs7861972 | 1.269 | -0.063 | -0.459 | -0.946 | -0.02 | -0.081 | -0.081 | 0 | 0.15 | 0.23 | 0.08 | 0 | 0.09 | 0.09 |
| 9 | 6774846 | rs12345872 | 0.47 | 0.091 | -0.451 | -0.878 | 0.008 | -0.12 | -0.12 | 0 | 0.19 | 0.32 | 0.16 | 0 | 0.2 | 0.13 |
| 9 | 6779505 | rs2779736 | -1.312 | -0.039 | -0.843 | -0.814 | -0.058 | 0.002 | 0.002 | 0 | 0.82 | 0.98 | 0.87 | 0.82 | 0.78 | 0.84 |
| 9 | 6782225 | rs4742263 | -0.692 | -0.196 | -0.639 | -0.813 | -0.026 | -0.026 | -0.026 | 2.759 | 0.66 | 0.74 | 0.6 | 0.39 | 0.6 | 0.55 |
| 9 | 6785243 | rs11794756 | 0.966 | -0.115 | -0.454 | -0.748 | 0.005 | -0.01 | -0.01 | 0 | 0.11 | 0.06 | 0.06 | 0.2 | 0.07 | 0.13 |
| 9 | 6786734 | rs7031834 | 0.177 | -0.125 | -0.42 | -0.769 | -0.007 | -0.001 | -0.001 | 0 | 0.25 | 0.24 | 0.15 | 0.2 | 0.16 | 0.22 |
| 9 | 6788085 | rs13297144 | 2.069 | -0.098 | -0.466 | -0.871 | -0.052 | 0.092 | 0.092 | 1.204 | 0.22 | 0.05 | 0.17 | 0 | 0.3 | 0.06 |
| 9 | 6790972 | rs10975830 | 1.338 | -0.141 | -0.408 | -0.698 | -0.006 | -0.039 | -0.039 | 0 | 0.14 | 0.14 | 0.08 | 0 | 0.09 | 0.09 |
| 9 | 6794938 | rs6477109 | -0.456 | 0.198 | -0.734 | -0.392 | -0.006 | -0.013 | -0.013 | 0 | 0.64 | 0.66 | 0.6 | 0.4 | 0.62 | 0.54 |
| 9 | 6811507 | rs11998950 | 0.223 | 0.14 | 0.082 | 0.006 | 0.043 | -0.03 | -0.03 | 0 | 0.06 | 0.19 | 0.09 | 0 | 0.11 | 0.04 |
| 9 | 6818299 | rs4740861 | -1.362 | 0.078 | 0.076 | 0.057 | -0.027 | 0.01 | 0.01 | 0 | 0.55 | 0.73 | 0.57 | 0.63 | 0.48 | 0.56 |
| 9 | 6820190 | rs7048549 | -1.397 | 0.073 | 0.083 | 0.054 | -0.033 | 0.007 | 0.007 | 0 | 0.55 | 0.76 | 0.57 | 0.63 | 0.48 | 0.54 |
| 9 | 6827316 | rs7861898 | -0.977 | 0.065 | 0.117 | 0.028 | -0.002 | -0.001 | -0.001 | 0 | 0.49 | 0.41 | 0.49 | 0.63 | 0.37 | 0.52 |
| 9 | 6830208 | rs7038349 | -1.51 | 0.014 | 0.088 | 0.049 | -0.007 | 0.008 | 0.008 | 0 | 0.54 | 0.62 | 0.56 | 0.63 | 0.48 | 0.56 |
| 9 | 6838348 | rs12553351 | 0.854 | -0.345 | 0.077 | 0.165 | -0.014 | 0.009 | 0.009 | 0 | 0.12 | 0.12 | 0.21 | 0.16 | 0.05 | 0.17 |
| 9 | 6838545 | rs10733530 | -1.556 | -0.318 | 0.066 | 0.138 | 0 | 0.031 | 0.031 | 0 | 0.52 | 0.72 | 0.55 | 0.63 | 0.45 | 0.55 |
| 9 | 6838925 | rs10975841 | -0.732 | -0.306 | 0.082 | 0.187 | 0.014 | 0.062 | 0.062 | 0 | 0.25 | 0.41 | 0.4 | 0.42 | 0.26 | 0.34 |
| 9 | 6841480 | rs7846927 | -0.215 | -0.39 | -0.297 | 0.19 | 0.001 | 0 | 0 | 0 | 0.3 | 0.26 | 0.32 | 0.21 | 0.23 | 0.27 |
| 9 | 6845517 | rs16924883 | 1.392 | -0.388 | -0.224 | 0.313 | -0.004 | 0.002 | 0.002 | 0 | 0.08 | 0.05 | 0.05 | 0.15 | 0.03 | 0.13 |
| 9 | 6845561 | rs6477117 | -0.176 | -0.406 | -0.436 | 0.364 | -0.009 | -0.008 | -0.008 | 0.595 | 0.29 | 0.3 | 0.31 | 0.2 | 0.21 | 0.26 |
| 9 | 6846877 | rs10975851 | 0.996 | 0.209 | -0.09 | 0.967 | -0.024 | -0.028 | -0.028 | 0 | 0.07 | 0.02 | 0.1 | 0.27 | 0.09 | 0.14 |
| 9 | 6847072 | rs4465021 | -1.413 | 0.085 | -0.019 | 0.72 | 0 | 0.003 | 0.003 | 0 | 0.32 | 0.65 | 0.33 | 0.63 | 0.3 | 0.41 |
| 9 | 6849317 | rs4742269 | 0.066 | -0.442 | 0.386 | 0.779 | -0.03 | -0.005 | -0.005 | 0 | 0.13 | 0.22 | 0.2 | 0.12 | 0.04 | 0.17 |
| 9 | 6849405 | rs10975854 | 0.444 | -0.22 | 0.049 | 0.956 | 0.022 | 0.106 | 0.106 | 0.013 | 0.09 | 0.22 | 0.08 | 0.2 | 0.08 | 0.1 |
| 9 | 6850376 | rs7018851 | -0.667 | -0.187 | -0.671 | 0.92 | -0.007 | 0.02 | 0.02 | 0 | 0.25 | 0.64 | 0.22 | 0.33 | 0.21 | 0.27 |
| 9 | 6851118 | rs10118127 | -0.252 | -0.05 | -0.818 | 1.139 | 0.026 | -0.015 | -0.015 | 0 | 0.31 | 0.11 | 0.27 | 0.26 | 0.25 | 0.36 |
| 9 | 6855838 | rs12352882 | -0.758 | 0.623 | 0.166 | 1.343 | 0.043 | 0.157 | 0.157 | 0.294 | 0.09 | 0.3 | 0.09 | 0.2 | 0.08 | 0.11 |
| 9 | 6858156 | rs10975858 | -0.234 | 0.373 | -0.748 | 1.027 | 0.027 | 0.025 | 0.025 | 0 | 0.45 | 0.28 | 0.47 | 0.42 | 0.41 | 0.46 |
| 9 | 6876538 | rs10815473 | 0.217 | 0.498 | -0.003 | 1.888 | -0.008 | -0.009 | -0.009 | 0.357 | 0.62 | 0.7 | 0.45 | 0.51 | 0.53 | 0.59 |
| 9 | 6878215 | rs35732562 | 0.371 | 0.398 | -0.659 | 1.311 | -0.02 | -0.066 | -0.066 | 0.002 | 0.21 | 0.04 | 0.15 | 0.23 | 0.18 | 0.35 |
| 9 | 6882169 | rs2381512 | -0.531 | 0.187 | -0.148 | 1.16 | 0.021 | -0.087 | -0.087 | 0 | 0.7 | 0.52 | 0.66 | 0.89 | 0.62 | 0.83 |
| 9 | 6883802 | rs1417292 | -0.177 | 0.297 | -0.192 | 1.651 | 0.015 | -0.008 | -0.008 | 0 | 0.64 | 0.52 | 0.43 | 0.7 | 0.56 | 0.65 |
| 9 | 6888611 | rs10117958 | 0.17 | 0.101 | -0.123 | 1.465 | -0.004 | 0.176 | 0.176 | 2.259 | 0.05 | 0.17 | 0.07 | 0.27 | 0.03 | 0.18 |
| 9 | 6892618 | rs7036701 | 0.057 | 0.089 | 0.003 | 1.427 | 0 | 0.001 | 0.001 | 0.022 | 0.59 | 0.49 | 0.4 | 0.47 | 0.53 | 0.57 |
| 9 | 6895374 | rs6477122 | 0.35 | 0.095 | -0.119 | 1.66 | 0.006 | 0.018 | 0.018 | 0.023 | 0.55 | 0.37 | 0.34 | 0.34 | 0.52 | 0.42 |
| 9 | 6900255 | rs12380608 | 1.536 | -0.253 | -0.113 | 1.403 | -0.062 | 0.189 | 0.189 | 1.866 | 0.31 | 0.05 | 0.18 | 0 | 0.3 | 0.05 |
| 9 | 6905429 | rs12349786 | -0.341 | -0.285 | 0.172 | 1.568 | 0.009 | 0.144 | 0.144 | 0.447 | 0.05 | 0.26 | 0.06 | 0.12 | 0.01 | 0.15 |
| 9 | 6908557 | rs10120163 | -0.468 | -0.285 | 0.15 | 1.565 | 0.028 | 0.172 | 0.172 | 0.906 | 0.05 | 0.3 | 0.06 | 0.12 | 0.01 | 0.15 |
| 9 | 6915393 | rs7018886 | 0.469 | 0.239 | -0.126 | 1.374 | 0.053 | -0.003 | -0.003 | 2.364 | 0.55 | 0.32 | 0.38 | 0.39 | 0.63 | 0.42 |
| 9 | 6917477 | rs1578324 | -0.036 | 0.281 | -0.214 | 1.399 | -0.039 | -0.059 | -0.059 | 0 | 0.28 | 0.17 | 0.21 | 0.35 | 0.33 | 0.41 |
| 9 | 6917615 | rs10975897 | 0.337 | 0.484 | -0.014 | 1.668 | -0.009 | 0.002 | 0.002 | 0 | 0.05 | 0.01 | 0.28 | 0.24 | 0.06 | 0.13 |
| 9 | 6925663 | rs11792447 | 0.382 | 0.483 | -0.066 | 1.564 | -0.053 | -0.005 | -0.005 | 0 | 0.27 | 0.08 | 0.17 | 0.19 | 0.3 | 0.38 |
| 9 | 6926738 | rs7043262 | -0.187 | 0.28 | 0.27 | 1.53 | -0.007 | 0.032 | 0.032 | 0.002 | 0.68 | 0.63 | 0.44 | 0.42 | 0.68 | 0.66 |
| 9 | 6928064 | rs4740862 | -0.256 | 0.27 | 0.719 | 1.555 | 0.005 | 0.144 | 0.144 | 0 | 0.94 | 0.87 | 0.7 | 0.73 | 0.94 | 0.86 |
| 9 | 6932013 | rs7031406 | 0.028 | 0.236 | 0.791 | 1.384 | -0.006 | -0.001 | -0.001 | 0 | 0.65 | 0.53 | 0.4 | 0.28 | 0.64 | 0.58 |
| 9 | 6943805 | rs11790720 | 0.315 | 0.398 | 0.464 | 0.578 | -0.047 | -0.018 | -0.018 | 0 | 0.28 | 0.09 | 0.18 | 0.22 | 0.31 | 0.44 |
| 9 | 6944415 | rs12683541 | 0.207 | 0.409 | 0.45 | 0.584 | -0.028 | -0.005 | -0.005 | 0 | 0.28 | 0.17 | 0.19 | 0.22 | 0.31 | 0.44 |
| 9 | 6946719 | rs4742272 | -1.028 | 0.392 | 0.49 | 0.336 | 0.011 | 0.024 | 0.024 | 2.698 | 0.59 | 0.39 | 0.74 | 0.93 | 0.65 | 0.85 |
| 9 | 6953554 | rs7019108 | 0.37 | 0.491 | -0.135 | 0.664 | -0.01 | -0.1 | -0.1 | 0 | 0.29 | 0.12 | 0.54 | 0.56 | 0.31 | 0.38 |
| 9 | 6956850 | rs12352785 | -1.208 | 0.515 | -0.319 | -0.04 | -0.003 | 0.021 | 0.021 | 2.017 | 0.66 | 0.79 | 0.78 | 1 | 0.67 | 0.94 |
| 9 | 6967846 | rs7865491 | 0.228 | 0.251 | -1.052 | 0.207 | 0.007 | -0.014 | -0.014 | 0.06 | 0.29 | 0.18 | 0.53 | 0.69 | 0.28 | 0.47 |
| 9 | 6969634 | rs702279 | -0.414 | 0.573 | -1.012 | 0.213 | 0.01 | 0.055 | 0.055 | 2.119 | 0.32 | 0.16 | 0.21 | 0.16 | 0.3 | 0.4 |
| 9 | 6974101 | rs6477134 | 0.04 | 0.889 | -0.814 | -0.119 | 0.001 | -0.013 | -0.013 | 0 | 0.55 | 0.5 | 0.36 | 0.75 | 0.58 | 0.49 |
| 9 | 6975623 | rs2792232 | 0.266 | 1.452 | -0.432 | 0.251 | 0.011 | 0.002 | 0.002 | 0 | 0.74 | 0.72 | 0.59 | 0.31 | 0.7 | 0.36 |
| 9 | 6978180 | rs11791347 | -0.081 | 1.649 | -0.274 | 1.079 | 0.011 | 0.024 | 0.024 | 0.247 | 0.19 | 0.2 | 0.33 | 0.43 | 0.25 | 0.5 |
| 9 | 6978321 | rs16925027 | 0.337 | 2.086 | 0.02 | 0.784 | 0.008 | -0.019 | -0.019 | 4.175 | 0.15 | 0.21 | 0.13 | 0.08 | 0.21 | 0.1 |
| 9 | 6978691 | rs1323410 | -0.157 | 1.951 | -0.152 | 0.894 | 0.021 | 0.042 | 0.042 | 0.224 | 0.81 | 0.78 | 0.67 | 0.57 | 0.75 | 0.5 |
| 9 | 6983331 | rs818909 | 0.036 | 1.346 | -0.314 | 0.387 | -0.002 | -0.037 | -0.037 | 0.608 | 0.37 | 0.53 | 0.38 | 0.25 | 0.39 | 0.24 |
| 9 | 6984172 | rs10815499 | 1.034 | 1.336 | -0.452 | 0.632 | -0.01 | -0.012 | -0.012 | 0 | 0.06 | 0 | 0.05 | 0.24 | 0.04 | 0.15 |
| 9 | 6984236 | rs2296067 | 0.362 | 1.458 | -0.415 | 0.576 | 0.012 | 0.033 | 0.033 | 0.022 | 0.19 | 0.21 | 0.32 | 0.43 | 0.22 | 0.5 |
| 9 | 6985602 | rs2381526 | -0.268 | 1.466 | -0.37 | 0.179 | -0.001 | -0.016 | -0.016 | 1.985 | 0.62 | 0.44 | 0.59 | 0.73 | 0.6 | 0.76 |

| | | | | | | | | | | | | | | | | |
|---|---|---|---|---|---|---|---|---|---|---|---|---|---|---|---|---|
| 9 | 6986190 | rs2381530 | 1.113 | 1.242 | -0.52 | 0.075 | 0.035 | 0.098 | 0.098 | 0 | 0.38 | 0.23 | 0.21 | 0.06 | 0.34 | 0.11 |
| 9 | 6987663 | rs1323398 | 0.316 | 0.619 | -0.491 | -0.157 | -0.018 | -0.041 | -0.041 | 0.693 | 0.39 | 0.48 | 0.32 | 0.19 | 0.35 | 0.12 |
| 9 | 6988475 | rs2760650 | -0.404 | 0.798 | 0.264 | 0.58 | -0.012 | -0.009 | -0.009 | 0 | 0.85 | 0.63 | 0.86 | 0.89 | 0.89 | 0.88 |
| 9 | 6990364 | rs2760657 | 1.375 | 0.92 | 0.033 | 0.577 | -0.018 | 0.006 | 0.006 | 0.746 | 0.2 | 0.15 | 0.28 | 0.18 | 0.3 | 0.14 |
| 9 | 6992425 | rs752646 | 1.426 | 0.92 | 0.023 | 0.563 | -0.023 | 0.004 | 0.004 | 0.991 | 0.2 | 0.12 | 0.27 | 0.18 | 0.3 | 0.12 |
| 9 | 6993811 | rs7848022 | 0.309 | 0.926 | 0.352 | 0.48 | 0.005 | 0.014 | 0.014 | 0 | 0.55 | 0.59 | 0.32 | 0.16 | 0.44 | 0.23 |
| 9 | 6993848 | rs2760660 | 0.13 | 0.926 | 0.491 | 0.785 | 0.008 | 0.019 | 0.019 | 0 | 0.71 | 0.64 | 0.59 | 0.51 | 0.69 | 0.45 |
| 9 | 6994542 | rs10815504 | -0.143 | 0.799 | 0.47 | 0.352 | 0 | -0.002 | -0.002 | 0.621 | 0.63 | 0.39 | 0.58 | 0.71 | 0.59 | 0.76 |
| 9 | 6996653 | rs10975955 | 0.161 | 0.101 | 0.379 | 0.247 | -0.014 | -0.047 | -0.047 | 0 | 0.17 | 0.23 | 0.11 | 0.02 | 0.12 | 0.11 |
| 9 | 6996934 | rs7847191 | 1.262 | 0.593 | 0.621 | 0.333 | -0.001 | 0.112 | 0.112 | 0 | 0.42 | 0.22 | 0.26 | 0.11 | 0.47 | 0.13 |
| 9 | 6997099 | rs818891 | 1.489 | 0.619 | 0.743 | 0.415 | 0 | 0 | 0 | 1.454 | 0.2 | 0.21 | 0.27 | 0.19 | 0.19 | 0.12 |
| 9 | 6997639 | rs946929 | 0.593 | 0.619 | 0.681 | 0.59 | 0.011 | 0.086 | 0.086 | 0 | 0.49 | 0.32 | 0.29 | 0.2 | 0.5 | 0.26 |
| 9 | 6998948 | rs11790877 | 1.268 | 0.63 | 0.651 | 0.568 | -0.005 | 0.116 | 0.116 | 0 | 0.42 | 0.22 | 0.26 | 0.12 | 0.47 | 0.14 |
| 9 | 6999846 | rs10122718 | -0.081 | 0.727 | 0.458 | 0.86 | 0.001 | -0.014 | -0.014 | 1.329 | 0.25 | 0.21 | 0.4 | 0.35 | 0.24 | 0.55 |
| 9 | 7000206 | rs10815506 | -0.141 | 0.667 | 0.469 | 0.866 | -0.007 | 0.006 | 0.006 | 0 | 0.08 | 0.1 | 0.03 | 0.09 | 0.04 | 0.12 |
| 9 | 7000595 | rs2792240 | 1.725 | 0.806 | 0.526 | 0.897 | 0.001 | 0.005 | 0.005 | 1.795 | 0.19 | 0.14 | 0.24 | 0.16 | 0.18 | 0.11 |
| 9 | 7000769 | rs2792238 | -0.937 | 0.822 | 0.561 | 0.872 | 0.005 | 0.027 | 0.027 | 0.232 | 0.51 | 0.42 | 0.67 | 0.6 | 0.46 | 0.8 |
| 9 | 7002389 | rs818887 | 0.774 | 0.891 | 0.623 | 0.977 | 0 | 0.003 | 0.003 | 0.156 | 0.24 | 0.26 | 0.29 | 0.44 | 0.25 | 0.19 |
| 9 | 7004340 | rs10975965 | 0.753 | 0.674 | 0.46 | 0.88 | -0.001 | -0.001 | -0.001 | 0 | 0.12 | 0.13 | 0.13 | 0.09 | 0.09 | 0.1 |
| 9 | 7006539 | rs1093707 | 0.485 | 0.72 | 0.407 | 0.947 | -0.003 | 0 | 0 | 1.323 | 0.26 | 0.29 | 0.27 | 0.26 | 0.22 | 0.24 |
| 9 | 7007046 | rs818876 | -0.249 | 0.649 | 0.28 | 0.818 | 0.001 | -0.001 | -0.001 | 0 | 0.4 | 0.45 | 0.58 | 0.66 | 0.38 | 0.65 |
| 9 | 7008732 | rs2890733 | 0.602 | 0.627 | 0.355 | 0.829 | 0.012 | 0.029 | 0.029 | 0 | 0.54 | 0.28 | 0.54 | 0.38 | 0.62 | 0.6 |
| 9 | 7012612 | rs818879 | 1.627 | 0.696 | -0.078 | 0.714 | -0.002 | 0.102 | 0.102 | 3.179 | 0.18 | 0.03 | 0.08 | 0 | 0.13 | 0.1 |
| 9 | 7015330 | rs818882 | 0.332 | 0.736 | 0.333 | 1.038 | 0.007 | 0.066 | 0.066 | 0 | 0.82 | 0.69 | 0.83 | 0.64 | 0.84 | 0.79 |
| 9 | 7017391 | rs7853037 | 0.455 | 0.733 | 0.458 | 1.087 | 0.004 | 0.006 | 0.006 | 0.003 | 0.4 | 0.32 | 0.39 | 0.36 | 0.38 | 0.53 |
| 9 | 7022554 | rs7852014 | -0.607 | 0.699 | 0.409 | 1.046 | -0.029 | 0.116 | 0.116 | 0.94 | 0.55 | 0.83 | 0.72 | 0.74 | 0.51 | 0.74 |
| 9 | 7026860 | rs12342947 | -0.007 | 0.724 | 0.599 | 1.099 | 0.005 | 0.056 | 0.056 | 0 | 0.05 | 0.21 | 0.1 | 0.08 | 0.04 | 0.09 |
| 9 | 7027194 | rs7045222 | 1.449 | 0.684 | 0.369 | 0.956 | 0.003 | -0.006 | -0.006 | 0 | 0.25 | 0.21 | 0.1 | 0.07 | 0.2 | 0.07 |
| 9 | 7030268 | rs10758821 | 0.017 | 0.774 | 0.428 | 1.133 | 0.005 | -0.001 | -0.001 | 0.823 | 0.45 | 0.38 | 0.49 | 0.46 | 0.42 | 0.61 |
| 9 | 7034123 | rs7865020 | -0.944 | 0.744 | 1.125 | 1.965 | 0.044 | 0.236 | 0.236 | 0.045 | 0.08 | 0.32 | 0.22 | 0.27 | 0.1 | 0.1 |
| 9 | 7036325 | rs7858552 | 0.258 | 0.77 | 1.031 | 1.915 | 0 | 0 | 0 | 0 | 0.58 | 0.54 | 0.69 | 0.55 | 0.68 | 0.58 |
| 9 | 7046917 | rs1417290 | -0.198 | 0.751 | 0.792 | 1.669 | 0 | 0.005 | 0.005 | 0 | 0.05 | 0.09 | 0.11 | 0.1 | 0.04 | 0.1 |
| 9 | 7050939 | rs7031625 | -0.301 | 0.742 | 0.899 | 1.665 | 0.002 | 0.012 | 0.012 | 0 | 0.66 | 0.51 | 0.66 | 0.58 | 0.68 | 0.75 |
| 9 | 7051826 | rs7019027 | 0.776 | 0.86 | 0.63 | 1.656 | -0.037 | -0.063 | -0.063 | 0.964 | 0.14 | 0.03 | 0.27 | 0.32 | 0.17 | 0.44 |
| 9 | 7051859 | rs7019042 | 0.495 | 0.785 | 0.777 | 1.577 | 0.006 | 0.007 | 0.007 | 0 | 0.55 | 0.27 | 0.51 | 0.42 | 0.6 | 0.57 |
| 9 | 7057701 | rs2381541 | -0.767 | 0.749 | 0.826 | 1.614 | -0.005 | 0.019 | 0.019 | 0 | 0.7 | 0.83 | 0.85 | 0.76 | 0.74 | 0.76 |
| 9 | 7058108 | rs10491779 | 0.267 | 0.77 | 0.489 | 1.487 | -0.007 | -0.001 | -0.001 | 0.102 | 0.58 | 0.48 | 0.68 | 0.6 | 0.67 | 0.58 |
| 9 | 7060854 | rs10976014 | -0.413 | 0.89 | -0.107 | 1.52 | -0.01 | -0.047 | -0.047 | 0.187 | 0.13 | 0.09 | 0.13 | 0.37 | 0.1 | 0.16 |
| 9 | 7064934 | rs4742298 | -0.654 | 1.409 | 0.092 | 1.492 | -0.014 | 0.01 | 0.01 | 0.02 | 0.72 | 0.88 | 0.86 | 0.77 | 0.74 | 0.79 |
| 9 | 7065141 | rs4742299 | 0.205 | 1.361 | -0.004 | 1.359 | -0.014 | -0.041 | -0.041 | 0 | 0.17 | 0.07 | 0.29 | 0.32 | 0.18 | 0.46 |
| 9 | 7065665 | rs4742301 | -1.093 | 1.408 | 0.067 | 1.216 | -0.004 | 0.137 | 0.137 | 0.514 | 0.75 | 0.94 | 0.87 | 1 | 0.78 | 0.84 |
| 9 | 7076463 | rs1570512 | -0.154 | 1.116 | 0.795 | 1.545 | 0.009 | 0.018 | 0.018 | 0 | 0.62 | 0.49 | 0.63 | 0.53 | 0.63 | 0.65 |
| 9 | 7081886 | rs7863122 | 0.472 | 1.003 | 0.466 | 1.05 | 0.014 | 0.079 | 0.079 | 0 | 0.44 | 0.28 | 0.32 | 0.19 | 0.45 | 0.2 |
| 9 | 7084213 | rs10429452 | -1.515 | 1.193 | 1.179 | 1.558 | 0.097 | 0.264 | 0.264 | 1.609 | 0.06 | 0.37 | 0.18 | 0.19 | 0.06 | 0.02 |
| 9 | 7091794 | rs10118608 | -0.902 | 1.877 | 0.682 | 1.383 | 0.078 | 0.03 | 0.03 | 0 | 0.26 | 0.41 | 0.45 | 0.39 | 0.37 | 0.55 |
| 9 | 7092759 | rs7033001 | 1.151 | 2.734 | 0.785 | 1.596 | 0.022 | 0.004 | 0.004 | 0.323 | 0.67 | 0.57 | 0.6 | 0.37 | 0.56 | 0.44 |
| 9 | 7093871 | rs4742308 | 1.273 | 2.715 | 0.934 | 1.562 | 0.03 | 0.003 | 0.003 | 0.767 | 0.66 | 0.55 | 0.59 | 0.36 | 0.56 | 0.44 |
| 9 | 7096061 | rs17456897 | -0.61 | 2.747 | 0.885 | 1.429 | 0.021 | 0.002 | 0.002 | 0 | 0.17 | 0.25 | 0.1 | 0.21 | 0.2 | 0.36 |
| 9 | 7098109 | rs13297081 | -0.613 | 2.694 | 0.868 | 1.412 | -0.024 | -0.004 | -0.004 | 0 | 0.16 | 0.06 | 0.1 | 0.21 | 0.2 | 0.36 |
| 9 | 7099205 | rs4742309 | -0.329 | 2.758 | 0.896 | 2.083 | -0.042 | -0.038 | -0.038 | 0.025 | 0.84 | 0.94 | 0.88 | 0.8 | 0.77 | 0.81 |
| 9 | 7101393 | rs16925350 | -0.304 | 2.671 | 0.877 | 1.923 | -0.007 | -0.011 | -0.011 | 2.723 | 0.13 | 0.06 | 0.09 | 0.18 | 0.14 | 0.31 |
| 9 | 7109651 | rs7860449 | 0.91 | 2.502 | 0.792 | 1.839 | -0.023 | -0.002 | -0.002 | 0 | 0.83 | 0.93 | 0.9 | 0.79 | 0.8 | 0.63 |
| 9 | 7113967 | rs7037866 | -1.073 | 2.467 | 1.079 | 1.772 | 0.055 | -0.02 | -0.02 | 0 | 0.29 | 0.44 | 0.43 | 0.7 | 0.41 | 0.47 |
| 9 | 7118990 | rs3802401 | -0.236 | 2.537 | 1.149 | 2 | 0.013 | 0.024 | 0.024 | 0.021 | 0.11 | 0.12 | 0.13 | 0.27 | 0.17 | 0.26 |
| 9 | 7121748 | rs1323409 | 0.464 | 2.582 | 1.038 | 1.937 | -0.016 | -0.008 | -0.008 | 0 | 0.85 | 0.93 | 0.92 | 0.79 | 0.83 | 0.69 |
| 9 | 7122722 | rs10758834 | 1.301 | 2.565 | 1.522 | 2.052 | 0.012 | 0.004 | 0.004 | 0 | 0.67 | 0.64 | 0.77 | 0.6 | 0.59 | 0.5 |
| 9 | 7122872 | rs10976050 | 0.264 | 2.552 | 1.52 | 2.054 | 0.047 | 0.027 | 0.027 | 0.382 | 0.17 | 0.27 | 0.15 | 0.2 | 0.24 | 0.18 |
| 9 | 7124297 | rs10976051 | 0.319 | 2.552 | 1.474 | 1.663 | 0.01 | 0.045 | 0.045 | 0.233 | 0.17 | 0.22 | 0.23 | 0.43 | 0.24 | 0.2 |
| 9 | 7124815 | rs4742314 | 1.379 | 2.54 | 1.528 | 1.619 | 0.03 | -0.001 | -0.001 | 0 | 0.66 | 0.54 | 0.67 | 0.36 | 0.59 | 0.48 |
| 9 | 7129452 | rs7041376 | 1.469 | 2.488 | 1.659 | 1.489 | 0.026 | -0.011 | -0.011 | 0 | 0.66 | 0.43 | 0.67 | 0.35 | 0.59 | 0.48 |
| 9 | 7130475 | rs7033490 | 1.718 | 2.151 | 1.594 | 1.559 | 0.017 | 0.017 | 0.017 | 0 | 0.72 | 0.65 | 0.72 | 0.53 | 0.66 | 0.42 |
| 9 | 7130628 | rs10976056 | -0.594 | 2.195 | 1.393 | 1.558 | 0.033 | 0.043 | 0.043 | 0.252 | 0.12 | 0.2 | 0.2 | 0.26 | 0.17 | 0.26 |
| 9 | 7132826 | rs1535625 | 1.107 | 1.568 | 1.155 | 1.134 | 0.003 | -0.005 | -0.005 | 1.325 | 0.83 | 0.7 | 0.73 | 0.79 | 0.83 | 0.68 |
| 9 | 7135196 | rs2890741 | -0.228 | -0.302 | 0.466 | -0.043 | -0.049 | -0.078 | -0.078 | 0.437 | 0.29 | 0.38 | 0.09 | 0.04 | 0.1 | 0.27 |
| 9 | 7135405 | rs2381567 | -0.976 | -0.248 | 0.159 | -0.338 | 0.045 | 0.085 | 0.085 | 0.051 | 0.48 | 0.7 | 0.58 | 0.68 | 0.65 | 0.52 |
| 9 | 7135540 | rs10758836 | -0.081 | -0.37 | 0.224 | -0.31 | 0 | 0.036 | 0.036 | 0 | 0.42 | 0.32 | 0.49 | 0.62 | 0.53 | 0.48 |
| 9 | 7135716 | rs12005620 | 0.336 | 0.093 | 0.274 | 0.092 | 0.005 | 0.019 | 0.019 | 0.07 | 0.37 | 0.52 | 0.34 | 0.48 | 0.35 | 0.28 |
| 9 | 7136345 | rs10739116 | -0.294 | 0.205 | -0.334 | 0.158 | -0.017 | -0.019 | -0.019 | 0 | 0.88 | 0.95 | 0.76 | 0.79 | 0.84 | 0.69 |
| 9 | 7137500 | rs2224762 | 0.057 | 0.37 | -0.722 | 0.588 | -0.006 | -0.013 | -0.013 | 0.446 | 0.41 | 0.16 | 0.38 | 0.39 | 0.55 | 0.37 |

| | | | | | | | | | | | | | | | | |
|---|---|---|---|---|---|---|---|---|---|---|---|---|---|---|---|---|
| 9 | 7138349 | rs10976062 | 0.082 | 0.73 | -0.48 | 0.549 | 0.004 | -0.001 | -0.001 | 0 | 0.08 | 0.1 | 0.16 | 0.15 | 0.18 | 0.19 |
| 9 | 7143431 | rs1885975 | -0.234 | 0.406 | -0.341 | 0.332 | 0.004 | 0.012 | 0.012 | 0 | 0.46 | 0.45 | 0.53 | 0.65 | 0.57 | 0.48 |
| 9 | 7145264 | rs12335888 | -0.063 | -0.063 | -0.594 | 0.476 | -0.01 | 0.005 | 0.005 | 1.101 | 0.06 | 0.09 | 0.03 | 0.15 | 0.02 | 0.15 |
| 9 | 7146888 | rs722628 | 0.387 | 0.161 | -0.58 | 0.598 | -0.01 | -0.028 | -0.028 | 5.352 | 0.36 | 0.22 | 0.3 | 0.51 | 0.36 | 0.29 |
| 9 | 7150975 | rs10976070 | 0.12 | -0.635 | -0.259 | 0.872 | 0.015 | 0.011 | 0.011 | 0.023 | 0.44 | 0.34 | 0.44 | 0.39 | 0.58 | 0.44 |
| 9 | 7150997 | rs913581 | 0.142 | -0.385 | -0.494 | 0.839 | -0.005 | -0.005 | -0.005 | 0.034 | 0.87 | 0.94 | 0.76 | 0.79 | 0.83 | 0.7 |
| 9 | 7155356 | rs12001158 | 0.711 | -0.76 | -0.325 | 0.758 | 0.016 | 0.002 | 0.002 | 1.813 | 0.34 | 0.28 | 0.25 | 0.26 | 0.23 | 0.11 |
| 9 | 7163530 | rs7032509 | 0.308 | -0.965 | -0.291 | 0.692 | 0.006 | 0.002 | 0.002 | 1.68 | 0.39 | 0.37 | 0.27 | 0.29 | 0.24 | 0.2 |
| 9 | 7163686 | rs7046542 | 0.946 | -1.014 | -0.333 | 0.745 | 0.01 | 0.005 | 0.005 | 2.502 | 0.33 | 0.28 | 0.23 | 0.18 | 0.2 | 0.09 |
| 9 | 7164061 | rs10815532 | 0.687 | -0.613 | -0.213 | 0.856 | 0.004 | 0.008 | 0.008 | 0 | 0.41 | 0.34 | 0.45 | 0.34 | 0.58 | 0.4 |
| 9 | 7164382 | rs10976080 | 0.593 | -0.581 | -0.448 | 0.879 | 0.055 | -0.009 | -0.009 | 0.017 | 0.34 | 0.22 | 0.23 | 0.33 | 0.22 | 0.2 |
| 9 | 7166090 | rs10511462 | -0.058 | -0.124 | -0.403 | 1.308 | 0.007 | 0.015 | 0.015 | 0 | 0.51 | 0.59 | 0.54 | 0.59 | 0.58 | 0.49 |
| 9 | 7167481 | rs1887870 | -0.493 | -0.306 | -0.552 | 1.24 | -0.019 | -0.017 | -0.017 | 0.496 | 0.79 | 0.93 | 0.68 | 0.75 | 0.79 | 0.73 |
| 9 | 7169311 | rs10115707 | -0.294 | -0.493 | -0.468 | 1.229 | -0.035 | 0.022 | 0.022 | 0 | 0.07 | 0.13 | 0.02 | 0.15 | 0.02 | 0.15 |
| 9 | 7170165 | rs3829103 | 0.224 | -0.403 | -0.546 | 1.262 | 0 | 0 | 0 | 0 | 0.22 | 0.19 | 0.32 | 0.25 | 0.18 | 0.28 |
| 9 | 7170742 | rs7040123 | -0.195 | -0.547 | -0.474 | 1.246 | -0.013 | -0.014 | -0.014 | 0 | 0.08 | 0.12 | 0.04 | 0.04 | 0.04 | 0.11 |
| 9 | 7173500 | rs7022054 | 0.064 | -0.415 | -0.529 | 1.143 | 0.002 | 0.009 | 0.009 | 0.101 | 0.48 | 0.56 | 0.53 | 0.6 | 0.53 | 0.48 |
| 9 | 7173752 | rs7022345 | -0.836 | -0.404 | -0.565 | 1.141 | -0.01 | -0.019 | -0.019 | 0 | 0.8 | 0.94 | 0.72 | 0.75 | 0.83 | 0.71 |
| 9 | 7174673 | rs913588 | 1.3 | -0.448 | -0.47 | 1.323 | 0.007 | -0.024 | -0.024 | 0.062 | 0.35 | 0.32 | 0.39 | 0.12 | 0.48 | 0.31 |
| 9 | 7174773 | rs913589 | -0.012 | -0.298 | -0.539 | 1.443 | 0.012 | 0.05 | 0.05 | 0.204 | 0.49 | 0.64 | 0.54 | 0.59 | 0.52 | 0.47 |
| 9 | 7174898 | rs913590 | 0.1 | -0.298 | -0.548 | 1.476 | 0.004 | 0.037 | 0.037 | 0.12 | 0.48 | 0.64 | 0.53 | 0.58 | 0.51 | 0.47 |
| 9 | 7175734 | rs7045956 | -1.546 | -0.351 | -0.559 | 1.512 | -0.027 | 0.31 | 0.31 | 0.555 | 0.06 | 0.23 | 0.11 | 0.43 | 0.01 | 0.05 |
| 9 | 7176125 | rs2820934 | 0.018 | -0.324 | -0.755 | 1.313 | 0.007 | -0.024 | -0.024 | 0.12 | 0.72 | 0.84 | 0.68 | 0.59 | 0.77 | 0.57 |
| 9 | 7177936 | rs10976092 | 1.467 | -0.234 | -0.791 | 1.21 | 0.001 | -0.031 | -0.031 | 0.361 | 0.22 | 0.2 | 0.15 | 0 | 0.25 | 0.09 |
| 9 | 7192313 | rs2820917 | -0.949 | -0.1 | -0.64 | 1.885 | -0.05 | -0.043 | -0.043 | 2.029 | 0.83 | 0.99 | 0.82 | 0.79 | 0.84 | 0.89 |
| 9 | 7194521 | rs10511463 | 0.879 | 0.007 | -0.329 | 1.736 | 0 | -0.001 | -0.001 | 0 | 0.28 | 0.34 | 0.24 | 0.34 | 0.24 | 0.2 |
| 9 | 7195408 | rs2820914 | 0.471 | 0.064 | -0.314 | 1.502 | 0.013 | 0.014 | 0.014 | 0 | 0.32 | 0.32 | 0.18 | 0.2 | 0.26 | 0.28 |
| 9 | 7197023 | rs2770759 | 0.182 | 0.119 | -0.303 | 1.575 | 0.005 | 0.001 | 0.001 | 0 | 0.37 | 0.39 | 0.22 | 0.41 | 0.36 | 0.34 |
| 9 | 7197796 | rs2820910 | -0.343 | 0.11 | -0.193 | 1.244 | 0 | 0 | 0 | 0 | 0.76 | 0.79 | 0.67 | 0.76 | 0.77 | 0.64 |
| 9 | 7201988 | rs10976120 | 0.728 | 0.013 | -0.306 | 1.043 | -0.001 | 0 | 0 | 0 | 0.34 | 0.26 | 0.33 | 0.16 | 0.34 | 0.28 |
| 9 | 7203306 | rs2183118 | -0.23 | -0.294 | -0.233 | 1.293 | -0.002 | -0.005 | -0.005 | 0 | 0.73 | 0.59 | 0.73 | 0.76 | 0.76 | 0.64 |
| 9 | 7205742 | rs2770765 | 0.457 | -0.429 | -0.156 | 1.615 | -0.006 | 0.005 | 0.005 | 0 | 0.3 | 0.39 | 0.26 | 0.39 | 0.26 | 0.24 |
| 9 | 7209423 | rs12375454 | 0.68 | -0.271 | -0.168 | 1.412 | -0.02 | 0.027 | 0.027 | 0 | 0.38 | 0.25 | 0.44 | 0.25 | 0.45 | 0.34 |
| 9 | 7212446 | rs13301012 | 1.101 | -0.213 | -0.168 | 1.246 | 0.002 | 0.002 | 0.002 | 0 | 0.25 | 0.23 | 0.15 | 0.09 | 0.24 | 0.12 |
| 9 | 7213187 | rs10815554 | 1.112 | -0.211 | -0.25 | 1.229 | 0.002 | 0.003 | 0.003 | 0 | 0.25 | 0.23 | 0.15 | 0.08 | 0.24 | 0.12 |
| 9 | 7214749 | rs2183121 | 0.071 | -0.363 | 0.018 | 1.351 | 0.008 | 0.032 | 0.032 | 0 | 0.06 | 0.24 | 0.04 | 0.08 | 0.03 | 0.07 |
| 9 | 7214782 | rs913578 | -0.153 | -0.357 | -0.022 | 1.533 | -0.014 | 0.114 | 0.114 | 0 | 0.05 | 0.13 | 0.11 | 0.3 | 0.03 | 0.12 |
| 9 | 7216180 | rs913579 | -0.025 | -0.304 | -0.124 | 1.472 | 0.002 | -0.006 | -0.006 | 0 | 0.47 | 0.52 | 0.57 | 0.61 | 0.49 | 0.48 |
| 9 | 7222495 | rs10976129 | 0.476 | -0.218 | -0.161 | 1.316 | -0.012 | 0.012 | 0.012 | 0 | 0.39 | 0.29 | 0.44 | 0.31 | 0.46 | 0.34 |
| 9 | 7222742 | rs2149538 | -0.034 | 0.171 | -0.609 | 1.552 | -0.002 | 0.062 | 0.062 | 0 | 0.52 | 0.73 | 0.6 | 0.67 | 0.52 | 0.55 |
| 9 | 7225030 | rs2820957 | -1.142 | 0.157 | -0.004 | 1.636 | -0.019 | -0.046 | -0.046 | 0 | 0.45 | 0.87 | 0.58 | 0.5 | 0.46 | 0.43 |
| 9 | 7226519 | rs10976131 | 0.386 | 0.126 | 0.085 | 1.469 | 0.014 | 0.037 | 0.037 | 0 | 0.25 | 0.13 | 0.15 | 0.09 | 0.23 | 0.12 |
| 9 | 7226794 | rs2770701 | -0.042 | 0.366 | -0.194 | 1.616 | -0.038 | -0.011 | -0.011 | 0.169 | 0.31 | 0.53 | 0.27 | 0.22 | 0.25 | 0.19 |
| 9 | 7226997 | rs2820956 | -1.089 | 0.383 | -0.257 | 1.656 | -0.019 | -0.023 | -0.023 | 0 | 0.45 | 0.87 | 0.58 | 0.52 | 0.45 | 0.43 |
| 9 | 7227918 | rs10815558 | -0.958 | 1.143 | 0.761 | 2.279 | 0.132 | 0.169 | 0.169 | 0.408 | 0.14 | 0.4 | 0.31 | 0.3 | 0.2 | 0.24 |
| 9 | 7231559 | rs10976135 | -0.879 | 1.096 | 0.147 | 1.801 | 0.089 | 0.261 | 0.261 | 0.735 | 0.16 | 0.42 | 0.36 | 0.42 | 0.23 | 0.29 |
| 9 | 7234651 | rs7019840 | -0.371 | 0.45 | -0.624 | 1.156 | 0.006 | -0.014 | -0.014 | 1.038 | 0.16 | 0.12 | 0.09 | 0.21 | 0.1 | 0.06 |
| 9 | 7236712 | rs1018654 | -0.543 | 0.908 | -1.885 | 0.186 | -0.098 | 0.101 | 0.101 | 0 | 0.73 | 0.96 | 0.84 | 0.87 | 0.65 | 0.82 |
| 9 | 7238701 | rs2770705 | -0.523 | 1.066 | -1.993 | 0.192 | -0.112 | 0.068 | 0.068 | 0 | 0.73 | 0.96 | 0.83 | 0.84 | 0.64 | 0.81 |
| 9 | 7239198 | rs2381573 | -0.282 | 0.879 | -0.667 | 0.175 | -0.002 | 0.01 | 0.01 | 0.101 | 0.26 | 0.3 | 0.18 | 0.29 | 0.22 | 0.13 |
| 9 | 7239370 | rs10815565 | 0.131 | 0.069 | -0.747 | -0.596 | 0.014 | 0.037 | 0.037 | 1.463 | 0.43 | 0.33 | 0.42 | 0.19 | 0.34 | 0.33 |
| 9 | 7240918 | rs10976140 | 1.162 | 0.022 | -0.753 | -0.625 | 0.003 | -0.015 | -0.015 | 0 | 0.27 | 0.22 | 0.32 | 0.02 | 0.24 | 0.28 |
| 9 | 7241270 | rs16925635 | 0.086 | 0.056 | -0.207 | -0.718 | 0.018 | 0.003 | 0.003 | 0.154 | 0.14 | 0.21 | 0.1 | 0.1 | 0.2 | 0.1 |
| 9 | 7242537 | rs16925637 | 1.461 | 0.102 | -0.206 | -0.613 | 0.031 | 0.003 | 0.003 | 3.851 | 0.16 | 0.07 | 0.09 | 0.15 | 0.08 | 0.02 |
| 9 | 7244151 | rs11794497 | 0.124 | -0.151 | -0.141 | -0.784 | -0.023 | 0.079 | 0.079 | 1.551 | 0.39 | 0.24 | 0.41 | 0.12 | 0.43 | 0.38 |
| 9 | 7244823 | rs2820962 | 0.487 | -0.128 | -0.008 | -0.706 | 0.032 | 0.086 | 0.086 | 1.958 | 0.42 | 0.19 | 0.41 | 0.24 | 0.33 | 0.36 |
| 9 | 7245009 | rs7028940 | 0.781 | 0.418 | 0.102 | -0.646 | -0.008 | 0.128 | 0.128 | 0.186 | 0.18 | 0.32 | 0.32 | 0.53 | 0.17 | 0.37 |
| 9 | 7245314 | rs2770710 | -0.611 | 0.461 | 0.034 | -0.71 | -0.038 | 0.032 | 0.032 | 0.645 | 0.65 | 0.76 | 0.74 | 0.78 | 0.5 | 0.73 |
| 9 | 7248916 | rs12551734 | 0.337 | 0.318 | -0.368 | -0.732 | 0.065 | 0.172 | 0.172 | 3.899 | 0.46 | 0.12 | 0.43 | 0.19 | 0.35 | 0.32 |
| 9 | 7250826 | rs7856490 | -0.325 | 0.439 | -0.307 | -0.625 | -0.011 | 0.032 | 0.032 | 0.504 | 0.58 | 0.29 | 0.53 | 0.27 | 0.56 | 0.43 |
| 9 | 7255930 | rs4742321 | 0.155 | 0.088 | -0.492 | -0.88 | -0.02 | 0.041 | 0.041 | 0 | 0.24 | 0.34 | 0.32 | 0.59 | 0.19 | 0.38 |
| 9 | 7257213 | rs966015 | -1.64 | 0.296 | -0.78 | -0.926 | -0.116 | 0.08 | 0.08 | 0 | 0.74 | 0.96 | 0.82 | 0.85 | 0.64 | 0.82 |
| 9 | 7257340 | rs10976157 | 0.528 | 0.287 | -0.733 | -0.946 | -0.023 | 0.004 | 0.004 | 0.092 | 0.13 | 0.09 | 0.1 | 0.08 | 0.2 | 0.1 |
| 9 | 7258526 | rs2494244 | 0.144 | -0.025 | -0.084 | -1.068 | -0.044 | -0.033 | -0.033 | 0 | 0.34 | 0.4 | 0.2 | 0.21 | 0.2 | 0.12 |
| 9 | 7259614 | rs10976161 | 1.168 | 0.099 | -0.312 | -1.109 | -0.001 | -0.02 | -0.02 | 0.193 | 0.17 | 0.14 | 0.27 | 0.02 | 0.17 | 0.27 |
| 9 | 7260390 | rs2820950 | -1.891 | 0.195 | -0.549 | -1.018 | -0.072 | 0.069 | 0.069 | 0.004 | 0.78 | 0.95 | 0.85 | 0.87 | 0.7 | 0.83 |
| 9 | 7263472 | rs2770719 | 0.638 | -0.108 | -0.093 | -0.974 | 0.017 | 0.001 | 0.001 | 6.152 | 0.28 | 0.21 | 0.15 | 0.23 | 0.16 | 0.1 |
| 9 | 7264558 | rs2820949 | -1.054 | 0.012 | 0.116 | -0.915 | -0.024 | 0.056 | 0.056 | 0 | 0.71 | 0.84 | 0.79 | 0.85 | 0.64 | 0.8 |
| 9 | 7266283 | rs10976166 | 0.85 | -0.133 | 0.514 | -0.769 | 0.004 | 0.015 | 0.015 | 0 | 0.14 | 0.12 | 0.27 | 0.51 | 0.13 | 0.36 |

| | | | | | | | | | | | | | | | | |
|---|---|---|---|---|---|---|---|---|---|---|---|---|---|---|---|---|
| 9 | 7267176 | rs4740874 | -0.32 | -0.175 | 1.109 | -0.886 | 0.006 | 0.049 | 0.049 | 0.516 | 0.27 | 0.48 | 0.37 | 0.6 | 0.31 | 0.46 |
| 9 | 7267702 | rs10733533 | 0.428 | -0.755 | 0.885 | -1.317 | 0.001 | -0.003 | -0.003 | 0.935 | 0.41 | 0.4 | 0.37 | 0.18 | 0.27 | 0.29 |
| 9 | 7270158 | rs1928199 | -1.202 | -0.067 | 0.504 | -1.501 | -0.009 | -0.021 | -0.021 | 0 | 0.71 | 0.77 | 0.59 | 0.35 | 0.65 | 0.51 |
| 9 | 7270363 | rs1928200 | 1.253 | -0.575 | -0.624 | -1.789 | 0.011 | 0.046 | 0.046 | 0 | 0.25 | 0.14 | 0.28 | 0.01 | 0.2 | 0.25 |
| 9 | 7272542 | rs2820944 | 0.661 | -0.456 | -0.318 | -1.594 | -0.001 | 0 | 0 | 4.085 | 0.19 | 0.17 | 0.12 | 0.24 | 0.19 | 0.1 |
| 9 | 7272552 | rs7859003 | -0.399 | -0.542 | -0.28 | -1.638 | 0.001 | -0.002 | -0.002 | 0.322 | 0.18 | 0.18 | 0.11 | 0.1 | 0.19 | 0.11 |
| 9 | 7273950 | rs10815578 | 1.056 | -0.706 | -0.23 | -1.513 | -0.006 | 0.045 | 0.045 | 0.014 | 0.16 | 0.19 | 0.26 | 0.52 | 0.13 | 0.37 |
| 9 | 7274696 | rs1887867 | -0.162 | -0.628 | -0.064 | -1.771 | -0.005 | -0.019 | -0.019 | 0 | 0.35 | 0.4 | 0.38 | 0.77 | 0.32 | 0.47 |
| 9 | 7275995 | rs17528044 | 1.616 | -0.481 | -0.373 | -1.822 | -0.04 | -0.019 | -0.019 | 0 | 0.1 | 0.04 | 0.09 | 0.09 | 0.18 | 0.09 |
| 9 | 7276387 | rs4486259 | -1.076 | -0.555 | 0.129 | -1.806 | 0.055 | -0.012 | -0.012 | 0 | 0.76 | 0.67 | 0.81 | 0.79 | 0.64 | 0.79 |
| 9 | 7276798 | rs7034160 | 0.524 | -0.612 | -0.338 | -1.842 | -0.018 | 0.123 | 0.123 | 0 | 0.38 | 0.17 | 0.38 | 0.11 | 0.39 | 0.36 |
| 9 | 7284763 | rs10815582 | 0.669 | -0.587 | -0.335 | -1.901 | -0.033 | 0.125 | 0.125 | 0 | 0.37 | 0.15 | 0.37 | 0.11 | 0.38 | 0.35 |
| 9 | 7285207 | rs10758855 | 0.29 | -0.764 | -0.141 | -1.828 | -0.006 | 0.059 | 0.059 | 0 | 0.44 | 0.3 | 0.41 | 0.19 | 0.5 | 0.41 |
| 9 | 7285885 | rs2770725 | -1.911 | -0.413 | -0.358 | -1.563 | -0.01 | 0.009 | 0.009 | 0 | 0.84 | 0.87 | 0.84 | 0.88 | 0.77 | 0.86 |
| 9 | 7286273 | rs10976178 | 0.226 | -0.746 | -0.24 | -1.26 | 0 | -0.002 | -0.002 | 0 | 0.28 | 0.25 | 0.35 | 0.69 | 0.19 | 0.4 |
| 9 | 7286599 | rs4742322 | 0.27 | -0.74 | -0.236 | -1.264 | 0 | -0.002 | -0.002 | 0 | 0.28 | 0.25 | 0.35 | 0.68 | 0.2 | 0.4 |
| 9 | 7291256 | rs10976182 | -0.013 | 0.547 | -0.13 | -0.73 | 0.013 | 0.011 | 0.011 | 0 | 0.67 | 0.4 | 0.64 | 0.62 | 0.62 | 0.72 |
| 9 | 7291584 | rs10976183 | 0.844 | 0.865 | 0.5 | -1.996 | -0.004 | -0.028 | -0.028 | 0 | 0.17 | 0.19 | 0.17 | 0.06 | 0.16 | 0.08 |
| 9 | 7291611 | rs10758857 | -1.798 | 0.877 | 0.629 | -2.15 | 0.04 | 0.004 | 0.004 | 0 | 0.89 | 0.78 | 0.9 | 0.89 | 0.86 | 0.98 |
| 9 | 7295783 | rs11794291 | 1.183 | 0.188 | -0.102 | -2.501 | -0.026 | 0.155 | 0.155 | 0 | 0.36 | 0.1 | 0.32 | 0.12 | 0.39 | 0.21 |
| 9 | 7296644 | rs10815583 | 0.167 | 0.322 | 0.162 | -2.492 | 0.015 | 0.035 | 0.035 | 0 | 0.39 | 0.32 | 0.39 | 0.82 | 0.31 | 0.65 |
| 9 | 7301900 | rs16925803 | 0.414 | 0.191 | 0.136 | -2.473 | 0.025 | 0.077 | 0.077 | 0.778 | 0.45 | 0.29 | 0.36 | 0.12 | 0.42 | 0.27 |
| 9 | 7304762 | rs7039269 | 0.121 | 0.347 | 0.137 | -2.497 | -0.006 | 0.125 | 0.125 | 0.105 | 0.48 | 0.29 | 0.43 | 0.12 | 0.53 | 0.29 |
| 9 | 7309750 | rs10815590 | 0.151 | 0.34 | 0.125 | -2.499 | -0.006 | 0.097 | 0.097 | 0.119 | 0.48 | 0.3 | 0.43 | 0.12 | 0.53 | 0.29 |
| 9 | 7313773 | rs7034937 | 1.317 | 0.17 | 0.128 | -2.961 | -0.015 | -0.015 | -0.015 | 0 | 0.07 | 0.07 | 0.03 | 0.03 | 0.02 | 0.07 |
| 9 | 7313822 | rs12380136 | 0.022 | 0.156 | 0.437 | -2.978 | 0.001 | -0.005 | -0.005 | 0.068 | 0.5 | 0.46 | 0.44 | 0.1 | 0.53 | 0.28 |
| 9 | 7316085 | rs12555938 | -0.302 | 0.238 | 0.465 | -2.927 | 0 | -0.031 | -0.031 | 1.462 | 0.57 | 0.53 | 0.46 | 0.13 | 0.55 | 0.35 |
| 9 | 7316589 | rs1926393 | 0.043 | 0.035 | 0.345 | -2.911 | 0.015 | 0.026 | 0.026 | 1.398 | 0.52 | 0.41 | 0.38 | 0.13 | 0.43 | 0.33 |
| 9 | 7319116 | rs7028391 | -1.276 | 0.135 | 0.367 | -2.827 | 0.002 | -0.024 | -0.024 | 0.424 | 0.82 | 0.81 | 0.81 | 0.94 | 0.82 | 0.88 |
| 9 | 7320871 | rs10976233 | -0.111 | 0.056 | 0.258 | -2.899 | 0.008 | -0.082 | -0.082 | 0 | 0.19 | 0.28 | 0.14 | 0.03 | 0.17 | 0.12 |
| 9 | 7325093 | rs2148348 | 0.367 | 0.283 | -0.084 | -2.763 | -0.002 | 0.115 | 0.115 | 0.591 | 0.48 | 0.3 | 0.3 | 0.1 | 0.48 | 0.26 |
| 9 | 7328880 | rs1926387 | 0.993 | 0.15 | -0.35 | -2.337 | -0.001 | 0.005 | 0.005 | 0.642 | 0.25 | 0.25 | 0.35 | 0.81 | 0.26 | 0.56 |
| 9 | 7335597 | rs10976248 | 0.176 | -0.035 | -0.895 | -2.595 | 0.003 | 0.114 | 0.114 | 0.09 | 0.36 | 0.21 | 0.33 | 0.02 | 0.34 | 0.26 |
| 9 | 7335949 | rs731253 | 0.385 | -0.325 | -0.412 | -2.84 | -0.007 | -0.042 | -0.042 | 0 | 0.37 | 0.48 | 0.39 | 0.78 | 0.32 | 0.58 |
| 9 | 7336443 | rs12347615 | 1.102 | -0.834 | -0.489 | -3.341 | -0.012 | -0.02 | -0.02 | 0.756 | 0.16 | 0.22 | 0.17 | 0.04 | 0.09 | 0.09 |
| 9 | 7338345 | rs7035555 | -0.63 | -0.825 | -0.758 | -3.533 | 0.01 | 0.019 | 0.019 | 0 | 0.76 | 0.83 | 0.79 | 0.94 | 0.77 | 0.8 |
| 9 | 7338432 | rs12351222 | 1.806 | -1.182 | -0.899 | -3.781 | -0.009 | -0.041 | -0.041 | 0 | 0.13 | 0.2 | 0.03 | 0 | 0.06 | 0.06 |
| 9 | 7338609 | rs10976253 | 2.42 | -1.163 | -0.86 | -3.659 | -0.012 | -0.012 | -0.012 | 0 | 0.21 | 0.11 | 0.33 | 0.7 | 0.25 | 0.5 |
| 9 | 7339425 | rs1926381 | -1.17 | -1.013 | -0.913 | -3.573 | -0.019 | 0.079 | 0.079 | 0 | 0.79 | 0.97 | 0.8 | 0.94 | 0.78 | 0.8 |
| 9 | 7340826 | rs10815597 | 1.557 | -1.162 | -0.828 | -3.644 | -0.004 | 0.005 | 0.005 | 0.017 | 0.27 | 0.15 | 0.34 | 0.73 | 0.27 | 0.55 |
| 9 | 7341123 | rs6477187 | -0.026 | -1.133 | -0.771 | -3.737 | 0.001 | 0.173 | 0.173 | 0 | 0.52 | 0.73 | 0.63 | 0.84 | 0.5 | 0.67 |
| 9 | 7341260 | rs12005581 | 1.871 | -1.174 | -0.721 | -3.654 | -0.002 | -0.023 | -0.023 | 0 | 0.12 | 0.16 | 0.03 | 0 | 0.05 | 0.07 |
| 9 | 7342059 | rs7856288 | 1.472 | -1.063 | -0.789 | -3.555 | -0.003 | 0.004 | 0.004 | 0 | 0.27 | 0.15 | 0.35 | 0.73 | 0.27 | 0.56 |
| 9 | 7342797 | rs17601822 | 2.146 | -1.119 | -0.907 | -3.401 | -0.036 | -0.022 | -0.022 | 0 | 0.14 | 0.04 | 0.26 | 0.16 | 0.22 | 0.07 |
| 9 | 7343188 | rs1952178 | 0.451 | -1.24 | -0.597 | -3.467 | -0.002 | -0.004 | -0.004 | 0 | 0.36 | 0.34 | 0.44 | 0.22 | 0.32 | 0.15 |
| 9 | 7343325 | rs1926380 | 1.575 | -1.444 | -0.94 | -3.333 | -0.004 | -0.012 | -0.012 | 0 | 0.26 | 0.15 | 0.2 | 0.71 | 0.25 | 0.51 |
| 9 | 7344034 | rs1926379 | 1.186 | -1.402 | -0.914 | -3.244 | 0.008 | -0.051 | -0.051 | 0 | 0.1 | 0.25 | 0.05 | 0.01 | 0.07 | 0.1 |
| 9 | 7346029 | rs10511465 | 1.659 | -1.485 | -1.176 | -3.301 | 0.028 | 0.092 | 0.092 | 0 | 0.15 | 0.01 | 0.05 | 0 | 0.1 | 0.11 |
| 9 | 7347596 | rs746084 | -0.748 | -1.236 | -0.987 | -3.199 | 0.003 | -0.01 | -0.01 | 0 | 0.85 | 0.81 | 0.81 | 0.94 | 0.82 | 0.87 |
| 9 | 7348292 | rs9695318 | -1.534 | -1.172 | -0.582 | -3.286 | -0.022 | 0.005 | 0.005 | 0.734 | 0.69 | 0.86 | 0.66 | 0.3 | 0.64 | 0.45 |
| 9 | 7348662 | rs7024562 | 0.027 | -0.451 | -0.899 | -2.963 | 0.016 | -0.038 | -0.038 | 1.997 | 0.25 | 0.38 | 0.26 | 0.11 | 0.27 | 0.2 |
| 9 | 7351874 | rs10976259 | -0.485 | -0.473 | -0.985 | -3.003 | 0.006 | -0.048 | -0.048 | 1.679 | 0.31 | 0.42 | 0.28 | 0.11 | 0.3 | 0.26 |
| 9 | 7355129 | rs1926374 | 0.378 | -0.223 | -1.187 | -2.699 | -0.046 | 0.037 | 0.037 | 1.967 | 0.29 | 0.02 | 0.31 | 0.19 | 0.31 | 0.15 |
| 9 | 7357693 | rs12554101 | 0.14 | -0.187 | -1.066 | -2.711 | 0.004 | 0.018 | 0.018 | 0.841 | 0.3 | 0.22 | 0.31 | 0.2 | 0.31 | 0.14 |
| 9 | 7360902 | rs4742336 | -0.601 | -0.125 | -1.009 | -2.75 | 0.011 | 0.09 | 0.09 | 0 | 0.48 | 0.3 | 0.56 | 0.23 | 0.56 | 0.34 |
| 9 | 7361202 | rs10976265 | -0.668 | 0.409 | -0.476 | -2.39 | -0.004 | -0.013 | -0.013 | 0 | 0.23 | 0.35 | 0.27 | 0.14 | 0.16 | 0.17 |
| 9 | 7366297 | rs7853490 | -0.995 | 0.331 | -0.56 | -2.334 | 0.004 | 0.025 | 0.025 | 0 | 0.33 | 0.3 | 0.27 | 0.2 | 0.32 | 0.21 |
| 9 | 7366709 | rs7869986 | 1.022 | 0.359 | -0.706 | -2.307 | -0.012 | 0.007 | 0.007 | 0.095 | 0.15 | 0.08 | 0.13 | 0.01 | 0.21 | 0.03 |
| 9 | 7370679 | rs7849548 | -0.461 | 0.413 | -0.771 | -2.273 | 0.011 | 0.079 | 0.079 | 1.593 | 0.3 | 0.06 | 0.3 | 0.18 | 0.27 | 0.14 |
| 9 | 7370794 | rs11792571 | 0.81 | 0.788 | -0.847 | -2.319 | -0.052 | 0.022 | 0.022 | 0 | 0.14 | 0.02 | 0.19 | 0.11 | 0.17 | 0.15 |
| 9 | 7371646 | rs1926363 | -0.436 | 0.923 | -0.546 | -2.337 | 0.006 | -0.014 | -0.014 | 0.084 | 0.25 | 0.28 | 0.27 | 0.13 | 0.28 | 0.25 |
| 9 | 7372732 | rs1926361 | -0.256 | 0.881 | -0.664 | -2.298 | 0.077 | -0.063 | -0.063 | 0.045 | 0.14 | 0.48 | 0.11 | 0.03 | 0.13 | 0.12 |
| 9 | 7373020 | rs4740882 | 0.712 | 0.925 | -0.825 | -2.239 | -0.024 | 0.037 | 0.037 | 1.132 | 0.2 | 0.03 | 0.27 | 0.12 | 0.23 | 0.07 |
| 9 | 7373868 | rs4740884 | -1.378 | 0.823 | -0.622 | -1.925 | 0.002 | 0.01 | 0.01 | 0 | 0.58 | 0.64 | 0.59 | 0.32 | 0.58 | 0.4 |
| 9 | 7374332 | rs7865621 | -1.142 | 1 | -0.233 | -1.452 | -0.002 | -0.024 | -0.024 | 0 | 0.43 | 0.52 | 0.39 | 0.19 | 0.39 | 0.24 |
| 9 | 7374476 | rs7865746 | -0.284 | 1.535 | -0.401 | -0.945 | 0.049 | -0.025 | -0.025 | 0 | 0.26 | 0.33 | 0.39 | 0.13 | 0.44 | 0.19 |
| 9 | 7375169 | rs7855413 | -0.477 | 0.846 | -0.551 | -0.559 | 0 | -0.019 | -0.019 | 0 | 0.41 | 0.41 | 0.41 | 0.23 | 0.41 | 0.21 |
| 9 | 7375925 | rs1926359 | -0.038 | 0.521 | -0.665 | -0.855 | 0.003 | -0.007 | -0.007 | 0 | 0.47 | 0.46 | 0.45 | 0.13 | 0.4 | 0.25 |
| 9 | 7378095 | rs4360345 | -0.206 | 0.558 | -0.787 | 0.139 | -0.06 | -0.034 | -0.034 | 2.263 | 0.83 | 0.95 | 0.76 | 0.76 | 0.73 | 0.79 |

| | | | | | | | | | | | | | | | | |
|---|---|---|---|---|---|---|---|---|---|---|---|---|---|---|---|---|
| 9 | 7379168 | rs7871958 | 0.878 | -0.135 | -0.19 | 0.484 | 0.001 | 0 | 0 | 0.965 | 0.66 | 0.68 | 0.55 | 0.32 | 0.63 | 0.48 |
| 9 | 7379273 | rs7872098 | -0.262 | 0.129 | -0.326 | 1.009 | -0.064 | -0.079 | -0.079 | 1.191 | 0.82 | 0.95 | 0.75 | 0.58 | 0.72 | 0.62 |
| 9 | 7381341 | rs16925989 | -0.643 | -0.136 | 0.43 | 0.197 | -0.024 | 0.029 | 0.029 | 0.006 | 0.08 | 0.25 | 0.04 | 0.13 | 0.04 | 0.09 |
| 9 | 7381520 | rs6477190 | 0.006 | -0.506 | 0.233 | -0.056 | -0.014 | 0.031 | 0.031 | 1.412 | 0.61 | 0.83 | 0.57 | 0.3 | 0.62 | 0.49 |
| 9 | 7382274 | rs6477191 | -1.435 | -0.653 | 1.065 | 0.182 | -0.05 | 0.032 | 0.032 | 0.357 | 0.12 | 0.46 | 0.06 | 0.14 | 0.04 | 0.13 |
| 9 | 7382363 | rs7862635 | -1.114 | -0.662 | 1.051 | 0.182 | -0.034 | 0.027 | 0.027 | 0 | 0.1 | 0.32 | 0.04 | 0.12 | 0.04 | 0.09 |
| 9 | 7387537 | rs7041418 | -0.153 | 0.228 | -0.344 | -0.233 | 0.001 | 0.005 | 0.005 | 0 | 0.53 | 0.46 | 0.52 | 0.78 | 0.49 | 0.48 |
| 9 | 7387603 | rs7041546 | -0.317 | 0.82 | -0.954 | -0.626 | -0.112 | 0.033 | 0.033 | 0.548 | 0.83 | 0.99 | 0.71 | 0.94 | 0.72 | 0.79 |
| 9 | 7387954 | rs10815605 | 0.506 | 0.307 | -0.606 | -0.827 | 0.029 | 0.056 | 0.056 | 0 | 0.3 | 0.18 | 0.17 | 0.14 | 0.23 | 0.28 |
| 9 | 7388656 | rs4742341 | 0.19 | 0.138 | -0.475 | -0.487 | -0.005 | 0.02 | 0.02 | 0.498 | 0.56 | 0.64 | 0.51 | 0.66 | 0.51 | 0.47 |
| 9 | 7390508 | rs10815606 | -0.363 | 0.277 | -0.643 | -0.468 | -0.07 | 0.037 | 0.037 | 0.684 | 0.81 | 0.94 | 0.61 | 0.92 | 0.69 | 0.77 |
| 9 | 7391634 | rs7862381 | 0.593 | -0.098 | -0.395 | -0.503 | 0.001 | -0.001 | -0.001 | 0 | 0.23 | 0.22 | 0.2 | 0.27 | 0.2 | 0.33 |
| 9 | 7393934 | rs7033457 | 0.261 | -0.008 | -0.419 | -0.457 | -0.035 | 0.002 | 0.002 | 0 | 0.28 | 0.5 | 0.23 | 0.3 | 0.21 | 0.38 |
| 9 | 7395038 | rs2148356 | 0.438 | 0.637 | -0.673 | -0.133 | -0.002 | -0.048 | -0.048 | 0 | 0.26 | 0.16 | 0.31 | 0.48 | 0.29 | 0.17 |
| 9 | 7396956 | rs4740888 | 0.11 | 1.069 | -0.69 | -0.16 | -0.028 | -0.061 | -0.061 | 0 | 0.28 | 0.16 | 0.37 | 0.5 | 0.35 | 0.18 |
| 9 | 7399175 | rs943174 | 0.22 | 0.69 | -0.535 | -0.152 | -0.003 | -0.027 | -0.027 | 0 | 0.61 | 0.73 | 0.38 | 0.5 | 0.43 | 0.62 |
| 9 | 7400754 | rs4620331 | -0.606 | 0.656 | -0.447 | -0.141 | 0.016 | 0.032 | 0.032 | 0.038 | 0.33 | 0.54 | 0.5 | 0.5 | 0.38 | 0.4 |
| 9 | 7402208 | rs7467398 | 0.966 | 0.029 | -1.646 | -0.627 | -0.016 | 0.038 | 0.038 | 0 | 0.28 | 0.02 | 0.19 | 0.16 | 0.29 | 0.18 |
| 9 | 7403305 | rs943172 | -0.001 | 0.627 | -1.195 | -0.552 | -0.006 | -0.008 | -0.008 | 0.051 | 0.64 | 0.36 | 0.69 | 0.51 | 0.61 | 0.67 |
| 9 | 7405366 | rs2997547 | -0.003 | 0.657 | -1.092 | -0.375 | -0.002 | -0.004 | -0.004 | 0.008 | 0.64 | 0.39 | 0.69 | 0.57 | 0.61 | 0.67 |
| 9 | 7405911 | rs2997552 | 0.856 | 0.487 | -1.594 | -0.56 | -0.081 | 0.073 | 0.073 | 0 | 0.41 | 0.12 | 0.48 | 0.26 | 0.45 | 0.44 |
| 9 | 7408763 | rs2997578 | 0.428 | 0.112 | -0.947 | -0.714 | -0.003 | 0.127 | 0.127 | 0 | 0.48 | 0.25 | 0.45 | 0.22 | 0.48 | 0.5 |
| 9 | 7411160 | rs2997584 | 0.599 | 0.132 | -0.229 | -0.76 | 0.018 | 0.067 | 0.067 | 0 | 0.46 | 0.38 | 0.43 | 0.19 | 0.43 | 0.46 |
| 9 | 7413841 | rs1926415 | -0.839 | 0.732 | -0.527 | -0.345 | 0.006 | 0.054 | 0.054 | 0 | 0.84 | 0.85 | 0.86 | 0.56 | 0.81 | 0.8 |
| 9 | 7415094 | rs1926410 | 1.332 | 0.269 | -0.051 | -0.374 | -0.001 | 0.01 | 0.01 | 0 | 0.17 | 0.23 | 0.2 | 0.26 | 0.18 | 0.12 |
| 9 | 7415340 | rs2997544 | 0.358 | 0.28 | -0.34 | -0.423 | 0.005 | 0.01 | 0.01 | 0 | 0.66 | 0.68 | 0.71 | 0.54 | 0.61 | 0.66 |
| 9 | 7426221 | rs1926403 | 0.959 | 0.151 | -0.813 | -0.663 | -0.047 | 0.114 | 0.114 | 0 | 0.4 | 0.09 | 0.44 | 0.2 | 0.42 | 0.44 |
| 9 | 7436205 | rs13294039 | 1.559 | 0.157 | -0.384 | -0.534 | -0.002 | -0.009 | -0.009 | 0 | 0.18 | 0.07 | 0.19 | 0.27 | 0.19 | 0.15 |
| 9 | 7437070 | rs7857046 | 0.214 | 0.098 | -0.195 | -0.533 | -0.026 | -0.011 | -0.011 | 0 | 0.24 | 0.58 | 0.23 | 0.28 | 0.2 | 0.19 |
| 9 | 7437154 | rs7875445 | 0.196 | 0.086 | -0.202 | -0.508 | -0.027 | -0.007 | -0.007 | 0 | 0.24 | 0.58 | 0.23 | 0.28 | 0.2 | 0.19 |
| 9 | 7446748 | rs2997550 | 1.366 | 0.255 | -0.492 | 0.163 | -0.007 | 0.036 | 0.036 | 0.668 | 0.52 | 0.17 | 0.59 | 0.4 | 0.56 | 0.55 |
| 9 | 7448042 | rs16926110 | 0.832 | 0.529 | -0.413 | 0.264 | -0.017 | -0.011 | -0.011 | 0.146 | 0.06 | 0.01 | 0.13 | 0.23 | 0.1 | 0.13 |
| 9 | 7454698 | rs2997564 | -0.208 | 0.578 | 0.164 | 0.249 | 0.024 | 0.048 | 0.048 | 0 | 0.77 | 0.7 | 0.76 | 0.6 | 0.68 | 0.78 |
| 9 | 7454719 | rs2997565 | 0.38 | 0.623 | 0.099 | 0.206 | 0 | 0.005 | 0.005 | 0.698 | 0.71 | 0.31 | 0.73 | 0.6 | 0.68 | 0.76 |
| 9 | 7462598 | rs2997573 | 1.063 | 0.582 | 0.172 | 0.238 | -0.001 | 0.007 | 0.007 | 0.09 | 0.57 | 0.31 | 0.62 | 0.37 | 0.6 | 0.6 |
| 9 | 7465205 | rs2803469 | -0.307 | 0.672 | 0.217 | 0.562 | -0.013 | -0.021 | -0.021 | 0 | 0.73 | 0.82 | 0.7 | 0.6 | 0.66 | 0.76 |
| 9 | 7466024 | rs13298501 | 0.204 | 0.682 | 0.079 | 0.553 | 0.011 | 0.002 | 0.002 | 0.208 | 0.08 | 0.08 | 0.09 | 0.12 | 0.11 | 0.18 |
| 9 | 7467366 | rs1926389 | 2.328 | 0.739 | -0.224 | 0.291 | 0.016 | 0.037 | 0.037 | 0 | 0.31 | 0.17 | 0.3 | 0.14 | 0.33 | 0.18 |
| 9 | 7467934 | rs2986261 | 1.931 | 1.027 | 0.167 | 0.748 | 0.011 | 0.112 | 0.112 | 0 | 0.49 | 0.23 | 0.46 | 0.19 | 0.47 | 0.42 |
| 9 | 7468664 | rs943170 | 0.183 | 0.926 | 0.209 | 0.793 | -0.001 | 0.001 | 0.001 | 1.157 | 0.13 | 0.16 | 0.13 | 0.28 | 0.06 | 0.17 |
| 9 | 7468720 | rs2986262 | 0.545 | 0.992 | 0.351 | 0.776 | 0.02 | 0.028 | 0.028 | 0 | 0.65 | 0.51 | 0.67 | 0.53 | 0.58 | 0.61 |
| 9 | 7470753 | rs1926371 | 0.204 | 1.038 | 0.244 | 0.762 | 0.032 | 0.039 | 0.039 | 0 | 0.7 | 0.53 | 0.73 | 0.63 | 0.63 | 0.79 |
| 9 | 7472554 | rs1887084 | 0.034 | 1.076 | -0.231 | 0.559 | 0.005 | -0.021 | -0.021 | 0.484 | 0.14 | 0.12 | 0.13 | 0.26 | 0.1 | 0.18 |
| 9 | 7472661 | rs2381609 | 1.014 | 1.055 | -0.17 | 0.56 | -0.004 | -0.005 | -0.005 | 0 | 0.62 | 0.37 | 0.59 | 0.46 | 0.56 | 0.62 |
| 9 | 7473692 | rs2986267 | -0.287 | 1.001 | -0.216 | 0.879 | 0.001 | 0.009 | 0.009 | 0 | 0.76 | 0.75 | 0.7 | 0.58 | 0.66 | 0.81 |
| 9 | 7474066 | rs10124650 | -0.168 | 1.11 | -0.255 | 1.017 | -0.003 | -0.011 | -0.011 | 0 | 0.14 | 0.11 | 0.15 | 0.29 | 0.12 | 0.21 |
| 9 | 7478453 | rs13297172 | -0.089 | 0.974 | -0.176 | 0.978 | 0.001 | -0.001 | -0.001 | 0 | 0.08 | 0.06 | 0.07 | 0.11 | 0.07 | 0.15 |
| 9 | 7478718 | rs2986276 | -0.618 | 1.41 | 0.147 | 1.342 | 0.072 | 0.211 | 0.211 | 1.869 | 0.92 | 0.77 | 0.82 | 0.7 | 0.85 | 0.92 |
| 9 | 7482618 | rs3955044 | -1.904 | 1.246 | -0.192 | 0.757 | 0.032 | 0.088 | 0.088 | 0 | 0.53 | 0.75 | 0.67 | 0.8 | 0.59 | 0.6 |
| 9 | 7485142 | rs10976314 | 1.381 | 1.277 | -0.199 | 0.72 | -0.001 | 0.075 | 0.075 | 0 | 0.53 | 0.29 | 0.4 | 0.22 | 0.51 | 0.47 |
| 9 | 7487847 | rs16926236 | 0.372 | 0.953 | -0.158 | 0.743 | 0.001 | 0.001 | 0.001 | 0 | 0.08 | 0.05 | 0.06 | 0.09 | 0.07 | 0.14 |
| 9 | 7487974 | rs10976317 | 0.784 | 0.952 | 0.043 | 0.775 | -0.006 | -0.013 | -0.013 | 0 | 0.21 | 0.2 | 0.1 | 0.13 | 0.14 | 0.14 |
| 9 | 7489047 | rs10815617 | 0.886 | 1.034 | -0.05 | 0.904 | -0.001 | 0 | 0 | 0.104 | 0.59 | 0.42 | 0.59 | 0.46 | 0.54 | 0.63 |
| 9 | 7489720 | rs4742351 | 1.546 | 1.091 | -0.182 | 0.778 | 0.023 | 0.115 | 0.115 | 0 | 0.47 | 0.23 | 0.44 | 0.2 | 0.42 | 0.43 |
| 9 | 7491136 | rs16926241 | 0.025 | 1.11 | -0.055 | 0.992 | -0.002 | -0.002 | -0.002 | 0 | 0.08 | 0.04 | 0.07 | 0.12 | 0.11 | 0.15 |
| 9 | 7493198 | rs10119703 | -0.408 | 0.999 | 0.108 | 1.252 | 0.044 | 0.177 | 0.177 | 0 | 0.9 | 0.8 | 0.84 | 0.67 | 0.87 | 0.9 |
| 9 | 7496902 | rs2381616 | 1.258 | -0.08 | -0.281 | 0.451 | -0.01 | 0.071 | 0.071 | 0 | 0.34 | 0.16 | 0.31 | 0.15 | 0.41 | 0.37 |
| 9 | 7497427 | rs10119450 | -0.166 | -0.192 | -0.247 | 0.478 | 0.001 | 0 | 0 | 0 | 0.67 | 0.71 | 0.54 | 0.46 | 0.64 | 0.61 |
| 9 | 7497932 | rs7029335 | -0.768 | -0.13 | 0.417 | 0.788 | -0.006 | -0.004 | -0.004 | 0.715 | 0.32 | 0.38 | 0.21 | 0.33 | 0.27 | 0.26 |
| 9 | 7500955 | rs7019330 | -0.275 | -0.022 | 0.311 | 0.827 | 0.008 | -0.02 | -0.02 | 0.121 | 0.27 | 0.25 | 0.22 | 0.38 | 0.28 | 0.29 |
| 9 | 7501258 | rs10815619 | 0.233 | -0.04 | 0.302 | 0.835 | 0.012 | -0.031 | -0.031 | 0 | 0.22 | 0.14 | 0.15 | 0.37 | 0.18 | 0.28 |
| 9 | 7503994 | rs7021631 | -1.188 | -0.131 | 0.266 | 0.788 | 0.008 | -0.01 | -0.01 | 3.577 | 0.34 | 0.35 | 0.26 | 0.48 | 0.31 | 0.35 |
| 9 | 7504790 | rs7853724 | -0.051 | 0.208 | 0.601 | 0.896 | 0.037 | 0.023 | 0.023 | 0 | 0.11 | 0.24 | 0.11 | 0.12 | 0.12 | 0.15 |
| 9 | 7505902 | rs1986733 | -0.072 | 0.175 | 0.379 | 0.911 | 0.01 | -0.013 | -0.013 | 0 | 0.23 | 0.26 | 0.18 | 0.4 | 0.18 | 0.3 |
| 9 | 7506212 | rs12237671 | 1.201 | 0.151 | 0.325 | 0.938 | 0 | -0.012 | -0.012 | 0.022 | 0.14 | 0.02 | 0.06 | 0.17 | 0.11 | 0.11 |
| 9 | 7507691 | rs4742355 | -0.675 | 0.174 | 0.356 | 0.919 | 0.004 | -0.005 | -0.005 | 0 | 0.28 | 0.31 | 0.23 | 0.41 | 0.27 | 0.31 |
| 9 | 7508229 | rs4740890 | -0.922 | 0.401 | 0.216 | 0.739 | 0.033 | -0.006 | -0.006 | 0.84 | 0.16 | 0.3 | 0.17 | 0.13 | 0.22 | 0.16 |
| 9 | 7510496 | rs17613887 | 0.578 | 0.061 | -0.162 | 0.722 | -0.006 | 0.027 | 0.027 | 0 | 0.07 | 0.04 | 0.09 | 0 | 0.14 | 0.02 |
| 9 | 7510851 | rs7019104 | -1.047 | 0 | 0.015 | 0.826 | -0.001 | -0.001 | -0.001 | 0 | 0.38 | 0.59 | 0.29 | 0.35 | 0.35 | 0.28 |

| | | | | | | | | | | | | | | | | |
|---|---|---|---|---|---|---|---|---|---|---|---|---|---|---|---|---|
| 9 | 7511166 | rs10511474 | -0.966 | 0.435 | -0.181 | 0.83 | -0.011 | 0.01 | 0.01 | 0 | 0.13 | 0.24 | 0.12 | 0.18 | 0.1 | 0.15 |
| 9 | 7512186 | rs7873050 | -0.278 | 0.774 | -0.55 | 0.876 | -0.001 | 0.003 | 0.003 | 0.35 | 0.08 | 0.09 | 0.09 | 0.25 | 0.09 | 0.21 |
| 9 | 7512409 | rs2381623 | -0.777 | 1.041 | -0.46 | 0.333 | 0.016 | -0.009 | -0.009 | 0 | 0.33 | 0.38 | 0.44 | 0.27 | 0.36 | 0.38 |
| 9 | 7512610 | rs10511476 | -0.776 | 1.16 | -0.399 | 0.585 | 0.028 | 0.068 | 0.068 | 0 | 0.08 | 0.2 | 0.17 | 0.17 | 0.08 | 0.16 |
| 9 | 7512820 | rs17542133 | -0.617 | 1.158 | -1.028 | 0.024 | -0.046 | 0.017 | 0.017 | 0 | 0.27 | 0.16 | 0.41 | 0.26 | 0.36 | 0.22 |
| 9 | 7513175 | rs2381624 | -0.036 | 0.545 | -0.345 | -0.439 | -0.019 | 0.012 | 0.012 | 0 | 0.86 | 0.79 | 0.94 | 0.86 | 0.91 | 0.73 |
| 9 | 7513262 | rs10758869 | 0.898 | 0.126 | -0.757 | -0.925 | 0 | -0.003 | -0.003 | 0 | 0.33 | 0.28 | 0.37 | 0.62 | 0.32 | 0.3 |
| 9 | 7514442 | rs10511478 | -0.024 | -0.217 | -0.634 | -1.025 | 0.006 | 0 | 0 | 0.246 | 0.26 | 0.2 | 0.17 | 0.11 | 0.23 | 0.18 |
| 9 | 7514825 | rs10511479 | 0.755 | 0.562 | -1.022 | -1.022 | -0.01 | 0.013 | 0.013 | 0 | 0.65 | 0.71 | 0.63 | 0.77 | 0.65 | 0.52 |
| 9 | 7515370 | rs4623493 | 0.034 | 0.049 | -0.081 | -0.606 | -0.017 | 0.011 | 0.011 | 0.069 | 0.87 | 0.74 | 0.93 | 0.87 | 0.91 | 0.78 |
| 9 | 7517429 | rs2381626 | 0.63 | 0.26 | -0.405 | -0.976 | 0.007 | -0.002 | -0.002 | 0 | 0.81 | 0.84 | 0.88 | 1 | 0.8 | 0.9 |
| 9 | 7518656 | rs2381628 | 0.835 | 0.248 | 0.377 | -1.079 | -0.018 | -0.089 | -0.089 | 0.077 | 0.79 | 0.6 | 0.83 | 0.95 | 0.79 | 0.87 |
| 9 | 7519263 | rs7466961 | 1.047 | 0.394 | 0.601 | -1.315 | 0.001 | -0.007 | -0.007 | 0.194 | 0.71 | 0.71 | 0.82 | 0.94 | 0.7 | 0.84 |
| 9 | 7520035 | rs7865752 | 1.201 | 0.278 | 1.147 | -1.293 | -0.015 | -0.084 | -0.084 | 0 | 0.71 | 0.45 | 0.74 | 0.92 | 0.71 | 0.84 |
| 9 | 7521177 | rs7870059 | 0.713 | 0.26 | 1.217 | -1.261 | -0.024 | -0.091 | -0.091 | 0.831 | 0.79 | 0.51 | 0.84 | 0.94 | 0.8 | 0.87 |
| 9 | 7521577 | rs4606115 | 0.46 | 0.139 | 1.275 | -1.319 | -0.055 | -0.116 | -0.116 | 1.84 | 0.82 | 0.52 | 0.81 | 0.98 | 0.85 | 0.96 |
| 9 | 7530883 | rs10976357 | 0.148 | 0.599 | 0.576 | -1.753 | -0.063 | 0.029 | 0.029 | 0.235 | 0.19 | 0.06 | 0.23 | 0.12 | 0.24 | 0.26 |
| 9 | 7532386 | rs7025720 | -0.64 | 1.228 | 0.976 | -1.461 | 0.049 | 0 | 0 | 0 | 0.3 | 0.43 | 0.42 | 0.33 | 0.43 | 0.38 |
| 9 | 7533050 | rs4740893 | 1.838 | 1.15 | 0.886 | -1.551 | 0.045 | 0.073 | 0.073 | 0 | 0.44 | 0.16 | 0.43 | 0.67 | 0.34 | 0.56 |
| 9 | 7533529 | rs4742366 | 0.086 | 1.135 | 0.994 | -1.53 | -0.018 | -0.096 | -0.096 | 3.33 | 0.75 | 0.66 | 0.85 | 1 | 0.77 | 0.94 |
| 9 | 7534603 | rs6477214 | 0.342 | 1.119 | 1.002 | -1.531 | -0.002 | -0.053 | -0.053 | 0 | 0.75 | 0.75 | 0.86 | 1 | 0.77 | 0.94 |
| 9 | 7535872 | rs10815631 | 1.141 | 1.238 | 0.877 | -1.327 | 0.012 | 0.046 | 0.046 | 0 | 0.53 | 0.33 | 0.48 | 0.65 | 0.42 | 0.57 |
| 9 | 7539001 | rs2890755 | 0.894 | 1.289 | 0.931 | -1.387 | 0.001 | 0.005 | 0.005 | 0 | 0.57 | 0.39 | 0.57 | 0.78 | 0.58 | 0.65 |
| 9 | 7539567 | rs2381635 | 1.168 | 1.078 | 0.811 | -1.38 | -0.001 | -0.007 | -0.007 | 0 | 0.49 | 0.46 | 0.54 | 0.76 | 0.5 | 0.62 |
| 9 | 7540496 | rs12349302 | 0.631 | 1.08 | 0.849 | -1.289 | 0.018 | -0.06 | -0.06 | 0.365 | 0.1 | 0.23 | 0.06 | 0 | 0.09 | 0.02 |
| 9 | 7541338 | rs7849878 | -0.613 | 0.982 | 0.66 | -1.438 | 0.008 | -0.007 | -0.007 | 0 | 0.86 | 0.86 | 0.89 | 1 | 0.85 | 0.97 |
| 9 | 7541473 | rs10976380 | -0.013 | 1.317 | 0.825 | -1.431 | 0.005 | 0.001 | 0.001 | 0.352 | 0.19 | 0.25 | 0.3 | 0.21 | 0.25 | 0.31 |
| 9 | 7542487 | rs12348699 | -0.9 | 0.727 | 0.475 | -1.62 | 0.009 | 0.022 | 0.022 | 0.008 | 0.23 | 0.19 | 0.14 | 0.03 | 0.15 | 0.1 |
| 9 | 7542967 | rs10976384 | 1.725 | 0.719 | 0.538 | -1.591 | 0.001 | 0.001 | 0.001 | 0 | 0.4 | 0.33 | 0.47 | 0.69 | 0.31 | 0.52 |
| 9 | 7543336 | rs12353094 | -0.125 | 0.655 | 0.461 | -1.388 | -0.002 | -0.026 | -0.026 | 0 | 0.17 | 0.19 | 0.07 | 0.02 | 0.11 | 0.06 |
| 9 | 7544922 | rs10739135 | 0.403 | 0.73 | 0.169 | -1.414 | 0.004 | 0.043 | 0.043 | 0 | 0.73 | 0.81 | 0.85 | 0.98 | 0.74 | 0.91 |
| 9 | 7546436 | rs4742374 | -0.951 | 0.676 | 0.459 | -1.146 | -0.004 | -0.029 | -0.029 | 0 | 0.89 | 0.89 | 0.91 | 1 | 0.86 | 0.98 |
| 9 | 7547443 | rs2381634 | 1.051 | 0.535 | 0.234 | -1.335 | 0.004 | 0.006 | 0.006 | 0 | 0.39 | 0.3 | 0.47 | 0.65 | 0.31 | 0.52 |
| 9 | 7557664 | rs4740896 | 1.114 | 0.505 | 0.508 | -1.063 | 0.024 | -0.001 | -0.001 | 0 | 0.37 | 0.21 | 0.42 | 0.64 | 0.27 | 0.51 |
| 9 | 7558182 | rs7855527 | 0.689 | 0.385 | 1.258 | -1.158 | -0.012 | -0.018 | -0.018 | 0 | 0.43 | 0.54 | 0.48 | 0.72 | 0.31 | 0.58 |
| 9 | 7558261 | rs4400445 | 0.602 | 0.744 | 1.245 | -1.314 | 0.007 | -0.01 | -0.01 | 0.224 | 0.16 | 0.16 | 0.13 | 0.1 | 0.18 | 0.12 |
| 9 | 7560568 | rs12376869 | 0.56 | 0.801 | 1.138 | -1.351 | 0.001 | 0 | 0 | 0 | 0.23 | 0.18 | 0.29 | 0.18 | 0.36 | 0.26 |
| 9 | 7560779 | rs2381640 | 1.34 | 0.74 | 1.17 | -1.181 | 0.026 | 0.017 | 0.017 | 0 | 0.37 | 0.21 | 0.43 | 0.69 | 0.27 | 0.52 |
| 9 | 7561271 | rs4740897 | -0.232 | 0.558 | 1.063 | -1.431 | 0.024 | -0.019 | -0.019 | 0.018 | 0.09 | 0.27 | 0.14 | 0.03 | 0.09 | 0.1 |
| 9 | 7561756 | rs10117858 | -0.23 | 0.448 | 1.242 | -1.335 | 0.051 | 0.064 | 0.064 | 0 | 0.05 | 0.3 | 0.04 | 0.03 | 0.02 | 0.04 |
| 9 | 7563692 | rs2381642 | 0.896 | 0.597 | 1.212 | -1.06 | 0 | 0.006 | 0.006 | 0 | 0.42 | 0.38 | 0.49 | 0.72 | 0.37 | 0.58 |
| 9 | 7570610 | rs725988 | 0.363 | 0.537 | 0.977 | -1.155 | 0.009 | 0.029 | 0.029 | 0.117 | 0.54 | 0.42 | 0.61 | 0.76 | 0.52 | 0.62 |
| 9 | 7578222 | rs4742378 | 1.205 | 0.654 | 1.044 | -1.131 | 0.004 | 0.028 | 0.028 | 0.025 | 0.42 | 0.26 | 0.48 | 0.7 | 0.37 | 0.52 |
| 9 | 7579004 | rs1485184 | -0.28 | 0.567 | 1.156 | -1.166 | 0 | -0.021 | -0.021 | 0 | 0.65 | 0.61 | 0.65 | 0.8 | 0.58 | 0.69 |
| 9 | 7580951 | rs2381645 | -0.109 | 0.559 | 1.181 | -1.168 | 0 | -0.004 | -0.004 | 0 | 0.65 | 0.66 | 0.66 | 0.8 | 0.58 | 0.69 |
| 9 | 7581533 | rs4742380 | 0.909 | 0.599 | 1.206 | -1.191 | 0.001 | 0.003 | 0.003 | 0.645 | 0.45 | 0.43 | 0.52 | 0.71 | 0.39 | 0.55 |
| 9 | 7582234 | rs10976420 | -0.168 | 0.552 | 1.221 | -1.233 | 0.001 | -0.047 | -0.047 | 0 | 0.66 | 0.62 | 0.67 | 0.94 | 0.6 | 0.8 |
| 9 | 7582632 | rs7873807 | 1.274 | 0.519 | 0.892 | -1.183 | 0.009 | 0.04 | 0.04 | 0.032 | 0.41 | 0.25 | 0.45 | 0.7 | 0.35 | 0.5 |
| 9 | 7582673 | rs7873817 | 0.57 | 0.058 | 0.996 | -1.032 | -0.001 | -0.004 | -0.004 | 0 | 0.63 | 0.39 | 0.69 | 0.72 | 0.62 | 0.68 |
| 9 | 7583375 | rs7036891 | 1.037 | -0.075 | 0.791 | -0.29 | 0.024 | 0.001 | 0.001 | 0 | 0.43 | 0.26 | 0.37 | 0.6 | 0.37 | 0.51 |
| 9 | 7584003 | rs4740900 | -0.184 | -0.063 | 1.054 | -0.325 | 0.014 | -0.042 | -0.042 | 0 | 0.64 | 0.55 | 0.54 | 0.81 | 0.59 | 0.78 |
| 9 | 7585382 | rs10125595 | 0.471 | -0.059 | 1.065 | -0.281 | 0.008 | -0.005 | -0.005 | 0 | 0.2 | 0.32 | 0.17 | 0.19 | 0.21 | 0.27 |
| 9 | 7585409 | rs1385447 | -0.203 | -0.042 | 1.08 | -0.19 | 0.011 | -0.03 | -0.03 | 0 | 0.65 | 0.58 | 0.54 | 0.81 | 0.58 | 0.77 |
| 9 | 7587282 | rs10976421 | 1.032 | -0.045 | 0.732 | -0.181 | 0.025 | 0.001 | 0.001 | 0 | 0.43 | 0.26 | 0.37 | 0.6 | 0.38 | 0.5 |
| 9 | 7587716 | rs4740901 | -0.157 | 0.092 | 0.76 | -0.132 | 0.006 | -0.025 | -0.025 | 0 | 0.68 | 0.64 | 0.56 | 0.84 | 0.61 | 0.81 |
| 9 | 7591465 | rs10758888 | -0.956 | -0.102 | 0.562 | 0.061 | -0.026 | -0.001 | -0.001 | 0 | 0.87 | 0.75 | 0.82 | 0.89 | 0.87 | 0.98 |
| 9 | 7592407 | rs3852399 | -1.125 | -0.082 | 0.394 | 0.075 | -0.019 | -0.001 | -0.001 | 0 | 0.88 | 0.8 | 0.83 | 0.89 | 0.87 | 0.98 |
| 9 | 7601812 | rs16926462 | 1.605 | -0.378 | -0.546 | 0.504 | 0.019 | -0.066 | -0.066 | 0 | 0.2 | 0.03 | 0.17 | 0.28 | 0.17 | 0.29 |
| 9 | 7604410 | rs10815645 | 0.817 | 0.085 | -0.021 | 0.278 | 0.013 | -0.031 | -0.031 | 2.445 | 0.26 | 0.28 | 0.3 | 0.09 | 0.29 | 0.23 |
| 9 | 7607425 | rs10815647 | -0.494 | -0.339 | 0.254 | 0.525 | -0.041 | 0.015 | 0.015 | 0.069 | 0.65 | 0.51 | 0.66 | 0.63 | 0.68 | 0.66 |
| 9 | 7612143 | rs10976450 | 0.233 | -0.156 | -0.37 | 0.282 | -0.053 | -0.073 | -0.073 | 0.045 | 0.37 | 0.14 | 0.35 | 0.44 | 0.41 | 0.37 |
| 9 | 7613834 | rs10976453 | 1.074 | -0.082 | -0.377 | 0.076 | -0.008 | 0.033 | 0.033 | 0 | 0.2 | 0.14 | 0.14 | 0.11 | 0.21 | 0.18 |
| 9 | 7617978 | rs10815648 | 1.115 | -0.102 | -0.298 | 0.114 | -0.013 | 0.026 | 0.026 | 0 | 0.23 | 0.14 | 0.31 | 0.11 | 0.23 | 0.22 |
| 9 | 7625060 | rs3912454 | -0.013 | -0.124 | -0.051 | 0.246 | 0 | -0.001 | -0.001 | 0 | 0.54 | 0.57 | 0.56 | 0.43 | 0.51 | 0.57 |
| 9 | 7633155 | rs7867163 | 1.502 | 0.057 | -0.521 | 0.113 | -0.035 | 0.055 | 0.055 | 0.107 | 0.2 | 0.06 | 0.28 | 0.1 | 0.21 | 0.17 |
| 9 | 7636050 | rs10976463 | 2.051 | -0.55 | -0.746 | 0.127 | -0.008 | -0.076 | -0.076 | 0 | 0.16 | 0.02 | 0.13 | 0.29 | 0.19 | 0.26 |
| 9 | 7636071 | rs7865284 | 1.506 | -0.385 | -0.723 | 0.036 | -0.042 | 0.057 | 0.057 | 0 | 0.22 | 0.06 | 0.31 | 0.1 | 0.24 | 0.21 |
| 9 | 7638453 | rs1564845 | 0.686 | -0.409 | -0.281 | 0.166 | 0.006 | 0.045 | 0.045 | 0 | 0.25 | 0.32 | 0.35 | 0.42 | 0.26 | 0.24 |
| 9 | 7641251 | rs10976470 | 2.165 | -0.425 | -0.515 | 0.186 | -0.023 | -0.07 | -0.07 | 0.033 | 0.15 | 0.02 | 0.13 | 0.26 | 0.19 | 0.24 |

| | | | | | | | | | | | | | | | | |
|---|---|---|---|---|---|---|---|---|---|---|---|---|---|---|---|---|
| 9 | 7641620 | rs7864806 | -0.261 | -0.555 | -0.285 | 0.059 | 0.001 | 0.003 | 0.003 | 0 | 0.59 | 0.58 | 0.55 | 0.51 | 0.57 | 0.6 |
| 9 | 7642033 | rs3005752 | 0.917 | -0.479 | -0.658 | 0.241 | -0.015 | -0.052 | -0.052 | 4.905 | 0.21 | 0.13 | 0.26 | 0.38 | 0.24 | 0.19 |
| 9 | 7646850 | rs10976473 | 1.239 | -0.574 | -0.858 | 0.166 | 0.016 | -0.081 | -0.081 | 0.361 | 0.22 | 0.03 | 0.14 | 0.3 | 0.2 | 0.27 |
| 9 | 7649442 | rs2930713 | -0.677 | -0.55 | -0.547 | -0.02 | -0.018 | 0.041 | 0.041 | 0 | 0.59 | 0.72 | 0.56 | 0.78 | 0.56 | 0.61 |
| 9 | 7656826 | rs2930727 | 0.176 | -0.743 | -0.344 | 0.085 | 0.011 | -0.012 | -0.012 | 4.413 | 0.44 | 0.28 | 0.62 | 0.47 | 0.51 | 0.43 |
| 9 | 7658953 | rs2930732 | -0.081 | -0.907 | -0.076 | -0.021 | -0.007 | -0.017 | -0.017 | 0 | 0.62 | 0.45 | 0.77 | 0.83 | 0.69 | 0.68 |
| 9 | 7662501 | rs2930736 | -1.142 | -0.89 | -0.049 | -0.012 | 0 | 0 | 0 | 0 | 0.79 | 0.8 | 0.88 | 0.87 | 0.81 | 0.79 |
| 9 | 7662660 | rs10976483 | 1.466 | -0.942 | 0.183 | 0.048 | 0.013 | -0.018 | -0.018 | 0.007 | 0.2 | 0.12 | 0.15 | 0.33 | 0.18 | 0.26 |
| 9 | 7664428 | rs11792985 | 1.277 | -0.596 | 0.255 | -0.045 | -0.003 | 0.001 | 0.001 | 0 | 0.22 | 0.25 | 0.34 | 0.18 | 0.28 | 0.25 |
| 9 | 7665194 | rs10976486 | 1.423 | -0.67 | 0.19 | -0.034 | 0.031 | -0.032 | -0.032 | 0 | 0.23 | 0.09 | 0.14 | 0.31 | 0.19 | 0.28 |
| 9 | 7665856 | rs9942919 | 1.262 | -0.546 | 0.241 | -0.015 | 0.016 | -0.008 | -0.008 | 0 | 0.22 | 0.25 | 0.35 | 0.19 | 0.28 | 0.28 |
| 9 | 7666387 | rs3005761 | -1.182 | -0.863 | 0.222 | -0.098 | -0.001 | -0.006 | -0.006 | 0 | 0.8 | 0.82 | 0.88 | 0.88 | 0.81 | 0.79 |
| 9 | 7668843 | rs3124028 | 0.339 | -0.806 | 0.136 | 0.006 | -0.004 | -0.006 | -0.006 | 1.286 | 0.39 | 0.36 | 0.58 | 0.54 | 0.5 | 0.45 |
| 9 | 7672125 | rs10815654 | 0.751 | -0.79 | 0.077 | 0.024 | 0.018 | -0.011 | -0.011 | 0.927 | 0.26 | 0.15 | 0.2 | 0.29 | 0.21 | 0.31 |
| 9 | 7673654 | rs10976492 | 1.226 | -0.89 | 0.037 | 0.043 | 0.003 | -0.003 | -0.003 | 0 | 0.21 | 0.16 | 0.12 | 0.26 | 0.15 | 0.26 |
| 9 | 7677811 | rs7046958 | 1.097 | -1.035 | -0.406 | -0.255 | 0.022 | 0.098 | 0.098 | 0.011 | 0.18 | 0.04 | 0.07 | 0 | 0.12 | 0.02 |
| 9 | 7682063 | rs10976498 | 1.602 | -0.745 | 0.164 | 0.184 | 0.018 | -0.013 | -0.013 | 0.28 | 0.18 | 0.11 | 0.11 | 0.25 | 0.14 | 0.24 |
| 9 | 7682601 | rs842304 | 0.275 | -0.426 | 0.627 | 0.221 | -0.003 | 0.005 | 0.005 | 0.121 | 0.33 | 0.5 | 0.36 | 0.64 | 0.33 | 0.46 |
| 9 | 7682860 | rs4740904 | 0.369 | -0.44 | 0.779 | 0.097 | -0.003 | -0.015 | -0.015 | 0 | 0.42 | 0.49 | 0.48 | 0.35 | 0.42 | 0.49 |
| 9 | 7684244 | rs328847 | 0.394 | -0.185 | 1.006 | 0.403 | 0.059 | 0.218 | 0.218 | 0 | 0.16 | 0.49 | 0.25 | 0.43 | 0.19 | 0.24 |
| 9 | 7685789 | rs12376595 | 1.647 | -0.625 | 0.768 | 0.2 | -0.001 | -0.002 | -0.002 | 0 | 0.18 | 0.13 | 0.11 | 0.24 | 0.14 | 0.26 |
| 9 | 7688633 | rs842302 | 0.476 | -0.384 | 0.765 | 0.163 | -0.005 | -0.001 | -0.001 | 0 | 0.31 | 0.22 | 0.33 | 0.68 | 0.35 | 0.58 |
| 9 | 7689651 | rs1332911 | 0.997 | -0.645 | 0.403 | -0.175 | -0.012 | 0.026 | 0.026 | 0 | 0.26 | 0.19 | 0.38 | 0.11 | 0.29 | 0.24 |
| 9 | 7690568 | rs7046363 | 1.071 | -0.659 | 0.178 | -0.175 | -0.063 | 0.08 | 0.08 | 0 | 0.25 | 0.06 | 0.37 | 0.11 | 0.29 | 0.31 |
| 9 | 7694465 | rs12002314 | 1.859 | -0.727 | 0.229 | -0.101 | 0.007 | -0.005 | -0.005 | 0 | 0.19 | 0.12 | 0.11 | 0.24 | 0.15 | 0.26 |
| 9 | 7694484 | rs842301 | -1.104 | -0.748 | 0.374 | -0.247 | 0.006 | 0.056 | 0.056 | 0 | 0.67 | 0.82 | 0.83 | 0.81 | 0.67 | 0.9 |
| 9 | 7695593 | rs12553833 | 0.496 | -0.314 | 0.517 | 0.048 | -0.04 | 0.076 | 0.076 | 0 | 0.26 | 0.12 | 0.37 | 0.09 | 0.3 | 0.31 |
| 9 | 7703280 | rs328838 | -0.93 | -0.204 | 0.619 | 0.254 | 0.004 | 0.004 | 0.004 | 0.108 | 0.44 | 0.71 | 0.7 | 0.55 | 0.51 | 0.63 |
| 9 | 7706762 | rs1332905 | -0.616 | -0.098 | 0.69 | 0.338 | 0 | -0.01 | -0.01 | 0 | 0.35 | 0.37 | 0.47 | 0.16 | 0.35 | 0.36 |
| 9 | 7707372 | rs170620 | 0.793 | 0.016 | 0.351 | 0.3 | -0.016 | -0.013 | -0.013 | 0.065 | 0.14 | 0.1 | 0.24 | 0.45 | 0.21 | 0.32 |
| 9 | 7711440 | rs328864 | -0.99 | -0.7 | 0.355 | 0.152 | 0.031 | 0.023 | 0.023 | 0.818 | 0.56 | 0.76 | 0.77 | 0.62 | 0.61 | 0.65 |
| 9 | 7711546 | rs328863 | -0.744 | -0.055 | 0.776 | 0.626 | 0.004 | 0.011 | 0.011 | 0.155 | 0.39 | 0.51 | 0.62 | 0.51 | 0.51 | 0.57 |
| 9 | 7711758 | rs4742386 | -1.037 | -0.008 | 0.804 | 0.645 | -0.001 | 0 | 0 | 0 | 0.43 | 0.63 | 0.7 | 0.55 | 0.52 | 0.62 |
| 9 | 7712116 | rs1905761 | -0.511 | 0.012 | 0.524 | 0.658 | 0.002 | 0.003 | 0.003 | 0.053 | 0.37 | 0.4 | 0.61 | 0.52 | 0.49 | 0.57 |
| 9 | 7720636 | rs842296 | -0.673 | 0.034 | -0.091 | 0.623 | -0.058 | -0.068 | -0.068 | 0.175 | 0.39 | 0.21 | 0.61 | 0.49 | 0.45 | 0.55 |
| 9 | 7725346 | rs186629 | 0.327 | 0.403 | 0.178 | 0.53 | 0.018 | 0.053 | 0.053 | 0.129 | 0.15 | 0.25 | 0.28 | 0.45 | 0.23 | 0.31 |
| 9 | 7725409 | rs328862 | 0.268 | 0.335 | 0.433 | 0.494 | -0.007 | 0.007 | 0.007 | 0 | 0.94 | 0.88 | 0.92 | 0.95 | 0.96 | 0.93 |
| 9 | 7725912 | rs328860 | -0.711 | 0.339 | 0.235 | 0.555 | -0.039 | -0.031 | -0.031 | 2.601 | 0.36 | 0.26 | 0.54 | 0.45 | 0.49 | 0.49 |
| 9 | 7726450 | rs1332900 | 0.82 | 0.325 | 0.737 | 0.144 | 0.056 | 0.127 | 0.127 | 0.878 | 0.76 | 0.47 | 0.61 | 0.48 | 0.72 | 0.62 |
| 9 | 7726901 | rs328857 | 1.073 | 0.293 | 0.482 | 0.142 | -0.002 | 0.019 | 0.019 | 0.113 | 0.91 | 0.76 | 0.9 | 0.93 | 0.95 | 0.92 |
| 9 | 7727062 | rs702283 | 0.121 | 0.305 | 0.272 | 0.149 | -0.007 | 0.006 | 0.006 | 0 | 0.94 | 0.9 | 0.92 | 0.95 | 0.96 | 0.94 |
| 9 | 7727088 | rs842300 | 0.798 | 0.356 | 0.141 | 0.167 | -0.072 | -0.087 | -0.087 | 0 | 0.13 | 0.06 | 0.26 | 0.45 | 0.23 | 0.31 |
| 9 | 7729573 | rs17546550 | 0.888 | 0.229 | 0.104 | 0.098 | -0.069 | 0.055 | 0.055 | 0 | 0.17 | 0.01 | 0.29 | 0.05 | 0.23 | 0.19 |
| 9 | 7730729 | rs7854298 | 0.975 | 0.164 | 0.382 | 0.074 | -0.005 | 0 | 0 | 0 | 0.58 | 0.34 | 0.31 | 0.43 | 0.5 | 0.42 |
| 9 | 7734480 | rs12344462 | 0.189 | 0.277 | -0.119 | 0.102 | -0.016 | 0.058 | 0.058 | 0 | 0.22 | 0.13 | 0.33 | 0.05 | 0.26 | 0.19 |
| 9 | 7735283 | rs17629093 | 0.01 | 0.279 | -0.113 | 0.101 | -0.011 | 0.036 | 0.036 | 0 | 0.24 | 0.17 | 0.34 | 0.05 | 0.28 | 0.22 |
| 9 | 7736266 | rs1412770 | 0.549 | 0.639 | -0.176 | 0.375 | -0.047 | -0.057 | -0.057 | 0 | 0.14 | 0.1 | 0.27 | 0.45 | 0.23 | 0.31 |
| 9 | 7742476 | rs1332906 | -0.333 | 0.47 | -0.136 | 0.137 | -0.005 | 0.001 | 0.001 | 0 | 0.95 | 0.95 | 0.93 | 0.95 | 0.99 | 0.96 |
| 9 | 7755443 | rs1332924 | 0.911 | -0.036 | -0.361 | -0.121 | 0 | 0.003 | 0.003 | 0 | 0.4 | 0.36 | 0.26 | 0.27 | 0.41 | 0.35 |
| 9 | 7757657 | rs7021164 | 0.085 | 0.067 | -0.434 | -0.027 | 0.01 | 0.028 | 0.028 | 0.006 | 0.77 | 0.72 | 0.63 | 0.47 | 0.74 | 0.61 |
| 9 | 7757891 | rs7863770 | 0.664 | -0.207 | -0.513 | 0.147 | 0 | -0.006 | -0.006 | 0 | 0.37 | 0.37 | 0.25 | 0.26 | 0.39 | 0.34 |
| 9 | 7758579 | rs10815672 | -0.403 | 0.65 | -0.848 | 0.568 | -0.004 | -0.004 | -0.004 | 0.678 | 0.32 | 0.28 | 0.58 | 0.52 | 0.44 | 0.54 |
| 9 | 7759020 | rs11791458 | 0.036 | 0.205 | -1.383 | -0.012 | 0.024 | 0.034 | 0.034 | 3.129 | 0.35 | 0.12 | 0.26 | 0.24 | 0.28 | 0.23 |
| 9 | 7759331 | rs10119431 | 0.836 | 0.59 | -0.386 | 0.401 | 0 | 0.099 | 0.099 | 0 | 0.79 | 0.6 | 0.57 | 0.5 | 0.83 | 0.77 |
| 9 | 7759917 | rs16926737 | -0.793 | 0.538 | -0.759 | -0.599 | 0.005 | -0.002 | -0.002 | 0 | 0.09 | 0.09 | 0.03 | 0 | 0.06 | 0.09 |
| 9 | 7764226 | rs1576939 | -0.318 | 0.877 | -0.708 | -1.27 | -0.001 | -0.023 | -0.023 | 0 | 0.59 | 0.65 | 0.65 | 0.89 | 0.56 | 0.65 |
| 9 | 7764377 | rs1576938 | 0.645 | 1.042 | -0.608 | -1.84 | 0.004 | 0.026 | 0.026 | 0 | 0.29 | 0.28 | 0.5 | 0.85 | 0.28 | 0.41 |
| 9 | 7769626 | rs1360585 | 0.668 | 1.078 | -0.563 | -1.844 | 0.012 | 0.046 | 0.046 | 0 | 0.29 | 0.22 | 0.49 | 0.85 | 0.27 | 0.41 |
| 9 | 7770233 | rs9407382 | -0.294 | 1.036 | -0.32 | -2.011 | 0.025 | -0.088 | -0.088 | 0 | 0.19 | 0.27 | 0.12 | 0.01 | 0.21 | 0.14 |
| 9 | 7770942 | rs10758900 | -0.255 | 0.592 | -0.18 | -1.695 | -0.008 | -0.04 | -0.04 | 0 | 0.15 | 0.22 | 0.06 | 0.03 | 0.11 | 0.13 |
| 9 | 7773856 | rs7044932 | -0.393 | 1.114 | 0.028 | -1.563 | 0.002 | 0.005 | 0.005 | 0 | 0.66 | 0.73 | 0.73 | 0.89 | 0.67 | 0.7 |
| 9 | 7774180 | rs4742409 | -0.36 | 1.057 | -0.511 | -1.516 | 0.029 | 0.072 | 0.072 | 0.121 | 0.63 | 0.33 | 0.71 | 0.89 | 0.68 | 0.71 |
| 9 | 7775651 | rs9407384 | -0.792 | 1.556 | 0.093 | 0.281 | 0.025 | 0.111 | 0.111 | 0.045 | 0.12 | 0.16 | 0.37 | 0.42 | 0.23 | 0.14 |
| 9 | 7780159 | rs7863054 | 0.374 | 1.421 | -0.904 | -0.051 | -0.036 | -0.002 | -0.002 | 0 | 0.36 | 0.08 | 0.2 | 0.26 | 0.34 | 0.32 |
| 9 | 7782594 | rs327752 | -0.69 | 1.485 | -0.13 | -0.018 | 0.041 | -0.046 | -0.046 | 0 | 0.17 | 0.25 | 0.36 | 0.06 | 0.26 | 0.12 |
| 9 | 7783210 | rs182753 | -0.748 | 1.495 | -0.388 | 0.385 | 0.045 | 0.042 | 0.042 | 0 | 0.17 | 0.32 | 0.48 | 0.29 | 0.25 | 0.25 |
| 9 | 7784512 | rs3904482 | -0.158 | 0.823 | -1.066 | 0.806 | -0.078 | -0.082 | -0.082 | 0 | 0.39 | 0.04 | 0.22 | 0.33 | 0.37 | 0.35 |
| 9 | 7787532 | rs7035953 | -0.105 | 0.774 | -0.355 | 0.584 | -0.007 | 0.003 | 0.003 | 0.147 | 0.41 | 0.48 | 0.33 | 0.57 | 0.35 | 0.47 |
| 9 | 7789577 | rs12347887 | -0.13 | 0.713 | -0.286 | 0.629 | -0.004 | -0.024 | -0.024 | 0 | 0.31 | 0.38 | 0.27 | 0.26 | 0.34 | 0.24 |

| | | | | | | | | | | | | | | | | |
|---|---|---|---|---|---|---|---|---|---|---|---|---|---|---|---|---|
| 9 | 7790322 | rs6477238 | -0.64 | 1.004 | -0.475 | 1.085 | 0.004 | 0.01 | 0.01 | 4.684 | 0.53 | 0.64 | 0.62 | 0.58 | 0.55 | 0.56 |
| 9 | 7790940 | rs327756 | -0.188 | 0.946 | -0.931 | 1.281 | -0.002 | -0.021 | -0.021 | 0 | 0.24 | 0.19 | 0.26 | 0.34 | 0.26 | 0.31 |
| 9 | 7791704 | rs4740914 | 0.14 | 0.995 | -1.091 | 1.149 | -0.016 | -0.009 | -0.009 | 0.213 | 0.56 | 0.68 | 0.61 | 0.58 | 0.55 | 0.53 |
| 9 | 7792318 | rs10815682 | 0.615 | 0.517 | -0.413 | 0.805 | 0.004 | 0.019 | 0.019 | 0 | 0.43 | 0.32 | 0.35 | 0.33 | 0.44 | 0.34 |
| 9 | 7792599 | rs327755 | -0.042 | 0.448 | -0.002 | 0.729 | 0.003 | 0.016 | 0.016 | 0 | 0.67 | 0.67 | 0.6 | 0.55 | 0.68 | 0.61 |
| 9 | 7792930 | rs327754 | -0.507 | 0.911 | 0.275 | 0.506 | 0.008 | 0.006 | 0.006 | 0 | 0.24 | 0.35 | 0.32 | 0.28 | 0.33 | 0.29 |
| 9 | 7793309 | rs327753 | -1.223 | 0.525 | 0.338 | 0.463 | -0.002 | -0.009 | -0.009 | 0 | 0.33 | 0.49 | 0.34 | 0.28 | 0.36 | 0.32 |
| 9 | 7796237 | rs10976580 | 0.938 | 0.196 | 0.185 | 0.712 | 0.011 | 0.051 | 0.051 | 0.281 | 0.46 | 0.31 | 0.42 | 0.28 | 0.45 | 0.33 |
| 9 | 7796329 | rs2990600 | -0.422 | 0.43 | 0.028 | 0.91 | 0.002 | 0.009 | 0.009 | 0.194 | 0.19 | 0.21 | 0.27 | 0.28 | 0.25 | 0.26 |
| 9 | 7802252 | rs9407389 | -0.358 | 0.379 | -0.015 | 1.076 | -0.009 | -0.002 | -0.002 | 0.403 | 0.77 | 0.82 | 0.77 | 0.75 | 0.73 | 0.66 |
| 9 | 7804131 | rs10123827 | -0.526 | 0.299 | 0.369 | 1.239 | -0.007 | 0.061 | 0.061 | 0.059 | 0.07 | 0.18 | 0.05 | 0.17 | 0.04 | 0.11 |
| 9 | 7805650 | rs327759 | -0.268 | 0.122 | -0.014 | 1.13 | -0.023 | 0.047 | 0.047 | 0 | 0.29 | 0.46 | 0.19 | 0.35 | 0.23 | 0.25 |
| 9 | 7806697 | rs4740916 | 0.091 | 0.265 | 0.006 | 0.784 | 0 | 0 | 0 | 0 | 0.54 | 0.52 | 0.63 | 0.58 | 0.54 | 0.59 |
| 9 | 7812484 | rs12057040 | 0.951 | 0.404 | -0.724 | 0.712 | -0.005 | 0.02 | 0.02 | 0 | 0.39 | 0.21 | 0.53 | 0.37 | 0.43 | 0.42 |
| 9 | 7813034 | rs13297065 | 1.713 | 0.456 | -1.722 | 0.722 | -0.089 | -0.051 | -0.051 | 0.02 | 0.29 | 0.02 | 0.3 | 0.28 | 0.31 | 0.29 |
| 9 | 7814465 | rs1332916 | -0.97 | 0.872 | -1.104 | 1.296 | -0.005 | -0.005 | -0.005 | 0 | 0.76 | 0.85 | 0.79 | 0.72 | 0.74 | 0.73 |
| 9 | 7815105 | rs1360583 | 0.78 | 0.65 | -0.861 | 1.392 | -0.004 | 0.042 | 0.042 | 0 | 0.45 | 0.23 | 0.54 | 0.4 | 0.48 | 0.44 |
| 9 | 7815375 | rs7849315 | 0.444 | 0.821 | -1.55 | 1.388 | -0.009 | -0.017 | -0.017 | 0.429 | 0.57 | 0.7 | 0.63 | 0.57 | 0.55 | 0.58 |
| 9 | 7815985 | rs9407390 | -0.46 | 1.161 | -1.297 | 0.847 | -0.014 | -0.006 | -0.006 | 0.827 | 0.74 | 0.84 | 0.76 | 0.7 | 0.69 | 0.72 |
| 9 | 7818641 | rs10123429 | -0.11 | 1.145 | -0.9 | 0.782 | -0.003 | 0.018 | 0.018 | 0 | 0.48 | 0.67 | 0.51 | 0.46 | 0.46 | 0.6 |
| 9 | 7820080 | rs10976603 | -0.794 | 1.848 | -0.468 | 0.881 | 0.03 | 0.007 | 0.007 | 0 | 0.12 | 0.2 | 0.22 | 0.15 | 0.2 | 0.08 |
| 9 | 7820479 | rs9408714 | -0.064 | 1.276 | -0.374 | 0.911 | 0 | 0.002 | 0.002 | 0 | 0.51 | 0.5 | 0.48 | 0.58 | 0.48 | 0.66 |
| 9 | 7822486 | rs2184034 | 0.737 | 1.574 | -0.093 | 1.146 | -0.017 | -0.008 | -0.008 | 0.833 | 0.17 | 0.22 | 0.11 | 0.18 | 0.12 | 0.13 |
| 9 | 7824326 | rs928459 | -0.449 | 1.45 | 0.125 | 0.706 | -0.007 | 0.006 | 0.006 | 0 | 0.46 | 0.65 | 0.45 | 0.62 | 0.44 | 0.6 |
| 9 | 7824725 | rs9408716 | -0.522 | 1.407 | 0.374 | 0.658 | 0.008 | 0.047 | 0.047 | 0 | 0.46 | 0.74 | 0.57 | 0.74 | 0.49 | 0.63 |
| 9 | 7827205 | rs1360582 | -0.972 | 1.211 | 0.592 | 0.689 | 0.031 | 0.091 | 0.091 | 0 | 0.52 | 0.83 | 0.6 | 0.74 | 0.53 | 0.65 |
| 9 | 7829623 | rs9299057 | -0.43 | 1.216 | 0.686 | 0.689 | 0.005 | 0.041 | 0.041 | 0 | 0.46 | 0.7 | 0.57 | 0.73 | 0.49 | 0.63 |
| 9 | 7833215 | rs3847266 | 0.699 | 1.62 | 0.738 | 1.046 | -0.009 | -0.013 | -0.013 | 2.076 | 0.18 | 0.13 | 0.28 | 0.3 | 0.24 | 0.34 |
| 9 | 7842130 | rs10976614 | 1.135 | 1.217 | 0.138 | 0.61 | -0.012 | 0.155 | 0.155 | 0.72 | 0.34 | 0.04 | 0.25 | 0.04 | 0.35 | 0.2 |
| 9 | 7842694 | rs10976617 | 0.394 | 1.375 | 0.704 | 0.987 | -0.007 | 0.04 | 0.04 | 0 | 0.54 | 0.35 | 0.42 | 0.27 | 0.5 | 0.36 |
| 9 | 7852471 | rs7042604 | -0.342 | 1.591 | 0.756 | 1.121 | -0.016 | -0.007 | -0.007 | 0.18 | 0.69 | 0.68 | 0.85 | 0.74 | 0.79 | 0.72 |
| 9 | 7856992 | rs7847381 | -0.507 | 1.461 | 0.64 | 1.097 | -0.022 | -0.001 | -0.001 | 0.266 | 0.71 | 0.69 | 0.87 | 0.72 | 0.82 | 0.73 |
| 9 | 7871163 | rs10815701 | 1.157 | 1.55 | 0.632 | 1.194 | 0.003 | 0.006 | 0.006 | 0 | 0.16 | 0.15 | 0.29 | 0.32 | 0.23 | 0.34 |
| 9 | 7871717 | rs10758911 | 0.042 | 1.485 | 0.596 | 1.386 | 0.01 | -0.02 | -0.02 | 0.462 | 0.64 | 0.27 | 0.69 | 0.41 | 0.71 | 0.6 |
| 9 | 7874526 | rs7874427 | 0.544 | 1.372 | 0.514 | 1.096 | 0.01 | 0.018 | 0.018 | 0 | 0.19 | 0.19 | 0.44 | 0.45 | 0.32 | 0.39 |
| 9 | 7887951 | rs10758913 | -0.269 | 1.386 | 0.517 | 0.943 | -0.019 | 0.022 | 0.022 | 0 | 0.69 | 0.57 | 0.74 | 0.59 | 0.77 | 0.7 |
| 9 | 7891408 | rs17630650 | 0.878 | 1.366 | -0.495 | 0.864 | -0.004 | 0.08 | 0.08 | 0 | 0.46 | 0.16 | 0.41 | 0.27 | 0.47 | 0.29 |
| 9 | 7895438 | rs10511487 | 0.681 | 0.831 | -0.48 | 0.848 | -0.011 | -0.002 | -0.002 | 3.988 | 0.17 | 0.11 | 0.3 | 0.2 | 0.23 | 0.37 |
| 9 | 7897865 | rs10758914 | -0.035 | 0.145 | -0.136 | 0.891 | -0.043 | -0.006 | -0.006 | 0.036 | 0.71 | 0.68 | 0.89 | 0.74 | 0.83 | 0.75 |
| 9 | 7898810 | rs10732363 | -0.476 | 0.398 | -0.973 | 0.649 | 0.007 | 0.029 | 0.029 | 1.371 | 0.74 | 0.87 | 0.78 | 0.77 | 0.79 | 0.79 |
| 9 | 7898998 | rs10122520 | 0.157 | 0.677 | -0.997 | 0.913 | -0.02 | -0.009 | -0.009 | 0 | 0.4 | 0.23 | 0.69 | 0.54 | 0.53 | 0.47 |
| 9 | 7901308 | rs4302898 | -0.462 | 0.666 | -0.922 | 1.526 | 0.016 | -0.075 | -0.075 | 0 | 0.23 | 0.08 | 0.16 | 0.41 | 0.15 | 0.39 |
| 9 | 7902733 | rs10976654 | -1.224 | 1.025 | -0.127 | 1.51 | 0.001 | -0.012 | -0.012 | 0 | 0.33 | 0.38 | 0.2 | 0.51 | 0.27 | 0.43 |
| 9 | 7906830 | rs10815710 | -0.897 | 1.215 | 0.381 | 1.204 | 0.041 | 0.058 | 0.058 | 0.63 | 0.14 | 0.42 | 0.19 | 0.17 | 0.17 | 0.1 |
| 9 | 7906933 | rs12347060 | -0.311 | 1.217 | 0.509 | 1.171 | 0.02 | -0.018 | -0.018 | 0 | 0.12 | 0.3 | 0.07 | 0.03 | 0.12 | 0.03 |
| 9 | 7907459 | rs10815712 | -0.998 | 1.192 | 0.39 | 1.18 | 0.038 | 0.058 | 0.058 | 0.084 | 0.15 | 0.42 | 0.19 | 0.17 | 0.17 | 0.1 |
| 9 | 7908874 | rs16926845 | 0.161 | 1.172 | -0.007 | 0.879 | -0.007 | 0.006 | 0.006 | 1.451 | 0.09 | 0.06 | 0.05 | 0.03 | 0.12 | 0.03 |
| 9 | 7909429 | rs10976659 | -0.047 | 1.461 | -0.02 | 0.989 | 0.016 | 0.014 | 0.014 | 0.076 | 0.08 | 0.14 | 0.11 | 0.14 | 0.12 | 0.09 |
| 9 | 7909576 | rs7849795 | -1.72 | 1.153 | -1.034 | 0.287 | 0.017 | 0.142 | 0.142 | 0.07 | 0.68 | 0.97 | 0.77 | 0.93 | 0.74 | 0.84 |
| 9 | 7910358 | rs13297029 | 0.957 | 1.344 | -1.319 | 0.715 | -0.05 | -0.116 | -0.116 | 0 | 0.21 | 0.02 | 0.26 | 0.37 | 0.22 | 0.34 |
| 9 | 7911344 | rs10815713 | -0.267 | 1.405 | -0.871 | 0.416 | -0.001 | 0 | 0 | 0 | 0.28 | 0.7 | 0.37 | 0.38 | 0.27 | 0.26 |
| 9 | 7912161 | rs10976664 | -0.059 | 1.689 | -0.88 | 0.45 | -0.071 | -0.067 | -0.067 | 0 | 0.33 | 0.18 | 0.36 | 0.53 | 0.43 | 0.44 |
| 9 | 7913952 | rs2890768 | -0.151 | 1.657 | -0.843 | 0.411 | -0.044 | -0.039 | -0.039 | 0 | 0.34 | 0.22 | 0.37 | 0.54 | 0.43 | 0.45 |
| 9 | 7914008 | rs10815714 | 1.03 | 1.672 | -0.725 | 0.424 | 0.008 | 0.008 | 0.008 | 0.024 | 0.08 | 0.11 | 0.11 | 0.14 | 0.1 | 0.09 |
| 9 | 7915771 | rs1998371 | -0.062 | 1.512 | -0.514 | 0.377 | -0.023 | -0.008 | -0.008 | 0 | 0.25 | 0.5 | 0.28 | 0.19 | 0.22 | 0.29 |
| 9 | 7918996 | rs2224601 | -2.185 | 1.567 | -0.662 | 0.29 | 0.048 | 0.167 | 0.167 | 0.225 | 0.68 | 0.95 | 0.81 | 0.95 | 0.76 | 0.83 |
| 9 | 7919153 | rs2208742 | -2.148 | 1.554 | -0.668 | 0.284 | 0.05 | 0.17 | 0.17 | 2.209 | 0.67 | 0.95 | 0.81 | 0.95 | 0.76 | 0.83 |
| 9 | 7922071 | rs7868745 | -1.987 | 1.638 | -0.637 | 0.32 | 0.034 | 0.082 | 0.082 | 1.632 | 0.66 | 0.92 | 0.78 | 0.82 | 0.72 | 0.81 |
| 9 | 7924047 | rs4740922 | 0.119 | 1.629 | -0.209 | 0.311 | 0.003 | -0.001 | -0.001 | 0 | 0.23 | 0.36 | 0.31 | 0.24 | 0.25 | 0.22 |
| 9 | 7925381 | rs1535480 | -0.182 | 1.748 | -0.273 | 0.354 | 0 | 0 | 0 | 0 | 0.39 | 0.37 | 0.54 | 0.68 | 0.38 | 0.52 |
| 9 | 7925745 | rs11999820 | 1.065 | 1.866 | -0.237 | 0.394 | -0.012 | 0.006 | 0.006 | 0.713 | 0.09 | 0.08 | 0.06 | 0.03 | 0.14 | 0.05 |
| 9 | 7929372 | rs2093483 | 0.16 | 1.515 | -0.301 | 0.421 | -0.016 | -0.061 | -0.061 | 0.198 | 0.3 | 0.16 | 0.36 | 0.56 | 0.32 | 0.43 |
| 9 | 7930092 | rs875917 | 0.46 | 1.676 | -0.039 | 0.347 | 0.004 | -0.001 | -0.001 | 0 | 0.12 | 0.14 | 0.1 | 0.12 | 0.2 | 0.1 |
| 9 | 7932619 | rs10976676 | -0.025 | 1.462 | -0.016 | 0.259 | 0.011 | -0.023 | -0.023 | 0 | 0.09 | 0.29 | 0.06 | 0.02 | 0.07 | 0.16 |
| 9 | 7935389 | rs10815718 | 0.297 | 1.607 | -0.347 | 0.344 | 0.002 | -0.001 | -0.001 | 0 | 0.14 | 0.16 | 0.1 | 0.07 | 0.18 | 0.11 |
| 9 | 7938795 | rs10758915 | -0.232 | 0.743 | -0.582 | 0.348 | 0.036 | 0.128 | 0.128 | 0.083 | 0.1 | 0.37 | 0.12 | 0.21 | 0.1 | 0.18 |
| 9 | 7945146 | rs10815721 | -0.828 | 0.755 | -0.671 | 0.269 | 0.016 | 0.026 | 0.026 | 0.013 | 0.17 | 0.38 | 0.2 | 0.22 | 0.19 | 0.23 |
| 9 | 7947001 | rs7868963 | 0.815 | 0.271 | -1.216 | 0.473 | -0.045 | -0.139 | -0.139 | 3.678 | 0.22 | 0.02 | 0.29 | 0.43 | 0.22 | 0.35 |
| 9 | 7949546 | rs1322468 | -1.626 | 0.512 | -1.061 | 0.389 | 0.005 | 0.179 | 0.179 | 0.431 | 0.69 | 0.99 | 0.81 | 0.96 | 0.75 | 0.84 |

| | | | | | | | | | | | | | | | | |
|---|---|---|---|---|---|---|---|---|---|---|---|---|---|---|---|---|
| 9 | 7951024 | rs10815723 | -0.715 | 0.302 | -0.024 | 1.169 | -0.004 | -0.002 | -0.002 | 0 | 0.36 | 0.57 | 0.49 | 0.67 | 0.34 | 0.54 |
| 9 | 7951684 | rs10815725 | -0.646 | -0.005 | -0.283 | 0.637 | -0.017 | 0.033 | 0.033 | 0 | 0.34 | 0.44 | 0.4 | 0.47 | 0.29 | 0.5 |
| 9 | 7955178 | rs4742424 | -0.675 | 0.001 | -0.306 | 0.632 | -0.017 | 0.031 | 0.031 | 0 | 0.35 | 0.43 | 0.4 | 0.47 | 0.29 | 0.5 |
| 9 | 7962765 | rs10976692 | -0.604 | -0.4 | -0.187 | 0.14 | -0.051 | -0.046 | -0.046 | 0 | 0.21 | 0.37 | 0.09 | 0.12 | 0.09 | 0.09 |
| 9 | 7966280 | rs2065119 | 0.223 | -0.446 | 0.026 | 0.346 | 0 | 0.049 | 0.049 | 0.729 | 0.15 | 0.32 | 0.18 | 0.26 | 0.14 | 0.16 |
| 9 | 7968906 | rs1407467 | 0.584 | 0.091 | -0.875 | -0.421 | 0.005 | -0.027 | -0.027 | 0.776 | 0.27 | 0.29 | 0.3 | 0.08 | 0.39 | 0.3 |
| 9 | 7978514 | rs10514819 | 0.089 | -0.139 | -0.703 | -0.485 | -0.036 | -0.044 | -0.044 | 0.066 | 0.19 | 0.35 | 0.09 | 0.11 | 0.11 | 0.09 |
| 9 | 7979082 | rs2025192 | 0.115 | 0.016 | -1.191 | -0.66 | -0.022 | 0.008 | 0.008 | 0.117 | 0.4 | 0.3 | 0.62 | 0.59 | 0.51 | 0.56 |
| 9 | 7979694 | rs10976702 | 0.966 | 0.022 | -1.074 | -0.657 | -0.003 | 0.014 | 0.014 | 0.113 | 0.16 | 0.24 | 0.16 | 0.26 | 0.15 | 0.15 |
| 9 | 7979832 | rs1322473 | -2.091 | 0.122 | -1.399 | -0.63 | -0.044 | 0.132 | 0.132 | 2.297 | 0.76 | 0.99 | 0.88 | 0.96 | 0.76 | 0.8 |
| 9 | 7980064 | rs7875477 | -0.405 | 0.085 | -1.261 | -0.526 | -0.008 | -0.002 | -0.002 | 0 | 0.35 | 0.63 | 0.26 | 0.38 | 0.25 | 0.25 |
| 9 | 7983044 | rs7871884 | -1.601 | 0.037 | -1.627 | -0.392 | -0.046 | 0.097 | 0.097 | 2.356 | 0.75 | 0.99 | 0.82 | 0.92 | 0.76 | 0.79 |
| 9 | 7985651 | rs1407469 | 0.635 | 0.064 | -1.953 | -0.409 | -0.056 | 0.001 | 0.001 | 0 | 0.21 | 0.03 | 0.18 | 0.14 | 0.23 | 0.18 |
| 9 | 7988643 | rs1322475 | -2.127 | -0.026 | -1.887 | -0.41 | -0.015 | 0.099 | 0.099 | 2.43 | 0.79 | 0.96 | 0.87 | 0.96 | 0.77 | 0.82 |
| 9 | 7994225 | rs764310 | 0.907 | 0.347 | -0.528 | 0.163 | 0.063 | -0.005 | -0.005 | 0 | 0.15 | 0.31 | 0.2 | 0.11 | 0.22 | 0.22 |
| 9 | 7994349 | rs764308 | -0.12 | -0.202 | -1.11 | 0.51 | 0 | -0.142 | -0.142 | 0 | 0.19 | 0.03 | 0.32 | 0.44 | 0.14 | 0.3 |
| 9 | 7994891 | rs10976710 | -0.061 | -0.185 | -1.102 | 0.518 | -0.001 | -0.14 | -0.14 | 0 | 0.19 | 0.03 | 0.32 | 0.44 | 0.15 | 0.29 |
| 9 | 7995103 | rs2025193 | 0.272 | -0.006 | -0.367 | 0.17 | -0.009 | 0 | 0 | 0.046 | 0.09 | 0.11 | 0.06 | 0.1 | 0.05 | 0.07 |
| 9 | 7999201 | rs12348891 | -0.054 | -0.011 | -0.341 | 0.164 | -0.012 | -0.003 | -0.003 | 0.579 | 0.1 | 0.18 | 0.08 | 0.1 | 0.06 | 0.07 |
| 9 | 8009428 | rs9408721 | -0.812 | 0.038 | -0.297 | -0.132 | 0.001 | -0.004 | -0.004 | 0 | 0.62 | 0.69 | 0.76 | 0.79 | 0.67 | 0.75 |
| 9 | 8009807 | rs12337889 | 0.004 | 0.193 | -0.377 | 0.133 | 0 | -0.003 | -0.003 | 0 | 0.07 | 0.09 | 0.08 | 0.04 | 0.06 | 0.13 |
| 9 | 8010899 | rs7030273 | -0.159 | -0.076 | -1.008 | 0.394 | -0.012 | -0.096 | -0.096 | 0 | 0.26 | 0.06 | 0.28 | 0.35 | 0.26 | 0.23 |
| 9 | 8011107 | rs4740924 | 0.962 | 0.246 | -0.59 | -0.103 | 0.026 | -0.024 | -0.024 | 0.087 | 0.13 | 0.18 | 0.17 | 0.09 | 0.18 | 0.2 |
| 9 | 8011265 | rs10758920 | -1.116 | -0.047 | -0.395 | -0.139 | 0 | 0 | 0 | 0 | 0.63 | 0.39 | 0.74 | 0.72 | 0.72 | 0.65 |
| 9 | 8012608 | rs12685956 | -0.678 | 0.175 | -0.223 | 0.27 | 0 | -0.051 | -0.051 | 0 | 0.37 | 0.19 | 0.46 | 0.53 | 0.36 | 0.47 |
| 9 | 8012773 | rs10976719 | 1.073 | 0.464 | 0.091 | 0.169 | 0.034 | -0.012 | -0.012 | 2.754 | 0.11 | 0.15 | 0.15 | 0.07 | 0.18 | 0.17 |
| 9 | 8014674 | rs9299059 | 0.601 | 0.215 | 0.216 | 0.125 | -0.009 | -0.067 | -0.067 | 0 | 0.28 | 0.47 | 0.22 | 0.12 | 0.26 | 0.19 |
| 9 | 8021344 | rs1322458 | 0.756 | 0.582 | 0.138 | 0.305 | 0.013 | 0.001 | 0.001 | 0 | 0.12 | 0.16 | 0.17 | 0.13 | 0.18 | 0.18 |
| 9 | 8023123 | rs1407461 | -1.803 | 0.631 | 0.228 | 0.069 | -0.042 | -0.129 | -0.129 | 0 | 0.7 | 0.55 | 0.83 | 0.95 | 0.79 | 0.83 |
| 9 | 8025789 | rs7849676 | 0.068 | 0.584 | -0.07 | 0.294 | 0.001 | -0.044 | -0.044 | 0 | 0.28 | 0.15 | 0.38 | 0.33 | 0.26 | 0.31 |
| 9 | 8029327 | rs7024333 | -2.148 | 0.554 | 0.207 | 0.336 | -0.001 | -0.001 | -0.001 | 0 | 0.72 | 0.67 | 0.83 | 0.73 | 0.79 | 0.82 |
| 9 | 8029507 | rs7027896 | 0.288 | 0.597 | 0.159 | 0.208 | -0.007 | -0.067 | -0.067 | 0 | 0.27 | 0.12 | 0.38 | 0.35 | 0.25 | 0.29 |
| 9 | 8040076 | rs10976737 | -1.11 | 0.569 | 0.259 | -0.898 | -0.001 | -0.058 | -0.058 | 2.147 | 0.6 | 0.52 | 0.66 | 0.88 | 0.62 | 0.64 |
| 9 | 8045606 | rs2146180 | -1.069 | 0.557 | 0.271 | -0.929 | -0.001 | -0.042 | -0.042 | 0 | 0.59 | 0.52 | 0.66 | 0.82 | 0.61 | 0.66 |
| 9 | 8052825 | rs10976744 | 0.593 | 0.633 | 0.346 | -0.905 | -0.006 | 0.007 | 0.007 | 0 | 0.31 | 0.26 | 0.26 | 0.28 | 0.35 | 0.33 |
| 9 | 8059266 | rs7357637 | -0.244 | 0.672 | 0.302 | -0.898 | -0.001 | 0.001 | 0.001 | 0 | 0.43 | 0.51 | 0.44 | 0.36 | 0.53 | 0.5 |
| 9 | 8063984 | rs1322467 | -2.292 | 0.738 | 0.026 | -0.858 | 0.014 | 0.108 | 0.108 | 0.091 | 0.76 | 0.94 | 0.86 | 0.96 | 0.8 | 0.82 |
| 9 | 8065584 | rs6477256 | -2.064 | 1.03 | -0.041 | -0.74 | 0.002 | -0.022 | -0.022 | 0 | 0.72 | 0.69 | 0.84 | 0.96 | 0.8 | 0.82 |
| 9 | 8074897 | rs10815746 | 1.379 | 1.131 | -0.589 | -0.647 | -0.051 | -0.001 | -0.001 | 1.154 | 0.09 | 0.02 | 0.15 | 0.07 | 0.18 | 0.17 |
| 9 | 8075119 | rs2381700 | 1.071 | 1.16 | -0.554 | -0.622 | 0.038 | 0.008 | 0.008 | 0 | 0.11 | 0.16 | 0.17 | 0.13 | 0.18 | 0.17 |
| 9 | 8079286 | rs7846713 | 0.353 | 0.821 | -0.511 | -1.164 | -0.002 | -0.004 | -0.004 | 0.556 | 0.21 | 0.18 | 0.38 | 0.75 | 0.22 | 0.4 |
| 9 | 8087483 | rs10976758 | -0.18 | 0.466 | -0.59 | -1.254 | 0 | -0.022 | -0.022 | 0 | 0.45 | 0.42 | 0.64 | 0.89 | 0.53 | 0.63 |
| 9 | 8087860 | rs975299 | -0.721 | 0.435 | -0.431 | -0.508 | -0.019 | -0.077 | -0.077 | 0 | 0.72 | 0.63 | 0.84 | 0.96 | 0.8 | 0.83 |
| 9 | 8090327 | rs7020343 | -0.524 | 1.156 | -0.548 | 0.431 | -0.001 | 0.004 | 0.004 | 0 | 0.42 | 0.64 | 0.42 | 0.38 | 0.42 | 0.47 |
| 9 | 8090430 | rs7020466 | -0.398 | 1.153 | 0.033 | 0.71 | -0.038 | 0.001 | 0.001 | 0.056 | 0.31 | 0.53 | 0.14 | 0.31 | 0.21 | 0.22 |
| 9 | 8092506 | rs10976762 | -1.442 | 1.232 | 0.282 | 0.92 | 0.01 | 0.158 | 0.158 | 0 | 0.05 | 0.15 | 0.06 | 0.34 | 0.05 | 0.19 |
| 9 | 8093826 | rs10976763 | 0.16 | 0.884 | 0.908 | 1.705 | -0.009 | 0.008 | 0.008 | 0 | 0.2 | 0.19 | 0.1 | 0.17 | 0.17 | 0.17 |
| 9 | 8093954 | rs10815753 | 1.174 | 0.373 | 0.737 | 1.372 | 0.041 | 0.074 | 0.074 | 0.428 | 0.54 | 0.3 | 0.65 | 0.3 | 0.64 | 0.46 |
| 9 | 8094390 | rs11999343 | 0.635 | 0.357 | 0.665 | 1.538 | -0.009 | -0.003 | -0.003 | 0 | 0.2 | 0.19 | 0.1 | 0.14 | 0.17 | 0.17 |
| 9 | 8095638 | rs6477258 | 0.303 | 0.293 | 0.92 | 1.267 | 0.002 | -0.001 | -0.001 | 0 | 0.63 | 0.72 | 0.75 | 0.68 | 0.71 | 0.66 |
| 9 | 8102543 | rs10815759 | -0.951 | 0.324 | 0.852 | 1.283 | 0.01 | 0.016 | 0.016 | 0 | 0.84 | 0.91 | 0.86 | 0.88 | 0.89 | 0.83 |
| 9 | 8106282 | rs7862526 | 0.464 | 0.303 | 1.072 | 1.302 | -0.023 | 0.023 | 0.023 | 0 | 0.22 | 0.3 | 0.14 | 0.19 | 0.17 | 0.16 |
| 9 | 8108790 | rs9650711 | 1.578 | 0.695 | 0.727 | 1.217 | 0.003 | 0.145 | 0.145 | 0.518 | 0.5 | 0.13 | 0.43 | 0.3 | 0.57 | 0.41 |
| 9 | 8114588 | rs10758936 | 1.149 | 0.567 | 0.742 | 1.282 | 0.065 | 0.063 | 0.063 | 0.94 | 0.54 | 0.1 | 0.62 | 0.33 | 0.68 | 0.61 |
| 9 | 8115492 | rs989726 | -1.037 | 0.564 | 0.795 | 1.295 | 0.004 | 0.017 | 0.017 | 0 | 0.84 | 0.92 | 0.85 | 0.85 | 0.88 | 0.83 |
| 9 | 8116662 | rs12336338 | 0.218 | 0.593 | 1.014 | 1.371 | -0.001 | 0.003 | 0.003 | 0.007 | 0.23 | 0.19 | 0.33 | 0.23 | 0.24 | 0.21 |
| 9 | 8119459 | rs7048333 | 0.554 | 0.508 | 1.282 | 1.395 | 0.008 | -0.016 | -0.016 | 2.33 | 0.21 | 0.13 | 0.15 | 0.23 | 0.18 | 0.16 |
| 9 | 8121487 | rs7034109 | -0.924 | 0.507 | 1.608 | 1.376 | -0.01 | -0.014 | -0.014 | 0.985 | 0.84 | 0.82 | 0.84 | 0.85 | 0.88 | 0.84 |
| 9 | 8121725 | rs10976786 | -0.224 | 0.475 | 1.716 | 1.387 | -0.026 | 0.065 | 0.065 | 0 | 0.26 | 0.56 | 0.21 | 0.52 | 0.21 | 0.23 |
| 9 | 8125336 | rs10441691 | 0.569 | 0.483 | 1.666 | 1.389 | -0.008 | 0.008 | 0.008 | 0 | 0.21 | 0.19 | 0.15 | 0.21 | 0.17 | 0.16 |
| 9 | 8125410 | rs10758939 | 1.265 | 0.536 | 1.572 | 1.34 | 0.066 | 0.045 | 0.045 | 0.952 | 0.55 | 0.19 | 0.63 | 0.39 | 0.68 | 0.61 |
| 9 | 8131068 | rs1578118 | -0.429 | 0.563 | 1.657 | 1.398 | 0 | 0.049 | 0.049 | 0 | 0.29 | 0.56 | 0.39 | 0.49 | 0.29 | 0.27 |
| 9 | 8132822 | rs7855517 | 0.037 | 0.397 | 1.704 | 1.378 | 0.037 | 0.126 | 0.126 | 0 | 0.22 | 0.38 | 0.36 | 0.49 | 0.26 | 0.26 |
| 9 | 8133055 | rs1361317 | -1.058 | 0.368 | 1.647 | 1.337 | 0.006 | 0.013 | 0.013 | 0 | 0.84 | 0.9 | 0.85 | 0.88 | 0.88 | 0.83 |
| 9 | 8133859 | rs911763 | 0.365 | 0.354 | 1.836 | 1.416 | 0.02 | 0.047 | 0.047 | 0 | 0.19 | 0.27 | 0.33 | 0.27 | 0.25 | 0.2 |
| 9 | 8136132 | rs941904 | 1.096 | 0.365 | 1.569 | 1.464 | 0.002 | 0.009 | 0.009 | 0.022 | 0.59 | 0.27 | 0.47 | 0.43 | 0.62 | 0.57 |
| 9 | 8138580 | rs16927158 | 0.92 | 0.321 | 1.261 | 1.407 | 0.001 | 0.011 | 0.011 | 0 | 0.15 | 0.08 | 0.16 | 0.14 | 0.12 | 0.17 |
| 9 | 8139013 | rs964222 | 0.044 | 0.315 | 1.354 | 1.453 | -0.009 | 0.072 | 0.072 | 0 | 0.19 | 0.39 | 0.21 | 0.33 | 0.13 | 0.24 |
| 9 | 8143737 | rs7466234 | 0.879 | 0.314 | 0.08 | 1.186 | 0 | 0.002 | 0.002 | 0 | 0.15 | 0.12 | 0.16 | 0.12 | 0.13 | 0.17 |

| | | | | | | | | | | | | | | | | |
|---|---|---|---|---|---|---|---|---|---|---|---|---|---|---|---|---|
| 9 | 8144973 | rs10976799 | 1.108 | 0.398 | -0.148 | 1.283 | -0.017 | 0.029 | 0.029 | 0.294 | 0.55 | 0.1 | 0.43 | 0.4 | 0.6 | 0.56 |
| 9 | 8147857 | rs10976802 | 0.449 | 0.454 | -0.903 | 0.665 | -0.055 | -0.017 | -0.017 | 0 | 0.3 | 0.09 | 0.28 | 0.33 | 0.35 | 0.46 |
| 9 | 8156271 | rs1361315 | 1.888 | -0.234 | -0.738 | 0.293 | -0.022 | 0.118 | 0.118 | 0 | 0.26 | 0.04 | 0.17 | 0.08 | 0.27 | 0.11 |
| 9 | 8157022 | rs9696734 | 0.217 | -0.144 | -0.753 | 0.39 | -0.035 | -0.018 | -0.018 | 0 | 0.65 | 0.22 | 0.55 | 0.65 | 0.56 | 0.66 |
| 9 | 8157313 | rs10815774 | 0.602 | -0.121 | -0.78 | 0.4 | 0.038 | -0.028 | -0.028 | 0 | 0.49 | 0.16 | 0.39 | 0.54 | 0.44 | 0.51 |
| 9 | 8160714 | rs13287364 | 0.819 | -0.095 | -0.617 | 0.367 | -0.062 | -0.045 | -0.045 | 0 | 0.27 | 0.12 | 0.39 | 0.31 | 0.42 | 0.34 |
| 9 | 8161572 | rs10976812 | -0.157 | -0.047 | -0.938 | 0.438 | 0.023 | 0.094 | 0.094 | 0 | 0.38 | 0.2 | 0.3 | 0.21 | 0.29 | 0.48 |
| 9 | 8165056 | rs17631170 | 1.39 | -0.12 | -1.097 | 0.348 | 0.012 | 0.056 | 0.056 | 0 | 0.17 | 0.03 | 0.14 | 0.08 | 0.11 | 0.15 |
| 9 | 8165192 | rs12555677 | 0.227 | 0.056 | -0.833 | 0.288 | 0.017 | -0.004 | -0.004 | 0.532 | 0.56 | 0.6 | 0.62 | 0.39 | 0.7 | 0.44 |
| 9 | 8165446 | rs13299501 | 0.27 | 0.107 | -0.885 | 0.287 | 0.049 | 0.046 | 0.046 | 0 | 0.3 | 0.52 | 0.44 | 0.38 | 0.43 | 0.34 |
| 9 | 8169035 | rs11794801 | 1.555 | 0.386 | -1.51 | -0.294 | -0.02 | 0.133 | 0.133 | 0 | 0.25 | 0.04 | 0.17 | 0.04 | 0.27 | 0.11 |
| 9 | 8169707 | rs10976815 | 0.144 | 0.555 | -1.073 | 0.153 | 0.006 | 0.026 | 0.026 | 0 | 0.2 | 0.12 | 0.15 | 0.12 | 0.17 | 0.32 |
| 9 | 8170945 | rs13292463 | 0.973 | 1.069 | -0.977 | 0.472 | -0.025 | 0.034 | 0.034 | 0 | 0.14 | 0.02 | 0.07 | 0.05 | 0.14 | 0.1 |
| 9 | 8171386 | rs10739149 | 0.466 | 1.237 | -0.569 | 0.605 | -0.003 | 0.01 | 0.01 | 0 | 0.56 | 0.35 | 0.43 | 0.4 | 0.5 | 0.54 |
| 9 | 8172046 | rs12237835 | -0.478 | 0.937 | -0.114 | 1.287 | 0.026 | 0.103 | 0.103 | 0.02 | 0.06 | 0.19 | 0.09 | 0.2 | 0.07 | 0.15 |
| 9 | 8174284 | rs1538518 | 0.614 | 1.15 | 0.534 | 1.633 | 0 | 0.001 | 0.001 | 0 | 0.56 | 0.39 | 0.55 | 0.41 | 0.6 | 0.54 |
| 9 | 8176226 | rs10739150 | 0.334 | 1.058 | 0.372 | 1.411 | -0.008 | 0.034 | 0.034 | 0.051 | 0.73 | 0.55 | 0.59 | 0.61 | 0.79 | 0.71 |
| 9 | 8176622 | rs10733543 | 0.637 | 0.949 | 0.375 | 1.363 | 0.015 | -0.034 | -0.034 | 0 | 0.71 | 0.23 | 0.56 | 0.56 | 0.79 | 0.7 |
| 9 | 8179325 | rs4130580 | 0.811 | 0.985 | 0.336 | 1.324 | 0.003 | -0.007 | -0.007 | 0 | 0.17 | 0.18 | 0.17 | 0.18 | 0.16 | 0.15 |
| 9 | 8180646 | rs10976822 | 0.482 | 0.92 | 0.215 | 1.637 | -0.055 | -0.07 | -0.07 | 0 | 0.32 | 0.03 | 0.16 | 0.29 | 0.34 | 0.31 |
| 9 | 8180709 | rs10976823 | -0.671 | 0.868 | 0.26 | 1.613 | -0.003 | 0.013 | 0.013 | 0 | 0.51 | 0.3 | 0.34 | 0.42 | 0.5 | 0.47 |
| 9 | 8181494 | rs10976826 | 0.533 | 0.883 | 0.349 | 1.666 | 0.001 | 0.159 | 0.159 | 0.076 | 0.06 | 0.2 | 0.24 | 0.22 | 0.05 | 0.15 |
| 9 | 8181848 | rs10739153 | 0.917 | 0.881 | 0.226 | 1.604 | -0.031 | 0.184 | 0.184 | 0.928 | 0.87 | 0.37 | 0.7 | 0.64 | 0.94 | 0.84 |
| 9 | 8182214 | rs12003225 | 0.887 | 0.894 | 0.179 | 1.509 | 0.002 | -0.013 | -0.013 | 0.854 | 0.17 | 0.16 | 0.17 | 0.13 | 0.16 | 0.14 |
| 9 | 8184461 | rs10976830 | 0.495 | 0.899 | 0.057 | 1.511 | -0.055 | -0.123 | -0.123 | 0 | 0.32 | 0.03 | 0.19 | 0.34 | 0.35 | 0.32 |
| 9 | 8185170 | rs10976831 | 1.541 | 0.935 | 0.088 | 1.486 | -0.114 | 0.099 | 0.099 | 0 | 0.36 | 0.07 | 0.36 | 0.22 | 0.44 | 0.37 |
| 9 | 8193646 | rs10976839 | 0.807 | 0.811 | 0.156 | 1.591 | 0.019 | -0.039 | -0.039 | 2.541 | 0.7 | 0.19 | 0.56 | 0.57 | 0.79 | 0.69 |
| 9 | 8196511 | rs7044355 | -1.164 | 0.866 | -0.028 | 1.241 | -0.016 | 0.061 | 0.061 | 0 | 0.55 | 0.76 | 0.52 | 0.65 | 0.53 | 0.53 |
| 9 | 8206576 | rs1781276 | -0.159 | 0.635 | 0.323 | 1.703 | -0.009 | 0.003 | 0.003 | 0 | 0.31 | 0.4 | 0.32 | 0.33 | 0.24 | 0.31 |
| 9 | 8206769 | rs1538523 | 1.006 | 0.666 | 0.413 | 1.784 | 0.006 | 0.002 | 0.002 | 0 | 0.51 | 0.4 | 0.48 | 0.49 | 0.59 | 0.53 |
| 9 | 8208014 | rs2185233 | 0.669 | 0.753 | 0.41 | 1.415 | 0.006 | 0.039 | 0.039 | 0 | 0.13 | 0.14 | 0.13 | 0.31 | 0.18 | 0.15 |
| 9 | 8208407 | rs7869678 | 1.037 | 0.744 | 0.428 | 1.407 | 0.031 | 0.016 | 0.016 | 0 | 0.5 | 0.28 | 0.5 | 0.54 | 0.6 | 0.53 |
| 9 | 8208999 | rs9299064 | -0.848 | 0.434 | 0.533 | 1.377 | 0.013 | 0.042 | 0.042 | 0.222 | 0.45 | 0.73 | 0.48 | 0.62 | 0.42 | 0.46 |
| 9 | 8209809 | rs10815790 | 1.976 | 0.527 | 0.453 | 1.386 | -0.083 | 0.208 | 0.208 | 0 | 0.35 | 0.05 | 0.34 | 0.05 | 0.4 | 0.25 |
| 9 | 8213656 | rs4008257 | -1.042 | 0.406 | 0.521 | 1.471 | 0.015 | 0.075 | 0.075 | 0.335 | 0.47 | 0.86 | 0.49 | 0.62 | 0.42 | 0.46 |
| 9 | 8215609 | rs10815791 | -0.114 | 0.374 | 0.592 | 1.711 | -0.01 | 0.131 | 0.131 | 0 | 0.08 | 0.25 | 0.22 | 0.16 | 0.05 | 0.15 |
| 9 | 8219852 | rs3847274 | -0.255 | 0.413 | 0.154 | 1.316 | 0 | 0.019 | 0.019 | 0 | 0.37 | 0.38 | 0.26 | 0.46 | 0.38 | 0.31 |
| 9 | 8220067 | rs4127335 | 1.96 | 0.415 | 0.13 | 1.308 | -0.094 | 0.08 | 0.08 | 0 | 0.36 | 0.06 | 0.35 | 0.22 | 0.41 | 0.38 |
| 9 | 8221247 | rs4742469 | 0.422 | 0.03 | 0.349 | 1.333 | -0.023 | 0.014 | 0.014 | 0.061 | 0.72 | 0.44 | 0.61 | 0.69 | 0.79 | 0.69 |
| 9 | 8223541 | rs7867627 | -0.472 | 0.27 | 0.025 | 1.046 | -0.001 | 0.003 | 0.003 | 0.258 | 0.82 | 0.85 | 0.84 | 0.87 | 0.84 | 0.84 |
| 9 | 8227101 | rs10976860 | 1.523 | 0.589 | -0.17 | 1.124 | -0.085 | 0.123 | 0.123 | 0 | 0.39 | 0.07 | 0.38 | 0.24 | 0.45 | 0.39 |
| 9 | 8227693 | rs10815796 | 1.487 | 0.59 | -0.075 | 1.123 | -0.053 | 0.114 | 0.114 | 0 | 0.39 | 0.13 | 0.39 | 0.24 | 0.45 | 0.39 |
| 9 | 8229283 | rs13287379 | 0.526 | 1.067 | 0.079 | 1.632 | -0.049 | -0.089 | -0.089 | 0 | 0.14 | 0.01 | 0.1 | 0.27 | 0.17 | 0.15 |
| 9 | 8230268 | rs1781263 | 0.235 | 1.063 | 0.421 | 1.608 | 0.002 | 0.024 | 0.024 | 0 | 0.15 | 0.14 | 0.11 | 0.27 | 0.17 | 0.16 |
| 9 | 8230433 | rs1984216 | 1.06 | 1.037 | 0.643 | 1.672 | 0.002 | -0.005 | -0.005 | 0 | 0.54 | 0.55 | 0.66 | 0.47 | 0.54 | 0.57 |
| 9 | 8231940 | rs7038843 | 0.88 | 0.259 | 0.805 | 1.292 | -0.006 | -0.006 | -0.006 | 0.046 | 0.15 | 0.22 | 0.07 | 0.11 | 0.13 | 0.1 |
| 9 | 8233804 | rs1339287 | 0.749 | 0.372 | 0.429 | 1.354 | 0.008 | 0.028 | 0.028 | 0 | 0.67 | 0.59 | 0.72 | 0.6 | 0.67 | 0.67 |
| 9 | 8234235 | rs1768574 | -0.317 | 0.557 | -0.367 | 0.851 | 0.003 | -0.003 | -0.003 | 0 | 0.84 | 0.86 | 0.85 | 0.85 | 0.84 | 0.82 |
| 9 | 8235513 | rs1768575 | -0.419 | 0.59 | -0.517 | 0.862 | -0.004 | 0.001 | 0.001 | 0.036 | 0.84 | 0.91 | 0.85 | 0.85 | 0.84 | 0.81 |
| 9 | 8235633 | rs10815798 | 1.019 | 0.489 | -0.64 | 0.969 | -0.022 | 0.047 | 0.047 | 0 | 0.46 | 0.22 | 0.6 | 0.39 | 0.51 | 0.56 |
| 9 | 8235673 | rs6477284 | 0.37 | 0.448 | -0.343 | 0.695 | 0 | -0.004 | -0.004 | 0 | 0.7 | 0.68 | 0.78 | 0.74 | 0.71 | 0.72 |
| 9 | 8239748 | rs9407406 | -0.967 | -0.162 | -0.241 | 0.639 | -0.009 | -0.003 | -0.003 | 0 | 0.24 | 0.71 | 0.2 | 0.19 | 0.18 | 0.23 |
| 9 | 8241238 | rs10976869 | 0.876 | -0.303 | -0.89 | 0.724 | 0.059 | -0.012 | -0.012 | 0 | 0.54 | 0.23 | 0.58 | 0.45 | 0.59 | 0.61 |
| 9 | 8241736 | rs17550688 | 0.228 | -0.184 | -0.918 | 0.722 | 0.004 | -0.003 | -0.003 | 0.426 | 0.06 | 0.08 | 0.19 | 0.04 | 0.09 | 0.09 |
| 9 | 8241845 | rs7043322 | -0.007 | -0.312 | 0.193 | 0.883 | -0.047 | 0.051 | 0.051 | 0 | 0.16 | 0.33 | 0.07 | 0.23 | 0.09 | 0.15 |
| 9 | 8241971 | rs12238640 | -0.075 | -0.319 | 0.364 | 0.879 | -0.034 | 0.033 | 0.033 | 0 | 0.15 | 0.23 | 0.07 | 0.23 | 0.09 | 0.15 |
| 9 | 8242654 | rs3842929 | -0.479 | -0.412 | -0.268 | 0.903 | -0.072 | -0.077 | -0.077 | 8.944 | 0.75 | 0.99 | 0.7 | 0.7 | 0.68 | 0.76 |
| 9 | 8243576 | rs3847275 | -1.415 | -0.383 | 0.367 | 0.828 | 0.002 | -0.001 | -0.001 | 0 | 0.24 | 0.75 | 0.2 | 0.27 | 0.18 | 0.25 |
| 9 | 8247628 | rs1416592 | -0.428 | -0.343 | 0.108 | 0.94 | 0.003 | -0.005 | -0.005 | 0.046 | 0.14 | 0.12 | 0.06 | 0.23 | 0.09 | 0.15 |
| 9 | 8253019 | rs1344183 | 0.067 | 0.043 | 0.005 | 0.283 | 0.004 | 0.038 | 0.038 | 1.685 | 0.41 | 0.28 | 0.4 | 0.77 | 0.4 | 0.49 |
| 9 | 8255046 | rs7861890 | 0.947 | 0.214 | -0.128 | -0.085 | -0.019 | 0.076 | 0.076 | 2.736 | 0.36 | 0.22 | 0.29 | 0.07 | 0.37 | 0.26 |
| 9 | 8255691 | rs10733544 | 0.069 | 0.107 | 0.203 | -0.035 | -0.006 | -0.031 | -0.031 | 0 | 0.45 | 0.47 | 0.45 | 0.79 | 0.44 | 0.52 |
| 9 | 8257725 | rs10815803 | 0.426 | 1.087 | -0.237 | 0.524 | 0.058 | 0.066 | 0.066 | 0.484 | 0.89 | 0.77 | 0.62 | 0.72 | 0.77 | 0.75 |
| 9 | 8258063 | rs10815806 | 0.37 | 0.51 | -0.809 | -0.102 | -0.014 | 0.137 | 0.137 | 2.516 | 0.34 | 0.11 | 0.23 | 0.06 | 0.32 | 0.25 |
| 9 | 8258921 | rs7048696 | -0.474 | 0.485 | -0.109 | 0.909 | -0.005 | 0.057 | 0.057 | 0.223 | 0.27 | 0.42 | 0.36 | 0.6 | 0.22 | 0.3 |
| 9 | 8260565 | rs10976889 | -0.858 | 0.849 | 0.14 | 0.206 | 0.032 | -0.063 | -0.063 | 0.198 | 0.18 | 0.36 | 0.17 | 0.04 | 0.2 | 0.14 |
| 9 | 8260602 | rs10815807 | -0.687 | 0.766 | 0.422 | 1.179 | 0.011 | 0.231 | 0.231 | 0 | 0.08 | 0.2 | 0.23 | 0.4 | 0.08 | 0.2 |
| 9 | 8260902 | rs884540 | 0.207 | 0.309 | 0.025 | 0.444 | -0.017 | 0.03 | 0.03 | 0 | 0.27 | 0.3 | 0.34 | 0.41 | 0.24 | 0.43 |
| 9 | 8261021 | rs3858055 | -0.948 | 0.247 | -0.366 | 0.383 | 0.004 | -0.013 | -0.013 | 0 | 0.46 | 0.78 | 0.52 | 0.45 | 0.45 | 0.57 |

| | | | | | | | | | | | | | | | | |
|---|---|---|---|---|---|---|---|---|---|---|---|---|---|---|---|---|
| 9 | 8261551 | rs3847279 | 0.224 | 0.018 | -0.827 | 0.456 | -0.046 | -0.153 | -0.153 | 0 | 0.26 | 0.01 | 0.31 | 0.43 | 0.28 | 0.46 |
| 9 | 8261820 | rs3847280 | -0.861 | -0.551 | -0.468 | 0.034 | -0.013 | -0.025 | -0.025 | 0 | 0.28 | 0.63 | 0.31 | 0.12 | 0.18 | 0.17 |
| 9 | 8261910 | rs3847281 | -0.772 | -0.567 | -1.019 | 0.317 | -0.068 | -0.034 | -0.034 | 0.06 | 0.56 | 0.92 | 0.61 | 0.59 | 0.51 | 0.64 |
| 9 | 8262469 | rs7045605 | -1.309 | -0.747 | 0.397 | 0.762 | -0.005 | 0.033 | 0.033 | 0.083 | 0.2 | 0.27 | 0.38 | 0.41 | 0.22 | 0.43 |
| 9 | 8264662 | rs10976896 | -0.129 | -0.688 | 0.236 | 0.571 | 0.001 | 0.005 | 0.005 | 0 | 0.09 | 0.03 | 0.03 | 0.04 | 0.05 | 0.06 |
| 9 | 8268343 | rs10976907 | -0.978 | -0.785 | 1.2 | 0.532 | 0.004 | -0.001 | -0.001 | 0 | 0.11 | 0.45 | 0.07 | 0.04 | 0.06 | 0.06 |
| 9 | 8269073 | rs7025712 | -1.445 | -0.951 | 1.299 | 0.699 | 0.002 | 0.15 | 0.15 | 0.319 | 0.19 | 0.44 | 0.37 | 0.39 | 0.18 | 0.37 |
| 9 | 8269791 | rs3847282 | -1.666 | -0.943 | 1.502 | 0.697 | 0.001 | 0.117 | 0.117 | 0 | 0.19 | 0.38 | 0.37 | 0.4 | 0.19 | 0.42 |
| 9 | 8270002 | rs12005381 | -0.606 | -1.033 | 1.53 | 0.7 | -0.025 | -0.007 | -0.007 | 0.048 | 0.1 | 0.27 | 0.05 | 0.05 | 0.03 | 0.06 |
| 9 | 8270830 | rs10815816 | -1.933 | -1.034 | 1.504 | 0.706 | -0.013 | 0.026 | 0.026 | 0.12 | 0.27 | 0.61 | 0.41 | 0.42 | 0.21 | 0.4 |
| 9 | 8277169 | rs7864066 | -0.93 | -1.037 | 1.586 | 0.693 | 0.019 | 0.118 | 0.118 | 0 | 0.14 | 0.29 | 0.29 | 0.34 | 0.15 | 0.34 |
| 9 | 8279622 | rs1500304 | -2.346 | -1.08 | 1.475 | 0.563 | 0.01 | -0.014 | -0.014 | 0 | 0.3 | 0.71 | 0.42 | 0.41 | 0.22 | 0.42 |
| 9 | 8283676 | rs12343846 | -1.895 | -1.049 | 1.459 | 0.485 | -0.018 | -0.006 | -0.006 | 0 | 0.05 | 0.16 | 0.04 | 0.02 | 0.04 | 0.02 |
| 9 | 8284560 | rs1472535 | -2.125 | -1.082 | 1.549 | 0.502 | -0.025 | 0.049 | 0.049 | 0 | 0.27 | 0.54 | 0.42 | 0.41 | 0.22 | 0.45 |
| 9 | 8284965 | rs7872172 | -1.972 | -1.07 | 1.504 | 0.509 | -0.007 | 0.01 | 0.01 | 0.36 | 0.26 | 0.71 | 0.4 | 0.41 | 0.18 | 0.45 |
| 9 | 8285325 | rs7862136 | -1.531 | -1.021 | 1.262 | 0.535 | -0.013 | 0.108 | 0.108 | 0 | 0.22 | 0.38 | 0.37 | 0.41 | 0.18 | 0.46 |
| 9 | 8296344 | rs1500309 | -0.726 | -0.977 | 0.955 | 0.329 | -0.03 | -0.053 | -0.053 | 0.29 | 0.17 | 0.28 | 0.11 | 0.04 | 0.11 | 0.09 |
| 9 | 8299968 | rs970347 | -1.901 | -0.978 | 0.436 | 0.38 | 0.035 | -0.074 | -0.074 | 0.667 | 0.32 | 0.9 | 0.46 | 0.41 | 0.25 | 0.4 |
| 9 | 8301561 | rs10976929 | -0.08 | -0.957 | 0.727 | 0.224 | -0.012 | -0.03 | -0.03 | 0 | 0.13 | 0.22 | 0.06 | 0.04 | 0.08 | 0.06 |
| 9 | 8302687 | rs12336333 | -1.121 | -0.928 | 0.918 | 0.225 | -0.031 | -0.083 | -0.083 | 0 | 0.21 | 0.52 | 0.14 | 0.07 | 0.13 | 0.16 |
| 9 | 8310575 | rs10976939 | 0.134 | -1.027 | 0.839 | 0.258 | -0.003 | -0.009 | -0.009 | 4.833 | 0.12 | 0.14 | 0.07 | 0.04 | 0.08 | 0.06 |
| 9 | 8312169 | rs1155271 | -0.908 | -1.043 | 0.978 | 0.274 | -0.003 | -0.002 | -0.002 | 0 | 0.08 | 0.38 | 0.06 | 0.03 | 0.03 | 0.1 |
| 9 | 8318851 | rs7867924 | -1.835 | -0.98 | 0.37 | 0.356 | 0.014 | -0.072 | -0.072 | 0 | 0.34 | 0.89 | 0.44 | 0.41 | 0.26 | 0.48 |
| 9 | 8318948 | rs1353983 | -2.233 | -2.274 | -0.505 | 0.031 | -0.014 | -0.097 | -0.097 | 0.176 | 0.57 | 0.98 | 0.66 | 0.57 | 0.38 | 0.69 |
| 9 | 8322288 | rs7848626 | -1.524 | -1.809 | -0.13 | 0.122 | 0.02 | -0.085 | -0.085 | 2.036 | 0.35 | 0.91 | 0.46 | 0.44 | 0.28 | 0.49 |
| 9 | 8330652 | rs10739155 | -1.559 | -1.838 | 0.341 | 0.234 | 0.006 | 0.001 | 0.001 | 0 | 0.43 | 0.41 | 0.46 | 0.38 | 0.32 | 0.59 |
| 9 | 8332307 | rs16927547 | 0.676 | -1.854 | 0.22 | 0.042 | -0.005 | 0.029 | 0.029 | 0 | 0.07 | 0.05 | 0.27 | 0.35 | 0.13 | 0.22 |
| 9 | 8337939 | rs7032880 | 1.558 | -1.253 | 0.37 | 0.005 | 0.004 | -0.003 | -0.003 | 0.291 | 0.55 | 0.39 | 0.51 | 0.51 | 0.63 | 0.41 |
| 9 | 8338690 | rs3780336 | 0.669 | -1.017 | 0.313 | 0.131 | 0.007 | 0.071 | 0.071 | 0 | 0.07 | 0.09 | 0.27 | 0.34 | 0.13 | 0.23 |
| 9 | 8338878 | rs3847286 | -0.075 | -1.108 | 0.266 | -0.01 | -0.02 | -0.069 | -0.069 | 1.363 | 0.16 | 0.41 | 0.09 | 0.04 | 0.09 | 0.07 |
| 9 | 8344187 | rs10815825 | 0.617 | -0.901 | -0.335 | 0.027 | -0.001 | 0.022 | 0.022 | 0 | 0.13 | 0.07 | 0.07 | 0.04 | 0.1 | 0.07 |
| 9 | 8345923 | rs10511491 | 1.582 | -0.976 | -0.398 | 0.069 | 0.025 | -0.064 | -0.064 | 0 | 0.53 | 0.12 | 0.49 | 0.51 | 0.64 | 0.41 |
| 9 | 8346265 | rs7848705 | 0.192 | -0.951 | -0.23 | 0.122 | 0.01 | 0.094 | 0.094 | 0 | 0.09 | 0.12 | 0.28 | 0.34 | 0.12 | 0.23 |
| 9 | 8348602 | rs10815828 | 1.553 | -0.957 | -0.507 | 0.052 | 0.036 | -0.019 | -0.019 | 0 | 0.52 | 0.19 | 0.39 | 0.51 | 0.62 | 0.41 |
| 9 | 8366920 | rs10977002 | 0.56 | -0.899 | -0.765 | 0.017 | 0.002 | 0.013 | 0.013 | 0.058 | 0.12 | 0.12 | 0.06 | 0.04 | 0.09 | 0.06 |
| 9 | 8367795 | rs1500321 | 1.688 | -0.716 | -0.959 | 0.071 | 0.023 | -0.072 | -0.072 | 0 | 0.51 | 0.06 | 0.39 | 0.49 | 0.63 | 0.41 |
| 9 | 8368253 | rs12554953 | 0.398 | -0.796 | -0.825 | 0.191 | -0.005 | 0.012 | 0.012 | 0 | 0.08 | 0.05 | 0.36 | 0.34 | 0.12 | 0.22 |
| 9 | 8370763 | rs1500323 | 1.151 | -0.984 | -0.761 | 0.125 | 0.024 | -0.042 | -0.042 | 0 | 0.66 | 0.22 | 0.46 | 0.54 | 0.73 | 0.48 |
| 9 | 8371405 | rs4742488 | -0.62 | -1.088 | -0.824 | 0.107 | -0.037 | 0.053 | 0.053 | 0.568 | 0.25 | 0.62 | 0.47 | 0.43 | 0.17 | 0.34 |
| 9 | 8373105 | rs10977013 | 0.427 | -1.041 | -1.08 | 0.035 | 0.01 | 0.059 | 0.059 | 2.966 | 0.16 | 0.03 | 0.07 | 0.04 | 0.11 | 0.07 |
| 9 | 8373735 | rs7038767 | 0.53 | -0.755 | -0.929 | 0.216 | -0.012 | 0.072 | 0.072 | 0.572 | 0.76 | 0.43 | 0.52 | 0.58 | 0.79 | 0.68 |
| 9 | 8375333 | rs12342923 | -1.017 | -0.864 | -1.662 | 0.234 | 0.11 | -0.065 | -0.065 | 0.005 | 0.49 | 0.93 | 0.6 | 0.49 | 0.32 | 0.56 |
| 9 | 8376287 | rs1039336 | -0.842 | -0.369 | -0.829 | 0.482 | -0.01 | -0.008 | -0.008 | 0 | 0.39 | 0.57 | 0.55 | 0.64 | 0.31 | 0.54 |
| 9 | 8379313 | rs1392521 | -0.587 | 0.151 | -1.133 | 1.28 | -0.023 | -0.053 | -0.053 | 0 | 0.11 | 0.01 | 0.29 | 0.47 | 0.08 | 0.43 |
| 9 | 8379838 | rs1500325 | -0.685 | 0.443 | -2.13 | 0.502 | -0.036 | 0.076 | 0.076 | 0.285 | 0.7 | 0.93 | 0.83 | 0.84 | 0.67 | 0.82 |
| 9 | 8382718 | rs12375761 | 0.707 | 0.992 | -2.185 | 0.447 | -0.001 | 0.06 | 0.06 | 0 | 0.28 | 0.07 | 0.15 | 0.15 | 0.26 | 0.18 |
| 9 | 8382720 | rs1875238 | 0.06 | 1.204 | -1.247 | 0.03 | 0.016 | -0.053 | -0.053 | 0.244 | 0.16 | 0.38 | 0.1 | 0.05 | 0.16 | 0.09 |
| 9 | 8382966 | rs1500326 | 1.085 | 1.36 | -1.462 | -0.158 | -0.023 | 0.042 | 0.042 | 0 | 0.37 | 0.1 | 0.33 | 0.28 | 0.39 | 0.31 |
| 9 | 8385838 | rs10977022 | 0.202 | 0.974 | -1.012 | -0.369 | 0.008 | 0.093 | 0.093 | 0 | 0.36 | 0.14 | 0.22 | 0.13 | 0.36 | 0.18 |
| 9 | 8385969 | rs989813 | -1.188 | 1.082 | -0.346 | -0.366 | -0.016 | 0.034 | 0.034 | 0 | 0.57 | 0.64 | 0.57 | 0.64 | 0.49 | 0.64 |
| 9 | 8390467 | rs1500327 | -1.101 | 1.138 | 0.007 | -0.296 | -0.001 | 0.006 | 0.006 | 0 | 0.55 | 0.61 | 0.6 | 0.63 | 0.53 | 0.6 |
| 9 | 8390505 | rs1500328 | -0.256 | 0.884 | 0.041 | -0.241 | -0.009 | -0.049 | -0.049 | 0.278 | 0.19 | 0.08 | 0.3 | 0.54 | 0.17 | 0.48 |
| 9 | 8393528 | rs10977035 | 1.324 | 0.854 | 0.01 | -0.369 | 0 | 0.07 | 0.07 | 0 | 0.23 | 0.08 | 0.12 | 0.05 | 0.26 | 0.08 |
| 9 | 8395913 | rs1021719 | -0.76 | 0.812 | 0.475 | -0.316 | -0.017 | 0.068 | 0.068 | 0 | 0.27 | 0.41 | 0.35 | 0.57 | 0.2 | 0.5 |
| 9 | 8396388 | rs1909756 | 0.709 | 0.599 | 0.459 | -0.519 | -0.01 | -0.023 | -0.023 | 0 | 0.29 | 0.37 | 0.16 | 0.14 | 0.26 | 0.17 |
| 9 | 8396543 | rs1500334 | 0.665 | 0.549 | 0.332 | -0.5 | -0.009 | -0.019 | -0.019 | 1.281 | 0.29 | 0.35 | 0.14 | 0.08 | 0.25 | 0.11 |
| 9 | 8397775 | rs10116650 | 0.572 | 0.61 | 0.328 | -0.452 | -0.008 | -0.038 | -0.038 | 0.169 | 0.29 | 0.28 | 0.2 | 0.11 | 0.28 | 0.13 |
| 9 | 8399766 | rs10124689 | -0.248 | 0.602 | 0.202 | -0.454 | 0.003 | 0.005 | 0.005 | 0.076 | 0.39 | 0.54 | 0.26 | 0.15 | 0.36 | 0.16 |
| 9 | 8401031 | rs10815854 | 1.123 | 0.761 | -0.035 | -0.47 | 0.001 | 0.005 | 0.005 | 7.425 | 0.71 | 0.3 | 0.76 | 0.82 | 0.74 | 0.81 |
| 9 | 8403640 | rs7876037 | -0.359 | 0.742 | -0.116 | -0.469 | -0.018 | 0.066 | 0.066 | 0.928 | 0.64 | 0.87 | 0.77 | 0.8 | 0.64 | 0.77 |
| 9 | 8404095 | rs7042145 | 0.98 | 0.799 | -0.07 | -0.418 | 0.006 | 0.057 | 0.057 | 0 | 0.51 | 0.41 | 0.45 | 0.32 | 0.57 | 0.35 |
| 9 | 8404987 | rs3739577 | 1.004 | 0.882 | -0.441 | 0.145 | -0.018 | -0.049 | -0.049 | 0.541 | 0.1 | 0.01 | 0.25 | 0.39 | 0.08 | 0.41 |
| 9 | 8405543 | rs10977054 | 0.996 | 0.875 | -0.426 | -0.167 | -0.034 | -0.007 | -0.007 | 0 | 0.09 | 0.01 | 0.09 | 0.06 | 0.11 | 0.05 |
| 9 | 8406003 | rs1500336 | -2.116 | 0.863 | -0.391 | -0.158 | -0.007 | -0.004 | -0.004 | 0 | 0.93 | 0.95 | 0.95 | 0.91 | 0.92 | 0.89 |
| 9 | 8408453 | rs10815855 | 1.317 | 0.841 | -0.515 | -0.255 | 0.011 | 0.068 | 0.068 | 0 | 0.23 | 0.08 | 0.11 | 0.06 | 0.21 | 0.08 |
| 9 | 8408724 | rs12156512 | 0.701 | 0.847 | -0.545 | -0.247 | -0.032 | -0.003 | -0.003 | 0 | 0.1 | 0.01 | 0.1 | 0.06 | 0.1 | 0.06 |
| 9 | 8408803 | rs3903826 | -0.361 | 0.855 | -0.543 | -0.203 | -0.036 | -0.089 | -0.089 | 0 | 0.84 | 0.97 | 0.66 | 0.42 | 0.84 | 0.47 |
| 9 | 8409329 | rs10977059 | 1.802 | 0.741 | -0.549 | -0.291 | 0.016 | 0.053 | 0.053 | 0 | 0.19 | 0.07 | 0.08 | 0.05 | 0.18 | 0.07 |
| 9 | 8411328 | rs13292038 | 1.483 | 0.667 | -0.493 | -0.303 | -0.052 | 0.08 | 0.08 | 0.061 | 0.41 | 0.19 | 0.39 | 0.28 | 0.48 | 0.31 |

| | | | | | | | | | | | | | | | | |
|---|---|---|---|---|---|---|---|---|---|---|---|---|---|---|---|---|
| 9 | 8411848 | rs7856797 | 0.011 | 0.712 | -0.274 | -0.304 | -0.004 | -0.038 | -0.038 | 3.485 | 0.34 | 0.4 | 0.19 | 0.09 | 0.34 | 0.13 |
| 9 | 8416196 | rs10511494 | 1.37 | 0.484 | -0.39 | -0.309 | 0.009 | 0.056 | 0.056 | 0 | 0.21 | 0.08 | 0.09 | 0.06 | 0.22 | 0.08 |
| 9 | 8423755 | rs10977081 | 1.222 | 0.603 | -0.194 | -0.134 | -0.01 | 0.001 | 0.001 | 4.87 | 0.07 | 0.13 | 0.04 | 0.09 | 0.04 | 0.08 |
| 9 | 8428249 | rs10815867 | 0.068 | 0.712 | -0.139 | -0.162 | 0.001 | -0.001 | -0.001 | 0 | 0.39 | 0.38 | 0.45 | 0.53 | 0.33 | 0.47 |
| 9 | 8428332 | rs12348387 | 1.515 | 0.684 | -0.09 | -0.14 | -0.01 | 0.034 | 0.034 | 7.172 | 0.07 | 0.15 | 0.04 | 0.13 | 0.04 | 0.09 |
| 9 | 8428492 | rs10815868 | -0.497 | 0.72 | -0.188 | -0.176 | 0.011 | 0.014 | 0.014 | 0 | 0.47 | 0.38 | 0.54 | 0.59 | 0.42 | 0.53 |
| 9 | 8429048 | rs9792404 | 0.069 | 0.747 | -0.19 | -0.151 | 0.003 | -0.014 | -0.014 | 0 | 0.15 | 0.12 | 0.13 | 0.18 | 0.13 | 0.14 |
| 9 | 8430033 | rs4742502 | -1.669 | 0.675 | 0.124 | -0.232 | 0.008 | -0.01 | -0.01 | 0 | 0.64 | 0.65 | 0.64 | 0.73 | 0.55 | 0.75 |
| 9 | 8430742 | rs942176 | -0.251 | 0.516 | -0.078 | -0.167 | 0.006 | 0.004 | 0.004 | 0 | 0.45 | 0.39 | 0.53 | 0.59 | 0.37 | 0.51 |
| 9 | 8431163 | rs2381798 | -0.226 | 0.605 | -0.045 | 0.296 | 0.006 | -0.003 | -0.003 | 0 | 0.46 | 0.4 | 0.55 | 0.47 | 0.42 | 0.52 |
| 9 | 8432073 | rs7866424 | -0.108 | 0.595 | -0.058 | 0.271 | 0.006 | 0.061 | 0.061 | 0 | 0.08 | 0.35 | 0.05 | 0.14 | 0.04 | 0.07 |
| 9 | 8432349 | rs16927848 | 0.906 | 0.612 | -0.265 | 0.279 | -0.005 | 0.021 | 0.021 | 0 | 0.05 | 0.12 | 0.04 | 0.14 | 0.04 | 0.07 |
| 9 | 8432653 | rs7040015 | 0.299 | 0.608 | -1.104 | -0.276 | 0.015 | 0.001 | 0.001 | 0.245 | 0.35 | 0.35 | 0.28 | 0.41 | 0.32 | 0.22 |
| 9 | 8432782 | rs10977093 | 1.003 | 0.823 | -1.767 | -0.354 | -0.011 | 0.105 | 0.105 | 0 | 0.29 | 0.09 | 0.2 | 0.07 | 0.29 | 0.14 |
| 9 | 8433160 | rs7027844 | -0.653 | 1.236 | -1.626 | -0.067 | -0.11 | 0.07 | 0.07 | 0 | 0.62 | 0.92 | 0.67 | 0.73 | 0.52 | 0.76 |
| 9 | 8433698 | rs7047641 | 0.347 | 1.097 | -1.591 | -0.01 | -0.063 | 0.069 | 0.069 | 0 | 0.38 | 0.08 | 0.32 | 0.27 | 0.45 | 0.24 |
| 9 | 8435110 | rs2381800 | -0.118 | 0.552 | -0.976 | 0.399 | -0.009 | 0.012 | 0.012 | 0 | 0.4 | 0.46 | 0.43 | 0.43 | 0.33 | 0.55 |
| 9 | 8436194 | rs10815873 | -0.302 | 0.679 | -1.666 | 0.625 | 0.021 | 0.009 | 0.009 | 0.025 | 0.5 | 0.46 | 0.48 | 0.43 | 0.56 | 0.55 |
| 9 | 8436963 | rs10977103 | -0.371 | 0.708 | -2.087 | 0.874 | -0.028 | -0.045 | -0.045 | 1.776 | 0.08 | 0.01 | 0.19 | 0.24 | 0.09 | 0.27 |
| 9 | 8441203 | rs10977109 | 0.13 | 0.62 | -1.988 | 0.909 | 0 | 0.024 | 0.024 | 0 | 0.06 | 0.04 | 0.16 | 0.26 | 0.06 | 0.25 |
| 9 | 8441646 | rs10815875 | -0.036 | 0.544 | -2.3 | 0.719 | -0.01 | -0.079 | -0.079 | 0 | 0.15 | 0.02 | 0.22 | 0.26 | 0.15 | 0.27 |
| 9 | 8445288 | rs10815876 | -0.669 | 0.712 | -3.016 | 0.487 | -0.091 | -0.053 | -0.053 | 1.254 | 0.48 | 0.08 | 0.62 | 0.6 | 0.51 | 0.56 |
| 9 | 8445864 | rs7855100 | -0.774 | 0.697 | -2.817 | 0.356 | -0.01 | 0.088 | 0.088 | 0 | 0.52 | 0.27 | 0.71 | 0.79 | 0.53 | 0.7 |
| 9 | 8447732 | rs2890795 | -0.839 | 0.631 | -2.837 | 0.185 | 0.012 | 0.127 | 0.127 | 0 | 0.53 | 0.26 | 0.74 | 0.85 | 0.58 | 0.71 |
| 9 | 8448775 | rs16927892 | 1.885 | 0.981 | -2.63 | 0.441 | -0.026 | -0.034 | -0.034 | 0.007 | 0.06 | 0 | 0.07 | 0.11 | 0.07 | 0.1 |
| 9 | 8450441 | rs10815877 | 0.065 | 0.79 | -2.772 | 0.426 | -0.052 | -0.091 | -0.091 | 0.035 | 0.35 | 0.11 | 0.46 | 0.57 | 0.41 | 0.36 |
| 9 | 8451191 | rs10815878 | -0.415 | 0.757 | -2.696 | 0.458 | -0.017 | -0.006 | -0.006 | 0 | 0.41 | 0.25 | 0.51 | 0.64 | 0.44 | 0.43 |
| 9 | 8451755 | rs10977117 | -0.278 | 0.603 | -2.409 | 0.437 | -0.005 | -0.033 | -0.033 | 0.145 | 0.43 | 0.23 | 0.49 | 0.57 | 0.46 | 0.36 |
| 9 | 8452953 | rs12375674 | 0.706 | 1.405 | -2.169 | 0.496 | 0 | 0.015 | 0.015 | 0.245 | 0.27 | 0.26 | 0.35 | 0.52 | 0.28 | 0.25 |
| 9 | 8453781 | rs1323588 | 0.926 | 1.314 | -1.886 | 0.491 | -0.007 | 0.004 | 0.004 | 0 | 0.47 | 0.29 | 0.45 | 0.6 | 0.5 | 0.36 |
| 9 | 8454566 | rs7031287 | -1.073 | 0.942 | -1.345 | 0.682 | 0.003 | 0.002 | 0.002 | 0.744 | 0.43 | 0.68 | 0.48 | 0.37 | 0.37 | 0.55 |
| 9 | 8454921 | rs7870456 | -0.495 | 0.422 | -0.896 | 0.69 | -0.027 | -0.013 | -0.013 | 0 | 0.32 | 0.58 | 0.33 | 0.24 | 0.22 | 0.41 |
| 9 | 8458811 | rs10122691 | -0.928 | 0.898 | -0.023 | 0.99 | 0.022 | 0.071 | 0.071 | 0 | 0.22 | 0.41 | 0.3 | 0.35 | 0.25 | 0.2 |
| 9 | 8460178 | rs2296097 | -1.445 | 0.584 | 0.244 | 0.913 | 0 | 0.073 | 0.073 | 0 | 0.25 | 0.51 | 0.29 | 0.38 | 0.25 | 0.21 |
| 9 | 8465598 | rs2281747 | -0.34 | 0.593 | 0.482 | 1.036 | -0.003 | 0.017 | 0.017 | 0 | 0.15 | 0.16 | 0.19 | 0.31 | 0.16 | 0.08 |
| 9 | 8468567 | rs12340424 | -1.113 | 0.532 | 0.575 | 1.003 | -0.001 | 0.025 | 0.025 | 0 | 0.18 | 0.24 | 0.21 | 0.32 | 0.2 | 0.16 |
| 9 | 8469173 | rs10977131 | -0.3 | 0.493 | 0.555 | 0.987 | -0.01 | 0.032 | 0.032 | 0 | 0.14 | 0.19 | 0.17 | 0.26 | 0.15 | 0.06 |
| 9 | 8470572 | rs7862199 | -0.107 | 0.517 | 0.564 | 1 | -0.006 | 0.02 | 0.02 | 0 | 0.13 | 0.16 | 0.17 | 0.26 | 0.14 | 0.06 |
| 9 | 8470974 | rs1359120 | -1.395 | 0.567 | 0.351 | 0.879 | 0.003 | 0 | 0 | 0 | 0.05 | 0.06 | 0.04 | 0.05 | 0.06 | 0.1 |
| 9 | 8471712 | rs12346891 | -2.423 | 0.529 | 0.519 | 0.859 | 0.02 | 0.001 | 0.001 | 0.406 | 0.1 | 0.33 | 0.07 | 0.07 | 0.06 | 0.11 |
| 9 | 8474233 | rs10977144 | -2.292 | 0.47 | 0.82 | 0.935 | 0.023 | 0.028 | 0.028 | 0 | 0.1 | 0.3 | 0.12 | 0.1 | 0.07 | 0.14 |
| 9 | 8478068 | rs2381804 | -2.06 | 0.995 | 0.691 | 0.803 | 0.005 | 0.008 | 0.008 | 0 | 0.32 | 0.47 | 0.34 | 0.39 | 0.33 | 0.31 |
| 9 | 8480144 | rs10815884 | 0.628 | 1.134 | 0.074 | 0.993 | -0.046 | -0.077 | -0.077 | 0.224 | 0.17 | 0.03 | 0.24 | 0.22 | 0.19 | 0.35 |
| 9 | 8480428 | rs10815885 | -1.584 | 0.74 | 0.699 | 1.133 | -0.004 | -0.005 | -0.005 | 0 | 0.29 | 0.28 | 0.31 | 0.37 | 0.31 | 0.26 |
| 9 | 8480519 | rs13294270 | -0.603 | 0.768 | 0.625 | 1.151 | -0.009 | -0.009 | -0.009 | 0.007 | 0.05 | 0.02 | 0.09 | 0.08 | 0.07 | 0.14 |
| 9 | 8482408 | rs10117986 | -1.559 | 0.767 | 0.671 | 1.125 | 0.008 | 0.017 | 0.017 | 0 | 0.27 | 0.43 | 0.32 | 0.37 | 0.29 | 0.25 |
| 9 | 8484039 | rs16928002 | -1.295 | 0.768 | 0.7 | 1.096 | 0.043 | 0.036 | 0.036 | 0 | 0.07 | 0.23 | 0.1 | 0.08 | 0.08 | 0.14 |
| 9 | 8489006 | rs2296101 | -1.72 | 0.742 | 0.75 | 1.098 | 0.007 | 0.013 | 0.013 | 0 | 0.28 | 0.5 | 0.31 | 0.32 | 0.29 | 0.24 |
| 9 | 8494221 | rs7858815 | -0.579 | 0.757 | 0.657 | 1.129 | 0 | -0.005 | -0.005 | 0 | 0.2 | 0.22 | 0.21 | 0.25 | 0.21 | 0.12 |
| 9 | 8495756 | rs7025468 | -0.931 | 0.79 | 0.645 | 1.142 | -0.002 | -0.005 | -0.005 | 0 | 0.22 | 0.23 | 0.22 | 0.3 | 0.23 | 0.14 |
| 9 | 8499295 | rs10124277 | -1.692 | 0.778 | 0.684 | 1.068 | 0.023 | 0.003 | 0.003 | 0 | 0.1 | 0.25 | 0.1 | 0.07 | 0.1 | 0.13 |
| 9 | 8500472 | rs7035558 | -2.001 | 0.755 | 0.614 | 1.283 | -0.001 | 0.042 | 0.042 | 0 | 0.31 | 0.48 | 0.32 | 0.45 | 0.31 | 0.27 |
| 9 | 8501698 | rs6477312 | -2.241 | 0.808 | 0.624 | 1.284 | 0.01 | 0.039 | 0.039 | 0 | 0.33 | 0.49 | 0.35 | 0.49 | 0.34 | 0.29 |
| 9 | 8502071 | rs13296910 | -1.626 | 0.851 | 0.571 | 1.246 | 0.002 | 0.01 | 0.01 | 0 | 0.09 | 0.1 | 0.1 | 0.18 | 0.11 | 0.14 |
| 9 | 8505087 | rs7868790 | -0.543 | 0.756 | 0.534 | 1.216 | -0.021 | -0.024 | -0.024 | 0 | 0.19 | 0.1 | 0.21 | 0.26 | 0.23 | 0.13 |
| 9 | 8508148 | rs1998516 | -1.795 | 0.681 | 0.594 | 1.156 | 0.004 | 0.011 | 0.011 | 0 | 0.28 | 0.44 | 0.32 | 0.34 | 0.29 | 0.23 |
| 9 | 8509378 | rs1535678 | -1.932 | 1.245 | 0.54 | 1.131 | 0.013 | 0.016 | 0.016 | 0 | 0.1 | 0.16 | 0.12 | 0.19 | 0.15 | 0.13 |
| 9 | 8509880 | rs16928072 | -1.933 | 1.215 | 0.537 | 1.155 | 0 | 0.012 | 0.012 | 0 | 0.11 | 0.16 | 0.12 | 0.19 | 0.11 | 0.12 |
| 9 | 8512133 | rs1998517 | -2.022 | 1.342 | 0.622 | 1.242 | 0.029 | 0.062 | 0.062 | 0 | 0.29 | 0.45 | 0.34 | 0.49 | 0.35 | 0.29 |
| 9 | 8513123 | rs1323578 | -1.651 | 1.305 | 0.615 | 1.134 | 0 | 0.001 | 0.001 | 0 | 0.09 | 0.15 | 0.11 | 0.09 | 0.09 | 0.12 |
| 9 | 8515987 | rs10124872 | -2.005 | 1.365 | 0.655 | 1.146 | 0.025 | 0.022 | 0.022 | 0 | 0.29 | 0.46 | 0.32 | 0.37 | 0.33 | 0.28 |
| 9 | 8517383 | rs10815897 | -2.336 | 1.495 | 1.582 | 1.257 | 0.035 | 0.062 | 0.062 | 0 | 0.22 | 0.55 | 0.29 | 0.38 | 0.25 | 0.27 |
| 9 | 8517545 | rs10120190 | -1.054 | 1.394 | 1.516 | 1.298 | 0.016 | 0.086 | 0.086 | 0 | 0.12 | 0.25 | 0.16 | 0.29 | 0.11 | 0.15 |
| 9 | 8518143 | rs7026388 | -1.671 | 1.213 | 1.37 | 1.106 | 0.01 | 0.068 | 0.068 | 0 | 0.22 | 0.44 | 0.27 | 0.39 | 0.16 | 0.21 |
| 9 | 8518395 | rs3763653 | -0.889 | 1.206 | 1.229 | 1.144 | 0.013 | 0.076 | 0.076 | 0 | 0.11 | 0.21 | 0.16 | 0.28 | 0.1 | 0.12 |
| 9 | 8520287 | rs7039690 | -1.882 | 0.974 | 1.167 | 0.965 | -0.024 | -0.028 | -0.028 | 0 | 0.2 | 0.47 | 0.17 | 0.12 | 0.13 | 0.22 |
| 9 | 8521121 | rs10511499 | 0.103 | 1.039 | 1.014 | 1.001 | 0.032 | 0.132 | 0.132 | 0 | 0.06 | 0.15 | 0.13 | 0.29 | 0.07 | 0.07 |
| 9 | 8522326 | rs6477314 | -1.391 | 0.999 | 1.055 | 1.045 | -0.014 | 0.028 | 0.028 | 0 | 0.16 | 0.32 | 0.15 | 0.21 | 0.11 | 0.22 |
| 9 | 8524448 | rs10977176 | 0.868 | 0.684 | 0.331 | 0.429 | -0.095 | 0.096 | 0.096 | 0.257 | 0.33 | 0.07 | 0.37 | 0.1 | 0.43 | 0.35 |

| | | | | | | | | | | | | | | | | |
|---|---|---|---|---|---|---|---|---|---|---|---|---|---|---|---|---|
| 9 | 8524854 | rs2273998 | -0.059 | 0.785 | 1.33 | 1.021 | 0.009 | 0.055 | 0.055 | 0.124 | 0.07 | 0.09 | 0.14 | 0.29 | 0.1 | 0.08 |
| 9 | 8533412 | rs10758982 | -1.903 | 0.788 | 1.369 | 0.981 | 0.039 | 0.107 | 0.107 | 0.356 | 0.21 | 0.45 | 0.26 | 0.38 | 0.25 | 0.3 |
| 9 | 8534385 | rs7046918 | -1.45 | 0.714 | 0.709 | 0.728 | -0.001 | -0.002 | -0.002 | 0 | 0.21 | 0.23 | 0.14 | 0.11 | 0.15 | 0.23 |
| 9 | 8538919 | rs2184978 | -1.523 | 0.794 | 0.258 | 1.03 | 0.004 | -0.023 | -0.023 | 0 | 0.19 | 0.49 | 0.13 | 0.11 | 0.17 | 0.21 |
| 9 | 8546095 | rs7856850 | -1.194 | 0.193 | -0.217 | 1.089 | -0.008 | 0.037 | 0.037 | 0.082 | 0.24 | 0.58 | 0.26 | 0.39 | 0.21 | 0.25 |
| 9 | 8554938 | rs10217447 | -1.017 | -0.006 | -0.926 | 1.2 | 0.024 | -0.032 | -0.032 | 0 | 0.22 | 0.87 | 0.23 | 0.26 | 0.16 | 0.19 |
| 9 | 8558331 | rs7026937 | -1.308 | -0.108 | 0.282 | 1.042 | 0.017 | 0.04 | 0.04 | 1.786 | 0.06 | 0.58 | 0.06 | 0.05 | 0.01 | 0.02 |
| 9 | 8562080 | rs13294631 | -0.05 | -0.826 | -1.432 | 0.67 | -0.007 | 0.017 | 0.017 | 0 | 0.32 | 0.01 | 0.14 | 0.16 | 0.23 | 0.24 |
| 9 | 8569186 | rs10977207 | 1.464 | 0.221 | -1.993 | 1.197 | -0.091 | -0.111 | -0.111 | 0 | 0.18 | 0.02 | 0.34 | 0.42 | 0.3 | 0.36 |
| 9 | 8574577 | rs1337811 | -0.727 | -0.265 | -1.128 | 0.966 | 0.017 | 0.02 | 0.02 | 0 | 0.49 | 0.71 | 0.44 | 0.36 | 0.41 | 0.41 |
| 9 | 8578261 | rs10977210 | -0.918 | -0.361 | -0.838 | 0.99 | 0.017 | 0.004 | 0.004 | 0 | 0.51 | 0.64 | 0.46 | 0.43 | 0.41 | 0.42 |
| 9 | 8580675 | rs12683419 | 0.08 | -0.468 | -0.279 | 1.056 | 0.039 | 0.043 | 0.043 | 0 | 0.36 | 0.16 | 0.18 | 0.21 | 0.26 | 0.26 |
| 9 | 8584312 | rs13293506 | -0.155 | -0.651 | -0.319 | 0.749 | 0.003 | 0.002 | 0.002 | 0 | 0.38 | 0.36 | 0.19 | 0.27 | 0.25 | 0.28 |
| 9 | 8585188 | rs7861140 | -0.019 | -0.552 | -0.46 | 0.668 | 0.025 | 0.006 | 0.006 | 0 | 0.14 | 0.23 | 0.15 | 0.2 | 0.15 | 0.14 |
| 9 | 8595338 | rs7020239 | 0.041 | -0.859 | -0.503 | 0.47 | 0.008 | 0.007 | 0.007 | 0 | 0.34 | 0.27 | 0.15 | 0.19 | 0.24 | 0.25 |
| 9 | 8598737 | rs9792470 | 0.484 | -1.183 | -1.036 | 0.553 | -0.017 | -0.083 | -0.083 | 0.179 | 0.14 | 0.01 | 0.09 | 0.23 | 0.13 | 0.1 |
| 9 | 8603606 | rs9792461 | 0.231 | -1.36 | -0.974 | 0.136 | -0.003 | -0.01 | -0.01 | 0.846 | 0.13 | 0.08 | 0.08 | 0.18 | 0.15 | 0.06 |
| 9 | 8608349 | rs16928196 | -0.362 | -1.916 | -1.025 | -0.195 | 0.038 | 0.048 | 0.048 | 0 | 0.3 | 0.25 | 0.15 | 0.09 | 0.17 | 0.33 |
| 9 | 8610513 | rs10977230 | -0.812 | -1.976 | -0.911 | 0.021 | -0.03 | 0.046 | 0.046 | 0 | 0.23 | 0.55 | 0.24 | 0.33 | 0.16 | 0.09 |
| 9 | 8615045 | rs1361117 | 0.238 | -1.801 | -1.418 | 0.121 | 0.085 | -0.011 | -0.011 | 0 | 0.24 | 0.02 | 0.09 | 0.18 | 0.13 | 0.17 |
| 9 | 8619879 | rs16928221 | -1.024 | -1.87 | -1.03 | 0.112 | -0.041 | -0.011 | -0.011 | 0 | 0.39 | 0.45 | 0.29 | 0.32 | 0.23 | 0.35 |
| 9 | 8620382 | rs10758985 | 0.594 | -1.579 | -1.453 | -0.418 | 0.002 | -0.017 | -0.017 | 0.235 | 0.4 | 0.56 | 0.57 | 0.71 | 0.51 | 0.59 |
| 9 | 8635810 | rs1512381 | -0.21 | -1.606 | -1.555 | -0.286 | 0.047 | 0.019 | 0.019 | 0.229 | 0.32 | 0.07 | 0.23 | 0.22 | 0.22 | 0.26 |
| 9 | 8645448 | rs16928234 | 0.294 | -1.717 | -1.351 | -0.276 | -0.006 | 0.001 | 0.001 | 0.025 | 0.25 | 0.2 | 0.15 | 0.19 | 0.1 | 0.16 |
| 9 | 8649429 | rs7041298 | -1.406 | -1.365 | -1.593 | -0.516 | -0.008 | 0.048 | 0.048 | 0 | 0.77 | 0.89 | 0.84 | 0.94 | 0.75 | 0.87 |
| 9 | 8659757 | rs1028018 | 0.528 | -1.325 | -1.442 | -0.403 | 0 | 0.003 | 0.003 | 0.16 | 0.43 | 0.63 | 0.59 | 0.68 | 0.55 | 0.58 |
| 9 | 8664838 | rs1025547 | -0.426 | -1.34 | -1.289 | -0.309 | 0.008 | -0.059 | -0.059 | 0.607 | 0.63 | 0.55 | 0.71 | 0.84 | 0.63 | 0.72 |
| 9 | 8669129 | rs12683865 | 0.331 | -1.714 | -1.722 | -0.348 | 0.06 | -0.041 | -0.041 | 0.562 | 0.18 | 0.06 | 0.17 | 0.23 | 0.1 | 0.13 |
| 9 | 8669273 | rs10815923 | 0.696 | -1.765 | -1.543 | -0.28 | 0.003 | 0 | 0 | 0 | 0.43 | 0.54 | 0.54 | 0.62 | 0.52 | 0.58 |
| 9 | 8670737 | rs7859849 | 0.71 | -1.748 | -1.534 | -0.423 | 0.004 | -0.004 | -0.004 | 0 | 0.43 | 0.53 | 0.54 | 0.63 | 0.52 | 0.58 |
| 9 | 8672042 | rs10977243 | 0.914 | -1.212 | -1.567 | 0.222 | -0.027 | -0.028 | -0.028 | 0.264 | 0.06 | 0 | 0.08 | 0.13 | 0.09 | 0.15 |
| 9 | 8672923 | rs16924692 | 0.601 | -1.589 | -1.362 | -0.1 | 0.039 | -0.043 | -0.043 | 0.107 | 0.14 | 0.02 | 0.14 | 0.17 | 0.1 | 0.11 |
| 9 | 8682672 | rs10511506 | 0.532 | -1.342 | -1.364 | -0.242 | -0.03 | 0.033 | 0.033 | 0 | 0.13 | 0.02 | 0.1 | 0.05 | 0.13 | 0.16 |
| 9 | 8692722 | rs7021243 | -0.492 | -1.522 | -0.922 | -0.21 | -0.022 | -0.009 | -0.009 | 0 | 0.27 | 0.41 | 0.24 | 0.22 | 0.2 | 0.24 |
| 9 | 8693517 | rs1434264 | -0.536 | -1.547 | -0.841 | -0.174 | -0.026 | -0.003 | -0.003 | 0 | 0.27 | 0.43 | 0.3 | 0.26 | 0.2 | 0.25 |
| 9 | 8694330 | rs10125358 | -1.191 | -1.735 | -0.794 | -0.243 | -0.024 | -0.005 | -0.005 | 0 | 0.33 | 0.52 | 0.34 | 0.32 | 0.21 | 0.27 |
| 9 | 8695233 | rs16928280 | 0.119 | -1.639 | -0.789 | -0.219 | -0.019 | -0.001 | -0.001 | 0.58 | 0.06 | 0.06 | 0.1 | 0.1 | 0.01 | 0.04 |
| 9 | 8695439 | rs12685924 | -1.261 | -1.769 | -0.901 | -0.307 | -0.022 | -0.008 | -0.008 | 0 | 0.38 | 0.5 | 0.34 | 0.35 | 0.23 | 0.29 |
| 9 | 8696227 | rs1368981 | -1.131 | -1.762 | -0.882 | -0.324 | 0.014 | 0.004 | 0.004 | 0 | 0.35 | 0.27 | 0.33 | 0.32 | 0.22 | 0.29 |
| 9 | 8700193 | rs4442186 | -1.112 | -1.724 | -1.27 | -0.301 | -0.031 | -0.012 | -0.012 | 0.007 | 0.35 | 0.43 | 0.27 | 0.26 | 0.23 | 0.26 |
| 9 | 8700396 | rs10481623 | 1.269 | -0.88 | -2.396 | 0.025 | -0.06 | -0.038 | -0.038 | 0 | 0.25 | 0.12 | 0.38 | 0.32 | 0.34 | 0.33 |
| 9 | 8702964 | rs10758987 | -1.042 | -1.439 | -2.185 | 0.351 | 0.007 | -0.032 | -0.032 | 0.788 | 0.36 | 0.15 | 0.31 | 0.48 | 0.31 | 0.47 |
| 9 | 8703025 | rs1543083 | 1.299 | -1.384 | -2.329 | 0.243 | -0.038 | -0.013 | -0.013 | 3.602 | 0.58 | 0.84 | 0.65 | 0.54 | 0.58 | 0.51 |
| 9 | 8704192 | rs1550002 | -0.151 | -1.302 | -2.351 | 0.363 | -0.019 | -0.074 | -0.074 | 0.009 | 0.21 | 0.04 | 0.19 | 0.34 | 0.2 | 0.32 |
| 9 | 8706540 | rs4742531 | 1.235 | -1.508 | -2.457 | 0.317 | 0.035 | -0.005 | -0.005 | 0 | 0.53 | 0.1 | 0.56 | 0.44 | 0.6 | 0.51 |
| 9 | 8710180 | rs17653604 | -0.092 | -0.897 | -1.361 | 0.543 | 0.001 | 0.019 | 0.019 | 0.013 | 0.17 | 0.07 | 0.13 | 0.14 | 0.11 | 0.15 |
| 9 | 8712365 | rs10758988 | 1.169 | -0.971 | -0.92 | 0.523 | -0.001 | 0.001 | 0.001 | 0 | 0.65 | 0.41 | 0.62 | 0.45 | 0.72 | 0.53 |
| 9 | 8715082 | rs1821429 | 0.939 | -0.949 | -1.162 | 0.335 | -0.026 | 0.058 | 0.058 | 0.124 | 0.54 | 0.7 | 0.45 | 0.3 | 0.5 | 0.43 |
| 9 | 8718703 | rs17575409 | 0.613 | -1.027 | -1.288 | 0.58 | -0.032 | -0.026 | -0.026 | 0 | 0.12 | 0 | 0.08 | 0.11 | 0.1 | 0.12 |
| 9 | 8719261 | rs7866922 | 0.957 | -1.117 | -0.64 | 0.383 | 0 | 0.001 | 0.001 | 0 | 0.54 | 0.43 | 0.45 | 0.34 | 0.54 | 0.47 |
| 9 | 8722635 | rs10977261 | -0.402 | -0.757 | -0.558 | 0.517 | 0.009 | 0.098 | 0.098 | 1.097 | 0.07 | 0.38 | 0.05 | 0.06 | 0.01 | 0.08 |
| 9 | 8726153 | rs1473822 | 0.712 | -0.324 | -1.533 | 0.669 | -0.054 | -0.015 | -0.015 | 0 | 0.6 | 0.86 | 0.5 | 0.38 | 0.55 | 0.47 |
| 9 | 8736813 | rs1434250 | -0.677 | -0.628 | -1.023 | 0.925 | 0.017 | -0.076 | -0.076 | 0 | 0.27 | 0.19 | 0.27 | 0.53 | 0.22 | 0.38 |
| 9 | 8737316 | rs6477333 | 0.962 | -0.868 | -0.498 | 0.857 | 0.002 | -0.006 | -0.006 | 0 | 0.63 | 0.67 | 0.56 | 0.39 | 0.65 | 0.53 |
| 9 | 8738267 | rs4742533 | 0.696 | -0.67 | -0.276 | 0.635 | -0.001 | 0.002 | 0.002 | 0 | 0.78 | 0.74 | 0.72 | 0.54 | 0.74 | 0.77 |
| 9 | 8739061 | rs10465099 | 1.765 | -1.244 | -0.817 | 0.287 | -0.025 | -0.069 | -0.069 | 0 | 0.48 | 0.04 | 0.49 | 0.42 | 0.55 | 0.57 |
| 9 | 8741389 | rs2033554 | -0.236 | -0.778 | -0.07 | 0.511 | 0.007 | 0.013 | 0.013 | 0 | 0.19 | 0.14 | 0.18 | 0.05 | 0.16 | 0.1 |
| 9 | 8743598 | rs10977272 | 1.467 | -0.526 | 0.761 | 0.637 | 0 | 0.015 | 0.015 | 0 | 0.68 | 0.37 | 0.63 | 0.48 | 0.68 | 0.72 |
| 9 | 8754549 | rs10733548 | 0.664 | -0.322 | 0.635 | 0.724 | 0.017 | 0.042 | 0.042 | 0 | 0.77 | 0.67 | 0.69 | 0.54 | 0.71 | 0.76 |
| 9 | 8774025 | rs1036327 | 0.949 | -0.406 | 0.655 | 0.767 | 0.014 | 0.112 | 0.112 | 0 | 0.73 | 0.51 | 0.68 | 0.5 | 0.7 | 0.73 |
| 9 | 8774381 | rs1036331 | 0.765 | -0.101 | 0.038 | 0.653 | 0.013 | 0.044 | 0.044 | 0 | 0.77 | 0.66 | 0.69 | 0.54 | 0.71 | 0.75 |
| 9 | 8777779 | rs9299072 | -1.288 | -1.17 | -0.341 | -0.282 | -0.014 | -0.041 | -0.041 | 0 | 0.3 | 0.37 | 0.27 | 0.06 | 0.19 | 0.15 |
| 9 | 8778606 | rs10977298 | -0.594 | -0.572 | -0.753 | -0.013 | 0.018 | 0.067 | 0.067 | 0 | 0.18 | 0.08 | 0.14 | 0.04 | 0.15 | 0.08 |
| 9 | 8784855 | rs10977299 | 0.424 | 0.174 | -0.464 | 0.723 | -0.003 | -0.001 | -0.001 | 0 | 0.58 | 0.37 | 0.54 | 0.4 | 0.6 | 0.61 |
| 9 | 8786827 | rs13283979 | 0.509 | -0.259 | -0.35 | 0.786 | 0.021 | 0.036 | 0.036 | 0.133 | 0.53 | 0.3 | 0.57 | 0.57 | 0.61 | 0.58 |
| 9 | 8787733 | rs3802396 | 0.093 | -0.232 | -0.386 | 1.041 | -0.002 | 0.131 | 0.131 | 0 | 0.11 | 0.22 | 0.17 | 0.4 | 0.09 | 0.11 |
| 9 | 8791925 | rs7862354 | 1.094 | -0.28 | -0.281 | 0.93 | -0.014 | 0.112 | 0.112 | 0 | 0.44 | 0.26 | 0.42 | 0.17 | 0.51 | 0.47 |
| 9 | 8791981 | rs12348514 | -0.198 | -0.34 | 0.186 | 1.058 | -0.031 | 0.037 | 0.037 | 0 | 0.16 | 0.44 | 0.17 | 0.2 | 0.1 | 0.14 |
| 9 | 8794314 | rs2082633 | 0.007 | -0.291 | 0.013 | 0.963 | 0 | -0.001 | -0.001 | 0.428 | 0.2 | 0.16 | 0.25 | 0.54 | 0.17 | 0.24 |

| | | | | | | | | | | | | | | | | |
|---|---|---|---|---|---|---|---|---|---|---|---|---|---|---|---|---|
| 9 | 8794775 | rs765800 | 0.211 | -0.029 | -0.447 | 0.44 | 0 | -0.009 | -0.009 | 0 | 0.41 | 0.47 | 0.53 | 0.62 | 0.42 | 0.45 |
| 9 | 8795072 | rs7032107 | -0.721 | -0.175 | -0.63 | 0.358 | -0.057 | 0.047 | 0.047 | 0 | 0.71 | 0.95 | 0.75 | 0.77 | 0.7 | 0.72 |
| 9 | 8796449 | rs1550004 | 0.077 | -0.148 | -0.565 | 0.442 | 0 | -0.004 | -0.004 | 0 | 0.43 | 0.54 | 0.55 | 0.62 | 0.43 | 0.45 |
| 9 | 8798347 | rs9299075 | -0.213 | -0.065 | 0.216 | 0.787 | 0.005 | -0.018 | -0.018 | 0 | 0.75 | 0.69 | 0.75 | 0.77 | 0.73 | 0.73 |
| 9 | 8805462 | rs4562389 | 0.357 | -0.013 | -0.216 | 0.78 | -0.003 | -0.01 | -0.01 | 0.027 | 0.7 | 0.38 | 0.7 | 0.75 | 0.71 | 0.73 |
| 9 | 8809042 | rs6477355 | -0.034 | 0.01 | -0.92 | 1.159 | 0.011 | -0.042 | -0.042 | 0 | 0.73 | 0.83 | 0.78 | 0.64 | 0.75 | 0.76 |
| 9 | 8812415 | rs10733549 | 0.82 | 0.094 | -1.447 | 0.948 | 0.008 | 0.033 | 0.033 | 0.447 | 0.52 | 0.31 | 0.55 | 0.48 | 0.57 | 0.51 |
| 9 | 8812871 | rs7855709 | -0.476 | 0.025 | -1.854 | 1.154 | -0.029 | -0.126 | -0.126 | 0 | 0.3 | 0.03 | 0.25 | 0.4 | 0.26 | 0.3 |
| 9 | 8813497 | rs10815954 | -0.685 | -0.919 | -1.561 | 0.377 | 0.007 | 0.009 | 0.009 | 0 | 0.25 | 0.21 | 0.15 | 0.18 | 0.15 | 0.28 |
| 9 | 8815247 | rs10815957 | 0.316 | -0.568 | -1.838 | 0.729 | 0.006 | -0.032 | -0.032 | 0.322 | 0.25 | 0.21 | 0.21 | 0.44 | 0.18 | 0.36 |
| 9 | 8816479 | rs10815959 | 0.084 | -0.183 | -2.484 | 0.52 | -0.017 | -0.047 | -0.047 | 0 | 0.69 | 0.75 | 0.63 | 0.5 | 0.67 | 0.61 |
| 9 | 8817060 | rs1434273 | 0.603 | -0.286 | -2.204 | 0.388 | 0.014 | 0.089 | 0.089 | 1.94 | 0.51 | 0.65 | 0.46 | 0.24 | 0.51 | 0.44 |
| 9 | 8817244 | rs1434274 | -0.369 | -0.233 | -2.2 | 0.508 | -0.037 | -0.062 | -0.062 | 0 | 0.76 | 0.86 | 0.65 | 0.6 | 0.69 | 0.67 |
| 9 | 8826108 | rs10815964 | -0.141 | -0.272 | -2.179 | 0.39 | -0.035 | 0.071 | 0.071 | 0.277 | 0.6 | 0.81 | 0.49 | 0.3 | 0.54 | 0.5 |
| 9 | 8828693 | rs10815966 | 1.879 | -0.242 | -2.163 | 0.475 | -0.029 | -0.076 | -0.076 | 0 | 0.16 | 0.03 | 0.16 | 0.3 | 0.15 | 0.16 |
| 9 | 8828849 | rs10977313 | 1.042 | -0.757 | -2.611 | 0.138 | 0.026 | -0.024 | -0.024 | 0 | 0.15 | 0.01 | 0.13 | 0.13 | 0.1 | 0.18 |
| 9 | 8829390 | rs10815967 | 2.044 | -0.42 | -2.397 | 0.319 | -0.032 | -0.08 | -0.08 | 0 | 0.16 | 0.03 | 0.16 | 0.3 | 0.15 | 0.16 |
| 9 | 8829567 | rs10758997 | 0.302 | 0.021 | -2.448 | 0.137 | 0.052 | 0.115 | 0.115 | 1.283 | 0.52 | 0.72 | 0.45 | 0.24 | 0.45 | 0.42 |
| 9 | 8830796 | rs1434256 | 0.34 | -0.014 | -2.17 | 0.135 | 0.033 | 0.064 | 0.064 | 1.617 | 0.52 | 0.63 | 0.44 | 0.24 | 0.45 | 0.42 |
| 9 | 8831995 | rs7030557 | 1.28 | 0.022 | -2.201 | 0.142 | 0.008 | -0.015 | -0.015 | 0 | 0.08 | 0.13 | 0.05 | 0.06 | 0.09 | 0.09 |
| 9 | 8832025 | rs11791648 | 1.086 | 0.517 | -2.397 | 0.327 | -0.07 | -0.083 | -0.083 | 0 | 0.13 | 0.01 | 0.22 | 0.25 | 0.19 | 0.13 |
| 9 | 8832038 | rs7048621 | -0.115 | 0.442 | -2.355 | 0.132 | -0.033 | 0.066 | 0.066 | 0.024 | 0.6 | 0.79 | 0.49 | 0.3 | 0.54 | 0.5 |
| 9 | 8840840 | rs2570299 | 0.237 | 0.597 | -2.394 | 0.085 | 0.019 | 0.084 | 0.084 | 1.185 | 0.55 | 0.72 | 0.47 | 0.24 | 0.48 | 0.43 |
| 9 | 8845429 | rs2570961 | 0.829 | 0.22 | -1.945 | -0.024 | -0.014 | 0.011 | 0.011 | 0.302 | 0.28 | 0.61 | 0.26 | 0.3 | 0.19 | 0.16 |
| 9 | 8845598 | rs1865343 | -0.346 | 0.415 | -1.853 | 0.118 | 0.013 | 0.007 | 0.007 | 0 | 0.81 | 0.77 | 0.83 | 0.77 | 0.79 | 0.71 |
| 9 | 8845911 | rs1865341 | -0.641 | 0.431 | -1.923 | 0.112 | -0.004 | 0.003 | 0.003 | 0.053 | 0.83 | 0.89 | 0.86 | 0.85 | 0.8 | 0.81 |
| 9 | 8846052 | rs2570960 | -1.413 | 0.287 | -1.908 | 0.11 | 0.002 | -0.001 | -0.001 | 0.86 | 0.88 | 0.9 | 0.9 | 0.9 | 0.88 | 0.88 |
| 9 | 8846955 | rs1975197 | 1.199 | 0.33 | -1.841 | 0.246 | -0.019 | -0.021 | -0.021 | 0 | 0.11 | 0.09 | 0.23 | 0.3 | 0.17 | 0.23 |
| 9 | 8847013 | rs2117100 | 0.661 | 0.088 | -2.143 | -0.227 | 0.025 | 0.042 | 0.042 | 0.377 | 0.33 | 0.76 | 0.24 | 0.11 | 0.27 | 0.18 |
| 9 | 8847233 | rs2117099 | 0.536 | 0.172 | -1.742 | -0.037 | 0.003 | -0.05 | -0.05 | 0 | 0.1 | 0.54 | 0.09 | 0 | 0.03 | 0 |
| 9 | 8850237 | rs10123580 | 0.945 | 0.276 | -2.181 | 0 | 0.005 | -0.062 | -0.062 | 0 | 0.33 | 0.08 | 0.3 | 0.34 | 0.27 | 0.34 |
| 9 | 8851012 | rs10759003 | 1.561 | 0.326 | -2.205 | -0.037 | -0.026 | 0.034 | 0.034 | 0 | 0.17 | 0 | 0.14 | 0.05 | 0.14 | 0.19 |
| 9 | 8852226 | rs2570301 | -0.113 | 0.45 | -2.238 | -0.06 | -0.01 | -0.004 | -0.004 | 0 | 0.52 | 0.83 | 0.51 | 0.48 | 0.5 | 0.5 |
| 9 | 8852668 | rs10977321 | 0.215 | 0.476 | -2.247 | -0.031 | -0.017 | 0.032 | 0.032 | 0 | 0.32 | 0.14 | 0.34 | 0.22 | 0.35 | 0.34 |
| 9 | 8853690 | rs3891649 | -0.555 | 0.442 | -1.736 | -0.09 | -0.001 | -0.004 | -0.004 | 2.178 | 0.58 | 0.62 | 0.58 | 0.52 | 0.53 | 0.51 |
| 9 | 8854069 | rs7466280 | 0.821 | 0.364 | -1.98 | -0.511 | -0.044 | -0.028 | -0.028 | 0 | 0.14 | 0.02 | 0.19 | 0.18 | 0.21 | 0.16 |
| 9 | 8855603 | rs10759006 | -0.274 | 0.317 | -1.861 | -0.774 | 0.056 | 0.113 | 0.113 | 0.126 | 0.58 | 0.15 | 0.7 | 0.76 | 0.67 | 0.64 |
| 9 | 8856908 | rs751372 | 0.463 | 0.748 | -1.043 | 0.024 | -0.002 | 0.006 | 0.006 | 0 | 0.08 | 0.13 | 0.06 | 0.13 | 0.07 | 0.17 |
| 9 | 8857079 | rs13284801 | 0.966 | 1.074 | -1.174 | 0.109 | -0.027 | 0.038 | 0.038 | 0 | 0.1 | 0.02 | 0.12 | 0.03 | 0.14 | 0.1 |
| 9 | 8860355 | rs10815977 | 0.961 | 1.023 | -0.859 | 0.14 | -0.019 | 0.037 | 0.037 | 0 | 0.57 | 0.25 | 0.52 | 0.67 | 0.53 | 0.53 |
| 9 | 8865925 | rs10977327 | 0.24 | 1.039 | -1.196 | 0.441 | -0.073 | -0.102 | -0.102 | 0 | 0.21 | 0.01 | 0.22 | 0.31 | 0.23 | 0.26 |
| 9 | 8867519 | rs723145 | 0.407 | 0.689 | -0.328 | 0.314 | -0.002 | -0.021 | -0.021 | 2.806 | 0.81 | 0.36 | 0.78 | 0.9 | 0.84 | 0.81 |
| 9 | 8868177 | rs2217711 | -0.897 | 0.783 | -0.298 | 0.052 | 0.015 | -0.047 | -0.047 | 0.431 | 0.89 | 0.81 | 0.87 | 0.97 | 0.87 | 0.91 |
| 9 | 8868774 | rs10815979 | -0.08 | 0.63 | -0.659 | 0.205 | -0.003 | -0.036 | -0.036 | 0.109 | 0.23 | 0.13 | 0.23 | 0.3 | 0.22 | 0.26 |
| 9 | 8870553 | rs12002381 | -0.067 | 0.142 | 0.115 | -0.237 | 0.027 | -0.017 | -0.017 | 7.256 | 0.07 | 0.29 | 0.06 | 0 | 0.02 | 0 |
| 9 | 8873348 | rs10759008 | 0.199 | 0.205 | -0.824 | -0.669 | -0.029 | 0.084 | 0.084 | 0 | 0.27 | 0.11 | 0.2 | 0.1 | 0.33 | 0.29 |
| 9 | 8873449 | rs10759009 | -0.031 | 0.095 | -0.761 | -0.723 | -0.01 | 0.067 | 0.067 | 0 | 0.3 | 0.17 | 0.2 | 0.1 | 0.33 | 0.29 |
| 9 | 8875954 | rs10759013 | 0.931 | -0.04 | 0.027 | -0.585 | -0.009 | -0.074 | -0.074 | 0 | 0.86 | 0.32 | 0.86 | 0.99 | 0.9 | 0.95 |
| 9 | 8877814 | rs12000930 | -0.497 | 0.064 | 0.006 | 0.278 | 0.031 | 0.185 | 0.185 | 0 | 0.08 | 0.24 | 0.21 | 0.29 | 0.09 | 0.06 |
| 9 | 8879118 | rs17584499 | -0.039 | -0.107 | -1.031 | 0.028 | -0.001 | 0.088 | 0.088 | 0 | 0.25 | 0.03 | 0.21 | 0.14 | 0.18 | 0.24 |
| 9 | 8879472 | rs7023424 | -1.294 | 0.48 | -0.202 | 0.821 | 0.005 | 0.133 | 0.133 | 0 | 0.08 | 0.15 | 0.18 | 0.36 | 0.09 | 0.14 |
| 9 | 8881741 | rs10815983 | -0.502 | 0.542 | -0.388 | 0.208 | 0.001 | 0 | 0 | 0 | 0.48 | 0.51 | 0.43 | 0.58 | 0.53 | 0.64 |
| 9 | 8884015 | rs13287579 | 1.177 | 0.424 | -0.97 | -0.165 | 0.003 | 0.108 | 0.108 | 0 | 0.31 | 0.06 | 0.29 | 0.11 | 0.27 | 0.17 |
| 9 | 8884844 | rs10815984 | 1.421 | 0.931 | -1.269 | -0.194 | -0.062 | 0.029 | 0.029 | 0.019 | 0.13 | 0.02 | 0.12 | 0.05 | 0.24 | 0.16 |
| 9 | 8885169 | rs10815986 | 1.421 | 0.931 | -1.269 | -0.194 | -0.062 | 0.029 | 0.029 | 0.019 | 0.13 | 0.02 | 0.12 | 0.05 | 0.24 | 0.16 |
| 9 | 8885476 | rs10759014 | 0.237 | 0.81 | -0.92 | -0.116 | 0.003 | 0.002 | 0.002 | 0.533 | 0.27 | 0.27 | 0.32 | 0.62 | 0.26 | 0.46 |
| 9 | 8889419 | rs10511512 | 0.664 | 1.206 | -1.309 | 0.274 | -0.031 | -0.034 | -0.034 | 0 | 0.08 | 0.02 | 0.16 | 0.22 | 0.12 | 0.09 |
| 9 | 8890725 | rs2890841 | -0.031 | 0.729 | -0.76 | 0.58 | 0.057 | 0.346 | 0.346 | 0 | 0.06 | 0.26 | 0.21 | 0.32 | 0.09 | 0.07 |
| 9 | 8890946 | rs7867067 | -0.918 | 0.619 | -1.428 | -0.086 | 0.015 | 0.03 | 0.03 | 0 | 0.68 | 0.78 | 0.7 | 0.89 | 0.73 | 0.83 |
| 9 | 8891463 | rs10815989 | 0.147 | 1.153 | -0.696 | 0.473 | 0.017 | -0.015 | -0.015 | 0 | 0.75 | 0.65 | 0.7 | 0.73 | 0.63 | 0.73 |
| 9 | 8893834 | rs7850165 | -0.343 | 0.937 | -1.165 | 0.845 | -0.008 | -0.013 | -0.013 | 0 | 0.15 | 0.11 | 0.1 | 0.21 | 0.14 | 0.26 |
| 9 | 8894070 | rs10977337 | -0.206 | 0.548 | -1.383 | 0.773 | 0.007 | -0.035 | -0.035 | 0 | 0.3 | 0.2 | 0.3 | 0.51 | 0.29 | 0.46 |
| 9 | 8895687 | rs1368680 | -0.493 | 0.42 | -1.546 | 0.134 | -0.002 | 0.11 | 0.11 | 0 | 0.55 | 0.81 | 0.64 | 0.82 | 0.57 | 0.65 |
| 9 | 8897215 | rs7866753 | 0.549 | 0.49 | -1.499 | 0.145 | 0 | -0.001 | -0.001 | 0 | 0.18 | 0.17 | 0.15 | 0.19 | 0.26 | 0.23 |
| 9 | 8897230 | rs6477360 | -1.131 | 0.515 | -1.46 | 0.099 | -0.053 | 0.146 | 0.146 | 0 | 0.56 | 0.91 | 0.65 | 0.82 | 0.57 | 0.62 |
| 9 | 8897676 | rs11794550 | 0.094 | 0.167 | -1.721 | 0.587 | -0.001 | 0.007 | 0.007 | 0 | 0.19 | 0.17 | 0.24 | 0.52 | 0.2 | 0.15 |
| 9 | 8900067 | rs10511511 | 0.671 | -0.084 | -1.678 | 0.495 | 0.006 | -0.072 | -0.072 | 0 | 0.13 | 0.02 | 0.19 | 0.28 | 0.12 | 0.27 |
| 9 | 8901169 | rs12554855 | 0.393 | -0.05 | -1.387 | 0.525 | -0.028 | 0.092 | 0.092 | 0 | 0.15 | 0.33 | 0.22 | 0.29 | 0.12 | 0.29 |
| 9 | 8901682 | rs1433540 | 0.706 | 0.137 | -2.392 | 0.538 | 0.012 | -0.077 | -0.077 | 0 | 0.6 | 0.09 | 0.58 | 0.53 | 0.61 | 0.45 |

| | | | | | | | | | | | | | | | | |
|---|---|---|---|---|---|---|---|---|---|---|---|---|---|---|---|---|
| 9 | 8906136 | rs10759016 | 0.709 | 0.141 | -2.389 | 0.487 | 0.013 | -0.094 | -0.094 | 0.707 | 0.6 | 0.09 | 0.58 | 0.52 | 0.62 | 0.46 |
| 9 | 8915142 | rs10815996 | -0.027 | 0.778 | -1.975 | 0.055 | -0.104 | 0.154 | 0.154 | 0.396 | 0.24 | 0.02 | 0.15 | 0.02 | 0.31 | 0.24 |
| 9 | 8916449 | rs10046838 | 0.373 | 0.342 | -1.399 | 0.49 | 0 | -0.004 | -0.004 | 0 | 0.55 | 0.75 | 0.57 | 0.47 | 0.49 | 0.58 |
| 9 | 8921426 | rs16928621 | -0.612 | -0.263 | -1.436 | 0.592 | 0.016 | -0.013 | -0.013 | 0 | 0.21 | 0.13 | 0.17 | 0.25 | 0.15 | 0.29 |
| 9 | 8922286 | rs4740969 | 0.107 | -0.242 | -0.949 | 0.524 | 0.01 | 0.065 | 0.065 | 0 | 0.75 | 0.65 | 0.72 | 0.64 | 0.75 | 0.75 |
| 9 | 8922467 | rs10977362 | 1.584 | -0.204 | -1.444 | 0.117 | -0.034 | 0.04 | 0.04 | 0 | 0.31 | 0.09 | 0.31 | 0.17 | 0.37 | 0.18 |
| 9 | 8923010 | rs7858751 | 0.185 | -0.833 | -1.098 | -0.133 | -0.008 | 0.089 | 0.089 | 0.176 | 0.62 | 0.78 | 0.73 | 0.86 | 0.65 | 0.58 |
| 9 | 8923351 | rs7858252 | 0.185 | -0.808 | -1.098 | -0.106 | -0.01 | 0.087 | 0.087 | 0.176 | 0.62 | 0.78 | 0.73 | 0.86 | 0.65 | 0.58 |
| 9 | 8923381 | rs16928627 | 1.139 | -0.737 | -1.011 | -0.022 | -0.023 | 0.021 | 0.021 | 0 | 0.07 | 0.11 | 0.1 | 0.19 | 0.01 | 0.08 |
| 9 | 8923715 | rs7020371 | 1.018 | -0.49 | -0.818 | -0.1 | -0.024 | 0.025 | 0.025 | 0 | 0.14 | 0.08 | 0.17 | 0.07 | 0.17 | 0.17 |
| 9 | 8924538 | rs1368684 | 0.221 | -0.654 | -0.422 | 0.088 | -0.004 | 0.077 | 0.077 | 0.013 | 0.57 | 0.73 | 0.71 | 0.73 | 0.6 | 0.47 |
| 9 | 8925157 | rs9644893 | 0.27 | -0.648 | -0.332 | 0.09 | 0 | 0.062 | 0.062 | 0.004 | 0.56 | 0.7 | 0.7 | 0.73 | 0.6 | 0.45 |
| 9 | 8926071 | rs10977363 | 0.946 | -0.88 | -0.03 | 0.039 | 0.005 | 0.001 | 0.001 | 0.876 | 0.39 | 0.38 | 0.39 | 0.36 | 0.34 | 0.24 |
| 9 | 8926956 | rs1897676 | 0.403 | -0.635 | -0.266 | -0.291 | 0.005 | -0.001 | -0.001 | 0.492 | 0.49 | 0.54 | 0.69 | 0.79 | 0.44 | 0.45 |
| 9 | 8936967 | rs10759020 | -0.442 | -0.725 | -0.1 | 0.125 | -0.017 | 0.04 | 0.04 | 0 | 0.18 | 0.24 | 0.33 | 0.52 | 0.13 | 0.26 |
| 9 | 8937116 | rs1433548 | -0.002 | -0.623 | -0.017 | -0.724 | -0.011 | 0.017 | 0.017 | 0 | 0.63 | 0.71 | 0.8 | 0.89 | 0.64 | 0.59 |
| 9 | 8937478 | rs10511514 | -0.11 | -0.648 | -0.557 | -0.741 | -0.132 | 0.218 | 0.218 | 0.016 | 0.34 | 0.03 | 0.16 | 0.11 | 0.35 | 0.41 |
| 9 | 8939271 | rs1368687 | 1.48 | -1.013 | -0.516 | -0.686 | -0.012 | 0.081 | 0.081 | 0.774 | 0.33 | 0.09 | 0.29 | 0.16 | 0.35 | 0.17 |
| 9 | 8940141 | rs7047447 | -1.09 | -0.861 | 0.219 | -0.855 | 0.002 | 0 | 0 | 0 | 0.5 | 0.47 | 0.58 | 0.75 | 0.47 | 0.62 |
| 9 | 8941696 | rs10759021 | 1.88 | -0.969 | 0.475 | -0.881 | 0.002 | 0.04 | 0.04 | 0.644 | 0.33 | 0.23 | 0.31 | 0.15 | 0.34 | 0.17 |
| 9 | 8950179 | rs9696786 | -0.293 | -1.014 | 0.633 | -1.001 | 0.015 | 0.027 | 0.027 | 0 | 0.34 | 0.26 | 0.47 | 0.74 | 0.26 | 0.46 |
| 9 | 8951749 | rs10816018 | 1.164 | -1.007 | 0.62 | -1.03 | -0.018 | 0.028 | 0.028 | 0 | 0.14 | 0.08 | 0.07 | 0.1 | 0.16 | 0.18 |
| 9 | 8953425 | rs7848109 | -0.82 | -1.008 | 0.593 | -1.048 | -0.009 | 0.078 | 0.078 | 0 | 0.42 | 0.28 | 0.56 | 0.75 | 0.47 | 0.62 |
| 9 | 8955010 | rs7852703 | 1.669 | -1.009 | 0.659 | -1.018 | -0.035 | 0.085 | 0.085 | 0.001 | 0.4 | 0.24 | 0.39 | 0.15 | 0.54 | 0.31 |
| 9 | 8955509 | rs10977400 | 0.312 | -1.205 | 0.582 | -0.201 | 0 | 0.004 | 0.004 | 0 | 0.25 | 0.25 | 0.34 | 0.48 | 0.23 | 0.24 |
| 9 | 8956534 | rs4526401 | 0.522 | -1.24 | 0.721 | -0.201 | -0.047 | 0.059 | 0.059 | 0 | 0.87 | 0.74 | 0.83 | 0.74 | 0.95 | 0.78 |
| 9 | 8960011 | rs7037483 | 0.92 | -1.259 | 0.501 | 0.381 | -0.002 | 0.123 | 0.123 | 0 | 0.11 | 0.18 | 0.19 | 0.32 | 0.12 | 0.11 |
| 9 | 8960329 | rs7851829 | 0.909 | -1.337 | 0.473 | 0.125 | 0.005 | -0.024 | -0.024 | 0 | 0.14 | 0.18 | 0.08 | 0.1 | 0.16 | 0.2 |
| 9 | 8967563 | rs1433552 | -0.541 | -1.221 | 0.26 | 0.173 | -0.025 | -0.018 | -0.018 | 0.468 | 0.33 | 0.24 | 0.43 | 0.49 | 0.44 | 0.4 |
| 9 | 8970493 | rs10816022 | -2.67 | -1.258 | 0.308 | 0.147 | 0 | 0.045 | 0.045 | 1.351 | 0.67 | 0.8 | 0.69 | 0.85 | 0.65 | 0.83 |
| 9 | 8971666 | rs13289308 | 1.052 | -1.29 | 0.393 | 0.435 | 0.003 | -0.001 | -0.001 | 0 | 0.1 | 0.06 | 0.07 | 0.16 | 0.1 | 0.15 |
| 9 | 8972570 | rs1368676 | -2.567 | -0.83 | -0.162 | -0.276 | 0.006 | 0.045 | 0.045 | 0.957 | 0.69 | 0.84 | 0.71 | 0.86 | 0.68 | 0.84 |
| 9 | 8976241 | rs7022702 | -1.109 | -0.998 | 0.053 | -0.333 | -0.029 | -0.025 | -0.025 | 0 | 0.34 | 0.47 | 0.25 | 0.36 | 0.21 | 0.43 |
| 9 | 8977444 | rs17586734 | 0.803 | -0.812 | -0.121 | -0.34 | 0.028 | -0.038 | -0.038 | 0 | 0.1 | 0.13 | 0.11 | 0 | 0.23 | 0.16 |
| 9 | 8980881 | rs12554851 | 0.894 | -0.905 | -0.338 | -0.287 | 0.018 | -0.013 | -0.013 | 0 | 0.06 | 0.01 | 0.13 | 0.25 | 0.02 | 0.06 |
| 9 | 8981644 | rs1036278 | -1.013 | -0.967 | -0.244 | -0.461 | 0.017 | -0.102 | -0.102 | 1.344 | 0.34 | 0.45 | 0.21 | 0.11 | 0.42 | 0.36 |
| 9 | 8982308 | rs10511516 | 0.03 | -0.913 | -0.275 | -0.309 | -0.023 | 0.01 | 0.01 | 0 | 0.31 | 0.39 | 0.38 | 0.56 | 0.2 | 0.28 |
| 9 | 8982620 | rs10816027 | -0.164 | -0.836 | -0.314 | -0.398 | -0.027 | -0.009 | -0.009 | 0 | 0.34 | 0.43 | 0.38 | 0.62 | 0.21 | 0.41 |
| 9 | 8984469 | rs4742554 | -2.417 | -0.705 | -0.926 | -0.368 | -0.017 | 0.021 | 0.021 | 0.844 | 0.68 | 0.86 | 0.6 | 0.74 | 0.63 | 0.77 |
| 9 | 8985404 | rs10977424 | 0.884 | -0.297 | -0.752 | -0.176 | -0.018 | 0.034 | 0.034 | 0 | 0.13 | 0.08 | 0.07 | 0.11 | 0.15 | 0.19 |
| 9 | 8987930 | rs9299083 | 0.068 | -0.353 | -0.477 | 0.239 | 0.029 | -0.047 | -0.047 | 3.218 | 0.11 | 0.16 | 0.1 | 0 | 0.23 | 0.16 |
| 9 | 8988375 | rs2039331 | -0.948 | -0.334 | -0.683 | 0.425 | -0.004 | -0.002 | -0.002 | 0 | 0.37 | 0.64 | 0.41 | 0.64 | 0.23 | 0.45 |
| 9 | 8993027 | rs9299084 | -0.686 | 0.34 | -1.372 | 0.644 | -0.016 | -0.031 | -0.031 | 0 | 0.34 | 0.26 | 0.33 | 0.38 | 0.42 | 0.39 |
| 9 | 8996113 | rs1331665 | -0.619 | 0.904 | -2.054 | 0.3 | -0.027 | 0.068 | 0.068 | 0 | 0.39 | 0.15 | 0.26 | 0.28 | 0.4 | 0.47 |
| 9 | 8996408 | rs10977429 | 0.282 | 0.625 | -2.042 | 0.133 | 0.001 | -0.009 | -0.009 | 0 | 0.09 | 0.05 | 0.06 | 0 | 0.07 | 0.05 |
| 9 | 8999697 | rs10977432 | 1.158 | 0.426 | -2.839 | 0.342 | 0.007 | 0.001 | 0.001 | 1.155 | 0.34 | 0.05 | 0.38 | 0.24 | 0.27 | 0.31 |
| 9 | 9000033 | rs973117 | 0.349 | 0.532 | -2.107 | 0.323 | 0.056 | 0.067 | 0.067 | 0.056 | 0.5 | 0.69 | 0.6 | 0.66 | 0.34 | 0.44 |
| 9 | 9000709 | rs7871755 | 0.4 | 0.809 | -1.898 | 0.394 | -0.001 | -0.005 | -0.005 | 0 | 0.67 | 0.33 | 0.46 | 0.47 | 0.64 | 0.75 |
| 9 | 9001207 | rs10511519 | 1.386 | 0.68 | -2.506 | 0.032 | -0.064 | 0.117 | 0.117 | 0.451 | 0.35 | 0.06 | 0.22 | 0.11 | 0.35 | 0.25 |
| 9 | 9001239 | rs10816028 | 0.828 | 0.635 | -2.435 | 0.191 | -0.003 | 0.049 | 0.049 | 0.599 | 0.5 | 0.27 | 0.34 | 0.32 | 0.47 | 0.48 |
| 9 | 9001538 | rs10511520 | 0.425 | 0.954 | -2.275 | 0.538 | -0.04 | 0.005 | 0.005 | 0 | 0.1 | 0.02 | 0.14 | 0.08 | 0.16 | 0.07 |
| 9 | 9003758 | rs12006331 | 0.815 | 0.896 | -1.919 | 0.363 | -0.004 | -0.041 | -0.041 | 0 | 0.07 | 0.15 | 0.06 | 0 | 0.06 | 0.05 |
| 9 | 9004122 | rs17587560 | -0.569 | 0.871 | -0.774 | 0.548 | -0.005 | 0.017 | 0.017 | 0 | 0.27 | 0.42 | 0.27 | 0.3 | 0.28 | 0.28 |
| 9 | 9010662 | rs1331662 | 0.61 | 1.078 | 0.109 | 1.089 | 0.083 | 0.206 | 0.206 | 0.951 | 0.91 | 0.57 | 0.86 | 0.72 | 0.86 | 0.77 |
| 9 | 9011209 | rs7046257 | 0.363 | 1.078 | -0.367 | 0.832 | 0.003 | 0.026 | 0.026 | 3.001 | 0.67 | 0.44 | 0.49 | 0.41 | 0.66 | 0.6 |
| 9 | 9014274 | rs613613 | 0.58 | 0.828 | -0.136 | 0.038 | -0.001 | 0.001 | 0.001 | 0 | 0.25 | 0.23 | 0.19 | 0.11 | 0.29 | 0.17 |
| 9 | 9016071 | rs440238 | -0.559 | 0.506 | -0.043 | -0.26 | 0.001 | -0.01 | -0.01 | 0.856 | 0.43 | 0.42 | 0.28 | 0.17 | 0.4 | 0.31 |
| 9 | 9018656 | rs7029807 | 0.048 | 0.457 | 0.167 | 0.074 | -0.003 | 0.038 | 0.038 | 0 | 0.19 | 0.11 | 0.1 | 0 | 0.2 | 0.18 |
| 9 | 9020223 | rs378363 | 1.185 | 0.696 | 0.391 | 0.268 | 0.002 | -0.019 | -0.019 | 0 | 0.2 | 0.27 | 0.19 | 0.2 | 0.18 | 0.18 |
| 9 | 9022772 | rs1041506 | 1.69 | 1.028 | 0.03 | 0.416 | -0.008 | -0.072 | -0.072 | 0.299 | 0.15 | 0.04 | 0.2 | 0.33 | 0.14 | 0.17 |
| 9 | 9023522 | rs7875273 | 0.077 | 1.148 | 0.202 | 0.387 | -0.003 | -0.002 | -0.002 | 0 | 0.4 | 0.39 | 0.33 | 0.4 | 0.43 | 0.45 |
| 9 | 9026157 | rs540259 | -0.537 | 1.245 | 0.153 | -0.035 | 0.005 | 0.047 | 0.047 | 0.254 | 0.52 | 0.64 | 0.4 | 0.37 | 0.54 | 0.48 |
| 9 | 9026854 | rs167785 | 0.052 | 1.255 | 0.241 | -0.069 | -0.001 | 0.016 | 0.016 | 0.716 | 0.46 | 0.4 | 0.31 | 0.17 | 0.47 | 0.31 |
| 9 | 9027495 | rs1348641 | -0.969 | 0.884 | 0.322 | 0.359 | 0.049 | 0.084 | 0.084 | 0.012 | 0.07 | 0.22 | 0.09 | 0.2 | 0.07 | 0.15 |
| 9 | 9029463 | rs324495 | 0.453 | 0.455 | 0.183 | 0.199 | 0.007 | 0.001 | 0.001 | 0 | 0.67 | 0.32 | 0.52 | 0.4 | 0.73 | 0.66 |
| 9 | 9030732 | rs368381 | 0.267 | 0.469 | 0.409 | 0.122 | 0.01 | 0.012 | 0.012 | 0 | 0.48 | 0.33 | 0.54 | 0.64 | 0.45 | 0.55 |
| 9 | 9030773 | rs10120940 | 0.325 | 0.395 | 0.262 | -0.078 | 0.007 | 0.04 | 0.04 | 0 | 0.06 | 0.11 | 0.05 | 0.12 | 0.09 | 0.06 |
| 9 | 9031563 | rs382364 | 0.83 | 0.444 | 0.085 | -0.102 | 0.011 | 0.025 | 0.025 | 0.298 | 0.34 | 0.23 | 0.26 | 0.32 | 0.3 | 0.37 |
| 9 | 9031977 | rs1889817 | 0.553 | 0.589 | -0.269 | -0.072 | -0.069 | 0.104 | 0.104 | 0 | 0.35 | 0.14 | 0.26 | 0.13 | 0.43 | 0.31 |

| | | | | | | | | | | | | | | | | |
|---|---|---|---|---|---|---|---|---|---|---|---|---|---|---|---|---|
| 9 | 9032105 | rs182719 | -1.147 | 0.381 | 0.755 | 0.793 | -0.004 | 0.057 | 0.057 | 0.002 | 0.85 | 0.86 | 0.64 | 0.58 | 0.88 | 0.79 |
| 9 | 9032872 | rs172862 | 0.675 | 0.575 | 0.794 | 0.011 | 0.003 | -0.013 | -0.013 | 0 | 0.45 | 0.48 | 0.37 | 0.36 | 0.47 | 0.43 |
| 9 | 9033824 | rs1999740 | -0.203 | -0.046 | 0.972 | 0.052 | 0.006 | 0.029 | 0.029 | 0 | 0.08 | 0.19 | 0.07 | 0.12 | 0.08 | 0.22 |
| 9 | 9037625 | rs324517 | 1.136 | 0.013 | 1.158 | -0.066 | -0.006 | 0.024 | 0.024 | 0.733 | 0.38 | 0.31 | 0.28 | 0.19 | 0.39 | 0.26 |
| 9 | 9038430 | rs324519 | 0.542 | -0.078 | 1.151 | -0.131 | 0.005 | 0.003 | 0.003 | 1.374 | 0.69 | 0.49 | 0.72 | 0.69 | 0.64 | 0.57 |
| 9 | 9039222 | rs1331660 | 0.081 | -0.1 | 1.077 | -0.118 | 0.009 | 0.022 | 0.022 | 0 | 0.08 | 0.19 | 0.07 | 0.12 | 0.08 | 0.2 |
| 9 | 9040946 | rs324524 | 1.67 | -0.067 | 0.802 | -0.175 | -0.014 | 0.094 | 0.094 | 1.374 | 0.34 | 0.16 | 0.23 | 0.15 | 0.34 | 0.22 |
| 9 | 9041579 | rs10739183 | 1.58 | -0.153 | 0.502 | -0.002 | 0.011 | -0.082 | -0.082 | 0.367 | 0.17 | 0.05 | 0.24 | 0.44 | 0.14 | 0.19 |
| 9 | 9042075 | rs324526 | 0.906 | -0.143 | 0.529 | -0.037 | 0.042 | 0.062 | 0.062 | 0 | 0.55 | 0.17 | 0.44 | 0.38 | 0.64 | 0.46 |
| 9 | 9042311 | rs324527 | 0.119 | -0.038 | 0.696 | -0.066 | 0.008 | 0.011 | 0.011 | 0.423 | 0.72 | 0.58 | 0.74 | 0.7 | 0.67 | 0.6 |
| 9 | 9042383 | rs7034007 | 0.27 | -0.215 | 0.443 | -0.342 | 0.004 | 0.025 | 0.025 | 0 | 0.08 | 0.2 | 0.06 | 0.1 | 0.07 | 0.19 |
| 9 | 9042840 | rs324529 | 1.04 | -0.256 | 0.16 | -0.348 | 0.032 | -0.02 | -0.02 | 0 | 0.49 | 0.14 | 0.56 | 0.48 | 0.57 | 0.48 |
| 9 | 9043421 | rs88818 | 0.111 | -0.228 | 0.225 | -0.332 | 0.018 | -0.028 | -0.028 | 0.037 | 0.68 | 0.22 | 0.69 | 0.51 | 0.76 | 0.61 |
| 9 | 9045011 | rs324533 | 0.962 | -0.218 | 0.093 | -0.351 | -0.049 | 0.163 | 0.163 | 3.314 | 0.45 | 0.13 | 0.3 | 0.21 | 0.46 | 0.3 |
| 9 | 9047488 | rs1118100 | -0.32 | -0.08 | 0.035 | -0.245 | -0.088 | -0.061 | -0.061 | 0 | 0.29 | 0.08 | 0.39 | 0.33 | 0.41 | 0.39 |
| 9 | 9048128 | rs13285004 | 0.691 | -0.033 | 0.106 | -0.194 | -0.046 | -0.048 | -0.048 | 0.313 | 0.19 | 0.08 | 0.34 | 0.29 | 0.31 | 0.3 |
| 9 | 9049813 | rs7035296 | 1.09 | -0.141 | 0.271 | -0.214 | -0.017 | 0.049 | 0.049 | 0 | 0.05 | 0.18 | 0.03 | 0.09 | 0.02 | 0.16 |
| 9 | 9052619 | rs10759035 | 0.144 | -0.18 | -0.007 | -0.321 | -0.067 | 0.038 | 0.038 | 1.221 | 0.35 | 0.11 | 0.56 | 0.71 | 0.42 | 0.49 |
| 9 | 9052964 | rs12685303 | 0.507 | -0.15 | 0.371 | -0.083 | -0.025 | 0.057 | 0.057 | 0 | 0.07 | 0.24 | 0.03 | 0.11 | 0.02 | 0.16 |
| 9 | 9054075 | rs12005189 | -0.376 | -0.252 | 0.438 | -0.177 | -0.011 | -0.035 | -0.035 | 0 | 0.09 | 0.16 | 0.05 | 0 | 0.05 | 0.02 |
| 9 | 9054313 | rs2225482 | -0.375 | -0.154 | 0.394 | -0.179 | 0 | 0.077 | 0.077 | 0 | 0.48 | 0.36 | 0.63 | 0.81 | 0.5 | 0.68 |
| 9 | 9054913 | rs1412096 | 1.154 | -0.124 | 0.137 | -0.104 | -0.088 | -0.074 | -0.074 | 0.053 | 0.22 | 0.03 | 0.27 | 0.23 | 0.31 | 0.27 |
| 9 | 9055544 | rs10977457 | 1.477 | -0.157 | 0.227 | -0.187 | 0.019 | -0.081 | -0.081 | 0.02 | 0.16 | 0.05 | 0.19 | 0.38 | 0.12 | 0.19 |
| 9 | 9057632 | rs10816040 | 1.354 | -0.146 | 0.242 | -0.235 | 0.023 | -0.07 | -0.07 | 0.199 | 0.17 | 0.07 | 0.2 | 0.37 | 0.12 | 0.19 |
| 9 | 9059182 | rs17665194 | 0.592 | 0.429 | 0.94 | -1.015 | -0.008 | 0.004 | 0.004 | 0 | 0.06 | 0.01 | 0.04 | 0.04 | 0.08 | 0.08 |
| 9 | 9059545 | rs324498 | 1.425 | 0.331 | 1.025 | -1.069 | -0.001 | 0.001 | 0.001 | 1.058 | 0.77 | 0.29 | 0.79 | 0.76 | 0.86 | 0.77 |
| 9 | 9059937 | rs1412102 | -0.297 | 0.324 | 0.558 | -1.059 | -0.001 | 0.004 | 0.004 | 0 | 0.13 | 0.11 | 0.11 | 0.07 | 0.15 | 0.12 |
| 9 | 9060444 | rs12555043 | -0.759 | 0.293 | 0.716 | -1.032 | 0.001 | 0.072 | 0.072 | 0 | 0.06 | 0.26 | 0.08 | 0.13 | 0.01 | 0.01 |
| 9 | 9061331 | rs12006155 | -0.159 | 0.494 | 0.61 | -0.987 | 0.024 | 0.032 | 0.032 | 0 | 0.09 | 0.26 | 0.07 | 0.11 | 0.09 | 0.18 |
| 9 | 9061819 | rs324501 | 0.739 | 0.6 | 0.064 | -1.04 | -0.032 | 0.094 | 0.094 | 1.897 | 0.33 | 0.1 | 0.24 | 0.2 | 0.34 | 0.22 |
| 9 | 9064311 | rs10977464 | -0.11 | 0.801 | -0.332 | -0.875 | 0.01 | 0.001 | 0.001 | 0 | 0.15 | 0.26 | 0.12 | 0.15 | 0.18 | 0.27 |
| 9 | 9064330 | rs10759037 | 0.928 | 0.621 | -0.564 | -0.949 | -0.012 | 0.031 | 0.031 | 0.01 | 0.38 | 0.1 | 0.52 | 0.67 | 0.37 | 0.48 |
| 9 | 9065478 | rs7026475 | -0.8 | 0.231 | -0.414 | -0.446 | -0.009 | -0.013 | -0.013 | 1.256 | 0.21 | 0.32 | 0.16 | 0.16 | 0.2 | 0.3 |
| 9 | 9065718 | rs16928863 | -0.088 | 0.403 | -0.997 | -0.554 | -0.005 | -0.033 | -0.033 | 0 | 0.08 | 0.1 | 0.06 | 0 | 0.08 | 0.04 |
| 9 | 9066755 | rs324506 | -0.199 | 0.626 | -1.224 | -0.09 | -0.018 | 0.073 | 0.073 | 1.282 | 0.31 | 0.17 | 0.21 | 0.15 | 0.31 | 0.2 |
| 9 | 9069839 | rs408960 | 0.131 | 0.096 | -0.571 | 0.707 | -0.019 | 0.026 | 0.026 | 0 | 0.68 | 0.6 | 0.72 | 0.52 | 0.76 | 0.55 |
| 9 | 9070768 | rs324513 | -0.021 | 0.028 | -0.29 | 0.969 | -0.003 | 0.001 | 0.001 | 0 | 0.7 | 0.71 | 0.79 | 0.59 | 0.78 | 0.56 |
| 9 | 9072276 | rs634921 | 0.684 | 0.247 | 0.008 | 0.973 | 0.047 | 0.056 | 0.056 | 0 | 0.52 | 0.31 | 0.48 | 0.31 | 0.62 | 0.43 |
| 9 | 9073241 | rs324482 | 0.018 | 0.207 | 0.075 | 1.064 | 0.011 | -0.009 | -0.009 | 0.834 | 0.62 | 0.37 | 0.65 | 0.49 | 0.71 | 0.54 |
| 9 | 9074042 | rs324484 | 0.406 | 0.205 | 0.347 | 1.086 | -0.012 | -0.004 | -0.004 | 0 | 0.55 | 0.44 | 0.54 | 0.36 | 0.65 | 0.48 |
| 9 | 9074830 | rs324486 | 0.044 | 0.195 | 0.371 | 1.094 | 0.02 | -0.01 | -0.01 | 0.308 | 0.62 | 0.35 | 0.66 | 0.47 | 0.72 | 0.53 |
| 9 | 9077607 | rs324475 | -0.349 | 0.199 | 0.437 | 1.078 | -0.004 | 0.003 | 0.003 | 0 | 0.67 | 0.66 | 0.72 | 0.54 | 0.74 | 0.56 |
| 9 | 9079771 | rs324479 | -0.349 | 0.34 | 0.535 | 0.969 | -0.016 | 0.006 | 0.006 | 0 | 0.64 | 0.56 | 0.71 | 0.57 | 0.7 | 0.54 |
| 9 | 9079956 | rs324480 | 0.642 | 0.323 | 0.488 | 0.962 | 0.031 | 0.034 | 0.034 | 0 | 0.49 | 0.39 | 0.49 | 0.35 | 0.58 | 0.41 |
| 9 | 9086923 | rs369166 | -0.14 | 0.219 | 0.464 | 0.932 | 0 | -0.001 | -0.001 | 0 | 0.61 | 0.44 | 0.7 | 0.58 | 0.69 | 0.53 |
| 9 | 9092671 | rs10977480 | -0.491 | 0.124 | 0.305 | 0.882 | 0.002 | -0.03 | -0.03 | 2.014 | 0.13 | 0.08 | 0.21 | 0.23 | 0.11 | 0.13 |
| 9 | 9093111 | rs10977482 | -0.578 | 0.144 | 0.45 | 0.892 | -0.004 | 0.024 | 0.024 | 0 | 0.13 | 0.17 | 0.22 | 0.23 | 0.11 | 0.13 |
| 9 | 9094805 | rs472324 | 0.786 | 0.163 | 0.493 | 0.978 | 0.035 | 0.06 | 0.06 | 0 | 0.48 | 0.32 | 0.41 | 0.29 | 0.58 | 0.39 |
| 9 | 9102409 | rs324542 | -0.469 | 0.141 | 0.546 | 0.999 | 0.009 | -0.007 | -0.007 | 0 | 0.64 | 0.72 | 0.73 | 0.56 | 0.71 | 0.52 |
| 9 | 9104869 | rs10977497 | -0.559 | 0.136 | 0.357 | 0.993 | -0.002 | 0.028 | 0.028 | 0 | 0.11 | 0.16 | 0.22 | 0.23 | 0.09 | 0.13 |
| 9 | 9108336 | rs324544 | 0.573 | 0.165 | 0.206 | 0.949 | 0.035 | 0.042 | 0.042 | 0 | 0.51 | 0.35 | 0.47 | 0.32 | 0.61 | 0.42 |
| 9 | 9109538 | rs9299087 | 0.026 | 0.16 | -0.305 | 1.064 | -0.007 | -0.027 | -0.027 | 0 | 0.08 | 0.04 | 0.16 | 0.17 | 0.07 | 0.1 |
| 9 | 9109843 | rs10816047 | 0.026 | 0.16 | -0.305 | 1.064 | -0.007 | -0.027 | -0.027 | 0 | 0.08 | 0.04 | 0.16 | 0.17 | 0.07 | 0.1 |
| 9 | 9119229 | rs17589981 | 0.559 | 0.904 | -0.403 | 0.584 | -0.062 | 0.082 | 0.082 | 0.233 | 0.35 | 0.06 | 0.27 | 0.18 | 0.37 | 0.19 |
| 9 | 9139360 | rs9299088 | -0.411 | 0.712 | -0.173 | 0.683 | -0.003 | -0.005 | -0.005 | 1.056 | 0.64 | 0.65 | 0.58 | 0.44 | 0.6 | 0.6 |
| 9 | 9142828 | rs957155 | -0.605 | 0.874 | 0.15 | 1.172 | 0.007 | 0.023 | 0.023 | 0 | 0.11 | 0.12 | 0.07 | 0.24 | 0.11 | 0.15 |
| 9 | 9144085 | rs10816049 | -0.29 | -0.144 | 0.294 | 1.324 | -0.022 | 0.006 | 0.006 | 0.831 | 0.27 | 0.38 | 0.29 | 0.26 | 0.19 | 0.39 |
| 9 | 9145908 | rs1992672 | -0.508 | -0.528 | 0.069 | 1.245 | 0.02 | -0.004 | -0.004 | 0 | 0.51 | 0.72 | 0.45 | 0.47 | 0.56 | 0.43 |
| 9 | 9146300 | rs10121402 | 1.129 | -0.626 | -0.053 | 1.129 | -0.072 | 0.054 | 0.054 | 0.113 | 0.27 | 0.11 | 0.23 | 0.12 | 0.42 | 0.29 |
| 9 | 9149128 | rs1999741 | -1.916 | -0.628 | -0.126 | 1.198 | 0.005 | 0.009 | 0.009 | 0.685 | 0.68 | 0.75 | 0.69 | 0.7 | 0.71 | 0.74 |
| 9 | 9149546 | rs10977530 | 0.81 | -0.544 | -0.147 | 1.224 | -0.008 | -0.038 | -0.038 | 0.641 | 0.15 | 0.03 | 0.19 | 0.18 | 0.13 | 0.27 |
| 9 | 9150790 | rs2381968 | 0.136 | -0.573 | 0.104 | 1.161 | 0 | -0.001 | -0.001 | 0 | 0.19 | 0.18 | 0.23 | 0.19 | 0.17 | 0.3 |
| 9 | 9151280 | rs7861959 | -0.214 | -0.644 | 0.095 | 1.172 | 0.006 | -0.008 | -0.008 | 0 | 0.23 | 0.17 | 0.33 | 0.24 | 0.23 | 0.33 |
| 9 | 9151640 | rs12003392 | 0.247 | -0.664 | 0.302 | 1.256 | -0.018 | -0.004 | -0.004 | 0.001 | 0.09 | 0.22 | 0.1 | 0.08 | 0.06 | 0.1 |
| 9 | 9153346 | rs7854497 | -0.418 | -0.675 | 0.235 | 1.309 | -0.033 | 0.114 | 0.114 | 0.312 | 0.15 | 0.37 | 0.11 | 0.29 | 0.07 | 0.1 |
| 9 | 9155409 | rs6477369 | -0.437 | -0.563 | 0.167 | 1.213 | -0.004 | 0.001 | 0.001 | 0.001 | 0.83 | 0.83 | 0.8 | 0.81 | 0.88 | 0.73 |
| 9 | 9155721 | rs7036801 | -0.631 | -0.553 | -0.353 | 1.249 | -0.057 | 0.077 | 0.077 | 0.689 | 0.23 | 0.53 | 0.23 | 0.35 | 0.14 | 0.14 |
| 9 | 9156055 | rs7040160 | 1.07 | -0.521 | -0.344 | 1.216 | -0.012 | 0.001 | 0.001 | 0.784 | 0.61 | 0.44 | 0.58 | 0.46 | 0.74 | 0.58 |
| 9 | 9158221 | rs12375909 | 1.357 | -0.343 | -0.432 | 1.114 | -0.001 | 0.049 | 0.049 | 0.52 | 0.35 | 0.18 | 0.32 | 0.21 | 0.34 | 0.29 |

| | | | | | | | | | | | | | | | | |
|---|---|---|---|---|---|---|---|---|---|---|---|---|---|---|---|---|
| 9 | 9161679 | rs12003567 | 0.088 | -0.533 | -0.267 | 0.929 | -0.051 | -0.003 | -0.003 | 1.866 | 0.16 | 0.36 | 0.19 | 0.12 | 0.1 | 0.1 |
| 9 | 9161898 | rs2890854 | 1.137 | -0.44 | -0.424 | 0.969 | -0.033 | 0.027 | 0.027 | 0 | 0.28 | 0.23 | 0.23 | 0.13 | 0.42 | 0.29 |
| 9 | 9162042 | rs10816054 | -0.396 | -0.407 | -0.5 | 0.886 | 0.02 | -0.007 | -0.007 | 0 | 0.51 | 0.73 | 0.45 | 0.46 | 0.55 | 0.43 |
| 9 | 9164473 | rs1445206 | 0.526 | -0.433 | -0.066 | 0.861 | -0.025 | 0.01 | 0.01 | 2.018 | 0.71 | 0.62 | 0.62 | 0.7 | 0.78 | 0.65 |
| 9 | 9169458 | rs10977545 | -0.178 | 0.418 | -0.437 | 0.455 | 0 | -0.003 | -0.003 | 0 | 0.24 | 0.23 | 0.32 | 0.13 | 0.21 | 0.19 |
| 9 | 9173304 | rs968079 | 0.499 | -0.047 | 0.318 | 1.171 | -0.008 | 0.212 | 0.212 | 0.171 | 0.87 | 0.46 | 0.85 | 0.67 | 0.86 | 0.96 |
| 9 | 9177013 | rs1968589 | 0.54 | -0.614 | -0.272 | 0.171 | -0.043 | -0.019 | -0.019 | 0 | 0.37 | 0.22 | 0.45 | 0.6 | 0.47 | 0.67 |
| 9 | 9178137 | rs1348644 | 0.915 | -0.417 | -0.375 | 0.569 | -0.013 | -0.011 | -0.011 | 0.095 | 0.24 | 0.17 | 0.24 | 0.4 | 0.37 | 0.41 |
| 9 | 9180827 | rs2381970 | -1.454 | -0.483 | -0.213 | 0.854 | 0.002 | -0.002 | -0.002 | 0.379 | 0.93 | 0.93 | 0.96 | 0.85 | 0.93 | 0.98 |
| 9 | 9181361 | rs1889820 | -0.553 | -0.583 | 0.079 | 0.753 | -0.018 | -0.008 | -0.008 | 1.684 | 0.64 | 0.56 | 0.74 | 0.7 | 0.65 | 0.86 |
| 9 | 9182216 | rs1445213 | -0.248 | -0.937 | -0.145 | 0.477 | 0 | 0 | 0 | 0 | 0.44 | 0.52 | 0.46 | 0.55 | 0.52 | 0.69 |
| 9 | 9183618 | rs1445212 | -0.971 | -0.454 | -0.12 | 0.479 | 0 | 0 | 0 | 0.094 | 0.73 | 0.71 | 0.63 | 0.72 | 0.74 | 0.8 |
| 9 | 9184911 | rs10816060 | 1.222 | -0.336 | -0.695 | 0.466 | -0.063 | 0.034 | 0.034 | 1.131 | 0.23 | 0.02 | 0.29 | 0.1 | 0.2 | 0.18 |
| 9 | 9186346 | rs945464 | 0.527 | 0.02 | -0.603 | 0.543 | 0.016 | 0.005 | 0.005 | 0.031 | 0.26 | 0.32 | 0.23 | 0.33 | 0.38 | 0.39 |
| 9 | 9187539 | rs10977555 | 0.663 | 0.038 | -0.767 | 0.434 | -0.054 | 0.005 | 0.005 | 1.111 | 0.24 | 0.02 | 0.33 | 0.14 | 0.21 | 0.19 |
| 9 | 9188865 | rs1959404 | -0.213 | -0.115 | -0.553 | 0.317 | -0.001 | 0.002 | 0.002 | 0.289 | 0.61 | 0.66 | 0.5 | 0.67 | 0.61 | 0.75 |
| 9 | 9195240 | rs12000334 | -0.304 | 0.032 | -0.714 | -0.308 | 0.003 | 0.001 | 0.001 | 0 | 0.48 | 0.45 | 0.53 | 0.78 | 0.38 | 0.71 |
| 9 | 9195415 | rs2028779 | -0.299 | 0.328 | -0.65 | -0.349 | 0 | 0 | 0 | 0 | 0.29 | 0.29 | 0.35 | 0.73 | 0.33 | 0.49 |
| 9 | 9195670 | rs2028780 | -0.518 | 0.437 | -0.534 | -1.248 | 0 | -0.05 | -0.05 | 1.956 | 0.78 | 0.79 | 0.8 | 1 | 0.73 | 0.86 |
| 9 | 9196661 | rs10977564 | 0.502 | 0.63 | -0.238 | -0.413 | 0.009 | 0.096 | 0.096 | 0.158 | 0.1 | 0.21 | 0.18 | 0.35 | 0.16 | 0.23 |
| 9 | 9197160 | rs13440065 | 0.622 | 0.455 | -0.18 | -1.058 | -0.003 | -0.006 | -0.006 | 2.241 | 0.13 | 0.12 | 0.08 | 0.02 | 0.15 | 0.09 |
| 9 | 9197298 | rs10435819 | 0.587 | 0.406 | -0.071 | -0.963 | 0.009 | 0.066 | 0.066 | 0.024 | 0.11 | 0.22 | 0.16 | 0.3 | 0.16 | 0.22 |
| 9 | 9197812 | rs1157067 | -0.984 | -0.023 | -0.295 | -1.486 | -0.001 | -0.002 | -0.002 | 0 | 0.43 | 0.41 | 0.43 | 0.37 | 0.43 | 0.48 |
| 9 | 9202754 | rs9987697 | 0.928 | 0.142 | -0.146 | -1.238 | -0.016 | 0.026 | 0.026 | 0 | 0.12 | 0.05 | 0.07 | 0.02 | 0.14 | 0.09 |
| 9 | 9202872 | rs10977567 | 1.804 | 0.354 | 0.118 | -1.377 | 0.026 | -0.025 | -0.025 | 0.181 | 0.14 | 0.24 | 0.13 | 0.1 | 0.16 | 0.09 |
| 9 | 9204531 | rs1982319 | -0.807 | -0.265 | 0.019 | -1.311 | -0.001 | -0.001 | -0.001 | 0 | 0.66 | 0.64 | 0.62 | 0.47 | 0.75 | 0.64 |
| 9 | 9214833 | rs4596705 | 0.403 | 0.394 | -0.625 | -1.548 | -0.034 | 0.215 | 0.215 | 0.206 | 0.43 | 0.1 | 0.34 | 0.09 | 0.44 | 0.33 |
| 9 | 9215776 | rs13299658 | 0.974 | 0.578 | -0.22 | -1.547 | 0.016 | -0.041 | -0.041 | 0 | 0.13 | 0.13 | 0.09 | 0.02 | 0.17 | 0.08 |
| 9 | 9216473 | rs7034934 | 1.796 | 0.277 | -0.081 | -1.732 | -0.005 | -0.021 | -0.021 | 0 | 0.22 | 0.21 | 0.2 | 0.08 | 0.21 | 0.17 |
| 9 | 9221800 | rs4620334 | -0.822 | 0.293 | -0.15 | -1.755 | 0.001 | 0.027 | 0.027 | 0.064 | 0.73 | 0.64 | 0.68 | 0.48 | 0.74 | 0.66 |
| 9 | 9222258 | rs13294494 | 1.107 | 0.294 | -0.186 | -1.907 | 0.005 | 0.078 | 0.078 | 1.401 | 0.31 | 0.13 | 0.27 | 0.07 | 0.3 | 0.24 |
| 9 | 9223809 | rs4357351 | 1.424 | 0.316 | -0.08 | -1.873 | 0.011 | -0.009 | -0.009 | 0 | 0.28 | 0.24 | 0.29 | 0.32 | 0.27 | 0.32 |
| 9 | 9225919 | rs7035832 | 0.083 | 0.255 | 0.347 | -1.636 | -0.005 | 0.01 | 0.01 | 0 | 0.44 | 0.63 | 0.45 | 0.41 | 0.48 | 0.47 |
| 9 | 9228094 | rs4621866 | 2.165 | 0.342 | 0.213 | -1.666 | -0.003 | 0.003 | 0.003 | 0.69 | 0.15 | 0.13 | 0.12 | 0.1 | 0.16 | 0.1 |
| 9 | 9228107 | rs12336488 | 1.177 | 0.286 | 0.228 | -1.744 | -0.013 | 0.064 | 0.064 | 0 | 0.32 | 0.14 | 0.23 | 0.02 | 0.32 | 0.18 |
| 9 | 9229566 | rs17667884 | 0.049 | 0.22 | 0.296 | -1.669 | 0 | -0.011 | -0.011 | 0 | 0.49 | 0.39 | 0.43 | 0.16 | 0.52 | 0.42 |
| 9 | 9230415 | rs7018892 | -1.269 | 0.216 | 0.217 | -1.534 | -0.004 | 0.127 | 0.127 | 0 | 0.91 | 0.79 | 0.81 | 0.5 | 0.94 | 0.82 |
| 9 | 9230674 | rs6477375 | 0.277 | 0.077 | -0.16 | -1.798 | 0.005 | 0.037 | 0.037 | 0 | 0.51 | 0.39 | 0.54 | 0.39 | 0.53 | 0.46 |
| 9 | 9231566 | rs10977596 | 2.073 | 0.299 | -0.36 | -1.725 | 0.002 | -0.006 | -0.006 | 0 | 0.15 | 0.14 | 0.12 | 0.09 | 0.17 | 0.1 |
| 9 | 9235200 | rs4742569 | 0.514 | 0.139 | -0.708 | -1.721 | 0.018 | 0.131 | 0.131 | 1.275 | 0.31 | 0.05 | 0.25 | 0.06 | 0.28 | 0.24 |
| 9 | 9235372 | rs4742571 | 0.497 | 0.13 | -0.597 | -1.75 | 0.024 | 0.085 | 0.085 | 1.37 | 0.31 | 0.12 | 0.25 | 0.06 | 0.28 | 0.24 |
| 9 | 9238929 | rs4742578 | 0.577 | 0.169 | -0.671 | -1.878 | 0.018 | 0.129 | 0.129 | 1.697 | 0.3 | 0.05 | 0.25 | 0.06 | 0.28 | 0.24 |
| 9 | 9241520 | rs10816071 | 1.413 | 0.251 | -0.239 | -1.634 | 0.018 | 0.17 | 0.17 | 0 | 0.09 | 0.28 | 0.16 | 0.34 | 0.1 | 0.2 |
| 9 | 9246647 | rs4568657 | 0.372 | 0.234 | -0.763 | -1.986 | -0.006 | 0.011 | 0.011 | 0 | 0.42 | 0.61 | 0.44 | 0.41 | 0.47 | 0.46 |
| 9 | 9247105 | rs10977614 | 2.255 | 0.278 | -0.678 | -1.84 | -0.055 | 0.076 | 0.076 | 0.534 | 0.14 | 0.03 | 0.09 | 0.02 | 0.17 | 0.09 |
| 9 | 9249795 | rs11791447 | 1.308 | 0.177 | -0.678 | -1.927 | 0.004 | -0.008 | -0.008 | 0 | 0.2 | 0.13 | 0.1 | 0.02 | 0.2 | 0.18 |
| 9 | 9249912 | rs7032620 | -0.588 | 0.367 | -0.678 | -1.489 | -0.011 | -0.038 | -0.038 | 0 | 0.74 | 0.76 | 0.7 | 0.47 | 0.76 | 0.71 |
| 9 | 9252865 | rs6477382 | 0.206 | 0.484 | -0.421 | -1.371 | 0.017 | 0.02 | 0.02 | 0.039 | 0.29 | 0.59 | 0.33 | 0.42 | 0.35 | 0.4 |
| 9 | 9258729 | rs4560852 | 0.382 | 0.388 | -0.75 | -1.178 | -0.017 | -0.04 | -0.04 | 0 | 0.09 | 0.01 | 0.19 | 0.52 | 0.08 | 0.21 |
| 9 | 9258963 | rs6477383 | -0.013 | 0.122 | -0.046 | -1.733 | -0.006 | 0.038 | 0.038 | 0 | 0.23 | 0.56 | 0.35 | 0.39 | 0.24 | 0.32 |
| 9 | 9260044 | rs17605087 | 1.643 | 0.044 | -0.799 | -2.052 | 0.016 | 0.136 | 0.136 | 0 | 0.23 | 0.03 | 0.15 | 0 | 0.18 | 0.13 |
| 9 | 9261737 | rs4626664 | 1.053 | 0.414 | 0.198 | -1.632 | 0.07 | 0.239 | 0.239 | 0 | 0.08 | 0.29 | 0.26 | 0.39 | 0.14 | 0.24 |
| 9 | 9261826 | rs11792302 | 1.222 | 0.297 | 0.265 | -1.713 | 0.003 | 0 | 0 | 0 | 0.13 | 0.09 | 0.05 | 0.02 | 0.1 | 0.08 |
| 9 | 9262323 | rs4626665 | 0.185 | 0.349 | 0.293 | -1.691 | 0.01 | 0.073 | 0.073 | 0 | 0.21 | 0.38 | 0.33 | 0.4 | 0.24 | 0.32 |
| 9 | 9263980 | rs7863177 | -0.675 | 0.455 | 0.158 | -1.926 | 0.014 | 0.005 | 0.005 | 0 | 0.41 | 0.47 | 0.48 | 0.49 | 0.49 | 0.44 |
| 9 | 9264439 | rs10816077 | 2.112 | 0.577 | 0.291 | -1.713 | 0.032 | 0.072 | 0.072 | 0 | 0.05 | 0.13 | 0.19 | 0.25 | 0.12 | 0.18 |
| 9 | 9264932 | rs9299090 | 0.544 | 0.761 | 0.182 | -1.798 | 0.104 | 0.124 | 0.124 | 0.031 | 0.13 | 0.29 | 0.34 | 0.4 | 0.27 | 0.25 |
| 9 | 9269284 | rs4363255 | -0.623 | 0.541 | 0.38 | -1.871 | 0.003 | 0.002 | 0.002 | 0 | 0.32 | 0.62 | 0.42 | 0.44 | 0.4 | 0.38 |
| 9 | 9270436 | rs10816081 | -1.004 | 0.082 | 0.121 | -2.084 | 0.05 | 0.005 | 0.005 | 0 | 0.54 | 0.66 | 0.57 | 0.5 | 0.66 | 0.53 |
| 9 | 9271454 | rs4289870 | 2.189 | 0.329 | 0.031 | -1.82 | 0.03 | 0.079 | 0.079 | 0 | 0.05 | 0.13 | 0.2 | 0.27 | 0.12 | 0.18 |
| 9 | 9272114 | rs16929125 | 1.434 | 0.272 | -0.083 | -1.877 | 0.007 | 0.031 | 0.031 | 0 | 0.13 | 0.02 | 0.07 | 0.05 | 0.09 | 0.07 |
| 9 | 9272841 | rs4302899 | 1.294 | 0.494 | 0.186 | -1.752 | 0.054 | 0.134 | 0.134 | 0 | 0.08 | 0.2 | 0.24 | 0.34 | 0.15 | 0.23 |
| 9 | 9277320 | rs4563945 | -0.298 | 0.233 | 0.237 | -1.854 | -0.002 | 0.003 | 0.003 | 0 | 0.4 | 0.48 | 0.5 | 0.5 | 0.54 | 0.46 |
| 9 | 9282982 | rs4492438 | -0.265 | 0.435 | 0.627 | -1.492 | 0.049 | 0.046 | 0.046 | 0.129 | 0.19 | 0.35 | 0.32 | 0.35 | 0.27 | 0.28 |
| 9 | 9283557 | rs10122609 | 1.418 | 0.473 | 0.234 | -1.513 | -0.014 | 0.01 | 0.01 | 0 | 0.09 | 0.05 | 0.07 | 0.02 | 0.14 | 0.06 |
| 9 | 9287321 | rs4146134 | 0.207 | 0.095 | 0.25 | -1.744 | -0.001 | 0.003 | 0.003 | 0.76 | 0.77 | 0.73 | 0.72 | 0.45 | 0.76 | 0.7 |
| 9 | 9287406 | rs4146135 | 0.959 | 0.172 | 0.108 | -1.631 | 0.037 | 0.151 | 0.151 | 0 | 0.09 | 0.24 | 0.25 | 0.35 | 0.13 | 0.22 |
| 9 | 9290343 | rs12350625 | 0.903 | 0.207 | 0.109 | -1.643 | 0.049 | 0.174 | 0.174 | 0 | 0.09 | 0.24 | 0.25 | 0.36 | 0.13 | 0.22 |
| 9 | 9292430 | rs11795233 | 1.39 | 0.044 | -0.546 | -1.933 | 0.004 | 0.002 | 0.002 | 0 | 0.12 | 0.1 | 0.08 | 0.11 | 0.11 | 0.12 |

| | | | | | | | | | | | | | | | | |
|---|---|---|---|---|---|---|---|---|---|---|---|---|---|---|---|---|
| 9 | 9294332 | rs1412886 | 1.995 | -0.109 | -0.918 | -2.206 | 0.008 | 0.106 | 0.106 | 0 | 0.21 | 0.04 | 0.16 | 0 | 0.17 | 0.13 |
| 9 | 9295875 | rs2151324 | 1.976 | -0.087 | -1.07 | -2.298 | -0.007 | 0.077 | 0.077 | 1.346 | 0.19 | 0.02 | 0.11 | 0.04 | 0.16 | 0.17 |
| 9 | 9297697 | rs1981049 | -0.609 | -0.121 | -0.707 | -1.953 | 0.055 | -0.076 | -0.076 | 0 | 0.66 | 0.8 | 0.64 | 0.52 | 0.73 | 0.66 |
| 9 | 9300084 | rs1157904 | 0.494 | -0.087 | -0.614 | -1.918 | -0.002 | 0.028 | 0.028 | 0.479 | 0.14 | 0.2 | 0.22 | 0.5 | 0.09 | 0.24 |
| 9 | 9301290 | rs12156461 | 1.966 | 0.085 | -0.714 | -1.97 | -0.048 | 0.051 | 0.051 | 0 | 0.13 | 0.03 | 0.1 | 0.02 | 0.2 | 0.09 |
| 9 | 9303510 | rs1333103 | -1.69 | -0.194 | -0.642 | -1.867 | 0.008 | 0.131 | 0.131 | 0 | 0.78 | 0.97 | 0.85 | 1 | 0.82 | 0.87 |
| 9 | 9310461 | rs1547288 | -0.31 | -0.207 | -0.658 | -1.76 | 0.015 | -0.043 | -0.043 | 0 | 0.62 | 0.79 | 0.6 | 0.49 | 0.63 | 0.58 |
| 9 | 9311838 | rs10977651 | 0.919 | 0.301 | -0.071 | -1.231 | 0.048 | 0.173 | 0.173 | 0 | 0.09 | 0.25 | 0.26 | 0.34 | 0.14 | 0.22 |
| 9 | 9321951 | rs9408754 | -0.953 | 0.137 | -0.676 | -1.434 | 0.014 | 0.026 | 0.026 | 0 | 0.66 | 0.75 | 0.77 | 0.93 | 0.71 | 0.78 |
| 9 | 9322503 | rs7039887 | -0.85 | 0.153 | -0.672 | -1.402 | 0.014 | 0.023 | 0.023 | 0 | 0.66 | 0.74 | 0.76 | 0.93 | 0.72 | 0.75 |
| 9 | 9326943 | rs4562383 | -0.762 | 0.199 | -0.497 | -1.231 | -0.01 | 0.104 | 0.104 | 0 | 0.53 | 0.71 | 0.68 | 0.9 | 0.54 | 0.69 |
| 9 | 9328048 | rs2780069 | -0.217 | 0.502 | -0.594 | -1.081 | -0.025 | 0.01 | 0.01 | 0 | 0.42 | 0.77 | 0.49 | 0.42 | 0.45 | 0.46 |
| 9 | 9337454 | rs1332800 | 0.444 | 1.003 | 0.066 | -0.608 | 0.028 | 0.104 | 0.104 | 0 | 0.09 | 0.23 | 0.26 | 0.36 | 0.14 | 0.22 |
| 9 | 9338113 | rs1332797 | -0.538 | 0.989 | -0.058 | -1.032 | -0.028 | -0.009 | -0.009 | 0 | 0.29 | 0.83 | 0.38 | 0.37 | 0.33 | 0.3 |
| 9 | 9340617 | rs4742585 | 0.566 | 0.926 | 0.519 | -0.814 | 0.074 | 0.188 | 0.188 | 0 | 0.09 | 0.28 | 0.26 | 0.34 | 0.14 | 0.22 |
| 9 | 9348102 | rs1332802 | -0.528 | 0.398 | 0.292 | -1.358 | 0 | -0.001 | -0.001 | 0.94 | 0.69 | 0.66 | 0.56 | 0.65 | 0.65 | 0.65 |
| 9 | 9348853 | rs10816097 | 1.514 | 0.784 | 0.169 | -1.56 | -0.031 | 0.023 | 0.023 | 0 | 0.17 | 0.1 | 0.14 | 0.03 | 0.3 | 0.1 |
| 9 | 9349832 | rs7858684 | 0.377 | 0.719 | 0.252 | -1.616 | 0.078 | 0.064 | 0.064 | 0 | 0.46 | 0.2 | 0.38 | 0.61 | 0.35 | 0.56 |
| 9 | 9350608 | rs10977673 | 1.469 | 0.824 | 0.16 | -1.595 | -0.063 | 0.057 | 0.057 | 0 | 0.18 | 0.06 | 0.14 | 0.03 | 0.3 | 0.1 |
| 9 | 9351184 | rs2151325 | -0.462 | 0.786 | 0.374 | -1.518 | 0 | -0.006 | -0.006 | 0.391 | 0.68 | 0.56 | 0.56 | 0.65 | 0.65 | 0.66 |
| 9 | 9352861 | rs10977679 | 0.301 | 0.605 | -0.226 | -1.542 | 0.011 | 0.048 | 0.048 | 0.145 | 0.53 | 0.31 | 0.32 | 0.12 | 0.55 | 0.41 |
| 9 | 9357065 | rs1889107 | -1.303 | 1.048 | -0.755 | -1.399 | -0.011 | 0.119 | 0.119 | 0 | 0.78 | 0.96 | 0.82 | 1 | 0.78 | 0.88 |
| 9 | 9357877 | rs10125869 | 0.422 | 1.163 | 0.128 | -0.477 | 0.068 | 0.233 | 0.233 | 0 | 0.1 | 0.36 | 0.26 | 0.36 | 0.14 | 0.22 |
| 9 | 9363616 | rs1855217 | -0.007 | 0.836 | -0.84 | -0.447 | 0.049 | 0.008 | 0.008 | 0 | 0.29 | 0.53 | 0.42 | 0.39 | 0.43 | 0.32 |
| 9 | 9374664 | rs17606608 | 0.595 | 0.643 | -1.554 | -0.515 | -0.037 | 0.078 | 0.078 | 0.826 | 0.18 | 0.02 | 0.11 | 0.04 | 0.19 | 0.18 |
| 9 | 9377679 | rs2780078 | -0.122 | 0.283 | -1.4 | 0.17 | 0.014 | -0.006 | -0.006 | 0 | 0.55 | 0.77 | 0.5 | 0.49 | 0.56 | 0.56 |
| 9 | 9380781 | rs10435830 | 0.297 | -0.041 | -1.18 | -0.151 | 0.008 | 0.014 | 0.014 | 0 | 0.29 | 0.23 | 0.24 | 0.16 | 0.27 | 0.22 |
| 9 | 9381760 | rs10977690 | 0.208 | -0.103 | -0.563 | -0.183 | -0.011 | -0.018 | -0.018 | 0 | 0.3 | 0.37 | 0.25 | 0.16 | 0.27 | 0.22 |
| 9 | 9385640 | rs10977709 | 0.332 | -0.069 | -0.811 | -0.161 | 0.007 | 0.014 | 0.014 | 0 | 0.29 | 0.23 | 0.24 | 0.16 | 0.27 | 0.22 |
| 9 | 9387487 | rs4742590 | -0.391 | -0.177 | -0.412 | 0.648 | -0.016 | 0.053 | 0.053 | 0 | 0.19 | 0.36 | 0.35 | 0.36 | 0.18 | 0.26 |
| 9 | 9389169 | rs10977713 | 0.102 | -0.196 | -1.159 | -0.529 | 0.009 | 0.011 | 0.011 | 0.192 | 0.29 | 0.26 | 0.14 | 0.06 | 0.27 | 0.21 |
| 9 | 9389992 | rs10816114 | 1.024 | 0.028 | -0.99 | -0.905 | 0.004 | 0.007 | 0.007 | 0 | 0.36 | 0.28 | 0.24 | 0.14 | 0.3 | 0.29 |
| 9 | 9392168 | rs10816119 | 1.108 | 0.594 | -0.967 | -1.052 | 0.001 | -0.001 | -0.001 | 1.102 | 0.1 | 0.12 | 0.12 | 0.05 | 0.15 | 0.07 |
| 9 | 9392387 | rs2382004 | -1.269 | 0.738 | -0.817 | -0.481 | 0.006 | 0.035 | 0.035 | 0 | 0.89 | 0.87 | 0.85 | 0.7 | 0.88 | 0.84 |
| 9 | 9393275 | rs7857734 | 1.312 | 0.661 | -0.862 | -0.516 | 0.005 | 0.012 | 0.012 | 0 | 0.37 | 0.28 | 0.23 | 0.16 | 0.3 | 0.29 |
| 9 | 9393744 | rs11795240 | 1.374 | 0.682 | -0.868 | -0.515 | -0.004 | -0.001 | -0.001 | 0.266 | 0.09 | 0.09 | 0.11 | 0.05 | 0.15 | 0.07 |
| 9 | 9394684 | rs2151322 | 0.676 | 0.428 | -0.941 | -0.879 | -0.011 | -0.057 | -0.057 | 0 | 0.31 | 0.15 | 0.41 | 0.46 | 0.34 | 0.3 |
| 9 | 9394902 | rs16929280 | 1.095 | 0.441 | -0.942 | -0.892 | -0.015 | -0.038 | -0.038 | 0 | 0.12 | 0.31 | 0.09 | 0.06 | 0.09 | 0.18 |
| 9 | 9395077 | rs2184077 | 0.657 | 0.45 | -1.039 | -0.864 | -0.032 | -0.098 | -0.098 | 0 | 0.31 | 0.08 | 0.4 | 0.46 | 0.34 | 0.3 |
| 9 | 9397312 | rs7848297 | 0.743 | 0.498 | -0.966 | -0.831 | 0.006 | 0.021 | 0.021 | 0 | 0.68 | 0.38 | 0.67 | 0.59 | 0.64 | 0.6 |
| 9 | 9399018 | rs1332798 | 0.609 | 0.481 | -0.998 | -0.965 | 0.003 | -0.017 | -0.017 | 0 | 0.29 | 0.29 | 0.15 | 0.06 | 0.27 | 0.19 |
| 9 | 9400177 | rs1412722 | 0.202 | 0.534 | -0.989 | -0.901 | -0.002 | 0.01 | 0.01 | 0.418 | 0.48 | 0.39 | 0.57 | 0.57 | 0.48 | 0.46 |
| 9 | 9404813 | rs10759063 | 1.073 | 0.731 | -0.661 | -0.974 | -0.021 | 0.027 | 0.027 | 1.725 | 0.23 | 0.16 | 0.18 | 0.09 | 0.31 | 0.15 |
| 9 | 9405003 | rs10759064 | -0.255 | 0.555 | -0.688 | -1.007 | 0.013 | -0.001 | -0.001 | 0 | 0.53 | 0.39 | 0.61 | 0.54 | 0.61 | 0.42 |
| 9 | 9405431 | rs10122096 | -0.376 | 0.501 | -0.721 | -0.869 | 0 | -0.004 | -0.004 | 0 | 0.85 | 0.88 | 0.83 | 0.67 | 0.88 | 0.83 |
| 9 | 9409606 | rs10977735 | 1.526 | 0.534 | -0.696 | -0.967 | -0.004 | 0.012 | 0.012 | 0.529 | 0.19 | 0.14 | 0.14 | 0.09 | 0.22 | 0.14 |
| 9 | 9412518 | rs2065514 | 1.349 | 0.555 | -0.38 | -0.784 | -0.01 | -0.053 | -0.053 | 6.36 | 0.26 | 0.31 | 0.13 | 0.06 | 0.2 | 0.29 |
| 9 | 9416962 | rs1333111 | 0.646 | 0.261 | -0.462 | -0.564 | 0 | -0.026 | -0.026 | 0 | 0.69 | 0.63 | 0.45 | 0.15 | 0.68 | 0.5 |
| 9 | 9417343 | rs928476 | -0.371 | 0.332 | -0.482 | -0.32 | 0.004 | 0.062 | 0.062 | 0 | 0.9 | 0.89 | 0.83 | 0.63 | 0.89 | 0.8 |
| 9 | 9417430 | rs16929315 | 0.933 | 0.327 | -0.487 | -0.306 | -0.003 | -0.006 | -0.006 | 0 | 0.09 | 0.1 | 0.06 | 0.02 | 0.06 | 0.07 |
| 9 | 9422159 | rs4342663 | 0.214 | 0.31 | -0.447 | -0.388 | 0.017 | 0 | 0 | 0 | 0.52 | 0.45 | 0.64 | 0.55 | 0.61 | 0.43 |
| 9 | 9423523 | rs1412727 | 0.666 | 0.936 | -0.123 | 0.588 | 0.003 | 0.033 | 0.033 | 0 | 0.84 | 0.82 | 0.78 | 0.67 | 0.83 | 0.78 |
| 9 | 9423558 | rs11792435 | 0.452 | 0.5 | -0.377 | 0.499 | 0.001 | 0.001 | 0.001 | 0.004 | 0.08 | 0.07 | 0.05 | 0.02 | 0.06 | 0.07 |
| 9 | 9425605 | rs3858066 | 0.044 | 0.716 | -0.4 | 0.744 | 0 | 0.042 | 0.042 | 0 | 0.12 | 0.14 | 0.32 | 0.46 | 0.16 | 0.26 |
| 9 | 9437369 | rs1412877 | -0.119 | 0.704 | -0.192 | 0.391 | 0.004 | -0.012 | -0.012 | 0 | 0.29 | 0.32 | 0.3 | 0.21 | 0.31 | 0.22 |
| 9 | 9437436 | rs1412875 | -0.24 | 0.32 | 0.39 | 0.258 | -0.041 | 0.122 | 0.122 | 0 | 0.21 | 0.44 | 0.3 | 0.44 | 0.18 | 0.29 |
| 9 | 9437674 | rs1412873 | -0.311 | 0.794 | 0.15 | 0.32 | -0.007 | 0.038 | 0.038 | 0 | 0.92 | 0.94 | 0.85 | 0.64 | 0.89 | 0.87 |
| 9 | 9439700 | rs4142435 | 0.388 | 0.742 | 0.218 | 0.268 | 0.008 | 0.011 | 0.011 | 0.247 | 0.65 | 0.64 | 0.58 | 0.55 | 0.58 | 0.66 |
| 9 | 9440057 | rs10977747 | 0.905 | 0.909 | -0.014 | 0.169 | -0.023 | 0.034 | 0.034 | 0.317 | 0.19 | 0.09 | 0.23 | 0.08 | 0.23 | 0.19 |
| 9 | 9440664 | rs16929395 | 0.192 | 0.618 | 0.482 | 0.138 | 0.004 | 0.13 | 0.13 | 0 | 0.13 | 0.3 | 0.27 | 0.44 | 0.15 | 0.21 |
| 9 | 9440925 | rs1360592 | 0.388 | 0.57 | 0.802 | 0.347 | 0.023 | 0.029 | 0.029 | 0.162 | 0.64 | 0.48 | 0.56 | 0.51 | 0.58 | 0.66 |
| 9 | 9441003 | rs1360591 | 0.936 | 0.652 | 0.619 | 0.063 | 0.002 | -0.001 | -0.001 | 0 | 0.55 | 0.5 | 0.5 | 0.48 | 0.51 | 0.55 |
| 9 | 9442794 | rs12376687 | -0.433 | 0.571 | 0.118 | -0.857 | 0.01 | 0.025 | 0.025 | 0 | 0.2 | 0.14 | 0.1 | 0.01 | 0.17 | 0.13 |
| 9 | 9443985 | rs1332211 | -0.899 | 0.956 | -0.025 | -0.62 | -0.031 | -0.029 | -0.029 | 1.387 | 0.95 | 0.99 | 0.87 | 0.82 | 0.86 | 0.9 |
| 9 | 9454258 | rs1332810 | -0.55 | 0.802 | 0.41 | -0.763 | -0.009 | -0.059 | -0.059 | 0 | 0.21 | 0.25 | 0.11 | 0.01 | 0.16 | 0.13 |
| 9 | 9455222 | rs10977771 | -0.435 | 0.797 | 0.44 | -0.765 | 0 | -0.028 | -0.028 | 0 | 0.2 | 0.2 | 0.1 | 0.01 | 0.16 | 0.13 |
| 9 | 9456615 | rs987353 | 1.416 | 0.809 | 0.68 | -0.98 | 0.004 | -0.03 | -0.03 | 2.676 | 0.63 | 0.45 | 0.63 | 0.68 | 0.6 | 0.63 |
| 9 | 9459099 | rs1333108 | 0.66 | 0.756 | 0.664 | -1.212 | -0.002 | 0 | 0 | 0 | 0.73 | 0.78 | 0.7 | 0.7 | 0.68 | 0.69 |
| 9 | 9461717 | rs10759075 | 0.593 | 0.717 | 0.742 | -1.242 | 0.013 | -0.015 | -0.015 | 0 | 0.74 | 0.7 | 0.71 | 0.7 | 0.69 | 0.71 |

| | | | | | | | | | | | | | | | | |
|---|---|---|---|---|---|---|---|---|---|---|---|---|---|---|---|---|
| 9 | 9464561 | rs10491613 | -0.913 | 0.641 | 0.745 | -1.321 | -0.003 | -0.06 | -0.06 | 0 | 0.31 | 0.33 | 0.15 | 0.01 | 0.23 | 0.17 |
| 9 | 9465911 | rs12338165 | 1.134 | 0.601 | 0.638 | -1.261 | -0.002 | -0.027 | -0.027 | 0.611 | 0.11 | 0.11 | 0.05 | 0 | 0.07 | 0.05 |
| 9 | 9469877 | rs10816135 | 0.267 | 0.66 | 0.656 | -1.248 | 0.012 | -0.026 | -0.026 | 0 | 0.74 | 0.65 | 0.73 | 0.72 | 0.69 | 0.76 |
| 9 | 9472132 | rs7027963 | 0.521 | 0.555 | 0.616 | -1.302 | 0.009 | -0.04 | -0.04 | 0.012 | 0.72 | 0.59 | 0.72 | 0.72 | 0.69 | 0.76 |
| 9 | 9472883 | rs2026984 | 1.841 | 0.51 | 0.556 | -1.367 | -0.005 | 0.07 | 0.07 | 0 | 0.52 | 0.37 | 0.58 | 0.7 | 0.52 | 0.57 |
| 9 | 9474970 | rs16929560 | 1.114 | 0.541 | 0.578 | -1.34 | 0.004 | -0.03 | -0.03 | 0 | 0.63 | 0.45 | 0.64 | 0.7 | 0.6 | 0.62 |
| 9 | 9475644 | rs10977801 | 1.554 | 0.624 | -0.083 | -1.263 | 0.002 | 0.009 | 0.009 | 0 | 0.1 | 0.05 | 0.04 | 0 | 0.07 | 0.04 |
| 9 | 9476195 | rs10759077 | 0.48 | 0.694 | 0.023 | -1.148 | 0.003 | -0.022 | -0.022 | 0 | 0.72 | 0.64 | 0.73 | 0.71 | 0.7 | 0.75 |
| 9 | 9476543 | rs13287051 | 0.346 | 0.796 | -0.623 | -1.034 | -0.031 | 0.045 | 0.045 | 0.423 | 0.1 | 0.01 | 0.09 | 0.02 | 0.11 | 0.16 |
| 9 | 9483287 | rs10977816 | 0.438 | 0.711 | -0.605 | -1.088 | 0.002 | -0.006 | -0.006 | 0 | 0.12 | 0.08 | 0.05 | 0.01 | 0.11 | 0.1 |
| 9 | 9483573 | rs2382017 | -0.15 | 0.629 | -0.606 | -1.355 | -0.001 | -0.002 | -0.002 | 0 | 0.21 | 0.18 | 0.14 | 0.04 | 0.19 | 0.18 |
| 9 | 9483973 | rs1332195 | 1.524 | 0.681 | -0.541 | -1.258 | -0.006 | 0.048 | 0.048 | 0 | 0.53 | 0.59 | 0.61 | 0.7 | 0.51 | 0.61 |
| 9 | 9488626 | rs12339036 | 0.846 | 0.489 | -0.463 | -1.05 | -0.006 | -0.048 | -0.048 | 0 | 0.12 | 0.15 | 0.06 | 0 | 0.09 | 0.05 |
| 9 | 9489220 | rs10977821 | 0.703 | 0.26 | -1.468 | -0.682 | 0.03 | -0.033 | -0.033 | 0.061 | 0.3 | 0.16 | 0.4 | 0.35 | 0.21 | 0.31 |
| 9 | 9494729 | rs6477398 | -0.7 | 0.88 | -1.732 | -1.236 | -0.078 | 0.031 | 0.031 | 0 | 0.72 | 0.9 | 0.7 | 0.69 | 0.6 | 0.84 |
| 9 | 9495226 | rs4551426 | 0.136 | 0.784 | -1.676 | -1.275 | -0.047 | 0.085 | 0.085 | 0.051 | 0.6 | 0.8 | 0.65 | 0.68 | 0.49 | 0.72 |
| 9 | 9499126 | rs2382019 | -0.498 | 0.815 | -1.632 | -1.12 | -0.028 | 0.028 | 0.028 | 0 | 0.72 | 0.87 | 0.72 | 0.69 | 0.66 | 0.88 |
| 9 | 9502055 | rs10977825 | -0.414 | 0.339 | -1.001 | -1.449 | -0.023 | -0.072 | -0.072 | 0 | 0.28 | 0.35 | 0.15 | 0.07 | 0.2 | 0.17 |
| 9 | 9503477 | rs1332199 | -0.414 | 1.554 | -1.376 | -1.267 | -0.039 | -0.034 | -0.034 | 0 | 0.94 | 0.99 | 0.89 | 0.85 | 0.84 | 0.92 |
| 9 | 9503941 | rs4742605 | 0.91 | 1.083 | -1.188 | -1.819 | -0.001 | 0.001 | 0.001 | 0 | 0.27 | 0.29 | 0.46 | 0.76 | 0.3 | 0.5 |
| 9 | 9504430 | rs12337005 | 0.355 | 0.882 | -1.444 | -1.853 | 0.016 | 0.017 | 0.017 | 0 | 0.21 | 0.14 | 0.09 | 0 | 0.16 | 0.11 |
| 9 | 9506433 | rs16929628 | 0.318 | 1.218 | -0.904 | -1.73 | 0 | -0.018 | -0.018 | 0 | 0.06 | 0.08 | 0.05 | 0.01 | 0.07 | 0.05 |
| 9 | 9516348 | rs668205 | 0.064 | 1.293 | -0.854 | -2.02 | -0.014 | 0.016 | 0.016 | 0.784 | 0.3 | 0.22 | 0.26 | 0.21 | 0.38 | 0.14 |
| 9 | 9517332 | rs16929647 | 0.213 | 1.237 | -0.982 | -2.149 | -0.001 | 0.007 | 0.007 | 0 | 0.11 | 0.1 | 0.08 | 0.07 | 0.1 | 0.17 |
| 9 | 9518199 | rs828836 | 0.621 | 1.161 | -0.733 | -1.994 | -0.06 | 0.011 | 0.011 | 0 | 0.23 | 0.06 | 0.25 | 0.2 | 0.31 | 0.12 |
| 9 | 9519701 | rs7044063 | -1.046 | 0.688 | 0.141 | -2.051 | 0.008 | -0.085 | -0.085 | 0 | 0.13 | 0.39 | 0.06 | 0 | 0.09 | 0.05 |
| 9 | 9525866 | rs632770 | 0.623 | 0.477 | 0.396 | -2.102 | -0.008 | -0.092 | -0.092 | 0 | 0.85 | 0.67 | 0.93 | 1 | 0.88 | 0.95 |
| 9 | 9526811 | rs694959 | 0.327 | 0.231 | 0.56 | -2.238 | -0.039 | -0.132 | -0.132 | 0.898 | 0.83 | 0.56 | 0.93 | 1 | 0.9 | 0.97 |
| 9 | 9530755 | rs7049205 | 1.085 | 0.79 | 0.129 | -2.078 | 0 | -0.015 | -0.015 | 1.163 | 0.17 | 0.16 | 0.14 | 0.04 | 0.18 | 0.04 |
| 9 | 9533062 | rs2038963 | 0.541 | 0.841 | 0.06 | -2.144 | -0.001 | -0.003 | -0.003 | 0.158 | 0.19 | 0.15 | 0.15 | 0.04 | 0.21 | 0.06 |
| 9 | 9534643 | rs7047252 | 1.011 | 0.815 | 0.064 | -2.14 | 0 | -0.013 | -0.013 | 0.005 | 0.17 | 0.15 | 0.13 | 0.04 | 0.18 | 0.04 |
| 9 | 9538589 | rs625165 | -0.42 | 1.229 | 0.133 | -2.136 | 0.024 | -0.035 | -0.035 | 0.175 | 0.28 | 0.39 | 0.32 | 0.21 | 0.37 | 0.18 |
| 9 | 9539114 | rs626804 | -0.329 | 1.252 | 0.129 | -2.108 | 0.079 | 0.078 | 0.078 | 0 | 0.06 | 0.24 | 0.13 | 0.17 | 0.12 | 0.07 |
| 9 | 9540141 | rs16929713 | 0.112 | 1.304 | -0.24 | -2.147 | 0 | -0.037 | -0.037 | 2.396 | 0.23 | 0.19 | 0.2 | 0.04 | 0.25 | 0.11 |
| 9 | 9553037 | rs17179053 | 0.067 | 0.885 | -0.382 | -1.925 | 0.012 | 0.03 | 0.03 | 0 | 0.1 | 0.04 | 0.05 | 0 | 0.08 | 0.09 |
| 9 | 9556992 | rs828828 | -0.892 | 1.241 | 0.096 | -1.571 | 0.057 | 0.057 | 0.057 | 0 | 0.07 | 0.16 | 0.13 | 0.15 | 0.12 | 0.1 |
| 9 | 9568362 | rs647190 | 0.259 | 1.364 | -0.486 | -1.565 | -0.051 | -0.021 | -0.021 | 0 | 0.13 | 0.03 | 0.19 | 0.18 | 0.23 | 0.08 |
| 9 | 9581738 | rs679640 | 0.036 | 1.204 | -0.143 | -1.522 | 0.069 | -0.024 | -0.024 | 1.021 | 0.2 | 0.4 | 0.22 | 0.18 | 0.3 | 0.08 |
| 9 | 9588336 | rs610789 | 0.293 | 1.089 | -0.345 | -1.498 | 0.007 | -0.006 | -0.006 | 0 | 0.23 | 0.23 | 0.23 | 0.19 | 0.33 | 0.13 |
| 9 | 9599213 | rs1412398 | -0.827 | 0.63 | -0.964 | -2.11 | -0.02 | 0.121 | 0.121 | 0 | 0.45 | 0.26 | 0.34 | 0.22 | 0.5 | 0.22 |
| 9 | 9600239 | rs664869 | -0.444 | 0.445 | -0.787 | -2.219 | 0.013 | -0.062 | -0.062 | 1.581 | 0.36 | 0.52 | 0.31 | 0.19 | 0.42 | 0.17 |
| 9 | 9600700 | rs16929890 | -0.124 | 0.428 | -0.841 | -2.169 | -0.014 | -0.069 | -0.069 | 0 | 0.24 | 0.27 | 0.12 | 0 | 0.14 | 0.09 |
| 9 | 9602686 | rs1326772 | -1.021 | 0.431 | -0.975 | -2.216 | -0.008 | 0.072 | 0.072 | 0 | 0.45 | 0.29 | 0.32 | 0.24 | 0.49 | 0.22 |
| 9 | 9604060 | rs10977869 | -0.666 | 0.536 | -0.979 | -2.258 | 0.012 | 0.048 | 0.048 | 0 | 0.43 | 0.3 | 0.28 | 0.1 | 0.41 | 0.19 |
| 9 | 9605832 | rs7029761 | -1.034 | 0.557 | -1.125 | -2.253 | 0.01 | 0.044 | 0.044 | 0.075 | 0.47 | 0.37 | 0.31 | 0.1 | 0.44 | 0.2 |
| 9 | 9609283 | rs10977874 | 0.938 | 0.664 | -1.334 | -2.203 | 0 | -0.022 | -0.022 | 0.294 | 0.22 | 0.25 | 0.19 | 0.11 | 0.27 | 0.1 |
| 9 | 9616646 | rs10491608 | 0.656 | 0.691 | -1.414 | -2.203 | -0.06 | 0.1 | 0.1 | 0 | 0.24 | 0.03 | 0.21 | 0.09 | 0.31 | 0.12 |
| 9 | 9620141 | rs17786597 | 0.493 | 1.068 | -2.008 | -2.528 | -0.084 | -0.006 | -0.006 | 0.002 | 0.15 | 0.03 | 0.16 | 0.16 | 0.25 | 0.24 |
| 9 | 9621379 | rs16929917 | 0.408 | 0.931 | -1.602 | -2.28 | -0.006 | 0.01 | 0.01 | 0 | 0.26 | 0.23 | 0.23 | 0.11 | 0.32 | 0.12 |
| 9 | 9621487 | rs16929919 | 0.293 | 0.933 | -1.6 | -2.279 | -0.004 | 0.002 | 0.002 | 0 | 0.26 | 0.25 | 0.23 | 0.11 | 0.32 | 0.12 |
| 9 | 9624597 | rs1831033 | -0.453 | 0.791 | -1.759 | -2.322 | 0.004 | 0.14 | 0.14 | 0 | 0.38 | 0.19 | 0.26 | 0.1 | 0.39 | 0.19 |
| 9 | 9630855 | rs10977896 | 0.288 | 1.06 | -1.925 | -2.131 | -0.047 | 0.087 | 0.087 | 0 | 0.25 | 0.05 | 0.21 | 0.13 | 0.32 | 0.14 |
| 9 | 9632921 | rs10977897 | 0.186 | 0.934 | -1.595 | -2.229 | -0.019 | -0.105 | -0.105 | 0 | 0.2 | 0.29 | 0.15 | 0 | 0.17 | 0.08 |
| 9 | 9638432 | rs10816151 | -0.062 | 0.918 | -1.675 | -2.258 | -0.021 | -0.12 | -0.12 | 0.244 | 0.22 | 0.38 | 0.15 | 0 | 0.18 | 0.08 |
| 9 | 9640335 | rs7864331 | -1.249 | 1.209 | -1.786 | -1.581 | 0 | -0.002 | -0.002 | 0 | 0.49 | 0.52 | 0.44 | 0.32 | 0.54 | 0.32 |
| 9 | 9643972 | rs10816156 | -0.218 | 0.584 | -2.118 | -1.024 | -0.001 | 0 | 0 | 0 | 0.3 | 0.33 | 0.49 | 0.33 | 0.31 | 0.28 |
| 9 | 9647916 | rs10759083 | 0.848 | 0.823 | -1.922 | -0.878 | -0.01 | 0.02 | 0.02 | 0 | 0.19 | 0.25 | 0.38 | 0.29 | 0.19 | 0.2 |
| 9 | 9649668 | rs17183242 | 1.179 | 1.304 | -1.98 | -0.846 | -0.064 | 0.031 | 0.031 | 0 | 0.19 | 0.04 | 0.18 | 0.2 | 0.31 | 0.14 |
| 9 | 9667071 | rs7854287 | -1.466 | 1.119 | -1.842 | -0.875 | -0.015 | 0.019 | 0.019 | 0 | 0.63 | 0.53 | 0.72 | 0.52 | 0.73 | 0.45 |
| 9 | 9673320 | rs2821477 | -0.42 | 1.249 | -1.897 | -0.935 | 0.007 | -0.053 | -0.053 | 0 | 0.42 | 0.48 | 0.35 | 0.2 | 0.49 | 0.21 |
| 9 | 9683443 | rs7039700 | -0.378 | 1.111 | -1.931 | -0.917 | 0.014 | 0.024 | 0.024 | 0.016 | 0.39 | 0.25 | 0.5 | 0.29 | 0.39 | 0.29 |
| 9 | 9684129 | rs1934269 | -1.234 | 1.013 | -1.932 | -0.892 | -0.002 | 0.004 | 0.004 | 0 | 0.59 | 0.51 | 0.7 | 0.52 | 0.68 | 0.4 |
| 9 | 9685304 | rs7856030 | 0.426 | 1.161 | -1.587 | -0.175 | -0.026 | -0.094 | -0.094 | 0 | 0.2 | 0.31 | 0.1 | 0 | 0.12 | 0.06 |
| 9 | 9687487 | rs4742610 | -0.279 | 1.35 | -1.486 | 0.087 | 0.033 | 0.014 | 0.014 | 0.038 | 0.37 | 0.25 | 0.45 | 0.33 | 0.31 | 0.24 |
| 9 | 9701254 | rs9657637 | -0.407 | 1.402 | -1.4 | 0.278 | -0.002 | 0.017 | 0.017 | 0.226 | 0.39 | 0.13 | 0.49 | 0.35 | 0.41 | 0.33 |
| 9 | 9701354 | rs9657638 | -1.123 | 1.532 | -1.31 | 0.39 | 0 | 0.014 | 0.014 | 0 | 0.5 | 0.44 | 0.65 | 0.73 | 0.52 | 0.62 |
| 9 | 9701768 | rs10511526 | -0.327 | 1.676 | -1.196 | 0.924 | -0.032 | 0.052 | 0.052 | 0 | 0.05 | 0.01 | 0.11 | 0.38 | 0.1 | 0.16 |
| 9 | 9708107 | rs4628308 | -0.979 | 0.763 | -1.007 | 0.963 | -0.011 | 0.186 | 0.186 | 0 | 0.05 | 0.12 | 0.08 | 0.37 | 0.02 | 0.13 |
| 9 | 9709405 | rs10733553 | -0.574 | 0.647 | -1.268 | 0.728 | 0.003 | 0.058 | 0.058 | 0 | 0.41 | 0.21 | 0.56 | 0.68 | 0.44 | 0.46 |

| | | | | | | | | | | | | | | | | |
|---|---|---|---|---|---|---|---|---|---|---|---|---|---|---|---|---|
| 9 | 9711904 | rs4391483 | -0.369 | 0.64 | -1.097 | 0.571 | 0 | 0.006 | 0.006 | 0 | 0.38 | 0.32 | 0.5 | 0.31 | 0.42 | 0.32 |
| 9 | 9714838 | rs16930096 | 0.853 | 0.483 | -1.32 | 0.037 | 0.022 | 0.014 | 0.014 | 0 | 0.15 | 0.07 | 0.06 | 0 | 0.1 | 0.09 |
| 9 | 9718066 | rs10511528 | 1.226 | 0.804 | -1.494 | -0.207 | 0.05 | 0.196 | 0.196 | 0 | 0.47 | 0.76 | 0.31 | 0.07 | 0.35 | 0.32 |
| 9 | 9724976 | rs4273904 | 0.436 | 0.99 | -1.606 | -0.071 | -0.095 | 0.139 | 0.139 | 0 | 0.66 | 0.84 | 0.41 | 0.09 | 0.51 | 0.45 |
| 9 | 9729134 | rs10816169 | 0.783 | 1.054 | -1.671 | 0.223 | -0.096 | -0.075 | -0.075 | 0 | 0.22 | 0.02 | 0.29 | 0.28 | 0.32 | 0.3 |
| 9 | 9730366 | rs1934268 | 0.248 | 1.144 | -1.644 | 0.364 | -0.044 | -0.047 | -0.047 | 0.544 | 0.91 | 0.96 | 0.91 | 0.77 | 0.84 | 0.86 |
| 9 | 9733974 | rs2761762 | 0.821 | 1.134 | -1.639 | 0.397 | -0.033 | -0.052 | -0.052 | 0 | 0.88 | 0.94 | 0.71 | 0.35 | 0.83 | 0.74 |
| 9 | 9734267 | rs10977936 | 0.436 | 0.952 | -1.464 | 0.09 | -0.017 | -0.076 | -0.076 | 1.07 | 0.2 | 0.29 | 0.12 | 0.02 | 0.15 | 0.14 |
| 9 | 9736054 | rs1323800 | 2.028 | 0.753 | -1.451 | -0.027 | -0.026 | -0.123 | -0.123 | 0 | 0.37 | 0.41 | 0.21 | 0.05 | 0.24 | 0.14 |
| 9 | 9736252 | rs1408123 | 0.391 | 0.813 | -1.529 | 0.107 | -0.079 | 0.203 | 0.203 | 0.467 | 0.64 | 0.91 | 0.38 | 0.07 | 0.42 | 0.39 |
| 9 | 9740949 | rs1768885 | 0.245 | 0.786 | -1.525 | 0.126 | -0.108 | -0.126 | -0.126 | 0 | 0.67 | 0.93 | 0.57 | 0.48 | 0.45 | 0.5 |
| 9 | 9750139 | rs1359176 | 1.212 | 0.851 | -1.348 | 0.016 | 0.003 | -0.02 | -0.02 | 3.083 | 0.46 | 0.54 | 0.31 | 0.07 | 0.39 | 0.33 |
| 9 | 9751345 | rs1768891 | 0.556 | 0.985 | -1.501 | 0.093 | -0.107 | 0.179 | 0.179 | 0.018 | 0.65 | 0.88 | 0.39 | 0.07 | 0.46 | 0.39 |
| 9 | 9756868 | rs1408122 | -0.659 | 0.731 | 0.067 | 1.291 | -0.076 | 0.076 | 0.076 | 0.026 | 0.23 | 0.4 | 0.17 | 0.39 | 0.08 | 0.2 |
| 9 | 9763352 | rs1174581 | 1.374 | 0.975 | -0.033 | 1.365 | -0.072 | -0.064 | -0.064 | 1.644 | 0.89 | 0.99 | 0.76 | 0.75 | 0.77 | 0.81 |
| 9 | 9770741 | rs768225 | -2.087 | 0.842 | 0.056 | 1.354 | -0.019 | -0.002 | -0.002 | 0 | 0.34 | 0.48 | 0.44 | 0.67 | 0.3 | 0.39 |
| 9 | 9772397 | rs10977968 | -1.378 | 0.843 | 0.127 | 1.527 | -0.046 | -0.054 | -0.054 | 0 | 0.09 | 0.01 | 0.24 | 0.26 | 0.17 | 0.18 |
| 9 | 9777072 | rs2821498 | -2.275 | 0.761 | 0.3 | 1.518 | -0.019 | -0.018 | -0.018 | 0 | 0.39 | 0.54 | 0.49 | 0.7 | 0.36 | 0.57 |
| 9 | 9778653 | rs2761723 | -2.02 | 0.801 | 0.307 | 1.423 | -0.012 | -0.02 | -0.02 | 0 | 0.37 | 0.5 | 0.39 | 0.3 | 0.35 | 0.44 |
| 9 | 9782303 | rs1746793 | -2.341 | 0.742 | 0.22 | 0.83 | 0.015 | 0.021 | 0.021 | 0 | 0.42 | 0.68 | 0.51 | 0.72 | 0.38 | 0.57 |
| 9 | 9782351 | rs1746794 | -1.353 | 0.763 | 0.302 | 0.917 | -0.035 | -0.06 | -0.06 | 0 | 0.29 | 0.53 | 0.19 | 0.15 | 0.22 | 0.35 |
| 9 | 9783693 | rs4740988 | -1.862 | 0.823 | 0.204 | 0.824 | -0.002 | -0.018 | -0.018 | 0 | 0.21 | 0.17 | 0.39 | 0.7 | 0.26 | 0.47 |
| 9 | 9785685 | rs2382056 | 2.154 | 0.877 | 0.196 | 0.806 | -0.008 | -0.014 | -0.014 | 0 | 0.77 | 0.81 | 0.59 | 0.29 | 0.68 | 0.5 |
| 9 | 9787587 | rs10977976 | 1.965 | 0.936 | 0.266 | 0.772 | 0.005 | 0.023 | 0.023 | 0 | 0.58 | 0.31 | 0.47 | 0.29 | 0.57 | 0.43 |
| 9 | 9789086 | rs7870880 | -2.373 | 1.433 | -0.19 | 1.522 | 0 | -0.005 | -0.005 | 0.149 | 0.23 | 0.24 | 0.39 | 0.71 | 0.27 | 0.49 |
| 9 | 9789202 | rs17232558 | -1.384 | 1.102 | -0.309 | 1.363 | 0.007 | -0.004 | -0.004 | 2.346 | 0.12 | 0.05 | 0.09 | 0.12 | 0.11 | 0.27 |
| 9 | 9793342 | rs2821487 | 1.321 | 1.177 | -0.446 | 1.135 | -0.003 | -0.039 | -0.039 | 0 | 0.82 | 0.89 | 0.66 | 0.33 | 0.78 | 0.59 |
| 9 | 9796116 | rs2761735 | -0.852 | 0.983 | 0.496 | 0.498 | -0.028 | -0.019 | -0.019 | 0 | 0.24 | 0.49 | 0.22 | 0.17 | 0.18 | 0.24 |
| 9 | 9800624 | rs1746799 | -1.946 | 1.936 | 0.039 | 0.11 | -0.05 | 0.242 | 0.242 | 0 | 0.54 | 0.94 | 0.71 | 0.87 | 0.52 | 0.52 |
| 9 | 9801153 | rs1323802 | -0.225 | 2.338 | 0.647 | 0.28 | -0.001 | -0.002 | -0.002 | 0 | 0.14 | 0.12 | 0.13 | 0.09 | 0.19 | 0.12 |
| 9 | 9803954 | rs2296170 | -0.182 | 2.38 | 0.828 | 0.279 | 0.008 | -0.013 | -0.013 | 0 | 0.13 | 0.15 | 0.13 | 0.09 | 0.19 | 0.12 |
| 9 | 9805134 | rs4740990 | -1.446 | 2.005 | 1.268 | 0.337 | 0.038 | 0.017 | 0.017 | 0 | 0.29 | 0.49 | 0.49 | 0.71 | 0.34 | 0.28 |
| 9 | 9805193 | rs4742622 | -0.875 | 2.001 | 1.492 | 0.756 | -0.001 | -0.004 | -0.004 | 0 | 0.16 | 0.16 | 0.15 | 0.09 | 0.24 | 0.13 |
| 9 | 9812870 | rs1746805 | -1.922 | 1.883 | 1.328 | 0.108 | -0.01 | 0.148 | 0.148 | 0 | 0.4 | 0.89 | 0.36 | 0.17 | 0.37 | 0.39 |
| 9 | 9816063 | rs2761741 | -0.444 | 1.82 | 1.747 | 0.051 | -0.023 | -0.045 | -0.045 | 0.442 | 0.21 | 0.36 | 0.15 | 0.07 | 0.13 | 0.25 |
| 9 | 9817393 | rs2250434 | -1.873 | 1.908 | 1.486 | 0.101 | -0.002 | 0.113 | 0.113 | 0 | 0.4 | 0.83 | 0.35 | 0.16 | 0.37 | 0.39 |
| 9 | 9831068 | rs1570591 | -1.3 | 2.043 | 1.3 | 0.346 | 0.012 | 0.053 | 0.053 | 0 | 0.07 | 0.09 | 0.33 | 0.72 | 0.11 | 0.15 |
| 9 | 9842176 | rs7851513 | -0.527 | 1.603 | 1.431 | 0.292 | -0.003 | 0 | 0 | 0.025 | 0.13 | 0.12 | 0.13 | 0.09 | 0.19 | 0.13 |
| 9 | 9851382 | rs4740993 | -1.17 | 1.614 | 1.15 | 0.346 | -0.013 | -0.006 | -0.006 | 0 | 0.17 | 0.1 | 0.43 | 0.8 | 0.25 | 0.29 |
| 9 | 9853357 | rs7037023 | -0.542 | 1.455 | 1.182 | -0.039 | -0.063 | 0.036 | 0.036 | 0.03 | 0.13 | 0.01 | 0.13 | 0.08 | 0.19 | 0.13 |
| 9 | 9853397 | rs7033762 | 0.594 | 1.2 | 1.594 | -0.144 | -0.038 | -0.083 | -0.083 | 0 | 0.79 | 0.51 | 0.85 | 0.93 | 0.87 | 0.75 |
| 9 | 9860402 | rs10978043 | -1.013 | 1.429 | 1.342 | 0.286 | -0.017 | 0.029 | 0.029 | 0 | 0.13 | 0.05 | 0.14 | 0.07 | 0.19 | 0.13 |
| 9 | 9886403 | rs7873669 | -1.144 | 1.097 | 1.756 | 0.127 | -0.026 | -0.087 | -0.087 | 0.846 | 0.17 | 0.42 | 0.15 | 0.06 | 0.13 | 0.15 |
| 9 | 9893071 | rs10978066 | -1.133 | 1.26 | 1.181 | 0.143 | 0.01 | -0.086 | -0.086 | 0.007 | 0.19 | 0.39 | 0.2 | 0.06 | 0.2 | 0.16 |
| 9 | 9895814 | rs7874380 | 1.257 | 1.136 | 1.275 | 0.428 | 0.071 | 0.002 | 0.002 | 0 | 0.95 | 0.78 | 0.95 | 0.99 | 0.94 | 0.98 |
| 9 | 9896971 | rs10491914 | 0.489 | 0.543 | 0.898 | 0.686 | 0.035 | -0.027 | -0.027 | 0 | 0.58 | 0.24 | 0.74 | 0.53 | 0.72 | 0.7 |
| 9 | 9898870 | rs10117170 | -0.261 | 0.472 | 1.313 | 0.906 | -0.052 | -0.066 | -0.066 | 0.981 | 0.34 | 0.47 | 0.14 | 0.15 | 0.18 | 0.22 |
| 9 | 9899274 | rs10117851 | -0.177 | 0.556 | 1.455 | 1.037 | -0.01 | -0.012 | -0.012 | 0 | 0.33 | 0.33 | 0.13 | 0.19 | 0.18 | 0.21 |
| 9 | 9904274 | rs16930522 | -1.046 | 0.603 | 1.428 | 0.979 | 0.018 | -0.025 | -0.025 | 0 | 0.06 | 0.15 | 0.07 | 0.01 | 0.08 | 0.01 |
| 9 | 9904498 | rs13284014 | 0.53 | 0.513 | 1.337 | 1.023 | 0.012 | -0.014 | -0.014 | 0 | 0.59 | 0.34 | 0.76 | 0.47 | 0.71 | 0.69 |
| 9 | 9905809 | rs11999385 | 0.265 | 0.184 | 0.533 | 1.128 | -0.003 | 0.001 | 0.001 | 0 | 0.66 | 0.68 | 0.83 | 0.44 | 0.79 | 0.7 |
| 9 | 9906970 | rs10816223 | 0.029 | 0.326 | 0.006 | 0.776 | 0.002 | 0.006 | 0.006 | 0 | 0.51 | 0.69 | 0.62 | 0.62 | 0.54 | 0.6 |
| 9 | 9907987 | rs4740998 | 0.903 | -0.083 | 0.44 | 0.483 | -0.008 | -0.03 | -0.03 | 0 | 0.93 | 0.87 | 0.98 | 1 | 0.97 | 0.97 |
| 9 | 9908296 | rs10978077 | -0.244 | -0.403 | 0.887 | 0.668 | -0.007 | 0.032 | 0.032 | 0.41 | 0.18 | 0.22 | 0.32 | 0.38 | 0.2 | 0.22 |
| 9 | 9908722 | rs10759100 | -0.703 | -0.467 | 1.556 | 0.513 | -0.008 | 0.03 | 0.03 | 0.149 | 0.22 | 0.38 | 0.35 | 0.38 | 0.24 | 0.23 |
| 9 | 9908821 | rs10978078 | 0.373 | -0.836 | 1.328 | 0.203 | 0.006 | 0.013 | 0.013 | 0 | 0.17 | 0.11 | 0.09 | 0.08 | 0.15 | 0.22 |
| 9 | 9909726 | rs10759101 | -0.309 | -0.841 | 1.447 | 0.206 | -0.005 | 0.046 | 0.046 | 0.179 | 0.21 | 0.39 | 0.34 | 0.38 | 0.24 | 0.23 |
| 9 | 9909795 | rs10978079 | 0.316 | -0.837 | 1.075 | 0.21 | -0.001 | 0.003 | 0.003 | 0.454 | 0.17 | 0.16 | 0.31 | 0.38 | 0.19 | 0.22 |
| 9 | 9910028 | rs12348066 | 1.709 | -0.958 | 1.131 | 0.288 | 0.027 | -0.026 | -0.026 | 0.006 | 0.48 | 0.24 | 0.5 | 0.43 | 0.53 | 0.41 |
| 9 | 9910993 | rs10978080 | 1.548 | -1.213 | 1.145 | 0.312 | 0.083 | -0.014 | -0.014 | 2.967 | 0.5 | 0.27 | 0.53 | 0.43 | 0.59 | 0.45 |
| 9 | 9911283 | rs16930559 | 0.46 | -1.228 | 1.144 | 0.282 | 0.007 | 0.009 | 0.009 | 0 | 0.17 | 0.12 | 0.07 | 0.08 | 0.12 | 0.21 |
| 9 | 9912338 | rs10816226 | 0.725 | -1.437 | 1.123 | 0.369 | -0.001 | -0.001 | -0.001 | 0 | 0.14 | 0.12 | 0.23 | 0.15 | 0.16 | 0.16 |
| 9 | 9912827 | rs10816229 | 0.447 | -1.422 | 0.601 | 0.402 | -0.034 | 0.013 | 0.013 | 0 | 0.74 | 0.45 | 0.64 | 0.63 | 0.8 | 0.74 |
| 9 | 9913373 | rs10978082 | 0.421 | -1.362 | 0.489 | 0.378 | -0.003 | -0.008 | -0.008 | 0 | 0.16 | 0.45 | 0.29 | 0.15 | 0.17 | 0.16 |
| 9 | 9914428 | rs10978083 | -0.623 | -1.397 | 0.325 | 0.36 | -0.041 | -0.047 | -0.047 | 0 | 0.33 | 0.49 | 0.33 | 0.22 | 0.27 | 0.38 |
| 9 | 9915186 | rs4461956 | 1.352 | -0.799 | 0.09 | 0.791 | 0.025 | -0.012 | -0.012 | 0.091 | 0.64 | 0.31 | 0.59 | 0.5 | 0.7 | 0.63 |
| 9 | 9916467 | rs7867164 | 1.933 | -0.364 | -0.201 | 1.217 | 0.013 | -0.023 | -0.023 | 0 | 0.84 | 0.92 | 0.89 | 0.67 | 0.9 | 0.81 |
| 9 | 9918634 | rs2382062 | 1.168 | -0.403 | -0.226 | 1.212 | 0.017 | -0.021 | -0.021 | 0.764 | 0.91 | 0.95 | 0.93 | 0.78 | 0.96 | 0.9 |
| 9 | 9918976 | rs4742640 | 1.267 | -0.609 | 0.347 | 1.048 | -0.052 | 0.024 | 0.024 | 0 | 0.75 | 0.57 | 0.87 | 0.71 | 0.87 | 0.79 |

| | | | | | | | | | | | | | | | | |
|---|---|---|---|---|---|---|---|---|---|---|---|---|---|---|---|---|
| 9 | 9920242 | rs10816235 | -1.839 | -0.518 | 1.093 | 1.85 | 0.026 | 0.308 | 0.308 | 0 | 0.06 | 0.31 | 0.27 | 0.29 | 0.02 | 0.15 |
| 9 | 9920345 | rs10816236 | -0.357 | -0.431 | 0.1 | 1.909 | -0.001 | 0.001 | 0.001 | 0 | 0.41 | 0.35 | 0.27 | 0.58 | 0.4 | 0.35 |
| 9 | 9929916 | rs12683380 | -0.041 | -0.254 | 0.069 | 1.92 | -0.014 | -0.092 | -0.092 | 3.567 | 0.34 | 0.12 | 0.21 | 0.43 | 0.33 | 0.31 |
| 9 | 9937753 | rs2382082 | -0.45 | -0.343 | 0.986 | 2.142 | 0.008 | 0.015 | 0.015 | 0 | 0.39 | 0.36 | 0.49 | 0.74 | 0.35 | 0.51 |
| 9 | 9939766 | rs7849423 | -0.428 | -0.612 | 1.48 | 2.042 | 0.005 | 0.007 | 0.007 | 0.012 | 0.39 | 0.62 | 0.28 | 0.61 | 0.35 | 0.36 |
| 9 | 9940329 | rs4636255 | -2.023 | -0.61 | 1.537 | 2.037 | 0.061 | 0.278 | 0.278 | 3.685 | 0.06 | 0.54 | 0.06 | 0.18 | 0.02 | 0.02 |
| 9 | 9940521 | rs7467612 | -0.388 | -0.616 | 1.704 | 2.02 | 0.012 | 0.053 | 0.053 | 0 | 0.05 | 0.06 | 0.05 | 0.18 | 0.07 | 0.04 |
| 9 | 9945265 | rs10759112 | -0.697 | -0.735 | 1.766 | 2.824 | 0 | 0.013 | 0.013 | 1.376 | 0.43 | 0.64 | 0.33 | 0.62 | 0.4 | 0.44 |
| 9 | 9954379 | rs16930664 | 0.07 | -0.642 | 1.282 | 2.582 | -0.028 | -0.101 | -0.101 | 0 | 0.34 | 0.11 | 0.23 | 0.49 | 0.33 | 0.35 |
| 9 | 9957301 | rs7860466 | 0.415 | -0.673 | 1.444 | 1.228 | -0.016 | -0.086 | -0.086 | 0 | 0.15 | 0.28 | 0.11 | 0 | 0.14 | 0.09 |
| 9 | 9960401 | rs7851920 | -1.356 | -0.897 | 1.433 | 1.511 | 0.015 | 0.253 | 0.253 | 0.401 | 0.07 | 0.22 | 0.05 | 0.37 | 0.04 | 0.04 |
| 9 | 9960537 | rs2382087 | -0.338 | -0.88 | 1.328 | 1.283 | 0 | -0.014 | -0.014 | 0 | 0.41 | 0.39 | 0.5 | 0.92 | 0.37 | 0.56 |
| 9 | 9961495 | rs10978137 | 0.884 | -0.857 | 1.19 | 1.286 | -0.011 | -0.09 | -0.09 | 0 | 0.29 | 0.1 | 0.41 | 0.54 | 0.28 | 0.43 |
| 9 | 9961522 | rs7027476 | -1.502 | -0.681 | 1.106 | 1.079 | -0.071 | 0.223 | 0.223 | 0.374 | 0.6 | 0.9 | 0.66 | 0.92 | 0.55 | 0.7 |
| 9 | 9962204 | rs7860014 | -0.73 | -0.638 | 1.135 | 1.145 | 0.035 | 0.312 | 0.312 | 0.227 | 0.06 | 0.24 | 0.06 | 0.37 | 0.04 | 0.04 |
| 9 | 9962381 | rs7470838 | 0.683 | -0.651 | 0.88 | 1.128 | -0.012 | -0.074 | -0.074 | 0 | 0.3 | 0.12 | 0.42 | 0.55 | 0.3 | 0.43 |
| 9 | 9963943 | rs10491907 | 0.212 | -0.664 | 0.961 | 0.269 | -0.001 | -0.037 | -0.037 | 0.61 | 0.18 | 0.2 | 0.13 | 0 | 0.18 | 0.15 |
| 9 | 9965824 | rs7873262 | -0.477 | -0.531 | 0.696 | 0.44 | 0.042 | -0.024 | -0.024 | 0 | 0.08 | 0.32 | 0.06 | 0 | 0.05 | 0.09 |
| 9 | 9968038 | rs10739206 | -1.16 | -0.454 | -0.272 | 0.493 | -0.085 | 0.238 | 0.238 | 0.117 | 0.6 | 0.92 | 0.66 | 0.92 | 0.53 | 0.72 |
| 9 | 9970747 | rs7870939 | 0.274 | -0.553 | -0.496 | 0.982 | 0.005 | -0.063 | -0.063 | 0.753 | 0.86 | 0.97 | 0.7 | 0.43 | 0.85 | 0.78 |
| 9 | 9972321 | rs2382089 | 0.901 | 0.013 | -0.822 | 0.336 | -0.071 | -0.005 | -0.005 | 0 | 0.25 | 0.03 | 0.18 | 0.21 | 0.28 | 0.31 |
| 9 | 9972784 | rs10115147 | -0.506 | -0.087 | -0.051 | 0.198 | 0 | -0.004 | -0.004 | 4.222 | 0.47 | 0.57 | 0.33 | 0.24 | 0.46 | 0.48 |
| 9 | 9974152 | rs7873526 | -0.252 | -0.447 | 0.293 | 0.039 | -0.032 | -0.141 | -0.141 | 1.261 | 0.22 | 0.53 | 0.17 | 0.03 | 0.19 | 0.15 |
| 9 | 9977113 | rs6477436 | 0.217 | -0.37 | 0.473 | 0.745 | -0.009 | 0.223 | 0.223 | 0 | 0.88 | 0.75 | 0.75 | 0.51 | 0.88 | 0.78 |
| 9 | 9979990 | rs9299106 | 0.93 | 0.28 | 0.505 | 0.703 | 0.013 | 0.114 | 0.114 | 0 | 0.83 | 0.67 | 0.71 | 0.37 | 0.82 | 0.68 |
| 9 | 10000548 | rs10978155 | 0.778 | 0.266 | -0.84 | -0.113 | -0.107 | -0.008 | -0.008 | 0 | 0.27 | 0.01 | 0.2 | 0.23 | 0.32 | 0.32 |
| 9 | 10000931 | rs10978156 | -0.259 | 0.226 | -0.13 | 0.113 | 0.034 | 0.068 | 0.068 | 0 | 0.06 | 0.16 | 0.07 | 0.12 | 0.07 | 0.08 |
| 9 | 10004946 | rs10816275 | 1.145 | -0.451 | -1.036 | -0.621 | 0.062 | 0.024 | 0.024 | 1.159 | 0.57 | 0.09 | 0.45 | 0.37 | 0.65 | 0.6 |
| 9 | 10008726 | rs2382104 | 0.511 | -0.585 | -0.471 | -0.708 | -0.027 | 0.222 | 0.222 | 0 | 0.82 | 0.59 | 0.63 | 0.48 | 0.86 | 0.76 |
| 9 | 10010180 | rs10809004 | 1.309 | -0.605 | -0.622 | -0.728 | 0.061 | 0.039 | 0.039 | 1.123 | 0.57 | 0.09 | 0.45 | 0.32 | 0.65 | 0.6 |
| 9 | 10018121 | rs4333667 | 0.488 | -0.392 | -0.708 | -0.796 | -0.021 | 0.047 | 0.047 | 0 | 0.08 | 0.01 | 0.04 | 0 | 0.09 | 0.03 |
| 9 | 10023441 | rs10809012 | 0.697 | -0.321 | -0.543 | -0.762 | 0.006 | 0.158 | 0.158 | 0 | 0.79 | 0.58 | 0.59 | 0.49 | 0.77 | 0.74 |
| 9 | 10025975 | rs727756 | 0.872 | -0.028 | -1.261 | -0.687 | 0.002 | 0.033 | 0.033 | 0.203 | 0.54 | 0.09 | 0.41 | 0.37 | 0.56 | 0.59 |
| 9 | 10030499 | rs1441400 | 0.373 | -0.032 | -0.552 | -0.27 | -0.11 | -0.012 | -0.012 | 0 | 0.26 | 0.01 | 0.19 | 0.2 | 0.32 | 0.32 |
| 9 | 10044092 | rs291267 | -0.498 | -0.065 | 0.804 | -0.202 | 0.011 | 0.005 | 0.005 | 0 | 0.31 | 0.51 | 0.33 | 0.35 | 0.33 | 0.34 |
| 9 | 10044783 | rs7868769 | -0.479 | -0.111 | 0.645 | -0.191 | -0.001 | -0.001 | -0.001 | 0 | 0.31 | 0.33 | 0.23 | 0.35 | 0.32 | 0.33 |
| 9 | 10047844 | rs2136530 | 0.174 | -0.1 | 0.76 | -0.077 | -0.014 | 0.119 | 0.119 | 0 | 0.15 | 0.27 | 0.28 | 0.51 | 0.14 | 0.22 |
| 9 | 10050872 | rs1995845 | -0.077 | -0.38 | 0.7 | -0.228 | -0.009 | 0.039 | 0.039 | 0 | 0.33 | 0.16 | 0.23 | 0.25 | 0.33 | 0.31 |
| 9 | 10054740 | rs442100 | -1.716 | -0.197 | 0.551 | -0.218 | 0.003 | 0.095 | 0.095 | 0 | 0.55 | 0.75 | 0.7 | 0.86 | 0.57 | 0.6 |
| 9 | 10054994 | rs447910 | -1.17 | 0.393 | 0.92 | 0.009 | -0.011 | 0.016 | 0.016 | 0 | 0.43 | 0.61 | 0.43 | 0.38 | 0.42 | 0.38 |
| 9 | 10056666 | rs6474531 | -1.173 | 0.71 | 0.638 | 0.149 | 0.054 | 0.05 | 0.05 | 0 | 0.09 | 0.28 | 0.18 | 0.12 | 0.1 | 0.12 |
| 9 | 10057995 | rs428721 | -1.135 | 0.162 | -0.174 | -0.023 | -0.009 | 0.02 | 0.02 | 0 | 0.39 | 0.81 | 0.29 | 0.34 | 0.35 | 0.42 |
| 9 | 10061106 | rs399780 | -1.291 | 0.103 | -0.122 | 0.046 | -0.021 | 0.016 | 0.016 | 0 | 0.4 | 0.81 | 0.35 | 0.37 | 0.35 | 0.43 |
| 9 | 10061712 | rs417403 | -1.175 | -0.004 | 0.899 | 0.161 | -0.035 | 0.079 | 0.079 | 0.169 | 0.12 | 0.54 | 0.09 | 0.16 | 0.04 | 0.1 |
| 9 | 10063148 | rs368297 | -1.44 | -0.021 | 0.632 | 0.153 | -0.028 | 0.012 | 0.012 | 1.248 | 0.13 | 0.72 | 0.11 | 0.16 | 0.04 | 0.12 |
| 9 | 10064950 | rs565315 | -1.732 | -0.039 | 0.485 | 0.124 | 0.009 | -0.008 | -0.008 | 0 | 0.16 | 0.79 | 0.21 | 0.23 | 0.08 | 0.14 |
| 9 | 10077180 | rs7023147 | -0.699 | 0.113 | 1.176 | 0.247 | -0.001 | 0.015 | 0.015 | 0 | 0.29 | 0.52 | 0.19 | 0.36 | 0.28 | 0.34 |
| 9 | 10077962 | rs10511531 | -0.122 | 0.352 | 0.606 | 0.078 | -0.015 | 0.028 | 0.028 | 0 | 0.24 | 0.13 | 0.16 | 0.15 | 0.28 | 0.32 |
| 9 | 10087429 | rs10809027 | -0.397 | 0.201 | 0.633 | 0.049 | -0.002 | 0.011 | 0.011 | 0 | 0.24 | 0.18 | 0.16 | 0.15 | 0.23 | 0.37 |
| 9 | 10101169 | rs403114 | -1.512 | 0.079 | 0.59 | -0.04 | 0.003 | 0.038 | 0.038 | 0 | 0.55 | 0.71 | 0.67 | 0.7 | 0.56 | 0.65 |
| 9 | 10101415 | rs440834 | -0.405 | -0.441 | 1.133 | 0.19 | -0.009 | 0.068 | 0.068 | 0 | 0.28 | 0.47 | 0.4 | 0.55 | 0.29 | 0.24 |
| 9 | 10101925 | rs450485 | -0.78 | -0.444 | 1.152 | 0.183 | 0.002 | 0.061 | 0.061 | 0 | 0.31 | 0.55 | 0.52 | 0.55 | 0.32 | 0.28 |
| 9 | 10112766 | rs2890901 | 0.589 | -0.331 | 0.959 | 0.092 | 0.012 | 0.001 | 0.001 | 0 | 0.15 | 0.08 | 0.07 | 0.14 | 0.11 | 0.29 |
| 9 | 10113900 | rs291296 | 1.66 | -0.296 | 1.004 | 0.116 | 0.017 | 0.08 | 0.08 | 0 | 0.5 | 0.28 | 0.38 | 0.29 | 0.54 | 0.39 |
| 9 | 10114943 | rs725261 | -0.281 | -0.275 | 1.033 | 0.121 | -0.002 | -0.004 | -0.004 | 0 | 0.19 | 0.24 | 0.2 | 0.15 | 0.15 | 0.34 |
| 9 | 10119157 | rs10118866 | -1.267 | -0.465 | 1.205 | 0.532 | -0.004 | -0.002 | -0.002 | 0 | 0.32 | 0.32 | 0.41 | 0.3 | 0.22 | 0.46 |
| 9 | 10124920 | rs294868 | -0.306 | -0.624 | 1.331 | 0.687 | 0.033 | 0.147 | 0.147 | 0 | 0.84 | 0.58 | 0.75 | 0.6 | 0.77 | 0.88 |
| 9 | 10127876 | rs7043462 | 0.192 | -0.673 | 1.231 | 0.643 | 0.004 | 0.002 | 0.002 | 0 | 0.17 | 0.15 | 0.2 | 0.15 | 0.12 | 0.29 |
| 9 | 10128645 | rs150409 | 1.897 | -0.603 | 1.1 | 0.527 | 0.014 | 0.088 | 0.088 | 0 | 0.48 | 0.21 | 0.32 | 0.3 | 0.53 | 0.35 |
| 9 | 10128701 | rs10958899 | 0.034 | -0.6 | 1.038 | 0.576 | -0.02 | -0.01 | -0.01 | 0.018 | 0.18 | 0.28 | 0.22 | 0.15 | 0.12 | 0.29 |
| 9 | 10129732 | rs10958904 | 0.519 | -0.539 | 0.755 | 0.607 | 0.011 | 0.003 | 0.003 | 0 | 0.16 | 0.1 | 0.19 | 0.15 | 0.12 | 0.29 |
| 9 | 10130219 | rs291304 | -0.352 | -0.535 | 1.065 | 0.629 | 0.001 | 0.022 | 0.022 | 0 | 0.84 | 0.8 | 0.76 | 0.6 | 0.77 | 0.88 |
| 9 | 10130546 | rs10119277 | 0.65 | -0.548 | 0.84 | 0.562 | 0.012 | -0.007 | -0.007 | 0 | 0.12 | 0.06 | 0.1 | 0.15 | 0.07 | 0.13 |
| 9 | 10132437 | rs10958908 | 0.5 | -0.546 | 0.827 | 0.563 | -0.008 | 0.002 | 0.002 | 0 | 0.13 | 0.15 | 0.11 | 0.15 | 0.07 | 0.12 |
| 9 | 10139153 | rs11525900 | -0.828 | -0.517 | 0.853 | 0.607 | 0.009 | -0.001 | -0.001 | 0.057 | 0.29 | 0.28 | 0.3 | 0.3 | 0.19 | 0.44 |
| 9 | 10139580 | rs294845 | -0.136 | -0.353 | 1.09 | 0.74 | 0.027 | 0.133 | 0.133 | 0 | 0.83 | 0.61 | 0.81 | 0.61 | 0.78 | 0.88 |
| 9 | 10145099 | rs9721595 | -0.456 | -0.211 | 0.982 | 0.792 | 0.027 | 0.107 | 0.107 | 0 | 0.06 | 0.18 | 0.21 | 0.17 | 0.08 | 0.05 |
| 9 | 10145186 | rs16931167 | -0.309 | -0.445 | 0.743 | 0.52 | 0.042 | 0.053 | 0.053 | 0.024 | 0.26 | 0.11 | 0.11 | 0.16 | 0.15 | 0.38 |
| 9 | 10145460 | rs291288 | -0.092 | -0.233 | 0.865 | 0.709 | 0.004 | 0.018 | 0.018 | 0 | 0.82 | 0.77 | 0.74 | 0.65 | 0.75 | 0.89 |

| | | | | | | | | | | | | | | | | |
|---|---|---|---|---|---|---|---|---|---|---|---|---|---|---|---|---|
| 9 | 10146521 | rs16931176 | -0.619 | -0.56 | 1.017 | 0.853 | 0.023 | 0.034 | 0.034 | 0 | 0.06 | 0.18 | 0.19 | 0.1 | 0.08 | 0.05 |
| 9 | 10148779 | rs291289 | 1.939 | -1.152 | 0.758 | 0.757 | 0.072 | -0.01 | -0.01 | 0 | 0.52 | 0.26 | 0.53 | 0.5 | 0.68 | 0.41 |
| 9 | 10158866 | rs13297654 | -1.067 | -1.21 | 0.698 | 0.742 | -0.001 | -0.001 | -0.001 | 0 | 0.16 | 0.17 | 0.32 | 0.16 | 0.16 | 0.23 |
| 9 | 10162452 | rs1304641 | 0.433 | -1.262 | 0.514 | 0.767 | -0.02 | -0.015 | -0.015 | 0 | 0.16 | 0.01 | 0.06 | 0.08 | 0.12 | 0.22 |
| 9 | 10167327 | rs7032659 | -1.081 | -1.336 | 1.025 | 0.758 | -0.015 | 0.012 | 0.012 | 0 | 0.3 | 0.25 | 0.23 | 0.4 | 0.18 | 0.36 |
| 9 | 10171341 | rs10809034 | -2.114 | -1.297 | 0.681 | 0.704 | 0.052 | -0.009 | -0.009 | 0 | 0.46 | 0.69 | 0.47 | 0.46 | 0.3 | 0.57 |
| 9 | 10173689 | rs10119783 | -1.361 | -1.008 | 0.616 | 0.569 | 0.001 | -0.005 | -0.005 | 0 | 0.3 | 0.35 | 0.34 | 0.2 | 0.29 | 0.4 |
| 9 | 10175819 | rs16931225 | -1.171 | -1 | 0.492 | 0.597 | 0 | -0.001 | -0.001 | 0 | 0.29 | 0.33 | 0.33 | 0.22 | 0.29 | 0.39 |
| 9 | 10175938 | rs17620334 | -0.881 | -0.999 | 0.441 | 0.598 | 0 | -0.004 | -0.004 | 0 | 0.24 | 0.2 | 0.19 | 0.22 | 0.23 | 0.3 |
| 9 | 10178560 | rs10117980 | -0.753 | -1.054 | 0.565 | 0.734 | 0.001 | -0.001 | -0.001 | 0.095 | 0.26 | 0.33 | 0.26 | 0.23 | 0.25 | 0.39 |
| 9 | 10178870 | rs10958936 | -1.697 | -1.129 | 0.578 | 0.764 | 0.007 | -0.001 | -0.001 | 0 | 0.39 | 0.59 | 0.45 | 0.36 | 0.24 | 0.5 |
| 9 | 10184746 | rs294848 | -0.993 | -0.611 | 0.097 | 0.582 | -0.001 | -0.002 | -0.002 | 0 | 0.29 | 0.34 | 0.3 | 0.2 | 0.26 | 0.36 |
| 9 | 10185596 | rs294852 | 0.798 | -0.633 | -0.587 | 0.641 | 0.017 | -0.018 | -0.018 | 0 | 0.68 | 0.31 | 0.58 | 0.55 | 0.76 | 0.7 |
| 9 | 10187598 | rs1322163 | 0.04 | -0.233 | -0.461 | 0.396 | -0.009 | 0.007 | 0.007 | 0 | 0.11 | 0.08 | 0.06 | 0.08 | 0.16 | 0.13 |
| 9 | 10192290 | rs294856 | -1.657 | -0.719 | -1.283 | 0.521 | -0.058 | -0.023 | -0.023 | 0 | 0.55 | 0.87 | 0.39 | 0.57 | 0.44 | 0.52 |
| 9 | 10192858 | rs294857 | -0.681 | -0.974 | -0.655 | 0.374 | 0.001 | 0.009 | 0.009 | 0 | 0.27 | 0.31 | 0.29 | 0.55 | 0.28 | 0.41 |
| 9 | 10196809 | rs294830 | -1.558 | -1.115 | -0.478 | 0.293 | 0.002 | 0.017 | 0.017 | 0 | 0.35 | 0.47 | 0.34 | 0.49 | 0.33 | 0.46 |
| 9 | 10197122 | rs294829 | -0.112 | -1.204 | -0.379 | 0.205 | -0.051 | -0.13 | -0.13 | 1.694 | 0.79 | 0.58 | 0.9 | 0.97 | 0.89 | 0.94 |
| 9 | 10202765 | rs294814 | -1.461 | -1.049 | -0.255 | 0.332 | 0.019 | 0.004 | 0.004 | 0.379 | 0.43 | 0.6 | 0.32 | 0.44 | 0.29 | 0.46 |
| 9 | 10204129 | rs410601 | 0.342 | -1.058 | -0.511 | 0.487 | -0.002 | 0.047 | 0.047 | 0 | 0.15 | 0.21 | 0.14 | 0.41 | 0.13 | 0.34 |
| 9 | 10205835 | rs1322133 | -1.233 | -1.224 | -0.593 | 0.142 | 0.035 | 0.121 | 0.121 | 0.442 | 0.43 | 0.76 | 0.22 | 0.11 | 0.32 | 0.27 |
| 9 | 10211759 | rs2498596 | -0.348 | -0.867 | 0.287 | 0.905 | 0.004 | 0.15 | 0.15 | 0 | 0.17 | 0.44 | 0.17 | 0.43 | 0.15 | 0.35 |
| 9 | 10212865 | rs911692 | -0.763 | -0.708 | 0.242 | 0.729 | 0.002 | -0.004 | -0.004 | 0.008 | 0.9 | 0.9 | 0.93 | 0.93 | 0.86 | 0.9 |
| 9 | 10214007 | rs2498598 | 0.142 | -0.71 | 0.435 | 0.804 | 0.023 | -0.004 | -0.004 | 0 | 0.08 | 0.13 | 0.13 | 0.07 | 0.13 | 0.15 |
| 9 | 10215130 | rs2475324 | -1.201 | -0.8 | 0.514 | 0.845 | 0.034 | 0.082 | 0.082 | 0 | 0.27 | 0.55 | 0.25 | 0.48 | 0.28 | 0.44 |
| 9 | 10225500 | rs7846730 | 1.74 | -1.22 | -0.47 | -0.039 | -0.064 | -0.059 | -0.059 | 0 | 0.37 | 0.13 | 0.5 | 0.43 | 0.47 | 0.4 |
| 9 | 10225533 | rs1885431 | 1.458 | -1.447 | -0.523 | -0.219 | 0.011 | 0.026 | 0.026 | 0.072 | 0.5 | 0.21 | 0.6 | 0.44 | 0.59 | 0.49 |
| 9 | 10227095 | rs16925550 | 3.491 | -1.525 | -0.388 | -0.139 | -0.007 | 0.059 | 0.059 | 0.567 | 0.06 | 0.07 | 0.02 | 0.33 | 0.02 | 0.11 |
| 9 | 10228203 | rs4143460 | 0.854 | -1.565 | -0.281 | -0.269 | 0.016 | -0.003 | -0.003 | 0 | 0.61 | 0.34 | 0.71 | 0.51 | 0.72 | 0.6 |
| 9 | 10229902 | rs10958962 | 0.215 | -1.53 | -0.514 | -0.302 | -0.012 | 0.063 | 0.063 | 0 | 0.24 | 0.15 | 0.17 | 0.08 | 0.25 | 0.19 |
| 9 | 10230744 | rs1322137 | 0.879 | -1.423 | -0.469 | -0.326 | 0.028 | 0.017 | 0.017 | 0.521 | 0.55 | 0.34 | 0.6 | 0.54 | 0.63 | 0.59 |
| 9 | 10233188 | rs7871439 | 0.272 | -1.641 | -0.376 | -0.221 | -0.025 | 0.04 | 0.04 | 1.35 | 0.15 | 0.22 | 0.07 | 0.38 | 0.07 | 0.19 |
| 9 | 10233361 | rs1322142 | 2.027 | -1.523 | -0.542 | -0.223 | -0.072 | -0.069 | -0.069 | 0 | 0.36 | 0.13 | 0.54 | 0.43 | 0.48 | 0.37 |
| 9 | 10237682 | rs10738156 | 2.007 | -1.829 | -0.572 | -0.211 | -0.083 | -0.085 | -0.085 | 0 | 0.35 | 0.15 | 0.36 | 0.41 | 0.48 | 0.34 |
| 9 | 10238391 | rs12353282 | 2.093 | -1.84 | -0.633 | -0.22 | -0.13 | -0.128 | -0.128 | 0.043 | 0.34 | 0.07 | 0.36 | 0.41 | 0.47 | 0.31 |
| 9 | 10238994 | rs10217306 | -0.152 | -2.008 | -0.106 | -0.357 | 0 | -0.085 | -0.085 | 0 | 0.76 | 0.63 | 0.81 | 0.89 | 0.79 | 0.78 |
| 9 | 10239309 | rs2382198 | 1.532 | -2.061 | -0.286 | -0.369 | 0.043 | 0.001 | 0.001 | 2.033 | 0.46 | 0.21 | 0.44 | 0.49 | 0.61 | 0.44 |
| 9 | 10240296 | rs2145990 | 1.646 | -2.03 | -0.254 | -0.421 | 0.018 | 0.033 | 0.033 | 0.226 | 0.14 | 0.07 | 0.11 | 0.04 | 0.11 | 0.09 |
| 9 | 10240428 | rs1570267 | -0.011 | -1.973 | -0.19 | -0.387 | -0.005 | -0.095 | -0.095 | 0 | 0.75 | 0.58 | 0.81 | 0.88 | 0.79 | 0.78 |
| 9 | 10246635 | rs9298652 | 1.277 | -2.119 | -0.638 | -0.353 | 0.053 | 0.083 | 0.083 | 0.229 | 0.52 | 0.26 | 0.56 | 0.71 | 0.61 | 0.62 |
| 9 | 10251603 | rs2498608 | 0.857 | -1.907 | -0.384 | -0.132 | 0.003 | -0.025 | -0.025 | 0 | 0.13 | 0.14 | 0.13 | 0.05 | 0.15 | 0.1 |
| 9 | 10252022 | rs12236941 | 1.469 | -2.011 | -0.647 | -0.092 | 0.084 | -0.009 | -0.009 | 0 | 0.48 | 0.08 | 0.49 | 0.5 | 0.65 | 0.5 |
| 9 | 10254449 | rs4741024 | 1.044 | -1.946 | -0.502 | -0.074 | 0.048 | 0.068 | 0.068 | 0 | 0.57 | 0.26 | 0.54 | 0.74 | 0.68 | 0.64 |
| 9 | 10257609 | rs16925616 | 1.983 | -1.79 | -0.473 | 0.035 | 0.006 | -0.03 | -0.03 | 0.157 | 0.1 | 0.03 | 0.06 | 0.28 | 0.09 | 0.26 |
| 9 | 10258144 | rs13294419 | 0.457 | -1.61 | -0.359 | -0.005 | -0.002 | 0.003 | 0.003 | 0 | 0.15 | 0.15 | 0.09 | 0.11 | 0.18 | 0.11 |
| 9 | 10260263 | rs2475335 | 0.177 | -1.745 | -0.154 | -0.015 | -0.051 | -0.023 | -0.023 | 0 | 0.7 | 0.49 | 0.67 | 0.8 | 0.81 | 0.71 |
| 9 | 10262484 | rs2475339 | -0.358 | -1.822 | -0.023 | -0.074 | -0.035 | -0.082 | -0.082 | 0 | 0.74 | 0.64 | 0.82 | 0.93 | 0.82 | 0.8 |
| 9 | 10263407 | rs2498627 | 0.827 | -1.791 | -0.182 | -0.071 | 0.012 | -0.039 | -0.039 | 0 | 0.13 | 0.21 | 0.13 | 0.06 | 0.15 | 0.09 |
| 9 | 10265282 | rs13285525 | 1.177 | -1.792 | -0.24 | -0.06 | -0.047 | -0.003 | -0.003 | 0.002 | 0.12 | 0.02 | 0.07 | 0.1 | 0.15 | 0.11 |
| 9 | 10266608 | rs2475348 | 0.809 | -1.722 | -0.155 | -0.017 | 0.012 | -0.043 | -0.043 | 0.05 | 0.13 | 0.21 | 0.13 | 0.06 | 0.15 | 0.09 |
| 9 | 10268794 | rs2475354 | 0.12 | -1.743 | -0.257 | 0.099 | -0.002 | 0 | 0 | 0 | 0.19 | 0.25 | 0.16 | 0.27 | 0.17 | 0.23 |
| 9 | 10270339 | rs2475357 | 0.883 | -1.545 | -0.827 | 0.271 | 0.067 | 0.031 | 0.031 | 0.101 | 0.52 | 0.27 | 0.49 | 0.53 | 0.66 | 0.52 |
| 9 | 10270636 | rs1322155 | 0.974 | -1.532 | -0.821 | 0.316 | 0.072 | 0.015 | 0.015 | 0.231 | 0.52 | 0.26 | 0.47 | 0.49 | 0.65 | 0.49 |
| 9 | 10273068 | rs1323505 | 0.441 | -1.402 | -0.778 | 0.237 | -0.002 | 0 | 0 | 0 | 0.14 | 0.14 | 0.09 | 0.12 | 0.15 | 0.13 |
| 9 | 10273124 | rs1323504 | 0.315 | -1.401 | -0.764 | 0.181 | 0.002 | -0.005 | -0.005 | 0 | 0.14 | 0.2 | 0.11 | 0.13 | 0.15 | 0.17 |
| 9 | 10273253 | rs7861410 | 1.603 | -1.495 | -0.819 | 0.373 | -0.053 | -0.002 | -0.002 | 0.232 | 0.42 | 0.15 | 0.4 | 0.55 | 0.49 | 0.49 |
| 9 | 10274080 | rs7865468 | 0.644 | -1.465 | -0.639 | 0.406 | 0.017 | 0.017 | 0.017 | 0 | 0.58 | 0.37 | 0.52 | 0.68 | 0.68 | 0.65 |
| 9 | 10274655 | rs12002909 | 0.352 | -1.133 | -0.671 | -0.095 | 0.004 | -0.011 | -0.011 | 0 | 0.13 | 0.21 | 0.09 | 0.1 | 0.15 | 0.11 |
| 9 | 10276100 | rs1359095 | 1.52 | -1.055 | -0.66 | -0.02 | -0.029 | 0.013 | 0.013 | 0 | 0.41 | 0.25 | 0.58 | 0.5 | 0.52 | 0.47 |
| 9 | 10277054 | rs10959002 | 0.764 | -1.127 | -0.85 | 0.193 | 0.041 | 0.065 | 0.065 | 0.938 | 0.12 | 0.03 | 0.08 | 0.01 | 0.06 | 0.05 |
| 9 | 10277319 | rs10959007 | -0.829 | -1.079 | -0.761 | 0.221 | 0.018 | 0.024 | 0.024 | 0.668 | 0.33 | 0.28 | 0.21 | 0.31 | 0.27 | 0.3 |
| 9 | 10277602 | rs16925679 | -0.335 | -0.199 | -0.147 | 0.806 | 0 | -0.001 | -0.001 | 0 | 0.21 | 0.22 | 0.12 | 0.31 | 0.21 | 0.25 |
| 9 | 10278722 | rs10738159 | 1.18 | 0.11 | 0.439 | 1.255 | -0.009 | 0.026 | 0.026 | 0.234 | 0.73 | 0.42 | 0.7 | 0.63 | 0.77 | 0.59 |
| 9 | 10282750 | rs10959013 | -1.511 | -0.247 | 0.793 | 1.519 | -0.008 | 0.045 | 0.045 | 0 | 0.05 | 0.08 | 0.1 | 0.14 | 0.03 | 0.2 |
| 9 | 10284628 | rs10959018 | -2.252 | -0.296 | 1.186 | 1.545 | -0.018 | 0.093 | 0.093 | 0 | 0.07 | 0.19 | 0.1 | 0.18 | 0.03 | 0.2 |
| 9 | 10285562 | rs7875861 | -0.855 | -0.427 | 1.511 | 1.501 | -0.013 | 0.016 | 0.016 | 0 | 0.07 | 0.08 | 0.06 | 0.16 | 0.04 | 0.15 |
| 9 | 10286986 | rs12344926 | -2.855 | -0.47 | 1.855 | 1.53 | 0.001 | -0.002 | -0.002 | 0 | 0.3 | 0.66 | 0.48 | 0.39 | 0.2 | 0.56 |
| 9 | 10287901 | rs1323499 | -2.094 | -0.485 | 1.961 | 1.356 | -0.033 | 0.017 | 0.017 | 0 | 0.23 | 0.43 | 0.37 | 0.25 | 0.17 | 0.36 |
| 9 | 10298444 | rs10809062 | -1.937 | -0.465 | 2.196 | 1.46 | 0.009 | 0.062 | 0.062 | 1.167 | 0.14 | 0.38 | 0.16 | 0.15 | 0.14 | 0.25 |

| | | | | | | | | | | | | | | | | |
|---|---|---|---|---|---|---|---|---|---|---|---|---|---|---|---|---|
| 9 | 10298970 | rs10809063 | -0.761 | -0.35 | 2.097 | 1.333 | 0.008 | -0.009 | -0.009 | 0.015 | 0.09 | 0.16 | 0.06 | 0.04 | 0.11 | 0.06 |
| 9 | 10299084 | rs661198 | -1.545 | -0.393 | 2.288 | 1.385 | -0.023 | 0.01 | 0.01 | 0 | 0.11 | 0.15 | 0.08 | 0.17 | 0.07 | 0.19 |
| 9 | 10309001 | rs12238349 | -0.318 | -0.389 | 2.213 | 1.383 | 0.005 | -0.001 | -0.001 | 0.112 | 0.06 | 0.05 | 0.06 | 0.11 | 0.04 | 0.12 |
| 9 | 10309897 | rs589991 | -2.224 | -0.375 | 2.322 | 1.345 | -0.017 | -0.002 | -0.002 | 0 | 0.2 | 0.34 | 0.17 | 0.21 | 0.18 | 0.27 |
| 9 | 10311000 | rs605318 | -3.04 | -0.43 | 2.309 | 1.372 | -0.041 | 0.011 | 0.011 | 0.19 | 0.25 | 0.51 | 0.39 | 0.25 | 0.18 | 0.35 |
| 9 | 10315193 | rs16925800 | -0.28 | -0.422 | 2.253 | 1.371 | 0.005 | 0.002 | 0.002 | 0 | 0.06 | 0.04 | 0.06 | 0.07 | 0.05 | 0.12 |
| 9 | 10315723 | rs635580 | -2.222 | -0.331 | 2.258 | 1.393 | -0.011 | 0.011 | 0.011 | 0 | 0.19 | 0.35 | 0.17 | 0.21 | 0.17 | 0.27 |
| 9 | 10317212 | rs603924 | -2.423 | -0.35 | 2.279 | 1.413 | -0.025 | 0.023 | 0.023 | 0 | 0.23 | 0.36 | 0.37 | 0.26 | 0.19 | 0.33 |
| 9 | 10318490 | rs587791 | -2.198 | -0.358 | 2.23 | 1.321 | -0.034 | -0.008 | -0.008 | 0 | 0.21 | 0.54 | 0.39 | 0.17 | 0.16 | 0.27 |
| 9 | 10330850 | rs10959051 | -2.496 | -0.127 | 1.435 | 1.324 | -0.007 | 0.028 | 0.028 | 0.296 | 0.27 | 0.57 | 0.43 | 0.33 | 0.25 | 0.36 |
| 9 | 10333382 | rs10959053 | -0.257 | -0.082 | 1.341 | 1.318 | -0.06 | -0.043 | -0.043 | 0.353 | 0.1 | 0.01 | 0.1 | 0.11 | 0.17 | 0.07 |
| 9 | 10335382 | rs1590347 | -2.289 | -0.211 | 1.476 | 1.261 | -0.068 | 0.053 | 0.053 | 0.144 | 0.17 | 0.47 | 0.32 | 0.2 | 0.07 | 0.28 |
| 9 | 10341531 | rs10959055 | -0.38 | -0.189 | 1.192 | 1.247 | -0.002 | 0.005 | 0.005 | 2.62 | 0.11 | 0.11 | 0.07 | 0.05 | 0.13 | 0.07 |
| 9 | 10341540 | rs10809071 | -2.307 | -0.103 | 1.072 | 1.003 | -0.005 | -0.013 | -0.013 | 0.416 | 0.33 | 0.62 | 0.49 | 0.2 | 0.28 | 0.36 |
| 9 | 10342442 | rs7871892 | -2.674 | -0.133 | 0.82 | 0.999 | 0.016 | 0.019 | 0.019 | 0.101 | 0.38 | 0.67 | 0.51 | 0.29 | 0.31 | 0.45 |
| 9 | 10342841 | rs12552756 | -0.938 | 0.009 | 0.536 | 0.944 | 0.003 | 0.001 | 0.001 | 1.598 | 0.14 | 0.12 | 0.28 | 0.1 | 0.13 | 0.13 |
| 9 | 10346862 | rs10959061 | -1.424 | 0.098 | 0.568 | 0.999 | -0.018 | 0 | 0 | 0.624 | 0.25 | 0.2 | 0.42 | 0.2 | 0.32 | 0.24 |
| 9 | 10348659 | rs7852334 | -2.338 | 0.162 | -0.259 | 0.939 | -0.065 | -0.084 | -0.084 | 0 | 0.56 | 0.92 | 0.71 | 0.51 | 0.53 | 0.75 |
| 9 | 10352639 | rs10809075 | -1.327 | -0.567 | 0.176 | 0.395 | -0.038 | -0.042 | -0.042 | 0 | 0.25 | 0.53 | 0.24 | 0.19 | 0.17 | 0.32 |
| 9 | 10354430 | rs17703834 | -0.02 | -0.554 | -0.002 | 0.396 | -0.026 | 0.022 | 0.022 | 0 | 0.14 | 0.03 | 0.16 | 0.09 | 0.14 | 0.21 |
| 9 | 10354770 | rs16925863 | -0.785 | -0.52 | 0.346 | 0.46 | -0.042 | 0.025 | 0.025 | 0.341 | 0.1 | 0.4 | 0.07 | 0.09 | 0.03 | 0.14 |
| 9 | 10355941 | rs10125156 | -0.257 | 0.182 | -0.331 | 0.438 | 0.002 | 0.001 | 0.001 | 0 | 0.2 | 0.23 | 0.21 | 0.26 | 0.21 | 0.3 |
| 9 | 10357348 | rs9298656 | -0.174 | 0.271 | -0.44 | 0.489 | 0.002 | 0.003 | 0.003 | 0 | 0.21 | 0.27 | 0.22 | 0.27 | 0.21 | 0.31 |
| 9 | 10358672 | rs10756029 | -0.933 | 0.209 | -1.161 | 0.652 | -0.023 | 0.082 | 0.082 | 3.879 | 0.51 | 0.85 | 0.5 | 0.35 | 0.48 | 0.59 |
| 9 | 10362221 | rs10959075 | 1.016 | -0.218 | -0.588 | 0.751 | 0.006 | 0.021 | 0.021 | 0 | 0.65 | 0.74 | 0.49 | 0.27 | 0.68 | 0.55 |
| 9 | 10362315 | rs10959076 | 1.016 | -0.218 | -0.523 | 0.733 | 0.006 | 0.023 | 0.023 | 0 | 0.65 | 0.72 | 0.49 | 0.26 | 0.68 | 0.55 |
| 9 | 10364655 | rs7033338 | -0.207 | -0.197 | -0.284 | 0.633 | 0 | -0.004 | -0.004 | 0.515 | 0.05 | 0.09 | 0.03 | 0.06 | 0.04 | 0.08 |
| 9 | 10365636 | rs16925876 | -0.726 | -0.407 | 0.607 | 0.455 | 0.006 | 0.044 | 0.044 | 0 | 0.09 | 0.28 | 0.1 | 0.1 | 0.06 | 0.19 |
| 9 | 10366812 | rs12551331 | -0.14 | -0.694 | -0.052 | 1.27 | -0.004 | -0.122 | -0.122 | 0.57 | 0.16 | 0.01 | 0.28 | 0.47 | 0.1 | 0.14 |
| 9 | 10367106 | rs1359091 | 2.137 | -0.974 | -0.265 | 1.249 | -0.012 | 0.152 | 0.152 | 0.372 | 0.49 | 0.04 | 0.37 | 0.2 | 0.61 | 0.32 |
| 9 | 10368596 | rs7034193 | 1.049 | -0.985 | -0.008 | 1.238 | 0.026 | -0.056 | -0.056 | 0 | 0.75 | 0.86 | 0.61 | 0.45 | 0.84 | 0.78 |
| 9 | 10369161 | rs10738162 | -0.113 | -0.885 | -0.084 | 1.239 | -0.01 | -0.117 | -0.117 | 0.087 | 0.15 | 0.01 | 0.31 | 0.55 | 0.11 | 0.14 |
| 9 | 10369718 | rs4595175 | 1.239 | -0.98 | 1.138 | 0.985 | -0.027 | -0.052 | -0.052 | 0 | 0.73 | 0.46 | 0.77 | 0.82 | 0.76 | 0.61 |
| 9 | 10371281 | rs1535658 | 0.928 | -0.88 | 1.37 | 0.99 | -0.007 | -0.027 | -0.027 | 0 | 0.73 | 0.6 | 0.76 | 0.83 | 0.74 | 0.68 |
| 9 | 10372488 | rs16925900 | -1.29 | -1.517 | 1.46 | 1.469 | -0.004 | 0.014 | 0.014 | 0 | 0.09 | 0.09 | 0.08 | 0.14 | 0.06 | 0.19 |
| 9 | 10373600 | rs10959080 | -0.76 | -1.641 | 1.338 | 1.406 | -0.006 | -0.071 | -0.071 | 0 | 0.13 | 0.04 | 0.34 | 0.64 | 0.1 | 0.26 |
| 9 | 10374386 | rs1570524 | 0.909 | -1.685 | 1.933 | 1.324 | -0.064 | -0.064 | -0.064 | 1.406 | 0.78 | 0.48 | 0.79 | 0.89 | 0.86 | 0.78 |
| 9 | 10374705 | rs6474539 | 2.169 | -1.735 | 2.116 | 1.463 | 0.048 | 0.038 | 0.038 | 0 | 0.61 | 0.26 | 0.46 | 0.28 | 0.72 | 0.42 |
| 9 | 10381819 | rs10959084 | -1.149 | -1.677 | 2.212 | 1.536 | -0.004 | 0.044 | 0.044 | 0 | 0.16 | 0.21 | 0.3 | 0.63 | 0.12 | 0.23 |
| 9 | 10387436 | rs10959087 | 0.138 | -1.669 | 2.233 | 1.54 | -0.014 | -0.035 | -0.035 | 0 | 0.09 | 0.01 | 0.25 | 0.47 | 0.08 | 0.14 |
| 9 | 10390416 | rs10959088 | 1.929 | -1.785 | 2.344 | 0.909 | -0.001 | 0 | 0 | 0 | 0.63 | 0.38 | 0.47 | 0.23 | 0.76 | 0.46 |
| 9 | 10391459 | rs10809084 | 0.616 | -1.697 | 2.136 | 0.897 | 0.006 | 0.006 | 0.006 | 0 | 0.81 | 0.85 | 0.84 | 0.88 | 0.86 | 0.78 |
| 9 | 10391604 | rs10756031 | 0.503 | -1.695 | 2.006 | 0.898 | 0.03 | 0.03 | 0.03 | 0 | 0.82 | 0.93 | 0.85 | 0.88 | 0.86 | 0.78 |
| 9 | 10395912 | rs10959090 | 2.244 | -1.621 | 2.007 | 0.883 | 0.041 | 0.054 | 0.054 | 0 | 0.57 | 0.32 | 0.44 | 0.18 | 0.71 | 0.38 |
| 9 | 10404336 | rs2382208 | 0.882 | -1.404 | 1.974 | 1.047 | -0.031 | -0.033 | -0.033 | 0 | 0.8 | 0.72 | 0.83 | 0.88 | 0.86 | 0.78 |
| 9 | 10409555 | rs7855950 | 0.829 | -1.404 | 1.824 | 1.09 | 0.012 | 0.01 | 0.01 | 0 | 0.81 | 0.87 | 0.69 | 0.88 | 0.86 | 0.77 |
| 9 | 10421687 | rs10809092 | -0.35 | -1.007 | 2.005 | 1.664 | -0.014 | -0.038 | -0.038 | 0.029 | 0.09 | 0.01 | 0.13 | 0.57 | 0.08 | 0.15 |
| 9 | 10423799 | rs12344987 | -0.35 | -1.013 | 2.037 | 1.695 | -0.014 | -0.038 | -0.038 | 0.033 | 0.09 | 0.01 | 0.13 | 0.57 | 0.08 | 0.15 |
| 9 | 10424164 | rs12346029 | -1.449 | -1.036 | 2.202 | 1.654 | -0.007 | 0.114 | 0.114 | 0.327 | 0.14 | 0.32 | 0.23 | 0.66 | 0.1 | 0.24 |
| 9 | 10432920 | rs619126 | -1.603 | -1.013 | 1.703 | 1.558 | -0.003 | 0.023 | 0.023 | 0.554 | 0.18 | 0.17 | 0.15 | 0.57 | 0.13 | 0.21 |
| 9 | 10436451 | rs618288 | -1.356 | -0.857 | 1.252 | 1.677 | -0.014 | 0.102 | 0.102 | 0.964 | 0.16 | 0.26 | 0.08 | 0.49 | 0.11 | 0.15 |
| 9 | 10436921 | rs620558 | -1.165 | -0.733 | 1.339 | 0.977 | -0.024 | 0.046 | 0.046 | 0.239 | 0.12 | 0.41 | 0.25 | 0.12 | 0.05 | 0.13 |
| 9 | 10448624 | rs1565898 | -0.038 | 0.369 | 0.157 | 0.084 | 0.016 | -0.057 | -0.057 | 2.735 | 0.15 | 0.16 | 0.17 | 0.02 | 0.24 | 0.13 |
| 9 | 10457788 | rs13287491 | -0.276 | -0.394 | -0.582 | 0.149 | -0.004 | 0.02 | 0.02 | 0 | 0.16 | 0.01 | 0.1 | 0.07 | 0.13 | 0.1 |
| 9 | 10459930 | rs13297915 | -0.26 | -0.367 | -0.582 | 0.149 | -0.009 | 0.019 | 0.019 | 0 | 0.16 | 0.01 | 0.09 | 0.07 | 0.13 | 0.1 |
| 9 | 10462105 | rs7850565 | 0.357 | -0.559 | -0.294 | -0.077 | -0.006 | -0.016 | -0.016 | 0.473 | 0.82 | 0.83 | 0.91 | 0.95 | 0.85 | 0.94 |
| 9 | 10463261 | rs10124165 | 0.561 | -0.646 | -0.251 | 0.556 | -0.011 | 0.015 | 0.015 | 1.457 | 0.65 | 0.39 | 0.72 | 0.55 | 0.7 | 0.63 |
| 9 | 10464815 | rs1537812 | -0.298 | -0.866 | -0.96 | 0.912 | 0.045 | -0.078 | -0.078 | 0 | 0.24 | 0.13 | 0.34 | 0.46 | 0.17 | 0.24 |
| 9 | 10465127 | rs4740452 | -0.116 | -0.508 | -0.759 | 1.186 | 0.002 | 0.061 | 0.061 | 0 | 0.08 | 0.14 | 0.11 | 0.35 | 0.09 | 0.28 |
| 9 | 10465615 | rs4740453 | -0.162 | -0.863 | -0.915 | 1.133 | 0.027 | -0.074 | -0.074 | 1.254 | 0.18 | 0.06 | 0.3 | 0.44 | 0.13 | 0.19 |
| 9 | 10468666 | rs12343415 | -0.863 | -0.52 | -0.734 | 1.368 | 0 | -0.019 | -0.019 | 0 | 0.1 | 0.03 | 0.05 | 0.3 | 0.08 | 0.13 |
| 9 | 10470220 | rs12352981 | -0.241 | 0.057 | -0.595 | 0.41 | -0.014 | 0.013 | 0.013 | 0 | 0.85 | 0.91 | 0.92 | 1 | 0.84 | 0.93 |
| 9 | 10478387 | rs2784624 | 0.033 | 0.007 | -0.086 | 0.55 | 0.004 | -0.01 | -0.01 | 0 | 0.84 | 0.81 | 0.89 | 0.91 | 0.84 | 0.88 |
| 9 | 10479816 | rs2784623 | -0.148 | -0.318 | 0.108 | 0.748 | -0.003 | 0.06 | 0.06 | 0 | 0.29 | 0.4 | 0.47 | 0.48 | 0.25 | 0.23 |
| 9 | 10479902 | rs10756041 | 0.659 | -0.163 | -0.692 | -0.04 | -0.064 | -0.102 | -0.102 | 0 | 0.36 | 0.14 | 0.59 | 0.55 | 0.48 | 0.48 |
| 9 | 10480030 | rs1855850 | -0.22 | -0.321 | -0.377 | -0.156 | 0.002 | 0.001 | 0.001 | 0 | 0.56 | 0.57 | 0.76 | 0.64 | 0.67 | 0.55 |
| 9 | 10481115 | rs1853231 | -0.067 | -0.379 | -0.666 | -0.136 | 0.044 | 0.012 | 0.012 | 0 | 0.45 | 0.23 | 0.24 | 0.39 | 0.33 | 0.44 |
| 9 | 10481205 | rs1853232 | 0.75 | -0.588 | -0.618 | -0.028 | 0.005 | 0.007 | 0.007 | 0 | 0.18 | 0.13 | 0.12 | 0.13 | 0.13 | 0.22 |
| 9 | 10481639 | rs7870466 | 0.079 | -0.235 | -0.318 | 0.282 | -0.004 | 0.002 | 0.002 | 0.01 | 0.43 | 0.39 | 0.61 | 0.59 | 0.5 | 0.53 |

| | | | | | | | | | | | | | | | | |
|---|---|---|---|---|---|---|---|---|---|---|---|---|---|---|---|---|
| 9 | 10484849 | rs2784607 | -0.881 | 0.319 | 0.166 | 0.989 | 0.019 | 0.079 | 0.079 | 0.273 | 0.13 | 0.22 | 0.32 | 0.33 | 0.2 | 0.27 |
| 9 | 10490752 | rs833408 | -0.886 | 0.099 | 0.288 | 0.836 | 0.026 | 0.127 | 0.127 | 0.054 | 0.19 | 0.43 | 0.34 | 0.32 | 0.21 | 0.3 |
| 9 | 10497345 | rs1555483 | 0.854 | 0.044 | -0.009 | 0.856 | -0.004 | 0.052 | 0.052 | 0.544 | 0.79 | 0.4 | 0.63 | 0.66 | 0.79 | 0.69 |
| 9 | 10501953 | rs2181156 | 0.016 | 0.036 | -1.136 | 0.21 | 0.015 | 0.056 | 0.056 | 0.696 | 0.37 | 0.14 | 0.27 | 0.24 | 0.36 | 0.24 |
| 9 | 10505161 | rs833417 | -0.47 | -0.237 | -1.109 | -0.285 | -0.003 | -0.01 | -0.01 | 0 | 0.64 | 0.63 | 0.74 | 0.76 | 0.68 | 0.79 |
| 9 | 10506167 | rs10959130 | 0.62 | 0.476 | -1.382 | 0.506 | -0.011 | -0.014 | -0.014 | 0 | 0.33 | 0.32 | 0.34 | 0.4 | 0.34 | 0.5 |
| 9 | 10509682 | rs2784589 | 0.477 | 0.911 | -1.577 | 0.641 | -0.019 | -0.011 | -0.011 | 0 | 0.1 | 0.05 | 0.33 | 0.16 | 0.24 | 0.19 |
| 9 | 10512117 | rs12379948 | 0.647 | 0.716 | -1.595 | 0.508 | 0.013 | 0.018 | 0.018 | 0 | 0.29 | 0.12 | 0.21 | 0.24 | 0.27 | 0.22 |
| 9 | 10512307 | rs2784598 | -0.136 | 1.26 | -1.178 | 0.806 | -0.018 | -0.015 | -0.015 | 0 | 0.1 | 0.05 | 0.34 | 0.17 | 0.25 | 0.2 |
| 9 | 10516888 | rs10115302 | 0.36 | 1.025 | -1.343 | 0.835 | -0.025 | -0.052 | -0.052 | 0 | 0.25 | 0.2 | 0.32 | 0.46 | 0.27 | 0.32 |
| 9 | 10523334 | rs833403 | -0.495 | 1.113 | -1.082 | 0.206 | -0.006 | 0.034 | 0.034 | 0 | 0.52 | 0.7 | 0.66 | 0.68 | 0.48 | 0.61 |
| 9 | 10523685 | rs4461952 | 1.261 | 0.672 | -1.228 | -0.124 | 0.006 | 0.049 | 0.049 | 0 | 0.21 | 0.09 | 0.1 | 0 | 0.18 | 0.1 |
| 9 | 10526845 | rs10959147 | 1.144 | 0.999 | -1.212 | 0.013 | -0.015 | 0.004 | 0.004 | 0 | 0.09 | 0.05 | 0.28 | 0.07 | 0.21 | 0.08 |
| 9 | 10527045 | rs833425 | 0.257 | 0.576 | -1.099 | 0.013 | 0 | 0.049 | 0.049 | 0 | 0.15 | 0.22 | 0.26 | 0.53 | 0.11 | 0.4 |
| 9 | 10530844 | rs702130 | -1.087 | 0.118 | -0.644 | -0.545 | -0.008 | -0.094 | -0.094 | 0.044 | 0.66 | 0.58 | 0.8 | 0.9 | 0.73 | 0.85 |
| 9 | 10530953 | rs10959149 | 0.439 | -0.313 | -0.362 | -0.67 | -0.001 | -0.055 | -0.055 | 0.383 | 0.34 | 0.37 | 0.38 | 0.08 | 0.34 | 0.16 |
| 9 | 10531641 | rs10756060 | 1.577 | -0.301 | -0.479 | -0.715 | -0.044 | -0.024 | -0.024 | 0 | 0.25 | 0.05 | 0.21 | 0.21 | 0.3 | 0.26 |
| 9 | 10532820 | rs833432 | -1.73 | -0.259 | -0.295 | -0.654 | 0.005 | -0.033 | -0.033 | 3.59 | 0.78 | 0.79 | 0.86 | 0.91 | 0.79 | 0.86 |
| 9 | 10534642 | rs833435 | -0.071 | -0.351 | -0.109 | -0.368 | -0.014 | 0.131 | 0.131 | 0 | 0.17 | 0.37 | 0.27 | 0.54 | 0.12 | 0.41 |
| 9 | 10538067 | rs2784609 | -2.119 | -0.29 | -0.106 | -0.752 | 0.02 | -0.026 | -0.026 | 0.776 | 0.86 | 0.66 | 0.9 | 0.91 | 0.9 | 0.92 |
| 9 | 10543899 | rs10959161 | 1.547 | -0.235 | -0.587 | -0.922 | -0.008 | -0.024 | -0.024 | 0 | 0.17 | 0.06 | 0.16 | 0.2 | 0.2 | 0.24 |
| 9 | 10546257 | rs2181158 | 0.926 | -0.05 | -0.382 | -0.774 | -0.024 | 0.122 | 0.122 | 0 | 0.46 | 0.14 | 0.32 | 0.21 | 0.48 | 0.34 |
| 9 | 10546814 | rs1008981 | -0.571 | -0.043 | -0.181 | -0.735 | -0.012 | 0.05 | 0.05 | 0 | 0.71 | 0.49 | 0.65 | 0.36 | 0.81 | 0.55 |
| 9 | 10547481 | rs833455 | -0.761 | -0.052 | -0.217 | -0.786 | -0.033 | -0.03 | -0.03 | 0 | 0.7 | 0.4 | 0.85 | 0.82 | 0.8 | 0.81 |
| 9 | 10548301 | rs10959163 | 1.504 | 0.158 | -0.255 | -0.399 | -0.012 | -0.008 | -0.008 | 0 | 0.06 | 0.03 | 0.05 | 0.07 | 0.06 | 0.11 |
| 9 | 10551203 | rs2784611 | -0.776 | 0.143 | -0.192 | -0.436 | -0.005 | 0.001 | 0.001 | 0 | 0.35 | 0.34 | 0.57 | 0.69 | 0.4 | 0.54 |
| 9 | 10552890 | rs1885508 | -0.28 | 0.162 | -0.094 | -0.105 | 0.001 | 0.008 | 0.008 | 0 | 0.14 | 0.15 | 0.25 | 0.54 | 0.09 | 0.38 |
| 9 | 10555471 | rs833412 | 0.113 | 0.053 | 0.003 | -0.282 | 0.002 | 0.001 | 0.001 | 0 | 0.5 | 0.42 | 0.66 | 0.58 | 0.52 | 0.46 |
| 9 | 10558102 | rs833416 | 0.22 | 0.343 | -0.109 | -0.245 | -0.004 | 0.043 | 0.043 | 0 | 0.17 | 0.18 | 0.26 | 0.54 | 0.12 | 0.4 |
| 9 | 10560924 | rs2757873 | 0.365 | 0.312 | -0.088 | -0.263 | -0.007 | 0.071 | 0.071 | 0 | 0.16 | 0.19 | 0.26 | 0.52 | 0.12 | 0.4 |
| 9 | 10561180 | rs10756061 | 1.148 | 0.126 | -0.343 | -0.365 | -0.014 | -0.027 | -0.027 | 0 | 0.27 | 0.05 | 0.21 | 0.23 | 0.31 | 0.26 |
| 9 | 10561818 | rs2038592 | 0.624 | 0.086 | -0.097 | -0.382 | -0.001 | -0.021 | -0.021 | 0 | 0.31 | 0.25 | 0.23 | 0.33 | 0.35 | 0.31 |
| 9 | 10562824 | rs1322310 | 0.992 | 0.092 | -0.199 | -0.382 | -0.013 | 0 | 0 | 0 | 0.29 | 0.09 | 0.21 | 0.22 | 0.32 | 0.29 |
| 9 | 10565658 | rs7465708 | 1.811 | -0.082 | -0.078 | -0.505 | 0.005 | 0.003 | 0.003 | 0.852 | 0.11 | 0.08 | 0.04 | 0.09 | 0.06 | 0.11 |
| 9 | 10569790 | rs10959172 | 1.059 | 0.106 | -0.001 | -0.536 | 0 | -0.001 | -0.001 | 0 | 0.29 | 0.15 | 0.21 | 0.22 | 0.3 | 0.29 |
| 9 | 10572234 | rs988954 | -1.716 | 0.545 | 0.037 | -0.628 | 0.036 | -0.002 | -0.002 | 0 | 0.93 | 0.93 | 0.72 | 0.93 | 0.8 | 0.91 |
| 9 | 10572433 | rs17203683 | 2.505 | 0.387 | 0.109 | -0.811 | -0.005 | -0.044 | -0.044 | 0 | 0.12 | 0.15 | 0.06 | 0 | 0.07 | 0.02 |
| 9 | 10572534 | rs7025985 | 1.276 | 0.239 | -0.003 | -0.906 | 0.006 | -0.118 | -0.118 | 0.606 | 0.29 | 0.46 | 0.18 | 0 | 0.25 | 0.09 |
| 9 | 10574347 | rs1322314 | 0.081 | 0.321 | -0.13 | -0.733 | 0.002 | 0.001 | 0.001 | 0 | 0.5 | 0.52 | 0.43 | 0.5 | 0.37 | 0.49 |
| 9 | 10575273 | rs4741037 | -0.11 | 0.399 | -0.175 | -0.718 | 0 | 0 | 0 | 2.356 | 0.54 | 0.52 | 0.49 | 0.53 | 0.47 | 0.5 |
| 9 | 10577775 | rs10756063 | 1.294 | 0.288 | -0.265 | -0.647 | 0 | -0.001 | -0.001 | 0 | 0.26 | 0.24 | 0.19 | 0.32 | 0.25 | 0.34 |
| 9 | 10580326 | rs10738168 | -0.828 | 0.35 | -0.285 | -0.589 | 0.003 | -0.025 | -0.025 | 0 | 0.82 | 0.77 | 0.68 | 0.93 | 0.75 | 0.86 |
| 9 | 10581485 | rs10959176 | 0.392 | 0.389 | -0.731 | -0.632 | 0.014 | -0.039 | -0.039 | 0 | 0.26 | 0.16 | 0.32 | 0.53 | 0.22 | 0.41 |
| 9 | 10582246 | rs1322280 | -1.553 | 0.751 | -0.602 | -0.572 | 0.033 | -0.002 | -0.002 | 0.092 | 0.93 | 0.92 | 0.71 | 0.93 | 0.79 | 0.91 |
| 9 | 10583223 | rs12683583 | 2.117 | 0.471 | -1.152 | -0.586 | -0.044 | 0.003 | 0.003 | 0 | 0.16 | 0.06 | 0.1 | 0.09 | 0.21 | 0.09 |
| 9 | 10585399 | rs1160259 | 0.035 | 0.488 | -0.941 | -0.246 | -0.012 | 0.051 | 0.051 | 0 | 0.21 | 0.25 | 0.27 | 0.55 | 0.14 | 0.25 |
| 9 | 10586527 | rs13298283 | 1.114 | 0.623 | -1.005 | -0.396 | -0.003 | -0.006 | -0.006 | 0 | 0.22 | 0.07 | 0.14 | 0.19 | 0.21 | 0.25 |
| 9 | 10589257 | rs10115631 | 0.487 | 0.823 | -0.931 | -0.084 | 0.016 | 0.176 | 0.176 | 0 | 0.09 | 0.18 | 0.24 | 0.6 | 0.1 | 0.19 |
| 9 | 10592492 | rs1322291 | -0.181 | 0.825 | -0.926 | -0.148 | 0.019 | 0.178 | 0.178 | 0 | 0.12 | 0.24 | 0.24 | 0.58 | 0.11 | 0.21 |
| 9 | 10595208 | rs10756066 | 0.177 | 1.005 | -0.731 | -0.328 | 0.014 | -0.006 | -0.006 | 0 | 0.23 | 0.31 | 0.33 | 0.18 | 0.26 | 0.13 |
| 9 | 10596853 | rs1407398 | 0.371 | 1.286 | -1.087 | -0.03 | -0.011 | -0.037 | -0.037 | 0 | 0.23 | 0.15 | 0.35 | 0.52 | 0.3 | 0.44 |
| 9 | 10598429 | rs10809130 | 1.055 | 1.043 | -1.006 | 0.206 | -0.035 | -0.006 | -0.006 | 0 | 0.06 | 0.01 | 0.23 | 0.05 | 0.13 | 0.03 |
| 9 | 10602787 | rs977124 | 1.049 | -0.067 | -1.326 | 0.051 | 0.013 | -0.058 | -0.058 | 0.015 | 0.16 | 0.03 | 0.07 | 0.19 | 0.13 | 0.22 |
| 9 | 10603102 | rs2382220 | -0.894 | 0.25 | -0.521 | -0.196 | 0.005 | 0.004 | 0.004 | 0 | 0.52 | 0.41 | 0.5 | 0.55 | 0.46 | 0.55 |
| 9 | 10603845 | rs985563 | -0.738 | 1.256 | -0.198 | 0.711 | 0.006 | 0.023 | 0.023 | 0.103 | 0.3 | 0.36 | 0.43 | 0.55 | 0.36 | 0.49 |
| 9 | 10620453 | rs7863685 | -0.708 | 1.574 | 0.049 | 0.436 | 0.095 | 0.008 | 0.008 | 0 | 0.1 | 0.25 | 0.28 | 0.05 | 0.21 | 0.1 |
| 9 | 10620520 | rs1407402 | 0.959 | 1.429 | -0.187 | -0.084 | 0.026 | -0.027 | -0.027 | 0.028 | 0.73 | 0.64 | 0.61 | 0.89 | 0.63 | 0.77 |
| 9 | 10626531 | rs10756070 | -0.76 | 1.241 | 0.16 | 0.869 | 0.044 | 0.115 | 0.115 | 0 | 0.12 | 0.2 | 0.48 | 0.48 | 0.24 | 0.19 |
| 9 | 10635719 | rs7048007 | -0.31 | -0.015 | -0.001 | 0.331 | -0.026 | -0.002 | -0.002 | 0 | 0.31 | 0.39 | 0.11 | 0.33 | 0.14 | 0.34 |
| 9 | 10637778 | rs1040910 | 0.494 | -0.275 | -0.583 | -0.135 | -0.106 | -0.125 | -0.125 | 0.858 | 0.28 | 0.07 | 0.56 | 0.48 | 0.41 | 0.38 |
| 9 | 10642041 | rs7040199 | 1.016 | -0.426 | -0.411 | -0.071 | 0.012 | 0.086 | 0.086 | 0 | 0.06 | 0.24 | 0.03 | 0.11 | 0.02 | 0.09 |
| 9 | 10649660 | rs2208707 | 0.513 | -0.237 | -0.574 | -0.187 | -0.101 | -0.125 | -0.125 | 0 | 0.27 | 0.07 | 0.55 | 0.47 | 0.4 | 0.38 |
| 9 | 10651724 | rs7868009 | 0.354 | -0.479 | -0.353 | -0.208 | -0.086 | 0.018 | 0.018 | 0.529 | 0.25 | 0.57 | 0.11 | 0.29 | 0.09 | 0.33 |
| 9 | 10653484 | rs4131202 | 0.57 | -0.531 | -0.356 | -0.264 | 0.068 | 0.032 | 0.032 | 0 | 0.6 | 0.19 | 0.76 | 0.69 | 0.78 | 0.57 |
| 9 | 10655642 | rs10756076 | -0.316 | -0.366 | -0.504 | -0.323 | -0.003 | -0.049 | -0.049 | 0 | 0.81 | 0.73 | 0.87 | 0.94 | 0.84 | 0.86 |
| 9 | 10656930 | rs4372056 | 0.757 | -0.396 | -0.41 | -0.28 | -0.075 | 0.069 | 0.069 | 0.378 | 0.19 | 0.37 | 0.09 | 0.3 | 0.07 | 0.33 |
| 9 | 10661677 | rs10809150 | 0.848 | 0.724 | -0.78 | -0.823 | -0.04 | -0.03 | -0.03 | 0.059 | 0.07 | 0 | 0.26 | 0.06 | 0.2 | 0.07 |
| 9 | 10663815 | rs4427223 | 0.269 | -0.278 | -0.417 | -1.145 | -0.002 | 0.076 | 0.076 | 0 | 0.51 | 0.66 | 0.67 | 0.77 | 0.49 | 0.71 |
| 9 | 10668198 | rs7045753 | 1.287 | -0.392 | 0.466 | -0.584 | -0.006 | -0.087 | -0.087 | 0.491 | 0.8 | 0.61 | 0.86 | 0.94 | 0.84 | 0.86 |

| | | | | | | | | | | | | | | | | |
|---|---|---|---|---|---|---|---|---|---|---|---|---|---|---|---|---|
| 9 | 10673850 | rs7029106 | -0.067 | -0.118 | 0.512 | -0.434 | -0.01 | -0.062 | -0.062 | 0 | 0.11 | 0.26 | 0.06 | 0 | 0.06 | 0.02 |
| 9 | 10674573 | rs4443711 | 0.632 | 0.239 | -0.066 | 0.057 | 0.022 | -0.006 | -0.006 | 0.433 | 0.92 | 0.82 | 0.91 | 0.94 | 0.9 | 0.9 |
| 9 | 10678556 | rs4992817 | -0.265 | -0.628 | -0.244 | -0.354 | -0.059 | -0.069 | -0.069 | 1.123 | 0.32 | 0.43 | 0.23 | 0.13 | 0.2 | 0.34 |
| 9 | 10679889 | rs10738180 | 0.388 | 0.163 | -0.625 | -0.042 | -0.001 | -0.001 | -0.001 | 1.068 | 0.27 | 0.24 | 0.2 | 0.41 | 0.31 | 0.3 |
| 9 | 10679912 | rs7852800 | -0.261 | 0.72 | -0.761 | -0.617 | -0.019 | 0.027 | 0.027 | 0 | 0.76 | 0.82 | 0.85 | 0.94 | 0.68 | 0.9 |
| 9 | 10681522 | rs947215 | 0.574 | -0.291 | -0.23 | -0.631 | -0.024 | -0.083 | -0.083 | 0 | 0.79 | 0.39 | 0.89 | 1 | 0.89 | 0.96 |
| 9 | 10683110 | rs4113604 | 0.69 | -0.152 | -0.128 | -0.194 | -0.018 | -0.028 | -0.028 | 0 | 0.67 | 0.65 | 0.66 | 0.79 | 0.74 | 0.89 |
| 9 | 10684803 | rs7023148 | 0.676 | -0.24 | 0.376 | -0.438 | -0.028 | -0.081 | -0.081 | 0.843 | 0.85 | 0.72 | 0.92 | 0.98 | 0.92 | 0.83 |
| 9 | 10684841 | rs13285976 | 0.23 | 0.356 | -0.651 | -0.605 | -0.074 | 0.039 | 0.039 | 0 | 0.34 | 0.11 | 0.29 | 0.23 | 0.43 | 0.23 |
| 9 | 10686202 | rs10511552 | 0.492 | -0.101 | -0.136 | -0.039 | 0.008 | 0.081 | 0.081 | 0 | 0.19 | 0.36 | 0.39 | 0.35 | 0.18 | 0.26 |
| 9 | 10690633 | rs10809156 | -0.375 | 0.325 | -0.01 | -0.014 | 0.019 | 0.004 | 0.004 | 0 | 0.54 | 0.61 | 0.7 | 0.58 | 0.65 | 0.52 |
| 9 | 10691050 | rs7857207 | -0.02 | 0.193 | -0.251 | 0.081 | 0.019 | 0.021 | 0.021 | 0 | 0.5 | 0.41 | 0.4 | 0.34 | 0.64 | 0.34 |
| 9 | 10692461 | rs10756099 | 0.792 | -0.025 | -1.035 | 0.052 | -0.02 | -0.094 | -0.094 | 0 | 0.35 | 0.06 | 0.25 | 0.41 | 0.29 | 0.46 |
| 9 | 10696313 | rs11787634 | 0.602 | 0.279 | -0.866 | 0.144 | -0.069 | 0.015 | 0.015 | 0 | 0.34 | 0.13 | 0.28 | 0.25 | 0.45 | 0.26 |
| 9 | 10699855 | rs3923120 | -0.425 | -0.223 | 0.138 | 0.409 | -0.036 | 0.13 | 0.13 | 0 | 0.13 | 0.45 | 0.34 | 0.24 | 0.05 | 0.14 |
| 9 | 10714842 | rs10465037 | 0.318 | -0.078 | -1.239 | 0.181 | -0.137 | 0.007 | 0.007 | 0.398 | 0.33 | 0.09 | 0.29 | 0.25 | 0.5 | 0.41 |
| 9 | 10723877 | rs10756104 | 0.499 | -1.151 | -0.752 | 0.107 | 0.005 | -0.006 | -0.006 | 0 | 0.36 | 0.37 | 0.28 | 0.45 | 0.26 | 0.35 |
| 9 | 10727372 | rs16926356 | 0.785 | -1.334 | -0.763 | 0.035 | 0.048 | -0.098 | -0.098 | 0 | 0.32 | 0.08 | 0.26 | 0.47 | 0.25 | 0.35 |
| 9 | 10736580 | rs10959258 | 0.212 | -1.461 | -0.723 | -0.022 | 0.147 | 0.13 | 0.13 | 0.178 | 0.49 | 0.1 | 0.37 | 0.29 | 0.67 | 0.44 |
| 9 | 10737038 | rs1154843 | 0.036 | -1.26 | -0.458 | 0.005 | -0.044 | 0.094 | 0.094 | 0 | 0.86 | 0.47 | 0.65 | 0.76 | 0.93 | 0.79 |
| 9 | 10749312 | rs10809183 | -0.074 | -1.044 | -1.267 | -0.003 | 0.164 | 0.129 | 0.129 | 0.494 | 0.5 | 0.1 | 0.36 | 0.28 | 0.67 | 0.47 |
| 9 | 10752338 | rs1831295 | 0.521 | -1.07 | -0.942 | 0.017 | 0.063 | -0.025 | -0.025 | 0 | 0.42 | 0.25 | 0.31 | 0.53 | 0.32 | 0.35 |
| 9 | 10753219 | rs10809185 | 0.991 | -0.969 | -1.144 | -0.089 | -0.019 | 0.066 | 0.066 | 0 | 0.17 | 0.02 | 0.08 | 0.02 | 0.19 | 0.08 |
| 9 | 10753557 | rs10511554 | 0.676 | -0.922 | -1.241 | -0.06 | 0.068 | -0.034 | -0.034 | 0.064 | 0.4 | 0.23 | 0.3 | 0.53 | 0.3 | 0.32 |
| 9 | 10754695 | rs13283737 | -0.044 | 0.085 | -1.048 | 0.074 | -0.107 | 0.012 | 0.012 | 0.877 | 0.36 | 0.08 | 0.29 | 0.26 | 0.5 | 0.42 |
| 9 | 10758044 | rs7023802 | -0.271 | -0.163 | -0.107 | 0.546 | 0.005 | -0.002 | -0.002 | 0.279 | 0.9 | 0.97 | 0.95 | 0.92 | 0.93 | 0.89 |
| 9 | 10770620 | rs10959282 | 1.571 | -0.068 | 0.732 | 0.826 | 0.003 | 0.085 | 0.085 | 0.022 | 0.92 | 0.76 | 0.7 | 0.76 | 0.93 | 0.83 |
| 9 | 10773858 | rs16926439 | -0.683 | -0.185 | 0.996 | 0.512 | 0.001 | -0.049 | -0.049 | 0 | 0.13 | 0.19 | 0.1 | 0 | 0.13 | 0.04 |
| 9 | 10777503 | rs10959288 | 1.803 | -0.487 | 1.019 | 0.443 | -0.009 | -0.032 | -0.032 | 0 | 0.59 | 0.49 | 0.38 | 0.15 | 0.61 | 0.49 |
| 9 | 10784500 | rs10809190 | -0.38 | -0.287 | 0.668 | 0.491 | 0.026 | -0.069 | -0.069 | 0 | 0.21 | 0.07 | 0.24 | 0.6 | 0.19 | 0.3 |
| 9 | 10790580 | rs10117531 | -1.962 | -0.316 | 0.881 | 0.512 | 0.014 | 0 | 0 | 0.278 | 0.4 | 0.67 | 0.34 | 0.61 | 0.38 | 0.39 |
| 9 | 10792826 | rs10959301 | -0.552 | -0.247 | 0.497 | 0.576 | 0.024 | -0.115 | -0.115 | 0 | 0.22 | 0.04 | 0.23 | 0.6 | 0.19 | 0.3 |
| 9 | 10810659 | rs10959310 | -1.428 | -0.217 | 0.798 | 0.546 | 0.073 | 0.016 | 0.016 | 0 | 0.06 | 0.31 | 0.1 | 0.01 | 0.07 | 0.06 |
| 9 | 10830012 | rs7022854 | -1.04 | -0.194 | 0.555 | 0.662 | 0.004 | -0.008 | -0.008 | 0.01 | 0.24 | 0.17 | 0.3 | 0.6 | 0.24 | 0.36 |
| 9 | 10834971 | rs10809209 | -0.546 | -0.225 | -0.02 | 0.703 | 0.014 | -0.129 | -0.129 | 0.029 | 0.2 | 0.03 | 0.21 | 0.59 | 0.17 | 0.3 |
| 9 | 10856892 | rs3904270 | -1.893 | 0.414 | 0.791 | 1.228 | -0.001 | 0.051 | 0.051 | 0 | 0.18 | 0.28 | 0.26 | 0.6 | 0.17 | 0.3 |
| 9 | 10858265 | rs3899613 | -0.763 | 0.401 | 0.774 | 1.022 | 0.019 | -0.055 | -0.055 | 1.081 | 0.11 | 0.42 | 0.08 | 0 | 0.08 | 0.05 |
| 9 | 10861806 | rs2890930 | 0.164 | 1.069 | 0.137 | 1.119 | -0.038 | -0.035 | -0.035 | 0 | 0.06 | 0 | 0.04 | 0.18 | 0.11 | 0.24 |
| 9 | 10862544 | rs3858019 | -0.769 | 0.321 | 0.471 | 1.125 | 0.003 | -0.012 | -0.012 | 0 | 0.07 | 0.1 | 0.12 | 0.03 | 0.09 | 0.06 |
| 9 | 10867150 | rs7850706 | -0.337 | 0.254 | 0.317 | 1.12 | 0 | -0.036 | -0.036 | 0 | 0.08 | 0.16 | 0.06 | 0 | 0.08 | 0.05 |
| 9 | 10868263 | rs10809233 | -1.708 | 0.242 | -0.079 | 1.256 | -0.007 | 0.02 | 0.02 | 0 | 0.28 | 0.68 | 0.34 | 0.61 | 0.25 | 0.33 |
| 9 | 10878677 | rs3858023 | -0.772 | 0.766 | -0.835 | 1.601 | -0.026 | -0.053 | -0.053 | 1.981 | 0.1 | 0.04 | 0.19 | 0.61 | 0.12 | 0.22 |
| 9 | 10878948 | rs2382245 | -1.023 | 0.622 | -0.728 | 0.688 | 0 | 0 | 0 | 0 | 0.34 | 0.41 | 0.52 | 0.79 | 0.3 | 0.44 |
| 9 | 10880887 | rs954492 | -0.368 | 1.03 | -1.248 | 0.891 | -0.067 | -0.068 | -0.068 | 0 | 0.15 | 0.01 | 0.2 | 0.29 | 0.21 | 0.18 |
| 9 | 10883851 | rs7873741 | -0.682 | 0.29 | -0.394 | 0.224 | -0.024 | -0.056 | -0.056 | 0.796 | 0.18 | 0.4 | 0.12 | 0.05 | 0.15 | 0.11 |
| 9 | 10884836 | rs12376782 | -1.342 | -0.196 | -0.327 | -0.077 | -0.036 | -0.058 | -0.058 | 0.126 | 0.3 | 0.57 | 0.18 | 0.12 | 0.21 | 0.24 |
| 9 | 10894863 | rs7867915 | -0.804 | -0.188 | -0.117 | -0.102 | 0.002 | -0.006 | -0.006 | 1.181 | 0.1 | 0.14 | 0.09 | 0.07 | 0.1 | 0.16 |
| 9 | 10904815 | rs16926616 | 0.666 | -0.086 | -0.96 | -0.316 | -0.032 | 0.011 | 0.011 | 0.099 | 0.33 | 0.1 | 0.25 | 0.26 | 0.37 | 0.2 |
| 9 | 10909144 | rs10809238 | 0.765 | -0.232 | -0.545 | -0.315 | -0.005 | -0.004 | -0.004 | 1.254 | 0.12 | 0.15 | 0.07 | 0.05 | 0.09 | 0.07 |
| 9 | 10921838 | rs985337 | -1.081 | -0.441 | -0.009 | -0.396 | -0.011 | -0.01 | -0.01 | 0.85 | 0.61 | 0.68 | 0.42 | 0.39 | 0.57 | 0.46 |
| 9 | 10923290 | rs1536997 | -0.921 | -0.357 | 0.055 | -0.404 | -0.001 | -0.001 | -0.001 | 0 | 0.59 | 0.59 | 0.42 | 0.4 | 0.58 | 0.46 |
| 9 | 10923969 | rs10959370 | 0.641 | -0.385 | 0.115 | -0.428 | -0.012 | -0.009 | -0.009 | 0 | 0.12 | 0.19 | 0.07 | 0.05 | 0.08 | 0.1 |
| 9 | 10924493 | rs10959372 | 0.566 | -0.452 | 0.287 | -0.436 | -0.031 | -0.018 | -0.018 | 0 | 0.11 | 0.22 | 0.07 | 0.05 | 0.06 | 0.07 |
| 9 | 10925989 | rs7466612 | -0.937 | -0.254 | 0.027 | -0.364 | -0.009 | -0.012 | -0.012 | 1.135 | 0.61 | 0.67 | 0.42 | 0.4 | 0.56 | 0.46 |
| 9 | 10928314 | rs7021524 | -0.459 | -0.544 | 0.007 | -0.551 | 0.007 | 0.027 | 0.027 | 0 | 0.5 | 0.64 | 0.36 | 0.36 | 0.49 | 0.31 |
| 9 | 10928755 | rs7863959 | -0.847 | -0.546 | 0.004 | -0.699 | -0.006 | -0.001 | -0.001 | 0 | 0.56 | 0.62 | 0.38 | 0.38 | 0.53 | 0.42 |
| 9 | 10929786 | rs4740464 | 1.146 | -0.588 | 0.323 | -0.741 | -0.01 | -0.039 | -0.039 | 0 | 0.85 | 0.76 | 0.91 | 0.93 | 0.89 | 0.86 |
| 9 | 10931501 | rs10756144 | -0.415 | -0.573 | 0.301 | -0.722 | 0.004 | 0.025 | 0.025 | 0 | 0.49 | 0.64 | 0.4 | 0.32 | 0.51 | 0.3 |
| 9 | 10931959 | rs10738194 | -0.042 | -0.561 | 0.308 | -0.715 | 0.006 | 0.011 | 0.011 | 0 | 0.45 | 0.63 | 0.38 | 0.32 | 0.46 | 0.29 |
| 9 | 10940127 | rs4378061 | -0.77 | -0.592 | 0.363 | -0.906 | -0.005 | 0.211 | 0.211 | 0 | 0.52 | 0.83 | 0.34 | 0.12 | 0.49 | 0.31 |
| 9 | 10942628 | rs7848707 | -1.373 | -0.586 | 0.265 | -0.826 | -0.081 | -0.05 | -0.05 | 0 | 0.6 | 0.89 | 0.42 | 0.38 | 0.56 | 0.41 |
| 9 | 10943698 | rs10959388 | 0.781 | -0.579 | 0.928 | -1.094 | 0.007 | -0.105 | -0.105 | 0 | 0.29 | 0.33 | 0.22 | 0 | 0.36 | 0.12 |
| 9 | 10948272 | rs1888189 | 0.136 | -0.558 | 0.88 | -1.059 | 0.015 | -0.165 | -0.165 | 1.174 | 0.34 | 0.47 | 0.26 | 0 | 0.41 | 0.15 |
| 9 | 10957549 | rs10809261 | -0.245 | -0.582 | 0.408 | -0.985 | -0.014 | 0.03 | 0.03 | 0 | 0.39 | 0.73 | 0.34 | 0.26 | 0.44 | 0.21 |
| 9 | 10962971 | rs7858983 | -0.875 | -0.638 | 0.117 | -1.079 | 0.003 | 0.166 | 0.166 | 0 | 0.5 | 0.84 | 0.35 | 0.22 | 0.45 | 0.31 |
| 9 | 10970165 | rs16926678 | -0.303 | -0.091 | 0.524 | -1.015 | 0.046 | 0.01 | 0.01 | 0 | 0.09 | 0.43 | 0.07 | 0.05 | 0.06 | 0.12 |
| 9 | 10972504 | rs10959419 | 0.472 | 0.008 | -0.052 | -1.114 | -0.031 | 0.059 | 0.059 | 0 | 0.28 | 0.14 | 0.2 | 0 | 0.36 | 0.12 |
| 9 | 10975246 | rs10448153 | 0.492 | 0.013 | -0.064 | -1.112 | -0.03 | 0.054 | 0.054 | 0 | 0.28 | 0.15 | 0.2 | 0 | 0.35 | 0.12 |
| 9 | 10990046 | rs1330257 | 0.563 | -0.271 | -0.306 | -0.974 | -0.009 | 0.011 | 0.011 | 0 | 0.35 | 0.24 | 0.59 | 0.66 | 0.39 | 0.63 |

| | | | | | | | | | | | | | | | | |
|---|---|---|---|---|---|---|---|---|---|---|---|---|---|---|---|---|
| 9 | 10996025 | rs913923 | 0.474 | -0.325 | 0.366 | -1.148 | 0.008 | -0.083 | -0.083 | 0 | 0.3 | 0.37 | 0.23 | 0 | 0.38 | 0.14 |
| 9 | 10998674 | rs7853830 | 0.64 | -0.334 | 0.458 | -1.161 | -0.002 | -0.042 | -0.042 | 0 | 0.29 | 0.27 | 0.21 | 0 | 0.35 | 0.14 |
| 9 | 11004888 | rs4381046 | -0.146 | -0.438 | 0.77 | -1.224 | 0.015 | -0.099 | -0.099 | 0 | 0.38 | 0.49 | 0.28 | 0.05 | 0.44 | 0.26 |
| 9 | 11012926 | rs10959456 | -1.391 | -0.329 | 0.641 | -0.591 | 0 | 0.16 | 0.16 | 0.72 | 0.92 | 0.82 | 0.72 | 0.75 | 0.94 | 0.71 |
| 9 | 11013198 | rs11791663 | -0.192 | -0.42 | 0.768 | -1.681 | 0.005 | -0.055 | -0.055 | 0 | 0.37 | 0.39 | 0.26 | 0.05 | 0.43 | 0.26 |
| 9 | 11018077 | rs12346562 | 0.036 | -0.425 | 0.531 | -1.718 | 0.014 | 0.077 | 0.077 | 0 | 0.32 | 0.18 | 0.15 | 0.05 | 0.28 | 0.19 |
| 9 | 11021845 | rs10756163 | -0.515 | -0.51 | 0.589 | -1.885 | 0.021 | -0.139 | -0.139 | 0 | 0.69 | 0.54 | 0.76 | 1 | 0.63 | 0.86 |
| 9 | 11022051 | rs10959464 | 0.931 | -0.563 | 0.487 | -1.773 | -0.001 | 0.008 | 0.008 | 0 | 0.35 | 0.26 | 0.56 | 0.7 | 0.35 | 0.62 |
| 9 | 11022581 | rs10959466 | 1.471 | -0.353 | 0.161 | -1.812 | -0.019 | -0.062 | -0.062 | 3.165 | 0.2 | 0.02 | 0.12 | 0.25 | 0.2 | 0.09 |
| 9 | 11024400 | rs10959468 | 0.736 | -0.19 | 0.504 | -1.799 | -0.047 | 0.065 | 0.065 | 0 | 0.26 | 0.12 | 0.2 | 0 | 0.37 | 0.14 |
| 9 | 11028959 | rs10959470 | 0.678 | -0.262 | 0.595 | -1.81 | 0.015 | 0.069 | 0.069 | 0 | 0.29 | 0.13 | 0.11 | 0 | 0.25 | 0.08 |
| 9 | 11036869 | rs10809302 | 0.075 | -0.246 | 0.68 | -1.815 | 0.007 | -0.086 | -0.086 | 0.681 | 0.63 | 0.39 | 0.72 | 0.95 | 0.6 | 0.77 |
| 9 | 11039639 | rs7044282 | 1.001 | -0.221 | 0.712 | -1.589 | -0.001 | 0 | 0 | 0 | 0.33 | 0.26 | 0.56 | 0.67 | 0.35 | 0.64 |
| 9 | 11042142 | rs10959478 | 0.705 | 0.013 | 0.637 | -1.617 | -0.05 | 0.05 | 0.05 | 0 | 0.2 | 0.09 | 0.2 | 0 | 0.34 | 0.11 |
| 9 | 11063519 | rs4097990 | -0.284 | -0.238 | 0.958 | -1.307 | 0.046 | 0.009 | 0.009 | 0 | 0.92 | 0.66 | 0.93 | 0.96 | 0.94 | 0.9 |
| 9 | 11070804 | rs6474617 | -0.565 | -0.633 | 0.539 | -1.519 | 0.054 | 0.044 | 0.044 | 0.417 | 0.65 | 0.74 | 0.84 | 0.75 | 0.78 | 0.89 |
| 9 | 11073873 | rs10959509 | 0.331 | -0.612 | 0.516 | -1.53 | 0.052 | 0.046 | 0.046 | 0 | 0.34 | 0.21 | 0.16 | 0.23 | 0.23 | 0.12 |
| 9 | 11073967 | rs10809325 | 1.293 | -0.345 | 0.523 | -1.682 | 0.009 | -0.012 | -0.012 | 0 | 0.34 | 0.38 | 0.6 | 0.71 | 0.37 | 0.64 |
| 9 | 11075596 | rs10809326 | 0.658 | -0.321 | 0.1 | -1.716 | -0.036 | 0.034 | 0.034 | 0 | 0.22 | 0.11 | 0.18 | 0 | 0.34 | 0.11 |
| 9 | 11075772 | rs7858670 | 0.377 | -0.587 | 0.113 | -1.564 | 0.065 | 0.042 | 0.042 | 0 | 0.34 | 0.15 | 0.15 | 0.25 | 0.22 | 0.13 |
| 9 | 11083426 | rs1334045 | 0.542 | 0.02 | 0.823 | -1.058 | -0.002 | 0 | 0 | 0 | 0.67 | 0.66 | 0.74 | 0.69 | 0.63 | 0.7 |
| 9 | 11083979 | rs10121358 | 0.526 | 0.067 | 0.741 | -1.05 | 0.002 | -0.003 | -0.003 | 0 | 0.65 | 0.38 | 0.7 | 0.69 | 0.62 | 0.7 |
| 9 | 11091254 | rs1931401 | -0.26 | 0.093 | 0.396 | -1.059 | -0.002 | 0.003 | 0.003 | 0 | 0.29 | 0.3 | 0.23 | 0.24 | 0.31 | 0.2 |
| 9 | 11101956 | rs7044787 | -0.846 | -0.102 | 0.742 | -1.273 | -0.009 | -0.048 | -0.048 | 0 | 0.16 | 0.26 | 0.07 | 0.05 | 0.11 | 0.12 |
| 9 | 11105401 | rs1334038 | 1.329 | -0.038 | 0.931 | -1.529 | -0.001 | -0.132 | -0.132 | 0 | 0.68 | 0.43 | 0.83 | 0.98 | 0.74 | 0.84 |
| 9 | 11108517 | rs10959531 | 1.303 | -0.028 | 0.964 | -1.517 | -0.001 | -0.119 | -0.119 | 0 | 0.68 | 0.41 | 0.83 | 0.98 | 0.74 | 0.84 |
| 9 | 11111096 | rs7042371 | -0.102 | -0.141 | 1.051 | -1.454 | 0.002 | -0.035 | -0.035 | 0 | 0.07 | 0.14 | 0.04 | 0 | 0.06 | 0.03 |
| 9 | 11114436 | rs7868584 | -0.826 | -0.13 | 1.279 | -1.554 | 0.015 | -0.096 | -0.096 | 0 | 0.22 | 0.33 | 0.13 | 0 | 0.21 | 0.09 |
| 9 | 11114781 | rs10809343 | 0.841 | -0.263 | 1.277 | -1.448 | -0.037 | -0.045 | -0.045 | 0 | 0.9 | 0.66 | 0.95 | 0.98 | 0.95 | 0.94 |
| 9 | 11116389 | rs12377914 | 1.449 | -0.107 | 0.441 | -1.847 | -0.056 | 0.116 | 0.116 | 0 | 0.19 | 0.02 | 0.1 | 0 | 0.25 | 0.05 |
| 9 | 11122042 | rs7863726 | -0.254 | -0.38 | 0.478 | -1.751 | 0 | 0.005 | 0.005 | 0 | 0.15 | 0.13 | 0.09 | 0.07 | 0.13 | 0.09 |
| 9 | 11127388 | rs10809352 | -0.084 | -0.425 | 0.754 | -1.749 | 0.003 | 0.001 | 0.001 | 0 | 0.94 | 0.77 | 0.97 | 0.97 | 0.96 | 0.94 |
| 9 | 11129139 | rs10809353 | -0.334 | -0.652 | 0.601 | -1.197 | 0.016 | -0.03 | -0.03 | 0.077 | 0.17 | 0.12 | 0.18 | 0.47 | 0.1 | 0.18 |
| 9 | 11129613 | rs1931405 | 0.955 | -0.693 | 1.118 | -1.22 | -0.027 | 0.038 | 0.038 | 0.451 | 0.73 | 0.39 | 0.75 | 0.51 | 0.82 | 0.73 |
| 9 | 11131170 | rs2184258 | 1.635 | -0.561 | 0.789 | -1.289 | -0.002 | 0.032 | 0.032 | 0.05 | 0.55 | 0.17 | 0.63 | 0.56 | 0.62 | 0.65 |
| 9 | 11135805 | rs9298665 | 0.594 | -0.384 | 1.183 | -0.854 | 0.004 | 0.022 | 0.022 | 0 | 0.1 | 0.06 | 0.09 | 0.43 | 0.04 | 0.17 |
| 9 | 11136285 | rs7870863 | 0.259 | -0.395 | 1.358 | -0.776 | -0.003 | 0.044 | 0.044 | 0 | 0.11 | 0.13 | 0.15 | 0.49 | 0.04 | 0.17 |
| 9 | 11140883 | rs7866090 | 0.447 | -0.229 | 1.536 | -0.867 | -0.007 | -0.016 | -0.016 | 0 | 0.14 | 0.09 | 0.09 | 0 | 0.19 | 0.07 |
| 9 | 11141099 | rs10959563 | 0.054 | -0.274 | 1.843 | -0.859 | -0.014 | 0.071 | 0.071 | 0 | 0.13 | 0.18 | 0.17 | 0.49 | 0.07 | 0.17 |
| 9 | 11141424 | rs10809356 | -1.107 | -0.405 | 1.872 | -1.139 | -0.003 | -0.063 | -0.063 | 0.524 | 0.76 | 0.7 | 0.86 | 1 | 0.75 | 0.95 |
| 9 | 11143413 | rs10959565 | 1.258 | -0.41 | 1.89 | -1.123 | -0.013 | -0.013 | -0.013 | 0 | 0.45 | 0.22 | 0.58 | 0.51 | 0.49 | 0.67 |
| 9 | 11143915 | rs10116927 | 1.093 | -0.447 | 1.878 | -1.128 | -0.001 | 0.001 | 0.001 | 0.019 | 0.47 | 0.37 | 0.59 | 0.52 | 0.49 | 0.66 |
| 9 | 11144684 | rs944997 | -1.118 | -0.449 | 1.891 | -1.15 | -0.004 | -0.078 | -0.078 | 0.073 | 0.76 | 0.66 | 0.85 | 1 | 0.75 | 0.95 |
| 9 | 11144781 | rs10959567 | 0.855 | -0.44 | 1.663 | -1.071 | 0.002 | 0.005 | 0.005 | 0.151 | 0.48 | 0.36 | 0.59 | 0.52 | 0.49 | 0.69 |
| 9 | 11155468 | rs10959573 | 0.147 | -0.523 | 1.457 | -1.335 | -0.008 | -0.009 | -0.009 | 0 | 0.59 | 0.32 | 0.75 | 1 | 0.57 | 0.89 |
| 9 | 11156238 | rs11999125 | 1.123 | -0.401 | 1.259 | -1.157 | -0.031 | -0.002 | -0.002 | 0 | 0.45 | 0.18 | 0.58 | 0.57 | 0.5 | 0.69 |
| 9 | 11167194 | rs6474635 | -0.082 | -0.325 | 1.194 | -0.978 | 0.012 | -0.012 | -0.012 | 0 | 0.14 | 0.13 | 0.16 | 0.44 | 0.07 | 0.2 |
| 9 | 11173836 | rs7037473 | -0.45 | -0.435 | 1.37 | -1.26 | 0.01 | 0.002 | 0.002 | 0.217 | 0.41 | 0.62 | 0.27 | 0.37 | 0.31 | 0.27 |
| 9 | 11174994 | rs10959581 | 0.572 | -0.323 | 1.383 | -1.335 | -0.002 | -0.022 | -0.022 | 1.135 | 0.14 | 0.1 | 0.1 | 0 | 0.19 | 0.04 |
| 9 | 11178159 | rs10959584 | -0.918 | -0.388 | 1.588 | -1.347 | 0 | 0 | 0 | 0 | 0.3 | 0.26 | 0.21 | 0.37 | 0.26 | 0.26 |
| 9 | 11178672 | rs12375980 | -1.357 | -0.186 | 1.633 | -1.249 | -0.007 | 0.046 | 0.046 | 0 | 0.52 | 0.72 | 0.37 | 0.37 | 0.5 | 0.31 |
| 9 | 11181745 | rs10756199 | 0.468 | 0.025 | 1.731 | -1.132 | 0.006 | 0.008 | 0.008 | 0 | 0.75 | 0.73 | 0.79 | 0.63 | 0.73 | 0.74 |
| 9 | 11187028 | rs2200787 | 0.899 | 0.034 | 1.847 | -1.167 | -0.018 | -0.155 | -0.155 | 0 | 0.24 | 0.42 | 0.13 | 0 | 0.23 | 0.07 |
| 9 | 11191795 | rs10809375 | -0.43 | 0.034 | 1.646 | -1.145 | -0.005 | -0.001 | -0.001 | 0.236 | 0.39 | 0.54 | 0.23 | 0.37 | 0.3 | 0.26 |
| 9 | 11194115 | rs10125755 | 0.9 | 0.085 | 1.359 | -1.156 | 0.001 | 0.064 | 0.064 | 0 | 0.49 | 0.27 | 0.65 | 0.63 | 0.49 | 0.69 |
| 9 | 11194132 | rs7465994 | 0.236 | -0.047 | 1.33 | -1.112 | 0.021 | 0.003 | 0.003 | 0.1 | 0.61 | 0.38 | 0.76 | 0.63 | 0.7 | 0.74 |
| 9 | 11203149 | rs1931388 | 0.04 | 0.151 | 0.722 | -1.086 | -0.013 | -0.023 | -0.023 | 0 | 0.32 | 0.28 | 0.2 | 0 | 0.4 | 0.09 |
| 9 | 11218042 | rs10511566 | 1.308 | 0.093 | 0.229 | -1.026 | 0.009 | 0.019 | 0.019 | 0 | 0.2 | 0.11 | 0.08 | 0 | 0.19 | 0.04 |
| 9 | 11218170 | rs11788082 | 0.156 | 0.248 | 0.364 | -1.032 | 0.045 | -0.057 | -0.057 | 0 | 0.13 | 0.19 | 0.14 | 0 | 0.22 | 0.05 |
| 9 | 11235661 | rs10809396 | -0.215 | -0.311 | -0.228 | -0.794 | 0.046 | -0.022 | -0.022 | 0 | 0.34 | 0.16 | 0.24 | 0.39 | 0.26 | 0.23 |
| 9 | 11246297 | rs10119093 | -0.268 | -0.191 | -0.401 | -0.378 | -0.001 | 0.008 | 0.008 | 0 | 0.16 | 0.16 | 0.18 | 0.39 | 0.08 | 0.19 |
| 9 | 11254070 | rs1387292 | 0.363 | 0.076 | -0.457 | -0.658 | -0.002 | 0.028 | 0.028 | 0 | 0.49 | 0.39 | 0.6 | 0.61 | 0.51 | 0.7 |
| 9 | 11257448 | rs10959672 | -0.589 | -0.122 | -1.287 | -0.939 | -0.072 | -0.048 | -0.048 | 0 | 0.29 | 0.07 | 0.31 | 0.36 | 0.32 | 0.28 |
| 9 | 11262339 | rs10756216 | -0.813 | -0.188 | -0.865 | -1.163 | 0.002 | -0.041 | -0.041 | 0 | 0.72 | 0.31 | 0.84 | 1 | 0.76 | 0.94 |
| 9 | 11270533 | rs1352519 | 1.556 | -0.162 | -1.475 | -0.271 | -0.032 | -0.066 | -0.066 | 0 | 0.1 | 0 | 0.25 | 0.27 | 0.13 | 0.37 |
| 9 | 11274798 | rs10959676 | 0.587 | -0.582 | -1.715 | -0.246 | 0.046 | -0.124 | -0.124 | 0 | 0.16 | 0 | 0.22 | 0.42 | 0.06 | 0.19 |
| 9 | 11275227 | rs1489177 | -1.411 | -0.798 | -1.141 | -0.579 | -0.023 | -0.138 | -0.138 | 0 | 0.74 | 0.47 | 0.87 | 1 | 0.78 | 0.96 |
| 9 | 11277879 | rs10959682 | 0.472 | -0.563 | -1.229 | -0.62 | -0.017 | -0.006 | -0.006 | 3.256 | 0.47 | 0.21 | 0.54 | 0.55 | 0.53 | 0.72 |
| 9 | 11278380 | rs1489175 | -1.765 | -0.68 | -0.994 | -0.689 | -0.028 | -0.144 | -0.144 | 0.548 | 0.78 | 0.6 | 0.9 | 1 | 0.82 | 0.97 |

| | | | | | | | | | | | | | | | | |
|---|---|---|---|---|---|---|---|---|---|---|---|---|---|---|---|---|
| 9 | 11305052 | rs11496296 | -0.5 | -0.705 | -0.965 | -0.619 | -0.024 | 0.006 | 0.006 | 0 | 0.61 | 0.45 | 0.69 | 0.58 | 0.75 | 0.8 |
| 9 | 11316481 | rs10756234 | -0.413 | -0.734 | -0.524 | -0.586 | -0.077 | 0.2 | 0.2 | 0.322 | 0.82 | 0.52 | 0.77 | 0.58 | 0.93 | 0.8 |
| 9 | 11327345 | rs10959723 | 0.578 | -0.515 | -0.575 | -0.628 | 0.01 | -0.019 | -0.019 | 0.423 | 0.12 | 0.11 | 0.13 | 0 | 0.22 | 0.08 |
| 9 | 11329927 | rs10116684 | 0.284 | -0.608 | -0.568 | -0.665 | 0.004 | 0.019 | 0.019 | 0 | 0.5 | 0.28 | 0.56 | 0.57 | 0.54 | 0.69 |
| 9 | 11337156 | rs12376216 | 1.044 | -0.553 | -0.557 | -0.674 | 0.011 | -0.005 | -0.005 | 0 | 0.1 | 0.06 | 0.13 | 0.01 | 0.2 | 0.08 |
| 9 | 11339109 | rs4741148 | -0.327 | -0.759 | -1.169 | -0.803 | 0.05 | -0.028 | -0.028 | 0.555 | 0.67 | 0.6 | 0.68 | 0.73 | 0.54 | 0.74 |
| 9 | 11343385 | rs10959734 | 2.357 | -0.674 | -1.411 | -0.69 | -0.036 | -0.058 | -0.058 | 0.802 | 0.09 | 0 | 0.26 | 0.23 | 0.13 | 0.34 |
| 9 | 11345138 | rs12552533 | 0.765 | -0.626 | -1.144 | -0.651 | 0.006 | -0.009 | -0.009 | 0.046 | 0.29 | 0.18 | 0.39 | 0.31 | 0.28 | 0.51 |
| 9 | 11359656 | rs1414202 | 1.07 | -0.687 | -1.199 | -0.605 | -0.023 | 0.174 | 0.174 | 0 | 0.4 | 0.15 | 0.24 | 0.11 | 0.42 | 0.2 |
| 9 | 11359686 | rs1335445 | 0.478 | -0.582 | -1.196 | -0.457 | 0.002 | -0.006 | -0.006 | 0.609 | 0.31 | 0.28 | 0.4 | 0.5 | 0.29 | 0.53 |
| 9 | 11366549 | rs10809441 | 0.537 | -0.425 | -1.302 | -0.573 | -0.095 | 0.078 | 0.078 | 0 | 0.29 | 0.08 | 0.29 | 0.12 | 0.45 | 0.24 |
| 9 | 11369213 | rs12378446 | 1.026 | -0.212 | -1.04 | -0.49 | 0.012 | -0.013 | -0.013 | 0 | 0.11 | 0.07 | 0.13 | 0.01 | 0.21 | 0.09 |
| 9 | 11374073 | rs16927457 | 0.675 | -0.413 | -1.126 | -0.364 | 0 | -0.003 | -0.003 | 0 | 0.28 | 0.25 | 0.38 | 0.32 | 0.29 | 0.49 |
| 9 | 11396601 | rs1335477 | 0.098 | -0.63 | -1.364 | -0.459 | 0.049 | -0.044 | -0.044 | 0 | 0.37 | 0.2 | 0.37 | 0.54 | 0.31 | 0.39 |
| 9 | 11398305 | rs1335475 | 1.943 | -0.552 | -1.346 | -0.61 | 0.004 | 0.003 | 0.003 | 0 | 0.21 | 0.2 | 0.08 | 0 | 0.18 | 0.03 |
| 9 | 11402634 | rs4483255 | 0.17 | -0.614 | -1.439 | -0.588 | -0.017 | -0.018 | -0.018 | 0.657 | 0.57 | 0.25 | 0.44 | 0.55 | 0.5 | 0.45 |
| 9 | 11403465 | rs10738232 | -0.51 | -0.497 | -1.44 | -0.674 | 0.01 | -0.025 | -0.025 | 0.337 | 0.68 | 0.29 | 0.57 | 0.55 | 0.71 | 0.55 |
| 9 | 11408052 | rs10959776 | 0.613 | -0.46 | -1.149 | -0.731 | -0.034 | 0.035 | 0.035 | 0 | 0.34 | 0.27 | 0.22 | 0.01 | 0.4 | 0.14 |
| 9 | 11414180 | rs1577436 | 2.159 | -0.454 | -1.168 | -0.703 | 0.002 | 0.046 | 0.046 | 0 | 0.2 | 0.13 | 0.08 | 0 | 0.18 | 0.03 |
| 9 | 11419061 | rs1335467 | -0.109 | -0.664 | -1.307 | -0.657 | -0.016 | -0.009 | -0.009 | 1.392 | 0.48 | 0.2 | 0.41 | 0.44 | 0.46 | 0.34 |
| 9 | 11427927 | rs1414213 | 0.155 | -0.794 | -1.134 | -0.566 | -0.111 | 0.157 | 0.157 | 0.519 | 0.2 | 0.5 | 0.23 | 0.43 | 0.03 | 0.21 |
| 9 | 11431990 | rs10959797 | 1.064 | -0.604 | -1.182 | -0.676 | -0.025 | 0.019 | 0.019 | 0.005 | 0.12 | 0.04 | 0.14 | 0.01 | 0.21 | 0.1 |
| 9 | 11434678 | rs10959800 | -0.034 | -0.711 | -1.236 | -0.715 | 0.049 | -0.082 | -0.082 | 0 | 0.34 | 0.4 | 0.32 | 0.12 | 0.47 | 0.27 |
| 9 | 11439075 | rs4524893 | 0 | -0.744 | -1.392 | -0.753 | 0.028 | 0.056 | 0.056 | 0 | 0.56 | 0.64 | 0.41 | 0.12 | 0.65 | 0.31 |
| 9 | 11440848 | rs10809476 | 0.975 | -0.633 | -1.105 | -0.705 | 0.027 | -0.032 | -0.032 | 0 | 0.13 | 0.11 | 0.14 | 0.01 | 0.21 | 0.1 |
| 9 | 11440880 | rs1414207 | 0.255 | -0.854 | -1.148 | -0.323 | -0.044 | 0.001 | 0.001 | 0 | 0.4 | 0.52 | 0.23 | 0.4 | 0.21 | 0.24 |
| 9 | 11441728 | rs10809477 | 1.039 | -0.876 | -1.223 | -0.19 | 0.08 | -0.103 | -0.103 | 0 | 0.16 | 0.02 | 0.12 | 0.4 | 0.03 | 0.2 |
| 9 | 11444251 | rs7848864 | -0.251 | -0.526 | -1.189 | -0.088 | 0.002 | -0.014 | -0.014 | 0 | 0.65 | 0.66 | 0.57 | 0.47 | 0.68 | 0.41 |
| 9 | 11448773 | rs7038987 | 0.988 | -0.537 | -0.728 | -0.161 | -0.001 | -0.015 | -0.015 | 0 | 0.22 | 0.25 | 0.17 | 0.08 | 0.25 | 0.15 |
| 9 | 11463799 | rs9406428 | 0.966 | -0.569 | -0.424 | -0.043 | -0.028 | -0.034 | -0.034 | 0.295 | 0.24 | 0.64 | 0.12 | 0.15 | 0.16 | 0.05 |
| 9 | 11465606 | rs950868 | 2.298 | -0.546 | -0.523 | -0.032 | -0.001 | 0.009 | 0.009 | 0 | 0.17 | 0.06 | 0.06 | 0.16 | 0.15 | 0.05 |
| 9 | 11467111 | rs10809488 | -0.207 | -0.479 | -0.566 | 0.208 | 0.004 | 0.017 | 0.017 | 0 | 0.47 | 0.37 | 0.33 | 0.41 | 0.4 | 0.31 |
| 9 | 11469190 | rs10959841 | 1.125 | -0.483 | -0.655 | 0.14 | -0.106 | 0.139 | 0.139 | 0 | 0.28 | 0.08 | 0.19 | 0.01 | 0.37 | 0.13 |
| 9 | 11471766 | rs10756254 | -0.728 | -0.409 | -0.544 | 0.264 | -0.001 | 0.038 | 0.038 | 0 | 0.67 | 0.49 | 0.49 | 0.48 | 0.64 | 0.46 |
| 9 | 11487579 | rs10738236 | -0.004 | -0.443 | -0.539 | 0.227 | 0.004 | -0.001 | -0.001 | 0.027 | 0.29 | 0.34 | 0.27 | 0.26 | 0.24 | 0.27 |
| 9 | 11488348 | rs725910 | 2.432 | -0.428 | -0.924 | 0.115 | -0.012 | 0.096 | 0.096 | 0 | 0.17 | 0.05 | 0.06 | 0 | 0.15 | 0.04 |
| 9 | 11490767 | rs7030303 | 1.777 | -0.406 | -0.937 | 0.153 | -0.061 | 0.023 | 0.023 | 0 | 0.18 | 0.01 | 0.14 | 0.08 | 0.25 | 0.15 |
| 9 | 11494328 | rs10959862 | -1.15 | -0.405 | -0.82 | 0.254 | -0.033 | -0.109 | -0.109 | 0.139 | 0.71 | 0.87 | 0.52 | 0.47 | 0.65 | 0.46 |
| 9 | 11499438 | rs1596341 | 0.762 | -0.337 | -0.573 | 0.146 | 0.003 | 0.088 | 0.088 | 0 | 0.38 | 0.29 | 0.24 | 0.07 | 0.4 | 0.19 |
| 9 | 11514645 | rs968995 | -0.947 | -0.299 | -0.751 | 0.181 | -0.014 | -0.081 | -0.081 | 0 | 0.69 | 0.84 | 0.53 | 0.47 | 0.65 | 0.46 |
| 9 | 11519008 | rs10114729 | -0.765 | -0.389 | -0.835 | 0.093 | -0.005 | 0.051 | 0.051 | 0 | 0.52 | 0.77 | 0.45 | 0.34 | 0.5 | 0.42 |
| 9 | 11525541 | rs7871636 | 0.641 | -0.441 | -0.769 | 0.023 | 0.005 | 0.03 | 0.03 | 0 | 0.4 | 0.38 | 0.25 | 0.21 | 0.4 | 0.2 |
| 9 | 11530473 | rs10809510 | -0.191 | -0.345 | -0.555 | 0.132 | 0.003 | 0 | 0 | 0 | 0.3 | 0.58 | 0.28 | 0.26 | 0.25 | 0.27 |
| 9 | 11549994 | rs10959919 | 1.325 | -0.232 | -0.749 | 0.034 | -0.032 | 0.027 | 0.027 | 0.018 | 0.11 | 0.03 | 0.13 | 0 | 0.21 | 0.1 |
| 9 | 11551772 | rs11792400 | 2.369 | -0.436 | -0.878 | 0.004 | -0.013 | 0.021 | 0.021 | 0 | 0.17 | 0.05 | 0.06 | 0.13 | 0.15 | 0.05 |
| 9 | 11552356 | rs10959923 | 1.284 | -0.581 | -0.873 | 0.15 | 0.073 | -0.08 | -0.08 | 0 | 0.16 | 0.01 | 0.1 | 0.24 | 0.03 | 0.17 |
| 9 | 11553641 | rs12349156 | 0.817 | -0.677 | -0.829 | 0.102 | -0.017 | 0.002 | 0.002 | 1.115 | 0.15 | 0.17 | 0.11 | 0.22 | 0.03 | 0.03 |
| 9 | 11558301 | rs12341457 | 0.609 | -0.585 | -0.877 | -0.025 | -0.014 | -0.013 | -0.013 | 0 | 0.16 | 0.19 | 0.12 | 0.06 | 0.07 | 0.03 |
| 9 | 11558362 | rs10809521 | -1.063 | -0.476 | -0.689 | -0.104 | -0.002 | 0.027 | 0.027 | 0 | 0.35 | 0.69 | 0.31 | 0.23 | 0.29 | 0.28 |
| 9 | 11560184 | rs1348384 | -0.654 | -0.561 | -0.549 | -0.046 | 0 | -0.01 | -0.01 | 0 | 0.31 | 0.41 | 0.29 | 0.23 | 0.25 | 0.27 |
| 9 | 11561438 | rs7047379 | -1.21 | -0.567 | -0.731 | -0.041 | -0.016 | -0.082 | -0.082 | 0.179 | 0.7 | 0.87 | 0.52 | 0.47 | 0.66 | 0.47 |
| 9 | 11574204 | rs10959951 | 0.289 | -0.518 | -0.703 | -0.058 | -0.001 | 0 | 0 | 0 | 0.33 | 0.33 | 0.32 | 0.25 | 0.48 | 0.26 |
| 9 | 11574930 | rs10119906 | -1.247 | -0.489 | -0.604 | -0.068 | -0.001 | -0.011 | -0.011 | 0 | 0.54 | 0.72 | 0.47 | 0.47 | 0.52 | 0.44 |
| 9 | 11578233 | rs10809526 | 0.062 | -0.392 | -0.563 | -0.093 | 0.056 | -0.091 | -0.091 | 0 | 0.17 | 0.62 | 0.19 | 0 | 0.22 | 0.1 |
| 9 | 11586357 | rs10959970 | 0.315 | -0.458 | -0.628 | 0.02 | 0.04 | -0.047 | -0.047 | 0 | 0.36 | 0.2 | 0.28 | 0.47 | 0.29 | 0.33 |
| 9 | 11590844 | rs1116978 | -0.728 | -0.354 | -0.66 | 0.026 | 0.012 | 0.003 | 0.003 | 0 | 0.5 | 0.64 | 0.46 | 0.47 | 0.51 | 0.43 |
| 9 | 11602294 | rs7026627 | -1.141 | -0.202 | -0.523 | 0.158 | -0.022 | -0.052 | -0.052 | 0 | 0.7 | 0.84 | 0.52 | 0.41 | 0.64 | 0.44 |
| 9 | 11605039 | rs10117313 | 0.936 | -0.256 | -0.493 | 0.003 | 0.001 | 0.038 | 0.038 | 0 | 0.4 | 0.32 | 0.26 | 0.25 | 0.43 | 0.2 |
| 9 | 11609469 | rs1838132 | 0.726 | -0.302 | -0.326 | -0.054 | 0.007 | 0.023 | 0.023 | 0 | 0.43 | 0.66 | 0.28 | 0.25 | 0.41 | 0.2 |
| 9 | 11609873 | rs4741169 | -0.975 | -0.332 | -0.33 | 0.074 | -0.003 | -0.043 | -0.043 | 0 | 0.7 | 0.81 | 0.54 | 0.47 | 0.66 | 0.47 |
| 9 | 11613259 | rs2821181 | -1.014 | -0.339 | -0.718 | 0.071 | -0.017 | -0.093 | -0.093 | 0 | 0.7 | 0.87 | 0.55 | 0.47 | 0.67 | 0.47 |
| 9 | 11617265 | rs1822387 | -1.934 | -0.478 | -1.286 | 0.213 | -0.05 | -0.099 | -0.099 | 2.459 | 0.87 | 0.99 | 0.65 | 0.55 | 0.81 | 0.64 |
| 9 | 11618203 | rs12236939 | 1.025 | -0.356 | -1.235 | 0.093 | 0.037 | -0.048 | -0.048 | 0 | 0.16 | 0.03 | 0.12 | 0.21 | 0.05 | 0.19 |
| 9 | 11645495 | rs10960058 | 0.622 | -0.324 | -1.159 | 0.046 | 0.046 | -0.015 | -0.015 | 0 | 0.51 | 0.23 | 0.35 | 0.47 | 0.44 | 0.37 |
| 9 | 11649257 | rs4741174 | 0.415 | -0.387 | -0.975 | 0.014 | 0.015 | -0.011 | -0.011 | 0.178 | 0.55 | 0.38 | 0.37 | 0.47 | 0.45 | 0.37 |
| 9 | 11656743 | rs10738244 | -0.082 | -0.431 | -1.005 | -0.101 | 0.006 | -0.036 | -0.036 | 0 | 0.59 | 0.8 | 0.4 | 0.45 | 0.44 | 0.37 |
| 9 | 11676507 | rs958842 | 1.982 | -0.443 | -1.373 | -0.16 | -0.059 | -0.068 | -0.068 | 0 | 0.19 | 0.01 | 0.16 | 0.25 | 0.25 | 0.17 |
| 9 | 11683177 | rs7033859 | 1.787 | 0.155 | -1.178 | -0.182 | -0.02 | -0.008 | -0.008 | 0 | 0.2 | 0.09 | 0.18 | 0.24 | 0.3 | 0.16 |
| 9 | 11683901 | rs2821204 | -1.118 | -0.243 | -1.194 | -0.24 | -0.038 | -0.023 | -0.023 | 0 | 0.65 | 0.43 | 0.78 | 0.79 | 0.81 | 0.79 |

| | | | | | | | | | | | | | | | | |
|---|---|---|---|---|---|---|---|---|---|---|---|---|---|---|---|---|
| 9 | 11683979 | rs2821203 | 1.208 | -0.037 | -1.899 | -0.313 | -0.032 | -0.111 | -0.111 | 0.14 | 0.18 | 0.03 | 0.37 | 0.47 | 0.2 | 0.39 |
| 9 | 11684502 | rs12378383 | 0.562 | 0.118 | -1.706 | -0.424 | -0.052 | 0.049 | 0.049 | 0.001 | 0.25 | 0.16 | 0.23 | 0.08 | 0.36 | 0.26 |
| 9 | 11693400 | rs2586335 | 0.599 | -0.041 | -1.973 | -0.323 | 0.037 | 0.057 | 0.057 | 0 | 0.45 | 0.13 | 0.53 | 0.67 | 0.37 | 0.54 |
| 9 | 11695108 | rs1470308 | 1.431 | -0.115 | -2.096 | -0.276 | -0.064 | -0.127 | -0.127 | 0.047 | 0.3 | 0.1 | 0.42 | 0.46 | 0.35 | 0.38 |
| 9 | 11695785 | rs7018532 | 1.932 | -0.247 | -2.023 | -0.371 | -0.008 | -0.016 | -0.016 | 0 | 0.2 | 0.08 | 0.16 | 0.25 | 0.25 | 0.16 |
| 9 | 11700487 | rs13296683 | 1.356 | -0.143 | -2.043 | -0.285 | -0.021 | -0.095 | -0.095 | 0.154 | 0.3 | 0.1 | 0.28 | 0.44 | 0.32 | 0.38 |
| 9 | 11711459 | rs10756267 | 1.714 | -0.148 | -2.064 | -0.218 | -0.021 | -0.076 | -0.076 | 0 | 0.14 | 0.03 | 0.34 | 0.42 | 0.15 | 0.33 |
| 9 | 11714572 | rs455131 | 0.462 | -0.188 | -1.804 | -0.433 | 0.056 | 0.059 | 0.059 | 0 | 0.45 | 0.18 | 0.53 | 0.66 | 0.34 | 0.54 |
| 9 | 11716862 | rs1486803 | -0.12 | -0.191 | -1.402 | -0.527 | 0.046 | -0.088 | -0.088 | 0 | 0.68 | 0.51 | 0.74 | 0.91 | 0.62 | 0.71 |
| 9 | 11725843 | rs4741178 | 0.313 | -0.215 | -1.456 | -0.387 | 0.053 | 0.061 | 0.061 | 0 | 0.47 | 0.32 | 0.55 | 0.66 | 0.38 | 0.55 |
| 9 | 11727020 | rs4741179 | -0.428 | 0.088 | -1.623 | -0.383 | -0.015 | 0.022 | 0.022 | 0 | 0.4 | 0.23 | 0.62 | 0.65 | 0.42 | 0.62 |
| 9 | 11743752 | rs10809575 | 0.005 | 0.372 | -1.54 | -0.013 | -0.023 | -0.003 | -0.003 | 0 | 0.19 | 0.12 | 0.3 | 0.16 | 0.22 | 0.25 |
| 9 | 11743846 | rs10809576 | -0.084 | 0.343 | -1.765 | -0.122 | -0.078 | -0.038 | -0.038 | 0 | 0.38 | 0.15 | 0.47 | 0.4 | 0.48 | 0.42 |
| 9 | 11747149 | rs1564999 | 1.54 | 0.606 | -1.59 | -0.135 | -0.013 | -0.008 | -0.008 | 0.58 | 0.06 | 0.03 | 0.19 | 0.16 | 0.08 | 0.16 |
| 9 | 11747626 | rs1486806 | 1.293 | 0.177 | -1.658 | -0.216 | -0.014 | 0.069 | 0.069 | 0 | 0.5 | 0.22 | 0.46 | 0.66 | 0.6 | 0.54 |
| 9 | 11747876 | rs10960164 | 0.977 | 0.474 | -1.634 | -0.273 | -0.023 | 0.014 | 0.014 | 0 | 0.13 | 0.1 | 0.14 | 0 | 0.2 | 0.1 |
| 9 | 11749911 | rs956095 | 0.591 | 0.259 | -1.491 | -0.187 | 0.014 | 0.029 | 0.029 | 0 | 0.25 | 0.12 | 0.18 | 0.18 | 0.18 | 0.21 |
| 9 | 11759484 | rs1979548 | 0.569 | 0.275 | -1.448 | -0.179 | 0.012 | 0.027 | 0.027 | 0 | 0.26 | 0.13 | 0.18 | 0.18 | 0.18 | 0.2 |
| 9 | 11775361 | rs10429595 | -0.654 | 0.479 | -0.889 | -0.2 | 0.125 | -0.107 | -0.107 | 1.451 | 0.85 | 0.58 | 0.83 | 1 | 0.8 | 0.9 |
| 9 | 11780023 | rs12376679 | 0.305 | 0.335 | -1.202 | -0.17 | -0.006 | 0.011 | 0.011 | 0.03 | 0.45 | 0.13 | 0.34 | 0.43 | 0.43 | 0.38 |
| 9 | 11782345 | rs10809590 | 0.32 | 0.328 | -1.206 | -0.174 | -0.007 | 0.009 | 0.009 | 0.015 | 0.45 | 0.13 | 0.34 | 0.42 | 0.43 | 0.38 |
| 9 | 11787410 | rs1825739 | 1.003 | 0.242 | -1.185 | -0.07 | 0 | 0 | 0 | 0.046 | 0.35 | 0.41 | 0.34 | 0.41 | 0.36 | 0.38 |
| 9 | 11787966 | rs10960221 | 0.288 | -0.014 | -1.283 | -0.022 | -0.006 | 0.011 | 0.011 | 0.034 | 0.45 | 0.13 | 0.34 | 0.43 | 0.43 | 0.38 |
| 9 | 11790536 | rs7867311 | -0.065 | 0.092 | -1.068 | -0.159 | 0.033 | 0.029 | 0.029 | 0.024 | 0.5 | 0.33 | 0.51 | 0.58 | 0.44 | 0.52 |
| 9 | 11798994 | rs1160534 | -0.351 | 0.371 | -0.844 | -0.392 | 0.037 | -0.097 | -0.097 | 0 | 0.84 | 0.49 | 0.83 | 1 | 0.84 | 0.9 |
| 9 | 11813213 | rs11999415 | 0.311 | -0.169 | -1.506 | -0.242 | 0.036 | 0.034 | 0.034 | 0 | 0.23 | 0.11 | 0.17 | 0.16 | 0.14 | 0.22 |
| 9 | 11813745 | rs1601615 | -0.218 | -0.075 | -1.773 | -0.251 | 0.031 | 0.033 | 0.033 | 0 | 0.44 | 0.25 | 0.38 | 0.39 | 0.43 | 0.4 |
| 9 | 11814915 | rs2129385 | 0.734 | 0.109 | -2.095 | -0.611 | 0.021 | 0.194 | 0.194 | 0 | 0.49 | 0.1 | 0.36 | 0.23 | 0.61 | 0.28 |
| 9 | 11820083 | rs10960278 | 2.146 | 0.195 | -1.935 | -0.394 | -0.065 | -0.058 | -0.058 | 0 | 0.2 | 0.01 | 0.18 | 0.23 | 0.28 | 0.17 |
| 9 | 11827745 | rs16928154 | 0.384 | -0.047 | -1.919 | -0.519 | 0.009 | 0.008 | 0.008 | 0 | 0.26 | 0.25 | 0.2 | 0.16 | 0.15 | 0.16 |
| 9 | 11829232 | rs10960295 | 2.442 | 0.034 | -1.969 | -0.472 | -0.066 | -0.06 | -0.06 | 0 | 0.18 | 0.01 | 0.17 | 0.23 | 0.26 | 0.18 |
| 9 | 11835471 | rs10732328 | -0.621 | -0.055 | -1.855 | -0.326 | -0.009 | 0.013 | 0.013 | 0 | 0.59 | 0.4 | 0.43 | 0.28 | 0.51 | 0.46 |
| 9 | 11838399 | rs10756295 | 1.289 | -0.05 | -1.938 | -0.394 | -0.063 | 0.064 | 0.064 | 0 | 0.3 | 0.1 | 0.19 | 0.11 | 0.33 | 0.26 |
| 9 | 11848378 | rs10960316 | 0.515 | -0.098 | -1.641 | 0.104 | 0.004 | 0.005 | 0.005 | 0 | 0.25 | 0.26 | 0.2 | 0.17 | 0.15 | 0.19 |
| 9 | 11850283 | rs16928182 | 1.849 | 0.032 | -1.643 | 0.192 | -0.001 | -0.001 | -0.001 | 0 | 0.06 | 0.06 | 0.03 | 0.08 | 0.06 | 0.01 |
| 9 | 11852043 | rs1948080 | 1.105 | 0.009 | -1.717 | 0.139 | -0.068 | 0.079 | 0.079 | 0 | 0.32 | 0.1 | 0.21 | 0.11 | 0.36 | 0.25 |
| 9 | 11857464 | rs6474675 | -0.294 | -0.015 | -1.696 | 0.142 | -0.009 | 0.013 | 0.013 | 0 | 0.57 | 0.43 | 0.4 | 0.28 | 0.49 | 0.45 |
| 9 | 11860446 | rs11999029 | 0.517 | -0.039 | -1.689 | 0.143 | 0.006 | 0.006 | 0.006 | 0 | 0.26 | 0.26 | 0.21 | 0.17 | 0.15 | 0.2 |
| 9 | 11861962 | rs973330 | -0.858 | 0.073 | -1.657 | 0.222 | 0.002 | 0.002 | 0.002 | 0.018 | 0.63 | 0.49 | 0.43 | 0.37 | 0.54 | 0.46 |
| 9 | 11869293 | rs4425849 | -1.882 | 0.127 | -1.372 | 0.329 | 0.032 | 0.216 | 0.216 | 1.01 | 0.84 | 0.65 | 0.64 | 0.6 | 0.83 | 0.66 |
| 9 | 11880552 | rs7049164 | -0.344 | 0.066 | -1.582 | 0.252 | 0.009 | 0.011 | 0.011 | 0.239 | 0.52 | 0.41 | 0.4 | 0.48 | 0.47 | 0.41 |
| 9 | 11883299 | rs10809637 | 0.953 | 0.103 | -1.74 | 0.219 | -0.066 | 0.109 | 0.109 | 0 | 0.32 | 0.09 | 0.21 | 0.08 | 0.36 | 0.25 |
| 9 | 11884852 | rs12006517 | 0.814 | -0.612 | -1.828 | 0.071 | 0.002 | 0.002 | 0.002 | 0.042 | 0.18 | 0.16 | 0.15 | 0.16 | 0.09 | 0.07 |
| 9 | 11885258 | rs1486802 | 0.314 | -0.559 | -1.738 | 0.204 | -0.02 | 0.08 | 0.08 | 0 | 0.44 | 0.26 | 0.27 | 0.17 | 0.43 | 0.41 |
| 9 | 11886098 | rs10960350 | 1.421 | -0.502 | -1.688 | 0.342 | -0.04 | -0.025 | -0.025 | 0 | 0.23 | 0.11 | 0.19 | 0.26 | 0.31 | 0.2 |
| 9 | 11887764 | rs17832803 | 2.072 | -0.458 | -1.889 | 0.111 | -0.045 | 0.11 | 0.11 | 0 | 0.19 | 0.04 | 0.1 | 0 | 0.22 | 0.05 |
| 9 | 11890045 | rs4401971 | -1.259 | -0.546 | -1.634 | 0.236 | 0.01 | 0.033 | 0.033 | 0.471 | 0.68 | 0.66 | 0.48 | 0.42 | 0.59 | 0.47 |
| 9 | 11891381 | rs12553961 | 1.435 | -0.89 | -1.8 | 0.092 | -0.005 | -0.002 | -0.002 | 0 | 0.14 | 0.13 | 0.11 | 0.16 | 0.03 | 0.02 |
| 9 | 11893047 | rs11532849 | 0.529 | -0.879 | -1.813 | 0.084 | -0.015 | 0.026 | 0.026 | 0 | 0.09 | 0.16 | 0.05 | 0.16 | 0.04 | 0.17 |
| 9 | 11903938 | rs12683024 | 2.667 | -0.775 | -1.929 | 0.08 | -0.052 | -0.013 | -0.013 | 0 | 0.17 | 0.02 | 0.15 | 0.17 | 0.25 | 0.21 |
| 9 | 11919732 | rs1999377 | -0.313 | -1.026 | -1.761 | 0.022 | -0.067 | 0.03 | 0.03 | 0.505 | 0.23 | 0.36 | 0.19 | 0.32 | 0.07 | 0.18 |
| 9 | 11929890 | rs10960376 | -0.275 | -0.95 | -1.979 | 0.046 | -0.005 | 0.001 | 0.001 | 0.057 | 0.36 | 0.32 | 0.25 | 0.28 | 0.37 | 0.38 |
| 9 | 11936751 | rs4642743 | 1.462 | -0.983 | -1.952 | 0.084 | -0.041 | 0.004 | 0.004 | 0 | 0.13 | 0.21 | 0.13 | 0.16 | 0.03 | 0.02 |
| 9 | 11937390 | rs10960385 | -0.364 | -0.859 | -2.101 | 0.058 | -0.006 | 0 | 0 | 0.316 | 0.37 | 0.34 | 0.25 | 0.29 | 0.38 | 0.39 |
| 9 | 11949207 | rs10960393 | 2.78 | -0.865 | -2.414 | 0.126 | -0.061 | -0.038 | -0.038 | 0.325 | 0.17 | 0.01 | 0.19 | 0.2 | 0.25 | 0.19 |
| 9 | 11951222 | rs1929412 | 1.725 | -1.063 | -2.183 | -0.174 | -0.014 | -0.001 | -0.001 | 0 | 0.12 | 0.14 | 0.12 | 0.17 | 0.03 | 0.02 |
| 9 | 11952643 | rs1929409 | -0.033 | -0.982 | -2.225 | -0.228 | -0.002 | 0.013 | 0.013 | 0 | 0.32 | 0.25 | 0.22 | 0.2 | 0.31 | 0.38 |
| 9 | 11953279 | rs10960397 | 1.621 | -0.752 | -2.435 | -0.149 | -0.108 | -0.173 | -0.173 | 0 | 0.3 | 0.03 | 0.49 | 0.55 | 0.41 | 0.54 |
| 9 | 11958106 | rs10809655 | 1.867 | -0.653 | -1.666 | 0.534 | -0.061 | 0.104 | 0.104 | 0 | 0.17 | 0.01 | 0.09 | 0 | 0.19 | 0.04 |
| 9 | 11960277 | rs7037332 | -0.918 | -0.805 | -0.734 | 1.418 | -0.017 | 0.039 | 0.039 | 0 | 0.23 | 0.35 | 0.18 | 0.42 | 0.11 | 0.17 |
| 9 | 11967114 | rs10960403 | 0.973 | -0.556 | -1.155 | 1.544 | 0.005 | 0.014 | 0.014 | 0 | 0.06 | 0.03 | 0.03 | 0.16 | 0.01 | 0.15 |
| 9 | 11976830 | rs16928298 | 0.169 | -0.093 | -0.918 | 1.034 | -0.002 | 0.007 | 0.007 | 0 | 0.15 | 0.22 | 0.14 | 0.25 | 0.08 | 0.03 |
| 9 | 11995297 | rs6474681 | 0.916 | -0.006 | -0.902 | 1.091 | 0.002 | -0.006 | -0.006 | 0 | 0.11 | 0.12 | 0.26 | 0.27 | 0.06 | 0.05 |
| 9 | 11996831 | rs4512479 | 1.076 | 0.19 | -0.827 | 1.049 | -0.008 | 0.011 | 0.011 | 0 | 0.4 | 0.31 | 0.29 | 0.33 | 0.43 | 0.49 |
| 9 | 12000099 | rs10809663 | 1.563 | 0.228 | -1.692 | 0.948 | -0.091 | 0.137 | 0.137 | 0 | 0.35 | 0.04 | 0.23 | 0.17 | 0.41 | 0.38 |
| 9 | 12003252 | rs10809666 | 0.421 | 1.083 | -1.524 | 1.009 | -0.08 | -0.118 | -0.118 | 0 | 0.2 | 0.03 | 0.22 | 0.38 | 0.25 | 0.31 |
| 9 | 12005070 | rs7048594 | 1.071 | 0.964 | -1.392 | 1.014 | -0.004 | 0.04 | 0.04 | 0.007 | 0.11 | 0.19 | 0.27 | 0.27 | 0.06 | 0.06 |
| 9 | 12007658 | rs12001461 | 0.472 | 0.927 | -1.431 | 0.929 | 0.004 | 0.071 | 0.071 | 0 | 0.24 | 0.07 | 0.15 | 0.01 | 0.26 | 0.14 |
| 9 | 12013711 | rs10756313 | -0.363 | 1.228 | -0.733 | 2.23 | -0.08 | -0.117 | -0.117 | 0.018 | 0.19 | 0.03 | 0.22 | 0.4 | 0.25 | 0.31 |

| | | | | | | | | | | | | | | | | |
|---|---|---|---|---|---|---|---|---|---|---|---|---|---|---|---|---|
| 9 | 12020873 | rs1446255 | 0.005 | 0.863 | 0.174 | 1.941 | -0.004 | 0.046 | 0.046 | 0 | 0.11 | 0.15 | 0.27 | 0.27 | 0.06 | 0.05 |
| 9 | 12046842 | rs12339814 | -0.769 | 1.402 | -0.391 | 1.806 | -0.001 | 0.036 | 0.036 | 0 | 0.1 | 0.11 | 0.33 | 0.37 | 0.14 | 0.26 |
| 9 | 12048206 | rs12238357 | -1.66 | 0.284 | 0.075 | 1.554 | -0.019 | 0.016 | 0.016 | 0 | 0.23 | 0.38 | 0.37 | 0.65 | 0.18 | 0.39 |
| 9 | 12051998 | rs9886865 | 0.24 | 0.248 | 0.579 | 1.944 | 0.02 | 0.157 | 0.157 | 0 | 0.91 | 0.88 | 0.65 | 0.54 | 0.87 | 0.76 |
| 9 | 12054659 | rs9969851 | 1.736 | 0.077 | 0.676 | 1.852 | -0.041 | 0.016 | 0.016 | 0 | 0.74 | 0.5 | 0.57 | 0.32 | 0.81 | 0.58 |
| 9 | 12056279 | rs7040466 | -1.026 | -0.031 | 0.55 | 1.781 | -0.009 | -0.004 | -0.004 | 0 | 0.15 | 0.18 | 0.06 | 0.17 | 0.07 | 0.14 |
| 9 | 12058197 | rs10960455 | -0.423 | 0.073 | 0.523 | 1.88 | -0.016 | 0.025 | 0.025 | 0 | 0.09 | 0.04 | 0.32 | 0.44 | 0.13 | 0.25 |
| 9 | 12065319 | rs1374177 | 1.561 | 0.263 | 0.364 | 1.561 | -0.021 | 0.017 | 0.017 | 0 | 0.75 | 0.59 | 0.57 | 0.36 | 0.82 | 0.58 |
| 9 | 12067552 | rs12686721 | 0.129 | 0.23 | -0.044 | 0.992 | -0.008 | -0.006 | -0.006 | 0.409 | 0.14 | 0.07 | 0.18 | 0.18 | 0.15 | 0.07 |
| 9 | 12067914 | rs7863023 | -0.795 | -0.222 | 0.011 | 0.948 | 0 | -0.008 | -0.008 | 0 | 0.13 | 0.14 | 0.05 | 0.17 | 0.05 | 0.16 |
| 9 | 12078499 | rs4449884 | -0.855 | 0.226 | 0.054 | 0.659 | 0 | -0.02 | -0.02 | 0 | 0.07 | 0.11 | 0.04 | 0 | 0.07 | 0.03 |
| 9 | 12082380 | rs10960462 | -1.544 | -0.211 | -0.028 | -0.006 | -0.001 | -0.004 | -0.004 | 0.043 | 0.54 | 0.58 | 0.39 | 0.29 | 0.51 | 0.42 |
| 9 | 12084080 | rs7859230 | -1.911 | -0.282 | 0.064 | 0.062 | -0.013 | -0.013 | -0.013 | 0 | 0.62 | 0.75 | 0.43 | 0.43 | 0.54 | 0.54 |
| 9 | 12085717 | rs7036903 | 0.92 | -0.346 | -0.262 | -0.201 | -0.002 | -0.005 | -0.005 | 0.703 | 0.71 | 0.72 | 0.53 | 0.41 | 0.72 | 0.64 |
| 9 | 12092193 | rs10960466 | 1.814 | -0.327 | -0.345 | -0.44 | -0.043 | 0.037 | 0.037 | 0 | 0.07 | 0 | 0.05 | 0 | 0.12 | 0.08 |
| 9 | 12092895 | rs10809694 | 0.722 | -0.288 | -0.262 | -0.374 | -0.031 | 0.012 | 0.012 | 0 | 0.16 | 0.04 | 0.11 | 0.1 | 0.17 | 0.14 |
| 9 | 12093072 | rs13297287 | 0.684 | -0.416 | -0.362 | -0.519 | -0.003 | 0.06 | 0.06 | 10.873 | 0.16 | 0.03 | 0.1 | 0.04 | 0.12 | 0.13 |
| 9 | 12095022 | rs10738261 | 0.281 | -0.43 | 0.077 | -0.525 | -0.02 | -0.037 | -0.037 | 0 | 0.64 | 0.5 | 0.73 | 0.78 | 0.7 | 0.74 |
| 9 | 12098000 | rs1446268 | -0.888 | -0.532 | 0.601 | -0.582 | -0.012 | -0.078 | -0.078 | 0 | 0.83 | 0.63 | 0.88 | 0.97 | 0.83 | 0.93 |
| 9 | 12099118 | rs10960477 | 0.279 | -0.499 | 0.489 | -0.577 | -0.039 | 0.038 | 0.038 | 0 | 0.2 | 0.06 | 0.13 | 0.1 | 0.22 | 0.14 |
| 9 | 12100880 | rs2382310 | 3.586 | -0.533 | 0.499 | -0.646 | -0.052 | 0.101 | 0.101 | 0 | 0.27 | 0.07 | 0.23 | 0.08 | 0.38 | 0.25 |
| 9 | 12101436 | rs1348742 | 1.634 | -0.522 | 0.573 | -0.591 | -0.008 | 0.024 | 0.024 | 0 | 0.41 | 0.33 | 0.3 | 0.22 | 0.47 | 0.4 |
| 9 | 12106293 | rs12000494 | 0.337 | -0.528 | 0.51 | -0.574 | -0.041 | 0.038 | 0.038 | 0 | 0.2 | 0.06 | 0.13 | 0.1 | 0.22 | 0.15 |
| 9 | 12107758 | rs11515183 | 1.53 | -0.366 | 0.914 | -0.218 | -0.013 | 0.036 | 0.036 | 0 | 0.35 | 0.2 | 0.26 | 0.22 | 0.42 | 0.37 |
| 9 | 12108916 | rs12351590 | -0.239 | -0.407 | 0.984 | -0.136 | -0.001 | 0.23 | 0.23 | 0 | 0.92 | 0.8 | 0.69 | 0.52 | 0.94 | 0.78 |
| 9 | 12112566 | rs7468068 | -0.311 | -0.402 | 0.99 | -0.167 | -0.002 | 0.024 | 0.024 | 0.151 | 0.41 | 0.31 | 0.31 | 0.25 | 0.35 | 0.34 |
| 9 | 12113011 | rs10435753 | -0.761 | -0.369 | 0.952 | -0.062 | -0.01 | 0.051 | 0.051 | 0 | 0.13 | 0.2 | 0.34 | 0.63 | 0.09 | 0.34 |
| 9 | 12124410 | rs10960488 | 0.141 | -0.343 | 1.001 | -0.04 | -0.003 | 0.19 | 0.19 | 0 | 0.08 | 0.18 | 0.34 | 0.5 | 0.06 | 0.22 |
| 9 | 12131415 | rs10960493 | 1.008 | -0.352 | 1.034 | -0.155 | -0.002 | -0.001 | -0.001 | 0 | 0.42 | 0.43 | 0.31 | 0.25 | 0.48 | 0.4 |
| 9 | 12133321 | rs1114756 | 2.396 | -0.327 | 0.991 | -0.212 | -0.027 | 0.06 | 0.06 | 0 | 0.29 | 0.14 | 0.24 | 0.07 | 0.38 | 0.26 |
| 9 | 12139870 | rs791773 | -0.554 | -0.282 | 0.91 | -0.165 | 0 | -0.01 | -0.01 | 0 | 0.48 | 0.44 | 0.33 | 0.15 | 0.45 | 0.33 |
| 9 | 12144717 | rs10960499 | 0.113 | -0.36 | 0.787 | -0.167 | -0.003 | 0.052 | 0.052 | 1.697 | 0.34 | 0.21 | 0.26 | 0.15 | 0.29 | 0.26 |
| 9 | 12149367 | rs7048056 | 0.63 | -0.158 | 0.422 | -0.097 | -0.001 | -0.028 | -0.028 | 0.068 | 0.15 | 0.14 | 0.12 | 0 | 0.19 | 0.08 |
| 9 | 12152990 | rs1446264 | 0.243 | -0.225 | 0.35 | -0.03 | -0.028 | 0.006 | 0.006 | 0.319 | 0.33 | 0.08 | 0.27 | 0.27 | 0.3 | 0.31 |
| 9 | 12156819 | rs10809701 | 1.401 | -0.167 | 0.438 | 0.125 | -0.015 | 0.037 | 0.037 | 0 | 0.36 | 0.13 | 0.25 | 0.28 | 0.4 | 0.38 |
| 9 | 12161870 | rs10117456 | 2.009 | -0.213 | 0.408 | 0.125 | -0.032 | 0.07 | 0.07 | 0 | 0.31 | 0.1 | 0.23 | 0.15 | 0.39 | 0.28 |
| 9 | 12163583 | rs10115948 | 2.068 | -0.195 | 0.411 | 0.1 | -0.037 | 0.073 | 0.073 | 0 | 0.3 | 0.08 | 0.22 | 0.14 | 0.38 | 0.28 |
| 9 | 12165072 | rs10960502 | -0.151 | -0.193 | 0.522 | 0.11 | -0.013 | 0.1 | 0.1 | 0.178 | 0.46 | 0.27 | 0.33 | 0.16 | 0.47 | 0.31 |
| 9 | 12165758 | rs7025259 | 0.393 | -0.051 | 0.448 | 0.262 | -0.003 | 0.181 | 0.181 | 0 | 0.91 | 0.81 | 0.65 | 0.45 | 0.93 | 0.71 |
| 9 | 12168247 | rs10809708 | 1.903 | 0.031 | -0.055 | 0.155 | -0.03 | 0.068 | 0.068 | 0 | 0.3 | 0.1 | 0.22 | 0.15 | 0.38 | 0.27 |
| 9 | 12185185 | rs4285563 | -0.689 | 0.154 | -0.009 | 0.181 | -0.005 | 0.077 | 0.077 | 0 | 0.94 | 0.96 | 0.68 | 0.53 | 0.93 | 0.8 |
| 9 | 12192428 | rs10809716 | 1.788 | 0.041 | -0.121 | -0.042 | -0.038 | 0.117 | 0.117 | 0 | 0.31 | 0.08 | 0.23 | 0.06 | 0.38 | 0.27 |
| 9 | 12194296 | rs10738266 | 0.46 | 0.012 | 0.13 | -0.111 | 0.007 | 0.081 | 0.081 | 0 | 0.74 | 0.59 | 0.54 | 0.51 | 0.71 | 0.71 |
| 9 | 12194855 | rs2066096 | 0.731 | -0.017 | -0.064 | -0.095 | 0.006 | -0.004 | -0.004 | 0.226 | 0.05 | 0.03 | 0.02 | 0.16 | 0.03 | 0.12 |
| 9 | 12197936 | rs1339272 | 0.068 | -0.002 | -0.005 | -0.114 | -0.005 | 0.076 | 0.076 | 0 | 0.79 | 0.71 | 0.57 | 0.52 | 0.77 | 0.72 |
| 9 | 12200957 | rs7852109 | -0.663 | 0.058 | -0.048 | -0.437 | -0.009 | -0.018 | -0.018 | 0 | 0.26 | 0.7 | 0.2 | 0.15 | 0.24 | 0.16 |
| 9 | 12201384 | rs7870825 | -0.514 | 0.172 | -0.103 | -0.488 | -0.035 | -0.026 | -0.026 | 0 | 0.64 | 0.9 | 0.48 | 0.37 | 0.65 | 0.56 |
| 9 | 12201866 | rs16928550 | -1.279 | -0.026 | 0.031 | -0.353 | 0.001 | -0.001 | -0.001 | 0.521 | 0.31 | 0.74 | 0.24 | 0.34 | 0.28 | 0.29 |
| 9 | 12213414 | rs1010663 | 0.199 | 0.316 | 0.192 | 0.007 | -0.004 | 0.008 | 0.008 | 0 | 0.46 | 0.64 | 0.54 | 0.43 | 0.48 | 0.47 |
| 9 | 12213879 | rs16928562 | -1.588 | 0.488 | 0.876 | 0.6 | 0.04 | 0.132 | 0.132 | 0.047 | 0.1 | 0.41 | 0.27 | 0.21 | 0.08 | 0.09 |
| 9 | 12216024 | rs10733241 | 0.894 | 0.417 | 0.193 | 0.495 | -0.002 | 0.008 | 0.008 | 0.053 | 0.49 | 0.32 | 0.44 | 0.53 | 0.53 | 0.53 |
| 9 | 12216756 | rs7875540 | -0.789 | 0.345 | -0.21 | 0.62 | -0.04 | 0.01 | 0.01 | 0.623 | 0.42 | 0.2 | 0.32 | 0.61 | 0.4 | 0.65 |
| 9 | 12224275 | rs10960535 | 0.381 | 0.403 | -0.329 | 0.48 | -0.033 | 0.029 | 0.029 | 0.556 | 0.14 | 0.03 | 0.06 | 0.07 | 0.17 | 0.11 |
| 9 | 12234642 | rs10116548 | 1.316 | 0.587 | 0.255 | 0.707 | -0.077 | 0.197 | 0.197 | 0 | 0.43 | 0.09 | 0.34 | 0.16 | 0.5 | 0.27 |
| 9 | 12234969 | rs946452 | -0.104 | 0.568 | 0.419 | 0.724 | 0.008 | -0.041 | -0.041 | 0 | 0.37 | 0.25 | 0.33 | 0.5 | 0.33 | 0.53 |
| 9 | 12237417 | rs1339284 | 1.47 | 0.558 | 0.43 | 0.654 | -0.006 | 0.043 | 0.043 | 0 | 0.8 | 0.34 | 0.67 | 0.67 | 0.84 | 0.8 |
| 9 | 12239604 | rs10960541 | 1.43 | 0.573 | 0.392 | 0.63 | -0.096 | 0.208 | 0.208 | 0 | 0.43 | 0.07 | 0.35 | 0.16 | 0.52 | 0.27 |
| 9 | 12240518 | rs2382337 | 0.556 | 0.549 | 0.459 | 0.628 | -0.007 | 0.031 | 0.031 | 0 | 0.87 | 0.58 | 0.91 | 0.9 | 0.88 | 0.91 |
| 9 | 12241585 | rs10809740 | -0.171 | 0.626 | 0.183 | 0.674 | 0.017 | -0.039 | -0.039 | 0.011 | 0.37 | 0.25 | 0.33 | 0.51 | 0.32 | 0.54 |
| 9 | 12243007 | rs4741216 | 0.576 | 0.63 | 0.411 | 0.64 | 0 | 0 | 0 | 0 | 0.53 | 0.53 | 0.59 | 0.39 | 0.57 | 0.38 |
| 9 | 12247973 | rs4477124 | 0.838 | 0.7 | 0.193 | 0.673 | 0 | 0.003 | 0.003 | 0 | 0.51 | 0.44 | 0.36 | 0.27 | 0.51 | 0.37 |
| 9 | 12248419 | rs4008045 | -0.005 | 0.702 | 0.187 | 0.697 | 0.011 | 0.02 | 0.02 | 0 | 0.35 | 0.31 | 0.4 | 0.27 | 0.28 | 0.26 |
| 9 | 12255238 | rs7860853 | -0.517 | 0.263 | 0.016 | 0.719 | 0.002 | -0.026 | -0.026 | 0.066 | 0.42 | 0.46 | 0.25 | 0.24 | 0.37 | 0.37 |
| 9 | 12256058 | rs12340485 | 0.123 | 0.296 | -0.057 | 0.72 | -0.04 | 0.134 | 0.134 | 0.499 | 0.34 | 0.09 | 0.2 | 0.14 | 0.32 | 0.27 |
| 9 | 12256453 | rs11791609 | 1.146 | 0.625 | -0.269 | 0.906 | -0.048 | 0.185 | 0.185 | 0.49 | 0.42 | 0.07 | 0.32 | 0.17 | 0.45 | 0.25 |
| 9 | 12261388 | rs4740521 | 0.979 | 0.591 | -0.231 | 0.872 | -0.009 | 0.079 | 0.079 | 0.946 | 0.45 | 0.27 | 0.54 | 0.3 | 0.47 | 0.26 |
| 9 | 12263833 | rs1416576 | 0.208 | 0.592 | -0.286 | 0.945 | 0.002 | 0.147 | 0.147 | 0 | 0.87 | 0.76 | 0.78 | 0.54 | 0.82 | 0.63 |
| 9 | 12267378 | rs1538515 | -0.707 | 0.77 | -0.752 | -0.361 | -0.001 | -0.037 | -0.037 | 0.157 | 0.4 | 0.47 | 0.23 | 0.26 | 0.34 | 0.35 |
| 9 | 12272744 | rs7874304 | -0.434 | 0.571 | -1.324 | -0.687 | -0.001 | -0.028 | -0.028 | 0.066 | 0.37 | 0.38 | 0.23 | 0.14 | 0.35 | 0.24 |

| | | | | | | | | | | | | | | | |
|---|---|---|---|---|---|---|---|---|---|---|---|---|---|---|---|
| 9 | 12276668 | rs17262250 | -0.051 | 0.371 | -2.267 | -0.603 | -0.009 | 0.217 | 0.217 | 0 | 0.37 | 0.02 | 0.23 | 0.07 | 0.37 | 0.32 |
| 9 | 12277388 | rs10809747 | -0.893 | 1.04 | -1.235 | 0.919 | 0.018 | 0.101 | 0.101 | 0 | 0.12 | 0.21 | 0.21 | 0.58 | 0.14 | 0.24 |
| 9 | 12278150 | rs951176 | -0.67 | 0.47 | -1.581 | 1.61 | 0.003 | -0.045 | -0.045 | 1.563 | 0.09 | 0.02 | 0.27 | 0.36 | 0.06 | 0.2 |
| 9 | 12284611 | rs10809748 | -0.623 | 0.254 | -1.392 | 1.372 | 0 | -0.001 | -0.001 | 0.366 | 0.3 | 0.31 | 0.15 | 0.2 | 0.24 | 0.3 |
| 9 | 12285170 | rs10283912 | -1.404 | 0.751 | -1.084 | 0.992 | 0.015 | 0.113 | 0.113 | 0 | 0.52 | 0.8 | 0.66 | 0.8 | 0.49 | 0.67 |
| 9 | 12288602 | rs7873941 | -0.262 | 1.113 | -0.432 | 0.672 | 0.045 | 0.022 | 0.022 | 0 | 0.13 | 0.39 | 0.12 | 0.12 | 0.11 | 0.14 |
| 9 | 12290837 | rs11794979 | -0.11 | 0.896 | -0.756 | 0.502 | -0.074 | 0.096 | 0.096 | 0 | 0.3 | 0.04 | 0.16 | 0.16 | 0.25 | 0.3 |
| 9 | 12297444 | rs7031915 | -1.616 | 0.897 | 0.44 | 0.506 | 0.071 | 0.021 | 0.021 | 1.088 | 0.07 | 0.52 | 0.07 | 0 | 0.06 | 0.04 |
| 9 | 12305664 | rs7043787 | 0.65 | 0.954 | 0.808 | 0.49 | -0.007 | 0.005 | 0.005 | 0 | 0.84 | 0.89 | 0.83 | 0.9 | 0.81 | 0.79 |
| 9 | 12317717 | rs10809751 | -0.741 | 0.673 | 0.917 | 0.273 | 0 | 0.039 | 0.039 | 0 | 0.48 | 0.37 | 0.28 | 0.27 | 0.41 | 0.42 |
| 9 | 12318093 | rs7040701 | -1.286 | 0.66 | 1.169 | 0.267 | -0.011 | 0.072 | 0.072 | 0 | 0.54 | 0.74 | 0.34 | 0.3 | 0.47 | 0.45 |
| 9 | 12319300 | rs10960592 | -0.026 | 0.679 | 1.045 | 0.235 | 0.005 | 0.055 | 0.055 | 0 | 0.37 | 0.27 | 0.2 | 0.16 | 0.31 | 0.31 |
| 9 | 12333231 | rs6474701 | -1.159 | 0.626 | 0.85 | 0.179 | -0.005 | 0.062 | 0.062 | 0 | 0.53 | 0.69 | 0.3 | 0.27 | 0.47 | 0.44 |
| 9 | 12350564 | rs1928873 | 1.481 | 0.288 | 0.361 | 0.156 | -0.099 | 0.001 | 0.001 | 0 | 0.33 | 0.02 | 0.26 | 0.22 | 0.41 | 0.36 |
| 9 | 12350993 | rs7869172 | -0.089 | 0.25 | 0.604 | 0.091 | 0.032 | 0.058 | 0.058 | 0 | 0.12 | 0.33 | 0.1 | 0.14 | 0.11 | 0.12 |
| 9 | 12351048 | rs4741220 | -0.795 | 0.261 | -0.26 | -0.298 | 0.001 | -0.034 | -0.034 | 4.835 | 0.88 | 0.83 | 0.85 | 1 | 0.87 | 0.88 |
| 9 | 12352035 | rs10960614 | 0.268 | 0.268 | -0.328 | -0.293 | -0.004 | 0.158 | 0.158 | 0.01 | 0.37 | 0.17 | 0.15 | 0.13 | 0.31 | 0.31 |
| 9 | 12352439 | rs10960615 | 1.725 | 0.305 | -0.334 | -0.294 | -0.095 | 0.002 | 0.002 | 0 | 0.33 | 0.02 | 0.26 | 0.22 | 0.41 | 0.39 |
| 9 | 12352971 | rs10960616 | 0.043 | 0.222 | 0.028 | -0.634 | -0.004 | 0.158 | 0.158 | 0.01 | 0.37 | 0.17 | 0.15 | 0.13 | 0.31 | 0.31 |
| 9 | 12352992 | rs7863513 | 0.489 | 0.239 | 0.061 | -0.534 | 0.002 | 0.075 | 0.075 | 0 | 0.5 | 0.61 | 0.68 | 0.86 | 0.56 | 0.59 |
| 9 | 12356017 | rs1331350 | -0.08 | 0.266 | 0.264 | -0.21 | 0.004 | -0.059 | -0.059 | 0 | 0.87 | 0.8 | 0.84 | 1 | 0.87 | 0.89 |
| 9 | 12359563 | rs4628342 | -0.826 | 0.29 | 0.268 | -0.196 | 0.004 | 0 | 0 | 0 | 0.55 | 0.4 | 0.67 | 0.59 | 0.59 | 0.42 |
| 9 | 12363134 | rs1412074 | -0.829 | 0.614 | 0.811 | -0.371 | -0.004 | 0.002 | 0.002 | 0 | 0.55 | 0.62 | 0.66 | 0.58 | 0.48 | 0.38 |
| 9 | 12363456 | rs1331359 | 0.46 | 0.568 | 0.638 | -0.401 | -0.005 | -0.05 | -0.05 | 0.388 | 0.14 | 0.05 | 0.42 | 0.53 | 0.13 | 0.16 |
| 9 | 12364718 | rs12338989 | 0.795 | 0.487 | 0.909 | -0.393 | -0.032 | -0.015 | -0.015 | 0 | 0.44 | 0.21 | 0.32 | 0.42 | 0.53 | 0.57 |
| 9 | 12366893 | rs12686127 | 0.598 | 0.252 | 0.902 | -0.334 | -0.014 | 0.045 | 0.045 | 0 | 0.11 | 0.21 | 0.05 | 0.27 | 0.07 | 0.22 |
| 9 | 12367988 | rs1929425 | -0.829 | 0.259 | 0.637 | -0.377 | 0.043 | 0.007 | 0.007 | 0.072 | 0.55 | 0.27 | 0.66 | 0.58 | 0.59 | 0.43 |
| 9 | 12370395 | rs10809762 | 1.315 | 0.208 | 0.703 | -0.354 | -0.033 | 0.085 | 0.085 | 0 | 0.36 | 0.15 | 0.29 | 0.15 | 0.45 | 0.4 |
| 9 | 12381912 | rs1331367 | 0.685 | 0.19 | 0.715 | -0.308 | -0.039 | 0.067 | 0.067 | 1.421 | 0.85 | 0.3 | 0.9 | 0.74 | 0.92 | 0.77 |
| 9 | 12382539 | rs1952692 | 0.117 | 0.299 | 0.509 | -0.467 | 0.001 | 0.06 | 0.06 | 0 | 0.38 | 0.23 | 0.2 | 0.05 | 0.33 | 0.22 |
| 9 | 12397405 | rs1360461 | 0.595 | 0.391 | 0.521 | -0.354 | 0.009 | 0.058 | 0.058 | 0 | 0.47 | 0.31 | 0.73 | 0.69 | 0.59 | 0.54 |
| 9 | 12397876 | rs2777007 | 0.559 | 0.375 | 0.647 | -0.321 | 0.003 | 0.115 | 0.115 | 0 | 0.16 | 0.28 | 0.46 | 0.55 | 0.14 | 0.22 |
| 9 | 12401666 | rs10756369 | 1.688 | 0.346 | 0.498 | -0.45 | -0.143 | 0.062 | 0.062 | 0 | 0.31 | 0.02 | 0.27 | 0.13 | 0.45 | 0.33 |
| 9 | 12404591 | rs1331346 | -0.573 | 0.344 | 0.666 | -0.365 | 0.012 | 0.211 | 0.211 | 0 | 0.84 | 0.63 | 0.53 | 0.46 | 0.86 | 0.78 |
| 9 | 12407578 | rs10116714 | 0.564 | 0.347 | 0.653 | -0.364 | 0.008 | 0.167 | 0.167 | 0 | 0.16 | 0.33 | 0.47 | 0.55 | 0.14 | 0.22 |
| 9 | 12421476 | rs7871312 | -1.666 | 0.32 | 0.511 | -0.369 | -0.113 | 0.042 | 0.042 | 0 | 0.67 | 0.97 | 0.72 | 0.83 | 0.55 | 0.64 |
| 9 | 12421929 | rs12683218 | 0.059 | 0.362 | 0.382 | -0.698 | -0.002 | 0 | 0 | 0.051 | 0.29 | 0.34 | 0.16 | 0.32 | 0.25 | 0.38 |
| 9 | 12425767 | rs10491747 | 0.685 | 0.345 | 0.27 | -0.81 | 0.013 | 0.095 | 0.095 | 0 | 0.54 | 0.26 | 0.39 | 0.13 | 0.61 | 0.41 |
| 9 | 12433463 | rs7856061 | 0.265 | 0.381 | 0.288 | -0.833 | -0.005 | -0.082 | -0.082 | 0 | 0.19 | 0.25 | 0.1 | 0 | 0.13 | 0.06 |
| 9 | 12442019 | rs1999380 | 0.519 | 0.266 | 0.348 | -0.049 | 0.003 | 0.003 | 0.003 | 0 | 0.55 | 0.62 | 0.66 | 0.44 | 0.58 | 0.43 |
| 9 | 12443586 | rs7860071 | -0.176 | 0.225 | 0.391 | -0.209 | -0.011 | 0.015 | 0.015 | 0.251 | 0.67 | 0.79 | 0.75 | 0.74 | 0.69 | 0.67 |
| 9 | 12445143 | rs1537181 | 0.008 | 0.23 | 0.296 | -0.307 | 0.003 | -0.06 | -0.06 | 0 | 0.23 | 0.24 | 0.14 | 0 | 0.23 | 0.13 |
| 9 | 12448743 | rs10756372 | -0.223 | 0.085 | 0.232 | -0.245 | -0.01 | 0.015 | 0.015 | 0.244 | 0.67 | 0.78 | 0.75 | 0.73 | 0.7 | 0.67 |
| 9 | 12452348 | rs10809779 | 0.692 | 0.16 | 0.326 | -0.905 | 0.007 | 0.047 | 0.047 | 0 | 0.53 | 0.35 | 0.39 | 0.12 | 0.59 | 0.42 |
| 9 | 12453240 | rs4740523 | -1.109 | 0.235 | 0.173 | -0.879 | -0.001 | -0.014 | -0.014 | 0.077 | 0.8 | 0.82 | 0.9 | 0.96 | 0.83 | 0.84 |
| 9 | 12456328 | rs2065367 | 1.086 | 0.328 | -0.113 | -0.694 | 0.006 | 0.031 | 0.031 | 0.091 | 0.12 | 0.04 | 0.06 | 0 | 0.09 | 0.02 |
| 9 | 12456576 | rs10960672 | 0.157 | 0.192 | -0.047 | -0.576 | 0.008 | 0.065 | 0.065 | 0 | 0.53 | 0.59 | 0.41 | 0.16 | 0.59 | 0.41 |
| 9 | 12460256 | rs4008031 | 0.994 | -0.122 | 0.036 | 0.185 | -0.004 | -0.025 | -0.025 | 0 | 0.3 | 0.34 | 0.27 | 0.15 | 0.34 | 0.28 |
| 9 | 12463411 | rs2026326 | 0.819 | -0.077 | 0.082 | 0.762 | -0.049 | 0.146 | 0.146 | 1.131 | 0.83 | 0.57 | 0.9 | 0.66 | 0.89 | 0.77 |
| 9 | 12469893 | rs11787776 | 0.227 | 0.441 | -1.056 | -0.008 | -0.041 | 0.066 | 0.066 | 0.013 | 0.33 | 0.16 | 0.27 | 0.11 | 0.47 | 0.36 |
| 9 | 12470048 | rs10756373 | 0.556 | 0.065 | -0.718 | 0.474 | -0.006 | 0.013 | 0.013 | 0.015 | 0.79 | 0.71 | 0.83 | 0.7 | 0.84 | 0.72 |
| 9 | 12476727 | rs7045858 | 1.307 | 0.054 | -0.787 | 0.519 | -0.001 | 0.001 | 0.001 | 0 | 0.79 | 0.72 | 0.55 | 0.19 | 0.82 | 0.66 |
| 9 | 12493565 | rs791658 | -0.949 | -0.253 | -0.75 | 1.149 | -0.032 | 0.283 | 0.283 | 0.268 | 0.07 | 0.23 | 0.33 | 0.57 | 0.01 | 0.09 |
| 9 | 12494501 | rs791660 | -0.58 | -0.262 | -0.695 | 1.192 | -0.029 | 0.026 | 0.026 | 0.075 | 0.06 | 0.19 | 0.07 | 0.07 | 0.01 | 0.03 |
| 9 | 12495309 | rs7020220 | 0.345 | -0.333 | -1.439 | 0.264 | -0.063 | 0.098 | 0.098 | 0.586 | 0.4 | 0.12 | 0.3 | 0.14 | 0.5 | 0.38 |
| 9 | 12497036 | rs7467390 | -1.114 | -0.375 | -1.107 | 0.336 | -0.015 | 0.038 | 0.038 | 0 | 0.17 | 0.26 | 0.38 | 0.64 | 0.09 | 0.17 |
| 9 | 12499211 | rs2021997 | 0.449 | 0.051 | -1.599 | -0.156 | -0.1 | 0.102 | 0.102 | 0 | 0.33 | 0.05 | 0.26 | 0.11 | 0.49 | 0.34 |
| 9 | 12501498 | rs791662 | -1.08 | 0.003 | -1.582 | -0.58 | -0.09 | 0.067 | 0.067 | 0 | 0.79 | 0.93 | 0.71 | 0.99 | 0.54 | 0.87 |
| 9 | 12502531 | rs4741226 | -0.549 | -0.765 | -1.066 | -1.115 | -0.014 | 0.004 | 0.004 | 0 | 0.67 | 0.74 | 0.67 | 0.98 | 0.47 | 0.8 |
| 9 | 12509667 | rs16929075 | 0.687 | -1.777 | -0.822 | -0.831 | -0.004 | 0.005 | 0.005 | 0.078 | 0.23 | 0.24 | 0.42 | 0.63 | 0.14 | 0.16 |
| 9 | 12510515 | rs791667 | -0.57 | -2.2 | -0.773 | -0.875 | -0.002 | -0.001 | -0.001 | 0 | 0.25 | 0.32 | 0.16 | 0.24 | 0.2 | 0.41 |
| 9 | 12512973 | rs1325129 | -0.565 | -2.279 | -0.349 | -1.138 | 0.008 | 0.007 | 0.007 | 0 | 0.39 | 0.43 | 0.55 | 0.87 | 0.28 | 0.49 |
| 9 | 12513587 | rs1041102 | -0.511 | -2.269 | -0.386 | -1.131 | 0.027 | 0.038 | 0.038 | 0.035 | 0.39 | 0.37 | 0.54 | 0.87 | 0.26 | 0.47 |
| 9 | 12515876 | rs791669 | -0.683 | -2.116 | 0.368 | -0.572 | 0.04 | 0.127 | 0.127 | 1.163 | 0.06 | 0.42 | 0.1 | 0.07 | 0.04 | 0.02 |
| 9 | 12521469 | rs10960694 | 0.562 | -2.462 | -0.2 | -0.676 | 0.001 | 0.001 | 0.001 | 0.057 | 0.21 | 0.25 | 0.14 | 0.09 | 0.12 | 0.1 |
| 9 | 12522037 | rs791681 | -1.833 | -2.792 | -0.742 | -1.186 | -0.051 | 0.076 | 0.076 | 0 | 0.69 | 0.89 | 0.69 | 0.98 | 0.48 | 0.8 |
| 9 | 12525999 | rs1325132 | 0.646 | -3.162 | -0.708 | -0.934 | 0 | -0.037 | -0.037 | 0 | 0.11 | 0.05 | 0.38 | 0.78 | 0.1 | 0.3 |
| 9 | 12526192 | rs702131 | -0.506 | -3.353 | -0.479 | -1.097 | 0.038 | 0.045 | 0.045 | 1.456 | 0.5 | 0.6 | 0.26 | 0.2 | 0.3 | 0.48 |
| 9 | 12527414 | rs791688 | -0.856 | -3.269 | -0.025 | -0.991 | -0.036 | -0.046 | -0.046 | 0.581 | 0.34 | 0.54 | 0.55 | 0.86 | 0.26 | 0.4 |

| | | | | | | | | | | | | | | | | |
|---|---|---|---|---|---|---|---|---|---|---|---|---|---|---|---|---|
| 9 | 12529952 | rs791695 | 0.115 | -3.186 | -0.095 | -1.409 | -0.056 | -0.068 | -0.068 | 0 | 0.24 | 0.62 | 0.18 | 0.08 | 0.15 | 0.1 |
| 9 | 12531493 | rs16929115 | 0.626 | -3.193 | -0.057 | -1.45 | -0.002 | -0.056 | -0.056 | 0 | 0.1 | 0.24 | 0.06 | 0 | 0.07 | 0.06 |
| 9 | 12532584 | rs791700 | 0.059 | -3.24 | -0.181 | -1.465 | -0.04 | -0.077 | -0.077 | 0 | 0.27 | 0.6 | 0.18 | 0.08 | 0.14 | 0.14 |
| 9 | 12537279 | rs1570781 | -1.457 | -3.221 | -0.464 | -1.759 | 0.017 | 0.144 | 0.144 | 0 | 0.6 | 0.84 | 0.66 | 0.98 | 0.41 | 0.73 |
| 9 | 12540358 | rs12235745 | 0.377 | -3.452 | -0.53 | -1.49 | 0.02 | -0.008 | -0.008 | 0 | 0.26 | 0.17 | 0.13 | 0.26 | 0.18 | 0.53 |
| 9 | 12541971 | rs12000969 | 1.004 | -3.182 | -0.313 | -1.406 | -0.005 | -0.027 | -0.027 | 0 | 0.09 | 0.16 | 0.06 | 0 | 0.08 | 0.01 |
| 9 | 12564122 | rs10756376 | 0.179 | -3.257 | -0.231 | -1.285 | -0.037 | 0.03 | 0.03 | 0 | 0.24 | 0.6 | 0.48 | 0.73 | 0.14 | 0.18 |
| 9 | 12570945 | rs1325149 | 0.468 | -3.216 | -0.162 | -1.358 | -0.037 | 0.053 | 0.053 | 0 | 0.22 | 0.46 | 0.47 | 0.73 | 0.14 | 0.19 |
| 9 | 12574724 | rs16923032 | 0.561 | -3.109 | -0.667 | -1.293 | 0.008 | -0.027 | -0.027 | 0 | 0.08 | 0.03 | 0.06 | 0.18 | 0.06 | 0.25 |
| 9 | 12578438 | rs10960708 | 1.772 | -3.108 | -0.869 | -1.729 | -0.044 | 0.099 | 0.099 | 0 | 0.35 | 0.15 | 0.27 | 0.01 | 0.54 | 0.2 |
| 9 | 12581270 | rs10809797 | -0.242 | -4.185 | -0.68 | -1.496 | 0.024 | -0.006 | -0.006 | 0.129 | 0.74 | 0.82 | 0.84 | 0.75 | 0.87 | 0.46 |
| 9 | 12582565 | rs1325154 | -0.422 | -4.112 | -0.722 | -1.387 | -0.006 | -0.029 | -0.029 | 0 | 0.91 | 0.96 | 0.89 | 0.82 | 0.9 | 0.74 |
| 9 | 12582787 | rs10429629 | 1.663 | -4.156 | -0.726 | -1.518 | 0.04 | 0.196 | 0.196 | 0 | 0.43 | 0.15 | 0.3 | 0.02 | 0.62 | 0.3 |
| 9 | 12585186 | rs10124604 | 0.333 | -3.806 | -0.223 | -1.223 | -0.016 | 0.078 | 0.078 | 0 | 0.22 | 0.5 | 0.49 | 0.73 | 0.17 | 0.17 |
| 9 | 12589981 | rs10809800 | -0.409 | -3.971 | -0.347 | -1.27 | -0.012 | 0.004 | 0.004 | 0 | 0.28 | 0.62 | 0.53 | 0.73 | 0.22 | 0.18 |
| 9 | 12593124 | rs4428755 | 0.749 | -4.801 | -0.081 | -1.521 | -0.011 | -0.004 | -0.004 | 0.317 | 0.67 | 0.68 | 0.8 | 0.75 | 0.8 | 0.42 |
| 9 | 12594030 | rs1359442 | -0.562 | -5.004 | 0.047 | -1.638 | -0.054 | -0.03 | -0.03 | 0 | 0.4 | 0.49 | 0.51 | 0.78 | 0.19 | 0.45 |
| 9 | 12594967 | rs10756380 | -0.788 | -4.897 | 0.032 | -1.581 | -0.003 | -0.022 | -0.022 | 1.452 | 0.91 | 0.96 | 0.89 | 0.86 | 0.92 | 0.73 |
| 9 | 12599561 | rs13283146 | 1.837 | -5.007 | 0.09 | -1.71 | 0.017 | -0.007 | -0.007 | 0 | 0.43 | 0.41 | 0.35 | 0.03 | 0.66 | 0.26 |
| 9 | 12599803 | rs10809802 | 0.54 | -5.025 | 0.083 | -1.692 | -0.003 | 0.001 | 0.001 | 0 | 0.22 | 0.22 | 0.45 | 0.72 | 0.13 | 0.16 |
| 9 | 12602681 | rs1408790 | 2.05 | -5.143 | 0.016 | -1.892 | 0.007 | 0.003 | 0.003 | 0 | 0.37 | 0.34 | 0.33 | 0.02 | 0.64 | 0.22 |
| 9 | 12606092 | rs10960718 | 1.117 | -5.091 | -0.129 | -1.845 | 0.05 | 0.029 | 0.029 | 0 | 0.49 | 0.42 | 0.37 | 0.03 | 0.73 | 0.27 |
| 9 | 12608432 | rs1325116 | -0.528 | -4.987 | -0.011 | -1.743 | 0.006 | 0.041 | 0.041 | 0.245 | 0.9 | 0.78 | 0.88 | 0.83 | 0.91 | 0.72 |
| 9 | 12610410 | rs7874576 | 0.801 | -5.034 | -0.302 | -1.79 | -0.018 | -0.037 | -0.037 | 0 | 0.1 | 0.23 | 0.05 | 0.02 | 0.05 | 0.05 |
| 9 | 12611123 | rs10809806 | 1.079 | -5.095 | -0.358 | -1.818 | -0.021 | -0.064 | -0.064 | 0.464 | 0.59 | 0.52 | 0.4 | 0.1 | 0.73 | 0.54 |
| 9 | 12612897 | rs1359444 | -0.622 | -5.053 | -0.333 | -1.707 | -0.042 | -0.011 | -0.011 | 0 | 0.33 | 0.46 | 0.5 | 0.73 | 0.18 | 0.2 |
| 9 | 12619787 | rs12350416 | 0.941 | -5.154 | -0.396 | -1.725 | -0.07 | -0.032 | -0.032 | 0 | 0.2 | 0.26 | 0.07 | 0.08 | 0.05 | 0.3 |
| 9 | 12619795 | rs10738286 | -0.855 | -5.085 | -0.574 | -1.694 | -0.012 | -0.038 | -0.038 | 0 | 0.91 | 0.98 | 0.9 | 0.83 | 0.91 | 0.72 |
| 9 | 12622878 | rs10960723 | 2.466 | -5.101 | -0.563 | -1.747 | -0.022 | 0.154 | 0.154 | 0.106 | 0.33 | 0.1 | 0.29 | 0.02 | 0.62 | 0.21 |
| 9 | 12628213 | rs1408793 | -0.125 | -5.061 | -0.452 | -1.654 | 0.005 | -0.003 | -0.003 | 0 | 0.27 | 0.25 | 0.48 | 0.71 | 0.17 | 0.21 |
| 9 | 12637846 | rs10124166 | 0.491 | -4.835 | -0.225 | -1.56 | -0.078 | -0.066 | -0.066 | 2.92 | 0.23 | 0.38 | 0.1 | 0.08 | 0.11 | 0.33 |
| 9 | 12641987 | rs10960736 | 0.898 | -4.815 | -0.186 | -1.555 | -0.102 | -0.064 | -0.064 | 0.049 | 0.2 | 0.34 | 0.07 | 0.07 | 0.05 | 0.32 |
| 9 | 12642366 | rs2031454 | 1.391 | -4.939 | -0.526 | -1.363 | 0.025 | -0.04 | -0.04 | 0 | 0.16 | 0.08 | 0.36 | 0.62 | 0.07 | 0.08 |
| 9 | 12643558 | rs6474720 | -0.976 | -4.868 | -0.648 | -1.624 | -0.009 | -0.031 | -0.031 | 1.104 | 0.92 | 0.97 | 0.89 | 0.81 | 0.91 | 0.75 |
| 9 | 12647332 | rs1155509 | 0.012 | -4.925 | -0.605 | -1.664 | 0.002 | -0.002 | -0.002 | 0.4 | 0.27 | 0.25 | 0.46 | 0.71 | 0.16 | 0.18 |
| 9 | 12650288 | rs9657586 | -1.484 | -5.019 | -0.764 | -1.717 | 0.162 | 0.14 | 0.14 | 4.668 | 0.56 | 0.89 | 0.57 | 0.79 | 0.23 | 0.51 |
| 9 | 12651340 | rs1408794 | 2.436 | -4.967 | -0.774 | -1.757 | -0.05 | 0.194 | 0.194 | 0.248 | 0.33 | 0.07 | 0.29 | 0.02 | 0.61 | 0.22 |
| 9 | 12651413 | rs1408795 | 2.168 | -4.972 | -0.773 | -1.756 | -0.001 | 0.209 | 0.209 | 0.401 | 0.35 | 0.08 | 0.31 | 0.02 | 0.66 | 0.23 |
| 9 | 12652983 | rs996697 | 0.664 | -4.986 | -0.657 | -1.737 | 0.008 | 0.007 | 0.007 | 0 | 0.61 | 0.38 | 0.76 | 0.73 | 0.8 | 0.39 |
| 9 | 12657603 | rs10435754 | -0.287 | -4.82 | -0.663 | -1.61 | 0.005 | -0.003 | -0.003 | 0 | 0.29 | 0.26 | 0.48 | 0.71 | 0.21 | 0.19 |
| 9 | 12668328 | rs1325127 | 1.84 | -4.626 | -0.318 | -1.38 | 0.026 | 0.03 | 0.03 | 0 | 0.4 | 0.27 | 0.66 | 0.71 | 0.65 | 0.32 |
| 9 | 12669346 | rs9298679 | 0.942 | -4.646 | -0.292 | -1.442 | 0 | 0 | 0 | 0 | 0.76 | 0.73 | 0.81 | 0.78 | 0.77 | 0.63 |
| 9 | 12671059 | rs1041105 | -0.538 | -4.633 | -0.319 | -1.431 | -0.016 | -0.032 | -0.032 | 0 | 0.89 | 0.96 | 0.88 | 0.8 | 0.88 | 0.69 |
| 9 | 12671856 | rs1408798 | -0.404 | -4.632 | -0.273 | -1.391 | 0 | -0.006 | -0.006 | 0 | 0.91 | 0.93 | 0.92 | 0.8 | 0.91 | 0.71 |
| 9 | 12672097 | rs1408799 | 1.589 | -4.8 | -0.293 | -1.513 | 0.074 | 0.188 | 0.188 | 0 | 0.47 | 0.19 | 0.36 | 0.02 | 0.65 | 0.22 |
| 9 | 12672320 | rs1408801 | -0.163 | -4.736 | -0.208 | -1.453 | -0.073 | -0.013 | -0.013 | 0 | 0.29 | 0.56 | 0.46 | 0.77 | 0.12 | 0.41 |
| 9 | 12674124 | rs16929340 | 1.188 | -4.419 | -0.261 | -1.579 | 0.08 | 0.094 | 0.094 | 0.073 | 0.46 | 0.31 | 0.4 | 0.03 | 0.73 | 0.28 |
| 9 | 12675264 | rs10960751 | 2.206 | -4.356 | -0.351 | -1.552 | -0.061 | 0.166 | 0.166 | 0.049 | 0.32 | 0.08 | 0.29 | 0.02 | 0.6 | 0.2 |
| 9 | 12683639 | rs7847593 | 0.511 | -4.21 | -0.114 | -1.387 | -0.004 | -0.039 | -0.039 | 0 | 0.12 | 0.22 | 0.09 | 0.02 | 0.08 | 0.08 |
| 9 | 12698363 | rs2075508 | 0.144 | -4.149 | -0.167 | -1.239 | -0.074 | -0.051 | -0.051 | 0.162 | 0.23 | 0.32 | 0.1 | 0.12 | 0.08 | 0.32 |
| 9 | 12699776 | rs2762462 | 0.516 | -4.177 | -0.212 | -1.361 | -0.019 | -0.07 | -0.07 | 1.1 | 0.58 | 0.52 | 0.5 | 0.14 | 0.73 | 0.53 |
| 9 | 12706236 | rs2209277 | 1.66 | -4.037 | -0.463 | -1.328 | 0.003 | 0.084 | 0.084 | 0 | 0.36 | 0.2 | 0.41 | 0.02 | 0.65 | 0.21 |
| 9 | 12707861 | rs10809828 | -0.637 | -4.097 | -0.501 | -1.572 | -0.034 | -0.04 | -0.04 | 0.043 | 0.4 | 0.54 | 0.48 | 0.75 | 0.23 | 0.32 |
| 9 | 12709305 | rs683 | 1.606 | -3.367 | -1.174 | -1.36 | -0.038 | 0.158 | 0.158 | 0 | 0.33 | 0.11 | 0.38 | 0.01 | 0.63 | 0.21 |
| 9 | 12710035 | rs910 | 1.353 | -3.361 | -1.201 | -1.358 | -0.01 | 0.193 | 0.193 | 0 | 0.36 | 0.1 | 0.41 | 0.02 | 0.65 | 0.21 |
| 9 | 12710090 | rs1063380 | 1.317 | -3.351 | -1.192 | -1.353 | 0.002 | 0.152 | 0.152 | 0 | 0.37 | 0.14 | 0.42 | 0.02 | 0.65 | 0.21 |
| 9 | 12712157 | rs1137134 | 1.907 | -3.283 | -1.174 | -1.322 | -0.105 | 0.193 | 0.193 | 0 | 0.31 | 0.05 | 0.35 | 0.02 | 0.58 | 0.2 |
| 9 | 12722357 | rs7871257 | -1.236 | -3.025 | -0.799 | -1.069 | 0.107 | 0.169 | 0.169 | 1.243 | 0.54 | 0.82 | 0.52 | 0.83 | 0.26 | 0.52 |
| 9 | 12734504 | rs2382364 | -0.058 | -2.015 | -0.538 | -0.387 | 0.008 | -0.028 | -0.028 | 0.772 | 0.89 | 0.97 | 0.91 | 0.82 | 0.9 | 0.75 |
| 9 | 12738690 | rs970946 | -1.114 | -2.511 | -0.135 | -0.737 | 0.029 | 0.041 | 0.041 | 0 | 0.48 | 0.57 | 0.51 | 0.81 | 0.31 | 0.54 |
| 9 | 12748633 | rs7019486 | -0.867 | -2.531 | -0.052 | -0.764 | -0.003 | -0.013 | -0.013 | 0 | 0.45 | 0.52 | 0.49 | 0.81 | 0.25 | 0.5 |
| 9 | 12757058 | rs10125771 | -0.466 | -2.833 | 0.058 | -0.733 | -0.031 | -0.014 | -0.014 | 0 | 0.3 | 0.41 | 0.42 | 0.68 | 0.17 | 0.28 |
| 9 | 12758881 | rs10960779 | 2.082 | -2.878 | 0.027 | -0.876 | 0.004 | -0.048 | -0.048 | 1.527 | 0.42 | 0.42 | 0.42 | 0.01 | 0.6 | 0.21 |
| 9 | 12759352 | rs10809837 | -0.505 | -3.065 | 0.241 | -0.516 | -0.039 | 0.049 | 0.049 | 0.019 | 0.26 | 0.35 | 0.32 | 0.62 | 0.14 | 0.31 |
| 9 | 12763001 | rs1041174 | 0.214 | -2.957 | 0.281 | -0.421 | -0.025 | 0.14 | 0.14 | 0.736 | 0.14 | 0.26 | 0.26 | 0.54 | 0.07 | 0.08 |
| 9 | 12765117 | rs10119113 | 0.706 | -2.85 | 0.109 | -0.429 | 0.003 | -0.005 | -0.005 | 0.683 | 0.72 | 0.74 | 0.75 | 0.66 | 0.8 | 0.51 |
| 9 | 12770108 | rs1326795 | -0.759 | -2.945 | 0.357 | -0.469 | -0.04 | 0.02 | 0.02 | 0 | 0.23 | 0.39 | 0.19 | 0.18 | 0.12 | 0.22 |
| 9 | 12770868 | rs1887175 | -0.262 | -2.868 | 0.205 | -0.567 | -0.002 | 0.048 | 0.048 | 0.245 | 0.9 | 0.73 | 0.91 | 0.88 | 0.92 | 0.75 |
| 9 | 12771222 | rs1887177 | 0.659 | -2.89 | 0.198 | -0.584 | -0.007 | 0.031 | 0.031 | 0 | 0.86 | 0.66 | 0.87 | 0.82 | 0.87 | 0.72 |

| | | | | | | | | | | | | | | | | |
|---|---|---|---|---|---|---|---|---|---|---|---|---|---|---|---|---|
| 9 | 12772498 | rs2209273 | 1.11 | -2.906 | 0.161 | -0.606 | -0.004 | 0.002 | 0.002 | 0.306 | 0.83 | 0.59 | 0.84 | 0.82 | 0.82 | 0.73 |
| 9 | 12773263 | rs7855624 | 1.193 | -2.957 | 0.033 | -0.592 | -0.009 | 0.003 | 0.003 | 0 | 0.68 | 0.41 | 0.7 | 0.73 | 0.73 | 0.55 |
| 9 | 12774019 | rs1326796 | 0.275 | -2.376 | -0.407 | -0.455 | 0 | -0.003 | -0.003 | 0 | 0.88 | 0.85 | 0.88 | 0.81 | 0.86 | 0.75 |
| 9 | 12775488 | rs10491742 | 1.646 | -2.518 | -0.636 | -0.632 | -0.044 | 0.299 | 0.299 | 0 | 0.4 | 0.05 | 0.39 | 0.01 | 0.61 | 0.21 |
| 9 | 12776516 | rs3750502 | -0.675 | -2.23 | 0.195 | 0.143 | -0.007 | -0.001 | -0.001 | 0.033 | 0.29 | 0.29 | 0.21 | 0.25 | 0.15 | 0.37 |
| 9 | 12778224 | rs1326800 | 0.9 | -1.463 | 0.389 | 0.681 | 0.025 | -0.033 | -0.033 | 0 | 0.82 | 0.86 | 0.83 | 0.72 | 0.88 | 0.63 |
| 9 | 12780670 | rs10809843 | 1.071 | -1.603 | 0.149 | 0.579 | -0.076 | 0.027 | 0.027 | 0 | 0.75 | 0.56 | 0.79 | 0.75 | 0.88 | 0.6 |
| 9 | 12785073 | rs7042370 | 0.322 | 0.176 | -0.026 | 0.495 | 0.016 | 0.007 | 0.007 | 0 | 0.7 | 0.67 | 0.45 | 0.34 | 0.58 | 0.56 |
| 9 | 12791031 | rs7029119 | -0.684 | 0.004 | 0.002 | 0.161 | 0.015 | -0.028 | -0.028 | 0.147 | 0.1 | 0.19 | 0.07 | 0.06 | 0.14 | 0.09 |
| 9 | 12794833 | rs1887174 | -0.472 | -1.084 | -0.473 | 0.185 | 0.07 | 0.037 | 0.037 | 0 | 0.43 | 0.69 | 0.41 | 0.43 | 0.24 | 0.29 |
| 9 | 12795698 | rs1041177 | 0.181 | -1.638 | -0.733 | 0.152 | 0.092 | 0.006 | 0.006 | 0 | 0.23 | 0.06 | 0.21 | 0.21 | 0.09 | 0.18 |
| 9 | 12796718 | rs16929574 | 0.008 | -1.726 | 0.213 | 0.093 | -0.009 | -0.006 | -0.006 | 0 | 0.24 | 0.21 | 0.1 | 0.1 | 0.09 | 0.24 |
| 9 | 12801148 | rs9298684 | -0.094 | -1.594 | 0.631 | 0.274 | -0.003 | 0.021 | 0.021 | 0 | 0.14 | 0.14 | 0.27 | 0.32 | 0.11 | 0.09 |
| 9 | 12813037 | rs4111024 | -0.968 | -1.402 | 1.303 | 0.218 | -0.023 | -0.004 | -0.004 | 0 | 0.38 | 0.42 | 0.25 | 0.43 | 0.23 | 0.33 |
| 9 | 12815091 | rs6474732 | -0.208 | -1.43 | 1.35 | 0.251 | -0.011 | 0.055 | 0.055 | 0 | 0.15 | 0.25 | 0.15 | 0.36 | 0.11 | 0.14 |
| 9 | 12818455 | rs6474734 | -1.167 | -1.303 | 1.844 | 0.15 | -0.015 | -0.006 | -0.006 | 0 | 0.39 | 0.52 | 0.26 | 0.42 | 0.26 | 0.34 |
| 9 | 12818692 | rs7846724 | -0.203 | -1.297 | 1.68 | 0.178 | 0.008 | 0.044 | 0.044 | 0 | 0.15 | 0.3 | 0.16 | 0.31 | 0.14 | 0.09 |
| 9 | 12822458 | rs16929629 | -1.049 | -1.197 | 0.905 | 0.061 | -0.001 | 0 | 0 | 0 | 0.38 | 0.6 | 0.27 | 0.42 | 0.23 | 0.33 |
| 9 | 12822678 | rs16929630 | 0.32 | -1.782 | -0.441 | -0.467 | 0.092 | 0.08 | 0.08 | 0 | 0.22 | 0.04 | 0.08 | 0.09 | 0.09 | 0.19 |
| 9 | 12830070 | rs617572 | -0.202 | -0.51 | -0.116 | 0.085 | 0.003 | 0.027 | 0.027 | 0 | 0.32 | 0.46 | 0.28 | 0.51 | 0.3 | 0.33 |
| 9 | 12839484 | rs652278 | -0.467 | -0.502 | 0.395 | 0.415 | 0.007 | -0.008 | -0.008 | 0 | 0.36 | 0.45 | 0.33 | 0.72 | 0.36 | 0.4 |
| 9 | 12842241 | rs10960816 | 1.948 | -0.384 | 0.154 | 0.756 | -0.012 | -0.02 | -0.02 | 0 | 0.05 | 0.02 | 0.06 | 0.29 | 0.06 | 0.02 |
| 9 | 12844891 | rs620255 | -0.772 | -0.46 | 0.457 | 0.772 | 0.042 | 0.116 | 0.116 | 0.34 | 0.87 | 0.79 | 0.83 | 0.61 | 0.8 | 0.65 |
| 9 | 12846053 | rs16923061 | 0.555 | -1.126 | -0.321 | -0.296 | 0.073 | 0.081 | 0.081 | 0 | 0.32 | 0.14 | 0.14 | 0.17 | 0.18 | 0.27 |
| 9 | 12846491 | rs483855 | -1.561 | -1.266 | -0.449 | -0.436 | -0.13 | 0.082 | 0.082 | 0.783 | 0.73 | 0.96 | 0.56 | 0.89 | 0.57 | 0.8 |
| 9 | 12852488 | rs645121 | -0.119 | -1.323 | -0.229 | -0.409 | 0.003 | -0.003 | -0.003 | 0.303 | 0.33 | 0.38 | 0.3 | 0.69 | 0.33 | 0.4 |
| 9 | 12855260 | rs7868380 | 0.898 | -1.563 | -0.305 | -0.505 | -0.024 | -0.02 | -0.02 | 0 | 0.24 | 0.32 | 0.12 | 0.15 | 0.11 | 0.22 |
| 9 | 12856773 | rs662135 | -0.98 | -0.716 | -0.409 | -0.467 | -0.002 | 0 | 0 | 0 | 0.77 | 0.82 | 0.53 | 0.82 | 0.61 | 0.75 |
| 9 | 12860574 | rs4406510 | -0.74 | -1.026 | 0.02 | -0.405 | 0.046 | 0.153 | 0.153 | 0.581 | 0.06 | 0.42 | 0.09 | 0.06 | 0.01 | 0.12 |
| 9 | 12861280 | rs13291898 | 0.545 | -0.91 | -0.665 | -0.561 | 0.054 | 0.088 | 0.088 | 0 | 0.33 | 0.15 | 0.14 | 0.14 | 0.19 | 0.26 |
| 9 | 12875204 | rs558579 | -0.109 | -0.137 | -0.321 | -0.408 | -0.013 | -0.012 | -0.012 | 0 | 0.89 | 0.79 | 0.9 | 0.93 | 0.93 | 0.83 |
| 9 | 12886899 | rs6474741 | -0.054 | -0.371 | -0.263 | -0.562 | 0.017 | 0.02 | 0.02 | 0 | 0.43 | 0.62 | 0.25 | 0.22 | 0.25 | 0.43 |
| 9 | 12890233 | rs1929918 | -0.269 | 0.264 | -0.157 | -0.267 | 0.008 | 0 | 0 | 0.165 | 0.73 | 0.5 | 0.77 | 0.75 | 0.73 | 0.6 |
| 9 | 12897342 | rs2183864 | 0.069 | 0.617 | -0.087 | -0.274 | 0.047 | 0.013 | 0.013 | 0.869 | 0.77 | 0.62 | 0.8 | 0.75 | 0.71 | 0.58 |
| 9 | 12899790 | rs7032650 | -0.833 | 0.518 | -0.12 | -0.578 | 0.006 | 0.014 | 0.014 | 0 | 0.47 | 0.55 | 0.51 | 0.17 | 0.41 | 0.34 |
| 9 | 12900789 | rs1332083 | -1.058 | 0.586 | 0.045 | 0.191 | -0.008 | -0.016 | -0.016 | 1.314 | 0.12 | 0.24 | 0.11 | 0.07 | 0.08 | 0.16 |
| 9 | 12904064 | rs13295077 | -0.645 | 0.921 | -0.341 | 0.275 | 0.038 | -0.015 | -0.015 | 0 | 0.11 | 0.24 | 0.12 | 0.05 | 0.16 | 0.2 |
| 9 | 12904223 | rs1332085 | 0.613 | 0.471 | -1.117 | 1.193 | 0.013 | -0.01 | -0.01 | 0.014 | 0.09 | 0.05 | 0.15 | 0.43 | 0.08 | 0.02 |
| 9 | 12905153 | rs10960828 | 0.134 | 0.533 | -1.552 | 0.291 | -0.015 | 0.034 | 0.034 | 0.184 | 0.25 | 0.05 | 0.18 | 0.1 | 0.23 | 0.08 |
| 9 | 12906962 | rs10435758 | -0.285 | 0.705 | -0.99 | 0.222 | -0.003 | -0.002 | -0.002 | 0 | 0.36 | 0.42 | 0.57 | 0.52 | 0.39 | 0.54 |
| 9 | 12907048 | rs13292327 | -0.741 | 0.973 | -0.836 | -0.139 | 0.01 | -0.008 | -0.008 | 0 | 0.1 | 0.09 | 0.07 | 0.12 | 0.15 | 0.16 |
| 9 | 12907902 | rs13293601 | 0.123 | 0.763 | -0.741 | -0.403 | 0.006 | -0.026 | -0.026 | 0.07 | 0.17 | 0.25 | 0.08 | 0.07 | 0.19 | 0.13 |
| 9 | 12908447 | rs10756424 | 0.168 | 0.654 | -1.547 | -0.574 | -0.075 | -0.039 | -0.039 | 0 | 0.38 | 0.12 | 0.47 | 0.55 | 0.49 | 0.6 |
| 9 | 12910124 | rs1332053 | -1.912 | 0.124 | -1.405 | -1.067 | -0.001 | -0.01 | -0.01 | 0.021 | 0.89 | 0.91 | 0.95 | 0.97 | 0.92 | 0.97 |
| 9 | 12910295 | rs10448182 | 1.214 | 0.319 | -1.746 | -1.231 | -0.069 | 0.075 | 0.075 | 0 | 0.2 | 0.01 | 0.14 | 0.06 | 0.27 | 0.15 |
| 9 | 12913298 | rs10960834 | -1.522 | 0.309 | -1.124 | -0.78 | -0.011 | -0.031 | -0.031 | 0.759 | 0.88 | 0.88 | 0.94 | 0.97 | 0.91 | 0.97 |
| 9 | 12914345 | rs10809860 | 0.664 | 1.028 | -1.198 | -0.59 | -0.055 | 0.129 | 0.129 | 0.733 | 0.26 | 0.05 | 0.19 | 0.09 | 0.34 | 0.23 |
| 9 | 12917048 | rs1984050 | 0.21 | 0.28 | -0.514 | -0.585 | 0.014 | -0.057 | -0.057 | 3.553 | 0.2 | 0.36 | 0.11 | 0.05 | 0.21 | 0.11 |
| 9 | 12920004 | rs10125252 | 0.208 | 0.242 | -0.755 | -0.583 | 0.002 | -0.026 | -0.026 | 0 | 0.21 | 0.3 | 0.11 | 0.05 | 0.2 | 0.14 |
| 9 | 12921644 | rs1888859 | 0.506 | 0.393 | -1.495 | -0.315 | -0.102 | -0.058 | -0.058 | 0 | 0.35 | 0.1 | 0.51 | 0.62 | 0.5 | 0.61 |
| 9 | 12922330 | rs2065423 | -0.516 | -0.025 | -1.033 | -0.014 | 0.053 | -0.012 | -0.012 | 0 | 0.58 | 0.78 | 0.63 | 0.56 | 0.7 | 0.77 |
| 9 | 12924111 | rs10511575 | -0.529 | -0.375 | -0.121 | -0.761 | 0 | -0.069 | -0.069 | 0 | 0.17 | 0.4 | 0.09 | 0.04 | 0.13 | 0.09 |
| 9 | 12924582 | rs7020025 | -0.757 | -0.354 | -1.017 | -0.52 | 0.003 | -0.017 | -0.017 | 0 | 0.3 | 0.21 | 0.27 | 0.28 | 0.28 | 0.23 |
| 9 | 12924863 | rs7020076 | -0.05 | -0.187 | -1.034 | -1.29 | -0.007 | 0.078 | 0.078 | 0 | 0.71 | 0.79 | 0.82 | 0.94 | 0.69 | 0.95 |
| 9 | 12925921 | rs10511576 | 0.155 | -0.204 | -0.643 | -0.88 | -0.013 | 0.061 | 0.061 | 0 | 0.26 | 0.36 | 0.39 | 0.54 | 0.21 | 0.45 |
| 9 | 12926360 | rs1412294 | 1.118 | -0.158 | -0.77 | -1.138 | -0.044 | -0.054 | -0.054 | 0 | 0.27 | 0.12 | 0.31 | 0.36 | 0.31 | 0.42 |
| 9 | 12926571 | rs1412296 | 0.551 | -0.281 | -0.286 | -1.839 | 0.002 | -0.006 | -0.006 | 1.476 | 0.52 | 0.45 | 0.7 | 0.88 | 0.51 | 0.86 |
| 9 | 12928475 | rs1332057 | 0.729 | -0.507 | 0.491 | -0.616 | -0.032 | -0.058 | -0.058 | 0 | 0.77 | 0.52 | 0.85 | 0.89 | 0.8 | 0.95 |
| 9 | 12931403 | rs16929942 | -0.525 | -0.025 | -0.144 | -0.274 | -0.025 | -0.044 | -0.044 | 0 | 0.26 | 0.37 | 0.13 | 0.12 | 0.19 | 0.07 |
| 9 | 12954308 | rs4740537 | 0.678 | 0.416 | 0.114 | -0.119 | 0.003 | -0.027 | -0.027 | 0 | 0.94 | 0.83 | 0.97 | 1 | 0.96 | 0.99 |
| 9 | 12956312 | rs10809863 | 0.243 | 0.164 | -0.459 | -0.016 | 0.027 | 0.01 | 0.01 | 0 | 0.5 | 0.18 | 0.49 | 0.51 | 0.58 | 0.33 |
| 9 | 12957884 | rs10448183 | 0.168 | -0.01 | 0.612 | -0.099 | 0 | -0.001 | -0.001 | 0.326 | 0.41 | 0.43 | 0.45 | 0.49 | 0.37 | 0.61 |
| 9 | 12959188 | rs10809865 | -0.129 | -0.005 | 0.547 | -0.098 | 0.006 | -0.003 | -0.003 | 4.326 | 0.44 | 0.63 | 0.47 | 0.49 | 0.38 | 0.65 |
| 9 | 12960088 | rs4741264 | -0.051 | 0.063 | 0.53 | -0.178 | 0.002 | -0.028 | -0.028 | 0 | 0.95 | 0.92 | 0.98 | 1 | 0.96 | 0.99 |
| 9 | 12971966 | rs1536791 | 1.139 | -0.15 | 0.901 | -0.27 | -0.015 | -0.053 | -0.053 | 0.556 | 0.9 | 0.75 | 0.89 | 0.99 | 0.95 | 0.97 |
| 9 | 12972382 | rs1536792 | 1.653 | -0.793 | 1.099 | 0.131 | -0.06 | 0.024 | 0.024 | 0 | 0.77 | 0.49 | 0.86 | 0.76 | 0.9 | 0.72 |
| 9 | 12982583 | rs13289590 | -0.231 | -0.831 | 0.934 | -0.125 | 0.01 | 0.019 | 0.019 | 0 | 0.07 | 0.03 | 0.02 | 0 | 0.04 | 0.01 |
| 9 | 12984009 | rs10491615 | 1.051 | -1.27 | 0.432 | -0.859 | 0.011 | 0.111 | 0.111 | 0.014 | 0.46 | 0.27 | 0.65 | 0.82 | 0.55 | 0.38 |
| 9 | 12985487 | rs1412301 | 1.435 | -1.227 | 0.782 | -0.756 | -0.038 | -0.049 | -0.049 | 0 | 0.75 | 0.37 | 0.85 | 0.87 | 0.89 | 0.74 |

| | | | | | | | | | | | | | | | | |
|---|---|---|---|---|---|---|---|---|---|---|---|---|---|---|---|---|
| 9 | 12988126 | rs1332071 | -0.033 | -0.514 | 0.607 | -0.673 | 0.051 | 0.114 | 0.114 | 0.012 | 0.17 | 0.34 | 0.35 | 0.32 | 0.27 | 0.09 |
| 9 | 12995112 | rs1887884 | 1.505 | -0.895 | 0.673 | -1.274 | -0.05 | -0.036 | -0.036 | 0 | 0.76 | 0.39 | 0.8 | 0.85 | 0.89 | 0.72 |
| 9 | 13007129 | rs1360517 | 0.63 | -0.798 | 0.473 | -1.259 | -0.006 | -0.002 | -0.002 | 0.015 | 0.85 | 0.79 | 0.94 | 0.87 | 0.92 | 0.76 |
| 9 | 13014232 | rs10756439 | 1.323 | -0.816 | 0.021 | -1.374 | -0.046 | -0.055 | -0.055 | 0 | 0.73 | 0.48 | 0.74 | 0.86 | 0.86 | 0.62 |
| 9 | 13020010 | rs12351740 | 0.138 | -0.508 | 0.021 | -1.118 | 0.006 | -0.019 | -0.019 | 0 | 0.08 | 0.04 | 0.03 | 0.13 | 0.06 | 0.23 |
| 9 | 13025284 | rs10514822 | 0.476 | -0.613 | -0.442 | -1.643 | 0.001 | -0.035 | -0.035 | 0 | 0.24 | 0.25 | 0.15 | 0.03 | 0.21 | 0.03 |
| 9 | 13027530 | rs10809876 | -0.182 | -0.616 | -0.909 | -2.111 | 0.01 | -0.125 | -0.125 | 0 | 0.4 | 0.46 | 0.38 | 0.05 | 0.4 | 0.3 |
| 9 | 13028726 | rs4741270 | 0.802 | -0.467 | -1.976 | -1.64 | -0.125 | 0.087 | 0.087 | 0 | 0.39 | 0.05 | 0.49 | 0.79 | 0.5 | 0.42 |
| 9 | 13029816 | rs9792560 | -0.106 | -0.195 | -1.749 | -1.664 | -0.001 | -0.018 | -0.018 | 0 | 0.53 | 0.58 | 0.62 | 0.82 | 0.55 | 0.46 |
| 9 | 13030676 | rs7848632 | 0.865 | -0.084 | -1.789 | -1.553 | -0.044 | 0.067 | 0.067 | 0.699 | 0.14 | 0.04 | 0.2 | 0.04 | 0.18 | 0.27 |
| 9 | 13032490 | rs7029019 | 0.011 | -0.34 | -1.516 | -1.231 | -0.011 | 0.001 | 0.001 | 0.194 | 0.11 | 0.5 | 0.12 | 0.03 | 0.04 | 0.02 |
| 9 | 13033268 | rs10435759 | 0.299 | 0.272 | -2.724 | -0.894 | -0.132 | 0.081 | 0.081 | 0 | 0.38 | 0.05 | 0.49 | 0.79 | 0.51 | 0.43 |
| 9 | 13044407 | rs17542206 | 0.136 | 1.031 | -2.507 | -0.979 | -0.13 | 0.122 | 0.122 | 0 | 0.2 | 0.01 | 0.21 | 0.04 | 0.29 | 0.27 |
| 9 | 13046042 | rs10960908 | -0.616 | 0.061 | -2.505 | -0.758 | -0.037 | 0.03 | 0.03 | 0 | 0.59 | 0.82 | 0.44 | 0.33 | 0.48 | 0.61 |
| 9 | 13048141 | rs1360059 | 0.751 | 0.024 | -2.705 | -0.76 | -0.124 | -0.042 | -0.042 | 0 | 0.35 | 0.05 | 0.53 | 0.67 | 0.49 | 0.38 |
| 9 | 13052570 | rs10960916 | 0.257 | -0.628 | -2.655 | -1.112 | 0.001 | 0.059 | 0.059 | 0 | 0.22 | 0.06 | 0.14 | 0 | 0.2 | 0.08 |
| 9 | 13055896 | rs4741277 | 0.858 | 0.034 | -3.365 | -1.183 | -0.111 | -0.035 | -0.035 | 0 | 0.37 | 0.07 | 0.52 | 0.66 | 0.48 | 0.3 |
| 9 | 13063892 | rs7861153 | 0.447 | 0.254 | -2.118 | -1.571 | 0.012 | -0.041 | -0.041 | 3.143 | 0.3 | 0.44 | 0.31 | 0.15 | 0.29 | 0.5 |
| 9 | 13064793 | rs999648 | 0.824 | -0.022 | -2.463 | -1.486 | -0.074 | 0.16 | 0.16 | 0.6 | 0.43 | 0.12 | 0.55 | 0.85 | 0.5 | 0.44 |
| 9 | 13071072 | rs10809894 | 0.502 | -0.103 | -2.442 | -0.735 | -0.054 | -0.075 | -0.075 | 0.191 | 0.48 | 0.07 | 0.57 | 0.5 | 0.59 | 0.32 |
| 9 | 13077072 | rs10960928 | -0.17 | -0.159 | -1.328 | -0.375 | -0.009 | 0.015 | 0.015 | 1.383 | 0.69 | 0.41 | 0.7 | 0.46 | 0.77 | 0.37 |
| 9 | 13088211 | rs10738320 | 0.157 | -0.214 | -1.504 | -0.522 | 0.038 | 0.027 | 0.027 | 0.857 | 0.6 | 0.25 | 0.72 | 0.65 | 0.72 | 0.69 |
| 9 | 13092076 | rs12346832 | 0.696 | 0.007 | -1.281 | 0.243 | -0.001 | -0.005 | -0.005 | 1.759 | 0.26 | 0.25 | 0.18 | 0.3 | 0.24 | 0.39 |
| 9 | 13093345 | rs10960942 | 0.548 | -0.248 | -1.236 | 0.153 | -0.002 | -0.03 | -0.03 | 0 | 0.28 | 0.17 | 0.18 | 0.3 | 0.25 | 0.4 |
| 9 | 13095067 | rs10960943 | -0.934 | -0.296 | -0.428 | 0.092 | -0.007 | 0.021 | 0.021 | 0 | 0.66 | 0.61 | 0.53 | 0.52 | 0.62 | 0.61 |
| 9 | 13095872 | rs10756449 | -0.034 | -0.601 | -0.082 | 0.246 | 0.05 | 0.211 | 0.211 | 0.198 | 0.95 | 0.67 | 0.88 | 0.87 | 0.98 | 0.87 |
| 9 | 13099797 | rs10514824 | 0.506 | -0.778 | -0.038 | -0.066 | 0 | -0.003 | -0.003 | 0 | 0.24 | 0.27 | 0.24 | 0.14 | 0.25 | 0.17 |
| 9 | 13105638 | rs10733253 | -0.319 | -0.889 | 0.325 | -0.021 | -0.049 | 0.018 | 0.018 | 0 | 0.85 | 0.64 | 0.84 | 0.85 | 0.94 | 0.83 |
| 9 | 13106335 | rs722651 | 1.08 | -0.589 | 0.229 | -0.102 | 0 | -0.004 | -0.004 | 0.006 | 0.37 | 0.36 | 0.45 | 0.49 | 0.36 | 0.39 |
| 9 | 13106598 | rs3264 | -0.451 | -0.333 | -0.043 | -0.25 | -0.003 | 0.005 | 0.005 | 0 | 0.35 | 0.33 | 0.36 | 0.16 | 0.39 | 0.23 |
| 9 | 13107679 | rs10217351 | -0.291 | -0.376 | -0.224 | 0.099 | -0.003 | -0.001 | -0.001 | 0 | 0.52 | 0.5 | 0.5 | 0.5 | 0.61 | 0.55 |
| 9 | 13108231 | rs7871226 | -0.086 | -0.976 | -0.579 | 0.124 | -0.048 | 0.002 | 0.002 | 0 | 0.84 | 0.69 | 0.85 | 0.85 | 0.94 | 0.83 |
| 9 | 13108794 | rs3765550 | 0.894 | -0.314 | -1.161 | -0.005 | -0.059 | -0.017 | -0.017 | 0 | 0.33 | 0.12 | 0.41 | 0.37 | 0.45 | 0.3 |
| 9 | 13115422 | rs2274647 | -0.534 | -1.032 | -0.831 | 0.072 | -0.002 | 0.003 | 0.003 | 0.96 | 0.33 | 0.38 | 0.28 | 0.63 | 0.29 | 0.55 |
| 9 | 13120380 | rs10960954 | 0.894 | -0.903 | -1.328 | -0.109 | -0.111 | -0.1 | -0.1 | 0 | 0.45 | 0.05 | 0.47 | 0.44 | 0.54 | 0.36 |
| 9 | 13124767 | rs9886797 | 0.473 | -1.4 | -0.312 | -0.167 | -0.003 | 0.004 | 0.004 | 0 | 0.64 | 0.39 | 0.66 | 0.51 | 0.69 | 0.63 |
| 9 | 13133479 | rs10809906 | -0.701 | -1.443 | -0.12 | -0.21 | 0 | 0.041 | 0.041 | 0 | 0.16 | 0.24 | 0.09 | 0.48 | 0.14 | 0.34 |
| 9 | 13147764 | rs10809911 | 0.202 | -1.543 | 0.341 | -0.412 | -0.011 | -0.134 | -0.134 | 0 | 0.24 | 0.43 | 0.26 | 0.01 | 0.2 | 0.06 |
| 9 | 13150531 | rs10756457 | -0.97 | -1.598 | 0.121 | -0.297 | 0.008 | -0.005 | -0.005 | 0 | 0.41 | 0.7 | 0.35 | 0.5 | 0.33 | 0.4 |
| 9 | 13158958 | rs1331670 | -1.408 | -1.741 | -0.028 | -0.389 | 0.036 | -0.017 | -0.017 | 0 | 0.44 | 0.84 | 0.36 | 0.5 | 0.32 | 0.35 |
| 9 | 13164434 | rs2225483 | -1.721 | -1.923 | -0.139 | -0.471 | 0.081 | 0.007 | 0.007 | 0.456 | 0.5 | 0.9 | 0.38 | 0.51 | 0.34 | 0.45 |
| 9 | 13171410 | rs12351444 | 0.911 | -1.73 | 0.842 | -0.26 | -0.012 | 0.005 | 0.005 | 0 | 0.61 | 0.46 | 0.68 | 0.5 | 0.74 | 0.59 |
| 9 | 13171577 | rs1331673 | 0.911 | -1.727 | 0.842 | -0.26 | -0.011 | 0.005 | 0.005 | 0 | 0.61 | 0.46 | 0.67 | 0.51 | 0.74 | 0.59 |
| 9 | 13176312 | rs2274856 | -0.709 | -1.712 | 0.413 | -0.239 | -0.015 | 0.035 | 0.035 | 0 | 0.33 | 0.53 | 0.3 | 0.5 | 0.26 | 0.31 |
| 9 | 13177389 | rs1331675 | 0.881 | -1.626 | -0.693 | -0.21 | 0.009 | -0.08 | -0.08 | 0.441 | 0.58 | 0.07 | 0.64 | 0.51 | 0.7 | 0.57 |
| 9 | 13185149 | rs10756461 | 1.217 | -1.522 | -0.737 | -0.205 | 0.017 | -0.112 | -0.112 | 0 | 0.52 | 0.03 | 0.61 | 0.41 | 0.66 | 0.53 |
| 9 | 13209807 | rs10809915 | 1.115 | -0.895 | -0.155 | -0.104 | -0.025 | 0.056 | 0.056 | 0.519 | 0.11 | 0 | 0.08 | 0 | 0.12 | 0.17 |
| 9 | 13217885 | rs1412112 | -0.521 | -1.048 | 0.392 | -0.164 | -0.008 | -0.102 | -0.102 | 0.923 | 0.83 | 0.69 | 0.87 | 0.99 | 0.85 | 0.89 |
| 9 | 13265893 | rs11793993 | 1.392 | -0.994 | -0.039 | 0.195 | -0.067 | -0.111 | -0.111 | 2.047 | 0.47 | 0.04 | 0.64 | 0.41 | 0.55 | 0.4 |
| 9 | 13267800 | rs10809923 | -0.355 | -1.095 | 0.412 | 0.103 | -0.016 | -0.129 | -0.129 | 1.271 | 0.81 | 0.64 | 0.84 | 1 | 0.83 | 0.89 |
| 9 | 13269740 | rs10809924 | -0.981 | -1.058 | 0.516 | 0.241 | -0.009 | 0.046 | 0.046 | 1.216 | 0.34 | 0.55 | 0.21 | 0.58 | 0.29 | 0.47 |
| 9 | 13300367 | rs10809933 | 1.192 | -0.999 | 0.288 | 0.591 | -0.018 | -0.125 | -0.125 | 0 | 0.58 | 0.04 | 0.68 | 0.39 | 0.64 | 0.59 |
| 9 | 13300442 | rs13439896 | -0.102 | -1.084 | 0.676 | 0.575 | -0.035 | 0.189 | 0.189 | 1.051 | 0.17 | 0.5 | 0.13 | 0.42 | 0.11 | 0.17 |
| 9 | 13315466 | rs13291386 | 0.263 | -1.063 | 0.719 | 0.525 | -0.009 | 0.009 | 0.009 | 0 | 0.2 | 0.12 | 0.15 | 0 | 0.24 | 0.14 |
| 9 | 13317775 | rs7036858 | -0.13 | -1.141 | 0.91 | 0.622 | -0.033 | 0.167 | 0.167 | 0.009 | 0.17 | 0.43 | 0.13 | 0.4 | 0.11 | 0.14 |
| 9 | 13319020 | rs10756472 | 0.618 | -1.141 | 0.783 | 0.626 | -0.018 | 0.027 | 0.027 | 0.812 | 0.8 | 0.3 | 0.82 | 0.58 | 0.88 | 0.84 |
| 9 | 13330233 | rs7032201 | -0.211 | -0.942 | 0.307 | 0.406 | -0.054 | 0.128 | 0.128 | 1.25 | 0.21 | 0.56 | 0.14 | 0.62 | 0.11 | 0.27 |
| 9 | 13345642 | rs11794329 | 0.16 | -0.824 | 0.41 | 0.235 | 0.01 | -0.092 | -0.092 | 0 | 0.2 | 0.31 | 0.14 | 0.01 | 0.23 | 0.16 |
| 9 | 13348599 | rs10961028 | 0.242 | -1.232 | 0.423 | 0.457 | -0.07 | 0.253 | 0.253 | 1.058 | 0.18 | 0.47 | 0.12 | 0.59 | 0.08 | 0.21 |
| 9 | 13350478 | rs1169239 | -0.039 | -1.172 | -0.072 | 0.501 | -0.06 | 0.102 | 0.102 | 1.623 | 0.22 | 0.63 | 0.16 | 0.59 | 0.1 | 0.24 |
| 9 | 13352297 | rs1147779 | 0.17 | -1.163 | -0.086 | -0.157 | -0.066 | 0.066 | 0.066 | 1.634 | 0.2 | 0.58 | 0.15 | 0.46 | 0.09 | 0.25 |
| 9 | 13352941 | rs1147782 | -1.009 | -1.134 | -0.119 | -0.18 | 0.037 | 0.019 | 0.019 | 0.38 | 0.4 | 0.72 | 0.29 | 0.46 | 0.31 | 0.39 |
| 9 | 13354300 | rs1811647 | 1.145 | -0.842 | -0.601 | -0.303 | -0.042 | 0.063 | 0.063 | 1.013 | 0.11 | 0 | 0.08 | 0 | 0.14 | 0.18 |
| 9 | 13357550 | rs1186041 | -0.373 | -1.142 | -0.011 | 0.235 | -0.034 | 0.028 | 0.028 | 0.783 | 0.23 | 0.65 | 0.17 | 0.47 | 0.12 | 0.28 |
| 9 | 13362974 | rs10511579 | -0.53 | -0.367 | -0.124 | -0.237 | 0.007 | 0.031 | 0.031 | 0.483 | 0.2 | 0.1 | 0.13 | 0.01 | 0.21 | 0.12 |
| 9 | 13365816 | rs7038474 | -0.409 | -0.24 | -0.305 | 0.553 | 0.014 | -0.006 | -0.006 | 0 | 0.39 | 0.3 | 0.32 | 0.57 | 0.24 | 0.36 |
| 9 | 13375758 | rs12341870 | -1.134 | -0.102 | 0.43 | 0.288 | 0.022 | 0.003 | 0.003 | 0.075 | 0.07 | 0.36 | 0.05 | 0 | 0.02 | 0.04 |
| 9 | 13380289 | rs928174 | 0.463 | 0.701 | -0.459 | 0.979 | -0.072 | -0.093 | -0.093 | 0 | 0.87 | 0.99 | 0.73 | 0.71 | 0.8 | 0.78 |
| 9 | 13380507 | rs928175 | -0.051 | 0.444 | -0.658 | 0.026 | -0.009 | 0.033 | 0.033 | 0 | 0.33 | 0.23 | 0.21 | 0.16 | 0.39 | 0.34 |

| | | | | | | | | | | | | | | | | |
|---|---|---|---|---|---|---|---|---|---|---|---|---|---|---|---|---|
| 9 | 13388038 | rs7045869 | -0.178 | -0.132 | 0.095 | -0.105 | -0.011 | -0.005 | -0.005 | 0.367 | 0.18 | 0.3 | 0.17 | 0.16 | 0.17 | 0.11 |
| 9 | 13396958 | rs10961039 | -0.141 | -0.533 | -0.585 | -0.12 | -0.073 | -0.029 | -0.029 | 0 | 0.18 | 0 | 0.1 | 0.15 | 0.16 | 0.21 |
| 9 | 13405961 | rs7027951 | -0.086 | -0.629 | 0.118 | -0.259 | -0.01 | -0.006 | -0.006 | 0.006 | 0.18 | 0.31 | 0.17 | 0.15 | 0.18 | 0.09 |
| 9 | 13413469 | rs10809945 | 0.591 | -0.575 | -0.293 | -0.516 | -0.015 | 0.05 | 0.05 | 0.825 | 0.09 | 0 | 0.03 | 0 | 0.08 | 0.14 |
| 9 | 13422337 | rs12000439 | -0.394 | -0.609 | 0.411 | -0.276 | -0.037 | -0.032 | -0.032 | 0.529 | 0.22 | 0.51 | 0.19 | 0.15 | 0.17 | 0.14 |
| 9 | 13444932 | rs10491751 | 0.606 | -0.436 | 0.046 | 0.018 | -0.005 | 0 | 0 | 0 | 0.65 | 0.27 | 0.48 | 0.32 | 0.67 | 0.62 |
| 9 | 13451243 | rs2184230 | 0.167 | -0.405 | -0.511 | 0.124 | -0.071 | -0.109 | -0.109 | 0.662 | 0.77 | 0.91 | 0.59 | 0.45 | 0.72 | 0.69 |
| 9 | 13453984 | rs1556577 | 0.626 | -0.325 | -0.531 | -0.15 | 0.009 | 0.059 | 0.059 | 0 | 0.54 | 0.23 | 0.44 | 0.29 | 0.61 | 0.53 |
| 9 | 13466310 | rs10756475 | 0.31 | -0.172 | -0.017 | 0.199 | -0.003 | 0.036 | 0.036 | 0.018 | 0.73 | 0.56 | 0.59 | 0.64 | 0.73 | 0.75 |
| 9 | 13466329 | rs12238169 | 0.465 | -0.19 | -0.795 | -0.371 | -0.048 | 0.155 | 0.155 | 0.001 | 0.33 | 0.08 | 0.18 | 0.07 | 0.35 | 0.31 |
| 9 | 13467026 | rs983618 | -0.696 | -0.159 | -0.63 | -0.405 | -0.005 | -0.041 | -0.041 | 0 | 0.82 | 0.89 | 0.63 | 0.65 | 0.8 | 0.88 |
| 9 | 13468616 | rs10809954 | 0.895 | -0.163 | -0.626 | -0.416 | 0 | -0.033 | -0.033 | 0.017 | 0.08 | 0.14 | 0.03 | 0 | 0.08 | 0.11 |
| 9 | 13469565 | rs12237974 | 0.394 | -0.216 | -0.987 | -0.431 | -0.008 | 0.056 | 0.056 | 0 | 0.34 | 0.22 | 0.27 | 0.12 | 0.35 | 0.31 |
| 9 | 13469705 | rs2182906 | 0.79 | -0.195 | -1.261 | -0.364 | 0 | -0.013 | -0.013 | 0.469 | 0.39 | 0.34 | 0.32 | 0.51 | 0.38 | 0.43 |
| 9 | 13491120 | rs2890969 | 0.174 | -0.123 | -1.71 | -1.239 | -0.005 | 0.038 | 0.038 | 0 | 0.34 | 0.28 | 0.28 | 0.11 | 0.35 | 0.34 |
| 9 | 13492623 | rs1889297 | -0.595 | -0.19 | -2.106 | -1.296 | -0.089 | 0.067 | 0.067 | 0.168 | 0.5 | 0.96 | 0.37 | 0.31 | 0.47 | 0.53 |
| 9 | 13503997 | rs10738335 | -0.092 | 0.688 | -2.018 | -2.207 | 0.002 | -0.113 | -0.113 | 0 | 0.63 | 0.54 | 0.71 | 0.96 | 0.65 | 0.7 |
| 9 | 13504924 | rs1333996 | -0.254 | 0.816 | -2.455 | -1.313 | -0.046 | 0.112 | 0.112 | 0 | 0.28 | 0.09 | 0.17 | 0.06 | 0.31 | 0.26 |
| 9 | 13507764 | rs10809958 | -0.306 | 0.898 | -2.382 | -1.376 | -0.075 | 0.135 | 0.135 | 0.104 | 0.32 | 0.12 | 0.27 | 0.04 | 0.46 | 0.24 |
| 9 | 13508282 | rs3737150 | -0.49 | -0.398 | -2.742 | -1.415 | 0.09 | 0.209 | 0.209 | 0.294 | 0.44 | 0.9 | 0.47 | 0.89 | 0.26 | 0.48 |
| 9 | 13510394 | rs12684544 | -0.266 | -0.676 | -1.935 | -0.75 | -0.001 | 0.003 | 0.003 | 3.527 | 0.24 | 0.28 | 0.33 | 0.68 | 0.19 | 0.37 |
| 9 | 13511067 | rs4741307 | -0.761 | -1.117 | -2.761 | -1.7 | -0.001 | 0.282 | 0.282 | 0 | 0.57 | 0.92 | 0.55 | 0.96 | 0.34 | 0.69 |
| 9 | 13520190 | rs1543713 | -0.672 | -1.145 | -1.644 | -1.917 | 0.003 | -0.066 | -0.066 | 0 | 0.49 | 0.52 | 0.61 | 0.99 | 0.42 | 0.69 |
| 9 | 13526525 | rs10961109 | -0.204 | -1.302 | -1.832 | -0.684 | 0.029 | -0.067 | -0.067 | 0 | 0.11 | 0.01 | 0.27 | 0.57 | 0.06 | 0.2 |
| 9 | 13528833 | rs10961110 | -0.351 | -0.87 | -1.447 | -0.822 | 0.04 | 0.015 | 0.015 | 2.704 | 0.11 | 0.14 | 0.09 | 0.15 | 0.17 | 0.1 |
| 9 | 13530711 | rs933034 | -1.302 | -1.435 | -0.762 | -1.109 | -0.002 | 0.039 | 0.039 | 0 | 0.57 | 0.67 | 0.59 | 0.87 | 0.43 | 0.66 |
| 9 | 13531732 | rs974350 | -0.809 | -1.973 | -0.057 | -0.912 | -0.027 | 0.018 | 0.018 | 0.192 | 0.24 | 0.36 | 0.32 | 0.63 | 0.14 | 0.23 |
| 9 | 13532846 | rs2018555 | -0.858 | -2.565 | 0.411 | -0.458 | 0.023 | 0.028 | 0.028 | 0 | 0.51 | 0.6 | 0.38 | 0.8 | 0.28 | 0.52 |
| 9 | 13536161 | rs7039074 | -1.048 | -2.561 | 0.506 | -0.581 | 0.039 | 0.111 | 0.111 | 0 | 0.57 | 0.82 | 0.38 | 0.81 | 0.32 | 0.6 |
| 9 | 13546652 | rs1333989 | 1.379 | -2.987 | 0.382 | -0.966 | -0.022 | 0.08 | 0.08 | 0.043 | 0.39 | 0.14 | 0.58 | 0.18 | 0.64 | 0.39 |
| 9 | 13548060 | rs10491754 | 0.013 | -3.212 | 0.907 | -0.348 | 0.019 | 0.017 | 0.017 | 0 | 0.33 | 0.29 | 0.14 | 0.21 | 0.18 | 0.29 |
| 9 | 13549596 | rs7042381 | -0.345 | -3.2 | 0.978 | -0.345 | -0.052 | 0.098 | 0.098 | 0.132 | 0.25 | 0.51 | 0.24 | 0.6 | 0.14 | 0.31 |
| 9 | 13553285 | rs10961135 | 0.341 | -3.166 | 1.208 | -0.317 | -0.009 | 0.212 | 0.212 | 0 | 0.1 | 0.24 | 0.18 | 0.6 | 0.05 | 0.28 |
| 9 | 13555912 | rs11999938 | 0.451 | -3.055 | 1.079 | -0.356 | 0.014 | 0.028 | 0.028 | 0 | 0.3 | 0.22 | 0.13 | 0.11 | 0.19 | 0.25 |
| 9 | 13560266 | rs10491757 | -1.552 | -3.014 | 0.769 | -0.189 | 0.026 | 0.107 | 0.107 | 0 | 0.57 | 0.79 | 0.38 | 0.8 | 0.33 | 0.59 |
| 9 | 13560795 | rs10961141 | 0.026 | -3.054 | 1.056 | -0.068 | -0.019 | 0.072 | 0.072 | 0.154 | 0.21 | 0.28 | 0.22 | 0.58 | 0.13 | 0.29 |
| 9 | 13564273 | rs10491758 | 0.227 | -2.989 | 0.994 | -0.096 | -0.004 | 0.216 | 0.216 | 0 | 0.1 | 0.25 | 0.18 | 0.69 | 0.05 | 0.29 |
| 9 | 13565582 | rs10961147 | -1.52 | -3.074 | 0.637 | -0.2 | 0.016 | 0.128 | 0.128 | 0 | 0.56 | 0.89 | 0.4 | 0.8 | 0.33 | 0.59 |
| 9 | 13568063 | rs1324186 | -0.472 | -3.056 | 0.891 | -0.253 | 0.027 | 0.041 | 0.041 | 0 | 0.42 | 0.33 | 0.19 | 0.15 | 0.28 | 0.3 |
| 9 | 13577494 | rs7847125 | -0.535 | -2.835 | 0.89 | -0.386 | -0.045 | -0.162 | -0.162 | 0 | 0.28 | 0.55 | 0.18 | 0.01 | 0.22 | 0.16 |
| 9 | 13586599 | rs7018881 | -1.141 | -2.734 | -0.024 | 0.021 | 0.048 | -0.043 | -0.043 | 0 | 0.37 | 0.79 | 0.33 | 0.53 | 0.27 | 0.45 |
| 9 | 13590418 | rs10961156 | 0.086 | -2.916 | 0.412 | 0.42 | -0.02 | 0.215 | 0.215 | 0.076 | 0.05 | 0.14 | 0.16 | 0.6 | 0.02 | 0.23 |
| 9 | 13590443 | rs10756485 | 0.423 | -3.441 | 0.343 | 0.012 | 0.016 | 0.015 | 0.015 | 0.09 | 0.56 | 0.59 | 0.8 | 0.9 | 0.75 | 0.72 |
| 9 | 13590687 | rs10961157 | 0.987 | -3.485 | 0.448 | 0.01 | 0.024 | 0.024 | 0.024 | 0 | 0.5 | 0.38 | 0.64 | 0.3 | 0.74 | 0.49 |
| 9 | 13591906 | rs13299116 | 0.549 | -3.591 | 0.452 | -0.115 | -0.017 | -0.051 | -0.051 | 0 | 0.17 | 0.3 | 0.07 | 0.02 | 0.12 | 0.05 |
| 9 | 13592377 | rs1029139 | 0.444 | -3.572 | 0.028 | -0.092 | 0.045 | 0.062 | 0.062 | 0.022 | 0.57 | 0.69 | 0.81 | 0.9 | 0.76 | 0.72 |
| 9 | 13592766 | rs10123222 | -0.043 | -3.524 | 0.055 | 0.071 | 0.003 | 0.002 | 0.002 | 0.134 | 0.68 | 0.71 | 0.83 | 0.8 | 0.85 | 0.72 |
| 9 | 13592808 | rs10123240 | 0.768 | -3.386 | -0.115 | -0.364 | 0.002 | 0.002 | 0.002 | 0 | 0.51 | 0.48 | 0.65 | 0.3 | 0.74 | 0.49 |
| 9 | 13593125 | rs10123887 | -0.089 | -2.943 | -0.083 | -0.088 | 0.033 | 0.038 | 0.038 | 0 | 0.73 | 0.85 | 0.88 | 0.92 | 0.88 | 0.77 |
| 9 | 13598413 | rs1408320 | 0.788 | -2.274 | 0.226 | 0.325 | -0.049 | -0.095 | -0.095 | 0.611 | 0.72 | 0.55 | 0.83 | 0.92 | 0.83 | 0.81 |
| 9 | 13600898 | rs1590408 | 0.567 | -2.12 | -0.72 | 0.377 | 0.016 | 0.019 | 0.019 | 0 | 0.74 | 0.75 | 0.85 | 0.92 | 0.84 | 0.82 |
| 9 | 13603327 | rs10961168 | -0.113 | -0.886 | -0.983 | 0.394 | -0.023 | -0.021 | -0.021 | 2.89 | 0.07 | 0.02 | 0.13 | 0.05 | 0.12 | 0.11 |
| 9 | 13606721 | rs10809998 | 0.404 | -1.595 | -1.032 | -0.177 | 0.039 | 0.061 | 0.061 | 0 | 0.74 | 0.9 | 0.86 | 0.95 | 0.87 | 0.81 |
| 9 | 13609119 | rs12336782 | -1.457 | -1.556 | -0.524 | 0.093 | -0.008 | -0.001 | -0.001 | 0 | 0.12 | 0.21 | 0.11 | 0.09 | 0.08 | 0.2 |
| 9 | 13609898 | rs1359253 | 0.46 | -2.509 | -0.198 | 0.536 | 0.003 | -0.003 | -0.003 | 0 | 0.63 | 0.6 | 0.85 | 0.44 | 0.83 | 0.64 |
| 9 | 13609910 | rs4278236 | 1.186 | -2.317 | -0.31 | 0.206 | -0.002 | -0.035 | -0.035 | 0.654 | 0.47 | 0.44 | 0.5 | 0.11 | 0.66 | 0.43 |
| 9 | 13610778 | rs10738339 | -0.165 | -2.556 | -0.226 | 0.665 | 0.008 | -0.003 | -0.003 | 0.953 | 0.7 | 0.72 | 0.87 | 0.62 | 0.84 | 0.64 |
| 9 | 13612345 | rs1324192 | 1.301 | -2.747 | -0.275 | 0.527 | 0.021 | 0.043 | 0.043 | 0 | 0.48 | 0.59 | 0.51 | 0.18 | 0.65 | 0.36 |
| 9 | 13614384 | rs7019262 | 1.956 | -2.662 | 0.013 | 0.451 | 0.004 | 0.154 | 0.154 | 0 | 0.41 | 0.2 | 0.42 | 0.07 | 0.63 | 0.32 |
| 9 | 13614760 | rs16930557 | 1.564 | -2.602 | -0.253 | 0.575 | 0.004 | -0.032 | -0.032 | 0 | 0.06 | 0 | 0.02 | 0.1 | 0.03 | 0.04 |
| 9 | 13616003 | rs2382402 | -0.763 | -2.574 | -0.223 | 0.442 | 0 | -0.072 | -0.072 | 0 | 0.24 | 0.12 | 0.37 | 0.44 | 0.23 | 0.31 |
| 9 | 13616063 | rs1324193 | -1.107 | -2.657 | 0.07 | 0.47 | -0.005 | 0.064 | 0.064 | 0.946 | 0.12 | 0.49 | 0.13 | 0.12 | 0.06 | 0.16 |
| 9 | 13616431 | rs1535836 | -0.107 | -2.48 | 0.096 | 0.581 | -0.009 | 0.001 | 0.001 | 0 | 0.69 | 0.68 | 0.85 | 0.62 | 0.84 | 0.64 |
| 9 | 13617635 | rs10810000 | -0.351 | -1.873 | -0.034 | 0.845 | 0.009 | -0.04 | -0.04 | 0 | 0.17 | 0.09 | 0.29 | 0.4 | 0.15 | 0.17 |
| 9 | 13619784 | rs12341613 | -0.925 | -1.629 | 0.073 | 0.352 | -0.002 | -0.008 | -0.008 | 0.29 | 0.08 | 0.1 | 0.06 | 0.04 | 0.06 | 0.21 |
| 9 | 13630162 | rs1853609 | -1.472 | -2.291 | 0.376 | 0.56 | 0.007 | -0.001 | -0.001 | 0.142 | 0.32 | 0.65 | 0.22 | 0.31 | 0.18 | 0.37 |
| 9 | 13631992 | rs13440249 | -1.152 | -2.351 | 1.006 | 0.539 | -0.007 | 0.002 | 0.002 | 0 | 0.28 | 0.27 | 0.19 | 0.31 | 0.18 | 0.37 |
| 9 | 13635181 | rs12352645 | -1.389 | -2.433 | 1.226 | 0.447 | -0.014 | -0.083 | -0.083 | 0 | 0.13 | 0.4 | 0.06 | 0.01 | 0.05 | 0.13 |
| 9 | 13639416 | rs16930601 | -0.848 | -2.389 | 1.193 | 0.557 | -0.06 | 0.029 | 0.029 | 0 | 0.24 | 0.53 | 0.16 | 0.28 | 0.1 | 0.16 |

| | | | | | | | | | | | | | | | | |
|---|---|---|---|---|---|---|---|---|---|---|---|---|---|---|---|---|
| 9 | 13648463 | rs10810001 | -0.656 | -2.69 | 1.187 | 0.144 | -0.01 | -0.037 | -0.037 | 0.362 | 0.83 | 0.75 | 0.92 | 0.97 | 0.9 | 0.91 |
| 9 | 13657571 | rs10961187 | 2.031 | -2.851 | 0.834 | 0.472 | 0 | -0.001 | -0.001 | 0.063 | 0.4 | 0.36 | 0.48 | 0.34 | 0.61 | 0.39 |
| 9 | 13660128 | rs8192964 | 1.835 | -2.853 | 0.89 | 0.438 | 0.005 | -0.012 | -0.012 | 0 | 0.4 | 0.46 | 0.49 | 0.33 | 0.58 | 0.39 |
| 9 | 13671296 | rs642806 | -0.385 | -2.982 | 0.852 | 0.333 | 0.009 | -0.029 | -0.029 | 0 | 0.91 | 0.75 | 0.93 | 0.96 | 0.9 | 0.92 |
| 9 | 13673697 | rs7872836 | 2.437 | -2.983 | 0.916 | 0.297 | -0.001 | 0 | 0 | 0 | 0.37 | 0.32 | 0.47 | 0.32 | 0.58 | 0.38 |
| 9 | 13674291 | rs571485 | 1.33 | -3.276 | 0.755 | 0.297 | 0.024 | -0.007 | -0.007 | 0 | 0.59 | 0.64 | 0.64 | 0.56 | 0.7 | 0.64 |
| 9 | 13677557 | rs688703 | 0.818 | -3.272 | 0.774 | 0.381 | -0.006 | 0.004 | 0.004 | 0.622 | 0.72 | 0.65 | 0.69 | 0.65 | 0.79 | 0.78 |
| 9 | 13682827 | rs613444 | -2.12 | -3.283 | 0.64 | 0.53 | 0.05 | 0.023 | 0.023 | 0 | 0.49 | 0.57 | 0.41 | 0.58 | 0.29 | 0.48 |
| 9 | 13682981 | rs522594 | -0.368 | -3.189 | 0.501 | 0.604 | 0.043 | -0.009 | -0.009 | 0 | 0.3 | 0.1 | 0.18 | 0.29 | 0.17 | 0.28 |
| 9 | 13688437 | rs4740554 | 0.286 | -3.123 | 0.557 | 0.667 | -0.018 | 0.099 | 0.099 | 0 | 0.16 | 0.31 | 0.14 | 0.27 | 0.09 | 0.15 |
| 9 | 13689066 | rs518054 | -0.237 | -3.115 | 0.062 | 0.726 | 0.016 | -0.049 | -0.049 | 0.526 | 0.28 | 0.05 | 0.26 | 0.33 | 0.2 | 0.33 |
| 9 | 13690235 | rs571221 | -0.149 | -3.105 | 0.065 | 0.686 | 0.019 | -0.052 | -0.052 | 0.014 | 0.27 | 0.04 | 0.31 | 0.32 | 0.2 | 0.33 |
| 9 | 13697152 | rs1590404 | 2.504 | -3.143 | 0.016 | 0.605 | -0.086 | -0.036 | -0.036 | 0 | 0.4 | 0.05 | 0.46 | 0.35 | 0.59 | 0.44 |
| 9 | 13703826 | rs562328 | 1.397 | -3.248 | 0.211 | 0.606 | -0.014 | 0.002 | 0.002 | 0 | 0.59 | 0.44 | 0.62 | 0.58 | 0.7 | 0.64 |
| 9 | 13704539 | rs1998584 | 2.677 | -3.235 | 0.341 | 0.606 | -0.004 | 0.006 | 0.006 | 0 | 0.41 | 0.29 | 0.47 | 0.35 | 0.59 | 0.38 |
| 9 | 13705679 | rs626778 | 1.451 | -3.244 | 0.403 | 0.627 | 0.032 | 0.006 | 0.006 | 0 | 0.59 | 0.24 | 0.54 | 0.39 | 0.7 | 0.62 |
| 9 | 13705882 | rs992457 | 1.797 | -3.227 | 0.319 | 0.413 | 0.003 | 0.001 | 0.001 | 0 | 0.49 | 0.43 | 0.56 | 0.57 | 0.62 | 0.5 |
| 9 | 13706483 | rs612828 | 1.678 | -3.138 | -0.38 | 0.416 | 0.005 | 0.031 | 0.031 | 0 | 0.11 | 0.03 | 0.06 | 0.01 | 0.08 | 0.13 |
| 9 | 13706973 | rs610593 | -0.57 | -3.033 | -0.275 | 0.456 | 0.026 | -0.023 | -0.023 | 0.072 | 0.9 | 0.72 | 0.89 | 0.96 | 0.88 | 0.96 |
| 9 | 13708607 | rs524126 | 1.182 | -2.94 | -0.744 | 0.486 | 0.029 | -0.018 | -0.018 | 0 | 0.6 | 0.25 | 0.6 | 0.58 | 0.7 | 0.64 |
| 9 | 13711595 | rs504659 | 0.937 | -2.806 | -0.808 | 0.447 | 0.021 | 0.024 | 0.024 | 0.013 | 0.16 | 0.07 | 0.06 | 0.01 | 0.08 | 0.13 |
| 9 | 13730047 | rs10120807 | 1.094 | -2.922 | -0.82 | 0.554 | -0.003 | 0.001 | 0.001 | 0 | 0.62 | 0.37 | 0.61 | 0.58 | 0.7 | 0.65 |
| 9 | 13730340 | rs4741325 | 1.125 | -2.88 | -1.119 | 0.636 | -0.023 | -0.076 | -0.076 | 0 | 0.12 | 0.01 | 0.22 | 0.3 | 0.11 | 0.19 |
| 9 | 13730665 | rs979203 | -0.858 | -2.818 | -0.902 | 0.417 | 0.001 | -0.017 | -0.017 | 0 | 0.91 | 0.8 | 0.95 | 0.96 | 0.92 | 0.97 |
| 9 | 13732363 | rs12339356 | 1.17 | -2.151 | -1.605 | 0.406 | -0.02 | 0.077 | 0.077 | 0 | 0.14 | 0.01 | 0.07 | 0.01 | 0.11 | 0.09 |
| 9 | 13734430 | rs2382407 | 1.743 | -2.203 | -1.674 | 0.646 | -0.116 | -0.012 | -0.012 | 0.886 | 0.4 | 0.05 | 0.45 | 0.32 | 0.57 | 0.43 |
| 9 | 13736662 | rs1556491 | 1.929 | -2.086 | -1.742 | 0.365 | -0.147 | 0.178 | 0.178 | 0 | 0.38 | 0.05 | 0.4 | 0.08 | 0.55 | 0.2 |
| 9 | 13736965 | rs717605 | 0.388 | -1.401 | -1.741 | 0.38 | 0.003 | 0.001 | 0.001 | 0 | 0.54 | 0.42 | 0.58 | 0.53 | 0.64 | 0.53 |
| 9 | 13737420 | rs7040897 | 0.202 | 0.734 | -2.935 | -0.041 | -0.102 | 0.027 | 0.027 | 0 | 0.19 | 0.03 | 0.27 | 0.07 | 0.34 | 0.14 |
| 9 | 13737975 | rs4741326 | 0.09 | 0.199 | -1.974 | 0.668 | 0 | -0.01 | -0.01 | 0 | 0.47 | 0.14 | 0.6 | 0.41 | 0.64 | 0.41 |
| 9 | 13738362 | rs10810012 | 0.595 | 0.264 | -2.01 | -0.236 | -0.104 | 0.167 | 0.167 | 0.678 | 0.37 | 0.07 | 0.36 | 0.08 | 0.49 | 0.15 |
| 9 | 13743830 | rs11792535 | -0.41 | 0.937 | -1.224 | 0.45 | 0.011 | 0.03 | 0.03 | 0 | 0.12 | 0.18 | 0.21 | 0.32 | 0.2 | 0.23 |
| 9 | 13744658 | rs1887456 | 0.968 | 0.695 | -1.839 | -0.148 | -0.142 | 0.157 | 0.157 | 0.335 | 0.33 | 0.03 | 0.37 | 0.07 | 0.44 | 0.17 |
| 9 | 13745502 | rs10961216 | -1.086 | 0.671 | -1.597 | -0.145 | -0.012 | 0.127 | 0.127 | 0.376 | 0.63 | 0.91 | 0.64 | 0.9 | 0.61 | 0.82 |
| 9 | 13746595 | rs10810013 | -0.145 | 0.433 | -1.707 | -0.144 | -0.024 | 0.045 | 0.045 | 0.422 | 0.43 | 0.25 | 0.35 | 0.69 | 0.44 | 0.53 |
| 9 | 13749977 | rs930543 | 1.919 | 0.626 | -1.695 | -0.256 | -0.051 | 0.073 | 0.073 | 0.081 | 0.26 | 0.02 | 0.3 | 0.08 | 0.28 | 0.15 |
| 9 | 13751127 | rs10738341 | -0.606 | 0.669 | -1.483 | -0.248 | 0.018 | 0.142 | 0.142 | 0.676 | 0.54 | 0.27 | 0.56 | 0.9 | 0.55 | 0.78 |
| 9 | 13752127 | rs10810020 | -1.579 | 0.702 | -1.607 | -0.168 | -0.03 | 0.106 | 0.106 | 0.105 | 0.69 | 0.94 | 0.67 | 0.9 | 0.67 | 0.84 |
| 9 | 13759474 | rs12347123 | 0.759 | 0.553 | -1.027 | 0.119 | 0.002 | 0.073 | 0.073 | 0 | 0.05 | 0.09 | 0.18 | 0.22 | 0.07 | 0.22 |
| 9 | 13759929 | rs10756494 | -0.281 | 0.556 | -1.033 | -0.051 | 0.013 | 0.088 | 0.088 | 0.307 | 0.5 | 0.21 | 0.37 | 0.68 | 0.5 | 0.57 |
| 9 | 13762898 | rs10961227 | 0.565 | 0.497 | -1.126 | 0.08 | -0.033 | -0.064 | -0.064 | 1.23 | 0.38 | 0.04 | 0.18 | 0.41 | 0.3 | 0.32 |
| 9 | 13763734 | rs10756497 | -0.244 | 0.578 | -1.045 | 0.024 | 0.026 | 0.09 | 0.09 | 0.217 | 0.5 | 0.19 | 0.38 | 0.68 | 0.51 | 0.57 |
| 9 | 13765292 | rs966992 | -1.404 | 0.516 | -0.48 | -0.08 | -0.018 | -0.078 | -0.078 | 0 | 0.67 | 0.52 | 0.64 | 0.91 | 0.69 | 0.85 |
| 9 | 13765759 | rs10810025 | -0.412 | 0.268 | -0.575 | -0.136 | -0.021 | -0.013 | -0.013 | 0 | 0.24 | 0.18 | 0.26 | 0.33 | 0.32 | 0.31 |
| 9 | 13768750 | rs10961229 | 1.565 | 0.458 | -0.608 | -0.255 | -0.037 | 0.112 | 0.112 | 0 | 0.3 | 0.09 | 0.31 | 0.07 | 0.36 | 0.18 |
| 9 | 13770466 | rs1034167 | -1.91 | 0.241 | -0.494 | -0.13 | 0.003 | 0.063 | 0.063 | 0.05 | 0.72 | 0.87 | 0.7 | 0.91 | 0.7 | 0.86 |
| 9 | 13771744 | rs1034164 | -0.17 | 0.221 | -0.582 | -0.111 | 0.009 | 0.003 | 0.003 | 0.224 | 0.26 | 0.33 | 0.26 | 0.34 | 0.33 | 0.39 |
| 9 | 13772055 | rs10756500 | -1.119 | 0.254 | -0.523 | -0.066 | -0.005 | -0.005 | -0.005 | 0 | 0.65 | 0.63 | 0.5 | 0.7 | 0.62 | 0.69 |
| 9 | 13774329 | rs1562045 | 0.269 | 0.292 | -0.48 | -0.046 | 0.019 | 0.05 | 0.05 | 0.042 | 0.2 | 0.42 | 0.21 | 0.34 | 0.22 | 0.28 |
| 9 | 13774683 | rs1359848 | -0.797 | 0.315 | -0.535 | -0.025 | 0.021 | 0.046 | 0.046 | 0.706 | 0.32 | 0.48 | 0.45 | 0.55 | 0.39 | 0.56 |
| 9 | 13775193 | rs10810029 | 0.909 | 0.357 | -0.88 | -0.112 | -0.033 | -0.048 | -0.048 | 0.016 | 0.16 | 0.09 | 0.18 | 0.34 | 0.21 | 0.28 |
| 9 | 13777299 | rs7030346 | 0.627 | 0.595 | -0.961 | -0.081 | 0.01 | -0.038 | -0.038 | 0 | 0.09 | 0.15 | 0.07 | 0 | 0.11 | 0.06 |
| 9 | 13779856 | rs10756504 | 0.251 | 0.279 | -0.807 | 0.202 | -0.016 | 0.023 | 0.023 | 0.001 | 0.44 | 0.29 | 0.3 | 0.35 | 0.42 | 0.35 |
| 9 | 13780058 | rs10756505 | -0.182 | -0.072 | -0.845 | 0.053 | -0.007 | -0.004 | -0.004 | 1.118 | 0.64 | 0.62 | 0.47 | 0.7 | 0.63 | 0.6 |
| 9 | 13787648 | rs10810035 | 0.861 | 0.21 | -1.509 | -0.037 | -0.057 | 0.105 | 0.105 | 0.009 | 0.26 | 0.02 | 0.31 | 0.06 | 0.29 | 0.11 |
| 9 | 13790202 | rs10756511 | 0.08 | 0.142 | -1.267 | 0.088 | -0.027 | -0.042 | -0.042 | 0 | 0.21 | 0.36 | 0.13 | 0.1 | 0.14 | 0.05 |
| 9 | 13791531 | rs1854163 | 0.031 | 0.647 | -1.93 | 0.509 | -0.005 | 0.003 | 0.003 | 0 | 0.68 | 0.88 | 0.5 | 0.72 | 0.64 | 0.63 |
| 9 | 13798727 | rs7858704 | -0.194 | 0.798 | -1.93 | 0.501 | -0.017 | 0.033 | 0.033 | 1.552 | 0.47 | 0.78 | 0.65 | 0.64 | 0.52 | 0.68 |
| 9 | 13799421 | rs10961259 | -0.783 | 0.644 | -1.226 | -0.11 | -0.004 | -0.049 | -0.049 | 0.179 | 0.21 | 0.38 | 0.14 | 0.07 | 0.19 | 0.1 |
| 9 | 13799717 | rs10756514 | -0.004 | 0.643 | -1.867 | -0.04 | -0.049 | -0.006 | -0.006 | 0 | 0.61 | 0.84 | 0.61 | 0.53 | 0.61 | 0.42 |
| 9 | 13802044 | rs1610343 | 0.049 | 0.654 | -1.738 | -0.016 | -0.032 | -0.004 | -0.004 | 0 | 0.6 | 0.79 | 0.57 | 0.48 | 0.6 | 0.41 |
| 9 | 13803088 | rs1410035 | 0.57 | 0.675 | -1.759 | 0.053 | 0.001 | 0.003 | 0.003 | 0 | 0.49 | 0.48 | 0.52 | 0.5 | 0.51 | 0.37 |
| 9 | 13806091 | rs4741336 | -0.103 | 0.692 | -1.908 | 0.281 | -0.039 | 0.023 | 0.023 | 0 | 0.61 | 0.72 | 0.57 | 0.23 | 0.6 | 0.39 |
| 9 | 13809735 | rs894822 | 0.192 | 0.767 | -1.316 | -0.085 | 0.006 | 0.013 | 0.013 | 0 | 0.53 | 0.37 | 0.54 | 0.39 | 0.57 | 0.42 |
| 9 | 13812037 | rs987416 | -0.968 | 0.343 | -1.021 | -0.967 | 0.002 | -0.022 | -0.022 | 0 | 0.19 | 0.2 | 0.13 | 0.01 | 0.14 | 0.11 |
| 9 | 13812685 | rs7866165 | -1.403 | 0.551 | -0.791 | -1.342 | -0.002 | -0.102 | -0.102 | 0 | 0.3 | 0.4 | 0.23 | 0.01 | 0.26 | 0.17 |
| 9 | 13815721 | rs986974 | -0.975 | 0.539 | -0.635 | -1.334 | 0.005 | -0.107 | -0.107 | 3.179 | 0.22 | 0.38 | 0.17 | 0 | 0.18 | 0.13 |
| 9 | 13825565 | rs11788150 | -1.469 | 0.65 | -1.116 | -1.62 | -0.035 | 0.108 | 0.108 | 0 | 0.36 | 0.21 | 0.23 | 0.02 | 0.35 | 0.24 |
| 9 | 13828594 | rs2078639 | 0.328 | 0.692 | -1.152 | -1.519 | -0.029 | 0.055 | 0.055 | 0 | 0.11 | 0.01 | 0.06 | 0.01 | 0.12 | 0.09 |

| | | | | | | | | | | | | | | | | |
|---|---|---|---|---|---|---|---|---|---|---|---|---|---|---|---|---|
| 9 | 13829452 | rs17818373 | -0.777 | 0.699 | -0.981 | -1.549 | 0.001 | -0.034 | -0.034 | 0.003 | 0.18 | 0.2 | 0.12 | 0.01 | 0.2 | 0.13 |
| 9 | 13830766 | rs1410040 | -1.074 | 0.378 | -0.93 | -1.674 | 0.021 | 0.067 | 0.067 | 0 | 0.25 | 0.13 | 0.14 | 0.02 | 0.2 | 0.19 |
| 9 | 13832491 | rs726353 | -2.42 | -0.183 | -0.257 | -1.899 | -0.001 | -0.037 | -0.037 | 2.244 | 0.46 | 0.51 | 0.27 | 0.13 | 0.36 | 0.29 |
| 9 | 13832587 | rs9407545 | -0.849 | -0.17 | -0.267 | -1.873 | 0.016 | 0.026 | 0.026 | 0 | 0.25 | 0.17 | 0.14 | 0.01 | 0.21 | 0.19 |
| 9 | 13836332 | rs17219480 | -0.554 | -0.227 | -0.285 | -1.873 | 0.006 | -0.01 | -0.01 | 0 | 0.14 | 0.12 | 0.07 | 0 | 0.08 | 0.11 |
| 9 | 13839436 | rs10961271 | -0.54 | -0.118 | -0.141 | -1.942 | 0.01 | -0.023 | -0.023 | 1.582 | 0.21 | 0.31 | 0.14 | 0.12 | 0.19 | 0.1 |
| 9 | 13844771 | rs12003992 | -2.442 | -0.09 | 0.023 | -1.962 | 0.003 | 0.004 | 0.004 | 0 | 0.47 | 0.63 | 0.27 | 0.13 | 0.36 | 0.31 |
| 9 | 13846800 | rs10491766 | -0.414 | -0.134 | 0.029 | -1.981 | 0.008 | 0.049 | 0.049 | 0.255 | 0.07 | 0.2 | 0.04 | 0.1 | 0.05 | 0.03 |
| 9 | 13848797 | rs7019634 | -2.423 | -0.146 | -0.053 | -1.988 | -0.001 | -0.04 | -0.04 | 0 | 0.46 | 0.56 | 0.29 | 0.12 | 0.39 | 0.28 |
| 9 | 13851999 | rs9407554 | -0.471 | -0.161 | -0.238 | -2.023 | 0.029 | 0.063 | 0.063 | 0.485 | 0.14 | 0.05 | 0.07 | 0 | 0.09 | 0.1 |
| 9 | 13852782 | rs10961283 | -0.348 | -0.155 | -0.246 | -2.317 | -0.009 | 0.017 | 0.017 | 0 | 0.17 | 0.14 | 0.1 | 0.09 | 0.14 | 0.09 |
| 9 | 13854370 | rs7036600 | -1.334 | -0.274 | 0.111 | -2.45 | -0.023 | -0.149 | -0.149 | 0 | 0.29 | 0.54 | 0.19 | 0.03 | 0.21 | 0.15 |
| 9 | 13854965 | rs12237041 | -1.53 | -0.32 | -0.072 | -2.543 | 0.009 | 0.081 | 0.081 | 0 | 0.34 | 0.19 | 0.19 | 0.1 | 0.28 | 0.21 |
| 9 | 13856937 | rs13295592 | -0.845 | -0.266 | 0.129 | -2.436 | 0.061 | 0.03 | 0.03 | 0.928 | 0.06 | 0.37 | 0.04 | 0.02 | 0.02 | 0.02 |
| 9 | 13857272 | rs17221355 | -0.594 | -0.157 | -0.131 | -2.462 | -0.023 | 0.041 | 0.041 | 0 | 0.2 | 0.06 | 0.11 | 0.08 | 0.17 | 0.1 |
| 9 | 13857339 | rs13300184 | -2.341 | -0.106 | -0.117 | -2.582 | 0.008 | 0.074 | 0.074 | 0 | 0.49 | 0.67 | 0.31 | 0.11 | 0.41 | 0.26 |
| 9 | 13857405 | rs16931028 | -0.59 | 0.235 | 0.121 | -2.394 | 0.006 | -0.146 | -0.146 | 2.994 | 0.2 | 0.48 | 0.13 | 0.03 | 0.16 | 0.07 |
| 9 | 13859052 | rs12341720 | -0.5 | 0.473 | -0.554 | -2.429 | -0.039 | 0.096 | 0.096 | 0 | 0.25 | 0.09 | 0.17 | 0.01 | 0.26 | 0.13 |
| 9 | 13859987 | rs7028624 | -1.579 | 0.319 | 0.068 | -2.084 | -0.009 | -0.065 | -0.065 | 0 | 0.19 | 0.55 | 0.14 | 0.09 | 0.14 | 0.14 |
| 9 | 13861427 | rs16931045 | -0.547 | 1.11 | -0.216 | -1.988 | 0.003 | -0.018 | -0.018 | 0.01 | 0.08 | 0.13 | 0.07 | 0 | 0.12 | 0.09 |
| 9 | 13864265 | rs10117920 | -0.222 | 1.155 | -0.496 | -2.426 | -0.032 | 0.077 | 0.077 | 0 | 0.24 | 0.07 | 0.16 | 0.01 | 0.25 | 0.12 |
| 9 | 13864770 | rs12341611 | -0.222 | 1.229 | -0.561 | -2.488 | -0.013 | 0.008 | 0.008 | 0 | 0.12 | 0.05 | 0.06 | 0 | 0.13 | 0.07 |
| 9 | 13865260 | rs7858214 | 0.714 | 1.167 | -0.201 | -2.468 | -0.002 | -0.041 | -0.041 | 0 | 0.14 | 0.21 | 0.12 | 0.01 | 0.13 | 0.05 |
| 9 | 13867306 | rs2224553 | -1.018 | 1.205 | 0.325 | -2.547 | 0.032 | -0.129 | -0.129 | 0 | 0.24 | 0.46 | 0.22 | 0.03 | 0.28 | 0.15 |
| 9 | 13868750 | rs10810049 | 0.111 | 1.221 | 0.455 | -2.555 | 0.04 | -0.104 | -0.104 | 0 | 0.18 | 0.43 | 0.17 | 0.02 | 0.2 | 0.1 |
| 9 | 13871187 | rs12006148 | -2.153 | 1.193 | 0.582 | -2.639 | 0.011 | -0.094 | -0.094 | 1.732 | 0.37 | 0.56 | 0.28 | 0.03 | 0.4 | 0.22 |
| 9 | 13873487 | rs2382423 | 0.97 | 1.159 | 0.489 | -2.583 | -0.009 | 0.032 | 0.032 | 0 | 0.13 | 0.08 | 0.11 | 0.01 | 0.15 | 0.05 |
| 9 | 13876562 | rs10961291 | -0.142 | 1.149 | 0.506 | -2.608 | -0.026 | 0.052 | 0.052 | 0 | 0.21 | 0.14 | 0.18 | 0.01 | 0.26 | 0.1 |
| 9 | 13879572 | rs12344958 | -2.071 | 1.07 | 0.542 | -2.671 | 0.006 | -0.093 | -0.093 | 0.007 | 0.35 | 0.63 | 0.25 | 0.03 | 0.35 | 0.21 |
| 9 | 13879604 | rs9407563 | 1.335 | 1.054 | -0.807 | -2.26 | -0.097 | -0.163 | -0.163 | 0 | 0.26 | 0.02 | 0.28 | 0.59 | 0.29 | 0.41 |
| 9 | 13880787 | rs7036035 | -2.094 | 0.449 | -0.78 | -2.691 | -0.005 | 0.101 | 0.101 | 0 | 0.48 | 0.7 | 0.36 | 0.11 | 0.46 | 0.3 |
| 9 | 13887476 | rs1576066 | 0.189 | 0.566 | -0.613 | -2.642 | 0.011 | -0.056 | -0.056 | 0 | 0.11 | 0.25 | 0.11 | 0.01 | 0.1 | 0.05 |
| 9 | 13891525 | rs9969743 | 1.974 | 0.603 | -0.828 | -2.671 | -0.042 | 0.069 | 0.069 | 0 | 0.13 | 0.02 | 0.12 | 0.01 | 0.17 | 0.06 |
| 9 | 13892287 | rs1831087 | -1.216 | 0.447 | -0.709 | -2.694 | 0.027 | -0.124 | -0.124 | 0 | 0.18 | 0.47 | 0.16 | 0.02 | 0.18 | 0.13 |
| 9 | 13896279 | rs7034760 | -1.026 | 0.511 | -0.913 | -2.687 | 0.023 | -0.15 | -0.15 | 0.896 | 0.24 | 0.5 | 0.18 | 0.02 | 0.21 | 0.13 |
| 9 | 13898481 | rs13292371 | 2.239 | 0.555 | -0.955 | -2.683 | -0.027 | 0.07 | 0.07 | 0 | 0.12 | 0 | 0.1 | 0.01 | 0.13 | 0.05 |
| 9 | 13898615 | rs10121526 | 0.808 | 0.6 | -0.847 | -2.591 | -0.026 | 0.043 | 0.043 | 0 | 0.12 | 0.03 | 0.05 | 0 | 0.11 | 0.08 |
| 9 | 13898956 | rs1923352 | -1.217 | 0.548 | -0.812 | -2.538 | -0.001 | -0.115 | -0.115 | 3.487 | 0.4 | 0.54 | 0.29 | 0.03 | 0.35 | 0.21 |
| 9 | 13902048 | rs10217667 | 2.184 | 0.139 | -1.318 | -2.553 | 0.044 | -0.018 | -0.018 | 0 | 0.26 | 0.01 | 0.3 | 0.17 | 0.19 | 0.28 |
| 9 | 13904860 | rs10961296 | -1.687 | 0.791 | -0.487 | -2.178 | -0.009 | 0.079 | 0.079 | 0 | 0.46 | 0.7 | 0.35 | 0.14 | 0.44 | 0.31 |
| 9 | 13912195 | rs12335400 | 0.326 | 0.984 | -0.555 | -2.016 | -0.012 | 0.01 | 0.01 | 0 | 0.13 | 0.1 | 0.07 | 0 | 0.13 | 0.08 |
| 9 | 13912482 | rs10113975 | 0.217 | 0.972 | -0.544 | -2.022 | -0.009 | 0.005 | 0.005 | 0 | 0.13 | 0.11 | 0.07 | 0 | 0.13 | 0.08 |
| 9 | 13917482 | rs1969938 | -1.698 | 0.576 | -0.468 | -2.131 | 0 | 0.047 | 0.047 | 0 | 0.49 | 0.65 | 0.35 | 0.1 | 0.47 | 0.35 |
| 9 | 13921272 | rs10961300 | -1.366 | 0.484 | -0.288 | -2.324 | -0.002 | -0.062 | -0.062 | 0 | 0.45 | 0.61 | 0.34 | 0.03 | 0.45 | 0.31 |
| 9 | 13921788 | rs7850903 | -1.285 | 0.507 | -0.291 | -2.315 | 0.007 | -0.122 | -0.122 | 0.157 | 0.42 | 0.46 | 0.33 | 0.03 | 0.45 | 0.25 |
| 9 | 13926247 | rs10961303 | -1.553 | 0.672 | -0.534 | -2.182 | -0.002 | 0.017 | 0.017 | 0 | 0.39 | 0.71 | 0.3 | 0.08 | 0.36 | 0.3 |
| 9 | 13928028 | rs2382427 | -1.766 | 0.816 | -0.566 | -2.19 | -0.004 | -0.005 | -0.005 | 0.001 | 0.42 | 0.64 | 0.3 | 0.06 | 0.39 | 0.35 |
| 9 | 13929064 | rs10123937 | -1.713 | 0.838 | -0.545 | -2.251 | -0.009 | -0.118 | -0.118 | 2.197 | 0.26 | 0.55 | 0.16 | 0.03 | 0.18 | 0.2 |
| 9 | 13930857 | rs10117560 | -1.246 | 0.996 | -0.473 | -2.238 | -0.002 | -0.125 | -0.125 | 0 | 0.34 | 0.52 | 0.26 | 0.05 | 0.29 | 0.2 |
| 9 | 13933371 | rs17277138 | 0.484 | 0.906 | -0.472 | -2.172 | -0.03 | 0.035 | 0.035 | 0.071 | 0.11 | 0.04 | 0.05 | 0 | 0.13 | 0.08 |
| 9 | 13937099 | rs16931157 | -1.86 | 0.971 | -0.539 | -2.299 | -0.001 | -0.039 | -0.039 | 5.528 | 0.42 | 0.6 | 0.3 | 0.07 | 0.43 | 0.38 |
| 9 | 13937519 | rs13285663 | -0.019 | 0.987 | -0.534 | -2.296 | -0.064 | 0.118 | 0.118 | 0.044 | 0.26 | 0.06 | 0.18 | 0.02 | 0.31 | 0.15 |
| 9 | 13943319 | rs16931161 | -1.127 | 0.838 | -0.426 | -2.221 | 0.012 | 0.019 | 0.019 | 0.005 | 0.08 | 0.5 | 0.05 | 0.01 | 0.02 | 0.03 |
| 9 | 13950593 | rs17277593 | 1.487 | 0.941 | -0.601 | -2.229 | -0.041 | 0.096 | 0.096 | 0.009 | 0.15 | 0.01 | 0.12 | 0.02 | 0.18 | 0.08 |
| 9 | 13953898 | rs16931165 | -1.873 | 0.954 | -0.602 | -2.246 | -0.001 | -0.038 | -0.038 | 0 | 0.44 | 0.59 | 0.27 | 0.06 | 0.43 | 0.37 |
| 9 | 13959282 | rs13290672 | 1.652 | 0.894 | -0.613 | -2.153 | -0.051 | 0.064 | 0.064 | 0 | 0.13 | 0.01 | 0.14 | 0.03 | 0.2 | 0.05 |
| 9 | 13960309 | rs12335962 | -0.939 | 0.854 | -0.01 | -2.196 | 0.004 | -0.073 | -0.073 | 1.549 | 0.33 | 0.38 | 0.24 | 0.03 | 0.35 | 0.18 |
| 9 | 13961462 | rs7856131 | -1.434 | 1.271 | 0.356 | -2.337 | 0 | -0.054 | -0.054 | 0 | 0.4 | 0.42 | 0.31 | 0.05 | 0.5 | 0.31 |
| 9 | 13968369 | rs7855714 | -1.361 | 1.274 | 0.279 | -2.197 | 0.005 | -0.076 | -0.076 | 0 | 0.39 | 0.58 | 0.26 | 0.05 | 0.41 | 0.35 |
| 9 | 13970315 | rs1590263 | -1.077 | 1.448 | 0.181 | -2.124 | 0.018 | -0.125 | -0.125 | 0.429 | 0.36 | 0.53 | 0.26 | 0.05 | 0.38 | 0.23 |
| 9 | 13974077 | rs16923127 | -0.888 | 0.648 | 0.113 | -1.687 | 0.014 | -0.11 | -0.11 | 1.939 | 0.2 | 0.37 | 0.11 | 0.01 | 0.18 | 0.18 |
| 9 | 13979872 | rs7037099 | -0.711 | -0.207 | -0.298 | -0.768 | -0.004 | -0.053 | -0.053 | 1.432 | 0.18 | 0.24 | 0.07 | 0.02 | 0.13 | 0.11 |
| 9 | 13980085 | rs7048074 | -0.883 | -1.052 | -0.588 | -1.078 | 0 | -0.001 | -0.001 | 0 | 0.44 | 0.42 | 0.39 | 0.27 | 0.35 | 0.54 |
| 9 | 13982131 | rs10810064 | 0.77 | -1.054 | -0.614 | -1.062 | 0.005 | 0.023 | 0.023 | 0 | 0.27 | 0.16 | 0.33 | 0.2 | 0.24 | 0.41 |
| 9 | 13986066 | rs7870444 | -0.26 | -0.648 | -0.34 | -0.837 | -0.018 | -0.059 | -0.059 | 0 | 0.18 | 0.26 | 0.06 | 0.02 | 0.1 | 0.1 |
| 9 | 13986757 | rs755267 | 1.075 | -0.786 | -0.287 | -1.161 | -0.008 | -0.028 | -0.028 | 1.293 | 0.55 | 0.44 | 0.57 | 0.25 | 0.61 | 0.55 |
| 9 | 13989084 | rs7025591 | -0.887 | -0.948 | -0.042 | -0.838 | -0.015 | -0.057 | -0.057 | 0.015 | 0.34 | 0.5 | 0.36 | 0.22 | 0.29 | 0.46 |
| 9 | 13992283 | rs10810073 | 0.787 | -0.795 | -0.29 | -0.745 | -0.04 | -0.002 | -0.002 | 0 | 0.2 | 0.04 | 0.29 | 0.2 | 0.22 | 0.38 |
| 9 | 14003679 | rs1322130 | 0.568 | -0.738 | -0.111 | -0.858 | -0.007 | 0.146 | 0.146 | 0.017 | 0.56 | 0.75 | 0.38 | 0.05 | 0.51 | 0.32 |

| | | | | | | | | | | | | | | | | |
|---|---|---|---|---|---|---|---|---|---|---|---|---|---|---|---|---|
| 9 | 14004053 | rs12005778 | -0.724 | -0.81 | 0.129 | -0.889 | -0.016 | -0.066 | -0.066 | 0 | 0.12 | 0.38 | 0.07 | 0.02 | 0.08 | 0.08 |
| 9 | 14005900 | rs7019866 | 1.488 | -0.828 | -0.16 | -0.949 | -0.002 | 0.007 | 0.007 | 0 | 0.43 | 0.34 | 0.28 | 0.03 | 0.4 | 0.2 |
| 9 | 14010270 | rs2181073 | -1.656 | -0.895 | -0.91 | -0.419 | 0.019 | 0.027 | 0.027 | 0 | 0.33 | 0.75 | 0.19 | 0.16 | 0.23 | 0.29 |
| 9 | 14010321 | rs10961326 | 1.343 | -0.842 | -0.949 | 0.163 | -0.029 | 0.03 | 0.03 | 0 | 0.06 | 0.01 | 0.26 | 0.48 | 0.14 | 0.19 |
| 9 | 14012093 | rs10961329 | 1.47 | -0.818 | -0.951 | 0.227 | -0.031 | 0.042 | 0.042 | 0 | 0.06 | 0.01 | 0.25 | 0.47 | 0.14 | 0.17 |
| 9 | 14012301 | rs16931224 | -1.349 | -0.939 | -0.924 | 0.01 | 0.006 | 0.008 | 0.008 | 0 | 0.31 | 0.71 | 0.18 | 0.13 | 0.22 | 0.23 |
| 9 | 14012596 | rs7848517 | -0.977 | -0.936 | -1.025 | 0.026 | -0.03 | -0.02 | -0.02 | 0 | 0.77 | 0.97 | 0.73 | 0.71 | 0.79 | 0.68 |
| 9 | 14013955 | rs7039809 | -0.107 | -0.982 | -0.902 | -0.18 | -0.002 | 0.015 | 0.015 | 0 | 0.68 | 0.77 | 0.45 | 0.15 | 0.62 | 0.42 |
| 9 | 14014614 | rs12001724 | -0.865 | -1.087 | -0.494 | 0.028 | -0.018 | -0.075 | -0.075 | 0 | 0.12 | 0.38 | 0.07 | 0.01 | 0.08 | 0.08 |
| 9 | 14014968 | rs7857245 | -1.704 | -1.071 | -0.703 | 0.039 | 0.01 | 0.013 | 0.013 | 0 | 0.33 | 0.73 | 0.19 | 0.15 | 0.22 | 0.29 |
| 9 | 14016560 | rs16931233 | -0.09 | -0.769 | -0.64 | 0.173 | -0.02 | -0.021 | -0.021 | 0 | 0.17 | 0.34 | 0.1 | 0.09 | 0.12 | 0.1 |
| 9 | 14020982 | rs1853195 | -0.316 | -0.388 | -1.081 | 0.32 | -0.003 | -0.076 | -0.076 | 0 | 0.75 | 0.97 | 0.68 | 0.62 | 0.79 | 0.6 |
| 9 | 14021990 | rs1977235 | 0.626 | -0.051 | -0.793 | 0.437 | -0.009 | -0.031 | -0.031 | 0 | 0.18 | 0.15 | 0.29 | 0.49 | 0.25 | 0.25 |
| 9 | 14023577 | rs12351326 | 1.405 | -0.109 | -0.902 | 0.018 | -0.052 | 0.137 | 0.137 | 0 | 0.4 | 0.15 | 0.31 | 0.05 | 0.44 | 0.21 |
| 9 | 14025101 | rs7022407 | -0.473 | -0.075 | -1.171 | 0.084 | -0.026 | 0.005 | 0.005 | 0 | 0.77 | 0.92 | 0.76 | 0.75 | 0.78 | 0.64 |
| 9 | 14025508 | rs12550926 | 0.504 | -0.142 | -1.013 | -0.103 | 0.002 | 0.003 | 0.003 | 0 | 0.62 | 0.39 | 0.67 | 0.71 | 0.69 | 0.58 |
| 9 | 14026089 | rs10810080 | 0.647 | -0.01 | -1.738 | 0.133 | -0.004 | 0.005 | 0.005 | 0 | 0.47 | 0.38 | 0.33 | 0.41 | 0.48 | 0.24 |
| 9 | 14026510 | rs16931256 | -0.301 | -0.299 | -0.909 | -0.344 | -0.02 | 0.026 | 0.026 | 0 | 0.13 | 0.35 | 0.07 | 0.11 | 0.08 | 0.08 |
| 9 | 14028713 | rs16931261 | 0.119 | -0.173 | -1.003 | -0.345 | 0.001 | 0.032 | 0.032 | 0 | 0.11 | 0.35 | 0.06 | 0.1 | 0.09 | 0.06 |
| 9 | 14029188 | rs6474805 | 0.175 | 0.074 | -2.093 | -0.587 | 0.004 | 0.029 | 0.029 | 0 | 0.77 | 0.97 | 0.77 | 0.77 | 0.82 | 0.67 |
| 9 | 14030100 | rs10961344 | 0.163 | 0.422 | -2.164 | -1.127 | -0.076 | 0.318 | 0.318 | 0 | 0.43 | 0.05 | 0.39 | 0.03 | 0.52 | 0.26 |
| 9 | 14030444 | rs10961346 | -1.239 | 1.152 | -0.463 | -0.355 | 0.021 | -0.042 | -0.042 | 0 | 0.24 | 0.47 | 0.31 | 0.11 | 0.22 | 0.21 |
| 9 | 14030829 | rs10810082 | 0.955 | 0.696 | -0.371 | 0.097 | 0.024 | 0.048 | 0.048 | 0 | 0.66 | 0.52 | 0.42 | 0.51 | 0.48 | 0.52 |
| 9 | 14031216 | rs10961347 | -0.548 | 0.952 | -0.429 | -0.457 | 0 | -0.073 | -0.073 | 0.088 | 0.16 | 0.48 | 0.12 | 0.06 | 0.12 | 0.17 |
| 9 | 14032023 | rs3899674 | -0.797 | 1.052 | -1.766 | -0.282 | -0.122 | 0.104 | 0.104 | 0 | 0.36 | 0.91 | 0.51 | 0.15 | 0.51 | 0.31 |
| 9 | 14036242 | rs3920568 | 0.168 | 0.545 | -0.758 | 0.265 | 0.003 | 0.002 | 0.002 | 0.63 | 0.55 | 0.44 | 0.34 | 0.61 | 0.42 | 0.52 |
| 9 | 14037831 | rs1853197 | -0.361 | 0.832 | -0.332 | -0.382 | 0.01 | -0.017 | -0.017 | 0.004 | 0.48 | 0.44 | 0.42 | 0.93 | 0.53 | 0.62 |
| 9 | 14043162 | rs10756524 | -0.385 | 0.785 | -0.036 | -0.397 | 0 | -0.068 | -0.068 | 2.606 | 0.48 | 0.54 | 0.44 | 0.95 | 0.51 | 0.66 |
| 9 | 14044604 | rs10756525 | -0.274 | 1.412 | -0.756 | 0.371 | 0.002 | 0.199 | 0.199 | 0.039 | 0.44 | 0.21 | 0.39 | 0.93 | 0.5 | 0.63 |
| 9 | 14045245 | rs10756526 | -0.079 | 1.023 | -0.72 | 0.468 | -0.122 | -0.095 | -0.095 | 0.968 | 0.41 | 0.11 | 0.27 | 0.57 | 0.43 | 0.58 |
| 9 | 14045745 | rs7874668 | -0.217 | 1.081 | -0.639 | 0.41 | -0.022 | -0.004 | -0.004 | 0 | 0.43 | 0.31 | 0.31 | 0.59 | 0.43 | 0.6 |
| 9 | 14058402 | rs1156572 | 0.196 | 0.967 | -0.833 | -0.327 | -0.016 | 0.017 | 0.017 | 0 | 0.81 | 0.88 | 0.86 | 1 | 0.79 | 0.87 |
| 9 | 14058548 | rs10116586 | 0.05 | 0.589 | -1.184 | -0.391 | -0.029 | 0.014 | 0.014 | 0.016 | 0.33 | 0.3 | 0.24 | 0.25 | 0.48 | 0.34 |
| 9 | 14063510 | rs1407345 | -0.304 | -0.332 | -1.55 | -0.456 | 0 | 0.067 | 0.067 | 0.054 | 0.67 | 0.84 | 0.71 | 0.89 | 0.68 | 0.74 |
| 9 | 14064741 | rs7039300 | 0.404 | -1.425 | -1.165 | -0.266 | 0.031 | 0.022 | 0.022 | 0 | 0.34 | 0.22 | 0.35 | 0.26 | 0.2 | 0.42 |
| 9 | 14064891 | rs10283831 | 1.727 | -1.597 | -1.281 | -0.431 | 0.003 | -0.02 | -0.02 | 0 | 0.17 | 0.14 | 0.11 | 0 | 0.13 | 0.07 |
| 9 | 14065522 | rs564854 | 0.774 | -1.583 | -1.331 | -0.422 | -0.054 | 0.105 | 0.105 | 0 | 0.18 | 0.03 | 0.12 | 0 | 0.21 | 0.14 |
| 9 | 14067281 | rs512477 | -0.53 | -2.229 | -0.797 | -0.518 | -0.006 | 0.018 | 0.018 | 0.286 | 0.62 | 0.64 | 0.41 | 0.18 | 0.45 | 0.4 |
| 9 | 14070422 | rs10961361 | -0.564 | -2.695 | -0.295 | 0.211 | -0.023 | 0.109 | 0.109 | 0 | 0.14 | 0.28 | 0.37 | 0.3 | 0.08 | 0.38 |
| 9 | 14072370 | rs569114 | -1.453 | -3.026 | -0.494 | 0.777 | 0.015 | 0.069 | 0.069 | 0 | 0.57 | 0.87 | 0.62 | 0.73 | 0.39 | 0.78 |
| 9 | 14074022 | rs516325 | -1.532 | -3.386 | 0.065 | 1.078 | 0.099 | 0.027 | 0.027 | 0 | 0.54 | 0.75 | 0.57 | 0.38 | 0.35 | 0.66 |
| 9 | 14074099 | rs10961364 | -0.786 | -3.61 | 0.192 | 0.978 | -0.086 | 0.011 | 0.011 | 0.052 | 0.36 | 0.48 | 0.45 | 0.38 | 0.16 | 0.5 |
| 9 | 14078878 | rs16931322 | 0.596 | -3.822 | 0.38 | 0.876 | -0.037 | -0.043 | -0.043 | 0 | 0.19 | 0.26 | 0.07 | 0.06 | 0.08 | 0.07 |
| 9 | 14079067 | rs480700 | -2.18 | -4.116 | 0.168 | 1.034 | 0.04 | -0.01 | -0.01 | 0 | 0.6 | 0.92 | 0.62 | 0.65 | 0.38 | 0.76 |
| 9 | 14085583 | rs484555 | -2.024 | -3.589 | 0.726 | 1.469 | 0.083 | 0.005 | 0.005 | 0.043 | 0.48 | 0.72 | 0.57 | 0.57 | 0.35 | 0.73 |
| 9 | 14087769 | rs7858 | 0.956 | -3.536 | 0.867 | 1.4 | 0.009 | 0.029 | 0.029 | 0 | 0.16 | 0.08 | 0.1 | 0.04 | 0.17 | 0.24 |
| 9 | 14088789 | rs548824 | 0.764 | -3.528 | 0.878 | 1.425 | 0.007 | 0.028 | 0.028 | 0 | 0.17 | 0.08 | 0.12 | 0.05 | 0.18 | 0.26 |
| 9 | 14098867 | rs542733 | -1.671 | -3.63 | 0.929 | 1.853 | 0.041 | -0.022 | -0.022 | 0 | 0.39 | 0.71 | 0.32 | 0.47 | 0.3 | 0.45 |
| 9 | 14107096 | rs508940 | -2.152 | -3.678 | 1.217 | 1.922 | 0.128 | -0.02 | -0.02 | 0.024 | 0.49 | 0.78 | 0.57 | 0.5 | 0.34 | 0.71 |
| 9 | 14109456 | rs510970 | 0.402 | -3.648 | 1.245 | 1.938 | 0.009 | 0.01 | 0.01 | 0 | 0.19 | 0.08 | 0.21 | 0.11 | 0.2 | 0.27 |
| 9 | 14110242 | rs571693 | 0.336 | -3.643 | 1.259 | 1.948 | 0.011 | 0.002 | 0.002 | 0 | 0.19 | 0.09 | 0.21 | 0.14 | 0.2 | 0.27 |
| 9 | 14113903 | rs573057 | -0.532 | -3.685 | 1.749 | 1.945 | -0.037 | -0.007 | -0.007 | 1.923 | 0.26 | 0.5 | 0.25 | 0.18 | 0.22 | 0.4 |
| 9 | 14116484 | rs498010 | 0.016 | -3.588 | 1.814 | 1.792 | -0.019 | 0.047 | 0.047 | 0 | 0.21 | 0.35 | 0.23 | 0.32 | 0.2 | 0.36 |
| 9 | 14117221 | rs580501 | -2.604 | -3.623 | 1.551 | 1.73 | 0.122 | 0.007 | 0.007 | 0.05 | 0.52 | 0.87 | 0.59 | 0.59 | 0.36 | 0.76 |
| 9 | 14119583 | rs10810092 | 0.146 | -3.374 | 1.741 | 1.578 | 0 | 0 | 0 | 0 | 0.19 | 0.18 | 0.18 | 0.11 | 0.17 | 0.33 |
| 9 | 14120803 | rs2307079 | -0.098 | -3.358 | 1.738 | 1.587 | -0.029 | -0.009 | -0.009 | 0 | 0.21 | 0.37 | 0.24 | 0.13 | 0.19 | 0.34 |
| 9 | 14136495 | rs3780582 | 0.008 | -3.448 | 1.274 | 1.797 | -0.013 | 0.029 | 0.029 | 0 | 0.21 | 0.26 | 0.23 | 0.33 | 0.19 | 0.34 |
| 9 | 14139828 | rs2382437 | -1.49 | -3.449 | 1.16 | 1.161 | -0.058 | -0.042 | -0.042 | 0 | 0.29 | 0.53 | 0.34 | 0.22 | 0.15 | 0.39 |
| 9 | 14144821 | rs7042365 | -1.559 | -3.162 | 1.197 | 1.526 | -0.065 | -0.015 | -0.015 | 0.041 | 0.26 | 0.45 | 0.35 | 0.28 | 0.15 | 0.37 |
| 9 | 14145487 | rs7031735 | -1.83 | -3.268 | 1.097 | 1.495 | -0.062 | -0.02 | -0.02 | 0 | 0.29 | 0.46 | 0.35 | 0.3 | 0.17 | 0.38 |
| 9 | 14152436 | rs10756531 | 2.448 | -2.983 | 0.766 | 1.823 | 0.077 | -0.04 | -0.04 | 0.198 | 0.54 | 0.03 | 0.52 | 0.48 | 0.66 | 0.43 |
| 9 | 14154231 | rs723968 | -1.197 | -2.902 | 1.064 | 1.894 | -0.003 | 0.001 | 0.001 | 0 | 0.2 | 0.21 | 0.3 | 0.28 | 0.14 | 0.34 |
| 9 | 14157468 | rs7044402 | 1.427 | -2.854 | 1.035 | 1.703 | -0.041 | -0.021 | -0.021 | 0 | 0.75 | 0.58 | 0.68 | 0.76 | 0.85 | 0.69 |
| 9 | 14159460 | rs2382443 | 1.389 | -2.806 | 1.005 | 1.397 | -0.01 | -0.027 | -0.027 | 0 | 0.64 | 0.56 | 0.61 | 0.76 | 0.68 | 0.51 |
| 9 | 14161927 | rs10810099 | 2.508 | -1.612 | 1.229 | 2.014 | 0.018 | -0.018 | -0.018 | 0.672 | 0.69 | 0.14 | 0.53 | 0.57 | 0.72 | 0.6 |
| 9 | 14171645 | rs12377502 | -0.004 | -1.246 | 1.243 | 1.831 | -0.005 | 0.032 | 0.032 | 0 | 0.07 | 0.06 | 0.13 | 0.36 | 0.14 | 0.05 |
| 9 | 14172048 | rs6474821 | -0.954 | -1.118 | 1.329 | 1.414 | 0.027 | 0.033 | 0.033 | 0 | 0.13 | 0.08 | 0.06 | 0.03 | 0.09 | 0.04 |
| 9 | 14185939 | rs16931436 | -1.133 | -0.909 | 1.127 | 1.594 | 0.001 | 0.001 | 0.001 | 0 | 0.2 | 0.18 | 0.29 | 0.17 | 0.13 | 0.29 |
| 9 | 14196340 | rs2382454 | -1.011 | 0.001 | 0.166 | 2.174 | 0.041 | 0.118 | 0.118 | 0 | 0.06 | 0.13 | 0.13 | 0.33 | 0.15 | 0.05 |

| | | | | | | | | | | | | | | | | |
|---|---|---|---|---|---|---|---|---|---|---|---|---|---|---|---|---|
| 9 | 14197498 | rs10738355 | 1.094 | -0.038 | 0.016 | 2.175 | -0.002 | 0.043 | 0.043 | 1.163 | 0.94 | 0.95 | 0.88 | 0.65 | 0.86 | 0.95 |
| 9 | 14199422 | rs10810110 | 0.867 | 0.033 | 0.16 | 2.322 | 0.041 | 0.153 | 0.153 | 0.357 | 0.95 | 0.91 | 0.86 | 0.59 | 0.85 | 0.94 |
| 9 | 14200005 | rs10810113 | 1.416 | -0.127 | 0.287 | 2.336 | 0.064 | 0.209 | 0.209 | 0 | 0.94 | 0.8 | 0.83 | 0.62 | 0.87 | 0.96 |
| 9 | 14205169 | rs12551619 | -1.593 | -1.028 | -0.426 | 1.675 | 0.088 | 0.009 | 0.009 | 0.125 | 0.41 | 0.79 | 0.52 | 0.36 | 0.31 | 0.58 |
| 9 | 14206835 | rs12339675 | -1.706 | -1.046 | -0.315 | 1.531 | 0.067 | 0.004 | 0.004 | 0 | 0.42 | 0.76 | 0.49 | 0.36 | 0.33 | 0.58 |
| 9 | 14208724 | rs10961424 | -0.844 | -1.136 | -0.242 | 1.579 | 0.002 | 0 | 0 | 0 | 0.26 | 0.23 | 0.34 | 0.24 | 0.25 | 0.42 |
| 9 | 14210653 | rs12340584 | -1.153 | -1.116 | -0.193 | 1.63 | -0.019 | 0.015 | 0.015 | 0.095 | 0.18 | 0.24 | 0.28 | 0.24 | 0.13 | 0.39 |
| 9 | 14211843 | rs10961426 | -0.692 | -0.7 | -0.185 | 1.603 | -0.002 | 0.031 | 0.031 | 0.138 | 0.16 | 0.24 | 0.28 | 0.25 | 0.13 | 0.37 |
| 9 | 14213974 | rs12682686 | -0.753 | -0.977 | -0.149 | 1.611 | -0.026 | 0.001 | 0.001 | 0 | 0.17 | 0.25 | 0.26 | 0.2 | 0.11 | 0.37 |
| 9 | 14215878 | rs10511593 | -0.996 | -0.695 | -0.706 | 1.693 | -0.055 | -0.075 | -0.075 | 0 | 0.16 | 0.01 | 0.21 | 0.18 | 0.2 | 0.16 |
| 9 | 14221591 | rs2382444 | 1.401 | -0.573 | -0.197 | 1.079 | 0.001 | -0.002 | -0.002 | 0 | 0.57 | 0.58 | 0.37 | 0.2 | 0.54 | 0.38 |
| 9 | 14226139 | rs10810119 | -0.875 | -0.506 | 0.016 | 1.608 | -0.003 | -0.017 | -0.017 | 0 | 0.16 | 0.12 | 0.25 | 0.6 | 0.19 | 0.24 |
| 9 | 14229179 | rs10961436 | -0.727 | -0.6 | 0.499 | 1.557 | -0.018 | 0.171 | 0.171 | 0 | 0.15 | 0.37 | 0.22 | 0.56 | 0.12 | 0.23 |
| 9 | 14234315 | rs10961440 | -1.173 | -0.802 | 0.055 | 1.225 | 0.002 | 0.002 | 0.002 | 0 | 0.23 | 0.22 | 0.18 | 0.14 | 0.19 | 0.21 |
| 9 | 14237649 | rs10756542 | 0.991 | 0.135 | -0.454 | 1.003 | -0.027 | 0.014 | 0.014 | 0.004 | 0.58 | 0.8 | 0.66 | 0.68 | 0.45 | 0.6 |
| 9 | 14241817 | rs1555553 | 0.355 | 0.219 | -0.235 | 0.934 | 0.002 | 0 | 0 | 0.04 | 0.7 | 0.33 | 0.58 | 0.65 | 0.6 | 0.51 |
| 9 | 14245551 | rs7030457 | -0.889 | 0.125 | -0.274 | 0.543 | -0.009 | 0.012 | 0.012 | 0.477 | 0.21 | 0.27 | 0.39 | 0.81 | 0.18 | 0.44 |
| 9 | 14247488 | rs10961450 | -0.214 | 0.709 | -0.71 | 0.707 | -0.005 | -0.002 | -0.002 | 0 | 0.18 | 0.08 | 0.19 | 0.14 | 0.19 | 0.21 |
| 9 | 14247788 | rs7866589 | -0.017 | 1.157 | -0.14 | 0.68 | -0.03 | 0.027 | 0.027 | 0 | 0.81 | 0.91 | 0.87 | 0.94 | 0.72 | 0.81 |
| 9 | 14248274 | rs12346458 | -1.342 | 0.459 | 0.69 | 1.258 | 0.01 | 0.108 | 0.108 | 0.579 | 0.16 | 0.47 | 0.29 | 0.26 | 0.12 | 0.39 |
| 9 | 14248934 | rs7027899 | 1.25 | 0.675 | 0.148 | 1.397 | 0.037 | 0.161 | 0.161 | 0.875 | 0.82 | 0.51 | 0.67 | 0.66 | 0.83 | 0.57 |
| 9 | 14249488 | rs7874591 | -0.132 | 0.643 | -0.987 | 0.562 | 0.003 | 0.015 | 0.015 | 0 | 0.81 | 0.89 | 0.88 | 1 | 0.84 | 0.89 |
| 9 | 14250285 | rs3761668 | 0.259 | 0.561 | -0.929 | 1.753 | 0.039 | -0.058 | -0.058 | 0 | 0.74 | 0.84 | 0.73 | 0.61 | 0.84 | 0.71 |
| 9 | 14262699 | rs1322990 | 1.526 | 1.998 | 0.93 | 2.319 | 0.164 | 0.329 | 0.329 | 0.204 | 0.92 | 0.61 | 0.76 | 0.69 | 0.87 | 0.88 |
| 9 | 14267601 | rs7846889 | 1.764 | 1.582 | 1.711 | 2.348 | 0.119 | 0.465 | 0.465 | 0.363 | 0.94 | 0.56 | 0.81 | 0.7 | 0.96 | 0.91 |
| 9 | 14269556 | rs1061820 | 0.733 | 1.253 | 1.25 | 2.294 | -0.006 | -0.021 | -0.021 | 0 | 0.6 | 0.44 | 0.46 | 0.27 | 0.62 | 0.52 |
| 9 | 14270342 | rs1322991 | 1.403 | 1.037 | 0.561 | 2.283 | -0.002 | 0.017 | 0.017 | 0 | 0.89 | 0.89 | 0.81 | 0.73 | 0.92 | 0.87 |
| 9 | 14274950 | rs1033837 | 1.275 | 0.97 | 0.652 | 2.357 | -0.014 | 0.036 | 0.036 | 0 | 0.91 | 0.82 | 0.81 | 0.77 | 0.94 | 0.94 |
| 9 | 14281896 | rs4740567 | -0.141 | 0.989 | 0.366 | 2.391 | -0.054 | -0.034 | -0.034 | 3.178 | 0.26 | 0.07 | 0.31 | 0.29 | 0.29 | 0.27 |
| 9 | 14286126 | rs6474827 | 1.623 | 0.721 | 0.702 | 2.528 | -0.005 | 0.065 | 0.065 | 0 | 0.9 | 0.82 | 0.81 | 0.73 | 0.93 | 0.9 |
| 9 | 14286997 | rs10738358 | 1.532 | 0.663 | 0.945 | 2.526 | -0.009 | 0.115 | 0.115 | 0 | 0.9 | 0.74 | 0.82 | 0.74 | 0.95 | 0.91 |
| 9 | 14290978 | rs10810124 | 0.356 | 0.55 | -0.291 | 2.02 | -0.003 | 0.008 | 0.008 | 0 | 0.41 | 0.28 | 0.36 | 0.37 | 0.38 | 0.28 |
| 9 | 14292207 | rs10810126 | 0.41 | 0.343 | -0.913 | 2.344 | -0.009 | -0.047 | -0.047 | 0 | 0.4 | 0.18 | 0.39 | 0.49 | 0.37 | 0.34 |
| 9 | 14292880 | rs10961466 | 0.442 | 0.468 | -1.007 | 2.289 | -0.002 | 0.023 | 0.023 | 0 | 0.37 | 0.07 | 0.29 | 0.28 | 0.31 | 0.3 |
| 9 | 14294350 | rs10810127 | 0.07 | 0.177 | -1.672 | 1.178 | 0.011 | 0.081 | 0.081 | 1.592 | 0.52 | 0.1 | 0.5 | 0.31 | 0.6 | 0.63 |
| 9 | 14294980 | rs17213362 | -0.325 | -0.763 | -1.454 | 0.546 | 0.049 | 0.026 | 0.026 | 0 | 0.4 | 0.16 | 0.29 | 0.25 | 0.29 | 0.29 |
| 9 | 14297978 | rs10429540 | 1.095 | -0.8 | 0.129 | 0.674 | -0.064 | 0.052 | 0.052 | 0.01 | 0.86 | 0.29 | 0.85 | 0.77 | 0.97 | 0.87 |
| 9 | 14300901 | rs11999144 | -1.548 | -0.864 | 0.546 | 0.804 | -0.027 | 0.151 | 0.151 | 0 | 0.07 | 0.21 | 0.15 | 0.22 | 0.04 | 0.03 |
| 9 | 14303061 | rs2382468 | 1.069 | -1.008 | 0.648 | 0.104 | -0.056 | -0.032 | -0.032 | 0 | 0.88 | 0.3 | 0.94 | 0.9 | 0.97 | 0.87 |
| 9 | 14303488 | rs7021322 | 0.838 | -0.742 | 1.162 | 0.102 | -0.009 | -0.004 | -0.004 | 0.668 | 0.89 | 0.51 | 0.95 | 0.89 | 0.96 | 0.88 |
| 9 | 14304973 | rs10810130 | 2.21 | -0.625 | 0.833 | -0.021 | 0.022 | 0.031 | 0.031 | 0.453 | 0.8 | 0.13 | 0.87 | 0.89 | 0.87 | 0.84 |
| 9 | 14310504 | rs10810131 | 0.683 | -0.46 | 0.265 | -0.134 | -0.011 | -0.016 | -0.016 | 0 | 0.21 | 0.12 | 0.29 | 0.27 | 0.27 | 0.1 |
| 9 | 14312724 | rs10961473 | 0.179 | -0.285 | 0.632 | -0.213 | -0.009 | -0.007 | -0.007 | 0 | 0.64 | 0.55 | 0.63 | 0.62 | 0.66 | 0.75 |
| 9 | 14313712 | rs2273933 | 1.214 | 0.079 | 1.122 | -0.363 | -0.004 | -0.036 | -0.036 | 0 | 0.87 | 0.36 | 0.93 | 0.9 | 0.91 | 0.85 |
| 9 | 14316570 | rs7874736 | -1.245 | 0.373 | -0.042 | -0.515 | -0.009 | 0.051 | 0.051 | 0 | 0.32 | 0.21 | 0.19 | 0.19 | 0.36 | 0.48 |
| 9 | 14323346 | rs7040685 | 0.921 | 0.14 | 0.235 | 0.028 | -0.039 | 0 | 0 | 0.151 | 0.86 | 0.38 | 0.89 | 0.89 | 0.94 | 0.92 |
| 9 | 14334138 | rs7048960 | 0.881 | 0.464 | -0.339 | -0.38 | -0.017 | -0.078 | -0.078 | 0.663 | 0.7 | 0.46 | 0.73 | 0.89 | 0.72 | 0.68 |
| 9 | 14338470 | rs10810132 | -1.139 | 0.922 | -1.011 | 0.049 | -0.021 | 0.043 | 0.043 | 0 | 0.16 | 0.03 | 0.12 | 0.09 | 0.18 | 0.17 |
| 9 | 14343670 | rs598296 | 1.18 | 1.518 | -0.381 | 0.907 | 0.031 | 0.097 | 0.097 | 0 | 0.82 | 0.7 | 0.73 | 0.56 | 0.78 | 0.71 |
| 9 | 14347811 | rs877571 | -0.401 | 0.44 | -0.809 | -0.542 | 0.028 | 0.041 | 0.041 | 0 | 0.37 | 0.3 | 0.22 | 0.13 | 0.25 | 0.28 |
| 9 | 14348563 | rs877569 | -1.172 | 0.599 | 0.201 | -0.575 | 0.012 | -0.05 | -0.05 | 0.122 | 0.09 | 0.33 | 0.07 | 0.01 | 0.07 | 0.04 |
| 9 | 14352457 | rs10961483 | -1.307 | 0.364 | 0.383 | -0.464 | -0.017 | 0.008 | 0.008 | 0 | 0.07 | 0.17 | 0.08 | 0.1 | 0.05 | 0.1 |
| 9 | 14356795 | rs7864996 | -1.328 | 0.583 | 0.706 | -1.022 | -0.004 | -0.076 | -0.076 | 0.379 | 0.23 | 0.36 | 0.11 | 0.13 | 0.17 | 0.29 |
| 9 | 14358681 | rs13296301 | -0.161 | 1.087 | 0.646 | -1.284 | 0.01 | -0.102 | -0.102 | 0.096 | 0.28 | 0.52 | 0.23 | 0.06 | 0.32 | 0.2 |
| 9 | 14359000 | rs12379369 | -0.602 | 0.884 | -0.051 | -1.041 | -0.011 | -0.014 | -0.014 | 0.904 | 0.08 | 0.02 | 0.04 | 0.11 | 0.05 | 0.14 |
| 9 | 14364590 | rs10756546 | -0.037 | 0.893 | -0.016 | -0.663 | 0.003 | -0.004 | -0.004 | 0 | 0.6 | 0.49 | 0.44 | 0.44 | 0.57 | 0.58 |
| 9 | 14365269 | rs683474 | -0.217 | 0.67 | -0.573 | -0.838 | -0.051 | 0.023 | 0.023 | 0.013 | 0.66 | 0.77 | 0.46 | 0.32 | 0.56 | 0.53 |
| 9 | 14365307 | rs2021306 | 0.463 | 0.902 | -1.331 | -0.701 | -0.064 | -0.019 | -0.019 | 0.092 | 0.25 | 0.02 | 0.17 | 0.19 | 0.23 | 0.22 |
| 9 | 14367505 | rs471088 | -0.171 | 0.818 | 0.041 | -0.184 | 0.02 | 0.042 | 0.042 | 0 | 0.09 | 0.3 | 0.09 | 0.14 | 0.06 | 0.13 |
| 9 | 14370172 | rs10961490 | -0.547 | 0.925 | -0.558 | -0.37 | 0 | -0.045 | -0.045 | 0 | 0.15 | 0.17 | 0.13 | 0.01 | 0.19 | 0.13 |
| 9 | 14376009 | rs4741365 | 0.361 | 1.834 | -0.675 | 0.639 | 0.015 | 0.022 | 0.022 | 0.061 | 0.93 | 0.9 | 0.9 | 0.8 | 0.84 | 0.93 |
| 9 | 14376038 | rs4741366 | -0.108 | 1.171 | -1.941 | 0.669 | 0.043 | 0.043 | 0.043 | 0 | 0.45 | 0.03 | 0.34 | 0.35 | 0.32 | 0.43 |
| 9 | 14378712 | rs2382469 | 1.1 | 1.034 | -0.837 | 1.025 | 0.044 | 0.111 | 0.111 | 0 | 0.76 | 0.52 | 0.67 | 0.49 | 0.67 | 0.55 |
| 9 | 14378776 | rs509609 | -0.019 | 1.05 | -0.948 | 1.186 | -0.028 | -0.045 | -0.045 | 0 | 0.87 | 0.93 | 0.89 | 0.77 | 0.82 | 0.88 |
| 9 | 14380528 | rs568008 | -0.23 | 0.728 | -0.364 | 0.876 | -0.007 | -0.008 | -0.008 | 0.021 | 0.43 | 0.37 | 0.42 | 0.3 | 0.33 | 0.5 |
| 9 | 14380945 | rs630973 | -0.96 | 0.699 | -0.161 | 1.023 | -0.003 | 0.095 | 0.095 | 0 | 0.1 | 0.23 | 0.18 | 0.23 | 0.08 | 0.21 |
| 9 | 14383296 | rs7867427 | 0.843 | 1.16 | -0.858 | 0.841 | -0.018 | 0.033 | 0.033 | 0 | 0.36 | 0.25 | 0.4 | 0.31 | 0.43 | 0.28 |
| 9 | 14384950 | rs679613 | -0.486 | 0.728 | -0.788 | 0.895 | 0.023 | 0.022 | 0.022 | 0 | 0.5 | 0.67 | 0.64 | 0.51 | 0.47 | 0.56 |
| 9 | 14384976 | rs10810134 | 0.013 | 0.074 | -0.496 | 0.5 | -0.009 | -0.01 | -0.01 | 0 | 0.32 | 0.31 | 0.22 | 0.13 | 0.25 | 0.25 |

| | | | | | | | | | | | | | | | | |
|---|---|---|---|---|---|---|---|---|---|---|---|---|---|---|---|---|
| 9 | 14387817 | rs2382470 | -0.131 | -0.466 | -1.275 | 0.078 | 0.081 | 0.08 | 0.08 | 0 | 0.63 | 0.78 | 0.76 | 0.78 | 0.75 | 0.74 |
| 9 | 14389011 | rs717932 | -0.005 | -0.294 | -1.174 | 0.147 | 0 | 0.001 | 0.001 | 0 | 0.36 | 0.55 | 0.33 | 0.52 | 0.51 | 0.42 |
| 9 | 14392785 | rs2208919 | 0.248 | 0.486 | -0.818 | 0.029 | 0.025 | -0.011 | -0.011 | 1.267 | 0.19 | 0.17 | 0.2 | 0.05 | 0.35 | 0.18 |
| 9 | 14393854 | rs10810136 | -0.77 | -0.549 | -1.163 | -0.379 | 0.031 | 0.067 | 0.067 | 0 | 0.5 | 0.63 | 0.37 | 0.71 | 0.55 | 0.51 |
| 9 | 14394325 | rs6474832 | 0.303 | -0.975 | -1.217 | -0.552 | 0.012 | 0.02 | 0.02 | 0 | 0.45 | 0.37 | 0.67 | 0.31 | 0.6 | 0.51 |
| 9 | 14395035 | rs2076872 | -0.009 | -1.061 | -0.546 | -0.485 | 0.02 | -0.009 | -0.009 | 0 | 0.32 | 0.31 | 0.16 | 0.65 | 0.2 | 0.33 |
| 9 | 14395595 | rs1807429 | 1.043 | -0.885 | -0.658 | -0.603 | 0.05 | -0.027 | -0.027 | 1.373 | 0.32 | 0.25 | 0.46 | 0.46 | 0.22 | 0.41 |
| 9 | 14396833 | rs10481545 | 0.398 | 0.275 | -0.448 | -0.318 | 0.003 | 0.028 | 0.028 | 0 | 0.5 | 0.35 | 0.62 | 0.44 | 0.53 | 0.5 |
| 9 | 14398201 | rs10810137 | 0.715 | -0.023 | -0.407 | -0.022 | 0.008 | 0.004 | 0.004 | 0 | 0.07 | 0.1 | 0.05 | 0.12 | 0.12 | 0.11 |
| 9 | 14401281 | rs17709614 | 0.666 | -0.217 | -0.72 | -0.097 | 0.033 | -0.006 | -0.006 | 0.225 | 0.52 | 0.27 | 0.66 | 0.56 | 0.59 | 0.53 |
| 9 | 14401932 | rs1323349 | -0.66 | -0.019 | -0.054 | 0.219 | -0.005 | 0.03 | 0.03 | 0.277 | 0.24 | 0.69 | 0.19 | 0.4 | 0.2 | 0.28 |
| 9 | 14402446 | rs7873382 | -1.035 | -0.09 | -0.124 | 0.144 | -0.001 | 0.007 | 0.007 | 2.284 | 0.29 | 0.69 | 0.21 | 0.4 | 0.23 | 0.32 |
| 9 | 14405158 | rs7042750 | -0.313 | 0.135 | 0.255 | 0.194 | 0.005 | 0.197 | 0.197 | 0.143 | 0.18 | 0.54 | 0.14 | 0.43 | 0.16 | 0.22 |
| 9 | 14405297 | rs1323350 | -0.432 | 0.137 | 0.271 | 0.19 | 0.029 | 0.19 | 0.19 | 0.132 | 0.19 | 0.54 | 0.15 | 0.43 | 0.18 | 0.23 |
| 9 | 14405555 | rs11788403 | 1.015 | 0.35 | -0.417 | 0.195 | -0.015 | 0.009 | 0.009 | 0 | 0.07 | 0.01 | 0.05 | 0.03 | 0.06 | 0.08 |
| 9 | 14405801 | rs1570499 | 0.26 | -0.046 | -0.33 | 0.056 | 0.008 | -0.029 | -0.029 | 0 | 0.69 | 0.75 | 0.84 | 0.56 | 0.73 | 0.66 |
| 9 | 14410591 | rs10961505 | 0.177 | 1.031 | -0.708 | -0.179 | -0.025 | 0.02 | 0.02 | 0 | 0.13 | 0.09 | 0.12 | 0.01 | 0.25 | 0.11 |
| 9 | 14411546 | rs4740570 | 0.719 | 0.431 | -0.673 | -0.359 | 0.006 | 0.01 | 0.01 | 0 | 0.48 | 0.55 | 0.69 | 0.58 | 0.47 | 0.55 |
| 9 | 14412287 | rs4741374 | 0.223 | -0.693 | -0.374 | 0.141 | -0.028 | 0.006 | 0.006 | 0 | 0.64 | 0.52 | 0.81 | 0.59 | 0.76 | 0.69 |
| 9 | 14419053 | rs10756549 | 0.19 | -0.796 | -0.713 | 0.193 | 0.058 | 0.001 | 0.001 | 0.106 | 0.64 | 0.8 | 0.84 | 0.61 | 0.78 | 0.63 |
| 9 | 14419092 | rs10810142 | 0.576 | -0.947 | -0.913 | 0.242 | 0.027 | -0.062 | -0.062 | 0.65 | 0.15 | 0.05 | 0.09 | 0.35 | 0.1 | 0.09 |
| 9 | 14419894 | rs10738360 | 0.519 | -0.829 | -0.1 | 0.364 | -0.015 | 0 | 0 | 0.012 | 0.6 | 0.64 | 0.82 | 0.52 | 0.76 | 0.56 |
| 9 | 14422035 | rs7875762 | 1.284 | 0.015 | -0.452 | 0.381 | -0.046 | -0.016 | -0.016 | 0 | 0.33 | 0.17 | 0.57 | 0.36 | 0.46 | 0.33 |
| 9 | 14422868 | rs2025319 | -0.344 | 0.347 | 0.399 | 0.3 | 0.063 | 0.039 | 0.039 | 0 | 0.11 | 0.31 | 0.09 | 0.12 | 0.11 | 0.21 |
| 9 | 14424929 | rs2031195 | -0.859 | 0.243 | 0.151 | 0.236 | -0.003 | 0.018 | 0.018 | 0.05 | 0.85 | 0.89 | 0.87 | 0.95 | 0.84 | 0.94 |
| 9 | 14426289 | rs1028704 | 0.555 | 0.189 | 0.537 | 0.239 | -0.001 | -0.015 | -0.015 | 0 | 0.71 | 0.56 | 0.76 | 0.81 | 0.73 | 0.8 |
| 9 | 14427560 | rs990938 | 0.49 | 0.129 | 0.478 | 0.423 | -0.001 | 0 | 0 | 0.047 | 0.1 | 0.13 | 0.06 | 0.19 | 0.07 | 0.11 |
| 9 | 14429307 | rs4741377 | 1.478 | 0.762 | 0.513 | 0.289 | -0.003 | -0.006 | -0.006 | 0 | 0.34 | 0.28 | 0.57 | 0.4 | 0.41 | 0.28 |
| 9 | 14430286 | rs373034 | 0.171 | 0.793 | 0.518 | 0.309 | -0.012 | 0 | 0 | 0 | 0.38 | 0.52 | 0.21 | 0.43 | 0.27 | 0.52 |
| 9 | 14430328 | rs303737 | -0.176 | 0.817 | 0.491 | 0.263 | 0.001 | -0.001 | -0.001 | 0 | 0.43 | 0.57 | 0.26 | 0.44 | 0.37 | 0.56 |
| 9 | 14432097 | rs991115 | 0.51 | 0.714 | 0.574 | 0.27 | -0.002 | -0.001 | -0.001 | 0.017 | 0.47 | 0.45 | 0.66 | 0.56 | 0.56 | 0.57 |
| 9 | 14433910 | rs501360 | 0.546 | 0.742 | 0.245 | 0.293 | -0.044 | -0.003 | -0.003 | 0 | 0.33 | 0.41 | 0.2 | 0.36 | 0.22 | 0.33 |
| 9 | 14434284 | rs958957 | 1.405 | 0.891 | -0.34 | 0.42 | -0.018 | -0.031 | -0.031 | 0 | 0.32 | 0.14 | 0.52 | 0.4 | 0.37 | 0.28 |
| 9 | 14437476 | rs508259 | -1.172 | 0.734 | -0.37 | 0.182 | -0.004 | 0.01 | 0.01 | 0.032 | 0.88 | 0.93 | 0.88 | 0.95 | 0.87 | 0.95 |
| 9 | 14437943 | rs303733 | 0.041 | 0.471 | -0.204 | 0.327 | 0.018 | 0.008 | 0.008 | 0 | 0.4 | 0.66 | 0.24 | 0.36 | 0.3 | 0.39 |
| 9 | 14440552 | rs303727 | -1.022 | 0.501 | -0.434 | 0.299 | -0.038 | -0.051 | -0.051 | 0.923 | 0.58 | 0.83 | 0.31 | 0.5 | 0.43 | 0.65 |
| 9 | 14440739 | rs215498 | 0.012 | 0.465 | -0.076 | 0.339 | 0.011 | 0.007 | 0.007 | 0 | 0.45 | 0.54 | 0.25 | 0.38 | 0.33 | 0.49 |
| 9 | 14442265 | rs1323351 | 0.624 | 0.486 | -0.051 | 0.286 | -0.008 | -0.009 | -0.009 | 0 | 0.09 | 0.22 | 0.03 | 0.09 | 0.06 | 0.09 |
| 9 | 14443468 | rs551112 | 0.024 | 0.272 | 0.051 | 0.127 | -0.02 | -0.019 | -0.019 | 0.027 | 0.13 | 0.31 | 0.07 | 0.1 | 0.1 | 0.1 |
| 9 | 14443784 | rs10491775 | 0.902 | -0.078 | -0.241 | 0.052 | 0.016 | -0.055 | -0.055 | 0 | 0.16 | 0.04 | 0.12 | 0.3 | 0.11 | 0.06 |
| 9 | 14444316 | rs1323339 | 2.053 | 0.473 | -0.261 | 0.107 | -0.055 | -0.067 | -0.067 | 3.974 | 0.26 | 0.12 | 0.53 | 0.45 | 0.36 | 0.28 |
| 9 | 14446001 | rs1556032 | 1.093 | 0.399 | -0.333 | -0.026 | -0.064 | -0.052 | -0.052 | 0 | 0.35 | 0.16 | 0.6 | 0.47 | 0.47 | 0.35 |
| 9 | 14447233 | rs567977 | 0.558 | -0.041 | -0.004 | -0.199 | -0.002 | -0.001 | -0.001 | 0.084 | 0.51 | 0.58 | 0.66 | 0.62 | 0.59 | 0.55 |
| 9 | 14447766 | rs10810148 | -0.215 | -0.029 | -0.313 | -0.181 | 0.011 | 0.001 | 0.001 | 0 | 0.42 | 0.6 | 0.21 | 0.43 | 0.26 | 0.43 |
| 9 | 14448497 | rs535076 | -0.301 | 0.402 | -0.513 | 0.026 | -0.019 | 0.021 | 0.021 | 0 | 0.84 | 0.93 | 0.84 | 0.94 | 0.79 | 0.9 |
| 9 | 14452729 | rs10810150 | 0.496 | 0.149 | -0.194 | 0.082 | 0.008 | -0.004 | -0.004 | 0 | 0.28 | 0.21 | 0.14 | 0.33 | 0.16 | 0.32 |
| 9 | 14453076 | rs17761983 | 2.297 | 1.034 | -0.841 | 0.072 | -0.06 | 0.102 | 0.102 | 0 | 0.27 | 0.02 | 0.24 | 0.07 | 0.26 | 0.08 |
| 9 | 14456012 | rs7021022 | -0.594 | 0.894 | -0.291 | -0.038 | -0.004 | -0.002 | -0.002 | 0 | 0.46 | 0.56 | 0.25 | 0.41 | 0.32 | 0.41 |
| 9 | 14456738 | rs4741378 | 1.739 | 1.161 | -0.475 | 0.142 | -0.028 | -0.037 | -0.037 | 2.277 | 0.31 | 0.16 | 0.55 | 0.46 | 0.38 | 0.33 |
| 9 | 14457263 | rs527093 | -1.939 | 1.173 | -0.428 | 0.209 | -0.016 | -0.022 | -0.022 | 0.623 | 0.93 | 1 | 0.9 | 0.95 | 0.88 | 0.96 |
| 9 | 14460872 | rs6474837 | 0.846 | 1.15 | -0.087 | 0.224 | 0.006 | 0.004 | 0.004 | 0.014 | 0.09 | 0.22 | 0.04 | 0.1 | 0.07 | 0.09 |
| 9 | 14464744 | rs913416 | 0.571 | 1 | -0.117 | 0.141 | -0.001 | 0.002 | 0.002 | 0 | 0.46 | 0.51 | 0.65 | 0.56 | 0.53 | 0.42 |
| 9 | 14465710 | rs913412 | -0.53 | 1.681 | -0.133 | 0.101 | 0.026 | -0.04 | -0.04 | 0 | 0.18 | 0.35 | 0.17 | 0.1 | 0.19 | 0.11 |
| 9 | 14465961 | rs877451 | 0.757 | 1.725 | -0.163 | 0.177 | 0.021 | 0 | 0 | 3.43 | 0.09 | 0.19 | 0.1 | 0.1 | 0.11 | 0.08 |
| 9 | 14468137 | rs303724 | 0.682 | 1.476 | -0.356 | 0.153 | 0.008 | -0.018 | -0.018 | 0.029 | 0.6 | 0.56 | 0.64 | 0.78 | 0.5 | 0.62 |
| 9 | 14468321 | rs1998495 | -0.077 | 1.591 | -0.431 | 0.108 | 0.028 | -0.037 | -0.037 | 1.512 | 0.2 | 0.33 | 0.2 | 0.12 | 0.25 | 0.17 |
| 9 | 14469146 | rs303723 | -0.316 | 1.433 | -0.812 | 0.581 | -0.007 | 0.015 | 0.015 | 0 | 0.8 | 0.9 | 0.84 | 0.9 | 0.75 | 0.78 |
| 9 | 14470759 | rs516641 | 0.198 | 1.032 | -0.539 | 0.591 | 0.008 | 0.02 | 0.02 | 0 | 0.66 | 0.78 | 0.81 | 0.85 | 0.67 | 0.5 |
| 9 | 14470833 | rs10961534 | -0.197 | 0.858 | -0.472 | 0.999 | -0.003 | 0.014 | 0.014 | 0 | 0.14 | 0.21 | 0.14 | 0.31 | 0.1 | 0.08 |
| 9 | 14473378 | rs1323344 | 0.474 | 0.92 | -0.078 | 0.743 | 0.012 | 0.001 | 0.001 | 0 | 0.6 | 0.53 | 0.39 | 0.34 | 0.48 | 0.44 |
| 9 | 14473556 | rs303721 | 0.191 | 0.982 | 0.018 | 0.893 | -0.071 | -0.061 | -0.061 | 0 | 0.64 | 0.87 | 0.54 | 0.46 | 0.53 | 0.49 |
| 9 | 14475352 | rs501463 | -0.452 | 1.075 | 0.328 | 0.945 | 0.085 | 0.127 | 0.127 | 0.149 | 0.1 | 0.4 | 0.17 | 0.13 | 0.1 | 0.09 |
| 9 | 14478588 | rs4141905 | 0.722 | 0.981 | 0.309 | 0.895 | 0.007 | 0.001 | 0.001 | 0 | 0.61 | 0.46 | 0.39 | 0.35 | 0.5 | 0.46 |
| 9 | 14479228 | rs487674 | -0.893 | 0.753 | 0.665 | 0.91 | -0.001 | 0.002 | 0.002 | 0 | 0.28 | 0.66 | 0.34 | 0.33 | 0.25 | 0.35 |
| 9 | 14479361 | rs186228 | -0.551 | 1.078 | 0.88 | 0.925 | 0.141 | 0.24 | 0.24 | 0 | 0.05 | 0.35 | 0.16 | 0.12 | 0.06 | 0.06 |
| 9 | 14480141 | rs392125 | -0.779 | 1.077 | 0.692 | 0.924 | 0.166 | 0.274 | 0.274 | 0.011 | 0.06 | 0.43 | 0.16 | 0.12 | 0.06 | 0.06 |
| 9 | 14491347 | rs439269 | -1.624 | 1.502 | 0.789 | 0.819 | 0.154 | 0.146 | 0.146 | 0.028 | 0.09 | 0.5 | 0.22 | 0.15 | 0.16 | 0.04 |
| 9 | 14493251 | rs10961542 | 1.884 | 0.776 | -0.371 | 0.236 | 0.006 | 0.112 | 0.112 | 0.451 | 0.27 | 0.01 | 0.14 | 0.08 | 0.18 | 0.09 |
| 9 | 14494388 | rs1323348 | -0.752 | 0.991 | -0.319 | 0.737 | -0.002 | 0.006 | 0.006 | 0.099 | 0.32 | 0.65 | 0.53 | 0.48 | 0.3 | 0.41 |

| | | | | | | | | | | | | | | | | |
|---|---|---|---|---|---|---|---|---|---|---|---|---|---|---|---|---|
| 9 | 14494845 | rs7875078 | -1.273 | 1.173 | -0.557 | 0.726 | 0.007 | 0.023 | 0.023 | 0.154 | 0.49 | 0.69 | 0.66 | 0.6 | 0.48 | 0.68 |
| 9 | 14496808 | rs4741384 | -0.232 | 1.539 | -0.281 | 0.89 | 0.01 | 0.032 | 0.032 | 0 | 0.24 | 0.56 | 0.34 | 0.27 | 0.26 | 0.28 |
| 9 | 14498336 | rs191006 | -0.005 | 1.86 | -0.572 | 0.84 | -0.072 | -0.048 | -0.048 | 0 | 0.8 | 0.93 | 0.85 | 0.77 | 0.69 | 0.87 |
| 9 | 14511645 | rs7023433 | 0.445 | 1.491 | 0.494 | 0.809 | 0.001 | -0.004 | -0.004 | 0 | 0.6 | 0.46 | 0.42 | 0.36 | 0.58 | 0.32 |
| 9 | 14514366 | rs4741388 | 0.184 | 1.566 | 0.735 | 0.931 | -0.007 | 0.008 | 0.008 | 0 | 0.63 | 0.57 | 0.55 | 0.48 | 0.69 | 0.34 |
| 9 | 14515364 | rs1318729 | -0.275 | 1.525 | 0.723 | 1.048 | 0.001 | 0.04 | 0.04 | 0 | 0.79 | 0.65 | 0.78 | 0.73 | 0.78 | 0.68 |
| 9 | 14518646 | rs10756570 | 0.06 | 1.685 | 0.807 | 1.11 | -0.004 | -0.036 | -0.036 | 0.698 | 0.24 | 0.16 | 0.39 | 0.4 | 0.24 | 0.37 |
| 9 | 14518829 | rs873553 | 1.131 | 1.558 | 0.993 | 0.891 | -0.001 | 0.08 | 0.08 | 0 | 0.51 | 0.33 | 0.37 | 0.32 | 0.52 | 0.31 |
| 9 | 14518985 | rs7851867 | 0.19 | 1.546 | 0.898 | 0.904 | -0.005 | -0.035 | -0.035 | 0 | 0.23 | 0.16 | 0.39 | 0.4 | 0.24 | 0.36 |
| 9 | 14519301 | rs1106726 | -0.201 | 1.558 | 1.036 | 0.815 | 0.002 | 0 | 0 | 0 | 0.74 | 0.64 | 0.76 | 0.75 | 0.73 | 0.7 |
| 9 | 14520560 | rs13288173 | 0.374 | 1.53 | 0.994 | 0.823 | -0.002 | 0.01 | 0.01 | 0 | 0.22 | 0.3 | 0.2 | 0.25 | 0.21 | 0.24 |
| 9 | 14520611 | rs913417 | -0.102 | 1.535 | 0.987 | 0.826 | 0 | 0.029 | 0.029 | 0 | 0.76 | 0.57 | 0.77 | 0.72 | 0.76 | 0.71 |
| 9 | 14524438 | rs927969 | 0.616 | 1.524 | 1.047 | 0.807 | 0.078 | 0.049 | 0.049 | 0.092 | 0.94 | 0.73 | 0.91 | 0.97 | 0.92 | 0.92 |
| 9 | 14524784 | rs4741390 | 0.293 | 1.519 | 0.368 | 0.845 | -0.006 | -0.056 | -0.056 | 0 | 0.22 | 0.13 | 0.35 | 0.42 | 0.2 | 0.37 |
| 9 | 14525980 | rs913418 | 0.591 | 1.546 | 0.335 | 0.646 | 0.001 | 0.004 | 0.004 | 0 | 0.55 | 0.49 | 0.45 | 0.33 | 0.59 | 0.37 |
| 9 | 14527083 | rs10961553 | 0.706 | 1.541 | 0.252 | 0.592 | -0.001 | 0.005 | 0.005 | 0 | 0.53 | 0.49 | 0.42 | 0.31 | 0.54 | 0.3 |
| 9 | 14529623 | rs10810172 | 1.668 | 1.167 | -0.456 | 0.456 | 0.022 | 0.07 | 0.07 | 0 | 0.31 | 0.17 | 0.2 | 0.13 | 0.25 | 0.13 |
| 9 | 14529836 | rs12376530 | -0.522 | 1.656 | 0.132 | 1.18 | -0.001 | 0.043 | 0.043 | 0 | 0.26 | 0.36 | 0.43 | 0.46 | 0.26 | 0.44 |
| 9 | 14533251 | rs7042232 | -1.245 | 2.304 | -0.012 | 1.121 | -0.024 | -0.007 | -0.007 | 0 | 0.29 | 0.22 | 0.48 | 0.72 | 0.37 | 0.44 |
| 9 | 14533865 | rs6474842 | 1.785 | 2.451 | 0.048 | 0.622 | 0.045 | 0.117 | 0.117 | 0.005 | 0.44 | 0.24 | 0.32 | 0.17 | 0.31 | 0.28 |
| 9 | 14534103 | rs7046317 | 1.003 | 2.638 | 0.323 | 1.134 | 0.005 | -0.006 | -0.006 | 0 | 0.58 | 0.44 | 0.58 | 0.54 | 0.6 | 0.43 |
| 9 | 14534505 | rs10756571 | 1.199 | 2.563 | 0.547 | 1.152 | 0.111 | 0.135 | 0.135 | 0 | 0.82 | 0.45 | 0.7 | 0.62 | 0.68 | 0.82 |
| 9 | 14534704 | rs1556028 | 0.905 | 2.706 | 0.988 | 1.299 | 0.01 | 0.088 | 0.088 | 0 | 0.76 | 0.6 | 0.58 | 0.52 | 0.73 | 0.65 |
| 9 | 14535957 | rs10810175 | 2.025 | 2.896 | 0.805 | 0.988 | 0.08 | 0.129 | 0.129 | 0.001 | 0.45 | 0.25 | 0.3 | 0.17 | 0.29 | 0.28 |
| 9 | 14536315 | rs12379841 | -0.873 | 2.903 | 0.957 | 1.217 | -0.003 | 0.04 | 0.04 | 0.439 | 0.33 | 0.41 | 0.3 | 0.45 | 0.32 | 0.51 |
| 9 | 14536379 | rs2039051 | 1.416 | 2.733 | 0.949 | 1.14 | -0.007 | 0.021 | 0.021 | 0.039 | 0.56 | 0.41 | 0.54 | 0.32 | 0.49 | 0.37 |
| 9 | 14538577 | rs10961558 | -0.14 | 2.463 | 1.607 | 1.796 | -0.006 | 0.107 | 0.107 | 0 | 0.11 | 0.22 | 0.18 | 0.33 | 0.07 | 0.24 |
| 9 | 14539758 | rs10961560 | 0.29 | 2.388 | 1.337 | 1.735 | 0.012 | 0.01 | 0.01 | 0.209 | 0.1 | 0.16 | 0.06 | 0.12 | 0.11 | 0.24 |
| 9 | 14540362 | rs4741397 | -1.644 | 2.567 | 1.259 | 1.708 | 0.015 | 0.044 | 0.044 | 0.256 | 0.28 | 0.4 | 0.4 | 0.59 | 0.34 | 0.56 |
| 9 | 14540539 | rs7847421 | 0.195 | 2.911 | 0.842 | 1.745 | 0.04 | 0.08 | 0.08 | 0 | 0.92 | 0.83 | 0.93 | 0.78 | 0.86 | 0.91 |
| 9 | 14540818 | rs7847828 | 1.333 | 2.701 | 0.792 | 1.428 | 0.011 | 0.023 | 0.023 | 0.445 | 0.72 | 0.39 | 0.7 | 0.52 | 0.61 | 0.57 |
| 9 | 14541236 | rs7850989 | 0.951 | 2.473 | 0.598 | 1.711 | -0.002 | 0.041 | 0.041 | 0 | 0.83 | 0.77 | 0.7 | 0.65 | 0.84 | 0.66 |
| 9 | 14543216 | rs11789001 | -1.262 | 2.885 | 0.178 | 1.629 | -0.045 | -0.047 | -0.047 | 0 | 0.12 | 0.08 | 0.18 | 0.28 | 0.22 | 0.07 |
| 9 | 14543548 | rs927971 | -1.902 | 2.355 | 0.951 | 1.63 | 0.118 | 0.202 | 0.202 | 0 | 0.13 | 0.45 | 0.19 | 0.3 | 0.19 | 0.06 |
| 9 | 14544555 | rs2149104 | -1.003 | 2.205 | 0.473 | 1.617 | -0.011 | -0.037 | -0.037 | 0 | 0.15 | 0.07 | 0.27 | 0.62 | 0.17 | 0.37 |
| 9 | 14547556 | rs10810178 | -0.421 | 2.159 | 0.637 | 1.713 | -0.004 | 0.03 | 0.03 | 0 | 0.11 | 0.08 | 0.16 | 0.39 | 0.07 | 0.34 |
| 9 | 14550138 | rs10491772 | 1.392 | 2.094 | -0.387 | 0.463 | -0.098 | 0.085 | 0.085 | 0 | 0.28 | 0.02 | 0.3 | 0.09 | 0.31 | 0.13 |
| 9 | 14552292 | rs1536647 | 1.726 | 2.445 | 1.052 | 1.111 | 0.11 | 0.016 | 0.016 | 0 | 0.94 | 0.76 | 0.86 | 0.93 | 0.87 | 0.97 |
| 9 | 14555896 | rs7047109 | -0.992 | 2.059 | 1.08 | 1.07 | 0.037 | -0.037 | -0.037 | 1.73 | 0.08 | 0.26 | 0.16 | 0.03 | 0.12 | 0.02 |
| 9 | 14556659 | rs10756576 | -2.205 | 1.862 | 1.154 | 1.182 | 0.003 | 0.004 | 0.004 | 0 | 0.31 | 0.52 | 0.42 | 0.68 | 0.34 | 0.47 |
| 9 | 14557806 | rs10961571 | -0.394 | 1.056 | 0.752 | 1.352 | 0.016 | -0.001 | -0.001 | 0 | 0.1 | 0.04 | 0.16 | 0.43 | 0.06 | 0.34 |
| 9 | 14560916 | rs4741405 | -1.407 | 1.336 | 1.382 | 0.478 | 0.089 | -0.005 | -0.005 | 0 | 0.09 | 0.4 | 0.13 | 0.05 | 0.1 | 0.06 |
| 9 | 14562630 | rs1556029 | 1.452 | 1.104 | 1.066 | 0.131 | -0.01 | -0.098 | -0.098 | 0 | 0.79 | 0.45 | 0.8 | 0.93 | 0.8 | 0.83 |
| 9 | 14564090 | rs10491771 | 1.971 | 1.493 | 0.789 | 0.643 | -0.013 | -0.004 | -0.004 | 0 | 0.63 | 0.26 | 0.49 | 0.36 | 0.61 | 0.54 |
| 9 | 14566116 | rs7867360 | -0.465 | 1.686 | 1.017 | 0.996 | 0 | -0.021 | -0.021 | 0.074 | 0.09 | 0.12 | 0.06 | 0.02 | 0.11 | 0.09 |
| 9 | 14566406 | rs7040652 | 1.617 | 1.447 | 0.525 | 0.833 | -0.005 | 0.005 | 0.005 | 0 | 0.68 | 0.35 | 0.69 | 0.73 | 0.64 | 0.77 |
| 9 | 14573137 | rs12352208 | -0.495 | 1.306 | 0.35 | 1.189 | -0.005 | 0.045 | 0.045 | 0 | 0.12 | 0.15 | 0.26 | 0.4 | 0.07 | 0.25 |
| 9 | 14573613 | rs10116154 | -1.429 | 1.505 | 0.811 | 0.724 | 0.051 | 0.022 | 0.022 | 0 | 0.22 | 0.42 | 0.21 | 0.25 | 0.3 | 0.22 |
| 9 | 14585052 | rs7029906 | 0.7 | 1.327 | -0.029 | -0.009 | 0.009 | 0.065 | 0.065 | 0.129 | 0.49 | 0.3 | 0.35 | 0.36 | 0.53 | 0.48 |
| 9 | 14588049 | rs12238310 | 1.751 | 1.631 | -0.512 | -0.368 | 0.015 | 0.029 | 0.029 | 0 | 0.32 | 0.23 | 0.21 | 0.21 | 0.3 | 0.24 |
| 9 | 14589694 | rs10961601 | 0.333 | 1.513 | -0.298 | -0.322 | 0.006 | 0.025 | 0.025 | 0.17 | 0.5 | 0.44 | 0.37 | 0.31 | 0.54 | 0.42 |
| 9 | 14590805 | rs7857143 | 0.517 | 1.275 | -0.628 | -0.155 | -0.071 | -0.071 | -0.071 | 0.186 | 0.27 | 0.05 | 0.3 | 0.31 | 0.32 | 0.16 |
| 9 | 14592180 | rs2225163 | 0.47 | 1.368 | -0.489 | -0.278 | -0.002 | -0.005 | -0.005 | 0 | 0.54 | 0.48 | 0.49 | 0.32 | 0.62 | 0.45 |
| 9 | 14594350 | rs10756583 | 0.466 | 1.47 | -0.12 | -0.313 | 0.014 | 0.077 | 0.077 | 0.007 | 0.54 | 0.7 | 0.48 | 0.32 | 0.58 | 0.45 |
| 9 | 14594501 | rs11794863 | 0.967 | 1.667 | -0.311 | -0.402 | 0.006 | -0.017 | -0.017 | 0 | 0.08 | 0.09 | 0.06 | 0.03 | 0.1 | 0.11 |
| 9 | 14604569 | rs4741409 | 1.1 | 1.366 | -0.214 | -0.348 | -0.004 | 0.016 | 0.016 | 0 | 0.45 | 0.44 | 0.41 | 0.28 | 0.48 | 0.34 |
| 9 | 14604822 | rs7850166 | 0.083 | 1.478 | -0.232 | -0.329 | 0.026 | -0.032 | -0.032 | 0 | 0.11 | 0.2 | 0.11 | 0.03 | 0.16 | 0.11 |
| 9 | 14610866 | rs1343705 | 0.338 | 1.313 | -0.613 | -0.29 | -0.059 | -0.012 | -0.012 | 0.55 | 0.61 | 0.86 | 0.67 | 0.62 | 0.56 | 0.69 |
| 9 | 14613104 | rs10810184 | 1.654 | 1.231 | -0.402 | -0.345 | -0.001 | 0.019 | 0.019 | 0 | 0.41 | 0.36 | 0.27 | 0.26 | 0.4 | 0.3 |
| 9 | 14615350 | rs10738372 | 0.284 | 1.244 | -0.433 | -0.33 | -0.057 | -0.017 | -0.017 | 0.364 | 0.61 | 0.86 | 0.67 | 0.62 | 0.55 | 0.69 |
| 9 | 14616590 | rs12236525 | 1.672 | 1.378 | -0.389 | -0.334 | -0.016 | 0.042 | 0.042 | 0 | 0.41 | 0.31 | 0.28 | 0.26 | 0.44 | 0.31 |
| 9 | 14616757 | rs10961620 | 2.122 | 1.25 | -0.443 | -0.417 | 0.002 | 0.023 | 0.023 | 0 | 0.35 | 0.31 | 0.24 | 0.17 | 0.36 | 0.22 |
| 9 | 14616829 | rs7044386 | 0.594 | 1.168 | -0.377 | -0.353 | -0.001 | -0.007 | -0.007 | 0 | 0.09 | 0.08 | 0.09 | 0.02 | 0.15 | 0.11 |
| 9 | 14617069 | rs10810186 | 0.757 | 1.121 | -0.308 | -0.339 | 0.015 | -0.019 | -0.019 | 0 | 0.14 | 0.09 | 0.26 | 0.35 | 0.08 | 0.36 |
| 9 | 14618856 | rs3802482 | 0.615 | 1.146 | -0.276 | -0.332 | 0.004 | -0.005 | -0.005 | 0.012 | 0.58 | 0.54 | 0.66 | 0.61 | 0.56 | 0.69 |
| 9 | 14620540 | rs10961621 | 0.475 | 0.711 | -0.484 | -0.125 | 0.002 | 0.011 | 0.011 | 0 | 0.14 | 0.12 | 0.26 | 0.35 | 0.08 | 0.36 |
| 9 | 14622952 | rs7874535 | -0.23 | 0.92 | -0.497 | -0.118 | -0.044 | -0.098 | -0.098 | 0 | 0.7 | 0.94 | 0.77 | 0.63 | 0.71 | 0.8 |
| 9 | 14645813 | rs10120588 | -0.392 | 0.785 | -0.823 | -0.349 | -0.041 | -0.038 | -0.038 | 0 | 0.34 | 0.11 | 0.36 | 0.35 | 0.37 | 0.22 |
| 9 | 14649728 | rs10961636 | 0.791 | 0.563 | -0.612 | -0.354 | -0.013 | 0.044 | 0.044 | 0 | 0.18 | 0.23 | 0.16 | 0.33 | 0.16 | 0.36 |

| | | | | | | | | | | | | | | | | |
|---|---|---|---|---|---|---|---|---|---|---|---|---|---|---|---|---|
| 9 | 14650493 | rs13284901 | 1.44 | 0.557 | -0.376 | -0.454 | 0.007 | -0.035 | -0.035 | 0 | 0.39 | 0.48 | 0.36 | 0.22 | 0.43 | 0.34 |
| 9 | 14650684 | rs7851730 | 0.567 | 0.845 | -0.212 | -0.297 | -0.025 | 0.058 | 0.058 | 0 | 0.2 | 0.29 | 0.17 | 0.34 | 0.16 | 0.4 |
| 9 | 14651283 | rs11790280 | 1.684 | 0.837 | -0.384 | -0.441 | 0.002 | -0.046 | -0.046 | 0 | 0.37 | 0.57 | 0.27 | 0.16 | 0.38 | 0.26 |
| 9 | 14656681 | rs10119411 | -0.818 | 0.934 | -0.448 | -0.466 | 0 | 0.05 | 0.05 | 0.417 | 0.48 | 0.32 | 0.41 | 0.68 | 0.47 | 0.62 |
| 9 | 14657139 | rs10756597 | 0.302 | 0.813 | -0.562 | -0.462 | 0.003 | -0.001 | -0.001 | 0 | 0.91 | 0.94 | 0.87 | 0.92 | 0.94 | 0.96 |
| 9 | 14660700 | rs10481503 | 0.286 | 1.028 | -0.379 | -0.28 | -0.041 | -0.055 | -0.055 | 0 | 0.27 | 0.09 | 0.25 | 0.35 | 0.3 | 0.2 |
| 9 | 14662459 | rs6474850 | 0.964 | 0.701 | -0.159 | -0.265 | -0.002 | -0.002 | -0.002 | 0.714 | 0.84 | 0.85 | 0.68 | 0.85 | 0.85 | 0.85 |
| 9 | 14670949 | rs10961649 | 1.937 | 0.788 | -0.067 | -0.375 | 0.004 | 0.045 | 0.045 | 0 | 0.33 | 0.23 | 0.23 | 0.13 | 0.34 | 0.24 |
| 9 | 14672267 | rs2890992 | 1.151 | 0.411 | 0.123 | -0.196 | -0.003 | 0 | 0 | 0 | 0.89 | 0.79 | 0.78 | 0.9 | 0.89 | 0.95 |
| 9 | 14676870 | rs10810211 | -0.195 | 0.42 | 0.08 | -0.183 | -0.003 | 0.005 | 0.005 | 0 | 0.95 | 0.89 | 0.9 | 0.9 | 0.97 | 0.97 |
| 9 | 14677538 | rs10810212 | 1.449 | 0.461 | 0.035 | -0.183 | -0.017 | 0.084 | 0.084 | 0.003 | 0.41 | 0.26 | 0.34 | 0.19 | 0.43 | 0.35 |
| 9 | 14679206 | rs10810213 | 0.074 | 0.452 | -0.254 | -0.125 | -0.03 | -0.029 | -0.029 | 0 | 0.29 | 0.14 | 0.35 | 0.62 | 0.36 | 0.45 |
| 9 | 14686308 | rs17288444 | 0.766 | 0.426 | -0.156 | -0.273 | -0.006 | 0.053 | 0.053 | 0.112 | 0.47 | 0.6 | 0.47 | 0.23 | 0.5 | 0.37 |
| 9 | 14689088 | rs10810220 | 0.311 | 0.624 | -0.341 | 0.142 | -0.018 | -0.045 | -0.045 | 0 | 0.21 | 0.09 | 0.22 | 0.33 | 0.25 | 0.4 |
| 9 | 14689191 | rs10756603 | 0.628 | 0.579 | -0.175 | 0.128 | 0.004 | 0.057 | 0.057 | 0 | 0.95 | 0.79 | 0.91 | 0.93 | 0.98 | 0.97 |
| 9 | 14692676 | rs10756604 | 0.328 | 0.643 | -0.434 | 0.157 | -0.022 | -0.051 | -0.051 | 0 | 0.21 | 0.08 | 0.22 | 0.34 | 0.25 | 0.4 |
| 9 | 14694211 | rs2890989 | -0.069 | 0.386 | -0.467 | 0.113 | -0.052 | -0.078 | -0.078 | 0 | 0.25 | 0.07 | 0.44 | 0.39 | 0.35 | 0.43 |
| 9 | 14694302 | rs2382479 | -1.256 | 0.333 | -0.283 | -0.125 | -0.003 | 0.028 | 0.028 | 0 | 0.58 | 0.61 | 0.74 | 0.85 | 0.56 | 0.66 |
| 9 | 14694536 | rs4326470 | 0.09 | -0.069 | 0.051 | 0.37 | -0.011 | 0.042 | 0.042 | 4.518 | 0.22 | 0.3 | 0.18 | 0.36 | 0.18 | 0.11 |
| 9 | 14696674 | rs1858633 | -0.409 | -0.23 | 0.203 | -0.173 | -0.023 | 0.019 | 0.019 | 0 | 0.05 | 0.12 | 0.08 | 0.1 | 0.01 | 0.06 |
| 9 | 14701230 | rs7021169 | -0.02 | -0.184 | 0.099 | -0.526 | 0.011 | 0.02 | 0.02 | 0.284 | 0.47 | 0.6 | 0.33 | 0.21 | 0.4 | 0.38 |
| 9 | 14701616 | rs12685826 | -0.98 | 0.025 | -0.212 | -0.521 | 0.006 | -0.008 | -0.008 | 0 | 0.21 | 0.31 | 0.26 | 0.2 | 0.21 | 0.35 |
| 9 | 14712257 | rs7024505 | 0.141 | 0.574 | -0.35 | -0.202 | 0.008 | 0.042 | 0.042 | 0 | 0.43 | 0.32 | 0.27 | 0.26 | 0.39 | 0.36 |
| 9 | 14722616 | rs10115703 | -0.8 | 0.874 | 0.034 | 0.071 | -0.001 | -0.009 | -0.009 | 1.763 | 0.12 | 0.11 | 0.13 | 0.02 | 0.08 | 0.13 |
| 9 | 14723637 | rs7035643 | -1.781 | 0.794 | 0.678 | -0.153 | -0.005 | -0.064 | -0.064 | 0 | 0.18 | 0.72 | 0.2 | 0.02 | 0.12 | 0.16 |
| 9 | 14724502 | rs10810226 | -2.12 | 1.548 | 0.226 | -0.421 | -0.039 | 0.102 | 0.102 | 0.285 | 0.28 | 0.89 | 0.28 | 0.13 | 0.25 | 0.26 |
| 9 | 14726304 | rs11794846 | -0.24 | 1.616 | 0.27 | -0.341 | -0.039 | 0.038 | 0.038 | 0 | 0.07 | 0.01 | 0.06 | 0 | 0.12 | 0.07 |
| 9 | 14728730 | rs12345917 | -1.033 | 2.092 | 0.805 | -0.133 | 0.051 | -0.018 | -0.018 | 0 | 0.2 | 0.35 | 0.32 | 0.17 | 0.33 | 0.25 |
| 9 | 14730124 | rs1494338 | 1.813 | 2.136 | 0.885 | 0.667 | 0.025 | 0.023 | 0.023 | 0.41 | 0.63 | 0.43 | 0.54 | 0.54 | 0.54 | 0.44 |
| 9 | 14730234 | rs10511595 | -0.439 | 1.921 | 0.765 | 0.666 | -0.005 | -0.003 | -0.003 | 0 | 0.13 | 0.11 | 0.24 | 0.17 | 0.2 | 0.24 |
| 9 | 14733642 | rs13287172 | 1.992 | 1.824 | 0.947 | 0.613 | 0.022 | 0.022 | 0.022 | 0.147 | 0.65 | 0.59 | 0.56 | 0.54 | 0.57 | 0.48 |
| 9 | 14733912 | rs1494341 | 1.354 | 1.598 | 0.942 | 0.644 | 0.025 | 0.043 | 0.043 | 0 | 0.77 | 0.62 | 0.8 | 0.71 | 0.78 | 0.7 |
| 9 | 14735053 | rs1048070 | 1.926 | 1.459 | 0.547 | 0.301 | -0.002 | -0.031 | -0.031 | 0 | 0.55 | 0.25 | 0.44 | 0.54 | 0.57 | 0.44 |
| 9 | 14735187 | rs7047712 | -0.396 | 1.969 | 0.756 | 0.391 | 0.025 | -0.047 | -0.047 | 0.657 | 0.1 | 0.14 | 0.1 | 0 | 0.15 | 0.04 |
| 9 | 14735610 | rs10756607 | 0.079 | 1.912 | 0.458 | 0.719 | -0.051 | -0.048 | -0.048 | 0 | 0.1 | 0.01 | 0.22 | 0.17 | 0.18 | 0.16 |
| 9 | 14737506 | rs10961689 | 0.935 | 1.901 | 0.834 | 0.77 | 0.001 | 0 | 0 | 0.044 | 0.8 | 0.76 | 0.74 | 0.78 | 0.77 | 0.62 |
| 9 | 14740682 | rs8181217 | -1.498 | 1.728 | 0.564 | 0.891 | 0 | 0.013 | 0.013 | 0 | 0.08 | 0.11 | 0.08 | 0.21 | 0.05 | 0.05 |
| 9 | 14747133 | rs2270529 | -1.185 | 2.039 | 1.283 | 0.853 | 0.024 | 0.013 | 0.013 | 0 | 0.23 | 0.36 | 0.34 | 0.28 | 0.3 | 0.4 |
| 9 | 14747412 | rs4741426 | 0.196 | 1.986 | 0.865 | 0.922 | -0.052 | -0.051 | -0.051 | 1.017 | 0.1 | 0.01 | 0.22 | 0.18 | 0.18 | 0.16 |
| 9 | 14749434 | rs10117516 | -1.394 | 2.02 | 1.217 | 0.778 | 0.028 | -0.015 | -0.015 | 0 | 0.32 | 0.55 | 0.4 | 0.27 | 0.4 | 0.44 |
| 9 | 14751872 | rs2890993 | -1.353 | 1.877 | 0.973 | 1.147 | 0.006 | 0.142 | 0.142 | 0.755 | 0.16 | 0.4 | 0.22 | 0.36 | 0.16 | 0.17 |
| 9 | 14754010 | rs2035987 | 0.583 | 1.424 | -0.622 | 0.686 | -0.035 | 0.035 | 0.035 | 1.206 | 0.5 | 0.13 | 0.52 | 0.59 | 0.51 | 0.47 |
| 9 | 14757744 | rs10738377 | 0.234 | 1.581 | -1.038 | 0.539 | -0.095 | -0.056 | -0.056 | 0.231 | 0.18 | 0.01 | 0.13 | 0.22 | 0.24 | 0.26 |
| 9 | 14757877 | rs7864052 | -0.919 | 1.46 | 0.091 | 0.305 | 0.039 | -0.023 | -0.023 | 0 | 0.3 | 0.53 | 0.24 | 0.28 | 0.4 | 0.44 |
| 9 | 14761002 | rs13294547 | 2.216 | 1.133 | -0.411 | 0.154 | 0.035 | 0.121 | 0.121 | 0 | 0.28 | 0.02 | 0.16 | 0.1 | 0.23 | 0.22 |
| 9 | 14761226 | rs1724 | -1.183 | 1.327 | -0.88 | 0.143 | 0.02 | -0.009 | -0.009 | 0 | 0.88 | 0.84 | 0.74 | 0.9 | 0.81 | 0.9 |
| 9 | 14761260 | rs10124839 | 1.262 | 1.276 | -0.81 | 0.123 | 0.006 | -0.006 | -0.006 | 0.053 | 0.06 | 0.07 | 0.07 | 0.06 | 0.14 | 0.14 |
| 9 | 14761788 | rs7858690 | 0.965 | 1.512 | -0.933 | 0.128 | -0.07 | -0.059 | -0.059 | 0 | 0.18 | 0.01 | 0.12 | 0.22 | 0.22 | 0.26 |
| 9 | 14762700 | rs12685742 | 1.363 | 1.214 | -0.477 | 0.044 | -0.027 | 0.12 | 0.12 | 0.003 | 0.43 | 0.18 | 0.43 | 0.2 | 0.46 | 0.37 |
| 9 | 14763056 | rs16932272 | 2.131 | 0.359 | -0.793 | 0.17 | -0.002 | 0.011 | 0.011 | 21.641 | 0.1 | 0.07 | 0.06 | 0.07 | 0.07 | 0.07 |
| 9 | 14763855 | rs7039708 | 0.249 | 0.333 | -0.547 | 0.529 | -0.002 | 0.009 | 0.009 | 0 | 0.82 | 0.82 | 0.75 | 0.65 | 0.9 | 0.87 |
| 9 | 14765030 | rs923926 | 1.964 | 0.422 | -0.678 | 0.394 | 0.05 | 0.086 | 0.086 | 1.775 | 0.4 | 0.14 | 0.24 | 0.26 | 0.32 | 0.33 |
| 9 | 14767916 | rs6474855 | 0.421 | 0.829 | -0.216 | 0.623 | 0.025 | 0.023 | 0.023 | 0.604 | 0.72 | 0.69 | 0.49 | 0.53 | 0.58 | 0.68 |
| 9 | 14770658 | rs17219005 | -0.544 | 0.827 | 0.014 | 0.352 | 0.004 | -0.028 | -0.028 | 2.557 | 0.24 | 0.27 | 0.27 | 0.08 | 0.41 | 0.24 |
| 9 | 14770825 | rs7860370 | 1.199 | 0.71 | -0.198 | 0.523 | -0.054 | -0.072 | -0.072 | 0.244 | 0.17 | 0.01 | 0.15 | 0.22 | 0.17 | 0.25 |
| 9 | 14771575 | rs954116 | -1.446 | 0.57 | -0.013 | 0.377 | 0.002 | 0.002 | 0.002 | 15.554 | 0.85 | 0.79 | 0.87 | 0.85 | 0.88 | 0.87 |
| 9 | 14771864 | rs7849041 | -1.165 | 0.526 | -0.003 | 0.432 | -0.005 | -0.004 | -0.004 | 0 | 0.31 | 0.35 | 0.42 | 0.39 | 0.42 | 0.27 |
| 9 | 14773370 | rs7028801 | -1.36 | 0.448 | -0.019 | 0.43 | 0.001 | 0.001 | 0.001 | 15.301 | 0.85 | 0.79 | 0.87 | 0.85 | 0.88 | 0.86 |
| 9 | 14774967 | rs6474858 | -1.36 | 0.448 | -0.034 | 0.43 | 0.001 | 0.001 | 0.001 | 15.301 | 0.85 | 0.79 | 0.87 | 0.85 | 0.88 | 0.86 |
| 9 | 14775779 | rs1874108 | 2.814 | 0.316 | -0.068 | 0.386 | 0.03 | 0.07 | 0.07 | 0 | 0.31 | 0.05 | 0.19 | 0.16 | 0.27 | 0.29 |
| 9 | 14775853 | rs1032474 | -1.397 | 0.528 | -0.014 | 0.404 | 0.001 | 0.001 | 0.001 | 15.534 | 0.85 | 0.79 | 0.87 | 0.85 | 0.88 | 0.87 |
| 9 | 14775859 | rs10733289 | -1.37 | 0.769 | 0.234 | 0.744 | 0.007 | 0.035 | 0.035 | 0.097 | 0.41 | 0.29 | 0.44 | 0.33 | 0.58 | 0.49 |
| 9 | 14782497 | rs1494354 | 0.311 | 0.781 | 0.562 | 0.868 | -0.001 | 0.017 | 0.017 | 0.02 | 0.2 | 0.17 | 0.19 | 0.33 | 0.2 | 0.33 |
| 9 | 14782934 | rs10961704 | 1.605 | 1.208 | 0.428 | 0.788 | 0.009 | 0.001 | 0.001 | 0 | 0.06 | 0.04 | 0.07 | 0.06 | 0.16 | 0.13 |
| 9 | 14783623 | rs12683551 | 1.12 | 1.263 | 0.607 | 0.708 | 0.036 | -0.042 | -0.042 | 0.528 | 0.11 | 0.18 | 0.11 | 0.02 | 0.19 | 0.08 |
| 9 | 14784665 | rs10810239 | -1.128 | 1.066 | 0.59 | 0.904 | 0.005 | 0.041 | 0.041 | 0 | 0.21 | 0.33 | 0.33 | 0.35 | 0.23 | 0.19 |
| 9 | 14784778 | rs10810240 | -1.851 | 1.052 | 0.657 | 0.934 | 0.003 | 0.048 | 0.048 | 0 | 0.26 | 0.36 | 0.36 | 0.42 | 0.27 | 0.27 |
| 9 | 14785224 | rs7027322 | 0.044 | 0.76 | 0.554 | 0.928 | 0.006 | 0.055 | 0.055 | 0 | 0.78 | 0.71 | 0.8 | 0.68 | 0.78 | 0.67 |
| 9 | 14788407 | rs7025956 | 2.294 | 0.597 | -0.391 | 0.843 | 0.018 | 0.126 | 0.126 | 1.269 | 0.44 | 0.09 | 0.29 | 0.26 | 0.36 | 0.38 |

| | | | | | | | | | | | | | | | | |
|---|---|---|---|---|---|---|---|---|---|---|---|---|---|---|---|---|
| 9 | 14788886 | rs16932282 | -0.054 | 0.626 | -0.384 | 1.007 | 0.025 | -0.043 | -0.043 | 0.492 | 0.15 | 0.1 | 0.25 | 0.27 | 0.09 | 0.06 |
| 9 | 14789400 | rs10810242 | 1.309 | 0.66 | -0.42 | 0.92 | -0.021 | -0.078 | -0.078 | 0 | 0.14 | 0.01 | 0.15 | 0.24 | 0.14 | 0.23 |
| 9 | 14790363 | rs1494348 | 2.307 | 0.575 | -0.266 | 0.819 | 0.03 | 0.107 | 0.107 | 1.358 | 0.42 | 0.19 | 0.29 | 0.26 | 0.35 | 0.36 |
| 9 | 14791007 | rs2291680 | 1.548 | 0.874 | 0.114 | 0.952 | 0.041 | 0.034 | 0.034 | 0.204 | 0.66 | 0.52 | 0.54 | 0.57 | 0.54 | 0.71 |
| 9 | 14795265 | rs12338615 | -0.148 | -0.037 | -0.407 | 0.82 | 0.013 | -0.008 | -0.008 | 8.743 | 0.4 | 0.4 | 0.39 | 0.44 | 0.27 | 0.26 |
| 9 | 14795358 | rs10961709 | 1.426 | 0.247 | -0.31 | 0.751 | 0.019 | 0.01 | 0.01 | 0 | 0.05 | 0.06 | 0.05 | 0.09 | 0.09 | 0.11 |
| 9 | 14797143 | rs1908303 | 0.201 | 0.078 | -0.408 | 0.802 | 0.001 | 0.001 | 0.001 | 1.992 | 0.35 | 0.37 | 0.25 | 0.2 | 0.26 | 0.23 |
| 9 | 14797880 | rs9886892 | 0.805 | 0.411 | -1.029 | 0.792 | -0.028 | -0.026 | -0.026 | 0.09 | 0.14 | 0.06 | 0.19 | 0.2 | 0.23 | 0.3 |
| 9 | 14798176 | rs10124784 | -0.357 | 0.319 | -1.166 | 0.587 | -0.021 | -0.059 | -0.059 | 0.548 | 0.13 | 0.22 | 0.06 | 0 | 0.06 | 0.04 |
| 9 | 14800779 | rs10738379 | 0.829 | 0.728 | -1.654 | 1.016 | -0.083 | -0.083 | -0.083 | 0 | 0.14 | 0.01 | 0.2 | 0.23 | 0.23 | 0.3 |
| 9 | 14801687 | rs10738380 | 0.84 | 0.734 | -1.625 | 1.018 | -0.069 | -0.069 | -0.069 | 0 | 0.14 | 0.03 | 0.19 | 0.23 | 0.23 | 0.25 |
| 9 | 14802948 | rs12350382 | 0.518 | 0.496 | -0.911 | 0.844 | -0.001 | 0.001 | 0.001 | 0 | 0.1 | 0.09 | 0.07 | 0.12 | 0.1 | 0.13 |
| 9 | 14804141 | rs17220118 | 0.761 | 0.478 | -1.198 | 0.792 | -0.06 | 0.043 | 0.043 | 0.491 | 0.33 | 0.07 | 0.42 | 0.27 | 0.4 | 0.37 |
| 9 | 14804733 | rs10810248 | -0.728 | 0.208 | -0.371 | 1.046 | -0.005 | 0.003 | 0.003 | 0.061 | 0.19 | 0.21 | 0.1 | 0.22 | 0.15 | 0.24 |
| 9 | 14808325 | rs10810251 | 1.544 | 0.446 | -1.006 | 1.088 | -0.051 | 0.004 | 0.004 | 0.219 | 0.21 | 0.06 | 0.19 | 0.14 | 0.3 | 0.22 |
| 9 | 14810847 | rs12347393 | -0.176 | 0.243 | -1.127 | 1.002 | -0.036 | -0.019 | -0.019 | 1.358 | 0.14 | 0.02 | 0.14 | 0.11 | 0.13 | 0.17 |
| 9 | 14813976 | rs10961726 | 0.057 | 0.477 | -0.486 | 1.117 | -0.026 | 0.005 | 0.005 | 0 | 0.06 | 0.01 | 0.09 | 0.07 | 0.08 | 0.08 |
| 9 | 14819370 | rs7023244 | 1.007 | 0.267 | -0.664 | 0.891 | 0.001 | 0.149 | 0.149 | 9.005 | 0.33 | 0.08 | 0.26 | 0.08 | 0.25 | 0.21 |
| 9 | 14819813 | rs10961730 | 0.582 | 0.302 | -0.415 | 0.97 | -0.001 | -0.001 | -0.001 | 0.313 | 0.14 | 0.1 | 0.09 | 0.16 | 0.14 | 0.16 |
| 9 | 14820223 | rs2779502 | 0.492 | 0.328 | 0.063 | 1.044 | 0.007 | -0.01 | -0.01 | 1.119 | 0.39 | 0.35 | 0.48 | 0.59 | 0.33 | 0.38 |
| 9 | 14821333 | rs7025001 | 1.075 | 0.326 | -0.11 | 0.997 | 0.019 | 0.126 | 0.126 | 8.678 | 0.33 | 0.12 | 0.26 | 0.08 | 0.25 | 0.21 |
| 9 | 14821819 | rs2779503 | 0.714 | 0.913 | 0.52 | 1.302 | 0.012 | -0.03 | -0.03 | 0 | 0.84 | 0.77 | 0.81 | 0.93 | 0.79 | 0.85 |
| 9 | 14824799 | rs12236053 | -0.045 | 0.516 | 0.773 | 1.024 | -0.007 | 0.012 | 0.012 | 0 | 0.14 | 0.22 | 0.08 | 0.17 | 0.1 | 0.15 |
| 9 | 14826616 | rs2818939 | 0.741 | 0.716 | 0.29 | 1.154 | -0.002 | 0.08 | 0.08 | 0.171 | 0.05 | 0.05 | 0.22 | 0.45 | 0.08 | 0.19 |
| 9 | 14830821 | rs7868862 | 0.289 | 0.551 | 0.484 | 0.875 | -0.002 | -0.011 | -0.011 | 0 | 0.2 | 0.34 | 0.11 | 0.14 | 0.17 | 0.17 |
| 9 | 14830989 | rs10810258 | -0.166 | 0.463 | 0.367 | 0.767 | -0.003 | -0.013 | -0.013 | 0 | 0.46 | 0.44 | 0.36 | 0.23 | 0.38 | 0.32 |
| 9 | 14831233 | rs2642406 | 0.533 | 0.455 | -0.327 | 0.954 | -0.008 | 0.01 | 0.01 | 0.62 | 0.08 | 0.05 | 0.23 | 0.42 | 0.12 | 0.2 |
| 9 | 14832349 | rs9792613 | 0.388 | 0.507 | -0.271 | 0.921 | 0.031 | 0.108 | 0.108 | 1.644 | 0.85 | 0.67 | 0.68 | 0.56 | 0.77 | 0.76 |
| 9 | 14833003 | rs10046797 | 0.09 | 0.231 | -0.773 | 0.444 | 0.001 | 0.062 | 0.062 | 0 | 0.3 | 0.53 | 0.36 | 0.56 | 0.3 | 0.37 |
| 9 | 14833488 | rs2779508 | -0.691 | 0.298 | -1.455 | 0.405 | -0.02 | 0.01 | 0.01 | 0.287 | 0.58 | 0.74 | 0.64 | 0.66 | 0.52 | 0.53 |
| 9 | 14834172 | rs12555189 | 0.09 | -0.11 | -0.561 | 0.069 | 0 | 0.062 | 0.062 | 2.846 | 0.26 | 0.12 | 0.27 | 0.1 | 0.22 | 0.15 |
| 9 | 14835055 | rs1855139 | 0.209 | -0.077 | -0.539 | 0.04 | 0.037 | -0.011 | -0.011 | 5.608 | 0.93 | 0.8 | 0.92 | 0.98 | 0.89 | 0.95 |
| 9 | 14835081 | rs10961746 | -0.089 | -0.455 | -1.228 | 0.172 | -0.008 | 0.078 | 0.078 | 2.856 | 0.27 | 0.1 | 0.27 | 0.09 | 0.23 | 0.15 |
| 9 | 14838860 | rs2779509 | 1.183 | -1.215 | -0.06 | 1.242 | -0.085 | 0.149 | 0.149 | 0.008 | 0.82 | 0.38 | 0.83 | 0.54 | 0.93 | 0.87 |
| 9 | 14842557 | rs1353223 | -0.605 | -0.978 | -0.132 | 0.801 | 0.002 | -0.038 | -0.038 | 0 | 0.08 | 0.15 | 0.05 | 0 | 0.08 | 0.02 |
| 9 | 14844777 | rs10810269 | -0.534 | -1.388 | -0.456 | 1.086 | 0.011 | -0.043 | -0.043 | 0 | 0.08 | 0.01 | 0.14 | 0.24 | 0.05 | 0.09 |
| 9 | 14848892 | rs1021493 | -0.201 | -1.67 | -1.218 | 0.147 | -0.025 | 0.11 | 0.11 | 0.828 | 0.79 | 0.97 | 0.85 | 0.98 | 0.8 | 0.91 |
| 9 | 14849216 | rs12236064 | -1.296 | -1.806 | -1.31 | 0.207 | 0.011 | -0.013 | -0.013 | 0 | 0.08 | 0.01 | 0.06 | 0.14 | 0.05 | 0.08 |
| 9 | 14850256 | rs16932354 | -1.011 | -1.973 | -0.919 | 0.43 | 0.005 | 0.008 | 0.008 | 0 | 0.14 | 0.15 | 0.24 | 0.56 | 0.11 | 0.14 |
| 9 | 14850816 | rs2779499 | 2.083 | -1.857 | -1.507 | 0.05 | -0.154 | 0.206 | 0.206 | 2.235 | 0.39 | 0.02 | 0.26 | 0.06 | 0.47 | 0.37 |
| 9 | 14851617 | rs2818940 | -0.191 | -1.793 | -0.521 | 0.282 | -0.037 | -0.09 | -0.09 | 0.114 | 0.74 | 0.49 | 0.8 | 0.96 | 0.81 | 0.89 |
| 9 | 14851703 | rs729492 | 0.691 | -1.761 | -0.485 | 0.307 | 0.023 | 0.027 | 0.027 | 2.143 | 0.64 | 0.32 | 0.73 | 0.82 | 0.76 | 0.84 |
| 9 | 14851841 | rs729491 | -0.466 | -1.735 | -0.478 | 0.34 | -0.001 | 0.042 | 0.042 | 0 | 0.12 | 0.13 | 0.23 | 0.54 | 0.12 | 0.11 |
| 9 | 14856284 | rs1889053 | -1.91 | -1.709 | -0.47 | 0.314 | 0.116 | 0.107 | 0.107 | 2.215 | 0.5 | 0.8 | 0.51 | 0.71 | 0.38 | 0.29 |
| 9 | 14858083 | rs10511599 | -1.338 | -1.415 | 0.534 | 0.25 | 0.009 | 0 | 0 | 0 | 0.09 | 0.07 | 0.06 | 0.14 | 0.05 | 0.04 |
| 9 | 14864321 | rs1889051 | -2.103 | -1.411 | 0.409 | 0.475 | 0.048 | 0.064 | 0.064 | 0.326 | 0.5 | 0.93 | 0.5 | 0.7 | 0.42 | 0.35 |
| 9 | 14866251 | rs16932379 | -1.608 | -1.374 | 0.753 | 0.502 | 0.004 | -0.011 | -0.011 | 0 | 0.2 | 0.14 | 0.25 | 0.56 | 0.17 | 0.21 |
| 9 | 14867958 | rs16932386 | -1.549 | -1.352 | 0.991 | 0.456 | 0.005 | 0 | 0 | 0 | 0.11 | 0.08 | 0.07 | 0.16 | 0.08 | 0.07 |
| 9 | 14869861 | rs6474865 | -1.8 | -1.363 | 1.6 | 0.43 | 0.031 | 0.033 | 0.033 | 0 | 0.47 | 0.65 | 0.46 | 0.7 | 0.41 | 0.34 |
| 9 | 14878094 | rs10810279 | -0.71 | -1.214 | 1.558 | 0.521 | 0.016 | 0.047 | 0.047 | 0 | 0.06 | 0.21 | 0.07 | 0.14 | 0.05 | 0.04 |
| 9 | 14878376 | rs10810280 | -0.779 | -1.215 | 1.556 | 0.52 | 0.015 | 0.046 | 0.046 | 0 | 0.06 | 0.22 | 0.07 | 0.15 | 0.05 | 0.04 |
| 9 | 14880667 | rs7862716 | -1.957 | -1.29 | 1.034 | 0.473 | -0.007 | 0.004 | 0.004 | 0.193 | 0.28 | 0.65 | 0.27 | 0.71 | 0.23 | 0.24 |
| 9 | 14884447 | rs10810283 | -1.196 | -1.186 | 0.444 | 0.725 | -0.014 | 0.003 | 0.003 | 0 | 0.41 | 0.46 | 0.4 | 0.64 | 0.37 | 0.32 |
| 9 | 14890268 | rs1996893 | -1.873 | -0.708 | -0.342 | 0.79 | -0.073 | 0.217 | 0.217 | 0.165 | 0.54 | 0.95 | 0.64 | 0.88 | 0.49 | 0.59 |
| 9 | 14891670 | rs10810285 | -1.397 | -0.292 | 0.488 | 1.459 | -0.001 | 0.025 | 0.025 | 0 | 0.14 | 0.11 | 0.16 | 0.44 | 0.15 | 0.14 |
| 9 | 14892957 | rs9776139 | -0.802 | -0.242 | 0.337 | 1.49 | 0.011 | -0.059 | -0.059 | 0.141 | 0.28 | 0.15 | 0.29 | 0.44 | 0.24 | 0.19 |
| 9 | 14898161 | rs10961780 | -1.511 | 0.545 | 0.925 | 1.788 | 0.076 | 0.307 | 0.307 | 0 | 0.11 | 0.45 | 0.12 | 0.36 | 0.1 | 0.11 |
| 9 | 14899708 | rs10810289 | -1.225 | 1.337 | 0.368 | 1.63 | 0.001 | 0.001 | 0.001 | 0.373 | 0.28 | 0.3 | 0.23 | 0.46 | 0.35 | 0.3 |
| 9 | 14900356 | rs4740593 | 0.857 | 0.956 | -1.342 | 0.472 | -0.041 | 0.156 | 0.156 | 0 | 0.36 | 0.03 | 0.36 | 0.12 | 0.33 | 0.37 |
| 9 | 14900565 | rs4741443 | -0.278 | 1.026 | -0.244 | 0.991 | 0.002 | 0.03 | 0.03 | 0 | 0.7 | 0.6 | 0.65 | 0.46 | 0.72 | 0.61 |
| 9 | 14901322 | rs1984063 | 1.224 | 1.061 | -0.612 | 0.821 | -0.124 | 0.166 | 0.166 | 0.151 | 0.42 | 0.02 | 0.38 | 0.16 | 0.47 | 0.36 |
| 9 | 14903275 | rs12337423 | 0.453 | 0.864 | -0.415 | 0.477 | 0.01 | 0.03 | 0.03 | 1.728 | 0.22 | 0.11 | 0.16 | 0 | 0.2 | 0.05 |
| 9 | 14903800 | rs10961785 | 0.92 | 0.944 | 0.315 | 0.828 | 0.005 | 0.012 | 0.012 | 0.104 | 0.62 | 0.43 | 0.5 | 0.58 | 0.61 | 0.58 |
| 9 | 14906868 | rs940120 | 0.005 | 1.005 | 0.267 | 0.836 | 0.004 | 0.046 | 0.046 | 0.448 | 0.87 | 0.86 | 0.65 | 0.59 | 0.84 | 0.65 |
| 9 | 14908056 | rs7034380 | 0.757 | 0.929 | 0.384 | 0.84 | -0.008 | -0.026 | -0.026 | 0.189 | 0.66 | 0.73 | 0.49 | 0.6 | 0.65 | 0.59 |
| 9 | 14919539 | rs3850442 | 1.047 | 0.288 | 0.375 | 1.097 | -0.004 | -0.004 | -0.004 | 0 | 0.78 | 0.81 | 0.61 | 0.58 | 0.75 | 0.57 |
| 9 | 14925706 | rs1916314 | 0.357 | 0.065 | 0.323 | 1.086 | -0.002 | 0.011 | 0.011 | 0.35 | 0.83 | 0.85 | 0.69 | 0.62 | 0.82 | 0.63 |
| 9 | 14929835 | rs10961795 | 1.188 | -0.099 | 0.256 | 0.96 | 0.001 | -0.016 | -0.016 | 0.287 | 0.58 | 0.27 | 0.44 | 0.5 | 0.61 | 0.54 |
| 9 | 14941672 | rs12348103 | 0.499 | 0.194 | 0.312 | 1.216 | -0.013 | -0.021 | -0.021 | 6.148 | 0.81 | 0.85 | 0.63 | 0.63 | 0.79 | 0.61 |

| | | | | | | | | | | | | | | | | |
|---|---|---|---|---|---|---|---|---|---|---|---|---|---|---|---|---|
| 9 | 14947472 | rs3861706 | 1.77 | -0.469 | -0.023 | 0.777 | -0.061 | 0.13 | 0.13 | 0 | 0.42 | 0.1 | 0.38 | 0.25 | 0.51 | 0.34 |
| 9 | 14949190 | rs9969815 | 0.904 | -0.472 | 0.054 | 0.775 | -0.032 | -0.001 | -0.001 | 0 | 0.7 | 0.82 | 0.59 | 0.33 | 0.68 | 0.43 |
| 9 | 14964952 | rs6415721 | 0.911 | -0.502 | 0.081 | 0.785 | -0.016 | -0.004 | -0.004 | 0.073 | 0.7 | 0.79 | 0.59 | 0.36 | 0.67 | 0.44 |
| 9 | 14989877 | rs10810310 | 2.085 | -0.573 | -0.198 | 0.732 | -0.156 | 0.043 | 0.043 | 0 | 0.42 | 0.02 | 0.39 | 0.34 | 0.51 | 0.37 |
| 9 | 14993419 | rs4548277 | 1.005 | -0.526 | 0.57 | 0.824 | 0.009 | 0.009 | 0.009 | 3.573 | 0.7 | 0.7 | 0.57 | 0.46 | 0.67 | 0.43 |
| 9 | 15005594 | rs1916309 | 0.001 | -0.435 | 0.375 | 0.853 | -0.001 | 0.007 | 0.007 | 0.323 | 0.83 | 0.8 | 0.74 | 0.76 | 0.82 | 0.67 |
| 9 | 15011825 | rs1403561 | 0.559 | -0.539 | 0.692 | 0.568 | -0.012 | -0.063 | -0.063 | 0 | 0.2 | 0.19 | 0.12 | 0 | 0.13 | 0.03 |
| 9 | 15017840 | rs16932518 | -0.906 | -0.409 | 0.604 | 0.849 | 0.063 | 0.005 | 0.005 | 0.053 | 0.09 | 0.32 | 0.18 | 0.08 | 0.07 | 0.08 |
| 9 | 15021481 | rs10511605 | 0.217 | -0.356 | -0.386 | 1.11 | -0.022 | -0.052 | -0.052 | 0.196 | 0.82 | 0.91 | 0.77 | 0.73 | 0.82 | 0.62 |
| 9 | 15047588 | rs4741446 | 0.413 | -0.378 | -0.371 | 0.952 | -0.019 | -0.045 | -0.045 | 0 | 0.81 | 0.87 | 0.72 | 0.63 | 0.79 | 0.61 |
| 9 | 15055607 | rs4445325 | -0.847 | -0.727 | -0.319 | 0.58 | 0.011 | 0.022 | 0.022 | 0 | 0.4 | 0.63 | 0.4 | 0.23 | 0.32 | 0.28 |
| 9 | 15057624 | rs7031476 | -0.95 | -0.675 | -0.297 | 0.652 | -0.032 | 0.024 | 0.024 | 0.174 | 0.16 | 0.63 | 0.18 | 0.21 | 0.14 | 0.19 |
| 9 | 15059821 | rs7040444 | -0.118 | -0.44 | -0.299 | 0.721 | -0.013 | -0.027 | -0.027 | 3.458 | 0.32 | 0.61 | 0.44 | 0.19 | 0.24 | 0.16 |
| 9 | 15060291 | rs6474884 | 0.954 | -0.516 | 0.11 | 1.112 | -0.006 | 0.012 | 0.012 | 0 | 0.63 | 0.57 | 0.8 | 0.57 | 0.7 | 0.49 |
| 9 | 15066819 | rs4146765 | 0.672 | -0.427 | -0.089 | 1.203 | 0 | 0.009 | 0.009 | 0.582 | 0.74 | 0.69 | 0.73 | 0.55 | 0.75 | 0.54 |
| 9 | 15066888 | rs4146764 | 0.59 | -0.397 | -0.081 | 1.132 | 0.001 | -0.006 | -0.006 | 0 | 0.75 | 0.8 | 0.8 | 0.68 | 0.83 | 0.61 |
| 9 | 15067831 | rs7849283 | -1.381 | -0.142 | -0.004 | 0.867 | 0.015 | 0.016 | 0.016 | 0 | 0.12 | 0.2 | 0.07 | 0.16 | 0.12 | 0.22 |
| 9 | 15075185 | rs7859597 | -1.042 | -0.292 | -0.281 | 0.909 | -0.001 | 0.006 | 0.006 | 0.137 | 0.28 | 0.27 | 0.41 | 0.54 | 0.26 | 0.44 |
| 9 | 15076859 | rs10756644 | 0.148 | -0.467 | -1.405 | 0.234 | 0.009 | 0.085 | 0.085 | 3.417 | 0.53 | 0.26 | 0.49 | 0.79 | 0.64 | 0.48 |
| 9 | 15078013 | rs10961867 | -0.101 | -0.59 | -1.36 | 0.153 | -0.013 | 0.007 | 0.007 | 0 | 0.19 | 0.19 | 0.14 | 0.28 | 0.11 | 0.16 |
| 9 | 15078522 | rs1936718 | -0.494 | -0.63 | -0.891 | 0.111 | -0.016 | -0.119 | -0.119 | 2.764 | 0.63 | 0.48 | 0.76 | 0.98 | 0.77 | 0.73 |
| 9 | 15078990 | rs10511608 | -0.99 | -0.895 | -0.538 | -0.106 | -0.017 | -0.064 | -0.064 | 0 | 0.79 | 0.71 | 0.83 | 0.99 | 0.86 | 0.83 |
| 9 | 15079847 | rs10961870 | -0.316 | -0.468 | -0.884 | 0.363 | -0.053 | 0.037 | 0.037 | 0 | 0.19 | 0.37 | 0.14 | 0.28 | 0.1 | 0.15 |
| 9 | 15081438 | rs10738386 | 0.326 | -0.253 | -0.85 | 0.214 | -0.002 | 0.041 | 0.041 | 1.997 | 0.51 | 0.36 | 0.43 | 0.73 | 0.59 | 0.43 |
| 9 | 15082933 | rs16932587 | -0.437 | 0.133 | -0.434 | 1.151 | -0.041 | 0.023 | 0.023 | 0 | 0.09 | 0.16 | 0.04 | 0.14 | 0.01 | 0.09 |
| 9 | 15084766 | rs4741452 | 0.756 | 0.88 | -0.811 | 1.149 | 0.001 | 0.013 | 0.013 | 0 | 0.89 | 0.85 | 0.73 | 0.75 | 0.84 | 0.73 |
| 9 | 15085796 | rs4364728 | -0.65 | 0.995 | -1.126 | 0.481 | 0 | 0.01 | 0.01 | 0 | 0.36 | 0.65 | 0.31 | 0.72 | 0.32 | 0.29 |
| 9 | 15086021 | rs10961876 | -0.349 | 1.455 | -1.931 | 0.613 | -0.064 | -0.047 | -0.047 | 2.845 | 0.24 | 0.06 | 0.25 | 0.19 | 0.31 | 0.18 |
| 9 | 15086653 | rs10124119 | 0.124 | 0.919 | -1.98 | -0.24 | 0.004 | 0.06 | 0.06 | 2.647 | 0.25 | 0.16 | 0.2 | 0.04 | 0.27 | 0.21 |
| 9 | 15087382 | rs7047010 | 0.414 | 0.871 | -1.622 | -0.942 | -0.003 | -0.102 | -0.102 | 0.039 | 0.27 | 0.45 | 0.21 | 0.08 | 0.25 | 0.16 |
| 9 | 15090559 | rs4741454 | 0.739 | 1.127 | -2.148 | -0.682 | -0.067 | 0.054 | 0.054 | 0.333 | 0.33 | 0.07 | 0.46 | 0.72 | 0.39 | 0.5 |
| 9 | 15103918 | rs10481554 | 0.209 | 0.853 | -1.257 | -0.082 | 0 | -0.011 | -0.011 | 0 | 0.33 | 0.35 | 0.37 | 0.25 | 0.36 | 0.33 |
| 9 | 15111167 | rs7871542 | 0.299 | 0.594 | -1.08 | -0.228 | 0.002 | 0.007 | 0.007 | 0 | 0.58 | 0.37 | 0.6 | 0.72 | 0.62 | 0.61 |
| 9 | 15115717 | rs1780155 | 0.092 | 0.724 | -0.825 | -0.231 | 0.003 | -0.037 | -0.037 | 0 | 0.34 | 0.43 | 0.38 | 0.26 | 0.36 | 0.39 |
| 9 | 15116228 | rs2800605 | 0.121 | 0.72 | -0.891 | -0.233 | 0.001 | -0.015 | -0.015 | 0.583 | 0.33 | 0.36 | 0.37 | 0.26 | 0.36 | 0.39 |
| 9 | 15122969 | rs2761137 | 0.834 | 0.644 | -0.852 | 0.112 | -0.003 | 0.068 | 0.068 | 0.344 | 0.85 | 0.51 | 0.8 | 0.74 | 0.9 | 0.81 |
| 9 | 15124737 | rs1323594 | 0.756 | 0.58 | -0.813 | -0.162 | -0.042 | -0.059 | -0.059 | 0 | 0.34 | 0.07 | 0.33 | 0.28 | 0.33 | 0.24 |
| 9 | 15127943 | rs1780160 | 0.151 | 0.465 | -0.542 | -0.347 | 0.013 | -0.007 | -0.007 | 0 | 0.92 | 0.55 | 0.96 | 0.97 | 0.98 | 0.98 |
| 9 | 15131912 | rs1780157 | -0.093 | 0.536 | -0.704 | -0.11 | 0 | 0.001 | 0.001 | 0.469 | 0.54 | 0.4 | 0.52 | 0.67 | 0.58 | 0.61 |
| 9 | 15133642 | rs7860820 | 1.085 | 0.694 | -0.384 | 0.284 | -0.005 | -0.005 | -0.005 | 0 | 0.06 | 0.05 | 0.11 | 0.12 | 0.08 | 0.15 |
| 9 | 15138120 | rs2761112 | 0.025 | 0.636 | -0.126 | 0.253 | 0.012 | 0.034 | 0.034 | 0.109 | 0.09 | 0.32 | 0.13 | 0.11 | 0.07 | 0.1 |
| 9 | 15141249 | rs1590349 | 0.114 | 0.643 | -0.253 | 0.207 | 0.002 | 0.021 | 0.021 | 0 | 0.29 | 0.3 | 0.29 | 0.62 | 0.29 | 0.41 |
| 9 | 15145385 | rs2800575 | -0.641 | 0.713 | -0.094 | -0.276 | 0.002 | -0.01 | -0.01 | 0 | 0.34 | 0.65 | 0.19 | 0.19 | 0.35 | 0.35 |
| 9 | 15153943 | rs12551441 | 0.035 | 0.829 | 0.252 | -0.337 | 0.004 | 0.008 | 0.008 | 0 | 0.3 | 0.25 | 0.19 | 0.16 | 0.22 | 0.18 |
| 9 | 15157746 | rs2096366 | -0.881 | 0.99 | 0.202 | -0.596 | -0.016 | 0.057 | 0.057 | 0.967 | 0.68 | 0.8 | 0.7 | 0.89 | 0.63 | 0.76 |
| 9 | 15159378 | rs9298726 | 0.467 | 0.587 | 0.107 | -1.203 | 0.008 | 0.009 | 0.009 | 0.103 | 0.28 | 0.22 | 0.18 | 0.17 | 0.23 | 0.24 |
| 9 | 15159928 | rs932712 | -0.884 | 0.59 | 0.069 | -1.235 | -0.011 | -0.132 | -0.132 | 0.229 | 0.68 | 0.5 | 0.71 | 0.96 | 0.7 | 0.82 |
| 9 | 15169211 | rs10961905 | 0.763 | 0.497 | -0.322 | -1.162 | 0.019 | 0.026 | 0.026 | 0 | 0.24 | 0.12 | 0.15 | 0.15 | 0.19 | 0.16 |
| 9 | 15170552 | rs1746331 | -1.289 | 0.795 | -0.738 | -1.299 | -0.008 | 0.083 | 0.083 | 4.069 | 0.74 | 0.86 | 0.75 | 0.93 | 0.72 | 0.81 |
| 9 | 15175798 | rs971835 | -0.408 | 0.848 | -0.676 | -1.403 | 0 | -0.055 | -0.055 | 0 | 0.67 | 0.62 | 0.69 | 0.95 | 0.66 | 0.76 |
| 9 | 15179840 | rs10738389 | -0.88 | 0.595 | -0.652 | -1.609 | -0.012 | -0.072 | -0.072 | 0 | 0.71 | 0.61 | 0.72 | 0.95 | 0.76 | 0.8 |
| 9 | 15186030 | rs2761134 | -0.285 | 0.881 | -0.853 | -1.64 | -0.002 | -0.007 | -0.007 | 0 | 0.11 | 0.11 | 0.1 | 0.02 | 0.17 | 0.07 |
| 9 | 15188106 | rs1407977 | -0.545 | 0.721 | -0.914 | -1.687 | -0.046 | 0.139 | 0.139 | 0.842 | 0.65 | 0.85 | 0.71 | 0.92 | 0.6 | 0.73 |
| 9 | 15188743 | rs10810360 | 1.006 | 0.623 | -1.048 | -1.592 | -0.034 | -0.021 | -0.021 | 0 | 0.32 | 0.07 | 0.33 | 0.24 | 0.31 | 0.28 |
| 9 | 15191599 | rs12554907 | 0.583 | 0.609 | -0.865 | -1.495 | -0.018 | 0.066 | 0.066 | 0 | 0.28 | 0.46 | 0.34 | 0.64 | 0.27 | 0.45 |
| 9 | 15193713 | rs7860410 | -1.213 | 0.481 | -1.01 | -1.716 | 0.007 | 0.053 | 0.053 | 1.487 | 0.74 | 0.84 | 0.8 | 0.91 | 0.76 | 0.8 |
| 9 | 15199085 | rs10733291 | -0.138 | 0.604 | -0.766 | -1.675 | 0 | 0.012 | 0.012 | 0.089 | 0.63 | 0.67 | 0.69 | 0.87 | 0.62 | 0.73 |
| 9 | 15201410 | rs517928 | -0.314 | 0.732 | -0.608 | -1.696 | 0.02 | -0.056 | -0.056 | 0 | 0.14 | 0.26 | 0.11 | 0.03 | 0.16 | 0.07 |
| 9 | 15201720 | rs527705 | -0.834 | 0.689 | -0.806 | -1.709 | 0.011 | 0.063 | 0.063 | 0.04 | 0.5 | 0.81 | 0.43 | 0.71 | 0.51 | 0.49 |
| 9 | 15202417 | rs693196 | 1.057 | 0.716 | -0.386 | -1.39 | 0 | 0.002 | 0.002 | 0.143 | 0.24 | 0.29 | 0.37 | 0.19 | 0.26 | 0.33 |
| 9 | 15205014 | rs569078 | -0.464 | 0.76 | -0.409 | -1.1 | 0.009 | -0.007 | -0.007 | 0 | 0.37 | 0.71 | 0.32 | 0.68 | 0.34 | 0.4 |
| 9 | 15207485 | rs1215177 | -1.129 | 0.697 | -0.383 | -1.494 | 0.009 | -0.003 | -0.003 | 0 | 0.76 | 0.82 | 0.84 | 0.96 | 0.84 | 0.81 |
| 9 | 15212216 | rs12343442 | -0.18 | 1.236 | 1.03 | -0.725 | -0.013 | -0.064 | -0.064 | 2.572 | 0.26 | 0.34 | 0.14 | 0.1 | 0.24 | 0.16 |
| 9 | 15215002 | rs472622 | -0.333 | 1.497 | 1.46 | -0.503 | 0.053 | 0.026 | 0.026 | 0 | 0.06 | 0.29 | 0.09 | 0.03 | 0.06 | 0.07 |
| 9 | 15216292 | rs11791131 | 0.368 | 1.77 | 1.008 | -0.515 | -0.042 | 0.01 | 0.01 | 0 | 0.07 | 0 | 0.07 | 0.05 | 0.13 | 0.06 |
| 9 | 15218947 | rs7870690 | -0.874 | 1.321 | 1.475 | -0.531 | -0.021 | -0.043 | -0.043 | 0 | 0.13 | 0.43 | 0.11 | 0.04 | 0.07 | 0.09 |
| 9 | 15229622 | rs475957 | -0.436 | 1.232 | 1.378 | -0.231 | 0.069 | 0.349 | 0.349 | 0 | 0.09 | 0.32 | 0.14 | 0.59 | 0.08 | 0.19 |
| 9 | 15229766 | rs16932848 | 0.456 | 1.265 | 1.005 | -0.215 | 0.009 | 0.129 | 0.129 | 0 | 0.06 | 0.04 | 0.12 | 0.58 | 0.08 | 0.18 |
| 9 | 15235893 | rs591647 | 0.169 | 1.243 | 0.482 | -0.682 | 0.003 | -0.021 | -0.021 | 0 | 0.18 | 0.22 | 0.12 | 0.08 | 0.22 | 0.14 |

| | | | | | | | | | | | | | | | | |
|---|---|---|---|---|---|---|---|---|---|---|---|---|---|---|---|---|
| 9 | 15240033 | rs513270 | 0.304 | 1.271 | 0.359 | -0.679 | 0.001 | -0.01 | -0.01 | 0 | 0.17 | 0.19 | 0.11 | 0.08 | 0.22 | 0.13 |
| 9 | 15240454 | rs619105 | -1.962 | 1.395 | 0.093 | -0.693 | -0.028 | 0.114 | 0.114 | 0.502 | 0.46 | 0.84 | 0.41 | 0.78 | 0.51 | 0.49 |
| 9 | 15240912 | rs12377170 | -0.083 | 1.056 | 0.443 | -1.182 | -0.007 | 0.018 | 0.018 | 0 | 0.18 | 0.11 | 0.12 | 0.08 | 0.23 | 0.14 |
| 9 | 15243411 | rs1215128 | -0.973 | 1.108 | 0.235 | -1.009 | 0.024 | -0.071 | -0.071 | 0 | 0.3 | 0.57 | 0.26 | 0.15 | 0.4 | 0.28 |
| 9 | 15246652 | rs519664 | -1.375 | 0.639 | -0.026 | -0.56 | 0.006 | 0.029 | 0.029 | 1.791 | 0.25 | 0.81 | 0.22 | 0.16 | 0.21 | 0.23 |
| 9 | 15258381 | rs10961940 | -1.346 | 0.51 | 0.872 | -0.452 | 0.016 | -0.065 | -0.065 | 0.238 | 0.15 | 0.51 | 0.14 | 0.03 | 0.13 | 0.13 |
| 9 | 15259432 | rs7864179 | -1.802 | 0.506 | 1.454 | -0.502 | 0.039 | -0.012 | -0.012 | 0 | 0.1 | 0.25 | 0.12 | 0.1 | 0.12 | 0.15 |
| 9 | 15264197 | rs10961946 | -2.176 | 0.302 | 1.464 | -0.464 | -0.016 | -0.028 | -0.028 | 0 | 0.2 | 0.75 | 0.16 | 0.11 | 0.15 | 0.17 |
| 9 | 15270247 | rs10810367 | 0.454 | 0.231 | 0.589 | -0.361 | -0.032 | -0.053 | -0.053 | 0 | 0.22 | 0.03 | 0.4 | 0.22 | 0.25 | 0.27 |
| 9 | 15270875 | rs9632874 | -0.076 | -0.311 | 1.528 | -0.585 | -0.032 | -0.049 | -0.049 | 0.015 | 0.11 | 0.25 | 0.06 | 0.02 | 0.06 | 0.07 |
| 9 | 15275697 | rs7873299 | -1.768 | -0.343 | 0.889 | -0.541 | -0.015 | -0.019 | -0.019 | 0 | 0.23 | 0.76 | 0.14 | 0.09 | 0.15 | 0.19 |
| 9 | 15279214 | rs10122040 | -1.011 | -0.269 | 0.981 | -0.411 | -0.048 | -0.03 | -0.03 | 0 | 0.16 | 0.6 | 0.11 | 0.07 | 0.09 | 0.1 |
| 9 | 15284100 | rs641579 | 0.394 | -0.876 | 0.394 | -1.061 | -0.051 | -0.101 | -0.101 | 0 | 0.65 | 0.48 | 0.8 | 0.89 | 0.76 | 0.73 |
| 9 | 15286744 | rs10810368 | 0.944 | -0.257 | 0.101 | -1.134 | -0.003 | 0.002 | 0.002 | 0 | 0.39 | 0.37 | 0.48 | 0.73 | 0.42 | 0.46 |
| 9 | 15289941 | rs1122677 | 0.64 | -0.282 | 0.06 | -1.181 | -0.067 | 0.019 | 0.019 | 0.31 | 0.23 | 0.04 | 0.31 | 0.16 | 0.33 | 0.25 |
| 9 | 15290012 | rs638491 | -0.468 | -0.761 | 0.618 | -1.384 | -0.026 | -0.04 | -0.04 | 2.137 | 0.76 | 0.7 | 0.84 | 0.88 | 0.84 | 0.78 |
| 9 | 15290174 | rs7875255 | -0.413 | -1.311 | -0.107 | -1.699 | -0.055 | -0.102 | -0.102 | 2.723 | 0.22 | 0.38 | 0.1 | 0.05 | 0.14 | 0.22 |
| 9 | 15291691 | rs7022687 | 0.924 | -0.946 | -0.129 | -1.424 | -0.059 | 0.018 | 0.018 | 0.248 | 0.23 | 0.04 | 0.31 | 0.14 | 0.31 | 0.23 |
| 9 | 15293513 | rs585002 | -0.331 | -0.987 | 0.157 | -1.507 | -0.054 | -0.105 | -0.105 | 1.426 | 0.72 | 0.46 | 0.82 | 0.89 | 0.81 | 0.78 |
| 9 | 15295073 | rs10114494 | 0.886 | -0.788 | -0.084 | -1.095 | -0.028 | -0.011 | -0.011 | 0.047 | 0.24 | 0.11 | 0.32 | 0.2 | 0.3 | 0.23 |
| 9 | 15295505 | rs570474 | 0.406 | -0.818 | 0.504 | -1.112 | 0.009 | 0.061 | 0.061 | 0 | 0.49 | 0.34 | 0.5 | 0.68 | 0.5 | 0.55 |
| 9 | 15296119 | rs4741484 | 1.016 | -0.864 | 0.479 | -1.087 | -0.111 | -0.022 | -0.022 | 0 | 0.38 | 0.08 | 0.45 | 0.61 | 0.45 | 0.41 |
| 9 | 15299590 | rs11792305 | 0.53 | -1.287 | 0.554 | -1.309 | 0.003 | 0.007 | 0.007 | 0.158 | 0.5 | 0.48 | 0.55 | 0.71 | 0.55 | 0.6 |
| 9 | 15299715 | rs583204 | 0.451 | -1.25 | 0.449 | -1.405 | -0.009 | -0.031 | -0.031 | 0 | 0.13 | 0.21 | 0.08 | 0.06 | 0.12 | 0.14 |
| 9 | 15300116 | rs3866794 | -0.032 | -1.237 | 0.376 | -1.397 | -0.029 | -0.034 | -0.034 | 2.62 | 0.12 | 0.44 | 0.07 | 0.08 | 0.07 | 0.18 |
| 9 | 15300968 | rs4008004 | 0.936 | -1.098 | 0.293 | -1.308 | -0.004 | -0.008 | -0.008 | 0 | 0.22 | 0.13 | 0.3 | 0.2 | 0.23 | 0.23 |
| 9 | 15301428 | rs2849049 | -0.181 | -1.108 | 0.238 | -1.301 | -0.003 | -0.006 | -0.006 | 0 | 0.35 | 0.41 | 0.38 | 0.26 | 0.35 | 0.37 |
| 9 | 15304782 | rs686030 | -0.788 | -0.662 | 0.047 | -0.771 | 0.007 | 0.034 | 0.034 | 13.065 | 0.85 | 0.91 | 0.89 | 0.97 | 0.87 | 0.96 |
| 9 | 15308154 | rs491322 | 0.248 | -0.271 | -0.081 | -0.869 | -0.019 | 0.043 | 0.043 | 0 | 0.59 | 0.8 | 0.53 | 0.75 | 0.59 | 0.68 |
| 9 | 15309928 | rs2245641 | 0.759 | -0.236 | 0.422 | -0.407 | -0.014 | -0.044 | -0.044 | 0 | 0.74 | 0.59 | 0.77 | 0.85 | 0.79 | 0.71 |
| 9 | 15310704 | rs2616752 | -1.393 | -0.189 | 0.348 | 0.008 | -0.006 | -0.004 | -0.004 | 0 | 0.38 | 0.47 | 0.46 | 0.33 | 0.37 | 0.46 |
| 9 | 15319412 | rs682553 | -0.158 | -0.126 | 0.421 | 0.049 | 0.022 | -0.023 | -0.023 | 0 | 0.07 | 0.15 | 0.11 | 0.02 | 0.13 | 0.03 |
| 9 | 15319506 | rs682092 | 0.574 | -0.095 | 0.117 | 0.063 | -0.032 | 0.018 | 0.018 | 0 | 0.05 | 0.01 | 0.1 | 0.02 | 0.13 | 0.03 |
| 9 | 15321542 | rs652185 | -2.151 | -0.039 | 0.085 | 0.897 | 0.009 | 0.026 | 0.026 | 0 | 0.52 | 0.68 | 0.45 | 0.63 | 0.48 | 0.69 |
| 9 | 15330631 | rs12554359 | -0.09 | -0.181 | 0.095 | 0.814 | 0.034 | 0.038 | 0.038 | 1.115 | 0.24 | 0.08 | 0.22 | 0.13 | 0.19 | 0.16 |
| 9 | 15333006 | rs10124733 | -0.519 | 0.031 | 0.31 | 1.029 | 0.003 | 0.001 | 0.001 | 0.005 | 0.34 | 0.33 | 0.26 | 0.28 | 0.26 | 0.43 |
| 9 | 15333788 | rs2798748 | -0.566 | -0.197 | -0.318 | 1.169 | -0.002 | -0.004 | -0.004 | 0 | 0.39 | 0.62 | 0.37 | 0.29 | 0.39 | 0.45 |
| 9 | 15335853 | rs688254 | -1.579 | 0.076 | -0.348 | 1.345 | -0.008 | 0.009 | 0.009 | 0.055 | 0.54 | 0.64 | 0.46 | 0.62 | 0.53 | 0.72 |
| 9 | 15335914 | rs16933006 | -1.77 | -0.012 | -0.379 | 1.553 | -0.005 | -0.013 | -0.013 | 0.019 | 0.13 | 0.07 | 0.09 | 0.33 | 0.14 | 0.26 |
| 9 | 15335996 | rs10511609 | -0.378 | -0.284 | -0.475 | 1.575 | -0.013 | -0.029 | -0.029 | 0.619 | 0.06 | 0 | 0.05 | 0.26 | 0.05 | 0.08 |
| 9 | 15337857 | rs12554331 | -0.934 | -0.462 | 0.088 | 1.484 | -0.05 | 0.081 | 0.081 | 0 | 0.26 | 0.47 | 0.17 | 0.46 | 0.2 | 0.39 |
| 9 | 15337952 | rs540457 | 0.977 | -0.529 | -0.027 | 0.879 | -0.042 | -0.022 | -0.022 | 0 | 0.76 | 0.39 | 0.78 | 0.78 | 0.84 | 0.73 |
| 9 | 15339722 | rs10961970 | 0.363 | 0.192 | -0.791 | 0.93 | 0.012 | -0.006 | -0.006 | 0 | 0.52 | 0.26 | 0.53 | 0.46 | 0.44 | 0.44 |
| 9 | 15340686 | rs10756657 | -0.07 | 0.404 | -0.838 | 0.84 | -0.019 | 0.005 | 0.005 | 0.429 | 0.17 | 0.06 | 0.18 | 0.15 | 0.21 | 0.14 |
| 9 | 15342231 | rs518519 | -0.254 | 0.454 | 0.235 | 1.104 | 0.021 | 0.081 | 0.081 | 0 | 0.9 | 0.73 | 0.83 | 0.77 | 0.86 | 0.8 |
| 9 | 15343613 | rs1215129 | 1.076 | 0.131 | 0.355 | 1.153 | 0.003 | 0.001 | 0.001 | 0.268 | 0.56 | 0.46 | 0.63 | 0.46 | 0.63 | 0.46 |
| 9 | 15347022 | rs498404 | 0.411 | -0.015 | 0.599 | 1.182 | -0.005 | 0.111 | 0.111 | 0 | 0.81 | 0.59 | 0.8 | 0.63 | 0.83 | 0.73 |
| 9 | 15347518 | rs1215134 | 1.234 | 0.018 | 0.494 | 1.127 | 0.03 | 0.022 | 0.022 | 0.23 | 0.54 | 0.34 | 0.62 | 0.45 | 0.63 | 0.46 |
| 9 | 15349934 | rs473315 | -1.05 | 0.556 | 0.898 | 0.41 | 0 | -0.003 | -0.003 | 0 | 0.42 | 0.52 | 0.27 | 0.21 | 0.43 | 0.49 |
| 9 | 15350556 | rs7870630 | -0.323 | 0.506 | 1.309 | 0.381 | -0.017 | 0 | 0 | 0 | 0.25 | 0.36 | 0.15 | 0.16 | 0.19 | 0.28 |
| 9 | 15351505 | rs681093 | 0.819 | 0.581 | 1.103 | 0.354 | -0.043 | 0.055 | 0.055 | 4.14 | 0.14 | 0.05 | 0.09 | 0.04 | 0.22 | 0.21 |
| 9 | 15355480 | rs632317 | -0.36 | 0.119 | 0.952 | 0.256 | 0.003 | -0.001 | -0.001 | 0.073 | 0.26 | 0.27 | 0.34 | 0.28 | 0.3 | 0.28 |
| 9 | 15357041 | rs519761 | 0.76 | 0.568 | 0.717 | 0.347 | -0.054 | 0.068 | 0.068 | 3.425 | 0.16 | 0.05 | 0.11 | 0.04 | 0.25 | 0.22 |
| 9 | 15362057 | rs1942476 | 0.987 | 0.329 | 0.669 | 0.421 | -0.015 | -0.011 | -0.011 | 0 | 0.53 | 0.32 | 0.5 | 0.56 | 0.5 | 0.46 |
| 9 | 15367759 | rs10810383 | 1.694 | 0.596 | 0.302 | 0.446 | 0.025 | -0.047 | -0.047 | 2.455 | 0.36 | 0.14 | 0.39 | 0.42 | 0.29 | 0.26 |
| 9 | 15370575 | rs1105191 | 1.087 | 0.68 | 0.527 | 0.575 | 0.01 | -0.01 | -0.01 | 0 | 0.57 | 0.32 | 0.68 | 0.58 | 0.61 | 0.49 |
| 9 | 15370832 | rs10961996 | 1.554 | 0.701 | 0.359 | 0.539 | 0.026 | -0.015 | -0.015 | 0 | 0.39 | 0.19 | 0.41 | 0.41 | 0.33 | 0.26 |
| 9 | 15371675 | rs10961997 | 0.01 | 0.609 | 0.188 | 0.349 | -0.024 | 0.028 | 0.028 | 0 | 0.17 | 0.08 | 0.12 | 0.14 | 0.21 | 0.2 |
| 9 | 15378449 | rs2804982 | 0.105 | 0.499 | 0.701 | 0.446 | -0.028 | 0.087 | 0.087 | 0.016 | 0.79 | 0.56 | 0.85 | 0.71 | 0.85 | 0.7 |
| 9 | 15380657 | rs1891847 | -0.589 | 0.506 | 0.214 | 0.435 | -0.01 | -0.016 | -0.016 | 0.277 | 0.35 | 0.2 | 0.24 | 0.31 | 0.35 | 0.49 |
| 9 | 15382572 | rs2798750 | 0.483 | 0.776 | -0.36 | 0.452 | -0.093 | -0.025 | -0.025 | 0.095 | 0.18 | 0.01 | 0.32 | 0.19 | 0.28 | 0.22 |
| 9 | 15384135 | rs4741495 | 0.816 | 0.431 | 0.124 | 0.59 | 0.008 | -0.034 | -0.034 | 0 | 0.4 | 0.13 | 0.36 | 0.39 | 0.35 | 0.27 |
| 9 | 15386433 | rs12375629 | -1.008 | 0.959 | 0.672 | 0.808 | 0.008 | 0.052 | 0.052 | 0 | 0.23 | 0.35 | 0.25 | 0.31 | 0.24 | 0.31 |
| 9 | 15386490 | rs10962014 | -0.779 | 0.873 | 0.42 | 0.921 | -0.001 | 0.091 | 0.091 | 0.517 | 0.21 | 0.56 | 0.23 | 0.31 | 0.21 | 0.31 |
| 9 | 15387949 | rs10962015 | -0.543 | 1.375 | -0.317 | 0.522 | -0.05 | 0.005 | 0.005 | 0 | 0.14 | 0.04 | 0.2 | 0.14 | 0.21 | 0.2 |
| 9 | 15389610 | rs2490855 | 0.12 | 0.686 | 0.47 | 0.178 | -0.028 | -0.031 | -0.031 | 0.621 | 0.9 | 0.68 | 0.95 | 0.97 | 0.95 | 0.97 |
| 9 | 15393381 | rs2040071 | 0.94 | 0.396 | 0.505 | -0.066 | -0.049 | -0.042 | -0.042 | 0 | 0.74 | 0.54 | 0.91 | 0.85 | 0.86 | 0.68 |
| 9 | 15397969 | rs2798716 | 0.852 | 0.343 | 0.623 | -0.043 | -0.062 | 0.005 | 0.005 | 7.076 | 0.82 | 0.64 | 0.93 | 0.85 | 0.92 | 0.76 |
| 9 | 15415461 | rs6474915 | -1.845 | 0.471 | 0.768 | 0.079 | -0.004 | -0.003 | -0.003 | 0.414 | 0.34 | 0.59 | 0.28 | 0.32 | 0.29 | 0.44 |

| | | | | | | | | | | | | | | | | |
|---|---|---|---|---|---|---|---|---|---|---|---|---|---|---|---|---|
| 9 | 15420805 | rs10117968 | -2.133 | 0.18 | 0.948 | -0.168 | 0.001 | 0.008 | 0.008 | 0 | 0.41 | 0.63 | 0.25 | 0.28 | 0.36 | 0.52 |
| 9 | 15421423 | rs2175259 | -1.564 | 0.228 | 1.055 | -0.128 | -0.009 | -0.025 | -0.025 | 0.009 | 0.33 | 0.52 | 0.21 | 0.28 | 0.3 | 0.43 |
| 9 | 15429539 | rs10738395 | 1.349 | -0.036 | 0.868 | -0.292 | -0.033 | -0.008 | -0.008 | 0 | 0.71 | 0.39 | 0.89 | 0.76 | 0.87 | 0.62 |
| 9 | 15431326 | rs1123038 | 1.069 | -0.088 | 1.287 | -0.228 | -0.04 | 0.022 | 0.022 | 1.843 | 0.79 | 0.61 | 0.93 | 0.75 | 0.91 | 0.64 |
| 9 | 15443430 | rs907076 | 2.051 | -0.073 | 1.252 | -0.252 | -0.006 | -0.003 | -0.003 | 0.633 | 0.64 | 0.36 | 0.87 | 0.72 | 0.81 | 0.56 |
| 9 | 15459821 | rs3087653 | 0.636 | -0.024 | 1.145 | -0.243 | -0.005 | 0.007 | 0.007 | 0.003 | 0.07 | 0.04 | 0.02 | 0 | 0.06 | 0.06 |
| 9 | 15461506 | rs12093 | 1.521 | -0.04 | 1.186 | -0.226 | -0.028 | -0.01 | -0.01 | 0.005 | 0.73 | 0.57 | 0.91 | 0.76 | 0.85 | 0.58 |
| 9 | 15467175 | rs909323 | 0.165 | 0.053 | 1.443 | -0.304 | -0.02 | -0.036 | -0.036 | 0.917 | 0.1 | 0.32 | 0.06 | 0.03 | 0.05 | 0.02 |
| 9 | 15482976 | rs7862428 | -1.621 | 0.01 | 1.384 | -0.256 | -0.03 | -0.032 | -0.032 | 0 | 0.23 | 0.66 | 0.12 | 0.14 | 0.12 | 0.22 |
| 9 | 15483211 | rs1033056 | 2.039 | -0.024 | 1.694 | -0.189 | 0.024 | 0.026 | 0.026 | 0.054 | 0.64 | 0.28 | 0.85 | 0.75 | 0.81 | 0.6 |
| 9 | 15516375 | rs7857933 | -0.944 | 0.254 | 1.347 | -0.19 | -0.006 | -0.012 | -0.012 | 0 | 0.12 | 0.2 | 0.05 | 0.1 | 0.09 | 0.13 |
| 9 | 15517742 | rs10125713 | 0.76 | 0.237 | 1.406 | -0.227 | -0.013 | -0.041 | -0.041 | 0 | 0.08 | 0.17 | 0.05 | 0.01 | 0.05 | 0.03 |
| 9 | 15521362 | rs7856710 | -0.527 | 0.287 | 1.322 | -0.196 | -0.005 | -0.009 | -0.009 | 0.062 | 0.11 | 0.17 | 0.05 | 0.09 | 0.07 | 0.13 |
| 9 | 15524911 | rs1397213 | 1.279 | 0.246 | 1.338 | 0.421 | 0 | -0.001 | -0.001 | 0.274 | 0.74 | 0.7 | 0.79 | 0.64 | 0.84 | 0.61 |
| 9 | 15530283 | rs7871956 | 1.141 | 0.331 | 0.973 | -0.154 | -0.034 | 0.011 | 0.011 | 2.999 | 0.59 | 0.76 | 0.63 | 0.62 | 0.6 | 0.43 |
| 9 | 15533394 | rs2457637 | 0.473 | -0.207 | 0.376 | -1.122 | -0.004 | 0.006 | 0.006 | 0.804 | 0.56 | 0.59 | 0.67 | 0.81 | 0.53 | 0.53 |
| 9 | 15533612 | rs2457638 | -0.271 | -0.222 | 0.548 | -1.341 | -0.028 | -0.013 | -0.013 | 0 | 0.14 | 0.18 | 0.08 | 0.11 | 0.07 | 0.12 |
| 9 | 15536429 | rs10962059 | 0.591 | -0.107 | 0.629 | -1.309 | -0.018 | -0.053 | -0.053 | 3.577 | 0.1 | 0.32 | 0.06 | 0.01 | 0.05 | 0.03 |
| 9 | 15552399 | rs10119257 | 0.706 | -0.089 | 0.453 | -1.328 | -0.021 | -0.055 | -0.055 | 0 | 0.09 | 0.21 | 0.05 | 0 | 0.05 | 0.03 |
| 9 | 15564904 | rs2821545 | 0.435 | -0.151 | 0.355 | -1.406 | -0.001 | -0.008 | -0.008 | 0 | 0.55 | 0.56 | 0.54 | 0.81 | 0.52 | 0.57 |
| 9 | 15574853 | rs4740615 | 1.11 | -0.191 | 0.325 | -1.053 | -0.041 | 0.103 | 0.103 | 3.11 | 0.44 | 0.16 | 0.6 | 0.73 | 0.48 | 0.54 |
| 9 | 15585395 | rs192466 | 0.498 | -0.2 | 0.317 | -1.06 | 0.002 | -0.045 | -0.045 | 3.454 | 0.54 | 0.5 | 0.55 | 0.83 | 0.52 | 0.58 |
| 9 | 15591372 | rs4741510 | 1.771 | -0.76 | -0.217 | -1.264 | -0.162 | -0.1 | -0.1 | 0 | 0.34 | 0.02 | 0.44 | 0.59 | 0.45 | 0.4 |
| 9 | 15602101 | rs2055772 | 0.704 | -0.546 | 0.193 | -1.237 | 0 | 0.02 | 0.02 | 1.591 | 0.46 | 0.4 | 0.61 | 0.74 | 0.47 | 0.54 |
| 9 | 15609608 | rs3129709 | -0.856 | -0.528 | -0.046 | -1.614 | -0.025 | 0.005 | 0.005 | 5.306 | 0.87 | 0.99 | 0.96 | 0.9 | 0.9 | 0.75 |
| 9 | 15612830 | rs276438 | 1.381 | -0.547 | 0.219 | -1.486 | -0.008 | -0.026 | -0.026 | 0 | 0.08 | 0.14 | 0.04 | 0 | 0.05 | 0.03 |
| 9 | 15623462 | rs405352 | 0.116 | -0.157 | 0.687 | -1.282 | -0.002 | -0.003 | -0.003 | 0 | 0.54 | 0.49 | 0.49 | 0.57 | 0.42 | 0.45 |
| 9 | 15634326 | rs4740619 | 0.558 | -0.662 | 0.37 | -2.269 | 0 | -0.009 | -0.009 | 0 | 0.47 | 0.44 | 0.63 | 0.75 | 0.48 | 0.55 |
| 9 | 15645713 | rs12340969 | 0.655 | -0.877 | 0.165 | -2.373 | -0.002 | 0.019 | 0.019 | 0.345 | 0.44 | 0.4 | 0.61 | 0.75 | 0.47 | 0.56 |
| 9 | 15656806 | rs1553727 | -1.496 | -0.867 | 0.298 | -2.409 | -0.016 | -0.038 | -0.038 | 0.591 | 0.91 | 0.86 | 0.96 | 1 | 0.95 | 0.97 |
| 9 | 15660497 | rs10756688 | 0.487 | -0.888 | 0.271 | -2.434 | 0 | 0.007 | 0.007 | 1.476 | 0.47 | 0.44 | 0.62 | 0.75 | 0.48 | 0.55 |
| 9 | 15661859 | rs6474941 | -1.867 | -0.877 | 0.286 | -2.415 | -0.015 | -0.038 | -0.038 | 0.064 | 0.92 | 0.87 | 0.96 | 1 | 0.95 | 0.98 |
| 9 | 15681694 | rs1355171 | 0.097 | -0.891 | 0.027 | -2.429 | -0.002 | 0.003 | 0.003 | 1.818 | 0.54 | 0.48 | 0.65 | 0.74 | 0.52 | 0.58 |
| 9 | 15681901 | rs10733295 | 0.128 | -0.873 | -0.021 | -2.421 | -0.001 | -0.003 | -0.003 | 0.376 | 0.55 | 0.57 | 0.66 | 0.75 | 0.53 | 0.58 |
| 9 | 15696281 | rs2175083 | -1.311 | -0.814 | -0.023 | -2.376 | -0.016 | -0.037 | -0.037 | 0 | 0.91 | 0.85 | 0.95 | 1 | 0.95 | 0.98 |
| 9 | 15705395 | rs1396707 | 0.777 | -0.858 | -0.071 | -2.466 | -0.004 | 0.029 | 0.029 | 2.588 | 0.43 | 0.36 | 0.6 | 0.76 | 0.47 | 0.56 |
| 9 | 15715215 | rs4741534 | 0.462 | -0.832 | -0.08 | -2.475 | 0 | -0.014 | -0.014 | 0 | 0.47 | 0.47 | 0.62 | 0.76 | 0.47 | 0.56 |
| 9 | 15726057 | rs10962128 | 0.914 | -0.522 | -0.496 | -2.585 | -0.077 | 0.024 | 0.024 | 0 | 0.12 | 0.01 | 0.15 | 0.06 | 0.23 | 0.12 |
| 9 | 15735854 | rs6474957 | -1.667 | -0.635 | -0.308 | -2.553 | -0.016 | -0.038 | -0.038 | 0 | 0.92 | 0.87 | 0.96 | 1 | 0.95 | 0.98 |
| 9 | 15746302 | rs7467207 | -0.445 | -0.69 | -0.271 | -2.642 | 0 | -0.016 | -0.016 | 0 | 0.46 | 0.51 | 0.35 | 0.24 | 0.47 | 0.42 |
| 9 | 15760988 | rs6474967 | -0.95 | -0.92 | -0.684 | -2.715 | -0.031 | 0.113 | 0.113 | 0.212 | 0.51 | 0.89 | 0.38 | 0.36 | 0.45 | 0.52 |
| 9 | 15765856 | rs1592099 | 1.062 | -0.921 | -0.69 | -2.714 | -0.06 | 0.056 | 0.056 | 0 | 0.42 | 0.12 | 0.59 | 0.63 | 0.49 | 0.45 |
| 9 | 15776138 | rs1539171 | -0.469 | -0.856 | -0.29 | -2.663 | -0.001 | -0.002 | -0.002 | 0.066 | 0.46 | 0.43 | 0.33 | 0.24 | 0.46 | 0.41 |
| 9 | 15776290 | rs1970498 | 0.972 | -0.789 | -0.041 | -2.94 | 0.03 | -0.027 | -0.027 | 0 | 0.12 | 0.19 | 0.13 | 0.06 | 0.18 | 0.12 |
| 9 | 15784631 | rs1539172 | -1.035 | -0.782 | -0.073 | -2.779 | -0.002 | 0.004 | 0.004 | 1.196 | 0.54 | 0.54 | 0.37 | 0.37 | 0.47 | 0.57 |
| 9 | 15788170 | rs955193 | -1.866 | -0.782 | 0.042 | -2.758 | -0.006 | -0.026 | -0.026 | 0 | 0.92 | 0.89 | 0.96 | 1 | 0.95 | 0.98 |
| 9 | 15799343 | rs2105317 | 1.085 | -0.794 | 0.001 | -2.75 | 0.005 | -0.005 | -0.005 | 0 | 0.38 | 0.39 | 0.6 | 0.61 | 0.47 | 0.41 |
| 9 | 15815246 | rs9406537 | -0.687 | -0.75 | -0.05 | -2.807 | 0.005 | 0.01 | 0.01 | 1.77 | 0.51 | 0.51 | 0.33 | 0.24 | 0.44 | 0.43 |
| 9 | 15823667 | rs7019205 | -0.641 | -0.726 | -0.181 | -2.807 | 0.001 | -0.008 | -0.008 | 0.166 | 0.5 | 0.5 | 0.34 | 0.24 | 0.45 | 0.43 |
| 9 | 15830506 | rs4741545 | 1.227 | -0.587 | -0.522 | -2.759 | -0.172 | -0.029 | -0.029 | 0 | 0.37 | 0.02 | 0.59 | 0.64 | 0.5 | 0.42 |
| 9 | 15841937 | rs2066292 | -0.825 | -0.809 | -0.553 | -3.21 | 0 | -0.004 | -0.004 | 0.014 | 0.44 | 0.43 | 0.32 | 0.24 | 0.44 | 0.48 |
| 9 | 15849304 | rs7852250 | -1.03 | -0.975 | -0.258 | -3.109 | -0.023 | 0.03 | 0.03 | 0 | 0.88 | 0.58 | 0.92 | 0.87 | 0.94 | 0.85 |
| 9 | 15850604 | rs1341740 | 0.772 | -0.775 | -0.239 | -3.092 | -0.036 | 0.024 | 0.024 | 0 | 0.15 | 0 | 0.06 | 0.07 | 0.13 | 0.22 |
| 9 | 15855317 | rs1578558 | -0.807 | -0.851 | 0.184 | -2.766 | -0.061 | -0.062 | -0.062 | 0.022 | 0.76 | 0.52 | 0.85 | 0.87 | 0.85 | 0.82 |
| 9 | 15869845 | rs10810456 | 0.236 | -0.767 | 0.074 | -2.83 | -0.003 | -0.006 | -0.006 | 0 | 0.22 | 0.21 | 0.17 | 0.08 | 0.23 | 0.2 |
| 9 | 15870465 | rs770204 | -0.772 | -0.785 | 0.044 | -2.628 | -0.044 | -0.04 | -0.04 | 0.026 | 0.77 | 0.57 | 0.84 | 0.84 | 0.85 | 0.81 |
| 9 | 15872875 | rs7867018 | 1.095 | -0.852 | 0.05 | -2.626 | -0.03 | 0.006 | 0.006 | 0 | 0.09 | 0.13 | 0.04 | 0.13 | 0.01 | 0.15 |
| 9 | 15876247 | rs1169470 | -0.492 | -0.788 | -0.026 | -2.598 | -0.043 | -0.038 | -0.038 | 0.003 | 0.71 | 0.58 | 0.84 | 0.82 | 0.85 | 0.79 |
| 9 | 15889411 | rs9406556 | 1.18 | -0.424 | -0.36 | -2.451 | -0.032 | -0.018 | -0.018 | 0.625 | 0.07 | 0.01 | 0.09 | 0.08 | 0.14 | 0.08 |
| 9 | 15895539 | rs1169478 | -0.654 | -0.45 | -0.161 | -2.674 | 0.004 | -0.009 | -0.009 | 0 | 0.33 | 0.38 | 0.39 | 0.23 | 0.4 | 0.4 |
| 9 | 15896393 | rs1773062 | -1.578 | -0.599 | -0.11 | -2.81 | 0.011 | 0.002 | 0.002 | 0 | 0.8 | 0.84 | 0.88 | 0.95 | 0.86 | 0.95 |
| 9 | 15898228 | rs1341729 | 0.321 | -0.597 | 0.115 | -2.825 | -0.004 | 0.004 | 0.004 | 0.92 | 0.27 | 0.32 | 0.29 | 0.15 | 0.26 | 0.31 |
| 9 | 15900348 | rs1891210 | 0.468 | -0.635 | 0.139 | -2.833 | 0.004 | 0.018 | 0.018 | 1.4 | 0.52 | 0.39 | 0.57 | 0.8 | 0.59 | 0.64 |
| 9 | 15909450 | rs3008702 | 0.461 | -0.761 | -0.085 | -2.734 | 0.043 | 0.056 | 0.056 | 0 | 0.61 | 0.35 | 0.73 | 0.75 | 0.7 | 0.72 |
| 9 | 15922137 | rs1328285 | -1.343 | -0.317 | -0.04 | -2.488 | -0.004 | -0.031 | -0.031 | 1.342 | 0.81 | 0.81 | 0.88 | 0.95 | 0.86 | 0.95 |
| 9 | 15923770 | rs3008683 | -1.26 | -0.273 | -0.042 | -2.488 | -0.014 | -0.062 | -0.062 | 0 | 0.8 | 0.75 | 0.87 | 0.95 | 0.84 | 0.94 |
| 9 | 15925200 | rs16933805 | 1.214 | -0.409 | -0.136 | -2.445 | 0.043 | -0.036 | -0.036 | 0 | 0.08 | 0.01 | 0.03 | 0.13 | 0.01 | 0.15 |
| 9 | 15937691 | rs7469281 | 0.515 | -0.338 | -0.105 | -2.51 | 0 | -0.031 | -0.031 | 0.811 | 0.24 | 0.21 | 0.19 | 0.05 | 0.26 | 0.08 |
| 9 | 15942554 | rs12684785 | 0.874 | -0.372 | 0.061 | -2.38 | -0.035 | 0.002 | 0.002 | 0 | 0.09 | 0.16 | 0.04 | 0.12 | 0.01 | 0.13 |

| | | | | | | | | | | | | | | | | |
|---|---|---|---|---|---|---|---|---|---|---|---|---|---|---|---|---|
| 9 | 15942765 | rs2987070 | -0.679 | -0.125 | -0.022 | -2.45 | 0.016 | -0.041 | -0.041 | 0 | 0.37 | 0.53 | 0.3 | 0.23 | 0.42 | 0.43 |
| 9 | 15959700 | rs4741552 | 2.033 | -0.145 | -0.607 | -2.412 | -0.136 | -0.079 | -0.079 | 0 | 0.33 | 0.02 | 0.41 | 0.59 | 0.41 | 0.38 |
| 9 | 15967745 | rs3008751 | -0.816 | -0.351 | -0.118 | -2.496 | -0.014 | -0.003 | -0.003 | 0.312 | 0.27 | 0.41 | 0.18 | 0.3 | 0.21 | 0.38 |
| 9 | 15967923 | rs3008750 | 0.06 | -0.303 | -0.502 | -2.486 | -0.031 | 0.049 | 0.049 | 3.095 | 0.26 | 0.15 | 0.18 | 0.15 | 0.27 | 0.35 |
| 9 | 15973288 | rs10756728 | 1.956 | -0.342 | -0.701 | -2.365 | -0.155 | -0.066 | -0.066 | 0.015 | 0.34 | 0.02 | 0.55 | 0.59 | 0.43 | 0.39 |
| 9 | 15975069 | rs10756730 | 1.96 | -0.317 | -0.698 | -2.362 | -0.166 | -0.066 | -0.066 | 3.24 | 0.34 | 0.02 | 0.56 | 0.59 | 0.45 | 0.39 |
| 9 | 15986487 | rs1927699 | -1.303 | -0.451 | -0.474 | -2.43 | 0.005 | 0.071 | 0.071 | 2.596 | 0.46 | 0.76 | 0.34 | 0.36 | 0.41 | 0.56 |
| 9 | 15997006 | rs2987036 | -1.056 | -0.467 | -0.327 | -2.469 | -0.031 | -0.104 | -0.104 | 0 | 0.78 | 0.46 | 0.87 | 0.95 | 0.84 | 0.94 |
| 9 | 16003441 | rs3008731 | 0.702 | -0.527 | -0.319 | -2.532 | -0.019 | 0.189 | 0.189 | 1.142 | 0.53 | 0.21 | 0.62 | 0.79 | 0.49 | 0.62 |
| 9 | 16009322 | rs3008725 | 0.165 | -0.481 | -0.328 | -2.422 | 0.016 | -0.001 | -0.001 | 0 | 0.81 | 0.83 | 0.93 | 0.79 | 0.93 | 0.73 |
| 9 | 16012046 | rs2987081 | 1.343 | -0.48 | -0.303 | -2.447 | -0.001 | -0.004 | -0.004 | 0 | 0.56 | 0.48 | 0.68 | 0.64 | 0.59 | 0.47 |
| 9 | 16012225 | rs10962251 | -0.067 | -0.299 | -0.958 | -2.368 | 0.013 | -0.028 | -0.028 | 0 | 0.25 | 0.34 | 0.25 | 0.15 | 0.35 | 0.26 |
| 9 | 16022910 | rs16933967 | 0.921 | -0.82 | -0.91 | -2.105 | 0.043 | -0.033 | -0.033 | 0 | 0.08 | 0 | 0.03 | 0.11 | 0.01 | 0.11 |
| 9 | 16026444 | rs10756734 | 1.89 | -0.756 | -1.112 | -2.262 | -0.106 | 0.013 | 0.013 | 1.224 | 0.36 | 0.08 | 0.57 | 0.62 | 0.46 | 0.41 |
| 9 | 16026664 | rs10810517 | 1.716 | -0.042 | -0.68 | -2.337 | -0.087 | 0.144 | 0.144 | 1.617 | 0.43 | 0.08 | 0.6 | 0.71 | 0.47 | 0.51 |
| 9 | 16028693 | rs7848880 | -0.161 | -0.058 | -0.535 | -2.359 | -0.003 | 0.002 | 0.002 | 5.094 | 0.25 | 0.28 | 0.24 | 0.16 | 0.31 | 0.25 |
| 9 | 16033109 | rs1887667 | 1.254 | -0.134 | -0.75 | -2.161 | -0.039 | 0.137 | 0.137 | 2.455 | 0.44 | 0.16 | 0.61 | 0.71 | 0.47 | 0.52 |
| 9 | 16042193 | rs1410454 | 1.209 | -0.138 | -0.764 | -2.168 | -0.023 | 0.113 | 0.113 | 2.781 | 0.44 | 0.2 | 0.61 | 0.7 | 0.47 | 0.52 |
| 9 | 16046371 | rs1328297 | -0.702 | -0.539 | -1.49 | -2.115 | -0.005 | 0.005 | 0.005 | 0 | 0.91 | 0.95 | 0.97 | 0.92 | 0.95 | 0.85 |
| 9 | 16046463 | rs3008694 | 0.807 | -0.447 | -1.493 | -2.139 | 0.002 | 0.031 | 0.031 | 0 | 0.25 | 0.17 | 0.19 | 0.13 | 0.29 | 0.13 |
| 9 | 16046983 | rs10962263 | 0.594 | -0.397 | -1.486 | -2.119 | -0.001 | -0.012 | -0.012 | 0 | 0.16 | 0.18 | 0.14 | 0.08 | 0.16 | 0.18 |
| 9 | 16047293 | rs1328299 | -0.023 | -0.145 | -1.992 | -1.925 | 0.024 | 0.073 | 0.073 | 2.91 | 0.69 | 0.17 | 0.79 | 0.84 | 0.79 | 0.68 |
| 9 | 16051490 | rs10733307 | 1.069 | -0.068 | -2.062 | -2.143 | -0.154 | 0.182 | 0.182 | 2.16 | 0.43 | 0.02 | 0.6 | 0.75 | 0.49 | 0.53 |
| 9 | 16053710 | rs1556045 | -0.091 | -0.091 | -1.961 | -2.183 | 0.015 | 0.036 | 0.036 | 2.65 | 0.69 | 0.24 | 0.79 | 0.88 | 0.78 | 0.66 |
| 9 | 16053792 | rs1014308 | -0.095 | 1.117 | -1.228 | -1.38 | -0.001 | 0.053 | 0.053 | 0 | 0.26 | 0.15 | 0.19 | 0.11 | 0.3 | 0.14 |
| 9 | 16056836 | rs2987089 | -0.765 | 0.468 | -0.381 | -0.988 | -0.014 | -0.044 | -0.044 | 0 | 0.11 | 0.2 | 0.05 | 0.02 | 0.07 | 0.03 |
| 9 | 16057378 | rs3008709 | -0.497 | 0.37 | -0.252 | -0.735 | 0 | 0 | 0 | 0 | 0.4 | 0.56 | 0.43 | 0.53 | 0.45 | 0.42 |
| 9 | 16060417 | rs2987093 | -1.031 | 0.312 | -0.149 | -1.159 | -0.001 | -0.005 | -0.005 | 0 | 0.37 | 0.64 | 0.27 | 0.24 | 0.29 | 0.4 |
| 9 | 16060746 | rs2026662 | 0.421 | 0.3 | -0.689 | -1.104 | -0.107 | -0.186 | -0.186 | 0.349 | 0.25 | 0.01 | 0.41 | 0.43 | 0.33 | 0.3 |
| 9 | 16061329 | rs10114770 | -0.281 | 0.05 | 0.875 | -1.619 | -0.002 | -0.013 | -0.013 | 0.001 | 0.13 | 0.16 | 0.15 | 0.05 | 0.08 | 0.15 |
| 9 | 16061960 | rs9886829 | -0.785 | 0.03 | 0.83 | -1.626 | -0.018 | 0.037 | 0.037 | 0 | 0.14 | 0.03 | 0.23 | 0.12 | 0.14 | 0.19 |
| 9 | 16063191 | rs9298745 | -1.494 | -0.098 | 1.174 | -1.66 | -0.001 | 0.004 | 0.004 | 0 | 0.29 | 0.25 | 0.3 | 0.25 | 0.27 | 0.3 |
| 9 | 16064646 | rs17784100 | 0.223 | -0.041 | 1.214 | -1.774 | -0.001 | -0.001 | -0.001 | 0 | 0.14 | 0.15 | 0.06 | 0.12 | 0.12 | 0.12 |
| 9 | 16066169 | rs10511612 | -0.632 | -0.026 | 1.335 | -1.877 | 0.001 | -0.019 | -0.019 | 1.487 | 0.14 | 0.16 | 0.24 | 0.1 | 0.13 | 0.17 |
| 9 | 16069629 | rs6475004 | -2.228 | -0.147 | 1.525 | -1.947 | 0.008 | 0.017 | 0.017 | 0.126 | 0.4 | 0.69 | 0.33 | 0.24 | 0.28 | 0.38 |
| 9 | 16070094 | rs4961643 | -0.676 | 0.161 | 1.905 | -1.839 | 0.003 | -0.029 | -0.029 | 1.726 | 0.14 | 0.2 | 0.23 | 0.1 | 0.13 | 0.15 |
| 9 | 16073639 | rs11999147 | -1.663 | 0.308 | 2.209 | -1.8 | 0.033 | -0.059 | -0.059 | 0 | 0.32 | 0.46 | 0.34 | 0.21 | 0.33 | 0.34 |
| 9 | 16074726 | rs10962269 | -1.465 | 0.341 | 2.374 | -1.784 | 0.049 | -0.072 | -0.072 | 0 | 0.2 | 0.52 | 0.29 | 0.12 | 0.21 | 0.28 |
| 9 | 16079296 | rs1887677 | -2.344 | 0.171 | 2.187 | -1.799 | -0.012 | 0.094 | 0.094 | 0 | 0.39 | 0.82 | 0.37 | 0.23 | 0.34 | 0.38 |
| 9 | 16081108 | rs11789353 | -0.21 | 0.117 | 2.419 | -1.854 | -0.004 | -0.014 | -0.014 | 0 | 0.15 | 0.21 | 0.06 | 0.09 | 0.11 | 0.12 |
| 9 | 16082467 | rs11789951 | -0.065 | 0.118 | 2.375 | -1.854 | -0.001 | -0.007 | -0.007 | 0 | 0.14 | 0.18 | 0.06 | 0.09 | 0.11 | 0.12 |
| 9 | 16083133 | rs10511613 | -1.896 | 0.166 | 2.519 | -1.836 | 0.007 | -0.07 | -0.07 | 0 | 0.24 | 0.59 | 0.27 | 0.12 | 0.25 | 0.26 |
| 9 | 16084769 | rs10962272 | -0.637 | 0.181 | 2.381 | -1.85 | 0.001 | -0.006 | -0.006 | 0 | 0.15 | 0.16 | 0.23 | 0.09 | 0.22 | 0.18 |
| 9 | 16085262 | rs7023221 | -2.247 | 0.018 | 2.089 | -1.849 | -0.015 | 0.093 | 0.093 | 0 | 0.38 | 0.82 | 0.32 | 0.21 | 0.36 | 0.4 |
| 9 | 16092087 | rs2891006 | -1.489 | -0.075 | 2.19 | -1.771 | 0.005 | -0.014 | -0.014 | 0 | 0.29 | 0.28 | 0.25 | 0.18 | 0.34 | 0.33 |
| 9 | 16092485 | rs9695323 | -0.483 | -0.007 | 2.078 | -1.727 | 0.031 | -0.018 | -0.018 | 0 | 0.16 | 0.33 | 0.1 | 0.13 | 0.16 | 0.14 |
| 9 | 16096547 | rs943784 | -1.677 | -0.055 | 1.246 | -1.704 | -0.003 | 0.007 | 0.007 | 0 | 0.36 | 0.61 | 0.28 | 0.19 | 0.37 | 0.32 |
| 9 | 16098105 | rs12005226 | -1.198 | 0.125 | 1.074 | -1.588 | 0.034 | -0.068 | -0.068 | 1.974 | 0.3 | 0.43 | 0.25 | 0.16 | 0.34 | 0.3 |
| 9 | 16103333 | rs867780 | -0.936 | 0.162 | 0.356 | -1.644 | 0.021 | -0.063 | -0.063 | 0.793 | 0.29 | 0.61 | 0.29 | 0.1 | 0.37 | 0.32 |
| 9 | 16106801 | rs765509 | -0.55 | -0.774 | 0.058 | -1.293 | 0.04 | 0.162 | 0.162 | 0 | 0.41 | 0.85 | 0.33 | 0.02 | 0.34 | 0.27 |
| 9 | 16110623 | rs16934049 | -2.08 | -0.796 | 1.224 | 0.036 | -0.016 | 0.017 | 0.017 | 0.137 | 0.12 | 0.28 | 0.05 | 0.17 | 0.07 | 0.1 |
| 9 | 16118758 | rs7873155 | -1.983 | -0.901 | 1.586 | 0.116 | -0.006 | 0.09 | 0.09 | 0 | 0.09 | 0.4 | 0.04 | 0.12 | 0.02 | 0.07 |
| 9 | 16123596 | rs7028308 | -2.506 | -0.898 | 0.497 | 0.097 | -0.06 | 0.005 | 0.005 | 0.938 | 0.12 | 0.39 | 0.04 | 0.13 | 0.02 | 0.19 |
| 9 | 16125443 | rs7036825 | -1.611 | -0.506 | -0.242 | 0.298 | -0.014 | 0.008 | 0.008 | 0 | 0.08 | 0.13 | 0.02 | 0.14 | 0.02 | 0.08 |
| 9 | 16130937 | rs7863615 | -0.924 | -0.699 | -0.405 | 0.019 | 0.006 | 0.006 | 0.006 | 1.762 | 0.16 | 0.12 | 0.07 | 0.07 | 0.12 | 0.06 |
| 9 | 16131805 | rs12553725 | -1.247 | -1.018 | -0.735 | 0.164 | 0.056 | 0.05 | 0.05 | 1.034 | 0.45 | 0.74 | 0.43 | 0.63 | 0.34 | 0.49 |
| 9 | 16133853 | rs10962290 | -1.313 | -1.173 | -0.59 | 0.249 | 0.052 | 0.029 | 0.029 | 0.129 | 0.46 | 0.73 | 0.42 | 0.61 | 0.34 | 0.47 |
| 9 | 16135924 | rs12341724 | -0.768 | -1.295 | -0.043 | 0.022 | 0.003 | -0.028 | -0.028 | 0 | 0.19 | 0.17 | 0.07 | 0 | 0.12 | 0.13 |
| 9 | 16136054 | rs10738431 | -1.579 | -1.14 | 0.252 | 0.402 | 0.02 | 0.004 | 0.004 | 0.398 | 0.43 | 0.65 | 0.38 | 0.57 | 0.34 | 0.43 |
| 9 | 16145299 | rs10810530 | -1.045 | -1.199 | 0.423 | 0.403 | -0.024 | -0.005 | -0.005 | 0 | 0.23 | 0.5 | 0.24 | 0.25 | 0.17 | 0.24 |
| 9 | 16147687 | rs1410448 | -1.822 | -1.352 | -0.623 | 0.41 | 0.02 | 0.092 | 0.092 | 1.082 | 0.5 | 0.97 | 0.51 | 0.71 | 0.38 | 0.56 |
| 9 | 16150498 | rs7872685 | -0.838 | -0.717 | -0.398 | 0.183 | 0.003 | 0.015 | 0.015 | 0 | 0.36 | 0.33 | 0.26 | 0.27 | 0.32 | 0.42 |
| 9 | 16151623 | rs2067789 | -0.386 | -0.921 | -0.329 | 0.369 | 0.009 | -0.015 | -0.015 | 0 | 0.17 | 0.14 | 0.14 | 0.46 | 0.13 | 0.26 |
| 9 | 16151659 | rs2067790 | -1.21 | -0.154 | -0.519 | 0.151 | -0.007 | -0.056 | -0.056 | 0.005 | 0.25 | 0.17 | 0.21 | 0.48 | 0.26 | 0.3 |
| 9 | 16152308 | rs7023487 | -1.149 | 0.053 | -0.111 | 0.021 | -0.023 | 0.076 | 0.076 | 0.894 | 0.41 | 0.72 | 0.35 | 0.19 | 0.43 | 0.37 |
| 9 | 16152635 | rs7851897 | 0.19 | -0.103 | 0.826 | -0.024 | 0.044 | -0.005 | -0.005 | 0 | 0.22 | 0.39 | 0.24 | 0.19 | 0.26 | 0.3 |
| 9 | 16155473 | rs7022227 | 1.03 | -0.337 | 0.94 | 0.001 | -0.002 | -0.011 | -0.011 | 0 | 0.09 | 0.1 | 0.02 | 0 | 0.04 | 0.04 |
| 9 | 16155596 | rs729119 | -0.076 | -0.237 | 1.18 | 0.019 | 0.039 | -0.035 | -0.035 | 0.243 | 0.07 | 0.11 | 0.07 | 0 | 0.12 | 0.02 |
| 9 | 16160638 | rs10962302 | -1.79 | -0.479 | 1.629 | 0.063 | -0.004 | 0.024 | 0.024 | 0.016 | 0.29 | 0.62 | 0.24 | 0.5 | 0.25 | 0.26 |

| | | | | | | | | | | | | | | | | |
|---|---|---|---|---|---|---|---|---|---|---|---|---|---|---|---|---|
| 9 | 16161217 | rs11999148 | -0.973 | -0.543 | 1.825 | 0.017 | -0.006 | -0.125 | -0.125 | 0.588 | 0.19 | 0.43 | 0.11 | 0 | 0.17 | 0.07 |
| 9 | 16161233 | rs4961664 | -2.422 | -0.624 | 1.381 | -0.077 | 0.053 | -0.068 | -0.068 | 0 | 0.4 | 0.84 | 0.28 | 0.49 | 0.31 | 0.29 |
| 9 | 16162546 | rs10962304 | -0.47 | -0.569 | 1.29 | 0.036 | 0.018 | -0.035 | -0.035 | 2.178 | 0.18 | 0.16 | 0.14 | 0.44 | 0.12 | 0.22 |
| 9 | 16163344 | rs10962305 | -1.42 | -0.516 | 1.322 | 0.008 | 0 | 0 | 0 | 0 | 0.27 | 0.33 | 0.23 | 0.5 | 0.28 | 0.26 |
| 9 | 16168116 | rs9298746 | -1.983 | -0.523 | 1.348 | 0.086 | -0.016 | 0.021 | 0.021 | 0 | 0.36 | 0.57 | 0.27 | 0.5 | 0.32 | 0.28 |
| 9 | 16172580 | rs1328293 | 0.3 | -0.492 | 0.962 | 0.797 | 0.004 | -0.003 | -0.003 | 0 | 0.13 | 0.14 | 0.1 | 0.28 | 0.09 | 0.16 |
| 9 | 16177206 | rs12238080 | -1.283 | -0.586 | 0.987 | 0.766 | 0.01 | -0.027 | -0.027 | 0 | 0.31 | 0.24 | 0.23 | 0.48 | 0.27 | 0.23 |
| 9 | 16178112 | rs7029038 | 0.277 | -0.574 | 0.928 | 0.732 | -0.009 | -0.019 | -0.019 | 0 | 0.13 | 0.04 | 0.12 | 0.2 | 0.14 | 0.05 |
| 9 | 16185090 | rs16934192 | -0.46 | -0.422 | 1.17 | 0.934 | -0.013 | 0.106 | 0.106 | 0 | 0.18 | 0.28 | 0.18 | 0.44 | 0.14 | 0.22 |
| 9 | 16185621 | rs10113994 | 0.474 | -0.455 | 0.968 | 0.895 | -0.008 | -0.001 | -0.001 | 0.022 | 0.13 | 0.03 | 0.12 | 0.11 | 0.15 | 0.04 |
| 9 | 16186417 | rs16934196 | -0.367 | -0.45 | 1.139 | 0.885 | -0.012 | 0.076 | 0.076 | 0 | 0.17 | 0.21 | 0.15 | 0.51 | 0.13 | 0.21 |
| 9 | 16186705 | rs1999472 | -2.514 | -0.071 | 0.506 | 0.837 | -0.084 | 0.074 | 0.074 | 0 | 0.62 | 0.98 | 0.62 | 0.78 | 0.61 | 0.61 |
| 9 | 16189987 | rs7852086 | -1.338 | 0.002 | 1.434 | 1.049 | -0.023 | 0.056 | 0.056 | 0 | 0.31 | 0.43 | 0.24 | 0.47 | 0.28 | 0.25 |
| 9 | 16190403 | rs10962321 | -0.37 | 0 | 1.154 | 1.048 | 0.004 | -0.009 | -0.009 | 1.984 | 0.17 | 0.15 | 0.15 | 0.47 | 0.14 | 0.2 |
| 9 | 16191258 | rs4961665 | -1.946 | -0.028 | 1.386 | 1.053 | 0.001 | -0.003 | -0.003 | 0 | 0.34 | 0.66 | 0.31 | 0.53 | 0.29 | 0.27 |
| 9 | 16191512 | rs16934202 | -0.761 | -0.002 | 1.144 | 1.18 | -0.011 | 0.034 | 0.034 | 0 | 0.19 | 0.27 | 0.12 | 0.29 | 0.14 | 0.19 |
| 9 | 16195381 | rs16934205 | -0.781 | -0.012 | 0.987 | 1.145 | -0.009 | 0.033 | 0.033 | 0 | 0.19 | 0.27 | 0.17 | 0.35 | 0.14 | 0.21 |
| 9 | 16196717 | rs12002003 | -0.749 | 0.007 | 0.972 | 1.05 | -0.003 | 0.024 | 0.024 | 0 | 0.19 | 0.25 | 0.21 | 0.56 | 0.14 | 0.22 |
| 9 | 16201094 | rs7856829 | -0.521 | -0.094 | 1.109 | 1.233 | -0.033 | 0.091 | 0.091 | 0 | 0.17 | 0.32 | 0.14 | 0.41 | 0.09 | 0.24 |
| 9 | 16201392 | rs7872577 | 0.132 | 0.363 | 1.059 | 1.223 | -0.014 | 0.003 | 0.003 | 0 | 0.13 | 0.05 | 0.11 | 0.12 | 0.13 | 0.05 |
| 9 | 16201612 | rs16934229 | -1.314 | 0.317 | 1.341 | 1.177 | 0.09 | 0.042 | 0.042 | 0 | 0.05 | 0.33 | 0.04 | 0 | 0.04 | 0.03 |
| 9 | 16202988 | rs7034501 | 0.153 | 0.326 | 1.048 | 1.201 | 0.002 | 0.006 | 0.006 | 0 | 0.13 | 0.11 | 0.11 | 0.11 | 0.13 | 0.04 |
| 9 | 16205165 | rs2151061 | -2.351 | 0.856 | 0.84 | 1.288 | 0.019 | -0.072 | -0.072 | 0 | 0.41 | 0.85 | 0.33 | 0.56 | 0.32 | 0.37 |
| 9 | 16206138 | rs10122810 | 0.173 | 0.852 | 0.899 | 1.245 | 0.001 | 0.004 | 0.004 | 0 | 0.13 | 0.11 | 0.11 | 0.1 | 0.13 | 0.04 |
| 9 | 16206695 | rs4961669 | -1.166 | 0.832 | 1.049 | 1.322 | -0.045 | 0.072 | 0.072 | 0 | 0.23 | 0.52 | 0.14 | 0.34 | 0.15 | 0.29 |
| 9 | 16207279 | rs1332488 | -0.776 | 0.872 | 1.121 | 1.374 | -0.06 | 0.121 | 0.121 | 0 | 0.2 | 0.49 | 0.12 | 0.34 | 0.11 | 0.26 |
| 9 | 16207913 | rs12350437 | -0.405 | 1.074 | 1.01 | 1.216 | 0.001 | -0.004 | -0.004 | 0 | 0.16 | 0.16 | 0.14 | 0.12 | 0.16 | 0.05 |
| 9 | 16209181 | rs1115245 | -1.51 | 0.805 | 0.95 | 1.239 | 0.015 | -0.017 | -0.017 | 0 | 0.31 | 0.7 | 0.27 | 0.57 | 0.21 | 0.33 |
| 9 | 16210163 | rs1412539 | -1.483 | 0.946 | 0.846 | 1.085 | 0.005 | -0.072 | -0.072 | 0 | 0.3 | 0.18 | 0.21 | 0.48 | 0.25 | 0.46 |
| 9 | 16210295 | rs1412538 | -0.465 | 1.027 | 0.73 | 1.091 | 0.009 | 0 | 0 | 0 | 0.84 | 0.79 | 0.85 | 0.86 | 0.82 | 0.84 |
| 9 | 16210330 | rs1412537 | -1.186 | 0.228 | 1.486 | 1.356 | 0.003 | -0.008 | -0.008 | 0 | 0.24 | 0.22 | 0.18 | 0.47 | 0.2 | 0.45 |
| 9 | 16210610 | rs10962333 | -1.053 | 0.268 | 1.292 | 1.516 | -0.01 | 0.022 | 0.022 | 0 | 0.06 | 0.04 | 0.08 | 0.36 | 0.06 | 0.07 |
| 9 | 16211137 | rs7044238 | -1.962 | 0.347 | 1.312 | 1.532 | 0.004 | -0.005 | -0.005 | 0 | 0.39 | 0.7 | 0.4 | 0.63 | 0.37 | 0.48 |
| 9 | 16211712 | rs12554091 | -0.351 | 0.499 | 0.756 | 1.614 | 0.133 | 0.126 | 0.126 | 0 | 0.06 | 0.21 | 0.17 | 0.17 | 0.15 | 0.02 |
| 9 | 16212766 | rs10511616 | -0.619 | 0.398 | 0.118 | 1.233 | -0.019 | 0.071 | 0.071 | 0 | 0.22 | 0.05 | 0.14 | 0.11 | 0.2 | 0.38 |
| 9 | 16212992 | rs10125444 | -1.252 | 0.306 | 0.411 | 1.248 | -0.001 | 0 | 0 | 0 | 0.43 | 0.44 | 0.4 | 0.64 | 0.43 | 0.49 |
| 9 | 16215744 | rs16934264 | -1.752 | -0.429 | 1.061 | 1.461 | -0.054 | 0.159 | 0.159 | 1.351 | 0.13 | 0.53 | 0.08 | 0.34 | 0.05 | 0.24 |
| 9 | 16216102 | rs17725257 | -1.229 | -0.192 | 0.633 | 1.103 | -0.015 | 0.041 | 0.041 | 0 | 0.08 | 0.03 | 0.05 | 0.01 | 0.08 | 0.14 |
| 9 | 16216437 | rs1412536 | -0.82 | -0.413 | 0.891 | 1.055 | 0.002 | 0.004 | 0.004 | 0 | 0.24 | 0.21 | 0.2 | 0.21 | 0.2 | 0.28 |
| 9 | 16217587 | rs7040148 | -2.233 | -0.019 | 1.664 | 1.558 | -0.011 | 0.171 | 0.171 | 0.674 | 0.13 | 0.35 | 0.12 | 0.48 | 0.1 | 0.19 |
| 9 | 16217690 | rs7023845 | -1.635 | 0.006 | 1.674 | 1.59 | 0.001 | 0.026 | 0.026 | 1.006 | 0.08 | 0.25 | 0.05 | 0.08 | 0.08 | 0.13 |
| 9 | 16218382 | rs16934273 | -0.049 | -0.027 | 1.399 | 1.498 | -0.007 | -0.012 | -0.012 | 0 | 0.11 | 0.16 | 0.07 | 0.11 | 0.07 | 0.2 |
| 9 | 16221343 | rs12004126 | -2.156 | -0.016 | 1.603 | 1.258 | 0.01 | 0.035 | 0.035 | 0 | 0.37 | 0.73 | 0.25 | 0.25 | 0.29 | 0.53 |
| 9 | 16221771 | rs12004220 | -2.249 | -0.107 | 1.632 | 1.326 | 0.045 | -0.064 | -0.064 | 0 | 0.37 | 0.85 | 0.28 | 0.59 | 0.24 | 0.58 |
| 9 | 16221932 | rs11793155 | -0.527 | 0.063 | 1.565 | 1.27 | 0.001 | 0.059 | 0.059 | 0 | 0.16 | 0.1 | 0.04 | 0.01 | 0.1 | 0.19 |
| 9 | 16222474 | rs7038757 | -2.363 | 0.132 | 1.757 | 1.369 | 0.002 | -0.01 | -0.01 | 1.281 | 0.42 | 0.78 | 0.35 | 0.64 | 0.36 | 0.6 |
| 9 | 16230184 | rs16934286 | -0.554 | -0.097 | 1.804 | 1.322 | -0.002 | -0.016 | -0.016 | 0.913 | 0.15 | 0.17 | 0.11 | 0.11 | 0.13 | 0.26 |
| 9 | 16230349 | rs7861430 | -2.578 | -0.021 | 1.969 | 1.508 | 0.01 | 0.022 | 0.022 | 0 | 0.45 | 0.75 | 0.36 | 0.68 | 0.37 | 0.64 |
| 9 | 16231174 | rs9407727 | -1.181 | -0.129 | 1.908 | 1.362 | -0.006 | 0.033 | 0.033 | 0 | 0.18 | 0.23 | 0.16 | 0.47 | 0.14 | 0.3 |
| 9 | 16231861 | rs9407728 | -0.722 | -0.085 | 2.045 | 1.032 | -0.03 | -0.051 | -0.051 | 0.38 | 0.16 | 0.38 | 0.1 | 0.11 | 0.09 | 0.23 |
| 9 | 16233085 | rs10810534 | -1.888 | -0.03 | 1.58 | 1.028 | -0.013 | 0.013 | 0.013 | 0 | 0.4 | 0.55 | 0.26 | 0.52 | 0.31 | 0.59 |
| 9 | 16234356 | rs1576840 | -2.195 | -0.012 | 0.988 | 0.674 | 0.025 | 0.011 | 0.011 | 0.043 | 0.45 | 0.82 | 0.39 | 0.63 | 0.35 | 0.62 |
| 9 | 16236676 | rs4961681 | -0.572 | -0.093 | 0.81 | 0.494 | -0.02 | -0.052 | -0.052 | 0 | 0.15 | 0.35 | 0.1 | 0.09 | 0.09 | 0.2 |
| 9 | 16237614 | rs10810537 | -1.285 | 0.142 | 0.277 | 0.781 | -0.008 | 0.019 | 0.019 | 0 | 0.29 | 0.43 | 0.27 | 0.56 | 0.23 | 0.42 |
| 9 | 16241198 | rs16934302 | -0.817 | 0.732 | -0.158 | 0.17 | -0.005 | -0.002 | -0.002 | 0 | 0.23 | 0.21 | 0.19 | 0.11 | 0.25 | 0.3 |
| 9 | 16242088 | rs7875247 | -0.605 | 0.349 | -0.263 | 0.448 | 0 | -0.004 | -0.004 | 0 | 0.33 | 0.67 | 0.36 | 0.41 | 0.33 | 0.23 |
| 9 | 16243361 | rs7864095 | -0.712 | -0.235 | 0.529 | -0.388 | -0.042 | -0.065 | -0.065 | 0.02 | 0.17 | 0.4 | 0.12 | 0.06 | 0.1 | 0.12 |
| 9 | 16243514 | rs7850689 | 0.331 | -0.76 | 0.8 | -0.961 | -0.02 | -0.046 | -0.046 | 0 | 0.85 | 0.67 | 0.86 | 0.94 | 0.89 | 0.88 |
| 9 | 16244120 | rs1412535 | 0.229 | -0.472 | 0.322 | -2.267 | 0.001 | -0.02 | -0.02 | 0.295 | 0.6 | 0.59 | 0.58 | 0.95 | 0.6 | 0.61 |
| 9 | 16244199 | rs2021322 | 0.387 | -0.429 | 0.352 | -2.283 | 0.002 | -0.017 | -0.017 | 0.422 | 0.58 | 0.59 | 0.56 | 0.95 | 0.59 | 0.61 |
| 9 | 16245170 | rs10738433 | 0.417 | -0.326 | 0.377 | -2.354 | 0.001 | -0.031 | -0.031 | 0 | 0.58 | 0.59 | 0.56 | 0.95 | 0.56 | 0.61 |
| 9 | 16245716 | rs2151065 | 0.74 | -0.535 | 0.36 | -2.513 | -0.003 | 0.23 | 0.23 | 0 | 0.51 | 0.21 | 0.5 | 0.93 | 0.51 | 0.54 |
| 9 | 16247466 | rs1571907 | 0.275 | -0.456 | -0.043 | -2.9 | 0.002 | 0.003 | 0.003 | 0 | 0.14 | 0.12 | 0.08 | 0.03 | 0.11 | 0.14 |
| 9 | 16251695 | rs2095074 | 0.987 | -0.883 | 0.147 | -3.267 | 0 | -0.082 | -0.082 | 0 | 0.56 | 0.44 | 0.71 | 0.98 | 0.6 | 0.61 |
| 9 | 16252410 | rs885386 | -0.167 | -1.115 | -0.175 | -3.258 | -0.005 | -0.041 | -0.041 | 0 | 0.22 | 0.29 | 0.1 | 0.01 | 0.15 | 0.27 |
| 9 | 16252807 | rs12683780 | 0.869 | -0.57 | -1.173 | -3.341 | -0.048 | 0.07 | 0.07 | 0.455 | 0.34 | 0.04 | 0.25 | 0.19 | 0.35 | 0.3 |
| 9 | 16253904 | rs7027156 | -0.246 | -0.753 | -0.827 | -3.339 | -0.015 | -0.101 | -0.101 | 0 | 0.79 | 0.42 | 0.7 | 0.97 | 0.79 | 0.86 |
| 9 | 16255704 | rs3898525 | -1.95 | -0.705 | -0.795 | -3.278 | 0.003 | 0.018 | 0.018 | 0 | 0.88 | 0.94 | 0.91 | 0.99 | 0.87 | 0.94 |
| 9 | 16256275 | rs10810541 | 2.021 | -0.582 | -1.281 | -3.281 | -0.047 | -0.035 | -0.035 | 0.114 | 0.17 | 0.08 | 0.32 | 0.76 | 0.28 | 0.32 |

| | | | | | | | | | | | | | | | | |
|---|---|---|---|---|---|---|---|---|---|---|---|---|---|---|---|---|
| 9 | 16256964 | rs7040587 | -0.339 | -0.864 | -0.818 | -3.217 | 0.002 | 0.002 | 0.002 | 0.01 | 0.3 | 0.32 | 0.17 | 0.03 | 0.25 | 0.28 |
| 9 | 16258118 | rs1888952 | 1.134 | -0.843 | -1.042 | -3.202 | 0.012 | 0.277 | 0.277 | 0 | 0.53 | 0.14 | 0.56 | 0.94 | 0.58 | 0.64 |
| 9 | 16260837 | rs1888951 | -1.526 | -0.943 | -0.698 | -2.717 | 0.002 | -0.01 | -0.01 | 0.867 | 0.88 | 0.93 | 0.91 | 1 | 0.88 | 0.94 |
| 9 | 16261940 | rs9406600 | -0.162 | -0.868 | -1.219 | -2.21 | -0.011 | -0.024 | -0.024 | 0 | 0.18 | 0.24 | 0.07 | 0.03 | 0.07 | 0.2 |
| 9 | 16263887 | rs9298751 | -0.259 | -0.796 | -1.442 | -2.272 | 0.017 | 0.019 | 0.019 | 0 | 0.24 | 0.22 | 0.17 | 0.12 | 0.16 | 0.18 |
| 9 | 16265063 | rs12005048 | -0.258 | -0.605 | -1.93 | -2.521 | 0.024 | 0.081 | 0.081 | 1.818 | 0.32 | 0.16 | 0.21 | 0.13 | 0.26 | 0.18 |
| 9 | 16266732 | rs7857514 | 0.467 | -0.631 | -1.79 | -2.21 | 0.003 | 0.007 | 0.007 | 0 | 0.12 | 0.09 | 0.2 | 0.01 | 0.1 | 0.06 |
| 9 | 16267557 | rs7033852 | -0.836 | -0.97 | -2.119 | -1.895 | 0.09 | 0.135 | 0.135 | 0.743 | 0.43 | 0.83 | 0.43 | 0.17 | 0.3 | 0.35 |
| 9 | 16271779 | rs7856337 | -0.378 | -0.255 | -2.12 | -0.826 | 0.016 | 0.086 | 0.086 | 0.8 | 0.46 | 0.82 | 0.46 | 0.65 | 0.45 | 0.52 |
| 9 | 16273781 | rs4335227 | -0.828 | 0.065 | -2.27 | -0.582 | -0.071 | 0.087 | 0.087 | 0.098 | 0.55 | 0.93 | 0.64 | 0.65 | 0.5 | 0.55 |
| 9 | 16275107 | rs6475020 | -0.251 | -0.777 | -1.841 | -0.926 | 0.104 | 0.142 | 0.142 | 1.469 | 0.4 | 0.79 | 0.38 | 0.1 | 0.25 | 0.35 |
| 9 | 16277713 | rs9406606 | -0.12 | -0.901 | -1.588 | -1.155 | -0.016 | 0.011 | 0.011 | 0 | 0.4 | 0.25 | 0.43 | 0.59 | 0.46 | 0.38 |
| 9 | 16278130 | rs1332478 | -0.372 | -1.15 | -1.103 | -1.315 | -0.001 | 0.015 | 0.015 | 0 | 0.49 | 0.56 | 0.57 | 0.63 | 0.49 | 0.46 |
| 9 | 16280194 | rs1105177 | -1.314 | -0.907 | -0.923 | -1.523 | 0.003 | -0.001 | -0.001 | 0 | 0.76 | 0.79 | 0.84 | 0.65 | 0.75 | 0.75 |
| 9 | 16283279 | rs7026127 | 0.679 | -0.659 | -0.625 | -1.431 | -0.005 | -0.018 | -0.018 | 2.82 | 0.18 | 0.2 | 0.14 | 0.07 | 0.15 | 0.17 |
| 9 | 16284231 | rs7858291 | -0.151 | -0.93 | -0.791 | -1.585 | 0.017 | 0.057 | 0.057 | 0 | 0.49 | 0.33 | 0.64 | 0.59 | 0.57 | 0.53 |
| 9 | 16286587 | rs10962366 | 0.832 | -0.937 | -0.802 | -1.292 | 0.026 | 0.055 | 0.055 | 0 | 0.25 | 0.38 | 0.29 | 0.47 | 0.32 | 0.22 |
| 9 | 16287839 | rs9406607 | -1.087 | -1.048 | -0.685 | -1.395 | 0.001 | 0.009 | 0.009 | 0 | 0.64 | 0.74 | 0.7 | 0.6 | 0.68 | 0.69 |
| 9 | 16287981 | rs9407735 | -0.332 | -1.123 | -0.724 | -1.439 | 0.002 | 0.056 | 0.056 | 0 | 0.5 | 0.71 | 0.5 | 0.54 | 0.54 | 0.48 |
| 9 | 16289428 | rs6475025 | -0.209 | -1.12 | -0.66 | -1.399 | 0.01 | 0.016 | 0.016 | 0 | 0.4 | 0.32 | 0.42 | 0.12 | 0.35 | 0.44 |
| 9 | 16289910 | rs7850784 | 1.068 | -0.897 | -0.443 | -1.253 | 0.049 | 0.101 | 0.101 | 0 | 0.24 | 0.4 | 0.29 | 0.46 | 0.32 | 0.22 |
| 9 | 16291478 | rs9406609 | 1.232 | -1.08 | -0.589 | -1.303 | -0.024 | -0.015 | -0.015 | 0.447 | 0.14 | 0.23 | 0.18 | 0.07 | 0.11 | 0.13 |
| 9 | 16291556 | rs9407736 | 0.041 | -1.163 | -0.609 | -1.447 | -0.008 | -0.026 | -0.026 | 0 | 0.29 | 0.35 | 0.4 | 0.11 | 0.27 | 0.3 |
| 9 | 16292669 | rs1360549 | -0.167 | -0.835 | -0.568 | -1.462 | 0.019 | 0.04 | 0.04 | 0 | 0.5 | 0.69 | 0.47 | 0.55 | 0.48 | 0.47 |
| 9 | 16296851 | rs9406613 | -0.786 | -1.515 | -0.654 | -1.247 | -0.008 | 0.023 | 0.023 | 0 | 0.66 | 0.85 | 0.8 | 0.64 | 0.69 | 0.7 |
| 9 | 16299066 | rs9407746 | 0.217 | -1.47 | -0.557 | -1.415 | 0.022 | 0.057 | 0.057 | 0 | 0.48 | 0.33 | 0.65 | 0.62 | 0.59 | 0.51 |
| 9 | 16303495 | rs7036684 | 0.774 | -1.211 | -0.367 | -1.269 | -0.005 | -0.034 | -0.034 | 0.029 | 0.13 | 0.11 | 0.05 | 0 | 0.11 | 0.11 |
| 9 | 16305615 | rs10962371 | -0.044 | -1.096 | -0.408 | -1.214 | 0.004 | -0.028 | -0.028 | 0 | 0.09 | 0.18 | 0.09 | 0.02 | 0.09 | 0.04 |
| 9 | 16306523 | rs7018975 | -0.667 | -1.534 | -0.55 | -1.109 | 0.004 | -0.014 | -0.014 | 0 | 0.71 | 0.79 | 0.71 | 0.65 | 0.72 | 0.79 |
| 9 | 16306543 | rs6475034 | -0.06 | -1.83 | -0.493 | -1.435 | -0.01 | -0.09 | -0.09 | 1.52 | 0.32 | 0.41 | 0.27 | 0.07 | 0.28 | 0.33 |
| 9 | 16308000 | rs7043435 | -0.797 | -1.877 | -0.326 | -1.081 | -0.001 | 0 | 0 | 0 | 0.72 | 0.7 | 0.7 | 0.64 | 0.72 | 0.82 |
| 9 | 16308805 | rs12376143 | 0.81 | -2.048 | -0.434 | -1.151 | 0.004 | 0.005 | 0.005 | 0 | 0.4 | 0.41 | 0.34 | 0.49 | 0.44 | 0.45 |
| 9 | 16309527 | rs9406618 | 0.994 | -1.996 | -0.304 | -0.988 | -0.023 | -0.012 | -0.012 | 1.627 | 0.13 | 0.21 | 0.15 | 0.09 | 0.11 | 0.11 |
| 9 | 16310723 | rs9407751 | 0.596 | -2.053 | -0.307 | -1.003 | -0.017 | -0.013 | -0.013 | 0 | 0.14 | 0.23 | 0.15 | 0.1 | 0.13 | 0.16 |
| 9 | 16313409 | rs6475036 | -0.298 | -2.318 | -0.234 | -1.072 | -0.025 | -0.041 | -0.041 | 0 | 0.32 | 0.53 | 0.38 | 0.2 | 0.28 | 0.42 |
| 9 | 16314909 | rs7867771 | -0.169 | -2.495 | -0.504 | -1.088 | -0.009 | 0 | 0 | 0 | 0.64 | 0.59 | 0.7 | 0.62 | 0.69 | 0.7 |
| 9 | 16316899 | rs7034935 | -1.459 | -2.361 | -0.726 | -0.961 | -0.04 | -0.071 | -0.071 | 0.879 | 0.84 | 0.96 | 0.88 | 0.69 | 0.81 | 0.86 |
| 9 | 16318055 | rs7853960 | 0.187 | -2.896 | -0.435 | -1.217 | -0.085 | -0.046 | -0.046 | 0 | 0.23 | 0.57 | 0.17 | 0.13 | 0.11 | 0.17 |
| 9 | 16318314 | rs7026479 | -0.207 | -2.582 | -0.685 | -1.28 | -0.011 | 0.013 | 0.013 | 0 | 0.68 | 0.61 | 0.66 | 0.57 | 0.71 | 0.71 |
| 9 | 16318869 | rs12552591 | 0.709 | -2.629 | -1.43 | -1.238 | -0.032 | 0.032 | 0.032 | 0 | 0.2 | 0.02 | 0.22 | 0.11 | 0.18 | 0.35 |
| 9 | 16321008 | rs914656 | 1.618 | -2.65 | -1.603 | -1.366 | -0.194 | -0.167 | -0.167 | 0 | 0.32 | 0.01 | 0.31 | 0.42 | 0.44 | 0.28 |
| 9 | 16321578 | rs16934439 | 0.496 | -2.498 | -1.053 | -1.464 | -0.001 | -0.027 | -0.027 | 0 | 0.2 | 0.18 | 0.12 | 0.02 | 0.12 | 0.1 |
| 9 | 16324236 | rs12686344 | 1.713 | -2.436 | -1.021 | -1.041 | -0.07 | -0.058 | -0.058 | 2.402 | 0.29 | 0.12 | 0.29 | 0.39 | 0.4 | 0.26 |
| 9 | 16324261 | rs7857417 | -0.726 | -2.697 | -0.808 | -1.199 | -0.005 | -0.039 | -0.039 | 0 | 0.45 | 0.53 | 0.38 | 0.16 | 0.34 | 0.46 |
| 9 | 16324374 | rs7872913 | -0.565 | -2.294 | -0.42 | -0.837 | -0.005 | 0.034 | 0.034 | 0 | 0.74 | 0.67 | 0.67 | 0.55 | 0.74 | 0.72 |
| 9 | 16326380 | rs12002852 | -0.008 | -2.135 | -0.942 | -0.963 | -0.008 | -0.035 | -0.035 | 1.248 | 0.1 | 0.17 | 0.12 | 0.01 | 0.06 | 0.16 |
| 9 | 16328103 | rs11788833 | 1.033 | -2.291 | -1.229 | -1.099 | -0.008 | 0.079 | 0.079 | 0 | 0.14 | 0.03 | 0.1 | 0.02 | 0.14 | 0.1 |
| 9 | 16328605 | rs4961697 | 0.529 | -2.558 | -1.098 | -1.079 | 0.009 | -0.005 | -0.005 | 0 | 0.57 | 0.4 | 0.63 | 0.5 | 0.6 | 0.61 |
| 9 | 16328688 | rs12347671 | 1.32 | -2.487 | -1.616 | -0.974 | -0.007 | -0.048 | -0.048 | 0 | 0.43 | 0.17 | 0.51 | 0.49 | 0.54 | 0.45 |
| 9 | 16329537 | rs1888955 | 0.501 | -2.466 | -1.544 | -0.968 | 0.074 | -0.033 | -0.033 | 0.935 | 0.57 | 0.19 | 0.64 | 0.51 | 0.69 | 0.55 |
| 9 | 16331304 | rs9407756 | -0.471 | -1.873 | -1.302 | -0.61 | 0.094 | -0.025 | -0.025 | 0 | 0.49 | 0.81 | 0.44 | 0.52 | 0.38 | 0.54 |
| 9 | 16334412 | rs7019421 | 0.42 | -1.335 | -0.805 | -0.279 | -0.016 | -0.032 | -0.032 | 0.125 | 0.4 | 0.2 | 0.43 | 0.46 | 0.5 | 0.43 |
| 9 | 16334761 | rs12683609 | 0.427 | -1.174 | -1.065 | -0.199 | -0.018 | -0.036 | -0.036 | 0 | 0.4 | 0.25 | 0.44 | 0.46 | 0.5 | 0.43 |
| 9 | 16336250 | rs10810557 | 0.261 | -1.217 | -0.623 | -0.104 | 0.031 | 0.006 | 0.006 | 0 | 0.36 | 0.54 | 0.35 | 0.59 | 0.43 | 0.44 |
| 9 | 16339369 | rs4961699 | 0.118 | -1.244 | -1.211 | -0.065 | -0.061 | -0.005 | -0.005 | 0 | 0.47 | 0.12 | 0.46 | 0.53 | 0.5 | 0.48 |
| 9 | 16339512 | rs4961701 | -0.017 | -1.383 | -1.309 | -0.006 | 0.046 | 0.03 | 0.03 | 0 | 0.17 | 0.05 | 0.19 | 0.1 | 0.11 | 0.17 |
| 9 | 16340357 | rs4961702 | 0.482 | -1.256 | -1.034 | 0.006 | -0.002 | 0.005 | 0.005 | 0 | 0.77 | 0.28 | 0.79 | 0.62 | 0.78 | 0.78 |
| 9 | 16342065 | rs11794667 | 1.001 | -1.408 | -1.202 | -0.144 | -0.057 | 0.194 | 0.194 | 0 | 0.32 | 0.02 | 0.28 | 0.05 | 0.3 | 0.21 |
| 9 | 16342994 | rs13298313 | 0.416 | -0.469 | -0.809 | 0.553 | 0.043 | 0.095 | 0.095 | 0 | 0.46 | 0.16 | 0.57 | 0.28 | 0.4 | 0.44 |
| 9 | 16343135 | rs7035494 | 0.514 | -0.422 | -0.638 | 0.03 | -0.005 | 0.035 | 0.035 | 0 | 0.79 | 0.33 | 0.8 | 0.63 | 0.8 | 0.78 |
| 9 | 16345371 | rs10962386 | 0.065 | -0.705 | -0.544 | 0.058 | 0.041 | 0.194 | 0.194 | 0 | 0.47 | 0.19 | 0.48 | 0.16 | 0.42 | 0.4 |
| 9 | 16348639 | rs12380074 | 0.344 | -0.476 | -0.555 | 0.312 | 0.041 | 0.162 | 0.162 | 0 | 0.39 | 0.13 | 0.4 | 0.14 | 0.32 | 0.34 |
| 9 | 16349135 | rs7030722 | -0.858 | -0.289 | -0.456 | 0.519 | -0.067 | -0.086 | -0.086 | 0.145 | 0.58 | 0.88 | 0.57 | 0.56 | 0.46 | 0.66 |
| 9 | 16349953 | rs4961706 | 0.197 | -0.304 | -0.253 | 0.516 | 0.023 | 0.042 | 0.042 | 0 | 0.45 | 0.36 | 0.44 | 0.19 | 0.34 | 0.47 |
| 9 | 16350160 | rs4961707 | -0.555 | -0.275 | -0.293 | 0.504 | -0.01 | 0.169 | 0.169 | 0 | 0.15 | 0.53 | 0.16 | 0.39 | 0.11 | 0.2 |
| 9 | 16350474 | rs4961708 | -0.765 | -0.102 | -0.288 | 0.632 | -0.114 | -0.138 | -0.138 | 0.395 | 0.59 | 0.96 | 0.58 | 0.56 | 0.46 | 0.66 |
| 9 | 16353534 | rs4526456 | -0.604 | -0.325 | 0.159 | 0.865 | -0.008 | 0.194 | 0.194 | 1.242 | 0.13 | 0.56 | 0.13 | 0.37 | 0.09 | 0.19 |
| 9 | 16354862 | rs4961479 | -0.355 | 0.437 | -0.626 | 0.654 | -0.061 | -0.075 | -0.075 | 0 | 0.6 | 0.84 | 0.61 | 0.58 | 0.5 | 0.69 |
| 9 | 16354904 | rs4961480 | -0.149 | 0.384 | -0.202 | 0.654 | -0.005 | -0.004 | -0.004 | 0 | 0.57 | 0.58 | 0.6 | 0.59 | 0.52 | 0.68 |

| | | | | | | | | | | | | | | | |
|---|---|---|---|---|---|---|---|---|---|---|---|---|---|---|---|
| 9 | 16355626 | rs7861586 | 0.338 | -0.163 | -0.357 | 0.127 | -0.006 | -0.012 | -0.012 | 0 | 0.41 | 0.4 | 0.36 | 0.24 | 0.28 | 0.52 |
| 9 | 16355870 | rs6475044 | -0.321 | 0.43 | -1.16 | 0.606 | -0.09 | -0.077 | -0.077 | 0 | 0.58 | 0.93 | 0.68 | 0.61 | 0.49 | 0.73 |
| 9 | 16358278 | rs16934512 | 0.209 | 0.483 | -1.187 | 0.627 | -0.1 | -0.102 | -0.102 | 0.277 | 0.38 | 0.04 | 0.28 | 0.38 | 0.42 | 0.28 |
| 9 | 16360889 | rs4961485 | -0.16 | 0.551 | -0.22 | 0.886 | 0.075 | 0.255 | 0.255 | 0 | 0.07 | 0.34 | 0.18 | 0.24 | 0.05 | 0.2 |
| 9 | 16360934 | rs4961712 | -0.181 | 0.479 | -0.567 | 0.342 | -0.01 | 0.059 | 0.059 | 0 | 0.47 | 0.72 | 0.48 | 0.63 | 0.47 | 0.44 |
| 9 | 16363460 | rs9407760 | -0.389 | 0.395 | -0.704 | 0.294 | -0.01 | 0.095 | 0.095 | 0 | 0.52 | 0.75 | 0.51 | 0.65 | 0.52 | 0.44 |
| 9 | 16364351 | rs7019169 | 0.087 | 0.145 | -1.03 | 0.054 | 0.031 | 0.018 | 0.018 | 0.008 | 0.53 | 0.71 | 0.53 | 0.4 | 0.37 | 0.59 |
| 9 | 16364612 | rs9407761 | 0.395 | 0.047 | -0.554 | -0.181 | 0.028 | 0.058 | 0.058 | 0.678 | 0.48 | 0.35 | 0.44 | 0.22 | 0.38 | 0.38 |
| 9 | 16364727 | rs12000507 | -0.193 | 0.007 | 0.021 | 0.046 | 0.018 | 0.024 | 0.024 | 0 | 0.06 | 0.2 | 0.04 | 0.07 | 0.03 | 0.13 |
| 9 | 16364925 | rs10962397 | 0.671 | 0.285 | -0.439 | -0.153 | -0.079 | -0.124 | -0.124 | 0.042 | 0.24 | 0.06 | 0.32 | 0.48 | 0.35 | 0.2 |
| 9 | 16366266 | rs10511618 | 0.233 | -0.762 | -0.241 | -0.636 | -0.024 | -0.115 | -0.115 | 0 | 0.2 | 0.37 | 0.16 | 0 | 0.15 | 0.13 |
| 9 | 16368732 | rs7864648 | 0.449 | -0.776 | -0.604 | -0.632 | 0.01 | 0.012 | 0.012 | 0.152 | 0.42 | 0.33 | 0.26 | 0.32 | 0.31 | 0.48 |
| 9 | 16369067 | rs10123982 | -0.776 | -0.872 | -0.509 | -0.717 | -0.008 | -0.009 | -0.009 | 0 | 0.32 | 0.37 | 0.28 | 0.21 | 0.25 | 0.36 |
| 9 | 16373545 | rs10123044 | -0.169 | -0.755 | -0.421 | -0.695 | -0.028 | -0.084 | -0.084 | 0.312 | 0.23 | 0.47 | 0.23 | 0.15 | 0.18 | 0.19 |
| 9 | 16375403 | rs9298759 | -0.991 | -0.752 | -0.448 | -0.525 | -0.015 | 0.011 | 0.011 | 0 | 0.33 | 0.47 | 0.36 | 0.38 | 0.27 | 0.42 |
| 9 | 16376437 | rs9298760 | -0.61 | -0.702 | -0.42 | -0.599 | -0.053 | 0.004 | 0.004 | 0.242 | 0.63 | 0.77 | 0.47 | 0.36 | 0.49 | 0.62 |
| 9 | 16377360 | rs9407764 | 0.185 | -0.685 | -0.514 | -0.065 | 0.001 | -0.001 | -0.001 | 0 | 0.5 | 0.46 | 0.38 | 0.49 | 0.4 | 0.53 |
| 9 | 16384212 | rs4620378 | -0.39 | -0.428 | -0.377 | 0.067 | 0.006 | 0.017 | 0.017 | 0.069 | 0.65 | 0.63 | 0.66 | 0.53 | 0.59 | 0.67 |
| 9 | 16388447 | rs4612476 | -1.051 | -0.114 | -0.955 | 0.131 | -0.12 | -0.131 | -0.131 | 0.111 | 0.74 | 0.97 | 0.68 | 0.66 | 0.64 | 0.76 |
| 9 | 16388491 | rs4495533 | 0.484 | -0.889 | -0.604 | -0.484 | 0.001 | -0.002 | -0.002 | 1.817 | 0.45 | 0.38 | 0.49 | 0.53 | 0.43 | 0.52 |
| 9 | 16390697 | rs7033140 | -0.207 | -1 | -0.26 | -0.624 | -0.027 | -0.104 | -0.104 | 0.397 | 0.29 | 0.57 | 0.19 | 0.11 | 0.2 | 0.24 |
| 9 | 16391294 | rs4442246 | 0.218 | -1.048 | -0.412 | -0.579 | 0.02 | 0.013 | 0.013 | 0.38 | 0.5 | 0.34 | 0.49 | 0.56 | 0.45 | 0.52 |
| 9 | 16392810 | rs7041706 | -0.6 | -0.983 | -0.319 | -0.691 | -0.024 | -0.12 | -0.12 | 0 | 0.32 | 0.57 | 0.21 | 0.12 | 0.24 | 0.25 |
| 9 | 16395766 | rs7023798 | -0.53 | -0.836 | 0.335 | -0.138 | 0.057 | -0.068 | -0.068 | 0 | 0.11 | 0.32 | 0.13 | 0 | 0.13 | 0.08 |
| 9 | 16397614 | rs10962417 | 0.852 | -1.002 | -0.099 | 0.01 | 0 | 0.001 | 0.001 | 0 | 0.68 | 0.68 | 0.56 | 0.59 | 0.67 | 0.59 |
| 9 | 16398267 | rs4287030 | 0.696 | -1.048 | -0.06 | 0.197 | 0 | 0.008 | 0.008 | 0 | 0.75 | 0.74 | 0.62 | 0.57 | 0.76 | 0.6 |
| 9 | 16398279 | rs7036631 | 0.313 | -0.746 | 0.139 | -0.027 | -0.002 | -0.022 | -0.022 | 0 | 0.28 | 0.26 | 0.15 | 0.13 | 0.21 | 0.25 |
| 9 | 16399927 | rs17797890 | -0.519 | -0.121 | 0.546 | 0.617 | 0.024 | -0.028 | -0.028 | 0 | 0.05 | 0.11 | 0.05 | 0 | 0.08 | 0.02 |
| 9 | 16402878 | rs2209867 | -1.58 | 0.037 | 0.445 | 1.201 | 0.004 | 0.004 | 0.004 | 0 | 0.18 | 0.17 | 0.36 | 0.36 | 0.28 | 0.31 |
| 9 | 16404794 | rs9407773 | -0.217 | -0.304 | 0.825 | 1.327 | -0.013 | 0.031 | 0.031 | 0 | 0.1 | 0.22 | 0.15 | 0.15 | 0.07 | 0.14 |
| 9 | 16405573 | rs9406646 | -1.335 | -0.728 | 1.165 | 1.385 | -0.064 | 0.014 | 0.014 | 0 | 0.25 | 0.55 | 0.22 | 0.3 | 0.14 | 0.22 |
| 9 | 16406897 | rs2182901 | -1.875 | -0.702 | 1.08 | 1.315 | 0.011 | 0.026 | 0.026 | 0.611 | 0.29 | 0.36 | 0.49 | 0.52 | 0.33 | 0.55 |
| 9 | 16407821 | rs9406647 | -1.238 | -0.75 | 1.009 | 1.313 | -0.001 | 0.003 | 0.003 | 0.544 | 0.22 | 0.22 | 0.42 | 0.5 | 0.26 | 0.53 |
| 9 | 16408042 | rs7025750 | -0.155 | -0.784 | 1.094 | 1.323 | -0.051 | 0.003 | 0.003 | 0 | 0.16 | 0.33 | 0.06 | 0.15 | 0.07 | 0.08 |
| 9 | 16408826 | rs717267 | -2.537 | -0.839 | 0.985 | 1.272 | -0.044 | -0.044 | -0.044 | 0 | 0.46 | 0.95 | 0.55 | 0.66 | 0.4 | 0.63 |
| 9 | 16411463 | rs7853747 | -2.536 | -0.841 | 1.136 | 1.295 | -0.005 | 0.01 | 0.01 | 0 | 0.46 | 0.91 | 0.56 | 0.68 | 0.4 | 0.63 |
| 9 | 16414756 | rs16934621 | -0.357 | -0.906 | 1.373 | 1.236 | -0.06 | -0.017 | -0.017 | 0 | 0.16 | 0.36 | 0.07 | 0.14 | 0.07 | 0.08 |
| 9 | 16415459 | rs2296864 | -0.357 | -0.906 | 1.373 | 1.236 | -0.06 | -0.017 | -0.017 | 0 | 0.16 | 0.35 | 0.07 | 0.14 | 0.07 | 0.08 |
| 9 | 16423164 | rs10511621 | -0.926 | -0.98 | 1.1 | 1.22 | -0.068 | -0.031 | -0.031 | 0.005 | 0.2 | 0.59 | 0.09 | 0.14 | 0.08 | 0.1 |
| 9 | 16424985 | rs10511619 | -1.178 | -0.84 | 1.044 | 1.363 | -0.011 | -0.01 | -0.01 | 0 | 0.18 | 0.15 | 0.37 | 0.4 | 0.28 | 0.32 |
| 9 | 16426951 | rs16934644 | 0.517 | -0.894 | 0.902 | 1.284 | 0.008 | 0.016 | 0.016 | 0 | 0.09 | 0.05 | 0.05 | 0.01 | 0.07 | 0.05 |
| 9 | 16430599 | rs7859226 | -1.978 | -0.957 | 0.942 | 1.342 | -0.001 | 0.005 | 0.005 | 0.551 | 0.33 | 0.35 | 0.47 | 0.53 | 0.33 | 0.58 |
| 9 | 16432118 | rs6475053 | -2.468 | -0.936 | 1.022 | 1.256 | 0.003 | -0.038 | -0.038 | 0 | 0.4 | 0.74 | 0.47 | 0.55 | 0.36 | 0.52 |
| 9 | 16435714 | rs3739715 | -0.059 | -0.674 | 1.147 | 1.267 | -0.011 | -0.018 | -0.018 | 0 | 0.11 | 0.16 | 0.05 | 0.03 | 0.07 | 0.05 |
| 9 | 16449788 | rs12338207 | -2.22 | -0.337 | 1.357 | 1.635 | 0.002 | 0.003 | 0.003 | 0.047 | 0.34 | 0.39 | 0.5 | 0.58 | 0.37 | 0.6 |
| 9 | 16450274 | rs10810564 | -0.624 | -0.343 | 1.352 | 1.58 | -0.013 | 0.011 | 0.011 | 0.014 | 0.11 | 0.21 | 0.1 | 0.15 | 0.07 | 0.24 |
| 9 | 16456759 | rs7864652 | -1.606 | -0.319 | 1.539 | 1.419 | 0.024 | -0.07 | -0.07 | 0 | 0.11 | 0.34 | 0.08 | 0 | 0.09 | 0.04 |
| 9 | 16458889 | rs10115759 | -1.338 | -0.447 | 1.755 | 1.554 | -0.03 | 0.07 | 0.07 | 0.366 | 0.14 | 0.37 | 0.11 | 0.27 | 0.07 | 0.28 |
| 9 | 16459175 | rs1887620 | -1.647 | -0.239 | 1.395 | 1.704 | 0.017 | 0.035 | 0.035 | 0 | 0.23 | 0.29 | 0.39 | 0.38 | 0.28 | 0.33 |
| 9 | 16464980 | rs10733310 | -1.955 | -0.059 | 1.084 | 1.354 | 0.002 | 0 | 0 | 0.509 | 0.35 | 0.37 | 0.52 | 0.61 | 0.41 | 0.6 |
| 9 | 16473118 | rs2274557 | -2.191 | -0.458 | 0.894 | 1.02 | 0.007 | -0.026 | -0.026 | 0 | 0.28 | 0.36 | 0.27 | 0.16 | 0.27 | 0.3 |
| 9 | 16476144 | rs7046884 | -0.867 | -0.492 | 0.979 | 0.977 | -0.014 | -0.06 | -0.06 | 0 | 0.14 | 0.22 | 0.07 | 0 | 0.07 | 0.04 |
| 9 | 16480296 | rs7868285 | -2.035 | -0.624 | 1.3 | 1.559 | -0.023 | 0.109 | 0.109 | 0.795 | 0.18 | 0.61 | 0.27 | 0.31 | 0.12 | 0.2 |
| 9 | 16482083 | rs4961719 | -0.183 | -0.486 | 1.113 | 1.7 | -0.043 | -0.044 | -0.044 | 3.045 | 0.08 | 0 | 0.19 | 0.26 | 0.13 | 0.12 |
| 9 | 16484731 | rs997693 | -1.395 | -0.506 | 1.168 | 1.576 | -0.034 | -0.026 | -0.026 | 0 | 0.13 | 0.06 | 0.2 | 0.16 | 0.2 | 0.28 |
| 9 | 16484870 | rs997694 | -2.589 | -0.586 | 1.303 | 1.561 | -0.003 | -0.014 | -0.014 | 0.345 | 0.35 | 0.55 | 0.38 | 0.24 | 0.32 | 0.36 |
| 9 | 16486142 | rs943750 | -0.792 | -0.767 | 1.679 | 1.299 | -0.014 | 0.016 | 0.016 | 0 | 0.1 | 0.29 | 0.11 | 0.07 | 0.06 | 0.07 |
| 9 | 16488167 | rs7872124 | -1.59 | -0.756 | 1.308 | 1.256 | 0.045 | 0.052 | 0.052 | 0 | 0.05 | 0.33 | 0.05 | 0.03 | 0 | 0.16 |
| 9 | 16489981 | rs1927633 | -1.629 | -0.634 | 0.958 | 1.323 | -0.026 | 0.034 | 0.034 | 0 | 0.31 | 0.54 | 0.36 | 0.38 | 0.25 | 0.33 |
| 9 | 16492516 | rs12056994 | -1.354 | 0.128 | 0.964 | 1.738 | 0.007 | 0.039 | 0.039 | 0.457 | 0.11 | 0.14 | 0.2 | 0.29 | 0.14 | 0.14 |
| 9 | 16500789 | rs1927631 | -1.587 | 0.248 | 1.102 | 1.393 | 0.034 | 0.056 | 0.056 | 0.01 | 0.11 | 0.17 | 0.15 | 0.26 | 0.16 | 0.17 |
| 9 | 16505902 | rs7043485 | -2.366 | -0.072 | 1.451 | 1.28 | 0.009 | 0.04 | 0.04 | 0 | 0.22 | 0.66 | 0.24 | 0.32 | 0.18 | 0.4 |
| 9 | 16507737 | rs7849139 | -2.297 | -0.248 | 1.516 | 1.279 | -0.052 | 0.085 | 0.085 | 0.046 | 0.16 | 0.62 | 0.17 | 0.27 | 0.05 | 0.29 |
| 9 | 16520681 | rs10962472 | -0.555 | 0.897 | 1.479 | 1.549 | -0.025 | -0.018 | -0.018 | 0 | 0.08 | 0.04 | 0.21 | 0.24 | 0.15 | 0.24 |
| 9 | 16520958 | rs4961492 | -1.449 | 1.223 | 1.422 | 1.496 | -0.064 | -0.051 | -0.051 | 0 | 0.13 | 0.04 | 0.27 | 0.23 | 0.28 | 0.35 |
| 9 | 16525841 | rs7031782 | -1.001 | 1.052 | 1.76 | 1.313 | 0.031 | 0.007 | 0.007 | 0.463 | 0.07 | 0.33 | 0.03 | 0 | 0.01 | 0.06 |
| 9 | 16526910 | rs16934786 | -1.429 | 0.924 | 1.36 | 1.36 | -0.07 | -0.07 | -0.07 | 1.968 | 0.16 | 0.03 | 0.16 | 0.2 | 0.21 | 0.25 |
| 9 | 16527027 | rs16934787 | -3.306 | 0.348 | 0.977 | 0.954 | -0.057 | -0.082 | -0.082 | 0 | 0.49 | 0.94 | 0.57 | 0.46 | 0.51 | 0.76 |
| 9 | 16527354 | rs3814111 | -1.749 | 0.568 | 1.424 | 1.096 | -0.006 | -0.007 | -0.007 | 0 | 0.18 | 0.2 | 0.19 | 0.26 | 0.22 | 0.31 |

| | | | | | | | | | | | | | | | | |
|---|---|---|---|---|---|---|---|---|---|---|---|---|---|---|---|---|
| 9 | 16529174 | rs4961722 | -1.683 | 0.554 | 1.655 | 0.974 | -0.026 | -0.001 | -0.001 | 0 | 0.22 | 0.13 | 0.32 | 0.21 | 0.37 | 0.39 |
| 9 | 16529554 | rs10962474 | -2.668 | 0.372 | 1.671 | 0.871 | -0.067 | 0.075 | 0.075 | 0.107 | 0.18 | 0.52 | 0.18 | 0.22 | 0.06 | 0.31 |
| 9 | 16532764 | rs7866317 | -0.937 | 0.738 | 1.549 | 1.036 | 0.015 | 0.006 | 0.006 | 0.001 | 0.11 | 0.12 | 0.23 | 0.14 | 0.21 | 0.21 |
| 9 | 16538942 | rs10962477 | -1.783 | 0.542 | 1.408 | 1.065 | 0.006 | 0.006 | 0.006 | 1.207 | 0.23 | 0.37 | 0.22 | 0.19 | 0.25 | 0.33 |
| 9 | 16540987 | rs2891071 | -0.88 | 0.123 | 1.045 | 1.124 | -0.1 | -0.021 | -0.021 | 0.005 | 0.19 | 0.06 | 0.31 | 0.17 | 0.33 | 0.39 |
| 9 | 16547743 | rs2149157 | -0.986 | 0.059 | 1.056 | 1.066 | -0.112 | -0.045 | -0.045 | 1.767 | 0.2 | 0.06 | 0.31 | 0.22 | 0.38 | 0.4 |
| 9 | 16547755 | rs2149158 | -1.838 | -0.07 | 1.315 | 0.972 | 0.016 | 0.017 | 0.017 | 3.753 | 0.36 | 0.64 | 0.51 | 0.44 | 0.47 | 0.57 |
| 9 | 16554405 | rs6475059 | -1.916 | -0.694 | 1.245 | 0.562 | -0.098 | -0.149 | -0.149 | 0 | 0.2 | 0.49 | 0.13 | 0.02 | 0.08 | 0.29 |
| 9 | 16572137 | rs10962493 | -1.757 | -0.708 | 1.52 | 0.558 | -0.091 | -0.129 | -0.129 | 0 | 0.18 | 0.54 | 0.13 | 0.02 | 0.07 | 0.22 |
| 9 | 16572826 | rs1927618 | -2.149 | -0.77 | 1.486 | 0.879 | -0.09 | 0.054 | 0.054 | 0.002 | 0.22 | 0.54 | 0.22 | 0.28 | 0.09 | 0.32 |
| 9 | 16573685 | rs12352543 | -1.452 | -0.728 | 1.642 | 0.769 | -0.058 | -0.088 | -0.088 | 0 | 0.16 | 0.32 | 0.12 | 0.02 | 0.07 | 0.22 |
| 9 | 16578434 | rs10123558 | -1.549 | -0.692 | 1.384 | 0.864 | 0.008 | -0.008 | -0.008 | 0 | 0.15 | 0.17 | 0.16 | 0.19 | 0.07 | 0.25 |
| 9 | 16587648 | rs1415476 | 0.52 | -0.625 | 1.413 | 0.537 | -0.028 | -0.064 | -0.064 | 1.185 | 0.89 | 0.66 | 0.93 | 1 | 0.94 | 0.95 |
| 9 | 16596071 | rs10511622 | -1.506 | -0.654 | 1.376 | 0.525 | -0.064 | -0.095 | -0.095 | 0 | 0.15 | 0.44 | 0.12 | 0.02 | 0.06 | 0.19 |
| 9 | 16609109 | rs16934855 | -0.419 | -0.593 | 0.679 | 0.563 | -0.035 | -0.045 | -0.045 | 0.983 | 0.09 | 0.23 | 0.06 | 0.01 | 0.06 | 0.1 |
| 9 | 16611692 | rs1337391 | 1.424 | -0.599 | 0.113 | 0.526 | -0.083 | -0.129 | -0.129 | 0.607 | 0.81 | 0.46 | 0.84 | 0.97 | 0.91 | 0.72 |
| 9 | 16619529 | rs13290470 | -0.81 | -0.035 | -0.467 | 1.074 | -0.088 | -0.061 | -0.061 | 0 | 0.19 | 0.05 | 0.3 | 0.28 | 0.35 | 0.44 |
| 9 | 16628401 | rs1337387 | -2.361 | -0.445 | -0.585 | 1.07 | -0.04 | 0.046 | 0.046 | 3.348 | 0.49 | 0.89 | 0.58 | 0.56 | 0.48 | 0.79 |
| 9 | 16628828 | rs17739929 | 1.036 | -1.122 | -1.303 | 0.345 | 0.001 | 0.07 | 0.07 | 0.028 | 0.12 | 0.01 | 0.04 | 0 | 0.08 | 0.03 |
| 9 | 16629923 | rs10810580 | -1.18 | -0.723 | -0.737 | 0.98 | 0.026 | 0.031 | 0.031 | 2.618 | 0.28 | 0.35 | 0.4 | 0.47 | 0.4 | 0.57 |
| 9 | 16645242 | rs7039706 | -2.149 | -0.688 | -0.822 | 0.773 | -0.033 | 0.054 | 0.054 | 4.362 | 0.47 | 0.88 | 0.57 | 0.58 | 0.48 | 0.75 |
| 9 | 16656587 | rs10962519 | -0.607 | -0.377 | -0.377 | 0.605 | 0.108 | 0.07 | 0.07 | 0.779 | 0.17 | 0.33 | 0.32 | 0.29 | 0.35 | 0.39 |
| 9 | 16656653 | rs1415471 | 1.514 | -0.364 | -0.533 | 0.645 | -0.036 | 0.024 | 0.024 | 2.303 | 0.62 | 0.15 | 0.56 | 0.69 | 0.57 | 0.34 |
| 9 | 16658392 | rs10756776 | 1.095 | 0.337 | -0.48 | 0.598 | 0.045 | -0.041 | -0.041 | 0 | 0.75 | 0.66 | 0.76 | 0.86 | 0.65 | 0.58 |
| 9 | 16658915 | rs10810584 | -0.862 | -0.198 | -0.906 | 0.775 | 0.002 | -0.003 | -0.003 | 0.596 | 0.31 | 0.7 | 0.4 | 0.57 | 0.3 | 0.45 |
| 9 | 16659259 | rs10115913 | 0.128 | -0.487 | -1.183 | 0.742 | 0.002 | 0.015 | 0.015 | 0.115 | 0.49 | 0.56 | 0.48 | 0.6 | 0.52 | 0.39 |
| 9 | 16661210 | rs7860104 | -1.114 | -2.089 | -1.203 | 1.305 | 0.04 | -0.054 | -0.054 | 0.613 | 0.19 | 0.11 | 0.23 | 0.61 | 0.08 | 0.47 |
| 9 | 16664759 | rs7046154 | 1.249 | -2.58 | -1.605 | 0.305 | -0.003 | 0.101 | 0.101 | 0 | 0.37 | 0.21 | 0.33 | 0.1 | 0.61 | 0.14 |
| 9 | 16669114 | rs12344619 | 0.022 | -2.728 | -1.677 | 0.232 | 0.005 | -0.006 | -0.006 | 0 | 0.06 | 0.03 | 0.05 | 0.1 | 0.06 | 0.03 |
| 9 | 16669912 | rs1337395 | -0.142 | -3.47 | -1.436 | 0.27 | 0.013 | 0.011 | 0.011 | 0 | 0.68 | 0.79 | 0.52 | 0.24 | 0.76 | 0.44 |
| 9 | 16671045 | rs10810585 | 0.125 | -3.748 | -1.38 | 0.49 | 0.039 | -0.029 | -0.029 | 0 | 0.71 | 0.85 | 0.65 | 0.73 | 0.77 | 0.76 |
| 9 | 16673676 | rs10810586 | -0.866 | -4.074 | -0.606 | 0.997 | 0.022 | -0.009 | -0.009 | 0 | 0.22 | 0.15 | 0.16 | 0.23 | 0.14 | 0.25 |
| 9 | 16676486 | rs7871223 | 0.024 | -3.56 | -0.365 | 0.986 | 0.003 | 0.018 | 0.018 | 0.022 | 0.1 | 0.15 | 0.07 | 0.18 | 0.09 | 0.17 |
| 9 | 16676601 | rs4961729 | -1.784 | -3.824 | 0.179 | 0.758 | -0.029 | -0.028 | -0.028 | 0 | 0.39 | 0.6 | 0.4 | 0.83 | 0.25 | 0.68 |
| 9 | 16677221 | rs7875154 | -0.311 | -3.14 | 0.062 | 0.827 | 0.002 | 0.005 | 0.005 | 0 | 0.11 | 0.16 | 0.07 | 0.12 | 0.09 | 0.17 |
| 9 | 16680138 | rs10738445 | 1.835 | -4.002 | 0.228 | 0.958 | 0.059 | -0.005 | -0.005 | 0 | 0.6 | 0.26 | 0.63 | 0.36 | 0.79 | 0.28 |
| 9 | 16682689 | rs10962540 | -0.087 | -3.69 | 0.014 | 0.939 | -0.005 | -0.009 | -0.009 | 0.002 | 0.09 | 0.01 | 0.19 | 0.11 | 0.07 | 0.25 |
| 9 | 16683543 | rs4961731 | 0.663 | -3.726 | 0.375 | 1.084 | -0.018 | 0.004 | 0.004 | 0 | 0.9 | 0.88 | 0.89 | 0.85 | 0.92 | 0.67 |
| 9 | 16699832 | rs12375507 | -0.58 | -3.46 | 0.506 | 1.005 | -0.01 | -0.007 | -0.007 | 0.067 | 0.12 | 0.17 | 0.22 | 0.11 | 0.08 | 0.26 |
| 9 | 16702394 | rs16934920 | -1.484 | -3.185 | 0.473 | 1.069 | -0.008 | -0.018 | -0.018 | 0 | 0.1 | 0.15 | 0.04 | 0.01 | 0.06 | 0.15 |
| 9 | 16708269 | rs7038172 | -1.114 | -3.002 | 0.413 | 1.415 | -0.015 | 0.123 | 0.123 | 0 | 0.1 | 0.17 | 0.08 | 0.46 | 0.06 | 0.14 |
| 9 | 16713666 | rs10738451 | -1.302 | -3.626 | 0.088 | 0.331 | 0.103 | 0.134 | 0.134 | 0.031 | 0.49 | 0.74 | 0.56 | 0.87 | 0.29 | 0.58 |
| 9 | 16715826 | rs1330303 | 1.715 | -3.987 | 0.702 | 0.02 | 0.038 | 0.052 | 0.052 | 0.822 | 0.42 | 0.24 | 0.32 | 0 | 0.63 | 0.16 |
| 9 | 16719445 | rs10962549 | 0.104 | -4.391 | 0.864 | 0.472 | 0.008 | -0.006 | -0.006 | 0 | 0.24 | 0.19 | 0.21 | 0.25 | 0.16 | 0.14 |
| 9 | 16721067 | rs4961737 | -0.507 | -4.35 | 1.087 | 0.521 | -0.027 | 0.047 | 0.047 | 0 | 0.2 | 0.29 | 0.33 | 0.61 | 0.11 | 0.43 |
| 9 | 16722304 | rs10810597 | -1.605 | -4.292 | 0.944 | 0.306 | 0.105 | 0.144 | 0.144 | 0.135 | 0.48 | 0.78 | 0.56 | 0.9 | 0.28 | 0.57 |
| 9 | 16727619 | rs2282071 | -1.092 | -4.169 | 1.218 | 0.674 | -0.02 | 0.086 | 0.086 | 0 | 0.12 | 0.19 | 0.08 | 0.52 | 0.07 | 0.3 |
| 9 | 16730653 | rs1856132 | 0.274 | -4.09 | 1.177 | 0.386 | -0.004 | -0.001 | -0.001 | 0 | 0.11 | 0.09 | 0.25 | 0.14 | 0.07 | 0.16 |
| 9 | 16730753 | rs1330302 | 0.69 | -4.108 | 1.008 | 0.127 | -0.016 | -0.007 | -0.007 | 0 | 0.87 | 0.88 | 0.88 | 0.9 | 0.91 | 0.74 |
| 9 | 16738889 | rs1970285 | 0.625 | -4.106 | 0.998 | 0.077 | -0.01 | -0.007 | -0.007 | 0.433 | 0.87 | 0.9 | 0.88 | 0.91 | 0.91 | 0.76 |
| 9 | 16744613 | rs7034961 | 0.972 | -4.154 | 0.99 | 0.048 | -0.038 | -0.027 | -0.027 | 0 | 0.85 | 0.71 | 0.87 | 0.91 | 0.91 | 0.76 |
| 9 | 16755121 | rs1330298 | -1.142 | -4.097 | 0.866 | -0.029 | -0.028 | -0.072 | -0.072 | 0.469 | 0.43 | 0.57 | 0.52 | 0.91 | 0.22 | 0.59 |
| 9 | 16761648 | rs10810618 | 1.059 | -3.684 | 1.146 | -0.039 | -0.011 | -0.1 | -0.1 | 0 | 0.86 | 0.6 | 0.87 | 0.99 | 0.92 | 0.83 |
| 9 | 16762537 | rs1330300 | 0.226 | -3.687 | 0.832 | -0.032 | -0.004 | -0.028 | -0.028 | 0.548 | 0.89 | 0.88 | 0.89 | 0.99 | 0.93 | 0.82 |
| 9 | 16764989 | rs10511626 | 0.396 | -4.112 | 0.36 | 0.274 | -0.003 | -0.102 | -0.102 | 0 | 0.22 | 0.01 | 0.18 | 0.24 | 0.12 | 0.23 |
| 9 | 16766118 | rs7850168 | 0.163 | -3.74 | 0.713 | 0.211 | 0.005 | -0.008 | -0.008 | 0 | 0.89 | 0.93 | 0.9 | 0.99 | 0.93 | 0.84 |
| 9 | 16768707 | rs7859111 | 0.021 | -3.713 | 1.019 | 0.225 | -0.015 | -0.052 | -0.052 | 0 | 0.9 | 0.81 | 0.89 | 0.99 | 0.93 | 0.86 |
| 9 | 16776500 | rs10810621 | -0.808 | -3.835 | 1.469 | 0.519 | 0.06 | 0.057 | 0.057 | 0 | 0.42 | 0.34 | 0.5 | 0.8 | 0.23 | 0.65 |
| 9 | 16778780 | rs10123118 | 0.235 | -3.737 | 1.168 | 0.734 | -0.002 | 0 | 0 | 0 | 0.11 | 0.1 | 0.24 | 0.14 | 0.07 | 0.18 |
| 9 | 16783380 | rs10962589 | -0.133 | -3.823 | 1.316 | 0.701 | 0.016 | 0.007 | 0.007 | 0 | 0.32 | 0.31 | 0.26 | 0.7 | 0.15 | 0.46 |
| 9 | 16783482 | rs10122736 | 0.053 | -3.773 | 1.47 | 0.698 | -0.017 | 0.056 | 0.056 | 0 | 0.12 | 0.15 | 0.25 | 0.29 | 0.08 | 0.23 |
| 9 | 16786071 | rs10810627 | -0.198 | -3.798 | 1.395 | 0.749 | 0.018 | 0.004 | 0.004 | 0 | 0.33 | 0.31 | 0.27 | 0.68 | 0.15 | 0.47 |
| 9 | 16786898 | rs7860932 | 0.313 | -3.789 | 1.002 | 0.614 | -0.02 | 0.036 | 0.036 | 0.193 | 0.88 | 0.85 | 0.88 | 0.81 | 0.92 | 0.8 |
| 9 | 16787406 | rs10810628 | 1.425 | -3.548 | 0.814 | 0.484 | 0.068 | 0.233 | 0.233 | 0.132 | 0.42 | 0.1 | 0.35 | 0 | 0.7 | 0.17 |
| 9 | 16787670 | rs3927536 | 0.904 | -3.473 | 0.741 | 0.581 | 0.187 | 0.312 | 0.312 | 0.955 | 0.51 | 0.12 | 0.58 | 0.15 | 0.77 | 0.34 |
| 9 | 16789024 | rs10810632 | 0.539 | -3.58 | 1.492 | 1.269 | -0.016 | 0.18 | 0.18 | 1.093 | 0.89 | 0.72 | 0.85 | 0.74 | 0.94 | 0.81 |
| 9 | 16789081 | rs10756815 | 1.306 | -3.688 | 1.469 | 1.381 | -0.049 | 0.125 | 0.125 | 0.25 | 0.78 | 0.57 | 0.57 | 0.57 | 0.85 | 0.62 |
| 9 | 16789878 | rs4961760 | 0.47 | -3.653 | 1.391 | 1.372 | -0.024 | 0.03 | 0.03 | 0.85 | 0.89 | 0.8 | 0.89 | 0.88 | 0.94 | 0.81 |
| 9 | 16791743 | rs10962594 | -0.162 | -3.946 | 1.022 | 1.004 | 0.029 | 0.004 | 0.004 | 0 | 0.36 | 0.33 | 0.35 | 0.63 | 0.14 | 0.45 |

| | | | | | | | | | | | | | | | | |
|---|---|---|---|---|---|---|---|---|---|---|---|---|---|---|---|---|
| 9 | 16793104 | rs10120562 | 0.319 | -3.826 | 0.786 | 0.663 | 0.005 | -0.007 | -0.007 | 0 | 0.1 | 0.06 | 0.27 | 0.17 | 0.08 | 0.19 |
| 9 | 16795241 | rs10810635 | 0.318 | -3.837 | 0.346 | 1.054 | 0.011 | -0.014 | -0.014 | 0.092 | 0.09 | 0.06 | 0.3 | 0.29 | 0.06 | 0.18 |
| 9 | 16795790 | rs16935073 | 0.419 | -3.877 | 0.286 | 1.027 | 0.008 | -0.001 | -0.001 | 0.229 | 0.09 | 0.07 | 0.3 | 0.31 | 0.06 | 0.18 |
| 9 | 16797461 | rs4961497 | -0.114 | -3.868 | 0.373 | 0.879 | -0.008 | 0.004 | 0.004 | 0 | 0.35 | 0.39 | 0.26 | 0.76 | 0.13 | 0.44 |
| 9 | 16799569 | rs10962604 | -0.015 | -3.906 | 0.354 | 0.865 | -0.018 | 0.001 | 0.001 | 0 | 0.35 | 0.4 | 0.26 | 0.76 | 0.13 | 0.46 |
| 9 | 16803158 | rs12236540 | 1.295 | -3.042 | 0.243 | 1.357 | -0.013 | 0 | 0 | 0.874 | 0.06 | 0.04 | 0.01 | 0.15 | 0.01 | 0.05 |
| 9 | 16804447 | rs7861591 | 0.084 | -2.763 | 0.227 | 1.27 | 0.006 | -0.012 | -0.012 | 0 | 0.1 | 0.05 | 0.26 | 0.15 | 0.08 | 0.18 |
| 9 | 16806521 | rs12351269 | 0.045 | -2.762 | 0.236 | 1.165 | 0.005 | -0.007 | -0.007 | 0 | 0.1 | 0.06 | 0.31 | 0.32 | 0.09 | 0.19 |
| 9 | 16810347 | rs11788776 | 0.818 | -1.499 | 0.364 | 0.013 | 0.019 | -0.063 | -0.063 | 0 | 0.32 | 0.34 | 0.27 | 0.08 | 0.53 | 0.21 |
| 9 | 16812767 | rs7852779 | -0.125 | -1.185 | 0.631 | 0.613 | 0.142 | 0.281 | 0.281 | 0 | 0.94 | 0.79 | 0.89 | 0.69 | 0.86 | 0.9 |
| 9 | 16815861 | rs10962619 | 0.234 | -1.769 | -0.307 | -0.05 | 0.036 | -0.018 | -0.018 | 0 | 0.33 | 0.28 | 0.43 | 0.48 | 0.19 | 0.37 |
| 9 | 16824430 | rs10810639 | -0.526 | -2.175 | 0.027 | 0.454 | 0.023 | -0.015 | -0.015 | 0 | 0.39 | 0.35 | 0.5 | 0.56 | 0.25 | 0.48 |
| 9 | 16825110 | rs7043062 | -0.203 | -1.817 | -0.145 | 0.794 | -0.032 | 0.027 | 0.027 | 0 | 0.33 | 0.48 | 0.43 | 0.54 | 0.19 | 0.36 |
| 9 | 16831748 | rs7030528 | -0.347 | -1.273 | -0.651 | -0.165 | 0 | -0.009 | -0.009 | 0.838 | 0.42 | 0.63 | 0.45 | 0.89 | 0.2 | 0.55 |
| 9 | 16838211 | rs4995398 | -0.579 | -0.223 | -0.68 | 0.677 | -0.001 | 0.001 | 0.001 | 0.011 | 0.26 | 0.24 | 0.23 | 0.4 | 0.27 | 0.42 |
| 9 | 16841515 | rs10810642 | 0.747 | -0.637 | -1.094 | 0.17 | -0.043 | 0.023 | 0.023 | 0 | 0.27 | 0.18 | 0.34 | 0.17 | 0.47 | 0.14 |
| 9 | 16841628 | rs10810643 | -0.057 | -0.482 | -1.007 | 0.496 | -0.02 | -0.011 | -0.011 | 1.244 | 0.12 | 0.07 | 0.16 | 0.36 | 0.21 | 0.36 |
| 9 | 16844949 | rs4961499 | -1.028 | -1.954 | -0.522 | -0.3 | -0.01 | -0.051 | -0.051 | 0.454 | 0.88 | 0.81 | 0.93 | 1 | 0.89 | 0.95 |
| 9 | 16846323 | rs1339548 | -0.593 | -2.744 | -0.318 | -0.346 | 0.028 | 0.021 | 0.021 | 0 | 0.51 | 0.4 | 0.32 | 0.65 | 0.34 | 0.5 |
| 9 | 16854839 | rs1339550 | -0.398 | -3.169 | -0.584 | -0.586 | 0.009 | -0.041 | -0.041 | 0.502 | 0.87 | 0.44 | 0.91 | 0.91 | 0.92 | 0.97 |
| 9 | 16858084 | rs10756819 | 0.95 | -3.634 | -1.196 | -0.835 | 0.027 | -0.004 | -0.004 | 0.686 | 0.54 | 0.08 | 0.56 | 0.56 | 0.68 | 0.55 |
| 9 | 16862280 | rs7861573 | -0.279 | -3.603 | -0.597 | -0.739 | -0.013 | -0.048 | -0.048 | 0.078 | 0.73 | 0.62 | 0.69 | 0.92 | 0.77 | 0.72 |
| 9 | 16862691 | rs12683636 | 0.572 | -3.6 | -0.592 | -0.726 | -0.029 | -0.017 | -0.017 | 0 | 0.14 | 0.26 | 0.23 | 0.08 | 0.07 | 0.24 |
| 9 | 16864521 | rs2153271 | 1.71 | -3.664 | -0.933 | -0.694 | -0.048 | 0.055 | 0.055 | 0 | 0.32 | 0.15 | 0.36 | 0.23 | 0.58 | 0.19 |
| 9 | 16869240 | rs7019402 | -0.251 | -3.746 | -0.809 | -0.464 | 0.081 | -0.037 | -0.037 | 0 | 0.7 | 0.83 | 0.76 | 0.43 | 0.89 | 0.5 |
| 9 | 16870421 | rs10810649 | -0.362 | -3.828 | -0.85 | -0.268 | 0.011 | -0.007 | -0.007 | 0 | 0.85 | 0.88 | 0.86 | 0.52 | 0.89 | 0.66 |
| 9 | 16877758 | rs10810656 | 0.702 | -3.756 | -0.68 | -0.225 | 0.025 | 0.018 | 0.018 | 0 | 0.56 | 0.59 | 0.52 | 0.36 | 0.81 | 0.26 |
| 9 | 16881914 | rs7868212 | 0.007 | -3.836 | -0.823 | -0.281 | 0.061 | 0.039 | 0.039 | 0.012 | 0.3 | 0.19 | 0.34 | 0.16 | 0.09 | 0.39 |
| 9 | 16882186 | rs1890640 | 1.305 | -3.807 | -0.88 | -0.302 | 0.118 | 0.132 | 0.132 | 0.731 | 0.44 | 0.15 | 0.43 | 0.24 | 0.73 | 0.23 |
| 9 | 16887073 | rs12006360 | 0.515 | -3.787 | -0.812 | -0.15 | 0.03 | -0.048 | -0.048 | 0 | 0.13 | 0.08 | 0.14 | 0.48 | 0.11 | 0.34 |
| 9 | 16894953 | rs4436220 | -0.114 | -3.849 | -0.615 | -0.192 | -0.037 | -0.007 | -0.007 | 1.146 | 0.71 | 0.66 | 0.58 | 0.8 | 0.91 | 0.62 |
| 9 | 16895610 | rs10733325 | -0.556 | -3.396 | -0.484 | -0.089 | -0.032 | 0.01 | 0.01 | 2.883 | 0.89 | 0.52 | 0.94 | 0.89 | 0.97 | 0.98 |
| 9 | 16903446 | rs10733333 | 0.303 | -3.419 | -0.698 | -0.039 | -0.052 | 0.032 | 0.032 | 0.05 | 0.74 | 0.39 | 0.8 | 0.4 | 0.86 | 0.61 |
| 9 | 16904846 | rs7033354 | 0.671 | -3.198 | -1.126 | -0.328 | 0.039 | 0.049 | 0.049 | 0 | 0.54 | 0.65 | 0.51 | 0.36 | 0.65 | 0.33 |
| 9 | 16911666 | rs10810666 | 0.961 | -2.052 | -1.204 | -0.456 | -0.046 | 0.047 | 0.047 | 0 | 0.12 | 0.03 | 0.1 | 0 | 0.18 | 0.05 |
| 9 | 16915021 | rs3814113 | -0.49 | -1.993 | -1.041 | -0.158 | 0.054 | 0.079 | 0.079 | 2.431 | 0.5 | 0.62 | 0.57 | 0.27 | 0.31 | 0.44 |
| 9 | 16920749 | rs6475097 | 0.24 | -2.1 | -1.247 | 0.434 | 0.038 | -0.039 | -0.039 | 0.507 | 0.74 | 0.83 | 0.78 | 0.43 | 0.84 | 0.58 |
| 9 | 16929803 | rs10810681 | 0.723 | -1.05 | -2.203 | -0.339 | -0.107 | 0.183 | 0.183 | 0.259 | 0.38 | 0.03 | 0.52 | 0.09 | 0.57 | 0.42 |
| 9 | 16931342 | rs10962718 | 0.705 | -0.671 | -2.032 | -0.47 | -0.028 | 0.195 | 0.195 | 0.115 | 0.39 | 0.08 | 0.55 | 0.08 | 0.62 | 0.41 |
| 9 | 16933037 | rs10962725 | -1.044 | -0.486 | -0.233 | 0.109 | 0.119 | 0.308 | 0.308 | 2.422 | 0.07 | 0.39 | 0.12 | 0.33 | 0.08 | 0.05 |
| 9 | 16936082 | rs7019530 | 0.479 | -1.018 | 0.036 | -0.011 | 0.005 | 0.054 | 0.054 | 0.034 | 0.81 | 0.61 | 0.79 | 0.29 | 0.81 | 0.7 |
| 9 | 16936283 | rs7035981 | 0.793 | -1.774 | -0.215 | -0.619 | 0.063 | 0.073 | 0.073 | 0.001 | 0.59 | 0.26 | 0.67 | 0.24 | 0.72 | 0.5 |
| 9 | 16937341 | rs10810685 | 0.733 | -1.776 | -0.013 | -0.502 | 0.028 | 0.003 | 0.003 | 0 | 0.6 | 0.33 | 0.67 | 0.4 | 0.73 | 0.51 |
| 9 | 16941006 | rs7875833 | -0.596 | -1.806 | 0.179 | -0.311 | -0.012 | -0.033 | -0.033 | 0 | 0.75 | 0.68 | 0.84 | 0.9 | 0.84 | 0.77 |
| 9 | 16942901 | rs1538589 | -0.879 | -2.018 | -0.258 | -0.602 | 0 | -0.002 | -0.002 | 0 | 0.41 | 0.6 | 0.33 | 0.52 | 0.38 | 0.53 |
| 9 | 16943046 | rs1538590 | 0.556 | -2.023 | -0.366 | -0.472 | -0.039 | 0.211 | 0.211 | 0.016 | 0.15 | 0.3 | 0.17 | 0.5 | 0.11 | 0.25 |
| 9 | 16948004 | rs6475098 | -0.177 | -2.058 | -0.343 | -0.685 | -0.02 | -0.043 | -0.043 | 0 | 0.72 | 0.47 | 0.84 | 0.83 | 0.79 | 0.73 |
| 9 | 16948706 | rs10756848 | -0.375 | -1.954 | 0.122 | -0.378 | -0.013 | 0.009 | 0.009 | 0.81 | 0.39 | 0.51 | 0.28 | 0.5 | 0.31 | 0.52 |
| 9 | 16950860 | rs10962763 | 1.884 | -1.798 | -0.13 | -0.505 | -0.145 | 0.019 | 0.019 | 0 | 0.3 | 0.02 | 0.49 | 0.23 | 0.43 | 0.19 |
| 9 | 16951644 | rs4961428 | -0.804 | -1.886 | 0.078 | -0.42 | 0.013 | 0.044 | 0.044 | 0 | 0.77 | 0.89 | 0.85 | 0.88 | 0.83 | 0.76 |
| 9 | 16959184 | rs7042688 | -0.439 | -1.667 | -0.363 | -0.163 | -0.019 | 0.016 | 0.016 | 2.49 | 0.39 | 0.41 | 0.29 | 0.46 | 0.31 | 0.44 |
| 9 | 16961911 | rs16935195 | 1.271 | -0.045 | -1.363 | -0.375 | -0.082 | -0.088 | -0.088 | 0 | 0.18 | 0.01 | 0.36 | 0.28 | 0.21 | 0.23 |
| 9 | 16963863 | rs11791614 | 0.661 | -0.155 | -1.312 | -0.957 | -0.007 | 0.011 | 0.011 | 0 | 0.21 | 0.14 | 0.13 | 0.01 | 0.24 | 0.18 |
| 9 | 16969197 | rs10511629 | 0.836 | -0.46 | -1.65 | -0.802 | 0.012 | -0.02 | -0.02 | 0 | 0.09 | 0.05 | 0.03 | 0.18 | 0.06 | 0.1 |
| 9 | 16969505 | rs10511628 | 0.803 | -0.466 | -1.585 | -0.805 | 0.009 | -0.012 | -0.012 | 0 | 0.09 | 0.07 | 0.04 | 0.18 | 0.06 | 0.1 |
| 9 | 16971165 | rs4961513 | 0.84 | -0.758 | -1.377 | -0.743 | -0.004 | 0.01 | 0.01 | 0 | 0.29 | 0.3 | 0.25 | 0.43 | 0.29 | 0.27 |
| 9 | 16974716 | rs953586 | 0.764 | -0.804 | -1.095 | -0.823 | 0.007 | 0.091 | 0.091 | 0 | 0.51 | 0.33 | 0.69 | 0.8 | 0.51 | 0.59 |
| 9 | 16979089 | rs3933182 | 0.654 | -0.819 | -0.973 | -0.863 | -0.003 | -0.019 | -0.019 | 0.036 | 0.23 | 0.2 | 0.16 | 0.02 | 0.31 | 0.21 |
| 9 | 16982584 | rs1032217 | -0.407 | -1.366 | -0.8 | -0.844 | -0.024 | -0.048 | -0.048 | 0 | 0.75 | 0.56 | 0.86 | 0.82 | 0.83 | 0.81 |
| 9 | 16983321 | rs10810702 | 0.536 | -0.953 | -0.726 | -0.557 | 0 | -0.039 | -0.039 | 3.908 | 0.21 | 0.2 | 0.12 | 0.01 | 0.23 | 0.19 |
| 9 | 16984995 | rs10962794 | 0.433 | -0.956 | -0.791 | -0.533 | 0 | -0.041 | -0.041 | 0 | 0.22 | 0.2 | 0.12 | 0.01 | 0.23 | 0.19 |
| 9 | 16986264 | rs1487511 | 0.088 | -0.967 | -0.685 | 0.443 | -0.034 | 0.113 | 0.113 | 0.004 | 0.16 | 0.27 | 0.16 | 0.47 | 0.11 | 0.24 |
| 9 | 16986415 | rs1487509 | 0.147 | -0.932 | -0.665 | 0.446 | -0.035 | 0.128 | 0.128 | 0 | 0.16 | 0.28 | 0.16 | 0.47 | 0.1 | 0.24 |
| 9 | 16987379 | rs10962795 | -0.008 | -0.647 | -1.257 | -0.32 | -0.035 | 0.118 | 0.118 | 0.835 | 0.2 | 0.06 | 0.13 | 0 | 0.22 | 0.11 |
| 9 | 16997535 | rs406268 | -0.19 | -0.788 | -0.238 | -0.062 | -0.004 | -0.007 | -0.007 | 0 | 0.88 | 0.65 | 0.89 | 0.95 | 0.92 | 0.95 |
| 9 | 16998247 | rs374785 | 0.849 | -0.806 | -0.041 | -0.097 | 0.016 | 0.058 | 0.058 | 0.157 | 0.85 | 0.52 | 0.8 | 0.84 | 0.86 | 0.9 |
| 9 | 16998460 | rs448034 | 1.754 | -1.663 | -0.191 | -0.791 | 0.052 | 0.133 | 0.133 | 0 | 0.54 | 0.24 | 0.65 | 0.8 | 0.64 | 0.65 |
| 9 | 17000855 | rs1487504 | 0.497 | -1.362 | -0.054 | -0.571 | 0.129 | 0.13 | 0.13 | 1.176 | 0.94 | 0.64 | 0.89 | 0.94 | 0.92 | 0.98 |
| 9 | 17002275 | rs10962808 | -0.544 | -1.387 | 0.119 | -0.603 | 0.008 | -0.005 | -0.005 | 0 | 0.13 | 0.28 | 0.12 | 0.13 | 0.14 | 0.16 |

| | | | | | | | | | | | | | | | | |
|---|---|---|---|---|---|---|---|---|---|---|---|---|---|---|---|---|
| 9 | 17004418 | rs10810705 | 1.371 | -1.255 | -0.483 | -0.043 | -0.001 | -0.02 | -0.02 | 0 | 0.31 | 0.22 | 0.32 | 0.54 | 0.28 | 0.43 |
| 9 | 17008137 | rs263628 | -1.75 | -1.425 | -0.759 | 0.083 | -0.117 | -0.133 | -0.133 | 0 | 0.74 | 0.99 | 0.65 | 0.7 | 0.63 | 0.72 |
| 9 | 17009481 | rs263655 | -0.511 | -1.236 | -0.118 | 0.568 | -0.001 | -0.013 | -0.013 | 0 | 0.53 | 0.73 | 0.57 | 0.53 | 0.53 | 0.49 |
| 9 | 17010201 | rs10962814 | 0.202 | -1.284 | -0.237 | 0.518 | -0.006 | 0.032 | 0.032 | 1.364 | 0.17 | 0.01 | 0.13 | 0.1 | 0.16 | 0.1 |
| 9 | 17010426 | rs7469906 | -0.024 | -1.266 | -0.176 | 0.622 | 0.005 | 0.013 | 0.013 | 0 | 0.48 | 0.41 | 0.46 | 0.47 | 0.45 | 0.47 |
| 9 | 17011346 | rs1386570 | -0.065 | -1.254 | -0.173 | 0.61 | 0.009 | 0.013 | 0.013 | 0 | 0.49 | 0.42 | 0.46 | 0.48 | 0.45 | 0.47 |
| 9 | 17012877 | rs1487500 | 1.336 | -1.249 | -0.706 | 0.634 | -0.05 | -0.014 | -0.014 | 0 | 0.31 | 0.13 | 0.34 | 0.37 | 0.36 | 0.4 |
| 9 | 17012980 | rs263650 | -1.289 | -1.347 | -0.722 | 0.607 | -0.026 | -0.035 | -0.035 | 0.349 | 0.66 | 0.89 | 0.64 | 0.68 | 0.62 | 0.62 |
| 9 | 17014026 | rs1487499 | 0.159 | -1.22 | -0.494 | 0.66 | -0.002 | 0.029 | 0.029 | 1.112 | 0.16 | 0.02 | 0.13 | 0.1 | 0.16 | 0.07 |
| 9 | 17015692 | rs13299954 | 1.116 | -0.982 | -0.157 | 0.918 | -0.026 | -0.018 | -0.018 | 5.124 | 0.29 | 0.1 | 0.32 | 0.37 | 0.29 | 0.37 |
| 9 | 17015867 | rs263648 | 0.274 | -0.993 | -0.138 | 0.982 | -0.011 | -0.013 | -0.013 | 0 | 0.36 | 0.72 | 0.44 | 0.45 | 0.35 | 0.39 |
| 9 | 17019121 | rs263640 | -1.907 | -0.924 | 0.014 | 0.912 | 0.05 | 0.067 | 0.067 | 4.165 | 0.08 | 0.69 | 0.13 | 0.09 | 0.07 | 0.08 |
| 9 | 17022423 | rs263636 | -1.672 | -1.03 | -0.195 | 1.221 | -0.007 | 0.019 | 0.019 | 3.55 | 0.22 | 0.72 | 0.35 | 0.32 | 0.16 | 0.39 |
| 9 | 17024496 | rs450114 | -0.517 | -1.006 | -0.032 | 1.204 | 0.047 | 0.158 | 0.158 | 1.878 | 0.07 | 0.47 | 0.21 | 0.14 | 0.02 | 0.18 |
| 9 | 17030146 | rs263569 | -1.095 | -0.988 | -0.032 | 1.232 | -0.015 | 0.172 | 0.172 | 3.299 | 0.2 | 0.54 | 0.34 | 0.49 | 0.15 | 0.32 |
| 9 | 17034019 | rs263576 | -0.606 | -1.087 | -0.319 | 1.052 | 0.001 | 0.011 | 0.011 | 0 | 0.47 | 0.73 | 0.52 | 0.55 | 0.42 | 0.51 |
| 9 | 17041557 | rs263585 | -1.019 | -1.523 | -0.201 | 1.096 | -0.066 | 0.101 | 0.101 | 1.751 | 0.18 | 0.45 | 0.33 | 0.28 | 0.09 | 0.3 |
| 9 | 17053484 | rs366216 | 0.386 | -1.274 | -0.438 | 0.797 | -0.001 | -0.018 | -0.018 | 0 | 0.32 | 0.42 | 0.34 | 0.29 | 0.28 | 0.3 |
| 9 | 17058576 | rs2492870 | -0.053 | -1.514 | -0.81 | 0.704 | 0 | -0.003 | -0.003 | 0 | 0.44 | 0.47 | 0.35 | 0.3 | 0.43 | 0.18 |
| 9 | 17071162 | rs2129626 | 1.446 | -1.523 | -0.914 | 0.675 | -0.069 | -0.015 | -0.015 | 0.777 | 0.24 | 0.03 | 0.16 | 0.23 | 0.25 | 0.12 |
| 9 | 17078922 | rs263561 | -0.075 | -1.709 | -0.796 | 0.831 | 0.002 | 0 | 0 | 0 | 0.44 | 0.5 | 0.36 | 0.44 | 0.44 | 0.28 |
| 9 | 17082040 | rs10962860 | 0.894 | -1.66 | -0.934 | 0.806 | -0.03 | 0.006 | 0.006 | 0.032 | 0.16 | 0.01 | 0.15 | 0.06 | 0.18 | 0.07 |
| 9 | 17088668 | rs10962862 | 0.918 | -1.692 | -0.934 | 0.807 | -0.028 | 0.006 | 0.006 | 0.033 | 0.16 | 0.01 | 0.15 | 0.06 | 0.18 | 0.08 |
| 9 | 17105586 | rs2200599 | -0.131 | -1.958 | -0.839 | 0.868 | 0.001 | 0 | 0 | 0 | 0.45 | 0.49 | 0.45 | 0.58 | 0.4 | 0.5 |
| 9 | 17117875 | rs2029247 | -0.428 | -1.77 | -0.93 | 0.97 | -0.03 | -0.06 | -0.06 | 0 | 0.29 | 0.07 | 0.26 | 0.19 | 0.3 | 0.18 |
| 9 | 17118007 | rs16935340 | 0.35 | -1.666 | -0.862 | 0.897 | -0.006 | -0.033 | -0.033 | 0 | 0.12 | 0.04 | 0.1 | 0.13 | 0.12 | 0.11 |
| 9 | 17120621 | rs972464 | -0.629 | -1.605 | -0.657 | 0.83 | -0.001 | -0.003 | -0.003 | 0 | 0.57 | 0.55 | 0.44 | 0.56 | 0.56 | 0.38 |
| 9 | 17130360 | rs411167 | -0.031 | -1.57 | -0.397 | 0.97 | -0.009 | 0 | 0 | 0 | 0.42 | 0.53 | 0.3 | 0.53 | 0.32 | 0.35 |
| 9 | 17134245 | rs4493995 | 0.368 | -1.3 | -1.144 | 0.658 | -0.014 | 0.019 | 0.019 | 0 | 0.37 | 0.07 | 0.26 | 0.21 | 0.31 | 0.24 |
| 9 | 17147361 | rs2779771 | -0.696 | -1.709 | -0.13 | 1.203 | -0.008 | 0.017 | 0.017 | 0 | 0.26 | 0.26 | 0.22 | 0.43 | 0.18 | 0.37 |
| 9 | 17147702 | rs2815179 | -1.665 | -2.073 | -0.097 | 1.023 | -0.008 | -0.003 | -0.003 | 0 | 0.37 | 0.58 | 0.26 | 0.58 | 0.21 | 0.43 |
| 9 | 17158245 | rs9407787 | -0.433 | -2.125 | -0.081 | 1.232 | -0.011 | 0.004 | 0.004 | 0 | 0.26 | 0.26 | 0.21 | 0.19 | 0.18 | 0.28 |
| 9 | 17161527 | rs10511633 | -0.344 | -2.076 | -0.056 | 1.256 | -0.01 | 0.007 | 0.007 | 0 | 0.24 | 0.24 | 0.21 | 0.18 | 0.18 | 0.28 |
| 9 | 17162413 | rs6475111 | -0.601 | -2.273 | -0.147 | 0.969 | -0.001 | 0 | 0 | 0 | 0.35 | 0.35 | 0.22 | 0.48 | 0.26 | 0.48 |
| 9 | 17165698 | rs2815175 | 0.055 | -2.105 | -0.055 | 1.166 | 0.02 | 0.049 | 0.049 | 0 | 0.12 | 0.26 | 0.11 | 0.13 | 0.11 | 0.12 |
| 9 | 17167963 | rs7037191 | 0.507 | -1.985 | -0.019 | 1.152 | -0.001 | 0.001 | 0.001 | 0 | 0.09 | 0.07 | 0.02 | 0.16 | 0.03 | 0.06 |
| 9 | 17181174 | rs2815173 | -0.44 | -2.025 | -0.001 | 1.137 | -0.034 | 0.025 | 0.025 | 0 | 0.25 | 0.39 | 0.22 | 0.18 | 0.17 | 0.27 |
| 9 | 17181689 | rs2779761 | -1.452 | -2.034 | -0.019 | 1.162 | 0 | -0.002 | -0.002 | 0 | 0.37 | 0.61 | 0.26 | 0.37 | 0.21 | 0.4 |
| 9 | 17208589 | rs4961534 | -0.355 | -1.962 | 0.006 | 1.086 | -0.036 | 0.022 | 0.022 | 0 | 0.24 | 0.35 | 0.21 | 0.18 | 0.16 | 0.27 |
| 9 | 17229028 | rs16935412 | 0.209 | -1.953 | 0.131 | 1.125 | -0.013 | 0.023 | 0.023 | 0 | 0.1 | 0.16 | 0.04 | 0.16 | 0.05 | 0.06 |
| 9 | 17230848 | rs4961535 | -0.358 | -1.973 | 0.154 | 1.178 | -0.026 | 0.16 | 0.16 | 0 | 0.12 | 0.29 | 0.07 | 0.42 | 0.05 | 0.12 |
| 9 | 17233530 | rs4458955 | -1.505 | -1.991 | 0.054 | 1.137 | -0.005 | -0.001 | -0.001 | 0 | 0.35 | 0.62 | 0.25 | 0.59 | 0.17 | 0.4 |
| 9 | 17243746 | rs1582029 | -0.551 | -1.961 | 0.032 | 1.118 | -0.044 | 0.02 | 0.02 | 0 | 0.25 | 0.55 | 0.21 | 0.18 | 0.13 | 0.28 |
| 9 | 17250170 | rs6475125 | -0.408 | -1.967 | 0.11 | 1.115 | -0.05 | 0.034 | 0.034 | 0 | 0.24 | 0.41 | 0.2 | 0.18 | 0.13 | 0.28 |
| 9 | 17261209 | rs1591138 | -1.852 | -2.088 | 0.121 | 0.924 | 0.001 | 0.004 | 0.004 | 0.362 | 0.6 | 0.6 | 0.42 | 0.43 | 0.44 | 0.72 |
| 9 | 17269437 | rs10962988 | -1.949 | -2.109 | 0.116 | 0.952 | -0.015 | 0.032 | 0.032 | 0.881 | 0.61 | 0.74 | 0.45 | 0.68 | 0.45 | 0.76 |
| 9 | 17271858 | rs654521 | -1.455 | -1.921 | 0.2 | 0.893 | 0.015 | 0.025 | 0.025 | 0.392 | 0.52 | 0.63 | 0.41 | 0.26 | 0.41 | 0.65 |
| 9 | 17273731 | rs3808795 | 0.106 | -1.924 | 0.16 | 0.892 | 0 | 0 | 0 | 0 | 0.29 | 0.33 | 0.25 | 0.35 | 0.29 | 0.44 |
| 9 | 17276104 | rs2027014 | 0.026 | -1.9 | 0.178 | 0.904 | -0.002 | -0.001 | -0.001 | 6.631 | 0.3 | 0.37 | 0.25 | 0.35 | 0.28 | 0.44 |
| 9 | 17278584 | rs2779773 | -1.596 | -1.893 | 0.104 | 0.861 | 0.024 | 0.011 | 0.011 | 0.884 | 0.53 | 0.72 | 0.43 | 0.53 | 0.42 | 0.71 |
| 9 | 17278726 | rs13292550 | 0.335 | -1.73 | -0.155 | 0.572 | 0 | 0.056 | 0.056 | 0.305 | 0.27 | 0.18 | 0.2 | 0.09 | 0.28 | 0.38 |
| 9 | 17280616 | rs870270 | -0.338 | -1.569 | 0.03 | 0.919 | -0.02 | 0.012 | 0.012 | 0 | 0.23 | 0.26 | 0.18 | 0.18 | 0.13 | 0.27 |
| 9 | 17296630 | rs601960 | -0.354 | -1.564 | 0.029 | 0.928 | -0.018 | 0.012 | 0.012 | 0 | 0.23 | 0.26 | 0.18 | 0.17 | 0.13 | 0.27 |
| 9 | 17298726 | rs679666 | -0.583 | -1.603 | 0.118 | 0.918 | -0.042 | 0.023 | 0.023 | 0 | 0.24 | 0.39 | 0.19 | 0.17 | 0.13 | 0.27 |
| 9 | 17300938 | rs646025 | -0.311 | -1.608 | 0.115 | 0.915 | -0.02 | 0.013 | 0.013 | 0 | 0.22 | 0.26 | 0.18 | 0.17 | 0.13 | 0.27 |
| 9 | 17305351 | rs2780204 | -0.568 | -1.6 | 0.075 | 0.92 | -0.042 | 0.023 | 0.023 | 0 | 0.24 | 0.39 | 0.19 | 0.17 | 0.13 | 0.27 |
| 9 | 17307204 | rs1759458 | -0.461 | -1.6 | 0.083 | 0.92 | -0.033 | 0.02 | 0.02 | 0 | 0.23 | 0.32 | 0.18 | 0.17 | 0.13 | 0.27 |
| 9 | 17309813 | rs6475128 | 0.521 | -1.607 | 0.064 | 1.106 | -0.042 | 0.049 | 0.049 | 0 | 0.76 | 0.61 | 0.79 | 0.76 | 0.87 | 0.73 |
| 9 | 17318681 | rs4621917 | 0.419 | -1.591 | 0.005 | 0.893 | -0.033 | 0.021 | 0.021 | 0 | 0.77 | 0.68 | 0.82 | 0.83 | 0.87 | 0.73 |
| 9 | 17322582 | rs10963033 | 0.969 | -1.649 | -0.156 | 0.904 | 0.035 | -0.023 | -0.023 | 0 | 0.12 | 0 | 0.09 | 0.05 | 0.06 | 0.16 |
| 9 | 17334075 | rs10963040 | 0.208 | -1.845 | -0.192 | 0.602 | 0.005 | 0.021 | 0.021 | 0.145 | 0.29 | 0.23 | 0.2 | 0.09 | 0.28 | 0.38 |
| 9 | 17340864 | rs3808782 | 0.699 | -1.901 | -0.245 | 0.889 | -0.029 | -0.042 | -0.042 | 0 | 0.07 | 0.01 | 0.07 | 0.13 | 0.06 | 0.11 |
| 9 | 17345018 | rs10963050 | -0.537 | -1.928 | -0.225 | 0.857 | 0.047 | 0.104 | 0.104 | 0 | 0.42 | 0.25 | 0.3 | 0.14 | 0.34 | 0.54 |
| 9 | 17349864 | rs2044119 | 1.034 | -1.907 | -0.117 | 0.627 | 0 | 0 | 0 | 0.172 | 0.49 | 0.53 | 0.62 | 0.74 | 0.59 | 0.34 |
| 9 | 17378776 | rs10963072 | 0.365 | -1.894 | -0.284 | 0.5 | 0.011 | 0.059 | 0.059 | 0.04 | 0.27 | 0.17 | 0.2 | 0.09 | 0.25 | 0.35 |
| 9 | 17384923 | rs10121841 | 1.314 | -1.942 | -0.081 | 0.63 | 0.016 | 0.009 | 0.009 | 0.773 | 0.45 | 0.27 | 0.55 | 0.45 | 0.59 | 0.28 |
| 9 | 17391531 | rs1442522 | 0.118 | -1.965 | 0.016 | 0.646 | -0.031 | 0.03 | 0.03 | 0 | 0.79 | 0.67 | 0.82 | 0.83 | 0.87 | 0.75 |
| 9 | 17394536 | rs7035276 | 0.118 | -1.965 | 0.016 | 0.658 | -0.031 | 0.032 | 0.032 | 0 | 0.79 | 0.67 | 0.83 | 0.83 | 0.87 | 0.75 |
| 9 | 17394996 | rs3739489 | -0.285 | -1.972 | 0.061 | 0.656 | 0.015 | 0.151 | 0.151 | 0 | 0.9 | 0.75 | 0.85 | 0.83 | 0.92 | 0.89 |

| | | | | | | | | | | | | | | | | |
|---|---|---|---|---|---|---|---|---|---|---|---|---|---|---|---|---|
| 9 | 17398404 | rs4961557 | 1.333 | -2.007 | -0.096 | 0.642 | 0.019 | 0.004 | 0.004 | 0.714 | 0.45 | 0.27 | 0.54 | 0.46 | 0.58 | 0.28 |
| 9 | 17400701 | rs4961559 | -0.134 | -1.912 | -0.141 | 0.561 | 0.005 | 0.003 | 0.003 | 1.024 | 0.32 | 0.31 | 0.27 | 0.37 | 0.29 | 0.45 |
| 9 | 17419319 | rs1442514 | 0.999 | -1.666 | -0.013 | 0.681 | -0.038 | 0.023 | 0.023 | 0.087 | 0.8 | 0.37 | 0.79 | 0.39 | 0.92 | 0.75 |
| 9 | 17429714 | rs1372698 | 0.407 | -1.579 | 0.493 | 0.853 | -0.009 | 0.12 | 0.12 | 0 | 0.83 | 0.74 | 0.84 | 0.68 | 0.89 | 0.83 |
| 9 | 17430610 | rs1953105 | 1.512 | -1.732 | 0.146 | 0.611 | -0.069 | 0.063 | 0.063 | 0.709 | 0.38 | 0.14 | 0.51 | 0.3 | 0.54 | 0.24 |
| 9 | 17436662 | rs1967357 | 0.923 | -1.242 | 1.021 | 1.038 | -0.004 | 0.072 | 0.072 | 0 | 0.7 | 0.56 | 0.73 | 0.66 | 0.71 | 0.79 |
| 9 | 17442000 | rs11792612 | 0.243 | -1.206 | 0.854 | 1.064 | -0.021 | -0.011 | -0.011 | 0.035 | 0.17 | 0.03 | 0.11 | 0.16 | 0.21 | 0.07 |
| 9 | 17444294 | rs1160340 | -0.335 | -1.29 | 0.894 | 0.992 | 0.001 | -0.004 | -0.004 | 0 | 0.82 | 0.82 | 0.87 | 0.91 | 0.9 | 0.69 |
| 9 | 17451550 | rs2593397 | -0.049 | -0.779 | -0.102 | 0.663 | -0.002 | 0.102 | 0.102 | 0 | 0.89 | 0.77 | 0.86 | 0.83 | 0.92 | 0.86 |
| 9 | 17454799 | rs2593382 | 0.684 | -0.535 | -0.947 | -0.302 | 0.03 | 0.169 | 0.169 | 1.682 | 0.49 | 0.14 | 0.55 | 0.29 | 0.6 | 0.35 |
| 9 | 17457854 | rs2197152 | 0.586 | 0.213 | 0.364 | -0.399 | 0.02 | -0.123 | -0.123 | 3.612 | 0.77 | 0.52 | 0.62 | 0.94 | 0.74 | 0.69 |
| 9 | 17463615 | rs10756881 | 0.972 | 0.064 | 0.776 | 0.093 | -0.018 | -0.019 | -0.019 | 0 | 0.83 | 0.41 | 0.89 | 0.88 | 0.9 | 0.79 |
| 9 | 17466468 | rs1837944 | 0.796 | 0.012 | 0.872 | 0.085 | -0.011 | -0.011 | -0.011 | 0 | 0.84 | 0.5 | 0.9 | 0.89 | 0.9 | 0.79 |
| 9 | 17480669 | rs10756885 | 1.121 | 0.046 | -1.509 | 0.13 | -0.016 | -0.012 | -0.012 | 0 | 0.81 | 0.28 | 0.89 | 0.87 | 0.9 | 0.78 |
| 9 | 17485598 | rs2383022 | 0.237 | 0.74 | -1.919 | 0.316 | -0.033 | -0.033 | -0.033 | 0.129 | 0.32 | 0.18 | 0.45 | 0.41 | 0.36 | 0.2 |
| 9 | 17486802 | rs3780256 | 0.407 | 0.679 | -2.274 | 0.049 | -0.041 | -0.013 | -0.013 | 2.096 | 0.49 | 0.13 | 0.37 | 0.45 | 0.52 | 0.58 |
| 9 | 17488111 | rs2243877 | -0.934 | 0.53 | -1.355 | -0.04 | 0.005 | 0.027 | 0.027 | 0 | 0.11 | 0.07 | 0.07 | 0 | 0.08 | 0.18 |
| 9 | 17493937 | rs10738478 | -0.527 | 0.79 | -1.684 | 0.229 | -0.023 | -0.029 | -0.029 | 0 | 0.33 | 0.16 | 0.5 | 0.56 | 0.38 | 0.31 |
| 9 | 17496114 | rs4466495 | -0.453 | 0.631 | -0.781 | 0.146 | -0.011 | -0.01 | -0.01 | 1.288 | 0.31 | 0.24 | 0.52 | 0.58 | 0.37 | 0.29 |
| 9 | 17502205 | rs2248136 | 0.578 | 0.191 | 0.187 | -0.255 | -0.011 | -0.005 | -0.005 | 0 | 0.93 | 0.31 | 0.94 | 0.95 | 0.97 | 0.96 |
| 9 | 17505228 | rs12377369 | 0.767 | -0.044 | -1.459 | -1.281 | -0.019 | 0.145 | 0.145 | 0.639 | 0.26 | 0.06 | 0.16 | 0.01 | 0.24 | 0.13 |
| 9 | 17507277 | rs10963119 | 0.792 | -0.036 | -1.43 | -1.27 | -0.017 | 0.123 | 0.123 | 1.206 | 0.24 | 0.06 | 0.16 | 0.01 | 0.23 | 0.11 |
| 9 | 17509083 | rs12337784 | 0.412 | 0.114 | -1.591 | -0.674 | -0.088 | -0.167 | -0.167 | 0 | 0.31 | 0.05 | 0.47 | 0.42 | 0.38 | 0.3 |
| 9 | 17511064 | rs2593362 | 0.468 | 0.06 | -1.458 | -0.51 | -0.033 | -0.07 | -0.07 | 0 | 0.37 | 0.11 | 0.31 | 0.52 | 0.37 | 0.55 |
| 9 | 17524038 | rs7350278 | 0.282 | -0.136 | -2.068 | -1.396 | -0.067 | 0.185 | 0.185 | 0.101 | 0.28 | 0.02 | 0.15 | 0.01 | 0.26 | 0.16 |
| 9 | 17531968 | rs2593371 | 0.437 | -0.528 | -2.116 | -1.708 | 0.032 | 0.256 | 0.256 | 0 | 0.56 | 0.13 | 0.74 | 0.94 | 0.67 | 0.78 |
| 9 | 17539320 | rs7866802 | -0.074 | -0.37 | -1.463 | -1.664 | 0.002 | -0.013 | -0.013 | 0 | 0.39 | 0.63 | 0.23 | 0.01 | 0.31 | 0.19 |
| 9 | 17564267 | rs1885170 | 0.687 | -0.543 | -1.26 | -1.552 | -0.011 | -0.021 | -0.021 | 0 | 0.93 | 0.75 | 0.93 | 0.95 | 0.96 | 0.96 |
| 9 | 17572069 | rs2754341 | 0.042 | -0.322 | -2.152 | -0.807 | -0.014 | -0.025 | -0.025 | 1.271 | 0.76 | 0.79 | 0.52 | 0.47 | 0.68 | 0.63 |
| 9 | 17581039 | rs7029216 | 1.601 | -0.456 | -2.428 | -1.193 | -0.092 | 0.182 | 0.182 | 0.034 | 0.3 | 0.06 | 0.18 | 0.01 | 0.33 | 0.15 |
| 9 | 17585896 | rs7047611 | -0.535 | -0.43 | -2.553 | -0.482 | 0.007 | -0.158 | -0.158 | 1.35 | 0.64 | 0.94 | 0.71 | 0.52 | 0.68 | 0.51 |
| 9 | 17586101 | rs10810812 | -0.161 | -0.417 | -1.847 | 0.024 | -0.076 | -0.106 | -0.106 | 7.753 | 0.29 | 0.1 | 0.55 | 0.52 | 0.4 | 0.42 |
| 9 | 17598287 | rs3808772 | 0.082 | -0.25 | -1.497 | 0.233 | 0.026 | -0.049 | -0.049 | 0 | 0.67 | 0.75 | 0.76 | 0.52 | 0.71 | 0.56 |
| 9 | 17599212 | rs2208485 | -0.111 | -0.022 | -1.787 | 0.923 | -0.083 | -0.103 | -0.103 | 5.061 | 0.25 | 0.09 | 0.56 | 0.58 | 0.39 | 0.41 |
| 9 | 17599511 | rs1010547 | -0.005 | -0.97 | -0.372 | 1.378 | -0.08 | -0.002 | -0.002 | 0 | 0.74 | 0.52 | 0.89 | 0.8 | 0.85 | 0.67 |
| 9 | 17600659 | rs3808769 | -0.031 | -0.548 | -0.809 | 1.43 | 0.017 | -0.036 | -0.036 | 0 | 0.77 | 0.93 | 0.89 | 0.75 | 0.82 | 0.67 |
| 9 | 17602001 | rs1535346 | 0.972 | -0.755 | -0.8 | 1.552 | 0.061 | -0.062 | -0.062 | 0 | 0.64 | 0.18 | 0.71 | 0.49 | 0.72 | 0.54 |
| 9 | 17603496 | rs10810813 | -0.126 | -0.624 | -0.841 | 1.444 | -0.047 | -0.035 | -0.035 | 0.255 | 0.12 | 0.01 | 0.15 | 0.18 | 0.17 | 0.16 |
| 9 | 17604934 | rs1543479 | 0.615 | -0.724 | -0.37 | 1.363 | 0.008 | 0.001 | 0.001 | 0.081 | 0.71 | 0.31 | 0.81 | 0.74 | 0.84 | 0.66 |
| 9 | 17605449 | rs10115064 | 0.524 | -0.711 | -0.286 | 1.37 | -0.017 | -0.002 | -0.002 | 0 | 0.71 | 0.35 | 0.82 | 0.74 | 0.84 | 0.67 |
| 9 | 17609045 | rs3808762 | 0.862 | -0.705 | -0.6 | 1.39 | 0.071 | -0.069 | -0.069 | 0 | 0.66 | 0.17 | 0.76 | 0.51 | 0.78 | 0.57 |
| 9 | 17609302 | rs2145659 | 1.123 | -0.615 | -0.076 | 1.448 | -0.009 | 0.084 | 0.084 | 0 | 0.81 | 0.54 | 0.86 | 0.71 | 0.85 | 0.67 |
| 9 | 17610211 | rs1885166 | -0.668 | 0.013 | -0.703 | 1.471 | -0.02 | -0.089 | -0.089 | 0.761 | 0.35 | 0.12 | 0.32 | 0.51 | 0.35 | 0.43 |
| 9 | 17613834 | rs3808756 | -0.804 | -0.056 | -0.639 | 1.127 | 0.004 | -0.019 | -0.019 | 0 | 0.37 | 0.15 | 0.3 | 0.56 | 0.34 | 0.54 |
| 9 | 17616193 | rs3808755 | -1.178 | 0.242 | -0.328 | 0.415 | 0.001 | -0.047 | -0.047 | 0 | 0.66 | 0.43 | 0.69 | 0.89 | 0.64 | 0.86 |
| 9 | 17618048 | rs3780247 | -0.102 | 0.195 | -0.52 | 0.655 | 0.039 | 0.063 | 0.063 | 0 | 0.47 | 0.37 | 0.46 | 0.64 | 0.37 | 0.56 |
| 9 | 17621965 | rs1555422 | -0.385 | 0.412 | -0.942 | 0.771 | -0.037 | -0.034 | -0.034 | 0 | 0.18 | 0.07 | 0.2 | 0.3 | 0.26 | 0.29 |
| 9 | 17622213 | rs3808750 | -0.036 | 0.441 | -0.603 | 0.964 | -0.006 | 0 | 0 | 0.152 | 0.47 | 0.55 | 0.45 | 0.46 | 0.34 | 0.47 |
| 9 | 17622897 | rs9407817 | 0.086 | 0.526 | -0.806 | 0.712 | -0.031 | -0.02 | -0.02 | 0.004 | 0.16 | 0.1 | 0.19 | 0.23 | 0.26 | 0.25 |
| 9 | 17624902 | rs3824388 | 0.847 | 0.278 | -1.071 | 0.56 | 0.004 | -0.093 | -0.093 | 0 | 0.21 | 0.09 | 0.31 | 0.41 | 0.17 | 0.28 |
| 9 | 17625002 | rs2208494 | 0.268 | 0.184 | -0.632 | 0.187 | 0.007 | 0.122 | 0.122 | 0 | 0.51 | 0.26 | 0.36 | 0.19 | 0.48 | 0.3 |
| 9 | 17626616 | rs3808747 | -0.122 | -0.104 | 0.364 | 1.072 | 0.004 | 0.003 | 0.003 | 0 | 0.23 | 0.22 | 0.12 | 0.12 | 0.15 | 0.18 |
| 9 | 17627988 | rs3808745 | 0.947 | 0.822 | 0.847 | 1.385 | 0.061 | 0.167 | 0.167 | 0 | 0.88 | 0.73 | 0.85 | 0.71 | 0.85 | 0.73 |
| 9 | 17630500 | rs4504735 | 1.582 | 0.236 | 1.076 | 0.618 | -0.028 | -0.097 | -0.097 | 1.729 | 0.87 | 0.5 | 0.94 | 1 | 0.94 | 0.94 |
| 9 | 17630591 | rs3808742 | -0.896 | 0.484 | 0.946 | 0.749 | -0.004 | -0.005 | -0.005 | 0 | 0.07 | 0.05 | 0.1 | 0.1 | 0.11 | 0.11 |
| 9 | 17631186 | rs975697 | 1.371 | 0.458 | 1.244 | 0.729 | -0.001 | -0.048 | -0.048 | 0.449 | 0.9 | 0.59 | 0.95 | 1 | 0.97 | 0.97 |
| 9 | 17631552 | rs1998247 | -0.867 | 0.176 | 0.318 | 1.002 | -0.013 | -0.136 | -0.136 | 0.032 | 0.3 | 0.09 | 0.33 | 0.45 | 0.27 | 0.49 |
| 9 | 17633023 | rs2224448 | 2.072 | 0.066 | 0.76 | 0.977 | 0.039 | 0.037 | 0.037 | 0 | 0.53 | 0.26 | 0.52 | 0.45 | 0.59 | 0.35 |
| 9 | 17633059 | rs3808734 | 0.15 | -0.094 | 0.586 | 1.335 | -0.04 | -0.099 | -0.099 | 0 | 0.15 | 0.01 | 0.16 | 0.23 | 0.19 | 0.29 |
| 9 | 17633232 | rs2208496 | 2.798 | 0.249 | 0.559 | 1.204 | 0.013 | 0.048 | 0.048 | 0 | 0.51 | 0.25 | 0.43 | 0.45 | 0.55 | 0.3 |
| 9 | 17634332 | rs4961580 | 2.315 | 0.21 | 1.069 | 1.24 | -0.001 | 0 | 0 | 0.101 | 0.57 | 0.47 | 0.54 | 0.46 | 0.61 | 0.37 |
| 9 | 17635712 | rs6475158 | 2.112 | 0.195 | 1.206 | 1.248 | -0.002 | 0 | 0 | 0 | 0.59 | 0.47 | 0.54 | 0.47 | 0.61 | 0.39 |
| 9 | 17637196 | rs3808731 | -1.468 | 0.139 | 0.731 | 1.264 | -0.021 | -0.124 | -0.124 | 0 | 0.3 | 0.09 | 0.3 | 0.43 | 0.28 | 0.48 |
| 9 | 17640022 | rs2038521 | -1.1 | 0.092 | 1.075 | 1.181 | -0.037 | -0.049 | -0.049 | 0 | 0.27 | 0.17 | 0.43 | 0.35 | 0.38 | 0.47 |
| 9 | 17640642 | rs10963179 | -1.586 | 0.114 | 1.072 | 1.025 | -0.001 | -0.001 | -0.001 | 1.125 | 0.15 | 0.18 | 0.12 | 0.2 | 0.08 | 0.2 |
| 9 | 17649842 | rs3808727 | -1.368 | 0.192 | 1.003 | 0.875 | -0.039 | -0.065 | -0.065 | 0 | 0.3 | 0.18 | 0.41 | 0.53 | 0.36 | 0.52 |
| 9 | 17650027 | rs7039627 | 1.04 | 0.148 | 1.491 | 0.944 | 0.076 | 0.122 | 0.122 | 1.939 | 0.95 | 0.58 | 0.97 | 0.98 | 1 | 0.97 |
| 9 | 17651870 | rs7350202 | -2.156 | 0.41 | 0.83 | 0.861 | -0.021 | -0.002 | -0.002 | 0.196 | 0.43 | 0.75 | 0.56 | 0.54 | 0.46 | 0.65 |
| 9 | 17655744 | rs1536065 | -1.35 | 0.466 | 0.683 | 0.887 | -0.058 | -0.085 | -0.085 | 0.219 | 0.31 | 0.12 | 0.59 | 0.5 | 0.4 | 0.58 |

| | | | | | | | | | | | | | | | | |
|---|---|---|---|---|---|---|---|---|---|---|---|---|---|---|---|---|
| 9 | 17657019 | rs10810817 | -1.504 | 0.4 | 1.097 | 0.854 | -0.015 | -0.027 | -0.027 | 0.121 | 0.33 | 0.24 | 0.59 | 0.52 | 0.41 | 0.6 |
| 9 | 17657363 | rs7866898 | -2.458 | 0.315 | 1.54 | 1.033 | -0.011 | 0.05 | 0.05 | 0 | 0.45 | 0.77 | 0.65 | 0.65 | 0.45 | 0.66 |
| 9 | 17662411 | rs10963190 | -0.81 | 0.377 | 1.448 | 0.99 | -0.086 | -0.151 | -0.151 | 0 | 0.24 | 0.05 | 0.38 | 0.49 | 0.33 | 0.47 |
| 9 | 17664191 | rs3808720 | -1.557 | 0.366 | 1.568 | 0.971 | -0.021 | -0.039 | -0.039 | 0 | 0.32 | 0.21 | 0.61 | 0.5 | 0.39 | 0.59 |
| 9 | 17673601 | rs7047514 | -2.084 | 0.34 | 2.151 | 0.896 | 0.004 | -0.091 | -0.091 | 0 | 0.17 | 0.48 | 0.27 | 0.04 | 0.11 | 0.16 |
| 9 | 17690052 | rs10118522 | -1.312 | 0.472 | 2.015 | 1.095 | -0.023 | -0.043 | -0.043 | 0 | 0.29 | 0.23 | 0.4 | 0.51 | 0.35 | 0.41 |
| 9 | 17690836 | rs3808713 | -1.812 | 0.458 | 2.027 | 1.084 | -0.005 | -0.006 | -0.006 | 0 | 0.37 | 0.35 | 0.63 | 0.52 | 0.42 | 0.54 |
| 9 | 17694913 | rs3808707 | -1.576 | 0.459 | 1.914 | 1.08 | -0.015 | -0.033 | -0.033 | 0 | 0.33 | 0.24 | 0.6 | 0.51 | 0.38 | 0.5 |
| 9 | 17699981 | rs717372 | -0.178 | 0.709 | 1.784 | 1.245 | -0.082 | -0.11 | -0.11 | 0 | 0.17 | 0.02 | 0.24 | 0.35 | 0.25 | 0.25 |
| 9 | 17700366 | rs998599 | -2.375 | 0.616 | 2.083 | 1.169 | -0.027 | -0.008 | -0.008 | 0 | 0.4 | 0.7 | 0.65 | 0.52 | 0.45 | 0.53 |
| 9 | 17703386 | rs16935912 | -1.824 | 0.582 | 2.254 | 1.138 | 0.019 | 0.042 | 0.042 | 0 | 0.15 | 0.37 | 0.38 | 0.15 | 0.16 | 0.24 |
| 9 | 17708814 | rs10491539 | -1.97 | 0.407 | 2.227 | 1.115 | -0.019 | 0.036 | 0.036 | 0.027 | 0.17 | 0.58 | 0.4 | 0.24 | 0.15 | 0.26 |
| 9 | 17710876 | rs10756902 | -0.25 | 0.661 | 1.85 | 1.179 | -0.088 | -0.109 | -0.109 | 1.084 | 0.18 | 0.02 | 0.25 | 0.35 | 0.26 | 0.31 |
| 9 | 17717690 | rs6475168 | -2.665 | 0.534 | 2.196 | 0.998 | 0 | 0 | 0 | 0 | 0.24 | 0.69 | 0.4 | 0.26 | 0.19 | 0.31 |
| 9 | 17718918 | rs7865066 | -0.439 | 0.541 | 2.158 | 1.002 | 0.002 | -0.033 | -0.033 | 0 | 0.07 | 0.1 | 0.22 | 0.01 | 0.06 | 0.11 |
| 9 | 17720884 | rs10114474 | -2.298 | 0.525 | 1.943 | 1.024 | -0.024 | 0.073 | 0.073 | 0.522 | 0.14 | 0.58 | 0.2 | 0.26 | 0.11 | 0.2 |
| 9 | 17726844 | rs10756904 | -0.05 | 0.671 | 1.455 | 1.105 | -0.045 | -0.061 | -0.061 | 0 | 0.15 | 0.04 | 0.25 | 0.37 | 0.23 | 0.26 |
| 9 | 17732138 | rs1536078 | -2.202 | 0.634 | 1.746 | 1.155 | -0.004 | 0.042 | 0.042 | 0.375 | 0.21 | 0.52 | 0.38 | 0.28 | 0.19 | 0.31 |
| 9 | 17735707 | rs2383044 | -0.801 | 0.547 | 1.59 | 1.176 | -0.053 | -0.088 | -0.088 | 0 | 0.19 | 0.04 | 0.36 | 0.48 | 0.31 | 0.39 |
| 9 | 17735905 | rs1536066 | -0.324 | 0.018 | 0.834 | 0.267 | 0.013 | 0.018 | 0.018 | 0 | 0.48 | 0.64 | 0.55 | 0.51 | 0.55 | 0.48 |
| 9 | 17736342 | rs2209440 | 0.089 | 0.097 | 0.535 | 0.215 | 0.006 | 0.001 | 0.001 | 0.274 | 0.42 | 0.5 | 0.42 | 0.41 | 0.48 | 0.37 |
| 9 | 17738857 | rs720268 | 0.578 | 0.037 | -0.436 | -0.182 | 0.009 | 0.056 | 0.056 | 0 | 0.39 | 0.19 | 0.2 | 0.28 | 0.36 | 0.35 |
| 9 | 17745053 | rs4324999 | -0.767 | 0.206 | 0.556 | -0.054 | 0.022 | 0.043 | 0.043 | 0 | 0.09 | 0.3 | 0.32 | 0.08 | 0.11 | 0.13 |
| 9 | 17746447 | rs4338205 | 0.077 | 0.215 | 0.553 | -0.05 | -0.002 | -0.003 | -0.003 | 0 | 0.06 | 0.1 | 0.3 | 0.08 | 0.11 | 0.08 |
| 9 | 17747173 | rs3780226 | -0.196 | 0.118 | 0.583 | -0.613 | 0.007 | -0.034 | -0.034 | 0 | 0.38 | 0.54 | 0.52 | 0.15 | 0.44 | 0.29 |
| 9 | 17750886 | rs3808681 | 1.049 | -0.265 | 0.564 | -0.817 | -0.051 | 0.105 | 0.105 | 0 | 0.43 | 0.17 | 0.33 | 0.8 | 0.47 | 0.44 |
| 9 | 17751559 | rs2383047 | 1.038 | -0.339 | 0.164 | -0.949 | -0.005 | -0.014 | -0.014 | 0 | 0.8 | 0.8 | 0.81 | 0.91 | 0.83 | 0.71 |
| 9 | 17757573 | rs3824371 | -1.055 | -0.067 | 0.222 | -1.003 | -0.001 | 0.002 | 0.002 | 0 | 0.15 | 0.14 | 0.09 | 0.07 | 0.15 | 0.14 |
| 9 | 17758945 | rs3824370 | 1.902 | -0.584 | -0.168 | -1.287 | -0.023 | 0.023 | 0.023 | 0.409 | 0.28 | 0.05 | 0.13 | 0.21 | 0.26 | 0.13 |
| 9 | 17759497 | rs10810848 | 1.602 | -0.525 | -0.252 | -1.142 | -0.064 | -0.068 | -0.068 | 0 | 0.11 | 0.01 | 0.25 | 0.59 | 0.2 | 0.29 |
| 9 | 17760260 | rs9407838 | -0.321 | -0.988 | 0.254 | -1.811 | 0.001 | -0.002 | -0.002 | 0.433 | 0.11 | 0.14 | 0.08 | 0.07 | 0.13 | 0.12 |
| 9 | 17761379 | rs1886589 | 0.94 | -1.044 | 0.529 | -1.851 | -0.002 | 0.021 | 0.021 | 0 | 0.46 | 0.45 | 0.44 | 0.88 | 0.48 | 0.48 |
| 9 | 17762217 | rs7036989 | -0.237 | -1.114 | 0.408 | -1.778 | -0.001 | 0.027 | 0.027 | 0 | 0.62 | 0.75 | 0.53 | 0.96 | 0.62 | 0.67 |
| 9 | 17765633 | rs9406699 | 0.552 | -1.183 | 0.364 | -1.736 | -0.027 | 0.078 | 0.078 | 0 | 0.3 | 0.16 | 0.21 | 0.08 | 0.33 | 0.17 |
| 9 | 17770090 | rs4961593 | 1.026 | -1.177 | 0.422 | -1.707 | -0.005 | -0.03 | -0.03 | 0.075 | 0.75 | 0.69 | 0.6 | 0.89 | 0.73 | 0.63 |
| 9 | 17771828 | rs2224954 | 0.892 | -1.19 | 0.656 | -1.679 | -0.018 | 0.105 | 0.105 | 0 | 0.47 | 0.25 | 0.34 | 0.8 | 0.5 | 0.45 |
| 9 | 17772029 | rs9407852 | 0.006 | -1.184 | 0.637 | -1.691 | 0.002 | 0 | 0 | 0 | 0.09 | 0.11 | 0.08 | 0.08 | 0.11 | 0.12 |
| 9 | 17772909 | rs4302941 | 0.689 | -1.181 | 0.714 | -1.671 | -0.003 | 0.068 | 0.068 | 0 | 0.49 | 0.38 | 0.44 | 0.88 | 0.5 | 0.51 |
| 9 | 17774662 | rs10738489 | 1.346 | -1.199 | 0.38 | -1.661 | -0.084 | 0.064 | 0.064 | 0 | 0.41 | 0.12 | 0.33 | 0.74 | 0.49 | 0.44 |
| 9 | 17775367 | rs2209429 | 0.275 | -1.096 | 0.515 | -1.74 | -0.011 | -0.006 | -0.006 | 1.183 | 0.84 | 0.8 | 0.71 | 0.9 | 0.85 | 0.75 |
| 9 | 17777485 | rs9407863 | -0.197 | -1.052 | 0.35 | -1.738 | 0.002 | 0 | 0 | 2.618 | 0.1 | 0.11 | 0.2 | 0.08 | 0.12 | 0.13 |
| 9 | 17788028 | rs9407874 | 0.448 | -1.191 | 0.54 | -1.784 | 0.006 | 0.034 | 0.034 | 0 | 0.56 | 0.36 | 0.41 | 0.88 | 0.6 | 0.57 |
| 9 | 17792845 | rs9407881 | -0.941 | -1.045 | 0.76 | -1.808 | 0.025 | 0.022 | 0.022 | 1.074 | 0.06 | 0.29 | 0.04 | 0.04 | 0.04 | 0.09 |
| 9 | 17793818 | rs10963272 | 1.122 | -1.158 | 0.583 | -1.957 | -0.04 | 0.16 | 0.16 | 0 | 0.25 | 0.03 | 0.15 | 0 | 0.24 | 0.06 |
| 9 | 17793903 | rs9406736 | -0.419 | -1.191 | 0.81 | -1.982 | -0.003 | -0.052 | -0.052 | 0 | 0.39 | 0.6 | 0.22 | 0.07 | 0.3 | 0.27 |
| 9 | 17794877 | rs12552857 | 0.78 | -1.145 | 0.69 | -1.938 | -0.01 | -0.009 | -0.009 | 0 | 0.06 | 0.03 | 0.3 | 0.07 | 0.09 | 0.08 |
| 9 | 17795169 | rs7860861 | 0.74 | -1.163 | 0.683 | -1.927 | 0.001 | -0.026 | -0.026 | 0 | 0.51 | 0.51 | 0.4 | 0.89 | 0.59 | 0.58 |
| 9 | 17796668 | rs1049430 | 0.49 | -1.121 | 0.299 | -1.893 | 0.012 | 0.115 | 0.115 | 0 | 0.54 | 0.74 | 0.4 | 0.87 | 0.58 | 0.55 |
| 9 | 17801555 | rs11791520 | -0.442 | -1.102 | 0.365 | -1.814 | -0.009 | -0.019 | -0.019 | 5.208 | 0.17 | 0.2 | 0.42 | 0.09 | 0.14 | 0.27 |
| 9 | 17810983 | rs4961598 | -0.803 | -1.095 | 0.439 | -1.74 | -0.024 | -0.018 | -0.018 | 0.101 | 0.23 | 0.37 | 0.46 | 0.16 | 0.16 | 0.36 |
| 9 | 17814272 | rs10810853 | 1.093 | -1.179 | 0.543 | -2.004 | -0.008 | 0.116 | 0.116 | 0 | 0.45 | 0.28 | 0.37 | 0.84 | 0.55 | 0.48 |
| 9 | 17814761 | rs7847005 | -0.715 | -1.182 | 0.55 | -2.005 | -0.019 | -0.035 | -0.035 | 1.91 | 0.22 | 0.37 | 0.46 | 0.13 | 0.18 | 0.36 |
| 9 | 17815641 | rs4961449 | 0.915 | -1.149 | 0.447 | -2.021 | 0.009 | 0.179 | 0.179 | 0 | 0.47 | 0.22 | 0.36 | 0.84 | 0.56 | 0.53 |
| 9 | 17818646 | rs7863227 | -0.534 | -1.126 | 0.531 | -1.978 | -0.008 | -0.017 | -0.017 | 0 | 0.21 | 0.26 | 0.45 | 0.13 | 0.17 | 0.34 |
| 9 | 17819429 | rs10120630 | 1.539 | -1.137 | 0.415 | -1.95 | 0.031 | 0.042 | 0.042 | 1.064 | 0.69 | 0.24 | 0.5 | 0.84 | 0.8 | 0.55 |
| 9 | 17825216 | rs7869721 | 0.039 | -1.022 | 0.465 | -1.652 | -0.001 | 0.065 | 0.065 | 0 | 0.42 | 0.26 | 0.6 | 0.13 | 0.43 | 0.41 |
| 9 | 17829258 | rs10756917 | 0.887 | -0.964 | -0.005 | -1.599 | -0.045 | 0.165 | 0.165 | 0 | 0.43 | 0.18 | 0.35 | 0.84 | 0.54 | 0.48 |
| 9 | 17829334 | rs7029727 | -0.139 | -0.966 | 0.079 | -1.597 | -0.001 | -0.025 | -0.025 | 0 | 0.44 | 0.44 | 0.61 | 0.13 | 0.44 | 0.4 |
| 9 | 17833057 | rs10963299 | 0.747 | -0.885 | 0.114 | -1.555 | 0.027 | 0.018 | 0.018 | 0 | 0.92 | 0.63 | 0.95 | 0.97 | 0.97 | 0.94 |
| 9 | 17836749 | rs12682702 | 0.821 | -0.742 | -0.377 | -1.524 | -0.054 | 0.164 | 0.164 | 0 | 0.43 | 0.18 | 0.34 | 0.84 | 0.53 | 0.48 |
| 9 | 17842652 | rs6475180 | 0.858 | -0.763 | 0.489 | -0.906 | -0.037 | -0.044 | -0.044 | 2.44 | 0.89 | 0.71 | 0.96 | 0.97 | 0.97 | 0.93 |
| 9 | 17849996 | rs5022716 | -1.179 | -0.638 | 0.215 | -1.152 | -0.013 | -0.025 | -0.025 | 0 | 0.34 | 0.63 | 0.31 | 0.23 | 0.29 | 0.39 |
| 9 | 17857109 | rs6475184 | -0.415 | -0.877 | -0.482 | -1.48 | 0.012 | -0.005 | -0.005 | 0 | 0.16 | 0.18 | 0.11 | 0.22 | 0.1 | 0.1 |
| 9 | 17869637 | rs7035861 | 0.448 | -0.523 | -0.496 | -1.963 | 0.003 | 0.015 | 0.015 | 0 | 0.2 | 0.16 | 0.11 | 0.03 | 0.18 | 0.07 |
| 9 | 17874093 | rs4474113 | 0.063 | -0.11 | 0.307 | -1.9 | 0.102 | -0.009 | -0.009 | 0.103 | 0.93 | 0.72 | 0.92 | 0.99 | 0.9 | 0.95 |
| 9 | 17878879 | rs4335230 | -0.343 | -0.843 | -0.447 | -2.449 | -0.029 | 0.194 | 0.194 | 0.321 | 0.33 | 0.1 | 0.2 | 0.02 | 0.3 | 0.25 |
| 9 | 17880109 | rs10963315 | -0.48 | -0.918 | -0.084 | -2.446 | 0.008 | 0.076 | 0.076 | 0 | 0.35 | 0.21 | 0.21 | 0.03 | 0.31 | 0.26 |
| 9 | 17880914 | rs10963317 | -1.13 | -1.018 | -0.018 | -2.465 | -0.001 | -0.012 | -0.012 | 0.059 | 0.46 | 0.62 | 0.31 | 0.04 | 0.42 | 0.35 |
| 9 | 17889538 | rs4524895 | -1.475 | -1.125 | -0.08 | -2.338 | 0.002 | 0.056 | 0.056 | 0 | 0.51 | 0.62 | 0.42 | 0.16 | 0.44 | 0.48 |

| | | | | | | | | | | | | | | | | |
|---|---|---|---|---|---|---|---|---|---|---|---|---|---|---|---|---|
| 9 | 17892368 | rs4961604 | 0.297 | -1.354 | 0.085 | -2.305 | -0.025 | -0.128 | -0.128 | 0 | 0.77 | 0.58 | 0.87 | 1 | 0.85 | 0.82 |
| 9 | 17902936 | rs10810866 | 0.696 | -0.897 | -0.378 | -1.767 | -0.031 | 0.051 | 0.051 | 0 | 0.16 | 0.01 | 0.12 | 0.04 | 0.16 | 0.1 |
| 9 | 17904496 | rs16936077 | 0.041 | -0.772 | 0.304 | -1.512 | 0.013 | -0.002 | -0.002 | 0.184 | 0.05 | 0.22 | 0.04 | 0.01 | 0.03 | 0.03 |
| 9 | 17916555 | rs2383053 | 0.423 | -0.401 | -0.156 | -1.072 | 0.004 | -0.01 | -0.01 | 0 | 0.56 | 0.47 | 0.61 | 0.68 | 0.5 | 0.51 |
| 9 | 17918806 | rs2891114 | 0.429 | -0.648 | -0.066 | -0.861 | 0.009 | 0.007 | 0.007 | 0 | 0.62 | 0.29 | 0.69 | 0.65 | 0.72 | 0.59 |
| 9 | 17931364 | rs4325000 | 0.637 | -0.741 | -0.151 | -0.944 | 0.017 | 0.015 | 0.015 | 0 | 0.59 | 0.29 | 0.61 | 0.63 | 0.69 | 0.52 |
| 9 | 17936628 | rs1755288 | 0.027 | -0.461 | -0.033 | -1.197 | 0 | -0.04 | -0.04 | 0.687 | 0.48 | 0.47 | 0.65 | 0.89 | 0.52 | 0.59 |
| 9 | 17937036 | rs12684916 | 0.027 | -0.447 | 0.066 | -0.972 | 0.002 | 0.084 | 0.084 | 0 | 0.19 | 0.37 | 0.22 | 0.34 | 0.18 | 0.17 |
| 9 | 17937168 | rs12684923 | 0.021 | -0.443 | -0.011 | -0.849 | 0.003 | 0.105 | 0.105 | 0 | 0.2 | 0.39 | 0.22 | 0.38 | 0.18 | 0.18 |
| 9 | 17937391 | rs13287394 | 0.231 | -0.442 | -0.08 | -1.168 | 0.004 | 0.037 | 0.037 | 0 | 0.18 | 0.31 | 0.21 | 0.25 | 0.17 | 0.17 |
| 9 | 17938351 | rs1755289 | 0.565 | -0.386 | -0.498 | -1.364 | 0.026 | 0.018 | 0.018 | 0.004 | 0.54 | 0.25 | 0.63 | 0.6 | 0.63 | 0.55 |
| 9 | 17943442 | rs4517213 | 1.067 | -0.05 | -0.184 | -1.482 | -0.002 | -0.022 | -0.022 | 0.469 | 0.25 | 0.17 | 0.2 | 0 | 0.28 | 0.12 |
| 9 | 17948600 | rs884097 | 0.343 | -0.066 | -0.314 | -1.515 | 0.009 | 0.042 | 0.042 | 0.812 | 0.31 | 0.17 | 0.23 | 0.02 | 0.28 | 0.19 |
| 9 | 17952583 | rs12552709 | 0.93 | -0.033 | -0.312 | -1.477 | -0.006 | 0.024 | 0.024 | 0.709 | 0.25 | 0.14 | 0.19 | 0 | 0.27 | 0.1 |
| 9 | 17961642 | rs1620986 | 0.231 | -0.199 | -0.015 | -1.611 | 0.001 | -0.038 | -0.038 | 0 | 0.48 | 0.49 | 0.66 | 0.91 | 0.58 | 0.58 |
| 9 | 17962477 | rs1329917 | 1.485 | -0.031 | -0.19 | -1.375 | -0.034 | -0.102 | -0.102 | 0 | 0.3 | 0.11 | 0.4 | 0.52 | 0.32 | 0.39 |
| 9 | 17963004 | rs2772686 | 0.091 | -0.16 | 0.075 | -1.665 | 0.006 | -0.015 | -0.015 | 0 | 0.49 | 0.59 | 0.67 | 0.92 | 0.58 | 0.59 |
| 9 | 17964514 | rs2811806 | -0.061 | -0.359 | 0.304 | -1.153 | -0.021 | -0.102 | -0.102 | 0 | 0.75 | 0.45 | 0.83 | 0.93 | 0.78 | 0.75 |
| 9 | 17967886 | rs1778170 | -0.203 | -0.501 | -0.249 | -0.998 | -0.035 | -0.109 | -0.109 | 2.873 | 0.78 | 0.59 | 0.87 | 0.94 | 0.86 | 0.74 |
| 9 | 17972059 | rs2248505 | 0.879 | -0.214 | -1.326 | -0.835 | -0.029 | -0.078 | -0.078 | 0.686 | 0.3 | 0.1 | 0.41 | 0.57 | 0.32 | 0.4 |
| 9 | 17976625 | rs2840787 | 0.286 | -0.398 | -0.617 | -0.977 | -0.008 | -0.038 | -0.038 | 0 | 0.8 | 0.75 | 0.89 | 0.94 | 0.86 | 0.76 |
| 9 | 17985166 | rs2772695 | 0.338 | -0.298 | -0.633 | -0.923 | -0.006 | -0.023 | -0.023 | 0 | 0.74 | 0.72 | 0.86 | 0.92 | 0.84 | 0.7 |
| 9 | 17987408 | rs2891117 | -0.205 | 0.17 | -0.787 | -0.878 | 0.001 | 0.015 | 0.015 | 0 | 0.5 | 0.55 | 0.35 | 0.16 | 0.51 | 0.42 |
| 9 | 17988492 | rs2253259 | -0.005 | 0.193 | -1.017 | -0.03 | -0.051 | -0.069 | -0.069 | 0 | 0.13 | 0.01 | 0.21 | 0.25 | 0.19 | 0.14 |
| 9 | 17990785 | rs2026764 | -0.321 | -0.399 | -0.433 | -0.485 | -0.028 | -0.03 | -0.03 | 0 | 0.16 | 0.2 | 0.05 | 0.03 | 0.06 | 0.09 |
| 9 | 17999021 | rs10963386 | -0.01 | 0.44 | -0.184 | -0.773 | -0.002 | -0.056 | -0.056 | 0 | 0.25 | 0.31 | 0.35 | 0.11 | 0.26 | 0.34 |
| 9 | 18002205 | rs10963391 | -0.762 | 0.351 | 0.046 | -0.651 | -0.006 | -0.054 | -0.054 | 0.483 | 0.36 | 0.49 | 0.44 | 0.18 | 0.34 | 0.42 |
| 9 | 18003633 | rs12346508 | -0.741 | 0.499 | 0.113 | 0.293 | -0.003 | -0.01 | -0.01 | 0 | 0.36 | 0.52 | 0.46 | 0.38 | 0.36 | 0.44 |
| 9 | 18005075 | rs10810880 | -0.008 | 0.416 | 0.344 | 0.288 | 0.003 | -0.006 | -0.006 | 0 | 0.26 | 0.45 | 0.32 | 0.32 | 0.29 | 0.33 |
| 9 | 18005255 | rs10511650 | -0.742 | -0.069 | 0.181 | -0.192 | -0.016 | -0.038 | -0.038 | 0 | 0.13 | 0.17 | 0.03 | 0.01 | 0.05 | 0.11 |
| 9 | 18009671 | rs10810882 | -0.021 | 0.235 | -0.082 | -0.634 | -0.011 | 0.033 | 0.033 | 0 | 0.37 | 0.29 | 0.36 | 0.09 | 0.42 | 0.34 |
| 9 | 18010619 | rs953923 | 1.486 | 0.47 | -0.235 | -0.175 | -0.031 | -0.029 | -0.029 | 0 | 0.05 | 0.01 | 0.22 | 0.17 | 0.13 | 0.08 |
| 9 | 18013733 | rs7041139 | -0.123 | -0.179 | 0.074 | -0.048 | -0.011 | 0.012 | 0.012 | 1.508 | 0.3 | 0.52 | 0.33 | 0.4 | 0.3 | 0.37 |
| 9 | 18016234 | rs1329931 | 0.336 | -0.201 | 0.053 | -0.104 | -0.002 | -0.047 | -0.047 | 0 | 0.25 | 0.31 | 0.3 | 0.1 | 0.26 | 0.31 |
| 9 | 18017007 | rs1411250 | -1.282 | -0.447 | -0.097 | -0.075 | -0.003 | -0.002 | -0.002 | 0 | 0.44 | 0.52 | 0.41 | 0.46 | 0.4 | 0.62 |
| 9 | 18017562 | rs7027669 | -1.182 | -0.576 | 0.386 | -0.133 | 0.047 | 0.036 | 0.036 | 0.35 | 0.43 | 0.84 | 0.37 | 0.4 | 0.34 | 0.51 |
| 9 | 18018207 | rs10810886 | 0.113 | -0.706 | 0.52 | -0.365 | -0.007 | 0.023 | 0.023 | 0.038 | 0.22 | 0.21 | 0.27 | 0.07 | 0.23 | 0.32 |
| 9 | 18023123 | rs947923 | -0.641 | -1.012 | 1.108 | -0.019 | 0.001 | -0.076 | -0.076 | 1.772 | 0.44 | 0.47 | 0.36 | 0.11 | 0.44 | 0.54 |
| 9 | 18023977 | rs10810888 | -1.572 | -1.004 | 1.118 | 0.341 | 0.002 | -0.008 | -0.008 | 0 | 0.57 | 0.61 | 0.51 | 0.49 | 0.59 | 0.66 |
| 9 | 18028358 | rs11789500 | -1.133 | -0.933 | 1.089 | 0.38 | 0.021 | 0.217 | 0.217 | 0 | 0.09 | 0.2 | 0.05 | 0.37 | 0.09 | 0.1 |
| 9 | 18032367 | rs10963401 | -0.839 | -1.094 | 1.364 | 0.278 | -0.001 | 0.007 | 0.007 | 0 | 0.47 | 0.52 | 0.41 | 0.46 | 0.52 | 0.48 |
| 9 | 18032732 | rs10963402 | -1.436 | -1.107 | 1.4 | 0.32 | 0.013 | -0.005 | -0.005 | 0 | 0.53 | 0.57 | 0.49 | 0.51 | 0.57 | 0.56 |
| 9 | 18034266 | rs10756924 | 0.009 | -1.136 | 1.409 | 0.3 | -0.042 | -0.051 | -0.051 | 0 | 0.36 | 0.19 | 0.43 | 0.49 | 0.49 | 0.45 |
| 9 | 18036892 | rs7034015 | -0.38 | -1.095 | 1.483 | 0.262 | -0.056 | -0.073 | -0.073 | 0 | 0.14 | 0.31 | 0.04 | 0.03 | 0.07 | 0.1 |
| 9 | 18038013 | rs1360235 | -0.883 | -1.094 | 1.365 | 0.291 | -0.004 | 0.027 | 0.027 | 0 | 0.47 | 0.61 | 0.41 | 0.41 | 0.51 | 0.52 |
| 9 | 18044104 | rs10810892 | -1.439 | -1.128 | 1.406 | 0.252 | -0.001 | 0.008 | 0.008 | 0 | 0.52 | 0.67 | 0.44 | 0.5 | 0.53 | 0.54 |
| 9 | 18044802 | rs10963408 | -0.009 | -1.032 | 1.292 | 0.328 | -0.002 | 0.117 | 0.117 | 0.11 | 0.06 | 0.09 | 0.04 | 0.35 | 0.05 | 0.08 |
| 9 | 18046709 | rs10810894 | -1.318 | -1.057 | 1.355 | 0.288 | -0.023 | 0.17 | 0.17 | 1.507 | 0.11 | 0.28 | 0.06 | 0.38 | 0.06 | 0.14 |
| 9 | 18051005 | rs10511649 | 0.474 | -0.982 | 1.441 | 0.076 | -0.036 | 0.066 | 0.066 | 0 | 0.3 | 0.21 | 0.34 | 0.08 | 0.38 | 0.34 |
| 9 | 18054030 | rs12002358 | -0.53 | -1.065 | 1.398 | 0.153 | 0.003 | 0.017 | 0.017 | 0 | 0.14 | 0.08 | 0.09 | 0.09 | 0.09 | 0.26 |
| 9 | 18055440 | rs7866191 | -1.43 | -1.013 | 1.71 | 0.33 | 0.011 | -0.003 | -0.003 | 0.126 | 0.56 | 0.62 | 0.51 | 0.5 | 0.6 | 0.71 |
| 9 | 18058912 | rs16936281 | 0.381 | -1.033 | 1.934 | 0.269 | -0.013 | -0.038 | -0.038 | 1.283 | 0.08 | 0.24 | 0.05 | 0 | 0.05 | 0.01 |
| 9 | 18059000 | rs10963421 | -0.423 | -1.05 | 1.886 | 0.26 | -0.01 | 0.04 | 0.04 | 0 | 0.41 | 0.34 | 0.46 | 0.15 | 0.5 | 0.53 |
| 9 | 18061140 | rs6475196 | -1.517 | -0.926 | 1.79 | 0.327 | 0.005 | 0.046 | 0.046 | 0 | 0.55 | 0.63 | 0.49 | 0.17 | 0.56 | 0.64 |
| 9 | 18066074 | rs7860145 | -0.779 | -0.862 | 1.824 | 0.435 | -0.001 | -0.003 | -0.003 | 0 | 0.46 | 0.45 | 0.43 | 0.44 | 0.53 | 0.38 |
| 9 | 18070475 | rs768703 | -1.073 | -0.843 | 1.804 | 0.464 | 0.011 | 0.019 | 0.019 | 0 | 0.49 | 0.61 | 0.44 | 0.47 | 0.54 | 0.49 |
| 9 | 18071787 | rs7860016 | -1.107 | -0.81 | 1.804 | 0.505 | -0.008 | 0.104 | 0.104 | 0.035 | 0.12 | 0.27 | 0.06 | 0.33 | 0.09 | 0.12 |
| 9 | 18073936 | rs7041361 | -1.358 | -0.861 | 1.874 | 0.275 | 0.006 | 0.013 | 0.013 | 0 | 0.53 | 0.55 | 0.48 | 0.16 | 0.55 | 0.61 |
| 9 | 18074369 | rs10738497 | -1.255 | -0.866 | 1.814 | 0.296 | 0.006 | 0.002 | 0.002 | 0 | 0.52 | 0.49 | 0.5 | 0.5 | 0.58 | 0.54 |
| 9 | 18075710 | rs12000388 | -0.78 | -0.771 | 1.396 | 0.211 | -0.029 | -0.02 | -0.02 | 2.21 | 0.14 | 0.3 | 0.1 | 0.09 | 0.08 | 0.25 |
| 9 | 18080592 | rs12156463 | 0.575 | -0.714 | 0.891 | 0.139 | -0.046 | 0.045 | 0.045 | 0 | 0.28 | 0.19 | 0.34 | 0.07 | 0.4 | 0.35 |
| 9 | 18082567 | rs1329924 | -1.408 | -0.958 | 1.205 | 0.274 | -0.064 | -0.064 | -0.064 | 0 | 0.27 | 0.53 | 0.16 | 0.1 | 0.15 | 0.28 |
| 9 | 18082704 | rs1329923 | -1.622 | -0.942 | 1.161 | 0.467 | -0.042 | 0.058 | 0.058 | 0 | 0.28 | 0.56 | 0.18 | 0.47 | 0.18 | 0.31 |
| 9 | 18083153 | rs10756929 | -1.502 | -1.08 | 1.329 | 0.602 | -0.071 | 0.097 | 0.097 | 0 | 0.23 | 0.56 | 0.16 | 0.44 | 0.13 | 0.24 |
| 9 | 18090973 | rs10963442 | 0.198 | -0.817 | 1.158 | 0.67 | -0.007 | 0.146 | 0.146 | 0 | 0.06 | 0.14 | 0.03 | 0.36 | 0.05 | 0.11 |
| 9 | 18091440 | rs10963443 | -0.222 | -0.822 | 1.373 | 0.386 | -0.001 | -0.009 | -0.009 | 0 | 0.36 | 0.35 | 0.44 | 0.16 | 0.45 | 0.45 |
| 9 | 18093377 | rs12348651 | -0.158 | -0.593 | 1.408 | 0.617 | -0.021 | -0.006 | -0.006 | 0.353 | 0.09 | 0.14 | 0.1 | 0.07 | 0.07 | 0.11 |
| 9 | 18101081 | rs7031350 | -1.223 | -0.554 | 1.593 | 1.116 | -0.013 | 0.222 | 0.222 | 0 | 0.14 | 0.51 | 0.08 | 0.37 | 0.08 | 0.14 |
| 9 | 18106940 | rs1411799 | -0.951 | -0.542 | 0.53 | 1.073 | -0.005 | -0.015 | -0.015 | 0 | 0.48 | 0.8 | 0.44 | 0.54 | 0.52 | 0.51 |

| | | | | | | | | | | | | | | | | |
|---|---|---|---|---|---|---|---|---|---|---|---|---|---|---|---|---|
| 9 | 18109831 | rs473309 | 0.594 | -0.238 | 0.388 | 0.028 | -0.004 | 0.036 | 0.036 | 0.268 | 0.52 | 0.42 | 0.65 | 0.81 | 0.53 | 0.56 |
| 9 | 18110037 | rs549861 | 0.702 | -0.446 | 0.12 | 0.307 | -0.002 | -0.004 | -0.004 | 0 | 0.12 | 0.13 | 0.14 | 0.22 | 0.12 | 0.08 |
| 9 | 18111646 | rs569644 | -0.79 | -0.192 | 0.102 | -0.155 | -0.001 | -0.027 | -0.027 | 0 | 0.7 | 0.74 | 0.75 | 0.89 | 0.7 | 0.69 |
| 9 | 18112889 | rs492960 | 0.69 | -0.162 | -0.655 | -0.07 | -0.01 | -0.013 | -0.013 | 0 | 0.15 | 0.06 | 0.14 | 0.22 | 0.15 | 0.11 |
| 9 | 18115934 | rs1556352 | 1.012 | -0.063 | -0.485 | -0.017 | -0.017 | -0.011 | -0.011 | 0 | 0.38 | 0.27 | 0.53 | 0.55 | 0.45 | 0.49 |
| 9 | 18119661 | rs4618799 | 0.922 | -0.274 | -0.554 | -0.06 | -0.029 | -0.002 | -0.002 | 0 | 0.38 | 0.23 | 0.55 | 0.58 | 0.48 | 0.47 |
| 9 | 18121492 | rs565342 | -0.308 | -0.204 | -0.352 | -0.153 | -0.007 | -0.084 | -0.084 | 0 | 0.68 | 0.54 | 0.73 | 0.88 | 0.7 | 0.71 |
| 9 | 18122560 | rs7852679 | -0.49 | -0.149 | -0.647 | 0.401 | 0.008 | -0.001 | -0.001 | 0 | 0.1 | 0.09 | 0.02 | 0.09 | 0.01 | 0.08 |
| 9 | 18123222 | rs16935857 | -0.833 | -0.091 | -0.741 | 0.5 | 0.037 | -0.018 | -0.018 | 0.007 | 0.12 | 0.05 | 0.04 | 0.14 | 0.05 | 0.09 |
| 9 | 18125480 | rs7864916 | 0.585 | 0.224 | -0.133 | 0.344 | -0.009 | -0.021 | -0.021 | 0 | 0.73 | 0.53 | 0.85 | 0.77 | 0.78 | 0.85 |
| 9 | 18127068 | rs10810904 | 0.768 | 0.461 | 0.345 | 0.338 | 0.019 | -0.005 | -0.005 | 0 | 0.75 | 0.52 | 0.83 | 0.75 | 0.75 | 0.79 |
| 9 | 18130036 | rs10963462 | 0.112 | 0.728 | 0.075 | 0.323 | -0.066 | -0.041 | -0.041 | 0.648 | 0.14 | 0.03 | 0.13 | 0.2 | 0.21 | 0.13 |
| 9 | 18130563 | rs10963464 | 0.944 | 1.18 | -0.574 | 0.68 | -0.048 | -0.095 | -0.095 | 0 | 0.13 | 0.01 | 0.2 | 0.33 | 0.19 | 0.18 |
| 9 | 18130574 | rs2156684 | 0.47 | 0.957 | -0.721 | 0.026 | -0.03 | -0.033 | -0.033 | 0 | 0.45 | 0.14 | 0.39 | 0.44 | 0.48 | 0.47 |
| 9 | 18132972 | rs1360413 | -1.195 | 0.891 | -0.042 | 0.027 | -0.012 | 0.015 | 0.015 | 0 | 0.7 | 0.56 | 0.56 | 0.65 | 0.73 | 0.66 |
| 9 | 18134914 | rs763976 | 0.807 | 0.951 | -0.302 | 0.058 | -0.067 | -0.039 | -0.039 | 0.083 | 0.16 | 0.04 | 0.13 | 0.21 | 0.22 | 0.2 |
| 9 | 18138042 | rs4961625 | 1.338 | 0.796 | -0.243 | 0.075 | 0.004 | 0.004 | 0.004 | 0 | 0.27 | 0.22 | 0.4 | 0.25 | 0.24 | 0.31 |
| 9 | 18139364 | rs1360412 | 0.385 | 0.724 | -0.167 | 0.074 | -0.014 | -0.012 | -0.012 | 0.204 | 0.47 | 0.17 | 0.4 | 0.44 | 0.48 | 0.48 |
| 9 | 18140836 | rs983230 | 0.458 | 0.749 | -0.102 | -0.005 | -0.026 | -0.005 | -0.005 | 0 | 0.4 | 0.28 | 0.52 | 0.45 | 0.45 | 0.47 |
| 9 | 18142416 | rs7034973 | 1.333 | 1.119 | -0.177 | -0.117 | 0.033 | 0.113 | 0.113 | 0 | 0.3 | 0.09 | 0.17 | 0.11 | 0.23 | 0.28 |
| 9 | 18142898 | rs7849137 | 0.087 | 1.329 | -0.031 | -0.008 | -0.034 | 0.09 | 0.09 | 0 | 0.77 | 0.47 | 0.76 | 0.66 | 0.8 | 0.8 |
| 9 | 18149174 | rs1411792 | -0.523 | 1.369 | 0.156 | 0.015 | 0.003 | 0.22 | 0.22 | 0 | 0.82 | 0.55 | 0.74 | 0.61 | 0.81 | 0.81 |
| 9 | 18151987 | rs1537136 | -0.275 | 1.379 | 0.154 | 0.013 | 0.011 | 0.193 | 0.193 | 0 | 0.8 | 0.53 | 0.73 | 0.62 | 0.76 | 0.79 |
| 9 | 18154099 | rs1411790 | 0.219 | 1.328 | -0.098 | -0.085 | 0.011 | -0.002 | -0.002 | 0 | 0.47 | 0.36 | 0.57 | 0.55 | 0.52 | 0.52 |
| 9 | 18155747 | rs12378327 | 0.621 | 1.313 | 0.096 | -0.146 | 0.006 | -0.012 | -0.012 | 0.018 | 0.36 | 0.31 | 0.19 | 0.11 | 0.28 | 0.29 |
| 9 | 18161098 | rs10810913 | 0.523 | 1.433 | -0.101 | -0.017 | -0.018 | 0.006 | 0.006 | 0 | 0.42 | 0.31 | 0.53 | 0.44 | 0.45 | 0.48 |
| 9 | 18163165 | rs2183645 | 1.319 | 1.587 | -0.128 | -0.193 | -0.011 | 0.02 | 0.02 | 0 | 0.29 | 0.22 | 0.43 | 0.22 | 0.32 | 0.31 |
| 9 | 18163621 | rs10963480 | 0.975 | 1.522 | -0.164 | -0.202 | -0.009 | 0.139 | 0.139 | 0 | 0.33 | 0.11 | 0.19 | 0.12 | 0.29 | 0.32 |
| 9 | 18164545 | rs10810914 | -0.125 | 1.657 | -0.099 | -0.181 | -0.03 | 0.13 | 0.13 | 0 | 0.77 | 0.48 | 0.76 | 0.58 | 0.81 | 0.8 |
| 9 | 18166112 | rs960232 | 0.91 | 1.618 | -0.105 | -0.169 | -0.006 | 0.005 | 0.005 | 0 | 0.32 | 0.3 | 0.45 | 0.28 | 0.35 | 0.34 |
| 9 | 18166899 | rs12006002 | 1.292 | 1.56 | -0.112 | -0.194 | -0.002 | 0.024 | 0.024 | 0 | 0.29 | 0.22 | 0.41 | 0.15 | 0.29 | 0.33 |
| 9 | 18166934 | rs10963486 | 1.287 | 1.429 | -0.139 | -0.257 | 0.009 | 0.111 | 0.111 | 0 | 0.3 | 0.12 | 0.21 | 0.12 | 0.28 | 0.3 |
| 9 | 18170133 | rs10810918 | 0.141 | 1.451 | -0.145 | -0.203 | 0.02 | -0.001 | -0.001 | 0.087 | 0.47 | 0.36 | 0.55 | 0.5 | 0.54 | 0.51 |
| 9 | 18174349 | rs1411795 | -0.364 | 1.44 | 0.009 | -0.16 | -0.037 | 0.175 | 0.175 | 0 | 0.79 | 0.61 | 0.78 | 0.61 | 0.81 | 0.81 |
| 9 | 18177395 | rs7025808 | -0.166 | 1.186 | -0.275 | -0.182 | -0.004 | 0.001 | 0.001 | 0 | 0.54 | 0.54 | 0.61 | 0.51 | 0.57 | 0.55 |
| 9 | 18177598 | rs10756945 | 1.099 | 1.381 | -0.217 | -0.092 | -0.005 | -0.016 | -0.016 | 0 | 0.16 | 0.11 | 0.24 | 0.32 | 0.23 | 0.19 |
| 9 | 18182792 | rs11789984 | 0.397 | 1.503 | -0.106 | -0.246 | -0.026 | -0.009 | -0.009 | 0 | 0.19 | 0.17 | 0.17 | 0.24 | 0.28 | 0.2 |
| 9 | 18185683 | rs10810920 | 0.618 | 1.2 | -0.163 | -0.461 | -0.003 | -0.021 | -0.021 | 0 | 0.3 | 0.21 | 0.33 | 0.42 | 0.36 | 0.44 |
| 9 | 18187403 | rs7031744 | 0.275 | 1.21 | -0.175 | -0.467 | -0.011 | 0.034 | 0.034 | 0 | 0.33 | 0.38 | 0.34 | 0.46 | 0.32 | 0.44 |
| 9 | 18187805 | rs7032289 | -0.56 | 1.527 | -0.147 | -0.309 | 0.042 | 0.013 | 0.013 | 0 | 0.81 | 0.68 | 0.83 | 0.76 | 0.74 | 0.82 |
| 9 | 18190434 | rs4961627 | 0.869 | 1.374 | -0.421 | -0.383 | 0.003 | -0.002 | -0.002 | 0 | 0.33 | 0.25 | 0.43 | 0.34 | 0.29 | 0.33 |
| 9 | 18193796 | rs2187455 | 0.691 | 1.445 | -0.372 | -0.54 | -0.033 | -0.022 | -0.022 | 0 | 0.17 | 0.1 | 0.14 | 0.24 | 0.21 | 0.2 |
| 9 | 18195685 | rs7027648 | 0.535 | 1.447 | -0.326 | -0.433 | -0.016 | -0.008 | -0.008 | 0 | 0.42 | 0.3 | 0.53 | 0.48 | 0.43 | 0.47 |
| 9 | 18199327 | rs12378891 | -0.056 | 1.491 | -0.507 | -0.442 | 0.022 | 0.016 | 0.016 | 0 | 0.47 | 0.3 | 0.35 | 0.55 | 0.42 | 0.47 |
| 9 | 18205105 | rs10963511 | 0.959 | 1.691 | -0.678 | -0.268 | -0.065 | -0.048 | -0.048 | 0 | 0.15 | 0.04 | 0.12 | 0.24 | 0.21 | 0.17 |
| 9 | 18208852 | rs10963518 | 0.515 | 1.939 | -0.666 | -0.369 | -0.029 | -0.025 | -0.025 | 0 | 0.15 | 0.12 | 0.12 | 0.25 | 0.21 | 0.18 |
| 9 | 18212306 | rs10963523 | 1.144 | 1.981 | -0.55 | -0.468 | -0.005 | 0.003 | 0.003 | 0 | 0.31 | 0.26 | 0.45 | 0.29 | 0.33 | 0.34 |
| 9 | 18214878 | rs4424360 | -0.434 | 1.741 | -0.428 | -0.663 | -0.002 | 0.004 | 0.004 | 0.173 | 0.66 | 0.65 | 0.65 | 0.46 | 0.65 | 0.54 |
| 9 | 18216312 | rs1944765 | -1.346 | 1.712 | -0.295 | -0.72 | -0.017 | -0.04 | -0.04 | 0 | 0.92 | 0.83 | 0.96 | 1 | 0.95 | 0.93 |
| 9 | 18216477 | rs1944764 | 1.355 | 1.833 | -0.504 | -0.703 | -0.044 | -0.048 | -0.048 | 0 | 0.12 | 0.05 | 0.11 | 0.24 | 0.17 | 0.17 |
| 9 | 18221377 | rs10963526 | 0.464 | 1.607 | -0.423 | -0.697 | -0.034 | -0.016 | -0.016 | 0.899 | 0.18 | 0.12 | 0.16 | 0.24 | 0.26 | 0.2 |
| 9 | 18224660 | rs10756949 | 0.313 | 1.521 | -0.439 | -0.749 | 0.029 | -0.04 | -0.04 | 0 | 0.42 | 0.21 | 0.32 | 0.47 | 0.38 | 0.42 |
| 9 | 18224772 | rs1944751 | 0.107 | 1.449 | -0.243 | -0.7 | 0.001 | -0.002 | -0.002 | 0 | 0.74 | 0.64 | 0.79 | 0.76 | 0.69 | 0.75 |
| 9 | 18231265 | rs1944752 | -0.312 | 1.312 | -0.337 | -0.88 | -0.004 | -0.013 | -0.013 | 0.375 | 0.46 | 0.5 | 0.47 | 0.74 | 0.59 | 0.62 |
| 9 | 18233494 | rs7048005 | 0.997 | 0.961 | -0.375 | -0.863 | 0.021 | 0.023 | 0.023 | 0.353 | 0.2 | 0.11 | 0.09 | 0 | 0.15 | 0.05 |
| 9 | 18238314 | rs7038985 | -0.09 | 1.153 | -0.295 | -0.851 | -0.008 | 0.027 | 0.027 | 0.044 | 0.42 | 0.34 | 0.45 | 0.69 | 0.54 | 0.57 |
| 9 | 18243696 | rs10963534 | 1.104 | 1.289 | -0.242 | -0.769 | -0.001 | 0.015 | 0.015 | 0 | 0.26 | 0.25 | 0.31 | 0.45 | 0.3 | 0.37 |
| 9 | 18244652 | rs1944745 | 0.322 | 1.098 | -0.385 | -0.926 | -0.016 | -0.096 | -0.096 | 0 | 0.27 | 0.31 | 0.14 | 0 | 0.21 | 0.06 |
| 9 | 18246545 | rs2891126 | 1.253 | 1.317 | -0.651 | -0.689 | -0.003 | 0.004 | 0.004 | 0 | 0.3 | 0.25 | 0.43 | 0.29 | 0.3 | 0.36 |
| 9 | 18248172 | rs11790387 | 0.497 | 1.07 | -0.789 | -0.733 | -0.006 | -0.034 | -0.034 | 0 | 0.07 | 0.16 | 0.04 | 0 | 0.05 | 0.01 |
| 9 | 18250459 | rs17199255 | 1.001 | 1.138 | -1.072 | -0.631 | -0.001 | -0.022 | -0.022 | 0 | 0.07 | 0.07 | 0.06 | 0 | 0.08 | 0.04 |
| 9 | 18251855 | rs10810935 | 0.059 | 1.176 | -1.281 | -0.619 | -0.08 | -0.011 | -0.011 | 0 | 0.29 | 0.04 | 0.16 | 0.25 | 0.28 | 0.21 |
| 9 | 18256588 | rs10511645 | -0.476 | 1.141 | -1.038 | -0.652 | -0.007 | 0.083 | 0.083 | 0.11 | 0.38 | 0.18 | 0.22 | 0.15 | 0.37 | 0.29 |
| 9 | 18258312 | rs7846812 | -0.944 | 1.161 | -0.668 | -0.584 | 0.003 | -0.019 | -0.019 | 0.048 | 0.34 | 0.42 | 0.21 | 0.22 | 0.34 | 0.29 |
| 9 | 18259760 | rs2244327 | -0.232 | 0.788 | -0.834 | -0.697 | 0.041 | 0.104 | 0.104 | 0 | 0.48 | 0.67 | 0.35 | 0.27 | 0.44 | 0.42 |
| 9 | 18261124 | rs2819204 | -0.258 | 0.69 | -0.795 | -0.158 | -0.032 | -0.012 | -0.012 | 0 | 0.63 | 0.78 | 0.76 | 0.62 | 0.55 | 0.75 |
| 9 | 18261485 | rs2809854 | -0.323 | 0.66 | -0.006 | -0.108 | -0.016 | 0.015 | 0.015 | 0 | 0.32 | 0.55 | 0.56 | 0.36 | 0.28 | 0.32 |
| 9 | 18268581 | rs13286417 | 0.547 | 1.043 | 0.269 | -0.36 | -0.007 | 0.067 | 0.067 | 0 | 0.47 | 0.25 | 0.3 | 0.26 | 0.51 | 0.52 |
| 9 | 18270369 | rs7852708 | 0.536 | 1.118 | 0.306 | -0.369 | -0.003 | -0.016 | -0.016 | 0 | 0.47 | 0.45 | 0.28 | 0.28 | 0.44 | 0.5 |

| | | | | | | | | | | | | | | | | |
|---|---|---|---|---|---|---|---|---|---|---|---|---|---|---|---|---|
| 9 | 18272700 | rs7861151 | 0.346 | 0.849 | -0.068 | -0.307 | 0.015 | 0.106 | 0.106 | 0 | 0.49 | 0.23 | 0.29 | 0.29 | 0.46 | 0.51 |
| 9 | 18277618 | rs7024439 | -0.242 | 0.884 | 0.013 | 0.04 | -0.003 | 0.015 | 0.015 | 0 | 0.62 | 0.47 | 0.77 | 0.58 | 0.62 | 0.75 |
| 9 | 18280143 | rs10810942 | 0.556 | 0.589 | -0.438 | 0.046 | 0 | 0.001 | 0.001 | 0 | 0.17 | 0.25 | 0.5 | 0.27 | 0.17 | 0.25 |
| 9 | 18290410 | rs7862129 | 0.843 | 0.701 | -0.536 | -0.354 | -0.009 | 0.073 | 0.073 | 0 | 0.16 | 0.06 | 0.07 | 0 | 0.17 | 0.07 |
| 9 | 18291313 | rs10963559 | 0.501 | 0.787 | -0.501 | -0.229 | -0.003 | 0.035 | 0.035 | 0 | 0.54 | 0.3 | 0.7 | 0.63 | 0.47 | 0.71 |
| 9 | 18294234 | rs7849559 | -0.525 | 0.531 | -0.722 | -0.374 | -0.012 | -0.014 | -0.014 | 1.144 | 0.68 | 0.33 | 0.78 | 0.62 | 0.66 | 0.72 |
| 9 | 18294429 | rs7865006 | 0.986 | 0.386 | -0.646 | -0.365 | 0.002 | -0.003 | -0.003 | 0 | 0.17 | 0.13 | 0.08 | 0 | 0.17 | 0.07 |
| 9 | 18295768 | rs2891122 | -1.08 | 0.433 | -0.145 | -0.039 | 0.017 | 0.029 | 0.029 | 0 | 0.76 | 0.69 | 0.82 | 0.64 | 0.71 | 0.82 |
| 9 | 18296065 | rs10124717 | 1.088 | 0.356 | -0.42 | -0.031 | 0.007 | -0.023 | -0.023 | 0 | 0.15 | 0.14 | 0.46 | 0.35 | 0.11 | 0.23 |
| 9 | 18298313 | rs7865911 | -0.603 | 0.565 | -0.128 | -0.036 | 0.012 | 0.028 | 0.028 | 0 | 0.71 | 0.61 | 0.81 | 0.62 | 0.69 | 0.75 |
| 9 | 18298447 | rs7865940 | -0.676 | 0.61 | -0.111 | -0.025 | 0.018 | 0.03 | 0.03 | 0 | 0.72 | 0.61 | 0.81 | 0.63 | 0.68 | 0.75 |
| 9 | 18300778 | rs10963560 | 0.294 | 0.438 | -0.288 | 0.069 | 0.005 | 0.025 | 0.025 | 0.541 | 0.11 | 0.17 | 0.08 | 0.21 | 0.11 | 0.21 |
| 9 | 18301116 | rs4610856 | -0.581 | -0.086 | -0.747 | -0.447 | 0.001 | 0.002 | 0.002 | 0.026 | 0.52 | 0.49 | 0.69 | 0.62 | 0.42 | 0.66 |
| 9 | 18301195 | rs10810948 | 0.799 | -0.058 | -0.647 | -0.168 | -0.009 | -0.022 | -0.022 | 0 | 0.1 | 0.07 | 0.08 | 0.24 | 0.12 | 0.19 |
| 9 | 18301446 | rs4546750 | -1.1 | 0.288 | -0.73 | -0.3 | 0.013 | 0.015 | 0.015 | 0 | 0.74 | 0.67 | 0.79 | 0.66 | 0.65 | 0.69 |
| 9 | 18302439 | rs10963564 | 0.467 | 0.288 | -0.299 | -0.125 | 0.047 | 0.164 | 0.164 | 0.277 | 0.09 | 0.29 | 0.09 | 0.26 | 0.1 | 0.19 |
| 9 | 18303691 | rs12002789 | 1.192 | 0.099 | -0.845 | -0.469 | 0.006 | -0.005 | -0.005 | 0.121 | 0.13 | 0.12 | 0.03 | 0 | 0.07 | 0.05 |
| 9 | 18317731 | rs7023546 | 0.221 | 0.112 | -0.73 | -0.175 | 0.007 | -0.017 | -0.017 | 0.002 | 0.36 | 0.29 | 0.59 | 0.38 | 0.32 | 0.46 |
| 9 | 18323073 | rs4436218 | -0.577 | 0.171 | -0.77 | -0.362 | -0.006 | 0 | 0 | 0 | 0.54 | 0.59 | 0.69 | 0.39 | 0.47 | 0.5 |
| 9 | 18323742 | rs10756956 | 1.362 | 0.374 | -0.702 | -0.199 | -0.022 | -0.011 | -0.011 | 0 | 0.06 | 0.03 | 0.06 | 0.26 | 0.1 | 0.17 |
| 9 | 18327215 | rs4129608 | 0.837 | -0.073 | -0.715 | -0.487 | -0.024 | -0.165 | -0.165 | 0 | 0.23 | 0.44 | 0.12 | 0 | 0.16 | 0.08 |
| 9 | 18332000 | rs10810957 | 0.95 | 0.055 | -1.231 | -0.17 | -0.022 | -0.059 | -0.059 | 0 | 0.2 | 0.13 | 0.5 | 0.36 | 0.21 | 0.26 |
| 9 | 18333039 | rs16936603 | 0.129 | 0.006 | -1.363 | -0.091 | -0.019 | -0.089 | -0.089 | 0 | 0.29 | 0.15 | 0.54 | 0.51 | 0.26 | 0.47 |
| 9 | 18334961 | rs7021177 | -0.906 | -0.041 | -0.561 | -0.228 | 0.004 | 0.022 | 0.022 | 0 | 0.52 | 0.59 | 0.68 | 0.63 | 0.49 | 0.64 |
| 9 | 18343434 | rs4961638 | -1.381 | -0.195 | -0.461 | -0.321 | 0.001 | 0.02 | 0.02 | 0 | 0.6 | 0.68 | 0.76 | 0.65 | 0.57 | 0.67 |
| 9 | 18343563 | rs10156465 | 1.401 | 0.222 | -0.55 | -0.044 | -0.017 | -0.011 | -0.011 | 0 | 0.07 | 0.03 | 0.06 | 0.26 | 0.1 | 0.18 |
| 9 | 18349993 | rs4007665 | 0.775 | 0.065 | -0.273 | -0.112 | -0.002 | -0.001 | -0.001 | 0 | 0.21 | 0.23 | 0.51 | 0.4 | 0.21 | 0.31 |
| 9 | 18358128 | rs7047314 | -0.018 | -0.153 | -0.243 | -0.457 | -0.001 | 0.002 | 0.002 | 1.626 | 0.52 | 0.53 | 0.42 | 0.6 | 0.53 | 0.52 |
| 9 | 18359014 | rs9407897 | -0.417 | -0.171 | -0.41 | -0.44 | -0.001 | 0 | 0 | 0 | 0.57 | 0.41 | 0.44 | 0.62 | 0.6 | 0.54 |
| 9 | 18362865 | rs10511652 | -0.383 | -0.11 | -0.448 | -0.393 | 0 | 0.001 | 0.001 | 0.044 | 0.54 | 0.45 | 0.44 | 0.65 | 0.59 | 0.54 |
| 9 | 18363377 | rs1411153 | -2.444 | -0.151 | -0.424 | -0.475 | 0 | -0.019 | -0.019 | 0.211 | 0.84 | 0.86 | 0.93 | 0.99 | 0.87 | 0.95 |
| 9 | 18364503 | rs7020284 | 0.709 | -0.132 | -0.547 | -0.435 | -0.001 | 0.03 | 0.03 | 1.835 | 0.3 | 0.41 | 0.49 | 0.34 | 0.28 | 0.4 |
| 9 | 18365045 | rs9406744 | 0.347 | 0.224 | -1.117 | -0.084 | -0.003 | 0.033 | 0.033 | 0.002 | 0.45 | 0.32 | 0.37 | 0.37 | 0.49 | 0.35 |
| 9 | 18368546 | rs13284884 | 1.68 | 0.164 | -1.042 | -0.08 | -0.009 | -0.032 | -0.032 | 0 | 0.08 | 0.1 | 0.06 | 0 | 0.07 | 0.01 |
| 9 | 18371966 | rs10810961 | 1.062 | 0.129 | -1.214 | 0.082 | -0.016 | -0.057 | -0.057 | 0.716 | 0.11 | 0.03 | 0.08 | 0.28 | 0.11 | 0.22 |
| 9 | 18372052 | rs2809862 | 0.58 | 0.029 | -1.31 | -0.103 | -0.011 | 0.019 | 0.019 | 0 | 0.4 | 0.31 | 0.36 | 0.34 | 0.48 | 0.24 |
| 9 | 18374849 | rs2791443 | 0.883 | 0.102 | -1.278 | -0.036 | -0.006 | 0.006 | 0.006 | 1.419 | 0.36 | 0.3 | 0.32 | 0.33 | 0.4 | 0.22 |
| 9 | 18375279 | rs12337847 | 1.425 | -0.177 | -1.376 | -0.073 | -0.014 | -0.041 | -0.041 | 0 | 0.09 | 0.03 | 0.07 | 0.22 | 0.1 | 0.19 |
| 9 | 18380943 | rs9298792 | 0.177 | -0.152 | -1.122 | -0.471 | 0.008 | 0.039 | 0.039 | 0.001 | 0.48 | 0.25 | 0.4 | 0.65 | 0.55 | 0.43 |
| 9 | 18383380 | rs9407902 | 0.721 | -0.142 | -1.093 | -0.475 | -0.029 | -0.008 | -0.008 | 0 | 0.43 | 0.25 | 0.37 | 0.6 | 0.5 | 0.4 |
| 9 | 18386757 | rs10120545 | 0.334 | -0.377 | -1.143 | -0.755 | 0 | -0.003 | -0.003 | 0 | 0.15 | 0.16 | 0.41 | 0.23 | 0.14 | 0.19 |
| 9 | 18387361 | rs1329719 | 0.176 | -0.29 | -1.092 | -1.293 | 0.001 | -0.007 | -0.007 | 0 | 0.61 | 0.61 | 0.75 | 0.89 | 0.63 | 0.63 |
| 9 | 18389998 | rs7390284 | 0.009 | -0.427 | -0.732 | -0.541 | 0.032 | 0.129 | 0.129 | 0.034 | 0.6 | 0.25 | 0.78 | 0.91 | 0.68 | 0.64 |
| 9 | 18390077 | rs7389852 | -0.341 | 0.144 | 0.095 | 0.401 | 0.001 | 0.003 | 0.003 | 0.272 | 0.11 | 0.12 | 0.4 | 0.27 | 0.1 | 0.18 |
| 9 | 18392900 | rs2791446 | 0.024 | 0.203 | -0.474 | -0.308 | -0.032 | 0.013 | 0.013 | 0 | 0.11 | 0.01 | 0.12 | 0.05 | 0.13 | 0.25 |
| 9 | 18393875 | rs2809861 | -0.827 | -0.043 | 0.165 | -0.622 | -0.002 | -0.051 | -0.051 | 0 | 0.31 | 0.36 | 0.3 | 0.13 | 0.3 | 0.35 |
| 9 | 18395376 | rs4961644 | -0.904 | 0.081 | 0.818 | -0.634 | 0.007 | 0.007 | 0.007 | 0 | 0.25 | 0.41 | 0.26 | 0.24 | 0.27 | 0.34 |
| 9 | 18401585 | rs1329718 | -1.126 | 0.132 | 0.165 | -1.073 | -0.007 | -0.021 | -0.021 | 0 | 0.41 | 0.53 | 0.47 | 0.26 | 0.36 | 0.59 |
| 9 | 18403928 | rs2153417 | -1.689 | 0.125 | -0.562 | -1.053 | -0.006 | 0.081 | 0.081 | 0.12 | 0.52 | 0.7 | 0.55 | 0.27 | 0.51 | 0.63 |
| 9 | 18405209 | rs17203690 | 0.361 | 0.192 | -0.561 | -0.716 | -0.015 | 0.002 | 0.002 | 0 | 0.11 | 0 | 0.12 | 0.07 | 0.09 | 0.21 |
| 9 | 18407521 | rs10810964 | 0.702 | 0.314 | -0.179 | -0.868 | 0 | -0.012 | -0.012 | 0.098 | 0.67 | 0.37 | 0.65 | 0.84 | 0.65 | 0.61 |
| 9 | 18408771 | rs4961646 | -0.145 | 0.468 | -0.446 | -0.572 | -0.003 | 0.007 | 0.007 | 0 | 0.17 | 0.08 | 0.19 | 0.1 | 0.17 | 0.24 |
| 9 | 18408970 | rs2809856 | -0.452 | 0.455 | 0.111 | -0.537 | 0.018 | -0.061 | -0.061 | 0 | 0.21 | 0.41 | 0.27 | 0.12 | 0.21 | 0.35 |
| 9 | 18411779 | rs1591928 | 0.584 | 0.426 | -0.195 | -0.664 | -0.002 | 0.006 | 0.006 | 0.694 | 0.55 | 0.41 | 0.54 | 0.79 | 0.51 | 0.47 |
| 9 | 18420347 | rs10810967 | 0.203 | 0.421 | -0.788 | -0.345 | -0.035 | -0.063 | -0.063 | 1.158 | 0.36 | 0.13 | 0.38 | 0.46 | 0.41 | 0.35 |
| 9 | 18424272 | rs12554111 | 0.316 | -0.042 | -0.831 | -0.68 | 0.022 | -0.061 | -0.061 | 0 | 0.27 | 0.13 | 0.29 | 0.42 | 0.26 | 0.3 |
| 9 | 18428324 | rs7873681 | -0.968 | 0.032 | -0.09 | -0.867 | 0.015 | -0.039 | -0.039 | 0.131 | 0.7 | 0.63 | 0.72 | 0.86 | 0.68 | 0.73 |
| 9 | 18430297 | rs2066151 | -1.128 | 0.064 | -0.237 | -0.973 | 0.003 | -0.026 | -0.026 | 0 | 0.72 | 0.7 | 0.73 | 0.91 | 0.69 | 0.76 |
| 9 | 18430609 | rs1329725 | 0.391 | 0.083 | -0.391 | -0.637 | -0.009 | -0.051 | -0.051 | 0.078 | 0.35 | 0.17 | 0.41 | 0.47 | 0.37 | 0.36 |
| 9 | 18438196 | rs11789537 | 1.012 | 0.272 | -0.2 | -0.876 | -0.037 | 0.078 | 0.078 | 0 | 0.27 | 0.11 | 0.24 | 0.05 | 0.3 | 0.16 |
| 9 | 18442710 | rs5026813 | -0.06 | 0.055 | 0.038 | -1.072 | 0.003 | -0.069 | -0.069 | 0 | 0.4 | 0.46 | 0.33 | 0.07 | 0.42 | 0.21 |
| 9 | 18444138 | rs6475216 | -0.469 | -0.413 | -0.164 | -0.902 | -0.011 | 0.002 | 0.002 | 0 | 0.52 | 0.31 | 0.51 | 0.55 | 0.55 | 0.46 |
| 9 | 18446300 | rs17205381 | 2.093 | -0.36 | -0.243 | -0.934 | -0.025 | 0.036 | 0.036 | 0 | 0.08 | 0 | 0.05 | 0 | 0.12 | 0.02 |
| 9 | 18446682 | rs1888065 | 0.58 | -0.501 | -0.079 | -0.93 | 0.003 | -0.089 | -0.089 | 0 | 0.33 | 0.45 | 0.28 | 0.08 | 0.3 | 0.23 |
| 9 | 18458068 | rs1340043 | -0.951 | 0.049 | 0.223 | -0.704 | 0 | 0.004 | 0.004 | 0 | 0.66 | 0.65 | 0.62 | 0.57 | 0.68 | 0.6 |
| 9 | 18463476 | rs4961649 | 0.115 | 0.089 | 0.302 | -0.536 | 0 | -0.002 | -0.002 | 0 | 0.43 | 0.46 | 0.45 | 0.35 | 0.45 | 0.46 |
| 9 | 18465657 | rs10123481 | -0.774 | 0.041 | 0.217 | -0.545 | 0.002 | 0 | 0 | 0 | 0.56 | 0.56 | 0.56 | 0.41 | 0.64 | 0.51 |
| 9 | 18470996 | rs1888067 | 0.373 | 0.098 | -0.051 | -0.446 | -0.068 | -0.005 | -0.005 | 1.04 | 0.4 | 0.15 | 0.43 | 0.35 | 0.43 | 0.38 |
| 9 | 18472546 | rs10511638 | 1.357 | 0.215 | -0.006 | -0.461 | -0.001 | -0.009 | -0.009 | 0 | 0.08 | 0.06 | 0.05 | 0 | 0.12 | 0.02 |

| | | | | | | | | | | | | | | | | |
|---|---|---|---|---|---|---|---|---|---|---|---|---|---|---|---|---|
| 9 | 18485434 | rs4961652 | -0.794 | 0.11 | -0.015 | -0.484 | 0.002 | -0.003 | -0.003 | 0.175 | 0.56 | 0.56 | 0.53 | 0.52 | 0.57 | 0.6 |
| 9 | 18486761 | rs17207339 | 0.805 | 0.288 | -0.071 | -0.426 | -0.011 | -0.002 | -0.002 | 0 | 0.1 | 0.07 | 0.09 | 0.1 | 0.12 | 0.16 |
| 9 | 18494532 | rs4961653 | 0.379 | 0.371 | -0.037 | -0.522 | -0.064 | -0.026 | -0.026 | 1.249 | 0.39 | 0.15 | 0.42 | 0.39 | 0.42 | 0.46 |
| 9 | 18498954 | rs1538999 | -0.664 | 0.43 | -0.424 | -0.436 | 0.02 | 0.006 | 0.006 | 0.669 | 0.54 | 0.68 | 0.53 | 0.43 | 0.58 | 0.57 |
| 9 | 18512570 | rs10810974 | -0.532 | 0.789 | -0.503 | -0.23 | -0.006 | 0.002 | 0.002 | 1.324 | 0.57 | 0.57 | 0.56 | 0.44 | 0.63 | 0.59 |
| 9 | 18513871 | rs12002347 | 0.6 | 0.926 | -0.68 | -0.363 | 0.007 | -0.037 | -0.037 | 0 | 0.09 | 0.15 | 0.07 | 0 | 0.1 | 0.02 |
| 9 | 18516637 | rs10963622 | 0.03 | 0.921 | -1.344 | -0.305 | 0.041 | 0.066 | 0.066 | 0 | 0.42 | 0.18 | 0.54 | 0.73 | 0.36 | 0.47 |
| 9 | 18516996 | rs13298611 | -0.033 | 1.519 | -1.307 | -0.482 | -0.022 | 0.024 | 0.024 | 0 | 0.19 | 0.13 | 0.14 | 0.09 | 0.26 | 0.1 |
| 9 | 18519699 | rs1341061 | -0.459 | 1.085 | -1.546 | -0.332 | 0.006 | 0.083 | 0.083 | 0.006 | 0.49 | 0.23 | 0.59 | 0.74 | 0.54 | 0.48 |
| 9 | 18527831 | rs1928535 | 0.407 | 1.011 | -1.499 | -0.357 | 0.006 | -0.015 | -0.015 | 0.271 | 0.31 | 0.23 | 0.24 | 0.65 | 0.28 | 0.24 |
| 9 | 18528775 | rs1341059 | -1.203 | 0.933 | -1.354 | -0.424 | 0.019 | 0.052 | 0.052 | 0 | 0.53 | 0.31 | 0.61 | 0.74 | 0.57 | 0.49 |
| 9 | 18530339 | rs10963625 | 2.012 | 0.865 | -1.897 | -0.858 | -0.031 | 0.009 | 0.009 | 0 | 0.18 | 0.02 | 0.13 | 0.11 | 0.19 | 0.22 |
| 9 | 18531748 | rs7863071 | -1.167 | 0.863 | -0.939 | -0.871 | -0.003 | -0.011 | -0.011 | 0 | 0.55 | 0.52 | 0.62 | 0.79 | 0.6 | 0.51 |
| 9 | 18533731 | rs10810981 | 1.8 | 0.656 | -1.144 | -0.404 | -0.048 | -0.143 | -0.143 | 0 | 0.19 | 0.01 | 0.16 | 0.55 | 0.21 | 0.14 |
| 9 | 18537444 | rs13295269 | 1.052 | 0.879 | -0.881 | -0.765 | -0.057 | 0.058 | 0.058 | 0 | 0.21 | 0.08 | 0.38 | 0.1 | 0.31 | 0.26 |
| 9 | 18538877 | rs6475227 | -1.153 | 0.826 | -0.9 | -0.763 | 0.003 | 0.008 | 0.008 | 0 | 0.57 | 0.58 | 0.64 | 0.8 | 0.61 | 0.52 |
| 9 | 18540040 | rs7388822 | 0.619 | 0.814 | -0.844 | -0.765 | 0.042 | -0.039 | -0.039 | 0 | 0.23 | 0.35 | 0.4 | 0.15 | 0.32 | 0.28 |
| 9 | 18541710 | rs6475228 | 0.193 | 0.619 | -1.269 | -0.454 | 0.005 | -0.022 | -0.022 | 0.164 | 0.33 | 0.22 | 0.25 | 0.63 | 0.3 | 0.24 |
| 9 | 18542869 | rs12376289 | 1.939 | 0.746 | -1.565 | -0.395 | -0.016 | 0.011 | 0.011 | 0 | 0.18 | 0.02 | 0.13 | 0.11 | 0.17 | 0.21 |
| 9 | 18545209 | rs7042159 | 0.164 | 0.843 | -0.876 | 0.087 | 0.002 | -0.022 | -0.022 | 0 | 0.31 | 0.22 | 0.24 | 0.63 | 0.29 | 0.2 |
| 9 | 18572900 | rs13294657 | 0.215 | 0.802 | -0.926 | -0.529 | -0.009 | 0.045 | 0.045 | 0 | 0.27 | 0.18 | 0.38 | 0.11 | 0.31 | 0.27 |
| 9 | 18574806 | rs10756976 | 0.599 | 0.599 | -1.072 | -0.441 | -0.001 | -0.007 | -0.007 | 0 | 0.25 | 0.22 | 0.29 | 0.66 | 0.24 | 0.28 |
| 9 | 18575898 | rs10732338 | -1.517 | 0.351 | -0.535 | -0.368 | 0.025 | 0.062 | 0.062 | 0 | 0.61 | 0.76 | 0.72 | 0.83 | 0.64 | 0.6 |
| 9 | 18576556 | rs10119114 | -0.68 | 0.288 | -0.036 | -0.322 | 0.047 | 0.106 | 0.106 | 0.31 | 0.05 | 0.35 | 0.03 | 0.07 | 0.01 | 0.02 |
| 9 | 18580905 | rs6475232 | -1.205 | 0.515 | -0.701 | -0.322 | 0.019 | 0.042 | 0.042 | 0 | 0.6 | 0.71 | 0.71 | 0.83 | 0.63 | 0.56 |
| 9 | 18582065 | rs7875881 | -0.216 | 0.73 | -0.461 | -0.5 | 0 | 0.004 | 0.004 | 0.47 | 0.19 | 0.23 | 0.31 | 0.17 | 0.2 | 0.3 |
| 9 | 18583822 | rs7040072 | 0.301 | 0.613 | -0.718 | -0.779 | -0.013 | 0.065 | 0.065 | 0.507 | 0.16 | 0.03 | 0.28 | 0.06 | 0.16 | 0.24 |
| 9 | 18590952 | rs1576897 | 0.172 | 0.039 | -0.875 | -0.828 | 0.055 | -0.028 | -0.028 | 2.488 | 0.43 | 0.19 | 0.39 | 0.53 | 0.31 | 0.44 |
| 9 | 18592374 | rs776794 | 0.242 | -0.129 | -0.599 | -0.852 | 0.014 | -0.012 | -0.012 | 1.117 | 0.39 | 0.28 | 0.43 | 0.54 | 0.31 | 0.43 |
| 9 | 18595548 | rs949682 | -0.977 | 0.677 | -0.318 | -0.882 | 0.002 | -0.002 | -0.002 | 0 | 0.7 | 0.64 | 0.65 | 0.73 | 0.62 | 0.67 |
| 9 | 18598470 | rs4433237 | -0.446 | 0.755 | -0.346 | -0.891 | 0.006 | -0.004 | -0.004 | 0.236 | 0.63 | 0.55 | 0.58 | 0.69 | 0.53 | 0.57 |
| 9 | 18599029 | rs10810992 | 1.574 | 0.634 | -0.711 | -0.947 | -0.011 | -0.114 | -0.114 | 0 | 0.28 | 0.05 | 0.25 | 0.47 | 0.24 | 0.22 |
| 9 | 18601460 | rs10756981 | 0.05 | 1.472 | 0.207 | -0.998 | 0.003 | -0.004 | -0.004 | 0 | 0.65 | 0.45 | 0.6 | 0.65 | 0.66 | 0.67 |
| 9 | 18601564 | rs2184002 | 0.159 | 1.492 | 0.329 | -1.024 | -0.021 | 0.012 | 0.012 | 1.086 | 0.61 | 0.33 | 0.57 | 0.63 | 0.54 | 0.65 |
| 9 | 18602403 | rs1539000 | 0.879 | 1.517 | 0.309 | -1.042 | 0.042 | -0.01 | -0.01 | 0 | 0.5 | 0.18 | 0.44 | 0.52 | 0.43 | 0.47 |
| 9 | 18606825 | rs1889007 | -0.187 | 1.457 | 0.594 | -1.183 | 0.017 | -0.054 | -0.054 | 0 | 0.63 | 0.5 | 0.6 | 0.79 | 0.56 | 0.75 |
| 9 | 18613798 | rs10963668 | 0.023 | 1.368 | 0.437 | -1.043 | -0.01 | 0.078 | 0.078 | 0 | 0.06 | 0.09 | 0.07 | 0.22 | 0.02 | 0.1 |
| 9 | 18614351 | rs16936905 | 1.712 | 1.356 | 0.353 | -1.127 | 0.071 | -0.093 | -0.093 | 0 | 0.37 | 0.1 | 0.34 | 0.49 | 0.25 | 0.46 |
| 9 | 18614724 | rs10810995 | -0.478 | 1.478 | 0.467 | -1.148 | 0.017 | -0.048 | -0.048 | 0 | 0.68 | 0.55 | 0.64 | 0.78 | 0.62 | 0.74 |
| 9 | 18615434 | rs7868185 | 1.211 | 1.347 | 0.024 | -1.127 | 0.001 | -0.02 | -0.02 | 0 | 0.09 | 0.07 | 0.08 | 0 | 0.13 | 0.01 |
| 9 | 18617420 | rs17229415 | 0.401 | 1.369 | 0.033 | -1.117 | -0.008 | 0.01 | 0.01 | 0 | 0.1 | 0.05 | 0.07 | 0.06 | 0.13 | 0.08 |
| 9 | 18617617 | rs1889005 | -0.41 | 1.408 | 0.066 | -1.126 | 0.026 | -0.05 | -0.05 | 0.04 | 0.67 | 0.48 | 0.63 | 0.78 | 0.61 | 0.74 |
| 9 | 18617804 | rs1889006 | 1.537 | 1.227 | -0.042 | -1.001 | 0.051 | -0.045 | -0.045 | 0.236 | 0.37 | 0.21 | 0.34 | 0.49 | 0.24 | 0.45 |
| 9 | 18619044 | rs10963675 | 0.191 | 1.285 | -0.103 | -0.946 | -0.003 | 0.022 | 0.022 | 1.62 | 0.56 | 0.38 | 0.56 | 0.72 | 0.48 | 0.65 |
| 9 | 18622043 | rs10963676 | -0.22 | 1.156 | -0.564 | -0.507 | -0.002 | -0.014 | -0.014 | 0 | 0.14 | 0.09 | 0.13 | 0.23 | 0.15 | 0.11 |
| 9 | 18629283 | rs10963680 | 0.634 | 0.63 | -0.691 | -0.419 | 0.059 | 0.008 | 0.008 | 1.518 | 0.38 | 0.16 | 0.28 | 0.62 | 0.24 | 0.42 |
| 9 | 18629792 | rs1576900 | 0.358 | 0.72 | -0.747 | -0.504 | 0.062 | 0.038 | 0.038 | 0.453 | 0.42 | 0.17 | 0.32 | 0.62 | 0.3 | 0.51 |
| 9 | 18631044 | rs10963684 | -0.403 | 1.658 | 0.213 | -0.11 | -0.007 | 0.004 | 0.004 | 1.534 | 0.09 | 0.08 | 0.22 | 0.12 | 0.16 | 0.11 |
| 9 | 18632108 | rs10481564 | -1.628 | 1.621 | 0.248 | -0.099 | -0.007 | -0.004 | -0.004 | 0.689 | 0.19 | 0.17 | 0.28 | 0.25 | 0.25 | 0.25 |
| 9 | 18632128 | rs1332712 | -0.989 | 1.431 | 0.154 | -0.084 | 0.034 | 0.013 | 0.013 | 1.184 | 0.25 | 0.51 | 0.36 | 0.31 | 0.29 | 0.26 |
| 9 | 18636286 | rs10963691 | -1.182 | 0.805 | -0.032 | 0.227 | -0.001 | 0.03 | 0.03 | 0.002 | 0.11 | 0.22 | 0.07 | 0.13 | 0.09 | 0.15 |
| 9 | 18636986 | rs17229989 | -0.025 | 0.62 | -0.574 | -0.502 | 0.011 | 0.039 | 0.039 | 0 | 0.22 | 0.13 | 0.11 | 0.04 | 0.2 | 0.21 |
| 9 | 18637737 | rs10811002 | -1.048 | 0.923 | 0.014 | -1.043 | 0.008 | 0.009 | 0.009 | 0 | 0.56 | 0.38 | 0.52 | 0.32 | 0.64 | 0.51 |
| 9 | 18638143 | rs1412671 | -0.757 | 1.482 | -0.23 | -1.248 | -0.004 | 0.017 | 0.017 | 0 | 0.45 | 0.36 | 0.38 | 0.21 | 0.53 | 0.41 |
| 9 | 18638517 | rs12352788 | 0.564 | 1.454 | -0.274 | -1.259 | -0.034 | 0.04 | 0.04 | 0 | 0.11 | 0.01 | 0.14 | 0.1 | 0.11 | 0.1 |
| 9 | 18640364 | rs10963695 | 0.061 | 1.645 | 0.475 | -1.367 | 0.001 | -0.003 | -0.003 | 0 | 0.08 | 0.07 | 0.12 | 0.1 | 0.08 | 0.14 |
| 9 | 18640431 | rs11795307 | 0.374 | 2.1 | 0.65 | -1.498 | -0.001 | 0.014 | 0.014 | 0 | 0.29 | 0.23 | 0.14 | 0.12 | 0.29 | 0.26 |
| 9 | 18640590 | rs7863100 | -0.534 | 1.96 | 0.56 | -1.418 | -0.009 | 0.006 | 0.006 | 0 | 0.16 | 0.12 | 0.24 | 0.1 | 0.24 | 0.15 |
| 9 | 18641667 | rs776754 | 1.161 | 1.937 | 0.631 | -1.421 | -0.022 | 0.032 | 0.032 | 0.93 | 0.69 | 0.75 | 0.59 | 0.79 | 0.58 | 0.65 |
| 9 | 18651444 | rs776785 | 0.387 | 1.32 | 0.366 | -1.713 | 0.025 | 0.078 | 0.078 | 0 | 0.5 | 0.74 | 0.57 | 0.67 | 0.56 | 0.57 |
| 9 | 18653324 | rs702203 | -0.02 | 1.279 | 0.402 | -1.906 | 0.007 | 0.072 | 0.072 | 0 | 0.58 | 0.7 | 0.67 | 0.8 | 0.57 | 0.55 |
| 9 | 18653380 | rs947144 | -0.061 | 1.371 | 0.346 | -1.906 | -0.023 | 0.025 | 0.025 | 0 | 0.18 | 0.06 | 0.22 | 0.1 | 0.21 | 0.08 |
| 9 | 18653773 | rs776782 | -0.939 | 1.405 | 0.626 | -1.938 | 0.027 | -0.052 | -0.052 | 0 | 0.26 | 0.43 | 0.31 | 0.1 | 0.32 | 0.16 |
| 9 | 18655961 | rs4977520 | 0.435 | 1.362 | 0.473 | -1.886 | 0.002 | 0.085 | 0.085 | 0 | 0.48 | 0.26 | 0.57 | 0.8 | 0.47 | 0.47 |
| 9 | 18656065 | rs4977522 | 1.783 | 1.358 | 0.412 | -1.851 | 0.006 | 0.005 | 0.005 | 0 | 0.3 | 0.19 | 0.35 | 0.71 | 0.24 | 0.39 |
| 9 | 18657019 | rs776778 | -1.096 | 1.082 | 0.2 | -1.781 | 0.002 | 0.018 | 0.018 | 0 | 0.73 | 0.8 | 0.83 | 0.91 | 0.72 | 0.75 |
| 9 | 18657061 | rs16936951 | 0.602 | 0.83 | -0.112 | -1.522 | 0.022 | -0.005 | -0.005 | 0 | 0.11 | 0.04 | 0.12 | 0.1 | 0.05 | 0.07 |
| 9 | 18657487 | rs776777 | 0.301 | 0.827 | -0.02 | -1.523 | 0 | 0.001 | 0.001 | 0 | 0.51 | 0.48 | 0.57 | 0.8 | 0.48 | 0.58 |
| 9 | 18657558 | rs776776 | -0.27 | 1.024 | -0.273 | -1.322 | 0.002 | -0.048 | -0.048 | 0 | 0.61 | 0.42 | 0.68 | 0.9 | 0.54 | 0.65 |

| | | | | | | | | | | | | | | | | |
|---|---|---|---|---|---|---|---|---|---|---|---|---|---|---|---|---|
| 9 | 18659794 | rs776773 | 0.428 | 1.116 | -0.224 | -0.691 | -0.002 | 0.002 | 0.002 | 0 | 0.34 | 0.3 | 0.4 | 0.69 | 0.31 | 0.4 |
| 9 | 18662292 | rs776771 | -0.677 | 0.985 | -0.176 | 0.045 | -0.003 | 0.026 | 0.026 | 0 | 0.29 | 0.36 | 0.15 | 0.43 | 0.27 | 0.49 |
| 9 | 18663543 | rs776770 | -0.409 | 1.095 | -0.122 | 0.022 | 0.06 | 0.238 | 0.238 | 0 | 0.12 | 0.29 | 0.08 | 0.4 | 0.14 | 0.29 |
| 9 | 18664041 | rs12352272 | -0.812 | 1.021 | -0.269 | -0.032 | 0.005 | 0.012 | 0.012 | 0 | 0.33 | 0.28 | 0.27 | 0.24 | 0.34 | 0.34 |
| 9 | 18664420 | rs702202 | -1.558 | 0.981 | 0.187 | 0.415 | -0.001 | 0 | 0 | 0 | 0.43 | 0.55 | 0.34 | 0.51 | 0.46 | 0.6 |
| 9 | 18666780 | rs1332708 | -0.226 | 1.151 | 0.499 | 0.62 | 0.014 | 0.076 | 0.076 | 0 | 0.11 | 0.22 | 0.07 | 0.28 | 0.11 | 0.27 |
| 9 | 18671467 | rs10811008 | -0.579 | 1.711 | -0.062 | 0.458 | 0.027 | 0.002 | 0.002 | 0 | 0.28 | 0.35 | 0.47 | 0.7 | 0.38 | 0.43 |
| 9 | 18672095 | rs1332706 | 0.881 | 1.509 | 0.244 | 0.289 | 0.022 | 0.001 | 0.001 | 0 | 0.66 | 0.39 | 0.33 | 0.44 | 0.51 | 0.67 |
| 9 | 18675325 | rs1360567 | 0.12 | 0.751 | 0.183 | -0.29 | 0.003 | -0.014 | -0.014 | 0 | 0.37 | 0.46 | 0.34 | 0.31 | 0.42 | 0.27 |
| 9 | 18676496 | rs2487477 | 0.471 | 1.019 | 0.151 | -0.123 | 0.009 | 0.014 | 0.014 | 0 | 0.13 | 0.16 | 0.08 | 0.21 | 0.16 | 0.22 |
| 9 | 18676945 | rs7045779 | 0.654 | 0.792 | 0.11 | -0.101 | -0.003 | -0.001 | -0.001 | 1.917 | 0.58 | 0.62 | 0.44 | 0.43 | 0.55 | 0.49 |
| 9 | 18679376 | rs10963718 | 2.235 | 0.551 | 0.047 | -0.207 | -0.002 | -0.004 | -0.004 | 0 | 0.22 | 0.2 | 0.11 | 0.14 | 0.15 | 0.23 |
| 9 | 18680771 | rs776761 | 1.598 | 0.596 | -0.062 | 0.088 | 0.023 | 0.051 | 0.051 | 0 | 0.08 | 0.15 | 0.04 | 0.18 | 0.1 | 0.19 |
| 9 | 18682318 | rs2811802 | -0.259 | 0.471 | -0.352 | -0.053 | -0.011 | -0.068 | -0.068 | 3.921 | 0.77 | 0.91 | 0.6 | 0.66 | 0.76 | 0.8 |
| 9 | 18683172 | rs6475237 | 0.286 | 0.166 | -0.356 | -0.011 | -0.008 | 0.005 | 0.005 | 0 | 0.37 | 0.31 | 0.35 | 0.36 | 0.47 | 0.32 |
| 9 | 18683680 | rs2482972 | 0.942 | 0.437 | -0.543 | 0.127 | -0.043 | -0.096 | -0.096 | 0 | 0.15 | 0.01 | 0.26 | 0.29 | 0.18 | 0.13 |
| 9 | 18689243 | rs944720 | -0.065 | 0.503 | -0.092 | 0.082 | -0.009 | 0.019 | 0.019 | 0 | 0.26 | 0.32 | 0.36 | 0.34 | 0.24 | 0.23 |
| 9 | 18700566 | rs2570569 | -0.635 | 0.316 | -0.411 | -0.092 | -0.001 | 0 | 0 | 0.658 | 0.46 | 0.4 | 0.58 | 0.52 | 0.47 | 0.52 |
| 9 | 18708175 | rs521911 | -1.658 | 0.153 | -0.368 | 0.109 | -0.016 | -0.015 | -0.015 | 0 | 0.79 | 0.71 | 0.9 | 0.84 | 0.86 | 0.81 |
| 9 | 18709694 | rs518607 | -1.321 | 0.097 | -0.177 | 0.023 | -0.025 | -0.046 | -0.046 | 0 | 0.75 | 0.39 | 0.8 | 0.86 | 0.85 | 0.78 |
| 9 | 18709767 | rs517764 | 0.474 | 0 | -0.65 | -0.121 | -0.03 | 0.005 | 0.005 | 0 | 0.24 | 0.15 | 0.27 | 0.19 | 0.34 | 0.19 |
| 9 | 18717903 | rs530120 | -1.856 | -0.026 | -0.569 | -0.133 | -0.038 | -0.041 | -0.041 | 0.776 | 0.77 | 0.63 | 0.9 | 0.86 | 0.88 | 0.8 |
| 9 | 18718086 | rs4977338 | 0.976 | -0.065 | -0.789 | -0.04 | 0.01 | -0.044 | -0.044 | 0 | 0.21 | 0.13 | 0.27 | 0.34 | 0.17 | 0.18 |
| 9 | 18720104 | rs482009 | -1.901 | 0.174 | -0.586 | 0.062 | -0.041 | -0.043 | -0.043 | 0.002 | 0.77 | 0.64 | 0.9 | 0.87 | 0.9 | 0.81 |
| 9 | 18723698 | rs578602 | -0.046 | 0.251 | -0.716 | 0.078 | 0.034 | -0.021 | -0.021 | 0 | 0.28 | 0.43 | 0.36 | 0.2 | 0.38 | 0.23 |
| 9 | 18724225 | rs574090 | -2.453 | 0.223 | -0.801 | 0.038 | -0.015 | -0.012 | -0.012 | 0 | 0.8 | 0.74 | 0.91 | 0.84 | 0.89 | 0.82 |
| 9 | 18727082 | rs944716 | 1.099 | 0.091 | -1.041 | 0.106 | -0.054 | -0.066 | -0.066 | 0.001 | 0.25 | 0.04 | 0.25 | 0.33 | 0.29 | 0.4 |
| 9 | 18727548 | rs532543 | -1.405 | 0.08 | -0.903 | -0.007 | -0.018 | -0.032 | -0.032 | 0 | 0.71 | 0.41 | 0.81 | 0.84 | 0.82 | 0.76 |
| 9 | 18728689 | rs563805 | -2.007 | 0.071 | -1.322 | 0.1 | -0.009 | -0.01 | -0.01 | 0.235 | 0.77 | 0.74 | 0.9 | 0.84 | 0.85 | 0.81 |
| 9 | 18730826 | rs476104 | 0.635 | 0.424 | -1.646 | 0.34 | -0.045 | 0.035 | 0.035 | 0 | 0.22 | 0.11 | 0.25 | 0.14 | 0.34 | 0.17 |
| 9 | 18731160 | rs748108 | 0.589 | 0.234 | -1.696 | 0.437 | 0.019 | -0.071 | -0.071 | 3.308 | 0.24 | 0.13 | 0.29 | 0.41 | 0.19 | 0.19 |
| 9 | 18732114 | rs944717 | -0.331 | 0.623 | -1.573 | 0.215 | -0.063 | 0.009 | 0.009 | 0 | 0.77 | 0.89 | 0.73 | 0.81 | 0.63 | 0.81 |
| 9 | 18733462 | rs576026 | 0.054 | 0.493 | -1.313 | 0.418 | -0.005 | 0.039 | 0.039 | 0 | 0.52 | 0.73 | 0.41 | 0.42 | 0.51 | 0.42 |
| 9 | 18738576 | rs568116 | -1.119 | 0.434 | -1.36 | 0.664 | 0.003 | -0.038 | -0.038 | 1.157 | 0.8 | 0.87 | 0.72 | 0.69 | 0.82 | 0.83 |
| 9 | 18739843 | rs2482970 | 0.398 | 0.475 | -1.915 | 0.454 | -0.007 | 0.01 | 0.01 | 0 | 0.22 | 0.12 | 0.24 | 0.16 | 0.27 | 0.16 |
| 9 | 18742434 | rs564014 | 0.265 | 0.449 | -1.911 | 0.308 | 0 | 0.049 | 0.049 | 0 | 0.23 | 0.15 | 0.24 | 0.11 | 0.27 | 0.14 |
| 9 | 18742990 | rs10963743 | 1.764 | 0.509 | -1.697 | 0.22 | 0.005 | 0.009 | 0.009 | 0 | 0.23 | 0.13 | 0.11 | 0.04 | 0.16 | 0.15 |
| 9 | 18744966 | rs1332704 | 0.322 | 0.578 | -1.649 | 0.317 | 0.008 | 0.027 | 0.027 | 0 | 0.5 | 0.39 | 0.42 | 0.46 | 0.5 | 0.6 |
| 9 | 18746561 | rs1549989 | 0.767 | 0.647 | -2.032 | 0.24 | -0.048 | -0.014 | -0.014 | 0 | 0.23 | 0.07 | 0.29 | 0.29 | 0.3 | 0.39 |
| 9 | 18751126 | rs1433808 | 0.037 | 0.28 | -1.684 | 0.277 | -0.008 | 0.053 | 0.053 | 0 | 0.21 | 0.11 | 0.22 | 0.12 | 0.26 | 0.14 |
| 9 | 18751638 | rs7043157 | 0.667 | 0.255 | -1.58 | 0.567 | 0.001 | 0.022 | 0.022 | 0 | 0.49 | 0.31 | 0.44 | 0.44 | 0.5 | 0.55 |
| 9 | 18752860 | rs2507122 | -0.419 | 0.314 | -1.647 | 0.471 | -0.008 | 0.008 | 0.008 | 0 | 0.21 | 0.1 | 0.22 | 0.16 | 0.26 | 0.15 |
| 9 | 18754211 | rs10733356 | 0.436 | -0.229 | -1.498 | 0.101 | 0.034 | 0.039 | 0.039 | 0 | 0.85 | 0.92 | 0.94 | 0.96 | 0.9 | 0.88 |
| 9 | 18758205 | rs2164000 | -0.034 | -0.207 | -1.319 | 0.054 | -0.011 | -0.019 | -0.019 | 0 | 0.61 | 0.6 | 0.76 | 0.77 | 0.72 | 0.72 |
| 9 | 18759822 | rs10429480 | 0.168 | 0.124 | -1.906 | 0.338 | 0.001 | -0.007 | -0.007 | 0.42 | 0.59 | 0.32 | 0.54 | 0.54 | 0.6 | 0.57 |
| 9 | 18760031 | rs4246839 | -0.142 | 0.439 | -2.001 | 0.301 | 0.01 | 0.018 | 0.018 | 0 | 0.87 | 0.93 | 0.95 | 0.91 | 0.92 | 0.85 |
| 9 | 18760665 | rs11794380 | 0.585 | 0.51 | -1.803 | 0.376 | 0.006 | 0.005 | 0.005 | 0 | 0.24 | 0.12 | 0.18 | 0.17 | 0.2 | 0.15 |
| 9 | 18761099 | rs1978746 | -0.418 | 1.451 | -1.767 | 0.648 | -0.037 | -0.037 | -0.037 | 0 | 0.08 | 0.02 | 0.25 | 0.15 | 0.2 | 0.24 |
| 9 | 18767301 | rs1433810 | 0.207 | 0.587 | -1.844 | -0.135 | -0.033 | 0.143 | 0.143 | 0 | 0.71 | 0.91 | 0.8 | 0.96 | 0.7 | 0.78 |
| 9 | 18767830 | rs10118309 | -0.028 | 0.325 | -1.289 | 0.477 | -0.012 | 0.122 | 0.122 | 0 | 0.11 | 0.31 | 0.16 | 0.31 | 0.08 | 0.18 |
| 9 | 18768201 | rs1991883 | 0.306 | 0.607 | -1.741 | -0.302 | -0.027 | 0.118 | 0.118 | 0 | 0.76 | 0.93 | 0.83 | 0.96 | 0.75 | 0.81 |
| 9 | 18772370 | rs7869627 | -0.084 | 0.412 | -1.181 | 0.27 | 0.011 | 0.015 | 0.015 | 0 | 0.5 | 0.38 | 0.6 | 0.56 | 0.51 | 0.55 |
| 9 | 18776116 | rs4295757 | 0.215 | 0.027 | -1.738 | -0.527 | 0.048 | 0.184 | 0.184 | 0 | 0.36 | 0.15 | 0.2 | 0.05 | 0.29 | 0.35 |
| 9 | 18778319 | rs2289006 | 0.995 | 0.151 | -1.396 | -0.922 | -0.013 | 0.032 | 0.032 | 0.121 | 0.33 | 0.24 | 0.22 | 0.13 | 0.36 | 0.18 |
| 9 | 18778885 | rs11789625 | 1.123 | -0.001 | -1.44 | -1.008 | 0.007 | 0.187 | 0.187 | 0.04 | 0.3 | 0.02 | 0.18 | 0.05 | 0.26 | 0.25 |
| 9 | 18779938 | rs12377348 | 1.771 | 0.388 | -1.387 | -1.093 | -0.001 | -0.047 | -0.047 | 0 | 0.21 | 0.21 | 0.13 | 0 | 0.18 | 0.04 |
| 9 | 18781154 | rs1549988 | -0.188 | 0.61 | -0.639 | -0.33 | 0.008 | 0.034 | 0.034 | 0.096 | 0.74 | 0.72 | 0.54 | 0.45 | 0.7 | 0.57 |
| 9 | 18782512 | rs7047163 | -1.037 | 0.544 | -0.682 | -0.175 | -0.015 | 0.045 | 0.045 | 0 | 0.84 | 0.95 | 0.9 | 0.94 | 0.86 | 0.93 |
| 9 | 18782704 | rs10811032 | 0.787 | 0.576 | -0.381 | 0.089 | 0.015 | 0.141 | 0.141 | 0 | 0.06 | 0.16 | 0.11 | 0.27 | 0.07 | 0.1 |
| 9 | 18786181 | rs715212 | 0.135 | 0.098 | -0.62 | -0.556 | 0.001 | 0.01 | 0.01 | 0 | 0.29 | 0.31 | 0.18 | 0.12 | 0.27 | 0.24 |
| 9 | 18786812 | rs12379407 | -0.053 | 0.104 | -0.535 | -0.257 | 0.028 | 0.247 | 0.247 | 0 | 0.09 | 0.32 | 0.12 | 0.3 | 0.08 | 0.12 |
| 9 | 18787794 | rs10963760 | 0.569 | -0.043 | -0.934 | -0.341 | -0.036 | -0.002 | -0.002 | 0 | 0.24 | 0.08 | 0.4 | 0.26 | 0.31 | 0.34 |
| 9 | 18789128 | rs4977432 | 1.29 | -0.151 | -0.914 | -0.261 | -0.027 | -0.006 | -0.006 | 0 | 0.13 | 0.07 | 0.11 | 0.13 | 0.17 | 0.13 |
| 9 | 18789352 | rs2033541 | 1.058 | -0.674 | -0.523 | -0.303 | 0.01 | 0.005 | 0.005 | 0 | 0.36 | 0.29 | 0.24 | 0.25 | 0.25 | 0.24 |
| 9 | 18791355 | rs10963764 | -0.414 | -0.444 | -0.189 | -0.007 | -0.03 | 0.15 | 0.15 | 0.183 | 0.11 | 0.32 | 0.11 | 0.3 | 0.06 | 0.14 |
| 9 | 18795058 | rs10963766 | -0.47 | -0.519 | -0.333 | -0.287 | -0.031 | 0.053 | 0.053 | 0 | 0.3 | 0.55 | 0.44 | 0.55 | 0.27 | 0.38 |
| 9 | 18795304 | rs2289005 | 0.038 | -0.527 | -0.2 | -0.267 | -0.007 | -0.004 | -0.004 | 0 | 0.24 | 0.28 | 0.35 | 0.24 | 0.24 | 0.27 |
| 9 | 18795679 | rs7847591 | 1.987 | -0.608 | -0.452 | -0.304 | 0.002 | 0.013 | 0.013 | 0 | 0.2 | 0.11 | 0.13 | 0 | 0.17 | 0.04 |
| 9 | 18796886 | rs1433825 | -0.613 | -0.372 | -0.56 | 0.297 | -0.012 | 0.117 | 0.117 | 0 | 0.09 | 0.28 | 0.05 | 0.21 | 0.05 | 0.12 |

| | | | | | | | | | | | | | | | | |
|---|---|---|---|---|---|---|---|---|---|---|---|---|---|---|---|---|
| 9 | 18797256 | rs2289003 | 0.4 | -0.179 | -1.544 | 0.214 | -0.027 | -0.027 | -0.027 | 0.345 | 0.57 | 0.84 | 0.42 | 0.5 | 0.45 | 0.47 |
| 9 | 18797920 | rs10963767 | -0.04 | 0.275 | -1.102 | 0.695 | -0.033 | -0.032 | -0.032 | 0 | 0.22 | 0.13 | 0.39 | 0.37 | 0.36 | 0.36 |
| 9 | 18798603 | rs1433819 | -0.19 | 0.086 | -0.873 | 0.673 | -0.007 | -0.031 | -0.031 | 0 | 0.06 | 0.01 | 0.04 | 0.2 | 0.06 | 0.12 |
| 9 | 18801305 | rs10963769 | 0.569 | -0.085 | -0.631 | 0.031 | 0.001 | 0.008 | 0.008 | 0 | 0.24 | 0.16 | 0.17 | 0.03 | 0.24 | 0.15 |
| 9 | 18804130 | rs7034696 | -0.104 | 0.115 | -0.648 | 0.454 | -0.006 | -0.007 | -0.007 | 0.024 | 0.21 | 0.22 | 0.3 | 0.36 | 0.27 | 0.34 |
| 9 | 18804418 | rs4977444 | 0.111 | -0.231 | -0.256 | 0.689 | -0.008 | 0.027 | 0.027 | 0 | 0.79 | 0.76 | 0.75 | 0.63 | 0.83 | 0.75 |
| 9 | 18805813 | rs10963773 | 1.084 | -0.342 | 0.189 | 0.096 | -0.005 | -0.015 | -0.015 | 0 | 0.32 | 0.38 | 0.27 | 0.21 | 0.27 | 0.26 |
| 9 | 18806093 | rs7870115 | -0.204 | -0.324 | 0.285 | 0.352 | -0.001 | -0.009 | -0.009 | 0 | 0.55 | 0.47 | 0.56 | 0.7 | 0.57 | 0.63 |
| 9 | 18806927 | rs4977446 | -0.894 | -0.251 | 0.372 | 0.451 | -0.002 | -0.004 | -0.004 | 0 | 0.29 | 0.3 | 0.28 | 0.49 | 0.33 | 0.4 |
| 9 | 18806999 | rs4977447 | 0.346 | -0.136 | 0.152 | 0.815 | -0.077 | -0.109 | -0.109 | 0 | 0.18 | 0.03 | 0.22 | 0.32 | 0.26 | 0.18 |
| 9 | 18807071 | rs4977448 | -0.314 | -0.193 | 0.183 | 0.777 | -0.02 | -0.042 | -0.042 | 0 | 0.23 | 0.17 | 0.26 | 0.47 | 0.32 | 0.37 |
| 9 | 18809482 | rs7854121 | 0.772 | -0.366 | 0.345 | 0.547 | 0.009 | 0.006 | 0.006 | 0 | 0.62 | 0.45 | 0.51 | 0.6 | 0.57 | 0.62 |
| 9 | 18810029 | rs2148847 | -0.1 | -0.545 | 0.064 | 0.211 | 0.015 | 0.053 | 0.053 | 0 | 0.52 | 0.75 | 0.51 | 0.69 | 0.56 | 0.6 |
| 9 | 18811047 | rs10963779 | 0.737 | -0.28 | 0.004 | 0.36 | -0.027 | -0.023 | -0.023 | 0 | 0.6 | 0.66 | 0.42 | 0.45 | 0.5 | 0.5 |
| 9 | 18817246 | rs1433809 | -1.22 | -0.269 | 0.039 | 0.137 | 0.007 | 0.011 | 0.011 | 0 | 0.86 | 0.91 | 0.79 | 0.95 | 0.91 | 0.91 |
| 9 | 18817594 | rs7874108 | 1.736 | -0.426 | 0.145 | -0.104 | -0.002 | -0.112 | -0.112 | 0.012 | 0.45 | 0.41 | 0.32 | 0.01 | 0.4 | 0.21 |
| 9 | 18818890 | rs10156461 | -0.264 | 0 | 0.303 | 0.469 | 0.031 | 0.145 | 0.145 | 0 | 0.1 | 0.27 | 0.1 | 0.28 | 0.1 | 0.27 |
| 9 | 18819568 | rs1980867 | -0.93 | -0.139 | 0.204 | 0.64 | -0.01 | 0.066 | 0.066 | 0 | 0.15 | 0.2 | 0.1 | 0.48 | 0.1 | 0.26 |
| 9 | 18823505 | rs7019448 | -0.125 | -0.15 | 0.075 | 0.533 | 0.023 | -0.018 | -0.018 | 0 | 0.81 | 0.91 | 0.76 | 0.74 | 0.86 | 0.82 |
| 9 | 18826500 | rs934469 | 1.09 | -0.095 | 0.392 | 0.569 | -0.01 | 0.049 | 0.049 | 0 | 0.7 | 0.48 | 0.66 | 0.47 | 0.77 | 0.56 |
| 9 | 18826610 | rs10963786 | 0.581 | -0.06 | 0.277 | 0.694 | -0.004 | 0.08 | 0.08 | 0 | 0.08 | 0.1 | 0.03 | 0.26 | 0.05 | 0.15 |
| 9 | 18827034 | rs718452 | 0.274 | -0.071 | 0.291 | 0.677 | 0.001 | 0.03 | 0.03 | 0 | 0.77 | 0.7 | 0.69 | 0.67 | 0.81 | 0.66 |
| 9 | 18830444 | rs16937127 | 0.49 | -0.045 | -0.18 | 0.615 | 0.007 | -0.033 | -0.033 | 2.195 | 0.06 | 0.01 | 0.02 | 0.2 | 0.04 | 0.1 |
| 9 | 18831022 | rs7858256 | 0.684 | -0.153 | -0.119 | 0.285 | 0.004 | 0.025 | 0.025 | 0 | 0.35 | 0.25 | 0.32 | 0.23 | 0.35 | 0.36 |
| 9 | 18831340 | rs7858859 | -0.247 | -0.201 | 0.133 | 0.408 | -0.003 | -0.006 | -0.006 | 0 | 0.45 | 0.6 | 0.4 | 0.51 | 0.46 | 0.6 |
| 9 | 18831444 | rs7859001 | 0.39 | -0.351 | 0.107 | 0.422 | -0.017 | 0.053 | 0.053 | 0.05 | 0.12 | 0.23 | 0.19 | 0.22 | 0.11 | 0.21 |
| 9 | 18832027 | rs10122919 | 0.853 | -0.309 | -0.03 | 0.459 | -0.002 | 0.008 | 0.008 | 0 | 0.1 | 0.15 | 0.2 | 0.24 | 0.11 | 0.21 |
| 9 | 18832659 | rs10811043 | 0.579 | -0.244 | -0.078 | 0.46 | -0.005 | 0.131 | 0.131 | 0 | 0.08 | 0.16 | 0.04 | 0.27 | 0.06 | 0.15 |
| 9 | 18833014 | rs7039188 | 0.827 | -0.283 | -0.092 | 0.533 | -0.036 | 0.04 | 0.04 | 0 | 0.69 | 0.59 | 0.52 | 0.51 | 0.75 | 0.61 |
| 9 | 18833547 | rs10963788 | 0.851 | 0.357 | -0.93 | 0.488 | -0.07 | -0.073 | -0.073 | 1.086 | 0.14 | 0.01 | 0.2 | 0.23 | 0.22 | 0.14 |
| 9 | 18835308 | rs1347265 | 0.473 | -0.04 | -0.431 | 0.759 | 0 | 0.003 | 0.003 | 0.01 | 0.35 | 0.35 | 0.19 | 0.47 | 0.32 | 0.39 |
| 9 | 18835570 | rs10756991 | -0.046 | -0.004 | -0.519 | 0.519 | 0.001 | 0.029 | 0.029 | 0 | 0.44 | 0.34 | 0.36 | 0.7 | 0.43 | 0.6 |
| 9 | 18847325 | rs7859917 | 0.112 | 0.032 | -0.824 | 0.655 | 0.001 | -0.008 | -0.008 | 0 | 0.39 | 0.66 | 0.31 | 0.55 | 0.33 | 0.33 |
| 9 | 18848066 | rs1368771 | -0.743 | -0.129 | -1.766 | 0.581 | -0.02 | 0.022 | 0.022 | 0 | 0.63 | 0.91 | 0.57 | 0.72 | 0.64 | 0.77 |
| 9 | 18848686 | rs1001379 | -0.568 | -0.244 | -1.07 | 0.997 | -0.001 | 0.181 | 0.181 | 3.595 | 0.09 | 0.59 | 0.06 | 0.23 | 0.05 | 0.24 |
| 9 | 18853257 | rs1814370 | 0.97 | -0.299 | -1.678 | 0.768 | -0.069 | 0.081 | 0.081 | 0 | 0.42 | 0.12 | 0.32 | 0.27 | 0.45 | 0.29 |
| 9 | 18853327 | rs11791292 | 0.18 | -0.302 | -1.701 | 0.785 | -0.01 | -0.019 | -0.019 | 0.414 | 0.11 | 0.08 | 0.19 | 0.25 | 0.13 | 0.24 |
| 9 | 18857005 | rs10963806 | -0.165 | -0.424 | -1.401 | 0.658 | -0.024 | 0.143 | 0.143 | 1.893 | 0.1 | 0.54 | 0.05 | 0.18 | 0.03 | 0.15 |
| 9 | 18861021 | rs10811051 | -0.058 | -0.341 | -1.627 | 0.624 | 0.038 | -0.013 | -0.013 | 0 | 0.52 | 0.24 | 0.51 | 0.52 | 0.57 | 0.54 |
| 9 | 18863560 | rs7039790 | -0.764 | -0.579 | -1.133 | 0.718 | -0.053 | 0.071 | 0.071 | 2.336 | 0.14 | 0.55 | 0.07 | 0.15 | 0.05 | 0.17 |
| 9 | 18865803 | rs12685549 | 0.308 | -0.146 | -1.376 | 1.013 | -0.034 | -0.054 | -0.054 | 0 | 0.09 | 0.02 | 0.11 | 0.29 | 0.12 | 0.1 |
| 9 | 18866563 | rs1897725 | 0.988 | -0.391 | -1.259 | 0.569 | 0.008 | 0.011 | 0.011 | 0 | 0.66 | 0.54 | 0.39 | 0.44 | 0.61 | 0.45 |
| 9 | 18872273 | rs10963814 | 1.433 | -0.41 | -1.185 | 0.88 | 0 | 0.013 | 0.013 | 0 | 0.05 | 0.04 | 0.16 | 0.17 | 0.06 | 0.09 |
| 9 | 18877987 | rs7047294 | 0.405 | -0.318 | -1.206 | 0.613 | -0.002 | -0.001 | -0.001 | 0 | 0.62 | 0.63 | 0.45 | 0.47 | 0.55 | 0.5 |
| 9 | 18878842 | rs2082580 | 0.316 | -0.459 | -1.408 | 0.376 | -0.004 | -0.016 | -0.016 | 0 | 0.4 | 0.48 | 0.26 | 0.29 | 0.38 | 0.26 |
| 9 | 18882562 | rs1368772 | -0.456 | -0.36 | -1.378 | 0.416 | 0 | 0 | 0 | 1.504 | 0.7 | 0.72 | 0.64 | 0.64 | 0.64 | 0.75 |
| 9 | 18883301 | rs2209778 | 1.269 | -0.359 | -1.422 | 0.539 | 0 | 0.011 | 0.011 | 0 | 0.07 | 0.05 | 0.18 | 0.21 | 0.1 | 0.11 |
| 9 | 18884432 | rs2099392 | 0.218 | -0.387 | -1.357 | 0.522 | 0.001 | -0.001 | -0.001 | 0.015 | 0.62 | 0.62 | 0.39 | 0.46 | 0.55 | 0.51 |
| 9 | 18885598 | rs4977265 | 1.13 | -0.459 | -1.822 | 0.044 | -0.055 | -0.016 | -0.016 | 0 | 0.28 | 0.06 | 0.37 | 0.23 | 0.34 | 0.28 |
| 9 | 18886370 | rs2082581 | 2.721 | -0.535 | -1.296 | 0.252 | 0.011 | 0.012 | 0.012 | 0.064 | 0.17 | 0.09 | 0.09 | 0 | 0.12 | 0.04 |
| 9 | 18888920 | rs3780223 | 0.112 | -0.431 | -1.104 | 0.436 | -0.004 | 0.078 | 0.078 | 0.076 | 0.09 | 0.21 | 0.19 | 0.17 | 0.09 | 0.11 |
| 9 | 18891040 | rs7030565 | -1.287 | -0.616 | -1.256 | 0.361 | -0.026 | 0.02 | 0.02 | 0.642 | 0.22 | 0.68 | 0.26 | 0.3 | 0.17 | 0.23 |
| 9 | 18891487 | rs934480 | 2.25 | -0.701 | -1.446 | 0.125 | 0.012 | 0.016 | 0.016 | 0.354 | 0.2 | 0.11 | 0.12 | 0 | 0.16 | 0.2 |
| 9 | 18891793 | rs2164001 | 0.253 | -0.866 | -1.545 | 0.137 | -0.019 | -0.022 | -0.022 | 0 | 0.34 | 0.26 | 0.42 | 0.51 | 0.46 | 0.45 |
| 9 | 18891910 | rs7031396 | 0.513 | -0.954 | -1.397 | 0.085 | 0.007 | 0.005 | 0.005 | 0.565 | 0.33 | 0.65 | 0.17 | 0.13 | 0.17 | 0.16 |
| 9 | 18894714 | rs10811055 | 0.878 | -0.812 | -1.282 | 0.399 | 0 | 0.004 | 0.004 | 0.033 | 0.08 | 0.08 | 0.21 | 0.19 | 0.13 | 0.13 |
| 9 | 18896061 | rs2082582 | -0.466 | -0.862 | -1.111 | 0.248 | -0.074 | 0.007 | 0.007 | 1.905 | 0.15 | 0.49 | 0.06 | 0.12 | 0.04 | 0.11 |
| 9 | 18903330 | rs1327504 | -0.594 | -0.419 | -1.301 | 0.232 | -0.005 | 0.009 | 0.009 | 0.962 | 0.48 | 0.65 | 0.55 | 0.62 | 0.55 | 0.63 |
| 9 | 18903625 | rs7467533 | -0.361 | -0.407 | -1.312 | 0.223 | -0.004 | 0.002 | 0.002 | 0.954 | 0.46 | 0.65 | 0.52 | 0.62 | 0.51 | 0.47 |
| 9 | 18905289 | rs7864878 | 2.34 | -0.447 | -1.387 | -0.123 | -0.007 | -0.051 | -0.051 | 0 | 0.19 | 0.19 | 0.11 | 0.01 | 0.13 | 0.15 |
| 9 | 18905525 | rs13285304 | -0.211 | -0.366 | -1.475 | 0.513 | -0.038 | -0.021 | -0.021 | 0.206 | 0.64 | 0.82 | 0.42 | 0.45 | 0.56 | 0.66 |
| 9 | 18907551 | rs945440 | -0.005 | -0.488 | -0.939 | 0.427 | 0.003 | 0.012 | 0.012 | 0.092 | 0.1 | 0.13 | 0.31 | 0.17 | 0.16 | 0.14 |
| 9 | 18907769 | rs945441 | -1.178 | -0.87 | -0.538 | 0.218 | -0.065 | -0.048 | -0.048 | 0 | 0.19 | 0.54 | 0.17 | 0.08 | 0.1 | 0.33 |
| 9 | 18909597 | rs16937177 | -0.484 | -0.5 | -0.295 | 0.239 | -0.011 | 0.011 | 0.011 | 0 | 0.12 | 0.33 | 0.16 | 0.1 | 0.05 | 0.06 |
| 9 | 18909703 | rs10117672 | 1.181 | -0.264 | -0.685 | 0.087 | -0.017 | 0.047 | 0.047 | 0 | 0.37 | 0.28 | 0.28 | 0.16 | 0.42 | 0.26 |
| 9 | 18910581 | rs11794832 | -0.463 | -0.266 | -0.011 | 0.656 | 0.001 | 0.002 | 0.002 | 0 | 0.13 | 0.16 | 0.29 | 0.21 | 0.21 | 0.3 |
| 9 | 18911029 | rs16937178 | -0.696 | -0.415 | 0.126 | 0.646 | -0.016 | 0.017 | 0.017 | 0 | 0.12 | 0.34 | 0.16 | 0.11 | 0.06 | 0.08 |
| 9 | 18913052 | rs17808244 | 0.263 | -0.186 | -0.673 | 0.569 | -0.016 | -0.01 | -0.01 | 0 | 0.08 | 0 | 0.07 | 0.08 | 0.07 | 0.06 |
| 9 | 18913080 | rs10811060 | 0.81 | -0.089 | -0.774 | 0.577 | -0.023 | -0.003 | -0.003 | 0 | 0.23 | 0.07 | 0.25 | 0.24 | 0.26 | 0.16 |

| | | | | | | | | | | | | | | | | |
|---|---|---|---|---|---|---|---|---|---|---|---|---|---|---|---|---|
| 9 | 18914103 | rs10811061 | -0.385 | -0.176 | -0.563 | 0.634 | 0.011 | -0.011 | -0.011 | 0.193 | 0.34 | 0.68 | 0.34 | 0.53 | 0.24 | 0.46 |
| 9 | 18914729 | rs7860467 | -0.943 | 0.085 | -0.478 | 0.561 | -0.043 | 0.012 | 0.012 | 0 | 0.59 | 0.84 | 0.6 | 0.63 | 0.5 | 0.55 |
| 9 | 18916311 | rs12345835 | -0.354 | 0.454 | 0.442 | 0.757 | 0.101 | 0.005 | 0.005 | 0.132 | 0.14 | 0.58 | 0.2 | 0.12 | 0.18 | 0.12 |
| 9 | 18917251 | rs1932458 | -1.063 | 0.475 | -0.125 | 0.788 | 0.001 | 0.035 | 0.035 | 0 | 0.58 | 0.81 | 0.62 | 0.68 | 0.59 | 0.51 |
| 9 | 18917347 | rs2182729 | -0.193 | 0.437 | 0.307 | 0.772 | 0.02 | 0.012 | 0.012 | 0 | 0.46 | 0.39 | 0.48 | 0.57 | 0.41 | 0.45 |
| 9 | 18917483 | rs2148850 | -0.794 | 0.501 | -0.535 | 0.913 | 0.004 | 0.015 | 0.015 | 0 | 0.57 | 0.67 | 0.62 | 0.62 | 0.59 | 0.52 |
| 9 | 18918845 | rs4540491 | -0.448 | 0.982 | -0.69 | 1.64 | -0.017 | -0.036 | -0.036 | 0 | 0.94 | 1 | 0.93 | 0.86 | 0.91 | 0.9 |
| 9 | 18921386 | rs10811063 | -0.647 | 0.912 | -0.526 | 1.6 | -0.022 | -0.105 | -0.105 | 3.788 | 0.24 | 0.08 | 0.31 | 0.54 | 0.25 | 0.28 |
| 9 | 18924136 | rs731529 | 0.269 | 0.571 | -0.237 | 1.311 | -0.003 | -0.062 | -0.062 | 0 | 0.15 | 0.03 | 0.14 | 0.24 | 0.11 | 0.09 |
| 9 | 18924463 | rs7022140 | -0.912 | 0.639 | 0.183 | 1.205 | 0.028 | -0.011 | -0.011 | 0.247 | 0.19 | 0.59 | 0.22 | 0.16 | 0.21 | 0.25 |
| 9 | 18925343 | rs4977266 | -1.211 | 0.562 | -0.445 | 1.028 | -0.003 | -0.021 | -0.021 | 0 | 0.37 | 0.71 | 0.37 | 0.38 | 0.33 | 0.35 |
| 9 | 18926997 | rs7031603 | -0.632 | 0.009 | 0.357 | 1.273 | -0.055 | 0.072 | 0.072 | 0.01 | 0.2 | 0.38 | 0.2 | 0.27 | 0.1 | 0.11 |
| 9 | 18927177 | rs4977467 | -0.708 | 0.124 | 0.414 | 1.251 | -0.028 | 0.034 | 0.034 | 0.634 | 0.22 | 0.35 | 0.16 | 0.28 | 0.14 | 0.2 |
| 9 | 18927768 | rs945442 | 0.875 | 0.825 | 0.499 | 1.533 | 0.023 | 0.062 | 0.062 | 0.004 | 0.79 | 0.6 | 0.78 | 0.69 | 0.77 | 0.67 |
| 9 | 18928039 | rs1052448 | -0.015 | 0.596 | -0.162 | 1.516 | -0.021 | -0.099 | -0.099 | 0 | 0.2 | 0.01 | 0.14 | 0.23 | 0.16 | 0.12 |
| 9 | 18929692 | rs7871217 | -0.618 | 1.476 | 0.392 | 0.979 | 0.091 | -0.061 | -0.061 | 0.21 | 0.11 | 0.3 | 0.14 | 0.01 | 0.16 | 0.12 |
| 9 | 18930880 | rs4977469 | -0.235 | 1.549 | 0.492 | 1.132 | 0.022 | 0.124 | 0.124 | 0 | 0.9 | 0.72 | 0.81 | 0.71 | 0.84 | 0.84 |
| 9 | 18932805 | rs1932456 | -0.804 | 1.552 | 0.504 | 0.838 | 0.098 | -0.062 | -0.062 | 0 | 0.11 | 0.32 | 0.14 | 0.01 | 0.16 | 0.17 |
| 9 | 18934303 | rs4977474 | -0.388 | 0.303 | -0.425 | 1.279 | -0.009 | -0.052 | -0.052 | 0 | 0.23 | 0.01 | 0.13 | 0.19 | 0.17 | 0.21 |
| 9 | 18935680 | rs10963844 | -1.489 | 0.415 | -0.144 | 1.434 | 0.029 | -0.001 | -0.001 | 0 | 0.43 | 0.77 | 0.43 | 0.53 | 0.35 | 0.41 |
| 9 | 18937624 | rs1977445 | 0.74 | 0.683 | -0.029 | 1.597 | -0.038 | -0.062 | -0.062 | 0 | 0.71 | 0.87 | 0.66 | 0.48 | 0.62 | 0.65 |
| 9 | 18939948 | rs945445 | 0.997 | 0.835 | 0.759 | 1.379 | 0.006 | 0.019 | 0.019 | 0 | 0.73 | 0.66 | 0.64 | 0.6 | 0.72 | 0.65 |
| 9 | 18940950 | rs10757004 | 1.623 | 0.608 | 0.381 | 1.063 | -0.044 | -0.089 | -0.089 | 0 | 0.61 | 0.09 | 0.56 | 0.53 | 0.61 | 0.6 |
| 9 | 18945868 | rs7864679 | -0.438 | 0.48 | 1.08 | 1.006 | -0.031 | -0.087 | -0.087 | 0 | 0.13 | 0.39 | 0.05 | 0.03 | 0.05 | 0.16 |
| 9 | 18947253 | rs7045203 | 0.168 | 0.573 | 0.977 | 1.121 | -0.039 | -0.06 | -0.06 | 0.61 | 0.85 | 0.95 | 0.69 | 0.57 | 0.77 | 0.79 |
| 9 | 18948086 | rs4977476 | 1.086 | 0.534 | 1.234 | 1.089 | 0.011 | 0.042 | 0.042 | 0 | 0.77 | 0.77 | 0.64 | 0.54 | 0.73 | 0.7 |
| 9 | 18952553 | rs12115446 | 0.441 | 0.454 | 1.332 | 1.097 | -0.002 | -0.021 | -0.021 | 0.032 | 0.07 | 0.18 | 0.05 | 0.03 | 0.05 | 0.02 |
| 9 | 18953625 | rs4977477 | 1.47 | 0.448 | 0.931 | 1.097 | 0.006 | -0.025 | -0.025 | 0 | 0.65 | 0.29 | 0.61 | 0.52 | 0.71 | 0.61 |
| 9 | 18955861 | rs2039010 | 0.247 | 0.501 | 0.97 | 1.167 | -0.001 | 0.019 | 0.019 | 0 | 0.85 | 0.88 | 0.7 | 0.57 | 0.8 | 0.8 |
| 9 | 18956317 | rs1006120 | 0.055 | 0.505 | 0.945 | 1.169 | -0.029 | -0.05 | -0.05 | 1.36 | 0.86 | 0.95 | 0.72 | 0.57 | 0.8 | 0.8 |
| 9 | 18958107 | rs6475276 | 0.307 | 0.586 | 1.063 | 1.171 | 0.034 | 0.128 | 0.128 | 0 | 0.84 | 0.7 | 0.69 | 0.57 | 0.78 | 0.79 |
| 9 | 18969422 | rs10511666 | -0.711 | 0.484 | 0.939 | 1.161 | -0.043 | -0.046 | -0.046 | 0 | 0.15 | 0.65 | 0.07 | 0.09 | 0.05 | 0.15 |
| 9 | 18973653 | rs7856285 | 1.717 | 0.456 | 0.809 | 1.212 | 0.004 | -0.098 | -0.098 | 0.04 | 0.63 | 0.12 | 0.61 | 0.46 | 0.72 | 0.58 |
| 9 | 18973918 | rs6475294 | 0.218 | 0.685 | 0.829 | 1.198 | -0.006 | 0.005 | 0.005 | 0 | 0.85 | 0.91 | 0.72 | 0.57 | 0.81 | 0.77 |
| 9 | 18974072 | rs6475295 | 0.95 | 0.54 | 0.805 | 1.177 | 0.005 | 0.079 | 0.079 | 0 | 0.78 | 0.67 | 0.67 | 0.55 | 0.78 | 0.74 |
| 9 | 18978625 | rs6475298 | 0.023 | 0.67 | 0.381 | 1.262 | -0.065 | -0.108 | -0.108 | 1.865 | 0.86 | 0.99 | 0.74 | 0.58 | 0.82 | 0.76 |
| 9 | 18984007 | rs7037837 | 1.648 | 0.071 | 0.337 | 0.669 | 0.041 | -0.005 | -0.005 | 0 | 0.53 | 0.25 | 0.51 | 0.52 | 0.65 | 0.51 |
| 9 | 18986580 | rs12552916 | 0.631 | 0.026 | 0.373 | 0.629 | 0.008 | 0.037 | 0.037 | 0 | 0.11 | 0.03 | 0.11 | 0.01 | 0.07 | 0.1 |
| 9 | 18989696 | rs10511667 | 0.231 | -0.038 | 0.337 | 0.67 | 0.001 | 0.047 | 0.047 | 0 | 0.13 | 0.03 | 0.14 | 0.03 | 0.1 | 0.11 |
| 9 | 18990027 | rs4506313 | 1.151 | -0.065 | 0.95 | 0.667 | -0.019 | 0.033 | 0.033 | 0 | 0.67 | 0.57 | 0.66 | 0.55 | 0.74 | 0.67 |
| 9 | 18995323 | rs10963898 | 1.548 | -0.021 | 1.033 | 0.715 | 0.028 | -0.007 | -0.007 | 1.106 | 0.54 | 0.3 | 0.5 | 0.52 | 0.63 | 0.56 |
| 9 | 19007561 | rs7858618 | -0.363 | -0.008 | 1.003 | 0.714 | -0.014 | -0.039 | -0.039 | 0 | 0.86 | 0.93 | 0.75 | 0.61 | 0.83 | 0.77 |
| 9 | 19009041 | rs10811094 | 2.135 | 0.056 | 0.656 | 0.61 | -0.061 | -0.021 | -0.021 | 1.705 | 0.39 | 0.17 | 0.4 | 0.39 | 0.42 | 0.4 |
| 9 | 19010440 | rs10811095 | -0.71 | 0.031 | 0.719 | 0.578 | -0.034 | -0.1 | -0.1 | 0.955 | 0.3 | 0.4 | 0.23 | 0.09 | 0.19 | 0.21 |
| 9 | 19010574 | rs17236081 | 1.624 | 0.051 | 0.704 | 0.594 | 0.03 | -0.004 | -0.004 | 0 | 0.52 | 0.26 | 0.47 | 0.51 | 0.55 | 0.47 |
| 9 | 19010867 | rs10811097 | -0.703 | 0.003 | 0.727 | 0.591 | -0.039 | -0.112 | -0.112 | 0 | 0.3 | 0.45 | 0.22 | 0.09 | 0.19 | 0.21 |
| 9 | 19011300 | rs16937267 | 0.652 | 0.02 | 0.655 | 0.607 | -0.019 | -0.027 | -0.027 | 0.031 | 0.12 | 0.01 | 0.07 | 0.12 | 0.13 | 0.07 |
| 9 | 19017723 | rs10811099 | 1.308 | 0.107 | 0.714 | 0.579 | 0.017 | -0.002 | -0.002 | 0 | 0.54 | 0.36 | 0.51 | 0.52 | 0.63 | 0.56 |
| 9 | 19017868 | rs10963906 | 0.543 | 0.207 | 0.59 | 0.59 | -0.027 | -0.034 | -0.034 | 0 | 0.13 | 0.01 | 0.07 | 0.13 | 0.14 | 0.1 |
| 9 | 19019820 | rs12349207 | -0.486 | 0.122 | 0.668 | 0.612 | 0.008 | 0.061 | 0.061 | 0 | 0.85 | 0.8 | 0.73 | 0.59 | 0.81 | 0.77 |
| 9 | 19022470 | rs10511668 | 2.182 | 0.083 | 0.561 | 0.598 | -0.083 | -0.037 | -0.037 | 0 | 0.38 | 0.11 | 0.4 | 0.38 | 0.42 | 0.38 |
| 9 | 19022995 | rs10963912 | 1.651 | 0.056 | 0.596 | 0.584 | 0.02 | -0.009 | -0.009 | 0.409 | 0.51 | 0.18 | 0.48 | 0.51 | 0.55 | 0.49 |
| 9 | 19026350 | rs7028384 | -0.791 | 0.165 | 0.645 | 0.391 | -0.032 | -0.13 | -0.13 | 0 | 0.32 | 0.46 | 0.24 | 0.08 | 0.2 | 0.2 |
| 9 | 19030014 | rs10757024 | -1.681 | 0.155 | 0.635 | 0.393 | 0.05 | 0.14 | 0.14 | 0 | 0.48 | 0.75 | 0.35 | 0.21 | 0.4 | 0.37 |
| 9 | 19031231 | rs10963919 | 0.574 | 0.161 | 0.611 | 0.399 | 0.005 | -0.003 | -0.003 | 0 | 0.13 | 0.07 | 0.07 | 0.13 | 0.13 | 0.09 |
| 9 | 19032907 | rs17810415 | 0.071 | 0.152 | 0.915 | 0.377 | 0 | -0.03 | -0.03 | 0 | 0.17 | 0.23 | 0.19 | 0.03 | 0.18 | 0.14 |
| 9 | 19033006 | rs10963921 | -1.537 | 0.228 | 0.387 | 0.427 | 0.03 | 0.076 | 0.076 | 0 | 0.47 | 0.66 | 0.33 | 0.21 | 0.4 | 0.37 |
| 9 | 19033058 | rs10963922 | 2.218 | 0.248 | 0.323 | 0.729 | -0.047 | -0.066 | -0.066 | 0 | 0.37 | 0.16 | 0.45 | 0.49 | 0.42 | 0.4 |
| 9 | 19045502 | rs10125646 | -1.615 | 0.419 | 0.453 | 0.223 | 0.025 | 0.066 | 0.066 | 0 | 0.47 | 0.66 | 0.35 | 0.21 | 0.41 | 0.38 |
| 9 | 19051208 | rs7022613 | 2.017 | 0.088 | 0.392 | 0.375 | -0.054 | -0.003 | -0.003 | 0 | 0.39 | 0.17 | 0.4 | 0.37 | 0.44 | 0.43 |
| 9 | 19053247 | rs7031392 | 0.634 | 0.006 | 0.36 | 0.253 | -0.002 | -0.005 | -0.005 | 0 | 0.11 | 0.09 | 0.06 | 0.13 | 0.16 | 0.11 |
| 9 | 19058821 | rs10511671 | 1.134 | -0.035 | 0.372 | 0.284 | 0.004 | -0.009 | -0.009 | 0 | 0.1 | 0.07 | 0.05 | 0.13 | 0.09 | 0.05 |
| 9 | 19059678 | rs4977494 | 0.747 | -0.022 | 0.325 | 0.281 | -0.008 | 0.051 | 0.051 | 0 | 0.13 | 0.03 | 0.15 | 0.03 | 0.1 | 0.11 |
| 9 | 19065295 | rs7467618 | -1.567 | -0.022 | 0.39 | 0.348 | 0.043 | 0.076 | 0.076 | 0 | 0.47 | 0.69 | 0.37 | 0.24 | 0.36 | 0.34 |
| 9 | 19069911 | rs10811110 | -0.972 | -0.034 | 0.431 | 0.342 | -0.022 | -0.034 | -0.034 | 0.568 | 0.31 | 0.57 | 0.16 | 0.17 | 0.26 | 0.18 |
| 9 | 19074740 | rs17238021 | 0.113 | 0.015 | 0.243 | 0.324 | -0.001 | 0.026 | 0.026 | 0 | 0.17 | 0.12 | 0.22 | 0.07 | 0.17 | 0.2 |
| 9 | 19074820 | rs10963938 | 1.89 | 0.097 | 0.293 | 0.473 | -0.042 | 0.026 | 0.026 | 0.003 | 0.4 | 0.19 | 0.4 | 0.33 | 0.45 | 0.42 |
| 9 | 19084444 | rs10963949 | 1.304 | 0.09 | 0.48 | 0.388 | -0.002 | -0.006 | -0.006 | 0 | 0.54 | 0.5 | 0.47 | 0.4 | 0.58 | 0.47 |
| 9 | 19086979 | rs2171957 | -0.223 | 0.075 | 0.527 | 0.36 | -0.006 | -0.021 | -0.021 | 0 | 0.22 | 0.24 | 0.25 | 0.09 | 0.15 | 0.15 |

| | | | | | | | | | | | | | | | | |
|---|---|---|---|---|---|---|---|---|---|---|---|---|---|---|---|---|
| 9 | 19087429 | rs7860122 | -0.751 | 0.077 | 0.395 | 0.352 | -0.019 | -0.076 | -0.076 | 0 | 0.29 | 0.42 | 0.27 | 0.11 | 0.23 | 0.24 |
| 9 | 19091387 | rs7467137 | -0.712 | 0.014 | 0.308 | 0.415 | -0.028 | -0.037 | -0.037 | 0 | 0.26 | 0.39 | 0.11 | 0.16 | 0.22 | 0.22 |
| 9 | 19098712 | rs6415731 | -0.551 | -0.455 | -0.125 | 0.622 | -0.019 | -0.033 | -0.033 | 0 | 0.86 | 0.92 | 0.78 | 0.61 | 0.81 | 0.75 |
| 9 | 19102828 | rs7865126 | 0.417 | -0.072 | -0.231 | 0.734 | 0.002 | -0.004 | -0.004 | 0 | 0.12 | 0.1 | 0.05 | 0.13 | 0.14 | 0.06 |
| 9 | 19108827 | rs12554335 | 0.389 | -0.024 | -0.283 | 0.706 | -0.009 | 0.028 | 0.028 | 0 | 0.15 | 0.04 | 0.21 | 0.06 | 0.12 | 0.12 |
| 9 | 19111637 | rs12338361 | 0.801 | -0.089 | -0.291 | 0.712 | 0.035 | 0.002 | 0.002 | 0 | 0.57 | 0.37 | 0.66 | 0.42 | 0.69 | 0.62 |
| 9 | 19115088 | rs12237614 | 0.704 | 0.204 | -0.625 | 0.6 | -0.001 | -0.003 | -0.003 | 0 | 0.1 | 0.09 | 0.04 | 0.14 | 0.05 | 0.04 |
| 9 | 19126476 | rs3802335 | -0.307 | 0.484 | -0.753 | 0.923 | -0.009 | 0.032 | 0.032 | 0 | 0.75 | 0.69 | 0.7 | 0.53 | 0.78 | 0.7 |
| 9 | 19126565 | rs3824369 | 0.061 | 1.351 | -1.319 | 1.018 | -0.028 | 0.022 | 0.022 | 0 | 0.41 | 0.26 | 0.42 | 0.37 | 0.48 | 0.47 |
| 9 | 19127877 | rs3808657 | 0.147 | 1.178 | -1.139 | 0.751 | -0.03 | 0.035 | 0.035 | 0.245 | 0.34 | 0.21 | 0.35 | 0.24 | 0.37 | 0.37 |
| 9 | 19132051 | rs2046841 | 0.983 | 1.074 | -0.726 | 1.364 | 0.003 | 0.047 | 0.047 | 0 | 0.87 | 0.86 | 0.79 | 0.58 | 0.84 | 0.83 |
| 9 | 19133732 | rs10757028 | -0.066 | 0.333 | -0.803 | 0.183 | 0.016 | 0.018 | 0.018 | 0 | 0.48 | 0.4 | 0.73 | 0.69 | 0.62 | 0.63 |
| 9 | 19142014 | rs10963987 | 0.077 | -0.674 | -0.854 | -0.408 | 0.01 | 0.001 | 0.001 | 0 | 0.19 | 0.12 | 0.06 | 0.01 | 0.08 | 0.11 |
| 9 | 19144191 | rs10963994 | -0.102 | -0.732 | -0.133 | -0.437 | -0.067 | -0.106 | -0.106 | 0 | 0.2 | 0.36 | 0.08 | 0.01 | 0.08 | 0.11 |
| 9 | 19145435 | rs4977499 | -0.405 | -0.897 | 0.126 | -0.439 | 0.019 | 0.013 | 0.013 | 0 | 0.67 | 0.74 | 0.8 | 0.7 | 0.7 | 0.73 |
| 9 | 19149599 | rs882881 | 0.657 | 0.332 | 0.203 | -0.739 | 0.102 | -0.033 | -0.033 | 5.261 | 0.07 | 0.2 | 0.21 | 0 | 0.19 | 0.05 |
| 9 | 19151845 | rs4977502 | -0.756 | 0.209 | 0.068 | -0.855 | -0.031 | -0.01 | -0.01 | 0 | 0.69 | 0.58 | 0.82 | 0.7 | 0.75 | 0.77 |
| 9 | 19154572 | rs10964006 | -0.845 | 0.401 | 0.116 | -1.376 | -0.001 | -0.106 | -0.106 | 0 | 0.29 | 0.51 | 0.3 | 0.02 | 0.31 | 0.19 |
| 9 | 19155134 | rs10964007 | -0.179 | -0.156 | -0.749 | -1.235 | 0.005 | -0.009 | -0.009 | 0.691 | 0.21 | 0.17 | 0.08 | 0.01 | 0.12 | 0.12 |
| 9 | 19156784 | rs3004297 | 0.26 | -0.26 | -1.975 | -1.239 | 0.047 | 0.07 | 0.07 | 0 | 0.37 | 0.19 | 0.35 | 0.77 | 0.31 | 0.5 |
| 9 | 19161239 | rs10964015 | -0.08 | 0.552 | -1.92 | -1.249 | -0.055 | 0.048 | 0.048 | 0 | 0.23 | 0.02 | 0.24 | 0.13 | 0.23 | 0.21 |
| 9 | 19162016 | rs3003715 | 0.482 | 1.172 | -1.239 | -1.01 | -0.041 | 0.038 | 0.038 | 0 | 0.68 | 0.26 | 0.6 | 0.85 | 0.54 | 0.73 |
| 9 | 19163207 | rs2503349 | 0.611 | 0.784 | -0.874 | -0.654 | 0.015 | -0.03 | -0.03 | 0.081 | 0.94 | 0.87 | 0.93 | 1 | 0.91 | 0.97 |
| 9 | 19171247 | rs13295225 | -0.333 | 0.979 | -1.475 | -1.116 | -0.016 | 0.106 | 0.106 | 0 | 0.31 | 0.15 | 0.24 | 0.02 | 0.33 | 0.23 |
| 9 | 19174259 | rs10964024 | 0.61 | 1.111 | -1.006 | -0.717 | 0.012 | 0.021 | 0.021 | 0 | 0.21 | 0.32 | 0.28 | 0.37 | 0.22 | 0.25 |
| 9 | 19176406 | rs10964025 | 0.694 | 1.079 | -0.894 | -0.733 | 0.001 | 0.003 | 0.003 | 0 | 0.2 | 0.21 | 0.27 | 0.38 | 0.22 | 0.26 |
| 9 | 19182893 | rs2503351 | -0.665 | 1.051 | -0.899 | -0.634 | 0.005 | 0.001 | 0.001 | 0 | 0.89 | 0.88 | 0.85 | 0.88 | 0.82 | 0.82 |
| 9 | 19184484 | rs13299148 | 1.122 | 1.097 | -1.159 | -0.903 | -0.044 | 0.14 | 0.14 | 0 | 0.26 | 0.01 | 0.19 | 0.02 | 0.26 | 0.18 |
| 9 | 19186670 | rs2451649 | -0.529 | 0.516 | -0.841 | -1.122 | -0.002 | -0.002 | -0.002 | 0 | 0.71 | 0.72 | 0.6 | 0.4 | 0.66 | 0.53 |
| 9 | 19191447 | rs13288647 | 0.947 | 0.116 | -0.871 | -1.324 | -0.045 | 0.137 | 0.137 | 0 | 0.3 | 0.1 | 0.25 | 0.02 | 0.34 | 0.25 |
| 9 | 19192694 | rs4977286 | -0.399 | 0.1 | -0.605 | -1.167 | 0.001 | 0.003 | 0.003 | 0 | 0.69 | 0.64 | 0.6 | 0.42 | 0.65 | 0.49 |
| 9 | 19201738 | rs457062 | 0.733 | 0.147 | -0.585 | -1.23 | 0.027 | 0.041 | 0.041 | 0.021 | 0.06 | 0.14 | 0.1 | 0.12 | 0.11 | 0.19 |
| 9 | 19205996 | rs4262389 | 0.806 | -0.242 | -0.745 | -1.65 | -0.012 | -0.044 | -0.044 | 0.485 | 0.18 | 0.17 | 0.07 | 0 | 0.09 | 0.02 |
| 9 | 19206930 | rs461567 | -0.413 | 0.092 | -1.128 | -1.3 | -0.03 | 0.11 | 0.11 | 0 | 0.69 | 0.84 | 0.75 | 0.91 | 0.67 | 0.74 |
| 9 | 19211852 | rs4083651 | -0.2 | -0.49 | -0.732 | -1.14 | -0.004 | -0.003 | -0.003 | 0.108 | 0.68 | 0.68 | 0.79 | 0.82 | 0.78 | 0.76 |
| 9 | 19223304 | rs10757033 | 0.502 | 0.304 | -0.314 | -1.214 | 0.015 | 0.058 | 0.058 | 0.067 | 0.5 | 0.27 | 0.52 | 0.58 | 0.55 | 0.65 |
| 9 | 19231643 | rs10123714 | 0.573 | 0.283 | -0.556 | -1.514 | -0.016 | 0.014 | 0.014 | 0.555 | 0.43 | 0.24 | 0.5 | 0.57 | 0.48 | 0.56 |
| 9 | 19234830 | rs4977291 | -1.451 | -0.229 | -0.142 | -1.524 | 0.001 | -0.001 | -0.001 | 0 | 0.81 | 0.83 | 0.9 | 0.79 | 0.88 | 0.88 |
| 9 | 19234908 | rs1329731 | 0.377 | -0.244 | 0.122 | -1.554 | 0.008 | -0.009 | -0.009 | 0 | 0.26 | 0.27 | 0.36 | 0.22 | 0.32 | 0.27 |
| 9 | 19235270 | rs1854551 | -0.878 | -0.286 | 0.135 | -1.551 | -0.035 | -0.01 | -0.01 | 0.542 | 0.77 | 0.66 | 0.88 | 0.79 | 0.87 | 0.88 |
| 9 | 19236489 | rs16937415 | 1.456 | -0.286 | 0.283 | -1.561 | 0.013 | 0.016 | 0.016 | 0 | 0.14 | 0.08 | 0.1 | 0.02 | 0.09 | 0.17 |
| 9 | 19239168 | rs4977517 | -0.848 | -0.275 | 0.319 | -1.552 | -0.041 | -0.012 | -0.012 | 0 | 0.76 | 0.65 | 0.87 | 0.79 | 0.87 | 0.88 |
| 9 | 19244788 | rs7027485 | 1.027 | -0.137 | 0.305 | -1.574 | 0.059 | 0.02 | 0.02 | 0 | 0.11 | 0.19 | 0.26 | 0.2 | 0.22 | 0.09 |
| 9 | 19253268 | rs10811150 | 0.747 | -0.298 | 0.28 | -1.433 | 0.008 | 0.021 | 0.021 | 0 | 0.51 | 0.22 | 0.51 | 0.54 | 0.53 | 0.69 |
| 9 | 19258183 | rs2383089 | -0.06 | -0.322 | 0.33 | -1.453 | 0.006 | 0.022 | 0.022 | 0 | 0.67 | 0.47 | 0.63 | 0.56 | 0.64 | 0.82 |
| 9 | 19281501 | rs10757049 | 0.659 | -0.298 | 0.326 | -1.302 | 0.005 | 0.011 | 0.011 | 0 | 0.51 | 0.4 | 0.52 | 0.55 | 0.52 | 0.71 |
| 9 | 19290143 | rs10122709 | -0.616 | -0.498 | 0.328 | -1.478 | 0.008 | -0.017 | -0.017 | 0 | 0.68 | 0.74 | 0.84 | 0.98 | 0.84 | 0.82 |
| 9 | 19298581 | rs7851902 | -0.527 | -0.288 | 0.389 | -1.448 | -0.062 | -0.132 | -0.132 | 0.049 | 0.67 | 0.44 | 0.82 | 0.98 | 0.81 | 0.8 |
| 9 | 19303086 | rs7022273 | -1.703 | -0.3 | 0.316 | -1.459 | -0.005 | -0.034 | -0.034 | 0 | 0.82 | 0.84 | 0.93 | 1 | 0.91 | 0.98 |
| 9 | 19308145 | rs10757052 | 2.011 | 0.07 | -0.403 | -1.886 | -0.071 | 0.115 | 0.115 | 0 | 0.21 | 0.03 | 0.17 | 0.02 | 0.27 | 0.13 |
| 9 | 19326053 | rs2256093 | 0.535 | 0.196 | 0.092 | -1.453 | -0.101 | -0.155 | -0.155 | 2.388 | 0.32 | 0.53 | 0.17 | 0.02 | 0.15 | 0.19 |
| 9 | 19331571 | rs17818670 | 0.431 | 0.492 | 0.098 | -1.207 | 0.042 | 0.066 | 0.066 | 0 | 0.14 | 0.19 | 0.31 | 0.42 | 0.26 | 0.14 |
| 9 | 19343291 | rs2666797 | -0.127 | -0.116 | -0.155 | -2.216 | 0.003 | -0.054 | -0.054 | 0 | 0.42 | 0.38 | 0.21 | 0.02 | 0.28 | 0.2 |
| 9 | 19359195 | rs4578040 | -0.575 | -0.185 | -0.093 | -2.104 | 0.018 | 0.109 | 0.109 | 0 | 0.51 | 0.76 | 0.48 | 0.34 | 0.45 | 0.37 |
| 9 | 19364042 | rs7863119 | 0.859 | -0.251 | -0.193 | -2.078 | -0.028 | 0.023 | 0.023 | 0.094 | 0.41 | 0.22 | 0.47 | 0.58 | 0.47 | 0.52 |
| 9 | 19370108 | rs10114375 | 0.556 | -0.211 | 0.394 | -2.079 | -0.032 | -0.076 | -0.076 | 0 | 0.18 | 0.28 | 0.12 | 0.02 | 0.11 | 0.19 |
| 9 | 19370343 | rs10117525 | 0.613 | -0.205 | 0.402 | -2.076 | -0.026 | -0.064 | -0.064 | 0 | 0.18 | 0.24 | 0.12 | 0.02 | 0.11 | 0.18 |
| 9 | 19376144 | rs10119465 | 0.869 | -0.132 | 0.214 | -2.006 | -0.049 | -0.101 | -0.101 | 1.122 | 0.17 | 0.54 | 0.13 | 0.02 | 0.09 | 0.18 |
| 9 | 19386565 | rs6475326 | -0.537 | 0.063 | -0.102 | -1.87 | 0.01 | 0.071 | 0.071 | 0 | 0.51 | 0.7 | 0.47 | 0.36 | 0.46 | 0.38 |
| 9 | 19397040 | rs7048650 | -0.028 | 0.213 | -0.352 | -1.705 | 0.003 | 0.025 | 0.025 | 0 | 0.3 | 0.21 | 0.3 | 0.2 | 0.26 | 0.09 |
| 9 | 19403973 | rs4418423 | -0.403 | -0.511 | -0.369 | -2.047 | 0.069 | 0.093 | 0.093 | 0.015 | 0.38 | 0.79 | 0.23 | 0.15 | 0.24 | 0.35 |
| 9 | 19419528 | rs10757058 | 0.794 | -0.152 | -0.038 | -1.855 | -0.001 | -0.001 | -0.001 | 0 | 0.18 | 0.14 | 0.12 | 0.15 | 0.18 | 0.26 |
| 9 | 19420040 | rs10118089 | 0.85 | -0.165 | -0.174 | -1.858 | 0.007 | 0.009 | 0.009 | 0 | 0.17 | 0.08 | 0.12 | 0.15 | 0.17 | 0.26 |
| 9 | 19426464 | rs7039309 | -0.88 | 0.063 | -0.655 | -2.134 | -0.004 | -0.017 | -0.017 | 0 | 0.72 | 0.88 | 0.68 | 0.37 | 0.71 | 0.6 |
| 9 | 19426904 | rs7023819 | 0.27 | 0.152 | 0.379 | -2.241 | -0.018 | -0.049 | -0.049 | 1.615 | 0.23 | 0.47 | 0.17 | 0.16 | 0.2 | 0.32 |
| 9 | 19428721 | rs10125228 | 0.93 | 0.179 | -0.338 | -2.193 | 0.001 | 0.008 | 0.008 | 0 | 0.18 | 0.09 | 0.12 | 0.16 | 0.19 | 0.28 |
| 9 | 19438279 | rs10811184 | 0.164 | 0.712 | -0.164 | -2.278 | 0.031 | -0.103 | -0.103 | 0 | 0.27 | 0.42 | 0.22 | 0.02 | 0.34 | 0.18 |
| 9 | 19442558 | rs10964139 | 0.045 | 0.676 | -1.238 | -2.067 | 0.017 | -0.088 | -0.088 | 0.266 | 0.22 | 0.7 | 0.24 | 0.02 | 0.26 | 0.14 |
| 9 | 19446663 | rs10757059 | -0.623 | 0.658 | -2.013 | -2.054 | 0.004 | 0.135 | 0.135 | 0 | 0.44 | 0.74 | 0.47 | 0.18 | 0.43 | 0.24 |

| | | | | | | | | | | | | | | | | |
|---|---|---|---|---|---|---|---|---|---|---|---|---|---|---|---|---|
| 9 | 19451807 | rs12343276 | 0.605 | -0.131 | -1.632 | -1.64 | 0.062 | 0.063 | 0.063 | 0.803 | 0.13 | 0.02 | 0.06 | 0 | 0.05 | 0.01 |
| 9 | 19452990 | rs7022373 | -0.357 | 0.519 | -1.392 | -1.319 | 0.017 | 0.106 | 0.106 | 0 | 0.58 | 0.74 | 0.6 | 0.2 | 0.64 | 0.31 |
| 9 | 19454906 | rs7045369 | -0.384 | 0.752 | -0.532 | -0.287 | 0.024 | -0.028 | -0.028 | 0.431 | 0.06 | 0.13 | 0.04 | 0.01 | 0.07 | 0 |
| 9 | 19457585 | rs7027520 | -0.507 | 0.469 | -1.209 | 0 | 0.01 | 0.013 | 0.013 | 0 | 0.31 | 0.2 | 0.4 | 0.76 | 0.29 | 0.37 |
| 9 | 19458939 | rs960908 | -0.273 | 0.555 | -1.158 | 0.769 | 0 | -0.019 | -0.019 | 0 | 0.27 | 0.29 | 0.29 | 0.45 | 0.26 | 0.16 |
| 9 | 19465900 | rs12336251 | -0.729 | 0.406 | -1.069 | 0.158 | 0 | 0.024 | 0.024 | 1.147 | 0.34 | 0.26 | 0.28 | 0.16 | 0.34 | 0.34 |
| 9 | 19468530 | rs7021552 | -0.879 | 0.324 | -1.106 | 0.037 | 0.003 | 0.068 | 0.068 | 0.169 | 0.35 | 0.22 | 0.22 | 0.11 | 0.35 | 0.3 |
| 9 | 19468696 | rs10964151 | -0.303 | 0.432 | -1.056 | 0.197 | -0.009 | 0.061 | 0.061 | 0.736 | 0.29 | 0.13 | 0.18 | 0.12 | 0.31 | 0.29 |
| 9 | 19469713 | rs10964155 | -0.274 | 0.348 | -1.339 | 0.251 | -0.035 | 0.12 | 0.12 | 0.824 | 0.27 | 0.02 | 0.19 | 0.09 | 0.27 | 0.25 |
| 9 | 19469846 | rs11790131 | -0.188 | 0.278 | -1.384 | 0.223 | 0 | 0.109 | 0.109 | 0.913 | 0.26 | 0.02 | 0.16 | 0.09 | 0.21 | 0.25 |
| 9 | 19470804 | rs12335928 | -0.331 | 0.239 | -1.271 | 0.776 | -0.019 | 0.1 | 0.1 | 0.899 | 0.26 | 0.02 | 0.19 | 0.09 | 0.26 | 0.25 |
| 9 | 19472460 | rs11792752 | -0.642 | -0.242 | -0.366 | 1.549 | -0.029 | 0.052 | 0.052 | 1.955 | 0.26 | 0.41 | 0.31 | 0.49 | 0.26 | 0.38 |
| 9 | 19472784 | rs10964160 | -0.511 | -0.404 | -0.165 | 1.73 | 0.008 | -0.015 | -0.015 | 0 | 0.21 | 0.15 | 0.27 | 0.39 | 0.22 | 0.33 |
| 9 | 19476532 | rs17566007 | -1.276 | -0.534 | 0.1 | 1.493 | -0.001 | -0.005 | -0.005 | 0.158 | 0.14 | 0.15 | 0.08 | 0.09 | 0.12 | 0.19 |
| 9 | 19477947 | rs6475332 | -0.39 | -0.004 | 0.086 | 0.726 | -0.008 | 0.019 | 0.019 | 0 | 0.71 | 0.76 | 0.73 | 0.83 | 0.71 | 0.77 |
| 9 | 19478972 | rs12346257 | -0.721 | 0.018 | 0.149 | 0.757 | 0.007 | 0.015 | 0.015 | 0.258 | 0.14 | 0.09 | 0.07 | 0.08 | 0.1 | 0.19 |
| 9 | 19479033 | rs7045446 | 0.468 | 0.129 | 0.358 | 1.027 | 0.001 | 0.007 | 0.007 | 0 | 0.49 | 0.58 | 0.44 | 0.38 | 0.47 | 0.4 |
| 9 | 19479536 | rs7046383 | 0.725 | -0.004 | 0.43 | 0.773 | -0.016 | 0.068 | 0.068 | 0 | 0.46 | 0.27 | 0.36 | 0.27 | 0.46 | 0.39 |
| 9 | 19484869 | rs7858547 | -1.074 | -0.566 | 0.761 | 0.701 | -0.061 | 0.004 | 0.004 | 0 | 0.14 | 0.32 | 0.04 | 0.15 | 0.03 | 0.17 |
| 9 | 19486059 | rs10964178 | 1.555 | -0.11 | 0.491 | 0.741 | 0.003 | 0.006 | 0.006 | 0.098 | 0.35 | 0.28 | 0.33 | 0.28 | 0.35 | 0.23 |
| 9 | 19492973 | rs10117909 | 0.064 | -0.121 | 0.474 | 0.792 | 0 | 0.069 | 0.069 | 0.731 | 0.23 | 0.3 | 0.41 | 0.53 | 0.26 | 0.38 |
| 9 | 19509905 | rs4977544 | 0.283 | -0.283 | -0.077 | 0.173 | -0.058 | 0.099 | 0.099 | 0 | 0.65 | 0.82 | 0.71 | 0.91 | 0.58 | 0.7 |
| 9 | 19510856 | rs7864646 | -0.064 | -0.725 | -0.609 | -0.272 | -0.159 | 0.168 | 0.168 | 0 | 0.68 | 0.95 | 0.71 | 0.93 | 0.53 | 0.77 |
| 9 | 19511096 | rs7867513 | 2.556 | -0.806 | -0.161 | -0.007 | 0.001 | -0.012 | -0.012 | 0 | 0.27 | 0.22 | 0.25 | 0.31 | 0.28 | 0.27 |
| 9 | 19512746 | rs4977545 | 0.388 | -0.539 | -0.045 | -0.143 | 0.031 | -0.05 | -0.05 | 0.047 | 0.66 | 0.56 | 0.73 | 0.79 | 0.61 | 0.64 |
| 9 | 19514307 | rs7022987 | 2.213 | -0.356 | -0.173 | 0.228 | -0.003 | -0.014 | -0.014 | 1.877 | 0.27 | 0.24 | 0.21 | 0.33 | 0.29 | 0.27 |
| 9 | 19517795 | rs2225143 | -0.044 | -0.204 | 0.19 | 0.54 | -0.04 | 0.053 | 0.053 | 0.787 | 0.31 | 0.41 | 0.48 | 0.49 | 0.22 | 0.38 |
| 9 | 19523984 | rs4977303 | 0.874 | 0.233 | 0.136 | 0.498 | -0.001 | 0 | 0 | 0 | 0.43 | 0.44 | 0.36 | 0.5 | 0.39 | 0.27 |
| 9 | 19526674 | rs4977547 | -0.08 | -0.049 | 0.173 | 0.361 | -0.013 | 0.018 | 0.018 | 4.651 | 0.34 | 0.35 | 0.49 | 0.56 | 0.26 | 0.56 |
| 9 | 19526712 | rs4977548 | 0.006 | -0.052 | 0.175 | 0.36 | 0.008 | -0.015 | -0.015 | 0 | 0.33 | 0.22 | 0.48 | 0.56 | 0.26 | 0.56 |
| 9 | 19528751 | rs2209802 | 0.576 | 0.349 | 0.29 | -0.383 | -0.016 | 0.01 | 0.01 | 0 | 0.67 | 0.65 | 0.71 | 0.92 | 0.56 | 0.79 |
| 9 | 19531262 | rs3780218 | 1.23 | 0.364 | 0.447 | -0.342 | 0.053 | -0.096 | -0.096 | 5.801 | 0.78 | 0.53 | 0.78 | 0.92 | 0.74 | 0.9 |
| 9 | 19532826 | rs10811199 | 0.736 | 0.108 | -0.059 | -0.55 | 0.013 | -0.076 | -0.076 | 0 | 0.82 | 0.66 | 0.85 | 1 | 0.79 | 0.95 |
| 9 | 19551434 | rs10738542 | 1.093 | -0.006 | -0.308 | -0.657 | -0.023 | 0.057 | 0.057 | 0 | 0.8 | 0.86 | 0.86 | 1 | 0.78 | 0.93 |
| 9 | 19551907 | rs1006240 | 1.929 | -0.141 | -0.721 | -0.938 | -0.046 | 0.146 | 0.146 | 1.85 | 0.27 | 0.03 | 0.23 | 0.04 | 0.29 | 0.2 |
| 9 | 19554109 | rs4977557 | 1.784 | -0.419 | 0.859 | -0.111 | 0.007 | -0.114 | -0.114 | 0 | 0.77 | 0.38 | 0.82 | 1 | 0.79 | 0.94 |
| 9 | 19556699 | rs12340099 | -0.496 | -0.121 | 0.759 | 0.007 | 0.054 | -0.066 | -0.066 | 0 | 0.11 | 0.43 | 0.11 | 0 | 0.12 | 0.03 |
| 9 | 19556835 | rs12349316 | 0.085 | -0.111 | 0.731 | 0.012 | 0.032 | -0.051 | -0.051 | 2.322 | 0.09 | 0.17 | 0.09 | 0 | 0.12 | 0.03 |
| 9 | 19570345 | rs7036497 | 1.505 | -0.054 | 0.703 | 0.025 | -0.002 | -0.014 | -0.014 | 0.177 | 0.9 | 0.89 | 0.93 | 1 | 0.91 | 0.97 |
| 9 | 19572704 | rs731407 | -0.62 | -0.128 | 0.799 | 0.569 | 0.072 | 0.285 | 0.285 | 0 | 0.12 | 0.41 | 0.15 | 0.34 | 0.15 | 0.13 |
| 9 | 19576362 | rs11790824 | 0.397 | -0.096 | 0.564 | 0.562 | 0.037 | -0.046 | -0.046 | 0.501 | 0.08 | 0.14 | 0.08 | 0 | 0.12 | 0.03 |
| 9 | 19576949 | rs2383101 | 1.334 | -0.079 | 0.552 | 0.559 | 0.027 | -0.097 | -0.097 | 1.573 | 0.82 | 0.78 | 0.85 | 1 | 0.78 | 0.94 |
| 9 | 19579427 | rs3780215 | -1.619 | -0.064 | 0.52 | 0.635 | 0.034 | 0.115 | 0.115 | 0 | 0.18 | 0.49 | 0.22 | 0.33 | 0.21 | 0.15 |
| 9 | 19582883 | rs7848995 | -1.446 | 0.121 | -0.215 | 0.599 | 0.03 | 0.028 | 0.028 | 0 | 0.17 | 0.32 | 0.19 | 0.23 | 0.2 | 0.17 |
| 9 | 19586463 | rs10811214 | -1.154 | 0.122 | -0.363 | 0.467 | 0.036 | 0.079 | 0.079 | 0 | 0.18 | 0.43 | 0.22 | 0.3 | 0.2 | 0.18 |
| 9 | 19591153 | rs2383103 | -0.672 | 0.226 | -0.546 | 0.476 | 0.014 | 0.102 | 0.102 | 0 | 0.14 | 0.29 | 0.17 | 0.32 | 0.17 | 0.18 |
| 9 | 19591691 | rs2209797 | 0.495 | 0.086 | -0.649 | 0.374 | 0.01 | -0.054 | -0.054 | 0 | 0.88 | 0.95 | 0.92 | 0.67 | 0.92 | 0.82 |
| 9 | 19593235 | rs9792664 | 1.334 | 0.081 | -0.627 | 0.382 | 0.002 | 0.012 | 0.012 | 0 | 0.72 | 0.59 | 0.76 | 0.67 | 0.75 | 0.77 |
| 9 | 19598020 | rs1885219 | 0.249 | 0.17 | -0.65 | 0.423 | -0.004 | -0.023 | -0.023 | 0 | 0.86 | 0.85 | 0.88 | 0.69 | 0.88 | 0.86 |
| 9 | 19598186 | rs12000488 | -0.15 | 0.085 | -0.384 | 0.201 | 0.009 | -0.059 | -0.059 | 0 | 0.11 | 0.14 | 0.1 | 0 | 0.13 | 0.04 |
| 9 | 19599114 | rs10964233 | -0.344 | 0.107 | -0.585 | 0.212 | 0.021 | -0.08 | -0.08 | 0 | 0.12 | 0.28 | 0.11 | 0.01 | 0.14 | 0.04 |
| 9 | 19600362 | rs10964234 | -0.352 | -0.041 | -1.63 | 0.699 | 0.006 | -0.144 | -0.144 | 0.041 | 0.21 | 0.01 | 0.22 | 0.41 | 0.19 | 0.38 |
| 9 | 19604078 | rs2180940 | -0.218 | 0.348 | -0.976 | -0.489 | 0 | 0.032 | 0.032 | 0.175 | 0.5 | 0.57 | 0.62 | 0.78 | 0.5 | 0.61 |
| 9 | 19606810 | rs2180939 | -0.348 | 0.046 | -0.765 | -0.741 | -0.004 | 0.096 | 0.096 | 1.727 | 0.53 | 0.69 | 0.66 | 0.84 | 0.51 | 0.63 |
| 9 | 19608329 | rs10964236 | 0.011 | -0.061 | -0.668 | -0.611 | -0.001 | 0.01 | 0.01 | 0.666 | 0.49 | 0.5 | 0.61 | 0.74 | 0.49 | 0.58 |
| 9 | 19613009 | rs4977230 | -0.046 | -0.284 | -1.023 | -1.21 | 0.003 | 0.005 | 0.005 | 0.374 | 0.59 | 0.57 | 0.76 | 0.77 | 0.68 | 0.73 |
| 9 | 19614952 | rs2208552 | -0.521 | -0.308 | -0.668 | -1.249 | 0.005 | 0.017 | 0.017 | 0.046 | 0.66 | 0.72 | 0.82 | 0.87 | 0.72 | 0.77 |
| 9 | 19622268 | rs4977308 | -0.579 | -0.295 | -0.559 | -1.239 | 0.009 | 0.034 | 0.034 | 0.048 | 0.66 | 0.75 | 0.82 | 0.87 | 0.72 | 0.77 |
| 9 | 19623126 | rs10757082 | -0.56 | -0.292 | -0.558 | -1.235 | 0.009 | 0.035 | 0.035 | 0.026 | 0.66 | 0.75 | 0.81 | 0.86 | 0.72 | 0.78 |
| 9 | 19626709 | rs7020568 | -0.484 | -0.314 | -0.367 | -1.24 | 0.012 | 0.049 | 0.049 | 0.081 | 0.66 | 0.8 | 0.82 | 0.86 | 0.72 | 0.77 |
| 9 | 19628633 | rs718623 | -0.132 | -0.373 | 0.248 | -1.308 | -0.012 | -0.028 | -0.028 | 0 | 0.6 | 0.48 | 0.75 | 0.76 | 0.7 | 0.72 |
| 9 | 19637920 | rs10811219 | -0.474 | -0.353 | 0.314 | -1.314 | -0.021 | -0.081 | -0.081 | 0.628 | 0.63 | 0.53 | 0.8 | 0.85 | 0.71 | 0.77 |
| 9 | 19645715 | rs2208553 | 0.304 | -0.337 | 0.639 | -1.34 | -0.002 | 0.032 | 0.032 | 0.366 | 0.48 | 0.43 | 0.59 | 0.72 | 0.49 | 0.59 |
| 9 | 19646058 | rs2208554 | 0.061 | -0.331 | 0.579 | -1.353 | -0.005 | 0.061 | 0.061 | 0.705 | 0.52 | 0.66 | 0.64 | 0.83 | 0.51 | 0.63 |
| 9 | 19649738 | rs7870850 | -0.583 | -0.369 | 0.489 | -1.308 | 0.01 | 0.042 | 0.042 | 0.103 | 0.66 | 0.75 | 0.82 | 0.85 | 0.72 | 0.77 |
| 9 | 19652765 | rs16937678 | -0.165 | -0.583 | 0.368 | -1.464 | 0.001 | -0.011 | -0.011 | 0 | 0.08 | 0.11 | 0.04 | 0 | 0.06 | 0.02 |
| 9 | 19652826 | rs16937679 | -0.648 | -0.594 | 0.4 | -1.379 | 0.002 | -0.002 | -0.002 | 0.068 | 0.1 | 0.12 | 0.07 | 0.12 | 0.06 | 0.05 |
| 9 | 19655267 | rs17492143 | 0.649 | 0.189 | 0.396 | -1.447 | 0.059 | -0.058 | -0.058 | 0.692 | 0.17 | 0.3 | 0.22 | 0.04 | 0.29 | 0.15 |
| 9 | 19656396 | rs16937690 | -0.363 | 0.183 | 0.383 | -1.447 | 0.018 | -0.076 | -0.076 | 0 | 0.25 | 0.35 | 0.25 | 0.04 | 0.34 | 0.17 |

| | | | | | | | | | | | | | | | | |
|---|---|---|---|---|---|---|---|---|---|---|---|---|---|---|---|---|
| 9 | 19657975 | rs10738546 | 0.525 | 0.2 | 0.493 | -1.405 | -0.006 | -0.019 | -0.019 | 0 | 0.43 | 0.49 | 0.57 | 0.71 | 0.39 | 0.6 |
| 9 | 19659564 | rs4977232 | 0.167 | -0.296 | 0.532 | -1.819 | -0.005 | -0.021 | -0.021 | 4.501 | 0.68 | 0.65 | 0.84 | 0.88 | 0.75 | 0.71 |
| 9 | 19659652 | rs4977233 | 0.686 | -0.149 | 0.611 | -1.846 | -0.004 | 0.103 | 0.103 | 0 | 0.53 | 0.32 | 0.64 | 0.85 | 0.49 | 0.64 |
| 9 | 19660406 | rs766576 | 0.894 | -0.185 | 0.597 | -1.855 | 0.001 | 0.109 | 0.109 | 0 | 0.51 | 0.26 | 0.61 | 0.76 | 0.46 | 0.59 |
| 9 | 19661573 | rs12686758 | -0.274 | -0.442 | 0.59 | -1.863 | -0.005 | 0.002 | 0.002 | 0.103 | 0.08 | 0.11 | 0.06 | 0.09 | 0.06 | 0.07 |
| 9 | 19661935 | rs2208557 | -0.848 | -0.37 | 0.692 | -1.903 | -0.006 | -0.02 | -0.02 | 0.017 | 0.79 | 0.74 | 0.86 | 0.88 | 0.82 | 0.82 |
| 9 | 19666510 | rs7040903 | 0.746 | 0.112 | 0.238 | -1.877 | 0.001 | 0.159 | 0.159 | 0 | 0.52 | 0.27 | 0.62 | 0.85 | 0.45 | 0.64 |
| 9 | 19668550 | rs7035054 | 0.45 | -0.501 | 0.251 | -1.736 | 0.002 | -0.011 | -0.011 | 0 | 0.2 | 0.22 | 0.2 | 0.1 | 0.23 | 0.15 |
| 9 | 19678269 | rs2383113 | 0.463 | -0.751 | 0.077 | -2.006 | -0.003 | -0.043 | -0.043 | 0 | 0.68 | 0.37 | 0.8 | 0.94 | 0.7 | 0.71 |
| 9 | 19682676 | rs13299310 | 0.28 | -0.337 | 0.156 | -1.608 | -0.013 | 0.029 | 0.029 | 0 | 0.2 | 0.09 | 0.2 | 0.09 | 0.22 | 0.15 |
| 9 | 19690739 | rs1885233 | 0.487 | -0.296 | 0.16 | -1.663 | -0.011 | 0.033 | 0.033 | 0 | 0.18 | 0.09 | 0.17 | 0.05 | 0.2 | 0.11 |
| 9 | 19692697 | rs13287081 | 0.284 | -0.301 | 0.177 | -1.553 | -0.008 | 0.001 | 0.001 | 0 | 0.19 | 0.11 | 0.22 | 0.14 | 0.22 | 0.13 |
| 9 | 19698751 | rs7862478 | -0.954 | -0.461 | 0.65 | -1.671 | -0.003 | -0.013 | -0.013 | 0 | 0.31 | 0.64 | 0.27 | 0.15 | 0.29 | 0.21 |
| 9 | 19700975 | rs12000690 | -0.008 | -0.748 | 0.35 | -1.857 | 0.003 | -0.011 | -0.011 | 0 | 0.14 | 0.11 | 0.04 | 0 | 0.07 | 0.06 |
| 9 | 19702659 | rs7857042 | -0.222 | -0.544 | 0.525 | -1.733 | -0.006 | -0.056 | -0.056 | 0 | 0.1 | 0.31 | 0.04 | 0 | 0.07 | 0.09 |
| 9 | 19705396 | rs7869055 | -0.778 | -0.51 | 0.387 | -1.726 | -0.005 | -0.073 | -0.073 | 0.087 | 0.3 | 0.46 | 0.26 | 0.11 | 0.29 | 0.2 |
| 9 | 19712170 | rs7869360 | -0.036 | -0.461 | 0.344 | -1.631 | -0.012 | -0.043 | -0.043 | 0.006 | 0.09 | 0.18 | 0.03 | 0 | 0.07 | 0.08 |
| 9 | 19712378 | rs7855757 | -0.397 | -0.483 | 0.497 | -1.642 | -0.007 | -0.058 | -0.058 | 0 | 0.11 | 0.35 | 0.05 | 0 | 0.07 | 0.08 |
| 9 | 19713500 | rs12000054 | -1.412 | -0.577 | -0.26 | -1.658 | -0.013 | -0.041 | -0.041 | 0 | 0.43 | 0.5 | 0.28 | 0.15 | 0.36 | 0.27 |
| 9 | 19728716 | rs2208549 | 0.675 | 0.215 | -0.217 | -1.097 | -0.047 | 0.003 | 0.003 | 0 | 0.15 | 0.02 | 0.15 | 0.09 | 0.21 | 0.06 |
| 9 | 19729664 | rs1182653 | 0.512 | 0.222 | -0.175 | -1.089 | -0.002 | 0.008 | 0.008 | 0 | 0.55 | 0.57 | 0.66 | 0.8 | 0.5 | 0.7 |
| 9 | 19731896 | rs1182654 | 0.544 | 0.274 | -0.248 | -1.143 | -0.002 | 0.005 | 0.005 | 0 | 0.56 | 0.56 | 0.65 | 0.85 | 0.5 | 0.71 |
| 9 | 19733052 | rs1182657 | 0.47 | 0.706 | -0.505 | -0.869 | 0 | -0.004 | -0.004 | 0.155 | 0.58 | 0.41 | 0.61 | 0.75 | 0.51 | 0.7 |
| 9 | 19736871 | rs1182665 | 0.364 | 1.27 | -0.938 | -0.477 | -0.018 | 0.005 | 0.005 | 0 | 0.78 | 0.77 | 0.79 | 0.83 | 0.65 | 0.79 |
| 9 | 19746160 | rs957910 | -0.529 | 1.166 | -1.257 | 0.598 | -0.035 | 0.056 | 0.056 | 0 | 0.19 | 0.1 | 0.17 | 0.05 | 0.26 | 0.15 |
| 9 | 19751159 | rs11790276 | -0.587 | 0.747 | -1.06 | 1.255 | 0.001 | -0.04 | -0.04 | 0 | 0.27 | 0.21 | 0.25 | 0.53 | 0.26 | 0.31 |
| 9 | 19751357 | rs1182672 | 0.341 | 0.79 | -0.797 | 0.738 | -0.012 | -0.055 | -0.055 | 0 | 0.35 | 0.45 | 0.24 | 0.19 | 0.29 | 0.3 |
| 9 | 19752860 | rs1327407 | 0.521 | 0.512 | -1.03 | 0.687 | -0.05 | 0.08 | 0.08 | 0.093 | 0.24 | 0.01 | 0.15 | 0.06 | 0.23 | 0.25 |
| 9 | 19754476 | rs10514825 | 0.498 | 0.569 | -0.966 | 0.68 | -0.002 | 0.1 | 0.1 | 0.024 | 0.24 | 0.04 | 0.15 | 0.05 | 0.21 | 0.25 |
| 9 | 19755034 | rs11788522 | 0.41 | 0.554 | -0.527 | 0.777 | 0.009 | 0.037 | 0.037 | 0 | 0.28 | 0.21 | 0.19 | 0.07 | 0.25 | 0.29 |
| 9 | 19756086 | rs16937762 | -0.495 | 0.643 | -0.045 | 1.377 | -0.005 | 0.017 | 0.017 | 0 | 0.05 | 0.1 | 0.02 | 0.12 | 0.04 | 0.07 |
| 9 | 19758207 | rs1007910 | 0.638 | 0.794 | -0.183 | 1.472 | -0.076 | 0.048 | 0.048 | 0.08 | 0.33 | 0.14 | 0.53 | 0.2 | 0.47 | 0.28 |
| 9 | 19760153 | rs1007907 | -0.866 | 0.617 | -0.049 | 1.483 | -0.007 | -0.028 | -0.028 | 0 | 0.75 | 0.67 | 0.82 | 0.8 | 0.77 | 0.78 |
| 9 | 19769000 | rs6475366 | 0.014 | 0.787 | -0.375 | 1.567 | 0.007 | -0.019 | -0.019 | 0 | 0.4 | 0.54 | 0.62 | 0.74 | 0.46 | 0.42 |
| 9 | 19771890 | rs1327409 | -0.299 | 0.448 | -0.289 | 1.484 | -0.016 | -0.003 | -0.003 | 0 | 0.68 | 0.48 | 0.78 | 0.67 | 0.71 | 0.63 |
| 9 | 19779172 | rs4977324 | -0.382 | 1.124 | -0.055 | 2.251 | -0.043 | -0.007 | -0.007 | 0.887 | 0.38 | 0.29 | 0.57 | 0.37 | 0.49 | 0.41 |
| 9 | 19787182 | rs1409954 | -0.271 | 1.2 | 0.028 | 2.226 | -0.03 | 0.017 | 0.017 | 0 | 0.35 | 0.28 | 0.55 | 0.24 | 0.44 | 0.32 |
| 9 | 19787952 | rs2383122 | -0.156 | 1.246 | -0.147 | 2.295 | -0.056 | 0.014 | 0.014 | 0 | 0.34 | 0.2 | 0.55 | 0.24 | 0.46 | 0.34 |
| 9 | 19792882 | rs1831083 | 1.266 | 0.944 | 1.071 | 2.806 | 0.098 | 0.447 | 0.447 | 1.276 | 0.92 | 0.66 | 0.82 | 0.56 | 0.92 | 0.8 |
| 9 | 19797850 | rs1327413 | 2.271 | 0.735 | 1.327 | 2.858 | -0.012 | 0.099 | 0.099 | 0.366 | 0.86 | 0.37 | 0.86 | 0.66 | 0.92 | 0.78 |
| 9 | 19801918 | rs10811244 | 1.741 | 0.727 | 1.284 | 2.457 | 0.026 | 0.29 | 0.29 | 0 | 0.88 | 0.66 | 0.81 | 0.52 | 0.92 | 0.78 |
| 9 | 19803724 | rs10811245 | 1.978 | 0.719 | 1.149 | 2.418 | 0.024 | 0.338 | 0.338 | 0.041 | 0.87 | 0.54 | 0.8 | 0.51 | 0.92 | 0.78 |
| 9 | 19840697 | rs7032894 | -0.721 | 0.743 | 0.486 | 1.637 | -0.02 | -0.031 | -0.031 | 3.641 | 0.09 | 0.16 | 0.04 | 0.01 | 0.07 | 0.04 |
| 9 | 19849403 | rs4977334 | -0.112 | 0.513 | -0.094 | 1.306 | 0.024 | 0.076 | 0.076 | 0 | 0.53 | 0.41 | 0.59 | 0.21 | 0.62 | 0.36 |
| 9 | 19857154 | rs7849437 | -0.099 | 0.348 | 0.459 | 1.208 | 0 | -0.006 | -0.006 | 1.067 | 0.56 | 0.48 | 0.61 | 0.24 | 0.64 | 0.38 |
| 9 | 19867614 | rs10964321 | 0.034 | -0.136 | 0.176 | 1.202 | 0.082 | 0.046 | 0.046 | 0 | 0.54 | 0.21 | 0.61 | 0.38 | 0.7 | 0.51 |
| 9 | 19868092 | rs1576043 | 0.683 | -0.191 | -0.142 | 1.507 | 0.01 | 0.015 | 0.015 | 0.015 | 0.32 | 0.01 | 0.22 | 0.2 | 0.22 | 0.3 |
| 9 | 19877230 | rs1327404 | -0.333 | 0.035 | 0.133 | 1.147 | 0.014 | 0.005 | 0.005 | 0 | 0.54 | 0.66 | 0.63 | 0.4 | 0.68 | 0.51 |
| 9 | 19881264 | rs7025940 | -0.876 | 0.279 | 0.196 | 0.708 | 0.034 | 0.019 | 0.019 | 0.016 | 0.05 | 0.26 | 0.04 | 0.03 | 0.04 | 0.07 |
| 9 | 19881812 | rs7026700 | 0.559 | 0.647 | -0.699 | 0.188 | 0.008 | 0.061 | 0.061 | 0.761 | 0.24 | 0.15 | 0.17 | 0.06 | 0.19 | 0.08 |
| 9 | 19881986 | rs12377344 | 0.416 | 0.971 | -0.761 | 0.1 | -0.031 | 0.025 | 0.025 | 0 | 0.22 | 0.12 | 0.3 | 0.09 | 0.36 | 0.24 |
| 9 | 19882045 | rs7030078 | 0.521 | 0.509 | -0.564 | 0.311 | 0.012 | 0.072 | 0.072 | 0.935 | 0.23 | 0.15 | 0.19 | 0.06 | 0.18 | 0.08 |
| 9 | 19882156 | rs10811261 | -0.095 | 0.327 | -0.291 | 0.749 | -0.014 | 0.067 | 0.067 | 0 | 0.69 | 0.47 | 0.69 | 0.48 | 0.75 | 0.76 |
| 9 | 19882454 | rs1857438 | 0.832 | 0.245 | 0.032 | 0.389 | 0.023 | -0.021 | -0.021 | 0 | 0.94 | 0.8 | 0.96 | 0.99 | 0.97 | 0.94 |
| 9 | 19883421 | rs10964323 | -0.073 | -0.17 | -0.724 | 0.339 | 0.004 | 0.004 | 0.004 | 2.912 | 0.47 | 0.55 | 0.27 | 0.33 | 0.34 | 0.35 |
| 9 | 19889326 | rs2051134 | 0.382 | -0.519 | -1.897 | -0.209 | 0.085 | 0.198 | 0.198 | 0 | 0.5 | 0.16 | 0.55 | 0.18 | 0.62 | 0.5 |
| 9 | 19897119 | rs1539683 | -0.136 | -0.75 | -1.396 | 0.082 | -0.057 | 0.083 | 0.083 | 0 | 0.82 | 0.58 | 0.87 | 0.75 | 0.88 | 0.86 |
| 9 | 19897264 | rs4977348 | 1.181 | -0.651 | -1.443 | 0.048 | -0.04 | 0.034 | 0.034 | 0 | 0.25 | 0.16 | 0.36 | 0.09 | 0.39 | 0.26 |
| 9 | 19898792 | rs10811278 | 0.134 | -0.844 | -0.91 | 0.514 | -0.004 | 0.066 | 0.066 | 0.09 | 0.29 | 0.41 | 0.33 | 0.58 | 0.26 | 0.37 |
| 9 | 19899443 | rs4977349 | 0.347 | -0.83 | -1.17 | 0.611 | 0.042 | -0.029 | -0.029 | 0 | 0.61 | 0.21 | 0.59 | 0.37 | 0.72 | 0.6 |
| 9 | 19901220 | rs10757104 | 0.822 | -0.494 | -0.519 | 0.867 | -0.038 | 0.06 | 0.06 | 0 | 0.8 | 0.62 | 0.87 | 0.75 | 0.86 | 0.86 |
| 9 | 19902822 | rs2383131 | -0.108 | -0.229 | -0.632 | 1.012 | -0.002 | -0.006 | -0.006 | 0.879 | 0.36 | 0.33 | 0.58 | 0.38 | 0.55 | 0.34 |
| 9 | 19903133 | rs1887393 | 0.271 | 0.09 | -0.879 | 0.999 | -0.019 | 0.014 | 0.014 | 0 | 0.32 | 0.27 | 0.39 | 0.19 | 0.45 | 0.25 |
| 9 | 19904103 | rs10738554 | 0.08 | -0.322 | -0.13 | 1.425 | -0.002 | -0.002 | -0.002 | 0 | 0.61 | 0.36 | 0.46 | 0.53 | 0.5 | 0.63 |
| 9 | 19912414 | rs16937827 | 0.3 | -0.509 | -0.227 | 1.542 | -0.013 | -0.018 | -0.018 | 0 | 0.06 | 0 | 0.1 | 0.22 | 0.04 | 0.11 |
| 9 | 19913075 | rs10811281 | 0.335 | -0.494 | -0.07 | 1.502 | -0.041 | 0.015 | 0.015 | 0 | 0.33 | 0.28 | 0.41 | 0.2 | 0.51 | 0.32 |
| 9 | 19914792 | rs1419258 | 0.322 | -0.574 | 0.419 | 1.48 | -0.004 | -0.013 | -0.013 | 0 | 0.58 | 0.31 | 0.45 | 0.58 | 0.45 | 0.57 |
| 9 | 19918297 | rs10964340 | 1.359 | -1.388 | -0.375 | 0.921 | -0.024 | 0.171 | 0.171 | 0.071 | 0.33 | 0.02 | 0.21 | 0.1 | 0.27 | 0.26 |
| 9 | 19919331 | rs7852773 | 0.529 | -1.382 | -0.157 | 0.962 | -0.013 | 0.005 | 0.005 | 0 | 0.36 | 0.33 | 0.44 | 0.26 | 0.55 | 0.33 |

| | | | | | | | | | | | | | | | | |
|---|---|---|---|---|---|---|---|---|---|---|---|---|---|---|---|---|
| 9 | 19932591 | rs17832714 | 0.672 | -1.24 | -0.463 | 1.064 | 0.006 | 0.013 | 0.013 | 0.559 | 0.39 | 0.07 | 0.31 | 0.33 | 0.32 | 0.37 |
| 9 | 19937440 | rs10811285 | 0.722 | -1.243 | -0.226 | 1.029 | 0.044 | -0.013 | -0.013 | 0 | 0.35 | 0.52 | 0.39 | 0.25 | 0.49 | 0.27 |
| 9 | 19939119 | rs10811286 | 1.332 | -1.31 | -0.314 | 1.066 | -0.026 | 0.152 | 0.152 | 0 | 0.33 | 0.03 | 0.25 | 0.12 | 0.28 | 0.28 |
| 9 | 19939469 | rs10811288 | 0.534 | -1.28 | 0.136 | 1.026 | 0.019 | -0.014 | -0.014 | 0 | 0.37 | 0.48 | 0.39 | 0.2 | 0.5 | 0.28 |
| 9 | 19942311 | rs1573224 | 0.871 | -1.267 | 0.055 | 1.049 | -0.046 | -0.031 | -0.031 | 0.033 | 0.38 | 0.03 | 0.34 | 0.33 | 0.33 | 0.38 |
| 9 | 19942824 | rs9774948 | 0.843 | -1.285 | 0.093 | 1.051 | 0.029 | -0.011 | -0.011 | 0.694 | 0.33 | 0.48 | 0.39 | 0.2 | 0.49 | 0.25 |
| 9 | 19946627 | rs7869796 | -0.649 | -1.36 | 0.173 | 1.053 | -0.012 | 0.027 | 0.027 | 0 | 0.8 | 0.76 | 0.79 | 0.58 | 0.88 | 0.67 |
| 9 | 19949350 | rs7039172 | 0.77 | -1.196 | 0.185 | 1.017 | 0.028 | -0.005 | -0.005 | 0.26 | 0.34 | 0.49 | 0.39 | 0.25 | 0.49 | 0.24 |
| 9 | 19950818 | rs7021102 | 0.872 | -1.11 | -0.032 | 0.949 | 0.017 | 0.028 | 0.028 | 0.101 | 0.38 | 0.16 | 0.35 | 0.31 | 0.32 | 0.38 |
| 9 | 19951575 | rs10964353 | 0.501 | -1.083 | -0.082 | 0.953 | 0.016 | 0.052 | 0.052 | 0.602 | 0.43 | 0.12 | 0.37 | 0.32 | 0.37 | 0.44 |
| 9 | 19958389 | rs9407950 | -0.471 | -1.191 | 0.166 | 0.847 | -0.003 | 0.008 | 0.008 | 0.574 | 0.79 | 0.79 | 0.8 | 0.57 | 0.86 | 0.66 |
| 9 | 19959158 | rs9407951 | 0.495 | -1.017 | 0.254 | 0.817 | -0.004 | 0.016 | 0.016 | 0 | 0.42 | 0.43 | 0.34 | 0.15 | 0.54 | 0.27 |
| 9 | 19969904 | rs7863041 | -0.488 | -1.037 | 0.279 | 0.863 | -0.016 | 0.032 | 0.032 | 0.634 | 0.79 | 0.76 | 0.8 | 0.57 | 0.86 | 0.67 |
| 9 | 19971287 | rs9406760 | 0.101 | -1.057 | 0.27 | 0.819 | 0.003 | 0.012 | 0.012 | 0 | 0.47 | 0.59 | 0.36 | 0.18 | 0.58 | 0.32 |
| 9 | 19979174 | rs9406764 | 0.613 | -1.114 | 0.105 | 0.704 | 0.002 | -0.019 | -0.019 | 0.016 | 0.41 | 0.49 | 0.34 | 0.11 | 0.54 | 0.25 |
| 9 | 19979623 | rs2154091 | 0.815 | -1.139 | -0.03 | 0.777 | 0.001 | -0.079 | -0.079 | 0 | 0.34 | 0.08 | 0.42 | 0.42 | 0.28 | 0.37 |
| 9 | 19987053 | rs1936451 | 0.111 | -1.168 | 0.067 | 0.836 | -0.035 | 0.059 | 0.059 | 0.614 | 0.74 | 0.61 | 0.76 | 0.56 | 0.82 | 0.63 |
| 9 | 19989693 | rs7872451 | -0.38 | -1.159 | 0.013 | 0.844 | -0.014 | 0.044 | 0.044 | 0.462 | 0.79 | 0.72 | 0.78 | 0.6 | 0.83 | 0.66 |
| 9 | 19994895 | rs9406768 | 0.266 | -1.099 | -0.247 | 0.598 | -0.003 | 0.014 | 0.014 | 0 | 0.44 | 0.45 | 0.33 | 0.13 | 0.54 | 0.3 |
| 9 | 20000600 | rs12057077 | 0.785 | -1.127 | -0.506 | 0.783 | -0.005 | -0.101 | -0.101 | 0 | 0.33 | 0.06 | 0.46 | 0.44 | 0.28 | 0.36 |
| 9 | 20001782 | rs7862804 | 0.506 | -1.128 | -0.233 | 0.444 | -0.005 | 0.018 | 0.018 | 0 | 0.41 | 0.4 | 0.29 | 0.12 | 0.54 | 0.29 |
| 9 | 20005312 | rs7037807 | 0.833 | -0.384 | -0.276 | 0.106 | 0.004 | -0.045 | -0.045 | 0.023 | 0.33 | 0.39 | 0.25 | 0.06 | 0.45 | 0.29 |
| 9 | 20005735 | rs6475389 | -0.208 | -0.501 | -0.533 | 0.359 | 0.017 | -0.02 | -0.02 | 0 | 0.7 | 0.79 | 0.77 | 0.61 | 0.78 | 0.68 |
| 9 | 20006704 | rs1112017 | 0.378 | -0.234 | -0.487 | 0.608 | -0.005 | -0.024 | -0.024 | 1.59 | 0.35 | 0.21 | 0.5 | 0.51 | 0.33 | 0.26 |
| 9 | 20014872 | rs1892014 | 0.22 | -0.338 | -0.537 | 0.083 | 0.005 | 0.002 | 0.002 | 0 | 0.66 | 0.67 | 0.82 | 0.66 | 0.75 | 0.67 |
| 9 | 20016698 | rs4977370 | 0.039 | -0.322 | -0.525 | 0.026 | 0.002 | 0.001 | 0.001 | 0 | 0.65 | 0.66 | 0.8 | 0.66 | 0.72 | 0.66 |
| 9 | 20016778 | rs4977371 | 0.821 | 0.009 | -0.869 | -0.251 | -0.04 | -0.057 | -0.057 | 0 | 0.22 | 0.08 | 0.52 | 0.57 | 0.25 | 0.35 |
| 9 | 20021047 | rs7860695 | 1.058 | -0.107 | -0.354 | -0.691 | -0.011 | 0.037 | 0.037 | 0 | 0.38 | 0.34 | 0.27 | 0.09 | 0.48 | 0.27 |
| 9 | 20022425 | rs10964389 | 0.141 | 0.001 | -0.065 | -0.131 | 0.002 | 0.286 | 0.286 | 0 | 0.1 | 0.29 | 0.17 | 0.43 | 0.06 | 0.22 |
| 9 | 20022680 | rs10811299 | -0.03 | 0.034 | -0.161 | -0.201 | 0.032 | 0.054 | 0.054 | 0 | 0.23 | 0.37 | 0.55 | 0.64 | 0.22 | 0.35 |
| 9 | 20025247 | rs10964392 | 1.005 | 0.08 | -0.253 | -0.239 | -0.006 | -0.007 | -0.007 | 0.71 | 0.17 | 0.12 | 0.37 | 0.22 | 0.17 | 0.17 |
| 9 | 20027231 | rs10964393 | -0.14 | -0.036 | -0.249 | -0.814 | 0.011 | 0.016 | 0.016 | 0 | 0.4 | 0.35 | 0.22 | 0.33 | 0.34 | 0.36 |
| 9 | 20030553 | rs10964396 | 0.251 | -0.444 | -0.29 | -0.886 | -0.001 | 0.008 | 0.008 | 0 | 0.22 | 0.33 | 0.12 | 0.24 | 0.19 | 0.14 |
| 9 | 20031656 | rs7851611 | -0.975 | -0.583 | -0.356 | -0.978 | 0.014 | 0.036 | 0.036 | 0 | 0.49 | 0.62 | 0.68 | 0.73 | 0.38 | 0.64 |
| 9 | 20032728 | rs12341973 | 0.63 | -0.547 | -0.459 | -0.947 | 0 | -0.001 | -0.001 | 0 | 0.2 | 0.14 | 0.1 | 0.23 | 0.18 | 0.12 |
| 9 | 20033087 | rs4246835 | -2.202 | -0.571 | -0.458 | -1.368 | -0.021 | 0.044 | 0.044 | 0 | 0.68 | 0.76 | 0.78 | 0.96 | 0.57 | 0.75 |
| 9 | 20033977 | rs6475394 | 0.511 | -0.87 | -0.109 | -1.073 | -0.037 | -0.055 | -0.055 | 0 | 0.24 | 0.38 | 0.13 | 0.12 | 0.15 | 0.26 |
| 9 | 20035279 | rs9792640 | -0.696 | -0.628 | 0.232 | -0.917 | 0.006 | -0.039 | -0.039 | 0.046 | 0.29 | 0.35 | 0.57 | 0.84 | 0.27 | 0.41 |
| 9 | 20035364 | rs9792484 | -1.37 | -0.481 | 0.053 | -1.014 | -0.003 | -0.015 | -0.015 | 0 | 0.63 | 0.64 | 0.78 | 0.92 | 0.55 | 0.7 |
| 9 | 20038436 | rs10964399 | 1.203 | -0.426 | -0.031 | -0.829 | 0.001 | 0.012 | 0.012 | 0 | 0.11 | 0.14 | 0.06 | 0.16 | 0.11 | 0.06 |
| 9 | 20039441 | rs7041787 | 0.542 | -0.53 | -0.057 | -0.92 | 0.009 | -0.001 | -0.001 | 0 | 0.36 | 0.35 | 0.21 | 0.1 | 0.29 | 0.32 |
| 9 | 20045332 | rs10964401 | -0.638 | -0.456 | -0.129 | -0.73 | 0.008 | 0.076 | 0.076 | 0 | 0.52 | 0.64 | 0.73 | 0.79 | 0.45 | 0.66 |
| 9 | 20046681 | rs12348353 | -0.198 | -0.417 | -0.128 | -0.508 | -0.01 | 0.002 | 0.002 | 0.041 | 0.19 | 0.26 | 0.1 | 0.23 | 0.14 | 0.15 |
| 9 | 20058939 | rs12684532 | 0.8 | -0.244 | -0.793 | -0.42 | -0.001 | 0 | 0 | 0 | 0.13 | 0.15 | 0.07 | 0.16 | 0.12 | 0.07 |
| 9 | 20059283 | rs10964409 | 1.25 | -0.163 | -0.911 | 0.085 | -0.013 | 0.018 | 0.018 | 0.713 | 0.07 | 0.01 | 0.18 | 0.47 | 0.06 | 0.2 |
| 9 | 20068518 | rs7037514 | 1.276 | -0.407 | -0.754 | -0.639 | -0.029 | -0.097 | -0.097 | 0 | 0.27 | 0.46 | 0.19 | 0.08 | 0.23 | 0.21 |
| 9 | 20072686 | rs4977378 | 0.261 | -0.084 | -0.905 | -0.23 | -0.025 | -0.059 | -0.059 | 0.001 | 0.16 | 0.05 | 0.49 | 0.64 | 0.16 | 0.34 |
| 9 | 20072967 | rs4246838 | -0.959 | -0.244 | -0.982 | -0.59 | -0.022 | -0.011 | -0.011 | 0 | 0.32 | 0.52 | 0.58 | 0.72 | 0.24 | 0.46 |
| 9 | 20083157 | rs3933328 | -1.155 | 0.095 | -0.762 | -0.382 | -0.019 | 0.036 | 0.036 | 0 | 0.63 | 0.74 | 0.78 | 0.94 | 0.53 | 0.71 |
| 9 | 20084210 | rs12342499 | 0.517 | 0.064 | -1.088 | -0.366 | -0.005 | -0.022 | -0.022 | 0 | 0.4 | 0.46 | 0.27 | 0.22 | 0.35 | 0.28 |
| 9 | 20084264 | rs4977384 | -0.577 | 0.252 | -1.875 | -0.489 | -0.047 | 0.113 | 0.113 | 0 | 0.63 | 0.82 | 0.78 | 0.95 | 0.53 | 0.71 |
| 9 | 20093557 | rs2383134 | -0.239 | -0.047 | -1.679 | 0.502 | -0.006 | -0.034 | -0.034 | 0.166 | 0.15 | 0.07 | 0.39 | 0.19 | 0.12 | 0.16 |
| 9 | 20095103 | rs10811316 | -0.434 | -1.099 | -2.226 | 0.139 | 0.001 | 0.041 | 0.041 | 0 | 0.39 | 0.06 | 0.53 | 0.22 | 0.33 | 0.34 |
| 9 | 20096390 | rs7871772 | -1.271 | -2.044 | -0.94 | 0.715 | -0.011 | 0.011 | 0.011 | 0.511 | 0.32 | 0.36 | 0.33 | 0.6 | 0.22 | 0.4 |
| 9 | 20097117 | rs5025178 | -0.492 | -1.982 | -0.838 | 0.806 | -0.005 | -0.033 | -0.033 | 0 | 0.17 | 0.21 | 0.12 | 0.02 | 0.12 | 0.11 |
| 9 | 20101645 | rs951303 | -0.114 | -2.125 | -0.882 | 1.042 | -0.006 | -0.006 | -0.006 | 0 | 0.08 | 0.02 | 0.18 | 0.51 | 0.08 | 0.2 |
| 9 | 20111554 | rs1333787 | -0.752 | -2.594 | -1.166 | 0.647 | 0.028 | 0.04 | 0.04 | 0 | 0.27 | 0.16 | 0.14 | 0.03 | 0.15 | 0.12 |
| 9 | 20123538 | rs10811319 | -1.056 | -2.31 | -0.977 | 0.879 | 0.014 | -0.03 | -0.03 | 0 | 0.24 | 0.17 | 0.3 | 0.56 | 0.17 | 0.33 |
| 9 | 20124467 | rs950020 | -1.309 | -2.168 | -0.694 | 0.972 | -0.004 | 0.009 | 0.009 | 1.122 | 0.25 | 0.3 | 0.26 | 0.55 | 0.18 | 0.34 |
| 9 | 20131746 | rs10964441 | 0.209 | -1.769 | -0.549 | 1.521 | -0.003 | 0.05 | 0.05 | 1.465 | 0.06 | 0.01 | 0.17 | 0.43 | 0.05 | 0.21 |
| 9 | 20136893 | rs7855796 | -0.139 | -1.753 | -0.24 | 1.163 | 0.002 | 0.218 | 0.218 | 1.841 | 0.06 | 0.21 | 0.06 | 0.31 | 0.04 | 0.18 |
| 9 | 20144936 | rs4977246 | 0.407 | -2.402 | -0.064 | 0.6 | -0.055 | -0.042 | -0.042 | 1.259 | 0.78 | 0.64 | 0.9 | 0.91 | 0.88 | 0.87 |
| 9 | 20145988 | rs2039461 | 1.123 | -2.463 | -1.155 | -0.043 | 0.081 | 0.338 | 0.338 | 0 | 0.5 | 0.1 | 0.33 | 0.11 | 0.62 | 0.45 |
| 9 | 20153762 | rs994975 | 0.609 | -2.191 | -0.103 | 0.843 | -0.061 | 0.245 | 0.245 | 0 | 0.85 | 0.51 | 0.86 | 0.61 | 0.92 | 0.69 |
| 9 | 20159208 | rs7019466 | -0.736 | -2.175 | -0.535 | -0.5 | 0.006 | 0.001 | 0.001 | 0 | 0.41 | 0.57 | 0.54 | 0.84 | 0.29 | 0.41 |
| 9 | 20168739 | rs7863754 | -0.282 | -0.725 | -0.591 | 0.013 | -0.02 | -0.116 | -0.116 | 0.026 | 0.23 | 0.48 | 0.18 | 0.07 | 0.17 | 0.18 |
| 9 | 20171664 | rs11562206 | -1.078 | -0.234 | -1.212 | 1.057 | -0.028 | -0.076 | -0.076 | 0 | 0.14 | 0.02 | 0.45 | 0.69 | 0.17 | 0.29 |
| 9 | 20178680 | rs1413254 | -0.802 | -0.07 | -0.788 | 1.603 | -0.001 | 0.114 | 0.114 | 0.392 | 0.08 | 0.09 | 0.25 | 0.44 | 0.07 | 0.29 |
| 9 | 20179356 | rs869149 | 0.292 | -0.972 | -0.908 | 0.564 | 0.008 | -0.016 | -0.016 | 0 | 0.21 | 0.12 | 0.11 | 0.02 | 0.1 | 0.17 |

| | | | | | | | | | | | | | | | |
|---|---|---|---|---|---|---|---|---|---|---|---|---|---|---|---|
| 9 | 20180241 | rs10738560 | 0.33 | -0.441 | -1.511 | -0.376 | -0.039 | -0.043 | -0.043 | 0 | 0.36 | 0.17 | 0.43 | 0.5 | 0.43 | 0.45 |
| 9 | 20180613 | rs1889245 | 0.494 | -0.147 | -0.864 | -0.229 | 0.023 | 0.027 | 0.027 | 0 | 0.13 | 0.21 | 0.16 | 0.27 | 0.22 | 0.17 |
| 9 | 20180680 | rs1889246 | -0.434 | -0.62 | -0.501 | -0.65 | -0.016 | 0.001 | 0.001 | 0 | 0.81 | 0.72 | 0.83 | 0.82 | 0.89 | 0.92 |
| 9 | 20186459 | rs13284316 | -0.164 | -0.348 | -0.397 | -0.707 | -0.001 | -0.024 | -0.024 | 0.368 | 0.11 | 0.14 | 0.11 | 0.01 | 0.12 | 0.08 |
| 9 | 20187462 | rs321205 | -0.478 | -0.516 | -0.123 | -0.815 | 0 | 0 | 0 | 3.651 | 0.54 | 0.56 | 0.51 | 0.31 | 0.58 | 0.26 |
| 9 | 20187574 | rs1889247 | -0.14 | -0.799 | 0.048 | -0.544 | -0.028 | 0.01 | 0.01 | 0 | 0.16 | 0.27 | 0.19 | 0.17 | 0.1 | 0.04 |
| 9 | 20190838 | rs4556171 | -0.299 | -0.342 | -0.428 | -0.683 | -0.01 | 0.028 | 0.028 | 0 | 0.09 | 0.03 | 0.06 | 0.01 | 0.09 | 0.08 |
| 9 | 20196466 | rs1456959 | -1.507 | -0.759 | -0.394 | -0.908 | 0.013 | 0.043 | 0.043 | 0 | 0.44 | 0.68 | 0.6 | 0.68 | 0.4 | 0.69 |
| 9 | 20209017 | rs403468 | 0.666 | -0.969 | -0.56 | -1.43 | 0.003 | 0.045 | 0.045 | 0 | 0.74 | 0.83 | 0.79 | 0.97 | 0.76 | 0.88 |
| 9 | 20209879 | rs1333790 | 0.247 | -0.887 | -0.325 | -0.939 | 0.005 | 0.008 | 0.008 | 0 | 0.08 | 0.13 | 0.07 | 0.14 | 0.1 | 0.14 |
| 9 | 20219315 | rs2034875 | -1.115 | -1.093 | -0.323 | -0.831 | 0.022 | -0.003 | -0.003 | 0.156 | 0.26 | 0.17 | 0.48 | 0.68 | 0.21 | 0.53 |
| 9 | 20238152 | rs10511678 | -0.72 | -0.915 | -0.607 | -1.049 | -0.013 | 0.094 | 0.094 | 0 | 0.18 | 0.01 | 0.14 | 0.02 | 0.16 | 0.1 |
| 9 | 20239894 | rs10964494 | -0.418 | -0.912 | -0.574 | -0.954 | -0.013 | 0.03 | 0.03 | 0 | 0.16 | 0.04 | 0.21 | 0.08 | 0.15 | 0.11 |
| 9 | 20241069 | rs7867635 | -0.092 | -1.196 | -0.651 | -1.574 | 0.003 | 0.153 | 0.153 | 1.22 | 0.57 | 0.8 | 0.56 | 0.9 | 0.43 | 0.85 |
| 9 | 20241907 | rs7853226 | -0.098 | -1.51 | -0.535 | -1.528 | 0.09 | 0.16 | 0.16 | 0 | 0.53 | 0.75 | 0.33 | 0.24 | 0.35 | 0.33 |
| 9 | 20244920 | rs1379216 | -0.185 | -1.615 | -0.199 | -1.601 | 0 | -0.06 | -0.06 | 0 | 0.17 | 0.34 | 0.19 | 0.06 | 0.19 | 0.08 |
| 9 | 20248264 | rs10964496 | 1.743 | -1.846 | -0.178 | -1.715 | 0.05 | 0.063 | 0.063 | 0.339 | 0.27 | 0.17 | 0.1 | 0.13 | 0.15 | 0.29 |
| 9 | 20250993 | rs1456962 | -1.651 | -1.737 | -0.404 | -1.82 | -0.18 | 0.188 | 0.188 | 1.387 | 0.68 | 0.96 | 0.69 | 0.96 | 0.52 | 0.92 |
| 9 | 20252271 | rs7049230 | 0.148 | -1.834 | 0.087 | -1.858 | 0.013 | 0.003 | 0.003 | 1.885 | 0.47 | 0.57 | 0.5 | 0.9 | 0.34 | 0.83 |
| 9 | 20253425 | rs7039444 | 0.081 | -1.827 | 0.098 | -1.853 | 0.016 | 0.008 | 0.008 | 1.591 | 0.48 | 0.57 | 0.5 | 0.9 | 0.34 | 0.85 |
| 9 | 20255355 | rs1551534 | -0.198 | -1.781 | -0.052 | -1.25 | -0.008 | 0.024 | 0.024 | 0 | 0.13 | 0.19 | 0.07 | 0.22 | 0.09 | 0.18 |
| 9 | 20255922 | rs4977400 | -1.924 | -1.701 | -0.456 | -1.384 | -0.177 | 0.162 | 0.162 | 1.271 | 0.71 | 0.96 | 0.7 | 0.96 | 0.53 | 0.92 |
| 9 | 20259528 | rs7018634 | 0.349 | -1.695 | -0.261 | -1.258 | -0.008 | 0.035 | 0.035 | 0 | 0.14 | 0.05 | 0.13 | 0.05 | 0.17 | 0.08 |
| 9 | 20262063 | rs12345147 | 1.705 | -1.899 | -0.099 | -1.282 | 0.035 | 0.042 | 0.042 | 0.3 | 0.26 | 0.16 | 0.11 | 0.13 | 0.15 | 0.24 |
| 9 | 20267556 | rs4977406 | 1.276 | -1.684 | -0.027 | -0.986 | 0.004 | 0.036 | 0.036 | 0.065 | 0.07 | 0.04 | 0.3 | 0.51 | 0.06 | 0.4 |
| 9 | 20269201 | rs13287033 | 0.204 | -1.665 | 0.527 | -1.195 | -0.012 | -0.008 | -0.008 | 0 | 0.45 | 0.55 | 0.19 | 0.39 | 0.29 | 0.43 |
| 9 | 20270444 | rs1551535 | 1.361 | -1.626 | 0.376 | -1.144 | 0.003 | 0.055 | 0.055 | 0.142 | 0.07 | 0.05 | 0.3 | 0.51 | 0.06 | 0.37 |
| 9 | 20270528 | rs4977407 | -0.694 | -1.665 | 0.714 | -1.261 | 0.002 | -0.004 | -0.004 | 0 | 0.58 | 0.55 | 0.32 | 0.44 | 0.45 | 0.51 |
| 9 | 20272411 | rs1031602 | 0.444 | -1.646 | 0.192 | -1.258 | 0.036 | 0.028 | 0.028 | 0 | 0.43 | 0.35 | 0.17 | 0.39 | 0.28 | 0.42 |
| 9 | 20272442 | rs13440226 | 0.271 | -1.616 | 0.212 | -1.274 | 0 | 0.012 | 0.012 | 0 | 0.15 | 0.11 | 0.14 | 0.05 | 0.17 | 0.08 |
| 9 | 20273783 | rs1840789 | 0.22 | -1.195 | 0.095 | -1.296 | 0.042 | 0.046 | 0.046 | 0 | 0.46 | 0.36 | 0.21 | 0.38 | 0.33 | 0.43 |
| 9 | 20273793 | rs7019257 | 0.49 | -0.923 | -0.253 | -1.082 | 0 | 0.006 | 0.006 | 0 | 0.12 | 0.08 | 0.1 | 0.05 | 0.12 | 0.06 |
| 9 | 20274450 | rs10811339 | -2.308 | 0.231 | -0.58 | -1.3 | -0.039 | 0.037 | 0.037 | 0 | 0.92 | 0.97 | 0.92 | 1 | 0.82 | 0.97 |
| 9 | 20274698 | rs4146635 | 1.451 | -1.012 | -0.289 | -0.945 | 0 | 0.052 | 0.052 | 0.078 | 0.07 | 0.06 | 0.31 | 0.52 | 0.07 | 0.41 |
| 9 | 20275143 | rs7868882 | 0.656 | -0.981 | -0.341 | -1.029 | 0.001 | 0.005 | 0.005 | 0 | 0.11 | 0.08 | 0.1 | 0.05 | 0.11 | 0.07 |
| 9 | 20282519 | rs12683533 | 0.8 | -1.14 | -0.618 | -1.089 | 0.076 | 0.117 | 0.117 | 0.515 | 0.37 | 0.17 | 0.16 | 0.18 | 0.25 | 0.28 |
| 9 | 20283406 | rs10811340 | 1.581 | -1.182 | -0.778 | -1.076 | 0.081 | 0.111 | 0.111 | 0.113 | 0.29 | 0.1 | 0.1 | 0.13 | 0.16 | 0.22 |
| 9 | 20285162 | rs10811342 | -0.69 | -0.933 | -0.385 | -0.994 | -0.003 | 0.013 | 0.013 | 0.276 | 0.57 | 0.68 | 0.35 | 0.42 | 0.46 | 0.51 |
| 9 | 20286678 | rs12353526 | 0.815 | -0.994 | -0.286 | -0.952 | 0.045 | 0.036 | 0.036 | 0 | 0.37 | 0.27 | 0.15 | 0.31 | 0.24 | 0.39 |
| 9 | 20287139 | rs717081 | -0.235 | -1.025 | 0.002 | -0.661 | 0.026 | 0.041 | 0.041 | 2.458 | 0.5 | 0.58 | 0.58 | 0.7 | 0.41 | 0.66 |
| 9 | 20287282 | rs717082 | 0.132 | -1.115 | -0.405 | -1.061 | 0.008 | 0.002 | 0.002 | 0.427 | 0.43 | 0.42 | 0.27 | 0.19 | 0.34 | 0.29 |
| 9 | 20287612 | rs7867670 | 0.544 | -0.873 | -1.274 | -0.735 | 0.063 | 0.093 | 0.093 | 0.013 | 0.34 | 0.16 | 0.17 | 0.16 | 0.24 | 0.27 |
| 9 | 20287948 | rs7026582 | -0.226 | -0.288 | -0.806 | -0.127 | -0.074 | 0.094 | 0.094 | 0.107 | 0.67 | 0.83 | 0.71 | 0.93 | 0.5 | 0.84 |
| 9 | 20289153 | rs882123 | -0.961 | -0.81 | 0.526 | 1.284 | -0.031 | 0.082 | 0.082 | 0 | 0.15 | 0.32 | 0.22 | 0.27 | 0.07 | 0.09 |
| 9 | 20292245 | rs10757131 | -1.05 | -0.799 | 0.473 | 1.255 | -0.015 | 0.119 | 0.119 | 0 | 0.17 | 0.43 | 0.23 | 0.28 | 0.1 | 0.12 |
| 9 | 20301704 | rs12341095 | 0.469 | -0.703 | -0.465 | 1.057 | 0.001 | 0.089 | 0.089 | 0.471 | 0.4 | 0.03 | 0.2 | 0.24 | 0.28 | 0.52 |
| 9 | 20304873 | rs17262780 | 0.533 | -0.627 | 0.258 | 0.861 | -0.006 | 0.002 | 0.002 | 0.068 | 0.37 | 0.32 | 0.25 | 0.07 | 0.39 | 0.15 |
| 9 | 20308956 | rs16938009 | 0.044 | -0.628 | 0.824 | 1.389 | -0.008 | 0.302 | 0.302 | 0.259 | 0.07 | 0.32 | 0.18 | 0.29 | 0.01 | 0.04 |
| 9 | 20309831 | rs224025 | -0.069 | -0.361 | 0.246 | 1.573 | 0.029 | 0.006 | 0.006 | 1.619 | 0.46 | 0.83 | 0.44 | 0.42 | 0.39 | 0.19 |
| 9 | 20313099 | rs224022 | -0.345 | -0.278 | 0.391 | 1.341 | 0.004 | 0.016 | 0.016 | 1.086 | 0.51 | 0.58 | 0.44 | 0.37 | 0.44 | 0.22 |
| 9 | 20318797 | rs7023422 | 0.718 | -0.493 | -0.068 | 1.004 | 0.045 | 0.059 | 0.059 | 0 | 0.4 | 0.26 | 0.23 | 0.25 | 0.28 | 0.54 |
| 9 | 20319623 | rs7044062 | 0.593 | -0.521 | -0.528 | 0.963 | 0.089 | 0.116 | 0.116 | 0 | 0.44 | 0.22 | 0.22 | 0.23 | 0.26 | 0.53 |
| 9 | 20322257 | rs224016 | -0.286 | -0.407 | -0.731 | 1.516 | -0.035 | -0.066 | -0.066 | 0 | 0.43 | 0.91 | 0.45 | 0.46 | 0.38 | 0.24 |
| 9 | 20322932 | rs7048991 | -0.907 | -0.622 | 0.961 | 2.478 | -0.034 | 0.236 | 0.236 | 0 | 0.11 | 0.45 | 0.18 | 0.29 | 0.02 | 0.05 |
| 9 | 20323424 | rs224015 | 1.103 | -0.155 | 0.144 | 2.372 | 0.089 | 0.235 | 0.235 | 0.048 | 0.92 | 0.81 | 0.73 | 0.58 | 0.77 | 0.81 |
| 9 | 20327754 | rs10964534 | -1.505 | -1.851 | -0.409 | 1.959 | 0.141 | -0.035 | -0.035 | 0 | 0.32 | 0.87 | 0.4 | 0.67 | 0.1 | 0.35 |
| 9 | 20331681 | rs3739934 | -1.09 | -1.81 | 0.142 | 2.195 | -0.04 | 0.009 | 0.009 | 0.785 | 0.2 | 0.74 | 0.18 | 0.26 | 0.03 | 0.11 |
| 9 | 20333293 | rs10435776 | -0.976 | -1.7 | 0.196 | 2.25 | -0.047 | 0.037 | 0.037 | 1.002 | 0.15 | 0.75 | 0.18 | 0.25 | 0.03 | 0.11 |
| 9 | 20337131 | rs7856579 | -1.134 | -1.683 | 0.308 | 2.283 | -0.045 | 0.037 | 0.037 | 0.172 | 0.16 | 0.74 | 0.21 | 0.27 | 0.04 | 0.15 |
| 9 | 20341702 | rs3824576 | -2.193 | -2.057 | 0.511 | 2.421 | -0.038 | -0.001 | -0.001 | 0 | 0.3 | 0.67 | 0.38 | 0.71 | 0.09 | 0.39 |
| 9 | 20341734 | rs4977421 | -2.277 | -2.064 | 0.422 | 2.425 | 0.078 | 0 | 0 | 0 | 0.31 | 0.82 | 0.39 | 0.71 | 0.1 | 0.39 |
| 9 | 20342044 | rs10964538 | -0.523 | -1.282 | 0.369 | 2.162 | 0.023 | 0 | 0 | 0.002 | 0.06 | 0.06 | 0.1 | 0.06 | 0.13 | 0.16 |
| 9 | 20347240 | rs7862166 | -2.32 | -1.335 | 0.31 | 2.26 | 0.023 | -0.094 | -0.094 | 0.762 | 0.3 | 0.92 | 0.33 | 0.39 | 0.19 | 0.32 |
| 9 | 20354575 | rs2301548 | -1.411 | -1.43 | 0.619 | 2.005 | -0.054 | 0.011 | 0.011 | 0.006 | 0.17 | 0.7 | 0.18 | 0.22 | 0.03 | 0.14 |
| 9 | 20354796 | rs2301550 | -1.632 | -1.437 | 0.51 | 2.024 | 0.044 | -0.022 | -0.022 | 0.01 | 0.18 | 0.85 | 0.22 | 0.24 | 0.03 | 0.15 |
| 9 | 20357805 | rs7866741 | -1.47 | -1.41 | 0.574 | 2.003 | -0.017 | -0.001 | -0.001 | 0 | 0.2 | 0.78 | 0.21 | 0.24 | 0.03 | 0.16 |
| 9 | 20379884 | rs10453171 | -1.38 | -1.228 | 1.584 | 1.993 | 0.038 | 0.218 | 0.218 | 0 | 0.06 | 0.3 | 0.1 | 0.18 | 0.01 | 0.04 |
| 9 | 20391116 | rs1413554 | -2.177 | -1.244 | 1.724 | 2.003 | 0.012 | 0.206 | 0.206 | 0.016 | 0.08 | 0.55 | 0.13 | 0.19 | 0.02 | 0.06 |
| 9 | 20405630 | rs10964552 | -1.234 | -1.092 | 1.502 | 1.967 | -0.001 | -0.031 | -0.031 | 0 | 0.18 | 0.24 | 0.14 | 0.07 | 0.15 | 0.2 |

| | | | | | | | | | | | | | | | | |
|---|---|---|---|---|---|---|---|---|---|---|---|---|---|---|---|---|
| 9 | 20411354 | rs16938058 | -1.206 | -0.946 | 1.302 | 2.455 | 0.014 | 0.172 | 0.172 | 0 | 0.05 | 0.23 | 0.09 | 0.18 | 0.01 | 0.04 |
| 9 | 20418323 | rs10811349 | -1.803 | -0.989 | 1.07 | 2.253 | -0.053 | 0.155 | 0.155 | 0 | 0.13 | 0.32 | 0.3 | 0.68 | 0.05 | 0.27 |
| 9 | 20464018 | rs4977256 | -0.305 | -0.6 | -0.414 | 0.646 | 0.012 | 0.02 | 0.02 | 0 | 0.36 | 0.67 | 0.41 | 0.16 | 0.28 | 0.37 |
| 9 | 20464362 | rs3904577 | -0.625 | 0.096 | -0.866 | 0.599 | -0.006 | -0.008 | -0.008 | 0 | 0.64 | 0.68 | 0.5 | 0.56 | 0.57 | 0.7 |
| 9 | 20464928 | rs6475431 | -0.491 | -0.302 | -0.845 | 0.647 | 0.002 | 0.002 | 0.002 | 0 | 0.36 | 0.33 | 0.19 | 0.31 | 0.28 | 0.39 |
| 9 | 20465928 | rs10964566 | -0.07 | -0.221 | -0.905 | 0.762 | 0.009 | -0.044 | -0.044 | 0.04 | 0.13 | 0.05 | 0.05 | 0.32 | 0.07 | 0.27 |
| 9 | 20469336 | rs12344417 | 0.353 | 0.011 | -0.614 | 0.252 | -0.008 | -0.094 | -0.094 | 1.028 | 0.22 | 0.27 | 0.14 | 0 | 0.2 | 0.11 |
| 9 | 20471312 | rs3934325 | 0.379 | 1.372 | -1.668 | 0.752 | -0.141 | -0.101 | -0.101 | 0.144 | 0.78 | 0.94 | 0.53 | 0.71 | 0.57 | 0.84 |
| 9 | 20475176 | rs12350051 | -0.081 | 1.082 | -1.87 | 0.275 | -0.021 | 0.033 | 0.033 | 0 | 0.24 | 0.16 | 0.17 | 0.13 | 0.25 | 0.22 |
| 9 | 20478188 | rs3893395 | -0.098 | 0.977 | -1.55 | 0.223 | -0.02 | 0.039 | 0.039 | 0 | 0.24 | 0.16 | 0.17 | 0.12 | 0.25 | 0.21 |
| 9 | 20482380 | rs6475434 | 0.086 | 1.47 | -1.294 | 0.671 | -0.016 | -0.016 | -0.016 | 0 | 0.85 | 0.87 | 0.67 | 0.7 | 0.72 | 0.77 |
| 9 | 20484509 | rs6475438 | 0.173 | 1.379 | -0.747 | 0.53 | 0.033 | 0.017 | 0.017 | 0.565 | 0.84 | 0.79 | 0.7 | 0.8 | 0.73 | 0.78 |
| 9 | 20484604 | rs6475439 | 0.202 | 1.381 | -0.725 | 0.528 | 0.034 | 0.019 | 0.019 | 0.508 | 0.84 | 0.78 | 0.7 | 0.78 | 0.73 | 0.77 |
| 9 | 20497142 | rs6475448 | -0.053 | 0.63 | -1.348 | 0.047 | -0.011 | 0.081 | 0.081 | 0 | 0.65 | 0.87 | 0.69 | 0.77 | 0.63 | 0.74 |
| 9 | 20498930 | rs1411726 | 0.281 | 0.093 | -1.012 | 0.07 | -0.001 | 0 | 0 | 1.008 | 0.28 | 0.26 | 0.21 | 0.34 | 0.19 | 0.21 |
| 9 | 20498974 | rs1411727 | 0.029 | -0.091 | -0.292 | -0.02 | -0.03 | 0.009 | 0.009 | 2.041 | 0.3 | 0.48 | 0.25 | 0.34 | 0.19 | 0.21 |
| 9 | 20508503 | rs10964583 | 0.101 | 0.301 | 0.131 | 0.249 | 0.036 | -0.021 | -0.021 | 6.533 | 0.2 | 0.45 | 0.18 | 0.18 | 0.19 | 0.19 |
| 9 | 20514400 | rs10811359 | 0.628 | 0.23 | 0.252 | 0.223 | 0.001 | 0.004 | 0.004 | 0.163 | 0.46 | 0.54 | 0.53 | 0.63 | 0.45 | 0.55 |
| 9 | 20519521 | rs7868240 | 0.508 | 0.047 | 0.229 | 0.178 | 0.01 | 0 | 0 | 0 | 0.48 | 0.42 | 0.48 | 0.49 | 0.45 | 0.55 |
| 9 | 20521543 | rs7854782 | 0.696 | 0.095 | 0.257 | -0.139 | 0.007 | 0.01 | 0.01 | 0.076 | 0.46 | 0.37 | 0.47 | 0.51 | 0.45 | 0.55 |
| 9 | 20527695 | rs10757135 | -0.407 | 0.339 | 0.21 | -0.141 | -0.003 | 0.062 | 0.062 | 0 | 0.68 | 0.79 | 0.71 | 0.83 | 0.66 | 0.75 |
| 9 | 20533643 | rs10117696 | 0.849 | 0.556 | -0.242 | -0.163 | 0.018 | 0.015 | 0.015 | 0 | 0.46 | 0.3 | 0.46 | 0.51 | 0.43 | 0.52 |
| 9 | 20537343 | rs6475457 | 0.608 | 0.251 | -0.236 | -0.083 | 0.003 | 0.014 | 0.014 | 0.146 | 0.49 | 0.54 | 0.57 | 0.63 | 0.48 | 0.55 |
| 9 | 20549744 | rs10964600 | 1.118 | 0.194 | -0.305 | -0.161 | -0.004 | -0.017 | -0.017 | 0.751 | 0.42 | 0.19 | 0.44 | 0.47 | 0.42 | 0.52 |
| 9 | 20553571 | rs1111766 | 1.02 | 0.852 | 0.377 | 0.786 | 0.016 | 0.033 | 0.033 | 0 | 0.7 | 0.44 | 0.71 | 0.51 | 0.69 | 0.62 |
| 9 | 20563799 | rs1360378 | 0.039 | 0.899 | -0.906 | 0.548 | -0.003 | 0.01 | 0.01 | 0 | 0.9 | 0.95 | 0.9 | 0.87 | 0.88 | 0.82 |
| 9 | 20578041 | rs12380857 | 0.59 | 1.355 | -0.961 | 1.06 | -0.041 | -0.036 | -0.036 | 0 | 0.07 | 0.01 | 0.06 | 0.22 | 0.14 | 0.07 |
| 9 | 20581841 | rs10811371 | -0.131 | 0.728 | -0.882 | -0.168 | -0.001 | 0.058 | 0.058 | 1.01 | 0.51 | 0.37 | 0.34 | 0.22 | 0.47 | 0.34 |
| 9 | 20581910 | rs10811372 | -0.187 | 0.67 | -0.677 | 0.399 | -0.002 | 0.019 | 0.019 | 0.837 | 0.52 | 0.39 | 0.39 | 0.36 | 0.47 | 0.35 |
| 9 | 20590209 | rs7025895 | -0.172 | 0.957 | -0.196 | 1.156 | 0.016 | 0.073 | 0.073 | 0.133 | 0.89 | 0.82 | 0.72 | 0.75 | 0.82 | 0.81 |
| 9 | 20597576 | rs6475464 | 0.598 | 0.72 | 0.062 | 1.214 | -0.009 | -0.014 | -0.014 | 0 | 0.36 | 0.16 | 0.28 | 0.38 | 0.35 | 0.47 |
| 9 | 20599426 | rs636100 | -0.149 | 0.765 | 0.248 | 1.085 | 0.047 | 0.144 | 0.144 | 0.183 | 0.89 | 0.71 | 0.72 | 0.75 | 0.82 | 0.82 |
| 9 | 20601177 | rs668703 | 0.361 | 0.754 | 0.126 | 1.014 | -0.005 | 0.126 | 0.126 | 0.936 | 0.51 | 0.26 | 0.37 | 0.26 | 0.47 | 0.36 |
| 9 | 20603364 | rs2775264 | 0.296 | 0.748 | 0.111 | 1.148 | -0.009 | 0.079 | 0.079 | 0.839 | 0.51 | 0.28 | 0.36 | 0.29 | 0.47 | 0.37 |
| 9 | 20606428 | rs623828 | 0.383 | 0.454 | -0.022 | 0.998 | -0.013 | 0.023 | 0.023 | 0.014 | 0.29 | 0.22 | 0.17 | 0.2 | 0.31 | 0.28 |
| 9 | 20608377 | rs2780840 | -0.128 | 0.524 | 0.342 | 1.245 | 0.007 | 0.009 | 0.009 | 0 | 0.87 | 0.83 | 0.68 | 0.74 | 0.81 | 0.82 |
| 9 | 20608912 | rs2188230 | 0.514 | 0.437 | 0.367 | 0.962 | 0.012 | 0.009 | 0.009 | 0 | 0.6 | 0.53 | 0.49 | 0.46 | 0.53 | 0.54 |
| 9 | 20610914 | rs2780839 | -0.269 | 0.459 | -0.2 | 0.977 | -0.005 | -0.003 | -0.003 | 0 | 0.88 | 0.93 | 0.69 | 0.68 | 0.81 | 0.81 |
| 9 | 20617114 | rs10811376 | 0.531 | 0.272 | -0.081 | 0.79 | -0.031 | 0.024 | 0.024 | 0 | 0.57 | 0.24 | 0.42 | 0.3 | 0.51 | 0.52 |
| 9 | 20623446 | rs2073854 | 0.326 | -0.298 | -0.135 | 0.583 | -0.079 | -0.062 | -0.062 | 0 | 0.22 | 0.39 | 0.15 | 0.1 | 0.11 | 0.09 |
| 9 | 20625228 | rs2519454 | 0.193 | -0.005 | -0.538 | 0.482 | -0.039 | 0.027 | 0.027 | 0.198 | 0.26 | 0.22 | 0.34 | 0.19 | 0.34 | 0.35 |
| 9 | 20625875 | rs2188229 | 0.826 | 0.153 | -0.655 | 0.346 | -0.005 | 0.075 | 0.075 | 0.662 | 0.42 | 0.25 | 0.32 | 0.17 | 0.43 | 0.46 |
| 9 | 20629336 | rs2780838 | -0.97 | -0.027 | -0.555 | 0.466 | -0.015 | 0.11 | 0.11 | 1.041 | 0.54 | 0.73 | 0.65 | 0.84 | 0.52 | 0.51 |
| 9 | 20629553 | rs1548378 | -0.704 | 0.199 | -0.401 | 0.616 | -0.008 | 0.127 | 0.127 | 0.167 | 0.49 | 0.73 | 0.64 | 0.84 | 0.5 | 0.49 |
| 9 | 20631633 | rs1548377 | -0.51 | 0.363 | -0.528 | 0.729 | -0.001 | 0.011 | 0.011 | 0.296 | 0.47 | 0.42 | 0.55 | 0.67 | 0.47 | 0.48 |
| 9 | 20638248 | rs6475467 | -0.613 | 0.066 | -1.086 | 0.538 | 0.02 | 0.005 | 0.005 | 1.539 | 0.44 | 0.7 | 0.48 | 0.47 | 0.38 | 0.29 |
| 9 | 20641181 | rs2383141 | -0.891 | -0.146 | -1.329 | 0.347 | 0.031 | 0.06 | 0.06 | 0.725 | 0.51 | 0.7 | 0.55 | 0.45 | 0.43 | 0.35 |
| 9 | 20641792 | rs10811380 | -0.997 | -0.223 | -1.351 | 0.078 | 0.032 | 0.076 | 0.076 | 0.38 | 0.51 | 0.72 | 0.52 | 0.39 | 0.42 | 0.3 |
| 9 | 20642438 | rs4129311 | -0.691 | 0.64 | -1.297 | 0.389 | -0.018 | 0.002 | 0.002 | 0.001 | 0.28 | 0.18 | 0.37 | 0.31 | 0.35 | 0.27 |
| 9 | 20652174 | rs10811383 | -0.166 | 0.408 | -1.428 | 0.933 | -0.017 | -0.067 | -0.067 | 0 | 0.11 | 0.01 | 0.25 | 0.27 | 0.13 | 0.12 |
| 9 | 20665071 | rs7859426 | 0.269 | 0.619 | -1.19 | 0.782 | -0.001 | -0.004 | -0.004 | 0 | 0.81 | 0.78 | 0.84 | 0.72 | 0.81 | 0.88 |
| 9 | 20670187 | rs4596728 | 0.412 | 0.488 | -1.291 | 0.801 | 0.001 | -0.001 | -0.001 | 0 | 0.17 | 0.17 | 0.14 | 0.15 | 0.16 | 0.11 |
| 9 | 20670573 | rs7872857 | -0.147 | 0.45 | -1.287 | 0.918 | -0.004 | -0.019 | -0.019 | 0 | 0.8 | 0.81 | 0.83 | 0.7 | 0.8 | 0.87 |
| 9 | 20676454 | rs7867822 | 0.443 | 0.307 | -0.722 | 0.808 | 0.009 | 0.038 | 0.038 | 0 | 0.71 | 0.56 | 0.75 | 0.55 | 0.69 | 0.76 |
| 9 | 20700838 | rs10964665 | 0.181 | 0.461 | -1.135 | -0.256 | 0.001 | 0.024 | 0.024 | 0 | 0.5 | 0.46 | 0.4 | 0.22 | 0.51 | 0.5 |
| 9 | 20701305 | rs1332311 | 1.602 | 0.425 | -1.608 | 0.23 | -0.079 | -0.062 | -0.062 | 0 | 0.16 | 0.01 | 0.36 | 0.25 | 0.21 | 0.12 |
| 9 | 20703928 | rs1332312 | 0.097 | 1.235 | -1.55 | 0.042 | 0.036 | 0.04 | 0.04 | 0 | 0.34 | 0.38 | 0.49 | 0.48 | 0.42 | 0.37 |
| 9 | 20704320 | rs10811394 | 0.566 | 1.09 | -1.728 | -0.066 | -0.043 | 0.156 | 0.156 | 0.18 | 0.47 | 0.13 | 0.38 | 0.22 | 0.5 | 0.49 |
| 9 | 20704772 | rs17832026 | 0.58 | 1.156 | -0.74 | -0.049 | -0.001 | 0.001 | 0.001 | 0 | 0.3 | 0.25 | 0.43 | 0.48 | 0.29 | 0.36 |
| 9 | 20710798 | rs1930050 | 0.311 | 1.171 | -0.644 | -0.045 | 0.027 | 0.03 | 0.03 | 0 | 0.31 | 0.37 | 0.48 | 0.47 | 0.41 | 0.35 |
| 9 | 20710896 | rs12376200 | 1.734 | 1.074 | -0.742 | -0.007 | -0.017 | -0.041 | -0.041 | 0.643 | 0.16 | 0.01 | 0.13 | 0.15 | 0.13 | 0.11 |
| 9 | 20714014 | rs2039391 | -1.43 | 1.225 | -0.841 | 0.148 | -0.002 | 0.023 | 0.023 | 0 | 0.91 | 0.87 | 0.91 | 0.68 | 0.86 | 0.85 |
| 9 | 20716880 | rs12684948 | 0.314 | 1.077 | -0.914 | -0.504 | 0.008 | 0.086 | 0.086 | 0 | 0.47 | 0.31 | 0.37 | 0.19 | 0.46 | 0.49 |
| 9 | 20717020 | rs12683110 | 0.455 | 0.618 | -1.143 | -0.458 | 0.044 | 0.131 | 0.131 | 0.006 | 0.44 | 0.16 | 0.32 | 0.19 | 0.36 | 0.49 |
| 9 | 20723087 | rs7046563 | 0.293 | 1.085 | -0.586 | -0.665 | 0.004 | -0.002 | -0.002 | 0 | 0.52 | 0.52 | 0.39 | 0.18 | 0.41 | 0.46 |
| 9 | 20726571 | rs1888920 | 0.069 | 1.353 | -0.716 | -0.396 | -0.005 | -0.037 | -0.037 | 0.734 | 0.3 | 0.12 | 0.27 | 0.38 | 0.29 | 0.3 |
| 9 | 20735234 | rs1556476 | -0.496 | 1.767 | -0.364 | -0.322 | 0.008 | 0.008 | 0.008 | 0 | 0.38 | 0.47 | 0.54 | 0.51 | 0.41 | 0.35 |
| 9 | 20737847 | rs10511689 | 1.043 | 1.656 | -0.486 | -0.38 | 0.007 | 0.024 | 0.024 | 0 | 0.21 | 0.06 | 0.18 | 0.16 | 0.2 | 0.14 |
| 9 | 20739042 | rs7031527 | -0.926 | 1.547 | -0.157 | -0.507 | 0.001 | 0.003 | 0.003 | 0 | 0.56 | 0.51 | 0.7 | 0.69 | 0.6 | 0.49 |

| | | | | | | | | | | | | | | | |
|---|---|---|---|---|---|---|---|---|---|---|---|---|---|---|---|
| 9 | 20740671 | rs7030093 | 1.263 | 1.96 | 0.156 | -0.605 | -0.003 | -0.015 | -0.015 | 0 | 0.35 | 0.28 | 0.28 | 0.04 | 0.4 | 0.38 |
| 9 | 20742770 | rs883601 | 0.912 | 1.974 | 0.161 | -0.527 | 0.024 | 0.152 | 0.152 | 0 | 0.52 | 0.22 | 0.41 | 0.22 | 0.47 | 0.51 |
| 9 | 20747070 | rs7043296 | -0.579 | 1.958 | 0.247 | -0.401 | 0.011 | 0.036 | 0.036 | 0 | 0.84 | 0.66 | 0.91 | 0.74 | 0.83 | 0.83 |
| 9 | 20749871 | rs1332322 | -0.298 | 2.102 | -0.007 | -0.448 | -0.009 | 0.035 | 0.035 | 0 | 0.34 | 0.4 | 0.45 | 0.52 | 0.26 | 0.33 |
| 9 | 20754419 | rs17685780 | 1.626 | 1.861 | -0.214 | -0.57 | 0.036 | 0.134 | 0.134 | 0 | 0.31 | 0.11 | 0.21 | 0.04 | 0.24 | 0.36 |
| 9 | 20756547 | rs1475660 | -0.153 | 1.792 | -0.049 | -0.285 | 0 | 0.015 | 0.015 | 0 | 0.68 | 0.6 | 0.57 | 0.47 | 0.7 | 0.72 |
| 9 | 20762926 | rs17832189 | 1.062 | 2.003 | -0.043 | -0.333 | 0.005 | 0.027 | 0.027 | 0 | 0.22 | 0.1 | 0.19 | 0.16 | 0.23 | 0.16 |
| 9 | 20763840 | rs16938162 | 0.965 | 2.105 | 0.126 | -0.478 | -0.005 | 0 | 0 | 0 | 0.34 | 0.26 | 0.28 | 0.04 | 0.39 | 0.38 |
| 9 | 20764870 | rs10511687 | 0.894 | 2.056 | 0.137 | -0.475 | 0 | -0.047 | -0.047 | 0 | 0.34 | 0.31 | 0.28 | 0.04 | 0.38 | 0.39 |
| 9 | 20766254 | rs10811399 | -0.281 | 2.171 | 0.258 | -0.281 | 0.006 | 0.006 | 0.006 | 0 | 0.38 | 0.53 | 0.55 | 0.51 | 0.42 | 0.31 |
| 9 | 20776743 | rs12551126 | -0.003 | 2.085 | 0.257 | -0.289 | 0.014 | 0.032 | 0.032 | 0 | 0.34 | 0.38 | 0.4 | 0.49 | 0.35 | 0.32 |
| 9 | 20779843 | rs4978125 | 0.026 | 2.099 | 0.072 | -0.251 | -0.008 | 0.016 | 0.016 | 0 | 0.34 | 0.36 | 0.39 | 0.51 | 0.25 | 0.3 |
| 9 | 20782100 | rs7027599 | -0.543 | 1.515 | -0.492 | -0.187 | -0.032 | -0.067 | -0.067 | 0.109 | 0.7 | 0.79 | 0.69 | 0.56 | 0.66 | 0.69 |
| 9 | 20786691 | rs17685984 | 1.175 | 1.196 | -0.599 | -0.477 | 0.022 | 0.14 | 0.14 | 0 | 0.31 | 0.09 | 0.22 | 0.05 | 0.25 | 0.38 |
| 9 | 20789869 | rs10964702 | 1.021 | 1.17 | -0.625 | -0.442 | -0.022 | 0.025 | 0.025 | 0 | 0.22 | 0.02 | 0.19 | 0.18 | 0.23 | 0.16 |
| 9 | 20793269 | rs7036612 | 1.206 | 1.169 | -0.532 | -0.467 | 0.034 | 0.067 | 0.067 | 0 | 0.31 | 0.18 | 0.2 | 0.04 | 0.22 | 0.37 |
| 9 | 20801654 | rs10964709 | 0.835 | 1.137 | -0.407 | -0.132 | 0.004 | 0.029 | 0.029 | 0 | 0.22 | 0.07 | 0.19 | 0.2 | 0.23 | 0.16 |
| 9 | 20811421 | rs10511682 | 0.808 | 1.144 | -0.627 | -0.108 | -0.004 | 0.016 | 0.016 | 0 | 0.22 | 0.05 | 0.19 | 0.19 | 0.23 | 0.16 |
| 9 | 20821287 | rs1556475 | -1.132 | 1.524 | -0.467 | 0.212 | -0.009 | -0.006 | -0.006 | 0 | 0.9 | 0.89 | 0.92 | 0.76 | 0.84 | 0.82 |
| 9 | 20822078 | rs3898946 | 0.161 | 1.44 | -0.173 | -0.046 | 0.003 | 0.018 | 0.018 | 0 | 0.33 | 0.32 | 0.45 | 0.47 | 0.35 | 0.3 |
| 9 | 20823314 | rs17759424 | 1.707 | 1.338 | -0.382 | 0.126 | -0.014 | -0.067 | -0.067 | 0 | 0.1 | 0.01 | 0.28 | 0.26 | 0.1 | 0.13 |
| 9 | 20825682 | rs7851807 | 1.075 | 1.398 | -0.041 | -0.064 | -0.005 | -0.049 | -0.049 | 0.017 | 0.33 | 0.33 | 0.27 | 0.06 | 0.25 | 0.36 |
| 9 | 20834837 | rs2026994 | -0.735 | 1.292 | 0.004 | -0.063 | 0.001 | 0.009 | 0.009 | 0 | 0.61 | 0.6 | 0.7 | 0.68 | 0.64 | 0.48 |
| 9 | 20840838 | rs7870627 | -0.354 | 1.023 | 0.117 | -0.036 | -0.006 | 0.019 | 0.019 | 0 | 0.54 | 0.54 | 0.61 | 0.67 | 0.55 | 0.44 |
| 9 | 20862515 | rs4977780 | -0.323 | 1.024 | 0.182 | -0.053 | 0.001 | -0.006 | -0.006 | 0 | 0.55 | 0.48 | 0.61 | 0.67 | 0.55 | 0.45 |
| 9 | 20867809 | rs2383154 | -1.379 | 1.133 | 0.178 | -0.038 | 0.037 | 0.118 | 0.118 | 0 | 0.91 | 0.76 | 0.92 | 0.73 | 0.86 | 0.82 |
| 9 | 20868172 | rs2383155 | -1.868 | 1.138 | 0.138 | -0.026 | -0.019 | -0.008 | -0.008 | 0.604 | 0.92 | 0.95 | 0.93 | 0.77 | 0.86 | 0.84 |
| 9 | 20872842 | rs11789503 | 1.608 | 0.907 | 0.311 | -0.094 | -0.003 | -0.003 | -0.003 | 0 | 0.14 | 0.12 | 0.11 | 0.18 | 0.19 | 0.13 |
| 9 | 20876259 | rs1986633 | -0.197 | 1.038 | 0.162 | -0.278 | -0.001 | 0 | 0 | 0 | 0.53 | 0.42 | 0.6 | 0.67 | 0.54 | 0.44 |
| 9 | 20876513 | rs1986632 | -0.641 | 1.072 | 0.037 | -0.275 | -0.008 | 0.008 | 0.008 | 0 | 0.59 | 0.59 | 0.66 | 0.68 | 0.58 | 0.46 |
| 9 | 20883241 | rs2891154 | 1.326 | 1.114 | -0.059 | -0.297 | -0.037 | 0.17 | 0.17 | 0 | 0.31 | 0.06 | 0.25 | 0.06 | 0.26 | 0.36 |
| 9 | 20886882 | rs7044578 | 1.057 | 1.127 | -0.035 | -0.186 | -0.003 | 0.016 | 0.016 | 0 | 0.22 | 0.08 | 0.17 | 0.19 | 0.22 | 0.17 |
| 9 | 20890466 | rs10811422 | -0.365 | 0.959 | 0.19 | -0.151 | -0.006 | -0.019 | -0.019 | 0.027 | 0.67 | 0.67 | 0.74 | 0.53 | 0.62 | 0.64 |
| 9 | 20891171 | rs7853915 | 0.694 | 0.883 | 0.057 | -0.251 | 0 | -0.003 | -0.003 | 0.152 | 0.35 | 0.3 | 0.3 | 0.06 | 0.3 | 0.36 |
| 9 | 20898233 | rs12236288 | -0.345 | 1.015 | -0.216 | -0.029 | -0.02 | -0.045 | -0.045 | 0 | 0.68 | 0.73 | 0.72 | 0.53 | 0.59 | 0.64 |
| 9 | 20908374 | rs4288451 | 0.146 | 0.914 | 0.045 | 0.019 | -0.007 | 0.035 | 0.035 | 0 | 0.33 | 0.49 | 0.42 | 0.47 | 0.31 | 0.29 |
| 9 | 20923579 | rs7044452 | -0.828 | 0.99 | 0.153 | -0.311 | 0 | -0.038 | -0.038 | 1.085 | 0.63 | 0.63 | 0.69 | 0.94 | 0.69 | 0.64 |
| 9 | 20928323 | rs9406776 | 0.376 | 1.141 | -0.003 | -0.099 | -0.022 | -0.046 | -0.046 | 0 | 0.31 | 0.21 | 0.25 | 0.42 | 0.37 | 0.34 |
| 9 | 20929977 | rs9407968 | -0.274 | 1.103 | 0.104 | -0.114 | -0.002 | 0.009 | 0.009 | 0 | 0.64 | 0.47 | 0.52 | 0.52 | 0.64 | 0.71 |
| 9 | 20932868 | rs2383171 | 0.701 | 1.107 | 0.114 | -0.156 | 0.005 | 0.061 | 0.061 | 0 | 0.36 | 0.26 | 0.31 | 0.1 | 0.29 | 0.38 |
| 9 | 20944558 | rs12552707 | 0.694 | 1.143 | 0.223 | -0.156 | 0.003 | 0.021 | 0.021 | 0 | 0.36 | 0.3 | 0.31 | 0.1 | 0.29 | 0.37 |
| 9 | 20953049 | rs3206852 | 0.309 | 1.158 | 0.197 | -0.016 | -0.018 | -0.039 | -0.039 | 1.877 | 0.32 | 0.24 | 0.27 | 0.44 | 0.4 | 0.34 |
| 9 | 20953750 | rs4977848 | 0.52 | 1.132 | 0.152 | -0.029 | -0.064 | -0.112 | -0.112 | 2.373 | 0.3 | 0.08 | 0.25 | 0.43 | 0.37 | 0.34 |
| 9 | 20963374 | rs10964769 | 0.417 | 1.08 | 0.561 | -0.032 | -0.002 | 0.004 | 0.004 | 0 | 0.32 | 0.32 | 0.42 | 0.47 | 0.3 | 0.28 |
| 9 | 20966250 | rs7873691 | 0.34 | 1.071 | 0.486 | -0.018 | -0.013 | -0.029 | -0.029 | 0 | 0.31 | 0.26 | 0.26 | 0.43 | 0.38 | 0.34 |
| 9 | 20975121 | rs4977863 | -0.894 | 1.063 | 0.414 | -0.118 | 0.001 | -0.005 | -0.005 | 0 | 0.65 | 0.72 | 0.7 | 0.94 | 0.71 | 0.65 |
| 9 | 20994176 | rs10738571 | 0.187 | 1.056 | 0.475 | 0.049 | -0.004 | -0.011 | -0.011 | 0 | 0.33 | 0.34 | 0.26 | 0.48 | 0.38 | 0.36 |
| 9 | 20998354 | rs10757171 | -0.939 | 0.965 | 0.303 | -0.117 | 0.001 | 0.048 | 0.048 | 0 | 0.66 | 0.79 | 0.71 | 0.94 | 0.71 | 0.65 |
| 9 | 20999646 | rs4278230 | -0.195 | 1.051 | 0.316 | 0.014 | 0.002 | -0.011 | -0.011 | 0.096 | 0.43 | 0.57 | 0.53 | 0.7 | 0.5 | 0.47 |
| 9 | 21002225 | rs12004236 | 0.78 | 1.086 | 0.143 | -0.104 | -0.006 | 0.11 | 0.11 | 0 | 0.36 | 0.18 | 0.29 | 0.04 | 0.3 | 0.36 |
| 9 | 21003372 | rs3884620 | 1.42 | 1.039 | -0.214 | 0.158 | -0.029 | -0.071 | -0.071 | 0.976 | 0.11 | 0.01 | 0.31 | 0.27 | 0.13 | 0.14 |
| 9 | 21005744 | rs4129714 | 0.959 | 1.066 | -0.081 | -0.087 | -0.008 | 0.116 | 0.116 | 0 | 0.36 | 0.16 | 0.28 | 0.04 | 0.3 | 0.35 |
| 9 | 21006934 | rs1134090 | 1.315 | 1.029 | -0.216 | -0.139 | -0.043 | 0.173 | 0.173 | 0 | 0.31 | 0.07 | 0.25 | 0.05 | 0.27 | 0.34 |
| 9 | 21007935 | rs7041474 | -1.08 | 1.183 | 0.118 | -0.25 | 0 | -0.039 | -0.039 | 0 | 0.66 | 0.72 | 0.7 | 0.96 | 0.69 | 0.67 |
| 9 | 21015651 | rs7873640 | 0.077 | 1.127 | 0.295 | 0.088 | -0.007 | -0.007 | -0.007 | 0 | 0.35 | 0.35 | 0.3 | 0.42 | 0.41 | 0.36 |
| 9 | 21017743 | rs2275886 | -0.948 | 0.97 | 0.259 | 0.156 | -0.001 | -0.005 | -0.005 | 0 | 0.75 | 0.79 | 0.76 | 0.78 | 0.73 | 0.83 |
| 9 | 21017828 | rs2275887 | 0.851 | 0.737 | 0.508 | 0.215 | 0.01 | -0.017 | -0.017 | 0 | 0.16 | 0.11 | 0.29 | 0.34 | 0.1 | 0.14 |
| 9 | 21019817 | rs7874876 | -0.091 | 0.87 | 0.491 | -0.01 | -0.001 | 0.001 | 0.001 | 0 | 0.65 | 0.43 | 0.55 | 0.5 | 0.68 | 0.7 |
| 9 | 21029330 | rs2298260 | 0.262 | 0.755 | 0.247 | 0.068 | 0.001 | -0.012 | -0.012 | 4.366 | 0.3 | 0.28 | 0.39 | 0.5 | 0.29 | 0.3 |
| 9 | 21037771 | rs10738578 | 0.021 | 0.937 | 0.153 | -0.033 | -0.002 | -0.004 | -0.004 | 0 | 0.33 | 0.24 | 0.22 | 0.4 | 0.32 | 0.34 |
| 9 | 21046482 | rs10811457 | 1.039 | 0.456 | -0.368 | -0.09 | 0.002 | -0.025 | -0.025 | 0 | 0.22 | 0.09 | 0.43 | 0.26 | 0.16 | 0.15 |
| 9 | 21046674 | rs10757178 | -0.265 | 0.426 | -0.313 | -0.23 | -0.002 | -0.001 | -0.001 | 0 | 0.55 | 0.38 | 0.65 | 0.52 | 0.48 | 0.6 |
| 9 | 21046790 | rs17692145 | 0.918 | 0.47 | -0.19 | -0.194 | -0.006 | -0.006 | -0.006 | 0.108 | 0.27 | 0.27 | 0.22 | 0.27 | 0.38 | 0.23 |
| 9 | 21051966 | rs7033704 | 0.92 | 0.43 | -0.166 | -0.17 | 0.007 | 0.047 | 0.047 | 0 | 0.14 | 0.19 | 0.09 | 0.25 | 0.12 | 0.21 |
| 9 | 21056085 | rs10964817 | 0.134 | 0.355 | -0.217 | -0.32 | 0 | 0.002 | 0.002 | 0 | 0.49 | 0.57 | 0.66 | 0.56 | 0.49 | 0.41 |
| 9 | 21058908 | rs1539741 | -0.586 | 0.345 | -0.431 | -0.425 | -0.001 | -0.006 | -0.006 | 0.274 | 0.72 | 0.74 | 0.79 | 0.81 | 0.66 | 0.63 |
| 9 | 21059531 | rs1539743 | -1.431 | 0.286 | -0.289 | -0.412 | 0.006 | -0.013 | -0.013 | 0 | 0.88 | 0.85 | 0.91 | 0.86 | 0.84 | 0.85 |
| 9 | 21060974 | rs1857651 | 1.052 | 0.135 | -0.22 | -0.369 | -0.001 | 0.002 | 0.002 | 0 | 0.32 | 0.41 | 0.26 | 0.29 | 0.36 | 0.23 |
| 9 | 21061642 | rs2383176 | -0.441 | 0.225 | -0.298 | -0.402 | -0.005 | 0.004 | 0.004 | 0.498 | 0.7 | 0.75 | 0.78 | 0.85 | 0.63 | 0.64 |

| | | | | | | | | | | | | | | | | |
|---|---|---|---|---|---|---|---|---|---|---|---|---|---|---|---|---|
| 9 | 21062144 | rs10811460 | 0.981 | 0.24 | -0.262 | -0.375 | -0.005 | 0.004 | 0.004 | 0 | 0.32 | 0.49 | 0.27 | 0.29 | 0.36 | 0.23 |
| 9 | 21065299 | rs1539744 | -0.426 | 0.262 | -0.356 | -0.462 | -0.017 | 0.047 | 0.047 | 0.584 | 0.7 | 0.81 | 0.79 | 0.85 | 0.63 | 0.63 |
| 9 | 21073945 | rs1424860 | 1.243 | 0.096 | -0.423 | -0.343 | -0.001 | 0 | 0 | 0 | 0.16 | 0.12 | 0.08 | 0.2 | 0.11 | 0.19 |
| 9 | 21077716 | rs1051922 | 1.092 | 0.075 | -0.479 | -0.226 | -0.001 | 0.003 | 0.003 | 0 | 0.32 | 0.4 | 0.26 | 0.32 | 0.36 | 0.26 |
| 9 | 21078952 | rs9333358 | 0.702 | -0.392 | -0.655 | 0.012 | 0.003 | -0.004 | -0.004 | 0 | 0.23 | 0.19 | 0.44 | 0.27 | 0.17 | 0.2 |
| 9 | 21080518 | rs10811465 | 0.899 | -0.3 | -0.921 | 0.057 | 0.002 | -0.013 | -0.013 | 0.889 | 0.22 | 0.13 | 0.44 | 0.27 | 0.17 | 0.19 |
| 9 | 21082814 | rs10811466 | 0.808 | 0.009 | -0.7 | 0.04 | -0.012 | -0.01 | -0.01 | 0.321 | 0.12 | 0.06 | 0.08 | 0.2 | 0.16 | 0.15 |
| 9 | 21093083 | rs10757181 | -0.136 | -0.185 | -0.399 | 0.222 | -0.001 | 0.001 | 0.001 | 0 | 0.29 | 0.29 | 0.17 | 0.38 | 0.27 | 0.33 |
| 9 | 21094151 | rs4977948 | 0.189 | -0.514 | -0.557 | 0.121 | 0.003 | -0.023 | -0.023 | 0 | 0.33 | 0.27 | 0.5 | 0.58 | 0.25 | 0.32 |
| 9 | 21095633 | rs10964832 | 1.728 | -0.388 | -0.296 | 0.035 | 0 | 0 | 0 | 0 | 0.15 | 0.14 | 0.14 | 0.12 | 0.2 | 0.12 |
| 9 | 21097135 | rs10757182 | 0.674 | -0.477 | -0.233 | -0.084 | 0.002 | 0 | 0 | 0 | 0.3 | 0.33 | 0.23 | 0.27 | 0.27 | 0.3 |
| 9 | 21098843 | rs12349659 | 1.797 | -0.469 | -0.179 | 0.303 | 0.008 | -0.002 | -0.002 | 0 | 0.14 | 0.16 | 0.07 | 0.14 | 0.07 | 0.18 |
| 9 | 21099471 | rs10757183 | -0.575 | -0.229 | -0.207 | 0.38 | -0.003 | 0.032 | 0.032 | 0 | 0.46 | 0.34 | 0.6 | 0.69 | 0.42 | 0.5 |
| 9 | 21102280 | rs1892130 | -0.642 | 0.111 | -0.363 | 0.333 | -0.005 | -0.019 | -0.019 | 0 | 0.83 | 0.77 | 0.89 | 0.88 | 0.84 | 0.81 |
| 9 | 21105078 | rs10757184 | -0.218 | -0.037 | -0.815 | 0.384 | 0.025 | 0.057 | 0.057 | 0 | 0.49 | 0.26 | 0.62 | 0.62 | 0.38 | 0.58 |
| 9 | 21105154 | rs4977959 | -0.084 | 0.088 | -1.121 | 0.166 | 0.04 | 0.023 | 0.023 | 0 | 0.52 | 0.29 | 0.7 | 0.51 | 0.56 | 0.6 |
| 9 | 21106901 | rs11795225 | 1.534 | -0.186 | -0.466 | 0.156 | -0.002 | -0.013 | -0.013 | 0 | 0.21 | 0.26 | 0.17 | 0.11 | 0.24 | 0.07 |
| 9 | 21108817 | rs2113718 | 0.613 | -0.223 | -0.515 | 0.265 | 0.011 | -0.033 | -0.033 | 0 | 0.35 | 0.23 | 0.54 | 0.46 | 0.25 | 0.42 |
| 9 | 21114105 | rs2032061 | -1.811 | -0.205 | 0.094 | 0.347 | -0.031 | 0.035 | 0.035 | 0 | 0.86 | 0.76 | 0.9 | 0.82 | 0.92 | 0.83 |
| 9 | 21114237 | rs10811474 | 0.219 | 0.021 | 0.072 | 0.352 | 0.001 | 0.01 | 0.01 | 0 | 0.48 | 0.43 | 0.62 | 0.62 | 0.39 | 0.56 |
| 9 | 21115002 | rs1364614 | 1.457 | 0.063 | -0.116 | -0.036 | -0.002 | -0.013 | -0.013 | 0 | 0.2 | 0.25 | 0.16 | 0.11 | 0.24 | 0.06 |
| 9 | 21117059 | rs4255228 | 0.191 | -0.02 | -0.096 | 0.094 | 0 | 0.011 | 0.011 | 0 | 0.48 | 0.45 | 0.62 | 0.69 | 0.4 | 0.56 |
| 9 | 21121917 | rs9886796 | -0.424 | 0.093 | -0.018 | 0.028 | 0.021 | -0.043 | -0.043 | 0 | 0.64 | 0.53 | 0.71 | 0.8 | 0.49 | 0.71 |
| 9 | 21122681 | rs1016129 | -1.126 | 0.221 | -0.118 | 0.061 | 0.026 | -0.011 | -0.011 | 4.178 | 0.84 | 0.8 | 0.88 | 0.91 | 0.72 | 0.78 |
| 9 | 21129246 | rs10964854 | 0.311 | 0.074 | -0.263 | 0.296 | -0.003 | 0.026 | 0.026 | 0 | 0.46 | 0.24 | 0.57 | 0.58 | 0.38 | 0.53 |
| 9 | 21129719 | rs1593270 | 1.16 | -0.271 | -0.17 | 0.689 | 0.02 | -0.013 | -0.013 | 0 | 0.15 | 0.09 | 0.13 | 0.21 | 0.11 | 0.18 |
| 9 | 21132664 | rs7854237 | -0.184 | 0.04 | 0.117 | 0.698 | -0.017 | -0.002 | -0.002 | 0 | 0.65 | 0.48 | 0.68 | 0.63 | 0.65 | 0.76 |
| 9 | 21142604 | rs10757189 | 0.152 | 0.558 | -0.325 | 0.867 | -0.008 | -0.017 | -0.017 | 0 | 0.28 | 0.21 | 0.37 | 0.35 | 0.27 | 0.27 |
| 9 | 21149963 | rs7046606 | 0.212 | 0.541 | -0.327 | 0.877 | -0.007 | -0.023 | -0.023 | 0.046 | 0.28 | 0.18 | 0.37 | 0.36 | 0.25 | 0.27 |
| 9 | 21150650 | rs7855438 | -0.628 | 0.371 | -0.178 | 0.671 | -0.011 | 0.038 | 0.038 | 0 | 0.67 | 0.48 | 0.69 | 0.52 | 0.65 | 0.67 |
| 9 | 21151553 | rs10964862 | -0.27 | 0.339 | -0.13 | 0.055 | 0.002 | -0.016 | -0.016 | 0 | 0.33 | 0.41 | 0.29 | 0.16 | 0.36 | 0.38 |
| 9 | 21153985 | rs10811482 | -0.251 | 0.846 | -0.27 | 0.112 | 0.009 | -0.038 | -0.038 | 4.048 | 0.15 | 0.12 | 0.1 | 0 | 0.26 | 0.21 |
| 9 | 21157373 | rs10757193 | -0.908 | 0.051 | -0.134 | 0.425 | 0.029 | 0.061 | 0.061 | 0 | 0.5 | 0.28 | 0.4 | 0.35 | 0.44 | 0.61 |
| 9 | 21157695 | rs10283629 | -1.591 | 0.222 | 0.183 | 0.702 | 0.01 | -0.045 | -0.045 | 0.008 | 0.61 | 0.52 | 0.65 | 0.86 | 0.59 | 0.77 |
| 9 | 21163761 | rs10964871 | 0.322 | 0.255 | 0.277 | 1.08 | 0.012 | 0.104 | 0.104 | 0 | 0.1 | 0.15 | 0.23 | 0.42 | 0.15 | 0.08 |
| 9 | 21166004 | rs2939 | 0.297 | 0.255 | 0.277 | 1.08 | 0.012 | 0.102 | 0.102 | 0 | 0.1 | 0.15 | 0.23 | 0.42 | 0.15 | 0.08 |
| 9 | 21170595 | rs7037868 | -1.785 | 0.177 | 0.516 | 1.022 | -0.003 | -0.004 | -0.004 | 0 | 0.63 | 0.62 | 0.67 | 0.88 | 0.59 | 0.76 |
| 9 | 21177609 | rs7022884 | -0.394 | 0.115 | 0.796 | 1.099 | 0.045 | 0.231 | 0.231 | 0.044 | 0.12 | 0.3 | 0.24 | 0.41 | 0.16 | 0.08 |
| 9 | 21178332 | rs1477478 | 0.051 | 0.17 | 0.651 | 1.126 | 0.025 | 0.151 | 0.151 | 0 | 0.11 | 0.22 | 0.24 | 0.4 | 0.16 | 0.08 |
| 9 | 21179174 | rs2019226 | -0.758 | 0.137 | 0.621 | 1.12 | 0.034 | 0.062 | 0.062 | 0 | 0.5 | 0.28 | 0.4 | 0.36 | 0.44 | 0.6 |
| 9 | 21196356 | rs2007448 | -0.115 | 0.292 | 0.26 | 1.352 | 0.033 | 0.184 | 0.184 | 0 | 0.11 | 0.27 | 0.24 | 0.41 | 0.16 | 0.09 |
| 9 | 21198384 | rs12348971 | -0.692 | 0.284 | 0.274 | 1.374 | 0.042 | 0.31 | 0.31 | 0 | 0.13 | 0.34 | 0.26 | 0.52 | 0.16 | 0.16 |
| 9 | 21199680 | rs4977686 | 1.917 | -0.081 | -0.049 | 1.111 | -0.155 | 0.119 | 0.119 | 0.206 | 0.4 | 0.02 | 0.34 | 0.14 | 0.45 | 0.25 |
| 9 | 21214877 | rs2015345 | -0.156 | 0.283 | 0.683 | 1.209 | 0.004 | 0.061 | 0.061 | 0 | 0.11 | 0.16 | 0.23 | 0.4 | 0.16 | 0.08 |
| 9 | 21216728 | rs3919593 | 0.997 | 0.319 | 0.606 | 1.097 | -0.02 | 0.011 | 0.011 | 0 | 0.09 | 0.18 | 0.04 | 0.1 | 0.04 | 0.09 |
| 9 | 21234721 | rs7033839 | -0.468 | 0.377 | 0.48 | 1.203 | 0.034 | 0.179 | 0.179 | 0.241 | 0.12 | 0.34 | 0.24 | 0.38 | 0.15 | 0.08 |
| 9 | 21255150 | rs1330320 | -1.447 | 0.339 | 0.377 | 1.072 | 0.01 | 0.027 | 0.027 | 0 | 0.32 | 0.4 | 0.38 | 0.51 | 0.37 | 0.32 |
| 9 | 21285389 | rs1475547 | -0.906 | 0.156 | 0.659 | 1.261 | -0.008 | 0.076 | 0.076 | 0.139 | 0.2 | 0.34 | 0.28 | 0.47 | 0.2 | 0.17 |
| 9 | 21289069 | rs1330311 | -1.014 | 0.157 | 0.659 | 1.261 | -0.007 | 0.083 | 0.083 | 0.048 | 0.2 | 0.34 | 0.28 | 0.47 | 0.2 | 0.13 |
| 9 | 21294921 | rs4978097 | -0.793 | 0.171 | 0.681 | 1.268 | -0.003 | 0.073 | 0.073 | 0 | 0.19 | 0.29 | 0.27 | 0.47 | 0.2 | 0.12 |
| 9 | 21295318 | rs10120117 | -0.354 | 0.114 | 0.608 | 1.343 | 0.007 | 0.07 | 0.07 | 0.075 | 0.11 | 0.2 | 0.23 | 0.35 | 0.15 | 0.09 |
| 9 | 21295828 | rs1330312 | -0.331 | 0.127 | 0.614 | 1.349 | 0.008 | 0.072 | 0.072 | 0.065 | 0.11 | 0.2 | 0.23 | 0.35 | 0.15 | 0.09 |
| 9 | 21300864 | rs4978107 | -0.198 | 0.126 | 0.627 | 1.348 | 0.004 | 0.048 | 0.048 | 0.033 | 0.11 | 0.16 | 0.23 | 0.36 | 0.15 | 0.09 |
| 9 | 21304057 | rs7021906 | -0.949 | 0.126 | 0.624 | 1.344 | 0.005 | 0.13 | 0.13 | 0.199 | 0.13 | 0.27 | 0.26 | 0.47 | 0.15 | 0.12 |
| 9 | 21306241 | rs12156640 | 0.392 | 0.119 | 0.491 | 1.295 | -0.009 | 0.028 | 0.028 | 0 | 0.11 | 0.02 | 0.07 | 0.06 | 0.08 | 0.24 |
| 9 | 21306318 | rs7031048 | -1.024 | 0.102 | 0.661 | 1.392 | 0.014 | 0.198 | 0.198 | 0.229 | 0.13 | 0.32 | 0.27 | 0.47 | 0.15 | 0.12 |
| 9 | 21323089 | rs10117202 | -0.26 | 0.11 | 0.631 | 1.222 | 0.012 | 0.085 | 0.085 | 0.093 | 0.11 | 0.19 | 0.24 | 0.35 | 0.15 | 0.09 |
| 9 | 21327141 | rs913931 | -0.573 | 0.146 | 0.704 | 1.062 | -0.001 | -0.001 | -0.001 | 0 | 0.15 | 0.13 | 0.11 | 0.1 | 0.14 | 0.26 |
| 9 | 21335404 | rs7863385 | 1.332 | 0.101 | 0.864 | 1.244 | -0.015 | -0.04 | -0.04 | 0.035 | 0.08 | 0.14 | 0.03 | 0 | 0.04 | 0.01 |
| 9 | 21340786 | rs10119678 | -1.169 | 0.116 | 0.831 | 1.411 | -0.014 | 0.055 | 0.055 | 0.039 | 0.19 | 0.31 | 0.26 | 0.37 | 0.15 | 0.22 |
| 9 | 21352862 | rs614541 | -0.403 | 0.221 | 0.584 | 1.443 | 0.007 | 0.048 | 0.048 | 0.239 | 0.11 | 0.16 | 0.23 | 0.29 | 0.15 | 0.08 |
| 9 | 21366894 | rs2480927 | -1.615 | -0.243 | 1.085 | 1.444 | -0.012 | 0.108 | 0.108 | 0 | 0.2 | 0.48 | 0.29 | 0.37 | 0.19 | 0.22 |
| 9 | 21370864 | rs596180 | -0.963 | -0.017 | 0.661 | 1.506 | 0.01 | 0.055 | 0.055 | 0 | 0.09 | 0.14 | 0.24 | 0.31 | 0.15 | 0.09 |
| 9 | 21381286 | rs647167 | -0.508 | -0.04 | 0.536 | 1.301 | 0.001 | 0.003 | 0.003 | 0 | 0.08 | 0.09 | 0.19 | 0.15 | 0.14 | 0.05 |
| 9 | 21383734 | rs615544 | 0.004 | -0.03 | 0.427 | 1.3 | -0.005 | -0.018 | -0.018 | 0 | 0.07 | 0.05 | 0.08 | 0.14 | 0.11 | 0.04 |
| 9 | 21384363 | rs10120977 | 0.366 | -0.312 | -0.065 | 0.738 | -0.044 | 0.137 | 0.137 | 0 | 0.22 | 0.02 | 0.13 | 0 | 0.22 | 0.21 |
| 9 | 21395611 | rs591537 | -1.682 | -0.351 | 0.743 | 1.599 | -0.009 | 0.031 | 0.031 | 0 | 0.28 | 0.65 | 0.33 | 0.43 | 0.24 | 0.32 |
| 9 | 21397773 | rs10121687 | 1.468 | -0.277 | 0.443 | 1.499 | -0.001 | -0.015 | -0.015 | 0 | 0.07 | 0.07 | 0.02 | 0 | 0.04 | 0.01 |
| 9 | 21398147 | rs10757218 | -1.06 | -0.246 | 0.785 | 1.676 | 0.013 | 0.044 | 0.044 | 0 | 0.14 | 0.26 | 0.23 | 0.33 | 0.17 | 0.19 |

| | | | | | | | | | | | | | | | | |
|---|---|---|---|---|---|---|---|---|---|---|---|---|---|---|---|---|
| 9 | 21401106 | rs4978113 | -0.829 | -0.216 | 0.531 | 1.528 | -0.018 | -0.026 | -0.026 | 2.551 | 0.13 | 0.09 | 0.22 | 0.31 | 0.17 | 0.19 |
| 9 | 21403636 | rs16938392 | 0.329 | -0.175 | 0.37 | 1.374 | -0.031 | -0.031 | -0.031 | 0 | 0.06 | 0.03 | 0.09 | 0.14 | 0.11 | 0.04 |
| 9 | 21405363 | rs10757219 | -1.406 | -0.313 | 0.636 | 1.421 | -0.004 | 0.008 | 0.008 | 0 | 0.27 | 0.67 | 0.36 | 0.63 | 0.23 | 0.36 |
| 9 | 21411631 | rs9333353 | -0.784 | -0.571 | 0.307 | 1.13 | -0.006 | -0.003 | -0.003 | 0 | 0.44 | 0.4 | 0.43 | 0.55 | 0.43 | 0.42 |
| 9 | 21413085 | rs1332180 | 0.802 | -0.216 | 0.116 | 0.735 | -0.002 | 0 | 0 | 0.013 | 0.06 | 0.1 | 0.05 | 0.04 | 0.05 | 0.11 |
| 9 | 21415687 | rs7025006 | -0.66 | -0.497 | 0.13 | 0.668 | 0.02 | 0.026 | 0.026 | 0 | 0.5 | 0.66 | 0.48 | 0.56 | 0.42 | 0.53 |
| 9 | 21419723 | rs2104880 | 0.407 | -0.454 | 0.183 | 0.105 | -0.003 | -0.025 | -0.025 | 0 | 0.1 | 0.14 | 0.03 | 0 | 0.05 | 0.01 |
| 9 | 21420677 | rs1332179 | -1.149 | -0.642 | 0.615 | 0.26 | -0.044 | 0.02 | 0.02 | 0 | 0.21 | 0.56 | 0.13 | 0.27 | 0.1 | 0.26 |
| 9 | 21423520 | rs1591032 | -0.774 | -0.326 | 0.49 | 0.091 | 0.013 | 0.005 | 0.005 | 0.128 | 0.16 | 0.2 | 0.25 | 0.14 | 0.18 | 0.17 |
| 9 | 21428770 | rs7043990 | -0.301 | -0.287 | 0.367 | 0.087 | 0.014 | 0.024 | 0.024 | 1.633 | 0.09 | 0.14 | 0.15 | 0.12 | 0.13 | 0.11 |
| 9 | 21438684 | rs1332190 | -1.676 | -0.586 | 0.033 | -0.051 | -0.004 | -0.003 | -0.003 | 0 | 0.36 | 0.58 | 0.36 | 0.34 | 0.29 | 0.42 |
| 9 | 21456776 | rs10491569 | 1.225 | -0.727 | -0.148 | -0.564 | 0.012 | 0.035 | 0.035 | 1.841 | 0.29 | 0.15 | 0.19 | 0.16 | 0.31 | 0.22 |
| 9 | 21470997 | rs7861480 | -1.513 | -0.383 | 0.271 | -0.291 | -0.053 | 0.026 | 0.026 | 0.102 | 0.21 | 0.55 | 0.19 | 0.28 | 0.1 | 0.33 |
| 9 | 21471726 | rs2383192 | 0.763 | -0.26 | 0.149 | -0.314 | 0.003 | 0.041 | 0.041 | 0 | 0.51 | 0.38 | 0.53 | 0.69 | 0.51 | 0.49 |
| 9 | 21485974 | rs10757222 | -0.012 | 0.234 | -0.138 | -0.06 | -0.021 | 0.026 | 0.026 | 0.133 | 0.64 | 0.69 | 0.62 | 0.73 | 0.58 | 0.57 |
| 9 | 21494300 | rs4978121 | 0.301 | 0.375 | -0.165 | -0.092 | -0.045 | 0.118 | 0.118 | 2.95 | 0.59 | 0.81 | 0.59 | 0.83 | 0.5 | 0.69 |
| 9 | 21497120 | rs2383194 | -0.433 | 0.184 | 0.584 | -0.37 | 0.045 | -0.096 | -0.096 | 0 | 0.7 | 0.54 | 0.71 | 0.92 | 0.64 | 0.81 |
| 9 | 21499793 | rs7849575 | -0.575 | 0.209 | 0.597 | -0.4 | 0.008 | -0.029 | -0.029 | 0 | 0.71 | 0.63 | 0.72 | 0.93 | 0.65 | 0.81 |
| 9 | 21503812 | rs4977726 | -0.71 | 0.327 | 0.587 | -0.405 | -0.003 | -0.006 | -0.006 | 0 | 0.72 | 0.7 | 0.74 | 0.93 | 0.69 | 0.81 |
| 9 | 21503821 | rs17729561 | -0.879 | 0.297 | 0.553 | -0.419 | 0.004 | -0.004 | -0.004 | 0.258 | 0.07 | 0.08 | 0.04 | 0 | 0.06 | 0.05 |
| 9 | 21504203 | rs1332184 | 0.634 | 0.629 | 0.669 | -0.29 | -0.011 | 0.035 | 0.035 | 0.451 | 0.57 | 0.62 | 0.62 | 0.73 | 0.6 | 0.54 |
| 9 | 21504973 | rs7848909 | 0.915 | 0.923 | 0.778 | -0.115 | 0.002 | -0.013 | -0.013 | 0 | 0.58 | 0.43 | 0.59 | 0.74 | 0.57 | 0.5 |
| 9 | 21506356 | rs1537276 | 0.047 | 1.002 | 0.137 | -0.083 | -0.037 | 0.076 | 0.076 | 0.791 | 0.69 | 0.81 | 0.71 | 0.95 | 0.63 | 0.78 |
| 9 | 21507589 | rs1537275 | 0.798 | 1.321 | 0.15 | -0.078 | 0.007 | -0.019 | -0.019 | 0 | 0.59 | 0.5 | 0.58 | 0.74 | 0.54 | 0.5 |
| 9 | 21516772 | rs10491566 | -0.742 | 1.193 | -0.064 | 0.006 | -0.011 | 0.013 | 0.013 | 0.096 | 0.74 | 0.79 | 0.76 | 0.9 | 0.68 | 0.82 |
| 9 | 21519921 | rs7026731 | -0.925 | 1.072 | -0.02 | 0.092 | -0.009 | 0.018 | 0.018 | 0 | 0.1 | 0.22 | 0.12 | 0.17 | 0.06 | 0.28 |
| 9 | 21530609 | rs10811561 | 0.546 | 1.083 | -0.081 | 0.145 | -0.012 | 0.005 | 0.005 | 0.69 | 0.6 | 0.68 | 0.58 | 0.67 | 0.54 | 0.52 |
| 9 | 21531629 | rs2025327 | -0.203 | 1.114 | -0.067 | 0.151 | -0.009 | 0.01 | 0.01 | 0.608 | 0.69 | 0.74 | 0.69 | 0.89 | 0.59 | 0.8 |
| 9 | 21536423 | rs12379721 | -1.047 | 0.35 | -0.09 | 0.299 | 0.01 | -0.021 | -0.021 | 0.379 | 0.1 | 0.06 | 0.11 | 0.23 | 0.06 | 0.28 |
| 9 | 21543444 | rs10811568 | -1.175 | 0.604 | 0.015 | 0.416 | 0 | -0.029 | -0.029 | 0 | 0.11 | 0.08 | 0.12 | 0.23 | 0.08 | 0.3 |
| 9 | 21548586 | rs7863623 | 0.174 | 0.833 | -0.454 | -0.078 | -0.003 | 0.009 | 0.009 | 0.73 | 0.27 | 0.21 | 0.26 | 0.1 | 0.29 | 0.17 |
| 9 | 21549638 | rs10965063 | -0.718 | 0.676 | -0.252 | 0.533 | -0.002 | -0.011 | -0.011 | 0 | 0.07 | 0.04 | 0.06 | 0.09 | 0.06 | 0.09 |
| 9 | 21551513 | rs991794 | -1.21 | 0.625 | 0.298 | -0.005 | -0.002 | -0.111 | -0.111 | 0 | 0.22 | 0.48 | 0.13 | 0.04 | 0.21 | 0.15 |
| 9 | 21551647 | rs991792 | -0.492 | 0.403 | 0.479 | -0.046 | -0.01 | -0.039 | -0.039 | 0.049 | 0.08 | 0.15 | 0.05 | 0 | 0.04 | 0.01 |
| 9 | 21552654 | rs1983810 | 0.979 | 0.534 | 0.397 | 0.359 | 0 | 0 | 0 | 0 | 0.7 | 0.33 | 0.68 | 0.73 | 0.73 | 0.75 |
| 9 | 21553004 | rs1983811 | 1.481 | 0.255 | 0.582 | 0.673 | 0.019 | -0.021 | -0.021 | 0 | 0.63 | 0.17 | 0.63 | 0.61 | 0.7 | 0.57 |
| 9 | 21555283 | rs9298816 | 1.522 | 0.178 | 0.96 | 0.728 | 0.003 | -0.017 | -0.017 | 0.047 | 0.66 | 0.34 | 0.67 | 0.97 | 0.73 | 0.8 |
| 9 | 21555445 | rs10757225 | 1.666 | 0.178 | 0.743 | 0.848 | 0.012 | -0.072 | -0.072 | 5.074 | 0.61 | 0.08 | 0.62 | 0.58 | 0.71 | 0.56 |
| 9 | 21565674 | rs7033598 | 1.641 | 0.306 | 0.652 | 0.628 | 0.016 | 0.079 | 0.079 | 0 | 0.6 | 0.21 | 0.62 | 0.84 | 0.68 | 0.58 |
| 9 | 21566730 | rs7865428 | 1.005 | 0.469 | -0.45 | 0.547 | -0.057 | -0.029 | -0.029 | 0 | 0.2 | 0.02 | 0.23 | 0.16 | 0.22 | 0.17 |
| 9 | 21573016 | rs1452653 | 0.386 | 0.15 | 0.822 | 0.51 | 0.002 | -0.103 | -0.103 | 0 | 0.77 | 0.54 | 0.72 | 0.96 | 0.79 | 0.84 |
| 9 | 21580763 | rs12685827 | -1.065 | 0.012 | 0.996 | 0.608 | -0.014 | -0.011 | -0.011 | 0.228 | 0.16 | 0.29 | 0.09 | 0.12 | 0.1 | 0.25 |
| 9 | 21581712 | rs7026552 | -0.006 | 0.058 | 0.999 | 0.538 | 0.005 | -0.035 | -0.035 | 0 | 0.07 | 0.11 | 0.04 | 0 | 0.08 | 0.06 |
| 9 | 21588969 | rs1542302 | 0.465 | 0.253 | 0.941 | 0.595 | 0.005 | -0.017 | -0.017 | 3.141 | 0.81 | 0.54 | 0.86 | 0.88 | 0.82 | 0.85 |
| 9 | 21589041 | rs2123340 | 1.368 | 0.274 | 0.395 | 0.642 | 0.021 | -0.031 | -0.031 | 0 | 0.61 | 0.2 | 0.74 | 0.54 | 0.68 | 0.55 |
| 9 | 21598941 | rs7032413 | -0.982 | -0.166 | 0.695 | 0.456 | -0.049 | 0 | 0 | 0 | 0.26 | 0.65 | 0.15 | 0.23 | 0.1 | 0.38 |
| 9 | 21600424 | rs12343168 | -1.005 | -0.07 | 1.034 | 0.443 | -0.034 | 0 | 0 | 0.149 | 0.26 | 0.6 | 0.15 | 0.23 | 0.14 | 0.4 |
| 9 | 21604067 | rs16938481 | -0.309 | -0.135 | 1.203 | 0.399 | -0.016 | -0.023 | -0.023 | 0.088 | 0.17 | 0.26 | 0.09 | 0.11 | 0.11 | 0.19 |
| 9 | 21604969 | rs10965079 | 0.98 | 0.1 | -0.029 | 0.155 | -0.118 | -0.013 | -0.013 | 0 | 0.37 | 0.02 | 0.45 | 0.27 | 0.41 | 0.36 |
| 9 | 21606566 | rs12115747 | 1.302 | -0.387 | 0.814 | 0.404 | -0.006 | -0.02 | -0.02 | 0.009 | 0.05 | 0.08 | 0.04 | 0 | 0.03 | 0 |
| 9 | 21611256 | rs7848640 | -0.848 | -0.359 | 0.717 | 0.468 | -0.041 | 0.003 | 0.003 | 0 | 0.26 | 0.56 | 0.16 | 0.24 | 0.15 | 0.4 |
| 9 | 21620510 | rs10123003 | -0.849 | -0.502 | 0.587 | 0.222 | -0.064 | -0.033 | -0.033 | 0.166 | 0.27 | 0.59 | 0.15 | 0.13 | 0.11 | 0.37 |
| 9 | 21626200 | rs10491562 | 0.069 | -0.545 | 0.304 | 0.113 | 0.003 | -0.008 | -0.008 | 0 | 0.17 | 0.16 | 0.08 | 0.02 | 0.11 | 0.17 |
| 9 | 21626573 | rs754819 | 0.475 | -0.327 | -0.127 | 0.365 | -0.047 | -0.07 | -0.07 | 1.912 | 0.11 | 0.01 | 0.31 | 0.24 | 0.17 | 0.13 |
| 9 | 21626953 | rs13297146 | -0.159 | -0.316 | 0.435 | 0.152 | 0.046 | -0.014 | -0.014 | 0 | 0.06 | 0.26 | 0.05 | 0 | 0.07 | 0.01 |
| 9 | 21638846 | rs10115623 | 0.603 | -0.396 | 0.488 | 0.194 | -0.022 | 0.016 | 0.016 | 0.001 | 0.07 | 0.1 | 0.03 | 0.09 | 0.01 | 0.22 |
| 9 | 21638902 | rs13294794 | 0.313 | -0.351 | 0.468 | 0.151 | 0.019 | -0.028 | -0.028 | 0 | 0.05 | 0.17 | 0.04 | 0 | 0.07 | 0.01 |
| 9 | 21641325 | rs10757227 | -0.137 | -0.275 | -0.151 | -0.023 | 0.016 | 0.002 | 0.002 | 0 | 0.44 | 0.63 | 0.42 | 0.62 | 0.34 | 0.51 |
| 9 | 21642590 | rs10757229 | 1.209 | -0.248 | -0.689 | 0.027 | -0.002 | -0.115 | -0.115 | 0.062 | 0.3 | 0.11 | 0.34 | 0.53 | 0.26 | 0.28 |
| 9 | 21651620 | rs11788139 | -1.015 | -0.147 | -0.025 | 0.076 | -0.029 | -0.028 | -0.028 | 0.156 | 0.28 | 0.33 | 0.12 | 0.12 | 0.14 | 0.41 |
| 9 | 21653928 | rs16938501 | 0.659 | -0.1 | 0.113 | 0.366 | -0.012 | 0.007 | 0.007 | 0 | 0.06 | 0.05 | 0.02 | 0.09 | 0.01 | 0.22 |
| 9 | 21658652 | rs13287480 | -0.575 | -0.029 | -0.081 | 0.297 | -0.002 | -0.028 | -0.028 | 0 | 0.1 | 0.17 | 0.05 | 0 | 0.08 | 0.01 |
| 9 | 21663040 | rs4512473 | -1.041 | -0.076 | -0.152 | 0.332 | -0.051 | -0.067 | -0.067 | 0 | 0.3 | 0.49 | 0.14 | 0.12 | 0.19 | 0.4 |
| 9 | 21670461 | rs12342723 | -0.512 | 0.218 | -0.56 | 0.128 | -0.022 | -0.089 | -0.089 | 1.63 | 0.23 | 0.39 | 0.12 | 0.02 | 0.18 | 0.17 |
| 9 | 21672769 | rs10120643 | -0.548 | 0.236 | -0.55 | 0.137 | -0.023 | -0.089 | -0.089 | 0 | 0.24 | 0.4 | 0.12 | 0.02 | 0.18 | 0.17 |
| 9 | 21673461 | rs16938525 | -0.304 | 0.701 | -1.243 | 0.986 | -0.047 | -0.059 | -0.059 | 0 | 0.1 | 0.01 | 0.31 | 0.2 | 0.15 | 0.12 |
| 9 | 21674425 | rs10965097 | -0.528 | 0.11 | -1.063 | 0.661 | 0.032 | 0.02 | 0.02 | 0 | 0.41 | 0.66 | 0.26 | 0.38 | 0.27 | 0.43 |
| 9 | 21675373 | rs2039585 | -0.942 | 0.153 | -0.422 | 1.205 | -0.018 | 0.196 | 0.196 | 0.153 | 0.18 | 0.51 | 0.21 | 0.36 | 0.15 | 0.23 |
| 9 | 21675545 | rs1377195 | -1.269 | 0.396 | -1.483 | 0.741 | -0.028 | 0.155 | 0.155 | 0.996 | 0.48 | 0.88 | 0.63 | 0.73 | 0.47 | 0.43 |

| | | | | | | | | | | | | | | | | |
|---|---|---|---|---|---|---|---|---|---|---|---|---|---|---|---|---|
| 9 | 21682759 | rs10965099 | 0.33 | 0.358 | -1.363 | 0.757 | -0.016 | -0.022 | -0.022 | 0 | 0.21 | 0.13 | 0.38 | 0.33 | 0.27 | 0.18 |
| 9 | 21683198 | rs16938544 | 0.587 | 0.28 | -1.16 | 0.801 | 0 | -0.001 | -0.001 | 0 | 0.09 | 0.08 | 0.02 | 0.04 | 0.04 | 0.06 |
| 9 | 21683289 | rs7860632 | -0.367 | 0.253 | -1.084 | 0.777 | -0.015 | 0.188 | 0.188 | 0.472 | 0.17 | 0.53 | 0.2 | 0.37 | 0.15 | 0.2 |
| 9 | 21684722 | rs7865502 | -0.826 | 0.3 | -1.205 | 0.789 | -0.015 | 0.03 | 0.03 | 0.595 | 0.27 | 0.66 | 0.22 | 0.4 | 0.19 | 0.26 |
| 9 | 21690958 | rs1869204 | -0.039 | 0.557 | -1.92 | 0.218 | -0.052 | 0.055 | 0.055 | 0 | 0.13 | 0.02 | 0.14 | 0.02 | 0.18 | 0.18 |
| 9 | 21695893 | rs12380505 | -1.044 | 0.453 | -1.747 | 0.473 | -0.054 | 0.103 | 0.103 | 0.69 | 0.47 | 0.86 | 0.44 | 0.69 | 0.54 | 0.42 |
| 9 | 21701432 | rs7848524 | -1.031 | 0.442 | -1.669 | 0.612 | -0.04 | 0.074 | 0.074 | 0.437 | 0.47 | 0.82 | 0.43 | 0.67 | 0.54 | 0.41 |
| 9 | 21701942 | rs10811586 | 0.049 | 0.431 | -1.279 | 0.864 | -0.018 | -0.005 | -0.005 | 0.369 | 0.25 | 0.16 | 0.22 | 0.29 | 0.4 | 0.24 |
| 9 | 21704498 | rs7874607 | -0.158 | 0.347 | -1.064 | 0.912 | -0.031 | 0.214 | 0.214 | 0.895 | 0.14 | 0.57 | 0.19 | 0.36 | 0.08 | 0.16 |
| 9 | 21706571 | rs896655 | -0.079 | 0.712 | -0.847 | 1.01 | -0.018 | -0.005 | -0.005 | 0 | 0.25 | 0.17 | 0.22 | 0.29 | 0.4 | 0.24 |
| 9 | 21721615 | rs2210779 | -0.703 | 0.493 | -0.414 | 1.06 | -0.021 | 0.142 | 0.142 | 0.367 | 0.18 | 0.54 | 0.19 | 0.38 | 0.1 | 0.1 |
| 9 | 21721939 | rs10123637 | -0.034 | 0.691 | -0.577 | 1.016 | 0.009 | 0.003 | 0.003 | 0 | 0.24 | 0.27 | 0.44 | 0.31 | 0.39 | 0.25 |
| 9 | 21728683 | rs6475564 | -1.334 | 0.684 | -0.569 | 0.72 | -0.065 | 0.04 | 0.04 | 0.952 | 0.42 | 0.83 | 0.62 | 0.69 | 0.5 | 0.35 |
| 9 | 21737822 | rs1335503 | 0.15 | 0.675 | -0.476 | 1.097 | -0.015 | -0.018 | -0.018 | 0 | 0.23 | 0.16 | 0.42 | 0.43 | 0.39 | 0.26 |
| 9 | 21745756 | rs1345026 | -2.054 | 0.185 | -0.936 | 0.711 | -0.058 | 0.027 | 0.027 | 0.442 | 0.59 | 0.94 | 0.75 | 0.71 | 0.58 | 0.56 |
| 9 | 21747803 | rs4636294 | -1.282 | 0.618 | -0.474 | 1.108 | -0.07 | 0.034 | 0.034 | 0.441 | 0.42 | 0.82 | 0.62 | 0.68 | 0.5 | 0.35 |
| 9 | 21760639 | rs7866787 | -1.284 | 0.667 | -0.327 | 1.09 | -0.05 | 0.041 | 0.041 | 0.525 | 0.42 | 0.79 | 0.62 | 0.69 | 0.5 | 0.36 |
| 9 | 21764086 | rs1414259 | -0.701 | 0.652 | -0.223 | 1.122 | -0.021 | 0.047 | 0.047 | 0.554 | 0.18 | 0.58 | 0.19 | 0.27 | 0.11 | 0.1 |
| 9 | 21772037 | rs7037577 | -1.328 | 0.638 | -0.273 | 1.116 | -0.073 | 0.045 | 0.045 | 0.784 | 0.42 | 0.83 | 0.62 | 0.69 | 0.5 | 0.37 |
| 9 | 21772064 | rs7037277 | -0.124 | 0.585 | -0.14 | 0.894 | 0.061 | 0.277 | 0.277 | 0.902 | 0.08 | 0.5 | 0.16 | 0.23 | 0.05 | 0.07 |
| 9 | 21782164 | rs10811615 | -1.561 | 0.653 | -0.366 | 0.883 | -0.06 | 0.087 | 0.087 | 0.583 | 0.44 | 0.84 | 0.61 | 0.7 | 0.47 | 0.4 |
| 9 | 21795661 | rs12380373 | -1.172 | 0.474 | -0.127 | 0.877 | -0.022 | 0.007 | 0.007 | 0.13 | 0.21 | 0.64 | 0.2 | 0.26 | 0.11 | 0.11 |
| 9 | 21810531 | rs7871477 | -1.493 | 0.209 | 0.854 | 0.873 | -0.021 | -0.001 | -0.001 | 0.156 | 0.25 | 0.64 | 0.21 | 0.26 | 0.12 | 0.17 |
| 9 | 21811027 | rs6475584 | -1.435 | 0.201 | 0.921 | 0.869 | -0.034 | 0.001 | 0.001 | 0.012 | 0.24 | 0.59 | 0.21 | 0.26 | 0.11 | 0.16 |
| 9 | 21812720 | rs10811624 | 0.312 | 0.205 | 0.78 | 0.869 | 0.006 | 0.036 | 0.036 | 0.307 | 0.11 | 0.02 | 0.02 | 0.03 | 0.06 | 0.04 |
| 9 | 21815166 | rs2039971 | -0.81 | 0.211 | 0.849 | 0.878 | -0.017 | 0.083 | 0.083 | 0 | 0.1 | 0.25 | 0.16 | 0.23 | 0.05 | 0.12 |
| 9 | 21816528 | rs7023329 | -1.324 | 0.381 | 0.74 | 0.867 | -0.001 | -0.002 | -0.002 | 1.141 | 0.41 | 0.51 | 0.5 | 0.47 | 0.48 | 0.27 |
| 9 | 21816758 | rs7023954 | -0.226 | 0.39 | 0.733 | 0.867 | 0.06 | 0.109 | 0.109 | 0.231 | 0.27 | 0.43 | 0.44 | 0.44 | 0.39 | 0.23 |
| 9 | 21818310 | rs3928894 | -2.176 | 0.351 | 0.626 | 0.869 | -0.032 | 0.128 | 0.128 | 0.114 | 0.52 | 0.84 | 0.66 | 0.7 | 0.53 | 0.4 |
| 9 | 21825996 | rs7867176 | -1.416 | 0.298 | 0.975 | 1.294 | -0.035 | 0.043 | 0.043 | 0 | 0.22 | 0.38 | 0.2 | 0.32 | 0.12 | 0.17 |
| 9 | 21826387 | rs7029287 | -1.402 | 0.3 | 0.963 | 1.253 | -0.034 | 0.019 | 0.019 | 0 | 0.21 | 0.38 | 0.2 | 0.26 | 0.11 | 0.17 |
| 9 | 21830479 | rs1544195 | 0.026 | 0.44 | 0.593 | 1.14 | 0.005 | 0.012 | 0.012 | 0 | 0.11 | 0.06 | 0.03 | 0.03 | 0.06 | 0.04 |
| 9 | 21840298 | rs10811629 | -0.275 | 0.969 | 0.63 | 1.288 | -0.029 | -0.034 | -0.034 | 0.378 | 0.24 | 0.17 | 0.5 | 0.44 | 0.41 | 0.29 |
| 9 | 21846285 | rs7850937 | -0.062 | 0.413 | 0.701 | 1.37 | 0.003 | 0.018 | 0.018 | 0 | 0.18 | 0.17 | 0.43 | 0.42 | 0.25 | 0.25 |
| 9 | 21846327 | rs7851133 | 0.018 | 0.439 | 0.694 | 1.369 | -0.001 | 0.012 | 0.012 | 0 | 0.18 | 0.16 | 0.43 | 0.42 | 0.23 | 0.24 |
| 9 | 21853221 | rs4478653 | -2.181 | 0.221 | 0.933 | 1.369 | -0.013 | 0.097 | 0.097 | 0.181 | 0.44 | 0.74 | 0.64 | 0.73 | 0.41 | 0.48 |
| 9 | 21854199 | rs4636295 | -0.154 | 0.23 | 0.919 | 1.335 | -0.007 | -0.008 | -0.008 | 0 | 0.2 | 0.16 | 0.43 | 0.42 | 0.25 | 0.29 |
| 9 | 21855038 | rs3802393 | -1.439 | 0.148 | 1.337 | 1.372 | -0.033 | 0.018 | 0.018 | 0 | 0.22 | 0.45 | 0.19 | 0.25 | 0.15 | 0.17 |
| 9 | 21856777 | rs7871127 | -0.408 | 0.135 | 1.713 | 1.64 | 0.001 | 0.055 | 0.055 | 0 | 0.07 | 0.14 | 0.15 | 0.19 | 0.08 | 0.1 |
| 9 | 21857244 | rs7047648 | -1.978 | 0.079 | 1.631 | 1.828 | -0.001 | 0.001 | 0.001 | 0 | 0.42 | 0.58 | 0.64 | 0.55 | 0.41 | 0.44 |
| 9 | 21857303 | rs7047899 | -1.809 | -0.283 | 2.221 | 2.086 | -0.016 | 0.078 | 0.078 | 0.481 | 0.12 | 0.55 | 0.18 | 0.19 | 0.08 | 0.17 |
| 9 | 21864098 | rs10117507 | -0.806 | -0.185 | 2.098 | 2.204 | 0.001 | 0.029 | 0.029 | 0 | 0.22 | 0.23 | 0.44 | 0.39 | 0.19 | 0.24 |
| 9 | 21864535 | rs3802394 | -0.597 | -0.175 | 1.609 | 2.21 | -0.003 | -0.019 | -0.019 | 0 | 0.21 | 0.14 | 0.43 | 0.39 | 0.19 | 0.25 |
| 9 | 21865842 | rs6475592 | -1.884 | -0.183 | 1.302 | 2.112 | 0.01 | -0.005 | -0.005 | 0.258 | 0.34 | 0.71 | 0.62 | 0.61 | 0.27 | 0.42 |
| 9 | 21881338 | rs4607709 | -1.874 | -0.195 | 1.271 | 2.058 | 0.009 | 0.001 | 0.001 | 0 | 0.34 | 0.7 | 0.62 | 0.62 | 0.27 | 0.42 |
| 9 | 21902354 | rs756641 | -0.739 | -0.099 | 1.355 | 2.092 | 0.001 | 0.011 | 0.011 | 0 | 0.22 | 0.23 | 0.43 | 0.37 | 0.2 | 0.24 |
| 9 | 21909979 | rs16938590 | -1.348 | -0.754 | 1.43 | 2.018 | -0.011 | 0.217 | 0.217 | 0 | 0.12 | 0.45 | 0.17 | 0.27 | 0.07 | 0.18 |
| 9 | 21910698 | rs10811635 | -0.678 | -0.802 | 0.834 | 1.886 | -0.001 | 0.007 | 0.007 | 0 | 0.2 | 0.19 | 0.36 | 0.35 | 0.18 | 0.22 |
| 9 | 21920346 | rs4977746 | -1.532 | -0.311 | 0.567 | 1.367 | 0.026 | -0.033 | -0.033 | 0 | 0.3 | 0.74 | 0.48 | 0.54 | 0.19 | 0.36 |
| 9 | 21925855 | rs7852128 | -1.496 | -0.287 | 0.565 | 1.204 | 0.024 | -0.037 | -0.037 | 0 | 0.3 | 0.73 | 0.48 | 0.48 | 0.19 | 0.34 |
| 9 | 21930505 | rs10965187 | -1.743 | 0.152 | 1.715 | 0.501 | 0.061 | 0.057 | 0.057 | 3.704 | 0.06 | 0.48 | 0.04 | 0 | 0 | 0 |
| 9 | 21936381 | rs7875199 | -2.077 | 0.061 | 0.476 | 0.553 | -0.033 | -0.005 | -0.005 | 0 | 0.09 | 0.8 | 0.06 | 0.05 | 0.01 | 0.13 |
| 9 | 21944317 | rs4977750 | 1.803 | 0.671 | 0.87 | 0.176 | 0.01 | -0.062 | -0.062 | 0.075 | 0.86 | 0.35 | 0.89 | 0.99 | 0.86 | 0.91 |
| 9 | 21944818 | rs4382559 | 1.379 | 0.709 | 2.005 | 0.177 | 0.058 | -0.067 | -0.067 | 0.082 | 0.89 | 0.69 | 0.93 | 1 | 0.87 | 0.97 |
| 9 | 21948666 | rs10965197 | 1.152 | 1.635 | 1.537 | -0.163 | 0.04 | -0.043 | -0.043 | 0.487 | 0.7 | 0.58 | 0.5 | 0.85 | 0.59 | 0.69 |
| 9 | 21952926 | rs2518722 | -1.639 | 1.106 | 1.719 | -0.54 | -0.013 | -0.117 | -0.117 | 0 | 0.26 | 0.56 | 0.2 | 0.02 | 0.27 | 0.08 |
| 9 | 21961866 | rs7041637 | 1.849 | 0.997 | 0.185 | -0.304 | 0.052 | -0.09 | -0.09 | 0.829 | 0.4 | 0.08 | 0.27 | 0.5 | 0.28 | 0.4 |
| 9 | 21966221 | rs3731257 | 1.936 | 0.734 | -0.083 | -0.362 | 0.052 | -0.077 | -0.077 | 0 | 0.37 | 0.08 | 0.26 | 0.55 | 0.25 | 0.38 |
| 9 | 21974218 | rs3731239 | 0.319 | 1.251 | -0.364 | -0.347 | -0.149 | 0.085 | 0.085 | 0 | 0.24 | 0.01 | 0.18 | 0.11 | 0.38 | 0.24 |
| 9 | 21981583 | rs4074785 | 0.981 | 1.295 | 0.121 | 0.002 | 0.005 | 0.009 | 0.009 | 0 | 0.1 | 0.15 | 0.33 | 0.13 | 0.08 | 0.18 |
| 9 | 21984661 | rs3731217 | -0.119 | 1.208 | 0.034 | 0.06 | -0.003 | -0.003 | -0.003 | 2.67 | 0.11 | 0.09 | 0.09 | 0.19 | 0.15 | 0.1 |
| 9 | 21990457 | rs7036656 | 1.21 | 1.204 | 0.214 | 0.008 | 0.001 | 0 | 0 | 0 | 0.78 | 0.75 | 0.83 | 0.79 | 0.71 | 0.85 |
| 9 | 21998035 | rs2811712 | 0.706 | 1.093 | 0.562 | 0.179 | -0.001 | 0.081 | 0.081 | 0 | 0.9 | 0.74 | 0.94 | 0.79 | 0.9 | 0.89 |
| 9 | 22000247 | rs3218005 | -0.827 | 1.093 | 0.563 | 0.179 | -0.01 | 0.079 | 0.079 | 0 | 0.1 | 0.26 | 0.06 | 0.21 | 0.08 | 0.11 |
| 9 | 22003223 | rs3217992 | 1.648 | 1.294 | 0.127 | 0.463 | 0.07 | -0.027 | -0.027 | 0.003 | 0.52 | 0.13 | 0.37 | 0.52 | 0.4 | 0.39 |
| 9 | 22003367 | rs1063192 | -0.375 | 1.382 | -0.049 | 0.429 | -0.172 | -0.033 | -0.033 | 0 | 0.78 | 0.99 | 0.8 | 0.83 | 0.57 | 0.73 |
| 9 | 22005330 | rs3217986 | 1.613 | 1.333 | -0.023 | 0.591 | -0.033 | -0.032 | -0.032 | 0 | 0.09 | 0.01 | 0.32 | 0.1 | 0.08 | 0.17 |
| 9 | 22012051 | rs13298881 | 0.37 | 1.278 | -0.22 | 0.448 | -0.017 | 0.024 | 0.024 | 0 | 0.08 | 0.04 | 0.07 | 0 | 0.12 | 0.02 |
| 9 | 22012422 | rs545226 | -1.928 | 1.624 | -0.257 | 0.507 | 0.01 | -0.011 | -0.011 | 0 | 0.49 | 0.93 | 0.62 | 0.4 | 0.57 | 0.61 |

| | | | | | | | | | | | | | | | | |
|---|---|---|---|---|---|---|---|---|---|---|---|---|---|---|---|---|
| 9 | 22028801 | rs7049105 | 0.79 | 1.289 | 0.08 | 0.681 | -0.034 | 0.004 | 0.004 | 0.303 | 0.68 | 0.75 | 0.75 | 0.7 | 0.49 | 0.61 |
| 9 | 22029547 | rs564398 | 0.198 | 1.324 | 0.035 | 0.628 | -0.164 | 0.086 | 0.086 | 0 | 0.22 | 0.01 | 0.19 | 0.09 | 0.41 | 0.27 |
| 9 | 22044059 | rs1333035 | 0.425 | 1.168 | 0.159 | 0.706 | -0.007 | 0.057 | 0.057 | 0 | 0.89 | 0.75 | 0.94 | 0.82 | 0.9 | 0.89 |
| 9 | 22044122 | rs1333034 | -0.335 | 1.182 | 0.125 | 0.658 | 0 | -0.009 | -0.009 | 1.291 | 0.92 | 0.89 | 0.95 | 0.99 | 0.91 | 0.96 |
| 9 | 22053591 | rs11790231 | 0.215 | 0.515 | -0.221 | 0.686 | 0.009 | -0.033 | -0.033 | 0.219 | 0.12 | 0.04 | 0.08 | 0.19 | 0.14 | 0.12 |
| 9 | 22056499 | rs10120688 | 1.16 | 0.929 | 0.578 | 0.558 | 0.019 | -0.02 | -0.02 | 0.008 | 0.63 | 0.42 | 0.72 | 0.72 | 0.5 | 0.61 |
| 9 | 22062134 | rs1011970 | -0.349 | 0.987 | 0.509 | 0.297 | 0.008 | -0.039 | -0.039 | 0.988 | 0.19 | 0.34 | 0.39 | 0.11 | 0.16 | 0.26 |
| 9 | 22067593 | rs10811650 | 1.745 | 1.031 | 0.482 | 0.356 | 0.04 | -0.026 | -0.026 | 0 | 0.53 | 0.2 | 0.38 | 0.53 | 0.44 | 0.4 |
| 9 | 22068652 | rs4977756 | 0.237 | 1.166 | 0.128 | 0.033 | 0.037 | -0.008 | -0.008 | 0 | 0.76 | 0.68 | 0.79 | 0.79 | 0.6 | 0.73 |
| 9 | 22072638 | rs9632885 | 0.83 | 1.062 | 0.154 | -0.075 | -0.02 | -0.024 | -0.024 | 0 | 0.65 | 0.75 | 0.48 | 0.55 | 0.49 | 0.54 |
| 9 | 22072719 | rs10757270 | 1.655 | 0.988 | 0.263 | -0.111 | 0.011 | -0.004 | -0.004 | 0 | 0.55 | 0.33 | 0.4 | 0.55 | 0.44 | 0.53 |
| 9 | 22074793 | rs7855162 | -1.306 | 1.191 | 0.641 | 0.213 | -0.006 | 0.164 | 0.164 | 0 | 0.07 | 0.4 | 0.05 | 0.17 | 0.01 | 0.08 |
| 9 | 22077543 | rs1412832 | -0.599 | 1.493 | 0.544 | 0.077 | -0.106 | -0.098 | -0.098 | 0 | 0.83 | 0.99 | 0.87 | 0.72 | 0.68 | 0.75 |
| 9 | 22078305 | rs7855660 | -1.594 | 0.485 | 0.764 | 0.185 | 0.042 | 0.039 | 0.039 | 2.725 | 0.05 | 0.33 | 0.02 | 0 | 0 | 0.05 |
| 9 | 22082375 | rs1547705 | -0.921 | 0.4 | 0.723 | 0.168 | -0.005 | -0.059 | -0.059 | 0 | 0.13 | 0.25 | 0.11 | 0 | 0.14 | 0.03 |
| 9 | 22083404 | rs1333040 | 0.211 | 0.313 | 0.723 | 0.568 | 0.003 | 0 | 0 | 0 | 0.69 | 0.64 | 0.51 | 0.7 | 0.57 | 0.61 |
| 9 | 22084310 | rs1537370 | 0.176 | -0.217 | 0.568 | 0.673 | -0.026 | 0.004 | 0.004 | 0 | 0.65 | 0.72 | 0.46 | 0.7 | 0.48 | 0.58 |
| 9 | 22088260 | rs10757272 | 0.947 | -0.24 | 0.592 | 0.657 | -0.021 | 0.038 | 0.038 | 1.151 | 0.56 | 0.18 | 0.42 | 0.69 | 0.49 | 0.56 |
| 9 | 22096055 | rs10757274 | 0.995 | -0.199 | 0.584 | 0.747 | -0.021 | -0.037 | -0.037 | 7.177 | 0.55 | 0.18 | 0.42 | 0.53 | 0.49 | 0.48 |
| 9 | 22098574 | rs4977574 | 0.894 | -0.191 | 0.447 | 0.751 | -0.033 | -0.05 | -0.05 | 0 | 0.55 | 0.14 | 0.42 | 0.53 | 0.49 | 0.48 |
| 9 | 22098619 | rs2891168 | 0.892 | -0.189 | 0.478 | 0.752 | -0.019 | -0.035 | -0.035 | 0 | 0.55 | 0.17 | 0.42 | 0.53 | 0.49 | 0.48 |
| 9 | 22103813 | rs1333042 | -0.256 | -0.15 | -0.192 | 0.779 | -0.159 | -0.093 | -0.093 | 1.109 | 0.7 | 0.97 | 0.5 | 0.7 | 0.5 | 0.59 |
| 9 | 22114495 | rs10738609 | 0.929 | -0.189 | -0.156 | 0.801 | -0.035 | -0.041 | -0.041 | 0 | 0.55 | 0.17 | 0.46 | 0.53 | 0.5 | 0.49 |
| 9 | 22115026 | rs2383206 | 0.49 | 0.65 | -0.128 | 1.214 | 0.02 | 0.022 | 0.022 | 0 | 0.64 | 0.41 | 0.5 | 0.53 | 0.51 | 0.52 |
| 9 | 22115105 | rs10965235 | -0.974 | 0.649 | -0.127 | 1.214 | 0.05 | 0.268 | 0.268 | 6.375 | 0.06 | 0.55 | 0.04 | 0.17 | 0 | 0.08 |
| 9 | 22115959 | rs2383207 | 0.16 | 1.136 | -1.56 | 0.762 | -0.165 | -0.093 | -0.093 | 0 | 0.7 | 0.97 | 0.55 | 0.69 | 0.51 | 0.6 |
| 9 | 22127613 | rs10757281 | -0.889 | 0.485 | -0.158 | 0.722 | -0.015 | -0.02 | -0.02 | 0.46 | 0.21 | 0.49 | 0.13 | 0.18 | 0.14 | 0.18 |
| 9 | 22128600 | rs10811658 | -0.925 | 1.034 | -0.246 | 1.143 | 0.018 | 0.054 | 0.054 | 0 | 0.27 | 0.48 | 0.24 | 0.5 | 0.31 | 0.27 |
| 9 | 22130065 | rs10965243 | -1.078 | 1.19 | -0.432 | 1.394 | 0.015 | 0.116 | 0.116 | 0.354 | 0.07 | 0.21 | 0.1 | 0.4 | 0.09 | 0.13 |
| 9 | 22133251 | rs7045889 | -0.963 | 1.012 | -0.603 | 0.601 | -0.001 | -0.008 | -0.008 | 0.629 | 0.61 | 0.51 | 0.55 | 0.72 | 0.61 | 0.6 |
| 9 | 22133645 | rs10217762 | 0.084 | 0.947 | -1.185 | 0.673 | -0.033 | 0.012 | 0.012 | 0 | 0.4 | 0.2 | 0.44 | 0.64 | 0.44 | 0.44 |
| 9 | 22133716 | rs10811659 | 0.545 | 0.986 | -1.032 | 0.67 | -0.019 | 0.027 | 0.027 | 1.812 | 0.17 | 0.06 | 0.11 | 0.1 | 0.24 | 0.17 |
| 9 | 22133984 | rs10757282 | -0.136 | 1.014 | -1.074 | 0.85 | -0.027 | 0.018 | 0.018 | 0 | 0.4 | 0.23 | 0.44 | 0.64 | 0.44 | 0.44 |
| 9 | 22137685 | rs7018475 | 0.283 | 0.644 | -0.959 | 0.904 | 0.005 | -0.016 | -0.016 | 0.353 | 0.3 | 0.26 | 0.33 | 0.4 | 0.27 | 0.35 |
| 9 | 22138762 | rs4977761 | -0.469 | 0.481 | -0.582 | 0.696 | -0.009 | 0.003 | 0.003 | 0 | 0.61 | 0.48 | 0.54 | 0.6 | 0.67 | 0.61 |
| 9 | 22140224 | rs2065501 | 0.322 | 0.79 | -0.578 | -0.054 | -0.001 | -0.004 | -0.004 | 0 | 0.69 | 0.7 | 0.66 | 0.27 | 0.68 | 0.81 |
| 9 | 22141034 | rs7854629 | -0.593 | 1.044 | -1.101 | -1.251 | -0.022 | 0.106 | 0.106 | 0 | 0.62 | 0.77 | 0.77 | 0.97 | 0.58 | 0.75 |
| 9 | 22141269 | rs7026735 | 0.326 | 1.425 | -0.321 | -1.386 | 0.018 | -0.011 | -0.011 | 0.037 | 0.25 | 0.31 | 0.4 | 0.24 | 0.29 | 0.39 |
| 9 | 22141552 | rs2065504 | -0.807 | 1.184 | -0.365 | -1.57 | -0.002 | 0.065 | 0.065 | 0 | 0.48 | 0.62 | 0.51 | 0.33 | 0.48 | 0.68 |
| 9 | 22141790 | rs2065505 | -1.535 | 1.254 | -0.261 | -1.721 | -0.006 | 0.076 | 0.076 | 0 | 0.66 | 0.76 | 0.78 | 0.97 | 0.64 | 0.81 |
| 9 | 22142907 | rs10811664 | 0.976 | 1.157 | -0.184 | -1.644 | -0.003 | 0.035 | 0.035 | 0 | 0.16 | 0.24 | 0.31 | 0.63 | 0.17 | 0.05 |
| 9 | 22144530 | rs7853656 | -0.021 | 1.113 | -0.192 | -1.705 | -0.011 | -0.047 | -0.047 | 0 | 0.23 | 0.3 | 0.1 | 0.09 | 0.18 | 0.29 |
| 9 | 22145694 | rs2065500 | 1.035 | 0.963 | -0.336 | -1.599 | 0 | -0.032 | -0.032 | 0.014 | 0.15 | 0.09 | 0.29 | 0.64 | 0.17 | 0.06 |
| 9 | 22151465 | rs10965256 | 0.414 | 0.846 | 0.111 | -1.687 | -0.009 | -0.003 | -0.003 | 2.917 | 0.15 | 0.14 | 0.04 | 0.09 | 0.06 | 0.17 |
| 9 | 22153360 | rs7853123 | -1.333 | 0.915 | 0.019 | -1.811 | -0.073 | 0.184 | 0.184 | 0 | 0.6 | 0.84 | 0.76 | 0.97 | 0.52 | 0.71 |
| 9 | 22155709 | rs944802 | 1.404 | 0.761 | 0.042 | -1.701 | 0.008 | -0.033 | -0.033 | 0 | 0.14 | 0.1 | 0.12 | 0.61 | 0.13 | 0.05 |
| 9 | 22157908 | rs1333052 | 0.497 | 0.822 | 0.404 | -1.598 | 0.009 | -0.024 | -0.024 | 0 | 0.3 | 0.4 | 0.43 | 0.28 | 0.31 | 0.47 |
| 9 | 22159416 | rs10757288 | -1.41 | 0.82 | 0.237 | -1.779 | -0.062 | 0.16 | 0.16 | 0 | 0.61 | 0.84 | 0.75 | 0.95 | 0.53 | 0.72 |
| 9 | 22159982 | rs10811667 | -1.569 | 0.897 | 0.057 | -1.798 | -0.072 | 0.189 | 0.189 | 0 | 0.63 | 0.88 | 0.77 | 0.95 | 0.58 | 0.76 |
| 9 | 22161828 | rs10965267 | 0.274 | 0.82 | 0.128 | -1.784 | -0.029 | -0.023 | -0.023 | 0.002 | 0.16 | 0.22 | 0.05 | 0.08 | 0.08 | 0.16 |
| 9 | 22162061 | rs10965269 | 1.12 | 0.872 | 0.094 | -1.757 | 0.006 | -0.019 | -0.019 | 0 | 0.16 | 0.14 | 0.16 | 0.62 | 0.16 | 0.06 |
| 9 | 22164991 | rs10811668 | -0.2 | 0.932 | 0.052 | -1.809 | -0.02 | -0.012 | -0.012 | 0.103 | 0.31 | 0.37 | 0.32 | 0.75 | 0.21 | 0.28 |
| 9 | 22166769 | rs2779748 | 1.056 | 1.002 | -0.658 | -1.858 | -0.074 | 0.187 | 0.187 | 0 | 0.36 | 0.12 | 0.22 | 0.05 | 0.42 | 0.24 |
| 9 | 22176961 | rs10757292 | -0.05 | 0.967 | 0.099 | -1.357 | -0.039 | -0.061 | -0.061 | 0 | 0.14 | 0.29 | 0.05 | 0.05 | 0.06 | 0.17 |
| 9 | 22184674 | rs4977764 | 0.494 | 0.952 | 0.164 | -1.141 | -0.003 | 0.036 | 0.036 | 0 | 0.06 | 0.07 | 0.15 | 0.22 | 0.04 | 0.2 |
| 9 | 22184997 | rs2767409 | -0.769 | 1.525 | 0.018 | -1.343 | -0.005 | -0.02 | -0.02 | 0 | 0.4 | 0.49 | 0.45 | 0.32 | 0.35 | 0.48 |
| 9 | 22189716 | rs1095898 | -0.429 | 1.495 | -0.061 | -1.551 | -0.001 | -0.001 | -0.001 | 0.05 | 0.34 | 0.33 | 0.29 | 0.07 | 0.33 | 0.37 |
| 9 | 22195321 | rs1679029 | -1.619 | 1.727 | -0.676 | -1.689 | -0.005 | 0.032 | 0.032 | 0 | 0.62 | 0.72 | 0.78 | 0.98 | 0.61 | 0.74 |
| 9 | 22195820 | rs2219849 | 1.457 | 2.118 | -0.813 | -0.395 | -0.002 | 0 | 0 | 0 | 0.56 | 0.58 | 0.38 | 0.58 | 0.5 | 0.46 |
| 9 | 22201586 | rs12350824 | 0.706 | 2.257 | -1.166 | -1.291 | -0.004 | 0.006 | 0.006 | 0 | 0.17 | 0.12 | 0.2 | 0 | 0.16 | 0.1 |
| 9 | 22202151 | rs2025796 | 0.278 | 2.11 | -0.852 | -1.052 | -0.036 | 0.051 | 0.051 | 0 | 0.63 | 0.74 | 0.55 | 0.94 | 0.54 | 0.7 |
| 9 | 22203800 | rs1409772 | -0.475 | 1.071 | -1.221 | -0.64 | 0.009 | 0.025 | 0.025 | 0 | 0.38 | 0.28 | 0.41 | 0.28 | 0.32 | 0.45 |
| 9 | 22206559 | rs9696092 | -0.64 | 1.301 | -1.565 | -0.977 | -0.086 | 0.136 | 0.136 | 0 | 0.32 | 0.13 | 0.21 | 0.06 | 0.44 | 0.18 |
| 9 | 22206987 | rs1679013 | 0.677 | 0.826 | -1.132 | -0.877 | -0.015 | 0.056 | 0.056 | 0.06 | 0.58 | 0.3 | 0.37 | 0.79 | 0.48 | 0.64 |
| 9 | 22228560 | rs12003808 | -0.917 | 0.469 | 0.507 | -0.491 | 0.007 | 0.014 | 0.014 | 0.4 | 0.08 | 0.44 | 0.15 | 0.04 | 0.03 | 0.12 |
| 9 | 22228756 | rs10811678 | 0.278 | -0.16 | -0.341 | -0.181 | -0.009 | -0.048 | -0.048 | 0 | 0.22 | 0.01 | 0.11 | 0.22 | 0.15 | 0.22 |
| 9 | 22237616 | rs7859353 | -0.892 | 0.169 | 0.802 | 0.033 | -0.001 | -0.004 | -0.004 | 0.491 | 0.07 | 0.37 | 0.05 | 0.03 | 0.03 | 0.12 |
| 9 | 22245423 | rs1679020 | 1.506 | 0.108 | 1.213 | 0.044 | -0.013 | -0.02 | -0.02 | 0 | 0.88 | 0.32 | 0.89 | 0.92 | 0.92 | 0.84 |
| 9 | 22246794 | rs1679022 | 1.429 | 0.105 | 1.211 | 0.043 | -0.012 | -0.015 | -0.015 | 0.063 | 0.89 | 0.33 | 0.89 | 0.93 | 0.93 | 0.84 |

| | | | | | | | | | | | | | | | | |
|---|---|---|---|---|---|---|---|---|---|---|---|---|---|---|---|---|
| 9 | 22272866 | rs1679026 | 1.208 | 0.135 | 1.315 | 0.065 | 0.033 | 0 | 0 | 0.007 | 0.9 | 0.47 | 0.91 | 0.94 | 0.92 | 0.83 |
| 9 | 22291931 | rs1609591 | 1.478 | 0.074 | 1.649 | 0.035 | 0.004 | -0.01 | -0.01 | 0 | 0.89 | 0.4 | 0.9 | 0.93 | 0.93 | 0.83 |
| 9 | 22301053 | rs1575971 | 0.593 | 0.219 | 1.315 | 0.362 | 0.022 | 0 | 0 | 0 | 0.93 | 0.68 | 0.94 | 0.98 | 0.96 | 0.9 |
| 9 | 22321234 | rs1327053 | -1.203 | 0.222 | 1.169 | 0.364 | 0.021 | -0.01 | -0.01 | 0.721 | 0.09 | 0.37 | 0.08 | 0.04 | 0.07 | 0.16 |
| 9 | 22332703 | rs943394 | 1.158 | 1.014 | 0.453 | 0.312 | 0.034 | -0.016 | -0.016 | 0.067 | 0.72 | 0.31 | 0.52 | 0.83 | 0.5 | 0.67 |
| 9 | 22333578 | rs7856683 | 0.923 | -0.123 | -0.798 | 0.216 | 0.001 | 0.042 | 0.042 | 0.328 | 0.38 | 0.01 | 0.17 | 0.28 | 0.27 | 0.31 |
| 9 | 22336434 | rs716580 | 0.125 | 0.297 | -0.432 | 0.118 | -0.089 | 0.011 | 0.011 | 3.287 | 0.82 | 0.92 | 0.6 | 0.89 | 0.56 | 0.83 |
| 9 | 22336954 | rs1359741 | 0.92 | 0.157 | 0.035 | 0.073 | 0.054 | -0.026 | -0.026 | 0 | 0.72 | 0.58 | 0.51 | 0.85 | 0.49 | 0.66 |
| 9 | 22340159 | rs9406787 | 0.021 | 0.26 | -0.398 | 0.129 | -0.137 | -0.007 | -0.007 | 0.021 | 0.82 | 0.97 | 0.61 | 0.88 | 0.55 | 0.8 |
| 9 | 22364080 | rs10811697 | 0.687 | 0.187 | -0.201 | 0.191 | 0.048 | -0.023 | -0.023 | 0.044 | 0.72 | 0.36 | 0.51 | 0.85 | 0.49 | 0.67 |
| 9 | 22379219 | rs10757321 | 0.657 | -0.511 | -0.893 | 0.486 | 0.036 | -0.011 | -0.011 | 0 | 0.42 | 0.05 | 0.2 | 0.36 | 0.29 | 0.48 |
| 9 | 22380028 | rs944033 | 0.491 | 0.728 | -0.016 | 0.405 | 0.019 | -0.027 | -0.027 | 0 | 0.77 | 0.56 | 0.61 | 0.88 | 0.67 | 0.74 |
| 9 | 22382449 | rs944037 | 0.183 | 0.69 | 0.085 | 0.386 | -0.031 | 0.025 | 0.025 | 0.018 | 0.79 | 0.8 | 0.63 | 0.88 | 0.67 | 0.74 |
| 9 | 22383816 | rs4452893 | 0.62 | 0.688 | 0.122 | 0.601 | 0.02 | -0.013 | -0.013 | 0 | 0.77 | 0.54 | 0.61 | 0.82 | 0.67 | 0.74 |
| 9 | 22392359 | rs7024019 | 0.254 | 0.333 | 0.454 | 0.551 | -0.001 | 0.029 | 0.029 | 2.057 | 0.38 | 0.27 | 0.23 | 0.15 | 0.35 | 0.38 |
| 9 | 22395664 | rs9696718 | -0.484 | 0.142 | 0.478 | 0.465 | -0.002 | -0.009 | -0.009 | 0 | 0.55 | 0.58 | 0.71 | 0.76 | 0.6 | 0.63 |
| 9 | 22399693 | rs7024096 | -0.13 | 0.28 | 0.436 | 0.12 | 0.022 | -0.005 | -0.005 | 0 | 0.2 | 0.4 | 0.38 | 0.19 | 0.29 | 0.26 |
| 9 | 22412570 | rs10965342 | -0.769 | 0.165 | 0.471 | 0.044 | 0.028 | 0.042 | 0.042 | 0.788 | 0.58 | 0.8 | 0.64 | 0.32 | 0.64 | 0.56 |
| 9 | 22417016 | rs7875648 | 0.231 | 0.122 | 0.55 | -0.228 | 0.006 | 0.029 | 0.029 | 3.76 | 0.43 | 0.63 | 0.28 | 0.16 | 0.36 | 0.37 |
| 9 | 22417552 | rs7846900 | 0.367 | 0.082 | 0.582 | -0.25 | -0.003 | -0.046 | -0.046 | 2.206 | 0.42 | 0.48 | 0.27 | 0.15 | 0.37 | 0.37 |
| 9 | 22430804 | rs16906084 | 0.046 | 0.493 | 0.929 | -0.083 | 0.006 | -0.009 | -0.009 | 9.526 | 0.14 | 0.26 | 0.29 | 0.09 | 0.15 | 0.18 |
| 9 | 22434723 | rs676417 | -0.617 | 0.46 | 0.957 | -0.186 | 0.022 | 0.026 | 0.026 | 0.43 | 0.56 | 0.71 | 0.62 | 0.32 | 0.62 | 0.53 |
| 9 | 22442343 | rs1576334 | 0.42 | 0.48 | 0.823 | -0.174 | -0.007 | 0.004 | 0.004 | 0 | 0.13 | 0.08 | 0.28 | 0.09 | 0.15 | 0.18 |
| 9 | 22442888 | rs596543 | 0.392 | 0.494 | 0.83 | -0.172 | 0.006 | -0.019 | -0.019 | 0 | 0.42 | 0.47 | 0.33 | 0.23 | 0.47 | 0.36 |
| 9 | 22446297 | rs693370 | 0.383 | 0.7 | 0.854 | -0.223 | 0.069 | 0.009 | 0.009 | 0.507 | 0.08 | 0.26 | 0.08 | 0.05 | 0.12 | 0.07 |
| 9 | 22446828 | rs655497 | -0.628 | 0.702 | 0.847 | -0.227 | 0.038 | 0.017 | 0.017 | 0.021 | 0.2 | 0.33 | 0.4 | 0.23 | 0.3 | 0.26 |
| 9 | 22451031 | rs563802 | -0.862 | 0.31 | 0.728 | -0.225 | 0.003 | 0.053 | 0.053 | 0.213 | 0.19 | 0.52 | 0.36 | 0.23 | 0.18 | 0.25 |
| 9 | 22454125 | rs17835669 | 0.749 | 0.357 | 0.645 | -0.212 | -0.021 | -0.006 | -0.006 | 0.005 | 0.11 | 0.06 | 0.28 | 0.11 | 0.18 | 0.17 |
| 9 | 22454150 | rs6475646 | -0.997 | -0.149 | 0.1 | -0.821 | -0.007 | -0.037 | -0.037 | 5.224 | 0.77 | 0.73 | 0.85 | 0.95 | 0.82 | 0.85 |
| 9 | 22455518 | rs10733377 | -0.603 | -0.049 | 0.023 | -0.896 | -0.001 | -0.006 | -0.006 | 0 | 0.39 | 0.57 | 0.56 | 0.75 | 0.38 | 0.52 |
| 9 | 22456296 | rs10738628 | -0.165 | 0.031 | 0.064 | -0.834 | -0.001 | -0.016 | -0.016 | 0.318 | 0.66 | 0.62 | 0.58 | 0.86 | 0.68 | 0.71 |
| 9 | 22460095 | rs665908 | -0.01 | 0.066 | 0.009 | -0.664 | 0.002 | 0.072 | 0.072 | 0.421 | 0.17 | 0.47 | 0.33 | 0.2 | 0.15 | 0.2 |
| 9 | 22463201 | rs7868400 | -0.168 | 0.185 | -0.12 | -0.634 | -0.008 | 0.038 | 0.038 | 0.715 | 0.46 | 0.28 | 0.49 | 0.64 | 0.46 | 0.45 |
| 9 | 22471418 | rs630871 | 0.807 | -0.092 | 0.196 | -0.466 | 0.005 | 0.071 | 0.071 | 0.798 | 0.06 | 0.25 | 0.03 | 0.05 | 0 | 0.07 |
| 9 | 22471832 | rs7039553 | 0.1 | 0.032 | -0.199 | -0.687 | 0.001 | 0.004 | 0.004 | 0.168 | 0.33 | 0.35 | 0.35 | 0.21 | 0.32 | 0.25 |
| 9 | 22472068 | rs16906189 | 1.413 | 0.018 | -0.206 | -0.78 | 0.013 | 0.028 | 0.028 | 2.053 | 0.2 | 0.13 | 0.12 | 0 | 0.16 | 0.08 |
| 9 | 22472289 | rs7027027 | 0.806 | 0.01 | -0.118 | -0.65 | -0.001 | -0.001 | -0.001 | 0 | 0.13 | 0.13 | 0.2 | 0.12 | 0.16 | 0.15 |
| 9 | 22481286 | rs556958 | 0.035 | -0.088 | -0.283 | -0.723 | -0.02 | 0.004 | 0.004 | 0 | 0.29 | 0.45 | 0.38 | 0.7 | 0.26 | 0.4 |
| 9 | 22487177 | rs667115 | -0.125 | -0.143 | -0.295 | -0.979 | -0.023 | -0.029 | -0.029 | 0.992 | 0.3 | 0.45 | 0.42 | 0.76 | 0.26 | 0.41 |
| 9 | 22488678 | rs586716 | -0.808 | -0.129 | -0.518 | -1.048 | -0.013 | 0.183 | 0.183 | 0.116 | 0.52 | 0.9 | 0.57 | 0.77 | 0.44 | 0.57 |
| 9 | 22508807 | rs7022740 | 0.617 | -0.206 | -0.297 | -0.91 | -0.002 | -0.011 | -0.011 | 0 | 0.23 | 0.19 | 0.37 | 0.71 | 0.26 | 0.35 |
| 9 | 22510754 | rs1392084 | -0.163 | -0.217 | -0.233 | -0.918 | -0.017 | -0.024 | -0.024 | 0 | 0.71 | 0.53 | 0.56 | 0.24 | 0.74 | 0.58 |
| 9 | 22513603 | rs17694987 | 0.829 | -0.144 | -0.461 | -0.901 | -0.034 | -0.013 | -0.013 | 0 | 0.13 | 0.05 | 0.2 | 0.12 | 0.16 | 0.14 |
| 9 | 22518195 | rs13284006 | 1.303 | -0.21 | -0.448 | -0.981 | 0.007 | 0.066 | 0.066 | 2.024 | 0.2 | 0.12 | 0.11 | 0 | 0.18 | 0.08 |
| 9 | 22520468 | rs2221132 | -0.792 | -0.31 | -0.267 | -1 | -0.022 | -0.065 | -0.065 | 0.986 | 0.76 | 0.64 | 0.84 | 0.95 | 0.82 | 0.86 |
| 9 | 22523533 | rs12000593 | -0.174 | -0.309 | -0.238 | -1.021 | -0.006 | -0.081 | -0.081 | 0 | 0.35 | 0.53 | 0.34 | 0.12 | 0.33 | 0.29 |
| 9 | 22525505 | rs10811719 | 0.549 | -0.225 | -0.562 | -1.029 | -0.003 | -0.009 | -0.009 | 0 | 0.24 | 0.18 | 0.39 | 0.72 | 0.26 | 0.36 |
| 9 | 22527750 | rs10965372 | 1.361 | -0.23 | -0.643 | -1.108 | -0.066 | 0.159 | 0.159 | 0 | 0.36 | 0.1 | 0.24 | 0.12 | 0.4 | 0.35 |
| 9 | 22529933 | rs7859900 | -1.551 | -0.264 | -0.586 | -1.129 | -0.002 | -0.059 | -0.059 | 0.774 | 0.81 | 0.76 | 0.88 | 0.99 | 0.83 | 0.84 |
| 9 | 22531190 | rs10738632 | -1.535 | -0.346 | -0.62 | -1.162 | -0.01 | 0.042 | 0.042 | 0 | 0.63 | 0.73 | 0.75 | 0.88 | 0.59 | 0.65 |
| 9 | 22532249 | rs10811721 | 1.439 | -0.349 | -0.8 | -0.593 | -0.051 | -0.054 | -0.054 | 0 | 0.1 | 0.01 | 0.08 | 0.22 | 0.13 | 0.1 |
| 9 | 22535511 | rs2891171 | 0.1 | -0.861 | -0.414 | -0.396 | -0.027 | -0.005 | -0.005 | 0 | 0.8 | 0.42 | 0.73 | 0.83 | 0.87 | 0.75 |
| 9 | 22542285 | rs10811723 | 0.175 | -0.981 | -0.512 | 0.08 | 0.017 | -0.053 | -0.053 | 0 | 0.67 | 0.21 | 0.56 | 0.54 | 0.69 | 0.73 |
| 9 | 22544607 | rs1948678 | 1.066 | -1.211 | -0.823 | -0.496 | -0.017 | 0.031 | 0.031 | 0 | 0.17 | 0.01 | 0.13 | 0.12 | 0.15 | 0.24 |
| 9 | 22547508 | rs4977586 | -0.676 | -1.253 | -1.197 | -0.454 | 0.032 | 0.048 | 0.048 | 0.642 | 0.5 | 0.72 | 0.59 | 0.59 | 0.45 | 0.52 |
| 9 | 22551240 | rs13289947 | 0.828 | -1.073 | -1.394 | -0.467 | 0.016 | 0.077 | 0.077 | 1.274 | 0.28 | 0.15 | 0.14 | 0.04 | 0.21 | 0.24 |
| 9 | 22554601 | rs1158382 | 0.544 | -0.538 | -2.085 | -0.042 | 0.018 | 0.26 | 0.26 | 0 | 0.05 | 0.14 | 0.27 | 0.43 | 0.12 | 0.12 |
| 9 | 22565072 | rs9644863 | 0.766 | -0.484 | -2.813 | -0.121 | -0.089 | 0.03 | 0.03 | 0 | 0.36 | 0.14 | 0.37 | 0.35 | 0.48 | 0.28 |
| 9 | 22583181 | rs1499558 | 0.748 | -0.97 | -2.923 | -0.92 | -0.021 | 0.052 | 0.052 | 0 | 0.23 | 0.02 | 0.15 | 0.12 | 0.17 | 0.29 |
| 9 | 22583561 | rs10811743 | 0.078 | -0.031 | -2.211 | -0.339 | -0.039 | 0.072 | 0.072 | 0 | 0.4 | 0.11 | 0.64 | 0.78 | 0.63 | 0.35 |
| 9 | 22596808 | rs7469285 | 0.505 | 0.056 | -1.472 | -0.194 | 0.002 | 0.091 | 0.091 | 0.119 | 0.45 | 0.25 | 0.56 | 0.21 | 0.46 | 0.34 |
| 9 | 22596867 | rs10965398 | 0.152 | -0.004 | -1.555 | 0.252 | -0.037 | -0.075 | -0.075 | 0 | 0.27 | 0.13 | 0.31 | 0.5 | 0.35 | 0.39 |
| 9 | 22598741 | rs10217416 | -0.241 | -0.047 | -0.473 | 0.243 | -0.021 | -0.005 | -0.005 | 0 | 0.72 | 0.52 | 0.86 | 0.71 | 0.81 | 0.71 |
| 9 | 22599344 | rs10757346 | -0.514 | 0.091 | -0.674 | 0.51 | 0.031 | 0.134 | 0.134 | 0.525 | 0.86 | 0.63 | 0.8 | 0.65 | 0.79 | 0.79 |
| 9 | 22607894 | rs10965400 | 0.946 | 0.097 | -0.912 | 0.363 | -0.004 | -0.01 | -0.01 | 0 | 0.65 | 0.27 | 0.71 | 0.37 | 0.63 | 0.64 |
| 9 | 22608275 | rs11793406 | 1.69 | -0.377 | -1.41 | -0.221 | -0.063 | 0.105 | 0.105 | 0.113 | 0.31 | 0.1 | 0.52 | 0.14 | 0.43 | 0.25 |
| 9 | 22612584 | rs12350224 | 0.806 | -0.381 | -0.925 | -0.003 | 0.003 | 0.024 | 0.024 | 0 | 0.18 | 0.21 | 0.1 | 0.28 | 0.15 | 0.12 |
| 9 | 22613796 | rs10118516 | 0.72 | -0.892 | -0.98 | -0.229 | 0.043 | 0.003 | 0.003 | 1.878 | 0.66 | 0.22 | 0.83 | 0.64 | 0.81 | 0.68 |
| 9 | 22614985 | rs10757350 | -0.093 | -1.028 | -0.731 | -0.021 | -0.002 | -0.006 | -0.006 | 0.15 | 0.29 | 0.27 | 0.31 | 0.49 | 0.35 | 0.42 |

| | | | | | | | | | | | | | | | | |
|---|---|---|---|---|---|---|---|---|---|---|---|---|---|---|---|---|
| 9 | 22616426 | rs2065846 | 1.098 | -1.021 | -0.67 | -0.018 | 0.004 | 0.025 | 0.025 | 0 | 0.18 | 0.17 | 0.1 | 0.29 | 0.15 | 0.14 |
| 9 | 22617009 | rs2065848 | -0.88 | -0.871 | -0.655 | 0.097 | 0.007 | -0.003 | -0.003 | 0 | 0.45 | 0.55 | 0.24 | 0.49 | 0.32 | 0.48 |
| 9 | 22617269 | rs1414696 | 1.141 | -0.861 | -0.688 | 0.088 | -0.011 | 0.125 | 0.125 | 1.069 | 0.49 | 0.2 | 0.57 | 0.22 | 0.5 | 0.37 |
| 9 | 22619647 | rs7024672 | -0.598 | -0.687 | -0.579 | 0.202 | 0.084 | 0.16 | 0.16 | 0 | 0.94 | 0.78 | 0.81 | 0.71 | 0.82 | 0.85 |
| 9 | 22622698 | rs10965404 | 1.58 | -0.74 | -0.734 | 0.16 | -0.037 | -0.045 | -0.045 | 0 | 0.13 | 0.01 | 0.09 | 0.13 | 0.13 | 0.24 |
| 9 | 22624610 | rs10965407 | 1.535 | -0.71 | -0.735 | 0.16 | -0.039 | -0.044 | -0.044 | 0 | 0.13 | 0.01 | 0.09 | 0.13 | 0.13 | 0.24 |
| 9 | 22625034 | rs1932605 | 0.274 | -0.995 | -0.821 | 0.06 | 0.005 | -0.012 | -0.012 | 0 | 0.59 | 0.28 | 0.66 | 0.41 | 0.63 | 0.7 |
| 9 | 22625247 | rs13290572 | 0.875 | -0.969 | -0.768 | 0.044 | -0.004 | 0.064 | 0.064 | 0.034 | 0.46 | 0.29 | 0.56 | 0.28 | 0.5 | 0.44 |
| 9 | 22625577 | rs7027641 | -0.901 | -0.741 | -0.876 | 0.21 | 0.003 | 0.048 | 0.048 | 0 | 0.8 | 0.6 | 0.76 | 0.69 | 0.79 | 0.82 |
| 9 | 22628385 | rs635827 | -1.044 | -0.619 | -0.89 | 0.215 | -0.023 | 0.067 | 0.067 | 0 | 0.81 | 0.59 | 0.78 | 0.69 | 0.85 | 0.83 |
| 9 | 22634275 | rs17695326 | 1.444 | -0.688 | -1.191 | -0.276 | -0.146 | 0.024 | 0.024 | 5.68 | 0.34 | 0.04 | 0.51 | 0.2 | 0.47 | 0.32 |
| 9 | 22638990 | rs1999760 | 0.461 | -0.716 | -1.016 | -0.306 | -0.02 | 0.003 | 0.003 | 0 | 0.28 | 0.4 | 0.15 | 0.29 | 0.2 | 0.19 |
| 9 | 22639410 | rs6475671 | 0.038 | -0.693 | -0.968 | -0.268 | 0 | 0.037 | 0.037 | 0 | 0.65 | 0.48 | 0.66 | 0.5 | 0.65 | 0.51 |
| 9 | 22639567 | rs10965408 | 1.933 | -0.736 | -0.942 | -0.263 | -0.035 | -0.017 | -0.017 | 0 | 0.15 | 0.2 | 0.05 | 0.08 | 0.03 | 0.08 |
| 9 | 22641126 | rs670126 | -0.932 | -0.571 | -1.06 | -0.198 | -0.077 | -0.104 | -0.104 | 2.002 | 0.82 | 0.93 | 0.74 | 0.59 | 0.69 | 0.66 |
| 9 | 22641633 | rs1448792 | -0.009 | -0.612 | -1.122 | -0.289 | 0 | -0.002 | -0.002 | 0.077 | 0.72 | 0.66 | 0.74 | 0.59 | 0.72 | 0.55 |
| 9 | 22644110 | rs1448789 | 1.306 | -0.904 | -1.66 | -0.237 | -0.136 | 0.023 | 0.023 | 8.038 | 0.35 | 0.04 | 0.52 | 0.22 | 0.47 | 0.34 |
| 9 | 22644806 | rs842837 | -1.103 | -0.705 | -1.508 | -0.131 | -0.076 | -0.097 | -0.097 | 0 | 0.82 | 0.93 | 0.73 | 0.59 | 0.68 | 0.68 |
| 9 | 22648562 | rs12553349 | 1.064 | -0.632 | -1.518 | -0.267 | -0.1 | 0.048 | 0.048 | 4.987 | 0.36 | 0.05 | 0.52 | 0.21 | 0.45 | 0.35 |
| 9 | 22653899 | rs10965412 | 1.517 | -0.843 | -1.101 | -0.311 | -0.046 | -0.029 | -0.029 | 0.771 | 0.16 | 0.3 | 0.07 | 0.08 | 0.04 | 0.16 |
| 9 | 22658001 | rs17695493 | 1.121 | -0.701 | -1.351 | -0.293 | -0.098 | 0.051 | 0.051 | 4.83 | 0.36 | 0.04 | 0.52 | 0.21 | 0.45 | 0.33 |
| 9 | 22664213 | rs578315 | -1.209 | -0.566 | -1.298 | -0.131 | -0.043 | -0.071 | -0.071 | 0 | 0.83 | 0.9 | 0.76 | 0.59 | 0.72 | 0.7 |
| 9 | 22668988 | rs12551699 | 2.095 | -0.619 | -1.201 | -0.198 | -0.07 | -0.02 | -0.02 | 0.715 | 0.28 | 0.08 | 0.53 | 0.23 | 0.45 | 0.28 |
| 9 | 22669166 | rs2210865 | 2.105 | -0.616 | -1.2 | -0.196 | -0.07 | -0.02 | -0.02 | 0 | 0.28 | 0.08 | 0.53 | 0.23 | 0.45 | 0.28 |
| 9 | 22673444 | rs842834 | -0.532 | -0.915 | -0.847 | -0.229 | 0.034 | 0.025 | 0.025 | 3.39 | 0.44 | 0.62 | 0.2 | 0.36 | 0.23 | 0.35 |
| 9 | 22690913 | rs12350664 | 1.675 | -0.795 | -0.751 | -0.277 | -0.01 | -0.017 | -0.017 | 1.416 | 0.15 | 0.16 | 0.05 | 0.07 | 0.05 | 0.09 |
| 9 | 22699397 | rs1619363 | 0.423 | -0.845 | -0.541 | -0.215 | 0.002 | 0.023 | 0.023 | 0 | 0.21 | 0.33 | 0.11 | 0.28 | 0.17 | 0.15 |
| 9 | 22700554 | rs7859540 | 0.387 | -0.855 | -0.54 | -0.224 | 0.001 | 0.025 | 0.025 | 0 | 0.21 | 0.32 | 0.11 | 0.29 | 0.17 | 0.15 |
| 9 | 22706144 | rs1375290 | 2.112 | -0.491 | -0.963 | -0.344 | -0.024 | -0.011 | -0.011 | 0 | 0.27 | 0.14 | 0.51 | 0.24 | 0.39 | 0.24 |
| 9 | 22711948 | rs474342 | -0.56 | -0.489 | -0.931 | -0.39 | -0.017 | -0.028 | -0.028 | 0.566 | 0.78 | 0.79 | 0.72 | 0.6 | 0.66 | 0.67 |
| 9 | 22724188 | rs565595 | -0.564 | -0.468 | -0.978 | -0.392 | -0.071 | -0.073 | -0.073 | 0 | 0.77 | 0.87 | 0.72 | 0.59 | 0.63 | 0.64 |
| 9 | 22725295 | rs688581 | -0.692 | -0.699 | -1.074 | -0.466 | 0.085 | 0.076 | 0.076 | 1.209 | 0.49 | 0.71 | 0.24 | 0.37 | 0.25 | 0.36 |
| 9 | 22730877 | rs593909 | -0.991 | -0.654 | -1.129 | -0.443 | -0.077 | -0.08 | -0.08 | 0 | 0.8 | 0.9 | 0.73 | 0.6 | 0.64 | 0.64 |
| 9 | 22731935 | rs4133767 | 1.219 | -0.801 | -0.864 | -0.507 | -0.026 | 0.033 | 0.033 | 1.328 | 0.39 | 0.16 | 0.55 | 0.24 | 0.44 | 0.29 |
| 9 | 22739631 | rs10125443 | 1.78 | -0.841 | -0.661 | -0.369 | -0.016 | 0.006 | 0.006 | 0.342 | 0.15 | 0.17 | 0.05 | 0.13 | 0.05 | 0.1 |
| 9 | 22742136 | rs640648 | -0.159 | -0.811 | -0.488 | -0.469 | -0.027 | 0.012 | 0.012 | 0 | 0.27 | 0.42 | 0.13 | 0.32 | 0.17 | 0.23 |
| 9 | 22746734 | rs842845 | 0.766 | -0.853 | -0.432 | -0.47 | -0.004 | 0.062 | 0.062 | 0 | 0.18 | 0.3 | 0.1 | 0.32 | 0.14 | 0.09 |
| 9 | 22747274 | rs842846 | 0.207 | -0.898 | -0.581 | -0.499 | 0 | 0 | 0 | 0.606 | 0.52 | 0.57 | 0.63 | 0.46 | 0.56 | 0.47 |
| 9 | 22748083 | rs842847 | 0.164 | -0.802 | -0.652 | -0.462 | 0 | -0.001 | -0.001 | 0.021 | 0.53 | 0.52 | 0.63 | 0.45 | 0.56 | 0.47 |
| 9 | 22748466 | rs527427 | 0.146 | -0.798 | -0.65 | -0.46 | 0 | -0.001 | -0.001 | 0 | 0.53 | 0.51 | 0.63 | 0.46 | 0.56 | 0.47 |
| 9 | 22749869 | rs644917 | -0.392 | -0.991 | -0.188 | -0.148 | -0.025 | 0.012 | 0.012 | 0 | 0.27 | 0.42 | 0.13 | 0.32 | 0.17 | 0.23 |
| 9 | 22752087 | rs535119 | 0.146 | -1.013 | -0.071 | -0.089 | -0.024 | 0.041 | 0.041 | 0 | 0.18 | 0.24 | 0.07 | 0.3 | 0.11 | 0.14 |
| 9 | 22754513 | rs10123107 | 1.107 | -1.219 | -0.115 | -0.103 | -0.037 | 0.047 | 0.047 | 0.834 | 0.15 | 0.3 | 0.09 | 0.17 | 0.05 | 0.13 |
| 9 | 22755025 | rs16906524 | 1.231 | -1.473 | -0.099 | -0.296 | -0.028 | -0.058 | -0.058 | 0.68 | 0.13 | 0.21 | 0.04 | 0 | 0.04 | 0.04 |
| 9 | 22756708 | rs495897 | -0.746 | -0.797 | -0.507 | -0.512 | -0.129 | -0.098 | -0.098 | 0.006 | 0.78 | 0.93 | 0.67 | 0.6 | 0.52 | 0.7 |
| 9 | 22765106 | rs4607705 | 0.814 | -0.881 | -0.432 | -0.564 | 0.037 | 0.055 | 0.055 | 0.272 | 0.5 | 0.57 | 0.56 | 0.29 | 0.37 | 0.46 |
| 9 | 22767164 | rs505061 | -0.033 | -0.685 | -0.472 | -0.543 | 0.011 | 0.005 | 0.005 | 0 | 0.63 | 0.64 | 0.58 | 0.42 | 0.47 | 0.57 |
| 9 | 22767236 | rs1349151 | 2.304 | -0.805 | -0.725 | -0.688 | -0.032 | 0.083 | 0.083 | 1.236 | 0.28 | 0.1 | 0.44 | 0.13 | 0.29 | 0.26 |
| 9 | 22767738 | rs1375292 | 0.828 | -0.877 | -0.49 | -0.706 | 0 | -0.017 | -0.017 | 0 | 0.51 | 0.49 | 0.53 | 0.14 | 0.39 | 0.34 |
| 9 | 22768289 | rs643958 | -0.658 | -0.245 | -0.209 | -0.252 | 0.009 | 0.004 | 0.004 | 0 | 0.8 | 0.79 | 0.71 | 0.63 | 0.59 | 0.73 |
| 9 | 22768666 | rs10965447 | 1.279 | -0.48 | -0.028 | -0.235 | -0.036 | 0.045 | 0.045 | 1.263 | 0.15 | 0.24 | 0.08 | 0.2 | 0.06 | 0.13 |
| 9 | 22770044 | rs10115398 | 1.54 | -0.442 | -0.035 | -0.216 | -0.026 | 0.04 | 0.04 | 1.321 | 0.14 | 0.18 | 0.08 | 0.21 | 0.05 | 0.13 |
| 9 | 22774651 | rs593899 | 0.599 | -0.491 | -0.156 | -0.214 | 0.008 | -0.007 | -0.007 | 0 | 0.6 | 0.59 | 0.61 | 0.4 | 0.49 | 0.5 |
| 9 | 22776141 | rs6475682 | 0.329 | -0.698 | -0.145 | -0.222 | -0.038 | -0.004 | -0.004 | 0 | 0.25 | 0.53 | 0.14 | 0.2 | 0.09 | 0.24 |
| 9 | 22780930 | rs1098977 | -0.763 | -0.79 | -0.194 | -0.226 | 0.025 | 0.012 | 0.012 | 0.534 | 0.4 | 0.64 | 0.21 | 0.34 | 0.24 | 0.23 |
| 9 | 22781709 | rs547279 | 0.146 | -0.641 | -0.202 | -0.247 | -0.024 | -0.02 | -0.02 | 0.155 | 0.72 | 0.77 | 0.68 | 0.6 | 0.54 | 0.68 |
| 9 | 22784110 | rs501290 | -1.662 | -0.189 | -0.263 | -0.109 | -0.032 | 0.013 | 0.013 | 0 | 0.88 | 0.96 | 0.87 | 0.9 | 0.81 | 0.87 |
| 9 | 22790288 | rs7852316 | 0.517 | -0.703 | 0.118 | -0.134 | -0.057 | -0.134 | -0.134 | 0 | 0.24 | 0.4 | 0.11 | 0.04 | 0.11 | 0.09 |
| 9 | 22791520 | rs10811773 | 1.475 | -0.601 | 0.123 | -0.022 | -0.028 | -0.062 | -0.062 | 0.86 | 0.14 | 0.33 | 0.07 | 0.04 | 0.05 | 0.04 |
| 9 | 22796540 | rs586554 | -1.043 | -0.148 | -0.125 | 0.199 | -0.064 | -0.064 | -0.064 | 0 | 0.82 | 0.92 | 0.72 | 0.63 | 0.6 | 0.75 |
| 9 | 22799090 | rs7032336 | -0.294 | -0.25 | -0.196 | 0.098 | -0.023 | -0.02 | -0.02 | 0 | 0.73 | 0.78 | 0.67 | 0.59 | 0.53 | 0.71 |
| 9 | 22799213 | rs12555613 | 1.281 | -0.374 | -0.221 | 0.023 | 0.005 | 0.017 | 0.017 | 0.858 | 0.54 | 0.53 | 0.57 | 0.3 | 0.4 | 0.57 |
| 9 | 22801570 | rs842853 | 0.493 | -0.161 | -0.579 | 0.282 | 0.002 | -0.016 | -0.016 | 0 | 0.16 | 0.17 | 0.1 | 0.31 | 0.14 | 0.15 |
| 9 | 22805644 | rs10125975 | 1.488 | -0.269 | -0.647 | 0.13 | 0.026 | 0.039 | 0.039 | 0.235 | 0.43 | 0.24 | 0.5 | 0.31 | 0.32 | 0.51 |
| 9 | 22811336 | rs9298841 | 1.161 | -0.17 | -0.539 | 0.205 | 0.029 | 0.042 | 0.042 | 0.027 | 0.47 | 0.3 | 0.53 | 0.34 | 0.38 | 0.53 |
| 9 | 22818289 | rs7874413 | 0.562 | -0.171 | -0.179 | 0.178 | 0 | -0.007 | -0.007 | 0 | 0.55 | 0.57 | 0.58 | 0.44 | 0.5 | 0.59 |
| 9 | 22819064 | rs4571809 | -1.743 | -0.173 | -0.748 | -0.049 | -0.046 | 0.179 | 0.179 | 3.036 | 0.62 | 0.85 | 0.54 | 0.87 | 0.64 | 0.56 |
| 9 | 22819407 | rs7865016 | -1.073 | -0.237 | -0.457 | 0.084 | 0.007 | 0.044 | 0.044 | 0 | 0.48 | 0.62 | 0.47 | 0.7 | 0.6 | 0.43 |
| 9 | 22829231 | rs6475687 | 0.746 | -0.27 | -0.556 | 0.303 | 0.044 | -0.023 | -0.023 | 0 | 0.62 | 0.35 | 0.78 | 0.46 | 0.78 | 0.65 |

| | | | | | | | | | | | | | | | | |
|---|---|---|---|---|---|---|---|---|---|---|---|---|---|---|---|---|
| 9 | 22834082 | rs4369069 | -0.364 | -0.109 | -0.342 | 0.424 | -0.043 | 0 | 0 | 0 | 0.3 | 0.21 | 0.23 | 0.25 | 0.47 | 0.27 |
| 9 | 22836549 | rs7846858 | 0.046 | 0.147 | 0.517 | 0.857 | 0.018 | 0.135 | 0.135 | 0 | 0.86 | 0.68 | 0.89 | 0.66 | 0.88 | 0.85 |
| 9 | 22841425 | rs4602981 | 0.657 | 0.124 | 0.568 | 0.82 | 0.036 | 0.157 | 0.157 | 0 | 0.83 | 0.51 | 0.87 | 0.66 | 0.86 | 0.78 |
| 9 | 22847961 | rs4446815 | 1.143 | 0.069 | 0.346 | 0.717 | 0.004 | 0.017 | 0.017 | 0 | 0.55 | 0.34 | 0.67 | 0.41 | 0.45 | 0.53 |
| 9 | 22854439 | rs4977797 | 0.105 | 0.074 | 0.41 | 0.783 | 0.037 | 0.135 | 0.135 | 0 | 0.85 | 0.63 | 0.87 | 0.67 | 0.86 | 0.84 |
| 9 | 22854786 | rs10965476 | 0.294 | 0.403 | 0.174 | 0.698 | -0.019 | -0.004 | -0.004 | 0 | 0.28 | 0.16 | 0.17 | 0.23 | 0.33 | 0.25 |
| 9 | 22856899 | rs7857348 | 0.499 | 0.359 | 0.26 | 0.688 | -0.016 | -0.003 | -0.003 | 0 | 0.27 | 0.2 | 0.17 | 0.23 | 0.34 | 0.25 |
| 9 | 22857606 | rs9969855 | 0.022 | 0.378 | 0.309 | 0.708 | 0.025 | 0.164 | 0.164 | 0 | 0.85 | 0.59 | 0.87 | 0.67 | 0.88 | 0.84 |
| 9 | 22859129 | rs7041327 | 1.177 | 0.258 | -0.134 | 0.564 | -0.002 | 0.006 | 0.006 | 0 | 0.71 | 0.36 | 0.8 | 0.66 | 0.75 | 0.7 |
| 9 | 22862721 | rs13297845 | 1.893 | -0.145 | -0.001 | 0.718 | 0.016 | 0.002 | 0.002 | 0.03 | 0.39 | 0.2 | 0.61 | 0.41 | 0.37 | 0.43 |
| 9 | 22866134 | rs12004142 | -0.202 | -0.118 | -0.057 | 0.295 | -0.052 | 0.031 | 0.031 | 0.103 | 0.38 | 0.16 | 0.23 | 0.25 | 0.47 | 0.35 |
| 9 | 22870294 | rs4977798 | 1.966 | -0.073 | -0.154 | 0.369 | 0.012 | 0.057 | 0.057 | 0 | 0.37 | 0.18 | 0.55 | 0.22 | 0.34 | 0.29 |
| 9 | 22889619 | rs10811794 | 0.359 | 0.183 | 0.24 | 0.685 | 0.011 | 0.136 | 0.136 | 0 | 0.84 | 0.7 | 0.86 | 0.46 | 0.86 | 0.71 |
| 9 | 22890635 | rs2086011 | 0.891 | 0.035 | 0.152 | 0.471 | 0.013 | 0.088 | 0.088 | 0.063 | 0.49 | 0.35 | 0.64 | 0.23 | 0.42 | 0.4 |
| 9 | 22894794 | rs922364 | 0.362 | 0.055 | 0.183 | 0.973 | -0.014 | 0.129 | 0.129 | 0 | 0.85 | 0.73 | 0.87 | 0.49 | 0.89 | 0.75 |
| 9 | 22905600 | rs1381924 | -0.041 | -0.128 | 0.106 | 1.008 | -0.018 | 0.109 | 0.109 | 0 | 0.88 | 0.79 | 0.85 | 0.45 | 0.91 | 0.77 |
| 9 | 22906349 | rs996367 | 0.625 | 0.054 | 0.149 | 0.956 | 0.006 | 0.155 | 0.155 | 0 | 0.84 | 0.67 | 0.82 | 0.42 | 0.86 | 0.7 |
| 9 | 22906729 | rs10811800 | 1.575 | -0.209 | -1.178 | 0.524 | 0.023 | 0.237 | 0.237 | 0.246 | 0.46 | 0.13 | 0.56 | 0.17 | 0.4 | 0.34 |
| 9 | 22909348 | rs1381923 | -0.363 | -0.272 | -0.994 | 0.504 | -0.052 | 0.015 | 0.015 | 0.442 | 0.36 | 0.22 | 0.24 | 0.27 | 0.48 | 0.37 |
| 9 | 22909498 | rs1381922 | 1.31 | -0.245 | -1.1 | 0.506 | -0.013 | 0.2 | 0.2 | 0.339 | 0.47 | 0.09 | 0.64 | 0.21 | 0.45 | 0.37 |
| 9 | 22909814 | rs1381921 | 0.936 | -0.223 | -1.023 | 0.572 | -0.064 | 0.174 | 0.174 | 0.005 | 0.54 | 0.13 | 0.67 | 0.22 | 0.5 | 0.41 |
| 9 | 22911500 | rs10965517 | -0.175 | -0.04 | -0.939 | 0.57 | -0.045 | 0.01 | 0.01 | 0.065 | 0.33 | 0.16 | 0.2 | 0.24 | 0.42 | 0.32 |
| 9 | 22913604 | rs1954543 | 1.185 | 0.223 | -0.349 | 0.896 | -0.001 | 0.035 | 0.035 | 0 | 0.55 | 0.26 | 0.65 | 0.4 | 0.47 | 0.56 |
| 9 | 22913891 | rs1937037 | -0.412 | 0 | -0.76 | 0.996 | -0.066 | -0.027 | -0.027 | 0 | 0.31 | 0.12 | 0.26 | 0.61 | 0.46 | 0.35 |
| 9 | 22915652 | rs12343679 | 0.216 | -0.083 | -0.842 | 0.508 | 0.004 | -0.008 | -0.008 | 0 | 0.13 | 0.11 | 0.06 | 0.01 | 0.07 | 0.07 |
| 9 | 22916961 | rs10965520 | 0.205 | 0.124 | -1.246 | 0.908 | -0.029 | -0.067 | -0.067 | 0 | 0.1 | 0.01 | 0.09 | 0.32 | 0.1 | 0.14 |
| 9 | 22917331 | rs1463006 | 1.188 | 0.048 | -0.996 | 0.266 | -0.004 | 0.147 | 0.147 | 0 | 0.5 | 0.25 | 0.65 | 0.22 | 0.48 | 0.38 |
| 9 | 22917801 | rs10757373 | -0.52 | 0.726 | -1.413 | 0.765 | -0.053 | 0.009 | 0.009 | 0 | 0.33 | 0.13 | 0.29 | 0.65 | 0.45 | 0.37 |
| 9 | 22919588 | rs1463033 | -0.924 | 0.743 | -1.506 | 0.747 | -0.075 | 0.047 | 0.047 | 0.068 | 0.37 | 0.17 | 0.32 | 0.68 | 0.51 | 0.47 |
| 9 | 22925116 | rs2000341 | -0.334 | 0.761 | -1.251 | 0.962 | -0.046 | -0.012 | -0.012 | 0 | 0.3 | 0.17 | 0.25 | 0.62 | 0.43 | 0.38 |
| 9 | 22934328 | rs1463019 | 0.05 | 0.791 | -0.656 | 1.636 | -0.002 | 0.053 | 0.053 | 0 | 0.08 | 0.06 | 0.11 | 0.35 | 0.07 | 0.1 |
| 9 | 22937343 | rs1381919 | 0.954 | 0.531 | -0.419 | 1.371 | -0.037 | 0.031 | 0.031 | 0.193 | 0.74 | 0.39 | 0.75 | 0.61 | 0.84 | 0.74 |
| 9 | 22943509 | rs10965529 | 0.007 | 0.857 | -1.271 | 1.285 | -0.024 | -0.035 | -0.035 | 0 | 0.23 | 0.17 | 0.32 | 0.37 | 0.36 | 0.32 |
| 9 | 22947192 | rs12341716 | -1.321 | 0.235 | -0.308 | 1.943 | -0.001 | -0.01 | -0.01 | 0 | 0.22 | 0.15 | 0.22 | 0.33 | 0.21 | 0.25 |
| 9 | 22947998 | rs10511719 | -0.068 | 0.19 | -0.936 | 1.947 | -0.03 | -0.002 | -0.002 | 0 | 0.06 | 0.01 | 0.09 | 0.26 | 0.12 | 0.12 |
| 9 | 22956519 | rs1599698 | -0.275 | -0.349 | -0.504 | 1.276 | 0.001 | -0.001 | -0.001 | 0 | 0.92 | 0.89 | 0.93 | 0.96 | 0.93 | 0.93 |
| 9 | 22961073 | rs10965536 | -0.505 | -0.155 | -0.458 | 1.431 | -0.035 | -0.064 | -0.064 | 0.298 | 0.1 | 0.01 | 0.12 | 0.28 | 0.11 | 0.16 |
| 9 | 22966185 | rs10965542 | 0.679 | -0.374 | 0.059 | 0.667 | 0 | -0.001 | -0.001 | 0.267 | 0.73 | 0.27 | 0.75 | 0.68 | 0.83 | 0.58 |
| 9 | 22967726 | rs10757377 | 0.604 | -0.31 | 0.058 | 0.692 | -0.013 | 0.002 | 0.002 | 1.668 | 0.73 | 0.34 | 0.77 | 0.68 | 0.84 | 0.58 |
| 9 | 22972837 | rs7020651 | -1.014 | 0.17 | 0.409 | 0.662 | 0.019 | -0.004 | -0.004 | 0.456 | 0.06 | 0.18 | 0.06 | 0.07 | 0.09 | 0.05 |
| 9 | 22982002 | rs7867193 | 0.074 | 0.061 | 0.012 | 0.102 | -0.004 | -0.033 | -0.033 | 0.654 | 0.58 | 0.45 | 0.47 | 0.21 | 0.63 | 0.44 |
| 9 | 22985180 | rs10811812 | -0.839 | -0.33 | -0.226 | -0.335 | -0.011 | -0.029 | -0.029 | 0 | 0.38 | 0.51 | 0.4 | 0.8 | 0.35 | 0.65 |
| 9 | 22986069 | rs10965553 | 0.636 | -0.223 | -0.74 | -0.308 | 0.006 | 0.024 | 0.024 | 0 | 0.16 | 0.08 | 0.08 | 0.03 | 0.13 | 0.16 |
| 9 | 22987003 | rs958562 | 0.671 | -0.247 | -0.906 | 0.173 | -0.018 | -0.129 | -0.129 | 0.301 | 0.16 | 0.01 | 0.25 | 0.51 | 0.19 | 0.3 |
| 9 | 22987088 | rs10811815 | 0.367 | -0.267 | -0.858 | -0.018 | -0.002 | -0.047 | -0.047 | 1.715 | 0.18 | 0.29 | 0.11 | 0.1 | 0.18 | 0.14 |
| 9 | 22987182 | rs1350996 | 0.817 | 0 | -1.51 | -0.032 | -0.102 | 0.096 | 0.096 | 0 | 0.34 | 0.04 | 0.32 | 0.11 | 0.41 | 0.2 |
| 9 | 22988063 | rs931026 | 0.28 | 0.23 | -0.413 | 0.369 | 0.001 | -0.052 | -0.052 | 0.509 | 0.7 | 0.59 | 0.75 | 0.91 | 0.77 | 0.82 |
| 9 | 23003005 | rs1023913 | -1.19 | -0.32 | 0.334 | 0.681 | -0.023 | -0.086 | -0.086 | 0.092 | 0.13 | 0.57 | 0.07 | 0 | 0.05 | 0.07 |
| 9 | 23012833 | rs1463009 | 1.625 | 0.642 | 0.518 | 0.216 | 0.006 | -0.062 | -0.062 | 0 | 0.78 | 0.24 | 0.81 | 0.98 | 0.76 | 0.81 |
| 9 | 23017610 | rs10123865 | 1.697 | 0.575 | 0.376 | 0.071 | -0.008 | 0.01 | 0.01 | 1.566 | 0.74 | 0.19 | 0.79 | 0.98 | 0.73 | 0.8 |
| 9 | 23018731 | rs1576676 | 2.143 | 0.497 | 0.189 | 0.337 | -0.003 | -0.002 | -0.002 | 0.963 | 0.71 | 0.22 | 0.75 | 0.73 | 0.69 | 0.78 |
| 9 | 23023866 | rs11999590 | -0.762 | 0.474 | 0.941 | 0.258 | -0.018 | -0.051 | -0.051 | 0.016 | 0.09 | 0.28 | 0.05 | 0 | 0.04 | 0.06 |
| 9 | 23024202 | rs7859974 | 0.937 | 0.251 | -0.706 | -0.083 | -0.163 | -0.18 | -0.18 | 0 | 0.34 | 0.03 | 0.49 | 0.53 | 0.42 | 0.39 |
| 9 | 23039269 | rs4977816 | 1.498 | 0.572 | -0.959 | -1.23 | -0.081 | -0.164 | -0.164 | 0 | 0.31 | 0.03 | 0.43 | 0.54 | 0.33 | 0.41 |
| 9 | 23042805 | rs10965572 | 0.045 | 0.252 | -0.618 | -2.808 | -0.004 | -0.051 | -0.051 | 0 | 0.32 | 0.35 | 0.17 | 0.03 | 0.26 | 0.21 |
| 9 | 23050746 | rs1331597 | 0.424 | 0.241 | -0.126 | -2.683 | 0 | -0.002 | -0.002 | 0 | 0.33 | 0.28 | 0.28 | 0.05 | 0.34 | 0.22 |
| 9 | 23051397 | rs7019344 | 0.066 | 0.438 | -0.296 | -2.503 | 0.012 | -0.04 | -0.04 | 0 | 0.86 | 0.82 | 0.86 | 0.99 | 0.77 | 0.88 |
| 9 | 23055362 | rs2383251 | -0.44 | 0.085 | -0.037 | -2.29 | 0.01 | 0.014 | 0.014 | 0.002 | 0.43 | 0.6 | 0.33 | 0.08 | 0.36 | 0.34 |
| 9 | 23059458 | rs12375567 | -1.378 | 0.735 | -0.141 | -1.686 | -0.003 | 0.011 | 0.011 | 0 | 0.13 | 0.09 | 0.09 | 0.07 | 0.14 | 0.2 |
| 9 | 23059968 | rs10965577 | -0.013 | 0.773 | -0.225 | -1.684 | -0.038 | -0.011 | -0.011 | 0 | 0.06 | 0.01 | 0.06 | 0.06 | 0.14 | 0.09 |
| 9 | 23060945 | rs10965580 | -1.023 | 0.724 | 0.127 | -1.673 | 0.006 | -0.029 | -0.029 | 0 | 0.15 | 0.23 | 0.11 | 0.08 | 0.16 | 0.2 |
| 9 | 23062503 | rs10965583 | -0.998 | 0.659 | 0.206 | -1.681 | -0.016 | -0.051 | -0.051 | 0.015 | 0.19 | 0.36 | 0.13 | 0.08 | 0.14 | 0.2 |
| 9 | 23064815 | rs12375524 | -0.979 | 0.664 | 0.189 | -1.687 | -0.015 | -0.055 | -0.055 | 0 | 0.19 | 0.36 | 0.13 | 0.07 | 0.14 | 0.21 |
| 9 | 23073339 | rs1416505 | 0.63 | 0.553 | -0.88 | -2.257 | 0.001 | 0.096 | 0.096 | 0.231 | 0.44 | 0.3 | 0.63 | 0.84 | 0.46 | 0.71 |
| 9 | 23073969 | rs10511716 | 0.376 | 0.463 | -0.837 | -2.454 | -0.002 | 0.1 | 0.1 | 0 | 0.33 | 0.12 | 0.21 | 0.12 | 0.28 | 0.18 |
| 9 | 23074207 | rs1416503 | 0.605 | 0.393 | -0.277 | -2.241 | 0.017 | -0.094 | -0.094 | 0.205 | 0.77 | 0.48 | 0.84 | 0.96 | 0.74 | 0.91 |
| 9 | 23074432 | rs2225965 | 1.2 | 0.472 | -0.691 | -2.22 | 0 | -0.018 | -0.018 | 0 | 0.7 | 0.31 | 0.68 | 0.9 | 0.65 | 0.65 |
| 9 | 23078402 | rs4977596 | 1.405 | 0.267 | -0.65 | -2.058 | 0.008 | 0.096 | 0.096 | 0 | 0.69 | 0.16 | 0.78 | 0.97 | 0.69 | 0.75 |
| 9 | 23081217 | rs10811830 | 0.635 | 0.627 | 0.125 | -1.627 | 0.059 | -0.082 | -0.082 | 0.145 | 0.82 | 0.49 | 0.83 | 0.97 | 0.75 | 0.89 |

| | | | | | | | | | | | | | | | | |
|---|---|---|---|---|---|---|---|---|---|---|---|---|---|---|---|---|
| 9 | 23081882 | rs12352937 | -0.541 | 0.052 | -0.227 | -1.397 | 0.016 | 0.04 | 0.04 | 0 | 0.45 | 0.32 | 0.32 | 0.13 | 0.36 | 0.49 |
| 9 | 23082438 | rs10811831 | -0.183 | 0.161 | -1.129 | -1.416 | 0.01 | 0.212 | 0.212 | 0.234 | 0.4 | 0.1 | 0.28 | 0.08 | 0.34 | 0.32 |
| 9 | 23087926 | rs9792672 | 1.415 | 0.149 | 0.384 | -0.396 | -0.026 | 0.039 | 0.039 | 1.784 | 0.9 | 0.65 | 0.9 | 0.89 | 0.97 | 0.89 |
| 9 | 23103266 | rs1934115 | 1.573 | 0.384 | 1.087 | -1.089 | -0.017 | -0.137 | -0.137 | 0 | 0.8 | 0.57 | 0.84 | 0.98 | 0.79 | 0.88 |
| 9 | 23121911 | rs4977824 | 2.09 | -0.059 | 0.855 | -1.29 | 0.028 | 0.032 | 0.032 | 2.667 | 0.8 | 0.17 | 0.84 | 0.9 | 0.92 | 0.83 |
| 9 | 23127310 | rs1538492 | 0.742 | 0.009 | 1.719 | -1.254 | 0.032 | 0.023 | 0.023 | 0 | 0.93 | 0.7 | 0.96 | 0.99 | 0.97 | 0.94 |
| 9 | 23127562 | rs4977826 | 2.375 | -0.105 | 1.501 | -1.274 | 0.038 | 0.035 | 0.035 | 3.324 | 0.81 | 0.13 | 0.85 | 0.9 | 0.93 | 0.85 |
| 9 | 23127648 | rs10757391 | 2.417 | -0.052 | 1.59 | -1.33 | 0.008 | 0.05 | 0.05 | 0.57 | 0.76 | 0.18 | 0.76 | 0.89 | 0.76 | 0.82 |
| 9 | 23128281 | rs10757392 | 2.194 | -0.039 | 1.702 | -1.358 | 0.002 | -0.01 | -0.01 | 0.981 | 0.78 | 0.27 | 0.84 | 0.98 | 0.76 | 0.86 |
| 9 | 23132263 | rs10965655 | 1.427 | 0.259 | 0.279 | -2.312 | -0.043 | 0.251 | 0.251 | 0.246 | 0.48 | 0.11 | 0.62 | 0.83 | 0.46 | 0.47 |
| 9 | 23132496 | rs10811853 | 1.588 | 0.081 | 0.848 | -2.361 | -0.005 | -0.057 | -0.057 | 0 | 0.77 | 0.38 | 0.84 | 0.96 | 0.76 | 0.86 |
| 9 | 23132614 | rs10757394 | 0.249 | 0.054 | 0.974 | -2.341 | 0.016 | 0.012 | 0.012 | 0.029 | 0.93 | 0.71 | 0.96 | 0.98 | 0.97 | 0.93 |
| 9 | 23137966 | rs10965663 | 0.41 | 0.431 | 0.842 | -2.359 | -0.081 | 0.131 | 0.131 | 0 | 0.26 | 0.02 | 0.13 | 0.05 | 0.26 | 0.32 |
| 9 | 23138534 | rs10811855 | 1.172 | 0.398 | 0.982 | -2.339 | -0.038 | 0.021 | 0.021 | 0.659 | 0.87 | 0.6 | 0.88 | 0.89 | 0.94 | 0.84 |
| 9 | 23141807 | rs10441708 | 0.374 | 0.407 | 1.035 | -2.338 | -0.001 | -0.004 | -0.004 | 0.139 | 0.92 | 0.71 | 0.96 | 0.98 | 0.97 | 0.92 |
| 9 | 23142396 | rs7873939 | 1.623 | -0.088 | 0.278 | -2.668 | -0.066 | 0.2 | 0.2 | 0.265 | 0.45 | 0.18 | 0.63 | 0.85 | 0.5 | 0.49 |
| 9 | 23156922 | rs4977602 | 2.067 | 0.38 | 0.295 | -2.155 | 0.081 | 0.073 | 0.073 | 5.427 | 0.76 | 0.15 | 0.84 | 0.9 | 0.93 | 0.82 |
| 9 | 23161023 | rs4007613 | -1.71 | 0.491 | 0.761 | -2.232 | 0.04 | -0.128 | -0.128 | 0.048 | 0.17 | 0.45 | 0.15 | 0.01 | 0.21 | 0.08 |
| 9 | 23161189 | rs2789476 | 0.021 | 0.622 | 0.642 | -2.327 | -0.009 | -0.057 | -0.057 | 0 | 0.65 | 0.66 | 0.84 | 0.93 | 0.76 | 0.65 |
| 9 | 23162207 | rs1591842 | 1.324 | 0.59 | 0.619 | -2.321 | 0.094 | 0.278 | 0.278 | 1.366 | 0.51 | 0.12 | 0.74 | 0.86 | 0.72 | 0.55 |
| 9 | 23166743 | rs7871386 | 0.073 | 0.708 | 0.647 | -2.288 | -0.003 | -0.056 | -0.056 | 0.197 | 0.65 | 0.65 | 0.82 | 0.93 | 0.73 | 0.65 |
| 9 | 23171045 | rs7869437 | -1.362 | 0.555 | 0.551 | -2.029 | 0.062 | -0.054 | -0.054 | 1.019 | 0.09 | 0.27 | 0.13 | 0.01 | 0.2 | 0.07 |
| 9 | 23172096 | rs1111819 | 1.25 | 0.294 | -0.398 | -2.039 | 0.071 | 0.215 | 0.215 | 1.344 | 0.56 | 0.13 | 0.73 | 0.84 | 0.72 | 0.59 |
| 9 | 23176101 | rs7033378 | 1.162 | 0.283 | 0.022 | -2.379 | 0.054 | 0.119 | 0.119 | 0.138 | 0.56 | 0.31 | 0.64 | 0.83 | 0.7 | 0.56 |
| 9 | 23181445 | rs1227466 | 0.741 | 0.77 | 1.068 | -2.584 | 0.01 | -0.102 | -0.102 | 0 | 0.63 | 0.52 | 0.62 | 0.89 | 0.6 | 0.64 |
| 9 | 23182825 | rs1227463 | 0.19 | 0.688 | 1.088 | -2.628 | 0.006 | -0.007 | -0.007 | 0 | 0.72 | 0.76 | 0.74 | 0.9 | 0.81 | 0.68 |
| 9 | 23183078 | rs4610838 | -0.68 | 0.577 | 1.144 | -2.632 | -0.016 | 0.02 | 0.02 | 3.634 | 0.1 | 0.43 | 0.04 | 0.08 | 0.01 | 0.1 |
| 9 | 23184130 | rs1227458 | 1.157 | 0.629 | 1.061 | -2.628 | 0.016 | 0.057 | 0.057 | 0.003 | 0.61 | 0.35 | 0.66 | 0.83 | 0.72 | 0.56 |
| 9 | 23184533 | rs1227456 | 0.892 | 0.594 | 1.065 | -2.625 | 0.034 | 0.04 | 0.04 | 0.204 | 0.64 | 0.36 | 0.7 | 0.85 | 0.81 | 0.59 |
| 9 | 23186453 | rs10757399 | -0.132 | 0.649 | 1.165 | -2.546 | 0.074 | -0.036 | -0.036 | 0 | 0.92 | 0.75 | 0.92 | 0.99 | 0.88 | 0.97 |
| 9 | 23191038 | rs10811866 | -1.589 | -0.017 | 0.819 | -2.782 | -0.004 | -0.046 | -0.046 | 0 | 0.45 | 0.42 | 0.35 | 0.15 | 0.31 | 0.45 |
| 9 | 23198470 | rs7849833 | -1.721 | -0.117 | 0.787 | -2.839 | 0 | -0.039 | -0.039 | 0 | 0.48 | 0.48 | 0.39 | 0.17 | 0.38 | 0.48 |
| 9 | 23199445 | rs10811867 | -2.355 | -0.116 | 0.855 | -2.862 | -0.001 | 0.16 | 0.16 | 0 | 0.54 | 0.73 | 0.48 | 0.19 | 0.47 | 0.55 |
| 9 | 23200026 | rs1339160 | -1.495 | -0.133 | 0.817 | -2.84 | -0.007 | -0.055 | -0.055 | 0.635 | 0.46 | 0.47 | 0.28 | 0.17 | 0.35 | 0.45 |
| 9 | 23200174 | rs10811869 | -0.842 | -0.156 | 0.822 | -2.852 | 0.001 | -0.01 | -0.01 | 0 | 0.36 | 0.33 | 0.27 | 0.15 | 0.34 | 0.43 |
| 9 | 23204090 | rs10811872 | -1.067 | -0.242 | 0.84 | -2.848 | 0 | -0.022 | -0.022 | 0 | 0.38 | 0.38 | 0.29 | 0.16 | 0.37 | 0.46 |
| 9 | 23207852 | rs10120394 | -1.617 | -0.255 | 0.56 | -2.851 | 0.103 | 0.155 | 0.155 | 1.126 | 0.47 | 0.71 | 0.27 | 0.18 | 0.27 | 0.39 |
| 9 | 23216803 | rs10965704 | -1.569 | -0.387 | 0.595 | -2.847 | -0.007 | -0.049 | -0.049 | 0 | 0.44 | 0.46 | 0.27 | 0.15 | 0.31 | 0.48 |
| 9 | 23222186 | rs1339140 | -1.135 | -0.674 | 0.19 | -2.703 | 0.018 | 0.012 | 0.012 | 0 | 0.38 | 0.61 | 0.24 | 0.15 | 0.25 | 0.45 |
| 9 | 23226243 | rs9298846 | -1.458 | -0.604 | 0.159 | -2.568 | 0.024 | 0.069 | 0.069 | 0 | 0.43 | 0.65 | 0.33 | 0.15 | 0.33 | 0.48 |
| 9 | 23242643 | rs1578104 | -1.976 | -0.952 | -0.255 | -2.541 | 0.063 | 0.157 | 0.157 | 3.547 | 0.41 | 0.82 | 0.31 | 0.13 | 0.31 | 0.4 |
| 9 | 23245863 | rs17769769 | -0.813 | -1.117 | -0.258 | -2.64 | -0.001 | 0.145 | 0.145 | 0.451 | 0.25 | 0.04 | 0.19 | 0.05 | 0.21 | 0.23 |
| 9 | 23246526 | rs1934110 | -1.844 | -1.093 | -0.273 | -2.538 | 0.083 | 0.114 | 0.114 | 4.19 | 0.37 | 0.8 | 0.23 | 0.13 | 0.21 | 0.37 |
| 9 | 23254622 | rs7871692 | -2.023 | -1.117 | -0.506 | -2.507 | 0.086 | 0.216 | 0.216 | 3.181 | 0.39 | 0.89 | 0.29 | 0.13 | 0.26 | 0.38 |
| 9 | 23259295 | rs7848673 | -1.155 | -1.134 | 0.519 | -2.376 | -0.079 | -0.083 | -0.083 | 0 | 0.13 | 0.48 | 0.04 | 0.02 | 0.02 | 0.1 |
| 9 | 23262261 | rs7868316 | -0.988 | -1.137 | 0.754 | -2.377 | -0.069 | -0.072 | -0.072 | 1.198 | 0.12 | 0.37 | 0.03 | 0.02 | 0.02 | 0.1 |
| 9 | 23262705 | rs6475722 | -2.119 | -1.192 | 0.866 | -2.463 | -0.016 | -0.027 | -0.027 | 0 | 0.18 | 0.77 | 0.06 | 0.02 | 0.03 | 0.12 |
| 9 | 23267111 | rs2225964 | 1.802 | -1.439 | 0.499 | -2.63 | -0.015 | -0.084 | -0.084 | 0 | 0.61 | 0.45 | 0.73 | 0.84 | 0.73 | 0.64 |
| 9 | 23274023 | rs7022904 | 1.674 | -1.486 | 0.668 | -2.642 | -0.031 | -0.092 | -0.092 | 0.003 | 0.66 | 0.44 | 0.77 | 0.84 | 0.79 | 0.69 |
| 9 | 23277704 | rs7864583 | -0.708 | -1.488 | 0.698 | -2.67 | -0.007 | -0.057 | -0.057 | 0.061 | 0.2 | 0.28 | 0.16 | 0.03 | 0.12 | 0.14 |
| 9 | 23285981 | rs6475727 | -1.231 | -1.404 | 0.411 | -2.454 | -0.005 | -0.039 | -0.039 | 0 | 0.31 | 0.33 | 0.22 | 0.11 | 0.22 | 0.23 |
| 9 | 23288364 | rs10757408 | 1.87 | -1.511 | -0.006 | -2.433 | 0.082 | 0.213 | 0.213 | 0.295 | 0.55 | 0.11 | 0.73 | 0.8 | 0.75 | 0.6 |
| 9 | 23289501 | rs10757409 | 1.289 | -1.351 | 0.178 | -2.395 | -0.015 | -0.082 | -0.082 | 0 | 0.62 | 0.51 | 0.73 | 0.84 | 0.74 | 0.69 |
| 9 | 23290141 | rs1934107 | 1.073 | -1.349 | 0.185 | -2.325 | -0.019 | -0.073 | -0.073 | 0.816 | 0.65 | 0.53 | 0.75 | 0.84 | 0.76 | 0.72 |
| 9 | 23297808 | rs7047280 | 1.504 | -1.162 | -0.546 | -1.949 | 0.017 | 0.108 | 0.108 | 0 | 0.48 | 0.23 | 0.51 | 0.7 | 0.63 | 0.47 |
| 9 | 23301612 | rs10757411 | 1.531 | -1.211 | -0.633 | -2.042 | 0.087 | 0.174 | 0.174 | 0.016 | 0.51 | 0.17 | 0.54 | 0.73 | 0.69 | 0.49 |
| 9 | 23307283 | rs10738660 | 1.699 | -1.192 | -0.599 | -2.014 | 0.01 | 0.127 | 0.127 | 0 | 0.47 | 0.17 | 0.53 | 0.71 | 0.65 | 0.47 |
| 9 | 23307377 | rs1544161 | 0.201 | -1.282 | -0.638 | -2.048 | -0.048 | 0.129 | 0.129 | 0 | 0.71 | 0.95 | 0.76 | 0.87 | 0.75 | 0.71 |
| 9 | 23313019 | rs10965762 | -0.862 | -1.601 | -0.081 | -2.109 | -0.087 | 0.032 | 0.032 | 0 | 0.17 | 0.47 | 0.2 | 0.16 | 0.06 | 0.14 |
| 9 | 23316148 | rs10965764 | -0.72 | -1.553 | -0.195 | -2.092 | -0.076 | 0.031 | 0.031 | 0 | 0.16 | 0.48 | 0.2 | 0.15 | 0.07 | 0.14 |
| 9 | 23320406 | rs10965766 | -0.877 | -1.543 | -0.217 | -2.088 | -0.071 | 0.013 | 0.013 | 0 | 0.17 | 0.53 | 0.2 | 0.16 | 0.07 | 0.16 |
| 9 | 23322467 | rs10811886 | -1.129 | -1.544 | -0.303 | -2.17 | -0.056 | -0.012 | -0.012 | 0.407 | 0.19 | 0.64 | 0.21 | 0.14 | 0.07 | 0.16 |
| 9 | 23325027 | rs7873743 | -1.165 | -1.464 | -0.397 | -2.15 | -0.029 | -0.017 | -0.017 | 0.712 | 0.22 | 0.64 | 0.21 | 0.14 | 0.09 | 0.2 |
| 9 | 23325084 | rs10811888 | 1.08 | -0.994 | -0.388 | -1.928 | 0.051 | 0.145 | 0.145 | 0 | 0.48 | 0.22 | 0.54 | 0.73 | 0.67 | 0.49 |
| 9 | 23341848 | rs16907121 | -1.287 | -0.275 | 0.198 | -1.853 | 0.046 | -0.047 | -0.047 | 0 | 0.09 | 0.29 | 0.11 | 0.01 | 0.12 | 0.05 |
| 9 | 23342211 | rs17780168 | 0.654 | -0.264 | -0.328 | -1.064 | -0.014 | 0.01 | 0.01 | 0 | 0.19 | 0.2 | 0.12 | 0.3 | 0.14 | 0.09 |
| 9 | 23343995 | rs1929171 | -0.454 | -0.498 | -0.348 | -1.339 | 0.005 | 0.007 | 0.007 | 0 | 0.66 | 0.71 | 0.74 | 0.85 | 0.75 | 0.63 |
| 9 | 23345347 | rs11794152 | 1.75 | -0.521 | -0.411 | -1.078 | -0.079 | -0.117 | -0.117 | 0.279 | 0.31 | 0.09 | 0.36 | 0.43 | 0.43 | 0.41 |
| 9 | 23353893 | rs7466754 | 0.312 | -0.568 | -0.195 | -1.265 | -0.054 | 0.027 | 0.027 | 0 | 0.25 | 0.37 | 0.29 | 0.34 | 0.18 | 0.1 |

| | | | | | | | | | | | | | | | | |
|---|---|---|---|---|---|---|---|---|---|---|---|---|---|---|---|---|
| 9 | 23354494 | rs10757418 | 0.028 | -0.409 | -0.206 | -1.313 | 0.003 | -0.073 | -0.073 | 0 | 0.61 | 0.54 | 0.66 | 0.84 | 0.63 | 0.59 |
| 9 | 23358448 | rs7029374 | -0.577 | -0.861 | -0.462 | -1.175 | 0.002 | -0.013 | -0.013 | 0 | 0.69 | 0.7 | 0.76 | 0.85 | 0.8 | 0.68 |
| 9 | 23378488 | rs1360380 | 0.295 | -0.152 | -0.233 | -0.793 | -0.001 | -0.004 | -0.004 | 0 | 0.4 | 0.37 | 0.46 | 0.47 | 0.59 | 0.49 |
| 9 | 23379719 | rs1360382 | 0.959 | 0.879 | -0.886 | -1.102 | -0.003 | -0.039 | -0.039 | 0 | 0.32 | 0.12 | 0.32 | 0.52 | 0.32 | 0.43 |
| 9 | 23384589 | rs1330924 | -0.938 | 0.757 | -0.86 | -1.532 | -0.009 | -0.016 | -0.016 | 0 | 0.59 | 0.61 | 0.68 | 0.66 | 0.68 | 0.63 |
| 9 | 23386537 | rs1888540 | -0.606 | 0.929 | -0.76 | -1.547 | 0.005 | 0.012 | 0.012 | 0 | 0.57 | 0.46 | 0.65 | 0.65 | 0.63 | 0.59 |
| 9 | 23386738 | rs1330159 | -1.198 | 1.056 | -0.865 | -1.598 | -0.02 | 0.029 | 0.029 | 0 | 0.29 | 0.28 | 0.34 | 0.09 | 0.34 | 0.2 |
| 9 | 23387019 | rs1330164 | -0.241 | 1.158 | -0.835 | -1.587 | 0.019 | -0.013 | -0.013 | 0.52 | 0.14 | 0.2 | 0.15 | 0.05 | 0.27 | 0.16 |
| 9 | 23388570 | rs17836155 | 1.856 | 1.06 | -1.028 | -1.502 | -0.007 | -0.049 | -0.049 | 0.54 | 0.26 | 0.08 | 0.3 | 0.6 | 0.3 | 0.39 |
| 9 | 23400606 | rs10965837 | 1.424 | 0.982 | -0.968 | -1.443 | -0.01 | -0.039 | -0.039 | 2.109 | 0.3 | 0.09 | 0.33 | 0.59 | 0.33 | 0.39 |
| 9 | 23402710 | rs4409483 | -0.479 | 0.922 | -0.953 | -1.588 | 0.027 | 0.064 | 0.064 | 0 | 0.52 | 0.42 | 0.63 | 0.64 | 0.62 | 0.58 |
| 9 | 23403315 | rs10811922 | 0.574 | 1.045 | -1.019 | -1.575 | 0.014 | 0.025 | 0.025 | 0 | 0.37 | 0.26 | 0.49 | 0.6 | 0.35 | 0.42 |
| 9 | 23404131 | rs10811923 | -0.522 | 0.92 | -0.822 | -1.578 | -0.004 | -0.015 | -0.015 | 0.053 | 0.52 | 0.56 | 0.63 | 0.64 | 0.61 | 0.56 |
| 9 | 23407386 | rs7035046 | 0.171 | 0.856 | -0.96 | -1.626 | 0.037 | 0.064 | 0.064 | 1.394 | 0.43 | 0.27 | 0.51 | 0.63 | 0.66 | 0.59 |
| 9 | 23407435 | rs1581979 | -0.384 | 0.869 | -0.995 | -1.668 | 0.068 | 0.124 | 0.124 | 0.399 | 0.51 | 0.31 | 0.66 | 0.65 | 0.66 | 0.58 |
| 9 | 23408637 | rs4617240 | 0.459 | 1.113 | -0.869 | -1.49 | -0.001 | -0.001 | -0.001 | 0 | 0.38 | 0.4 | 0.37 | 0.61 | 0.38 | 0.44 |
| 9 | 23416920 | rs10511722 | -0.874 | 1.374 | -0.783 | -1.384 | 0.092 | -0.051 | -0.051 | 0 | 0.17 | 0.39 | 0.27 | 0.05 | 0.3 | 0.2 |
| 9 | 23420202 | rs10511724 | 0.678 | 1.44 | -1.428 | -1.384 | -0.041 | 0.038 | 0.038 | 0 | 0.08 | 0.04 | 0.21 | 0.02 | 0.22 | 0.06 |
| 9 | 23422031 | rs2210293 | 0.552 | 0.478 | -1.623 | -1.497 | -0.013 | 0.064 | 0.064 | 0.119 | 0.41 | 0.11 | 0.39 | 0.63 | 0.42 | 0.54 |
| 9 | 23423892 | rs10811927 | -0.038 | -0.228 | -0.785 | -1.195 | -0.009 | -0.014 | -0.014 | 0.099 | 0.58 | 0.58 | 0.72 | 0.66 | 0.71 | 0.68 |
| 9 | 23425710 | rs7873308 | -0.56 | -0.142 | -0.948 | -1.079 | 0.01 | 0.04 | 0.04 | 0.093 | 0.53 | 0.64 | 0.49 | 0.26 | 0.56 | 0.48 |
| 9 | 23427158 | rs7023393 | 0.587 | -0.225 | -0.182 | -0.938 | 0.011 | -0.075 | -0.075 | 0 | 0.27 | 0.31 | 0.24 | 0.03 | 0.36 | 0.34 |
| 9 | 23430020 | rs1541106 | -0.542 | -0.359 | -0.229 | -0.902 | 0 | -0.005 | -0.005 | 0 | 0.4 | 0.58 | 0.38 | 0.25 | 0.4 | 0.4 |
| 9 | 23431906 | rs1330927 | 0.546 | -0.27 | -0.226 | -0.955 | 0.007 | -0.071 | -0.071 | 0.792 | 0.29 | 0.31 | 0.23 | 0.03 | 0.36 | 0.31 |
| 9 | 23437895 | rs7030403 | -0.795 | -0.717 | 0.371 | 0.052 | -0.054 | 0.092 | 0.092 | 0 | 0.16 | 0.43 | 0.18 | 0.29 | 0.05 | 0.14 |
| 9 | 23442387 | rs7043141 | -0.118 | -0.806 | -0.073 | 0.228 | -0.051 | 0.031 | 0.031 | 0.127 | 0.2 | 0.39 | 0.21 | 0.26 | 0.11 | 0.2 |
| 9 | 23442605 | rs10738675 | 0.046 | 0.45 | 0.19 | 1.324 | -0.025 | -0.043 | -0.043 | 0 | 0.83 | 0.89 | 0.7 | 0.58 | 0.74 | 0.64 |
| 9 | 23463060 | rs10115057 | -0.366 | -0.021 | 0.676 | 0.463 | 0.006 | -0.025 | -0.025 | 0 | 0.08 | 0.4 | 0.05 | 0 | 0.03 | 0.04 |
| 9 | 23479588 | rs404479 | 0.138 | 0.079 | -0.261 | 0.46 | -0.112 | -0.077 | -0.077 | 1.154 | 0.7 | 0.92 | 0.62 | 0.4 | 0.59 | 0.56 |
| 9 | 23485158 | rs9696502 | 0.435 | 0.047 | 0.443 | 0.602 | -0.011 | -0.032 | -0.032 | 0 | 0.06 | 0.14 | 0.02 | 0 | 0.03 | 0.04 |
| 9 | 23488018 | rs10811933 | 0.719 | 0.033 | 0.194 | 0.482 | -0.002 | -0.02 | -0.02 | 0 | 0.44 | 0.5 | 0.42 | 0.34 | 0.43 | 0.44 |
| 9 | 23498139 | rs12338776 | 0.043 | -0.1 | -0.11 | 1.507 | -0.021 | -0.094 | -0.094 | 0 | 0.17 | 0.02 | 0.09 | 0.33 | 0.2 | 0.32 |
| 9 | 23513235 | rs274915 | 0.292 | -0.041 | -0.147 | 1.461 | -0.029 | -0.112 | -0.112 | 1.269 | 0.64 | 0.94 | 0.64 | 0.5 | 0.66 | 0.57 |
| 9 | 23516235 | rs17836259 | -0.197 | 0.089 | 0.576 | 1.407 | -0.041 | -0.13 | -0.13 | 1.297 | 0.16 | 0.02 | 0.17 | 0.42 | 0.2 | 0.33 |
| 9 | 23516288 | rs10965890 | -0.431 | -0.078 | 1.393 | 0.592 | -0.013 | 0.046 | 0.046 | 0 | 0.08 | 0.16 | 0.17 | 0.17 | 0.05 | 0.16 |
| 9 | 23517331 | rs6475745 | -1.297 | -0.045 | 1.723 | 0.557 | 0.001 | 0.095 | 0.095 | 0 | 0.11 | 0.44 | 0.19 | 0.16 | 0.06 | 0.18 |
| 9 | 23523491 | rs175631 | -1.43 | -0.103 | 1.31 | 0.547 | -0.018 | 0.075 | 0.075 | 0 | 0.12 | 0.48 | 0.21 | 0.16 | 0.07 | 0.17 |
| 9 | 23529626 | rs156396 | -0.525 | -0.015 | 0.607 | 0.521 | -0.005 | 0.024 | 0.024 | 0.382 | 0.44 | 0.85 | 0.45 | 0.36 | 0.44 | 0.38 |
| 9 | 23530791 | rs381813 | -0.339 | 0.097 | 0.573 | 0.545 | -0.001 | -0.002 | -0.002 | 0 | 0.42 | 0.61 | 0.43 | 0.3 | 0.43 | 0.37 |
| 9 | 23532809 | rs7865248 | -0.508 | 0.301 | 0.286 | 0.817 | -0.002 | 0.016 | 0.016 | 0 | 0.08 | 0.1 | 0.18 | 0.17 | 0.05 | 0.14 |
| 9 | 23542707 | rs655258 | -0.377 | 0.122 | -0.062 | 0.68 | 0.002 | 0.014 | 0.014 | 0.019 | 0.5 | 0.45 | 0.51 | 0.4 | 0.55 | 0.46 |
| 9 | 23545229 | rs274958 | -1.044 | -0.01 | 0.197 | 1.075 | -0.017 | 0.062 | 0.062 | 0 | 0.08 | 0.18 | 0.2 | 0.18 | 0.06 | 0.12 |
| 9 | 23557134 | rs11789575 | -0.817 | -0.124 | 0.083 | 0.436 | -0.028 | -0.049 | -0.049 | 0.507 | 0.16 | 0.28 | 0.09 | 0.04 | 0.1 | 0.15 |
| 9 | 23557227 | rs10511729 | -0.829 | 0.155 | 0.068 | 0.781 | -0.005 | 0.012 | 0.012 | 0 | 0.34 | 0.48 | 0.25 | 0.61 | 0.31 | 0.49 |
| 9 | 23569119 | rs12552015 | -0.476 | 0.365 | -0.249 | -0.144 | 0.004 | 0.091 | 0.091 | 0.404 | 0.37 | 0.2 | 0.28 | 0.09 | 0.33 | 0.37 |
| 9 | 23569870 | rs17698074 | -0.402 | 0.388 | -0.26 | -0.131 | 0.016 | 0.087 | 0.087 | 0.403 | 0.36 | 0.18 | 0.27 | 0.09 | 0.3 | 0.37 |
| 9 | 23570688 | rs10116397 | 0.238 | -0.133 | -0.176 | 0.319 | 0.002 | -0.002 | -0.002 | 0.234 | 0.11 | 0.09 | 0.07 | 0.19 | 0.06 | 0.16 |
| 9 | 23575805 | rs7852159 | -0.503 | 0.417 | -0.258 | -0.373 | -0.001 | -0.045 | -0.045 | 0 | 0.53 | 0.49 | 0.35 | 0.13 | 0.41 | 0.45 |
| 9 | 23585350 | rs274946 | -0.192 | 0.481 | -0.239 | -0.135 | 0 | -0.014 | -0.014 | 0 | 0.52 | 0.45 | 0.39 | 0.16 | 0.51 | 0.47 |
| 9 | 23599604 | rs12348953 | -1.306 | 0.248 | -0.094 | 1.029 | -0.023 | 0.015 | 0.015 | 0.096 | 0.13 | 0.18 | 0.09 | 0.16 | 0.09 | 0.09 |
| 9 | 23602407 | rs10811936 | -0.741 | 0.588 | -0.256 | 2.233 | 0 | 0.131 | 0.131 | 0 | 0.05 | 0.06 | 0.16 | 0.54 | 0.09 | 0.15 |
| 9 | 23603120 | rs4977617 | -0.996 | 0.594 | -0.252 | 2.234 | 0 | 0.121 | 0.121 | 0 | 0.06 | 0.07 | 0.16 | 0.54 | 0.09 | 0.15 |
| 9 | 23612594 | rs7357751 | -1.133 | 0.743 | -0.125 | 3.165 | 0.005 | 0.144 | 0.144 | 0 | 0.06 | 0.08 | 0.19 | 0.53 | 0.12 | 0.14 |
| 9 | 23613593 | rs1041350 | 0.084 | 0.338 | -0.24 | 2.603 | 0.02 | 0.029 | 0.029 | 0 | 0.34 | 0.28 | 0.1 | 0.15 | 0.18 | 0.24 |
| 9 | 23613815 | rs10511731 | -0.576 | 0.723 | -0.146 | 2.985 | -0.001 | 0.028 | 0.028 | 0 | 0.05 | 0.05 | 0.19 | 0.17 | 0.1 | 0.07 |
| 9 | 23619821 | rs7867905 | -1.493 | 0.555 | -0.06 | 2.832 | -0.003 | 0.064 | 0.064 | 0 | 0.13 | 0.17 | 0.21 | 0.54 | 0.15 | 0.21 |
| 9 | 23622640 | rs10965977 | -1.604 | 0.523 | -0.067 | 2.826 | -0.003 | 0.05 | 0.05 | 0 | 0.13 | 0.17 | 0.25 | 0.56 | 0.16 | 0.21 |
| 9 | 23634624 | rs1411241 | -0.599 | 1.359 | -0.347 | 2.454 | -0.046 | -0.036 | -0.036 | 0 | 0.11 | 0.03 | 0.2 | 0.13 | 0.16 | 0.24 |
| 9 | 23643682 | rs4977871 | -0.23 | 1.273 | 0.147 | 1.757 | -0.002 | 0.006 | 0.006 | 0.12 | 0.32 | 0.41 | 0.6 | 0.29 | 0.4 | 0.24 |
| 9 | 23645189 | rs9886753 | -0.372 | 1.467 | 0.315 | 2.004 | 0.04 | 0.126 | 0.126 | 0 | 0.05 | 0.18 | 0.2 | 0.14 | 0.1 | 0.02 |
| 9 | 23646551 | rs10966007 | -1.122 | 1.769 | 0.042 | 1.704 | 0.018 | 0.032 | 0.032 | 0 | 0.17 | 0.22 | 0.25 | 0.45 | 0.24 | 0.37 |
| 9 | 23651429 | rs10966009 | -1.162 | 1.449 | 0.079 | 1.134 | 0.009 | 0.001 | 0.001 | 0 | 0.11 | 0.18 | 0.18 | 0.11 | 0.12 | 0.19 |
| 9 | 23653899 | rs16907478 | -0.489 | 1.3 | 0.271 | 0.436 | -0.005 | -0.056 | -0.056 | 0.232 | 0.15 | 0.2 | 0.09 | 0 | 0.11 | 0.06 |
| 9 | 23655747 | rs10757442 | -0.354 | 1.174 | -0.716 | -0.242 | -0.027 | 0.056 | 0.056 | 0.161 | 0.39 | 0.25 | 0.39 | 0.22 | 0.44 | 0.37 |
| 9 | 23656678 | rs1329035 | -0.519 | 1.074 | -0.886 | -0.427 | -0.004 | 0.012 | 0.012 | 0 | 0.84 | 0.88 | 0.9 | 0.9 | 0.82 | 0.89 |
| 9 | 23657690 | rs6475751 | 0.77 | 1.049 | -0.25 | -0.386 | 0.005 | -0.002 | -0.002 | 0 | 0.7 | 0.64 | 0.68 | 0.75 | 0.65 | 0.69 |
| 9 | 23658011 | rs6475752 | 0.567 | 0.62 | -0.004 | -0.384 | 0 | -0.001 | -0.001 | 0 | 0.72 | 0.66 | 0.74 | 0.75 | 0.71 | 0.72 |
| 9 | 23659183 | rs4977875 | -1.381 | 0.58 | -0.278 | -0.385 | -0.007 | 0.034 | 0.034 | 0 | 0.89 | 0.95 | 0.95 | 1 | 0.89 | 0.92 |
| 9 | 23660419 | rs7855153 | 0.586 | 0.298 | 0.104 | 0.343 | -0.021 | 0.009 | 0.009 | 0 | 0.63 | 0.49 | 0.75 | 0.58 | 0.69 | 0.71 |

| | | | | | | | | | | | | | | | | |
|---|---|---|---|---|---|---|---|---|---|---|---|---|---|---|---|---|
| 9 | 23661974 | rs7873735 | 0.141 | 0.181 | 0.687 | 0.246 | -0.02 | 0.07 | 0.07 | 0 | 0.2 | 0.38 | 0.17 | 0.38 | 0.16 | 0.17 |
| 9 | 23662907 | rs10448166 | 1.036 | 0.043 | 0.203 | 0.069 | 0.012 | 0.033 | 0.033 | 0 | 0.48 | 0.27 | 0.43 | 0.38 | 0.52 | 0.48 |
| 9 | 23663533 | rs4977622 | -0.225 | 0.343 | 0.451 | 0.275 | -0.007 | 0.006 | 0.006 | 0 | 0.68 | 0.53 | 0.77 | 0.63 | 0.72 | 0.73 |
| 9 | 23664696 | rs7871964 | 0.647 | 0.323 | 0.246 | 0.238 | 0.004 | 0.05 | 0.05 | 0 | 0.09 | 0.13 | 0.3 | 0.24 | 0.14 | 0.2 |
| 9 | 23665940 | rs10733391 | -0.374 | 0.336 | 0.339 | 0.309 | -0.017 | 0.018 | 0.018 | 0 | 0.7 | 0.54 | 0.78 | 0.61 | 0.75 | 0.75 |
| 9 | 23668205 | rs7858686 | -0.394 | 0.287 | 0.367 | 0.342 | -0.002 | 0.009 | 0.009 | 0.004 | 0.69 | 0.57 | 0.77 | 0.61 | 0.71 | 0.73 |
| 9 | 23668266 | rs10732211 | 0.258 | 0.365 | -0.275 | -0.071 | -0.002 | 0.021 | 0.021 | 0.083 | 0.57 | 0.23 | 0.46 | 0.36 | 0.57 | 0.53 |
| 9 | 23670123 | rs1410844 | 0.174 | 0.518 | 0.151 | 0.311 | 0 | 0 | 0 | 0.036 | 0.25 | 0.22 | 0.26 | 0.24 | 0.28 | 0.36 |
| 9 | 23677432 | rs1036105 | 0.427 | 0.211 | -0.187 | -0.019 | 0.006 | 0.059 | 0.059 | 0 | 0.79 | 0.91 | 0.88 | 0.91 | 0.82 | 0.84 |
| 9 | 23677964 | rs1036107 | 0.971 | 0.702 | -0.103 | -0.227 | -0.001 | -0.058 | -0.058 | 0.139 | 0.68 | 0.45 | 0.8 | 0.89 | 0.72 | 0.8 |
| 9 | 23679836 | rs17468071 | -0.263 | 0.899 | -0.18 | -0.038 | 0.007 | -0.018 | -0.018 | 0 | 0.06 | 0.09 | 0.05 | 0 | 0.1 | 0.03 |
| 9 | 23680743 | rs16907598 | -0.603 | 0.536 | 0.578 | -0.104 | 0.001 | -0.006 | -0.006 | 0.021 | 0.06 | 0.28 | 0.02 | 0 | 0 | 0.03 |
| 9 | 23681361 | rs2383268 | -0.391 | 0.903 | -0.515 | -0.724 | -0.042 | 0.134 | 0.134 | 0.623 | 0.57 | 0.81 | 0.5 | 0.8 | 0.47 | 0.47 |
| 9 | 23685144 | rs10966021 | -0.953 | 1.69 | -0.56 | -0.988 | 0.054 | -0.056 | -0.056 | 0 | 0.23 | 0.41 | 0.32 | 0.11 | 0.31 | 0.15 |
| 9 | 23685236 | rs10738679 | -1.118 | 1.829 | -1.157 | -0.906 | 0.017 | 0.094 | 0.094 | 0 | 0.69 | 0.86 | 0.77 | 0.9 | 0.77 | 0.58 |
| 9 | 23686123 | rs932883 | -0.173 | 1.416 | -1.591 | -0.575 | 0.002 | 0.032 | 0.032 | 0.091 | 0.39 | 0.76 | 0.25 | 0.32 | 0.32 | 0.25 |
| 9 | 23687455 | rs2116227 | 1.127 | 1.439 | -1.358 | -0.455 | 0 | -0.003 | -0.003 | 0 | 0.25 | 0.23 | 0.14 | 0.09 | 0.24 | 0.2 |
| 9 | 23687696 | rs4977884 | -1.251 | 1.617 | -1.701 | -0.045 | 0.02 | 0.087 | 0.087 | 0 | 0.71 | 0.88 | 0.78 | 0.91 | 0.79 | 0.62 |
| 9 | 23689717 | rs4977886 | -0.242 | 2.002 | -1.267 | 0.271 | -0.03 | 0.004 | 0.004 | 0.436 | 0.12 | 0.07 | 0.28 | 0.11 | 0.29 | 0.11 |
| 9 | 23694188 | rs10733392 | 0.187 | 1.343 | -1.445 | 0.693 | -0.015 | -0.076 | -0.076 | 0.426 | 0.78 | 0.89 | 0.64 | 0.5 | 0.74 | 0.71 |
| 9 | 23699886 | rs1854442 | 1.063 | 0.786 | -2.096 | 0.51 | 0.041 | 0.015 | 0.015 | 0 | 0.29 | 0.1 | 0.29 | 0.22 | 0.2 | 0.41 |
| 9 | 23702349 | rs1410840 | -0.424 | 1.584 | -0.985 | 0.603 | 0.082 | 0.027 | 0.027 | 0.013 | 0.15 | 0.44 | 0.14 | 0.16 | 0.19 | 0.11 |
| 9 | 23704310 | rs1023704 | -0.534 | 1.501 | -0.787 | 0.676 | 0.01 | 0.041 | 0.041 | 0 | 0.29 | 0.63 | 0.39 | 0.54 | 0.3 | 0.31 |
| 9 | 23705108 | rs10966032 | 0.271 | 1.477 | -0.719 | 0.589 | 0.113 | 0.02 | 0.02 | 0.073 | 0.08 | 0.28 | 0.08 | 0.06 | 0.14 | 0.05 |
| 9 | 23706348 | rs1329024 | 0.891 | 1.77 | -1.462 | 0.505 | -0.023 | -0.007 | -0.007 | 0 | 0.07 | 0 | 0.06 | 0.06 | 0.08 | 0.04 |
| 9 | 23713284 | rs1360226 | -0.705 | 1.89 | -1.214 | 0.575 | 0.005 | 0.001 | 0.001 | 0 | 0.14 | 0.13 | 0.16 | 0.22 | 0.25 | 0.11 |
| 9 | 23713680 | rs3829088 | 0.517 | 1.56 | -1.232 | 0.674 | 0.022 | 0.026 | 0.026 | 0 | 0.07 | 0.09 | 0.1 | 0.16 | 0.15 | 0.05 |
| 9 | 23718363 | rs7847989 | 0.956 | 0.399 | -1.788 | 0.164 | 0.057 | 0.031 | 0.031 | 0 | 0.33 | 0.05 | 0.29 | 0.22 | 0.2 | 0.39 |
| 9 | 23718520 | rs4977887 | 0.448 | 0.89 | -1.291 | 0.426 | -0.016 | 0.023 | 0.023 | 1.296 | 0.59 | 0.26 | 0.5 | 0.38 | 0.53 | 0.54 |
| 9 | 23720904 | rs1329041 | -0.457 | -0.083 | -1.368 | 0.94 | -0.025 | 0.003 | 0.003 | 0 | 0.38 | 0.46 | 0.4 | 0.38 | 0.28 | 0.37 |
| 9 | 23726069 | rs12347232 | 0.088 | -0.305 | -1.314 | 1.153 | -0.061 | 0.067 | 0.067 | 0 | 0.18 | 0.33 | 0.12 | 0.31 | 0.08 | 0.11 |
| 9 | 23732000 | rs7861646 | -0.649 | -0.117 | -1.518 | 1.023 | -0.024 | 0.014 | 0.014 | 0 | 0.39 | 0.46 | 0.44 | 0.5 | 0.3 | 0.42 |
| 9 | 23737627 | rs3793577 | 0.254 | 0.095 | -1.414 | 1.102 | 0 | -0.001 | -0.001 | 0 | 0.49 | 0.5 | 0.46 | 0.52 | 0.51 | 0.48 |
| 9 | 23738335 | rs3793578 | 0.106 | -0.075 | -1.438 | 1.363 | 0.034 | -0.068 | -0.068 | 0.429 | 0.13 | 0.01 | 0.08 | 0.23 | 0.07 | 0.08 |
| 9 | 23751700 | rs12236195 | -0.764 | 0.154 | -1.209 | 1.225 | 0.001 | 0.024 | 0.024 | 0.008 | 0.17 | 0.19 | 0.26 | 0.32 | 0.18 | 0.33 |
| 9 | 23753106 | rs16907695 | -0.704 | -0.32 | -1.458 | 0.89 | 0.053 | 0.037 | 0.037 | 0 | 0.21 | 0.08 | 0.13 | 0.09 | 0.1 | 0.12 |
| 9 | 23759368 | rs12057071 | -0.869 | -0.341 | -1.49 | 1.136 | 0.047 | -0.002 | -0.002 | 0 | 0.22 | 0.14 | 0.16 | 0.23 | 0.12 | 0.15 |
| 9 | 23765511 | rs9298850 | 0.173 | -0.085 | -2.224 | 0.482 | -0.01 | 0.147 | 0.147 | 0.065 | 0.47 | 0.84 | 0.66 | 0.73 | 0.53 | 0.49 |
| 9 | 23785967 | rs2195731 | 0.114 | -0.395 | -0.974 | 1.18 | -0.02 | -0.002 | -0.002 | 0 | 0.15 | 0.14 | 0.24 | 0.18 | 0.07 | 0.08 |
| 9 | 23787897 | rs992548 | -0.256 | -0.342 | -1.135 | 1.357 | -0.051 | -0.063 | -0.063 | 2.086 | 0.12 | 0.04 | 0.2 | 0.26 | 0.19 | 0.32 |
| 9 | 23788214 | rs16907731 | -1.173 | -0.356 | -1.118 | 1.35 | 0.007 | -0.019 | -0.019 | 0 | 0.28 | 0.19 | 0.44 | 0.44 | 0.27 | 0.4 |
| 9 | 23788680 | rs1410846 | -1.795 | -0.457 | -0.996 | 1.39 | -0.036 | 0.035 | 0.035 | 0.514 | 0.36 | 0.48 | 0.54 | 0.54 | 0.28 | 0.46 |
| 9 | 23790800 | rs1653081 | -0.92 | -0.581 | -0.531 | 1.024 | -0.107 | 0.015 | 0.015 | 0.172 | 0.24 | 0.44 | 0.34 | 0.29 | 0.09 | 0.14 |
| 9 | 23790801 | rs1728256 | -0.936 | -0.58 | -0.535 | 1.024 | -0.109 | 0.015 | 0.015 | 0.231 | 0.24 | 0.45 | 0.34 | 0.29 | 0.09 | 0.14 |
| 9 | 23802442 | rs16907782 | -0.676 | -0.389 | -1.012 | 0.987 | 0 | 0.001 | 0.001 | 0 | 0.22 | 0.15 | 0.32 | 0.2 | 0.09 | 0.11 |
| 9 | 23811127 | rs3793595 | -0.342 | 0.435 | -1.221 | 0.98 | -0.012 | -0.008 | -0.008 | 1.228 | 0.21 | 0.07 | 0.31 | 0.23 | 0.22 | 0.15 |
| 9 | 23813840 | rs702223 | -1.371 | 0.681 | -1.009 | 1.38 | 0.001 | 0.022 | 0.022 | 1.424 | 0.33 | 0.38 | 0.49 | 0.53 | 0.36 | 0.45 |
| 9 | 23816554 | rs4977892 | -0.307 | 0.578 | -1.185 | 1.192 | 0.005 | -0.012 | -0.012 | 0 | 0.21 | 0.07 | 0.3 | 0.25 | 0.19 | 0.14 |
| 9 | 23817548 | rs17551174 | -0.157 | 0.508 | -1.236 | 1.055 | -0.007 | 0.011 | 0.011 | 1.257 | 0.19 | 0.02 | 0.1 | 0.14 | 0.17 | 0.13 |
| 9 | 23829728 | rs7870728 | -0.712 | 0.736 | -1.049 | 0.73 | 0.001 | 0.075 | 0.075 | 0.047 | 0.35 | 0.16 | 0.22 | 0.23 | 0.38 | 0.21 |
| 9 | 23829824 | rs12341793 | 1.23 | 0.5 | -0.884 | 0.72 | -0.023 | -0.003 | -0.003 | 0.113 | 0.28 | 0.39 | 0.22 | 0.21 | 0.17 | 0.17 |
| 9 | 23831031 | rs7869806 | 1.292 | 0.571 | -0.971 | 0.155 | -0.009 | -0.02 | -0.02 | 0.299 | 0.09 | 0.03 | 0.08 | 0.16 | 0.11 | 0.08 |
| 9 | 23831141 | rs4142149 | -0.79 | 0.567 | -1.029 | 0.237 | 0.002 | 0.004 | 0.004 | 0.275 | 0.38 | 0.17 | 0.26 | 0.37 | 0.39 | 0.22 |
| 9 | 23831902 | rs10811973 | -0.117 | 0.623 | -1.046 | -0.839 | -0.006 | 0.055 | 0.055 | 4.334 | 0.25 | 0.09 | 0.17 | 0.14 | 0.27 | 0.16 |
| 9 | 23832717 | rs10966093 | 0.492 | 0.608 | -1.037 | -0.892 | 0.006 | 0.027 | 0.027 | 3.415 | 0.17 | 0.08 | 0.09 | 0 | 0.16 | 0.08 |
| 9 | 23832770 | rs783455 | -1.53 | 0.79 | -0.936 | -0.841 | -0.006 | 0.064 | 0.064 | 0 | 0.72 | 0.88 | 0.78 | 0.91 | 0.71 | 0.6 |
| 9 | 23837464 | rs13286215 | 1.353 | 0.553 | -0.739 | -0.794 | -0.004 | -0.049 | -0.049 | 0 | 0.24 | 0.32 | 0.14 | 0.04 | 0.17 | 0.14 |
| 9 | 23840388 | rs10966097 | -0.146 | 0.692 | -1.196 | -0.448 | 0.001 | 0.053 | 0.053 | 4.217 | 0.26 | 0.11 | 0.17 | 0.14 | 0.27 | 0.16 |
| 9 | 23842612 | rs10811978 | -0.319 | 0.691 | -1.181 | -0.376 | 0.004 | 0.014 | 0.014 | 3.48 | 0.27 | 0.13 | 0.37 | 0.25 | 0.28 | 0.18 |
| 9 | 23844194 | rs9650694 | -0.075 | 0.763 | -1.297 | -0.448 | -0.006 | 0.058 | 0.058 | 4.334 | 0.25 | 0.09 | 0.17 | 0.14 | 0.27 | 0.16 |
| 9 | 23852266 | rs10811980 | -0.816 | 0.918 | -1.225 | -0.893 | -0.019 | 0.134 | 0.134 | 0.485 | 0.45 | 0.15 | 0.43 | 0.76 | 0.51 | 0.47 |
| 9 | 23859906 | rs10811984 | -0.824 | 0.764 | -1.255 | -0.936 | -0.009 | 0.154 | 0.154 | 0.52 | 0.47 | 0.17 | 0.47 | 0.77 | 0.58 | 0.51 |
| 9 | 23880959 | rs7847255 | 0.436 | 1.005 | -0.186 | -0.949 | 0.021 | 0.041 | 0.041 | 0 | 0.46 | 0.4 | 0.24 | 0.13 | 0.35 | 0.24 |
| 9 | 23892846 | rs1528049 | 0.279 | 1.201 | -0.942 | -0.807 | -0.017 | -0.019 | -0.019 | 0 | 0.18 | 0.12 | 0.3 | 0.64 | 0.3 | 0.34 |
| 9 | 23898021 | rs542527 | 1.373 | 0.872 | -0.921 | -0.922 | -0.064 | 0.125 | 0.125 | 1.066 | 0.23 | 0.02 | 0.11 | 0.04 | 0.23 | 0.17 |
| 9 | 23901927 | rs2383296 | -0.716 | 1.285 | 0.069 | -0.017 | -0.014 | -0.008 | -0.008 | 0 | 0.25 | 0.2 | 0.35 | 0.66 | 0.38 | 0.4 |
| 9 | 23940420 | rs10966141 | -0.067 | 1.014 | -0.042 | -0.022 | -0.032 | 0.004 | 0.004 | 0.031 | 0.15 | 0.01 | 0.08 | 0.12 | 0.14 | 0.1 |
| 9 | 23972006 | rs4977903 | -0.693 | 0.748 | 0.59 | -0.182 | -0.015 | 0.047 | 0.047 | 0 | 0.43 | 0.28 | 0.45 | 0.83 | 0.48 | 0.62 |
| 9 | 23978172 | rs7858484 | 0.609 | 0.896 | -0.213 | -0.088 | -0.004 | -0.03 | -0.03 | 1.451 | 0.56 | 0.64 | 0.34 | 0.03 | 0.5 | 0.33 |

| | | | | | | | | | | | | | | | | |
|---|---|---|---|---|---|---|---|---|---|---|---|---|---|---|---|---|
| 9 | 23983817 | rs1901155 | 0.489 | 1.037 | -0.722 | -0.013 | -0.026 | 0.095 | 0.095 | 0 | 0.58 | 0.77 | 0.54 | 0.16 | 0.51 | 0.37 |
| 9 | 24007751 | rs10812016 | -1.284 | 1.163 | -0.193 | 1.68 | 0.025 | 0.237 | 0.237 | 0 | 0.06 | 0.18 | 0.13 | 0.43 | 0.1 | 0.13 |
| 9 | 24013674 | rs10966173 | -0.153 | 1.286 | -0.709 | 1.509 | -0.037 | -0.046 | -0.046 | 0.091 | 0.11 | 0 | 0.04 | 0.17 | 0.09 | 0.07 |
| 9 | 24032281 | rs4615671 | -0.021 | 1.202 | 0.292 | 1.636 | -0.036 | -0.01 | -0.01 | 0.097 | 0.1 | 0 | 0.04 | 0.09 | 0.08 | 0.06 |
| 9 | 24034072 | rs4350058 | -1.187 | 1.108 | 0.428 | 2.205 | 0.002 | 0.142 | 0.142 | 0 | 0.06 | 0.07 | 0.1 | 0.49 | 0.1 | 0.15 |
| 9 | 24035011 | rs4548269 | -0.722 | 1.17 | 0.405 | 1.28 | 0.023 | 0.061 | 0.061 | 0 | 0.52 | 0.69 | 0.42 | 0.68 | 0.54 | 0.67 |
| 9 | 24052569 | rs11999150 | -1.334 | 1.172 | 0.639 | 2.137 | 0.01 | 0.159 | 0.159 | 0 | 0.06 | 0.11 | 0.11 | 0.57 | 0.1 | 0.15 |
| 9 | 24060455 | rs7862410 | -2.129 | 1.116 | 0.581 | 2.072 | 0.031 | 0.327 | 0.327 | 0.141 | 0.1 | 0.33 | 0.14 | 0.5 | 0.11 | 0.24 |
| 9 | 24063404 | rs12683447 | -1.585 | 1.019 | 0.355 | 1.895 | 0.022 | 0.267 | 0.267 | 0 | 0.06 | 0.22 | 0.21 | 0.57 | 0.06 | 0.17 |
| 9 | 24065751 | rs10966191 | -1.141 | 1.054 | 0.137 | 1.861 | 0.002 | 0.112 | 0.112 | 0 | 0.05 | 0.08 | 0.27 | 0.63 | 0.06 | 0.17 |
| 9 | 24072205 | rs10812022 | -0.842 | 1.288 | 0.113 | 2.36 | 0.014 | 0.19 | 0.19 | 0 | 0.05 | 0.08 | 0.27 | 0.47 | 0.12 | 0.19 |
| 9 | 24075368 | rs12379069 | -0.381 | 1.386 | -0.177 | 0.748 | -0.011 | 0 | 0 | 1.752 | 0.25 | 0.2 | 0.24 | 0.03 | 0.32 | 0.2 |
| 9 | 24091818 | rs4977912 | -0.393 | 0.866 | -0.548 | 0.519 | 0.006 | -0.018 | -0.018 | 0 | 0.22 | 0.16 | 0.31 | 0.59 | 0.21 | 0.23 |
| 9 | 24095294 | rs4416909 | -1.584 | 0.872 | -0.337 | 0.49 | 0 | -0.009 | -0.009 | 0.045 | 0.48 | 0.51 | 0.62 | 0.71 | 0.53 | 0.44 |
| 9 | 24097219 | rs10966207 | -1.596 | 0.876 | -0.308 | 0.471 | 0 | -0.002 | -0.002 | 0 | 0.48 | 0.54 | 0.63 | 0.68 | 0.53 | 0.43 |
| 9 | 24102399 | rs1333337 | -1.522 | 0.913 | -0.606 | 0.442 | -0.002 | 0.008 | 0.008 | 0 | 0.47 | 0.47 | 0.56 | 0.62 | 0.53 | 0.42 |
| 9 | 24105508 | rs4977917 | -1.514 | 0.884 | -0.608 | 0.42 | -0.001 | 0.008 | 0.008 | 0 | 0.47 | 0.47 | 0.56 | 0.65 | 0.53 | 0.42 |
| 9 | 24111280 | rs2225614 | -0.556 | 0.974 | -0.253 | 0.909 | 0.002 | -0.017 | -0.017 | 0 | 0.19 | 0.14 | 0.3 | 0.61 | 0.2 | 0.23 |
| 9 | 24113158 | rs10966220 | -0.811 | 1.392 | 0.048 | 0.424 | 0.004 | -0.028 | -0.028 | 0 | 0.16 | 0.18 | 0.15 | 0.03 | 0.24 | 0.11 |
| 9 | 24113936 | rs10812027 | -1.059 | 1.382 | 0.064 | 0.419 | 0.038 | -0.082 | -0.082 | 0 | 0.17 | 0.29 | 0.15 | 0.03 | 0.24 | 0.11 |
| 9 | 24121196 | rs12376503 | -1.281 | 1.534 | -0.503 | 0.378 | -0.012 | 0.026 | 0.026 | 0 | 0.14 | 0.06 | 0.12 | 0 | 0.18 | 0.04 |
| 9 | 24125018 | rs7848340 | -2.047 | 1.513 | -0.244 | 0.496 | 0.015 | -0.088 | -0.088 | 0 | 0.18 | 0.31 | 0.11 | 0.02 | 0.2 | 0.1 |
| 9 | 24125062 | rs7862252 | -0.567 | 1.53 | -0.213 | 0.517 | 0.022 | -0.042 | -0.042 | 0 | 0.08 | 0.15 | 0.05 | 0 | 0.11 | 0.03 |
| 9 | 24133577 | rs12554249 | -1.169 | 1.47 | -0.324 | 0.657 | 0.002 | 0.005 | 0.005 | 0 | 0.1 | 0.13 | 0.1 | 0.24 | 0.11 | 0.11 |
| 9 | 24133646 | rs10966238 | -2.033 | 1.489 | -0.348 | 0.62 | 0.008 | -0.051 | -0.051 | 0.031 | 0.2 | 0.31 | 0.12 | 0.13 | 0.23 | 0.14 |
| 9 | 24146203 | rs10812030 | -1.487 | 0.805 | -0.177 | 0.633 | 0.003 | 0.004 | 0.004 | 0 | 0.12 | 0.19 | 0.11 | 0.21 | 0.13 | 0.11 |
| 9 | 24146436 | rs2298620 | -1.574 | 0.802 | -0.175 | 0.632 | 0.006 | 0.008 | 0.008 | 0 | 0.12 | 0.23 | 0.11 | 0.21 | 0.13 | 0.11 |
| 9 | 24152518 | rs7850169 | -1.446 | 0.857 | -0.071 | 0.277 | -0.008 | -0.077 | -0.077 | 1.687 | 0.14 | 0.29 | 0.09 | 0.02 | 0.14 | 0.08 |
| 9 | 24156484 | rs17176112 | -1.066 | 1.498 | 0.074 | 0.63 | 0.04 | 0.012 | 0.012 | 0 | 0.23 | 0.33 | 0.41 | 0.23 | 0.29 | 0.18 |
| 9 | 24156917 | rs16923681 | -1.293 | 1.34 | 0.375 | 0.817 | 0.046 | 0.045 | 0.045 | 0 | 0.24 | 0.39 | 0.41 | 0.3 | 0.28 | 0.21 |
| 9 | 24164332 | rs7848321 | -1.296 | 1.127 | 0.372 | 0.496 | -0.019 | -0.028 | -0.028 | 2.228 | 0.11 | 0.27 | 0.13 | 0.05 | 0.07 | 0.11 |
| 9 | 24175234 | rs10124866 | -1.687 | 1.123 | 0.687 | 0.503 | 0.025 | 0.022 | 0.022 | 0 | 0.23 | 0.33 | 0.36 | 0.26 | 0.25 | 0.21 |
| 9 | 24189914 | rs7850234 | -1.612 | 1.173 | 0.683 | 0.54 | 0.067 | 0.051 | 0.051 | 0 | 0.2 | 0.34 | 0.36 | 0.25 | 0.26 | 0.17 |
| 9 | 24190635 | rs4977646 | -0.682 | 1.166 | 0.731 | 0.536 | -0.017 | -0.015 | -0.015 | 0 | 0.14 | 0.11 | 0.25 | 0.21 | 0.22 | 0.1 |
| 9 | 24190647 | rs7033936 | -0.551 | 1.163 | 0.741 | 0.56 | -0.002 | 0.015 | 0.015 | 0 | 0.06 | 0.16 | 0.1 | 0.05 | 0.04 | 0.07 |
| 9 | 24192164 | rs17771522 | -0.207 | 1.322 | 0.706 | 0.519 | -0.015 | -0.014 | -0.014 | 0 | 0.1 | 0.07 | 0.23 | 0.21 | 0.17 | 0.07 |
| 9 | 24192964 | rs7028478 | -1.902 | 1.312 | 0.705 | 0.507 | 0.005 | 0.004 | 0.004 | 0 | 0.21 | 0.19 | 0.36 | 0.27 | 0.27 | 0.2 |
| 9 | 24193119 | rs7031817 | -1.989 | 1.313 | 0.682 | 0.507 | 0.057 | 0.045 | 0.045 | 0 | 0.21 | 0.35 | 0.37 | 0.27 | 0.27 | 0.2 |
| 9 | 24201121 | rs4977922 | -0.243 | 1.344 | 0.504 | 0.332 | -0.034 | -0.022 | -0.022 | 0 | 0.1 | 0.03 | 0.23 | 0.15 | 0.17 | 0.05 |
| 9 | 24205004 | rs7871735 | -2.172 | 0.599 | 0.264 | 0.179 | -0.001 | 0.001 | 0.001 | 0 | 0.36 | 0.31 | 0.46 | 0.3 | 0.32 | 0.33 |
| 9 | 24212352 | rs17195961 | -0.15 | 0.789 | 0.397 | 0.161 | -0.001 | -0.011 | -0.011 | 0 | 0.08 | 0.08 | 0.03 | 0 | 0.05 | 0.06 |
| 9 | 24231246 | rs480502 | -0.152 | 0.887 | -0.172 | -0.19 | -0.04 | 0.017 | 0.017 | 0.389 | 0.8 | 0.92 | 0.67 | 0.81 | 0.72 | 0.76 |
| 9 | 24233393 | rs718339 | -1.548 | 0.718 | -0.159 | -0.144 | 0.005 | -0.002 | -0.002 | 0.683 | 0.24 | 0.25 | 0.35 | 0.22 | 0.25 | 0.19 |
| 9 | 24234492 | rs4977927 | -0.644 | 0.856 | -0.121 | -0.039 | -0.009 | -0.007 | -0.007 | 0 | 0.16 | 0.1 | 0.32 | 0.22 | 0.21 | 0.14 |
| 9 | 24254082 | rs10812040 | 0.589 | 0.849 | -0.546 | -0.075 | -0.001 | -0.067 | -0.067 | 0 | 0.1 | 0.01 | 0.15 | 0.29 | 0.11 | 0.17 |
| 9 | 24262091 | rs17771619 | -0.87 | 1.008 | -0.284 | -0.016 | 0.021 | 0.006 | 0.006 | 0 | 0.17 | 0.2 | 0.32 | 0.19 | 0.21 | 0.14 |
| 9 | 24272883 | rs1410964 | -0.804 | 0.949 | -0.345 | -0.051 | -0.008 | -0.003 | -0.003 | 0 | 0.19 | 0.15 | 0.34 | 0.2 | 0.23 | 0.14 |
| 9 | 24290018 | rs7031702 | -0.148 | 0.767 | -0.394 | -0.732 | 0.021 | 0.044 | 0.044 | 0 | 0.13 | 0.06 | 0.05 | 0 | 0.08 | 0.09 |
| 9 | 24292583 | rs7868331 | -1.585 | 0.769 | -0.358 | -0.721 | -0.007 | 0.018 | 0.018 | 0 | 0.31 | 0.24 | 0.38 | 0.13 | 0.31 | 0.22 |
| 9 | 24298508 | rs6475790 | -2.074 | 0.608 | -0.4 | -0.791 | 0.026 | -0.03 | -0.03 | 0.755 | 0.62 | 0.89 | 0.79 | 0.46 | 0.65 | 0.58 |
| 9 | 24322545 | rs17198062 | -1.857 | 0.956 | 0.081 | -0.564 | -0.016 | 0.066 | 0.066 | 0 | 0.31 | 0.19 | 0.38 | 0.11 | 0.3 | 0.21 |
| 9 | 24332121 | rs12345327 | -1.359 | 1.462 | 0.778 | -0.001 | 0.04 | -0.051 | -0.051 | 0 | 0.2 | 0.37 | 0.34 | 0.1 | 0.23 | 0.11 |
| 9 | 24335841 | rs16908250 | -1.033 | 0.454 | 1.145 | 0.435 | 0.004 | 0.029 | 0.029 | 0 | 0.06 | 0.18 | 0.11 | 0.06 | 0.04 | 0.05 |
| 9 | 24346050 | rs10491801 | -1.549 | 0.843 | 1.088 | 0.35 | 0.086 | 0.029 | 0.029 | 0.743 | 0.07 | 0.37 | 0.17 | 0.03 | 0.05 | 0.02 |
| 9 | 24375619 | rs16908321 | -1.298 | 0.69 | 0.934 | 0.309 | 0.048 | 0.02 | 0.02 | 0.612 | 0.06 | 0.32 | 0.16 | 0.03 | 0.03 | 0.02 |
| 9 | 24460541 | rs13294480 | -0.824 | 0.608 | -0.186 | 0.507 | -0.021 | -0.038 | -0.038 | 0 | 0.1 | 0.01 | 0.07 | 0.17 | 0.12 | 0.14 |
| 9 | 24467698 | rs4618784 | -1.526 | 0.718 | 0.419 | 0.27 | 0.048 | -0.014 | -0.014 | 0.063 | 0.06 | 0.23 | 0.04 | 0 | 0.05 | 0.05 |
| 9 | 24471843 | rs7852101 | -1.67 | 0.678 | 0.366 | 0.252 | 0.024 | -0.03 | -0.03 | 0 | 0.07 | 0.19 | 0.04 | 0 | 0.05 | 0.05 |
| 9 | 24479050 | rs10757476 | -1.795 | 0.459 | 0.076 | 0.443 | 0 | 0 | 0 | 0 | 0.19 | 0.15 | 0.21 | 0.25 | 0.21 | 0.2 |
| 9 | 24482743 | rs10812053 | -1.576 | 0.471 | 0.165 | 0.523 | -0.002 | 0.005 | 0.005 | 0.019 | 0.17 | 0.17 | 0.18 | 0.25 | 0.18 | 0.19 |
| 9 | 24493958 | rs11788362 | -0.948 | 1.12 | -0.122 | 0.242 | -0.005 | 0.003 | 0.003 | 0 | 0.13 | 0.09 | 0.25 | 0.12 | 0.24 | 0.11 |
| 9 | 24495241 | rs10812054 | -0.518 | 0.748 | -0.647 | -0.073 | -0.014 | 0.051 | 0.051 | 0 | 0.47 | 0.3 | 0.44 | 0.35 | 0.53 | 0.47 |
| 9 | 24527357 | rs2257210 | -1.173 | 0.382 | -0.624 | -0.067 | 0.002 | 0 | 0 | 0 | 0.25 | 0.26 | 0.3 | 0.24 | 0.33 | 0.23 |
| 9 | 24536825 | rs10966331 | 0.962 | -0.147 | -1.672 | -0.084 | 0.004 | -0.043 | -0.043 | 0 | 0.32 | 0.06 | 0.36 | 0.24 | 0.24 | 0.43 |
| 9 | 24539632 | rs1888109 | -0.786 | -0.147 | -1.077 | -0.246 | 0.005 | -0.014 | -0.014 | 0 | 0.27 | 0.34 | 0.33 | 0.21 | 0.34 | 0.25 |
| 9 | 24540910 | rs16908477 | -0.013 | -0.19 | -1.036 | -0.494 | -0.013 | -0.042 | -0.042 | 1.509 | 0.08 | 0.25 | 0.04 | 0 | 0.05 | 0.01 |
| 9 | 24545513 | rs6475797 | 0.289 | -0.529 | -1.95 | -1.009 | -0.143 | 0.094 | 0.094 | 0 | 0.64 | 0.95 | 0.57 | 0.81 | 0.51 | 0.73 |
| 9 | 24573717 | rs2149913 | 0.079 | -0.432 | -1.236 | -0.968 | 0.01 | -0.083 | -0.083 | 0 | 0.74 | 0.66 | 0.72 | 0.95 | 0.67 | 0.82 |
| 9 | 24587252 | rs1759039 | 0.639 | -0.356 | -1.63 | -1.239 | -0.029 | 0.071 | 0.071 | 0 | 0.13 | 0.01 | 0.11 | 0.01 | 0.12 | 0.08 |

| | | | | | | | | | | | | | | | | |
|---|---|---|---|---|---|---|---|---|---|---|---|---|---|---|---|---|
| 9 | 24595311 | rs7018906 | 0.387 | -0.763 | -1.463 | -0.981 | -0.002 | -0.023 | -0.023 | 0 | 0.69 | 0.59 | 0.6 | 0.69 | 0.63 | 0.67 |
| 9 | 24605821 | rs1162608 | -0.1 | -1.326 | -2.081 | -1.555 | 0.005 | 0.098 | 0.098 | 0 | 0.29 | 0.09 | 0.23 | 0.14 | 0.24 | 0.13 |
| 9 | 24611322 | rs13284068 | 1.217 | -0.926 | -2.432 | -1.572 | -0.004 | 0.062 | 0.062 | 0.024 | 0.18 | 0.02 | 0.26 | 0.04 | 0.12 | 0.13 |
| 9 | 24613381 | rs1333490 | 0.117 | -0.548 | -2.341 | -1.153 | -0.034 | -0.018 | -0.018 | 0 | 0.13 | 0.02 | 0.31 | 0.16 | 0.14 | 0.12 |
| 9 | 24616032 | rs10812061 | 0.743 | -0.716 | -2.277 | -1.345 | 0.001 | -0.013 | -0.013 | 0.55 | 0.66 | 0.65 | 0.44 | 0.82 | 0.55 | 0.71 |
| 9 | 24625115 | rs1162610 | -0.524 | -0.695 | -2.215 | -1.297 | -0.005 | 0.003 | 0.003 | 0 | 0.82 | 0.84 | 0.71 | 0.82 | 0.69 | 0.78 |
| 9 | 24625154 | rs12341864 | 1.696 | -0.762 | -2.79 | -1.681 | -0.036 | 0.063 | 0.063 | 0 | 0.19 | 0.08 | 0.15 | 0.03 | 0.23 | 0.14 |
| 9 | 24626089 | rs1329851 | -0.247 | -0.595 | -2.363 | -0.352 | -0.035 | -0.047 | -0.047 | 0.216 | 0.42 | 0.12 | 0.49 | 0.31 | 0.36 | 0.52 |
| 9 | 24631812 | rs2772571 | 0.796 | -0.493 | -1.862 | -0.126 | 0.05 | -0.057 | -0.057 | 0 | 0.25 | 0.08 | 0.31 | 0.24 | 0.15 | 0.4 |
| 9 | 24635445 | rs7048683 | 0.611 | 0.684 | -0.875 | -0.206 | -0.026 | -0.027 | -0.027 | 0 | 0.05 | 0.12 | 0.03 | 0 | 0.03 | 0.01 |
| 9 | 24651946 | rs1360563 | 0.274 | 0.896 | -0.846 | -0.214 | -0.002 | 0 | 0 | 0 | 0.79 | 0.82 | 0.72 | 0.84 | 0.66 | 0.8 |
| 9 | 24658725 | rs6475805 | 1.252 | 0.791 | -0.845 | -0.167 | 0.005 | -0.018 | -0.018 | 0.229 | 0.6 | 0.57 | 0.52 | 0.69 | 0.5 | 0.67 |
| 9 | 24670982 | rs10966380 | -0.831 | 1.572 | -0.516 | 0.064 | -0.044 | 0.033 | 0.033 | 0 | 0.23 | 0.16 | 0.27 | 0.07 | 0.43 | 0.27 |
| 9 | 24671257 | rs10966381 | -0.775 | 1.469 | -0.511 | 0.259 | -0.063 | -0.005 | -0.005 | 0 | 0.29 | 0.17 | 0.33 | 0.35 | 0.46 | 0.28 |
| 9 | 24671602 | rs1332677 | -1.131 | 1.49 | -0.51 | 0.228 | 0.027 | -0.011 | -0.011 | 0 | 0.35 | 0.42 | 0.38 | 0.34 | 0.5 | 0.3 |
| 9 | 24672959 | rs10966384 | -0.969 | 1.233 | -0.721 | 0.479 | 0.008 | -0.001 | -0.001 | 0.002 | 0.07 | 0.09 | 0.07 | 0.05 | 0.1 | 0.11 |
| 9 | 24677263 | rs10122018 | -0.837 | 1.259 | -0.675 | 0.461 | 0.022 | -0.007 | -0.007 | 0.151 | 0.07 | 0.14 | 0.06 | 0.03 | 0.09 | 0.06 |
| 9 | 24683161 | rs12004244 | -1.338 | 1.292 | -0.873 | 0.479 | 0.007 | -0.032 | -0.032 | 0.51 | 0.1 | 0.14 | 0.08 | 0.01 | 0.12 | 0.05 |
| 9 | 24687157 | rs10966398 | -0.548 | 1.602 | -1.164 | 0.679 | -0.009 | 0.008 | 0.008 | 0 | 0.05 | 0.03 | 0.05 | 0.01 | 0.09 | 0.04 |
| 9 | 24689992 | rs11792697 | -0.605 | 1.643 | -1.155 | 0.699 | -0.013 | 0 | 0 | 0 | 0.05 | 0.02 | 0.05 | 0.04 | 0.1 | 0.09 |
| 9 | 24696280 | rs1332678 | 0.173 | 1.624 | -1.161 | 0.727 | -0.001 | -0.001 | -0.001 | 0 | 0.76 | 0.7 | 0.7 | 0.64 | 0.67 | 0.72 |
| 9 | 24698546 | rs10966405 | -0.761 | 1.263 | -1.243 | 0.477 | -0.016 | -0.012 | -0.012 | 1.493 | 0.05 | 0.03 | 0.05 | 0.07 | 0.1 | 0.05 |
| 9 | 24708079 | rs10966415 | -0.819 | 1.223 | -1.231 | 0.128 | -0.007 | 0.006 | 0.006 | 0 | 0.06 | 0.04 | 0.05 | 0.02 | 0.1 | 0.05 |
| 9 | 24716478 | rs10966430 | -0.891 | 1.249 | -1.479 | 0.231 | -0.014 | -0.008 | -0.008 | 0.514 | 0.07 | 0.04 | 0.1 | 0.07 | 0.1 | 0.08 |
| 9 | 24721911 | rs10812089 | -0.082 | 0.238 | -1.963 | -0.386 | 0.044 | 0.107 | 0.107 | 0 | 0.28 | 0.05 | 0.32 | 0.06 | 0.18 | 0.25 |
| 9 | 24724747 | rs11789683 | -0.341 | 1.702 | -1.767 | -0.592 | -0.021 | 0.01 | 0.01 | 0 | 0.13 | 0.11 | 0.17 | 0 | 0.31 | 0.08 |
| 9 | 24725459 | rs7870157 | -0.24 | 0.548 | -1.39 | -1.055 | -0.021 | -0.048 | -0.048 | 0 | 0.37 | 0.48 | 0.55 | 0.87 | 0.27 | 0.58 |
| 9 | 24727103 | rs16908770 | 0.168 | 0.604 | -0.9 | -0.72 | -0.002 | 0.021 | 0.021 | 0 | 0.08 | 0.24 | 0.09 | 0.14 | 0.07 | 0.19 |
| 9 | 24731623 | rs2056416 | -0.048 | 0.346 | -1.137 | -0.795 | -0.002 | 0.004 | 0.004 | 0 | 0.21 | 0.25 | 0.46 | 0.68 | 0.19 | 0.37 |
| 9 | 24740388 | rs7859091 | 0.236 | 0.166 | -1.04 | 0.325 | -0.006 | 0.084 | 0.084 | 0 | 0.08 | 0.09 | 0.27 | 0.3 | 0.04 | 0.22 |
| 9 | 24740580 | rs7873039 | -0.548 | -0.442 | -1.519 | -0.729 | 0.044 | 0.099 | 0.099 | 0 | 0.47 | 0.29 | 0.54 | 0.72 | 0.36 | 0.58 |
| 9 | 24741297 | rs4977954 | -0.939 | 0.127 | -0.963 | -0.057 | 0.004 | 0.002 | 0.002 | 0 | 0.25 | 0.24 | 0.42 | 0.73 | 0.17 | 0.46 |
| 9 | 24744456 | rs10966438 | -0.661 | 0.58 | -1.189 | 0.135 | 0.019 | -0.035 | -0.035 | 0.009 | 0.24 | 0.15 | 0.23 | 0.6 | 0.17 | 0.33 |
| 9 | 24745655 | rs10114014 | 0.122 | 1.943 | -0.968 | -0.24 | -0.006 | -0.003 | -0.003 | 0 | 0.12 | 0.11 | 0.16 | 0 | 0.29 | 0.07 |
| 9 | 24754015 | rs7865352 | 0.072 | 0.828 | -0.644 | 0.636 | -0.001 | 0.041 | 0.041 | 0.446 | 0.77 | 0.73 | 0.88 | 0.57 | 0.82 | 0.77 |
| 9 | 24756027 | rs10124916 | 0.759 | 2.031 | -0.828 | 0.291 | 0.004 | 0.013 | 0.013 | 1.039 | 0.61 | 0.55 | 0.57 | 0.53 | 0.48 | 0.63 |
| 9 | 24756335 | rs13295819 | -0.172 | 1.718 | -1.045 | 0.114 | -0.068 | 0.059 | 0.059 | 0 | 0.14 | 0.05 | 0.3 | 0 | 0.32 | 0.13 |
| 9 | 24757755 | rs7468674 | -0.843 | 0.371 | -0.162 | 0.714 | 0.007 | 0.029 | 0.029 | 0 | 0.14 | 0.25 | 0.29 | 0.17 | 0.15 | 0.24 |
| 9 | 24768179 | rs4319186 | 1.004 | -0.126 | -0.918 | 0.233 | -0.101 | 0.045 | 0.045 | 0.079 | 0.37 | 0.09 | 0.45 | 0.24 | 0.46 | 0.39 |
| 9 | 24770615 | rs4284108 | 0.142 | -0.94 | -0.194 | 0.225 | -0.075 | 0.136 | 0.136 | 0.019 | 0.77 | 0.45 | 0.81 | 0.44 | 0.89 | 0.72 |
| 9 | 24770659 | rs10966450 | 0.63 | -1.067 | 0.083 | 0.194 | -0.004 | 0.044 | 0.044 | 0 | 0.12 | 0.16 | 0.15 | 0.59 | 0.08 | 0.17 |
| 9 | 24771927 | rs10116739 | 0.661 | -1.163 | 0.227 | -0.326 | -0.006 | -0.003 | -0.003 | 0 | 0.59 | 0.38 | 0.66 | 0.37 | 0.69 | 0.63 |
| 9 | 24775586 | rs4145038 | -0.274 | -0.93 | 0.303 | -0.48 | 0.007 | 0 | 0 | 0 | 0.09 | 0.12 | 0.17 | 0.12 | 0.16 | 0.13 |
| 9 | 24781165 | rs4625115 | 2.276 | -1.069 | -0.012 | -0.564 | -0.149 | 0.056 | 0.056 | 0 | 0.33 | 0.02 | 0.4 | 0.16 | 0.38 | 0.32 |
| 9 | 24781300 | rs10123842 | 0.307 | -1.495 | 0.442 | -0.361 | -0.031 | 0.013 | 0.013 | 0 | 0.13 | 0.18 | 0.05 | 0.15 | 0.07 | 0.18 |
| 9 | 24782317 | rs12337557 | 0.176 | -1.429 | 0.22 | -0.466 | 0.005 | 0.008 | 0.008 | 0 | 0.22 | 0.23 | 0.17 | 0.07 | 0.19 | 0.08 |
| 9 | 24783531 | rs9298853 | -1.036 | -1.243 | 0.012 | -0.395 | 0 | -0.002 | -0.002 | 0 | 0.42 | 0.64 | 0.37 | 0.22 | 0.43 | 0.42 |
| 9 | 24785247 | rs10812095 | -0.174 | -1.289 | 0.148 | -0.179 | 0.042 | -0.048 | -0.048 | 0 | 0.22 | 0.14 | 0.19 | 0.61 | 0.13 | 0.25 |
| 9 | 24788179 | rs10732345 | -2.208 | -1.182 | 0.026 | -0.498 | -0.017 | 0.031 | 0.031 | 0 | 0.62 | 0.77 | 0.54 | 0.82 | 0.53 | 0.65 |
| 9 | 24788197 | rs10732346 | -2.447 | -1.189 | -0.063 | -0.539 | -0.12 | 0.111 | 0.111 | 0.004 | 0.64 | 0.93 | 0.56 | 0.82 | 0.55 | 0.66 |
| 9 | 24795767 | rs12551729 | -0.115 | -1.226 | 0.102 | -0.169 | 0.037 | -0.049 | -0.049 | 0 | 0.2 | 0.14 | 0.19 | 0.62 | 0.12 | 0.21 |
| 9 | 24797747 | rs7025209 | 0.339 | -1.239 | -0.168 | -0.195 | 0.013 | 0.048 | 0.048 | 0 | 0.19 | 0.08 | 0.13 | 0.03 | 0.17 | 0.08 |
| 9 | 24799884 | rs9406815 | -0.486 | -0.391 | 0.205 | 0.372 | -0.033 | 0.047 | 0.047 | 0 | 0.22 | 0.28 | 0.2 | 0.62 | 0.13 | 0.26 |
| 9 | 24805161 | rs11795390 | -0.997 | 0.657 | -0.131 | 0.139 | 0.038 | 0.008 | 0.008 | 0 | 0.07 | 0.1 | 0.16 | 0.1 | 0.18 | 0.05 |
| 9 | 24805660 | rs10966479 | -0.513 | 0.547 | -0.119 | 0.057 | 0 | -0.005 | -0.005 | 0 | 0.47 | 0.53 | 0.36 | 0.23 | 0.46 | 0.45 |
| 9 | 24821064 | rs9408006 | -0.236 | 0.612 | -0.002 | 0.079 | 0.001 | -0.015 | -0.015 | 0 | 0.44 | 0.44 | 0.21 | 0.14 | 0.31 | 0.46 |
| 9 | 24821079 | rs17772576 | -0.999 | 0.732 | 0.028 | 0.139 | 0.091 | 0.024 | 0.024 | 0 | 0.07 | 0.14 | 0.16 | 0.1 | 0.19 | 0.04 |
| 9 | 24821877 | rs7039242 | -1.315 | 0.663 | 0.04 | 0.137 | 0.096 | 0.025 | 0.025 | 0 | 0.07 | 0.17 | 0.16 | 0.1 | 0.19 | 0.1 |
| 9 | 24823390 | rs10115985 | -0.02 | -0.245 | -0.185 | 0.393 | -0.035 | 0.173 | 0.173 | 0.015 | 0.16 | 0.37 | 0.18 | 0.57 | 0.09 | 0.2 |
| 9 | 24823420 | rs10966490 | -0.067 | -0.239 | -0.196 | 0.25 | -0.035 | -0.065 | -0.065 | 0 | 0.42 | 0.45 | 0.19 | 0.12 | 0.27 | 0.27 |
| 9 | 24833789 | rs2026590 | 0.157 | -0.052 | -0.322 | 0.338 | 0.008 | -0.001 | -0.001 | 0 | 0.39 | 0.27 | 0.16 | 0.12 | 0.25 | 0.27 |
| 9 | 24837092 | rs9406825 | -0.006 | -0.141 | -0.461 | 0.455 | 0.015 | 0.02 | 0.02 | 0 | 0.42 | 0.32 | 0.29 | 0.24 | 0.35 | 0.34 |
| 9 | 24837608 | rs7867489 | -0.224 | -0.142 | -0.489 | 0.455 | 0.005 | 0.008 | 0.008 | 0 | 0.45 | 0.38 | 0.33 | 0.28 | 0.42 | 0.37 |
| 9 | 24838852 | rs2038965 | -1.095 | -0.357 | -0.555 | 0.448 | -0.044 | 0.131 | 0.131 | 0 | 0.6 | 0.93 | 0.48 | 0.77 | 0.44 | 0.59 |
| 9 | 24848892 | rs973778 | -0.309 | -0.414 | -0.435 | 0.359 | 0.018 | 0.026 | 0.026 | 0 | 0.48 | 0.57 | 0.35 | 0.28 | 0.4 | 0.48 |
| 9 | 24850287 | rs7861757 | 0.141 | -0.28 | -1.335 | -0.133 | 0.012 | 0.013 | 0.013 | 0 | 0.4 | 0.29 | 0.29 | 0.12 | 0.34 | 0.27 |
| 9 | 24855304 | rs7024994 | -0.424 | 0.61 | -2.039 | 0.069 | -0.141 | 0.128 | 0.128 | 0.047 | 0.62 | 0.97 | 0.54 | 0.81 | 0.51 | 0.63 |
| 9 | 24868290 | rs7871116 | -0.769 | 0.082 | -0.233 | 0.561 | -0.005 | -0.038 | -0.038 | 0.014 | 0.16 | 0.58 | 0.14 | 0.09 | 0.16 | 0.16 |
| 9 | 24875821 | rs7863590 | -0.742 | 0.074 | 0.603 | 0.564 | 0.016 | -0.009 | -0.009 | 0 | 0.13 | 0.43 | 0.12 | 0.1 | 0.14 | 0.14 |

| | | | | | | | | | | | | | | | | |
|---|---|---|---|---|---|---|---|---|---|---|---|---|---|---|---|---|
| 9 | 24878417 | rs7875618 | -0.659 | -0.382 | -0.2 | 0.739 | -0.003 | 0 | 0 | 0 | 0.55 | 0.44 | 0.44 | 0.46 | 0.45 | 0.44 |
| 9 | 24881879 | rs7866172 | -0.137 | -0.921 | -0.214 | 0.946 | -0.024 | 0.046 | 0.046 | 0 | 0.07 | 0.15 | 0.12 | 0.15 | 0.01 | 0.02 |
| 9 | 24883179 | rs7873820 | -0.957 | -1.06 | -0.24 | 1.092 | 0.029 | -0.016 | -0.016 | 0 | 0.2 | 0.88 | 0.15 | 0.22 | 0.17 | 0.19 |
| 9 | 24883440 | rs9408015 | -0.053 | -1.05 | -0.226 | 1.073 | 0.002 | -0.023 | -0.023 | 0 | 0.15 | 0.49 | 0.12 | 0.1 | 0.15 | 0.16 |
| 9 | 24884969 | rs10812114 | -0.002 | -1.145 | -0.288 | 1.128 | 0.085 | -0.044 | -0.044 | 0 | 0.43 | 0.11 | 0.35 | 0.42 | 0.32 | 0.32 |
| 9 | 24895221 | rs10118854 | -0.091 | -1.056 | -0.079 | 1.108 | -0.008 | -0.027 | -0.027 | 0.528 | 0.15 | 0.53 | 0.12 | 0.09 | 0.15 | 0.16 |
| 9 | 24895412 | rs7045754 | -0.135 | -1.056 | -0.097 | 1.108 | 0.003 | -0.02 | -0.02 | 0 | 0.15 | 0.44 | 0.12 | 0.09 | 0.15 | 0.16 |
| 9 | 24895992 | rs7046259 | -0.578 | -1.064 | -0.142 | 1.104 | -0.01 | -0.02 | -0.02 | 0.132 | 0.17 | 0.74 | 0.14 | 0.09 | 0.16 | 0.17 |
| 9 | 24902466 | rs7022130 | -1.057 | -0.016 | 0.499 | 1.071 | -0.006 | 0.008 | 0.008 | 0.37 | 0.59 | 0.45 | 0.46 | 0.54 | 0.47 | 0.46 |
| 9 | 24905439 | rs12686755 | 1.189 | 0.157 | 0.306 | 1.247 | -0.162 | -0.051 | -0.051 | 0 | 0.35 | 0.04 | 0.43 | 0.33 | 0.42 | 0.46 |
| 9 | 24906802 | rs4316218 | 0.046 | -0.003 | 0.255 | 1.18 | 0.103 | -0.031 | -0.031 | 0.308 | 0.43 | 0.09 | 0.35 | 0.41 | 0.31 | 0.29 |
| 9 | 24911674 | rs1318995 | -1.424 | 0.017 | 0.403 | 1.149 | -0.021 | 0.007 | 0.007 | 1.075 | 0.63 | 0.82 | 0.51 | 0.66 | 0.5 | 0.49 |
| 9 | 24912753 | rs7849819 | -0.598 | -0.015 | 0.653 | 1.091 | -0.01 | -0.041 | -0.041 | 0 | 0.18 | 0.62 | 0.15 | 0.1 | 0.19 | 0.19 |
| 9 | 24920900 | rs10812137 | -0.051 | 0 | 0.972 | 1.091 | 0 | -0.022 | -0.022 | 0 | 0.14 | 0.27 | 0.11 | 0.1 | 0.16 | 0.17 |
| 9 | 24922485 | rs2126176 | -0.951 | -0.827 | 0.379 | 0.494 | -0.023 | -0.074 | -0.074 | 0.416 | 0.36 | 0.55 | 0.26 | 0.1 | 0.34 | 0.28 |
| 9 | 24924449 | rs10966571 | -1.034 | -0.803 | 0.399 | 0.717 | 0.002 | 0.001 | 0.001 | 0 | 0.48 | 0.47 | 0.4 | 0.58 | 0.34 | 0.31 |
| 9 | 24932377 | rs10966576 | 0.331 | -0.844 | 0.182 | 0.586 | -0.003 | -0.038 | -0.038 | 0.837 | 0.18 | 0.21 | 0.14 | 0 | 0.17 | 0.11 |
| 9 | 24942014 | rs10812142 | -0.1 | -0.844 | 0.299 | 0.685 | 0.033 | -0.037 | -0.037 | 0.005 | 0.29 | 0.2 | 0.25 | 0.55 | 0.18 | 0.22 |
| 9 | 24946349 | rs1461333 | 0.215 | -0.732 | 0.341 | 0.642 | 0.076 | -0.037 | -0.037 | 0 | 0.83 | 0.48 | 0.84 | 0.9 | 0.77 | 0.8 |
| 9 | 24956623 | rs7856326 | 0.515 | -0.86 | 0.211 | 0.886 | 0.027 | -0.024 | -0.024 | 0 | 0.23 | 0.1 | 0.15 | 0.24 | 0.16 | 0.2 |
| 9 | 24960893 | rs7851190 | 0.636 | -0.701 | 0.367 | 0.874 | 0.02 | 0.006 | 0.006 | 1.691 | 0.78 | 0.32 | 0.8 | 0.76 | 0.72 | 0.77 |
| 9 | 24973797 | rs10738700 | 2.009 | -0.8 | 0.235 | 0.923 | -0.178 | -0.077 | -0.077 | 0 | 0.36 | 0.03 | 0.45 | 0.37 | 0.44 | 0.5 |
| 9 | 24976322 | rs7038176 | 0.745 | -0.809 | 0.381 | 0.923 | 0.001 | 0 | 0 | 0 | 0.55 | 0.55 | 0.59 | 0.47 | 0.61 | 0.65 |
| 9 | 24977201 | rs1461338 | -0.346 | -0.811 | 0.447 | 1.065 | -0.016 | 0.128 | 0.128 | 1.489 | 0.19 | 0.47 | 0.21 | 0.39 | 0.18 | 0.18 |
| 9 | 24979796 | rs7021001 | 0.4 | -0.861 | 0.251 | 0.967 | 0.023 | -0.031 | -0.031 | 0.483 | 0.23 | 0.17 | 0.23 | 0.4 | 0.14 | 0.21 |
| 9 | 24984658 | rs10757505 | 1.253 | -0.865 | 0.191 | 0.871 | -0.025 | 0.1 | 0.1 | 0.8 | 0.59 | 0.14 | 0.65 | 0.77 | 0.56 | 0.69 |
| 9 | 24985359 | rs10812151 | 0.328 | -0.894 | 0.181 | 0.889 | 0.034 | -0.048 | -0.048 | 0 | 0.23 | 0.11 | 0.22 | 0.4 | 0.14 | 0.21 |
| 9 | 24985660 | rs1461340 | 0.328 | -0.894 | 0.181 | 0.76 | 0.037 | -0.038 | -0.038 | 0 | 0.23 | 0.11 | 0.22 | 0.33 | 0.13 | 0.2 |
| 9 | 24985964 | rs10757506 | 1.813 | -0.885 | 0.137 | 0.748 | -0.105 | -0.059 | -0.059 | 0 | 0.37 | 0.07 | 0.44 | 0.43 | 0.43 | 0.49 |
| 9 | 24988020 | rs10812152 | 1.821 | -0.857 | 0.018 | 0.767 | -0.147 | -0.097 | -0.097 | 0 | 0.37 | 0.04 | 0.44 | 0.44 | 0.43 | 0.51 |
| 9 | 24996421 | rs1461331 | -0.322 | -0.899 | 0.032 | 0.584 | 0.035 | 0.011 | 0.011 | 0 | 0.19 | 0.12 | 0.1 | 0.14 | 0.11 | 0.11 |
| 9 | 24998099 | rs10966629 | 0.426 | -0.852 | 0.702 | 0.295 | 0.098 | -0.064 | -0.064 | 0.122 | 0.79 | 0.49 | 0.79 | 0.89 | 0.69 | 0.81 |
| 9 | 24998156 | rs10812156 | -0.354 | -0.957 | 0.338 | 0.495 | 0.011 | 0.072 | 0.072 | 1.058 | 0.74 | 0.49 | 0.69 | 0.55 | 0.76 | 0.75 |
| 9 | 24999327 | rs1125516 | 0.605 | -0.707 | -0.159 | 0.649 | 0.041 | -0.012 | -0.012 | 0 | 0.55 | 0.27 | 0.57 | 0.55 | 0.67 | 0.65 |
| 9 | 25003295 | rs10812158 | -0.905 | -0.3 | 0.347 | 0.322 | 0.003 | -0.063 | -0.063 | 0.457 | 0.19 | 0.42 | 0.15 | 0.09 | 0.22 | 0.17 |
| 9 | 25004127 | rs7857156 | -1.126 | -0.363 | -0.46 | 0.184 | 0.016 | 0.043 | 0.043 | 0 | 0.24 | 0.16 | 0.15 | 0.06 | 0.22 | 0.18 |
| 9 | 25004901 | rs6475831 | -1.153 | 0.386 | -1.27 | 0.398 | -0.061 | -0.067 | -0.067 | 0.093 | 0.57 | 0.9 | 0.56 | 0.55 | 0.48 | 0.51 |
| 9 | 25005553 | rs964084 | -0.883 | 0.962 | -1.259 | -0.015 | -0.019 | -0.032 | -0.032 | 0 | 0.23 | 0.15 | 0.37 | 0.57 | 0.31 | 0.27 |
| 9 | 25008242 | rs7046929 | -0.493 | 0.127 | -1.692 | 0.156 | 0.011 | -0.102 | -0.102 | 0.563 | 0.14 | 0.02 | 0.25 | 0.54 | 0.09 | 0.19 |
| 9 | 25010710 | rs10124126 | -0.382 | 0.414 | -0.843 | 0.132 | 0.087 | 0.112 | 0.112 | 0.131 | 0.07 | 0.53 | 0.23 | 0.09 | 0.04 | 0.08 |
| 9 | 25027430 | rs1563950 | -1.56 | 0.583 | -2.537 | 0.013 | -0.05 | 0.022 | 0.022 | 0 | 0.55 | 0.9 | 0.65 | 0.69 | 0.47 | 0.59 |
| 9 | 25029338 | rs13286283 | -0.485 | 0.177 | -1.944 | -0.702 | 0.029 | 0.157 | 0.157 | 0 | 0.28 | 0.04 | 0.11 | 0.04 | 0.25 | 0.26 |
| 9 | 25035251 | rs9298861 | 0.585 | 0.597 | -1.593 | -0.124 | -0.002 | 0.012 | 0.012 | 0 | 0.69 | 0.82 | 0.63 | 0.34 | 0.68 | 0.55 |
| 9 | 25049507 | rs754636 | -0.832 | 1.536 | -1.354 | -0.434 | -0.012 | 0.029 | 0.029 | 0 | 0.31 | 0.21 | 0.44 | 0.74 | 0.35 | 0.49 |
| 9 | 25068476 | rs10812169 | -1.357 | 1.366 | -0.257 | -0.01 | -0.015 | 0.065 | 0.065 | 0 | 0.09 | 0.25 | 0.04 | 0.12 | 0.04 | 0.15 |
| 9 | 25090139 | rs10966726 | -0.55 | 1.423 | -0.379 | -0.044 | -0.095 | -0.104 | -0.104 | 0.588 | 0.28 | 0.05 | 0.41 | 0.62 | 0.35 | 0.38 |
| 9 | 25093183 | rs10812183 | -1.345 | 0.8 | 0.287 | -0.483 | -0.045 | 0.068 | 0.068 | 0 | 0.2 | 0.47 | 0.13 | 0.26 | 0.12 | 0.3 |
| 9 | 25100472 | rs7467005 | -0.463 | 1.198 | 0.21 | -0.368 | -0.155 | -0.102 | -0.102 | 0.171 | 0.29 | 0.05 | 0.44 | 0.63 | 0.43 | 0.39 |
| 9 | 25117372 | rs1925478 | -0.461 | 0.978 | 1.056 | -0.528 | -0.106 | -0.061 | -0.061 | 0 | 0.29 | 0.08 | 0.44 | 0.65 | 0.42 | 0.38 |
| 9 | 25122214 | rs1886709 | -0.807 | 0.934 | 0.867 | -0.527 | -0.058 | 0.002 | 0.002 | 0 | 0.35 | 0.17 | 0.48 | 0.69 | 0.48 | 0.45 |
| 9 | 25127661 | rs3126961 | -1.749 | 0.377 | 0.4 | -1.249 | 0.013 | 0.057 | 0.057 | 1.404 | 0.52 | 0.58 | 0.56 | 0.9 | 0.56 | 0.65 |
| 9 | 25139818 | rs3118280 | -1.663 | 0.35 | 0.831 | -1.057 | -0.007 | -0.034 | -0.034 | 0 | 0.3 | 0.55 | 0.17 | 0.22 | 0.23 | 0.26 |
| 9 | 25142445 | rs3126975 | -1.065 | 0.327 | 1.007 | -1.093 | -0.003 | -0.002 | -0.002 | 0 | 0.22 | 0.24 | 0.1 | 0.13 | 0.12 | 0.2 |
| 9 | 25143480 | rs3118292 | -1.886 | 0.354 | 1.082 | -1.007 | 0.014 | -0.081 | -0.081 | 0 | 0.19 | 0.47 | 0.13 | 0.14 | 0.17 | 0.16 |
| 9 | 25157678 | rs3118318 | -1.029 | 0.346 | 1.038 | -0.991 | -0.018 | 0.009 | 0.009 | 0.002 | 0.12 | 0.25 | 0.07 | 0.13 | 0.08 | 0.19 |
| 9 | 25163771 | rs1952053 | -2.316 | 0.312 | 1.119 | -1.033 | -0.003 | 0.017 | 0.017 | 3.078 | 0.23 | 0.86 | 0.16 | 0.22 | 0.19 | 0.26 |
| 9 | 25176251 | rs7034635 | -1.929 | 0.452 | 1.301 | -1.01 | 0.065 | -0.027 | -0.027 | 3.704 | 0.1 | 0.61 | 0.09 | 0.09 | 0.12 | 0.12 |
| 9 | 25178084 | rs12337176 | -1.078 | 0.458 | 1.195 | -1.006 | 0.127 | 0.02 | 0.02 | 2.304 | 0.07 | 0.29 | 0.06 | 0.09 | 0.11 | 0.07 |
| 9 | 25178540 | rs1886716 | -2.535 | 0.447 | 0.824 | -1.009 | 0.009 | -0.015 | -0.015 | 0 | 0.13 | 0.74 | 0.13 | 0.14 | 0.15 | 0.19 |
| 9 | 25188506 | rs7033077 | 2.23 | 0.31 | 0.719 | -0.92 | 0.003 | 0.03 | 0.03 | 3.557 | 0.8 | 0.16 | 0.86 | 0.85 | 0.84 | 0.8 |
| 9 | 25193401 | rs9406859 | 1.789 | 0.188 | 0.335 | -0.738 | 0 | 0.225 | 0.225 | 0.048 | 0.48 | 0.07 | 0.44 | 0.21 | 0.44 | 0.41 |
| 9 | 25195023 | rs7039724 | -1.624 | 0.037 | 0.957 | -0.566 | 0.004 | -0.045 | -0.045 | 0.109 | 0.17 | 0.63 | 0.09 | 0.11 | 0.13 | 0.2 |
| 9 | 25196181 | rs7871597 | 0.959 | 0.033 | 0.989 | -0.579 | -0.033 | -0.045 | -0.045 | 0.204 | 0.89 | 0.73 | 0.94 | 0.95 | 0.94 | 0.81 |
| 9 | 25212649 | rs1412517 | 1.823 | 0.073 | 0.871 | -0.411 | 0.015 | 0.021 | 0.021 | 4.101 | 0.69 | 0.15 | 0.83 | 0.72 | 0.78 | 0.62 |
| 9 | 25218359 | rs1332437 | 1.547 | 0.238 | 1.261 | -0.352 | 0.142 | 0.045 | 0.045 | 2.883 | 0.91 | 0.58 | 0.9 | 0.87 | 0.88 | 0.87 |
| 9 | 25227233 | rs9408080 | -1.054 | 0.246 | 1.255 | -0.351 | 0.119 | 0.018 | 0.018 | 2.184 | 0.07 | 0.28 | 0.06 | 0.08 | 0.1 | 0.05 |
| 9 | 25227443 | rs9408081 | -1.054 | 0.246 | 1.255 | -0.351 | 0.119 | 0.018 | 0.018 | 2.184 | 0.07 | 0.28 | 0.06 | 0.08 | 0.1 | 0.05 |
| 9 | 25227846 | rs10966805 | -0.545 | 0.307 | 0.761 | -0.802 | 0.027 | 0.004 | 0.004 | 0 | 0.35 | 0.4 | 0.49 | 0.63 | 0.5 | 0.29 |
| 9 | 25233460 | rs10966810 | 0.58 | 0.294 | 0.679 | -0.78 | 0.007 | -0.003 | -0.003 | 1.299 | 0.62 | 0.45 | 0.47 | 0.32 | 0.47 | 0.52 |

| | | | | | | | | | | | | | | | | |
|---|---|---|---|---|---|---|---|---|---|---|---|---|---|---|---|---|
| 9 | 25233484 | rs10966811 | 0.402 | 0.387 | 0.202 | -0.727 | -0.096 | -0.115 | -0.115 | 0.378 | 0.27 | 0.05 | 0.42 | 0.56 | 0.39 | 0.23 |
| 9 | 25241082 | rs7027321 | 1.523 | 0.252 | 0.181 | -0.746 | -0.003 | 0.117 | 0.117 | 0 | 0.4 | 0.14 | 0.4 | 0.19 | 0.37 | 0.36 |
| 9 | 25262246 | rs10738710 | 1.322 | 0.704 | 0.756 | -0.318 | 0.143 | 0.02 | 0.02 | 2.524 | 0.92 | 0.57 | 0.93 | 0.94 | 0.9 | 0.82 |
| 9 | 25263760 | rs1625700 | -0.415 | 0.809 | 0.539 | -0.25 | 0.122 | 0.022 | 0.022 | 0.263 | 0.05 | 0.25 | 0.06 | 0.06 | 0.1 | 0.06 |
| 9 | 25263918 | rs1758734 | 0.141 | 0.633 | -0.352 | -0.171 | -0.058 | -0.057 | -0.057 | 0.317 | 0.68 | 0.86 | 0.52 | 0.45 | 0.55 | 0.76 |
| 9 | 25275213 | rs10812209 | -0.362 | 0.817 | 0.216 | 0.541 | -0.005 | -0.014 | -0.014 | 0 | 0.33 | 0.27 | 0.35 | 0.39 | 0.4 | 0.21 |
| 9 | 25275893 | rs7857330 | 0.132 | 0.582 | 0.319 | 0.422 | -0.01 | -0.037 | -0.037 | 0 | 0.1 | 0.21 | 0.08 | 0.04 | 0.08 | 0.06 |
| 9 | 25276559 | rs1332438 | -0.119 | 0.969 | -0.927 | 0.915 | -0.053 | -0.076 | -0.076 | 0 | 0.13 | 0.01 | 0.26 | 0.25 | 0.18 | 0.22 |
| 9 | 25281072 | rs10966831 | 0.056 | 0.797 | -1.013 | 0.982 | -0.009 | -0.074 | -0.074 | 0.69 | 0.11 | 0.01 | 0.06 | 0.28 | 0.08 | 0.15 |
| 9 | 25281912 | rs10966833 | 0.444 | 1.408 | -1.29 | 0.935 | -0.122 | -0.136 | -0.136 | 0 | 0.26 | 0.03 | 0.39 | 0.4 | 0.36 | 0.2 |
| 9 | 25283201 | rs10757517 | -0.184 | 1.03 | -0.271 | 0.756 | 0 | 0 | 0 | 0.039 | 0.4 | 0.4 | 0.55 | 0.5 | 0.56 | 0.3 |
| 9 | 25287180 | rs1799615 | 0.946 | 0.865 | -0.847 | 0.28 | -0.009 | 0.132 | 0.132 | 0 | 0.29 | 0.07 | 0.19 | 0.04 | 0.26 | 0.18 |
| 9 | 25296601 | rs10966845 | 0.012 | 1.13 | -0.586 | 1.681 | -0.034 | -0.128 | -0.128 | 0.309 | 0.22 | 0.01 | 0.11 | 0.32 | 0.18 | 0.13 |
| 9 | 25298264 | rs1668962 | 0.47 | 1.453 | 1.183 | 2.026 | 0.014 | 0.132 | 0.132 | 3.245 | 0.85 | 0.56 | 0.86 | 0.69 | 0.86 | 0.77 |
| 9 | 25299770 | rs17782830 | 0.027 | 1.18 | 1.026 | 1.856 | 0.003 | -0.015 | -0.015 | 1.655 | 0.11 | 0.07 | 0.04 | 0.26 | 0.1 | 0.17 |
| 9 | 25301050 | rs1332440 | -0.737 | 1.427 | 1.288 | 1.995 | 0.001 | 0.005 | 0.005 | 0 | 0.23 | 0.24 | 0.38 | 0.33 | 0.26 | 0.39 |
| 9 | 25303763 | rs17176169 | 0.52 | 1.386 | 1.221 | 2.003 | -0.016 | -0.066 | -0.066 | 0.264 | 0.11 | 0 | 0.04 | 0.27 | 0.1 | 0.19 |
| 9 | 25306889 | rs2084743 | 1.753 | 1.328 | 2.006 | 1.917 | -0.002 | 0.003 | 0.003 | 1.057 | 0.6 | 0.39 | 0.59 | 0.4 | 0.62 | 0.4 |
| 9 | 25307811 | rs1751817 | -1.764 | 1.428 | 2.244 | 1.968 | 0.004 | -0.002 | -0.002 | 0.441 | 0.38 | 0.53 | 0.4 | 0.6 | 0.38 | 0.6 |
| 9 | 25314595 | rs1751813 | -0.661 | 1.465 | 2.35 | 2.053 | 0.004 | 0.009 | 0.009 | 0 | 0.22 | 0.23 | 0.32 | 0.32 | 0.26 | 0.37 |
| 9 | 25316440 | rs1855074 | 1.562 | 1.53 | 2.454 | 2.039 | 0.001 | -0.002 | -0.002 | 0 | 0.63 | 0.41 | 0.61 | 0.42 | 0.66 | 0.42 |
| 9 | 25323165 | rs1372875 | -1.611 | 1.483 | 2.477 | 2.074 | 0.01 | 0.018 | 0.018 | 1.174 | 0.34 | 0.49 | 0.38 | 0.58 | 0.35 | 0.55 |
| 9 | 25325829 | rs1372873 | -1.59 | 1.484 | 2.495 | 2.199 | 0.013 | 0.043 | 0.043 | 1.307 | 0.34 | 0.43 | 0.38 | 0.45 | 0.35 | 0.55 |
| 9 | 25325895 | rs1372872 | -0.613 | 1.494 | 2.405 | 2.15 | -0.002 | -0.002 | -0.002 | 0 | 0.22 | 0.21 | 0.32 | 0.32 | 0.25 | 0.37 |
| 9 | 25330810 | rs1327385 | -0.339 | 1.538 | 2.461 | 2.034 | 0.001 | 0.073 | 0.073 | 3.845 | 0.88 | 0.78 | 0.95 | 0.73 | 0.9 | 0.82 |
| 9 | 25334987 | rs1751811 | -0.737 | 1.569 | 2.206 | 1.846 | -0.006 | -0.013 | -0.013 | 0 | 0.23 | 0.2 | 0.32 | 0.33 | 0.26 | 0.38 |
| 9 | 25336447 | rs1668986 | -0.481 | 1.572 | 2.165 | 1.853 | -0.002 | 0.002 | 0.002 | 0.794 | 0.17 | 0.17 | 0.3 | 0.29 | 0.22 | 0.37 |
| 9 | 25352201 | rs1758726 | -0.764 | 1.892 | 2.076 | 0.905 | -0.001 | -0.003 | -0.003 | 0 | 0.22 | 0.22 | 0.2 | 0.04 | 0.21 | 0.13 |
| 9 | 25353887 | rs980462 | -2.12 | 1.856 | 2.222 | 0.95 | -0.016 | 0.018 | 0.018 | 0.955 | 0.42 | 0.69 | 0.44 | 0.37 | 0.46 | 0.36 |
| 9 | 25359869 | rs7862611 | -1.007 | 1.155 | 2.168 | 1.038 | -0.005 | 0.092 | 0.092 | 2.261 | 0.2 | 0.39 | 0.2 | 0.43 | 0.19 | 0.22 |
| 9 | 25363830 | rs967671 | -2.054 | 1.085 | 2.119 | 1.026 | 0.001 | -0.006 | -0.006 | 0 | 0.42 | 0.67 | 0.41 | 0.47 | 0.4 | 0.35 |
| 9 | 25368388 | rs10122848 | -0.333 | 1.05 | 1.675 | 1.06 | -0.001 | 0.076 | 0.076 | 2.885 | 0.16 | 0.28 | 0.1 | 0.4 | 0.17 | 0.19 |
| 9 | 25377010 | rs3849934 | 0.902 | 0.995 | 1.699 | 0.838 | -0.003 | -0.009 | -0.009 | 0 | 0.73 | 0.6 | 0.69 | 0.8 | 0.76 | 0.83 |
| 9 | 25384487 | rs1576033 | 1.609 | 1.108 | 1.335 | 0.841 | -0.004 | 0.041 | 0.041 | 0.295 | 0.59 | 0.31 | 0.66 | 0.8 | 0.59 | 0.67 |
| 9 | 25385317 | rs10757521 | 1.839 | 1.137 | 1.317 | 0.742 | -0.001 | -0.01 | -0.01 | 0.945 | 0.57 | 0.3 | 0.57 | 0.55 | 0.58 | 0.64 |
| 9 | 25389512 | rs1576031 | 1.487 | 1.149 | 1.28 | 0.738 | 0.007 | -0.026 | -0.026 | 0 | 0.61 | 0.5 | 0.68 | 0.71 | 0.59 | 0.67 |
| 9 | 25392322 | rs10733401 | 1.478 | 1.167 | 1.252 | 0.747 | 0.007 | -0.025 | -0.025 | 0 | 0.61 | 0.5 | 0.68 | 0.71 | 0.59 | 0.67 |
| 9 | 25396789 | rs17701704 | 0.071 | 1.175 | 1.17 | 0.742 | -0.01 | -0.036 | -0.036 | 7.675 | 0.13 | 0.04 | 0.08 | 0.26 | 0.16 | 0.19 |
| 9 | 25398069 | rs945811 | 1.746 | 1.22 | 1.175 | 0.516 | -0.003 | -0.012 | -0.012 | 0.381 | 0.57 | 0.28 | 0.56 | 0.56 | 0.58 | 0.63 |
| 9 | 25412587 | rs2031735 | 0.914 | 1.182 | 1.393 | -0.175 | 0.022 | -0.105 | -0.105 | 0 | 0.8 | 0.69 | 0.78 | 1 | 0.77 | 0.86 |
| 9 | 25416556 | rs1831078 | 0.824 | 1.176 | 0.92 | -0.125 | 0.016 | -0.112 | -0.112 | 0.764 | 0.74 | 0.56 | 0.76 | 0.96 | 0.75 | 0.86 |
| 9 | 25426760 | rs1831077 | -0.973 | 0.756 | -0.354 | 0.345 | -0.008 | 0.035 | 0.035 | 0.625 | 0.2 | 0.08 | 0.21 | 0.09 | 0.23 | 0.17 |
| 9 | 25427205 | rs4977977 | -1.116 | 0.511 | -0.193 | 0.384 | 0.01 | 0.02 | 0.02 | 0.734 | 0.19 | 0.06 | 0.18 | 0.14 | 0.18 | 0.17 |
| 9 | 25440150 | rs12380757 | -1.475 | 0.834 | 0.497 | 0.343 | -0.003 | -0.009 | -0.009 | 0.675 | 0.23 | 0.26 | 0.31 | 0.16 | 0.22 | 0.18 |
| 9 | 25440872 | rs7035858 | -1.185 | 0.871 | 0.451 | 0.318 | 0.007 | 0.017 | 0.017 | 0 | 0.09 | 0.13 | 0.14 | 0.11 | 0.1 | 0.05 |
| 9 | 25442538 | rs10966910 | -1.266 | 0.845 | 0.464 | 0.3 | -0.001 | -0.044 | -0.044 | 0.083 | 0.18 | 0.25 | 0.29 | 0.06 | 0.17 | 0.14 |
| 9 | 25451196 | rs7856650 | -0.374 | 0.847 | 0.819 | 0.294 | 0.003 | 0.032 | 0.032 | 0.295 | 0.13 | 0.05 | 0.16 | 0.06 | 0.12 | 0.13 |
| 9 | 25452963 | rs7861656 | -1.902 | 0.804 | 1.541 | 0.25 | 0.007 | -0.029 | -0.029 | 0.047 | 0.24 | 0.44 | 0.33 | 0.17 | 0.24 | 0.18 |
| 9 | 25457130 | rs10966921 | -1.466 | 0.836 | 1.565 | 0.27 | 0.03 | 0.024 | 0.024 | 0 | 0.1 | 0.3 | 0.06 | 0.1 | 0.12 | 0.05 |
| 9 | 25461099 | rs10812251 | -1.305 | 0.647 | 0.856 | 0.521 | 0.004 | 0.026 | 0.026 | 0.318 | 0.26 | 0.4 | 0.16 | 0.33 | 0.24 | 0.22 |
| 9 | 25462188 | rs10812252 | -0.267 | 0.116 | 0.402 | 0.236 | 0.019 | -0.007 | -0.007 | 0 | 0.22 | 0.18 | 0.15 | 0.23 | 0.14 | 0.2 |
| 9 | 25463339 | rs1576026 | 1.625 | -0.078 | 1.045 | 0.137 | 0.025 | 0.022 | 0.022 | 0.129 | 0.52 | 0.32 | 0.62 | 0.61 | 0.63 | 0.63 |
| 9 | 25471466 | rs12345447 | -0.665 | -0.112 | 1.325 | 0.31 | -0.001 | 0.01 | 0.01 | 0 | 0.24 | 0.35 | 0.14 | 0.28 | 0.2 | 0.19 |
| 9 | 25472081 | rs7028841 | -0.662 | -0.107 | 1.24 | 0.313 | -0.001 | 0.004 | 0.004 | 0 | 0.24 | 0.31 | 0.16 | 0.29 | 0.2 | 0.19 |
| 9 | 25481661 | rs7857140 | -0.367 | 0.694 | -0.138 | 0.883 | -0.03 | -0.064 | -0.064 | 0 | 0.22 | 0.07 | 0.32 | 0.34 | 0.25 | 0.19 |
| 9 | 25485676 | rs10966932 | -0.266 | -0.053 | -0.436 | 0.416 | 0.028 | 0.021 | 0.021 | 0.186 | 0.31 | 0.21 | 0.23 | 0.23 | 0.24 | 0.14 |
| 9 | 25487360 | rs7024522 | 1.191 | 0.143 | -0.698 | 0.308 | -0.059 | 0.009 | 0.009 | 0.695 | 0.37 | 0.2 | 0.39 | 0.39 | 0.48 | 0.55 |
| 9 | 25491653 | rs10757544 | -0.06 | 0.212 | -0.388 | 0.415 | -0.008 | -0.044 | -0.044 | 0.031 | 0.28 | 0.13 | 0.34 | 0.38 | 0.27 | 0.28 |
| 9 | 25496361 | rs10966934 | 0.42 | 0.161 | 0.063 | 0.398 | 0.003 | 0.018 | 0.018 | 0 | 0.67 | 0.48 | 0.63 | 0.63 | 0.7 | 0.69 |
| 9 | 25498831 | rs1591029 | 0.391 | 0.155 | 0.075 | 0.369 | 0 | 0.004 | 0.004 | 0 | 0.69 | 0.38 | 0.63 | 0.63 | 0.72 | 0.72 |
| 9 | 25499528 | rs1360523 | 0.2 | 0.161 | 0.175 | 0.374 | 0.006 | 0.041 | 0.041 | 0 | 0.7 | 0.53 | 0.64 | 0.63 | 0.73 | 0.73 |
| 9 | 25500238 | rs1332173 | 0.15 | 0.173 | 0.119 | 0.392 | -0.006 | -0.036 | -0.036 | 0 | 0.25 | 0.15 | 0.32 | 0.37 | 0.24 | 0.27 |
| 9 | 25502753 | rs1576786 | 0.831 | -0.179 | -0.134 | 0.198 | 0 | -0.002 | -0.002 | 0 | 0.56 | 0.5 | 0.53 | 0.44 | 0.61 | 0.58 |
| 9 | 25503991 | rs16909495 | 0.89 | -0.398 | -0.343 | 0.003 | -0.013 | -0.032 | -0.032 | 4.595 | 0.18 | 0.2 | 0.1 | 0.03 | 0.13 | 0.04 |
| 9 | 25509545 | rs4275297 | -0.441 | -0.219 | -0.509 | 0.25 | -0.001 | -0.004 | -0.004 | 0.406 | 0.31 | 0.31 | 0.25 | 0.36 | 0.3 | 0.17 |
| 9 | 25512096 | rs1576789 | -1.327 | -0.044 | -0.533 | 0.279 | 0.025 | 0.048 | 0.048 | 0 | 0.54 | 0.35 | 0.52 | 0.66 | 0.55 | 0.42 |
| 9 | 25512140 | rs10812261 | 0.483 | 0.011 | -0.535 | 0.32 | -0.06 | -0.113 | -0.113 | 0 | 0.25 | 0.05 | 0.32 | 0.37 | 0.27 | 0.27 |
| 9 | 25512506 | rs6475847 | -0.424 | 0.095 | -0.263 | 0.127 | -0.009 | 0.018 | 0.018 | 0 | 0.4 | 0.3 | 0.46 | 0.64 | 0.37 | 0.38 |
| 9 | 25514397 | rs2383542 | -1.585 | 0.055 | -0.259 | 0.201 | 0.003 | -0.009 | -0.009 | 0.704 | 0.61 | 0.57 | 0.52 | 0.63 | 0.55 | 0.46 |

| | | | | | | | | | | | | | | | | |
|---|---|---|---|---|---|---|---|---|---|---|---|---|---|---|---|---|
| 9 | 25517233 | rs6475848 | -0.059 | 0.186 | 0.292 | 0.116 | 0.014 | -0.116 | -0.116 | 0.975 | 0.19 | 0.38 | 0.13 | 0.01 | 0.19 | 0.06 |
| 9 | 25522087 | rs10966943 | 0.973 | 0.642 | -0.132 | 0.327 | 0.012 | 0.063 | 0.063 | 0 | 0.55 | 0.21 | 0.54 | 0.69 | 0.61 | 0.71 |
| 9 | 25523107 | rs17774840 | 0.663 | 0.533 | -0.227 | 0.42 | -0.041 | 0.013 | 0.013 | 0 | 0.17 | 0.06 | 0.16 | 0.16 | 0.18 | 0.2 |
| 9 | 25527621 | rs12003047 | 0.088 | 0.381 | 0.242 | 0.345 | 0.007 | -0.005 | -0.005 | 0 | 0.22 | 0.21 | 0.16 | 0.03 | 0.19 | 0.08 |
| 9 | 25532512 | rs4977676 | 1.045 | 0.501 | -0.071 | 0.759 | -0.039 | -0.017 | -0.017 | 0.008 | 0.54 | 0.11 | 0.45 | 0.43 | 0.51 | 0.57 |
| 9 | 25539330 | rs12004657 | -0.447 | 0.326 | -0.052 | 0.723 | -0.001 | -0.003 | -0.003 | 0 | 0.28 | 0.29 | 0.24 | 0.29 | 0.3 | 0.22 |
| 9 | 25540603 | rs12002354 | -0.555 | 0.315 | -0.049 | 0.718 | 0 | -0.002 | -0.002 | 0.033 | 0.29 | 0.31 | 0.24 | 0.29 | 0.3 | 0.22 |
| 9 | 25545690 | rs4631560 | -0.677 | 0.097 | -0.079 | 1.026 | 0.018 | -0.032 | -0.032 | 0 | 0.32 | 0.25 | 0.25 | 0.45 | 0.29 | 0.28 |
| 9 | 25547130 | rs12686619 | 1.445 | 0.09 | -0.144 | 0.97 | 0.027 | 0.011 | 0.011 | 1.516 | 0.55 | 0.37 | 0.59 | 0.41 | 0.66 | 0.7 |
| 9 | 25548004 | rs1412378 | 0.762 | -0.005 | -0.227 | 0.72 | -0.005 | 0.001 | 0.001 | 2.645 | 0.63 | 0.38 | 0.62 | 0.57 | 0.73 | 0.76 |
| 9 | 25548319 | rs17774954 | 0.616 | 0.107 | -0.669 | 0.659 | -0.069 | 0.04 | 0.04 | 0 | 0.18 | 0.05 | 0.16 | 0.13 | 0.2 | 0.2 |
| 9 | 25548513 | rs4977992 | 1.616 | 0.151 | -0.551 | 0.725 | -0.042 | -0.016 | -0.016 | 0 | 0.43 | 0.13 | 0.43 | 0.44 | 0.49 | 0.54 |
| 9 | 25551426 | rs10812270 | 0.584 | 0.408 | -0.419 | 0.613 | -0.016 | 0.037 | 0.037 | 0 | 0.6 | 0.3 | 0.54 | 0.75 | 0.58 | 0.71 |
| 9 | 25551627 | rs7028018 | -0.386 | 0.487 | -0.469 | 0.56 | 0 | -0.017 | -0.017 | 0 | 0.32 | 0.42 | 0.42 | 0.26 | 0.33 | 0.23 |
| 9 | 25552106 | rs17783673 | 0.883 | 0.514 | -0.772 | 0.756 | -0.01 | -0.069 | -0.069 | 0 | 0.11 | 0.01 | 0.07 | 0.25 | 0.09 | 0.14 |
| 9 | 25552702 | rs1332160 | -0.137 | 0.538 | -0.907 | 0.275 | 0.001 | -0.009 | -0.009 | 0 | 0.64 | 0.63 | 0.66 | 0.76 | 0.6 | 0.7 |
| 9 | 25557615 | rs2498725 | 1.242 | 1.026 | -1.058 | 0.466 | -0.019 | -0.009 | -0.009 | 0.011 | 0.16 | 0.11 | 0.17 | 0.19 | 0.2 | 0.2 |
| 9 | 25561285 | rs1929973 | -1.089 | 0.827 | -1.097 | 0.436 | -0.008 | -0.009 | -0.009 | 0 | 0.58 | 0.35 | 0.42 | 0.5 | 0.54 | 0.48 |
| 9 | 25562454 | rs1412385 | -0.725 | 0.521 | -1.259 | 0.426 | 0.019 | -0.012 | -0.012 | 0 | 0.44 | 0.35 | 0.38 | 0.46 | 0.37 | 0.29 |
| 9 | 25562784 | rs2498738 | -0.761 | 0.658 | -1.893 | 0.269 | -0.004 | -0.016 | -0.016 | 0 | 0.57 | 0.29 | 0.42 | 0.5 | 0.54 | 0.49 |
| 9 | 25564648 | rs2482527 | -0.064 | 0.793 | -1.882 | -0.445 | -0.046 | 0.049 | 0.049 | 0.21 | 0.31 | 0.14 | 0.21 | 0.2 | 0.3 | 0.3 |
| 9 | 25564892 | rs2498716 | -0.41 | 0.879 | -1.266 | -0.44 | 0.003 | 0.014 | 0.014 | 1.575 | 0.7 | 0.76 | 0.65 | 0.83 | 0.68 | 0.77 |
| 9 | 25565442 | rs7848711 | -0.631 | 1.228 | -1.311 | -0.145 | -0.005 | 0.002 | 0.002 | 0 | 0.25 | 0.25 | 0.2 | 0.17 | 0.36 | 0.22 |
| 9 | 25576240 | rs10757553 | 0.192 | -0.076 | -1.761 | -0.739 | 0.035 | 0.117 | 0.117 | 0.217 | 0.64 | 0.16 | 0.54 | 0.91 | 0.71 | 0.82 |
| 9 | 25576545 | rs10757554 | 0.62 | -0.498 | -2.055 | -0.4 | 0.073 | 0.133 | 0.133 | 0.333 | 0.57 | 0.13 | 0.41 | 0.75 | 0.68 | 0.75 |
| 9 | 25587832 | rs12340663 | 0.025 | -0.793 | -1.202 | -0.42 | -0.043 | -0.048 | -0.048 | 1.022 | 0.23 | 0.68 | 0.34 | 0.1 | 0.1 | 0.05 |
| 9 | 25594961 | rs12684364 | 0.672 | -0.375 | -1.84 | 0.211 | -0.011 | -0.054 | -0.054 | 0 | 0.26 | 0.1 | 0.12 | 0.35 | 0.23 | 0.33 |
| 9 | 25598014 | rs10966975 | 0.237 | -0.867 | -1.886 | -0.186 | 0.01 | 0.009 | 0.009 | 0 | 0.24 | 0.2 | 0.26 | 0.16 | 0.14 | 0.3 |
| 9 | 25598674 | rs4451410 | 0.58 | -0.902 | -0.985 | -0.188 | -0.003 | -0.002 | -0.002 | 0 | 0.56 | 0.45 | 0.36 | 0.57 | 0.63 | 0.52 |
| 9 | 25599995 | rs4246861 | -0.421 | -0.974 | -0.718 | -0.206 | -0.03 | 0.003 | 0.003 | 0 | 0.34 | 0.44 | 0.55 | 0.42 | 0.21 | 0.38 |
| 9 | 25604885 | rs10123020 | -1.018 | -0.565 | -0.614 | -0.053 | 0.008 | -0.031 | -0.031 | 0 | 0.9 | 0.88 | 0.91 | 0.99 | 0.83 | 0.89 |
| 9 | 25606107 | rs10966982 | 0.987 | -0.54 | -0.755 | 0.036 | 0.006 | -0.108 | -0.108 | 0 | 0.26 | 0.08 | 0.11 | 0.45 | 0.23 | 0.35 |
| 9 | 25607913 | rs4433235 | 0.151 | -0.423 | -0.278 | 0.248 | 0.008 | 0.124 | 0.124 | 0 | 0.12 | 0.33 | 0.3 | 0.3 | 0.08 | 0.09 |
| 9 | 25610272 | rs7032627 | -0.706 | -0.191 | -0.279 | 0.067 | 0.007 | -0.028 | -0.028 | 0 | 0.89 | 0.88 | 0.9 | 0.99 | 0.8 | 0.89 |
| 9 | 25612202 | rs6475857 | -0.274 | -0.416 | -0.329 | 0.336 | 0.126 | 0.092 | 0.092 | 1.665 | 0.05 | 0.32 | 0.18 | 0.08 | 0.05 | 0.03 |
| 9 | 25612704 | rs10511761 | -0.327 | -0.635 | -1.184 | 0.324 | -0.018 | -0.026 | -0.026 | 0 | 0.59 | 0.28 | 0.72 | 0.53 | 0.56 | 0.54 |
| 9 | 25616984 | rs10966996 | 0.21 | -1.272 | -1.377 | 0.22 | 0.065 | -0.04 | -0.04 | 0.064 | 0.15 | 0.02 | 0.15 | 0.18 | 0.05 | 0.07 |
| 9 | 25618052 | rs7036377 | -0.644 | -1.894 | -1.06 | -0.199 | -0.027 | -0.037 | -0.037 | 0 | 0.32 | 0.55 | 0.27 | 0.16 | 0.15 | 0.24 |
| 9 | 25618261 | rs7036435 | -0.17 | -1.447 | -1.154 | -0.054 | -0.02 | -0.012 | -0.012 | 0 | 0.42 | 0.46 | 0.48 | 0.64 | 0.32 | 0.52 |
| 9 | 25627885 | rs10967005 | 0.812 | -0.203 | -1.228 | 0.739 | 0.013 | 0.033 | 0.033 | 0 | 0.39 | 0.22 | 0.25 | 0.31 | 0.28 | 0.36 |
| 9 | 25628210 | rs10967006 | 0.375 | -0.177 | -1.422 | 0.695 | 0.02 | 0.034 | 0.034 | 0 | 0.47 | 0.33 | 0.29 | 0.4 | 0.34 | 0.43 |
| 9 | 25628819 | rs4478657 | 0.7 | -0.045 | -1.314 | 0.82 | 0.01 | 0.009 | 0.009 | 0 | 0.42 | 0.34 | 0.4 | 0.37 | 0.32 | 0.38 |
| 9 | 25629550 | rs10812286 | 0.789 | 0.05 | -1.262 | 0.556 | 0.006 | 0.015 | 0.015 | 0 | 0.39 | 0.33 | 0.27 | 0.27 | 0.28 | 0.33 |
| 9 | 25630951 | rs17177480 | -0.387 | 0.393 | -1.457 | 1.109 | 0.01 | -0.034 | -0.034 | 0 | 0.11 | 0.07 | 0.04 | 0.27 | 0.07 | 0.17 |
| 9 | 25631130 | rs10511762 | -0.314 | 1.152 | -1.637 | 0.61 | -0.072 | 0.024 | 0.024 | 0 | 0.27 | 0.09 | 0.4 | 0.22 | 0.42 | 0.31 |
| 9 | 25636098 | rs4531139 | -0.314 | 0.671 | -0.622 | 0.671 | -0.001 | -0.001 | -0.001 | 1.071 | 0.13 | 0.13 | 0.11 | 0.12 | 0.11 | 0.23 |
| 9 | 25640498 | rs4431675 | -0.424 | 0.583 | 0.574 | 1.016 | -0.016 | 0.125 | 0.125 | 0 | 0.15 | 0.34 | 0.11 | 0.37 | 0.11 | 0.24 |
| 9 | 25643882 | rs4512475 | -0.125 | 0.593 | 0.617 | 0.967 | 0.011 | 0.014 | 0.014 | 0 | 0.14 | 0.31 | 0.12 | 0.14 | 0.12 | 0.25 |
| 9 | 25646816 | rs7871606 | -1.173 | 0.678 | 0.751 | 0.975 | -0.005 | 0.053 | 0.053 | 0 | 0.28 | 0.38 | 0.22 | 0.5 | 0.24 | 0.51 |
| 9 | 25648199 | rs7036161 | 0.598 | 1.276 | 0.414 | -0.305 | 0.007 | -0.022 | -0.022 | 0 | 0.74 | 0.7 | 0.8 | 0.85 | 0.69 | 0.64 |
| 9 | 25648783 | rs10967022 | 0.433 | 1.539 | -0.077 | -0.594 | -0.048 | 0.023 | 0.023 | 0 | 0.22 | 0.08 | 0.35 | 0.16 | 0.32 | 0.14 |
| 9 | 25648883 | rs4335219 | 1.793 | 1.657 | 0.257 | -0.53 | 0.032 | 0.007 | 0.007 | 0 | 0.37 | 0.25 | 0.33 | 0.63 | 0.23 | 0.38 |
| 9 | 25651865 | rs4361838 | 1.108 | 1.732 | 0.284 | -0.524 | -0.004 | -0.009 | -0.009 | 0 | 0.55 | 0.52 | 0.6 | 0.7 | 0.37 | 0.53 |
| 9 | 25652508 | rs4534200 | 0.349 | 1.966 | -0.167 | -0.436 | -0.063 | 0.047 | 0.047 | 0 | 0.71 | 0.81 | 0.72 | 0.85 | 0.49 | 0.8 |
| 9 | 25652937 | rs4144933 | 0.534 | 1.89 | -0.137 | -0.445 | -0.007 | 0.001 | 0.001 | 0 | 0.18 | 0.18 | 0.21 | 0.16 | 0.28 | 0.1 |
| 9 | 25658015 | rs10117752 | -0.244 | 1.839 | -0.107 | -0.422 | -0.008 | 0.003 | 0.003 | 0 | 0.3 | 0.27 | 0.28 | 0.29 | 0.36 | 0.36 |
| 9 | 25658076 | rs4978003 | 1.577 | 1.294 | 0.137 | -0.297 | 0.015 | 0.009 | 0.009 | 0 | 0.41 | 0.32 | 0.35 | 0.63 | 0.28 | 0.36 |
| 9 | 25658472 | rs12001336 | -0.722 | 1.291 | 0.203 | -0.131 | -0.024 | -0.066 | -0.066 | 0 | 0.19 | 0.37 | 0.27 | 0.08 | 0.15 | 0.19 |
| 9 | 25659418 | rs10967030 | 0.735 | 1.672 | 0.213 | 0.253 | 0.003 | -0.024 | -0.024 | 0 | 0.59 | 0.57 | 0.61 | 0.77 | 0.41 | 0.65 |
| 9 | 25659637 | rs10124907 | 0.689 | 1.372 | 0.067 | 0.097 | 0.001 | -0.015 | -0.015 | 0 | 0.59 | 0.59 | 0.62 | 0.79 | 0.43 | 0.66 |
| 9 | 25661720 | rs4978005 | 0.596 | 1.083 | 0.052 | 0.842 | 0.001 | 0.001 | 0.001 | 0 | 0.58 | 0.57 | 0.51 | 0.59 | 0.42 | 0.64 |
| 9 | 25666644 | rs13285063 | -0.248 | 1.291 | -0.511 | 0.772 | -0.031 | 0.025 | 0.025 | 0 | 0.08 | 0.02 | 0.09 | 0.04 | 0.18 | 0.08 |
| 9 | 25667662 | rs11999484 | -0.45 | 1.283 | -0.418 | 0.769 | 0.011 | -0.017 | -0.017 | 0 | 0.09 | 0.11 | 0.1 | 0.04 | 0.18 | 0.08 |
| 9 | 25667695 | rs4260970 | 0.091 | 1.169 | -0.667 | 0.892 | -0.036 | -0.056 | -0.056 | 0 | 0.64 | 0.83 | 0.63 | 0.55 | 0.6 | 0.7 |
| 9 | 25677215 | rs12348 | 0.733 | 0.936 | -0.198 | 0.687 | 0.001 | 0.002 | 0.002 | 0 | 0.51 | 0.45 | 0.49 | 0.66 | 0.38 | 0.67 |
| 9 | 25677349 | rs1128953 | 0.22 | 1.057 | 0.119 | 0.63 | -0.021 | 0.003 | 0.003 | 0 | 0.6 | 0.71 | 0.6 | 0.67 | 0.55 | 0.74 |
| 9 | 25679242 | rs11999257 | -0.668 | 1.11 | 0.498 | 0.172 | 0.04 | -0.041 | -0.041 | 0 | 0.08 | 0.24 | 0.09 | 0 | 0.15 | 0.05 |
| 9 | 25683220 | rs4574937 | -0.871 | 1.151 | 0.005 | 0.309 | 0.004 | -0.011 | -0.011 | 1.891 | 0.4 | 0.6 | 0.33 | 0.21 | 0.53 | 0.26 |
| 9 | 25683697 | rs13301551 | -0.856 | 1.082 | -0.139 | 0.366 | 0.013 | -0.04 | -0.04 | 0 | 0.08 | 0.15 | 0.09 | 0 | 0.15 | 0.05 |

| | | | | | | | | | | | | | | | | |
|---|---|---|---|---|---|---|---|---|---|---|---|---|---|---|---|---|
| 9 | 25689954 | rs10967037 | -0.325 | 0.814 | -0.636 | 0 | -0.001 | 0.054 | 0.054 | 0 | 0.41 | 0.28 | 0.25 | 0.21 | 0.41 | 0.3 |
| 9 | 25690311 | rs4461986 | -0.764 | 0.7 | -0.091 | 0.658 | -0.007 | -0.011 | -0.011 | 0.034 | 0.59 | 0.66 | 0.48 | 0.51 | 0.6 | 0.38 |
| 9 | 25691032 | rs7031899 | -0.905 | 0.735 | -0.222 | 0.795 | 0.003 | -0.011 | -0.011 | 0 | 0.65 | 0.77 | 0.53 | 0.58 | 0.67 | 0.55 |
| 9 | 25692824 | rs10757564 | 0.141 | 0.163 | 0.386 | 0.966 | -0.005 | 0.074 | 0.074 | 0.907 | 0.83 | 0.58 | 0.88 | 0.74 | 0.84 | 0.71 |
| 9 | 25694413 | rs4131608 | -0.936 | -0.356 | 0.073 | 1.326 | 0.008 | -0.013 | -0.013 | 0 | 0.3 | 0.7 | 0.33 | 0.5 | 0.26 | 0.32 |
| 9 | 25695333 | rs7033322 | -0.917 | -0.377 | 0.047 | 1.321 | 0.028 | -0.036 | -0.036 | 0.368 | 0.3 | 0.73 | 0.32 | 0.5 | 0.24 | 0.32 |
| 9 | 25700484 | rs12554383 | -0.339 | -0.526 | 0.429 | 1.486 | 0.001 | -0.003 | -0.003 | 0.792 | 0.23 | 0.18 | 0.28 | 0.32 | 0.23 | 0.27 |
| 9 | 25700669 | rs1937940 | -0.865 | -0.535 | -0.231 | 1.496 | 0.022 | -0.027 | -0.027 | 0 | 0.28 | 0.71 | 0.31 | 0.44 | 0.22 | 0.24 |
| 9 | 25717386 | rs1231330 | -0.212 | -0.041 | -0.423 | 0.64 | 0.002 | 0.041 | 0.041 | 0 | 0.58 | 0.65 | 0.57 | 0.72 | 0.58 | 0.66 |
| 9 | 25717537 | rs12352086 | -1.009 | -0.177 | 0.204 | 1.135 | -0.019 | 0.02 | 0.02 | 0 | 0.3 | 0.55 | 0.27 | 0.32 | 0.27 | 0.37 |
| 9 | 25719019 | rs13286578 | -0.72 | -0.279 | 0.802 | 1.242 | -0.02 | 0.049 | 0.049 | 0.053 | 0.25 | 0.57 | 0.24 | 0.32 | 0.21 | 0.24 |
| 9 | 25722702 | rs7857572 | -0.172 | -0.27 | 0.896 | 1.29 | -0.024 | 0.033 | 0.033 | 0 | 0.19 | 0.25 | 0.21 | 0.25 | 0.15 | 0.25 |
| 9 | 25727225 | rs1231336 | 0.211 | -0.221 | 0.094 | 1.126 | -0.001 | 0 | 0 | 0 | 0.81 | 0.79 | 0.77 | 0.76 | 0.84 | 0.76 |
| 9 | 25728739 | rs4335220 | -1.068 | -0.103 | 0.335 | 0.997 | -0.001 | 0.001 | 0.001 | 0.018 | 0.28 | 0.63 | 0.25 | 0.32 | 0.3 | 0.32 |
| 9 | 25730285 | rs10812304 | -0.666 | -0.076 | 0.102 | 1.095 | -0.004 | 0.041 | 0.041 | 0.02 | 0.25 | 0.58 | 0.24 | 0.32 | 0.26 | 0.28 |
| 9 | 25737567 | rs757099 | 0.232 | -0.242 | -0.957 | 0.983 | -0.033 | 0.069 | 0.069 | 0 | 0.49 | 0.2 | 0.4 | 0.38 | 0.44 | 0.51 |
| 9 | 25737868 | rs17039 | 0.146 | -0.348 | -0.364 | 1.175 | 0.002 | 0.047 | 0.047 | 0 | 0.5 | 0.43 | 0.54 | 0.74 | 0.42 | 0.64 |
| 9 | 25740863 | rs1231339 | -0.113 | -0.153 | -0.562 | 1.417 | -0.003 | -0.004 | -0.004 | 0.758 | 0.61 | 0.56 | 0.46 | 0.54 | 0.52 | 0.57 |
| 9 | 25743020 | rs10491876 | -0.582 | 0.045 | -1.105 | 1.721 | -0.012 | -0.043 | -0.043 | 0 | 0.14 | 0.08 | 0.2 | 0.33 | 0.12 | 0.19 |
| 9 | 25745122 | rs1231345 | 0.689 | -1.016 | -1.657 | 1.317 | 0.1 | -0.069 | -0.069 | 0 | 0.62 | 0.13 | 0.51 | 0.39 | 0.79 | 0.58 |
| 9 | 25745702 | rs1231346 | 1.663 | -0.683 | -2.192 | 1.003 | -0.151 | -0.025 | -0.025 | 0.011 | 0.4 | 0.05 | 0.29 | 0.3 | 0.42 | 0.45 |
| 9 | 25748799 | rs1231353 | 0.184 | -1.002 | -1.324 | 1.388 | -0.023 | -0.001 | -0.001 | 3.351 | 0.64 | 0.49 | 0.63 | 0.62 | 0.8 | 0.65 |
| 9 | 25750661 | rs1231360 | 1.788 | -0.983 | -1.383 | 1.333 | -0.099 | 0.024 | 0.024 | 0.878 | 0.42 | 0.09 | 0.29 | 0.3 | 0.43 | 0.47 |
| 9 | 25751461 | rs17290832 | 0.65 | -1.008 | -1.281 | 1.339 | -0.028 | 0.02 | 0.02 | 0 | 0.19 | 0.14 | 0.22 | 0.11 | 0.36 | 0.13 |
| 9 | 25752310 | rs1231362 | 1.96 | -1.028 | -1.302 | 1.366 | -0.141 | -0.002 | -0.002 | 0 | 0.4 | 0.05 | 0.28 | 0.29 | 0.42 | 0.45 |
| 9 | 25752734 | rs1231365 | 0.082 | -1.351 | -1.185 | 1.296 | -0.038 | 0.03 | 0.03 | 0 | 0.71 | 0.49 | 0.66 | 0.6 | 0.87 | 0.66 |
| 9 | 25755422 | rs1231373 | 0.501 | -1.279 | -1.129 | 1.307 | -0.018 | -0.003 | -0.003 | 0 | 0.65 | 0.48 | 0.62 | 0.65 | 0.8 | 0.64 |
| 9 | 25755952 | rs1231375 | 0.514 | -1.138 | -1.536 | 1.062 | 0.069 | -0.054 | -0.054 | 0 | 0.67 | 0.23 | 0.54 | 0.43 | 0.86 | 0.61 |
| 9 | 25772172 | rs1634404 | -0.322 | -0.647 | -1.373 | 0.608 | -0.04 | 0.08 | 0.08 | 1.087 | 0.29 | 0.15 | 0.27 | 0.1 | 0.45 | 0.18 |
| 9 | 25775262 | rs10967072 | 0.005 | -0.64 | -0.7 | 0.808 | 0.013 | 0.008 | 0.008 | 0 | 0.18 | 0.37 | 0.27 | 0.22 | 0.25 | 0.15 |
| 9 | 25775447 | rs1147808 | -2.093 | -1.252 | -0.42 | 1.005 | 0.06 | 0.02 | 0.02 | 0.028 | 0.5 | 0.76 | 0.53 | 0.6 | 0.44 | 0.51 |
| 9 | 25783111 | rs2522032 | 0.811 | -1.607 | -0.412 | 0.975 | -0.073 | 0.024 | 0.024 | 0.016 | 0.71 | 0.54 | 0.76 | 0.66 | 0.88 | 0.68 |
| 9 | 25787040 | rs1220090 | -2.378 | -1.52 | -0.517 | 0.874 | -0.046 | -0.02 | -0.02 | 1.794 | 0.57 | 0.88 | 0.5 | 0.62 | 0.52 | 0.53 |
| 9 | 25787431 | rs4084056 | 0.037 | -1.515 | -0.485 | 0.846 | -0.002 | 0.007 | 0.007 | 0 | 0.29 | 0.32 | 0.25 | 0.24 | 0.37 | 0.17 |
| 9 | 25788721 | rs1220087 | -2.537 | -1.542 | -0.517 | 0.793 | -0.011 | 0.005 | 0.005 | 0.714 | 0.53 | 0.87 | 0.49 | 0.62 | 0.49 | 0.52 |
| 9 | 25792486 | rs1220081 | -2.008 | -1.549 | -0.502 | 0.796 | 0.05 | -0.02 | -0.02 | 0.379 | 0.38 | 0.69 | 0.41 | 0.46 | 0.23 | 0.45 |
| 9 | 25794090 | rs1220078 | 0.412 | -1.591 | -0.756 | 0.432 | -0.095 | 0.151 | 0.151 | 0 | 0.26 | 0.03 | 0.23 | 0.06 | 0.37 | 0.14 |
| 9 | 25795808 | rs1220091 | -1.616 | -1.676 | -0.644 | 0.624 | -0.05 | -0.013 | -0.013 | 0 | 0.33 | 0.52 | 0.26 | 0.3 | 0.19 | 0.37 |
| 9 | 25800000 | rs1235054 | -0.104 | -1.659 | -0.734 | 0.499 | -0.103 | 0.194 | 0.194 | 0 | 0.31 | 0.04 | 0.26 | 0.07 | 0.45 | 0.15 |
| 9 | 25805162 | rs7849755 | -0.261 | -1.676 | -0.548 | 0.701 | -0.037 | 0 | 0 | 0.199 | 0.25 | 0.59 | 0.33 | 0.26 | 0.13 | 0.3 |
| 9 | 25808485 | rs1782030 | -1.053 | -1.633 | -0.387 | 0.704 | -0.062 | -0.006 | -0.006 | 0 | 0.3 | 0.46 | 0.25 | 0.3 | 0.16 | 0.34 |
| 9 | 25812031 | rs1220073 | 0.451 | -1.53 | -0.693 | 0.731 | -0.074 | 0.093 | 0.093 | 0 | 0.25 | 0.03 | 0.32 | 0.14 | 0.36 | 0.15 |
| 9 | 25815074 | rs1220071 | -1.426 | -1.456 | -0.236 | 0.896 | -0.044 | -0.016 | -0.016 | 0 | 0.33 | 0.53 | 0.38 | 0.3 | 0.21 | 0.36 |
| 9 | 25815700 | rs989053 | -1.337 | -1.539 | -0.59 | 1.003 | -0.027 | -0.022 | -0.022 | 0 | 0.49 | 0.95 | 0.67 | 0.65 | 0.49 | 0.49 |
| 9 | 25816219 | rs7025024 | -1.045 | -1.263 | -0.35 | 0.433 | 0.018 | 0.046 | 0.046 | 0.354 | 0.46 | 0.67 | 0.64 | 0.4 | 0.48 | 0.44 |
| 9 | 25816366 | rs10967086 | 1.203 | -1.185 | -0.402 | 0.441 | -0.032 | 0.05 | 0.05 | 0 | 0.19 | 0.09 | 0.3 | 0.12 | 0.3 | 0.1 |
| 9 | 25816707 | rs975346 | -1.644 | -2.711 | 0.255 | 1.416 | -0.028 | -0.005 | -0.005 | 0 | 0.31 | 0.56 | 0.37 | 0.3 | 0.2 | 0.35 |
| 9 | 25819089 | rs1892406 | -1.585 | -2.852 | 0.954 | 1.55 | -0.035 | -0.008 | -0.008 | 8.211 | 0.31 | 0.55 | 0.36 | 0.28 | 0.16 | 0.36 |
| 9 | 25825595 | rs1220064 | 0.608 | -2.865 | 1.213 | 1.445 | 0.001 | 0.006 | 0.006 | 0 | 0.47 | 0.42 | 0.44 | 0.33 | 0.52 | 0.44 |
| 9 | 25830763 | rs1220052 | 0.163 | -2.936 | 1.789 | 1.452 | 0.012 | -0.039 | -0.039 | 0 | 0.56 | 0.8 | 0.65 | 0.46 | 0.61 | 0.55 |
| 9 | 25833408 | rs9408099 | 0.564 | -3.065 | 1.442 | 1.349 | 0.039 | 0.053 | 0.053 | 0.016 | 0.49 | 0.2 | 0.37 | 0.36 | 0.58 | 0.5 |
| 9 | 25837224 | rs1220074 | 0.588 | -3.09 | 0.897 | 1.41 | -0.037 | -0.012 | -0.012 | 0.512 | 0.48 | 0.05 | 0.37 | 0.37 | 0.56 | 0.52 |
| 9 | 25839353 | rs1220044 | 0.45 | -3.147 | 1.035 | 1.416 | 0.043 | 0.034 | 0.034 | 0 | 0.52 | 0.2 | 0.39 | 0.38 | 0.57 | 0.51 |
| 9 | 25845171 | rs1843463 | -0.344 | -3.076 | 0.445 | 1.23 | -0.03 | 0.193 | 0.193 | 0 | 0.63 | 0.9 | 0.79 | 0.92 | 0.62 | 0.7 |
| 9 | 25859709 | rs1147813 | 0.053 | -2.977 | 0.666 | 1.431 | 0.01 | 0.185 | 0.185 | 1.244 | 0.55 | 0.81 | 0.77 | 0.9 | 0.59 | 0.65 |
| 9 | 25889362 | rs2171099 | 0.514 | -2.419 | 0.563 | 1.825 | -0.048 | 0.008 | 0.008 | 0.083 | 0.47 | 0.05 | 0.36 | 0.32 | 0.55 | 0.47 |
| 9 | 25890226 | rs993233 | -0.12 | -2.581 | 1.188 | 1.809 | 0.01 | -0.001 | -0.001 | 0.176 | 0.6 | 0.4 | 0.6 | 0.56 | 0.64 | 0.68 |
| 9 | 25890597 | rs993232 | -0.382 | -2.721 | 0.868 | 1.881 | -0.012 | 0.007 | 0.007 | 0 | 0.63 | 0.61 | 0.71 | 0.56 | 0.68 | 0.68 |
| 9 | 25897335 | rs836954 | -1.968 | -2.989 | 1.41 | 2.524 | 0.039 | -0.022 | -0.022 | 0.788 | 0.36 | 0.76 | 0.3 | 0.43 | 0.16 | 0.54 |
| 9 | 25899227 | rs16909867 | -1.748 | -3.061 | 1.847 | 2.901 | -0.034 | 0.172 | 0.172 | 0.503 | 0.1 | 0.21 | 0.05 | 0.29 | 0.04 | 0.26 |
| 9 | 25901396 | rs1219937 | -1.515 | -3.073 | 2.314 | 2.718 | -0.072 | 0.023 | 0.023 | 6.046 | 0.26 | 0.54 | 0.22 | 0.31 | 0.13 | 0.45 |
| 9 | 25901711 | rs1219936 | -1.056 | -3.08 | 2.209 | 2.765 | -0.054 | 0.085 | 0.085 | 2.347 | 0.21 | 0.36 | 0.14 | 0.36 | 0.1 | 0.37 |
| 9 | 25902763 | rs1219935 | -0.849 | -3.076 | 2.141 | 2.72 | -0.035 | 0.032 | 0.032 | 1.879 | 0.18 | 0.21 | 0.1 | 0.28 | 0.08 | 0.36 |
| 9 | 25917668 | rs1219938 | -1.597 | -2.944 | 1.127 | 2.282 | -0.069 | 0.035 | 0.035 | 2.464 | 0.32 | 0.42 | 0.22 | 0.56 | 0.15 | 0.55 |
| 9 | 25919851 | rs1219942 | -1.805 | -2.904 | -0.443 | 2.239 | 0.087 | -0.041 | -0.041 | 0 | 0.37 | 0.8 | 0.33 | 0.46 | 0.18 | 0.55 |
| 9 | 25923262 | rs666778 | -0.588 | -2.535 | -0.46 | 2.355 | 0.029 | -0.04 | -0.04 | 0.006 | 0.21 | 0.15 | 0.13 | 0.4 | 0.11 | 0.42 |
| 9 | 25929148 | rs658858 | -0.707 | -2.153 | -0.83 | 2.336 | 0.084 | -0.056 | -0.056 | 5.819 | 0.31 | 0.82 | 0.29 | 0.41 | 0.16 | 0.5 |
| 9 | 25934932 | rs541805 | -0.705 | -1.938 | -1.031 | 2.037 | 0.138 | -0.101 | -0.101 | 0.365 | 0.36 | 0.88 | 0.35 | 0.53 | 0.18 | 0.55 |
| 9 | 25953815 | rs6475872 | -1.351 | -0.645 | 0.325 | 2.357 | -0.013 | 0.347 | 0.347 | 0 | 0.06 | 0.29 | 0.19 | 0.39 | 0.01 | 0.01 |

| | | | | | | | | | | | | | | | | |
|---|---|---|---|---|---|---|---|---|---|---|---|---|---|---|---|---|
| 9 | 25954098 | rs1855027 | -1.449 | -0.668 | 0.314 | 2.345 | -0.016 | 0.281 | 0.281 | 0 | 0.06 | 0.29 | 0.19 | 0.35 | 0.01 | 0.01 |
| 9 | 25955007 | rs9886671 | -0.66 | -0.729 | 0.341 | 1.067 | -0.01 | -0.054 | -0.054 | 0 | 0.28 | 0.39 | 0.23 | 0.11 | 0.24 | 0.17 |
| 9 | 25956931 | rs1332339 | -0.779 | -0.683 | 0.044 | 1.39 | -0.001 | -0.01 | -0.01 | 0.715 | 0.1 | 0.07 | 0.3 | 0.67 | 0.11 | 0.31 |
| 9 | 25966740 | rs10453196 | -0.527 | -1.91 | 0.911 | 0.718 | -0.073 | 0.03 | 0.03 | 1.228 | 0.14 | 0.42 | 0.07 | 0.17 | 0.04 | 0.25 |
| 9 | 25971497 | rs844356 | 1.137 | -1.955 | 1.478 | 0.542 | -0.066 | 0 | 0 | 3.19 | 0.81 | 0.37 | 0.88 | 0.78 | 0.93 | 0.71 |
| 9 | 25973166 | rs473107 | 0.671 | -1.517 | 0.295 | -0.835 | -0.065 | 0.113 | 0.113 | 0 | 0.27 | 0.07 | 0.25 | 0 | 0.35 | 0.14 |
| 9 | 25973292 | rs1332348 | 1.282 | -1.337 | -0.556 | -1.775 | -0.023 | 0.143 | 0.143 | 0 | 0.3 | 0.09 | 0.2 | 0.03 | 0.3 | 0.19 |
| 9 | 25979072 | rs651998 | -0.078 | -1.32 | -0.05 | -1.747 | 0.002 | -0.005 | -0.005 | 0 | 0.71 | 0.7 | 0.53 | 0.22 | 0.68 | 0.59 |
| 9 | 25979514 | rs475033 | 0.38 | -1.028 | 0.084 | -2.051 | 0.002 | -0.02 | -0.02 | 2.88 | 0.62 | 0.6 | 0.42 | 0.22 | 0.6 | 0.58 |
| 9 | 25988943 | rs2778457 | 0.241 | -1.067 | -0.033 | -2.035 | -0.008 | -0.004 | -0.004 | 0 | 0.72 | 0.3 | 0.78 | 0.78 | 0.84 | 0.68 |
| 9 | 26005487 | rs1332340 | 0.452 | -0.903 | -0.179 | -2.021 | -0.033 | 0.018 | 0.018 | 0 | 0.25 | 0.1 | 0.44 | 0.76 | 0.28 | 0.42 |
| 9 | 26014600 | rs2383609 | 1.588 | -0.786 | -0.404 | -2.161 | -0.062 | 0.104 | 0.104 | 0 | 0.25 | 0.07 | 0.25 | 0 | 0.35 | 0.14 |
| 9 | 26020806 | rs10738742 | 0.34 | -0.625 | -0.27 | -2.047 | -0.014 | 0.02 | 0.02 | 0 | 0.26 | 0.17 | 0.44 | 0.79 | 0.28 | 0.44 |
| 9 | 26022050 | rs9942876 | -0.756 | -0.604 | -0.292 | -1.931 | 0 | -0.005 | -0.005 | 0 | 0.92 | 0.92 | 0.89 | 1 | 0.91 | 0.99 |
| 9 | 26028352 | rs877755 | 0.54 | -0.619 | -0.378 | -1.941 | -0.009 | 0.027 | 0.027 | 0.216 | 0.08 | 0.02 | 0.12 | 0 | 0.09 | 0.01 |
| 9 | 26034790 | rs12551101 | 0.459 | -0.967 | -0.234 | -2.209 | -0.013 | -0.024 | -0.024 | 0 | 0.36 | 0.43 | 0.17 | 0.2 | 0.27 | 0.42 |
| 9 | 26035315 | rs17494475 | 1.541 | -1.029 | -0.977 | -2.248 | -0.056 | 0.103 | 0.103 | 0.673 | 0.21 | 0.01 | 0.1 | 0.04 | 0.21 | 0.16 |
| 9 | 26038217 | rs10757585 | 0.027 | -0.72 | -0.636 | -2.148 | 0.035 | 0.094 | 0.094 | 0 | 0.55 | 0.27 | 0.68 | 0.8 | 0.64 | 0.56 |
| 9 | 26038747 | rs10757587 | 0.027 | -0.325 | -0.786 | -1.804 | -0.024 | 0.014 | 0.014 | 0 | 0.25 | 0.11 | 0.42 | 0.77 | 0.28 | 0.43 |
| 9 | 26044500 | rs13293014 | 0.213 | -0.281 | -0.569 | -1.571 | -0.008 | 0.008 | 0.008 | 0.01 | 0.08 | 0.03 | 0.11 | 0 | 0.09 | 0.01 |
| 9 | 26058260 | rs7019491 | -0.195 | -0.918 | -1.544 | -2.168 | 0.048 | 0.17 | 0.17 | 0 | 0.67 | 0.91 | 0.87 | 0.98 | 0.75 | 0.72 |
| 9 | 26058823 | rs10967180 | -0.407 | -0.371 | -1.479 | -1.641 | 0 | 0 | 0 | 0 | 0.09 | 0.07 | 0.05 | 0.14 | 0.07 | 0.08 |
| 9 | 26059028 | rs957252 | 1.149 | -0.503 | -1.502 | -1.893 | -0.037 | 0.046 | 0.046 | 0 | 0.44 | 0.18 | 0.57 | 0.7 | 0.53 | 0.49 |
| 9 | 26062582 | rs9406901 | -0.2 | -0.345 | -1.098 | -1.857 | 0.028 | 0.135 | 0.135 | 0.014 | 0.61 | 0.83 | 0.73 | 0.98 | 0.66 | 0.71 |
| 9 | 26062842 | rs16910121 | 0.813 | -0.275 | -1.044 | -1.787 | 0.002 | -0.018 | -0.018 | 0 | 0.06 | 0.07 | 0.13 | 0 | 0.09 | 0.01 |
| 9 | 26068261 | rs2383614 | -0.147 | -0.339 | 0.121 | -2.064 | -0.009 | -0.01 | -0.01 | 0 | 0.54 | 0.48 | 0.53 | 0.28 | 0.63 | 0.44 |
| 9 | 26069415 | rs946035 | 1.738 | -0.267 | 0.007 | -2.024 | -0.139 | 0.052 | 0.052 | 0.015 | 0.31 | 0.08 | 0.29 | 0.14 | 0.47 | 0.23 |
| 9 | 26078764 | rs4978040 | 0.668 | -0.27 | 0.584 | -1.962 | 0.02 | 0.063 | 0.063 | 0 | 0.45 | 0.34 | 0.62 | 0.84 | 0.6 | 0.55 |
| 9 | 26086274 | rs10812346 | 0.652 | -0.071 | 0.499 | -1.882 | 0.006 | 0.024 | 0.024 | 0.015 | 0.05 | 0.07 | 0.06 | 0.13 | 0.08 | 0.11 |
| 9 | 26087714 | rs10967194 | 1.58 | -0.275 | 0.603 | -1.997 | -0.042 | 0.018 | 0.018 | 0 | 0.12 | 0.18 | 0.04 | 0.15 | 0.04 | 0.16 |
| 9 | 26087907 | rs7046442 | -0.688 | -0.297 | 0.625 | -2.008 | -0.005 | -0.016 | -0.016 | 0.601 | 0.55 | 0.57 | 0.81 | 0.82 | 0.71 | 0.58 |
| 9 | 26110570 | rs4977692 | 0.131 | 0.035 | 0.608 | -1.887 | 0.009 | -0.032 | -0.032 | 0 | 0.09 | 0.09 | 0.16 | 0 | 0.13 | 0.02 |
| 9 | 26117767 | rs16910176 | 1.487 | -0.034 | 0.84 | -1.857 | -0.014 | -0.001 | -0.001 | 0.289 | 0.11 | 0.12 | 0.04 | 0.12 | 0.04 | 0.17 |
| 9 | 26121103 | rs6475884 | -1.575 | 0.132 | 0.985 | -1.321 | -0.009 | 0.003 | 0.003 | 0 | 0.63 | 0.67 | 0.48 | 0.35 | 0.56 | 0.59 |
| 9 | 26125099 | rs7035912 | 1.819 | 0.16 | 1.153 | -1.255 | -0.024 | -0.01 | -0.01 | 1.537 | 0.09 | 0.2 | 0.03 | 0.06 | 0.04 | 0.16 |
| 9 | 26127824 | rs7030771 | 1.368 | 0.216 | 1.058 | -1.282 | -0.004 | -0.003 | -0.003 | 0 | 0.06 | 0.05 | 0.13 | 0 | 0.09 | 0.01 |
| 9 | 26128568 | rs1434863 | -1.143 | 0.139 | 1.308 | -1.284 | 0.016 | -0.016 | -0.016 | 3.873 | 0.26 | 0.45 | 0.28 | 0.2 | 0.25 | 0.3 |
| 9 | 26133821 | rs10812354 | -1.946 | 0.139 | 1.29 | -1.253 | 0.047 | 0.019 | 0.019 | 0 | 0.18 | 0.38 | 0.25 | 0.17 | 0.19 | 0.13 |
| 9 | 26134368 | rs1434880 | -1.26 | 0.155 | 1.044 | -1.199 | 0.005 | -0.003 | -0.003 | 0 | 0.27 | 0.54 | 0.28 | 0.25 | 0.23 | 0.3 |
| 9 | 26136235 | rs10757596 | 0.31 | 0.211 | 0.458 | -1.195 | -0.001 | 0.008 | 0.008 | 0.111 | 0.06 | 0.08 | 0.08 | 0.14 | 0.09 | 0.12 |
| 9 | 26136894 | rs2250791 | -2.553 | 0.33 | 0.162 | -1.472 | -0.129 | 0.184 | 0.184 | 0.08 | 0.74 | 0.98 | 0.76 | 1 | 0.6 | 0.88 |
| 9 | 26151961 | rs2787165 | 0.982 | 0.251 | 0.311 | -1.366 | 0.034 | 0.201 | 0.201 | 0 | 0.36 | 0.1 | 0.18 | 0.07 | 0.27 | 0.31 |
| 9 | 26153793 | rs2571525 | 0.878 | 0.227 | 0.323 | -1.334 | 0.001 | 0.204 | 0.204 | 0 | 0.37 | 0.11 | 0.21 | 0.07 | 0.32 | 0.33 |
| 9 | 26155175 | rs1434876 | 0.824 | 0.234 | 0.329 | -1.282 | 0 | 0.179 | 0.179 | 0 | 0.37 | 0.11 | 0.22 | 0.1 | 0.33 | 0.33 |
| 9 | 26178732 | rs10757598 | -0.989 | 0.222 | 0.779 | -1.235 | 0.008 | -0.033 | -0.033 | 0 | 0.57 | 0.51 | 0.67 | 0.75 | 0.5 | 0.61 |
| 9 | 26179065 | rs1434860 | -1.867 | 0.166 | 0.746 | -1.275 | -0.104 | 0.086 | 0.086 | 0.003 | 0.68 | 0.91 | 0.72 | 0.87 | 0.54 | 0.77 |
| 9 | 26180249 | rs1836463 | 1.908 | 0.131 | 0.966 | -1.26 | -0.026 | -0.016 | -0.016 | 0 | 0.09 | 0.19 | 0.03 | 0.06 | 0.04 | 0.16 |
| 9 | 26181177 | rs10812357 | -0.597 | 0.143 | 1.05 | -1.301 | 0.003 | 0.005 | 0.005 | 0 | 0.51 | 0.55 | 0.56 | 0.75 | 0.44 | 0.6 |
| 9 | 26184917 | rs2571519 | 1.495 | 0.107 | 1.116 | -1.23 | -0.034 | -0.02 | -0.02 | 0 | 0.11 | 0.31 | 0.04 | 0.06 | 0.04 | 0.16 |
| 9 | 26187635 | rs1821553 | -0.716 | 0.24 | 1.101 | -1.185 | 0.003 | 0.016 | 0.016 | 0 | 0.52 | 0.43 | 0.64 | 0.74 | 0.44 | 0.57 |
| 9 | 26189592 | rs1933276 | -1.307 | 0.203 | 0.889 | -1.16 | -0.048 | 0.119 | 0.119 | 0 | 0.59 | 0.88 | 0.73 | 0.8 | 0.54 | 0.6 |
| 9 | 26190234 | rs10967221 | 0.855 | 0.106 | 0.794 | -1.092 | -0.019 | -0.042 | -0.042 | 0 | 0.08 | 0.01 | 0.31 | 0.66 | 0.08 | 0.27 |
| 9 | 26191012 | rs10757601 | -0.552 | 0.087 | 0.963 | -1.137 | 0.017 | 0.059 | 0.059 | 0 | 0.5 | 0.6 | 0.58 | 0.8 | 0.42 | 0.58 |
| 9 | 26194166 | rs2787180 | -1.33 | 0.036 | 1.024 | -1.128 | -0.019 | 0.085 | 0.085 | 0 | 0.58 | 0.79 | 0.68 | 0.75 | 0.53 | 0.56 |
| 9 | 26195613 | rs1369198 | -0.244 | 0.041 | 0.929 | -1.22 | 0.063 | 0.044 | 0.044 | 0 | 0.06 | 0.27 | 0.04 | 0.06 | 0.05 | 0.02 |
| 9 | 26196672 | rs2152969 | -1.735 | 0.376 | 0.622 | -1.371 | -0.045 | 0.145 | 0.145 | 0 | 0.66 | 0.87 | 0.73 | 0.88 | 0.61 | 0.69 |
| 9 | 26200057 | rs2068369 | 1.014 | 0.432 | 0.716 | -1.255 | -0.005 | 0.184 | 0.184 | 0 | 0.34 | 0.06 | 0.18 | 0.04 | 0.31 | 0.26 |
| 9 | 26201458 | rs13293826 | 0.525 | 0.398 | 0.701 | -1.253 | -0.084 | 0.24 | 0.24 | 0 | 0.39 | 0.06 | 0.24 | 0.05 | 0.43 | 0.31 |
| 9 | 26203244 | rs10511766 | 1.336 | 0.673 | 0.861 | -0.94 | 0.003 | -0.012 | -0.012 | 0 | 0.05 | 0.05 | 0.12 | 0 | 0.06 | 0.01 |
| 9 | 26205604 | rs12684405 | -0.323 | 0.844 | 1.259 | -0.559 | 0.03 | 0.186 | 0.186 | 0 | 0.11 | 0.25 | 0.28 | 0.62 | 0.09 | 0.28 |
| 9 | 26205694 | rs2217796 | 0.915 | 0.649 | 0.943 | -0.867 | -0.006 | 0.011 | 0.011 | 0 | 0.43 | 0.41 | 0.37 | 0.21 | 0.53 | 0.29 |
| 9 | 26207267 | rs1434883 | -0.197 | 0.576 | 0.89 | -0.911 | 0.033 | 0.082 | 0.082 | 0 | 0.58 | 0.74 | 0.74 | 0.88 | 0.64 | 0.56 |
| 9 | 26210349 | rs10511767 | -0.896 | 0.544 | 0.873 | -0.989 | 0 | 0.032 | 0.032 | 0 | 0.64 | 0.75 | 0.8 | 0.89 | 0.65 | 0.73 |
| 9 | 26210446 | rs10967234 | 1.768 | 0.602 | 0.984 | -0.932 | -0.012 | 0.082 | 0.082 | 0 | 0.34 | 0.17 | 0.24 | 0.11 | 0.4 | 0.22 |
| 9 | 26211820 | rs4978054 | -0.8 | 0.689 | 1.145 | -0.585 | 0.012 | 0.078 | 0.078 | 0 | 0.14 | 0.22 | 0.4 | 0.67 | 0.13 | 0.28 |
| 9 | 26226392 | rs717704 | -0.491 | 0.398 | 1.118 | -0.584 | -0.001 | 0.056 | 0.056 | 0 | 0.09 | 0.29 | 0.08 | 0.15 | 0.07 | 0.06 |
| 9 | 26230579 | rs13300363 | 0.747 | 0.154 | 0.588 | -0.434 | -0.066 | 0.03 | 0.03 | 0 | 0.3 | 0.03 | 0.18 | 0.16 | 0.32 | 0.27 |
| 9 | 26233664 | rs10967253 | 0.277 | 0.202 | 0.564 | -0.581 | -0.075 | 0.17 | 0.17 | 0 | 0.33 | 0.03 | 0.18 | 0.06 | 0.34 | 0.26 |
| 9 | 26241359 | rs1434857 | -0.399 | 0.182 | 0.942 | -0.411 | 0.005 | 0.01 | 0.01 | 0 | 0.42 | 0.38 | 0.45 | 0.82 | 0.33 | 0.55 |

| | | | | | | | | | | | | | | | | |
|---|---|---|---|---|---|---|---|---|---|---|---|---|---|---|---|---|
| 9 | 26248035 | rs1932713 | 0.352 | 0.2 | 1.158 | -0.224 | -0.004 | 0.143 | 0.143 | 0 | 0.85 | 0.64 | 0.69 | 0.37 | 0.88 | 0.71 |
| 9 | 26257238 | rs623893 | -0.446 | -0.006 | 0.909 | -0.443 | -0.021 | -0.04 | -0.04 | 0 | 0.83 | 0.74 | 0.89 | 0.94 | 0.87 | 0.77 |
| 9 | 26261340 | rs443014 | -0.543 | 0.039 | 0.197 | -0.652 | 0.001 | -0.008 | -0.008 | 0 | 0.4 | 0.56 | 0.59 | 0.84 | 0.43 | 0.58 |
| 9 | 26275374 | rs514616 | 0.489 | -0.016 | 0.243 | -0.46 | -0.012 | -0.096 | -0.096 | 4.631 | 0.7 | 0.6 | 0.73 | 0.96 | 0.76 | 0.76 |
| 9 | 26275493 | rs693087 | 0.368 | -0.211 | 0.178 | -0.402 | -0.034 | 0.085 | 0.085 | 0.799 | 0.75 | 0.58 | 0.67 | 0.47 | 0.83 | 0.7 |
| 9 | 26275964 | rs1897888 | -1.03 | -0.145 | 0.045 | -1.079 | -0.004 | -0.062 | -0.062 | 0 | 0.86 | 0.77 | 0.93 | 1 | 0.88 | 0.96 |
| 9 | 26276367 | rs7470949 | 2.148 | -0.132 | -0.065 | -1.101 | -0.047 | 0.083 | 0.083 | 0 | 0.32 | 0.08 | 0.25 | 0.12 | 0.38 | 0.21 |
| 9 | 26289217 | rs353436 | -1.196 | -0.371 | 0.115 | -1.006 | -0.016 | -0.06 | -0.06 | 1.033 | 0.86 | 0.8 | 0.94 | 1 | 0.91 | 0.97 |
| 9 | 26292267 | rs2571510 | -0.708 | -0.422 | 0.135 | -1.077 | -0.011 | -0.08 | -0.08 | 0.665 | 0.31 | 0.34 | 0.31 | 0.04 | 0.25 | 0.24 |
| 9 | 26294087 | rs10967292 | 0.061 | -0.397 | 0.183 | -0.968 | -0.011 | 0.138 | 0.138 | 0 | 0.12 | 0.22 | 0.31 | 0.63 | 0.08 | 0.28 |
| 9 | 26296319 | rs589340 | 0.096 | -0.411 | -0.015 | -1.129 | -0.027 | 0.054 | 0.054 | 0.099 | 0.29 | 0.17 | 0.31 | 0.06 | 0.35 | 0.29 |
| 9 | 26297312 | rs17776101 | 1.961 | -0.317 | 0.036 | -1.069 | -0.06 | 0.098 | 0.098 | 0 | 0.34 | 0.14 | 0.26 | 0.12 | 0.44 | 0.22 |
| 9 | 26297693 | rs367832 | -0.257 | -0.3 | 0.03 | -1.087 | -0.008 | -0.028 | -0.028 | 0 | 0.77 | 0.76 | 0.87 | 0.96 | 0.86 | 0.82 |
| 9 | 26298970 | rs2618032 | 0.845 | -0.158 | -0.307 | -1.021 | -0.089 | 0.102 | 0.102 | 0.041 | 0.25 | 0.02 | 0.16 | 0.06 | 0.3 | 0.25 |
| 9 | 26302964 | rs17719825 | -0.873 | -0.126 | -0.115 | -0.92 | 0.012 | -0.032 | -0.032 | 0.742 | 0.07 | 0.1 | 0.16 | 0 | 0.08 | 0.05 |
| 9 | 26307529 | rs11793655 | 1.214 | 0.197 | -0.232 | -0.894 | -0.02 | 0.128 | 0.128 | 0 | 0.43 | 0.11 | 0.33 | 0.16 | 0.46 | 0.37 |
| 9 | 26313753 | rs10967301 | 0.934 | 0.729 | 0.02 | -0.633 | -0.019 | 0.044 | 0.044 | 0 | 0.39 | 0.26 | 0.39 | 0.13 | 0.47 | 0.26 |
| 9 | 26313814 | rs2618012 | 0.536 | 0.203 | 0.122 | -0.543 | -0.027 | -0.024 | -0.024 | 0.547 | 0.26 | 0.4 | 0.19 | 0.17 | 0.17 | 0.24 |
| 9 | 26316310 | rs10967303 | 1.121 | 0.013 | -0.068 | -0.246 | -0.053 | 0.094 | 0.094 | 0 | 0.33 | 0.11 | 0.25 | 0.12 | 0.41 | 0.22 |
| 9 | 26317242 | rs989494 | -0.414 | 0.575 | 0.792 | 1.012 | -0.018 | 0.194 | 0.194 | 0 | 0.09 | 0.19 | 0.25 | 0.62 | 0.04 | 0.24 |
| 9 | 26317973 | rs2618014 | 0.603 | 0.971 | -0.5 | -0.08 | -0.013 | 0.012 | 0.012 | 0.104 | 0.72 | 0.8 | 0.8 | 0.93 | 0.66 | 0.77 |
| 9 | 26318151 | rs2785103 | 1.135 | 0.304 | -0.908 | 0.321 | -0.043 | 0.048 | 0.048 | 0.569 | 0.8 | 0.97 | 0.88 | 0.89 | 0.79 | 0.79 |
| 9 | 26318333 | rs2785102 | 1.285 | 0.415 | -0.266 | 0.85 | 0.002 | 0.012 | 0.012 | 0.723 | 0.87 | 0.8 | 0.93 | 0.78 | 0.88 | 0.86 |
| 9 | 26324013 | rs1414924 | 1.405 | -0.308 | 0.734 | 1.039 | -0.011 | 0.077 | 0.077 | 1.149 | 0.88 | 0.39 | 0.9 | 0.83 | 0.92 | 0.85 |
| 9 | 26328954 | rs1336461 | 1.527 | -0.345 | 0.738 | 1.36 | -0.006 | 0.135 | 0.135 | 1.655 | 0.89 | 0.38 | 0.9 | 0.72 | 0.92 | 0.85 |
| 9 | 26329422 | rs2618011 | -1.724 | -0.612 | 1.21 | 1.39 | -0.006 | 0.18 | 0.18 | 6.149 | 0.08 | 0.53 | 0.05 | 0.21 | 0.02 | 0.08 |
| 9 | 26333942 | rs1336466 | 0.047 | -1.348 | 0.151 | 1.784 | 0.002 | -0.112 | -0.112 | 0 | 0.23 | 0.02 | 0.33 | 0.34 | 0.2 | 0.29 |
| 9 | 26359485 | rs915508 | 1.925 | -1.849 | 1.232 | 1.946 | -0.053 | 0.144 | 0.144 | 0.736 | 0.79 | 0.52 | 0.87 | 0.56 | 0.86 | 0.72 |
| 9 | 26364526 | rs2484918 | 1.593 | -1.826 | 1.259 | 1.919 | 0.025 | 0.025 | 0.025 | 0 | 0.9 | 0.61 | 0.92 | 0.91 | 0.92 | 0.93 |
| 9 | 26371680 | rs406058 | 0.541 | -1.82 | 1.178 | 1.958 | -0.012 | -0.011 | -0.011 | 0 | 0.94 | 0.98 | 0.94 | 0.81 | 0.92 | 0.93 |
| 9 | 26376870 | rs958843 | 2.439 | -1.925 | 1.212 | 1.804 | -0.009 | 0.003 | 0.003 | 3.91 | 0.78 | 0.32 | 0.86 | 0.68 | 0.87 | 0.72 |
| 9 | 26381874 | rs2769970 | -1.447 | -1.984 | 1.234 | 1.756 | -0.076 | 0.072 | 0.072 | 2.911 | 0.15 | 0.43 | 0.06 | 0.24 | 0.06 | 0.2 |
| 9 | 26382205 | rs400528 | -1.667 | -1.978 | 1.32 | 1.766 | -0.075 | 0.051 | 0.051 | 8.093 | 0.16 | 0.58 | 0.18 | 0.26 | 0.07 | 0.22 |
| 9 | 26384371 | rs694150 | -1.058 | -1.934 | 1.247 | 1.782 | 0.036 | 0.252 | 0.252 | 3.722 | 0.07 | 0.42 | 0.04 | 0.21 | 0.02 | 0.1 |
| 9 | 26391495 | rs2618022 | 1.371 | -1.925 | 1.286 | 1.774 | 0.032 | 0.047 | 0.047 | 0 | 0.91 | 0.65 | 0.93 | 0.91 | 0.92 | 0.92 |
| 9 | 26392058 | rs2152634 | 1.666 | -1.621 | 1.229 | 1.75 | -0.006 | -0.018 | -0.018 | 0 | 0.83 | 0.51 | 0.88 | 0.88 | 0.86 | 0.8 |
| 9 | 26394003 | rs1887720 | -1.525 | -1.655 | 1.19 | 1.726 | -0.039 | -0.062 | -0.062 | 0 | 0.16 | 0.53 | 0.18 | 0.04 | 0.08 | 0.15 |
| 9 | 26397837 | rs3896192 | 1.669 | -1.858 | 1.104 | 1.781 | -0.028 | 0.044 | 0.044 | 0.06 | 0.84 | 0.7 | 0.9 | 0.73 | 0.89 | 0.85 |
| 9 | 26403617 | rs16910426 | 1.143 | -2.392 | 0.226 | 1.153 | 0.01 | 0.009 | 0.009 | 1.597 | 0.11 | 0.06 | 0.06 | 0 | 0.07 | 0.01 |
| 9 | 26409664 | rs367129 | 0.237 | -2.261 | 0.436 | 1.497 | -0.018 | -0.022 | -0.022 | 0 | 0.94 | 0.99 | 0.95 | 0.91 | 0.94 | 0.93 |
| 9 | 26410031 | rs10812392 | 1.959 | -2.319 | 0.494 | 1.518 | -0.029 | 0.012 | 0.012 | 2.869 | 0.77 | 0.35 | 0.77 | 0.67 | 0.87 | 0.71 |
| 9 | 26414780 | rs1855980 | 0.384 | -2.253 | 0.019 | 1.799 | 0.018 | -0.137 | -0.137 | 0 | 0.23 | 0.02 | 0.32 | 0.4 | 0.2 | 0.29 |
| 9 | 26425046 | rs454086 | 0.649 | -1.685 | 0.587 | 1.887 | -0.011 | -0.008 | -0.008 | 0.004 | 0.94 | 0.98 | 0.95 | 0.79 | 0.93 | 0.92 |
| 9 | 26433194 | rs10967335 | -0.569 | -1.734 | 0.565 | 1.398 | -0.016 | -0.02 | -0.02 | 0.47 | 0.06 | 0.01 | 0.05 | 0.09 | 0.07 | 0.07 |
| 9 | 26433751 | rs10812394 | 1.663 | -2.04 | 0.941 | 1.258 | -0.07 | 0.005 | 0.005 | 6.383 | 0.8 | 0.36 | 0.82 | 0.75 | 0.94 | 0.76 |
| 9 | 26440477 | rs1571363 | -0.825 | -1.967 | 1.102 | 1.22 | -0.078 | -0.043 | -0.043 | 0 | 0.14 | 0.34 | 0.14 | 0.04 | 0.04 | 0.12 |
| 9 | 26444583 | rs7864971 | -1.491 | -2.024 | 0.901 | 1.279 | -0.115 | 0.032 | 0.032 | 3.477 | 0.19 | 0.49 | 0.16 | 0.25 | 0.05 | 0.21 |
| 9 | 26449874 | rs10967346 | -1.554 | -2.011 | 1.004 | 1.154 | -0.026 | 0.098 | 0.098 | 4.397 | 0.1 | 0.57 | 0.07 | 0.18 | 0.04 | 0.12 |
| 9 | 26453356 | rs6475895 | -1.79 | -2.418 | 0.967 | 1.125 | -0.095 | -0.002 | -0.002 | 0 | 0.24 | 0.54 | 0.16 | 0.24 | 0.09 | 0.32 |
| 9 | 26453723 | rs6475897 | -2.003 | -2.28 | 0.929 | 1.314 | -0.021 | 0.075 | 0.075 | 2.107 | 0.15 | 0.65 | 0.18 | 0.26 | 0.1 | 0.21 |
| 9 | 26457170 | rs1328425 | -1.516 | -2.342 | 0.711 | 0.95 | 0 | -0.001 | -0.001 | 0 | 0.53 | 0.5 | 0.53 | 0.71 | 0.38 | 0.65 |
| 9 | 26463378 | rs4576502 | -0.713 | -2.117 | 1.123 | 1.166 | -0.015 | 0.001 | 0.001 | 1.53 | 0.35 | 0.47 | 0.37 | 0.58 | 0.3 | 0.44 |
| 9 | 26486471 | rs2094537 | 0.397 | -1.903 | 0.829 | -0.03 | -0.002 | -0.064 | -0.064 | 0.129 | 0.6 | 0.5 | 0.45 | 0.15 | 0.57 | 0.58 |
| 9 | 26489225 | rs12554403 | 0.425 | -1.902 | 0.901 | -0.043 | -0.001 | -0.041 | -0.041 | 0.071 | 0.6 | 0.46 | 0.45 | 0.15 | 0.57 | 0.57 |
| 9 | 26506732 | rs1854313 | 0.292 | -2.059 | 1.048 | -0.153 | -0.002 | -0.04 | -0.04 | 0 | 0.62 | 0.46 | 0.45 | 0.15 | 0.61 | 0.61 |
| 9 | 26507211 | rs12339423 | 0.905 | -1.753 | 0.261 | -1.121 | -0.004 | -0.02 | -0.02 | 0 | 0.42 | 0.38 | 0.37 | 0.03 | 0.47 | 0.42 |
| 9 | 26511000 | rs17753877 | 0.584 | -1.753 | 0.224 | -1.142 | 0.021 | 0.21 | 0.21 | 1.09 | 0.53 | 0.21 | 0.41 | 0.1 | 0.54 | 0.49 |
| 9 | 26511835 | rs7867285 | 1.003 | -1.767 | 0.225 | -1.157 | -0.006 | -0.003 | -0.003 | 0 | 0.4 | 0.32 | 0.36 | 0.03 | 0.43 | 0.38 |
| 9 | 26515052 | rs1359967 | 0.534 | -1.761 | 0.351 | -1.118 | 0.007 | 0.029 | 0.029 | 0.986 | 0.53 | 0.37 | 0.42 | 0.1 | 0.54 | 0.48 |
| 9 | 26515617 | rs7867988 | 0.378 | -1.928 | 0.694 | -0.964 | 0.001 | 0.018 | 0.018 | 0.206 | 0.58 | 0.59 | 0.47 | 0.17 | 0.55 | 0.5 |
| 9 | 26516013 | rs4333705 | 0.639 | -1.904 | 0.513 | -1.338 | 0.006 | 0.015 | 0.015 | 0.807 | 0.54 | 0.38 | 0.42 | 0.09 | 0.54 | 0.48 |
| 9 | 26518883 | rs10511789 | 0.921 | -2.068 | 0.229 | -0.978 | 0.042 | -0.05 | -0.05 | 0 | 0.19 | 0.05 | 0.3 | 0.67 | 0.13 | 0.26 |
| 9 | 26523911 | rs12000866 | 1.566 | -1.262 | -0.022 | -1.557 | -0.004 | 0.048 | 0.048 | 0 | 0.32 | 0.23 | 0.3 | 0.03 | 0.34 | 0.36 |
| 9 | 26524416 | rs4111166 | 0.282 | -1.327 | 0.113 | -1.61 | 0.001 | -0.061 | -0.061 | 0 | 0.57 | 0.5 | 0.41 | 0.15 | 0.52 | 0.57 |
| 9 | 26526309 | rs12002423 | 1.82 | -1.507 | 0.071 | -1.663 | -0.034 | 0.05 | 0.05 | 0 | 0.15 | 0.03 | 0.07 | 0.08 | 0.13 | 0.11 |
| 9 | 26526453 | rs12004473 | 1.035 | -1.517 | 0.037 | -1.727 | -0.018 | 0.156 | 0.156 | 0.161 | 0.47 | 0.25 | 0.38 | 0.1 | 0.48 | 0.4 |
| 9 | 26528575 | rs2225078 | 0.388 | -1.462 | -0.02 | -1.645 | 0.003 | 0.039 | 0.039 | 0.447 | 0.23 | 0.12 | 0.09 | 0.12 | 0.17 | 0.2 |
| 9 | 26530126 | rs10429615 | -0.119 | -1.457 | 0.053 | -1.677 | -0.006 | 0.061 | 0.061 | 0 | 0.65 | 0.79 | 0.49 | 0.15 | 0.61 | 0.53 |
| 9 | 26530632 | rs10967384 | 0.044 | -1.394 | 0.124 | -1.653 | -0.001 | -0.058 | -0.058 | 0.793 | 0.62 | 0.44 | 0.46 | 0.15 | 0.58 | 0.51 |

| | | | | | | | | | | | | | | | |
|---|---|---|---|---|---|---|---|---|---|---|---|---|---|---|---|
| 9 | 26531228 | rs4606151 | 0.621 | -1.365 | 0.078 | -1.636 | -0.001 | 0.035 | 0.035 | 0 | 0.45 | 0.37 | 0.39 | 0.08 | 0.44 | 0.38 |
| 9 | 26538805 | rs994390 | -0.135 | -1.522 | -0.232 | -1.568 | -0.006 | 0.043 | 0.043 | 0 | 0.64 | 0.73 | 0.58 | 0.15 | 0.59 | 0.55 |
| 9 | 26546431 | rs13294038 | 0.531 | -1.571 | -0.225 | -1.607 | -0.011 | 0.078 | 0.078 | 0 | 0.47 | 0.37 | 0.52 | 0.08 | 0.49 | 0.41 |
| 9 | 26547991 | rs10429492 | 0.37 | -1.568 | -0.169 | -1.555 | -0.008 | -0.028 | -0.028 | 0 | 0.09 | 0.19 | 0.03 | 0.05 | 0.06 | 0.11 |
| 9 | 26551026 | rs17754132 | 0.567 | -1.66 | -0.287 | -1.59 | 0.007 | 0.052 | 0.052 | 0.352 | 0.23 | 0.1 | 0.1 | 0.12 | 0.17 | 0.21 |
| 9 | 26551726 | rs10757612 | -0.584 | -1.549 | -0.131 | -1.046 | -0.013 | 0.006 | 0.006 | 0 | 0.7 | 0.81 | 0.64 | 0.29 | 0.64 | 0.56 |
| 9 | 26555763 | rs2383698 | 0.81 | -1.403 | -0.08 | -0.866 | -0.004 | 0.007 | 0.007 | 0 | 0.58 | 0.44 | 0.62 | 0.73 | 0.53 | 0.65 |
| 9 | 26556311 | rs1926757 | 0.435 | -1.337 | -0.548 | -0.795 | -0.039 | -0.017 | -0.017 | 0 | 0.61 | 0.8 | 0.65 | 0.59 | 0.53 | 0.65 |
| 9 | 26561000 | rs1983901 | -0.486 | -1.162 | -0.322 | -0.706 | 0.015 | 0.004 | 0.004 | 0 | 0.85 | 0.84 | 0.78 | 0.81 | 0.75 | 0.79 |
| 9 | 26565214 | rs1328426 | 1.007 | -1.341 | -0.399 | -0.801 | -0.004 | 0.006 | 0.006 | 0.02 | 0.72 | 0.26 | 0.76 | 0.77 | 0.69 | 0.68 |
| 9 | 26565913 | rs1887221 | -1.05 | -1.595 | -0.593 | -0.877 | -0.013 | 0.009 | 0.009 | 0 | 0.45 | 0.67 | 0.37 | 0.4 | 0.46 | 0.39 |
| 9 | 26569013 | rs17754201 | 1.087 | -2.049 | -0.463 | -1.265 | 0.037 | 0.044 | 0.044 | 0 | 0.13 | 0.01 | 0.02 | 0.05 | 0.06 | 0.11 |
| 9 | 26569055 | rs12550891 | 1.082 | -2.014 | -0.584 | -1.402 | 0.017 | 0.034 | 0.034 | 0 | 0.63 | 0.27 | 0.72 | 0.77 | 0.68 | 0.65 |
| 9 | 26570613 | rs7847445 | -0.201 | -1.816 | -0.126 | -1.28 | 0.037 | 0.033 | 0.033 | 0.235 | 0.15 | 0.38 | 0.18 | 0.18 | 0.2 | 0.23 |
| 9 | 26571285 | rs12379692 | 0.843 | -1.803 | -0.593 | -1.243 | -0.024 | -0.025 | -0.025 | 0.571 | 0.1 | 0.07 | 0.15 | 0.17 | 0.18 | 0.19 |
| 9 | 26572110 | rs2148601 | 0.13 | -2.399 | -0.965 | -1.611 | -0.016 | -0.014 | -0.014 | 0 | 0.19 | 0.07 | 0.16 | 0.18 | 0.21 | 0.24 |
| 9 | 26572537 | rs10118162 | -0.705 | -1.861 | -0.48 | -1.29 | -0.016 | -0.028 | -0.028 | 0 | 0.3 | 0.57 | 0.26 | 0.23 | 0.28 | 0.3 |
| 9 | 26576086 | rs4978082 | 0.691 | -1.71 | -0.842 | -0.942 | -0.002 | -0.001 | -0.001 | 0.046 | 0.75 | 0.29 | 0.78 | 0.77 | 0.76 | 0.68 |
| 9 | 26576117 | rs964855 | 0.546 | -1.297 | -0.843 | -0.735 | -0.015 | -0.015 | -0.015 | 0 | 0.08 | 0.07 | 0.15 | 0.18 | 0.19 | 0.21 |
| 9 | 26580067 | rs996414 | -0.075 | -1.345 | -0.82 | -1.095 | -0.016 | -0.112 | -0.112 | 0 | 0.75 | 0.55 | 0.88 | 0.91 | 0.83 | 0.81 |
| 9 | 26582104 | rs1926771 | 0.944 | -1.572 | -1.302 | -1.324 | 0.013 | 0.052 | 0.052 | 0.86 | 0.62 | 0.24 | 0.53 | 0.75 | 0.6 | 0.65 |
| 9 | 26582312 | rs10511787 | 1.639 | -1.466 | -1.256 | -1.282 | 0.001 | 0.006 | 0.006 | 0 | 0.07 | 0.04 | 0.21 | 0.02 | 0.09 | 0.05 |
| 9 | 26593785 | rs10812417 | -0.271 | -1.588 | -1.083 | -1.065 | 0.026 | 0.03 | 0.03 | 0.018 | 0.71 | 0.65 | 0.76 | 0.63 | 0.67 | 0.67 |
| 9 | 26595257 | rs12379432 | 0.877 | -1.731 | -1.083 | -1.179 | 0.002 | 0.006 | 0.006 | 0.097 | 0.53 | 0.49 | 0.68 | 0.58 | 0.56 | 0.58 |
| 9 | 26595380 | rs10812418 | -0.456 | -1.592 | -0.998 | -0.97 | 0.006 | -0.002 | -0.002 | 0 | 0.27 | 0.26 | 0.3 | 0.08 | 0.22 | 0.17 |
| 9 | 26597540 | rs17692544 | 0.899 | -1.558 | -1.058 | -0.941 | 0.027 | 0.067 | 0.067 | 0 | 0.15 | 0.03 | 0.07 | 0.05 | 0.11 | 0.09 |
| 9 | 26598313 | rs4316219 | -0.454 | -1.546 | -0.952 | -0.939 | -0.025 | -0.06 | -0.06 | 0.212 | 0.27 | 0.36 | 0.31 | 0.08 | 0.22 | 0.18 |
| 9 | 26600187 | rs10812420 | 0.363 | -1.31 | -1.058 | -0.742 | 0.014 | 0.06 | 0.06 | 0 | 0.1 | 0.13 | 0.15 | 0.3 | 0.19 | 0.19 |
| 9 | 26601230 | rs10967431 | -0.617 | -1.898 | -1.353 | -0.794 | 0.038 | -0.034 | -0.034 | 0 | 0.29 | 0.25 | 0.33 | 0.57 | 0.17 | 0.37 |
| 9 | 26603243 | rs913161 | 0.694 | -1.272 | -1.061 | -0.769 | 0.001 | 0.004 | 0.004 | 0 | 0.1 | 0.09 | 0.22 | 0.03 | 0.11 | 0.09 |
| 9 | 26605757 | rs10812423 | -0.825 | -1.066 | -1.088 | -0.748 | -0.022 | -0.007 | -0.007 | 0.292 | 0.2 | 0.16 | 0.36 | 0.2 | 0.32 | 0.27 |
| 9 | 26608337 | rs1926760 | -1.017 | -1.322 | -0.916 | -1 | -0.004 | -0.076 | -0.076 | 0 | 0.83 | 0.69 | 0.91 | 0.92 | 0.88 | 0.89 |
| 9 | 26609613 | rs10812426 | 0.055 | -1.186 | -0.716 | -0.832 | 0.081 | 0.053 | 0.053 | 0 | 0.12 | 0.31 | 0.17 | 0.17 | 0.2 | 0.18 |
| 9 | 26612557 | rs7869769 | -1.651 | -1.497 | -0.989 | -0.853 | -0.001 | -0.001 | -0.001 | 0 | 0.4 | 0.45 | 0.46 | 0.36 | 0.43 | 0.36 |
| 9 | 26612670 | rs10967440 | -0.494 | -1.198 | -0.819 | -0.757 | 0.018 | 0.002 | 0.002 | 0 | 0.2 | 0.31 | 0.37 | 0.2 | 0.32 | 0.27 |
| 9 | 26614847 | rs10812428 | -1.286 | -1.265 | -0.789 | -0.792 | 0.026 | 0.022 | 0.022 | 0 | 0.3 | 0.47 | 0.45 | 0.33 | 0.4 | 0.29 |
| 9 | 26618389 | rs7849701 | -0.771 | -1.34 | -0.602 | -0.828 | -0.025 | -0.049 | -0.049 | 0 | 0.87 | 0.57 | 0.94 | 0.95 | 0.93 | 0.92 |
| 9 | 26623297 | rs10757619 | -0.048 | -1.431 | -0.996 | -0.925 | -0.02 | -0.019 | -0.019 | 0 | 0.28 | 0.23 | 0.26 | 0.37 | 0.33 | 0.35 |
| 9 | 26624871 | rs7868631 | -1.179 | -1.495 | -1.023 | -0.963 | -0.005 | 0.06 | 0.06 | 0 | 0.46 | 0.32 | 0.52 | 0.28 | 0.52 | 0.45 |
| 9 | 26626081 | rs1998909 | 1.383 | -1.423 | -1.675 | -0.939 | -0.012 | 0.051 | 0.051 | 0 | 0.51 | 0.24 | 0.46 | 0.63 | 0.51 | 0.56 |
| 9 | 26628498 | rs10967447 | 1.758 | -1.454 | -1.679 | -0.916 | -0.027 | -0.007 | -0.007 | 0 | 0.43 | 0.24 | 0.44 | 0.58 | 0.46 | 0.49 |
| 9 | 26629083 | rs1926775 | 0.737 | -1.428 | -1.62 | -0.972 | -0.001 | 0 | 0 | 0 | 0.62 | 0.42 | 0.69 | 0.66 | 0.63 | 0.66 |
| 9 | 26629382 | rs1926773 | 0.83 | -1.157 | -1.921 | -1.226 | -0.001 | 0.05 | 0.05 | 0 | 0.62 | 0.25 | 0.53 | 0.75 | 0.63 | 0.65 |
| 9 | 26631112 | rs913163 | 0.307 | -1.018 | -1.897 | -1.103 | -0.077 | 0.002 | 0.002 | 0.216 | 0.23 | 0.06 | 0.22 | 0.22 | 0.28 | 0.26 |
| 9 | 26633841 | rs10812432 | 0.642 | -0.866 | -1.64 | -0.928 | 0.014 | 0.114 | 0.114 | 0.927 | 0.53 | 0.24 | 0.52 | 0.71 | 0.55 | 0.59 |
| 9 | 26636413 | rs7851334 | 0.653 | -1.073 | -1.853 | -1.011 | 0.019 | 0.068 | 0.068 | 0.029 | 0.28 | 0.1 | 0.19 | 0.13 | 0.26 | 0.31 |
| 9 | 26637253 | rs2164097 | -0.297 | -0.119 | -1.817 | -1.328 | -0.032 | 0.052 | 0.052 | 0 | 0.71 | 0.87 | 0.68 | 0.86 | 0.67 | 0.78 |
| 9 | 26641557 | rs10967453 | 0.421 | 0.18 | -2.143 | -1.69 | -0.006 | 0.083 | 0.083 | 0 | 0.23 | 0.08 | 0.14 | 0.09 | 0.2 | 0.17 |
| 9 | 26642033 | rs1369052 | 0.197 | 0.142 | -2.057 | -1.632 | -0.037 | 0.126 | 0.126 | 0 | 0.6 | 0.85 | 0.58 | 0.86 | 0.55 | 0.73 |
| 9 | 26649956 | rs10511783 | -1.39 | -0.118 | -1.472 | -1.536 | 0.004 | 0.004 | 0.004 | 0 | 0.7 | 0.75 | 0.81 | 0.87 | 0.75 | 0.71 |
| 9 | 26650017 | rs989947 | 0.522 | -0.219 | -1.383 | -1.741 | -0.028 | 0.025 | 0.025 | 0 | 0.35 | 0.18 | 0.42 | 0.69 | 0.38 | 0.41 |
| 9 | 26651364 | rs4978089 | -1.805 | -0.228 | -1.401 | -1.723 | 0.003 | 0.002 | 0.002 | 0 | 0.71 | 0.76 | 0.81 | 0.87 | 0.77 | 0.71 |
| 9 | 26652538 | rs10967458 | 0.214 | 0.026 | -1.448 | -0.77 | -0.039 | -0.075 | -0.075 | 0 | 0.34 | 0.14 | 0.42 | 0.57 | 0.38 | 0.4 |
| 9 | 26653154 | rs10812438 | 0.597 | 0.122 | -0.728 | -0.69 | 0.003 | 0.003 | 0.003 | 0 | 0.26 | 0.31 | 0.35 | 0.65 | 0.3 | 0.31 |
| 9 | 26657127 | rs10967459 | 0.342 | -0.382 | -1.057 | -0.926 | -0.037 | 0.025 | 0.025 | 1.058 | 0.13 | 0.01 | 0.06 | 0.1 | 0.12 | 0.14 |
| 9 | 26658407 | rs10121064 | -1.112 | -0.718 | -0.781 | -0.896 | -0.003 | -0.001 | -0.001 | 1.42 | 0.64 | 0.64 | 0.82 | 0.72 | 0.74 | 0.69 |
| 9 | 26658658 | rs10967460 | 1.412 | -0.379 | -0.673 | -1.445 | 0.035 | 0.003 | 0.003 | 0 | 0.09 | 0.11 | 0.24 | 0.06 | 0.17 | 0.08 |
| 9 | 26659562 | rs1033991 | -1.172 | -0.65 | -0.505 | -1.407 | 0.028 | 0.041 | 0.041 | 0 | 0.65 | 0.75 | 0.83 | 0.88 | 0.74 | 0.7 |
| 9 | 26665482 | rs10812442 | 0.285 | -0.611 | -0.274 | -1.522 | -0.002 | -0.06 | -0.06 | 0 | 0.16 | 0.34 | 0.1 | 0.1 | 0.12 | 0.14 |
| 9 | 26668917 | rs1434479 | 0.017 | -0.696 | -0.57 | -1.659 | 0.001 | 0.099 | 0.099 | 0 | 0.42 | 0.21 | 0.66 | 0.78 | 0.57 | 0.5 |
| 9 | 26673735 | rs12351975 | 0.616 | -0.77 | -0.429 | -1.702 | -0.024 | -0.107 | -0.107 | 0 | 0.25 | 0.45 | 0.16 | 0.1 | 0.17 | 0.16 |
| 9 | 26674986 | rs980876 | -1.686 | -0.803 | -0.545 | -1.648 | 0.007 | 0.008 | 0.008 | 0 | 0.69 | 0.71 | 0.81 | 0.85 | 0.75 | 0.7 |
| 9 | 26678360 | rs10738754 | -1.257 | -0.835 | -0.593 | -1.674 | 0.033 | 0.05 | 0.05 | 0 | 0.63 | 0.75 | 0.8 | 0.84 | 0.72 | 0.66 |
| 9 | 26681345 | rs10967475 | -0.031 | -0.788 | -0.347 | -1.701 | 0 | -0.006 | -0.006 | 1.234 | 0.41 | 0.36 | 0.66 | 0.76 | 0.56 | 0.52 |
| 9 | 26681499 | rs10812443 | 0.09 | -0.787 | -0.373 | -1.674 | -0.014 | 0.054 | 0.054 | 0 | 0.39 | 0.2 | 0.64 | 0.72 | 0.55 | 0.47 |
| 9 | 26684083 | rs10812444 | -0.818 | -0.55 | -0.415 | -1.8 | -0.003 | -0.043 | -0.043 | 0.473 | 0.42 | 0.46 | 0.43 | 0.2 | 0.39 | 0.39 |
| 9 | 26684890 | rs13289242 | 1.265 | -0.327 | -0.364 | -1.716 | -0.016 | 0.023 | 0.023 | 0 | 0.12 | 0.05 | 0.25 | 0.05 | 0.17 | 0.1 |
| 9 | 26684901 | rs12553807 | 1.07 | -0.335 | -0.212 | -1.675 | -0.004 | -0.001 | -0.001 | 0 | 0.27 | 0.29 | 0.39 | 0.66 | 0.36 | 0.29 |
| 9 | 26685529 | rs10812445 | 0.529 | -0.335 | -0.142 | -1.554 | -0.004 | -0.019 | -0.019 | 0 | 0.15 | 0.41 | 0.1 | 0.14 | 0.1 | 0.24 |

| | | | | | | | | | | | | | | | | |
|---|---|---|---|---|---|---|---|---|---|---|---|---|---|---|---|---|
| 9 | 26687936 | rs10812447 | 0.627 | -0.272 | -0.365 | -1.599 | 0 | -0.035 | -0.035 | 0.196 | 0.14 | 0.23 | 0.07 | 0.1 | 0.11 | 0.12 |
| 9 | 26689442 | rs13295742 | 1.072 | -0.206 | -0.711 | -1.524 | -0.014 | 0.021 | 0.021 | 0 | 0.12 | 0.05 | 0.25 | 0.05 | 0.17 | 0.1 |
| 9 | 26694184 | rs7871815 | -1.358 | -0.366 | -0.773 | -1.668 | 0.031 | 0.052 | 0.052 | 0 | 0.63 | 0.77 | 0.8 | 0.84 | 0.71 | 0.65 |
| 9 | 26698331 | rs13296414 | 0.975 | -0.139 | -0.629 | -1.595 | 0.001 | -0.007 | -0.007 | 0.563 | 0.12 | 0.08 | 0.25 | 0.06 | 0.18 | 0.1 |
| 9 | 26698644 | rs12708266 | -0.551 | -0.21 | -0.557 | -1.753 | -0.003 | -0.056 | -0.056 | 0 | 0.37 | 0.51 | 0.41 | 0.15 | 0.35 | 0.25 |
| 9 | 26699111 | rs10967482 | 0.363 | -0.449 | -0.482 | -1.561 | -0.022 | -0.104 | -0.104 | 5.19 | 0.25 | 0.43 | 0.16 | 0.1 | 0.17 | 0.15 |
| 9 | 26705815 | rs12115390 | 1.346 | -0.345 | -1.058 | -1.748 | -0.035 | 0.099 | 0.099 | 0 | 0.16 | 0.01 | 0.11 | 0 | 0.18 | 0.04 |
| 9 | 26706278 | rs10115168 | 0.146 | -0.621 | -0.795 | -1.781 | -0.002 | -0.018 | -0.018 | 0 | 0.29 | 0.36 | 0.16 | 0.13 | 0.21 | 0.22 |
| 9 | 26707333 | rs12376450 | 3.065 | -0.126 | -0.959 | -0.9 | -0.012 | 0.007 | 0.007 | 0 | 0.07 | 0 | 0.09 | 0.4 | 0.06 | 0.24 |
| 9 | 26708668 | rs1434482 | -0.051 | -0.599 | -0.84 | -2.29 | -0.002 | -0.017 | -0.017 | 0 | 0.8 | 0.7 | 0.82 | 0.85 | 0.79 | 0.76 |
| 9 | 26709294 | rs1865368 | -0.202 | -0.64 | -0.834 | -2.273 | -0.003 | -0.011 | -0.011 | 0 | 0.81 | 0.71 | 0.83 | 0.85 | 0.81 | 0.8 |
| 9 | 26714614 | rs2060409 | 0.118 | -0.811 | -1.149 | -2.408 | 0.032 | 0.108 | 0.108 | 0.161 | 0.43 | 0.2 | 0.27 | 0.28 | 0.35 | 0.34 |
| 9 | 26717062 | rs10967491 | 0.7 | -0.939 | -1.246 | -2.494 | 0.068 | 0.193 | 0.193 | 0 | 0.35 | 0.08 | 0.12 | 0.1 | 0.21 | 0.27 |
| 9 | 26719332 | rs10117522 | 0.091 | -0.383 | -0.91 | -2.457 | 0.074 | -0.012 | -0.012 | 0 | 0.22 | 0.47 | 0.42 | 0.21 | 0.34 | 0.15 |
| 9 | 26722736 | rs7045601 | -1.251 | -0.036 | -0.978 | -2.508 | -0.015 | 0.003 | 0.003 | 0.075 | 0.74 | 0.81 | 0.64 | 0.3 | 0.68 | 0.57 |
| 9 | 26728455 | rs7868048 | -1.091 | -0.048 | -0.968 | -2.801 | -0.008 | 0.002 | 0.002 | 0 | 0.76 | 0.77 | 0.61 | 0.28 | 0.66 | 0.56 |
| 9 | 26730057 | rs10967496 | 0.861 | -0.263 | -1.59 | -3.016 | -0.012 | 0.081 | 0.081 | 0 | 0.19 | 0.02 | 0.08 | 0.08 | 0.13 | 0.2 |
| 9 | 26734671 | rs1918350 | -0.953 | 0.362 | -0.748 | -2.604 | 0.023 | -0.017 | -0.017 | 0 | 0.73 | 0.61 | 0.62 | 0.28 | 0.67 | 0.56 |
| 9 | 26735409 | rs12347998 | 1.76 | 0.034 | -0.482 | -2.506 | -0.01 | -0.077 | -0.077 | 0.928 | 0.15 | 0.3 | 0.11 | 0.01 | 0.13 | 0.13 |
| 9 | 26736162 | rs10812452 | 0.663 | 0.126 | -0.959 | -2.136 | 0.049 | 0.024 | 0.024 | 0 | 0.08 | 0.21 | 0.13 | 0.14 | 0.14 | 0.04 |
| 9 | 26742892 | rs10967507 | 0.533 | -0.841 | -1.718 | -2.243 | 0.068 | 0.194 | 0.194 | 0 | 0.36 | 0.06 | 0.13 | 0.11 | 0.21 | 0.31 |
| 9 | 26749094 | rs1434478 | -0.621 | 0.339 | -0.529 | -1.957 | 0.009 | -0.038 | -0.038 | 0 | 0.33 | 0.42 | 0.48 | 0.2 | 0.51 | 0.38 |
| 9 | 26750142 | rs10757627 | -0.145 | 0.379 | -0.335 | -1.941 | 0.004 | -0.042 | -0.042 | 0 | 0.83 | 0.79 | 0.89 | 0.97 | 0.83 | 0.75 |
| 9 | 26750721 | rs9694997 | 0.419 | 0.227 | -0.56 | -2.293 | -0.047 | 0.036 | 0.036 | 0.192 | 0.27 | 0.22 | 0.44 | 0.15 | 0.44 | 0.11 |
| 9 | 26751013 | rs4978100 | 0.193 | -0.989 | -0.752 | -2.682 | 0.057 | 0.11 | 0.11 | 0.247 | 0.53 | 0.33 | 0.76 | 0.9 | 0.68 | 0.55 |
| 9 | 26751543 | rs7026561 | -0.736 | -0.748 | -0.507 | -2.575 | -0.001 | -0.022 | -0.022 | 0 | 0.84 | 0.83 | 0.89 | 0.97 | 0.85 | 0.75 |
| 9 | 26751666 | rs7041124 | 0.029 | -1.387 | 0.15 | -2.621 | -0.012 | -0.046 | -0.046 | 5.695 | 0.56 | 0.49 | 0.44 | 0.82 | 0.39 | 0.64 |
| 9 | 26755930 | rs10283761 | -0.237 | -1.176 | 0.911 | -2.546 | -0.032 | -0.12 | -0.12 | 0 | 0.24 | 0.49 | 0.17 | 0.06 | 0.16 | 0.38 |
| 9 | 26760477 | rs1369050 | -1.248 | -0.709 | 0.599 | -2.666 | 0.034 | 0.165 | 0.165 | 0 | 0.5 | 0.75 | 0.52 | 0.14 | 0.53 | 0.57 |
| 9 | 26766179 | rs1469773 | 1.415 | -0.28 | 0.751 | -2.535 | 0.03 | -0.03 | -0.03 | 0 | 0.11 | 0.1 | 0.28 | 0 | 0.24 | 0.06 |
| 9 | 26767510 | rs7853898 | -0.047 | -0.346 | 1.102 | -2.598 | 0.088 | -0.114 | -0.114 | 3.723 | 0.2 | 0.58 | 0.39 | 0.03 | 0.31 | 0.29 |
| 9 | 26767842 | rs12350711 | 0.52 | -0.475 | 1.004 | -2.67 | -0.028 | -0.048 | -0.048 | 7.195 | 0.2 | 0.27 | 0.11 | 0.05 | 0.12 | 0.33 |
| 9 | 26767853 | rs12555089 | 1.633 | -0.314 | 0.414 | -2.557 | 0.063 | -0.049 | -0.049 | 0 | 0.11 | 0.14 | 0.29 | 0 | 0.24 | 0.06 |
| 9 | 26770030 | rs7046166 | -0.04 | -0.377 | 0.348 | -2.556 | 0.015 | -0.04 | -0.04 | 0 | 0.27 | 0.32 | 0.37 | 0.07 | 0.36 | 0.2 |
| 9 | 26772805 | rs1880947 | 0.289 | -0.504 | -0.218 | -2.451 | 0.051 | 0.129 | 0.129 | 0 | 0.28 | 0.02 | 0.07 | 0.06 | 0.13 | 0.23 |
| 9 | 26777330 | rs2383716 | 0.368 | 0.074 | 0.733 | -2.349 | 0.115 | -0.102 | -0.102 | 2.021 | 0.76 | 0.52 | 0.56 | 0.93 | 0.61 | 0.67 |
| 9 | 26783996 | rs6475937 | 1.029 | 0.331 | 0.612 | -2.256 | 0.077 | -0.059 | -0.059 | 0.051 | 0.12 | 0.2 | 0.29 | 0 | 0.24 | 0.06 |
| 9 | 26788498 | rs10511775 | 0.889 | -0.343 | 0.174 | -2.041 | 0.007 | 0.087 | 0.087 | 0 | 0.42 | 0.25 | 0.27 | 0.17 | 0.39 | 0.26 |
| 9 | 26796160 | rs10116852 | 0.732 | -0.484 | 0.047 | -1.995 | 0.007 | 0.072 | 0.072 | 0 | 0.43 | 0.3 | 0.28 | 0.17 | 0.4 | 0.26 |
| 9 | 26797676 | rs10967540 | 1.098 | -0.695 | -0.016 | -1.64 | 0.004 | 0.028 | 0.028 | 0 | 0.07 | 0.02 | 0.04 | 0.01 | 0.07 | 0.22 |
| 9 | 26800359 | rs12338262 | 0.079 | -0.98 | 0.392 | -1.817 | -0.027 | -0.054 | -0.054 | 0 | 0.28 | 0.38 | 0.11 | 0.07 | 0.15 | 0.25 |
| 9 | 26804802 | rs12339392 | 0.719 | -1.088 | 0.346 | -1.909 | -0.038 | -0.054 | -0.054 | 0 | 0.17 | 0.25 | 0.07 | 0.03 | 0.1 | 0.11 |
| 9 | 26808023 | rs1336336 | 1.065 | -0.984 | 0.103 | -2.165 | 0.001 | 0.071 | 0.071 | 0.05 | 0.46 | 0.33 | 0.46 | 0.17 | 0.5 | 0.28 |
| 9 | 26834139 | rs10812476 | 0.937 | -0.801 | 0.06 | -1.401 | -0.001 | 0 | 0 | 0 | 0.08 | 0.09 | 0.08 | 0.13 | 0.09 | 0.24 |
| 9 | 26838839 | rs7872881 | 0.193 | -1.007 | 0.147 | -1.655 | 0.009 | 0.002 | 0.002 | 0 | 0.27 | 0.26 | 0.09 | 0.06 | 0.14 | 0.25 |
| 9 | 26839219 | rs10967557 | 1.191 | -0.656 | -0.095 | -1.523 | -0.051 | 0.024 | 0.024 | 0.118 | 0.09 | 0.02 | 0.13 | 0.05 | 0.19 | 0.05 |
| 9 | 26843342 | rs1336332 | 1.159 | -0.893 | -0.018 | -1.464 | -0.015 | 0.051 | 0.051 | 0 | 0.13 | 0.01 | 0.04 | 0.03 | 0.08 | 0.14 |
| 9 | 26851480 | rs10967566 | 1.304 | -0.924 | 0.204 | -1.305 | -0.01 | 0.081 | 0.081 | 0 | 0.06 | 0.07 | 0.08 | 0.7 | 0.07 | 0.18 |
| 9 | 26866642 | rs7036871 | 0.994 | -0.913 | 0.201 | -1.277 | 0.008 | 0.029 | 0.029 | 0 | 0.14 | 0.11 | 0.06 | 0.03 | 0.09 | 0.15 |
| 9 | 26869532 | rs7018943 | 0.767 | -0.834 | 0.472 | -1.094 | -0.012 | 0.133 | 0.133 | 0 | 0.07 | 0.15 | 0.21 | 0.69 | 0.08 | 0.18 |
| 9 | 26893662 | rs7038314 | -0.807 | -0.895 | 0.347 | -1.096 | -0.004 | 0.003 | 0.003 | 0 | 0.88 | 0.9 | 0.86 | 0.93 | 0.81 | 0.95 |
| 9 | 26893731 | rs7038218 | -0.539 | -1.618 | 0.405 | -1.221 | -0.038 | -0.014 | -0.014 | 0 | 0.35 | 0.47 | 0.31 | 0.7 | 0.23 | 0.41 |
| 9 | 26899690 | rs7018962 | 1.032 | -1.55 | 0.295 | -1.238 | 0.009 | 0.022 | 0.022 | 0 | 0.14 | 0.11 | 0.06 | 0.01 | 0.09 | 0.12 |
| 9 | 26902593 | rs10114508 | 1.161 | -1.39 | 0.23 | -1.704 | 0.039 | 0.065 | 0.065 | 1.03 | 0.38 | 0.24 | 0.19 | 0.18 | 0.27 | 0.24 |
| 9 | 26904699 | rs16910873 | 1.272 | -1.285 | 0.234 | -1.565 | 0.009 | 0.022 | 0.022 | 0 | 0.14 | 0.11 | 0.06 | 0.01 | 0.09 | 0.12 |
| 9 | 26917718 | rs7871286 | 0.072 | -1.303 | 0.4 | -1.437 | -0.014 | -0.063 | -0.063 | 0 | 0.31 | 0.38 | 0.13 | 0.04 | 0.19 | 0.24 |
| 9 | 26919607 | rs10120342 | 1.119 | -1.29 | 0.297 | -1.441 | 0.046 | 0.065 | 0.065 | 1.065 | 0.38 | 0.24 | 0.19 | 0.19 | 0.26 | 0.24 |
| 9 | 26931255 | rs10757635 | -0.463 | -1.611 | 0.616 | -0.994 | -0.051 | -0.026 | -0.026 | 0 | 0.36 | 0.51 | 0.32 | 0.7 | 0.24 | 0.41 |
| 9 | 26940723 | rs12003612 | 1.071 | -1.426 | 0.496 | -0.924 | -0.034 | -0.052 | -0.052 | 0 | 0.16 | 0.22 | 0.07 | 0.03 | 0.1 | 0.11 |
| 9 | 26960085 | rs10812503 | -0.37 | -1.777 | 0.342 | -1.041 | -0.038 | -0.023 | -0.023 | 0 | 0.39 | 0.52 | 0.32 | 0.7 | 0.25 | 0.44 |
| 9 | 27006575 | rs10967657 | 0.971 | -1.325 | -0.041 | -0.995 | -0.032 | 0.095 | 0.095 | 0.898 | 0.43 | 0.25 | 0.46 | 0.18 | 0.48 | 0.26 |
| 9 | 27009738 | rs17756299 | 1.325 | -1.621 | -0.302 | -1.153 | 0.018 | 0.053 | 0.053 | 0 | 0.14 | 0.04 | 0.04 | 0.01 | 0.09 | 0.12 |
| 9 | 27032385 | rs7857669 | 0.139 | -1.539 | 0.123 | -1.067 | 0.039 | 0.017 | 0.017 | 0 | 0.35 | 0.29 | 0.27 | 0.7 | 0.19 | 0.43 |
| 9 | 27047246 | rs10812520 | -0.699 | -1.426 | 0.3 | -1.179 | 0.035 | -0.015 | -0.015 | 0.78 | 0.84 | 0.77 | 0.79 | 0.88 | 0.74 | 0.72 |
| 9 | 27047460 | rs16910953 | 1.724 | -2.084 | -0.351 | -1.561 | 0.023 | 0.042 | 0.042 | 0 | 0.18 | 0.08 | 0.1 | 0.05 | 0.13 | 0.08 |
| 9 | 27050593 | rs1413824 | 1.687 | -1.771 | -0.488 | -1.392 | -0.001 | -0.008 | -0.008 | 0.115 | 0.07 | 0.09 | 0.08 | 0.05 | 0.08 | 0.23 |
| 9 | 27062721 | rs3429 | 0.732 | -1.949 | -0.759 | -1.32 | 0.11 | 0.182 | 0.182 | 0 | 0.3 | 0.04 | 0.07 | 0.03 | 0.14 | 0.25 |
| 9 | 27074233 | rs620090 | -1.166 | -1.559 | -0.362 | -1.049 | 0.021 | -0.004 | -0.004 | 0.494 | 0.88 | 0.84 | 0.85 | 0.87 | 0.82 | 0.86 |
| 9 | 27081879 | rs16910976 | 0.627 | -2.168 | -0.646 | -1.144 | 0.035 | 0.057 | 0.057 | 0 | 0.24 | 0.1 | 0.12 | 0.06 | 0.16 | 0.13 |

| | | | | | | | | | | | | | | | | |
|---|---|---|---|---|---|---|---|---|---|---|---|---|---|---|---|---|
| 9 | 27084462 | rs10812525 | -0.308 | -1.712 | 0.097 | -0.628 | 0.015 | 0.01 | 0.01 | 0 | 0.36 | 0.35 | 0.28 | 0.72 | 0.22 | 0.46 |
| 9 | 27098940 | rs1161603 | -0.155 | -1.756 | -0.014 | -0.91 | 0.005 | -0.026 | -0.026 | 0 | 0.69 | 0.72 | 0.9 | 0.96 | 0.83 | 0.74 |
| 9 | 27099244 | rs596427 | -0.534 | -1.391 | 0.27 | -0.64 | -0.012 | -0.039 | -0.039 | 1.854 | 0.76 | 0.74 | 0.93 | 0.98 | 0.88 | 0.79 |
| 9 | 27107349 | rs10967716 | 0.546 | -0.309 | 0.881 | -0.256 | -0.007 | 0.006 | 0.006 | 0 | 0.14 | 0.11 | 0.16 | 0.05 | 0.21 | 0.08 |
| 9 | 27108433 | rs10967717 | 0.097 | -0.396 | 0.935 | 0.192 | -0.005 | 0.187 | 0.187 | 0 | 0.05 | 0.09 | 0.19 | 0.62 | 0.05 | 0.22 |
| 9 | 27108975 | rs657867 | -0.387 | -0.33 | 0.941 | 0.031 | 0.057 | -0.059 | -0.059 | 1.963 | 0.83 | 0.65 | 0.8 | 0.95 | 0.79 | 0.92 |
| 9 | 27112311 | rs615375 | 0.357 | -0.509 | 0.631 | 0.213 | 0.027 | 0.071 | 0.071 | 0 | 0.76 | 0.44 | 0.61 | 0.44 | 0.74 | 0.74 |
| 9 | 27115768 | rs10738763 | -0.352 | -0.325 | 0.551 | -0.425 | 0.046 | -0.07 | -0.07 | 0 | 0.79 | 0.56 | 0.67 | 0.92 | 0.77 | 0.88 |
| 9 | 27118807 | rs621786 | 0.231 | 0.159 | 0.065 | -0.376 | 0.01 | 0.014 | 0.014 | 0 | 0.42 | 0.37 | 0.31 | 0.68 | 0.39 | 0.63 |
| 9 | 27120369 | rs666812 | -0.496 | 0.015 | -0.149 | -0.765 | -0.004 | -0.074 | -0.074 | 1.13 | 0.6 | 0.56 | 0.65 | 0.91 | 0.67 | 0.72 |
| 9 | 27123260 | rs603085 | 0.572 | 0.07 | -0.362 | 0.04 | -0.005 | 0.034 | 0.034 | 0.168 | 0.17 | 0.16 | 0.17 | 0.53 | 0.14 | 0.49 |
| 9 | 27124745 | rs4242698 | 0.737 | 0.474 | -0.424 | -0.613 | -0.012 | -0.01 | -0.01 | 0 | 0.16 | 0.14 | 0.32 | 0 | 0.29 | 0.04 |
| 9 | 27132514 | rs7044842 | 0.04 | 0.202 | -0.241 | -1.298 | 0 | -0.086 | -0.086 | 0 | 0.54 | 0.54 | 0.47 | 0.03 | 0.55 | 0.28 |
| 9 | 27133016 | rs16911047 | 1.879 | -0.016 | -0.05 | -1.541 | -0.002 | -0.047 | -0.047 | 0 | 0.09 | 0.15 | 0.06 | 0 | 0.1 | 0.03 |
| 9 | 27133385 | rs7032817 | 0.301 | -0.174 | 0.023 | -1.663 | 0.057 | -0.134 | -0.134 | 1.397 | 0.23 | 0.42 | 0.37 | 0 | 0.37 | 0.07 |
| 9 | 27134868 | rs10511799 | 1.651 | -0.125 | 0.006 | -1.605 | -0.029 | 0.02 | 0.02 | 0 | 0.13 | 0.08 | 0.29 | 0 | 0.27 | 0.05 |
| 9 | 27135831 | rs7042119 | 0.622 | -0.175 | -0.004 | -1.608 | -0.019 | -0.103 | -0.103 | 0.084 | 0.41 | 0.46 | 0.18 | 0.03 | 0.28 | 0.23 |
| 9 | 27137242 | rs669441 | -0.826 | -0.156 | -0.02 | -1.103 | 0 | 0 | 0 | 0 | 0.7 | 0.7 | 0.66 | 0.65 | 0.67 | 0.76 |
| 9 | 27144863 | rs12005388 | 0.316 | -0.08 | -0.816 | -1.206 | 0 | 0.014 | 0.014 | 0 | 0.45 | 0.4 | 0.42 | 0.03 | 0.45 | 0.25 |
| 9 | 27149669 | rs4878149 | 1.252 | -0.057 | -1.194 | -1.21 | -0.1 | 0.095 | 0.095 | 0 | 0.2 | 0.01 | 0.25 | 0.04 | 0.25 | 0.11 |
| 9 | 27151394 | rs7034505 | -0.014 | -0.239 | -0.785 | -0.473 | -0.002 | 0.008 | 0.008 | 0 | 0.51 | 0.61 | 0.55 | 0.58 | 0.5 | 0.65 |
| 9 | 27151645 | rs7862217 | 0.86 | 0.222 | -0.389 | -0.958 | 0.102 | -0.041 | -0.041 | 0 | 0.07 | 0.19 | 0.27 | 0 | 0.2 | 0.03 |
| 9 | 27151706 | rs7035021 | -0.033 | -0.156 | -0.586 | -0.995 | 0.017 | 0.02 | 0.02 | 0 | 0.41 | 0.31 | 0.26 | 0.33 | 0.28 | 0.51 |
| 9 | 27152292 | rs4601425 | -0.048 | 0.055 | -0.446 | -0.844 | -0.004 | 0.039 | 0.039 | 0 | 0.34 | 0.28 | 0.34 | 0.03 | 0.33 | 0.23 |
| 9 | 27152308 | rs664461 | -0.737 | -0.018 | -0.389 | -0.613 | 0.002 | 0.004 | 0.004 | 0 | 0.49 | 0.54 | 0.49 | 0.37 | 0.4 | 0.55 |
| 9 | 27153866 | rs600728 | 0.561 | -0.815 | -0.723 | -0.465 | -0.034 | 0.06 | 0.06 | 0.342 | 0.15 | 0.26 | 0.14 | 0.31 | 0.08 | 0.33 |
| 9 | 27153874 | rs507441 | -0.102 | -0.447 | -0.586 | -0.272 | 0.016 | 0.011 | 0.011 | 0 | 0.42 | 0.36 | 0.23 | 0.34 | 0.24 | 0.53 |
| 9 | 27161811 | rs1413825 | 0.078 | 0.146 | -0.518 | -0.063 | 0.005 | 0.021 | 0.021 | 0 | 0.59 | 0.68 | 0.76 | 0.34 | 0.64 | 0.64 |
| 9 | 27163395 | rs506685 | -0.208 | 0.041 | -0.568 | -0.043 | -0.004 | -0.004 | -0.004 | 0 | 0.76 | 0.76 | 0.82 | 0.38 | 0.71 | 0.74 |
| 9 | 27164220 | rs7036743 | 0.333 | 0.575 | -0.039 | -0.012 | 0.029 | 0.053 | 0.053 | 0 | 0.36 | 0.26 | 0.35 | 0.16 | 0.3 | 0.44 |
| 9 | 27164846 | rs1107774 | 0.106 | 1.966 | 0.743 | 0.535 | 0.016 | -0.02 | -0.02 | 0 | 0.23 | 0.24 | 0.4 | 0.1 | 0.35 | 0.28 |
| 9 | 27165030 | rs671804 | -0.006 | 1.933 | 0.885 | 0.801 | 0.064 | 0.134 | 0.134 | 0 | 0.91 | 0.81 | 0.82 | 0.59 | 0.82 | 0.88 |
| 9 | 27166159 | rs10967750 | 0.312 | 1.892 | 0.752 | 0.254 | 0.018 | -0.066 | -0.066 | 0 | 0.22 | 0.24 | 0.37 | 0.02 | 0.32 | 0.16 |
| 9 | 27169124 | rs3818283 | 0.861 | 1.731 | 1.007 | 1.326 | 0.164 | 0.201 | 0.201 | 0.017 | 0.9 | 0.64 | 0.81 | 0.68 | 0.79 | 0.88 |
| 9 | 27170934 | rs565586 | 1.042 | 1.471 | 0.87 | 1.404 | 0.006 | 0.026 | 0.026 | 0 | 0.86 | 0.85 | 0.78 | 0.63 | 0.77 | 0.87 |
| 9 | 27171373 | rs10812529 | 0.788 | 1.792 | 0.734 | 1.419 | 0.165 | 0.238 | 0.238 | 0.059 | 0.89 | 0.58 | 0.81 | 0.68 | 0.82 | 0.89 |
| 9 | 27172838 | rs3739542 | 0.688 | 1.659 | 0.071 | 1.256 | 0.054 | 0.079 | 0.079 | 1.014 | 0.79 | 0.71 | 0.77 | 0.58 | 0.68 | 0.78 |
| 9 | 27173062 | rs3824410 | -0.032 | 0.704 | -0.631 | 0.996 | 0.02 | -0.03 | -0.03 | 0 | 0.25 | 0.2 | 0.25 | 0.44 | 0.2 | 0.39 |
| 9 | 27173180 | rs666478 | 0.22 | 0.486 | -0.816 | 0.178 | -0.006 | 0.031 | 0.031 | 0 | 0.55 | 0.56 | 0.55 | 0.29 | 0.52 | 0.41 |
| 9 | 27173704 | rs12554799 | 0.071 | 0.812 | -0.81 | -0.239 | -0.004 | 0.008 | 0.008 | 0 | 0.1 | 0.07 | 0.16 | 0.01 | 0.13 | 0.15 |
| 9 | 27176554 | rs2208637 | 0.429 | 1.548 | -0.613 | -0.668 | 0.011 | -0.036 | -0.036 | 0 | 0.11 | 0.13 | 0.26 | 0 | 0.14 | 0.04 |
| 9 | 27176886 | rs2224530 | 0.508 | 1.888 | -0.907 | -1.078 | -0.009 | 0.009 | 0.009 | 0 | 0.17 | 0.14 | 0.16 | 0.12 | 0.21 | 0.17 |
| 9 | 27179075 | rs7047856 | -0.023 | 1.359 | -0.773 | -1.094 | -0.015 | -0.012 | -0.012 | 0 | 0.3 | 0.46 | 0.32 | 0.27 | 0.3 | 0.39 |
| 9 | 27182802 | rs669102 | -0.934 | 0.779 | -1.189 | -1.386 | -0.03 | -0.038 | -0.038 | 0 | 0.68 | 0.75 | 0.54 | 0.47 | 0.59 | 0.76 |
| 9 | 27182825 | rs10967760 | 1.128 | 0.447 | -0.747 | -1.559 | -0.023 | 0.012 | 0.012 | 0 | 0.22 | 0.26 | 0.17 | 0.27 | 0.21 | 0.25 |
| 9 | 27184714 | rs16911136 | 2.042 | 0.164 | -0.909 | -1.806 | -0.019 | -0.051 | -0.051 | 0 | 0.13 | 0.18 | 0.05 | 0.01 | 0.05 | 0.13 |
| 9 | 27185762 | rs17834811 | 0.262 | -0.029 | -1.017 | -1.795 | -0.027 | 0.007 | 0.007 | 0 | 0.34 | 0.37 | 0.22 | 0.35 | 0.26 | 0.28 |
| 9 | 27187422 | rs581724 | -0.15 | 0.278 | -0.999 | -1.845 | -0.001 | 0.006 | 0.006 | 0 | 0.46 | 0.44 | 0.59 | 0.12 | 0.47 | 0.51 |
| 9 | 27188862 | rs2152066 | 0.097 | 0.303 | -0.998 | -1.781 | -0.012 | 0 | 0 | 0 | 0.38 | 0.38 | 0.34 | 0.34 | 0.33 | 0.43 |
| 9 | 27190655 | rs1334811 | 2.232 | 0.109 | -1.06 | -1.627 | 0.007 | -0.003 | -0.003 | 0 | 0.12 | 0.12 | 0.04 | 0.01 | 0.05 | 0.15 |
| 9 | 27190890 | rs481294 | -1.901 | 0.527 | -1.165 | -1.538 | -0.007 | -0.047 | -0.047 | 0.79 | 0.79 | 0.84 | 0.82 | 0.45 | 0.76 | 0.8 |
| 9 | 27192036 | rs2224529 | 0.318 | 0.473 | -0.79 | -1.645 | -0.009 | -0.014 | -0.014 | 0 | 0.34 | 0.4 | 0.43 | 0.28 | 0.33 | 0.26 |
| 9 | 27193503 | rs4879250 | 0.009 | 0.778 | -0.864 | -1.769 | -0.006 | -0.115 | -0.115 | 0 | 0.34 | 0.46 | 0.48 | 0.01 | 0.32 | 0.34 |
| 9 | 27193541 | rs491892 | -0.227 | 1.408 | -0.856 | -1.237 | -0.001 | -0.002 | -0.002 | 0 | 0.45 | 0.37 | 0.35 | 0.44 | 0.44 | 0.45 |
| 9 | 27193847 | rs578327 | 1.407 | 1.418 | -1.082 | -1.254 | -0.033 | 0.016 | 0.016 | 0 | 0.16 | 0.08 | 0.16 | 0.11 | 0.23 | 0.18 |
| 9 | 27196290 | rs3780315 | 0.048 | 1.823 | -0.99 | -1.251 | -0.001 | 0.003 | 0.003 | 0 | 0.32 | 0.31 | 0.4 | 0.32 | 0.31 | 0.29 |
| 9 | 27197282 | rs10812532 | 2.042 | 1.829 | -0.956 | -1.231 | 0.002 | -0.006 | -0.006 | 0 | 0.1 | 0.12 | 0.07 | 0.04 | 0.09 | 0.15 |
| 9 | 27199373 | rs7027984 | 0.859 | 1.821 | -1.057 | -1.23 | 0 | 0.011 | 0.011 | 0 | 0.22 | 0.26 | 0.35 | 0.26 | 0.23 | 0.23 |
| 9 | 27200738 | rs4242670 | 1.836 | 1.83 | -1.169 | -1.347 | 0.043 | -0.115 | -0.115 | 0.88 | 0.29 | 0.11 | 0.17 | 0.53 | 0.22 | 0.27 |
| 9 | 27201830 | rs692946 | -0.489 | 2.421 | -0.908 | -0.984 | 0.002 | 0 | 0 | 0 | 0.39 | 0.36 | 0.53 | 0.4 | 0.52 | 0.42 |
| 9 | 27201968 | rs1413829 | 0.568 | 2.457 | -1.097 | -0.964 | 0.007 | 0.011 | 0.011 | 0 | 0.23 | 0.26 | 0.36 | 0.28 | 0.27 | 0.24 |
| 9 | 27202663 | rs7876024 | 0.669 | 2.464 | -1.1 | -0.966 | 0.001 | 0.014 | 0.014 | 0 | 0.22 | 0.26 | 0.35 | 0.27 | 0.23 | 0.23 |
| 9 | 27202870 | rs639225 | -0.41 | 2.49 | -1.103 | -0.97 | -0.003 | -0.001 | -0.001 | 0 | 0.39 | 0.33 | 0.51 | 0.4 | 0.48 | 0.4 |
| 9 | 27203134 | rs638203 | -0.41 | 2.49 | -1.1 | -0.97 | -0.001 | -0.001 | -0.001 | 0 | 0.39 | 0.34 | 0.51 | 0.4 | 0.48 | 0.4 |
| 9 | 27203625 | rs2756900 | 1.183 | 2.709 | -0.992 | -0.853 | -0.041 | 0.013 | 0.013 | 0 | 0.16 | 0.08 | 0.16 | 0.12 | 0.25 | 0.17 |
| 9 | 27205021 | rs542913 | 1.183 | 2.709 | -0.992 | -0.853 | -0.041 | 0.013 | 0.013 | 0 | 0.16 | 0.08 | 0.16 | 0.12 | 0.25 | 0.17 |
| 9 | 27206307 | rs510101 | -0.642 | 2.479 | -1.044 | -0.872 | 0.007 | -0.005 | -0.005 | 0 | 0.46 | 0.35 | 0.54 | 0.46 | 0.55 | 0.44 |
| 9 | 27208857 | rs641848 | -0.182 | 2.717 | -0.433 | -0.791 | 0.023 | 0.006 | 0.006 | 0 | 0.26 | 0.32 | 0.41 | 0.27 | 0.32 | 0.26 |
| 9 | 27212360 | rs10757641 | -0.705 | 1.931 | -0.97 | -0.701 | -0.037 | 0.013 | 0.013 | 0 | 0.83 | 0.93 | 0.87 | 0.87 | 0.77 | 0.82 |

| | | | | | | | | | | | | | | | | |
|---|---|---|---|---|---|---|---|---|---|---|---|---|---|---|---|---|
| 9 | 27212966 | rs1111782 | 0.976 | 1.322 | -0.991 | -0.794 | 0.005 | 0.002 | 0.002 | 1.36 | 0.48 | 0.5 | 0.39 | 0.59 | 0.33 | 0.47 |
| 9 | 27213580 | rs3737188 | 0.549 | 1.635 | -1.083 | -0.493 | -0.009 | -0.004 | -0.004 | 0 | 0.18 | 0.07 | 0.18 | 0.19 | 0.2 | 0.17 |
| 9 | 27213772 | rs3737187 | 0.501 | 1.147 | -0.818 | -0.374 | 0.035 | 0.013 | 0.013 | 0 | 0.52 | 0.44 | 0.4 | 0.59 | 0.38 | 0.47 |
| 9 | 27214719 | rs534129 | -0.095 | 1.506 | -0.631 | -0.336 | 0.043 | -0.011 | -0.011 | 0 | 0.3 | 0.43 | 0.44 | 0.26 | 0.38 | 0.34 |
| 9 | 27214972 | rs536078 | -0.305 | 0.778 | -1.063 | -0.322 | 0.011 | 0.024 | 0.024 | 0 | 0.88 | 0.93 | 0.82 | 1 | 0.88 | 0.86 |
| 9 | 27218053 | rs2756901 | 0.995 | 1.123 | -0.97 | -0.154 | -0.008 | 0.029 | 0.029 | 0.018 | 0.66 | 0.33 | 0.54 | 0.81 | 0.6 | 0.66 |
| 9 | 27219058 | rs3736819 | 0.969 | 0.613 | -1.247 | 0.472 | -0.015 | -0.069 | -0.069 | 0.02 | 0.11 | 0 | 0.09 | 0.18 | 0.09 | 0.12 |
| 9 | 27220066 | rs2273719 | 0.065 | 0.77 | -0.536 | 0.291 | 0.012 | 0.031 | 0.031 | 0 | 0.15 | 0.39 | 0.16 | 0.17 | 0.13 | 0.19 |
| 9 | 27226947 | rs7024846 | -1.3 | 1.142 | -0.753 | 0.478 | 0.016 | -0.015 | -0.015 | 0 | 0.33 | 0.52 | 0.61 | 0.33 | 0.43 | 0.43 |
| 9 | 27228022 | rs2273717 | -0.741 | 0.979 | -0.776 | 0.622 | 0.007 | -0.018 | -0.018 | 0 | 0.86 | 0.88 | 0.89 | 0.78 | 0.91 | 0.88 |
| 9 | 27228564 | rs10812540 | 0.077 | 0.881 | -0.712 | 0.607 | -0.003 | -0.003 | -0.003 | 0 | 0.05 | 0.05 | 0.21 | 0.13 | 0.09 | 0.12 |
| 9 | 27228754 | rs1555454 | -0.058 | 0.867 | -0.83 | 0.407 | 0 | -0.008 | -0.008 | 0.04 | 0.8 | 0.77 | 0.82 | 0.78 | 0.82 | 0.87 |
| 9 | 27229141 | rs2273715 | 0.354 | 0.892 | -0.691 | 0.49 | 0.004 | 0.027 | 0.027 | 0 | 0.15 | 0.38 | 0.16 | 0.19 | 0.13 | 0.2 |
| 9 | 27230152 | rs9987817 | 0.271 | 0.894 | -0.722 | 0.478 | 0.01 | -0.008 | -0.008 | 0.005 | 0.15 | 0.38 | 0.16 | 0.15 | 0.13 | 0.12 |
| 9 | 27232748 | rs1322048 | -0.872 | 0.836 | -0.727 | 0.448 | -0.005 | -0.037 | -0.037 | 0 | 0.3 | 0.55 | 0.39 | 0.15 | 0.32 | 0.23 |
| 9 | 27233157 | rs7858722 | 1.749 | 0.842 | -0.733 | 0.452 | -0.028 | -0.027 | -0.027 | 0.651 | 0.52 | 0.19 | 0.27 | 0.49 | 0.48 | 0.53 |
| 9 | 27233367 | rs4878368 | 0.008 | 0.862 | -0.693 | 0.518 | 0.001 | 0.016 | 0.016 | 0 | 0.82 | 0.74 | 0.66 | 0.64 | 0.8 | 0.76 |
| 9 | 27234029 | rs7035592 | 1.012 | 0.857 | -0.681 | 0.499 | 0 | 0.006 | 0.006 | 0 | 0.67 | 0.36 | 0.5 | 0.5 | 0.67 | 0.64 |
| 9 | 27235123 | rs10967797 | 0.067 | 0.77 | -0.451 | 0.271 | 0.008 | 0.036 | 0.036 | 0 | 0.15 | 0.38 | 0.16 | 0.19 | 0.13 | 0.2 |
| 9 | 27247314 | rs2057362 | -1.343 | 0.809 | -0.677 | 0.232 | -0.003 | 0.023 | 0.023 | 0.438 | 0.36 | 0.7 | 0.61 | 0.33 | 0.41 | 0.43 |
| 9 | 27248177 | rs7042745 | -0.732 | 0.854 | -0.612 | 0.226 | 0.005 | -0.017 | -0.017 | 0.087 | 0.19 | 0.26 | 0.43 | 0.13 | 0.27 | 0.23 |
| 9 | 27253049 | rs10120928 | -0.038 | 0.814 | -0.856 | 0.241 | 0.003 | 0.011 | 0.011 | 0 | 0.16 | 0.12 | 0.24 | 0.13 | 0.21 | 0.12 |
| 9 | 27258216 | rs997675 | -1.134 | 0.697 | -0.862 | 0.27 | -0.002 | 0.016 | 0.016 | 0 | 0.36 | 0.62 | 0.59 | 0.23 | 0.38 | 0.41 |
| 9 | 27260594 | rs12348317 | 0.469 | 0.614 | -0.711 | 0.549 | -0.018 | 0.112 | 0.112 | 0 | 0.18 | 0.53 | 0.2 | 0.33 | 0.14 | 0.2 |
| 9 | 27261030 | rs1853186 | 0.589 | 0.554 | -1.087 | 0.451 | 0.002 | 0.01 | 0.01 | 0 | 0.69 | 0.7 | 0.66 | 0.78 | 0.67 | 0.75 |
| 9 | 27261943 | rs744121 | 0.679 | 0.699 | -0.678 | 0.672 | -0.02 | 0.159 | 0.159 | 0 | 0.17 | 0.42 | 0.19 | 0.33 | 0.13 | 0.19 |
| 9 | 27263515 | rs967144 | 0.632 | 0.71 | -0.806 | 0.294 | -0.002 | 0.044 | 0.044 | 0.05 | 0.7 | 0.79 | 0.68 | 0.9 | 0.7 | 0.77 |
| 9 | 27264215 | rs10812546 | 0.664 | 0.71 | -0.799 | 0.293 | 0 | 0.029 | 0.029 | 0.057 | 0.7 | 0.73 | 0.67 | 0.9 | 0.7 | 0.77 |
| 9 | 27264539 | rs10967828 | 1.322 | 0.758 | -0.829 | 0.302 | 0.005 | -0.014 | -0.014 | 0 | 0.65 | 0.53 | 0.47 | 0.77 | 0.63 | 0.63 |
| 9 | 27271360 | rs12683092 | -0.068 | 0.618 | -0.913 | 0.354 | -0.003 | -0.014 | -0.014 | 0 | 0.79 | 0.73 | 0.86 | 0.9 | 0.83 | 0.88 |
| 9 | 27274428 | rs13288748 | -0.046 | 0.715 | -0.877 | 0.596 | -0.001 | 0.02 | 0.02 | 0 | 0.22 | 0.67 | 0.39 | 0.34 | 0.18 | 0.29 |
| 9 | 27278084 | rs10967849 | 0.253 | 0.798 | -0.963 | 0.482 | -0.005 | 0.007 | 0.007 | 0 | 0.1 | 0.08 | 0.2 | 0 | 0.14 | 0.11 |
| 9 | 27280676 | rs4878384 | -1.246 | 0.721 | -0.912 | 0.502 | 0 | 0.007 | 0.007 | 0 | 0.33 | 0.61 | 0.59 | 0.45 | 0.34 | 0.42 |
| 9 | 27282942 | rs17695487 | 0.003 | 0.684 | -0.786 | 0.498 | -0.001 | 0.003 | 0.003 | 0 | 0.11 | 0.13 | 0.2 | 0.07 | 0.15 | 0.11 |
| 9 | 27283020 | rs10967856 | -1.394 | 0.692 | -0.796 | 0.484 | 0 | 0.003 | 0.003 | 0 | 0.34 | 0.6 | 0.61 | 0.44 | 0.36 | 0.42 |
| 9 | 27284363 | rs1126107 | -0.008 | 0.702 | -0.8 | 0.448 | -0.001 | -0.013 | -0.013 | 0 | 0.12 | 0.2 | 0.22 | 0.04 | 0.15 | 0.12 |
| 9 | 27288113 | rs2038571 | 0.042 | 0.703 | -0.787 | 0.501 | 0.014 | 0.094 | 0.094 | 0 | 0.22 | 0.49 | 0.39 | 0.38 | 0.21 | 0.29 |
| 9 | 27288895 | rs2383722 | -1.31 | 0.559 | -0.864 | 0.448 | 0 | -0.003 | -0.003 | 0 | 0.36 | 0.6 | 0.61 | 0.42 | 0.37 | 0.42 |
| 9 | 27289110 | rs10967862 | 1.873 | 0.382 | -1.103 | 0.344 | -0.016 | 0.019 | 0.019 | 0.192 | 0.5 | 0.19 | 0.29 | 0.47 | 0.51 | 0.46 |
| 9 | 27289812 | rs1980661 | 1.167 | 0.458 | -0.995 | 0.487 | -0.006 | -0.016 | -0.016 | 0 | 0.63 | 0.36 | 0.51 | 0.51 | 0.66 | 0.59 |
| 9 | 27290984 | rs2383724 | -0.465 | 0.374 | -1.002 | 0.496 | -0.001 | -0.001 | -0.001 | 0.694 | 0.86 | 0.86 | 0.9 | 0.89 | 0.87 | 0.87 |
| 9 | 27292279 | rs3739534 | 0.501 | 0.467 | -0.818 | 0.7 | -0.003 | 0.144 | 0.144 | 0 | 0.18 | 0.43 | 0.19 | 0.34 | 0.15 | 0.21 |
| 9 | 27293009 | rs7873922 | -0.672 | 0.403 | -0.989 | 0.589 | 0.01 | 0.042 | 0.042 | 0 | 0.28 | 0.58 | 0.46 | 0.44 | 0.28 | 0.39 |
| 9 | 27296705 | rs3824406 | 1.332 | -1.067 | -2.042 | -0.042 | 0.014 | 0.082 | 0.082 | 0 | 0.25 | 0.01 | 0.08 | 0.2 | 0.19 | 0.15 |
| 9 | 27297301 | rs7875191 | 0.367 | -0.908 | -1.806 | -0.156 | -0.001 | 0.005 | 0.005 | 0 | 0.85 | 0.86 | 0.75 | 0.99 | 0.85 | 0.81 |
| 9 | 27297876 | rs10967868 | -0.118 | -1.442 | -1.413 | -0.143 | -0.003 | 0.007 | 0.007 | 0 | 0.12 | 0.05 | 0.24 | 0.01 | 0.15 | 0.21 |
| 9 | 27298500 | rs10217315 | -0.819 | -1.716 | -1.722 | 0.496 | 0.024 | -0.013 | -0.013 | 0 | 0.82 | 0.91 | 0.72 | 0.78 | 0.88 | 0.83 |
| 9 | 27299620 | rs17778819 | 0.437 | -1.733 | -1.617 | 0.984 | -0.023 | -0.136 | -0.136 | 0.171 | 0.33 | 0.05 | 0.31 | 0.39 | 0.31 | 0.39 |
| 9 | 27307857 | rs6475988 | -0.731 | -1.772 | -0.415 | 0.751 | 0 | 0.009 | 0.009 | 0 | 0.55 | 0.65 | 0.49 | 0.64 | 0.47 | 0.53 |
| 9 | 27309659 | rs10967875 | 0.78 | -1.856 | -0.607 | 0.682 | -0.036 | 0.015 | 0.015 | 0 | 0.33 | 0.02 | 0.29 | 0.2 | 0.31 | 0.29 |
| 9 | 27315514 | rs10967882 | 0.627 | -1.861 | -0.423 | 0.674 | 0.002 | 0 | 0 | 0 | 0.45 | 0.31 | 0.59 | 0.47 | 0.55 | 0.5 |
| 9 | 27316971 | rs10217619 | 0.878 | -1.596 | -0.7 | -0.111 | -0.003 | -0.011 | -0.011 | 0.08 | 0.18 | 0.23 | 0.25 | 0.15 | 0.16 | 0.14 |
| 9 | 27326060 | rs3410 | -0.164 | -1.694 | -0.621 | -0.1 | -0.002 | 0.002 | 0.002 | 0 | 0.29 | 0.47 | 0.39 | 0.26 | 0.27 | 0.22 |
| 9 | 27326088 | rs3409 | 1.433 | -1.544 | -1.362 | -0.085 | 0.016 | 0.101 | 0.101 | 0 | 0.22 | 0.05 | 0.08 | 0.07 | 0.19 | 0.18 |
| 9 | 27326780 | rs1555453 | -0.503 | -1.075 | -0.733 | 0.45 | 0 | -0.018 | -0.018 | 2.66 | 0.86 | 0.93 | 0.77 | 0.86 | 0.87 | 0.88 |
| 9 | 27326904 | rs2181029 | 0.39 | -0.974 | -0.619 | 0.435 | -0.003 | -0.01 | -0.01 | 0 | 0.18 | 0.24 | 0.15 | 0.15 | 0.15 | 0.14 |
| 9 | 27328964 | rs3739531 | -0.217 | -0.575 | -1.144 | 0.501 | -0.006 | -0.007 | -0.007 | 0.493 | 0.84 | 0.86 | 0.72 | 0.84 | 0.81 | 0.82 |
| 9 | 27329792 | rs1984008 | 0.498 | -0.993 | -1.34 | 0.723 | 0.009 | -0.077 | -0.077 | 0 | 0.27 | 0.14 | 0.19 | 0.44 | 0.26 | 0.3 |
| 9 | 27330514 | rs3739530 | 0.409 | -1.129 | -0.924 | 0.693 | -0.037 | -0.054 | -0.054 | 0.02 | 0.23 | 0.35 | 0.12 | 0.08 | 0.14 | 0.15 |
| 9 | 27331324 | rs6475989 | 0.679 | -0.928 | -0.856 | 0.874 | -0.037 | 0.039 | 0.039 | 0 | 0.26 | 0.51 | 0.21 | 0.31 | 0.19 | 0.26 |
| 9 | 27332818 | rs10967894 | 1.085 | -0.823 | -1.217 | 0.945 | -0.027 | -0.006 | -0.006 | 0 | 0.23 | 0.15 | 0.38 | 0.31 | 0.33 | 0.36 |
| 9 | 27332892 | rs1571649 | 0.176 | -0.589 | -1.957 | 0.585 | -0.007 | -0.024 | -0.024 | 0.052 | 0.46 | 0.2 | 0.44 | 0.44 | 0.48 | 0.39 |
| 9 | 27333155 | rs10812569 | 0.536 | 0.199 | -1.413 | 1.027 | -0.005 | -0.039 | -0.039 | 2.123 | 0.23 | 0.15 | 0.18 | 0.31 | 0.24 | 0.28 |
| 9 | 27334289 | rs10738769 | 0.501 | 0.139 | -1.455 | 1.002 | -0.004 | -0.026 | -0.026 | 0 | 0.24 | 0.16 | 0.19 | 0.3 | 0.24 | 0.27 |
| 9 | 27335339 | rs10967899 | -0.227 | 0.611 | -0.853 | 1.149 | 0.065 | 0.051 | 0.051 | 0 | 0.27 | 0.47 | 0.44 | 0.45 | 0.35 | 0.38 |
| 9 | 27336207 | rs16911416 | -0.456 | 0.17 | -1.104 | 0.841 | 0.015 | -0.015 | -0.015 | 1.154 | 0.11 | 0.03 | 0.05 | 0.11 | 0.07 | 0.09 |
| 9 | 27338480 | rs10967901 | -0.434 | 0.571 | -1.119 | 0.765 | -0.061 | -0.008 | -0.008 | 0.073 | 0.63 | 0.77 | 0.53 | 0.63 | 0.56 | 0.56 |
| 9 | 27340967 | rs10812570 | -0.228 | 0.123 | -0.454 | 0.452 | 0.022 | 0.04 | 0.04 | 1.354 | 0.44 | 0.3 | 0.28 | 0.29 | 0.41 | 0.35 |
| 9 | 27341338 | rs7041639 | -1.065 | -0.17 | 0.628 | 0.64 | -0.043 | 0.024 | 0.024 | 0 | 0.14 | 0.28 | 0.06 | 0.17 | 0.08 | 0.15 |

| | | | | | | | | | | | | | | | | |
|---|---|---|---|---|---|---|---|---|---|---|---|---|---|---|---|---|
| 9 | 27344662 | rs7867777 | -1.829 | 0.047 | 2.075 | 0.741 | 0.079 | 0.197 | 0.197 | 2.244 | 0.06 | 0.44 | 0.05 | 0.08 | 0.02 | 0.01 |
| 9 | 27346374 | rs13298013 | -1.807 | 0.1 | 2.079 | 0.746 | 0.094 | 0.207 | 0.207 | 2.419 | 0.06 | 0.43 | 0.05 | 0.08 | 0.03 | 0.01 |
| 9 | 27349580 | rs12003803 | -1.892 | 0.12 | 2.542 | 0.738 | 0.088 | 0.2 | 0.2 | 2.196 | 0.06 | 0.43 | 0.05 | 0.08 | 0.03 | 0.01 |
| 9 | 27353460 | rs1888380 | -0.271 | 0.397 | 1.871 | 0.5 | -0.001 | 0 | 0 | 0.103 | 0.45 | 0.5 | 0.38 | 0.4 | 0.45 | 0.38 |
| 9 | 27354556 | rs7044577 | -0.406 | 0.378 | 1.846 | 0.49 | -0.001 | 0 | 0 | 0 | 0.45 | 0.5 | 0.38 | 0.4 | 0.45 | 0.38 |
| 9 | 27358115 | rs10812573 | -0.041 | 0.389 | 1.878 | 0.127 | -0.015 | 0.022 | 0.022 | 0 | 0.39 | 0.26 | 0.32 | 0.31 | 0.4 | 0.3 |
| 9 | 27361872 | rs7849799 | -1.722 | 0.66 | 2.384 | -0.135 | 0.066 | -0.052 | -0.052 | 0.256 | 0.1 | 0.47 | 0.1 | 0 | 0.09 | 0.05 |
| 9 | 27364824 | rs7034798 | -1.411 | 0.686 | 1.675 | -0.086 | 0.067 | -0.032 | -0.032 | 0 | 0.09 | 0.4 | 0.09 | 0 | 0.08 | 0.05 |
| 9 | 27367278 | rs1330921 | -1.478 | 0.713 | 1.441 | -0.085 | 0.053 | -0.043 | -0.043 | 0 | 0.09 | 0.39 | 0.09 | 0 | 0.08 | 0.05 |
| 9 | 27371123 | rs4483251 | -1.349 | 1.09 | 0.804 | -0.216 | 0.014 | -0.048 | -0.048 | 0 | 0.12 | 0.17 | 0.22 | 0.01 | 0.11 | 0.18 |
| 9 | 27372924 | rs1111764 | -0.888 | 0.884 | 0.426 | -0.25 | -0.001 | 0.005 | 0.005 | 0 | 0.51 | 0.56 | 0.5 | 0.37 | 0.49 | 0.54 |
| 9 | 27375843 | rs10967916 | -0.179 | 0.856 | 0.32 | -0.409 | 0.019 | -0.005 | -0.005 | 0.419 | 0.4 | 0.21 | 0.28 | 0.36 | 0.37 | 0.42 |
| 9 | 27378301 | rs1330920 | -0.761 | 0.664 | 0.363 | -0.378 | -0.008 | 0.021 | 0.021 | 0 | 0.28 | 0.21 | 0.36 | 0.15 | 0.28 | 0.34 |
| 9 | 27384204 | rs12346755 | -1.688 | 0.234 | 0.951 | -0.15 | 0.003 | 0.053 | 0.053 | 0 | 0.08 | 0.22 | 0.06 | 0.12 | 0.05 | 0.13 |
| 9 | 27386505 | rs1110155 | 0.072 | -0.169 | 0.052 | -1.023 | 0.017 | 0.019 | 0.019 | 0 | 0.39 | 0.36 | 0.42 | 0.77 | 0.26 | 0.58 |
| 9 | 27389824 | rs7021452 | -0.301 | -0.244 | 0.549 | -1.08 | -0.011 | -0.019 | -0.019 | 0 | 0.43 | 0.54 | 0.44 | 0.77 | 0.3 | 0.6 |
| 9 | 27398306 | rs11791484 | -0.957 | 0.056 | 0.666 | -0.912 | 0.003 | 0.009 | 0.009 | 0 | 0.06 | 0.06 | 0.09 | 0.11 | 0.08 | 0.13 |
| 9 | 27401297 | rs1888382 | -1.094 | 0.044 | 0.404 | -1.495 | 0.002 | 0.038 | 0.038 | 0 | 0.8 | 0.9 | 0.89 | 1 | 0.8 | 0.96 |
| 9 | 27402961 | rs2150336 | -0.183 | 0.03 | 0.476 | -1.362 | -0.013 | -0.028 | -0.028 | 0 | 0.41 | 0.54 | 0.47 | 0.77 | 0.33 | 0.6 |
| 9 | 27407914 | rs11787669 | 0.104 | 0.166 | 0.293 | -1.204 | -0.008 | -0.008 | -0.008 | 0 | 0.05 | 0.02 | 0.08 | 0.11 | 0.07 | 0.13 |
| 9 | 27412769 | rs11788670 | 1.096 | 0.266 | 0.332 | -1.399 | -0.012 | 0.012 | 0.012 | 0 | 0.3 | 0.23 | 0.22 | 0.2 | 0.38 | 0.25 |
| 9 | 27428550 | rs17696142 | 0.791 | 0.214 | 0.308 | -1.504 | 0.004 | 0.017 | 0.017 | 0.264 | 0.23 | 0.14 | 0.12 | 0 | 0.22 | 0.05 |
| 9 | 27430232 | rs1161680 | -0.173 | 0.017 | 0.335 | -0.773 | -0.003 | 0.004 | 0.004 | 0 | 0.58 | 0.61 | 0.39 | 0.36 | 0.6 | 0.36 |
| 9 | 27441537 | rs2589053 | -0.553 | -0.191 | 0.112 | -0.772 | 0 | -0.028 | -0.028 | 0 | 0.66 | 0.74 | 0.48 | 0.47 | 0.69 | 0.54 |
| 9 | 27443802 | rs2589054 | -0.569 | -0.276 | 0.137 | -0.707 | -0.011 | -0.004 | -0.004 | 0 | 0.38 | 0.56 | 0.27 | 0.32 | 0.32 | 0.3 |
| 9 | 27447089 | rs1058326 | 0.864 | -0.254 | 0.051 | -0.557 | -0.002 | 0.002 | 0.002 | 0.103 | 0.37 | 0.36 | 0.39 | 0.2 | 0.4 | 0.28 |
| 9 | 27447337 | rs10967939 | 1.695 | -0.073 | -0.118 | -0.385 | -0.027 | -0.148 | -0.148 | 0 | 0.21 | 0.02 | 0.3 | 0.49 | 0.19 | 0.33 |
| 9 | 27447434 | rs1822725 | 0.48 | -0.086 | -0.001 | -0.33 | 0.003 | 0.082 | 0.082 | 0 | 0.9 | 0.83 | 0.9 | 0.72 | 0.9 | 0.74 |
| 9 | 27450211 | rs944404 | 0.006 | 0.098 | -0.413 | -0.612 | -0.001 | -0.005 | -0.005 | 0 | 0.35 | 0.37 | 0.44 | 0.54 | 0.34 | 0.43 |
| 9 | 27451482 | rs4879507 | 0.975 | 0.191 | -0.863 | -0.659 | 0.002 | -0.017 | -0.017 | 0.107 | 0.23 | 0.23 | 0.12 | 0.04 | 0.19 | 0.06 |
| 9 | 27451960 | rs765709 | 0.799 | 0.218 | -0.893 | -0.614 | -0.042 | 0.027 | 0.027 | 0 | 0.08 | 0 | 0.07 | 0.03 | 0.11 | 0.08 |
| 9 | 27453327 | rs10812599 | 1.403 | 0.181 | -0.826 | -0.523 | 0.004 | -0.089 | -0.089 | 0.165 | 0.21 | 0.07 | 0.3 | 0.47 | 0.19 | 0.31 |
| 9 | 27455823 | rs10967945 | -0.076 | 0.662 | -0.06 | -0.851 | 0.029 | -0.042 | -0.042 | 0 | 0.11 | 0.18 | 0.11 | 0.03 | 0.15 | 0.09 |
| 9 | 27456928 | rs7021930 | -0.211 | 0.471 | -0.464 | -0.675 | 0.002 | -0.093 | -0.093 | 0.775 | 0.34 | 0.41 | 0.23 | 0.07 | 0.34 | 0.14 |
| 9 | 27458939 | rs725804 | -0.519 | 0.361 | -0.835 | -0.459 | 0.001 | -0.038 | -0.038 | 1.37 | 0.42 | 0.56 | 0.32 | 0.12 | 0.43 | 0.23 |
| 9 | 27467670 | rs10757663 | 0.331 | -0.33 | -1.596 | 0.755 | 0.047 | -0.072 | -0.072 | 0.912 | 0.73 | 0.85 | 0.63 | 0.44 | 0.83 | 0.63 |
| 9 | 27468461 | rs10511816 | 0.698 | 0.203 | -2.69 | -0.251 | -0.118 | 0.157 | 0.157 | 0.15 | 0.34 | 0.05 | 0.3 | 0.12 | 0.42 | 0.22 |
| 9 | 27473099 | rs10967951 | 0.822 | 0.271 | -2.483 | 0.644 | -0.041 | -0.095 | -0.095 | 0.879 | 0.15 | 0.01 | 0.21 | 0.33 | 0.17 | 0.24 |
| 9 | 27474214 | rs10967952 | 0.158 | 0.255 | -2.887 | 1.083 | -0.009 | -0.048 | -0.048 | 0 | 0.2 | 0.07 | 0.25 | 0.26 | 0.21 | 0.26 |
| 9 | 27474593 | rs7866248 | -0.028 | -0.238 | -2.036 | 0.962 | -0.005 | 0.008 | 0.008 | 1.089 | 0.76 | 0.27 | 0.7 | 0.58 | 0.82 | 0.68 |
| 9 | 27478052 | rs1822723 | -0.409 | 0.445 | -1.873 | 1.255 | -0.025 | -0.058 | -0.058 | 1.279 | 0.24 | 0.12 | 0.38 | 0.37 | 0.27 | 0.39 |
| 9 | 27483959 | rs895023 | 0.933 | 0.302 | -0.243 | 0.881 | 0.073 | 0.029 | 0.029 | 0.114 | 0.95 | 0.77 | 0.96 | 0.99 | 0.96 | 0.91 |
| 9 | 27488092 | rs17779457 | 0.887 | -0.265 | -0.371 | 1.03 | 0.019 | -0.015 | -0.015 | 0 | 0.33 | 0.24 | 0.31 | 0.33 | 0.24 | 0.27 |
| 9 | 27488906 | rs868857 | 0.891 | -0.272 | -0.351 | 1.026 | 0.019 | -0.015 | -0.015 | 0 | 0.33 | 0.24 | 0.32 | 0.32 | 0.23 | 0.27 |
| 9 | 27489251 | rs868856 | -0.535 | -0.355 | 0.35 | 0.983 | 0.002 | 0 | 0 | 0.243 | 0.61 | 0.45 | 0.63 | 0.67 | 0.72 | 0.62 |
| 9 | 27490967 | rs7046653 | -0.51 | -0.365 | 0.358 | 0.978 | 0.004 | 0 | 0 | 0.813 | 0.61 | 0.44 | 0.63 | 0.67 | 0.72 | 0.62 |
| 9 | 27502986 | rs1977661 | 0.477 | -0.66 | 0.656 | 1.27 | 0 | 0.001 | 0.001 | 0.58 | 0.15 | 0.16 | 0.21 | 0.32 | 0.11 | 0.19 |
| 9 | 27508687 | rs2166128 | 0.111 | -0.47 | 0.506 | 1.098 | 0.025 | 0.009 | 0.009 | 0.399 | 0.92 | 0.8 | 0.94 | 0.91 | 0.93 | 0.85 |
| 9 | 27510360 | rs10812605 | -0.517 | -0.509 | 0.699 | 1.072 | 0.009 | -0.002 | -0.002 | 0.016 | 0.57 | 0.41 | 0.6 | 0.58 | 0.66 | 0.52 |
| 9 | 27511284 | rs2484314 | 0.999 | -0.472 | 0.367 | 1.078 | 0.005 | -0.005 | -0.005 | 0 | 0.35 | 0.33 | 0.33 | 0.33 | 0.27 | 0.32 |
| 9 | 27516640 | rs774351 | 1.095 | -0.474 | 0.365 | 1.202 | 0.004 | 0.001 | 0.001 | 0 | 0.35 | 0.34 | 0.33 | 0.29 | 0.27 | 0.32 |
| 9 | 27519173 | rs11792285 | 1.067 | -0.412 | -0.125 | 0.998 | -0.101 | -0.011 | -0.011 | 0 | 0.27 | 0.07 | 0.27 | 0.23 | 0.38 | 0.21 |
| 9 | 27526047 | rs700782 | 1.303 | -0.247 | 0.332 | 0.661 | 0.014 | -0.002 | -0.002 | 0 | 0.32 | 0.23 | 0.31 | 0.28 | 0.24 | 0.27 |
| 9 | 27529316 | rs903603 | 0.253 | -0.193 | 0.096 | 0.617 | -0.101 | -0.017 | -0.017 | 0 | 0.44 | 0.09 | 0.49 | 0.61 | 0.51 | 0.46 |
| 9 | 27533452 | rs4879541 | 0.186 | -0.187 | 0.107 | 0.582 | -0.028 | 0.019 | 0.019 | 0 | 0.46 | 0.15 | 0.5 | 0.6 | 0.53 | 0.5 |
| 9 | 27536397 | rs2814707 | 1.56 | -0.247 | 0.091 | 0.314 | 0.008 | 0.043 | 0.043 | 0.894 | 0.3 | 0.22 | 0.15 | 0.07 | 0.24 | 0.16 |
| 9 | 27542261 | rs10812611 | 0.316 | -0.239 | -0.052 | 0.348 | -0.046 | -0.022 | -0.022 | 0 | 0.46 | 0.09 | 0.51 | 0.6 | 0.57 | 0.49 |
| 9 | 27543281 | rs3849942 | -1.636 | -0.238 | 0.139 | 0.407 | 0.007 | 0.025 | 0.025 | 0.783 | 0.69 | 0.75 | 0.85 | 0.91 | 0.77 | 0.84 |
| 9 | 27547313 | rs3739526 | 0.035 | -0.199 | -0.246 | 0.559 | 0.002 | -0.031 | -0.031 | 0 | 0.15 | 0.1 | 0.27 | 0.29 | 0.16 | 0.28 |
| 9 | 27553876 | rs12349820 | 0.956 | 0.001 | -0.083 | 0.474 | 0.006 | -0.002 | -0.002 | 0.071 | 0.15 | 0.11 | 0.14 | 0.12 | 0.25 | 0.2 |
| 9 | 27556780 | rs10122902 | -0.228 | -0.056 | -0.095 | 0.513 | -0.002 | 0.01 | 0.01 | 0.119 | 0.17 | 0.15 | 0.28 | 0.3 | 0.16 | 0.28 |
| 9 | 27559733 | rs1565948 | -0.935 | -0.012 | -0.022 | 0.474 | -0.01 | -0.008 | -0.008 | 0 | 0.34 | 0.26 | 0.45 | 0.43 | 0.48 | 0.53 |
| 9 | 27560594 | rs3849944 | -1.063 | -0.005 | -0.062 | 0.474 | 0.016 | 0.011 | 0.011 | 0.192 | 0.35 | 0.33 | 0.46 | 0.43 | 0.49 | 0.53 |
| 9 | 27560965 | rs12347222 | -0.288 | 0.01 | -0.099 | 0.481 | 0 | 0.011 | 0.011 | 0.075 | 0.17 | 0.17 | 0.3 | 0.3 | 0.21 | 0.29 |
| 9 | 27561049 | rs774359 | 1.465 | -0.096 | -0.409 | -0.313 | 0.005 | 0.061 | 0.061 | 0.36 | 0.32 | 0.25 | 0.16 | 0.1 | 0.25 | 0.2 |
| 9 | 27561628 | rs17769294 | 1.154 | -0.093 | -0.605 | -0.316 | -0.006 | 0.049 | 0.049 | 0 | 0.09 | 0.02 | 0.07 | 0 | 0.11 | 0.06 |
| 9 | 27561800 | rs2453554 | 1.749 | -0.075 | -0.524 | -0.226 | 0.003 | 0.089 | 0.089 | 0.557 | 0.3 | 0.19 | 0.15 | 0.09 | 0.24 | 0.16 |
| 9 | 27563868 | rs2453555 | 1.754 | -0.085 | -0.541 | -0.229 | 0.003 | 0.088 | 0.088 | 0.537 | 0.3 | 0.19 | 0.15 | 0.09 | 0.23 | 0.16 |
| 9 | 27567145 | rs10757668 | 0.919 | 0.102 | -0.629 | -0.034 | 0.012 | -0.005 | -0.005 | 0 | 0.15 | 0.14 | 0.15 | 0.12 | 0.25 | 0.19 |

| | | | | | | | | | | | | | | | | |
|---|---|---|---|---|---|---|---|---|---|---|---|---|---|---|---|---|
| 9 | 27570051 | rs4879572 | 1.532 | -0.201 | -1.042 | 0.276 | -0.003 | -0.038 | -0.038 | 0 | 0.07 | 0.01 | 0.14 | 0.15 | 0.06 | 0.12 |
| 9 | 27572255 | rs2282241 | -0.076 | -0.013 | -0.936 | 0.09 | -0.01 | 0.098 | 0.098 | 0 | 0.43 | 0.1 | 0.61 | 0.68 | 0.44 | 0.5 |
| 9 | 27575785 | rs1948522 | -0.246 | -0.016 | -0.781 | 0.094 | -0.015 | 0.025 | 0.025 | 0 | 0.15 | 0.06 | 0.14 | 0.11 | 0.19 | 0.11 |
| 9 | 27579560 | rs1982915 | -0.133 | 0.736 | -0.341 | 0.09 | 0 | -0.023 | -0.023 | 0 | 0.53 | 0.53 | 0.34 | 0.22 | 0.47 | 0.37 |
| 9 | 27580674 | rs12350076 | -0.199 | 0.689 | -0.275 | 0.454 | -0.002 | -0.001 | -0.001 | 0 | 0.53 | 0.47 | 0.49 | 0.44 | 0.47 | 0.48 |
| 9 | 27581239 | rs2783010 | -0.219 | 0.671 | -0.293 | 0.475 | -0.002 | 0 | 0 | 0 | 0.54 | 0.54 | 0.47 | 0.44 | 0.43 | 0.47 |
| 9 | 27581980 | rs10967992 | 0.665 | 0.77 | -0.33 | 0.415 | -0.015 | 0.01 | 0.01 | 0 | 0.16 | 0.08 | 0.15 | 0.11 | 0.27 | 0.2 |
| 9 | 27584530 | rs7868845 | 0.352 | 0.302 | -0.938 | 0.405 | -0.006 | -0.047 | -0.047 | 0 | 0.38 | 0.42 | 0.23 | 0.12 | 0.31 | 0.21 |
| 9 | 27586162 | rs2453556 | -0.054 | 0.509 | -0.739 | 0.854 | 0.003 | -0.002 | -0.002 | 0.752 | 0.43 | 0.6 | 0.41 | 0.44 | 0.36 | 0.39 |
| 9 | 27588731 | rs702231 | 0.762 | 1.069 | -0.3 | 1.097 | 0.042 | 0.16 | 0.16 | 0 | 0.81 | 0.56 | 0.62 | 0.66 | 0.78 | 0.71 |
| 9 | 27599202 | rs10757671 | -0.038 | 1.248 | -1.569 | 0.188 | -0.082 | 0.065 | 0.065 | 0 | 0.2 | 0.02 | 0.14 | 0.1 | 0.26 | 0.16 |
| 9 | 27599746 | rs2477518 | -0.777 | 0.668 | -0.79 | 0.591 | 0.004 | -0.008 | -0.008 | 0 | 0.22 | 0.32 | 0.32 | 0.15 | 0.25 | 0.18 |
| 9 | 27600645 | rs10968001 | -1.29 | -0.003 | -0.963 | 0.954 | -0.001 | 0 | 0 | 0 | 0.16 | 0.15 | 0.09 | 0.22 | 0.08 | 0.19 |
| 9 | 27601332 | rs480866 | -0.14 | 0.172 | -1.363 | 1.05 | 0.003 | -0.056 | -0.056 | 0 | 0.56 | 0.77 | 0.47 | 0.52 | 0.58 | 0.45 |
| 9 | 27602921 | rs633583 | -0.119 | 0.087 | -1.403 | 1.019 | 0.009 | -0.023 | -0.023 | 0.505 | 0.56 | 0.77 | 0.47 | 0.57 | 0.59 | 0.45 |
| 9 | 27606780 | rs615849 | 0.009 | 0.198 | -0.693 | 0.849 | -0.001 | -0.003 | -0.003 | 0 | 0.47 | 0.45 | 0.42 | 0.33 | 0.53 | 0.38 |
| 9 | 27609207 | rs813504 | 0.916 | 0.675 | -0.479 | 0.826 | 0.009 | 0.001 | 0.001 | 0 | 0.94 | 0.92 | 0.83 | 0.93 | 0.89 | 0.9 |
| 9 | 27609514 | rs810174 | 1.384 | 0.624 | 0.085 | 0.85 | 0.024 | 0.022 | 0.022 | 0 | 0.8 | 0.6 | 0.77 | 0.76 | 0.78 | 0.8 |
| 9 | 27610116 | rs17779794 | -0.281 | 0.8 | -0.706 | 1.196 | -0.025 | -0.049 | -0.049 | 0 | 0.09 | 0 | 0.12 | 0.18 | 0.11 | 0.1 |
| 9 | 27610657 | rs2889829 | -0.782 | 0.339 | -0.967 | 0.629 | 0.009 | 0.029 | 0.029 | 0 | 0.6 | 0.7 | 0.51 | 0.78 | 0.64 | 0.57 |
| 9 | 27615794 | rs504532 | -0.578 | 0.317 | -0.78 | 0.791 | -0.007 | -0.034 | -0.034 | 0 | 0.3 | 0.07 | 0.23 | 0.28 | 0.29 | 0.23 |
| 9 | 27620642 | rs10812622 | -0.687 | 0.244 | -0.685 | 0.79 | 0.011 | -0.006 | -0.006 | 0 | 0.31 | 0.17 | 0.28 | 0.3 | 0.29 | 0.23 |
| 9 | 27621563 | rs10968015 | -0.576 | 0.049 | -0.663 | 0.798 | 0.017 | -0.004 | -0.004 | 0 | 0.29 | 0.17 | 0.23 | 0.29 | 0.26 | 0.22 |
| 9 | 27627074 | rs10968017 | -0.907 | 0.13 | -0.737 | 0.878 | 0.007 | -0.046 | -0.046 | 0 | 0.33 | 0.12 | 0.28 | 0.39 | 0.29 | 0.33 |
| 9 | 27627362 | rs536635 | 0.621 | 0.143 | -0.452 | 0.812 | 0.001 | 0.005 | 0.005 | 0 | 0.65 | 0.71 | 0.65 | 0.74 | 0.64 | 0.67 |
| 9 | 27630050 | rs4879626 | 0.181 | -0.001 | -0.636 | 1.224 | -0.005 | 0.14 | 0.14 | 0 | 0.93 | 0.74 | 0.86 | 0.77 | 0.96 | 0.9 |
| 9 | 27630126 | rs10968022 | -0.574 | 0.227 | -1.401 | 0.762 | -0.028 | -0.016 | -0.016 | 0 | 0.28 | 0.05 | 0.15 | 0.25 | 0.26 | 0.23 |
| 9 | 27644077 | rs7847656 | -1.243 | 0.34 | -0.766 | 1.063 | 0 | 0 | 0 | 0 | 0.14 | 0.17 | 0.09 | 0.26 | 0.12 | 0.17 |
| 9 | 27652540 | rs2643801 | -0.534 | 0.229 | -0.198 | 0.746 | 0.006 | -0.038 | -0.038 | 0 | 0.15 | 0.22 | 0.32 | 0.07 | 0.18 | 0.27 |
| 9 | 27653714 | rs536246 | -0.098 | 0.454 | -0.695 | 0.204 | -0.001 | -0.004 | -0.004 | 0.608 | 0.59 | 0.37 | 0.7 | 0.7 | 0.67 | 0.68 |
| 9 | 27664162 | rs10511813 | 1.505 | 0.316 | -1.069 | -0.004 | -0.093 | 0.069 | 0.069 | 0.237 | 0.2 | 0.03 | 0.18 | 0.04 | 0.23 | 0.16 |
| 9 | 27670083 | rs3904255 | -0.997 | 0.516 | -0.441 | 0.787 | 0.006 | 0.032 | 0.032 | 1.756 | 0.35 | 0.47 | 0.51 | 0.54 | 0.42 | 0.55 |
| 9 | 27671898 | rs9695050 | 1.201 | 0.482 | -0.221 | 0.763 | -0.005 | -0.032 | -0.032 | 0 | 0.31 | 0.12 | 0.28 | 0.38 | 0.31 | 0.17 |
| 9 | 27674224 | rs1330204 | 0.176 | 0.423 | 0.118 | 0.949 | 0.008 | 0.157 | 0.157 | 0.317 | 0.08 | 0.26 | 0.04 | 0.31 | 0.06 | 0.07 |
| 9 | 27674777 | rs12235880 | 1.234 | 0.406 | -0.359 | 0.946 | -0.033 | -0.078 | -0.078 | 0 | 0.3 | 0.06 | 0.27 | 0.38 | 0.31 | 0.17 |
| 9 | 27678194 | rs7036816 | -0.659 | 0.462 | -0.344 | 0.948 | 0.033 | 0.203 | 0.203 | 0.029 | 0.18 | 0.52 | 0.19 | 0.5 | 0.22 | 0.25 |
| 9 | 27679922 | rs4436203 | 1.131 | 0.383 | -0.408 | 0.774 | -0.025 | -0.032 | -0.032 | 0 | 0.08 | 0.03 | 0.12 | 0.19 | 0.15 | 0.17 |
| 9 | 27683762 | rs6476005 | -0.807 | 0.301 | -0.285 | 0.638 | 0.003 | 0.027 | 0.027 | 1.813 | 0.36 | 0.5 | 0.51 | 0.53 | 0.42 | 0.55 |
| 9 | 27688290 | rs2150021 | 1.128 | 0.287 | -0.497 | 0.633 | -0.009 | -0.059 | -0.059 | 0.019 | 0.3 | 0.1 | 0.27 | 0.4 | 0.3 | 0.18 |
| 9 | 27688488 | rs10757678 | -0.855 | 0.44 | -0.519 | 0.389 | 0 | -0.008 | -0.008 | 0.043 | 0.66 | 0.61 | 0.78 | 0.93 | 0.72 | 0.72 |
| 9 | 27696928 | rs7039566 | 0.692 | 0.424 | -0.444 | 0.919 | 0.003 | -0.007 | -0.007 | 0 | 0.32 | 0.31 | 0.25 | 0.42 | 0.3 | 0.17 |
| 9 | 27702989 | rs13291888 | 0.423 | 0.228 | -1.039 | 1.083 | -0.029 | -0.042 | -0.042 | 0 | 0.09 | 0.03 | 0.13 | 0.25 | 0.15 | 0.2 |
| 9 | 27703432 | rs10968078 | -0.127 | 0.219 | -1.113 | 0.161 | -0.03 | 0.072 | 0.072 | 0.019 | 0.18 | 0.02 | 0.33 | 0.05 | 0.22 | 0.32 |
| 9 | 27704877 | rs944183 | -0.302 | 0.187 | -1.006 | 0.153 | -0.01 | 0.047 | 0.047 | 0 | 0.19 | 0.07 | 0.34 | 0.05 | 0.22 | 0.32 |
| 9 | 27706404 | rs6476012 | 0.361 | 0.142 | -0.989 | 0.012 | -0.012 | -0.02 | -0.02 | 0.845 | 0.39 | 0.44 | 0.37 | 0.72 | 0.37 | 0.32 |
| 9 | 27707461 | rs1411381 | -0.81 | 0.142 | -1.185 | 0.053 | -0.002 | -0.028 | -0.028 | 0 | 0.26 | 0.33 | 0.4 | 0.17 | 0.28 | 0.4 |
| 9 | 27707666 | rs1411379 | -0.124 | 0.261 | -1.34 | 0.103 | 0.011 | -0.05 | -0.05 | 0 | 0.18 | 0.28 | 0.36 | 0.05 | 0.22 | 0.31 |
| 9 | 27711260 | rs17697408 | 0.669 | 0.625 | -1.862 | 0.349 | -0.04 | 0.095 | 0.095 | 0 | 0.35 | 0.16 | 0.22 | 0.1 | 0.35 | 0.27 |
| 9 | 27711270 | rs12555932 | -0.425 | 0.556 | -1.518 | 0.805 | 0.001 | -0.006 | -0.006 | 0 | 0.05 | 0.04 | 0.04 | 0.12 | 0.05 | 0.09 |
| 9 | 27715940 | rs1360267 | -0.519 | 0.547 | -1.553 | 0.941 | 0.021 | 0.003 | 0.003 | 0 | 0.43 | 0.19 | 0.59 | 0.35 | 0.43 | 0.46 |
| 9 | 27722059 | rs951970 | -0.766 | 1.033 | -1.087 | 0.911 | -0.014 | -0.008 | -0.008 | 0.194 | 0.11 | 0.07 | 0.28 | 0.17 | 0.17 | 0.23 |
| 9 | 27723713 | rs2026147 | 0.373 | 0.455 | -1.161 | 0.797 | 0.035 | 0.024 | 0.024 | 0 | 0.53 | 0.71 | 0.48 | 0.4 | 0.61 | 0.39 |
| 9 | 27723858 | rs2026146 | -0.525 | 0.255 | -0.357 | 1.11 | -0.013 | 0.066 | 0.066 | 1.258 | 0.25 | 0.35 | 0.33 | 0.4 | 0.2 | 0.33 |
| 9 | 27724415 | rs10757683 | 0.244 | 0.833 | -0.643 | 0.881 | -0.009 | -0.014 | -0.014 | 0 | 0.78 | 0.8 | 0.66 | 0.67 | 0.66 | 0.67 |
| 9 | 27725305 | rs10732349 | 0.724 | -0.058 | 0.381 | 0.005 | -0.047 | -0.095 | -0.095 | 0 | 0.86 | 0.51 | 0.93 | 1 | 0.93 | 0.8 |
| 9 | 27726731 | rs4242689 | 0.89 | -0.153 | 0.449 | -0.042 | -0.052 | -0.093 | -0.093 | 0 | 0.86 | 0.44 | 0.92 | 1 | 0.93 | 0.8 |
| 9 | 27726758 | rs4559330 | 1.081 | 0.189 | -0.32 | -0.28 | -0.07 | 0.038 | 0.038 | 1.976 | 0.31 | 0.09 | 0.25 | 0.15 | 0.34 | 0.24 |
| 9 | 27729356 | rs4425847 | 0.649 | 0.062 | 0.162 | -0.157 | -0.061 | -0.105 | -0.105 | 0 | 0.85 | 0.43 | 0.93 | 1 | 0.93 | 0.8 |
| 9 | 27733472 | rs1857079 | 0.319 | 0.372 | -0.223 | 0.088 | -0.036 | -0.022 | -0.022 | 0 | 0.07 | 0 | 0.27 | 0.08 | 0.1 | 0.12 |
| 9 | 27745466 | rs10968089 | -0.135 | 0.198 | -0.093 | -0.163 | -0.005 | 0.056 | 0.056 | 0.117 | 0.57 | 0.32 | 0.67 | 0.85 | 0.59 | 0.56 |
| 9 | 27746785 | rs1411373 | 0.826 | -0.03 | 1.004 | 0.442 | -0.011 | 0.114 | 0.114 | 0 | 0.9 | 0.67 | 0.95 | 0.83 | 0.95 | 0.9 |
| 9 | 27749775 | rs1774037 | 0.01 | -0.088 | 0.463 | -0.205 | 0 | 0.092 | 0.092 | 0 | 0.53 | 0.33 | 0.66 | 0.76 | 0.47 | 0.63 |
| 9 | 27753894 | rs1536982 | -0.084 | -0.116 | 0.92 | -0.232 | -0.002 | 0.056 | 0.056 | 0 | 0.54 | 0.4 | 0.68 | 0.76 | 0.48 | 0.63 |
| 9 | 27756019 | rs10812662 | 0.584 | 0.147 | 0.633 | -0.697 | -0.132 | 0.142 | 0.142 | 0 | 0.33 | 0.09 | 0.26 | 0.08 | 0.49 | 0.25 |
| 9 | 27766802 | rs7851492 | -0.885 | 0.078 | 1.342 | -0.51 | 0.014 | -0.036 | -0.036 | 0.144 | 0.09 | 0.36 | 0.03 | 0 | 0.05 | 0.08 |
| 9 | 27792965 | rs10968110 | 0.61 | -0.036 | 0.386 | -0.767 | -0.137 | 0.169 | 0.169 | 0 | 0.37 | 0.08 | 0.36 | 0.08 | 0.5 | 0.29 |
| 9 | 27800007 | rs10968114 | 0.352 | -0.411 | 0.613 | -0.727 | 0.005 | 0.162 | 0.162 | 0 | 0.46 | 0.21 | 0.56 | 0.84 | 0.44 | 0.5 |
| 9 | 27801225 | rs7863376 | -1.11 | -0.442 | 1.072 | -0.722 | -0.009 | 0.024 | 0.024 | 0.056 | 0.09 | 0.29 | 0.04 | 0.08 | 0.06 | 0.15 |
| 9 | 27803120 | rs10968119 | 0.719 | -0.402 | 0.906 | -0.735 | -0.112 | 0.173 | 0.173 | 0 | 0.37 | 0.08 | 0.35 | 0.08 | 0.51 | 0.29 |

| | | | | | | | | | | | | | | | | |
|---|---|---|---|---|---|---|---|---|---|---|---|---|---|---|---|---|
| 9 | 27804540 | rs10968120 | 0.002 | -0.322 | 0.973 | -0.635 | 0.054 | 0.015 | 0.015 | 0.633 | 0.06 | 0.26 | 0.02 | 0 | 0.04 | 0.04 |
| 9 | 27808448 | rs10968122 | 0.112 | -0.119 | 0.546 | -0.734 | -0.015 | 0.213 | 0.213 | 0 | 0.49 | 0.19 | 0.58 | 0.84 | 0.49 | 0.52 |
| 9 | 27819540 | rs10812667 | -0.372 | 0.062 | 0.982 | -0.421 | 0.044 | 0.094 | 0.094 | 0.279 | 0.06 | 0.34 | 0.04 | 0.08 | 0.04 | 0.1 |
| 9 | 27821606 | rs17770626 | 0.353 | 0.074 | 0.008 | -0.51 | -0.006 | 0.001 | 0.001 | 0 | 0.13 | 0.13 | 0.18 | 0.09 | 0.19 | 0.15 |
| 9 | 27824832 | rs7851811 | 0.214 | -0.104 | 0.064 | -0.359 | 0.127 | 0.181 | 0.181 | 0 | 0.5 | 0.24 | 0.53 | 0.17 | 0.69 | 0.41 |
| 9 | 27836272 | rs10116939 | -0.434 | 0.391 | 1.058 | 0.068 | 0.059 | 0.02 | 0.02 | 0 | 0.06 | 0.26 | 0.02 | 0 | 0.04 | 0.04 |
| 9 | 27838705 | rs11999840 | -0.797 | 0.468 | 0.861 | 0.054 | -0.006 | -0.02 | -0.02 | 0 | 0.13 | 0.16 | 0.16 | 0.07 | 0.09 | 0.17 |
| 9 | 27839530 | rs7035013 | -0.399 | 1.05 | 0.345 | 0.459 | -0.01 | -0.004 | -0.004 | 0 | 0.51 | 0.68 | 0.44 | 0.43 | 0.48 | 0.51 |
| 9 | 27839795 | rs11789249 | 1.048 | 1.212 | 0.432 | 0.288 | 0.012 | 0.055 | 0.055 | 0 | 0.31 | 0.17 | 0.24 | 0.11 | 0.29 | 0.23 |
| 9 | 27840808 | rs12376417 | 0.643 | 1.249 | 0.787 | 0.359 | 0.019 | 0.066 | 0.066 | 0 | 0.36 | 0.13 | 0.26 | 0.13 | 0.33 | 0.29 |
| 9 | 27840948 | rs1342095 | -0.701 | 1.377 | 1.154 | 0.609 | 0.004 | 0.017 | 0.017 | 0.466 | 0.54 | 0.73 | 0.45 | 0.37 | 0.45 | 0.51 |
| 9 | 27845760 | rs7865479 | -1.181 | 1.039 | 1.308 | 0.64 | 0.007 | 0.174 | 0.174 | 0.226 | 0.08 | 0.41 | 0.13 | 0.22 | 0.02 | 0.11 |
| 9 | 27850239 | rs10812673 | 1.206 | 0.975 | 0.984 | 0.672 | -0.034 | 0.038 | 0.038 | 0 | 0.3 | 0.05 | 0.25 | 0.13 | 0.3 | 0.24 |
| 9 | 27856707 | rs947521 | 0.326 | 0.913 | 1.347 | 0.671 | -0.004 | -0.003 | -0.003 | 0 | 0.4 | 0.42 | 0.3 | 0.31 | 0.36 | 0.37 |
| 9 | 27860390 | rs4581139 | 0.273 | 0.944 | 1.133 | 0.707 | 0.007 | 0.004 | 0.004 | 0 | 0.39 | 0.32 | 0.28 | 0.31 | 0.36 | 0.37 |
| 9 | 27863830 | rs10968155 | -0.829 | 1.09 | 0.895 | 1.055 | -0.003 | 0.045 | 0.045 | 0 | 0.11 | 0.09 | 0.11 | 0.26 | 0.14 | 0.21 |
| 9 | 27865600 | rs1004158 | 0.07 | 1.28 | 1.05 | 0.719 | 0.005 | 0.023 | 0.023 | 0 | 0.52 | 0.36 | 0.41 | 0.16 | 0.45 | 0.38 |
| 9 | 27869510 | rs2383733 | 2.096 | 1.062 | 1.665 | 1.762 | 0.021 | 0.364 | 0.364 | 0.73 | 0.9 | 0.58 | 0.74 | 0.65 | 0.93 | 0.88 |
| 9 | 27872232 | rs7865942 | 1.841 | 1.505 | 2.143 | 1.694 | 0.124 | 0.074 | 0.074 | 0 | 0.88 | 0.54 | 0.8 | 0.88 | 0.84 | 0.88 |
| 9 | 27874679 | rs6476027 | -1.891 | 1.447 | 2.194 | 1.872 | 0.063 | 0.144 | 0.144 | 0 | 0.16 | 0.3 | 0.21 | 0.34 | 0.19 | 0.22 |
| 9 | 27875173 | rs2795345 | 0.547 | 1.208 | 1.868 | 1.769 | 0.032 | 0.046 | 0.046 | 0 | 0.46 | 0.4 | 0.43 | 0.21 | 0.37 | 0.45 |
| 9 | 27875247 | rs2795344 | -0.397 | 1.21 | 1.842 | 1.767 | -0.008 | 0.017 | 0.017 | 0 | 0.56 | 0.64 | 0.61 | 0.55 | 0.52 | 0.63 |
| 9 | 27878335 | rs2778430 | -1.61 | 1.298 | 1.944 | 1.702 | 0.129 | 0.078 | 0.078 | 0 | 0.11 | 0.32 | 0.2 | 0.13 | 0.17 | 0.12 |
| 9 | 27880815 | rs10481575 | -1.982 | 1.273 | 1.912 | 1.683 | 0.09 | 0.201 | 0.201 | 0 | 0.17 | 0.43 | 0.23 | 0.36 | 0.2 | 0.23 |
| 9 | 27885011 | rs2383741 | -1.996 | 1.277 | 1.793 | 1.685 | 0.066 | 0.155 | 0.155 | 0 | 0.17 | 0.53 | 0.24 | 0.36 | 0.2 | 0.22 |
| 9 | 27894033 | rs7469938 | 1.139 | 0.642 | 1.545 | 1.116 | 0.02 | 0.104 | 0.104 | 0 | 0.36 | 0.11 | 0.26 | 0.14 | 0.33 | 0.37 |
| 9 | 27894751 | rs1316921 | -1.711 | 0.799 | 2.057 | 1.252 | 0.04 | 0.114 | 0.114 | 0.024 | 0.14 | 0.33 | 0.17 | 0.24 | 0.12 | 0.2 |
| 9 | 27895649 | rs10491821 | -1.119 | 0.803 | 2.05 | 1.222 | 0.072 | -0.025 | -0.025 | 0.219 | 0.08 | 0.21 | 0.13 | 0 | 0.09 | 0.09 |
| 9 | 27899045 | rs1930023 | -1.514 | 0.786 | 1.851 | 1.258 | 0.036 | 0.104 | 0.104 | 0.156 | 0.15 | 0.38 | 0.17 | 0.23 | 0.12 | 0.2 |
| 9 | 27912332 | rs10812687 | 1.066 | 0.77 | 0.822 | 1.21 | 0.029 | -0.033 | -0.033 | 0 | 0.36 | 0.16 | 0.27 | 0.35 | 0.33 | 0.47 |
| 9 | 27914251 | rs7873614 | -1.323 | 0.781 | 1.184 | 1.161 | 0.014 | -0.036 | -0.036 | 0 | 0.13 | 0.22 | 0.16 | 0.02 | 0.12 | 0.13 |
| 9 | 27924051 | rs10968196 | -1.576 | 0.681 | 1.45 | 1.261 | 0.026 | 0.174 | 0.174 | 2.289 | 0.12 | 0.52 | 0.17 | 0.23 | 0.07 | 0.19 |
| 9 | 27926593 | rs7030798 | -0.75 | 0.662 | 1.528 | 1.342 | 0.01 | 0.087 | 0.087 | 0 | 0.09 | 0.21 | 0.14 | 0.24 | 0.07 | 0.19 |
| 9 | 27926612 | rs1930037 | 0.196 | 0.624 | 1.37 | 1.303 | -0.031 | -0.102 | -0.102 | 1.01 | 0.46 | 0.1 | 0.43 | 0.57 | 0.56 | 0.37 |
| 9 | 27927133 | rs10812695 | -0.653 | 0.312 | 1.398 | 1.293 | 0.058 | 0.09 | 0.09 | 0 | 0.55 | 0.28 | 0.57 | 0.81 | 0.63 | 0.56 |
| 9 | 27930953 | rs10491825 | 0.707 | 0.373 | 0.957 | 1.304 | 0.014 | -0.001 | -0.001 | 0 | 0.41 | 0.41 | 0.3 | 0.35 | 0.36 | 0.51 |
| 9 | 27931179 | rs10968202 | 0.721 | 0.367 | 0.991 | 1.306 | 0.038 | 0.104 | 0.104 | 0 | 0.39 | 0.19 | 0.28 | 0.14 | 0.36 | 0.41 |
| 9 | 27938780 | rs1930042 | 1.092 | 0.299 | 0.269 | 1.319 | 0.035 | -0.025 | -0.025 | 3.225 | 0.36 | 0.15 | 0.27 | 0.33 | 0.32 | 0.47 |
| 9 | 27945758 | rs9644872 | -0.071 | 0.615 | -0.024 | 1.249 | -0.043 | -0.101 | -0.101 | 0 | 0.44 | 0.08 | 0.5 | 0.59 | 0.57 | 0.52 |
| 9 | 27950863 | rs10757699 | -0.354 | 0.554 | 0.344 | 0.976 | 0.08 | 0.122 | 0.122 | 0.109 | 0.5 | 0.25 | 0.54 | 0.81 | 0.64 | 0.58 |
| 9 | 27973766 | rs10812712 | -1.015 | 1.048 | 1.419 | 1.141 | 0.003 | 0 | 0 | 4.608 | 0.34 | 0.29 | 0.34 | 0.32 | 0.39 | 0.29 |
| 9 | 27974789 | rs10968243 | -0.13 | 1.471 | 1.227 | 1.289 | -0.044 | -0.041 | -0.041 | 0 | 0.07 | 0.01 | 0.1 | 0.15 | 0.17 | 0.07 |
| 9 | 27977762 | rs10968246 | -1.154 | 1.333 | 1.369 | 1.184 | 0.006 | 0.006 | 0.006 | 0 | 0.36 | 0.3 | 0.36 | 0.51 | 0.42 | 0.43 |
| 9 | 27979665 | rs4142124 | -0.925 | 1.347 | 1.22 | 0.386 | -0.072 | 0.01 | 0.01 | 0 | 0.33 | 0.07 | 0.23 | 0.27 | 0.4 | 0.35 |
| 9 | 27992290 | rs10968258 | -1.458 | 1.121 | 1.577 | 0.805 | 0.098 | 0.129 | 0.129 | 0 | 0.13 | 0.4 | 0.24 | 0.19 | 0.17 | 0.1 |
| 9 | 27997217 | rs1930014 | -0.776 | 1.282 | 1.338 | 0.853 | 0.046 | 0.062 | 0.062 | 0 | 0.1 | 0.14 | 0.23 | 0.19 | 0.16 | 0.1 |
| 9 | 28001944 | rs1197944 | 0.206 | 0.78 | 0.744 | -0.123 | -0.006 | 0.126 | 0.126 | 0 | 0.47 | 0.71 | 0.57 | 0.83 | 0.51 | 0.48 |
| 9 | 28002851 | rs1197920 | 0.302 | 0.862 | 0.198 | -0.073 | -0.002 | 0.011 | 0.011 | 0 | 0.46 | 0.53 | 0.55 | 0.82 | 0.51 | 0.47 |
| 9 | 28004967 | rs10812720 | 0.745 | 0.613 | 0.38 | -0.05 | -0.016 | -0.04 | -0.04 | 0 | 0.67 | 0.55 | 0.82 | 0.84 | 0.75 | 0.71 |
| 9 | 28007069 | rs7028286 | 0.505 | 0.655 | 0.682 | -0.116 | -0.054 | -0.05 | -0.05 | 2.825 | 0.71 | 0.54 | 0.88 | 0.86 | 0.85 | 0.75 |
| 9 | 28011383 | rs4880001 | 0.815 | 1.215 | 0.161 | -0.758 | -0.037 | 0.049 | 0.049 | 0 | 0.19 | 0.09 | 0.31 | 0.03 | 0.35 | 0.19 |
| 9 | 28013298 | rs10968281 | 0.027 | 0.63 | 0.925 | -0.384 | 0.001 | 0.003 | 0.003 | 0 | 0.14 | 0.11 | 0.08 | 0.05 | 0.13 | 0.08 |
| 9 | 28020746 | rs1343453 | 0.753 | 0.309 | 1.084 | 0.568 | -0.027 | -0.047 | -0.047 | 0.705 | 0.75 | 0.56 | 0.83 | 0.84 | 0.84 | 0.7 |
| 9 | 28022473 | rs1197913 | -1.449 | 0.304 | 1.484 | 0.563 | -0.019 | 0.046 | 0.046 | 0 | 0.33 | 0.51 | 0.18 | 0.45 | 0.26 | 0.4 |
| 9 | 28023564 | rs1197918 | -1.352 | 0.25 | 1.705 | 0.561 | -0.013 | 0.036 | 0.036 | 0 | 0.32 | 0.44 | 0.17 | 0.45 | 0.26 | 0.4 |
| 9 | 28024647 | rs869994 | 1.75 | 0.697 | 1.581 | 0.884 | 0 | 0 | 0 | 0 | 0.59 | 0.4 | 0.72 | 0.42 | 0.64 | 0.49 |
| 9 | 28028549 | rs12684725 | -1.026 | 0.005 | 1.312 | 0.922 | 0.039 | 0.03 | 0.03 | 0 | 0.29 | 0.19 | 0.09 | 0.16 | 0.16 | 0.26 |
| 9 | 28034028 | rs10968293 | -0.886 | -0.083 | 1.614 | 1.107 | -0.028 | 0.076 | 0.076 | 0 | 0.2 | 0.32 | 0.12 | 0.44 | 0.1 | 0.45 |
| 9 | 28035209 | rs1343457 | 1.763 | -0.116 | 1.622 | 0.864 | 0.026 | -0.011 | -0.011 | 0 | 0.6 | 0.3 | 0.75 | 0.47 | 0.73 | 0.43 |
| 9 | 28035596 | rs10968295 | -0.057 | 0.033 | 1.211 | 0.813 | -0.01 | -0.003 | -0.003 | 0.167 | 0.15 | 0.21 | 0.06 | 0.13 | 0.08 | 0.29 |
| 9 | 28036634 | rs7875743 | 1.091 | 0.523 | 1.179 | 0.972 | 0.07 | -0.007 | -0.007 | 0.124 | 0.92 | 0.78 | 0.87 | 0.93 | 0.87 | 0.89 |
| 9 | 28036991 | rs1930018 | 0.503 | 0.396 | 1.191 | 1.091 | 0.002 | 0.101 | 0.101 | 0 | 0.81 | 0.65 | 0.87 | 0.65 | 0.84 | 0.8 |
| 9 | 28037082 | rs944646 | 0.756 | 0.36 | 1.043 | 1.026 | -0.008 | 0.077 | 0.077 | 0 | 0.74 | 0.59 | 0.78 | 0.6 | 0.8 | 0.69 |
| 9 | 28037611 | rs1452352 | 0.74 | 0.394 | 0.636 | 0.998 | -0.008 | 0.096 | 0.096 | 0 | 0.74 | 0.55 | 0.78 | 0.59 | 0.79 | 0.69 |
| 9 | 28040937 | rs1452353 | 0.848 | 0.479 | -0.247 | 0.731 | -0.009 | -0.006 | -0.006 | 0 | 0.07 | 0.02 | 0.08 | 0.08 | 0.07 | 0.02 |
| 9 | 28041638 | rs1198002 | -0.415 | -0.045 | 0.19 | 0.899 | -0.011 | 0.002 | 0.002 | 0 | 0.86 | 0.86 | 0.95 | 0.86 | 0.96 | 0.95 |
| 9 | 28043199 | rs16912396 | 0.979 | -0.021 | 0.13 | 0.97 | -0.005 | -0.013 | -0.013 | 0.015 | 0.07 | 0.02 | 0.07 | 0.11 | 0.07 | 0.05 |
| 9 | 28043247 | rs1198006 | -0.446 | -0.094 | 0.169 | 0.974 | 0.001 | -0.002 | -0.002 | 0 | 0.87 | 0.91 | 0.95 | 0.86 | 0.96 | 0.95 |
| 9 | 28044739 | rs1198009 | 0.855 | 0.313 | -0.604 | 1.027 | -0.022 | -0.069 | -0.069 | 0.654 | 0.18 | 0.06 | 0.18 | 0.43 | 0.28 | 0.11 |

| | | | | | | | | | | | | | | | | |
|---|---|---|---|---|---|---|---|---|---|---|---|---|---|---|---|---|
| 9 | 28045814 | rs12115635 | 2.204 | -0.1 | -1.271 | -0.019 | 0.009 | 0.105 | 0.105 | 0 | 0.18 | 0.03 | 0.21 | 0.01 | 0.14 | 0.11 |
| 9 | 28045924 | rs17511658 | 0.881 | 0.909 | -1.313 | 0.198 | -0.054 | -0.006 | -0.006 | 0.019 | 0.12 | 0.04 | 0.22 | 0.11 | 0.21 | 0.12 |
| 9 | 28049518 | rs1343456 | 0.552 | 1.001 | -1.048 | 0.171 | -0.018 | 0.003 | 0.003 | 0 | 0.13 | 0.07 | 0.26 | 0.12 | 0.24 | 0.13 |
| 9 | 28050777 | rs1147820 | 0.748 | 0.742 | -0.672 | 0.031 | 0.002 | -0.002 | -0.002 | 0 | 0.67 | 0.61 | 0.61 | 0.7 | 0.62 | 0.76 |
| 9 | 28053556 | rs10812732 | -0.112 | 0.204 | 0.044 | 0.208 | -0.049 | 0.015 | 0.015 | 0.403 | 0.21 | 0.61 | 0.15 | 0.25 | 0.08 | 0.2 |
| 9 | 28053878 | rs6476034 | -1.653 | 0.637 | -0.412 | 0.282 | -0.006 | -0.046 | -0.046 | 0 | 0.84 | 0.93 | 0.82 | 0.51 | 0.75 | 0.89 |
| 9 | 28055616 | rs10968314 | 0.652 | 0.363 | -0.936 | -0.277 | -0.042 | 0.012 | 0.012 | 0.019 | 0.38 | 0.1 | 0.46 | 0.31 | 0.4 | 0.39 |
| 9 | 28055877 | rs12553579 | 1.017 | 0.593 | -0.902 | -0.331 | -0.054 | -0.028 | -0.028 | 0 | 0.08 | 0.01 | 0.2 | 0.1 | 0.19 | 0.13 |
| 9 | 28055900 | rs1901565 | -0.483 | -0.082 | -0.945 | -0.787 | -0.032 | -0.017 | -0.017 | 1.09 | 0.33 | 0.23 | 0.56 | 0.4 | 0.41 | 0.51 |
| 9 | 28056067 | rs1901566 | -1.75 | -0.321 | -0.642 | -0.782 | -0.016 | 0.033 | 0.033 | 0 | 0.93 | 0.83 | 0.98 | 0.88 | 0.98 | 0.97 |
| 9 | 28056106 | rs10968316 | 2.36 | -0.343 | -1.03 | -0.964 | 0.013 | 0.117 | 0.117 | 0 | 0.2 | 0.01 | 0.09 | 0 | 0.13 | 0.05 |
| 9 | 28058180 | rs2167643 | 0.809 | -0.375 | -0.678 | -0.922 | 0.007 | 0.017 | 0.017 | 0 | 0.38 | 0.42 | 0.3 | 0.46 | 0.42 | 0.32 |
| 9 | 28058435 | rs2123204 | -1.117 | -0.411 | -0.531 | -0.68 | -0.009 | -0.024 | -0.024 | 6.628 | 0.71 | 0.63 | 0.81 | 0.85 | 0.76 | 0.82 |
| 9 | 28064690 | rs10812736 | 2.163 | -0.222 | -0.689 | -0.658 | -0.011 | 0.093 | 0.093 | 0.006 | 0.19 | 0.05 | 0.13 | 0.02 | 0.19 | 0.06 |
| 9 | 28064791 | rs2123205 | 1.534 | -0.054 | -0.243 | -0.39 | 0.008 | 0.076 | 0.076 | 0 | 0.17 | 0.21 | 0.18 | 0.43 | 0.24 | 0.09 |
| 9 | 28068081 | rs7853840 | 1.323 | -0.14 | -0.266 | -1.011 | -0.001 | -0.026 | -0.026 | 0 | 0.09 | 0.13 | 0.06 | 0.02 | 0.07 | 0.08 |
| 9 | 28069562 | rs7858580 | -0.834 | -0.299 | -0.215 | -1.045 | -0.025 | -0.063 | -0.063 | 0 | 0.89 | 0.77 | 0.8 | 0.99 | 0.92 | 0.92 |
| 9 | 28070283 | rs7042653 | -0.507 | -0.498 | -0.378 | -1.343 | -0.02 | -0.074 | -0.074 | 0 | 0.66 | 0.4 | 0.64 | 0.95 | 0.72 | 0.77 |
| 9 | 28072851 | rs10968329 | 2.291 | -0.402 | -0.613 | -1.266 | -0.03 | 0.055 | 0.055 | 0 | 0.18 | 0.03 | 0.13 | 0.03 | 0.19 | 0.07 |
| 9 | 28074530 | rs7857347 | -2.178 | -0.397 | 0.007 | -1.176 | 0.048 | 0.016 | 0.016 | 1.796 | 0.95 | 0.73 | 0.95 | 1 | 0.95 | 1 |
| 9 | 28075173 | rs7033571 | 2.006 | -0.182 | -0.237 | -1.337 | 0.011 | -0.019 | -0.019 | 0 | 0.12 | 0.17 | 0.06 | 0.03 | 0.12 | 0.13 |
| 9 | 28082935 | rs16912472 | 0.671 | -0.01 | -0.898 | -1.029 | 0 | -0.001 | -0.001 | 0 | 0.38 | 0.39 | 0.36 | 0.66 | 0.35 | 0.51 |
| 9 | 28083672 | rs1839967 | 1.095 | 0.166 | -0.968 | -1.183 | -0.024 | -0.026 | -0.026 | 0 | 0.18 | 0.09 | 0.23 | 0.27 | 0.23 | 0.19 |
| 9 | 28086281 | rs896585 | 0.259 | 0.295 | -1.001 | -1.109 | -0.004 | 0.019 | 0.019 | 0 | 0.43 | 0.33 | 0.43 | 0.69 | 0.47 | 0.57 |
| 9 | 28086344 | rs2167644 | 2.497 | 0.018 | -1.262 | -0.846 | -0.006 | 0.007 | 0.007 | 0 | 0.14 | 0.09 | 0.18 | 0.43 | 0.21 | 0.12 |
| 9 | 28086683 | rs10968337 | 1.769 | 0.175 | -1.131 | -0.932 | -0.009 | 0.045 | 0.045 | 0.025 | 0.09 | 0.18 | 0.06 | 0.2 | 0.06 | 0.03 |
| 9 | 28088526 | rs4305993 | 2.429 | 0.314 | -1.242 | -0.902 | 0.018 | -0.013 | -0.013 | 0 | 0.1 | 0.19 | 0.06 | 0.03 | 0.1 | 0.1 |
| 9 | 28093874 | rs7847672 | -0.732 | 0.177 | -1.628 | -0.664 | -0.005 | 0.01 | 0.01 | 0 | 0.64 | 0.58 | 0.64 | 0.36 | 0.65 | 0.54 |
| 9 | 28104987 | rs10968347 | 1.879 | -0.019 | -1.953 | -0.718 | 0.003 | 0.098 | 0.098 | 0 | 0.16 | 0.01 | 0.07 | 0 | 0.12 | 0.03 |
| 9 | 28106898 | rs10812741 | -0.164 | 0.377 | -1.906 | -0.331 | -0.052 | -0.015 | -0.015 | 2.073 | 0.29 | 0.1 | 0.42 | 0.32 | 0.33 | 0.52 |
| 9 | 28111873 | rs17771324 | -0.281 | 0.299 | -1.755 | -0.227 | -0.013 | -0.027 | -0.027 | 0 | 0.18 | 0.11 | 0.22 | 0.27 | 0.17 | 0.19 |
| 9 | 28119512 | rs7034231 | -1.402 | 0.324 | -1.596 | -0.241 | 0.001 | 0.008 | 0.008 | 0 | 0.84 | 0.82 | 0.78 | 0.96 | 0.83 | 0.92 |
| 9 | 28119660 | rs17698657 | 2.252 | 0.299 | -1.654 | -0.22 | 0.009 | 0.025 | 0.025 | 0 | 0.12 | 0.02 | 0.07 | 0.02 | 0.1 | 0.02 |
| 9 | 28120384 | rs1452333 | 0.824 | 0.231 | -1.053 | -0.497 | 0.002 | -0.003 | -0.003 | 0 | 0.27 | 0.24 | 0.15 | 0.08 | 0.19 | 0.22 |
| 9 | 28123402 | rs1452335 | -0.381 | 0.274 | -0.721 | -0.645 | -0.009 | -0.019 | -0.019 | 0.404 | 0.61 | 0.57 | 0.64 | 0.73 | 0.68 | 0.69 |
| 9 | 28127342 | rs1452338 | 0.58 | 0.322 | -0.684 | -0.636 | -0.082 | -0.027 | -0.027 | 0.062 | 0.33 | 0.14 | 0.47 | 0.65 | 0.5 | 0.48 |
| 9 | 28129815 | rs6476037 | 0.183 | 0.226 | -0.72 | -0.643 | 0.004 | 0.007 | 0.007 | 0 | 0.39 | 0.33 | 0.35 | 0.26 | 0.32 | 0.31 |
| 9 | 28136460 | rs1343452 | -0.029 | 0.239 | -1.266 | -0.623 | 0.011 | 0.011 | 0.011 | 0 | 0.4 | 0.33 | 0.23 | 0.28 | 0.27 | 0.3 |
| 9 | 28140215 | rs12552434 | 1.778 | 0.945 | -1.844 | -0.17 | -0.039 | -0.046 | -0.046 | 0 | 0.16 | 0.08 | 0.26 | 0.4 | 0.31 | 0.1 |
| 9 | 28141923 | rs4573372 | 0.386 | 0.748 | -1.709 | -0.525 | -0.049 | -0.011 | -0.011 | 0 | 0.35 | 0.2 | 0.6 | 0.64 | 0.52 | 0.51 |
| 9 | 28143059 | rs12684801 | 0.953 | 0.6 | -1.708 | -0.487 | 0.011 | -0.006 | -0.006 | 0.268 | 0.12 | 0.06 | 0.08 | 0.16 | 0.07 | 0.08 |
| 9 | 28145962 | rs2198292 | 0.696 | 0.759 | -1.601 | -0.491 | 0.004 | 0.014 | 0.014 | 0 | 0.28 | 0.26 | 0.18 | 0.11 | 0.23 | 0.24 |
| 9 | 28152658 | rs10812745 | 1.106 | 1.288 | -1.74 | 0.162 | 0.007 | -0.019 | -0.019 | 0.676 | 0.08 | 0.02 | 0.05 | 0.19 | 0.04 | 0.06 |
| 9 | 28158881 | rs10812748 | 0.107 | 1.67 | -1.648 | 0.203 | -0.036 | -0.019 | -0.019 | 0 | 0.32 | 0.21 | 0.45 | 0.63 | 0.45 | 0.45 |
| 9 | 28164645 | rs2044961 | -0.318 | 1.023 | -1.586 | 0.162 | -0.017 | -0.002 | -0.002 | 0 | 0.39 | 0.23 | 0.61 | 0.64 | 0.56 | 0.52 |
| 9 | 28173138 | rs2044962 | 0.073 | 1.032 | -1.465 | 0.505 | -0.034 | -0.059 | -0.059 | 0 | 0.31 | 0.16 | 0.37 | 0.49 | 0.4 | 0.39 |
| 9 | 28176868 | rs1452348 | 0.429 | 0.75 | -1.345 | 0.562 | -0.013 | -0.024 | -0.024 | 0 | 0.1 | 0.03 | 0.15 | 0.18 | 0.1 | 0.1 |
| 9 | 28181085 | rs1319535 | -0.178 | 0.318 | -1.048 | 0.347 | 0.013 | -0.006 | -0.006 | 0 | 0.55 | 0.61 | 0.68 | 0.52 | 0.61 | 0.39 |
| 9 | 28185536 | rs6476041 | 0.771 | -0.114 | -0.833 | 0.145 | -0.009 | -0.011 | -0.011 | 0.516 | 0.07 | 0.08 | 0.02 | 0.01 | 0.04 | 0.05 |
| 9 | 28186091 | rs7040482 | -0.788 | -0.225 | -0.783 | 0.09 | 0 | 0.001 | 0.001 | 0 | 0.71 | 0.74 | 0.77 | 0.75 | 0.73 | 0.65 |
| 9 | 28187972 | rs7045929 | -0.702 | -0.279 | -0.351 | 0.063 | -0.003 | -0.01 | -0.01 | 0.002 | 0.7 | 0.65 | 0.76 | 0.75 | 0.73 | 0.65 |
| 9 | 28188444 | rs4878817 | -0.261 | -0.341 | -0.337 | -0.038 | 0.016 | 0.028 | 0.028 | 0 | 0.5 | 0.44 | 0.66 | 0.69 | 0.59 | 0.53 |
| 9 | 28194546 | rs16912561 | 0.952 | -0.369 | -0.329 | -0.039 | 0.008 | 0.001 | 0.001 | 0 | 0.13 | 0.2 | 0.15 | 0.12 | 0.11 | 0.16 |
| 9 | 28195206 | rs10968379 | 1.028 | -0.699 | -0.212 | -0.192 | -0.003 | -0.006 | -0.006 | 0 | 0.19 | 0.23 | 0.12 | 0.14 | 0.18 | 0.29 |
| 9 | 28198828 | rs10968384 | 1.482 | 0.148 | -0.67 | -0.047 | -0.077 | -0.061 | -0.061 | 0.07 | 0.14 | 0.02 | 0.17 | 0.2 | 0.25 | 0.2 |
| 9 | 28199845 | rs17771757 | 1.173 | 0.371 | -0.722 | -0.02 | -0.009 | 0 | 0 | 0 | 0.05 | 0.03 | 0.14 | 0.04 | 0.08 | 0.15 |
| 9 | 28202675 | rs10812754 | 1.055 | 0.13 | -0.853 | 0.153 | -0.094 | -0.113 | -0.113 | 0 | 0.26 | 0.08 | 0.36 | 0.46 | 0.39 | 0.25 |
| 9 | 28205162 | rs10511824 | 1.663 | -0.185 | -0.995 | -0.055 | -0.005 | -0.046 | -0.046 | 2.322 | 0.11 | 0.01 | 0.12 | 0.21 | 0.09 | 0.09 |
| 9 | 28205326 | rs10968386 | 1.291 | -0.114 | -0.835 | -0.045 | -0.002 | -0.004 | -0.004 | 0 | 0.22 | 0.25 | 0.13 | 0.14 | 0.19 | 0.29 |
| 9 | 28207637 | rs12682967 | 1.016 | -0.157 | -0.779 | -0.045 | 0.009 | -0.004 | -0.004 | 0 | 0.18 | 0.12 | 0.14 | 0.22 | 0.14 | 0.14 |
| 9 | 28208463 | rs1562972 | -0.299 | -0.146 | -0.723 | 0.031 | -0.015 | 0.052 | 0.052 | 0 | 0.47 | 0.25 | 0.52 | 0.65 | 0.53 | 0.39 |
| 9 | 28209655 | rs7861219 | 2.233 | -0.202 | -0.602 | 0.141 | -0.002 | 0.001 | 0.001 | 0 | 0.13 | 0.11 | 0.2 | 0.27 | 0.14 | 0.05 |
| 9 | 28210699 | rs10968388 | 0.501 | -0.573 | -0.419 | -0.161 | 0.024 | 0.006 | 0.006 | 0 | 0.32 | 0.21 | 0.21 | 0.28 | 0.24 | 0.16 |
| 9 | 28215694 | rs4604546 | -1.369 | -0.219 | -0.008 | -0.343 | 0.005 | 0.007 | 0.007 | 0 | 0.87 | 0.74 | 0.86 | 0.88 | 0.9 | 0.85 |
| 9 | 28215883 | rs13296178 | 1.936 | -0.223 | -0.109 | -0.338 | -0.013 | -0.014 | -0.014 | 0 | 0.14 | 0.18 | 0.07 | 0.06 | 0.11 | 0.02 |
| 9 | 28216153 | rs10968391 | 0.634 | 0.025 | -0.665 | -0.192 | 0.015 | 0.027 | 0.027 | 0.896 | 0.22 | 0.12 | 0.11 | 0.14 | 0.19 | 0.28 |
| 9 | 28218512 | rs4574938 | -1.01 | 0.161 | -0.601 | 0.017 | -0.013 | -0.013 | -0.013 | 0.026 | 0.74 | 0.63 | 0.79 | 0.82 | 0.79 | 0.83 |
| 9 | 28221514 | rs10511827 | -0.014 | 0.55 | -0.947 | 0.175 | -0.001 | -0.004 | -0.004 | 3.082 | 0.25 | 0.25 | 0.25 | 0.15 | 0.26 | 0.34 |
| 9 | 28222606 | rs10511825 | -0.142 | 0.556 | -1.32 | 0.868 | -0.045 | -0.121 | -0.121 | 0 | 0.25 | 0.06 | 0.34 | 0.5 | 0.26 | 0.18 |

| | | | | | | | | | | | | | | | | |
|---|---|---|---|---|---|---|---|---|---|---|---|---|---|---|---|---|
| 9 | 28222934 | rs4399024 | -0.661 | 0.529 | -0.74 | 0.804 | 0.001 | -0.003 | -0.003 | 0.086 | 0.4 | 0.59 | 0.44 | 0.63 | 0.41 | 0.3 |
| 9 | 28227167 | rs7357773 | 0.007 | 0.353 | -0.506 | 0.879 | -0.006 | 0.023 | 0.023 | 0 | 0.2 | 0.3 | 0.19 | 0.28 | 0.19 | 0.07 |
| 9 | 28231047 | rs7865194 | -0.001 | 0.352 | -0.507 | 0.861 | -0.006 | 0.02 | 0.02 | 0 | 0.2 | 0.3 | 0.19 | 0.28 | 0.19 | 0.07 |
| 9 | 28236666 | rs7857732 | -0.107 | 0.46 | -0.816 | 0.625 | 0 | -0.001 | -0.001 | 0 | 0.2 | 0.21 | 0.19 | 0.13 | 0.17 | 0.1 |
| 9 | 28237928 | rs10968404 | -0.09 | 0.885 | -1.455 | 1.155 | -0.035 | -0.081 | -0.081 | 0.197 | 0.22 | 0.07 | 0.21 | 0.34 | 0.23 | 0.19 |
| 9 | 28250940 | rs7044982 | -0.184 | 1.738 | -1.074 | 1.176 | -0.02 | -0.016 | -0.016 | 0.668 | 0.16 | 0.12 | 0.25 | 0.25 | 0.26 | 0.21 |
| 9 | 28251718 | rs13440274 | -0.193 | 1.652 | -0.983 | 1.133 | -0.012 | -0.01 | -0.01 | 0.528 | 0.14 | 0.12 | 0.23 | 0.24 | 0.22 | 0.19 |
| 9 | 28252086 | rs7038123 | -0.608 | 0.893 | -0.772 | 1.199 | 0.005 | 0.03 | 0.03 | 0 | 0.24 | 0.29 | 0.25 | 0.41 | 0.28 | 0.24 |
| 9 | 28254742 | rs4279684 | -0.062 | -0.096 | -0.833 | 0.813 | 0.023 | -0.014 | -0.014 | 0 | 0.54 | 0.29 | 0.35 | 0.46 | 0.37 | 0.41 |
| 9 | 28256245 | rs4554572 | 0.231 | -0.225 | -1 | 0.707 | -0.093 | 0.072 | 0.072 | 0 | 0.32 | 0.14 | 0.34 | 0.2 | 0.52 | 0.26 |
| 9 | 28257148 | rs4559331 | 0.604 | -1.094 | -1.073 | 0.602 | 0.019 | 0.016 | 0.016 | 0 | 0.23 | 0.13 | 0.24 | 0.18 | 0.17 | 0.28 |
| 9 | 28263705 | rs10812768 | 0.946 | -1.416 | -0.382 | 0.723 | 0.006 | -0.045 | -0.045 | 0 | 0.24 | 0.11 | 0.19 | 0.26 | 0.19 | 0.25 |
| 9 | 28272794 | rs13298436 | 0.026 | -0.527 | 0.366 | 0.706 | 0.029 | 0.131 | 0.131 | 0.002 | 0.23 | 0.43 | 0.29 | 0.4 | 0.31 | 0.17 |
| 9 | 28273920 | rs10968427 | 1.814 | -0.533 | 0.173 | 0.621 | -0.033 | 0.029 | 0.029 | 0 | 0.2 | 0.11 | 0.19 | 0.16 | 0.27 | 0.2 |
| 9 | 28276698 | rs13301122 | 1.4 | -0.388 | 0.193 | 0.675 | -0.054 | 0.046 | 0.046 | 0 | 0.08 | 0.01 | 0.14 | 0 | 0.23 | 0.05 |
| 9 | 28280654 | rs7871547 | -0.131 | -0.445 | 0.293 | 0.68 | 0.001 | 0.003 | 0.003 | 0 | 0.46 | 0.58 | 0.48 | 0.56 | 0.59 | 0.37 |
| 9 | 28294231 | rs10812774 | 0.034 | -0.51 | 0.448 | 0.718 | 0.001 | 0.002 | 0.002 | 0 | 0.59 | 0.47 | 0.44 | 0.45 | 0.55 | 0.5 |
| 9 | 28299438 | rs7855717 | 1.441 | -0.61 | 0.091 | 0.706 | -0.042 | 0.038 | 0.038 | 0 | 0.23 | 0.1 | 0.2 | 0.17 | 0.28 | 0.2 |
| 9 | 28301145 | rs10812775 | -0.637 | -0.923 | 0.325 | 0.718 | 0.008 | 0.003 | 0.003 | 0 | 0.71 | 0.77 | 0.64 | 0.69 | 0.78 | 0.62 |
| 9 | 28305521 | rs1953003 | 0.546 | -0.971 | -0.267 | 0.711 | 0.02 | -0.021 | -0.021 | 0.066 | 0.17 | 0.12 | 0.08 | 0.22 | 0.1 | 0.13 |
| 9 | 28309788 | rs13299028 | 0.19 | -0.719 | -0.512 | 0.669 | -0.036 | 0 | 0 | 0 | 0.33 | 0.15 | 0.37 | 0.34 | 0.4 | 0.32 |
| 9 | 28321635 | rs7860817 | 0.678 | -0.691 | -0.666 | 0.276 | -0.068 | 0.113 | 0.113 | 0 | 0.4 | 0.16 | 0.32 | 0.22 | 0.45 | 0.3 |
| 9 | 28327563 | rs6476056 | -0.261 | -0.76 | -0.546 | 0.304 | 0.001 | -0.004 | -0.004 | 0 | 0.48 | 0.63 | 0.48 | 0.44 | 0.54 | 0.41 |
| 9 | 28328038 | rs1332286 | -0.444 | -1.008 | -0.35 | 0.344 | 0.005 | -0.005 | -0.005 | 0 | 0.57 | 0.38 | 0.63 | 0.48 | 0.63 | 0.55 |
| 9 | 28329088 | rs1999423 | 0.451 | -0.908 | -0.568 | 0.435 | 0.019 | -0.004 | -0.004 | 0 | 0.29 | 0.19 | 0.34 | 0.31 | 0.2 | 0.33 |
| 9 | 28334137 | rs7847504 | -0.031 | -0.814 | -0.864 | -0.088 | 0.026 | 0.028 | 0.028 | 0 | 0.44 | 0.38 | 0.41 | 0.28 | 0.31 | 0.37 |
| 9 | 28336185 | rs13291498 | 0.815 | -0.454 | -1.147 | 0.209 | -0.02 | 0.011 | 0.011 | 0 | 0.1 | 0.06 | 0.15 | 0 | 0.23 | 0.05 |
| 9 | 28337698 | rs13298550 | 1.006 | -0.434 | -1.274 | 0.228 | -0.046 | 0.044 | 0.044 | 0 | 0.09 | 0.03 | 0.15 | 0 | 0.23 | 0.05 |
| 9 | 28338911 | rs9792406 | 0.864 | -0.764 | -1.337 | 0.289 | 0.029 | -0.024 | -0.024 | 0 | 0.13 | 0.05 | 0.06 | 0.18 | 0.07 | 0.1 |
| 9 | 28339748 | rs10491897 | 0.654 | -0.273 | -1.244 | 0.489 | -0.062 | -0.017 | -0.017 | 0.606 | 0.15 | 0.05 | 0.25 | 0.16 | 0.27 | 0.15 |
| 9 | 28340158 | rs1360507 | -0.249 | -0.604 | -1.021 | 0.174 | 0.024 | 0.04 | 0.04 | 0 | 0.66 | 0.53 | 0.59 | 0.51 | 0.56 | 0.59 |
| 9 | 28340971 | rs10968521 | -0.283 | -1.088 | -0.567 | 0.277 | -0.067 | 0.015 | 0.015 | 0 | 0.27 | 0.34 | 0.27 | 0.3 | 0.12 | 0.25 |
| 9 | 28341892 | rs10968522 | 0.26 | 0.034 | -0.976 | -0.144 | -0.017 | 0.049 | 0.049 | 0 | 0.2 | 0.08 | 0.18 | 0.01 | 0.2 | 0.09 |
| 9 | 28342014 | rs12347591 | -0.813 | -0.165 | -0.284 | 0.847 | -0.036 | 0.104 | 0.104 | 0 | 0.19 | 0.34 | 0.31 | 0.53 | 0.13 | 0.26 |
| 9 | 28342179 | rs1331885 | -1.027 | 0.721 | -0.712 | 0.699 | -0.02 | -0.027 | -0.027 | 0 | 0.25 | 0.23 | 0.42 | 0.4 | 0.4 | 0.28 |
| 9 | 28355201 | rs10491895 | -0.156 | 0.515 | -0.737 | 0.288 | -0.03 | 0.052 | 0.052 | 0 | 0.09 | 0.01 | 0.17 | 0 | 0.14 | 0.05 |
| 9 | 28358436 | rs7032657 | -0.53 | 0.133 | -0.429 | 0.108 | 0 | -0.003 | -0.003 | 0 | 0.57 | 0.54 | 0.56 | 0.54 | 0.52 | 0.62 |
| 9 | 28359979 | rs3842979 | -0.096 | 0.131 | -0.019 | 0.079 | 0.009 | 0.011 | 0.011 | 0 | 0.52 | 0.61 | 0.42 | 0.63 | 0.42 | 0.59 |
| 9 | 28360213 | rs3843937 | -0.151 | 0.173 | -0.008 | 0.091 | 0.005 | 0.007 | 0.007 | 0 | 0.53 | 0.6 | 0.43 | 0.63 | 0.43 | 0.59 |
| 9 | 28360711 | rs3849956 | 1.509 | 0.199 | -0.351 | 0.126 | -0.108 | -0.004 | -0.004 | 0 | 0.27 | 0.07 | 0.34 | 0.25 | 0.39 | 0.32 |
| 9 | 28365308 | rs4879240 | 0.775 | 0.106 | -0.271 | 0.216 | -0.108 | 0.002 | 0.002 | 0 | 0.33 | 0.1 | 0.37 | 0.26 | 0.4 | 0.33 |
| 9 | 28366398 | rs4278232 | 0.687 | -0.513 | 0.212 | 0.471 | 0.008 | -0.008 | -0.008 | 0.297 | 0.33 | 0.32 | 0.3 | 0.36 | 0.3 | 0.47 |
| 9 | 28366643 | rs7856324 | 1.134 | -0.616 | 0.51 | 0.476 | -0.011 | 0.018 | 0.018 | 0 | 0.12 | 0.17 | 0.06 | 0.17 | 0.11 | 0.1 |
| 9 | 28367812 | rs1331891 | 0.423 | -0.608 | 0.385 | 0.459 | 0.017 | -0.016 | -0.016 | 0.281 | 0.38 | 0.33 | 0.33 | 0.45 | 0.33 | 0.49 |
| 9 | 28368404 | rs6476060 | -0.279 | -0.605 | 0.67 | 0.439 | -0.012 | 0.051 | 0.051 | 0 | 0.2 | 0.32 | 0.25 | 0.27 | 0.21 | 0.18 |
| 9 | 28370761 | rs16912763 | 1.07 | -0.633 | 0.498 | 0.422 | -0.001 | -0.002 | -0.002 | 1.105 | 0.08 | 0.08 | 0.18 | 0.11 | 0.1 | 0.08 |
| 9 | 28371227 | rs12684059 | 0.269 | -0.655 | 0.511 | 0.393 | 0.029 | 0.041 | 0.041 | 0 | 0.48 | 0.29 | 0.58 | 0.37 | 0.56 | 0.41 |
| 9 | 28372519 | rs2383769 | 0.112 | -0.669 | 0.602 | 0.412 | -0.038 | 0.054 | 0.054 | 0 | 0.81 | 0.55 | 0.87 | 0.73 | 0.86 | 0.89 |
| 9 | 28376487 | rs10491894 | 0.885 | -0.655 | 0.514 | 0.384 | -0.052 | 0.047 | 0.047 | 0 | 0.39 | 0.21 | 0.4 | 0.27 | 0.45 | 0.33 |
| 9 | 28377370 | rs1412423 | 0.172 | -0.714 | 0.608 | 0.414 | -0.039 | 0.055 | 0.055 | 0 | 0.81 | 0.56 | 0.87 | 0.74 | 0.86 | 0.88 |
| 9 | 28378229 | rs10757736 | 0.206 | -0.575 | 0.313 | 0.329 | 0.029 | 0.039 | 0.039 | 0 | 0.48 | 0.29 | 0.58 | 0.37 | 0.55 | 0.41 |
| 9 | 28380814 | rs2104859 | 0.71 | -1.051 | 0.266 | -0.078 | -0.01 | 0.04 | 0.04 | 0 | 0.29 | 0.22 | 0.21 | 0.17 | 0.28 | 0.25 |
| 9 | 28381681 | rs1571846 | 1.062 | -1.085 | 0.244 | -0.101 | -0.019 | 0.071 | 0.071 | 0 | 0.28 | 0.16 | 0.2 | 0.07 | 0.27 | 0.21 |
| 9 | 28381857 | rs1331879 | 1.618 | -1.093 | -0.004 | -0.074 | -0.072 | 0.011 | 0.011 | 0 | 0.33 | 0.13 | 0.36 | 0.26 | 0.39 | 0.33 |
| 9 | 28382153 | rs944594 | -0.951 | -1.169 | 0.457 | 0.398 | -0.012 | 0.031 | 0.031 | 0 | 0.86 | 0.82 | 0.91 | 0.82 | 0.88 | 0.89 |
| 9 | 28382826 | rs1331880 | 0.878 | -1.174 | 0.568 | 0.289 | -0.048 | 0.042 | 0.042 | 0 | 0.69 | 0.49 | 0.75 | 0.44 | 0.76 | 0.61 |
| 9 | 28383008 | rs2150862 | 0.299 | -1.18 | 0.575 | 0.296 | -0.064 | 0.087 | 0.087 | 0 | 0.75 | 0.49 | 0.77 | 0.44 | 0.81 | 0.62 |
| 9 | 28385612 | rs1412226 | 1.133 | -1.135 | 0.654 | 0.193 | -0.037 | -0.015 | -0.015 | 0 | 0.66 | 0.56 | 0.71 | 0.27 | 0.73 | 0.57 |
| 9 | 28390177 | rs10812795 | 0.935 | -1.093 | 0.62 | 0.226 | -0.035 | 0.021 | 0.021 | 0 | 0.68 | 0.6 | 0.71 | 0.37 | 0.74 | 0.6 |
| 9 | 28391010 | rs10968554 | 1.489 | -1.037 | 0.468 | 0.345 | -0.003 | 0.018 | 0.018 | 0 | 0.11 | 0.1 | 0.07 | 0.18 | 0.12 | 0.11 |
| 9 | 28391518 | rs10968555 | 1.522 | -1.047 | 0.485 | 0.629 | 0.007 | -0.008 | -0.008 | 0 | 0.08 | 0.06 | 0.12 | 0.17 | 0.04 | 0.17 |
| 9 | 28394358 | rs7019284 | -0.18 | -1.023 | 0.458 | 0.475 | -0.009 | -0.021 | -0.021 | 0 | 0.4 | 0.43 | 0.33 | 0.26 | 0.34 | 0.41 |
| 9 | 28395585 | rs10812802 | -0.066 | -1.013 | 0.43 | 0.473 | -0.005 | -0.03 | -0.03 | 0.041 | 0.39 | 0.4 | 0.33 | 0.16 | 0.34 | 0.37 |
| 9 | 28395773 | rs10968556 | 1.36 | -1.033 | 0.462 | 0.446 | -0.002 | -0.035 | -0.035 | 0.738 | 0.24 | 0.22 | 0.17 | 0.01 | 0.24 | 0.19 |
| 9 | 28396747 | rs7028515 | 1.321 | -0.68 | 0.112 | 0.88 | -0.003 | 0.015 | 0.015 | 0 | 0.11 | 0.1 | 0.07 | 0.19 | 0.12 | 0.11 |
| 9 | 28398636 | rs1331903 | 0.389 | -0.662 | 0.11 | 0.809 | -0.004 | -0.03 | -0.03 | 0 | 0.39 | 0.4 | 0.33 | 0.16 | 0.35 | 0.36 |
| 9 | 28398726 | rs4878360 | 1.223 | -0.639 | 0.027 | 0.797 | -0.034 | 0.018 | 0.018 | 0 | 0.34 | 0.29 | 0.36 | 0.26 | 0.39 | 0.33 |
| 9 | 28399683 | rs2183828 | 2.056 | -0.659 | -0.184 | 0.736 | -0.075 | -0.015 | -0.015 | 0 | 0.28 | 0.1 | 0.32 | 0.26 | 0.36 | 0.32 |
| 9 | 28400166 | rs1009216 | 1.145 | -0.334 | -0.095 | 0.134 | -0.003 | 0.015 | 0.015 | 0 | 0.11 | 0.1 | 0.07 | 0.2 | 0.12 | 0.11 |

| | | | | | | | | | | | | | | | | |
|---|---|---|---|---|---|---|---|---|---|---|---|---|---|---|---|---|
| 9 | 28400172 | rs1002406 | -0.151 | -0.243 | -0.174 | -0.514 | -0.007 | -0.043 | -0.043 | 0 | 0.4 | 0.47 | 0.33 | 0.17 | 0.34 | 0.36 |
| 9 | 28400551 | rs10812804 | -0.244 | -0.164 | -0.363 | -0.314 | 0.001 | 0.002 | 0.002 | 0 | 0.46 | 0.41 | 0.5 | 0.18 | 0.45 | 0.4 |
| 9 | 28402387 | rs1929807 | -0.374 | 0.235 | -0.581 | -0.253 | 0 | 0.009 | 0.009 | 0 | 0.44 | 0.46 | 0.4 | 0.76 | 0.41 | 0.65 |
| 9 | 28402960 | rs10125103 | -0.354 | -0.266 | -0.536 | -0.374 | 0 | 0 | 0 | 0 | 0.88 | 0.88 | 0.88 | 0.89 | 0.9 | 0.96 |
| 9 | 28403443 | rs7020846 | 0.733 | -1.183 | -0.166 | -0.333 | -0.045 | -0.006 | -0.006 | 0 | 0.7 | 0.58 | 0.78 | 0.68 | 0.81 | 0.73 |
| 9 | 28403813 | rs6476065 | 0.023 | -1.077 | -0.744 | -1.138 | 0.003 | 0.019 | 0.019 | 0 | 0.56 | 0.57 | 0.56 | 0.79 | 0.55 | 0.69 |
| 9 | 28407379 | rs1331909 | -1.058 | -1.075 | -0.794 | -0.897 | -0.005 | -0.036 | -0.036 | 0.078 | 0.89 | 0.86 | 0.94 | 1 | 0.91 | 0.98 |
| 9 | 28407568 | rs1331910 | -1.058 | -1.075 | -0.794 | -0.897 | -0.005 | -0.036 | -0.036 | 0 | 0.89 | 0.86 | 0.94 | 1 | 0.91 | 0.98 |
| 9 | 28409300 | rs7858089 | 0.648 | -1.023 | -1.028 | -0.882 | 0.093 | 0.156 | 0.156 | 0.277 | 0.51 | 0.25 | 0.61 | 0.32 | 0.65 | 0.43 |
| 9 | 28410944 | rs10968572 | 1.572 | -0.833 | 0.063 | -0.313 | 0.003 | -0.008 | -0.008 | 0 | 0.17 | 0.14 | 0.14 | 0.34 | 0.13 | 0.21 |
| 9 | 28412375 | rs2183825 | 1.041 | -0.76 | 0.26 | -0.462 | 0.051 | 0.038 | 0.038 | 1.013 | 0.2 | 0.29 | 0.26 | 0.27 | 0.31 | 0.23 |
| 9 | 28412544 | rs2150855 | 0.261 | -0.779 | 0.259 | -0.467 | 0.021 | -0.027 | -0.027 | 0 | 0.38 | 0.46 | 0.44 | 0.7 | 0.44 | 0.47 |
| 9 | 28413315 | rs10968574 | 0.308 | -0.904 | 0.213 | -0.546 | -0.041 | 0.03 | 0.03 | 0 | 0.17 | 0.32 | 0.14 | 0.23 | 0.09 | 0.22 |
| 9 | 28414339 | rs10968576 | 1.696 | -0.902 | 0.3 | -0.514 | 0.011 | 0.001 | 0.001 | 1.309 | 0.17 | 0.14 | 0.25 | 0.18 | 0.3 | 0.17 |
| 9 | 28415512 | rs10968577 | 1.136 | -0.901 | 0.309 | -0.557 | 0.007 | 0.004 | 0.004 | 1.129 | 0.19 | 0.16 | 0.26 | 0.27 | 0.31 | 0.23 |
| 9 | 28415970 | rs10757740 | 0.416 | -0.906 | 0.307 | -0.573 | -0.005 | 0.004 | 0.004 | 0 | 0.37 | 0.33 | 0.44 | 0.7 | 0.44 | 0.47 |
| 9 | 28417572 | rs7849276 | 1.03 | -0.968 | 0.168 | -0.726 | -0.003 | -0.032 | -0.032 | 0 | 0.17 | 0.27 | 0.23 | 0.02 | 0.13 | 0.09 |
| 9 | 28419193 | rs2183826 | 0.244 | -0.725 | 0.491 | -0.415 | -0.026 | 0.014 | 0.014 | 0 | 0.17 | 0.19 | 0.13 | 0.23 | 0.09 | 0.21 |
| 9 | 28420020 | rs7041128 | 0.309 | -0.288 | 0.467 | -0.147 | 0.005 | 0.014 | 0.014 | 0 | 0.52 | 0.47 | 0.5 | 0.29 | 0.46 | 0.52 |
| 9 | 28426078 | rs874183 | 1.381 | -0.143 | 0.24 | -0.073 | 0.002 | 0.003 | 0.003 | 0.084 | 0.18 | 0.17 | 0.18 | 0.44 | 0.13 | 0.23 |
| 9 | 28427576 | rs7855030 | 0.275 | -0.137 | 0.248 | -0.129 | -0.003 | -0.001 | -0.001 | 0 | 0.34 | 0.32 | 0.43 | 0.62 | 0.43 | 0.41 |
| 9 | 28437141 | rs10812826 | 0.911 | 0.035 | -0.161 | -0.509 | -0.069 | 0.225 | 0.225 | 0 | 0.38 | 0.11 | 0.4 | 0.06 | 0.45 | 0.28 |
| 9 | 28440394 | rs7021018 | 1.187 | -0.22 | -0.168 | -0.518 | -0.024 | 0.198 | 0.198 | 0 | 0.35 | 0.11 | 0.36 | 0.06 | 0.38 | 0.27 |
| 9 | 28442343 | rs10491888 | 1.265 | -0.436 | -0.097 | -0.358 | 0.005 | -0.017 | -0.017 | 0.496 | 0.19 | 0.15 | 0.15 | 0.37 | 0.13 | 0.24 |
| 9 | 28445047 | rs1331919 | 0.89 | -0.413 | -0.123 | -0.38 | -0.03 | 0.234 | 0.234 | 0.142 | 0.39 | 0.1 | 0.39 | 0.06 | 0.43 | 0.3 |
| 9 | 28450213 | rs10812831 | 0.871 | -0.32 | 0.177 | -0.274 | 0.003 | 0.01 | 0.01 | 0 | 0.13 | 0.12 | 0.15 | 0.2 | 0.13 | 0.26 |
| 9 | 28458823 | rs10968607 | -0.313 | -0.339 | -0.099 | -0.218 | -0.003 | 0.002 | 0.002 | 0.008 | 0.72 | 0.81 | 0.73 | 0.83 | 0.65 | 0.84 |
| 9 | 28463701 | rs7044907 | 1.111 | -0.047 | -0.178 | -0.066 | 0.003 | -0.018 | -0.018 | 0 | 0.19 | 0.15 | 0.17 | 0.38 | 0.15 | 0.22 |
| 9 | 28470551 | rs1888814 | 0.407 | 0.503 | 0.445 | 0.532 | -0.002 | 0.01 | 0.01 | 0 | 0.68 | 0.5 | 0.7 | 0.64 | 0.7 | 0.75 |
| 9 | 28477082 | rs10757744 | -0.598 | 0.936 | 0.421 | 0.744 | -0.009 | -0.084 | -0.084 | 0 | 0.83 | 0.8 | 0.9 | 1 | 0.86 | 0.97 |
| 9 | 28480661 | rs979525 | 0.783 | 0.846 | 0.518 | 0.841 | -0.031 | -0.019 | -0.019 | 0 | 0.15 | 0.06 | 0.18 | 0.17 | 0.21 | 0.12 |
| 9 | 28487096 | rs7469495 | -0.594 | 0.45 | 1.233 | 0.732 | -0.013 | -0.08 | -0.08 | 0.556 | 0.86 | 0.8 | 0.92 | 1 | 0.9 | 0.98 |
| 9 | 28499876 | rs10968622 | -0.627 | 0.91 | -0.156 | 1.186 | 0.047 | 0.081 | 0.081 | 0 | 0.13 | 0.3 | 0.22 | 0.3 | 0.18 | 0.28 |
| 9 | 28500104 | rs10968623 | 0.108 | 0.794 | -0.009 | 1.344 | 0.052 | 0.075 | 0.075 | 0 | 0.09 | 0.23 | 0.15 | 0.19 | 0.12 | 0.24 |
| 9 | 28503578 | rs7034070 | 0.105 | 0.202 | -1.149 | 0.767 | -0.004 | 0.06 | 0.06 | 0 | 0.24 | 0.12 | 0.15 | 0.13 | 0.26 | 0.21 |
| 9 | 28509666 | rs6476078 | 0.247 | 0.723 | -0.857 | 1.09 | 0.015 | 0.036 | 0.036 | 0 | 0.94 | 0.89 | 0.94 | 0.88 | 0.91 | 0.93 |
| 9 | 28509969 | rs7857851 | -0.224 | 0.381 | -0.851 | 0.865 | 0.014 | 0.004 | 0.004 | 0 | 0.44 | 0.65 | 0.39 | 0.43 | 0.41 | 0.35 |
| 9 | 28517001 | rs10491886 | 0.368 | 0.303 | -0.537 | 1.074 | -0.015 | 0.038 | 0.038 | 0.009 | 0.67 | 0.47 | 0.58 | 0.54 | 0.75 | 0.64 |
| 9 | 28520208 | rs10757754 | -0.047 | 0.101 | -0.685 | 1.311 | 0.012 | -0.014 | -0.014 | 0 | 0.41 | 0.26 | 0.33 | 0.48 | 0.39 | 0.47 |
| 9 | 28521811 | rs10757755 | 0.071 | 0.092 | -0.698 | 1.421 | 0.011 | -0.048 | -0.048 | 0 | 0.37 | 0.15 | 0.32 | 0.54 | 0.37 | 0.45 |
| 9 | 28539581 | rs12005012 | -1.068 | -0.018 | 0.364 | 0.992 | 0.014 | 0.089 | 0.089 | 0.537 | 0.05 | 0.21 | 0.07 | 0.09 | 0.03 | 0.02 |
| 9 | 28550930 | rs17832109 | 0.106 | 0.686 | -0.416 | 0.275 | -0.065 | 0.124 | 0.124 | 0 | 0.2 | 0.02 | 0.32 | 0.02 | 0.23 | 0.16 |
| 9 | 28562149 | rs10491884 | 0.525 | 1.4 | 0.608 | 0.828 | 0.067 | 0.123 | 0.123 | 0.044 | 0.87 | 0.64 | 0.72 | 0.71 | 0.82 | 0.69 |
| 9 | 28562256 | rs10491883 | -0.747 | 1.691 | 0.897 | 0.546 | 0.024 | -0.012 | -0.012 | 5.247 | 0.15 | 0.27 | 0.13 | 0.12 | 0.16 | 0.15 |
| 9 | 28566849 | rs10117773 | -1.65 | 1.593 | 1.31 | 0.565 | 0.022 | 0.117 | 0.117 | 0.22 | 0.19 | 0.51 | 0.21 | 0.35 | 0.2 | 0.33 |
| 9 | 28568013 | rs10812844 | -0.468 | 1.43 | 1.272 | 0.644 | 0 | 0.071 | 0.071 | 0 | 0.08 | 0.12 | 0.09 | 0.26 | 0.11 | 0.2 |
| 9 | 28569575 | rs7038354 | 1.224 | 1.163 | 1.335 | 0.211 | -0.009 | 0.023 | 0.023 | 0 | 0.56 | 0.4 | 0.36 | 0.62 | 0.5 | 0.49 |
| 9 | 28570652 | rs1521731 | -1.314 | 0.755 | 1.634 | 0.456 | 0.016 | 0.138 | 0.138 | 0.021 | 0.17 | 0.52 | 0.19 | 0.35 | 0.17 | 0.32 |
| 9 | 28571014 | rs10812846 | -1.075 | 0.744 | 1.493 | 0.459 | 0.015 | 0.16 | 0.16 | 0 | 0.15 | 0.39 | 0.17 | 0.35 | 0.16 | 0.3 |
| 9 | 28572086 | rs7855874 | 0.462 | 0.788 | 1.251 | 0.256 | 0.002 | 0.009 | 0.009 | 0.004 | 0.69 | 0.74 | 0.54 | 0.91 | 0.7 | 0.83 |
| 9 | 28577961 | rs1115012 | 1.246 | 0.636 | 1.423 | 0.395 | -0.004 | -0.004 | -0.004 | 0 | 0.62 | 0.48 | 0.45 | 0.69 | 0.66 | 0.64 |
| 9 | 28578198 | rs2383772 | 1.246 | 0.636 | 1.423 | 0.395 | -0.004 | -0.004 | -0.004 | 0.857 | 0.62 | 0.48 | 0.45 | 0.69 | 0.66 | 0.64 |
| 9 | 28587832 | rs2204076 | -0.723 | 0.651 | 1.026 | 0.369 | 0 | 0 | 0 | 0.087 | 0.29 | 0.25 | 0.46 | 0.12 | 0.26 | 0.21 |
| 9 | 28587994 | rs2891316 | 0.305 | 0.678 | 0.509 | 0.23 | -0.015 | 0.076 | 0.076 | 0.271 | 0.19 | 0.05 | 0.29 | 0.02 | 0.18 | 0.13 |
| 9 | 28595770 | rs16913363 | 0.144 | 0.701 | 0.587 | 0.228 | -0.001 | -0.01 | -0.01 | 0.023 | 0.2 | 0.12 | 0.3 | 0.02 | 0.18 | 0.13 |
| 9 | 28599107 | rs11998883 | 0.134 | 0.691 | 0.586 | 0.223 | -0.001 | 0.018 | 0.018 | 0.206 | 0.2 | 0.1 | 0.3 | 0.02 | 0.18 | 0.14 |
| 9 | 28601794 | rs7852443 | -0.441 | 0.716 | 0.575 | 0.271 | -0.002 | 0.007 | 0.007 | 0 | 0.26 | 0.2 | 0.36 | 0.2 | 0.26 | 0.29 |
| 9 | 28607069 | rs10968668 | 0.519 | 0.741 | 0.527 | 0.323 | -0.019 | -0.021 | -0.021 | 0.033 | 0.05 | 0.02 | 0.05 | 0.18 | 0.08 | 0.15 |
| 9 | 28609758 | rs10968671 | -0.88 | 0.722 | 0.592 | 0.308 | 0 | 0.001 | 0.001 | 0 | 0.3 | 0.24 | 0.41 | 0.22 | 0.28 | 0.33 |
| 9 | 28613702 | rs12347952 | -0.209 | 0.787 | 0.919 | 0.714 | -0.006 | -0.023 | -0.023 | 0 | 0.09 | 0.13 | 0.06 | 0.04 | 0.07 | 0.04 |
| 9 | 28616672 | rs10812853 | -1.355 | 0.899 | 1.295 | 0.985 | 0.061 | 0.21 | 0.21 | 0 | 0.08 | 0.3 | 0.14 | 0.22 | 0.1 | 0.17 |
| 9 | 28619450 | rs10968679 | -1.089 | 1.01 | 1.228 | 0.976 | 0.051 | 0.133 | 0.133 | 0 | 0.08 | 0.2 | 0.17 | 0.22 | 0.11 | 0.18 |
| 9 | 28628500 | rs10511828 | -1.799 | 0.986 | 1.146 | 1.167 | 0.004 | 0.043 | 0.043 | 0 | 0.07 | 0.11 | 0.28 | 0.14 | 0.07 | 0.11 |
| 9 | 28645019 | rs893496 | -2.464 | 0.872 | 1.384 | 1.031 | 0.013 | 0.066 | 0.066 | 0 | 0.28 | 0.48 | 0.5 | 0.38 | 0.27 | 0.33 |
| 9 | 28645626 | rs6476084 | -2.062 | 0.862 | 1.406 | 1.034 | 0.036 | 0.054 | 0.054 | 0 | 0.13 | 0.2 | 0.36 | 0.27 | 0.17 | 0.26 |
| 9 | 28647749 | rs1155414 | -2.043 | 0.838 | 1.404 | 1.025 | -0.005 | 0.082 | 0.082 | 0 | 0.24 | 0.51 | 0.31 | 0.33 | 0.21 | 0.24 |
| 9 | 28649685 | rs7855149 | -2.143 | 0.843 | 1.401 | 1.025 | -0.009 | 0.064 | 0.064 | 0 | 0.24 | 0.56 | 0.31 | 0.33 | 0.21 | 0.24 |
| 9 | 28649864 | rs10968702 | -0.736 | 0.837 | 1.35 | 0.987 | -0.019 | -0.058 | -0.058 | 0 | 0.15 | 0.25 | 0.13 | 0.04 | 0.1 | 0.07 |
| 9 | 28659143 | rs7046685 | -1.099 | 0.915 | 1.34 | 0.984 | -0.029 | -0.114 | -0.114 | 1.345 | 0.17 | 0.45 | 0.15 | 0.04 | 0.1 | 0.07 |

| | | | | | | | | | | | | | | | | |
|---|---|---|---|---|---|---|---|---|---|---|---|---|---|---|---|---|
| 9 | 28663042 | rs12341535 | -1.086 | 0.914 | 1.342 | 0.983 | -0.029 | -0.107 | -0.107 | 10.524 | 0.17 | 0.43 | 0.14 | 0.04 | 0.1 | 0.07 |
| 9 | 28677376 | rs11793079 | 1.691 | 0.764 | 0.671 | 1.065 | -0.105 | 0.071 | 0.071 | 1.205 | 0.42 | 0.03 | 0.29 | 0.26 | 0.42 | 0.28 |
| 9 | 28680131 | rs12345065 | -0.641 | 0.741 | 2.023 | 1.316 | -0.006 | -0.032 | -0.032 | 0 | 0.14 | 0.16 | 0.12 | 0.05 | 0.1 | 0.07 |
| 9 | 28687561 | rs16913525 | -1.084 | 0.769 | 1.968 | 1.331 | 0.005 | 0.004 | 0.004 | 0 | 0.05 | 0.11 | 0.26 | 0.07 | 0.04 | 0.11 |
| 9 | 28693005 | rs10968724 | -0.114 | 0.779 | 1.958 | 1.356 | 0.007 | 0.03 | 0.03 | 0 | 0.05 | 0.05 | 0.09 | 0.19 | 0.12 | 0.15 |
| 9 | 28697679 | rs723979 | -2.103 | 0.755 | 1.981 | 1.355 | -0.005 | 0.087 | 0.087 | 0 | 0.24 | 0.48 | 0.26 | 0.33 | 0.22 | 0.23 |
| 9 | 28705607 | rs4879261 | -1.062 | 0.774 | 1.807 | 1.221 | -0.029 | -0.082 | -0.082 | 0 | 0.16 | 0.32 | 0.14 | 0.06 | 0.1 | 0.07 |
| 9 | 28705864 | rs1583667 | -1.582 | 0.839 | 1.604 | 1.255 | -0.003 | 0.003 | 0.003 | 0 | 0.07 | 0.06 | 0.2 | 0.04 | 0.06 | 0.09 |
| 9 | 28715676 | rs7024629 | -2.467 | 0.758 | 1.882 | 1.492 | 0 | 0.036 | 0.036 | 0 | 0.28 | 0.58 | 0.51 | 0.4 | 0.25 | 0.34 |
| 9 | 28719213 | rs1438478 | -0.378 | 0.798 | 1.899 | 1.52 | -0.009 | 0 | 0 | 0 | 0.06 | 0.03 | 0.09 | 0.18 | 0.11 | 0.16 |
| 9 | 28722712 | rs10968740 | -1.899 | 0.788 | 1.982 | 1.494 | 0.004 | 0.038 | 0.038 | 0 | 0.27 | 0.59 | 0.51 | 0.39 | 0.25 | 0.33 |
| 9 | 28728183 | rs7041270 | -1.693 | 0.844 | 1.633 | 1.417 | 0.002 | 0.012 | 0.012 | 0 | 0.25 | 0.3 | 0.49 | 0.38 | 0.25 | 0.32 |
| 9 | 28732830 | rs4879262 | -1.188 | 0.361 | 1.593 | 1.282 | 0.001 | -0.011 | -0.011 | 0 | 0.19 | 0.12 | 0.32 | 0.28 | 0.16 | 0.27 |
| 9 | 28734729 | rs321747 | 1.122 | 0.563 | 1.684 | 1.18 | -0.004 | 0.002 | 0.002 | 0.048 | 0.71 | 0.37 | 0.52 | 0.68 | 0.78 | 0.69 |
| 9 | 28736988 | rs7851298 | -0.946 | 0.642 | 0.857 | 1.191 | 0 | -0.002 | -0.002 | 0 | 0.26 | 0.33 | 0.44 | 0.3 | 0.22 | 0.3 |
| 9 | 28752971 | rs16913617 | -1.134 | 0.918 | -0.124 | 1.559 | 0.005 | 0.037 | 0.037 | 0 | 0.08 | 0.11 | 0.08 | 0.21 | 0.07 | 0.13 |
| 9 | 28753619 | rs321736 | -1.113 | 0.81 | 0.128 | 1.331 | 0.013 | 0.042 | 0.042 | 0 | 0.16 | 0.36 | 0.14 | 0.24 | 0.13 | 0.17 |
| 9 | 28754535 | rs321738 | 0.357 | 0.226 | -0.298 | 1.113 | -0.024 | 0.004 | 0.004 | 0.682 | 0.6 | 0.55 | 0.74 | 0.45 | 0.71 | 0.49 |
| 9 | 28756883 | rs321741 | 0.154 | 0.339 | -0.371 | 1.161 | -0.017 | 0.011 | 0.011 | 0 | 0.92 | 0.87 | 0.98 | 0.87 | 0.97 | 0.86 |
| 9 | 28757900 | rs10812871 | -0.496 | 0.93 | -0.715 | 0.687 | 0.001 | 0.006 | 0.006 | 0 | 0.46 | 0.47 | 0.48 | 0.71 | 0.56 | 0.54 |
| 9 | 28758532 | rs10968754 | -0.76 | 1.395 | -0.961 | 1.167 | -0.027 | -0.025 | -0.025 | 0 | 0.13 | 0.04 | 0.13 | 0.15 | 0.18 | 0.09 |
| 9 | 28772999 | rs824249 | 1.013 | 1.647 | -0.162 | 1.016 | 0.014 | -0.022 | -0.022 | 0 | 0.88 | 0.75 | 0.87 | 0.94 | 0.91 | 0.89 |
| 9 | 28773458 | rs824251 | -1.569 | 1.914 | -0.689 | 1.561 | -0.047 | -0.084 | -0.084 | 0 | 0.33 | 0.84 | 0.42 | 0.48 | 0.34 | 0.38 |
| 9 | 28773602 | rs824252 | -0.068 | 1.903 | -0.25 | 1.649 | -0.001 | 0.02 | 0.02 | 0 | 0.1 | 0.07 | 0.07 | 0.32 | 0.1 | 0.16 |
| 9 | 28779376 | rs866630 | -2.143 | 1.765 | 0.384 | 1.576 | -0.048 | -0.067 | -0.067 | 1.324 | 0.38 | 0.83 | 0.42 | 0.5 | 0.4 | 0.4 |
| 9 | 28781231 | rs10812875 | -1.055 | 1.729 | 0.924 | 1.666 | 0.02 | -0.047 | -0.047 | 0 | 0.21 | 0.36 | 0.2 | 0.09 | 0.24 | 0.13 |
| 9 | 28782175 | rs824264 | -1.323 | 1.528 | 1.076 | 1.781 | 0.02 | 0.165 | 0.165 | 0 | 0.17 | 0.42 | 0.21 | 0.42 | 0.16 | 0.26 |
| 9 | 28784343 | rs824265 | -0.952 | 1.469 | 1.17 | 1.969 | 0.01 | 0.158 | 0.158 | 0 | 0.14 | 0.3 | 0.11 | 0.37 | 0.12 | 0.22 |
| 9 | 28787049 | rs824242 | -0.328 | 1.501 | 0.809 | 1.995 | -0.004 | -0.003 | -0.003 | 0 | 0.11 | 0.07 | 0.09 | 0.36 | 0.12 | 0.22 |
| 9 | 28793773 | rs1389729 | 0.973 | 1.822 | 0.722 | 1.704 | -0.009 | 0.012 | 0.012 | 0 | 0.81 | 0.81 | 0.8 | 0.88 | 0.79 | 0.84 |
| 9 | 28795700 | rs10968773 | 2.138 | 1.366 | -0.114 | 1.445 | -0.074 | 0.048 | 0.048 | 0 | 0.5 | 0.05 | 0.47 | 0.35 | 0.47 | 0.44 |
| 9 | 28796648 | rs1494337 | 1.89 | 1.356 | 0.988 | 1.5 | 0.008 | 0.011 | 0.011 | 0 | 0.61 | 0.48 | 0.58 | 0.45 | 0.59 | 0.53 |
| 9 | 28799706 | rs824222 | 0.784 | 1.318 | 0.891 | 1.549 | -0.012 | 0.015 | 0.015 | 0 | 0.77 | 0.79 | 0.71 | 0.82 | 0.76 | 0.79 |
| 9 | 28804098 | rs824230 | -0.864 | 1.295 | 1.176 | 1.591 | 0 | 0.029 | 0.029 | 0 | 0.16 | 0.22 | 0.12 | 0.35 | 0.16 | 0.27 |
| 9 | 28805269 | rs7038242 | 2.094 | 1.279 | 1.658 | 1.558 | 0 | 0 | 0 | 0 | 0.6 | 0.43 | 0.58 | 0.45 | 0.59 | 0.52 |
| 9 | 28812865 | rs7852834 | 2.087 | 1.276 | 1.633 | 1.557 | -0.007 | -0.011 | -0.011 | 0 | 0.59 | 0.34 | 0.58 | 0.45 | 0.59 | 0.53 |
| 9 | 28816121 | rs824236 | 0.302 | 1.298 | 1.644 | 1.485 | -0.005 | 0.004 | 0.004 | 0 | 0.81 | 0.8 | 0.8 | 0.88 | 0.79 | 0.84 |
| 9 | 28817957 | rs1389728 | -0.189 | 1.261 | 1.931 | 1.507 | 0.007 | 0.025 | 0.025 | 0.051 | 0.07 | 0.15 | 0.09 | 0.1 | 0.11 | 0.09 |
| 9 | 28819784 | rs1327362 | 2.033 | 1.247 | 1.862 | 1.505 | -0.005 | -0.006 | -0.006 | 0.337 | 0.6 | 0.37 | 0.58 | 0.46 | 0.57 | 0.53 |
| 9 | 28826452 | rs17374075 | -0.333 | 1.26 | 1.875 | 1.499 | -0.025 | 0.027 | 0.027 | 0 | 0.19 | 0.14 | 0.19 | 0.12 | 0.22 | 0.17 |
| 9 | 28827591 | rs824270 | -0.633 | 1.243 | 1.914 | 1.516 | -0.007 | -0.02 | -0.02 | 0 | 0.14 | 0.08 | 0.11 | 0.35 | 0.16 | 0.27 |
| 9 | 28839438 | rs702246 | 0.287 | 1.262 | 1.91 | 1.487 | -0.012 | 0.012 | 0.012 | 0 | 0.81 | 0.81 | 0.81 | 0.87 | 0.78 | 0.84 |
| 9 | 28842326 | rs702248 | 0.388 | 1.242 | 1.91 | 1.475 | -0.01 | 0.012 | 0.012 | 0 | 0.81 | 0.8 | 0.81 | 0.88 | 0.78 | 0.84 |
| 9 | 28843891 | rs13293352 | 2.002 | 1.224 | 1.966 | 1.538 | -0.001 | -0.001 | -0.001 | 0.074 | 0.6 | 0.41 | 0.57 | 0.46 | 0.57 | 0.53 |
| 9 | 28849378 | rs1952277 | -1.883 | 1.194 | 2.119 | 1.506 | -0.006 | 0.068 | 0.068 | 0.098 | 0.14 | 0.51 | 0.21 | 0.17 | 0.14 | 0.15 |
| 9 | 28853125 | rs16913751 | -1.092 | 1.217 | 2.175 | 1.447 | 0.043 | 0.1 | 0.1 | 0 | 0.09 | 0.35 | 0.11 | 0.1 | 0.1 | 0.09 |
| 9 | 28855071 | rs866632 | 1.393 | 1.13 | 2.038 | 1.372 | 0.01 | -0.015 | -0.015 | 1.07 | 0.71 | 0.4 | 0.7 | 0.77 | 0.68 | 0.75 |
| 9 | 28867478 | rs914822 | -1.548 | 0.831 | 2.085 | 1.56 | -0.005 | 0.038 | 0.038 | 0 | 0.11 | 0.54 | 0.13 | 0.1 | 0.1 | 0.09 |
| 9 | 28869800 | rs320947 | -1.507 | 0.866 | 2.007 | 1.706 | 0.01 | 0.109 | 0.109 | 0 | 0.2 | 0.31 | 0.23 | 0.4 | 0.2 | 0.3 |
| 9 | 28879500 | rs320955 | -2.279 | 0.867 | 1.819 | 1.697 | 0.002 | -0.015 | -0.015 | 0 | 0.29 | 0.65 | 0.34 | 0.52 | 0.3 | 0.39 |
| 9 | 28885610 | rs7854994 | 1.11 | 0.939 | 1.637 | 1.544 | -0.012 | 0.002 | 0.002 | 0 | 0.84 | 0.8 | 0.82 | 0.81 | 0.89 | 0.75 |
| 9 | 28887332 | rs4471128 | 1.747 | 0.926 | 1.787 | 1.571 | -0.021 | 0.047 | 0.047 | 0 | 0.79 | 0.66 | 0.75 | 0.7 | 0.81 | 0.69 |
| 9 | 28895042 | rs7046727 | 1.747 | 0.926 | 1.787 | 1.571 | -0.021 | 0.047 | 0.047 | 0 | 0.79 | 0.66 | 0.75 | 0.7 | 0.81 | 0.69 |
| 9 | 28895878 | rs1368992 | 1.107 | 0.917 | 1.726 | 1.572 | -0.012 | 0.002 | 0.002 | 0 | 0.84 | 0.8 | 0.82 | 0.81 | 0.89 | 0.76 |
| 9 | 28905235 | rs10757769 | 2.075 | 0.972 | 1.631 | 1.449 | -0.008 | 0.011 | 0.011 | 0 | 0.75 | 0.71 | 0.72 | 0.69 | 0.76 | 0.58 |
| 9 | 28911017 | rs13300441 | -1.416 | 0.973 | 1.62 | 1.44 | 0.004 | -0.001 | -0.001 | 0 | 0.13 | 0.11 | 0.19 | 0.12 | 0.17 | 0.21 |
| 9 | 28927193 | rs2995889 | 0.609 | 0.962 | 1.62 | 1.446 | -0.021 | -0.001 | -0.001 | 0 | 0.88 | 0.82 | 0.91 | 0.86 | 0.93 | 0.8 |
| 9 | 28934898 | rs1964389 | 0.296 | 0.96 | 1.614 | 1.448 | -0.003 | -0.001 | -0.001 | 0.228 | 0.89 | 0.9 | 0.92 | 0.86 | 0.93 | 0.8 |
| 9 | 28940111 | rs7871600 | 1.901 | 0.952 | 1.602 | 1.443 | -0.011 | 0.007 | 0.007 | 0 | 0.78 | 0.71 | 0.81 | 0.75 | 0.8 | 0.63 |
| 9 | 28942538 | rs13285474 | -1.57 | 0.963 | 1.573 | 1.443 | 0.005 | 0.008 | 0.008 | 0 | 0.14 | 0.13 | 0.2 | 0.17 | 0.17 | 0.22 |
| 9 | 28942608 | rs10491632 | -1.57 | 0.963 | 1.573 | 1.443 | 0.005 | 0.008 | 0.008 | 0 | 0.14 | 0.13 | 0.2 | 0.17 | 0.17 | 0.22 |
| 9 | 28951731 | rs7039434 | 0.559 | 0.924 | 1.512 | 1.372 | -0.01 | -0.001 | -0.001 | 0 | 0.88 | 0.86 | 0.92 | 0.87 | 0.93 | 0.8 |
| 9 | 28959079 | rs4879290 | 2.312 | 0.781 | 1.305 | 1.215 | 0.021 | 0.028 | 0.028 | 0 | 0.56 | 0.3 | 0.59 | 0.45 | 0.6 | 0.44 |
| 9 | 28960723 | rs13287598 | -0.889 | 0.786 | 1.348 | 1.152 | 0.011 | 0.008 | 0.008 | 0 | 0.1 | 0.13 | 0.1 | 0.11 | 0.12 | 0.19 |
| 9 | 28970237 | rs10968843 | 0.229 | 0.742 | 1.213 | 1.154 | -0.013 | -0.004 | -0.004 | 0.009 | 0.88 | 0.85 | 0.92 | 0.9 | 0.93 | 0.81 |
| 9 | 28975706 | rs1434302 | 0.124 | 0.749 | 1.186 | 1.157 | -0.001 | 0 | 0 | 0.053 | 0.89 | 0.91 | 0.92 | 0.9 | 0.93 | 0.81 |
| 9 | 28977705 | rs10968854 | 2.932 | 1.026 | 0.822 | 1.101 | -0.09 | 0.011 | 0.011 | 0.052 | 0.4 | 0.05 | 0.36 | 0.36 | 0.43 | 0.34 |
| 9 | 28979255 | rs10812910 | 1.261 | 1.035 | 1.152 | 1.126 | -0.024 | 0.046 | 0.046 | 0 | 0.66 | 0.47 | 0.7 | 0.58 | 0.72 | 0.63 |
| 9 | 28981481 | rs4879297 | 0.013 | 1.111 | 0.866 | 0.941 | 0.008 | -0.02 | -0.02 | 0 | 0.15 | 0.24 | 0.17 | 0.09 | 0.16 | 0.1 |

| | | | | | | | | | | | | | | | | |
|---|---|---|---|---|---|---|---|---|---|---|---|---|---|---|---|---|
| 9 | 29013529 | rs16914202 | 0.384 | 0.917 | 1.065 | 1.223 | 0 | 0.033 | 0.033 | 0 | 0.17 | 0.2 | 0.12 | 0.29 | 0.16 | 0.14 |
| 9 | 29015933 | rs12683606 | -0.086 | 0.917 | 1.035 | 1.19 | 0 | 0.01 | 0.01 | 0 | 0.2 | 0.2 | 0.21 | 0.34 | 0.21 | 0.21 |
| 9 | 29017258 | rs6476118 | 2.508 | 0.999 | 1.015 | 1.197 | 0.006 | 0.042 | 0.042 | 0 | 0.51 | 0.38 | 0.51 | 0.43 | 0.53 | 0.41 |
| 9 | 29020053 | rs10968884 | -0.443 | 1.168 | 0.851 | 1.023 | -0.03 | -0.024 | -0.024 | 0 | 0.1 | 0.02 | 0.14 | 0.14 | 0.14 | 0.1 |
| 9 | 29023122 | rs10812933 | -1.796 | 1.082 | 1.002 | 1.003 | -0.012 | -0.003 | -0.003 | 0 | 0.37 | 0.29 | 0.39 | 0.54 | 0.41 | 0.41 |
| 9 | 29024733 | rs12376696 | -1.63 | 1.087 | 1.117 | 1.022 | -0.007 | 0.003 | 0.003 | 0 | 0.2 | 0.17 | 0.28 | 0.19 | 0.24 | 0.27 |
| 9 | 29024894 | rs7847631 | -2.045 | 1.079 | 1.23 | 1.011 | 0.004 | -0.001 | -0.001 | 0 | 0.39 | 0.54 | 0.41 | 0.54 | 0.41 | 0.41 |
| 9 | 29032248 | rs10968899 | -1.006 | 1.069 | 1.013 | 1.002 | -0.021 | 0.008 | 0.008 | 0.064 | 0.16 | 0.08 | 0.18 | 0.14 | 0.2 | 0.23 |
| 9 | 29042229 | rs10968907 | 0.031 | 0.928 | 1.035 | 0.975 | -0.001 | 0.005 | 0.005 | 0.165 | 0.19 | 0.16 | 0.11 | 0.36 | 0.17 | 0.15 |
| 9 | 29052442 | rs10812937 | -1.746 | 0.883 | 1.074 | 0.986 | 0 | 0.001 | 0.001 | 0 | 0.35 | 0.34 | 0.3 | 0.5 | 0.37 | 0.37 |
| 9 | 29054853 | rs7468614 | -1.746 | 0.883 | 1.074 | 0.983 | 0 | 0.001 | 0.001 | 0 | 0.35 | 0.34 | 0.3 | 0.5 | 0.37 | 0.37 |
| 9 | 29074499 | rs13293280 | -1.395 | 0.776 | 1.163 | 0.899 | -0.001 | -0.01 | -0.01 | 0 | 0.18 | 0.28 | 0.2 | 0.14 | 0.18 | 0.25 |
| 9 | 29084708 | rs1418344 | -2.18 | 0.804 | 1.221 | 0.91 | 0 | 0.003 | 0.003 | 0 | 0.41 | 0.67 | 0.42 | 0.55 | 0.4 | 0.45 |
| 9 | 29088460 | rs13302466 | -1.463 | 0.782 | 1.095 | 0.869 | -0.001 | -0.003 | -0.003 | 0 | 0.19 | 0.24 | 0.2 | 0.14 | 0.19 | 0.26 |
| 9 | 29088547 | rs1954280 | -1.752 | 0.773 | 1.161 | 0.865 | -0.008 | -0.03 | -0.03 | 0 | 0.21 | 0.39 | 0.21 | 0.14 | 0.19 | 0.26 |
| 9 | 29092556 | rs10968935 | -0.408 | 0.871 | 0.962 | 0.994 | 0 | 0.051 | 0.051 | 0.018 | 0.22 | 0.27 | 0.21 | 0.41 | 0.21 | 0.2 |
| 9 | 29112301 | rs7019031 | -0.464 | 0.862 | 0.964 | 1.002 | 0 | 0.057 | 0.057 | 0.09 | 0.23 | 0.3 | 0.21 | 0.4 | 0.21 | 0.2 |
| 9 | 29119535 | rs7847094 | 2.394 | 1.343 | 0.897 | 0.993 | -0.009 | 0.114 | 0.114 | 0 | 0.51 | 0.32 | 0.52 | 0.33 | 0.52 | 0.41 |
| 9 | 29126329 | rs1361708 | 0.109 | 1.331 | 0.665 | 1.095 | 0.004 | 0.063 | 0.063 | 0.19 | 0.18 | 0.24 | 0.12 | 0.35 | 0.17 | 0.15 |
| 9 | 29132164 | rs1418358 | -1.776 | 1.382 | 0.625 | 0.917 | 0.004 | -0.008 | -0.008 | 0 | 0.38 | 0.49 | 0.39 | 0.58 | 0.39 | 0.39 |
| 9 | 29133769 | rs10812990 | 0.308 | 1.359 | 0.493 | 1.033 | -0.005 | -0.016 | -0.016 | 0 | 0.17 | 0.14 | 0.11 | 0.39 | 0.18 | 0.16 |
| 9 | 29150822 | rs1418352 | -1.561 | 1.59 | 0.573 | 0.731 | 0.003 | -0.006 | -0.006 | 0 | 0.41 | 0.54 | 0.4 | 0.52 | 0.41 | 0.4 |
| 9 | 29153469 | rs7870900 | 1.287 | 1.582 | 0.607 | 0.744 | 0.005 | 0.008 | 0.008 | 1.698 | 0.6 | 0.49 | 0.59 | 0.53 | 0.59 | 0.63 |
| 9 | 29161218 | rs10123671 | -0.995 | 1.529 | 0.612 | 0.66 | 0.002 | 0.001 | 0.001 | 0.803 | 0.16 | 0.13 | 0.15 | 0.18 | 0.17 | 0.23 |
| 9 | 29166390 | rs10968988 | 0.535 | 1.169 | 0.104 | 0.7 | -0.004 | -0.011 | -0.011 | 0 | 0.17 | 0.15 | 0.11 | 0.4 | 0.18 | 0.15 |
| 9 | 29174711 | rs17527789 | 0.797 | 0.198 | -1.515 | -0.73 | -0.115 | 0.01 | 0.01 | 0 | 0.36 | 0.04 | 0.54 | 0.35 | 0.42 | 0.42 |
| 9 | 29192125 | rs10757817 | -0.026 | 0.348 | -1.151 | -0.566 | -0.003 | 0.047 | 0.047 | 0.209 | 0.25 | 0.35 | 0.23 | 0.45 | 0.22 | 0.2 |
| 9 | 29198885 | rs10491658 | 1.093 | 0.364 | -0.726 | -0.76 | 0.018 | 0.028 | 0.028 | 0 | 0.06 | 0.21 | 0.12 | 0.05 | 0.05 | 0.05 |
| 9 | 29198980 | rs7024949 | -0.192 | 0.087 | -0.89 | -0.996 | -0.008 | -0.052 | -0.052 | 0 | 0.66 | 0.57 | 0.81 | 0.81 | 0.7 | 0.62 |
| 9 | 29203366 | rs2066346 | -0.107 | 0.305 | -0.755 | -0.547 | 0 | -0.006 | -0.006 | 0.259 | 0.15 | 0.14 | 0.19 | 0.02 | 0.15 | 0.09 |
| 9 | 29214289 | rs1928658 | 0.712 | 0.45 | -0.641 | -0.416 | 0.002 | -0.028 | -0.028 | 0 | 0.09 | 0.14 | 0.07 | 0 | 0.09 | 0.01 |
| 9 | 29218907 | rs1010587 | 1.23 | 0.419 | -0.895 | -0.312 | 0 | -0.004 | -0.004 | 0 | 0.27 | 0.2 | 0.36 | 0.33 | 0.28 | 0.4 |
| 9 | 29226705 | rs4879321 | -0.403 | 0.071 | -0.126 | -0.52 | -0.063 | -0.053 | -0.053 | 0 | 0.8 | 0.68 | 0.94 | 0.91 | 0.92 | 0.86 |
| 9 | 29226761 | rs16914502 | 1.423 | 0.131 | 0.073 | -0.598 | 0.007 | -0.034 | -0.034 | 0 | 0.08 | 0.14 | 0.07 | 0 | 0.1 | 0.01 |
| 9 | 29227533 | rs10757820 | 0.148 | 0.119 | 0.063 | -0.603 | -0.048 | -0.106 | -0.106 | 0 | 0.72 | 0.55 | 0.86 | 0.91 | 0.83 | 0.85 |
| 9 | 29230848 | rs10757821 | -0.029 | 0.137 | -0.016 | -0.521 | -0.048 | -0.032 | -0.032 | 0.624 | 0.77 | 0.69 | 0.84 | 0.86 | 0.88 | 0.74 |
| 9 | 29231781 | rs16914523 | 0.867 | 0.144 | 0.196 | -0.548 | 0.039 | -0.058 | -0.058 | 1.715 | 0.1 | 0.33 | 0.08 | 0 | 0.1 | 0.01 |
| 9 | 29233374 | rs10813031 | -0.016 | 0.263 | 0.287 | -0.436 | 0.008 | 0.003 | 0.003 | 3.305 | 0.7 | 0.33 | 0.85 | 0.92 | 0.83 | 0.85 |
| 9 | 29236498 | rs7045302 | -0.086 | 0.752 | 0.038 | -0.364 | -0.02 | -0.053 | -0.053 | 0 | 0.71 | 0.68 | 0.78 | 0.88 | 0.79 | 0.74 |
| 9 | 29244165 | rs10491659 | 0.554 | 1.13 | -0.872 | -0.35 | -0.037 | 0.006 | 0.006 | 0 | 0.18 | 0.03 | 0.12 | 0.11 | 0.2 | 0.21 |
| 9 | 29244767 | rs10969027 | 1.631 | 1.104 | -1.13 | 0.115 | -0.021 | -0.098 | -0.098 | 0 | 0.35 | 0.09 | 0.38 | 0.42 | 0.33 | 0.37 |
| 9 | 29247354 | rs1928691 | -0.019 | 0.902 | -0.689 | -0.139 | 0.007 | 0.037 | 0.037 | 0.499 | 0.69 | 0.27 | 0.73 | 0.81 | 0.74 | 0.72 |
| 9 | 29249698 | rs2891324 | -0.005 | 0.905 | -0.865 | -0.133 | 0.008 | 0.042 | 0.042 | 0.609 | 0.69 | 0.26 | 0.73 | 0.82 | 0.74 | 0.71 |
| 9 | 29258978 | rs10969037 | 1.439 | 1.103 | -1.295 | 0.221 | -0.08 | -0.151 | -0.151 | 0 | 0.33 | 0.03 | 0.39 | 0.43 | 0.36 | 0.35 |
| 9 | 29261935 | rs7852004 | 0.596 | 1.122 | -0.437 | 0.042 | -0.012 | -0.035 | -0.035 | 6.655 | 0.21 | 0.39 | 0.15 | 0.08 | 0.17 | 0.21 |
| 9 | 29262510 | rs1928676 | -1.209 | 1.157 | -0.267 | -0.11 | 0.03 | 0.043 | 0.043 | 0.123 | 0.78 | 0.87 | 0.93 | 0.94 | 0.84 | 0.82 |
| 9 | 29263492 | rs10813035 | 1.892 | 1.267 | -0.271 | -0.06 | -0.005 | -0.05 | -0.05 | 0.474 | 0.34 | 0.12 | 0.4 | 0.43 | 0.36 | 0.35 |
| 9 | 29264328 | rs10119913 | 0.388 | 1.256 | 0.128 | -0.078 | -0.005 | -0.02 | -0.02 | 2.61 | 0.56 | 0.52 | 0.78 | 0.86 | 0.67 | 0.61 |
| 9 | 29266639 | rs10813039 | 1.168 | 1.168 | 0.611 | 0.155 | -0.015 | 0.007 | 0.007 | 0 | 0.53 | 0.3 | 0.53 | 0.51 | 0.53 | 0.56 |
| 9 | 29269462 | rs1928669 | -0.606 | 1.324 | 0.692 | 0.136 | -0.036 | -0.079 | -0.079 | 0 | 0.76 | 0.68 | 0.91 | 0.94 | 0.84 | 0.82 |
| 9 | 29270224 | rs10757826 | -0.588 | 1.307 | 0.614 | 0.146 | 0.054 | 0.067 | 0.067 | 0.007 | 0.23 | 0.39 | 0.38 | 0.43 | 0.31 | 0.26 |
| 9 | 29276550 | rs7020596 | -0.488 | 1.124 | 0.564 | 0.169 | -0.046 | -0.093 | -0.093 | 0.111 | 0.76 | 0.64 | 0.91 | 0.94 | 0.84 | 0.82 |
| 9 | 29282192 | rs10757831 | -0.471 | 1.915 | 0.565 | 1.051 | 0.009 | 0.02 | 0.02 | 0 | 0.16 | 0.2 | 0.23 | 0.29 | 0.19 | 0.15 |
| 9 | 29285982 | rs7864608 | -0.609 | 1.922 | 0.56 | 0.945 | 0.007 | 0.03 | 0.03 | 0 | 0.17 | 0.2 | 0.23 | 0.36 | 0.19 | 0.16 |
| 9 | 29291279 | rs16914634 | -0.808 | 2.03 | 0.95 | 0.693 | 0.127 | 0.086 | 0.086 | 0 | 0.12 | 0.39 | 0.19 | 0.22 | 0.18 | 0.09 |
| 9 | 29296753 | rs10491661 | -0.799 | 2.16 | 0.876 | 0.528 | 0.067 | 0 | 0 | 0 | 0.09 | 0.21 | 0.18 | 0.11 | 0.16 | 0.08 |
| 9 | 29302701 | rs1928663 | -0.131 | 2.682 | 0.579 | 0.106 | -0.009 | 0.022 | 0.022 | 0 | 0.25 | 0.2 | 0.21 | 0.1 | 0.29 | 0.25 |
| 9 | 29309713 | rs1928661 | -1.127 | 2.472 | 0.482 | 0.383 | -0.02 | 0.027 | 0.027 | 0 | 0.37 | 0.78 | 0.39 | 0.25 | 0.4 | 0.5 |
| 9 | 29313166 | rs12682678 | -1.147 | 2.612 | 0.419 | 0.295 | -0.013 | 0.009 | 0.009 | 0 | 0.32 | 0.78 | 0.36 | 0.25 | 0.38 | 0.46 |
| 9 | 29315618 | rs10491662 | 0.187 | 2.662 | 0.837 | 0.267 | -0.002 | -0.013 | -0.013 | 0 | 0.2 | 0.25 | 0.19 | 0.04 | 0.27 | 0.19 |
| 9 | 29317769 | rs1169758 | 0.529 | 2.696 | 0.823 | 0.352 | 0.01 | -0.007 | -0.007 | 0 | 0.71 | 0.62 | 0.65 | 0.79 | 0.57 | 0.7 |
| 9 | 29318382 | rs1169757 | 1.57 | 2.899 | 0.735 | 0.331 | -0.051 | -0.043 | -0.043 | 0 | 0.6 | 0.12 | 0.51 | 0.57 | 0.5 | 0.44 |
| 9 | 29318695 | rs12378755 | -0.244 | 2.702 | 1.064 | 0.429 | 0.029 | 0.012 | 0.012 | 0 | 0.1 | 0.22 | 0.14 | 0.13 | 0.11 | 0.22 |
| 9 | 29325150 | rs1169753 | 0.04 | 2.774 | 0.448 | 0.401 | -0.004 | -0.002 | -0.002 | 0 | 0.85 | 0.88 | 0.81 | 0.82 | 0.8 | 0.89 |
| 9 | 29325412 | rs1928652 | -1.403 | 2.735 | 0.39 | 0.313 | -0.009 | 0.005 | 0.005 | 0 | 0.31 | 0.71 | 0.36 | 0.25 | 0.36 | 0.5 |
| 9 | 29326375 | rs7023568 | 0.311 | 2.889 | 0.283 | 0.266 | -0.015 | 0.015 | 0.015 | 0 | 0.13 | 0.14 | 0.15 | 0.04 | 0.23 | 0.18 |
| 9 | 29331710 | rs1169751 | 0.272 | 2.44 | 0.306 | 0.505 | -0.004 | -0.002 | -0.002 | 0 | 0.85 | 0.85 | 0.81 | 0.82 | 0.8 | 0.89 |
| 9 | 29335831 | rs7040052 | -0.262 | 1.988 | 0.139 | 0.12 | 0 | -0.002 | -0.002 | 0 | 0.33 | 0.34 | 0.39 | 0.31 | 0.39 | 0.28 |
| 9 | 29336303 | rs16914695 | 0.184 | 2.218 | 0.451 | -0.207 | 0.009 | -0.061 | -0.061 | 0 | 0.18 | 0.3 | 0.19 | 0 | 0.2 | 0.16 |

| | | | | | | | | | | | | | | | | |
|---|---|---|---|---|---|---|---|---|---|---|---|---|---|---|---|---|
| 9 | 29336679 | rs4529540 | -1.678 | 1.968 | 0.163 | -0.237 | 0.006 | -0.009 | -0.009 | 3.776 | 0.63 | 0.78 | 0.68 | 0.46 | 0.7 | 0.6 |
| 9 | 29349856 | rs9695393 | -0.414 | 1.777 | -0.063 | -0.404 | 0 | -0.041 | -0.041 | 0 | 0.29 | 0.38 | 0.28 | 0.05 | 0.29 | 0.26 |
| 9 | 29350462 | rs10757841 | 0.253 | 1.82 | 0.067 | -0.366 | -0.002 | -0.004 | -0.004 | 0 | 0.3 | 0.3 | 0.32 | 0.15 | 0.35 | 0.23 |
| 9 | 29350850 | rs1004617 | -1.266 | 1.444 | -0.073 | -0.234 | 0.014 | 0.024 | 0.024 | 0 | 0.56 | 0.74 | 0.58 | 0.29 | 0.64 | 0.51 |
| 9 | 29363582 | rs1929831 | -0.107 | 1.643 | -0.097 | -0.273 | -0.002 | 0.002 | 0.002 | 0 | 0.25 | 0.31 | 0.23 | 0.14 | 0.27 | 0.27 |
| 9 | 29364847 | rs2150865 | 0.01 | 1.475 | -0.305 | -0.586 | 0.001 | -0.029 | -0.029 | 0 | 0.33 | 0.37 | 0.36 | 0.11 | 0.37 | 0.23 |
| 9 | 29368665 | rs10969108 | 0.702 | 1.357 | -0.352 | -0.324 | -0.046 | 0.062 | 0.062 | 0 | 0.13 | 0.03 | 0.11 | 0 | 0.17 | 0.12 |
| 9 | 29374906 | rs4879335 | -0.226 | 1.235 | -0.233 | -0.284 | -0.004 | 0.023 | 0.023 | 0 | 0.39 | 0.35 | 0.41 | 0.16 | 0.42 | 0.35 |
| 9 | 29393359 | rs12686399 | 1.407 | 0.986 | -0.157 | -0.236 | -0.003 | 0.001 | 0.001 | 0 | 0.31 | 0.27 | 0.33 | 0.62 | 0.28 | 0.35 |
| 9 | 29398399 | rs4487868 | -1.823 | 0.993 | -0.133 | -0.315 | 0.002 | -0.022 | -0.022 | 0 | 0.89 | 0.83 | 0.94 | 0.97 | 0.9 | 0.87 |
| 9 | 29402291 | rs17533687 | 0.574 | 1.109 | -0.099 | -0.342 | -0.01 | 0.129 | 0.129 | 0 | 0.35 | 0.14 | 0.3 | 0.07 | 0.38 | 0.3 |
| 9 | 29403851 | rs12379537 | 1.058 | 1.164 | -0.049 | -0.297 | -0.012 | 0.013 | 0.013 | 0 | 0.14 | 0.11 | 0.12 | 0 | 0.16 | 0.11 |
| 9 | 29404372 | rs16914740 | 1.613 | 0.966 | -0.233 | -0.44 | 0 | -0.01 | -0.01 | 0.395 | 0.11 | 0.07 | 0.06 | 0 | 0.13 | 0.01 |
| 9 | 29408855 | rs13292178 | -1.735 | 0.971 | -0.056 | 0.137 | 0.002 | -0.022 | -0.022 | 0 | 0.89 | 0.83 | 0.94 | 0.97 | 0.89 | 0.87 |
| 9 | 29411568 | rs10491665 | 0.95 | 1.004 | -0.084 | 0.09 | -0.008 | 0 | 0 | 0 | 0.14 | 0.13 | 0.12 | 0 | 0.16 | 0.11 |
| 9 | 29412843 | rs10757851 | 0.213 | 0.966 | -0.061 | 0.071 | -0.001 | -0.001 | -0.001 | 0 | 0.41 | 0.36 | 0.46 | 0.37 | 0.44 | 0.44 |
| 9 | 29419802 | rs10813080 | -0.44 | 0.922 | -0.085 | 0.093 | -0.005 | -0.02 | -0.02 | 0 | 0.55 | 0.56 | 0.58 | 0.25 | 0.59 | 0.5 |
| 9 | 29426093 | rs1408512 | -1.64 | 0.96 | -0.104 | -0.039 | 0.001 | -0.022 | -0.022 | 0 | 0.89 | 0.82 | 0.93 | 0.97 | 0.9 | 0.87 |
| 9 | 29426335 | rs2891328 | 1.464 | 0.893 | -0.145 | 0.065 | 0.002 | -0.006 | -0.006 | 0 | 0.3 | 0.21 | 0.32 | 0.49 | 0.27 | 0.3 |
| 9 | 29431179 | rs1575599 | 0.148 | 1.127 | -0.617 | -0.362 | 0.007 | -0.026 | -0.026 | 0 | 0.72 | 0.63 | 0.78 | 0.87 | 0.7 | 0.74 |
| 9 | 29435030 | rs1999426 | -0.474 | 1.091 | -0.313 | 0.322 | 0.067 | 0.43 | 0.43 | 0 | 0.09 | 0.36 | 0.23 | 0.54 | 0.11 | 0.12 |
| 9 | 29435969 | rs13287338 | 0.383 | 0.873 | -0.625 | -0.062 | 0.002 | 0.031 | 0.031 | 0 | 0.35 | 0.23 | 0.3 | 0.11 | 0.37 | 0.32 |
| 9 | 29436763 | rs7043013 | -1.272 | 0.865 | -0.704 | -0.16 | 0.01 | 0.065 | 0.065 | 0 | 0.59 | 0.71 | 0.64 | 0.75 | 0.66 | 0.62 |
| 9 | 29436823 | rs13291586 | 0.41 | 0.834 | -0.754 | -0.219 | 0.003 | 0.031 | 0.031 | 0 | 0.35 | 0.23 | 0.3 | 0.11 | 0.37 | 0.32 |
| 9 | 29440074 | rs12379607 | 0.768 | 0.94 | -0.747 | -0.244 | 0.004 | -0.031 | -0.031 | 0 | 0.13 | 0.18 | 0.12 | 0 | 0.18 | 0.09 |
| 9 | 29440562 | rs7022573 | -0.205 | 0.898 | -0.747 | -0.114 | 0.003 | 0.02 | 0.02 | 0 | 0.46 | 0.6 | 0.54 | 0.65 | 0.48 | 0.45 |
| 9 | 29445609 | rs7856020 | 1.063 | 0.762 | -0.774 | -0.129 | 0.003 | 0.261 | 0.261 | 0 | 0.12 | 0.32 | 0.22 | 0.45 | 0.1 | 0.09 |
| 9 | 29445887 | rs17535735 | 1.964 | 0.56 | -1.182 | -0.447 | -0.042 | -0.068 | -0.068 | 0 | 0.2 | 0.02 | 0.24 | 0.24 | 0.2 | 0.22 |
| 9 | 29447139 | rs7849929 | -0.285 | 0.591 | -1.042 | -0.534 | -0.006 | 0.026 | 0.026 | 1.445 | 0.66 | 0.69 | 0.79 | 0.89 | 0.67 | 0.69 |
| 9 | 29448530 | rs7041148 | -1.565 | 0.397 | -1.099 | -0.483 | 0.005 | 0.033 | 0.033 | 1.329 | 0.81 | 0.88 | 0.9 | 0.96 | 0.86 | 0.86 |
| 9 | 29457656 | rs4879352 | -0.252 | 0.477 | -1.06 | -0.476 | 0 | 0.037 | 0.037 | 0 | 0.49 | 0.63 | 0.61 | 0.78 | 0.51 | 0.43 |
| 9 | 29465624 | rs13285271 | 0.618 | 0.474 | -1.089 | -0.418 | -0.001 | 0.01 | 0.01 | 0 | 0.31 | 0.23 | 0.29 | 0.11 | 0.35 | 0.34 |
| 9 | 29470565 | rs12554780 | 1.866 | 0.565 | -1.15 | -0.376 | -0.011 | -0.021 | -0.021 | 0 | 0.2 | 0.09 | 0.24 | 0.24 | 0.21 | 0.23 |
| 9 | 29472650 | rs13295051 | 0.057 | 0.655 | -0.886 | -0.191 | -0.001 | 0.008 | 0.008 | 0 | 0.35 | 0.25 | 0.33 | 0.27 | 0.38 | 0.46 |
| 9 | 29473498 | rs10491670 | 0.105 | 0.664 | -0.888 | -0.222 | -0.002 | 0.017 | 0.017 | 0 | 0.34 | 0.25 | 0.33 | 0.19 | 0.38 | 0.37 |
| 9 | 29476774 | rs13440118 | 1.219 | 0.326 | -0.457 | -0.058 | 0.008 | 0.272 | 0.272 | 0 | 0.11 | 0.26 | 0.21 | 0.52 | 0.1 | 0.1 |
| 9 | 29476855 | rs10813104 | 0.341 | 0.357 | -0.691 | -0.481 | -0.002 | -0.011 | -0.011 | 0.055 | 0.24 | 0.23 | 0.19 | 0.12 | 0.22 | 0.22 |
| 9 | 29483651 | rs7040084 | 0.815 | 0.17 | -0.626 | -0.433 | -0.008 | -0.035 | -0.035 | 0 | 0.21 | 0.23 | 0.15 | 0.04 | 0.18 | 0.2 |
| 9 | 29486705 | rs1590444 | 0.509 | 0.143 | -0.717 | -0.538 | -0.006 | 0.026 | 0.026 | 0 | 0.21 | 0.17 | 0.16 | 0.09 | 0.22 | 0.13 |
| 9 | 29489987 | rs10738817 | 1.3 | 0.198 | -0.565 | -0.251 | 0.008 | 0.272 | 0.272 | 0 | 0.11 | 0.26 | 0.22 | 0.55 | 0.09 | 0.1 |
| 9 | 29500531 | rs7864125 | 0.736 | 0.339 | -0.877 | -0.67 | -0.006 | -0.032 | -0.032 | 0 | 0.2 | 0.2 | 0.15 | 0.04 | 0.18 | 0.2 |
| 9 | 29501683 | rs12057038 | -0.007 | 0.556 | -0.989 | -0.222 | -0.009 | 0.03 | 0.03 | 0.02 | 0.2 | 0.17 | 0.16 | 0.09 | 0.22 | 0.13 |
| 9 | 29508543 | rs7026384 | -0.056 | 0.606 | -1.041 | -0.04 | -0.004 | -0.006 | -0.006 | 0.522 | 0.2 | 0.2 | 0.2 | 0.24 | 0.21 | 0.23 |
| 9 | 29516586 | rs571210 | 0.192 | 0.491 | -1.031 | -0.7 | 0.001 | 0.019 | 0.019 | 0 | 0.6 | 0.59 | 0.69 | 0.78 | 0.65 | 0.61 |
| 9 | 29516728 | rs11536732 | 0.531 | 0.526 | -1.148 | -0.778 | -0.007 | 0.001 | 0.001 | 0.25 | 0.17 | 0.17 | 0.18 | 0.16 | 0.19 | 0.11 |
| 9 | 29520544 | rs603216 | 0.245 | 0.343 | -1.072 | -0.786 | 0 | 0.027 | 0.027 | 0.617 | 0.67 | 0.71 | 0.77 | 0.78 | 0.72 | 0.76 |
| 9 | 29533691 | rs12380193 | 2.278 | 0.812 | -1.606 | -0.999 | -0.058 | -0.018 | -0.018 | 0 | 0.12 | 0.02 | 0.18 | 0.08 | 0.17 | 0.14 |
| 9 | 29535541 | rs621211 | 0.607 | 0.702 | -1.228 | -1.289 | 0.003 | 0.009 | 0.009 | 0.015 | 0.44 | 0.54 | 0.51 | 0.68 | 0.42 | 0.5 |
| 9 | 29536618 | rs640850 | 0.945 | 0.802 | -1.443 | -1.252 | -0.012 | 0.006 | 0.006 | 0 | 0.36 | 0.52 | 0.39 | 0.6 | 0.25 | 0.4 |
| 9 | 29539147 | rs681870 | -1.475 | 0.775 | -1.506 | -1.282 | 0.016 | 0.068 | 0.068 | 0.194 | 0.77 | 0.89 | 0.89 | 0.94 | 0.84 | 0.83 |
| 9 | 29539183 | rs10733440 | 1.759 | 0.447 | -1.102 | -1.334 | -0.026 | 0.09 | 0.09 | 0 | 0.2 | 0.26 | 0.23 | 0.55 | 0.13 | 0.17 |
| 9 | 29539919 | rs1889055 | 0.234 | 0.562 | -1.391 | -1.44 | -0.017 | 0.013 | 0.013 | 0 | 0.32 | 0.25 | 0.39 | 0.18 | 0.47 | 0.37 |
| 9 | 29540105 | rs10738821 | 0.919 | 0.574 | -1.211 | -1.381 | -0.011 | 0.016 | 0.016 | 0 | 0.3 | 0.54 | 0.36 | 0.71 | 0.25 | 0.24 |
| 9 | 29544132 | rs665477 | -0.359 | 0.776 | -1.208 | -1.256 | -0.007 | 0.11 | 0.11 | 0 | 0.55 | 0.75 | 0.64 | 0.84 | 0.54 | 0.58 |
| 9 | 29545875 | rs13290799 | 1.2 | 0.821 | -1.203 | -1.274 | -0.007 | -0.001 | -0.001 | 0.847 | 0.09 | 0.07 | 0.14 | 0.08 | 0.18 | 0.1 |
| 9 | 29546888 | rs683471 | 0.367 | 0.807 | -1.187 | -1.258 | 0.025 | 0.049 | 0.049 | 0 | 0.45 | 0.68 | 0.51 | 0.76 | 0.36 | 0.48 |
| 9 | 29548583 | rs482067 | -0.379 | 0.806 | -1.22 | -1.293 | -0.012 | 0.047 | 0.047 | 0.009 | 0.67 | 0.77 | 0.74 | 0.84 | 0.66 | 0.61 |
| 9 | 29556315 | rs10969232 | -0.427 | 0.852 | -1.228 | -1.394 | -0.009 | 0.04 | 0.04 | 0 | 0.68 | 0.77 | 0.75 | 0.88 | 0.65 | 0.63 |
| 9 | 29556577 | rs10813121 | 0.379 | 0.805 | -1.231 | -1.408 | 0.015 | 0.029 | 0.029 | 0 | 0.46 | 0.62 | 0.5 | 0.8 | 0.35 | 0.48 |
| 9 | 29556765 | rs646439 | -1.354 | 0.78 | -1.303 | -1.304 | 0.006 | 0.007 | 0.007 | 0.694 | 0.78 | 0.84 | 0.88 | 0.98 | 0.83 | 0.77 |
| 9 | 29565060 | rs490511 | -0.628 | 0.968 | -1.46 | -1.18 | -0.016 | 0.096 | 0.096 | 0 | 0.69 | 0.84 | 0.77 | 0.91 | 0.68 | 0.7 |
| 9 | 29569571 | rs10969259 | 1.21 | 1.024 | -1.492 | -1.214 | -0.016 | 0.022 | 0.022 | 0 | 0.21 | 0.14 | 0.25 | 0.09 | 0.3 | 0.15 |
| 9 | 29572017 | rs489383 | 0.104 | 1.062 | -1.317 | -1.16 | 0.006 | -0.004 | -0.004 | 0 | 0.31 | 0.38 | 0.37 | 0.24 | 0.4 | 0.21 |
| 9 | 29572075 | rs488550 | 1.432 | 1.065 | -1.456 | -1.084 | 0.002 | 0.006 | 0.006 | 0 | 0.09 | 0.17 | 0.11 | 0.15 | 0.1 | 0.07 |
| 9 | 29574928 | rs10491680 | 1.747 | 0.867 | -1.516 | -1.136 | 0.006 | -0.001 | -0.001 | 0 | 0.14 | 0.09 | 0.07 | 0.11 | 0.11 | 0.26 |
| 9 | 29575467 | rs483312 | 1.458 | 0.889 | -1.491 | -1.139 | 0.002 | 0.005 | 0.005 | 0 | 0.09 | 0.16 | 0.11 | 0.15 | 0.1 | 0.07 |
| 9 | 29584528 | rs7856040 | 0.079 | 0.819 | -1.568 | -1.267 | 0.003 | 0.031 | 0.031 | 0 | 0.45 | 0.54 | 0.53 | 0.76 | 0.45 | 0.62 |
| 9 | 29586738 | rs7864533 | 0.639 | 0.801 | -1.562 | -1.254 | 0.001 | 0.024 | 0.024 | 0 | 0.31 | 0.46 | 0.46 | 0.65 | 0.33 | 0.37 |
| 9 | 29587032 | rs12238051 | 0.223 | 1.455 | -1.45 | -1.148 | -0.002 | -0.01 | -0.01 | 0.166 | 0.23 | 0.23 | 0.17 | 0.26 | 0.22 | 0.32 |

| | | | | | | | | | | | | | | | | |
|---|---|---|---|---|---|---|---|---|---|---|---|---|---|---|---|---|
| 9 | 29588006 | rs2184023 | -0.549 | 1.427 | -1.459 | -1.136 | -0.005 | 0.013 | 0.013 | 0 | 0.45 | 0.39 | 0.43 | 0.35 | 0.51 | 0.47 |
| 9 | 29591047 | rs510290 | 1.159 | 1.418 | -1.371 | -1.054 | 0 | 0.01 | 0.01 | 0 | 0.08 | 0.16 | 0.1 | 0.15 | 0.08 | 0.06 |
| 9 | 29592836 | rs560764 | -0.527 | 1.521 | -1.421 | -1.098 | -0.015 | 0.003 | 0.003 | 0 | 0.38 | 0.33 | 0.47 | 0.33 | 0.55 | 0.35 |
| 9 | 29593098 | rs558097 | -0.519 | 1.531 | -1.349 | -1.034 | -0.027 | -0.008 | -0.008 | 0 | 0.25 | 0.21 | 0.29 | 0.27 | 0.38 | 0.28 |
| 9 | 29594380 | rs1482071 | 0.947 | 1.184 | -1.269 | -0.947 | -0.022 | -0.013 | -0.013 | 0 | 0.08 | 0.04 | 0.12 | 0.11 | 0.17 | 0.21 |
| 9 | 29594461 | rs503350 | -0.307 | 0.411 | -1.013 | -0.962 | 0.029 | -0.005 | -0.005 | 0 | 0.53 | 0.43 | 0.54 | 0.46 | 0.67 | 0.68 |
| 9 | 29595350 | rs670205 | 0.076 | 0.322 | -0.859 | -0.901 | 0.005 | -0.005 | -0.005 | 0 | 0.45 | 0.45 | 0.5 | 0.43 | 0.59 | 0.63 |
| 9 | 29595826 | rs580215 | 0.138 | 0.197 | -0.526 | -1.444 | -0.001 | -0.076 | -0.076 | 0 | 0.76 | 0.71 | 0.78 | 1 | 0.8 | 0.81 |
| 9 | 29596265 | rs1482069 | 0.018 | -0.263 | -0.199 | -1.299 | -0.009 | -0.017 | -0.017 | 0 | 0.4 | 0.44 | 0.34 | 0.74 | 0.4 | 0.32 |
| 9 | 29599001 | rs10757893 | 1.128 | 0.139 | 0.184 | -1.068 | -0.027 | 0.024 | 0.024 | 0 | 0.41 | 0.19 | 0.46 | 0.67 | 0.42 | 0.36 |
| 9 | 29599326 | rs4879368 | -0.199 | -0.231 | 0.521 | -1.229 | -0.001 | -0.026 | -0.026 | 0.886 | 0.92 | 0.83 | 0.88 | 1 | 0.94 | 0.87 |
| 9 | 29601893 | rs12345603 | -0.524 | -0.221 | 0.768 | -1.197 | 0.002 | 0.008 | 0.008 | 0 | 0.44 | 0.52 | 0.36 | 0.24 | 0.43 | 0.48 |
| 9 | 29614435 | rs10969287 | 0.407 | -0.235 | 0.269 | -1.167 | -0.031 | -0.05 | -0.05 | 0 | 0.26 | 0.31 | 0.14 | 0.11 | 0.18 | 0.23 |
| 9 | 29616943 | rs10757894 | 1.019 | -0.168 | 0.144 | -1.182 | -0.004 | 0.085 | 0.085 | 0 | 0.47 | 0.3 | 0.52 | 0.76 | 0.51 | 0.4 |
| 9 | 29618472 | rs10969289 | -0.114 | -0.452 | 0.025 | -0.703 | -0.05 | -0.059 | -0.059 | 1.517 | 0.26 | 0.4 | 0.19 | 0.14 | 0.18 | 0.29 |
| 9 | 29626137 | rs1930395 | 0.517 | 0.12 | -0.177 | -0.495 | -0.002 | -0.032 | -0.032 | 0 | 0.92 | 0.8 | 0.88 | 1 | 0.94 | 0.88 |
| 9 | 29630811 | rs7044847 | 0.006 | 0.205 | 0.308 | -0.658 | 0.013 | -0.011 | -0.011 | 0 | 0.2 | 0.33 | 0.23 | 0.17 | 0.26 | 0.21 |
| 9 | 29632142 | rs601162 | -0.254 | -0.066 | 0.143 | -1.101 | -0.019 | 0.078 | 0.078 | 0 | 0.79 | 0.91 | 0.79 | 0.92 | 0.81 | 0.72 |
| 9 | 29636965 | rs2128184 | 0.281 | 0.037 | 0.468 | -0.843 | 0.026 | 0.01 | 0.01 | 0.028 | 0.19 | 0.4 | 0.24 | 0.18 | 0.27 | 0.2 |
| 9 | 29639093 | rs12552131 | 0.705 | 0.006 | 0.592 | -0.855 | -0.002 | 0.001 | 0.001 | 0 | 0.16 | 0.16 | 0.16 | 0.12 | 0.23 | 0.12 |
| 9 | 29639398 | rs12379623 | 1.205 | -0.149 | 0.078 | -0.898 | -0.108 | 0.016 | 0.016 | 0 | 0.44 | 0.04 | 0.45 | 0.66 | 0.45 | 0.45 |
| 9 | 29640181 | rs7847579 | 0.246 | -0.146 | 0.243 | -0.954 | 0.014 | -0.013 | -0.013 | 0 | 0.2 | 0.37 | 0.24 | 0.17 | 0.27 | 0.21 |
| 9 | 29647032 | rs10813149 | 1.137 | 0.017 | -0.52 | -0.929 | -0.096 | 0.082 | 0.082 | 0 | 0.43 | 0.06 | 0.46 | 0.71 | 0.44 | 0.42 |
| 9 | 29653136 | rs7863931 | 0.662 | 0.32 | -0.129 | -0.65 | -0.011 | 0.007 | 0.007 | 0 | 0.16 | 0.12 | 0.16 | 0.12 | 0.24 | 0.12 |
| 9 | 29659560 | rs10969313 | 0.068 | 0.018 | -0.686 | -0.75 | 0.026 | 0.072 | 0.072 | 0 | 0.53 | 0.24 | 0.63 | 0.36 | 0.6 | 0.45 |
| 9 | 29661276 | rs7048366 | 0.046 | -0.003 | -0.697 | -0.765 | 0.027 | 0.072 | 0.072 | 0 | 0.53 | 0.23 | 0.62 | 0.36 | 0.6 | 0.44 |
| 9 | 29663089 | rs10813150 | 0.283 | 0.027 | -0.759 | -0.832 | 0.013 | 0.107 | 0.107 | 0 | 0.49 | 0.2 | 0.6 | 0.34 | 0.56 | 0.4 |
| 9 | 29664382 | rs7047143 | -0.946 | -0.256 | -0.473 | -0.927 | -0.001 | -0.038 | -0.038 | 0 | 0.79 | 0.89 | 0.86 | 0.46 | 0.8 | 0.8 |
| 9 | 29666098 | rs2086573 | 0.778 | -0.358 | -0.77 | -1.072 | -0.042 | 0.07 | 0.07 | 0 | 0.24 | 0.08 | 0.26 | 0.11 | 0.3 | 0.16 |
| 9 | 29675907 | rs10117742 | -0.633 | -0.339 | 0.226 | -0.91 | -0.006 | 0.045 | 0.045 | 0 | 0.77 | 0.69 | 0.83 | 0.46 | 0.77 | 0.74 |
| 9 | 29678786 | rs16915158 | -1.417 | -0.317 | 0.565 | -1.051 | 0.052 | -0.059 | -0.059 | 0 | 0.12 | 0.49 | 0.07 | 0.04 | 0.09 | 0.19 |
| 9 | 29679114 | rs10114966 | 0.624 | -0.34 | 0.466 | -1.057 | 0.002 | -0.018 | -0.018 | 0.955 | 0.65 | 0.21 | 0.76 | 0.42 | 0.68 | 0.59 |
| 9 | 29685572 | rs6476148 | 0.048 | -0.313 | 0.634 | -1.048 | -0.009 | -0.1 | -0.1 | 1.255 | 0.28 | 0.5 | 0.24 | 0.11 | 0.22 | 0.25 |
| 9 | 29691094 | rs12553141 | 0.534 | -0.332 | 0.827 | -0.955 | -0.004 | -0.029 | -0.029 | 0.471 | 0.23 | 0.27 | 0.2 | 0.08 | 0.18 | 0.25 |
| 9 | 29699426 | rs4574931 | -0.907 | -0.445 | 0.705 | -0.822 | 0 | 0.007 | 0.007 | 0 | 0.79 | 0.84 | 0.87 | 0.47 | 0.81 | 0.76 |
| 9 | 29700159 | rs1541132 | 0.432 | -0.449 | 0.72 | -0.847 | 0.012 | 0.023 | 0.023 | 0 | 0.56 | 0.39 | 0.66 | 0.37 | 0.64 | 0.47 |
| 9 | 29706139 | rs12555325 | 0.222 | -0.185 | 0.892 | -0.677 | -0.02 | -0.083 | -0.083 | 0 | 0.24 | 0.46 | 0.22 | 0.08 | 0.17 | 0.21 |
| 9 | 29713272 | rs2383834 | -0.646 | -0.219 | 0.452 | 0.119 | -0.004 | -0.001 | -0.001 | 0 | 0.54 | 0.52 | 0.64 | 0.53 | 0.57 | 0.44 |
| 9 | 29715833 | rs12375805 | 1.49 | -0.146 | 0.111 | 0.265 | -0.004 | 0.016 | 0.016 | 0 | 0.05 | 0 | 0.14 | 0.31 | 0.04 | 0.07 |
| 9 | 29718934 | rs13285631 | 1.16 | 0.043 | 0.262 | 0.201 | 0.028 | -0.035 | -0.035 | 0 | 0.36 | 0.1 | 0.28 | 0.41 | 0.31 | 0.36 |
| 9 | 29729921 | rs7859217 | 0.246 | 0.095 | 0.24 | 0.114 | -0.04 | -0.066 | -0.066 | 1.682 | 0.34 | 0.16 | 0.42 | 0.44 | 0.38 | 0.25 |
| 9 | 29730581 | rs6476151 | -0.72 | 0.075 | 0.764 | 0.086 | -0.006 | -0.002 | -0.002 | 0.021 | 0.57 | 0.57 | 0.65 | 0.55 | 0.58 | 0.56 |
| 9 | 29732140 | rs10757901 | 0.375 | 0.07 | 0.509 | 0.55 | -0.05 | 0.005 | 0.005 | 0 | 0.33 | 0.14 | 0.43 | 0.27 | 0.37 | 0.2 |
| 9 | 29736012 | rs12344997 | 1.12 | 0.146 | 0.939 | 0.432 | -0.01 | -0.01 | -0.01 | 0 | 0.18 | 0.28 | 0.18 | 0.12 | 0.13 | 0.3 |
| 9 | 29737005 | rs10757903 | -0.436 | 0.176 | 0.994 | 0.538 | 0 | -0.002 | -0.002 | 0 | 0.52 | 0.52 | 0.6 | 0.36 | 0.5 | 0.42 |
| 9 | 29739205 | rs928636 | 1.303 | 0.192 | 0.829 | 0.686 | 0.016 | -0.004 | -0.004 | 0 | 0.34 | 0.14 | 0.28 | 0.36 | 0.3 | 0.33 |
| 9 | 29740463 | rs1335172 | -0.485 | 0.361 | 1.458 | 0.307 | -0.001 | 0.008 | 0.008 | 0 | 0.51 | 0.57 | 0.6 | 0.38 | 0.49 | 0.52 |
| 9 | 29748884 | rs1932131 | -0.464 | 0.363 | 1.506 | 0.308 | 0 | 0.005 | 0.005 | 0 | 0.5 | 0.56 | 0.59 | 0.38 | 0.48 | 0.52 |
| 9 | 29750167 | rs1932133 | -0.926 | 0.376 | 1.464 | 0.307 | 0.007 | 0.005 | 0.005 | 0 | 0.54 | 0.68 | 0.65 | 0.38 | 0.55 | 0.55 |
| 9 | 29751705 | rs10813161 | 0.627 | 0.344 | 1.374 | 0.304 | -0.05 | -0.007 | -0.007 | 0 | 0.29 | 0.1 | 0.44 | 0.26 | 0.35 | 0.18 |
| 9 | 29753393 | rs4437734 | 0.858 | 1.011 | 1.618 | 0.303 | 0.01 | -0.014 | -0.014 | 0 | 0.19 | 0.25 | 0.19 | 0.1 | 0.2 | 0.21 |
| 9 | 29753415 | rs4471130 | 0.946 | 0.392 | 1.652 | 0.328 | -0.009 | -0.016 | -0.016 | 0 | 0.18 | 0.28 | 0.15 | 0.1 | 0.14 | 0.21 |
| 9 | 29758063 | rs972133 | -0.32 | 0.572 | 1.131 | 0.726 | 0.012 | 0.003 | 0.003 | 0 | 0.56 | 0.72 | 0.65 | 0.41 | 0.56 | 0.57 |
| 9 | 29761393 | rs1467825 | -0.394 | 1.031 | 1.068 | 0.592 | 0.008 | -0.012 | -0.012 | 0 | 0.05 | 0.1 | 0.05 | 0.01 | 0.08 | 0.04 |
| 9 | 29783989 | rs10969375 | -1.494 | 0.977 | 1.39 | 0.839 | 0.054 | 0.067 | 0.067 | 1.356 | 0.1 | 0.27 | 0.15 | 0.15 | 0.16 | 0.18 |
| 9 | 29785304 | rs10969376 | 0.396 | 0.832 | 1.056 | 0.79 | -0.036 | -0.022 | -0.022 | 0.94 | 0.43 | 0.12 | 0.39 | 0.46 | 0.47 | 0.47 |
| 9 | 29788364 | rs10813168 | -1.019 | 0.859 | 1.12 | 0.808 | 0.059 | 0.072 | 0.072 | 0.437 | 0.09 | 0.24 | 0.15 | 0.15 | 0.16 | 0.18 |
| 9 | 29788387 | rs10738825 | 0.221 | 0.801 | 1.72 | 0.715 | 0.036 | 0.013 | 0.013 | 0 | 0.92 | 0.65 | 0.94 | 0.98 | 0.95 | 0.97 |
| 9 | 29789814 | rs950316 | 0.242 | 0.757 | 1.698 | 0.665 | -0.004 | -0.001 | -0.001 | 0 | 0.46 | 0.4 | 0.55 | 0.48 | 0.49 | 0.32 |
| 9 | 29791410 | rs4879382 | 0.525 | 0.738 | 1.317 | 0.675 | -0.006 | -0.007 | -0.007 | 0 | 0.43 | 0.16 | 0.39 | 0.47 | 0.45 | 0.47 |
| 9 | 29791863 | rs2184513 | 0.881 | 0.104 | 1.357 | 0.749 | 0 | -0.003 | -0.003 | 1.032 | 0.36 | 0.12 | 0.4 | 0.34 | 0.33 | 0.16 |
| 9 | 29800646 | rs10969382 | 0.958 | 0.021 | 1.442 | 0.782 | 0.007 | -0.016 | -0.016 | 1.191 | 0.35 | 0.12 | 0.39 | 0.35 | 0.31 | 0.14 |
| 9 | 29803895 | rs10969385 | -1.463 | -0.436 | 2.429 | 0.584 | 0.035 | 0.222 | 0.222 | 2.32 | 0.05 | 0.35 | 0.05 | 0.14 | 0.02 | 0.11 |
| 9 | 29813230 | rs1360798 | 2.69 | -0.515 | 2.289 | 0.515 | -0.037 | 0.018 | 0.018 | 1.241 | 0.82 | 0.3 | 0.9 | 0.81 | 0.91 | 0.7 |
| 9 | 29815766 | rs7022877 | -2.535 | -0.517 | 2.389 | 0.516 | -0.021 | 0.124 | 0.124 | 2.868 | 0.11 | 0.6 | 0.06 | 0.18 | 0.04 | 0.27 |
| 9 | 29816084 | rs10117882 | -1.491 | -0.506 | 2.27 | 0.524 | 0.025 | 0.219 | 0.219 | 3.345 | 0.06 | 0.4 | 0.05 | 0.15 | 0.02 | 0.26 |
| 9 | 29816465 | rs1414029 | 2.111 | -0.485 | 1.572 | 0.522 | -0.037 | 0.018 | 0.018 | 1.241 | 0.82 | 0.3 | 0.9 | 0.81 | 0.91 | 0.7 |
| 9 | 29823067 | rs12352814 | -1.679 | -0.413 | 1.831 | 0.35 | -0.01 | -0.066 | -0.066 | 0.307 | 0.13 | 0.37 | 0.09 | 0.05 | 0.08 | 0.08 |
| 9 | 29824057 | rs10738827 | -0.205 | -0.387 | 1.774 | 0.266 | 0.004 | 0.005 | 0.005 | 0 | 0.57 | 0.65 | 0.61 | 0.61 | 0.6 | 0.5 |

| | | | | | | | | | | | | | | | | |
|---|---|---|---|---|---|---|---|---|---|---|---|---|---|---|---|---|
| 9 | 29825925 | rs10969394 | 1.085 | -0.42 | 1.825 | 0.254 | -0.069 | -0.03 | -0.03 | 0 | 0.42 | 0.1 | 0.51 | 0.55 | 0.51 | 0.26 |
| 9 | 29831400 | rs1543729 | -0.238 | -0.49 | 2.23 | 0.225 | 0.006 | 0.009 | 0.009 | 0 | 0.57 | 0.7 | 0.61 | 0.61 | 0.59 | 0.49 |
| 9 | 29833969 | rs12345817 | 0.158 | -0.462 | 1.861 | 0.186 | -0.007 | -0.022 | -0.022 | 0 | 0.06 | 0.11 | 0.03 | 0.01 | 0.05 | 0.03 |
| 9 | 29845253 | rs2152210 | 0.891 | -0.452 | 1.706 | 0.199 | -0.015 | 0.006 | 0.006 | 0 | 0.43 | 0.27 | 0.52 | 0.56 | 0.52 | 0.28 |
| 9 | 29849587 | rs7025492 | -0.623 | -0.602 | 1.212 | 0.032 | -0.03 | 0.064 | 0.064 | 0 | 0.67 | 0.89 | 0.66 | 0.76 | 0.65 | 0.63 |
| 9 | 29861353 | rs12347202 | -0.116 | -0.494 | 1.549 | 0.277 | -0.011 | -0.032 | -0.032 | 0.396 | 0.07 | 0.15 | 0.03 | 0 | 0.05 | 0.02 |
| 9 | 29869252 | rs10813187 | 1.241 | -0.775 | 1.209 | 0.155 | -0.018 | -0.005 | -0.005 | 0 | 0.45 | 0.11 | 0.56 | 0.6 | 0.56 | 0.3 |
| 9 | 29881379 | rs10813194 | 1.247 | -0.807 | 1.276 | 0.161 | 0 | 0.012 | 0.012 | 0 | 0.45 | 0.16 | 0.56 | 0.6 | 0.56 | 0.3 |
| 9 | 29894841 | rs7871951 | -0.521 | -0.826 | 1.833 | 0.174 | 0.002 | 0.001 | 0.001 | 0 | 0.43 | 0.4 | 0.34 | 0.38 | 0.36 | 0.44 |
| 9 | 29900225 | rs10969441 | -1.32 | -0.408 | 2.376 | 0.541 | -0.028 | -0.063 | -0.063 | 0 | 0.1 | 0.29 | 0.04 | 0 | 0.05 | 0.03 |
| 9 | 29903903 | rs6476171 | -2.254 | -0.411 | 2.217 | 0.388 | 0.022 | 0.022 | 0.022 | 0 | 0.44 | 0.79 | 0.41 | 0.4 | 0.39 | 0.67 |
| 9 | 29904043 | rs7048031 | -1.17 | -0.398 | 2.19 | 0.412 | 0.002 | 0.008 | 0.008 | 0 | 0.31 | 0.38 | 0.34 | 0.38 | 0.32 | 0.45 |
| 9 | 29911013 | rs6476173 | -0.783 | -0.267 | 1.77 | 0.169 | -0.022 | -0.019 | -0.019 | 0 | 0.05 | 0.09 | 0.03 | 0.02 | 0.01 | 0.21 |
| 9 | 29919239 | rs9408158 | -0.505 | -0.135 | 1.492 | 0.264 | -0.05 | 0.008 | 0.008 | 0 | 0.23 | 0.07 | 0.28 | 0.24 | 0.3 | 0.34 |
| 9 | 29928182 | rs12345491 | -0.267 | -0.182 | 1.768 | 0.343 | -0.009 | -0.026 | -0.026 | 0 | 0.06 | 0.11 | 0.02 | 0 | 0.05 | 0.01 |
| 9 | 29938824 | rs2492624 | -1.463 | -0.185 | 1 | 0.996 | 0.009 | -0.007 | -0.007 | 0.061 | 0.39 | 0.69 | 0.42 | 0.62 | 0.34 | 0.62 |
| 9 | 29938935 | rs2502106 | -0.666 | -0.039 | 1.305 | 1.217 | -0.001 | -0.01 | -0.01 | 0 | 0.2 | 0.17 | 0.25 | 0.39 | 0.18 | 0.28 |
| 9 | 29939556 | rs2492626 | -1.829 | -0.155 | 1.539 | 1.127 | 0.002 | -0.006 | -0.006 | 0.771 | 0.37 | 0.58 | 0.48 | 0.53 | 0.32 | 0.57 |
| 9 | 29940394 | rs4415403 | -1.243 | -0.077 | 1.715 | 1.151 | 0.035 | 0.123 | 0.123 | 0 | 0.16 | 0.35 | 0.14 | 0.25 | 0.18 | 0.34 |
| 9 | 29942472 | rs10813204 | -0.842 | -0.1 | 1.45 | 1.181 | 0.01 | 0.061 | 0.061 | 0 | 0.12 | 0.18 | 0.1 | 0.24 | 0.16 | 0.3 |
| 9 | 29943307 | rs9408166 | -0.479 | -0.138 | 1.04 | 1.177 | 0.011 | 0.012 | 0.012 | 0 | 0.23 | 0.13 | 0.35 | 0.27 | 0.2 | 0.22 |
| 9 | 29943652 | rs10813205 | 1.98 | 0.269 | 0.574 | 1.049 | -0.053 | -0.063 | -0.063 | 0 | 0.51 | 0.1 | 0.43 | 0.47 | 0.49 | 0.37 |
| 9 | 29950775 | rs7020357 | -0.608 | 0.414 | 0.827 | 0.833 | 0.002 | 0.016 | 0.016 | 0 | 0.93 | 0.92 | 0.95 | 0.88 | 0.9 | 0.9 |
| 9 | 29951544 | rs10969475 | 1.681 | 0.422 | 0.479 | 0.853 | -0.052 | 0.059 | 0.059 | 0 | 0.53 | 0.12 | 0.45 | 0.64 | 0.51 | 0.58 |
| 9 | 29958595 | rs10511835 | -1.297 | 0.547 | 0.81 | 0.722 | -0.001 | -0.001 | -0.001 | 0 | 0.35 | 0.34 | 0.45 | 0.24 | 0.38 | 0.31 |
| 9 | 29963072 | rs16915458 | -0.848 | 0.447 | 1.126 | 0.424 | 0 | -0.052 | -0.052 | 0 | 0.17 | 0.22 | 0.11 | 0 | 0.15 | 0.09 |
| 9 | 29964976 | rs7027471 | -0.697 | 0.456 | 1.058 | 0.43 | 0 | -0.039 | -0.039 | 0 | 0.15 | 0.21 | 0.11 | 0 | 0.15 | 0.07 |
| 9 | 29976426 | rs10813211 | 1.861 | 0.665 | 0.71 | 0.744 | -0.035 | -0.038 | -0.038 | 0 | 0.57 | 0.18 | 0.45 | 0.46 | 0.51 | 0.39 |
| 9 | 29978410 | rs12236822 | 0.094 | 0.688 | 0.734 | 0.926 | -0.03 | -0.042 | -0.042 | 0 | 0.17 | 0.07 | 0.34 | 0.34 | 0.25 | 0.27 |
| 9 | 29985815 | rs6476179 | -0.892 | 0.473 | 1.698 | 0.871 | 0.003 | -0.007 | -0.007 | 0 | 0.16 | 0.4 | 0.13 | 0.17 | 0.16 | 0.23 |
| 9 | 29985893 | rs9298912 | -0.14 | 0.456 | 2.264 | 0.864 | -0.003 | -0.001 | -0.001 | 0 | 0.07 | 0.14 | 0.04 | 0.03 | 0.06 | 0.09 |
| 9 | 29986250 | rs2813927 | -1.308 | 0.443 | 2.736 | 0.859 | 0.005 | -0.003 | -0.003 | 0.119 | 0.21 | 0.37 | 0.16 | 0.2 | 0.21 | 0.32 |
| 9 | 29989597 | rs2360192 | -1.326 | 0.422 | 2.73 | 0.835 | 0.005 | -0.075 | -0.075 | 0.369 | 0.21 | 0.44 | 0.16 | 0.03 | 0.2 | 0.15 |
| 9 | 29991337 | rs10121528 | -1.788 | 0.416 | 2.753 | 0.863 | -0.003 | -0.031 | -0.031 | 0 | 0.23 | 0.54 | 0.18 | 0.19 | 0.22 | 0.32 |
| 9 | 29991403 | rs16915477 | 0.104 | 0.438 | 2.65 | 0.821 | -0.013 | 0.012 | 0.012 | 0 | 0.11 | 0.08 | 0.09 | 0 | 0.14 | 0.06 |
| 9 | 29993527 | rs10969497 | -1.229 | 0.439 | 2.481 | 0.831 | 0.005 | -0.075 | -0.075 | 0 | 0.21 | 0.44 | 0.16 | 0.03 | 0.2 | 0.15 |
| 9 | 30002855 | rs9408174 | -1.177 | 0.485 | 2.627 | 1.03 | 0.009 | -0.07 | -0.07 | 0 | 0.2 | 0.41 | 0.16 | 0.03 | 0.2 | 0.15 |
| 9 | 30006830 | rs9298914 | -2.126 | 0.494 | 2.463 | 1.059 | -0.011 | -0.014 | -0.014 | 0 | 0.43 | 0.7 | 0.56 | 0.55 | 0.5 | 0.6 |
| 9 | 30009444 | rs1962543 | -1.88 | 0.554 | 2.314 | 1.008 | 0.016 | -0.003 | -0.003 | 0 | 0.4 | 0.53 | 0.53 | 0.39 | 0.48 | 0.43 |
| 9 | 30010493 | rs10969511 | 0 | 0.607 | 1.852 | 1.087 | -0.051 | -0.056 | -0.056 | 0 | 0.19 | 0.07 | 0.38 | 0.35 | 0.28 | 0.28 |
| 9 | 30022860 | rs9406958 | 0.511 | 0.558 | 2.034 | 1.183 | 0.031 | 0.063 | 0.063 | 0 | 0.74 | 0.51 | 0.59 | 0.58 | 0.7 | 0.71 |
| 9 | 30030188 | rs7032287 | 1.601 | 0.387 | 1.928 | 1.422 | -0.021 | -0.014 | -0.014 | 0 | 0.59 | 0.26 | 0.4 | 0.38 | 0.54 | 0.44 |
| 9 | 30038860 | rs12375503 | -0.169 | 0.394 | 1.993 | 1.615 | -0.013 | -0.017 | -0.017 | 0 | 0.22 | 0.21 | 0.39 | 0.36 | 0.29 | 0.25 |
| 9 | 30043590 | rs2804511 | 1.216 | 0.303 | 1.925 | 1.602 | 0.023 | 0.258 | 0.258 | 0 | 0.95 | 0.73 | 0.91 | 0.74 | 0.96 | 0.74 |
| 9 | 30057012 | rs2813941 | -0.986 | 0.276 | 1.834 | 1.594 | -0.011 | 0.089 | 0.089 | 0 | 0.09 | 0.24 | 0.11 | 0.23 | 0.09 | 0.31 |
| 9 | 30064199 | rs13299989 | 1.969 | -0.034 | 0.481 | 1.125 | 0.043 | 0.029 | 0.029 | 0.252 | 0.3 | 0.04 | 0.15 | 0.2 | 0.19 | 0.21 |
| 9 | 30064376 | rs9406965 | -0.53 | -0.017 | 0.566 | 1.026 | -0.014 | -0.013 | -0.013 | 0 | 0.1 | 0.22 | 0.04 | 0.03 | 0.07 | 0.09 |
| 9 | 30064499 | rs10969535 | -0.057 | 0.03 | 0.589 | 1.049 | -0.008 | -0.007 | -0.007 | 0 | 0.08 | 0.22 | 0.04 | 0.03 | 0.07 | 0.09 |
| 9 | 30067745 | rs10969539 | 0.613 | 0.461 | 0.109 | 1.167 | -0.031 | -0.057 | -0.057 | 0.689 | 0.19 | 0.07 | 0.36 | 0.35 | 0.24 | 0.23 |
| 9 | 30068009 | rs2360145 | -1.207 | 0.425 | 0.243 | 1.063 | 0.004 | -0.005 | -0.005 | 0.215 | 0.75 | 0.77 | 0.77 | 0.79 | 0.7 | 0.83 |
| 9 | 30075086 | rs10969547 | 0.268 | 0.486 | 0.162 | 1.236 | -0.05 | -0.058 | -0.058 | 0 | 0.2 | 0.07 | 0.38 | 0.35 | 0.28 | 0.25 |
| 9 | 30078773 | rs2360149 | 0.039 | 0.012 | 0.495 | 1.075 | 0.007 | 0.016 | 0.016 | 0 | 0.58 | 0.78 | 0.69 | 0.67 | 0.63 | 0.55 |
| 9 | 30080099 | rs1984299 | 0.226 | -0.151 | 0.78 | 1.039 | 0.002 | 0.003 | 0.003 | 0 | 0.55 | 0.62 | 0.68 | 0.67 | 0.62 | 0.55 |
| 9 | 30087981 | rs10969554 | -2.075 | -0.164 | 0.875 | 1.324 | -0.006 | 0.234 | 0.234 | 1.821 | 0.07 | 0.38 | 0.1 | 0.24 | 0.02 | 0.17 |
| 9 | 30089859 | rs1780063 | -0.06 | -0.105 | 0.366 | 1.07 | 0.007 | 0.015 | 0.015 | 0 | 0.59 | 0.8 | 0.69 | 0.67 | 0.64 | 0.57 |
| 9 | 30094419 | rs1633490 | 1.666 | 0.452 | 0.797 | 1.593 | 0.003 | 0.009 | 0.009 | 0.371 | 0.67 | 0.39 | 0.65 | 0.41 | 0.65 | 0.44 |
| 9 | 30101419 | rs2804505 | 0.113 | 0.442 | 0.962 | 1.548 | 0.016 | 0.022 | 0.022 | 0 | 0.93 | 0.87 | 0.88 | 0.89 | 0.91 | 0.86 |
| 9 | 30103515 | rs2245665 | 0.891 | 0.348 | 0.973 | 1.211 | 0.007 | 0.016 | 0.016 | 0.3 | 0.72 | 0.66 | 0.68 | 0.4 | 0.7 | 0.46 |
| 9 | 30106229 | rs1780055 | 1.194 | 0.393 | 0.677 | 1.233 | 0.004 | 0.016 | 0.016 | 0.287 | 0.7 | 0.4 | 0.66 | 0.4 | 0.7 | 0.46 |
| 9 | 30115050 | rs1633498 | 0.746 | 0.85 | -0.405 | 0.981 | -0.006 | -0.008 | -0.008 | 0 | 0.77 | 0.81 | 0.76 | 0.63 | 0.71 | 0.61 |
| 9 | 30122786 | rs1510909 | -0.874 | 0.677 | -0.303 | 0.1 | 0.001 | 0.062 | 0.062 | 0.098 | 0.43 | 0.72 | 0.4 | 0.16 | 0.4 | 0.41 |
| 9 | 30138908 | rs4878424 | 0.045 | 0.394 | 0.748 | -0.073 | -0.053 | -0.032 | -0.032 | 0.579 | 0.19 | 0.37 | 0.15 | 0.12 | 0.09 | 0.22 |
| 9 | 30143230 | rs1018254 | -0.382 | 0.494 | 0.953 | -0.171 | -0.001 | -0.013 | -0.013 | 0 | 0.24 | 0.26 | 0.25 | 0.05 | 0.32 | 0.21 |
| 9 | 30158784 | rs7872835 | -0.534 | 0.217 | 1.32 | -0.191 | -0.007 | -0.072 | -0.072 | 0 | 0.29 | 0.34 | 0.18 | 0.03 | 0.23 | 0.11 |
| 9 | 30175188 | rs13290342 | 0.433 | 0.196 | 1.33 | -0.2 | 0.008 | -0.06 | -0.06 | 0.108 | 0.15 | 0.26 | 0.14 | 0.03 | 0.16 | 0.05 |
| 9 | 30181745 | rs2800343 | 0.99 | 0.013 | 1.118 | 0.289 | 0.052 | 0.027 | 0.027 | 0.321 | 0.63 | 0.18 | 0.7 | 0.74 | 0.76 | 0.69 |
| 9 | 30194560 | rs1419375 | -0.688 | 0.167 | 0.697 | 0.345 | 0.018 | 0.007 | 0.007 | 0 | 0.86 | 0.95 | 0.88 | 0.89 | 0.92 | 0.8 |
| 9 | 30207721 | rs2800328 | -0.039 | 0.097 | 1.115 | 0.32 | -0.056 | -0.03 | -0.03 | 0 | 0.82 | 0.62 | 0.86 | 0.89 | 0.92 | 0.79 |
| 9 | 30207872 | rs1510913 | 1.722 | 0.263 | 0.294 | 0.334 | -0.036 | -0.114 | -0.114 | 0 | 0.31 | 0.05 | 0.38 | 0.5 | 0.34 | 0.31 |

| | | | | | | | | | | | | | | | | |
|---|---|---|---|---|---|---|---|---|---|---|---|---|---|---|---|---|
| 9 | 30212447 | rs2769635 | 0.187 | 0.285 | 0.803 | 0.483 | -0.056 | -0.03 | -0.03 | 0 | 0.18 | 0.38 | 0.14 | 0.11 | 0.09 | 0.21 |
| 9 | 30219494 | rs2769643 | -0.773 | 0.347 | 0.967 | 0.337 | -0.001 | -0.01 | -0.01 | 0.002 | 0.33 | 0.6 | 0.28 | 0.14 | 0.24 | 0.25 |
| 9 | 30222006 | rs1543047 | -0.232 | 0.386 | 1.387 | 0.383 | 0.009 | -0.012 | -0.012 | 0 | 0.82 | 0.61 | 0.78 | 0.86 | 0.84 | 0.81 |
| 9 | 30240159 | rs9696659 | 0.457 | 0.404 | 1.478 | 0.432 | 0.003 | -0.04 | -0.04 | 0.708 | 0.16 | 0.22 | 0.15 | 0.03 | 0.17 | 0.05 |
| 9 | 30247932 | rs12339642 | -0.388 | 0.336 | 1.605 | 0.427 | -0.054 | 0.018 | 0.018 | 0.349 | 0.23 | 0.58 | 0.17 | 0.22 | 0.1 | 0.26 |
| 9 | 30253168 | rs7021842 | 0.633 | 0.424 | 1.557 | 0.42 | 0.003 | -0.005 | -0.005 | 0 | 0.16 | 0.22 | 0.21 | 0.14 | 0.16 | 0.19 |
| 9 | 30255328 | rs1328165 | -1.21 | 0.419 | 1.562 | 0.416 | 0.045 | 0.038 | 0.038 | 0.383 | 0.38 | 0.83 | 0.3 | 0.25 | 0.26 | 0.31 |
| 9 | 30257784 | rs10813235 | 0.664 | 0.644 | 1.413 | 0.336 | -0.05 | 0.015 | 0.015 | 0 | 0.08 | 0.01 | 0.11 | 0.02 | 0.16 | 0.14 |
| 9 | 30257830 | rs7867533 | 0.665 | 0.358 | 1.584 | 0.392 | 0.004 | -0.042 | -0.042 | 0 | 0.14 | 0.22 | 0.14 | 0.03 | 0.15 | 0.05 |
| 9 | 30260156 | rs1576160 | 0.665 | 0.358 | 1.584 | 0.392 | 0.004 | -0.042 | -0.042 | 0 | 0.14 | 0.22 | 0.14 | 0.03 | 0.15 | 0.05 |
| 9 | 30276852 | rs12338157 | 0.552 | 0.348 | 1.709 | 0.415 | 0.002 | -0.004 | -0.004 | 0 | 0.16 | 0.25 | 0.21 | 0.14 | 0.16 | 0.19 |
| 9 | 30280905 | rs12336564 | -0.421 | 0.366 | 1.509 | 0.403 | -0.016 | -0.139 | -0.139 | 0.137 | 0.31 | 0.43 | 0.2 | 0.03 | 0.24 | 0.11 |
| 9 | 30282776 | rs2149165 | -1.042 | 0.341 | 1.542 | 0.431 | 0.026 | 0.018 | 0.018 | 0.003 | 0.37 | 0.75 | 0.3 | 0.26 | 0.26 | 0.3 |
| 9 | 30286600 | rs10511857 | -0.501 | 0.417 | 1.418 | 0.226 | -0.017 | -0.144 | -0.144 | 0 | 0.31 | 0.44 | 0.2 | 0.03 | 0.25 | 0.11 |
| 9 | 30297393 | rs10124607 | -0.95 | 0.203 | 1.363 | 0.693 | 0.009 | 0.001 | 0.001 | 0.283 | 0.36 | 0.69 | 0.29 | 0.28 | 0.25 | 0.33 |
| 9 | 30303562 | rs10125266 | -0.445 | 0.186 | 1.475 | 0.742 | -0.016 | -0.078 | -0.078 | 0 | 0.3 | 0.43 | 0.22 | 0.08 | 0.24 | 0.11 |
| 9 | 30310098 | rs2225172 | -0.886 | 0.131 | 1.488 | 0.731 | -0.023 | -0.002 | -0.002 | 0 | 0.35 | 0.58 | 0.28 | 0.28 | 0.25 | 0.34 |
| 9 | 30318865 | rs9298919 | -0.474 | 0.134 | 1.466 | 0.712 | -0.017 | -0.081 | -0.081 | 0 | 0.31 | 0.43 | 0.22 | 0.08 | 0.24 | 0.11 |
| 9 | 30326044 | rs16915873 | 0.827 | 0.164 | 1.263 | 0.695 | 0.011 | 0.084 | 0.084 | 0 | 0.14 | 0.01 | 0.06 | 0 | 0.09 | 0.06 |
| 9 | 30326115 | rs3853264 | 0.678 | 0.166 | 1.416 | 0.695 | 0.004 | -0.015 | -0.015 | 0 | 0.14 | 0.21 | 0.16 | 0.08 | 0.14 | 0.04 |
| 9 | 30329054 | rs3853265 | -0.9 | 0.182 | 1.327 | 0.746 | -0.023 | -0.014 | -0.014 | 0 | 0.34 | 0.58 | 0.23 | 0.23 | 0.25 | 0.17 |
| 9 | 30338229 | rs10125259 | -1.086 | 0.178 | 1.314 | 0.711 | -0.021 | -0.001 | -0.001 | 0.493 | 0.35 | 0.6 | 0.28 | 0.29 | 0.25 | 0.34 |
| 9 | 30360841 | rs7029121 | -1.091 | 0.166 | 1.239 | 0.61 | 0.033 | 0.078 | 0.078 | 0.116 | 0.35 | 0.78 | 0.25 | 0.09 | 0.24 | 0.1 |
| 9 | 30362055 | rs16915923 | 0.562 | 0.15 | 1.321 | 0.608 | -0.013 | -0.055 | -0.055 | 0.14 | 0.16 | 0.23 | 0.07 | 0 | 0.09 | 0.06 |
| 9 | 30366782 | rs10969641 | -1.081 | 0.27 | 1.162 | 0.611 | 0.024 | 0.07 | 0.07 | 0.095 | 0.35 | 0.77 | 0.25 | 0.08 | 0.25 | 0.1 |
| 9 | 30375986 | rs3904444 | -1.117 | 0.261 | 1.147 | 0.624 | 0.033 | 0.08 | 0.08 | 0.337 | 0.36 | 0.78 | 0.3 | 0.13 | 0.26 | 0.28 |
| 9 | 30378901 | rs1502888 | 0.733 | 0.632 | 0.922 | 0.667 | 0.007 | -0.008 | -0.008 | 0.046 | 0.72 | 0.35 | 0.67 | 0.79 | 0.69 | 0.86 |
| 9 | 30381063 | rs10511860 | 1.179 | 0.62 | 0.778 | 0.66 | 0.003 | 0.083 | 0.083 | 0 | 0.14 | 0.01 | 0.06 | 0 | 0.09 | 0.06 |
| 9 | 30399295 | rs10813243 | 1.438 | 0.668 | 0.793 | 0.764 | 0.006 | -0.05 | -0.05 | 0.108 | 0.54 | 0.12 | 0.55 | 0.59 | 0.59 | 0.56 |
| 9 | 30401266 | rs6476201 | -0.965 | 0.353 | 0.961 | 0.686 | 0.032 | 0.005 | 0.005 | 0.111 | 0.36 | 0.78 | 0.3 | 0.28 | 0.27 | 0.34 |
| 9 | 30403952 | rs10813246 | -0.228 | 0.432 | 0.678 | 0.721 | -0.006 | 0.081 | 0.081 | 0.019 | 0.2 | 0.54 | 0.23 | 0.28 | 0.17 | 0.28 |
| 9 | 30409103 | rs3853269 | 0.657 | 0.426 | 0.299 | 0.748 | 0.039 | 0.003 | 0.003 | 0 | 0.63 | 0.21 | 0.68 | 0.72 | 0.72 | 0.65 |
| 9 | 30430337 | rs10118094 | 0.12 | 0.285 | 0.615 | 1.017 | -0.061 | 0.073 | 0.073 | 0.125 | 0.77 | 0.45 | 0.8 | 0.67 | 0.85 | 0.74 |
| 9 | 30438383 | rs10969671 | 1.742 | 0.577 | 0.232 | 0.995 | -0.016 | 0.139 | 0.139 | 0 | 0.42 | 0.12 | 0.37 | 0.18 | 0.41 | 0.33 |
| 9 | 30443903 | rs3853275 | 0.01 | 0.207 | 0.583 | 1.09 | -0.067 | 0.104 | 0.104 | 0.002 | 0.78 | 0.51 | 0.8 | 0.67 | 0.86 | 0.74 |
| 9 | 30447342 | rs10757953 | 0.625 | 0.295 | 0.231 | 1.102 | -0.022 | 0.017 | 0.017 | 0 | 0.63 | 0.43 | 0.68 | 0.59 | 0.72 | 0.7 |
| 9 | 30448896 | rs10969681 | 0.434 | 0.453 | -0.461 | 1.051 | -0.004 | -0.019 | -0.019 | 0 | 0.16 | 0.22 | 0.14 | 0.08 | 0.14 | 0.04 |
| 9 | 30453151 | rs3866499 | -0.082 | 0.494 | -1.166 | 1.117 | -0.043 | -0.036 | -0.036 | 0 | 0.08 | 0.01 | 0.14 | 0.11 | 0.11 | 0.11 |
| 9 | 30458347 | rs4879437 | 0.394 | -0.224 | -1.04 | 1.014 | 0.049 | -0.034 | -0.034 | 0.75 | 0.67 | 0.21 | 0.73 | 0.6 | 0.81 | 0.73 |
| 9 | 30467137 | rs10813259 | 0.114 | 0.204 | -1.345 | 0.848 | -0.022 | -0.008 | -0.008 | 0 | 0.26 | 0.14 | 0.22 | 0.34 | 0.29 | 0.27 |
| 9 | 30467975 | rs10813260 | -0.072 | 0.155 | -0.848 | 1.388 | 0.001 | 0.007 | 0.007 | 0 | 0.28 | 0.27 | 0.28 | 0.57 | 0.31 | 0.29 |
| 9 | 30468541 | rs10738838 | -0.163 | 0.118 | -0.795 | 1.472 | 0.002 | 0.013 | 0.013 | 0 | 0.28 | 0.28 | 0.32 | 0.62 | 0.32 | 0.37 |
| 9 | 30471240 | rs7020697 | 0.512 | -0.372 | -0.175 | 1.217 | -0.071 | 0.027 | 0.027 | 2.014 | 0.15 | 0.42 | 0.07 | 0.11 | 0.05 | 0.07 |
| 9 | 30484037 | rs10813262 | -0.05 | -0.358 | -0.304 | 1.186 | -0.053 | 0.004 | 0.004 | 0.996 | 0.81 | 0.3 | 0.84 | 0.85 | 0.91 | 0.83 |
| 9 | 30493749 | rs4532690 | -0.479 | -0.371 | -0.169 | 1.18 | -0.064 | 0.004 | 0.004 | 1.018 | 0.84 | 0.4 | 0.89 | 0.89 | 0.91 | 0.92 |
| 9 | 30500252 | rs4878432 | -0.089 | -0.365 | -0.215 | 1.182 | -0.05 | 0.004 | 0.004 | 1.008 | 0.82 | 0.3 | 0.84 | 0.85 | 0.91 | 0.83 |
| 9 | 30504349 | rs17775810 | 0.731 | -0.144 | -0.755 | 0.921 | -0.06 | 0.023 | 0.023 | 0.059 | 0.2 | 0.01 | 0.2 | 0.09 | 0.21 | 0.18 |
| 9 | 30504425 | rs10969704 | 1.694 | -0.071 | -0.908 | 0.837 | -0.081 | 0.034 | 0.034 | 0 | 0.3 | 0.04 | 0.34 | 0.17 | 0.35 | 0.26 |
| 9 | 30521752 | rs10813280 | -0.035 | -0.197 | -1.202 | 0.413 | -0.013 | -0.01 | -0.01 | 0.078 | 0.76 | 0.3 | 0.84 | 0.88 | 0.84 | 0.91 |
| 9 | 30524250 | rs10813281 | 1.078 | -0.269 | -1.124 | 0.684 | -0.025 | -0.024 | -0.024 | 0 | 0.21 | 0.1 | 0.19 | 0.34 | 0.22 | 0.26 |
| 9 | 30555391 | rs10117990 | 1.023 | -0.28 | -0.73 | 0.641 | -0.042 | 0.011 | 0.011 | 0.24 | 0.15 | 0.32 | 0.08 | 0.12 | 0.09 | 0.08 |
| 9 | 30567014 | rs10813300 | 0.512 | -0.275 | -0.934 | 0.648 | -0.002 | 0.036 | 0.036 | 0.276 | 0.56 | 0.06 | 0.58 | 0.27 | 0.64 | 0.55 |
| 9 | 30584575 | rs10969757 | 0.475 | -0.273 | -0.871 | 0.65 | 0.028 | 0.055 | 0.055 | 0.205 | 0.56 | 0.11 | 0.59 | 0.27 | 0.65 | 0.56 |
| 9 | 30591737 | rs1104787 | 0.714 | -0.285 | -0.796 | 0.7 | -0.006 | -0.004 | -0.004 | 0 | 0.25 | 0.2 | 0.2 | 0.34 | 0.24 | 0.26 |
| 9 | 30598864 | rs10757981 | -0.248 | -0.288 | -0.749 | 0.692 | -0.001 | 0.04 | 0.04 | 0.021 | 0.71 | 0.4 | 0.66 | 0.39 | 0.73 | 0.64 |
| 9 | 30604262 | rs10969777 | 0.454 | -0.256 | -1.059 | 0.639 | 0.018 | 0.04 | 0.04 | 0.296 | 0.56 | 0.07 | 0.59 | 0.27 | 0.64 | 0.56 |
| 9 | 30613669 | rs12380109 | 0.502 | -0.266 | -1.107 | 0.629 | 0.011 | 0.039 | 0.039 | 0.297 | 0.56 | 0.07 | 0.58 | 0.27 | 0.64 | 0.56 |
| 9 | 30619070 | rs10969796 | 0.567 | -0.195 | -1.013 | 0.675 | -0.013 | -0.013 | -0.013 | 0 | 0.25 | 0.14 | 0.19 | 0.35 | 0.24 | 0.26 |
| 9 | 30621116 | rs10813324 | 1.007 | -0.128 | -1.217 | 0.476 | -0.077 | 0.045 | 0.045 | 0 | 0.35 | 0.04 | 0.36 | 0.18 | 0.37 | 0.31 |
| 9 | 30641824 | rs12004245 | -0.706 | -0.249 | -0.3 | 0.827 | 0.023 | 0.21 | 0.21 | 0 | 0.05 | 0.24 | 0.08 | 0.24 | 0.05 | 0.02 |
| 9 | 30645432 | rs12344302 | -0.939 | -0.26 | -0.306 | 0.86 | 0.015 | 0.225 | 0.225 | 0 | 0.06 | 0.24 | 0.13 | 0.27 | 0.06 | 0.1 |
| 9 | 30645756 | rs13302787 | 2.065 | -0.384 | -0.659 | 0.615 | -0.027 | 0.033 | 0.033 | 0 | 0.11 | 0.01 | 0.21 | 0.04 | 0.13 | 0.08 |
| 9 | 30646764 | rs7867594 | -0.562 | -0.358 | -0.264 | 0.887 | 0.021 | 0.207 | 0.207 | 0 | 0.05 | 0.24 | 0.08 | 0.24 | 0.05 | 0.02 |
| 9 | 30646780 | rs10969808 | 0.694 | -0.443 | -0.269 | 0.825 | -0.001 | -0.002 | -0.002 | 0 | 0.24 | 0.28 | 0.21 | 0.35 | 0.21 | 0.26 |
| 9 | 30647718 | rs1033216 | 0.111 | -0.528 | -0.416 | 0.795 | 0.011 | -0.009 | -0.009 | 0 | 0.61 | 0.3 | 0.71 | 0.53 | 0.69 | 0.65 |
| 9 | 30656199 | rs7857459 | -0.71 | -0.394 | -0.281 | 0.863 | 0.012 | 0.218 | 0.218 | 0 | 0.06 | 0.24 | 0.13 | 0.27 | 0.06 | 0.1 |
| 9 | 30666675 | rs10125968 | 0.104 | -0.427 | -0.329 | 0.845 | -0.056 | 0.098 | 0.098 | 0.028 | 0.21 | 0.59 | 0.2 | 0.38 | 0.15 | 0.18 |
| 9 | 30668911 | rs9650700 | 0.599 | -0.362 | -0.44 | 0.795 | -0.003 | 0.039 | 0.039 | 0.331 | 0.56 | 0.05 | 0.58 | 0.27 | 0.64 | 0.56 |
| 9 | 30669429 | rs10738851 | 0.433 | -0.452 | -0.32 | 0.809 | -0.053 | 0.009 | 0.009 | 0 | 0.19 | 0.42 | 0.14 | 0.14 | 0.12 | 0.17 |

| | | | | | | | | | | | | | | | | |
|---|---|---|---|---|---|---|---|---|---|---|---|---|---|---|---|---|
| 9 | 30677333 | rs12337598 | 0.177 | -0.464 | -0.379 | 0.759 | -0.055 | 0.105 | 0.105 | 0.044 | 0.21 | 0.59 | 0.2 | 0.38 | 0.15 | 0.19 |
| 9 | 30679704 | rs3849892 | -0.949 | -0.441 | -0.4 | 0.753 | 0.013 | 0.019 | 0.019 | 0 | 0.77 | 0.65 | 0.79 | 0.65 | 0.78 | 0.74 |
| 9 | 30695029 | rs7872163 | 0.296 | -0.411 | -0.411 | 0.788 | -0.06 | 0.169 | 0.169 | 0 | 0.19 | 0.5 | 0.19 | 0.38 | 0.14 | 0.18 |
| 9 | 30698142 | rs7027645 | 0.734 | -0.063 | -0.425 | 0.862 | 0 | 0.001 | 0.001 | 0 | 0.23 | 0.28 | 0.21 | 0.35 | 0.22 | 0.26 |
| 9 | 30713637 | rs10969834 | 1.15 | -0.074 | -0.507 | 0.841 | -0.032 | -0.038 | -0.038 | 0.092 | 0.2 | 0.07 | 0.18 | 0.35 | 0.22 | 0.26 |
| 9 | 30753998 | rs10969854 | 1.229 | -0.013 | -0.915 | 0.602 | -0.103 | 0.053 | 0.053 | 0 | 0.41 | 0.04 | 0.47 | 0.21 | 0.46 | 0.41 |
| 9 | 30770699 | rs2791599 | 0.627 | 0.612 | -0.252 | 0.805 | 0.007 | 0.061 | 0.061 | 0 | 0.19 | 0.35 | 0.23 | 0.33 | 0.2 | 0.24 |
| 9 | 30782465 | rs2249529 | 1.165 | 0.647 | -0.336 | 0.802 | -0.005 | 0.012 | 0.012 | 0 | 0.16 | 0.08 | 0.16 | 0.07 | 0.17 | 0.21 |
| 9 | 30795437 | rs1888892 | 1.005 | 0.657 | -0.402 | 0.677 | 0.003 | 0.02 | 0.02 | 0 | 0.17 | 0.15 | 0.22 | 0.33 | 0.2 | 0.19 |
| 9 | 30797625 | rs10969868 | 0.338 | 0.368 | -0.338 | 0.961 | -0.01 | 0.005 | 0.005 | 0 | 0.55 | 0.37 | 0.55 | 0.33 | 0.55 | 0.5 |
| 9 | 30804872 | rs2502921 | 0.743 | 0.48 | -0.316 | 1.053 | 0.009 | 0.085 | 0.085 | 0.217 | 0.9 | 0.81 | 0.88 | 0.66 | 0.9 | 0.78 |
| 9 | 30822433 | rs10969875 | -0.484 | 0.478 | -0.634 | 0.417 | -0.001 | -0.018 | -0.018 | 0 | 0.1 | 0.12 | 0.06 | 0 | 0.1 | 0.09 |
| 9 | 30825690 | rs1115506 | 0.073 | 0.314 | -1.033 | 0.352 | -0.016 | 0.017 | 0.017 | 0 | 0.43 | 0.29 | 0.43 | 0.38 | 0.5 | 0.23 |
| 9 | 30825999 | rs1115507 | 0.328 | 0.371 | -1.207 | 0.281 | 0 | 0.053 | 0.053 | 0 | 0.14 | 0.19 | 0.13 | 0.32 | 0.15 | 0.04 |
| 9 | 30840471 | rs10969885 | -1.343 | -0.019 | -0.618 | 0.526 | 0.005 | 0.027 | 0.027 | 0.11 | 0.6 | 0.75 | 0.65 | 0.76 | 0.59 | 0.6 |
| 9 | 30864788 | rs17776734 | 0.942 | 0.03 | -1.086 | 0.594 | -0.102 | 0.131 | 0.131 | 0 | 0.3 | 0.04 | 0.28 | 0.08 | 0.35 | 0.26 |
| 9 | 30872374 | rs7863381 | -0.446 | 0.27 | -0.359 | 0.87 | 0.001 | 0.047 | 0.047 | 0.054 | 0.26 | 0.42 | 0.35 | 0.67 | 0.26 | 0.38 |
| 9 | 30883856 | rs2210441 | 0.596 | 0.196 | -0.725 | 0.775 | -0.068 | 0.132 | 0.132 | 0 | 0.3 | 0.1 | 0.29 | 0.08 | 0.33 | 0.21 |
| 9 | 30907219 | rs10969963 | 0.315 | 0.369 | -0.585 | 0.96 | -0.005 | -0.011 | -0.011 | 0 | 0.21 | 0.17 | 0.33 | 0.4 | 0.28 | 0.24 |
| 9 | 30907279 | rs10969964 | 0.315 | 0.369 | -0.585 | 0.96 | -0.006 | -0.014 | -0.014 | 0 | 0.21 | 0.17 | 0.32 | 0.4 | 0.28 | 0.24 |
| 9 | 30909009 | rs10969969 | 0.367 | 0.35 | -0.639 | 0.967 | -0.034 | -0.078 | -0.078 | 0.502 | 0.21 | 0.1 | 0.33 | 0.4 | 0.28 | 0.25 |
| 9 | 30915973 | rs1332240 | 0.928 | 0.341 | -0.656 | 0.946 | -0.073 | 0.038 | 0.038 | 0 | 0.18 | 0.02 | 0.23 | 0.06 | 0.26 | 0.11 |
| 9 | 30918764 | rs10813381 | 0.323 | 0.344 | -0.614 | 0.934 | -0.01 | -0.025 | -0.025 | 0.227 | 0.21 | 0.15 | 0.32 | 0.4 | 0.28 | 0.25 |
| 9 | 30925753 | rs952695 | 0.188 | 0.185 | -0.157 | 0.881 | 0.007 | 0.035 | 0.035 | 0 | 0.21 | 0.3 | 0.34 | 0.4 | 0.29 | 0.24 |
| 9 | 30934859 | rs10969982 | 1.214 | 0.321 | -0.91 | 0.518 | -0.075 | -0.085 | -0.085 | 0 | 0.18 | 0.03 | 0.33 | 0.24 | 0.25 | 0.26 |
| 9 | 30953341 | rs1332234 | -0.505 | -0.729 | -0.337 | -0.635 | 0.007 | -0.003 | -0.003 | 0 | 0.6 | 0.42 | 0.59 | 0.39 | 0.58 | 0.53 |
| 9 | 30955393 | rs7021156 | -0.698 | -0.793 | -0.293 | -0.641 | -0.006 | -0.002 | -0.002 | 0 | 0.67 | 0.69 | 0.52 | 0.4 | 0.63 | 0.59 |
| 9 | 30970211 | rs7867525 | -0.619 | -0.752 | -0.38 | -0.629 | 0.007 | -0.001 | -0.001 | 0 | 0.64 | 0.4 | 0.47 | 0.38 | 0.63 | 0.47 |
| 9 | 30974923 | rs11790297 | 0.094 | -0.764 | 0.05 | -0.71 | 0.023 | 0.067 | 0.067 | 0 | 0.43 | 0.21 | 0.24 | 0.32 | 0.38 | 0.36 |
| 9 | 30975578 | rs10970001 | -0.405 | -0.614 | 0.102 | -0.618 | 0.014 | 0.021 | 0.021 | 0 | 0.49 | 0.39 | 0.4 | 0.34 | 0.39 | 0.44 |
| 9 | 30979640 | rs2888998 | 0.258 | -0.384 | 0.48 | -0.372 | -0.01 | 0.005 | 0.005 | 0 | 0.22 | 0.23 | 0.33 | 0.73 | 0.16 | 0.25 |
| 9 | 30981578 | rs4626692 | 0.467 | -0.381 | 0.555 | -0.557 | 0.033 | 0.065 | 0.065 | 0 | 0.5 | 0.34 | 0.58 | 0.32 | 0.52 | 0.67 |
| 9 | 30982465 | rs7024400 | -1.979 | -0.383 | 0.623 | -0.605 | -0.001 | -0.003 | -0.003 | 0.078 | 0.87 | 0.87 | 0.83 | 0.97 | 0.79 | 0.91 |
| 9 | 30982620 | rs7024630 | 0.424 | -0.383 | 0.729 | -0.57 | 0.014 | 0.026 | 0.026 | 0 | 0.51 | 0.42 | 0.59 | 0.32 | 0.52 | 0.67 |
| 9 | 30988320 | rs10813392 | 0.86 | -0.19 | 0.632 | -0.54 | -0.022 | 0.09 | 0.09 | 0 | 0.45 | 0.25 | 0.36 | 0.22 | 0.46 | 0.52 |
| 9 | 30990917 | rs10813395 | 0.627 | -0.185 | 0.579 | -0.527 | -0.001 | 0.055 | 0.055 | 0 | 0.47 | 0.31 | 0.54 | 0.3 | 0.48 | 0.56 |
| 9 | 30995741 | rs10156409 | -0.808 | -0.209 | 0.55 | -0.485 | 0.005 | 0.01 | 0.01 | 0 | 0.47 | 0.51 | 0.52 | 0.75 | 0.42 | 0.43 |
| 9 | 30997203 | rs10813399 | -0.707 | -0.123 | 0.488 | -0.499 | -0.002 | -0.007 | -0.007 | 0 | 0.46 | 0.44 | 0.51 | 0.75 | 0.42 | 0.46 |
| 9 | 31004169 | rs10813415 | 0.9 | -0.238 | 0.492 | -0.426 | -0.012 | 0.023 | 0.023 | 0 | 0.18 | 0.19 | 0.17 | 0.65 | 0.13 | 0.2 |
| 9 | 31018578 | rs13290998 | 1.232 | -0.167 | 0.396 | -0.549 | 0.001 | -0.023 | -0.023 | 0.65 | 0.16 | 0.13 | 0.09 | 0 | 0.14 | 0.05 |
| 9 | 31021219 | rs10970058 | 0.73 | -0.186 | 0.465 | -0.298 | -0.006 | 0.021 | 0.021 | 0 | 0.19 | 0.18 | 0.3 | 0.58 | 0.14 | 0.19 |
| 9 | 31021567 | rs2778413 | 0.058 | -0.016 | 0.591 | -0.406 | 0.024 | -0.01 | -0.01 | 0 | 0.24 | 0.31 | 0.28 | 0.2 | 0.35 | 0.31 |
| 9 | 31021638 | rs12343404 | 1.065 | 0.056 | 0.539 | -0.355 | 0 | -0.001 | -0.001 | 0 | 0.1 | 0.09 | 0.09 | 0.08 | 0.12 | 0.13 |
| 9 | 31022187 | rs7036584 | -0.632 | 0.021 | 0.553 | -0.389 | -0.003 | -0.003 | -0.003 | 0 | 0.45 | 0.43 | 0.48 | 0.66 | 0.4 | 0.37 |
| 9 | 31023908 | rs10813420 | 0.207 | 0.072 | 0.543 | -0.422 | -0.008 | 0.003 | 0.003 | 0 | 0.65 | 0.45 | 0.51 | 0.33 | 0.64 | 0.59 |
| 9 | 31024144 | rs10757998 | -0.868 | 0.013 | 0.647 | -0.177 | -0.043 | 0.056 | 0.056 | 0 | 0.81 | 0.61 | 0.75 | 0.68 | 0.89 | 0.72 |
| 9 | 31025113 | rs7049218 | 0.196 | -0.006 | 0.646 | -0.231 | 0 | 0.001 | 0.001 | 0 | 0.31 | 0.33 | 0.32 | 0.36 | 0.39 | 0.17 |
| 9 | 31027115 | rs2992117 | -1.319 | -0.04 | 0.694 | -0.235 | -0.028 | 0.029 | 0.029 | 0 | 0.83 | 0.71 | 0.77 | 0.7 | 0.9 | 0.83 |
| 9 | 31027597 | rs10757999 | 0.447 | -0.02 | 0.593 | -0.22 | 0.017 | 0.058 | 0.058 | 0 | 0.5 | 0.28 | 0.42 | 0.32 | 0.5 | 0.54 |
| 9 | 31027880 | rs16916442 | 0.975 | 0.08 | 0.455 | -0.184 | -0.002 | 0.003 | 0.003 | 0 | 0.18 | 0.2 | 0.15 | 0.28 | 0.16 | 0.07 |
| 9 | 31029867 | rs17720893 | 1.526 | 0.237 | 0.333 | -0.045 | -0.011 | 0.004 | 0.004 | 0 | 0.13 | 0.08 | 0.17 | 0.07 | 0.23 | 0.1 |
| 9 | 31030917 | rs17720922 | 1.502 | 0.385 | 0.334 | -0.061 | -0.005 | 0.002 | 0.002 | 0 | 0.13 | 0.1 | 0.17 | 0.07 | 0.23 | 0.1 |
| 9 | 31037099 | rs12338719 | -0.041 | 0.294 | 0.249 | -0.211 | -0.015 | -0.022 | -0.022 | 0.178 | 0.31 | 0.21 | 0.36 | 0.43 | 0.42 | 0.34 |
| 9 | 31038399 | rs12000796 | 0.561 | 0.165 | 0.348 | -0.291 | -0.002 | 0.038 | 0.038 | 0 | 0.53 | 0.39 | 0.52 | 0.25 | 0.49 | 0.41 |
| 9 | 31050747 | rs13293430 | 0.818 | 0.062 | 0.376 | -0.233 | 0 | -0.014 | -0.014 | 0 | 0.2 | 0.19 | 0.24 | 0.03 | 0.17 | 0.09 |
| 9 | 31055720 | rs7862915 | 1.727 | 0.029 | 0.502 | -0.179 | -0.025 | 0.058 | 0.058 | 0 | 0.14 | 0.26 | 0.09 | 0.29 | 0.08 | 0.13 |
| 9 | 31063444 | rs10970083 | 1.053 | 0.1 | 0.352 | -0.375 | -0.056 | 0.069 | 0.069 | 0.53 | 0.4 | 0.15 | 0.32 | 0.24 | 0.42 | 0.38 |
| 9 | 31064657 | rs12552052 | 0.542 | 0.101 | 0.248 | -0.382 | 0.013 | 0.09 | 0.09 | 0.492 | 0.55 | 0.22 | 0.53 | 0.23 | 0.54 | 0.46 |
| 9 | 31066983 | rs3936927 | -0.7 | 0.453 | 0.552 | 0.039 | -0.039 | 0.202 | 0.202 | 0.37 | 0.82 | 0.54 | 0.83 | 0.4 | 0.88 | 0.73 |
| 9 | 31078653 | rs17777591 | 1.913 | 0.6 | 0.275 | 0.002 | -0.017 | 0.02 | 0.02 | 0 | 0.05 | 0 | 0.04 | 0 | 0.05 | 0.08 |
| 9 | 31084302 | rs10813447 | 0.389 | 0.431 | 0.529 | 0.083 | 0.007 | -0.006 | -0.006 | 0 | 0.62 | 0.35 | 0.59 | 0.39 | 0.71 | 0.66 |
| 9 | 31085518 | rs13301262 | 0.95 | 0.535 | 0.588 | -0.04 | 0.002 | -0.012 | -0.012 | 0.014 | 0.17 | 0.1 | 0.09 | 0.01 | 0.15 | 0.04 |
| 9 | 31089681 | rs6476259 | -0.531 | 0.651 | 0.457 | 0.085 | -0.01 | -0.093 | -0.093 | 0 | 0.77 | 0.69 | 0.76 | 0.95 | 0.81 | 0.83 |
| 9 | 31100668 | rs13285414 | 0.757 | 0.704 | 0.061 | 0.118 | -0.006 | -0.065 | -0.065 | 0.995 | 0.21 | 0.22 | 0.23 | 0.01 | 0.18 | 0.08 |
| 9 | 31101278 | rs6476261 | -0.518 | 0.816 | -0.449 | 0.157 | -0.01 | -0.12 | -0.12 | 3.336 | 0.76 | 0.61 | 0.74 | 0.94 | 0.78 | 0.77 |
| 9 | 31102874 | rs12347592 | 0.902 | 0.823 | -0.621 | 0.159 | 0.01 | -0.004 | -0.004 | 0 | 0.22 | 0.25 | 0.26 | 0.15 | 0.29 | 0.26 |
| 9 | 31105113 | rs1360797 | -0.753 | 0.892 | -0.895 | 0.353 | -0.023 | 0.026 | 0.026 | 0 | 0.76 | 0.57 | 0.68 | 0.68 | 0.8 | 0.8 |
| 9 | 31107451 | rs1475793 | -0.753 | 0.892 | -0.895 | 0.353 | -0.023 | 0.026 | 0.026 | 0 | 0.76 | 0.57 | 0.69 | 0.68 | 0.8 | 0.8 |
| 9 | 31108083 | rs10970110 | 0.626 | 0.816 | -0.874 | 0.661 | 0.01 | -0.003 | -0.003 | 0 | 0.22 | 0.25 | 0.26 | 0.15 | 0.29 | 0.27 |

| | | | | | | | | | | | | | | | | |
|---|---|---|---|---|---|---|---|---|---|---|---|---|---|---|---|---|
| 9 | 31111743 | rs975177 | -0.575 | 0.435 | -1.478 | 0.754 | 0.016 | -0.013 | -0.013 | 0 | 0.45 | 0.31 | 0.37 | 0.52 | 0.38 | 0.44 |
| 9 | 31116343 | rs2184505 | -0.257 | 1.044 | -1.17 | 0.149 | -0.005 | 0.024 | 0.024 | 0.099 | 0.32 | 0.22 | 0.28 | 0.22 | 0.3 | 0.3 |
| 9 | 31116823 | rs12553076 | -0.169 | 0.56 | -1.107 | 0.091 | 0.017 | 0.028 | 0.028 | 0.198 | 0.31 | 0.22 | 0.25 | 0.2 | 0.24 | 0.3 |
| 9 | 31121425 | rs2225740 | 0.323 | 0.978 | -0.878 | 0.202 | -0.02 | 0.055 | 0.055 | 0 | 0.55 | 0.25 | 0.34 | 0.68 | 0.46 | 0.55 |
| 9 | 31123932 | rs1335157 | 0.288 | 1.025 | -0.872 | 0.755 | -0.045 | -0.056 | -0.056 | 0 | 0.55 | 0.12 | 0.35 | 0.47 | 0.52 | 0.52 |
| 9 | 31136986 | rs16916555 | 0.15 | 1.01 | 0.055 | 0.672 | 0 | -0.015 | -0.015 | 0.302 | 0.58 | 0.52 | 0.37 | 0.66 | 0.45 | 0.52 |
| 9 | 31146393 | rs10813477 | 0.452 | 0.995 | 0.077 | 0.721 | 0.012 | -0.023 | -0.023 | 0 | 0.54 | 0.19 | 0.33 | 0.47 | 0.44 | 0.5 |
| 9 | 31152444 | rs1335160 | 0.144 | 0.998 | 0.307 | 0.721 | -0.001 | -0.002 | -0.002 | 0.605 | 0.58 | 0.58 | 0.37 | 0.69 | 0.45 | 0.52 |
| 9 | 31163494 | rs10758029 | 0.118 | 1.069 | 0.281 | 0.701 | 0 | 0.001 | 0.001 | 0 | 0.59 | 0.57 | 0.37 | 0.47 | 0.48 | 0.5 |
| 9 | 31174646 | rs10758031 | -0.095 | 1.312 | 0.028 | 0.502 | 0.038 | 0.03 | 0.03 | 0.48 | 0.81 | 0.64 | 0.65 | 0.74 | 0.79 | 0.65 |
| 9 | 31178755 | rs10970183 | 0.916 | 1.282 | -0.388 | 0.431 | -0.045 | -0.08 | -0.08 | 0 | 0.55 | 0.07 | 0.31 | 0.47 | 0.47 | 0.5 |
| 9 | 31182134 | rs11793357 | 0.307 | 1.299 | -0.149 | 0.387 | -0.041 | 0.076 | 0.076 | 0.14 | 0.22 | 0.09 | 0.26 | 0.05 | 0.3 | 0.13 |
| 9 | 31182384 | rs4878464 | -0.043 | 1.304 | 0.232 | 0.55 | 0.012 | 0.005 | 0.005 | 0 | 0.24 | 0.32 | 0.31 | 0.29 | 0.32 | 0.15 |
| 9 | 31189546 | rs7862164 | 0.302 | 0.934 | 0.206 | 0.797 | 0.002 | 0.067 | 0.067 | 0.609 | 0.75 | 0.43 | 0.6 | 0.5 | 0.76 | 0.62 |
| 9 | 31193338 | rs10465068 | 0.418 | 0.932 | 0.342 | 0.713 | 0.014 | -0.008 | -0.008 | 0.274 | 0.76 | 0.53 | 0.63 | 0.76 | 0.77 | 0.65 |
| 9 | 31222420 | rs10813516 | 0.285 | 0.893 | 0.167 | 0.586 | 0 | -0.005 | -0.005 | 0 | 0.75 | 0.35 | 0.62 | 0.76 | 0.77 | 0.65 |
| 9 | 31233122 | rs13291610 | 0.448 | 1.038 | -0.171 | 0.341 | -0.086 | 0.107 | 0.107 | 0.179 | 0.23 | 0.06 | 0.29 | 0.07 | 0.34 | 0.13 |
| 9 | 31233220 | rs13292646 | 0.314 | 1.029 | -0.14 | 0.336 | -0.049 | 0.085 | 0.085 | 0.255 | 0.24 | 0.14 | 0.3 | 0.07 | 0.34 | 0.13 |
| 9 | 31236422 | rs1360665 | 0.435 | 1.027 | -0.15 | 0.336 | -0.086 | 0.108 | 0.108 | 0.225 | 0.23 | 0.07 | 0.29 | 0.07 | 0.34 | 0.13 |
| 9 | 31241652 | rs2065579 | 0.314 | 1.027 | -0.145 | 0.335 | -0.054 | 0.091 | 0.091 | 0.35 | 0.24 | 0.13 | 0.3 | 0.07 | 0.34 | 0.13 |
| 9 | 31242351 | rs2050239 | 0.225 | 0.994 | -0.058 | 0.367 | 0 | -0.002 | -0.002 | 0 | 0.75 | 0.33 | 0.62 | 0.76 | 0.77 | 0.65 |
| 9 | 31248679 | rs7025164 | 0.944 | 0.893 | -0.106 | 0.423 | 0.011 | 0.117 | 0.117 | 0 | 0.52 | 0.19 | 0.34 | 0.67 | 0.46 | 0.52 |
| 9 | 31253291 | rs12344989 | -0.628 | 0.456 | 0.253 | 0.256 | -0.008 | 0.08 | 0.08 | 0 | 0.1 | 0.25 | 0.08 | 0.23 | 0.07 | 0.07 |
| 9 | 31254719 | rs12337164 | -0.517 | 0.052 | -0.156 | -0.211 | -0.001 | 0.003 | 0.003 | 1.286 | 0.54 | 0.61 | 0.49 | 0.34 | 0.56 | 0.36 |
| 9 | 31255726 | rs10970246 | -0.088 | -0.152 | 0.501 | -0.343 | 0.003 | 0.037 | 0.037 | 1.882 | 0.49 | 0.4 | 0.41 | 0.11 | 0.51 | 0.29 |
| 9 | 31262671 | rs10758046 | -0.228 | -0.136 | 0.517 | -0.32 | 0.002 | 0.044 | 0.044 | 2.796 | 0.52 | 0.39 | 0.46 | 0.35 | 0.56 | 0.36 |
| 9 | 31263902 | rs13293735 | 0.447 | -0.216 | 0.39 | -0.373 | -0.027 | 0.114 | 0.114 | 3.237 | 0.41 | 0.32 | 0.4 | 0.12 | 0.49 | 0.27 |
| 9 | 31269291 | rs1158566 | -0.595 | -0.163 | 0.332 | -0.303 | -0.002 | 0.006 | 0.006 | 1.279 | 0.57 | 0.59 | 0.49 | 0.34 | 0.58 | 0.34 |
| 9 | 31271770 | rs11791890 | 0.055 | -0.007 | -0.135 | -0.494 | -0.026 | 0.166 | 0.166 | 1.481 | 0.45 | 0.19 | 0.41 | 0.12 | 0.51 | 0.28 |
| 9 | 31282032 | rs12555645 | -0.521 | -0.393 | 0.191 | 0.284 | 0.001 | 0.009 | 0.009 | 0 | 0.55 | 0.63 | 0.51 | 0.61 | 0.41 | 0.68 |
| 9 | 31287949 | rs10465074 | 1.144 | -0.915 | -0.333 | 0.418 | -0.042 | -0.152 | -0.152 | 0.083 | 0.24 | 0.01 | 0.15 | 0.36 | 0.19 | 0.3 |
| 9 | 31288508 | rs10813525 | -0.701 | -0.691 | 0.366 | 0.385 | -0.011 | 0.102 | 0.102 | 0 | 0.57 | 0.78 | 0.54 | 0.88 | 0.43 | 0.7 |
| 9 | 31291138 | rs10970260 | -0.09 | -0.77 | 0.533 | 0.31 | 0.003 | 0.003 | 0.003 | 0 | 0.48 | 0.53 | 0.47 | 0.62 | 0.36 | 0.67 |
| 9 | 31307968 | rs10758050 | 0.539 | -0.776 | 0.519 | 0.253 | -0.006 | 0.026 | 0.026 | 0 | 0.43 | 0.26 | 0.48 | 0.39 | 0.58 | 0.32 |
| 9 | 31314587 | rs7020872 | 0.046 | -0.779 | 0.531 | 0.252 | -0.005 | -0.048 | -0.048 | 0 | 0.91 | 0.81 | 0.95 | 1 | 0.94 | 0.99 |
| 9 | 31318420 | rs7038347 | 0.296 | -0.769 | 0.705 | 0.311 | -0.001 | 0 | 0 | 0 | 0.45 | 0.44 | 0.5 | 0.39 | 0.59 | 0.33 |
| 9 | 31324237 | rs10970286 | 0.439 | -0.04 | 0.297 | 0.16 | -0.082 | 0.144 | 0.144 | 0 | 0.42 | 0.12 | 0.47 | 0.14 | 0.5 | 0.32 |
| 9 | 31325092 | rs10813534 | 0.227 | -0.21 | 0.601 | 0.32 | 0 | 0.02 | 0.02 | 0 | 0.43 | 0.26 | 0.42 | 0.56 | 0.4 | 0.61 |
| 9 | 31340536 | rs10125602 | -0.954 | -0.405 | 0.814 | 0.252 | -0.018 | 0.028 | 0.028 | 0 | 0.08 | 0.2 | 0.05 | 0.23 | 0.06 | 0.03 |
| 9 | 31347460 | rs6476284 | -0.636 | -0.299 | 0.747 | 0.421 | -0.013 | 0.088 | 0.088 | 0.008 | 0.55 | 0.73 | 0.5 | 0.79 | 0.48 | 0.66 |
| 9 | 31351194 | rs10813550 | 0.288 | -0.158 | 0.376 | 0.57 | -0.001 | 0.005 | 0.005 | 0 | 0.43 | 0.26 | 0.43 | 0.54 | 0.41 | 0.63 |
| 9 | 31359237 | rs10758055 | 0.242 | -0.151 | 0.336 | 0.589 | 0 | 0.004 | 0.004 | 0 | 0.43 | 0.32 | 0.43 | 0.54 | 0.42 | 0.61 |
| 9 | 31372120 | rs1888329 | 0.46 | -0.09 | -0.023 | 0.502 | -0.035 | 0.055 | 0.055 | 0 | 0.44 | 0.13 | 0.59 | 0.57 | 0.45 | 0.72 |
| 9 | 31381021 | rs10970317 | -1.493 | -0.195 | 0.897 | 0.437 | -0.053 | -0.007 | -0.007 | 0 | 0.16 | 0.55 | 0.08 | 0.19 | 0.06 | 0.04 |
| 9 | 31394025 | rs7857709 | -0.584 | -0.114 | 0.696 | 0.431 | -0.02 | 0.044 | 0.044 | 0 | 0.6 | 0.73 | 0.67 | 0.8 | 0.51 | 0.76 |
| 9 | 31394143 | rs13296212 | 0.884 | -0.07 | 0.637 | 0.384 | -0.063 | 0.042 | 0.042 | 0 | 0.42 | 0.05 | 0.57 | 0.62 | 0.42 | 0.69 |
| 9 | 31394275 | rs7873690 | -0.648 | -0.068 | 0.639 | 0.378 | -0.023 | 0.044 | 0.044 | 0 | 0.6 | 0.73 | 0.68 | 0.8 | 0.51 | 0.76 |
| 9 | 31396488 | rs1382291 | 0.882 | 0.02 | 0.531 | 0.399 | -0.059 | 0.02 | 0.02 | 0 | 0.42 | 0.05 | 0.57 | 0.58 | 0.42 | 0.69 |
| 9 | 31397676 | rs7857257 | -0.967 | 0.045 | 1.1 | 0.314 | -0.025 | -0.082 | -0.082 | 0.775 | 0.13 | 0.27 | 0.04 | 0 | 0.06 | 0.05 |
| 9 | 31405730 | rs7025296 | -0.459 | 0.275 | 0.849 | 0.337 | -0.019 | 0.038 | 0.038 | 0 | 0.6 | 0.73 | 0.66 | 0.8 | 0.51 | 0.76 |
| 9 | 31408125 | rs11793707 | 0.629 | 0.373 | 0.506 | 0.246 | -0.086 | 0.166 | 0.166 | 1.646 | 0.3 | 0.02 | 0.17 | 0.05 | 0.3 | 0.06 |
| 9 | 31424013 | rs7866477 | -2.743 | 0.243 | 2.225 | 1.772 | -0.01 | 0.05 | 0.05 | 0.065 | 0.24 | 0.63 | 0.32 | 0.39 | 0.19 | 0.36 |
| 9 | 31439249 | rs10758063 | -0.316 | 0.363 | 2.163 | 1.936 | 0 | 0.006 | 0.006 | 0.004 | 0.07 | 0.05 | 0.11 | 0.15 | 0.11 | 0.16 |
| 9 | 31446639 | rs10813582 | -2.029 | 0.261 | 2.479 | 1.84 | -0.001 | 0.03 | 0.03 | 0.017 | 0.17 | 0.18 | 0.27 | 0.27 | 0.16 | 0.3 |
| 9 | 31446664 | rs7856016 | -2.758 | 0.255 | 2.529 | 1.817 | -0.011 | 0.131 | 0.131 | 0.055 | 0.23 | 0.5 | 0.31 | 0.38 | 0.19 | 0.36 |
| 9 | 31454538 | rs10813591 | -2.341 | 0.36 | 2.544 | 1.847 | 0.002 | 0.078 | 0.078 | 0.002 | 0.18 | 0.31 | 0.27 | 0.27 | 0.16 | 0.28 |
| 9 | 31470393 | rs10970374 | -0.88 | 0.365 | 2.394 | 1.847 | -0.006 | 0.017 | 0.017 | 0.414 | 0.09 | 0.05 | 0.23 | 0.27 | 0.12 | 0.26 |
| 9 | 31471679 | rs10120052 | -2.601 | 0.359 | 2.459 | 1.843 | 0.003 | 0.115 | 0.115 | 0.272 | 0.19 | 0.42 | 0.28 | 0.27 | 0.16 | 0.28 |
| 9 | 31472633 | rs10813601 | -1.832 | 0.403 | 2.341 | 1.879 | 0.017 | 0.118 | 0.118 | 0.192 | 0.12 | 0.23 | 0.24 | 0.26 | 0.13 | 0.25 |
| 9 | 31473208 | rs2447721 | 1.847 | 0.392 | 2.286 | 1.848 | 0.014 | 0.11 | 0.11 | 0.292 | 0.88 | 0.72 | 0.75 | 0.74 | 0.87 | 0.73 |
| 9 | 31479215 | rs10970378 | -1.804 | 0.402 | 2.257 | 1.89 | 0.015 | 0.164 | 0.164 | 0.622 | 0.12 | 0.25 | 0.24 | 0.38 | 0.13 | 0.31 |
| 9 | 31494101 | rs1929104 | -1.493 | 0.371 | 2.414 | 1.508 | -0.014 | -0.048 | -0.048 | 1.361 | 0.09 | 0.28 | 0.04 | 0 | 0.04 | 0.02 |
| 9 | 31500197 | rs13297113 | -2.142 | 0.549 | 2.291 | 1.537 | 0.022 | 0.059 | 0.059 | 0.162 | 0.06 | 0.21 | 0.08 | 0.12 | 0.06 | 0.06 |
| 9 | 31524219 | rs10465076 | -0.007 | 0.568 | 2.095 | 1.337 | -0.012 | -0.007 | -0.007 | 0 | 0.06 | 0.02 | 0.1 | 0.14 | 0.11 | 0.14 |
| 9 | 31532212 | rs16917077 | -0.841 | 0.512 | 2.261 | 1.105 | -0.004 | -0.031 | -0.031 | 0 | 0.07 | 0.2 | 0.04 | 0 | 0.04 | 0.02 |
| 9 | 31533362 | rs2502176 | 2.421 | 0.496 | 2.473 | 1.056 | 0.027 | -0.04 | -0.04 | 0.569 | 0.91 | 0.59 | 0.89 | 1 | 0.94 | 0.97 |
| 9 | 31538491 | rs12342878 | -1.408 | 0.503 | 2.201 | 1.104 | -0.006 | -0.048 | -0.048 | 0 | 0.09 | 0.24 | 0.06 | 0 | 0.05 | 0.02 |
| 9 | 31540903 | rs10511880 | -0.577 | 0.581 | 2.136 | 1.424 | 0.007 | 0.055 | 0.055 | 0.727 | 0.1 | 0.13 | 0.25 | 0.26 | 0.14 | 0.21 |
| 9 | 31553846 | rs7034481 | 2.7 | 0.495 | 2.454 | 1.361 | -0.011 | -0.02 | -0.02 | 0.114 | 0.86 | 0.42 | 0.87 | 0.92 | 0.91 | 0.84 |

| | | | | | | | | | | | | | | | | |
|---|---|---|---|---|---|---|---|---|---|---|---|---|---|---|---|---|
| 9 | 31555977 | rs12337314 | -1.373 | 0.515 | 2.403 | 1.365 | 0.01 | -0.019 | -0.019 | 1.29 | 0.07 | 0.27 | 0.04 | 0 | 0.03 | 0.06 |
| 9 | 31568046 | rs16917154 | -0.906 | 0.529 | 1.816 | 1.387 | 0.002 | -0.028 | -0.028 | 0 | 0.06 | 0.2 | 0.03 | 0 | 0.03 | 0.05 |
| 9 | 31573759 | rs10970427 | -1.403 | 0.497 | 1.654 | 1.406 | 0.01 | 0.045 | 0.045 | 0 | 0.08 | 0.29 | 0.05 | 0.08 | 0.05 | 0.13 |
| 9 | 31586876 | rs4570254 | 1.383 | 0.572 | 0.723 | 1.393 | -0.001 | 0.011 | 0.011 | 0.618 | 0.75 | 0.24 | 0.62 | 0.68 | 0.78 | 0.61 |
| 9 | 31591132 | rs4242684 | 0.382 | 0.458 | 0.164 | 0.885 | 0.001 | 0 | 0 | 0 | 0.62 | 0.31 | 0.52 | 0.66 | 0.64 | 0.67 |
| 9 | 31602656 | rs16917200 | -0.7 | 0.328 | 0.81 | 1.041 | 0.012 | 0.153 | 0.153 | 0 | 0.05 | 0.27 | 0.1 | 0.11 | 0.02 | 0.12 |
| 9 | 31608609 | rs7035270 | 0.681 | 0.846 | 0.522 | 0.422 | 0.013 | -0.046 | -0.046 | 0 | 0.13 | 0.15 | 0.08 | 0 | 0.15 | 0.09 |
| 9 | 31612603 | rs1412339 | 0.021 | 0.529 | 0.699 | 0.477 | 0.025 | 0.075 | 0.075 | 0 | 0.1 | 0.21 | 0.11 | 0.23 | 0.11 | 0.14 |
| 9 | 31614031 | rs16924096 | -0.086 | 0.482 | 0.719 | 0.168 | -0.015 | -0.029 | -0.029 | 0 | 0.25 | 0.41 | 0.36 | 0.11 | 0.25 | 0.17 |
| 9 | 31619571 | rs1412341 | 0.324 | 0.179 | -0.148 | 0.181 | -0.024 | -0.014 | -0.014 | 0 | 0.39 | 0.51 | 0.3 | 0.67 | 0.26 | 0.46 |
| 9 | 31620128 | rs11791319 | 0.312 | 0.219 | -0.164 | -0.054 | 0.001 | 0.011 | 0.011 | 0 | 0.2 | 0.14 | 0.27 | 0 | 0.22 | 0.05 |
| 9 | 31628475 | rs7851583 | -0.212 | 0.235 | -0.072 | 0.011 | -0.005 | -0.012 | -0.012 | 0 | 0.25 | 0.36 | 0.35 | 0.11 | 0.25 | 0.14 |
| 9 | 31629459 | rs17179862 | -0.287 | 0.317 | -0.215 | -0.237 | -0.001 | -0.024 | -0.024 | 0.161 | 0.11 | 0.13 | 0.25 | 0 | 0.16 | 0.07 |
| 9 | 31629602 | rs781098 | 0.738 | 0.493 | -0.597 | -0.075 | -0.015 | -0.017 | -0.017 | 0 | 0.78 | 0.89 | 0.74 | 0.74 | 0.69 | 0.85 |
| 9 | 31629682 | rs781099 | 1.404 | 0.385 | -0.429 | -0.432 | 0 | -0.014 | -0.014 | 0 | 0.65 | 0.63 | 0.56 | 0.85 | 0.59 | 0.74 |
| 9 | 31629777 | rs781101 | 0.872 | 0.338 | -0.29 | -0.475 | -0.004 | -0.099 | -0.099 | 1.471 | 0.79 | 0.66 | 0.69 | 0.96 | 0.77 | 0.89 |
| 9 | 31636424 | rs17265305 | 0.29 | 0.359 | -0.34 | -0.397 | 0.01 | -0.055 | -0.055 | 0 | 0.1 | 0.15 | 0.24 | 0 | 0.14 | 0.05 |
| 9 | 31636880 | rs4879530 | -0.104 | 0.354 | -0.191 | -0.325 | 0.015 | 0.015 | 0.015 | 0 | 0.11 | 0.23 | 0.27 | 0.15 | 0.16 | 0.11 |
| 9 | 31643058 | rs431008 | 1.169 | 0.399 | -0.1 | -0.168 | 0.002 | 0 | 0 | 0 | 0.55 | 0.48 | 0.36 | 0.43 | 0.45 | 0.58 |
| 9 | 31645637 | rs4879531 | 0.797 | 0.234 | -0.116 | -0.185 | 0.004 | -0.034 | -0.034 | 0 | 0.09 | 0.1 | 0.24 | 0 | 0.14 | 0.05 |
| 9 | 31656498 | rs895288 | -0.421 | 0.219 | -0.339 | -0.378 | 0 | -0.009 | -0.009 | 0 | 0.22 | 0.24 | 0.36 | 0.13 | 0.26 | 0.14 |
| 9 | 31669518 | rs2044401 | -0.176 | 0.191 | 0.002 | -0.042 | 0.001 | -0.011 | -0.011 | 0 | 0.17 | 0.26 | 0.32 | 0.13 | 0.22 | 0.13 |
| 9 | 31673588 | rs410684 | 1.551 | 0.145 | 0.157 | -0.065 | 0.002 | 0.001 | 0.001 | 0 | 0.47 | 0.43 | 0.34 | 0.42 | 0.41 | 0.52 |
| 9 | 31682902 | rs11790434 | -0.183 | 0.297 | -0.099 | -0.056 | 0 | -0.008 | -0.008 | 0.021 | 0.17 | 0.22 | 0.32 | 0.13 | 0.22 | 0.13 |
| 9 | 31704278 | rs13297624 | -0.342 | 0.274 | 0.03 | -0.07 | 0.002 | -0.016 | -0.016 | 0.091 | 0.18 | 0.28 | 0.34 | 0.12 | 0.22 | 0.13 |
| 9 | 31705959 | rs12552096 | 0.342 | 0.378 | -0.627 | 0.051 | -0.019 | -0.021 | -0.021 | 0 | 0.08 | 0.03 | 0.11 | 0.14 | 0.13 | 0.14 |
| 9 | 31713085 | rs12552555 | 0.435 | 0.39 | -0.675 | -0.009 | -0.029 | -0.001 | -0.001 | 0 | 0.08 | 0.01 | 0.1 | 0.07 | 0.13 | 0.13 |
| 9 | 31714812 | rs10970502 | 0.594 | 0.034 | -0.553 | 0.059 | 0.005 | -0.002 | -0.002 | 0 | 0.11 | 0.08 | 0.1 | 0.31 | 0.06 | 0.13 |
| 9 | 31729270 | rs288901 | 0.683 | 0.046 | -0.395 | -0.07 | 0.007 | -0.006 | -0.006 | 1.722 | 0.71 | 0.52 | 0.57 | 0.73 | 0.64 | 0.75 |
| 9 | 31745109 | rs10758078 | 1.917 | -0.147 | -0.58 | -0.256 | -0.005 | -0.006 | -0.006 | 0 | 0.38 | 0.23 | 0.38 | 0.41 | 0.38 | 0.52 |
| 9 | 31745845 | rs1445648 | 0.593 | 0.05 | -0.715 | -0.254 | -0.004 | -0.001 | -0.001 | 1.12 | 0.11 | 0.09 | 0.25 | 0.01 | 0.16 | 0.05 |
| 9 | 31748036 | rs288884 | -0.5 | -0.017 | -0.792 | -0.016 | 0.004 | -0.002 | -0.002 | 0.255 | 0.27 | 0.19 | 0.17 | 0.31 | 0.24 | 0.21 |
| 9 | 31791211 | rs10970527 | 0.243 | 0.213 | -0.493 | 0.565 | 0.002 | 0.003 | 0.003 | 0 | 0.1 | 0.08 | 0.05 | 0.31 | 0.06 | 0.13 |
| 9 | 31811400 | rs10813645 | 0.801 | 0.565 | -0.415 | 0.319 | 0.003 | -0.013 | -0.013 | 0 | 0.57 | 0.26 | 0.49 | 0.54 | 0.55 | 0.57 |
| 9 | 31811669 | rs1928091 | 0.155 | 0.356 | 0.002 | 0.278 | -0.005 | -0.01 | -0.01 | 0 | 0.7 | 0.72 | 0.61 | 0.64 | 0.64 | 0.76 |
| 9 | 31817321 | rs928031 | 1.501 | 0.469 | -0.144 | 0.174 | 0.002 | 0.009 | 0.009 | 0 | 0.49 | 0.24 | 0.41 | 0.48 | 0.44 | 0.45 |
| 9 | 31819858 | rs780750 | 0.047 | 0.505 | -0.138 | 0.193 | -0.005 | -0.01 | -0.01 | 0 | 0.7 | 0.72 | 0.61 | 0.63 | 0.64 | 0.76 |
| 9 | 31820755 | rs10970532 | 0.4 | 0.591 | -0.175 | 0.231 | 0.001 | 0.034 | 0.034 | 0 | 0.13 | 0.17 | 0.27 | 0.28 | 0.17 | 0.14 |
| 9 | 31822916 | rs7026194 | -0.015 | 0.531 | -0.026 | 0.201 | 0.017 | 0.083 | 0.083 | 3.138 | 0.06 | 0.39 | 0.05 | 0.1 | 0.01 | 0.18 |
| 9 | 31825468 | rs780757 | 1.231 | 0.582 | -0.175 | 0.178 | 0.001 | -0.001 | -0.001 | 0.005 | 0.07 | 0.06 | 0.07 | 0 | 0.08 | 0.01 |
| 9 | 31825662 | rs780758 | -0.749 | 0.611 | -0.068 | 0.48 | -0.016 | 0.025 | 0.025 | 0 | 0.83 | 0.91 | 0.88 | 0.91 | 0.81 | 0.89 |
| 9 | 31825872 | rs1328740 | 0.483 | 0.684 | -0.294 | 0.517 | -0.017 | 0.007 | 0.007 | 0 | 0.08 | 0.02 | 0.08 | 0.05 | 0.12 | 0.11 |
| 9 | 31832821 | rs10813647 | -0.038 | 0.415 | -0.044 | 0.683 | 0.002 | 0.035 | 0.035 | 0 | 0.13 | 0.17 | 0.27 | 0.27 | 0.17 | 0.14 |
| 9 | 31836370 | rs10738875 | 0.257 | 0.367 | -0.261 | 0.651 | 0.01 | 0.043 | 0.043 | 0 | 0.69 | 0.52 | 0.59 | 0.64 | 0.64 | 0.78 |
| 9 | 31836555 | rs10511886 | 0.281 | 0.498 | -0.666 | 0.599 | -0.002 | 0.01 | 0.01 | 0.764 | 0.65 | 0.34 | 0.56 | 0.54 | 0.64 | 0.61 |
| 9 | 31837334 | rs10511887 | -0.814 | 0.507 | -1.401 | 0.476 | -0.025 | -0.013 | -0.013 | 0 | 0.23 | 0.05 | 0.35 | 0.28 | 0.28 | 0.28 |
| 9 | 31842496 | rs997609 | -0.25 | -0.25 | -0.75 | 0.695 | -0.017 | 0.025 | 0.025 | 0 | 0.13 | 0.22 | 0.1 | 0.21 | 0.08 | 0.15 |
| 9 | 31849082 | rs1576269 | 0.217 | -0.3 | -0.838 | 0.661 | -0.004 | -0.004 | -0.004 | 0 | 0.1 | 0.15 | 0.06 | 0.09 | 0.07 | 0.14 |
| 9 | 31849851 | rs10970559 | -1.277 | -0.359 | -1.234 | -0.097 | -0.038 | 0.131 | 0.131 | 0 | 0.57 | 0.71 | 0.54 | 0.9 | 0.53 | 0.8 |
| 9 | 31856797 | rs7870196 | 0.848 | -0.359 | -1.07 | 0.012 | -0.004 | -0.004 | -0.004 | 0 | 0.1 | 0.13 | 0.06 | 0.09 | 0.07 | 0.14 |
| 9 | 31868612 | rs10813650 | -1.227 | -0.22 | -1.289 | -0.038 | 0.013 | 0.143 | 0.143 | 0 | 0.53 | 0.65 | 0.51 | 0.9 | 0.5 | 0.79 |
| 9 | 31871335 | rs10970568 | 0.3 | -0.103 | -1.473 | 0.629 | 0.004 | 0.013 | 0.013 | 0 | 0.06 | 0.02 | 0.06 | 0.24 | 0.05 | 0.2 |
| 9 | 31872062 | rs10511888 | 0.112 | -0.201 | -1.38 | 0.527 | 0.005 | -0.062 | -0.062 | 0 | 0.28 | 0.11 | 0.29 | 0.47 | 0.31 | 0.38 |
| 9 | 31872624 | rs943927 | -1.174 | 0.189 | -1.471 | -0.358 | -0.06 | 0.188 | 0.188 | 0 | 0.57 | 0.77 | 0.65 | 0.99 | 0.55 | 0.78 |
| 9 | 31879919 | rs6476305 | -1.143 | -0.386 | -1.163 | 0.386 | -0.029 | 0.095 | 0.095 | 0 | 0.55 | 0.71 | 0.53 | 0.8 | 0.51 | 0.75 |
| 9 | 31892857 | rs17191070 | 0.973 | -0.626 | -1.005 | 0.118 | 0.004 | -0.011 | -0.011 | 0.595 | 0.13 | 0.09 | 0.08 | 0 | 0.08 | 0.03 |
| 9 | 31894436 | rs2375725 | 2.169 | -0.635 | -1.447 | -0.273 | -0.099 | 0.075 | 0.075 | 0.767 | 0.23 | 0.05 | 0.35 | 0.05 | 0.32 | 0.19 |
| 9 | 31894764 | rs7024321 | -0.304 | -0.77 | -0.724 | 0.609 | 0.002 | 0.032 | 0.032 | 0.54 | 0.22 | 0.14 | 0.15 | 0.13 | 0.18 | 0.12 |
| 9 | 31912519 | rs10125694 | 1.797 | -0.927 | 0.187 | 1.271 | 0.025 | 0.311 | 0.311 | 0 | 0.06 | 0.25 | 0.08 | 0.28 | 0.03 | 0.01 |
| 9 | 31915424 | rs10813661 | -0.062 | -0.905 | 0.142 | 1.252 | -0.033 | 0.066 | 0.066 | 0 | 0.18 | 0.33 | 0.16 | 0.28 | 0.11 | 0.05 |
| 9 | 31925048 | rs1887874 | -1.268 | -0.856 | 0.248 | 1.203 | -0.001 | 0 | 0 | 0 | 0.34 | 0.3 | 0.33 | 0.51 | 0.34 | 0.67 |
| 9 | 31930090 | rs4879543 | -0.319 | -0.77 | 0.484 | 1.477 | -0.005 | 0.001 | 0.001 | 0 | 0.23 | 0.21 | 0.25 | 0.25 | 0.22 | 0.35 |
| 9 | 31938495 | rs1602261 | -1.387 | -0.757 | 0.531 | 1.461 | -0.008 | 0.01 | 0.01 | 0.1 | 0.32 | 0.44 | 0.36 | 0.66 | 0.33 | 0.63 |
| 9 | 31951536 | rs2200964 | -2.181 | -0.576 | 0.843 | 1.321 | 0.022 | 0.038 | 0.038 | 0 | 0.48 | 0.6 | 0.47 | 0.68 | 0.41 | 0.7 |
| 9 | 31955694 | rs13284518 | -0.831 | -0.578 | 0.9 | 1.386 | -0.019 | 0.053 | 0.053 | 0 | 0.25 | 0.34 | 0.3 | 0.38 | 0.23 | 0.32 |
| 9 | 31970850 | rs7025945 | -1.143 | -0.446 | 0.848 | 1.292 | -0.026 | -0.035 | -0.035 | 0 | 0.39 | 0.44 | 0.34 | 0.26 | 0.3 | 0.35 |
| 9 | 31977623 | rs1038268 | -1.787 | -0.418 | 0.717 | 1.315 | 0.021 | -0.005 | -0.005 | 0 | 0.44 | 0.58 | 0.44 | 0.56 | 0.38 | 0.64 |
| 9 | 31989998 | rs2375811 | -1.778 | -0.418 | 0.689 | 1.315 | 0.02 | -0.005 | -0.005 | 1.086 | 0.44 | 0.57 | 0.44 | 0.56 | 0.38 | 0.64 |
| 9 | 32002358 | rs10813695 | 1.07 | -0.435 | 0.535 | 1.339 | -0.033 | -0.026 | -0.026 | 0.114 | 0.08 | 0.01 | 0.07 | 0.12 | 0.1 | 0.09 |

| | | | | | | | | | | | | | | | | |
|---|---|---|---|---|---|---|---|---|---|---|---|---|---|---|---|---|
| 9 | 32009054 | rs10970672 | 1.034 | -0.439 | 0.533 | 1.337 | -0.033 | -0.025 | -0.025 | 0.006 | 0.08 | 0.01 | 0.07 | 0.12 | 0.1 | 0.09 |
| 9 | 32041861 | rs6476322 | -1.03 | -0.445 | 0.36 | 1.301 | 0.001 | 0.001 | 0.001 | 0 | 0.37 | 0.28 | 0.33 | 0.26 | 0.3 | 0.35 |
| 9 | 32061188 | rs385280 | -1.654 | -0.44 | 0.383 | 1.314 | 0 | 0 | 0 | 0.132 | 0.46 | 0.4 | 0.41 | 0.38 | 0.4 | 0.44 |
| 9 | 32068378 | rs10813717 | 0.335 | -0.549 | 0.442 | 1.243 | -0.005 | -0.002 | -0.002 | 0 | 0.73 | 0.76 | 0.74 | 0.75 | 0.77 | 0.66 |
| 9 | 32071702 | rs1038270 | 0.587 | -0.568 | 0.551 | 1.327 | -0.035 | 0.064 | 0.064 | 0 | 0.71 | 0.6 | 0.69 | 0.56 | 0.76 | 0.67 |
| 9 | 32078175 | rs1508318 | -0.704 | -0.519 | 0.736 | 1.315 | 0 | -0.001 | -0.001 | 0 | 0.23 | 0.2 | 0.19 | 0.25 | 0.24 | 0.4 |
| 9 | 32081117 | rs10435795 | 1.691 | -0.531 | 0.716 | 1.326 | -0.036 | -0.003 | -0.003 | 0 | 0.07 | 0 | 0.04 | 0.07 | 0.1 | 0.08 |
| 9 | 32081823 | rs7048642 | 0.461 | -0.606 | 0.707 | 1.283 | -0.005 | 0.021 | 0.021 | 0 | 0.08 | 0.03 | 0.11 | 0.03 | 0.11 | 0.08 |
| 9 | 32081832 | rs10758102 | 1.007 | -0.62 | 0.817 | 1.287 | -0.029 | 0.051 | 0.051 | 0 | 0.71 | 0.58 | 0.69 | 0.57 | 0.76 | 0.67 |
| 9 | 32082616 | rs7038038 | 2.508 | -0.62 | 0.817 | 1.288 | 0.006 | 0.043 | 0.043 | 0 | 0.48 | 0.38 | 0.5 | 0.32 | 0.52 | 0.27 |
| 9 | 32087232 | rs9886775 | 1.003 | -0.621 | 0.811 | 1.288 | -0.027 | 0.047 | 0.047 | 0 | 0.71 | 0.61 | 0.69 | 0.57 | 0.76 | 0.67 |
| 9 | 32094230 | rs10738877 | -0.537 | -0.606 | 0.769 | 1.275 | -0.005 | -0.001 | -0.001 | 0.231 | 0.22 | 0.15 | 0.18 | 0.25 | 0.24 | 0.4 |
| 9 | 32104829 | rs1508310 | 0.488 | -0.595 | 0.769 | 1.374 | -0.004 | -0.001 | -0.001 | 0 | 0.08 | 0.03 | 0.17 | 0.05 | 0.11 | 0.08 |
| 9 | 32105254 | rs10970780 | -0.646 | -0.581 | 0.745 | 1.371 | 0 | -0.001 | -0.001 | 0 | 0.23 | 0.2 | 0.19 | 0.25 | 0.24 | 0.4 |
| 9 | 32105784 | rs6476341 | 0.969 | -0.545 | 0.817 | 1.447 | -0.029 | 0.052 | 0.052 | 0 | 0.71 | 0.58 | 0.69 | 0.57 | 0.76 | 0.67 |
| 9 | 32110777 | rs1842636 | 1.202 | -0.45 | 0.671 | 1.542 | -0.002 | 0.002 | 0.002 | 0 | 0.72 | 0.72 | 0.73 | 0.65 | 0.74 | 0.4 |
| 9 | 32116520 | rs16917898 | -0.315 | -0.465 | 0.713 | 1.536 | 0.012 | 0.001 | 0.001 | 0 | 0.17 | 0.07 | 0.07 | 0.19 | 0.11 | 0.25 |
| 9 | 32117175 | rs7849297 | 1.23 | -0.472 | 0.784 | 1.435 | -0.008 | 0.011 | 0.011 | 0 | 0.82 | 0.74 | 0.79 | 0.8 | 0.85 | 0.57 |
| 9 | 32117710 | rs7038301 | 1.553 | -0.45 | 0.77 | 1.451 | -0.001 | 0.001 | 0.001 | 0 | 0.64 | 0.66 | 0.65 | 0.52 | 0.63 | 0.31 |
| 9 | 32117729 | rs12000809 | -1.008 | -0.332 | 0.809 | 1.147 | 0 | -0.015 | -0.015 | 0.114 | 0.25 | 0.13 | 0.23 | 0.34 | 0.26 | 0.36 |
| 9 | 32119934 | rs11790716 | -0.963 | -0.316 | 0.577 | 1.157 | -0.007 | -0.037 | -0.037 | 0 | 0.25 | 0.09 | 0.23 | 0.34 | 0.26 | 0.36 |
| 9 | 32120093 | rs2495643 | 1.622 | -0.278 | 0.609 | 1.157 | -0.003 | -0.001 | -0.001 | 0 | 0.66 | 0.7 | 0.66 | 0.63 | 0.64 | 0.57 |
| 9 | 32121610 | rs10758117 | 0.188 | -0.352 | 0.578 | 1.163 | -0.019 | -0.081 | -0.081 | 0 | 0.16 | 0.03 | 0.16 | 0.31 | 0.15 | 0.23 |
| 9 | 32122888 | rs2174915 | -0.864 | -0.351 | 0.608 | 1.11 | 0 | -0.009 | -0.009 | 0.766 | 0.15 | 0.14 | 0.2 | 0.09 | 0.15 | 0.18 |
| 9 | 32125244 | rs10970789 | 0.179 | -0.377 | 0.495 | 1.141 | -0.019 | -0.064 | -0.064 | 0 | 0.16 | 0.03 | 0.1 | 0.28 | 0.15 | 0.22 |
| 9 | 32125843 | rs2819851 | -0.399 | -0.334 | 0.557 | 1.151 | 0.03 | -0.011 | -0.011 | 0.019 | 0.11 | 0.22 | 0.21 | 0.05 | 0.17 | 0.11 |
| 9 | 32127249 | rs10970792 | -0.609 | -0.599 | 0.178 | 1.257 | 0.014 | -0.021 | -0.021 | 0 | 0.22 | 0.09 | 0.13 | 0.31 | 0.19 | 0.32 |
| 9 | 32128828 | rs10970796 | -0.074 | -0.378 | 0.505 | 1.287 | -0.022 | 0.045 | 0.045 | 0 | 0.13 | 0.25 | 0.13 | 0.19 | 0.08 | 0.02 |
| 9 | 32134018 | rs2843575 | -0.975 | -0.014 | 0.622 | 1.003 | 0.014 | 0.016 | 0.016 | 0 | 0.3 | 0.44 | 0.36 | 0.37 | 0.33 | 0.35 |
| 9 | 32137678 | rs7859569 | -0.702 | -0.188 | 0.768 | 1.035 | 0.007 | -0.002 | -0.002 | 0 | 0.25 | 0.15 | 0.13 | 0.25 | 0.2 | 0.26 |
| 9 | 32138539 | rs2819846 | -0.458 | -0.338 | 1.01 | 0.869 | -0.003 | -0.013 | -0.013 | 0 | 0.69 | 0.56 | 0.73 | 0.71 | 0.72 | 0.63 |
| 9 | 32143671 | rs906585 | 1.012 | -0.351 | 1.103 | 0.969 | -0.015 | -0.002 | -0.002 | 2.448 | 0.43 | 0.32 | 0.58 | 0.4 | 0.52 | 0.36 |
| 9 | 32163518 | rs10970821 | 2.097 | -0.464 | 0.75 | 0.823 | -0.067 | 0.157 | 0.157 | 0.709 | 0.29 | 0.08 | 0.3 | 0.01 | 0.34 | 0.21 |
| 9 | 32165673 | rs10758121 | 0.32 | -0.495 | 0.799 | 0.815 | -0.005 | 0.015 | 0.015 | 0 | 0.54 | 0.25 | 0.45 | 0.41 | 0.52 | 0.71 |
| 9 | 32169502 | rs1567256 | 0.988 | -0.459 | 0.877 | 0.758 | -0.009 | -0.005 | -0.005 | 0 | 0.1 | 0.11 | 0.2 | 0.07 | 0.09 | 0.11 |
| 9 | 32169647 | rs1567255 | -0.482 | -0.466 | 0.832 | 0.761 | -0.009 | 0.017 | 0.017 | 0 | 0.68 | 0.75 | 0.72 | 0.81 | 0.64 | 0.83 |
| 9 | 32169982 | rs1567253 | 0.697 | -0.469 | 0.892 | 0.809 | -0.011 | 0.143 | 0.143 | 0 | 0.11 | 0.2 | 0.24 | 0.41 | 0.12 | 0.11 |
| 9 | 32171850 | rs7854381 | -0.175 | -0.442 | 0.901 | 0.789 | -0.006 | 0.004 | 0.004 | 0 | 0.66 | 0.73 | 0.7 | 0.66 | 0.64 | 0.55 |
| 9 | 32173751 | rs2819842 | -0.795 | -0.365 | 0.47 | 0.763 | -0.011 | 0.031 | 0.031 | 0 | 0.71 | 0.83 | 0.77 | 0.81 | 0.7 | 0.86 |
| 9 | 32176843 | rs16917960 | 0.558 | -0.396 | 0.467 | 0.767 | -0.012 | 0.138 | 0.138 | 0 | 0.12 | 0.2 | 0.24 | 0.41 | 0.12 | 0.11 |
| 9 | 32176938 | rs10970826 | 0.15 | -0.408 | 0.442 | 0.56 | 0.003 | -0.002 | -0.002 | 0.507 | 0.18 | 0.12 | 0.1 | 0.21 | 0.15 | 0.22 |
| 9 | 32181180 | rs10970828 | -0.2 | -0.316 | 0.463 | 0.592 | 0.007 | 0.002 | 0.002 | 0.174 | 0.23 | 0.15 | 0.11 | 0.21 | 0.15 | 0.23 |
| 9 | 32181573 | rs2843582 | -0.492 | -0.243 | 0.323 | 0.451 | -0.027 | 0.04 | 0.04 | 0 | 0.7 | 0.83 | 0.73 | 0.81 | 0.64 | 0.83 |
| 9 | 32191942 | rs1331250 | 1.024 | -0.241 | 0.394 | 0.526 | -0.01 | 0.005 | 0.005 | 0 | 0.1 | 0.15 | 0.21 | 0.1 | 0.11 | 0.11 |
| 9 | 32192407 | rs1929351 | -0.685 | -0.239 | -0.023 | 0.465 | -0.01 | 0.024 | 0.024 | 0 | 0.72 | 0.79 | 0.77 | 0.83 | 0.7 | 0.86 |
| 9 | 32203380 | rs1556395 | -0.254 | -0.413 | -0.057 | 0.556 | 0 | 0.015 | 0.015 | 1.477 | 0.62 | 0.82 | 0.72 | 0.65 | 0.64 | 0.77 |
| 9 | 32205704 | rs1331240 | 1.601 | -0.283 | -0.04 | 0.637 | -0.009 | 0.021 | 0.021 | 0.814 | 0.3 | 0.27 | 0.34 | 0.07 | 0.35 | 0.22 |
| 9 | 32211402 | rs7853053 | 0.257 | -0.235 | -0.603 | 0.726 | 0 | 0.014 | 0.014 | 0.778 | 0.5 | 0.65 | 0.48 | 0.58 | 0.47 | 0.62 |
| 9 | 32223204 | rs12379725 | 0.9 | 0.094 | -1.716 | 0.632 | -0.06 | 0.102 | 0.102 | 1.523 | 0.37 | 0.12 | 0.38 | 0.16 | 0.44 | 0.35 |
| 9 | 32231421 | rs13301455 | 1.035 | 0.209 | -1.595 | 0.722 | 0.003 | -0.001 | -0.001 | 0 | 0.08 | 0.09 | 0.07 | 0.08 | 0.1 | 0.12 |
| 9 | 32234067 | rs1331248 | -0.041 | 0.23 | -1.94 | 0.452 | -0.053 | -0.044 | -0.044 | 1.052 | 0.66 | 0.92 | 0.74 | 0.63 | 0.62 | 0.75 |
| 9 | 32239114 | rs1331249 | 1.29 | 0.337 | -1.894 | 0.149 | -0.013 | 0.065 | 0.065 | 0.888 | 0.31 | 0.22 | 0.33 | 0.09 | 0.34 | 0.23 |
| 9 | 32239645 | rs10970862 | 0.073 | -0.035 | -1.389 | 0.537 | 0.004 | -0.041 | -0.041 | 0 | 0.1 | 0.01 | 0.04 | 0.19 | 0.08 | 0.17 |
| 9 | 32246791 | rs1999368 | -0.731 | 0.04 | -0.454 | 0.461 | -0.041 | -0.049 | -0.049 | 0.652 | 0.24 | 0.44 | 0.29 | 0.16 | 0.18 | 0.3 |
| 9 | 32255125 | rs10970869 | -0.534 | 1.037 | -1.144 | 0.913 | -0.055 | -0.069 | -0.069 | 0 | 0.15 | 0.03 | 0.16 | 0.22 | 0.19 | 0.21 |
| 9 | 32257426 | rs10511898 | -0.995 | 0.566 | -0.322 | 0.577 | -0.006 | -0.013 | -0.013 | 0.017 | 0.11 | 0.21 | 0.1 | 0.08 | 0.08 | 0.14 |
| 9 | 32257573 | rs10114714 | 1.159 | 0.997 | -0.501 | 0.598 | -0.005 | -0.002 | -0.002 | 0 | 0.95 | 0.92 | 0.84 | 0.94 | 0.9 | 0.89 |
| 9 | 32258150 | rs12346810 | -1.647 | 0.334 | 0.421 | 0.042 | -0.051 | -0.118 | -0.118 | 0 | 0.17 | 0.35 | 0.1 | 0.01 | 0.08 | 0.04 |
| 9 | 32258600 | rs1411872 | 0.537 | -0.085 | 0.249 | 0.312 | 0.007 | -0.003 | -0.003 | 0 | 0.55 | 0.42 | 0.58 | 0.56 | 0.63 | 0.79 |
| 9 | 32259801 | rs10116526 | -0.317 | -0.435 | 1.035 | 0.65 | -0.018 | 0.065 | 0.065 | 0.165 | 0.2 | 0.35 | 0.2 | 0.35 | 0.17 | 0.11 |
| 9 | 32262917 | rs12336397 | -0.794 | -0.487 | 1.269 | 0.902 | 0.026 | 0.111 | 0.111 | 2.451 | 0.06 | 0.22 | 0.07 | 0.14 | 0.03 | 0.05 |
| 9 | 32267867 | rs1537156 | -0.493 | -0.961 | 1.699 | 1.233 | -0.012 | 0.004 | 0.004 | 0 | 0.27 | 0.35 | 0.26 | 0.25 | 0.2 | 0.19 |
| 9 | 32270454 | rs7848402 | -0.244 | -0.281 | 0.861 | 0.672 | -0.006 | -0.091 | -0.091 | 1.47 | 0.32 | 0.14 | 0.3 | 0.53 | 0.36 | 0.43 |
| 9 | 32271795 | rs7024902 | 0.445 | 0.109 | 0.783 | 0.519 | -0.001 | -0.002 | -0.002 | 0.485 | 0.61 | 0.61 | 0.65 | 0.47 | 0.57 | 0.54 |
| 9 | 32274310 | rs7037573 | -0.52 | 0.148 | 0.829 | 0.539 | -0.006 | 0.023 | 0.023 | 0.299 | 0.33 | 0.56 | 0.39 | 0.39 | 0.28 | 0.24 |
| 9 | 32275256 | rs1547699 | -1.285 | 0.056 | 0.497 | 0.316 | -0.001 | 0.031 | 0.031 | 0 | 0.93 | 0.99 | 0.97 | 1 | 0.94 | 0.97 |
| 9 | 32276752 | rs10970892 | 0.828 | 0.163 | 0.513 | 0.429 | -0.023 | -0.032 | -0.032 | 1.817 | 0.18 | 0.08 | 0.24 | 0.32 | 0.27 | 0.36 |
| 9 | 32278891 | rs3928808 | 2.141 | 0.049 | 0.75 | 0.374 | -0.004 | 0.004 | 0.004 | 0 | 0.32 | 0.29 | 0.36 | 0.06 | 0.34 | 0.31 |
| 9 | 32282978 | rs10970899 | 0.241 | 0.07 | 0.694 | 0.401 | -0.049 | -0.033 | -0.033 | 0 | 0.23 | 0.1 | 0.27 | 0.32 | 0.33 | 0.39 |

| | | | | | | | | | | | | | | | | |
|---|---|---|---|---|---|---|---|---|---|---|---|---|---|---|---|---|
| 9 | 32283644 | rs7039600 | 2.33 | 0.153 | 0.863 | 0.361 | 0 | 0.041 | 0.041 | 0.039 | 0.3 | 0.2 | 0.27 | 0.07 | 0.29 | 0.3 |
| 9 | 32292317 | rs1331226 | 0.708 | 0.021 | 0.844 | 0.364 | 0.008 | 0.061 | 0.061 | 0 | 0.1 | 0.14 | 0.21 | 0.26 | 0.1 | 0.15 |
| 9 | 32295707 | rs10511900 | 1.583 | -0.037 | 0.783 | 0.367 | 0.014 | -0.065 | -0.065 | 0 | 0.13 | 0.01 | 0.05 | 0.2 | 0.09 | 0.06 |
| 9 | 32302837 | rs11792665 | 2.825 | -0.039 | 0.791 | 0.318 | -0.034 | 0.138 | 0.138 | 0.131 | 0.29 | 0.09 | 0.27 | 0.07 | 0.29 | 0.3 |
| 9 | 32305904 | rs1331231 | -1.118 | 0.029 | 0.796 | 0.386 | 0.001 | 0.067 | 0.067 | 0 | 0.48 | 0.39 | 0.54 | 0.8 | 0.53 | 0.61 |
| 9 | 32310726 | rs992710 | 1.877 | 0.023 | 0.842 | 0.38 | 0.003 | -0.027 | -0.027 | 0 | 0.36 | 0.55 | 0.41 | 0.2 | 0.37 | 0.36 |
| 9 | 32314084 | rs1121130 | -0.775 | 0.106 | 0.474 | 0.43 | 0.02 | 0.027 | 0.027 | 0 | 0.85 | 0.93 | 0.95 | 1 | 0.9 | 0.96 |
| 9 | 32316967 | rs1411866 | -0.795 | 0.106 | 0.473 | 0.416 | 0.021 | 0.028 | 0.028 | 0 | 0.85 | 0.94 | 0.95 | 1 | 0.9 | 0.96 |
| 9 | 32319302 | rs7027564 | 0.568 | 0.168 | 0.515 | 0.517 | 0.007 | 0.064 | 0.064 | 0 | 0.1 | 0.14 | 0.21 | 0.27 | 0.1 | 0.15 |
| 9 | 32319634 | rs2094703 | -0.444 | 0.202 | 0.56 | 0.517 | 0.046 | 0.049 | 0.049 | 0 | 0.29 | 0.43 | 0.4 | 0.45 | 0.42 | 0.44 |
| 9 | 32328019 | rs7856471 | 2.072 | 0.199 | 0.339 | 0.456 | 0.002 | -0.012 | -0.012 | 0 | 0.34 | 0.35 | 0.27 | 0.07 | 0.29 | 0.31 |
| 9 | 32331414 | rs10970923 | -0.27 | 0.236 | 0.388 | 0.734 | 0.081 | 0.094 | 0.094 | 0.204 | 0.09 | 0.31 | 0.1 | 0.14 | 0.1 | 0.08 |
| 9 | 32332855 | rs10117454 | 0.066 | 0.024 | -0.266 | 0.615 | 0.052 | 0.016 | 0.016 | 0.159 | 0.71 | 0.93 | 0.88 | 0.79 | 0.8 | 0.89 |
| 9 | 32333058 | rs11999411 | 0.236 | 0.247 | -0.103 | 0.878 | 0.026 | 0.078 | 0.078 | 0 | 0.1 | 0.15 | 0.28 | 0.27 | 0.15 | 0.14 |
| 9 | 32333630 | rs16918170 | 1.038 | -0.114 | -0.523 | 0.908 | 0.013 | -0.06 | -0.06 | 0 | 0.13 | 0.01 | 0.05 | 0.2 | 0.09 | 0.07 |
| 9 | 32341611 | rs10970932 | 0.842 | 0.183 | -0.089 | 0.875 | 0.01 | -0.001 | -0.001 | 0 | 0.49 | 0.55 | 0.38 | 0.48 | 0.39 | 0.4 |
| 9 | 32343182 | rs4598348 | -1.04 | 0.41 | -0.195 | 1.385 | 0 | 0.003 | 0.003 | 0 | 0.22 | 0.22 | 0.27 | 0.46 | 0.26 | 0.36 |
| 9 | 32344707 | rs10970934 | -1.236 | 0.241 | -0.043 | 1.159 | -0.004 | 0.05 | 0.05 | 0.801 | 0.05 | 0.17 | 0.04 | 0.14 | 0.02 | 0.06 |
| 9 | 32344825 | rs10511903 | -0.7 | 0.417 | -0.427 | 1.465 | -0.007 | -0.038 | -0.038 | 0 | 0.12 | 0.03 | 0.13 | 0.25 | 0.15 | 0.13 |
| 9 | 32359091 | rs10813796 | -0.72 | 0.401 | -0.484 | 1.086 | 0.01 | 0.006 | 0.006 | 0 | 0.51 | 0.41 | 0.53 | 0.51 | 0.54 | 0.51 |
| 9 | 32361569 | rs10970941 | 0.496 | 0.482 | -0.355 | 1.016 | 0.001 | 0.006 | 0.006 | 0 | 0.11 | 0.18 | 0.08 | 0.19 | 0.13 | 0.07 |
| 9 | 32372280 | rs10970945 | 0.435 | 0.5 | -0.496 | 1.029 | -0.043 | 0 | 0 | 0 | 0.28 | 0.21 | 0.41 | 0.21 | 0.39 | 0.33 |
| 9 | 32372462 | rs7855483 | -2.15 | -0.032 | -0.106 | 0.894 | 0.009 | -0.003 | -0.003 | 0 | 0.73 | 0.82 | 0.73 | 0.72 | 0.81 | 0.62 |
| 9 | 32374666 | rs9792540 | 1.349 | -0.398 | -0.046 | 0.701 | 0.01 | 0.011 | 0.011 | 1.077 | 0.21 | 0.15 | 0.09 | 0.19 | 0.17 | 0.07 |
| 9 | 32376618 | rs1023087 | -1.962 | -0.395 | 0.235 | 0.742 | 0 | -0.001 | -0.001 | 0 | 0.7 | 0.74 | 0.71 | 0.68 | 0.78 | 0.6 |
| 9 | 32378956 | rs10813801 | -0.187 | -0.41 | 0.392 | 0.693 | 0.03 | -0.024 | -0.024 | 0 | 0.36 | 0.53 | 0.53 | 0.21 | 0.51 | 0.35 |
| 9 | 32381785 | rs1023085 | -2.247 | -0.477 | 0.411 | 0.702 | -0.002 | 0 | 0 | 0 | 0.72 | 0.74 | 0.71 | 0.67 | 0.81 | 0.61 |
| 9 | 32381910 | rs10813803 | 1.315 | -0.436 | 0.415 | 0.735 | 0.008 | 0.009 | 0.009 | 1.034 | 0.21 | 0.15 | 0.09 | 0.19 | 0.18 | 0.07 |
| 9 | 32383708 | rs867469 | -0.447 | -0.088 | 0.631 | 0.535 | 0 | -0.001 | -0.001 | 0 | 0.36 | 0.45 | 0.53 | 0.21 | 0.54 | 0.36 |
| 9 | 32392611 | rs10813805 | -0.005 | 0.087 | 0.576 | 0.535 | -0.004 | 0.001 | 0.001 | 0 | 0.33 | 0.34 | 0.49 | 0.21 | 0.46 | 0.33 |
| 9 | 32394422 | rs1041321 | 0.176 | -0.042 | 0.497 | 0.45 | 0.026 | 0.033 | 0.033 | 1.711 | 0.54 | 0.37 | 0.71 | 0.25 | 0.65 | 0.61 |
| 9 | 32395013 | rs7847742 | 0.153 | 0.025 | 0.477 | 0.481 | -0.004 | 0.001 | 0.001 | 0 | 0.33 | 0.34 | 0.49 | 0.21 | 0.46 | 0.33 |
| 9 | 32397920 | rs10813808 | 0.469 | -0.002 | 0.57 | 0.593 | 0.02 | -0.022 | -0.022 | 0 | 0.38 | 0.16 | 0.2 | 0.48 | 0.32 | 0.26 |
| 9 | 32398234 | rs7033149 | 0.463 | 0.121 | 0.19 | 0.674 | 0.045 | 0.154 | 0.154 | 0.131 | 0.91 | 0.7 | 0.84 | 0.73 | 0.87 | 0.85 |
| 9 | 32401610 | rs6476361 | -1.207 | 0.176 | -0.299 | 0.685 | -0.007 | -0.006 | -0.006 | 0 | 0.67 | 0.51 | 0.62 | 0.69 | 0.68 | 0.57 |
| 9 | 32403290 | rs10970961 | 0.676 | 0.728 | -0.855 | 0.568 | -0.052 | -0.016 | -0.016 | 1.1 | 0.26 | 0.1 | 0.37 | 0.21 | 0.33 | 0.31 |
| 9 | 32410850 | rs4442231 | 0.173 | 0.654 | -0.213 | 0.357 | 0.033 | -0.033 | -0.033 | 0 | 0.94 | 0.82 | 0.85 | 1 | 0.9 | 0.97 |
| 9 | 32412622 | rs4879584 | 0.453 | 0.426 | -0.735 | 0.654 | -0.003 | -0.052 | -0.052 | 0.053 | 0.17 | 0.05 | 0.13 | 0.34 | 0.16 | 0.19 |
| 9 | 32416386 | rs10970969 | 0.2 | -0.082 | -1.473 | 0.068 | -0.02 | -0.011 | -0.011 | 0.485 | 0.4 | 0.07 | 0.36 | 0.37 | 0.36 | 0.47 |
| 9 | 32418237 | rs2026739 | 0.091 | 0.18 | -0.306 | 0.401 | 0.012 | 0.039 | 0.039 | 0 | 0.75 | 0.57 | 0.68 | 0.64 | 0.73 | 0.73 |
| 9 | 32424210 | rs10970972 | 0.852 | 0.285 | -0.164 | 0.032 | 0.035 | 0.13 | 0.13 | 0 | 0.47 | 0.26 | 0.32 | 0.1 | 0.39 | 0.43 |
| 9 | 32424893 | rs10813814 | 0.023 | 0.208 | 0.132 | 0.595 | -0.012 | 0.046 | 0.046 | 0.007 | 0.74 | 0.39 | 0.68 | 0.48 | 0.76 | 0.72 |
| 9 | 32425270 | rs13293491 | 0.478 | 0.448 | 0.063 | 0.576 | -0.004 | -0.005 | -0.005 | 0.204 | 0.26 | 0.22 | 0.33 | 0.54 | 0.33 | 0.29 |
| 9 | 32425910 | rs3780473 | 0.315 | 0.353 | -0.124 | 0.517 | -0.013 | -0.023 | -0.023 | 0.312 | 0.28 | 0.26 | 0.34 | 0.55 | 0.34 | 0.3 |
| 9 | 32427874 | rs4879586 | 0.108 | -0.073 | 0.012 | 0.683 | -0.005 | -0.013 | -0.013 | 0.353 | 0.18 | 0.15 | 0.27 | 0.36 | 0.22 | 0.24 |
| 9 | 32431306 | rs12555576 | 1.537 | 0.119 | 0.164 | 0.523 | -0.034 | 0.077 | 0.077 | 0.385 | 0.2 | 0.06 | 0.11 | 0.04 | 0.2 | 0.32 |
| 9 | 32432681 | rs1467712 | -0.097 | 0.234 | 0.557 | 0.539 | 0.012 | 0.022 | 0.022 | 0 | 0.29 | 0.32 | 0.35 | 0.5 | 0.36 | 0.25 |
| 9 | 32435676 | rs10758139 | -0.135 | 0.218 | 0.671 | 0.543 | -0.019 | -0.034 | -0.034 | 0.088 | 0.3 | 0.24 | 0.36 | 0.5 | 0.36 | 0.26 |
| 9 | 32437249 | rs10813816 | -0.204 | 0.211 | 0.805 | 0.534 | 0.008 | 0.018 | 0.018 | 0 | 0.31 | 0.34 | 0.37 | 0.5 | 0.36 | 0.26 |
| 9 | 32449244 | rs10813818 | -0.68 | 0.075 | 0.814 | 0.508 | -0.002 | -0.002 | -0.002 | 0 | 0.53 | 0.56 | 0.49 | 0.58 | 0.57 | 0.59 |
| 9 | 32450187 | rs12985 | -0.226 | 0.275 | 0.541 | 0.561 | 0.027 | 0.037 | 0.037 | 0 | 0.31 | 0.43 | 0.39 | 0.52 | 0.37 | 0.26 |
| 9 | 32451144 | rs7042042 | -0.959 | 0.213 | 0.55 | 0.59 | -0.006 | -0.003 | -0.003 | 0 | 0.55 | 0.56 | 0.6 | 0.57 | 0.63 | 0.63 |
| 9 | 32452627 | rs17288914 | 1.498 | 0.407 | 0.26 | 0.58 | 0 | 0.045 | 0.045 | 0.779 | 0.21 | 0.11 | 0.11 | 0.06 | 0.2 | 0.33 |
| 9 | 32453278 | rs10970986 | 0.652 | 0.659 | 0.143 | 0.954 | -0.11 | -0.087 | -0.087 | 0.783 | 0.2 | 0.01 | 0.27 | 0.28 | 0.28 | 0.19 |
| 9 | 32456017 | rs12006123 | 1.441 | 0.493 | 0.264 | 0.717 | -0.002 | -0.017 | -0.017 | 0.067 | 0.22 | 0.24 | 0.12 | 0.06 | 0.2 | 0.32 |
| 9 | 32456656 | rs12555727 | 1.66 | 0.517 | 0.041 | 0.724 | -0.007 | 0.06 | 0.06 | 0.405 | 0.2 | 0.05 | 0.1 | 0.05 | 0.2 | 0.32 |
| 9 | 32458053 | rs10738889 | -0.31 | 0.583 | 0.502 | 1.062 | -0.026 | -0.023 | -0.023 | 0 | 0.25 | 0.22 | 0.43 | 0.37 | 0.35 | 0.24 |
| 9 | 32458081 | rs10738890 | -0.237 | 0.548 | 0.416 | 1.073 | -0.037 | -0.031 | -0.031 | 0 | 0.24 | 0.16 | 0.42 | 0.37 | 0.35 | 0.24 |
| 9 | 32459450 | rs3205166 | 0.107 | 0.165 | 0.414 | 0.907 | -0.016 | -0.038 | -0.038 | 0.179 | 0.35 | 0.2 | 0.36 | 0.45 | 0.35 | 0.25 |
| 9 | 32461331 | rs9650702 | 0.606 | 0.659 | 0.416 | -0.008 | -0.029 | 0.011 | 0.011 | 0 | 0.15 | 0.07 | 0.12 | 0.13 | 0.21 | 0.12 |
| 9 | 32467129 | rs10813821 | 0.263 | 0.571 | 0.468 | -0.227 | -0.04 | 0.069 | 0.069 | 1.078 | 0.36 | 0.14 | 0.34 | 0.21 | 0.42 | 0.46 |
| 9 | 32474690 | rs4384073 | 0.735 | 0.636 | 1.005 | -0.102 | 0.02 | 0.06 | 0.06 | 0 | 0.9 | 0.68 | 0.83 | 0.84 | 0.88 | 0.86 |
| 9 | 32487010 | rs7037171 | -0.332 | 0.416 | 1.048 | -0.075 | 0.002 | -0.025 | -0.025 | 0 | 0.94 | 0.89 | 0.98 | 1 | 0.93 | 0.92 |
| 9 | 32488421 | rs10971001 | 0.038 | 0.54 | 1.14 | -0.064 | -0.028 | 0.058 | 0.058 | 0.413 | 0.36 | 0.22 | 0.35 | 0.21 | 0.42 | 0.47 |
| 9 | 32490675 | rs639949 | -0.989 | 0.418 | 1.241 | -0.176 | -0.001 | 0.001 | 0.001 | 0 | 0.61 | 0.62 | 0.5 | 0.26 | 0.56 | 0.57 |
| 9 | 32492352 | rs17289927 | 1.094 | 0.236 | 1.335 | -0.107 | 0.028 | 0.096 | 0.096 | 2.281 | 0.23 | 0.06 | 0.12 | 0.01 | 0.14 | 0.06 |
| 9 | 32492903 | rs605383 | -1.248 | 0.242 | 1.346 | -0.107 | 0 | -0.003 | -0.003 | 1.615 | 0.66 | 0.64 | 0.52 | 0.26 | 0.63 | 0.61 |
| 9 | 32497007 | rs10813826 | -0.074 | 0.274 | 1.343 | -0.095 | -0.055 | 0.092 | 0.092 | 0.771 | 0.39 | 0.16 | 0.36 | 0.2 | 0.49 | 0.49 |
| 9 | 32502525 | rs7026407 | -1.319 | 0.261 | 1.387 | -0.067 | 0.001 | 0.002 | 0.002 | 0 | 0.66 | 0.64 | 0.53 | 0.38 | 0.62 | 0.62 |

| | | | | | | | | | | | | | | | | |
|---|---|---|---|---|---|---|---|---|---|---|---|---|---|---|---|---|
| 9 | 32503440 | rs669260 | 0.94 | 0.302 | 1.255 | -0.039 | 0.026 | 0.048 | 0.048 | 2.644 | 0.23 | 0.14 | 0.12 | 0.01 | 0.14 | 0.06 |
| 9 | 32516864 | rs621277 | -1.009 | 0.313 | 1.292 | -0.025 | -0.001 | -0.001 | -0.001 | 0 | 0.64 | 0.41 | 0.51 | 0.38 | 0.63 | 0.63 |
| 9 | 32519154 | rs10738891 | -1.263 | 0.318 | 1.266 | -0.024 | 0.001 | 0.009 | 0.009 | 0 | 0.64 | 0.49 | 0.51 | 0.39 | 0.63 | 0.64 |
| 9 | 32523737 | rs11795343 | -0.492 | 0.412 | 0.768 | 0.014 | 0.001 | -0.014 | -0.014 | 0 | 0.43 | 0.48 | 0.27 | 0.25 | 0.42 | 0.27 |
| 9 | 32524298 | rs17290242 | 1.373 | 0.269 | 0.666 | -0.018 | 0.001 | 0.042 | 0.042 | 0.076 | 0.22 | 0.05 | 0.22 | 0.11 | 0.21 | 0.36 |
| 9 | 32532469 | rs626214 | -0.547 | 0.292 | 0.691 | -0.013 | -0.006 | 0.001 | 0.001 | 0.997 | 0.44 | 0.63 | 0.44 | 0.43 | 0.49 | 0.57 |
| 9 | 32533829 | rs7025666 | 0.549 | 0.448 | 0.801 | 0.002 | 0.005 | -0.003 | -0.003 | 2.496 | 0.19 | 0.23 | 0.17 | 0.14 | 0.28 | 0.16 |
| 9 | 32534851 | rs657454 | -1.326 | 0.484 | 0.706 | 0.039 | 0.006 | 0 | 0 | 0.759 | 0.67 | 0.62 | 0.52 | 0.34 | 0.62 | 0.63 |
| 9 | 32537141 | rs597131 | -0.248 | 0.488 | 0.594 | 0.016 | -0.022 | 0.043 | 0.043 | 0.99 | 0.41 | 0.3 | 0.37 | 0.25 | 0.49 | 0.52 |
| 9 | 32542204 | rs10971019 | 0.687 | 0.618 | 0.456 | 0.098 | 0.005 | 0.036 | 0.036 | 0.009 | 0.22 | 0.07 | 0.2 | 0.11 | 0.21 | 0.36 |
| 9 | 32555049 | rs6476369 | 0.232 | 1.168 | 1.4 | 0.766 | 0.009 | -0.017 | -0.017 | 0.166 | 0.64 | 0.62 | 0.51 | 0.22 | 0.59 | 0.56 |
| 9 | 32559114 | rs10813835 | 0.376 | 1.123 | 1.038 | 0.771 | -0.003 | 0.004 | 0.004 | 0 | 0.63 | 0.41 | 0.5 | 0.22 | 0.59 | 0.56 |
| 9 | 32569145 | rs637421 | 0.349 | 1.51 | -0.088 | 0.574 | -0.001 | -0.001 | -0.001 | 0 | 0.67 | 0.68 | 0.54 | 0.37 | 0.6 | 0.58 |
| 9 | 32586822 | rs12000260 | -1.443 | 1.48 | 0.281 | 0.216 | 0.008 | -0.012 | -0.012 | 0 | 0.14 | 0.19 | 0.18 | 0.08 | 0.18 | 0.16 |
| 9 | 32592639 | rs688958 | -1.564 | 1.93 | 0.631 | 0.041 | 0.017 | -0.01 | -0.01 | 0 | 0.06 | 0.11 | 0.08 | 0.03 | 0.1 | 0.08 |
| 9 | 32596952 | rs2050455 | -1.244 | 1.865 | 0.741 | -0.028 | -0.01 | 0.009 | 0.009 | 0 | 0.12 | 0.12 | 0.17 | 0.05 | 0.18 | 0.11 |
| 9 | 32601501 | rs540467 | -2.617 | 1.551 | 1.914 | -0.225 | 0.013 | -0.081 | -0.081 | 0 | 0.19 | 0.6 | 0.17 | 0.04 | 0.21 | 0.1 |
| 9 | 32611604 | rs2196271 | -1.161 | 1.475 | 1.89 | -0.154 | 0 | -0.003 | -0.003 | 0 | 0.1 | 0.11 | 0.09 | 0.04 | 0.14 | 0.08 |
| 9 | 32618055 | rs10813840 | -1.846 | 1.319 | 2.03 | 0.73 | 0.003 | 0.026 | 0.026 | 0.009 | 0.11 | 0.13 | 0.22 | 0.31 | 0.15 | 0.21 |
| 9 | 32619211 | rs10813841 | -0.844 | 1.316 | 2.032 | 0.864 | 0.012 | 0.099 | 0.099 | 0 | 0.07 | 0.12 | 0.12 | 0.31 | 0.1 | 0.16 |
| 9 | 32625622 | rs10813843 | -0.674 | 1.309 | 2.024 | 0.852 | 0.01 | 0.007 | 0.007 | 0 | 0.07 | 0.09 | 0.05 | 0.09 | 0.1 | 0.14 |
| 9 | 32631369 | rs10971047 | -0.913 | 1.321 | 2.022 | 0.861 | 0.01 | 0.094 | 0.094 | 0 | 0.07 | 0.11 | 0.12 | 0.31 | 0.1 | 0.17 |
| 9 | 32642224 | rs10971049 | -1.831 | 1.311 | 2.012 | 0.853 | -0.001 | 0.017 | 0.017 | 0 | 0.11 | 0.13 | 0.22 | 0.31 | 0.15 | 0.22 |
| 9 | 32668703 | rs16918411 | -1.485 | 1.092 | 2.32 | 0.742 | 0.034 | 0.048 | 0.048 | 0 | 0.09 | 0.24 | 0.07 | 0.12 | 0.1 | 0.14 |
| 9 | 32670310 | rs10813852 | -1.496 | 1.078 | 2.412 | 0.735 | 0.043 | 0.058 | 0.058 | 0 | 0.09 | 0.32 | 0.09 | 0.12 | 0.1 | 0.14 |
| 9 | 32675417 | rs2117523 | -2.912 | 0.884 | 2.802 | 0.734 | 0.035 | 0.293 | 0.293 | 0.948 | 0.08 | 0.35 | 0.15 | 0.28 | 0.04 | 0.1 |
| 9 | 32698601 | rs1329005 | 2.639 | 0.738 | 3.266 | 0.719 | 0.034 | 0.396 | 0.396 | 4.659 | 0.94 | 0.62 | 0.9 | 0.76 | 1 | 0.96 |
| 9 | 32701164 | rs7846828 | -0.313 | 0.432 | 2.443 | -0.184 | 0 | -0.028 | -0.028 | 0 | 0.33 | 0.3 | 0.19 | 0.06 | 0.28 | 0.21 |
| 9 | 32701552 | rs7862812 | -0.623 | 0.788 | 2.4 | -0.393 | -0.01 | 0.03 | 0.03 | 0 | 0.39 | 0.3 | 0.25 | 0.09 | 0.38 | 0.26 |
| 9 | 32717013 | rs1328994 | -1.63 | 0.786 | 2.064 | -0.539 | 0.003 | -0.016 | -0.016 | 0 | 0.58 | 0.78 | 0.44 | 0.42 | 0.56 | 0.41 |
| 9 | 32717956 | rs1326917 | 0.446 | 0.731 | 1.977 | -0.584 | -0.001 | 0.037 | 0.037 | 0 | 0.54 | 0.66 | 0.72 | 0.89 | 0.56 | 0.68 |
| 9 | 32720834 | rs1328996 | 0.316 | 0.779 | 1.725 | -0.542 | -0.008 | 0.126 | 0.126 | 1.165 | 0.31 | 0.13 | 0.18 | 0.06 | 0.27 | 0.2 |
| 9 | 32721882 | rs4879604 | 1.498 | 0.794 | 1.837 | -0.53 | -0.017 | -0.044 | -0.044 | 0.706 | 0.41 | 0.16 | 0.57 | 0.58 | 0.45 | 0.6 |
| 9 | 32722999 | rs12003519 | -0.265 | 0.698 | 2.799 | 0.074 | 0.004 | -0.004 | -0.004 | 0.67 | 0.07 | 0.09 | 0.06 | 0.03 | 0.11 | 0.05 |
| 9 | 32723639 | rs3849901 | 1.211 | 0.72 | 2.99 | 0.052 | 0.001 | -0.076 | -0.076 | 0.303 | 0.82 | 0.62 | 0.88 | 0.95 | 0.83 | 0.9 |
| 9 | 32724290 | rs10813864 | 2.179 | 0.587 | 2.614 | -0.048 | -0.001 | -0.038 | -0.038 | 0 | 0.67 | 0.39 | 0.78 | 0.92 | 0.7 | 0.84 |
| 9 | 32724790 | rs7021315 | 0.282 | 0.804 | 1.903 | 0.035 | -0.007 | -0.017 | -0.017 | 0 | 0.08 | 0.08 | 0.04 | 0 | 0.02 | 0 |
| 9 | 32725612 | rs10971105 | -0.847 | 0.863 | 1.617 | 0.073 | 0.005 | -0.045 | -0.045 | 2.569 | 0.16 | 0.29 | 0.11 | 0.05 | 0.17 | 0.1 |
| 9 | 32725773 | rs7025476 | -1.162 | 0.66 | 1.152 | 0.515 | -0.007 | 0.091 | 0.091 | 0.49 | 0.19 | 0.51 | 0.18 | 0.3 | 0.13 | 0.1 |
| 9 | 32738005 | rs10813868 | -1.146 | 0.674 | 1.198 | -0.431 | -0.017 | -0.119 | -0.119 | 0 | 0.21 | 0.43 | 0.2 | 0.01 | 0.16 | 0.18 |
| 9 | 32738041 | rs10813869 | -1.276 | 0.656 | 1.492 | -0.439 | -0.01 | -0.075 | -0.075 | 0 | 0.2 | 0.27 | 0.18 | 0.01 | 0.16 | 0.18 |
| 9 | 32739599 | rs960083 | 1.338 | 1.239 | 1.882 | -0.982 | 0.032 | -0.119 | -0.119 | 0 | 0.74 | 0.47 | 0.77 | 0.93 | 0.69 | 0.83 |
| 9 | 32741428 | rs1410831 | 2.37 | 1.465 | 1.877 | -1.157 | -0.001 | 0.034 | 0.034 | 4.103 | 0.78 | 0.23 | 0.69 | 0.99 | 0.81 | 0.87 |
| 9 | 32741593 | rs1410832 | 1.919 | 1.466 | 2.232 | -1.149 | 0.002 | -0.043 | -0.043 | 0.383 | 0.84 | 0.33 | 0.79 | 1 | 0.85 | 0.91 |
| 9 | 32748365 | rs12551778 | -1.248 | 1.629 | 2.519 | -1.227 | 0.005 | -0.025 | -0.025 | 0 | 0.11 | 0.13 | 0.13 | 0 | 0.17 | 0.08 |
| 9 | 32754309 | rs7848558 | -1.907 | 1.537 | 3.194 | -1.082 | 0.029 | 0.02 | 0.02 | 0 | 0.08 | 0.22 | 0.04 | 0.08 | 0.08 | 0.12 |
| 9 | 32754685 | rs7849269 | 2.886 | 2.077 | 3.482 | -1.354 | 0.011 | -0.037 | -0.037 | 0 | 0.75 | 0.35 | 0.7 | 0.92 | 0.72 | 0.8 |
| 9 | 32755028 | rs10758161 | 3.212 | 2.023 | 3.603 | -1.357 | 0.011 | -0.025 | -0.025 | 0.248 | 0.86 | 0.46 | 0.92 | 0.92 | 0.89 | 0.88 |
| 9 | 32759228 | rs10813873 | 3.603 | 2.076 | 3.551 | -1.372 | 0.003 | -0.056 | -0.056 | 0 | 0.82 | 0.47 | 0.84 | 0.93 | 0.86 | 0.83 |
| 9 | 32764431 | rs11999320 | 0.084 | 2.093 | 3.424 | -1.378 | -0.006 | 0.003 | 0.003 | 0 | 0.07 | 0.06 | 0.11 | 0 | 0.13 | 0.05 |
| 9 | 32765337 | rs11788663 | 0.536 | 2.121 | 2.125 | -1.219 | -0.015 | -0.046 | -0.046 | 0.404 | 0.27 | 0.09 | 0.25 | 0.34 | 0.31 | 0.18 |
| 9 | 32769523 | rs664241 | 2.119 | 2.132 | 2.636 | -1.163 | 0.022 | -0.012 | -0.012 | 0 | 0.88 | 0.69 | 0.88 | 0.93 | 0.9 | 0.83 |
| 9 | 32773035 | rs630276 | 2.249 | 2.111 | 2.634 | -1.051 | 0.041 | -0.049 | -0.049 | 0 | 0.81 | 0.62 | 0.77 | 0.92 | 0.77 | 0.77 |
| 9 | 32780525 | rs17312997 | -0.662 | 2.097 | 2.686 | -1.053 | 0.006 | -0.02 | -0.02 | 0 | 0.05 | 0.16 | 0.03 | 0 | 0.03 | 0 |
| 9 | 32780898 | rs640747 | 2.46 | 2.111 | 2.432 | -1.053 | 0.015 | -0.062 | -0.062 | 0 | 0.75 | 0.41 | 0.73 | 0.92 | 0.74 | 0.77 |
| 9 | 32784838 | rs1810807 | 0.763 | 2.138 | 1.878 | -1.226 | -0.019 | 0.026 | 0.026 | 4.848 | 0.28 | 0.06 | 0.23 | 0.22 | 0.28 | 0.14 |
| 9 | 32786653 | rs300637 | 2.542 | 2.878 | 2.097 | -0.929 | 0.021 | -0.062 | -0.062 | 0 | 0.74 | 0.41 | 0.73 | 0.92 | 0.72 | 0.77 |
| 9 | 32788611 | rs10813876 | -0.847 | 2.889 | 2.071 | -0.918 | -0.009 | 0.004 | 0.004 | 0 | 0.13 | 0.09 | 0.13 | 0.07 | 0.2 | 0.16 |
| 9 | 32797839 | rs300629 | 2.636 | 3.059 | 2.005 | 0.217 | 0.03 | 0.026 | 0.026 | 0 | 0.71 | 0.41 | 0.61 | 0.67 | 0.63 | 0.72 |
| 9 | 32802703 | rs2439760 | 2.555 | 2.785 | 2.064 | -0.26 | 0.027 | -0.071 | -0.071 | 0 | 0.75 | 0.42 | 0.7 | 0.92 | 0.73 | 0.77 |
| 9 | 32803092 | rs10971142 | -0.309 | 2.722 | 1.424 | -0.119 | 0.011 | -0.028 | -0.028 | 0 | 0.1 | 0.14 | 0.15 | 0.01 | 0.15 | 0.11 |
| 9 | 32821457 | rs606902 | 1.14 | 2.695 | 2.019 | -0.073 | 0.086 | -0.003 | -0.003 | 0 | 0.93 | 0.69 | 0.94 | 1 | 0.93 | 1 |
| 9 | 32824325 | rs10971156 | 0.882 | 2.533 | 1.588 | -0.178 | -0.002 | -0.009 | -0.009 | 1.486 | 0.07 | 0.06 | 0.08 | 0 | 0.08 | 0.03 |
| 9 | 32827302 | rs1822234 | -1.865 | 3.077 | 2.012 | 0.618 | 0.053 | 0.134 | 0.134 | 0 | 0.18 | 0.37 | 0.34 | 0.51 | 0.23 | 0.14 |
| 9 | 32830551 | rs10971159 | 0.675 | 2.687 | 1.83 | 0.21 | 0.008 | -0.031 | -0.031 | 0 | 0.07 | 0.09 | 0.08 | 0 | 0.08 | 0.03 |
| 9 | 32831188 | rs12552521 | -0.269 | 2.677 | 1.923 | 0.205 | 0.043 | -0.052 | -0.052 | 0.861 | 0.09 | 0.33 | 0.1 | 0 | 0.08 | 0.02 |
| 9 | 32831281 | rs12555111 | -1.015 | 2.843 | 1.739 | 0.213 | 0.04 | -0.017 | -0.017 | 0 | 0.13 | 0.26 | 0.11 | 0.1 | 0.16 | 0.12 |
| 9 | 32834922 | rs596443 | 1.877 | 2.843 | 1.535 | 0.228 | 0.021 | -0.033 | -0.033 | 0 | 0.77 | 0.37 | 0.8 | 0.87 | 0.77 | 0.85 |
| 9 | 32850506 | rs12684467 | -0.697 | 1.871 | 0.859 | 0.028 | -0.011 | -0.003 | -0.003 | 0.198 | 0.12 | 0.15 | 0.03 | 0.11 | 0.07 | 0.08 |

| | | | | | | | | | | | | | | | | |
|---|---|---|---|---|---|---|---|---|---|---|---|---|---|---|---|---|
| 9 | 32854057 | rs10971164 | 1.437 | 1.316 | -1.279 | 0.378 | -0.117 | -0.137 | -0.137 | 6.356 | 0.44 | 0.03 | 0.4 | 0.5 | 0.43 | 0.67 |
| 9 | 32855756 | rs1442286 | 0.202 | 1.165 | -0.616 | 0.293 | 0.033 | 0.051 | 0.051 | 0 | 0.5 | 0.6 | 0.35 | 0.36 | 0.38 | 0.68 |
| 9 | 32857801 | rs7875919 | -0.049 | 1.255 | -0.381 | 0.278 | 0.007 | -0.046 | -0.046 | 0.01 | 0.61 | 0.48 | 0.66 | 0.87 | 0.57 | 0.79 |
| 9 | 32859234 | rs10813878 | 0.9 | 0.832 | -1.211 | 0.159 | -0.135 | -0.124 | -0.124 | 0 | 0.4 | 0.02 | 0.31 | 0.44 | 0.44 | 0.62 |
| 9 | 32861992 | rs1359719 | -0.691 | 1.017 | -0.721 | -0.925 | -0.009 | -0.01 | -0.01 | 0 | 0.67 | 0.58 | 0.75 | 0.75 | 0.73 | 0.74 |
| 9 | 32862544 | rs4879619 | -0.625 | 0.776 | -0.883 | -1.141 | -0.012 | -0.002 | -0.002 | 0 | 0.8 | 0.71 | 0.88 | 0.82 | 0.9 | 0.8 |
| 9 | 32865017 | rs16918694 | 2.041 | 0.786 | -0.904 | -1.141 | 0 | 0.004 | 0.004 | 0.422 | 0.13 | 0.09 | 0.13 | 0.07 | 0.17 | 0.06 |
| 9 | 32866253 | rs10971169 | 1.329 | 0.855 | -0.935 | -1.164 | -0.017 | 0.026 | 0.026 | 0 | 0.19 | 0.1 | 0.22 | 0.02 | 0.24 | 0.08 |
| 9 | 32870406 | rs10118347 | -0.033 | 0.84 | -0.675 | -1.168 | -0.02 | -0.022 | -0.022 | 0 | 0.66 | 0.52 | 0.75 | 0.75 | 0.73 | 0.73 |
| 9 | 32870789 | rs10758165 | 0.915 | 0.848 | -0.66 | -1.124 | -0.002 | 0.011 | 0.011 | 0.917 | 0.46 | 0.4 | 0.53 | 0.73 | 0.49 | 0.65 |
| 9 | 32873495 | rs4879621 | 0.037 | 0.802 | -0.295 | -1.024 | 0.004 | 0.004 | 0.004 | 0.556 | 0.64 | 0.34 | 0.74 | 0.75 | 0.73 | 0.73 |
| 9 | 32876932 | rs7865934 | 2.848 | 1.226 | -0.198 | -0.901 | 0.025 | -0.03 | -0.03 | 0 | 0.06 | 0.08 | 0.08 | 0 | 0.11 | 0.02 |
| 9 | 32879341 | rs10813887 | -0.039 | 0.602 | -0.251 | -0.924 | -0.036 | -0.031 | -0.031 | 0 | 0.7 | 0.4 | 0.85 | 0.82 | 0.84 | 0.78 |
| 9 | 32880323 | rs10971175 | 1.282 | 0.914 | -0.496 | -0.709 | -0.035 | 0.039 | 0.039 | 0.059 | 0.06 | 0.02 | 0.28 | 0.49 | 0.11 | 0.08 |
| 9 | 32883747 | rs12683296 | 0.348 | 0.745 | -0.476 | -1.189 | 0.051 | 0.084 | 0.084 | 0.934 | 0.57 | 0.28 | 0.73 | 0.84 | 0.68 | 0.73 |
| 9 | 32884738 | rs7847699 | 1.619 | 0.711 | -0.402 | -1.193 | -0.007 | -0.024 | -0.024 | 0.871 | 0.15 | 0.19 | 0.12 | 0.07 | 0.17 | 0.05 |
| 9 | 32884983 | rs10122775 | 0.456 | 0.888 | -0.377 | -1.214 | 0.055 | 0.038 | 0.038 | 0 | 0.28 | 0.45 | 0.53 | 0.61 | 0.36 | 0.16 |
| 9 | 32886639 | rs1888871 | 1.114 | 0.782 | -0.256 | -1.076 | 0.036 | 0.135 | 0.135 | 0 | 0.51 | 0.25 | 0.46 | 0.26 | 0.57 | 0.64 |
| 9 | 32886717 | rs1348255 | -0.708 | 0.69 | -0.319 | -1.066 | 0.006 | -0.057 | -0.057 | 0.021 | 0.9 | 0.7 | 0.92 | 1 | 0.93 | 0.92 |
| 9 | 32889200 | rs10511913 | 1.272 | 0.935 | -0.294 | -0.847 | -0.033 | 0.024 | 0.024 | 0.013 | 0.06 | 0.02 | 0.27 | 0.58 | 0.11 | 0.08 |
| 9 | 32899807 | rs4879635 | -0.613 | 0.785 | -0.7 | -0.923 | -0.001 | -0.006 | -0.006 | 0.236 | 0.74 | 0.69 | 0.87 | 0.91 | 0.85 | 0.78 |
| 9 | 32903868 | rs1470217 | -0.294 | 0.97 | -0.397 | -1.138 | 0.003 | 0.007 | 0.007 | 0 | 0.61 | 0.59 | 0.57 | 0.33 | 0.7 | 0.69 |
| 9 | 32907608 | rs7042500 | 0.313 | 1.251 | 0.34 | -0.77 | -0.017 | -0.067 | -0.067 | 0 | 0.17 | 0.45 | 0.14 | 0.07 | 0.17 | 0.05 |
| 9 | 32909543 | rs10971207 | 0.148 | 1.735 | 0.168 | -0.426 | 0.013 | -0.069 | -0.069 | 0.249 | 0.2 | 0.25 | 0.26 | 0.02 | 0.24 | 0.08 |
| 9 | 32911250 | rs10971212 | 0.075 | 1.747 | -0.121 | -0.414 | -0.009 | 0.002 | 0.002 | 0 | 0.2 | 0.12 | 0.23 | 0.02 | 0.24 | 0.08 |
| 9 | 32917428 | rs13289434 | -0.916 | 1.965 | 0.196 | 0.133 | 0.01 | 0.01 | 0.01 | 0.221 | 0.22 | 0.3 | 0.2 | 0.27 | 0.23 | 0.25 |
| 9 | 32921928 | rs4879641 | -1.503 | 2.008 | 0.034 | -0.225 | -0.012 | 0.06 | 0.06 | 0.08 | 0.42 | 0.35 | 0.41 | 0.79 | 0.5 | 0.36 |
| 9 | 32923482 | rs16918866 | -0.083 | 2.138 | -0.013 | -0.424 | 0.045 | -0.109 | -0.109 | 3.089 | 0.22 | 0.37 | 0.22 | 0.03 | 0.28 | 0.09 |
| 9 | 32923966 | rs12555433 | 0.311 | 1.995 | -0.307 | -0.242 | -0.021 | -0.015 | -0.015 | 0.043 | 0.11 | 0.06 | 0.09 | 0.17 | 0.14 | 0.17 |
| 9 | 32925505 | rs12552198 | -0.868 | 1.455 | -0.174 | -0.557 | 0.005 | -0.064 | -0.064 | 0 | 0.41 | 0.49 | 0.42 | 0.87 | 0.5 | 0.36 |
| 9 | 32932194 | rs10733465 | 0.782 | 1.846 | -0.008 | -0.428 | 0.051 | -0.07 | -0.07 | 0.167 | 0.76 | 0.58 | 0.73 | 0.94 | 0.63 | 0.89 |
| 9 | 32970518 | rs4879653 | 0.175 | 1.764 | 0.299 | 0.271 | 0.026 | -0.054 | -0.054 | 0 | 0.9 | 0.73 | 0.93 | 1 | 0.9 | 0.96 |
| 9 | 32978640 | rs10813913 | 0.637 | 1.612 | 0.516 | 0.938 | -0.003 | 0.037 | 0.037 | 0 | 0.12 | 0.15 | 0.06 | 0.29 | 0.1 | 0.18 |
| 9 | 32978806 | rs10971259 | -0.395 | 1.247 | 0.59 | 1.15 | -0.01 | 0.062 | 0.062 | 0 | 0.17 | 0.21 | 0.15 | 0.4 | 0.13 | 0.25 |
| 9 | 32982687 | rs1016674 | 1.138 | 0.843 | 0.598 | 0.83 | 0.011 | -0.036 | -0.036 | 0.978 | 0.65 | 0.53 | 0.69 | 0.84 | 0.62 | 0.86 |
| 9 | 32984072 | rs3824457 | 1.526 | 0.814 | 0.711 | 0.823 | -0.004 | -0.027 | -0.027 | 2.232 | 0.1 | 0.04 | 0.05 | 0.29 | 0.1 | 0.18 |
| 9 | 32990132 | rs10813916 | 0.327 | 0.841 | 0.74 | 0.821 | -0.011 | 0.026 | 0.026 | 0.036 | 0.22 | 0.14 | 0.19 | 0.03 | 0.25 | 0.08 |
| 9 | 32994500 | rs2183870 | 2.029 | 0.764 | 0.636 | 0.79 | 0.005 | 0.006 | 0.006 | 0.472 | 0.47 | 0.31 | 0.53 | 0.44 | 0.49 | 0.57 |
| 9 | 33007411 | rs10124051 | 1.085 | 0.783 | 0.613 | 0.8 | -0.003 | 0.038 | 0.038 | 0 | 0.12 | 0.15 | 0.06 | 0.29 | 0.1 | 0.18 |
| 9 | 33023833 | rs10758179 | 0.971 | 0.746 | 0.62 | 0.788 | -0.004 | 0.059 | 0.059 | 0 | 0.12 | 0.18 | 0.06 | 0.28 | 0.1 | 0.18 |
| 9 | 33025113 | rs3818644 | -2.481 | 0.335 | -0.066 | 0.637 | -0.02 | 0.023 | 0.023 | 4.14 | 0.66 | 0.87 | 0.57 | 0.73 | 0.64 | 0.58 |
| 9 | 33026572 | rs20583 | -2.46 | 0.443 | 0.091 | 0.715 | -0.038 | -0.03 | -0.03 | 0 | 0.55 | 0.86 | 0.48 | 0.57 | 0.51 | 0.44 |
| 9 | 33026638 | rs2274769 | -0.526 | 0.471 | 0.038 | 0.697 | -0.001 | -0.001 | -0.001 | 0 | 0.3 | 0.34 | 0.3 | 0.16 | 0.38 | 0.14 |
| 9 | 33034051 | rs10448231 | 0.048 | 0.476 | -0.011 | 0.704 | -0.001 | 0 | 0 | 0 | 0.26 | 0.28 | 0.25 | 0.16 | 0.28 | 0.11 |
| 9 | 33034665 | rs1330692 | 1.98 | 0.159 | -0.952 | 0.279 | -0.002 | -0.044 | -0.044 | 0 | 0.18 | 0.02 | 0.16 | 0.21 | 0.15 | 0.28 |
| 9 | 33045147 | rs10758180 | -1.762 | 0.302 | 0.826 | 1.003 | 0.005 | -0.018 | -0.018 | 0.204 | 0.41 | 0.52 | 0.41 | 0.29 | 0.41 | 0.26 |
| 9 | 33050343 | rs10971368 | 0.662 | 0.465 | 0.131 | 1.145 | -0.003 | 0.046 | 0.046 | 0 | 0.12 | 0.15 | 0.06 | 0.3 | 0.1 | 0.18 |
| 9 | 33058162 | rs12379261 | -0.871 | 0.46 | -0.286 | 0.633 | -0.05 | -0.033 | -0.033 | 0 | 0.09 | 0.02 | 0.15 | 0.13 | 0.14 | 0.14 |
| 9 | 33082969 | rs12552290 | -1.363 | 0.427 | -0.205 | 0.582 | -0.003 | 0.01 | 0.01 | 0.036 | 0.33 | 0.27 | 0.35 | 0.28 | 0.33 | 0.22 |
| 9 | 33084384 | rs10758185 | -1.287 | 0.47 | -0.33 | 0.603 | -0.008 | 0.011 | 0.011 | 0 | 0.32 | 0.25 | 0.35 | 0.29 | 0.32 | 0.21 |
| 9 | 33092835 | rs10813942 | -0.489 | 0.715 | -0.307 | 0.508 | -0.048 | 0.07 | 0.07 | 0 | 0.26 | 0.14 | 0.23 | 0.03 | 0.34 | 0.11 |
| 9 | 33097996 | rs13300152 | -0.593 | 0.233 | -0.415 | 0.748 | -0.048 | 0.048 | 0.048 | 0 | 0.25 | 0.09 | 0.26 | 0.15 | 0.31 | 0.1 |
| 9 | 33099668 | rs10971406 | -0.46 | 0.244 | -0.41 | 0.752 | -0.047 | 0.05 | 0.05 | 0 | 0.25 | 0.09 | 0.26 | 0.16 | 0.31 | 0.1 |
| 9 | 33103173 | rs544723 | 1.442 | 0.195 | -0.224 | 0.979 | -0.016 | 0.092 | 0.092 | 2.7 | 0.49 | 0.18 | 0.56 | 0.36 | 0.54 | 0.58 |
| 9 | 33104524 | rs10813946 | -1.153 | 0.118 | 0.498 | 1.054 | 0 | 0.005 | 0.005 | 0.197 | 0.31 | 0.34 | 0.21 | 0.34 | 0.29 | 0.26 |
| 9 | 33104540 | rs1013217 | -1.382 | 0.161 | 0.525 | 1.048 | -0.007 | 0.025 | 0.025 | 0 | 0.36 | 0.44 | 0.27 | 0.47 | 0.31 | 0.36 |
| 9 | 33106619 | rs750592 | -1.119 | 0.128 | 0.468 | 0.3 | -0.009 | -0.013 | -0.013 | 0.057 | 0.24 | 0.3 | 0.2 | 0.15 | 0.19 | 0.15 |
| 9 | 33113322 | rs7019909 | 0.258 | 0.344 | 0.969 | 0.911 | 0 | 0.175 | 0.175 | 0 | 0.11 | 0.18 | 0.07 | 0.5 | 0.1 | 0.26 |
| 9 | 33117524 | rs2031902 | 1.686 | 0.203 | 0.77 | 0.382 | -0.008 | 0.061 | 0.061 | 0.003 | 0.56 | 0.14 | 0.63 | 0.31 | 0.58 | 0.57 |
| 9 | 33120706 | rs3818593 | 0.179 | 0.54 | 0.974 | 0.492 | -0.003 | 0.032 | 0.032 | 0.054 | 0.22 | 0.23 | 0.13 | 0.48 | 0.19 | 0.3 |
| 9 | 33124872 | rs12342831 | 0.176 | 0.527 | 1.074 | 0.248 | -0.002 | 0.025 | 0.025 | 0.07 | 0.26 | 0.28 | 0.16 | 0.54 | 0.25 | 0.32 |
| 9 | 33125279 | rs4879666 | 1.297 | 0.497 | 0.68 | 0.206 | -0.043 | 0.136 | 0.136 | 0.512 | 0.48 | 0.15 | 0.61 | 0.28 | 0.49 | 0.53 |
| 9 | 33131651 | rs10758192 | -0.608 | -0.156 | 0.991 | 0.329 | -0.013 | 0.001 | 0.001 | 0.653 | 0.36 | 0.55 | 0.26 | 0.61 | 0.32 | 0.35 |
| 9 | 33137303 | rs1411609 | -1.229 | -0.012 | 0.956 | 0.272 | 0.013 | 0.025 | 0.025 | 0.149 | 0.46 | 0.66 | 0.34 | 0.61 | 0.44 | 0.36 |
| 9 | 33165985 | rs10813958 | -1.44 | -0.029 | 0.823 | 0.258 | 0.023 | 0.079 | 0.079 | 0.319 | 0.48 | 0.69 | 0.36 | 0.73 | 0.45 | 0.45 |
| 9 | 33180228 | rs7854765 | -0.329 | 0.103 | 0.428 | 0.47 | 0.002 | 0.056 | 0.056 | 0 | 0.08 | 0.28 | 0.05 | 0.15 | 0.05 | 0.09 |
| 9 | 33180362 | rs10813960 | -0.224 | 0.397 | -0.316 | 0.53 | 0.005 | -0.036 | -0.036 | 6.747 | 0.29 | 0.16 | 0.2 | 0.54 | 0.27 | 0.31 |
| 9 | 33186156 | rs7038463 | -0.947 | 0.495 | 0.001 | 0.249 | 0.001 | -0.028 | -0.028 | 0 | 0.11 | 0.22 | 0.09 | 0.03 | 0.12 | 0.02 |
| 9 | 33192940 | rs1702956 | -0.146 | 0.443 | -0.573 | 0.218 | -0.001 | 0.037 | 0.037 | 2.215 | 0.47 | 0.39 | 0.4 | 0.66 | 0.49 | 0.4 |

| | | | | | | | | | | | | | | | | |
|---|---|---|---|---|---|---|---|---|---|---|---|---|---|---|---|---|
| 9 | 33193935 | rs1702962 | -0.063 | 1.039 | -0.549 | 0.771 | -0.01 | -0.03 | -0.03 | 0 | 0.44 | 0.2 | 0.35 | 0.51 | 0.43 | 0.33 |
| 9 | 33200956 | rs478689 | -0.232 | 1.011 | -0.21 | 0.471 | 0.012 | 0.045 | 0.045 | 1.128 | 0.47 | 0.35 | 0.4 | 0.66 | 0.5 | 0.4 |
| 9 | 33203602 | rs545575 | 0.229 | 0.374 | -0.306 | 0.074 | 0.005 | 0.004 | 0.004 | 0 | 0.15 | 0.11 | 0.07 | 0.08 | 0.13 | 0.22 |
| 9 | 33204104 | rs976164 | 0.431 | 0.202 | -0.752 | -0.149 | -0.031 | 0.01 | 0.01 | 0 | 0.39 | 0.27 | 0.43 | 0.65 | 0.44 | 0.41 |
| 9 | 33205183 | rs12351178 | 0.571 | -0.176 | -0.939 | -0.434 | 0.009 | 0.03 | 0.03 | 0 | 0.12 | 0.05 | 0.08 | 0.06 | 0.11 | 0.12 |
| 9 | 33205408 | rs4879670 | 0.719 | -0.116 | -0.811 | -0.353 | -0.031 | 0.006 | 0.006 | 0 | 0.39 | 0.26 | 0.42 | 0.62 | 0.44 | 0.41 |
| 9 | 33206375 | rs4879671 | 0.776 | -0.148 | -0.852 | -0.07 | -0.036 | -0.017 | -0.017 | 0 | 0.38 | 0.25 | 0.38 | 0.54 | 0.44 | 0.41 |
| 9 | 33210959 | rs10758198 | 0.84 | -0.032 | -0.957 | -0.13 | -0.057 | -0.026 | -0.026 | 0 | 0.37 | 0.22 | 0.36 | 0.54 | 0.41 | 0.39 |
| 9 | 33212335 | rs10813965 | 1.176 | -0.599 | -1.199 | -0.28 | -0.125 | -0.011 | -0.011 | 0 | 0.27 | 0.02 | 0.39 | 0.19 | 0.34 | 0.25 |
| 9 | 33214128 | rs1622785 | -1.224 | -0.606 | -0.636 | -0.284 | -0.045 | 0.028 | 0.028 | 0.002 | 0.72 | 0.88 | 0.6 | 0.81 | 0.66 | 0.75 |
| 9 | 33215662 | rs1702639 | -0.24 | -0.697 | -0.56 | -0.305 | -0.07 | 0.039 | 0.039 | 0.251 | 0.6 | 0.87 | 0.47 | 0.67 | 0.54 | 0.63 |
| 9 | 33218297 | rs1702645 | -1.301 | -1.092 | -0.109 | -0.272 | 0.016 | 0.013 | 0.013 | 0.098 | 0.35 | 0.65 | 0.24 | 0.27 | 0.24 | 0.36 |
| 9 | 33218668 | rs539826 | -0.661 | -0.895 | 0.246 | -0.231 | -0.045 | -0.023 | -0.023 | 0 | 0.23 | 0.65 | 0.11 | 0.13 | 0.12 | 0.24 |
| 9 | 33228189 | rs830574 | -0.177 | -0.811 | -0.154 | -0.309 | 0.032 | 0.026 | 0.026 | 0 | 0.19 | 0.09 | 0.11 | 0.14 | 0.17 | 0.22 |
| 9 | 33232374 | rs3866501 | 0.758 | -1.096 | 0.515 | -0.428 | 0.009 | 0.01 | 0.01 | 3.214 | 0.71 | 0.24 | 0.83 | 0.86 | 0.83 | 0.78 |
| 9 | 33240225 | rs706107 | -1.153 | -1.116 | 0.657 | -0.203 | 0.027 | 0.014 | 0.014 | 1.771 | 0.32 | 0.71 | 0.26 | 0.26 | 0.25 | 0.27 |
| 9 | 33245076 | rs480583 | -0.088 | -1.072 | 1.174 | -0.232 | -0.006 | -0.004 | -0.004 | 0 | 0.21 | 0.24 | 0.19 | 0.16 | 0.19 | 0.18 |
| 9 | 33250767 | rs12350735 | 0.568 | -1.084 | 1.653 | -0.224 | -0.016 | -0.019 | -0.019 | 8.104 | 0.07 | 0.3 | 0.03 | 0 | 0 | 0.04 |
| 9 | 33256179 | rs706118 | -1.492 | -1.509 | 1.527 | -0.379 | 0.087 | 0.042 | 0.042 | 8.456 | 0.37 | 0.84 | 0.27 | 0.26 | 0.25 | 0.27 |
| 9 | 33263208 | rs3758271 | 0.243 | -1.627 | 1.538 | -0.405 | 0.003 | 0.011 | 0.011 | 0 | 0.14 | 0.09 | 0.16 | 0.09 | 0.17 | 0.18 |
| 9 | 33270748 | rs831271 | 0.338 | -1.394 | 1.244 | -0.378 | -0.006 | -0.004 | -0.004 | 0 | 0.75 | 0.72 | 0.82 | 0.84 | 0.82 | 0.79 |
| 9 | 33290532 | rs830576 | 0.211 | -1.356 | 0.81 | -0.362 | 0.023 | 0.017 | 0.017 | 1.265 | 0.76 | 0.81 | 0.83 | 0.84 | 0.83 | 0.79 |
| 9 | 33303900 | rs706127 | 0.932 | -1.356 | 0.38 | -0.35 | 0.016 | 0.009 | 0.009 | 4.789 | 0.66 | 0.27 | 0.74 | 0.74 | 0.76 | 0.73 |
| 9 | 33309913 | rs10511908 | 1.201 | -1.236 | 0.101 | -0.297 | -0.022 | -0.024 | -0.024 | 0.124 | 0.06 | 0.21 | 0.02 | 0 | 0 | 0.04 |
| 9 | 33328812 | rs2292243 | 0.862 | -0.851 | -1.055 | -0.059 | -0.024 | -0.025 | -0.025 | 0.042 | 0.1 | 0.02 | 0.13 | 0.16 | 0.13 | 0.18 |
| 9 | 33343397 | rs4879686 | 0.556 | -0.599 | -0.893 | 0.043 | 0.049 | -0.029 | -0.029 | 0 | 0.53 | 0.05 | 0.6 | 0.58 | 0.62 | 0.55 |
| 9 | 33345616 | rs706134 | 0.425 | -0.501 | -0.705 | 0.1 | 0.001 | 0.001 | 0.001 | 1.007 | 0.56 | 0.39 | 0.63 | 0.58 | 0.64 | 0.55 |
| 9 | 33367088 | rs12237401 | 0.336 | 0.038 | -1.917 | 0.363 | -0.11 | -0.144 | -0.144 | 3.636 | 0.28 | 0.04 | 0.28 | 0.4 | 0.34 | 0.32 |
| 9 | 33398606 | rs4879696 | -0.582 | -0.232 | -1.352 | -0.302 | 0.021 | 0.028 | 0.028 | 0 | 0.19 | 0.07 | 0.1 | 0.15 | 0.13 | 0.07 |
| 9 | 33404808 | rs3758267 | -0.938 | -0.317 | -0.893 | -0.884 | -0.007 | -0.03 | -0.03 | 0 | 0.31 | 0.32 | 0.26 | 0.16 | 0.21 | 0.14 |
| 9 | 33406297 | rs7029953 | -0.422 | -0.532 | -0.688 | -1.009 | 0.003 | -0.077 | -0.077 | 0 | 0.12 | 0.31 | 0.19 | 0 | 0.09 | 0.08 |
| 9 | 33406907 | rs12351969 | -0.066 | -0.496 | -0.794 | -0.992 | 0.002 | -0.058 | -0.058 | 0 | 0.11 | 0.21 | 0.18 | 0 | 0.09 | 0.08 |
| 9 | 33407780 | rs10813971 | 1.29 | -0.375 | -1.705 | -0.539 | -0.006 | -0.055 | -0.055 | 0.196 | 0.25 | 0.18 | 0.35 | 0.48 | 0.27 | 0.28 |
| 9 | 33408019 | rs17337873 | 0.089 | -0.306 | -1.666 | -0.47 | 0.03 | 0.04 | 0.04 | 0.05 | 0.2 | 0.06 | 0.07 | 0.16 | 0.12 | 0.06 |
| 9 | 33414173 | rs2777738 | 0.096 | 0.675 | -1.554 | -0.454 | -0.053 | 0.058 | 0.058 | 0.303 | 0.65 | 0.86 | 0.64 | 0.76 | 0.58 | 0.55 |
| 9 | 33420008 | rs10758202 | -0.207 | 0.584 | -1.061 | 0.144 | 0.002 | 0.014 | 0.014 | 0 | 0.26 | 0.34 | 0.21 | 0.35 | 0.31 | 0.27 |
| 9 | 33420062 | rs7022937 | -1.037 | 0.483 | -1.128 | 0.025 | 0.001 | 0.001 | 0.001 | 0 | 0.51 | 0.54 | 0.54 | 0.53 | 0.57 | 0.53 |
| 9 | 33420204 | rs7022959 | -0.486 | 0.38 | -1.251 | -0.126 | 0 | 0.004 | 0.004 | 0 | 0.39 | 0.49 | 0.39 | 0.53 | 0.43 | 0.46 |
| 9 | 33420448 | rs10813974 | -1.536 | -0.135 | -0.744 | -0.097 | -0.011 | 0.012 | 0.012 | 0.002 | 0.77 | 0.59 | 0.76 | 0.76 | 0.81 | 0.83 |
| 9 | 33420467 | rs17253315 | 0.802 | -0.021 | -0.926 | -0.134 | 0.004 | 0.008 | 0.008 | 0.062 | 0.27 | 0.17 | 0.23 | 0.26 | 0.25 | 0.31 |
| 9 | 33425385 | rs4878538 | 0.112 | 0.122 | -0.959 | -0.081 | -0.024 | -0.001 | -0.001 | 0 | 0.38 | 0.31 | 0.35 | 0.32 | 0.44 | 0.32 |
| 9 | 33427091 | rs4878539 | -0.068 | -0.559 | -0.781 | -0.203 | 0.002 | -0.003 | -0.003 | 0 | 0.6 | 0.39 | 0.57 | 0.49 | 0.65 | 0.6 |
| 9 | 33430585 | rs3860985 | -0.425 | -0.373 | -0.881 | -0.094 | 0.002 | 0.01 | 0.01 | 0 | 0.41 | 0.37 | 0.48 | 0.64 | 0.41 | 0.47 |
| 9 | 33434026 | rs3849904 | 0.331 | -0.89 | -0.384 | 0.149 | -0.004 | 0.003 | 0.003 | 0.01 | 0.68 | 0.38 | 0.62 | 0.57 | 0.75 | 0.62 |
| 9 | 33434153 | rs6476410 | 0.168 | -0.832 | -0.48 | 0.177 | -0.013 | 0.04 | 0.04 | 0.082 | 0.27 | 0.35 | 0.35 | 0.42 | 0.24 | 0.37 |
| 9 | 33436612 | rs10813979 | 0.331 | -0.809 | -0.545 | 0.33 | -0.001 | -0.001 | -0.001 | 0 | 0.22 | 0.2 | 0.23 | 0.23 | 0.21 | 0.19 |
| 9 | 33437244 | rs34239839 | 0.99 | -0.665 | -0.975 | -0.035 | -0.003 | 0.051 | 0.051 | 0.11 | 0.42 | 0.23 | 0.32 | 0.35 | 0.41 | 0.46 |
| 9 | 33439148 | rs3892777 | 0.63 | -0.441 | -0.981 | -0.068 | -0.006 | 0.007 | 0.007 | 0 | 0.35 | 0.17 | 0.26 | 0.35 | 0.33 | 0.19 |
| 9 | 33439738 | rs12237858 | 1.585 | -0.446 | -0.412 | 0.457 | -0.016 | 0.031 | 0.031 | 0 | 0.1 | 0.18 | 0.15 | 0.19 | 0.08 | 0.2 |
| 9 | 33439829 | rs567691 | 0.498 | -0.206 | -0.15 | 0.426 | 0.007 | 0.066 | 0.066 | 0 | 0.14 | 0.31 | 0.22 | 0.23 | 0.15 | 0.18 |
| 9 | 33441826 | rs16919255 | 1.105 | -0.016 | -1.158 | 0.653 | -0.013 | -0.039 | -0.039 | 0 | 0.12 | 0.05 | 0.19 | 0.33 | 0.15 | 0.17 |
| 9 | 33442272 | rs2231235 | 1.421 | -0.274 | -1.238 | 0.38 | 0.007 | -0.016 | -0.016 | 0 | 0.09 | 0 | 0.09 | 0.04 | 0.07 | 0.08 |
| 9 | 33442952 | rs2228332 | -0.457 | -0.004 | -1.388 | 0.277 | 0.013 | 0.074 | 0.074 | 0 | 0.57 | 0.76 | 0.69 | 0.69 | 0.59 | 0.54 |
| 9 | 33444861 | rs34391490 | 1.072 | -0.025 | -0.786 | 0.553 | -0.005 | 0.029 | 0.029 | 0 | 0.16 | 0.28 | 0.2 | 0.22 | 0.14 | 0.19 |
| 9 | 33447547 | rs12555686 | 1.6 | 0.206 | -1.252 | 0.456 | 0.022 | -0.028 | -0.028 | 0 | 0.1 | 0.01 | 0.16 | 0.14 | 0.06 | 0.07 |
| 9 | 33451136 | rs2777753 | -1.267 | 0.082 | -1.087 | 0.598 | 0.009 | -0.001 | -0.001 | 0.765 | 0.58 | 0.64 | 0.69 | 0.56 | 0.65 | 0.44 |
| 9 | 33457653 | rs505698 | 0.968 | 0.155 | -1.564 | 0.618 | -0.011 | 0.031 | 0.031 | 0 | 0.13 | 0.25 | 0.11 | 0.22 | 0.13 | 0.17 |
| 9 | 33460141 | rs3860989 | 0.717 | 0.58 | -2.07 | 0.47 | -0.048 | -0.047 | -0.047 | 0 | 0.25 | 0.07 | 0.44 | 0.29 | 0.31 | 0.21 |
| 9 | 33460846 | rs10971520 | -0.278 | 0.565 | -2.038 | 0.424 | -0.007 | 0.001 | 0.001 | 0 | 0.33 | 0.27 | 0.49 | 0.31 | 0.37 | 0.22 |
| 9 | 33461299 | rs12683438 | 2.398 | 0.655 | -1.906 | 0.435 | 0.015 | -0.014 | -0.014 | 0.675 | 0.08 | 0.19 | 0.05 | 0.02 | 0.06 | 0.01 |
| 9 | 33461574 | rs1060037 | 0.783 | 0.621 | -1.918 | 0.554 | -0.01 | 0.03 | 0.03 | 0 | 0.14 | 0.25 | 0.15 | 0.23 | 0.14 | 0.18 |
| 9 | 33465182 | rs2785210 | -1.411 | 0.829 | -2.29 | 0.505 | 0.002 | -0.001 | -0.001 | 0 | 0.48 | 0.57 | 0.54 | 0.43 | 0.55 | 0.37 |
| 9 | 33465928 | rs3739680 | 2.381 | 0.916 | -2.625 | 0.265 | -0.035 | 0.023 | 0.023 | 1.234 | 0.19 | 0.02 | 0.1 | 0.14 | 0.17 | 0.23 |
| 9 | 33466219 | rs3739681 | 0.587 | 1.405 | -1.639 | 0.554 | -0.003 | 0.013 | 0.013 | 0 | 0.13 | 0.17 | 0.14 | 0.23 | 0.14 | 0.17 |
| 9 | 33467577 | rs2275228 | 0.767 | 1.461 | -1.653 | 0.524 | 0.009 | 0.003 | 0.003 | 0 | 0.13 | 0.22 | 0.26 | 0.15 | 0.2 | 0.12 |
| 9 | 33468682 | rs11795028 | 0.343 | 0.782 | -1.821 | 0.572 | -0.004 | 0.02 | 0.02 | 0 | 0.14 | 0.21 | 0.14 | 0.23 | 0.14 | 0.17 |
| 9 | 33469191 | rs2777747 | 1.64 | 0.701 | -1.794 | 0.533 | 0.024 | 0.002 | 0.002 | 2.096 | 0.08 | 0.24 | 0.06 | 0.03 | 0.06 | 0.01 |
| 9 | 33469406 | rs2777746 | 0.166 | 0.658 | -1.914 | 0.538 | 0.007 | -0.003 | -0.003 | 0 | 0.24 | 0.32 | 0.33 | 0.18 | 0.28 | 0.13 |
| 9 | 33473893 | rs2777744 | -1.76 | 0.751 | -1.761 | 0.517 | -0.002 | -0.002 | -0.002 | 10.467 | 0.58 | 0.58 | 0.71 | 0.59 | 0.64 | 0.44 |

| | | | | | | | | | | | | | | | | |
|---|---|---|---|---|---|---|---|---|---|---|---|---|---|---|---|---|
| 9 | 33487023 | rs7022355 | 0.307 | 0.871 | -1.691 | 0.55 | 0.007 | -0.017 | -0.017 | 0 | 0.14 | 0.25 | 0.09 | 0.07 | 0.13 | 0.09 |
| 9 | 33506670 | rs10813990 | 1.062 | 0.651 | -1.608 | 0.512 | 0.005 | -0.002 | -0.002 | 0 | 0.45 | 0.59 | 0.3 | 0.48 | 0.37 | 0.58 |
| 9 | 33518281 | rs10758211 | -0.462 | 0.377 | -1.376 | 0.667 | -0.037 | 0.002 | 0.002 | 0.006 | 0.85 | 0.65 | 0.81 | 0.85 | 0.91 | 0.93 |
| 9 | 33524775 | rs7866888 | 0.742 | 0.873 | -1.796 | 0.698 | 0.003 | -0.001 | -0.001 | 0 | 0.47 | 0.4 | 0.34 | 0.46 | 0.43 | 0.58 |
| 9 | 33525447 | rs4120127 | -0.288 | 1.493 | -2.01 | 0.738 | -0.062 | -0.033 | -0.033 | 0.345 | 0.14 | 0.03 | 0.25 | 0.16 | 0.21 | 0.12 |
| 9 | 33527793 | rs10120325 | -0.264 | 1.266 | -1.966 | 0.738 | -0.016 | -0.013 | -0.013 | 0 | 0.15 | 0.09 | 0.23 | 0.19 | 0.21 | 0.11 |
| 9 | 33554121 | rs2477414 | -0.654 | 0.821 | -1.83 | 0.541 | -0.009 | 0.013 | 0.013 | 0.006 | 0.08 | 0.08 | 0.09 | 0.06 | 0.1 | 0.11 |
| 9 | 33572347 | rs7853994 | 0.09 | 1.07 | -1.8 | 0.786 | -0.006 | -0.008 | -0.008 | 0 | 0.17 | 0.17 | 0.21 | 0.21 | 0.2 | 0.19 |
| 9 | 33573212 | rs12378940 | 0.053 | 1.027 | -2.039 | 0.38 | -0.13 | -0.026 | -0.026 | 5.601 | 0.23 | 0.02 | 0.31 | 0.13 | 0.38 | 0.27 |
| 9 | 33576579 | rs1853722 | -0.306 | -0.594 | -0.892 | 0.22 | -0.064 | -0.026 | -0.026 | 0 | 0.33 | 0.43 | 0.21 | 0.31 | 0.19 | 0.29 |
| 9 | 33609013 | rs855471 | -0.093 | -0.18 | -0.971 | -0.105 | -0.001 | -0.019 | -0.019 | 0 | 0.11 | 0.15 | 0.05 | 0 | 0.07 | 0.02 |
| 9 | 33611709 | rs10971602 | 0.263 | -0.2 | -1.016 | -0.115 | -0.017 | 0.049 | 0.049 | 0 | 0.09 | 0 | 0.03 | 0 | 0.07 | 0.02 |
| 9 | 33613660 | rs855495 | -1.011 | -0.49 | -1.117 | -0.61 | 0.011 | 0.051 | 0.051 | 0 | 0.19 | 0.11 | 0.1 | 0 | 0.15 | 0.09 |
| 9 | 33613835 | rs855498 | -1.011 | -0.49 | -1.117 | -0.61 | 0.011 | 0.051 | 0.051 | 0 | 0.19 | 0.12 | 0.11 | 0 | 0.15 | 0.11 |
| 9 | 33614214 | rs855500 | -0.678 | -0.449 | -1.755 | -0.533 | 0.05 | 0.005 | 0.005 | 0.317 | 0.53 | 0.24 | 0.58 | 0.51 | 0.62 | 0.65 |
| 9 | 33617310 | rs855508 | -0.09 | -0.487 | -1.742 | -0.515 | -0.029 | -0.036 | -0.036 | 0.316 | 0.43 | 0.17 | 0.52 | 0.52 | 0.54 | 0.57 |
| 9 | 33618936 | rs855514 | 0.233 | -1.443 | -1.152 | -0.883 | -0.04 | -0.035 | -0.035 | 0 | 0.66 | 0.45 | 0.77 | 0.84 | 0.79 | 0.73 |
| 9 | 33624429 | rs855523 | 0.41 | -1.73 | -0.663 | -1.128 | -0.007 | -0.013 | -0.013 | 0 | 0.16 | 0.22 | 0.06 | 0.06 | 0.08 | 0.17 |
| 9 | 33625360 | rs10971607 | 0.829 | -1.322 | -0.6 | -1.223 | -0.005 | 0.013 | 0.013 | 0 | 0.3 | 0.21 | 0.29 | 0.29 | 0.31 | 0.19 |
| 9 | 33629249 | rs10814017 | 0.219 | -1.208 | -0.479 | -1.103 | -0.002 | 0.025 | 0.025 | 0 | 0.39 | 0.29 | 0.33 | 0.3 | 0.38 | 0.21 |
| 9 | 33633838 | rs855533 | -1.149 | -1.107 | 0.011 | -1.095 | -0.014 | 0.007 | 0.007 | 0.675 | 0.65 | 0.74 | 0.43 | 0.35 | 0.55 | 0.45 |
| 9 | 33643569 | rs1101845 | -1.074 | -1.137 | -0.021 | -1.107 | 0.003 | -0.004 | -0.004 | 0 | 0.63 | 0.61 | 0.42 | 0.35 | 0.55 | 0.44 |
| 9 | 33661286 | rs1856202 | 0.652 | -1.154 | 0.176 | -1.123 | -0.049 | -0.039 | -0.039 | 0.047 | 0.74 | 0.47 | 0.8 | 0.85 | 0.82 | 0.71 |
| 9 | 33663511 | rs1977008 | 0.263 | -1.013 | 0.258 | -1.197 | -0.016 | -0.028 | -0.028 | 0 | 0.3 | 0.44 | 0.28 | 0.22 | 0.29 | 0.13 |
| 9 | 33676027 | rs855465 | 0.206 | -1.034 | 0.364 | -1.212 | -0.026 | -0.051 | -0.051 | 0.16 | 0.14 | 0.25 | 0.07 | 0.03 | 0.06 | 0.18 |
| 9 | 33699373 | rs1765372 | -0.661 | -1.053 | 0.41 | -1.231 | -0.045 | -0.106 | -0.106 | 0 | 0.21 | 0.45 | 0.12 | 0.03 | 0.13 | 0.23 |
| 9 | 33700901 | rs855434 | -1.15 | -0.572 | -0.007 | -1.09 | -0.006 | 0.002 | 0.002 | 0 | 0.73 | 0.81 | 0.69 | 0.8 | 0.74 | 0.68 |
| 9 | 33702339 | rs855436 | -0.451 | -0.591 | 0.04 | -1.363 | 0.029 | 0.049 | 0.049 | 0.678 | 0.43 | 0.7 | 0.34 | 0.24 | 0.35 | 0.31 |
| 9 | 33731480 | rs855441 | -0.445 | -0.513 | 0.041 | -1.36 | 0.027 | 0.047 | 0.047 | 0.77 | 0.43 | 0.7 | 0.34 | 0.24 | 0.36 | 0.31 |
| 9 | 33738727 | rs10814040 | 0.627 | -0.276 | 0.209 | -1.206 | -0.011 | -0.022 | -0.022 | 0 | 0.29 | 0.45 | 0.27 | 0.22 | 0.29 | 0.18 |
| 9 | 33739343 | rs10758224 | 1.261 | -0.23 | 0.055 | -1.157 | -0.003 | -0.001 | -0.001 | 0 | 0.23 | 0.25 | 0.23 | 0.22 | 0.23 | 0.13 |
| 9 | 33751865 | rs10971679 | 1.241 | -0.222 | 0.067 | -1.154 | -0.003 | -0.001 | -0.001 | 0 | 0.23 | 0.25 | 0.23 | 0.22 | 0.22 | 0.13 |
| 9 | 33761112 | rs10971687 | 0.861 | -0.095 | -0.414 | -0.959 | -0.041 | -0.045 | -0.045 | 0 | 0.3 | 0.11 | 0.4 | 0.6 | 0.39 | 0.37 |
| 9 | 33785963 | rs3925857 | -0.511 | -0.252 | -0.313 | -0.884 | 0.001 | 0.018 | 0.018 | 0 | 0.69 | 0.75 | 0.73 | 0.83 | 0.72 | 0.61 |
| 9 | 33792265 | rs760098 | -0.511 | -0.252 | -0.336 | -0.884 | 0.001 | 0.023 | 0.023 | 0 | 0.69 | 0.75 | 0.73 | 0.83 | 0.72 | 0.61 |
| 9 | 33799370 | rs216345 | -0.656 | 0.372 | -0.594 | -1.143 | -0.001 | -0.01 | -0.01 | 0 | 0.6 | 0.6 | 0.65 | 0.76 | 0.63 | 0.67 |
| 9 | 33804813 | rs10971709 | 1.043 | 0.484 | -0.868 | -1.139 | -0.05 | -0.147 | -0.147 | 0.037 | 0.2 | 0.01 | 0.32 | 0.58 | 0.24 | 0.22 |
| 9 | 33810791 | rs4602979 | 1.724 | 0.269 | -1.066 | -1.331 | -0.018 | -0.032 | -0.032 | 0 | 0.17 | 0.03 | 0.17 | 0.21 | 0.16 | 0.21 |
| 9 | 33818257 | rs216364 | 0.265 | 0.47 | -0.691 | -1.243 | -0.023 | -0.054 | -0.054 | 0 | 0.34 | 0.51 | 0.28 | 0.17 | 0.25 | 0.4 |
| 9 | 33845079 | rs2486675 | -0.07 | 0.573 | -0.716 | -1.272 | 0.001 | -0.05 | -0.05 | 0 | 0.39 | 0.52 | 0.32 | 0.17 | 0.38 | 0.45 |
| 9 | 33846422 | rs4515641 | 0.637 | 0.49 | -0.518 | -1.318 | 0.001 | -0.001 | -0.001 | 0 | 0.31 | 0.31 | 0.23 | 0.07 | 0.24 | 0.37 |
| 9 | 33880690 | rs2050789 | 1.443 | 0.817 | -0.773 | -0.768 | -0.026 | -0.089 | -0.089 | 0 | 0.12 | 0.01 | 0.22 | 0.49 | 0.15 | 0.13 |
| 9 | 33888000 | rs13302810 | 2.525 | 0.831 | -0.415 | -1.16 | 0 | -0.016 | -0.016 | 0 | 0.12 | 0.1 | 0.11 | 0.01 | 0.15 | 0.06 |
| 9 | 33915558 | rs10971789 | 0.702 | 0.81 | -0.71 | -0.915 | -0.018 | -0.098 | -0.098 | 0 | 0.21 | 0.04 | 0.32 | 0.61 | 0.24 | 0.26 |
| 9 | 33917498 | rs4817 | -0.006 | 0.815 | -0.625 | -0.992 | 0.001 | -0.05 | -0.05 | 0 | 0.39 | 0.52 | 0.32 | 0.17 | 0.38 | 0.45 |
| 9 | 33932702 | rs307649 | 1.081 | 0.829 | -0.882 | -1.198 | -0.008 | -0.003 | -0.003 | 0 | 0.19 | 0.25 | 0.18 | 0.16 | 0.16 | 0.33 |
| 9 | 33935736 | rs307652 | -1.082 | 0.731 | -0.832 | -1.09 | -0.001 | -0.005 | -0.005 | 0 | 0.6 | 0.61 | 0.6 | 0.73 | 0.62 | 0.69 |
| 9 | 33941759 | rs307658 | -0.157 | 0.769 | -0.926 | -1.275 | 0.001 | -0.044 | -0.044 | 0 | 0.39 | 0.51 | 0.27 | 0.17 | 0.38 | 0.45 |
| 9 | 33952629 | rs16935299 | 0.52 | 0.728 | -1.083 | -0.757 | -0.012 | -0.082 | -0.082 | 0 | 0.21 | 0.05 | 0.28 | 0.61 | 0.24 | 0.26 |
| 9 | 33956619 | rs7869836 | 1.116 | 0.67 | -0.729 | -0.974 | -0.005 | -0.068 | -0.068 | 0.485 | 0.15 | 0.25 | 0.05 | 0 | 0.09 | 0.06 |
| 9 | 33960152 | rs307661 | -1.067 | 0.724 | -0.723 | -1.377 | -0.001 | -0.007 | -0.007 | 0.019 | 0.6 | 0.61 | 0.6 | 0.78 | 0.62 | 0.71 |
| 9 | 33960219 | rs10113865 | 1.459 | 0.687 | -0.71 | -1.362 | -0.005 | -0.068 | -0.068 | 0 | 0.15 | 0.25 | 0.05 | 0 | 0.1 | 0.06 |
| 9 | 33971740 | rs3739690 | 0.858 | 0.72 | -0.825 | -1.354 | -0.003 | -0.049 | -0.049 | 0.867 | 0.21 | 0.09 | 0.28 | 0.61 | 0.24 | 0.26 |
| 9 | 33976715 | rs834029 | -0.129 | 0.744 | -0.744 | -1.305 | -0.02 | -0.051 | -0.051 | 0.12 | 0.34 | 0.5 | 0.22 | 0.17 | 0.25 | 0.4 |
| 9 | 34001455 | rs932969 | 1.631 | 0.643 | -0.87 | -1.206 | -0.005 | 0.012 | 0.012 | 0.399 | 0.16 | 0.05 | 0.13 | 0.07 | 0.14 | 0.31 |
| 9 | 34065659 | rs7853333 | -0.966 | 0.435 | -0.38 | -1.321 | -0.026 | -0.016 | -0.016 | 0.14 | 0.89 | 0.79 | 0.86 | 0.9 | 0.94 | 0.96 |
| 9 | 34071541 | rs7871764 | 0.247 | 0.705 | -0.452 | -1.402 | 0.003 | -0.071 | -0.071 | 1.131 | 0.68 | 0.42 | 0.73 | 0.89 | 0.7 | 0.83 |
| 9 | 34082144 | rs10441723 | -0.037 | 0.966 | -0.372 | -1.128 | -0.002 | -0.001 | -0.001 | 3.665 | 0.63 | 0.6 | 0.62 | 0.29 | 0.69 | 0.68 |
| 9 | 34085859 | rs7036030 | 1.963 | 1.072 | -0.582 | -1.482 | 0.002 | -0.072 | -0.072 | 0 | 0.17 | 0.24 | 0.13 | 0.01 | 0.18 | 0.07 |
| 9 | 34087360 | rs1052656 | -0.222 | 1.053 | -0.62 | -1.465 | -0.002 | -0.001 | -0.001 | 0 | 0.63 | 0.6 | 0.62 | 0.29 | 0.68 | 0.67 |
| 9 | 34088701 | rs11793082 | 0.779 | 0.931 | -0.785 | -1.451 | -0.034 | 0.023 | 0.023 | 0 | 0.31 | 0.17 | 0.34 | 0.22 | 0.37 | 0.29 |
| 9 | 34095717 | rs7853415 | -1.87 | 0.787 | -0.804 | -1.31 | 0 | 0.013 | 0.013 | 0 | 0.85 | 0.92 | 0.92 | 0.85 | 0.89 | 0.91 |
| 9 | 34104443 | rs4879741 | 1.332 | 0.948 | -0.636 | -1.215 | 0.006 | 0.009 | 0.009 | 0 | 0.26 | 0.16 | 0.27 | 0.21 | 0.21 | 0.23 |
| 9 | 34107310 | rs10511915 | 2.01 | 0.952 | -0.627 | -1.199 | 0 | 0.006 | 0.006 | 0.334 | 0.15 | 0.05 | 0.13 | 0.07 | 0.14 | 0.31 |
| 9 | 34124860 | rs2275003 | 0.131 | 0.854 | -0.537 | -1.178 | -0.009 | 0.027 | 0.027 | 0 | 0.46 | 0.36 | 0.49 | 0.29 | 0.51 | 0.61 |
| 9 | 34127763 | rs10971942 | -0.064 | 0.649 | -0.433 | -1.082 | 0.009 | -0.058 | -0.058 | 0 | 0.42 | 0.48 | 0.42 | 0.08 | 0.54 | 0.45 |
| 9 | 34159144 | rs12376762 | 0.882 | 1.048 | -0.601 | -0.736 | -0.054 | 0.03 | 0.03 | 0 | 0.08 | 0 | 0.14 | 0.01 | 0.19 | 0.05 |
| 9 | 34162975 | rs7042499 | -0.176 | 0.613 | -0.097 | -1.263 | 0.013 | -0.069 | -0.069 | 0 | 0.68 | 0.38 | 0.66 | 0.95 | 0.58 | 0.82 |
| 9 | 34167733 | rs7018487 | 0.621 | 0.365 | -0.515 | -0.864 | 0.049 | 0.035 | 0.035 | 1.117 | 0.42 | 0.25 | 0.4 | 0.74 | 0.31 | 0.53 |

| | | | | | | | | | | | | | | | | |
|---|---|---|---|---|---|---|---|---|---|---|---|---|---|---|---|---|
| 9 | 34170980 | rs2769710 | -1.693 | 0.366 | -0.156 | -0.884 | -0.004 | -0.114 | -0.114 | 0 | 0.79 | 0.57 | 0.81 | 0.96 | 0.77 | 0.87 |
| 9 | 34187354 | rs1041251 | 0.059 | 0.261 | -0.318 | -0.892 | -0.137 | 0.141 | 0.141 | 0 | 0.32 | 0.06 | 0.34 | 0.06 | 0.46 | 0.31 |
| 9 | 34197250 | rs10971988 | 0.428 | 0.211 | -0.218 | -0.929 | 0.03 | 0.042 | 0.042 | 0.872 | 0.44 | 0.33 | 0.46 | 0.9 | 0.31 | 0.56 |
| 9 | 34198916 | rs2094998 | 0.898 | 0.208 | -0.207 | -0.901 | 0.044 | 0.031 | 0.031 | 1.327 | 0.42 | 0.25 | 0.4 | 0.74 | 0.31 | 0.53 |
| 9 | 34208138 | rs4879760 | 0.89 | 0.205 | 0.068 | -0.902 | 0.03 | 0.022 | 0.022 | 1.283 | 0.42 | 0.28 | 0.4 | 0.74 | 0.31 | 0.53 |
| 9 | 34209859 | rs10971998 | 0.435 | 0.205 | 0.066 | -0.902 | 0.009 | 0.001 | 0.001 | 0.951 | 0.44 | 0.36 | 0.46 | 0.9 | 0.31 | 0.56 |
| 9 | 34210872 | rs4319185 | -1.773 | 0.21 | 0.094 | -0.948 | -0.006 | -0.091 | -0.091 | 0.424 | 0.78 | 0.62 | 0.81 | 0.96 | 0.77 | 0.87 |
| 9 | 34213548 | rs10972003 | 1.708 | -0.13 | -0.489 | -0.466 | 0.022 | -0.016 | -0.016 | 0 | 0.24 | 0.11 | 0.17 | 0.29 | 0.17 | 0.32 |
| 9 | 34237044 | rs10972013 | 0.11 | -0.028 | -0.505 | -0.535 | 0.029 | 0.041 | 0.041 | 0.98 | 0.44 | 0.33 | 0.46 | 0.9 | 0.31 | 0.56 |
| 9 | 34237480 | rs17258448 | 1.084 | 0.026 | -0.527 | -0.558 | -0.064 | 0.087 | 0.087 | 0 | 0.24 | 0.06 | 0.2 | 0.05 | 0.26 | 0.26 |
| 9 | 34244227 | rs4008745 | 0.105 | -0.011 | -0.625 | -0.522 | 0.029 | 0.041 | 0.041 | 0.98 | 0.44 | 0.33 | 0.46 | 0.9 | 0.31 | 0.56 |
| 9 | 34252127 | rs1049567 | -1.587 | 0.019 | -0.549 | -0.47 | -0.012 | -0.079 | -0.079 | 0.051 | 0.76 | 0.35 | 0.79 | 0.96 | 0.77 | 0.87 |
| 9 | 34253097 | rs17258783 | 2.055 | 0.251 | -0.699 | -0.453 | -0.044 | 0.017 | 0.017 | 0.35 | 0.13 | 0.04 | 0.17 | 0.05 | 0.21 | 0.24 |
| 9 | 34253488 | rs4574933 | 1.894 | 0.208 | -0.688 | -0.444 | -0.048 | 0.022 | 0.022 | 0.094 | 0.14 | 0.04 | 0.17 | 0.05 | 0.21 | 0.25 |
| 9 | 34256347 | rs10814083 | 0.489 | 0.039 | -0.569 | -0.328 | 0.035 | 0.024 | 0.024 | 1.264 | 0.42 | 0.27 | 0.4 | 0.74 | 0.31 | 0.53 |
| 9 | 34269079 | rs10972032 | 0.747 | -0.032 | -0.684 | -0.267 | -0.033 | 0.063 | 0.063 | 0.476 | 0.2 | 0.29 | 0.3 | 0.61 | 0.15 | 0.24 |
| 9 | 34277053 | rs2890545 | -1.625 | -0.006 | -0.704 | -0.201 | -0.016 | -0.076 | -0.076 | 0 | 0.76 | 0.39 | 0.8 | 0.93 | 0.77 | 0.84 |
| 9 | 34284944 | rs17350373 | 0.387 | 0.114 | -0.987 | -0.263 | -0.066 | 0.055 | 0.055 | 0 | 0.47 | 0.1 | 0.36 | 0.35 | 0.43 | 0.58 |
| 9 | 34290407 | rs11790773 | 1.68 | 0.007 | -1.002 | -0.243 | 0.008 | -0.043 | -0.043 | 0 | 0.23 | 0.04 | 0.16 | 0.27 | 0.17 | 0.32 |
| 9 | 34300938 | rs10972043 | -0.134 | 0.111 | -0.861 | -0.157 | 0.068 | 0.029 | 0.029 | 0 | 0.56 | 0.11 | 0.5 | 0.34 | 0.62 | 0.62 |
| 9 | 34310357 | rs10972047 | 1.533 | 0.014 | -0.785 | -0.113 | 0.014 | -0.032 | -0.032 | 0 | 0.16 | 0.08 | 0.22 | 0.46 | 0.14 | 0.22 |
| 9 | 34310927 | rs10972048 | -0.193 | 0.142 | -0.867 | -0.08 | 0.069 | 0.031 | 0.031 | 0 | 0.56 | 0.11 | 0.5 | 0.34 | 0.62 | 0.62 |
| 9 | 34333855 | rs11788425 | 0.4 | 0.253 | -1.06 | -0.06 | -0.061 | 0.069 | 0.069 | 0 | 0.47 | 0.1 | 0.36 | 0.33 | 0.43 | 0.57 |
| 9 | 34342634 | rs10972067 | 0.915 | -0.008 | -0.987 | -0.03 | -0.034 | 0.06 | 0.06 | 0.401 | 0.2 | 0.29 | 0.3 | 0.63 | 0.14 | 0.28 |
| 9 | 34348743 | rs10972072 | -1.156 | 0.58 | -0.475 | -0.199 | -0.014 | -0.118 | -0.118 | 0.018 | 0.76 | 0.41 | 0.8 | 0.96 | 0.77 | 0.9 |
| 9 | 34373041 | rs10972084 | 0.947 | 1.161 | -0.927 | -0.281 | -0.056 | 0.032 | 0.032 | 0 | 0.08 | 0 | 0.14 | 0.01 | 0.2 | 0.05 |
| 9 | 34379692 | rs3737242 | 1.299 | 1.153 | -1.018 | -0.269 | -0.046 | -0.017 | -0.017 | 0 | 0.14 | 0.04 | 0.17 | 0.09 | 0.21 | 0.24 |
| 9 | 34382726 | rs3808876 | -0.632 | 0.811 | -0.657 | -0.265 | 0.001 | -0.011 | -0.011 | 0 | 0.63 | 0.37 | 0.63 | 0.86 | 0.56 | 0.66 |
| 9 | 34386011 | rs10972090 | 1.92 | 0.244 | -0.807 | 0.31 | 0.009 | -0.027 | -0.027 | 0 | 0.1 | 0.03 | 0.09 | 0.11 | 0.06 | 0.03 |
| 9 | 34387524 | rs10758257 | -0.147 | 0.461 | -0.558 | 0.11 | 0.023 | 0.048 | 0.048 | 1.242 | 0.55 | 0.32 | 0.49 | 0.86 | 0.37 | 0.62 |
| 9 | 34390330 | rs4879784 | -1.176 | 0.652 | -0.247 | -0.175 | -0.003 | -0.089 | -0.089 | 0 | 0.73 | 0.61 | 0.77 | 0.96 | 0.71 | 0.81 |
| 9 | 34393422 | rs7043224 | -0.565 | 0.525 | -0.01 | 0.096 | 0.007 | 0.008 | 0.008 | 0.784 | 0.45 | 0.56 | 0.55 | 0.27 | 0.64 | 0.41 |
| 9 | 34395816 | rs3933343 | -1.012 | 0.005 | -0.141 | 0.64 | -0.039 | 0.044 | 0.044 | 0 | 0.78 | 0.63 | 0.74 | 0.64 | 0.86 | 0.71 |
| 9 | 34396346 | rs4879785 | -0.564 | -0.111 | -0.197 | 0.781 | -0.047 | 0.075 | 0.075 | 0 | 0.76 | 0.59 | 0.68 | 0.49 | 0.86 | 0.65 |
| 9 | 34399002 | rs1002352 | -0.656 | -0.141 | -0.071 | 0.765 | -0.062 | 0.098 | 0.098 | 0 | 0.75 | 0.51 | 0.67 | 0.5 | 0.86 | 0.65 |
| 9 | 34399338 | rs885048 | 1.252 | 0.447 | -0.302 | 0.472 | -0.052 | 0.029 | 0.029 | 0 | 0.08 | 0 | 0.14 | 0.01 | 0.2 | 0.05 |
| 9 | 34406384 | rs10115705 | 0.742 | 0.394 | 0.045 | 0.634 | -0.001 | -0.001 | -0.001 | 0 | 0.58 | 0.4 | 0.4 | 0.44 | 0.52 | 0.47 |
| 9 | 34408204 | rs946847 | -1.351 | 0.404 | 0.008 | 0.644 | 0.004 | -0.003 | -0.003 | 0.047 | 0.79 | 0.8 | 0.75 | 0.69 | 0.86 | 0.73 |
| 9 | 34415648 | rs11788070 | 0.785 | 0.451 | -0.41 | 0.614 | -0.011 | -0.008 | -0.008 | 0 | 0.57 | 0.29 | 0.39 | 0.44 | 0.52 | 0.47 |
| 9 | 34419581 | rs10972104 | 0.962 | 0.714 | -0.444 | 0.551 | -0.022 | 0.021 | 0.021 | 0 | 0.09 | 0.03 | 0.15 | 0.01 | 0.2 | 0.05 |
| 9 | 34421215 | rs11788370 | 1.531 | 0.254 | -0.501 | 0.688 | -0.006 | -0.078 | -0.078 | 0 | 0.19 | 0.02 | 0.1 | 0.27 | 0.16 | 0.25 |
| 9 | 34422185 | rs13297645 | 0.808 | 0.348 | -0.378 | 0.613 | 0.013 | 0.028 | 0.028 | 0 | 0.43 | 0.27 | 0.42 | 0.27 | 0.41 | 0.33 |
| 9 | 34423498 | rs4590517 | 0.807 | 0.713 | 0.186 | 0.668 | 0.031 | 0.017 | 0.017 | 0 | 0.11 | 0.22 | 0.14 | 0.1 | 0.13 | 0.12 |
| 9 | 34423806 | rs10814110 | 0.553 | 0.463 | -0.29 | 0.616 | -0.021 | -0.009 | -0.009 | 2.216 | 0.62 | 0.26 | 0.51 | 0.55 | 0.57 | 0.57 |
| 9 | 34431079 | rs4256660 | 0.808 | -0.298 | 0.438 | -0.026 | 0.028 | -0.095 | -0.095 | 0.045 | 0.17 | 0.35 | 0.15 | 0.01 | 0.17 | 0.15 |
| 9 | 34437469 | rs2077403 | 0.867 | -0.701 | 0.523 | -0.038 | -0.011 | 0.002 | 0.002 | 0 | 0.2 | 0.2 | 0.26 | 0.34 | 0.12 | 0.3 |
| 9 | 34464433 | rs10125115 | 1.187 | -0.699 | 0.663 | -0.069 | 0.015 | -0.067 | -0.067 | 0 | 0.16 | 0.22 | 0.15 | 0.01 | 0.17 | 0.15 |
| 9 | 34467119 | rs10972129 | 1.842 | -0.587 | 0.279 | -0.198 | -0.062 | -0.117 | -0.117 | 0.041 | 0.17 | 0.02 | 0.11 | 0.36 | 0.2 | 0.26 |
| 9 | 34469025 | rs4879795 | -0.874 | -0.595 | 0.447 | -0.194 | 0.025 | -0.007 | -0.007 | 0 | 0.43 | 0.68 | 0.52 | 0.47 | 0.35 | 0.51 |
| 9 | 34492213 | rs3793472 | -2.032 | -0.62 | 0.095 | -0.603 | -0.004 | 0.006 | 0.006 | 0.252 | 0.76 | 0.78 | 0.71 | 0.83 | 0.71 | 0.89 |
| 9 | 34500821 | rs11793196 | 1.1 | -0.782 | 0.58 | -0.496 | 0.02 | -0.096 | -0.096 | 0 | 0.17 | 0.36 | 0.15 | 0.01 | 0.16 | 0.14 |
| 9 | 34504634 | rs10511917 | 1.02 | -0.768 | 0.587 | -0.489 | 0.018 | -0.097 | -0.097 | 2.374 | 0.17 | 0.36 | 0.15 | 0.01 | 0.16 | 0.14 |
| 9 | 34509746 | rs4879801 | -0.83 | -0.792 | 0.547 | -0.483 | 0.005 | 0.011 | 0.011 | 0 | 0.41 | 0.62 | 0.46 | 0.36 | 0.36 | 0.49 |
| 9 | 34512893 | rs17353380 | 0.609 | -0.363 | 0.38 | -0.718 | 0.001 | -0.037 | -0.037 | 3.746 | 0.17 | 0.21 | 0.14 | 0.01 | 0.17 | 0.1 |
| 9 | 34515101 | rs12002889 | -0.127 | -0.365 | 0.288 | -0.571 | 0.001 | -0.08 | -0.08 | 0 | 0.29 | 0.39 | 0.21 | 0.03 | 0.27 | 0.27 |
| 9 | 34515328 | rs12001113 | -0.328 | -0.324 | 0.147 | -0.557 | 0.005 | -0.079 | -0.079 | 0 | 0.32 | 0.43 | 0.26 | 0.05 | 0.34 | 0.3 |
| 9 | 34536873 | rs10114507 | -0.506 | -0.622 | -0.165 | -0.215 | 0.001 | -0.013 | -0.013 | 0 | 0.34 | 0.54 | 0.39 | 0.27 | 0.34 | 0.32 |
| 9 | 34540065 | rs4878578 | 0.298 | -1.042 | -0.132 | 0.106 | -0.024 | -0.001 | -0.001 | 0 | 0.16 | 0.21 | 0.23 | 0.17 | 0.1 | 0.31 |
| 9 | 34545932 | rs4879803 | 0.409 | -0.556 | -0.29 | -0.007 | 0.006 | 0.029 | 0.029 | 0.025 | 0.67 | 0.58 | 0.69 | 0.55 | 0.66 | 0.61 |
| 9 | 34548009 | rs4395980 | 0.119 | -0.475 | -0.312 | 0.104 | 0.002 | 0.018 | 0.018 | 0 | 0.21 | 0.2 | 0.25 | 0.44 | 0.24 | 0.3 |
| 9 | 34550295 | rs10814119 | 0.633 | -1.545 | -0.13 | -0.562 | 0.048 | 0.001 | 0.001 | 0 | 0.55 | 0.32 | 0.54 | 0.56 | 0.7 | 0.57 |
| 9 | 34550830 | rs4879804 | 0.581 | -1.552 | -0.419 | -0.487 | 0.033 | 0.002 | 0.002 | 0 | 0.13 | 0.02 | 0.12 | 0.1 | 0.09 | 0.16 |
| 9 | 34553667 | rs4363285 | 0.503 | -1.741 | 0.215 | -0.614 | -0.067 | 0.062 | 0.062 | 0.079 | 0.74 | 0.56 | 0.73 | 0.62 | 0.85 | 0.73 |
| 9 | 34554634 | rs1077962 | 1.001 | -1.695 | 0.003 | -0.893 | -0.011 | 0.07 | 0.07 | 0 | 0.17 | 0.06 | 0.18 | 0.05 | 0.14 | 0.16 |
| 9 | 34554999 | rs7044318 | 0.097 | -1.765 | 0.288 | -0.821 | -0.008 | -0.021 | -0.021 | 0 | 0.75 | 0.67 | 0.7 | 0.85 | 0.82 | 0.81 |
| 9 | 34555481 | rs12552724 | 0.988 | -1.464 | -0.075 | -0.715 | -0.018 | 0.075 | 0.075 | 0 | 0.17 | 0.03 | 0.17 | 0.05 | 0.14 | 0.16 |
| 9 | 34557313 | rs12553466 | 0.816 | -1.451 | -0.073 | -0.709 | -0.042 | 0.072 | 0.072 | 0 | 0.18 | 0.03 | 0.2 | 0.06 | 0.18 | 0.16 |
| 9 | 34561160 | rs10972149 | -0.14 | -1.366 | 0.385 | -0.357 | -0.003 | -0.001 | -0.001 | 0 | 0.18 | 0.16 | 0.15 | 0.28 | 0.11 | 0.25 |
| 9 | 34561808 | rs10814123 | -0.14 | -1.366 | 0.385 | -0.357 | -0.003 | -0.001 | -0.001 | 0 | 0.18 | 0.16 | 0.15 | 0.28 | 0.11 | 0.24 |

| | | | | | | | | | | | | | | | | |
|---|---|---|---|---|---|---|---|---|---|---|---|---|---|---|---|---|
| 9 | 34564274 | rs4879805 | -0.134 | -1.369 | 0.383 | -0.359 | -0.003 | -0.001 | -0.001 | 0 | 0.18 | 0.16 | 0.15 | 0.28 | 0.11 | 0.25 |
| 9 | 34565781 | rs10738931 | 0.597 | -1.486 | 1.005 | -0.398 | -0.085 | 0.085 | 0.085 | 0.046 | 0.78 | 0.46 | 0.77 | 0.63 | 0.89 | 0.75 |
| 9 | 34572767 | rs10758268 | 0.977 | -1.51 | 0.522 | -0.746 | 0.041 | 0.008 | 0.008 | 0 | 0.47 | 0.2 | 0.49 | 0.58 | 0.61 | 0.41 |
| 9 | 34572815 | rs1571401 | 0.037 | -1.613 | 0.615 | -0.641 | -0.016 | 0.008 | 0.008 | 0 | 0.17 | 0.17 | 0.12 | 0.28 | 0.09 | 0.23 |
| 9 | 34581525 | rs1080750 | 1.881 | -0.781 | -0.051 | -1.228 | -0.123 | 0.137 | 0.137 | 0 | 0.21 | 0.01 | 0.23 | 0 | 0.32 | 0.04 |
| 9 | 34584105 | rs3763615 | 0.303 | -1.439 | 1.026 | -0.935 | 0.001 | -0.024 | -0.024 | 0 | 0.28 | 0.29 | 0.3 | 0.08 | 0.29 | 0.33 |
| 9 | 34584319 | rs3763613 | -0.746 | -1.626 | 1.297 | -0.712 | -0.009 | 0.001 | 0.001 | 0 | 0.28 | 0.27 | 0.24 | 0.31 | 0.19 | 0.3 |
| 9 | 34587485 | rs7037232 | -0.252 | -1.434 | 1.472 | -0.607 | 0.045 | 0.054 | 0.054 | 0.019 | 0.08 | 0.34 | 0.12 | 0.1 | 0.07 | 0.09 |
| 9 | 34592699 | rs7036351 | 0.051 | -1.524 | 1.31 | -0.619 | -0.006 | 0.008 | 0.008 | 0 | 0.19 | 0.2 | 0.21 | 0.31 | 0.15 | 0.2 |
| 9 | 34607766 | rs16931689 | 0.06 | -1.514 | 1.32 | -0.611 | -0.003 | 0.005 | 0.005 | 0 | 0.19 | 0.19 | 0.26 | 0.31 | 0.16 | 0.21 |
| 9 | 34619397 | rs10814127 | 1.066 | -1.527 | 1.6 | -0.717 | -0.053 | 0.073 | 0.073 | 0 | 0.22 | 0.01 | 0.19 | 0.06 | 0.21 | 0.25 |
| 9 | 34619621 | rs10972168 | -0.189 | -1.6 | 1.774 | -0.746 | 0.019 | 0.062 | 0.062 | 0 | 0.31 | 0.15 | 0.27 | 0.15 | 0.24 | 0.35 |
| 9 | 34623718 | rs10814129 | 0.217 | -1.621 | 1.519 | -0.718 | -0.004 | 0.008 | 0.008 | 0 | 0.18 | 0.19 | 0.25 | 0.31 | 0.15 | 0.2 |
| 9 | 34625409 | rs10972175 | -0.781 | -1.602 | 1.586 | -0.766 | -0.008 | -0.046 | -0.046 | 0 | 0.38 | 0.53 | 0.29 | 0.17 | 0.31 | 0.43 |
| 9 | 34627333 | rs12553321 | 0.121 | -1.578 | 1.316 | -0.752 | 0.017 | 0.039 | 0.039 | 0.182 | 0.08 | 0.02 | 0.02 | 0 | 0.03 | 0.1 |
| 9 | 34631159 | rs10046843 | -0.764 | -1.57 | 1.407 | -0.728 | -0.006 | -0.035 | -0.035 | 0.21 | 0.39 | 0.55 | 0.34 | 0.17 | 0.31 | 0.44 |
| 9 | 34633529 | rs11791806 | 0.018 | -1.46 | 1.218 | -0.606 | -0.002 | 0.002 | 0.002 | 0 | 0.19 | 0.2 | 0.25 | 0.32 | 0.15 | 0.2 |
| 9 | 34637991 | rs10814130 | 0.072 | -1.451 | 0.925 | -0.597 | 0.009 | -0.012 | -0.012 | 0 | 0.19 | 0.16 | 0.25 | 0.32 | 0.15 | 0.2 |
| 9 | 34648418 | rs2070075 | 0.299 | -1.391 | 0.665 | -0.673 | 0.012 | 0.029 | 0.029 | 0 | 0.06 | 0.01 | 0.01 | 0 | 0.03 | 0.1 |
| 9 | 34649442 | rs2070074 | -0.176 | -1.449 | 0.8 | -0.704 | -0.022 | 0.048 | 0.048 | 0 | 0.11 | 0.02 | 0.06 | 0.02 | 0.1 | 0.19 |
| 9 | 34652330 | rs1061758 | 0.274 | -1.443 | 1.011 | -0.669 | -0.035 | 0.093 | 0.093 | 0 | 0.77 | 0.61 | 0.66 | 0.59 | 0.84 | 0.79 |
| 9 | 34654142 | rs3808868 | 1.616 | -1.434 | 0.836 | -0.641 | -0.045 | -0.061 | -0.061 | 0.328 | 0.42 | 0.17 | 0.39 | 0.5 | 0.54 | 0.35 |
| 9 | 34655436 | rs2812357 | 0.618 | -1.294 | 0.74 | -0.607 | 0.003 | -0.002 | -0.002 | 0 | 0.56 | 0.62 | 0.48 | 0.53 | 0.64 | 0.54 |
| 9 | 34664488 | rs913835 | -0.567 | -1.07 | -0.225 | -1.339 | -0.005 | -0.012 | -0.012 | 0 | 0.77 | 0.79 | 0.74 | 0.85 | 0.8 | 0.72 |
| 9 | 34680853 | rs2772554 | 0.184 | -0.641 | -0.12 | -1.05 | -0.013 | -0.046 | -0.046 | 0 | 0.73 | 0.68 | 0.67 | 0.85 | 0.78 | 0.77 |
| 9 | 34686157 | rs2772559 | -0.608 | -0.264 | -0.686 | -0.648 | -0.004 | 0.047 | 0.047 | 0.021 | 0.57 | 0.71 | 0.57 | 0.24 | 0.44 | 0.61 |
| 9 | 34700716 | rs7032259 | -1.433 | -0.423 | -0.603 | -0.632 | 0 | -0.005 | -0.005 | 0.911 | 0.09 | 0.08 | 0.04 | 0.02 | 0.04 | 0.07 |
| 9 | 34707373 | rs10972201 | -0.543 | 0.542 | -1.448 | -1.209 | -0.116 | 0.11 | 0.11 | 2.74 | 0.24 | 0.04 | 0.26 | 0.07 | 0.31 | 0.35 |
| 9 | 34708899 | rs10972203 | -0.877 | 0.226 | -0.212 | -1.378 | 0.056 | 0.021 | 0.021 | 0 | 0.06 | 0.35 | 0.05 | 0.02 | 0.03 | 0.06 |
| 9 | 34710260 | rs2812378 | 0.596 | 0.064 | 0.009 | -1.486 | 0.039 | -0.115 | -0.115 | 0.42 | 0.73 | 0.57 | 0.7 | 0.93 | 0.68 | 0.65 |
| 9 | 34710338 | rs11574914 | -0.561 | 0.069 | -0.117 | -1.484 | -0.102 | 0.12 | 0.12 | 2.465 | 0.25 | 0.06 | 0.27 | 0.07 | 0.32 | 0.35 |
| 9 | 34710450 | rs2812377 | 1.225 | -0.329 | -0.523 | -1.144 | -0.015 | -0.039 | -0.039 | 0 | 0.3 | 0.17 | 0.38 | 0.3 | 0.35 | 0.26 |
| 9 | 34713129 | rs10124246 | 0.619 | -0.687 | -0.376 | -1.307 | -0.025 | -0.06 | -0.06 | 0 | 0.26 | 0.41 | 0.17 | 0.08 | 0.16 | 0.23 |
| 9 | 34723035 | rs10814138 | 0.998 | -0.572 | -0.601 | -1.352 | -0.113 | 0.061 | 0.061 | 0.737 | 0.18 | 0.01 | 0.21 | 0.05 | 0.3 | 0.21 |
| 9 | 34723181 | rs2297132 | 0.148 | -0.924 | -0.273 | -1.198 | -0.133 | -0.126 | -0.126 | 0 | 0.41 | 0.07 | 0.4 | 0.59 | 0.48 | 0.55 |
| 9 | 34727831 | rs7035198 | -1.13 | -0.856 | 0.062 | -1.141 | -0.001 | 0.008 | 0.008 | 0.42 | 0.68 | 0.48 | 0.59 | 0.7 | 0.66 | 0.78 |
| 9 | 34730015 | rs2812344 | 1.244 | -0.836 | 0.007 | -1.138 | -0.006 | -0.023 | -0.023 | 0.078 | 0.3 | 0.21 | 0.39 | 0.3 | 0.34 | 0.21 |
| 9 | 34730395 | rs7024727 | 0.791 | -0.89 | -0.075 | -1.526 | 0.039 | 0.071 | 0.071 | 0 | 0.22 | 0.08 | 0.13 | 0.06 | 0.14 | 0.18 |
| 9 | 34741752 | rs2812373 | -0.454 | -0.873 | 0.042 | -1.503 | -0.046 | -0.091 | -0.091 | 0 | 0.75 | 0.63 | 0.84 | 0.94 | 0.85 | 0.8 |
| 9 | 34771875 | rs277580 | -0.051 | -0.856 | 0.706 | -0.313 | -0.054 | 0.06 | 0.06 | 0 | 0.73 | 0.57 | 0.79 | 0.37 | 0.84 | 0.71 |
| 9 | 34772268 | rs277579 | 0.472 | -0.624 | 0.437 | -0.452 | 0.033 | 0.033 | 0.033 | 0 | 0.57 | 0.32 | 0.61 | 0.3 | 0.66 | 0.46 |
| 9 | 34783711 | rs277608 | 0.474 | -0.646 | 0.493 | -0.442 | 0.036 | 0.029 | 0.029 | 0 | 0.57 | 0.29 | 0.61 | 0.31 | 0.66 | 0.45 |
| 9 | 34784168 | rs277607 | 0.474 | -0.646 | 0.493 | -0.442 | 0.036 | 0.029 | 0.029 | 0 | 0.57 | 0.29 | 0.61 | 0.31 | 0.66 | 0.45 |
| 9 | 34785308 | rs277606 | 0.604 | -0.613 | 0.191 | -0.425 | 0.069 | 0.057 | 0.057 | 0 | 0.55 | 0.17 | 0.6 | 0.31 | 0.66 | 0.43 |
| 9 | 34795035 | rs4879828 | 0.43 | -0.618 | 1.113 | 0.083 | -0.031 | 0.055 | 0.055 | 0 | 0.91 | 0.69 | 0.9 | 0.85 | 0.99 | 0.88 |
| 9 | 34799535 | rs2890604 | 0.626 | -0.702 | 1.82 | 0.061 | -0.03 | -0.024 | -0.024 | 0 | 0.9 | 0.49 | 0.92 | 0.96 | 0.99 | 0.93 |
| 9 | 34803230 | rs1411207 | -0.213 | -0.741 | 1.528 | -0.071 | 0.003 | -0.043 | -0.043 | 0 | 0.35 | 0.17 | 0.29 | 0.54 | 0.36 | 0.47 |
| 9 | 34807277 | rs277558 | -0.575 | -0.747 | 1.746 | -0.074 | -0.024 | -0.022 | -0.022 | 0 | 0.93 | 0.85 | 0.95 | 0.96 | 0.99 | 0.93 |
| 9 | 34811270 | rs10814157 | 0.048 | -0.7 | 1.577 | 0.003 | -0.006 | -0.029 | -0.029 | 2.146 | 0.33 | 0.17 | 0.28 | 0.57 | 0.36 | 0.45 |
| 9 | 34812795 | rs7850527 | -0.185 | -0.759 | 1.524 | -0.116 | 0.008 | -0.065 | -0.065 | 0 | 0.34 | 0.13 | 0.27 | 0.54 | 0.33 | 0.47 |
| 9 | 34823809 | rs277583 | 0.381 | -0.715 | 1.704 | -0.171 | -0.007 | -0.006 | -0.006 | 0 | 0.59 | 0.58 | 0.64 | 0.31 | 0.64 | 0.44 |
| 9 | 34830432 | rs4879832 | 1.106 | -0.933 | 1.534 | -0.32 | 0.08 | 0.079 | 0.079 | 0.401 | 0.51 | 0.25 | 0.58 | 0.31 | 0.64 | 0.35 |
| 9 | 34836905 | rs3802426 | -1.09 | -0.924 | 1.672 | -0.31 | 0.034 | 0.034 | 0.034 | 0.102 | 0.48 | 0.65 | 0.41 | 0.67 | 0.38 | 0.63 |
| 9 | 34839291 | rs924565 | -0.881 | -0.981 | 1.963 | -0.324 | -0.006 | -0.027 | -0.027 | 2.153 | 0.07 | 0.29 | 0.04 | 0 | 0.02 | 0.06 |
| 9 | 34843372 | rs10814165 | -0.421 | -0.893 | 1.738 | -0.265 | 0.014 | 0.008 | 0.008 | 0 | 0.4 | 0.32 | 0.36 | 0.67 | 0.35 | 0.56 |
| 9 | 34866054 | rs1566598 | -0.501 | -0.923 | 1.61 | -0.319 | 0.002 | -0.018 | -0.018 | 0.623 | 0.83 | 0.69 | 0.76 | 0.86 | 0.83 | 0.67 |
| 9 | 34866989 | rs1566595 | 1.238 | -0.789 | 1.644 | -0.509 | 0.018 | 0.034 | 0.034 | 0.891 | 0.47 | 0.34 | 0.52 | 0.35 | 0.56 | 0.26 |
| 9 | 34867913 | rs1948745 | -0.154 | -0.609 | 1.411 | -0.347 | -0.004 | -0.039 | -0.039 | 0 | 0.79 | 0.65 | 0.72 | 0.86 | 0.79 | 0.65 |
| 9 | 34895598 | rs276671 | 0.751 | -0.544 | 1.205 | -0.149 | -0.005 | 0.054 | 0.054 | 0 | 0.2 | 0.21 | 0.15 | 0.47 | 0.2 | 0.22 |
| 9 | 34895767 | rs276672 | 0.751 | -0.544 | 1.205 | -0.149 | -0.005 | 0.054 | 0.054 | 0 | 0.2 | 0.21 | 0.15 | 0.47 | 0.2 | 0.22 |
| 9 | 34910344 | rs2782403 | 0.683 | -0.591 | 1.039 | -0.13 | 0 | 0.01 | 0.01 | 0 | 0.21 | 0.18 | 0.14 | 0.49 | 0.2 | 0.23 |
| 9 | 34911879 | rs7033016 | 0.376 | -0.561 | 1.201 | -0.413 | -0.009 | -0.054 | -0.054 | 0 | 0.2 | 0.23 | 0.22 | 0.03 | 0.19 | 0.28 |
| 9 | 34911949 | rs686400 | -0.67 | -0.529 | 0.992 | -0.382 | 0.001 | 0.006 | 0.006 | 0 | 0.92 | 0.9 | 0.9 | 0.88 | 0.94 | 0.88 |
| 9 | 34914379 | rs2050362 | 0.96 | -0.508 | 0.747 | -0.33 | 0.007 | -0.045 | -0.045 | 0 | 0.2 | 0.11 | 0.13 | 0.49 | 0.2 | 0.2 |
| 9 | 34918201 | rs4879836 | 1.343 | -0.745 | 0.748 | -0.55 | -0.017 | 0.018 | 0.018 | 0.256 | 0.41 | 0.3 | 0.48 | 0.35 | 0.52 | 0.26 |
| 9 | 34920429 | rs2173897 | 1.297 | -0.738 | 0.763 | -0.547 | -0.011 | 0.015 | 0.015 | 0.147 | 0.42 | 0.32 | 0.48 | 0.35 | 0.52 | 0.26 |
| 9 | 34935155 | rs10972262 | -0.839 | -0.231 | 1.037 | -0.434 | 0.001 | -0.069 | -0.069 | 5.289 | 0.31 | 0.47 | 0.37 | 0.19 | 0.31 | 0.45 |
| 9 | 34938198 | rs7041922 | 0.222 | -0.328 | 0.492 | -0.39 | 0.009 | 0.032 | 0.032 | 0.214 | 0.2 | 0.11 | 0.2 | 0.07 | 0.18 | 0.28 |
| 9 | 34942073 | rs10283808 | 0.156 | -0.337 | 0.42 | -0.457 | 0.012 | 0.041 | 0.041 | 0.12 | 0.21 | 0.11 | 0.2 | 0.07 | 0.18 | 0.28 |

| | | | | | | | | | | | | | | | | |
|---|---|---|---|---|---|---|---|---|---|---|---|---|---|---|---|---|
| 9 | 34957613 | rs7031962 | 0.363 | 0.252 | 0.262 | -0.462 | 0.001 | 0.003 | 0.003 | 0 | 0.31 | 0.37 | 0.37 | 0.35 | 0.33 | 0.28 |
| 9 | 34966220 | rs7874491 | 0.501 | -0.216 | 0.678 | -0.159 | -0.012 | 0.029 | 0.029 | 0 | 0.22 | 0.31 | 0.17 | 0.68 | 0.19 | 0.3 |
| 9 | 34978431 | rs3739873 | 0.424 | -0.404 | 0.402 | 0.04 | 0.008 | -0.024 | -0.024 | 0 | 0.25 | 0.18 | 0.18 | 0.63 | 0.22 | 0.27 |
| 9 | 34982343 | rs3088305 | -0.994 | -1.086 | 0.919 | -0.865 | 0.006 | -0.1 | -0.1 | 3.82 | 0.58 | 0.53 | 0.46 | 0.92 | 0.47 | 0.68 |
| 9 | 34983974 | rs4879847 | 0.096 | -0.937 | 0.762 | -0.746 | 0.037 | 0.005 | 0.005 | 0 | 0.27 | 0.17 | 0.19 | 0.25 | 0.17 | 0.3 |
| 9 | 34996014 | rs4879849 | -0.612 | -0.853 | 1.052 | -0.593 | -0.01 | 0.085 | 0.085 | 0 | 0.13 | 0.25 | 0.13 | 0.39 | 0.1 | 0.2 |
| 9 | 34998003 | rs9608 | 1.047 | -1.264 | 1.155 | -0.637 | 0.004 | -0.02 | -0.02 | 0 | 0.25 | 0.19 | 0.16 | 0.53 | 0.23 | 0.26 |
| 9 | 34999992 | rs10814192 | 0.33 | -1.378 | 1.245 | -0.746 | 0.062 | 0.144 | 0.144 | 0.246 | 0.59 | 0.22 | 0.7 | 0.06 | 0.69 | 0.51 |
| 9 | 35000289 | rs10972278 | 0.419 | -1.408 | 1.244 | -0.783 | 0.074 | 0.195 | 0.195 | 0.476 | 0.58 | 0.16 | 0.7 | 0.06 | 0.69 | 0.5 |
| 9 | 35010738 | rs10972281 | -0.972 | -1.389 | 1.416 | -0.774 | -0.011 | 0.031 | 0.031 | 0 | 0.72 | 0.62 | 0.81 | 0.45 | 0.75 | 0.73 |
| 9 | 35011305 | rs4879852 | -0.089 | -1.366 | 1.414 | -0.739 | -0.029 | 0.102 | 0.102 | 0 | 0.11 | 0.23 | 0.11 | 0.37 | 0.07 | 0.2 |
| 9 | 35031343 | rs10972291 | 0.335 | -1.434 | 1.201 | -0.875 | 0.006 | 0.023 | 0.023 | 0 | 0.25 | 0.2 | 0.2 | 0.08 | 0.2 | 0.27 |
| 9 | 35041604 | rs10972297 | 1.353 | -1.47 | 1.124 | -0.868 | 0.004 | -0.029 | -0.029 | 0 | 0.24 | 0.16 | 0.15 | 0.55 | 0.23 | 0.26 |
| 9 | 35043291 | rs615474 | -1.022 | -1.459 | 1.111 | -0.867 | -0.016 | 0.04 | 0.04 | 0 | 0.71 | 0.6 | 0.81 | 0.45 | 0.75 | 0.73 |
| 9 | 35049508 | rs7048389 | -0.348 | -1.486 | 1.206 | -0.906 | -0.036 | 0.195 | 0.195 | 0.009 | 0.13 | 0.39 | 0.12 | 0.37 | 0.07 | 0.2 |
| 9 | 35051269 | rs642296 | 0.749 | -1.506 | 1.182 | -0.92 | -0.092 | 0.044 | 0.044 | 0 | 0.43 | 0.06 | 0.59 | 0.32 | 0.54 | 0.37 |
| 9 | 35054586 | rs565070 | 0.59 | -1.511 | 1.184 | -0.849 | 0.001 | 0.049 | 0.049 | 0 | 0.46 | 0.28 | 0.61 | 0.38 | 0.55 | 0.46 |
| 9 | 35055669 | rs595429 | 0.515 | -1.496 | 1.193 | -0.864 | 0.002 | 0.015 | 0.015 | 0 | 0.46 | 0.42 | 0.62 | 0.38 | 0.55 | 0.46 |
| 9 | 35062972 | rs514492 | -1.547 | -1.493 | 1.185 | -0.871 | -0.002 | 0.012 | 0.012 | 0 | 0.75 | 0.74 | 0.84 | 0.45 | 0.77 | 0.74 |
| 9 | 35068201 | rs10972300 | 0.57 | -1.488 | 1.08 | -0.923 | 0.003 | 0.046 | 0.046 | 0 | 0.23 | 0.12 | 0.2 | 0.08 | 0.2 | 0.27 |
| 9 | 35074917 | rs587118 | 1.311 | -1.324 | 0.648 | -0.798 | -0.188 | 0.257 | 0.257 | 0.187 | 0.34 | 0.02 | 0.5 | 0 | 0.48 | 0.25 |
| 9 | 35088142 | rs527459 | 0.591 | -1.038 | 0.126 | -0.071 | -0.035 | -0.057 | -0.057 | 0 | 0.46 | 0.11 | 0.6 | 0.51 | 0.55 | 0.39 |
| 9 | 35098155 | rs527365 | -0.589 | -0.863 | 0.63 | 0.456 | -0.012 | 0.012 | 0.012 | 0 | 0.72 | 0.64 | 0.83 | 0.64 | 0.77 | 0.75 |
| 9 | 35100468 | rs666549 | -0.433 | -0.868 | 0.614 | 0.403 | -0.015 | 0.025 | 0.025 | 0 | 0.71 | 0.63 | 0.82 | 0.46 | 0.76 | 0.74 |
| 9 | 35100569 | rs586017 | -0.708 | -0.767 | 0.465 | 0.472 | -0.009 | 0.031 | 0.031 | 0 | 0.75 | 0.7 | 0.84 | 0.46 | 0.78 | 0.76 |
| 9 | 35106987 | rs10972316 | 0.035 | -0.653 | 0.12 | 0.417 | -0.004 | 0.018 | 0.018 | 0 | 0.08 | 0.15 | 0.09 | 0.26 | 0.05 | 0.11 |
| 9 | 35109949 | rs944700 | 0.375 | -0.277 | -0.251 | -0.398 | -0.032 | 0.002 | 0.002 | 0 | 0.43 | 0.27 | 0.58 | 0.57 | 0.49 | 0.44 |
| 9 | 35112406 | rs10972320 | -0.156 | -0.639 | -0.052 | -0.253 | -0.008 | 0.001 | 0.001 | 0 | 0.64 | 0.41 | 0.8 | 0.63 | 0.75 | 0.71 |
| 9 | 35114394 | rs7852999 | 0.051 | -0.439 | 0.034 | -1.112 | -0.001 | 0.051 | 0.051 | 0.013 | 0.42 | 0.32 | 0.32 | 0.93 | 0.39 | 0.61 |
| 9 | 35114614 | rs10123856 | 0.133 | -0.432 | -0.065 | -1.075 | -0.011 | 0.114 | 0.114 | 0 | 0.38 | 0.22 | 0.28 | 0.93 | 0.35 | 0.55 |
| 9 | 35116932 | rs10972321 | -0.303 | -0.437 | 0.859 | -0.382 | -0.024 | -0.004 | -0.004 | 0.108 | 0.24 | 0.39 | 0.2 | 0.15 | 0.23 | 0.19 |
| 9 | 35121795 | rs1571953 | 0.364 | -0.477 | 0.393 | -0.401 | 0.003 | 0.002 | 0.002 | 0 | 0.73 | 0.23 | 0.79 | 0.85 | 0.77 | 0.81 |
| 9 | 35123369 | rs2381298 | 1.296 | -0.035 | 0.704 | -0.374 | 0.095 | 0.072 | 0.072 | 2.268 | 0.95 | 0.44 | 0.97 | 0.99 | 0.99 | 1 |
| 9 | 35141508 | rs4879869 | 0.037 | 0.109 | -0.204 | -0.793 | -0.004 | 0.008 | 0.008 | 0.348 | 0.41 | 0.36 | 0.34 | 0.92 | 0.39 | 0.68 |
| 9 | 35141705 | rs10972341 | -0.264 | 0.098 | -0.183 | -0.228 | -0.007 | 0.018 | 0.018 | 0.343 | 0.6 | 0.66 | 0.7 | 0.21 | 0.62 | 0.33 |
| 9 | 35149993 | rs10972351 | 0.565 | 0.14 | -0.743 | -0.574 | -0.036 | -0.036 | -0.036 | 0 | 0.17 | 0.01 | 0.2 | 0.15 | 0.21 | 0.15 |
| 9 | 35166763 | rs13293564 | 0.418 | -0.317 | -0.549 | -0.93 | -0.002 | -0.031 | -0.031 | 0 | 0.37 | 0.64 | 0.49 | 0.01 | 0.4 | 0.19 |
| 9 | 35171905 | rs4879876 | 0.147 | -0.077 | -0.803 | -0.62 | -0.042 | -0.004 | -0.004 | 0 | 0.2 | 0.01 | 0.16 | 0.14 | 0.24 | 0.15 |
| 9 | 35177345 | rs7865406 | -0.806 | 0.003 | -0.205 | -1.013 | -0.01 | -0.06 | -0.06 | 0.038 | 0.83 | 0.75 | 0.68 | 0.98 | 0.87 | 0.84 |
| 9 | 35191713 | rs3904435 | 0.146 | -0.306 | 0.292 | -0.198 | -0.033 | 0.039 | 0.039 | 0 | 0.76 | 0.58 | 0.61 | 0.68 | 0.83 | 0.72 |
| 9 | 35192366 | rs11531752 | 0.361 | -0.304 | 0.335 | -0.192 | -0.015 | 0.15 | 0.15 | 0 | 0.06 | 0.12 | 0.07 | 0.3 | 0.03 | 0.12 |
| 9 | 35208145 | rs10972381 | 0.89 | -0.494 | 0.495 | -0.302 | 0.01 | 0.002 | 0.002 | 0 | 0.57 | 0.36 | 0.45 | 0.55 | 0.6 | 0.52 |
| 9 | 35231047 | rs10972390 | 1.148 | -0.486 | 0.598 | -0.368 | 0.003 | 0.004 | 0.004 | 0 | 0.16 | 0.1 | 0.31 | 0.01 | 0.13 | 0.14 |
| 9 | 35250666 | rs10118130 | 0.218 | -0.512 | 0.775 | -0.376 | -0.01 | -0.091 | -0.091 | 0 | 0.63 | 0.52 | 0.52 | 0.87 | 0.63 | 0.68 |
| 9 | 35271817 | rs10114937 | 0.419 | 0.135 | 0.755 | -0.328 | 0.003 | 0.043 | 0.043 | 0.558 | 0.71 | 0.44 | 0.75 | 0.55 | 0.69 | 0.62 |
| 9 | 35289544 | rs10758303 | 0.888 | 0.179 | 0.755 | -0.33 | 0 | 0.001 | 0.001 | 0.902 | 0.56 | 0.41 | 0.44 | 0.55 | 0.55 | 0.48 |
| 9 | 35290205 | rs4419897 | 0.386 | 0.321 | 0.907 | -0.12 | 0.001 | 0.022 | 0.022 | 1.374 | 0.2 | 0.15 | 0.16 | 0.14 | 0.23 | 0.2 |
| 9 | 35302950 | rs10972428 | 0.386 | 0.321 | 0.995 | -0.107 | 0.001 | 0.024 | 0.024 | 1.374 | 0.2 | 0.15 | 0.16 | 0.13 | 0.23 | 0.2 |
| 9 | 35307408 | rs572527 | -0.627 | 0.326 | 1.051 | -0.091 | 0.009 | 0.055 | 0.055 | 1.772 | 0.28 | 0.51 | 0.26 | 0.44 | 0.31 | 0.38 |
| 9 | 35343563 | rs2069008 | 0.698 | 0.466 | 0.91 | 0.037 | -0.004 | 0.043 | 0.043 | 0 | 0.06 | 0.04 | 0.09 | 0.3 | 0.08 | 0.15 |
| 9 | 35361044 | rs10814231 | 0.632 | 0.386 | 0.896 | -0.063 | -0.022 | 0.02 | 0.02 | 3.871 | 0.18 | 0.07 | 0.15 | 0.14 | 0.23 | 0.29 |
| 9 | 35369125 | rs12684897 | 1.53 | 0.411 | 0.447 | 0.17 | -0.003 | -0.003 | -0.003 | 0 | 0.16 | 0.13 | 0.15 | 0.15 | 0.18 | 0.11 |
| 9 | 35374982 | rs2149368 | 0.906 | 0.466 | 0.419 | 0.246 | -0.022 | 0.02 | 0.02 | 3.871 | 0.18 | 0.07 | 0.15 | 0.14 | 0.23 | 0.29 |
| 9 | 35381504 | rs2281999 | -0.226 | 0.466 | 0.523 | 0.263 | 0.01 | 0.046 | 0.046 | 3.347 | 0.28 | 0.58 | 0.26 | 0.45 | 0.31 | 0.47 |
| 9 | 35400272 | rs4878613 | -0.108 | 0.508 | 0.743 | 0.407 | 0.108 | 0.314 | 0.314 | 0 | 0.09 | 0.4 | 0.1 | 0.31 | 0.09 | 0.18 |
| 9 | 35410334 | rs12238748 | 1.581 | 0.452 | 0.234 | 0.628 | -0.013 | -0.022 | -0.022 | 0.06 | 0.06 | 0 | 0.04 | 0.22 | 0.05 | 0.07 |
| 9 | 35413694 | rs4878615 | 0.614 | 0.485 | 0.399 | 0.546 | 0.038 | 0.163 | 0.163 | 0 | 0.07 | 0.19 | 0.09 | 0.31 | 0.09 | 0.15 |
| 9 | 35427324 | rs10972462 | 0.057 | 0.376 | 0.372 | 0.662 | 0.011 | 0.058 | 0.058 | 3.441 | 0.28 | 0.59 | 0.27 | 0.53 | 0.32 | 0.47 |
| 9 | 35446386 | rs9298963 | 0.391 | 0.268 | -0.094 | 0.89 | -0.005 | -0.017 | -0.017 | 0.27 | 0.09 | 0.04 | 0.06 | 0.33 | 0.08 | 0.13 |
| 9 | 35460769 | rs1971100 | 1.305 | 0.409 | -0.338 | 0.447 | -0.002 | -0.005 | -0.005 | 0.054 | 0.15 | 0.13 | 0.15 | 0.15 | 0.17 | 0.11 |
| 9 | 35475391 | rs10972486 | 0.382 | 0.683 | -0.585 | 0.261 | -0.004 | -0.042 | -0.042 | 0 | 0.25 | 0.11 | 0.3 | 0.42 | 0.22 | 0.26 |
| 9 | 35480564 | rs7040391 | -0.635 | 0.815 | -0.542 | -0.014 | -0.017 | 0.025 | 0.025 | 0 | 0.63 | 0.51 | 0.68 | 0.51 | 0.71 | 0.62 |
| 9 | 35482075 | rs7875285 | -0.189 | 1.024 | -0.629 | -0.058 | -0.007 | 0.008 | 0.008 | 0.093 | 0.45 | 0.37 | 0.52 | 0.44 | 0.48 | 0.34 |
| 9 | 35482149 | rs10972487 | 1.203 | 0.85 | -0.691 | -0.088 | -0.059 | -0.105 | -0.105 | 0.038 | 0.23 | 0.02 | 0.21 | 0.39 | 0.28 | 0.17 |
| 9 | 35482764 | rs7018605 | -0.038 | 0.494 | -0.541 | 0.042 | 0.002 | 0.009 | 0.009 | 0.075 | 0.46 | 0.38 | 0.53 | 0.44 | 0.51 | 0.34 |
| 9 | 35497219 | rs7870832 | 0.72 | 0.447 | -0.757 | -0.347 | 0.002 | -0.01 | -0.01 | 0.196 | 0.18 | 0.16 | 0.29 | 0.03 | 0.16 | 0.13 |
| 9 | 35500545 | rs1890591 | -0.723 | 0.737 | -0.405 | -0.085 | -0.014 | 0.005 | 0.005 | 0.466 | 0.64 | 0.58 | 0.69 | 0.61 | 0.7 | 0.63 |
| 9 | 35514653 | rs10972498 | -0.111 | 0.553 | -0.87 | 0.254 | 0.005 | 0.001 | 0.001 | 0.794 | 0.35 | 0.21 | 0.4 | 0.35 | 0.38 | 0.4 |
| 9 | 35518997 | rs4879901 | 0.274 | 0.259 | -0.874 | 0.244 | 0 | 0.024 | 0.024 | 2.575 | 0.41 | 0.24 | 0.61 | 0.32 | 0.51 | 0.31 |

| | | | | | | | | | | | | | | | | |
|---|---|---|---|---|---|---|---|---|---|---|---|---|---|---|---|---|
| 9 | 35522318 | rs7047617 | -0.742 | 0.292 | -0.558 | 0.011 | -0.039 | -0.011 | -0.011 | 0 | 0.62 | 0.61 | 0.77 | 0.64 | 0.71 | 0.59 |
| 9 | 35526605 | rs2115983 | -0.601 | 0.291 | -0.59 | 0 | -0.041 | -0.01 | -0.01 | 0 | 0.61 | 0.58 | 0.77 | 0.63 | 0.71 | 0.58 |
| 9 | 35532053 | rs10972503 | -0.497 | 0.271 | -0.428 | -0.014 | 0.026 | 0.008 | 0.008 | 1.721 | 0.59 | 0.35 | 0.75 | 0.62 | 0.71 | 0.55 |
| 9 | 35532091 | rs10972504 | -0.561 | 0.269 | -0.405 | -0.009 | 0.003 | 0.001 | 0.001 | 1.898 | 0.6 | 0.41 | 0.75 | 0.62 | 0.71 | 0.57 |
| 9 | 35533348 | rs7041770 | -1.073 | -0.045 | -0.546 | -0.111 | -0.082 | 0.017 | 0.017 | 5.057 | 0.7 | 0.52 | 0.83 | 0.63 | 0.86 | 0.65 |
| 9 | 35551434 | rs10814252 | 0.292 | 0.363 | -0.566 | 0.137 | 0.003 | -0.017 | -0.017 | 0 | 0.19 | 0.15 | 0.15 | 0.32 | 0.21 | 0.25 |
| 9 | 35557696 | rs1890590 | 0.852 | 0.166 | -0.144 | -0.042 | -0.007 | 0 | 0 | 0 | 0.35 | 0.29 | 0.19 | 0.38 | 0.27 | 0.4 |
| 9 | 35559905 | rs12338773 | 0.628 | 0.887 | -0.304 | -0.092 | -0.018 | 0.027 | 0.027 | 2.843 | 0.39 | 0.23 | 0.6 | 0.29 | 0.48 | 0.29 |
| 9 | 35562549 | rs755774 | 0.878 | 0.825 | -0.485 | -0.217 | -0.054 | 0.009 | 0.009 | 2.521 | 0.33 | 0.21 | 0.58 | 0.28 | 0.44 | 0.26 |
| 9 | 35566752 | rs4879907 | -0.812 | 0.65 | -0.368 | -0.097 | 0.022 | -0.055 | -0.055 | 1.184 | 0.82 | 0.95 | 0.86 | 0.7 | 0.82 | 0.76 |
| 9 | 35578001 | rs10758312 | -0.6 | 0.812 | -0.347 | -0.158 | 0.022 | 0.013 | 0.013 | 1.544 | 0.57 | 0.39 | 0.75 | 0.61 | 0.67 | 0.53 |
| 9 | 35592324 | rs7849006 | -0.333 | 1.809 | 0.046 | -0.151 | 0 | 0 | 0 | 0 | 0.34 | 0.39 | 0.46 | 0.38 | 0.4 | 0.37 |
| 9 | 35601967 | rs10814259 | -0.755 | 1.773 | 0.082 | -0.156 | 0.014 | 0.001 | 0.001 | 0 | 0.39 | 0.53 | 0.49 | 0.39 | 0.47 | 0.43 |
| 9 | 35606884 | rs2275422 | -0.756 | 1.772 | 0.082 | -0.156 | 0.014 | 0.001 | 0.001 | 0 | 0.39 | 0.53 | 0.49 | 0.38 | 0.47 | 0.43 |
| 9 | 35610912 | rs4879913 | 0.127 | 1.967 | 0.029 | -0.235 | 0.016 | 0.021 | 0.021 | 0 | 0.65 | 0.47 | 0.5 | 0.55 | 0.59 | 0.63 |
| 9 | 35618523 | rs10814260 | 0.529 | 1.734 | -0.727 | -0.106 | -0.004 | -0.018 | -0.018 | 0.062 | 0.17 | 0.14 | 0.15 | 0.4 | 0.21 | 0.24 |
| 9 | 35628739 | rs10972530 | 0.013 | 1.649 | -0.848 | -0.168 | -0.004 | -0.011 | -0.011 | 0 | 0.64 | 0.39 | 0.52 | 0.56 | 0.62 | 0.62 |
| 9 | 35631423 | rs10972531 | 0.399 | 1.662 | -1.26 | -0.155 | -0.06 | -0.044 | -0.044 | 0.314 | 0.59 | 0.17 | 0.48 | 0.55 | 0.55 | 0.57 |
| 9 | 35635683 | rs1890592 | 0.096 | 1.741 | -1.36 | -0.327 | -0.053 | 0.033 | 0.033 | 0.816 | 0.31 | 0.24 | 0.34 | 0.2 | 0.42 | 0.21 |
| 9 | 35641859 | rs10972540 | 1.206 | 1.568 | -1.465 | -0.292 | -0.026 | 0.037 | 0.037 | 0 | 0.08 | 0.04 | 0.05 | 0 | 0.09 | 0.02 |
| 9 | 35643558 | rs17466775 | 1.14 | 1.483 | -1.353 | -0.116 | -0.082 | -0.052 | -0.052 | 0 | 0.2 | 0.01 | 0.26 | 0.18 | 0.24 | 0.16 |
| 9 | 35648950 | rs3138083 | -0.231 | 1.004 | -1.27 | -0.694 | -0.014 | 0.041 | 0.041 | 0.958 | 0.39 | 0.29 | 0.45 | 0.86 | 0.45 | 0.56 |
| 9 | 35649216 | rs3138082 | 0.941 | 0.451 | -1.755 | -0.959 | -0.012 | 0.193 | 0.193 | 1.91 | 0.28 | 0.02 | 0.11 | 0.02 | 0.2 | 0.23 |
| 9 | 35650458 | rs2297880 | -0.391 | 0.532 | -0.619 | -0.128 | -0.03 | 0.054 | 0.054 | 0 | 0.67 | 0.55 | 0.64 | 0.56 | 0.74 | 0.58 |
| 9 | 35659591 | rs10441725 | -0.343 | 0.961 | -0.318 | -0.351 | 0.063 | -0.036 | -0.036 | 0.031 | 0.11 | 0.33 | 0.09 | 0.04 | 0.13 | 0.08 |
| 9 | 35660901 | rs2275420 | -0.343 | 0.961 | -0.318 | -0.351 | 0.063 | -0.036 | -0.036 | 0 | 0.11 | 0.33 | 0.09 | 0.04 | 0.13 | 0.08 |
| 9 | 35660990 | rs1339374 | -0.945 | 0.906 | -0.297 | -0.644 | 0.025 | -0.124 | -0.124 | 0 | 0.81 | 0.6 | 0.87 | 0.98 | 0.82 | 0.89 |
| 9 | 35661379 | rs10972554 | 0.576 | 0.828 | -1.13 | -0.114 | 0.015 | -0.104 | -0.104 | 0 | 0.32 | 0.06 | 0.35 | 0.52 | 0.32 | 0.38 |
| 9 | 35662251 | rs2297879 | 0.562 | 0.826 | -1.144 | -0.151 | 0.016 | -0.1 | -0.1 | 0.025 | 0.33 | 0.06 | 0.35 | 0.5 | 0.33 | 0.38 |
| 9 | 35671627 | rs1934158 | -0.35 | 0.932 | -0.214 | 0.246 | 0.065 | -0.002 | -0.002 | 0 | 0.11 | 0.39 | 0.1 | 0.11 | 0.13 | 0.09 |
| 9 | 35673891 | rs2071675 | 0.575 | 0.73 | -1.027 | 0.63 | -0.004 | 0.021 | 0.021 | 0 | 0.07 | 0.06 | 0.15 | 0.2 | 0.06 | 0.16 |
| 9 | 35674053 | rs2071676 | 0.467 | 0.825 | -1.386 | 0.57 | -0.029 | -0.122 | -0.122 | 0.481 | 0.36 | 0.08 | 0.41 | 0.44 | 0.36 | 0.44 |
| 9 | 35679061 | rs2857802 | -0.281 | 0.838 | -0.753 | 0.443 | -0.001 | 0.007 | 0.007 | 0 | 0.58 | 0.64 | 0.56 | 0.39 | 0.61 | 0.51 |
| 9 | 35679251 | rs3829078 | 0.212 | 1.042 | -0.918 | 0.192 | -0.026 | 0.014 | 0.014 | 0.006 | 0.06 | 0.02 | 0.06 | 0.03 | 0.1 | 0.05 |
| 9 | 35680564 | rs3750433 | 0.968 | 0.796 | -1.376 | -0.221 | 0.036 | 0.164 | 0.164 | 0 | 0.31 | 0.08 | 0.32 | 0.05 | 0.24 | 0.3 |
| 9 | 35681299 | rs1538535 | 1.223 | 1.059 | -1.507 | -0.121 | -0.01 | -0.019 | -0.019 | 0 | 0.22 | 0.08 | 0.24 | 0.26 | 0.25 | 0.19 |
| 9 | 35681368 | rs4879924 | 0.372 | 1.079 | -0.688 | -0.508 | 0.033 | -0.048 | -0.048 | 0.006 | 0.11 | 0.24 | 0.09 | 0.03 | 0.14 | 0.08 |
| 9 | 35682585 | rs17259350 | 1.158 | 1.103 | -0.792 | -0.428 | -0.032 | 0.027 | 0.027 | 0.005 | 0.08 | 0.01 | 0.04 | 0.03 | 0.1 | 0.07 |
| 9 | 35686376 | rs760402 | -0.779 | 0.837 | 0.193 | 0.143 | 0.003 | 0.016 | 0.016 | 0 | 0.75 | 0.71 | 0.69 | 0.59 | 0.69 | 0.79 |
| 9 | 35689373 | rs2145925 | -0.148 | 0.655 | 0.459 | -0.006 | 0.007 | -0.002 | -0.002 | 0 | 0.42 | 0.45 | 0.35 | 0.62 | 0.45 | 0.5 |
| 9 | 35690310 | rs2756892 | 1.076 | 0.607 | 0.42 | -0.129 | 0.014 | 0.043 | 0.043 | 0 | 0.33 | 0.21 | 0.34 | 0.05 | 0.27 | 0.3 |
| 9 | 35700150 | rs2295794 | 0.119 | 0.629 | 0.415 | 0.049 | -0.012 | -0.02 | -0.02 | 0 | 0.38 | 0.25 | 0.33 | 0.54 | 0.42 | 0.51 |
| 9 | 35705086 | rs867194 | 0.954 | 0.687 | 0.185 | -0.01 | 0.004 | -0.007 | -0.007 | 0 | 0.34 | 0.29 | 0.35 | 0.05 | 0.27 | 0.28 |
| 9 | 35712003 | rs2295795 | 0.957 | 0.863 | 0.022 | 0.484 | -0.061 | -0.114 | -0.114 | 0 | 0.21 | 0.02 | 0.27 | 0.34 | 0.27 | 0.19 |
| 9 | 35731004 | rs3763630 | 0.968 | 0.616 | 0.127 | 0.544 | -0.014 | -0.055 | -0.055 | 0.462 | 0.13 | 0.02 | 0.19 | 0.33 | 0.17 | 0.15 |
| 9 | 35734956 | rs10814274 | 0.906 | 0.575 | 0.301 | 0.343 | 0.031 | 0.035 | 0.035 | 1.264 | 0.51 | 0.25 | 0.43 | 0.41 | 0.43 | 0.43 |
| 9 | 35748806 | rs1570246 | 0.906 | 0.58 | 0.263 | 0.515 | 0.031 | 0.056 | 0.056 | 1.264 | 0.51 | 0.25 | 0.43 | 0.4 | 0.43 | 0.43 |
| 9 | 35751221 | rs1570248 | -0.821 | 0.685 | 0.512 | 0.371 | 0 | -0.003 | -0.003 | 0.206 | 0.73 | 0.75 | 0.67 | 0.82 | 0.66 | 0.59 |
| 9 | 35752252 | rs1570249 | 1.065 | 0.68 | 0.474 | 0.381 | 0.031 | 0.032 | 0.032 | 1.264 | 0.51 | 0.26 | 0.43 | 0.42 | 0.43 | 0.43 |
| 9 | 35757075 | rs1951432 | -0.531 | 0.839 | 0.474 | 0.025 | -0.002 | 0 | 0 | 0 | 0.73 | 0.75 | 0.67 | 0.82 | 0.66 | 0.59 |
| 9 | 35758578 | rs10814275 | 0.664 | 0.896 | -0.034 | -0.242 | 0.024 | -0.01 | -0.01 | 0 | 0.07 | 0.21 | 0.23 | 0.05 | 0.1 | 0.06 |
| 9 | 35766561 | rs4878628 | 0.826 | 0.942 | -0.511 | -0.154 | -0.094 | 0.094 | 0.094 | 0 | 0.19 | 0.02 | 0.22 | 0.07 | 0.26 | 0.26 |
| 9 | 35778800 | rs7020142 | 0.769 | 0.101 | -0.008 | -0.129 | -0.022 | -0.063 | -0.063 | 0.349 | 0.13 | 0.22 | 0.06 | 0 | 0.11 | 0.01 |
| 9 | 35780220 | rs7847621 | -0.197 | 0.697 | 0.476 | 0.348 | -0.015 | 0.005 | 0.005 | 0 | 0.72 | 0.77 | 0.63 | 0.76 | 0.62 | 0.6 |
| 9 | 35789720 | rs12337802 | -0.338 | 0.654 | 0.812 | 0.633 | -0.003 | 0.09 | 0.09 | 0.253 | 0.21 | 0.58 | 0.24 | 0.38 | 0.2 | 0.16 |
| 9 | 35798710 | rs3793543 | -0.225 | 0.655 | 0.81 | 0.635 | 0.011 | 0.154 | 0.154 | 0 | 0.19 | 0.5 | 0.23 | 0.38 | 0.2 | 0.16 |
| 9 | 35804149 | rs10758325 | 1.19 | 0.6 | 0.402 | 0.543 | 0.033 | 0.069 | 0.069 | 0.571 | 0.52 | 0.18 | 0.39 | 0.38 | 0.41 | 0.45 |
| 9 | 35809328 | rs2281645 | 1.029 | 0.755 | 0.114 | 0.544 | -0.115 | 0.015 | 0.015 | 0 | 0.2 | 0.03 | 0.26 | 0.14 | 0.31 | 0.25 |
| 9 | 35812623 | rs878749 | 0.879 | 0.672 | 0.328 | 0.617 | -0.007 | -0.037 | -0.037 | 0 | 0.13 | 0.1 | 0.2 | 0.31 | 0.18 | 0.15 |
| 9 | 35815356 | rs4879929 | 1.31 | 0.583 | 0.787 | 0.496 | -0.004 | 0.084 | 0.084 | 3.287 | 0.86 | 0.35 | 0.86 | 0.76 | 0.89 | 0.84 |
| 9 | 35817579 | rs13292880 | 1.279 | 0.64 | 0.754 | 0.514 | 0.018 | 0.047 | 0.047 | 0.355 | 0.53 | 0.28 | 0.42 | 0.36 | 0.41 | 0.43 |
| 9 | 35818647 | rs2281646 | 0.41 | 0.597 | 1.087 | 0.585 | 0.02 | 0.164 | 0.164 | 0.217 | 0.9 | 0.58 | 0.88 | 0.83 | 0.92 | 0.86 |
| 9 | 35823009 | rs734158 | 0.114 | 0.642 | 0.827 | 0.605 | 0 | 0.018 | 0.018 | 0 | 0.16 | 0.17 | 0.2 | 0.38 | 0.2 | 0.16 |
| 9 | 35825413 | rs4879930 | 0.947 | 0.635 | 0.811 | 0.563 | -0.001 | 0.016 | 0.016 | 0.15 | 0.74 | 0.34 | 0.67 | 0.45 | 0.72 | 0.69 |
| 9 | 35833448 | rs10738948 | -0.029 | 0.673 | 0.753 | 0.5 | 0.032 | -0.013 | -0.013 | 0 | 0.7 | 0.56 | 0.61 | 0.75 | 0.61 | 0.56 |
| 9 | 35841071 | rs2236292 | 1.159 | 0.672 | 0.354 | 0.584 | -0.04 | -0.085 | -0.085 | 0 | 0.12 | 0.01 | 0.18 | 0.32 | 0.17 | 0.16 |
| 9 | 35841783 | rs2236293 | 1.326 | 0.61 | 0.503 | 0.418 | 0.039 | 0.085 | 0.085 | 0.78 | 0.51 | 0.17 | 0.4 | 0.38 | 0.41 | 0.39 |
| 9 | 35842854 | rs13284995 | 1.241 | 0.617 | 0.53 | 0.421 | 0.031 | 0.059 | 0.059 | 0.494 | 0.52 | 0.25 | 0.41 | 0.37 | 0.41 | 0.39 |
| 9 | 35854437 | rs7035913 | 0.744 | 0.65 | 0.037 | 0.352 | -0.031 | 0.035 | 0.035 | 0 | 0.22 | 0.13 | 0.27 | 0.07 | 0.31 | 0.3 |

| | | | | | | | | | | | | | | | | |
|---|---|---|---|---|---|---|---|---|---|---|---|---|---|---|---|---|
| 9 | 35854565 | rs17204908 | 0.906 | 0.676 | -0.425 | 0.366 | -0.113 | 0.09 | 0.09 | 0 | 0.2 | 0.03 | 0.26 | 0.07 | 0.31 | 0.3 |
| 9 | 35855218 | rs12345598 | 0.345 | 0.797 | -0.255 | 0.572 | -0.018 | -0.052 | -0.052 | 0 | 0.14 | 0.06 | 0.19 | 0.37 | 0.19 | 0.17 |
| 9 | 35860483 | rs10814287 | 0.892 | 0.673 | -0.285 | 0.461 | 0.047 | 0.093 | 0.093 | 1.053 | 0.5 | 0.17 | 0.39 | 0.37 | 0.39 | 0.39 |
| 9 | 35860942 | rs11791889 | 0.891 | 0.64 | -0.283 | 0.467 | 0.057 | 0.09 | 0.09 | 1.071 | 0.5 | 0.19 | 0.38 | 0.37 | 0.38 | 0.39 |
| 9 | 35867901 | rs10814288 | -0.305 | 1.039 | 0.005 | 1.113 | -0.001 | 0.003 | 0.003 | 0 | 0.17 | 0.16 | 0.26 | 0.36 | 0.22 | 0.17 |
| 9 | 35881628 | rs7039312 | -0.153 | 1.293 | 0.189 | 0.987 | 0.003 | 0.068 | 0.068 | 0 | 0.06 | 0.13 | 0.04 | 0.18 | 0.08 | 0.15 |
| 9 | 35887412 | rs4879938 | 0.165 | 1.812 | 0.111 | 0.319 | 0 | -0.007 | -0.007 | 2.478 | 0.21 | 0.21 | 0.32 | 0.07 | 0.3 | 0.25 |
| 9 | 35893292 | rs10972610 | -0.729 | 1.454 | 0.481 | 1.047 | 0.006 | 0.033 | 0.033 | 0 | 0.06 | 0.11 | 0.03 | 0.19 | 0.06 | 0.05 |
| 9 | 35894737 | rs6476493 | -0.12 | 1.21 | 0.556 | 1.125 | -0.003 | 0.003 | 0.003 | 0.117 | 0.73 | 0.78 | 0.61 | 0.54 | 0.69 | 0.71 |
| 9 | 35903317 | rs2890719 | -0.662 | 1.189 | -0.23 | 0.783 | 0.009 | 0.04 | 0.04 | 0 | 0.06 | 0.12 | 0.03 | 0.19 | 0.06 | 0.06 |
| 9 | 35903370 | rs1467577 | 0.566 | 1.054 | -0.043 | 0.651 | -0.002 | 0.008 | 0.008 | 0 | 0.65 | 0.4 | 0.54 | 0.43 | 0.57 | 0.55 |
| 9 | 35905966 | rs2025175 | 0.494 | 1.147 | -0.346 | 0.509 | 0.002 | -0.006 | -0.006 | 0.056 | 0.62 | 0.38 | 0.52 | 0.49 | 0.61 | 0.53 |
| 9 | 35918778 | rs4483242 | -0.034 | 1.412 | 0.039 | 0.09 | 0.015 | -0.055 | -0.055 | 0 | 0.1 | 0.2 | 0.08 | 0 | 0.15 | 0.04 |
| 9 | 35920856 | rs2025174 | -0.116 | 1.792 | 0.152 | 0.253 | 0.001 | 0.034 | 0.034 | 0.133 | 0.78 | 0.77 | 0.54 | 0.4 | 0.73 | 0.71 |
| 9 | 35923384 | rs7046469 | -0.367 | 2.469 | 0.512 | 0.036 | 0.053 | 0.014 | 0.014 | 0 | 0.07 | 0.25 | 0.11 | 0.07 | 0.11 | 0.11 |
| 9 | 35933323 | rs10814300 | -0.829 | 2.542 | 0.138 | -0.266 | -0.003 | -0.001 | -0.001 | 5.105 | 0.32 | 0.35 | 0.21 | 0.15 | 0.38 | 0.28 |
| 9 | 35935138 | rs10814301 | -0.572 | 2.381 | -0.287 | -0.123 | 0.069 | -0.058 | -0.058 | 0.731 | 0.17 | 0.47 | 0.18 | 0.07 | 0.25 | 0.15 |
| 9 | 35935963 | rs10814303 | -0.572 | 2.381 | -0.287 | -0.123 | 0.069 | -0.058 | -0.058 | 0.731 | 0.17 | 0.47 | 0.19 | 0.07 | 0.25 | 0.15 |
| 9 | 35952224 | rs723425 | -0.183 | 1.965 | -1.132 | 0.328 | -0.054 | -0.02 | -0.02 | 0 | 0.2 | 0.05 | 0.12 | 0.21 | 0.24 | 0.25 |
| 9 | 35953318 | rs7857956 | -0.678 | 1.843 | -0.894 | -0.017 | -0.02 | 0.04 | 0.04 | 0 | 0.4 | 0.76 | 0.37 | 0.31 | 0.49 | 0.55 |
| 9 | 35953983 | rs10738950 | 0.7 | 1.592 | -0.489 | 0.174 | -0.003 | -0.007 | -0.007 | 0.691 | 0.37 | 0.38 | 0.29 | 0.24 | 0.32 | 0.25 |
| 9 | 35954375 | rs7034414 | 0.041 | 1.53 | -0.124 | 0.116 | -0.008 | 0.006 | 0.006 | 0.189 | 0.35 | 0.35 | 0.3 | 0.32 | 0.39 | 0.5 |
| 9 | 35957355 | rs2233568 | 0.61 | 1.293 | 0.105 | 0.173 | -0.016 | -0.02 | -0.02 | 0 | 0.37 | 0.49 | 0.3 | 0.3 | 0.28 | 0.29 |
| 9 | 35961915 | rs732233 | 0.443 | 1.08 | 0.207 | 0.059 | -0.009 | -0.013 | -0.013 | 0 | 0.39 | 0.51 | 0.3 | 0.31 | 0.29 | 0.29 |
| 9 | 35962782 | rs4878636 | 0.429 | 1.053 | 0.311 | 0.065 | -0.019 | -0.027 | -0.027 | 0 | 0.39 | 0.45 | 0.29 | 0.29 | 0.28 | 0.25 |
| 9 | 35968663 | rs10758338 | 0.101 | 1.485 | 0.667 | 0.025 | -0.018 | -0.017 | -0.017 | 1.183 | 0.56 | 0.64 | 0.45 | 0.52 | 0.44 | 0.48 |
| 9 | 35970516 | rs13292355 | 0.742 | 1.285 | 0.463 | -0.047 | -0.04 | -0.053 | -0.053 | 0 | 0.18 | 0.03 | 0.16 | 0.23 | 0.17 | 0.24 |
| 9 | 35971579 | rs12352266 | 0.131 | 1.405 | 0.968 | -0.273 | 0.074 | -0.025 | -0.025 | 0 | 0.08 | 0.29 | 0.06 | 0 | 0.09 | 0.04 |
| 9 | 35975125 | rs7048672 | -0.085 | 1.715 | 0.821 | -0.374 | 0.023 | -0.057 | -0.057 | 0 | 0.16 | 0.29 | 0.1 | 0.07 | 0.18 | 0.17 |
| 9 | 35978258 | rs10972658 | 1.641 | 2.005 | 0.51 | -0.427 | 0.042 | -0.02 | -0.02 | 0 | 0.31 | 0.05 | 0.18 | 0.29 | 0.18 | 0.24 |
| 9 | 35978321 | rs10972659 | 0.962 | 2.034 | 0.72 | 0.3 | -0.003 | -0.043 | -0.043 | 0.112 | 0.19 | 0.1 | 0.17 | 0.37 | 0.17 | 0.2 |
| 9 | 35978632 | rs7874748 | -1.304 | 2.237 | 0.804 | 0.313 | -0.012 | -0.005 | -0.005 | 0 | 0.79 | 0.82 | 0.56 | 0.81 | 0.66 | 0.79 |
| 9 | 35983405 | rs10814308 | 0.075 | 2.077 | 0.979 | 0.306 | 0.026 | -0.071 | -0.071 | 0 | 0.16 | 0.35 | 0.1 | 0.07 | 0.17 | 0.17 |
| 9 | 35988418 | rs12004566 | 1.957 | 2.016 | 0.867 | 0.32 | 0.075 | -0.004 | -0.004 | 1.002 | 0.3 | 0.05 | 0.16 | 0.29 | 0.15 | 0.21 |
| 9 | 35991104 | rs16932816 | 0.653 | 2.178 | 1.194 | 0.251 | 0.003 | -0.009 | -0.009 | 0 | 0.12 | 0.21 | 0.1 | 0.08 | 0.13 | 0.18 |
| 9 | 35991115 | rs16932818 | 0.03 | 2.188 | 1.208 | 0.245 | 0.015 | -0.057 | -0.057 | 0 | 0.16 | 0.3 | 0.11 | 0.07 | 0.18 | 0.17 |
| 9 | 35992302 | rs1408466 | 0.05 | 2.207 | 1.164 | 0.313 | 0.028 | -0.01 | -0.01 | 0 | 0.64 | 0.5 | 0.44 | 0.74 | 0.49 | 0.58 |
| 9 | 35993376 | rs10738955 | 0.051 | 2.198 | 1.16 | 0.309 | 0.028 | -0.01 | -0.01 | 0 | 0.64 | 0.5 | 0.44 | 0.74 | 0.49 | 0.58 |
| 9 | 35994554 | rs1535912 | -0.773 | 2.207 | 1.031 | 0.358 | -0.008 | -0.001 | -0.001 | 6.032 | 0.77 | 0.76 | 0.54 | 0.82 | 0.62 | 0.79 |
| 9 | 35998220 | rs1324461 | -0.811 | 1.683 | 1.308 | 0.366 | 0.036 | -0.074 | -0.074 | 0.999 | 0.18 | 0.35 | 0.12 | 0.06 | 0.21 | 0.2 |
| 9 | 35998726 | rs1324459 | -0.213 | 1.698 | 1.087 | 0.366 | 0.01 | -0.039 | -0.039 | 0 | 0.15 | 0.22 | 0.1 | 0.06 | 0.18 | 0.17 |
| 9 | 36002881 | rs10972680 | -0.614 | 1.673 | 1.087 | 0.372 | 0.025 | -0.058 | -0.058 | 0 | 0.17 | 0.27 | 0.12 | 0.06 | 0.2 | 0.2 |
| 9 | 36005428 | rs942480 | -0.241 | 1.449 | 1.088 | 0.837 | 0 | -0.006 | -0.006 | 0 | 0.13 | 0.28 | 0.12 | 0.09 | 0.13 | 0.21 |
| 9 | 36013929 | rs2017241 | -1.035 | 1.23 | 1.043 | 0.967 | -0.011 | 0.038 | 0.038 | 0.019 | 0.3 | 0.57 | 0.28 | 0.53 | 0.24 | 0.47 |
| 9 | 36014850 | rs2077559 | -0.405 | 1.207 | 1.086 | 1.297 | -0.004 | 0.054 | 0.054 | 0 | 0.14 | 0.27 | 0.11 | 0.27 | 0.13 | 0.2 |
| 9 | 36018310 | rs7029221 | -0.401 | 1.202 | 1.14 | 1.303 | -0.004 | 0.034 | 0.034 | 0 | 0.14 | 0.23 | 0.11 | 0.27 | 0.13 | 0.17 |
| 9 | 36018752 | rs6476506 | -0.873 | 1.013 | 0.564 | 0.91 | -0.044 | 0.049 | 0.049 | 0.192 | 0.7 | 0.83 | 0.52 | 0.83 | 0.57 | 0.77 |
| 9 | 36023246 | rs7027417 | -0.494 | 0.89 | 1.233 | 1.444 | -0.003 | 0.03 | 0.03 | 0 | 0.14 | 0.26 | 0.08 | 0.21 | 0.13 | 0.13 |
| 9 | 36023728 | rs10814320 | -0.023 | 0.702 | 1.394 | 1.424 | -0.006 | 0.125 | 0.125 | 2.678 | 0.16 | 0.37 | 0.15 | 0.31 | 0.1 | 0.3 |
| 9 | 36025452 | rs7863810 | 0.132 | 0.792 | 0.683 | 1.373 | -0.001 | 0.009 | 0.009 | 5.944 | 0.14 | 0.14 | 0.15 | 0.27 | 0.1 | 0.26 |
| 9 | 36028399 | rs12339284 | -0.599 | 0.667 | 0.734 | 1.135 | -0.012 | 0.046 | 0.046 | 0 | 0.14 | 0.27 | 0.08 | 0.2 | 0.12 | 0.1 |
| 9 | 36031610 | rs10814323 | -1.186 | 0.87 | 0.745 | 1.032 | 0.013 | 0.028 | 0.028 | 0.056 | 0.19 | 0.39 | 0.12 | 0.19 | 0.21 | 0.17 |
| 9 | 36036003 | rs7873354 | -1.026 | 0.289 | 0.863 | 1.742 | -0.038 | 0.139 | 0.139 | 1.418 | 0.2 | 0.53 | 0.21 | 0.51 | 0.13 | 0.35 |
| 9 | 36038361 | rs7024649 | 0.051 | 0.217 | 0.9 | 2.191 | 0.005 | -0.005 | -0.005 | 0 | 0.11 | 0.08 | 0.04 | 0.17 | 0.05 | 0.22 |
| 9 | 36040713 | rs10972705 | -0.116 | 0.192 | 1.055 | 2.37 | 0.002 | 0.114 | 0.114 | 0 | 0.07 | 0.1 | 0.12 | 0.27 | 0.08 | 0.09 |
| 9 | 36046596 | rs6476514 | -0.548 | 0.171 | 0.244 | 1.864 | 0.023 | -0.037 | -0.037 | 0.122 | 0.67 | 0.58 | 0.47 | 0.8 | 0.53 | 0.71 |
| 9 | 36050506 | rs1324457 | -0.044 | 0.024 | -0.34 | 1.337 | 0.002 | 0.004 | 0.004 | 0 | 0.56 | 0.47 | 0.36 | 0.58 | 0.44 | 0.65 |
| 9 | 36060451 | rs7035468 | -0.274 | -0.078 | -0.259 | 1.402 | 0.012 | 0.008 | 0.008 | 0 | 0.61 | 0.49 | 0.35 | 0.53 | 0.45 | 0.62 |
| 9 | 36066385 | rs6476519 | 0.441 | -0.15 | -0.148 | 1.404 | -0.042 | 0.075 | 0.075 | 0.449 | 0.14 | 0.33 | 0.09 | 0.24 | 0.05 | 0.25 |
| 9 | 36077871 | rs1324465 | -1.137 | -0.208 | -0.02 | 1.379 | -0.055 | 0.06 | 0.06 | 0.062 | 0.7 | 0.86 | 0.48 | 0.83 | 0.53 | 0.76 |
| 9 | 36087876 | rs16932912 | 0.892 | -0.022 | -0.452 | 1.258 | 0.002 | 0.114 | 0.114 | 0 | 0.07 | 0.1 | 0.12 | 0.27 | 0.08 | 0.09 |
| 9 | 36095505 | rs4879952 | 1.624 | -0.042 | -0.733 | 1.016 | -0.064 | 0.01 | 0.01 | 0.428 | 0.32 | 0.02 | 0.18 | 0.29 | 0.26 | 0.3 |
| 9 | 36099871 | rs6476527 | -0.664 | 0.319 | -0.287 | 1.204 | 0.007 | -0.003 | -0.003 | 0 | 0.67 | 0.64 | 0.49 | 0.71 | 0.52 | 0.69 |
| 9 | 36101562 | rs7029444 | 0.931 | 0.135 | -0.349 | 1.173 | 0.003 | -0.003 | -0.003 | 0 | 0.11 | 0.09 | 0.07 | 0.15 | 0.07 | 0.21 |
| 9 | 36113712 | rs13283904 | 0.949 | 0.072 | -0.379 | 1.158 | -0.009 | 0.02 | 0.02 | 0 | 0.08 | 0.05 | 0.08 | 0.03 | 0.1 | 0.05 |
| 9 | 36128544 | rs7875409 | -0.341 | 0.18 | -0.055 | 1.362 | -0.002 | -0.003 | -0.003 | 0.826 | 0.66 | 0.37 | 0.4 | 0.49 | 0.55 | 0.63 |
| 9 | 36130245 | rs7033900 | -0.98 | 0.336 | -0.021 | 1.277 | 0.024 | 0.007 | 0.007 | 0 | 0.75 | 0.65 | 0.56 | 0.75 | 0.63 | 0.75 |
| 9 | 36132335 | rs7869658 | -0.132 | 0.669 | 0.096 | 0.861 | 0 | 0.006 | 0.006 | 0 | 0.14 | 0.26 | 0.11 | 0.15 | 0.1 | 0.24 |
| 9 | 36135424 | rs10972754 | -0.559 | 0.614 | -0.368 | 0.772 | -0.002 | -0.007 | -0.007 | 0 | 0.57 | 0.41 | 0.37 | 0.55 | 0.51 | 0.67 |

| | | | | | | | | | | | | | | | | |
|---|---|---|---|---|---|---|---|---|---|---|---|---|---|---|---|---|
| 9 | 36137346 | rs752353 | 0.103 | 0.799 | -0.734 | 0.653 | -0.027 | -0.026 | -0.026 | 0 | 0.21 | 0.11 | 0.18 | 0.26 | 0.23 | 0.29 |
| 9 | 36137513 | rs752351 | -0.288 | 0.677 | -0.601 | 0.646 | 0.014 | 0.002 | 0.002 | 0 | 0.3 | 0.55 | 0.29 | 0.29 | 0.33 | 0.36 |
| 9 | 36140115 | rs11791833 | 0.005 | 0.528 | -1.219 | 0.775 | 0.011 | -0.009 | -0.009 | 1.009 | 0.34 | 0.17 | 0.15 | 0.38 | 0.26 | 0.32 |
| 9 | 36141514 | rs17835038 | -0.833 | 0.561 | -1.19 | 0.654 | -0.003 | -0.001 | -0.001 | 0 | 0.08 | 0.04 | 0.05 | 0.11 | 0.07 | 0.11 |
| 9 | 36141635 | rs1571268 | 0.294 | 0.458 | -1.667 | 0.385 | -0.05 | 0.003 | 0.003 | 1.409 | 0.39 | 0.11 | 0.52 | 0.31 | 0.46 | 0.28 |
| 9 | 36147128 | rs7022784 | 0.189 | 0.416 | -0.767 | 0.325 | 0.012 | -0.002 | -0.002 | 0 | 0.42 | 0.26 | 0.5 | 0.47 | 0.43 | 0.52 |
| 9 | 36154678 | rs10814339 | 0.01 | 0.824 | -0.764 | -0.02 | -0.009 | 0.027 | 0.027 | 0 | 0.22 | 0.13 | 0.23 | 0.12 | 0.26 | 0.16 |
| 9 | 36159456 | rs10972763 | 0.295 | 0.752 | -0.436 | 0.139 | -0.002 | 0.008 | 0.008 | 0 | 0.7 | 0.62 | 0.84 | 0.67 | 0.75 | 0.68 |
| 9 | 36160040 | rs17778964 | 0.328 | 0.557 | -0.772 | 0.076 | 0.001 | 0.009 | 0.009 | 0.521 | 0.23 | 0.15 | 0.16 | 0.21 | 0.22 | 0.28 |
| 9 | 36168940 | rs7038327 | -0.627 | 0.204 | -0.201 | -0.105 | 0 | 0 | 0 | 0 | 0.48 | 0.45 | 0.28 | 0.5 | 0.47 | 0.58 |
| 9 | 36169187 | rs10972770 | 1.269 | 0.226 | -0.326 | -0.016 | -0.032 | -0.012 | -0.012 | 0 | 0.07 | 0.02 | 0.08 | 0.05 | 0.15 | 0.11 |
| 9 | 36169494 | rs3739610 | -0.446 | 0.308 | -0.329 | -0.031 | -0.022 | 0.002 | 0.002 | 0 | 0.4 | 0.43 | 0.19 | 0.45 | 0.29 | 0.47 |
| 9 | 36169598 | rs3739609 | -0.932 | 0.026 | 0.023 | -0.074 | 0.006 | -0.003 | -0.003 | 0.071 | 0.49 | 0.59 | 0.3 | 0.5 | 0.45 | 0.58 |
| 9 | 36173934 | rs12237915 | -0.207 | -0.222 | 0.145 | -0.021 | -0.007 | 0.003 | 0.003 | 0.419 | 0.23 | 0.26 | 0.1 | 0.27 | 0.2 | 0.28 |
| 9 | 36186264 | rs10972784 | -1.976 | -0.258 | -0.246 | -0.349 | -0.003 | 0.015 | 0.015 | 0.61 | 0.76 | 0.75 | 0.66 | 0.47 | 0.81 | 0.66 |
| 9 | 36190198 | rs1999005 | 0.712 | -0.47 | -0.364 | -0.459 | 0.01 | -0.007 | -0.007 | 0.811 | 0.15 | 0.1 | 0.06 | 0.2 | 0.1 | 0.12 |
| 9 | 36191004 | rs2274523 | 1.737 | -0.843 | -0.44 | -0.607 | -0.036 | 0.009 | 0.009 | 0 | 0.14 | 0.17 | 0.06 | 0.18 | 0.05 | 0.17 |
| 9 | 36200678 | rs13287406 | 0.854 | -0.603 | -0.546 | -0.701 | 0.01 | -0.019 | -0.019 | 0 | 0.06 | 0.09 | 0.08 | 0 | 0.14 | 0.07 |
| 9 | 36203718 | rs4879960 | 1.444 | -0.498 | -0.808 | -0.917 | -0.083 | 0.082 | 0.082 | 0 | 0.23 | 0.02 | 0.29 | 0.07 | 0.28 | 0.12 |
| 9 | 36204455 | rs2025879 | -0.827 | -0.63 | -0.295 | -0.775 | 0.004 | 0.001 | 0.001 | 0 | 0.39 | 0.64 | 0.22 | 0.38 | 0.3 | 0.37 |
| 9 | 36207967 | rs16933076 | 0.748 | -0.237 | -0.507 | -0.745 | 0.023 | -0.028 | -0.028 | 1.113 | 0.06 | 0.13 | 0.08 | 0 | 0.14 | 0.07 |
| 9 | 36211827 | rs1053414 | 0.524 | -0.535 | -0.555 | -0.715 | -0.007 | 0 | 0 | 1.541 | 0.17 | 0.2 | 0.07 | 0.19 | 0.11 | 0.13 |
| 9 | 36214971 | rs1043313 | -1.255 | -0.678 | -0.501 | -0.838 | 0.028 | 0.014 | 0.014 | 0.102 | 0.61 | 0.7 | 0.59 | 0.31 | 0.75 | 0.47 |
| 9 | 36220436 | rs623166 | -2.137 | -0.68 | -0.593 | -0.507 | 0.016 | -0.079 | -0.079 | 0 | 0.75 | 0.87 | 0.73 | 0.59 | 0.81 | 0.67 |
| 9 | 36223654 | rs10758350 | 0.457 | -0.582 | -0.726 | -1.108 | -0.015 | 0.081 | 0.081 | 0.112 | 0.35 | 0.11 | 0.34 | 0.23 | 0.37 | 0.21 |
| 9 | 36231483 | rs4879961 | -1.986 | -0.769 | -0.631 | -0.936 | 0.025 | -0.072 | -0.072 | 0 | 0.73 | 0.85 | 0.65 | 0.48 | 0.8 | 0.64 |
| 9 | 36232413 | rs10738958 | 0.468 | -0.844 | -0.315 | -0.986 | -0.038 | -0.013 | -0.013 | 0.173 | 0.68 | 0.51 | 0.58 | 0.32 | 0.77 | 0.42 |
| 9 | 36234439 | rs7019324 | 1.58 | -0.564 | -0.027 | -0.826 | -0.044 | -0.002 | -0.002 | 0 | 0.15 | 0.2 | 0.07 | 0.17 | 0.05 | 0.19 |
| 9 | 36245759 | rs7871216 | 1.019 | -0.449 | -0.287 | -0.881 | 0.002 | -0.001 | -0.001 | 1.823 | 0.11 | 0.09 | 0.06 | 0.17 | 0.09 | 0.09 |
| 9 | 36248825 | rs4878646 | -0.375 | -0.098 | -0.23 | -0.787 | 0.034 | 0.079 | 0.079 | 0 | 0.49 | 0.3 | 0.49 | 0.3 | 0.59 | 0.38 |
| 9 | 36250665 | rs10758352 | -1.883 | -0.086 | -0.121 | -0.65 | 0.007 | -0.016 | -0.016 | 0 | 0.72 | 0.75 | 0.64 | 0.5 | 0.78 | 0.62 |
| 9 | 36257992 | rs10814362 | -1.869 | -0.08 | -0.146 | -0.648 | 0.008 | -0.018 | -0.018 | 0 | 0.72 | 0.73 | 0.63 | 0.5 | 0.78 | 0.62 |
| 9 | 36261111 | rs7036270 | -1.879 | -0.076 | -0.394 | -0.645 | 0.015 | -0.034 | -0.034 | 0 | 0.72 | 0.79 | 0.64 | 0.5 | 0.78 | 0.62 |
| 9 | 36276024 | rs12379203 | 0.827 | 0.417 | -0.516 | -0.651 | 0.006 | 0 | 0 | 0 | 0.12 | 0.12 | 0.15 | 0.07 | 0.22 | 0.17 |
| 9 | 36291214 | rs4879980 | -0.026 | 0.057 | -0.448 | -0.637 | -0.004 | 0.057 | 0.057 | 0 | 0.39 | 0.24 | 0.38 | 0.24 | 0.43 | 0.21 |
| 9 | 36295756 | rs4879983 | -1 | 0.569 | -0.713 | -0.355 | -0.026 | 0.005 | 0.005 | 0 | 0.63 | 0.67 | 0.5 | 0.43 | 0.57 | 0.45 |
| 9 | 36306809 | rs10814373 | -1.736 | 0.267 | -0.506 | 0.028 | 0.025 | 0 | 0 | 0.44 | 0.54 | 0.74 | 0.57 | 0.48 | 0.55 | 0.56 |
| 9 | 36307043 | rs10123040 | -1.548 | 0.315 | 0.042 | 0.034 | 0 | 0 | 0 | 0.577 | 0.52 | 0.52 | 0.55 | 0.48 | 0.55 | 0.56 |
| 9 | 36310642 | rs10972843 | -0.483 | 0.374 | 0.035 | 0.057 | 0.035 | 0.118 | 0.118 | 0.653 | 0.11 | 0.43 | 0.25 | 0.12 | 0.1 | 0.17 |
| 9 | 36314530 | rs7038460 | -1.538 | 0.189 | -0.264 | 0.035 | 0.014 | 0.02 | 0.02 | 0 | 0.51 | 0.74 | 0.52 | 0.48 | 0.49 | 0.55 |
| 9 | 36315900 | rs7043543 | -0.089 | 1.285 | -0.553 | -0.344 | -0.052 | 0.039 | 0.039 | 0 | 0.17 | 0.08 | 0.2 | 0.05 | 0.35 | 0.14 |
| 9 | 36318240 | rs12379853 | -0.075 | 1.003 | -0.529 | -0.277 | 0.075 | -0.046 | -0.046 | 0.479 | 0.17 | 0.22 | 0.23 | 0.05 | 0.34 | 0.14 |
| 9 | 36321726 | rs12685361 | 0.024 | -0.364 | -1.024 | 0.083 | 0.031 | -0.039 | -0.039 | 0 | 0.17 | 0.05 | 0.2 | 0.2 | 0.09 | 0.24 |
| 9 | 36322621 | rs10972856 | 0.938 | -0.614 | -0.31 | -0.013 | -0.045 | -0.007 | -0.007 | 0.152 | 0.77 | 0.43 | 0.74 | 0.77 | 0.88 | 0.65 |
| 9 | 36343164 | rs10814376 | -0.891 | -1.244 | -0.483 | -0.238 | 0.015 | 0.017 | 0.017 | 0 | 0.47 | 0.33 | 0.44 | 0.4 | 0.61 | 0.54 |
| 9 | 36346527 | rs12347593 | 0.155 | -1.361 | -0.074 | -0.307 | -0.025 | 0.084 | 0.084 | 0 | 0.92 | 0.7 | 0.83 | 0.91 | 0.98 | 0.96 |
| 9 | 36351800 | rs10814377 | 0.941 | -1.417 | -0.09 | -0.388 | 0.027 | 0.02 | 0.02 | 0 | 0.51 | 0.46 | 0.54 | 0.52 | 0.38 | 0.43 |
| 9 | 36359376 | rs12376663 | 0.639 | -0.896 | -0.397 | -0.521 | -0.063 | 0.047 | 0.047 | 0 | 0.19 | 0.03 | 0.15 | 0.11 | 0.27 | 0.18 |
| 9 | 36367338 | rs10972872 | 0.699 | -0.879 | -0.483 | -0.515 | -0.061 | 0.043 | 0.043 | 0 | 0.19 | 0.03 | 0.15 | 0.11 | 0.27 | 0.18 |
| 9 | 36382790 | rs12376699 | 1.115 | -0.516 | -0.405 | -0.481 | -0.071 | 0.056 | 0.056 | 0 | 0.11 | 0.01 | 0.13 | 0.02 | 0.22 | 0.11 |
| 9 | 36408108 | rs10814384 | 0.07 | -0.612 | -0.246 | -0.452 | 0.02 | 0.008 | 0.008 | 0 | 0.26 | 0.29 | 0.3 | 0.37 | 0.35 | 0.35 |
| 9 | 36419529 | rs954779 | -2.067 | -1.254 | -0.802 | -0.54 | -0.003 | -0.004 | -0.004 | 2.554 | 0.71 | 0.72 | 0.7 | 0.68 | 0.72 | 0.72 |
| 9 | 36458793 | rs10511929 | 1.555 | -0.413 | -0.958 | -0.391 | 0.028 | 0.007 | 0.007 | 0 | 0.13 | 0.02 | 0.06 | 0.11 | 0.06 | 0.16 |
| 9 | 36463744 | rs10118394 | -1.859 | -0.307 | -0.934 | -0.258 | 0.011 | -0.006 | -0.006 | 0 | 0.61 | 0.64 | 0.64 | 0.6 | 0.67 | 0.66 |
| 9 | 36481391 | rs10814392 | 0.315 | 0.497 | -0.97 | -0.806 | -0.034 | -0.019 | -0.019 | 0 | 0.12 | 0.06 | 0.25 | 0.1 | 0.21 | 0.05 |
| 9 | 36488261 | rs13291966 | -0.727 | 0.217 | -0.636 | -0.89 | -0.001 | -0.008 | -0.008 | 0.033 | 0.36 | 0.57 | 0.3 | 0.23 | 0.37 | 0.44 |
| 9 | 36500619 | rs329376 | 0.924 | 0.483 | -0.81 | -0.541 | -0.014 | 0.055 | 0.055 | 0.037 | 0.58 | 0.33 | 0.49 | 0.73 | 0.5 | 0.6 |
| 9 | 36507639 | rs329366 | 0.669 | 0.758 | -0.332 | -0.297 | -0.005 | 0.003 | 0.003 | 0.386 | 0.65 | 0.38 | 0.53 | 0.76 | 0.55 | 0.62 |
| 9 | 36525772 | rs426361 | 0.265 | 0.624 | -0.223 | -0.246 | -0.005 | -0.027 | -0.027 | 0 | 0.58 | 0.55 | 0.74 | 0.82 | 0.63 | 0.65 |
| 9 | 36530955 | rs7029237 | -0.924 | 0.942 | 0.361 | 0.455 | 0.041 | 0.048 | 0.048 | 0.523 | 0.26 | 0.54 | 0.43 | 0.32 | 0.36 | 0.3 |
| 9 | 36539122 | rs7863577 | 0.214 | -0.205 | 1.189 | -0.617 | -0.069 | -0.089 | -0.089 | 0.03 | 0.71 | 0.44 | 0.89 | 0.9 | 0.9 | 0.78 |
| 9 | 36550835 | rs373510 | 0.317 | -0.288 | 1.506 | -0.724 | -0.016 | -0.036 | -0.036 | 0 | 0.2 | 0.21 | 0.08 | 0.08 | 0.1 | 0.09 |
| 9 | 36565438 | rs444543 | 0.409 | 0.262 | 1.551 | -0.485 | 0.008 | 0 | 0 | 0 | 0.88 | 0.88 | 0.81 | 0.81 | 0.75 | 0.75 |
| 9 | 36635409 | rs2265346 | -0.353 | -0.213 | 2.483 | -0.97 | -0.011 | -0.024 | -0.024 | 0 | 0.26 | 0.25 | 0.09 | 0.11 | 0.11 | 0.19 |
| 9 | 36639442 | rs3780350 | 0.585 | -0.347 | 2.095 | -1.183 | 0.001 | 0.011 | 0.011 | 0 | 0.16 | 0.07 | 0.19 | 0.12 | 0.18 | 0.14 |
| 9 | 36640907 | rs13298120 | 1.167 | -0.367 | 1.854 | -1.235 | -0.012 | 0.067 | 0.067 | 0 | 0.12 | 0.02 | 0.15 | 0 | 0.14 | 0.06 |
| 9 | 36642362 | rs2483641 | 0.843 | -0.446 | 1.96 | -1.148 | 0.003 | -0.001 | -0.001 | 1.576 | 0.18 | 0.12 | 0.07 | 0.08 | 0.1 | 0.08 |
| 9 | 36647786 | rs10814422 | 0.543 | -0.444 | 2.032 | -1.167 | -0.047 | -0.082 | -0.082 | 0 | 0.69 | 0.43 | 0.88 | 0.89 | 0.87 | 0.8 |
| 9 | 36655648 | rs10973013 | 0.08 | -0.47 | 2.145 | -1.056 | -0.044 | -0.07 | -0.07 | 0 | 0.73 | 0.61 | 0.9 | 0.89 | 0.89 | 0.81 |

| | | | | | | | | | | | | | | | | |
|---|---|---|---|---|---|---|---|---|---|---|---|---|---|---|---|---|
| 9 | 36674842 | rs2274473 | 0.32 | -0.464 | 1.985 | -1.054 | -0.056 | -0.084 | -0.084 | 0.044 | 0.71 | 0.42 | 0.89 | 0.89 | 0.89 | 0.81 |
| 9 | 36687106 | rs10046792 | 0.116 | -0.409 | 1.462 | -1.021 | -0.025 | -0.016 | -0.016 | 0 | 0.06 | 0.13 | 0.01 | 0.03 | 0.01 | 0.11 |
| 9 | 36691231 | rs4504731 | -0.68 | -0.362 | 1.374 | -1.003 | -0.016 | -0.04 | -0.04 | 0.447 | 0.81 | 0.79 | 0.92 | 0.92 | 0.9 | 0.92 |
| 9 | 36693098 | rs10814430 | -0.251 | -0.208 | 0.852 | -1.038 | 0.002 | -0.03 | -0.03 | 0 | 0.15 | 0.2 | 0.12 | 0.03 | 0.18 | 0.2 |
| 9 | 36697735 | rs10511931 | 0.413 | -0.295 | 0.208 | -0.957 | -0.022 | -0.076 | -0.076 | 0.611 | 0.68 | 0.42 | 0.84 | 0.89 | 0.78 | 0.73 |
| 9 | 36702096 | rs2094394 | 0.689 | 0.04 | -0.764 | -0.67 | 0.008 | 0.149 | 0.149 | 0.024 | 0.48 | 0.2 | 0.59 | 0.74 | 0.51 | 0.6 |
| 9 | 36702485 | rs4880002 | 0.184 | 0.022 | -0.545 | -0.173 | 0.019 | 0.021 | 0.021 | 0.911 | 0.82 | 0.91 | 0.95 | 0.92 | 0.93 | 0.92 |
| 9 | 36718368 | rs10511932 | -0.385 | 0.406 | -0.507 | 0.357 | -0.018 | -0.033 | -0.033 | 1.841 | 0.09 | 0.01 | 0.1 | 0.16 | 0.12 | 0.2 |
| 9 | 36718812 | rs7863614 | -0.863 | 0.257 | -0.899 | 0.222 | 0.005 | 0.021 | 0.021 | 0 | 0.51 | 0.7 | 0.56 | 0.53 | 0.47 | 0.54 |
| 9 | 36718844 | rs7848373 | -0.715 | 0.186 | -0.923 | 0.208 | 0.021 | 0.007 | 0.007 | 0 | 0.47 | 0.69 | 0.48 | 0.52 | 0.39 | 0.48 |
| 9 | 36719751 | rs12379827 | 0.488 | 0.196 | -1.603 | 0.184 | 0.011 | 0.019 | 0.019 | 0 | 0.14 | 0.06 | 0.05 | 0.06 | 0.08 | 0.06 |
| 9 | 36722345 | rs11792514 | -0.619 | 0.319 | -1.223 | 0.401 | -0.01 | -0.046 | -0.046 | 0 | 0.3 | 0.18 | 0.39 | 0.45 | 0.31 | 0.42 |
| 9 | 36722854 | rs16933492 | 0.146 | 0.383 | -1.027 | 0.422 | -0.006 | -0.019 | -0.019 | 0 | 0.2 | 0.13 | 0.28 | 0.29 | 0.18 | 0.21 |
| 9 | 36725914 | rs11794314 | -0.205 | 0.299 | -1.257 | -0.129 | 0.009 | 0.043 | 0.043 | 0 | 0.24 | 0.1 | 0.15 | 0.17 | 0.21 | 0.12 |
| 9 | 36727388 | rs12552072 | -0.376 | 0.235 | -0.739 | -0.12 | -0.002 | -0.003 | -0.003 | 0 | 0.21 | 0.21 | 0.23 | 0.18 | 0.23 | 0.31 |
| 9 | 36731326 | rs1887432 | -0.296 | -0.321 | -0.457 | -0.508 | -0.035 | -0.064 | -0.064 | 0 | 0.22 | 0.42 | 0.2 | 0.11 | 0.15 | 0.12 |
| 9 | 36732969 | rs10738975 | 0.638 | 0.376 | -0.345 | -0.619 | -0.002 | 0.016 | 0.016 | 0 | 0.65 | 0.34 | 0.57 | 0.83 | 0.67 | 0.68 |
| 9 | 36736811 | rs10738976 | 0.815 | 0.378 | -0.291 | -0.58 | 0.001 | -0.009 | -0.009 | 0 | 0.66 | 0.39 | 0.57 | 0.82 | 0.68 | 0.66 |
| 9 | 36738641 | rs7857754 | 0.992 | 0.83 | 0.264 | -0.283 | 0.002 | -0.012 | -0.012 | 0 | 0.66 | 0.42 | 0.54 | 0.8 | 0.62 | 0.52 |
| 9 | 36740957 | rs7873864 | -0.904 | 1.328 | 0.126 | 0.038 | 0.056 | 0.002 | 0.002 | 0 | 0.14 | 0.35 | 0.21 | 0.07 | 0.17 | 0.2 |
| 9 | 36741027 | rs10116484 | -0.834 | 1.369 | -0.128 | 0.058 | 0.044 | 0 | 0 | 0 | 0.13 | 0.3 | 0.21 | 0.07 | 0.16 | 0.21 |
| 9 | 36744729 | rs752358 | -0.385 | 1.427 | -0.815 | 0.416 | 0.001 | 0.001 | 0.001 | 0.018 | 0.27 | 0.23 | 0.23 | 0.3 | 0.28 | 0.26 |
| 9 | 36744752 | rs752357 | -0.759 | 0.977 | -1.216 | 0.129 | 0.058 | 0.013 | 0.013 | 0.027 | 0.52 | 0.28 | 0.61 | 0.43 | 0.68 | 0.54 |
| 9 | 36748641 | rs6476575 | 0.236 | 0.79 | -1.184 | 0.287 | 0.002 | -0.001 | -0.001 | 4.464 | 0.68 | 0.75 | 0.76 | 0.68 | 0.73 | 0.83 |
| 9 | 36754763 | rs10814443 | 0.862 | 1.051 | -0.943 | 0.183 | 0.011 | -0.014 | -0.014 | 0.018 | 0.44 | 0.32 | 0.44 | 0.45 | 0.41 | 0.57 |
| 9 | 36756622 | rs1536544 | -0.059 | 1.193 | -0.574 | 0.201 | 0 | 0 | 0 | 0 | 0.62 | 0.64 | 0.71 | 0.67 | 0.64 | 0.81 |
| 9 | 36757802 | rs1887438 | -0.097 | 1.177 | -0.59 | 0.223 | 0 | 0.002 | 0.002 | 0 | 0.63 | 0.65 | 0.71 | 0.64 | 0.64 | 0.81 |
| 9 | 36765045 | rs10973061 | 0.186 | 1.011 | -0.773 | -0.132 | -0.017 | -0.015 | -0.015 | 0 | 0.59 | 0.66 | 0.49 | 0.49 | 0.5 | 0.72 |
| 9 | 36776224 | rs10814447 | 1.043 | 0.408 | -0.678 | -0.396 | 0.034 | -0.015 | -0.015 | 0 | 0.46 | 0.22 | 0.42 | 0.5 | 0.39 | 0.6 |
| 9 | 36782013 | rs2381583 | 0.604 | 0.749 | -0.091 | -0.137 | 0.058 | 0.088 | 0.088 | 1.723 | 0.77 | 0.43 | 0.58 | 0.67 | 0.67 | 0.79 |
| 9 | 36782358 | rs2800263 | -0.07 | 0.738 | 0.087 | -0.142 | 0.053 | 0.072 | 0.072 | 0 | 0.83 | 0.69 | 0.61 | 0.67 | 0.67 | 0.89 |
| 9 | 36785184 | rs10973075 | 0.979 | 0.685 | -0.2 | -0.33 | -0.026 | 0.063 | 0.063 | 0 | 0.11 | 0.01 | 0.05 | 0 | 0.11 | 0.03 |
| 9 | 36788414 | rs10973078 | 1.42 | 0.843 | -0.02 | -0.475 | 0.031 | -0.009 | -0.009 | 0.008 | 0.48 | 0.27 | 0.41 | 0.49 | 0.39 | 0.6 |
| 9 | 36790042 | rs727653 | -0.586 | 0.755 | 0.009 | -0.369 | 0.065 | 0.094 | 0.094 | 1.245 | 0.23 | 0.55 | 0.43 | 0.35 | 0.34 | 0.22 |
| 9 | 36790902 | rs3849922 | 1.124 | 0.872 | -0.206 | -0.348 | -0.008 | 0.006 | 0.006 | 0 | 0.16 | 0.03 | 0.09 | 0.16 | 0.15 | 0.13 |
| 9 | 36791430 | rs10973080 | 0.385 | 0.722 | -0.243 | -0.387 | -0.007 | -0.008 | -0.008 | 0.166 | 0.07 | 0.09 | 0.03 | 0.03 | 0.04 | 0.07 |
| 9 | 36791532 | rs10973081 | -0.023 | 0.766 | -0.061 | -0.344 | 0 | 0 | 0 | 0 | 0.28 | 0.33 | 0.14 | 0.2 | 0.19 | 0.31 |
| 9 | 36793445 | rs2800247 | 0.193 | 0.088 | -0.57 | -0.16 | -0.009 | -0.033 | -0.033 | 0.092 | 0.35 | 0.27 | 0.51 | 0.49 | 0.43 | 0.34 |
| 9 | 36797328 | rs7025013 | 0.577 | 0.162 | -0.75 | -0.154 | -0.005 | -0.011 | -0.011 | 0 | 0.44 | 0.25 | 0.43 | 0.51 | 0.51 | 0.51 |
| 9 | 36798025 | rs13292929 | 0.653 | 0.412 | -0.672 | 0.146 | 0.012 | 0.001 | 0.001 | 0 | 0.26 | 0.16 | 0.25 | 0.22 | 0.25 | 0.29 |
| 9 | 36798742 | rs10758401 | 0.158 | 0.686 | -0.428 | 0.021 | -0.002 | 0.002 | 0.002 | 0 | 0.43 | 0.33 | 0.6 | 0.36 | 0.49 | 0.41 |
| 9 | 36808003 | rs10814455 | 0.377 | 0.479 | -0.621 | -0.023 | 0.014 | -0.011 | -0.011 | 0 | 0.3 | 0.18 | 0.26 | 0.31 | 0.25 | 0.3 |
| 9 | 36809712 | rs2851701 | -0.47 | 0.505 | -0.536 | 0.048 | 0.01 | -0.016 | -0.016 | 0.073 | 0.69 | 0.78 | 0.72 | 0.58 | 0.75 | 0.65 |
| 9 | 36812517 | rs7850874 | -0.351 | 1.39 | -0.628 | 0.54 | -0.014 | -0.011 | -0.011 | 0.019 | 0.1 | 0.09 | 0.15 | 0.16 | 0.16 | 0.08 |
| 9 | 36814647 | rs1176844 | 0.362 | 1.323 | -0.473 | 0.417 | 0.002 | -0.001 | -0.001 | 0 | 0.42 | 0.48 | 0.5 | 0.41 | 0.49 | 0.5 |
| 9 | 36815627 | rs11791373 | -0.476 | 1.431 | -0.479 | -0.069 | -0.008 | -0.005 | -0.005 | 0 | 0.11 | 0.12 | 0.16 | 0.16 | 0.18 | 0.08 |
| 9 | 36816714 | rs2800256 | -0.212 | 0.676 | -0.456 | -0.184 | -0.003 | -0.003 | -0.003 | 0 | 0.54 | 0.59 | 0.67 | 0.68 | 0.67 | 0.64 |
| 9 | 36819706 | rs2851696 | 0.544 | 1.252 | 0.041 | -0.441 | 0.004 | -0.076 | -0.076 | 0 | 0.67 | 0.53 | 0.73 | 0.84 | 0.64 | 0.67 |
| 9 | 36820262 | rs2851695 | 0.546 | 1.252 | 0.011 | -0.515 | 0.005 | -0.074 | -0.074 | 0 | 0.64 | 0.5 | 0.7 | 0.85 | 0.61 | 0.76 |
| 9 | 36820604 | rs2800266 | 0.523 | 1.012 | 0.071 | -0.376 | -0.001 | -0.051 | -0.051 | 0 | 0.64 | 0.46 | 0.7 | 0.8 | 0.6 | 0.75 |
| 9 | 36824086 | rs7848675 | -0.821 | 1.082 | 0.258 | -0.038 | 0.04 | -0.003 | -0.003 | 0 | 0.23 | 0.37 | 0.26 | 0.23 | 0.31 | 0.26 |
| 9 | 36830616 | rs2800252 | 0.915 | -0.479 | 0.271 | -0.423 | -0.07 | -0.058 | -0.058 | 0.055 | 0.79 | 0.39 | 0.84 | 0.93 | 0.94 | 0.87 |
| 9 | 36840446 | rs3739437 | -0.08 | -0.499 | -0.008 | -0.293 | 0.044 | -0.029 | -0.029 | 0.036 | 0.09 | 0.01 | 0.08 | 0.11 | 0.02 | 0.11 |
| 9 | 36843844 | rs4880018 | 0.184 | -0.546 | 1.132 | -0.347 | 0.029 | -0.011 | -0.011 | 0.041 | 0.94 | 0.75 | 0.96 | 1 | 0.97 | 0.99 |
| 9 | 36845973 | rs3780136 | -0.509 | -0.323 | 0.993 | -0.58 | -0.01 | 0.01 | 0.01 | 0.017 | 0.11 | 0.11 | 0.07 | 0.06 | 0.13 | 0.18 |
| 9 | 36847139 | rs7044118 | 1.913 | -0.494 | 1.178 | -0.65 | 0.011 | 0.017 | 0.017 | 2.341 | 0.7 | 0.28 | 0.78 | 0.83 | 0.82 | 0.7 |
| 9 | 36851358 | rs7852051 | -0.334 | -0.237 | 0.287 | -0.211 | -0.014 | 0.049 | 0.049 | 0 | 0.71 | 0.58 | 0.69 | 0.6 | 0.75 | 0.73 |
| 9 | 36852316 | rs17391260 | 0.236 | 0.083 | -0.412 | -0.548 | -0.055 | 0.076 | 0.076 | 0.061 | 0.18 | 0.06 | 0.12 | 0.03 | 0.23 | 0.2 |
| 9 | 36854541 | rs4510947 | -0.138 | -0.48 | -0.272 | -0.57 | -0.008 | 0 | 0 | 0 | 0.6 | 0.58 | 0.63 | 0.58 | 0.69 | 0.66 |
| 9 | 36855486 | rs4326462 | -0.049 | -0.014 | -0.405 | -0.488 | 0 | 0.002 | 0.002 | 0 | 0.51 | 0.41 | 0.54 | 0.6 | 0.5 | 0.43 |
| 9 | 36856268 | rs10511934 | 0.355 | 0.002 | -0.422 | -0.359 | 0.001 | 0.001 | 0.001 | 0 | 0.2 | 0.17 | 0.11 | 0.1 | 0.16 | 0.19 |
| 9 | 36856828 | rs2005154 | -0.634 | -0.023 | -0.976 | -0.267 | -0.001 | -0.001 | -0.001 | 0 | 0.89 | 0.87 | 0.83 | 0.87 | 0.88 | 0.9 |
| 9 | 36859519 | rs10814462 | 0.283 | 0.035 | -1.315 | -0.329 | 0.009 | 0.013 | 0.013 | 0 | 0.19 | 0.1 | 0.11 | 0.1 | 0.17 | 0.2 |
| 9 | 36861137 | rs10758407 | -0.008 | 0.007 | -1.46 | -0.427 | 0.009 | 0.037 | 0.037 | 0 | 0.24 | 0.12 | 0.13 | 0.1 | 0.19 | 0.19 |
| 9 | 36861407 | rs10973109 | 0.128 | 0.606 | -1.282 | -0.272 | -0.004 | 0.007 | 0.007 | 0 | 0.18 | 0.11 | 0.12 | 0.09 | 0.19 | 0.16 |
| 9 | 36863670 | rs4880020 | -0.114 | 0.847 | -1.422 | 0.418 | -0.058 | -0.108 | -0.108 | 0 | 0.23 | 0.08 | 0.41 | 0.46 | 0.31 | 0.29 |
| 9 | 36866070 | rs16933658 | -0.807 | -0.067 | -0.355 | -0.131 | -0.019 | 0.006 | 0.006 | 0 | 0.16 | 0.33 | 0.1 | 0.12 | 0.07 | 0.11 |
| 9 | 36872314 | rs7850607 | 0.434 | -0.311 | -0.052 | -0.595 | -0.03 | -0.135 | -0.135 | 0 | 0.79 | 0.53 | 0.92 | 0.96 | 0.89 | 0.82 |
| 9 | 36876541 | rs12684931 | 0.612 | -0.175 | -0.428 | -0.617 | -0.018 | -0.009 | -0.009 | 0.62 | 0.06 | 0 | 0.07 | 0.07 | 0.09 | 0.08 |

| | | | | | | | | | | | | | | | | |
|---|---|---|---|---|---|---|---|---|---|---|---|---|---|---|---|---|
| 9 | 36878774 | rs4074255 | 0.86 | -1.203 | -0.985 | -1.214 | 0.128 | 0.168 | 0.168 | 3.683 | 0.51 | 0.17 | 0.74 | 0.76 | 0.71 | 0.63 |
| 9 | 36879510 | rs10973115 | 0.465 | -0.834 | -0.741 | -1.013 | 0.001 | -0.002 | -0.002 | 0 | 0.07 | 0.05 | 0.05 | 0.1 | 0.04 | 0.04 |
| 9 | 36880930 | rs10758411 | 1.063 | -0.911 | -0.687 | -1.043 | -0.013 | -0.042 | -0.042 | 0 | 0.66 | 0.48 | 0.8 | 0.8 | 0.76 | 0.7 |
| 9 | 36882141 | rs2277143 | -0.336 | -0.708 | -1.214 | -0.852 | 0.006 | 0.015 | 0.015 | 0 | 0.13 | 0.04 | 0.07 | 0.1 | 0.07 | 0.05 |
| 9 | 36885450 | rs3780140 | 1.545 | 0.115 | -1.819 | -0.826 | -0.112 | 0.011 | 0.011 | 0 | 0.27 | 0.05 | 0.31 | 0.18 | 0.37 | 0.36 |
| 9 | 36885598 | rs3780142 | -0.137 | -0.1 | -1.379 | -0.743 | -0.005 | 0 | 0 | 0 | 0.91 | 0.87 | 0.94 | 0.9 | 0.96 | 0.95 |
| 9 | 36885746 | rs3824339 | 0.063 | -0.08 | -1.469 | -0.859 | -0.018 | -0.055 | -0.055 | 0 | 0.28 | 0.17 | 0.39 | 0.58 | 0.28 | 0.22 |
| 9 | 36885919 | rs3780143 | 0.873 | -0.045 | -1.494 | -0.842 | -0.079 | -0.165 | -0.165 | 0.112 | 0.2 | 0.02 | 0.34 | 0.49 | 0.25 | 0.2 |
| 9 | 36887241 | rs3780144 | 0.937 | -0.299 | -0.777 | -0.756 | 0.004 | 0 | 0 | 2.927 | 0.06 | 0.07 | 0.07 | 0.07 | 0.08 | 0.06 |
| 9 | 36887510 | rs4581141 | 0.222 | -0.475 | -0.311 | -0.803 | -0.015 | -0.029 | -0.029 | 0 | 0.78 | 0.61 | 0.89 | 0.85 | 0.87 | 0.76 |
| 9 | 36888148 | rs4268201 | 0.496 | -0.666 | -0.413 | -0.895 | -0.039 | -0.011 | -0.011 | 0 | 0.73 | 0.53 | 0.85 | 0.78 | 0.85 | 0.72 |
| 9 | 36888975 | rs9298973 | -0.069 | -0.659 | -0.39 | -0.895 | -0.044 | -0.016 | -0.016 | 0 | 0.8 | 0.55 | 0.9 | 0.88 | 0.9 | 0.77 |
| 9 | 36894151 | rs11999298 | -0.006 | -0.326 | -1.071 | -0.7 | 0.001 | 0 | 0 | 0 | 0.08 | 0.09 | 0.05 | 0.1 | 0.05 | 0.05 |
| 9 | 36895382 | rs10973120 | 0.871 | -0.139 | -1.435 | -0.786 | -0.048 | 0.041 | 0.041 | 0.609 | 0.46 | 0.06 | 0.64 | 0.68 | 0.58 | 0.55 |
| 9 | 36896143 | rs1573257 | 0.798 | -0.11 | -1.421 | -0.772 | -0.041 | 0.049 | 0.049 | 0.905 | 0.47 | 0.07 | 0.64 | 0.68 | 0.58 | 0.56 |
| 9 | 36901161 | rs1936637 | -0.408 | -0.066 | -0.77 | -0.725 | -0.001 | -0.008 | -0.008 | 0 | 0.41 | 0.33 | 0.42 | 0.58 | 0.33 | 0.44 |
| 9 | 36904177 | rs3824341 | 1.093 | -0.222 | -0.316 | -0.956 | -0.01 | -0.025 | -0.025 | 0.036 | 0.18 | 0.21 | 0.11 | 0.09 | 0.16 | 0.15 |
| 9 | 36904278 | rs10973122 | 0.672 | -0.342 | -0.018 | -1.078 | -0.019 | -0.06 | -0.06 | 1.168 | 0.18 | 0.25 | 0.08 | 0.04 | 0.09 | 0.21 |
| 9 | 36905150 | rs7034991 | 0.569 | -0.09 | -0.39 | -0.662 | -0.045 | -0.105 | -0.105 | 0.516 | 0.22 | 0.12 | 0.34 | 0.52 | 0.24 | 0.23 |
| 9 | 36906843 | rs10123925 | -0.148 | -0.373 | 0.009 | -0.78 | -0.001 | 0 | 0 | 0.506 | 0.35 | 0.61 | 0.28 | 0.2 | 0.24 | 0.41 |
| 9 | 36911573 | rs3758163 | -0.739 | 0.085 | 0.205 | -0.425 | 0 | -0.003 | -0.003 | 0 | 0.64 | 0.64 | 0.6 | 0.8 | 0.58 | 0.65 |
| 9 | 36913074 | rs7868521 | -0.836 | 0.094 | 0.065 | -0.413 | -0.013 | 0.04 | 0.04 | 0 | 0.67 | 0.78 | 0.72 | 0.86 | 0.59 | 0.74 |
| 9 | 36922489 | rs12553834 | 0.867 | 0.518 | -1.198 | -0.413 | -0.125 | 0.026 | 0.026 | 0 | 0.31 | 0.04 | 0.26 | 0.19 | 0.4 | 0.22 |
| 9 | 36926028 | rs3780151 | 0.052 | 0.747 | -0.63 | -0.6 | -0.001 | -0.008 | -0.008 | 0.591 | 0.54 | 0.51 | 0.56 | 0.75 | 0.54 | 0.48 |
| 9 | 36927364 | rs7032626 | 1.47 | 0.852 | -0.709 | -0.597 | -0.066 | 0.052 | 0.052 | 0 | 0.3 | 0.05 | 0.2 | 0.16 | 0.34 | 0.2 |
| 9 | 36932412 | rs4878670 | -1.36 | 0.52 | -0.569 | -0.947 | -0.009 | -0.043 | -0.043 | 0 | 0.87 | 0.84 | 0.95 | 1 | 0.92 | 0.78 |
| 9 | 36934760 | rs13288123 | 0.618 | 0.447 | -0.634 | -0.993 | -0.029 | 0.026 | 0.026 | 0 | 0.25 | 0.11 | 0.25 | 0.19 | 0.34 | 0.24 |
| 9 | 36936360 | rs3758165 | -0.432 | 0.383 | -0.35 | -0.879 | -0.038 | -0.043 | -0.043 | 4.201 | 0.79 | 0.68 | 0.81 | 0.93 | 0.88 | 0.72 |
| 9 | 36939356 | rs10973133 | 0.993 | 0.577 | -0.601 | -0.949 | -0.007 | 0.164 | 0.164 | 0 | 0.49 | 0.15 | 0.37 | 0.23 | 0.57 | 0.38 |
| 9 | 36941292 | rs7047533 | 0.697 | 0.508 | -0.301 | -0.904 | -0.004 | -0.013 | -0.013 | 0.6 | 0.21 | 0.19 | 0.33 | 0.56 | 0.2 | 0.27 |
| 9 | 36942624 | rs3780160 | -0.379 | 0.426 | -0.216 | -0.858 | -0.029 | -0.031 | -0.031 | 0 | 0.79 | 0.71 | 0.81 | 0.92 | 0.89 | 0.74 |
| 9 | 36943396 | rs7853360 | 0.206 | 0.434 | -0.165 | -0.86 | -0.01 | -0.015 | -0.015 | 0 | 0.73 | 0.62 | 0.73 | 0.82 | 0.77 | 0.68 |
| 9 | 36944882 | rs16933784 | 1.148 | 0.94 | -0.162 | -0.464 | -0.024 | -0.022 | -0.022 | 0 | 0.05 | 0.02 | 0.08 | 0.1 | 0.12 | 0.07 |
| 9 | 36950301 | rs4880042 | -0.908 | 0.745 | -0.31 | -0.726 | 0.007 | 0.039 | 0.039 | 0 | 0.49 | 0.61 | 0.6 | 0.77 | 0.55 | 0.57 |
| 9 | 36951035 | rs10511936 | 0.325 | 0.82 | -0.547 | -0.296 | -0.008 | 0.007 | 0.007 | 0.007 | 0.19 | 0.1 | 0.16 | 0.17 | 0.24 | 0.22 |
| 9 | 36952743 | rs4407980 | 0.189 | 0.289 | -0.33 | -0.162 | -0.026 | -0.002 | -0.002 | 0 | 0.72 | 0.57 | 0.73 | 0.82 | 0.75 | 0.82 |
| 9 | 36956031 | rs12351115 | 1.369 | 0.071 | -0.265 | -0.571 | -0.008 | -0.047 | -0.047 | 0 | 0.12 | 0.19 | 0.05 | 0.01 | 0.08 | 0.15 |
| 9 | 36957379 | rs11790931 | 0.544 | -0.103 | -0.036 | -0.595 | -0.009 | -0.004 | -0.004 | 0.067 | 0.19 | 0.28 | 0.1 | 0.12 | 0.19 | 0.07 |
| 9 | 36957447 | rs10973140 | 0.851 | 0.397 | -0.526 | 0.086 | -0.063 | -0.116 | -0.116 | 0 | 0.17 | 0.01 | 0.3 | 0.38 | 0.15 | 0.28 |
| 9 | 36957633 | rs7876045 | 0.012 | 0.121 | -0.132 | -0.13 | -0.014 | -0.021 | -0.021 | 3.415 | 0.8 | 0.69 | 0.78 | 0.93 | 0.85 | 0.73 |
| 9 | 36958016 | rs10814484 | 0.799 | 0.084 | -0.147 | -0.098 | -0.007 | 0 | 0 | 0 | 0.18 | 0.29 | 0.1 | 0.12 | 0.19 | 0.07 |
| 9 | 36958317 | rs1888016 | -0.736 | 0.122 | -0.16 | -0.06 | -0.017 | 0.004 | 0.004 | 0 | 0.92 | 0.82 | 0.83 | 0.94 | 0.94 | 0.89 |
| 9 | 36960765 | rs10758418 | -0.144 | 0.396 | -0.216 | -0.077 | -0.01 | 0.014 | 0.014 | 0 | 0.77 | 0.59 | 0.75 | 0.84 | 0.75 | 0.87 |
| 9 | 36962064 | rs7858508 | 1.101 | 0.38 | -0.302 | 0.002 | 0.001 | -0.002 | -0.002 | 0 | 0.16 | 0.19 | 0.1 | 0.04 | 0.14 | 0.19 |
| 9 | 36968384 | rs10973145 | 0.707 | 0.331 | 0.022 | -0.02 | 0.018 | 0.022 | 0.022 | 0 | 0.6 | 0.26 | 0.65 | 0.82 | 0.62 | 0.67 |
| 9 | 36968880 | rs10121384 | -0.161 | 0.344 | 0.219 | 0 | -0.01 | 0.007 | 0.007 | 0 | 0.77 | 0.6 | 0.75 | 0.85 | 0.76 | 0.86 |
| 9 | 36975108 | rs10814496 | 0.631 | 0.268 | 0.128 | -0.003 | 0.039 | 0.03 | 0.03 | 0 | 0.61 | 0.23 | 0.67 | 0.83 | 0.67 | 0.69 |
| 9 | 36977212 | rs13292007 | 1.983 | 0.316 | -0.563 | -0.442 | -0.058 | 0.084 | 0.084 | 0.09 | 0.24 | 0.02 | 0.15 | 0.1 | 0.22 | 0.14 |
| 9 | 36977798 | rs7043127 | -0.886 | 0.493 | -0.189 | 0.064 | -0.002 | 0.003 | 0.003 | 0 | 0.78 | 0.71 | 0.86 | 0.8 | 0.75 | 0.88 |
| 9 | 36979395 | rs3780169 | 0.128 | 0.579 | -0.126 | 0.004 | 0.012 | 0.022 | 0.022 | 0 | 0.29 | 0.31 | 0.46 | 0.62 | 0.33 | 0.49 |
| 9 | 36980920 | rs7019007 | 1.277 | 0.66 | 0.552 | 0.297 | -0.019 | -0.013 | -0.013 | 1.327 | 0.1 | 0.18 | 0.14 | 0.04 | 0.07 | 0.07 |
| 9 | 36983604 | rs7045954 | -0.646 | 0.604 | 0.43 | 0.3 | -0.003 | 0.005 | 0.005 | 0 | 0.78 | 0.69 | 0.85 | 0.81 | 0.75 | 0.87 |
| 9 | 36985143 | rs10814498 | -0.411 | 0.938 | 0.539 | 0.613 | -0.022 | -0.003 | -0.003 | 0.063 | 0.67 | 0.42 | 0.71 | 0.77 | 0.68 | 0.79 |
| 9 | 36989054 | rs4880048 | 0.505 | 0.69 | 0.172 | 0.663 | -0.004 | 0.012 | 0.012 | 0 | 0.46 | 0.3 | 0.55 | 0.69 | 0.48 | 0.51 |
| 9 | 36989746 | rs7868665 | -0.658 | 0.594 | 0.384 | 0.657 | -0.001 | -0.001 | -0.001 | 0 | 0.67 | 0.59 | 0.68 | 0.77 | 0.64 | 0.82 |
| 9 | 36993071 | rs10125775 | 0.118 | 0.649 | 0.225 | 1.05 | -0.006 | 0.007 | 0.007 | 0.053 | 0.45 | 0.34 | 0.55 | 0.69 | 0.46 | 0.52 |
| 9 | 36993868 | rs883966 | -0.345 | 1.183 | 0.865 | 1.954 | 0.003 | 0.032 | 0.032 | 0.252 | 0.11 | 0.11 | 0.29 | 0.34 | 0.11 | 0.24 |
| 9 | 36997417 | rs3758171 | -0.855 | 1.174 | 0.765 | 1.458 | -0.018 | -0.003 | -0.003 | 0 | 0.15 | 0.13 | 0.35 | 0.15 | 0.23 | 0.27 |
| 9 | 36998993 | rs1329573 | 0.4 | 1.028 | 1.284 | 1.533 | 0.009 | 0.067 | 0.067 | 0 | 0.89 | 0.7 | 0.94 | 0.79 | 0.93 | 0.75 |
| 9 | 36999024 | rs4880050 | 0.4 | 1.028 | 1.284 | 1.533 | 0.009 | 0.067 | 0.067 | 0 | 0.89 | 0.7 | 0.94 | 0.79 | 0.93 | 0.75 |
| 9 | 37002115 | rs7020413 | -1.017 | 1.071 | 1.355 | 1.548 | 0.031 | -0.008 | -0.008 | 0 | 0.2 | 0.31 | 0.38 | 0.18 | 0.25 | 0.29 |
| 9 | 37003075 | rs7852624 | -0.659 | 1.191 | 1.099 | 1.696 | -0.007 | -0.001 | -0.001 | 0 | 0.15 | 0.15 | 0.3 | 0.17 | 0.23 | 0.25 |
| 9 | 37005561 | rs6476606 | 1.144 | 0.978 | 1.471 | 1.747 | 0.018 | 0.005 | 0.005 | 0 | 0.77 | 0.74 | 0.55 | 0.74 | 0.66 | 0.6 |
| 9 | 37007475 | rs4880051 | 1.317 | 0.917 | 1.52 | 1.827 | 0.023 | 0.017 | 0.017 | 0 | 0.75 | 0.65 | 0.51 | 0.65 | 0.62 | 0.61 |
| 9 | 37011289 | rs959396 | 1.386 | 0.594 | 1.699 | 2.068 | 0.03 | 0.056 | 0.056 | 0 | 0.71 | 0.56 | 0.54 | 0.48 | 0.64 | 0.43 |
| 9 | 37017666 | rs2381595 | 1.757 | -0.433 | 2.069 | 1.936 | -0.078 | 0.062 | 0.062 | 0.135 | 0.88 | 0.57 | 0.95 | 0.82 | 1 | 0.74 |
| 9 | 37031876 | rs3013739 | 1.74 | -0.324 | 1.745 | 1.841 | -0.084 | 0.12 | 0.12 | 0.051 | 0.87 | 0.53 | 0.97 | 0.78 | 0.99 | 0.74 |
| 9 | 37036247 | rs2282079 | 0.059 | 0.114 | 1.712 | 1.786 | -0.001 | 0.024 | 0.024 | 0 | 0.05 | 0.08 | 0.06 | 0.11 | 0.04 | 0.02 |
| 9 | 37038258 | rs4256662 | 1.55 | 0.046 | 1.174 | 1.818 | -0.026 | 0.084 | 0.084 | 1.683 | 0.84 | 0.35 | 0.86 | 0.69 | 0.91 | 0.91 |

| | | | | | | | | | | | | | | | | |
|---|---|---|---|---|---|---|---|---|---|---|---|---|---|---|---|---|
| 9 | 37045825 | rs4483247 | -1.24 | 0.816 | 1.256 | 1.471 | 0.084 | 0.048 | 0.048 | 0 | 0.09 | 0.28 | 0.17 | 0.11 | 0.14 | 0.16 |
| 9 | 37046091 | rs1411060 | -0.042 | 0.332 | 0.266 | 0.666 | 0.015 | 0.075 | 0.075 | 0 | 0.3 | 0.15 | 0.16 | 0.15 | 0.26 | 0.21 |
| 9 | 37055109 | rs10973177 | 0.086 | 0.46 | 0.577 | 0.714 | 0.032 | 0.018 | 0.018 | 0 | 0.06 | 0.09 | 0.16 | 0.12 | 0.14 | 0.12 |
| 9 | 37056373 | rs10973179 | 0.015 | 0.457 | 0.607 | 0.713 | 0.036 | 0.02 | 0.02 | 0 | 0.06 | 0.1 | 0.16 | 0.12 | 0.14 | 0.12 |
| 9 | 37066231 | rs10973183 | -1.229 | -0.14 | 0.504 | 0.216 | 0.001 | 0.011 | 0.011 | 0 | 0.53 | 0.53 | 0.37 | 0.31 | 0.51 | 0.54 |
| 9 | 37070398 | rs7854002 | 0.379 | -0.22 | 0.695 | 0.147 | -0.016 | -0.024 | -0.024 | 0 | 0.21 | 0.27 | 0.05 | 0.08 | 0.13 | 0.22 |
| 9 | 37075859 | rs4132811 | -1.302 | -0.199 | 0.679 | 0.172 | 0.002 | 0.01 | 0.01 | 0 | 0.53 | 0.52 | 0.37 | 0.31 | 0.51 | 0.54 |
| 9 | 37081301 | rs7034554 | -0.289 | -0.351 | 0.684 | 0.099 | 0 | -0.001 | -0.001 | 0.199 | 0.4 | 0.35 | 0.19 | 0.14 | 0.36 | 0.32 |
| 9 | 37092433 | rs7872067 | -1.347 | -0.32 | 0.627 | 0.116 | 0.002 | 0.009 | 0.009 | 0 | 0.53 | 0.52 | 0.37 | 0.31 | 0.51 | 0.55 |
| 9 | 37097670 | rs7848138 | -1.364 | -0.316 | 0.629 | 0.118 | 0.001 | 0.009 | 0.009 | 0 | 0.53 | 0.52 | 0.37 | 0.31 | 0.51 | 0.55 |
| 9 | 37125911 | rs12376809 | -0.414 | -0.354 | 1.148 | 0.131 | 0.083 | 0.062 | 0.062 | 0.007 | 0.09 | 0.3 | 0.19 | 0.11 | 0.15 | 0.15 |
| 9 | 37139899 | rs10973229 | 0.335 | -0.355 | 1.04 | 0.13 | -0.011 | -0.011 | -0.011 | 10.692 | 0.07 | 0.03 | 0.17 | 0.11 | 0.15 | 0.14 |
| 9 | 37182050 | rs13285022 | 0.632 | -0.308 | 1.116 | 0.009 | -0.001 | 0.008 | 0.008 | 0 | 0.24 | 0.18 | 0.15 | 0.12 | 0.23 | 0.17 |
| 9 | 37193871 | rs10814515 | -1.571 | -0.38 | 1.065 | 0.041 | 0.004 | 0.059 | 0.059 | 0 | 0.54 | 0.74 | 0.38 | 0.31 | 0.51 | 0.55 |
| 9 | 37198176 | rs10973250 | 0.488 | -0.32 | 1.002 | 0.13 | -0.015 | -0.014 | -0.014 | 0.893 | 0.07 | 0.03 | 0.17 | 0.12 | 0.15 | 0.15 |
| 9 | 37201664 | rs7043643 | 1.399 | -0.374 | 1.038 | 0.117 | -0.022 | 0.026 | 0.026 | 0 | 0.05 | 0.09 | 0.01 | 0.06 | 0.01 | 0.08 |
| 9 | 37213151 | rs7848459 | 1.268 | -0.41 | 1.1 | 0.466 | -0.027 | 0.038 | 0.038 | 0 | 0.05 | 0.11 | 0.01 | 0.06 | 0 | 0.08 |
| 9 | 37218231 | rs7041839 | -0.407 | -0.553 | 1.087 | 0.344 | -0.011 | 0.026 | 0.026 | 0 | 0.35 | 0.25 | 0.32 | 0.23 | 0.38 | 0.32 |
| 9 | 37223751 | rs2029647 | 1.54 | -0.512 | 0.955 | 0.389 | -0.026 | 0.024 | 0.024 | 0 | 0.05 | 0.08 | 0.01 | 0.06 | 0 | 0.08 |
| 9 | 37254690 | rs10758428 | 2.184 | -0.707 | 0.501 | 0.981 | -0.113 | -0.074 | -0.074 | 0 | 0.3 | 0.03 | 0.47 | 0.29 | 0.38 | 0.32 |
| 9 | 37256393 | rs489657 | 1.734 | -0.693 | 0.565 | 0.943 | -0.026 | 0.015 | 0.015 | 0 | 0.05 | 0.08 | 0.01 | 0.05 | 0 | 0.08 |
| 9 | 37258237 | rs13296908 | -0.594 | -0.796 | 0.727 | 0.882 | -0.001 | -0.012 | -0.012 | 0 | 0.47 | 0.44 | 0.35 | 0.25 | 0.5 | 0.46 |
| 9 | 37259118 | rs10973274 | -0.099 | -0.76 | 0.579 | 0.718 | 0.058 | 0.041 | 0.041 | 0.039 | 0.1 | 0.29 | 0.19 | 0.12 | 0.15 | 0.15 |
| 9 | 37260487 | rs10973276 | 1.473 | -0.789 | 0.41 | 0.748 | 0.039 | -0.051 | -0.051 | 0 | 0.09 | 0.01 | 0.1 | 0.26 | 0.06 | 0.1 |
| 9 | 37277603 | rs308495 | -0.812 | -0.683 | 0.497 | 0.41 | 0.022 | 0.09 | 0.09 | 0 | 0.47 | 0.67 | 0.21 | 0.19 | 0.36 | 0.4 |
| 9 | 37277831 | rs308494 | 0.552 | -0.651 | 0.554 | 0.473 | -0.023 | 0.073 | 0.073 | 0.04 | 0.07 | 0.3 | 0.02 | 0.06 | 0 | 0.09 |
| 9 | 37302155 | rs10814518 | -0.078 | -0.639 | 0.349 | 0.371 | -0.045 | 0.054 | 0.054 | 0 | 0.31 | 0.13 | 0.31 | 0.22 | 0.37 | 0.35 |
| 9 | 37356403 | rs186299 | -0.741 | -0.643 | 0.388 | 0.518 | -0.003 | 0.011 | 0.011 | 0 | 0.85 | 0.84 | 0.87 | 0.62 | 0.9 | 0.88 |
| 9 | 37364094 | rs997318 | -0.5 | -0.497 | 0.514 | 0.278 | -0.001 | -0.004 | -0.004 | 0 | 0.4 | 0.35 | 0.27 | 0.22 | 0.37 | 0.31 |
| 9 | 37364675 | rs10814520 | -1.089 | -0.414 | 0.056 | 0.335 | 0.014 | 0.049 | 0.049 | 0 | 0.49 | 0.64 | 0.46 | 0.33 | 0.51 | 0.47 |
| 9 | 37370553 | rs308504 | -0.181 | -0.612 | 0.198 | 0.318 | -0.012 | -0.026 | -0.026 | 0 | 0.2 | 0.29 | 0.06 | 0.08 | 0.15 | 0.22 |
| 9 | 37375126 | rs927928 | 0.34 | -0.788 | 0.077 | 0.261 | 0.018 | 0.03 | 0.03 | 0.135 | 0.12 | 0.42 | 0.17 | 0.19 | 0.09 | 0.17 |
| 9 | 37376587 | rs3780335 | -0.035 | -0.991 | -0.367 | 0.051 | 0.003 | 0.055 | 0.055 | 0.001 | 0.32 | 0.18 | 0.13 | 0.09 | 0.27 | 0.26 |
| 9 | 37377959 | rs553850 | -0.494 | -1.18 | -0.164 | 0.47 | -0.05 | 0.038 | 0.038 | 0 | 0.2 | 0.54 | 0.19 | 0.25 | 0.11 | 0.25 |
| 9 | 37378628 | rs2768654 | -0.073 | -1.025 | -0.804 | 0.648 | 0.009 | -0.011 | -0.011 | 0 | 0.83 | 0.85 | 0.86 | 0.62 | 0.89 | 0.88 |
| 9 | 37382099 | rs503897 | -0.479 | -0.234 | -1.429 | -0.235 | -0.036 | -0.029 | -0.029 | 0 | 0.65 | 0.87 | 0.76 | 0.36 | 0.62 | 0.51 |
| 9 | 37384977 | rs7871498 | 0.456 | 0.057 | -1.174 | -0.505 | 0.001 | -0.002 | -0.002 | 0 | 0.16 | 0.26 | 0.37 | 0.18 | 0.16 | 0.19 |
| 9 | 37387177 | rs7025491 | 0.497 | 0.11 | -0.74 | -0.663 | -0.011 | 0.064 | 0.064 | 0 | 0.44 | 0.27 | 0.36 | 0.21 | 0.46 | 0.32 |
| 9 | 37387218 | rs7025809 | -0.668 | 0.052 | -0.757 | -0.533 | -0.003 | 0.017 | 0.017 | 0 | 0.62 | 0.58 | 0.74 | 0.44 | 0.62 | 0.6 |
| 9 | 37387456 | rs7026066 | 1.536 | 0.563 | -1.018 | -0.917 | 0.001 | -0.003 | -0.003 | 0 | 0.23 | 0.26 | 0.22 | 0.2 | 0.24 | 0.19 |
| 9 | 37390630 | rs10814529 | 0.258 | 0.555 | -1.032 | -0.924 | -0.014 | 0.06 | 0.06 | 0 | 0.42 | 0.28 | 0.34 | 0.17 | 0.46 | 0.28 |
| 9 | 37390964 | rs765567 | -0.034 | 0.46 | -0.977 | -0.483 | -0.001 | 0.003 | 0.003 | 0 | 0.41 | 0.62 | 0.61 | 0.4 | 0.39 | 0.39 |
| 9 | 37392087 | rs744985 | 0.047 | 0.407 | -0.87 | -0.355 | 0.02 | 0.052 | 0.052 | 0 | 0.49 | 0.59 | 0.4 | 0.21 | 0.57 | 0.32 |
| 9 | 37393048 | rs12555254 | 0.088 | 0.9 | -0.622 | 0.467 | -0.009 | 0.003 | 0.003 | 0 | 0.08 | 0.07 | 0.25 | 0.21 | 0.1 | 0.15 |
| 9 | 37394542 | rs4878685 | 0.236 | 0.687 | -0.945 | -0.238 | -0.033 | 0.087 | 0.087 | 0.82 | 0.6 | 0.25 | 0.57 | 0.8 | 0.55 | 0.61 |
| 9 | 37395039 | rs471579 | -0.151 | 0.563 | 0 | -0.329 | 0.02 | -0.045 | -0.045 | 0 | 0.64 | 0.53 | 0.61 | 0.83 | 0.56 | 0.66 |
| 9 | 37395138 | rs16934059 | 1.074 | 0.393 | -0.586 | -0.369 | 0.042 | -0.026 | -0.026 | 0.625 | 0.46 | 0.17 | 0.29 | 0.57 | 0.4 | 0.42 |
| 9 | 37397532 | rs7873371 | -0.006 | 0.496 | 0.542 | -0.729 | 0.03 | -0.049 | -0.049 | 1.046 | 0.65 | 0.55 | 0.62 | 0.83 | 0.55 | 0.68 |
| 9 | 37399384 | rs2768653 | 0.351 | 0.415 | 0.394 | -0.73 | 0.001 | -0.006 | -0.006 | 0 | 0.65 | 0.64 | 0.63 | 0.81 | 0.58 | 0.69 |
| 9 | 37404949 | rs2087358 | 0.367 | 0.992 | 0.353 | -0.889 | -0.013 | 0.048 | 0.048 | 0 | 0.24 | 0.13 | 0.16 | 0 | 0.25 | 0.13 |
| 9 | 37406391 | rs7047872 | -0.785 | 0.835 | 0.695 | -0.49 | 0.066 | 0.051 | 0.051 | 0.278 | 0.09 | 0.21 | 0.22 | 0.17 | 0.17 | 0.18 |
| 9 | 37411834 | rs1571231 | 1.561 | 0.603 | 0.744 | -0.511 | 0.061 | 0.038 | 0.038 | 0.033 | 0.86 | 0.47 | 0.76 | 0.82 | 0.84 | 0.8 |
| 9 | 37412704 | rs6476629 | -0.047 | 0.666 | 0.114 | -1.208 | 0.002 | -0.101 | -0.101 | 0.197 | 0.3 | 0.31 | 0.21 | 0.01 | 0.3 | 0.22 |
| 9 | 37413003 | rs17411255 | -0.581 | 0.72 | -0.406 | -0.421 | -0.032 | -0.021 | -0.021 | 0 | 0.05 | 0.01 | 0.13 | 0.07 | 0.09 | 0.11 |
| 9 | 37418935 | rs10814535 | -0.049 | 0.122 | -0.428 | -0.893 | 0.018 | 0.034 | 0.034 | 0 | 0.49 | 0.43 | 0.55 | 0.88 | 0.42 | 0.53 |
| 9 | 37421065 | rs7856656 | 0.46 | 0.125 | -0.982 | -0.031 | 0.011 | -0.026 | -0.026 | 2.92 | 0.37 | 0.26 | 0.48 | 0.45 | 0.35 | 0.32 |
| 9 | 37426152 | rs2768659 | 0.041 | 0.097 | -1.505 | -0.29 | 0.091 | -0.011 | -0.011 | 0 | 0.5 | 0.22 | 0.71 | 0.46 | 0.65 | 0.56 |
| 9 | 37429497 | rs309460 | -0.399 | 0.36 | -1.698 | -0.494 | 0.012 | 0.115 | 0.115 | 0 | 0.58 | 0.26 | 0.65 | 0.86 | 0.66 | 0.67 |
| 9 | 37429814 | rs309458 | -0.467 | 0.305 | -0.926 | -0.589 | 0.094 | 0.071 | 0.071 | 0 | 0.93 | 0.71 | 0.83 | 0.92 | 0.92 | 0.9 |
| 9 | 37432264 | rs309455 | -0.388 | 0.233 | -0.979 | -0.606 | 0.078 | 0.061 | 0.061 | 0 | 0.92 | 0.73 | 0.83 | 0.91 | 0.92 | 0.9 |
| 9 | 37433556 | rs309453 | -0.2 | 0.258 | -1.967 | -0.609 | 0.015 | 0.127 | 0.127 | 0 | 0.57 | 0.25 | 0.64 | 0.86 | 0.65 | 0.7 |
| 9 | 37436105 | rs7867764 | 1.735 | 0.051 | -2.123 | -0.723 | -0.011 | 0.144 | 0.144 | 0 | 0.25 | 0.04 | 0.11 | 0.01 | 0.21 | 0.12 |
| 9 | 37438526 | rs2840246 | 0.685 | 0.071 | -2.23 | -0.646 | 0.011 | -0.078 | -0.078 | 0 | 0.13 | 0.04 | 0.07 | 0.49 | 0.1 | 0.26 |
| 9 | 37439351 | rs7039333 | 1.163 | 0.383 | -1.777 | -0.317 | -0.001 | 0 | 0 | 0 | 0.06 | 0.07 | 0.05 | 0.04 | 0.05 | 0.08 |
| 9 | 37441650 | rs17502738 | 1.504 | 0.497 | -2.05 | -0.359 | -0.01 | 0.146 | 0.146 | 0 | 0.25 | 0.04 | 0.11 | 0.01 | 0.21 | 0.12 |
| 9 | 37444994 | rs10973340 | 0.45 | 1.28 | -1.212 | 0.399 | -0.054 | -0.075 | -0.075 | 3.817 | 0.12 | 0.02 | 0.37 | 0.36 | 0.21 | 0.2 |
| 9 | 37454874 | rs309444 | 0.774 | 0.77 | -0.657 | -0.209 | -0.001 | -0.006 | -0.006 | 0.005 | 0.83 | 0.38 | 0.79 | 0.88 | 0.87 | 0.84 |
| 9 | 37464909 | rs7049017 | -0.13 | 0.584 | -1.112 | -0.258 | -0.008 | 0.024 | 0.024 | 0 | 0.11 | 0.06 | 0.06 | 0.03 | 0.11 | 0.09 |
| 9 | 37483309 | rs913280 | -0.639 | 0.505 | -1.13 | -0.227 | -0.003 | 0.025 | 0.025 | 0 | 0.21 | 0.14 | 0.24 | 0.08 | 0.24 | 0.24 |

| | | | | | | | | | | | | | | | | |
|---|---|---|---|---|---|---|---|---|---|---|---|---|---|---|---|---|
| 9 | 37483388 | rs913279 | -1.211 | 0.409 | -1.333 | -0.027 | -0.004 | 0.02 | 0.02 | 0 | 0.4 | 0.8 | 0.55 | 0.3 | 0.42 | 0.4 |
| 9 | 37486383 | rs10973360 | 0.921 | -0.552 | -0.493 | -0.873 | -0.044 | -0.125 | -0.125 | 0.361 | 0.75 | 0.61 | 0.87 | 0.99 | 0.84 | 0.85 |
| 9 | 37486603 | rs1571234 | -0.182 | -0.48 | -0.294 | -0.781 | 0.008 | 0.011 | 0.011 | 0 | 0.47 | 0.57 | 0.46 | 0.29 | 0.54 | 0.5 |
| 9 | 37487667 | rs2296774 | 0.404 | -0.034 | 0.317 | -0.62 | 0.006 | -0.028 | -0.028 | 0 | 0.07 | 0.14 | 0.06 | 0.01 | 0.06 | 0.08 |
| 9 | 37488009 | rs10758432 | 1.647 | -0.033 | 0.165 | -0.147 | -0.038 | -0.035 | -0.035 | 0 | 0.07 | 0.01 | 0.25 | 0.23 | 0.14 | 0.12 |
| 9 | 37488725 | rs1887455 | -1.716 | 0.376 | 0.922 | -0.228 | -0.015 | -0.022 | -0.022 | 0.012 | 0.3 | 0.77 | 0.44 | 0.41 | 0.3 | 0.37 |
| 9 | 37489298 | rs3758223 | 0.156 | 0.514 | 0.833 | -0.197 | 0.007 | 0.009 | 0.009 | 0 | 0.14 | 0.17 | 0.31 | 0.24 | 0.2 | 0.19 |
| 9 | 37492679 | rs3739576 | 0.156 | 0.514 | 0.833 | -0.197 | 0.007 | 0.009 | 0.009 | 0 | 0.14 | 0.17 | 0.31 | 0.24 | 0.2 | 0.19 |
| 9 | 37492721 | rs3739575 | 1.133 | 0.548 | 0.8 | -0.205 | -0.033 | -0.031 | -0.031 | 0.005 | 0.05 | 0.01 | 0.07 | 0.16 | 0.09 | 0.14 |
| 9 | 37493548 | rs10814567 | -0.175 | 0.56 | 0.921 | -0.235 | 0.054 | 0.067 | 0.067 | 0 | 0.16 | 0.37 | 0.32 | 0.24 | 0.2 | 0.19 |
| 9 | 37493588 | rs10758434 | 0.17 | 0.56 | 0.908 | -0.235 | 0.007 | 0.009 | 0.009 | 0 | 0.14 | 0.17 | 0.31 | 0.24 | 0.2 | 0.19 |
| 9 | 37500895 | rs2277161 | -0.202 | 0.566 | 1.026 | -0.232 | 0.062 | 0.077 | 0.077 | 0 | 0.17 | 0.42 | 0.33 | 0.24 | 0.2 | 0.19 |
| 9 | 37503087 | rs10814571 | 0.168 | 0.609 | 0.546 | -0.206 | 0.007 | 0.009 | 0.009 | 0 | 0.14 | 0.17 | 0.31 | 0.24 | 0.2 | 0.19 |
| 9 | 37506966 | rs11794762 | 0.604 | 0.721 | 0.199 | -0.276 | -0.022 | -0.025 | -0.025 | 0 | 0.13 | 0.01 | 0.14 | 0.1 | 0.13 | 0.18 |
| 9 | 37509875 | rs1006763 | 0.952 | 1.026 | 0.701 | -0.414 | -0.032 | -0.03 | -0.03 | 0 | 0.05 | 0.01 | 0.07 | 0.16 | 0.09 | 0.14 |
| 9 | 37512808 | rs12237351 | -0.769 | 1.019 | 1.034 | -0.43 | 0.037 | 0.048 | 0.048 | 0 | 0.21 | 0.31 | 0.39 | 0.41 | 0.3 | 0.33 |
| 9 | 37522672 | rs10973394 | 1.856 | 0.912 | 0.661 | -1.012 | -0.007 | 0.005 | 0.005 | 0.043 | 0.69 | 0.28 | 0.56 | 0.7 | 0.7 | 0.62 |
| 9 | 37528404 | rs7027892 | -1.391 | 0.868 | 0.774 | -1.009 | 0.002 | -0.071 | -0.071 | 0 | 0.16 | 0.57 | 0.12 | 0.05 | 0.09 | 0.19 |
| 9 | 37537557 | rs7044153 | 1.624 | 0.872 | 0.684 | -0.947 | -0.004 | 0.001 | 0.001 | 0 | 0.7 | 0.31 | 0.54 | 0.7 | 0.66 | 0.62 |
| 9 | 37542954 | rs12337272 | -1.454 | 0.672 | 0.604 | -1.009 | -0.02 | -0.092 | -0.092 | 0 | 0.21 | 0.55 | 0.19 | 0.07 | 0.17 | 0.24 |
| 9 | 37547256 | rs10125441 | -0.546 | 0.473 | 0.807 | -0.901 | -0.025 | -0.066 | -0.066 | 0.007 | 0.12 | 0.34 | 0.08 | 0.01 | 0.05 | 0.09 |
| 9 | 37551276 | rs13291473 | 0.472 | 0.329 | 0.456 | -1.429 | 0.002 | -0.037 | -0.037 | 0 | 0.27 | 0.21 | 0.15 | 0.01 | 0.2 | 0.1 |
| 9 | 37552338 | rs10973409 | -1.309 | 0.547 | 0.301 | -1.459 | 0.004 | 0.065 | 0.065 | 0 | 0.48 | 0.66 | 0.54 | 0.3 | 0.49 | 0.42 |
| 9 | 37558780 | rs10118277 | -0.797 | 0.906 | 0.189 | -0.798 | 0.031 | 0.017 | 0.017 | 0 | 0.21 | 0.34 | 0.38 | 0.28 | 0.29 | 0.33 |
| 9 | 37565522 | rs10814578 | 0.132 | 1.372 | 0.016 | -0.621 | 0.001 | -0.001 | -0.001 | 0 | 0.12 | 0.13 | 0.29 | 0.24 | 0.23 | 0.15 |
| 9 | 37573330 | rs10758440 | -0.353 | 0.801 | -0.218 | -0.882 | -0.029 | 0.004 | 0.004 | 0 | 0.15 | 0.09 | 0.3 | 0.12 | 0.21 | 0.15 |
| 9 | 37583429 | rs9644884 | 1.183 | 0.932 | -0.089 | -0.876 | -0.034 | -0.026 | -0.026 | 0.391 | 0.07 | 0.02 | 0.25 | 0.12 | 0.16 | 0.12 |
| 9 | 37587742 | rs4621903 | 0.416 | 1.098 | 0.055 | -1.209 | -0.001 | -0.063 | -0.063 | 2.239 | 0.2 | 0.31 | 0.13 | 0 | 0.16 | 0.05 |
| 9 | 37589346 | rs13296887 | 1.048 | 1.063 | 0.021 | -1.226 | -0.019 | 0.042 | 0.042 | 1.155 | 0.16 | 0.1 | 0.11 | 0 | 0.16 | 0.03 |
| 9 | 37591236 | rs10758442 | -0.719 | 0.999 | 0.399 | -1.262 | 0.025 | -0.076 | -0.076 | 0 | 0.32 | 0.54 | 0.4 | 0.12 | 0.33 | 0.19 |
| 9 | 37592452 | rs7021977 | 0.136 | 0.532 | 0.303 | -1.001 | -0.001 | -0.063 | -0.063 | 0.057 | 0.2 | 0.31 | 0.13 | 0 | 0.16 | 0.05 |
| 9 | 37611238 | rs13283323 | 0.491 | 0.69 | 0.059 | -0.858 | 0 | -0.041 | -0.041 | 0 | 0.17 | 0.26 | 0.1 | 0 | 0.14 | 0.03 |
| 9 | 37626975 | rs12339112 | 0.566 | 0.52 | -0.094 | -0.908 | 0.001 | -0.021 | -0.021 | 0.244 | 0.21 | 0.23 | 0.17 | 0.12 | 0.24 | 0.24 |
| 9 | 37627872 | rs10123280 | -0.035 | 0.061 | -0.236 | -1.226 | -0.004 | -0.022 | -0.022 | 0.225 | 0.65 | 0.44 | 0.6 | 0.28 | 0.66 | 0.6 |
| 9 | 37629310 | rs17505776 | 1.175 | 0.025 | -0.686 | -0.985 | -0.042 | -0.011 | -0.011 | 0.306 | 0.17 | 0.04 | 0.19 | 0.16 | 0.16 | 0.16 |
| 9 | 37631583 | rs10973443 | 1.267 | 0.014 | -0.533 | -0.922 | -0.002 | 0.085 | 0.085 | 0 | 0.28 | 0.12 | 0.35 | 0.06 | 0.26 | 0.31 |
| 9 | 37635299 | rs999988 | 0.338 | -0.556 | 0.38 | -0.089 | 0.001 | 0.152 | 0.152 | 0 | 0.82 | 0.51 | 0.77 | 0.42 | 0.8 | 0.75 |
| 9 | 37638744 | rs10973446 | -0.371 | -0.577 | 0.266 | -0.133 | -0.005 | -0.001 | -0.001 | 0 | 0.36 | 0.45 | 0.38 | 0.6 | 0.4 | 0.45 |
| 9 | 37638972 | rs9919069 | -1.226 | -0.661 | 0.053 | 0.181 | -0.007 | 0.037 | 0.037 | 0 | 0.54 | 0.7 | 0.45 | 0.39 | 0.54 | 0.55 |
| 9 | 37641833 | rs2182318 | 0.677 | -0.701 | 0.193 | -0.129 | -0.002 | -0.002 | -0.002 | 0 | 0.18 | 0.19 | 0.15 | 0.17 | 0.13 | 0.18 |
| 9 | 37647741 | rs875780 | 0.468 | -0.716 | 0.355 | -0.175 | -0.005 | -0.045 | -0.045 | 0 | 0.24 | 0.3 | 0.17 | 0.09 | 0.25 | 0.25 |
| 9 | 37650203 | rs7021824 | 0.095 | -0.662 | 0.478 | 0.112 | 0.003 | 0.229 | 0.229 | 0 | 0.15 | 0.37 | 0.23 | 0.48 | 0.16 | 0.21 |
| 9 | 37651804 | rs7027222 | -0.27 | -1.255 | 0.149 | 0.175 | -0.016 | 0.017 | 0.017 | 0 | 0.34 | 0.42 | 0.31 | 0.51 | 0.31 | 0.39 |
| 9 | 37654930 | rs12684941 | -0.814 | -1.216 | -0.411 | 0.23 | 0.042 | 0.043 | 0.043 | 0 | 0.52 | 0.85 | 0.48 | 0.65 | 0.43 | 0.55 |
| 9 | 37656212 | rs1886909 | -1.674 | -1.193 | -0.554 | 0.077 | -0.05 | 0.036 | 0.036 | 0 | 0.64 | 0.83 | 0.55 | 0.76 | 0.53 | 0.7 |
| 9 | 37656636 | rs10973458 | 1.27 | -1.062 | -0.762 | 0.112 | -0.005 | 0.016 | 0.016 | 0 | 0.07 | 0.04 | 0.04 | 0 | 0.06 | 0.03 |
| 9 | 37657011 | rs6476655 | -0.24 | -1.147 | -0.733 | 0.168 | -0.032 | 0.164 | 0.164 | 0.048 | 0.15 | 0.3 | 0.14 | 0.5 | 0.07 | 0.27 |
| 9 | 37660696 | rs1325916 | -1.799 | -1.148 | -0.811 | 0.124 | -0.089 | 0.044 | 0.044 | 0 | 0.64 | 0.91 | 0.55 | 0.76 | 0.53 | 0.7 |
| 9 | 37663810 | rs6476656 | -0.808 | -1.107 | -0.596 | 0.148 | 0 | 0.01 | 0.01 | 0 | 0.55 | 0.63 | 0.51 | 0.76 | 0.47 | 0.67 |
| 9 | 37666361 | rs10973475 | 0.262 | -0.919 | -0.921 | 0.671 | -0.011 | -0.03 | -0.03 | 3.152 | 0.06 | 0.02 | 0.03 | 0.11 | 0.04 | 0.13 |
| 9 | 37666667 | rs10814594 | -0.355 | -0.78 | -0.839 | 0.255 | -0.034 | 0.092 | 0.092 | 0.114 | 0.24 | 0.53 | 0.19 | 0.5 | 0.14 | 0.3 |
| 9 | 37676472 | rs16934357 | -0.379 | 0.133 | -0.926 | -0.214 | 0 | -0.002 | -0.002 | 0 | 0.22 | 0.21 | 0.19 | 0.09 | 0.26 | 0.2 |
| 9 | 37684753 | rs7469365 | -0.647 | -0.071 | -1.156 | 0.224 | 0.008 | 0.011 | 0.011 | 0 | 0.2 | 0.13 | 0.1 | 0.17 | 0.18 | 0.18 |
| 9 | 37686417 | rs13298495 | -0.661 | 0.434 | -1.801 | 0.529 | -0.091 | -0.066 | -0.066 | 1.008 | 0.34 | 0.11 | 0.4 | 0.46 | 0.42 | 0.44 |
| 9 | 37689623 | rs10814601 | -1.096 | 0.181 | -1.222 | 0.182 | -0.004 | -0.07 | -0.07 | 0 | 0.66 | 0.57 | 0.69 | 0.94 | 0.71 | 0.7 |
| 9 | 37690945 | rs10973491 | 0.806 | -0.229 | -1.026 | 0.317 | -0.001 | -0.018 | -0.018 | 0 | 0.22 | 0.17 | 0.26 | 0.41 | 0.26 | 0.14 |
| 9 | 37692654 | rs3747539 | 0.582 | -0.109 | -0.983 | 0.357 | -0.003 | 0.005 | 0.005 | 0 | 0.2 | 0.1 | 0.09 | 0.16 | 0.19 | 0.17 |
| 9 | 37694070 | rs7028875 | -0.286 | -0.237 | -0.988 | 0.464 | -0.024 | 0.07 | 0.07 | 1.155 | 0.4 | 0.18 | 0.27 | 0.21 | 0.44 | 0.2 |
| 9 | 37702874 | rs10511942 | 0.785 | -0.186 | -1.342 | 0.495 | -0.025 | -0.01 | -0.01 | 0 | 0.19 | 0.04 | 0.09 | 0.17 | 0.18 | 0.17 |
| 9 | 37707274 | rs6476659 | 1.055 | -0.16 | -1.342 | 0.501 | -0.021 | 0.053 | 0.053 | 1.23 | 0.2 | 0.09 | 0.18 | 0.05 | 0.25 | 0.04 |
| 9 | 37708840 | rs6476660 | -1.688 | -0.225 | -1.146 | 0.431 | -0.006 | -0.082 | -0.082 | 0 | 0.67 | 0.59 | 0.7 | 0.95 | 0.7 | 0.69 |
| 9 | 37711594 | rs2296553 | -0.291 | -0.289 | -1.04 | 0.422 | -0.007 | 0.005 | 0.005 | 0.66 | 0.41 | 0.34 | 0.36 | 0.58 | 0.44 | 0.3 |
| 9 | 37712391 | rs7039767 | 0.862 | -0.231 | -1.036 | 0.48 | -0.018 | -0.018 | -0.018 | 0 | 0.16 | 0.12 | 0.31 | 0.32 | 0.23 | 0.31 |
| 9 | 37715724 | rs1952125 | -0.98 | -0.212 | -0.992 | 0.398 | 0 | -0.022 | -0.022 | 0 | 0.48 | 0.53 | 0.53 | 0.9 | 0.45 | 0.66 |
| 9 | 37719473 | rs10738986 | -1.424 | 0.34 | -1.274 | 0.465 | -0.009 | 0.03 | 0.03 | 0 | 0.64 | 0.51 | 0.62 | 0.58 | 0.69 | 0.57 |
| 9 | 37723330 | rs4878173 | 0.199 | 0.816 | -1.121 | 0.978 | -0.018 | -0.021 | -0.021 | 0 | 0.15 | 0.11 | 0.32 | 0.31 | 0.23 | 0.29 |
| 9 | 37724195 | rs10814609 | -0.461 | 0.627 | -0.98 | 0.979 | -0.01 | -0.003 | -0.003 | 2.145 | 0.36 | 0.32 | 0.42 | 0.46 | 0.41 | 0.47 |
| 9 | 37733013 | rs2026546 | -0.625 | 0.661 | -1.354 | 1.342 | -0.022 | -0.033 | -0.033 | 0 | 0.32 | 0.15 | 0.41 | 0.44 | 0.33 | 0.33 |
| 9 | 37733925 | rs750693 | -0.861 | 1.084 | -1.916 | 1.547 | -0.075 | -0.122 | -0.122 | 0.328 | 0.2 | 0.05 | 0.32 | 0.44 | 0.28 | 0.49 |

| | | | | | | | | | | | | | | | | |
|---|---|---|---|---|---|---|---|---|---|---|---|---|---|---|---|---|
| 9 | 37739932 | rs10758456 | -0.483 | 0.718 | -0.83 | 1.45 | 0.008 | 0.103 | 0.103 | 0.102 | 0.12 | 0.28 | 0.18 | 0.33 | 0.11 | 0.24 |
| 9 | 37740637 | rs2281872 | -1.278 | 0.7 | -0.735 | 0.492 | 0 | -0.001 | -0.001 | 0 | 0.6 | 0.63 | 0.65 | 0.54 | 0.61 | 0.68 |
| 9 | 37745413 | rs3747541 | -0.657 | 0.639 | -0.693 | 0.264 | -0.003 | -0.025 | -0.025 | 4.704 | 0.37 | 0.58 | 0.42 | 0.2 | 0.33 | 0.34 |
| 9 | 37747985 | rs10973517 | 0.267 | 0.557 | -0.046 | 0.157 | -0.027 | -0.094 | -0.094 | 0 | 0.25 | 0.42 | 0.26 | 0.1 | 0.18 | 0.11 |
| 9 | 37749213 | rs10511944 | -0.071 | 0.637 | 0.3 | 0.233 | 0.026 | 0.014 | 0.014 | 0 | 0.13 | 0.35 | 0.18 | 0.1 | 0.15 | 0.25 |
| 9 | 37751219 | rs10973520 | -1.262 | 0.67 | 0.466 | 0.146 | 0.001 | -0.007 | -0.007 | 0 | 0.48 | 0.68 | 0.61 | 0.52 | 0.49 | 0.6 |
| 9 | 37752907 | rs7873428 | -0.637 | 0.564 | 0.694 | -0.096 | 0.018 | -0.044 | -0.044 | 0 | 0.34 | 0.49 | 0.35 | 0.2 | 0.34 | 0.32 |
| 9 | 37752947 | rs7873440 | -1.225 | 0.633 | 0.689 | 0.018 | -0.001 | -0.005 | -0.005 | 0 | 0.46 | 0.58 | 0.6 | 0.54 | 0.49 | 0.59 |
| 9 | 37753058 | rs7861041 | 0.41 | 0.599 | 0.848 | -0.12 | 0.017 | 0.003 | 0.003 | 0.015 | 0.13 | 0.2 | 0.19 | 0.11 | 0.17 | 0.25 |
| 9 | 37755620 | rs10973524 | -0.622 | 0.59 | 0.861 | -0.169 | -0.015 | 0.034 | 0.034 | 0 | 0.32 | 0.28 | 0.34 | 0.19 | 0.33 | 0.32 |
| 9 | 37768742 | rs10115653 | -0.773 | -0.272 | 0.394 | -0.899 | -0.004 | -0.012 | -0.012 | 0 | 0.54 | 0.46 | 0.52 | 0.24 | 0.58 | 0.45 |
| 9 | 37771325 | rs7035295 | 0.696 | -0.588 | 0.177 | -1.054 | -0.008 | 0.004 | 0.004 | 0.033 | 0.17 | 0.14 | 0.21 | 0.12 | 0.17 | 0.28 |
| 9 | 37774893 | rs10973537 | 0.536 | -0.594 | 0.191 | -1.06 | -0.001 | 0 | 0 | 0 | 0.17 | 0.22 | 0.22 | 0.12 | 0.17 | 0.28 |
| 9 | 37775467 | rs4878174 | -0.532 | -0.62 | 0.233 | -1.058 | 0.005 | 0.006 | 0.006 | 0 | 0.5 | 0.46 | 0.49 | 0.22 | 0.57 | 0.42 |
| 9 | 37780558 | rs1059059 | 0.525 | -0.613 | 0.179 | -1.081 | -0.007 | -0.075 | -0.075 | 0 | 0.22 | 0.38 | 0.17 | 0.09 | 0.17 | 0.07 |
| 9 | 37788911 | rs10814620 | -0.226 | -0.906 | -0.016 | -1.294 | 0.006 | -0.038 | -0.038 | 0 | 0.41 | 0.5 | 0.4 | 0.15 | 0.42 | 0.41 |
| 9 | 37796342 | rs927631 | 1.759 | -0.96 | -0.151 | -1.32 | -0.049 | 0.049 | 0.049 | 0 | 0.15 | 0.01 | 0.12 | 0.07 | 0.16 | 0.04 |
| 9 | 37797412 | rs10120822 | 1.133 | -0.987 | 0.064 | -1.333 | 0.013 | -0.076 | -0.076 | 0 | 0.18 | 0.32 | 0.14 | 0.07 | 0.16 | 0.04 |
| 9 | 37806245 | rs872778 | 1.169 | -0.987 | 0.072 | -1.333 | 0.008 | -0.065 | -0.065 | 0 | 0.18 | 0.28 | 0.13 | 0.07 | 0.16 | 0.04 |
| 9 | 37814873 | rs2841954 | 0.734 | -1.047 | 0 | -1.383 | -0.018 | 0.029 | 0.029 | 0 | 0.34 | 0.29 | 0.35 | 0.13 | 0.41 | 0.39 |
| 9 | 37820416 | rs2841946 | 0.77 | -1.037 | -0.026 | -1.378 | -0.026 | 0.042 | 0.042 | 0 | 0.34 | 0.27 | 0.35 | 0.13 | 0.41 | 0.39 |
| 9 | 37861794 | rs10115457 | 0.682 | -0.847 | 0.116 | -1.169 | 0.012 | -0.077 | -0.077 | 0.061 | 0.18 | 0.32 | 0.14 | 0.07 | 0.16 | 0.04 |
| 9 | 37865943 | rs7859075 | 0.682 | -0.847 | 0.116 | -1.169 | 0.012 | -0.077 | -0.077 | 0 | 0.18 | 0.32 | 0.14 | 0.07 | 0.16 | 0.04 |
| 9 | 37866551 | rs10511954 | 1.089 | -0.831 | -0.018 | -1.161 | -0.029 | 0.058 | 0.058 | 0 | 0.16 | 0.05 | 0.12 | 0.07 | 0.16 | 0.04 |
| 9 | 37874090 | rs7873455 | 0.801 | -0.939 | -0.074 | -1.12 | 0.034 | -0.021 | -0.021 | 0 | 0.12 | 0.2 | 0.12 | 0.08 | 0.15 | 0.04 |
| 9 | 37886429 | rs13294843 | 0.675 | -0.951 | -0.211 | -0.842 | -0.023 | -0.064 | -0.064 | 0 | 0.13 | 0.03 | 0.25 | 0.34 | 0.15 | 0.25 |
| 9 | 37889241 | rs2025440 | -1.89 | -0.959 | -0.23 | -1.251 | -0.007 | 0.016 | 0.016 | 0.015 | 0.51 | 0.66 | 0.64 | 0.57 | 0.48 | 0.69 |
| 9 | 37889858 | rs2025439 | 0.601 | -0.38 | -0.539 | -0.872 | -0.014 | -0.049 | -0.049 | 0 | 0.13 | 0.04 | 0.25 | 0.34 | 0.15 | 0.25 |
| 9 | 37911531 | rs10973607 | 0.925 | -0.299 | -0.423 | -0.925 | -0.057 | -0.032 | -0.032 | 3.382 | 0.12 | 0.01 | 0.17 | 0.11 | 0.16 | 0.25 |
| 9 | 37916408 | rs3747547 | -0.369 | -0.367 | 0.278 | -1.037 | -0.001 | -0.005 | -0.005 | 0 | 0.14 | 0.3 | 0.09 | 0.1 | 0.1 | 0.13 |
| 9 | 37919743 | rs2281784 | -0.237 | -0.422 | -0.129 | -1.016 | -0.001 | -0.003 | -0.003 | 0 | 0.06 | 0.06 | 0.04 | 0.11 | 0.01 | 0.04 |
| 9 | 37932314 | rs1082865 | -1.102 | -0.544 | -0.001 | -1.052 | 0 | 0.026 | 0.026 | 0 | 0.76 | 0.73 | 0.76 | 0.57 | 0.71 | 0.84 |
| 9 | 37934140 | rs776023 | 0.031 | 0.39 | -0.085 | -1.247 | -0.003 | -0.026 | -0.026 | 0 | 0.58 | 0.62 | 0.4 | 0.07 | 0.51 | 0.51 |
| 9 | 37934780 | rs776022 | 2.01 | 0.233 | -0.122 | -1.239 | -0.001 | -0.079 | -0.079 | 2.95 | 0.31 | 0.36 | 0.22 | 0.01 | 0.28 | 0.18 |
| 9 | 37939673 | rs776019 | -0.229 | 0.034 | -0.213 | -0.913 | -0.001 | -0.011 | -0.011 | 3.842 | 0.61 | 0.62 | 0.49 | 0.22 | 0.56 | 0.58 |
| 9 | 37942152 | rs776015 | -0.195 | 0.059 | -0.236 | -0.915 | -0.001 | -0.016 | -0.016 | 0 | 0.6 | 0.6 | 0.49 | 0.22 | 0.56 | 0.59 |
| 9 | 37943571 | rs7027014 | -0.449 | -0.068 | -0.501 | -0.385 | 0.008 | 0.007 | 0.007 | 0 | 0.16 | 0.13 | 0.1 | 0.13 | 0.11 | 0.15 |
| 9 | 37944427 | rs735740 | -1.022 | -0.082 | -0.783 | -0.215 | 0.016 | -0.014 | -0.014 | 0 | 0.17 | 0.12 | 0.19 | 0.15 | 0.12 | 0.22 |
| 9 | 37949638 | rs7864512 | -0.645 | 0.01 | -0.543 | -0.147 | 0.009 | 0.006 | 0.006 | 0 | 0.16 | 0.18 | 0.11 | 0.14 | 0.11 | 0.15 |
| 9 | 37950936 | rs2103241 | 0.173 | -0.059 | -0.557 | -0.148 | 0.002 | -0.003 | -0.003 | 0 | 0.12 | 0.1 | 0.08 | 0.12 | 0.1 | 0.13 |
| 9 | 37951176 | rs2091415 | -0.655 | -0.058 | -0.489 | -0.147 | 0.004 | 0.003 | 0.003 | 0 | 0.16 | 0.21 | 0.11 | 0.14 | 0.11 | 0.15 |
| 9 | 37956271 | rs7044060 | -0.944 | -0.016 | -0.468 | -0.388 | -0.018 | 0.084 | 0.084 | 0 | 0.54 | 0.66 | 0.6 | 0.82 | 0.54 | 0.55 |
| 9 | 37974481 | rs3802414 | -0.781 | -0.072 | -0.516 | -0.378 | -0.028 | 0.107 | 0.107 | 0 | 0.53 | 0.72 | 0.51 | 0.78 | 0.53 | 0.5 |
| 9 | 37974743 | rs1138374 | -0.755 | -0.076 | -0.518 | -0.38 | -0.027 | 0.11 | 0.11 | 0 | 0.52 | 0.72 | 0.51 | 0.78 | 0.53 | 0.5 |
| 9 | 37990801 | rs2149547 | 0.222 | -0.172 | 0.012 | -0.275 | 0.003 | -0.005 | -0.005 | 1.25 | 0.37 | 0.39 | 0.42 | 0.71 | 0.42 | 0.37 |
| 9 | 37993303 | rs11789441 | 1.253 | -0.184 | -0.066 | -0.332 | -0.019 | 0.031 | 0.031 | 0.11 | 0.1 | 0.01 | 0.06 | 0.02 | 0.1 | 0.11 |
| 9 | 37995845 | rs10973634 | 0.563 | -0.157 | -0.038 | -0.314 | -0.025 | 0.104 | 0.104 | 0 | 0.48 | 0.29 | 0.49 | 0.23 | 0.47 | 0.5 |
| 9 | 37996724 | rs10973635 | 0.979 | -0.067 | 0.08 | 0.048 | -0.005 | -0.014 | -0.014 | 0 | 0.13 | 0.04 | 0.17 | 0.31 | 0.14 | 0.23 |
| 9 | 38003567 | rs12345885 | -0.426 | 0.201 | -0.118 | -0.222 | 0.003 | 0.006 | 0.006 | 0.621 | 0.22 | 0.33 | 0.33 | 0.22 | 0.22 | 0.31 |
| 9 | 38020544 | rs4878737 | -1.936 | 0.176 | 0 | -0.113 | 0.005 | 0.051 | 0.051 | 0 | 0.78 | 0.71 | 0.75 | 0.58 | 0.72 | 0.86 |
| 9 | 38023500 | rs7856790 | 1.358 | -0.004 | -0.307 | -0.38 | 0.018 | 0.042 | 0.042 | 0 | 0.36 | 0.35 | 0.3 | 0.15 | 0.29 | 0.24 |
| 9 | 38027644 | rs943936 | 1.173 | -0.298 | 0.317 | 0.167 | 0.003 | 0.003 | 0.003 | 0 | 0.76 | 0.73 | 0.77 | 0.82 | 0.82 | 0.73 |
| 9 | 38040095 | rs7025303 | 0.78 | -0.714 | 0.517 | 0.562 | 0.028 | 0.138 | 0.138 | 0 | 0.54 | 0.2 | 0.44 | 0.13 | 0.57 | 0.57 |
| 9 | 38058424 | rs4878743 | -0.737 | -0.557 | 0.85 | 0.954 | -0.003 | 0.032 | 0.032 | 0 | 0.72 | 0.64 | 0.63 | 0.46 | 0.7 | 0.81 |
| 9 | 38074715 | rs1928251 | -1.412 | -1.423 | 0.716 | 0.874 | -0.076 | 0.031 | 0.031 | 1.361 | 0.14 | 0.45 | 0.22 | 0.16 | 0.02 | 0.07 |
| 9 | 38081261 | rs4262388 | 0.073 | -0.65 | 0.714 | 1.049 | -0.004 | 0.034 | 0.034 | 0 | 0.69 | 0.64 | 0.63 | 0.37 | 0.66 | 0.66 |
| 9 | 38085784 | rs7037490 | -1.092 | -0.454 | 0.758 | 1.115 | -0.008 | 0.045 | 0.045 | 0 | 0.81 | 0.73 | 0.79 | 0.69 | 0.8 | 0.86 |
| 9 | 38089638 | rs1928233 | 1.185 | -0.535 | 0.511 | 0.895 | 0.007 | 0.062 | 0.062 | 0 | 0.49 | 0.28 | 0.51 | 0.34 | 0.52 | 0.53 |
| 9 | 38089969 | rs7024427 | 0.386 | -0.975 | 0.953 | 0.823 | -0.012 | 0.005 | 0.005 | 0 | 0.63 | 0.58 | 0.61 | 0.55 | 0.65 | 0.65 |
| 9 | 38090793 | rs7021766 | -1.107 | -0.871 | 0.676 | 0.946 | -0.004 | 0.003 | 0.003 | 0.044 | 0.17 | 0.23 | 0.09 | 0.17 | 0.12 | 0.16 |
| 9 | 38098363 | rs4878750 | 0.312 | -0.947 | 0.675 | 0.91 | -0.013 | -0.007 | -0.007 | 0.011 | 0.64 | 0.42 | 0.7 | 0.71 | 0.72 | 0.69 |
| 9 | 38114557 | rs12552637 | 1.663 | -0.906 | 0.52 | 0.96 | -0.011 | 0.003 | 0.003 | 0.189 | 0.46 | 0.13 | 0.5 | 0.37 | 0.54 | 0.52 |
| 9 | 38125272 | rs11791344 | -0.516 | -0.764 | 0.547 | 0.971 | -0.01 | -0.026 | -0.026 | 0.001 | 0.08 | 0.02 | 0.06 | 0.17 | 0.08 | 0.15 |
| 9 | 38128597 | rs4878753 | 0.042 | -0.761 | 0.751 | 0.98 | 0.002 | 0.016 | 0.016 | 0 | 0.17 | 0.18 | 0.2 | 0.31 | 0.18 | 0.15 |
| 9 | 38129670 | rs7039254 | 1.373 | -0.838 | 0.721 | 0.951 | 0.013 | 0.016 | 0.016 | 0.117 | 0.47 | 0.32 | 0.52 | 0.4 | 0.54 | 0.54 |
| 9 | 38135564 | rs885431 | 1.293 | -0.843 | 0.726 | 0.883 | 0.011 | 0.01 | 0.01 | 0.047 | 0.48 | 0.34 | 0.52 | 0.4 | 0.55 | 0.54 |
| 9 | 38140772 | rs10973681 | -0.087 | -0.793 | 0.42 | 0.926 | 0.002 | 0.02 | 0.02 | 0 | 0.18 | 0.22 | 0.19 | 0.31 | 0.19 | 0.15 |
| 9 | 38141369 | rs10814661 | 1.505 | -0.811 | -0.057 | 0.877 | -0.006 | 0.006 | 0.006 | 0.487 | 0.46 | 0.17 | 0.51 | 0.38 | 0.54 | 0.52 |
| 9 | 38144829 | rs1928242 | -0.833 | -0.373 | 0.058 | 0.947 | 0.013 | 0.074 | 0.074 | 0 | 0.23 | 0.34 | 0.25 | 0.46 | 0.25 | 0.27 |

| | | | | | | | | | | | | | | | | |
|---|---|---|---|---|---|---|---|---|---|---|---|---|---|---|---|---|
| 9 | 38150167 | rs943935 | -0.737 | -0.363 | 0.136 | -0.049 | -0.023 | -0.07 | -0.07 | 0 | 0.77 | 0.72 | 0.87 | 0.93 | 0.82 | 0.85 |
| 9 | 38151268 | rs10973683 | 0.412 | 0.062 | -0.701 | 0.415 | 0.037 | 0.069 | 0.069 | 0.054 | 0.52 | 0.29 | 0.65 | 0.66 | 0.58 | 0.6 |
| 9 | 38154137 | rs7859035 | 0.099 | -0.038 | -0.518 | -0.071 | 0.016 | 0.037 | 0.037 | 0 | 0.58 | 0.42 | 0.7 | 0.81 | 0.64 | 0.72 |
| 9 | 38159475 | rs12004825 | -0.966 | -0.152 | 0.009 | 0.377 | -0.025 | 0.023 | 0.023 | 0 | 0.06 | 0.18 | 0.04 | 0.08 | 0.01 | 0.05 |
| 9 | 38164364 | rs10441687 | 0.036 | -0.075 | -0.586 | -0.356 | 0.003 | 0.164 | 0.164 | 0.005 | 0.48 | 0.22 | 0.53 | 0.74 | 0.43 | 0.52 |
| 9 | 38165646 | rs2165776 | 0.619 | 0.343 | -0.434 | -0.974 | -0.019 | 0.049 | 0.049 | 1.049 | 0.26 | 0.14 | 0.15 | 0.01 | 0.3 | 0.22 |
| 9 | 38168321 | rs10758476 | 0.295 | 0.347 | 0.149 | -1.102 | -0.002 | -0.026 | -0.026 | 0 | 0.6 | 0.44 | 0.67 | 0.92 | 0.53 | 0.59 |
| 9 | 38176640 | rs12235873 | 0.71 | 0.23 | -0.337 | 0.087 | -0.002 | 0.017 | 0.017 | 0 | 0.07 | 0.03 | 0.05 | 0.32 | 0.05 | 0.14 |
| 9 | 38178570 | rs4242645 | 0.206 | 0.212 | 0.363 | -0.137 | 0.015 | -0.027 | -0.027 | 0.388 | 0.76 | 0.53 | 0.7 | 0.77 | 0.74 | 0.73 |
| 9 | 38179054 | rs10758477 | 0.784 | 0.095 | 0.244 | 0.009 | -0.003 | 0.102 | 0.102 | 0.005 | 0.45 | 0.21 | 0.5 | 0.67 | 0.42 | 0.47 |
| 9 | 38182929 | rs4878759 | 0.829 | 0.231 | -0.388 | -0.107 | -0.048 | -0.047 | -0.047 | 4.96 | 0.14 | 0.01 | 0.2 | 0.12 | 0.15 | 0.22 |
| 9 | 38186096 | rs4878761 | 0.2 | 0.222 | -0.144 | -0.25 | 0.01 | 0.075 | 0.075 | 0 | 0.5 | 0.39 | 0.55 | 0.78 | 0.44 | 0.51 |
| 9 | 38187008 | rs10973694 | 0 | 0.225 | 0.095 | -0.238 | 0.001 | 0.007 | 0.007 | 0 | 0.52 | 0.45 | 0.55 | 0.75 | 0.44 | 0.5 |
| 9 | 38189527 | rs894520 | -0.488 | 0.124 | 0.015 | -0.458 | -0.002 | -0.033 | -0.033 | 0 | 0.33 | 0.5 | 0.38 | 0.28 | 0.33 | 0.42 |
| 9 | 38190816 | rs4878763 | -1.656 | 0.513 | 0.14 | -0.405 | -0.011 | 0.004 | 0.004 | 0 | 0.83 | 0.9 | 0.88 | 1 | 0.77 | 0.88 |
| 9 | 38191117 | rs4878764 | -1.588 | 0.513 | 0.15 | -0.406 | 0.002 | -0.037 | -0.037 | 0 | 0.82 | 0.86 | 0.88 | 1 | 0.77 | 0.88 |
| 9 | 38196563 | rs10118655 | -0.995 | 0.563 | 0.233 | -0.087 | 0.001 | -0.002 | -0.002 | 0 | 0.44 | 0.48 | 0.55 | 0.35 | 0.46 | 0.63 |
| 9 | 38197384 | rs12352894 | 0.724 | 0.565 | 0.065 | -0.107 | -0.033 | -0.041 | -0.041 | 2.165 | 0.14 | 0.02 | 0.19 | 0.12 | 0.15 | 0.24 |
| 9 | 38198629 | rs10973703 | 0.324 | 0.494 | 0.23 | -0.114 | -0.024 | -0.043 | -0.043 | 1.058 | 0.24 | 0.08 | 0.31 | 0.3 | 0.22 | 0.31 |
| 9 | 38199821 | rs2810747 | -0.668 | 0.347 | 0.536 | 0.006 | 0 | -0.001 | -0.001 | 0.386 | 0.45 | 0.5 | 0.56 | 0.46 | 0.48 | 0.64 |
| 9 | 38205001 | rs2053553 | -1.211 | 0.553 | 0.685 | -0.353 | 0.027 | -0.091 | -0.091 | 0.446 | 0.85 | 0.72 | 0.88 | 1 | 0.8 | 0.92 |
| 9 | 38207733 | rs920706 | 0.339 | 0.484 | 0.634 | -0.314 | 0.002 | -0.039 | -0.039 | 0 | 0.68 | 0.62 | 0.78 | 0.82 | 0.64 | 0.59 |
| 9 | 38208693 | rs10118416 | 0.911 | 0.531 | 0.486 | 0.081 | -0.009 | 0 | 0 | 0 | 0.39 | 0.44 | 0.28 | 0.38 | 0.31 | 0.25 |
| 9 | 38209247 | rs2810729 | 0.522 | 0.697 | 0.367 | 0.298 | -0.007 | -0.004 | -0.004 | 0.768 | 0.59 | 0.65 | 0.42 | 0.44 | 0.54 | 0.36 |
| 9 | 38210621 | rs920707 | 0.153 | 0.957 | -0.165 | 0.265 | 0.019 | 0.035 | 0.035 | 0 | 0.31 | 0.42 | 0.49 | 0.39 | 0.31 | 0.38 |
| 9 | 38214047 | rs2810726 | -0.772 | 0.981 | -0.555 | 0.246 | -0.004 | 0.006 | 0.006 | 0 | 0.45 | 0.62 | 0.61 | 0.63 | 0.48 | 0.61 |
| 9 | 38225264 | rs13300331 | -0.467 | 0.724 | -0.977 | 0.367 | -0.01 | -0.017 | -0.017 | 0.901 | 0.12 | 0.06 | 0.09 | 0.23 | 0.15 | 0.27 |
| 9 | 38227021 | rs4878187 | -0.48 | 0.706 | -0.975 | 0.129 | -0.001 | -0.001 | -0.001 | 0 | 0.12 | 0.1 | 0.13 | 0.19 | 0.12 | 0.22 |
| 9 | 38232043 | rs4271056 | 0.608 | 0.758 | -0.771 | -0.126 | 0.001 | -0.045 | -0.045 | 4.822 | 0.17 | 0.21 | 0.14 | 0.05 | 0.2 | 0.14 |
| 9 | 38233891 | rs894525 | 0.305 | 1.001 | -1.227 | -0.11 | 0.002 | 0.028 | 0.028 | 0 | 0.33 | 0.52 | 0.42 | 0.37 | 0.31 | 0.37 |
| 9 | 38235659 | rs1006087 | -0.535 | 0.895 | -1.135 | -0.175 | -0.001 | 0.01 | 0.01 | 0 | 0.52 | 0.68 | 0.56 | 0.42 | 0.52 | 0.54 |
| 9 | 38244221 | rs2810723 | 0.195 | 0.725 | -0.677 | 0.188 | -0.005 | -0.042 | -0.042 | 0.293 | 0.69 | 0.61 | 0.74 | 0.8 | 0.7 | 0.67 |
| 9 | 38250950 | rs2585692 | 0.266 | 1.44 | -0.95 | 0.714 | -0.023 | -0.025 | -0.025 | 1.224 | 0.76 | 0.83 | 0.6 | 0.58 | 0.67 | 0.56 |
| 9 | 38251684 | rs10511947 | -0.105 | 0.565 | -0.671 | 0.413 | -0.002 | -0.002 | -0.002 | 0 | 0.22 | 0.21 | 0.14 | 0.13 | 0.2 | 0.23 |
| 9 | 38252526 | rs4388534 | 0.682 | 0.28 | -0.784 | -0.127 | 0.027 | 0.094 | 0.094 | 0.23 | 0.47 | 0.32 | 0.65 | 0.74 | 0.54 | 0.61 |
| 9 | 38254018 | rs7045845 | 0.526 | 0.233 | -0.663 | -0.39 | -0.006 | 0.002 | 0.002 | 0.016 | 0.19 | 0.14 | 0.14 | 0 | 0.24 | 0.12 |
| 9 | 38257360 | rs2810762 | 0.925 | 0.596 | 0.117 | -0.039 | 0.02 | 0.024 | 0.024 | 0.408 | 0.78 | 0.48 | 0.68 | 0.74 | 0.76 | 0.53 |
| 9 | 38260952 | rs2810760 | 1.09 | 0.755 | 0.576 | -0.155 | 0.003 | 0.014 | 0.014 | 1.197 | 0.46 | 0.4 | 0.46 | 0.68 | 0.41 | 0.55 |
| 9 | 38272467 | rs1923425 | 0.834 | 0.589 | 1.003 | 0.217 | 0.035 | 0.148 | 0.148 | 0 | 0.9 | 0.77 | 0.76 | 0.72 | 0.85 | 0.86 |
| 9 | 38273626 | rs570941 | 0.813 | 0.591 | 1.004 | 0.218 | 0.037 | 0.153 | 0.153 | 0 | 0.9 | 0.77 | 0.76 | 0.72 | 0.85 | 0.86 |
| 9 | 38280691 | rs189315 | 0.804 | 0.596 | 1.034 | 0.114 | 0.034 | 0.129 | 0.129 | 0 | 0.9 | 0.79 | 0.76 | 0.74 | 0.85 | 0.88 |
| 9 | 38282317 | rs17443417 | 0.965 | 0.363 | 0.809 | -0.432 | -0.01 | 0.026 | 0.026 | 0 | 0.55 | 0.57 | 0.49 | 0.69 | 0.46 | 0.74 |
| 9 | 38282692 | rs4878777 | 0.473 | 0.341 | 0.828 | -0.472 | 0.007 | -0.027 | -0.027 | 0 | 0.77 | 0.63 | 0.71 | 0.88 | 0.72 | 0.82 |
| 9 | 38284646 | rs12238450 | 0.248 | 0.308 | 0.518 | -0.585 | -0.005 | 0.046 | 0.046 | 0 | 0.06 | 0.08 | 0.07 | 0.22 | 0.04 | 0.08 |
| 9 | 38285290 | rs341485 | 0.522 | 0.27 | 0.596 | -1.169 | -0.04 | -0.128 | -0.128 | 0 | 0.77 | 0.54 | 0.83 | 0.96 | 0.82 | 0.76 |
| 9 | 38286537 | rs12551041 | -0.191 | 0.689 | -0.129 | -0.851 | 0.005 | 0.021 | 0.021 | 0 | 0.25 | 0.17 | 0.16 | 0.04 | 0.24 | 0.24 |
| 9 | 38289065 | rs10973730 | 0.944 | 0.634 | -0.293 | -1.053 | -0.012 | 0.074 | 0.074 | 0 | 0.57 | 0.72 | 0.6 | 0.86 | 0.47 | 0.6 |
| 9 | 38290172 | rs1348335 | -0.104 | 0.847 | -0.244 | -0.839 | 0.024 | 0.037 | 0.037 | 0 | 0.17 | 0.03 | 0.15 | 0.07 | 0.16 | 0.18 |
| 9 | 38290694 | rs4878191 | 1.509 | 0.762 | 0.268 | -0.482 | 0.009 | 0.002 | 0.002 | 0 | 0.45 | 0.39 | 0.5 | 0.52 | 0.43 | 0.48 |
| 9 | 38293885 | rs10973731 | -0.766 | 1.294 | 1.253 | 0.252 | 0.021 | -0.002 | -0.002 | 0 | 0.07 | 0.07 | 0.1 | 0.07 | 0.13 | 0.06 |
| 9 | 38297774 | rs2840227 | 1.729 | 1.162 | 1.443 | 0.199 | -0.001 | -0.002 | -0.002 | 0 | 0.58 | 0.57 | 0.6 | 0.53 | 0.55 | 0.49 |
| 9 | 38298233 | rs726363 | 1.157 | 1.146 | 1.442 | 0.309 | 0.001 | 0 | 0 | 0.828 | 0.67 | 0.65 | 0.71 | 0.72 | 0.7 | 0.56 |
| 9 | 38298279 | rs763935 | 2.025 | 1.126 | 1.354 | 0.333 | -0.011 | -0.015 | -0.015 | 0 | 0.55 | 0.25 | 0.58 | 0.53 | 0.54 | 0.48 |
| 9 | 38299419 | rs1442977 | 1.62 | 0.32 | 1.487 | 0.453 | -0.005 | 0.001 | 0.001 | 0 | 0.61 | 0.45 | 0.66 | 0.59 | 0.65 | 0.52 |
| 9 | 38303243 | rs4878782 | 1.034 | 0.427 | 1.191 | 0.558 | -0.004 | 0.031 | 0.031 | 0.671 | 0.86 | 0.69 | 0.87 | 0.76 | 0.87 | 0.75 |
| 9 | 38309332 | rs1885490 | -0.859 | 0.56 | 0.787 | 0.181 | -0.002 | -0.001 | -0.001 | 3.266 | 0.29 | 0.22 | 0.26 | 0.24 | 0.29 | 0.34 |
| 9 | 38309523 | rs1885492 | 0.52 | 0.547 | 0.84 | 0.243 | -0.001 | -0.009 | -0.009 | 0 | 0.88 | 0.82 | 0.9 | 0.87 | 0.88 | 0.8 |
| 9 | 38312365 | rs560512 | -0.475 | 0.57 | 0.496 | 0.308 | 0 | 0.002 | 0.002 | 0 | 0.22 | 0.26 | 0.17 | 0.2 | 0.2 | 0.25 |
| 9 | 38316331 | rs4878783 | 1.253 | 0.735 | 0.92 | 0.077 | -0.002 | -0.025 | -0.025 | 1.646 | 0.75 | 0.49 | 0.78 | 0.78 | 0.75 | 0.66 |
| 9 | 38318134 | rs341466 | -0.548 | 0.369 | 0.824 | 0.008 | -0.03 | -0.027 | -0.027 | 0 | 0.14 | 0.25 | 0.09 | 0.09 | 0.07 | 0.13 |
| 9 | 38328183 | rs4878784 | 1.594 | 0.919 | 0.378 | 0.333 | -0.033 | 0.032 | 0.032 | 0 | 0.45 | 0.22 | 0.51 | 0.44 | 0.5 | 0.34 |
| 9 | 38330114 | rs16935064 | -0.895 | 1.901 | 0.75 | 0.01 | 0.034 | -0.036 | -0.036 | 0 | 0.14 | 0.22 | 0.17 | 0.04 | 0.21 | 0.1 |
| 9 | 38333064 | rs11792523 | -1.294 | 1.83 | 0.725 | 0.03 | 0.025 | -0.013 | -0.013 | 0 | 0.18 | 0.23 | 0.25 | 0.16 | 0.27 | 0.24 |
| 9 | 38335032 | rs10973753 | -0.134 | 1.669 | 0.602 | 0.151 | 0.001 | -0.029 | -0.029 | 3.149 | 0.09 | 0.1 | 0.1 | 0 | 0.11 | 0.04 |
| 9 | 38335475 | rs1885493 | -1.141 | 1.852 | 1.077 | 0.408 | 0.123 | 0.02 | 0.02 | 0 | 0.09 | 0.29 | 0.12 | 0.07 | 0.15 | 0.13 |
| 9 | 38337291 | rs1555481 | 1.089 | 2.035 | 0.698 | 0.682 | 0.041 | 0.03 | 0.03 | 0.212 | 0.8 | 0.68 | 0.76 | 0.71 | 0.7 | 0.7 |
| 9 | 38342744 | rs10732359 | -1.318 | 1.223 | 0.44 | 0.586 | 0 | 0 | 0 | 0 | 0.53 | 0.44 | 0.4 | 0.51 | 0.46 | 0.63 |
| 9 | 38355452 | rs10814680 | 0.868 | 1.353 | 0.624 | 0.401 | 0.002 | 0.027 | 0.027 | 0.152 | 0.51 | 0.36 | 0.53 | 0.65 | 0.52 | 0.46 |
| 9 | 38357065 | rs4878790 | 1.359 | 1.447 | -0.367 | 0.665 | -0.015 | -0.071 | -0.071 | 0.165 | 0.38 | 0.12 | 0.43 | 0.51 | 0.39 | 0.32 |

| | | | | | | | | | | | | | | | | |
|---|---|---|---|---|---|---|---|---|---|---|---|---|---|---|---|---|
| 9 | 38360890 | rs10973759 | 0.268 | 1.232 | -0.487 | 0.42 | -0.011 | 0.017 | 0.017 | 0 | 0.17 | 0.12 | 0.25 | 0.12 | 0.19 | 0.23 |
| 9 | 38361379 | rs4878791 | 1.506 | 1.518 | -0.528 | 0.576 | -0.019 | -0.081 | -0.081 | 0.074 | 0.38 | 0.12 | 0.43 | 0.51 | 0.4 | 0.33 |
| 9 | 38362171 | rs1004604 | -0.406 | 1.229 | -0.306 | 0.349 | 0.011 | -0.017 | -0.017 | 0.053 | 0.28 | 0.51 | 0.28 | 0.15 | 0.29 | 0.36 |
| 9 | 38365696 | rs1033794 | -0.83 | 1.234 | -0.912 | 0.557 | -0.028 | 0.093 | 0.093 | 0.597 | 0.54 | 0.78 | 0.53 | 0.23 | 0.49 | 0.59 |
| 9 | 38371434 | rs911755 | 0.808 | 1.023 | -1.019 | 0.648 | -0.009 | -0.068 | -0.068 | 0 | 0.38 | 0.12 | 0.43 | 0.54 | 0.39 | 0.32 |
| 9 | 38372888 | rs6476690 | 0.524 | 0.873 | -0.652 | 0.403 | -0.003 | 0.005 | 0.005 | 0 | 0.41 | 0.37 | 0.5 | 0.23 | 0.45 | 0.24 |
| 9 | 38376173 | rs4878795 | -0.699 | 0.004 | -0.459 | 0.794 | -0.015 | 0.019 | 0.019 | 0 | 0.22 | 0.36 | 0.3 | 0.29 | 0.17 | 0.36 |
| 9 | 38380350 | rs4878797 | -0.131 | -0.088 | -1.28 | 0.977 | -0.042 | -0.028 | -0.028 | 0.745 | 0.27 | 0.09 | 0.22 | 0.23 | 0.28 | 0.24 |
| 9 | 38380389 | rs10973770 | 0.014 | -0.201 | -0.913 | 0.994 | -0.025 | 0.024 | 0.024 | 0 | 0.17 | 0.27 | 0.2 | 0.29 | 0.1 | 0.33 |
| 9 | 38381308 | rs10973772 | -0.9 | -0.071 | -0.573 | 0.779 | 0.006 | 0.001 | 0.001 | 0 | 0.47 | 0.49 | 0.43 | 0.52 | 0.38 | 0.57 |
| 9 | 38382340 | rs7848268 | -0.588 | -0.164 | -0.137 | 1.068 | -0.008 | 0.086 | 0.086 | 0 | 0.23 | 0.46 | 0.16 | 0.34 | 0.23 | 0.18 |
| 9 | 38384578 | rs11790480 | 0.147 | -0.027 | -0.753 | 1.119 | -0.005 | -0.05 | -0.05 | 0.25 | 0.17 | 0.05 | 0.27 | 0.32 | 0.17 | 0.22 |
| 9 | 38386351 | rs1555480 | 0.406 | -0.071 | -0.906 | 0.286 | 0.034 | 0.081 | 0.081 | 0.414 | 0.41 | 0.23 | 0.22 | 0.2 | 0.29 | 0.33 |
| 9 | 38390937 | rs10738997 | 0.758 | 0.111 | -0.028 | 0.427 | 0.007 | 0.036 | 0.036 | 0.711 | 0.77 | 0.42 | 0.67 | 0.75 | 0.79 | 0.69 |
| 9 | 38392256 | rs7040397 | 0.678 | 0.149 | 0.203 | 0.383 | -0.003 | 0.056 | 0.056 | 0.562 | 0.81 | 0.53 | 0.86 | 0.77 | 0.85 | 0.76 |
| 9 | 38392938 | rs4646769 | 0.405 | 0.151 | 0.302 | 0.423 | 0.002 | 0.078 | 0.078 | 0.213 | 0.84 | 0.48 | 0.87 | 0.78 | 0.87 | 0.77 |
| 9 | 38393171 | rs4646770 | 1.072 | 0.53 | -0.679 | 0.307 | -0.093 | -0.039 | -0.039 | 0 | 0.22 | 0.03 | 0.19 | 0.24 | 0.3 | 0.24 |
| 9 | 38393209 | rs4242647 | -0.025 | 0.366 | 0.142 | 0.229 | -0.001 | 0.048 | 0.048 | 0.142 | 0.85 | 0.65 | 0.88 | 0.77 | 0.87 | 0.77 |
| 9 | 38394563 | rs7846941 | 0.837 | 0.599 | 0.108 | 0.293 | -0.001 | 0 | 0 | 0.249 | 0.23 | 0.21 | 0.18 | 0.22 | 0.32 | 0.21 |
| 9 | 38394806 | rs4646771 | 0.151 | 0.386 | 0.082 | 0.673 | 0 | 0.001 | 0.001 | 0 | 0.24 | 0.22 | 0.32 | 0.41 | 0.2 | 0.35 |
| 9 | 38394961 | rs4646772 | 0.441 | 0.465 | -0.034 | 0.717 | -0.01 | -0.071 | -0.071 | 0 | 0.22 | 0.08 | 0.3 | 0.4 | 0.2 | 0.35 |
| 9 | 38395928 | rs2073477 | 0.604 | 0.005 | 0.146 | 0.678 | -0.01 | 0.019 | 0.019 | 0 | 0.23 | 0.23 | 0.3 | 0.39 | 0.14 | 0.31 |
| 9 | 38396002 | rs2228093 | 0.727 | 0.016 | -0.446 | 0.651 | 0.02 | -0.063 | -0.063 | 0 | 0.22 | 0.08 | 0.29 | 0.39 | 0.15 | 0.31 |
| 9 | 38396065 | rs2073478 | -0.346 | 0.425 | -0.568 | 0.055 | 0.028 | 0.099 | 0.099 | 0.15 | 0.52 | 0.26 | 0.36 | 0.3 | 0.61 | 0.48 |
| 9 | 38397915 | rs7859877 | -0.647 | 0.442 | -0.476 | 0.124 | -0.005 | -0.002 | -0.002 | 0 | 0.55 | 0.51 | 0.52 | 0.42 | 0.64 | 0.46 |
| 9 | 38398826 | rs4646774 | -0.57 | 0.464 | -0.309 | -0.458 | -0.011 | -0.008 | -0.008 | 0 | 0.26 | 0.36 | 0.27 | 0.2 | 0.21 | 0.29 |
| 9 | 38399721 | rs7853954 | -0.007 | 0.567 | -0.827 | -0.632 | 0.001 | 0.052 | 0.052 | 0 | 0.18 | 0.06 | 0.07 | 0.09 | 0.14 | 0.19 |
| 9 | 38404105 | rs4878805 | -0.043 | 0.508 | -0.273 | -1.06 | -0.004 | -0.027 | -0.027 | 0.052 | 0.36 | 0.41 | 0.37 | 0.21 | 0.35 | 0.26 |
| 9 | 38404260 | rs4878807 | -1.202 | 1.01 | -0.529 | -1.352 | -0.014 | -0.013 | -0.013 | 0 | 0.59 | 0.49 | 0.54 | 0.3 | 0.63 | 0.55 |
| 9 | 38404352 | rs7865905 | 0.292 | 1.019 | -0.609 | -1.379 | -0.057 | 0.075 | 0.075 | 0 | 0.23 | 0.09 | 0.16 | 0.08 | 0.28 | 0.28 |
| 9 | 38404579 | rs4878808 | 0.805 | 1.06 | -0.293 | -1.469 | 0 | 0.001 | 0.001 | 0 | 0.28 | 0.26 | 0.32 | 0.2 | 0.26 | 0.15 |
| 9 | 38405834 | rs4878809 | -0.572 | 0.909 | -0.2 | -1.437 | 0.002 | 0.002 | 0.002 | 0 | 0.15 | 0.15 | 0.29 | 0.18 | 0.17 | 0.25 |
| 9 | 38406458 | rs11791018 | 0.76 | 0.793 | -0.233 | -1.387 | -0.014 | 0.014 | 0.014 | 0 | 0.1 | 0.03 | 0.12 | 0.08 | 0.12 | 0.21 |
| 9 | 38406590 | rs11361 | 1.179 | 0.771 | -0.204 | -1.469 | 0.005 | 0.016 | 0.016 | 0 | 0.24 | 0.21 | 0.16 | 0.11 | 0.22 | 0.12 |
| 9 | 38407090 | rs1475102 | 0.378 | -0.147 | 0.039 | -1.149 | 0.005 | -0.006 | -0.006 | 0 | 0.77 | 0.64 | 0.75 | 0.73 | 0.7 | 0.65 |
| 9 | 38407767 | rs7021469 | 2.008 | -0.23 | 0.041 | -1.189 | 0.005 | -0.006 | -0.006 | 3.271 | 0.23 | 0.18 | 0.23 | 0.45 | 0.13 | 0.21 |
| 9 | 38407860 | rs1923424 | 0.476 | -0.033 | -0.178 | -1.467 | 0.004 | -0.008 | -0.008 | 0.251 | 0.77 | 0.64 | 0.75 | 0.73 | 0.7 | 0.66 |
| 9 | 38408592 | rs2038589 | 0.476 | -0.033 | -0.146 | -1.473 | 0.004 | -0.008 | -0.008 | 0 | 0.77 | 0.64 | 0.75 | 0.73 | 0.7 | 0.66 |
| 9 | 38409081 | rs7857243 | -0.927 | -0.048 | -0.056 | -1.48 | 0.009 | 0.023 | 0.023 | 0 | 0.88 | 0.7 | 0.86 | 0.8 | 0.82 | 0.86 |
| 9 | 38409232 | rs7029786 | 0.461 | -0.057 | -0.086 | -1.472 | 0.004 | -0.008 | -0.008 | 0.471 | 0.77 | 0.64 | 0.75 | 0.73 | 0.7 | 0.66 |
| 9 | 38417990 | rs4086036 | -1.171 | -0.215 | -0.152 | -1.378 | -0.002 | 0 | 0 | 0.118 | 0.66 | 0.67 | 0.65 | 0.39 | 0.7 | 0.66 |
| 9 | 38421975 | rs1467574 | 1.008 | 0.589 | -0.46 | -1.204 | -0.086 | 0.038 | 0.038 | 0 | 0.24 | 0.01 | 0.17 | 0.11 | 0.27 | 0.15 |
| 9 | 38426961 | rs2381871 | 0.579 | 0.586 | -0.204 | -1.181 | -0.025 | 0.013 | 0.013 | 0 | 0.29 | 0.11 | 0.31 | 0.25 | 0.32 | 0.23 |
| 9 | 38427185 | rs12684944 | 2.481 | 0.441 | -0.354 | -1.132 | 0.045 | -0.14 | -0.14 | 0.043 | 0.22 | 0.03 | 0.25 | 0.41 | 0.13 | 0.16 |
| 9 | 38427593 | rs998226 | 2.481 | 0.441 | -0.354 | -1.132 | 0.045 | -0.14 | -0.14 | 0.067 | 0.22 | 0.03 | 0.25 | 0.41 | 0.13 | 0.16 |
| 9 | 38430109 | rs7046458 | 0.156 | 0.503 | 0.009 | -1.387 | -0.008 | 0.039 | 0.039 | 0 | 0.38 | 0.31 | 0.22 | 0.12 | 0.38 | 0.27 |
| 9 | 38431667 | rs10973813 | 0.127 | 0.621 | 0.32 | -1.385 | 0.016 | -0.015 | -0.015 | 0 | 0.22 | 0.41 | 0.3 | 0.21 | 0.24 | 0.27 |
| 9 | 38436525 | rs10973819 | 0.574 | 0.655 | 0.262 | -1.438 | 0.005 | -0.018 | -0.018 | 0 | 0.19 | 0.22 | 0.14 | 0.08 | 0.23 | 0.25 |
| 9 | 38436992 | rs769153 | 0.035 | 0.614 | 0.164 | -1.176 | 0.01 | 0.008 | 0.008 | 0 | 0.82 | 0.8 | 0.75 | 0.73 | 0.75 | 0.7 |
| 9 | 38440543 | rs4298574 | -0.946 | 0.671 | 0.831 | -1.753 | 0.009 | 0.017 | 0.017 | 0 | 0.59 | 0.69 | 0.38 | 0.19 | 0.61 | 0.52 |
| 9 | 38445742 | rs12551371 | 0.75 | 0.871 | 0.42 | -1.651 | -0.026 | -0.009 | -0.009 | 0 | 0.06 | 0 | 0.14 | 0.07 | 0.11 | 0.21 |
| 9 | 38450984 | rs4424353 | -0.295 | 0.718 | 0.819 | -1.71 | 0.013 | -0.043 | -0.043 | 0 | 0.81 | 0.77 | 0.76 | 0.89 | 0.73 | 0.67 |
| 9 | 38456365 | rs1110183 | 1.367 | 0.606 | 0.234 | -1.784 | 0.03 | -0.056 | -0.056 | 0 | 0.33 | 0.12 | 0.28 | 0.56 | 0.23 | 0.27 |
| 9 | 38456840 | rs4394481 | -0.148 | 0.561 | 0.314 | -1.8 | -0.005 | 0.064 | 0.064 | 0 | 0.46 | 0.27 | 0.46 | 0.33 | 0.51 | 0.4 |
| 9 | 38457824 | rs10973834 | 1.581 | -0.134 | 0.275 | -1.197 | 0.026 | -0.134 | -0.134 | 0 | 0.26 | 0.02 | 0.24 | 0.56 | 0.17 | 0.21 |
| 9 | 38466944 | rs2799731 | -0.344 | 0.153 | 0.702 | -1.874 | -0.02 | 0.072 | 0.072 | 0 | 0.59 | 0.79 | 0.6 | 0.79 | 0.48 | 0.53 |
| 9 | 38478777 | rs2799740 | -1.192 | 0.846 | 1.112 | -1.402 | 0.012 | -0.034 | -0.034 | 0 | 0.3 | 0.4 | 0.32 | 0.21 | 0.3 | 0.29 |
| 9 | 38483399 | rs10973853 | -0.434 | 1.016 | 0.439 | -1.076 | -0.057 | 0.007 | 0.007 | 0 | 0.13 | 0.01 | 0.08 | 0.09 | 0.16 | 0.14 |
| 9 | 38505602 | rs7855435 | 0.255 | 0.704 | 0.586 | -1.245 | 0.008 | 0.078 | 0.078 | 0 | 0.61 | 0.22 | 0.79 | 0.95 | 0.66 | 0.77 |
| 9 | 38522055 | rs2996368 | -1.56 | 0.702 | 1.734 | -0.135 | -0.028 | -0.031 | -0.031 | 0 | 0.08 | 0.58 | 0.04 | 0 | 0.01 | 0.02 |
| 9 | 38522203 | rs2993164 | -1.362 | 0.704 | 1.768 | -0.134 | 0.001 | -0.003 | -0.003 | 0.934 | 0.08 | 0.5 | 0.03 | 0 | 0.01 | 0.02 |
| 9 | 38533004 | rs10973874 | -0.685 | 1.211 | 0.033 | 0.262 | -0.095 | -0.077 | -0.077 | 0 | 0.16 | 0.01 | 0.19 | 0.22 | 0.26 | 0.24 |
| 9 | 38541855 | rs10973883 | -1.211 | 0.347 | 0.902 | 0.062 | 0.014 | 0.01 | 0.01 | 0.005 | 0.07 | 0.38 | 0.02 | 0 | 0.01 | 0.02 |
| 9 | 38563171 | rs2993129 | -0.511 | 0.86 | -0.068 | 0.234 | 0 | -0.012 | -0.012 | 3.155 | 0.34 | 0.59 | 0.25 | 0.26 | 0.32 | 0.23 |
| 9 | 38572216 | rs10814720 | -0.64 | 1.412 | -0.514 | 0.949 | -0.025 | -0.055 | -0.055 | 0 | 0.19 | 0.1 | 0.28 | 0.46 | 0.23 | 0.19 |
| 9 | 38615175 | rs2381858 | -1.23 | 1.901 | 0.095 | 0.599 | 0.077 | 0.012 | 0.012 | 0 | 0.05 | 0.14 | 0.08 | 0.07 | 0.12 | 0.04 |
| 9 | 38626309 | rs10973939 | -1.295 | 2.199 | 0.175 | 0.612 | 0.112 | 0.02 | 0.02 | 0 | 0.05 | 0.14 | 0.09 | 0.07 | 0.15 | 0.05 |
| 9 | 38633652 | rs10973948 | -2.027 | 1.297 | 0.87 | 0.146 | 0.003 | -0.039 | -0.039 | 0 | 0.18 | 0.65 | 0.15 | 0.08 | 0.17 | 0.17 |
| 9 | 38636463 | rs10973951 | -1.998 | 1.214 | 1.096 | 0.098 | 0.026 | -0.038 | -0.038 | 0.044 | 0.14 | 0.23 | 0.12 | 0.07 | 0.17 | 0.15 |

| | | | | | | | | | | | | | | | | |
|---|---|---|---|---|---|---|---|---|---|---|---|---|---|---|---|---|
| 9 | 38639318 | rs12340088 | -1.79 | 1.015 | 0.851 | -0.04 | 0.014 | -0.039 | -0.039 | 4.257 | 0.14 | 0.24 | 0.13 | 0.03 | 0.17 | 0.15 |
| 9 | 38639853 | rs10973953 | -1.87 | 1.316 | 0.085 | -0.101 | -0.016 | 0.069 | 0.069 | 2.916 | 0.28 | 0.83 | 0.26 | 0.09 | 0.26 | 0.33 |
| 9 | 38641599 | rs12708274 | -0.877 | 1.152 | 0.335 | 0.608 | 0.015 | 0.069 | 0.069 | 0.004 | 0.3 | 0.53 | 0.5 | 0.5 | 0.36 | 0.36 |
| 9 | 38642154 | rs883954 | -0.445 | 0.244 | 0.364 | -0.255 | -0.042 | -0.116 | -0.116 | 0 | 0.21 | 0.36 | 0.13 | 0.03 | 0.1 | 0.15 |
| 9 | 38646995 | rs11790597 | -0.203 | 0.446 | -0.375 | 0.744 | -0.011 | -0.024 | -0.024 | 0.063 | 0.13 | 0.08 | 0.31 | 0.37 | 0.17 | 0.17 |
| 9 | 38648775 | rs1890112 | -0.341 | -0.138 | -0.332 | 0.144 | -0.006 | -0.022 | -0.022 | 0 | 0.39 | 0.35 | 0.36 | 0.11 | 0.26 | 0.35 |
| 9 | 38650244 | rs10758511 | 0.216 | 0.561 | -0.682 | -0.026 | -0.016 | -0.017 | -0.017 | 0 | 0.36 | 0.29 | 0.48 | 0.5 | 0.51 | 0.33 |
| 9 | 38652915 | rs10758513 | -0.117 | 0.725 | -0.612 | -0.063 | 0.043 | 0.101 | 0.101 | 0 | 0.48 | 0.3 | 0.41 | 0.26 | 0.39 | 0.45 |
| 9 | 38652961 | rs11792859 | 1.318 | 0.225 | -1.109 | -0.196 | 0.047 | 0.125 | 0.125 | 0.037 | 0.25 | 0.02 | 0.25 | 0.05 | 0.16 | 0.21 |
| 9 | 38654738 | rs16931661 | 1.481 | 0.209 | -1.166 | -0.183 | 0.046 | 0.09 | 0.09 | 0.019 | 0.21 | 0.01 | 0.22 | 0.04 | 0.11 | 0.19 |
| 9 | 38654814 | rs9407070 | -0.654 | 0.458 | -0.321 | -0.148 | 0 | -0.003 | -0.003 | 0 | 0.64 | 0.6 | 0.75 | 0.76 | 0.64 | 0.54 |
| 9 | 38655153 | rs6476727 | -0.789 | 0.28 | -0.062 | 0.275 | 0.012 | 0.015 | 0.015 | 0 | 0.7 | 0.54 | 0.75 | 0.67 | 0.71 | 0.65 |
| 9 | 38655269 | rs1577869 | -0.841 | 0.389 | -0.084 | 0.347 | 0.047 | 0.037 | 0.037 | 0 | 0.73 | 0.54 | 0.74 | 0.68 | 0.69 | 0.66 |
| 9 | 38655438 | rs1577868 | 0.689 | 0.527 | 0.057 | 0.206 | 0.026 | 0.038 | 0.038 | 0 | 0.09 | 0.25 | 0.1 | 0.12 | 0.1 | 0.11 |
| 9 | 38655835 | rs9298981 | 0.13 | 0.301 | 0.116 | 0.07 | 0.007 | -0.009 | -0.009 | 0 | 0.42 | 0.48 | 0.54 | 0.66 | 0.54 | 0.46 |
| 9 | 38657104 | rs3124041 | -0.192 | 0.321 | 0.136 | 0.091 | 0 | -0.006 | -0.006 | 0 | 0.46 | 0.48 | 0.56 | 0.66 | 0.6 | 0.49 |
| 9 | 38657144 | rs17456345 | 1.165 | 0.406 | -0.211 | 0.251 | -0.016 | 0.01 | 0.01 | 0 | 0.24 | 0.14 | 0.16 | 0.18 | 0.34 | 0.22 |
| 9 | 38660694 | rs3124044 | 0.374 | 0.465 | -0.062 | 0.339 | 0.003 | 0.025 | 0.025 | 0 | 0.21 | 0.23 | 0.39 | 0.48 | 0.26 | 0.28 |
| 9 | 38661416 | rs7031991 | -0.941 | 0.386 | -0.084 | 0.22 | -0.017 | -0.011 | -0.011 | 0.064 | 0.7 | 0.72 | 0.55 | 0.4 | 0.6 | 0.59 |
| 9 | 38667363 | rs12337426 | 0.814 | 0.132 | -0.146 | 0.096 | -0.008 | -0.006 | -0.006 | 0 | 0.28 | 0.24 | 0.18 | 0.31 | 0.37 | 0.26 |
| 9 | 38668070 | rs12555495 | 1.287 | -0.093 | -0.139 | -0.1 | 0.059 | 0.063 | 0.063 | 0 | 0.29 | 0.13 | 0.26 | 0.04 | 0.13 | 0.22 |
| 9 | 38668792 | rs12352401 | 1.22 | 0.149 | -0.548 | 0.06 | -0.017 | 0.019 | 0.019 | 0 | 0.24 | 0.13 | 0.16 | 0.17 | 0.34 | 0.22 |
| 9 | 38670778 | rs7048582 | 0.505 | 0.312 | -0.491 | 0.182 | -0.03 | -0.029 | -0.029 | 5.065 | 0.36 | 0.21 | 0.48 | 0.56 | 0.5 | 0.42 |
| 9 | 38671889 | rs4074217 | -1.967 | 0.393 | -0.584 | 0.135 | -0.014 | -0.002 | -0.002 | 0 | 0.81 | 0.83 | 0.84 | 0.83 | 0.76 | 0.77 |
| 9 | 38673236 | rs13294686 | 1.396 | 0.196 | -0.775 | 0.365 | 0.016 | -0.013 | -0.013 | 0 | 0.26 | 0.18 | 0.27 | 0.31 | 0.19 | 0.24 |
| 9 | 38673327 | rs12686391 | 1.978 | 0.398 | -0.984 | 0.465 | -0.005 | -0.019 | -0.019 | 0 | 0.12 | 0.06 | 0.29 | 0.34 | 0.13 | 0.24 |
| 9 | 38673606 | rs7873962 | 1.348 | 0.496 | -1.152 | 0.253 | -0.018 | 0.013 | 0.013 | 0 | 0.23 | 0.14 | 0.18 | 0.18 | 0.36 | 0.18 |
| 9 | 38673912 | rs16931494 | 0.629 | 0.447 | -1.231 | 0.216 | 0.023 | 0.001 | 0.001 | 0.008 | 0.33 | 0.19 | 0.32 | 0.3 | 0.26 | 0.29 |
| 9 | 38675465 | rs7039377 | -0.204 | 0.332 | -1.363 | 0.112 | 0.006 | 0.009 | 0.009 | 0 | 0.44 | 0.68 | 0.47 | 0.48 | 0.5 | 0.49 |
| 9 | 38677071 | rs10973967 | 1.347 | 0.572 | -1.447 | 0.155 | -0.002 | 0.009 | 0.009 | 0 | 0.16 | 0.16 | 0.12 | 0.31 | 0.17 | 0.23 |
| 9 | 38677136 | rs10814734 | -0.802 | 0.369 | -2.219 | 0.005 | 0.026 | 0.116 | 0.116 | 0.021 | 0.68 | 0.87 | 0.69 | 0.93 | 0.78 | 0.76 |
| 9 | 38677513 | rs10973968 | 0.926 | 0.725 | -1.923 | 0.324 | -0.033 | -0.008 | -0.008 | 0 | 0.24 | 0.12 | 0.21 | 0.29 | 0.38 | 0.2 |
| 9 | 38677737 | rs7875312 | 0.284 | 0.731 | -1.367 | 0.424 | -0.009 | -0.005 | -0.005 | 0 | 0.48 | 0.34 | 0.5 | 0.62 | 0.57 | 0.52 |
| 9 | 38678740 | rs7849958 | -0.548 | 0.674 | -1.471 | 0.044 | -0.028 | 0.082 | 0.082 | 0.344 | 0.78 | 0.94 | 0.72 | 0.93 | 0.8 | 0.8 |
| 9 | 38679722 | rs4641154 | -0.255 | 0.621 | -0.714 | 0.587 | -0.001 | 0 | 0 | 2.277 | 0.37 | 0.46 | 0.45 | 0.35 | 0.35 | 0.42 |
| 9 | 38680826 | rs4242651 | -0.555 | 0.779 | -0.752 | 0.732 | 0.022 | 0.018 | 0.018 | 0 | 0.33 | 0.45 | 0.39 | 0.42 | 0.38 | 0.4 |
| 9 | 38683660 | rs4878845 | -0.657 | 1.397 | -0.845 | 1.13 | 0.069 | 0.143 | 0.143 | 0 | 0.1 | 0.24 | 0.25 | 0.29 | 0.16 | 0.25 |
| 9 | 38685261 | rs13292644 | -0.02 | 1.543 | -1.587 | 1.198 | 0.01 | 0.042 | 0.042 | 0 | 0.06 | 0.04 | 0.23 | 0.28 | 0.15 | 0.22 |
| 9 | 38685643 | rs4593645 | -0.32 | 0.598 | -2.275 | 0.701 | -0.043 | 0.069 | 0.069 | 0.047 | 0.52 | 0.87 | 0.49 | 0.67 | 0.53 | 0.72 |
| 9 | 38687510 | rs10739004 | -0.548 | 0.04 | -2.257 | 0.788 | 0.062 | 0.016 | 0.016 | 1.004 | 0.62 | 0.91 | 0.78 | 0.7 | 0.69 | 0.77 |
| 9 | 38693047 | rs4878219 | -0.574 | 0.384 | -1.461 | 1.413 | 0.003 | -0.006 | -0.006 | 0 | 0.18 | 0.13 | 0.33 | 0.34 | 0.15 | 0.22 |
| 9 | 38699239 | rs10973981 | -0.008 | 0.819 | -1.093 | 0.874 | 0.011 | -0.041 | -0.041 | 0.94 | 0.09 | 0.13 | 0.07 | 0 | 0.12 | 0.1 |
| 9 | 38708759 | rs16935357 | 0.172 | 0.561 | -2.065 | 0.971 | -0.018 | -0.021 | -0.021 | 0 | 0.14 | 0.07 | 0.09 | 0.21 | 0.17 | 0.18 |
| 9 | 38715865 | rs10511953 | 0.095 | 0.114 | -1.284 | 1.038 | 0.022 | -0.001 | -0.001 | 0.056 | 0.64 | 0.74 | 0.67 | 0.4 | 0.69 | 0.62 |
| 9 | 71233266 | rs7851511 | -0.985 | 0.783 | 0.699 | 1.05 | 0.023 | 0.004 | 0.004 | 0.04 | 0.05 | 0.2 | 0.03 | 0.01 | 0.04 | 0.05 |
| 9 | 71241117 | rs944550 | -1.682 | 1.135 | 0.454 | 0.732 | -0.007 | -0.071 | -0.071 | 0 | 0.16 | 0.6 | 0.09 | 0.01 | 0.11 | 0.05 |
| 9 | 71241343 | rs10869160 | -0.037 | 1.086 | 0.063 | 0.913 | -0.024 | -0.022 | -0.022 | 0 | 0.2 | 0.09 | 0.3 | 0.2 | 0.25 | 0.3 |
| 9 | 71244025 | rs4744669 | 1.357 | 0.874 | 0.617 | 1.228 | 0.046 | 0.296 | 0.296 | 0 | 0.92 | 0.68 | 0.89 | 0.69 | 0.93 | 0.95 |
| 9 | 71247375 | rs10869167 | -0.001 | 0.876 | 0.618 | 0.935 | -0.121 | 0.091 | 0.091 | 3.643 | 0.2 | 0.01 | 0.31 | 0.03 | 0.3 | 0.15 |
| 9 | 71250999 | rs4347049 | 0.778 | 0.92 | 1.09 | 1.307 | 0.027 | 0.046 | 0.046 | 2.508 | 0.69 | 0.54 | 0.63 | 0.5 | 0.64 | 0.64 |
| 9 | 71301557 | rs1014237 | -1.916 | 0.741 | 0.595 | 0.424 | 0.009 | -0.061 | -0.061 | 0 | 0.19 | 0.55 | 0.14 | 0.07 | 0.2 | 0.09 |
| 9 | 71305001 | rs12338099 | -1.593 | 0.637 | 0.799 | 0.519 | 0.018 | -0.007 | -0.007 | 0 | 0.13 | 0.45 | 0.09 | 0.07 | 0.1 | 0.07 |
| 9 | 71309707 | rs10869200 | -0.261 | 1.047 | -0.378 | 0.695 | -0.101 | 0.017 | 0.017 | 2.725 | 0.72 | 0.94 | 0.66 | 0.79 | 0.66 | 0.74 |
| 9 | 71313742 | rs10735616 | -0.229 | 1.061 | -0.384 | 0.89 | -0.103 | -0.034 | -0.034 | 0 | 0.72 | 0.94 | 0.61 | 0.76 | 0.66 | 0.73 |
| 9 | 71318128 | rs7023009 | 0.873 | 0.564 | -0.187 | 0.474 | -0.004 | 0.041 | 0.041 | 0.992 | 0.52 | 0.59 | 0.53 | 0.65 | 0.48 | 0.66 |
| 9 | 71318817 | rs1107108 | 0.875 | 0.562 | -0.189 | 0.472 | -0.004 | 0.041 | 0.041 | 0.946 | 0.52 | 0.59 | 0.53 | 0.65 | 0.48 | 0.66 |
| 9 | 71324475 | rs2151418 | 0.679 | 0.458 | -0.146 | 0.418 | 0.005 | 0.036 | 0.036 | 0.003 | 0.52 | 0.6 | 0.56 | 0.7 | 0.52 | 0.69 |
| 9 | 71325908 | rs4534183 | 0.585 | 0.441 | -0.113 | 0.409 | -0.002 | 0.091 | 0.091 | 0.11 | 0.53 | 0.73 | 0.57 | 0.7 | 0.53 | 0.69 |
| 9 | 71328378 | rs10121993 | -0.009 | 0.437 | -0.204 | 0.345 | -0.021 | 0.038 | 0.038 | 0.075 | 0.65 | 0.85 | 0.64 | 0.73 | 0.63 | 0.7 |
| 9 | 71334809 | rs4745223 | 0.738 | 0.51 | -0.294 | 0.369 | 0.008 | 0.041 | 0.041 | 0.001 | 0.52 | 0.6 | 0.57 | 0.7 | 0.53 | 0.69 |
| 9 | 71345123 | rs4745230 | -0.133 | 0.593 | -0.763 | 0.458 | -0.081 | 0.001 | 0.001 | 0.328 | 0.67 | 0.93 | 0.65 | 0.73 | 0.64 | 0.72 |
| 9 | 71346288 | rs11143568 | 0.37 | 0.632 | -0.812 | 0.393 | -0.083 | -0.012 | -0.012 | 0 | 0.29 | 0.02 | 0.16 | 0.21 | 0.32 | 0.25 |
| 9 | 71346717 | rs4745231 | 0.697 | 0.815 | -0.451 | 0.713 | 0.001 | 0.01 | 0.01 | 0.808 | 0.52 | 0.69 | 0.53 | 0.43 | 0.45 | 0.49 |
| 9 | 71347362 | rs17057092 | -0.363 | 0.661 | -0.33 | 0.978 | -0.003 | 0.178 | 0.178 | 0.048 | 0.06 | 0.18 | 0.02 | 0.17 | 0.02 | 0.05 |
| 9 | 71347988 | rs13296679 | -0.309 | 0.782 | -0.411 | 0.845 | 0.011 | 0.046 | 0.046 | 0.817 | 0.18 | 0.22 | 0.15 | 0.33 | 0.23 | 0.23 |
| 9 | 71352493 | rs7044422 | 1.564 | 0.655 | -0.831 | 0.876 | 0.006 | -0.017 | -0.017 | 0 | 0.33 | 0.28 | 0.35 | 0.41 | 0.27 | 0.43 |
| 9 | 71353056 | rs10781142 | -0.572 | 0.652 | -0.913 | 0.826 | -0.096 | 0.029 | 0.029 | 0.442 | 0.67 | 0.95 | 0.66 | 0.76 | 0.64 | 0.69 |
| 9 | 71354184 | rs10735624 | 0.233 | 0.553 | -0.571 | 1.085 | 0 | 0.025 | 0.025 | 0 | 0.95 | 0.93 | 0.96 | 0.79 | 0.95 | 0.88 |
| 9 | 71355133 | rs7044045 | -1.074 | 0.603 | -0.443 | 0.735 | 0 | -0.003 | -0.003 | 1.204 | 0.35 | 0.67 | 0.27 | 0.35 | 0.37 | 0.25 |

| | | | | | | | | | | | | | | | | |
|---|---|---|---|---|---|---|---|---|---|---|---|---|---|---|---|---|
| 9 | 71357605 | rs7866673 | -1.068 | 0.634 | -0.488 | 0.72 | 0 | -0.002 | -0.002 | 0 | 0.35 | 0.62 | 0.32 | 0.36 | 0.38 | 0.26 |
| 9 | 71364769 | rs10869263 | -0.058 | 0.467 | -0.532 | 0.877 | -0.001 | -0.025 | -0.025 | 0 | 0.38 | 0.27 | 0.4 | 0.42 | 0.38 | 0.27 |
| 9 | 71365013 | rs4744689 | -0.052 | 0.497 | -0.22 | 0.968 | -0.002 | 0.003 | 0.003 | 0.431 | 0.37 | 0.37 | 0.48 | 0.31 | 0.4 | 0.28 |
| 9 | 71370174 | rs10735625 | -0.385 | 0.01 | -0.218 | 0.622 | -0.013 | 0.03 | 0.03 | 0 | 0.65 | 0.82 | 0.63 | 0.79 | 0.65 | 0.66 |
| 9 | 71376291 | rs1336492 | 0.047 | -0.183 | 0.362 | 0.665 | 0 | 0.003 | 0.003 | 0 | 0.58 | 0.53 | 0.55 | 0.6 | 0.62 | 0.63 |
| 9 | 71383888 | rs945573 | -0.201 | -0.216 | 0.236 | 0.378 | 0.007 | 0.044 | 0.044 | 0 | 0.5 | 0.36 | 0.53 | 0.3 | 0.52 | 0.45 |
| 9 | 71384758 | rs7031263 | 0.541 | -0.209 | 0.204 | 0.306 | -0.018 | 0.046 | 0.046 | 0 | 0.35 | 0.15 | 0.37 | 0.19 | 0.35 | 0.34 |
| 9 | 71386041 | rs7035163 | -1.457 | -0.214 | 0.189 | 0.159 | -0.003 | 0.025 | 0.025 | 0 | 0.68 | 0.59 | 0.65 | 0.39 | 0.7 | 0.51 |
| 9 | 71387764 | rs10511964 | -0.128 | -0.145 | 0.099 | 0.074 | 0.005 | 0.064 | 0.064 | 0 | 0.5 | 0.34 | 0.48 | 0.26 | 0.51 | 0.45 |
| 9 | 71396426 | rs12349968 | 1.111 | 0.029 | -0.028 | 0.262 | 0.001 | 0.013 | 0.013 | 0 | 0.15 | 0.09 | 0.1 | 0 | 0.17 | 0.11 |
| 9 | 71396746 | rs975123 | -1.509 | 0.031 | -0.024 | 0.631 | 0.001 | 0.036 | 0.036 | 0 | 0.94 | 0.9 | 0.97 | 0.79 | 0.96 | 0.88 |
| 9 | 71397734 | rs1412988 | -1.047 | 0.02 | -0.006 | 0.598 | 0.005 | 0.056 | 0.056 | 0 | 0.93 | 0.86 | 0.96 | 0.79 | 0.95 | 0.87 |
| 9 | 71397747 | rs2017 | 0.696 | 0.018 | 0.076 | 0.627 | -0.002 | 0.014 | 0.014 | 0 | 0.61 | 0.52 | 0.75 | 0.55 | 0.62 | 0.66 |
| 9 | 71421343 | rs17391596 | 1.001 | -0.02 | -0.356 | 0.477 | -0.034 | 0.051 | 0.051 | 0 | 0.31 | 0.11 | 0.35 | 0.15 | 0.33 | 0.31 |
| 9 | 71421359 | rs10511963 | 1.96 | -0.248 | -0.699 | 0.626 | -0.082 | -0.097 | -0.097 | 0.016 | 0.22 | 0.02 | 0.36 | 0.34 | 0.27 | 0.32 |
| 9 | 71425223 | rs11143793 | 0.106 | -0.182 | -0.322 | 0.359 | -0.016 | 0.117 | 0.117 | 0 | 0.45 | 0.23 | 0.44 | 0.16 | 0.47 | 0.34 |
| 9 | 71433212 | rs1556751 | 0.464 | -0.257 | 0.078 | 0.52 | -0.002 | -0.002 | -0.002 | 0 | 0.63 | 0.6 | 0.77 | 0.64 | 0.65 | 0.72 |
| 9 | 71443540 | rs1889150 | 0.532 | -0.324 | 0.098 | 0.427 | 0 | -0.001 | -0.001 | 0 | 0.66 | 0.65 | 0.78 | 0.66 | 0.65 | 0.7 |
| 9 | 71453443 | rs4745321 | -0.464 | -0.335 | 0.11 | 0.387 | -0.004 | -0.003 | -0.003 | 0.509 | 0.78 | 0.79 | 0.87 | 0.78 | 0.8 | 0.77 |
| 9 | 71454316 | rs7871676 | 0.446 | -0.324 | 0.232 | 0.402 | 0 | 0.009 | 0.009 | 0 | 0.39 | 0.37 | 0.39 | 0.26 | 0.35 | 0.33 |
| 9 | 71457602 | rs10869400 | 1.226 | -0.3 | 0.265 | 0.475 | 0.003 | 0.047 | 0.047 | 0 | 0.29 | 0.39 | 0.41 | 0.52 | 0.31 | 0.42 |
| 9 | 71460428 | rs7874714 | 1.468 | -0.297 | 0.261 | 0.493 | 0.002 | 0.014 | 0.014 | 0 | 0.27 | 0.29 | 0.39 | 0.39 | 0.29 | 0.38 |
| 9 | 71464932 | rs10781232 | -0.643 | -0.33 | 0.283 | 0.484 | 0 | -0.001 | -0.001 | 0 | 0.8 | 0.79 | 0.87 | 0.78 | 0.79 | 0.77 |
| 9 | 71470111 | rs7041004 | -0.119 | -0.409 | 0.418 | 0.475 | 0.005 | 0.174 | 0.174 | 0.483 | 0.88 | 0.63 | 0.93 | 0.67 | 0.93 | 0.86 |
| 9 | 71471912 | rs6560397 | 0.113 | -0.333 | 0.34 | 0.396 | -0.005 | 0.08 | 0.08 | 0 | 0.49 | 0.28 | 0.47 | 0.27 | 0.5 | 0.43 |
| 9 | 71472782 | rs7043727 | 0.376 | -0.315 | 0.376 | 0.449 | 0.003 | 0.251 | 0.251 | 0.453 | 0.86 | 0.59 | 0.92 | 0.54 | 0.92 | 0.82 |
| 9 | 71473803 | rs7863342 | 0.349 | -0.282 | 0.22 | 0.383 | -0.001 | -0.003 | -0.003 | 0 | 0.39 | 0.36 | 0.48 | 0.5 | 0.44 | 0.47 |
| 9 | 71475381 | rs11144027 | 0.134 | -0.188 | 0.025 | 0.1 | -0.002 | 0.012 | 0.012 | 0 | 0.25 | 0.18 | 0.17 | 0.01 | 0.29 | 0.13 |
| 9 | 71482288 | rs2152647 | -0.522 | -0.106 | 0.461 | 0.515 | 0.003 | 0.019 | 0.019 | 0 | 0.51 | 0.43 | 0.48 | 0.37 | 0.51 | 0.37 |
| 9 | 71484028 | rs4744726 | 0.848 | -0.123 | 0.472 | 0.642 | -0.003 | -0.019 | -0.019 | 0 | 0.41 | 0.29 | 0.47 | 0.49 | 0.45 | 0.51 |
| 9 | 71509599 | rs953036 | 0.771 | -0.421 | 0.707 | 0.821 | -0.004 | 0.043 | 0.043 | 0.204 | 0.05 | 0.15 | 0.03 | 0.11 | 0.02 | 0.01 |
| 9 | 71512668 | rs6560420 | 0.656 | -0.493 | 0.725 | 0.775 | -0.003 | 0.023 | 0.023 | 0 | 0.78 | 0.68 | 0.86 | 0.67 | 0.8 | 0.84 |
| 9 | 71514547 | rs883951 | 0.824 | -0.598 | 0.51 | 0.243 | -0.023 | 0.058 | 0.058 | 0 | 0.28 | 0.13 | 0.35 | 0.14 | 0.29 | 0.16 |
| 9 | 71514651 | rs883952 | 1.006 | -0.611 | 0.659 | 0.305 | -0.004 | 0.018 | 0.018 | 0 | 0.47 | 0.35 | 0.5 | 0.39 | 0.5 | 0.6 |
| 9 | 71517082 | rs7855908 | 0.636 | -0.729 | 1.429 | 0.815 | 0.018 | 0.161 | 0.161 | 0.381 | 0.92 | 0.7 | 0.95 | 0.78 | 0.95 | 0.95 |
| 9 | 71517261 | rs7865397 | -0.546 | -0.785 | 1.319 | 0.756 | -0.002 | -0.002 | -0.002 | 0 | 0.16 | 0.22 | 0.1 | 0.13 | 0.16 | 0.08 |
| 9 | 71519136 | rs10781264 | 1.583 | -0.781 | 1.27 | 0.758 | -0.016 | 0.019 | 0.019 | 0.396 | 0.7 | 0.4 | 0.82 | 0.4 | 0.78 | 0.71 |
| 9 | 71521810 | rs7024695 | -0.948 | -0.764 | 1.204 | 0.302 | -0.011 | -0.018 | -0.018 | 0 | 0.21 | 0.43 | 0.11 | 0.12 | 0.17 | 0.05 |
| 9 | 71527774 | rs17058884 | 0.661 | -0.77 | 1.012 | 0.411 | 0.001 | 0.056 | 0.056 | 0 | 0.07 | 0.05 | 0.04 | 0.37 | 0.04 | 0.21 |
| 9 | 71533367 | rs7859638 | 0.517 | -0.77 | 1.17 | 0.326 | -0.019 | 0.253 | 0.253 | 0 | 0.88 | 0.68 | 0.93 | 0.59 | 0.95 | 0.74 |
| 9 | 71539731 | rs10869538 | -0.634 | -0.72 | 1.012 | 0.166 | -0.006 | -0.02 | -0.02 | 0 | 0.17 | 0.34 | 0.11 | 0.11 | 0.17 | 0.04 |
| 9 | 71540231 | rs7875472 | 0.463 | -0.714 | 1.547 | 0.21 | 0.042 | 0.123 | 0.123 | 2.165 | 0.92 | 0.5 | 0.96 | 0.9 | 0.97 | 0.99 |
| 9 | 71546444 | rs10781283 | 1.294 | -0.798 | 1.195 | 0.244 | 0.029 | -0.062 | -0.062 | 0 | 0.7 | 0.21 | 0.82 | 0.49 | 0.78 | 0.71 |
| 9 | 71554159 | rs4745402 | 1.264 | -0.886 | 0.439 | -0.273 | -0.134 | -0.136 | -0.136 | 0 | 0.33 | 0.02 | 0.42 | 0.44 | 0.38 | 0.5 |
| 9 | 71555522 | rs10746974 | 0.551 | -0.666 | 1.596 | 0.541 | -0.025 | 0.15 | 0.15 | 0 | 0.82 | 0.53 | 0.89 | 0.6 | 0.86 | 0.76 |
| 9 | 71557341 | rs3829061 | 0.424 | -0.599 | 1.639 | 0.464 | -0.007 | 0.041 | 0.041 | 0.291 | 0.05 | 0.17 | 0.03 | 0.1 | 0.02 | 0.01 |
| 9 | 71562055 | rs11144442 | -1.258 | -0.538 | 1.341 | 0.482 | -0.005 | 0.052 | 0.052 | 0.492 | 0.12 | 0.34 | 0.08 | 0.15 | 0.08 | 0.06 |
| 9 | 71574757 | rs10114872 | -0.655 | -0.6 | 0.774 | 0.181 | -0.016 | 0.097 | 0.097 | 0 | 0.2 | 0.36 | 0.13 | 0.51 | 0.16 | 0.24 |
| 9 | 71576640 | rs10124169 | -0.93 | -0.578 | 0.587 | 0.189 | -0.017 | 0.078 | 0.078 | 0 | 0.25 | 0.44 | 0.16 | 0.51 | 0.21 | 0.25 |
| 9 | 71577792 | rs7020192 | -1.033 | -0.578 | 0.519 | 0.189 | -0.023 | 0.085 | 0.085 | 0 | 0.26 | 0.51 | 0.16 | 0.51 | 0.21 | 0.25 |
| 9 | 71591393 | rs10781306 | -0.293 | -0.57 | -0.468 | -0.446 | -0.006 | -0.144 | -0.144 | 0 | 0.72 | 0.65 | 0.84 | 0.99 | 0.74 | 0.84 |
| 9 | 71595965 | rs7048825 | -0.54 | -0.589 | -1.076 | 0.602 | 0.007 | -0.041 | -0.041 | 0 | 0.16 | 0.08 | 0.1 | 0.3 | 0.14 | 0.16 |
| 9 | 71597840 | rs12378665 | 0.166 | -0.529 | -1.126 | 0.642 | 0.006 | -0.028 | -0.028 | 0.497 | 0.12 | 0.06 | 0.07 | 0.29 | 0.09 | 0.15 |
| 9 | 71598339 | rs7031059 | -0.83 | -0.465 | -1.105 | 0.365 | -0.011 | 0.013 | 0.013 | 0.086 | 0.3 | 0.69 | 0.2 | 0.54 | 0.21 | 0.24 |
| 9 | 71604123 | rs4745466 | 0.078 | 0.124 | -1.985 | -0.251 | 0.016 | 0.11 | 0.11 | 0.138 | 0.5 | 0.27 | 0.56 | 0.67 | 0.55 | 0.48 |
| 9 | 71606315 | rs2297231 | 0.017 | 0.453 | -2.222 | -0.065 | -0.063 | -0.04 | -0.04 | 1.091 | 0.39 | 0.13 | 0.52 | 0.55 | 0.52 | 0.45 |
| 9 | 71612891 | rs883751 | -0.026 | 0.916 | -1.863 | -0.644 | -0.014 | 0.018 | 0.018 | 0 | 0.22 | 0.16 | 0.25 | 0 | 0.23 | 0.18 |
| 9 | 71615766 | rs2871223 | 0.038 | 0.543 | -2.42 | -0.31 | 0.008 | -0.015 | -0.015 | 0 | 0.29 | 0.17 | 0.37 | 0.33 | 0.3 | 0.31 |
| 9 | 71618245 | rs10781329 | 1.024 | 0.514 | -1.978 | 0.147 | -0.047 | -0.065 | -0.065 | 0.053 | 0.24 | 0.1 | 0.23 | 0.31 | 0.33 | 0.23 |
| 9 | 71620614 | rs10869686 | 0.758 | 0.396 | -1.262 | 0.163 | -0.004 | 0.029 | 0.029 | 0 | 0.15 | 0.13 | 0.09 | 0.33 | 0.11 | 0.15 |
| 9 | 71621276 | rs7019661 | -0.05 | 0.395 | -0.851 | 0.036 | -0.005 | 0.043 | 0.043 | 0 | 0.2 | 0.33 | 0.12 | 0.35 | 0.14 | 0.22 |
| 9 | 71621922 | rs1541083 | -0.409 | 0.084 | -1.58 | -0.433 | -0.049 | 0.053 | 0.053 | 0 | 0.57 | 0.83 | 0.65 | 0.34 | 0.49 | 0.54 |
| 9 | 71624695 | rs872077 | 0.155 | -0.403 | -1.575 | 0.04 | 0.015 | -0.056 | -0.056 | 0.43 | 0.77 | 0.89 | 0.81 | 0.5 | 0.85 | 0.67 |
| 9 | 71626508 | rs28461568 | 1.333 | -0.112 | -1.993 | -0.427 | -0.082 | -0.06 | -0.06 | 0.645 | 0.33 | 0.15 | 0.3 | 0.39 | 0.47 | 0.34 |
| 9 | 71627283 | rs11144762 | 0.438 | -0.696 | -1.556 | -0.997 | 0 | -0.017 | -0.017 | 0 | 0.23 | 0.23 | 0.32 | 0.03 | 0.25 | 0.14 |
| 9 | 71627525 | rs12345886 | -0.203 | -1.059 | -1.344 | -1.176 | -0.005 | -0.049 | -0.049 | 0.051 | 0.26 | 0.34 | 0.38 | 0.05 | 0.22 | 0.19 |
| 9 | 71628207 | rs3730386 | -0.076 | -1.088 | -0.917 | -0.653 | -0.021 | -0.007 | -0.007 | 0 | 0.33 | 0.38 | 0.18 | 0.31 | 0.27 | 0.41 |
| 9 | 71629914 | rs4745514 | -1.027 | -1.121 | -0.902 | -0.656 | 0.007 | 0.006 | 0.006 | 0 | 0.35 | 0.38 | 0.49 | 0.28 | 0.25 | 0.3 |
| 9 | 71630576 | rs34539730 | 1.579 | -1.066 | -1.005 | -0.668 | 0.001 | 0.001 | 0.001 | 0.157 | 0.05 | 0.06 | 0.14 | 0.04 | 0.06 | 0.1 |

| | | | | | | | | | | | | | | | | |
|---|---|---|---|---|---|---|---|---|---|---|---|---|---|---|---|---|
| 9 | 71630663 | rs4745517 | 0.583 | -1.064 | -1.032 | -0.562 | 0.018 | -0.022 | -0.022 | 1.539 | 0.14 | 0.1 | 0.15 | 0.25 | 0.06 | 0.12 |
| 9 | 71630841 | rs10869721 | 0.493 | -0.952 | -0.899 | -0.581 | -0.004 | 0 | 0 | 0.37 | 0.29 | 0.25 | 0.16 | 0.31 | 0.27 | 0.36 |
| 9 | 71631733 | rs4745520 | 1.981 | -0.61 | -1.205 | -0.482 | -0.079 | -0.057 | -0.057 | 0 | 0.34 | 0.17 | 0.35 | 0.4 | 0.48 | 0.35 |
| 9 | 71642499 | rs10869745 | 0.874 | -0.784 | -0.945 | -0.588 | 0.012 | -0.007 | -0.007 | 0 | 0.57 | 0.43 | 0.67 | 0.48 | 0.65 | 0.49 |
| 9 | 71643131 | rs9314845 | -0.143 | -0.676 | -0.686 | -0.5 | -0.013 | 0.012 | 0.012 | 0 | 0.72 | 0.57 | 0.82 | 0.68 | 0.72 | 0.62 |
| 9 | 71647375 | rs3763608 | 1.807 | -0.301 | -0.766 | -0.661 | -0.054 | -0.068 | -0.068 | 0 | 0.31 | 0.14 | 0.42 | 0.43 | 0.43 | 0.35 |
| 9 | 71647410 | rs3763609 | 0.578 | -0.573 | -0.534 | -0.832 | -0.003 | 0 | 0 | 0 | 0.59 | 0.53 | 0.57 | 0.51 | 0.7 | 0.56 |
| 9 | 71648083 | rs7046232 | 0.901 | -0.607 | -0.605 | -0.878 | 0.024 | -0.005 | -0.005 | 0.137 | 0.55 | 0.6 | 0.57 | 0.5 | 0.67 | 0.55 |
| 9 | 71649464 | rs12000125 | -0.864 | -0.555 | -0.525 | -0.733 | 0.036 | -0.024 | -0.024 | 0 | 0.4 | 0.35 | 0.41 | 0.5 | 0.3 | 0.44 |
| 9 | 71651605 | rs1800651 | 0.27 | -0.153 | -0.169 | -0.799 | 0.008 | -0.051 | -0.051 | 0 | 0.24 | 0.26 | 0.14 | 0.08 | 0.26 | 0.22 |
| 9 | 71659280 | rs3793451 | 0.174 | -0.117 | -0.335 | -0.324 | 0.007 | -0.001 | -0.001 | 0.503 | 0.06 | 0.05 | 0.21 | 0.25 | 0.05 | 0.09 |
| 9 | 71668197 | rs3829062 | -1.384 | -0.015 | 0.164 | -0.525 | -0.013 | -0.004 | -0.004 | 0 | 0.59 | 0.4 | 0.55 | 0.56 | 0.53 | 0.51 |
| 9 | 71671116 | rs2309394 | 1.184 | -0.025 | 0.369 | -0.53 | 0.016 | 0.007 | 0.007 | 0.274 | 0.39 | 0.4 | 0.43 | 0.44 | 0.47 | 0.48 |
| 9 | 71679438 | rs3793465 | -1.779 | 0.416 | 0.714 | -0.319 | -0.002 | 0.004 | 0.004 | 0 | 0.6 | 0.76 | 0.68 | 0.65 | 0.59 | 0.64 |
| 9 | 71686181 | rs9314854 | -1.14 | 0.571 | 0.762 | -0.288 | -0.015 | 0.011 | 0.011 | 0 | 0.43 | 0.71 | 0.58 | 0.64 | 0.49 | 0.55 |
| 9 | 71686476 | rs7870295 | -0.825 | 0.587 | 0.866 | -0.31 | -0.004 | -0.008 | -0.008 | 0 | 0.39 | 0.37 | 0.54 | 0.57 | 0.43 | 0.43 |
| 9 | 71687924 | rs10890 | -1.059 | 0.587 | 0.918 | -0.31 | -0.002 | 0 | 0 | 0 | 0.42 | 0.57 | 0.57 | 0.64 | 0.49 | 0.55 |
| 9 | 71688300 | rs4744807 | -0.883 | 0.586 | 0.922 | -0.31 | -0.004 | -0.007 | -0.007 | 0.092 | 0.39 | 0.4 | 0.54 | 0.57 | 0.43 | 0.43 |
| 9 | 71688311 | rs4744808 | -1.06 | 0.586 | 0.922 | -0.31 | -0.002 | 0 | 0 | 0 | 0.42 | 0.57 | 0.57 | 0.64 | 0.49 | 0.55 |
| 9 | 71688476 | rs11145043 | 1.684 | 0.533 | 0.561 | -0.323 | 0.002 | 0.016 | 0.016 | 0 | 0.43 | 0.21 | 0.33 | 0.35 | 0.42 | 0.36 |
| 9 | 71689756 | rs4745580 | -1.014 | 0.389 | 0.373 | -0.124 | -0.004 | -0.008 | -0.008 | 0 | 0.39 | 0.4 | 0.54 | 0.57 | 0.43 | 0.43 |
| 9 | 71698202 | rs6560551 | -0.975 | 0.407 | 0.333 | -0.159 | -0.004 | -0.008 | -0.008 | 0 | 0.39 | 0.37 | 0.54 | 0.56 | 0.43 | 0.44 |
| 9 | 71698581 | rs11145103 | -0.74 | 0.62 | 0.52 | -0.583 | 0.007 | 0.026 | 0.026 | 0 | 0.13 | 0.19 | 0.14 | 0.31 | 0.18 | 0.26 |
| 9 | 71707902 | rs7027499 | 0.937 | 0.666 | 0.455 | -0.927 | 0.005 | 0.047 | 0.047 | 0 | 0.49 | 0.45 | 0.69 | 0.66 | 0.5 | 0.46 |
| 9 | 71709115 | rs7851458 | -0.944 | 0.536 | 0.656 | -1.097 | -0.005 | -0.038 | -0.038 | 0.122 | 0.88 | 0.92 | 0.94 | 1 | 0.92 | 0.92 |
| 9 | 71712224 | rs11790543 | 1.154 | 0.525 | 0.677 | -1.07 | 0.004 | 0.044 | 0.044 | 0 | 0.49 | 0.46 | 0.69 | 0.67 | 0.5 | 0.46 |
| 9 | 71712736 | rs11145193 | 0.705 | 0.492 | 0.807 | -1.217 | 0.003 | -0.012 | -0.012 | 0 | 0.21 | 0.16 | 0.12 | 0.01 | 0.2 | 0.1 |
| 9 | 71712866 | rs11145194 | 0.939 | 0.536 | 0.89 | -1.073 | 0.038 | -0.004 | -0.004 | 0 | 0.08 | 0.12 | 0.06 | 0.08 | 0.12 | 0.19 |
| 9 | 71719618 | rs4744826 | 0.28 | 0.615 | 0.801 | -1.106 | -0.003 | -0.003 | -0.003 | 0 | 0.81 | 0.87 | 0.83 | 0.93 | 0.8 | 0.82 |
| 9 | 71724126 | rs7870657 | 1.332 | 0.614 | 1.009 | -0.922 | 0.011 | 0.018 | 0.018 | 0 | 0.64 | 0.49 | 0.66 | 0.51 | 0.61 | 0.48 |
| 9 | 71733603 | rs11145330 | 1.008 | 0.688 | 0.972 | -1.317 | 0.034 | -0.004 | -0.004 | 0 | 0.08 | 0.1 | 0.05 | 0.08 | 0.12 | 0.19 |
| 9 | 71747395 | rs6560585 | 1.618 | 0.649 | 0.868 | -1.449 | 0.003 | 0.006 | 0.006 | 0 | 0.44 | 0.44 | 0.6 | 0.6 | 0.4 | 0.35 |
| 9 | 71753270 | rs7875573 | -0.928 | 0.662 | 0.77 | -1.568 | 0 | 0.029 | 0.029 | 0 | 0.87 | 0.95 | 0.92 | 1 | 0.87 | 0.92 |
| 9 | 71755629 | rs17061543 | 0.814 | 0.628 | 0.711 | -1.574 | 0.006 | -0.042 | -0.042 | 2.368 | 0.08 | 0.02 | 0.06 | 0.24 | 0.06 | 0.15 |
| 9 | 71755705 | rs11145422 | 0.688 | 0.702 | 0.802 | -1.609 | 0.001 | 0.016 | 0.016 | 0 | 0.21 | 0.16 | 0.11 | 0.01 | 0.21 | 0.12 |
| 9 | 71759542 | rs13286409 | 0.466 | 0.686 | 0.729 | -1.595 | -0.005 | -0.002 | -0.002 | 0 | 0.05 | 0.06 | 0.09 | 0.07 | 0.08 | 0.09 |
| 9 | 71765617 | rs5006364 | -0.79 | 0.582 | 0.978 | -1.591 | 0.018 | -0.056 | -0.056 | 0 | 0.35 | 0.41 | 0.27 | 0.13 | 0.42 | 0.41 |
| 9 | 71769323 | rs4745646 | 0.407 | 0.639 | 0.971 | -1.511 | -0.007 | 0 | 0 | 0 | 0.78 | 0.83 | 0.86 | 0.76 | 0.82 | 0.76 |
| 9 | 71787927 | rs1416738 | 1.624 | 0.719 | 0.538 | -1.511 | 0.019 | 0.038 | 0.038 | 0 | 0.44 | 0.32 | 0.59 | 0.62 | 0.4 | 0.36 |
| 9 | 71791043 | rs17061953 | 0.995 | 0.782 | 0.502 | -1.505 | 0.018 | -0.01 | -0.01 | 0 | 0.08 | 0.07 | 0.04 | 0.05 | 0.13 | 0.19 |
| 9 | 71791547 | rs1538583 | 0.605 | 0.764 | -0.033 | -1.564 | -0.001 | 0.001 | 0.001 | 0 | 0.22 | 0.19 | 0.12 | 0.01 | 0.21 | 0.13 |
| 9 | 71802285 | rs3002378 | -1.023 | 0.602 | -0.404 | -1.321 | 0 | 0.014 | 0.014 | 0 | 0.5 | 0.45 | 0.31 | 0.31 | 0.52 | 0.55 |
| 9 | 71813571 | rs7043149 | 0.726 | 0.666 | -0.328 | -1.189 | -0.013 | 0.039 | 0.039 | 0 | 0.55 | 0.34 | 0.66 | 0.62 | 0.52 | 0.43 |
| 9 | 71820740 | rs4338173 | -0.58 | 0.427 | -0.418 | -1.107 | 0.004 | -0.031 | -0.031 | 0 | 0.38 | 0.42 | 0.24 | 0.3 | 0.4 | 0.48 |
| 9 | 71838080 | rs11137572 | -0.341 | 0.508 | -1.303 | -1.699 | 0.003 | 0.055 | 0.055 | 0 | 0.39 | 0.33 | 0.19 | 0.03 | 0.38 | 0.3 |
| 9 | 71838985 | rs2486448 | -1.058 | 0.322 | -1.767 | -1.343 | -0.068 | 0.17 | 0.17 | 0.41 | 0.63 | 0.94 | 0.42 | 0.16 | 0.6 | 0.55 |
| 9 | 71840396 | rs17062682 | -1.046 | 0.135 | -0.532 | -0.526 | -0.003 | -0.008 | -0.008 | 0 | 0.17 | 0.28 | 0.18 | 0.12 | 0.14 | 0.19 |
| 9 | 71841018 | rs17062695 | -0.785 | 0.123 | -0.478 | -0.34 | -0.002 | 0.007 | 0.007 | 0 | 0.07 | 0.16 | 0.11 | 0.12 | 0.06 | 0.1 |
| 9 | 71842391 | rs11137588 | -0.382 | 0.211 | -0.754 | -0.174 | 0.001 | -0.003 | -0.003 | 0 | 0.13 | 0.07 | 0.07 | 0.16 | 0.14 | 0.17 |
| 9 | 71843023 | rs2309428 | 0.967 | 0.261 | -0.442 | -0.273 | -0.01 | -0.025 | -0.025 | 0 | 0.79 | 0.64 | 0.82 | 0.87 | 0.83 | 0.75 |
| 9 | 71846783 | rs955792 | -0.851 | 0.533 | -0.475 | 0.231 | 0.001 | 0.019 | 0.019 | 0 | 0.05 | 0.12 | 0.1 | 0.15 | 0.05 | 0.12 |
| 9 | 71849766 | rs10867187 | -0.606 | 0.789 | -0.146 | 0.941 | 0.009 | 0.069 | 0.069 | 0 | 0.16 | 0.22 | 0.44 | 0.43 | 0.19 | 0.22 |
| 9 | 71850141 | rs10746573 | -0.601 | 0.807 | -0.478 | 0.212 | -0.001 | -0.001 | -0.001 | 0.582 | 0.56 | 0.54 | 0.72 | 0.62 | 0.63 | 0.69 |
| 9 | 71850853 | rs2275431 | 1.082 | 0.773 | -0.746 | -0.014 | -0.035 | 0.067 | 0.067 | 0 | 0.28 | 0.08 | 0.2 | 0.17 | 0.33 | 0.4 |
| 9 | 71852760 | rs2309421 | 0.065 | 0.874 | -0.68 | 0.448 | -0.007 | -0.01 | -0.01 | 0.004 | 0.17 | 0.16 | 0.45 | 0.46 | 0.22 | 0.23 |
| 9 | 71855357 | rs2498421 | 0.027 | 0.532 | -0.768 | 0.059 | 0.003 | -0.021 | -0.021 | 0 | 0.59 | 0.53 | 0.41 | 0.22 | 0.54 | 0.63 |
| 9 | 71858517 | rs2486456 | -0.372 | 0.524 | -0.886 | 0.121 | 0.012 | 0.003 | 0.003 | 0 | 0.58 | 0.68 | 0.74 | 0.65 | 0.64 | 0.71 |
| 9 | 71862129 | rs3750551 | 1.288 | 0.579 | -0.945 | 0.594 | -0.009 | -0.026 | -0.026 | 0 | 0.08 | 0.02 | 0.33 | 0.27 | 0.11 | 0.11 |
| 9 | 71869473 | rs3812536 | 0.144 | 0.731 | -1.167 | 0.034 | -0.027 | -0.028 | -0.028 | 0 | 0.38 | 0.25 | 0.6 | 0.61 | 0.45 | 0.49 |
| 9 | 71873762 | rs2039785 | 0.192 | 0.246 | -0.456 | -0.06 | -0.002 | 0.003 | 0.003 | 0 | 0.48 | 0.46 | 0.58 | 0.6 | 0.49 | 0.7 |
| 9 | 71875458 | rs2039787 | -1.597 | 0.09 | -0.997 | -0.023 | -0.052 | -0.093 | -0.093 | 1.742 | 0.84 | 0.99 | 0.86 | 0.74 | 0.82 | 0.96 |
| 9 | 71877579 | rs11789945 | 0.558 | 0.061 | -0.95 | -0.231 | 0.002 | 0.033 | 0.033 | 0.097 | 0.2 | 0.06 | 0.12 | 0.1 | 0.14 | 0.16 |
| 9 | 71883224 | rs1538579 | 0.312 | -0.144 | -0.459 | 0.018 | 0.005 | -0.003 | -0.003 | 0 | 0.6 | 0.43 | 0.75 | 0.52 | 0.68 | 0.8 |
| 9 | 71886979 | rs12350141 | 0.093 | -0.073 | -0.243 | -0.112 | -0.028 | -0.134 | -0.134 | 1.721 | 0.25 | 0.45 | 0.14 | 0.02 | 0.16 | 0.21 |
| 9 | 71887136 | rs2095875 | -1.712 | -0.097 | -0.495 | 0.052 | -0.052 | -0.135 | -0.135 | 0 | 0.82 | 0.99 | 0.88 | 0.64 | 0.81 | 0.95 |
| 9 | 71888362 | rs7868867 | -1.755 | -0.094 | -0.494 | 0.118 | -0.052 | -0.099 | -0.099 | 0 | 0.83 | 0.99 | 0.88 | 0.74 | 0.81 | 0.95 |
| 9 | 71891756 | rs1538581 | 0.585 | -0.115 | -0.011 | -0.062 | 0.002 | -0.015 | -0.015 | 0 | 0.22 | 0.17 | 0.11 | 0.02 | 0.16 | 0.21 |
| 9 | 71898527 | rs17063341 | 1.382 | 0.151 | 0.068 | -0.011 | 0.001 | -0.047 | -0.047 | 0 | 0.11 | 0.22 | 0.05 | 0.01 | 0.11 | 0.09 |
| 9 | 71901883 | rs6559428 | -0.102 | 0.167 | 0.358 | 0.517 | -0.005 | 0.003 | 0.003 | 0.296 | 0.58 | 0.62 | 0.75 | 0.62 | 0.65 | 0.75 |

| | | | | | | | | | | | | | | | | |
|---|---|---|---|---|---|---|---|---|---|---|---|---|---|---|---|---|
| 9 | 71907945 | rs12001191 | 0.821 | 0.239 | -0.068 | 0.399 | -0.02 | -0.022 | -0.022 | 0 | 0.43 | 0.36 | 0.69 | 0.58 | 0.54 | 0.64 |
| 9 | 71908061 | rs7034960 | 0.669 | 0.116 | -0.095 | 0.308 | 0.01 | 0.012 | 0.012 | 0 | 0.13 | 0.12 | 0.04 | 0.05 | 0.08 | 0.08 |
| 9 | 71916472 | rs11138012 | 0.521 | 0.51 | -0.005 | 0.521 | -0.018 | -0.018 | -0.018 | 0 | 0.43 | 0.35 | 0.69 | 0.56 | 0.55 | 0.64 |
| 9 | 71917575 | rs4744873 | 0.445 | 0.603 | -0.238 | 0.566 | -0.006 | -0.007 | -0.007 | 0 | 0.44 | 0.4 | 0.69 | 0.56 | 0.55 | 0.64 |
| 9 | 71926658 | rs6559453 | -1.021 | 0.764 | -0.642 | 1.333 | -0.008 | -0.061 | -0.061 | 0.474 | 0.86 | 0.95 | 0.85 | 0.68 | 0.85 | 0.95 |
| 9 | 71926960 | rs1572762 | -1.019 | 0.772 | -0.642 | 1.354 | -0.006 | -0.061 | -0.061 | 0 | 0.86 | 0.95 | 0.85 | 0.68 | 0.85 | 0.95 |
| 9 | 71927067 | rs1962383 | 0.64 | 0.558 | -0.306 | 0.826 | -0.018 | 0.011 | 0.011 | 0 | 0.63 | 0.49 | 0.74 | 0.62 | 0.71 | 0.77 |
| 9 | 71935577 | rs10780297 | 1.255 | 0.763 | -0.436 | 0.863 | -0.003 | 0 | 0 | 0 | 0.56 | 0.47 | 0.7 | 0.49 | 0.59 | 0.66 |
| 9 | 71939431 | rs10867357 | 0.446 | 0.726 | -0.223 | 1.167 | -0.02 | 0.012 | 0.012 | 0 | 0.71 | 0.55 | 0.76 | 0.7 | 0.75 | 0.82 |
| 9 | 71950492 | rs1334310 | 0.871 | 1.315 | -0.862 | 1.328 | -0.032 | -0.05 | -0.05 | 0 | 0.66 | 0.8 | 0.62 | 0.54 | 0.6 | 0.66 |
| 9 | 71953195 | rs7041728 | -0.582 | 1.271 | -0.9 | 1.292 | -0.006 | 0.004 | 0.004 | 0 | 0.18 | 0.18 | 0.24 | 0.16 | 0.23 | 0.22 |
| 9 | 71953719 | rs1970162 | 0.514 | 1.322 | -0.84 | 1.444 | -0.027 | -0.051 | -0.051 | 0 | 0.67 | 0.8 | 0.62 | 0.55 | 0.62 | 0.74 |
| 9 | 71954614 | rs10780308 | -0.395 | 1.261 | -0.83 | 1.268 | 0.009 | -0.03 | -0.03 | 0 | 0.78 | 0.85 | 0.8 | 0.7 | 0.78 | 0.9 |
| 9 | 71962528 | rs7033857 | -0.88 | 1.55 | -0.166 | 1.367 | 0.004 | 0.01 | 0.01 | 0 | 0.2 | 0.21 | 0.21 | 0.35 | 0.25 | 0.31 |
| 9 | 71973203 | rs7466730 | -0.107 | 1.552 | -0.896 | 1.141 | -0.062 | -0.062 | -0.062 | 0.147 | 0.11 | 0.01 | 0.16 | 0.17 | 0.18 | 0.18 |
| 9 | 71973435 | rs4421413 | 0.089 | 0.902 | -0.725 | 0.635 | 0 | -0.001 | -0.001 | 1.947 | 0.45 | 0.58 | 0.35 | 0.3 | 0.41 | 0.51 |
| 9 | 71985151 | rs11138342 | 1.046 | 0.725 | -0.974 | 0.871 | -0.01 | -0.035 | -0.035 | 0 | 0.07 | 0 | 0.13 | 0.12 | 0.08 | 0.12 |
| 9 | 71986805 | rs10780321 | -0.247 | 0.397 | -1.08 | 0.724 | -0.004 | -0.003 | -0.003 | 8.433 | 0.4 | 0.6 | 0.34 | 0.38 | 0.39 | 0.49 |
| 9 | 71990645 | rs4351475 | 0.631 | 0.461 | -1.45 | 0.738 | -0.08 | -0.054 | -0.054 | 0 | 0.35 | 0.07 | 0.45 | 0.36 | 0.41 | 0.39 |
| 9 | 71991799 | rs10867423 | -0.574 | 0.41 | -1.433 | 0.705 | -0.008 | 0 | 0 | 0.286 | 0.75 | 0.69 | 0.79 | 0.74 | 0.79 | 0.88 |
| 9 | 71998391 | rs10780333 | 0.802 | 0.25 | -0.673 | 0.177 | -0.01 | -0.017 | -0.017 | 0.088 | 0.29 | 0.38 | 0.21 | 0.24 | 0.23 | 0.37 |
| 9 | 72008141 | rs7033674 | -0.424 | 0.072 | -0.829 | 0.147 | 0.001 | -0.014 | -0.014 | 0 | 0.72 | 0.76 | 0.73 | 0.6 | 0.73 | 0.8 |
| 9 | 72011075 | rs11138435 | 0.515 | -0.021 | 0.056 | 0.167 | 0.012 | 0.021 | 0.021 | 0 | 0.26 | 0.16 | 0.19 | 0.23 | 0.22 | 0.35 |
| 9 | 72016960 | rs10867453 | 0.464 | -0.02 | 0.113 | 0.168 | 0.009 | 0.015 | 0.015 | 0 | 0.26 | 0.19 | 0.2 | 0.23 | 0.22 | 0.35 |
| 9 | 72027018 | rs2253885 | -0.022 | -0.144 | 0.3 | 0.197 | 0.023 | 0.037 | 0.037 | 1.209 | 0.32 | 0.24 | 0.29 | 0.2 | 0.26 | 0.43 |
| 9 | 72028323 | rs2777860 | -1.57 | -0.094 | 0.344 | 0.315 | -0.073 | -0.033 | -0.033 | 0 | 0.54 | 0.91 | 0.66 | 0.6 | 0.56 | 0.66 |
| 9 | 72036369 | rs11138566 | -0.436 | -0.122 | 0.7 | 0.606 | 0.024 | 0.224 | 0.224 | 1.383 | 0.12 | 0.53 | 0.23 | 0.32 | 0.09 | 0.11 |
| 9 | 72042096 | rs1757957 | 0.245 | -0.132 | 0.473 | 0.454 | 0.016 | 0.022 | 0.022 | 0 | 0.3 | 0.24 | 0.29 | 0.23 | 0.26 | 0.42 |
| 9 | 72047599 | rs1217339 | 0.72 | -0.162 | 0.499 | 0.434 | 0.01 | 0.012 | 0.012 | 0.12 | 0.26 | 0.22 | 0.13 | 0.19 | 0.19 | 0.31 |
| 9 | 72056397 | rs2781530 | -0.084 | -0.15 | 0.831 | 0.561 | -0.009 | 0.009 | 0.009 | 0 | 0.32 | 0.35 | 0.29 | 0.49 | 0.25 | 0.4 |
| 9 | 72058006 | rs1806760 | 0.74 | -0.151 | 0.796 | 0.53 | 0.004 | 0.004 | 0.004 | 0.114 | 0.26 | 0.23 | 0.13 | 0.19 | 0.19 | 0.32 |
| 9 | 72067942 | rs7869375 | -0.587 | -0.097 | 0.911 | 0.476 | -0.022 | 0.018 | 0.018 | 0 | 0.39 | 0.47 | 0.52 | 0.53 | 0.35 | 0.51 |
| 9 | 72076547 | rs4744899 | -1.112 | -0.068 | 0.94 | 0.434 | 0.023 | 0 | 0 | 0 | 0.47 | 0.81 | 0.56 | 0.59 | 0.44 | 0.6 |
| 9 | 72077866 | rs2781536 | -0.36 | -0.109 | 1.085 | 0.413 | -0.01 | -0.019 | -0.019 | 0 | 0.35 | 0.43 | 0.38 | 0.29 | 0.36 | 0.48 |
| 9 | 72080317 | rs1761285 | 0.739 | -0.113 | 1.129 | 0.417 | 0.014 | 0.012 | 0.012 | 0.453 | 0.23 | 0.18 | 0.12 | 0.19 | 0.18 | 0.29 |
| 9 | 72080464 | rs1631780 | -1.323 | -0.097 | 1.418 | 0.431 | 0.019 | -0.004 | -0.004 | 1.239 | 0.45 | 0.75 | 0.55 | 0.58 | 0.43 | 0.58 |
| 9 | 72081462 | rs870694 | -0.607 | 0.035 | 1.499 | 0.523 | 0.019 | -0.001 | -0.001 | 0 | 0.12 | 0.19 | 0.26 | 0.1 | 0.19 | 0.19 |
| 9 | 72082970 | rs10867613 | -0.565 | 0.036 | 1.49 | 0.524 | 0.011 | -0.001 | -0.001 | 0 | 0.12 | 0.15 | 0.25 | 0.1 | 0.19 | 0.19 |
| 9 | 72087236 | rs4744901 | 0.824 | 0.042 | 1.452 | 0.559 | 0.005 | 0.104 | 0.104 | 0.031 | 0.06 | 0.11 | 0.16 | 0.29 | 0.06 | 0.09 |
| 9 | 72090480 | rs2781541 | -1.109 | -0.048 | 1.413 | 0.502 | 0.022 | 0.006 | 0.006 | 0.598 | 0.47 | 0.73 | 0.56 | 0.59 | 0.45 | 0.61 |
| 9 | 72091104 | rs2777862 | 0.667 | -0.071 | 1.515 | 0.438 | 0.009 | 0.01 | 0.01 | 0.403 | 0.27 | 0.23 | 0.14 | 0.19 | 0.2 | 0.33 |
| 9 | 72101672 | rs7857219 | -0.614 | 0.058 | 1.582 | 0.419 | 0.014 | 0 | 0 | 0.11 | 0.12 | 0.19 | 0.25 | 0.1 | 0.18 | 0.19 |
| 9 | 72106629 | rs10867647 | -0.103 | 0.096 | 1.608 | 0.477 | 0.036 | 0.202 | 0.202 | 0.625 | 0.09 | 0.32 | 0.18 | 0.29 | 0.07 | 0.09 |
| 9 | 72107335 | rs11138927 | 0.639 | 0.111 | 1.506 | 0.485 | 0.009 | 0.095 | 0.095 | 0.088 | 0.07 | 0.11 | 0.17 | 0.29 | 0.07 | 0.09 |
| 9 | 72115231 | rs2150878 | -0.893 | 0.01 | 1.455 | 0.422 | 0.022 | 0.026 | 0.026 | 0.393 | 0.73 | 0.82 | 0.87 | 0.81 | 0.8 | 0.68 |
| 9 | 72115498 | rs2309481 | -1.033 | 0.073 | 1.566 | 0.591 | 0.028 | 0.097 | 0.097 | 0.024 | 0.17 | 0.26 | 0.42 | 0.42 | 0.25 | 0.28 |
| 9 | 72132199 | rs11139063 | 0.368 | -0.001 | 1.599 | 0.536 | 0.01 | 0.007 | 0.007 | 0.453 | 0.59 | 0.29 | 0.65 | 0.58 | 0.67 | 0.53 |
| 9 | 72134376 | rs10481751 | -0.293 | -0.016 | 1.694 | 0.518 | 0.016 | 0.013 | 0.013 | 0.052 | 0.68 | 0.69 | 0.83 | 0.88 | 0.74 | 0.62 |
| 9 | 72136263 | rs12378318 | 1.285 | 0.03 | 1.509 | 0.53 | -0.016 | 0.022 | 0.022 | 2.075 | 0.48 | 0.13 | 0.39 | 0.39 | 0.46 | 0.35 |
| 9 | 72147700 | rs1412243 | -0.93 | 0.01 | 1.637 | 0.533 | 0.033 | 0.075 | 0.075 | 0.101 | 0.13 | 0.33 | 0.27 | 0.19 | 0.21 | 0.19 |
| 9 | 72148492 | rs7851858 | 0.301 | -0.035 | 1.592 | 0.37 | -0.016 | -0.01 | -0.01 | 0 | 0.61 | 0.45 | 0.65 | 0.58 | 0.67 | 0.54 |
| 9 | 72152776 | rs1029064 | 0.14 | 0.021 | 1.523 | 0.425 | 0.009 | 0.104 | 0.104 | 0.306 | 0.08 | 0.21 | 0.18 | 0.3 | 0.07 | 0.09 |
| 9 | 72160174 | rs7858171 | -0.932 | -0.035 | 1.289 | 0.332 | 0.034 | 0.066 | 0.066 | 0.443 | 0.73 | 0.85 | 0.87 | 0.88 | 0.8 | 0.68 |
| 9 | 72163388 | rs1330327 | 0.429 | 0.012 | 1.219 | 0.4 | 0.013 | 0.011 | 0.011 | 0.581 | 0.58 | 0.31 | 0.62 | 0.59 | 0.65 | 0.54 |
| 9 | 72166621 | rs11139300 | 1.2 | 0.045 | 0.738 | -0.221 | -0.051 | 0.036 | 0.036 | 1.704 | 0.36 | 0.08 | 0.32 | 0.24 | 0.38 | 0.24 |
| 9 | 72167347 | rs7864493 | -1.242 | -0.194 | 1.476 | -0.981 | 0.017 | -0.035 | -0.035 | 0.733 | 0.24 | 0.51 | 0.2 | 0.13 | 0.29 | 0.26 |
| 9 | 72174392 | rs1970017 | 0.706 | -0.308 | 1.607 | -0.753 | 0.005 | 0.015 | 0.015 | 0.998 | 0.59 | 0.34 | 0.7 | 0.79 | 0.58 | 0.52 |
| 9 | 72174528 | rs11139338 | -0.38 | -0.302 | 1.481 | -0.754 | 0.004 | 0.036 | 0.036 | 0.576 | 0.33 | 0.22 | 0.24 | 0.21 | 0.36 | 0.43 |
| 9 | 72181521 | rs12004586 | -0.218 | -0.165 | 1.618 | -0.638 | 0.024 | -0.02 | -0.02 | 0.063 | 0.07 | 0.29 | 0.06 | 0 | 0.08 | 0.06 |
| 9 | 72204608 | rs1330334 | 0.304 | -0.164 | 1.186 | -0.646 | -0.005 | -0.016 | -0.016 | 0 | 0.68 | 0.64 | 0.8 | 0.79 | 0.7 | 0.61 |
| 9 | 72209932 | rs12115441 | 0.492 | -0.107 | 0.882 | -0.624 | -0.002 | 0.004 | 0.004 | 0 | 0.05 | 0.05 | 0.08 | 0.16 | 0.12 | 0.12 |
| 9 | 72213799 | rs7853648 | 0.374 | -0.216 | 0.735 | -0.932 | 0.023 | 0.041 | 0.041 | 0.047 | 0.25 | 0.16 | 0.12 | 0.05 | 0.18 | 0.28 |
| 9 | 72230645 | rs10746682 | 0.399 | -0.172 | 0.56 | -0.872 | 0.015 | 0.04 | 0.04 | 1.18 | 0.6 | 0.26 | 0.73 | 0.83 | 0.62 | 0.56 |
| 9 | 72233182 | rs13288998 | 0.631 | 0.007 | -0.032 | -0.399 | -0.028 | -0.013 | -0.013 | 0 | 0.11 | 0.02 | 0.06 | 0.09 | 0.09 | 0.08 |
| 9 | 72243177 | rs975645 | 0.76 | -0.33 | 0.76 | -0.447 | -0.01 | -0.017 | -0.017 | 0 | 0.7 | 0.4 | 0.66 | 0.73 | 0.74 | 0.89 |
| 9 | 72252269 | rs10867845 | -0.852 | -0.493 | 0.668 | -0.971 | -0.012 | -0.051 | -0.051 | 0 | 0.27 | 0.4 | 0.21 | 0.14 | 0.25 | 0.16 |
| 9 | 72252915 | rs4629937 | 1.747 | -0.632 | 0.641 | -0.961 | 0.004 | -0.033 | -0.033 | 0 | 0.68 | 0.25 | 0.82 | 0.97 | 0.75 | 0.71 |
| 9 | 72258604 | rs7851932 | 0.021 | 0.072 | -0.023 | -0.506 | -0.01 | -0.022 | -0.022 | 0 | 0.11 | 0.11 | 0.04 | 0.03 | 0.05 | 0.11 |
| 9 | 72262481 | rs7868291 | -1.375 | 0.057 | 0.066 | -0.524 | -0.007 | -0.068 | -0.068 | 0 | 0.22 | 0.31 | 0.13 | 0.03 | 0.21 | 0.27 |

| | | | | | | | | | | | | | | | | |
|---|---|---|---|---|---|---|---|---|---|---|---|---|---|---|---|---|
| 9 | 72273898 | rs11139651 | -0.921 | 0.529 | -0.171 | -0.132 | 0.003 | -0.038 | -0.038 | 0 | 0.11 | 0.19 | 0.09 | 0 | 0.16 | 0.16 |
| 9 | 72291978 | rs10867879 | 0.19 | 0.703 | -0.332 | 0.426 | 0 | 0.006 | 0.006 | 0.035 | 0.62 | 0.55 | 0.69 | 0.58 | 0.59 | 0.53 |
| 9 | 72303882 | rs10867916 | -0.474 | 0.509 | 0.363 | 0.925 | 0.01 | 0.097 | 0.097 | 0.618 | 0.17 | 0.46 | 0.14 | 0.21 | 0.14 | 0.18 |
| 9 | 72322433 | rs10867932 | 0.482 | 1.31 | 0.506 | 0.842 | 0.013 | -0.003 | -0.003 | 2.679 | 0.48 | 0.31 | 0.38 | 0.56 | 0.45 | 0.37 |
| 9 | 72328727 | rs7857137 | 1.1 | 1.119 | 0.108 | 0.903 | -0.012 | -0.007 | -0.007 | 0 | 0.33 | 0.09 | 0.29 | 0.35 | 0.33 | 0.22 |
| 9 | 72338610 | rs11139997 | 1.045 | 1.13 | 0.193 | 1.008 | -0.074 | 0.084 | 0.084 | 0 | 0.32 | 0.03 | 0.45 | 0.14 | 0.35 | 0.4 |
| 9 | 72342930 | rs1411411 | 1.077 | 1.13 | 0.194 | 0.911 | -0.075 | 0.125 | 0.125 | 0 | 0.31 | 0.03 | 0.4 | 0.11 | 0.35 | 0.4 |
| 9 | 72362158 | rs10511969 | 1.874 | 0.986 | -0.256 | 0.943 | 0 | -0.015 | -0.015 | 0 | 0.11 | 0.06 | 0.08 | 0.18 | 0.11 | 0.09 |
| 9 | 72363992 | rs11793130 | 1.156 | 1.135 | -0.144 | 0.806 | -0.032 | 0.116 | 0.116 | 0 | 0.32 | 0.1 | 0.46 | 0.11 | 0.35 | 0.4 |
| 9 | 72373346 | rs2297172 | 1.151 | 1.122 | -0.02 | 0.884 | 0.002 | 0.019 | 0.019 | 0 | 0.13 | 0.14 | 0.14 | 0.23 | 0.14 | 0.18 |
| 9 | 72377357 | rs11140189 | 1.214 | 1.176 | -0.035 | 0.906 | -0.064 | 0.126 | 0.126 | 0 | 0.31 | 0.04 | 0.43 | 0.12 | 0.34 | 0.4 |
| 9 | 72383476 | rs1536993 | -0.429 | 1.262 | 0.138 | 0.966 | 0.001 | -0.015 | -0.015 | 0 | 0.6 | 0.42 | 0.53 | 0.78 | 0.59 | 0.54 |
| 9 | 72398515 | rs11521712 | 1.251 | 1.407 | 0.204 | 1.499 | -0.004 | 0.012 | 0.012 | 0 | 0.08 | 0.07 | 0.12 | 0.23 | 0.1 | 0.17 |
| 9 | 72403792 | rs6559749 | -0.091 | 1.213 | 0.321 | 1.122 | 0.073 | 0.109 | 0.109 | 0.485 | 0.95 | 0.74 | 0.97 | 0.89 | 0.94 | 0.94 |
| 9 | 72405717 | rs1819414 | -0.601 | 1.419 | -0.068 | 1.118 | 0.02 | 0.039 | 0.039 | 0 | 0.82 | 0.71 | 0.77 | 0.72 | 0.76 | 0.83 |
| 9 | 72408691 | rs10868073 | 0.919 | -0.471 | -1.074 | -1.074 | -0.03 | 0.054 | 0.054 | 0 | 0.2 | 0.01 | 0.25 | 0.1 | 0.21 | 0.15 |
| 9 | 72410323 | rs17517599 | 1.132 | -0.633 | -1.604 | -1.317 | -0.006 | 0.065 | 0.065 | 0 | 0.11 | 0.02 | 0.09 | 0 | 0.11 | 0.18 |
| 9 | 72413613 | rs7022623 | 1.783 | -1.043 | -0.57 | -1.351 | 0.037 | 0.084 | 0.084 | 0 | 0.17 | 0.26 | 0.19 | 0.3 | 0.18 | 0.14 |
| 9 | 72417239 | rs9314727 | 0.21 | -1.02 | -0.74 | -1.365 | 0.008 | 0.005 | 0.005 | 0 | 0.33 | 0.52 | 0.33 | 0.31 | 0.32 | 0.32 |
| 9 | 72422919 | rs11140368 | 1.602 | -1.151 | -0.705 | -1.287 | -0.003 | 0.003 | 0.003 | 0 | 0.1 | 0.08 | 0.16 | 0.28 | 0.13 | 0.13 |
| 9 | 72424523 | rs2871266 | -0.434 | -1.189 | -0.863 | -1.416 | -0.005 | -0.035 | -0.035 | 0 | 0.64 | 0.61 | 0.82 | 0.82 | 0.69 | 0.62 |
| 9 | 72425204 | rs2183380 | 1.424 | -1.105 | -0.749 | -1.208 | -0.001 | 0.01 | 0.01 | 0 | 0.11 | 0.12 | 0.17 | 0.29 | 0.13 | 0.13 |
| 9 | 72427015 | rs11791776 | 1.476 | -1.095 | -0.763 | -1.203 | -0.003 | 0.006 | 0.006 | 0 | 0.1 | 0.08 | 0.16 | 0.29 | 0.13 | 0.13 |
| 9 | 72429750 | rs7863087 | 1.227 | -1.328 | -0.895 | -1.464 | 0.009 | 0.032 | 0.032 | 0 | 0.13 | 0.09 | 0.02 | 0.08 | 0.07 | 0.16 |
| 9 | 72432080 | rs10124812 | -0.708 | -1.109 | -0.461 | -1.248 | 0 | 0.019 | 0.019 | 0 | 0.64 | 0.73 | 0.73 | 0.68 | 0.65 | 0.59 |
| 9 | 72440451 | rs10868131 | -0.246 | -1.127 | -0.33 | -1.306 | 0.006 | -0.002 | -0.002 | 1.971 | 0.63 | 0.37 | 0.54 | 0.62 | 0.64 | 0.67 |
| 9 | 72441391 | rs1330239 | 0.702 | -1.156 | -0.429 | -1.347 | -0.033 | 0.036 | 0.036 | 0 | 0.34 | 0.11 | 0.24 | 0.31 | 0.33 | 0.36 |
| 9 | 72442540 | rs10511972 | 0.3 | -1.151 | -0.464 | -1.371 | 0.005 | 0.005 | 0.005 | 0 | 0.53 | 0.54 | 0.46 | 0.61 | 0.52 | 0.48 |
| 9 | 72446051 | rs4439190 | 0.256 | -1.117 | -1.012 | -1.391 | 0.023 | 0.153 | 0.153 | 0 | 0.33 | 0.1 | 0.35 | 0.1 | 0.33 | 0.36 |
| 9 | 72448995 | rs10521434 | 1.292 | -0.999 | -1.05 | -1.254 | -0.001 | 0.037 | 0.037 | 0 | 0.11 | 0.07 | 0.1 | 0.01 | 0.12 | 0.2 |
| 9 | 72456491 | rs12685383 | 0.48 | -1.326 | -0.903 | -1.045 | -0.044 | 0.029 | 0.029 | 0 | 0.34 | 0.1 | 0.24 | 0.31 | 0.33 | 0.34 |
| 9 | 72460608 | rs1493050 | -0.41 | -1.146 | -0.488 | -0.996 | 0.009 | -0.005 | -0.005 | 0 | 0.55 | 0.66 | 0.4 | 0.58 | 0.41 | 0.49 |
| 9 | 72464074 | rs1389128 | -0.42 | -1.128 | -0.479 | -0.987 | 0.008 | -0.005 | -0.005 | 0.167 | 0.56 | 0.66 | 0.39 | 0.58 | 0.41 | 0.49 |
| 9 | 72466275 | rs10511973 | 0.034 | -0.565 | -0.786 | -0.692 | -0.002 | -0.022 | -0.022 | 0 | 0.37 | 0.29 | 0.29 | 0.46 | 0.36 | 0.4 |
| 9 | 72472831 | rs10780682 | 1.69 | 0.051 | -0.084 | -0.44 | -0.04 | -0.054 | -0.054 | 1.502 | 0.39 | 0.09 | 0.29 | 0.41 | 0.33 | 0.29 |
| 9 | 72479046 | rs2035921 | -0.468 | -0.258 | 0.564 | -0.082 | 0.017 | 0.046 | 0.046 | 0 | 0.86 | 0.57 | 0.84 | 0.84 | 0.86 | 0.86 |
| 9 | 72479663 | rs10868224 | 0.574 | -0.271 | 0.611 | -0.113 | -0.001 | 0 | 0 | 0 | 0.69 | 0.66 | 0.58 | 0.84 | 0.57 | 0.63 |
| 9 | 72480440 | rs766003 | 0.766 | -0.265 | 0.495 | -0.11 | 0.016 | -0.055 | -0.055 | 0 | 0.67 | 0.46 | 0.57 | 0.84 | 0.56 | 0.63 |
| 9 | 72484697 | rs7869694 | 0.578 | -0.221 | 0.379 | -0.063 | 0.012 | 0.09 | 0.09 | 0.063 | 0.09 | 0.12 | 0.17 | 0.26 | 0.13 | 0.15 |
| 9 | 72486904 | rs12235696 | 1.607 | -0.117 | -0.416 | -0.006 | -0.038 | -0.013 | -0.013 | 0 | 0.46 | 0.19 | 0.34 | 0.5 | 0.41 | 0.4 |
| 9 | 72488417 | rs13284442 | 0.079 | 0.078 | -0.558 | -0.609 | -0.027 | 0.03 | 0.03 | 0 | 0.14 | 0.08 | 0.26 | 0.01 | 0.28 | 0.22 |
| 9 | 72488442 | rs17085936 | -0.117 | -0.369 | -0.22 | -0.065 | 0.052 | 0.017 | 0.017 | 0 | 0.1 | 0.02 | 0.05 | 0.08 | 0.04 | 0.08 |
| 9 | 72501591 | rs4237236 | 0.458 | 0.776 | 0.379 | 0.236 | 0.002 | 0.016 | 0.016 | 0 | 0.87 | 0.78 | 0.85 | 0.84 | 0.87 | 0.86 |
| 9 | 72512292 | rs12378305 | -0.431 | 1.085 | -0.943 | -0.438 | -0.017 | 0.038 | 0.038 | 0 | 0.23 | 0.2 | 0.3 | 0.01 | 0.34 | 0.25 |
| 9 | 72512898 | rs10217478 | 0.181 | 0.881 | -0.757 | 0.011 | -0.004 | 0.014 | 0.014 | 0 | 0.66 | 0.36 | 0.6 | 0.54 | 0.66 | 0.56 |
| 9 | 72513240 | rs1907965 | -1.518 | 0.812 | 0.012 | -0.262 | 0.016 | -0.028 | -0.028 | 0.267 | 0.08 | 0.28 | 0.19 | 0.02 | 0.07 | 0.16 |
| 9 | 72514177 | rs10868248 | 1.049 | 0.814 | 0.735 | 0.259 | -0.002 | 0.08 | 0.08 | 0.82 | 0.82 | 0.47 | 0.89 | 0.67 | 0.87 | 0.74 |
| 9 | 72519172 | rs11140842 | -1.179 | 0.96 | 1.549 | 0.234 | 0.032 | -0.026 | -0.026 | 0 | 0.06 | 0.14 | 0.06 | 0.01 | 0.08 | 0.07 |
| 9 | 72527688 | rs17520192 | -1.387 | 0.873 | 1.057 | 0.294 | 0.02 | -0.013 | -0.013 | 0 | 0.19 | 0.29 | 0.23 | 0.15 | 0.21 | 0.2 |
| 9 | 72530831 | rs10511975 | 1.357 | 0.682 | -0.1 | -0.626 | -0.059 | 0.161 | 0.161 | 0 | 0.36 | 0.16 | 0.26 | 0.06 | 0.42 | 0.26 |
| 9 | 72531390 | rs6559850 | -0.661 | 0.527 | 0.214 | -0.527 | -0.006 | 0.067 | 0.067 | 0.045 | 0.87 | 0.78 | 0.93 | 0.68 | 0.89 | 0.79 |
| 9 | 72532067 | rs7852214 | -0.581 | 0.129 | -0.067 | -0.592 | -0.046 | -0.081 | -0.081 | 0 | 0.33 | 0.09 | 0.39 | 0.51 | 0.38 | 0.35 |
| 9 | 72532478 | rs12684384 | 0.205 | 0.1 | -0.153 | -0.661 | -0.053 | -0.131 | -0.131 | 1.688 | 0.25 | 0.03 | 0.34 | 0.51 | 0.3 | 0.23 |
| 9 | 72535613 | rs10511976 | 0.154 | 0.048 | -0.094 | -0.695 | 0.003 | -0.012 | -0.012 | 0 | 0.05 | 0.07 | 0.05 | 0.01 | 0.08 | 0.07 |
| 9 | 72552407 | rs7870323 | -0.636 | -0.126 | -0.14 | -0.745 | -0.028 | -0.052 | -0.052 | 0.933 | 0.21 | 0.61 | 0.2 | 0.09 | 0.14 | 0.22 |
| 9 | 72562231 | rs7032845 | 1.323 | -0.08 | -0.206 | -0.791 | -0.001 | 0.089 | 0.089 | 0 | 0.5 | 0.65 | 0.41 | 0.15 | 0.49 | 0.41 |
| 9 | 72562779 | rs1493048 | 0.62 | -0.151 | 0.091 | -0.812 | -0.018 | -0.008 | -0.008 | 0 | 0.11 | 0.21 | 0.14 | 0.09 | 0.07 | 0.15 |
| 9 | 72565221 | rs10868315 | -0.405 | -0.098 | -0.081 | -0.694 | -0.036 | -0.139 | -0.139 | 0.386 | 0.31 | 0.03 | 0.46 | 0.49 | 0.32 | 0.31 |
| 9 | 72566736 | rs1389119 | 1.04 | -0.111 | -0.034 | -0.701 | 0.002 | 0.065 | 0.065 | 0 | 0.53 | 0.66 | 0.42 | 0.15 | 0.52 | 0.46 |
| 9 | 72569410 | rs7041366 | -1.232 | -0.42 | 0.09 | -0.602 | 0.01 | 0.003 | 0.003 | 0 | 0.43 | 0.57 | 0.5 | 0.57 | 0.38 | 0.44 |
| 9 | 72579310 | rs12236860 | -0.465 | -0.447 | 0.25 | -0.594 | -0.025 | 0.068 | 0.068 | 0.074 | 0.31 | 0.43 | 0.39 | 0.51 | 0.29 | 0.25 |
| 9 | 72581578 | rs4744581 | 1.298 | -0.342 | 0.052 | -0.632 | 0.02 | 0.259 | 0.259 | 0 | 0.47 | 0.21 | 0.43 | 0.07 | 0.55 | 0.41 |
| 9 | 72590450 | rs10511977 | 0.076 | -0.132 | 0.071 | -0.629 | 0.001 | -0.013 | -0.013 | 0 | 0.06 | 0.06 | 0.05 | 0 | 0.08 | 0.07 |
| 9 | 72591329 | rs11141099 | 2.353 | -0.132 | -0.003 | -0.677 | -0.148 | 0.253 | 0.253 | 0 | 0.4 | 0.07 | 0.37 | 0.07 | 0.46 | 0.34 |
| 9 | 72591525 | rs11141100 | 2.343 | -0.132 | -0.003 | -0.677 | -0.148 | 0.254 | 0.254 | 0 | 0.4 | 0.07 | 0.37 | 0.07 | 0.46 | 0.34 |
| 9 | 72593325 | rs11141106 | -0.77 | -0.19 | 0.073 | -0.614 | 0 | 0.002 | 0.002 | 0 | 0.25 | 0.25 | 0.34 | 0.23 | 0.26 | 0.31 |
| 9 | 72597565 | rs902964 | -1.006 | -0.192 | 0.167 | -0.576 | -0.015 | 0.091 | 0.091 | 0.081 | 0.87 | 0.79 | 0.94 | 0.64 | 0.91 | 0.81 |
| 9 | 72598602 | rs10868350 | 2.46 | -0.176 | 0.063 | -0.614 | -0.109 | 0.209 | 0.209 | 0 | 0.38 | 0.12 | 0.37 | 0.07 | 0.44 | 0.29 |
| 9 | 72609439 | rs10868355 | 1.434 | -0.214 | 0.462 | -0.581 | 0.039 | -0.026 | -0.026 | 0 | 0.57 | 0.25 | 0.57 | 0.41 | 0.63 | 0.45 |

| | | | | | | | | | | | | | | | | |
|---|---|---|---|---|---|---|---|---|---|---|---|---|---|---|---|---|
| 9 | 72609532 | rs1565691 | 1.246 | -0.206 | 0.487 | -0.577 | -0.011 | 0.01 | 0.01 | 0 | 0.6 | 0.51 | 0.59 | 0.41 | 0.64 | 0.48 |
| 9 | 72611680 | rs10868358 | 0.174 | -0.196 | 0.466 | -0.109 | -0.001 | 0.06 | 0.06 | 0 | 0.81 | 0.68 | 0.88 | 0.62 | 0.83 | 0.74 |
| 9 | 72621005 | rs10868365 | 0.425 | -0.178 | 0.171 | -0.099 | 0 | -0.026 | -0.026 | 0 | 0.19 | 0.13 | 0.21 | 0.33 | 0.19 | 0.16 |
| 9 | 72629662 | rs12554412 | -0.154 | -0.349 | 0.401 | 0.112 | 0.007 | -0.005 | -0.005 | 0 | 0.18 | 0.13 | 0.29 | 0.22 | 0.18 | 0.21 |
| 9 | 72633252 | rs12238553 | 0.63 | -0.379 | 0.039 | 0.099 | 0.009 | -0.026 | -0.026 | 0 | 0.14 | 0.07 | 0.17 | 0.33 | 0.11 | 0.15 |
| 9 | 72634148 | rs10868393 | 0.855 | -0.456 | 0.096 | 0.044 | 0.021 | -0.015 | -0.015 | 1.652 | 0.64 | 0.36 | 0.64 | 0.4 | 0.73 | 0.55 |
| 9 | 72637119 | rs4446788 | 1.59 | -0.324 | -0.063 | -0.299 | 0.077 | 0.216 | 0.216 | 0 | 0.49 | 0.26 | 0.47 | 0.07 | 0.6 | 0.4 |
| 9 | 72648582 | rs12376586 | 2.087 | -0.227 | -0.573 | -0.195 | -0.091 | 0.272 | 0.272 | 0 | 0.42 | 0.09 | 0.4 | 0.07 | 0.51 | 0.31 |
| 9 | 72648997 | rs2975926 | 0.057 | -0.177 | 0.27 | 0.51 | -0.02 | 0.097 | 0.097 | 0.053 | 0.13 | 0.22 | 0.06 | 0.37 | 0.09 | 0.19 |
| 9 | 72649864 | rs17444393 | -0.805 | 0.099 | -0.233 | 0.073 | 0.002 | 0.003 | 0.003 | 0 | 0.22 | 0.19 | 0.28 | 0.19 | 0.19 | 0.27 |
| 9 | 72658280 | rs11141404 | 1.664 | 0.276 | -0.618 | 0.039 | 0.004 | -0.008 | -0.008 | 0 | 0.56 | 0.16 | 0.62 | 0.44 | 0.59 | 0.44 |
| 9 | 72662689 | rs2997657 | -0.733 | 0.211 | 0.413 | 0.02 | -0.003 | -0.031 | -0.031 | 0.745 | 0.21 | 0.38 | 0.24 | 0.2 | 0.21 | 0.24 |
| 9 | 72667128 | rs12352964 | -0.288 | 0.32 | 0.74 | 0.152 | 0.004 | -0.001 | -0.001 | 0 | 0.13 | 0.17 | 0.18 | 0.19 | 0.15 | 0.19 |
| 9 | 72672537 | rs6559939 | 1.966 | 0.254 | 0.674 | 0.13 | -0.019 | -0.051 | -0.051 | 0.276 | 0.61 | 0.09 | 0.66 | 0.46 | 0.64 | 0.47 |
| 9 | 72676057 | rs2997664 | -1.772 | 0.227 | 1.249 | 0.21 | -0.017 | -0.007 | -0.007 | 0 | 0.3 | 0.59 | 0.27 | 0.4 | 0.27 | 0.45 |
| 9 | 72679678 | rs2997668 | -0.002 | 0.176 | 1.31 | 0.185 | -0.038 | 0.018 | 0.018 | 0 | 0.15 | 0.44 | 0.08 | 0.21 | 0.08 | 0.2 |
| 9 | 72683911 | rs2997674 | 0.476 | 0.307 | 1.308 | 0.244 | 0.07 | 0.074 | 0.074 | 0.017 | 0.91 | 0.73 | 0.92 | 0.87 | 0.89 | 0.91 |
| 9 | 72696962 | rs12686668 | 0.545 | 0.229 | 1.156 | 0.212 | 0.014 | -0.008 | -0.008 | 0.01 | 0.14 | 0.1 | 0.06 | 0.21 | 0.1 | 0.22 |
| 9 | 72707609 | rs2997685 | 0.897 | 0.192 | 1.362 | 0.253 | 0.026 | 0.06 | 0.06 | 1.176 | 0.9 | 0.69 | 0.92 | 0.87 | 0.9 | 0.91 |
| 9 | 72710563 | rs4237238 | -0.043 | 0.209 | 1.311 | 0.247 | 0.003 | 0 | 0 | 0.202 | 0.12 | 0.14 | 0.18 | 0.19 | 0.15 | 0.19 |
| 9 | 72722865 | rs1927103 | 2.046 | 0.177 | 1.521 | 0.224 | 0.001 | -0.024 | -0.024 | 0 | 0.61 | 0.15 | 0.66 | 0.47 | 0.63 | 0.47 |
| 9 | 72726597 | rs2078391 | 2.101 | 0.299 | 0.459 | -0.162 | -0.116 | 0.182 | 0.182 | 0 | 0.42 | 0.02 | 0.44 | 0.11 | 0.44 | 0.35 |
| 9 | 72728388 | rs3015224 | -0.7 | 0.235 | 0.92 | 0.047 | -0.01 | 0.002 | 0.002 | 0 | 0.26 | 0.35 | 0.22 | 0.41 | 0.24 | 0.38 |
| 9 | 72728822 | rs4744977 | 0.908 | 0.271 | 0.938 | 0.033 | -0.003 | -0.007 | -0.007 | 0.014 | 0.71 | 0.52 | 0.74 | 0.6 | 0.7 | 0.58 |
| 9 | 72738339 | rs10511980 | 1.655 | 0.162 | 0.647 | -0.011 | 0.015 | -0.031 | -0.031 | 0 | 0.63 | 0.12 | 0.68 | 0.5 | 0.67 | 0.57 |
| 9 | 72739478 | rs10511981 | 1.394 | 0.357 | 0.847 | 0.098 | 0.131 | 0.055 | 0.055 | 3.579 | 0.95 | 0.6 | 0.95 | 1 | 0.94 | 0.96 |
| 9 | 72749590 | rs4744982 | -0.144 | 0.308 | 0.37 | 0.152 | -0.006 | -0.001 | -0.001 | 0 | 0.83 | 0.82 | 0.93 | 0.79 | 0.86 | 0.77 |
| 9 | 72765006 | rs7856528 | 0.524 | 0.385 | 0.378 | 0.137 | -0.001 | -0.002 | -0.002 | 0 | 0.8 | 0.73 | 0.84 | 0.75 | 0.81 | 0.68 |
| 9 | 72766144 | rs10780851 | 0.524 | 0.385 | 0.378 | 0.137 | -0.001 | -0.002 | -0.002 | 0.12 | 0.8 | 0.73 | 0.84 | 0.75 | 0.81 | 0.68 |
| 9 | 72766414 | rs1927117 | 0.866 | 0.399 | 0.355 | 0.14 | 0.016 | -0.048 | -0.048 | 0.041 | 0.13 | 0.01 | 0.03 | 0.22 | 0.09 | 0.19 |
| 9 | 72767405 | rs10125454 | 0.255 | 0.579 | 0.535 | 0.201 | 0.082 | -0.008 | -0.008 | 1.447 | 0.93 | 0.75 | 0.87 | 0.96 | 0.89 | 0.88 |
| 9 | 72783608 | rs10780871 | 0.256 | 0.579 | 0.522 | 0.217 | 0.08 | -0.001 | -0.001 | 0 | 0.93 | 0.76 | 0.87 | 0.95 | 0.89 | 0.87 |
| 9 | 72788372 | rs1927133 | 0.213 | 0.586 | 0.464 | 0.219 | 0.082 | 0 | 0 | 1.622 | 0.94 | 0.76 | 0.87 | 0.95 | 0.89 | 0.87 |
| 9 | 72873252 | rs2282068 | 0.63 | 0.415 | -0.155 | 0.244 | -0.069 | -0.149 | -0.149 | 0.019 | 0.18 | 0.01 | 0.23 | 0.4 | 0.23 | 0.16 |
| 9 | 72885267 | rs1180109 | -0.417 | 0.284 | 0.419 | 0.151 | 0.008 | 0.028 | 0.028 | 0 | 0.9 | 0.77 | 0.9 | 0.85 | 0.91 | 0.88 |
| 9 | 72897434 | rs1180116 | -0.417 | 0.284 | 0.419 | 0.151 | 0.008 | 0.028 | 0.028 | 0 | 0.9 | 0.77 | 0.9 | 0.85 | 0.91 | 0.88 |
| 9 | 72897440 | rs1180117 | -0.352 | 0.206 | 0.475 | 0.043 | -0.001 | 0 | 0 | 0 | 0.2 | 0.27 | 0.15 | 0.27 | 0.19 | 0.32 |
| 9 | 72897970 | rs11142348 | 0.155 | 0.211 | 0.419 | 0.036 | 0.035 | 0.004 | 0.004 | 1.642 | 0.06 | 0.21 | 0.08 | 0.05 | 0.06 | 0.12 |
| 9 | 72902503 | rs10868798 | 0.733 | 0.167 | 0.099 | 0.04 | -0.002 | 0.008 | 0.008 | 0.006 | 0.08 | 0.08 | 0.09 | 0.15 | 0.09 | 0.12 |
| 9 | 72918167 | rs10125617 | 1.976 | -0.04 | -0.227 | -0.25 | -0.002 | -0.046 | -0.046 | 0.188 | 0.58 | 0.09 | 0.66 | 0.49 | 0.64 | 0.49 |
| 9 | 72951417 | rs7049118 | -0.135 | -0.021 | 0.056 | -0.229 | -0.01 | 0.005 | 0.005 | 0.492 | 0.12 | 0.3 | 0.08 | 0.1 | 0.08 | 0.07 |
| 9 | 72977345 | rs12684881 | 0.751 | 0.014 | -0.487 | -0.321 | -0.001 | 0.009 | 0.009 | 0.011 | 0.09 | 0.09 | 0.09 | 0.15 | 0.09 | 0.12 |
| 9 | 72978218 | rs10491653 | 1.131 | -0.003 | -1.088 | -0.299 | 0.016 | -0.048 | -0.048 | 0.025 | 0.13 | 0.01 | 0.03 | 0.22 | 0.09 | 0.2 |
| 9 | 72979121 | rs12347747 | 1.818 | 0.159 | -1.08 | -0.355 | -0.05 | 0.173 | 0.173 | 0.012 | 0.4 | 0.12 | 0.44 | 0.09 | 0.42 | 0.33 |
| 9 | 72979246 | rs10868811 | -0.371 | 0.1 | -0.519 | 0.08 | -0.008 | -0.053 | -0.053 | 0 | 0.28 | 0.14 | 0.29 | 0.5 | 0.29 | 0.23 |
| 9 | 72989832 | rs4745008 | -0.698 | 0.134 | -0.29 | 0.018 | 0 | 0.009 | 0.009 | 0.001 | 0.34 | 0.45 | 0.37 | 0.48 | 0.34 | 0.2 |
| 9 | 72993185 | rs10780926 | -0.483 | 0.101 | -0.178 | 0.011 | -0.016 | -0.059 | -0.059 | 0 | 0.31 | 0.17 | 0.35 | 0.47 | 0.34 | 0.2 |
| 9 | 72998332 | rs11142387 | 0.828 | 0.151 | -0.263 | -0.248 | -0.008 | 0.031 | 0.031 | 0.185 | 0.55 | 0.36 | 0.51 | 0.41 | 0.52 | 0.69 |
| 9 | 72998425 | rs11142388 | 1.883 | 0.039 | -0.78 | -0.481 | -0.08 | 0.141 | 0.141 | 0 | 0.36 | 0.08 | 0.41 | 0.09 | 0.41 | 0.33 |
| 9 | 73009913 | rs10780928 | -0.86 | 0.12 | 0.09 | -0.339 | -0.003 | 0.018 | 0.018 | 0.078 | 0.45 | 0.64 | 0.46 | 0.58 | 0.46 | 0.3 |
| 9 | 73017053 | rs7034715 | 0.084 | 0.06 | 0.417 | -0.476 | 0.01 | -0.016 | -0.016 | 0 | 0.14 | 0.28 | 0.11 | 0.08 | 0.12 | 0.1 |
| 9 | 73021070 | rs11142402 | 1.486 | -0.005 | -0.062 | -0.517 | -0.052 | 0.13 | 0.13 | 0 | 0.39 | 0.15 | 0.47 | 0.18 | 0.42 | 0.4 |
| 9 | 73028766 | rs1056448 | 1.467 | 0.009 | -0.103 | -0.536 | -0.055 | 0.133 | 0.133 | 0 | 0.39 | 0.12 | 0.44 | 0.18 | 0.42 | 0.4 |
| 9 | 73029831 | rs10868824 | 1.467 | 0.003 | -0.107 | -0.571 | -0.054 | 0.137 | 0.137 | 0 | 0.39 | 0.12 | 0.46 | 0.18 | 0.42 | 0.4 |
| 9 | 73038018 | rs10868828 | 0.155 | 0.147 | 0.123 | -0.274 | -0.001 | 0.037 | 0.037 | 0 | 0.75 | 0.64 | 0.66 | 0.39 | 0.78 | 0.69 |
| 9 | 73040468 | rs6560132 | 0.68 | -0.021 | 0.06 | -0.493 | -0.009 | -0.017 | -0.017 | 0 | 0.61 | 0.41 | 0.62 | 0.28 | 0.68 | 0.5 |
| 9 | 73065727 | rs915490 | 0.388 | 0.18 | -0.059 | -0.335 | 0.001 | 0 | 0 | 0 | 0.19 | 0.15 | 0.09 | 0.23 | 0.17 | 0.39 |
| 9 | 73070585 | rs13299473 | 2.021 | 0.092 | -0.175 | -0.546 | -0.03 | 0.081 | 0.081 | 0 | 0.36 | 0.23 | 0.45 | 0.2 | 0.4 | 0.38 |
| 9 | 73079531 | rs7853214 | 0.434 | 0.165 | -0.019 | -0.538 | 0 | 0 | 0 | 0 | 0.19 | 0.18 | 0.09 | 0.23 | 0.17 | 0.37 |
| 9 | 73080367 | rs7873409 | 0.769 | 0.08 | 0.124 | -0.689 | -0.002 | -0.007 | -0.007 | 0 | 0.54 | 0.57 | 0.56 | 0.42 | 0.57 | 0.75 |
| 9 | 73087603 | rs4745013 | -0.418 | -0.077 | 0.13 | -0.257 | 0.006 | -0.01 | -0.01 | 0 | 0.34 | 0.25 | 0.36 | 0.62 | 0.31 | 0.46 |
| 9 | 73094035 | rs2184754 | 0.275 | -0.095 | 0.07 | -0.286 | -0.011 | -0.039 | -0.039 | 0 | 0.69 | 0.56 | 0.82 | 0.81 | 0.72 | 0.84 |
| 9 | 73094295 | rs963266 | 0.908 | -0.014 | -0.228 | -0.198 | 0.015 | 0.054 | 0.054 | 0 | 0.5 | 0.29 | 0.72 | 0.58 | 0.55 | 0.46 |
| 9 | 73095886 | rs11142467 | 0.742 | -0.11 | -0.017 | 0.17 | 0.003 | -0.067 | -0.067 | 0.065 | 0.15 | 0.07 | 0.26 | 0.39 | 0.14 | 0.09 |
| 9 | 73101159 | rs7029925 | -1.341 | -0.36 | 0.448 | -0.031 | -0.024 | 0.062 | 0.062 | 1.624 | 0.06 | 0.21 | 0.03 | 0.11 | 0.01 | 0.05 |
| 9 | 73102824 | rs3923699 | 0.485 | 0.116 | 0.575 | -0.026 | -0.008 | -0.004 | -0.004 | 0 | 0.56 | 0.7 | 0.53 | 0.47 | 0.48 | 0.51 |
| 9 | 73103998 | rs17054772 | 0.52 | 0.086 | 0.512 | -0.13 | -0.002 | -0.021 | -0.021 | 3.108 | 0.21 | 0.23 | 0.11 | 0.14 | 0.17 | 0.35 |
| 9 | 73104819 | rs4548247 | -0.025 | 0.161 | 0.475 | -0.113 | -0.015 | -0.033 | -0.033 | 0.017 | 0.72 | 0.82 | 0.62 | 0.51 | 0.64 | 0.8 |
| 9 | 73108351 | rs11792173 | 1.255 | 0.055 | 0.722 | -0.15 | 0.002 | 0.047 | 0.047 | 0 | 0.48 | 0.35 | 0.5 | 0.36 | 0.47 | 0.46 |

| | | | | | | | | | | | | | | | | |
|---|---|---|---|---|---|---|---|---|---|---|---|---|---|---|---|---|
| 9 | 73109347 | rs7865838 | 1.905 | -0.459 | 0.363 | -0.436 | 0.005 | -0.028 | -0.028 | 0 | 0.33 | 0.45 | 0.45 | 0.17 | 0.35 | 0.35 |
| 9 | 73113806 | rs915488 | 2.378 | -0.446 | 0.203 | -0.417 | -0.067 | 0.051 | 0.051 | 0 | 0.31 | 0.1 | 0.42 | 0.17 | 0.35 | 0.35 |
| 9 | 73115774 | rs7856388 | 0.569 | -0.473 | 0.244 | -0.403 | -0.028 | -0.007 | -0.007 | 0 | 0.18 | 0.26 | 0.07 | 0.19 | 0.11 | 0.11 |
| 9 | 73120468 | rs11999226 | 0.839 | -0.351 | -0.086 | -0.374 | -0.031 | 0.039 | 0.039 | 0 | 0.2 | 0.03 | 0.1 | 0.14 | 0.17 | 0.35 |
| 9 | 73121298 | rs7037072 | -0.973 | -0.335 | 0.011 | -0.38 | 0.002 | 0.059 | 0.059 | 0 | 0.48 | 0.25 | 0.48 | 0.64 | 0.53 | 0.54 |
| 9 | 73121323 | rs10780936 | 0.57 | -0.215 | -0.202 | -0.236 | -0.11 | -0.153 | -0.153 | 0 | 0.23 | 0.02 | 0.36 | 0.39 | 0.35 | 0.15 |
| 9 | 73127939 | rs17054874 | -0.105 | -0.764 | -0.006 | -0.391 | 0 | -0.001 | -0.001 | 0 | 0.09 | 0.07 | 0.05 | 0.12 | 0.09 | 0.07 |
| 9 | 73131895 | rs10868848 | -0.119 | -0.9 | 0.304 | -0.539 | -0.055 | -0.097 | -0.097 | 0 | 0.79 | 0.48 | 0.91 | 0.92 | 0.88 | 0.9 |
| 9 | 73133551 | rs4436186 | 1.433 | -0.867 | -0.463 | -0.531 | -0.006 | 0.036 | 0.036 | 1.393 | 0.47 | 0.05 | 0.75 | 0.61 | 0.62 | 0.47 |
| 9 | 73138287 | rs7046672 | 0.277 | -0.825 | 0.095 | -0.877 | -0.005 | -0.062 | -0.062 | 0.453 | 0.19 | 0.22 | 0.09 | 0.02 | 0.13 | 0.32 |
| 9 | 73147528 | rs4744600 | 0.554 | -0.442 | -0.434 | -1.002 | -0.051 | -0.101 | -0.101 | 0.082 | 0.26 | 0.1 | 0.42 | 0.5 | 0.4 | 0.21 |
| 9 | 73149897 | rs7853385 | 1.035 | -0.291 | -0.32 | -0.943 | -0.014 | 0.067 | 0.067 | 0 | 0.16 | 0.04 | 0.07 | 0.02 | 0.13 | 0.32 |
| 9 | 73150873 | rs17535963 | 2.388 | -0.506 | -0.454 | -1.032 | -0.073 | 0.048 | 0.048 | 0 | 0.26 | 0.07 | 0.33 | 0.18 | 0.28 | 0.27 |
| 9 | 73151715 | rs7033976 | -1.03 | -0.495 | -0.035 | -1.025 | -0.001 | -0.034 | -0.034 | 0 | 0.82 | 0.86 | 0.92 | 0.98 | 0.87 | 0.68 |
| 9 | 73151970 | rs3739776 | -0.976 | -0.484 | 0.133 | -0.91 | -0.003 | 0.009 | 0.009 | 0 | 0.54 | 0.57 | 0.57 | 0.7 | 0.58 | 0.39 |
| 9 | 73161554 | rs12338410 | 0.978 | -0.551 | 0.279 | -0.982 | -0.013 | -0.039 | -0.039 | 0 | 0.07 | 0.19 | 0.03 | 0 | 0.06 | 0.08 |
| 9 | 73162236 | rs1414850 | -1.374 | -0.601 | 0.417 | -1.102 | 0.003 | -0.032 | -0.032 | 0 | 0.85 | 0.84 | 0.91 | 0.97 | 0.87 | 0.68 |
| 9 | 73164712 | rs1889915 | -0.566 | -0.585 | 0.387 | -0.988 | 0.008 | 0.063 | 0.063 | 0.154 | 0.5 | 0.39 | 0.54 | 0.8 | 0.52 | 0.32 |
| 9 | 73165078 | rs10780947 | 1.353 | -0.25 | -0.364 | -0.837 | -0.082 | -0.154 | -0.154 | 0 | 0.19 | 0.01 | 0.33 | 0.51 | 0.28 | 0.16 |
| 9 | 73168680 | rs6560143 | -0.741 | 0.047 | 0.426 | -0.6 | -0.022 | 0.066 | 0.066 | 0 | 0.58 | 0.76 | 0.59 | 0.8 | 0.57 | 0.4 |
| 9 | 73170295 | rs10511984 | 1.8 | 0.062 | 0.275 | -0.624 | -0.031 | 0.045 | 0.045 | 0 | 0.27 | 0.15 | 0.33 | 0.17 | 0.3 | 0.28 |
| 9 | 73170382 | rs10746847 | -0.458 | 0.042 | 0.388 | -0.621 | 0.001 | -0.012 | -0.012 | 0 | 0.56 | 0.49 | 0.57 | 0.7 | 0.57 | 0.39 |
| 9 | 73172801 | rs4352910 | 1.827 | 0.156 | -0.299 | -0.524 | -0.084 | 0.053 | 0.053 | 0 | 0.26 | 0.06 | 0.32 | 0.17 | 0.3 | 0.28 |
| 9 | 73183281 | rs10746850 | -0.796 | 0.024 | -0.41 | -0.499 | -0.011 | 0.07 | 0.07 | 0.004 | 0.57 | 0.74 | 0.58 | 0.8 | 0.54 | 0.39 |
| 9 | 73193371 | rs10868854 | -0.122 | -0.326 | -0.521 | -0.456 | -0.032 | -0.032 | -0.032 | 1.221 | 0.39 | 0.21 | 0.45 | 0.51 | 0.46 | 0.3 |
| 9 | 73193428 | rs10780950 | 0.133 | -0.907 | 0.369 | -0.411 | -0.028 | 0.006 | 0.006 | 0 | 0.75 | 0.63 | 0.83 | 0.77 | 0.83 | 0.71 |
| 9 | 73195703 | rs1317103 | 0.584 | -0.903 | 0.057 | -0.482 | -0.034 | -0.018 | -0.018 | 0 | 0.67 | 0.5 | 0.79 | 0.77 | 0.77 | 0.63 |
| 9 | 73196666 | rs17458750 | 0.929 | -0.896 | -1.049 | -0.762 | -0.068 | -0.018 | -0.018 | 0 | 0.42 | 0.17 | 0.6 | 0.55 | 0.48 | 0.37 |
| 9 | 73199483 | rs4744604 | 1.159 | -1.227 | -0.859 | -0.558 | -0.06 | -0.046 | -0.046 | 0 | 0.46 | 0.12 | 0.62 | 0.54 | 0.52 | 0.37 |
| 9 | 73200907 | rs11790957 | -0.515 | -1.234 | -0.823 | -0.564 | -0.013 | -0.03 | -0.03 | 0 | 0.24 | 0.06 | 0.17 | 0.25 | 0.22 | 0.17 |
| 9 | 73203612 | rs10735599 | -0.63 | -1.326 | -0.159 | -0.737 | 0 | -0.002 | -0.002 | 0 | 0.8 | 0.78 | 0.87 | 0.85 | 0.77 | 0.76 |
| 9 | 73204431 | rs7865858 | 1.076 | -1.373 | 0.181 | -0.755 | 0.003 | 0.008 | 0.008 | 0 | 0.56 | 0.74 | 0.68 | 0.56 | 0.55 | 0.49 |
| 9 | 73210302 | rs11142498 | -0.317 | -1.225 | 0.09 | -0.47 | 0.001 | -0.014 | -0.014 | 0.014 | 0.79 | 0.72 | 0.87 | 0.85 | 0.76 | 0.76 |
| 9 | 73213735 | rs4744607 | -0.424 | -1.193 | -0.566 | -0.454 | -0.004 | 0.009 | 0.009 | 0 | 0.8 | 0.84 | 0.87 | 0.85 | 0.76 | 0.76 |
| 9 | 73218656 | rs4615645 | 0.714 | -1.045 | -0.521 | -0.462 | 0.057 | 0.018 | 0.018 | 0 | 0.53 | 0.24 | 0.67 | 0.55 | 0.6 | 0.39 |
| 9 | 73224560 | rs10114679 | 0.362 | -1.078 | -0.353 | -0.54 | 0.005 | -0.011 | -0.011 | 0 | 0.12 | 0.11 | 0.05 | 0 | 0.06 | 0.04 |
| 9 | 73225802 | rs3763619 | 0.616 | -0.995 | -0.47 | -0.335 | 0.032 | 0.009 | 0.009 | 0 | 0.47 | 0.25 | 0.66 | 0.53 | 0.58 | 0.38 |
| 9 | 73231662 | rs11142508 | 0.573 | -0.89 | -0.547 | -0.297 | 0.031 | -0.008 | -0.008 | 0 | 0.47 | 0.21 | 0.65 | 0.52 | 0.58 | 0.39 |
| 9 | 73239062 | rs11142515 | -0.288 | -0.203 | 0.108 | -0.085 | 0.009 | -0.009 | -0.009 | 0 | 0.29 | 0.4 | 0.24 | 0.27 | 0.4 | 0.53 |
| 9 | 73239605 | rs10868861 | -0.656 | -0.844 | 0.674 | -0.587 | -0.033 | -0.114 | -0.114 | 0.219 | 0.22 | 0.48 | 0.15 | 0.08 | 0.17 | 0.21 |
| 9 | 73245200 | rs11142518 | 1.29 | -0.8 | -0.243 | -0.462 | -0.017 | -0.106 | -0.106 | 1.629 | 0.2 | 0.04 | 0.3 | 0.38 | 0.18 | 0.14 |
| 9 | 73248523 | rs11142521 | 0.332 | -1.028 | -0.358 | -0.698 | 0.017 | -0.009 | -0.009 | 0.085 | 0.35 | 0.19 | 0.41 | 0.62 | 0.27 | 0.25 |
| 9 | 73250037 | rs4322073 | 0.858 | -0.927 | -0.176 | -0.65 | -0.022 | 0.048 | 0.048 | 2.91 | 0.42 | 0.21 | 0.43 | 0.3 | 0.57 | 0.52 |
| 9 | 73261860 | rs7849603 | -0.312 | -1.046 | -0.071 | -0.627 | 0.001 | -0.001 | -0.001 | 0 | 0.42 | 0.55 | 0.4 | 0.46 | 0.35 | 0.38 |
| 9 | 73281358 | rs7027906 | 1.109 | -1.003 | -0.042 | -0.688 | 0.001 | 0 | 0 | 0.011 | 0.22 | 0.26 | 0.15 | 0.19 | 0.29 | 0.33 |
| 9 | 73284190 | rs7854748 | -0.141 | -1.032 | -0.061 | -0.633 | -0.005 | 0.001 | 0.001 | 1.682 | 0.41 | 0.52 | 0.39 | 0.45 | 0.35 | 0.39 |
| 9 | 73287076 | rs4465028 | -1.114 | -1.026 | -0.184 | -0.611 | -0.036 | -0.009 | -0.009 | 0 | 0.63 | 0.81 | 0.54 | 0.64 | 0.64 | 0.73 |
| 9 | 73288256 | rs1011308 | 1.059 | -1.108 | 0.257 | -0.601 | 0.008 | -0.002 | -0.002 | 0 | 0.23 | 0.33 | 0.15 | 0.18 | 0.29 | 0.33 |
| 9 | 73293499 | rs1934474 | -1.354 | -1.153 | 0.144 | -0.584 | 0.005 | -0.008 | -0.008 | 0.064 | 0.84 | 0.83 | 0.83 | 0.76 | 0.91 | 0.9 |
| 9 | 73294910 | rs12345213 | 0.984 | -1.072 | 0.214 | -0.585 | -0.005 | 0.007 | 0.007 | 0 | 0.07 | 0.05 | 0.09 | 0.06 | 0.1 | 0.16 |
| 9 | 73305723 | rs4617221 | -0.185 | -1.073 | 0.376 | -0.566 | -0.005 | 0.001 | 0.001 | 1.738 | 0.4 | 0.45 | 0.38 | 0.45 | 0.34 | 0.39 |
| 9 | 73307490 | rs10780959 | -0.193 | -1.072 | 0.386 | -0.143 | -0.004 | -0.004 | -0.004 | 0.744 | 0.41 | 0.45 | 0.38 | 0.38 | 0.34 | 0.39 |
| 9 | 73309867 | rs7023662 | -0.193 | -1.072 | 0.386 | -0.523 | -0.004 | 0 | 0 | 1.814 | 0.41 | 0.45 | 0.38 | 0.45 | 0.34 | 0.39 |
| 9 | 73312129 | rs7860377 | 0.066 | -1.062 | 0.431 | -0.635 | -0.016 | -0.041 | -0.041 | 0.275 | 0.35 | 0.48 | 0.2 | 0.2 | 0.35 | 0.39 |
| 9 | 73315772 | rs11142556 | 1.817 | -0.836 | -0.089 | -0.552 | -0.092 | 0.044 | 0.044 | 0.323 | 0.21 | 0.02 | 0.29 | 0.12 | 0.28 | 0.18 |
| 9 | 73357667 | rs11142561 | 1.786 | -0.823 | -0.084 | -0.553 | -0.094 | 0.048 | 0.048 | 0.388 | 0.21 | 0.03 | 0.29 | 0.12 | 0.28 | 0.18 |
| 9 | 73369501 | rs12335434 | 0.738 | -0.922 | 0.039 | -0.649 | 0.011 | 0.035 | 0.035 | 0 | 0.14 | 0.07 | 0.07 | 0.01 | 0.11 | 0.09 |
| 9 | 73369687 | rs7849151 | 0.685 | -1.104 | 0.6 | -0.601 | -0.031 | 0.075 | 0.075 | 0 | 0.18 | 0.44 | 0.18 | 0.25 | 0.1 | 0.14 |
| 9 | 73375005 | rs4745035 | 0.262 | -1.048 | 0.36 | -0.616 | -0.021 | -0.015 | -0.015 | 0.452 | 0.64 | 0.44 | 0.68 | 0.69 | 0.73 | 0.65 |
| 9 | 73378890 | rs10868882 | 0.962 | -0.924 | -0.326 | -0.532 | -0.121 | -0.141 | -0.141 | 2.197 | 0.41 | 0.04 | 0.54 | 0.5 | 0.47 | 0.3 |
| 9 | 73389934 | rs879857 | 1.339 | -0.868 | -0.195 | -0.524 | 0 | -0.001 | -0.001 | 0.261 | 0.18 | 0.15 | 0.11 | 0.19 | 0.23 | 0.27 |
| 9 | 73391084 | rs10868885 | 0.827 | -0.539 | -0.699 | -0.445 | -0.092 | -0.111 | -0.111 | 0.209 | 0.41 | 0.08 | 0.54 | 0.51 | 0.46 | 0.3 |
| 9 | 73392983 | rs1328148 | 0.094 | -0.553 | -0.562 | -0.434 | 0.01 | -0.011 | -0.011 | 1.696 | 0.39 | 0.61 | 0.31 | 0.43 | 0.37 | 0.47 |
| 9 | 73397090 | rs6560153 | 0.08 | -0.498 | -0.322 | -0.266 | -0.017 | -0.084 | -0.084 | 0 | 0.8 | 0.63 | 0.85 | 0.94 | 0.83 | 0.77 |
| 9 | 73398448 | rs10435960 | 0.388 | -0.06 | -1.037 | 0.307 | -0.026 | -0.101 | -0.101 | 1.231 | 0.2 | 0.04 | 0.25 | 0.38 | 0.19 | 0.12 |
| 9 | 73399750 | rs1831143 | 0.13 | -0.118 | -0.773 | -0.155 | 0.004 | -0.003 | -0.003 | 0.042 | 0.2 | 0.15 | 0.12 | 0.2 | 0.19 | 0.22 |
| 9 | 73402201 | rs7040905 | -0.484 | 0.273 | -0.952 | -0.535 | 0.002 | 0.009 | 0.009 | 0 | 0.5 | 0.5 | 0.38 | 0.34 | 0.53 | 0.56 |
| 9 | 73405244 | rs11142594 | 0.768 | 0.204 | -1.249 | -0.424 | 0.006 | 0 | 0 | 0 | 0.18 | 0.08 | 0.11 | 0.18 | 0.18 | 0.27 |
| 9 | 73405864 | rs10118380 | 0.043 | 0.329 | -0.943 | -0.694 | 0 | -0.001 | -0.001 | 0.065 | 0.44 | 0.4 | 0.54 | 0.54 | 0.46 | 0.42 |

| | | | | | | | | | | | | | | | | |
|---|---|---|---|---|---|---|---|---|---|---|---|---|---|---|---|---|
| 9 | 73406551 | rs1028879 | 1.049 | 0.73 | -1.4 | -0.527 | -0.076 | -0.11 | -0.11 | 0.209 | 0.22 | 0.07 | 0.46 | 0.53 | 0.31 | 0.33 |
| 9 | 73408518 | rs10868890 | 0.13 | 0.346 | -0.729 | -0.437 | 0.016 | 0.035 | 0.035 | 0 | 0.76 | 0.54 | 0.71 | 0.71 | 0.78 | 0.74 |
| 9 | 73408650 | rs11142598 | 1.194 | 0.568 | -0.977 | -0.465 | 0.002 | -0.106 | -0.106 | 0 | 0.22 | 0.07 | 0.37 | 0.49 | 0.18 | 0.27 |
| 9 | 73409336 | rs7048454 | -0.36 | 0.6 | -0.177 | -0.436 | -0.017 | 0.023 | 0.023 | 0.023 | 0.82 | 0.58 | 0.85 | 0.82 | 0.9 | 0.88 |
| 9 | 73416062 | rs1328153 | -0.187 | 0.687 | -0.209 | -0.524 | -0.016 | -0.028 | -0.028 | 0.139 | 0.7 | 0.61 | 0.79 | 0.82 | 0.79 | 0.83 |
| 9 | 73416797 | rs1410373 | -0.171 | 0.457 | -0.235 | -0.598 | -0.022 | -0.037 | -0.037 | 0 | 0.7 | 0.57 | 0.78 | 0.82 | 0.79 | 0.85 |
| 9 | 73417675 | rs13285335 | 0.473 | 0.982 | -0.234 | -0.531 | 0.005 | -0.026 | -0.026 | 0 | 0.08 | 0.09 | 0.07 | 0 | 0.12 | 0.03 |
| 9 | 73420493 | rs7873697 | -0.956 | 0.8 | -0.187 | -0.873 | -0.001 | -0.027 | -0.027 | 0 | 0.31 | 0.61 | 0.26 | 0.16 | 0.27 | 0.29 |
| 9 | 73421171 | rs7863403 | -1.101 | 0.432 | -0.088 | -0.928 | 0.012 | 0.023 | 0.023 | 0.029 | 0.32 | 0.72 | 0.25 | 0.14 | 0.22 | 0.23 |
| 9 | 73422925 | rs995903 | -0.691 | 0.422 | -0.025 | -0.936 | -0.029 | -0.059 | -0.059 | 0.493 | 0.28 | 0.56 | 0.21 | 0.14 | 0.17 | 0.22 |
| 9 | 73427233 | rs10868894 | 0.971 | 0.831 | -0.863 | -0.763 | -0.076 | 0.066 | 0.066 | 0 | 0.24 | 0.11 | 0.34 | 0.14 | 0.37 | 0.35 |
| 9 | 73436971 | rs4526420 | 0.892 | 0.487 | -0.728 | -0.512 | -0.058 | -0.048 | -0.048 | 0.194 | 0.53 | 0.08 | 0.4 | 0.56 | 0.51 | 0.51 |
| 9 | 73437824 | rs7038646 | -0.438 | 0.455 | 0.031 | -0.55 | 0.011 | 0.018 | 0.018 | 0 | 0.71 | 0.63 | 0.61 | 0.67 | 0.68 | 0.7 |
| 9 | 73440705 | rs7863095 | -0.697 | 0.059 | -0.272 | -0.925 | 0.021 | -0.006 | -0.006 | 0.559 | 0.38 | 0.64 | 0.36 | 0.4 | 0.26 | 0.3 |
| 9 | 73440813 | rs7862322 | -0.264 | 0.033 | -0.079 | -0.924 | -0.039 | 0 | 0 | 0 | 0.33 | 0.45 | 0.27 | 0.28 | 0.24 | 0.17 |
| 9 | 73441437 | rs10117842 | 0.389 | -0.01 | -0.582 | -1.266 | -0.021 | 0.041 | 0.041 | 0 | 0.38 | 0.31 | 0.44 | 0.14 | 0.52 | 0.4 |
| 9 | 73442413 | rs2011851 | 2.297 | 0.145 | -0.757 | -1.217 | -0.081 | 0.014 | 0.014 | 0.789 | 0.15 | 0.03 | 0.28 | 0.13 | 0.26 | 0.22 |
| 9 | 73442724 | rs1034538 | -2.305 | 0.116 | -0.776 | -1.154 | -0.034 | -0.143 | -0.143 | 0 | 0.76 | 0.96 | 0.8 | 0.54 | 0.78 | 0.7 |
| 9 | 73446373 | rs10081686 | -0.375 | -0.019 | -0.264 | -1.18 | -0.011 | -0.004 | -0.004 | 0.896 | 0.43 | 0.5 | 0.45 | 0.41 | 0.41 | 0.41 |
| 9 | 73453045 | rs1890017 | 1.749 | 0.147 | -0.211 | -1.146 | 0.01 | -0.045 | -0.045 | 0 | 0.09 | 0.16 | 0.07 | 0 | 0.09 | 0.02 |
| 9 | 73462784 | rs7022926 | -0.722 | 0.083 | -0.254 | -1.195 | 0.004 | 0.014 | 0.014 | 0 | 0.53 | 0.59 | 0.56 | 0.27 | 0.58 | 0.59 |
| 9 | 73470611 | rs11142623 | 2.495 | 0.111 | -0.426 | -1.105 | -0.006 | -0.063 | -0.063 | 0.605 | 0.1 | 0 | 0.15 | 0.27 | 0.1 | 0.09 |
| 9 | 73470756 | rs1337026 | 0.077 | 0.175 | -0.225 | -1.124 | 0.011 | -0.046 | -0.046 | 0 | 0.32 | 0.38 | 0.35 | 0.13 | 0.36 | 0.29 |
| 9 | 73472360 | rs10868916 | -0.098 | 0.058 | -0.224 | -1.181 | -0.007 | 0.007 | 0.007 | 0 | 0.4 | 0.35 | 0.4 | 0.15 | 0.47 | 0.41 |
| 9 | 73472818 | rs1415225 | -2.087 | 0.058 | -0.114 | -1.029 | 0.003 | -0.025 | -0.025 | 0 | 0.72 | 0.8 | 0.79 | 0.54 | 0.77 | 0.7 |
| 9 | 73473596 | rs17555916 | 1.59 | 0.479 | -0.337 | -1.01 | -0.006 | 0.031 | 0.031 | 0 | 0.19 | 0.12 | 0.15 | 0.01 | 0.22 | 0.17 |
| 9 | 73474633 | rs11142627 | 2.114 | 0.546 | -0.378 | -1.009 | -0.004 | 0.003 | 0.003 | 0.724 | 0.15 | 0.15 | 0.26 | 0.13 | 0.24 | 0.22 |
| 9 | 73476497 | rs10511988 | 0.944 | 0.368 | -0.304 | -0.89 | -0.002 | 0.007 | 0.007 | 0 | 0.19 | 0.2 | 0.22 | 0.27 | 0.19 | 0.11 |
| 9 | 73477141 | rs17556165 | 1.109 | 0.383 | -0.356 | -0.914 | -0.001 | 0.015 | 0.015 | 0 | 0.22 | 0.18 | 0.22 | 0.14 | 0.24 | 0.29 |
| 9 | 73480240 | rs12003687 | 0.78 | 0.294 | -0.035 | -0.655 | -0.035 | -0.053 | -0.053 | 0 | 0.13 | 0.2 | 0.04 | 0.01 | 0.05 | 0.07 |
| 9 | 73480626 | rs1337024 | 0.06 | 0.934 | -0.255 | -0.402 | 0 | 0.004 | 0.004 | 0 | 0.22 | 0.25 | 0.27 | 0.26 | 0.25 | 0.12 |
| 9 | 73480718 | rs1415221 | -0.282 | 0.975 | -0.286 | -0.401 | -0.003 | -0.002 | -0.002 | 0 | 0.37 | 0.37 | 0.52 | 0.44 | 0.48 | 0.35 |
| 9 | 73482440 | rs12378024 | -0.156 | 0.911 | -0.237 | -0.404 | 0.003 | 0.001 | 0.001 | 0.217 | 0.34 | 0.36 | 0.48 | 0.39 | 0.43 | 0.33 |
| 9 | 73485048 | rs3812530 | 0.057 | 0.675 | 0.014 | -0.457 | 0.005 | 0.036 | 0.036 | 0 | 0.23 | 0.32 | 0.17 | 0.4 | 0.27 | 0.18 |
| 9 | 73485446 | rs7856482 | 1.016 | 0.766 | -0.196 | -0.794 | -0.019 | 0.009 | 0.009 | 0 | 0.19 | 0.11 | 0.35 | 0.19 | 0.27 | 0.24 |
| 9 | 73488564 | rs13293998 | 0.348 | 0.642 | 0.294 | -0.694 | 0.003 | 0.022 | 0.022 | 0 | 0.22 | 0.29 | 0.17 | 0.35 | 0.27 | 0.18 |
| 9 | 73490171 | rs9792446 | 0.95 | 0.677 | 0.264 | -0.717 | 0 | -0.01 | -0.01 | 0 | 0.17 | 0.15 | 0.14 | 0.35 | 0.21 | 0.17 |
| 9 | 73490284 | rs9792690 | 0.326 | 0.689 | 0.235 | -0.714 | 0.004 | 0.023 | 0.023 | 0 | 0.22 | 0.29 | 0.17 | 0.35 | 0.27 | 0.18 |
| 9 | 73492507 | rs4532663 | 0.412 | 0.413 | 0.351 | -1.164 | -0.007 | -0.074 | -0.074 | 0.713 | 0.19 | 0.3 | 0.12 | 0.05 | 0.15 | 0.06 |
| 9 | 73493728 | rs1890016 | -1.386 | 0.329 | -0.361 | -1.415 | -0.017 | -0.082 | -0.082 | 0.107 | 0.72 | 0.88 | 0.65 | 0.6 | 0.72 | 0.59 |
| 9 | 73494438 | rs11142635 | -0.432 | 0.233 | -0.054 | -1.473 | 0.011 | 0.02 | 0.02 | 0 | 0.48 | 0.62 | 0.29 | 0.41 | 0.44 | 0.35 |
| 9 | 73494939 | rs13283806 | 0.219 | 0.375 | 0.001 | -1.181 | 0.002 | -0.003 | -0.003 | 0 | 0.33 | 0.3 | 0.2 | 0.36 | 0.33 | 0.31 |
| 9 | 73496967 | rs17471974 | 0.553 | 0.414 | -0.129 | -1.237 | 0 | 0.006 | 0.006 | 0 | 0.19 | 0.13 | 0.13 | 0.01 | 0.17 | 0.14 |
| 9 | 73497518 | rs7021176 | -0.395 | 0.509 | 0.111 | -1.022 | 0.01 | 0.092 | 0.092 | 0 | 0.72 | 0.55 | 0.67 | 0.54 | 0.75 | 0.61 |
| 9 | 73500481 | rs11142636 | -0.726 | 0.646 | 0.084 | -1.015 | -0.001 | 0.037 | 0.037 | 0 | 0.78 | 0.72 | 0.7 | 0.6 | 0.81 | 0.65 |
| 9 | 73500857 | rs7851915 | 0.435 | 0.578 | -0.142 | -1.161 | -0.001 | -0.002 | -0.002 | 0 | 0.56 | 0.61 | 0.55 | 0.54 | 0.6 | 0.48 |
| 9 | 73502424 | rs10780982 | 0.64 | 0.729 | -0.256 | -0.968 | -0.012 | 0.133 | 0.133 | 0 | 0.42 | 0.17 | 0.46 | 0.2 | 0.42 | 0.36 |
| 9 | 73502636 | rs1361028 | 0.66 | 0.632 | -0.273 | -0.979 | 0.015 | -0.053 | -0.053 | 3.482 | 0.26 | 0.13 | 0.17 | 0.36 | 0.26 | 0.21 |
| 9 | 73504145 | rs13285568 | 1.027 | 0.618 | -0.36 | -1.014 | 0.017 | -0.073 | -0.073 | 0 | 0.22 | 0.08 | 0.15 | 0.35 | 0.22 | 0.19 |
| 9 | 73507057 | rs1337033 | 0.857 | 0.684 | -0.306 | -1.064 | -0.051 | 0.125 | 0.125 | 0 | 0.41 | 0.1 | 0.47 | 0.25 | 0.43 | 0.35 |
| 9 | 73507096 | rs10511992 | 1.634 | 0.606 | -0.146 | -1.178 | -0.001 | -0.018 | -0.018 | 0 | 0.08 | 0.12 | 0.02 | 0 | 0.06 | 0.09 |
| 9 | 73508497 | rs11142639 | 1.064 | 0.645 | -0.198 | -1.194 | -0.043 | 0.126 | 0.126 | 0 | 0.41 | 0.1 | 0.47 | 0.25 | 0.43 | 0.34 |
| 9 | 73508552 | rs7046928 | -1.363 | 0.653 | 0.286 | -1.022 | 0.017 | 0.054 | 0.054 | 0 | 0.82 | 0.79 | 0.69 | 0.62 | 0.78 | 0.74 |
| 9 | 73508693 | rs13287493 | 0.942 | 0.659 | -0.011 | -0.985 | 0.015 | -0.061 | -0.061 | 0.619 | 0.26 | 0.12 | 0.15 | 0.37 | 0.25 | 0.22 |
| 9 | 73509677 | rs17472220 | 1.404 | 0.8 | 0.058 | -1.097 | 0 | -0.015 | -0.015 | 0.022 | 0.08 | 0.11 | 0.03 | 0 | 0.06 | 0.09 |
| 9 | 73510880 | rs2993013 | -0.215 | 0.762 | 0.563 | -1.092 | -0.001 | 0.013 | 0.013 | 0 | 0.55 | 0.61 | 0.53 | 0.25 | 0.52 | 0.48 |
| 9 | 73514958 | rs7868945 | 0.691 | 0.9 | 0.661 | -1.012 | 0.061 | 0.03 | 0.03 | 0 | 0.06 | 0.27 | 0.04 | 0.05 | 0.05 | 0.04 |
| 9 | 73515771 | rs10868926 | 0.535 | 0.78 | 0.062 | -1.01 | -0.02 | 0.172 | 0.172 | 0 | 0.46 | 0.16 | 0.48 | 0.21 | 0.46 | 0.43 |
| 9 | 73517105 | rs4143736 | 1.01 | 0.771 | 0.179 | -0.882 | 0.008 | -0.027 | -0.027 | 0 | 0.26 | 0.14 | 0.19 | 0.35 | 0.29 | 0.22 |
| 9 | 73525515 | rs10868928 | 0.925 | 0.837 | 0.142 | -0.998 | -0.026 | 0.113 | 0.113 | 0 | 0.37 | 0.1 | 0.43 | 0.19 | 0.39 | 0.34 |
| 9 | 73527458 | rs1337036 | -0.379 | 0.818 | 0.591 | -0.982 | 0 | -0.075 | -0.075 | 0.636 | 0.16 | 0.59 | 0.08 | 0.06 | 0.12 | 0.19 |
| 9 | 73532529 | rs9696174 | -0.645 | 1.004 | 0.328 | -1.127 | 0.005 | -0.085 | -0.085 | 0 | 0.29 | 0.62 | 0.18 | 0.06 | 0.27 | 0.29 |
| 9 | 73536416 | rs3010419 | -0.445 | 1.027 | 0.24 | -1.092 | -0.005 | 0.1 | 0.1 | 0 | 0.52 | 0.73 | 0.52 | 0.24 | 0.5 | 0.51 |
| 9 | 73538154 | rs7849064 | 1.475 | 1.003 | 0.226 | -1.026 | 0.002 | 0.036 | 0.036 | 0 | 0.27 | 0.12 | 0.35 | 0.19 | 0.27 | 0.25 |
| 9 | 73539365 | rs3010421 | 0.301 | 0.997 | 0.64 | -1.016 | 0.017 | -0.022 | -0.022 | 0.189 | 0.12 | 0.46 | 0.07 | 0.06 | 0.08 | 0.09 |
| 9 | 73540269 | rs1337009 | -1.422 | 1.017 | 0.267 | -0.94 | -0.016 | -0.054 | -0.054 | 1.807 | 0.79 | 0.9 | 0.69 | 0.57 | 0.77 | 0.73 |
| 9 | 73547108 | rs1415219 | 1.449 | 1.02 | 0.283 | -1.214 | 0 | 0.038 | 0.038 | 0 | 0.27 | 0.1 | 0.35 | 0.18 | 0.27 | 0.25 |
| 9 | 73547773 | rs1337013 | 1.076 | 1.024 | 0.673 | -1.129 | 0.006 | -0.046 | -0.046 | 0.082 | 0.08 | 0.28 | 0.04 | 0 | 0.05 | 0.08 |
| 9 | 73557169 | rs12347867 | 0.909 | 1.163 | 0.421 | -1.103 | 0 | -0.027 | -0.027 | 0 | 0.14 | 0.14 | 0.1 | 0 | 0.16 | 0.17 |

| | | | | | | | | | | | | | | | | |
|---|---|---|---|---|---|---|---|---|---|---|---|---|---|---|---|---|
| 9 | 73563097 | rs1981161 | 0.534 | 1.084 | 0.377 | -1.123 | 0.004 | 0.021 | 0.021 | 0 | 0.37 | 0.31 | 0.41 | 0.24 | 0.35 | 0.34 |
| 9 | 73564396 | rs12377705 | 1.093 | 1.169 | -0.063 | -0.741 | 0.001 | -0.01 | -0.01 | 0.259 | 0.22 | 0.15 | 0.16 | 0.27 | 0.24 | 0.2 |
| 9 | 73567734 | rs2993000 | 1.106 | 1.065 | -0.017 | -0.732 | 0.003 | -0.006 | -0.006 | 0.597 | 0.22 | 0.18 | 0.16 | 0.27 | 0.23 | 0.19 |
| 9 | 73567782 | rs2993001 | -0.737 | 1.078 | -0.079 | -0.425 | -0.004 | -0.03 | -0.03 | 0.103 | 0.77 | 0.8 | 0.69 | 0.52 | 0.77 | 0.73 |
| 9 | 73574860 | rs945688 | -0.911 | 1.067 | -0.206 | -0.375 | 0.001 | -0.056 | -0.056 | 0 | 0.78 | 0.87 | 0.7 | 0.53 | 0.78 | 0.74 |
| 9 | 73575662 | rs1108226 | -0.964 | 1.082 | -0.244 | -0.326 | 0.001 | -0.065 | -0.065 | 0.038 | 0.78 | 0.88 | 0.7 | 0.54 | 0.79 | 0.74 |
| 9 | 73577326 | rs2993003 | -0.244 | 1.017 | 0.076 | -0.578 | 0.002 | 0.013 | 0.013 | 0 | 0.51 | 0.55 | 0.46 | 0.27 | 0.52 | 0.51 |
| 9 | 73583539 | rs7863158 | 0.923 | 1.06 | 0.168 | -0.759 | 0 | 0.003 | 0.003 | 0 | 0.3 | 0.28 | 0.34 | 0.26 | 0.3 | 0.26 |
| 9 | 73586356 | rs2993008 | 1.332 | 1.119 | 0.26 | -0.887 | 0 | -0.05 | -0.05 | 0 | 0.1 | 0.18 | 0.03 | 0 | 0.08 | 0.11 |
| 9 | 73586559 | rs3010434 | 0.393 | 1.016 | 0.436 | -0.855 | 0.011 | -0.113 | -0.113 | 0.682 | 0.22 | 0.36 | 0.13 | 0 | 0.22 | 0.25 |
| 9 | 73587520 | rs7857794 | 0.531 | 1.066 | 0.198 | -0.383 | -0.001 | -0.004 | -0.004 | 0.111 | 0.31 | 0.28 | 0.35 | 0.34 | 0.32 | 0.26 |
| 9 | 73590123 | rs12351733 | 0.515 | 1.073 | 0.204 | -0.38 | -0.001 | -0.003 | -0.003 | 0 | 0.31 | 0.29 | 0.35 | 0.34 | 0.32 | 0.26 |
| 9 | 73590376 | rs10868934 | 0.906 | 1.095 | -0.012 | -0.336 | -0.013 | -0.082 | -0.082 | 0.015 | 0.23 | 0.1 | 0.31 | 0.46 | 0.23 | 0.31 |
| 9 | 73593866 | rs3010438 | -0.391 | 1.08 | 0.231 | -0.131 | 0.001 | 0.004 | 0.004 | 0 | 0.69 | 0.61 | 0.64 | 0.66 | 0.68 | 0.74 |
| 9 | 73598308 | rs1558924 | 0.398 | 1.102 | -0.074 | 0.03 | -0.002 | -0.008 | -0.008 | 0 | 0.29 | 0.25 | 0.34 | 0.43 | 0.31 | 0.28 |
| 9 | 73598428 | rs10868936 | 0.391 | 1.047 | -0.171 | -0.286 | 0.003 | -0.001 | -0.001 | 0 | 0.36 | 0.33 | 0.27 | 0.12 | 0.31 | 0.38 |
| 9 | 73598465 | rs1558926 | 0.49 | 1.294 | -0.334 | 0.391 | -0.017 | -0.085 | -0.085 | 0.11 | 0.23 | 0.09 | 0.25 | 0.41 | 0.24 | 0.25 |
| 9 | 73598950 | rs10868937 | 0.042 | 1.126 | -0.014 | 0.164 | 0 | -0.001 | -0.001 | 0 | 0.6 | 0.63 | 0.46 | 0.34 | 0.55 | 0.49 |
| 9 | 73599501 | rs7857162 | -0.113 | 1.099 | 0.34 | 0.449 | 0.004 | 0.048 | 0.048 | 0 | 0.12 | 0.22 | 0.11 | 0.23 | 0.11 | 0.2 |
| 9 | 73600430 | rs719788 | -0.167 | 1.086 | 0.531 | 0.467 | 0.008 | 0.067 | 0.067 | 0 | 0.12 | 0.25 | 0.12 | 0.23 | 0.11 | 0.2 |
| 9 | 73602628 | rs1558928 | 1.009 | 1.193 | 0.526 | 0.103 | 0.038 | 0.032 | 0.032 | 0 | 0.46 | 0.18 | 0.39 | 0.35 | 0.42 | 0.38 |
| 9 | 73614737 | rs11142667 | 1.04 | 1.15 | 0.538 | 0.096 | 0.038 | 0.028 | 0.028 | 0 | 0.45 | 0.15 | 0.38 | 0.35 | 0.41 | 0.38 |
| 9 | 73615374 | rs2909292 | 1.118 | 1.392 | 0.645 | 0.046 | -0.003 | -0.001 | -0.001 | 0 | 0.73 | 0.28 | 0.64 | 0.69 | 0.7 | 0.67 |
| 9 | 73632244 | rs978790 | 1.143 | 1.305 | 0.468 | 0.007 | 0.039 | 0.032 | 0.032 | 0 | 0.46 | 0.23 | 0.38 | 0.35 | 0.42 | 0.39 |
| 9 | 73632464 | rs495259 | -0.586 | 1.39 | 0.857 | -0.021 | 0.035 | -0.048 | -0.048 | 1.917 | 0.9 | 0.83 | 0.93 | 1 | 0.88 | 0.9 |
| 9 | 73633405 | rs12551768 | 0.285 | 1.363 | 0.792 | 0.036 | 0.008 | 0.018 | 0.018 | 0 | 0.12 | 0.26 | 0.11 | 0.14 | 0.11 | 0.2 |
| 9 | 73644816 | rs11142672 | 1.2 | 1.349 | 0.56 | 0.098 | 0.037 | 0.04 | 0.04 | 0 | 0.45 | 0.13 | 0.38 | 0.33 | 0.42 | 0.38 |
| 9 | 73655072 | rs1411164 | 1.041 | 1.346 | 0.577 | 0.065 | -0.002 | 0.004 | 0.004 | 0 | 0.08 | 0.05 | 0.05 | 0.02 | 0.09 | 0.08 |
| 9 | 73656077 | rs6560173 | -0.515 | 1.339 | 0.603 | 0.066 | 0.029 | 0.034 | 0.034 | 0 | 0.16 | 0.39 | 0.27 | 0.23 | 0.18 | 0.23 |
| 9 | 73674356 | rs672801 | 0.999 | 1.409 | 0.135 | 0.068 | -0.048 | -0.087 | -0.087 | 0.64 | 0.27 | 0.06 | 0.25 | 0.34 | 0.29 | 0.3 |
| 9 | 73680233 | rs523734 | -0.729 | 1.43 | 0.338 | 0.006 | 0.036 | -0.045 | -0.045 | 0 | 0.91 | 0.84 | 0.93 | 1 | 0.88 | 0.89 |
| 9 | 73691183 | rs11142684 | 1.871 | 1.044 | -0.603 | -0.211 | -0.034 | 0.009 | 0.009 | 0 | 0.16 | 0.02 | 0.2 | 0.09 | 0.22 | 0.11 |
| 9 | 73695760 | rs552849 | -1.326 | 1.049 | -0.222 | -0.154 | -0.053 | 0.033 | 0.033 | 0 | 0.54 | 0.84 | 0.54 | 0.64 | 0.49 | 0.55 |
| 9 | 73697886 | rs13285838 | -0.976 | 1.13 | -0.094 | -0.074 | 0.002 | 0.015 | 0.015 | 1.254 | 0.22 | 0.67 | 0.3 | 0.31 | 0.21 | 0.26 |
| 9 | 73702948 | rs656875 | 0.937 | 1.116 | -0.634 | -0.161 | -0.02 | -0.097 | -0.097 | 0 | 0.28 | 0.1 | 0.28 | 0.42 | 0.29 | 0.28 |
| 9 | 73709579 | rs667136 | 1.014 | 1.144 | -0.778 | -0.147 | -0.04 | -0.124 | -0.124 | 0 | 0.28 | 0.05 | 0.28 | 0.42 | 0.29 | 0.28 |
| 9 | 73711655 | rs7026563 | -0.459 | 1.002 | -0.281 | 0.149 | 0.026 | 0.032 | 0.032 | 0 | 0.16 | 0.33 | 0.27 | 0.23 | 0.18 | 0.23 |
| 9 | 73718807 | rs1329748 | 0.808 | 1.041 | -1.284 | -0.008 | 0.008 | -0.002 | -0.002 | 0 | 0.42 | 0.11 | 0.38 | 0.36 | 0.41 | 0.39 |
| 9 | 73722116 | rs972386 | 0.135 | 1.298 | -0.595 | 0.055 | -0.001 | 0 | 0 | 0 | 0.08 | 0.07 | 0.05 | 0.02 | 0.09 | 0.08 |
| 9 | 73742312 | rs662975 | 0.059 | 1.329 | -0.446 | 0.29 | 0.038 | 0.045 | 0.045 | 0 | 0.28 | 0.43 | 0.52 | 0.42 | 0.41 | 0.51 |
| 9 | 73746381 | rs2183301 | 0.995 | 0.689 | -1.71 | -0.128 | -0.057 | -0.071 | -0.071 | 0 | 0.29 | 0.01 | 0.18 | 0.31 | 0.26 | 0.14 |
| 9 | 73753420 | rs7046666 | -0.782 | 1.062 | -0.662 | -0.201 | -0.021 | -0.026 | -0.026 | 0 | 0.43 | 0.75 | 0.59 | 0.45 | 0.5 | 0.63 |
| 9 | 73753735 | rs10868959 | 1.018 | 0.978 | -0.751 | -0.248 | -0.012 | 0.02 | 0.02 | 0.699 | 0.51 | 0.08 | 0.39 | 0.39 | 0.46 | 0.35 |
| 9 | 73757570 | rs4744622 | 0.556 | 0.897 | -0.379 | -0.292 | -0.029 | -0.034 | -0.034 | 0 | 0.85 | 0.69 | 0.95 | 0.92 | 0.92 | 0.94 |
| 9 | 73759434 | rs10746858 | 1.594 | 1.528 | -0.09 | -0.182 | 0.004 | -0.007 | -0.007 | 0 | 0.77 | 0.27 | 0.82 | 0.8 | 0.78 | 0.72 |
| 9 | 73762488 | rs12349882 | 0.349 | 1.61 | -0.093 | -0.125 | -0.005 | -0.005 | -0.005 | 6.716 | 0.1 | 0.16 | 0.04 | 0.08 | 0.05 | 0.05 |
| 9 | 73765910 | rs12686625 | 1.138 | 1.623 | -0.194 | -0.137 | -0.05 | -0.09 | -0.09 | 1.793 | 0.19 | 0.04 | 0.37 | 0.3 | 0.23 | 0.28 |
| 9 | 73768548 | rs17056365 | 1.175 | 1.597 | 0.202 | -0.166 | 0.025 | 0.027 | 0.027 | 0 | 0.06 | 0.14 | 0.04 | 0.1 | 0.08 | 0.1 |
| 9 | 73769283 | rs10868962 | -1.678 | 1.521 | 0.249 | -0.18 | 0.005 | 0 | 0 | 0.17 | 0.24 | 0.71 | 0.2 | 0.3 | 0.28 | 0.37 |
| 9 | 73775105 | rs11142707 | -0.462 | 1.693 | 0.311 | -0.06 | 0.19 | 0.104 | 0.104 | 1.438 | 0.08 | 0.41 | 0.13 | 0.13 | 0.14 | 0.22 |
| 9 | 73775928 | rs2031976 | 0.877 | 1.14 | -0.172 | -0.083 | 0.016 | 0.014 | 0.014 | 0 | 0.06 | 0.1 | 0.04 | 0.1 | 0.08 | 0.09 |
| 9 | 73778502 | rs11142708 | 0.857 | 1.128 | -0.825 | -0.361 | -0.049 | 0.016 | 0.016 | 0 | 0.15 | 0.01 | 0.1 | 0.08 | 0.16 | 0.07 |
| 9 | 73782712 | rs6560182 | -0.555 | 1.404 | -0.197 | -0.361 | -0.007 | -0.016 | -0.016 | 1.734 | 0.36 | 0.31 | 0.49 | 0.51 | 0.39 | 0.5 |
| 9 | 73786741 | rs7040038 | -1.208 | 1.573 | 0.088 | -0.487 | 0.035 | -0.015 | -0.015 | 0 | 0.46 | 0.79 | 0.59 | 0.52 | 0.58 | 0.6 |
| 9 | 73796648 | rs10868969 | 0.161 | 1.763 | 0.062 | -0.418 | -0.013 | -0.047 | -0.047 | 0 | 0.28 | 0.17 | 0.41 | 0.34 | 0.29 | 0.29 |
| 9 | 73798371 | rs1329767 | -0.29 | 1.833 | 0.466 | -0.44 | 0.001 | 0.014 | 0.014 | 0 | 0.31 | 0.33 | 0.47 | 0.45 | 0.31 | 0.4 |
| 9 | 73801266 | rs10512009 | 0.721 | 1.609 | 0.294 | -0.578 | -0.014 | -0.039 | -0.039 | 0 | 0.22 | 0.15 | 0.38 | 0.34 | 0.24 | 0.27 |
| 9 | 73820503 | rs17056479 | 0.36 | 1.611 | 0.29 | -0.606 | 0.006 | 0.005 | 0.005 | 0 | 0.11 | 0.13 | 0.11 | 0.17 | 0.15 | 0.23 |
| 9 | 73828288 | rs7018911 | 1.327 | 1.602 | 0.315 | -0.591 | 0.081 | -0.04 | -0.04 | 0 | 0.85 | 0.42 | 0.88 | 0.91 | 0.79 | 0.89 |
| 9 | 73833825 | rs4745072 | -1.614 | 1.572 | 0.301 | -0.593 | 0.053 | 0.025 | 0.025 | 0 | 0.49 | 0.66 | 0.63 | 0.6 | 0.64 | 0.62 |
| 9 | 73836535 | rs6560191 | -0.75 | 1.584 | 0.223 | -0.582 | -0.04 | -0.051 | -0.051 | 0.029 | 0.38 | 0.18 | 0.51 | 0.51 | 0.44 | 0.52 |
| 9 | 73850169 | rs11142747 | 0.358 | 1.68 | 0.219 | -0.546 | 0.006 | 0.004 | 0.004 | 0 | 0.1 | 0.12 | 0.11 | 0.17 | 0.15 | 0.23 |
| 9 | 73856839 | rs11142750 | 0.807 | 1.667 | 0.375 | -0.577 | 0.022 | -0.008 | -0.008 | 0.057 | 0.06 | 0.06 | 0.08 | 0.01 | 0.19 | 0.09 |
| 9 | 73859086 | rs9785312 | -1.176 | 1.63 | 0.498 | -0.58 | 0.198 | 0.012 | 0.012 | 1.434 | 0.11 | 0.47 | 0.12 | 0.08 | 0.2 | 0.1 |
| 9 | 73861353 | rs4745077 | -1.726 | 1.548 | 0.4 | -0.603 | 0.094 | 0.043 | 0.043 | 0.207 | 0.5 | 0.78 | 0.63 | 0.6 | 0.64 | 0.62 |
| 9 | 73873692 | rs17056622 | -0.406 | 1.646 | 0.468 | -0.588 | 0.189 | -0.036 | -0.036 | 0 | 0.09 | 0.39 | 0.1 | 0.01 | 0.19 | 0.09 |
| 9 | 73879320 | rs10781005 | 0.761 | 1.544 | 0.376 | -0.593 | -0.002 | -0.005 | -0.005 | 0.364 | 0.57 | 0.6 | 0.48 | 0.49 | 0.55 | 0.48 |
| 9 | 73886505 | rs10781009 | 0.787 | 1.438 | 0.283 | -0.66 | -0.007 | -0.006 | -0.006 | 0 | 0.55 | 0.63 | 0.45 | 0.49 | 0.51 | 0.47 |
| 9 | 73888408 | rs11142771 | 2.073 | 1.385 | 0.177 | -0.666 | 0.005 | 0.035 | 0.035 | 0.093 | 0.07 | 0.02 | 0.03 | 0 | 0.05 | 0.02 |

| | | | | | | | | | | | | | | | | |
|---|---|---|---|---|---|---|---|---|---|---|---|---|---|---|---|---|
| 9 | 73894279 | rs7850256 | 1.744 | 1.418 | 0.199 | -0.589 | 0.079 | -0.008 | -0.008 | 2.544 | 0.43 | 0.12 | 0.34 | 0.41 | 0.31 | 0.37 |
| 9 | 73895545 | rs7022575 | 1.531 | 1.436 | 0.23 | -0.595 | 0.069 | -0.015 | -0.015 | 0 | 0.45 | 0.19 | 0.35 | 0.48 | 0.32 | 0.38 |
| 9 | 73896545 | rs10869010 | 0.04 | 1.644 | -0.021 | -0.573 | 0.002 | -0.008 | -0.008 | 0 | 0.83 | 0.78 | 0.85 | 0.84 | 0.8 | 0.75 |
| 9 | 73902470 | rs1504388 | -0.57 | 1.832 | 0.33 | -0.342 | 0.143 | -0.045 | -0.045 | 4.234 | 0.09 | 0.3 | 0.1 | 0.01 | 0.19 | 0.09 |
| 9 | 73903664 | rs1532803 | -1.206 | 1.753 | 0.403 | -0.337 | 0.181 | 0.01 | 0.01 | 2.395 | 0.12 | 0.43 | 0.16 | 0.1 | 0.22 | 0.21 |
| 9 | 73907625 | rs11142786 | -0.394 | 0.868 | 0.331 | -0.074 | -0.011 | -0.07 | -0.07 | 0 | 0.21 | 0.4 | 0.12 | 0.09 | 0.18 | 0.16 |
| 9 | 73909625 | rs1604856 | -0.394 | 0.868 | 0.305 | -0.061 | -0.011 | -0.068 | -0.068 | 0 | 0.21 | 0.4 | 0.12 | 0.09 | 0.19 | 0.16 |
| 9 | 73909781 | rs13300497 | -0.952 | 0.894 | 0.306 | -0.005 | 0.003 | 0.001 | 0.001 | 0 | 0.24 | 0.43 | 0.18 | 0.27 | 0.23 | 0.29 |
| 9 | 73909871 | rs17533283 | -0.435 | 0.783 | 0.287 | -0.301 | -0.005 | -0.033 | -0.033 | 0 | 0.19 | 0.27 | 0.12 | 0.09 | 0.18 | 0.16 |
| 9 | 73914256 | rs11142792 | -0.213 | 0.996 | -0.195 | -0.209 | 0.007 | -0.021 | -0.021 | 0 | 0.19 | 0.42 | 0.14 | 0.17 | 0.2 | 0.19 |
| 9 | 73914605 | rs11142794 | -0.242 | 1.116 | -0.363 | 0.06 | 0 | -0.022 | -0.022 | 0.026 | 0.19 | 0.15 | 0.3 | 0.26 | 0.17 | 0.13 |
| 9 | 73915883 | rs1566837 | 0.382 | 0.873 | -0.354 | 0.174 | 0.008 | -0.003 | -0.003 | 0.02 | 0.61 | 0.53 | 0.51 | 0.67 | 0.53 | 0.59 |
| 9 | 73920359 | rs7023772 | 1.022 | 0.406 | -0.25 | 0.243 | -0.001 | 0.005 | 0.005 | 0.075 | 0.86 | 0.79 | 0.83 | 0.83 | 0.85 | 0.86 |
| 9 | 73920739 | rs2134860 | 1.125 | 0.365 | 0.028 | 0.302 | -0.004 | 0.028 | 0.028 | 0.062 | 0.85 | 0.72 | 0.81 | 0.79 | 0.85 | 0.78 |
| 9 | 73923082 | rs1021879 | 0.833 | 0.345 | -0.016 | 0.308 | -0.002 | 0.002 | 0.002 | 0.179 | 0.87 | 0.8 | 0.87 | 0.91 | 0.87 | 0.93 |
| 9 | 73925831 | rs4142472 | 1.004 | 0.063 | -0.519 | -0.274 | 0.035 | 0.066 | 0.066 | 1.233 | 0.54 | 0.17 | 0.52 | 0.73 | 0.61 | 0.56 |
| 9 | 73929083 | rs1472730 | 0.613 | -0.324 | 0.726 | -0.118 | 0.047 | 0.226 | 0.226 | 0.731 | 0.93 | 0.64 | 0.92 | 0.87 | 0.98 | 0.94 |
| 9 | 73931778 | rs4744628 | 0.571 | -0.55 | 0.442 | -0.583 | -0.01 | -0.009 | -0.009 | 0 | 0.71 | 0.36 | 0.82 | 0.82 | 0.81 | 0.77 |
| 9 | 73935165 | rs17617904 | 0.257 | -0.403 | 0.085 | -0.725 | 0.002 | 0.029 | 0.029 | 0.287 | 0.22 | 0.16 | 0.25 | 0.04 | 0.19 | 0.16 |
| 9 | 73935754 | rs10781013 | 1.133 | -0.321 | -0.425 | -0.691 | 0.029 | 0.125 | 0.125 | 0.301 | 0.48 | 0.12 | 0.56 | 0.77 | 0.62 | 0.62 |
| 9 | 73936166 | rs17617922 | -0.469 | -0.207 | 0.263 | -0.313 | -0.005 | -0.067 | -0.067 | 0 | 0.21 | 0.26 | 0.1 | 0.03 | 0.17 | 0.16 |
| 9 | 73940190 | rs11792896 | -1.154 | 0.095 | 0.349 | 0.622 | -0.005 | 0.059 | 0.059 | 0 | 0.06 | 0.15 | 0.14 | 0.2 | 0.04 | 0.17 |
| 9 | 73953144 | rs1897258 | 0.862 | -0.277 | 0.92 | 0.201 | -0.044 | -0.047 | -0.047 | 0 | 0.89 | 0.7 | 0.95 | 0.98 | 0.97 | 0.94 |
| 9 | 73958026 | rs7852227 | 0.145 | -0.168 | 0.415 | -0.68 | -0.017 | 0.065 | 0.065 | 0 | 0.26 | 0.11 | 0.18 | 0.04 | 0.3 | 0.17 |
| 9 | 73967943 | rs11142806 | 1.317 | 0.059 | 0.771 | 0.161 | -0.003 | 0.038 | 0.038 | 0 | 0.52 | 0.27 | 0.55 | 0.58 | 0.52 | 0.45 |
| 9 | 73973798 | rs4744630 | -0.852 | 0.173 | 0.931 | 0.06 | 0.024 | 0.001 | 0.001 | 0 | 0.38 | 0.47 | 0.43 | 0.41 | 0.48 | 0.54 |
| 9 | 73995267 | rs10512007 | -0.985 | 0.137 | 1.331 | 0.155 | -0.007 | 0.001 | 0.001 | 0 | 0.4 | 0.38 | 0.4 | 0.41 | 0.44 | 0.54 |
| 9 | 74006667 | rs12352829 | -0.621 | 0.274 | 1.298 | 0.349 | 0.036 | 0.067 | 0.067 | 0 | 0.11 | 0.29 | 0.2 | 0.26 | 0.14 | 0.3 |
| 9 | 74016773 | rs1074670 | -0.924 | 0.166 | 1.02 | 0.378 | 0.003 | -0.008 | -0.008 | 0 | 0.41 | 0.48 | 0.41 | 0.33 | 0.44 | 0.53 |
| 9 | 74018496 | rs1891301 | 1.049 | 0.244 | 0.22 | 0.242 | -0.022 | 0.038 | 0.038 | 0 | 0.52 | 0.19 | 0.56 | 0.59 | 0.54 | 0.42 |
| 9 | 74036883 | rs7852943 | -0.096 | 0.306 | -0.156 | -0.069 | -0.009 | 0.044 | 0.044 | 0 | 0.27 | 0.16 | 0.19 | 0.04 | 0.3 | 0.23 |
| 9 | 74037537 | rs12344569 | 0.627 | -0.099 | -0.15 | 0.764 | 0 | 0.027 | 0.027 | 0 | 0.06 | 0.06 | 0.09 | 0.21 | 0.03 | 0.11 |
| 9 | 74039166 | rs1935338 | 1.703 | -0.607 | -0.75 | 1.119 | 0.055 | -0.115 | -0.115 | 0 | 0.65 | 0.06 | 0.61 | 0.48 | 0.76 | 0.58 |
| 9 | 74042233 | rs4469528 | 1.696 | -0.541 | -0.706 | 1.119 | 0.055 | -0.101 | -0.101 | 0.098 | 0.66 | 0.08 | 0.61 | 0.48 | 0.76 | 0.58 |
| 9 | 74042353 | rs10869019 | -0.609 | -0.613 | 0.399 | 1.21 | 0.024 | 0.098 | 0.098 | 0 | 0.1 | 0.3 | 0.15 | 0.18 | 0.09 | 0.17 |
| 9 | 74050819 | rs12375722 | 0.873 | -0.445 | -0.95 | 0.537 | -0.142 | 0.141 | 0.141 | 0 | 0.31 | 0.02 | 0.38 | 0.07 | 0.39 | 0.17 |
| 9 | 74052978 | rs10116204 | 0.134 | -1.038 | 0.672 | 1.437 | -0.052 | 0.038 | 0.038 | 1.685 | 0.87 | 0.71 | 0.88 | 0.83 | 0.96 | 0.95 |
| 9 | 74059199 | rs12238624 | -0.623 | -0.579 | 0.498 | 2.052 | -0.01 | -0.048 | -0.048 | 0 | 0.11 | 0.03 | 0.1 | 0.24 | 0.1 | 0.16 |
| 9 | 74061310 | rs4745087 | 2.044 | -0.716 | 1.294 | 2.327 | -0.052 | 0.142 | 0.142 | 0 | 0.77 | 0.48 | 0.78 | 0.48 | 0.88 | 0.76 |
| 9 | 74067610 | rs12002628 | 2.153 | -0.688 | 1.476 | 2.305 | -0.007 | 0.015 | 0.015 | 0.022 | 0.73 | 0.32 | 0.73 | 0.47 | 0.82 | 0.67 |
| 9 | 74070471 | rs1821070 | 1.374 | -0.709 | 1.756 | 2.374 | 0.011 | 0.132 | 0.132 | 3.237 | 0.94 | 0.62 | 0.97 | 0.9 | 1 | 0.99 |
| 9 | 74072841 | rs7034150 | 0.054 | -0.673 | 1.554 | 2.413 | -0.019 | 0.11 | 0.11 | 0 | 0.08 | 0.16 | 0.1 | 0.29 | 0.02 | 0.07 |
| 9 | 74080676 | rs10869025 | 1.908 | -0.657 | 1.402 | 2.231 | -0.013 | 0.032 | 0.032 | 0.007 | 0.74 | 0.35 | 0.74 | 0.47 | 0.83 | 0.66 |
| 9 | 74085486 | rs11507232 | -0.588 | -0.413 | 1.107 | 1.997 | 0.014 | 0.118 | 0.118 | 0 | 0.1 | 0.26 | 0.14 | 0.21 | 0.07 | 0.17 |
| 9 | 74096528 | rs11142845 | -0.95 | -0.53 | 0.586 | 1.764 | 0.002 | -0.005 | -0.005 | 0 | 0.09 | 0.07 | 0.16 | 0.16 | 0.11 | 0.08 |
| 9 | 74101337 | rs10869028 | -0.993 | -0.8 | 1.005 | 2.199 | -0.019 | 0.155 | 0.155 | 0 | 0.14 | 0.28 | 0.18 | 0.54 | 0.07 | 0.25 |
| 9 | 74103833 | rs10869029 | 1.933 | -0.907 | 1.336 | 1.97 | -0.023 | 0.009 | 0.009 | 0.077 | 0.73 | 0.33 | 0.73 | 0.33 | 0.82 | 0.67 |
| 9 | 74104808 | rs4549848 | 1.721 | -0.892 | 1.368 | 1.982 | -0.044 | 0.035 | 0.035 | 0 | 0.74 | 0.47 | 0.73 | 0.33 | 0.82 | 0.67 |
| 9 | 74115344 | rs2309966 | -0.479 | -0.573 | 0.132 | 1.303 | -0.02 | -0.015 | -0.015 | 0 | 0.55 | 0.77 | 0.69 | 0.49 | 0.57 | 0.51 |
| 9 | 74122340 | rs2309968 | 0.08 | -0.85 | 0.668 | 0.694 | -0.016 | -0.054 | -0.054 | 0 | 0.1 | 0.26 | 0.05 | 0.01 | 0.04 | 0.11 |
| 9 | 74130286 | rs10869031 | 0.567 | -0.82 | -0.015 | 0.71 | -0.018 | -0.008 | -0.008 | 0.405 | 0.49 | 0.19 | 0.43 | 0.44 | 0.49 | 0.53 |
| 9 | 74135468 | rs11142863 | -0.077 | -0.8 | -0.098 | 0.83 | -0.001 | -0.06 | -0.06 | 0 | 0.14 | 0.04 | 0.12 | 0.27 | 0.11 | 0.25 |
| 9 | 74136094 | rs11791628 | 1.649 | -0.768 | -0.097 | 0.749 | -0.021 | 0.109 | 0.109 | 0 | 0.35 | 0.14 | 0.3 | 0.17 | 0.37 | 0.27 |
| 9 | 74151806 | rs11142873 | 1.081 | -0.63 | 0.119 | 0.751 | -0.028 | 0.046 | 0.046 | 0 | 0.27 | 0.18 | 0.38 | 0.11 | 0.38 | 0.18 |
| 9 | 74152609 | rs2309974 | 0.173 | -0.692 | 0.311 | 0.807 | -0.001 | -0.001 | -0.001 | 0 | 0.57 | 0.52 | 0.46 | 0.44 | 0.51 | 0.58 |
| 9 | 74154710 | rs2309975 | -0.136 | -0.69 | 0.341 | 0.829 | -0.005 | 0.089 | 0.089 | 0 | 0.85 | 0.75 | 0.84 | 0.54 | 0.9 | 0.74 |
| 9 | 74157998 | rs10124132 | 1.123 | -0.732 | 0.425 | 0.797 | 0.008 | -0.018 | -0.018 | 0 | 0.07 | 0.24 | 0.03 | 0 | 0.03 | 0.05 |
| 9 | 74158641 | rs7042172 | 0.719 | -0.692 | 0.066 | 0.803 | -0.024 | 0.081 | 0.081 | 0.112 | 0.78 | 0.44 | 0.81 | 0.54 | 0.87 | 0.71 |
| 9 | 74158862 | rs17056975 | 0.952 | -0.687 | 0.025 | 0.766 | -0.018 | 0.033 | 0.033 | 0 | 0.29 | 0.23 | 0.38 | 0.11 | 0.38 | 0.18 |
| 9 | 74167402 | rs2309979 | 0.977 | -0.857 | 0.219 | 0.996 | -0.025 | 0.066 | 0.066 | 0.444 | 0.77 | 0.42 | 0.81 | 0.54 | 0.87 | 0.71 |
| 9 | 74171129 | rs10746878 | 0.592 | -0.827 | 0.19 | 1.011 | -0.028 | 0.144 | 0.144 | 0.089 | 0.79 | 0.53 | 0.81 | 0.54 | 0.87 | 0.72 |
| 9 | 74172458 | rs7876052 | -0.078 | -0.83 | -0.271 | 0.952 | -0.008 | -0.022 | -0.022 | 0 | 0.13 | 0.03 | 0.07 | 0.13 | 0.11 | 0.13 |
| 9 | 74174742 | rs2309982 | -0.055 | -0.586 | -0.188 | 0.71 | 0 | 0.005 | 0.005 | 0 | 0.37 | 0.37 | 0.39 | 0.21 | 0.39 | 0.25 |
| 9 | 74175647 | rs2309983 | 1.895 | -0.695 | -0.429 | 0.999 | -0.098 | 0.114 | 0.114 | 0.005 | 0.35 | 0.04 | 0.32 | 0.17 | 0.4 | 0.29 |
| 9 | 74176418 | rs2232139 | 0.633 | -0.708 | -0.725 | 0.618 | -0.008 | 0.021 | 0.021 | 1.61 | 0.22 | 0.18 | 0.27 | 0.07 | 0.23 | 0.21 |
| 9 | 74176852 | rs7043834 | 1.025 | -0.715 | -0.76 | 0.66 | 0.006 | -0.027 | -0.027 | 0.244 | 0.14 | 0.04 | 0.11 | 0.13 | 0.16 | 0.05 |
| 9 | 74179834 | rs10124877 | -0.386 | -0.706 | -0.441 | 0.636 | 0.001 | 0.037 | 0.037 | 0 | 0.85 | 0.82 | 0.84 | 0.55 | 0.86 | 0.78 |
| 9 | 74181734 | rs11142892 | 0.696 | -0.66 | -0.689 | 0.554 | -0.02 | 0.051 | 0.051 | 1.516 | 0.21 | 0.1 | 0.27 | 0.07 | 0.23 | 0.21 |
| 9 | 74186310 | rs4606122 | 0.988 | -0.701 | -0.65 | 0.623 | -0.001 | -0.007 | -0.007 | 0 | 0.55 | 0.35 | 0.53 | 0.49 | 0.6 | 0.5 |

| | | | | | | | | | | | | | | | | |
|---|---|---|---|---|---|---|---|---|---|---|---|---|---|---|---|---|
| 9 | 74192592 | rs11142895 | 1.314 | -0.705 | -0.841 | 0.595 | -0.012 | 0.091 | 0.091 | 0.208 | 0.51 | 0.12 | 0.47 | 0.31 | 0.6 | 0.4 |
| 9 | 74199338 | rs13287882 | 1.592 | -0.77 | -1.006 | 0.269 | -0.076 | 0.025 | 0.025 | 0 | 0.42 | 0.09 | 0.39 | 0.31 | 0.48 | 0.38 |
| 9 | 74199813 | rs9314788 | -0.963 | -0.423 | -0.705 | 0.292 | 0.006 | -0.025 | -0.025 | 0 | 0.95 | 0.91 | 0.96 | 1 | 0.94 | 0.98 |
| 9 | 74200674 | rs2210016 | 0.217 | -0.448 | -0.831 | 0.397 | 0.006 | -0.088 | -0.088 | 0.836 | 0.15 | 0.03 | 0.16 | 0.3 | 0.16 | 0.15 |
| 9 | 74202802 | rs11142904 | -0.27 | -0.428 | -0.781 | 0.398 | 0.009 | -0.046 | -0.046 | 0 | 0.18 | 0.12 | 0.21 | 0.3 | 0.2 | 0.21 |
| 9 | 74204367 | rs11142906 | 0.514 | -0.19 | -0.699 | 0.164 | -0.005 | 0.02 | 0.02 | 0.39 | 0.21 | 0.16 | 0.27 | 0.07 | 0.21 | 0.19 |
| 9 | 74205232 | rs10869044 | 0.214 | -0.173 | -0.601 | 0.2 | 0 | -0.021 | -0.021 | 0 | 0.24 | 0.26 | 0.28 | 0.08 | 0.23 | 0.21 |
| 9 | 74208282 | rs12353518 | 1.313 | -0.144 | -0.717 | 0.419 | 0.001 | 0.019 | 0.019 | 0 | 0.46 | 0.34 | 0.39 | 0.28 | 0.47 | 0.37 |
| 9 | 74228247 | rs4069914 | -0.688 | -0.028 | -0.58 | 0.636 | -0.001 | 0.058 | 0.058 | 0 | 0.89 | 0.88 | 0.86 | 0.52 | 0.91 | 0.79 |
| 9 | 74229859 | rs1329411 | 0.781 | 0.005 | -0.813 | -0.184 | -0.002 | 0.007 | 0.007 | 0.472 | 0.55 | 0.37 | 0.46 | 0.28 | 0.59 | 0.4 |
| 9 | 74244432 | rs868147 | 0.804 | 0.003 | -1.398 | -0.408 | 0.003 | -0.005 | -0.005 | 0.648 | 0.43 | 0.44 | 0.32 | 0.37 | 0.53 | 0.54 |
| 9 | 74244754 | rs11142927 | -0.013 | 0.093 | -1.698 | -0.716 | -0.015 | 0.082 | 0.082 | 0 | 0.31 | 0.16 | 0.3 | 0.09 | 0.31 | 0.21 |
| 9 | 74245097 | rs13293927 | 1.866 | 0.616 | -2.183 | -1.038 | -0.067 | 0.151 | 0.151 | 0 | 0.28 | 0.01 | 0.17 | 0.06 | 0.28 | 0.21 |
| 9 | 74245141 | rs10781030 | -0.116 | 0.796 | -1.125 | -0.254 | 0.001 | 0.065 | 0.065 | 0 | 0.13 | 0.18 | 0.27 | 0.54 | 0.12 | 0.22 |
| 9 | 74245268 | rs6560217 | 0.56 | 0.579 | -1.26 | -1.024 | 0 | 0.003 | 0.003 | 0 | 0.68 | 0.72 | 0.69 | 0.91 | 0.7 | 0.79 |
| 9 | 74246308 | rs4745104 | -0.748 | 0.784 | -1.557 | -0.891 | -0.012 | 0.112 | 0.112 | 0 | 0.34 | 0.14 | 0.32 | 0.09 | 0.36 | 0.22 |
| 9 | 74248170 | rs4745106 | 0.31 | 1.333 | -1.085 | -0.492 | 0.006 | -0.018 | -0.018 | 0 | 0.12 | 0.11 | 0.21 | 0.04 | 0.17 | 0.12 |
| 9 | 74250387 | rs1928423 | 0.466 | 0.738 | -1.097 | -0.316 | 0.001 | 0.005 | 0.005 | 0 | 0.57 | 0.6 | 0.48 | 0.78 | 0.58 | 0.7 |
| 9 | 74254313 | rs2201127 | -0.418 | 0.962 | -1.716 | -1.231 | -0.048 | 0.124 | 0.124 | 0 | 0.66 | 0.82 | 0.8 | 1 | 0.63 | 0.82 |
| 9 | 74257162 | rs2871401 | -0.039 | 0.343 | -0.852 | -0.439 | 0 | 0.002 | 0.002 | 0 | 0.47 | 0.46 | 0.55 | 0.41 | 0.49 | 0.49 |
| 9 | 74258824 | rs2087716 | 0.358 | -0.115 | -1.193 | -0.455 | -0.07 | 0.013 | 0.013 | 0 | 0.36 | 0.08 | 0.33 | 0.33 | 0.46 | 0.37 |
| 9 | 74260812 | rs10781032 | 0.229 | -0.525 | -0.181 | 0.393 | -0.02 | 0.177 | 0.177 | 0.146 | 0.15 | 0.57 | 0.32 | 0.4 | 0.11 | 0.28 |
| 9 | 74261572 | rs10746883 | 0.584 | -0.495 | 0.529 | 0.481 | 0.02 | 0.324 | 0.324 | 0 | 0.13 | 0.44 | 0.3 | 0.42 | 0.11 | 0.27 |
| 9 | 74262200 | rs2131039 | 0.198 | -0.028 | -0.648 | 0.226 | -0.074 | -0.027 | -0.027 | 0 | 0.25 | 0.03 | 0.2 | 0.22 | 0.31 | 0.22 |
| 9 | 74265535 | rs1410989 | -1.69 | -0.56 | -0.457 | -0.231 | -0.013 | 0.069 | 0.069 | 0 | 0.64 | 0.79 | 0.76 | 0.9 | 0.66 | 0.77 |
| 9 | 74266320 | rs11142947 | 0.883 | -0.246 | -0.39 | 0.053 | -0.001 | 0.001 | 0.001 | 0 | 0.12 | 0.13 | 0.1 | 0.18 | 0.12 | 0.11 |
| 9 | 74270232 | rs970480 | -0.516 | -0.532 | -0.457 | 0.232 | -0.012 | -0.021 | -0.021 | 0.04 | 0.42 | 0.32 | 0.45 | 0.48 | 0.48 | 0.45 |
| 9 | 74272001 | rs11142949 | 0.055 | -0.471 | -0.575 | 0.12 | -0.024 | -0.013 | -0.013 | 0 | 0.31 | 0.23 | 0.37 | 0.3 | 0.42 | 0.36 |
| 9 | 74277315 | rs7854239 | -0.333 | -0.725 | -0.218 | -0.047 | 0 | -0.001 | -0.001 | 0.008 | 0.45 | 0.54 | 0.67 | 0.55 | 0.55 | 0.5 |
| 9 | 74278480 | rs11142952 | -0.146 | -1.531 | -0.756 | -0.39 | -0.043 | 0.062 | 0.062 | 0 | 0.27 | 0.53 | 0.19 | 0.44 | 0.18 | 0.34 |
| 9 | 74279487 | rs4745109 | -1.543 | -1.215 | -1.182 | -1.017 | -0.001 | 0.101 | 0.101 | 0.63 | 0.72 | 0.9 | 0.84 | 1 | 0.73 | 0.84 |
| 9 | 74281371 | rs4745111 | -1.543 | -1.212 | -1.182 | -1.017 | -0.004 | 0.101 | 0.101 | 0.63 | 0.72 | 0.9 | 0.83 | 1 | 0.72 | 0.84 |
| 9 | 74283609 | rs11142955 | 0.033 | -0.981 | -1.26 | -0.931 | -0.009 | 0.004 | 0.004 | 0 | 0.23 | 0.23 | 0.22 | 0.06 | 0.37 | 0.17 |
| 9 | 74285850 | rs1887971 | 0.082 | -0.801 | -1.138 | -0.565 | -0.006 | -0.037 | -0.037 | 0 | 0.43 | 0.2 | 0.53 | 0.51 | 0.54 | 0.53 |
| 9 | 74292618 | rs4745112 | -0.054 | -0.049 | -0.751 | -0.043 | -0.007 | 0.003 | 0.003 | 0.63 | 0.57 | 0.48 | 0.59 | 0.57 | 0.61 | 0.59 |
| 9 | 74296497 | rs10511999 | 0.038 | 0.347 | -1.45 | -0.449 | -0.146 | 0.019 | 0.019 | 0 | 0.18 | 0.02 | 0.26 | 0.08 | 0.43 | 0.22 |
| 9 | 74297245 | rs10511998 | 0.562 | -0.556 | -1.021 | -0.262 | -0.049 | 0.142 | 0.142 | 0.791 | 0.22 | 0.02 | 0.13 | 0 | 0.23 | 0.16 |
| 9 | 74298720 | rs6560226 | -0.606 | -0.63 | -0.985 | -0.25 | -0.019 | 0.112 | 0.112 | 0.395 | 0.78 | 0.95 | 0.87 | 1 | 0.78 | 0.84 |
| 9 | 74300640 | rs2310021 | -0.451 | -0.639 | -1.069 | -0.18 | -0.032 | 0.161 | 0.161 | 0.008 | 0.76 | 0.96 | 0.85 | 1 | 0.75 | 0.84 |
| 9 | 74304506 | rs4745114 | 1.046 | -0.587 | -1.001 | 0.274 | -0.027 | -0.055 | -0.055 | 0 | 0.43 | 0.08 | 0.45 | 0.44 | 0.53 | 0.42 |
| 9 | 74305098 | rs2297089 | 0.568 | -0.085 | -0.192 | 0.704 | -0.032 | -0.007 | -0.007 | 0 | 0.09 | 0.25 | 0.03 | 0.08 | 0.02 | 0.05 |
| 9 | 74308389 | rs3780605 | -0.197 | 0.203 | -0.192 | 0.95 | -0.002 | 0.097 | 0.097 | 0 | 0.13 | 0.22 | 0.16 | 0.34 | 0.14 | 0.28 |
| 9 | 74311872 | rs4745116 | -0.49 | 0.632 | -0.796 | 0.406 | -0.003 | -0.008 | -0.008 | 2.554 | 0.42 | 0.32 | 0.44 | 0.49 | 0.42 | 0.36 |
| 9 | 74312747 | rs1293 | 0.422 | 0.177 | -1.191 | 0.558 | -0.003 | -0.001 | -0.001 | 0 | 0.4 | 0.43 | 0.38 | 0.43 | 0.41 | 0.47 |
| 9 | 74313429 | rs17057129 | -0.658 | -0.046 | -1.478 | 0.913 | 0.013 | -0.081 | -0.081 | 0.409 | 0.23 | 0.05 | 0.24 | 0.35 | 0.22 | 0.2 |
| 9 | 74318137 | rs11142963 | -0.964 | 0.053 | -0.64 | 0.433 | -0.008 | -0.012 | -0.012 | 0 | 0.18 | 0.26 | 0.11 | 0.13 | 0.13 | 0.32 |
| 9 | 74319677 | rs17057133 | -0.194 | -0.467 | -0.732 | 0.47 | 0.008 | -0.003 | -0.003 | 0 | 0.09 | 0.06 | 0.12 | 0.1 | 0.03 | 0.03 |
| 9 | 74321381 | rs12341685 | -1.286 | -0.867 | 0.095 | -0.005 | -0.022 | -0.032 | -0.032 | 1.786 | 0.17 | 0.34 | 0.17 | 0.12 | 0.09 | 0.1 |
| 9 | 74324149 | rs7046059 | -1.163 | -0.881 | 0.232 | -0.012 | 0.002 | 0.003 | 0.003 | 1.168 | 0.14 | 0.13 | 0.16 | 0.12 | 0.09 | 0.1 |
| 9 | 74325433 | rs1010434 | -0.042 | -0.868 | 0.213 | -0.015 | -0.025 | 0.027 | 0.027 | 0.526 | 0.05 | 0.01 | 0.04 | 0.02 | 0.06 | 0.07 |
| 9 | 74327419 | rs7037458 | -0.115 | -1.193 | -0.406 | -0.481 | 0.007 | 0.009 | 0.009 | 0 | 0.55 | 0.45 | 0.6 | 0.74 | 0.66 | 0.54 |
| 9 | 74332784 | rs2307005 | -0.283 | -1.149 | -0.554 | -0.453 | 0.004 | 0.007 | 0.007 | 1.601 | 0.14 | 0.09 | 0.16 | 0.12 | 0.09 | 0.11 |
| 9 | 74340910 | rs11142978 | 0.591 | -1.256 | -0.645 | -0.546 | 0.016 | -0.016 | -0.016 | 0.228 | 0.46 | 0.13 | 0.47 | 0.63 | 0.64 | 0.67 |
| 9 | 74345158 | rs25694 | -0.098 | -1.23 | -0.583 | -0.519 | -0.007 | 0.018 | 0.018 | 0.046 | 0.08 | 0.04 | 0.05 | 0.02 | 0.08 | 0.22 |
| 9 | 74348424 | rs7030493 | 0.369 | -1.894 | -0.485 | -0.913 | -0.008 | -0.003 | -0.003 | 0 | 0.17 | 0.19 | 0.12 | 0.1 | 0.09 | 0.12 |
| 9 | 74350002 | rs2274698 | 0.729 | -1.739 | -0.564 | -1.062 | -0.026 | -0.028 | -0.028 | 0.006 | 0.11 | 0.03 | 0.2 | 0.16 | 0.15 | 0.14 |
| 9 | 74355028 | rs25690 | 0.184 | -1.732 | -0.577 | -1.063 | -0.007 | 0.018 | 0.018 | 0.046 | 0.08 | 0.04 | 0.05 | 0.02 | 0.08 | 0.22 |
| 9 | 74355185 | rs3753006 | -0.339 | -1.794 | 0.054 | -1.089 | 0.046 | 0.055 | 0.055 | 0 | 0.14 | 0.36 | 0.23 | 0.16 | 0.15 | 0.14 |
| 9 | 74360234 | rs25688 | 0.136 | -1.769 | -0.15 | -1.102 | -0.007 | 0.018 | 0.018 | 0.06 | 0.08 | 0.04 | 0.05 | 0.02 | 0.08 | 0.22 |
| 9 | 74374194 | rs2310024 | 0.314 | -2.016 | 0.003 | -0.918 | -0.022 | -0.012 | -0.012 | 0.774 | 0.6 | 0.47 | 0.72 | 0.77 | 0.8 | 0.81 |
| 9 | 74378277 | rs7030211 | -0.705 | -1.594 | -0.37 | -0.898 | 0.001 | -0.001 | -0.001 | 1.048 | 0.19 | 0.17 | 0.28 | 0.18 | 0.25 | 0.35 |
| 9 | 74381599 | rs12555916 | 0.335 | -1.399 | -0.342 | -0.855 | -0.007 | -0.006 | -0.006 | 3.582 | 0.12 | 0.07 | 0.22 | 0.16 | 0.17 | 0.13 |
| 9 | 74384876 | rs1404195 | 1.298 | -1.248 | -0.451 | -0.902 | -0.107 | -0.062 | -0.062 | 0 | 0.39 | 0.12 | 0.42 | 0.57 | 0.54 | 0.44 |
| 9 | 74392687 | rs4745122 | 0.49 | -1.291 | 0.046 | -1.113 | -0.015 | -0.047 | -0.047 | 0 | 0.62 | 0.42 | 0.73 | 0.89 | 0.74 | 0.63 |
| 9 | 74393843 | rs4745123 | 0.28 | -1.191 | -0.057 | -1.32 | -0.019 | -0.093 | -0.093 | 0 | 0.63 | 0.41 | 0.79 | 0.97 | 0.76 | 0.66 |
| 9 | 74401123 | rs990449 | 1.331 | -0.134 | 0.22 | -0.723 | 0.031 | -0.017 | -0.017 | 0 | 0.86 | 0.59 | 0.9 | 0.93 | 0.87 | 0.92 |
| 9 | 74410755 | rs10511995 | -1.48 | -0.165 | 0.234 | -0.738 | 0.02 | -0.022 | -0.022 | 0.64 | 0.14 | 0.37 | 0.1 | 0.07 | 0.13 | 0.08 |
| 9 | 74414358 | rs11143006 | 0.531 | 1.124 | -0.381 | -0.433 | -0.041 | -0.037 | -0.037 | 0 | 0.1 | 0.03 | 0.22 | 0.18 | 0.16 | 0.07 |
| 9 | 74427419 | rs1464758 | -0.61 | 0.887 | -0.193 | -0.483 | 0.004 | -0.003 | -0.003 | 0.046 | 0.1 | 0.14 | 0.08 | 0.07 | 0.12 | 0.07 |

| | | | | | | | | | | | | | | | | |
|---|---|---|---|---|---|---|---|---|---|---|---|---|---|---|---|---|
| 9 | 74445353 | rs2310041 | -0.556 | 0.975 | -0.267 | -0.551 | -0.011 | 0.01 | 0.01 | 0.016 | 0.11 | 0.1 | 0.1 | 0.06 | 0.15 | 0.07 |
| 9 | 74463295 | rs7025473 | 1.042 | 0.959 | -0.227 | -0.565 | 0.021 | -0.029 | -0.029 | 0 | 0.87 | 0.73 | 0.89 | 0.94 | 0.85 | 0.93 |
| 9 | 74491379 | rs1404190 | 1.324 | 0.94 | -0.164 | -0.575 | 0.024 | -0.043 | -0.043 | 0 | 0.85 | 0.64 | 0.88 | 0.94 | 0.85 | 0.93 |
| 9 | 74532618 | rs13299567 | -0.486 | 0.957 | -0.248 | -0.554 | -0.001 | 0 | 0 | 0 | 0.1 | 0.11 | 0.09 | 0.06 | 0.14 | 0.07 |
| 9 | 74552234 | rs17491074 | -0.458 | 1.162 | -0.31 | -0.538 | -0.012 | 0.009 | 0.009 | 0.008 | 0.1 | 0.09 | 0.09 | 0.06 | 0.14 | 0.07 |
| 9 | 74575181 | rs11143040 | -0.108 | 0.873 | -0.686 | -0.434 | -0.063 | -0.038 | -0.038 | 0 | 0.13 | 0.03 | 0.21 | 0.15 | 0.18 | 0.08 |
| 9 | 74599651 | rs7030498 | 0.144 | 0.895 | -0.86 | 0.079 | 0.002 | -0.027 | -0.027 | 0.589 | 0.65 | 0.58 | 0.63 | 0.74 | 0.59 | 0.69 |
| 9 | 74600212 | rs7046123 | 0.803 | 0.65 | -1.351 | 0.315 | 0.017 | -0.038 | -0.038 | 0 | 0.37 | 0.19 | 0.27 | 0.44 | 0.33 | 0.51 |
| 9 | 74600517 | rs10869073 | -0.25 | 0.738 | -1.11 | -0.004 | 0.003 | -0.016 | -0.016 | 0.887 | 0.26 | 0.25 | 0.23 | 0.16 | 0.36 | 0.12 |
| 9 | 74602933 | rs2147241 | 0.548 | -0.254 | -1.462 | 0.486 | 0.068 | 0.041 | 0.041 | 0.083 | 0.54 | 0.15 | 0.51 | 0.4 | 0.64 | 0.43 |
| 9 | 74607151 | rs10869076 | 0.398 | -0.571 | -1.02 | 0.463 | -0.001 | 0.009 | 0.009 | 0.034 | 0.53 | 0.4 | 0.37 | 0.31 | 0.52 | 0.47 |
| 9 | 74610494 | rs942391 | -0.203 | -0.677 | -0.827 | 0.921 | -0.01 | 0.104 | 0.104 | 0.095 | 0.1 | 0.24 | 0.07 | 0.28 | 0.08 | 0.03 |
| 9 | 74612197 | rs11787649 | -0.1 | -0.568 | -0.874 | 0.781 | -0.001 | 0.002 | 0.002 | 0.233 | 0.64 | 0.52 | 0.56 | 0.59 | 0.63 | 0.56 |
| 9 | 74619041 | rs942390 | 1.063 | -0.003 | -1.66 | 0.488 | -0.09 | 0.139 | 0.139 | 0.03 | 0.38 | 0.09 | 0.34 | 0.14 | 0.47 | 0.24 |
| 9 | 74621568 | rs12682751 | -0.548 | -0.018 | -1.295 | 1.153 | -0.007 | 0.004 | 0.004 | 0 | 0.09 | 0.07 | 0.08 | 0.28 | 0.11 | 0.05 |
| 9 | 74622195 | rs13285788 | -0.586 | -0.102 | -1.228 | 1.156 | 0.007 | 0.068 | 0.068 | 0 | 0.09 | 0.15 | 0.08 | 0.28 | 0.11 | 0.06 |
| 9 | 74630070 | rs956090 | 0.684 | -0.47 | -1.975 | 0.205 | 0.019 | 0.203 | 0.203 | 2.424 | 0.51 | 0.1 | 0.43 | 0.18 | 0.63 | 0.35 |
| 9 | 74637129 | rs2147244 | 0.944 | -0.603 | -1.97 | 0.016 | -0.052 | 0.266 | 0.266 | 1.48 | 0.45 | 0.06 | 0.34 | 0.09 | 0.59 | 0.27 |
| 9 | 74642649 | rs4745150 | -0.08 | -0.569 | -0.664 | 0.936 | 0.048 | 0.185 | 0.185 | 1.983 | 0.12 | 0.4 | 0.37 | 0.36 | 0.13 | 0.29 |
| 9 | 74649339 | rs4745152 | 0.28 | -1.469 | -0.371 | 1.077 | -0.01 | 0.049 | 0.049 | 0.017 | 0.79 | 0.76 | 0.88 | 0.61 | 0.82 | 0.65 |
| 9 | 74653124 | rs9314790 | -0.906 | -1.524 | 0.238 | 1.223 | -0.043 | 0.041 | 0.041 | 2.31 | 0.29 | 0.53 | 0.19 | 0.37 | 0.21 | 0.15 |
| 9 | 74664333 | rs1806868 | -0.336 | -1.04 | 0.578 | 1.325 | -0.027 | 0.111 | 0.111 | 0.121 | 0.15 | 0.29 | 0.29 | 0.42 | 0.08 | 0.27 |
| 9 | 74667770 | rs1570638 | 1.227 | -0.757 | 0.565 | 1.361 | -0.009 | 0.028 | 0.028 | 2.722 | 0.69 | 0.61 | 0.63 | 0.5 | 0.69 | 0.57 |
| 9 | 74668310 | rs4744654 | 0.488 | -0.404 | 0.424 | 1.062 | -0.002 | 0.008 | 0.008 | 0 | 0.86 | 0.82 | 0.84 | 0.8 | 0.88 | 0.65 |
| 9 | 74674427 | rs7047632 | -0.754 | -0.613 | 0.694 | 1.024 | -0.028 | 0.099 | 0.099 | 2.755 | 0.22 | 0.45 | 0.29 | 0.47 | 0.18 | 0.32 |
| 9 | 74675604 | rs12345527 | -0.539 | -0.617 | 0.949 | 0.995 | -0.004 | 0.022 | 0.022 | 2.397 | 0.2 | 0.27 | 0.28 | 0.47 | 0.18 | 0.32 |
| 9 | 74679126 | rs10869092 | 0.141 | -0.508 | 0.927 | 0.658 | 0.004 | -0.022 | -0.022 | 1.137 | 0.79 | 0.75 | 0.82 | 0.9 | 0.79 | 0.76 |
| 9 | 74679454 | rs7030573 | -0.611 | -0.613 | 0.766 | 0.318 | -0.017 | 0.035 | 0.035 | 0.009 | 0.29 | 0.47 | 0.32 | 0.48 | 0.24 | 0.34 |
| 9 | 74683057 | rs10869094 | 1.357 | -0.29 | 0.064 | 0.226 | -0.049 | -0.042 | -0.042 | 0 | 0.44 | 0.15 | 0.45 | 0.42 | 0.51 | 0.41 |
| 9 | 74695444 | rs10781064 | 0.843 | -0.212 | 0.667 | 0.348 | 0.008 | -0.008 | -0.008 | 0 | 0.63 | 0.27 | 0.62 | 0.52 | 0.69 | 0.66 |
| 9 | 74698289 | rs10746902 | 0.442 | -0.296 | 1.188 | 0.3 | -0.013 | 0.054 | 0.054 | 0.228 | 0.72 | 0.49 | 0.67 | 0.52 | 0.75 | 0.67 |
| 9 | 74724989 | rs10735611 | 1.063 | -0.425 | 0.886 | 0.094 | 0.003 | 0.003 | 0.003 | 3.799 | 0.68 | 0.28 | 0.71 | 0.72 | 0.74 | 0.75 |
| 9 | 74728011 | rs10435966 | 1.108 | -0.373 | 0.923 | 0.055 | 0.014 | 0.027 | 0.027 | 3.171 | 0.68 | 0.2 | 0.7 | 0.72 | 0.73 | 0.76 |
| 9 | 74733695 | rs4416886 | 0.826 | -0.378 | 1.294 | 0.024 | -0.001 | -0.062 | -0.062 | 0 | 0.11 | 0.34 | 0.05 | 0 | 0.05 | 0.01 |
| 9 | 74733832 | rs4744660 | 1.798 | -0.522 | 1.03 | 0.137 | -0.009 | 0.158 | 0.158 | 2.042 | 0.47 | 0.15 | 0.54 | 0.23 | 0.57 | 0.4 |
| 9 | 74737437 | rs9314794 | 0.173 | -0.516 | 1.046 | 0.155 | 0 | 0.012 | 0.012 | 2.652 | 0.78 | 0.6 | 0.76 | 0.72 | 0.78 | 0.77 |
| 9 | 74739999 | rs4745160 | 0.035 | -0.483 | 0.519 | 0.193 | 0.006 | -0.105 | -0.105 | 0.958 | 0.21 | 0.05 | 0.16 | 0.5 | 0.16 | 0.36 |
| 9 | 74745434 | rs4745167 | 1.666 | -0.476 | 0.527 | 0.181 | 0.005 | 0.134 | 0.134 | 1.902 | 0.47 | 0.2 | 0.55 | 0.23 | 0.57 | 0.4 |
| 9 | 74747787 | rs10869127 | -0.014 | -0.562 | 0.477 | 0.283 | 0.006 | -0.106 | -0.106 | 0 | 0.21 | 0.06 | 0.16 | 0.5 | 0.16 | 0.36 |
| 9 | 74756542 | rs10781076 | 1.485 | -0.185 | 0.339 | 0.076 | -0.04 | 0.177 | 0.177 | 1.13 | 0.47 | 0.15 | 0.52 | 0.2 | 0.54 | 0.42 |
| 9 | 74760556 | rs10511997 | 0.913 | 0.358 | 0.079 | -0.073 | -0.032 | -0.021 | -0.021 | 0.025 | 0.06 | 0 | 0.1 | 0.08 | 0.1 | 0.06 |
| 9 | 74761921 | rs4745173 | 0.044 | 0.246 | 0.549 | -0.026 | 0.012 | 0.025 | 0.025 | 3.719 | 0.8 | 0.69 | 0.74 | 0.67 | 0.75 | 0.77 |
| 9 | 74763415 | rs7036479 | 0.044 | 0.23 | 0.55 | -0.41 | 0.002 | -0.045 | -0.045 | 0 | 0.17 | 0.31 | 0.15 | 0.08 | 0.15 | 0.07 |
| 9 | 74763426 | rs7021424 | 0.27 | 0.225 | 0.63 | -0.413 | 0 | -0.022 | -0.022 | 0 | 0.16 | 0.21 | 0.15 | 0.08 | 0.15 | 0.07 |
| 9 | 74773148 | rs4540461 | 1.597 | 0.157 | 0.262 | -0.516 | -0.044 | 0.194 | 0.194 | 0.29 | 0.41 | 0.12 | 0.42 | 0.11 | 0.44 | 0.37 |
| 9 | 74776701 | rs10869134 | 0.161 | 0.205 | 0.23 | -0.49 | 0.001 | -0.045 | -0.045 | 0 | 0.17 | 0.31 | 0.15 | 0.08 | 0.15 | 0.07 |
| 9 | 74779404 | rs10869137 | 0.468 | 0.25 | 0.212 | -0.347 | 0 | -0.044 | -0.044 | 0 | 0.59 | 0.51 | 0.58 | 0.18 | 0.59 | 0.44 |
| 9 | 74797309 | rs10869142 | -0.258 | 0.29 | 0.492 | -0.055 | 0.007 | -0.054 | -0.054 | 0 | 0.17 | 0.38 | 0.16 | 0.07 | 0.15 | 0.07 |
| 9 | 74800525 | rs4484759 | -0.327 | 0.484 | 0.127 | 0.385 | 0 | 0.018 | 0.018 | 0 | 0.19 | 0.16 | 0.18 | 0.49 | 0.17 | 0.3 |
| 9 | 74814485 | rs10781085 | -0.964 | 0.395 | 0.838 | 0.189 | 0.027 | 0.067 | 0.067 | 0 | 0.27 | 0.47 | 0.29 | 0.56 | 0.26 | 0.36 |
| 9 | 74821250 | rs11143168 | -1.444 | 0.013 | 0.158 | -0.291 | -0.001 | -0.003 | -0.003 | 0 | 0.4 | 0.46 | 0.35 | 0.57 | 0.35 | 0.46 |
| 9 | 74825707 | rs11143170 | -0.95 | -0.039 | 0.173 | -0.256 | 0.001 | 0.027 | 0.027 | 0.331 | 0.28 | 0.37 | 0.28 | 0.54 | 0.25 | 0.38 |
| 9 | 74827066 | rs7030035 | -0.165 | -0.025 | 0.172 | -0.288 | -0.028 | -0.086 | -0.086 | 0.918 | 0.2 | 0.05 | 0.23 | 0.54 | 0.21 | 0.38 |
| 9 | 74828339 | rs7861301 | -0.274 | -0.263 | 0.391 | -0.46 | 0.007 | -0.036 | -0.036 | 0.041 | 0.07 | 0.21 | 0.05 | 0 | 0.04 | 0 |
| 9 | 74828973 | rs17494333 | 2.266 | -0.221 | -0.117 | -0.878 | -0.056 | 0.103 | 0.103 | 0.351 | 0.28 | 0.07 | 0.45 | 0.13 | 0.33 | 0.27 |
| 9 | 74832555 | rs5025470 | 0.993 | -0.346 | 0.161 | -0.837 | -0.006 | -0.069 | -0.069 | 0.032 | 0.13 | 0.24 | 0.08 | 0 | 0.08 | 0.01 |
| 9 | 74834384 | rs10869148 | -0.097 | -0.299 | 0.13 | -0.759 | -0.014 | -0.025 | -0.025 | 0.227 | 0.26 | 0.17 | 0.31 | 0.54 | 0.31 | 0.4 |
| 9 | 74836247 | rs10869149 | -1.735 | -0.55 | 0.092 | -0.921 | 0.001 | -0.004 | -0.004 | 4.621 | 0.56 | 0.53 | 0.45 | 0.57 | 0.47 | 0.52 |
| 9 | 74842516 | rs13295564 | -0.032 | -0.631 | -0.052 | -0.929 | 0.021 | 0.029 | 0.029 | 0.18 | 0.27 | 0.21 | 0.13 | 0.03 | 0.15 | 0.12 |
| 9 | 74845989 | rs10869150 | 0.137 | -0.541 | -0.045 | -0.903 | -0.019 | -0.057 | -0.057 | 0 | 0.26 | 0.11 | 0.28 | 0.54 | 0.28 | 0.4 |
| 9 | 74852965 | rs13298009 | -0.116 | -0.476 | 0.086 | -0.875 | 0.023 | 0.031 | 0.031 | 0.404 | 0.28 | 0.21 | 0.13 | 0.03 | 0.15 | 0.11 |
| 9 | 74853935 | rs11143193 | -1.666 | -0.485 | 0.032 | -0.881 | 0 | 0.002 | 0.002 | 0.947 | 0.52 | 0.45 | 0.41 | 0.57 | 0.41 | 0.49 |
| 9 | 74858118 | rs4130866 | 0.093 | -0.362 | -0.112 | -0.839 | -0.026 | -0.067 | -0.067 | 0 | 0.24 | 0.07 | 0.28 | 0.54 | 0.27 | 0.38 |
| 9 | 74865077 | rs2208744 | -1.46 | -0.724 | -0.052 | -0.983 | -0.002 | 0 | 0 | 1.751 | 0.57 | 0.54 | 0.51 | 0.56 | 0.47 | 0.5 |
| 9 | 74870018 | rs7029217 | -0.525 | -0.793 | 0.043 | -1.025 | 0.005 | -0.002 | -0.002 | 0 | 0.41 | 0.31 | 0.39 | 0.54 | 0.32 | 0.39 |
| 9 | 74874951 | rs969460 | 0.698 | -0.74 | -0.118 | -1.096 | 0.008 | 0.012 | 0.012 | 0 | 0.16 | 0.12 | 0.09 | 0.02 | 0.1 | 0.11 |
| 9 | 74878937 | rs10869157 | -1.241 | -0.657 | -0.027 | -1.044 | -0.002 | 0.002 | 0.002 | 1.686 | 0.55 | 0.53 | 0.51 | 0.61 | 0.46 | 0.49 |
| 9 | 74879449 | rs1322480 | 0.276 | -0.526 | -0.207 | -0.893 | -0.003 | -0.013 | -0.013 | 0 | 0.22 | 0.17 | 0.32 | 0.59 | 0.23 | 0.36 |
| 9 | 74884103 | rs7863783 | -0.533 | -0.56 | -0.223 | -0.47 | 0.008 | -0.006 | -0.006 | 0.066 | 0.45 | 0.31 | 0.43 | 0.49 | 0.39 | 0.46 |

| | | | | | | | | | | | | | | | | |
|---|---|---|---|---|---|---|---|---|---|---|---|---|---|---|---|---|
| 9 | 74887703 | rs11143230 | 2.196 | -0.727 | -0.883 | -0.766 | -0.086 | -0.006 | -0.006 | 0.039 | 0.23 | 0.07 | 0.38 | 0.23 | 0.34 | 0.26 |
| 9 | 74888969 | rs7859318 | -0.004 | -0.695 | -0.795 | -0.722 | 0.041 | -0.037 | -0.037 | 0 | 0.42 | 0.18 | 0.36 | 0.49 | 0.32 | 0.46 |
| 9 | 74892709 | rs7875593 | 0.332 | -0.49 | -0.705 | -0.889 | 0.001 | -0.03 | -0.03 | 0.656 | 0.15 | 0.17 | 0.16 | 0.02 | 0.21 | 0.07 |
| 9 | 74892832 | rs881510 | -0.011 | -0.858 | -0.478 | -0.682 | 0.028 | -0.024 | -0.024 | 0 | 0.41 | 0.22 | 0.37 | 0.48 | 0.32 | 0.46 |
| 9 | 74893024 | rs2871427 | -0.53 | -0.857 | -0.56 | -0.685 | -0.001 | -0.001 | -0.001 | 0 | 0.84 | 0.84 | 0.93 | 0.74 | 0.85 | 0.79 |
| 9 | 74894172 | rs1998372 | -0.302 | -0.867 | -0.25 | -0.676 | 0.003 | -0.002 | -0.002 | 0 | 0.42 | 0.35 | 0.38 | 0.49 | 0.32 | 0.46 |
| 9 | 74897034 | rs11143233 | 0.209 | -0.464 | -0.816 | -1.028 | -0.005 | 0.035 | 0.035 | 0 | 0.68 | 0.81 | 0.59 | 0.24 | 0.68 | 0.41 |
| 9 | 74897271 | rs4997577 | 0.205 | -0.415 | -0.83 | -1.044 | -0.003 | 0.015 | 0.015 | 0 | 0.67 | 0.76 | 0.58 | 0.25 | 0.67 | 0.43 |
| 9 | 74907597 | rs4745182 | -0.327 | -0.124 | -0.988 | -0.988 | 0.018 | 0.048 | 0.048 | 0 | 0.42 | 0.24 | 0.46 | 0.72 | 0.4 | 0.66 |
| 9 | 74909843 | rs7846745 | 1.848 | 0.081 | -1.507 | -1.18 | -0.039 | -0.05 | -0.05 | 2.73 | 0.17 | 0.04 | 0.24 | 0.27 | 0.18 | 0.08 |
| 9 | 74913868 | rs12378224 | 0.501 | 0.101 | -1.259 | -1.277 | 0 | -0.008 | -0.008 | 0 | 0.07 | 0.1 | 0.05 | 0 | 0.06 | 0.08 |
| 9 | 74921643 | rs10746908 | 0.086 | -0.079 | -0.946 | -1.722 | 0.005 | 0.043 | 0.043 | 0 | 0.42 | 0.32 | 0.41 | 0.29 | 0.4 | 0.19 |
| 9 | 74921883 | rs911861 | -1.023 | -0.003 | -0.57 | -1.925 | 0.017 | -0.06 | -0.06 | 0.056 | 0.8 | 0.66 | 0.85 | 0.96 | 0.79 | 0.84 |
| 9 | 74923489 | rs1322484 | 1.836 | -0.048 | -0.589 | -1.934 | -0.019 | -0.021 | -0.021 | 0 | 0.22 | 0.09 | 0.27 | 0.28 | 0.24 | 0.09 |
| 9 | 74925179 | rs17057618 | 1.813 | -0.031 | -0.525 | -1.814 | -0.008 | -0.055 | -0.055 | 0.404 | 0.14 | 0.19 | 0.09 | 0 | 0.1 | 0.01 |
| 9 | 74927609 | rs4745183 | 0.41 | 0.243 | -0.616 | -1.641 | 0 | 0.003 | 0.003 | 0 | 0.39 | 0.33 | 0.45 | 0.67 | 0.39 | 0.66 |
| 9 | 74932819 | rs9411042 | 0.561 | 0.306 | -0.811 | -1.525 | 0.01 | 0.003 | 0.003 | 0.213 | 0.29 | 0.28 | 0.48 | 0.66 | 0.41 | 0.61 |
| 9 | 74938125 | rs2104188 | 0.647 | 0.05 | -1.036 | -1.854 | -0.018 | 0.061 | 0.061 | 1.504 | 0.2 | 0.06 | 0.13 | 0.03 | 0.2 | 0.17 |
| 9 | 74940364 | rs13439944 | 0.114 | -0.388 | -0.389 | -1.568 | -0.051 | -0.159 | -0.159 | 0 | 0.23 | 0.47 | 0.11 | 0 | 0.11 | 0.1 |
| 9 | 74944437 | rs9411047 | 0.388 | 0.198 | -1.031 | -0.711 | -0.025 | -0.03 | -0.03 | 0 | 0.21 | 0.03 | 0.16 | 0.28 | 0.23 | 0.46 |
| 9 | 74948556 | rs11143267 | -0.013 | 0.253 | -0.277 | -1.052 | 0.009 | 0.008 | 0.008 | 0 | 0.42 | 0.46 | 0.5 | 0.45 | 0.47 | 0.39 |
| 9 | 74956781 | rs7849626 | -0.045 | 0.144 | 0.392 | -1.26 | -0.004 | -0.01 | -0.01 | 0 | 0.57 | 0.37 | 0.53 | 0.56 | 0.57 | 0.6 |
| 9 | 74968193 | rs12349878 | 1.999 | -0.145 | -0.112 | -1.553 | 0.014 | 0.047 | 0.047 | 1.52 | 0.11 | 0.04 | 0.06 | 0 | 0.06 | 0.01 |
| 9 | 74969076 | rs3398 | 0.399 | -0.183 | 0.142 | -1.517 | -0.03 | -0.035 | -0.035 | 0 | 0.44 | 0.19 | 0.44 | 0.56 | 0.5 | 0.59 |
| 9 | 74969864 | rs969 | 0.291 | -0.163 | 0.546 | -1.49 | 0.005 | 0.01 | 0.01 | 0 | 0.4 | 0.46 | 0.48 | 0.44 | 0.43 | 0.39 |
| 9 | 74974635 | rs11143274 | 0.795 | 0.198 | 0.357 | -1.73 | -0.007 | 0.005 | 0.005 | 0 | 0.1 | 0.06 | 0.09 | 0.02 | 0.15 | 0.08 |
| 9 | 74989643 | rs2984524 | 0.124 | 0.058 | 0.195 | -1.604 | -0.039 | -0.033 | -0.033 | 0 | 0.54 | 0.17 | 0.47 | 0.57 | 0.52 | 0.6 |
| 9 | 74997987 | rs2984519 | 0.054 | 0.089 | 0.383 | -1.606 | -0.002 | -0.002 | -0.002 | 8.734 | 0.56 | 0.41 | 0.48 | 0.57 | 0.51 | 0.61 |
| 9 | 75012258 | rs3012511 | -0.152 | 0.083 | 0.378 | -1.609 | 0.008 | 0.008 | 0.008 | 0 | 0.58 | 0.5 | 0.49 | 0.57 | 0.51 | 0.61 |
| 9 | 75021841 | rs7027119 | -0.869 | 0.119 | -0.153 | -1.82 | -0.037 | -0.13 | -0.13 | 0.07 | 0.75 | 0.95 | 0.77 | 0.45 | 0.69 | 0.45 |
| 9 | 75027824 | rs7846808 | -0.74 | 0.388 | -0.129 | -1.687 | -0.065 | -0.063 | -0.063 | 0.018 | 0.55 | 0.91 | 0.53 | 0.43 | 0.48 | 0.39 |
| 9 | 75046564 | rs10781101 | -0.052 | 0.236 | -0.001 | -1.658 | -0.002 | 0.001 | 0.001 | 0 | 0.45 | 0.54 | 0.55 | 0.42 | 0.55 | 0.41 |
| 9 | 75059164 | rs6560275 | -0.849 | 0.39 | -0.509 | -1.595 | -0.012 | -0.068 | -0.068 | 1.011 | 0.58 | 0.91 | 0.6 | 0.43 | 0.57 | 0.43 |
| 9 | 75078002 | rs11143303 | 0.604 | 0.838 | -0.266 | -1.588 | -0.019 | 0.013 | 0.013 | 0 | 0.12 | 0.07 | 0.14 | 0.02 | 0.23 | 0.11 |
| 9 | 75081322 | rs10511996 | -0.447 | 0.404 | -0.401 | -1.602 | -0.002 | -0.016 | -0.016 | 1.048 | 0.46 | 0.75 | 0.56 | 0.43 | 0.53 | 0.43 |
| 9 | 75092054 | rs10869180 | -0.197 | 0.371 | -0.042 | -1.604 | 0.001 | 0.005 | 0.005 | 0 | 0.44 | 0.5 | 0.54 | 0.43 | 0.53 | 0.43 |
| 9 | 75096601 | rs7023338 | -0.461 | 0.378 | -0.064 | -1.621 | -0.002 | -0.016 | -0.016 | 0.058 | 0.46 | 0.75 | 0.56 | 0.43 | 0.53 | 0.43 |
| 9 | 75096610 | rs12005469 | -0.262 | 0.385 | -0.055 | -1.617 | 0 | 0.002 | 0.002 | 0 | 0.44 | 0.52 | 0.55 | 0.43 | 0.53 | 0.43 |
| 9 | 75112780 | rs7040392 | -0.803 | 0.369 | -0.424 | -1.63 | -0.006 | -0.037 | -0.037 | 0 | 0.56 | 0.84 | 0.59 | 0.43 | 0.56 | 0.44 |
| 9 | 75123453 | rs7857674 | -0.806 | 0.477 | -0.491 | -1.571 | -0.012 | -0.075 | -0.075 | 0 | 0.77 | 0.89 | 0.8 | 0.45 | 0.73 | 0.5 |
| 9 | 75135825 | rs10869183 | -0.414 | 0.227 | -0.146 | -1.315 | 0.003 | 0.004 | 0.004 | 0 | 0.41 | 0.55 | 0.53 | 0.41 | 0.46 | 0.43 |
| 9 | 75135912 | rs13291144 | 0.532 | -0.222 | -0.066 | -1.362 | 0.003 | -0.021 | -0.021 | 0 | 0.08 | 0.15 | 0.04 | 0.03 | 0.07 | 0.02 |
| 9 | 75136789 | rs7022441 | -1.144 | -0.197 | -0.318 | -1.326 | -0.013 | -0.001 | -0.001 | 0 | 0.54 | 0.68 | 0.5 | 0.41 | 0.49 | 0.42 |
| 9 | 75139296 | rs12353502 | 1.677 | -0.231 | -0.568 | -1.166 | -0.021 | -0.041 | -0.041 | 0.014 | 0.12 | 0.22 | 0.04 | 0 | 0.03 | 0.01 |
| 9 | 75172587 | rs7875995 | 0.791 | 0.203 | -0.776 | -1.222 | -0.025 | 0.029 | 0.029 | 0 | 0.11 | 0.05 | 0.1 | 0.02 | 0.17 | 0.07 |
| 9 | 75177017 | rs7856874 | -1.46 | 0.056 | -0.926 | -1.288 | -0.083 | -0.07 | -0.07 | 2.335 | 0.61 | 0.91 | 0.52 | 0.41 | 0.52 | 0.46 |
| 9 | 75188225 | rs7864744 | -0.088 | -0.14 | -0.527 | -1.41 | 0 | 0.004 | 0.004 | 0.069 | 0.36 | 0.57 | 0.42 | 0.39 | 0.4 | 0.38 |
| 9 | 75188446 | rs9942926 | 0.36 | -0.335 | -0.482 | -1.405 | -0.008 | -0.031 | -0.031 | 0 | 0.2 | 0.28 | 0.09 | 0.02 | 0.1 | 0.04 |
| 9 | 75193817 | rs12344850 | 0.447 | -0.267 | -0.558 | -1.393 | -0.001 | 0.001 | 0.001 | 3.274 | 0.09 | 0.09 | 0.04 | 0.02 | 0.07 | 0.04 |
| 9 | 75194978 | rs7026124 | -0.824 | -0.356 | -0.562 | -1.503 | 0.049 | -0.001 | -0.001 | 0.759 | 0.45 | 0.76 | 0.42 | 0.39 | 0.34 | 0.34 |
| 9 | 75200657 | rs7853366 | -1.409 | -0.271 | -0.668 | -1.58 | -0.045 | -0.034 | -0.034 | 1.884 | 0.57 | 0.86 | 0.5 | 0.41 | 0.5 | 0.41 |
| 9 | 75207329 | rs7029452 | -0.679 | -0.275 | -0.264 | -1.56 | 0.009 | -0.001 | -0.001 | 0 | 0.44 | 0.64 | 0.41 | 0.39 | 0.34 | 0.34 |
| 9 | 75210243 | rs7045535 | 0.84 | -0.278 | -0.207 | -1.548 | -0.02 | 0.078 | 0.078 | 0.647 | 0.25 | 0.45 | 0.33 | 0.36 | 0.24 | 0.29 |
| 9 | 75211603 | rs13285932 | -0.126 | -0.262 | -0.225 | -1.571 | 0.003 | 0.023 | 0.023 | 0.137 | 0.36 | 0.5 | 0.42 | 0.39 | 0.4 | 0.36 |
| 9 | 75218470 | rs1899937 | 0.495 | 0.012 | -0.2 | -1.453 | 0 | 0.035 | 0.035 | 1.214 | 0.2 | 0.15 | 0.12 | 0.03 | 0.19 | 0.08 |
| 9 | 75229248 | rs920901 | -0.277 | -0.228 | -0.117 | -1.312 | -0.009 | 0.014 | 0.014 | 0 | 0.37 | 0.55 | 0.38 | 0.41 | 0.34 | 0.35 |
| 9 | 75231217 | rs10217204 | -0.717 | -0.395 | 0.006 | -1.181 | 0.004 | -0.005 | -0.005 | 0 | 0.46 | 0.63 | 0.46 | 0.44 | 0.46 | 0.41 |
| 9 | 75231370 | rs7026304 | -1.427 | -0.547 | -0.041 | -1.26 | -0.008 | -0.021 | -0.021 | 0.03 | 0.58 | 0.78 | 0.48 | 0.44 | 0.48 | 0.43 |
| 9 | 75232433 | rs10156444 | -0.461 | -0.542 | 0.294 | -1.4 | 0.007 | -0.035 | -0.035 | 0.696 | 0.22 | 0.36 | 0.2 | 0.07 | 0.28 | 0.09 |
| 9 | 75262770 | rs17058062 | 1.12 | -0.534 | 0.43 | -1.395 | -0.009 | -0.036 | -0.036 | 0 | 0.08 | 0.19 | 0.09 | 0 | 0.1 | 0.02 |
| 9 | 75276898 | rs1663762 | 0.966 | -0.615 | 0.447 | -1.426 | -0.016 | 0.075 | 0.075 | 0 | 0.23 | 0.31 | 0.26 | 0.38 | 0.18 | 0.31 |
| 9 | 75284559 | rs12339514 | -0.534 | -0.601 | 0.369 | -1.461 | 0.001 | -0.011 | -0.011 | 0.116 | 0.22 | 0.31 | 0.2 | 0.07 | 0.28 | 0.1 |
| 9 | 75295593 | rs1417614 | -0.17 | -0.513 | 0.244 | -1.311 | -0.005 | 0.024 | 0.024 | 0 | 0.33 | 0.35 | 0.33 | 0.42 | 0.29 | 0.37 |
| 9 | 75307719 | rs1444832 | -1.199 | -0.746 | 0.02 | -1.416 | 0.036 | -0.016 | -0.016 | 0 | 0.56 | 0.77 | 0.45 | 0.42 | 0.41 | 0.4 |
| 9 | 75315438 | rs1796993 | 1.12 | -0.793 | 0.083 | -1.425 | -0.014 | 0.057 | 0.057 | 0 | 0.22 | 0.27 | 0.25 | 0.38 | 0.16 | 0.31 |
| 9 | 75316142 | rs12552056 | 2.443 | -0.312 | -0.687 | -1.801 | -0.063 | 0.133 | 0.133 | 0 | 0.22 | 0.03 | 0.22 | 0.02 | 0.23 | 0.07 |
| 9 | 75331221 | rs2793162 | -1.356 | -0.418 | -0.542 | -1.684 | -0.011 | -0.029 | -0.029 | 0 | 0.6 | 0.79 | 0.48 | 0.44 | 0.47 | 0.42 |
| 9 | 75343789 | rs2487464 | -1.442 | -0.368 | -0.618 | -1.67 | -0.035 | -0.069 | -0.069 | 0 | 0.6 | 0.87 | 0.49 | 0.44 | 0.48 | 0.42 |
| 9 | 75349586 | rs2589632 | 1.214 | -0.534 | -0.362 | -1.699 | -0.013 | 0.047 | 0.047 | 0 | 0.22 | 0.29 | 0.25 | 0.38 | 0.16 | 0.31 |

| | | | | | | | | | | | | | | | | |
|---|---|---|---|---|---|---|---|---|---|---|---|---|---|---|---|---|
| 9 | 75352123 | rs1341032 | -0.599 | 0.099 | -0.578 | -1.393 | 0.104 | 0.005 | 0.005 | 0 | 0.51 | 0.86 | 0.4 | 0.4 | 0.35 | 0.36 |
| 9 | 75354330 | rs2589628 | -1.626 | 0.772 | -0.567 | -0.825 | -0.038 | -0.019 | -0.019 | 1.004 | 0.93 | 1 | 0.89 | 0.64 | 0.84 | 0.8 |
| 9 | 75365926 | rs10481772 | -0.735 | 0.305 | -0.363 | -1.225 | -0.011 | -0.086 | -0.086 | 0 | 0.18 | 0.44 | 0.14 | 0.02 | 0.17 | 0.04 |
| 9 | 75375810 | rs1550760 | -1 | 0.31 | -0.894 | -1.102 | -0.044 | -0.085 | -0.085 | 0 | 0.6 | 0.89 | 0.5 | 0.43 | 0.48 | 0.42 |
| 9 | 75382598 | rs7863483 | 0.483 | 0.28 | -0.662 | -1.1 | 0.029 | -0.005 | -0.005 | 0 | 0.05 | 0.12 | 0.04 | 0.02 | 0.06 | 0.02 |
| 9 | 75391254 | rs1341033 | 0.163 | 0.166 | -0.545 | -1.15 | 0.008 | 0.002 | 0.002 | 0 | 0.46 | 0.58 | 0.37 | 0.37 | 0.3 | 0.35 |
| 9 | 75400088 | rs13285263 | 0.19 | 0.474 | -0.234 | -0.697 | 0.029 | -0.005 | -0.005 | 0 | 0.05 | 0.12 | 0.04 | 0.02 | 0.06 | 0.01 |
| 9 | 75414490 | rs2994693 | -0.622 | 0.831 | -0.301 | -0.653 | -0.103 | -0.144 | -0.144 | 0 | 0.83 | 0.99 | 0.74 | 0.45 | 0.72 | 0.52 |
| 9 | 75415331 | rs17641064 | -0.339 | -0.757 | 0.408 | -0.85 | 0.019 | -0.007 | -0.007 | 0.054 | 0.05 | 0.11 | 0.04 | 0.02 | 0.06 | 0.01 |
| 9 | 75444298 | rs11999588 | -0.696 | -0.789 | 0.344 | -0.797 | 0.021 | 0.001 | 0.001 | 0 | 0.05 | 0.21 | 0.04 | 0.02 | 0.04 | 0.01 |
| 9 | 75445904 | rs4330753 | -1.482 | -0.934 | 0.384 | -0.979 | -0.014 | -0.138 | -0.138 | 0 | 0.24 | 0.56 | 0.21 | 0.04 | 0.26 | 0.1 |
| 9 | 75459316 | rs7027182 | -0.867 | -0.453 | -0.128 | -0.544 | 0.014 | -0.041 | -0.041 | 2.323 | 0.13 | 0.26 | 0.11 | 0.04 | 0.16 | 0.08 |
| 9 | 75498014 | rs660519 | -1.082 | -0.427 | -0.24 | -0.504 | 0.022 | -0.075 | -0.075 | 0 | 0.18 | 0.4 | 0.14 | 0.03 | 0.22 | 0.08 |
| 9 | 75507770 | rs10869198 | 1.437 | -0.909 | -1.222 | -0.221 | -0.106 | -0.15 | -0.15 | 0 | 0.36 | 0.05 | 0.48 | 0.58 | 0.49 | 0.54 |
| 9 | 75510629 | rs618596 | -1.021 | -0.854 | -1.006 | -0.038 | 0.009 | -0.064 | -0.064 | 0 | 0.18 | 0.36 | 0.14 | 0.03 | 0.22 | 0.08 |
| 9 | 75510887 | rs617590 | -1.021 | -0.854 | -1.006 | -0.038 | 0.009 | -0.064 | -0.064 | 0 | 0.18 | 0.36 | 0.14 | 0.03 | 0.22 | 0.08 |
| 9 | 75512300 | rs348479 | -1.045 | -0.86 | -0.971 | -0.047 | 0.009 | -0.064 | -0.064 | 0 | 0.19 | 0.36 | 0.14 | 0.03 | 0.22 | 0.08 |
| 9 | 75513722 | rs348480 | 0.289 | -0.895 | -0.995 | -0.064 | -0.04 | 0.064 | 0.064 | 0 | 0.11 | 0.01 | 0.08 | 0.01 | 0.16 | 0.06 |
| 9 | 75514436 | rs348481 | -1.25 | -0.852 | -0.966 | -0.043 | 0.009 | -0.064 | -0.064 | 0.055 | 0.18 | 0.35 | 0.14 | 0.03 | 0.22 | 0.08 |
| 9 | 75515114 | rs348483 | 0.491 | -0.843 | -1.021 | -0.038 | -0.029 | 0.051 | 0.051 | 0 | 0.1 | 0.03 | 0.08 | 0.01 | 0.16 | 0.06 |
| 9 | 75515325 | rs348484 | 0.491 | -0.799 | -1.237 | -0.016 | -0.029 | 0.051 | 0.051 | 0 | 0.1 | 0.03 | 0.08 | 0.01 | 0.16 | 0.06 |
| 9 | 75515454 | rs348485 | -0.539 | -0.873 | -1.195 | 0.022 | 0.005 | -0.036 | -0.036 | 2.283 | 0.17 | 0.23 | 0.13 | 0.03 | 0.22 | 0.08 |
| 9 | 75517582 | rs4646547 | 0.613 | -0.966 | -0.574 | 0.5 | -0.009 | -0.045 | -0.045 | 0 | 0.39 | 0.16 | 0.49 | 0.58 | 0.5 | 0.54 |
| 9 | 75519975 | rs348471 | -0.472 | -1.01 | -0.03 | 0.113 | 0.011 | 0.012 | 0.012 | 0 | 0.5 | 0.53 | 0.55 | 0.52 | 0.5 | 0.39 |
| 9 | 75521060 | rs348472 | 0.426 | -1.182 | -0.054 | -0.056 | -0.032 | -0.083 | -0.083 | 0 | 0.86 | 0.61 | 0.94 | 0.99 | 0.95 | 0.91 |
| 9 | 75521938 | rs8187981 | 0.537 | -1.289 | 0.185 | 0.208 | 0.018 | 0.053 | 0.053 | 0 | 0.45 | 0.19 | 0.51 | 0.44 | 0.5 | 0.61 |
| 9 | 75522835 | rs17058224 | -0.005 | -1.27 | 0.223 | 0.147 | -0.009 | -0.033 | -0.033 | 0 | 0.11 | 0.14 | 0.05 | 0.01 | 0.04 | 0.09 |
| 9 | 75523857 | rs1888203 | -1.838 | -1.44 | 0.513 | 0.052 | -0.016 | -0.022 | -0.022 | 0 | 0.05 | 0.23 | 0.02 | 0 | 0.01 | 0.02 |
| 9 | 75525879 | rs348475 | -0.77 | -1.453 | 0.432 | 0.073 | 0.021 | 0.032 | 0.032 | 0 | 0.54 | 0.7 | 0.49 | 0.52 | 0.5 | 0.38 |
| 9 | 75525939 | rs8187976 | -0.001 | -1.52 | 0.475 | -0.331 | -0.005 | -0.026 | -0.026 | 0 | 0.11 | 0.13 | 0.05 | 0.01 | 0.05 | 0.09 |
| 9 | 75529888 | rs348458 | -0.231 | -1.61 | 0.67 | -0.184 | -0.006 | -0.005 | -0.005 | 0 | 0.47 | 0.45 | 0.45 | 0.53 | 0.44 | 0.32 |
| 9 | 75533324 | rs610529 | -0.18 | -1.711 | 0.667 | -0.236 | -0.004 | -0.004 | -0.004 | 0 | 0.47 | 0.43 | 0.45 | 0.54 | 0.44 | 0.32 |
| 9 | 75534760 | rs348460 | 0.655 | -1.774 | 0.688 | -0.286 | -0.007 | -0.031 | -0.031 | 0.02 | 0.86 | 0.85 | 0.94 | 0.98 | 0.95 | 0.87 |
| 9 | 75537298 | rs8187945 | -0.951 | -1.969 | 0.735 | -0.229 | 0.018 | 0.008 | 0.008 | 0 | 0.15 | 0.09 | 0.05 | 0.02 | 0.05 | 0.12 |
| 9 | 75537590 | rs348459 | 0.904 | -1.824 | 0.807 | -0.155 | -0.014 | -0.043 | -0.043 | 0.016 | 0.85 | 0.81 | 0.94 | 0.98 | 0.95 | 0.87 |
| 9 | 75539077 | rs3739962 | 1.432 | -1.73 | 0.689 | -0.296 | -0.032 | -0.003 | -0.003 | 0 | 0.36 | 0.18 | 0.49 | 0.45 | 0.5 | 0.55 |
| 9 | 75540187 | rs2161811 | 1.45 | -1.719 | 0.694 | -0.293 | -0.032 | -0.005 | -0.005 | 0.108 | 0.36 | 0.18 | 0.49 | 0.46 | 0.5 | 0.54 |
| 9 | 75540828 | rs8187928 | 0.53 | -1.738 | 0.621 | -0.827 | 0.013 | 0.007 | 0.007 | 0 | 0.08 | 0.03 | 0.02 | 0 | 0.03 | 0.06 |
| 9 | 75541942 | rs2303317 | 1.463 | -1.751 | 0.623 | -0.835 | -0.042 | -0.01 | -0.01 | 0.523 | 0.36 | 0.16 | 0.49 | 0.46 | 0.5 | 0.54 |
| 9 | 75541962 | rs8187925 | 0.538 | -1.753 | 0.624 | -0.836 | 0.013 | 0.007 | 0.007 | 0 | 0.08 | 0.03 | 0.02 | 0 | 0.02 | 0.06 |
| 9 | 75542895 | rs7043217 | 0.826 | -1.754 | 0.632 | -0.837 | 0.018 | 0.038 | 0.038 | 0 | 0.45 | 0.25 | 0.51 | 0.46 | 0.53 | 0.6 |
| 9 | 75543243 | rs8187924 | 1.474 | -1.753 | 0.632 | -0.836 | -0.042 | -0.01 | -0.01 | 0 | 0.36 | 0.17 | 0.49 | 0.46 | 0.5 | 0.54 |
| 9 | 75544391 | rs2017362 | -0.052 | -1.969 | 1.009 | -1.024 | 0.01 | 0.005 | 0.005 | 0.167 | 0.41 | 0.34 | 0.37 | 0.53 | 0.36 | 0.34 |
| 9 | 75544797 | rs8187919 | 0.383 | -1.896 | 0.908 | -0.978 | -0.002 | -0.015 | -0.015 | 0.657 | 0.09 | 0.08 | 0.02 | 0 | 0.02 | 0.06 |
| 9 | 75544932 | rs4237253 | 1.739 | -1.909 | 0.908 | -0.984 | -0.059 | -0.022 | -0.022 | 0 | 0.35 | 0.12 | 0.49 | 0.45 | 0.5 | 0.54 |
| 9 | 75545060 | rs4237254 | 0.94 | -1.913 | 0.9 | -0.987 | 0.017 | 0.037 | 0.037 | 0 | 0.45 | 0.25 | 0.51 | 0.46 | 0.53 | 0.6 |
| 9 | 75545140 | rs4237255 | 1.633 | -1.842 | 0.853 | -0.948 | -0.042 | -0.01 | -0.01 | 0 | 0.36 | 0.17 | 0.49 | 0.45 | 0.5 | 0.54 |
| 9 | 75545456 | rs4646543 | 1.628 | -1.815 | 0.87 | -0.931 | -0.042 | -0.01 | -0.01 | 0 | 0.36 | 0.17 | 0.49 | 0.46 | 0.5 | 0.54 |
| 9 | 75545638 | rs3815836 | 1.618 | -1.815 | 0.87 | -0.931 | -0.042 | -0.01 | -0.01 | 0 | 0.36 | 0.17 | 0.49 | 0.46 | 0.5 | 0.54 |
| 9 | 75545882 | rs13959 | 1.626 | -1.81 | 0.873 | -0.928 | -0.042 | -0.01 | -0.01 | 2.315 | 0.36 | 0.17 | 0.49 | 0.46 | 0.5 | 0.54 |
| 9 | 75547612 | rs348463 | -0.201 | -1.118 | 1.54 | -0.619 | 0.005 | 0.034 | 0.034 | 2.518 | 0.26 | 0.32 | 0.33 | 0.53 | 0.28 | 0.28 |
| 9 | 75549810 | rs8187910 | -0.863 | -1.191 | 1.609 | -0.652 | -0.012 | -0.04 | -0.04 | 0 | 0.1 | 0.15 | 0.08 | 0.03 | 0.11 | 0.08 |
| 9 | 75550675 | rs7851899 | 1.307 | -1.27 | 1.618 | -0.685 | -0.058 | -0.08 | -0.08 | 0 | 0.86 | 0.68 | 0.96 | 0.98 | 0.97 | 0.93 |
| 9 | 75551300 | rs2773806 | 1.291 | -1.259 | 1.64 | -0.68 | -0.058 | -0.08 | -0.08 | 0 | 0.86 | 0.68 | 0.96 | 0.98 | 0.97 | 0.93 |
| 9 | 75551910 | rs186997 | 1.291 | -1.259 | 1.637 | -0.68 | -0.058 | -0.08 | -0.08 | 0.095 | 0.86 | 0.68 | 0.96 | 0.98 | 0.97 | 0.93 |
| 9 | 75551996 | rs8187902 | 0.09 | -1.255 | 1.614 | -0.678 | 0.012 | 0.004 | 0.004 | 0 | 0.08 | 0.04 | 0.02 | 0 | 0.02 | 0.06 |
| 9 | 75553212 | rs348454 | 1.645 | -1.252 | 1.635 | -0.656 | -0.027 | -0.025 | -0.025 | 0 | 0.93 | 0.8 | 0.97 | 0.96 | 0.97 | 0.93 |
| 9 | 75554224 | rs348450 | 1.465 | -1.251 | 1.632 | -0.655 | -0.026 | -0.024 | -0.024 | 0 | 0.93 | 0.81 | 0.98 | 0.96 | 0.97 | 0.93 |
| 9 | 75554952 | rs7027604 | 2.093 | -1.28 | 1.617 | -0.664 | 0.017 | 0.053 | 0.053 | 1.383 | 0.47 | 0.14 | 0.53 | 0.41 | 0.56 | 0.56 |
| 9 | 75554982 | rs348449 | 1.447 | -1.24 | 1.638 | -0.644 | -0.026 | -0.024 | -0.024 | 0 | 0.93 | 0.81 | 0.98 | 0.96 | 0.97 | 0.93 |
| 9 | 75556909 | rs8187895 | 0.132 | -1.253 | 1.677 | -0.657 | 0.003 | -0.036 | -0.036 | 0 | 0.07 | 0.13 | 0.07 | 0 | 0.08 | 0.02 |
| 9 | 75557007 | rs1469167 | 1.917 | -1.291 | 1.674 | -0.667 | -0.074 | -0.076 | -0.076 | 0 | 0.84 | 0.68 | 0.95 | 0.96 | 0.96 | 0.88 |
| 9 | 75557728 | rs1330285 | 0.319 | -1.138 | 1.638 | -0.593 | 0.001 | -0.002 | -0.002 | 0.2 | 0.07 | 0.04 | 0.02 | 0 | 0.01 | 0.05 |
| 9 | 75558799 | rs8187893 | 0.319 | -1.138 | 1.638 | -0.593 | 0.001 | -0.002 | -0.002 | 1.597 | 0.07 | 0.04 | 0.02 | 0 | 0.01 | 0.05 |
| 9 | 75559818 | rs647880 | -0.446 | -1.127 | 1.714 | -0.593 | 0.005 | 0.018 | 0.018 | 0.535 | 0.28 | 0.32 | 0.34 | 0.53 | 0.3 | 0.29 |
| 9 | 75559849 | rs8187890 | -0.475 | -1.116 | 1.685 | -0.693 | -0.003 | -0.045 | -0.045 | 0.361 | 0.08 | 0.17 | 0.07 | 0 | 0.09 | 0.02 |
| 9 | 75561149 | rs1928848 | 0.359 | -1.422 | 1.684 | -0.639 | 0.001 | -0.003 | -0.003 | 0 | 0.07 | 0.03 | 0.02 | 0 | 0.01 | 0.05 |
| 9 | 75561786 | rs8187884 | 0.246 | -1.388 | 1.733 | -0.626 | 0.002 | -0.033 | -0.033 | 0 | 0.06 | 0.1 | 0.06 | 0 | 0.09 | 0.02 |
| 9 | 75563557 | rs1424482 | 1.748 | -1.345 | 1.848 | -0.593 | 0.035 | 0.04 | 0.04 | 2.629 | 0.55 | 0.26 | 0.6 | 0.39 | 0.64 | 0.6 |

| | | | | | | | | | | | | | | | | |
|---|---|---|---|---|---|---|---|---|---|---|---|---|---|---|---|---|
| 9 | 75564954 | rs8187876 | 0.165 | -1.521 | 1.946 | -0.656 | -0.016 | 0.167 | 0.167 | 4.731 | 0.13 | 0.27 | 0.22 | 0.49 | 0.07 | 0.23 |
| 9 | 75565990 | rs8187871 | -0.321 | -1.397 | 1.64 | -0.633 | -0.045 | -0.046 | -0.046 | 0 | 0.08 | 0.19 | 0.02 | 0 | 0.01 | 0.05 |
| 9 | 75568803 | rs348445 | 1.747 | -1.397 | 1.605 | -0.634 | -0.027 | -0.025 | -0.025 | 0.026 | 0.93 | 0.82 | 0.98 | 0.96 | 0.97 | 0.94 |
| 9 | 75569482 | rs8187864 | 0.371 | -1.401 | 1.591 | -0.631 | -0.004 | -0.001 | -0.001 | 0.111 | 0.07 | 0.03 | 0.01 | 0 | 0.01 | 0.05 |
| 9 | 75572106 | rs694564 | 1.814 | -1.391 | 1.614 | -0.653 | -0.029 | -0.027 | -0.027 | 0.015 | 0.93 | 0.8 | 0.98 | 0.96 | 0.97 | 0.94 |
| 9 | 75576214 | rs1330284 | 2.098 | -1.319 | 1.536 | -0.67 | 0.016 | 0.089 | 0.089 | 1.12 | 0.48 | 0.13 | 0.53 | 0.4 | 0.55 | 0.56 |
| 9 | 75590794 | rs7030453 | 0.579 | -1.276 | 1.593 | -0.653 | -0.005 | -0.027 | -0.027 | 0.607 | 0.9 | 0.9 | 0.92 | 0.97 | 0.88 | 0.93 |
| 9 | 75593837 | rs1364454 | 1.955 | -1.284 | 1.526 | -0.656 | -0.079 | -0.128 | -0.128 | 0 | 0.78 | 0.56 | 0.89 | 0.96 | 0.88 | 0.88 |
| 9 | 75594556 | rs7858367 | 2.107 | -1.292 | 1.478 | -0.661 | 0.01 | 0.089 | 0.089 | 0.976 | 0.48 | 0.11 | 0.52 | 0.4 | 0.55 | 0.55 |
| 9 | 75597830 | rs7849271 | -0.642 | -1.257 | 1.479 | -0.628 | 0.01 | 0.017 | 0.017 | 0.835 | 0.29 | 0.37 | 0.36 | 0.54 | 0.32 | 0.33 |
| 9 | 75602903 | rs7856728 | 1.086 | -1.222 | 1.27 | -0.639 | -0.023 | -0.063 | -0.063 | 0 | 0.87 | 0.74 | 0.92 | 0.96 | 0.88 | 0.93 |
| 9 | 75608626 | rs10869208 | -0.65 | -1.204 | 1.264 | -0.624 | 0.01 | 0.015 | 0.015 | 0.913 | 0.29 | 0.37 | 0.37 | 0.55 | 0.31 | 0.33 |
| 9 | 75615188 | rs1582620 | 1.748 | -1.221 | 1.218 | -0.683 | -0.061 | -0.11 | -0.11 | 0 | 0.8 | 0.64 | 0.9 | 0.96 | 0.87 | 0.88 |
| 9 | 75624687 | rs1424484 | -1.553 | -0.763 | 1.257 | -0.459 | -0.022 | -0.069 | -0.069 | 0.567 | 0.13 | 0.29 | 0.09 | 0.04 | 0.13 | 0.07 |
| 9 | 75633424 | rs1593197 | -0.922 | -0.507 | 1.16 | -0.125 | -0.001 | -0.006 | -0.006 | 0 | 0.09 | 0.09 | 0.06 | 0.03 | 0.08 | 0.12 |
| 9 | 75636267 | rs7028573 | -0.98 | -0.567 | 1.165 | -0.121 | -0.001 | 0 | 0 | 2.708 | 0.29 | 0.65 | 0.37 | 0.55 | 0.26 | 0.31 |
| 9 | 75637712 | rs2226111 | -0.008 | -0.543 | 1.008 | -0.142 | -0.009 | 0.015 | 0.015 | 0 | 0.95 | 0.99 | 0.99 | 0.98 | 0.95 | 0.97 |
| 9 | 75640105 | rs4745209 | -0.724 | -0.53 | 1.247 | -0.127 | 0.016 | 0.038 | 0.038 | 2.953 | 0.27 | 0.44 | 0.36 | 0.55 | 0.26 | 0.31 |
| 9 | 75649046 | rs11143438 | -0.502 | -0.655 | 1.223 | -0.064 | -0.001 | 0 | 0 | 0 | 0.08 | 0.07 | 0.06 | 0.05 | 0.08 | 0.12 |
| 9 | 75669943 | rs11143450 | -0.463 | -0.637 | 0.985 | -0.068 | 0 | 0 | 0 | 0 | 0.08 | 0.07 | 0.07 | 0.05 | 0.09 | 0.13 |
| 9 | 75675129 | rs10869215 | 1.444 | -0.573 | 0.743 | -0.087 | -0.002 | 0.075 | 0.075 | 1.256 | 0.56 | 0.12 | 0.59 | 0.4 | 0.6 | 0.55 |
| 9 | 75682680 | rs1961830 | 1.21 | -0.356 | 0.738 | 0.045 | -0.001 | -0.011 | -0.011 | 0 | 0.6 | 0.46 | 0.61 | 0.4 | 0.6 | 0.55 |
| 9 | 75703530 | rs7046288 | 0.888 | -0.23 | -0.146 | 1.027 | -0.004 | 0.041 | 0.041 | 0.356 | 0.85 | 0.84 | 0.83 | 0.51 | 0.89 | 0.81 |
| 9 | 75710330 | rs2871480 | -0.858 | -0.42 | -0.08 | 0.318 | -0.025 | -0.014 | -0.014 | 0 | 0.08 | 0.11 | 0.07 | 0.05 | 0.02 | 0.04 |
| 9 | 75715402 | rs4744679 | -0.152 | -0.297 | -0.243 | -0.464 | -0.003 | 0.074 | 0.074 | 0 | 0.52 | 0.65 | 0.5 | 0.29 | 0.47 | 0.24 |
| 9 | 75719701 | rs1342020 | -0.183 | -0.318 | -0.251 | -0.488 | -0.004 | 0.073 | 0.073 | 0 | 0.52 | 0.66 | 0.5 | 0.29 | 0.47 | 0.25 |
| 9 | 75721599 | rs4744681 | -0.31 | -0.527 | -0.435 | -0.75 | -0.013 | 0.047 | 0.047 | 0.007 | 0.58 | 0.7 | 0.52 | 0.31 | 0.52 | 0.34 |
| 9 | 75730317 | rs17058479 | -0.118 | -0.697 | 0.393 | -0.789 | -0.012 | -0.015 | -0.015 | 0 | 0.09 | 0.11 | 0.08 | 0.05 | 0.07 | 0.04 |
| 9 | 75730662 | rs7875996 | 0.689 | -1.042 | 0.25 | -1.162 | 0.004 | 0.097 | 0.097 | 0 | 0.55 | 0.71 | 0.52 | 0.24 | 0.57 | 0.28 |
| 9 | 75732302 | rs1935316 | -0.739 | -1.02 | 0.304 | -1.097 | -0.033 | 0.006 | 0.006 | 0 | 0.7 | 0.85 | 0.61 | 0.32 | 0.69 | 0.44 |
| 9 | 75735898 | rs11143481 | 0.364 | -1.005 | 0.26 | -1.094 | 0.013 | 0.022 | 0.022 | 0 | 0.06 | 0.01 | 0.02 | 0.03 | 0.05 | 0.1 |
| 9 | 75744524 | rs11143486 | 0.842 | -0.939 | 0.498 | -1.386 | 0.005 | 0.006 | 0.006 | 0 | 0.51 | 0.43 | 0.45 | 0.15 | 0.57 | 0.24 |
| 9 | 75757209 | rs10869227 | -0.857 | -0.913 | 0.6 | -1.259 | -0.025 | 0.005 | 0.005 | 0 | 0.7 | 0.82 | 0.61 | 0.3 | 0.69 | 0.41 |
| 9 | 75761043 | rs11143503 | 0.082 | -0.995 | 0.562 | -1.265 | -0.01 | -0.003 | -0.003 | 0 | 0.1 | 0.11 | 0.07 | 0.1 | 0.07 | 0.04 |
| 9 | 75763159 | rs2795107 | 0.478 | -1.059 | 1.246 | -1.382 | 0.013 | 0.103 | 0.103 | 0 | 0.53 | 0.29 | 0.44 | 0.16 | 0.59 | 0.31 |
| 9 | 75765478 | rs2795108 | -1.63 | -1.182 | 1.596 | -1.304 | -0.008 | 0.046 | 0.046 | 0.588 | 0.17 | 0.41 | 0.18 | 0.2 | 0.13 | 0.2 |
| 9 | 75773161 | rs2795116 | -1.137 | -1.158 | 1.291 | -1.292 | -0.014 | 0.01 | 0.01 | 0.169 | 0.13 | 0.23 | 0.09 | 0.15 | 0.1 | 0.14 |
| 9 | 75778370 | rs2795118 | 0.204 | -1.156 | 0.924 | -1.286 | -0.01 | -0.017 | -0.017 | 0 | 0.07 | 0.09 | 0.02 | 0.01 | 0.05 | 0.1 |
| 9 | 75782131 | rs1342018 | 0.779 | -0.795 | 0.57 | -1.416 | -0.052 | 0.28 | 0.28 | 0 | 0.46 | 0.13 | 0.44 | 0.07 | 0.56 | 0.34 |
| 9 | 75782510 | rs1342017 | -0.177 | -0.91 | 1.116 | -1.424 | 0.005 | -0.049 | -0.049 | 0 | 0.61 | 0.57 | 0.53 | 0.17 | 0.63 | 0.37 |
| 9 | 75784264 | rs2811226 | -0.868 | -0.953 | 1 | -1.345 | -0.027 | 0.002 | 0.002 | 0 | 0.68 | 0.84 | 0.64 | 0.3 | 0.67 | 0.41 |
| 9 | 75788656 | rs4745219 | 0.134 | -0.846 | 1.207 | -0.97 | -0.002 | -0.001 | -0.001 | 0 | 0.09 | 0.08 | 0.06 | 0.1 | 0.07 | 0.05 |
| 9 | 75789935 | rs7357731 | 0.22 | -0.641 | 1.367 | -0.814 | 0.055 | 0.138 | 0.138 | 0 | 0.93 | 0.73 | 0.89 | 0.87 | 0.96 | 0.96 |
| 9 | 75792301 | rs10781126 | 0.497 | -0.999 | 0.816 | -0.377 | -0.004 | 0.024 | 0.024 | 0 | 0.73 | 0.7 | 0.69 | 0.59 | 0.71 | 0.73 |
| 9 | 75792846 | rs11143520 | 0.714 | -1.206 | 0.85 | -0.013 | -0.003 | 0.036 | 0.036 | 0.111 | 0.18 | 0.25 | 0.07 | 0.34 | 0.13 | 0.38 |
| 9 | 75793444 | rs17058589 | 0.466 | -1.253 | 0.629 | -0.053 | -0.003 | 0.009 | 0.009 | 0 | 0.09 | 0.04 | 0.07 | 0.08 | 0.09 | 0.11 |
| 9 | 75797835 | rs4256646 | 0.525 | -1.615 | -0.554 | -0.124 | -0.002 | -0.001 | -0.001 | 0 | 0.51 | 0.47 | 0.47 | 0.45 | 0.54 | 0.57 |
| 9 | 75800805 | rs10869237 | 0.579 | -1.783 | -0.665 | -0.183 | -0.027 | 0.11 | 0.11 | 0.172 | 0.24 | 0.44 | 0.15 | 0.45 | 0.18 | 0.41 |
| 9 | 75810204 | rs4581120 | 0.555 | -1.978 | -0.735 | -0.325 | 0.008 | 0.018 | 0.018 | 0 | 0.55 | 0.44 | 0.62 | 0.25 | 0.6 | 0.35 |
| 9 | 75810296 | rs10746916 | 2.019 | -1.906 | -1.07 | -0.516 | -0.153 | 0.188 | 0.188 | 0 | 0.27 | 0.02 | 0.32 | 0.01 | 0.36 | 0.16 |
| 9 | 75810393 | rs4581121 | 0.099 | -1.922 | -0.795 | -0.279 | -0.005 | -0.03 | -0.03 | 0.763 | 0.79 | 0.89 | 0.77 | 0.69 | 0.77 | 0.75 |
| 9 | 75813393 | rs7848091 | -0.903 | -1.159 | -0.518 | -0.088 | 0 | -0.037 | -0.037 | 0 | 0.89 | 0.94 | 0.84 | 0.75 | 0.88 | 0.86 |
| 9 | 75816850 | rs17653504 | 1.526 | -0.041 | -0.667 | -0.164 | 0.015 | 0.029 | 0.029 | 0 | 0.26 | 0.09 | 0.16 | 0.15 | 0.2 | 0.14 |
| 9 | 75817746 | rs10869241 | 1.741 | 0.193 | -0.401 | -0.13 | 0.011 | 0.054 | 0.054 | 0.202 | 0.19 | 0.27 | 0.1 | 0.38 | 0.17 | 0.39 |
| 9 | 75819224 | rs7034411 | 1.64 | 0.205 | -0.592 | -0.093 | -0.063 | -0.05 | -0.05 | 0.057 | 0.15 | 0.07 | 0.43 | 0.21 | 0.29 | 0.19 |
| 9 | 75819755 | rs13290663 | 1.482 | 0.059 | -0.541 | -0.161 | 0.005 | 0.122 | 0.122 | 0 | 0.23 | 0.05 | 0.11 | 0.02 | 0.21 | 0.13 |
| 9 | 75819945 | rs10746917 | -1.383 | 0.076 | -0.214 | 0.025 | 0.002 | -0.024 | -0.024 | 0 | 0.89 | 0.93 | 0.84 | 0.75 | 0.88 | 0.86 |
| 9 | 75828161 | rs10735621 | -0.768 | 0.154 | -0.74 | 0.056 | 0.023 | -0.031 | -0.031 | 0.003 | 0.63 | 0.84 | 0.68 | 0.6 | 0.68 | 0.71 |
| 9 | 75828707 | rs4237258 | 1.496 | 0.135 | -0.722 | 0.057 | 0.016 | 0.028 | 0.028 | 0 | 0.26 | 0.1 | 0.16 | 0.15 | 0.2 | 0.14 |
| 9 | 75837087 | rs11143541 | 0.104 | 0.175 | -0.473 | 0.039 | -0.095 | 0.061 | 0.061 | 0 | 0.4 | 0.14 | 0.54 | 0.22 | 0.51 | 0.31 |
| 9 | 75840034 | rs9657736 | -1.456 | 0.116 | 0.314 | 0.021 | -0.02 | 0.065 | 0.065 | 0 | 0.92 | 0.87 | 0.86 | 0.75 | 0.95 | 0.9 |
| 9 | 75866968 | rs10429583 | -0.135 | 0.098 | 0.374 | -0.974 | 0.002 | -0.003 | -0.003 | 0.032 | 0.65 | 0.62 | 0.7 | 0.64 | 0.68 | 0.76 |
| 9 | 75872324 | rs12001114 | -0.016 | -0.159 | 0.508 | -1.118 | -0.051 | -0.065 | -0.065 | 0 | 0.14 | 0.4 | 0.05 | 0.04 | 0.05 | 0.09 |
| 9 | 75877116 | rs7018507 | 0.895 | -0.243 | 0.388 | -1.076 | 0.107 | 0.072 | 0.072 | 0 | 0.49 | 0.08 | 0.57 | 0.41 | 0.63 | 0.62 |
| 9 | 75895599 | rs17058682 | 1.252 | -0.171 | 0.636 | -1.077 | -0.027 | -0.028 | -0.028 | 0 | 0.08 | 0.17 | 0.02 | 0.02 | 0.04 | 0.07 |
| 9 | 75897972 | rs7860450 | -1.026 | -0.17 | 0.435 | -1.088 | 0.034 | -0.069 | -0.069 | 0 | 0.72 | 0.87 | 0.74 | 0.64 | 0.74 | 0.8 |
| 9 | 75900508 | rs12343428 | 0.449 | 0.114 | 0.444 | -1.245 | 0.061 | -0.026 | -0.026 | 0 | 0.06 | 0.25 | 0.04 | 0 | 0.08 | 0.04 |
| 9 | 75905148 | rs10781135 | 1.484 | -0.008 | 0.258 | -1.329 | -0.124 | -0.051 | -0.051 | 0 | 0.36 | 0.03 | 0.25 | 0.4 | 0.37 | 0.48 |
| 9 | 75921571 | rs17058729 | 1.535 | -0.151 | 0.399 | -1.284 | -0.009 | -0.008 | -0.008 | 0 | 0.08 | 0.08 | 0.02 | 0.02 | 0.04 | 0.06 |

| | | | | | | | | | | | | | | | | |
|---|---|---|---|---|---|---|---|---|---|---|---|---|---|---|---|---|
| 9 | 75922205 | rs17058735 | 1.535 | -0.134 | 0.431 | -1.39 | -0.011 | -0.011 | -0.011 | 0 | 0.08 | 0.08 | 0.02 | 0.01 | 0.04 | 0.06 |
| 9 | 75932178 | rs10465189 | 0.816 | -0.091 | 0.198 | -1.472 | 0.07 | 0.071 | 0.071 | 0 | 0.5 | 0.06 | 0.53 | 0.4 | 0.62 | 0.62 |
| 9 | 75956329 | rs954227 | -1.073 | -0.334 | 0.392 | -1.457 | -0.012 | -0.076 | -0.076 | 0 | 0.12 | 0.6 | 0.07 | 0.03 | 0.07 | 0.07 |
| 9 | 75970379 | rs1932202 | 1.576 | -0.184 | 0.032 | -1.505 | -0.079 | 0.078 | 0.078 | 0 | 0.13 | 0.02 | 0.32 | 0.01 | 0.25 | 0.14 |
| 9 | 75975477 | rs1414165 | -0.613 | -0.716 | 0.29 | -1.25 | 0.033 | -0.022 | -0.022 | 0.382 | 0.7 | 0.82 | 0.66 | 0.44 | 0.73 | 0.76 |
| 9 | 75978400 | rs7863241 | -0.833 | -0.74 | 0.29 | -1.259 | -0.003 | -0.022 | -0.022 | 0 | 0.21 | 0.74 | 0.09 | 0.04 | 0.11 | 0.13 |
| 9 | 75979673 | rs11143574 | 0.24 | -0.746 | 0.395 | -1.262 | -0.01 | -0.064 | -0.064 | 2.342 | 0.13 | 0.37 | 0.06 | 0.01 | 0.09 | 0.1 |
| 9 | 75987179 | rs1932201 | -0.872 | -0.746 | 0.394 | -1.262 | -0.003 | -0.022 | -0.022 | 0.258 | 0.21 | 0.74 | 0.09 | 0.04 | 0.11 | 0.13 |
| 9 | 75991893 | rs10735622 | -0.824 | -0.69 | 0.408 | -1.232 | 0.028 | -0.024 | -0.024 | 0 | 0.71 | 0.83 | 0.67 | 0.44 | 0.74 | 0.76 |
| 9 | 76002121 | rs10114105 | 0.039 | -0.489 | 0.696 | -1.08 | -0.017 | -0.07 | -0.07 | 0 | 0.14 | 0.37 | 0.06 | 0.01 | 0.09 | 0.1 |
| 9 | 76017507 | rs1335385 | -1.021 | -0.524 | 0.528 | -1.101 | 0.004 | -0.011 | -0.011 | 0 | 0.22 | 0.75 | 0.1 | 0.04 | 0.11 | 0.13 |
| 9 | 76038127 | rs17058849 | -1.048 | -0.483 | 0.723 | -1.091 | 0.016 | 0.001 | 0.001 | 1.344 | 0.07 | 0.37 | 0.03 | 0.03 | 0.01 | 0.03 |
| 9 | 76054140 | rs280563 | -0.445 | -0.967 | 0.643 | -1.045 | -0.032 | -0.083 | -0.083 | 0 | 0.18 | 0.67 | 0.08 | 0.01 | 0.1 | 0.1 |
| 9 | 76078682 | rs1908811 | 0.799 | -1.047 | 0.316 | -1.039 | 0.105 | 0.089 | 0.089 | 0 | 0.52 | 0.21 | 0.59 | 0.4 | 0.65 | 0.62 |
| 9 | 76089602 | rs278747 | -0.877 | -0.995 | 0.196 | -0.83 | 0.001 | 0.05 | 0.05 | 0 | 0.91 | 0.89 | 0.83 | 0.63 | 0.94 | 0.91 |
| 9 | 76098252 | rs278738 | 1.606 | -1.005 | 0.194 | -0.822 | -0.01 | 0.114 | 0.114 | 0 | 0.07 | 0.11 | 0.16 | 0.37 | 0.03 | 0.08 |
| 9 | 76105546 | rs278736 | 0.235 | -1.004 | 0.187 | -1.003 | 0.021 | -0.01 | -0.01 | 0.009 | 0.29 | 0.21 | 0.33 | 0.57 | 0.25 | 0.23 |
| 9 | 76115057 | rs11143600 | 1.623 | -0.91 | -0.359 | -1.394 | -0.021 | 0.056 | 0.056 | 0.049 | 0.23 | 0.02 | 0.17 | 0.1 | 0.19 | 0.16 |
| 9 | 76118698 | rs11794852 | 2.17 | -0.688 | -0.555 | -1.433 | -0.095 | -0.077 | -0.077 | 0.376 | 0.16 | 0.01 | 0.09 | 0.3 | 0.24 | 0.34 |
| 9 | 76121101 | rs182183 | -0.536 | -0.757 | 0.557 | -0.861 | -0.005 | 0 | 0 | 0.014 | 0.34 | 0.39 | 0.38 | 0.56 | 0.31 | 0.3 |
| 9 | 76125591 | rs1390947 | 1.842 | -0.743 | 0.246 | -0.837 | -0.098 | 0.004 | 0.004 | 0 | 0.4 | 0.08 | 0.28 | 0.4 | 0.44 | 0.51 |
| 9 | 76126194 | rs993350 | 1.297 | -0.723 | 0.239 | -0.697 | -0.006 | 0.103 | 0.103 | 0 | 0.16 | 0.01 | 0.08 | 0.01 | 0.18 | 0.08 |
| 9 | 76150496 | rs407979 | -0.205 | -0.74 | 0.435 | -0.548 | 0.016 | -0.022 | -0.022 | 0 | 0.29 | 0.23 | 0.34 | 0.37 | 0.27 | 0.22 |
| 9 | 76152918 | rs10781146 | 1.405 | -0.837 | 0.482 | -0.651 | 0.077 | 0.083 | 0.083 | 0 | 0.49 | 0.19 | 0.47 | 0.4 | 0.59 | 0.62 |
| 9 | 76155313 | rs395937 | -0.814 | -0.811 | 0.666 | -0.656 | -0.011 | 0.008 | 0.008 | 0.19 | 0.37 | 0.46 | 0.4 | 0.37 | 0.34 | 0.24 |
| 9 | 76157987 | rs12552642 | -0.524 | -1.833 | 1.051 | -0.067 | 0.006 | -0.033 | -0.033 | 0 | 0.08 | 0.13 | 0.15 | 0 | 0.11 | 0.08 |
| 9 | 76159447 | rs278717 | -0.545 | -1.842 | 1.268 | -0.041 | -0.026 | -0.045 | -0.045 | 0 | 0.88 | 0.83 | 0.97 | 0.97 | 0.95 | 0.92 |
| 9 | 76161901 | rs278727 | 0.18 | -1.859 | 1.367 | 0.001 | -0.067 | 0.101 | 0.101 | 0 | 0.85 | 0.58 | 0.87 | 0.77 | 0.94 | 0.87 |
| 9 | 76161967 | rs278729 | 0.823 | -1.86 | 1.375 | -0.005 | -0.048 | 0.052 | 0.052 | 0 | 0.82 | 0.46 | 0.85 | 0.77 | 0.91 | 0.86 |
| 9 | 76164687 | rs452974 | -0.458 | -2.137 | 0.957 | 0.048 | -0.028 | -0.047 | -0.047 | 0 | 0.88 | 0.82 | 0.96 | 0.97 | 0.95 | 0.92 |
| 9 | 76175652 | rs275528 | 0.402 | -2.187 | 0.972 | 0.004 | -0.03 | -0.107 | -0.107 | 0 | 0.7 | 0.63 | 0.87 | 0.96 | 0.76 | 0.82 |
| 9 | 76184436 | rs157117 | -1.323 | -1.974 | 0.417 | 0.192 | -0.002 | 0 | 0 | 0.17 | 0.37 | 0.63 | 0.49 | 0.57 | 0.33 | 0.3 |
| 9 | 76185813 | rs12339193 | 0.382 | -1.685 | -0.9 | 0.057 | 0 | -0.033 | -0.033 | 0.53 | 0.18 | 0.2 | 0.09 | 0.01 | 0.19 | 0.09 |
| 9 | 76187476 | rs17659570 | 0.281 | -1.568 | -1.356 | 0.17 | 0.001 | 0.066 | 0.066 | 0 | 0.21 | 0.09 | 0.19 | 0.01 | 0.24 | 0.15 |
| 9 | 76189863 | rs10512023 | 0.465 | -0.868 | -1.952 | 0.736 | 0.122 | -0.009 | -0.009 | 0.223 | 0.51 | 0.17 | 0.54 | 0.58 | 0.62 | 0.67 |
| 9 | 76191829 | rs10512024 | 0.53 | -0.748 | -2.119 | 0.6 | 0.132 | 0.094 | 0.094 | 0.014 | 0.5 | 0.1 | 0.45 | 0.39 | 0.63 | 0.64 |
| 9 | 76198452 | rs157109 | -0.642 | 0.181 | -1.08 | 1.209 | 0.009 | -0.007 | -0.007 | 0 | 0.1 | 0.04 | 0.19 | 0.36 | 0.07 | 0.1 |
| 9 | 76203946 | rs2492825 | 0.047 | 0.353 | -0.893 | -0.33 | -0.003 | -0.069 | -0.069 | 0 | 0.55 | 0.55 | 0.79 | 0.93 | 0.62 | 0.72 |
| 9 | 76204270 | rs10217425 | -0.539 | -0.199 | -1.3 | 0.532 | 0.022 | -0.052 | -0.052 | 0.261 | 0.27 | 0.16 | 0.29 | 0.46 | 0.2 | 0.49 |
| 9 | 76204511 | rs275515 | -0.873 | 0.238 | -0.721 | 0.522 | -0.006 | 0.009 | 0.009 | 0.095 | 0.39 | 0.49 | 0.43 | 0.44 | 0.35 | 0.53 |
| 9 | 76205632 | rs7870301 | -1.058 | 0.145 | -0.484 | 0.84 | 0.002 | 0.027 | 0.027 | 0 | 0.06 | 0.11 | 0.13 | 0.13 | 0.07 | 0.07 |
| 9 | 76207300 | rs9314809 | 0.503 | 0.065 | -0.459 | 0.507 | -0.001 | -0.001 | -0.001 | 0 | 0.58 | 0.45 | 0.71 | 0.62 | 0.66 | 0.47 |
| 9 | 76224560 | rs1505518 | 1.447 | -0.264 | -0.765 | -0.02 | -0.039 | -0.057 | -0.057 | 0 | 0.31 | 0.17 | 0.37 | 0.46 | 0.42 | 0.29 |
| 9 | 76228755 | rs17660640 | 0.362 | -0.898 | -0.591 | -0.158 | 0.012 | 0.076 | 0.076 | 0 | 0.2 | 0.03 | 0.18 | 0.03 | 0.17 | 0.11 |
| 9 | 76231486 | rs7857209 | -0.288 | -1.018 | -0.211 | -0.071 | -0.013 | -0.004 | -0.004 | 0 | 0.1 | 0.18 | 0.13 | 0.12 | 0.06 | 0.15 |
| 9 | 76232053 | rs156762 | -1.268 | -1.215 | 0.245 | -0.163 | -0.042 | -0.005 | -0.005 | 0.632 | 0.14 | 0.57 | 0.15 | 0.13 | 0.06 | 0.15 |
| 9 | 76237760 | rs276781 | -1.252 | -1.859 | -0.575 | -0.072 | 0.173 | 0.058 | 0.058 | 0.116 | 0.47 | 0.82 | 0.35 | 0.37 | 0.24 | 0.45 |
| 9 | 76242155 | rs1398307 | -1.427 | -0.899 | 0.262 | 0.041 | 0.047 | 0.055 | 0.055 | 1.314 | 0.32 | 0.78 | 0.3 | 0.15 | 0.19 | 0.22 |
| 9 | 76246287 | rs156768 | -1.518 | -0.033 | 0.195 | -0.028 | 0.009 | 0.077 | 0.077 | 1.341 | 0.35 | 0.79 | 0.33 | 0.16 | 0.3 | 0.25 |
| 9 | 76246845 | rs1606326 | 2.251 | 0.626 | 1.173 | 1.167 | 0.007 | 0 | 0 | 0 | 0.69 | 0.37 | 0.52 | 0.67 | 0.62 | 0.65 |
| 9 | 76249347 | rs156772 | -2.182 | 0.531 | 2.048 | 1.024 | 0.044 | -0.003 | -0.003 | 0 | 0.08 | 0.47 | 0.05 | 0 | 0.03 | 0.03 |
| 9 | 76255260 | rs1505517 | -0.429 | 0.593 | 1.9 | 1.399 | -0.012 | -0.008 | -0.008 | 3.085 | 0.06 | 0.04 | 0.16 | 0.2 | 0.15 | 0.12 |
| 9 | 76256736 | rs1352514 | 2.675 | 0.569 | 1.847 | 1.369 | -0.058 | -0.054 | -0.054 | 0 | 0.68 | 0.13 | 0.61 | 0.61 | 0.67 | 0.65 |
| 9 | 76258721 | rs2502788 | -2.668 | 0.338 | 1.999 | 1.131 | -0.031 | -0.059 | -0.059 | 0.414 | 0.1 | 0.73 | 0.06 | 0 | 0.03 | 0.03 |
| 9 | 76300013 | rs11143657 | -0.653 | 0.372 | 2.513 | 1.49 | -0.022 | -0.013 | -0.013 | 0 | 0.07 | 0.03 | 0.16 | 0.24 | 0.15 | 0.11 |
| 9 | 76313134 | rs11143663 | 2.823 | 0.285 | 2.03 | 1.291 | -0.109 | -0.073 | -0.073 | 0.054 | 0.55 | 0.07 | 0.38 | 0.52 | 0.53 | 0.54 |
| 9 | 76321504 | rs11143667 | -2.45 | 0.353 | 2.131 | 1.282 | 0.032 | 0.021 | 0.021 | 0.228 | 0.27 | 0.42 | 0.46 | 0.31 | 0.33 | 0.27 |
| 9 | 76321671 | rs1938287 | -0.602 | 0.363 | 1.772 | 1.279 | -0.021 | 0.042 | 0.042 | 0 | 0.14 | 0.05 | 0.28 | 0.05 | 0.17 | 0.13 |
| 9 | 76325130 | rs17059293 | -0.313 | 0.435 | 1.762 | 1.337 | -0.043 | -0.035 | -0.035 | 0 | 0.07 | 0.01 | 0.16 | 0.24 | 0.15 | 0.11 |
| 9 | 76334135 | rs13288902 | 2.897 | 0.387 | 1.778 | 1.082 | -0.107 | -0.038 | -0.038 | 2.148 | 0.56 | 0.09 | 0.38 | 0.55 | 0.53 | 0.54 |
| 9 | 76343174 | rs2171659 | -0.149 | -0.309 | 2.014 | 1.128 | -0.049 | -0.036 | -0.036 | 0 | 0.87 | 0.98 | 0.87 | 0.83 | 0.86 | 0.81 |
| 9 | 76353087 | rs1387289 | 1.97 | -0.287 | 1.951 | 1.302 | 0.012 | -0.039 | -0.039 | 0 | 0.69 | 0.51 | 0.57 | 0.76 | 0.69 | 0.65 |
| 9 | 76355713 | rs17605754 | -0.798 | -0.25 | 1.786 | 1.302 | -0.029 | 0.016 | 0.016 | 0 | 0.13 | 0.05 | 0.27 | 0.07 | 0.2 | 0.15 |
| 9 | 76364970 | rs1116198 | 2.274 | -0.263 | 1.822 | 1.345 | -0.041 | 0.054 | 0.054 | 1.365 | 0.64 | 0.08 | 0.54 | 0.74 | 0.69 | 0.65 |
| 9 | 76371678 | rs10869280 | -1.464 | -0.215 | 1.863 | 1.149 | -0.024 | 0.049 | 0.049 | 0 | 0.3 | 0.18 | 0.42 | 0.21 | 0.31 | 0.35 |
| 9 | 76375693 | rs2007888 | 0.963 | -0.181 | 1.769 | 1.107 | 0.032 | 0.128 | 0.128 | 0.408 | 0.85 | 0.52 | 0.76 | 0.72 | 0.84 | 0.8 |
| 9 | 76389514 | rs2125391 | -0.717 | -0.522 | 0.978 | 1.036 | -0.027 | 0.05 | 0.05 | 0.338 | 0.19 | 0.02 | 0.19 | 0.08 | 0.16 | 0.19 |
| 9 | 76392885 | rs4745257 | 1.08 | -0.515 | 1.231 | 1.229 | 0.039 | 0.314 | 0.314 | 1.543 | 0.92 | 0.56 | 0.86 | 0.73 | 0.95 | 0.83 |
| 9 | 76412152 | rs1458490 | 1.424 | -0.265 | -0.689 | 1.101 | 0.036 | -0.022 | -0.022 | 0 | 0.66 | 0.16 | 0.64 | 0.65 | 0.77 | 0.62 |

| | | | | | | | | | | | | | | | | |
|---|---|---|---|---|---|---|---|---|---|---|---|---|---|---|---|---|
| 9 | 76427979 | rs1458495 | 1.385 | -0.036 | -0.868 | 0.99 | 0.011 | -0.074 | -0.074 | 0 | 0.6 | 0.14 | 0.53 | 0.54 | 0.68 | 0.56 |
| 9 | 76433950 | rs11999227 | -0.528 | -0.064 | -0.045 | 0.692 | -0.02 | -0.049 | -0.049 | 0.218 | 0.11 | 0.19 | 0.18 | 0 | 0.08 | 0.07 |
| 9 | 76448339 | rs4745262 | -0.907 | -0.02 | -0.038 | 1.133 | -0.014 | -0.016 | -0.016 | 0 | 0.14 | 0.14 | 0.14 | 0.24 | 0.18 | 0.22 |
| 9 | 76480444 | rs10869292 | 1.872 | -0.865 | -0.168 | 0.781 | 0.055 | -0.084 | -0.084 | 0 | 0.64 | 0.11 | 0.45 | 0.55 | 0.74 | 0.58 |
| 9 | 76491589 | rs1993566 | -0.519 | -1.043 | 0.165 | 0.341 | -0.002 | -0.002 | -0.002 | 0.036 | 0.89 | 0.9 | 0.89 | 0.95 | 0.96 | 0.88 |
| 9 | 76510172 | rs898048 | -0.816 | -1.115 | 1.032 | 0.293 | -0.035 | -0.093 | -0.093 | 0 | 0.13 | 0.46 | 0.09 | 0 | 0.08 | 0.07 |
| 9 | 76517066 | rs1379907 | -0.39 | -1.091 | 1.009 | 0.529 | 0.002 | 0.063 | 0.063 | 0 | 0.12 | 0.15 | 0.23 | 0.46 | 0.14 | 0.22 |
| 9 | 76539678 | rs17059634 | -0.997 | -1.167 | 1.306 | 0.32 | -0.028 | -0.079 | -0.079 | 0.095 | 0.14 | 0.29 | 0.1 | 0 | 0.1 | 0.09 |
| 9 | 76560279 | rs10869299 | 1.267 | -1.114 | 0.941 | 0.406 | -0.061 | -0.092 | -0.092 | 2.077 | 0.76 | 0.44 | 0.8 | 0.95 | 0.88 | 0.78 |
| 9 | 76565484 | rs10781171 | 1.925 | -0.946 | 0.237 | 0.488 | 0.065 | -0.074 | -0.074 | 0.143 | 0.59 | 0.13 | 0.54 | 0.49 | 0.71 | 0.53 |
| 9 | 76576244 | rs2029574 | -0.516 | -0.982 | 0.309 | 0.346 | 0.005 | 0.005 | 0.005 | 0 | 0.89 | 0.91 | 0.89 | 0.95 | 0.96 | 0.88 |
| 9 | 76596401 | rs1491457 | 0.911 | -0.983 | 0.261 | 0.341 | -0.011 | -0.026 | -0.026 | 0.216 | 0.78 | 0.75 | 0.82 | 0.95 | 0.88 | 0.81 |
| 9 | 76622068 | rs11143754 | 1.766 | -0.69 | -0.794 | 0.241 | -0.162 | 0.035 | 0.035 | 0 | 0.31 | 0.02 | 0.22 | 0.21 | 0.42 | 0.21 |
| 9 | 76638581 | rs726900 | 0.009 | -0.306 | -0.648 | 0.619 | 0.004 | 0.062 | 0.062 | 0 | 0.07 | 0.06 | 0.19 | 0.36 | 0.14 | 0.15 |
| 9 | 76648938 | rs11143764 | 1.919 | -0.604 | -0.631 | 0.136 | -0.069 | 0.054 | 0.054 | 0 | 0.3 | 0.05 | 0.19 | 0.21 | 0.34 | 0.17 |
| 9 | 76659915 | rs3012803 | -0.307 | -1.031 | -0.008 | 0.473 | 0.001 | -0.008 | -0.008 | 0 | 0.1 | 0.1 | 0.07 | 0 | 0.07 | 0.11 |
| 9 | 76669299 | rs2046913 | -0.515 | -1.027 | 0.062 | 0.536 | 0.009 | 0.068 | 0.068 | 0 | 0.13 | 0.18 | 0.27 | 0.47 | 0.22 | 0.22 |
| 9 | 76670504 | rs1032366 | 0.46 | -0.969 | -0.665 | 0.569 | 0.011 | 0.008 | 0.008 | 0 | 0.1 | 0 | 0.1 | 0.06 | 0.04 | 0.12 |
| 9 | 76676071 | rs10114881 | -1.028 | -0.927 | -0.482 | 0.509 | 0.006 | -0.02 | -0.02 | 0 | 0.31 | 0.66 | 0.43 | 0.52 | 0.29 | 0.36 |
| 9 | 76679339 | rs3012844 | -1.315 | -0.697 | -0.689 | 0.638 | 0.037 | -0.044 | -0.044 | 4.043 | 0.41 | 0.76 | 0.5 | 0.53 | 0.36 | 0.47 |
| 9 | 76710284 | rs7020553 | 1.075 | -0.707 | -0.922 | 0.697 | 0.047 | -0.06 | -0.06 | 0 | 0.6 | 0.15 | 0.53 | 0.47 | 0.66 | 0.53 |
| 9 | 76726435 | rs4425827 | 0.874 | -1.076 | -0.041 | 0.607 | -0.063 | -0.05 | -0.05 | 0.101 | 0.77 | 0.5 | 0.68 | 0.9 | 0.86 | 0.77 |
| 9 | 76731402 | rs12551119 | 0.09 | -1.042 | -0.746 | 0.803 | -0.026 | 0.006 | 0.006 | 0.169 | 0.28 | 0.09 | 0.25 | 0.21 | 0.34 | 0.27 |
| 9 | 76732506 | rs4745283 | -0.16 | -1.275 | 0.141 | 0.684 | -0.001 | -0.036 | -0.036 | 0 | 0.93 | 0.83 | 0.85 | 1 | 0.94 | 0.93 |
| 9 | 76732765 | rs10781187 | 0.794 | -1.352 | 0.014 | 0.675 | -0.024 | 0.019 | 0.019 | 4.004 | 0.88 | 0.48 | 0.82 | 0.94 | 0.94 | 0.89 |
| 9 | 76740830 | rs4288440 | 0.885 | -1.906 | -0.945 | 0.576 | 0.133 | 0.057 | 0.057 | 0 | 0.48 | 0.21 | 0.49 | 0.42 | 0.64 | 0.34 |
| 9 | 76741048 | rs4745285 | 0.511 | -2.521 | -0.896 | 0.415 | -0.02 | -0.01 | -0.01 | 0 | 0.73 | 0.3 | 0.77 | 0.89 | 0.86 | 0.7 |
| 9 | 76741242 | rs4745286 | -0.044 | -2.038 | -1.07 | 0.577 | -0.001 | -0.182 | -0.182 | 1.398 | 0.25 | 0.02 | 0.38 | 0.47 | 0.23 | 0.36 |
| 9 | 76743899 | rs7847369 | -0.638 | -1.804 | -0.931 | 0.244 | 0.057 | -0.007 | -0.007 | 0 | 0.47 | 0.17 | 0.65 | 0.53 | 0.44 | 0.52 |
| 9 | 76744909 | rs10869330 | -0.71 | -1.824 | -0.747 | 0.251 | 0.052 | 0.017 | 0.017 | 3.048 | 0.48 | 0.26 | 0.65 | 0.57 | 0.44 | 0.52 |
| 9 | 76753271 | rs11143804 | -0.491 | -1.773 | -0.959 | 0.233 | 0.025 | -0.03 | -0.03 | 0 | 0.46 | 0.1 | 0.55 | 0.56 | 0.43 | 0.51 |
| 9 | 76755616 | rs10781189 | -0.423 | -1.797 | -0.875 | 0.302 | 0.034 | -0.029 | -0.029 | 0 | 0.45 | 0.17 | 0.55 | 0.52 | 0.43 | 0.5 |
| 9 | 76758064 | rs4483218 | 0.288 | -2.143 | -1.067 | 0.483 | -0.051 | 0.017 | 0.017 | 0.417 | 0.28 | 0.04 | 0.32 | 0.22 | 0.33 | 0.28 |
| 9 | 76764649 | rs11143810 | -0.576 | -1.643 | 0.17 | 0.377 | 0.001 | 0 | 0 | 0.461 | 0.49 | 0.47 | 0.58 | 0.58 | 0.43 | 0.55 |
| 9 | 76765949 | rs10781190 | -0.423 | -1.666 | 0.233 | 0.379 | 0.013 | 0.001 | 0.001 | 0.811 | 0.48 | 0.38 | 0.57 | 0.53 | 0.43 | 0.51 |
| 9 | 76766914 | rs11143814 | -0.16 | -1.645 | -0.176 | 0.389 | 0.033 | -0.031 | -0.031 | 0.416 | 0.45 | 0.17 | 0.55 | 0.52 | 0.43 | 0.5 |
| 9 | 76767313 | rs12553320 | 0.914 | -1.058 | -0.476 | 0.516 | -0.066 | 0.02 | 0.02 | 0 | 0.35 | 0.18 | 0.3 | 0.29 | 0.44 | 0.24 |
| 9 | 76786297 | rs10746936 | -0.414 | -0.673 | -0.474 | 0.439 | -0.001 | -0.002 | -0.002 | 0 | 0.54 | 0.44 | 0.64 | 0.58 | 0.5 | 0.57 |
| 9 | 76787591 | rs966266 | -0.437 | -1.02 | -0.505 | 0.213 | -0.01 | 0.011 | 0.011 | 0 | 0.32 | 0.35 | 0.41 | 0.55 | 0.27 | 0.5 |
| 9 | 76845371 | rs4745304 | -0.132 | -0.346 | -1.332 | -0.975 | 0.01 | 0.047 | 0.047 | 6.757 | 0.6 | 0.28 | 0.72 | 0.88 | 0.68 | 0.78 |
| 9 | 76861089 | rs17611237 | -0.551 | -0.284 | -0.854 | -1.099 | 0.031 | -0.026 | -0.026 | 0 | 0.13 | 0.28 | 0.15 | 0.12 | 0.16 | 0.14 |
| 9 | 76872628 | rs2889274 | 0.022 | -0.314 | -1.248 | -1.12 | 0.008 | 0.041 | 0.041 | 5.332 | 0.61 | 0.28 | 0.72 | 0.88 | 0.68 | 0.78 |
| 9 | 76881401 | rs670010 | -0.681 | -0.459 | -1.536 | -1.237 | -0.049 | 0.103 | 0.103 | 0.647 | 0.78 | 0.95 | 0.88 | 1 | 0.77 | 0.9 |
| 9 | 76886080 | rs7022880 | 0.348 | -0.334 | -1.383 | -1.163 | 0.001 | -0.01 | -0.01 | 0 | 0.63 | 0.33 | 0.72 | 0.88 | 0.69 | 0.82 |
| 9 | 76916282 | rs11143904 | -0.874 | -0.504 | -1.595 | -1.023 | -0.014 | -0.003 | -0.003 | 0 | 0.14 | 0.23 | 0.13 | 0.14 | 0.09 | 0.21 |
| 9 | 76917840 | rs10746953 | 1.055 | -0.236 | -2.115 | -1.133 | -0.029 | 0.057 | 0.057 | 1.442 | 0.41 | 0.2 | 0.58 | 0.71 | 0.49 | 0.51 |
| 9 | 76924650 | rs11143913 | 0.404 | -0.141 | -1.887 | -1.066 | 0.003 | 0.001 | 0.001 | 0 | 0.06 | 0.08 | 0.11 | 0.1 | 0.07 | 0.15 |
| 9 | 76926329 | rs10869386 | 0.167 | -0.281 | -2.025 | -1.187 | 0.044 | 0.083 | 0.083 | 0.2 | 0.61 | 0.25 | 0.81 | 0.96 | 0.75 | 0.76 |
| 9 | 76935726 | rs11143917 | 1.526 | 0.029 | -1.814 | -0.517 | -0.009 | 0.03 | 0.03 | 1.574 | 0.61 | 0.25 | 0.64 | 0.66 | 0.64 | 0.51 |
| 9 | 76936998 | rs2991751 | -0.682 | 0.13 | -1.829 | -0.499 | -0.013 | 0.017 | 0.017 | 0.564 | 0.91 | 0.97 | 0.95 | 1 | 0.91 | 0.92 |
| 9 | 76938649 | rs2991753 | 0.138 | 0.107 | -1.824 | -0.493 | -0.052 | 0.006 | 0.006 | 0.253 | 0.79 | 0.92 | 0.8 | 0.8 | 0.73 | 0.72 |
| 9 | 76944002 | rs4237265 | 1.407 | 0.052 | -1.751 | -0.534 | 0.002 | 0.018 | 0.018 | 0.7 | 0.66 | 0.27 | 0.74 | 0.76 | 0.71 | 0.59 |
| 9 | 76944989 | rs12002539 | -0.256 | -0.128 | -1.52 | -0.469 | -0.043 | -0.014 | -0.014 | 0 | 0.09 | 0.2 | 0.02 | 0.04 | 0.02 | 0.14 |
| 9 | 76949180 | rs10491923 | 0.644 | 0.053 | -1.565 | -0.362 | -0.058 | -0.051 | -0.051 | 0 | 0.16 | 0.05 | 0.26 | 0.28 | 0.24 | 0.29 |
| 9 | 76951346 | rs7034999 | 0.777 | -0.105 | -1.743 | -0.389 | 0.013 | -0.04 | -0.04 | 0 | 0.32 | 0.19 | 0.43 | 0.5 | 0.31 | 0.34 |
| 9 | 76955137 | rs7040408 | -0.215 | -0.154 | -1.669 | -0.172 | 0.021 | -0.025 | -0.025 | 0.19 | 0.56 | 0.28 | 0.48 | 0.5 | 0.6 | 0.51 |
| 9 | 76956283 | rs1953287 | 1.316 | -0.146 | -1.899 | -0.18 | -0.054 | 0.073 | 0.073 | 0 | 0.33 | 0.04 | 0.2 | 0.18 | 0.33 | 0.18 |
| 9 | 76960254 | rs4255204 | 0.25 | -0.191 | -1.897 | -0.172 | 0.025 | -0.021 | -0.021 | 0.844 | 0.48 | 0.09 | 0.46 | 0.46 | 0.58 | 0.47 |
| 9 | 76960437 | rs7863592 | -0.817 | -0.791 | -1.296 | -0.512 | -0.067 | -0.025 | -0.025 | 0 | 0.11 | 0.62 | 0.05 | 0.04 | 0.02 | 0.05 |
| 9 | 76966194 | rs11143935 | 0.008 | -0.252 | -2.366 | -0.267 | 0.017 | 0.076 | 0.076 | 0 | 0.38 | 0.17 | 0.22 | 0.22 | 0.35 | 0.22 |
| 9 | 76967832 | rs11143937 | -0.035 | -0.247 | -2.346 | -0.248 | 0.018 | 0.08 | 0.08 | 0 | 0.39 | 0.17 | 0.22 | 0.22 | 0.35 | 0.22 |
| 9 | 76969241 | rs17060197 | -0.644 | -0.198 | -0.992 | 0.402 | -0.03 | -0.002 | -0.002 | 0 | 0.06 | 0.13 | 0.02 | 0.04 | 0.02 | 0.04 |
| 9 | 76970791 | rs17277553 | -0.458 | 0.358 | -1.29 | 0.574 | -0.013 | 0 | 0 | 0 | 0.12 | 0.01 | 0.06 | 0.09 | 0.12 | 0.16 |
| 9 | 76974920 | rs3903098 | -0.938 | 0.472 | -0.646 | 1.242 | -0.002 | 0.071 | 0.071 | 0 | 0.16 | 0.24 | 0.24 | 0.43 | 0.14 | 0.27 |
| 9 | 76976022 | rs17060208 | -0.602 | 0.588 | 0.06 | 0.801 | 0.026 | -0.011 | -0.011 | 0 | 0.06 | 0.16 | 0.14 | 0.01 | 0.05 | 0.04 |
| 9 | 76980484 | rs11143944 | -0.573 | 0.331 | -1.057 | 0.291 | 0.011 | 0.08 | 0.08 | 0 | 0.43 | 0.28 | 0.38 | 0.76 | 0.36 | 0.5 |
| 9 | 76994882 | rs11143948 | -0.014 | 0.129 | -1.269 | 0.568 | 0.003 | -0.025 | -0.025 | 0 | 0.28 | 0.25 | 0.28 | 0.53 | 0.22 | 0.29 |
| 9 | 77004242 | rs11792875 | 1.111 | 0.697 | -1.732 | -0.707 | -0.05 | -0.007 | -0.007 | 0 | 0.19 | 0.06 | 0.15 | 0.18 | 0.32 | 0.32 |
| 9 | 77010681 | rs11143958 | 0.499 | 0.533 | -1.69 | -0.819 | -0.018 | 0.018 | 0.018 | 0.885 | 0.25 | 0.16 | 0.29 | 0.19 | 0.37 | 0.37 |

| | | | | | | | | | | | | | | | | |
|---|---|---|---|---|---|---|---|---|---|---|---|---|---|---|---|---|
| 9 | 77023144 | rs4554546 | 0.367 | 0.532 | -1.559 | -0.704 | -0.001 | -0.023 | -0.023 | 0 | 0.06 | 0.1 | 0.14 | 0.01 | 0.05 | 0.05 |
| 9 | 77034979 | rs10869398 | -0.159 | 0.486 | -1.528 | -0.685 | 0.011 | 0.07 | 0.07 | 0 | 0.43 | 0.28 | 0.37 | 0.73 | 0.37 | 0.5 |
| 9 | 77038959 | rs17060276 | 0.351 | 0.57 | -1.249 | -0.862 | -0.005 | -0.024 | -0.024 | 0 | 0.06 | 0.11 | 0.13 | 0.01 | 0.04 | 0.04 |
| 9 | 77051764 | rs915291 | 0.188 | 0.468 | -0.479 | -0.916 | 0.027 | -0.018 | -0.018 | 0.672 | 0.92 | 0.74 | 0.85 | 0.99 | 0.95 | 0.95 |
| 9 | 77114274 | rs10869406 | -0.353 | 0.369 | -0.298 | -1.159 | 0.006 | 0.029 | 0.029 | 0 | 0.45 | 0.4 | 0.41 | 0.82 | 0.38 | 0.51 |
| 9 | 77136979 | rs13291991 | 0.326 | 0.172 | -0.819 | -1.301 | -0.117 | 0.08 | 0.08 | 0 | 0.37 | 0.05 | 0.22 | 0.2 | 0.45 | 0.4 |
| 9 | 77138246 | rs1018584 | 0.927 | 0.134 | -0.5 | -1.291 | -0.001 | -0.017 | -0.017 | 0 | 0.35 | 0.28 | 0.46 | 0.07 | 0.32 | 0.18 |
| 9 | 77140951 | rs10869412 | 0.308 | 0.123 | -0.37 | -1.303 | -0.006 | 0.001 | 0.001 | 0 | 0.26 | 0.35 | 0.31 | 0.74 | 0.23 | 0.42 |
| 9 | 77146468 | rs7870498 | -0.324 | 0.054 | -0.244 | -1.302 | 0.004 | -0.022 | -0.022 | 0.402 | 0.59 | 0.54 | 0.76 | 0.8 | 0.52 | 0.58 |
| 9 | 77149837 | rs7042950 | 0.4 | 0.085 | -0.252 | -1.292 | -0.003 | 0 | 0 | 0 | 0.25 | 0.32 | 0.31 | 0.74 | 0.22 | 0.4 |
| 9 | 77150289 | rs7873840 | -1.043 | 0.212 | -0.325 | -1.172 | 0.02 | -0.056 | -0.056 | 0 | 0.78 | 0.63 | 0.85 | 0.88 | 0.67 | 0.67 |
| 9 | 77158236 | rs17611535 | 1.307 | 0.098 | -0.789 | -0.882 | -0.025 | 0.005 | 0.005 | 0.303 | 0.17 | 0.04 | 0.1 | 0.08 | 0.15 | 0.09 |
| 9 | 77159265 | rs10217594 | 0.554 | 0.388 | -0.539 | -1.109 | -0.001 | -0.036 | -0.036 | 0 | 0.34 | 0.3 | 0.42 | 0.07 | 0.25 | 0.16 |
| 9 | 77159715 | rs10781235 | 0.044 | 0.352 | -0.579 | -0.976 | 0 | 0 | 0 | 2.356 | 0.26 | 0.29 | 0.27 | 0.73 | 0.22 | 0.38 |
| 9 | 77164006 | rs4745340 | -0.022 | 0.138 | -0.732 | -1.63 | 0 | 0.16 | 0.16 | 0 | 0.5 | 0.75 | 0.55 | 0.24 | 0.54 | 0.48 |
| 9 | 77188885 | rs17612218 | 2.459 | 1.008 | -0.574 | -1.396 | -0.034 | 0.042 | 0.042 | 0 | 0.08 | 0 | 0.04 | 0 | 0.14 | 0.04 |
| 9 | 77192841 | rs1323354 | 0.297 | 0.849 | 0.048 | -1.344 | 0.034 | -0.024 | -0.024 | 0 | 0.11 | 0.28 | 0.24 | 0.06 | 0.13 | 0.17 |
| 9 | 77194528 | rs10512036 | -0.88 | 1.097 | 0.079 | -1.538 | 0.004 | -0.061 | -0.061 | 0 | 0.68 | 0.57 | 0.69 | 0.82 | 0.61 | 0.69 |
| 9 | 77194612 | rs10781238 | 0.836 | 1.14 | 0.153 | -1.424 | -0.002 | 0.001 | 0.001 | 0 | 0.1 | 0.08 | 0.22 | 0.07 | 0.14 | 0.18 |
| 9 | 77218345 | rs11144020 | -0.086 | 1.043 | 0.355 | -1.519 | -0.009 | 0.028 | 0.028 | 0.004 | 0.57 | 0.37 | 0.45 | 0.76 | 0.47 | 0.5 |
| 9 | 77224088 | rs10869428 | 0.027 | 1.049 | 0.303 | -1.458 | -0.029 | 0.111 | 0.111 | 0.076 | 0.56 | 0.21 | 0.45 | 0.75 | 0.47 | 0.5 |
| 9 | 77224424 | rs17613014 | 0.854 | 1.156 | 0.344 | -1.482 | -0.002 | 0.003 | 0.003 | 0 | 0.31 | 0.27 | 0.3 | 0.19 | 0.38 | 0.32 |
| 9 | 77229161 | rs10119947 | 0.587 | 1.201 | 0.33 | -1.392 | 0.017 | -0.019 | -0.019 | 1.351 | 0.11 | 0.22 | 0.24 | 0.06 | 0.14 | 0.17 |
| 9 | 77236263 | rs10869430 | 0.677 | 0.665 | -0.31 | -1 | -0.008 | 0.008 | 0.008 | 0 | 0.43 | 0.19 | 0.38 | 0.55 | 0.35 | 0.28 |
| 9 | 77238103 | rs10869431 | -0.443 | 0.854 | 0.079 | -1.022 | 0.058 | -0.024 | -0.024 | 0 | 0.88 | 0.58 | 0.75 | 0.94 | 0.86 | 0.82 |
| 9 | 77241777 | rs1831121 | -0.139 | 0.718 | -0.124 | -0.911 | -0.014 | 0.037 | 0.037 | 0 | 0.57 | 0.34 | 0.44 | 0.74 | 0.46 | 0.52 |
| 9 | 77245873 | rs17691614 | 1.362 | 0.4 | -0.432 | -0.995 | -0.001 | -0.003 | -0.003 | 0.134 | 0.25 | 0.23 | 0.23 | 0.19 | 0.27 | 0.29 |
| 9 | 77247597 | rs7032677 | -0.34 | 0.596 | -0.971 | -0.922 | -0.03 | 0.119 | 0.119 | 0.214 | 0.58 | 0.19 | 0.47 | 0.76 | 0.53 | 0.52 |
| 9 | 77254334 | rs11144032 | 0.377 | 0.092 | -1.325 | -0.46 | -0.071 | -0.101 | -0.101 | 0 | 0.41 | 0.06 | 0.36 | 0.52 | 0.38 | 0.25 |
| 9 | 77257346 | rs2273975 | 1.363 | 0.012 | -1.488 | -0.4 | 0.014 | 0.058 | 0.058 | 0.112 | 0.26 | 0.04 | 0.16 | 0.09 | 0.17 | 0.07 |
| 9 | 77258265 | rs11144033 | -0.454 | 0.904 | 0.061 | 0.211 | -0.001 | -0.003 | -0.003 | 4.838 | 0.25 | 0.25 | 0.38 | 0.09 | 0.4 | 0.21 |
| 9 | 77264546 | rs1319551 | 0.509 | 0.998 | -0.158 | 0.313 | -0.021 | -0.002 | -0.002 | 0 | 0.12 | 0.08 | 0.28 | 0.09 | 0.24 | 0.17 |
| 9 | 77267620 | rs12352112 | -0.037 | 0.301 | 0.382 | 0.325 | -0.011 | -0.049 | -0.049 | 0.746 | 0.09 | 0.22 | 0.05 | 0 | 0.04 | 0.04 |
| 9 | 77268666 | rs10869433 | -0.524 | 0.498 | 0.094 | 0.803 | -0.01 | -0.011 | -0.011 | 0 | 0.21 | 0.39 | 0.15 | 0.2 | 0.2 | 0.24 |
| 9 | 77269060 | rs11144039 | 0.053 | 0.887 | -0.381 | 1.175 | -0.009 | 0.021 | 0.021 | 0 | 0.08 | 0.04 | 0.13 | 0.43 | 0.09 | 0.17 |
| 9 | 77272296 | rs17060398 | 0.053 | 0.887 | -0.484 | 1.175 | -0.019 | -0.006 | -0.006 | 0 | 0.08 | 0.03 | 0.13 | 0.43 | 0.09 | 0.17 |
| 9 | 77275913 | rs7865407 | -1.173 | 0.458 | -0.091 | 0.205 | 0.023 | 0.079 | 0.079 | 0 | 0.52 | 0.67 | 0.61 | 0.88 | 0.59 | 0.46 |
| 9 | 77279806 | rs12001830 | -0.065 | 0.561 | -0.337 | 0.267 | 0.006 | 0.032 | 0.032 | 0.353 | 0.19 | 0.08 | 0.11 | 0.01 | 0.19 | 0.1 |
| 9 | 77280145 | rs10121918 | -1.802 | 0.506 | -0.057 | 0.246 | 0.014 | 0.187 | 0.187 | 0 | 0.52 | 0.82 | 0.59 | 0.86 | 0.58 | 0.46 |
| 9 | 77283889 | rs1327837 | -1.678 | -0.113 | 0.724 | 1.443 | -0.001 | -0.001 | -0.001 | 0.86 | 0.38 | 0.64 | 0.36 | 0.67 | 0.35 | 0.28 |
| 9 | 77288320 | rs1323368 | -0.256 | -0.078 | 0.748 | 1.446 | 0.008 | 0.025 | 0.025 | 0.353 | 0.19 | 0.1 | 0.11 | 0.01 | 0.19 | 0.1 |
| 9 | 77292289 | rs1327836 | -0.769 | -0.053 | 0.746 | 1.49 | 0.003 | 0.116 | 0.116 | 0.716 | 0.19 | 0.55 | 0.26 | 0.66 | 0.15 | 0.18 |
| 9 | 77299740 | rs11144062 | -0.579 | 0.376 | 0.445 | 0.999 | -0.001 | -0.028 | -0.028 | 0.596 | 0.2 | 0.16 | 0.12 | 0.01 | 0.19 | 0.1 |
| 9 | 77305480 | rs1410227 | 1.106 | 0.413 | 0.837 | 1.747 | -0.007 | 0.152 | 0.152 | 3.478 | 0.83 | 0.45 | 0.85 | 0.65 | 0.86 | 0.89 |
| 9 | 77305695 | rs1410226 | -0.55 | 0.357 | 0.286 | 1.787 | 0.011 | -0.022 | -0.022 | 0.341 | 0.13 | 0.09 | 0.15 | 0.25 | 0.09 | 0.09 |
| 9 | 77308710 | rs1410225 | 1.058 | 0.392 | 0.293 | 1.089 | -0.001 | 0 | 0 | 0 | 0.71 | 0.28 | 0.69 | 0.43 | 0.78 | 0.8 |
| 9 | 77312690 | rs499922 | -0.032 | 0.623 | 0.21 | 0.826 | -0.014 | 0.003 | 0.003 | 0 | 0.86 | 0.79 | 0.94 | 0.8 | 0.92 | 0.8 |
| 9 | 77313531 | rs525065 | 1.55 | 0.725 | 0.177 | 0.986 | 0.009 | -0.003 | -0.003 | 0 | 0.66 | 0.29 | 0.62 | 0.35 | 0.7 | 0.62 |
| 9 | 77315134 | rs577787 | 0.067 | 0.61 | 0.541 | 0.993 | -0.026 | 0.014 | 0.014 | 0 | 0.85 | 0.68 | 0.93 | 0.8 | 0.92 | 0.8 |
| 9 | 77323772 | rs519067 | 0.755 | 1.224 | -0.637 | 0.416 | -0.008 | 0.025 | 0.025 | 0 | 0.57 | 0.27 | 0.44 | 0.33 | 0.54 | 0.57 |
| 9 | 77324251 | rs2245644 | 0.61 | 1.047 | -0.501 | 0.462 | -0.006 | -0.006 | -0.006 | 0 | 0.6 | 0.31 | 0.46 | 0.36 | 0.59 | 0.6 |
| 9 | 77329367 | rs11144073 | -1.093 | 1.226 | -0.065 | -0.249 | 0.018 | -0.087 | -0.087 | 0.241 | 0.19 | 0.29 | 0.34 | 0.02 | 0.21 | 0.06 |
| 9 | 77332549 | rs546542 | 0.251 | 1.729 | -0.724 | -0.389 | -0.031 | 0.001 | 0.001 | 0 | 0.71 | 0.82 | 0.6 | 0.31 | 0.66 | 0.55 |
| 9 | 77332745 | rs1536613 | -0.22 | 0.532 | -0.736 | -1.268 | -0.009 | -0.022 | -0.022 | 0 | 0.27 | 0.58 | 0.33 | 0.14 | 0.22 | 0.24 |
| 9 | 77333181 | rs2209820 | 0.762 | 0.898 | -0.879 | -0.522 | -0.003 | 0.005 | 0.005 | 0 | 0.06 | 0.04 | 0.12 | 0.17 | 0.09 | 0.07 |
| 9 | 77336810 | rs1327831 | 0.841 | 0.775 | -0.93 | -0.623 | -0.041 | 0.03 | 0.03 | 0 | 0.07 | 0 | 0.22 | 0 | 0.14 | 0.04 |
| 9 | 77349845 | rs1759831 | 1.144 | 0.753 | -0.238 | -1.11 | 0.005 | 0.011 | 0.011 | 0.046 | 0.55 | 0.41 | 0.31 | 0.17 | 0.44 | 0.41 |
| 9 | 77357968 | rs877809 | 0.068 | 0.418 | 0.311 | -1.234 | -0.039 | -0.013 | -0.013 | 0 | 0.22 | 0.64 | 0.11 | 0.16 | 0.08 | 0.21 |
| 9 | 77358430 | rs2254229 | 1.163 | 0.49 | 0.178 | -1.198 | -0.038 | -0.004 | -0.004 | 0 | 0.17 | 0.22 | 0.09 | 0.16 | 0.08 | 0.2 |
| 9 | 77361384 | rs476673 | -0.352 | 0.641 | -0.015 | -1.151 | -0.047 | -0.006 | -0.006 | 0 | 0.72 | 0.88 | 0.6 | 0.32 | 0.66 | 0.54 |
| 9 | 77366199 | rs11787707 | 0.966 | 0.321 | 0.13 | -0.946 | -0.004 | 0.001 | 0.001 | 0 | 0.06 | 0.04 | 0.11 | 0.15 | 0.09 | 0.07 |
| 9 | 77373129 | rs12002738 | 1.29 | 0.169 | 0.224 | -0.96 | -0.003 | -0.001 | -0.001 | 0 | 0.15 | 0.11 | 0.08 | 0.16 | 0.08 | 0.2 |
| 9 | 77376633 | rs2274925 | 0.175 | 0.153 | 0.593 | -0.966 | 0.046 | 0.073 | 0.073 | 0.017 | 0.08 | 0.35 | 0.07 | 0.16 | 0.08 | 0.1 |
| 9 | 77376647 | rs2274924 | -0.119 | 0.143 | 0.529 | -0.968 | -0.04 | 0.024 | 0.024 | 0.034 | 0.23 | 0.45 | 0.15 | 0.32 | 0.16 | 0.32 |
| 9 | 77377410 | rs3750425 | 0.995 | 0.131 | 0.19 | -0.918 | -0.043 | -0.004 | -0.004 | 2.859 | 0.17 | 0.3 | 0.09 | 0.15 | 0.08 | 0.2 |
| 9 | 77393689 | rs11144082 | -0.266 | 0.735 | -0.357 | -0.902 | -0.102 | -0.102 | -0.102 | 0 | 0.71 | 0.93 | 0.49 | 0.47 | 0.59 | 0.65 |
| 9 | 77396655 | rs6560408 | 0.601 | 0.445 | -0.42 | -1.378 | 0.017 | 0.031 | 0.031 | 0 | 0.5 | 0.41 | 0.29 | 0.28 | 0.42 | 0.4 |
| 9 | 77403203 | rs11144083 | 0.026 | 0.267 | -0.304 | -1.174 | 0.002 | 0.01 | 0.01 | 0 | 0.18 | 0.14 | 0.13 | 0.13 | 0.19 | 0.15 |
| 9 | 77404435 | rs11144085 | -1.023 | 0.726 | -0.18 | -0.701 | 0.001 | -0.002 | -0.002 | 1.055 | 0.4 | 0.53 | 0.58 | 0.64 | 0.42 | 0.62 |

| | | | | | | | | | | | | | | | | |
|---|---|---|---|---|---|---|---|---|---|---|---|---|---|---|---|---|
| 9 | 77406281 | rs2151424 | -0.422 | 0.964 | -0.402 | -0.67 | -0.002 | -0.052 | -0.052 | 0 | 0.25 | 0.36 | 0.18 | 0.11 | 0.25 | 0.26 |
| 9 | 77407636 | rs4145894 | 0.343 | 0.939 | -1.002 | -0.146 | -0.075 | -0.108 | -0.108 | 0.278 | 0.14 | 0.01 | 0.39 | 0.51 | 0.18 | 0.36 |
| 9 | 77415284 | rs7859201 | -0.901 | 0.593 | -0.768 | 0.111 | 0.007 | 0.001 | 0.001 | 2.465 | 0.37 | 0.74 | 0.34 | 0.35 | 0.37 | 0.37 |
| 9 | 77417489 | rs7848706 | 0.207 | 0.58 | -0.699 | 0.084 | 0 | 0.007 | 0.007 | 0 | 0.12 | 0.11 | 0.12 | 0.23 | 0.15 | 0.14 |
| 9 | 77423730 | rs17060535 | -0.072 | 0.205 | -0.255 | -0.634 | 0 | -0.001 | -0.001 | 0 | 0.2 | 0.19 | 0.17 | 0.12 | 0.22 | 0.23 |
| 9 | 77424957 | rs12551151 | 0.296 | 0.244 | -0.919 | -0.513 | -0.085 | -0.065 | -0.065 | 0 | 0.23 | 0.05 | 0.41 | 0.71 | 0.32 | 0.46 |
| 9 | 77441137 | rs4745361 | 0.269 | 0.107 | -0.787 | -0.23 | -0.035 | -0.084 | -0.084 | 0 | 0.16 | 0.01 | 0.13 | 0.26 | 0.17 | 0.23 |
| 9 | 77442616 | rs7045949 | -1.074 | 0.05 | -0.241 | -0.413 | 0.003 | 0.066 | 0.066 | 1.184 | 0.51 | 0.67 | 0.63 | 0.81 | 0.55 | 0.7 |
| 9 | 77443424 | rs17060568 | 0.751 | 0.344 | -1.063 | -0.217 | -0.026 | -0.027 | -0.027 | 0 | 0.05 | 0.01 | 0.05 | 0.12 | 0.1 | 0.12 |
| 9 | 77472066 | rs12378991 | 1.103 | -0.041 | -1.486 | -0.435 | 0 | 0.075 | 0.075 | 3.041 | 0.12 | 0.01 | 0.08 | 0 | 0.07 | 0.02 |
| 9 | 77475588 | rs6560417 | -0.318 | 0.04 | -1.449 | -0.382 | 0.004 | 0.046 | 0.046 | 0 | 0.94 | 0.91 | 0.86 | 0.81 | 0.91 | 0.83 |
| 9 | 77476386 | rs2184118 | 0.354 | 0.112 | -1.617 | -0.716 | -0.066 | -0.091 | -0.091 | 0.669 | 0.28 | 0.09 | 0.45 | 0.54 | 0.39 | 0.46 |
| 9 | 77502160 | rs1333342 | 0.19 | 0.159 | -1.464 | -0.066 | -0.098 | -0.087 | -0.087 | 0 | 0.39 | 0.11 | 0.53 | 0.54 | 0.46 | 0.48 |
| 9 | 77502322 | rs1333343 | 0.528 | 0.091 | -0.317 | 1.209 | -0.012 | 0.05 | 0.05 | 0 | 0.79 | 0.69 | 0.76 | 0.55 | 0.82 | 0.66 |
| 9 | 77516005 | rs11144144 | -0.584 | 0.42 | -0.467 | 0.485 | -0.008 | 0.029 | 0.029 | 0.126 | 0.44 | 0.67 | 0.63 | 0.33 | 0.44 | 0.46 |
| 9 | 77516923 | rs2800249 | 0.194 | 0.561 | -0.378 | 0.184 | -0.008 | 0.021 | 0.021 | 0.019 | 0.6 | 0.41 | 0.46 | 0.81 | 0.57 | 0.62 |
| 9 | 77520324 | rs13298337 | -0.285 | 0.42 | -0.492 | 0.06 | 0.001 | 0.007 | 0.007 | 0 | 0.7 | 0.68 | 0.66 | 0.5 | 0.67 | 0.6 |
| 9 | 77520677 | rs2800265 | -0.195 | 0.441 | 0.101 | -0.096 | 0.002 | 0.006 | 0.006 | 0 | 0.35 | 0.4 | 0.38 | 0.51 | 0.39 | 0.49 |
| 9 | 77523690 | rs2769064 | -0.13 | 0.368 | -0.083 | -0.137 | 0.007 | -0.052 | -0.052 | 0 | 0.6 | 0.53 | 0.48 | 0.83 | 0.58 | 0.62 |
| 9 | 77530589 | rs2769050 | -0.277 | 0.221 | -0.061 | -0.249 | 0.007 | -0.085 | -0.085 | 0 | 0.7 | 0.51 | 0.63 | 0.89 | 0.69 | 0.78 |
| 9 | 77534116 | rs1176828 | -0.676 | 0.28 | -0.506 | -0.308 | -0.006 | -0.034 | -0.034 | 0.001 | 0.93 | 0.88 | 0.97 | 1 | 0.96 | 0.98 |
| 9 | 77535512 | rs7029929 | 0.423 | 0.376 | -0.423 | -0.004 | 0.062 | -0.001 | -0.001 | 0.566 | 0.06 | 0.26 | 0.05 | 0 | 0.06 | 0.09 |
| 9 | 77535672 | rs13299921 | 1.011 | 0.387 | -1.636 | 0.123 | 0.02 | -0.06 | -0.06 | 0.01 | 0.28 | 0.09 | 0.12 | 0.31 | 0.24 | 0.22 |
| 9 | 77536753 | rs1359879 | -0.097 | 0.79 | -1.288 | 0.54 | -0.022 | -0.018 | -0.018 | 0 | 0.6 | 0.73 | 0.51 | 0.44 | 0.56 | 0.44 |
| 9 | 77543548 | rs4745367 | -0.04 | 0.795 | -1.516 | 0.289 | -0.031 | 0.037 | 0.037 | 0.338 | 0.24 | 0.1 | 0.31 | 0.15 | 0.25 | 0.14 |
| 9 | 77545238 | rs10118224 | -0.423 | 0.819 | -1.249 | -0.275 | 0.004 | -0.025 | -0.025 | 0 | 0.18 | 0.25 | 0.2 | 0.09 | 0.2 | 0.14 |
| 9 | 77551617 | rs1176816 | 0.052 | 1.652 | -1.209 | -0.676 | -0.017 | 0.01 | 0.01 | 0 | 0.78 | 0.83 | 0.62 | 0.8 | 0.64 | 0.66 |
| 9 | 77563087 | rs2769058 | 1.073 | 0.326 | -1.635 | 0.005 | 0.003 | -0.008 | -0.008 | 0 | 0.38 | 0.24 | 0.35 | 0.4 | 0.34 | 0.32 |
| 9 | 77573159 | rs10869460 | 0.53 | 0.259 | -1.96 | 0.004 | -0.118 | 0.032 | 0.032 | 0.014 | 0.2 | 0.02 | 0.19 | 0.13 | 0.33 | 0.24 |
| 9 | 77585644 | rs7867200 | -0.546 | -0.143 | -1.506 | -0.033 | -0.015 | -0.035 | -0.035 | 0 | 0.33 | 0.26 | 0.4 | 0.48 | 0.39 | 0.49 |
| 9 | 77595402 | rs11144177 | 0.363 | -0.146 | -1.452 | -0.01 | -0.047 | 0.019 | 0.019 | 0 | 0.22 | 0.13 | 0.37 | 0.21 | 0.35 | 0.34 |
| 9 | 77597362 | rs10869464 | 0.363 | -0.146 | -1.452 | -0.01 | -0.047 | 0.021 | 0.021 | 0 | 0.22 | 0.13 | 0.37 | 0.21 | 0.35 | 0.34 |
| 9 | 77608145 | rs11144179 | 1.224 | -0.322 | -1.768 | -0.316 | -0.002 | 0.021 | 0.021 | 0.068 | 0.1 | 0.09 | 0.04 | 0.28 | 0.03 | 0.15 |
| 9 | 77618438 | rs10781258 | -0.724 | -0.453 | -1.67 | -0.476 | -0.007 | -0.017 | -0.017 | 0 | 0.42 | 0.36 | 0.44 | 0.48 | 0.44 | 0.52 |
| 9 | 77624761 | rs11144184 | -0.26 | -0.48 | -1.609 | -0.475 | -0.003 | 0 | 0 | 0 | 0.2 | 0.23 | 0.07 | 0.27 | 0.09 | 0.18 |
| 9 | 77626873 | rs11793714 | 1.214 | -0.449 | -1.638 | -0.52 | 0.009 | 0.017 | 0.017 | 0 | 0.09 | 0.05 | 0.04 | 0 | 0.06 | 0.03 |
| 9 | 77641451 | rs10869474 | 1.157 | -0.408 | -1.659 | -0.515 | -0.108 | 0.104 | 0.104 | 0.026 | 0.2 | 0.03 | 0.18 | 0.08 | 0.33 | 0.24 |
| 9 | 77650696 | rs10781262 | -0.179 | -0.438 | -1.428 | -0.392 | -0.013 | -0.007 | -0.007 | 0 | 0.32 | 0.27 | 0.23 | 0.33 | 0.37 | 0.4 |
| 9 | 77652455 | rs10869476 | -0.662 | -0.372 | -1.452 | -0.382 | -0.017 | 0.015 | 0.015 | 0 | 0.4 | 0.28 | 0.26 | 0.34 | 0.43 | 0.42 |
| 9 | 77653755 | rs7850614 | -0.053 | -0.496 | -1.18 | -0.844 | -0.002 | -0.003 | -0.003 | 0 | 0.7 | 0.36 | 0.56 | 0.73 | 0.71 | 0.65 |
| 9 | 77662999 | rs773139 | 0.474 | -0.318 | -1.288 | -0.887 | -0.005 | -0.005 | -0.005 | 0 | 0.78 | 0.71 | 0.73 | 0.86 | 0.86 | 0.79 |
| 9 | 77670081 | rs11144203 | 0.902 | 0.526 | -1.635 | -0.966 | -0.041 | -0.037 | -0.037 | 0.074 | 0.07 | 0.01 | 0.22 | 0.14 | 0.15 | 0.15 |
| 9 | 77675778 | rs11144207 | 1.244 | 0.682 | -1.763 | -1.079 | -0.037 | 0.057 | 0.057 | 0 | 0.13 | 0.02 | 0.14 | 0.08 | 0.16 | 0.16 |
| 9 | 77676285 | rs11519 | 1.244 | 0.707 | -1.763 | -1.079 | -0.039 | 0.057 | 0.057 | 0 | 0.13 | 0.02 | 0.14 | 0.08 | 0.16 | 0.16 |
| 9 | 77680812 | rs911737 | 1.314 | 1.238 | -2.07 | -1.217 | 0.013 | 0.015 | 0.015 | 0.215 | 0.45 | 0.63 | 0.48 | 0.57 | 0.44 | 0.38 |
| 9 | 77682437 | rs3780175 | 1.377 | 1.16 | -2.047 | -1.201 | -0.039 | 0.058 | 0.058 | 0 | 0.13 | 0.02 | 0.14 | 0.07 | 0.16 | 0.16 |
| 9 | 77683026 | rs7021664 | -1.258 | 1.074 | -1.876 | -1.065 | -0.027 | 0.046 | 0.046 | 0 | 0.92 | 0.99 | 0.95 | 1 | 0.92 | 0.9 |
| 9 | 77683225 | rs7021926 | 0.462 | 1.093 | -1.826 | -1.029 | 0.021 | 0.035 | 0.035 | 0.607 | 0.65 | 0.74 | 0.85 | 0.79 | 0.76 | 0.69 |
| 9 | 77684439 | rs13289688 | 0.433 | 1.055 | -2.124 | -1.094 | -0.082 | 0.09 | 0.09 | 0 | 0.64 | 0.85 | 0.57 | 0.76 | 0.56 | 0.58 |
| 9 | 77684690 | rs3752955 | 0.86 | 1.185 | -1.788 | -1.193 | -0.029 | 0.026 | 0.026 | 0.693 | 0.56 | 0.74 | 0.59 | 0.68 | 0.53 | 0.52 |
| 9 | 77690463 | rs7857408 | 0.214 | 1.226 | -1.697 | -1.259 | -0.054 | 0.048 | 0.048 | 0 | 0.77 | 0.92 | 0.71 | 0.86 | 0.71 | 0.72 |
| 9 | 77690749 | rs2376397 | 0.68 | 0.93 | -0.902 | -1.262 | -0.008 | -0.018 | -0.018 | 1.341 | 0.22 | 0.38 | 0.21 | 0.16 | 0.21 | 0.19 |
| 9 | 77693316 | rs4442208 | 0.873 | 1.103 | -0.961 | -1.174 | 0.001 | -0.021 | -0.021 | 0.411 | 0.16 | 0.16 | 0.09 | 0 | 0.13 | 0.08 |
| 9 | 77696677 | rs11144217 | 0.23 | 0.905 | -0.741 | -1.341 | -0.006 | -0.012 | -0.012 | 0 | 0.73 | 0.33 | 0.66 | 0.85 | 0.76 | 0.78 |
| 9 | 77696918 | rs11144218 | 1.122 | 1.058 | -0.887 | -1.344 | -0.017 | 0.013 | 0.013 | 1.589 | 0.63 | 0.25 | 0.34 | 0.63 | 0.53 | 0.5 |
| 9 | 77699899 | rs10869495 | 1.09 | 0.989 | -0.805 | -1.313 | 0.03 | -0.002 | -0.002 | 0 | 0.51 | 0.31 | 0.33 | 0.56 | 0.43 | 0.38 |
| 9 | 77700084 | rs10869496 | 1.483 | 0.979 | -0.819 | -1.319 | 0.038 | -0.041 | -0.041 | 0.309 | 0.46 | 0.13 | 0.23 | 0.45 | 0.37 | 0.27 |
| 9 | 77705570 | rs10869497 | 1.112 | 1.018 | -0.79 | -1.29 | 0.03 | -0.002 | -0.002 | 0 | 0.51 | 0.31 | 0.34 | 0.55 | 0.42 | 0.38 |
| 9 | 77706944 | rs2376398 | 0.903 | 1.085 | -0.702 | -1.293 | -0.004 | 0.001 | 0.001 | 0.297 | 0.67 | 0.34 | 0.4 | 0.72 | 0.54 | 0.57 |
| 9 | 77713279 | rs10123421 | -0.581 | 0.871 | 0.141 | -1.304 | 0.013 | -0.02 | -0.02 | 0.1 | 0.2 | 0.32 | 0.31 | 0.13 | 0.29 | 0.23 |
| 9 | 77716145 | rs11144228 | -0.021 | 0.867 | 0.328 | -1.285 | 0.08 | 0.023 | 0.023 | 0 | 0.13 | 0.33 | 0.27 | 0.13 | 0.23 | 0.19 |
| 9 | 77722267 | rs1033777 | 1.82 | 0.809 | 0.098 | -1.282 | 0.002 | 0.041 | 0.041 | 0 | 0.08 | 0.02 | 0.04 | 0 | 0.06 | 0.04 |
| 9 | 77726636 | rs10869504 | 0.447 | 0.88 | 0.179 | -1.283 | 0.05 | 0.012 | 0.012 | 0 | 0.11 | 0.18 | 0.26 | 0.13 | 0.23 | 0.19 |
| 9 | 77727884 | rs3780180 | 1.579 | 0.75 | -0.131 | -1.36 | 0.014 | -0.095 | -0.095 | 0 | 0.37 | 0.07 | 0.19 | 0.44 | 0.3 | 0.24 |
| 9 | 77737029 | rs10869507 | -1.337 | 0.797 | 0.038 | -1.326 | 0.025 | -0.055 | -0.055 | 0 | 0.29 | 0.54 | 0.39 | 0.13 | 0.37 | 0.28 |
| 9 | 77744416 | rs6560421 | 1.125 | 0.639 | -0.248 | -1.438 | 0.014 | -0.026 | -0.026 | 0 | 0.39 | 0.27 | 0.29 | 0.55 | 0.35 | 0.31 |
| 9 | 77745424 | rs2295861 | -1.517 | 0.773 | -0.491 | -1.282 | -0.014 | -0.036 | -0.036 | 0 | 0.72 | 0.86 | 0.68 | 0.68 | 0.73 | 0.63 |
| 9 | 77748151 | rs7040349 | 0.16 | 0.996 | -0.28 | -1.137 | 0.026 | -0.005 | -0.005 | 0 | 0.15 | 0.18 | 0.26 | 0.13 | 0.25 | 0.23 |
| 9 | 77748968 | rs7872448 | -0.47 | 0.952 | -0.693 | -1.191 | 0.021 | -0.052 | -0.052 | 0 | 0.24 | 0.45 | 0.36 | 0.13 | 0.3 | 0.27 |

| | | | | | | | | | | | | | | | | |
|---|---|---|---|---|---|---|---|---|---|---|---|---|---|---|---|---|
| 9 | 77749569 | rs10869509 | 0.935 | 1.098 | -1.022 | -1.122 | -0.03 | -0.115 | -0.115 | 0 | 0.16 | 0.39 | 0.07 | 0 | 0.08 | 0.11 |
| 9 | 77752967 | rs1009282 | -0.091 | 1.59 | -1.182 | -0.501 | -0.06 | -0.012 | -0.012 | 0 | 0.41 | 0.21 | 0.48 | 0.56 | 0.56 | 0.4 |
| 9 | 77755469 | rs2273770 | 0.282 | 1.629 | -1.199 | -0.476 | 0.002 | -0.007 | -0.007 | 0 | 0.36 | 0.35 | 0.27 | 0.43 | 0.38 | 0.22 |
| 9 | 77758636 | rs7862075 | -1.112 | 1.696 | -1.631 | -0.498 | -0.007 | -0.01 | -0.01 | 0 | 0.58 | 0.58 | 0.63 | 0.68 | 0.67 | 0.54 |
| 9 | 77760543 | rs4376557 | 0.877 | 2.247 | -2.553 | 0.13 | -0.001 | -0.08 | -0.08 | 0 | 0.26 | 0.09 | 0.16 | 0.43 | 0.29 | 0.19 |
| 9 | 77761642 | rs2146044 | -0.014 | 2.071 | -2.351 | 0.254 | -0.03 | 0.021 | 0.021 | 0 | 0.13 | 0.01 | 0.12 | 0.11 | 0.11 | 0.15 |
| 9 | 77762449 | rs1407376 | 0.038 | 2.202 | -1.779 | 0.622 | -0.013 | -0.014 | -0.014 | 1.313 | 0.06 | 0.07 | 0.23 | 0.13 | 0.18 | 0.18 |
| 9 | 77763300 | rs13290703 | 0.159 | 1.521 | -1.991 | 0.613 | -0.028 | -0.089 | -0.089 | 0 | 0.24 | 0.1 | 0.19 | 0.44 | 0.28 | 0.18 |
| 9 | 77763924 | rs7041918 | -0.527 | 1.238 | -2.439 | 0.388 | -0.054 | -0.08 | -0.08 | 0 | 0.4 | 0.11 | 0.54 | 0.46 | 0.45 | 0.33 |
| 9 | 77764011 | rs17704078 | -0.635 | 1.162 | -2.281 | -0.29 | -0.077 | 0.042 | 0.042 | 0 | 0.18 | 0.02 | 0.17 | 0.12 | 0.21 | 0.14 |
| 9 | 77767024 | rs7859631 | -0.805 | 0.752 | -1.432 | -0.366 | 0.01 | -0.013 | -0.013 | 0 | 0.27 | 0.37 | 0.24 | 0.2 | 0.27 | 0.25 |
| 9 | 77767571 | rs10869516 | -1.165 | 0.812 | -1.553 | -0.52 | 0.004 | -0.008 | -0.008 | 0 | 0.31 | 0.38 | 0.28 | 0.2 | 0.34 | 0.25 |
| 9 | 77770054 | rs773024 | -0.699 | -0.143 | -1.38 | -0.698 | -0.002 | -0.008 | -0.008 | 0 | 0.43 | 0.49 | 0.32 | 0.26 | 0.36 | 0.41 |
| 9 | 77780144 | rs1081875 | -0.829 | -0.532 | -0.804 | -1.129 | 0.003 | 0.038 | 0.038 | 0 | 0.55 | 0.65 | 0.37 | 0.24 | 0.41 | 0.44 |
| 9 | 77782517 | rs1081877 | -0.968 | -0.636 | -0.797 | -1.179 | -0.003 | 0.065 | 0.065 | 0 | 0.57 | 0.75 | 0.38 | 0.24 | 0.41 | 0.44 |
| 9 | 77785171 | rs11144271 | 0.904 | -0.386 | -1.021 | -1.254 | -0.043 | 0.039 | 0.039 | 0 | 0.07 | 0.01 | 0.11 | 0 | 0.11 | 0.05 |
| 9 | 77789607 | rs10283463 | -0.353 | -0.715 | -0.613 | -1.156 | 0.005 | 0.012 | 0.012 | 0 | 0.47 | 0.38 | 0.35 | 0.3 | 0.39 | 0.34 |
| 9 | 77789735 | rs10283465 | -0.825 | -0.536 | -0.569 | -1.045 | 0 | 0.003 | 0.003 | 0 | 0.55 | 0.55 | 0.37 | 0.3 | 0.41 | 0.43 |
| 9 | 77806643 | rs12341208 | -0.855 | -0.642 | -0.13 | -1.103 | -0.006 | -0.07 | -0.07 | 0 | 0.13 | 0.31 | 0.07 | 0 | 0.08 | 0.14 |
| 9 | 77808519 | rs17060946 | -0.165 | -0.894 | 0.433 | -1.168 | -0.031 | -0.092 | -0.092 | 0 | 0.15 | 0.32 | 0.08 | 0.01 | 0.1 | 0.06 |
| 9 | 77809042 | rs10781268 | 0.668 | -0.672 | -0.103 | -1.361 | -0.014 | 0.02 | 0.02 | 0.02 | 0.37 | 0.23 | 0.58 | 0.71 | 0.46 | 0.52 |
| 9 | 77809075 | rs7030420 | -0.75 | -0.746 | 0.313 | -1.011 | -0.059 | -0.133 | -0.133 | 0.91 | 0.18 | 0.47 | 0.09 | 0.01 | 0.1 | 0.07 |
| 9 | 77812173 | rs10781269 | -0.269 | -0.319 | 0.072 | -0.482 | -0.022 | 0.009 | 0.009 | 0 | 0.3 | 0.49 | 0.36 | 0.57 | 0.27 | 0.42 |
| 9 | 77813220 | rs10283844 | -0.16 | -0.196 | -0.21 | -0.968 | 0 | -0.002 | -0.002 | 0 | 0.27 | 0.26 | 0.13 | 0.12 | 0.24 | 0.31 |
| 9 | 77813584 | rs1999462 | 0.275 | -0.436 | 0.502 | 0.104 | 0 | 0.236 | 0.236 | 0 | 0.05 | 0.18 | 0.02 | 0.23 | 0.01 | 0.21 |
| 9 | 77813720 | rs10781270 | -0.858 | -0.498 | 0.499 | 0.06 | -0.021 | 0.005 | 0.005 | 0 | 0.34 | 0.48 | 0.16 | 0.35 | 0.26 | 0.51 |
| 9 | 77814984 | rs10124785 | -0.37 | -0.641 | 0.021 | -0.321 | -0.006 | -0.005 | -0.005 | 0 | 0.59 | 0.53 | 0.58 | 0.72 | 0.58 | 0.68 |
| 9 | 77817068 | rs13294304 | 1.584 | -0.465 | -0.177 | -0.276 | -0.071 | -0.022 | -0.022 | 0 | 0.27 | 0.06 | 0.4 | 0.34 | 0.31 | 0.17 |
| 9 | 77825733 | rs1332349 | 0.354 | -0.558 | -0.038 | -0.316 | -0.001 | -0.015 | -0.015 | 2.518 | 0.28 | 0.27 | 0.16 | 0.13 | 0.26 | 0.31 |
| 9 | 77830570 | rs12550903 | 1.315 | -0.791 | -0.117 | -0.415 | -0.001 | -0.066 | -0.066 | 0.66 | 0.18 | 0.09 | 0.29 | 0.28 | 0.19 | 0.16 |
| 9 | 77830922 | rs1930069 | -0.515 | -0.82 | 0 | -0.51 | -0.002 | -0.004 | -0.004 | 0 | 0.45 | 0.48 | 0.24 | 0.38 | 0.39 | 0.63 |
| 9 | 77832575 | rs1332366 | 0.656 | -0.607 | -0.103 | -0.354 | -0.007 | 0.039 | 0.039 | 0 | 0.17 | 0.2 | 0.09 | 0.25 | 0.13 | 0.32 |
| 9 | 77833663 | rs1412461 | -1.771 | -0.59 | -0.231 | -0.382 | -0.039 | -0.002 | -0.002 | 2.942 | 0.73 | 0.86 | 0.62 | 0.66 | 0.7 | 0.84 |
| 9 | 77835651 | rs11559396 | 1.016 | -0.513 | -0.246 | -0.2 | -0.003 | -0.04 | -0.04 | 2.851 | 0.14 | 0.05 | 0.07 | 0.23 | 0.12 | 0.16 |
| 9 | 77846356 | rs6560430 | -0.053 | -0.655 | -0.039 | -0.446 | -0.001 | 0 | 0 | 0 | 0.57 | 0.51 | 0.51 | 0.59 | 0.5 | 0.58 |
| 9 | 77848630 | rs4629935 | -0.1 | -0.689 | 0.077 | -0.434 | 0.002 | -0.007 | -0.007 | 0 | 0.58 | 0.36 | 0.54 | 0.57 | 0.57 | 0.46 |
| 9 | 77848711 | rs4342676 | -0.1 | -0.632 | 0.077 | -0.434 | 0.001 | -0.007 | -0.007 | 0 | 0.58 | 0.36 | 0.54 | 0.57 | 0.57 | 0.46 |
| 9 | 77848790 | rs4614064 | 1.759 | -0.426 | -0.539 | -0.738 | -0.124 | -0.058 | -0.058 | 0 | 0.26 | 0.01 | 0.37 | 0.33 | 0.3 | 0.16 |
| 9 | 77849725 | rs7032088 | 0.236 | -0.559 | -0.292 | -0.772 | 0.002 | 0.03 | 0.03 | 0 | 0.48 | 0.33 | 0.5 | 0.36 | 0.5 | 0.47 |
| 9 | 77851696 | rs7852833 | 0.283 | -0.535 | -0.314 | -0.768 | 0 | 0.047 | 0.047 | 0 | 0.47 | 0.3 | 0.5 | 0.36 | 0.5 | 0.46 |
| 9 | 77852292 | rs11144325 | 1.557 | -0.349 | -0.885 | -0.503 | -0.112 | -0.073 | -0.073 | 0 | 0.27 | 0.03 | 0.51 | 0.37 | 0.34 | 0.16 |
| 9 | 77852659 | rs2095060 | -0.579 | -0.497 | -0.363 | -0.395 | 0.002 | -0.01 | -0.01 | 0 | 0.79 | 0.72 | 0.86 | 0.82 | 0.77 | 0.74 |
| 9 | 77853236 | rs2095057 | 0.745 | -0.582 | -0.373 | -0.453 | -0.008 | -0.036 | -0.036 | 0.015 | 0.1 | 0.15 | 0.04 | 0 | 0.06 | 0.09 |
| 9 | 77854544 | rs10869533 | 0.247 | -0.613 | -0.408 | -0.537 | -0.001 | -0.031 | -0.031 | 0 | 0.59 | 0.55 | 0.74 | 0.79 | 0.58 | 0.46 |
| 9 | 77854711 | rs10869534 | 0.978 | -0.53 | -0.417 | -0.688 | -0.003 | 0 | 0 | 0 | 0.16 | 0.15 | 0.1 | 0.1 | 0.2 | 0.24 |
| 9 | 77857783 | rs12000663 | 1.214 | 0.064 | -0.713 | -0.446 | -0.043 | -0.032 | -0.032 | 0 | 0.3 | 0.09 | 0.51 | 0.36 | 0.29 | 0.17 |
| 9 | 77858306 | rs1537295 | -0.243 | -0.017 | -0.623 | -0.518 | 0 | 0.028 | 0.028 | 0 | 0.5 | 0.64 | 0.37 | 0.61 | 0.53 | 0.6 |
| 9 | 77858578 | rs10119763 | -0.257 | -0.195 | -0.177 | -0.441 | -0.008 | -0.064 | -0.064 | 0.007 | 0.8 | 0.73 | 0.88 | 0.97 | 0.82 | 0.77 |
| 9 | 77860670 | rs4626675 | 0.49 | -0.104 | -0.461 | -0.005 | 0.005 | -0.003 | -0.003 | 0 | 0.47 | 0.32 | 0.64 | 0.47 | 0.53 | 0.29 |
| 9 | 77877793 | rs7854525 | -1.017 | -0.31 | -0.042 | -0.315 | -0.013 | -0.011 | -0.011 | 0.15 | 0.27 | 0.54 | 0.28 | 0.24 | 0.25 | 0.37 |
| 9 | 77881317 | rs7037086 | -0.71 | -0.408 | 0.355 | -0.377 | 0.005 | 0.019 | 0.019 | 0 | 0.16 | 0.29 | 0.1 | 0.16 | 0.12 | 0.18 |
| 9 | 77895520 | rs10869542 | -0.476 | -0.376 | 0.347 | -0.41 | 0.004 | -0.049 | -0.049 | 0 | 0.12 | 0.21 | 0.08 | 0 | 0.11 | 0.1 |
| 9 | 77905427 | rs7861441 | -0.269 | -0.398 | 0.065 | -0.416 | 0.001 | 0.02 | 0.02 | 0 | 0.11 | 0.07 | 0.17 | 0.06 | 0.13 | 0.19 |
| 9 | 77933300 | rs7028291 | -1.996 | -0.444 | -0.199 | -0.68 | 0.026 | 0.074 | 0.074 | 0 | 0.5 | 0.76 | 0.41 | 0.36 | 0.45 | 0.5 |
| 9 | 77936388 | rs4332197 | -1.754 | -0.457 | -0.211 | -0.679 | 0.014 | 0.057 | 0.057 | 0 | 0.48 | 0.76 | 0.41 | 0.36 | 0.45 | 0.5 |
| 9 | 77965572 | rs10115058 | 0.606 | -0.38 | -0.169 | -0.823 | -0.004 | -0.063 | -0.063 | 0 | 0.14 | 0.21 | 0.07 | 0 | 0.12 | 0.07 |
| 9 | 77976700 | rs2376734 | 0.648 | -0.274 | -0.128 | -0.366 | -0.018 | -0.004 | -0.004 | 0 | 0.11 | 0.03 | 0.14 | 0.13 | 0.13 | 0.16 |
| 9 | 77981257 | rs1887372 | -1.195 | -0.401 | 0.179 | -0.485 | -0.001 | -0.001 | -0.001 | 0 | 0.27 | 0.28 | 0.19 | 0.18 | 0.25 | 0.29 |
| 9 | 77988254 | rs11144406 | 0.68 | -0.282 | -0.087 | -0.333 | -0.019 | -0.017 | -0.017 | 0.668 | 0.11 | 0.03 | 0.14 | 0.18 | 0.13 | 0.16 |
| 9 | 77988304 | rs11144407 | -0.546 | -0.436 | 0.072 | -0.295 | -0.035 | 0.011 | 0.011 | 0.771 | 0.14 | 0.3 | 0.08 | 0.12 | 0.09 | 0.11 |
| 9 | 77991404 | rs1409937 | -2.618 | -0.49 | 0.15 | -0.334 | 0.007 | -0.006 | -0.006 | 0.198 | 0.52 | 0.74 | 0.44 | 0.48 | 0.47 | 0.56 |
| 9 | 77992822 | rs12004565 | -0.572 | -0.067 | -0.241 | -0.921 | -0.031 | 0.01 | 0.01 | 0.417 | 0.14 | 0.25 | 0.08 | 0.12 | 0.09 | 0.11 |
| 9 | 77996459 | rs2094372 | -0.428 | -0.076 | -0.307 | -0.955 | -0.013 | 0.023 | 0.023 | 0 | 0.22 | 0.11 | 0.24 | 0.19 | 0.26 | 0.27 |
| 9 | 77996819 | rs12377787 | -1.306 | -0.083 | -0.308 | -0.995 | -0.029 | 0.068 | 0.068 | 0 | 0.32 | 0.17 | 0.31 | 0.19 | 0.37 | 0.36 |
| 9 | 77999354 | rs7867522 | -1.415 | -0.236 | -0.084 | -0.685 | -0.005 | -0.005 | -0.005 | 0 | 0.36 | 0.4 | 0.32 | 0.31 | 0.35 | 0.38 |
| 9 | 78012662 | rs1327377 | -1.621 | -0.452 | -0.053 | -0.689 | -0.009 | -0.022 | -0.022 | 0.163 | 0.42 | 0.49 | 0.34 | 0.28 | 0.38 | 0.45 |
| 9 | 78012950 | rs10781285 | -0.827 | -0.325 | -0.397 | -0.587 | -0.03 | 0.064 | 0.064 | 0 | 0.27 | 0.09 | 0.24 | 0.18 | 0.28 | 0.35 |
| 9 | 78019028 | rs7019425 | -0.601 | -0.476 | 0.442 | -0.092 | -0.02 | 0.083 | 0.083 | 0.614 | 0.07 | 0.23 | 0.02 | 0.11 | 0.02 | 0.05 |
| 9 | 78027655 | rs17712651 | -0.579 | -0.547 | 0.224 | -0.377 | -0.012 | 0.057 | 0.057 | 0 | 0.16 | 0.06 | 0.13 | 0.01 | 0.17 | 0.07 |

| | | | | | | | | | | | | | | | | |
|---|---|---|---|---|---|---|---|---|---|---|---|---|---|---|---|---|
| 9 | 78028346 | rs489332 | -1.246 | -0.69 | 0.819 | -0.444 | -0.008 | -0.058 | -0.058 | 0.273 | 0.21 | 0.31 | 0.16 | 0.06 | 0.18 | 0.13 |
| 9 | 78029345 | rs569406 | -0.666 | -0.639 | 0.747 | -0.293 | 0.009 | 0.048 | 0.048 | 0 | 0.12 | 0.2 | 0.17 | 0.19 | 0.12 | 0.15 |
| 9 | 78029927 | rs1327378 | -0.377 | -0.73 | 1.087 | -0.303 | -0.009 | 0.101 | 0.101 | 0.998 | 0.07 | 0.3 | 0.02 | 0.09 | 0.02 | 0.04 |
| 9 | 78030021 | rs1327379 | -0.032 | -0.718 | 1.075 | -0.297 | -0.016 | 0.083 | 0.083 | 0.43 | 0.07 | 0.23 | 0.02 | 0.09 | 0.02 | 0.04 |
| 9 | 78032390 | rs477651 | -2.398 | -0.671 | 0.688 | -0.319 | 0.017 | 0 | 0 | 0.05 | 0.38 | 0.7 | 0.35 | 0.35 | 0.32 | 0.32 |
| 9 | 78038069 | rs11144428 | -0.253 | -0.477 | 0.43 | -0.336 | -0.004 | -0.009 | -0.009 | 0 | 0.1 | 0.07 | 0.16 | 0.19 | 0.12 | 0.14 |
| 9 | 78042927 | rs473299 | -1.59 | -0.346 | 0.425 | -0.765 | -0.035 | -0.044 | -0.044 | 0 | 0.24 | 0.5 | 0.16 | 0.11 | 0.19 | 0.1 |
| 9 | 78045277 | rs551421 | -1.007 | -0.205 | 0.337 | -0.743 | 0 | -0.034 | -0.034 | 0.016 | 0.18 | 0.18 | 0.13 | 0.01 | 0.18 | 0.07 |
| 9 | 78052850 | rs7854515 | -0.25 | -0.348 | 0.228 | -0.642 | 0.016 | 0.164 | 0.164 | 0.964 | 0.07 | 0.32 | 0.03 | 0.1 | 0.02 | 0.04 |
| 9 | 78056969 | rs13287590 | -0.575 | 0.024 | -0.84 | -0.896 | 0.006 | 0.043 | 0.043 | 0 | 0.13 | 0.06 | 0.07 | 0.01 | 0.11 | 0.05 |
| 9 | 78061074 | rs17713515 | -0.685 | 0.264 | -0.702 | -0.489 | -0.046 | 0.008 | 0.008 | 0 | 0.18 | 0.05 | 0.3 | 0.12 | 0.25 | 0.1 |
| 9 | 78066726 | rs891370 | -0.263 | -0.153 | -0.174 | -0.861 | -0.03 | -0.09 | -0.09 | 2.498 | 0.75 | 0.64 | 0.84 | 0.94 | 0.83 | 0.89 |
| 9 | 78068052 | rs13301187 | -0.024 | 0.063 | 0.161 | -1.139 | -0.005 | -0.108 | -0.108 | 0 | 0.19 | 0.37 | 0.24 | 0.01 | 0.17 | 0.1 |
| 9 | 78068102 | rs500859 | 0.581 | 0.489 | 0.061 | -0.949 | 0.027 | 0.002 | 0.002 | 0 | 0.26 | 0.31 | 0.34 | 0.33 | 0.38 | 0.28 |
| 9 | 78068222 | rs9314833 | -0.076 | -0.426 | -0.089 | -1.222 | 0.001 | -0.005 | -0.005 | 0.01 | 0.18 | 0.21 | 0.1 | 0.06 | 0.11 | 0.34 |
| 9 | 78069403 | rs578023 | -0.598 | -0.878 | 0.017 | -0.922 | -0.015 | -0.001 | -0.001 | 0 | 0.58 | 0.58 | 0.69 | 0.46 | 0.68 | 0.75 |
| 9 | 78069679 | rs516609 | 1.502 | -0.663 | 0.298 | -1.019 | 0.005 | -0.07 | -0.07 | 0 | 0.12 | 0.31 | 0.21 | 0 | 0.13 | 0.08 |
| 9 | 78070574 | rs2078096 | 0.905 | -0.302 | -0.132 | -1.123 | -0.023 | 0 | 0 | 0 | 0.3 | 0.23 | 0.39 | 0.32 | 0.44 | 0.23 |
| 9 | 78071910 | rs10123838 | 0.201 | -0.838 | 0.01 | -1.009 | 0.007 | 0.022 | 0.022 | 0 | 0.44 | 0.41 | 0.64 | 0.32 | 0.61 | 0.38 |
| 9 | 78080515 | rs1359806 | 0.125 | -0.838 | -0.039 | -1.016 | -0.004 | -0.015 | -0.015 | 0 | 0.45 | 0.54 | 0.64 | 0.31 | 0.6 | 0.38 |
| 9 | 78084508 | rs4744750 | 0.669 | -0.76 | -0.103 | -1.029 | 0.006 | -0.004 | -0.004 | 0.133 | 0.34 | 0.37 | 0.44 | 0.3 | 0.47 | 0.29 |
| 9 | 78085490 | rs2009306 | 0.679 | -0.76 | -0.103 | -1.029 | 0.007 | -0.004 | -0.004 | 2.079 | 0.34 | 0.37 | 0.44 | 0.3 | 0.47 | 0.29 |
| 9 | 78089813 | rs524346 | -0.198 | -0.758 | -0.297 | -1.056 | 0.047 | 0.128 | 0.128 | 0.016 | 0.5 | 0.84 | 0.67 | 0.31 | 0.61 | 0.51 |
| 9 | 78091442 | rs11144436 | 1.532 | -0.769 | -0.472 | -1.173 | -0.141 | -0.054 | -0.054 | 0 | 0.24 | 0.02 | 0.36 | 0.29 | 0.39 | 0.27 |
| 9 | 78092155 | rs7042130 | 0.842 | -1.072 | -0.301 | -1.293 | -0.024 | 0.011 | 0.011 | 0 | 0.32 | 0.27 | 0.45 | 0.3 | 0.5 | 0.3 |
| 9 | 78092361 | rs7028332 | 0.548 | -1.037 | -0.29 | -1.29 | 0.008 | -0.007 | -0.007 | 0 | 0.36 | 0.56 | 0.48 | 0.3 | 0.51 | 0.3 |
| 9 | 78096565 | rs969312 | 0.004 | -1.111 | -0.375 | -1.244 | 0.058 | 0.127 | 0.127 | 0 | 0.5 | 0.75 | 0.68 | 0.3 | 0.66 | 0.38 |
| 9 | 78099292 | rs522560 | -0.358 | -1.099 | -0.508 | -0.928 | 0.051 | 0.06 | 0.06 | 0 | 0.54 | 0.83 | 0.7 | 0.35 | 0.68 | 0.48 |
| 9 | 78108869 | rs1327369 | 0.206 | -0.979 | -0.344 | -0.987 | 0.002 | 0.009 | 0.009 | 0 | 0.46 | 0.47 | 0.63 | 0.33 | 0.59 | 0.51 |
| 9 | 78113669 | rs2778899 | -1.643 | -1.514 | -0.75 | -0.825 | 0.023 | 0.027 | 0.027 | 0 | 0.76 | 0.96 | 0.87 | 0.84 | 0.85 | 0.86 |
| 9 | 78113689 | rs12551990 | 1.824 | -1.533 | -0.752 | -0.9 | -0.085 | 0.001 | 0.001 | 0 | 0.27 | 0.1 | 0.39 | 0.3 | 0.43 | 0.28 |
| 9 | 78119359 | rs10512042 | 1.923 | -1.411 | -0.782 | -0.856 | -0.002 | -0.029 | -0.029 | 0 | 0.14 | 0.18 | 0.21 | 0.01 | 0.15 | 0.08 |
| 9 | 78120328 | rs10512043 | 1.401 | -1.309 | -0.814 | -0.568 | -0.034 | -0.103 | -0.103 | 0 | 0.16 | 0.05 | 0.18 | 0.41 | 0.2 | 0.12 |
| 9 | 78123330 | rs576367 | -0.878 | -1.036 | -0.517 | -0.768 | 0.008 | 0.002 | 0.002 | 0 | 0.6 | 0.86 | 0.68 | 0.44 | 0.65 | 0.73 |
| 9 | 78127980 | rs12156469 | 0.527 | -0.238 | -0.721 | -0.523 | -0.019 | 0.009 | 0.009 | 0 | 0.31 | 0.27 | 0.41 | 0.28 | 0.45 | 0.43 |
| 9 | 78141623 | rs928423 | 1.181 | -0.204 | -0.963 | -0.444 | -0.065 | -0.008 | -0.008 | 0.156 | 0.27 | 0.13 | 0.38 | 0.29 | 0.43 | 0.22 |
| 9 | 78143128 | rs1332518 | 1.181 | -0.214 | -0.963 | -0.441 | -0.075 | -0.008 | -0.008 | 0 | 0.27 | 0.12 | 0.38 | 0.29 | 0.42 | 0.22 |
| 9 | 78144832 | rs12000133 | 0.59 | -0.631 | -1.191 | -0.73 | 0.021 | 0.038 | 0.038 | 0 | 0.13 | 0.05 | 0.04 | 0.07 | 0.06 | 0.45 |
| 9 | 78146907 | rs12002365 | 0.249 | -0.441 | -1.224 | -0.754 | -0.006 | 0.042 | 0.042 | 0.004 | 0.14 | 0.05 | 0.08 | 0.07 | 0.11 | 0.45 |
| 9 | 78147051 | rs4745416 | -1.008 | -1.258 | -0.664 | -1.007 | -0.011 | -0.072 | -0.072 | 0 | 0.82 | 0.73 | 0.79 | 0.99 | 0.83 | 0.9 |
| 9 | 78155439 | rs10512045 | 0.357 | -1.236 | -1.269 | -1.071 | -0.013 | 0.078 | 0.078 | 0 | 0.19 | 0.05 | 0.07 | 0.07 | 0.16 | 0.46 |
| 9 | 78157110 | rs10781292 | -0.734 | -1.16 | -1.29 | -1.013 | 0.003 | 0.013 | 0.013 | 0 | 0.55 | 0.59 | 0.4 | 0.7 | 0.4 | 0.67 |
| 9 | 78158285 | rs11144480 | 0.975 | -1.476 | -1.29 | -0.222 | -0.006 | -0.001 | -0.001 | 0 | 0.17 | 0.17 | 0.08 | 0.15 | 0.06 | 0.1 |
| 9 | 78158939 | rs1010568 | 0.338 | -0.244 | -1.166 | 0.247 | 0.003 | 0.015 | 0.015 | 2.465 | 0.16 | 0.08 | 0.2 | 0.01 | 0.17 | 0.1 |
| 9 | 78163305 | rs10116662 | -0.135 | -0.135 | -1.428 | 0.424 | 0.004 | -0.028 | -0.028 | 2.132 | 0.19 | 0.13 | 0.23 | 0.48 | 0.18 | 0.11 |
| 9 | 78165213 | rs4451372 | -0.478 | -0.149 | -1.391 | 0.372 | 0.016 | -0.027 | -0.027 | 0 | 0.35 | 0.24 | 0.44 | 0.5 | 0.35 | 0.21 |
| 9 | 78171186 | rs10869587 | -0.349 | -0.143 | -1.542 | 0.285 | 0.002 | -0.149 | -0.149 | 0 | 0.29 | 0.06 | 0.16 | 0.4 | 0.22 | 0.15 |
| 9 | 78183488 | rs10435971 | 0.35 | -0.128 | -1.064 | 0.414 | -0.014 | -0.001 | -0.001 | 0 | 0.05 | 0.06 | 0.02 | 0.03 | 0.01 | 0.05 |
| 9 | 78185039 | rs12340637 | -0.161 | 0.357 | -0.274 | 0.364 | 0.012 | -0.036 | -0.036 | 1.429 | 0.12 | 0.33 | 0.17 | 0.04 | 0.12 | 0.35 |
| 9 | 78191069 | rs10869590 | -0.124 | 0.271 | -0.282 | 0.428 | 0.008 | -0.032 | -0.032 | 0 | 0.19 | 0.14 | 0.12 | 0.35 | 0.15 | 0.08 |
| 9 | 78193142 | rs7872039 | -0.424 | 0.509 | 0.011 | 0.394 | 0.012 | 0.002 | 0.002 | 0 | 0.52 | 0.35 | 0.42 | 0.49 | 0.53 | 0.34 |
| 9 | 78206328 | rs10869593 | 0.258 | 0.608 | -0.24 | 0.459 | 0.015 | -0.033 | -0.033 | 0 | 0.17 | 0.12 | 0.09 | 0.36 | 0.11 | 0.08 |
| 9 | 78206459 | rs7045152 | -1.227 | 0.664 | -0.008 | 0.423 | 0 | -0.005 | -0.005 | 0 | 0.66 | 0.77 | 0.6 | 0.58 | 0.65 | 0.71 |
| 9 | 78208290 | rs7019700 | -0.4 | 0.544 | -1.043 | 0.478 | 0.007 | -0.028 | -0.028 | 0 | 0.27 | 0.22 | 0.25 | 0.42 | 0.23 | 0.44 |
| 9 | 78212428 | rs7864568 | -1.18 | 0.763 | -1.276 | 0.414 | -0.01 | 0.007 | 0.007 | 0 | 0.66 | 0.59 | 0.59 | 0.58 | 0.67 | 0.71 |
| 9 | 78212566 | rs10118286 | -0.121 | 1.39 | -1.865 | -0.268 | -0.041 | 0.065 | 0.065 | 0 | 0.38 | 0.22 | 0.34 | 0.19 | 0.44 | 0.26 |
| 9 | 78215299 | rs11144509 | -0.413 | 1.038 | -1.816 | 0.33 | 0.001 | -0.001 | -0.001 | 0 | 0.09 | 0.11 | 0.16 | 0.07 | 0.12 | 0.34 |
| 9 | 78215586 | rs10114227 | -0.27 | 1.277 | -2.262 | 0.144 | -0.052 | 0.091 | 0.091 | 0 | 0.4 | 0.18 | 0.3 | 0.23 | 0.43 | 0.27 |
| 9 | 78218231 | rs11144511 | -0.337 | 1.412 | -2.392 | 0.571 | -0.106 | -0.108 | -0.108 | 0 | 0.33 | 0.12 | 0.51 | 0.57 | 0.46 | 0.42 |
| 9 | 78222406 | rs13300641 | -0.05 | -0.136 | -1.722 | -0.339 | -0.061 | -0.065 | -0.065 | 0.069 | 0.67 | 0.45 | 0.72 | 0.87 | 0.81 | 0.84 |
| 9 | 78223506 | rs12683806 | -0.033 | -0.092 | -2.016 | -0.34 | -0.009 | -0.011 | -0.011 | 0 | 0.07 | 0 | 0.04 | 0.04 | 0.07 | 0.04 |
| 9 | 78224432 | rs1328556 | -0.46 | -0.193 | -1.86 | -0.409 | -0.006 | -0.004 | -0.004 | 0.804 | 0.18 | 0.15 | 0.25 | 0.19 | 0.24 | 0.23 |
| 9 | 78228430 | rs11144516 | -0.82 | -0.227 | -1.544 | -0.507 | 0.006 | 0.001 | 0.001 | 0.376 | 0.2 | 0.21 | 0.27 | 0.19 | 0.28 | 0.27 |
| 9 | 78230339 | rs914660 | -0.803 | -0.47 | -1.346 | -0.634 | -0.054 | -0.072 | -0.072 | 0 | 0.64 | 0.49 | 0.78 | 0.88 | 0.79 | 0.83 |
| 9 | 78230797 | rs10869601 | -0.591 | -0.176 | -1.751 | -0.805 | -0.001 | -0.006 | -0.006 | 0.005 | 0.33 | 0.33 | 0.32 | 0.28 | 0.33 | 0.34 |
| 9 | 78239754 | rs11144526 | -0.057 | -0.087 | -1.758 | -0.825 | -0.039 | -0.031 | -0.031 | 0 | 0.17 | 0.07 | 0.24 | 0.19 | 0.22 | 0.25 |
| 9 | 78240443 | rs13290264 | 1.102 | -0.142 | -1.857 | -0.843 | -0.188 | -0.156 | -0.156 | 1.502 | 0.29 | 0.03 | 0.38 | 0.58 | 0.47 | 0.44 |
| 9 | 78243639 | rs7026536 | 1.198 | -0.162 | -1.744 | -0.851 | -0.014 | -0.021 | -0.021 | 0 | 0.15 | 0.17 | 0.06 | 0.09 | 0.09 | 0.11 |
| 9 | 78243859 | rs10869608 | 0.413 | -0.339 | -1.67 | -0.79 | 0 | 0.009 | 0.009 | 0 | 0.45 | 0.39 | 0.52 | 0.68 | 0.52 | 0.54 |

| | | | | | | | | | | | | | | | | |
|---|---|---|---|---|---|---|---|---|---|---|---|---|---|---|---|---|
| 9 | 78249553 | rs11144540 | 0.961 | -0.155 | -1.69 | -0.736 | -0.053 | -0.044 | -0.044 | 0 | 0.1 | 0.03 | 0.19 | 0.16 | 0.17 | 0.2 |
| 9 | 78250722 | rs17645740 | -0.06 | -0.55 | -1.297 | -0.834 | 0 | -0.009 | -0.009 | 0 | 0.11 | 0.12 | 0.07 | 0.04 | 0.12 | 0.08 |
| 9 | 78251232 | rs1328570 | -0.029 | -0.488 | -1.153 | -0.85 | -0.002 | -0.003 | -0.003 | 0 | 0.43 | 0.48 | 0.4 | 0.28 | 0.34 | 0.37 |
| 9 | 78251528 | rs1328569 | 0.704 | -0.187 | -1.205 | -0.645 | -0.043 | -0.046 | -0.046 | 0 | 0.1 | 0.03 | 0.18 | 0.17 | 0.15 | 0.19 |
| 9 | 78259163 | rs11144543 | 0.671 | -0.187 | -1.209 | -0.626 | -0.037 | -0.054 | -0.054 | 0 | 0.1 | 0.03 | 0.27 | 0.19 | 0.14 | 0.2 |
| 9 | 78259536 | rs7030860 | 0.336 | -0.526 | -0.663 | -1.037 | 0.037 | -0.058 | -0.058 | 0 | 0.11 | 0.44 | 0.08 | 0 | 0.09 | 0.04 |
| 9 | 78260121 | rs7031657 | 0.633 | -0.322 | -1.133 | -1.204 | 0.006 | 0.059 | 0.059 | 1.522 | 0.46 | 0.24 | 0.48 | 0.7 | 0.53 | 0.44 |
| 9 | 78260618 | rs1359999 | 0.506 | -0.349 | -0.976 | -0.836 | -0.013 | 0.012 | 0.012 | 0 | 0.1 | 0.01 | 0.06 | 0.03 | 0.1 | 0.06 |
| 9 | 78260714 | rs943860 | 0.547 | -0.364 | -0.949 | -0.903 | -0.012 | -0.012 | -0.012 | 0 | 0.82 | 0.86 | 0.68 | 0.81 | 0.77 | 0.68 |
| 9 | 78262577 | rs6560465 | -0.942 | -1.083 | -0.752 | -1.241 | -0.007 | -0.036 | -0.036 | 0 | 0.65 | 0.69 | 0.83 | 0.91 | 0.77 | 0.76 |
| 9 | 78263056 | rs12344781 | 0.452 | -1.176 | -0.271 | -1.322 | -0.007 | -0.048 | -0.048 | 0.182 | 0.09 | 0.34 | 0.06 | 0 | 0.06 | 0.04 |
| 9 | 78263400 | rs2094583 | 1.444 | -0.841 | -0.617 | -1.173 | -0.114 | -0.115 | -0.115 | 0 | 0.24 | 0.09 | 0.39 | 0.59 | 0.41 | 0.3 |
| 9 | 78263565 | rs3939133 | 1.049 | -0.83 | -0.503 | -1.215 | -0.031 | -0.03 | -0.03 | 0.905 | 0.07 | 0.01 | 0.17 | 0.13 | 0.12 | 0.25 |
| 9 | 78266575 | rs1328565 | -0.108 | -1.036 | -0.332 | -1.302 | -0.057 | -0.039 | -0.039 | 0 | 0.23 | 0.49 | 0.12 | 0.12 | 0.14 | 0.15 |
| 9 | 78267055 | rs10869620 | 0.991 | -1.061 | -0.531 | -1.355 | -0.005 | 0.016 | 0.016 | 0 | 0.1 | 0.02 | 0.06 | 0.03 | 0.1 | 0.06 |
| 9 | 78267078 | rs7038116 | -0.465 | -1.127 | -0.331 | -1.385 | -0.041 | -0.09 | -0.09 | 0.073 | 0.21 | 0.41 | 0.14 | 0.03 | 0.16 | 0.1 |
| 9 | 78267235 | rs1887796 | -1.294 | -1.132 | -0.325 | -1.475 | 0.016 | 0.01 | 0.01 | 0 | 0.65 | 0.73 | 0.83 | 0.91 | 0.77 | 0.75 |
| 9 | 78268985 | rs9696683 | -0.381 | -0.981 | -0.409 | -1.37 | -0.024 | -0.064 | -0.064 | 0.215 | 0.2 | 0.35 | 0.13 | 0.03 | 0.17 | 0.1 |
| 9 | 78271704 | rs1927986 | -0.109 | -0.917 | -0.56 | -1.297 | -0.002 | -0.004 | -0.004 | 0 | 0.45 | 0.53 | 0.53 | 0.64 | 0.58 | 0.41 |
| 9 | 78272025 | rs4745430 | -0.976 | -0.625 | -0.243 | -1.043 | 0.053 | 0.096 | 0.096 | 0 | 0.55 | 0.72 | 0.76 | 0.88 | 0.67 | 0.7 |
| 9 | 78272095 | rs4745431 | -0.183 | -0.222 | -0.624 | -0.865 | 0 | 0 | 0 | 0 | 0.41 | 0.45 | 0.64 | 0.74 | 0.6 | 0.59 |
| 9 | 78273456 | rs4442209 | -0.043 | 0.064 | -1.157 | -0.539 | -0.051 | 0.033 | 0.033 | 0 | 0.39 | 0.18 | 0.44 | 0.71 | 0.49 | 0.4 |
| 9 | 78274889 | rs1014766 | -0.777 | -0.232 | -0.334 | -0.613 | -0.019 | 0.006 | 0.006 | 0 | 0.31 | 0.47 | 0.29 | 0.29 | 0.26 | 0.4 |
| 9 | 78276017 | rs11144561 | 0.3 | 0.238 | -0.577 | -0.712 | -0.027 | -0.034 | -0.034 | 0.301 | 0.1 | 0.02 | 0.2 | 0.18 | 0.13 | 0.26 |
| 9 | 78277767 | rs2094588 | 0.036 | -0.087 | 0.051 | -1.265 | -0.007 | -0.071 | -0.071 | 0.11 | 0.13 | 0.27 | 0.06 | 0.01 | 0.09 | 0.07 |
| 9 | 78279942 | rs7857624 | 0.203 | -0.076 | 0.078 | -1.261 | -0.006 | -0.061 | -0.061 | 0 | 0.12 | 0.24 | 0.06 | 0.01 | 0.09 | 0.07 |
| 9 | 78282011 | rs11144565 | 0.142 | -0.023 | -0.227 | -1.263 | -0.004 | 0.011 | 0.011 | 0.004 | 0.08 | 0.05 | 0.05 | 0.05 | 0.09 | 0.05 |
| 9 | 78282779 | rs1571413 | -1.478 | -0.295 | -0.437 | -2.103 | 0.003 | -0.029 | -0.029 | 0 | 0.79 | 0.83 | 0.91 | 1 | 0.84 | 0.96 |
| 9 | 78282994 | rs1571414 | -0.675 | -0.381 | -0.341 | -2.171 | 0.012 | -0.004 | -0.004 | 0 | 0.72 | 0.78 | 0.87 | 1 | 0.77 | 0.92 |
| 9 | 78284116 | rs11144566 | 0.992 | -0.131 | -0.659 | -2.124 | -0.037 | -0.033 | -0.033 | 0.05 | 0.11 | 0.03 | 0.2 | 0.18 | 0.15 | 0.25 |
| 9 | 78284173 | rs943864 | -1.199 | -0.408 | -0.086 | -2.322 | -0.014 | -0.091 | -0.091 | 0.803 | 0.76 | 0.73 | 0.88 | 1 | 0.82 | 0.95 |
| 9 | 78284451 | rs943865 | -0.363 | -0.77 | 0.018 | -2.543 | -0.019 | -0.116 | -0.116 | 0.625 | 0.63 | 0.49 | 0.84 | 0.99 | 0.75 | 0.88 |
| 9 | 78286685 | rs11144569 | 1.598 | -0.16 | -0.595 | -2.767 | -0.008 | 0.014 | 0.014 | 0 | 0.13 | 0.07 | 0.11 | 0.02 | 0.18 | 0.12 |
| 9 | 78286736 | rs1410579 | 1.911 | -0.327 | -0.786 | -2.572 | 0.018 | 0.004 | 0.004 | 0 | 0.11 | 0.02 | 0.05 | 0.09 | 0.07 | 0.15 |
| 9 | 78289296 | rs7026438 | 0.884 | -0.138 | -0.337 | -2.422 | 0.002 | 0.002 | 0.002 | 0 | 0.29 | 0.38 | 0.44 | 0.64 | 0.33 | 0.38 |
| 9 | 78289935 | rs7031238 | -0.377 | -0.337 | -0.362 | -2.477 | -0.011 | 0.026 | 0.026 | 0 | 0.6 | 0.6 | 0.36 | 0.18 | 0.52 | 0.37 |
| 9 | 78291861 | rs7866089 | -0.044 | -0.39 | -0.367 | -2.584 | 0.019 | 0.067 | 0.067 | 0.144 | 0.46 | 0.68 | 0.38 | 0.2 | 0.38 | 0.37 |
| 9 | 78294845 | rs10869631 | -0.925 | -0.49 | -0.247 | -2.602 | 0.002 | 0.015 | 0.015 | 0.563 | 0.65 | 0.64 | 0.68 | 0.82 | 0.68 | 0.69 |
| 9 | 78296308 | rs12001533 | 0.854 | -0.371 | -0.188 | -2.49 | -0.002 | -0.048 | -0.048 | 0 | 0.11 | 0.21 | 0.07 | 0.01 | 0.09 | 0.07 |
| 9 | 78301522 | rs7856004 | 0.815 | -0.089 | -1.021 | -2.599 | -0.026 | 0.044 | 0.044 | 0 | 0.33 | 0.12 | 0.46 | 0.77 | 0.34 | 0.46 |
| 9 | 78302645 | rs4745436 | 1.487 | -0.24 | -0.912 | -2.612 | -0.015 | -0.018 | -0.018 | 0 | 0.11 | 0.06 | 0.21 | 0.15 | 0.15 | 0.26 |
| 9 | 78306320 | rs1013792 | 0.033 | -0.365 | -0.668 | -2.74 | 0.001 | -0.035 | -0.035 | 0 | 0.45 | 0.43 | 0.54 | 0.83 | 0.44 | 0.54 |
| 9 | 78306554 | rs1328549 | 1.31 | -0.372 | -0.612 | -2.733 | -0.03 | -0.047 | -0.047 | 0 | 0.11 | 0.25 | 0.05 | 0.03 | 0.05 | 0.06 |
| 9 | 78306600 | rs1410569 | 2.395 | -0.192 | -1.22 | -2.614 | -0.102 | -0.101 | -0.101 | 0 | 0.16 | 0.02 | 0.38 | 0.65 | 0.27 | 0.25 |
| 9 | 78307440 | rs17061560 | 1.722 | -0.346 | -1.196 | -2.701 | 0.02 | 0.019 | 0.019 | 0 | 0.17 | 0.02 | 0.06 | 0.12 | 0.11 | 0.22 |
| 9 | 78308057 | rs4745437 | -0.191 | 0.104 | -0.772 | -2.405 | -0.006 | -0.122 | -0.122 | 0.46 | 0.55 | 0.45 | 0.75 | 0.98 | 0.58 | 0.79 |
| 9 | 78308605 | rs1359998 | -1.822 | 0.106 | -0.792 | -2.483 | 0.016 | 0.021 | 0.021 | 2.416 | 0.77 | 0.82 | 0.9 | 1 | 0.82 | 0.95 |
| 9 | 78308750 | rs7863282 | 0.372 | 0.095 | -0.56 | -2.127 | 0 | -0.048 | -0.048 | 0.27 | 0.2 | 0.23 | 0.14 | 0.02 | 0.23 | 0.16 |
| 9 | 78309287 | rs1328550 | -0.271 | 0.071 | -0.405 | -1.4 | -0.011 | -0.016 | -0.016 | 0 | 0.39 | 0.35 | 0.34 | 0.3 | 0.31 | 0.54 |
| 9 | 78313509 | rs10746980 | -0.326 | 0.436 | -0.486 | -0.872 | -0.002 | 0 | 0 | 0 | 0.59 | 0.66 | 0.5 | 0.39 | 0.56 | 0.57 |
| 9 | 78313939 | rs1328554 | -0.642 | 1.144 | -0.417 | -0.374 | 0.007 | -0.045 | -0.045 | 0 | 0.64 | 0.82 | 0.74 | 0.46 | 0.71 | 0.61 |
| 9 | 78314854 | rs7874982 | 1.331 | 1.122 | -0.786 | -0.681 | -0.018 | -0.052 | -0.052 | 0.01 | 0.11 | 0.02 | 0.08 | 0.18 | 0.11 | 0.21 |
| 9 | 78315752 | rs7044203 | 1.041 | 1.24 | -0.328 | -1.164 | 0.02 | -0.004 | -0.004 | 0.476 | 0.1 | 0.18 | 0.24 | 0.08 | 0.2 | 0.09 |
| 9 | 78322176 | rs518417 | -1.53 | 0.68 | -0.067 | -1.432 | -0.017 | -0.082 | -0.082 | 4.676 | 0.8 | 0.74 | 0.92 | 0.99 | 0.85 | 0.96 |
| 9 | 78325082 | rs693362 | -2.108 | 0.7 | -0.505 | -1.425 | 0.004 | 0.058 | 0.058 | 0.052 | 0.84 | 0.91 | 0.93 | 1 | 0.85 | 0.97 |
| 9 | 78328761 | rs2479837 | -0.385 | 0.428 | -0.113 | -1.392 | 0.013 | 0.086 | 0.086 | 0.08 | 0.55 | 0.67 | 0.64 | 0.85 | 0.57 | 0.73 |
| 9 | 78330608 | rs11144589 | 1.24 | 0.223 | -0.255 | -1.391 | 0.036 | 0.047 | 0.047 | 0.255 | 0.2 | 0.08 | 0.07 | 0.07 | 0.11 | 0.17 |
| 9 | 78331204 | rs795085 | -1.401 | 0.442 | -0.2 | -1.423 | -0.003 | 0.001 | 0.001 | 0 | 0.76 | 0.72 | 0.69 | 0.92 | 0.69 | 0.87 |
| 9 | 78331945 | rs473295 | 0.517 | 0.643 | 0.021 | -1.329 | -0.002 | -0.001 | -0.001 | 0 | 0.39 | 0.43 | 0.28 | 0.71 | 0.29 | 0.47 |
| 9 | 78332861 | rs4744765 | 1.127 | 0.736 | 0.23 | -1.321 | 0.088 | -0.02 | -0.02 | 0 | 0.06 | 0.15 | 0.26 | 0.01 | 0.18 | 0.05 |
| 9 | 78332998 | rs2986388 | 0.069 | 0.765 | -0.287 | -1.324 | -0.003 | 0.019 | 0.019 | 0.157 | 0.46 | 0.6 | 0.54 | 0.72 | 0.47 | 0.51 |
| 9 | 78333685 | rs489571 | 1.221 | 0.797 | -1.228 | -0.957 | -0.018 | -0.078 | -0.078 | 0 | 0.21 | 0.08 | 0.16 | 0.64 | 0.19 | 0.32 |
| 9 | 78336697 | rs17061627 | 0.112 | 0.761 | -0.507 | -0.689 | 0.003 | -0.007 | -0.007 | 0 | 0.14 | 0.1 | 0.1 | 0.13 | 0.13 | 0.21 |
| 9 | 78339768 | rs2149385 | 0.008 | 0.773 | -0.328 | -0.57 | -0.006 | 0.014 | 0.014 | 0 | 0.66 | 0.66 | 0.62 | 0.79 | 0.59 | 0.66 |
| 9 | 78342898 | rs487502 | -0.039 | 0.383 | -0.189 | -0.82 | -0.005 | -0.035 | -0.035 | 6.789 | 0.3 | 0.36 | 0.34 | 0.06 | 0.28 | 0.23 |
| 9 | 78347251 | rs2479843 | -0.266 | 0.483 | -0.173 | -0.645 | 0.027 | 0.026 | 0.026 | 0.198 | 0.36 | 0.21 | 0.32 | 0.3 | 0.32 | 0.37 |
| 9 | 78348570 | rs10869637 | 0.441 | 0.512 | -0.165 | -0.623 | -0.001 | -0.002 | -0.002 | 0 | 0.13 | 0.12 | 0.13 | 0.22 | 0.17 | 0.19 |
| 9 | 78350057 | rs7025497 | 0.745 | 0.751 | -0.282 | -0.808 | 0.011 | 0.002 | 0.002 | 1.384 | 0.41 | 0.65 | 0.33 | 0.63 | 0.32 | 0.47 |
| 9 | 78350484 | rs561412 | -1.143 | 0.821 | -0.381 | -0.82 | -0.027 | 0.042 | 0.042 | 0 | 0.8 | 0.9 | 0.74 | 0.94 | 0.74 | 0.88 |

| | | | | | | | | | | | | | | | | |
|---|---|---|---|---|---|---|---|---|---|---|---|---|---|---|---|---|
| 9 | 78351260 | rs11144603 | 1.247 | 0.733 | 0.215 | -0.859 | 0 | -0.003 | -0.003 | 0 | 0.35 | 0.29 | 0.28 | 0.66 | 0.29 | 0.46 |
| 9 | 78354076 | rs1359715 | 0.964 | 0.736 | 0.103 | -0.941 | 0.033 | 0.079 | 0.079 | 0.489 | 0.23 | 0.09 | 0.21 | 0.06 | 0.15 | 0.19 |
| 9 | 78355920 | rs1328560 | 0.912 | 0.755 | 0.212 | -0.932 | 0.031 | 0.056 | 0.056 | 0.409 | 0.23 | 0.14 | 0.22 | 0.06 | 0.15 | 0.19 |
| 9 | 78368000 | rs7041935 | -0.167 | 0.77 | 0.253 | -0.948 | 0.046 | -0.012 | -0.012 | 2.026 | 0.07 | 0.32 | 0.09 | 0 | 0.08 | 0.04 |
| 9 | 78369830 | rs1751250 | 0.029 | 0.825 | 0.131 | -0.92 | -0.002 | 0.018 | 0.018 | 0.159 | 0.36 | 0.25 | 0.38 | 0.29 | 0.36 | 0.39 |
| 9 | 78369864 | rs1753320 | 0.005 | 0.891 | 0.124 | -0.926 | 0.009 | 0.018 | 0.018 | 0.133 | 0.35 | 0.25 | 0.33 | 0.27 | 0.32 | 0.36 |
| 9 | 78369986 | rs1753321 | 0.042 | 0.828 | 0.14 | -0.914 | -0.002 | 0.018 | 0.018 | 0.159 | 0.36 | 0.25 | 0.38 | 0.3 | 0.36 | 0.39 |
| 9 | 78371410 | rs528408 | -0.272 | 0.924 | 0.209 | -0.279 | 0.015 | -0.028 | -0.028 | 0 | 0.73 | 0.58 | 0.65 | 0.83 | 0.66 | 0.83 |
| 9 | 78372907 | rs7860318 | 1.289 | 1 | -0.325 | -0.151 | -0.012 | -0.03 | -0.03 | 0 | 0.09 | 0.04 | 0.07 | 0.22 | 0.1 | 0.15 |
| 9 | 78391161 | rs1926728 | 0.417 | 0.802 | -0.048 | -0.16 | -0.003 | -0.02 | -0.02 | 0 | 0.6 | 0.29 | 0.56 | 0.53 | 0.55 | 0.59 |
| 9 | 78397226 | rs1359716 | -0.757 | 0.562 | -0.472 | -0.574 | 0.013 | 0.033 | 0.033 | 0 | 0.75 | 0.87 | 0.88 | 1 | 0.81 | 0.9 |
| 9 | 78407484 | rs4745444 | 0.835 | 0.816 | -0.389 | -0.338 | 0.014 | -0.009 | -0.009 | 0.006 | 0.49 | 0.2 | 0.45 | 0.53 | 0.42 | 0.53 |
| 9 | 78412423 | rs1409689 | 0.788 | 0.813 | -0.545 | -0.562 | 0.025 | -0.004 | -0.004 | 0 | 0.49 | 0.16 | 0.47 | 0.53 | 0.41 | 0.52 |
| 9 | 78416165 | rs7856261 | 0.743 | 0.869 | -0.272 | -0.408 | 0.012 | 0.072 | 0.072 | 0.251 | 0.09 | 0.21 | 0.15 | 0.18 | 0.09 | 0.17 |
| 9 | 78416519 | rs10869646 | 1.116 | 0.815 | -0.41 | -0.404 | -0.008 | -0.017 | -0.017 | 0 | 0.08 | 0.05 | 0.14 | 0.21 | 0.09 | 0.17 |
| 9 | 78416549 | rs11144624 | 1.516 | 0.741 | -0.582 | -0.444 | -0.031 | -0.099 | -0.099 | 0 | 0.26 | 0.07 | 0.27 | 0.45 | 0.26 | 0.33 |
| 9 | 78430918 | rs2025767 | -0.567 | 0.773 | -0.373 | -0.447 | 0 | -0.007 | -0.007 | 0.356 | 0.73 | 0.71 | 0.76 | 0.94 | 0.77 | 0.87 |
| 9 | 78438434 | rs1326914 | 0.216 | 0.741 | -0.337 | -0.279 | 0.005 | 0.032 | 0.032 | 0.202 | 0.11 | 0.18 | 0.27 | 0.27 | 0.13 | 0.22 |
| 9 | 78439746 | rs11793539 | 1.406 | 0.472 | -0.617 | -0.486 | 0.023 | 0.056 | 0.056 | 2.709 | 0.18 | 0.04 | 0.08 | 0.03 | 0.12 | 0.12 |
| 9 | 78450924 | rs6560469 | 0.018 | 0.883 | -0.373 | -0.03 | -0.004 | -0.006 | -0.006 | 0 | 0.78 | 0.77 | 0.65 | 0.76 | 0.76 | 0.73 |
| 9 | 78452816 | rs4745450 | 0.789 | 0.916 | -0.594 | 0.56 | -0.021 | -0.038 | -0.038 | 0 | 0.09 | 0.03 | 0.09 | 0.26 | 0.13 | 0.15 |
| 9 | 78454330 | rs10869656 | -0.695 | 0.907 | -0.559 | 0.539 | -0.011 | -0.035 | -0.035 | 1.054 | 0.21 | 0.21 | 0.36 | 0.51 | 0.26 | 0.38 |
| 9 | 78456306 | rs7027513 | -0.232 | 0.827 | -0.329 | 0.643 | 0.004 | 0.046 | 0.046 | 0.375 | 0.12 | 0.23 | 0.27 | 0.25 | 0.13 | 0.23 |
| 9 | 78458130 | rs10869657 | 0.589 | 0.836 | -0.587 | 0.554 | -0.002 | -0.012 | -0.012 | 0 | 0.31 | 0.27 | 0.29 | 0.43 | 0.29 | 0.36 |
| 9 | 78458935 | rs1831037 | -0.784 | 0.911 | -0.516 | 1.324 | 0 | 0.015 | 0.015 | 0.2 | 0.12 | 0.17 | 0.27 | 0.25 | 0.13 | 0.22 |
| 9 | 78465639 | rs2148588 | -0.252 | 0.673 | -0.442 | 1.005 | -0.001 | 0.008 | 0.008 | 0 | 0.11 | 0.14 | 0.09 | 0.26 | 0.12 | 0.15 |
| 9 | 78473260 | rs7030848 | -0.239 | 0.525 | 0.077 | 1.245 | -0.009 | 0.066 | 0.066 | 0 | 0.16 | 0.3 | 0.19 | 0.26 | 0.14 | 0.18 |
| 9 | 78479072 | rs11144655 | 0.641 | 0.364 | 0.143 | 0.283 | 0.009 | 0.016 | 0.016 | 0 | 0.44 | 0.27 | 0.35 | 0.4 | 0.42 | 0.43 |
| 9 | 78479975 | rs1326904 | -0.398 | 0.475 | 0.309 | 0.098 | -0.005 | 0.012 | 0.012 | 0 | 0.46 | 0.63 | 0.5 | 0.56 | 0.43 | 0.47 |
| 9 | 78484125 | rs7025650 | 0.567 | 0.077 | 0.38 | -0.525 | 0.006 | -0.034 | -0.034 | 0.051 | 0.32 | 0.28 | 0.13 | 0.02 | 0.22 | 0.15 |
| 9 | 78485374 | rs1934737 | -0.649 | 0.097 | 0.403 | -0.348 | -0.002 | -0.003 | -0.003 | 0 | 0.55 | 0.6 | 0.4 | 0.48 | 0.46 | 0.44 |
| 9 | 78485754 | rs1934735 | 0.421 | 0.159 | 0.407 | -0.234 | 0.001 | 0.038 | 0.038 | 0 | 0.23 | 0.26 | 0.27 | 0.46 | 0.23 | 0.29 |
| 9 | 78489484 | rs1544181 | 0.848 | 0.279 | 0.53 | -0.275 | 0.007 | 0.014 | 0.014 | 0 | 0.35 | 0.34 | 0.43 | 0.47 | 0.38 | 0.5 |
| 9 | 78496742 | rs10869664 | 0.211 | 0.309 | 0.046 | -0.218 | -0.003 | -0.019 | -0.019 | 0 | 0.22 | 0.19 | 0.32 | 0.43 | 0.23 | 0.28 |
| 9 | 78497134 | rs12156495 | 0.826 | 0.17 | -0.048 | -1.223 | -0.008 | 0.011 | 0.011 | 0 | 0.09 | 0.07 | 0.12 | 0.02 | 0.14 | 0.03 |
| 9 | 78497266 | rs7862766 | 0.488 | 0.125 | -0.013 | -1.235 | 0.007 | -0.039 | -0.039 | 0 | 0.1 | 0.19 | 0.11 | 0.02 | 0.13 | 0.03 |
| 9 | 78502734 | rs2377423 | -0.117 | -0.191 | -0.469 | -1.167 | 0.026 | 0.083 | 0.083 | 0 | 0.69 | 0.83 | 0.86 | 0.98 | 0.77 | 0.85 |
| 9 | 78504729 | rs3814115 | -0.283 | -0.761 | -0.58 | -1.028 | 0.058 | 0.055 | 0.055 | 0.547 | 0.48 | 0.65 | 0.37 | 0.33 | 0.32 | 0.44 |
| 9 | 78506187 | rs7040769 | 0.012 | -1.098 | -0.614 | -1.539 | -0.061 | -0.157 | -0.157 | 0 | 0.27 | 0.53 | 0.11 | 0 | 0.14 | 0.05 |
| 9 | 78509414 | rs2050833 | -0.401 | -0.249 | -1.23 | -1.237 | -0.016 | -0.003 | -0.003 | 0.013 | 0.76 | 0.85 | 0.76 | 0.22 | 0.78 | 0.51 |
| 9 | 78510036 | rs7027130 | 0.438 | -0.305 | -0.578 | -1.179 | -0.017 | -0.083 | -0.083 | 0.245 | 0.13 | 0.32 | 0.05 | 0 | 0.08 | 0.02 |
| 9 | 78511363 | rs7031971 | -0.371 | 0.706 | -1.158 | -1.247 | 0.015 | 0.236 | 0.236 | 0 | 0.53 | 0.84 | 0.43 | 0.11 | 0.44 | 0.29 |
| 9 | 78512019 | rs4745464 | -0.347 | 1.032 | -0.994 | -1.056 | -0.062 | 0.174 | 0.174 | 0 | 0.58 | 0.83 | 0.47 | 0.11 | 0.51 | 0.34 |
| 9 | 78515871 | rs4744774 | 0.986 | 0.965 | -0.876 | -1.16 | -0.003 | -0.046 | -0.046 | 0 | 0.14 | 0.15 | 0.21 | 0.02 | 0.12 | 0.15 |
| 9 | 78518859 | rs4744775 | -0.874 | 1.081 | -1.383 | -1.06 | -0.033 | 0.015 | 0.015 | 0 | 0.69 | 0.87 | 0.59 | 0.22 | 0.7 | 0.44 |
| 9 | 78518999 | rs10869665 | 0.845 | 1.122 | -1.215 | -1.144 | -0.014 | -0.064 | -0.064 | 0 | 0.36 | 0.37 | 0.18 | 0.06 | 0.28 | 0.15 |
| 9 | 78527549 | rs1006279 | -0.276 | 1.3 | -1.257 | -1.078 | -0.069 | 0.17 | 0.17 | 0 | 0.6 | 0.84 | 0.47 | 0.1 | 0.52 | 0.38 |
| 9 | 78529571 | rs1006280 | 0.486 | 0.886 | -1.102 | -0.822 | -0.012 | -0.07 | -0.07 | 0 | 0.38 | 0.39 | 0.19 | 0.05 | 0.29 | 0.15 |
| 9 | 78536089 | rs4554552 | 0.09 | 0.984 | -1.653 | 0.58 | -0.032 | 0.009 | 0.009 | 0 | 0.71 | 0.86 | 0.59 | 0.21 | 0.71 | 0.5 |
| 9 | 78546389 | rs2377426 | -0.639 | 0.805 | -0.671 | 1.221 | -0.001 | 0.036 | 0.036 | 2.311 | 0.28 | 0.32 | 0.24 | 0.51 | 0.25 | 0.4 |
| 9 | 78546411 | rs10120870 | -0.826 | 1.517 | -1.356 | 0.375 | -0.053 | 0.079 | 0.079 | 0 | 0.25 | 0.09 | 0.16 | 0.07 | 0.28 | 0.14 |
| 9 | 78548556 | rs10869666 | 0.057 | 1.15 | -1.095 | -0.161 | 0.003 | 0.005 | 0.005 | 0 | 0.24 | 0.28 | 0.23 | 0.33 | 0.25 | 0.29 |
| 9 | 78551222 | rs10869667 | -0.694 | 0.018 | -1.144 | -0.483 | 0.015 | 0.014 | 0.014 | 0 | 0.79 | 0.87 | 0.94 | 0.87 | 0.88 | 0.79 |
| 9 | 78556095 | rs11144690 | 1.17 | 0.148 | -1.12 | -0.393 | -0.002 | 0.023 | 0.023 | 0 | 0.07 | 0.08 | 0.05 | 0.23 | 0.03 | 0.12 |
| 9 | 78557137 | rs4584211 | 0.04 | -0.175 | -0.754 | -0.937 | -0.003 | -0.008 | -0.008 | 3.6 | 0.57 | 0.52 | 0.42 | 0.36 | 0.58 | 0.46 |
| 9 | 78562669 | rs10869672 | 0.361 | -0.523 | -0.601 | -1.34 | 0.019 | 0.019 | 0.019 | 0.744 | 0.51 | 0.46 | 0.77 | 0.85 | 0.65 | 0.67 |
| 9 | 78567264 | rs4504704 | -0.006 | -0.566 | -0.195 | -1.207 | -0.016 | -0.02 | -0.02 | 0 | 0.69 | 0.69 | 0.86 | 0.85 | 0.8 | 0.73 |
| 9 | 78570116 | rs17061821 | 0.748 | -0.576 | -0.19 | -1.162 | -0.029 | -0.045 | -0.045 | 0 | 0.11 | 0.23 | 0.05 | 0.01 | 0.05 | 0.05 |
| 9 | 78570960 | rs2066188 | 1.839 | -0.104 | -0.221 | -0.863 | 0.006 | 0.077 | 0.077 | 0.097 | 0.1 | 0.09 | 0.42 | 0.51 | 0.2 | 0.29 |
| 9 | 78571504 | rs1340503 | 1.57 | -0.316 | -0.256 | -1.228 | -0.003 | -0.001 | -0.001 | 0 | 0.31 | 0.25 | 0.21 | 0.32 | 0.29 | 0.31 |
| 9 | 78573139 | rs11144698 | 1.474 | -0.322 | -0.129 | -1.229 | 0.001 | 0 | 0 | 0 | 0.32 | 0.34 | 0.22 | 0.33 | 0.29 | 0.31 |
| 9 | 78575644 | rs4745478 | -1.257 | -0.406 | -0.073 | -1.077 | -0.004 | 0 | 0 | 0 | 0.84 | 0.8 | 0.93 | 0.83 | 0.9 | 0.81 |
| 9 | 78576644 | rs1340510 | 0.351 | -0.42 | -0.173 | -1.198 | -0.024 | -0.054 | -0.054 | 0 | 0.69 | 0.53 | 0.81 | 0.83 | 0.73 | 0.76 |
| 9 | 78579094 | rs10781320 | 1.057 | -0.301 | -0.229 | -1.123 | 0.015 | 0.058 | 0.058 | 0 | 0.46 | 0.36 | 0.67 | 0.82 | 0.54 | 0.62 |
| 9 | 78579268 | rs10781321 | 1.682 | -1.058 | -0.51 | -0.553 | 0.006 | 0.074 | 0.074 | 0 | 0.09 | 0.09 | 0.42 | 0.5 | 0.2 | 0.28 |
| 9 | 78580233 | rs10869673 | 0.44 | -1.149 | -0.503 | -0.639 | 0 | -0.002 | -0.002 | 2.865 | 0.61 | 0.52 | 0.41 | 0.33 | 0.57 | 0.44 |
| 9 | 78581439 | rs914256 | -0.163 | -0.962 | -0.878 | -0.617 | -0.002 | -0.003 | -0.003 | 0 | 0.72 | 0.65 | 0.83 | 0.82 | 0.77 | 0.73 |
| 9 | 78583187 | rs10512052 | 1.885 | -0.582 | -1.008 | -0.331 | -0.053 | -0.017 | -0.017 | 0 | 0.08 | 0.02 | 0.41 | 0.5 | 0.2 | 0.25 |
| 9 | 78583414 | rs11144702 | 0.343 | -1.635 | -0.646 | -1.441 | -0.002 | -0.007 | -0.007 | 2.191 | 0.6 | 0.56 | 0.79 | 0.82 | 0.71 | 0.67 |

| | | | | | | | | | | | | | | | | |
|---|---|---|---|---|---|---|---|---|---|---|---|---|---|---|---|---|
| 9 | 78590288 | rs11144705 | 0.192 | -1.583 | -0.733 | -1.258 | 0.017 | 0 | 0 | 0 | 0.17 | 0.12 | 0.08 | 0 | 0.1 | 0.09 |
| 9 | 78590669 | rs10781322 | -0.02 | -1.048 | -0.373 | -0.953 | 0.022 | 0.028 | 0.028 | 0 | 0.72 | 0.61 | 0.44 | 0.39 | 0.66 | 0.53 |
| 9 | 78594609 | rs2153496 | 0.364 | -1.338 | -0.344 | -1.747 | -0.005 | -0.01 | -0.01 | 0 | 0.53 | 0.5 | 0.73 | 0.72 | 0.64 | 0.62 |
| 9 | 78596109 | rs7847011 | 0.085 | -1.79 | -0.227 | -1.895 | -0.017 | -0.024 | -0.024 | 0 | 0.69 | 0.64 | 0.8 | 0.84 | 0.74 | 0.72 |
| 9 | 78598605 | rs1340508 | 1.076 | -1.747 | -0.833 | -2.11 | 0.01 | 0.014 | 0.014 | 1.149 | 0.2 | 0.2 | 0.52 | 0.66 | 0.29 | 0.36 |
| 9 | 78599133 | rs6560480 | 0.013 | -2.163 | -0.735 | -2.237 | -0.033 | -0.05 | -0.05 | 0 | 0.63 | 0.48 | 0.76 | 0.83 | 0.71 | 0.65 |
| 9 | 78608872 | rs2789610 | 1.298 | -1.904 | -0.722 | -2.102 | 0.001 | 0.008 | 0.008 | 0.361 | 0.17 | 0.16 | 0.48 | 0.66 | 0.23 | 0.33 |
| 9 | 78613185 | rs1416556 | -0.153 | -2.126 | -0.77 | -2.206 | -0.052 | -0.049 | -0.049 | 0 | 0.32 | 0.52 | 0.22 | 0.17 | 0.24 | 0.31 |
| 9 | 78615371 | rs6560483 | 0.37 | -1.56 | -0.609 | -1.972 | -0.002 | -0.002 | -0.002 | 0 | 0.41 | 0.39 | 0.27 | 0.18 | 0.45 | 0.26 |
| 9 | 78619829 | rs12000040 | 0.224 | -1.413 | -0.769 | -1.934 | -0.001 | 0.001 | 0.001 | 0 | 0.46 | 0.45 | 0.29 | 0.34 | 0.48 | 0.38 |
| 9 | 78621536 | rs2789607 | 0.398 | -0.43 | -0.735 | -0.833 | 0.006 | 0.017 | 0.017 | 0.171 | 0.18 | 0.17 | 0.48 | 0.66 | 0.22 | 0.34 |
| 9 | 78622348 | rs1416553 | -0.279 | -0.516 | -0.945 | -0.699 | 0.003 | 0.004 | 0.004 | 0 | 0.17 | 0.15 | 0.11 | 0.11 | 0.12 | 0.13 |
| 9 | 78626324 | rs17061934 | -0.034 | -0.325 | -1.21 | -0.973 | 0.016 | 0.025 | 0.025 | 0 | 0.16 | 0.09 | 0.06 | 0 | 0.09 | 0.07 |
| 9 | 78631286 | rs1538505 | -0.37 | 0.053 | -0.933 | -0.268 | -0.025 | 0.002 | 0.002 | 1.581 | 0.31 | 0.19 | 0.54 | 0.75 | 0.37 | 0.49 |
| 9 | 78636676 | rs1339251 | -0.426 | -0.46 | -0.385 | -0.256 | -0.025 | -0.047 | -0.047 | 0 | 0.1 | 0.13 | 0.02 | 0 | 0.03 | 0.02 |
| 9 | 78639196 | rs10781324 | -0.868 | -0.194 | -0.458 | 0.324 | 0 | 0.001 | 0.001 | 0.219 | 0.19 | 0.19 | 0.13 | 0.3 | 0.22 | 0.21 |
| 9 | 78640399 | rs10781325 | -1.037 | -0.407 | -0.386 | -0.175 | -0.006 | -0.009 | -0.009 | 0 | 0.35 | 0.56 | 0.23 | 0.27 | 0.31 | 0.28 |
| 9 | 78641311 | rs10781328 | -0.567 | 0.14 | -0.468 | 0.421 | 0.005 | 0.033 | 0.033 | 0 | 0.06 | 0.09 | 0.04 | 0.22 | 0.12 | 0.16 |
| 9 | 78642228 | rs12342299 | -0.557 | -0.124 | -0.649 | -0.031 | 0.005 | 0.007 | 0.007 | 0.124 | 0.13 | 0.08 | 0.09 | 0.03 | 0.07 | 0.13 |
| 9 | 78642567 | rs12554374 | -0.454 | -0.039 | -1.465 | 0.452 | -0.034 | -0.108 | -0.108 | 0.891 | 0.18 | 0.04 | 0.36 | 0.63 | 0.21 | 0.35 |
| 9 | 78649934 | rs11144723 | -0.19 | -0.319 | -0.519 | 0.232 | 0.006 | -0.018 | -0.018 | 0 | 0.18 | 0.09 | 0.17 | 0.23 | 0.15 | 0.31 |
| 9 | 78651290 | rs10869682 | -0.211 | -0.283 | -0.367 | 0.254 | 0.003 | -0.006 | -0.006 | 0.01 | 0.18 | 0.11 | 0.17 | 0.23 | 0.16 | 0.27 |
| 9 | 78651491 | rs10521467 | 0.375 | 0.612 | -0.659 | 0.002 | -0.021 | -0.013 | -0.013 | 0.074 | 0.08 | 0.05 | 0.09 | 0.12 | 0.15 | 0.1 |
| 9 | 78651988 | rs7033416 | 0.811 | 0.362 | -0.528 | -0.927 | -0.005 | -0.161 | -0.161 | 0.575 | 0.25 | 0.5 | 0.19 | 0 | 0.26 | 0.14 |
| 9 | 78653976 | rs1029037 | -0.229 | 0.25 | -0.666 | -0.508 | 0.016 | 0.045 | 0.045 | 0 | 0.45 | 0.71 | 0.36 | 0.35 | 0.42 | 0.47 |
| 9 | 78654101 | rs1023181 | 0.319 | -0.034 | -1.26 | -0.577 | -0.008 | -0.095 | -0.095 | 1.264 | 0.33 | 0.09 | 0.5 | 0.55 | 0.32 | 0.33 |
| 9 | 78655927 | rs6560488 | -0.251 | 0.008 | -1.118 | -0.364 | 0.024 | -0.006 | -0.006 | 0 | 0.53 | 0.79 | 0.46 | 0.46 | 0.59 | 0.59 |
| 9 | 78656338 | rs6560490 | -0.045 | 0.086 | -0.966 | -0.629 | 0.014 | 0.018 | 0.018 | 1.154 | 0.31 | 0.56 | 0.29 | 0.38 | 0.31 | 0.49 |
| 9 | 78657751 | rs2777020 | 0.875 | 0.308 | -1.754 | 0.037 | -0.02 | -0.059 | -0.059 | 0 | 0.22 | 0.1 | 0.33 | 0.37 | 0.27 | 0.16 |
| 9 | 78659547 | rs11144728 | -0.429 | 0.463 | -1.421 | 0.142 | -0.014 | 0.007 | 0.007 | 0 | 0.41 | 0.32 | 0.47 | 0.36 | 0.49 | 0.32 |
| 9 | 78659798 | rs2185230 | 0.614 | 0.508 | -1.553 | 0.039 | 0.009 | 0.006 | 0.006 | 0 | 0.26 | 0.15 | 0.31 | 0.24 | 0.28 | 0.23 |
| 9 | 78663605 | rs2153228 | 0.29 | 0.76 | -1.526 | 0.305 | 0.045 | -0.098 | -0.098 | 0 | 0.22 | 0.09 | 0.3 | 0.46 | 0.16 | 0.43 |
| 9 | 78664405 | rs9886813 | -0.644 | 1.832 | -0.728 | -0.132 | 0.016 | -0.005 | -0.005 | 0 | 0.15 | 0.19 | 0.15 | 0.12 | 0.21 | 0.09 |
| 9 | 78670516 | rs13291278 | -0.682 | 1.844 | -0.816 | -0.172 | 0.036 | -0.006 | -0.006 | 0 | 0.16 | 0.22 | 0.15 | 0.12 | 0.22 | 0.1 |
| 9 | 78673593 | rs1258094 | -0.484 | 1.494 | -1.307 | -0.64 | -0.026 | 0.043 | 0.043 | 0.126 | 0.23 | 0.11 | 0.24 | 0 | 0.34 | 0.1 |
| 9 | 78683749 | rs2153227 | -0.086 | 0.966 | -1.424 | -0.395 | -0.014 | 0.038 | 0.038 | 0 | 0.66 | 0.84 | 0.69 | 0.75 | 0.62 | 0.81 |
| 9 | 78684215 | rs10869685 | 0.387 | 1.33 | -1.509 | -0.557 | -0.027 | 0.014 | 0.014 | 0 | 0.18 | 0.13 | 0.17 | 0.11 | 0.26 | 0.34 |
| 9 | 78686818 | rs2297344 | 0.378 | 0.705 | -1.351 | -0.161 | 0 | 0.002 | 0.002 | 0.114 | 0.13 | 0.15 | 0.12 | 0.24 | 0.18 | 0.13 |
| 9 | 78687998 | rs871449 | 0.156 | 0.618 | -1.226 | -0.203 | 0 | 0.005 | 0.005 | 0.054 | 0.15 | 0.18 | 0.12 | 0.24 | 0.18 | 0.13 |
| 9 | 78688079 | rs10512053 | 1.625 | -0.009 | -1.47 | -0.165 | 0.041 | -0.075 | -0.075 | 0 | 0.2 | 0.05 | 0.27 | 0.33 | 0.09 | 0.21 |
| 9 | 78688289 | rs1339254 | 0.542 | -0.032 | -1.266 | -0.204 | -0.016 | 0.001 | 0.001 | 0 | 0.2 | 0.18 | 0.17 | 0.16 | 0.27 | 0.36 |
| 9 | 78689238 | rs10781331 | -1.558 | -0.091 | -1.293 | -0.265 | -0.013 | 0.04 | 0.04 | 0 | 0.66 | 0.85 | 0.69 | 0.74 | 0.63 | 0.8 |
| 9 | 78690471 | rs1475615 | -0.85 | -0.214 | -1.211 | -0.197 | 0.077 | 0.058 | 0.058 | 0 | 0.49 | 0.71 | 0.52 | 0.62 | 0.36 | 0.45 |
| 9 | 78690813 | rs10869688 | -0.826 | -0.177 | -1.073 | -0.187 | -0.006 | 0 | 0 | 0.895 | 0.28 | 0.66 | 0.26 | 0.28 | 0.26 | 0.25 |
| 9 | 78705681 | rs1029036 | 0.455 | -0.27 | 0.171 | -0.121 | -0.016 | 0.006 | 0.006 | 0.47 | 0.31 | 0.31 | 0.32 | 0.38 | 0.16 | 0.25 |
| 9 | 78706282 | rs1571792 | -0.625 | 0.059 | 0.287 | -0.311 | 0.037 | -0.004 | -0.004 | 0 | 0.34 | 0.44 | 0.36 | 0.35 | 0.46 | 0.44 |
| 9 | 78708179 | rs10869692 | 0.945 | 0.443 | 0.045 | -0.209 | -0.018 | 0.01 | 0.01 | 0 | 0.16 | 0.13 | 0.21 | 0.1 | 0.26 | 0.28 |
| 9 | 78713300 | rs10781332 | -0.952 | 0.303 | 0.266 | -0.126 | 0.006 | 0.017 | 0.017 | 0 | 0.59 | 0.7 | 0.67 | 0.71 | 0.59 | 0.77 |
| 9 | 78715080 | rs12001902 | 0.864 | 0.387 | 0.183 | 0.065 | 0.002 | 0.032 | 0.032 | 0 | 0.06 | 0.07 | 0.06 | 0.24 | 0.09 | 0.09 |
| 9 | 78718544 | rs1416548 | 0.437 | 0.558 | 0.117 | -0.28 | -0.008 | 0.011 | 0.011 | 0 | 0.2 | 0.16 | 0.16 | 0.08 | 0.23 | 0.31 |
| 9 | 78718893 | rs10869695 | -0.135 | 0.06 | 0.104 | -0.108 | -0.001 | 0.067 | 0.067 | 0 | 0.08 | 0.1 | 0.21 | 0.35 | 0.06 | 0.13 |
| 9 | 78719646 | rs1339230 | -1.555 | 0.045 | 0.284 | -0.381 | -0.004 | 0.005 | 0.005 | 0.148 | 0.67 | 0.76 | 0.67 | 0.72 | 0.61 | 0.74 |
| 9 | 78721540 | rs10869696 | 0.142 | 0.247 | 0.099 | -0.309 | 0.004 | 0.07 | 0.07 | 0 | 0.08 | 0.1 | 0.23 | 0.35 | 0.1 | 0.13 |
| 9 | 78721689 | rs7860650 | 1.07 | 0.027 | 0.207 | -0.49 | -0.004 | -0.041 | -0.041 | 0 | 0.24 | 0.25 | 0.14 | 0 | 0.12 | 0.11 |
| 9 | 78722328 | rs2297343 | 0.968 | 0.161 | 0.158 | -0.411 | -0.001 | -0.006 | -0.006 | 0.901 | 0.06 | 0.07 | 0.04 | 0.03 | 0.07 | 0.1 |
| 9 | 78723048 | rs4744780 | 0.366 | 0.176 | 0.115 | -0.332 | 0.005 | -0.005 | -0.005 | 0 | 0.32 | 0.31 | 0.37 | 0.35 | 0.22 | 0.24 |
| 9 | 78723206 | rs914366 | 0.241 | 0.394 | -0.215 | -0.288 | -0.011 | -0.005 | -0.005 | 0 | 0.27 | 0.25 | 0.28 | 0.35 | 0.36 | 0.4 |
| 9 | 78723307 | rs914365 | -0.813 | 0.752 | -0.452 | 0.064 | 0.001 | 0.006 | 0.006 | 0 | 0.63 | 0.71 | 0.66 | 0.69 | 0.58 | 0.65 |
| 9 | 78726242 | rs10869699 | 0.245 | 1.161 | -0.588 | 0.606 | -0.037 | 0.008 | 0.008 | 0 | 0.06 | 0.01 | 0.22 | 0.34 | 0.1 | 0.13 |
| 9 | 78729213 | rs1571791 | -0.537 | 0.927 | -0.315 | 0.287 | 0 | -0.003 | -0.003 | 0 | 0.61 | 0.67 | 0.59 | 0.47 | 0.59 | 0.59 |
| 9 | 78730136 | rs7046252 | -0.049 | 0.748 | -0.395 | -0.218 | -0.006 | 0.001 | 0.001 | 0 | 0.29 | 0.27 | 0.26 | 0.09 | 0.32 | 0.25 |
| 9 | 78738971 | rs990657 | -1.015 | 0.81 | 0.036 | -0.091 | -0.003 | -0.01 | -0.01 | 0.007 | 0.41 | 0.55 | 0.58 | 0.68 | 0.55 | 0.48 |
| 9 | 78740235 | rs11144756 | 0.06 | 1.016 | -0.145 | -0.02 | -0.018 | 0.042 | 0.042 | 0 | 0.28 | 0.2 | 0.25 | 0.09 | 0.32 | 0.23 |
| 9 | 78741027 | rs7037905 | 0.005 | 1.03 | -0.208 | -0.013 | -0.005 | -0.006 | -0.006 | 0.043 | 0.28 | 0.27 | 0.25 | 0.09 | 0.32 | 0.23 |
| 9 | 78742125 | rs1339236 | -0.548 | 0.713 | -0.265 | 0.329 | -0.004 | -0.003 | -0.003 | 0.769 | 0.36 | 0.35 | 0.52 | 0.42 | 0.5 | 0.42 |
| 9 | 78745177 | rs7035578 | 0.469 | 1.262 | -0.637 | 0.131 | -0.005 | -0.003 | -0.003 | 0 | 0.11 | 0.1 | 0.15 | 0 | 0.13 | 0.06 |
| 9 | 78745460 | rs6560502 | -1.093 | 0.717 | -0.742 | 0.288 | 0.006 | 0.008 | 0.008 | 0.325 | 0.67 | 0.92 | 0.65 | 0.71 | 0.65 | 0.65 |
| 9 | 78746781 | rs10746992 | -0.789 | 1.36 | -0.262 | 0.527 | 0.005 | 0.004 | 0.004 | 0 | 0.37 | 0.43 | 0.53 | 0.42 | 0.5 | 0.42 |
| 9 | 78751632 | rs4745507 | -1.021 | 1.806 | -0.634 | 0.584 | 0.003 | 0.024 | 0.024 | 0 | 0.13 | 0.12 | 0.42 | 0.58 | 0.26 | 0.29 |

| | | | | | | | | | | | | | | | | |
|---|---|---|---|---|---|---|---|---|---|---|---|---|---|---|---|---|
| 9 | 78751683 | rs1416552 | 0.386 | 1.556 | -0.907 | -0.286 | -0.009 | 0.018 | 0.018 | 0.132 | 0.22 | 0.13 | 0.15 | 0 | 0.25 | 0.06 |
| 9 | 78755492 | rs1416325 | -0.624 | 1.684 | -0.665 | 0.316 | 0.007 | 0.051 | 0.051 | 0 | 0.1 | 0.09 | 0.47 | 0.6 | 0.17 | 0.2 |
| 9 | 78757167 | rs6560506 | 0.242 | 0.285 | -0.394 | -0.354 | -0.001 | -0.001 | -0.001 | 0 | 0.35 | 0.35 | 0.56 | 0.61 | 0.29 | 0.42 |
| 9 | 78757975 | rs10869708 | -0.144 | -0.046 | -0.214 | -0.777 | -0.006 | -0.004 | -0.004 | 0.514 | 0.59 | 0.59 | 0.68 | 0.94 | 0.46 | 0.6 |
| 9 | 78759288 | rs1331384 | -0.242 | -0.055 | -0.194 | -0.776 | -0.009 | 0.015 | 0.015 | 0 | 0.6 | 0.63 | 0.68 | 0.93 | 0.48 | 0.6 |
| 9 | 78762118 | rs9314838 | 0.417 | 0.129 | 0.17 | -0.296 | 0.001 | 0 | 0 | 0 | 0.46 | 0.46 | 0.48 | 0.4 | 0.44 | 0.62 |
| 9 | 78762295 | rs4745511 | 0.052 | 0.096 | 0.321 | -0.274 | -0.001 | 0.029 | 0.029 | 0 | 0.49 | 0.57 | 0.63 | 0.69 | 0.48 | 0.71 |
| 9 | 78762796 | rs4391491 | 0.891 | 0.124 | 0.334 | -0.314 | -0.002 | 0.002 | 0.002 | 0.127 | 0.16 | 0.16 | 0.11 | 0.08 | 0.22 | 0.25 |
| 9 | 78768830 | rs17776673 | 1.309 | -0.09 | 0.325 | -0.238 | 0 | 0.011 | 0.011 | 0 | 0.3 | 0.3 | 0.37 | 0.34 | 0.22 | 0.39 |
| 9 | 78772523 | rs2377525 | -0.8 | 0.16 | 0.128 | -0.477 | -0.02 | 0.022 | 0.022 | 0 | 0.73 | 0.85 | 0.76 | 1 | 0.67 | 0.85 |
| 9 | 78773209 | rs12236697 | 1.74 | 0.018 | 0.137 | -0.38 | -0.009 | -0.025 | -0.025 | 0 | 0.26 | 0.19 | 0.36 | 0.33 | 0.22 | 0.35 |
| 9 | 78773953 | rs2297342 | 1.761 | 0.026 | 0.007 | -0.412 | -0.025 | -0.058 | -0.058 | 1.403 | 0.26 | 0.16 | 0.36 | 0.33 | 0.22 | 0.35 |
| 9 | 78775570 | rs1888705 | 0.267 | 0.145 | 0.182 | -0.449 | -0.021 | 0.058 | 0.058 | 0 | 0.58 | 0.69 | 0.66 | 0.89 | 0.48 | 0.62 |
| 9 | 78776057 | rs10869713 | 0.006 | 0.388 | 0.072 | -0.493 | -0.002 | 0.012 | 0.012 | 0 | 0.32 | 0.27 | 0.22 | 0.12 | 0.37 | 0.31 |
| 9 | 78777937 | rs1411956 | 0.629 | 0.453 | 0.102 | -0.422 | -0.003 | -0.006 | -0.006 | 3.086 | 0.17 | 0.22 | 0.1 | 0.11 | 0.19 | 0.23 |
| 9 | 78779832 | rs6560507 | -0.068 | 0.43 | 0.229 | -0.364 | 0.008 | 0.018 | 0.018 | 0 | 0.84 | 0.75 | 0.73 | 0.73 | 0.83 | 0.86 |
| 9 | 78780475 | rs10869715 | 0.518 | 0.665 | 0.307 | -0.882 | -0.004 | -0.014 | -0.014 | 0.337 | 0.17 | 0.3 | 0.1 | 0.11 | 0.19 | 0.22 |
| 9 | 78780578 | rs7037635 | 0.884 | 0.89 | -0.113 | -0.627 | -0.003 | -0.003 | -0.003 | 0.207 | 0.17 | 0.12 | 0.14 | 0 | 0.19 | 0.08 |
| 9 | 78781260 | rs1331386 | -0.035 | 0.842 | 0.057 | 0.139 | 0.02 | -0.042 | -0.042 | 1.372 | 0.91 | 0.86 | 0.89 | 1 | 0.88 | 0.97 |
| 9 | 78785098 | rs10124834 | 0.131 | 0.461 | -0.229 | 0.559 | -0.001 | 0.002 | 0.002 | 0 | 0.31 | 0.36 | 0.51 | 0.6 | 0.25 | 0.45 |
| 9 | 78788473 | rs10869720 | -1.22 | 0.216 | 0.075 | 1.073 | -0.013 | 0.041 | 0.041 | 0.008 | 0.16 | 0.27 | 0.37 | 0.36 | 0.16 | 0.24 |
| 9 | 78788543 | rs17062174 | -0.488 | 0.414 | -0.242 | 0.91 | -0.005 | 0.013 | 0.013 | 0 | 0.07 | 0.04 | 0.09 | 0.02 | 0.1 | 0.12 |
| 9 | 78791189 | rs2377527 | 1.56 | 0.255 | 0.207 | 0.95 | -0.003 | 0.032 | 0.032 | 0 | 0.79 | 0.63 | 0.53 | 0.65 | 0.78 | 0.71 |
| 9 | 78791312 | rs10781344 | -1.089 | 0.186 | 0.463 | 1.132 | 0.002 | 0.191 | 0.191 | 0 | 0.06 | 0.2 | 0.28 | 0.33 | 0.06 | 0.11 |
| 9 | 78792438 | rs12684735 | -0.335 | -0.124 | -0.34 | 0.989 | 0 | -0.046 | -0.046 | 0.049 | 0.3 | 0.18 | 0.4 | 0.56 | 0.29 | 0.33 |
| 9 | 78793274 | rs1571793 | 0.434 | -0.1 | -0.559 | 0.769 | 0 | -0.001 | -0.001 | 0.491 | 0.19 | 0.12 | 0.2 | 0.2 | 0.21 | 0.29 |
| 9 | 78793895 | rs2377528 | 0.986 | -0.12 | -0.802 | 0.655 | -0.045 | 0.088 | 0.088 | 0 | 0.15 | 0.01 | 0.18 | 0.02 | 0.17 | 0.15 |
| 9 | 78795051 | rs7045212 | 0.437 | 0.301 | -1.014 | 0.814 | -0.056 | -0.077 | -0.077 | 2.976 | 0.84 | 0.98 | 0.74 | 0.78 | 0.82 | 0.92 |
| 9 | 78796522 | rs2270570 | 0.279 | 0.717 | -0.913 | 0.264 | -0.024 | -0.002 | -0.002 | 0 | 0.12 | 0.08 | 0.14 | 0.14 | 0.21 | 0.2 |
| 9 | 78800114 | rs2279659 | 0.505 | 0.496 | -0.155 | 0.377 | 0.009 | 0.109 | 0.109 | 0 | 0.84 | 0.64 | 0.84 | 0.72 | 0.82 | 0.71 |
| 9 | 78800281 | rs17062197 | 0.388 | 0.629 | -0.443 | 0.082 | 0.018 | 0.004 | 0.004 | 0.048 | 0.1 | 0.18 | 0.11 | 0.1 | 0.15 | 0.18 |
| 9 | 78800989 | rs10869723 | 1.2 | 0.38 | -0.622 | -0.189 | 0.007 | 0.054 | 0.054 | 0 | 0.33 | 0.21 | 0.19 | 0.08 | 0.28 | 0.14 |
| 9 | 78802474 | rs7872680 | 1.087 | 0.129 | -1.176 | -0.322 | 0.013 | 0.006 | 0.006 | 0 | 0.21 | 0.15 | 0.12 | 0.16 | 0.17 | 0.13 |
| 9 | 78803466 | rs7046850 | -0.969 | 0.204 | -1.114 | -1.007 | -0.004 | -0.027 | -0.027 | 0 | 0.92 | 0.91 | 0.91 | 1 | 0.93 | 0.95 |
| 9 | 78803512 | rs10521468 | 1.986 | 0.454 | -1.308 | -0.985 | -0.023 | 0.023 | 0.023 | 0 | 0.15 | 0.07 | 0.13 | 0 | 0.18 | 0.07 |
| 9 | 78804704 | rs10869726 | 1.102 | 0.165 | -1.229 | -0.942 | -0.022 | -0.022 | -0.022 | 0 | 0.2 | 0.25 | 0.09 | 0.09 | 0.12 | 0.12 |
| 9 | 78804842 | rs10869727 | 0.776 | 0.057 | -1.281 | -0.949 | -0.023 | -0.017 | -0.017 | 0 | 0.23 | 0.26 | 0.09 | 0.13 | 0.12 | 0.12 |
| 9 | 78808493 | rs10869729 | 1.466 | 0.656 | -1.613 | -0.672 | -0.038 | -0.036 | -0.036 | 0 | 0.07 | 0.01 | 0.1 | 0.11 | 0.12 | 0.14 |
| 9 | 78810203 | rs11793906 | 1.666 | 0.609 | -1.46 | -0.739 | -0.031 | 0.04 | 0.04 | 0 | 0.15 | 0.05 | 0.13 | 0 | 0.2 | 0.07 |
| 9 | 78810344 | rs966097 | 0.027 | 0.083 | -1.165 | -0.307 | -0.003 | -0.003 | -0.003 | 2.678 | 0.32 | 0.27 | 0.44 | 0.63 | 0.32 | 0.33 |
| 9 | 78810887 | rs10869730 | -0.235 | -0.217 | -1.777 | -0.879 | 0.019 | -0.004 | -0.004 | 0 | 0.4 | 0.25 | 0.43 | 0.58 | 0.36 | 0.38 |
| 9 | 78811377 | rs10869731 | 0.061 | 0.074 | -1.881 | -2.216 | 0.007 | 0.084 | 0.084 | 0 | 0.28 | 0.14 | 0.39 | 0.86 | 0.25 | 0.43 |
| 9 | 78811935 | rs7031594 | 1.026 | 0.12 | -1.608 | -2.17 | 0.001 | -0.021 | -0.021 | 2.926 | 0.13 | 0.14 | 0.13 | 0 | 0.12 | 0.11 |
| 9 | 78815368 | rs10746995 | -0.169 | -0.12 | -1.517 | -2.204 | -0.032 | -0.044 | -0.044 | 0 | 0.69 | 0.49 | 0.69 | 0.79 | 0.76 | 0.78 |
| 9 | 78816016 | rs7026021 | 0.434 | 0.142 | -1.858 | -2.225 | -0.002 | -0.001 | -0.001 | 0 | 0.27 | 0.4 | 0.37 | 0.67 | 0.23 | 0.4 |
| 9 | 78816092 | rs11144794 | 0.691 | 0.61 | -2.345 | -2.05 | -0.051 | 0.099 | 0.099 | 0 | 0.19 | 0.04 | 0.13 | 0.02 | 0.21 | 0.12 |
| 9 | 78817288 | rs12379097 | 0.494 | 0.39 | -1.861 | -1.424 | -0.049 | 0.046 | 0.046 | 0 | 0.18 | 0.01 | 0.13 | 0.1 | 0.21 | 0.21 |
| 9 | 78820927 | rs4745528 | 0.037 | 0.301 | -1.356 | -1.488 | 0.046 | 0.063 | 0.063 | 0 | 0.53 | 0.24 | 0.69 | 0.66 | 0.56 | 0.48 |
| 9 | 78823996 | rs990559 | 0.784 | 0.855 | -1.755 | -1.683 | -0.047 | -0.059 | -0.059 | 0 | 0.26 | 0.06 | 0.44 | 0.66 | 0.3 | 0.37 |
| 9 | 78826932 | rs1039034 | 0.606 | 0.316 | -0.882 | -1.515 | 0.01 | 0.018 | 0.018 | 0 | 0.31 | 0.23 | 0.18 | 0.28 | 0.31 | 0.31 |
| 9 | 78828051 | rs12336492 | 0.047 | 0.24 | -0.244 | -1.598 | 0.005 | 0.004 | 0.004 | 0 | 0.37 | 0.63 | 0.22 | 0.37 | 0.36 | 0.41 |
| 9 | 78834472 | rs10781347 | 0.601 | 0.153 | -0.414 | -1.572 | 0.009 | 0.016 | 0.016 | 0 | 0.31 | 0.25 | 0.18 | 0.28 | 0.31 | 0.31 |
| 9 | 78836985 | rs10869736 | 0.642 | 0.563 | -0.454 | -1.367 | -0.042 | -0.049 | -0.049 | 0 | 0.27 | 0.09 | 0.44 | 0.64 | 0.31 | 0.24 |
| 9 | 78843635 | rs10781349 | -0.39 | 0.519 | 0.164 | -1.547 | -0.003 | 0.001 | 0.001 | 0 | 0.68 | 0.73 | 0.52 | 0.31 | 0.66 | 0.66 |
| 9 | 78845010 | rs1979737 | 1.031 | 0.519 | 0.096 | -1.862 | 0.008 | 0.074 | 0.074 | 0 | 0.46 | 0.3 | 0.27 | 0.24 | 0.43 | 0.35 |
| 9 | 78849238 | rs10869741 | -0.021 | 0.583 | 0.316 | -1.906 | 0.011 | -0.036 | -0.036 | 0 | 0.61 | 0.52 | 0.42 | 0.26 | 0.58 | 0.54 |
| 9 | 78852069 | rs13293615 | 0.239 | 0.466 | 0.276 | -1.861 | -0.006 | -0.048 | -0.048 | 3.197 | 0.13 | 0.21 | 0.13 | 0.01 | 0.12 | 0.17 |
| 9 | 78854312 | rs2007744 | 0.177 | 0.458 | 0.492 | -1.87 | -0.011 | -0.074 | -0.074 | 0.041 | 0.83 | 0.72 | 0.84 | 0.99 | 0.84 | 0.73 |
| 9 | 78856589 | rs17720899 | -0.058 | 0.696 | 0.013 | -1.588 | 0.007 | 0.002 | 0.002 | 0 | 0.06 | 0.09 | 0.11 | 0.06 | 0.11 | 0.13 |
| 9 | 78859359 | rs1908495 | 0.718 | 0.726 | -0.022 | -1.622 | 0.005 | 0.014 | 0.014 | 0 | 0.48 | 0.44 | 0.28 | 0.24 | 0.43 | 0.34 |
| 9 | 78863548 | rs2261557 | 0.436 | 0.566 | -0.104 | -1.491 | -0.001 | -0.006 | -0.006 | 0 | 0.52 | 0.48 | 0.45 | 0.25 | 0.47 | 0.54 |
| 9 | 78865300 | rs1587103 | 0.335 | 0.473 | -0.513 | -1.375 | 0.001 | -0.013 | -0.013 | 0 | 0.54 | 0.5 | 0.48 | 0.3 | 0.53 | 0.47 |
| 9 | 78867869 | rs2643328 | -0.014 | 0.074 | -1.379 | -1.214 | -0.036 | 0.005 | 0.005 | 0 | 0.69 | 0.84 | 0.63 | 0.28 | 0.65 | 0.66 |
| 9 | 78868816 | rs10781353 | 0.173 | -0.984 | -1.327 | -1.042 | -0.059 | 0.011 | 0.011 | 0 | 0.33 | 0.12 | 0.5 | 0.67 | 0.53 | 0.42 |
| 9 | 78869524 | rs1002420 | -0.081 | -1.442 | -0.686 | -0.928 | 0.003 | 0 | 0 | 0 | 0.38 | 0.35 | 0.49 | 0.59 | 0.59 | 0.43 |
| 9 | 78869581 | rs2261722 | 0.187 | -2.266 | -1.375 | -1.3 | 0.044 | 0.039 | 0.039 | 0 | 0.4 | 0.24 | 0.43 | 0.34 | 0.31 | 0.42 |
| 9 | 78872846 | rs2842472 | 0.638 | -2.321 | -1.28 | -1.341 | 0.016 | 0.021 | 0.021 | 0.12 | 0.3 | 0.22 | 0.19 | 0.26 | 0.27 | 0.11 |
| 9 | 78874769 | rs1353342 | 0.045 | -1.795 | -0.586 | -1.07 | -0.063 | -0.056 | -0.056 | 2.932 | 0.74 | 0.46 | 0.87 | 0.92 | 0.89 | 0.83 |
| 9 | 78885928 | rs2643322 | 0.404 | -1.356 | -1.542 | -0.654 | 0.083 | -0.017 | -0.017 | 0 | 0.54 | 0.19 | 0.51 | 0.54 | 0.69 | 0.41 |

| | | | | | | | | | | | | | | | | |
|---|---|---|---|---|---|---|---|---|---|---|---|---|---|---|---|---|
| 9 | 78887992 | rs2490588 | 0.387 | -0.762 | -1.033 | -1.034 | -0.009 | 0.134 | 0.134 | 0 | 0.43 | 0.22 | 0.36 | 0.18 | 0.43 | 0.47 |
| 9 | 78888137 | rs1994964 | 0.434 | -0.824 | -1.149 | -0.957 | -0.122 | -0.012 | -0.012 | 0 | 0.34 | 0.09 | 0.48 | 0.71 | 0.46 | 0.36 |
| 9 | 78889021 | rs10781356 | 0.443 | -0.797 | -1.136 | -0.944 | -0.122 | -0.016 | -0.016 | 0 | 0.34 | 0.09 | 0.49 | 0.71 | 0.46 | 0.36 |
| 9 | 78892526 | rs3915861 | -0.498 | -0.528 | -0.668 | -0.882 | -0.002 | 0.036 | 0.036 | 0 | 0.53 | 0.59 | 0.61 | 0.8 | 0.53 | 0.45 |
| 9 | 78899083 | rs9314843 | -0.471 | -0.76 | -0.114 | -0.519 | -0.034 | -0.008 | -0.008 | 0 | 0.16 | 0.27 | 0.11 | 0.08 | 0.08 | 0.08 |
| 9 | 78900161 | rs10115735 | -0.895 | -0.876 | -0.191 | -0.636 | -0.026 | -0.012 | -0.012 | 0 | 0.2 | 0.27 | 0.07 | 0.09 | 0.09 | 0.14 |
| 9 | 78901112 | rs1495257 | -0.743 | -0.34 | -0.212 | -1.206 | -0.004 | 0.025 | 0.025 | 0 | 0.54 | 0.58 | 0.41 | 0.19 | 0.49 | 0.43 |
| 9 | 78902949 | rs2260411 | -0.136 | 0.09 | -0.268 | -1.806 | -0.014 | 0.072 | 0.072 | 0 | 0.28 | 0.16 | 0.21 | 0.05 | 0.27 | 0.16 |
| 9 | 78903962 | rs2842486 | -0.388 | -0.126 | -0.44 | -1.846 | -0.032 | 0.119 | 0.119 | 0 | 0.3 | 0.12 | 0.24 | 0.09 | 0.27 | 0.16 |
| 9 | 78904272 | rs903549 | -0.187 | -0.101 | -0.361 | -1.831 | -0.031 | 0.114 | 0.114 | 0 | 0.29 | 0.13 | 0.21 | 0.07 | 0.27 | 0.16 |
| 9 | 78904508 | rs903547 | -0.232 | -0.11 | -0.329 | -1.835 | -0.027 | 0.116 | 0.116 | 0 | 0.29 | 0.16 | 0.22 | 0.08 | 0.27 | 0.16 |
| 9 | 78904950 | rs17721076 | -0.112 | -0.23 | 0.507 | -2.082 | 0.005 | -0.035 | -0.035 | 0 | 0.18 | 0.23 | 0.17 | 0.06 | 0.22 | 0.25 |
| 9 | 78906150 | rs2842483 | -0.18 | -0.024 | 0.955 | -2.061 | 0.002 | -0.005 | -0.005 | 0 | 0.31 | 0.34 | 0.23 | 0.08 | 0.26 | 0.16 |
| 9 | 78907594 | rs7021909 | -0.779 | -0.176 | 1.382 | -2.097 | -0.007 | -0.091 | -0.091 | 0.095 | 0.27 | 0.52 | 0.23 | 0.06 | 0.26 | 0.28 |
| 9 | 78908826 | rs2260649 | -1.713 | -0.227 | 1.294 | -2.132 | -0.006 | 0.033 | 0.033 | 0 | 0.54 | 0.63 | 0.42 | 0.14 | 0.5 | 0.43 |
| 9 | 78910401 | rs10869747 | -0.889 | -0.112 | 1.288 | -2.014 | -0.02 | -0.043 | -0.043 | 0 | 0.24 | 0.3 | 0.18 | 0.06 | 0.19 | 0.26 |
| 9 | 78912008 | rs7039902 | -0.977 | 0.335 | 1.508 | -1.876 | 0.064 | -0.047 | -0.047 | 0 | 0.09 | 0.33 | 0.06 | 0 | 0.09 | 0.05 |
| 9 | 78912017 | rs7025902 | -0.979 | 0.4 | 1.498 | -1.88 | 0.085 | -0.047 | -0.047 | 0 | 0.09 | 0.32 | 0.07 | 0 | 0.11 | 0.05 |
| 9 | 78913563 | rs2842478 | 0.74 | -0.295 | 1.182 | -2.59 | -0.029 | -0.127 | -0.127 | 0 | 0.65 | 0.45 | 0.78 | 0.95 | 0.76 | 0.75 |
| 9 | 78915454 | rs1078868 | 0.585 | -0.222 | 1.096 | -2.559 | -0.003 | -0.042 | -0.042 | 0 | 0.67 | 0.65 | 0.79 | 0.93 | 0.76 | 0.75 |
| 9 | 78917378 | rs11788518 | 0.333 | -0.482 | 0.083 | -3.115 | 0.065 | 0.151 | 0.151 | 0.289 | 0.32 | 0.1 | 0.15 | 0.07 | 0.2 | 0.28 |
| 9 | 78919387 | rs2490591 | 0.005 | 0.103 | 0.632 | -2.494 | 0.012 | 0.013 | 0.013 | 0 | 0.72 | 0.76 | 0.83 | 0.9 | 0.81 | 0.83 |
| 9 | 78928591 | rs976825 | -0.404 | 0.185 | 0.459 | -2.312 | -0.013 | -0.025 | -0.025 | 0 | 0.29 | 0.32 | 0.18 | 0.17 | 0.2 | 0.21 |
| 9 | 78929086 | rs2055164 | -0.772 | 0.199 | 0.302 | -2.391 | -0.01 | -0.032 | -0.032 | 0 | 0.34 | 0.36 | 0.21 | 0.17 | 0.26 | 0.23 |
| 9 | 78930965 | rs2258063 | -0.449 | 0.937 | 0.171 | -2.284 | 0.015 | -0.004 | -0.004 | 0 | 0.12 | 0.18 | 0.11 | 0.1 | 0.18 | 0.21 |
| 9 | 78931893 | rs2258131 | -0.684 | 0.986 | -0.536 | -2.275 | 0.006 | -0.006 | -0.006 | 0 | 0.87 | 0.99 | 0.9 | 0.85 | 0.92 | 0.89 |
| 9 | 78932693 | rs1027384 | -1.33 | 1.782 | 0.198 | -2.258 | -0.015 | 0.034 | 0.034 | 0.223 | 0.4 | 0.7 | 0.34 | 0.19 | 0.45 | 0.44 |
| 9 | 78933874 | rs11144827 | -0.634 | 1.628 | 0.578 | -2.083 | 0.161 | 0.067 | 0.067 | 0 | 0.08 | 0.32 | 0.14 | 0.09 | 0.16 | 0.08 |
| 9 | 78936358 | rs1110222 | -0.912 | 1.411 | 0.309 | -2.206 | 0.001 | -0.064 | -0.064 | 1.071 | 0.33 | 0.46 | 0.24 | 0.14 | 0.34 | 0.35 |
| 9 | 78936564 | rs11144829 | 0.003 | 2.131 | 0.286 | -2.173 | 0.031 | -0.038 | -0.038 | 0 | 0.15 | 0.25 | 0.2 | 0.07 | 0.2 | 0.15 |
| 9 | 78937619 | rs2842468 | 0.193 | 1.62 | -0.186 | -1.926 | 0.008 | 0.01 | 0.01 | 0 | 0.09 | 0.07 | 0.05 | 0.03 | 0.07 | 0.09 |
| 9 | 78938628 | rs2842467 | -1.114 | 1.398 | -0.263 | -1.973 | 0.022 | 0.079 | 0.079 | 0.033 | 0.49 | 0.69 | 0.43 | 0.21 | 0.55 | 0.49 |
| 9 | 78940026 | rs2803433 | 0.015 | 1.263 | -0.409 | -1.886 | 0.008 | -0.039 | -0.039 | 0.03 | 0.11 | 0.2 | 0.07 | 0.03 | 0.12 | 0.11 |
| 9 | 78942956 | rs2259969 | -1.039 | 1.159 | -0.562 | -1.884 | 0.053 | 0.127 | 0.127 | 0.104 | 0.5 | 0.72 | 0.45 | 0.21 | 0.6 | 0.5 |
| 9 | 78947200 | rs11144834 | -0.52 | 1.268 | -0.463 | -1.733 | 0.017 | -0.062 | -0.062 | 0 | 0.38 | 0.55 | 0.37 | 0.18 | 0.43 | 0.39 |
| 9 | 78949109 | rs7026762 | 0.136 | 0.315 | -1.078 | -2.186 | 0.037 | 0.081 | 0.081 | 0 | 0.34 | 0.15 | 0.17 | 0.08 | 0.27 | 0.28 |
| 9 | 78950203 | rs2803439 | 0.023 | 0.476 | -0.767 | -1.502 | -0.011 | 0.004 | 0.004 | 0.114 | 0.73 | 0.47 | 0.82 | 0.69 | 0.73 | 0.79 |
| 9 | 78956690 | rs1390143 | 1.178 | 1.221 | -1.16 | -1.78 | -0.092 | 0.083 | 0.083 | 0.62 | 0.15 | 0.02 | 0.18 | 0.02 | 0.27 | 0.17 |
| 9 | 78958599 | rs10781357 | 0.142 | 0.797 | -0.483 | -1.729 | 0.014 | -0.012 | -0.012 | 0 | 0.28 | 0.36 | 0.53 | 0.76 | 0.36 | 0.42 |
| 9 | 78959341 | rs10869756 | 1.284 | 0.689 | -0.584 | -1.658 | -0.028 | -0.083 | -0.083 | 0.604 | 0.18 | 0.07 | 0.44 | 0.55 | 0.21 | 0.33 |
| 9 | 78959679 | rs2842495 | 0.466 | 0.564 | -0.452 | -1.8 | -0.004 | -0.061 | -0.061 | 0 | 0.58 | 0.45 | 0.38 | 0.14 | 0.56 | 0.49 |
| 9 | 78961388 | rs11144848 | 1.095 | -0.15 | -0.457 | -1.674 | 0.042 | 0.074 | 0.074 | 0.337 | 0.41 | 0.22 | 0.19 | 0.12 | 0.29 | 0.3 |
| 9 | 78962285 | rs13300668 | 0.457 | -0.105 | -0.4 | -1.614 | 0.023 | 0.039 | 0.039 | 0.47 | 0.49 | 0.38 | 0.27 | 0.33 | 0.39 | 0.39 |
| 9 | 78962759 | rs2803405 | -0.095 | -0.532 | -0.034 | -1.648 | -0.07 | -0.065 | -0.065 | 5.134 | 0.82 | 0.51 | 0.89 | 0.9 | 0.87 | 0.87 |
| 9 | 78965993 | rs2257220 | -0.433 | -0.482 | -0.256 | -1.654 | -0.049 | -0.045 | -0.045 | 0.046 | 0.83 | 0.64 | 0.9 | 0.9 | 0.87 | 0.88 |
| 9 | 78969059 | rs7036921 | 0.599 | -0.375 | -0.702 | -1.717 | 0.029 | 0.045 | 0.045 | 0.397 | 0.41 | 0.26 | 0.19 | 0.09 | 0.29 | 0.32 |
| 9 | 78971232 | rs10869765 | -0.462 | -0.465 | -1.082 | -2.297 | -0.055 | 0.13 | 0.13 | 0 | 0.6 | 0.78 | 0.71 | 0.98 | 0.53 | 0.64 |
| 9 | 78971253 | rs7034265 | 0.946 | -1.139 | -1.337 | -2.007 | -0.019 | 0.002 | 0.002 | 0 | 0.23 | 0.26 | 0.15 | 0.22 | 0.18 | 0.21 |
| 9 | 78975569 | rs2377637 | 0.892 | -0.393 | -1.589 | -1.102 | -0.01 | -0.072 | -0.072 | 0.297 | 0.19 | 0.08 | 0.43 | 0.57 | 0.18 | 0.28 |
| 9 | 78976710 | rs13288756 | 0.513 | -0.419 | -1.11 | -1.089 | -0.008 | -0.032 | -0.032 | 0 | 0.13 | 0.06 | 0.1 | 0.19 | 0.13 | 0.15 |
| 9 | 78987999 | rs2501936 | -0.631 | -0.406 | -0.666 | -1.43 | -0.008 | -0.031 | -0.031 | 0 | 0.9 | 0.9 | 0.96 | 1 | 0.93 | 0.94 |
| 9 | 78989528 | rs2490573 | 0.368 | -0.817 | -0.722 | -1.65 | 0.022 | 0.03 | 0.03 | 0 | 0.49 | 0.39 | 0.62 | 0.63 | 0.59 | 0.64 |
| 9 | 78989956 | rs2490574 | 1.538 | -0.12 | -1.037 | -1.806 | -0.058 | -0.086 | -0.086 | 0 | 0.16 | 0.04 | 0.37 | 0.59 | 0.25 | 0.35 |
| 9 | 78993077 | rs1961995 | 1.062 | -0.132 | -1.07 | -1.864 | -0.08 | -0.098 | -0.098 | 0 | 0.18 | 0.04 | 0.39 | 0.6 | 0.3 | 0.37 |
| 9 | 78994056 | rs7038009 | 0.867 | -0.272 | -0.922 | -1.887 | 0.001 | -0.045 | -0.045 | 0 | 0.33 | 0.33 | 0.25 | 0.03 | 0.34 | 0.28 |
| 9 | 79004213 | rs11144870 | 2.111 | -0.243 | -1.166 | -1.923 | -0.082 | 0.026 | 0.026 | 1.107 | 0.12 | 0.01 | 0.16 | 0.03 | 0.24 | 0.17 |
| 9 | 79019066 | rs7035385 | 0.688 | -0.404 | -0.793 | -1.855 | 0.001 | -0.045 | -0.045 | 0.025 | 0.36 | 0.37 | 0.27 | 0.03 | 0.42 | 0.3 |
| 9 | 79020386 | rs2501920 | 1.52 | -0.424 | -0.805 | -1.812 | -0.008 | -0.008 | -0.008 | 0 | 0.16 | 0.12 | 0.37 | 0.61 | 0.23 | 0.34 |
| 9 | 79032236 | rs10869770 | 0.708 | -0.466 | -0.733 | -1.872 | -0.021 | 0.025 | 0.025 | 0.171 | 0.35 | 0.28 | 0.27 | 0.03 | 0.42 | 0.3 |
| 9 | 79032427 | rs7847608 | -1.416 | -0.081 | -0.891 | -1.474 | -0.01 | -0.011 | -0.011 | 0 | 0.84 | 0.88 | 0.63 | 0.41 | 0.77 | 0.65 |
| 9 | 79036059 | rs12216907 | 1.356 | -0.292 | -0.918 | -1.459 | 0.002 | 0.019 | 0.019 | 0 | 0.08 | 0.03 | 0.04 | 0 | 0.07 | 0.05 |
| 9 | 79046297 | rs11144898 | 0.06 | -0.432 | -0.583 | -1.413 | 0.013 | 0.003 | 0.003 | 0.122 | 0.37 | 0.31 | 0.31 | 0.38 | 0.29 | 0.29 |
| 9 | 79048714 | rs11144899 | -0.492 | -0.094 | -0.446 | -1.617 | -0.006 | -0.005 | -0.005 | 0 | 0.54 | 0.47 | 0.48 | 0.41 | 0.57 | 0.48 |
| 9 | 79055143 | rs10869779 | 0.176 | -0.337 | -0.464 | -1.729 | -0.013 | 0.011 | 0.011 | 0 | 0.36 | 0.33 | 0.37 | 0.22 | 0.43 | 0.42 |
| 9 | 79056003 | rs12378119 | 0.878 | -0.312 | -0.602 | -1.692 | 0.006 | 0.004 | 0.004 | 0 | 0.23 | 0.17 | 0.2 | 0.19 | 0.19 | 0.25 |
| 9 | 79065299 | rs11144905 | 0.382 | -0.243 | -0.081 | -1.701 | 0.004 | -0.025 | -0.025 | 0.057 | 0.23 | 0.25 | 0.22 | 0.03 | 0.3 | 0.22 |
| 9 | 79068627 | rs10114264 | -1.091 | -0.505 | -0.184 | -1.627 | 0.007 | 0.035 | 0.035 | 0 | 0.82 | 0.78 | 0.63 | 0.41 | 0.77 | 0.65 |
| 9 | 79078038 | rs10735646 | -1.339 | -0.554 | -0.643 | -1.855 | -0.045 | -0.097 | -0.097 | 0.524 | 0.84 | 0.96 | 0.64 | 0.42 | 0.78 | 0.64 |
| 9 | 79078703 | rs1998587 | 1.611 | -1.349 | -0.465 | -1.896 | 0.022 | 0.031 | 0.031 | 0.64 | 0.17 | 0.11 | 0.09 | 0 | 0.13 | 0.12 |

| | | | | | | | | | | | | | | | | |
|---|---|---|---|---|---|---|---|---|---|---|---|---|---|---|---|---|
| 9 | 79085839 | rs7032380 | 1.341 | -1.414 | -0.082 | -1.569 | -0.026 | 0.031 | 0.031 | 0.537 | 0.08 | 0.2 | 0.04 | 0.11 | 0.03 | 0.02 |
| 9 | 79093664 | rs785936 | 0.207 | -1.211 | -0.258 | -1.685 | -0.004 | -0.004 | -0.004 | 0 | 0.56 | 0.42 | 0.67 | 0.69 | 0.67 | 0.67 |
| 9 | 79097144 | rs4745552 | 0.176 | -0.938 | -0.808 | -1.486 | 0.016 | 0.003 | 0.003 | 0 | 0.32 | 0.23 | 0.28 | 0.31 | 0.27 | 0.28 |
| 9 | 79100981 | rs785915 | -1.558 | -0.168 | -0.843 | -1.668 | -0.058 | -0.014 | -0.014 | 0.015 | 0.88 | 0.96 | 0.84 | 0.92 | 0.77 | 0.84 |
| 9 | 79102496 | rs684271 | 0.667 | -0.905 | -0.386 | -2.133 | -0.033 | -0.067 | -0.067 | 0.181 | 0.29 | 0.4 | 0.16 | 0.08 | 0.18 | 0.16 |
| 9 | 79111675 | rs619429 | -0.458 | -0.932 | -0.49 | -2.304 | -0.027 | 0.057 | 0.057 | 0 | 0.58 | 0.72 | 0.59 | 0.72 | 0.54 | 0.57 |
| 9 | 79115844 | rs4532668 | -0.628 | -1.047 | -0.352 | -2.364 | -0.003 | 0.038 | 0.038 | 0 | 0.59 | 0.64 | 0.68 | 0.84 | 0.59 | 0.69 |
| 9 | 79116108 | rs1969980 | 0.696 | -1.16 | -0.334 | -2.43 | -0.053 | -0.053 | -0.053 | 1.99 | 0.26 | 0.07 | 0.42 | 0.64 | 0.35 | 0.42 |
| 9 | 79117751 | rs2282683 | 0.84 | -1.11 | -0.252 | -2.376 | 0.011 | 0.033 | 0.033 | 0.007 | 0.29 | 0.24 | 0.25 | 0.2 | 0.23 | 0.27 |
| 9 | 79118329 | rs1057406 | 0.84 | -1.11 | -0.252 | -2.376 | 0.011 | 0.033 | 0.033 | 0.007 | 0.29 | 0.24 | 0.25 | 0.2 | 0.23 | 0.27 |
| 9 | 79118903 | rs707739 | 1.107 | -1.139 | -0.23 | -2.391 | 0.008 | 0.057 | 0.057 | 0.672 | 0.28 | 0.15 | 0.24 | 0.2 | 0.23 | 0.27 |
| 9 | 79119752 | rs683969 | -0.362 | -1.439 | -0.291 | -2.615 | -0.079 | -0.099 | -0.099 | 0 | 0.24 | 0.44 | 0.12 | 0.12 | 0.16 | 0.12 |
| 9 | 79121085 | rs785907 | -1.609 | -1.178 | -0.147 | -2.453 | 0.011 | -0.001 | -0.001 | 0 | 0.85 | 0.81 | 0.82 | 0.92 | 0.77 | 0.85 |
| 9 | 79123201 | rs1547140 | 1.264 | -1.204 | 0.142 | -2.317 | 0.061 | 0.098 | 0.098 | 0.082 | 0.16 | 0.25 | 0.35 | 0.59 | 0.25 | 0.31 |
| 9 | 79123715 | rs11144933 | 0.884 | -1.215 | -0.056 | -2.301 | 0.01 | 0.031 | 0.031 | 0.362 | 0.29 | 0.23 | 0.25 | 0.19 | 0.23 | 0.27 |
| 9 | 79125260 | rs7026142 | 0.373 | -1.274 | -0.289 | -2.324 | 0 | -0.006 | -0.006 | 0 | 0.48 | 0.46 | 0.54 | 0.61 | 0.52 | 0.58 |
| 9 | 79126033 | rs17180166 | 1.173 | -1.222 | -0.817 | -1.863 | 0.01 | 0.019 | 0.019 | 0.653 | 0.18 | 0.07 | 0.18 | 0.17 | 0.14 | 0.15 |
| 9 | 79134925 | rs543677 | 0.919 | -1.26 | -0.683 | -1.835 | 0.045 | 0.048 | 0.048 | 0.957 | 0.36 | 0.23 | 0.19 | 0.08 | 0.26 | 0.2 |
| 9 | 79148376 | rs621293 | 0.484 | -0.153 | -0.17 | -1.159 | -0.005 | -0.021 | -0.021 | 0 | 0.61 | 0.51 | 0.42 | 0.27 | 0.44 | 0.48 |
| 9 | 79150662 | rs652061 | 0.338 | 0.311 | -0.118 | -0.702 | 0.11 | -0.019 | -0.019 | 0 | 0.77 | 0.54 | 0.72 | 0.85 | 0.69 | 0.76 |
| 9 | 79156769 | rs1886268 | -0.55 | -1.015 | -0.526 | -0.281 | -0.078 | -0.035 | -0.035 | 0 | 0.32 | 0.48 | 0.21 | 0.31 | 0.12 | 0.29 |
| 9 | 79159993 | rs927632 | -0.088 | -0.095 | -0.81 | -0.107 | -0.009 | 0.035 | 0.035 | 0 | 0.34 | 0.2 | 0.22 | 0.17 | 0.39 | 0.26 |
| 9 | 79160759 | rs2771026 | 0.404 | -0.618 | -0.285 | -0.032 | 0.003 | 0.007 | 0.007 | 0.081 | 0.51 | 0.52 | 0.69 | 0.63 | 0.53 | 0.58 |
| 9 | 79164646 | rs13290974 | 0.505 | -0.254 | -0.103 | 0.436 | 0.003 | 0.007 | 0.007 | 0 | 0.15 | 0.27 | 0.35 | 0.16 | 0.17 | 0.2 |
| 9 | 79174014 | rs13298775 | 1.025 | -0.252 | -0.027 | 0.477 | -0.004 | -0.004 | -0.004 | 0.006 | 0.11 | 0.11 | 0.21 | 0.11 | 0.14 | 0.11 |
| 9 | 79179861 | rs785896 | 0.966 | -0.353 | -0.099 | 0.325 | -0.036 | -0.06 | -0.06 | 0 | 0.37 | 0.17 | 0.36 | 0.48 | 0.4 | 0.32 |
| 9 | 79186052 | rs7857967 | 0.36 | -0.621 | 0.378 | 0.702 | 0.009 | 0.018 | 0.018 | 0 | 0.52 | 0.47 | 0.7 | 0.64 | 0.57 | 0.53 |
| 9 | 79188714 | rs7042923 | 0.247 | -0.593 | 0.645 | 0.703 | 0.001 | 0.002 | 0.002 | 0 | 0.54 | 0.61 | 0.71 | 0.64 | 0.57 | 0.54 |
| 9 | 79189090 | rs7043420 | 0.908 | -0.668 | 0.421 | 0.489 | -0.028 | -0.008 | -0.008 | 0 | 0.1 | 0.16 | 0.06 | 0.1 | 0.04 | 0.16 |
| 9 | 79190847 | rs2031908 | -0.191 | -0.568 | 0.356 | 0.522 | -0.001 | -0.002 | -0.002 | 0 | 0.62 | 0.64 | 0.76 | 0.7 | 0.61 | 0.69 |
| 9 | 79198957 | rs4237274 | -0.873 | -0.447 | -0.172 | 0.24 | -0.001 | 0 | 0 | 0 | 0.12 | 0.16 | 0.17 | 0.17 | 0.12 | 0.18 |
| 9 | 79199853 | rs4504705 | -1.047 | -0.387 | -0.336 | -0.152 | -0.02 | 0.016 | 0.016 | 0 | 0.79 | 0.87 | 0.81 | 0.92 | 0.68 | 0.9 |
| 9 | 79201074 | rs2149589 | -1.125 | -0.754 | -0.04 | 0.073 | -0.005 | 0.002 | 0.002 | 0 | 0.57 | 0.67 | 0.37 | 0.45 | 0.55 | 0.55 |
| 9 | 79203526 | rs13291560 | 0.551 | -0.263 | 0.391 | 0.497 | -0.038 | 0.065 | 0.065 | 0 | 0.24 | 0.09 | 0.14 | 0.08 | 0.28 | 0.11 |
| 9 | 79207324 | rs7873112 | -0.581 | -0.26 | 0.506 | 0.524 | 0.001 | 0.002 | 0.002 | 0 | 0.38 | 0.4 | 0.23 | 0.33 | 0.35 | 0.48 |
| 9 | 79212638 | rs2007778 | -1.435 | -0.161 | 0.097 | 0.544 | -0.024 | -0.04 | -0.04 | 1.536 | 0.54 | 0.9 | 0.34 | 0.45 | 0.46 | 0.63 |
| 9 | 79219334 | rs6420275 | -0.765 | 0.134 | -0.247 | -0.188 | 0.007 | 0.038 | 0.038 | 0 | 0.43 | 0.62 | 0.26 | 0.1 | 0.39 | 0.28 |
| 9 | 79221312 | rs7038278 | -1.144 | 0.031 | 0.164 | 0.665 | -0.022 | -0.009 | -0.009 | 0 | 0.22 | 0.27 | 0.13 | 0.21 | 0.18 | 0.4 |
| 9 | 79223252 | rs11144971 | -0.812 | 0.207 | -0.052 | 0.282 | 0.015 | 0.034 | 0.034 | 0 | 0.21 | 0.11 | 0.08 | 0.03 | 0.18 | 0.21 |
| 9 | 79225259 | rs812533 | 1.733 | 0.623 | 0.504 | 1.242 | 0 | -0.013 | -0.013 | 0.286 | 0.62 | 0.33 | 0.69 | 0.55 | 0.62 | 0.57 |
| 9 | 79225328 | rs11144978 | -1.117 | 0.578 | 0.439 | 0.909 | -0.002 | 0.075 | 0.075 | 0 | 0.27 | 0.11 | 0.14 | 0.04 | 0.27 | 0.25 |
| 9 | 79228352 | rs3087612 | -1.103 | 0.299 | 1.577 | 1.146 | 0.085 | -0.01 | -0.01 | 0 | 0.06 | 0.27 | 0.09 | 0 | 0.09 | 0.02 |
| 9 | 79230108 | rs7865529 | -2.196 | 0.23 | 1.578 | 1.844 | 0.022 | 0.139 | 0.139 | 2.146 | 0.12 | 0.59 | 0.11 | 0.22 | 0.1 | 0.15 |
| 9 | 79230242 | rs700800 | 1.303 | -0.025 | 1.784 | 2.324 | -0.039 | 0.039 | 0.039 | 0 | 0.76 | 0.67 | 0.86 | 0.54 | 0.91 | 0.64 |
| 9 | 79230395 | rs7865889 | -0.868 | -0.077 | 1.815 | 2.086 | -0.052 | 0.024 | 0.024 | 0 | 0.21 | 0.34 | 0.06 | 0.31 | 0.07 | 0.32 |
| 9 | 79230575 | rs10512062 | -0.395 | -0.048 | 1.599 | 1.856 | 0.014 | 0.015 | 0.015 | 0 | 0.17 | 0.12 | 0.03 | 0.09 | 0.06 | 0.21 |
| 9 | 79232203 | rs1329020 | 0.162 | -0.096 | 1.666 | 1.852 | 0.017 | 0.017 | 0.017 | 0 | 0.85 | 0.87 | 0.96 | 0.91 | 0.95 | 0.8 |
| 9 | 79233313 | rs786318 | 2.211 | -0.116 | 2.058 | 2.374 | 0.013 | 0.015 | 0.015 | 0.044 | 0.61 | 0.3 | 0.6 | 0.33 | 0.7 | 0.57 |
| 9 | 79233633 | rs786319 | 1.086 | -0.115 | 2.723 | 2.649 | -0.023 | -0.016 | -0.016 | 0 | 0.77 | 0.68 | 0.88 | 0.78 | 0.85 | 0.76 |
| 9 | 79234712 | rs700802 | 1.949 | -0.235 | 2.927 | 2.807 | -0.049 | 0.014 | 0.014 | 0 | 0.67 | 0.44 | 0.67 | 0.42 | 0.8 | 0.6 |
| 9 | 79234786 | rs890152 | -1.462 | -0.007 | 2.775 | 2.591 | -0.004 | 0.085 | 0.085 | 0 | 0.15 | 0.24 | 0.27 | 0.32 | 0.15 | 0.09 |
| 9 | 79236165 | rs786320 | 1.021 | -0.027 | 2.615 | 2.262 | -0.021 | -0.031 | -0.031 | 0 | 0.79 | 0.63 | 0.88 | 0.83 | 0.85 | 0.78 |
| 9 | 79236274 | rs1536821 | -0.251 | -0.136 | 2.359 | 2.205 | 0.023 | 0.021 | 0.021 | 0 | 0.87 | 0.89 | 0.97 | 0.93 | 0.96 | 0.82 |
| 9 | 79237318 | rs7867457 | -1.583 | 0.687 | 2.333 | 2.349 | 0.051 | 0.12 | 0.12 | 0 | 0.18 | 0.36 | 0.33 | 0.28 | 0.24 | 0.16 |
| 9 | 79240050 | rs700809 | -0.2 | 0.43 | 1.525 | 1.603 | 0.001 | 0.003 | 0.003 | 0 | 0.87 | 0.87 | 0.97 | 0.92 | 0.95 | 0.82 |
| 9 | 79243280 | rs7035759 | -1.702 | 1.241 | 2.277 | 2.031 | 0.058 | 0.006 | 0.006 | 0 | 0.18 | 0.34 | 0.16 | 0.17 | 0.25 | 0.14 |
| 9 | 79251229 | rs561450 | 1.388 | 1.439 | 2.326 | 2.43 | 0.011 | 0.033 | 0.033 | 0 | 0.85 | 0.84 | 0.82 | 0.77 | 0.79 | 0.84 |
| 9 | 79253347 | rs3739518 | -1.178 | 1.474 | 2.34 | 2.357 | 0.032 | 0.003 | 0.003 | 0.106 | 0.13 | 0.18 | 0.17 | 0.16 | 0.23 | 0.16 |
| 9 | 79254447 | rs605523 | 2.433 | 1.249 | 2.459 | 2.58 | -0.036 | -0.017 | -0.017 | 0 | 0.65 | 0.2 | 0.62 | 0.44 | 0.55 | 0.61 |
| 9 | 79259850 | rs512951 | 1.942 | 1.326 | 2.59 | 2.618 | 0.018 | 0.022 | 0.022 | 0 | 0.71 | 0.68 | 0.66 | 0.62 | 0.57 | 0.7 |
| 9 | 79272580 | rs7018789 | -1.73 | 1.128 | 1.857 | 2.239 | 0.014 | 0.129 | 0.129 | 0.096 | 0.2 | 0.63 | 0.22 | 0.41 | 0.22 | 0.25 |
| 9 | 79282119 | rs7862059 | -0.575 | 0.885 | 1.716 | 1.662 | 0.011 | -0.011 | -0.011 | 0 | 0.08 | 0.21 | 0.05 | 0.07 | 0.07 | 0.04 |
| 9 | 79282170 | rs17062868 | -1.18 | 0.914 | 1.975 | 1.522 | 0 | -0.025 | -0.025 | 0.689 | 0.12 | 0.15 | 0.11 | 0.05 | 0.11 | 0.04 |
| 9 | 79282589 | rs666721 | 0.92 | -0.261 | 1.062 | 1.331 | 0.091 | -0.007 | -0.007 | 0 | 0.53 | 0.14 | 0.61 | 0.55 | 0.69 | 0.71 |
| 9 | 79293941 | rs514208 | 0.316 | -0.374 | 1.983 | 1.415 | 0.002 | -0.001 | -0.001 | 0.165 | 0.61 | 0.45 | 0.67 | 0.63 | 0.76 | 0.75 |
| 9 | 79298133 | rs680949 | 0.767 | -0.317 | 1.922 | 1.387 | 0.002 | 0.091 | 0.091 | 1.426 | 0.87 | 0.5 | 0.92 | 0.82 | 0.9 | 0.84 |
| 9 | 79298723 | rs10123752 | -0.365 | -0.311 | 1.692 | 1.404 | 0.035 | 0.025 | 0.025 | 0.152 | 0.33 | 0.2 | 0.3 | 0.25 | 0.21 | 0.13 |
| 9 | 79305709 | rs678943 | 1.083 | 0.673 | 1.411 | 0.678 | 0.042 | -0.121 | -0.121 | 0 | 0.78 | 0.49 | 0.78 | 0.92 | 0.74 | 0.75 |
| 9 | 79318367 | rs4745571 | 0.848 | 0.73 | 0.677 | 0.509 | 0.019 | 0.022 | 0.022 | 1.053 | 0.25 | 0.03 | 0.27 | 0.17 | 0.17 | 0.09 |

| | | | | | | | | | | | | | | | |
|---|---|---|---|---|---|---|---|---|---|---|---|---|---|---|---|
| 9 | 79318471 | rs560094 | 0.633 | 1.046 | 1.823 | 0.677 | 0.029 | -0.059 | -0.059 | 0.395 | 0.83 | 0.69 | 0.82 | 0.91 | 0.77 | 0.76 |
| 9 | 79318998 | rs620552 | 0.633 | 1.046 | 1.823 | 0.677 | 0.029 | -0.059 | -0.059 | 0 | 0.83 | 0.69 | 0.82 | 0.91 | 0.77 | 0.76 |
| 9 | 79320095 | rs3739524 | -0.987 | 1.435 | 1.94 | 1.13 | 0.077 | 0.265 | 0.265 | 0 | 0.11 | 0.35 | 0.18 | 0.47 | 0.17 | 0.19 |
| 9 | 79320640 | rs512110 | 0.457 | 1.406 | 1.931 | 1.088 | 0.021 | -0.048 | -0.048 | 1.84 | 0.83 | 0.74 | 0.82 | 0.91 | 0.77 | 0.76 |
| 9 | 79321276 | rs517634 | 0.463 | 1.4 | 1.91 | 1.09 | 0.027 | -0.059 | -0.059 | 0 | 0.83 | 0.69 | 0.82 | 0.91 | 0.77 | 0.76 |
| 9 | 79322674 | rs680775 | 0.45 | 1.399 | 2.071 | 1.09 | 0.024 | -0.054 | -0.054 | 0 | 0.83 | 0.7 | 0.82 | 0.91 | 0.77 | 0.77 |
| 9 | 79322720 | rs680729 | 0.468 | 1.399 | 2.007 | 1.09 | 0.036 | -0.066 | -0.066 | 0 | 0.83 | 0.66 | 0.82 | 0.91 | 0.77 | 0.76 |
| 9 | 79324439 | rs12238013 | -0.646 | 1.438 | 2.085 | 1.144 | 0.031 | 0.179 | 0.179 | 0 | 0.1 | 0.21 | 0.16 | 0.46 | 0.16 | 0.18 |
| 9 | 79325615 | rs611461 | 0.479 | 1.426 | 2.058 | 1.103 | 0.027 | -0.056 | -0.056 | 0.014 | 0.83 | 0.69 | 0.82 | 0.92 | 0.77 | 0.76 |
| 9 | 79328461 | rs9785303 | -0.66 | 1.377 | 1.659 | 1.291 | 0.032 | 0.179 | 0.179 | 0 | 0.1 | 0.21 | 0.16 | 0.46 | 0.15 | 0.18 |
| 9 | 79333056 | rs890150 | 0.235 | 1.221 | 1.028 | 0.319 | -0.008 | 0.043 | 0.043 | 0 | 0.68 | 0.73 | 0.74 | 0.88 | 0.66 | 0.68 |
| 9 | 79335621 | rs10869801 | -1.18 | 1.239 | 1.313 | 0.75 | 0.043 | 0.136 | 0.136 | 0 | 0.12 | 0.25 | 0.18 | 0.57 | 0.18 | 0.24 |
| 9 | 79348193 | rs11145032 | -0.847 | 0.948 | 1.381 | 0.735 | 0.008 | 0.185 | 0.185 | 0 | 0.11 | 0.23 | 0.17 | 0.49 | 0.11 | 0.17 |
| 9 | 79349008 | rs7040264 | 1.482 | 0.693 | 1.171 | 0.249 | 0.004 | 0.073 | 0.073 | 0.035 | 0.48 | 0.21 | 0.53 | 0.32 | 0.45 | 0.4 |
| 9 | 79351237 | rs7856888 | 0.353 | 0.701 | 1.225 | 0.203 | -0.02 | 0.082 | 0.082 | 0 | 0.62 | 0.75 | 0.72 | 0.8 | 0.58 | 0.6 |
| 9 | 79353911 | rs1078986 | 0.131 | 0.677 | 1.026 | 0.206 | -0.063 | 0.149 | 0.149 | 0 | 0.64 | 0.89 | 0.74 | 0.8 | 0.61 | 0.61 |
| 9 | 79360937 | rs10869812 | -0.974 | 0.663 | 1.39 | 0.457 | 0.021 | 0.228 | 0.228 | 0 | 0.12 | 0.37 | 0.16 | 0.49 | 0.13 | 0.2 |
| 9 | 79361021 | rs11145038 | -1.236 | 0.662 | 1.456 | 0.456 | 0.034 | 0.296 | 0.296 | 0.002 | 0.13 | 0.44 | 0.17 | 0.49 | 0.13 | 0.2 |
| 9 | 79370447 | rs2225245 | 0.78 | 0.741 | 1.157 | 0.305 | -0.001 | 0.056 | 0.056 | 0 | 0.54 | 0.62 | 0.63 | 0.72 | 0.48 | 0.52 |
| 9 | 79371206 | rs9314856 | -0.576 | 0.416 | 1.439 | 0.614 | -0.006 | 0.316 | 0.316 | 0 | 0.1 | 0.31 | 0.13 | 0.41 | 0.07 | 0.13 |
| 9 | 79372502 | rs1425284 | 0.601 | 0.724 | 0.988 | 0.225 | -0.029 | 0.079 | 0.079 | 0 | 0.62 | 0.79 | 0.71 | 0.72 | 0.58 | 0.51 |
| 9 | 79375225 | rs4327916 | 0.746 | 0.657 | 1.045 | 0.181 | 0.001 | -0.038 | -0.038 | 0.021 | 0.62 | 0.46 | 0.72 | 0.72 | 0.62 | 0.52 |
| 9 | 79381463 | rs10521471 | 2.34 | 0.362 | -0.005 | -0.052 | 0.005 | 0.018 | 0.018 | 0 | 0.21 | 0.02 | 0.21 | 0.14 | 0.16 | 0.09 |
| 9 | 79384370 | rs566234 | -0.345 | 0.896 | 0.641 | 0.592 | -0.062 | 0.023 | 0.023 | 0 | 0.31 | 0.09 | 0.22 | 0.3 | 0.34 | 0.44 |
| 9 | 79385980 | rs627587 | -1.661 | 0.895 | 0.87 | 0.599 | 0.001 | 0.104 | 0.104 | 0.276 | 0.48 | 0.83 | 0.44 | 0.69 | 0.46 | 0.59 |
| 9 | 79386445 | rs615037 | -1.253 | 0.913 | 1.187 | 0.608 | -0.001 | -0.014 | -0.014 | 0 | 0.45 | 0.58 | 0.41 | 0.69 | 0.46 | 0.59 |
| 9 | 79390020 | rs1936283 | -1.503 | 0.972 | 1.04 | 0.634 | 0.001 | 0.093 | 0.093 | 0.273 | 0.48 | 0.82 | 0.43 | 0.69 | 0.46 | 0.58 |
| 9 | 79392249 | rs12379174 | 0.18 | 1.077 | 1.162 | 0.589 | -0.003 | -0.006 | -0.006 | 0.041 | 0.18 | 0.28 | 0.11 | 0.22 | 0.14 | 0.25 |
| 9 | 79395872 | rs7861307 | -1.082 | 1.313 | 0.756 | 1.124 | 0 | 0.012 | 0.012 | 0.081 | 0.18 | 0.24 | 0.24 | 0.3 | 0.2 | 0.26 |
| 9 | 79396072 | rs7864683 | -0.481 | 1.191 | 0.885 | 0.722 | -0.016 | -0.006 | -0.006 | 1.3 | 0.26 | 0.37 | 0.17 | 0.29 | 0.22 | 0.3 |
| 9 | 79400295 | rs954974 | -0.877 | 1.314 | 0.753 | 0.674 | 0.029 | -0.026 | -0.026 | 0 | 0.27 | 0.47 | 0.2 | 0.28 | 0.27 | 0.36 |
| 9 | 79402584 | rs3924629 | -0.266 | 1.171 | 0.804 | 0.652 | 0.032 | 0.028 | 0.028 | 3.357 | 0.1 | 0.36 | 0.06 | 0.08 | 0.08 | 0.06 |
| 9 | 79404589 | rs10869820 | -0.249 | 1.165 | 0.699 | 0.649 | 0.036 | 0.031 | 0.031 | 4.217 | 0.1 | 0.39 | 0.06 | 0.08 | 0.08 | 0.06 |
| 9 | 79404749 | rs7854027 | 2.308 | 0.923 | 0.199 | 0.445 | 0.017 | 0.022 | 0.022 | 1.409 | 0.35 | 0.27 | 0.28 | 0.21 | 0.25 | 0.23 |
| 9 | 79408144 | rs10781380 | -1.079 | 0.85 | 0.38 | 0.749 | -0.006 | -0.005 | -0.005 | 0.001 | 0.29 | 0.54 | 0.2 | 0.33 | 0.26 | 0.31 |
| 9 | 79412910 | rs10781381 | -0.406 | 0.727 | 0.36 | 0.99 | -0.006 | 0.055 | 0.055 | 0.003 | 0.21 | 0.3 | 0.26 | 0.35 | 0.2 | 0.19 |
| 9 | 79420249 | rs11145077 | -0.125 | 0.851 | -0.039 | 0.986 | 0 | 0.001 | 0.001 | 0 | 0.2 | 0.18 | 0.09 | 0.23 | 0.16 | 0.22 |
| 9 | 79424168 | rs1343396 | 0.399 | 1.071 | -0.301 | 0.805 | -0.003 | -0.011 | -0.011 | 0 | 0.7 | 0.76 | 0.56 | 0.66 | 0.64 | 0.69 |
| 9 | 79425692 | rs10521470 | 0.932 | 1.325 | -0.731 | 0.892 | -0.019 | -0.024 | -0.024 | 0.503 | 0.1 | 0.07 | 0.09 | 0.13 | 0.12 | 0.16 |
| 9 | 79429800 | rs10124357 | 1.365 | 1.147 | -0.453 | 0.632 | -0.003 | 0.01 | 0.01 | 1.323 | 0.52 | 0.41 | 0.42 | 0.48 | 0.49 | 0.51 |
| 9 | 79432780 | rs3905245 | 1.315 | 1.139 | -0.457 | 0.628 | -0.003 | 0.008 | 0.008 | 1.183 | 0.53 | 0.42 | 0.43 | 0.48 | 0.49 | 0.51 |
| 9 | 79435440 | rs1343394 | 2.065 | 1.117 | -0.479 | 0.625 | 0.002 | 0.006 | 0.006 | 1.949 | 0.39 | 0.34 | 0.35 | 0.25 | 0.35 | 0.29 |
| 9 | 79436617 | rs7867090 | -1.511 | 1.231 | -0.702 | 0.784 | -0.052 | -0.078 | -0.078 | 0 | 0.88 | 0.99 | 0.68 | 0.72 | 0.81 | 0.84 |
| 9 | 79437798 | rs10869828 | -0.154 | 0.796 | -0.279 | -0.035 | -0.051 | -0.025 | -0.025 | 0 | 0.62 | 0.81 | 0.48 | 0.5 | 0.52 | 0.51 |
| 9 | 79438813 | rs10869829 | 0.561 | 1.132 | -0.438 | 0.001 | -0.021 | -0.003 | -0.003 | 0.037 | 0.13 | 0.06 | 0.08 | 0.2 | 0.13 | 0.2 |
| 9 | 79442798 | rs4744812 | 1.645 | 1.223 | -0.434 | -0.082 | 0.005 | 0.007 | 0.007 | 1.703 | 0.36 | 0.32 | 0.32 | 0.22 | 0.28 | 0.23 |
| 9 | 79444014 | rs10869830 | 0.517 | 1.213 | -0.806 | 0.024 | -0.023 | -0.019 | -0.019 | 0 | 0.13 | 0.06 | 0.08 | 0.23 | 0.14 | 0.2 |
| 9 | 79445633 | rs10869832 | 0.143 | 1.108 | -0.735 | -0.041 | -0.013 | 0.006 | 0.006 | 2.477 | 0.57 | 0.66 | 0.44 | 0.47 | 0.49 | 0.49 |
| 9 | 79450930 | rs2000289 | 0.475 | 0.815 | -0.679 | -0.232 | -0.001 | -0.003 | -0.003 | 1.327 | 0.41 | 0.57 | 0.37 | 0.24 | 0.34 | 0.28 |
| 9 | 79451775 | rs10869838 | 0.198 | 0.282 | -1.174 | -0.146 | 0.009 | 0.063 | 0.063 | 2.255 | 0.47 | 0.36 | 0.42 | 0.28 | 0.44 | 0.25 |
| 9 | 79453107 | rs7853555 | 0.126 | -0.936 | -1.422 | -0.192 | -0.006 | 0.012 | 0.012 | 0.102 | 0.39 | 0.16 | 0.39 | 0.35 | 0.56 | 0.48 |
| 9 | 79455225 | rs10781386 | 0.431 | -1.329 | -0.968 | -0.754 | 0.016 | 0.028 | 0.028 | 0 | 0.75 | 0.87 | 0.76 | 0.86 | 0.77 | 0.84 |
| 9 | 79459024 | rs1936285 | -0.194 | -1.633 | 0.061 | -0.737 | -0.022 | -0.041 | -0.041 | 0.061 | 0.32 | 0.35 | 0.15 | 0.16 | 0.21 | 0.22 |
| 9 | 79459478 | rs12685440 | -0.907 | -1.792 | -0.212 | -0.714 | 0.094 | 0.091 | 0.091 | 0 | 0.43 | 0.78 | 0.22 | 0.31 | 0.28 | 0.35 |
| 9 | 79460804 | rs17785193 | -1.126 | -0.976 | -0.017 | -0.181 | 0.009 | -0.014 | -0.014 | 0.358 | 0.08 | 0.06 | 0.05 | 0.14 | 0.07 | 0.06 |
| 9 | 79467016 | rs6560554 | -1.11 | -0.989 | 0.415 | -0.172 | -0.017 | 0.027 | 0.027 | 0.28 | 0.08 | 0.22 | 0.06 | 0.14 | 0.06 | 0.09 |
| 9 | 79467204 | rs1361856 | -0.144 | -1.51 | 0.442 | -0.586 | -0.04 | -0.064 | -0.064 | 0.019 | 0.32 | 0.41 | 0.15 | 0.16 | 0.23 | 0.18 |
| 9 | 79467459 | rs1361857 | -0.621 | -1.429 | 0.46 | -0.512 | -0.026 | -0.019 | -0.019 | 0.102 | 0.41 | 0.49 | 0.2 | 0.33 | 0.29 | 0.28 |
| 9 | 79471562 | rs4745588 | -0.041 | -0.864 | 0.176 | -0.377 | 0.018 | 0.082 | 0.082 | 0.09 | 0.56 | 0.7 | 0.65 | 0.8 | 0.58 | 0.64 |
| 9 | 79471995 | rs12238848 | 0.592 | -0.758 | -0.307 | -0.299 | -0.03 | 0.035 | 0.035 | 0 | 0.43 | 0.24 | 0.59 | 0.61 | 0.49 | 0.47 |
| 9 | 79477693 | rs10116340 | -1.259 | -0.818 | 0.406 | 0.262 | -0.04 | -0.005 | -0.005 | 0 | 0.18 | 0.62 | 0.07 | 0.19 | 0.09 | 0.17 |
| 9 | 79500154 | rs1125469 | -0.095 | -0.258 | -0.311 | 0.628 | -0.012 | -0.02 | -0.02 | 0 | 0.65 | 0.69 | 0.63 | 0.49 | 0.55 | 0.66 |
| 9 | 79504068 | rs11145117 | -0.158 | -0.584 | -0.67 | 0.662 | 0.011 | -0.045 | -0.045 | 0 | 0.16 | 0.02 | 0.16 | 0.25 | 0.11 | 0.2 |
| 9 | 79505493 | rs944561 | 0.58 | -0.675 | -0.24 | 0.85 | 0 | 0.015 | 0.015 | 0 | 0.48 | 0.59 | 0.47 | 0.32 | 0.45 | 0.49 |
| 9 | 79505647 | rs944559 | -0.237 | -0.575 | -0.104 | 0.793 | -0.027 | -0.02 | -0.02 | 1.124 | 0.65 | 0.76 | 0.65 | 0.58 | 0.56 | 0.72 |
| 9 | 79508251 | rs2377811 | -0.279 | -0.614 | -0.059 | 0.763 | -0.001 | -0.001 | -0.001 | 0 | 0.66 | 0.68 | 0.65 | 0.58 | 0.59 | 0.74 |
| 9 | 79511499 | rs7031918 | -0.768 | -0.654 | 0.585 | 0.255 | -0.024 | -0.008 | -0.008 | 0.279 | 0.19 | 0.3 | 0.19 | 0.23 | 0.11 | 0.22 |
| 9 | 79515295 | rs2154044 | 0.495 | -0.938 | 0.603 | 0.011 | 0.002 | -0.043 | -0.043 | 0 | 0.43 | 0.54 | 0.41 | 0.09 | 0.45 | 0.36 |
| 9 | 79518215 | rs7035057 | -0.427 | -1.118 | 0.402 | 0.219 | -0.028 | 0.009 | 0.009 | 0 | 0.61 | 0.79 | 0.62 | 0.38 | 0.54 | 0.68 |

| | | | | | | | | | | | | | | | | |
|---|---|---|---|---|---|---|---|---|---|---|---|---|---|---|---|---|
| 9 | 79521736 | rs10869843 | 0.506 | -0.855 | 1.122 | 0.118 | 0.003 | -0.038 | -0.038 | 0.382 | 0.44 | 0.53 | 0.51 | 0.14 | 0.53 | 0.39 |
| 9 | 79524756 | rs3843581 | -0.047 | -0.668 | 1.462 | 0.252 | -0.003 | -0.003 | -0.003 | 0 | 0.57 | 0.46 | 0.65 | 0.41 | 0.63 | 0.65 |
| 9 | 79526812 | rs17787855 | 0.32 | -0.653 | 0.639 | 0.232 | -0.023 | -0.058 | -0.058 | 0.099 | 0.1 | 0.01 | 0.15 | 0.27 | 0.09 | 0.25 |
| 9 | 79528772 | rs10869844 | -0.289 | -0.645 | 0.748 | 0.211 | 0.003 | 0.007 | 0.007 | 0 | 0.59 | 0.68 | 0.57 | 0.41 | 0.62 | 0.64 |
| 9 | 79530722 | rs1929533 | -0.358 | -0.632 | 0.356 | 0.207 | -0.008 | 0.004 | 0.004 | 0 | 0.6 | 0.81 | 0.58 | 0.41 | 0.62 | 0.64 |
| 9 | 79537871 | rs10121560 | 0.136 | -0.447 | 0.114 | -0.235 | -0.001 | -0.008 | -0.008 | 0 | 0.48 | 0.46 | 0.5 | 0.14 | 0.55 | 0.4 |
| 9 | 79546523 | rs6560565 | -0.206 | -0.086 | -0.612 | 0.126 | -0.019 | -0.018 | -0.018 | 3.772 | 0.64 | 0.78 | 0.71 | 0.44 | 0.6 | 0.64 |
| 9 | 79548154 | rs10116164 | 0.088 | -0.281 | -0.662 | -0.669 | 0.004 | 0.009 | 0.009 | 0 | 0.2 | 0.15 | 0.24 | 0.05 | 0.22 | 0.1 |
| 9 | 79561569 | rs10217550 | -0.566 | -0.64 | -0.138 | -1.26 | -0.042 | -0.182 | -0.182 | 0 | 0.31 | 0.45 | 0.27 | 0.02 | 0.25 | 0.12 |
| 9 | 79566864 | rs13285464 | 0.19 | 0.092 | -0.857 | -0.56 | -0.008 | -0.012 | -0.012 | 0 | 0.62 | 0.67 | 0.7 | 0.39 | 0.62 | 0.7 |
| 9 | 79566889 | rs7035636 | -0.841 | 0.524 | -1.321 | -0.446 | -0.091 | -0.152 | -0.152 | 0.449 | 0.8 | 0.99 | 0.82 | 0.41 | 0.76 | 0.81 |
| 9 | 79569218 | rs7027830 | -0.052 | 0.089 | -0.588 | -0.88 | -0.002 | -0.045 | -0.045 | 0.333 | 0.12 | 0.22 | 0.09 | 0.02 | 0.1 | 0.08 |
| 9 | 79571921 | rs7857541 | 0.026 | 0.293 | -0.703 | -0.977 | -0.002 | -0.03 | -0.03 | 0 | 0.41 | 0.48 | 0.43 | 0.18 | 0.41 | 0.48 |
| 9 | 79571944 | rs11145148 | 0.193 | 0.783 | -0.514 | -0.805 | 0 | -0.012 | -0.012 | 0.011 | 0.09 | 0.07 | 0.12 | 0 | 0.17 | 0.05 |
| 9 | 79574979 | rs2183770 | 0.12 | 0.369 | -0.644 | -0.975 | -0.006 | 0.029 | 0.029 | 0 | 0.54 | 0.58 | 0.47 | 0.19 | 0.49 | 0.54 |
| 9 | 79576188 | rs7852521 | -0.083 | 0.546 | -0.493 | -0.852 | -0.012 | 0.023 | 0.023 | 0.016 | 0.59 | 0.63 | 0.49 | 0.21 | 0.51 | 0.57 |
| 9 | 79577007 | rs7856464 | -0.09 | 0.813 | -0.56 | -0.95 | -0.007 | 0.008 | 0.008 | 0.094 | 0.68 | 0.66 | 0.63 | 0.13 | 0.62 | 0.6 |
| 9 | 79578016 | rs1929540 | -0.45 | 0.438 | 0.124 | -0.656 | -0.01 | -0.074 | -0.074 | 0 | 0.16 | 0.32 | 0.18 | 0.02 | 0.13 | 0.09 |
| 9 | 79581742 | rs4745601 | -0.819 | 0.996 | -0.838 | -0.173 | -0.086 | -0.064 | -0.064 | 0.473 | 0.3 | 0.11 | 0.4 | 0.36 | 0.39 | 0.34 |
| 9 | 79582862 | rs1929539 | -1.653 | 0.473 | -0.208 | 0.013 | 0.001 | 0.003 | 0.003 | 0 | 0.47 | 0.4 | 0.57 | 0.61 | 0.52 | 0.48 |
| 9 | 79584585 | rs7856621 | 0.093 | 0.54 | -0.311 | 0.096 | 0.008 | -0.012 | -0.012 | 6.327 | 0.15 | 0.12 | 0.16 | 0.24 | 0.13 | 0.14 |
| 9 | 79589352 | rs4744815 | 0.194 | 0.817 | -0.369 | 0.028 | -0.127 | -0.139 | -0.139 | 0.745 | 0.23 | 0.02 | 0.33 | 0.36 | 0.31 | 0.28 |
| 9 | 79592158 | rs11145168 | 0.889 | 0.532 | -0.115 | -0.036 | -0.003 | -0.023 | -0.023 | 0.248 | 0.09 | 0.13 | 0.04 | 0.01 | 0.05 | 0.09 |
| 9 | 79592957 | rs11145170 | -1.043 | -0.142 | 0.256 | -0.423 | -0.015 | -0.018 | -0.018 | 0 | 0.3 | 0.37 | 0.23 | 0.23 | 0.2 | 0.21 |
| 9 | 79594546 | rs7033468 | -2.006 | -0.184 | 0.431 | -0.539 | -0.024 | 0.025 | 0.025 | 0 | 0.57 | 0.7 | 0.6 | 0.64 | 0.5 | 0.45 |
| 9 | 79599678 | rs10869856 | 0.17 | -0.359 | 0.783 | -0.506 | -0.039 | -0.105 | -0.105 | 0 | 0.19 | 0.34 | 0.09 | 0.02 | 0.1 | 0.12 |
| 9 | 79603916 | rs11145189 | -0.83 | -0.381 | 0.868 | -0.415 | -0.006 | -0.002 | -0.002 | 0 | 0.2 | 0.28 | 0.21 | 0.24 | 0.16 | 0.19 |
| 9 | 79604018 | rs1929538 | -1.973 | -0.099 | 0.744 | -0.489 | -0.019 | 0.017 | 0.017 | 0 | 0.58 | 0.66 | 0.61 | 0.65 | 0.52 | 0.46 |
| 9 | 79605912 | rs1929536 | -0.735 | -0.327 | 0.738 | -0.497 | -0.003 | 0.001 | 0.001 | 0 | 0.43 | 0.42 | 0.44 | 0.44 | 0.39 | 0.36 |
| 9 | 79607007 | rs7853173 | -2.005 | -0.166 | 0.703 | -0.485 | -0.022 | 0.023 | 0.023 | 0.058 | 0.58 | 0.68 | 0.61 | 0.66 | 0.51 | 0.45 |
| 9 | 79608487 | rs7048169 | -0.742 | -0.139 | 0.712 | -0.764 | 0.006 | -0.005 | -0.005 | 0.416 | 0.44 | 0.37 | 0.43 | 0.44 | 0.4 | 0.37 |
| 9 | 79611749 | rs7030774 | 0.403 | -0.381 | 0.468 | -0.64 | -0.054 | -0.102 | -0.102 | 0 | 0.27 | 0.1 | 0.38 | 0.41 | 0.37 | 0.25 |
| 9 | 79612186 | rs7858152 | -0.09 | -0.15 | 0.827 | -0.394 | -0.025 | -0.046 | -0.046 | 1.144 | 0.16 | 0.35 | 0.13 | 0.1 | 0.08 | 0.11 |
| 9 | 79615741 | rs4745607 | -1.436 | -0.096 | 0.416 | -0.354 | -0.023 | -0.015 | -0.015 | 0 | 0.58 | 0.68 | 0.58 | 0.51 | 0.51 | 0.44 |
| 9 | 79623442 | rs4745608 | -0.417 | -0.078 | 0.252 | -0.385 | -0.01 | -0.032 | -0.032 | 1.403 | 0.43 | 0.21 | 0.44 | 0.41 | 0.45 | 0.34 |
| 9 | 79629068 | rs7027412 | -0.196 | 0.072 | 1.317 | -0.579 | -0.002 | -0.03 | -0.03 | 0 | 0.08 | 0.31 | 0.04 | 0.03 | 0.03 | 0.02 |
| 9 | 79629718 | rs10869866 | 0.061 | 0.166 | 1.089 | -0.57 | -0.011 | -0.038 | -0.038 | 0 | 0.08 | 0.16 | 0.05 | 0.03 | 0.04 | 0.02 |
| 9 | 79636426 | rs7044545 | -0.992 | 0.022 | 1.17 | -0.624 | -0.024 | -0.083 | -0.083 | 1.838 | 0.13 | 0.39 | 0.08 | 0.03 | 0.05 | 0.09 |
| 9 | 79643960 | rs7864598 | -0.325 | 0.27 | -0.094 | -0.388 | -0.013 | 0.018 | 0.018 | 0 | 0.39 | 0.32 | 0.51 | 0.35 | 0.45 | 0.39 |
| 9 | 79646546 | rs1928481 | -1.181 | 0.382 | 0.114 | -0.779 | 0.002 | -0.021 | -0.021 | 0.395 | 0.81 | 0.83 | 0.84 | 0.92 | 0.81 | 0.78 |
| 9 | 79653957 | rs10118945 | -0.401 | 0.362 | 0.963 | -0.667 | 0.025 | -0.026 | -0.026 | 0 | 0.09 | 0.34 | 0.09 | 0.01 | 0.07 | 0.09 |
| 9 | 79658445 | rs4744818 | -1.368 | 0.02 | 0.381 | -1.107 | 0.008 | 0.067 | 0.067 | 0 | 0.69 | 0.86 | 0.81 | 0.99 | 0.77 | 0.74 |
| 9 | 79664633 | rs11145238 | -0.842 | 0.012 | 0.827 | -1.323 | -0.006 | -0.042 | -0.042 | 0 | 0.2 | 0.31 | 0.19 | 0.06 | 0.18 | 0.21 |
| 9 | 79664647 | rs11145239 | -0.817 | 0.392 | 0.824 | -0.887 | 0 | 0.025 | 0.025 | 0.171 | 0.35 | 0.57 | 0.44 | 0.44 | 0.36 | 0.4 |
| 9 | 79665627 | rs4012486 | 0.999 | 0.415 | 0.83 | -0.87 | 0.02 | 0.118 | 0.118 | 0 | 0.18 | 0.33 | 0.29 | 0.42 | 0.22 | 0.22 |
| 9 | 79666892 | rs11145242 | -0.352 | 0.02 | 0.746 | -1.11 | -0.006 | -0.039 | -0.039 | 0 | 0.16 | 0.24 | 0.15 | 0.02 | 0.14 | 0.18 |
| 9 | 79667495 | rs1556152 | -1.359 | 0.097 | 0.635 | -1.021 | 0.022 | 0.105 | 0.105 | 0 | 0.59 | 0.82 | 0.75 | 0.99 | 0.69 | 0.65 |
| 9 | 79669104 | rs10435941 | -1.655 | 0.045 | 0.748 | -0.484 | 0.022 | 0.055 | 0.055 | 1.307 | 0.14 | 0.48 | 0.14 | 0.14 | 0.13 | 0.16 |
| 9 | 79677419 | rs12352362 | 0.094 | 0.257 | -0.179 | -0.498 | 0.003 | 0.001 | 0.001 | 2.14 | 0.13 | 0.09 | 0.06 | 0.05 | 0.12 | 0.05 |
| 9 | 79687816 | rs13302493 | -0.228 | 0.651 | 0.021 | -0.14 | 0 | 0 | 0 | 0 | 0.08 | 0.1 | 0.08 | 0.07 | 0.11 | 0.08 |
| 9 | 79691690 | rs12345375 | 0.07 | 0.736 | 0.501 | 0.013 | 0.02 | 0.099 | 0.099 | 0 | 0.24 | 0.44 | 0.35 | 0.4 | 0.28 | 0.29 |
| 9 | 79693492 | rs10869887 | 0.521 | 0.737 | 0.399 | 0.143 | 0.019 | 0.053 | 0.053 | 0 | 0.2 | 0.34 | 0.35 | 0.4 | 0.26 | 0.19 |
| 9 | 79695482 | rs7029178 | -0.137 | 0.682 | 0.089 | -0.006 | 0.012 | 0.045 | 0.045 | 0 | 0.67 | 0.51 | 0.61 | 0.56 | 0.64 | 0.62 |
| 9 | 79699406 | rs7870404 | 0.739 | 0.668 | 0.058 | -0.102 | 0.007 | 0.025 | 0.025 | 0 | 0.22 | 0.3 | 0.35 | 0.38 | 0.26 | 0.2 |
| 9 | 79699486 | rs7857478 | -0.245 | 0.787 | 0.552 | 0.287 | 0.002 | 0.014 | 0.014 | 0 | 0.7 | 0.67 | 0.63 | 0.56 | 0.66 | 0.63 |
| 9 | 79700008 | rs10156664 | -0.137 | 0.673 | 0.674 | 0.384 | 0.008 | 0.018 | 0.018 | 0 | 0.72 | 0.59 | 0.7 | 0.69 | 0.68 | 0.72 |
| 9 | 79702675 | rs10781417 | 0.969 | 0.547 | 0.538 | 0.19 | 0 | 0 | 0 | 0.06 | 0.55 | 0.43 | 0.63 | 0.54 | 0.51 | 0.57 |
| 9 | 79706374 | rs10781419 | 0.303 | 0.431 | 0.078 | 0.176 | -0.001 | -0.001 | -0.001 | 0.004 | 0.66 | 0.67 | 0.7 | 0.7 | 0.62 | 0.67 |
| 9 | 79707908 | rs4361823 | -0.244 | 0.537 | 0.198 | 0.465 | 0.002 | 0.007 | 0.007 | 0 | 0.73 | 0.69 | 0.65 | 0.7 | 0.69 | 0.71 |
| 9 | 79714137 | rs10869896 | -1.087 | 0.354 | 0.466 | 0.51 | -0.004 | 0.049 | 0.049 | 0.111 | 0.14 | 0.33 | 0.06 | 0.15 | 0.09 | 0.13 |
| 9 | 79724774 | rs7020571 | -0.504 | 0.359 | -0.173 | 0.413 | 0.017 | 0.023 | 0.023 | 0 | 0.47 | 0.66 | 0.56 | 0.66 | 0.4 | 0.52 |
| 9 | 79726518 | rs4484766 | -0.212 | 0.219 | 0.405 | 0.683 | 0 | -0.001 | -0.001 | 0.067 | 0.38 | 0.42 | 0.51 | 0.46 | 0.36 | 0.45 |
| 9 | 79741569 | rs4634718 | 0.585 | 0.381 | 0.508 | 0.529 | -0.023 | -0.036 | -0.036 | 0 | 0.17 | 0.13 | 0.41 | 0.49 | 0.26 | 0.3 |
| 9 | 79747317 | rs1953018 | -0.19 | 0.255 | 0.63 | 0.349 | -0.004 | -0.004 | -0.004 | 0 | 0.27 | 0.32 | 0.46 | 0.53 | 0.35 | 0.33 |
| 9 | 79757559 | rs12683654 | 0.327 | 0.465 | 0.406 | -0.071 | 0.011 | 0.017 | 0.017 | 0 | 0.21 | 0.14 | 0.12 | 0.04 | 0.17 | 0.17 |
| 9 | 79761711 | rs12345013 | 1.183 | 0.522 | 0.44 | 0.061 | 0.001 | 0 | 0 | 0 | 0.07 | 0.11 | 0.05 | 0.04 | 0.09 | 0.04 |
| 9 | 79763338 | rs10118161 | -0.226 | 0.47 | 0.567 | 0.021 | 0.014 | -0.009 | -0.009 | 0 | 0.11 | 0.29 | 0.07 | 0.04 | 0.11 | 0.09 |
| 9 | 79763758 | rs4548251 | -0.73 | 0.468 | 0.355 | 0.21 | -0.007 | 0.016 | 0.016 | 0 | 0.46 | 0.39 | 0.58 | 0.67 | 0.47 | 0.53 |
| 9 | 79776332 | rs10512064 | 0.538 | 0.116 | -0.342 | -0.322 | -0.002 | 0.009 | 0.009 | 8.271 | 0.13 | 0.12 | 0.08 | 0.04 | 0.13 | 0.09 |

| | | | | | | | | | | | | | | | | |
|---|---|---|---|---|---|---|---|---|---|---|---|---|---|---|---|---|
| 9 | 79783751 | rs12156556 | -1.441 | 0.422 | 0.116 | 0.191 | 0.001 | 0.01 | 0.01 | 0.125 | 0.54 | 0.39 | 0.49 | 0.54 | 0.5 | 0.56 |
| 9 | 79786834 | rs11790663 | 1.276 | 0.493 | 0.024 | 0.546 | -0.062 | -0.086 | -0.086 | 0 | 0.15 | 0.04 | 0.38 | 0.41 | 0.22 | 0.29 |
| 9 | 79790425 | rs17785751 | 0.695 | 0.483 | 0.075 | 0.388 | -0.001 | 0.004 | 0.004 | 8.612 | 0.13 | 0.13 | 0.08 | 0.04 | 0.13 | 0.08 |
| 9 | 79791793 | rs1054368 | -1.563 | 0.313 | 0.493 | 0.328 | 0.002 | -0.01 | -0.01 | 0 | 0.57 | 0.55 | 0.52 | 0.63 | 0.51 | 0.57 |
| 9 | 79802986 | rs12338283 | -0.062 | 0.174 | 0.552 | 0.139 | -0.004 | -0.023 | -0.023 | 0.07 | 0.25 | 0.26 | 0.16 | 0.06 | 0.21 | 0.23 |
| 9 | 79818630 | rs12684856 | 2.437 | 0.148 | -0.14 | -0.624 | -0.023 | 0.051 | 0.051 | 0 | 0.14 | 0.04 | 0.11 | 0.05 | 0.16 | 0.15 |
| 9 | 79824002 | rs10869910 | 1.422 | 0.194 | 0.004 | -0.306 | -0.027 | -0.065 | -0.065 | 0 | 0.17 | 0.07 | 0.3 | 0.41 | 0.2 | 0.32 |
| 9 | 79835101 | rs3802497 | 0.506 | -0.032 | -0.08 | -0.408 | -0.001 | -0.007 | -0.007 | 1.441 | 0.12 | 0.04 | 0.06 | 0.11 | 0.1 | 0.06 |
| 9 | 79846516 | rs11145347 | 0.372 | -0.043 | -0.038 | -0.43 | 0.008 | 0.018 | 0.018 | 0.361 | 0.24 | 0.21 | 0.14 | 0.06 | 0.2 | 0.21 |
| 9 | 79866051 | rs2377903 | -0.727 | -0.231 | 0.046 | -0.414 | 0.006 | 0.014 | 0.014 | 0.016 | 0.37 | 0.3 | 0.22 | 0.19 | 0.31 | 0.29 |
| 9 | 79873690 | rs2150904 | -1.591 | -0.181 | 0.152 | -0.33 | -0.001 | -0.002 | -0.002 | 0 | 0.59 | 0.61 | 0.63 | 0.68 | 0.55 | 0.61 |
| 9 | 79888146 | rs17063468 | 1 | -0.228 | 0.101 | -0.335 | -0.005 | -0.015 | -0.015 | 0 | 0.11 | 0.04 | 0.06 | 0.12 | 0.1 | 0.06 |
| 9 | 79888932 | rs10781429 | -1.249 | -0.224 | 0.146 | -0.245 | 0 | -0.003 | -0.003 | 0 | 0.56 | 0.46 | 0.6 | 0.58 | 0.53 | 0.61 |
| 9 | 79897117 | rs11145366 | 1.679 | -0.227 | 0.085 | -0.305 | 0 | 0.008 | 0.008 | 9.288 | 0.11 | 0.12 | 0.07 | 0.03 | 0.12 | 0.07 |
| 9 | 79901703 | rs12342114 | -0.716 | -0.236 | 0.11 | -0.313 | 0.007 | 0.023 | 0.023 | 0.021 | 0.37 | 0.29 | 0.19 | 0.16 | 0.31 | 0.27 |
| 9 | 79935018 | rs9696759 | -1.421 | -0.258 | 0.072 | -0.414 | -0.001 | -0.016 | -0.016 | 1.216 | 0.57 | 0.51 | 0.62 | 0.68 | 0.54 | 0.61 |
| 9 | 79946961 | rs17424026 | 0.412 | -0.196 | -0.18 | -0.355 | -0.001 | -0.009 | -0.009 | 0 | 0.12 | 0.03 | 0.06 | 0.11 | 0.1 | 0.06 |
| 9 | 79954545 | rs7025532 | 0.291 | -0.211 | -0.128 | -0.411 | 0.001 | -0.01 | -0.01 | 1.922 | 0.24 | 0.23 | 0.12 | 0.03 | 0.2 | 0.2 |
| 9 | 79973725 | rs10781433 | -1.54 | -0.212 | 0.053 | -0.351 | 0 | 0 | 0 | 1.134 | 0.61 | 0.61 | 0.61 | 0.57 | 0.54 | 0.64 |
| 9 | 79978900 | rs7041310 | -0.575 | -0.217 | -0.085 | -0.399 | 0.002 | 0.002 | 0.002 | 0.013 | 0.37 | 0.33 | 0.22 | 0.13 | 0.31 | 0.29 |
| 9 | 79986057 | rs3737289 | -0.381 | -0.199 | -0.053 | -0.103 | -0.004 | 0.017 | 0.017 | 0.13 | 0.46 | 0.3 | 0.52 | 0.35 | 0.44 | 0.54 |
| 9 | 79989953 | rs2275548 | 0.098 | -0.224 | -0.094 | -0.122 | -0.011 | -0.018 | -0.018 | 0 | 0.33 | 0.22 | 0.45 | 0.35 | 0.35 | 0.39 |
| 9 | 79998341 | rs12349390 | -0.707 | -0.323 | -0.032 | -0.033 | -0.006 | -0.001 | -0.001 | 0 | 0.37 | 0.4 | 0.25 | 0.37 | 0.32 | 0.3 |
| 9 | 80002460 | rs7034531 | -1.203 | -0.298 | 0.005 | -0.027 | -0.001 | -0.001 | -0.001 | 0.112 | 0.55 | 0.43 | 0.51 | 0.44 | 0.51 | 0.59 |
| 9 | 80004213 | rs4012460 | -1.716 | -0.384 | 0.205 | -0.075 | -0.012 | 0.009 | 0.009 | 0 | 0.62 | 0.71 | 0.55 | 0.65 | 0.52 | 0.64 |
| 9 | 80010025 | rs4744831 | 0.274 | -0.168 | -0.328 | -0.005 | -0.04 | -0.102 | -0.102 | 0 | 0.29 | 0.11 | 0.37 | 0.45 | 0.3 | 0.38 |
| 9 | 80017018 | rs12347526 | 0.487 | 0.043 | -0.422 | -0.276 | -0.003 | 0.023 | 0.023 | 0.362 | 0.29 | 0.21 | 0.21 | 0.12 | 0.31 | 0.24 |
| 9 | 80031144 | rs12343797 | 0.374 | 0.03 | -0.393 | -0.224 | 0.007 | 0.025 | 0.025 | 0.273 | 0.25 | 0.2 | 0.13 | 0.03 | 0.22 | 0.22 |
| 9 | 80039658 | rs17063627 | 0.183 | 0.089 | -0.353 | 0.01 | -0.001 | -0.001 | -0.001 | 0 | 0.09 | 0.08 | 0.07 | 0.13 | 0.08 | 0.08 |
| 9 | 80041641 | rs7860724 | -0.125 | 0.057 | -0.251 | -0.216 | -0.001 | -0.003 | -0.003 | 0 | 0.43 | 0.42 | 0.32 | 0.3 | 0.4 | 0.41 |
| 9 | 80041884 | rs10747015 | -0.113 | 0.33 | -0.367 | -0.293 | 0 | 0.005 | 0.005 | 0 | 0.55 | 0.58 | 0.67 | 0.66 | 0.56 | 0.55 |
| 9 | 80044056 | rs2296781 | -0.089 | 0.432 | -0.319 | -0.395 | 0 | -0.006 | -0.006 | 0 | 0.45 | 0.45 | 0.34 | 0.29 | 0.45 | 0.42 |
| 9 | 80044563 | rs12348357 | 0.59 | 0.465 | -0.538 | -0.509 | -0.059 | 0.134 | 0.134 | 0.786 | 0.31 | 0.13 | 0.24 | 0.06 | 0.35 | 0.28 |
| 9 | 80049298 | rs1801258 | 0.988 | 0.364 | -0.829 | -0.257 | -0.025 | 0.066 | 0.066 | 0.074 | 0.27 | 0.51 | 0.33 | 0.35 | 0.29 | 0.25 |
| 9 | 80050810 | rs12683817 | 0.252 | 0.117 | -1.115 | -0.446 | -0.018 | 0.017 | 0.017 | 0 | 0.33 | 0.25 | 0.26 | 0.31 | 0.34 | 0.3 |
| 9 | 80053953 | rs1327546 | -0.765 | 0.119 | -1.141 | -0.535 | -0.003 | -0.01 | -0.01 | 0.017 | 0.34 | 0.41 | 0.21 | 0.3 | 0.3 | 0.27 |
| 9 | 80057559 | rs1036955 | -0.767 | -0.23 | -1.349 | -0.321 | -0.058 | 0.041 | 0.041 | 0.876 | 0.62 | 0.83 | 0.7 | 0.66 | 0.59 | 0.63 |
| 9 | 80057994 | rs1034014 | 0.993 | -0.207 | -0.919 | -0.122 | -0.018 | 0.147 | 0.147 | 0.425 | 0.19 | 0.33 | 0.3 | 0.5 | 0.15 | 0.25 |
| 9 | 80058577 | rs10781437 | -0.104 | -0.172 | -0.968 | -0.152 | -0.003 | -0.006 | -0.006 | 0 | 0.45 | 0.56 | 0.43 | 0.33 | 0.47 | 0.4 |
| 9 | 80059966 | rs11145426 | 1.249 | -0.038 | -0.949 | 0.154 | -0.011 | 0.077 | 0.077 | 1.225 | 0.16 | 0.22 | 0.27 | 0.33 | 0.12 | 0.22 |
| 9 | 80060071 | rs12553676 | 1.249 | -0.038 | -0.949 | 0.154 | -0.011 | 0.077 | 0.077 | 1.225 | 0.16 | 0.22 | 0.27 | 0.33 | 0.12 | 0.22 |
| 9 | 80060712 | rs7039129 | 0.589 | -0.06 | -1.587 | 0.111 | 0.022 | -0.029 | -0.029 | 0 | 0.26 | 0.11 | 0.26 | 0.33 | 0.18 | 0.34 |
| 9 | 80064749 | rs12342328 | 0.387 | 0.138 | -1.74 | -0.176 | 0 | -0.016 | -0.016 | 0 | 0.06 | 0.07 | 0.04 | 0 | 0.04 | 0.01 |
| 9 | 80066047 | rs12336835 | 1.005 | 0.051 | -2.044 | -0.087 | -0.032 | -0.06 | -0.06 | 0 | 0.26 | 0.17 | 0.31 | 0.38 | 0.34 | 0.17 |
| 9 | 80066070 | rs4745639 | 0.487 | -0.322 | -1.304 | 0.137 | -0.011 | 0.146 | 0.146 | 0.734 | 0.2 | 0.39 | 0.3 | 0.42 | 0.15 | 0.22 |
| 9 | 80066118 | rs12336869 | 0.341 | -0.296 | -2.752 | -0.197 | 0.027 | 0.045 | 0.045 | 0 | 0.14 | 0.05 | 0.05 | 0 | 0.09 | 0.15 |
| 9 | 80066813 | rs10869926 | 0.806 | 0.107 | -2.478 | 0.376 | -0.009 | -0.089 | -0.089 | 1.403 | 0.16 | 0.02 | 0.26 | 0.32 | 0.13 | 0.24 |
| 9 | 80070995 | rs12352752 | 0.118 | 0.469 | -2.359 | -0.326 | -0.023 | 0.056 | 0.056 | 0 | 0.18 | 0.07 | 0.1 | 0 | 0.18 | 0.16 |
| 9 | 80071833 | rs7032444 | -0.765 | 0.486 | -3.174 | -0.452 | -0.069 | 0.048 | 0.048 | 0.004 | 0.69 | 0.95 | 0.64 | 0.83 | 0.69 | 0.63 |
| 9 | 80075134 | rs1547190 | -0.735 | 0.594 | -2.699 | -0.075 | -0.022 | -0.04 | -0.04 | 0.871 | 0.43 | 0.8 | 0.4 | 0.53 | 0.55 | 0.37 |
| 9 | 80076247 | rs3780293 | -0.917 | 0.334 | -2.923 | -0.469 | -0.039 | 0.2 | 0.2 | 2.529 | 0.39 | 0.96 | 0.27 | 0.16 | 0.35 | 0.27 |
| 9 | 80076442 | rs3824405 | -0.622 | 0.483 | -2.578 | -0.355 | 0.007 | 0.081 | 0.081 | 3.292 | 0.34 | 0.85 | 0.24 | 0.18 | 0.32 | 0.24 |
| 9 | 80077675 | rs7862678 | 0.224 | 0.681 | -2.207 | -0.119 | 0.025 | 0.008 | 0.008 | 0 | 0.42 | 0.32 | 0.36 | 0.36 | 0.33 | 0.57 |
| 9 | 80078016 | rs734468 | -0.352 | 1.274 | -1.852 | -0.049 | -0.127 | 0.027 | 0.027 | 0 | 0.31 | 0.07 | 0.44 | 0.72 | 0.44 | 0.5 |
| 9 | 80080976 | rs11145434 | -0.111 | 1.248 | -1.532 | 0.058 | 0.002 | -0.132 | -0.132 | 0 | 0.34 | 0.08 | 0.21 | 0.49 | 0.28 | 0.4 |
| 9 | 80081202 | rs2165962 | -0.995 | 1.279 | -1.054 | -0.211 | -0.062 | 0.171 | 0.171 | 0.255 | 0.57 | 0.84 | 0.48 | 0.86 | 0.5 | 0.55 |
| 9 | 80086534 | rs990000 | 0.331 | 1.175 | -0.483 | 0.059 | 0.012 | 0.042 | 0.042 | 0 | 0.13 | 0.15 | 0.21 | 0.34 | 0.19 | 0.11 |
| 9 | 80087031 | rs994362 | -0.992 | 1.157 | -0.01 | -0.174 | -0.015 | 0.099 | 0.099 | 0.277 | 0.56 | 0.77 | 0.47 | 0.83 | 0.51 | 0.58 |
| 9 | 80109669 | rs7029884 | -0.465 | 0.976 | 0.386 | -0.35 | -0.005 | 0.015 | 0.015 | 0.02 | 0.44 | 0.36 | 0.43 | 0.84 | 0.48 | 0.53 |
| 9 | 80110333 | rs10781441 | -1.256 | 0.951 | 0.171 | -0.363 | -0.059 | 0.165 | 0.165 | 0.369 | 0.57 | 0.87 | 0.49 | 0.85 | 0.53 | 0.58 |
| 9 | 80112213 | rs2889774 | 0.144 | 1.077 | 0.049 | -0.153 | 0.002 | -0.036 | -0.036 | 0.652 | 0.33 | 0.22 | 0.22 | 0.47 | 0.3 | 0.41 |
| 9 | 80115740 | rs7047853 | -0.808 | 1.191 | -0.005 | -0.177 | 0.002 | 0.059 | 0.059 | 0.053 | 0.53 | 0.64 | 0.43 | 0.84 | 0.51 | 0.57 |
| 9 | 80122834 | rs1888980 | -0.343 | 1.275 | -0.169 | 0.243 | -0.004 | -0.003 | -0.003 | 0 | 0.2 | 0.15 | 0.24 | 0.39 | 0.28 | 0.14 |
| 9 | 80123488 | rs7039992 | 0.379 | 0.695 | 0.025 | 0.255 | 0.016 | -0.026 | -0.026 | 0 | 0.29 | 0.24 | 0.16 | 0.49 | 0.21 | 0.41 |
| 9 | 80123765 | rs11788546 | 0.427 | 0.69 | 0.027 | 0.253 | 0.021 | -0.038 | -0.038 | 0 | 0.28 | 0.2 | 0.16 | 0.48 | 0.2 | 0.42 |
| 9 | 80124780 | rs6560604 | -0.924 | 0.896 | 0.053 | -0.063 | 0.001 | 0.002 | 0.002 | 0 | 0.57 | 0.63 | 0.67 | 0.98 | 0.56 | 0.74 |
| 9 | 80124810 | rs7041488 | -0.579 | 1.028 | -0.47 | 0.006 | 0.039 | 0.095 | 0.095 | 0.33 | 0.49 | 0.85 | 0.51 | 0.63 | 0.41 | 0.66 |
| 9 | 80124903 | rs7852777 | -0.749 | 0.565 | -0.19 | 0.72 | 0.004 | 0.045 | 0.045 | 0.11 | 0.41 | 0.29 | 0.36 | 0.81 | 0.38 | 0.53 |
| 9 | 80125731 | rs11145450 | 0.12 | 0.68 | -0.993 | 0.741 | 0.012 | -0.024 | -0.024 | 0 | 0.26 | 0.21 | 0.15 | 0.43 | 0.19 | 0.4 |

| | | | | | | | | | | | | | | | | |
|---|---|---|---|---|---|---|---|---|---|---|---|---|---|---|---|---|
| 9 | 80126898 | rs11145452 | 0.152 | 0.487 | -0.963 | 0.691 | 0.003 | -0.006 | -0.006 | 0 | 0.26 | 0.23 | 0.14 | 0.41 | 0.18 | 0.39 |
| 9 | 80130546 | rs17195438 | -0.367 | 1.283 | -0.87 | 0.654 | -0.004 | -0.002 | -0.002 | 0 | 0.16 | 0.12 | 0.19 | 0.22 | 0.18 | 0.12 |
| 9 | 80134405 | rs636562 | -0.456 | 0.935 | -0.939 | 0.574 | 0.015 | 0.024 | 0.024 | 0 | 0.53 | 0.68 | 0.62 | 0.66 | 0.6 | 0.58 |
| 9 | 80137434 | rs3780296 | 0.232 | 1.129 | -0.614 | 0.792 | -0.001 | -0.001 | -0.001 | 0 | 0.63 | 0.34 | 0.66 | 0.54 | 0.62 | 0.6 |
| 9 | 80143110 | rs11145462 | 0.424 | 0.679 | -0.966 | 0.588 | 0.008 | 0.009 | 0.009 | 0.24 | 0.49 | 0.36 | 0.63 | 0.44 | 0.54 | 0.49 |
| 9 | 80145512 | rs4745641 | -0.252 | 0.664 | -0.38 | 0.525 | -0.031 | 0.039 | 0.039 | 0.52 | 0.68 | 0.46 | 0.77 | 0.55 | 0.8 | 0.66 |
| 9 | 80149172 | rs10781444 | -0.533 | 0.56 | 0.16 | 0.649 | -0.01 | 0.014 | 0.014 | 0.117 | 0.72 | 0.65 | 0.8 | 0.61 | 0.81 | 0.72 |
| 9 | 80149187 | rs10869935 | -0.491 | 0.75 | 0.45 | 0.464 | 0.023 | -0.059 | -0.059 | 0 | 0.23 | 0.43 | 0.22 | 0.13 | 0.25 | 0.22 |
| 9 | 80159865 | rs1443864 | -0.493 | 0.325 | 0.496 | 0.677 | -0.007 | -0.038 | -0.038 | 4.428 | 0.69 | 0.85 | 0.73 | 0.63 | 0.69 | 0.7 |
| 9 | 80160010 | rs1373254 | 0.789 | 0.282 | 1.041 | 0.578 | 0.008 | -0.015 | -0.015 | 0 | 0.48 | 0.23 | 0.58 | 0.48 | 0.53 | 0.45 |
| 9 | 80162570 | rs7046214 | -1.034 | 0.287 | 0.835 | 0.454 | 0.012 | 0.014 | 0.014 | 1.638 | 0.29 | 0.72 | 0.23 | 0.18 | 0.24 | 0.28 |
| 9 | 80163049 | rs6560606 | -0.124 | 0.285 | 1.268 | 0.474 | -0.04 | -0.044 | -0.044 | 0 | 0.2 | 0.46 | 0.15 | 0.13 | 0.16 | 0.23 |
| 9 | 80164601 | rs10869944 | 0.755 | 0.337 | 0.632 | 0.995 | -0.048 | -0.055 | -0.055 | 0.107 | 0.45 | 0.12 | 0.56 | 0.49 | 0.52 | 0.44 |
| 9 | 80165255 | rs7040864 | 0.395 | 0.351 | 1.048 | 0.97 | -0.012 | -0.011 | -0.011 | 0.278 | 0.16 | 0.17 | 0.1 | 0.13 | 0.12 | 0.2 |
| 9 | 80168128 | rs10869945 | 0.727 | 0.344 | 0.944 | 1.001 | -0.045 | -0.042 | -0.042 | 0 | 0.45 | 0.12 | 0.57 | 0.56 | 0.53 | 0.45 |
| 9 | 80169316 | rs7035476 | 0.569 | 0.26 | 0.961 | 0.859 | -0.009 | -0.004 | -0.004 | 0 | 0.16 | 0.16 | 0.1 | 0.15 | 0.11 | 0.19 |
| 9 | 80177345 | rs10781447 | -0.193 | 0.798 | 1.078 | 0.92 | -0.004 | -0.002 | -0.002 | 0 | 0.6 | 0.64 | 0.64 | 0.46 | 0.55 | 0.6 |
| 9 | 80177554 | rs10781448 | 0.89 | 0.672 | 1.028 | 0.895 | 0.023 | -0.005 | -0.005 | 0 | 0.44 | 0.17 | 0.53 | 0.44 | 0.4 | 0.5 |
| 9 | 80180741 | rs7860549 | -0.824 | 0.626 | 1.162 | 0.839 | -0.016 | -0.016 | -0.016 | 1.765 | 0.14 | 0.4 | 0.1 | 0.07 | 0.1 | 0.08 |
| 9 | 80181725 | rs10781449 | -0.448 | 0.688 | 0.813 | 1.007 | 0.001 | 0.001 | 0.001 | 0.513 | 0.62 | 0.64 | 0.66 | 0.58 | 0.58 | 0.64 |
| 9 | 80181751 | rs7849646 | -0.584 | 0.68 | 0.204 | 1.004 | -0.045 | -0.06 | -0.06 | 0 | 0.64 | 0.84 | 0.68 | 0.53 | 0.57 | 0.64 |
| 9 | 80185929 | rs961572 | -0.384 | 0.295 | 0.445 | 0.264 | -0.013 | -0.039 | -0.039 | 0 | 0.64 | 0.59 | 0.76 | 0.86 | 0.71 | 0.69 |
| 9 | 80186307 | rs2440229 | -0.713 | 0.294 | -0.116 | 0.252 | -0.005 | -0.04 | -0.04 | 0 | 0.69 | 0.67 | 0.77 | 0.94 | 0.73 | 0.72 |
| 9 | 80191419 | rs1443868 | 0.994 | 0.364 | -1.123 | 0.299 | -0.166 | -0.116 | -0.116 | 0 | 0.39 | 0.03 | 0.52 | 0.44 | 0.43 | 0.37 |
| 9 | 80194682 | rs1626834 | -0.642 | 0.151 | -0.314 | 0.113 | 0.009 | 0.08 | 0.08 | 0.131 | 0.71 | 0.88 | 0.78 | 0.93 | 0.74 | 0.72 |
| 9 | 80196611 | rs716080 | 0.811 | -0.175 | -0.146 | 0.201 | -0.013 | 0.217 | 0.217 | 0.705 | 0.51 | 0.1 | 0.64 | 0.82 | 0.58 | 0.59 |
| 9 | 80196839 | rs1772921 | -0.26 | -0.029 | -0.06 | 0.311 | 0.004 | 0.071 | 0.071 | 0.321 | 0.64 | 0.83 | 0.76 | 0.84 | 0.69 | 0.66 |
| 9 | 80198591 | rs755486 | 0.344 | 0.026 | -0.545 | 0.651 | -0.103 | 0.129 | 0.129 | 0 | 0.42 | 0.08 | 0.6 | 0.81 | 0.52 | 0.59 |
| 9 | 80201301 | rs1162211 | 1.295 | -0.299 | 0.671 | 1.367 | 0.003 | -0.025 | -0.025 | 0 | 0.84 | 0.88 | 0.65 | 0.56 | 0.83 | 0.62 |
| 9 | 80205948 | rs1144434 | 1.67 | -0.311 | 1.244 | 0.847 | 0 | 0.006 | 0.006 | 1.718 | 0.7 | 0.62 | 0.5 | 0.58 | 0.66 | 0.49 |
| 9 | 80209125 | rs1144430 | 1.697 | -0.768 | 1.194 | 0.341 | 0.032 | 0.042 | 0.042 | 0.194 | 0.64 | 0.32 | 0.7 | 0.84 | 0.76 | 0.66 |
| 9 | 80209604 | rs1147182 | 0.396 | -0.632 | 1.851 | 0.395 | -0.005 | -0.05 | -0.05 | 0 | 0.84 | 0.77 | 0.89 | 0.96 | 0.85 | 0.86 |
| 9 | 80210750 | rs10120776 | 0.834 | -0.6 | 1.858 | 0.47 | -0.017 | -0.062 | -0.062 | 4.245 | 0.75 | 0.64 | 0.75 | 0.91 | 0.78 | 0.67 |
| 9 | 80213743 | rs17788791 | -0.071 | -0.469 | 1.009 | 0.627 | -0.015 | -0.018 | -0.018 | 0 | 0.05 | 0.02 | 0.22 | 0.15 | 0.08 | 0.18 |
| 9 | 80217685 | rs1932074 | 1.426 | -0.851 | 1.311 | 0.143 | 0.034 | 0.039 | 0.039 | 0.194 | 0.64 | 0.29 | 0.69 | 0.81 | 0.76 | 0.65 |
| 9 | 80219681 | rs2184481 | 0.967 | -0.037 | 1.005 | 0.353 | -0.01 | -0.034 | -0.034 | 0 | 0.76 | 0.71 | 0.76 | 0.91 | 0.77 | 0.67 |
| 9 | 80223608 | rs2225741 | -1.841 | 0.097 | 1.68 | 0.819 | 0.072 | 0.118 | 0.118 | 4.905 | 0.07 | 0.39 | 0.05 | 0.11 | 0.07 | 0.02 |
| 9 | 80223823 | rs2210755 | -2.443 | 0.202 | 2.832 | 0.782 | 0.155 | 0.116 | 0.116 | 4.982 | 0.05 | 0.41 | 0.07 | 0.05 | 0.08 | 0.02 |
| 9 | 80226683 | rs3905108 | -2.626 | 0.228 | 3.541 | 0.726 | 0.084 | 0.047 | 0.047 | 1.354 | 0.07 | 0.38 | 0.07 | 0.06 | 0.06 | 0.05 |
| 9 | 80230357 | rs17063807 | -2.186 | -0.421 | 3.554 | 0.914 | -0.006 | -0.001 | -0.001 | 0.017 | 0.24 | 0.72 | 0.29 | 0.29 | 0.19 | 0.29 |
| 9 | 80230606 | rs12339543 | -3.113 | -0.418 | 3.745 | 0.909 | 0.081 | 0.012 | 0.012 | 3.816 | 0.09 | 0.57 | 0.1 | 0.06 | 0.09 | 0.05 |
| 9 | 80236438 | rs13288161 | -1.982 | -0.617 | 3.599 | 0.884 | -0.067 | 0.013 | 0.013 | 0.614 | 0.23 | 0.51 | 0.25 | 0.29 | 0.16 | 0.29 |
| 9 | 80248577 | rs7864216 | -1.184 | -0.721 | 3.37 | 0.86 | 0.005 | -0.003 | -0.003 | 0 | 0.17 | 0.13 | 0.18 | 0.23 | 0.1 | 0.24 |
| 9 | 80248895 | rs13289616 | -1.812 | -0.76 | 3.735 | 0.842 | -0.085 | 0.005 | 0.005 | 0.375 | 0.22 | 0.55 | 0.24 | 0.28 | 0.11 | 0.27 |
| 9 | 80259039 | rs11145503 | -2.325 | -0.846 | 3.546 | 1.934 | 0.017 | -0.022 | -0.022 | 0.162 | 0.28 | 0.74 | 0.48 | 0.49 | 0.16 | 0.43 |
| 9 | 80260083 | rs11145504 | -1.307 | -0.863 | 3.97 | 2.004 | -0.051 | 0.009 | 0.009 | 0 | 0.19 | 0.28 | 0.21 | 0.27 | 0.11 | 0.26 |
| 9 | 80263100 | rs1122838 | -1.344 | -0.759 | 3.924 | 1.953 | -0.032 | 0.005 | 0.005 | 0 | 0.19 | 0.28 | 0.23 | 0.27 | 0.14 | 0.28 |
| 9 | 80263849 | rs2236542 | -2.313 | -0.786 | 3.979 | 1.979 | -0.033 | 0.033 | 0.033 | 0.491 | 0.28 | 0.66 | 0.49 | 0.5 | 0.2 | 0.45 |
| 9 | 80277522 | rs35727218 | -1.444 | -0.564 | 3.467 | 1.692 | -0.013 | -0.001 | -0.001 | 1.744 | 0.19 | 0.27 | 0.23 | 0.26 | 0.14 | 0.32 |
| 9 | 80286142 | rs6560609 | -1.803 | -0.616 | 3.128 | 1.585 | -0.059 | -0.011 | -0.011 | 0 | 0.25 | 0.48 | 0.19 | 0.26 | 0.17 | 0.3 |
| 9 | 80287843 | rs11145517 | 0.545 | -0.119 | 1.565 | 1.554 | -0.054 | -0.048 | -0.048 | 0 | 0.15 | 0.01 | 0.16 | 0.1 | 0.17 | 0.14 |
| 9 | 80293657 | rs10869955 | 1.051 | 0.022 | 1.662 | 1.162 | -0.001 | -0.002 | -0.002 | 0.035 | 0.48 | 0.52 | 0.59 | 0.62 | 0.48 | 0.39 |
| 9 | 80294359 | rs34782998 | -0.636 | -0.1 | 1.518 | 1.017 | 0.005 | 0.003 | 0.003 | 0 | 0.19 | 0.14 | 0.15 | 0.19 | 0.17 | 0.26 |
| 9 | 80334981 | rs2026000 | 0.377 | -0.701 | 0.88 | 1.209 | 0.048 | -0.006 | -0.006 | 0.167 | 0.59 | 0.18 | 0.55 | 0.46 | 0.7 | 0.59 |
| 9 | 80350999 | rs2839899 | 0.387 | -0.759 | 1.206 | 1.741 | 0.062 | -0.002 | -0.002 | 0 | 0.59 | 0.16 | 0.55 | 0.44 | 0.7 | 0.6 |
| 9 | 80355515 | rs4744849 | 1.497 | -0.534 | 0.87 | 1.59 | -0.165 | 0.17 | 0.17 | 0 | 0.39 | 0.05 | 0.38 | 0.13 | 0.49 | 0.29 |
| 9 | 80364115 | rs10869963 | -1.17 | -0.728 | 1.206 | 1.679 | -0.017 | -0.051 | -0.051 | 0.551 | 0.78 | 0.96 | 0.89 | 0.78 | 0.79 | 0.83 |
| 9 | 80366259 | rs10869964 | 0.304 | -0.762 | 2.134 | 1.678 | -0.019 | 0.013 | 0.013 | 0.227 | 0.66 | 0.49 | 0.61 | 0.46 | 0.72 | 0.6 |
| 9 | 80371529 | rs10747022 | -0.977 | -0.74 | 1.947 | 1.653 | 0.001 | -0.021 | -0.021 | 0.057 | 0.77 | 0.89 | 0.89 | 0.78 | 0.79 | 0.83 |
| 9 | 80381887 | rs11145558 | 0.28 | -0.485 | 1.927 | 1.548 | -0.006 | 0.006 | 0.006 | 0.231 | 0.65 | 0.44 | 0.59 | 0.48 | 0.68 | 0.58 |
| 9 | 80389921 | rs1328531 | -1.24 | -0.472 | 2.007 | 1.692 | 0.003 | 0.014 | 0.014 | 0.157 | 0.08 | 0.2 | 0.06 | 0.11 | 0.08 | 0.22 |
| 9 | 80414763 | rs2118021 | 0.282 | -0.514 | 1.795 | 1.66 | -0.004 | 0.002 | 0.002 | 0.239 | 0.65 | 0.41 | 0.59 | 0.48 | 0.68 | 0.57 |
| 9 | 80444279 | rs1410551 | 0.512 | -0.549 | 1.487 | 1.627 | 0.025 | -0.062 | -0.062 | 0 | 0.61 | 0.17 | 0.55 | 0.54 | 0.68 | 0.58 |
| 9 | 80457276 | rs10869977 | 0.929 | -0.993 | 0.353 | 1.017 | -0.09 | 0.18 | 0.18 | 0 | 0.46 | 0.07 | 0.41 | 0.19 | 0.52 | 0.39 |
| 9 | 80469561 | rs7048503 | 0.427 | -1.11 | 0.555 | 1.281 | -0.002 | -0.012 | -0.012 | 0.021 | 0.16 | 0.1 | 0.1 | 0.24 | 0.17 | 0.18 |
| 9 | 80469747 | rs7048811 | 0.105 | -1.239 | 1.102 | 1.347 | -0.007 | 0.005 | 0.005 | 0 | 0.65 | 0.4 | 0.57 | 0.41 | 0.7 | 0.56 |
| 9 | 80472931 | rs11145589 | -0.285 | -1.225 | 1.133 | 1.393 | -0.019 | 0.05 | 0.05 | 0 | 0.69 | 0.47 | 0.59 | 0.54 | 0.75 | 0.79 |
| 9 | 80480997 | rs10735654 | -0.987 | -1.157 | 0.713 | 1.369 | 0 | -0.024 | -0.024 | 0.013 | 0.76 | 0.92 | 0.88 | 0.75 | 0.78 | 0.82 |
| 9 | 80485033 | rs12352936 | -0.236 | -1.65 | 0.819 | 0.731 | -0.023 | -0.064 | -0.064 | 0 | 0.15 | 0.18 | 0.06 | 0 | 0.11 | 0.09 |

| | | | | | | | | | | | | | | | | |
|---|---|---|---|---|---|---|---|---|---|---|---|---|---|---|---|---|
| 9 | 80485795 | rs1410556 | 0.58 | -1.633 | 0.968 | 0.993 | 0.048 | -0.029 | -0.029 | 0 | 0.54 | 0.21 | 0.48 | 0.52 | 0.62 | 0.65 |
| 9 | 80485900 | rs1410555 | -1.248 | -1.363 | 1.557 | 1.222 | 0 | 0.001 | 0.001 | 0 | 0.75 | 0.72 | 0.84 | 0.72 | 0.76 | 0.82 |
| 9 | 80487849 | rs2209911 | -1.312 | -1.358 | 1.53 | 1.226 | -0.003 | -0.01 | -0.01 | 0 | 0.76 | 0.82 | 0.85 | 0.72 | 0.76 | 0.84 |
| 9 | 80496768 | rs11145603 | -0.2 | -1.349 | 0.7 | 0.587 | 0.004 | -0.013 | -0.013 | 0 | 0.21 | 0.16 | 0.14 | 0.34 | 0.19 | 0.39 |
| 9 | 80500405 | rs10781468 | -0.966 | -1.309 | 0.813 | 0.591 | -0.005 | 0.058 | 0.058 | 0 | 0.71 | 0.42 | 0.69 | 0.52 | 0.74 | 0.77 |
| 9 | 80506587 | rs2296937 | 0.096 | -1.39 | 0.672 | 0.588 | 0.017 | -0.011 | -0.011 | 2.55 | 0.18 | 0.11 | 0.11 | 0.26 | 0.13 | 0.16 |
| 9 | 80515456 | rs10781471 | -0.306 | -1.318 | 0.754 | 0.564 | 0 | -0.025 | -0.025 | 0 | 0.66 | 0.3 | 0.65 | 0.51 | 0.68 | 0.72 |
| 9 | 80544252 | rs4641131 | -1.561 | -1.246 | 0.658 | 0.487 | 0.018 | 0.01 | 0.01 | 0 | 0.37 | 0.26 | 0.22 | 0.35 | 0.31 | 0.46 |
| 9 | 80561271 | rs11145626 | 0.982 | -1.243 | 0.586 | 0.502 | -0.019 | -0.065 | -0.065 | 0 | 0.14 | 0.01 | 0.1 | 0.24 | 0.13 | 0.15 |
| 9 | 80561672 | rs7035689 | -0.869 | -1.132 | 0.753 | 0.436 | 0.027 | -0.002 | -0.002 | 0 | 0.3 | 0.22 | 0.18 | 0.34 | 0.24 | 0.4 |
| 9 | 80580046 | rs10869995 | -0.767 | -1.146 | 0.732 | 0.426 | 0.04 | -0.01 | -0.01 | 0 | 0.3 | 0.14 | 0.17 | 0.34 | 0.24 | 0.42 |
| 9 | 80584749 | rs17064001 | 0.384 | -1.112 | 0.719 | 0.434 | -0.001 | -0.025 | -0.025 | 0.018 | 0.07 | 0.07 | 0.04 | 0.01 | 0.06 | 0.06 |
| 9 | 80589760 | rs17064005 | -1.333 | -1.069 | 0.69 | 0.482 | 0.005 | 0.003 | 0.003 | 0 | 0.37 | 0.32 | 0.22 | 0.34 | 0.3 | 0.48 |
| 9 | 80602561 | rs7846810 | -1.464 | -0.837 | 0.783 | 0.495 | -0.007 | -0.007 | -0.007 | 0 | 0.39 | 0.42 | 0.24 | 0.36 | 0.32 | 0.49 |
| 9 | 80613593 | rs1075369 | -1.749 | -1.102 | 0.901 | 0.492 | -0.005 | -0.044 | -0.044 | 0 | 0.75 | 0.83 | 0.72 | 0.51 | 0.74 | 0.77 |
| 9 | 80618079 | rs10870004 | -1.688 | -1.102 | 1.042 | 0.472 | -0.001 | 0 | 0 | 0 | 0.74 | 0.75 | 0.72 | 0.52 | 0.74 | 0.79 |
| 9 | 80618374 | rs4745686 | 1.706 | -1.129 | 1.181 | 0.44 | -0.017 | 0.021 | 0.021 | 0 | 0.34 | 0.28 | 0.48 | 0.18 | 0.44 | 0.3 |
| 9 | 80619811 | rs4745687 | -1.595 | -1.097 | 0.921 | 0.516 | 0.005 | 0.005 | 0.005 | 0 | 0.4 | 0.41 | 0.24 | 0.3 | 0.3 | 0.49 |
| 9 | 80628858 | rs11145654 | -1.371 | -1.1 | 0.947 | 0.451 | 0.011 | 0.007 | 0.007 | 0 | 0.37 | 0.32 | 0.23 | 0.35 | 0.3 | 0.48 |
| 9 | 80630123 | rs10870007 | -1.392 | -1.066 | 0.963 | 0.468 | 0.011 | 0.007 | 0.007 | 0 | 0.38 | 0.33 | 0.23 | 0.35 | 0.3 | 0.48 |
| 9 | 80638165 | rs1041584 | -1.447 | -1.035 | 0.987 | 0.587 | -0.002 | -0.001 | -0.001 | 0 | 0.38 | 0.4 | 0.24 | 0.37 | 0.3 | 0.45 |
| 9 | 80650658 | rs10870014 | -0.596 | -1.025 | 0.702 | 0.339 | 0.023 | -0.004 | -0.004 | 1.307 | 0.28 | 0.17 | 0.15 | 0.32 | 0.24 | 0.42 |
| 9 | 80665785 | rs4745694 | 0.693 | -1.116 | 0.678 | 0.287 | 0.013 | 0.028 | 0.028 | 0 | 0.12 | 0.05 | 0.06 | 0.01 | 0.11 | 0.09 |
| 9 | 80666802 | rs7850444 | -1.654 | -1.132 | 0.746 | 0.272 | 0.019 | 0.013 | 0.013 | 0 | 0.38 | 0.33 | 0.23 | 0.34 | 0.3 | 0.48 |
| 9 | 80687873 | rs10870028 | 0.605 | -1.067 | 0.632 | 0.329 | 0.012 | -0.021 | -0.021 | 0 | 0.17 | 0.12 | 0.11 | 0.33 | 0.13 | 0.33 |
| 9 | 80704731 | rs950141 | -1.604 | -1.036 | 0.691 | 0.337 | 0.001 | 0.001 | 0.001 | 0 | 0.39 | 0.4 | 0.24 | 0.34 | 0.3 | 0.48 |
| 9 | 80712566 | rs7037675 | -0.767 | -0.945 | 0.703 | 0.419 | -0.014 | 0.001 | 0.001 | 0 | 0.27 | 0.36 | 0.17 | 0.33 | 0.19 | 0.38 |
| 9 | 80717576 | rs10512069 | 1.472 | -0.815 | 0.721 | 0.467 | 0.019 | -0.015 | -0.015 | 0 | 0.36 | 0.46 | 0.5 | 0.24 | 0.45 | 0.29 |
| 9 | 80717949 | rs9314866 | -0.514 | -0.833 | 0.686 | 0.507 | 0.008 | -0.018 | -0.018 | 4.672 | 0.62 | 0.78 | 0.66 | 0.56 | 0.64 | 0.66 |
| 9 | 80718596 | rs10781492 | -0.907 | -0.718 | 0.718 | 0.624 | 0 | -0.001 | -0.001 | 0 | 0.25 | 0.28 | 0.16 | 0.33 | 0.19 | 0.38 |
| 9 | 80722684 | rs7039240 | -0.07 | -0.744 | 1.083 | 0.358 | -0.013 | -0.062 | -0.062 | 0 | 0.14 | 0.21 | 0.06 | 0.01 | 0.11 | 0.08 |
| 9 | 80723135 | rs7039739 | -1.931 | -0.6 | 1.233 | 0.743 | 0.022 | -0.001 | -0.001 | 0 | 0.39 | 0.73 | 0.38 | 0.49 | 0.34 | 0.49 |
| 9 | 80724067 | rs9969693 | -1.176 | -0.971 | 1.309 | 0.971 | -0.023 | 0.061 | 0.061 | 0.439 | 0.07 | 0.23 | 0.17 | 0.18 | 0.04 | 0.11 |
| 9 | 80727620 | rs11145699 | 0.205 | -0.894 | 1.014 | 0.937 | 0.001 | -0.003 | -0.003 | 1.187 | 0.16 | 0.13 | 0.12 | 0.3 | 0.15 | 0.3 |
| 9 | 80742806 | rs11145712 | -0.969 | -0.827 | 1.064 | 0.95 | -0.019 | 0.033 | 0.033 | 0.091 | 0.06 | 0.13 | 0.16 | 0.18 | 0.04 | 0.11 |
| 9 | 80746139 | rs10512071 | -0.106 | -0.965 | 1.545 | 0.754 | -0.021 | -0.077 | -0.077 | 0 | 0.14 | 0.32 | 0.08 | 0.01 | 0.11 | 0.08 |
| 9 | 80747897 | rs7854825 | -1.362 | -1.212 | 1.399 | 0.71 | -0.023 | -0.021 | -0.021 | 0 | 0.32 | 0.51 | 0.18 | 0.3 | 0.27 | 0.37 |
| 9 | 80747989 | rs7869891 | -1.943 | -1.235 | 0.903 | 0.709 | 0.088 | -0.025 | -0.025 | 0 | 0.4 | 0.84 | 0.37 | 0.48 | 0.29 | 0.48 |
| 9 | 80767173 | rs10870056 | -0.891 | -1.171 | 1.243 | 0.779 | -0.029 | 0.059 | 0.059 | 0 | 0.25 | 0.57 | 0.3 | 0.54 | 0.2 | 0.4 |
| 9 | 80772413 | rs7023397 | -1.699 | -1.168 | 1.067 | 0.735 | 0.059 | -0.035 | -0.035 | 0 | 0.38 | 0.76 | 0.36 | 0.55 | 0.3 | 0.5 |
| 9 | 80782539 | rs357631 | -0.812 | -1.014 | 0.574 | 0.671 | 0.025 | 0.073 | 0.073 | 0 | 0.47 | 0.69 | 0.42 | 0.79 | 0.41 | 0.54 |
| 9 | 80793672 | rs357642 | 1.073 | -1.025 | 0.336 | 0.841 | -0.01 | -0.052 | -0.052 | 0 | 0.23 | 0.12 | 0.12 | 0.26 | 0.2 | 0.15 |
| 9 | 80805624 | rs7047353 | -1.711 | -0.962 | 0.265 | 0.583 | -0.023 | 0.039 | 0.039 | 0.261 | 0.61 | 0.73 | 0.62 | 0.79 | 0.53 | 0.66 |
| 9 | 80813104 | rs10747028 | -1.63 | -0.954 | 0.354 | 0.718 | -0.013 | 0.017 | 0.017 | 0.028 | 0.6 | 0.7 | 0.55 | 0.7 | 0.53 | 0.65 |
| 9 | 80813860 | rs10870064 | 0.355 | -1.146 | 0.416 | 0.64 | -0.004 | -0.026 | -0.026 | 0 | 0.14 | 0.14 | 0.07 | 0.01 | 0.1 | 0.1 |
| 9 | 80827249 | rs12238895 | 0.272 | -1.014 | 0.284 | 1.023 | 0.001 | -0.02 | -0.02 | 0 | 0.29 | 0.32 | 0.44 | 0.41 | 0.27 | 0.26 |
| 9 | 80834899 | rs876400 | 0.617 | -1.041 | 0.326 | 1.022 | 0 | -0.006 | -0.006 | 4.452 | 0.26 | 0.29 | 0.27 | 0.46 | 0.24 | 0.2 |
| 9 | 80839235 | rs10217530 | -0.136 | -1.28 | 0.201 | 1.014 | 0.02 | -0.013 | -0.013 | 0 | 0.37 | 0.27 | 0.21 | 0.41 | 0.29 | 0.33 |
| 9 | 80842398 | rs1571869 | -0.084 | -0.879 | -0.392 | 0.858 | -0.096 | -0.038 | -0.038 | 0 | 0.72 | 0.99 | 0.83 | 0.72 | 0.68 | 0.69 |
| 9 | 80845906 | rs7861244 | 1.955 | -1 | 0.04 | 0.749 | -0.035 | 0.034 | 0.034 | 0.135 | 0.34 | 0.16 | 0.33 | 0.25 | 0.38 | 0.37 |
| 9 | 80847924 | rs4877498 | 1.255 | -0.863 | 0.371 | 0.785 | -0.001 | -0.009 | -0.009 | 0 | 0.59 | 0.5 | 0.48 | 0.66 | 0.58 | 0.62 |
| 9 | 80854608 | rs11137568 | 1.947 | -0.558 | 0.781 | 0.842 | -0.026 | 0.02 | 0.02 | 0.237 | 0.33 | 0.15 | 0.32 | 0.27 | 0.36 | 0.37 |
| 9 | 80863088 | rs1929402 | -0.389 | -0.781 | 0.637 | 0.821 | -0.011 | -0.027 | -0.027 | 0.095 | 0.75 | 0.86 | 0.86 | 0.64 | 0.75 | 0.69 |
| 9 | 80865783 | rs10867167 | -1.044 | -0.805 | 1.064 | 0.741 | 0.068 | 0.093 | 0.093 | 0.108 | 0.07 | 0.26 | 0.31 | 0.09 | 0.08 | 0.08 |
| 9 | 80874338 | rs12555718 | 0.358 | -0.833 | 0.962 | 0.719 | -0.024 | 0.012 | 0.012 | 0.017 | 0.57 | 0.73 | 0.73 | 0.38 | 0.54 | 0.54 |
| 9 | 80879149 | rs1057650 | 0.342 | -0.694 | 0.948 | 0.692 | -0.002 | -0.019 | -0.019 | 0 | 0.15 | 0.14 | 0.08 | 0.01 | 0.08 | 0.09 |
| 9 | 80901335 | rs11137586 | -1.102 | -0.441 | 1.084 | 0.871 | 0.01 | 0.093 | 0.093 | 1.071 | 0.06 | 0.3 | 0.17 | 0.09 | 0.05 | 0.07 |
| 9 | 80906946 | rs10429454 | -1.151 | -0.403 | 0.792 | 0.877 | 0.05 | 0.069 | 0.069 | 0.929 | 0.07 | 0.29 | 0.31 | 0.09 | 0.08 | 0.08 |
| 9 | 80911234 | rs17725890 | 0.052 | -0.467 | -0.068 | 0.833 | 0.006 | -0.004 | -0.004 | 0 | 0.32 | 0.22 | 0.19 | 0.3 | 0.3 | 0.25 |
| 9 | 80911421 | rs3824548 | 1.418 | 0.05 | -0.402 | 1.061 | -0.055 | -0.094 | -0.094 | 0 | 0.14 | 0.01 | 0.09 | 0.28 | 0.17 | 0.11 |
| 9 | 80916382 | rs4877550 | 1.587 | -0.224 | -0.252 | 0.562 | 0.001 | -0.006 | -0.006 | 0 | 0.34 | 0.33 | 0.38 | 0.22 | 0.35 | 0.35 |
| 9 | 80917070 | rs3739472 | 1.565 | -0.273 | -0.434 | 0.764 | -0.038 | -0.064 | -0.064 | 0.433 | 0.14 | 0.05 | 0.1 | 0.25 | 0.18 | 0.11 |
| 9 | 80918333 | rs3758200 | 0.91 | -0.675 | 0.378 | 0.815 | -0.007 | -0.01 | -0.01 | 0 | 0.54 | 0.5 | 0.73 | 0.67 | 0.62 | 0.66 |
| 9 | 80918420 | rs3758201 | 0.931 | -0.673 | 0.38 | 0.816 | -0.007 | -0.01 | -0.01 | 0.044 | 0.54 | 0.5 | 0.73 | 0.67 | 0.62 | 0.66 |
| 9 | 80918995 | rs7849813 | 0.284 | -0.146 | 0.147 | 1.136 | -0.006 | -0.012 | -0.012 | 0 | 0.81 | 0.89 | 0.88 | 0.69 | 0.79 | 0.82 |
| 9 | 80920757 | rs944514 | 0.509 | -0.388 | 0.323 | 0.929 | -0.001 | -0.003 | -0.003 | 0.402 | 0.76 | 0.71 | 0.83 | 0.68 | 0.74 | 0.79 |
| 9 | 80921732 | rs10116267 | 1.865 | -0.397 | -0.047 | 0.923 | -0.014 | 0.086 | 0.086 | 0 | 0.46 | 0.17 | 0.42 | 0.25 | 0.48 | 0.42 |
| 9 | 80923250 | rs3739476 | -1.307 | -0.35 | 0.079 | 1.06 | -0.004 | 0.119 | 0.119 | 0 | 0.11 | 0.2 | 0.29 | 0.42 | 0.11 | 0.23 |
| 9 | 80924008 | rs10867178 | -1.165 | -0.381 | 0.193 | 1.016 | -0.005 | 0.008 | 0.008 | 0 | 0.29 | 0.34 | 0.39 | 0.43 | 0.26 | 0.37 |

| | | | | | | | | | | | | | | | | |
|---|---|---|---|---|---|---|---|---|---|---|---|---|---|---|---|---|
| 9 | 80925219 | rs7874921 | -1.006 | -0.38 | 0.141 | 1.016 | 0.001 | -0.004 | -0.004 | 0 | 0.27 | 0.27 | 0.39 | 0.43 | 0.26 | 0.37 |
| 9 | 80927815 | rs17064306 | -0.088 | -0.364 | -0.26 | 0.014 | 0.011 | 0.022 | 0.022 | 0 | 0.18 | 0.15 | 0.11 | 0.01 | 0.15 | 0.15 |
| 9 | 80930439 | rs7858423 | 0.452 | -0.356 | -0.46 | 0.067 | 0.004 | 0.073 | 0.073 | 0 | 0.14 | 0.03 | 0.09 | 0.01 | 0.13 | 0.12 |
| 9 | 80932515 | rs3780193 | -0.898 | 0.032 | -0.088 | 0.509 | 0.008 | 0.019 | 0.019 | 0 | 0.16 | 0.17 | 0.21 | 0.32 | 0.22 | 0.26 |
| 9 | 80932887 | rs7043123 | 1.293 | -0.181 | -0.389 | 0.266 | -0.011 | -0.017 | -0.017 | 0 | 0.76 | 0.82 | 0.77 | 0.63 | 0.69 | 0.62 |
| 9 | 80933020 | rs10735532 | 1.293 | -0.181 | -0.389 | 0.266 | -0.011 | -0.017 | -0.017 | 0.193 | 0.76 | 0.82 | 0.77 | 0.63 | 0.69 | 0.62 |
| 9 | 80935799 | rs11137606 | -0.265 | -0.581 | 0.225 | 0.341 | -0.052 | 0.043 | 0.043 | 0.117 | 0.25 | 0.54 | 0.27 | 0.32 | 0.14 | 0.12 |
| 9 | 80938001 | rs11137607 | 1.39 | -0.53 | 0.55 | 0.133 | -0.001 | 0.026 | 0.026 | 0 | 0.56 | 0.34 | 0.51 | 0.3 | 0.55 | 0.5 |
| 9 | 80943271 | rs3758203 | -0.483 | -0.498 | 0.692 | 0.159 | 0.003 | -0.003 | -0.003 | 1.307 | 0.12 | 0.12 | 0.17 | 0.26 | 0.11 | 0.23 |
| 9 | 80943620 | rs10780231 | -1.194 | -0.618 | 1.221 | 0.146 | 0.005 | -0.003 | -0.003 | 0 | 0.36 | 0.64 | 0.45 | 0.61 | 0.28 | 0.41 |
| 9 | 80946291 | rs2776859 | 1.252 | -0.323 | 1.059 | -0.148 | -0.01 | -0.006 | -0.006 | 0 | 0.59 | 0.21 | 0.56 | 0.48 | 0.6 | 0.6 |
| 9 | 80958478 | rs7468433 | 1.296 | -1.407 | 1.207 | -0.066 | 0.009 | 0.033 | 0.033 | 0 | 0.14 | 0.04 | 0.06 | 0.05 | 0.09 | 0.12 |
| 9 | 80958634 | rs2776861 | 1.159 | -1.389 | 1.493 | -0.067 | 0.008 | -0.003 | -0.003 | 0.058 | 0.63 | 0.38 | 0.63 | 0.48 | 0.72 | 0.63 |
| 9 | 80958922 | rs2788867 | 0.407 | -1.367 | 1.754 | -0.09 | -0.016 | -0.045 | -0.045 | 0 | 0.85 | 0.75 | 0.8 | 0.91 | 0.86 | 0.86 |
| 9 | 80965822 | rs17790192 | 0.867 | -1.217 | 1.475 | -1.103 | 0 | -0.007 | -0.007 | 0.109 | 0.15 | 0.14 | 0.07 | 0.05 | 0.09 | 0.12 |
| 9 | 80968185 | rs4069725 | -0.435 | -1.566 | 1.499 | -1.006 | -0.006 | 0.148 | 0.148 | 0 | 0.12 | 0.28 | 0.18 | 0.33 | 0.11 | 0.23 |
| 9 | 80968877 | rs10867186 | -1.722 | -1.922 | 1.366 | -1.24 | 0.023 | -0.009 | -0.009 | 1.239 | 0.35 | 0.69 | 0.28 | 0.38 | 0.25 | 0.37 |
| 9 | 80972768 | rs11137621 | -0.604 | -1.802 | 1.345 | -1.663 | -0.028 | -0.071 | -0.071 | 0 | 0.32 | 0.38 | 0.13 | 0.1 | 0.2 | 0.22 |
| 9 | 80981110 | rs3010744 | 1.406 | -1.506 | 0.884 | -1.519 | -0.026 | 0.144 | 0.144 | 0 | 0.26 | 0.01 | 0.26 | 0.06 | 0.26 | 0.18 |
| 9 | 80983748 | rs2989577 | -0.439 | -1.453 | 1.467 | -1.51 | 0.004 | -0.041 | -0.041 | 0 | 0.49 | 0.52 | 0.37 | 0.14 | 0.38 | 0.34 |
| 9 | 80985484 | rs11137631 | -0.915 | -1.001 | 2.375 | -0.279 | -0.003 | 0.07 | 0.07 | 0 | 0.09 | 0.15 | 0.17 | 0.26 | 0.09 | 0.16 |
| 9 | 80985638 | rs10123128 | -1.184 | -0.949 | 2.523 | -0.443 | 0.022 | -0.013 | -0.013 | 2.642 | 0.07 | 0.3 | 0.05 | 0.03 | 0.05 | 0.08 |
| 9 | 80998764 | rs7024405 | -0.808 | -0.927 | 2.058 | -0.386 | -0.001 | 0.057 | 0.057 | 0 | 0.09 | 0.15 | 0.17 | 0.2 | 0.09 | 0.16 |
| 9 | 81004205 | rs10867190 | -0.808 | -0.919 | 1.851 | -0.386 | -0.001 | 0.062 | 0.062 | 0 | 0.09 | 0.15 | 0.17 | 0.2 | 0.09 | 0.16 |
| 9 | 81004253 | rs10867191 | -1.437 | -0.915 | 1.742 | -0.393 | 0 | 0.099 | 0.099 | 0 | 0.12 | 0.28 | 0.19 | 0.24 | 0.11 | 0.21 |
| 9 | 81012358 | rs7027964 | -1.623 | -1.001 | 0.857 | -0.471 | -0.008 | 0.002 | 0.002 | 0 | 0.25 | 0.25 | 0.19 | 0.29 | 0.19 | 0.29 |
| 9 | 81016144 | rs7849881 | -1.926 | -0.966 | 0.056 | -0.474 | 0.076 | 0.007 | 0.007 | 0.963 | 0.34 | 0.88 | 0.27 | 0.29 | 0.23 | 0.34 |
| 9 | 81038220 | rs7019259 | -0.753 | -0.704 | 0.353 | -0.17 | -0.003 | 0.045 | 0.045 | 0 | 0.08 | 0.09 | 0.13 | 0.23 | 0.08 | 0.2 |
| 9 | 81054215 | rs7848904 | -0.778 | -0.533 | 0.222 | -0.32 | -0.002 | -0.016 | -0.016 | 0 | 0.08 | 0.04 | 0.09 | 0.18 | 0.09 | 0.17 |
| 9 | 81057409 | rs4498630 | -1.108 | -0.885 | 0.066 | -0.53 | 0.033 | 0.015 | 0.015 | 4.929 | 0.28 | 0.81 | 0.23 | 0.21 | 0.15 | 0.32 |
| 9 | 81059643 | rs12376992 | -0.884 | -0.159 | 0.131 | -0.358 | 0.029 | 0.08 | 0.08 | 0.167 | 0.43 | 0.78 | 0.35 | 0.26 | 0.33 | 0.4 |
| 9 | 81064328 | rs10780236 | -0.857 | 0.21 | 0.682 | 0.903 | 0.023 | 0.137 | 0.137 | 0 | 0.06 | 0.2 | 0.13 | 0.18 | 0.06 | 0.16 |
| 9 | 81071842 | rs7863422 | -1.265 | 0.078 | 1.259 | 0.299 | 0.032 | -0.009 | -0.009 | 0 | 0.07 | 0.29 | 0.05 | 0.01 | 0.05 | 0.06 |
| 9 | 81078138 | rs3010671 | -1.805 | -0.441 | 1.282 | -0.032 | -0.024 | -0.057 | -0.057 | 0 | 0.16 | 0.73 | 0.12 | 0.02 | 0.08 | 0.07 |
| 9 | 81083897 | rs10125737 | 0.783 | -0.133 | 0.327 | 0.236 | -0.017 | -0.012 | -0.012 | 0.064 | 0.46 | 0.24 | 0.33 | 0.57 | 0.51 | 0.53 |
| 9 | 81089830 | rs3010681 | -0.682 | -0.462 | 1.093 | 0.077 | -0.038 | -0.063 | -0.063 | 0.944 | 0.26 | 0.4 | 0.18 | 0.09 | 0.17 | 0.23 |
| 9 | 81091143 | rs7855727 | 0.887 | 0.193 | 1.967 | 0.142 | 0.011 | 0.013 | 0.013 | 0.505 | 0.61 | 0.52 | 0.41 | 0.56 | 0.55 | 0.6 |
| 9 | 81094291 | rs3010690 | -0.218 | 0.167 | 1.998 | 0.201 | 0.013 | -0.025 | -0.025 | 0.011 | 0.77 | 0.71 | 0.55 | 0.91 | 0.67 | 0.84 |
| 9 | 81096795 | rs7872028 | -0.234 | 0.143 | 2.138 | 0.287 | -0.01 | -0.009 | -0.009 | 0 | 0.18 | 0.2 | 0.14 | 0.1 | 0.12 | 0.23 |
| 9 | 81098829 | rs2938889 | -0.53 | 0.154 | 2.071 | 0.337 | 0.025 | -0.024 | -0.024 | 0 | 0.79 | 0.71 | 0.55 | 0.88 | 0.68 | 0.85 |
| 9 | 81100971 | rs1394933 | 1.908 | 0.181 | 2.078 | 0.389 | 0.029 | 0.146 | 0.146 | 0.758 | 0.5 | 0.22 | 0.31 | 0.77 | 0.46 | 0.56 |
| 9 | 81101590 | rs2938900 | -0.494 | 0.283 | 2.195 | 0.484 | -0.008 | 0.004 | 0.004 | 0 | 0.79 | 0.77 | 0.56 | 0.89 | 0.68 | 0.85 |
| 9 | 81102028 | rs2998680 | -1.023 | -0.241 | 2.1 | 0.672 | -0.024 | -0.039 | -0.039 | 0 | 0.22 | 0.37 | 0.19 | 0.1 | 0.18 | 0.25 |
| 9 | 81106266 | rs2998683 | -1.589 | -0.369 | 2.231 | 0.785 | -0.006 | -0.005 | -0.005 | 0 | 0.12 | 0.45 | 0.08 | 0.07 | 0.05 | 0.03 |
| 9 | 81106476 | rs2135238 | -0.163 | -0.042 | 2.17 | 0.775 | 0.006 | -0.014 | -0.014 | 0.548 | 0.83 | 0.77 | 0.71 | 0.89 | 0.79 | 0.86 |
| 9 | 81111931 | rs2938923 | 0.002 | -0.053 | 2.127 | 0.754 | 0.016 | -0.033 | -0.033 | 0 | 0.82 | 0.74 | 0.7 | 0.89 | 0.79 | 0.83 |
| 9 | 81116860 | rs3010702 | -1.215 | -0.154 | 2.172 | 0.743 | -0.006 | -0.006 | -0.006 | 0 | 0.11 | 0.42 | 0.07 | 0.07 | 0.05 | 0.03 |
| 9 | 81119846 | rs10867212 | 1.287 | -0.151 | 2.123 | 0.742 | -0.001 | -0.016 | -0.016 | 0 | 0.71 | 0.31 | 0.63 | 0.85 | 0.74 | 0.81 |
| 9 | 81120990 | rs3010713 | -0.147 | -0.153 | 2.145 | 0.74 | 0.004 | -0.019 | -0.019 | 0 | 0.83 | 0.81 | 0.71 | 0.92 | 0.79 | 0.83 |
| 9 | 81121174 | rs3849862 | 1.33 | -0.151 | 2.019 | 0.74 | -0.001 | -0.011 | -0.011 | 0.371 | 0.71 | 0.3 | 0.63 | 0.85 | 0.74 | 0.8 |
| 9 | 81123772 | rs3010716 | 0.011 | -0.017 | 2.03 | 0.656 | 0.024 | -0.052 | -0.052 | 0.032 | 0.83 | 0.69 | 0.7 | 0.91 | 0.79 | 0.83 |
| 9 | 81132551 | rs10867213 | 1.629 | -0.141 | 1.722 | 0.92 | 0.014 | -0.006 | -0.006 | 0 | 0.58 | 0.3 | 0.44 | 0.5 | 0.62 | 0.55 |
| 9 | 81135377 | rs3913784 | -0.627 | -0.046 | 1.128 | 1.221 | -0.015 | 0.031 | 0.031 | 0 | 0.22 | 0.25 | 0.2 | 0.38 | 0.15 | 0.28 |
| 9 | 81148338 | rs1032889 | -1.6 | 0.073 | 0.757 | 1.002 | -0.004 | 0.002 | 0.002 | 0 | 0.4 | 0.57 | 0.54 | 0.51 | 0.37 | 0.45 |
| 9 | 81152970 | rs3010737 | 0.129 | 0.21 | -0.291 | 1.04 | -0.01 | 0.01 | 0.01 | 0 | 0.15 | 0.14 | 0.32 | 0.11 | 0.2 | 0.14 |
| 9 | 81154501 | rs1472773 | -0.737 | -0.35 | -0.389 | 1.402 | 0.021 | -0.033 | -0.033 | 0 | 0.2 | 0.14 | 0.2 | 0.4 | 0.12 | 0.27 |
| 9 | 81158113 | rs7027911 | 0.402 | -0.31 | -1.1 | 0.975 | 0.003 | 0.064 | 0.064 | 1.401 | 0.45 | 0.11 | 0.43 | 0.34 | 0.58 | 0.48 |
| 9 | 81158443 | rs3009696 | -0.545 | -0.497 | -0.35 | 1.204 | 0.021 | 0.053 | 0.053 | 0 | 0.2 | 0.36 | 0.37 | 0.33 | 0.27 | 0.21 |
| 9 | 81161599 | rs1566977 | -0.595 | -0.871 | -0.774 | 0.464 | 0.025 | 0.037 | 0.037 | 0 | 0.41 | 0.27 | 0.4 | 0.77 | 0.28 | 0.52 |
| 9 | 81162746 | rs1105025 | -0.98 | -0.294 | -0.707 | 0.979 | -0.004 | -0.014 | -0.014 | 0.161 | 0.16 | 0.11 | 0.28 | 0.69 | 0.18 | 0.27 |
| 9 | 81164099 | rs10867216 | -0.98 | -0.294 | -0.707 | 0.979 | -0.004 | -0.014 | -0.014 | 0 | 0.16 | 0.11 | 0.28 | 0.69 | 0.18 | 0.29 |
| 9 | 81167717 | rs3009698 | -0.456 | -0.325 | -0.775 | 1.47 | 0 | 0.071 | 0.071 | 0.02 | 0.95 | 0.97 | 0.93 | 0.57 | 0.96 | 0.97 |
| 9 | 81168218 | rs1505618 | -0.838 | 0.488 | -0.463 | 1.319 | 0.017 | -0.008 | -0.008 | 0 | 0.16 | 0.17 | 0.29 | 0.11 | 0.32 | 0.29 |
| 9 | 81169107 | rs1394926 | 0.58 | 0.511 | -0.558 | 0.601 | 0.02 | -0.043 | -0.043 | 0 | 0.33 | 0.55 | 0.34 | 0.23 | 0.37 | 0.36 |
| 9 | 81169950 | rs2047995 | 1.38 | 0.229 | -0.724 | 0.4 | 0.076 | 0.091 | 0.091 | 1.042 | 0.37 | 0.12 | 0.21 | 0.19 | 0.23 | 0.29 |
| 9 | 81171898 | rs1466959 | -0.462 | 0.79 | -0.523 | 0.73 | 0.019 | -0.002 | -0.002 | 0 | 0.15 | 0.17 | 0.31 | 0.17 | 0.32 | 0.29 |
| 9 | 81174574 | rs11792922 | 1.707 | 0.754 | -0.274 | 0.663 | 0.009 | 0.016 | 0.016 | 1.672 | 0.68 | 0.37 | 0.52 | 0.44 | 0.61 | 0.64 |
| 9 | 81178322 | rs870541 | 0.883 | 1.33 | -0.832 | 0.905 | -0.091 | 0.048 | 0.048 | 0.319 | 0.18 | 0.01 | 0.2 | 0.07 | 0.27 | 0.16 |
| 9 | 81178478 | rs870542 | -0.553 | 0.946 | 0.173 | 1.565 | 0.044 | 0.007 | 0.007 | 0 | 0.1 | 0.16 | 0.15 | 0.16 | 0.23 | 0.25 |

| | | | | | | | | | | | | | | | | |
|---|---|---|---|---|---|---|---|---|---|---|---|---|---|---|---|---|
| 9 | 81178617 | rs870543 | -0.562 | 0.948 | 0.086 | 1.566 | 0.067 | 0.014 | 0.014 | 0 | 0.1 | 0.21 | 0.15 | 0.16 | 0.23 | 0.25 |
| 9 | 81179936 | rs883883 | -0.158 | 0.905 | -0.224 | 1.636 | 0.001 | 0.054 | 0.054 | 0.043 | 0.95 | 0.97 | 0.93 | 0.63 | 0.96 | 0.97 |
| 9 | 81180304 | rs884585 | 0.912 | 0.937 | -0.172 | 1.222 | 0.038 | 0.093 | 0.093 | 1.124 | 0.77 | 0.59 | 0.7 | 0.46 | 0.69 | 0.69 |
| 9 | 81188738 | rs11137740 | 0.19 | -0.038 | -0.527 | 0.597 | -0.013 | -0.006 | -0.006 | 0 | 0.63 | 0.59 | 0.68 | 0.35 | 0.72 | 0.86 |
| 9 | 81189439 | rs11790951 | 0.017 | -0.122 | -0.537 | -0.069 | -0.001 | -0.012 | -0.012 | 0.145 | 0.34 | 0.32 | 0.38 | 0.11 | 0.3 | 0.29 |
| 9 | 81190597 | rs9695134 | -0.127 | 0.322 | -0.637 | 0.111 | -0.014 | -0.009 | -0.009 | 0 | 0.61 | 0.75 | 0.62 | 0.41 | 0.57 | 0.68 |
| 9 | 81191252 | rs12551231 | 0.71 | 0.053 | -0.87 | -0.1 | -0.062 | -0.065 | -0.065 | 2.777 | 0.25 | 0.05 | 0.27 | 0.29 | 0.32 | 0.38 |
| 9 | 81194641 | rs1505641 | -0.456 | 0.512 | -0.092 | 0.723 | -0.002 | 0.003 | 0.003 | 0 | 0.71 | 0.72 | 0.72 | 0.72 | 0.69 | 0.71 |
| 9 | 81196948 | rs1505637 | -0.541 | 0.585 | -0.16 | 1.052 | -0.018 | -0.02 | -0.02 | 0.594 | 0.71 | 0.89 | 0.72 | 0.65 | 0.7 | 0.71 |
| 9 | 81217496 | rs769290 | 0.593 | 0.432 | -0.387 | 1.028 | -0.015 | -0.089 | -0.089 | 2.404 | 0.14 | 0.01 | 0.18 | 0.29 | 0.14 | 0.13 |
| 9 | 81217634 | rs6559399 | 0.248 | 0.417 | -0.117 | 1.035 | -0.004 | -0.057 | -0.057 | 0 | 0.7 | 0.87 | 0.65 | 0.52 | 0.7 | 0.69 |
| 9 | 81223723 | rs11137759 | -0.152 | 0.204 | -0.305 | 0.96 | 0 | -0.008 | -0.008 | 2.097 | 0.27 | 0.26 | 0.27 | 0.39 | 0.24 | 0.24 |
| 9 | 81225091 | rs1859061 | 1.853 | -0.281 | -1.054 | 0.322 | -0.055 | 0.141 | 0.141 | 1.527 | 0.22 | 0.01 | 0.18 | 0.01 | 0.22 | 0.18 |
| 9 | 81226171 | rs234798 | 0.166 | -0.457 | -0.554 | 0.948 | -0.001 | 0.067 | 0.067 | 0 | 0.06 | 0.03 | 0.03 | 0.34 | 0.05 | 0.11 |
| 9 | 81226596 | rs7856547 | -0.482 | -0.551 | -0.401 | 0.857 | -0.005 | 0.006 | 0.006 | 5.333 | 0.31 | 0.39 | 0.25 | 0.39 | 0.24 | 0.26 |
| 9 | 81229089 | rs4877771 | 0.497 | -0.521 | -1.462 | 0.872 | -0.012 | -0.026 | -0.026 | 0 | 0.19 | 0.17 | 0.23 | 0.39 | 0.23 | 0.25 |
| 9 | 81231055 | rs234795 | -0.787 | -0.425 | -1.677 | 0.029 | 0.023 | 0.039 | 0.039 | 3.841 | 0.4 | 0.71 | 0.31 | 0.73 | 0.3 | 0.37 |
| 9 | 81240595 | rs234792 | -0.195 | -0.08 | -1.95 | -0.168 | 0.007 | 0.047 | 0.047 | 3.782 | 0.44 | 0.15 | 0.37 | 0.28 | 0.38 | 0.42 |
| 9 | 81250966 | rs2188902 | -0.02 | 0.047 | -1.972 | -0.012 | 0 | -0.044 | -0.044 | 0 | 0.49 | 0.9 | 0.61 | 0.51 | 0.56 | 0.5 |
| 9 | 81254538 | rs2106804 | -0.203 | 0.086 | -1.687 | -0.602 | 0.001 | 0.097 | 0.097 | 0 | 0.52 | 0.73 | 0.44 | 0.29 | 0.42 | 0.35 |
| 9 | 81259195 | rs10867228 | 0.967 | 0.145 | -1.651 | -0.448 | -0.022 | 0.014 | 0.014 | 0 | 0.19 | 0.11 | 0.26 | 0.2 | 0.18 | 0.11 |
| 9 | 81265380 | rs7870042 | 0.807 | 0.079 | -1.013 | -0.432 | 0.001 | -0.011 | -0.011 | 0 | 0.22 | 0.31 | 0.28 | 0.22 | 0.19 | 0.12 |
| 9 | 81272770 | rs3009691 | -0.588 | 0.412 | -1.522 | -0.368 | 0.029 | -0.067 | -0.067 | 1.265 | 0.66 | 0.92 | 0.83 | 0.69 | 0.7 | 0.51 |
| 9 | 81275274 | rs13297134 | 1.127 | 0.639 | -0.831 | -0.169 | 0.001 | -0.013 | -0.013 | 0 | 0.25 | 0.24 | 0.14 | 0.08 | 0.21 | 0.21 |
| 9 | 81284527 | rs2905478 | 0.715 | 0.886 | -0.839 | 0.431 | -0.025 | -0.023 | -0.023 | 0.023 | 0.14 | 0.05 | 0.39 | 0.4 | 0.3 | 0.17 |
| 9 | 81285865 | rs3008200 | 0.859 | 0.865 | -0.82 | 0.417 | -0.023 | -0.02 | -0.02 | 0 | 0.13 | 0.05 | 0.38 | 0.38 | 0.27 | 0.17 |
| 9 | 81301082 | rs2905481 | -0.678 | 0.132 | -1.651 | 0.308 | -0.006 | -0.036 | -0.036 | 1.655 | 0.87 | 0.98 | 0.95 | 0.85 | 0.88 | 0.75 |
| 9 | 81315526 | rs1757947 | 0.08 | 0.32 | -1.427 | 0.147 | 0.019 | 0.012 | 0.012 | 4.771 | 0.56 | 0.81 | 0.77 | 0.68 | 0.6 | 0.48 |
| 9 | 81328432 | rs2521904 | -0.334 | 0.435 | -0.866 | 0.009 | -0.002 | 0.004 | 0.004 | 0 | 0.59 | 0.59 | 0.33 | 0.71 | 0.46 | 0.62 |
| 9 | 81331339 | rs6559402 | -1.011 | 0.911 | -1.612 | 0.391 | -0.099 | -0.079 | -0.079 | 0 | 0.84 | 0.99 | 0.79 | 0.77 | 0.77 | 0.87 |
| 9 | 81333004 | rs1634349 | -0.77 | 0.967 | -1.319 | 0.25 | -0.035 | 0.041 | 0.041 | 0 | 0.63 | 0.77 | 0.35 | 0.7 | 0.5 | 0.64 |
| 9 | 81337236 | rs7868916 | 0.331 | 0.292 | -1.289 | -0.062 | 0.07 | -0.07 | -0.07 | 1.902 | 0.35 | 0.18 | 0.15 | 0.45 | 0.24 | 0.38 |
| 9 | 81339569 | rs1757945 | -0.277 | 0.269 | -0.721 | -0.027 | -0.01 | -0.004 | -0.004 | 0 | 0.74 | 0.79 | 0.72 | 0.73 | 0.73 | 0.71 |
| 9 | 81344701 | rs1929883 | 0.735 | 0.398 | -0.558 | -0.139 | -0.001 | 0.006 | 0.006 | 0.042 | 0.38 | 0.62 | 0.55 | 0.27 | 0.41 | 0.3 |
| 9 | 81348300 | rs1619442 | -1.347 | -0.474 | -1.066 | 0.081 | 0.023 | -0.023 | -0.023 | 4.386 | 0.82 | 0.99 | 0.92 | 0.88 | 0.84 | 0.77 |
| 9 | 81356984 | rs17788537 | 1.042 | -0.074 | -0.835 | 0.145 | 0.003 | 0.019 | 0.019 | 0 | 0.07 | 0.01 | 0.03 | 0.03 | 0.03 | 0.15 |
| 9 | 81359983 | rs11137821 | 0.734 | 0.004 | -0.444 | 0.145 | -0.01 | 0.024 | 0.024 | 0 | 0.39 | 0.34 | 0.66 | 0.29 | 0.53 | 0.37 |
| 9 | 81365721 | rs10867255 | 0.911 | -0.055 | -0.333 | 0.223 | 0.018 | -0.032 | -0.032 | 0 | 0.36 | 0.52 | 0.53 | 0.29 | 0.45 | 0.3 |
| 9 | 81370463 | rs10867258 | 1.137 | -0.024 | -1.055 | 0.547 | -0.017 | -0.038 | -0.038 | 0 | 0.2 | 0.11 | 0.12 | 0.39 | 0.2 | 0.13 |
| 9 | 81372519 | rs10512079 | 0.169 | -0.145 | -0.869 | 0.634 | -0.029 | -0.031 | -0.031 | 0 | 0.37 | 0.23 | 0.34 | 0.46 | 0.36 | 0.38 |
| 9 | 81372838 | rs10867263 | 0.078 | 0.916 | -0.851 | 0.544 | -0.029 | 0.029 | 0.029 | 0 | 0.33 | 0.23 | 0.52 | 0.28 | 0.44 | 0.3 |
| 9 | 81378381 | rs6559406 | -0.698 | 0.753 | -0.601 | 0.771 | 0.002 | 0.02 | 0.02 | 0 | 0.28 | 0.31 | 0.43 | 0.54 | 0.31 | 0.37 |
| 9 | 81381363 | rs12554407 | -0.249 | 0.744 | -1.038 | 0.292 | -0.014 | 0.023 | 0.023 | 0 | 0.19 | 0.07 | 0.23 | 0.18 | 0.2 | 0.18 |
| 9 | 81384173 | rs7030787 | -0.466 | 0.513 | -0.954 | -0.13 | 0 | 0.004 | 0.004 | 0.025 | 0.36 | 0.64 | 0.33 | 0.22 | 0.36 | 0.44 |
| 9 | 81384366 | rs1929889 | -0.895 | 0.521 | -1.55 | -0.215 | -0.096 | -0.086 | -0.086 | 0 | 0.56 | 0.92 | 0.43 | 0.45 | 0.5 | 0.5 |
| 9 | 81385545 | rs12347476 | 0.006 | 0.543 | -1.645 | -0.355 | -0.004 | -0.011 | -0.011 | 0 | 0.31 | 0.63 | 0.22 | 0.18 | 0.3 | 0.42 |
| 9 | 81387331 | rs11137830 | 0.673 | 0.475 | -1.548 | -0.347 | -0.01 | -0.013 | -0.013 | 0 | 0.19 | 0.06 | 0.08 | 0.23 | 0.14 | 0.06 |
| 9 | 81391164 | rs4529531 | 0.436 | 0.3 | -0.52 | -0.606 | 0.058 | 0.071 | 0.071 | 0.321 | 0.13 | 0.35 | 0.12 | 0.14 | 0.19 | 0.2 |
| 9 | 81394118 | rs7848869 | 0.835 | 0.351 | -0.257 | -0.629 | -0.005 | 0.032 | 0.032 | 0.018 | 0.09 | 0.17 | 0.1 | 0.14 | 0.11 | 0.15 |
| 9 | 81394862 | rs10867271 | -0.045 | 0.451 | -0.235 | -1.061 | -0.004 | -0.069 | -0.069 | 0.148 | 0.31 | 0.34 | 0.3 | 0.05 | 0.27 | 0.32 |
| 9 | 81394990 | rs10867272 | -0.037 | 0.468 | -0.265 | -0.983 | 0 | 0.008 | 0.008 | 0.174 | 0.15 | 0.17 | 0.21 | 0.18 | 0.17 | 0.18 |
| 9 | 81399854 | rs10117274 | 0.256 | 0.152 | -0.244 | -1.2 | -0.003 | -0.053 | -0.053 | 0 | 0.26 | 0.26 | 0.26 | 0.04 | 0.22 | 0.27 |
| 9 | 81402187 | rs10867274 | 0.308 | 0.38 | -0.481 | -1.024 | -0.029 | 0.015 | 0.015 | 0.293 | 0.27 | 0.14 | 0.23 | 0.24 | 0.33 | 0.16 |
| 9 | 81402625 | rs12349155 | 0.912 | 0.314 | -0.926 | -1.115 | -0.011 | 0.016 | 0.016 | 0 | 0.14 | 0.11 | 0.22 | 0.05 | 0.17 | 0.22 |
| 9 | 81403141 | rs7022062 | -1.05 | 0.205 | -0.899 | -1.108 | -0.008 | 0.035 | 0.035 | 0 | 0.51 | 0.69 | 0.39 | 0.38 | 0.48 | 0.36 |
| 9 | 81404818 | rs11137843 | 0.749 | 0.241 | -1.103 | -1.132 | -0.089 | -0.013 | -0.013 | 0.512 | 0.23 | 0.02 | 0.21 | 0.23 | 0.25 | 0.11 |
| 9 | 81404992 | rs1929871 | -0.774 | 0.037 | -0.96 | -0.915 | -0.011 | 0.036 | 0.036 | 0 | 0.52 | 0.72 | 0.39 | 0.39 | 0.48 | 0.36 |
| 9 | 81407851 | rs4300072 | -0.767 | 0.047 | -0.963 | -0.911 | -0.01 | 0.036 | 0.036 | 0 | 0.51 | 0.72 | 0.39 | 0.39 | 0.48 | 0.36 |
| 9 | 81409486 | rs3897787 | -0.812 | 0.068 | -1.249 | -0.9 | -0.031 | 0.039 | 0.039 | 0 | 0.52 | 0.77 | 0.39 | 0.39 | 0.48 | 0.36 |
| 9 | 81414687 | rs1929877 | -1.631 | 0.09 | -1.474 | -0.788 | -0.013 | -0.068 | -0.068 | 0 | 0.67 | 0.92 | 0.62 | 0.44 | 0.66 | 0.59 |
| 9 | 81420820 | rs7851210 | -0.293 | -0.225 | -0.936 | -0.691 | -0.005 | -0.001 | -0.001 | 0 | 0.23 | 0.25 | 0.15 | 0.15 | 0.18 | 0.23 |
| 9 | 81422195 | rs6559412 | -0.476 | -0.025 | -0.718 | -0.497 | -0.008 | -0.017 | -0.017 | 0 | 0.36 | 0.42 | 0.36 | 0.2 | 0.32 | 0.42 |
| 9 | 81426189 | rs880403 | -0.537 | 0.01 | -1.152 | -0.379 | 0.02 | 0.043 | 0.043 | 0.055 | 0.38 | 0.3 | 0.36 | 0.17 | 0.34 | 0.35 |
| 9 | 81429025 | rs12348374 | -0.57 | -0.141 | -2.02 | 0.068 | -0.07 | -0.005 | -0.005 | 0 | 0.56 | 0.93 | 0.43 | 0.64 | 0.43 | 0.42 |
| 9 | 81434850 | rs10780261 | 0.005 | -0.765 | -1.875 | 0.239 | 0.024 | -0.021 | -0.021 | 0 | 0.25 | 0.16 | 0.11 | 0.29 | 0.13 | 0.2 |
| 9 | 81435312 | rs7045330 | -1.337 | -0.686 | -1.391 | 0.34 | -0.005 | -0.019 | -0.019 | 0 | 0.55 | 0.83 | 0.42 | 0.52 | 0.43 | 0.41 |
| 9 | 81436632 | rs11137859 | 0.492 | -0.876 | -1.799 | 0.461 | 0.019 | 0.043 | 0.043 | 4.426 | 0.19 | 0.07 | 0.08 | 0.05 | 0.11 | 0.13 |
| 9 | 81436959 | rs7867205 | -1.258 | -0.987 | -1.799 | 0.477 | -0.039 | -0.058 | -0.058 | 0.469 | 0.55 | 0.91 | 0.43 | 0.52 | 0.43 | 0.42 |
| 9 | 81442685 | rs11137862 | 0.487 | -1.024 | -1.733 | 0.478 | 0.025 | 0.036 | 0.036 | 4.297 | 0.19 | 0.09 | 0.07 | 0.04 | 0.1 | 0.14 |

| | | | | | | | | | | | | | | | | |
|---|---|---|---|---|---|---|---|---|---|---|---|---|---|---|---|---|
| 9 | 81443855 | rs6559415 | -1.164 | -1.176 | -1.694 | 0.297 | 0.085 | -0.013 | -0.013 | 0 | 0.4 | 0.79 | 0.3 | 0.35 | 0.24 | 0.31 |
| 9 | 81447644 | rs1958925 | 0.067 | -1.243 | -1.551 | 0.305 | -0.029 | -0.041 | -0.041 | 0 | 0.14 | 0.62 | 0.13 | 0.06 | 0.09 | 0.09 |
| 9 | 81451697 | rs1543690 | 0.624 | -1.015 | -1.545 | 0.413 | -0.022 | 0.005 | 0.005 | 0.039 | 0.16 | 0.12 | 0.19 | 0.17 | 0.21 | 0.11 |
| 9 | 81460812 | rs10867289 | -0.161 | -1.055 | -1.508 | 0.443 | 0.028 | -0.026 | -0.026 | 0 | 0.25 | 0.16 | 0.17 | 0.29 | 0.13 | 0.2 |
| 9 | 81474084 | rs12378461 | 0.444 | -0.85 | -1.413 | -0.107 | 0.007 | 0.021 | 0.021 | 0.427 | 0.19 | 0.12 | 0.09 | 0.05 | 0.13 | 0.14 |
| 9 | 81476382 | rs11137879 | -0.602 | -0.822 | -1.416 | -0.097 | 0 | 0.048 | 0.048 | 0 | 0.35 | 0.25 | 0.27 | 0.22 | 0.31 | 0.24 |
| 9 | 81493973 | rs11137922 | 0.355 | -0.899 | -1.41 | -0.11 | 0.012 | 0.029 | 0.029 | 3.102 | 0.2 | 0.12 | 0.09 | 0.05 | 0.13 | 0.14 |
| 9 | 81497316 | rs11137930 | 0.725 | -0.86 | -1.345 | -0.119 | -0.003 | 0.001 | 0.001 | 0.023 | 0.15 | 0.16 | 0.18 | 0.16 | 0.17 | 0.1 |
| 9 | 81497488 | rs7028992 | -1.482 | -0.995 | -1.605 | -0.128 | 0.02 | 0.141 | 0.141 | 0.161 | 0.52 | 0.89 | 0.4 | 0.26 | 0.4 | 0.34 |
| 9 | 81500880 | rs10283740 | -0.502 | -0.991 | -1.418 | -0.111 | 0.011 | 0.019 | 0.019 | 0.195 | 0.31 | 0.73 | 0.31 | 0.22 | 0.27 | 0.2 |
| 9 | 81506587 | rs7028820 | -1.044 | -0.69 | -1.21 | 0.071 | 0.041 | 0.148 | 0.148 | 0.012 | 0.51 | 0.87 | 0.4 | 0.26 | 0.4 | 0.34 |
| 9 | 81508579 | rs4877311 | 0.313 | -0.5 | -1.105 | 0.314 | 0.009 | -0.041 | -0.041 | 0.105 | 0.31 | 0.19 | 0.32 | 0.44 | 0.34 | 0.53 |
| 9 | 81509121 | rs10867297 | -0.929 | -0.557 | -0.295 | -0.017 | 0.012 | -0.029 | -0.029 | 0 | 0.15 | 0.26 | 0.13 | 0.08 | 0.16 | 0.17 |
| 9 | 81518152 | rs7867093 | -0.315 | -0.726 | -0.418 | 0.312 | 0.004 | 0.014 | 0.014 | 0.549 | 0.28 | 0.36 | 0.35 | 0.32 | 0.33 | 0.34 |
| 9 | 81535490 | rs10125496 | -0.792 | -0.851 | -1.152 | -0.18 | -0.003 | 0.105 | 0.105 | 0.248 | 0.52 | 0.92 | 0.42 | 0.26 | 0.4 | 0.46 |
| 9 | 81535908 | rs10125637 | -0.564 | -0.863 | -0.779 | -0.183 | 0.044 | 0.1 | 0.1 | 0.263 | 0.5 | 0.74 | 0.41 | 0.26 | 0.4 | 0.46 |
| 9 | 81552231 | rs10780277 | -0.76 | -0.866 | -1.033 | -0.195 | 0.011 | 0.115 | 0.115 | 0.025 | 0.52 | 0.88 | 0.38 | 0.25 | 0.4 | 0.46 |
| 9 | 81553066 | rs11137960 | 0.823 | -1.099 | -0.502 | -0.388 | -0.004 | -0.014 | -0.014 | 0 | 0.2 | 0.18 | 0.1 | 0.05 | 0.08 | 0.14 |
| 9 | 81553828 | rs1438806 | 1.28 | -0.697 | -1.411 | -0.224 | -0.105 | -0.116 | -0.116 | 0.513 | 0.23 | 0.03 | 0.31 | 0.36 | 0.33 | 0.34 |
| 9 | 81554948 | rs10512076 | -1.106 | -1.285 | -0.662 | -0.673 | -0.002 | 0.087 | 0.087 | 0 | 0.54 | 0.92 | 0.38 | 0.26 | 0.4 | 0.5 |
| 9 | 81555504 | rs1438805 | 0.025 | -1.314 | -0.272 | -0.683 | -0.007 | -0.001 | -0.001 | 0 | 0.28 | 0.23 | 0.29 | 0.24 | 0.32 | 0.35 |
| 9 | 81560828 | rs1999497 | -1.088 | -1.226 | -0.272 | -0.695 | -0.014 | 0.066 | 0.066 | 0 | 0.56 | 0.84 | 0.38 | 0.26 | 0.42 | 0.54 |
| 9 | 81568457 | rs11137969 | 0.662 | -1.078 | -0.196 | -0.674 | -0.004 | 0 | 0 | 0 | 0.09 | 0.07 | 0.07 | 0.06 | 0.12 | 0.13 |
| 9 | 81569012 | rs7860309 | 0.316 | -1.255 | -0.009 | -0.763 | -0.071 | -0.11 | -0.11 | 0.044 | 0.27 | 0.43 | 0.12 | 0.05 | 0.1 | 0.21 |
| 9 | 81569998 | rs2218075 | -0.774 | -1.094 | -0.351 | -0.678 | 0.114 | 0.134 | 0.134 | 0 | 0.48 | 0.77 | 0.35 | 0.23 | 0.3 | 0.39 |
| 9 | 81572766 | rs10512074 | -0.52 | -0.704 | -0.264 | -0.538 | 0.016 | 0 | 0 | 0 | 0.29 | 0.44 | 0.3 | 0.24 | 0.32 | 0.34 |
| 9 | 81574729 | rs10867315 | 0.161 | -1.046 | -0.616 | -0.484 | 0.001 | -0.002 | -0.002 | 0.047 | 0.19 | 0.18 | 0.22 | 0.19 | 0.2 | 0.2 |
| 9 | 81576070 | rs7852123 | -0.379 | -0.668 | -1.006 | -0.313 | 0.031 | 0.035 | 0.035 | 0 | 0.44 | 0.62 | 0.3 | 0.2 | 0.28 | 0.35 |
| 9 | 81589458 | rs10118058 | -0.103 | -0.036 | -1.485 | 0.06 | 0.031 | 0.024 | 0.024 | 0 | 0.5 | 0.77 | 0.54 | 0.44 | 0.47 | 0.48 |
| 9 | 81591540 | rs10867325 | -0.562 | 0.071 | -1.187 | 0.244 | -0.002 | -0.003 | -0.003 | 0 | 0.13 | 0.1 | 0.34 | 0.23 | 0.13 | 0.23 |
| 9 | 81594461 | rs2033817 | -0.301 | 0.008 | -0.971 | -0.07 | -0.001 | 0 | 0 | 0 | 0.39 | 0.63 | 0.38 | 0.48 | 0.37 | 0.35 |
| 9 | 81596162 | rs4637922 | -0.086 | -0.076 | -0.362 | -0.146 | 0.002 | 0.008 | 0.008 | 0.144 | 0.36 | 0.39 | 0.37 | 0.49 | 0.37 | 0.36 |
| 9 | 81606411 | rs7852195 | -0.66 | -0.52 | 0.154 | 0.112 | -0.023 | -0.027 | -0.027 | 0.066 | 0.09 | 0.54 | 0.05 | 0 | 0.01 | 0 |
| 9 | 81620737 | rs10512072 | -0.573 | -0.503 | -0.62 | -0.176 | -0.011 | 0.084 | 0.084 | 7.83 | 0.8 | 0.96 | 0.89 | 0.96 | 0.84 | 0.81 |
| 9 | 81622277 | rs11137990 | 0.004 | -0.462 | -0.476 | 0.124 | -0.038 | 0.081 | 0.081 | 3.161 | 0.71 | 0.5 | 0.77 | 0.57 | 0.81 | 0.76 |
| 9 | 81626352 | rs11137996 | -0.256 | -0.351 | -0.967 | -0.344 | 0.005 | 0.001 | 0.001 | 0.176 | 0.24 | 0.06 | 0.31 | 0.21 | 0.18 | 0.34 |
| 9 | 81627699 | rs1341082 | 0.088 | -0.637 | -0.863 | -0.137 | 0.015 | -0.023 | -0.023 | 0 | 0.17 | 0.12 | 0.18 | 0.62 | 0.12 | 0.18 |
| 9 | 81628077 | rs8181181 | -0.97 | -0.398 | -0.142 | -0.573 | -0.005 | -0.031 | -0.031 | 0 | 0.41 | 0.39 | 0.52 | 0.87 | 0.33 | 0.52 |
| 9 | 81628944 | rs1036598 | -0.72 | 0.302 | -0.186 | -0.718 | 0.004 | -0.016 | -0.016 | 0.026 | 0.85 | 0.81 | 0.85 | 0.96 | 0.74 | 0.88 |
| 9 | 81629264 | rs12376398 | 1.235 | 0.174 | -0.075 | -0.815 | -0.008 | 0.071 | 0.071 | 0.138 | 0.45 | 0.31 | 0.35 | 0.11 | 0.47 | 0.44 |
| 9 | 81633695 | rs11137998 | 0.439 | 0.093 | 0.076 | -0.672 | -0.004 | -0.008 | -0.008 | 0 | 0.2 | 0.12 | 0.32 | 0.28 | 0.2 | 0.36 |
| 9 | 81634304 | rs1014331 | 1.126 | 0.085 | -0.147 | -0.557 | -0.003 | 0.041 | 0.041 | 0.004 | 0.47 | 0.32 | 0.44 | 0.64 | 0.44 | 0.46 |
| 9 | 81635735 | rs7859992 | 0.219 | -0.275 | 0.17 | -0.028 | 0.027 | -0.049 | -0.049 | 0.122 | 0.17 | 0.08 | 0.18 | 0.6 | 0.12 | 0.18 |
| 9 | 81642434 | rs10867331 | -0.484 | -0.246 | 0.562 | -0.147 | -0.03 | 0.007 | 0.007 | 0.269 | 0.24 | 0.44 | 0.34 | 0.25 | 0.15 | 0.28 |
| 9 | 81646444 | rs6559441 | -1.281 | -0.366 | 0.416 | -0.308 | -0.011 | -0.017 | -0.017 | 0.081 | 0.4 | 0.53 | 0.4 | 0.28 | 0.29 | 0.45 |
| 9 | 81647793 | rs6559443 | -1.694 | 0.514 | 0.905 | 0.215 | 0 | 0.002 | 0.002 | 2.182 | 0.35 | 0.67 | 0.41 | 0.28 | 0.3 | 0.48 |
| 9 | 81648357 | rs7874846 | 0.536 | 0.582 | 0.954 | 0.292 | 0.01 | 0.002 | 0.002 | 0 | 0.93 | 0.94 | 0.95 | 0.93 | 0.89 | 0.87 |
| 9 | 81651946 | rs10122426 | -0.534 | 0.941 | 0.489 | 0.563 | -0.063 | -0.02 | -0.02 | 0 | 0.9 | 0.99 | 0.88 | 0.91 | 0.86 | 0.93 |
| 9 | 81657195 | rs10512080 | 1.495 | 0.286 | 0.853 | 0.572 | -0.004 | -0.005 | -0.005 | 1.026 | 0.67 | 0.35 | 0.61 | 0.71 | 0.75 | 0.65 |
| 9 | 81659736 | rs1417653 | -0.172 | 0.266 | 0.893 | 0.399 | -0.001 | 0.005 | 0.005 | 0.07 | 0.2 | 0.11 | 0.31 | 0.2 | 0.17 | 0.3 |
| 9 | 81669140 | rs6559445 | 1.002 | 0.317 | 1.017 | 0.417 | -0.011 | -0.014 | -0.014 | 0 | 0.9 | 0.76 | 0.93 | 0.96 | 0.92 | 0.93 |
| 9 | 81671791 | rs4242641 | 1.901 | 0.33 | 1.133 | 0.445 | 0.008 | 0.03 | 0.03 | 2.839 | 0.61 | 0.3 | 0.53 | 0.2 | 0.67 | 0.45 |
| 9 | 81673370 | rs1341086 | 1.149 | 0.363 | 1.172 | 0.253 | 0.008 | -0.003 | -0.003 | 0 | 0.89 | 0.75 | 0.92 | 0.93 | 0.89 | 0.86 |
| 9 | 81675754 | rs7038375 | 1.205 | 0.387 | 0.81 | 0.283 | -0.003 | -0.006 | -0.006 | 0 | 0.68 | 0.37 | 0.6 | 0.75 | 0.73 | 0.62 |
| 9 | 81678250 | rs7031319 | -0.523 | 0.475 | 1.023 | 0.526 | 0.023 | -0.027 | -0.027 | 0.602 | 0.14 | 0.27 | 0.13 | 0.05 | 0.1 | 0.06 |
| 9 | 81679224 | rs17082093 | -1.287 | 0.322 | 1.188 | 0.449 | -0.008 | -0.005 | -0.005 | 1.64 | 0.27 | 0.66 | 0.38 | 0.25 | 0.19 | 0.31 |
| 9 | 81679875 | rs4877387 | -1.414 | 0.173 | 1.38 | 0.65 | 0.001 | -0.002 | -0.002 | 0.039 | 0.3 | 0.36 | 0.46 | 0.77 | 0.26 | 0.4 |
| 9 | 81680447 | rs17082096 | -0.77 | 0.27 | 1.665 | 0.837 | 0.002 | 0 | 0 | 0.476 | 0.18 | 0.17 | 0.31 | 0.21 | 0.14 | 0.27 |
| 9 | 81692440 | rs1361537 | -1.593 | 0.365 | 1.624 | 0.37 | 0.042 | 0.032 | 0.032 | 1.895 | 0.06 | 0.48 | 0.05 | 0 | 0.01 | 0 |
| 9 | 81696557 | rs11138049 | -1.508 | 0.485 | 0.923 | 0.423 | -0.003 | 0.006 | 0.006 | 0 | 0.1 | 0.13 | 0.13 | 0.09 | 0.09 | 0.14 |
| 9 | 81706625 | rs2066259 | -0.957 | 0.404 | 0.72 | 0.485 | -0.011 | 0 | 0 | 1.645 | 0.23 | 0.65 | 0.37 | 0.25 | 0.16 | 0.27 |
| 9 | 81709503 | rs7866466 | -1.309 | 0.156 | 0.282 | 0.437 | 0.001 | 0.001 | 0.001 | 1.638 | 0.34 | 0.63 | 0.46 | 0.21 | 0.24 | 0.34 |
| 9 | 81720891 | rs1544315 | -0.97 | -0.105 | 0.288 | 0.692 | -0.024 | 0.023 | 0.023 | 0.194 | 0.24 | 0.32 | 0.42 | 0.3 | 0.16 | 0.37 |
| 9 | 81725579 | rs10780291 | -0.527 | 0.216 | -0.124 | 0.582 | -0.007 | 0.007 | 0.007 | 0.053 | 0.65 | 0.53 | 0.65 | 0.33 | 0.67 | 0.64 |
| 9 | 81729460 | rs11138074 | -0.467 | 0.264 | -0.381 | 0.087 | 0.004 | 0.019 | 0.019 | 0 | 0.33 | 0.24 | 0.22 | 0.21 | 0.3 | 0.38 |
| 9 | 81738012 | rs12001272 | -0.582 | 0.717 | -0.185 | 0.215 | 0.013 | 0.03 | 0.03 | 0 | 0.08 | 0.15 | 0.13 | 0.16 | 0.13 | 0.16 |
| 9 | 81739976 | rs2236578 | -0.41 | 0.713 | -0.183 | 0.211 | -0.007 | -0.004 | -0.004 | 0 | 0.07 | 0.04 | 0.12 | 0.16 | 0.13 | 0.22 |
| 9 | 81743062 | rs11138084 | -0.27 | 0.168 | -0.283 | -0.158 | -0.005 | -0.019 | -0.019 | 0 | 0.22 | 0.34 | 0.16 | 0.14 | 0.19 | 0.33 |
| 9 | 81745376 | rs11138085 | -0.638 | 0.22 | -0.372 | -0.195 | -0.001 | -0.005 | -0.005 | 0.571 | 0.57 | 0.38 | 0.49 | 0.17 | 0.51 | 0.46 |

| | | | | | | | | | | | | | | | | |
|---|---|---|---|---|---|---|---|---|---|---|---|---|---|---|---|---|
| 9 | 81745871 | rs12552083 | 0.221 | 0.367 | -0.829 | -0.09 | -0.003 | 0.085 | 0.085 | 0 | 0.35 | 0.15 | 0.33 | 0.03 | 0.33 | 0.14 |
| 9 | 81747584 | rs10780292 | -0.033 | 0.552 | -0.504 | 0.345 | -0.001 | 0.014 | 0.014 | 0 | 0.57 | 0.6 | 0.49 | 0.81 | 0.58 | 0.75 |
| 9 | 81757988 | rs6559456 | -0.041 | 0.477 | -0.227 | 0.845 | 0.001 | -0.011 | -0.011 | 0.077 | 0.57 | 0.48 | 0.4 | 0.62 | 0.57 | 0.67 |
| 9 | 81758498 | rs11138089 | -0.28 | 0.908 | -0.442 | 0.899 | -0.019 | 0.004 | 0.004 | 0 | 0.13 | 0.08 | 0.16 | 0.09 | 0.28 | 0.21 |
| 9 | 81758706 | rs7023500 | -0.724 | 0.9 | -0.449 | 0.862 | -0.041 | 0.056 | 0.056 | 0 | 0.62 | 0.86 | 0.44 | 0.75 | 0.58 | 0.69 |
| 9 | 81758804 | rs7023856 | 0.858 | 0.833 | -0.467 | 1.289 | -0.005 | -0.001 | -0.001 | 0.054 | 0.42 | 0.41 | 0.23 | 0.36 | 0.29 | 0.33 |
| 9 | 81759693 | rs9314676 | -0.493 | 0.834 | -0.616 | 0.92 | -0.008 | 0.037 | 0.037 | 0 | 0.6 | 0.73 | 0.5 | 0.68 | 0.58 | 0.74 |
| 9 | 81765660 | rs11790699 | 0.135 | 1.229 | -0.749 | 0.997 | -0.041 | -0.03 | -0.03 | 0 | 0.07 | 0.01 | 0.11 | 0.09 | 0.16 | 0.14 |
| 9 | 81766500 | rs10732690 | -0.869 | 0.91 | -0.67 | 0.905 | -0.011 | 0 | 0 | 0 | 0.57 | 0.55 | 0.74 | 0.46 | 0.65 | 0.56 |
| 9 | 81768535 | rs12378835 | -0.182 | 1.258 | -1.134 | 0.811 | -0.04 | 0.001 | 0.001 | 0.021 | 0.12 | 0.06 | 0.17 | 0.09 | 0.28 | 0.21 |
| 9 | 81771012 | rs4294266 | 0.75 | 0.74 | -1.053 | 0.941 | 0.004 | -0.002 | -0.002 | 0 | 0.4 | 0.59 | 0.24 | 0.45 | 0.29 | 0.26 |
| 9 | 81771711 | rs1032404 | 0.045 | 0.649 | -1.386 | 0.989 | -0.003 | 0 | 0 | 0 | 0.39 | 0.11 | 0.54 | 0.34 | 0.36 | 0.41 |
| 9 | 81775281 | rs12682880 | -0.558 | 0.832 | -1.208 | 0.645 | 0 | 0.004 | 0.004 | 2.863 | 0.3 | 0.3 | 0.4 | 0.2 | 0.31 | 0.3 |
| 9 | 81775361 | rs10512082 | -0.127 | 0.794 | -1.307 | 0.564 | -0.015 | 0.064 | 0.064 | 2.528 | 0.26 | 0.11 | 0.39 | 0.08 | 0.3 | 0.27 |
| 9 | 81775520 | rs1341090 | 0.946 | 1.343 | -1.362 | 0.759 | -0.019 | 0.005 | 0.005 | 4.831 | 0.62 | 0.68 | 0.45 | 0.72 | 0.48 | 0.57 |
| 9 | 81776086 | rs1027263 | 0.338 | 0.982 | -1.324 | 1.022 | 0.005 | -0.005 | -0.005 | 0.051 | 0.92 | 0.93 | 0.85 | 0.9 | 0.79 | 0.87 |
| 9 | 81777861 | rs1873738 | -1.401 | 0.169 | -1.156 | 1.137 | -0.037 | 0.024 | 0.024 | 1.883 | 0.21 | 0.33 | 0.13 | 0.33 | 0.1 | 0.37 |
| 9 | 81778487 | rs12554475 | -0.305 | 0.454 | -1.462 | 0.82 | 0.01 | 0 | 0 | 0 | 0.12 | 0.07 | 0.1 | 0.09 | 0.07 | 0.16 |
| 9 | 81781384 | rs732027 | 1.186 | 0.309 | -2.06 | 0.555 | -0.023 | 0.006 | 0.006 | 2.718 | 0.58 | 0.73 | 0.68 | 0.61 | 0.55 | 0.3 |
| 9 | 81782185 | rs1361536 | 0.607 | 0.566 | -1.973 | 0.647 | -0.022 | 0.014 | 0.014 | 0.004 | 0.73 | 0.77 | 0.78 | 0.76 | 0.63 | 0.56 |
| 9 | 81784956 | rs2378309 | -0.651 | 0.691 | -2.127 | 0.471 | -0.064 | 0.048 | 0.048 | 0 | 0.84 | 0.97 | 0.82 | 0.92 | 0.75 | 0.77 |
| 9 | 81786242 | rs1341088 | -0.651 | 0.691 | -2.127 | 0.512 | -0.064 | -0.01 | -0.01 | 0.249 | 0.84 | 0.97 | 0.82 | 0.87 | 0.75 | 0.76 |
| 9 | 81791960 | rs7850379 | 1.337 | 0.35 | -2.056 | 0.555 | -0.023 | 0.002 | 0.002 | 1.147 | 0.6 | 0.72 | 0.66 | 0.62 | 0.53 | 0.37 |
| 9 | 81800073 | rs17791273 | 0.728 | 0.401 | -2.229 | 0.572 | 0.001 | -0.03 | -0.03 | 0 | 0.1 | 0 | 0.09 | 0.09 | 0.07 | 0.16 |
| 9 | 81801506 | rs7025892 | -0.087 | 0.356 | -2.258 | 0.541 | -0.086 | 0.018 | 0.018 | 0.186 | 0.79 | 0.95 | 0.78 | 0.87 | 0.64 | 0.68 |
| 9 | 81810992 | rs4877474 | 0.136 | 0.309 | -2.389 | 0.505 | -0.07 | 0.029 | 0.029 | 0 | 0.73 | 0.91 | 0.78 | 0.77 | 0.63 | 0.58 |
| 9 | 81811031 | rs4877475 | 1.221 | 0.259 | -2.41 | 0.547 | -0.037 | 0.036 | 0.036 | 0.225 | 0.59 | 0.79 | 0.67 | 0.66 | 0.53 | 0.39 |
| 9 | 81812343 | rs10122209 | 1.178 | 0.267 | -2.406 | 0.537 | -0.025 | -0.007 | -0.007 | 0 | 0.59 | 0.78 | 0.67 | 0.61 | 0.54 | 0.38 |
| 9 | 81814421 | rs902908 | -0.113 | 0.465 | -2.476 | 0.474 | -0.102 | 0.096 | 0.096 | 0 | 0.76 | 0.98 | 0.85 | 0.91 | 0.68 | 0.69 |
| 9 | 81816784 | rs7872379 | 0.013 | 0.273 | -2.335 | 0.493 | 0.012 | 0.001 | 0.001 | 0 | 0.14 | 0.08 | 0.11 | 0.11 | 0.1 | 0.2 |
| 9 | 81821091 | rs2493600 | -0.355 | 0.644 | -2.443 | 0.529 | -0.064 | -0.014 | -0.014 | 0.098 | 0.27 | 0.11 | 0.22 | 0.27 | 0.37 | 0.42 |
| 9 | 81821747 | rs11138128 | 0.726 | 0.079 | -2.907 | 0.66 | 0.035 | -0.097 | -0.097 | 0 | 0.23 | 0.09 | 0.38 | 0.38 | 0.2 | 0.11 |
| 9 | 81824781 | rs7846875 | -0.368 | 0.872 | -2.267 | 0.715 | 0.008 | -0.001 | -0.001 | 0 | 0.39 | 0.27 | 0.37 | 0.32 | 0.34 | 0.46 |
| 9 | 81829336 | rs273464 | 0.562 | 0.897 | -2.61 | 0.557 | -0.003 | 0.01 | 0.01 | 0.341 | 0.26 | 0.18 | 0.17 | 0.21 | 0.25 | 0.33 |
| 9 | 81834426 | rs17791387 | 0.78 | 0.805 | -2.644 | 0.593 | -0.019 | -0.036 | -0.036 | 0 | 0.14 | 0.01 | 0.15 | 0.11 | 0.14 | 0.22 |
| 9 | 81835737 | rs273465 | 0.825 | 0.836 | -2.623 | 0.577 | -0.001 | 0.016 | 0.016 | 0.739 | 0.23 | 0.14 | 0.15 | 0.1 | 0.21 | 0.21 |
| 9 | 81837587 | rs17791393 | 0.628 | 0.917 | -2.644 | 0.584 | -0.026 | -0.034 | -0.034 | 0 | 0.14 | 0.01 | 0.15 | 0.11 | 0.15 | 0.23 |
| 9 | 81839609 | rs2769957 | -0.604 | 1.539 | -2.43 | 0.514 | -0.076 | -0.078 | -0.078 | 0 | 0.86 | 0.98 | 0.84 | 0.69 | 0.76 | 0.87 |
| 9 | 81843768 | rs273453 | -0.04 | 0.829 | -1.791 | -0.099 | -0.002 | -0.003 | -0.003 | 0 | 0.28 | 0.31 | 0.18 | 0.23 | 0.23 | 0.34 |
| 9 | 81847769 | rs190640 | 0.002 | 0.919 | -1.783 | -0.056 | -0.003 | -0.006 | -0.006 | 0 | 0.28 | 0.32 | 0.18 | 0.23 | 0.23 | 0.34 |
| 9 | 81866882 | rs273432 | -0.466 | 1.412 | -1.616 | -0.24 | 0.008 | -0.029 | -0.029 | 0.02 | 0.26 | 0.39 | 0.15 | 0.13 | 0.29 | 0.37 |
| 9 | 81869419 | rs1887719 | -0.277 | 1.255 | -1.972 | 0.329 | -0.005 | -0.001 | -0.001 | 1.139 | 0.32 | 0.27 | 0.33 | 0.31 | 0.32 | 0.53 |
| 9 | 81870379 | rs11792810 | -0.555 | 1.23 | -2.281 | 0.322 | -0.001 | -0.004 | -0.004 | 0 | 0.57 | 0.39 | 0.5 | 0.43 | 0.57 | 0.72 |
| 9 | 81870811 | rs1753061 | -0.439 | 1.347 | -2.6 | 0.72 | -0.102 | -0.037 | -0.037 | 0.86 | 0.71 | 0.94 | 0.71 | 0.68 | 0.59 | 0.71 |
| 9 | 81871500 | rs113123 | 0.157 | 0.697 | -2.092 | 0.693 | 0.01 | 0 | 0 | 0.682 | 0.38 | 0.64 | 0.41 | 0.38 | 0.3 | 0.22 |
| 9 | 81872880 | rs10746589 | 0.584 | 1.361 | -2.177 | 1.139 | -0.052 | -0.071 | -0.071 | 0 | 0.89 | 0.98 | 0.87 | 0.71 | 0.84 | 0.88 |
| 9 | 81873321 | rs12351466 | 0.057 | 0.797 | -1.858 | 0.145 | -0.04 | -0.136 | -0.136 | 0.615 | 0.25 | 0.46 | 0.17 | 0.09 | 0.2 | 0.28 |
| 9 | 81874005 | rs273471 | 0.865 | 0.977 | -2.26 | 0.413 | -0.067 | -0.04 | -0.04 | 0.663 | 0.2 | 0.04 | 0.19 | 0.22 | 0.28 | 0.18 |
| 9 | 81876910 | rs273472 | 0.467 | 0.471 | -1.804 | 0.699 | 0.003 | -0.001 | -0.001 | 0 | 0.52 | 0.53 | 0.42 | 0.46 | 0.6 | 0.68 |
| 9 | 81877286 | rs273473 | 0.379 | 0.402 | -1.341 | 1.307 | -0.004 | 0.027 | 0.027 | 0 | 0.78 | 0.75 | 0.65 | 0.56 | 0.81 | 0.83 |
| 9 | 81881142 | rs4877480 | -0.331 | 0.298 | -1.611 | 1.115 | 0.013 | -0.035 | -0.035 | 0 | 0.19 | 0.07 | 0.13 | 0.27 | 0.14 | 0.3 |
| 9 | 81882407 | rs4486277 | -0.901 | 0.206 | -1.137 | 0.514 | 0.004 | 0.003 | 0.003 | 0.396 | 0.42 | 0.39 | 0.43 | 0.36 | 0.34 | 0.51 |
| 9 | 81890076 | rs1492611 | 0.134 | 0.404 | -1.276 | 0.457 | 0.019 | -0.024 | -0.024 | 0.205 | 0.41 | 0.2 | 0.35 | 0.49 | 0.34 | 0.43 |
| 9 | 81894224 | rs10780298 | 1.054 | 1.227 | -1.654 | 0.479 | -0.053 | -0.061 | -0.061 | 0 | 0.11 | 0.02 | 0.15 | 0.33 | 0.21 | 0.14 |
| 9 | 81896716 | rs273449 | 0.329 | 0.922 | -1.264 | 0.33 | 0.003 | -0.007 | -0.007 | 0 | 0.44 | 0.33 | 0.51 | 0.54 | 0.39 | 0.57 |
| 9 | 81897650 | rs273450 | -0.386 | 0.834 | -0.936 | 0.257 | -0.003 | -0.003 | -0.003 | 0.038 | 0.61 | 0.63 | 0.59 | 0.54 | 0.48 | 0.64 |
| 9 | 81904763 | rs17789673 | 0.948 | 0.686 | -0.88 | 0.106 | 0.003 | 0.016 | 0.016 | 0.538 | 0.14 | 0.05 | 0.14 | 0.05 | 0.1 | 0.16 |
| 9 | 81908179 | rs7027362 | 0.096 | 0.663 | -0.57 | 0.073 | -0.002 | -0.016 | -0.016 | 0 | 0.19 | 0.26 | 0.31 | 0.1 | 0.16 | 0.32 |
| 9 | 81911067 | rs1688964 | -1.467 | 1.024 | -0.616 | -0.15 | 0.035 | -0.081 | -0.081 | 2.86 | 0.85 | 0.71 | 0.86 | 0.98 | 0.78 | 0.88 |
| 9 | 81918369 | rs273438 | 0.028 | 1.103 | -0.814 | 0.117 | 0.011 | 0.001 | 0.001 | 1.71 | 0.66 | 0.63 | 0.67 | 0.63 | 0.53 | 0.77 |
| 9 | 81926600 | rs166648 | 0.682 | 0.663 | -0.636 | 0.057 | 0.009 | -0.003 | -0.003 | 0 | 0.52 | 0.37 | 0.36 | 0.5 | 0.43 | 0.46 |
| 9 | 81928572 | rs7026633 | -0.569 | 0.473 | -0.676 | -0.278 | 0 | -0.043 | -0.043 | 0.102 | 0.23 | 0.33 | 0.29 | 0.1 | 0.23 | 0.27 |
| 9 | 81929517 | rs12380408 | 0.234 | 0.516 | -0.774 | -0.284 | 0.001 | 0.032 | 0.032 | 0.231 | 0.17 | 0.09 | 0.11 | 0 | 0.16 | 0.11 |
| 9 | 81932554 | rs10867361 | 1.537 | 0.499 | -0.673 | -0.014 | -0.001 | 0 | 0 | 0 | 0.05 | 0.04 | 0.03 | 0.07 | 0.05 | 0.02 |
| 9 | 81936632 | rs4877483 | -0.02 | 0.409 | -0.651 | 0.105 | -0.026 | 0.027 | 0.027 | 0.068 | 0.61 | 0.74 | 0.64 | 0.69 | 0.52 | 0.72 |
| 9 | 81946034 | rs12006184 | 0.92 | 0.291 | -0.434 | 0.06 | 0.011 | -0.008 | -0.008 | 0 | 0.11 | 0.17 | 0.22 | 0.09 | 0.14 | 0.2 |
| 9 | 81946719 | rs4143031 | -0.637 | 0.44 | -0.697 | -0.01 | -0.001 | -0.004 | -0.004 | 0 | 0.74 | 0.74 | 0.8 | 0.89 | 0.71 | 0.81 |
| 9 | 81951578 | rs12683085 | 2.297 | -0.101 | -0.825 | 0.342 | 0.034 | -0.087 | -0.087 | 0 | 0.2 | 0.02 | 0.16 | 0.3 | 0.12 | 0.28 |
| 9 | 81951599 | rs12683067 | -0.219 | 0.206 | -0.453 | 0.671 | -0.007 | 0.015 | 0.015 | 1.271 | 0.55 | 0.67 | 0.6 | 0.58 | 0.47 | 0.7 |

| | | | | | | | | | | | | | | | | |
|---|---|---|---|---|---|---|---|---|---|---|---|---|---|---|---|---|
| 9 | 81953539 | rs7020903 | -0.56 | 0.114 | 0.1 | 0.42 | 0.001 | -0.042 | -0.042 | 0 | 0.17 | 0.35 | 0.24 | 0.09 | 0.15 | 0.2 |
| 9 | 81953661 | rs7021045 | 1.15 | 0.423 | -0.433 | 0.494 | -0.001 | -0.009 | -0.009 | 0.358 | 0.08 | 0.07 | 0.06 | 0 | 0.08 | 0.05 |
| 9 | 81953998 | rs7849719 | 1.05 | 0.451 | -0.372 | 0.808 | -0.006 | 0.002 | 0.002 | 0.021 | 0.28 | 0.29 | 0.22 | 0.3 | 0.21 | 0.3 |
| 9 | 81966076 | rs7043897 | -0.879 | 1.458 | 0.253 | 0.567 | 0.016 | -0.016 | -0.016 | 0 | 0.38 | 0.52 | 0.33 | 0.3 | 0.42 | 0.24 |
| 9 | 81969812 | rs12551528 | 0.579 | 1.963 | 0.039 | 0.474 | -0.039 | 0.014 | 0.014 | 0 | 0.1 | 0.02 | 0.11 | 0.06 | 0.14 | 0.06 |
| 9 | 81972841 | rs7026908 | -0.051 | 1.68 | 0.4 | 0.432 | -0.022 | 0.03 | 0.03 | 0 | 0.26 | 0.15 | 0.23 | 0.16 | 0.3 | 0.1 |
| 9 | 81974200 | rs7859622 | -0.051 | 1.68 | 0.42 | 0.432 | -0.021 | 0.028 | 0.028 | 0 | 0.26 | 0.15 | 0.23 | 0.16 | 0.3 | 0.1 |
| 9 | 81975178 | rs12344408 | 0.02 | 1.668 | 0.868 | 0.434 | 0.037 | 0.021 | 0.021 | 0.035 | 0.11 | 0.22 | 0.05 | 0.13 | 0.12 | 0.14 |
| 9 | 81975570 | rs10780304 | -2.017 | 1.417 | 0.806 | 0.556 | 0.084 | 0.011 | 0.011 | 0 | 0.57 | 0.85 | 0.65 | 0.64 | 0.7 | 0.54 |
| 9 | 81978708 | rs10780305 | -1.138 | 1.415 | 1.02 | 0.833 | -0.003 | -0.005 | -0.005 | 0.147 | 0.31 | 0.7 | 0.41 | 0.48 | 0.4 | 0.43 |
| 9 | 81983320 | rs11138168 | -1.031 | 1.301 | 0.904 | 0.92 | 0.007 | 0.025 | 0.025 | 0.754 | 0.29 | 0.61 | 0.25 | 0.4 | 0.33 | 0.35 |
| 9 | 81988322 | rs12338373 | -0.573 | 1.388 | 0.146 | 0.95 | 0.01 | 0.02 | 0.02 | 0 | 0.26 | 0.27 | 0.2 | 0.38 | 0.33 | 0.35 |
| 9 | 81999215 | rs12345803 | -0.607 | 1.384 | 0.411 | 0.892 | 0.037 | 0.049 | 0.049 | 0.369 | 0.12 | 0.33 | 0.09 | 0.15 | 0.11 | 0.14 |
| 9 | 82001689 | rs2378367 | -0.303 | 1.386 | 0.288 | 0.83 | 0.026 | 0.019 | 0.019 | 0.107 | 0.11 | 0.24 | 0.05 | 0.13 | 0.1 | 0.14 |
| 9 | 82004814 | rs10867378 | -0.053 | 1.022 | -0.198 | 1.015 | -0.006 | -0.087 | -0.087 | 2.347 | 0.24 | 0.1 | 0.18 | 0.38 | 0.27 | 0.29 |
| 9 | 82010265 | rs2378373 | 0.445 | 1.198 | -0.016 | 0.771 | 0 | -0.012 | -0.012 | 0 | 0.2 | 0.19 | 0.16 | 0.26 | 0.24 | 0.21 |
| 9 | 82012607 | rs10867379 | -0.703 | 0.843 | -0.224 | 0.48 | 0.023 | 0.013 | 0.013 | 0 | 0.52 | 0.72 | 0.49 | 0.43 | 0.59 | 0.32 |
| 9 | 82015733 | rs17800992 | -0.11 | 1.149 | 0.13 | 0.164 | 0.017 | 0.015 | 0.015 | 0 | 0.07 | 0.15 | 0.16 | 0.1 | 0.1 | 0.1 |
| 9 | 82015765 | rs17082349 | -0.685 | 1.095 | 0.64 | 0.175 | 0.048 | 0.06 | 0.06 | 0 | 0.08 | 0.31 | 0.2 | 0.11 | 0.1 | 0.1 |
| 9 | 82018917 | rs9314677 | -1.218 | 0.935 | -0.036 | 0.083 | -0.047 | -0.152 | -0.152 | 0.953 | 0.64 | 0.97 | 0.62 | 0.5 | 0.67 | 0.52 |
| 9 | 82019860 | rs2378377 | -2.14 | 1.524 | 1.664 | -1.062 | 0.058 | 0.011 | 0.011 | 0.005 | 0.15 | 0.57 | 0.24 | 0.12 | 0.15 | 0.12 |
| 9 | 82032352 | rs4556160 | -2.422 | 1.488 | 1.829 | -1.062 | 0.026 | 0.004 | 0.004 | 0 | 0.16 | 0.64 | 0.24 | 0.14 | 0.15 | 0.12 |
| 9 | 82032456 | rs2378380 | -2.609 | 1.683 | 1.674 | -1.16 | -0.013 | 0.016 | 0.016 | 0.249 | 0.3 | 0.78 | 0.39 | 0.23 | 0.31 | 0.25 |
| 9 | 82042491 | rs12554999 | -0.55 | 1.058 | 1.841 | -0.958 | 0.054 | -0.018 | -0.018 | 0.114 | 0.06 | 0.22 | 0.04 | 0 | 0.05 | 0.01 |
| 9 | 82060528 | rs11138220 | -1.631 | 1.077 | 1.635 | -0.923 | 0.015 | 0.027 | 0.027 | 0 | 0.06 | 0.15 | 0.17 | 0.13 | 0.1 | 0.11 |
| 9 | 82066164 | rs11138231 | -1.301 | 1.087 | 1.587 | -0.93 | 0.004 | 0.008 | 0.008 | 0 | 0.05 | 0.09 | 0.16 | 0.13 | 0.1 | 0.1 |
| 9 | 82080249 | rs10491790 | -1.352 | 0.982 | 1.979 | -0.986 | 0.04 | -0.032 | -0.032 | 0 | 0.11 | 0.49 | 0.07 | 0.01 | 0.05 | 0.01 |
| 9 | 82083763 | rs10867389 | -2.066 | 0.974 | 1.822 | -1.116 | 0.012 | -0.028 | -0.028 | 0 | 0.2 | 0.41 | 0.26 | 0.15 | 0.22 | 0.21 |
| 9 | 82086682 | rs10117770 | -1.671 | 0.976 | 1.952 | -1.117 | -0.007 | -0.088 | -0.088 | 6.523 | 0.14 | 0.35 | 0.11 | 0.02 | 0.12 | 0.1 |
| 9 | 82086816 | rs10117809 | -1.79 | 0.973 | 2.015 | -1.118 | -0.016 | -0.11 | -0.11 | 0 | 0.16 | 0.44 | 0.12 | 0.02 | 0.12 | 0.1 |
| 9 | 82087111 | rs11138252 | -1.365 | 0.986 | 1.905 | -1.106 | -0.025 | -0.022 | -0.022 | 0.712 | 0.08 | 0.06 | 0.16 | 0.13 | 0.13 | 0.13 |
| 9 | 82088036 | rs7030540 | -2.411 | 0.924 | 2.09 | -1.138 | -0.035 | -0.082 | -0.082 | 0.329 | 0.2 | 0.64 | 0.15 | 0.03 | 0.12 | 0.11 |
| 9 | 82096194 | rs10491791 | -1.118 | 1.048 | 2.03 | -1.114 | 0.019 | 0.033 | 0.033 | 0 | 0.05 | 0.1 | 0.16 | 0.13 | 0.1 | 0.12 |
| 9 | 82099407 | rs12336218 | -2.134 | 0.991 | 1.993 | -1.156 | 0.019 | -0.015 | -0.015 | 0.698 | 0.21 | 0.48 | 0.29 | 0.16 | 0.22 | 0.21 |
| 9 | 82116941 | rs7867387 | -1.218 | 0.922 | 1.918 | -1.174 | -0.018 | -0.074 | -0.074 | 0 | 0.13 | 0.28 | 0.09 | 0.02 | 0.1 | 0.1 |
| 9 | 82118185 | rs10867395 | -2.192 | 0.94 | 1.249 | -1.179 | -0.001 | -0.007 | -0.007 | 0 | 0.29 | 0.69 | 0.3 | 0.16 | 0.25 | 0.23 |
| 9 | 82121219 | rs7870740 | -1.493 | 1.201 | 1.295 | -0.996 | 0.036 | -0.026 | -0.026 | 0 | 0.09 | 0.3 | 0.06 | 0.03 | 0.09 | 0.1 |
| 9 | 82122285 | rs7859384 | -1.342 | 1.267 | 0.386 | -1.699 | 0.018 | -0.026 | -0.026 | 0 | 0.7 | 0.85 | 0.63 | 0.33 | 0.75 | 0.62 |
| 9 | 82122617 | rs10867396 | -2.314 | 1.437 | 1.891 | -1.071 | -0.008 | -0.097 | -0.097 | 0.161 | 0.19 | 0.63 | 0.11 | 0.04 | 0.14 | 0.14 |
| 9 | 82127183 | rs12347524 | -1.209 | 1.434 | 1.976 | -1.028 | 0.057 | 0.016 | 0.016 | 3.999 | 0.07 | 0.37 | 0.07 | 0.03 | 0.04 | 0.07 |
| 9 | 82130427 | rs10867398 | -1.25 | 1.471 | 2.125 | -1.033 | 0.044 | -0.006 | -0.006 | 0 | 0.07 | 0.28 | 0.12 | 0.03 | 0.06 | 0.08 |
| 9 | 82130861 | rs1417080 | -2.591 | 1.448 | 2.02 | -1.116 | -0.007 | -0.024 | -0.024 | 0.178 | 0.26 | 0.66 | 0.29 | 0.17 | 0.24 | 0.25 |
| 9 | 82132941 | rs11138277 | -1.205 | 1.472 | 1.965 | -1.099 | -0.003 | 0.004 | 0.004 | 0 | 0.13 | 0.09 | 0.09 | 0.01 | 0.15 | 0.07 |
| 9 | 82140447 | rs12351223 | -0.839 | 1.419 | 2.123 | -1.108 | 0.015 | -0.024 | -0.024 | 0 | 0.07 | 0.16 | 0.05 | 0.03 | 0.08 | 0.06 |
| 9 | 82144364 | rs1572803 | -2.166 | 1.405 | 2.234 | -1.116 | 0.046 | -0.036 | -0.036 | 0.185 | 0.11 | 0.48 | 0.18 | 0.04 | 0.1 | 0.09 |
| 9 | 82148988 | rs7870915 | -1.929 | 1.437 | 2.162 | -1.163 | 0.012 | -0.118 | -0.118 | 1.949 | 0.2 | 0.44 | 0.14 | 0.03 | 0.23 | 0.14 |
| 9 | 82152549 | rs1538752 | -2.399 | 1.427 | 2.092 | -1.173 | -0.001 | -0.11 | -0.11 | 0 | 0.25 | 0.61 | 0.24 | 0.04 | 0.25 | 0.17 |
| 9 | 82157404 | rs6559483 | -3.149 | 1.503 | 1.739 | -1.489 | -0.035 | 0.123 | 0.123 | 0.328 | 0.49 | 0.92 | 0.39 | 0.33 | 0.48 | 0.37 |
| 9 | 82159036 | rs7048062 | -3.203 | 1.45 | 1.877 | -1.477 | -0.008 | 0.043 | 0.043 | 0.898 | 0.42 | 0.73 | 0.36 | 0.33 | 0.44 | 0.29 |
| 9 | 82165005 | rs13291796 | -0.639 | 1.458 | 1.945 | -1.481 | -0.002 | 0 | 0 | 0 | 0.13 | 0.09 | 0.15 | 0.16 | 0.16 | 0.07 |
| 9 | 82167845 | rs12348335 | -0.706 | 1.464 | 1.878 | -1.483 | -0.003 | -0.034 | -0.034 | 0 | 0.1 | 0.17 | 0.05 | 0.03 | 0.08 | 0.06 |
| 9 | 82168805 | rs10491794 | -1.847 | 1.458 | 1.461 | -1.447 | -0.006 | -0.039 | -0.039 | 0 | 0.29 | 0.58 | 0.21 | 0.17 | 0.29 | 0.22 |
| 9 | 82169035 | rs7037685 | -2.094 | 1.386 | 0.925 | -1.36 | -0.007 | 0.044 | 0.044 | 0.885 | 0.43 | 0.73 | 0.38 | 0.33 | 0.44 | 0.29 |
| 9 | 82179241 | rs7855614 | -0.985 | 1.088 | 1.327 | -0.974 | -0.036 | -0.062 | -0.062 | 0 | 0.12 | 0.21 | 0.11 | 0.01 | 0.07 | 0.08 |
| 9 | 82179623 | rs7031185 | -1.679 | 1.287 | 1.404 | -0.905 | -0.01 | -0.034 | -0.034 | 0.002 | 0.15 | 0.25 | 0.14 | 0.1 | 0.13 | 0.11 |
| 9 | 82185354 | rs2807304 | -1.96 | 1.505 | 1.651 | -1.023 | 0.028 | -0.027 | -0.027 | 0 | 0.24 | 0.46 | 0.22 | 0.15 | 0.3 | 0.2 |
| 9 | 82197735 | rs2807302 | -2.041 | 1.487 | 1.659 | -1.01 | 0.032 | 0.008 | 0.008 | 0.556 | 0.19 | 0.58 | 0.19 | 0.15 | 0.22 | 0.15 |
| 9 | 82217647 | rs11999556 | -1.173 | 0.958 | 1.731 | -0.921 | -0.01 | -0.031 | -0.031 | 0 | 0.14 | 0.22 | 0.13 | 0.1 | 0.12 | 0.09 |
| 9 | 82225053 | rs2807310 | -1.872 | 0.949 | 1.775 | -0.925 | 0.034 | 0.014 | 0.014 | 0.252 | 0.19 | 0.48 | 0.17 | 0.15 | 0.2 | 0.16 |
| 9 | 82229981 | rs11138311 | -0.051 | 0.942 | 1.701 | -0.943 | -0.019 | 0.035 | 0.035 | 0.109 | 0.07 | 0.01 | 0.04 | 0 | 0.09 | 0.06 |
| 9 | 82233842 | rs7044519 | -1.62 | 0.828 | 2.279 | -0.648 | 0.001 | -0.003 | -0.003 | 0 | 0.16 | 0.15 | 0.09 | 0.02 | 0.17 | 0.11 |
| 9 | 82234395 | rs7029276 | -0.73 | 0.822 | 2.281 | -0.646 | -0.001 | -0.022 | -0.022 | 0 | 0.1 | 0.15 | 0.06 | 0.02 | 0.09 | 0.05 |
| 9 | 82237934 | rs4242612 | 2.658 | 0.862 | 2.195 | -0.666 | 0.014 | -0.032 | -0.032 | 0 | 0.72 | 0.41 | 0.77 | 0.84 | 0.67 | 0.76 |
| 9 | 82243701 | rs12337384 | -2.052 | 0.84 | 2.342 | -0.673 | -0.011 | -0.087 | -0.087 | 0.217 | 0.2 | 0.39 | 0.11 | 0.02 | 0.19 | 0.12 |
| 9 | 82246520 | rs13440432 | -0.321 | 0.841 | 2.33 | -0.644 | 0.005 | 0.003 | 0.003 | 0.416 | 0.08 | 0.21 | 0.1 | 0.1 | 0.07 | 0.04 |
| 9 | 82251896 | rs11138314 | -1.968 | 0.877 | 2.36 | -0.667 | -0.002 | -0.084 | -0.084 | 0 | 0.16 | 0.49 | 0.1 | 0.02 | 0.14 | 0.05 |
| 9 | 82263667 | rs12553469 | -0.102 | 0.884 | 1.897 | -0.662 | -0.024 | 0.022 | 0.022 | 0 | 0.07 | 0.02 | 0.04 | 0.02 | 0.1 | 0.05 |
| 9 | 82272674 | rs7022420 | -0.571 | 0.877 | 2.005 | -0.661 | -0.007 | -0.033 | -0.033 | 0 | 0.13 | 0.16 | 0.12 | 0.01 | 0.09 | 0.08 |
| 9 | 82275591 | rs2027005 | 2.519 | 0.885 | 1.978 | -0.642 | -0.003 | 0.024 | 0.024 | 1.149 | 0.67 | 0.19 | 0.68 | 0.74 | 0.66 | 0.74 |

| | | | | | | | | | | | | | | | | |
|---|---|---|---|---|---|---|---|---|---|---|---|---|---|---|---|---|
| 9 | 82280362 | rs4877146 | 2.553 | 0.664 | 1.858 | -0.523 | -0.012 | 0.027 | 0.027 | 1.991 | 0.63 | 0.15 | 0.66 | 0.71 | 0.62 | 0.73 |
| 9 | 82282697 | rs12342140 | -1.619 | 0.617 | 2.12 | -0.508 | 0.016 | -0.006 | -0.006 | 0 | 0.16 | 0.4 | 0.11 | 0.15 | 0.15 | 0.07 |
| 9 | 82298288 | rs7859802 | -1.685 | 0.654 | 1.989 | -0.655 | -0.027 | -0.129 | -0.129 | 0 | 0.25 | 0.44 | 0.18 | 0.02 | 0.2 | 0.14 |
| 9 | 82298324 | rs11138329 | 2.467 | 0.628 | 1.917 | -0.635 | -0.01 | 0.052 | 0.052 | 2.606 | 0.63 | 0.15 | 0.67 | 0.75 | 0.63 | 0.73 |
| 9 | 82302985 | rs12349682 | -0.646 | 0.645 | 1.933 | -0.618 | -0.013 | -0.044 | -0.044 | 0 | 0.16 | 0.32 | 0.14 | 0.1 | 0.13 | 0.11 |
| 9 | 82308670 | rs7025437 | 0.907 | 0.676 | 1.136 | -0.592 | 0.026 | 0.038 | 0.038 | 0 | 0.94 | 0.9 | 0.91 | 0.87 | 0.87 | 0.89 |
| 9 | 82312046 | rs7039267 | 2.125 | 0.671 | 0.494 | -0.516 | 0.004 | 0.014 | 0.014 | 3.628 | 0.64 | 0.15 | 0.66 | 0.7 | 0.64 | 0.73 |
| 9 | 82316246 | rs11138337 | -0.793 | 0.849 | 0.93 | -0.281 | 0.001 | -0.021 | -0.021 | 0 | 0.1 | 0.15 | 0.05 | 0.02 | 0.1 | 0.04 |
| 9 | 82329295 | rs11138340 | 2.565 | 0.872 | 0.953 | -0.26 | 0 | -0.003 | -0.003 | 0.449 | 0.73 | 0.33 | 0.71 | 0.81 | 0.69 | 0.76 |
| 9 | 82330095 | rs12345782 | -1.908 | 1.49 | -0.163 | 0.061 | 0 | -0.002 | -0.002 | 0.089 | 0.14 | 0.16 | 0.08 | 0.06 | 0.14 | 0.1 |
| 9 | 82333000 | rs7022923 | -0.99 | 1.759 | 0.041 | 0.142 | 0.009 | -0.03 | -0.03 | 0 | 0.1 | 0.19 | 0.06 | 0.02 | 0.11 | 0.04 |
| 9 | 82342503 | rs4355856 | 1.374 | 1.662 | -0.056 | 0.149 | 0.027 | 0.011 | 0.011 | 0 | 0.83 | 0.74 | 0.83 | 0.84 | 0.76 | 0.85 |
| 9 | 82345399 | rs2027007 | 1.11 | 1.353 | 0.024 | 0.225 | 0.023 | 0.019 | 0.019 | 0 | 0.85 | 0.74 | 0.84 | 0.84 | 0.79 | 0.85 |
| 9 | 82352046 | rs7023371 | 0.015 | 0.803 | -0.297 | -0.267 | -0.005 | -0.026 | -0.026 | 5.742 | 0.6 | 0.56 | 0.47 | 0.78 | 0.67 | 0.77 |
| 9 | 82352255 | rs6559497 | 0.119 | 0.707 | -0.413 | -0.623 | 0.024 | 0.151 | 0.151 | 0.785 | 0.6 | 0.82 | 0.74 | 0.89 | 0.65 | 0.78 |
| 9 | 82354524 | rs10867418 | 0.511 | 0.589 | -0.566 | -0.511 | -0.043 | 0.009 | 0.009 | 0.164 | 0.41 | 0.16 | 0.5 | 0.57 | 0.54 | 0.53 |
| 9 | 82354924 | rs2149882 | 0.583 | 0.659 | 0.072 | -0.461 | 0 | 0 | 0 | 0.003 | 0.15 | 0.16 | 0.08 | 0.17 | 0.13 | 0.27 |
| 9 | 82359588 | rs11138350 | 0.299 | -0.096 | -0.515 | -0.541 | -0.031 | -0.029 | -0.029 | 0 | 0.42 | 0.08 | 0.57 | 0.57 | 0.6 | 0.52 |
| 9 | 82361923 | rs4144534 | 0.494 | -0.001 | -0.632 | -0.627 | -0.102 | -0.071 | -0.071 | 0.623 | 0.37 | 0.06 | 0.5 | 0.54 | 0.53 | 0.51 |
| 9 | 82362167 | rs10867419 | -1.186 | -0.255 | -0.417 | -0.772 | 0.053 | 0.143 | 0.143 | 1.937 | 0.64 | 0.88 | 0.67 | 0.9 | 0.71 | 0.84 |
| 9 | 82364676 | rs1329743 | -0.771 | -0.403 | 0.223 | -0.865 | 0.027 | 0.034 | 0.034 | 1.427 | 0.59 | 0.7 | 0.64 | 0.88 | 0.7 | 0.8 |
| 9 | 82374355 | rs10746599 | -0.427 | -0.517 | 0.95 | -0.842 | -0.033 | -0.029 | -0.029 | 0 | 0.57 | 0.55 | 0.64 | 0.77 | 0.74 | 0.7 |
| 9 | 82377154 | rs1411160 | 1.01 | -0.501 | 1.316 | -0.809 | 0.001 | 0.002 | 0.002 | 0 | 0.13 | 0.16 | 0.06 | 0.21 | 0.12 | 0.17 |
| 9 | 82377443 | rs1888073 | -0.718 | -0.535 | 1.38 | -0.822 | -0.008 | -0.03 | -0.03 | 4.804 | 0.6 | 0.61 | 0.64 | 0.88 | 0.75 | 0.82 |
| 9 | 82383308 | rs10867421 | -1.114 | -0.293 | 1.011 | -0.859 | 0.056 | 0.172 | 0.172 | 0.956 | 0.65 | 0.89 | 0.67 | 0.9 | 0.75 | 0.84 |
| 9 | 82392719 | rs6559502 | 0.348 | -0.263 | 1.363 | -0.734 | 0.001 | 0.002 | 0.002 | 1.124 | 0.42 | 0.4 | 0.55 | 0.56 | 0.58 | 0.52 |
| 9 | 82395528 | rs4475583 | -0.41 | -0.223 | 0.907 | -0.805 | 0.055 | 0.141 | 0.141 | 0.471 | 0.5 | 0.75 | 0.6 | 0.68 | 0.61 | 0.65 |
| 9 | 82399186 | rs977219 | -0.295 | -0.217 | 1.038 | -0.8 | 0.036 | 0.102 | 0.102 | 1.251 | 0.48 | 0.7 | 0.59 | 0.68 | 0.6 | 0.62 |
| 9 | 82429923 | rs12345318 | 0.485 | 0.867 | 0.784 | 0.05 | -0.014 | -0.011 | -0.011 | 0.969 | 0.87 | 0.9 | 0.71 | 0.71 | 0.78 | 0.71 |
| 9 | 82430122 | rs7039148 | 0.477 | 0.834 | 0.747 | 0.034 | -0.023 | -0.026 | -0.026 | 0.119 | 0.87 | 0.9 | 0.71 | 0.71 | 0.78 | 0.71 |
| 9 | 82432302 | rs11138361 | 0.421 | 0.569 | 0.988 | -0.06 | -0.026 | 0.195 | 0.195 | 0.667 | 0.19 | 0.45 | 0.18 | 0.39 | 0.17 | 0.22 |
| 9 | 82433182 | rs7037957 | 0.793 | 0.731 | 0.725 | -0.443 | 0.044 | 0.196 | 0.196 | 0.661 | 0.49 | 0.2 | 0.42 | 0.3 | 0.42 | 0.35 |
| 9 | 82437835 | rs11138362 | 0.715 | 0.697 | 0.435 | -0.267 | -0.022 | 0.148 | 0.148 | 0.677 | 0.18 | 0.34 | 0.17 | 0.39 | 0.17 | 0.22 |
| 9 | 82438585 | rs4877152 | -0.079 | 0.702 | 0.444 | -0.291 | -0.07 | -0.037 | -0.037 | 0.514 | 0.69 | 0.81 | 0.6 | 0.69 | 0.57 | 0.58 |
| 9 | 82439058 | rs4273932 | 0.403 | 0.004 | 0.01 | -0.553 | 0.036 | 0.009 | 0.009 | 0 | 0.51 | 0.37 | 0.33 | 0.4 | 0.36 | 0.36 |
| 9 | 82443844 | rs11138364 | 0.017 | 0.131 | 0.684 | 0.201 | -0.023 | 0.171 | 0.171 | 0.686 | 0.18 | 0.37 | 0.17 | 0.4 | 0.17 | 0.23 |
| 9 | 82449195 | rs11138368 | 0.829 | 0.105 | 0.019 | -0.267 | -0.008 | 0.006 | 0.006 | 0 | 0.07 | 0.1 | 0.03 | 0.08 | 0.08 | 0.04 |
| 9 | 82455838 | rs7873885 | 0.579 | 0.02 | -1.929 | -0.53 | -0.018 | -0.068 | -0.068 | 0 | 0.56 | 0.15 | 0.53 | 0.48 | 0.57 | 0.52 |
| 9 | 82456587 | rs4877517 | 0.055 | -1.03 | -2.12 | -0.986 | -0.033 | -0.083 | -0.083 | 0 | 0.66 | 0.89 | 0.64 | 0.53 | 0.65 | 0.56 |
| 9 | 82457398 | rs7035737 | -0.661 | -0.762 | -1.982 | -0.759 | -0.073 | -0.102 | -0.102 | 0 | 0.77 | 0.95 | 0.77 | 0.59 | 0.71 | 0.65 |
| 9 | 82457493 | rs11138373 | 1.531 | -0.74 | -2.451 | -1.136 | -0.05 | -0.103 | -0.103 | 0 | 0.15 | 0.02 | 0.3 | 0.43 | 0.23 | 0.27 |
| 9 | 82462882 | rs12342683 | 0.973 | -0.826 | -1.787 | -1.181 | -0.025 | -0.029 | -0.029 | 0 | 0.13 | 0.51 | 0.1 | 0.07 | 0.11 | 0.06 |
| 9 | 82462958 | rs7858020 | -0.957 | -0.868 | -2.096 | -1.467 | 0 | 0 | 0 | 1.664 | 0.4 | 0.57 | 0.48 | 0.55 | 0.48 | 0.49 |
| 9 | 82465583 | rs10512094 | 1.554 | -0.903 | -2.143 | -1.503 | 0.026 | 0.145 | 0.145 | 5.625 | 0.32 | 0.08 | 0.26 | 0.04 | 0.22 | 0.16 |
| 9 | 82472582 | rs6559507 | 1.45 | -0.839 | -2.071 | -1.451 | -0.011 | -0.043 | -0.043 | 0.547 | 0.17 | 0.09 | 0.31 | 0.44 | 0.23 | 0.28 |
| 9 | 82473507 | rs11138385 | 0.289 | -0.6 | -1.835 | -1.373 | 0.009 | -0.087 | -0.087 | 0.062 | 0.23 | 0.43 | 0.17 | 0.11 | 0.25 | 0.21 |
| 9 | 82474979 | rs4877153 | -0.39 | -0.232 | -1.859 | -1.094 | 0.002 | 0.01 | 0.01 | 1.009 | 0.63 | 0.46 | 0.68 | 0.52 | 0.59 | 0.57 |
| 9 | 82478788 | rs7039360 | -0.032 | -0.71 | -1.535 | -1.46 | -0.002 | 0.01 | 0.01 | 0.935 | 0.57 | 0.58 | 0.43 | 0.15 | 0.46 | 0.37 |
| 9 | 82485633 | rs3894377 | 0.506 | -0.263 | -1.368 | -1.096 | 0.001 | 0.004 | 0.004 | 0 | 0.54 | 0.53 | 0.47 | 0.63 | 0.51 | 0.56 |
| 9 | 82488433 | rs12345503 | -0.094 | -0.389 | -1.936 | -1.386 | 0.019 | 0.137 | 0.137 | 4.28 | 0.47 | 0.3 | 0.42 | 0.07 | 0.39 | 0.26 |
| 9 | 82489613 | rs2274733 | -0.288 | -0.411 | -1.967 | -1.287 | 0.045 | -0.021 | -0.021 | 0 | 0.46 | 0.75 | 0.43 | 0.58 | 0.42 | 0.54 |
| 9 | 82489957 | rs4877518 | 1.503 | -0.401 | -1.956 | -1.353 | 0.005 | 0.07 | 0.07 | 0.929 | 0.28 | 0.19 | 0.33 | 0.03 | 0.24 | 0.13 |
| 9 | 82494427 | rs17728874 | 0.554 | -0.363 | -1.982 | -1.317 | -0.016 | 0.061 | 0.061 | 0.906 | 0.17 | 0.03 | 0.08 | 0.05 | 0.16 | 0.12 |
| 9 | 82495376 | rs6559511 | -0.884 | -0.299 | -1.883 | -1.217 | -0.031 | -0.04 | -0.04 | 0 | 0.73 | 0.81 | 0.74 | 0.61 | 0.66 | 0.68 |
| 9 | 82495768 | rs1483023 | 1.089 | 0.147 | -1.635 | -0.987 | -0.026 | 0.123 | 0.123 | 4.085 | 0.25 | 0.44 | 0.32 | 0.54 | 0.27 | 0.42 |
| 9 | 82503489 | rs267583 | -0.876 | 0.375 | -2.118 | -0.785 | -0.064 | -0.061 | -0.061 | 0.286 | 0.75 | 0.93 | 0.8 | 0.65 | 0.7 | 0.71 |
| 9 | 82510157 | rs4877521 | -0.027 | 0.013 | -1.226 | -1.218 | 0.005 | 0.101 | 0.101 | 0.893 | 0.28 | 0.13 | 0.34 | 0.06 | 0.23 | 0.14 |
| 9 | 82517423 | rs7021014 | -0.544 | -0.003 | -1.289 | -1.213 | 0.014 | 0.118 | 0.118 | 0.599 | 0.32 | 0.14 | 0.37 | 0.06 | 0.28 | 0.16 |
| 9 | 82523992 | rs12553223 | -0.173 | 0.53 | -1.685 | -1.033 | -0.02 | 0.153 | 0.153 | 0.759 | 0.27 | 0.04 | 0.33 | 0.03 | 0.24 | 0.12 |
| 9 | 82533026 | rs267571 | 0.773 | 1.175 | -0.47 | -0.773 | 0.031 | -0.09 | -0.09 | 0.737 | 0.76 | 0.47 | 0.8 | 0.92 | 0.71 | 0.87 |
| 9 | 82535883 | rs267574 | 0.912 | 1.389 | 0.209 | -1.091 | 0.008 | -0.034 | -0.034 | 0 | 0.71 | 0.72 | 0.73 | 0.92 | 0.61 | 0.85 |
| 9 | 82536522 | rs11138411 | 0.96 | 0.973 | -0.565 | -0.837 | -0.026 | 0.036 | 0.036 | 0 | 0.33 | 0.43 | 0.33 | 0.53 | 0.29 | 0.41 |
| 9 | 82536663 | rs10746608 | 0.871 | 1.002 | -0.667 | -0.871 | -0.008 | 0.009 | 0.009 | 0 | 0.34 | 0.36 | 0.33 | 0.53 | 0.29 | 0.41 |
| 9 | 82541282 | rs12238216 | 1.168 | 0.859 | -0.397 | -0.917 | 0.017 | -0.054 | -0.054 | 0 | 0.31 | 0.14 | 0.3 | 0.53 | 0.29 | 0.41 |
| 9 | 82542041 | rs388621 | 0.556 | 1.091 | -0.424 | -0.856 | -0.049 | -0.059 | -0.059 | 0 | 0.36 | 0.06 | 0.39 | 0.39 | 0.32 | 0.44 |
| 9 | 82547473 | rs267587 | 0.649 | 1.395 | 0.074 | -1.045 | 0.033 | -0.115 | -0.115 | 0 | 0.81 | 0.45 | 0.81 | 0.98 | 0.79 | 0.9 |
| 9 | 82551242 | rs10867442 | 0.278 | 1.15 | 0.032 | -0.944 | 0.006 | 0.016 | 0.016 | 0 | 0.44 | 0.37 | 0.42 | 0.59 | 0.46 | 0.47 |
| 9 | 82555561 | rs10867444 | -1.289 | 1.816 | 0.639 | -1.076 | 0.038 | -0.069 | -0.069 | 0 | 0.29 | 0.32 | 0.27 | 0.06 | 0.43 | 0.19 |
| 9 | 82559959 | rs1329579 | -1.066 | 1.956 | 0.679 | -0.799 | 0.075 | -0.022 | -0.022 | 0 | 0.24 | 0.43 | 0.45 | 0.22 | 0.32 | 0.21 |

| | | | | | | | | | | | | | | | | |
|---|---|---|---|---|---|---|---|---|---|---|---|---|---|---|---|---|
| 9 | 82562184 | rs17729039 | -0.158 | 1.844 | 0.74 | -0.825 | 0 | 0 | 0 | 0.875 | 0.1 | 0.09 | 0.08 | 0.03 | 0.14 | 0.07 |
| 9 | 82572208 | rs962883 | -0.95 | 1.425 | 0.608 | -0.504 | 0.024 | -0.016 | -0.016 | 0 | 0.67 | 0.82 | 0.83 | 0.61 | 0.75 | 0.66 |
| 9 | 82572623 | rs11138425 | 0.44 | 1.602 | 0.634 | -0.516 | 0.006 | -0.02 | -0.02 | 0.022 | 0.07 | 0.07 | 0.11 | 0 | 0.14 | 0.03 |
| 9 | 82579569 | rs7853567 | -0.219 | 1.212 | 0.467 | -0.477 | 0.068 | -0.071 | -0.071 | 0 | 0.19 | 0.23 | 0.19 | 0.03 | 0.29 | 0.1 |
| 9 | 82579689 | rs17801845 | 0.217 | 1.178 | 0.369 | -0.491 | 0.003 | 0.007 | 0.007 | 0 | 0.08 | 0.06 | 0.05 | 0.03 | 0.09 | 0.06 |
| 9 | 82582453 | rs12686796 | 1.296 | 1.148 | 0.392 | -0.457 | 0.021 | -0.069 | -0.069 | 0 | 0.33 | 0.11 | 0.29 | 0.38 | 0.3 | 0.39 |
| 9 | 82582993 | rs4877524 | -0.587 | 1.367 | 0.563 | -0.243 | 0.022 | 0.001 | 0.001 | 0 | 0.66 | 0.81 | 0.81 | 0.64 | 0.71 | 0.66 |
| 9 | 82586369 | rs1329580 | -0.482 | 1.729 | 0.391 | -0.428 | 0.07 | -0.071 | -0.071 | 0.227 | 0.18 | 0.25 | 0.19 | 0.03 | 0.28 | 0.1 |
| 9 | 82588226 | rs913757 | -0.247 | 0.56 | -0.14 | -0.15 | -0.002 | -0.002 | -0.002 | 0.007 | 0.64 | 0.63 | 0.8 | 0.66 | 0.7 | 0.65 |
| 9 | 82592333 | rs11138436 | 0.429 | 0.535 | -0.472 | 0.205 | 0.003 | -0.003 | -0.003 | 0.85 | 0.46 | 0.41 | 0.5 | 0.45 | 0.42 | 0.53 |
| 9 | 82596501 | rs7851008 | 0.143 | 0.207 | -1.36 | 0.435 | 0.02 | 0.004 | 0.004 | 0 | 0.62 | 0.73 | 0.47 | 0.38 | 0.74 | 0.66 |
| 9 | 82597404 | rs7024109 | -0.189 | -0.097 | -1.43 | 0.338 | -0.045 | 0.007 | 0.007 | 0 | 0.64 | 0.51 | 0.69 | 0.58 | 0.76 | 0.62 |
| 9 | 82598756 | rs13297228 | 1.638 | 0.21 | -1.348 | 0.422 | -0.059 | 0.049 | 0.049 | 0.377 | 0.33 | 0.03 | 0.21 | 0.21 | 0.34 | 0.37 |
| 9 | 82598985 | rs7859610 | -0.992 | 0.485 | -0.554 | 0.382 | 0.092 | -0.058 | -0.058 | 0 | 0.24 | 0.47 | 0.23 | 0.13 | 0.35 | 0.19 |
| 9 | 82602724 | rs10780347 | -1.097 | 0.771 | -0.528 | 0.584 | 0.073 | 0.079 | 0.079 | 0 | 0.27 | 0.47 | 0.44 | 0.39 | 0.37 | 0.25 |
| 9 | 82603623 | rs10867451 | -0.762 | 1.073 | -0.822 | 0.097 | -0.008 | 0.005 | 0.005 | 0 | 0.23 | 0.22 | 0.21 | 0.13 | 0.37 | 0.11 |
| 9 | 82607865 | rs7048183 | -0.849 | 0.821 | -0.739 | -0.016 | -0.004 | 0.065 | 0.065 | 0.028 | 0.52 | 0.66 | 0.32 | 0.28 | 0.43 | 0.38 |
| 9 | 82609825 | rs1329576 | -1.129 | 1.114 | 0.063 | -0.669 | 0.041 | -0.104 | -0.104 | 0 | 0.18 | 0.36 | 0.13 | 0.01 | 0.18 | 0.18 |
| 9 | 82613329 | rs10780348 | 0.088 | 0.502 | -0.058 | -1.244 | -0.033 | -0.165 | -0.165 | 0 | 0.73 | 0.5 | 0.86 | 0.99 | 0.82 | 0.73 |
| 9 | 82613770 | rs11138447 | 0.648 | 0.934 | -0.701 | -1.604 | -0.032 | 0.012 | 0.012 | 0 | 0.22 | 0.1 | 0.2 | 0.23 | 0.28 | 0.31 |
| 9 | 82616777 | rs10746614 | 0.025 | 0.809 | -0.356 | -1.901 | 0.002 | -0.041 | -0.041 | 0 | 0.69 | 0.34 | 0.83 | 0.98 | 0.76 | 0.72 |
| 9 | 82623065 | rs11138454 | 0.491 | 0.533 | -0.97 | -2.248 | 0.008 | 0.17 | 0.17 | 0 | 0.51 | 0.26 | 0.71 | 0.92 | 0.56 | 0.61 |
| 9 | 82636239 | rs1410499 | -2.113 | 0.44 | -1.218 | -2.418 | 0.002 | 0.082 | 0.082 | 0 | 0.86 | 1 | 0.95 | 0.99 | 0.9 | 0.82 |
| 9 | 82646539 | rs1328430 | 0.845 | 0.292 | -1.333 | -2.574 | -0.012 | 0.137 | 0.137 | 0 | 0.48 | 0.28 | 0.7 | 0.9 | 0.53 | 0.58 |
| 9 | 82654332 | rs1410502 | 1.258 | 0.344 | -1.275 | -2.4 | -0.066 | 0.04 | 0.04 | 0 | 0.18 | 0.02 | 0.11 | 0.07 | 0.21 | 0.11 |
| 9 | 82662587 | rs11138465 | 0.529 | 0.292 | -1.326 | -2.462 | -0.088 | 0.079 | 0.079 | 0 | 0.23 | 0.02 | 0.14 | 0.08 | 0.28 | 0.14 |
| 9 | 82663163 | rs10867455 | -0.542 | 0.347 | -1.225 | -2.458 | -0.008 | 0.022 | 0.022 | 1.357 | 0.33 | 0.71 | 0.23 | 0.09 | 0.35 | 0.18 |
| 9 | 82672704 | rs12347897 | -0.426 | 0.101 | 0.137 | -2.468 | 0 | -0.027 | -0.027 | 0 | 0.4 | 0.58 | 0.23 | 0.07 | 0.37 | 0.32 |
| 9 | 82675016 | rs7026368 | -1.116 | 0.486 | 1.015 | -1.832 | 0.078 | -0.014 | -0.014 | 0.796 | 0.09 | 0.42 | 0.07 | 0.01 | 0.07 | 0.06 |
| 9 | 82690543 | rs13299280 | -0.626 | 0.921 | 1.03 | -1.839 | 0.138 | -0.038 | -0.038 | 3.814 | 0.09 | 0.34 | 0.1 | 0 | 0.15 | 0.06 |
| 9 | 82695996 | rs10746617 | -0.832 | 0.524 | 0.051 | -2.314 | 0.009 | 0.119 | 0.119 | 0.892 | 0.68 | 0.92 | 0.81 | 0.91 | 0.72 | 0.79 |
| 9 | 82699351 | rs1887722 | 0.493 | 0.463 | 0.288 | -2.677 | 0.015 | 0.136 | 0.136 | 0.673 | 0.5 | 0.31 | 0.65 | 0.95 | 0.48 | 0.62 |
| 9 | 82703907 | rs11138484 | -0.12 | 0.523 | 0.695 | -2.686 | 0.078 | -0.051 | -0.051 | 0 | 0.09 | 0.23 | 0.09 | 0 | 0.15 | 0.06 |
| 9 | 82705241 | rs12005290 | 0.369 | 0.404 | 0.895 | -2.662 | 0.009 | -0.047 | -0.047 | 0 | 0.1 | 0.16 | 0.07 | 0.01 | 0.09 | 0.12 |
| 9 | 82706489 | rs4877158 | 0.737 | 0.34 | 0.209 | -2.958 | 0.006 | 0.025 | 0.025 | 0 | 0.33 | 0.27 | 0.42 | 0.9 | 0.29 | 0.37 |
| 9 | 82713213 | rs7865549 | -1.571 | 0.373 | 0.076 | -2.899 | 0.014 | 0.068 | 0.068 | 0.724 | 0.69 | 0.83 | 0.82 | 0.92 | 0.71 | 0.78 |
| 9 | 82716095 | rs12686238 | 1.908 | 0.67 | -0.434 | -2.61 | -0.028 | -0.036 | -0.036 | 0.017 | 0.07 | 0 | 0.09 | 0.1 | 0.09 | 0.11 |
| 9 | 82721529 | rs10867459 | -1.575 | 0.62 | 0.165 | -2.789 | -0.017 | -0.101 | -0.101 | 0 | 0.68 | 0.57 | 0.8 | 0.96 | 0.72 | 0.79 |
| 9 | 82726068 | rs11138492 | 0.385 | 0.72 | -0.487 | -2.674 | 0.01 | 0.054 | 0.054 | 0 | 0.32 | 0.21 | 0.44 | 0.9 | 0.28 | 0.39 |
| 9 | 82727367 | rs7850306 | 1.024 | 0.513 | -0.698 | -2.29 | -0.046 | 0.077 | 0.077 | 0.087 | 0.19 | 0.04 | 0.22 | 0.06 | 0.23 | 0.28 |
| 9 | 82750377 | rs7867554 | 0.631 | 0.226 | -0.102 | -2.328 | -0.006 | -0.08 | -0.08 | 0.197 | 0.37 | 0.48 | 0.22 | 0.04 | 0.34 | 0.3 |
| 9 | 82759463 | rs10114675 | 0.183 | 0.463 | 0.431 | -0.934 | 0 | 0.009 | 0.009 | 0 | 0.11 | 0.29 | 0.07 | 0.09 | 0.11 | 0.13 |
| 9 | 82759818 | rs7044051 | 0.335 | 0.431 | 0.292 | -0.492 | 0.009 | 0.019 | 0.019 | 0 | 0.65 | 0.48 | 0.67 | 0.5 | 0.64 | 0.58 |
| 9 | 82762576 | rs4607700 | -0.899 | 0.457 | 0.365 | -0.503 | 0.007 | 0.041 | 0.041 | 0 | 0.3 | 0.43 | 0.41 | 0.5 | 0.32 | 0.39 |
| 9 | 82768721 | rs11138518 | 0.209 | 0.574 | 0.242 | -0.473 | -0.008 | 0.007 | 0.007 | 0 | 0.08 | 0.06 | 0.07 | 0.04 | 0.13 | 0.2 |
| 9 | 82770941 | rs1034053 | -0.234 | 0.725 | 0.419 | -0.427 | 0.018 | 0.043 | 0.043 | 0 | 0.72 | 0.59 | 0.71 | 0.54 | 0.69 | 0.7 |
| 9 | 82772034 | rs11138521 | -0.167 | 0.761 | 0.324 | -0.509 | 0 | 0.018 | 0.018 | 0.629 | 0.78 | 0.8 | 0.78 | 0.53 | 0.77 | 0.65 |
| 9 | 82772815 | rs11138522 | 0.761 | 0.892 | -0.017 | -0.557 | -0.053 | 0.052 | 0.052 | 0 | 0.1 | 0.03 | 0.1 | 0 | 0.15 | 0.07 |
| 9 | 82773983 | rs11138523 | 0.308 | 0.865 | 0.045 | -0.486 | -0.009 | 0.009 | 0.009 | 0 | 0.08 | 0.06 | 0.07 | 0.03 | 0.13 | 0.18 |
| 9 | 82776693 | rs11138527 | 0.308 | 0.866 | 0.045 | -0.485 | -0.01 | 0.009 | 0.009 | 0 | 0.08 | 0.05 | 0.07 | 0.04 | 0.13 | 0.21 |
| 9 | 82780965 | rs17176689 | 2.02 | 0.628 | -0.243 | -0.658 | -0.001 | 0.238 | 0.238 | 0.48 | 0.39 | 0.1 | 0.3 | 0.06 | 0.37 | 0.3 |
| 9 | 82781519 | rs11788986 | 0.849 | 0.603 | -0.087 | -0.442 | -0.001 | -0.014 | -0.014 | 0 | 0.13 | 0.09 | 0.3 | 0.4 | 0.13 | 0.14 |
| 9 | 82782923 | rs7863707 | 0.206 | 0.595 | 0.127 | -0.479 | 0.025 | 0.034 | 0.034 | 0 | 0.7 | 0.53 | 0.71 | 0.59 | 0.68 | 0.69 |
| 9 | 82788836 | rs11787922 | 1.023 | 0.562 | 0.248 | -0.402 | -0.005 | 0.106 | 0.106 | 0.01 | 0.11 | 0.18 | 0.3 | 0.4 | 0.1 | 0.13 |
| 9 | 82789034 | rs11787929 | -0.418 | 0.548 | 0.245 | -0.439 | -0.014 | 0.097 | 0.097 | 0.536 | 0.22 | 0.39 | 0.36 | 0.48 | 0.22 | 0.26 |
| 9 | 82794251 | rs4877161 | 0.528 | 0.567 | -0.037 | -0.383 | -0.004 | -0.039 | -0.039 | 0 | 0.13 | 0.08 | 0.3 | 0.38 | 0.13 | 0.14 |
| 9 | 82796396 | rs10156438 | -0.143 | 0.473 | 0.243 | -0.441 | -0.013 | 0.076 | 0.076 | 0 | 0.77 | 0.65 | 0.8 | 0.54 | 0.81 | 0.73 |
| 9 | 82796707 | rs11138539 | 0.232 | 0.638 | -0.203 | -0.487 | -0.035 | -0.052 | -0.052 | 0 | 0.2 | 0.11 | 0.39 | 0.38 | 0.26 | 0.2 |
| 9 | 82805488 | rs10867476 | 1.447 | 0.562 | -0.345 | -0.631 | -0.054 | 0.307 | 0.307 | 0.493 | 0.47 | 0.11 | 0.37 | 0.1 | 0.5 | 0.51 |
| 9 | 82811242 | rs1329559 | -0.874 | 0.431 | 0.072 | -0.451 | -0.009 | 0.011 | 0.011 | 0 | 0.35 | 0.56 | 0.34 | 0.57 | 0.28 | 0.36 |
| 9 | 82815148 | rs13294474 | -0.042 | 0.445 | 0.114 | -0.478 | 0.011 | 0.029 | 0.029 | 0 | 0.22 | 0.23 | 0.39 | 0.4 | 0.26 | 0.21 |
| 9 | 82819611 | rs1329555 | -0.377 | 0.142 | 0.133 | -0.638 | -0.007 | 0.028 | 0.028 | 0 | 0.31 | 0.36 | 0.31 | 0.46 | 0.27 | 0.35 |
| 9 | 82820878 | rs1329554 | -2.59 | 0.305 | -0.053 | -0.735 | -0.021 | 0.231 | 0.231 | 0.573 | 0.64 | 0.92 | 0.78 | 0.95 | 0.65 | 0.67 |
| 9 | 82829118 | rs659968 | -1.216 | 0.272 | 0.099 | -0.601 | 0.008 | -0.003 | -0.003 | 0 | 0.42 | 0.65 | 0.39 | 0.58 | 0.39 | 0.47 |
| 9 | 82851561 | rs17082971 | 0.694 | 0.276 | 0.366 | -0.372 | -0.005 | 0.125 | 0.125 | 0.272 | 0.12 | 0.2 | 0.1 | 0.36 | 0.1 | 0.12 |
| 9 | 82853486 | rs552594 | 0.086 | 0.282 | 0.16 | -0.312 | 0 | -0.008 | -0.008 | 0 | 0.25 | 0.21 | 0.4 | 0.51 | 0.24 | 0.32 |
| 9 | 82855536 | rs12553692 | 0.904 | 0.322 | 0.063 | -0.349 | -0.001 | -0.009 | -0.009 | 0 | 0.05 | 0.09 | 0.15 | 0 | 0.1 | 0.1 |
| 9 | 82857894 | rs7033100 | 1.811 | 0.214 | -0.082 | -0.469 | -0.004 | 0.23 | 0.23 | 0.366 | 0.38 | 0.11 | 0.22 | 0.03 | 0.36 | 0.34 |
| 9 | 82862960 | rs10123588 | 1.624 | 0.224 | 0.266 | -0.124 | -0.015 | 0.077 | 0.077 | 0 | 0.07 | 0.2 | 0.03 | 0.09 | 0.03 | 0.05 |

| | | | | | | | | | | | | | | | | |
|---|---|---|---|---|---|---|---|---|---|---|---|---|---|---|---|---|
| 9 | 82864918 | rs517274 | 1.635 | 0.322 | 0.069 | 0.096 | 0 | 0.024 | 0.024 | 0 | 0.07 | 0.07 | 0.09 | 0.26 | 0.07 | 0.11 |
| 9 | 82873587 | rs589737 | -0.117 | 0.356 | -0.022 | -0.186 | 0 | -0.013 | -0.013 | 3.044 | 0.26 | 0.25 | 0.5 | 0.5 | 0.27 | 0.37 |
| 9 | 82874140 | rs587379 | -0.749 | 0.364 | 0.205 | -0.384 | 0.016 | -0.076 | -0.076 | 0 | 0.84 | 0.75 | 0.89 | 1 | 0.83 | 0.92 |
| 9 | 82875336 | rs10867485 | -0.003 | 0.303 | -0.082 | -0.48 | -0.001 | 0.056 | 0.056 | 0 | 0.2 | 0.25 | 0.14 | 0.46 | 0.19 | 0.22 |
| 9 | 82884577 | rs2695 | 0.027 | 0.289 | -0.97 | -0.415 | 0 | -0.013 | -0.013 | 0 | 0.26 | 0.25 | 0.51 | 0.5 | 0.27 | 0.37 |
| 9 | 82886345 | rs953506 | 1.829 | 0.371 | -1.314 | -0.779 | -0.019 | 0.238 | 0.238 | 0.588 | 0.37 | 0.08 | 0.22 | 0.03 | 0.36 | 0.32 |
| 9 | 82889273 | rs602630 | -0.697 | 0.697 | -0.83 | -0.326 | -0.035 | 0.05 | 0.05 | 0 | 0.86 | 0.94 | 0.89 | 1 | 0.83 | 0.92 |
| 9 | 82900888 | rs10780367 | 0.271 | 0.186 | -1.366 | -0.154 | 0.027 | -0.038 | -0.038 | 0.137 | 0.29 | 0.2 | 0.18 | 0.47 | 0.18 | 0.32 |
| 9 | 82903566 | rs11138590 | 0.705 | 0.466 | -1.798 | 0.032 | -0.038 | -0.016 | -0.016 | 0 | 0.42 | 0.24 | 0.53 | 0.44 | 0.51 | 0.44 |
| 9 | 82913245 | rs7034213 | 0.031 | 1.059 | -1.248 | 0.809 | -0.003 | 0.025 | 0.025 | 0 | 0.12 | 0.1 | 0.32 | 0.4 | 0.17 | 0.15 |
| 9 | 82916993 | rs6559541 | 0.639 | 0.994 | -1.252 | 0.664 | -0.015 | -0.02 | -0.02 | 0.064 | 0.68 | 0.76 | 0.56 | 0.51 | 0.63 | 0.72 |
| 9 | 82917653 | rs1333937 | 0.09 | 1.057 | -1.182 | 1.243 | -0.042 | -0.047 | -0.047 | 0 | 0.08 | 0 | 0.25 | 0.2 | 0.1 | 0.08 |
| 9 | 82922380 | rs11793341 | -0.166 | 0.765 | -1.081 | 0.95 | -0.007 | -0.03 | -0.03 | 0 | 0.08 | 0.03 | 0.07 | 0.13 | 0.08 | 0.14 |
| 9 | 82926278 | rs7044696 | -0.243 | 0.759 | -1.151 | 0.22 | 0.001 | 0.082 | 0.082 | 0.193 | 0.51 | 0.39 | 0.55 | 0.8 | 0.57 | 0.66 |
| 9 | 82929577 | rs10867496 | 0.018 | 0.782 | -0.811 | 0.647 | 0.016 | 0.066 | 0.066 | 0 | 0.13 | 0.27 | 0.11 | 0.24 | 0.16 | 0.18 |
| 9 | 82931943 | rs1572145 | -1.06 | 0.973 | -1.192 | -0.13 | -0.024 | 0.058 | 0.058 | 0 | 0.82 | 0.93 | 0.87 | 1 | 0.82 | 0.91 |
| 9 | 82932712 | rs915004 | 0.248 | 1.318 | -0.937 | 0.082 | 0 | 0 | 0 | 0.021 | 0.46 | 0.53 | 0.56 | 0.51 | 0.54 | 0.6 |
| 9 | 82934337 | rs1333936 | 0.301 | 1.13 | -0.598 | 0.039 | 0.002 | 0.001 | 0.001 | 0 | 0.48 | 0.43 | 0.54 | 0.51 | 0.54 | 0.63 |
| 9 | 82935097 | rs1333934 | -0.089 | 1.016 | -0.511 | -0.066 | -0.001 | -0.001 | -0.001 | 2.485 | 0.64 | 0.67 | 0.61 | 0.57 | 0.62 | 0.69 |
| 9 | 82940654 | rs10867499 | 1.863 | 0.879 | -0.569 | -0.09 | 0.015 | 0.034 | 0.034 | 0.168 | 0.16 | 0.11 | 0.06 | 0.08 | 0.08 | 0.05 |
| 9 | 82949474 | rs10122600 | -1.313 | 1.23 | -0.489 | -0.067 | -0.013 | 0.021 | 0.021 | 0 | 0.8 | 0.89 | 0.86 | 1 | 0.8 | 0.9 |
| 9 | 82949861 | rs1577101 | -0.59 | 0.751 | -0.291 | 0.183 | -0.018 | -0.001 | -0.001 | 0 | 0.69 | 0.78 | 0.63 | 0.66 | 0.67 | 0.75 |
| 9 | 82950716 | rs11138611 | 1.06 | 0.474 | -0.371 | 0.219 | 0.003 | 0.019 | 0.019 | 0 | 0.12 | 0.15 | 0.11 | 0.22 | 0.16 | 0.21 |
| 9 | 82954954 | rs7041680 | 0.048 | 0.127 | -0.58 | 0.002 | 0.046 | 0.003 | 0.003 | 2.186 | 0.39 | 0.27 | 0.37 | 0.32 | 0.27 | 0.28 |
| 9 | 82956688 | rs10746630 | -0.975 | 0.211 | -0.923 | 0.024 | 0.016 | 0.016 | 0.016 | 1.881 | 0.7 | 0.65 | 0.61 | 0.57 | 0.65 | 0.73 |
| 9 | 82962544 | rs11787758 | 0.376 | 0.277 | -1.38 | 0.496 | -0.002 | 0.013 | 0.013 | 0 | 0.3 | 0.25 | 0.22 | 0.25 | 0.37 | 0.44 |
| 9 | 82965051 | rs1889283 | -0.101 | -0.119 | -1.078 | 1.016 | 0.019 | -0.054 | -0.054 | 1.68 | 0.17 | 0.08 | 0.3 | 0.24 | 0.15 | 0.18 |
| 9 | 82966983 | rs7047495 | -0.372 | 0.172 | -0.639 | 0.801 | -0.001 | 0.036 | 0.036 | 0 | 0.13 | 0.2 | 0.11 | 0.25 | 0.16 | 0.25 |
| 9 | 82968379 | rs6559543 | -0.022 | -0.103 | -1.144 | 0.224 | 0.01 | 0.037 | 0.037 | 0 | 0.12 | 0.02 | 0.42 | 0.08 | 0.13 | 0.26 |
| 9 | 82971893 | rs1333940 | -0.204 | -0.429 | -1.091 | 0.121 | 0.045 | 0.009 | 0.009 | 0 | 0.49 | 0.68 | 0.68 | 0.59 | 0.45 | 0.52 |
| 9 | 82975319 | rs17083204 | -0.604 | 0.084 | -1.232 | -0.348 | -0.018 | -0.056 | -0.056 | 4.123 | 0.26 | 0.21 | 0.18 | 0.42 | 0.27 | 0.25 |
| 9 | 82976239 | rs11138628 | 0.656 | -0.155 | -1.401 | -0.36 | -0.089 | -0.083 | -0.083 | 0.425 | 0.45 | 0.09 | 0.62 | 0.61 | 0.52 | 0.48 |
| 9 | 82977316 | rs10491946 | 0.507 | -0.175 | -1.276 | -0.492 | 0.022 | -0.015 | -0.015 | 0 | 0.08 | 0.5 | 0.06 | 0 | 0.02 | 0.05 |
| 9 | 82980785 | rs928537 | 0.734 | -0.17 | -1.392 | -0.478 | -0.109 | -0.115 | -0.115 | 1.734 | 0.45 | 0.07 | 0.62 | 0.6 | 0.52 | 0.48 |
| 9 | 82981744 | rs7027955 | 0.157 | -0.057 | -1.303 | -0.39 | -0.001 | -0.005 | -0.005 | 0 | 0.52 | 0.65 | 0.68 | 0.6 | 0.54 | 0.53 |
| 9 | 82985544 | rs10746633 | -0.713 | 0.015 | -1.463 | -0.48 | -0.005 | 0.024 | 0.024 | 0.216 | 0.65 | 0.76 | 0.79 | 0.94 | 0.66 | 0.71 |
| 9 | 82988782 | rs7849222 | -0.458 | 0.029 | -1.615 | -0.537 | -0.018 | -0.029 | -0.029 | 2.272 | 0.77 | 0.86 | 0.78 | 0.72 | 0.72 | 0.78 |
| 9 | 82989464 | rs4877167 | 0.101 | 0.047 | -1.474 | -0.509 | -0.013 | 0.007 | 0.007 | 0.877 | 0.56 | 0.7 | 0.68 | 0.67 | 0.55 | 0.59 |
| 9 | 82990664 | rs11138638 | 1.021 | 0.243 | -1.849 | -0.438 | -0.062 | 0.005 | 0.005 | 1.508 | 0.11 | 0.01 | 0.07 | 0.08 | 0.14 | 0.07 |
| 9 | 82991662 | rs4877557 | 1.434 | 0.276 | -1.703 | -0.455 | 0.004 | -0.09 | -0.09 | 0.712 | 0.34 | 0.1 | 0.56 | 0.53 | 0.38 | 0.42 |
| 9 | 82993629 | rs4278224 | 1.312 | 0.765 | -1.38 | -0.516 | -0.055 | 0.006 | 0.006 | 0.007 | 0.09 | 0.01 | 0.27 | 0.06 | 0.14 | 0.08 |
| 9 | 82994364 | rs1544230 | -0.27 | 0.64 | -1.23 | -0.495 | -0.011 | 0.022 | 0.022 | 0 | 0.78 | 0.88 | 0.8 | 0.91 | 0.74 | 0.77 |
| 9 | 82996676 | rs2378475 | 0.919 | 0.306 | -1.252 | -0.383 | -0.02 | 0.04 | 0.04 | 0.306 | 0.51 | 0.18 | 0.6 | 0.65 | 0.54 | 0.47 |
| 9 | 82999854 | rs1541958 | 1.02 | 0.258 | -0.41 | -0.496 | 0.007 | -0.002 | -0.002 | 0 | 0.5 | 0.38 | 0.62 | 0.58 | 0.54 | 0.46 |
| 9 | 83000507 | rs7861030 | 0.88 | 0.183 | -0.508 | -0.562 | 0.014 | 0.062 | 0.062 | 2.605 | 0.48 | 0.17 | 0.41 | 0.67 | 0.51 | 0.58 |
| 9 | 83001846 | rs4877564 | -0.819 | 0.259 | -0.207 | -0.575 | 0.015 | -0.006 | -0.006 | 1.898 | 0.28 | 0.41 | 0.41 | 0.23 | 0.27 | 0.28 |
| 9 | 83010685 | rs17083258 | 0.948 | 0.282 | -0.448 | -0.56 | 0 | -0.023 | -0.023 | 0 | 0.09 | 0.05 | 0.29 | 0.18 | 0.1 | 0.16 |
| 9 | 83013727 | rs7864141 | -0.382 | 0.307 | -0.447 | -0.593 | -0.004 | 0 | 0 | 2.109 | 0.69 | 0.8 | 0.75 | 0.74 | 0.67 | 0.69 |
| 9 | 83016971 | rs10867523 | 1.428 | 0.381 | -0.551 | -0.656 | -0.017 | 0.014 | 0.014 | 0 | 0.1 | 0.07 | 0.07 | 0.04 | 0.14 | 0.05 |
| 9 | 83018572 | rs10867524 | 1.543 | 0.295 | -0.6 | -0.687 | -0.001 | -0.091 | -0.091 | 0.483 | 0.38 | 0.09 | 0.32 | 0.51 | 0.4 | 0.41 |
| 9 | 83021572 | rs11138651 | 1.099 | 0.453 | -0.402 | -0.661 | 0.023 | -0.025 | -0.025 | 0 | 0.11 | 0.18 | 0.08 | 0.04 | 0.15 | 0.05 |
| 9 | 83024270 | rs991677 | 0.648 | 0.446 | -0.253 | -0.664 | 0.056 | -0.05 | -0.05 | 0 | 0.12 | 0.31 | 0.09 | 0.04 | 0.15 | 0.05 |
| 9 | 83025449 | rs1347374 | -0.171 | 0.471 | -0.687 | -0.615 | -0.005 | 0.001 | 0.001 | 0 | 0.68 | 0.8 | 0.76 | 0.75 | 0.68 | 0.69 |
| 9 | 83027012 | rs7047256 | 0.877 | 0.435 | -0.929 | -0.656 | -0.041 | 0.112 | 0.112 | 0.626 | 0.56 | 0.16 | 0.44 | 0.77 | 0.56 | 0.62 |
| 9 | 83029869 | rs7029470 | 1.335 | 0.454 | -0.848 | -0.574 | 0.026 | -0.053 | -0.053 | 0 | 0.39 | 0.19 | 0.33 | 0.52 | 0.39 | 0.42 |
| 9 | 83039827 | rs11138655 | 0.744 | 0.341 | -0.489 | -0.603 | 0.004 | 0.03 | 0.03 | 1.212 | 0.53 | 0.66 | 0.67 | 0.71 | 0.53 | 0.64 |
| 9 | 83040540 | rs4877574 | 0.752 | 0.365 | -0.492 | -0.603 | 0.002 | 0.015 | 0.015 | 1.221 | 0.52 | 0.61 | 0.67 | 0.71 | 0.53 | 0.64 |
| 9 | 83041867 | rs1109201 | 0.776 | 0.521 | -0.767 | -0.496 | 0.013 | -0.039 | -0.039 | 5.143 | 0.14 | 0.03 | 0.11 | 0.2 | 0.1 | 0.05 |
| 9 | 83041919 | rs1109202 | 0.155 | 0.47 | -0.572 | -0.394 | -0.012 | 0.02 | 0.02 | 1.139 | 0.59 | 0.72 | 0.69 | 0.76 | 0.6 | 0.81 |
| 9 | 83043376 | rs892686 | 0.456 | 0.165 | -0.274 | -0.401 | 0.027 | 0.03 | 0.03 | 1.107 | 0.49 | 0.4 | 0.61 | 0.69 | 0.45 | 0.61 |
| 9 | 83044079 | rs4144881 | 1.027 | 0.452 | -0.303 | -0.479 | -0.012 | -0.037 | -0.037 | 0.081 | 0.12 | 0.03 | 0.12 | 0.23 | 0.11 | 0.08 |
| 9 | 83044102 | rs1419273 | -0.535 | 0.233 | -0.392 | -0.907 | 0 | -0.005 | -0.005 | 0 | 0.82 | 0.87 | 0.88 | 1 | 0.83 | 0.91 |
| 9 | 83046008 | rs1419274 | -0.784 | 0.373 | -0.196 | -0.839 | -0.001 | -0.03 | -0.03 | 0.105 | 0.84 | 0.82 | 0.88 | 1 | 0.84 | 0.94 |
| 9 | 83048003 | rs11138659 | 0.557 | 0.663 | 0.36 | -0.474 | -0.007 | -0.014 | -0.014 | 0.053 | 0.17 | 0.08 | 0.16 | 0.22 | 0.16 | 0.12 |
| 9 | 83048441 | rs10512102 | 0.998 | 0.744 | -0.031 | -0.639 | -0.039 | 0.042 | 0.042 | 2.589 | 0.13 | 0.01 | 0.09 | 0.04 | 0.14 | 0.07 |
| 9 | 83048528 | rs10512103 | -0.215 | 0.489 | 0.194 | -0.964 | -0.015 | -0.098 | -0.098 | 0 | 0.27 | 0.48 | 0.16 | 0.08 | 0.21 | 0.37 |
| 9 | 83065765 | rs1857469 | -1.409 | 0.594 | 0.047 | -0.92 | -0.016 | 0.057 | 0.057 | 0 | 0.52 | 0.81 | 0.36 | 0.36 | 0.47 | 0.64 |
| 9 | 83066545 | rs7872658 | -1.541 | 0.786 | -0.18 | -0.271 | -0.007 | -0.029 | -0.029 | 0.168 | 0.59 | 0.85 | 0.41 | 0.59 | 0.55 | 0.68 |
| 9 | 83069568 | rs12238292 | 0.079 | 0.883 | 0.018 | -0.232 | 0.024 | 0.029 | 0.029 | 0 | 0.24 | 0.32 | 0.2 | 0.28 | 0.25 | 0.28 |

| | | | | | | | | | | | | | | | | |
|---|---|---|---|---|---|---|---|---|---|---|---|---|---|---|---|---|
| 9 | 83070204 | rs11138672 | 0.052 | 0.893 | 0.22 | -0.014 | -0.009 | 0.016 | 0.016 | 0 | 0.2 | 0.37 | 0.11 | 0.28 | 0.16 | 0.34 |
| 9 | 83070605 | rs11138673 | 0.182 | 0.888 | 0.324 | -0.017 | -0.006 | 0.011 | 0.011 | 0 | 0.19 | 0.28 | 0.11 | 0.27 | 0.16 | 0.33 |
| 9 | 83070979 | rs7860480 | 1.576 | 0.955 | 0.247 | -0.008 | 0.017 | -0.002 | -0.002 | 0 | 0.05 | 0.07 | 0.02 | 0.05 | 0.08 | 0.16 |
| 9 | 83072844 | rs4877174 | -0.987 | 0.806 | 0.028 | 0.299 | 0.001 | 0.004 | 0.004 | 0 | 0.41 | 0.61 | 0.48 | 0.54 | 0.41 | 0.61 |
| 9 | 83073408 | rs2378479 | -0.337 | 0.531 | -0.145 | 0.509 | -0.007 | 0.01 | 0.01 | 0 | 0.19 | 0.29 | 0.1 | 0.27 | 0.18 | 0.36 |
| 9 | 83077768 | rs2164186 | 0.252 | 0.572 | -0.637 | 0.892 | 0.002 | 0.003 | 0.003 | 0 | 0.1 | 0.08 | 0.12 | 0.27 | 0.11 | 0.07 |
| 9 | 83081971 | rs2196244 | -0.408 | 1.187 | -0.003 | 1.081 | 0.077 | 0.119 | 0.119 | 0 | 0.93 | 0.78 | 0.68 | 0.83 | 0.89 | 0.9 |
| 9 | 83083647 | rs7862553 | -0.769 | 1.052 | -0.283 | 0.488 | -0.001 | -0.006 | -0.006 | 0.47 | 0.47 | 0.38 | 0.35 | 0.56 | 0.47 | 0.49 |
| 9 | 83085633 | rs7020677 | 1.06 | 0.995 | -0.267 | 0.464 | 0.004 | 0.009 | 0.009 | 0 | 0.46 | 0.41 | 0.34 | 0.28 | 0.43 | 0.41 |
| 9 | 83088437 | rs17083422 | 0.982 | 0.99 | -0.273 | 0.537 | 0.002 | 0.001 | 0.001 | 0 | 0.1 | 0.08 | 0.12 | 0.26 | 0.11 | 0.07 |
| 9 | 83089867 | rs4612444 | 0.064 | 1.128 | -0.338 | 0.341 | 0.02 | 0.009 | 0.009 | 0 | 0.8 | 0.72 | 0.63 | 0.78 | 0.78 | 0.6 |
| 9 | 83090265 | rs1550051 | 0.355 | 1.123 | -0.508 | 0.452 | 0 | 0.02 | 0.02 | 0 | 0.14 | 0.19 | 0.17 | 0.26 | 0.17 | 0.11 |
| 9 | 83095955 | rs11138688 | 1.273 | 1.085 | -0.568 | 0.421 | -0.001 | 0.005 | 0.005 | 0 | 0.11 | 0.16 | 0.06 | 0.22 | 0.1 | 0.04 |
| 9 | 83096573 | rs10867550 | -0.676 | 1.034 | -0.534 | 0.411 | -0.003 | -0.008 | -0.008 | 0 | 0.38 | 0.37 | 0.32 | 0.51 | 0.41 | 0.22 |
| 9 | 83096816 | rs11138689 | 1.366 | 1.073 | -0.94 | 0.244 | -0.023 | -0.059 | -0.059 | 0.08 | 0.12 | 0.02 | 0.13 | 0.22 | 0.12 | 0.09 |
| 9 | 83099363 | rs10867551 | -0.191 | 0.969 | -0.649 | 0.052 | 0.002 | -0.004 | -0.004 | 0.118 | 0.3 | 0.34 | 0.25 | 0.26 | 0.3 | 0.17 |
| 9 | 83099364 | rs4877578 | -0.319 | 1.019 | -0.61 | 0.066 | 0.01 | 0.015 | 0.015 | 0.133 | 0.84 | 0.78 | 0.89 | 0.8 | 0.83 | 0.64 |
| 9 | 83100591 | rs17256174 | 1.118 | 1.04 | -0.684 | 0.035 | 0.004 | 0.02 | 0.02 | 0 | 0.1 | 0.06 | 0.03 | 0.09 | 0.08 | 0.26 |
| 9 | 83101100 | rs10119230 | 0.798 | 1.034 | -0.347 | 0.153 | 0.005 | 0.004 | 0.004 | 0 | 0.6 | 0.47 | 0.45 | 0.5 | 0.58 | 0.52 |
| 9 | 83104358 | rs17256298 | 1.327 | 1.055 | -0.388 | 0.099 | 0.001 | 0.018 | 0.018 | 0 | 0.09 | 0.05 | 0.03 | 0.09 | 0.08 | 0.26 |
| 9 | 83105091 | rs978282 | -1.354 | 1.106 | -0.292 | 0.233 | 0.08 | 0.022 | 0.022 | 0 | 0.93 | 0.79 | 0.92 | 0.94 | 0.9 | 0.9 |
| 9 | 83107844 | rs10867555 | -0.082 | 0.998 | -0.386 | 0.209 | -0.035 | 0.003 | 0.003 | 0 | 0.3 | 0.12 | 0.18 | 0.33 | 0.31 | 0.37 |
| 9 | 83109040 | rs2378481 | 1.219 | 0.982 | -0.297 | 0.224 | -0.002 | 0.012 | 0.012 | 1.092 | 0.08 | 0.1 | 0.09 | 0.09 | 0.07 | 0.06 |
| 9 | 83110616 | rs10780407 | -0.708 | 0.987 | -0.289 | 0.22 | -0.004 | -0.001 | -0.001 | 0 | 0.37 | 0.31 | 0.28 | 0.42 | 0.37 | 0.43 |
| 9 | 83111466 | rs17341528 | 1.331 | 0.94 | -0.587 | 0.225 | 0.001 | 0.015 | 0.015 | 0 | 0.09 | 0.05 | 0.03 | 0.1 | 0.08 | 0.26 |
| 9 | 83113022 | rs11138696 | 2.425 | 0.652 | -0.77 | -0.383 | 0.02 | 0.055 | 0.055 | 0 | 0.23 | 0.11 | 0.19 | 0 | 0.21 | 0.15 |
| 9 | 83113481 | rs7034862 | 0.681 | 1.136 | -0.506 | -0.058 | -0.001 | -0.001 | -0.001 | 0 | 0.56 | 0.47 | 0.66 | 0.52 | 0.55 | 0.48 |
| 9 | 83115815 | rs10465167 | 0.302 | 0.945 | -1.291 | -0.208 | -0.014 | 0.006 | 0.006 | 0 | 0.29 | 0.18 | 0.18 | 0.33 | 0.3 | 0.37 |
| 9 | 83118945 | rs1972804 | 1.305 | 0.797 | -1.538 | -0.139 | 0.033 | 0.058 | 0.058 | 0 | 0.35 | 0.14 | 0.49 | 0.27 | 0.35 | 0.2 |
| 9 | 83120662 | rs11138702 | -0.091 | 0.88 | -1.261 | -0.127 | -0.033 | 0.006 | 0.006 | 0.174 | 0.31 | 0.12 | 0.23 | 0.34 | 0.32 | 0.39 |
| 9 | 83127968 | rs2164183 | 1.029 | 1.261 | -1.349 | -0.115 | -0.022 | 0.083 | 0.083 | 0 | 0.48 | 0.16 | 0.6 | 0.58 | 0.49 | 0.35 |
| 9 | 83128214 | rs7019895 | 0.66 | 1.621 | -0.471 | -0.181 | 0.058 | -0.022 | -0.022 | 0 | 0.08 | 0.19 | 0.12 | 0.03 | 0.12 | 0.06 |
| 9 | 83139208 | rs7851229 | 0.726 | 1.686 | 0.213 | -0.3 | 0.108 | -0.092 | -0.092 | 0 | 0.81 | 0.54 | 0.79 | 0.94 | 0.73 | 0.82 |
| 9 | 83141248 | rs4877582 | 0.685 | 1.697 | 0.219 | -0.293 | 0.11 | -0.09 | -0.09 | 0 | 0.82 | 0.54 | 0.79 | 0.94 | 0.73 | 0.83 |
| 9 | 83156867 | rs2149342 | -0.093 | 1.675 | 0.222 | -0.291 | 0.095 | -0.054 | -0.054 | 0.23 | 0.88 | 0.55 | 0.89 | 0.96 | 0.85 | 0.89 |
| 9 | 83162551 | rs4394477 | -0.284 | 1.699 | 0.413 | -0.274 | 0.129 | -0.02 | -0.02 | 0.706 | 0.9 | 0.68 | 0.84 | 0.94 | 0.79 | 0.84 |
| 9 | 83162993 | rs4590514 | 1.85 | 1.752 | -0.271 | -0.082 | 0.015 | -0.024 | -0.024 | 0 | 0.38 | 0.11 | 0.27 | 0.27 | 0.33 | 0.26 |
| 9 | 83163922 | rs12554921 | 1.165 | 1.778 | -0.006 | 0.101 | -0.006 | -0.01 | -0.01 | 2.113 | 0.12 | 0.1 | 0.35 | 0.3 | 0.17 | 0.1 |
| 9 | 83169108 | rs7022798 | -0.52 | 1.648 | 0.226 | -0.244 | 0.022 | 0.002 | 0.002 | 0 | 0.83 | 0.66 | 0.87 | 0.78 | 0.8 | 0.87 |
| 9 | 83169355 | rs2209907 | 1.105 | 1.629 | 0.254 | -0.245 | -0.004 | 0.001 | 0.001 | 0 | 0.13 | 0.18 | 0.09 | 0.19 | 0.11 | 0.04 |
| 9 | 83169672 | rs10780428 | 2.135 | 1.538 | 0.094 | -0.329 | 0.024 | -0.014 | -0.014 | 0.041 | 0.39 | 0.13 | 0.27 | 0.27 | 0.33 | 0.26 |
| 9 | 83169994 | rs10867574 | 1.207 | 1.78 | 0.572 | -0.127 | -0.004 | 0.002 | 0.002 | 0 | 0.69 | 0.34 | 0.72 | 0.75 | 0.62 | 0.79 |
| 9 | 83171575 | rs2296935 | 1.88 | 1.822 | 0.313 | -0.099 | -0.026 | -0.058 | -0.058 | 0 | 0.09 | 0.01 | 0.1 | 0.2 | 0.09 | 0.1 |
| 9 | 83172180 | rs1328500 | 0.417 | 1.703 | 0.933 | -0.194 | 0.075 | -0.013 | -0.013 | 0.809 | 0.77 | 0.5 | 0.76 | 0.75 | 0.67 | 0.81 |
| 9 | 83173323 | rs11791976 | 1.397 | 1.581 | 0.544 | -0.185 | 0.002 | 0.028 | 0.028 | 0 | 0.06 | 0.01 | 0.03 | 0 | 0.06 | 0.01 |
| 9 | 83174096 | rs10867575 | 1.366 | 1.686 | 0.668 | -0.184 | -0.003 | 0.002 | 0.002 | 0 | 0.69 | 0.34 | 0.72 | 0.75 | 0.62 | 0.79 |
| 9 | 83174331 | rs928012 | 0.555 | 1.341 | 0.758 | -0.072 | 0.005 | -0.011 | -0.011 | 0 | 0.77 | 0.66 | 0.83 | 0.78 | 0.74 | 0.85 |
| 9 | 83175451 | rs11138731 | 0.022 | 1.015 | 0.086 | 0.03 | 0.001 | 0.002 | 0.002 | 0 | 0.18 | 0.18 | 0.12 | 0.19 | 0.17 | 0.05 |
| 9 | 83197939 | rs10780435 | 1.362 | 0.878 | -0.248 | 0.179 | -0.004 | 0.073 | 0.073 | 0 | 0.45 | 0.13 | 0.28 | 0.25 | 0.4 | 0.23 |
| 9 | 83198003 | rs6559559 | -0.482 | 1.106 | 0.743 | 0.64 | 0.012 | 0.113 | 0.113 | 0 | 0.21 | 0.41 | 0.45 | 0.38 | 0.25 | 0.18 |
| 9 | 83199814 | rs1328521 | -0.752 | 1.396 | 0.489 | 0.694 | 0.005 | 0.001 | 0.001 | 0 | 0.86 | 0.83 | 0.85 | 0.79 | 0.81 | 0.87 |
| 9 | 83202032 | rs7849462 | -0.046 | 1.379 | 0.46 | 0.693 | 0.045 | 0.017 | 0.017 | 0 | 0.82 | 0.57 | 0.82 | 0.76 | 0.8 | 0.87 |
| 9 | 83205767 | rs17258867 | 1.136 | 1.399 | -0.119 | 0.639 | -0.031 | 0.016 | 0.016 | 0.12 | 0.07 | 0 | 0.09 | 0.02 | 0.12 | 0.07 |
| 9 | 83207932 | rs12551055 | 0.677 | 1.31 | 0.037 | 0.736 | -0.004 | -0.013 | -0.013 | 1.375 | 0.11 | 0.07 | 0.33 | 0.36 | 0.13 | 0.11 |
| 9 | 83210614 | rs1410550 | -0.087 | 1.235 | 0.175 | 0.351 | 0.021 | -0.002 | -0.002 | 0 | 0.83 | 0.67 | 0.83 | 0.79 | 0.8 | 0.87 |
| 9 | 83217377 | rs1410542 | 0.73 | 1.179 | -0.24 | 0.362 | -0.016 | -0.004 | -0.004 | 0 | 0.63 | 0.68 | 0.38 | 0.37 | 0.51 | 0.64 |
| 9 | 83221548 | rs11138755 | -0.442 | 1.093 | -0.147 | 0.365 | 0.003 | 0.035 | 0.035 | 0 | 0.21 | 0.22 | 0.46 | 0.53 | 0.29 | 0.46 |
| 9 | 83235735 | rs10123659 | -0.388 | 0.854 | 0.342 | 0.222 | 0.005 | 0.168 | 0.168 | 0.173 | 0.15 | 0.34 | 0.21 | 0.37 | 0.16 | 0.1 |
| 9 | 83238535 | rs724372 | 0.592 | 0.854 | 0.481 | 0.071 | 0.005 | 0.015 | 0.015 | 0 | 0.7 | 0.66 | 0.49 | 0.49 | 0.64 | 0.49 |
| 9 | 83241416 | rs1328515 | -1.277 | 0.885 | 0.48 | 0.11 | -0.017 | 0.008 | 0.008 | 0.012 | 0.88 | 0.92 | 0.89 | 0.99 | 0.82 | 0.88 |
| 9 | 83246314 | rs4877600 | 0.354 | 0.879 | 0.324 | 0.142 | 0.03 | 0.096 | 0.096 | 0 | 0.78 | 0.55 | 0.77 | 0.64 | 0.78 | 0.84 |
| 9 | 83247124 | rs7019006 | 0.566 | 0.871 | 0.062 | 0.138 | 0.002 | 0.015 | 0.015 | 0 | 0.76 | 0.38 | 0.75 | 0.64 | 0.77 | 0.84 |
| 9 | 83247711 | rs13284117 | -0.263 | 0.855 | 0.056 | 0.129 | 0.018 | 0.07 | 0.07 | 0.257 | 0.83 | 0.4 | 0.77 | 0.64 | 0.81 | 0.86 |
| 9 | 83249863 | rs10453249 | 0.596 | 0.857 | 0.056 | 0.13 | 0.002 | 0.015 | 0.015 | 0 | 0.76 | 0.38 | 0.75 | 0.64 | 0.77 | 0.84 |
| 9 | 83252173 | rs7040874 | -0.267 | 0.886 | 0.088 | 0.163 | 0.019 | 0.071 | 0.071 | 0.231 | 0.83 | 0.4 | 0.77 | 0.64 | 0.81 | 0.86 |
| 9 | 83252509 | rs10867599 | -0.267 | 0.886 | 0.088 | 0.163 | 0.019 | 0.071 | 0.071 | 0.231 | 0.83 | 0.4 | 0.77 | 0.64 | 0.81 | 0.86 |
| 9 | 83254196 | rs11793167 | 0.13 | 0.566 | -0.385 | -0.011 | 0.007 | 0.022 | 0.022 | 0 | 0.14 | 0.09 | 0.06 | 0.03 | 0.12 | 0.19 |
| 9 | 83259922 | rs10867602 | -0.819 | 0.769 | -0.328 | -0.279 | 0.001 | -0.011 | -0.011 | 0 | 0.27 | 0.28 | 0.49 | 0.55 | 0.32 | 0.59 |
| 9 | 83264458 | rs12336241 | -1.274 | 0.771 | -0.207 | -0.316 | 0 | 0.013 | 0.013 | 0 | 0.31 | 0.38 | 0.52 | 0.55 | 0.38 | 0.61 |

| | | | | | | | | | | | | | | | | |
|---|---|---|---|---|---|---|---|---|---|---|---|---|---|---|---|---|
| 9 | 83279833 | rs10867605 | 0.194 | 0.783 | -0.694 | -0.288 | -0.003 | 0.012 | 0.012 | 0 | 0.13 | 0.13 | 0.37 | 0.5 | 0.18 | 0.37 |
| 9 | 83284684 | rs4877611 | 0.758 | 0.785 | -0.692 | -0.263 | -0.001 | 0.044 | 0.044 | 0 | 0.11 | 0.12 | 0.34 | 0.42 | 0.16 | 0.14 |
| 9 | 83284737 | rs4877612 | -1.057 | 0.906 | -0.672 | -0.307 | -0.012 | -0.035 | -0.035 | 3.474 | 0.29 | 0.21 | 0.51 | 0.57 | 0.38 | 0.61 |
| 9 | 83290499 | rs11138807 | 1.514 | 0.79 | -1.071 | -0.697 | -0.025 | 0.004 | 0.004 | 0 | 0.14 | 0.01 | 0.1 | 0.1 | 0.14 | 0.06 |
| 9 | 83293714 | rs10123247 | 0.684 | 1.265 | -0.334 | -0.517 | 0.036 | -0.071 | -0.071 | 0 | 0.73 | 0.58 | 0.72 | 0.89 | 0.63 | 0.64 |
| 9 | 83306095 | rs9695862 | 0.792 | 1.043 | -0.725 | -0.823 | 0 | -0.022 | -0.022 | 0 | 0.61 | 0.59 | 0.66 | 0.87 | 0.5 | 0.55 |
| 9 | 83309521 | rs4287017 | 1.717 | 0.818 | -0.369 | -0.686 | -0.014 | -0.024 | -0.024 | 5.663 | 0.56 | 0.23 | 0.57 | 0.52 | 0.55 | 0.35 |
| 9 | 83311139 | rs7036247 | -0.029 | 0.949 | -0.426 | -0.932 | -0.007 | 0.005 | 0.005 | 0 | 0.8 | 0.79 | 0.82 | 0.9 | 0.73 | 0.7 |
| 9 | 83316951 | rs10746668 | 1.12 | 0.919 | -0.426 | -0.925 | 0.025 | -0.065 | -0.065 | 0.479 | 0.69 | 0.53 | 0.7 | 0.84 | 0.6 | 0.4 |
| 9 | 83318011 | rs17231264 | 1.014 | 0.877 | -0.257 | -0.855 | 0 | -0.016 | -0.016 | 0.302 | 0.1 | 0.06 | 0.03 | 0 | 0.05 | 0.03 |
| 9 | 83328730 | rs17083946 | -0.613 | 0.518 | -0.135 | -0.783 | 0 | -0.036 | -0.036 | 0 | 0.18 | 0.32 | 0.18 | 0.11 | 0.2 | 0.27 |
| 9 | 83333000 | rs10491932 | -1.208 | 0.426 | -0.124 | -0.82 | -0.025 | -0.074 | -0.074 | 0 | 0.28 | 0.39 | 0.2 | 0.11 | 0.24 | 0.29 |
| 9 | 83367779 | rs17083997 | 0.517 | 0.49 | -0.032 | -0.682 | -0.004 | -0.006 | -0.006 | 0 | 0.11 | 0.19 | 0.08 | 0.09 | 0.09 | 0.19 |
| 9 | 83373425 | rs7865140 | -0.654 | 0.397 | -0.26 | -0.658 | -0.046 | -0.047 | -0.047 | 0.313 | 0.25 | 0.38 | 0.14 | 0.14 | 0.16 | 0.23 |
| 9 | 83383795 | rs12002026 | -0.098 | 0.396 | -0.272 | -0.811 | -0.067 | -0.12 | -0.12 | 0 | 0.17 | 0.34 | 0.09 | 0 | 0.08 | 0.03 |
| 9 | 83387588 | rs10867619 | 1.652 | 0.583 | -0.393 | -0.729 | -0.005 | -0.018 | -0.018 | 0.248 | 0.54 | 0.25 | 0.56 | 0.52 | 0.53 | 0.34 |
| 9 | 83397162 | rs4268198 | -0.879 | 0.955 | 0.04 | -0.106 | -0.042 | -0.073 | -0.073 | 1.008 | 0.24 | 0.36 | 0.14 | 0.1 | 0.16 | 0.23 |
| 9 | 83397674 | rs6559588 | -0.246 | 0.972 | 0.071 | -0.127 | -0.017 | 0.005 | 0.005 | 0 | 0.84 | 0.86 | 0.81 | 0.98 | 0.72 | 0.81 |
| 9 | 83398603 | rs7019400 | 0.425 | 0.969 | 0.117 | -0.093 | -0.02 | -0.027 | -0.027 | 2.379 | 0.79 | 0.82 | 0.71 | 0.61 | 0.69 | 0.56 |
| 9 | 83399798 | rs1576757 | 0.242 | 0.958 | 0.206 | -0.393 | -0.002 | -0.002 | -0.002 | 0 | 0.1 | 0.11 | 0.07 | 0.1 | 0.09 | 0.19 |
| 9 | 83404085 | rs12346044 | 1.587 | 0.984 | 0.172 | -0.397 | 0.002 | -0.073 | -0.073 | 0.753 | 0.6 | 0.5 | 0.67 | 0.88 | 0.56 | 0.58 |
| 9 | 83408197 | rs10491936 | -0.564 | 0.84 | 0.518 | 0.162 | -0.028 | -0.052 | -0.052 | 0 | 0.21 | 0.29 | 0.12 | 0.1 | 0.15 | 0.23 |
| 9 | 83412886 | rs1359249 | 1.066 | 1.117 | 0.088 | 0.493 | -0.011 | -0.003 | -0.003 | 1.259 | 0.11 | 0.05 | 0.09 | 0.14 | 0.12 | 0.17 |
| 9 | 83429265 | rs10735555 | 1.925 | 1.041 | 0.219 | 0.783 | 0 | 0.012 | 0.012 | 0.688 | 0.51 | 0.37 | 0.55 | 0.43 | 0.5 | 0.29 |
| 9 | 83429615 | rs4284104 | -0.905 | 1.537 | -0.499 | -0.36 | 0.015 | -0.035 | -0.035 | 0 | 0.89 | 0.87 | 0.9 | 1 | 0.83 | 0.9 |
| 9 | 83439786 | rs4877622 | 1.896 | 1.578 | -0.457 | -0.362 | 0.001 | 0.01 | 0.01 | 1.555 | 0.54 | 0.58 | 0.63 | 0.72 | 0.46 | 0.3 |
| 9 | 83441767 | rs12000148 | -0.719 | 1.104 | -0.348 | -0.317 | -0.009 | -0.021 | -0.021 | 0.062 | 0.25 | 0.26 | 0.15 | 0.14 | 0.18 | 0.21 |
| 9 | 83445616 | rs11794073 | 0.385 | 0.813 | -0.223 | -0.318 | 0.031 | 0.033 | 0.033 | 0.434 | 0.12 | 0.05 | 0.05 | 0 | 0.06 | 0.04 |
| 9 | 83447633 | rs728242 | 1.353 | 0.886 | -0.264 | -0.259 | 0 | -0.007 | -0.007 | 0.608 | 0.55 | 0.54 | 0.6 | 0.69 | 0.51 | 0.32 |
| 9 | 83454689 | rs12002342 | -0.491 | 0.469 | 0.361 | -0.268 | -0.036 | -0.066 | -0.066 | 0 | 0.13 | 0.2 | 0.06 | 0 | 0.06 | 0.04 |
| 9 | 83455332 | rs7035022 | 0.442 | 1.46 | -0.007 | 0.47 | 0.013 | -0.035 | -0.035 | 0 | 0.89 | 0.87 | 0.9 | 1 | 0.83 | 0.89 |
| 9 | 83457320 | rs10435943 | -1.136 | 0.459 | 0.568 | 1.267 | -0.015 | 0.2 | 0.2 | 0 | 0.09 | 0.3 | 0.22 | 0.27 | 0.03 | 0.1 |
| 9 | 83462500 | rs1041039 | -1.543 | 0.932 | 0.171 | 1.213 | 0.007 | 0.064 | 0.064 | 0 | 0.23 | 0.33 | 0.46 | 0.39 | 0.26 | 0.18 |
| 9 | 83465888 | rs10746670 | -1.125 | 0.563 | -0.183 | 1.222 | -0.001 | -0.001 | -0.001 | 0 | 0.65 | 0.6 | 0.75 | 0.54 | 0.69 | 0.44 |
| 9 | 83485092 | rs10491940 | -1.123 | 0.407 | -0.022 | 1.085 | 0.011 | -0.012 | -0.012 | 0 | 0.61 | 0.7 | 0.73 | 0.54 | 0.66 | 0.43 |
| 9 | 83488864 | rs11138885 | -0.549 | 0.423 | 0.423 | 0.97 | -0.022 | 0.07 | 0.07 | 0 | 0.16 | 0.25 | 0.27 | 0.33 | 0.1 | 0.18 |
| 9 | 83499061 | rs12379326 | 0.563 | 0.389 | 0.531 | 0.871 | -0.026 | 0.06 | 0.06 | 0 | 0.31 | 0.16 | 0.21 | 0.14 | 0.33 | 0.18 |
| 9 | 83501287 | rs6559594 | -1.549 | 0.31 | 0.854 | 0.862 | 0 | -0.001 | -0.001 | 0 | 0.61 | 0.56 | 0.71 | 0.54 | 0.66 | 0.43 |
| 9 | 83503203 | rs11138895 | -0.629 | 0.32 | 1.037 | 0.949 | -0.024 | 0.083 | 0.083 | 0 | 0.15 | 0.25 | 0.25 | 0.32 | 0.1 | 0.17 |
| 9 | 83503893 | rs10780467 | -0.525 | 0.439 | 0.88 | 0.93 | 0 | 0 | 0 | 0 | 0.47 | 0.51 | 0.48 | 0.47 | 0.43 | 0.35 |
| 9 | 83513137 | rs7025679 | -0.951 | 0.433 | 0.879 | 0.714 | -0.001 | -0.001 | -0.001 | 0 | 0.52 | 0.5 | 0.69 | 0.53 | 0.58 | 0.35 |
| 9 | 83514230 | rs9314695 | -0.119 | 0.508 | 0.799 | 0.552 | 0.002 | -0.009 | -0.009 | 0.448 | 0.33 | 0.3 | 0.44 | 0.44 | 0.3 | 0.23 |
| 9 | 83524227 | rs10122814 | -0.618 | 0.719 | 1.055 | 0.879 | 0.032 | 0.066 | 0.066 | 0 | 0.21 | 0.36 | 0.47 | 0.39 | 0.31 | 0.15 |
| 9 | 83526643 | rs2209263 | 0.151 | 0.735 | 0.816 | 0.688 | -0.006 | 0.022 | 0.022 | 0 | 0.26 | 0.15 | 0.2 | 0.15 | 0.24 | 0.17 |
| 9 | 83539913 | rs12552136 | 1.775 | 0.721 | 1.129 | 0.767 | 0.013 | 0.01 | 0.01 | 0 | 0.42 | 0.26 | 0.22 | 0.38 | 0.36 | 0.62 |
| 9 | 83540744 | rs1853607 | 0.797 | 0.682 | 1.33 | 0.794 | -0.028 | 0.191 | 0.191 | 0 | 0.86 | 0.58 | 0.68 | 0.62 | 0.88 | 0.91 |
| 9 | 83541272 | rs11138911 | 1.763 | 0.685 | 1.346 | 0.774 | 0.003 | 0.003 | 0.003 | 0 | 0.43 | 0.38 | 0.23 | 0.38 | 0.36 | 0.63 |
| 9 | 83544183 | rs10780471 | 1.868 | 0.674 | 1.383 | 0.769 | 0.012 | 0.009 | 0.009 | 0 | 0.42 | 0.24 | 0.22 | 0.38 | 0.36 | 0.63 |
| 9 | 83549419 | rs10780472 | 0.947 | 0.775 | 1.445 | 0.788 | 0.012 | 0.046 | 0.046 | 0.052 | 0.72 | 0.56 | 0.51 | 0.56 | 0.65 | 0.85 |
| 9 | 83549681 | rs715976 | 0.947 | 0.775 | 1.446 | 0.788 | 0.012 | 0.048 | 0.048 | 0.052 | 0.72 | 0.56 | 0.51 | 0.56 | 0.65 | 0.85 |
| 9 | 83550453 | rs10867651 | -0.177 | 0.768 | 1.42 | 0.777 | -0.008 | 0.019 | 0.019 | 0 | 0.29 | 0.17 | 0.28 | 0.19 | 0.3 | 0.23 |
| 9 | 83553517 | rs10491944 | 0.183 | 0.772 | 1.378 | 0.788 | 0.012 | 0.021 | 0.021 | 0 | 0.1 | 0.03 | 0.06 | 0.03 | 0.09 | 0.06 |
| 9 | 83559923 | rs7859443 | 0.261 | 0.774 | 1.42 | 0.814 | -0.019 | -0.02 | -0.02 | 0 | 0.88 | 0.8 | 0.91 | 0.91 | 0.91 | 0.94 |
| 9 | 83562465 | rs10867654 | 0.967 | 0.762 | 1.412 | 0.792 | 0.014 | 0.057 | 0.057 | 0.019 | 0.72 | 0.51 | 0.5 | 0.56 | 0.65 | 0.82 |
| 9 | 83567044 | rs11138939 | 1.761 | 0.766 | 1.138 | 0.696 | 0.008 | 0.01 | 0.01 | 0 | 0.42 | 0.31 | 0.23 | 0.38 | 0.36 | 0.6 |
| 9 | 83579776 | rs727433 | 0.492 | 1.019 | 0.66 | 0.612 | -0.075 | 0.003 | 0.003 | 0 | 0.77 | 0.92 | 0.67 | 0.78 | 0.66 | 0.82 |
| 9 | 83583372 | rs9314697 | -0.236 | 1.013 | 0.663 | 0.62 | -0.005 | -0.021 | -0.021 | 0 | 0.9 | 0.93 | 0.83 | 0.81 | 0.88 | 0.87 |
| 9 | 83587426 | rs17245550 | 0.114 | 0.931 | 0.681 | 0.576 | 0.015 | 0.021 | 0.021 | 0 | 0.11 | 0.05 | 0.06 | 0.03 | 0.09 | 0.06 |
| 9 | 83591485 | rs10512111 | -0.326 | 0.928 | 0.897 | 0.545 | -0.042 | 0.068 | 0.068 | 0.081 | 0.28 | 0.13 | 0.23 | 0.07 | 0.36 | 0.15 |
| 9 | 83603221 | rs12344028 | -1.33 | 0.87 | 1.306 | 0.769 | -0.03 | 0.139 | 0.139 | 0 | 0.2 | 0.5 | 0.37 | 0.37 | 0.18 | 0.14 |
| 9 | 83603802 | rs1984064 | 0.692 | 0.926 | 1.232 | 0.743 | -0.069 | 0.065 | 0.065 | 0.07 | 0.13 | 0.01 | 0.15 | 0.03 | 0.22 | 0.06 |
| 9 | 83604324 | rs11138958 | 2.198 | 0.879 | 1.263 | 0.758 | 0.01 | 0.007 | 0.007 | 0 | 0.42 | 0.3 | 0.23 | 0.38 | 0.35 | 0.57 |
| 9 | 83615318 | rs2378554 | 0.804 | 0.927 | 1.171 | 0.779 | -0.059 | 0.06 | 0.06 | 0.208 | 0.62 | 0.82 | 0.6 | 0.75 | 0.53 | 0.72 |
| 9 | 83615409 | rs884065 | -0.173 | 0.902 | 1.25 | 0.848 | 0.018 | 0.173 | 0.173 | 0 | 0.07 | 0.2 | 0.3 | 0.34 | 0.1 | 0.09 |
| 9 | 83637462 | rs7038882 | 0.841 | 0.85 | 1.132 | 0.739 | -0.001 | 0.006 | 0.006 | 0 | 0.15 | 0.1 | 0.07 | 0.03 | 0.13 | 0.1 |
| 9 | 83637927 | rs1332854 | 0.773 | 0.893 | 1.177 | 0.706 | -0.005 | -0.028 | -0.028 | 2.13 | 0.64 | 0.56 | 0.66 | 0.77 | 0.66 | 0.75 |
| 9 | 83643130 | rs10867678 | 0.483 | 0.876 | 1.123 | 0.681 | 0.004 | -0.064 | -0.064 | 1.598 | 0.17 | 0.34 | 0.09 | 0.03 | 0.13 | 0.1 |
| 9 | 83643410 | rs1556447 | 2.371 | 0.943 | 1.083 | 0.724 | 0.014 | 0.007 | 0.007 | 0 | 0.41 | 0.23 | 0.22 | 0.38 | 0.35 | 0.57 |
| 9 | 83643525 | rs1572003 | 1.602 | 0.933 | 1.108 | 0.747 | 0.018 | 0.018 | 0.018 | 0 | 0.54 | 0.27 | 0.38 | 0.41 | 0.58 | 0.62 |

| | | | | | | | | | | | | | | | | |
|---|---|---|---|---|---|---|---|---|---|---|---|---|---|---|---|---|
| 9 | 83651543 | rs4877200 | 1.622 | 0.939 | 1.122 | 0.751 | 0.012 | 0.012 | 0.012 | 0 | 0.55 | 0.3 | 0.38 | 0.41 | 0.58 | 0.63 |
| 9 | 83653077 | rs12004880 | 0.007 | 0.964 | 1.148 | 0.863 | 0.009 | 0.153 | 0.153 | 0 | 0.07 | 0.16 | 0.29 | 0.33 | 0.08 | 0.08 |
| 9 | 83655465 | rs1332865 | -0.353 | 0.957 | 0.894 | 0.838 | -0.002 | -0.016 | -0.016 | 0 | 0.9 | 0.91 | 0.83 | 0.8 | 0.88 | 0.87 |
| 9 | 83661308 | rs2151225 | -1.1 | 1.14 | 1.069 | 0.656 | 0.115 | 0.214 | 0.214 | 0 | 0.12 | 0.41 | 0.39 | 0.36 | 0.22 | 0.12 |
| 9 | 83669782 | rs7863653 | 0.029 | 1.127 | 1.002 | 0.606 | 0.003 | -0.01 | -0.01 | 0 | 0.2 | 0.19 | 0.16 | 0.06 | 0.27 | 0.14 |
| 9 | 83670423 | rs1186145 | -1.628 | 1.027 | 0.786 | 0.823 | -0.027 | 0.042 | 0.042 | 0 | 0.28 | 0.49 | 0.45 | 0.35 | 0.26 | 0.16 |
| 9 | 83675496 | rs7022470 | -1.818 | 1.038 | 0.725 | 0.843 | 0.043 | 0.095 | 0.095 | 0 | 0.24 | 0.53 | 0.45 | 0.39 | 0.31 | 0.18 |
| 9 | 83686226 | rs1186462 | -0.773 | 1.096 | -0.09 | 0.523 | 0.018 | 0.078 | 0.078 | 0.114 | 0.19 | 0.04 | 0.14 | 0.03 | 0.18 | 0.08 |
| 9 | 83691262 | rs1537342 | 0.702 | 1.489 | 0.001 | 0.852 | -0.025 | -0.012 | -0.012 | 1.237 | 0.87 | 0.91 | 0.73 | 0.78 | 0.74 | 0.85 |
| 9 | 83691628 | rs1332879 | 0.696 | 1.481 | 0.014 | 0.848 | -0.022 | -0.011 | -0.011 | 0.109 | 0.87 | 0.91 | 0.73 | 0.77 | 0.74 | 0.85 |
| 9 | 83693384 | rs2779622 | -0.171 | 1.501 | -0.018 | 0.87 | -0.008 | -0.03 | -0.03 | 0 | 0.92 | 0.98 | 0.84 | 0.84 | 0.89 | 0.89 |
| 9 | 83693947 | rs7047671 | -0.854 | 1.41 | 0.004 | 0.702 | 0 | -0.007 | -0.007 | 0.013 | 0.16 | 0.22 | 0.14 | 0.03 | 0.17 | 0.1 |
| 9 | 83695176 | rs2225495 | 1.455 | 1.445 | 0.069 | 0.75 | -0.017 | 0.012 | 0.012 | 0 | 0.7 | 0.69 | 0.59 | 0.74 | 0.57 | 0.75 |
| 9 | 83695647 | rs2779625 | -0.272 | 1.297 | 0.138 | 1.024 | -0.019 | 0.098 | 0.098 | 0 | 0.12 | 0.29 | 0.3 | 0.27 | 0.1 | 0.05 |
| 9 | 83699819 | rs2779627 | 0.677 | 1.553 | 0.341 | 1.253 | -0.007 | 0.073 | 0.073 | 0 | 0.81 | 0.7 | 0.57 | 0.56 | 0.8 | 0.85 |
| 9 | 83701040 | rs10867692 | -1.124 | 1.637 | -0.339 | 1.11 | 0 | 0.002 | 0.002 | 0 | 0.13 | 0.15 | 0.1 | 0.03 | 0.12 | 0.1 |
| 9 | 83701931 | rs7856684 | -1.151 | 1.858 | -0.212 | 1.257 | 0.023 | -0.012 | -0.012 | 0.185 | 0.14 | 0.23 | 0.18 | 0.1 | 0.24 | 0.13 |
| 9 | 83714890 | rs4390020 | -0.49 | 1.935 | -1.02 | 0.888 | -0.032 | -0.019 | -0.019 | 0.269 | 0.27 | 0.23 | 0.31 | 0.38 | 0.41 | 0.54 |
| 9 | 83719971 | rs4877203 | 1.185 | 2.23 | -0.29 | 0.658 | 0.042 | 0.086 | 0.086 | 0 | 0.78 | 0.58 | 0.79 | 0.63 | 0.72 | 0.5 |
| 9 | 83720918 | rs1929715 | 1.069 | 2.303 | -0.248 | 0.398 | 0.037 | -0.061 | -0.061 | 0.309 | 0.72 | 0.62 | 0.73 | 0.91 | 0.57 | 0.73 |
| 9 | 83721188 | rs7862233 | 0.148 | 2.228 | -0.279 | 0.461 | -0.008 | -0.008 | -0.008 | 0.81 | 0.12 | 0.08 | 0.14 | 0.34 | 0.15 | 0.41 |
| 9 | 83721515 | rs10867699 | 1.01 | 0.857 | -0.722 | -0.929 | 0.028 | 0.06 | 0.06 | 0 | 0.21 | 0.07 | 0.12 | 0.03 | 0.15 | 0.04 |
| 9 | 83726045 | rs3927559 | 0.424 | 0.897 | -0.331 | -0.801 | 0.005 | -0.041 | -0.041 | 0 | 0.86 | 0.65 | 0.91 | 0.96 | 0.85 | 0.89 |
| 9 | 83726176 | rs3927561 | 1.172 | 0.904 | -0.285 | -0.826 | 0.01 | -0.078 | -0.078 | 2.248 | 0.82 | 0.54 | 0.87 | 0.96 | 0.81 | 0.87 |
| 9 | 83727201 | rs11139034 | 0.238 | 0.776 | -0.221 | -0.597 | 0 | 0.038 | 0.038 | 0 | 0.07 | 0.22 | 0.03 | 0.09 | 0.05 | 0.04 |
| 9 | 83733322 | rs957692 | -0.636 | 0.771 | -0.468 | -0.859 | 0.029 | -0.077 | -0.077 | 0 | 0.28 | 0.45 | 0.26 | 0.09 | 0.37 | 0.25 |
| 9 | 83738285 | rs7026961 | -0.965 | 0.794 | -0.445 | -0.865 | 0.035 | -0.093 | -0.093 | 0.006 | 0.31 | 0.49 | 0.29 | 0.09 | 0.4 | 0.26 |
| 9 | 83751479 | rs7860605 | -0.589 | 0.699 | -0.436 | -0.84 | 0.037 | -0.082 | -0.082 | 0 | 0.27 | 0.46 | 0.27 | 0.09 | 0.36 | 0.25 |
| 9 | 83751955 | rs17084584 | -0.769 | 0.691 | -0.342 | -0.852 | 0.01 | -0.033 | -0.033 | 0.208 | 0.12 | 0.33 | 0.09 | 0.04 | 0.13 | 0.09 |
| 9 | 83752021 | rs7470833 | 0.922 | 0.755 | -0.642 | -0.831 | -0.077 | 0.041 | 0.041 | 0 | 0.14 | 0.01 | 0.17 | 0.06 | 0.23 | 0.16 |
| 9 | 83759957 | rs11790515 | 1.056 | 0.289 | -0.881 | -1.097 | 0.006 | 0.015 | 0.015 | 0 | 0.14 | 0.11 | 0.1 | 0.03 | 0.11 | 0.03 |
| 9 | 83763114 | rs1999415 | 0.975 | 0.198 | -0.745 | -1.149 | 0.002 | -0.004 | -0.004 | 1.151 | 0.16 | 0.14 | 0.1 | 0.03 | 0.11 | 0.03 |
| 9 | 83763800 | rs2809849 | -0.884 | -0.036 | -0.297 | -1.304 | -0.002 | -0.015 | -0.015 | 0.145 | 0.42 | 0.63 | 0.4 | 0.12 | 0.47 | 0.28 |
| 9 | 83774420 | rs2777673 | 0.673 | -1.352 | -0.86 | -0.759 | -0.024 | -0.006 | -0.006 | 0 | 0.42 | 0.21 | 0.38 | 0.47 | 0.52 | 0.61 |
| 9 | 83774885 | rs2777674 | -0.217 | -1.711 | -0.705 | -0.796 | 0.002 | 0.106 | 0.106 | 0 | 0.59 | 0.77 | 0.65 | 0.89 | 0.59 | 0.75 |
| 9 | 83776258 | rs2809845 | -0.929 | -1.993 | -0.603 | -0.782 | 0.011 | 0.017 | 0.017 | 0 | 0.8 | 0.87 | 0.85 | 0.94 | 0.86 | 0.94 |
| 9 | 83790397 | rs2777678 | 0.525 | -2.285 | -0.498 | -0.857 | 0.042 | -0.027 | -0.027 | 0.02 | 0.61 | 0.67 | 0.57 | 0.55 | 0.75 | 0.8 |
| 9 | 83796167 | rs10780493 | 1.141 | -2.096 | -0.81 | -0.873 | -0.061 | 0.027 | 0.027 | 0 | 0.36 | 0.15 | 0.22 | 0.27 | 0.43 | 0.6 |
| 9 | 83811684 | rs11139068 | -0.591 | -1.882 | 0.326 | -0.478 | -0.032 | 0.125 | 0.125 | 0 | 0.09 | 0.33 | 0.19 | 0.25 | 0.04 | 0.03 |
| 9 | 83822268 | rs10114161 | 0.371 | -1.919 | 0.192 | -0.567 | 0.012 | -0.005 | -0.005 | 0 | 0.27 | 0.55 | 0.39 | 0.31 | 0.31 | 0.22 |
| 9 | 83838856 | rs7869606 | -0.535 | -2.14 | -0.093 | -0.585 | 0.022 | 0.049 | 0.049 | 0 | 0.45 | 0.74 | 0.64 | 0.71 | 0.44 | 0.36 |
| 9 | 83848710 | rs4614076 | 0.352 | -2.166 | -0.091 | -0.486 | -0.01 | -0.018 | -0.018 | 2.384 | 0.31 | 0.51 | 0.39 | 0.31 | 0.31 | 0.22 |
| 9 | 83850716 | rs7867933 | 1.054 | -1.964 | -0.184 | -0.107 | 0.016 | 0.019 | 0.019 | 0 | 0.07 | 0.06 | 0.2 | 0.39 | 0.04 | 0.1 |
| 9 | 83858558 | rs4484778 | 0.573 | -2.003 | 0.174 | -0.454 | -0.008 | 0.009 | 0.009 | 0.194 | 0.4 | 0.33 | 0.25 | 0.29 | 0.45 | 0.62 |
| 9 | 83861324 | rs4526433 | 0.445 | -1.223 | -0.035 | -0.787 | 0 | -0.025 | -0.025 | 0 | 0.33 | 0.55 | 0.32 | 0.28 | 0.29 | 0.27 |
| 9 | 83865218 | rs12000290 | 1.215 | -1.248 | 0.045 | -0.805 | 0.005 | -0.062 | -0.062 | 0.044 | 0.28 | 0.29 | 0.15 | 0.03 | 0.27 | 0.24 |
| 9 | 83867125 | rs10125662 | 0.287 | -1.117 | 0.137 | -0.645 | -0.01 | 0.145 | 0.145 | 0 | 0.07 | 0.31 | 0.17 | 0.26 | 0.03 | 0.06 |
| 9 | 83868297 | rs4517197 | 0.938 | -1.139 | -0.24 | -0.654 | 0.001 | -0.056 | -0.056 | 0 | 0.32 | 0.3 | 0.18 | 0.03 | 0.3 | 0.25 |
| 9 | 83872854 | rs9987634 | 0.268 | -0.917 | -0.197 | -0.366 | -0.011 | -0.019 | -0.019 | 0 | 0.32 | 0.47 | 0.34 | 0.3 | 0.31 | 0.25 |
| 9 | 83873695 | rs12377896 | 0.517 | -0.862 | -0.581 | -0.14 | -0.007 | 0.027 | 0.027 | 0 | 0.06 | 0.06 | 0.15 | 0.25 | 0.03 | 0.06 |
| 9 | 83874230 | rs10867720 | -0.273 | -0.855 | -0.622 | -0.209 | -0.001 | 0.016 | 0.016 | 0.021 | 0.19 | 0.22 | 0.26 | 0.53 | 0.23 | 0.44 |
| 9 | 83882277 | rs11139105 | 1.162 | -0.614 | -1.195 | -0.405 | -0.076 | 0.047 | 0.047 | 0.797 | 0.24 | 0.01 | 0.15 | 0.11 | 0.28 | 0.15 |
| 9 | 83887652 | rs17271965 | 1.161 | -0.613 | -1.182 | -0.395 | -0.072 | 0.058 | 0.058 | 0.989 | 0.23 | 0.01 | 0.15 | 0.09 | 0.28 | 0.15 |
| 9 | 83890416 | rs6559617 | -1.484 | -0.841 | -1.245 | -0.438 | -0.009 | -0.109 | -0.109 | 0.022 | 0.76 | 0.96 | 0.63 | 0.64 | 0.81 | 0.83 |
| 9 | 83893170 | rs10780502 | -0.204 | -0.592 | -0.17 | -0.325 | 0.004 | 0.079 | 0.079 | 0 | 0.19 | 0.23 | 0.26 | 0.47 | 0.22 | 0.44 |
| 9 | 83898946 | rs4428744 | -0.374 | -0.783 | -0.088 | -0.83 | 0.015 | 0 | 0 | 0 | 0.42 | 0.62 | 0.4 | 0.48 | 0.42 | 0.49 |
| 9 | 83909232 | rs13298109 | 1.237 | -0.23 | -0.029 | -1.421 | 0.014 | -0.04 | -0.04 | 0 | 0.19 | 0.25 | 0.13 | 0.02 | 0.25 | 0.21 |
| 9 | 83928428 | rs4639587 | 0.782 | -0.439 | -0.485 | -1.53 | -0.01 | -0.009 | -0.009 | 1.104 | 0.31 | 0.3 | 0.22 | 0.02 | 0.38 | 0.27 |
| 9 | 83928740 | rs7869128 | 0.995 | -0.525 | -1.189 | -1.563 | -0.089 | 0.154 | 0.154 | 0 | 0.3 | 0.1 | 0.22 | 0.02 | 0.39 | 0.27 |
| 9 | 83933198 | rs10867734 | 0.612 | -0.322 | -1.364 | -1.538 | -0.056 | 0.05 | 0.05 | 0 | 0.15 | 0.01 | 0.08 | 0.06 | 0.18 | 0.14 |
| 9 | 83941281 | rs10119867 | -1.125 | -0.624 | -1.514 | -1.345 | -0.084 | -0.034 | -0.034 | 0 | 0.67 | 0.97 | 0.43 | 0.29 | 0.68 | 0.71 |
| 9 | 83950452 | rs1888172 | 0.855 | -0.728 | -1.162 | -1.459 | -0.018 | 0.053 | 0.053 | 0 | 0.37 | 0.24 | 0.26 | 0.17 | 0.44 | 0.54 |
| 9 | 83950550 | rs1330210 | -0.644 | -0.271 | -1.393 | -0.934 | -0.041 | 0.038 | 0.038 | 0 | 0.67 | 0.91 | 0.43 | 0.26 | 0.68 | 0.7 |
| 9 | 83959411 | rs10780510 | -0.872 | 0.217 | -1.082 | -0.769 | -0.042 | -0.018 | -0.018 | 0 | 0.71 | 0.91 | 0.44 | 0.35 | 0.69 | 0.75 |
| 9 | 83963524 | rs10867740 | -0.672 | -0.138 | -0.6 | -1.213 | -0.018 | -0.062 | -0.062 | 0 | 0.27 | 0.51 | 0.15 | 0.14 | 0.24 | 0.16 |
| 9 | 83974745 | rs4452888 | 1.113 | 0.336 | -1.096 | -1.414 | 0.012 | 0.23 | 0.23 | 0.17 | 0.36 | 0.07 | 0.19 | 0.03 | 0.34 | 0.29 |
| 9 | 83975947 | rs10867741 | -0.084 | 0.469 | -0.867 | -1.399 | -0.001 | 0.203 | 0.203 | 0 | 0.54 | 0.16 | 0.34 | 0.22 | 0.63 | 0.67 |
| 9 | 83977478 | rs11139170 | 1.215 | 0.51 | -0.916 | -1.382 | -0.045 | 0.015 | 0.015 | 0.007 | 0.08 | 0.01 | 0.07 | 0.03 | 0.16 | 0.1 |
| 9 | 83978906 | rs11792869 | 0.702 | 0.483 | -0.911 | -1.405 | -0.057 | 0.218 | 0.218 | 0.036 | 0.43 | 0.08 | 0.27 | 0.19 | 0.47 | 0.58 |

| | | | | | | | | | | | | | | | | |
|---|---|---|---|---|---|---|---|---|---|---|---|---|---|---|---|---|
| 9 | 83984974 | rs2183369 | -0.628 | 0.166 | -0.34 | -1.353 | 0.003 | 0.021 | 0.021 | 0 | 0.62 | 0.39 | 0.39 | 0.29 | 0.67 | 0.72 |
| 9 | 83985310 | rs7043176 | -0.015 | 0.211 | -0.349 | -1.372 | 0.025 | 0.127 | 0.127 | 0 | 0.55 | 0.25 | 0.35 | 0.26 | 0.65 | 0.71 |
| 9 | 83985424 | rs11139174 | 1.145 | 0.292 | -0.461 | -1.418 | -0.012 | 0.18 | 0.18 | 0.302 | 0.38 | 0.07 | 0.22 | 0.18 | 0.36 | 0.53 |
| 9 | 83986104 | rs11139175 | 0.585 | 0.128 | -0.242 | -1.247 | -0.007 | 0.01 | 0.01 | 0 | 0.1 | 0.1 | 0.07 | 0.03 | 0.16 | 0.09 |
| 9 | 83988392 | rs7874548 | -0.164 | 0.094 | 0.199 | -1.204 | 0.013 | 0.072 | 0.072 | 0 | 0.58 | 0.33 | 0.36 | 0.26 | 0.65 | 0.71 |
| 9 | 83993434 | rs11139182 | 0.675 | 0.048 | 0.081 | -1.388 | -0.029 | 0.215 | 0.215 | 0 | 0.46 | 0.13 | 0.37 | 0.18 | 0.47 | 0.54 |
| 9 | 83998047 | rs1329387 | -0.848 | 0.155 | 0.158 | -0.943 | -0.02 | 0.001 | 0.001 | 0 | 0.68 | 0.82 | 0.51 | 0.38 | 0.66 | 0.69 |
| 9 | 83998114 | rs12376965 | 0.204 | 0.18 | 0.079 | -0.851 | -0.038 | -0.047 | -0.047 | 0 | 0.09 | 0.02 | 0.07 | 0.17 | 0.16 | 0.12 |
| 9 | 84006993 | rs3906153 | -0.49 | 0.069 | 0.3 | -0.885 | -0.001 | 0.002 | 0.002 | 0 | 0.61 | 0.67 | 0.48 | 0.4 | 0.61 | 0.67 |
| 9 | 84007124 | rs3860892 | 0.687 | 0.098 | 0.282 | -1.032 | -0.013 | 0.114 | 0.114 | 0 | 0.44 | 0.29 | 0.37 | 0.18 | 0.49 | 0.57 |
| 9 | 84007235 | rs3860894 | 0.656 | 0.358 | -0.031 | -0.741 | 0.002 | -0.009 | -0.009 | 0.41 | 0.47 | 0.56 | 0.38 | 0.16 | 0.43 | 0.54 |
| 9 | 84007864 | rs7869726 | 0.069 | 0.508 | -0.795 | -0.473 | -0.032 | 0.031 | 0.031 | 0 | 0.58 | 0.82 | 0.42 | 0.38 | 0.47 | 0.59 |
| 9 | 84008633 | rs10780512 | -0.417 | 0.351 | -0.647 | -0.251 | -0.031 | -0.026 | -0.026 | 0.021 | 0.68 | 0.92 | 0.51 | 0.39 | 0.65 | 0.71 |
| 9 | 84009625 | rs3908631 | -0.205 | 0.177 | -0.595 | -0.235 | -0.012 | 0.014 | 0.014 | 0 | 0.62 | 0.86 | 0.49 | 0.38 | 0.62 | 0.67 |
| 9 | 84011111 | rs7866986 | -0.745 | 0.198 | -0.489 | 0.469 | -0.003 | -0.004 | -0.004 | 0.185 | 0.57 | 0.75 | 0.62 | 0.61 | 0.58 | 0.49 |
| 9 | 84012054 | rs10780513 | -1.549 | -0.041 | 0.292 | 0.377 | -0.012 | -0.051 | -0.051 | 2.675 | 0.24 | 0.55 | 0.17 | 0.15 | 0.22 | 0.13 |
| 9 | 84013508 | rs10867751 | -0.679 | -0.032 | 0.263 | 0.362 | -0.007 | 0.016 | 0.016 | 0 | 0.16 | 0.09 | 0.11 | 0.13 | 0.2 | 0.1 |
| 9 | 84023250 | rs11139218 | -0.314 | 0.246 | 0.767 | 0.712 | -0.016 | 0.004 | 0.004 | 0 | 0.07 | 0.04 | 0.07 | 0.05 | 0.12 | 0.08 |
| 9 | 84023759 | rs11139219 | 0.407 | 0.269 | 0.407 | 1.267 | -0.058 | -0.175 | -0.175 | 0 | 0.23 | 0.02 | 0.42 | 0.44 | 0.22 | 0.32 |
| 9 | 84029173 | rs12345913 | -1.402 | 0.345 | 1.593 | 1.128 | 0.002 | 0.041 | 0.041 | 0 | 0.06 | 0.19 | 0.03 | 0.09 | 0.03 | 0.02 |
| 9 | 84029999 | rs10118881 | -2.809 | 0.338 | 2.027 | 1.09 | 0.007 | -0.028 | -0.028 | 0 | 0.18 | 0.55 | 0.14 | 0.13 | 0.17 | 0.12 |
| 9 | 84030732 | rs12339226 | -1.343 | 0.31 | 1.906 | 1.104 | 0.003 | 0.043 | 0.043 | 0 | 0.06 | 0.19 | 0.03 | 0.09 | 0.03 | 0.02 |
| 9 | 84033464 | rs7028565 | -1.009 | 0.313 | 1.795 | 1.111 | -0.006 | 0.02 | 0.02 | 0 | 0.05 | 0.1 | 0.03 | 0.09 | 0.03 | 0.02 |
| 9 | 84034924 | rs7036348 | -2.793 | 0.291 | 1.785 | 1.067 | -0.003 | -0.031 | -0.031 | 0 | 0.19 | 0.56 | 0.14 | 0.13 | 0.17 | 0.12 |
| 9 | 84035574 | rs11794168 | -1.472 | 0.308 | 1.48 | 1.046 | 0.021 | -0.034 | -0.034 | 0.042 | 0.11 | 0.26 | 0.1 | 0.05 | 0.14 | 0.1 |
| 9 | 84053760 | rs6559627 | -2.19 | -0.108 | 1.381 | 1.551 | -0.034 | 0.106 | 0.106 | 0.047 | 0.15 | 0.43 | 0.13 | 0.23 | 0.1 | 0.1 |
| 9 | 84055480 | rs12344650 | -1.522 | -0.078 | 1.367 | 1.388 | -0.009 | -0.023 | -0.023 | 0 | 0.09 | 0.27 | 0.07 | 0.04 | 0.07 | 0.06 |
| 9 | 84059393 | rs12348817 | -2.257 | -0.131 | 1.332 | 1.545 | -0.028 | 0.069 | 0.069 | 0 | 0.14 | 0.3 | 0.12 | 0.2 | 0.1 | 0.1 |
| 9 | 84070398 | rs11139263 | -1.757 | -0.079 | 1.368 | 1.542 | -0.013 | 0.011 | 0.011 | 0 | 0.11 | 0.22 | 0.09 | 0.13 | 0.09 | 0.08 |
| 9 | 84070426 | rs1329383 | -1.242 | -0.009 | 0.106 | 1.411 | -0.079 | 0.056 | 0.056 | 1.586 | 0.67 | 0.99 | 0.74 | 0.79 | 0.67 | 0.77 |
| 9 | 84073624 | rs11139265 | 0.335 | -0.293 | 0.234 | 1.218 | -0.01 | 0.001 | 0.001 | 0 | 0.06 | 0 | 0.06 | 0.04 | 0.07 | 0.04 |
| 9 | 84088631 | rs663838 | -1.074 | 0.052 | 0.144 | 1.728 | -0.049 | -0.009 | -0.009 | 0 | 0.65 | 0.91 | 0.64 | 0.67 | 0.63 | 0.75 |
| 9 | 84090670 | rs7871047 | -1.661 | -0.15 | 0.944 | 1.428 | -0.029 | 0.034 | 0.034 | 0 | 0.19 | 0.45 | 0.15 | 0.21 | 0.14 | 0.13 |
| 9 | 84102451 | rs10123329 | -1.602 | -0.205 | 0.446 | 1.558 | -0.041 | 0.042 | 0.042 | 0.551 | 0.21 | 0.56 | 0.16 | 0.28 | 0.14 | 0.14 |
| 9 | 84105189 | rs10512118 | 0.039 | 0.102 | -0.425 | 1.222 | -0.008 | -0.03 | -0.03 | 0 | 0.06 | 0 | 0.06 | 0.11 | 0.07 | 0.05 |
| 9 | 84113944 | rs584846 | -0.862 | -0.079 | -0.648 | 0.99 | -0.013 | 0.08 | 0.08 | 1.366 | 0.55 | 0.77 | 0.4 | 0.3 | 0.57 | 0.56 |
| 9 | 84115380 | rs602405 | -1.294 | -0.187 | -1.141 | 1.621 | -0.126 | -0.036 | -0.036 | 0 | 0.63 | 0.97 | 0.63 | 0.71 | 0.61 | 0.79 |
| 9 | 84116843 | rs495519 | 0.388 | -0.261 | -0.478 | 0.946 | -0.01 | 0.025 | 0.025 | 0 | 0.37 | 0.33 | 0.29 | 0.18 | 0.45 | 0.43 |
| 9 | 84117747 | rs524783 | -0.877 | -0.289 | -0.383 | 0.926 | 0.003 | 0.007 | 0.007 | 0 | 0.26 | 0.38 | 0.22 | 0.21 | 0.3 | 0.25 |
| 9 | 84124991 | rs594548 | -1.696 | -0.162 | -0.822 | 0.887 | -0.119 | 0.125 | 0.125 | 2.082 | 0.65 | 0.95 | 0.72 | 0.85 | 0.63 | 0.74 |
| 9 | 84127773 | rs530177 | -1.132 | -0.179 | -0.231 | 0.808 | 0.002 | 0.009 | 0.009 | 0.704 | 0.55 | 0.65 | 0.43 | 0.33 | 0.61 | 0.53 |
| 9 | 84140380 | rs10115264 | -0.571 | -0.571 | 0.187 | 0.549 | -0.002 | -0.011 | -0.011 | 0.005 | 0.46 | 0.53 | 0.27 | 0.27 | 0.4 | 0.4 |
| 9 | 84141525 | rs1576733 | -0.157 | -0.663 | 0.228 | 0.502 | -0.002 | -0.019 | -0.019 | 0 | 0.39 | 0.42 | 0.22 | 0.15 | 0.34 | 0.36 |
| 9 | 84144714 | rs11139298 | 0.453 | -0.473 | 0.124 | 0.566 | -0.003 | 0.027 | 0.027 | 0 | 0.07 | 0.16 | 0.06 | 0.12 | 0.06 | 0.04 |
| 9 | 84146693 | rs12376717 | 0.761 | -0.652 | -0.558 | 0.411 | -0.024 | 0.087 | 0.087 | 0.08 | 0.29 | 0.11 | 0.27 | 0.12 | 0.32 | 0.31 |
| 9 | 84148456 | rs11139301 | 1.03 | -0.351 | -0.511 | 0.751 | -0.011 | -0.031 | -0.031 | 2.288 | 0.06 | 0 | 0.04 | 0.12 | 0.06 | 0.04 |
| 9 | 84160842 | rs4877214 | 0.112 | -0.293 | -0.914 | 1.25 | 0.022 | -0.041 | -0.041 | 0 | 0.09 | 0.02 | 0.2 | 0.23 | 0.04 | 0.22 |
| 9 | 84164493 | rs7043518 | -0.522 | -0.184 | -0.726 | 0.994 | 0.03 | 0.013 | 0.013 | 0.979 | 0.51 | 0.68 | 0.56 | 0.48 | 0.42 | 0.58 |
| 9 | 84165251 | rs815863 | 0.502 | -0.07 | -0.238 | 1.37 | 0.027 | 0.067 | 0.067 | 0 | 0.79 | 0.66 | 0.69 | 0.66 | 0.7 | 0.72 |
| 9 | 84170326 | rs815866 | 0.505 | -0.862 | -0.114 | 1.447 | -0.024 | 0.05 | 0.05 | 0 | 0.88 | 0.82 | 0.79 | 0.71 | 0.94 | 0.75 |
| 9 | 84170701 | rs815867 | 0.331 | -1.29 | -0.488 | 0.75 | 0.005 | 0.011 | 0.011 | 0 | 0.46 | 0.58 | 0.39 | 0.1 | 0.58 | 0.34 |
| 9 | 84173486 | rs7029222 | 0.348 | -1.125 | -0.578 | 0.954 | 0 | -0.002 | -0.002 | 0.923 | 0.35 | 0.33 | 0.35 | 0.48 | 0.3 | 0.38 |
| 9 | 84181238 | rs11139315 | 0.202 | -0.978 | -0.579 | 0.691 | -0.004 | 0.097 | 0.097 | 0 | 0.4 | 0.18 | 0.3 | 0.26 | 0.38 | 0.41 |
| 9 | 84182228 | rs944591 | 1.146 | -1.18 | -0.25 | 0.811 | 0.011 | -0.002 | -0.002 | 0 | 0.77 | 0.77 | 0.63 | 0.24 | 0.86 | 0.68 |
| 9 | 84182613 | rs7045345 | -0.703 | -1.244 | 0.316 | 0.767 | -0.033 | -0.001 | -0.001 | 0 | 0.16 | 0.25 | 0.09 | 0.13 | 0.08 | 0.06 |
| 9 | 84183227 | rs10867775 | 0.983 | -1.315 | 0.158 | 0.632 | -0.013 | 0.125 | 0.125 | 0 | 0.34 | 0.11 | 0.26 | 0.14 | 0.33 | 0.44 |
| 9 | 84183462 | rs10780519 | -1.209 | -1.096 | 0.266 | 0.742 | -0.004 | 0.007 | 0.007 | 0 | 0.36 | 0.55 | 0.37 | 0.51 | 0.35 | 0.26 |
| 9 | 84185097 | rs12377054 | 1.431 | -0.966 | 0.191 | 0.398 | -0.009 | -0.033 | -0.033 | 0 | 0.07 | 0.01 | 0.04 | 0.1 | 0.06 | 0.03 |
| 9 | 84185365 | rs10780521 | 0.956 | -0.966 | 0.232 | 0.398 | 0.006 | -0.041 | -0.041 | 0 | 0.36 | 0.2 | 0.34 | 0.47 | 0.33 | 0.47 |
| 9 | 84187273 | rs11139320 | 0.825 | -0.964 | 0.35 | 0.399 | 0.001 | -0.003 | -0.003 | 0.53 | 0.37 | 0.35 | 0.47 | 0.47 | 0.33 | 0.47 |
| 9 | 84191049 | rs10780522 | 0.738 | -0.965 | 0.22 | 0.4 | -0.007 | 0.007 | 0.007 | 0 | 0.38 | 0.42 | 0.35 | 0.46 | 0.33 | 0.46 |
| 9 | 84191267 | rs7027019 | -1.595 | -1.003 | 0.186 | 0.394 | -0.006 | -0.009 | -0.009 | 0 | 0.7 | 0.71 | 0.64 | 0.61 | 0.68 | 0.7 |
| 9 | 84191525 | rs11139324 | 1.387 | -0.86 | 0.033 | 0.41 | -0.012 | -0.033 | -0.033 | 0.489 | 0.08 | 0.01 | 0.04 | 0.1 | 0.06 | 0.03 |
| 9 | 84192036 | rs1412224 | 0.631 | -0.816 | 0.117 | 0.364 | 0.002 | -0.013 | -0.013 | 0 | 0.17 | 0.2 | 0.21 | 0.02 | 0.27 | 0.17 |
| 9 | 84193897 | rs17084936 | 0.693 | -0.835 | -0.001 | 0.417 | 0.001 | -0.003 | -0.003 | 0 | 0.38 | 0.36 | 0.35 | 0.46 | 0.33 | 0.46 |
| 9 | 84195189 | rs4877217 | 1.489 | -0.765 | -0.703 | 0.262 | -0.039 | -0.057 | -0.057 | 0.014 | 0.28 | 0.05 | 0.29 | 0.32 | 0.26 | 0.46 |
| 9 | 84196490 | rs10481754 | 0.36 | -0.632 | -0.246 | 0.554 | -0.001 | 0.015 | 0.015 | 1.713 | 0.48 | 0.55 | 0.48 | 0.33 | 0.52 | 0.62 |
| 9 | 84197169 | rs11139328 | 0.382 | 0.071 | -0.568 | 0.333 | -0.037 | -0.023 | -0.023 | 0 | 0.11 | 0.01 | 0.1 | 0.1 | 0.13 | 0.04 |
| 9 | 84202226 | rs2796462 | 0.957 | 0.36 | -0.286 | 0.002 | 0 | -0.052 | -0.052 | 0 | 0.25 | 0.25 | 0.14 | 0 | 0.26 | 0.07 |

| | | | | | | | | | | | | | | | | |
|---|---|---|---|---|---|---|---|---|---|---|---|---|---|---|---|---|
| 9 | 84204332 | rs2777772 | 1.14 | 0.302 | -0.838 | 0.13 | -0.06 | 0.113 | 0.113 | 0 | 0.23 | 0.06 | 0.14 | 0.01 | 0.25 | 0.07 |
| 9 | 84204374 | rs911639 | -0.176 | 0.288 | -0.696 | 0.663 | -0.003 | 0.165 | 0.165 | 0 | 0.89 | 0.73 | 0.87 | 0.56 | 0.92 | 0.75 |
| 9 | 84204779 | rs2796466 | 0.263 | 0.59 | -0.417 | -0.167 | 0.015 | -0.064 | -0.064 | 0.1 | 0.36 | 0.36 | 0.3 | 0.03 | 0.46 | 0.27 |
| 9 | 84207360 | rs6559629 | -0.073 | 0.069 | -0.944 | 0.353 | 0 | -0.001 | -0.001 | 0 | 0.45 | 0.48 | 0.5 | 0.62 | 0.5 | 0.72 |
| 9 | 84210148 | rs7860224 | -0.199 | -0.505 | -1.382 | 0.034 | 0.004 | 0 | 0 | 0 | 0.15 | 0.14 | 0.08 | 0.02 | 0.09 | 0.08 |
| 9 | 84210209 | rs7860431 | -0.283 | -0.246 | -1.689 | 0.31 | 0.103 | 0.079 | 0.079 | 0.144 | 0.48 | 0.12 | 0.34 | 0.33 | 0.38 | 0.43 |
| 9 | 84210838 | rs7036755 | -1.117 | 0.416 | -0.736 | 0.544 | 0.016 | 0.118 | 0.118 | 0.227 | 0.14 | 0.31 | 0.26 | 0.29 | 0.14 | 0.09 |
| 9 | 84211820 | rs864179 | -0.429 | 0.372 | -1.972 | -0.427 | 0.01 | -0.051 | -0.051 | 0 | 0.67 | 0.94 | 0.73 | 0.71 | 0.73 | 0.61 |
| 9 | 84212536 | rs1573078 | -0.175 | 0.608 | -1.357 | -1.19 | 0.007 | -0.011 | -0.011 | 0.483 | 0.5 | 0.47 | 0.34 | 0.06 | 0.49 | 0.37 |
| 9 | 84214974 | rs7039820 | 0.272 | 0.886 | -2.381 | -1.237 | 0.003 | 0.055 | 0.055 | 0 | 0.16 | 0.07 | 0.15 | 0.02 | 0.18 | 0.21 |
| 9 | 84216090 | rs815845 | -0.037 | 0.959 | -1.892 | -0.958 | -0.009 | 0.126 | 0.126 | 0 | 0.33 | 0.14 | 0.19 | 0.04 | 0.33 | 0.14 |
| 9 | 84233892 | rs10867781 | -0.671 | 1.302 | -1.279 | 0.055 | -0.023 | -0.013 | -0.013 | 0.43 | 0.21 | 0.12 | 0.29 | 0.26 | 0.29 | 0.27 |
| 9 | 84241442 | rs11139346 | -0.943 | 0.62 | -0.723 | 0.771 | -0.009 | 0.077 | 0.077 | 0 | 0.14 | 0.19 | 0.44 | 0.57 | 0.14 | 0.19 |
| 9 | 84244131 | rs11139349 | -0.295 | 0.865 | -1.243 | 0.427 | -0.038 | -0.091 | -0.091 | 0.159 | 0.21 | 0.02 | 0.15 | 0.28 | 0.25 | 0.28 |
| 9 | 84245004 | rs12379295 | -0.49 | 0.767 | -0.078 | -0.037 | 0.017 | -0.035 | -0.035 | 0 | 0.16 | 0.49 | 0.1 | 0.11 | 0.14 | 0.13 |
| 9 | 84247426 | rs1342478 | -0.308 | 1.261 | -0.42 | -0.497 | 0.011 | -0.084 | -0.084 | 0.89 | 0.75 | 0.69 | 0.83 | 0.99 | 0.73 | 0.92 |
| 9 | 84247453 | rs10780524 | -0.512 | 0.723 | -0.514 | 0.783 | 0 | 0.009 | 0.009 | 0 | 0.18 | 0.19 | 0.42 | 0.5 | 0.19 | 0.14 |
| 9 | 84249963 | rs10867784 | -1.014 | 0.916 | -0.483 | 0.428 | 0 | -0.002 | -0.002 | 0 | 0.11 | 0.11 | 0.09 | 0.07 | 0.1 | 0.1 |
| 9 | 84252359 | rs11139351 | -0.393 | 0.412 | -0.552 | 1.011 | 0.009 | -0.024 | -0.024 | 0 | 0.21 | 0.15 | 0.29 | 0.49 | 0.18 | 0.45 |
| 9 | 84252766 | rs3814532 | 0.109 | 0.711 | -0.563 | 0.277 | -0.013 | -0.005 | -0.005 | 0 | 0.64 | 0.64 | 0.64 | 0.61 | 0.54 | 0.74 |
| 9 | 84254819 | rs10867787 | 0.489 | 0.982 | -1.065 | 0.299 | -0.048 | -0.03 | -0.03 | 0 | 0.11 | 0.01 | 0.14 | 0.15 | 0.2 | 0.11 |
| 9 | 84256984 | rs12551040 | 0.142 | 0.84 | -0.732 | 0.259 | 0.005 | -0.008 | -0.008 | 0 | 0.23 | 0.19 | 0.3 | 0.49 | 0.18 | 0.46 |
| 9 | 84261124 | rs11999426 | 1.524 | 0.709 | -0.062 | 0.204 | 0.002 | 0.064 | 0.064 | 1.389 | 0.42 | 0.33 | 0.35 | 0.18 | 0.36 | 0.33 |
| 9 | 84262485 | rs4877669 | -1.133 | 0.973 | -0.116 | 0.264 | -0.002 | -0.007 | -0.007 | 1.271 | 0.44 | 0.43 | 0.51 | 0.69 | 0.48 | 0.59 |
| 9 | 84262687 | rs10867791 | -0.394 | 1.405 | -0.087 | 0.214 | 0.013 | -0.008 | -0.008 | 0 | 0.21 | 0.26 | 0.22 | 0.21 | 0.29 | 0.13 |
| 9 | 84270400 | rs2796449 | -0.058 | 1.492 | -0.548 | 0.367 | -0.003 | -0.007 | -0.007 | 0 | 0.89 | 0.9 | 0.92 | 0.89 | 0.88 | 0.93 |
| 9 | 84275014 | rs2627002 | -0.132 | 1.529 | -0.699 | 0.385 | -0.009 | -0.017 | -0.017 | 0.443 | 0.89 | 0.92 | 0.92 | 0.89 | 0.88 | 0.91 |
| 9 | 84281312 | rs10119841 | 0.65 | 1.81 | -0.775 | 0.302 | -0.041 | 0.011 | 0.011 | 0.044 | 0.8 | 0.89 | 0.81 | 0.85 | 0.67 | 0.81 |
| 9 | 84288353 | rs2591 | 0.417 | 1.258 | -1.104 | 0.246 | -0.06 | 0.012 | 0.012 | 0.232 | 0.82 | 0.98 | 0.89 | 0.91 | 0.79 | 0.84 |
| 9 | 84289295 | rs3739581 | -0.181 | 0.621 | -1.595 | -0.074 | 0.063 | 0.043 | 0.043 | 0.113 | 0.39 | 0.76 | 0.23 | 0.23 | 0.24 | 0.17 |
| 9 | 84297696 | rs10867795 | 0.277 | 0.901 | -1.544 | 0.898 | -0.001 | -0.028 | -0.028 | 0 | 0.1 | 0.05 | 0.24 | 0.49 | 0.11 | 0.23 |
| 9 | 84304985 | rs3814535 | -0.476 | 1.864 | -0.593 | 0.581 | 0.108 | 0.04 | 0.04 | 0.957 | 0.14 | 0.32 | 0.23 | 0.23 | 0.25 | 0.16 |
| 9 | 84305732 | rs1484113 | 0.361 | 1.83 | -1.192 | 0.173 | -0.036 | 0.048 | 0.048 | 0.05 | 0.22 | 0.14 | 0.24 | 0.1 | 0.3 | 0.24 |
| 9 | 84308948 | rs2796441 | -0.063 | 1.898 | -0.992 | 0.168 | -0.05 | -0.064 | -0.064 | 1.889 | 0.33 | 0.15 | 0.49 | 0.59 | 0.42 | 0.45 |
| 9 | 84314437 | rs7018614 | 0.159 | 1.761 | -0.559 | 0.025 | 0 | 0 | 0 | 0 | 0.14 | 0.15 | 0.17 | 0.2 | 0.16 | 0.17 |
| 9 | 84323545 | rs11787711 | 0.551 | 1.649 | -0.021 | 0.121 | -0.014 | -0.023 | -0.023 | 0.296 | 0.13 | 0.07 | 0.17 | 0.25 | 0.17 | 0.16 |
| 9 | 84323900 | rs4877223 | 0.184 | 1.446 | -0.169 | 0.031 | -0.005 | -0.008 | -0.008 | 0 | 0.1 | 0.04 | 0.16 | 0.08 | 0.11 | 0.18 |
| 9 | 84327210 | rs2627013 | -0.516 | 1.447 | -0.051 | -0.032 | -0.027 | 0.021 | 0.021 | 0.005 | 0.46 | 0.28 | 0.57 | 0.61 | 0.54 | 0.59 |
| 9 | 84344542 | rs3860897 | -0.612 | 1.571 | -0.243 | 0.125 | -0.055 | -0.088 | -0.088 | 1.814 | 0.69 | 0.86 | 0.62 | 0.51 | 0.62 | 0.62 |
| 9 | 84348762 | rs4434691 | 0.747 | 1.395 | 0.02 | 0.14 | 0.017 | 0.021 | 0.021 | 0.377 | 0.51 | 0.32 | 0.4 | 0.4 | 0.44 | 0.39 |
| 9 | 84352443 | rs10512121 | 0.127 | 1.619 | -0.118 | 0.402 | -0.029 | -0.041 | -0.041 | 0 | 0.33 | 0.17 | 0.38 | 0.49 | 0.38 | 0.39 |
| 9 | 84353330 | rs11139391 | 0.104 | 1.631 | -0.062 | 0.395 | -0.022 | -0.031 | -0.031 | 0 | 0.33 | 0.21 | 0.38 | 0.49 | 0.38 | 0.39 |
| 9 | 84358072 | rs7041897 | 1.425 | 1.099 | 0.035 | 0.282 | 0.025 | 0.022 | 0.022 | 0.085 | 0.3 | 0.23 | 0.18 | 0.13 | 0.21 | 0.2 |
| 9 | 84365210 | rs12683003 | 0.459 | 1.191 | -0.292 | 0.272 | -0.001 | -0.002 | -0.002 | 0.089 | 0.14 | 0.13 | 0.08 | 0.05 | 0.13 | 0.08 |
| 9 | 84369450 | rs7864030 | 0.031 | 1.403 | -0.323 | 0.409 | -0.001 | -0.001 | -0.001 | 0 | 0.6 | 0.65 | 0.45 | 0.48 | 0.54 | 0.56 |
| 9 | 84370124 | rs13287994 | 0.508 | 1.38 | -0.442 | 0.416 | -0.001 | -0.012 | -0.012 | 0 | 0.14 | 0.14 | 0.09 | 0.05 | 0.13 | 0.08 |
| 9 | 84371927 | rs1007995 | 0.425 | 1.389 | -0.442 | 0.436 | 0.023 | 0.013 | 0.013 | 0 | 0.5 | 0.65 | 0.41 | 0.42 | 0.43 | 0.36 |
| 9 | 84373209 | rs11139400 | 0.825 | 0.924 | -0.867 | 0.055 | 0.021 | 0.045 | 0.045 | 0.96 | 0.4 | 0.24 | 0.26 | 0.18 | 0.34 | 0.27 |
| 9 | 84373428 | rs12377016 | 0.293 | 1.06 | -0.484 | 0.502 | -0.007 | 0.034 | 0.034 | 0.022 | 0.53 | 0.31 | 0.32 | 0.24 | 0.46 | 0.48 |
| 9 | 84386712 | rs1010745 | 0.313 | 1.2 | -0.465 | 0.873 | -0.012 | -0.009 | -0.009 | 0 | 0.59 | 0.7 | 0.44 | 0.5 | 0.52 | 0.57 |
| 9 | 84389168 | rs17085042 | -0.73 | 1.176 | -1.158 | 1.203 | -0.043 | -0.077 | -0.077 | 0 | 0.14 | 0.05 | 0.21 | 0.37 | 0.19 | 0.24 |
| 9 | 84389731 | rs7847362 | -0.419 | 0.138 | -0.8 | 0.279 | -0.023 | -0.038 | -0.038 | 0 | 0.19 | 0.27 | 0.1 | 0.03 | 0.14 | 0.18 |
| 9 | 84389914 | rs10867812 | -0.235 | 0.805 | -0.951 | 0.612 | 0.003 | 0.01 | 0.01 | 0 | 0.28 | 0.3 | 0.47 | 0.61 | 0.34 | 0.38 |
| 9 | 84391174 | rs10081631 | -0.261 | 1.041 | -1.19 | 0.519 | 0.001 | 0.016 | 0.016 | 0 | 0.26 | 0.31 | 0.25 | 0.45 | 0.24 | 0.35 |
| 9 | 84392044 | rs637736 | -0.088 | 1.072 | -1.718 | -0.128 | 0 | 0.002 | 0.002 | 0 | 0.65 | 0.41 | 0.57 | 0.51 | 0.58 | 0.67 |
| 9 | 84399614 | rs676113 | -0.135 | 0.78 | -2.039 | -0.99 | -0.012 | 0.042 | 0.042 | 0 | 0.81 | 0.86 | 0.86 | 0.96 | 0.8 | 0.89 |
| 9 | 84402241 | rs564171 | 0.653 | 0.755 | -2.13 | -0.742 | 0.03 | 0.111 | 0.111 | 0.049 | 0.29 | 0.05 | 0.24 | 0.07 | 0.21 | 0.24 |
| 9 | 84410212 | rs7030006 | -0.111 | 1.24 | -1.217 | -0.062 | -0.009 | 0.007 | 0.007 | 0 | 0.29 | 0.27 | 0.48 | 0.83 | 0.39 | 0.5 |
| 9 | 84414429 | rs630561 | -0.93 | 1.094 | -0.515 | 0.5 | 0.03 | 0.049 | 0.049 | 0 | 0.06 | 0.17 | 0.1 | 0.1 | 0.08 | 0.07 |
| 9 | 84419758 | rs669223 | 0.629 | 1.354 | -0.549 | 0.31 | 0.022 | -0.033 | -0.033 | 0 | 0.8 | 0.7 | 0.82 | 0.91 | 0.75 | 0.8 |
| 9 | 84431758 | rs7048179 | -1.446 | 0.697 | -0.878 | 1.072 | 0.007 | -0.016 | -0.016 | 2.14 | 0.2 | 0.76 | 0.19 | 0.41 | 0.17 | 0.23 |
| 9 | 84433130 | rs1889528 | 0.419 | 0.763 | -1.422 | 0.615 | -0.052 | 0.067 | 0.067 | 0 | 0.41 | 0.2 | 0.49 | 0.32 | 0.5 | 0.32 |
| 9 | 84441027 | rs10121323 | -0.679 | 0.527 | -0.233 | 0.62 | 0.008 | 0.166 | 0.166 | 0 | 0.16 | 0.35 | 0.18 | 0.43 | 0.16 | 0.33 |
| 9 | 84445414 | rs11139438 | 0.118 | 0.608 | -0.662 | -1.042 | 0.002 | 0.018 | 0.018 | 0.818 | 0.37 | 0.32 | 0.26 | 0.08 | 0.34 | 0.28 |
| 9 | 84451844 | rs2031757 | -0.826 | 0.488 | -0.962 | -1.154 | -0.038 | 0.018 | 0.018 | 0 | 0.72 | 0.86 | 0.84 | 0.85 | 0.7 | 0.74 |
| 9 | 84454027 | rs10867818 | 0.022 | 0.49 | -0.711 | -1.332 | 0.005 | 0.013 | 0.013 | 0 | 0.35 | 0.3 | 0.34 | 0.09 | 0.32 | 0.27 |
| 9 | 84456361 | rs10780536 | -0.009 | 0.604 | -0.736 | -1.224 | -0.001 | -0.031 | -0.031 | 0.74 | 0.42 | 0.37 | 0.4 | 0.1 | 0.45 | 0.35 |
| 9 | 84457989 | rs478787 | -0.316 | 0.519 | -0.948 | -1.34 | 0.022 | 0.112 | 0.112 | 0 | 0.52 | 0.64 | 0.38 | 0.17 | 0.55 | 0.48 |
| 9 | 84458091 | rs7867888 | 0.013 | 0.366 | -0.863 | -1.411 | -0.029 | -0.105 | -0.105 | 0 | 0.19 | 0.3 | 0.12 | 0 | 0.15 | 0.1 |

| | | | | | | | | | | | | | | | | |
|---|---|---|---|---|---|---|---|---|---|---|---|---|---|---|---|---|
| 9 | 84458774 | rs11139442 | 0.321 | 0.582 | -1.566 | -1.129 | 0.001 | 0.003 | 0.003 | 0 | 0.47 | 0.57 | 0.54 | 0.81 | 0.45 | 0.58 |
| 9 | 84459072 | rs478381 | 0.292 | 1.032 | -2.038 | -0.832 | -0.05 | 0.065 | 0.065 | 0 | 0.86 | 0.98 | 0.88 | 0.97 | 0.86 | 0.96 |
| 9 | 84460636 | rs1854158 | -0.05 | 1.187 | -2.217 | -0.623 | -0.061 | 0.03 | 0.03 | 0 | 0.73 | 0.91 | 0.63 | 0.8 | 0.69 | 0.71 |
| 9 | 84460855 | rs10867821 | 0.182 | 0.475 | -2.255 | -0.263 | 0.028 | -0.048 | -0.048 | 0.06 | 0.3 | 0.18 | 0.14 | 0.59 | 0.23 | 0.46 |
| 9 | 84461318 | rs17370182 | -0.613 | 1.027 | -1.723 | -0.215 | -0.019 | -0.024 | -0.024 | 0 | 0.05 | 0 | 0.05 | 0.08 | 0.08 | 0.06 |
| 9 | 84462943 | rs4999155 | -1.035 | 1.287 | -1.559 | -0.139 | -0.001 | 0.003 | 0.003 | 0 | 0.06 | 0.04 | 0.24 | 0.14 | 0.12 | 0.08 |
| 9 | 84473474 | rs7024973 | -0.687 | 0.682 | -1.325 | -0.675 | 0.025 | 0.059 | 0.059 | 0.179 | 0.4 | 0.68 | 0.45 | 0.19 | 0.29 | 0.23 |
| 9 | 84481989 | rs7037280 | -1.001 | 0.412 | -0.455 | -0.31 | -0.016 | -0.012 | -0.012 | 0 | 0.31 | 0.57 | 0.28 | 0.26 | 0.2 | 0.18 |
| 9 | 84495227 | rs2378592 | -0.723 | 0.659 | -0.006 | -0.359 | 0.003 | 0.008 | 0.008 | 10.549 | 0.27 | 0.2 | 0.26 | 0.18 | 0.24 | 0.14 |
| 9 | 84519154 | rs12351812 | -0.796 | 0.709 | 0.702 | -0.235 | 0.007 | -0.016 | -0.016 | 0 | 0.05 | 0.1 | 0.05 | 0.01 | 0.08 | 0.01 |
| 9 | 84528161 | rs10217389 | -1.989 | 0.686 | 0.978 | -0.272 | 0.004 | -0.038 | -0.038 | 0 | 0.09 | 0.23 | 0.06 | 0.01 | 0.1 | 0.03 |
| 9 | 84565837 | rs6559647 | -2.755 | 0.721 | 1.181 | -0.345 | 0.015 | -0.064 | -0.064 | 0 | 0.12 | 0.33 | 0.09 | 0.02 | 0.13 | 0.03 |
| 9 | 84589488 | rs11139464 | -2.247 | 0.746 | 1.515 | -0.656 | 0.004 | -0.035 | -0.035 | 0.02 | 0.33 | 0.69 | 0.2 | 0.03 | 0.27 | 0.14 |
| 9 | 84589869 | rs11139465 | -2.811 | 0.746 | 1.689 | -0.654 | 0.031 | -0.079 | -0.079 | 0.258 | 0.13 | 0.48 | 0.1 | 0.02 | 0.13 | 0.04 |
| 9 | 84600972 | rs4877228 | -2.728 | 0.864 | 1.523 | -0.031 | 0.038 | -0.001 | -0.001 | 0 | 0.45 | 0.78 | 0.39 | 0.58 | 0.41 | 0.35 |
| 9 | 84605293 | rs10118455 | -0.314 | 0.763 | 1.11 | -0.102 | 0.009 | -0.014 | -0.014 | 3.371 | 0.05 | 0.11 | 0.05 | 0.01 | 0.09 | 0.01 |
| 9 | 84618966 | rs10867827 | -2.621 | 0.753 | 1.103 | -0.067 | 0.026 | -0.069 | -0.069 | 0.592 | 0.13 | 0.47 | 0.1 | 0.03 | 0.13 | 0.03 |
| 9 | 84642933 | rs12344007 | -0.488 | 0.752 | 1.259 | -0.11 | 0.011 | -0.006 | -0.006 | 0 | 0.2 | 0.18 | 0.1 | 0.01 | 0.14 | 0.1 |
| 9 | 84672279 | rs12342760 | -2.412 | 0.604 | 1.075 | -0.011 | 0.01 | -0.063 | -0.063 | 0 | 0.12 | 0.31 | 0.08 | 0.02 | 0.13 | 0.03 |
| 9 | 84686091 | rs7867431 | -2.433 | 0.604 | 1.041 | -0.011 | 0.009 | -0.064 | -0.064 | 0 | 0.13 | 0.33 | 0.08 | 0.02 | 0.13 | 0.03 |
| 9 | 84736303 | rs1319017 | -1.785 | 0.58 | 0.645 | 0.315 | 0.043 | 0.034 | 0.034 | 0.458 | 0.42 | 0.87 | 0.38 | 0.68 | 0.35 | 0.36 |
| 9 | 84743408 | rs12338960 | -2.055 | 0.531 | 0.602 | 0.328 | 0.005 | 0.085 | 0.085 | 2.287 | 0.47 | 0.9 | 0.42 | 0.69 | 0.45 | 0.38 |
| 9 | 84763025 | rs11139500 | -1.895 | 0.173 | 0.574 | 0.324 | 0.046 | 0.033 | 0.033 | 0.144 | 0.43 | 0.87 | 0.38 | 0.67 | 0.36 | 0.33 |
| 9 | 84765612 | rs4877686 | -1.282 | 0.011 | 0.828 | 0.266 | -0.004 | 0.03 | 0.03 | 0.111 | 0.29 | 0.48 | 0.3 | 0.65 | 0.28 | 0.29 |
| 9 | 84789243 | rs2767719 | -1.087 | 0.187 | 1.086 | 0.403 | 0 | 0.026 | 0.026 | 0 | 0.27 | 0.39 | 0.28 | 0.66 | 0.26 | 0.29 |
| 9 | 84828677 | rs2209725 | -2.628 | 0.231 | 0.949 | 0.511 | -0.008 | 0.042 | 0.042 | 4.674 | 0.49 | 0.98 | 0.44 | 0.67 | 0.44 | 0.35 |
| 9 | 84843011 | rs2788099 | -2.381 | 0.179 | 1.112 | 0.563 | 0.064 | -0.065 | -0.065 | 0.031 | 0.43 | 0.88 | 0.38 | 0.53 | 0.34 | 0.34 |
| 9 | 84894926 | rs1933577 | 1.396 | 0.164 | 1.297 | 0.59 | 0.008 | -0.029 | -0.029 | 0 | 0.7 | 0.29 | 0.7 | 0.48 | 0.75 | 0.71 |
| 9 | 84929102 | rs11794308 | 0.909 | 0.183 | 1.31 | 0.61 | -0.005 | 0.095 | 0.095 | 0 | 0.73 | 0.49 | 0.72 | 0.5 | 0.76 | 0.73 |
| 9 | 84935895 | rs7859493 | 0.741 | 0.183 | 1.268 | 0.6 | -0.001 | 0.114 | 0.114 | 0 | 0.74 | 0.6 | 0.72 | 0.5 | 0.76 | 0.73 |
| 9 | 84950712 | rs10512124 | -0.02 | 0.266 | 0.675 | 0.602 | 0.003 | 0.063 | 0.063 | 2.554 | 0.11 | 0.01 | 0.06 | 0.01 | 0.09 | 0.02 |
| 9 | 84951226 | rs1933580 | 1.798 | 0.269 | 0.75 | 0.66 | 0.032 | -0.055 | -0.055 | 0 | 0.61 | 0.21 | 0.63 | 0.49 | 0.67 | 0.71 |
| 9 | 84982792 | rs6559659 | -2.09 | 0.332 | 0.465 | 0.688 | 0.045 | -0.06 | -0.06 | 0 | 0.41 | 0.81 | 0.38 | 0.53 | 0.34 | 0.32 |
| 9 | 85027979 | rs10123160 | -1.139 | 0.504 | 1.203 | 0.836 | 0.004 | 0.104 | 0.104 | 0 | 0.24 | 0.44 | 0.35 | 0.59 | 0.24 | 0.25 |
| 9 | 85036598 | rs12551702 | -0.926 | 0.555 | 1.214 | 0.928 | 0.006 | 0.071 | 0.071 | 0 | 0.22 | 0.3 | 0.33 | 0.52 | 0.24 | 0.24 |
| 9 | 85072253 | rs12684844 | -0.644 | 0.164 | 0.652 | 1.039 | -0.031 | -0.1 | -0.1 | 0.289 | 0.18 | 0.07 | 0.3 | 0.53 | 0.21 | 0.22 |
| 9 | 85080395 | rs12685656 | -0.926 | 0.144 | 0.675 | 1.028 | 0.001 | 0.029 | 0.029 | 0 | 0.21 | 0.27 | 0.32 | 0.53 | 0.22 | 0.25 |
| 9 | 85088990 | rs7046558 | -1.62 | 0.052 | 0.568 | 0.961 | -0.007 | 0.049 | 0.049 | 0.495 | 0.29 | 0.36 | 0.36 | 0.54 | 0.26 | 0.28 |
| 9 | 85097651 | rs1411661 | -1.729 | 0.049 | 0.542 | 0.992 | -0.011 | 0.06 | 0.06 | 0.553 | 0.3 | 0.41 | 0.36 | 0.55 | 0.27 | 0.28 |
| 9 | 85101768 | rs11139579 | -1.641 | -0.024 | 0.432 | 0.967 | -0.016 | 0.002 | 0.002 | 3.216 | 0.13 | 0.69 | 0.11 | 0.09 | 0.11 | 0.04 |
| 9 | 85106906 | rs1411664 | -2.486 | -0.034 | -0.231 | 0.981 | 0.01 | -0.021 | -0.021 | 0 | 0.41 | 0.91 | 0.46 | 0.63 | 0.37 | 0.32 |
| 9 | 85107147 | rs7875416 | -1.644 | 0.262 | 0.062 | 1.245 | -0.01 | 0.023 | 0.023 | 2.963 | 0.13 | 0.61 | 0.1 | 0.09 | 0.11 | 0.04 |
| 9 | 85117420 | rs955975 | -0.82 | 0.581 | -1.221 | 1.163 | -0.033 | -0.096 | -0.096 | 0 | 0.2 | 0.06 | 0.29 | 0.58 | 0.22 | 0.25 |
| 9 | 85128341 | rs928223 | -1.315 | 0.583 | -0.97 | 1.167 | -0.002 | 0.006 | 0.006 | 0 | 0.09 | 0.15 | 0.07 | 0.12 | 0.11 | 0.04 |
| 9 | 85134671 | rs1330828 | -1.11 | 0.717 | -1.401 | 1.051 | 0.004 | -0.045 | -0.045 | 0 | 0.35 | 0.86 | 0.41 | 0.6 | 0.35 | 0.33 |
| 9 | 85139211 | rs10780547 | -0.031 | 0.749 | -1.868 | 0.716 | -0.079 | -0.018 | -0.018 | 0 | 0.66 | 0.91 | 0.41 | 0.67 | 0.63 | 0.68 |
| 9 | 85147648 | rs6559670 | -0.302 | 0.239 | -1.484 | 0.35 | -0.022 | 0.048 | 0.048 | 0 | 0.26 | 0.62 | 0.14 | 0.58 | 0.2 | 0.4 |
| 9 | 85147773 | rs11139601 | 0.81 | 0.206 | -1.416 | 0.266 | 0.001 | 0.116 | 0.116 | 0 | 0.39 | 0.23 | 0.26 | 0.06 | 0.41 | 0.28 |
| 9 | 85150039 | rs2378640 | -0.533 | 0.161 | -1.061 | 0.421 | -0.004 | 0.004 | 0.004 | 0 | 0.27 | 0.69 | 0.14 | 0.63 | 0.21 | 0.41 |
| 9 | 85161570 | rs11139606 | -0.029 | 0.415 | -1.272 | 0.322 | 0 | -0.009 | -0.009 | 0 | 0.07 | 0.06 | 0.07 | 0.02 | 0.07 | 0.01 |
| 9 | 85163682 | rs10867852 | 0.15 | 0.297 | -1.018 | 1.088 | -0.001 | 0.056 | 0.056 | 0.146 | 0.07 | 0.08 | 0.05 | 0.37 | 0.07 | 0.27 |
| 9 | 85166081 | rs11139607 | -0.155 | 0.059 | 0.091 | 0.979 | -0.022 | 0.083 | 0.083 | 0 | 0.15 | 0.36 | 0.07 | 0.25 | 0.13 | 0.14 |
| 9 | 85167648 | rs10780548 | 0.462 | 0.503 | -0.375 | 0.858 | -0.114 | -0.027 | -0.027 | 0 | 0.55 | 0.04 | 0.29 | 0.6 | 0.54 | 0.57 |
| 9 | 85168377 | rs1009037 | 0.115 | 0.508 | 0.711 | 0.623 | -0.002 | 0.001 | 0.001 | 0 | 0.65 | 0.65 | 0.4 | 0.27 | 0.63 | 0.49 |
| 9 | 85168848 | rs7035264 | -0.707 | 0.834 | 0.745 | 0.416 | -0.004 | -0.014 | -0.014 | 1.035 | 0.11 | 0.09 | 0.08 | 0 | 0.15 | 0.07 |
| 9 | 85175417 | rs1933015 | 1.276 | 0.436 | 1.134 | 0.436 | 0.031 | 0.043 | 0.043 | 0 | 0.43 | 0.23 | 0.38 | 0.29 | 0.4 | 0.31 |
| 9 | 85182996 | rs1933016 | 0.656 | 0.507 | 1.451 | 0.735 | 0.026 | 0.097 | 0.097 | 0 | 0.52 | 0.34 | 0.42 | 0.75 | 0.48 | 0.63 |
| 9 | 85190433 | rs7019337 | 0.119 | 0.591 | 1.742 | 0.709 | -0.001 | -0.004 | -0.004 | 0 | 0.58 | 0.62 | 0.47 | 0.8 | 0.52 | 0.69 |
| 9 | 85197241 | rs11139631 | -0.029 | 0.358 | 1.668 | 0.912 | 0.002 | 0.051 | 0.051 | 0 | 0.08 | 0.09 | 0.05 | 0.46 | 0.07 | 0.33 |
| 9 | 85200045 | rs11139635 | -1.308 | 0.334 | 1.623 | 0.886 | -0.013 | 0.221 | 0.221 | 0 | 0.15 | 0.39 | 0.09 | 0.5 | 0.11 | 0.4 |
| 9 | 85209810 | rs7028728 | -0.485 | 0.482 | 1.509 | 0.745 | 0.009 | -0.036 | -0.036 | 3.484 | 0.11 | 0.13 | 0.09 | 0 | 0.14 | 0.06 |
| 9 | 85210881 | rs17085618 | 0.612 | 0.473 | 1.378 | 0.889 | 0.003 | 0.042 | 0.042 | 0 | 0.06 | 0.02 | 0.03 | 0.47 | 0.05 | 0.31 |
| 9 | 85216270 | rs11139649 | -0.791 | 0.435 | 1.737 | 0.879 | -0.003 | 0.201 | 0.201 | 0 | 0.12 | 0.33 | 0.07 | 0.5 | 0.09 | 0.36 |
| 9 | 85227767 | rs10512130 | -0.536 | 0.44 | 1.755 | 0.823 | 0.023 | -0.056 | -0.056 | 0 | 0.12 | 0.15 | 0.09 | 0 | 0.14 | 0.07 |
| 9 | 85228687 | rs17085644 | 0.626 | 0.43 | 1.742 | 0.993 | 0.005 | 0.042 | 0.042 | 0 | 0.06 | 0.03 | 0.03 | 0.41 | 0.05 | 0.3 |
| 9 | 85237252 | rs7873931 | 0.046 | 0.404 | 1.582 | 0.885 | -0.031 | 0.069 | 0.069 | 0 | 0.6 | 0.82 | 0.49 | 0.8 | 0.53 | 0.7 |
| 9 | 85256142 | rs7031058 | -1.856 | 0.422 | 1.391 | 0.871 | -0.001 | 0 | 0 | 0 | 0.27 | 0.7 | 0.19 | 0.51 | 0.24 | 0.46 |
| 9 | 85267179 | rs1537302 | 1.58 | 0.148 | 1.23 | 0.672 | 0.005 | 0.004 | 0.004 | 0 | 0.36 | 0.23 | 0.35 | 0.3 | 0.36 | 0.3 |
| 9 | 85272505 | rs12057032 | 0.208 | 0.197 | 1.194 | 0.715 | 0.019 | -0.046 | -0.046 | 0.002 | 0.15 | 0.08 | 0.07 | 0.4 | 0.12 | 0.34 |

| | | | | | | | | | | | | | | | | |
|---|---|---|---|---|---|---|---|---|---|---|---|---|---|---|---|---|
| 9 | 85282258 | rs17085684 | 0.198 | 0.24 | 1.251 | 0.645 | 0.012 | -0.019 | -0.019 | 0 | 0.15 | 0.11 | 0.07 | 0.42 | 0.12 | 0.34 |
| 9 | 85288781 | rs10867868 | 1.141 | 0.726 | 1.841 | 0.413 | 0.028 | 0.109 | 0.109 | 0 | 0.5 | 0.28 | 0.42 | 0.72 | 0.48 | 0.64 |
| 9 | 85290284 | rs7875440 | -0.103 | 0.758 | 2.039 | 0.392 | 0.001 | -0.018 | -0.018 | 0 | 0.67 | 0.58 | 0.54 | 0.81 | 0.66 | 0.79 |
| 9 | 85296585 | rs1571905 | -1.561 | 0.781 | 2.055 | 0.381 | 0.045 | -0.067 | -0.067 | 0.361 | 0.19 | 0.36 | 0.13 | 0.09 | 0.18 | 0.15 |
| 9 | 85297244 | rs7025180 | -0.66 | 0.777 | 2.066 | 0.371 | 0.077 | -0.034 | -0.034 | 0 | 0.08 | 0.26 | 0.07 | 0 | 0.11 | 0.06 |
| 9 | 85303653 | rs11139689 | -1.684 | 0.777 | 2.042 | 0.372 | 0.037 | -0.062 | -0.062 | 0 | 0.18 | 0.35 | 0.13 | 0.09 | 0.18 | 0.15 |
| 9 | 85311147 | rs668853 | 0.437 | 0.76 | 1.858 | 0.396 | 0 | -0.005 | -0.005 | 0 | 0.58 | 0.41 | 0.48 | 0.72 | 0.58 | 0.7 |
| 9 | 85321895 | rs493701 | -1.157 | 0.691 | 0.918 | 0.456 | 0.023 | 0.081 | 0.081 | 0 | 0.5 | 0.69 | 0.57 | 0.28 | 0.52 | 0.36 |
| 9 | 85326428 | rs509091 | -1.166 | 0.764 | 0.364 | 0.434 | 0.035 | 0.143 | 0.143 | 0 | 0.5 | 0.8 | 0.58 | 0.27 | 0.52 | 0.35 |
| 9 | 85329887 | rs12236312 | -0.272 | 0.699 | 0.287 | 0.523 | -0.011 | -0.05 | -0.05 | 0.8 | 0.12 | 0.03 | 0.06 | 0.24 | 0.09 | 0.11 |
| 9 | 85332687 | rs581729 | -1.253 | 0.709 | 0.248 | 0.553 | 0.031 | 0.014 | 0.014 | 0 | 0.49 | 0.74 | 0.57 | 0.43 | 0.52 | 0.34 |
| 9 | 85339429 | rs11139710 | -0.207 | 0.996 | 0.09 | 0.595 | 0.039 | -0.039 | -0.039 | 0 | 0.07 | 0.17 | 0.07 | 0 | 0.11 | 0.06 |
| 9 | 85347613 | rs609444 | -0.236 | 0.962 | -0.474 | 0.656 | 0.015 | -0.038 | -0.038 | 0 | 0.37 | 0.57 | 0.53 | 0.2 | 0.44 | 0.24 |
| 9 | 85353762 | rs11139720 | -0.199 | 0.965 | -0.507 | 0.64 | 0.018 | -0.043 | -0.043 | 0.417 | 0.37 | 0.56 | 0.52 | 0.2 | 0.44 | 0.24 |
| 9 | 85359511 | rs7043569 | -0.986 | 0.998 | -1.868 | 0.859 | -0.021 | -0.063 | -0.063 | 0 | 0.45 | 0.92 | 0.38 | 0.44 | 0.42 | 0.31 |
| 9 | 85365505 | rs11139732 | -0.466 | 1.459 | -0.934 | 1.587 | -0.011 | -0.06 | -0.06 | 0 | 0.16 | 0.05 | 0.18 | 0.42 | 0.21 | 0.31 |
| 9 | 85374993 | rs11139754 | -0.978 | 1.291 | -0.064 | 1.526 | -0.002 | 0.02 | 0.02 | 0 | 0.21 | 0.31 | 0.21 | 0.69 | 0.23 | 0.37 |
| 9 | 85390604 | rs11139766 | -0.942 | 1.288 | -0.021 | 1.11 | 0.004 | 0.062 | 0.062 | 0.668 | 0.23 | 0.35 | 0.25 | 0.43 | 0.28 | 0.34 |
| 9 | 85391724 | rs7859706 | -0.866 | 1.879 | -0.086 | 0.898 | 0.051 | -0.037 | -0.037 | 0 | 0.07 | 0.19 | 0.09 | 0 | 0.13 | 0.05 |
| 9 | 85391790 | rs7875501 | -1.676 | 1.867 | -0.045 | 1.023 | 0.072 | 0.177 | 0.177 | 0.812 | 0.09 | 0.28 | 0.11 | 0.27 | 0.16 | 0.1 |
| 9 | 85394979 | rs1570228 | -1.772 | 1.863 | -0.293 | 0.693 | 0.003 | 0.003 | 0.003 | 0.502 | 0.32 | 0.36 | 0.36 | 0.45 | 0.45 | 0.36 |
| 9 | 85395554 | rs11139775 | -1.237 | 1.825 | -0.4 | 0.715 | -0.04 | -0.054 | -0.054 | 0 | 0.13 | 0.05 | 0.12 | 0.27 | 0.19 | 0.1 |
| 9 | 85397699 | rs11788842 | -0.066 | 1.882 | -0.356 | 0.591 | 0.001 | -0.005 | -0.005 | 0 | 0.05 | 0.04 | 0.08 | 0 | 0.13 | 0.04 |
| 9 | 85400266 | rs4877709 | 0.028 | 1.86 | -0.399 | 0.667 | -0.025 | -0.013 | -0.013 | 0 | 0.17 | 0.05 | 0.21 | 0.21 | 0.24 | 0.26 |
| 9 | 85406002 | rs963727 | -2.337 | 1.456 | -0.412 | 0.261 | -0.01 | 0.187 | 0.187 | 0.481 | 0.56 | 0.89 | 0.56 | 0.85 | 0.63 | 0.7 |
| 9 | 85409526 | rs1998304 | 0.106 | 1.73 | -0.27 | 0.368 | -0.018 | 0.004 | 0.004 | 0 | 0.17 | 0.1 | 0.22 | 0.18 | 0.25 | 0.26 |
| 9 | 85410090 | rs10867884 | -2.286 | 1.248 | -0.12 | 0.203 | 0.008 | 0.113 | 0.113 | 0 | 0.57 | 0.78 | 0.56 | 0.85 | 0.63 | 0.68 |
| 9 | 85412760 | rs35433085 | 0.463 | 1.688 | -0.423 | 0.596 | -0.034 | -0.042 | -0.042 | 0 | 0.12 | 0.01 | 0.17 | 0.18 | 0.17 | 0.17 |
| 9 | 85415537 | rs1040870 | -0.623 | 1.473 | -0.269 | 0.854 | 0.017 | 0.033 | 0.033 | 0.098 | 0.26 | 0.35 | 0.33 | 0.5 | 0.41 | 0.35 |
| 9 | 85417267 | rs1005796 | -0.636 | 1.478 | -0.339 | 0.862 | 0.026 | 0.052 | 0.052 | 0.194 | 0.26 | 0.38 | 0.34 | 0.51 | 0.41 | 0.35 |
| 9 | 85421474 | rs12342232 | -0.48 | 1.012 | -0.674 | 1.425 | -0.007 | -0.076 | -0.076 | 0 | 0.17 | 0.04 | 0.15 | 0.34 | 0.18 | 0.32 |
| 9 | 85428983 | rs1885348 | -0.708 | 1.202 | 0.006 | -0.272 | 0.006 | -0.046 | -0.046 | 0 | 0.34 | 0.4 | 0.36 | 0.12 | 0.38 | 0.28 |
| 9 | 85430251 | rs10491545 | -0.212 | 1.297 | 0.112 | -0.327 | -0.006 | 0.068 | 0.068 | 0 | 0.28 | 0.15 | 0.31 | 0.06 | 0.3 | 0.24 |
| 9 | 85430415 | rs10491544 | -0.596 | 1.335 | 0.143 | -0.306 | 0.002 | -0.023 | -0.023 | 0 | 0.33 | 0.36 | 0.37 | 0.12 | 0.38 | 0.28 |
| 9 | 85430642 | rs2378645 | -0.335 | 1.352 | 0.146 | -0.307 | -0.001 | -0.026 | -0.026 | 1.699 | 0.31 | 0.32 | 0.34 | 0.06 | 0.35 | 0.25 |
| 9 | 85432032 | rs987371 | -0.595 | 1.145 | 0.221 | -0.3 | -0.002 | -0.054 | -0.054 | 0.971 | 0.33 | 0.43 | 0.33 | 0.12 | 0.33 | 0.27 |
| 9 | 85436498 | rs6559685 | -1.172 | 0.869 | -0.375 | -0.404 | 0.004 | 0.033 | 0.033 | 0 | 0.62 | 0.76 | 0.72 | 0.71 | 0.63 | 0.62 |
| 9 | 85437593 | rs7874804 | -0.166 | 0.707 | -0.215 | 0.268 | 0.004 | 0.029 | 0.029 | 0 | 0.3 | 0.44 | 0.39 | 0.59 | 0.29 | 0.34 |
| 9 | 85443579 | rs10867893 | -0.402 | -0.168 | -0.569 | -0.383 | 0.007 | 0.001 | 0.001 | 0 | 0.25 | 0.2 | 0.19 | 0.07 | 0.15 | 0.23 |
| 9 | 85444907 | rs2208619 | -0.219 | -0.257 | -0.563 | -0.433 | 0.019 | 0.02 | 0.02 | 0 | 0.2 | 0.12 | 0.15 | 0.07 | 0.09 | 0.22 |
| 9 | 85451313 | rs11139822 | -0.234 | -0.257 | -0.561 | -0.433 | 0.024 | 0.026 | 0.026 | 0 | 0.2 | 0.12 | 0.15 | 0.07 | 0.09 | 0.22 |
| 9 | 85451416 | rs10867903 | -0.411 | -0.771 | -0.337 | -0.063 | -0.011 | 0.077 | 0.077 | 0 | 0.1 | 0.16 | 0.09 | 0.67 | 0.05 | 0.33 |
| 9 | 85452097 | rs12349496 | 0.017 | -0.704 | -0.4 | -0.151 | 0.024 | 0.026 | 0.026 | 0 | 0.2 | 0.12 | 0.15 | 0.07 | 0.09 | 0.22 |
| 9 | 85453867 | rs13440407 | -0.589 | -0.674 | -0.341 | -0.143 | -0.003 | -0.011 | -0.011 | 0 | 0.07 | 0.11 | 0.05 | 0.03 | 0.06 | 0.02 |
| 9 | 85458060 | rs2224524 | 0.993 | -0.448 | 0.044 | -1.421 | -0.075 | 0.067 | 0.067 | 0.333 | 0.32 | 0.15 | 0.45 | 0.16 | 0.44 | 0.25 |
| 9 | 85459107 | rs2208622 | 0.371 | -0.155 | -0.223 | -2.174 | -0.013 | 0 | 0 | 0 | 0.22 | 0.13 | 0.43 | 0.25 | 0.34 | 0.26 |
| 9 | 85460807 | rs10867906 | -0.555 | -0.368 | 0.005 | -2.282 | 0 | -0.001 | -0.001 | 3.321 | 0.71 | 0.7 | 0.79 | 0.32 | 0.74 | 0.58 |
| 9 | 85478265 | rs12002556 | 0.979 | -0.094 | -0.302 | -2.209 | -0.001 | -0.001 | -0.001 | 0 | 0.15 | 0.12 | 0.16 | 0.09 | 0.2 | 0.12 |
| 9 | 85478756 | rs12684912 | 0.994 | -0.149 | -0.315 | -2.319 | 0.017 | -0.058 | -0.058 | 0.438 | 0.35 | 0.47 | 0.43 | 0.16 | 0.35 | 0.25 |
| 9 | 85481263 | rs12683498 | 0.449 | -0.136 | -0.341 | -2.275 | -0.003 | -0.003 | -0.003 | 0 | 0.21 | 0.13 | 0.39 | 0.25 | 0.25 | 0.25 |
| 9 | 85489244 | rs9987635 | 0.828 | -0.197 | -0.2 | -2.042 | 0.013 | -0.036 | -0.036 | 0 | 0.37 | 0.52 | 0.44 | 0.23 | 0.36 | 0.26 |
| 9 | 85490188 | rs10867909 | -0.385 | -0.251 | -0.253 | -2.089 | -0.002 | -0.002 | -0.002 | 0 | 0.71 | 0.7 | 0.75 | 0.36 | 0.65 | 0.57 |
| 9 | 85490520 | rs11139849 | 0.347 | -0.253 | -0.245 | -2.088 | 0.021 | 0.045 | 0.045 | 0.385 | 0.35 | 0.21 | 0.32 | 0.13 | 0.29 | 0.31 |
| 9 | 85492701 | rs12235637 | -0.044 | -0.256 | -0.28 | -2.119 | 0 | 0.013 | 0.013 | 0.111 | 0.16 | 0.19 | 0.17 | 0.64 | 0.21 | 0.35 |
| 9 | 85493025 | rs10867910 | -0.232 | -0.259 | -0.271 | -2.105 | -0.002 | -0.001 | -0.001 | 0 | 0.7 | 0.69 | 0.75 | 0.31 | 0.64 | 0.56 |
| 9 | 85495552 | rs4357374 | 0.696 | -0.125 | -0.271 | -1.996 | 0 | -0.004 | -0.004 | 0 | 0.38 | 0.63 | 0.44 | 0.24 | 0.36 | 0.31 |
| 9 | 85495968 | rs7875448 | 1.038 | -0.102 | -0.148 | -2.024 | 0.018 | -0.047 | -0.047 | 0 | 0.34 | 0.47 | 0.42 | 0.18 | 0.35 | 0.3 |
| 9 | 85506296 | rs7038698 | -0.893 | -0.306 | -0.687 | -2.154 | 0.046 | 0.116 | 0.116 | 0 | 0.63 | 0.83 | 0.75 | 0.96 | 0.75 | 0.78 |
| 9 | 85509355 | rs10867914 | -0.806 | 0.235 | -0.198 | -1.849 | -0.014 | -0.003 | -0.003 | 0.084 | 0.22 | 0.19 | 0.29 | 0.78 | 0.36 | 0.44 |
| 9 | 85511305 | rs10746692 | -1.606 | 0.257 | -0.374 | -1.912 | 0.001 | -0.006 | -0.006 | 0.235 | 0.84 | 0.88 | 0.9 | 1 | 0.85 | 0.92 |
| 9 | 85511504 | rs11139863 | -0.349 | 0.339 | -0.164 | -1.066 | 0.014 | 0.009 | 0.009 | 0 | 0.12 | 0.14 | 0.16 | 0.19 | 0.2 | 0.18 |
| 9 | 85521271 | rs7026976 | 0.433 | 0.569 | -0.535 | -1.232 | -0.05 | 0.021 | 0.021 | 0 | 0.26 | 0.14 | 0.23 | 0.2 | 0.42 | 0.23 |
| 9 | 85524530 | rs4554557 | -0.603 | 0.026 | 0.088 | -0.579 | 0.004 | 0.009 | 0.009 | 0 | 0.15 | 0.18 | 0.15 | 0.2 | 0.18 | 0.19 |
| 9 | 85524569 | rs4604527 | -0.998 | -0.437 | 0.003 | -0.82 | 0.039 | 0.038 | 0.038 | 0 | 0.45 | 0.65 | 0.39 | 0.29 | 0.34 | 0.44 |
| 9 | 85527035 | rs11139868 | -1.341 | 0.037 | 0.303 | -0.471 | -0.003 | 0.015 | 0.015 | 0.341 | 0.25 | 0.56 | 0.22 | 0.26 | 0.24 | 0.33 |
| 9 | 85527950 | rs3897456 | -1.059 | 0.01 | -0.615 | 0.554 | -0.054 | -0.115 | -0.115 | 0 | 0.84 | 0.97 | 0.82 | 0.51 | 0.76 | 0.75 |
| 9 | 85540229 | rs12340476 | -1.173 | -0.458 | 0.554 | 0.077 | -0.009 | 0.021 | 0.021 | 0 | 0.21 | 0.3 | 0.17 | 0.27 | 0.18 | 0.31 |
| 9 | 85547017 | rs7860062 | -1.415 | -0.716 | 0.369 | -0.256 | 0.065 | 0.083 | 0.083 | 0 | 0.47 | 0.79 | 0.4 | 0.32 | 0.36 | 0.5 |
| 9 | 85552173 | rs17745151 | 0.085 | -0.312 | 0.166 | -0.033 | 0.013 | 0.019 | 0.019 | 0 | 0.08 | 0.09 | 0.14 | 0.19 | 0.16 | 0.15 |

| | | | | | | | | | | | | | | | | |
|---|---|---|---|---|---|---|---|---|---|---|---|---|---|---|---|---|
| 9 | 85557505 | rs7032152 | -1.356 | -0.466 | 0.443 | -0.104 | 0.014 | -0.009 | -0.009 | 0.523 | 0.22 | 0.43 | 0.2 | 0.2 | 0.24 | 0.25 |
| 9 | 85559944 | rs7029717 | 0.501 | 0.079 | 0.074 | 0.003 | 0.045 | 0.066 | 0.066 | 0 | 0.19 | 0.06 | 0.17 | 0.05 | 0.11 | 0.12 |
| 9 | 85570577 | rs11139881 | -0.088 | 0.188 | 0.14 | -0.023 | 0.035 | -0.021 | -0.021 | 0 | 0.06 | 0.22 | 0.05 | 0.02 | 0.07 | 0.06 |
| 9 | 85574053 | rs10867918 | -0.755 | 0.307 | -0.045 | 0.099 | -0.009 | -0.018 | -0.018 | 0 | 0.37 | 0.52 | 0.37 | 0.27 | 0.34 | 0.35 |
| 9 | 85577007 | rs10780559 | 0.036 | -0.336 | -0.971 | -0.436 | 0.01 | -0.001 | -0.001 | 2.659 | 0.27 | 0.22 | 0.22 | 0.05 | 0.18 | 0.17 |
| 9 | 85579528 | rs4620356 | -0.52 | 0.272 | -1.425 | 0.221 | -0.042 | 0.013 | 0.013 | 1.858 | 0.58 | 0.92 | 0.65 | 0.35 | 0.56 | 0.55 |
| 9 | 85590972 | rs2999948 | -0.046 | 0.121 | -0.943 | -0.217 | -0.017 | 0.109 | 0.109 | 0.151 | 0.45 | 0.27 | 0.44 | 0.19 | 0.45 | 0.37 |
| 9 | 85596321 | rs10735559 | -0.648 | 0.244 | -0.557 | -0.351 | -0.038 | 0.107 | 0.107 | 0.416 | 0.58 | 0.79 | 0.54 | 0.84 | 0.55 | 0.68 |
| 9 | 85601796 | rs4385545 | 0.157 | 0.01 | 0.598 | -0.335 | 0.005 | 0.072 | 0.072 | 0 | 0.46 | 0.32 | 0.49 | 0.78 | 0.5 | 0.56 |
| 9 | 85601818 | rs4355855 | -0.62 | 0.026 | 1.418 | -0.336 | -0.003 | 0.001 | 0.001 | 0 | 0.55 | 0.5 | 0.53 | 0.84 | 0.55 | 0.68 |
| 9 | 85608152 | rs2279814 | 0.039 | -0.031 | 1.627 | 0.079 | 0 | -0.001 | -0.001 | 0 | 0.48 | 0.41 | 0.45 | 0.73 | 0.51 | 0.55 |
| 9 | 85609781 | rs10867921 | -0.005 | -0.084 | 1.854 | 0.073 | 0.001 | -0.009 | -0.009 | 0 | 0.06 | 0.12 | 0.04 | 0.03 | 0.07 | 0.05 |
| 9 | 85613043 | rs6559694 | -0.226 | -0.127 | 2.049 | 0.028 | 0.006 | 0.049 | 0.049 | 0 | 0.5 | 0.57 | 0.47 | 0.79 | 0.52 | 0.62 |
| 9 | 85623560 | rs4348583 | 0.014 | -0.121 | 2.011 | 0.064 | -0.001 | -0.003 | -0.003 | 0 | 0.47 | 0.52 | 0.46 | 0.73 | 0.51 | 0.53 |
| 9 | 85627408 | rs4146960 | 0.047 | -0.122 | 2.011 | 0.063 | -0.001 | -0.004 | -0.004 | 0 | 0.47 | 0.51 | 0.46 | 0.73 | 0.51 | 0.53 |
| 9 | 85639259 | rs7019227 | -0.088 | -0.128 | 2.168 | 0.09 | 0.006 | 0.066 | 0.066 | 0 | 0.48 | 0.63 | 0.48 | 0.73 | 0.51 | 0.53 |
| 9 | 85643603 | rs10867929 | 0.114 | -0.142 | 1.946 | 0.766 | -0.001 | -0.005 | -0.005 | 0 | 0.46 | 0.43 | 0.44 | 0.63 | 0.51 | 0.52 |
| 9 | 85650916 | rs1125915 | -0.753 | -0.02 | 1.762 | 0.855 | -0.058 | 0.045 | 0.045 | 0.78 | 0.57 | 0.81 | 0.5 | 0.69 | 0.56 | 0.67 |
| 9 | 85663674 | rs10116216 | -0.201 | -0.132 | 2.048 | 0.793 | 0 | 0.002 | 0.002 | 0 | 0.49 | 0.44 | 0.45 | 0.62 | 0.53 | 0.57 |
| 9 | 85680418 | rs1995423 | -0.797 | -0.092 | 1.521 | 0.734 | -0.071 | 0.156 | 0.156 | 0.962 | 0.57 | 0.83 | 0.52 | 0.79 | 0.56 | 0.68 |
| 9 | 85698493 | rs7024275 | -0.536 | -0.014 | 1.502 | 0.721 | 0.013 | 0.059 | 0.059 | 0.121 | 0.48 | 0.34 | 0.43 | 0.79 | 0.5 | 0.68 |
| 9 | 85702264 | rs1502687 | -0.537 | -0.009 | 1.517 | 0.72 | 0.009 | 0.044 | 0.044 | 0.415 | 0.48 | 0.37 | 0.44 | 0.79 | 0.5 | 0.68 |
| 9 | 85704334 | rs7855133 | -0.43 | 0.14 | 1.473 | 0.789 | 0.012 | 0.006 | 0.006 | 0.159 | 0.45 | 0.34 | 0.39 | 0.61 | 0.43 | 0.56 |
| 9 | 85704575 | rs11536913 | 0.816 | -0.09 | 0.525 | 1.285 | -0.015 | -0.082 | -0.082 | 0.101 | 0.15 | 0.04 | 0.23 | 0.42 | 0.15 | 0.24 |
| 9 | 85715039 | rs7020876 | 0.309 | 0.096 | 0.972 | 0.944 | 0.029 | -0.053 | -0.053 | 0 | 0.32 | 0.18 | 0.31 | 0.57 | 0.3 | 0.44 |
| 9 | 85715943 | rs6559697 | -0.16 | 0.212 | 1.248 | 1.025 | 0.028 | -0.003 | -0.003 | 0 | 0.41 | 0.29 | 0.37 | 0.6 | 0.39 | 0.54 |
| 9 | 85719699 | rs1995424 | -0.756 | 0.176 | 1.752 | 1 | 0.038 | 0.065 | 0.065 | 2.401 | 0.49 | 0.69 | 0.43 | 0.68 | 0.45 | 0.65 |
| 9 | 85721071 | rs4311721 | -0.252 | 0.222 | 1.847 | 0.603 | 0.008 | 0.01 | 0.01 | 0.704 | 0.43 | 0.39 | 0.39 | 0.7 | 0.38 | 0.55 |
| 9 | 85721119 | rs2378646 | -0.129 | 0.235 | 1.609 | 0.61 | 0.035 | 0.046 | 0.046 | 0.605 | 0.41 | 0.28 | 0.38 | 0.7 | 0.38 | 0.55 |
| 9 | 85725775 | rs11795101 | -0.687 | 0.353 | 1.199 | 0.659 | 0.011 | 0.029 | 0.029 | 0.738 | 0.05 | 0.14 | 0.04 | 0.09 | 0.07 | 0.11 |
| 9 | 85735656 | rs17307723 | -1.392 | 0.355 | 1.179 | 0.635 | 0.003 | 0.007 | 0.007 | 0 | 0.07 | 0.09 | 0.05 | 0.09 | 0.1 | 0.11 |
| 9 | 85736106 | rs7862814 | -0.295 | 0.278 | 1.134 | 0.56 | -0.013 | -0.023 | -0.023 | 0 | 0.43 | 0.52 | 0.41 | 0.71 | 0.41 | 0.56 |
| 9 | 85748119 | rs11139921 | -0.444 | 0.36 | 0.449 | 0.446 | -0.001 | -0.018 | -0.018 | 0 | 0.46 | 0.5 | 0.44 | 0.77 | 0.46 | 0.67 |
| 9 | 85756459 | rs1502680 | -0.548 | 1.101 | 0.009 | 1.017 | 0.012 | 0.068 | 0.068 | 0 | 0.51 | 0.67 | 0.53 | 0.68 | 0.49 | 0.5 |
| 9 | 85756528 | rs1502681 | 0.303 | 0.982 | 0.552 | 1.218 | -0.004 | 0.013 | 0.013 | 0 | 0.29 | 0.25 | 0.28 | 0.53 | 0.28 | 0.35 |
| 9 | 85764146 | rs17308295 | 1.28 | 1.54 | 0.253 | 1.115 | 0.005 | 0.194 | 0.194 | 0.449 | 0.37 | 0.09 | 0.34 | 0.04 | 0.3 | 0.28 |
| 9 | 85764196 | rs11139927 | 0.463 | 1.624 | 0.388 | 1.842 | 0.018 | 0.13 | 0.13 | 0 | 0.07 | 0.11 | 0.09 | 0.28 | 0.12 | 0.18 |
| 9 | 85765521 | rs7040878 | 0.915 | 1.628 | 0.151 | 1.794 | 0.028 | 0.066 | 0.066 | 0.114 | 0.44 | 0.31 | 0.38 | 0.23 | 0.35 | 0.37 |
| 9 | 85766721 | rs952744 | 1.037 | 1.4 | 0.032 | 1.628 | -0.034 | 0.004 | 0.004 | 1.536 | 0.19 | 0.06 | 0.22 | 0.15 | 0.2 | 0.15 |
| 9 | 85778241 | rs7043147 | -0.388 | 1.493 | 0.479 | 1.851 | 0.035 | 0.156 | 0.156 | 0 | 0.08 | 0.22 | 0.09 | 0.24 | 0.12 | 0.2 |
| 9 | 85781109 | rs7860585 | 1.055 | 1.433 | 0.298 | 1.689 | -0.035 | 0.017 | 0.017 | 0.359 | 0.19 | 0.07 | 0.22 | 0.14 | 0.21 | 0.15 |
| 9 | 85781709 | rs11139936 | 0.94 | 1.571 | 0.371 | 1.822 | 0.041 | 0.041 | 0.041 | 0.558 | 0.44 | 0.31 | 0.36 | 0.32 | 0.31 | 0.35 |
| 9 | 85787718 | rs4322086 | 0.315 | 1.553 | 0.481 | 1.804 | 0.012 | 0.004 | 0.004 | 0.447 | 0.63 | 0.47 | 0.58 | 0.46 | 0.52 | 0.49 |
| 9 | 85788822 | rs1575506 | 1.031 | 1.55 | 0.47 | 1.801 | 0.037 | 0.036 | 0.036 | 0.695 | 0.43 | 0.31 | 0.36 | 0.32 | 0.3 | 0.35 |
| 9 | 85788944 | rs12555032 | 1.117 | 1.544 | 0.374 | 1.782 | -0.032 | 0.019 | 0.019 | 0.409 | 0.19 | 0.08 | 0.22 | 0.14 | 0.21 | 0.15 |
| 9 | 85793511 | rs17308435 | -0.29 | 1.582 | 1.363 | 2.874 | 0 | 0 | 0 | 0 | 0.28 | 0.26 | 0.3 | 0.26 | 0.28 | 0.2 |
| 9 | 85794098 | rs17080551 | 0.994 | 1.644 | 1.32 | 2.91 | 0.129 | 0.196 | 0.196 | 0 | 0.93 | 0.81 | 0.89 | 0.72 | 0.81 | 0.7 |
| 9 | 85797104 | rs11139942 | -0.958 | 1.263 | 1.093 | 2.604 | -0.06 | 0.058 | 0.058 | 0.058 | 0.18 | 0.4 | 0.22 | 0.3 | 0.09 | 0.2 |
| 9 | 85797799 | rs13290606 | -0.181 | 1.327 | 0.837 | 2.661 | 0.001 | -0.004 | -0.004 | 0 | 0.28 | 0.26 | 0.3 | 0.3 | 0.29 | 0.2 |
| 9 | 85797854 | rs12336763 | 0.311 | 1.3 | 0.844 | 1.898 | -0.01 | -0.014 | -0.014 | 0 | 0.24 | 0.32 | 0.21 | 0.18 | 0.19 | 0.17 |
| 9 | 85798722 | rs11139944 | -0.916 | 1.545 | 0.334 | 1.948 | -0.007 | 0 | 0 | 0 | 0.42 | 0.56 | 0.43 | 0.48 | 0.29 | 0.38 |
| 9 | 85798771 | rs11139945 | 1.449 | 2.372 | 1.362 | 2.93 | 0.123 | 0.218 | 0.218 | 0 | 0.87 | 0.69 | 0.79 | 0.62 | 0.73 | 0.68 |
| 9 | 85799271 | rs1323763 | -0.849 | 0.931 | 0.572 | 1.931 | 0.002 | -0.003 | -0.003 | 0 | 0.1 | 0.08 | 0.17 | 0.17 | 0.05 | 0.14 |
| 9 | 85806577 | rs10867947 | -1.538 | 1 | 0.666 | 1.914 | -0.031 | 0.124 | 0.124 | 0.156 | 0.18 | 0.6 | 0.28 | 0.5 | 0.08 | 0.32 |
| 9 | 85808708 | rs4282648 | 0.686 | 1.647 | 0.32 | 1.308 | 0.011 | 0.011 | 0.011 | 0 | 0.85 | 0.82 | 0.9 | 0.73 | 0.79 | 0.87 |
| 9 | 85808790 | rs7046769 | 0.967 | 1.479 | 0.401 | 1.278 | 0.032 | 0.02 | 0.02 | 0 | 0.72 | 0.51 | 0.67 | 0.66 | 0.69 | 0.62 |
| 9 | 85810910 | rs10115371 | -1.277 | 1.012 | 0.382 | 1.347 | -0.003 | 0.027 | 0.027 | 0 | 0.2 | 0.28 | 0.21 | 0.71 | 0.21 | 0.29 |
| 9 | 85811540 | rs11139953 | 0.586 | 1.303 | 0.595 | 0.626 | 0.028 | -0.025 | -0.025 | 3.218 | 0.68 | 0.55 | 0.52 | 0.79 | 0.63 | 0.69 |
| 9 | 85811825 | rs1323766 | 0.81 | 1.173 | 1.04 | 0.562 | -0.003 | 0.002 | 0.002 | 0 | 0.66 | 0.34 | 0.5 | 0.79 | 0.62 | 0.69 |
| 9 | 85812142 | rs7864580 | 0.875 | 1.144 | 1 | 0.548 | -0.012 | 0.016 | 0.016 | 0 | 0.65 | 0.3 | 0.5 | 0.79 | 0.63 | 0.69 |
| 9 | 85812953 | rs17400257 | 0.308 | 1.374 | 0.613 | 0.431 | -0.03 | 0.023 | 0.023 | 0 | 0.12 | 0 | 0.22 | 0.04 | 0.11 | 0.14 |
| 9 | 85813385 | rs11139955 | -0.595 | 1.481 | 1.13 | 0.379 | -0.01 | 0.006 | 0.006 | 0 | 0.2 | 0.16 | 0.28 | 0.16 | 0.2 | 0.25 |
| 9 | 85813844 | rs6559711 | 0.051 | 1.317 | 1.048 | 1.049 | 0.068 | -0.024 | -0.024 | 0 | 0.87 | 0.66 | 0.8 | 0.95 | 0.83 | 0.94 |
| 9 | 85821621 | rs12682758 | -0.11 | 1.096 | 0.348 | 1.108 | -0.019 | -0.048 | -0.048 | 0.313 | 0.09 | 0.01 | 0.14 | 0.19 | 0.12 | 0.1 |
| 9 | 85833127 | rs1323771 | 0.382 | 1.551 | 0.737 | 0.756 | 0.019 | -0.071 | -0.071 | 0 | 0.83 | 0.79 | 0.85 | 1 | 0.76 | 0.94 |
| 9 | 85834349 | rs1408101 | 1.417 | 1.354 | 0.914 | 0.859 | 0.003 | -0.01 | -0.01 | 0.128 | 0.6 | 0.47 | 0.47 | 0.78 | 0.56 | 0.7 |
| 9 | 85835108 | rs10114830 | -0.11 | 1.418 | 0.226 | 0.736 | 0.024 | -0.042 | -0.042 | 0 | 0.85 | 0.75 | 0.79 | 0.95 | 0.81 | 0.95 |
| 9 | 85835157 | rs11139976 | -0.605 | 1.435 | -0.428 | 0.926 | -0.038 | 0.027 | 0.027 | 0 | 0.22 | 0.09 | 0.3 | 0.13 | 0.24 | 0.21 |
| 9 | 85845746 | rs10867954 | -0.914 | 1.4 | -0.466 | 0.879 | -0.046 | 0.029 | 0.029 | 0 | 0.24 | 0.11 | 0.37 | 0.15 | 0.3 | 0.28 |

| | | | | | | | | | | | | | | | | |
|---|---|---|---|---|---|---|---|---|---|---|---|---|---|---|---|---|
| 9 | 85847317 | rs10867956 | 1.468 | 1.266 | -0.401 | 0.817 | -0.042 | 0.097 | 0.097 | 0 | 0.59 | 0.22 | 0.45 | 0.8 | 0.57 | 0.69 |
| 9 | 85849458 | rs10780579 | 0.452 | 1.094 | -0.464 | 0.513 | -0.003 | -0.025 | -0.025 | 0 | 0.77 | 0.33 | 0.74 | 0.95 | 0.81 | 0.94 |
| 9 | 85851830 | rs4877740 | -1.404 | 1.213 | -0.53 | 0.508 | -0.032 | 0.009 | 0.009 | 0.286 | 0.32 | 0.21 | 0.45 | 0.75 | 0.4 | 0.49 |
| 9 | 85855600 | rs10117817 | 0.232 | 1.221 | -0.442 | 0.527 | -0.008 | -0.058 | -0.058 | 0.015 | 0.78 | 0.38 | 0.75 | 0.95 | 0.81 | 0.94 |
| 9 | 85859648 | rs1048228 | -0.628 | 1.228 | -0.764 | 0.637 | 0 | 0.016 | 0.016 | 0 | 0.11 | 0.12 | 0.15 | 0.6 | 0.15 | 0.22 |
| 9 | 85860018 | rs1048223 | 0.816 | 1.225 | -1.045 | 0.486 | 0.019 | -0.022 | -0.022 | 0 | 0.66 | 0.22 | 0.65 | 0.35 | 0.69 | 0.7 |
| 9 | 85863091 | rs10114696 | 1.398 | 1.282 | -1.17 | 0.338 | -0.003 | 0.113 | 0.113 | 0 | 0.42 | 0.14 | 0.28 | 0.2 | 0.39 | 0.44 |
| 9 | 85865631 | rs4877748 | -0.22 | 1.319 | -0.644 | 0.502 | -0.026 | -0.007 | -0.007 | 3.659 | 0.07 | 0.01 | 0.06 | 0.07 | 0.08 | 0.07 |
| 9 | 85872664 | rs4877750 | 0.6 | 1.196 | -0.307 | 0.202 | 0.001 | -0.022 | -0.022 | 0 | 0.8 | 0.82 | 0.77 | 0.95 | 0.8 | 0.91 |
| 9 | 85872940 | rs13294854 | 1.186 | 0.94 | -0.253 | 0.527 | -0.004 | 0.041 | 0.041 | 0 | 0.49 | 0.29 | 0.44 | 0.38 | 0.53 | 0.54 |
| 9 | 85874113 | rs2889953 | -1.237 | 0.647 | -0.481 | -0.068 | 0.004 | 0.026 | 0.026 | 0 | 0.41 | 0.66 | 0.42 | 0.28 | 0.38 | 0.27 |
| 9 | 85876399 | rs12339855 | 0.993 | 0.409 | -0.53 | -0.331 | 0.002 | 0.005 | 0.005 | 0.615 | 0.68 | 0.33 | 0.69 | 0.9 | 0.73 | 0.88 |
| 9 | 85876633 | rs10867967 | 1.852 | 0.19 | -0.542 | -0.372 | -0.003 | 0.056 | 0.056 | 0 | 0.57 | 0.19 | 0.56 | 0.7 | 0.58 | 0.78 |
| 9 | 85879636 | rs1323781 | 0.413 | 0.196 | -0.526 | -0.32 | -0.039 | -0.045 | -0.045 | 1.539 | 0.09 | 0.01 | 0.13 | 0.17 | 0.15 | 0.1 |
| 9 | 85881204 | rs9695210 | -0.755 | 0.136 | 0.813 | -0.363 | 0.019 | -0.037 | -0.037 | 0.134 | 0.92 | 0.74 | 0.93 | 1 | 0.93 | 1 |
| 9 | 85882323 | rs13296597 | 0.802 | 0.211 | 0.885 | -0.393 | 0.009 | 0.021 | 0.021 | 0 | 0.08 | 0.02 | 0.03 | 0 | 0.05 | 0.02 |
| 9 | 85883879 | rs10512138 | 0.128 | 0.181 | 0.918 | -0.402 | 0.022 | -0.007 | -0.007 | 0 | 0.16 | 0.39 | 0.21 | 0.1 | 0.16 | 0.09 |
| 9 | 85885299 | rs4877751 | 1.206 | 0.004 | 0.781 | -0.51 | -0.002 | 0.007 | 0.007 | 0 | 0.65 | 0.32 | 0.59 | 0.73 | 0.62 | 0.79 |
| 9 | 85891266 | rs13298532 | -0.916 | 0.071 | 1.064 | -0.512 | -0.012 | -0.063 | -0.063 | 0 | 0.26 | 0.48 | 0.24 | 0.1 | 0.2 | 0.07 |
| 9 | 85900401 | rs7856762 | 0.229 | 0.096 | 1.35 | -0.509 | 0.005 | -0.008 | -0.008 | 0 | 0.14 | 0.23 | 0.2 | 0.08 | 0.16 | 0.05 |
| 9 | 85904384 | rs2209186 | 1.836 | 0.016 | 1.153 | -0.331 | 0 | 0.052 | 0.052 | 0 | 0.55 | 0.21 | 0.55 | 0.69 | 0.58 | 0.77 |
| 9 | 85904778 | rs2378658 | 1.226 | 0.103 | 1.283 | -0.221 | -0.005 | 0.003 | 0.003 | 0 | 0.65 | 0.29 | 0.59 | 0.69 | 0.63 | 0.79 |
| 9 | 85905060 | rs1535753 | 1.226 | 0.103 | 1.283 | -0.221 | -0.005 | 0.003 | 0.003 | 0 | 0.65 | 0.29 | 0.59 | 0.69 | 0.62 | 0.79 |
| 9 | 85909882 | rs10117251 | 1.423 | -0.062 | 0.52 | -0.342 | 0.009 | 0.047 | 0.047 | 0 | 0.56 | 0.21 | 0.54 | 0.69 | 0.59 | 0.7 |
| 9 | 85910512 | rs1570582 | -0.75 | -0.082 | 1.114 | -0.391 | 0 | -0.05 | -0.05 | 0.015 | 0.91 | 0.78 | 0.94 | 1 | 0.93 | 1 |
| 9 | 85910642 | rs11140009 | -0.088 | -0.057 | 0.811 | -0.375 | -0.006 | -0.031 | -0.031 | 0.561 | 0.22 | 0.3 | 0.22 | 0.09 | 0.2 | 0.14 |
| 9 | 85910950 | rs1535750 | 1.531 | -0.07 | 0.766 | -0.373 | 0.009 | 0.047 | 0.047 | 0 | 0.56 | 0.21 | 0.54 | 0.69 | 0.59 | 0.7 |
| 9 | 85912228 | rs11140010 | 0.027 | -0.08 | 0.434 | -0.404 | -0.001 | -0.017 | -0.017 | 0 | 0.2 | 0.26 | 0.22 | 0.09 | 0.19 | 0.22 |
| 9 | 85914243 | rs1359168 | 1.274 | 0.01 | -0.101 | -0.041 | -0.012 | 0.004 | 0.004 | 0 | 0.57 | 0.14 | 0.54 | 0.65 | 0.6 | 0.61 |
| 9 | 85925309 | rs11140018 | 0.242 | 0.388 | -0.23 | 0.559 | -0.025 | -0.049 | -0.049 | 1.568 | 0.18 | 0.1 | 0.28 | 0.39 | 0.28 | 0.25 |
| 9 | 85927157 | rs2031261 | 0.379 | 0.324 | -0.039 | 0.327 | -0.001 | 0.003 | 0.003 | 0 | 0.19 | 0.16 | 0.12 | 0.06 | 0.16 | 0.21 |
| 9 | 85927600 | rs10867985 | 0.234 | 0.373 | -0.583 | 0.105 | -0.002 | -0.018 | -0.018 | 0.081 | 0.19 | 0.21 | 0.16 | 0.04 | 0.18 | 0.13 |
| 9 | 85929821 | rs1408106 | -1.985 | 0.333 | -0.48 | 0.494 | 0.002 | 0.016 | 0.016 | 0 | 0.64 | 0.75 | 0.63 | 0.73 | 0.69 | 0.67 |
| 9 | 85930972 | rs10123088 | 0.196 | 0.539 | -0.399 | 0.891 | 0 | 0.032 | 0.032 | 0 | 0.06 | 0.07 | 0.06 | 0.23 | 0.07 | 0.08 |
| 9 | 85932455 | rs17086147 | -0.325 | 0.553 | -0.58 | 0.881 | -0.002 | 0.005 | 0.005 | 0 | 0.27 | 0.39 | 0.19 | 0.29 | 0.23 | 0.29 |
| 9 | 85937637 | rs4361831 | -0.088 | 0.637 | -1.121 | 1.126 | -0.02 | -0.05 | -0.05 | 0.367 | 0.18 | 0.09 | 0.28 | 0.39 | 0.28 | 0.25 |
| 9 | 85942341 | rs3903842 | -0.273 | 0.415 | -1.01 | 1.19 | -0.001 | -0.005 | -0.005 | 0 | 0.35 | 0.63 | 0.37 | 0.47 | 0.41 | 0.33 |
| 9 | 85943575 | rs10867988 | 0.669 | 0.063 | -1.195 | 1.04 | 0.006 | 0.055 | 0.055 | 0 | 0.46 | 0.33 | 0.55 | 0.29 | 0.46 | 0.4 |
| 9 | 85944604 | rs10867991 | 0.652 | 0.075 | -1.112 | 1.033 | 0.005 | 0.057 | 0.057 | 0 | 0.46 | 0.32 | 0.54 | 0.28 | 0.46 | 0.4 |
| 9 | 85944803 | rs10867992 | 1.59 | 0.373 | -1.305 | 0.893 | -0.005 | 0.036 | 0.036 | 0 | 0.24 | 0.14 | 0.17 | 0.15 | 0.22 | 0.24 |
| 9 | 85945797 | rs17086156 | 0.757 | 0.46 | -1.371 | 1.212 | 0.004 | 0.067 | 0.067 | 0 | 0.07 | 0.12 | 0.06 | 0.23 | 0.07 | 0.08 |
| 9 | 85947759 | rs12555197 | 0.879 | 0.386 | -1.389 | 0.853 | 0.004 | -0.013 | -0.013 | 0.491 | 0.27 | 0.4 | 0.3 | 0.22 | 0.33 | 0.24 |
| 9 | 85949206 | rs7861973 | -0.208 | 0.407 | -1.672 | 0.833 | 0.048 | -0.029 | -0.029 | 0 | 0.81 | 0.9 | 0.91 | 0.75 | 0.87 | 0.74 |
| 9 | 85951471 | rs10780595 | -0.287 | 0.436 | -1.715 | 0.815 | 0.047 | -0.033 | -0.033 | 0.557 | 0.82 | 0.92 | 0.91 | 0.75 | 0.87 | 0.74 |
| 9 | 85954902 | rs2889243 | -0.281 | 0.439 | -1.715 | 0.808 | 0.042 | -0.024 | -0.024 | 0 | 0.82 | 0.9 | 0.91 | 0.75 | 0.87 | 0.74 |
| 9 | 85955438 | rs4242620 | -0.437 | 1.475 | -1.188 | 0.512 | -0.023 | 0.019 | 0.019 | 0 | 0.22 | 0.12 | 0.26 | 0.14 | 0.24 | 0.18 |
| 9 | 85959871 | rs3860902 | 0.073 | 1.066 | -0.844 | 0.823 | 0 | 0.007 | 0.007 | 0 | 0.72 | 0.71 | 0.78 | 0.55 | 0.72 | 0.58 |
| 9 | 85960840 | rs10780596 | 0.636 | 1.148 | -0.866 | -0.201 | -0.001 | -0.014 | -0.014 | 0 | 0.3 | 0.28 | 0.32 | 0.07 | 0.35 | 0.22 |
| 9 | 85964735 | rs2277175 | 0.698 | 1.481 | -0.967 | 0.044 | -0.01 | -0.017 | -0.017 | 0 | 0.13 | 0.09 | 0.18 | 0.35 | 0.14 | 0.18 |
| 9 | 85969893 | rs3889596 | 1.031 | 1.175 | -0.674 | -0.323 | -0.011 | 0.009 | 0.009 | 0 | 0.11 | 0.16 | 0.19 | 0.11 | 0.08 | 0.05 |
| 9 | 85970165 | rs11140046 | 1.262 | 1.071 | -0.34 | -0.392 | -0.007 | 0.072 | 0.072 | 0 | 0.3 | 0.15 | 0.32 | 0.07 | 0.35 | 0.21 |
| 9 | 85977276 | rs7035160 | -0.524 | 1.043 | -0.337 | -0.425 | -0.002 | 0.002 | 0.002 | 0 | 0.61 | 0.63 | 0.65 | 0.38 | 0.61 | 0.51 |
| 9 | 85980087 | rs10868000 | 1.193 | 1.027 | -0.336 | -0.326 | -0.008 | 0.069 | 0.069 | 0 | 0.3 | 0.16 | 0.32 | 0.11 | 0.35 | 0.21 |
| 9 | 85981967 | rs11140056 | 1.206 | 0.997 | -0.351 | -0.346 | -0.009 | 0.068 | 0.068 | 0 | 0.3 | 0.16 | 0.32 | 0.1 | 0.35 | 0.21 |
| 9 | 85986905 | rs1556149 | 0.678 | 0.886 | 0.048 | -0.374 | 0.001 | -0.015 | -0.015 | 0 | 0.36 | 0.4 | 0.38 | 0.3 | 0.39 | 0.29 |
| 9 | 85987935 | rs3739657 | -0.919 | 0.832 | 0.445 | -0.292 | -0.001 | -0.001 | -0.001 | 0 | 0.42 | 0.59 | 0.47 | 0.45 | 0.37 | 0.37 |
| 9 | 85988205 | rs3739656 | -0.339 | 0.813 | 0.541 | -0.298 | 0.001 | -0.005 | -0.005 | 0 | 0.32 | 0.44 | 0.27 | 0.32 | 0.29 | 0.33 |
| 9 | 85989187 | rs4877757 | -1.232 | 0.773 | 0.551 | -0.344 | -0.002 | 0.011 | 0.011 | 0 | 0.71 | 0.7 | 0.79 | 0.53 | 0.71 | 0.57 |
| 9 | 85997312 | rs10512140 | 1.323 | 1.179 | 0.435 | -0.372 | -0.038 | 0.08 | 0.08 | 0 | 0.3 | 0.12 | 0.35 | 0.1 | 0.38 | 0.21 |
| 9 | 85998802 | rs4084066 | 0.648 | 1.155 | 0.374 | -0.344 | -0.001 | 0.02 | 0.02 | 0 | 0.39 | 0.29 | 0.51 | 0.21 | 0.42 | 0.25 |
| 9 | 86001996 | rs7870557 | -0.276 | 1.109 | 0.441 | -0.306 | 0.001 | -0.002 | -0.002 | 0.042 | 0.31 | 0.4 | 0.27 | 0.33 | 0.29 | 0.32 |
| 9 | 86003591 | rs17086209 | 2.252 | 0.84 | -0.056 | -0.751 | 0.011 | 0.072 | 0.072 | 0 | 0.12 | 0.01 | 0.04 | 0 | 0.06 | 0.08 |
| 9 | 86004436 | rs3739655 | -0.059 | 1.078 | 0.196 | -0.61 | 0.002 | 0.005 | 0.005 | 0 | 0.25 | 0.32 | 0.25 | 0.33 | 0.23 | 0.31 |
| 9 | 86007105 | rs10117284 | -1.353 | 0.976 | 0.113 | -0.602 | 0.018 | -0.042 | -0.042 | 0 | 0.71 | 0.78 | 0.83 | 0.53 | 0.75 | 0.6 |
| 9 | 86007259 | rs11140065 | 0.362 | 1.016 | 0.187 | -0.714 | -0.012 | 0.028 | 0.028 | 0 | 0.42 | 0.32 | 0.55 | 0.22 | 0.48 | 0.26 |
| 9 | 86011428 | rs6559722 | 0.505 | 1.501 | 0.171 | -0.734 | 0 | -0.001 | -0.001 | 0 | 0.41 | 0.41 | 0.52 | 0.21 | 0.41 | 0.25 |
| 9 | 86014433 | rs4145729 | 0.031 | 1.411 | 0.198 | -0.518 | 0.007 | 0.053 | 0.053 | 0 | 0.5 | 0.29 | 0.56 | 0.59 | 0.48 | 0.38 |
| 9 | 86016834 | rs10868004 | 0.682 | 1.467 | 0.032 | -0.472 | -0.015 | -0.01 | -0.01 | 0 | 0.4 | 0.22 | 0.38 | 0.64 | 0.41 | 0.45 |
| 9 | 86020127 | rs17086223 | 1.39 | 1.242 | 0.202 | -0.239 | -0.022 | -0.034 | -0.034 | 0 | 0.11 | 0.04 | 0.16 | 0.14 | 0.11 | 0.07 |

| | | | | | | | | | | | | | | | | |
|---|---|---|---|---|---|---|---|---|---|---|---|---|---|---|---|---|
| 9 | 86021132 | rs7035287 | -0.9 | 1.259 | 0.348 | -0.328 | -0.001 | 0.006 | 0.006 | 0 | 0.4 | 0.61 | 0.48 | 0.26 | 0.38 | 0.31 |
| 9 | 86021400 | rs3860916 | -0.068 | 1.33 | 0.573 | -0.261 | 0.015 | 0.024 | 0.024 | 0 | 0.22 | 0.3 | 0.38 | 0.26 | 0.23 | 0.14 |
| 9 | 86029329 | rs7039438 | 0.105 | 1.325 | 0.575 | -0.258 | 0.014 | 0.011 | 0.011 | 0 | 0.08 | 0.12 | 0.19 | 0.12 | 0.14 | 0.07 |
| 9 | 86031577 | rs11140072 | 1.18 | 0.961 | 0.562 | -0.39 | 0.017 | 0.027 | 0.027 | 0 | 0.07 | 0.17 | 0.05 | 0.11 | 0.07 | 0.23 |
| 9 | 86038298 | rs3860918 | 0.446 | 0.976 | 0.541 | -0.362 | 0.015 | 0.043 | 0.043 | 0 | 0.1 | 0.18 | 0.07 | 0.26 | 0.11 | 0.33 |
| 9 | 86038847 | rs3889637 | 0.506 | 0.973 | 0.469 | -0.463 | -0.003 | 0.002 | 0.002 | 0.016 | 0.23 | 0.2 | 0.25 | 0.14 | 0.26 | 0.24 |
| 9 | 86041189 | rs11140081 | 1.048 | 0.851 | 0.619 | -0.49 | -0.018 | -0.005 | -0.005 | 0.021 | 0.15 | 0.31 | 0.21 | 0.12 | 0.09 | 0.07 |
| 9 | 86041302 | rs11140082 | -0.343 | 0.863 | 0.614 | -0.487 | -0.007 | 0.039 | 0.039 | 0 | 0.25 | 0.46 | 0.28 | 0.38 | 0.19 | 0.4 |
| 9 | 86042009 | rs10217669 | 1.326 | 0.857 | 0.593 | -0.483 | -0.011 | -0.003 | -0.003 | 1.67 | 0.14 | 0.2 | 0.2 | 0.12 | 0.09 | 0.07 |
| 9 | 86042288 | rs7019130 | -0.863 | 0.893 | 0.433 | -0.51 | -0.003 | 0.024 | 0.024 | 0 | 0.46 | 0.69 | 0.5 | 0.34 | 0.42 | 0.53 |
| 9 | 86044173 | rs4877761 | -1.275 | 0.903 | 0.424 | -0.544 | -0.004 | -0.007 | -0.007 | 0 | 0.5 | 0.79 | 0.55 | 0.51 | 0.49 | 0.64 |
| 9 | 86045746 | rs11140087 | -0.225 | 0.91 | 0.466 | -0.542 | 0.011 | -0.011 | -0.011 | 0 | 0.31 | 0.41 | 0.3 | 0.23 | 0.33 | 0.46 |
| 9 | 86048288 | rs11140088 | -1.119 | 0.92 | 0.176 | -0.532 | -0.001 | -0.004 | -0.004 | 0 | 0.49 | 0.71 | 0.55 | 0.51 | 0.48 | 0.64 |
| 9 | 86052275 | rs7875766 | 1.41 | 0.923 | 0.187 | -0.601 | 0.008 | 0.006 | 0.006 | 0 | 0.07 | 0.14 | 0.04 | 0.09 | 0.07 | 0.23 |
| 9 | 86055337 | rs2375919 | -1.1 | 0.948 | -0.288 | -0.569 | -0.003 | -0.006 | -0.006 | 0.59 | 0.5 | 0.79 | 0.55 | 0.51 | 0.49 | 0.64 |
| 9 | 86056670 | rs11999447 | 0.304 | 0.886 | -0.458 | -0.438 | -0.003 | 0.008 | 0.008 | 0.051 | 0.23 | 0.19 | 0.24 | 0.09 | 0.25 | 0.24 |
| 9 | 86057113 | rs11140090 | 0.117 | 0.84 | -0.661 | -0.473 | -0.004 | -0.003 | -0.003 | 0 | 0.26 | 0.24 | 0.27 | 0.31 | 0.29 | 0.34 |
| 9 | 86058964 | rs4145727 | -1.195 | 0.474 | -0.679 | -0.614 | 0.008 | 0.058 | 0.058 | 0.009 | 0.88 | 0.96 | 0.92 | 0.99 | 0.86 | 0.93 |
| 9 | 86069237 | rs4877767 | -0.329 | 0.64 | -0.09 | -0.477 | -0.002 | -0.038 | -0.038 | 0 | 0.23 | 0.45 | 0.26 | 0.17 | 0.17 | 0.3 |
| 9 | 86071342 | rs11140102 | 0.251 | 0.09 | -0.376 | -0.042 | 0.001 | -0.001 | -0.001 | 0 | 0.17 | 0.16 | 0.14 | 0.17 | 0.13 | 0.2 |
| 9 | 86075411 | rs4014024 | 0.214 | -0.098 | -0.657 | -0.47 | 0.001 | 0.047 | 0.047 | 0 | 0.45 | 0.6 | 0.49 | 0.25 | 0.55 | 0.42 |
| 9 | 86078128 | rs11140109 | 1.234 | -0.106 | -0.755 | -0.437 | -0.006 | -0.027 | -0.027 | 0 | 0.16 | 0.04 | 0.13 | 0.18 | 0.14 | 0.2 |
| 9 | 86080928 | rs7855690 | 0.203 | -0.125 | -0.52 | -0.416 | 0.001 | 0.067 | 0.067 | 0 | 0.46 | 0.28 | 0.38 | 0.26 | 0.45 | 0.37 |
| 9 | 86086600 | rs7850633 | 0.125 | -0.179 | 0.143 | -0.399 | 0.004 | 0.03 | 0.03 | 0 | 0.52 | 0.42 | 0.49 | 0.25 | 0.52 | 0.49 |
| 9 | 86088061 | rs12000033 | 1.208 | 0.016 | 0.086 | -0.366 | 0.068 | -0.033 | -0.033 | 0.026 | 0.07 | 0.23 | 0.09 | 0 | 0.08 | 0.02 |
| 9 | 86088871 | rs10746709 | -1.35 | -0.006 | -0.038 | -0.319 | -0.005 | -0.045 | -0.045 | 0.022 | 0.72 | 0.76 | 0.62 | 0.45 | 0.66 | 0.58 |
| 9 | 86089295 | rs7032223 | -1.913 | -0.043 | -0.036 | -0.309 | 0 | -0.034 | -0.034 | 0 | 0.74 | 0.76 | 0.66 | 0.63 | 0.74 | 0.7 |
| 9 | 86093237 | rs17086319 | 0.982 | -0.003 | 0.177 | -0.522 | 0.005 | -0.056 | -0.056 | 0 | 0.35 | 0.36 | 0.35 | 0.07 | 0.39 | 0.29 |
| 9 | 86093654 | rs5014093 | 0.593 | -0.013 | 0.156 | -0.503 | 0.011 | -0.058 | -0.058 | 0.023 | 0.39 | 0.45 | 0.41 | 0.25 | 0.46 | 0.4 |
| 9 | 86099092 | rs10868019 | -0.596 | 0.044 | 0.57 | -0.143 | 0.05 | 0.026 | 0.026 | 0.274 | 0.14 | 0.39 | 0.16 | 0.15 | 0.13 | 0.07 |
| 9 | 86100539 | rs10746712 | -0.596 | 0.044 | 0.564 | -0.143 | 0.049 | 0.026 | 0.026 | 0.274 | 0.14 | 0.39 | 0.16 | 0.15 | 0.13 | 0.07 |
| 9 | 86102933 | rs7873385 | -0.665 | -0.165 | -0.206 | -0.242 | 0.007 | -0.015 | -0.015 | 0.214 | 0.64 | 0.86 | 0.63 | 0.41 | 0.68 | 0.62 |
| 9 | 86103862 | rs6559728 | 0.349 | -0.22 | 0.042 | -0.326 | 0.013 | 0.033 | 0.033 | 0 | 0.53 | 0.54 | 0.5 | 0.08 | 0.56 | 0.45 |
| 9 | 86105137 | rs3860935 | -0.586 | -0.225 | 0.221 | -0.288 | 0.019 | 0.007 | 0.007 | 0 | 0.63 | 0.74 | 0.63 | 0.42 | 0.68 | 0.62 |
| 9 | 86108665 | rs3860941 | 0.578 | -0.38 | 0 | -0.491 | 0.001 | -0.029 | -0.029 | 0 | 0.48 | 0.49 | 0.49 | 0.08 | 0.51 | 0.42 |
| 9 | 86109124 | rs11140121 | -0.445 | -0.344 | -0.638 | -0.427 | -0.001 | 0.035 | 0.035 | 0 | 0.1 | 0.19 | 0.12 | 0.34 | 0.12 | 0.16 |
| 9 | 86111068 | rs11140123 | 1.726 | -0.454 | -0.754 | -0.425 | 0 | -0.001 | -0.001 | 0.081 | 0.1 | 0.09 | 0.07 | 0.24 | 0.07 | 0.08 |
| 9 | 86111242 | rs12551103 | 1.262 | -0.503 | -0.85 | -0.248 | 0.007 | -0.046 | -0.046 | 0.014 | 0.08 | 0 | 0.09 | 0.23 | 0.05 | 0.06 |
| 9 | 86112734 | rs7863524 | -0.532 | -0.517 | -0.816 | -0.308 | 0.016 | 0.013 | 0.013 | 0 | 0.63 | 0.75 | 0.62 | 0.41 | 0.65 | 0.61 |
| 9 | 86117401 | rs12552693 | 0.443 | -0.121 | -0.575 | -0.26 | -0.002 | 0.049 | 0.049 | 0 | 0.5 | 0.37 | 0.49 | 0.3 | 0.48 | 0.47 |
| 9 | 86127539 | rs7043959 | -0.921 | 0.005 | -1.508 | 0.106 | -0.028 | -0.112 | -0.112 | 0.294 | 0.75 | 0.91 | 0.73 | 0.64 | 0.74 | 0.68 |
| 9 | 86135188 | rs17086403 | 0.478 | 0.177 | -0.731 | 0.198 | -0.004 | 0.007 | 0.007 | 0.152 | 0.11 | 0.14 | 0.07 | 0.24 | 0.07 | 0.07 |
| 9 | 86149610 | rs7849075 | 0.605 | 0.679 | -1.053 | -0.033 | -0.022 | 0.095 | 0.095 | 0.076 | 0.44 | 0.29 | 0.44 | 0.16 | 0.46 | 0.65 |
| 9 | 86149786 | rs7863215 | -0.071 | 0.094 | -0.754 | -0.128 | -0.006 | -0.002 | -0.002 | 0 | 0.66 | 0.66 | 0.71 | 0.65 | 0.71 | 0.79 |
| 9 | 86155392 | rs1888746 | -0.273 | 0.366 | -1.194 | 0.078 | -0.038 | -0.035 | -0.035 | 0 | 0.69 | 0.97 | 0.74 | 0.7 | 0.7 | 0.83 |
| 9 | 86156348 | rs2210381 | 1.722 | 0.708 | -1.273 | -0.142 | -0.023 | 0.12 | 0.12 | 0 | 0.3 | 0.14 | 0.33 | 0.03 | 0.29 | 0.3 |
| 9 | 86164428 | rs13289150 | 0.082 | 0.933 | -1.443 | 0.066 | -0.017 | -0.057 | -0.057 | 15.119 | 0.35 | 0.1 | 0.33 | 0.42 | 0.38 | 0.25 |
| 9 | 86168692 | rs6559732 | 0.683 | 1.508 | -0.687 | 0.811 | 0.011 | 0.009 | 0.009 | 0.145 | 0.83 | 0.71 | 0.85 | 0.77 | 0.82 | 0.82 |
| 9 | 86177401 | rs3934902 | 0.392 | 1.447 | -0.733 | 0.592 | -0.003 | 0.003 | 0.003 | 0.084 | 0.15 | 0.14 | 0.17 | 0.3 | 0.15 | 0.27 |
| 9 | 86178646 | rs7025704 | -0.609 | 1.3 | -0.436 | 0.373 | 0.04 | -0.052 | -0.052 | 0 | 0.09 | 0.28 | 0.07 | 0.01 | 0.08 | 0.07 |
| 9 | 86179563 | rs4451390 | 0.378 | 1.181 | -1.367 | 0.768 | -0.065 | -0.11 | -0.11 | 0 | 0.78 | 0.97 | 0.74 | 0.58 | 0.75 | 0.79 |
| 9 | 86184504 | rs11793821 | -0.725 | 0.726 | -0.131 | 0.11 | 0.002 | -0.033 | -0.033 | 0.938 | 0.15 | 0.39 | 0.17 | 0.11 | 0.11 | 0.07 |
| 9 | 86185455 | rs4877258 | 0.658 | 1.291 | -0.403 | -0.222 | -0.003 | 0.002 | 0.002 | 0 | 0.77 | 0.81 | 0.58 | 0.25 | 0.71 | 0.69 |
| 9 | 86191111 | rs7032583 | -0.512 | 0.541 | 0.271 | -0.487 | -0.037 | -0.016 | -0.016 | 0 | 0.19 | 0.43 | 0.11 | 0.15 | 0.15 | 0.1 |
| 9 | 86199324 | rs2780991 | -1.661 | 0.099 | -0.738 | -1.783 | -0.002 | 0.166 | 0.166 | 0 | 0.74 | 0.95 | 0.9 | 1 | 0.82 | 0.8 |
| 9 | 86200547 | rs1572134 | 0.742 | 0.572 | -0.902 | -1.76 | -0.051 | 0.079 | 0.079 | 0 | 0.29 | 0.15 | 0.31 | 0.13 | 0.39 | 0.38 |
| 9 | 86203246 | rs12001341 | 0.425 | 0.524 | -0.54 | -1.628 | -0.042 | -0.053 | -0.053 | 0.524 | 0.18 | 0.41 | 0.09 | 0.11 | 0.11 | 0.1 |
| 9 | 86203301 | rs17086501 | 0.669 | 0.547 | -0.241 | -1.617 | -0.02 | -0.026 | -0.026 | 0 | 0.16 | 0.21 | 0.07 | 0.11 | 0.11 | 0.1 |
| 9 | 86206343 | rs17321848 | 0.518 | 1.318 | -0.925 | -1.658 | -0.007 | 0.001 | 0.001 | 0 | 0.12 | 0.12 | 0.12 | 0.11 | 0.19 | 0.22 |
| 9 | 86216008 | rs10113885 | 0.11 | 1.269 | -1.02 | -1.836 | -0.053 | 0.089 | 0.089 | 0 | 0.3 | 0.17 | 0.31 | 0.11 | 0.41 | 0.39 |
| 9 | 86216100 | rs2811897 | -0.864 | 0.507 | -0.586 | -1.844 | 0.031 | 0.04 | 0.04 | 0.03 | 0.53 | 0.41 | 0.77 | 0.84 | 0.67 | 0.69 |
| 9 | 86220040 | rs10868031 | 0.317 | 1.191 | -0.965 | -0.333 | -0.039 | -0.107 | -0.107 | 0 | 0.15 | 0.02 | 0.33 | 0.45 | 0.13 | 0.13 |
| 9 | 86221436 | rs11790364 | 0.878 | 0.976 | -1.138 | -0.894 | -0.006 | 0.07 | 0.07 | 3.8 | 0.13 | 0.01 | 0.05 | 0.03 | 0.09 | 0.13 |
| 9 | 86226119 | rs7865279 | -0.457 | 1.717 | -0.608 | -1.014 | 0.005 | -0.034 | -0.034 | 0.066 | 0.23 | 0.25 | 0.23 | 0.07 | 0.27 | 0.35 |
| 9 | 86229249 | rs7848656 | -0.145 | 1.748 | -0.488 | -0.934 | 0.047 | -0.019 | -0.019 | 0.396 | 0.09 | 0.17 | 0.16 | 0.03 | 0.16 | 0.21 |
| 9 | 86231370 | rs949864 | 0.009 | 1.644 | 0.087 | -0.613 | -0.001 | 0.046 | 0.046 | 0 | 0.18 | 0.2 | 0.37 | 0.49 | 0.15 | 0.18 |
| 9 | 86232809 | rs2780982 | -1.422 | 1.276 | -0.463 | -1.358 | -0.006 | 0.029 | 0.029 | 0 | 0.78 | 0.92 | 0.81 | 0.89 | 0.76 | 0.87 |
| 9 | 86234596 | rs2780980 | 0.481 | 1.071 | -0.518 | -1.559 | 0 | -0.014 | -0.014 | 0 | 0.47 | 0.39 | 0.37 | 0.05 | 0.53 | 0.51 |
| 9 | 86244359 | rs2811907 | 0.44 | 1.028 | -0.349 | -1.551 | 0.001 | -0.014 | -0.014 | 0 | 0.48 | 0.42 | 0.37 | 0.05 | 0.53 | 0.52 |

| | | | | | | | | | | | | | | | | |
|---|---|---|---|---|---|---|---|---|---|---|---|---|---|---|---|---|
| 9 | 86253482 | rs2811913 | 1.39 | 1.229 | -0.535 | -1.599 | -0.039 | 0.087 | 0.087 | 1.21 | 0.15 | 0.02 | 0.12 | 0.03 | 0.19 | 0.15 |
| 9 | 86258685 | rs1052690 | 1.39 | 1.234 | -0.532 | -1.597 | -0.039 | 0.087 | 0.087 | 1.888 | 0.15 | 0.02 | 0.13 | 0.03 | 0.19 | 0.15 |
| 9 | 86260234 | rs9410586 | 0.285 | 1.322 | -0.434 | -1.634 | -0.032 | 0.032 | 0.032 | 0.68 | 0.21 | 0.13 | 0.23 | 0.04 | 0.33 | 0.3 |
| 9 | 86265425 | rs2811921 | -0.663 | 0.777 | -0.46 | -1.526 | -0.017 | 0.127 | 0.127 | 0 | 0.41 | 0.19 | 0.54 | 0.84 | 0.41 | 0.5 |
| 9 | 86273487 | rs2781006 | 1.003 | 0.802 | -0.383 | -1.543 | 0.009 | -0.003 | -0.003 | 0.28 | 0.42 | 0.35 | 0.27 | 0.04 | 0.39 | 0.37 |
| 9 | 86278817 | rs7866234 | -0.151 | 0.941 | -0.614 | -1.471 | 0.004 | 0.023 | 0.023 | 0 | 0.27 | 0.2 | 0.42 | 0.81 | 0.22 | 0.34 |
| 9 | 86281195 | rs12344615 | 2.241 | 0.924 | -0.576 | -1.487 | 0.003 | -0.035 | -0.035 | 0 | 0.26 | 0.25 | 0.13 | 0 | 0.2 | 0.21 |
| 9 | 86281995 | rs2058912 | 0.678 | 1.485 | -0.86 | -0.966 | -0.03 | 0.034 | 0.034 | 0.08 | 0.27 | 0.14 | 0.29 | 0.12 | 0.38 | 0.27 |
| 9 | 86283423 | rs2811929 | -0.783 | 1.48 | -0.862 | -0.968 | -0.031 | 0.036 | 0.036 | 0.095 | 0.73 | 0.86 | 0.71 | 0.88 | 0.62 | 0.73 |
| 9 | 86284178 | rs1044175 | -0.854 | 1.522 | -0.839 | -0.947 | -0.03 | 0.034 | 0.034 | 0.08 | 0.73 | 0.86 | 0.71 | 0.88 | 0.62 | 0.73 |
| 9 | 86297669 | rs11140212 | -0.632 | 0.745 | -0.803 | -0.164 | -0.033 | 0.011 | 0.011 | 0 | 0.25 | 0.06 | 0.41 | 0.81 | 0.22 | 0.34 |
| 9 | 86304008 | rs4877796 | -0.759 | 0.766 | -0.804 | -0.194 | 0.003 | 0.03 | 0.03 | 0 | 0.27 | 0.2 | 0.42 | 0.85 | 0.23 | 0.36 |
| 9 | 86323309 | rs3814507 | 0.029 | 0.995 | -1.7 | 0.285 | -0.034 | -0.126 | -0.126 | 1.605 | 0.17 | 0.02 | 0.37 | 0.5 | 0.15 | 0.17 |
| 9 | 86347254 | rs7020313 | 0.514 | 0.582 | -1.321 | -0.349 | -0.038 | 0.03 | 0.03 | 0.295 | 0.23 | 0.03 | 0.17 | 0.11 | 0.24 | 0.16 |
| 9 | 86368105 | rs10868053 | -0.732 | 0.361 | -0.629 | -0.494 | 0 | -0.003 | -0.003 | 0 | 0.58 | 0.44 | 0.7 | 0.6 | 0.59 | 0.51 |
| 9 | 86368660 | rs7047907 | -1.193 | 0.666 | -0.401 | -0.607 | -0.017 | -0.008 | -0.008 | 1.63 | 0.65 | 0.54 | 0.79 | 0.71 | 0.72 | 0.62 |
| 9 | 86369356 | rs10868054 | -0.554 | 0.668 | -0.409 | -0.607 | 0 | -0.003 | -0.003 | 0.001 | 0.58 | 0.44 | 0.7 | 0.6 | 0.59 | 0.51 |
| 9 | 86381582 | rs7855177 | -2.153 | 0.409 | -0.074 | -0.815 | 0.002 | -0.037 | -0.037 | 0 | 0.74 | 0.75 | 0.83 | 1 | 0.77 | 0.8 |
| 9 | 86381814 | rs11140240 | 0.353 | 0.267 | 0.128 | -0.852 | -0.002 | -0.048 | -0.048 | 0 | 0.34 | 0.21 | 0.52 | 0.46 | 0.36 | 0.36 |
| 9 | 86399815 | rs7021357 | -0.475 | 0.409 | 0.314 | -0.732 | -0.001 | 0.005 | 0.005 | 1.289 | 0.57 | 0.37 | 0.69 | 0.6 | 0.59 | 0.51 |
| 9 | 86404584 | rs4297107 | 0.478 | 0.446 | 0.306 | -0.683 | -0.015 | -0.083 | -0.083 | 1.175 | 0.32 | 0.17 | 0.51 | 0.46 | 0.34 | 0.33 |
| 9 | 86426620 | rs9314723 | -0.593 | 0.402 | 0.285 | -0.704 | -0.013 | -0.082 | -0.082 | 0 | 0.67 | 0.83 | 0.49 | 0.54 | 0.66 | 0.65 |
| 9 | 86443840 | rs7021609 | -0.115 | 0.536 | 0.52 | -0.709 | 0.073 | -0.005 | -0.005 | 0 | 0.08 | 0.22 | 0.1 | 0.11 | 0.13 | 0.11 |
| 9 | 86448944 | rs7027121 | -0.115 | 0.536 | 0.52 | -0.709 | 0.073 | -0.005 | -0.005 | 0 | 0.08 | 0.22 | 0.1 | 0.11 | 0.13 | 0.11 |
| 9 | 86488586 | rs7853625 | -0.115 | 0.536 | 0.567 | -0.709 | 0.103 | 0.006 | 0.006 | 0 | 0.08 | 0.25 | 0.1 | 0.11 | 0.13 | 0.11 |
| 9 | 86497334 | rs13290882 | 0.907 | 0.465 | 0.64 | -0.718 | -0.024 | -0.013 | -0.013 | 0 | 0.36 | 0.51 | 0.21 | 0.29 | 0.28 | 0.37 |
| 9 | 86518796 | rs12001918 | 0.125 | 0.476 | 0.661 | -0.671 | 0.035 | -0.008 | -0.008 | 0.678 | 0.08 | 0.15 | 0.09 | 0.11 | 0.13 | 0.11 |
| 9 | 86534480 | rs10868072 | -0.103 | 0.479 | 0.663 | -0.667 | 0.108 | 0.022 | 0.022 | 0 | 0.08 | 0.25 | 0.1 | 0.14 | 0.15 | 0.11 |
| 9 | 86535730 | rs12552069 | 0.219 | 0.478 | 0.582 | -0.667 | 0.022 | -0.006 | -0.006 | 0 | 0.08 | 0.12 | 0.09 | 0.11 | 0.13 | 0.11 |
| 9 | 86539603 | rs4298563 | 0.192 | 0.463 | 0.386 | -0.695 | -0.009 | 0.009 | 0.009 | 3.883 | 0.43 | 0.73 | 0.34 | 0.43 | 0.42 | 0.49 |
| 9 | 86549939 | rs10124390 | -0.107 | 0.419 | 0.335 | -0.735 | -0.043 | -0.026 | -0.026 | 0 | 0.62 | 0.74 | 0.37 | 0.41 | 0.53 | 0.53 |
| 9 | 86553589 | rs3180887 | 0.619 | 0.397 | 0.395 | -0.698 | 0.031 | 0.018 | 0.018 | 3.597 | 0.37 | 0.63 | 0.22 | 0.28 | 0.27 | 0.35 |
| 9 | 86558270 | rs11140304 | 0.343 | 0.424 | 0.406 | -0.666 | 0.002 | 0.021 | 0.021 | 0 | 0.05 | 0.03 | 0.03 | 0.29 | 0.04 | 0.17 |
| 9 | 86571798 | rs1572031 | -0.866 | 0.212 | 0.01 | -0.817 | -0.019 | -0.068 | -0.068 | 0 | 0.73 | 0.83 | 0.51 | 0.53 | 0.69 | 0.65 |
| 9 | 86595070 | rs296887 | 0.867 | 0.136 | -0.451 | -0.64 | -0.011 | -0.003 | -0.003 | 0 | 0.34 | 0.43 | 0.21 | 0.29 | 0.28 | 0.37 |
| 9 | 86595498 | rs296888 | 0.92 | 0.133 | -0.435 | -0.642 | -0.008 | -0.002 | -0.002 | 0 | 0.33 | 0.39 | 0.21 | 0.29 | 0.28 | 0.37 |
| 9 | 86617265 | rs1982151 | -1.074 | 0.141 | -0.471 | -0.637 | -0.01 | -0.003 | -0.003 | 2.13 | 0.66 | 0.58 | 0.79 | 0.71 | 0.72 | 0.63 |
| 9 | 86623268 | rs4877815 | -0.445 | 0.076 | -0.527 | -0.654 | -0.004 | -0.001 | -0.001 | 0 | 0.43 | 0.54 | 0.64 | 0.61 | 0.48 | 0.48 |
| 9 | 86625243 | rs6559758 | 1 | 0.246 | -0.828 | -0.771 | -0.042 | 0.051 | 0.051 | 0.139 | 0.23 | 0.03 | 0.15 | 0.1 | 0.25 | 0.15 |
| 9 | 86646572 | rs10868083 | -1.88 | -0.126 | -0.593 | -0.759 | 0.002 | -0.021 | -0.021 | 0 | 0.75 | 0.74 | 0.85 | 0.97 | 0.79 | 0.83 |
| 9 | 86666444 | rs11793330 | 0.686 | -0.039 | -0.501 | -0.379 | -0.006 | 0.037 | 0.037 | 0 | 0.26 | 0.12 | 0.12 | 0.08 | 0.26 | 0.12 |
| 9 | 86667098 | rs973460 | -0.987 | 0.018 | -0.573 | -0.24 | -0.006 | -0.003 | -0.003 | 0.073 | 0.49 | 0.73 | 0.38 | 0.5 | 0.47 | 0.46 |
| 9 | 86669330 | rs11140345 | -0.473 | 0.028 | -0.343 | -0.155 | 0.012 | -0.01 | -0.01 | 0 | 0.4 | 0.32 | 0.19 | 0.39 | 0.36 | 0.33 |
| 9 | 86676854 | rs11140350 | -0.665 | 0.342 | -0.508 | 0.145 | -0.006 | 0.03 | 0.03 | 0.906 | 0.51 | 0.68 | 0.4 | 0.43 | 0.51 | 0.45 |
| 9 | 86679251 | rs10868091 | -0.78 | 0.856 | -0.39 | 0.542 | -0.004 | 0.07 | 0.07 | 0 | 0.14 | 0.2 | 0.14 | 0.38 | 0.11 | 0.19 |
| 9 | 86682315 | rs1898321 | -0.583 | 0.857 | -0.432 | 0.554 | -0.013 | 0.023 | 0.023 | 0 | 0.13 | 0.17 | 0.13 | 0.27 | 0.07 | 0.18 |
| 9 | 86691975 | rs7857500 | -0.068 | 0.153 | -0.726 | 0.605 | 0.003 | 0.004 | 0.004 | 0 | 0.09 | 0.09 | 0.11 | 0.29 | 0.06 | 0.17 |
| 9 | 86693470 | rs2118166 | 0.536 | -0.028 | -0.683 | 0.286 | 0.002 | -0.007 | -0.007 | 0 | 0.24 | 0.22 | 0.41 | 0.43 | 0.25 | 0.34 |
| 9 | 86694759 | rs10746732 | -1.51 | 0.075 | -0.427 | -0.023 | 0 | -0.033 | -0.033 | 0 | 0.74 | 0.77 | 0.86 | 1 | 0.77 | 0.8 |
| 9 | 86696287 | rs12339554 | -0.178 | -0.195 | -0.132 | 1.09 | 0.011 | -0.024 | -0.024 | 0 | 0.27 | 0.23 | 0.41 | 0.5 | 0.25 | 0.38 |
| 9 | 86696749 | rs10868094 | -0.575 | -0.011 | -0.521 | 0.703 | -0.026 | 0.004 | 0.004 | 0.716 | 0.36 | 0.51 | 0.47 | 0.6 | 0.31 | 0.44 |
| 9 | 86700593 | rs17086797 | -1.112 | 0.428 | -0.712 | 1.164 | 0.001 | -0.014 | -0.014 | 0.018 | 0.2 | 0.19 | 0.36 | 0.39 | 0.17 | 0.26 |
| 9 | 86700737 | rs4471123 | -1.323 | 0.472 | -2.048 | 0.488 | -0.131 | -0.136 | -0.136 | 0.874 | 0.62 | 0.93 | 0.66 | 0.51 | 0.51 | 0.46 |
| 9 | 86700760 | rs11788077 | -0.048 | 0.517 | -0.728 | 0.215 | -0.021 | -0.161 | -0.161 | 0 | 0.39 | 0.48 | 0.24 | 0.01 | 0.32 | 0.17 |
| 9 | 86707888 | rs7029403 | -0.135 | -0.112 | -0.584 | -0.043 | -0.047 | -0.142 | -0.142 | 1.248 | 0.29 | 0.51 | 0.15 | 0.01 | 0.15 | 0.15 |
| 9 | 86709041 | rs17086818 | -0.211 | 0.444 | -0.692 | 0.774 | -0.003 | 0.038 | 0.038 | 0.016 | 0.17 | 0.17 | 0.29 | 0.32 | 0.13 | 0.12 |
| 9 | 86713990 | rs7032296 | -1.327 | 0.533 | -0.432 | 0.759 | -0.001 | 0.004 | 0.004 | 0.028 | 0.49 | 0.66 | 0.59 | 0.43 | 0.45 | 0.37 |
| 9 | 86714002 | rs7032430 | -0.127 | 0.493 | -0.727 | 0.738 | -0.001 | -0.002 | -0.002 | 0 | 0.22 | 0.22 | 0.27 | 0.31 | 0.23 | 0.21 |
| 9 | 86715566 | rs7853970 | -2.127 | 0.498 | -1.325 | 0.657 | -0.12 | -0.092 | -0.092 | 0 | 0.59 | 0.97 | 0.68 | 0.55 | 0.53 | 0.5 |
| 9 | 86716873 | rs10868097 | -0.18 | 0.536 | -1.048 | 0.581 | -0.057 | 0.001 | 0.001 | 0 | 0.13 | 0.01 | 0.13 | 0.06 | 0.15 | 0.07 |
| 9 | 86718822 | rs893150 | -1.664 | 0.483 | -1.168 | 0.872 | -0.077 | -0.017 | -0.017 | 0 | 0.49 | 0.96 | 0.56 | 0.56 | 0.44 | 0.42 |
| 9 | 86719547 | rs1529952 | -1.391 | 0.428 | -0.246 | 0.799 | 0.005 | 0.005 | 0.005 | 0.026 | 0.32 | 0.65 | 0.49 | 0.56 | 0.37 | 0.34 |
| 9 | 86721015 | rs10868100 | -1.17 | 0.494 | 0.258 | 0.899 | 0.002 | 0.08 | 0.08 | 0.644 | 0.17 | 0.58 | 0.28 | 0.24 | 0.13 | 0.15 |
| 9 | 86721077 | rs11789466 | -0.477 | 0.301 | -0.095 | 0.737 | -0.059 | -0.049 | -0.049 | 0.191 | 0.16 | 0.01 | 0.2 | 0.2 | 0.23 | 0.16 |
| 9 | 86721929 | rs6559768 | -1.395 | 0.432 | 0.041 | 0.719 | -0.015 | -0.015 | -0.015 | 0 | 0.26 | 0.18 | 0.29 | 0.33 | 0.32 | 0.39 |
| 9 | 86724391 | rs11140381 | -1.058 | 0.416 | -0.164 | 0.599 | -0.074 | -0.004 | -0.004 | 0 | 0.22 | 0.02 | 0.25 | 0.19 | 0.28 | 0.24 |
| 9 | 86725070 | rs11140383 | -0.052 | 0.432 | -0.082 | 0.674 | -0.05 | -0.055 | -0.055 | 0 | 0.13 | 0.01 | 0.19 | 0.2 | 0.2 | 0.15 |
| 9 | 86735718 | rs7848501 | -1.911 | 0.403 | 0.22 | 0.618 | 0.008 | 0.006 | 0.006 | 0 | 0.58 | 0.69 | 0.62 | 0.64 | 0.63 | 0.61 |
| 9 | 86736004 | rs11140391 | 0.687 | 0.261 | 0.536 | 0.467 | -0.047 | -0.084 | -0.084 | 0.345 | 0.13 | 0.31 | 0.05 | 0 | 0.04 | 0.07 |

| | | | | | | | | | | | | | | | | |
|---|---|---|---|---|---|---|---|---|---|---|---|---|---|---|---|---|
| 9 | 86736068 | rs2889956 | -1.282 | 0.737 | 0.531 | 0.791 | 0.003 | 0.013 | 0.013 | 0.054 | 0.25 | 0.34 | 0.29 | 0.36 | 0.26 | 0.38 |
| 9 | 86742278 | rs2118169 | -0.414 | 0.598 | 0.601 | 0.743 | 0.001 | 0.046 | 0.046 | 0 | 0.09 | 0.19 | 0.16 | 0.21 | 0.08 | 0.08 |
| 9 | 86743347 | rs7028167 | -1.412 | 0.563 | 0.657 | 0.624 | -0.001 | -0.033 | -0.033 | 0 | 0.23 | 0.41 | 0.29 | 0.14 | 0.22 | 0.25 |
| 9 | 86743543 | rs13284104 | -0.124 | 1.147 | 0.672 | 0.777 | 0.064 | 0.055 | 0.055 | 0.35 | 0.24 | 0.43 | 0.29 | 0.37 | 0.34 | 0.13 |
| 9 | 86744106 | rs7357702 | 0.406 | 0.551 | 0.051 | 0.648 | -0.003 | 0.079 | 0.079 | 0 | 0.12 | 0.37 | 0.07 | 0.12 | 0.08 | 0.16 |
| 9 | 86744475 | rs1246285 | 0.331 | 0.438 | -0.356 | 0.404 | -0.087 | -0.083 | -0.083 | 0 | 0.27 | 0.1 | 0.33 | 0.38 | 0.4 | 0.26 |
| 9 | 86749003 | rs11140398 | 0.706 | 0.423 | -0.538 | 0.381 | -0.018 | -0.007 | -0.007 | 0 | 0.07 | 0.02 | 0.15 | 0.09 | 0.09 | 0.09 |
| 9 | 86750762 | rs2799850 | -0.962 | 0.343 | -0.495 | 0.424 | -0.023 | -0.009 | -0.009 | 0 | 0.43 | 0.24 | 0.45 | 0.42 | 0.52 | 0.44 |
| 9 | 86750781 | rs11140400 | -0.084 | 0.339 | -1.003 | -0.18 | 0.006 | 0.076 | 0.076 | 0 | 0.17 | 0.03 | 0.11 | 0.04 | 0.14 | 0.19 |
| 9 | 86751792 | rs10512144 | 0.077 | 0.34 | -0.992 | -0.174 | 0.011 | 0.028 | 0.028 | 0 | 0.15 | 0.08 | 0.11 | 0.04 | 0.13 | 0.18 |
| 9 | 86757023 | rs7029843 | -0.254 | 0.299 | -0.623 | 0.145 | -0.001 | -0.014 | -0.014 | 0 | 0.34 | 0.34 | 0.33 | 0.14 | 0.3 | 0.34 |
| 9 | 86758069 | rs1246267 | -0.302 | 0.489 | -1.087 | 0.374 | 0.016 | 0.015 | 0.015 | 0.659 | 0.33 | 0.41 | 0.36 | 0.55 | 0.42 | 0.33 |
| 9 | 86761781 | rs1246264 | -1.392 | 0.382 | -2.267 | 0.104 | 0.016 | -0.013 | -0.013 | 0 | 0.67 | 0.9 | 0.7 | 0.68 | 0.7 | 0.67 |
| 9 | 86763217 | rs1273061 | -0.338 | 0.911 | -1.512 | -0.063 | -0.102 | -0.003 | -0.003 | 0 | 0.26 | 0.05 | 0.3 | 0.24 | 0.35 | 0.15 |
| 9 | 86770003 | rs1246274 | 0.96 | -0.152 | -0.256 | -0.291 | -0.022 | -0.116 | -0.116 | 0 | 0.82 | 0.42 | 0.84 | 1 | 0.84 | 0.91 |
| 9 | 86770610 | rs7027670 | 0.145 | -0.487 | -0.579 | -0.648 | 0 | -0.003 | -0.003 | 0 | 0.17 | 0.16 | 0.11 | 0.04 | 0.14 | 0.22 |
| 9 | 86771168 | rs996503 | 1.052 | -1.149 | 0.226 | -0.757 | -0.008 | -0.019 | -0.019 | 0 | 0.64 | 0.4 | 0.59 | 0.82 | 0.68 | 0.63 |
| 9 | 86772993 | rs963761 | 1.275 | -1.772 | 0.018 | -0.952 | -0.046 | -0.055 | -0.055 | 0.241 | 0.69 | 0.48 | 0.65 | 0.81 | 0.75 | 0.83 |
| 9 | 86774339 | rs1147187 | 1.987 | -2.56 | -0.38 | -0.257 | -0.025 | -0.026 | -0.026 | 0 | 0.36 | 0.17 | 0.46 | 0.39 | 0.52 | 0.33 |
| 9 | 86774848 | rs1147189 | 0.067 | -2.472 | 0.054 | -0.239 | -0.025 | -0.028 | -0.028 | 0 | 0.81 | 0.59 | 0.7 | 0.86 | 0.81 | 0.84 |
| 9 | 86776381 | rs6559770 | -0.401 | -1.912 | -0.156 | 0.143 | -0.002 | 0 | 0 | 0 | 0.37 | 0.34 | 0.29 | 0.54 | 0.31 | 0.6 |
| 9 | 86777046 | rs2250159 | 1.601 | -1.943 | -0.238 | 0.125 | -0.018 | 0.031 | 0.031 | 0 | 0.39 | 0.22 | 0.38 | 0.28 | 0.46 | 0.23 |
| 9 | 86778137 | rs2250005 | 1.515 | -2.041 | -0.42 | 0.175 | 0.009 | 0.037 | 0.037 | 0 | 0.43 | 0.26 | 0.45 | 0.33 | 0.51 | 0.3 |
| 9 | 86778277 | rs2249998 | -0.16 | -1.589 | -0.323 | 0.401 | -0.021 | -0.021 | -0.021 | 0 | 0.87 | 0.75 | 0.87 | 0.92 | 0.87 | 0.91 |
| 9 | 86778415 | rs1932652 | -0.148 | -1.091 | -0.73 | 0.606 | 0.02 | -0.044 | -0.044 | 1.863 | 0.18 | 0.1 | 0.08 | 0.37 | 0.13 | 0.27 |
| 9 | 86786231 | rs2246348 | 0.182 | -1.323 | -0.445 | 0.2 | 0.007 | 0.012 | 0.012 | 0.145 | 0.69 | 0.75 | 0.72 | 0.86 | 0.71 | 0.58 |
| 9 | 86787001 | rs2246240 | 0.778 | -1.842 | -0.01 | 0.767 | 0.007 | -0.002 | -0.002 | 0 | 0.54 | 0.59 | 0.64 | 0.5 | 0.6 | 0.3 |
| 9 | 86788022 | rs10868107 | 0.18 | -1.924 | 0.017 | 0.768 | -0.003 | 0.02 | 0.02 | 0 | 0.15 | 0.15 | 0.08 | 0.36 | 0.12 | 0.29 |
| 9 | 86789200 | rs2245993 | 0.31 | -2.061 | 0.182 | 1.165 | -0.02 | 0.068 | 0.068 | 0 | 0.8 | 0.69 | 0.8 | 0.71 | 0.8 | 0.71 |
| 9 | 86789306 | rs11140416 | 0.427 | -1.12 | -0.077 | 1.289 | -0.061 | -0.076 | -0.076 | 1.843 | 0.21 | 0.05 | 0.37 | 0.31 | 0.24 | 0.27 |
| 9 | 86789462 | rs2245980 | -0.431 | -1.109 | -0.055 | 1.275 | -0.018 | 0.061 | 0.061 | 0 | 0.83 | 0.71 | 0.8 | 0.78 | 0.82 | 0.75 |
| 9 | 86791333 | rs7039156 | -1.445 | -1.319 | 0.227 | 0.913 | 0.001 | 0.004 | 0.004 | 0 | 0.48 | 0.58 | 0.55 | 0.71 | 0.45 | 0.66 |
| 9 | 86792058 | rs7042904 | -0.977 | -1.405 | 0.37 | 0.618 | -0.031 | -0.03 | -0.03 | 0 | 0.29 | 0.47 | 0.2 | 0.3 | 0.23 | 0.4 |
| 9 | 86793008 | rs10868111 | 0.664 | -1.355 | 0.216 | 0.631 | -0.003 | 0.003 | 0.003 | 0 | 0.15 | 0.17 | 0.09 | 0.28 | 0.12 | 0.28 |
| 9 | 86794631 | rs2256603 | 2.021 | -1.247 | 0.163 | 0.414 | -0.045 | 0.115 | 0.115 | 0 | 0.34 | 0.15 | 0.25 | 0.05 | 0.38 | 0.1 |
| 9 | 86795274 | rs17425371 | -0.03 | -1.16 | 0.172 | 0.541 | -0.002 | 0.03 | 0.03 | 0.003 | 0.09 | 0.05 | 0.08 | 0 | 0.08 | 0.09 |
| 9 | 86802590 | rs2265721 | 1.563 | -0.964 | 0.473 | 0.547 | -0.001 | -0.006 | -0.006 | 0 | 0.36 | 0.34 | 0.22 | 0.13 | 0.34 | 0.1 |
| 9 | 86806595 | rs1272566 | 0.456 | -1.481 | 0.392 | 0.813 | 0.004 | 0.006 | 0.006 | 0 | 0.68 | 0.69 | 0.65 | 0.74 | 0.74 | 0.65 |
| 9 | 86811147 | rs2251760 | 0.366 | -1.507 | 0.299 | 1.136 | 0.004 | -0.018 | -0.018 | 0 | 0.69 | 0.72 | 0.49 | 0.54 | 0.73 | 0.57 |
| 9 | 86812940 | rs2251530 | 0.307 | -1.485 | 0.308 | 1.06 | 0.003 | -0.012 | -0.012 | 1.851 | 0.69 | 0.73 | 0.49 | 0.49 | 0.73 | 0.57 |
| 9 | 86820581 | rs2484894 | 1.341 | -1.612 | 0.174 | 0.743 | 0.018 | 0.205 | 0.205 | 0 | 0.46 | 0.17 | 0.34 | 0.1 | 0.58 | 0.25 |
| 9 | 86824068 | rs7466124 | -0.384 | -1.826 | -0.554 | 0.696 | -0.08 | 0.095 | 0.095 | 0 | 0.19 | 0.57 | 0.3 | 0.25 | 0.09 | 0.22 |
| 9 | 86826872 | rs7861108 | -1.399 | -1.893 | -0.664 | 0.483 | 0.129 | -0.004 | -0.004 | 0 | 0.39 | 0.91 | 0.43 | 0.67 | 0.25 | 0.53 |
| 9 | 86830499 | rs7874528 | -0.101 | -1.764 | -0.424 | 0.48 | -0.079 | -0.132 | -0.132 | 0 | 0.18 | 0.53 | 0.12 | 0 | 0.08 | 0.14 |
| 9 | 86831936 | rs4877271 | 0.227 | -1.445 | -0.971 | 0.651 | -0.045 | -0.056 | -0.056 | 0.486 | 0.17 | 0.04 | 0.25 | 0.26 | 0.19 | 0.22 |
| 9 | 86834719 | rs7045243 | -0.651 | -1.569 | -0.869 | 0.615 | 0.042 | 0.021 | 0.021 | 0.644 | 0.34 | 0.72 | 0.23 | 0.34 | 0.22 | 0.44 |
| 9 | 86836666 | rs6559774 | -0.672 | -1.49 | -0.926 | 0.606 | 0.009 | 0.007 | 0.007 | 0.054 | 0.34 | 0.68 | 0.23 | 0.34 | 0.25 | 0.45 |
| 9 | 86838505 | rs4877823 | -1.128 | -1.35 | -0.737 | 0.591 | 0.055 | -0.002 | -0.002 | 0 | 0.5 | 0.74 | 0.48 | 0.6 | 0.41 | 0.65 |
| 9 | 86841402 | rs10780656 | -1.842 | -1.029 | -1.19 | 0.428 | -0.162 | 0.123 | 0.123 | 0 | 0.69 | 0.96 | 0.77 | 0.9 | 0.6 | 0.82 |
| 9 | 86853384 | rs7024751 | -0.262 | -0.971 | 0.177 | 1.373 | 0.028 | 0.032 | 0.032 | 0.013 | 0.07 | 0.18 | 0.08 | 0.1 | 0.1 | 0.04 |
| 9 | 86855859 | rs4877824 | -1.958 | -0.6 | -0.386 | 1.121 | -0.052 | 0.075 | 0.075 | 0 | 0.51 | 0.91 | 0.67 | 0.62 | 0.42 | 0.48 |
| 9 | 86857433 | rs4877826 | 0.136 | -0.699 | -0.368 | 1.17 | -0.059 | -0.014 | -0.014 | 0 | 0.14 | 0.04 | 0.24 | 0.16 | 0.16 | 0.1 |
| 9 | 86859971 | rs4877827 | -0.245 | -0.714 | -0.278 | 1.17 | -0.037 | -0.079 | -0.079 | 0 | 0.16 | 0.07 | 0.42 | 0.42 | 0.17 | 0.18 |
| 9 | 86867036 | rs883431 | -0.776 | -0.802 | 0.566 | 1.129 | 0.014 | 0.142 | 0.142 | 0 | 0.2 | 0.4 | 0.44 | 0.5 | 0.17 | 0.19 |
| 9 | 86868157 | rs1015951 | -1.526 | -0.877 | 0.822 | 0.806 | -0.004 | -0.027 | -0.027 | 0 | 0.41 | 0.46 | 0.4 | 0.26 | 0.31 | 0.37 |
| 9 | 86868447 | rs11140471 | -0.246 | -0.667 | 1.238 | 1.144 | 0.027 | 0.045 | 0.045 | 0.168 | 0.07 | 0.25 | 0.07 | 0.09 | 0.07 | 0.03 |
| 9 | 86872684 | rs11140476 | -0.129 | -0.742 | 1.097 | 0.975 | -0.005 | -0.019 | -0.019 | 0 | 0.21 | 0.26 | 0.13 | 0.06 | 0.13 | 0.25 |
| 9 | 86873532 | rs4877829 | -2.297 | -0.76 | 0.913 | 1.035 | -0.024 | 0.044 | 0.044 | 0 | 0.48 | 0.91 | 0.59 | 0.6 | 0.38 | 0.46 |
| 9 | 86878682 | rs7857113 | -1.399 | -0.705 | 1.322 | 1.151 | 0.015 | 0.002 | 0.002 | 0.329 | 0.25 | 0.54 | 0.29 | 0.27 | 0.22 | 0.14 |
| 9 | 86879344 | rs12348345 | -0.492 | -0.707 | 1.645 | 1.154 | 0.029 | -0.008 | -0.008 | 0 | 0.17 | 0.34 | 0.23 | 0.17 | 0.15 | 0.1 |
| 9 | 86880493 | rs11140482 | -0.977 | -0.717 | 1.665 | 1.166 | 0.013 | 0.153 | 0.153 | 0.011 | 0.2 | 0.54 | 0.36 | 0.48 | 0.16 | 0.19 |
| 9 | 86884890 | rs2184747 | -2.032 | -0.714 | 1.63 | 1.028 | -0.001 | -0.024 | -0.024 | 0 | 0.28 | 0.76 | 0.42 | 0.56 | 0.24 | 0.23 |
| 9 | 86886897 | rs11790651 | -0.313 | -0.7 | 1.594 | 1.032 | 0.005 | 0.026 | 0.026 | 0.311 | 0.08 | 0.19 | 0.06 | 0.1 | 0.07 | 0.04 |
| 9 | 86888818 | rs1332538 | -2.436 | -0.777 | 1.481 | 0.976 | -0.009 | 0.038 | 0.038 | 0 | 0.48 | 0.9 | 0.58 | 0.58 | 0.37 | 0.52 |
| 9 | 86889644 | rs12003423 | -0.369 | -0.757 | 1.581 | 1.014 | 0.027 | -0.012 | -0.012 | 0 | 0.17 | 0.31 | 0.22 | 0.16 | 0.15 | 0.1 |
| 9 | 86890758 | rs17426961 | -0.023 | -0.775 | 1.638 | 1.019 | 0.015 | 0.055 | 0.055 | 0 | 0.2 | 0.09 | 0.16 | 0.02 | 0.14 | 0.24 |
| 9 | 86893313 | rs11140490 | -0.513 | -0.75 | 1.646 | 0.848 | 0.024 | -0.015 | -0.015 | 0 | 0.17 | 0.34 | 0.23 | 0.16 | 0.15 | 0.1 |
| 9 | 86896586 | rs12686790 | -0.208 | -0.743 | 1.659 | 0.856 | 0.001 | 0.017 | 0.017 | 0 | 0.07 | 0.14 | 0.05 | 0.1 | 0.07 | 0.09 |
| 9 | 86900016 | rs4877833 | -0.481 | -0.736 | 1.647 | 0.836 | 0.025 | -0.014 | -0.014 | 0.161 | 0.17 | 0.34 | 0.23 | 0.16 | 0.15 | 0.1 |

| | | | | | | | | | | | | | | | | |
|---|---|---|---|---|---|---|---|---|---|---|---|---|---|---|---|---|
| 9 | 86900613 | rs7853066 | -0.316 | -0.735 | 1.654 | 0.845 | 0.022 | -0.008 | -0.008 | 0 | 0.16 | 0.34 | 0.22 | 0.16 | 0.13 | 0.1 |
| 9 | 86900926 | rs7853758 | -0.511 | -0.833 | 1.604 | 0.819 | 0.01 | -0.022 | -0.022 | 0 | 0.18 | 0.34 | 0.23 | 0.16 | 0.14 | 0.11 |
| 9 | 86904185 | rs4585823 | 0.038 | -0.794 | 1.662 | 0.863 | 0.008 | 0.012 | 0.012 | 0 | 0.19 | 0.15 | 0.17 | 0.03 | 0.14 | 0.24 |
| 9 | 86904754 | rs7042987 | -0.638 | 0.006 | 2.283 | 1.496 | 0.007 | 0.009 | 0.009 | 0.14 | 0.08 | 0.15 | 0.06 | 0.11 | 0.1 | 0.09 |
| 9 | 86905551 | rs4877834 | -0.639 | -0.002 | 2.291 | 1.484 | 0.019 | -0.013 | -0.013 | 0 | 0.17 | 0.34 | 0.22 | 0.16 | 0.13 | 0.1 |
| 9 | 86906008 | rs7047315 | -0.158 | -0.017 | 2.395 | 1.477 | -0.01 | 0 | 0 | 0 | 0.14 | 0.1 | 0.2 | 0.16 | 0.13 | 0.1 |
| 9 | 86906107 | rs6559779 | -2.259 | -0.036 | 2.437 | 1.611 | -0.021 | 0.16 | 0.16 | 4.595 | 0.15 | 0.66 | 0.22 | 0.42 | 0.11 | 0.18 |
| 9 | 86906932 | rs7864584 | -2.277 | -0.036 | 2.437 | 1.611 | -0.022 | 0.158 | 0.158 | 3.953 | 0.15 | 0.66 | 0.22 | 0.42 | 0.11 | 0.18 |
| 9 | 86908830 | rs12343928 | -2.047 | -0.03 | 2.434 | 1.618 | -0.004 | 0.272 | 0.272 | 2.512 | 0.14 | 0.51 | 0.21 | 0.38 | 0.11 | 0.17 |
| 9 | 86909082 | rs10868137 | -0.106 | -0.023 | 2.413 | 1.636 | -0.01 | -0.002 | -0.002 | 0 | 0.14 | 0.1 | 0.2 | 0.16 | 0.13 | 0.1 |
| 9 | 86909399 | rs7030202 | -0.665 | -0.025 | 2.433 | 1.646 | 0.006 | 0.009 | 0.009 | 0 | 0.08 | 0.15 | 0.07 | 0.11 | 0.1 | 0.09 |
| 9 | 86909435 | rs885003 | -1.929 | -0.025 | 2.414 | 1.648 | 0.007 | 0.317 | 0.317 | 2.344 | 0.13 | 0.46 | 0.2 | 0.38 | 0.11 | 0.17 |
| 9 | 86909550 | rs885004 | -0.091 | -0.011 | 2.204 | 1.644 | -0.009 | -0.003 | -0.003 | 0 | 0.13 | 0.1 | 0.2 | 0.16 | 0.13 | 0.1 |
| 9 | 86914268 | rs11791990 | -0.681 | -0.008 | 2.261 | 1.656 | 0.007 | 0.006 | 0.006 | 0 | 0.08 | 0.15 | 0.07 | 0.08 | 0.1 | 0.09 |
| 9 | 86914372 | rs17087049 | -1.091 | -0.022 | 2.345 | 1.651 | 0.035 | 0.422 | 0.422 | 1.767 | 0.06 | 0.34 | 0.14 | 0.27 | 0.01 | 0.08 |
| 9 | 86915542 | rs7861242 | -1.25 | -0.059 | 1.841 | 1.529 | 0.004 | -0.01 | -0.01 | 0 | 0.18 | 0.26 | 0.22 | 0.11 | 0.18 | 0.2 |
| 9 | 86916851 | rs4877835 | -0.058 | -0.054 | 1.832 | 1.482 | -0.01 | -0.004 | -0.004 | 0.266 | 0.14 | 0.1 | 0.2 | 0.18 | 0.13 | 0.1 |
| 9 | 86916910 | rs12342617 | -0.739 | -0.055 | 1.753 | 1.482 | 0.01 | 0.008 | 0.008 | 0 | 0.08 | 0.19 | 0.07 | 0.08 | 0.1 | 0.09 |
| 9 | 86917088 | rs4877836 | -0.037 | -0.052 | 1.739 | 1.484 | -0.017 | -0.008 | -0.008 | 0.056 | 0.13 | 0.08 | 0.2 | 0.18 | 0.13 | 0.1 |
| 9 | 86917382 | rs4877837 | -1.265 | -0.036 | 1.689 | 1.378 | 0.004 | -0.013 | -0.013 | 0 | 0.18 | 0.26 | 0.22 | 0.1 | 0.18 | 0.2 |
| 9 | 86920236 | rs7867504 | -2.72 | -0.05 | 1.645 | 1.394 | -0.014 | 0.005 | 0.005 | 0 | 0.39 | 0.84 | 0.58 | 0.6 | 0.32 | 0.39 |
| 9 | 86920632 | rs12685409 | -0.52 | -0.027 | 1.539 | 1.409 | 0.005 | 0.005 | 0.005 | 0.129 | 0.08 | 0.11 | 0.06 | 0.08 | 0.1 | 0.09 |
| 9 | 86920711 | rs4242626 | -2.511 | -0.012 | 1.475 | 1.42 | -0.009 | 0.006 | 0.006 | 0 | 0.38 | 0.82 | 0.57 | 0.6 | 0.32 | 0.39 |
| 9 | 86921615 | rs10868142 | -1.246 | 0.035 | 1.615 | 1.453 | 0.002 | -0.013 | -0.013 | 0.182 | 0.18 | 0.29 | 0.22 | 0.11 | 0.17 | 0.2 |
| 9 | 86922777 | rs13288052 | -0.443 | 0.199 | 1.245 | 1.305 | 0.023 | 0.057 | 0.057 | 0 | 0.07 | 0.14 | 0.2 | 0.19 | 0.1 | 0.11 |
| 9 | 86925655 | rs9792674 | -2.31 | -0.13 | 1.062 | 1.134 | 0.003 | 0.025 | 0.025 | 0 | 0.36 | 0.81 | 0.56 | 0.62 | 0.29 | 0.41 |
| 9 | 86925748 | rs11140507 | -1.898 | 0.172 | 1.721 | 1.375 | 0.004 | 0.055 | 0.055 | 0 | 0.19 | 0.27 | 0.36 | 0.54 | 0.19 | 0.26 |
| 9 | 86926892 | rs11140508 | -1.308 | 0.181 | 1.567 | 1.439 | -0.008 | -0.007 | -0.007 | 0 | 0.14 | 0.11 | 0.23 | 0.18 | 0.17 | 0.12 |
| 9 | 86928133 | rs11140510 | -1.429 | 0.185 | 1.569 | 1.441 | -0.009 | -0.007 | -0.007 | 0 | 0.14 | 0.11 | 0.23 | 0.18 | 0.17 | 0.12 |
| 9 | 86928905 | rs11140511 | -2.236 | 0.134 | 2.021 | 1.412 | 0.031 | -0.003 | -0.003 | 4.735 | 0.27 | 0.45 | 0.41 | 0.26 | 0.26 | 0.22 |
| 9 | 86929380 | rs4877842 | -2.337 | 0.116 | 2.035 | 1.27 | 0.022 | 0.002 | 0.002 | 0 | 0.28 | 0.46 | 0.52 | 0.63 | 0.28 | 0.36 |
| 9 | 86931003 | rs7853538 | -0.73 | 0.102 | 1.778 | 1.068 | 0.072 | 0.068 | 0.068 | 3.825 | 0.05 | 0.26 | 0.01 | 0 | 0 | 0 |
| 9 | 86932764 | rs11140519 | -0.167 | 0.153 | 1.241 | 1.122 | -0.015 | -0.016 | -0.016 | 0 | 0.08 | 0.06 | 0.17 | 0.13 | 0.1 | 0.1 |
| 9 | 86932859 | rs12347156 | -1.07 | 0.136 | 1.314 | 1.094 | 0.022 | 0.018 | 0.018 | 3.878 | 0.07 | 0.48 | 0.03 | 0 | 0 | 0.04 |
| 9 | 86933046 | rs17336552 | 0.074 | 0.138 | 1.309 | 1.115 | 0.002 | -0.001 | -0.001 | 1.017 | 0.08 | 0.11 | 0.17 | 0.1 | 0.1 | 0.09 |
| 9 | 86933405 | rs7035188 | -1.896 | 0.108 | 1.15 | 1.211 | -0.032 | 0.211 | 0.211 | 1.83 | 0.15 | 0.57 | 0.19 | 0.38 | 0.09 | 0.19 |
| 9 | 86933934 | rs10868145 | -2.173 | 0.11 | 1.047 | 1.207 | 0.011 | 0.011 | 0.011 | 0 | 0.27 | 0.41 | 0.51 | 0.61 | 0.29 | 0.34 |
| 9 | 86934074 | rs10868146 | -0.444 | -0.085 | 0.967 | 1.461 | 0 | 0.111 | 0.111 | 0.769 | 0.08 | 0.1 | 0.16 | 0.38 | 0.09 | 0.15 |
| 9 | 86934277 | rs10868147 | -0.444 | -0.085 | 0.857 | 1.461 | 0 | 0.088 | 0.088 | 0.769 | 0.08 | 0.08 | 0.16 | 0.38 | 0.09 | 0.15 |
| 9 | 86935306 | rs11140521 | -1.232 | -0.049 | 1.074 | 0.476 | 0.027 | 0.023 | 0.023 | 3.658 | 0.07 | 0.42 | 0.02 | 0 | 0 | 0.04 |
| 9 | 86935429 | rs11140522 | -1.108 | 0.294 | -0.285 | 0.642 | -0.005 | -0.005 | -0.005 | 0.028 | 0.1 | 0.08 | 0.17 | 0.13 | 0.1 | 0.1 |
| 9 | 86937315 | rs11140525 | -0.808 | 0.2 | -0.596 | 0.242 | 0.001 | 0.037 | 0.037 | 0.623 | 0.35 | 0.8 | 0.45 | 0.25 | 0.26 | 0.24 |
| 9 | 86939823 | rs4877845 | 0.085 | 0.639 | 0.081 | 0.544 | 0.026 | 0.011 | 0.011 | 0 | 0.73 | 0.46 | 0.71 | 0.73 | 0.75 | 0.73 |
| 9 | 86941443 | rs13283683 | 0.207 | 0.001 | 0.415 | 0.433 | 0 | 0 | 0 | 3.105 | 0.16 | 0.15 | 0.12 | 0.19 | 0.13 | 0.25 |
| 9 | 86944451 | rs4305983 | -0.23 | -0.262 | 0.169 | 0.294 | -0.026 | 0.048 | 0.048 | 0 | 0.71 | 0.85 | 0.82 | 0.83 | 0.72 | 0.79 |
| 9 | 86944921 | rs4242627 | 1.184 | -0.433 | 0.282 | 1.056 | -0.001 | 0 | 0 | 0 | 0.52 | 0.51 | 0.53 | 0.35 | 0.56 | 0.44 |
| 9 | 86946073 | rs17343066 | 1.598 | -0.475 | 0.244 | 1.037 | -0.092 | 0.07 | 0.07 | 0.326 | 0.46 | 0.09 | 0.49 | 0.31 | 0.54 | 0.41 |
| 9 | 86946417 | rs4877847 | 1.194 | -0.377 | 0.383 | 1.144 | -0.001 | 0.003 | 0.003 | 0 | 0.52 | 0.51 | 0.58 | 0.37 | 0.56 | 0.44 |
| 9 | 86946607 | rs980292 | -1.056 | -0.283 | 0.254 | 0.886 | -0.034 | 0.032 | 0.032 | 0 | 0.79 | 0.89 | 0.85 | 0.87 | 0.78 | 0.84 |
| 9 | 86948616 | rs4266723 | -1.133 | -0.286 | 0.253 | 0.885 | -0.034 | 0.03 | 0.03 | 0 | 0.8 | 0.9 | 0.86 | 0.87 | 0.79 | 0.84 |
| 9 | 86949847 | rs13283321 | 0.147 | -0.29 | 0.726 | 1.076 | -0.012 | -0.001 | -0.001 | 0 | 0.17 | 0.24 | 0.12 | 0.19 | 0.13 | 0.24 |
| 9 | 86949973 | rs10868149 | -0.806 | -0.287 | 1.131 | 1.155 | -0.019 | 0.032 | 0.032 | 0 | 0.26 | 0.37 | 0.28 | 0.37 | 0.22 | 0.39 |
| 9 | 86950870 | rs4877848 | -0.787 | -0.282 | 1.213 | 1.121 | -0.02 | -0.003 | -0.003 | 0 | 0.25 | 0.32 | 0.17 | 0.28 | 0.19 | 0.31 |
| 9 | 86951615 | rs4877850 | -0.48 | -0.003 | 1.416 | 1.032 | 0.03 | 0.024 | 0.024 | 0 | 0.81 | 0.68 | 0.75 | 0.74 | 0.79 | 0.79 |
| 9 | 86953674 | rs7043257 | -0.489 | -0.006 | 1.348 | 1.145 | 0.009 | 0.021 | 0.021 | 0 | 0.81 | 0.75 | 0.85 | 0.67 | 0.79 | 0.84 |
| 9 | 86954132 | rs4877851 | -0.517 | -0.165 | 1.384 | 1.053 | -0.016 | -0.009 | -0.009 | 0 | 0.22 | 0.64 | 0.13 | 0.2 | 0.14 | 0.26 |
| 9 | 86954466 | rs7027983 | 0.546 | -0.151 | 1.591 | 0.968 | 0.011 | 0.01 | 0.01 | 0 | 0.64 | 0.48 | 0.65 | 0.37 | 0.64 | 0.54 |
| 9 | 86955157 | rs3812509 | 0.6 | -0.147 | 1.359 | 0.979 | 0.015 | 0.022 | 0.022 | 0 | 0.66 | 0.54 | 0.65 | 0.41 | 0.64 | 0.55 |
| 9 | 86955682 | rs7035753 | 1.106 | -0.062 | 0.752 | 0.985 | -0.042 | -0.029 | -0.029 | 0 | 0.59 | 0.12 | 0.61 | 0.38 | 0.63 | 0.53 |
| 9 | 86960060 | rs10780664 | -1.288 | -0.196 | 1.884 | 1.249 | 0.001 | 0.001 | 0.001 | 1.624 | 0.26 | 0.72 | 0.26 | 0.47 | 0.2 | 0.36 |
| 9 | 86963692 | rs7860537 | -0.08 | -0.196 | 1.934 | 1.23 | 0.002 | 0.002 | 0.002 | 0 | 0.16 | 0.16 | 0.08 | 0.17 | 0.13 | 0.26 |
| 9 | 86964575 | rs13297855 | -0.04 | -0.196 | 1.934 | 1.23 | 0.001 | 0.002 | 0.002 | 0 | 0.16 | 0.15 | 0.08 | 0.17 | 0.13 | 0.26 |
| 9 | 86964656 | rs13298157 | -0.758 | -0.153 | 1.422 | 1.254 | -0.027 | -0.037 | -0.037 | 0 | 0.22 | 0.53 | 0.13 | 0.2 | 0.16 | 0.28 |
| 9 | 86966200 | rs11140543 | -1.072 | -0.132 | 1.077 | 1.238 | 0.005 | -0.011 | -0.011 | 0 | 0.26 | 0.75 | 0.26 | 0.47 | 0.2 | 0.36 |
| 9 | 86966759 | rs11140544 | -0.998 | -0.17 | 1.061 | 1.14 | 0 | -0.001 | -0.001 | 0 | 0.25 | 0.72 | 0.15 | 0.38 | 0.18 | 0.3 |
| 9 | 86969052 | rs17087148 | -1.762 | -0.048 | 1.173 | 1.35 | 0.029 | 0.303 | 0.303 | 0.043 | 0.05 | 0.26 | 0.15 | 0.27 | 0.04 | 0.09 |
| 9 | 86970543 | rs10868152 | -1.616 | -0.134 | 1.17 | 0.957 | 0.01 | -0.018 | -0.018 | 0 | 0.32 | 0.75 | 0.26 | 0.44 | 0.29 | 0.28 |
| 9 | 86971246 | rs4877854 | -0.986 | -0.149 | 1.676 | 0.813 | -0.025 | -0.007 | -0.007 | 0.371 | 0.2 | 0.49 | 0.14 | 0.18 | 0.15 | 0.18 |
| 9 | 86972086 | rs12552512 | -0.219 | -0.036 | 1.568 | 0.845 | -0.002 | 0.006 | 0.006 | 0 | 0.16 | 0.23 | 0.11 | 0.18 | 0.14 | 0.17 |

| | | | | | | | | | | | | | | | | |
|---|---|---|---|---|---|---|---|---|---|---|---|---|---|---|---|---|
| 9 | 86983368 | rs10868153 | -2.096 | -0.232 | 1.796 | 0.527 | -0.01 | -0.009 | -0.009 | 0 | 0.1 | 0.48 | 0.06 | 0.04 | 0.05 | 0.04 |
| 9 | 86983672 | rs12378361 | 0.039 | 0.099 | 1.129 | 0.442 | -0.044 | -0.048 | -0.048 | 0 | 0.14 | 0.03 | 0.12 | 0.19 | 0.19 | 0.11 |
| 9 | 86991225 | rs11140555 | -1.456 | 0.037 | 1.606 | 0.422 | -0.006 | -0.01 | -0.01 | 2.415 | 0.23 | 0.54 | 0.18 | 0.19 | 0.24 | 0.14 |
| 9 | 86994670 | rs7021071 | -1.271 | 0.025 | 1.659 | 0.416 | 0.002 | -0.003 | -0.003 | 0 | 0.22 | 0.45 | 0.17 | 0.19 | 0.24 | 0.13 |
| 9 | 86994945 | rs11140557 | -0.152 | 0.037 | 1.587 | 0.421 | -0.015 | -0.02 | -0.02 | 0 | 0.16 | 0.1 | 0.12 | 0.19 | 0.19 | 0.11 |
| 9 | 87000794 | rs12555868 | -1.032 | 0.07 | 1.877 | 0.418 | 0.005 | 0.002 | 0.002 | 0 | 0.2 | 0.28 | 0.15 | 0.19 | 0.23 | 0.13 |
| 9 | 87004182 | rs586046 | 0.9 | -0.209 | 0.777 | -0.054 | -0.027 | 0.066 | 0.066 | 0.471 | 0.28 | 0.02 | 0.28 | 0.15 | 0.2 | 0.37 |
| 9 | 87008860 | rs12379161 | -0.801 | -0.107 | 1.45 | 0.172 | 0.001 | 0.008 | 0.008 | 0 | 0.22 | 0.37 | 0.16 | 0.23 | 0.23 | 0.14 |
| 9 | 87013515 | rs10780674 | -1.285 | -0.111 | 1.464 | 0.172 | -0.014 | -0.005 | -0.005 | 2.5 | 0.26 | 0.6 | 0.17 | 0.23 | 0.22 | 0.14 |
| 9 | 87015151 | rs11140582 | -0.961 | -0.123 | 1.573 | 0.155 | -0.009 | 0.005 | 0.005 | 0 | 0.23 | 0.41 | 0.15 | 0.23 | 0.22 | 0.14 |
| 9 | 87026298 | rs7042275 | -1.371 | 0.119 | 1.376 | 0.228 | 0.057 | 0.049 | 0.049 | 2.988 | 0.06 | 0.26 | 0.08 | 0.06 | 0.06 | 0.11 |
| 9 | 87050410 | rs10780677 | 0.295 | 0.099 | 1.068 | 0.234 | -0.01 | -0.017 | -0.017 | 0.722 | 0.15 | 0.11 | 0.1 | 0.17 | 0.17 | 0.1 |
| 9 | 87050694 | rs7870219 | -1.604 | 0.019 | 1.535 | 0.201 | 0.002 | 0.002 | 0.002 | 0.947 | 0.28 | 0.73 | 0.23 | 0.24 | 0.28 | 0.23 |
| 9 | 87057020 | rs17087228 | -1.087 | -0.3 | 1.542 | 0.101 | -0.032 | -0.024 | -0.024 | 0.165 | 0.19 | 0.39 | 0.1 | 0.13 | 0.14 | 0.08 |
| 9 | 87062568 | rs7868516 | -0.527 | -0.268 | 1.292 | 0.132 | -0.009 | -0.002 | -0.002 | 0 | 0.15 | 0.23 | 0.08 | 0.13 | 0.1 | 0.06 |
| 9 | 87075137 | rs1332535 | -1.531 | -0.313 | 1.224 | 0.127 | -0.004 | -0.002 | -0.002 | 0 | 0.25 | 0.7 | 0.19 | 0.2 | 0.2 | 0.17 |
| 9 | 87082451 | rs7875211 | 0.094 | -0.228 | 0.816 | 0.19 | -0.012 | 0.023 | 0.023 | 0 | 0.47 | 0.33 | 0.46 | 0.56 | 0.5 | 0.33 |
| 9 | 87085182 | rs1326930 | 1.182 | -0.224 | 0.71 | 0.181 | -0.029 | -0.044 | -0.044 | 0 | 0.32 | 0.2 | 0.39 | 0.43 | 0.4 | 0.27 |
| 9 | 87086912 | rs2778944 | 0.981 | -0.172 | 0.189 | 0.015 | -0.072 | 0.057 | 0.057 | 0.288 | 0.31 | 0.03 | 0.31 | 0.16 | 0.27 | 0.37 |
| 9 | 87089546 | rs4877276 | -0.531 | 0.04 | 0.736 | 0.275 | 0 | 0.056 | 0.056 | 0.011 | 0.53 | 0.72 | 0.56 | 0.62 | 0.55 | 0.42 |
| 9 | 87097512 | rs4433222 | -1.437 | -0.037 | 1.52 | 0.236 | -0.007 | -0.019 | -0.019 | 0 | 0.15 | 0.72 | 0.16 | 0.09 | 0.13 | 0.13 |
| 9 | 87099638 | rs11140629 | -1.518 | 0.095 | 1.102 | 0.317 | -0.003 | -0.015 | -0.015 | 0 | 0.24 | 0.51 | 0.2 | 0.21 | 0.22 | 0.18 |
| 9 | 87117500 | rs12685438 | -0.867 | -0.284 | 0.839 | 0.887 | -0.003 | -0.031 | -0.031 | 0 | 0.13 | 0.02 | 0.06 | 0.15 | 0.09 | 0.05 |
| 9 | 87125877 | rs12685148 | -0.631 | -0.29 | 1.131 | 0.868 | -0.009 | -0.027 | -0.027 | 0.011 | 0.12 | 0.03 | 0.06 | 0.14 | 0.09 | 0.05 |
| 9 | 87127736 | rs10868171 | -1.409 | -0.424 | 2.026 | 0.789 | 0.092 | 0.198 | 0.198 | 1.358 | 0.05 | 0.46 | 0.06 | 0.08 | 0.01 | 0.03 |
| 9 | 87135644 | rs7871278 | -0.665 | -0.635 | 0.996 | 0.375 | 0 | 0 | 0 | 0 | 0.39 | 0.61 | 0.43 | 0.42 | 0.3 | 0.41 |
| 9 | 87143351 | rs1926746 | -0.569 | -0.67 | 1.038 | 0.449 | -0.015 | 0.005 | 0.005 | 0 | 0.32 | 0.4 | 0.24 | 0.36 | 0.21 | 0.38 |
| 9 | 87167205 | rs11140663 | 0.623 | -0.812 | 0.746 | 0.344 | -0.041 | -0.078 | -0.078 | 0 | 0.2 | 0.02 | 0.19 | 0.28 | 0.17 | 0.3 |
| 9 | 87173097 | rs1347857 | -0.995 | -0.785 | 0.458 | 0.27 | 0.043 | -0.022 | -0.022 | 0 | 0.36 | 0.79 | 0.42 | 0.42 | 0.25 | 0.4 |
| 9 | 87173333 | rs1347858 | -1.562 | -0.81 | 0.371 | 0.376 | 0.063 | -0.034 | -0.034 | 0 | 0.43 | 0.84 | 0.46 | 0.56 | 0.3 | 0.45 |
| 9 | 87177985 | rs12115546 | -0.703 | -0.742 | 0.592 | 0.329 | -0.017 | -0.014 | -0.014 | 0 | 0.09 | 0.18 | 0.18 | 0.05 | 0.06 | 0.05 |
| 9 | 87179264 | rs1866439 | -1.014 | -0.667 | -0.057 | 0.344 | 0.041 | -0.024 | -0.024 | 0 | 0.36 | 0.79 | 0.42 | 0.42 | 0.25 | 0.4 |
| 9 | 87190178 | rs11140676 | 0.116 | -0.152 | -0.276 | 0.429 | 0 | -0.008 | -0.008 | 0 | 0.27 | 0.25 | 0.25 | 0.37 | 0.25 | 0.36 |
| 9 | 87193728 | rs2118819 | -0.621 | -0.176 | -0.26 | 0.395 | 0.002 | -0.009 | -0.009 | 0 | 0.32 | 0.32 | 0.34 | 0.43 | 0.28 | 0.37 |
| 9 | 87200556 | rs2218090 | -1.74 | 0.048 | -0.812 | 0.2 | -0.042 | -0.039 | -0.039 | 0 | 0.54 | 0.88 | 0.5 | 0.55 | 0.47 | 0.47 |
| 9 | 87201325 | rs9886924 | 0.434 | 0.096 | -0.704 | 0.279 | -0.004 | -0.026 | -0.026 | 0 | 0.25 | 0.18 | 0.24 | 0.38 | 0.24 | 0.32 |
| 9 | 87203787 | rs11788721 | 0.503 | 0.214 | -0.69 | 0.107 | -0.001 | -0.002 | -0.002 | 0 | 0.15 | 0.14 | 0.1 | 0.12 | 0.15 | 0.08 |
| 9 | 87208419 | rs7021722 | -1.091 | 0.292 | -1.06 | 0.347 | -0.002 | 0.01 | 0.01 | 0 | 0.51 | 0.86 | 0.49 | 0.56 | 0.45 | 0.45 |
| 9 | 87216845 | rs715243 | -1.241 | 0.278 | -1.11 | 0.329 | -0.076 | 0.008 | 0.008 | 0 | 0.59 | 0.9 | 0.5 | 0.34 | 0.52 | 0.35 |
| 9 | 87217628 | rs4111912 | -0.259 | -0.075 | -0.108 | 0.184 | 0 | -0.003 | -0.003 | 0 | 0.38 | 0.38 | 0.27 | 0.3 | 0.35 | 0.32 |
| 9 | 87226088 | rs1020120 | -1.57 | -0.203 | -0.261 | 0.219 | -0.034 | -0.037 | -0.037 | 0.009 | 0.61 | 0.84 | 0.51 | 0.54 | 0.54 | 0.46 |
| 9 | 87231641 | rs4877869 | -0.417 | 0.18 | -0.114 | 0.185 | -0.014 | 0.026 | 0.026 | 0.035 | 0.37 | 0.22 | 0.26 | 0.3 | 0.37 | 0.33 |
| 9 | 87234111 | rs10868215 | -0.634 | 0.226 | -0.246 | 0.195 | -0.001 | 0.002 | 0.002 | 0 | 0.39 | 0.37 | 0.27 | 0.3 | 0.38 | 0.34 |
| 9 | 87236739 | rs12005792 | -0.843 | -0.387 | -0.411 | 0.751 | 0.006 | -0.052 | -0.052 | 0 | 0.29 | 0.04 | 0.17 | 0.3 | 0.21 | 0.31 |
| 9 | 87238610 | rs11789677 | -0.442 | -0.111 | -0.052 | 0.591 | -0.006 | 0.001 | 0.001 | 0 | 0.21 | 0.18 | 0.17 | 0.18 | 0.23 | 0.12 |
| 9 | 87239147 | rs11140693 | -0.223 | -0.145 | -0.075 | 0.869 | 0 | -0.016 | -0.016 | 0 | 0.06 | 0.03 | 0.06 | 0.16 | 0.06 | 0.05 |
| 9 | 87240130 | rs11140694 | -0.856 | -0.156 | -0.065 | 0.821 | -0.011 | -0.058 | -0.058 | 0.188 | 0.18 | 0.03 | 0.12 | 0.22 | 0.13 | 0.27 |
| 9 | 87240403 | rs11140695 | -1.318 | -0.262 | 0.148 | 0.813 | 0.012 | -0.041 | -0.041 | 0 | 0.24 | 0.09 | 0.17 | 0.27 | 0.17 | 0.29 |
| 9 | 87246051 | rs17087472 | -0.03 | -0.216 | 0.172 | 0.793 | 0.049 | -0.03 | -0.03 | 0.892 | 0.06 | 0.16 | 0.06 | 0 | 0.09 | 0.03 |
| 9 | 87248164 | rs11140704 | -1.314 | -0.275 | 0.108 | 0.804 | 0.003 | -0.071 | -0.071 | 0 | 0.24 | 0.05 | 0.17 | 0.27 | 0.17 | 0.29 |
| 9 | 87250190 | rs11140707 | -1.756 | -0.311 | 0.874 | 0.805 | 0.031 | -0.073 | -0.073 | 0.535 | 0.4 | 0.12 | 0.29 | 0.45 | 0.3 | 0.38 |
| 9 | 87251183 | rs985543 | -2.119 | -0.267 | 0.892 | 0.83 | 0.002 | 0.005 | 0.005 | 0 | 0.47 | 0.34 | 0.41 | 0.48 | 0.42 | 0.42 |
| 9 | 87251376 | rs10868220 | 0.656 | -0.218 | 0.923 | 0.806 | -0.003 | -0.048 | -0.048 | 0 | 0.16 | 0.02 | 0.11 | 0.18 | 0.13 | 0.09 |
| 9 | 87253419 | rs17087514 | -0.654 | -0.382 | 1.589 | 0.076 | 0.068 | -0.016 | -0.016 | 0.815 | 0.07 | 0.21 | 0.12 | 0.03 | 0.11 | 0.04 |
| 9 | 87266352 | rs1187333 | 0.686 | -0.372 | 1.57 | 0.075 | 0.059 | -0.021 | -0.021 | 1.482 | 0.92 | 0.79 | 0.87 | 0.97 | 0.88 | 0.95 |
| 9 | 87266700 | rs11140714 | -0.485 | -0.451 | 1.601 | 0.012 | 0.001 | 0.002 | 0.002 | 0 | 0.25 | 0.25 | 0.17 | 0.13 | 0.2 | 0.13 |
| 9 | 87267699 | rs1439054 | -0.61 | -0.538 | 1.465 | 0.071 | 0.029 | -0.044 | -0.044 | 0.009 | 0.16 | 0.06 | 0.11 | 0.26 | 0.09 | 0.25 |
| 9 | 87272824 | rs11140720 | -0.518 | -0.526 | 1.455 | 0.076 | 0.029 | -0.042 | -0.042 | 0.514 | 0.15 | 0.06 | 0.11 | 0.21 | 0.09 | 0.24 |
| 9 | 87275895 | rs1147199 | 1.541 | -0.578 | 1.526 | 0.035 | 0 | -0.001 | -0.001 | 0 | 0.77 | 0.78 | 0.82 | 0.79 | 0.82 | 0.73 |
| 9 | 87281307 | rs1307279 | 1.561 | -0.496 | 1.856 | 0.057 | -0.014 | 0.019 | 0.019 | 0.182 | 0.74 | 0.5 | 0.74 | 0.7 | 0.79 | 0.7 |
| 9 | 87282527 | rs1212171 | 1.963 | -0.477 | 1.593 | 0.013 | 0.013 | 0.024 | 0.024 | 0.005 | 0.49 | 0.28 | 0.57 | 0.58 | 0.59 | 0.57 |
| 9 | 87282615 | rs3758317 | 0.332 | -0.442 | 1.467 | 0.045 | -0.038 | 0.083 | 0.083 | 0.075 | 0.19 | 0.03 | 0.1 | 0.07 | 0.17 | 0.12 |
| 9 | 87288193 | rs1439050 | -1.634 | -0.367 | 1.751 | 0.133 | -0.009 | -0.002 | -0.002 | 0 | 0.44 | 0.54 | 0.37 | 0.42 | 0.32 | 0.4 |
| 9 | 87295237 | rs1187350 | 1.512 | -0.284 | 1.395 | 0.254 | 0 | 0.006 | 0.006 | 0 | 0.48 | 0.4 | 0.57 | 0.56 | 0.56 | 0.57 |
| 9 | 87302196 | rs1619120 | 1.187 | -0.268 | 1.414 | 0.213 | 0.001 | 0.003 | 0.003 | 0 | 0.55 | 0.57 | 0.7 | 0.63 | 0.62 | 0.62 |
| 9 | 87308783 | rs1187343 | 0.778 | -0.303 | 1.218 | 0.245 | -0.025 | 0.083 | 0.083 | 0 | 0.62 | 0.81 | 0.76 | 0.82 | 0.64 | 0.7 |
| 9 | 87316037 | rs1187337 | 1.348 | -0.324 | 1.34 | 0.278 | 0.011 | 0.055 | 0.055 | 0 | 0.52 | 0.68 | 0.68 | 0.7 | 0.57 | 0.6 |
| 9 | 87346919 | rs1419245 | -0.545 | -0.302 | 1.238 | 0.325 | 0.004 | -0.024 | -0.024 | 0 | 0.07 | 0.01 | 0.06 | 0.09 | 0.05 | 0.12 |
| 9 | 87352788 | rs3780632 | 1.878 | -0.515 | 1.788 | 0.101 | 0.008 | 0.002 | 0.002 | 0.418 | 0.65 | 0.34 | 0.66 | 0.68 | 0.69 | 0.68 |

| | | | | | | | | | | | | | | | | |
|---|---|---|---|---|---|---|---|---|---|---|---|---|---|---|---|---|
| 9 | 87355358 | rs11140745 | 1.89 | -0.516 | 1.738 | 0.105 | 0.016 | 0.006 | 0.006 | 0.341 | 0.64 | 0.31 | 0.65 | 0.68 | 0.69 | 0.68 |
| 9 | 87387622 | rs1573219 | 1.712 | -0.522 | 2.095 | 0.103 | -0.021 | -0.015 | -0.015 | 0 | 0.69 | 0.53 | 0.75 | 0.74 | 0.73 | 0.7 |
| 9 | 87392582 | rs10465180 | 2.107 | -0.532 | 2.128 | 0.099 | -0.012 | -0.003 | -0.003 | 0 | 0.66 | 0.44 | 0.67 | 0.68 | 0.69 | 0.67 |
| 9 | 87392708 | rs4877880 | 1.065 | -0.551 | 2.359 | 0.092 | 0 | 0 | 0 | 0 | 0.77 | 0.79 | 0.84 | 0.87 | 0.78 | 0.83 |
| 9 | 87395810 | rs1778970 | 2.053 | -0.549 | 2.352 | 0.096 | 0.016 | 0.004 | 0.004 | 0.272 | 0.64 | 0.3 | 0.65 | 0.68 | 0.7 | 0.67 |
| 9 | 87395876 | rs1662697 | 1.744 | -0.5 | 1.908 | 0.12 | -0.011 | -0.007 | -0.007 | 0 | 0.69 | 0.64 | 0.76 | 0.73 | 0.73 | 0.7 |
| 9 | 87405212 | rs1662695 | -0.507 | -0.205 | 0.715 | 0.095 | -0.022 | -0.03 | -0.03 | 0 | 0.92 | 0.99 | 0.93 | 0.87 | 0.9 | 0.89 |
| 9 | 87406168 | rs1443440 | 0.669 | -0.137 | 0.872 | 0.142 | -0.001 | -0.003 | -0.003 | 0 | 0.75 | 0.72 | 0.82 | 0.83 | 0.77 | 0.81 |
| 9 | 87409025 | rs1899640 | 1.182 | -0.151 | 0.742 | 0.174 | -0.005 | -0.003 | -0.003 | 0 | 0.68 | 0.65 | 0.68 | 0.72 | 0.69 | 0.66 |
| 9 | 87409862 | rs17087710 | -1.17 | -0.012 | 0.431 | 0.517 | -0.011 | -0.041 | -0.041 | 0 | 0.08 | 0.01 | 0.08 | 0.13 | 0.06 | 0.15 |
| 9 | 87414794 | rs1187287 | 0.41 | -0.442 | 0.925 | 0.064 | -0.029 | -0.054 | -0.054 | 0 | 0.71 | 0.53 | 0.82 | 0.82 | 0.8 | 0.8 |
| 9 | 87415028 | rs1187286 | 0.975 | -0.497 | 0.948 | 0.136 | -0.026 | -0.007 | -0.007 | 0.178 | 0.64 | 0.49 | 0.75 | 0.68 | 0.76 | 0.7 |
| 9 | 87416825 | rs1187280 | -0.229 | -0.527 | 0.785 | 0.042 | -0.013 | 0.055 | 0.055 | 0 | 0.79 | 0.89 | 0.86 | 0.91 | 0.8 | 0.89 |
| 9 | 87419117 | rs716893 | 1.467 | -0.262 | 0.947 | -0.139 | 0.009 | 0.017 | 0.017 | 0 | 0.61 | 0.3 | 0.56 | 0.71 | 0.66 | 0.7 |
| 9 | 87421631 | rs3739804 | -1.647 | -0.197 | 1.363 | -0.095 | 0.02 | 0.127 | 0.127 | 0.69 | 0.12 | 0.42 | 0.16 | 0.18 | 0.1 | 0.14 |
| 9 | 87425973 | rs1047896 | -0.236 | -0.177 | 1.266 | -0.174 | -0.01 | 0.046 | 0.046 | 0 | 0.78 | 0.87 | 0.84 | 0.92 | 0.8 | 0.88 |
| 9 | 87426298 | rs2013566 | -1.554 | -0.207 | 1.435 | -0.125 | 0.022 | 0.116 | 0.116 | 0.267 | 0.11 | 0.4 | 0.16 | 0.18 | 0.1 | 0.14 |
| 9 | 87429290 | rs1624327 | 0.472 | -0.142 | 0.828 | -0.125 | -0.002 | -0.038 | -0.038 | 0.022 | 0.71 | 0.66 | 0.69 | 0.87 | 0.72 | 0.84 |
| 9 | 87430491 | rs3654 | -0.505 | -0.07 | 0.018 | 0.014 | -0.011 | -0.025 | -0.025 | 0.269 | 0.06 | 0.01 | 0.1 | 0.17 | 0.06 | 0.09 |
| 9 | 87441475 | rs10512154 | 0.096 | -0.174 | 0.081 | -0.041 | -0.013 | 0.045 | 0.045 | 0 | 0.22 | 0.13 | 0.18 | 0.1 | 0.23 | 0.12 |
| 9 | 87442302 | rs10780687 | -0.988 | -0.029 | 0.124 | 0.021 | -0.012 | -0.003 | -0.003 | 0 | 0.35 | 0.49 | 0.4 | 0.32 | 0.3 | 0.33 |
| 9 | 87447045 | rs2083828 | -0.761 | 0.332 | -0.543 | -0.397 | -0.001 | 0.008 | 0.008 | 0.445 | 0.52 | 0.54 | 0.51 | 0.33 | 0.5 | 0.35 |
| 9 | 87450766 | rs1867283 | 0.888 | 0.174 | -1.745 | 0.121 | -0.068 | -0.093 | -0.093 | 0 | 0.4 | 0.12 | 0.46 | 0.47 | 0.5 | 0.58 |
| 9 | 87452058 | rs7855888 | -0.808 | -0.355 | -1.265 | 0.552 | 0.008 | -0.011 | -0.011 | 0 | 0.3 | 0.7 | 0.32 | 0.41 | 0.24 | 0.28 |
| 9 | 87454817 | rs11140783 | 0.479 | -0.548 | -0.994 | 0.687 | -0.016 | -0.047 | -0.047 | 0 | 0.08 | 0 | 0.04 | 0.19 | 0.08 | 0.06 |
| 9 | 87455347 | rs4486281 | -0.81 | -1.032 | -1.303 | 0.699 | 0.061 | 0.038 | 0.038 | 0.32 | 0.4 | 0.75 | 0.4 | 0.32 | 0.32 | 0.34 |
| 9 | 87465791 | rs11791654 | -0.153 | -0.878 | -0.307 | 1.064 | 0.003 | -0.015 | -0.015 | 0.313 | 0.11 | 0.13 | 0.19 | 0.06 | 0.14 | 0.13 |
| 9 | 87473009 | rs1838158 | 0.309 | -0.817 | -0.59 | 1.738 | -0.018 | -0.072 | -0.072 | 0 | 0.86 | 0.98 | 0.89 | 0.75 | 0.87 | 0.91 |
| 9 | 87474397 | rs10868231 | -1.065 | -1.036 | 0.229 | 1.779 | -0.006 | -0.001 | -0.001 | 0 | 0.3 | 0.32 | 0.34 | 0.3 | 0.29 | 0.22 |
| 9 | 87477274 | rs10116287 | 0.515 | -1.052 | 0.21 | 1.776 | 0.019 | -0.065 | -0.065 | 0 | 0.17 | 0.03 | 0.09 | 0.22 | 0.15 | 0.08 |
| 9 | 87478135 | rs7048015 | -1.397 | -0.899 | 1.159 | 1.917 | 0.008 | 0.035 | 0.035 | 0.574 | 0.19 | 0.49 | 0.38 | 0.26 | 0.18 | 0.31 |
| 9 | 87484613 | rs4877289 | 2.447 | -0.928 | 1.602 | 1.875 | 0.01 | 0.101 | 0.101 | 0 | 0.46 | 0.25 | 0.4 | 0.27 | 0.52 | 0.54 |
| 9 | 87488718 | rs3780634 | -0.277 | -0.954 | 1.478 | 1.874 | 0.013 | -0.037 | -0.037 | 0 | 0.13 | 0.25 | 0.26 | 0.08 | 0.14 | 0.15 |
| 9 | 87491253 | rs10780691 | 2.518 | -0.909 | 1.277 | 1.892 | -0.028 | 0.122 | 0.122 | 0 | 0.46 | 0.12 | 0.38 | 0.27 | 0.52 | 0.52 |
| 9 | 87492610 | rs1221 | -0.342 | -0.616 | 1.387 | 1.933 | 0.02 | -0.048 | -0.048 | 0 | 0.13 | 0.28 | 0.19 | 0.06 | 0.14 | 0.14 |
| 9 | 87499252 | rs9314736 | 0.12 | -0.76 | 1.917 | 2.598 | -0.014 | -0.049 | -0.049 | 0 | 0.85 | 0.91 | 0.87 | 0.75 | 0.84 | 0.93 |
| 9 | 87506932 | rs11140799 | -0.649 | -0.614 | 1.901 | 1.864 | 0.05 | 0.047 | 0.047 | 0 | 0.06 | 0.26 | 0.17 | 0.08 | 0.06 | 0.04 |
| 9 | 87512669 | rs11140803 | -0.569 | -0.674 | 1.667 | 1.881 | 0.005 | -0.003 | -0.003 | 0 | 0.2 | 0.18 | 0.12 | 0.22 | 0.18 | 0.09 |
| 9 | 87530511 | rs4304401 | -0.959 | -0.456 | 1.56 | 1.097 | 0.064 | 0.06 | 0.06 | 0.064 | 0.07 | 0.34 | 0.18 | 0.08 | 0.06 | 0.04 |
| 9 | 87530935 | rs10868238 | 1.6 | -0.374 | 0.432 | 1.074 | 0.041 | 0.085 | 0.085 | 0.661 | 0.54 | 0.24 | 0.49 | 0.28 | 0.58 | 0.65 |
| 9 | 87533389 | rs6559836 | 0.517 | 0.249 | 0.411 | 0.652 | -0.023 | 0.065 | 0.065 | 0.01 | 0.78 | 0.94 | 0.82 | 0.92 | 0.78 | 0.91 |
| 9 | 87542728 | rs7045900 | -1.535 | 0.723 | 1.69 | 1.092 | 0.047 | 0.109 | 0.109 | 0.799 | 0.08 | 0.34 | 0.2 | 0.13 | 0.08 | 0.07 |
| 9 | 87551964 | rs12340212 | -0.319 | 0.829 | 1.92 | 1.325 | 0.001 | 0.022 | 0.022 | 0 | 0.16 | 0.27 | 0.16 | 0.24 | 0.15 | 0.13 |
| 9 | 87553563 | rs6559838 | -1.642 | 0.822 | 1.905 | 1.316 | 0.005 | 0.056 | 0.056 | 0.769 | 0.23 | 0.61 | 0.35 | 0.37 | 0.23 | 0.23 |
| 9 | 87558294 | rs2808707 | -2.16 | 0.8 | 1.503 | 1.2 | -0.004 | -0.006 | -0.006 | 0 | 0.42 | 0.64 | 0.5 | 0.45 | 0.44 | 0.29 |
| 9 | 87563252 | rs1078947 | 0.155 | 0.927 | 1.068 | 1.193 | 0.01 | 0.04 | 0.04 | 0 | 0.85 | 0.74 | 0.83 | 0.73 | 0.83 | 0.86 |
| 9 | 87563562 | rs2289656 | -0.212 | 0.966 | 0.912 | 1.166 | -0.005 | 0.048 | 0.048 | 0 | 0.19 | 0.11 | 0.15 | 0.03 | 0.2 | 0.08 |
| 9 | 87566307 | rs1002261 | -0.822 | 1.307 | 1.367 | 2.031 | 0.07 | 0.11 | 0.11 | 0 | 0.06 | 0.24 | 0.18 | 0.14 | 0.07 | 0.08 |
| 9 | 87570912 | rs2586566 | 1.08 | 1.256 | 0.221 | 1.815 | 0.018 | 0.082 | 0.082 | 1.713 | 0.76 | 0.45 | 0.68 | 0.57 | 0.76 | 0.75 |
| 9 | 87573977 | rs2277192 | -0.707 | 1.332 | 0.191 | 1.874 | -0.038 | -0.04 | -0.04 | 6.206 | 0.09 | 0.02 | 0.11 | 0.38 | 0.13 | 0.1 |
| 9 | 87574009 | rs2277193 | -1.56 | 1.314 | 0.888 | 1.627 | -0.001 | -0.027 | -0.027 | 0.41 | 0.3 | 0.5 | 0.42 | 0.22 | 0.28 | 0.23 |
| 9 | 87575500 | rs6559840 | -1.674 | 1.239 | 1.448 | 1.663 | 0 | 0.034 | 0.034 | 0 | 0.31 | 0.39 | 0.36 | 0.46 | 0.26 | 0.29 |
| 9 | 87575920 | rs17087959 | -1.569 | 1.254 | 1.598 | 1.627 | -0.005 | 0.017 | 0.017 | 0.435 | 0.25 | 0.26 | 0.16 | 0.43 | 0.18 | 0.19 |
| 9 | 87576008 | rs1899637 | -0.541 | 1.527 | 1.552 | 1.507 | 0.006 | -0.001 | -0.001 | 0.024 | 0.11 | 0.07 | 0.08 | 0.08 | 0.08 | 0.04 |
| 9 | 87580367 | rs2034107 | -1.165 | 1.586 | 1.649 | 1.265 | 0.014 | -0.012 | -0.012 | 0.792 | 0.09 | 0.18 | 0.21 | 0.04 | 0.09 | 0.12 |
| 9 | 87582827 | rs984430 | 1.357 | 1.457 | 1.53 | 1.43 | 0.011 | 0.019 | 0.019 | 0 | 0.87 | 0.82 | 0.78 | 0.76 | 0.82 | 0.87 |
| 9 | 87585624 | rs3860945 | -1.317 | 1.331 | 1.584 | 1.21 | 0.002 | -0.033 | -0.033 | 0 | 0.17 | 0.4 | 0.31 | 0.08 | 0.14 | 0.16 |
| 9 | 87589165 | rs7026417 | -0.159 | 1.54 | 1.577 | 1.378 | -0.002 | 0.046 | 0.046 | 0 | 0.08 | 0.07 | 0.08 | 0.37 | 0.1 | 0.08 |
| 9 | 87590382 | rs4877894 | 2.549 | 1.502 | 1.625 | 0.592 | -0.003 | 0.001 | 0.001 | 0 | 0.59 | 0.41 | 0.44 | 0.29 | 0.57 | 0.57 |
| 9 | 87593028 | rs10868241 | 2.025 | 1.498 | 0.783 | 0.715 | -0.012 | -0.009 | -0.009 | 0 | 0.75 | 0.81 | 0.76 | 0.37 | 0.7 | 0.73 |
| 9 | 87595734 | rs4361832 | 0.964 | 1.503 | 0.524 | 0.561 | -0.061 | -0.051 | -0.051 | 0.713 | 0.85 | 0.97 | 0.89 | 0.8 | 0.81 | 0.82 |
| 9 | 87600078 | rs4504715 | 0.071 | 1.519 | 0.552 | 0.579 | -0.049 | -0.042 | -0.042 | 0 | 0.9 | 0.99 | 0.97 | 0.89 | 0.88 | 0.86 |
| 9 | 87608999 | rs10512159 | 1.232 | 0.909 | 0.335 | 0.265 | -0.046 | 0.077 | 0.077 | 0 | 0.13 | 0.01 | 0.09 | 0 | 0.18 | 0.06 |
| 9 | 87609384 | rs4419891 | 1.93 | 0.717 | 1.434 | 1.166 | 0.005 | -0.003 | -0.003 | 0 | 0.74 | 0.7 | 0.72 | 0.29 | 0.71 | 0.67 |
| 9 | 87616257 | rs1948308 | 2.752 | 0.597 | 1.599 | 1.057 | 0.005 | -0.032 | -0.032 | 0 | 0.57 | 0.49 | 0.45 | 0.26 | 0.56 | 0.55 |
| 9 | 87616532 | rs923559 | 1.694 | 0.598 | 1.546 | 1.064 | -0.019 | -0.02 | -0.02 | 0 | 0.78 | 0.86 | 0.79 | 0.38 | 0.71 | 0.72 |
| 9 | 87618012 | rs7036165 | 1.682 | 0.597 | 1.546 | 1.064 | -0.019 | -0.02 | -0.02 | 0.933 | 0.78 | 0.86 | 0.79 | 0.38 | 0.71 | 0.72 |
| 9 | 87618385 | rs4394478 | 1.682 | 0.599 | 1.547 | 1.065 | -0.019 | -0.02 | -0.02 | 0.029 | 0.78 | 0.86 | 0.79 | 0.38 | 0.71 | 0.72 |
| 9 | 87619071 | rs13440094 | -1.862 | 0.433 | 0.935 | 0.882 | -0.016 | -0.05 | -0.05 | 0 | 0.21 | 0.44 | 0.35 | 0.12 | 0.15 | 0.17 |

| | | | | | | | | | | | | | | | | |
|---|---|---|---|---|---|---|---|---|---|---|---|---|---|---|---|---|
| 9 | 87625848 | rs17418241 | -0.282 | 0.856 | 0.811 | 1.121 | 0.021 | -0.012 | -0.012 | 0 | 0.08 | 0.16 | 0.21 | 0.03 | 0.08 | 0.09 |
| 9 | 87632993 | rs1387924 | 1.281 | 0.919 | 0.527 | 1.298 | -0.04 | -0.057 | -0.057 | 0 | 0.83 | 0.98 | 0.93 | 0.81 | 0.81 | 0.82 |
| 9 | 87647100 | rs1073049 | -0.132 | 0.615 | 0.809 | 0.926 | 0.004 | -0.032 | -0.032 | 0 | 0.16 | 0.24 | 0.29 | 0.03 | 0.14 | 0.09 |
| 9 | 87647454 | rs11140831 | 1.971 | 0.518 | 0.4 | 0.87 | -0.081 | 0.214 | 0.214 | 1.099 | 0.5 | 0.04 | 0.38 | 0.26 | 0.53 | 0.48 |
| 9 | 87651621 | rs10435949 | 1.084 | 0.917 | 1.155 | 0.964 | 0.002 | -0.016 | -0.016 | 0.012 | 0.86 | 0.84 | 0.93 | 0.95 | 0.86 | 0.78 |
| 9 | 87654393 | rs10780697 | 2.075 | 0.943 | 1.046 | 1.02 | -0.001 | -0.001 | -0.001 | 0 | 0.66 | 0.39 | 0.6 | 0.7 | 0.71 | 0.66 |
| 9 | 87659399 | rs681329 | 0.423 | 1.295 | 0.367 | 0.101 | 0 | -0.004 | -0.004 | 0 | 0.89 | 0.88 | 0.95 | 0.97 | 0.87 | 0.84 |
| 9 | 87660797 | rs474899 | 1.593 | 1.311 | 0.093 | 0.124 | 0.011 | -0.062 | -0.062 | 0.191 | 0.7 | 0.54 | 0.64 | 0.86 | 0.72 | 0.67 |
| 9 | 87676586 | rs475335 | 0.913 | 1.291 | -0.893 | 0.394 | -0.023 | 0.031 | 0.031 | 0.249 | 0.88 | 0.96 | 0.95 | 0.95 | 0.86 | 0.79 |
| 9 | 87694716 | rs7042038 | -0.867 | 1.181 | -0.208 | 0.769 | 0.021 | 0.033 | 0.033 | 0 | 0.24 | 0.44 | 0.47 | 0.68 | 0.26 | 0.25 |
| 9 | 87696145 | rs17348377 | -1.09 | 0.936 | -0.152 | 0.834 | -0.001 | -0.001 | -0.001 | 0 | 0.16 | 0.16 | 0.25 | 0.19 | 0.14 | 0.14 |
| 9 | 87697154 | rs615189 | 0.507 | 0.509 | -0.943 | 0.65 | -0.011 | 0.004 | 0.004 | 0 | 0.41 | 0.66 | 0.37 | 0.32 | 0.46 | 0.38 |
| 9 | 87699138 | rs544862 | -0.423 | 0.35 | 0.058 | 0.539 | -0.003 | 0 | 0 | 0 | 0.53 | 0.54 | 0.61 | 0.51 | 0.61 | 0.51 |
| 9 | 87699646 | rs2780936 | 0.929 | -0.069 | 0.042 | 0.609 | -0.106 | 0.009 | 0.009 | 0 | 0.31 | 0.12 | 0.33 | 0.19 | 0.45 | 0.3 |
| 9 | 87701841 | rs11999991 | 0.517 | -0.642 | -0.586 | -0.071 | -0.017 | 0.081 | 0.081 | 0 | 0.14 | 0.02 | 0.08 | 0.02 | 0.13 | 0.07 |
| 9 | 87703126 | rs11140851 | -0.127 | -0.747 | 0.128 | 0.073 | 0.001 | 0.002 | 0.002 | 0 | 0.19 | 0.16 | 0.25 | 0.2 | 0.15 | 0.17 |
| 9 | 87704881 | rs630426 | -0.396 | -0.893 | 0.36 | 0.151 | -0.028 | 0.031 | 0.031 | 0.149 | 0.66 | 0.61 | 0.69 | 0.49 | 0.74 | 0.56 |
| 9 | 87705388 | rs1809699 | 0.346 | -1.188 | 0.225 | 0.034 | -0.057 | 0.068 | 0.068 | 0.075 | 0.29 | 0.4 | 0.26 | 0.51 | 0.17 | 0.4 |
| 9 | 87707585 | rs7044762 | -1.132 | -1.057 | 0.948 | -0.527 | -0.007 | -0.078 | -0.078 | 0.11 | 0.12 | 0.55 | 0.13 | 0 | 0.09 | 0.04 |
| 9 | 87710734 | rs7034839 | -0.282 | -0.95 | 0.476 | -1.212 | 0 | -0.012 | -0.012 | 0 | 0.54 | 0.46 | 0.54 | 0.13 | 0.58 | 0.27 |
| 9 | 87718566 | rs622445 | -0.618 | -1.152 | 0.084 | -0.301 | -0.009 | -0.043 | -0.043 | 2.033 | 0.9 | 0.99 | 0.89 | 0.48 | 0.91 | 0.66 |
| 9 | 87726012 | rs4877295 | 0.157 | -1.013 | 0.269 | -1.114 | -0.013 | -0.033 | -0.033 | 0 | 0.3 | 0.54 | 0.32 | 0.2 | 0.28 | 0.25 |
| 9 | 87728550 | rs7868560 | 0.338 | -1.138 | 0.325 | -1.335 | -0.012 | -0.155 | -0.155 | 1.83 | 0.3 | 0.42 | 0.18 | 0.01 | 0.26 | 0.05 |
| 9 | 87752417 | rs7852361 | 0.672 | -1.585 | 1.24 | -1.228 | -0.037 | -0.085 | -0.085 | 0.062 | 0.22 | 0.63 | 0.14 | 0.04 | 0.15 | 0.21 |
| 9 | 87761907 | rs10868257 | 0.651 | -1.476 | 0.748 | -1.218 | -0.049 | 0.072 | 0.072 | 0 | 0.34 | 0.15 | 0.42 | 0.18 | 0.44 | 0.15 |
| 9 | 87766276 | rs6559849 | -1.128 | -1.446 | 0.596 | -1.225 | -0.023 | 0.081 | 0.081 | 0 | 0.61 | 0.84 | 0.61 | 0.22 | 0.6 | 0.38 |
| 9 | 87769185 | rs1931101 | 1.742 | -1.478 | 0.808 | -1.215 | -0.026 | -0.074 | -0.074 | 0 | 0.17 | 0.35 | 0.07 | 0.05 | 0.11 | 0.18 |
| 9 | 87769748 | rs12004450 | 0.938 | -1.441 | 0.685 | -1.194 | -0.034 | -0.103 | -0.103 | 0.315 | 0.22 | 0.41 | 0.13 | 0.05 | 0.16 | 0.21 |
| 9 | 87771842 | rs496239 | -1.926 | -1.622 | 0.309 | -0.969 | -0.051 | -0.015 | -0.015 | 0.067 | 0.72 | 0.88 | 0.74 | 0.75 | 0.69 | 0.72 |
| 9 | 87772807 | rs526454 | 1.19 | -1.339 | 0.457 | -0.697 | 0.007 | -0.002 | -0.002 | 0 | 0.12 | 0.06 | 0.12 | 0.56 | 0.09 | 0.41 |
| 9 | 87778956 | rs620420 | -0.412 | -1.477 | 0.521 | -0.821 | 0 | 0.059 | 0.059 | 0 | 0.48 | 0.3 | 0.58 | 0.74 | 0.52 | 0.56 |
| 9 | 87782211 | rs665889 | -0.559 | -1.477 | 0.586 | -0.829 | 0.001 | 0.004 | 0.004 | 0 | 0.49 | 0.42 | 0.6 | 0.78 | 0.53 | 0.61 |
| 9 | 87784527 | rs4877909 | 0.797 | -1.157 | 0.704 | -0.831 | -0.002 | 0.001 | 0.001 | 0 | 0.39 | 0.33 | 0.48 | 0.17 | 0.42 | 0.15 |
| 9 | 87785672 | rs4877910 | 0.916 | -1.081 | 0.656 | -0.811 | -0.011 | 0.051 | 0.051 | 0 | 0.38 | 0.22 | 0.46 | 0.18 | 0.41 | 0.15 |
| 9 | 87788048 | rs1387927 | 0.73 | -1.066 | 0.488 | -0.8 | -0.002 | 0.006 | 0.006 | 0 | 0.4 | 0.34 | 0.47 | 0.21 | 0.43 | 0.2 |
| 9 | 87788660 | rs13289484 | 0.885 | -1.069 | 0.355 | -0.823 | -0.012 | 0.055 | 0.055 | 0 | 0.39 | 0.25 | 0.46 | 0.16 | 0.42 | 0.17 |
| 9 | 87795226 | rs329200 | -0.42 | -1.17 | 0.222 | -0.704 | 0.001 | 0 | 0 | 0 | 0.39 | 0.65 | 0.27 | 0.66 | 0.28 | 0.61 |
| 9 | 87796745 | rs1887618 | 0.924 | -1.128 | -0.014 | -0.736 | -0.005 | 0.041 | 0.041 | 0 | 0.39 | 0.25 | 0.45 | 0.15 | 0.39 | 0.17 |
| 9 | 87799182 | rs329202 | -1.672 | -1.098 | -0.248 | -0.694 | -0.057 | -0.01 | -0.01 | 2.248 | 0.78 | 0.89 | 0.73 | 0.81 | 0.67 | 0.78 |
| 9 | 87799412 | rs329204 | -1.643 | -1.098 | -0.248 | -0.637 | -0.058 | -0.034 | -0.034 | 0 | 0.78 | 0.89 | 0.73 | 0.74 | 0.67 | 0.75 |
| 9 | 87802989 | rs329219 | -1.057 | -1.678 | 0.617 | 0.075 | 0.025 | 0 | 0 | 0 | 0.39 | 0.71 | 0.28 | 0.64 | 0.28 | 0.61 |
| 9 | 87806205 | rs329206 | -0.31 | -1.43 | 0.187 | 0.44 | -0.019 | 0.014 | 0.014 | 0 | 0.3 | 0.38 | 0.2 | 0.65 | 0.21 | 0.58 |
| 9 | 87808147 | rs329209 | 0.179 | -1.467 | -0.03 | 0.484 | -0.005 | -0.091 | -0.091 | 0 | 0.13 | 0.01 | 0.11 | 0.56 | 0.1 | 0.41 |
| 9 | 87808253 | rs11140885 | 1.209 | -1.42 | 0.079 | 0.195 | -0.014 | -0.035 | -0.035 | 3.231 | 0.16 | 0.25 | 0.06 | 0.06 | 0.11 | 0.15 |
| 9 | 87811640 | rs752184 | -0.581 | -0.922 | 0.043 | 1.138 | 0.002 | 0.014 | 0.014 | 0 | 0.14 | 0.11 | 0.15 | 0.63 | 0.11 | 0.43 |
| 9 | 87820596 | rs7850866 | -1.013 | -1.045 | 0.25 | 0.664 | -0.013 | -0.026 | -0.026 | 0 | 0.39 | 0.58 | 0.23 | 0.18 | 0.29 | 0.15 |
| 9 | 87822415 | rs1931114 | -0.993 | -1.115 | 0.262 | 0.893 | -0.004 | -0.004 | -0.004 | 0 | 0.4 | 0.59 | 0.24 | 0.25 | 0.29 | 0.16 |
| 9 | 87832469 | rs1931093 | -0.865 | -1.243 | 0.617 | 0.824 | 0.028 | 0.028 | 0.028 | 0 | 0.37 | 0.31 | 0.22 | 0.26 | 0.29 | 0.16 |
| 9 | 87834342 | rs11140892 | -0.294 | -1.352 | 0.287 | 0.671 | -0.003 | -0.026 | -0.026 | 2.991 | 0.19 | 0.05 | 0.13 | 0.22 | 0.18 | 0.12 |
| 9 | 87835091 | rs7030902 | -2.573 | -1.119 | -0.291 | 0.898 | -0.126 | 0.16 | 0.16 | 0.14 | 0.58 | 0.98 | 0.64 | 0.87 | 0.49 | 0.62 |
| 9 | 87844016 | rs11140900 | -0.242 | -1.246 | 0.383 | 0.974 | 0.005 | -0.007 | -0.007 | 0 | 0.2 | 0.16 | 0.14 | 0.23 | 0.18 | 0.12 |
| 9 | 87844548 | rs1931094 | -2.034 | -1.15 | 0.857 | 1.655 | 0.05 | 0.094 | 0.094 | 0 | 0.5 | 0.82 | 0.55 | 0.67 | 0.42 | 0.44 |
| 9 | 87845035 | rs11140902 | -0.919 | -1.191 | 1.796 | 1.914 | -0.054 | -0.02 | -0.02 | 0 | 0.29 | 0.46 | 0.19 | 0.23 | 0.21 | 0.15 |
| 9 | 87847392 | rs766348 | -0.287 | -1.029 | 1.029 | 2.202 | -0.016 | -0.014 | -0.014 | 0 | 0.09 | 0.05 | 0.29 | 0.32 | 0.14 | 0.2 |
| 9 | 87855507 | rs7018725 | -1.473 | -1.093 | 0.922 | 1.469 | -0.001 | -0.001 | -0.001 | 0 | 0.39 | 0.62 | 0.52 | 0.69 | 0.35 | 0.4 |
| 9 | 87861650 | rs913444 | -1.866 | -1.45 | 0.023 | 1.374 | 0.092 | 0.047 | 0.047 | 0.522 | 0.44 | 0.92 | 0.52 | 0.68 | 0.28 | 0.43 |
| 9 | 87864354 | rs7862326 | -1.087 | -1.552 | 0.626 | 1.799 | -0.009 | 0.072 | 0.072 | 0 | 0.15 | 0.27 | 0.33 | 0.28 | 0.11 | 0.23 |
| 9 | 87875615 | rs10122948 | -2.047 | -1.535 | -0.207 | 1.644 | 0.107 | 0.064 | 0.064 | 0.371 | 0.44 | 0.88 | 0.51 | 0.69 | 0.28 | 0.43 |
| 9 | 87877165 | rs1927605 | -0.497 | -1.113 | 0.302 | 1.508 | 0.024 | -0.029 | -0.029 | 0 | 0.21 | 0.14 | 0.11 | 0.33 | 0.15 | 0.1 |
| 9 | 87890350 | rs2841502 | 1.346 | 0.064 | 0.462 | 0.837 | -0.031 | 0.002 | 0.002 | 0 | 0.68 | 0.86 | 0.55 | 0.3 | 0.67 | 0.71 |
| 9 | 87892071 | rs1927602 | -1.998 | 0.25 | 1.203 | 0.809 | 0 | 0.015 | 0.015 | 3.039 | 0.1 | 0.64 | 0.11 | 0.07 | 0.06 | 0.11 |
| 9 | 87897177 | rs2841445 | 2.191 | 0.163 | 1.454 | 0.679 | -0.024 | 0.162 | 0.162 | 0.12 | 0.56 | 0.1 | 0.44 | 0.23 | 0.6 | 0.59 |
| 9 | 87903467 | rs2841473 | 1.108 | 0.192 | 1.518 | 0.778 | -0.011 | 0.032 | 0.032 | 0 | 0.71 | 0.83 | 0.68 | 0.94 | 0.68 | 0.87 |
| 9 | 87909739 | rs2769612 | 2.395 | 0.202 | 1.687 | 0.787 | -0.023 | 0.14 | 0.14 | 0.015 | 0.56 | 0.15 | 0.43 | 0.23 | 0.58 | 0.59 |
| 9 | 87910924 | rs17088419 | -0.609 | 0.201 | 1.789 | 0.836 | -0.006 | 0.128 | 0.128 | 0 | 0.05 | 0.14 | 0.14 | 0.64 | 0.04 | 0.16 |
| 9 | 87917458 | rs1983944 | 1.615 | 0.2 | 1.963 | 0.825 | -0.001 | 0.001 | 0.001 | 0 | 0.65 | 0.66 | 0.53 | 0.3 | 0.64 | 0.71 |
| 9 | 87922664 | rs4877913 | -0.865 | 0.241 | 1.675 | 0.846 | -0.01 | 0.154 | 0.154 | 0 | 0.06 | 0.16 | 0.15 | 0.64 | 0.04 | 0.16 |
| 9 | 87923891 | rs2841484 | 2.327 | 0.235 | 1.754 | 0.821 | -0.011 | 0.085 | 0.085 | 0.036 | 0.57 | 0.23 | 0.44 | 0.23 | 0.58 | 0.6 |
| 9 | 87926046 | rs2841486 | 1.801 | 0.38 | 1.639 | 0.58 | 0.003 | -0.019 | -0.019 | 0 | 0.64 | 0.43 | 0.58 | 0.87 | 0.62 | 0.76 |

| | | | | | | | | | | | | | | | | |
|---|---|---|---|---|---|---|---|---|---|---|---|---|---|---|---|---|
| 9 | 87928035 | rs12551160 | -0.614 | 0.468 | 1.577 | 0.516 | -0.002 | 0.004 | 0.004 | 0 | 0.13 | 0.08 | 0.07 | 0 | 0.14 | 0.03 |
| 9 | 87930045 | rs2841498 | 0.234 | 0.465 | 1.573 | 0.531 | -0.047 | 0.025 | 0.025 | 0.042 | 0.86 | 0.97 | 0.75 | 0.94 | 0.83 | 0.9 |
| 9 | 87930294 | rs2841500 | 1.545 | 0.45 | 1.783 | 0.562 | -0.026 | 0.059 | 0.059 | 0 | 0.69 | 0.83 | 0.64 | 0.88 | 0.67 | 0.81 |
| 9 | 87937563 | rs7026075 | -1.672 | 0.398 | 1.918 | 0.657 | 0.124 | 0.081 | 0.081 | 0.692 | 0.05 | 0.4 | 0.06 | 0.02 | 0.04 | 0.04 |
| 9 | 87952694 | rs4439204 | -1.334 | 0.307 | 1.843 | 0.899 | -0.01 | 0.043 | 0.043 | 0 | 0.21 | 0.37 | 0.27 | 0.69 | 0.2 | 0.24 |
| 9 | 87960028 | rs4877920 | -1.963 | 0.202 | 2.378 | 0.823 | -0.008 | 0.018 | 0.018 | 0.673 | 0.25 | 0.53 | 0.31 | 0.71 | 0.23 | 0.29 |
| 9 | 87983336 | rs10122851 | -2.189 | 0.211 | 2.271 | 0.84 | 0.06 | 0.035 | 0.035 | 0.055 | 0.07 | 0.31 | 0.06 | 0.02 | 0.06 | 0.06 |
| 9 | 87992789 | rs2814734 | 1.491 | 0.145 | 1.911 | 0.569 | 0.005 | -0.032 | -0.032 | 3.659 | 0.76 | 0.68 | 0.65 | 0.91 | 0.76 | 0.85 |
| 9 | 87994507 | rs7860844 | -1.017 | -0.089 | 2.005 | 0.588 | -0.006 | 0.081 | 0.081 | 0.113 | 0.2 | 0.41 | 0.28 | 0.69 | 0.19 | 0.25 |
| 9 | 87995971 | rs2814737 | 2.159 | 0.009 | 1.296 | 0.542 | 0.006 | 0.049 | 0.049 | 0.288 | 0.58 | 0.35 | 0.44 | 0.22 | 0.62 | 0.62 |
| 9 | 87996304 | rs2841443 | 0.649 | 0.158 | 0.778 | 0.318 | -0.04 | 0.011 | 0.011 | 0 | 0.84 | 0.99 | 0.77 | 0.93 | 0.87 | 0.9 |
| 9 | 88000506 | rs2592991 | 0.477 | 0.449 | 0.863 | 0.402 | -0.044 | -0.01 | -0.01 | 0 | 0.87 | 0.97 | 0.72 | 0.93 | 0.86 | 0.9 |
| 9 | 88009912 | rs1867956 | 2.013 | 0.34 | 1.131 | 0.373 | -0.001 | -0.006 | -0.006 | 3.663 | 0.67 | 0.67 | 0.49 | 0.17 | 0.65 | 0.47 |
| 9 | 88010388 | rs1447029 | 1.883 | 0.345 | 0.775 | 0.373 | -0.015 | 0.063 | 0.063 | 0 | 0.68 | 0.85 | 0.51 | 0.17 | 0.67 | 0.52 |
| 9 | 88012713 | rs1374507 | -0.939 | 0.264 | 0.267 | 0.347 | 0.007 | -0.053 | -0.053 | 0.131 | 0.2 | 0.05 | 0.23 | 0.76 | 0.18 | 0.36 |
| 9 | 88013822 | rs2841451 | 1.68 | 0.253 | 0.59 | 0.135 | 0.028 | 0.031 | 0.031 | 0.116 | 0.78 | 0.44 | 0.66 | 0.82 | 0.78 | 0.64 |
| 9 | 88013998 | rs2593016 | 0.169 | 0.244 | 0.455 | 0.118 | -0.042 | -0.016 | -0.016 | 0.41 | 0.87 | 0.97 | 0.77 | 0.93 | 0.86 | 0.9 |
| 9 | 88014485 | rs2841452 | -0.923 | 0.22 | 0.473 | 0.377 | 0.01 | -0.08 | -0.08 | 0 | 0.2 | 0.05 | 0.23 | 0.65 | 0.18 | 0.35 |
| 9 | 88015543 | rs2841454 | 1.45 | 0.486 | 0.862 | 0.641 | 0.025 | 0.082 | 0.082 | 0.123 | 0.78 | 0.44 | 0.66 | 0.69 | 0.78 | 0.63 |
| 9 | 88017606 | rs12685505 | -0.908 | 0.267 | 0.598 | 0.683 | -0.002 | 0.071 | 0.071 | 0 | 0.06 | 0.08 | 0.11 | 0.67 | 0.06 | 0.33 |
| 9 | 88024130 | rs10512167 | -0.728 | 0.18 | 0.291 | 0.365 | -0.016 | -0.009 | -0.009 | 0.275 | 0.12 | 0.01 | 0.28 | 0.06 | 0.12 | 0.09 |
| 9 | 88031251 | rs2814732 | -0.899 | 0.853 | 0.858 | -0.019 | 0.045 | -0.071 | -0.071 | 0.027 | 0.12 | 0.21 | 0.08 | 0.01 | 0.15 | 0.07 |
| 9 | 88042156 | rs12236186 | -0.714 | 0.959 | 0.838 | -0.426 | -0.001 | -0.002 | -0.002 | 0.633 | 0.16 | 0.17 | 0.11 | 0.14 | 0.14 | 0.2 |
| 9 | 88043188 | rs13292809 | 2.969 | 0.808 | 0.809 | -0.642 | -0.033 | 0.173 | 0.173 | 0.017 | 0.44 | 0.18 | 0.37 | 0.12 | 0.48 | 0.21 |
| 9 | 88043805 | rs7028056 | -2.309 | 0.416 | 1.5 | -0.008 | -0.009 | -0.045 | -0.045 | 0 | 0.35 | 0.44 | 0.52 | 0.85 | 0.31 | 0.62 |
| 9 | 88044756 | rs10868292 | -1.798 | 0.428 | 1.044 | -0.067 | 0 | 0 | 0 | 0 | 0.29 | 0.28 | 0.35 | 0.2 | 0.25 | 0.3 |
| 9 | 88047238 | rs2256555 | -1.635 | 0.582 | 0.314 | 0.207 | -0.011 | 0.17 | 0.17 | 0.16 | 0.13 | 0.24 | 0.16 | 0.62 | 0.11 | 0.36 |
| 9 | 88059730 | rs11140970 | -0.398 | 0.615 | -0.079 | 0.045 | -0.005 | -0.015 | -0.015 | 0.828 | 0.06 | 0.02 | 0.05 | 0.09 | 0.04 | 0.11 |
| 9 | 88069314 | rs12553871 | -0.442 | 0.432 | -0.044 | 0.039 | -0.009 | -0.009 | -0.009 | 0 | 0.11 | 0.01 | 0.23 | 0.06 | 0.11 | 0.11 |
| 9 | 88087511 | rs11140985 | -1.028 | 0.648 | 0.383 | 0.23 | 0.002 | 0.002 | 0.002 | 0 | 0.15 | 0.06 | 0.09 | 0.12 | 0.13 | 0.19 |
| 9 | 88098058 | rs10868306 | -0.84 | 0.655 | 0.602 | 0.499 | 0 | 0.118 | 0.118 | 0 | 0.06 | 0.06 | 0.16 | 0.58 | 0.06 | 0.29 |
| 9 | 88099866 | rs1992812 | 1.155 | 0.592 | 0.509 | -0.987 | -0.013 | -0.051 | -0.051 | 0.23 | 0.91 | 0.83 | 0.94 | 0.99 | 0.94 | 0.94 |
| 9 | 88101673 | rs7042753 | -1.163 | 0.962 | 0.028 | -1.089 | -0.001 | -0.058 | -0.058 | 0.039 | 0.32 | 0.66 | 0.33 | 0.03 | 0.34 | 0.18 |
| 9 | 88104161 | rs1447025 | -0.473 | 1.279 | -0.067 | -0.881 | 0.024 | -0.095 | -0.095 | 0 | 0.23 | 0.32 | 0.39 | 0 | 0.34 | 0.12 |
| 9 | 88105301 | rs10114046 | 0.055 | 0.522 | -0.267 | 0.12 | -0.001 | -0.004 | -0.004 | 0 | 0.73 | 0.79 | 0.81 | 0.75 | 0.73 | 0.76 |
| 9 | 88111373 | rs9314742 | -0.626 | 0.373 | -0.244 | 0.168 | -0.014 | -0.014 | -0.014 | 0.212 | 0.34 | 0.55 | 0.26 | 0.23 | 0.27 | 0.28 |
| 9 | 88113727 | rs7041064 | -0.659 | 0.527 | -0.563 | 0.407 | 0 | 0 | 0 | 0 | 0.41 | 0.59 | 0.34 | 0.71 | 0.36 | 0.48 |
| 9 | 88116256 | rs10512168 | -0.69 | 0.649 | -1.579 | 0.637 | -0.052 | -0.101 | -0.101 | 0.319 | 0.22 | 0.04 | 0.31 | 0.6 | 0.3 | 0.38 |
| 9 | 88117073 | rs876927 | -0.139 | -0.16 | -1.5 | -0.259 | 0.02 | 0.008 | 0.008 | 0 | 0.33 | 0.29 | 0.33 | 0.31 | 0.19 | 0.38 |
| 9 | 88118172 | rs4588941 | -0.351 | 0.268 | -2.127 | -0.648 | -0.002 | 0.036 | 0.036 | 0 | 0.42 | 0.21 | 0.43 | 0.35 | 0.4 | 0.42 |
| 9 | 88118237 | rs935466 | -0.162 | 0.856 | -2.176 | -0.494 | -0.028 | 0.005 | 0.005 | 0 | 0.36 | 0.21 | 0.41 | 0.35 | 0.4 | 0.38 |
| 9 | 88122906 | rs7854574 | 0.095 | 0.545 | -2.081 | -0.704 | 0.007 | 0.018 | 0.018 | 0 | 0.13 | 0.05 | 0.07 | 0.08 | 0.12 | 0.15 |
| 9 | 88122937 | rs7043100 | 1.163 | 0.536 | -2.19 | -1.252 | -0.001 | -0.005 | -0.005 | 0 | 0.15 | 0.12 | 0.09 | 0.02 | 0.12 | 0.06 |
| 9 | 88123347 | rs3860946 | 0.609 | 0.663 | -1.974 | -0.789 | -0.017 | 0.003 | 0.003 | 0 | 0.61 | 0.72 | 0.56 | 0.65 | 0.47 | 0.67 |
| 9 | 88123838 | rs10115062 | 0.502 | 1.041 | -1.967 | -0.803 | -0.047 | -0.006 | -0.006 | 0 | 0.63 | 0.77 | 0.58 | 0.66 | 0.5 | 0.69 |
| 9 | 88128899 | rs10868312 | 1.17 | 0.592 | -0.98 | -0.624 | 0.014 | 0.252 | 0.252 | 0 | 0.11 | 0.26 | 0.12 | 0.5 | 0.1 | 0.2 |
| 9 | 88131557 | rs10746763 | -0.137 | 0.308 | -0.951 | -0.774 | 0.037 | 0.004 | 0.004 | 0 | 0.57 | 0.75 | 0.5 | 0.63 | 0.39 | 0.65 |
| 9 | 88133513 | rs12343187 | 0.191 | 0.228 | -0.943 | -0.801 | -0.015 | -0.029 | -0.029 | 0.495 | 0.12 | 0.13 | 0.03 | 0 | 0.03 | 0.04 |
| 9 | 88137639 | rs1250792 | 1.289 | 0.721 | -1.065 | -0.74 | -0.068 | -0.066 | -0.066 | 0 | 0.11 | 0.02 | 0.32 | 0.26 | 0.34 | 0.11 |
| 9 | 88138617 | rs11141010 | 2.436 | -0.073 | -1.202 | -0.916 | -0.009 | 0.06 | 0.06 | 1.934 | 0.28 | 0.02 | 0.33 | 0.14 | 0.25 | 0.36 |
| 9 | 88140497 | rs10780727 | 0.56 | -0.133 | -0.533 | -0.749 | -0.017 | 0.271 | 0.271 | 0.838 | 0.16 | 0.54 | 0.13 | 0.5 | 0.1 | 0.24 |
| 9 | 88141571 | rs7032476 | 0.773 | -0.127 | -0.292 | -0.746 | 0.009 | 0.339 | 0.339 | 0.535 | 0.14 | 0.51 | 0.12 | 0.5 | 0.1 | 0.24 |
| 9 | 88143119 | rs7850808 | -0.369 | -0.225 | -0.242 | -0.806 | -0.049 | 0.106 | 0.106 | 0 | 0.23 | 0.39 | 0.13 | 0.5 | 0.13 | 0.28 |
| 9 | 88149101 | rs2278111 | -1.119 | -0.206 | -0.086 | -0.84 | -0.021 | 0.023 | 0.023 | 0.642 | 0.29 | 0.65 | 0.17 | 0.51 | 0.16 | 0.3 |
| 9 | 88153227 | rs7863713 | -1.151 | -0.155 | -0.113 | -1.094 | 0.06 | -0.059 | -0.059 | 0 | 0.49 | 0.89 | 0.28 | 0.52 | 0.32 | 0.38 |
| 9 | 88154909 | rs17088766 | 1.955 | -0.178 | -0.022 | -1.093 | -0.001 | 0.023 | 0.023 | 1.281 | 0.19 | 0.14 | 0.11 | 0.02 | 0.16 | 0.08 |
| 9 | 88154941 | rs2059403 | -0.415 | 0.12 | 0.458 | -0.709 | -0.045 | 0.102 | 0.102 | 0 | 0.23 | 0.35 | 0.12 | 0.49 | 0.13 | 0.28 |
| 9 | 88160238 | rs17349624 | 2.138 | 0.153 | 0.401 | -0.759 | 0.034 | 0.181 | 0.181 | 0 | 0.46 | 0.15 | 0.43 | 0.15 | 0.38 | 0.43 |
| 9 | 88164213 | rs11141018 | -1.355 | 0.151 | 0.402 | -0.76 | 0.036 | -0.04 | -0.04 | 0.05 | 0.3 | 0.77 | 0.18 | 0.5 | 0.16 | 0.3 |
| 9 | 88167035 | rs10780728 | -1.263 | 0.159 | 0.393 | -0.756 | 0.012 | -0.014 | -0.014 | 0 | 0.29 | 0.71 | 0.17 | 0.5 | 0.16 | 0.29 |
| 9 | 88171820 | rs10120410 | -0.433 | 0.175 | 0.308 | -0.741 | -0.112 | -0.078 | -0.078 | 3.119 | 0.75 | 0.92 | 0.61 | 0.66 | 0.55 | 0.74 |
| 9 | 88180032 | rs10780729 | 1.227 | 0.163 | 0.309 | -0.726 | 0.009 | 0.253 | 0.253 | 0 | 0.11 | 0.27 | 0.11 | 0.53 | 0.1 | 0.23 |
| 9 | 88191184 | rs10115559 | -0.49 | 0.12 | 0.26 | -0.739 | -0.17 | -0.105 | -0.105 | 5.046 | 0.76 | 0.98 | 0.64 | 0.72 | 0.55 | 0.75 |
| 9 | 88204998 | rs4645620 | 1.173 | 0.145 | 0.307 | -0.705 | 0.008 | 0.253 | 0.253 | 0 | 0.12 | 0.27 | 0.11 | 0.53 | 0.1 | 0.23 |
| 9 | 88205819 | rs1469202 | -0.856 | 0.181 | 0.192 | -0.676 | -0.027 | 0.053 | 0.053 | 0.036 | 0.27 | 0.6 | 0.18 | 0.54 | 0.16 | 0.29 |
| 9 | 88215534 | rs1571119 | -0.378 | 0.264 | 0.025 | -0.61 | -0.104 | -0.062 | -0.062 | 0 | 0.75 | 0.9 | 0.63 | 0.71 | 0.55 | 0.75 |
| 9 | 88226038 | rs1819284 | -0.518 | 0.125 | 0.128 | -0.639 | -0.049 | 0.102 | 0.102 | 0 | 0.23 | 0.36 | 0.14 | 0.52 | 0.13 | 0.28 |
| 9 | 88234954 | rs618242 | 0.262 | 0.378 | -0.021 | -0.616 | -0.095 | -0.052 | -0.052 | 2.333 | 0.25 | 0.1 | 0.37 | 0.28 | 0.45 | 0.26 |
| 9 | 88242953 | rs10117805 | -1.268 | -0.124 | 0.35 | -0.414 | -0.004 | 0.039 | 0.039 | 1.707 | 0.27 | 0.59 | 0.35 | 0.57 | 0.24 | 0.31 |

| | | | | | | | | | | | | | | | | |
|---|---|---|---|---|---|---|---|---|---|---|---|---|---|---|---|---|
| 9 | 88244337 | rs590506 | -1.695 | -0.049 | 0.62 | -0.279 | 0.016 | -0.005 | -0.005 | 0 | 0.38 | 0.7 | 0.4 | 0.38 | 0.34 | 0.4 |
| 9 | 88277116 | rs7031647 | -0.782 | -0.012 | 0.629 | -0.169 | 0.02 | 0.19 | 0.19 | 0 | 0.19 | 0.37 | 0.3 | 0.53 | 0.2 | 0.23 |
| 9 | 88279039 | rs4877947 | -0.988 | 0.001 | 0.584 | -0.156 | 0.004 | 0.186 | 0.186 | 0 | 0.22 | 0.42 | 0.31 | 0.55 | 0.2 | 0.26 |
| 9 | 88279765 | rs9314744 | -1.976 | -0.066 | 0.279 | -0.189 | 0.027 | 0.153 | 0.153 | 0.228 | 0.45 | 0.82 | 0.49 | 0.84 | 0.38 | 0.56 |
| 9 | 88299460 | rs10868338 | 1.022 | 0.176 | 0.642 | 0.075 | 0.041 | 0.109 | 0.109 | 0 | 0.05 | 0.2 | 0.17 | 0.19 | 0.05 | 0.06 |
| 9 | 88308127 | rs7863248 | -1.048 | 0.178 | -0.09 | -0.322 | 0.001 | 0.094 | 0.094 | 0 | 0.25 | 0.5 | 0.34 | 0.54 | 0.24 | 0.28 |
| 9 | 88331840 | rs10780740 | -1.089 | -0.556 | 0.06 | -0.267 | -0.016 | 0.044 | 0.044 | 0.167 | 0.16 | 0.37 | 0.2 | 0.75 | 0.11 | 0.26 |
| 9 | 88340454 | rs4877950 | -1.514 | -0.478 | -0.496 | -0.78 | -0.069 | 0.069 | 0.069 | 0.056 | 0.7 | 0.99 | 0.64 | 0.85 | 0.68 | 0.6 |
| 9 | 88351591 | rs10868342 | -1.009 | -0.955 | 0.169 | -0.168 | -0.004 | 0.037 | 0.037 | 0.069 | 0.16 | 0.34 | 0.21 | 0.75 | 0.12 | 0.27 |
| 9 | 88353231 | rs3916184 | -1.465 | -1.13 | 0.205 | -0.168 | -0.037 | 0.04 | 0.04 | 1.287 | 0.19 | 0.5 | 0.21 | 0.75 | 0.11 | 0.27 |
| 9 | 88357495 | rs3812508 | 1.078 | -0.618 | -0.279 | 0.167 | -0.003 | 0.147 | 0.147 | 0 | 0.07 | 0.12 | 0.1 | 0.55 | 0.05 | 0.15 |
| 9 | 88371919 | rs10868345 | 0.22 | -0.72 | -0.301 | -0.836 | 0.023 | 0.042 | 0.042 | 0.225 | 0.47 | 0.33 | 0.43 | 0.09 | 0.63 | 0.3 |
| 9 | 88477869 | rs7018466 | 0.169 | -1.06 | 0.679 | -0.341 | -0.03 | 0.001 | 0.001 | 0 | 0.35 | 0.48 | 0.41 | 0.36 | 0.24 | 0.48 |
| 9 | 88518441 | rs7039482 | -1.094 | -1.131 | 0.011 | -0.375 | -0.015 | -0.031 | -0.031 | 0 | 0.17 | 0.23 | 0.1 | 0.05 | 0.12 | 0.12 |
| 9 | 88520203 | rs11141141 | -0.612 | -1.053 | -0.379 | -0.333 | 0.009 | 0.029 | 0.029 | 0 | 0.14 | 0.06 | 0.09 | 0.05 | 0.11 | 0.11 |
| 9 | 88521040 | rs11141146 | -0.263 | -1.02 | -0.361 | -0.316 | 0.002 | 0.019 | 0.019 | 0 | 0.12 | 0.05 | 0.09 | 0.05 | 0.11 | 0.11 |
| 9 | 88530006 | rs570516 | -0.211 | -1.473 | -0.526 | -0.416 | 0.026 | -0.033 | -0.033 | 0.033 | 0.39 | 0.17 | 0.46 | 0.41 | 0.33 | 0.55 |
| 9 | 88530616 | rs2770831 | -0.517 | -1.444 | -0.212 | -0.53 | 0.022 | 0.006 | 0.006 | 2.09 | 0.46 | 0.27 | 0.57 | 0.46 | 0.43 | 0.63 |
| 9 | 88533596 | rs645063 | 0.469 | -1.23 | -0.144 | -0.38 | 0.011 | -0.006 | -0.006 | 0 | 0.3 | 0.28 | 0.34 | 0.39 | 0.21 | 0.46 |
| 9 | 88533866 | rs12552584 | -0.126 | -0.634 | -0.115 | -0.523 | 0.013 | -0.022 | -0.022 | 0 | 0.08 | 0.08 | 0.16 | 0.01 | 0.14 | 0.06 |
| 9 | 88538024 | rs483334 | -0.571 | -1.522 | -0.109 | -0.36 | 0.006 | 0.024 | 0.024 | 0 | 0.56 | 0.71 | 0.62 | 0.62 | 0.45 | 0.67 |
| 9 | 88538154 | rs649213 | 0.872 | -2.032 | 0.069 | -0.591 | 0.007 | -0.006 | -0.006 | 0.07 | 0.3 | 0.27 | 0.36 | 0.59 | 0.22 | 0.53 |
| 9 | 88540410 | rs10512169 | -0.306 | -1.912 | 0.126 | -0.64 | -0.004 | -0.02 | -0.02 | 0 | 0.09 | 0.05 | 0.05 | 0 | 0.07 | 0.04 |
| 9 | 88560387 | rs11141156 | -1.388 | -1.921 | 0.082 | -0.165 | 0 | -0.005 | -0.005 | 0.253 | 0.2 | 0.19 | 0.22 | 0.02 | 0.21 | 0.11 |
| 9 | 88563064 | rs954124 | -2.151 | -2.066 | 0.339 | -0.228 | -0.002 | 0.002 | 0.002 | 0.677 | 0.28 | 0.73 | 0.32 | 0.37 | 0.22 | 0.23 |
| 9 | 88597262 | rs10868357 | -1.346 | -1.786 | -0.687 | 0.077 | 0.009 | -0.01 | -0.01 | 0.36 | 0.14 | 0.12 | 0.13 | 0.36 | 0.08 | 0.14 |
| 9 | 88605205 | rs4877955 | -1.261 | -1.554 | -0.567 | 0.211 | -0.011 | 0.037 | 0.037 | 0.024 | 0.15 | 0.09 | 0.21 | 0.02 | 0.18 | 0.11 |
| 9 | 88646703 | rs4877959 | -0.786 | -1.353 | -0.619 | -0.257 | -0.002 | -0.01 | -0.01 | 0 | 0.13 | 0.15 | 0.21 | 0.02 | 0.18 | 0.09 |
| 9 | 88650029 | rs7046093 | -0.903 | -1.674 | -0.857 | -0.02 | -0.027 | -0.048 | -0.048 | 0.23 | 0.75 | 0.99 | 0.78 | 0.75 | 0.78 | 0.57 |
| 9 | 88669475 | rs7031120 | -0.085 | -1.803 | -0.045 | -0.035 | -0.02 | -0.024 | -0.024 | 0 | 0.34 | 0.4 | 0.35 | 0.2 | 0.26 | 0.48 |
| 9 | 88671087 | rs10115014 | -0.557 | -1.622 | 0.48 | 0.829 | -0.017 | 0.149 | 0.149 | 0 | 0.1 | 0.18 | 0.11 | 0.46 | 0.08 | 0.15 |
| 9 | 88679834 | rs7865836 | -1.569 | -2.14 | 0.125 | 0.407 | 0.021 | -0.016 | -0.016 | 0 | 0.33 | 0.76 | 0.26 | 0.6 | 0.23 | 0.37 |
| 9 | 88679955 | rs7866281 | -1.281 | -1.999 | 0.058 | 0.504 | -0.031 | 0.048 | 0.048 | 0 | 0.28 | 0.44 | 0.22 | 0.57 | 0.23 | 0.34 |
| 9 | 88692838 | rs2183126 | 0.082 | -1.799 | -0.043 | 0.393 | 0.004 | 0.011 | 0.011 | 0 | 0.11 | 0.07 | 0.06 | 0.05 | 0.12 | 0.13 |
| 9 | 88693460 | rs7019241 | -0.643 | -1.79 | 0.276 | 0.649 | -0.023 | 0.222 | 0.222 | 0 | 0.14 | 0.37 | 0.15 | 0.56 | 0.09 | 0.24 |
| 9 | 88693763 | rs10125666 | -0.688 | -1.802 | 0.27 | 0.653 | -0.024 | 0.221 | 0.221 | 0 | 0.15 | 0.38 | 0.15 | 0.56 | 0.09 | 0.24 |
| 9 | 88698673 | rs957617 | 0.653 | -2.163 | -1.282 | -0.186 | 0.013 | 0.159 | 0.159 | 0.213 | 0.42 | 0.24 | 0.29 | 0.05 | 0.57 | 0.27 |
| 9 | 88698820 | rs1928228 | -0.287 | -2.376 | -1.131 | -0.035 | -0.031 | 0.145 | 0.145 | 0.235 | 0.17 | 0.6 | 0.17 | 0.54 | 0.09 | 0.24 |
| 9 | 88700060 | rs10868366 | -0.287 | -2.376 | -1.131 | -0.035 | -0.031 | 0.145 | 0.145 | 0.235 | 0.17 | 0.6 | 0.16 | 0.54 | 0.09 | 0.24 |
| 9 | 88707692 | rs10868372 | -0.803 | -2.503 | -1.239 | 0.161 | 0.058 | 0.095 | 0.095 | 0 | 0.62 | 0.87 | 0.54 | 0.84 | 0.69 | 0.65 |
| 9 | 88715004 | rs2225237 | -0.769 | -1.573 | -0.271 | 0.732 | -0.034 | 0.143 | 0.143 | 0.235 | 0.17 | 0.6 | 0.17 | 0.54 | 0.09 | 0.24 |
| 9 | 88729691 | rs7875522 | 0.42 | -1.333 | -0.859 | -0.568 | -0.003 | 0.093 | 0.093 | 0 | 0.4 | 0.27 | 0.4 | 0.07 | 0.62 | 0.27 |
| 9 | 88733933 | rs7870214 | -0.706 | -2.041 | -0.97 | -0.294 | 0.08 | 0.037 | 0.037 | 0 | 0.66 | 0.9 | 0.6 | 0.25 | 0.82 | 0.53 |
| 9 | 88738450 | rs2770811 | 0.659 | -2.227 | -1.029 | -0.883 | -0.027 | 0.01 | 0.01 | 0.079 | 0.36 | 0.69 | 0.35 | 0 | 0.48 | 0.14 |
| 9 | 88741540 | rs10115513 | -0.489 | -2.182 | 0.039 | -0.74 | 0.038 | 0.021 | 0.021 | 1.82 | 0.05 | 0.27 | 0.04 | 0 | 0.02 | 0.01 |
| 9 | 88743556 | rs2812003 | -0.927 | -2.284 | -0.237 | -0.715 | 0.052 | 0.038 | 0.038 | 0 | 0.67 | 0.89 | 0.62 | 0.26 | 0.79 | 0.51 |
| 9 | 88755951 | rs12378988 | 0.511 | -2.136 | 0.061 | -0.609 | 0 | -0.005 | -0.005 | 0 | 0.07 | 0.05 | 0.05 | 0.09 | 0.05 | 0.1 |
| 9 | 88756246 | rs11141262 | 1.113 | -2.574 | -0.185 | 0.141 | 0.025 | -0.037 | -0.037 | 0 | 0.3 | 0.09 | 0.34 | 0.34 | 0.21 | 0.4 |
| 9 | 88766713 | rs11507200 | -0.523 | -2.533 | 0.532 | 0.033 | -0.002 | -0.001 | -0.001 | 0.559 | 0.55 | 0.53 | 0.57 | 0.51 | 0.46 | 0.68 |
| 9 | 88778654 | rs12002764 | -0.778 | -2.457 | 0.558 | 0.073 | -0.032 | 0.195 | 0.195 | 0 | 0.14 | 0.3 | 0.12 | 0.5 | 0.06 | 0.19 |
| 9 | 88782362 | rs4620358 | -0.419 | -2.43 | 0.4 | 0.062 | 0.007 | 0.004 | 0.004 | 0 | 0.54 | 0.3 | 0.59 | 0.51 | 0.48 | 0.68 |
| 9 | 88784346 | rs11141283 | 0.411 | -2.253 | 0.396 | 0.275 | -0.001 | -0.006 | -0.006 | 0 | 0.07 | 0.05 | 0.05 | 0.09 | 0.05 | 0.1 |
| 9 | 88790268 | rs7855538 | -0.194 | -2.342 | 0.464 | 0.131 | 0 | 0 | 0 | 0.318 | 0.23 | 0.25 | 0.22 | 0.18 | 0.25 | 0.28 |
| 9 | 88794032 | rs4877963 | -0.412 | -2.406 | 0.574 | 0.171 | 0.008 | -0.014 | -0.014 | 0.223 | 0.29 | 0.25 | 0.21 | 0.57 | 0.26 | 0.4 |
| 9 | 88795219 | rs11141289 | -0.341 | -2.143 | 1.086 | 0.117 | 0.005 | 0.01 | 0.01 | 0 | 0.08 | 0.26 | 0.07 | 0.06 | 0.05 | 0.08 |
| 9 | 88800714 | rs1359170 | 0.316 | -1.405 | 0.745 | -0.325 | -0.007 | 0 | 0 | 0 | 0.31 | 0.47 | 0.46 | 0.33 | 0.29 | 0.37 |
| 9 | 88811119 | rs7847143 | -1.729 | -1.266 | 0.356 | -0.434 | -0.101 | 0.064 | 0.064 | 0.273 | 0.76 | 0.94 | 0.75 | 0.89 | 0.65 | 0.88 |
| 9 | 88824018 | rs10868394 | 0.242 | -1.277 | 0.571 | -0.417 | -0.002 | 0.008 | 0.008 | 0 | 0.31 | 0.51 | 0.49 | 0.32 | 0.31 | 0.38 |
| 9 | 88833465 | rs4316216 | 0.951 | -1.206 | 0.617 | -0.399 | -0.015 | -0.029 | -0.029 | 1.251 | 0.26 | 0.18 | 0.43 | 0.33 | 0.3 | 0.37 |
| 9 | 88839915 | rs1002960 | 0.471 | -1.429 | 0.554 | -0.703 | -0.018 | -0.041 | -0.041 | 0 | 0.26 | 0.32 | 0.21 | 0.11 | 0.2 | 0.2 |
| 9 | 88844634 | rs11141306 | -0.56 | -1.168 | 0.708 | -0.016 | -0.031 | 0.07 | 0.07 | 0 | 0.23 | 0.35 | 0.13 | 0.4 | 0.13 | 0.31 |
| 9 | 88847477 | rs942284 | -0.305 | -1.269 | 0.76 | -0.136 | -0.024 | 0.072 | 0.072 | 0 | 0.22 | 0.32 | 0.15 | 0.46 | 0.12 | 0.33 |
| 9 | 88854546 | rs10118760 | -0.955 | -1.28 | 0.225 | -0.464 | 0.042 | 0.193 | 0.193 | 0 | 0.5 | 0.78 | 0.55 | 0.9 | 0.4 | 0.73 |
| 9 | 88856534 | rs6559918 | 0.479 | -1.305 | 0.136 | -0.371 | 0 | 0.057 | 0.057 | 0 | 0.27 | 0.41 | 0.39 | 0.43 | 0.27 | 0.38 |
| 9 | 88859144 | rs7848944 | -0.152 | -0.852 | -0.171 | -1.022 | -0.001 | -0.031 | -0.031 | 0.125 | 0.46 | 0.53 | 0.54 | 0.86 | 0.39 | 0.67 |
| 9 | 88860820 | rs4484780 | -0.38 | -0.674 | -1.308 | -0.959 | 0.04 | 0.219 | 0.219 | 0 | 0.49 | 0.81 | 0.56 | 0.86 | 0.39 | 0.67 |
| 9 | 88869630 | rs11791066 | 0.281 | -0.803 | -1.442 | -0.789 | 0.02 | 0.081 | 0.081 | 0 | 0.22 | 0.03 | 0.17 | 0.06 | 0.18 | 0.13 |
| 9 | 88870898 | rs1547629 | -1.45 | -0.283 | -2.11 | -1.472 | -0.174 | 0.196 | 0.196 | 0.528 | 0.72 | 0.98 | 0.8 | 0.99 | 0.63 | 0.85 |
| 9 | 88874117 | rs1061946 | -0.262 | -0.62 | -1.273 | -1.094 | 0.016 | 0.001 | 0.001 | 0 | 0.5 | 0.46 | 0.5 | 0.52 | 0.45 | 0.47 |

| | | | | | | | | | | | | | | | | |
|---|---|---|---|---|---|---|---|---|---|---|---|---|---|---|---|---|
| 9 | 88877447 | rs4877966 | 0.174 | -0.632 | -1.274 | -1.095 | -0.013 | 0.141 | 0.141 | 0.575 | 0.22 | 0.53 | 0.3 | 0.47 | 0.18 | 0.38 |
| 9 | 88902165 | rs10780756 | -0.27 | -0.636 | -1.207 | -1.099 | 0.027 | 0.002 | 0.002 | 0 | 0.5 | 0.43 | 0.5 | 0.52 | 0.45 | 0.47 |
| 9 | 88910161 | rs4587414 | -0.224 | -0.663 | -1.218 | -1.111 | 0.024 | 0.001 | 0.001 | 0 | 0.51 | 0.43 | 0.51 | 0.52 | 0.45 | 0.47 |
| 9 | 88914554 | rs10115526 | 0.181 | -0.658 | -1.216 | -1.108 | -0.013 | 0.122 | 0.122 | 0.597 | 0.22 | 0.55 | 0.3 | 0.47 | 0.18 | 0.38 |
| 9 | 88926377 | rs7028494 | -0.183 | -1.015 | -0.849 | -0.959 | 0.062 | -0.04 | -0.04 | 0 | 0.47 | 0.17 | 0.47 | 0.47 | 0.45 | 0.43 |
| 9 | 88933672 | rs700759 | -0.733 | -1.069 | -0.683 | -0.97 | 0 | 0.002 | 0.002 | 0 | 0.76 | 0.71 | 0.69 | 0.59 | 0.73 | 0.7 |
| 9 | 88942674 | rs791313 | 0.229 | -1.076 | -0.825 | -0.98 | 0.027 | 0.002 | 0.002 | 0 | 0.5 | 0.57 | 0.49 | 0.48 | 0.55 | 0.53 |
| 9 | 88959938 | rs791323 | 0.072 | -1.264 | -0.85 | -0.976 | 0.068 | -0.036 | -0.036 | 0 | 0.52 | 0.83 | 0.53 | 0.53 | 0.54 | 0.57 |
| 9 | 88963008 | rs17461620 | 0.518 | -1.017 | -0.841 | -1.234 | 0.037 | 0.096 | 0.096 | 0 | 0.25 | 0.03 | 0.18 | 0.06 | 0.19 | 0.13 |
| 9 | 88985734 | rs4877971 | 0.528 | -0.19 | -0.74 | -0.96 | -0.014 | -0.032 | -0.032 | 0 | 0.33 | 0.26 | 0.27 | 0.01 | 0.42 | 0.19 |
| 9 | 89000219 | rs11141370 | -0.181 | -0.532 | -0.609 | 0.194 | 0.005 | -0.04 | -0.04 | 0.947 | 0.14 | 0.07 | 0.21 | 0.4 | 0.13 | 0.27 |
| 9 | 89001745 | rs10868424 | -0.145 | -0.554 | -0.496 | 0.183 | 0.004 | -0.007 | -0.007 | 3.061 | 0.15 | 0.1 | 0.22 | 0.4 | 0.13 | 0.28 |
| 9 | 89011328 | rs6559926 | -0.239 | -0.535 | -0.615 | 0.193 | 0.006 | -0.037 | -0.037 | 0 | 0.15 | 0.06 | 0.21 | 0.4 | 0.13 | 0.28 |
| 9 | 89022298 | rs13284082 | -0.63 | -0.505 | -0.042 | -0.059 | -0.018 | 0.087 | 0.087 | 1.209 | 0.25 | 0.46 | 0.32 | 0.42 | 0.19 | 0.35 |
| 9 | 89023508 | rs11141381 | -0.748 | -0.713 | -1.118 | -0.324 | 0.042 | 0.009 | 0.009 | 0.889 | 0.45 | 0.33 | 0.44 | 0.4 | 0.33 | 0.45 |
| 9 | 89025919 | rs10868434 | -0.551 | -0.339 | -1.02 | 0.12 | 0.004 | -0.023 | -0.023 | 0 | 0.23 | 0.19 | 0.3 | 0.41 | 0.2 | 0.36 |
| 9 | 89026453 | rs353806 | -0.186 | 0.146 | -1.55 | 0.229 | -0.057 | -0.109 | -0.109 | 0 | 0.86 | 0.99 | 0.71 | 0.41 | 0.77 | 0.77 |
| 9 | 89029703 | rs187117 | 0.204 | 0.079 | -1.565 | -0.135 | 0.017 | -0.008 | -0.008 | 0 | 0.41 | 0.72 | 0.37 | 0.53 | 0.28 | 0.61 |
| 9 | 89030832 | rs416513 | 1.734 | -0.123 | -2.008 | -0.462 | -0.04 | 0.101 | 0.101 | 0.121 | 0.2 | 0.04 | 0.14 | 0.05 | 0.18 | 0.21 |
| 9 | 89030908 | rs17051878 | -0.234 | -0.169 | -0.7 | -0.312 | -0.028 | 0.154 | 0.154 | 0 | 0.18 | 0.36 | 0.2 | 0.48 | 0.1 | 0.41 |
| 9 | 89031511 | rs2889968 | -0.32 | -0.017 | -1.137 | -0.502 | -0.032 | -0.07 | -0.07 | 2.545 | 0.8 | 0.93 | 0.69 | 0.59 | 0.74 | 0.8 |
| 9 | 89031729 | rs806269 | 0.355 | 0.004 | -0.596 | -0.479 | 0.014 | 0.179 | 0.179 | 0 | 0.81 | 0.6 | 0.76 | 0.51 | 0.83 | 0.6 |
| 9 | 89033309 | rs11792677 | 0.349 | -0.038 | -1.084 | -0.948 | -0.088 | -0.011 | -0.011 | 0.185 | 0.66 | 0.85 | 0.62 | 0.31 | 0.52 | 0.46 |
| 9 | 89035754 | rs1331844 | 0.353 | -0.069 | -0.638 | -0.939 | -0.088 | -0.013 | -0.013 | 0 | 0.66 | 0.85 | 0.62 | 0.31 | 0.52 | 0.46 |
| 9 | 89035877 | rs17051885 | 0.769 | -0.417 | -0.486 | -1.253 | -0.007 | 0 | 0 | 0 | 0.27 | 0.3 | 0.34 | 0.22 | 0.22 | 0.2 |
| 9 | 89036027 | rs354815 | -0.508 | -0.083 | -0.532 | -0.918 | -0.037 | 0.006 | 0.006 | 0 | 0.37 | 0.27 | 0.39 | 0.69 | 0.49 | 0.54 |
| 9 | 89040610 | rs1041526 | 0.214 | -0.202 | -0.372 | -0.976 | -0.015 | -0.016 | -0.016 | 0 | 0.39 | 0.46 | 0.41 | 0.23 | 0.32 | 0.22 |
| 9 | 89041277 | rs11141391 | 1.445 | 0.087 | -0.941 | -0.566 | -0.074 | -0.086 | -0.086 | 0.94 | 0.29 | 0.05 | 0.36 | 0.6 | 0.3 | 0.47 |
| 9 | 89042419 | rs1576728 | 0.823 | 0.009 | 0.181 | -0.48 | 0.007 | 0.24 | 0.24 | 0 | 0.1 | 0.25 | 0.17 | 0.64 | 0.08 | 0.26 |
| 9 | 89046282 | rs4877980 | 1.985 | 0.172 | -0.649 | -1.12 | -0.052 | 0.121 | 0.121 | 0.012 | 0.27 | 0.1 | 0.18 | 0.08 | 0.26 | 0.33 |
| 9 | 89051246 | rs184789 | -0.907 | 0.212 | -0.674 | -1.58 | -0.027 | 0.038 | 0.038 | 0 | 0.75 | 0.84 | 0.81 | 0.95 | 0.68 | 0.91 |
| 9 | 89057171 | rs189544 | 1.41 | 0.132 | -0.746 | -1.729 | -0.048 | 0.157 | 0.157 | 0 | 0.42 | 0.2 | 0.32 | 0.15 | 0.45 | 0.41 |
| 9 | 89058963 | rs410206 | 1.359 | 0.129 | -0.808 | -1.836 | -0.022 | 0.029 | 0.029 | 0 | 0.12 | 0.06 | 0.12 | 0.01 | 0.22 | 0.06 |
| 9 | 89059014 | rs9314753 | 1.233 | -0.153 | -0.069 | -1.802 | -0.025 | -0.007 | -0.007 | 0 | 0.22 | 0.25 | 0.32 | 0.16 | 0.16 | 0.22 |
| 9 | 89060516 | rs7851100 | 1.935 | 0.029 | -0.119 | -1.879 | -0.016 | 0.054 | 0.054 | 0 | 0.19 | 0.07 | 0.14 | 0.04 | 0.21 | 0.08 |
| 9 | 89062045 | rs7038873 | 1.923 | -0.017 | 0.093 | -1.735 | -0.008 | 0.114 | 0.114 | 0 | 0.1 | 0.21 | 0.16 | 0.71 | 0.07 | 0.28 |
| 9 | 89063984 | rs1929802 | 1.382 | 0.053 | -0.615 | -1.712 | 0.013 | 0 | 0 | 0 | 0.2 | 0.15 | 0.3 | 0.16 | 0.16 | 0.22 |
| 9 | 89066329 | rs353826 | 1.57 | 0.131 | -0.643 | -1.748 | -0.029 | 0.039 | 0.039 | 0 | 0.11 | 0.04 | 0.12 | 0.01 | 0.21 | 0.06 |
| 9 | 89066515 | rs4877981 | 2.296 | -0.2 | -0.478 | -1.488 | -0.018 | 0.128 | 0.128 | 1.212 | 0.28 | 0.09 | 0.19 | 0.08 | 0.23 | 0.31 |
| 9 | 89068546 | rs12346290 | 0.073 | 0.121 | 0.048 | -0.91 | -0.043 | -0.018 | -0.018 | 0 | 0.24 | 0.35 | 0.32 | 0.16 | 0.16 | 0.21 |
| 9 | 89071996 | rs11141409 | 0.124 | 1.309 | -1.101 | -2.021 | -0.084 | 0.078 | 0.078 | 0 | 0.28 | 0.05 | 0.22 | 0.1 | 0.39 | 0.16 |
| 9 | 89073689 | rs353834 | 1.201 | 0.827 | -1.004 | -2.184 | -0.023 | 0.128 | 0.128 | 0 | 0.55 | 0.24 | 0.65 | 0.8 | 0.48 | 0.8 |
| 9 | 89074684 | rs7026472 | -0.039 | 1.14 | -0.828 | -1.912 | -0.01 | 0.01 | 0.01 | 0 | 0.29 | 0.19 | 0.24 | 0.2 | 0.39 | 0.16 |
| 9 | 89081586 | rs460811 | 0.369 | 0.22 | -0.892 | -2.285 | -0.041 | 0.053 | 0.053 | 0 | 0.66 | 0.77 | 0.7 | 0.94 | 0.5 | 0.8 |
| 9 | 89086848 | rs432957 | 1.06 | -0.184 | -0.899 | -2.435 | -0.024 | 0.1 | 0.1 | 0 | 0.56 | 0.29 | 0.65 | 0.9 | 0.51 | 0.81 |
| 9 | 89102306 | rs653030 | -0.57 | 0.366 | -1.01 | -2.319 | 0.014 | -0.001 | -0.001 | 0 | 0.87 | 0.87 | 0.85 | 0.9 | 0.73 | 0.91 |
| 9 | 89107564 | rs168493 | 0.722 | -0.162 | -1.615 | -2.444 | 0.009 | -0.037 | -0.037 | 0.682 | 0.25 | 0.16 | 0.33 | 0.61 | 0.23 | 0.49 |
| 9 | 89115887 | rs6559931 | -0.506 | 0.143 | -1.368 | -3.07 | -0.069 | 0.076 | 0.076 | 0 | 0.74 | 0.9 | 0.74 | 0.97 | 0.56 | 0.88 |
| 9 | 89117014 | rs7341859 | -0.573 | -0.037 | -1.221 | -3.138 | -0.023 | 0.005 | 0.005 | 0 | 0.75 | 0.87 | 0.79 | 1 | 0.61 | 0.91 |
| 9 | 89134626 | rs4877322 | -0.214 | 0.106 | -1.619 | -3.062 | -0.024 | 0.019 | 0.019 | 0 | 0.71 | 0.86 | 0.78 | 0.98 | 0.6 | 0.84 |
| 9 | 89142905 | rs2148996 | 0.748 | -0.304 | -1.758 | -3.359 | -0.004 | 0.067 | 0.067 | 0 | 0.51 | 0.7 | 0.53 | 0.84 | 0.43 | 0.66 |
| 9 | 89148945 | rs11792704 | 0.633 | 0.604 | -1.576 | -2.965 | 0.024 | 0.148 | 0.148 | 0 | 0.38 | 0.11 | 0.27 | 0.15 | 0.31 | 0.32 |
| 9 | 89157066 | rs2065376 | 0.257 | 0.779 | -0.452 | -2.442 | 0.01 | -0.018 | -0.018 | 0 | 0.28 | 0.26 | 0.19 | 0 | 0.22 | 0.09 |
| 9 | 89160794 | rs1331433 | -0.614 | 0.877 | -0.389 | -2.453 | 0.031 | -0.06 | -0.06 | 0 | 0.86 | 0.86 | 0.88 | 1 | 0.75 | 0.93 |
| 9 | 89162181 | rs10491846 | 0.882 | 0.761 | -0.521 | -2.4 | -0.002 | 0.049 | 0.049 | 0 | 0.52 | 0.38 | 0.68 | 0.91 | 0.52 | 0.78 |
| 9 | 89162323 | rs10780767 | 0.827 | 0.928 | -0.548 | -2.223 | -0.002 | -0.008 | -0.008 | 0.982 | 0.55 | 0.42 | 0.68 | 0.92 | 0.53 | 0.82 |
| 9 | 89166936 | rs12551728 | 0.262 | 1.186 | -0.729 | -2.185 | 0.005 | -0.033 | -0.033 | 0 | 0.26 | 0.26 | 0.18 | 0.01 | 0.2 | 0.06 |
| 9 | 89167524 | rs962704 | -0.202 | 2.363 | -0.685 | -0.862 | -0.012 | 0.012 | 0.012 | 0 | 0.11 | 0.07 | 0.15 | 0.1 | 0.15 | 0.14 |
| 9 | 89168700 | rs1331436 | 0.817 | 2.001 | -0.312 | -0.515 | 0.049 | -0.074 | -0.074 | 0.962 | 0.85 | 0.82 | 0.88 | 1 | 0.75 | 0.93 |
| 9 | 89172482 | rs913357 | 0.099 | 1.12 | -0.59 | -0.004 | -0.003 | -0.003 | -0.003 | 0.021 | 0.52 | 0.45 | 0.48 | 0.38 | 0.49 | 0.47 |
| 9 | 89172667 | rs913356 | 0.189 | 0.935 | -0.655 | 0.024 | -0.034 | -0.007 | -0.007 | 0 | 0.56 | 0.25 | 0.5 | 0.41 | 0.57 | 0.44 |
| 9 | 89174807 | rs10868444 | 0.287 | 1.171 | -0.928 | -0.33 | -0.001 | 0.008 | 0.008 | 0 | 0.47 | 0.26 | 0.5 | 0.42 | 0.5 | 0.42 |
| 9 | 89174943 | rs17051989 | 0.593 | 1.019 | -0.526 | -0.578 | -0.007 | -0.049 | -0.049 | 1.451 | 0.09 | 0.21 | 0.06 | 0 | 0.05 | 0.04 |
| 9 | 89177743 | rs7850197 | 0.778 | 1.169 | -1.083 | -0.437 | -0.005 | -0.032 | -0.032 | 0.121 | 0.39 | 0.19 | 0.36 | 0.41 | 0.44 | 0.39 |
| 9 | 89178164 | rs1537190 | 0.433 | 1.244 | -0.904 | -0.29 | 0.016 | 0.01 | 0.01 | 0 | 0.47 | 0.25 | 0.41 | 0.42 | 0.48 | 0.4 |
| 9 | 89178987 | rs7026807 | 0.104 | 1.191 | -0.812 | -0.394 | 0.013 | -0.021 | -0.021 | 0.335 | 0.11 | 0.16 | 0.16 | 0.06 | 0.12 | 0.04 |
| 9 | 89188290 | rs2065378 | -0.032 | 1.107 | -0.847 | -0.095 | 0.001 | 0.004 | 0.004 | 0 | 0.58 | 0.42 | 0.57 | 0.47 | 0.6 | 0.43 |
| 9 | 89191784 | rs10780771 | 0.378 | 0.978 | -1.543 | -0.127 | -0.048 | -0.013 | -0.013 | 0.398 | 0.41 | 0.19 | 0.33 | 0.37 | 0.49 | 0.48 |
| 9 | 89193482 | rs10117819 | 0.09 | 0.637 | -2.409 | 0.164 | -0.136 | -0.139 | -0.139 | 0 | 0.66 | 0.94 | 0.4 | 0.42 | 0.49 | 0.61 |

| | | | | | | | | | | | | | | | | |
|---|---|---|---|---|---|---|---|---|---|---|---|---|---|---|---|---|
| 9 | 89199480 | rs2183731 | 0.295 | -0.224 | -2.033 | -0.506 | 0.048 | 0.074 | 0.074 | 0 | 0.45 | 0.25 | 0.25 | 0.24 | 0.32 | 0.39 |
| 9 | 89199498 | rs2183732 | 0.3 | -0.232 | -2 | -0.51 | 0.047 | 0.073 | 0.073 | 0 | 0.45 | 0.25 | 0.25 | 0.24 | 0.32 | 0.39 |
| 9 | 89207149 | rs7048627 | -0.464 | -0.075 | -1.222 | 0.029 | -0.003 | -0.014 | -0.014 | 0.236 | 0.17 | 0.23 | 0.12 | 0.06 | 0.15 | 0.23 |
| 9 | 89212523 | rs2860107 | -0.577 | -0.543 | -1.465 | -0.102 | 0.006 | 0.018 | 0.018 | 0 | 0.48 | 0.4 | 0.44 | 0.4 | 0.46 | 0.57 |
| 9 | 89213659 | rs7852670 | -0.011 | -0.296 | -1.238 | -0.117 | -0.012 | -0.023 | -0.023 | 0.768 | 0.14 | 0.2 | 0.1 | 0.05 | 0.12 | 0.21 |
| 9 | 89219411 | rs985863 | -0.514 | -0.6 | -0.926 | -0.029 | 0.001 | 0.003 | 0.003 | 0 | 0.48 | 0.42 | 0.47 | 0.41 | 0.46 | 0.59 |
| 9 | 89219781 | rs868823 | 0.275 | -0.589 | -1.256 | -0.017 | -0.018 | -0.012 | -0.012 | 0 | 0.34 | 0.18 | 0.37 | 0.35 | 0.34 | 0.38 |
| 9 | 89227561 | rs11141456 | -1.034 | -0.523 | -1.088 | 0.216 | -0.044 | -0.041 | -0.041 | 0 | 0.58 | 0.84 | 0.5 | 0.51 | 0.44 | 0.65 |
| 9 | 89227755 | rs10868451 | -0.845 | -0.551 | -1.042 | 0.234 | 0.005 | -0.012 | -0.012 | 0 | 0.55 | 0.8 | 0.47 | 0.52 | 0.41 | 0.63 |
| 9 | 89230485 | rs12343626 | 0.446 | -0.567 | -0.971 | 0.227 | 0.006 | 0.007 | 0.007 | 0 | 0.08 | 0.08 | 0.05 | 0.05 | 0.05 | 0.19 |
| 9 | 89230779 | rs9696070 | 0.041 | -0.582 | -0.56 | 0.228 | -0.006 | 0.004 | 0.004 | 0 | 0.44 | 0.47 | 0.4 | 0.44 | 0.37 | 0.46 |
| 9 | 89236894 | rs9695351 | -0.08 | 0.261 | -0.947 | 0.227 | 0.011 | 0 | 0 | 0 | 0.43 | 0.64 | 0.46 | 0.4 | 0.33 | 0.47 |
| 9 | 89240136 | rs4492454 | -0.299 | 0.874 | -1.068 | 0.057 | 0.022 | 0.027 | 0.027 | 0 | 0.49 | 0.65 | 0.59 | 0.38 | 0.55 | 0.49 |
| 9 | 89240189 | rs1888713 | 0.496 | 0.784 | -1.403 | 0.068 | 0 | 0.001 | 0.001 | 0 | 0.12 | 0.06 | 0.16 | 0.12 | 0.11 | 0.2 |
| 9 | 89240203 | rs1888714 | -0.856 | 0.704 | -1.35 | 0.158 | 0.035 | -0.053 | -0.053 | 0 | 0.61 | 0.78 | 0.75 | 0.5 | 0.66 | 0.69 |
| 9 | 89245979 | rs914394 | -0.264 | 1.268 | -1.309 | 0.165 | -0.021 | 0.05 | 0.05 | 0 | 0.44 | 0.77 | 0.57 | 0.25 | 0.47 | 0.49 |
| 9 | 89246924 | rs7851559 | -0.722 | 0.442 | -1.851 | 0.168 | 0.09 | -0.076 | -0.076 | 0 | 0.59 | 0.91 | 0.77 | 0.38 | 0.69 | 0.63 |
| 9 | 89248754 | rs1331453 | -0.771 | 1.075 | -0.571 | -0.141 | 0.004 | -0.062 | -0.062 | 3.156 | 0.11 | 0.24 | 0.11 | 0 | 0.1 | 0.08 |
| 9 | 89252706 | rs10512176 | -0.568 | 1.475 | -0.691 | -0.074 | -0.019 | 0.017 | 0.017 | 0 | 0.24 | 0.15 | 0.27 | 0.18 | 0.34 | 0.16 |
| 9 | 89269078 | rs1411987 | -1.127 | 0.894 | -0.708 | 0.058 | 0.024 | -0.013 | -0.013 | 0 | 0.58 | 0.74 | 0.73 | 0.48 | 0.67 | 0.54 |
| 9 | 89269261 | rs11141473 | 0.556 | 0.911 | 0.143 | -0.031 | 0.003 | -0.029 | -0.029 | 0 | 0.23 | 0.38 | 0.19 | 0.17 | 0.23 | 0.33 |
| 9 | 89271532 | rs4877995 | -0.121 | 0.875 | 0.326 | 0.048 | -0.021 | 0.003 | 0.003 | 0 | 0.28 | 0.18 | 0.38 | 0.25 | 0.4 | 0.17 |
| 9 | 89271591 | rs2183730 | -1.124 | 0.718 | 0.276 | -0.169 | -0.007 | 0 | 0 | 0 | 0.55 | 0.55 | 0.58 | 0.48 | 0.69 | 0.5 |
| 9 | 89275786 | rs7869597 | -0.148 | 0.675 | 0.454 | 0.032 | -0.034 | 0.001 | 0.001 | 0 | 0.28 | 0.14 | 0.37 | 0.25 | 0.4 | 0.17 |
| 9 | 89278227 | rs4877329 | -1.419 | 0.54 | 0.85 | -0.029 | 0.025 | -0.005 | -0.005 | 0 | 0.54 | 0.81 | 0.59 | 0.48 | 0.63 | 0.49 |
| 9 | 89279055 | rs7853196 | 0.727 | 0.604 | 0.915 | 0.014 | -0.002 | 0.001 | 0.001 | 0 | 0.21 | 0.23 | 0.17 | 0.21 | 0.22 | 0.32 |
| 9 | 89280867 | rs932956 | -0.718 | 0.7 | 0.552 | 0.009 | -0.083 | -0.079 | -0.079 | 0.11 | 0.84 | 0.98 | 0.72 | 0.71 | 0.71 | 0.86 |
| 9 | 89285932 | rs7860045 | 0.047 | 0.829 | 1.477 | 0.067 | 0.004 | -0.022 | -0.022 | 0 | 0.25 | 0.49 | 0.19 | 0.21 | 0.24 | 0.32 |
| 9 | 89286335 | rs11141482 | -0.737 | 0.736 | 1.639 | -0.021 | -0.038 | -0.112 | -0.112 | 1.124 | 0.18 | 0.33 | 0.25 | 0.01 | 0.12 | 0.06 |
| 9 | 89287374 | rs6559946 | -1.417 | 0.797 | 1.134 | 0.086 | 0.032 | 0.125 | 0.125 | 0 | 0.43 | 0.82 | 0.45 | 0.22 | 0.37 | 0.4 |
| 9 | 89290300 | rs1929463 | 0.491 | 0.896 | 1.219 | 0.294 | -0.009 | 0 | 0 | 0 | 0.22 | 0.17 | 0.18 | 0.21 | 0.24 | 0.34 |
| 9 | 89293393 | rs11141486 | 0.14 | 0.799 | 1.211 | 0.222 | -0.016 | 0.012 | 0.012 | 0 | 0.23 | 0.17 | 0.33 | 0.19 | 0.26 | 0.35 |
| 9 | 89296599 | rs10868464 | 0.003 | 0.772 | 1.401 | 0.223 | 0.007 | -0.019 | -0.019 | 0 | 0.26 | 0.36 | 0.35 | 0.19 | 0.27 | 0.36 |
| 9 | 89297754 | rs10868467 | -0.25 | 0.721 | 1.342 | 0.21 | 0.007 | -0.042 | -0.042 | 0 | 0.27 | 0.51 | 0.35 | 0.19 | 0.25 | 0.35 |
| 9 | 89302560 | rs10868469 | 1.363 | 0.688 | 0.577 | 0.342 | 0.002 | -0.083 | -0.083 | 0.501 | 0.45 | 0.1 | 0.26 | 0.51 | 0.35 | 0.47 |
| 9 | 89305431 | rs1475617 | 0.154 | 0.595 | 1.065 | 0.648 | 0.023 | 0.002 | 0.002 | 0 | 0.72 | 0.65 | 0.57 | 0.75 | 0.53 | 0.79 |
| 9 | 89306305 | rs11141494 | 0.528 | 0.572 | 0.924 | 0.767 | -0.018 | -0.018 | -0.018 | 1.884 | 0.19 | 0.09 | 0.14 | 0.28 | 0.19 | 0.2 |
| 9 | 89308609 | rs10780781 | 0.571 | 0.51 | 0.927 | 0.771 | 0.002 | -0.001 | -0.001 | 0.014 | 0.68 | 0.66 | 0.4 | 0.71 | 0.5 | 0.68 |
| 9 | 89309429 | rs4878002 | -0.306 | 0.562 | 0.748 | 0.833 | -0.001 | -0.001 | -0.001 | 0 | 0.79 | 0.75 | 0.85 | 0.71 | 0.8 | 0.8 |
| 9 | 89315229 | rs1331443 | -0.376 | 0.325 | 1.027 | 0.521 | 0.01 | -0.001 | -0.001 | 0 | 0.32 | 0.33 | 0.36 | 0.32 | 0.45 | 0.29 |
| 9 | 89316235 | rs7032797 | 1.335 | 0.204 | 0.976 | 0.653 | 0.02 | 0.007 | 0.007 | 0 | 0.44 | 0.34 | 0.28 | 0.39 | 0.32 | 0.43 |
| 9 | 89317479 | rs10746782 | 0.98 | 0.203 | 0.965 | 0.652 | 0.036 | 0.014 | 0.014 | 0 | 0.48 | 0.36 | 0.45 | 0.43 | 0.35 | 0.54 |
| 9 | 89321913 | rs1927642 | 1.309 | 0.194 | 0.936 | 0.659 | 0.038 | 0.013 | 0.013 | 0 | 0.44 | 0.34 | 0.28 | 0.39 | 0.32 | 0.43 |
| 9 | 89323257 | rs755056 | -0.593 | 0.283 | 0.564 | 0.678 | 0.001 | 0.001 | 0.001 | 0 | 0.8 | 0.74 | 0.81 | 0.75 | 0.81 | 0.82 |
| 9 | 89327965 | rs7047160 | 0.719 | 0.333 | 0.77 | 0.707 | 0.052 | 0.022 | 0.022 | 0 | 0.49 | 0.33 | 0.45 | 0.43 | 0.37 | 0.54 |
| 9 | 89331137 | rs2378715 | -1.264 | 0.866 | 0.882 | 0.819 | -0.025 | -0.008 | -0.008 | 0 | 0.34 | 0.7 | 0.55 | 0.39 | 0.46 | 0.38 |
| 9 | 89336987 | rs4878010 | -0.378 | 1.019 | 0.928 | 0.824 | 0.019 | 0.012 | 0.012 | 0 | 0.24 | 0.27 | 0.32 | 0.36 | 0.39 | 0.29 |
| 9 | 89347214 | rs7864375 | -1.151 | 0.803 | 1.424 | 0.721 | 0.05 | 0.03 | 0.03 | 1.047 | 0.06 | 0.34 | 0.04 | 0 | 0.03 | 0 |
| 9 | 89351232 | rs10046817 | 0.551 | 0.836 | 1.498 | 0.721 | 0.041 | 0.013 | 0.013 | 0 | 0.71 | 0.6 | 0.49 | 0.65 | 0.58 | 0.62 |
| 9 | 89351788 | rs932858 | 1.202 | 0.845 | 1.033 | 0.724 | -0.018 | 0.001 | 0.001 | 0 | 0.66 | 0.33 | 0.46 | 0.65 | 0.54 | 0.62 |
| 9 | 89356423 | rs1571333 | 0.273 | 1.108 | 0.387 | 0.937 | -0.011 | -0.009 | -0.009 | 0 | 0.76 | 0.77 | 0.68 | 0.65 | 0.6 | 0.72 |
| 9 | 89365622 | rs11141530 | 0.783 | 1.448 | -0.3 | 1.38 | -0.05 | -0.053 | -0.053 | 0 | 0.1 | 0.01 | 0.13 | 0.19 | 0.13 | 0.1 |
| 9 | 89368334 | rs10514832 | -0.339 | 1.268 | -0.259 | 1.304 | -0.024 | -0.019 | -0.019 | 0.02 | 0.18 | 0.12 | 0.22 | 0.26 | 0.21 | 0.17 |
| 9 | 89371743 | rs2274558 | 0.062 | 1.25 | -0.007 | 0.928 | 0.08 | -0.023 | -0.023 | 0 | 0.74 | 0.62 | 0.55 | 0.78 | 0.55 | 0.71 |
| 9 | 89372582 | rs2274559 | 0.943 | -0.064 | -1.181 | 0.574 | 0.113 | 0.085 | 0.085 | 0.323 | 0.46 | 0.05 | 0.32 | 0.3 | 0.33 | 0.46 |
| 9 | 89372790 | rs2274561 | 0.318 | -0.42 | -1.278 | 0.54 | -0.006 | -0.032 | -0.032 | 0 | 0.16 | 0.06 | 0.13 | 0.19 | 0.17 | 0.19 |
| 9 | 89375314 | rs994029 | -0.833 | -0.183 | -1.441 | -0.119 | 0.04 | 0.186 | 0.186 | 0 | 0.58 | 0.89 | 0.66 | 0.85 | 0.62 | 0.59 |
| 9 | 89377344 | rs10868482 | 0.084 | -0.171 | -0.564 | 0.171 | 0.018 | 0.013 | 0.013 | 0 | 0.35 | 0.41 | 0.46 | 0.54 | 0.44 | 0.35 |
| 9 | 89378267 | rs10746786 | -0.879 | -0.307 | -0.063 | 0.208 | 0.006 | 0 | 0 | 0 | 0.31 | 0.72 | 0.27 | 0.32 | 0.25 | 0.3 |
| 9 | 89378977 | rs11141538 | -0.882 | -0.307 | -0.091 | 0.208 | 0.014 | 0 | 0 | 1.244 | 0.31 | 0.73 | 0.27 | 0.32 | 0.25 | 0.3 |
| 9 | 89382631 | rs12351577 | -1.067 | -0.344 | 0.291 | -0.104 | -0.041 | -0.072 | -0.072 | 0.053 | 0.11 | 0.6 | 0.07 | 0 | 0.04 | 0.01 |
| 9 | 89382759 | rs12338073 | -0.762 | -0.351 | 0.663 | -0.107 | 0.008 | -0.031 | -0.031 | 0 | 0.09 | 0.49 | 0.06 | 0 | 0.04 | 0.01 |
| 9 | 89394222 | rs7043314 | -1.099 | -0.283 | 0.467 | -0.036 | -0.013 | -0.013 | -0.013 | 0 | 0.32 | 0.6 | 0.27 | 0.27 | 0.26 | 0.3 |
| 9 | 89402003 | rs7852002 | -0.361 | -0.255 | 0.916 | -0.06 | -0.005 | -0.022 | -0.022 | 0 | 0.07 | 0.23 | 0.03 | 0 | 0.02 | 0 |
| 9 | 89404665 | rs7036971 | -1.103 | -0.312 | 0.269 | -0.006 | -0.007 | -0.005 | -0.005 | 0.011 | 0.31 | 0.63 | 0.27 | 0.28 | 0.26 | 0.29 |
| 9 | 89408927 | rs7851591 | 0.108 | -0.255 | 0.157 | 0.063 | -0.002 | -0.022 | -0.022 | 0 | 0.21 | 0.14 | 0.21 | 0.28 | 0.21 | 0.27 |
| 9 | 89415661 | rs10514834 | 0.22 | -0.058 | 0.128 | 0.058 | -0.009 | -0.014 | -0.014 | 0.614 | 0.2 | 0.13 | 0.22 | 0.27 | 0.24 | 0.24 |
| 9 | 89419847 | rs7048294 | -1.413 | -0.509 | -0.87 | 0.013 | 0.018 | 0.002 | 0.002 | 0 | 0.66 | 0.98 | 0.73 | 0.76 | 0.76 | 0.58 |
| 9 | 89422169 | rs374503 | -0.723 | -0.335 | -0.092 | 0.176 | -0.001 | -0.001 | -0.001 | 0 | 0.31 | 0.66 | 0.29 | 0.28 | 0.28 | 0.25 |

| | | | | | | | | | | | | | | | | |
|---|---|---|---|---|---|---|---|---|---|---|---|---|---|---|---|---|
| 9 | 89429681 | rs4877337 | -0.031 | 0.336 | -0.213 | -0.087 | 0.019 | -0.049 | -0.049 | 0 | 0.16 | 0.37 | 0.22 | 0.11 | 0.18 | 0.14 |
| 9 | 89429897 | rs12236362 | 0.499 | 0.282 | -1.012 | 0.187 | -0.012 | -0.025 | -0.025 | 0 | 0.2 | 0.13 | 0.3 | 0.32 | 0.25 | 0.21 |
| 9 | 89432027 | rs452723 | -0.377 | 0.412 | -1.19 | 0.406 | -0.004 | 0.007 | 0.007 | 0 | 0.41 | 0.69 | 0.55 | 0.42 | 0.44 | 0.37 |
| 9 | 89436746 | rs11141559 | -0.02 | 1.38 | -1.621 | 0.85 | -0.101 | -0.126 | -0.126 | 0.86 | 0.17 | 0.02 | 0.25 | 0.37 | 0.29 | 0.29 |
| 9 | 89437343 | rs10868488 | -0.2 | 0.138 | -1.321 | -0.059 | 0.011 | 0.019 | 0.019 | 0.682 | 0.74 | 0.84 | 0.8 | 0.87 | 0.76 | 0.71 |
| 9 | 89439307 | rs4878016 | -0.008 | 0.462 | -0.784 | 0.473 | 0.005 | 0.03 | 0.03 | 0 | 0.78 | 0.77 | 0.78 | 0.61 | 0.77 | 0.75 |
| 9 | 89440597 | rs4242637 | 0.363 | 0.37 | -1.29 | -0.047 | -0.037 | -0.035 | -0.035 | 1.304 | 0.31 | 0.2 | 0.44 | 0.42 | 0.34 | 0.34 |
| 9 | 89440695 | rs10868490 | 0.557 | -0.145 | -0.927 | -0.567 | 0.029 | 0.093 | 0.093 | 0.029 | 0.41 | 0.26 | 0.29 | 0.17 | 0.41 | 0.39 |
| 9 | 89443026 | rs10780795 | 0.347 | -0.136 | -0.431 | -0.468 | 0 | 0.002 | 0.002 | 0 | 0.43 | 0.51 | 0.53 | 0.46 | 0.41 | 0.27 |
| 9 | 89443078 | rs11141567 | 0.608 | -0.373 | -0.741 | -0.534 | 0.035 | 0.074 | 0.074 | 0 | 0.29 | 0.1 | 0.17 | 0.15 | 0.26 | 0.28 |
| 9 | 89444725 | rs1028880 | 0.911 | -0.43 | -0.239 | -0.614 | -0.005 | -0.015 | -0.015 | 0 | 0.37 | 0.28 | 0.49 | 0.46 | 0.36 | 0.25 |
| 9 | 89445581 | rs1977459 | 0.197 | -0.302 | 0.041 | -0.514 | 0.014 | 0.011 | 0.011 | 0.948 | 0.66 | 0.41 | 0.67 | 0.61 | 0.61 | 0.54 |
| 9 | 89448000 | rs10780796 | 0.041 | -0.546 | -0.198 | -0.514 | -0.001 | -0.001 | -0.001 | 0 | 0.66 | 0.63 | 0.7 | 0.61 | 0.62 | 0.52 |
| 9 | 89453847 | rs7021048 | 0.739 | -0.645 | -0.24 | -0.631 | -0.003 | -0.005 | -0.005 | 1.791 | 0.36 | 0.31 | 0.46 | 0.41 | 0.39 | 0.21 |
| 9 | 89456443 | rs10746789 | -0.02 | -0.395 | -0.053 | -0.617 | 0.005 | 0.008 | 0.008 | 1.994 | 0.64 | 0.58 | 0.63 | 0.57 | 0.58 | 0.51 |
| 9 | 89468654 | rs7032556 | 0.624 | -0.62 | 0.061 | -0.751 | 0.028 | -0.002 | -0.002 | 0.009 | 0.07 | 0.31 | 0.04 | 0 | 0.03 | 0.02 |
| 9 | 89472439 | rs13302123 | 1.221 | -0.754 | -0.614 | -0.838 | -0.035 | 0.027 | 0.027 | 0.002 | 0.21 | 0.01 | 0.12 | 0.1 | 0.17 | 0.22 |
| 9 | 89474289 | rs7048161 | 1.149 | -0.509 | 0.302 | -0.331 | 0.003 | 0.006 | 0.006 | 2.933 | 0.57 | 0.26 | 0.62 | 0.59 | 0.55 | 0.5 |
| 9 | 89478860 | rs10868498 | 0.148 | -0.401 | -0.357 | -0.151 | -0.088 | -0.107 | -0.107 | 0.093 | 0.77 | 0.97 | 0.75 | 0.68 | 0.71 | 0.6 |
| 9 | 89481996 | rs1169848 | 0.326 | -0.244 | 0.355 | 0.321 | -0.027 | -0.041 | -0.041 | 0.41 | 0.08 | 0.01 | 0.08 | 0.25 | 0.1 | 0.25 |
| 9 | 89488447 | rs10868499 | -1.254 | -0.267 | 0.574 | -0.215 | -0.018 | -0.028 | -0.028 | 1.098 | 0.11 | 0.27 | 0.06 | 0.1 | 0.11 | 0.07 |
| 9 | 89501366 | rs7018734 | -0.584 | -0.04 | 0.268 | -0.577 | -0.001 | -0.034 | -0.034 | 0 | 0.28 | 0.23 | 0.21 | 0.03 | 0.33 | 0.24 |
| 9 | 89505042 | rs10868500 | -1.322 | -0.6 | 0.362 | -0.513 | 0.025 | 0.099 | 0.099 | 0.155 | 0.46 | 0.69 | 0.43 | 0.21 | 0.42 | 0.32 |
| 9 | 89505232 | rs12352609 | -1.322 | -0.6 | 0.564 | -0.513 | 0.018 | 0.063 | 0.063 | 0.155 | 0.46 | 0.67 | 0.43 | 0.21 | 0.42 | 0.32 |
| 9 | 89509444 | rs10780800 | -1.318 | -0.597 | 0.464 | -0.494 | 0.027 | 0.108 | 0.108 | 0.154 | 0.46 | 0.71 | 0.43 | 0.22 | 0.42 | 0.32 |
| 9 | 89512335 | rs10868502 | -0.932 | -0.801 | 0.83 | -0.026 | -0.027 | -0.013 | -0.013 | 2.198 | 0.35 | 0.5 | 0.37 | 0.22 | 0.29 | 0.35 |
| 9 | 89514067 | rs7021050 | -1.121 | -0.621 | 0.725 | 0.2 | 0.019 | -0.013 | -0.013 | 0 | 0.37 | 0.65 | 0.42 | 0.34 | 0.31 | 0.37 |
| 9 | 89519033 | rs10512177 | -0.36 | -0.662 | 0.58 | -0.307 | 0.012 | 0.064 | 0.064 | 4.006 | 0.28 | 0.16 | 0.19 | 0.04 | 0.27 | 0.25 |
| 9 | 89521299 | rs10780802 | -0.144 | -0.699 | 0.698 | -0.163 | 0.01 | 0.031 | 0.031 | 4.434 | 0.23 | 0.16 | 0.17 | 0.04 | 0.21 | 0.19 |
| 9 | 89525066 | rs11141591 | -0.386 | -0.698 | 0.895 | -0.163 | -0.01 | -0.035 | -0.035 | 0 | 0.25 | 0.29 | 0.19 | 0.04 | 0.21 | 0.19 |
| 9 | 89525449 | rs11141592 | -0.808 | -0.706 | 0.67 | -0.138 | -0.027 | -0.046 | -0.046 | 0 | 0.31 | 0.57 | 0.35 | 0.09 | 0.21 | 0.2 |
| 9 | 89554097 | rs7036455 | -1.275 | -0.142 | -0.907 | -0.036 | -0.042 | -0.032 | -0.032 | 0 | 0.56 | 0.87 | 0.55 | 0.45 | 0.48 | 0.48 |
| 9 | 89558064 | rs12350534 | -0.214 | 0.164 | -0.346 | -0.129 | 0.018 | -0.026 | -0.026 | 0 | 0.05 | 0.08 | 0.05 | 0 | 0.06 | 0.04 |
| 9 | 89567590 | rs1927081 | -0.68 | -0.268 | -0.818 | -0.573 | 0 | 0.053 | 0.053 | 0 | 0.24 | 0.1 | 0.13 | 0 | 0.24 | 0.15 |
| 9 | 89570731 | rs10868508 | 0.753 | -0.4 | -0.716 | -0.644 | -0.005 | -0.021 | -0.021 | 0 | 0.05 | 0.09 | 0.08 | 0 | 0.03 | 0.05 |
| 9 | 89573083 | rs9792548 | -2.165 | -0.74 | -1.024 | -0.582 | 0.034 | 0.002 | 0.002 | 0.584 | 0.5 | 0.9 | 0.5 | 0.45 | 0.38 | 0.4 |
| 9 | 89574128 | rs12555168 | 0.146 | -0.842 | -0.618 | -0.392 | 0.003 | -0.004 | -0.004 | 0.019 | 0.1 | 0.11 | 0.07 | 0.15 | 0.06 | 0.08 |
| 9 | 89574902 | rs7034424 | -1.773 | -0.862 | 0.007 | -0.438 | 0.008 | 0.001 | 0.001 | 0 | 0.4 | 0.67 | 0.27 | 0.41 | 0.34 | 0.35 |
| 9 | 89575478 | rs1927084 | -2.076 | -0.878 | 0.192 | -0.373 | 0.031 | -0.01 | -0.01 | 0.349 | 0.41 | 0.74 | 0.39 | 0.45 | 0.32 | 0.32 |
| 9 | 89575529 | rs7862522 | -0.792 | -0.952 | 0.393 | -0.371 | -0.004 | -0.014 | -0.014 | 0 | 0.26 | 0.51 | 0.18 | 0.25 | 0.25 | 0.24 |
| 9 | 89576961 | rs10119026 | -1.688 | -0.972 | 0.456 | -0.604 | -0.05 | 0.011 | 0.011 | 0.103 | 0.21 | 0.39 | 0.31 | 0.19 | 0.1 | 0.14 |
| 9 | 89579246 | rs7849517 | -0.652 | -0.838 | 0.338 | -0.659 | 0.004 | -0.028 | -0.028 | 0 | 0.23 | 0.2 | 0.15 | 0 | 0.19 | 0.14 |
| 9 | 89579576 | rs11141634 | -1.679 | -0.85 | 0.326 | -0.642 | 0.016 | 0.017 | 0.017 | 0 | 0.36 | 0.32 | 0.38 | 0.19 | 0.28 | 0.21 |
| 9 | 89584572 | rs12555841 | -0.287 | -0.995 | 0.045 | -0.895 | 0.031 | 0.109 | 0.109 | 0 | 0.26 | 0.08 | 0.27 | 0.04 | 0.22 | 0.13 |
| 9 | 89589210 | rs10448319 | -1.06 | -0.905 | 0.273 | -0.786 | -0.018 | 0.012 | 0.012 | 0 | 0.17 | 0.3 | 0.16 | 0.17 | 0.12 | 0.14 |
| 9 | 89593328 | rs12379195 | -1.437 | -0.902 | 0.27 | -0.789 | -0.019 | 0.012 | 0.012 | 0 | 0.2 | 0.3 | 0.31 | 0.21 | 0.13 | 0.15 |
| 9 | 89593967 | rs11141638 | -0.683 | -0.894 | 0.278 | -0.786 | 0 | -0.001 | -0.001 | 0 | 0.15 | 0.15 | 0.14 | 0.17 | 0.11 | 0.14 |
| 9 | 89594294 | rs4878029 | -1.754 | -0.918 | 0.174 | -0.886 | 0.013 | 0.018 | 0.018 | 0 | 0.43 | 0.38 | 0.42 | 0.21 | 0.34 | 0.27 |
| 9 | 89597085 | rs10114955 | -2.177 | -0.211 | 0.494 | -1.036 | -0.02 | 0.086 | 0.086 | 0.471 | 0.55 | 0.72 | 0.57 | 0.22 | 0.48 | 0.3 |
| 9 | 89600651 | rs7047338 | -1.573 | 0.28 | 1.088 | -0.129 | -0.013 | 0.037 | 0.037 | 0 | 0.17 | 0.3 | 0.29 | 0.22 | 0.12 | 0.13 |
| 9 | 89605153 | rs6559958 | -1.54 | 0.817 | 1.343 | 0.221 | 0.03 | 0.22 | 0.22 | 0.348 | 0.11 | 0.32 | 0.25 | 0.41 | 0.12 | 0.17 |
| 9 | 89606711 | rs1927070 | 1.256 | 1.373 | 1.333 | 0.167 | -0.01 | 0.009 | 0.009 | 0 | 0.88 | 0.88 | 0.84 | 0.95 | 0.81 | 0.77 |
| 9 | 89606981 | rs13296748 | 1.371 | 1.347 | 1.155 | 0.029 | -0.022 | 0.06 | 0.06 | 0 | 0.76 | 0.86 | 0.79 | 0.95 | 0.73 | 0.71 |
| 9 | 89607073 | rs10868521 | 1.371 | 1.347 | 1.142 | 0.029 | -0.023 | 0.063 | 0.063 | 0 | 0.76 | 0.86 | 0.79 | 0.95 | 0.73 | 0.71 |
| 9 | 89608864 | rs4493983 | -1.341 | 1.33 | 1.181 | 0.103 | 0.037 | 0.22 | 0.22 | 0.401 | 0.11 | 0.33 | 0.25 | 0.42 | 0.13 | 0.17 |
| 9 | 89614276 | rs7863740 | -1.379 | 1.326 | 1.191 | 0.117 | 0.057 | 0.362 | 0.362 | 1.165 | 0.07 | 0.31 | 0.17 | 0.41 | 0.08 | 0.11 |
| 9 | 89619991 | rs7031306 | 2.293 | 1.292 | 1.576 | 0.06 | -0.021 | -0.031 | -0.031 | 0 | 0.62 | 0.23 | 0.51 | 0.55 | 0.61 | 0.55 |
| 9 | 89620854 | rs1927087 | 1.329 | 1.373 | 1.484 | -0.019 | -0.002 | 0.002 | 0.002 | 0 | 0.89 | 0.88 | 0.85 | 0.95 | 0.83 | 0.78 |
| 9 | 89627369 | rs7022771 | 2.168 | 1.281 | 1.523 | 0.078 | -0.01 | -0.021 | -0.021 | 0.055 | 0.63 | 0.27 | 0.49 | 0.55 | 0.62 | 0.54 |
| 9 | 89633112 | rs12553476 | -0.445 | 1.324 | 1.433 | 0.111 | -0.005 | 0.065 | 0.065 | 0 | 0.17 | 0.3 | 0.21 | 0.4 | 0.16 | 0.18 |
| 9 | 89650543 | rs7873337 | 0.789 | 1.338 | 1.464 | 0.143 | -0.013 | 0.013 | 0.013 | 0 | 0.86 | 0.89 | 0.82 | 0.95 | 0.81 | 0.76 |
| 9 | 89652307 | rs6559962 | 1.604 | 1.328 | 1.453 | 0.174 | 0.016 | 0.026 | 0.026 | 0 | 0.65 | 0.44 | 0.52 | 0.56 | 0.62 | 0.54 |
| 9 | 89655363 | rs7023578 | -0.561 | 0.667 | 1.133 | 0.324 | -0.002 | 0.023 | 0.023 | 0 | 0.21 | 0.3 | 0.29 | 0.39 | 0.19 | 0.22 |
| 9 | 89655704 | rs7870183 | -0.546 | 0.67 | 1.135 | 0.326 | -0.002 | 0.023 | 0.023 | 0 | 0.2 | 0.3 | 0.29 | 0.39 | 0.19 | 0.22 |
| 9 | 89670005 | rs4242638 | 1.537 | 0.983 | 1.405 | -0.281 | 0.049 | -0.061 | -0.061 | 0 | 0.85 | 0.56 | 0.82 | 0.94 | 0.84 | 0.76 |
| 9 | 89670188 | rs4878036 | 1.553 | 0.737 | 1.11 | -0.239 | 0.007 | 0.057 | 0.057 | 0 | 0.53 | 0.35 | 0.62 | 0.7 | 0.55 | 0.54 |
| 9 | 89673000 | rs13302497 | -0.411 | 0.304 | 1.025 | -0.268 | 0.021 | 0.029 | 0.029 | 0 | 0.31 | 0.11 | 0.21 | 0.24 | 0.29 | 0.24 |
| 9 | 89687854 | rs4878043 | 1.584 | 1.099 | 1.507 | 0.032 | 0.015 | 0.058 | 0.058 | 0.104 | 0.54 | 0.34 | 0.63 | 0.69 | 0.57 | 0.54 |
| 9 | 89688153 | rs1028863 | 0.164 | 1.208 | 1.497 | 0.073 | 0.036 | -0.01 | -0.01 | 0.019 | 0.91 | 0.84 | 0.87 | 0.93 | 0.86 | 0.8 |

| | | | | | | | | | | | | | | | | |
|---|---|---|---|---|---|---|---|---|---|---|---|---|---|---|---|---|
| 9 | 89704509 | rs13292789 | -0.534 | 0.584 | 1.276 | -0.025 | 0.053 | 0.046 | 0.046 | 0 | 0.32 | 0.11 | 0.21 | 0.23 | 0.28 | 0.24 |
| 9 | 89709884 | rs1012593 | -1.405 | 0.533 | 1.729 | -0.092 | 0.029 | 0.031 | 0.031 | 0.015 | 0.41 | 0.61 | 0.31 | 0.24 | 0.31 | 0.27 |
| 9 | 89711103 | rs7863730 | -1.662 | 0.611 | 1.833 | 0.083 | 0.016 | -0.006 | -0.006 | 0 | 0.07 | 0.29 | 0.04 | 0 | 0.02 | 0.03 |
| 9 | 89722325 | rs7858361 | -0.607 | 0.688 | 1.405 | 0.083 | 0.006 | 0.011 | 0.011 | 0 | 0.33 | 0.3 | 0.26 | 0.21 | 0.31 | 0.25 |
| 9 | 89726511 | rs7874988 | -0.837 | 0.648 | 1.495 | 0.182 | -0.011 | -0.019 | -0.019 | 0.003 | 0.37 | 0.44 | 0.28 | 0.26 | 0.33 | 0.27 |
| 9 | 89738445 | rs10115421 | -1.053 | 0.68 | 1.561 | 0.157 | -0.024 | -0.042 | -0.042 | 0 | 0.39 | 0.55 | 0.29 | 0.26 | 0.33 | 0.26 |
| 9 | 89745271 | rs277742 | 1.665 | 0.961 | 1.21 | 0.297 | -0.019 | 0.097 | 0.097 | 0.515 | 0.51 | 0.19 | 0.56 | 0.66 | 0.5 | 0.52 |
| 9 | 89753471 | rs277736 | 0.121 | 1.117 | 1.229 | 0.314 | 0.02 | -0.019 | -0.019 | 4.24 | 0.9 | 0.8 | 0.86 | 0.96 | 0.84 | 0.78 |
| 9 | 89754133 | rs277733 | 0.794 | 0.972 | 1.319 | 0.173 | 0.008 | -0.037 | -0.037 | 2.203 | 0.75 | 0.65 | 0.71 | 0.92 | 0.71 | 0.68 |
| 9 | 89763278 | rs13295659 | -1.111 | 0.764 | 1.148 | -0.07 | 0.017 | 0.024 | 0.024 | 0.161 | 0.49 | 0.59 | 0.33 | 0.27 | 0.38 | 0.34 |
| 9 | 89765887 | rs166750 | -0.032 | 0.774 | 1.167 | 0.454 | 0.048 | -0.018 | -0.018 | 2.17 | 0.92 | 0.87 | 0.88 | 0.96 | 0.84 | 0.81 |
| 9 | 89766465 | rs277763 | 1.704 | 0.686 | 0.887 | 1.213 | -0.064 | 0.053 | 0.053 | 0.078 | 0.48 | 0.07 | 0.54 | 0.37 | 0.48 | 0.42 |
| 9 | 89772284 | rs463064 | -0.448 | 0.936 | 1.056 | 0.893 | 0 | 0.005 | 0.005 | 0 | 0.93 | 0.95 | 0.89 | 0.97 | 0.84 | 0.83 |
| 9 | 89772413 | rs277755 | 1.221 | 0.781 | 1.772 | 0.939 | 0.001 | -0.017 | -0.017 | 0 | 0.57 | 0.42 | 0.62 | 0.7 | 0.55 | 0.58 |
| 9 | 89773016 | rs277753 | 1.139 | 0.857 | 1.788 | 0.939 | 0 | -0.008 | -0.008 | 0 | 0.58 | 0.49 | 0.62 | 0.7 | 0.55 | 0.58 |
| 9 | 89775429 | rs277750 | 1.108 | 0.864 | 1.793 | 0.943 | 0 | -0.006 | -0.006 | 1.382 | 0.59 | 0.51 | 0.63 | 0.7 | 0.55 | 0.58 |
| 9 | 89777497 | rs7868122 | -0.834 | 0.854 | 1.805 | 0.939 | 0.002 | 0.003 | 0.003 | 0 | 0.33 | 0.35 | 0.25 | 0.27 | 0.29 | 0.23 |
| 9 | 89782806 | rs277747 | 0.908 | 0.774 | 1.797 | 0.982 | -0.007 | 0.021 | 0.021 | 0 | 0.6 | 0.64 | 0.64 | 0.7 | 0.55 | 0.58 |
| 9 | 89801126 | rs277722 | 1.027 | 0.845 | 2.234 | 1.525 | 0.001 | 0 | 0 | 0 | 0.59 | 0.42 | 0.62 | 0.56 | 0.55 | 0.57 |
| 9 | 89803351 | rs277714 | 1.424 | 0.775 | 2.27 | 1.632 | -0.007 | -0.023 | -0.023 | 0 | 0.54 | 0.26 | 0.6 | 0.5 | 0.53 | 0.52 |
| 9 | 89809139 | rs12554839 | -0.971 | 0.973 | 2.428 | 1.649 | 0.006 | -0.015 | -0.015 | 0 | 0.2 | 0.28 | 0.21 | 0.13 | 0.22 | 0.27 |
| 9 | 89817082 | rs1885782 | -0.929 | 0.976 | 2.499 | 1.643 | 0.013 | -0.027 | -0.027 | 0 | 0.21 | 0.35 | 0.22 | 0.13 | 0.23 | 0.27 |
| 9 | 89820895 | rs427361 | -0.155 | 1.004 | 2.55 | 1.759 | 0 | 0.004 | 0.004 | 0 | 0.2 | 0.16 | 0.15 | 0.38 | 0.25 | 0.18 |
| 9 | 89821219 | rs373546 | -0.629 | 0.977 | 2.941 | 1.734 | 0.022 | 0.065 | 0.065 | 0 | 0.24 | 0.38 | 0.17 | 0.39 | 0.27 | 0.21 |
| 9 | 89827518 | rs12235447 | -0.126 | 1.006 | 2.606 | 1.762 | 0.003 | -0.011 | -0.011 | 0 | 0.2 | 0.12 | 0.1 | 0.39 | 0.19 | 0.18 |
| 9 | 89833267 | rs278709 | 0.902 | 1.244 | 2.473 | 1.516 | 0 | -0.005 | -0.005 | 0 | 0.77 | 0.68 | 0.76 | 0.88 | 0.74 | 0.7 |
| 9 | 89836173 | rs7863624 | 0.252 | 1.27 | 2.469 | 1.526 | 0.012 | 0.007 | 0.007 | 0 | 0.08 | 0.11 | 0.09 | 0.12 | 0.14 | 0.21 |
| 9 | 89843781 | rs10113878 | 0.037 | 1.194 | 2.331 | 1.595 | -0.006 | 0.023 | 0.023 | 0 | 0.2 | 0.19 | 0.11 | 0.38 | 0.19 | 0.18 |
| 9 | 89844462 | rs11141716 | 0.071 | 1.253 | 2.173 | 1.526 | 0.005 | 0.005 | 0.005 | 0 | 0.2 | 0.2 | 0.14 | 0.29 | 0.25 | 0.15 |
| 9 | 89844907 | rs2031126 | 0.147 | 1.024 | 2.032 | 1.496 | 0.002 | -0.014 | -0.014 | 0 | 0.86 | 0.83 | 0.86 | 1 | 0.88 | 0.91 |
| 9 | 89849237 | rs10512182 | 0.064 | 1.121 | 1.712 | 1.335 | 0 | -0.002 | -0.002 | 0 | 0.19 | 0.15 | 0.1 | 0.31 | 0.2 | 0.14 |
| 9 | 89854486 | rs11141719 | -0.484 | 1.14 | 1.894 | 1.312 | 0.033 | 0.017 | 0.017 | 0 | 0.23 | 0.43 | 0.17 | 0.31 | 0.28 | 0.17 |
| 9 | 89854595 | rs7862236 | 0.975 | 1.143 | 1.195 | 1.313 | -0.023 | -0.024 | -0.024 | 0 | 0.52 | 0.12 | 0.57 | 0.48 | 0.45 | 0.5 |
| 9 | 89858267 | rs10122994 | 0.667 | 1.07 | 1.167 | 1.32 | -0.001 | -0.001 | -0.001 | 1.022 | 0.57 | 0.42 | 0.65 | 0.49 | 0.53 | 0.53 |
| 9 | 89862258 | rs7026586 | 0.833 | 0.687 | 0.42 | 1.293 | 0.01 | 0.024 | 0.024 | 0.065 | 0.52 | 0.19 | 0.56 | 0.5 | 0.42 | 0.5 |
| 9 | 89869968 | rs12348439 | -1.562 | 0.508 | 1.575 | 1.747 | 0.04 | 0.195 | 0.195 | 0 | 0.06 | 0.22 | 0.06 | 0.27 | 0.05 | 0.03 |
| 9 | 89874518 | rs7025731 | -0.847 | 0.412 | 1.689 | 1.676 | 0.008 | 0 | 0 | 0 | 0.19 | 0.23 | 0.17 | 0.18 | 0.27 | 0.23 |
| 9 | 89874893 | rs1969763 | 1.902 | 0.259 | 2.457 | 1.902 | 0.012 | 0.037 | 0.037 | 0 | 0.68 | 0.46 | 0.69 | 0.45 | 0.61 | 0.51 |
| 9 | 89874932 | rs1969762 | 1.402 | 0.209 | 2.273 | 1.735 | 0.004 | 0.141 | 0.141 | 0 | 0.87 | 0.72 | 0.87 | 0.62 | 0.88 | 0.73 |
| 9 | 89878104 | rs1570421 | 1.456 | 0.143 | 2.233 | 1.691 | -0.003 | 0.098 | 0.098 | 0 | 0.86 | 0.78 | 0.87 | 0.63 | 0.88 | 0.73 |
| 9 | 89880234 | rs4877347 | 1.053 | -0.303 | 1.832 | 1.354 | -0.01 | 0.009 | 0.009 | 0 | 0.76 | 0.78 | 0.85 | 0.63 | 0.84 | 0.72 |
| 9 | 89888137 | rs7859923 | 1.57 | -0.194 | 2.199 | 1.234 | -0.027 | 0.015 | 0.015 | 0 | 0.67 | 0.47 | 0.55 | 0.56 | 0.73 | 0.66 |
| 9 | 89890786 | rs2889975 | 1.843 | -0.348 | 2.19 | 1.3 | 0.092 | -0.099 | -0.099 | 0.988 | 0.61 | 0.15 | 0.51 | 0.47 | 0.73 | 0.61 |
| 9 | 89893787 | rs10125155 | -0.294 | -0.262 | 2.664 | 1.35 | -0.034 | -0.076 | -0.076 | 0 | 0.12 | 0.33 | 0.09 | 0 | 0.09 | 0.07 |
| 9 | 89898642 | rs7041849 | -2.444 | -0.27 | 2.851 | 1.363 | 0.046 | 0.184 | 0.184 | 7.595 | 0.06 | 0.46 | 0.05 | 0.1 | 0.01 | 0.05 |
| 9 | 89903208 | rs1923797 | 2.422 | -0.164 | 2.485 | 1.425 | -0.024 | 0.05 | 0.05 | 0 | 0.54 | 0.13 | 0.42 | 0.3 | 0.55 | 0.45 |
| 9 | 89903221 | rs7020618 | -0.48 | -0.109 | 2.308 | 1.437 | 0.057 | 0.074 | 0.074 | 0 | 0.12 | 0.24 | 0.13 | 0.27 | 0.19 | 0.21 |
| 9 | 89905542 | rs4878051 | 2.648 | -0.069 | 1.51 | 1.385 | -0.1 | 0.048 | 0.048 | 0 | 0.51 | 0.06 | 0.39 | 0.3 | 0.5 | 0.45 |
| 9 | 89905888 | rs4878052 | 0.552 | 0.037 | 1.625 | 1.49 | 0.004 | -0.014 | -0.014 | 0 | 0.8 | 0.87 | 0.65 | 0.57 | 0.83 | 0.73 |
| 9 | 89910813 | rs10746796 | 1.008 | 0.135 | 1.603 | 1.565 | 0.001 | -0.002 | -0.002 | 1.729 | 0.7 | 0.72 | 0.58 | 0.57 | 0.75 | 0.67 |
| 9 | 89917347 | rs17472787 | 0.145 | 0.854 | 1.484 | 1.915 | -0.053 | -0.044 | -0.044 | 0.654 | 0.08 | 0.02 | 0.1 | 0.17 | 0.2 | 0.17 |
| 9 | 89922395 | rs944655 | 1.449 | 0.385 | 1.885 | 1.942 | -0.006 | 0.05 | 0.05 | 0 | 0.83 | 0.71 | 0.65 | 0.57 | 0.86 | 0.81 |
| 9 | 89924306 | rs10868554 | -0.43 | 0.642 | 2.271 | 1.857 | 0.082 | 0.016 | 0.016 | 0 | 0.13 | 0.3 | 0.15 | 0.15 | 0.24 | 0.14 |
| 9 | 89924898 | rs10868555 | -0.105 | 0.739 | 2.143 | 1.896 | 0.008 | 0 | 0 | 0 | 0.12 | 0.14 | 0.14 | 0.15 | 0.25 | 0.14 |
| 9 | 89931500 | rs10780823 | 1.44 | 0.485 | 2.093 | 1.905 | -0.006 | 0.078 | 0.078 | 0 | 0.83 | 0.69 | 0.66 | 0.59 | 0.86 | 0.81 |
| 9 | 89934837 | rs1923789 | -0.487 | 0.491 | 1.981 | 1.961 | -0.002 | 0.025 | 0.025 | 0 | 0.1 | 0.15 | 0.29 | 0.29 | 0.08 | 0.11 |
| 9 | 89936504 | rs4878057 | 0.719 | 0.272 | 1.805 | 2.034 | -0.021 | 0.036 | 0.036 | 0 | 0.85 | 0.78 | 0.88 | 0.72 | 0.92 | 0.76 |
| 9 | 89939154 | rs1923786 | 0.934 | 0.306 | 1.773 | 1.981 | -0.001 | 0.011 | 0.011 | 0 | 0.94 | 0.89 | 0.95 | 0.89 | 0.94 | 0.92 |
| 9 | 89940123 | rs6559979 | 1.487 | 0.298 | 1.573 | 1.896 | -0.029 | 0.068 | 0.068 | 0 | 0.74 | 0.63 | 0.61 | 0.44 | 0.84 | 0.65 |
| 9 | 89941523 | rs1923783 | 0.726 | 0.308 | 0.873 | 1.065 | 0.007 | -0.03 | -0.03 | 0 | 0.94 | 0.88 | 0.96 | 1 | 0.95 | 0.97 |
| 9 | 89944405 | rs10868558 | -0.64 | 0.379 | 0.537 | 1.075 | 0.039 | 0.018 | 0.018 | 0 | 0.31 | 0.54 | 0.49 | 0.45 | 0.43 | 0.38 |
| 9 | 89946647 | rs7026815 | 0.85 | 0.673 | -0.448 | 0.72 | -0.027 | 0.013 | 0.013 | 0 | 0.53 | 0.18 | 0.39 | 0.38 | 0.49 | 0.43 |
| 9 | 89951180 | rs7874448 | -0.624 | 0.781 | -0.22 | 0.744 | -0.001 | -0.002 | -0.002 | 0 | 0.38 | 0.4 | 0.51 | 0.58 | 0.4 | 0.51 |
| 9 | 89951915 | rs1888924 | 0.653 | 0.947 | -0.195 | 0.098 | 0.009 | -0.024 | -0.024 | 0 | 0.93 | 0.91 | 0.95 | 1 | 0.9 | 0.98 |
| 9 | 89954522 | rs11141752 | -0.113 | 0.764 | -1.29 | -0.7 | -0.006 | 0.125 | 0.125 | 0.527 | 0.35 | 0.09 | 0.32 | 0.17 | 0.32 | 0.45 |
| 9 | 89955458 | rs4878060 | -0.641 | 1.089 | -0.837 | -1.123 | 0 | -0.014 | -0.014 | 0 | 0.59 | 0.45 | 0.56 | 0.24 | 0.63 | 0.59 |
| 9 | 89958743 | rs11141754 | 0.758 | 0.701 | -1.396 | -1.641 | 0.023 | 0.076 | 0.076 | 0 | 0.29 | 0.03 | 0.24 | 0.17 | 0.22 | 0.35 |
| 9 | 89959135 | rs1923782 | -0.321 | 1.353 | -1.094 | -1.944 | 0.019 | -0.025 | -0.025 | 0 | 0.23 | 0.29 | 0.24 | 0.09 | 0.4 | 0.16 |
| 9 | 89959906 | rs4877351 | -1.254 | 1.2 | -0.415 | -2.037 | 0.005 | -0.011 | -0.011 | 0 | 0.34 | 0.62 | 0.35 | 0.2 | 0.47 | 0.42 |

| | | | | | | | | | | | | | | | | |
|---|---|---|---|---|---|---|---|---|---|---|---|---|---|---|---|---|
| 9 | 89961029 | rs4878063 | -0.457 | 1.228 | -0.272 | -2.048 | 0.037 | -0.077 | -0.077 | 0 | 0.26 | 0.57 | 0.3 | 0.09 | 0.37 | 0.3 |
| 9 | 89961776 | rs912067 | -0.436 | 1.2 | 0.05 | -2.083 | 0.005 | -0.003 | -0.003 | 0.14 | 0.25 | 0.28 | 0.27 | 0.19 | 0.35 | 0.4 |
| 9 | 89963474 | rs1923800 | -0.682 | 1.162 | 0.098 | -2.063 | 0.009 | -0.04 | -0.04 | 0 | 0.29 | 0.35 | 0.28 | 0.12 | 0.39 | 0.35 |
| 9 | 89965089 | rs4878066 | -1.379 | 1.052 | -0.281 | -1.98 | -0.003 | -0.001 | -0.001 | 0 | 0.66 | 0.67 | 0.59 | 0.39 | 0.73 | 0.78 |
| 9 | 89967335 | rs10746798 | -1.326 | 1.012 | 0.058 | -2.002 | -0.028 | -0.032 | -0.032 | 6.372 | 0.64 | 0.56 | 0.58 | 0.31 | 0.72 | 0.71 |
| 9 | 89967771 | rs10868564 | 0.751 | 0.742 | -0.573 | -2.006 | -0.067 | 0.102 | 0.102 | 0 | 0.34 | 0.08 | 0.29 | 0.2 | 0.34 | 0.36 |
| 9 | 89970895 | rs1930056 | 2.23 | 0.18 | -0.743 | -2.017 | 0.077 | -0.101 | -0.101 | 0 | 0.21 | 0.03 | 0.29 | 0.55 | 0.12 | 0.19 |
| 9 | 89979884 | rs1930057 | 0.554 | 1.125 | 0.139 | -1.839 | 0.076 | -0.021 | -0.021 | 0.034 | 0.82 | 0.66 | 0.84 | 0.84 | 0.76 | 0.89 |
| 9 | 89985014 | rs2378733 | -0.722 | 1.052 | 0.214 | -1.82 | 0.009 | -0.026 | -0.026 | 0 | 0.89 | 0.82 | 0.9 | 0.93 | 0.87 | 0.97 |
| 9 | 89985250 | rs7861579 | 0.117 | 0.881 | 0.044 | -1.911 | -0.003 | 0.02 | 0.02 | 0 | 0.54 | 0.61 | 0.6 | 0.67 | 0.54 | 0.53 |
| 9 | 89991119 | rs11141771 | 1.526 | 0.702 | -0.524 | -1.778 | 0.082 | -0.08 | -0.08 | 0 | 0.34 | 0.02 | 0.39 | 0.58 | 0.21 | 0.3 |
| 9 | 89996566 | rs1958938 | 0.158 | 1.591 | -0.082 | -1.699 | -0.009 | 0.006 | 0.006 | 0 | 0.81 | 0.85 | 0.8 | 0.91 | 0.67 | 0.86 |
| 9 | 89997130 | rs10125807 | -0.674 | 1.631 | 0.089 | -1.648 | 0.1 | -0.022 | -0.022 | 0.141 | 0.14 | 0.35 | 0.11 | 0.08 | 0.22 | 0.14 |
| 9 | 90001560 | rs1930055 | -0.015 | 1.429 | -0.151 | -1.503 | 0.097 | 0.035 | 0.035 | 0 | 0.06 | 0.19 | 0.06 | 0.08 | 0.11 | 0.11 |
| 9 | 90002875 | rs10868569 | 0.84 | 1.384 | -0.097 | -1.643 | 0.034 | -0.056 | -0.056 | 1.511 | 0.68 | 0.44 | 0.7 | 0.83 | 0.53 | 0.77 |
| 9 | 90002911 | rs12376350 | 1.54 | 0.302 | -0.154 | -1.674 | 0.055 | -0.003 | -0.003 | 1.112 | 0.36 | 0.19 | 0.44 | 0.55 | 0.21 | 0.29 |
| 9 | 90004329 | rs4878073 | 0.232 | 0.491 | -0.189 | -1.69 | 0.015 | -0.09 | -0.09 | 0 | 0.76 | 0.47 | 0.81 | 0.9 | 0.74 | 0.89 |
| 9 | 90006563 | rs4144392 | 0.83 | -1.171 | -0.281 | -1.578 | 0.022 | 0.014 | 0.014 | 0.122 | 0.28 | 0.15 | 0.21 | 0.27 | 0.21 | 0.47 |
| 9 | 90007371 | rs10868572 | 1.02 | -1.165 | -0.282 | -1.586 | 0.022 | 0.042 | 0.042 | 0.088 | 0.26 | 0.14 | 0.2 | 0.2 | 0.2 | 0.37 |
| 9 | 90007462 | rs1958937 | 0.385 | -0.833 | -0.083 | -1.246 | 0.001 | -0.017 | -0.017 | 0.377 | 0.64 | 0.4 | 0.63 | 0.79 | 0.43 | 0.76 |
| 9 | 90012840 | rs12344505 | 1.441 | -0.781 | -1.074 | -1.311 | 0.076 | -0.053 | -0.053 | 0 | 0.32 | 0.09 | 0.43 | 0.59 | 0.2 | 0.34 |
| 9 | 90013672 | rs10746805 | 1.389 | -0.737 | -1.029 | -1.287 | 0.067 | -0.027 | -0.027 | 0 | 0.33 | 0.17 | 0.44 | 0.59 | 0.21 | 0.37 |
| 9 | 90013698 | rs10780833 | 0.941 | -0.622 | -1.332 | -1.367 | 0.017 | 0.042 | 0.042 | 0.023 | 0.27 | 0.05 | 0.2 | 0.2 | 0.22 | 0.3 |
| 9 | 90018842 | rs4878075 | 0.457 | -0.372 | -0.821 | -1.299 | 0.002 | -0.045 | -0.045 | 0 | 0.63 | 0.47 | 0.67 | 0.86 | 0.41 | 0.77 |
| 9 | 90020128 | rs7039493 | -0.042 | 0.059 | -0.566 | -1.075 | -0.013 | -0.012 | -0.012 | 0.318 | 0.36 | 0.39 | 0.26 | 0.31 | 0.26 | 0.49 |
| 9 | 90034195 | rs10746806 | 0.636 | 1.319 | -0.205 | -0.589 | 0.045 | -0.031 | -0.031 | 5.549 | 0.73 | 0.65 | 0.76 | 0.85 | 0.51 | 0.8 |
| 9 | 90034729 | rs2183939 | 0.786 | 1.884 | -1.049 | -0.596 | -0.062 | 0.064 | 0.064 | 0.082 | 0.77 | 0.93 | 0.83 | 0.9 | 0.66 | 0.87 |
| 9 | 90036219 | rs2314 | 1.66 | 1.171 | -1.205 | -0.59 | 0.093 | 0.042 | 0.042 | 0 | 0.42 | 0.11 | 0.56 | 0.61 | 0.29 | 0.42 |
| 9 | 90041160 | rs4579610 | 0.021 | 1.556 | -0.854 | -0.283 | -0.041 | 0.018 | 0.018 | 0 | 0.87 | 0.95 | 0.89 | 0.92 | 0.75 | 0.9 |
| 9 | 90043880 | rs7870463 | -1.242 | 1.056 | -0.115 | -0.03 | -0.023 | -0.011 | -0.011 | 0 | 0.34 | 0.52 | 0.3 | 0.32 | 0.24 | 0.36 |
| 9 | 90045801 | rs6559987 | -0.581 | 0.806 | 0.171 | -0.12 | -0.027 | -0.022 | -0.022 | 0.162 | 0.12 | 0.35 | 0.07 | 0.06 | 0.06 | 0.05 |
| 9 | 90046983 | rs7025941 | -0.012 | 0.906 | -0.223 | 0.002 | -0.002 | 0.002 | 0.002 | 0 | 0.14 | 0.14 | 0.14 | 0.13 | 0.14 | 0.21 |
| 9 | 90049204 | rs4878077 | -0.069 | 1.555 | -0.249 | -0.044 | -0.077 | 0.014 | 0.014 | 0 | 0.86 | 0.97 | 0.86 | 0.92 | 0.7 | 0.9 |
| 9 | 90051391 | rs10512185 | 1.361 | 1.474 | -0.091 | -0.059 | -0.008 | 0.003 | 0.003 | 0.628 | 0.75 | 0.73 | 0.74 | 0.82 | 0.64 | 0.79 |
| 9 | 90052125 | rs10868585 | 0.487 | 1.483 | -0.346 | -0.05 | 0.005 | 0.006 | 0.006 | 7.403 | 0.07 | 0.03 | 0.04 | 0 | 0.04 | 0.02 |
| 9 | 90052537 | rs2149809 | 1.173 | 1.408 | 0.118 | -0.047 | 0.015 | -0.045 | -0.045 | 0 | 0.68 | 0.45 | 0.74 | 0.84 | 0.61 | 0.84 |
| 9 | 90054710 | rs10780837 | 1.19 | 1.386 | 0.133 | -0.074 | 0.021 | -0.058 | -0.058 | 0.834 | 0.68 | 0.5 | 0.74 | 0.83 | 0.61 | 0.84 |
| 9 | 90063194 | rs11141813 | 1.409 | 1.429 | 0.196 | 0.058 | 0.015 | -0.021 | -0.021 | 2.217 | 0.66 | 0.45 | 0.68 | 0.73 | 0.61 | 0.79 |
| 9 | 90066956 | rs10868592 | 1.384 | 1.37 | 0.197 | 0.059 | 0.014 | -0.021 | -0.021 | 0 | 0.66 | 0.45 | 0.68 | 0.73 | 0.61 | 0.79 |
| 9 | 90067655 | rs17475756 | 0.189 | 1.394 | -0.229 | 0.074 | -0.048 | -0.018 | -0.018 | 0.192 | 0.13 | 0.01 | 0.11 | 0.13 | 0.14 | 0.22 |
| 9 | 90068509 | rs10124291 | 0.785 | 1.324 | -0.029 | 0.082 | 0.011 | -0.028 | -0.028 | 0 | 0.71 | 0.58 | 0.78 | 0.82 | 0.65 | 0.86 |
| 9 | 90073584 | rs7849703 | 1.749 | 1.246 | -0.687 | 0.065 | 0.033 | 0.003 | 0.003 | 0.036 | 0.46 | 0.08 | 0.5 | 0.59 | 0.38 | 0.51 |
| 9 | 90079395 | rs1928474 | -1.623 | 1.283 | 0.161 | 1.038 | -0.001 | 0.02 | 0.02 | 3.034 | 0.29 | 0.67 | 0.42 | 0.43 | 0.27 | 0.42 |
| 9 | 90081038 | rs4457416 | 0.261 | 1.424 | 0.052 | 1.225 | -0.025 | 0.025 | 0.025 | 1.531 | 0.79 | 0.89 | 0.85 | 0.88 | 0.68 | 0.91 |
| 9 | 90085205 | rs1411090 | -0.023 | 1.263 | -0.073 | 1.465 | -0.034 | -0.079 | -0.079 | 0 | 0.13 | 0.01 | 0.25 | 0.28 | 0.14 | 0.26 |
| 9 | 90095024 | rs10780842 | 1.802 | 1.275 | 0.491 | 1.505 | 0.001 | -0.001 | -0.001 | 0 | 0.54 | 0.43 | 0.49 | 0.46 | 0.43 | 0.5 |
| 9 | 90095233 | rs10868605 | 1.978 | 1.283 | 0.332 | 1.562 | 0.025 | 0.029 | 0.029 | 0.247 | 0.49 | 0.21 | 0.47 | 0.39 | 0.42 | 0.45 |
| 9 | 90098265 | rs2094707 | 0.524 | 1.565 | 1.019 | 1.526 | 0.011 | -0.009 | -0.009 | 0.653 | 0.78 | 0.77 | 0.81 | 0.87 | 0.63 | 0.9 |
| 9 | 90101096 | rs11141844 | 1.847 | 1.414 | 1.255 | 1.986 | 0.019 | -0.008 | -0.008 | 0 | 0.61 | 0.5 | 0.62 | 0.66 | 0.49 | 0.58 |
| 9 | 90101620 | rs7043393 | 1.882 | 1.412 | 1.253 | 2.061 | 0.016 | 0.004 | 0.004 | 0.384 | 0.6 | 0.5 | 0.61 | 0.59 | 0.48 | 0.53 |
| 9 | 90110124 | rs3758308 | 1.809 | 1.394 | 1.248 | 1.674 | 0.014 | 0.006 | 0.006 | 0.393 | 0.6 | 0.5 | 0.5 | 0.44 | 0.47 | 0.5 |
| 9 | 90114746 | rs2058882 | -1.287 | 1.463 | 1.126 | 1.638 | -0.001 | 0.151 | 0.151 | 0.82 | 0.22 | 0.46 | 0.36 | 0.43 | 0.2 | 0.42 |
| 9 | 90125129 | rs10125534 | -0.884 | 1.373 | 0.847 | 1.361 | -0.001 | 0.042 | 0.042 | 0 | 0.19 | 0.23 | 0.32 | 0.44 | 0.18 | 0.37 |
| 9 | 90125146 | rs10122321 | -0.657 | 1.485 | 0.532 | 1.375 | -0.001 | -0.001 | -0.001 | 0.023 | 0.18 | 0.14 | 0.32 | 0.43 | 0.18 | 0.29 |
| 9 | 90132563 | rs1475524 | 1.16 | 1.532 | 0.648 | 1.129 | 0.013 | 0.008 | 0.008 | 1.329 | 0.75 | 0.64 | 0.77 | 0.69 | 0.61 | 0.87 |
| 9 | 90133673 | rs1556 | 1.148 | 1.56 | 0.28 | 1.109 | 0.03 | 0.014 | 0.014 | 2.934 | 0.72 | 0.51 | 0.76 | 0.69 | 0.6 | 0.87 |
| 9 | 90135027 | rs17399090 | -0.837 | 1.755 | -0.622 | 0.983 | -0.042 | -0.019 | -0.019 | 0 | 0.15 | 0.06 | 0.14 | 0.14 | 0.27 | 0.07 |
| 9 | 90136246 | rs1105384 | 0.769 | 0.888 | -1.373 | 0.522 | -0.054 | -0.149 | -0.149 | 0 | 0.34 | 0.05 | 0.33 | 0.39 | 0.31 | 0.44 |
| 9 | 90137616 | rs871495 | 0.595 | 1.267 | -0.419 | 0.111 | 0.002 | 0.02 | 0.02 | 0 | 0.66 | 0.61 | 0.41 | 0.47 | 0.62 | 0.53 |
| 9 | 90137909 | rs10780849 | 1.194 | 1.162 | 0.172 | -0.026 | 0 | 0 | 0 | 0 | 0.4 | 0.41 | 0.23 | 0.41 | 0.3 | 0.4 |
| 9 | 90138130 | rs10746816 | 0.027 | 1.126 | 0.089 | 0.087 | 0.013 | -0.018 | -0.018 | 0 | 0.73 | 0.65 | 0.6 | 0.81 | 0.71 | 0.74 |
| 9 | 90139441 | rs11141862 | 0.423 | 0.726 | -0.077 | 0.014 | -0.072 | 0.098 | 0.098 | 0 | 0.25 | 0.07 | 0.17 | 0.06 | 0.33 | 0.14 |
| 9 | 90140505 | rs17477673 | 0.469 | 0.671 | 0.571 | -0.085 | -0.012 | 0.027 | 0.027 | 0.952 | 0.6 | 0.37 | 0.67 | 0.87 | 0.54 | 0.78 |
| 9 | 90142383 | rs11793890 | 0.873 | 0.717 | 0.54 | -0.113 | -0.068 | 0.077 | 0.077 | 0 | 0.22 | 0.07 | 0.17 | 0.06 | 0.32 | 0.12 |
| 9 | 90143575 | rs4878089 | 0.976 | 0.668 | 0.599 | -0.064 | -0.001 | 0.019 | 0.019 | 0.269 | 0.53 | 0.33 | 0.49 | 0.54 | 0.46 | 0.57 |
| 9 | 90145381 | rs10512186 | 0.903 | 0.614 | 0.865 | -0.067 | -0.001 | 0.01 | 0.01 | 0.116 | 0.53 | 0.4 | 0.5 | 0.54 | 0.46 | 0.58 |
| 9 | 90146658 | rs7040394 | -0.734 | 0.597 | 1.186 | -0.077 | 0 | -0.003 | -0.003 | 0 | 0.87 | 0.86 | 0.89 | 0.95 | 0.86 | 0.91 |
| 9 | 90148464 | rs12686192 | -0.042 | 0.647 | 0.844 | -0.072 | 0 | 0 | 0 | 0 | 0.16 | 0.14 | 0.21 | 0.13 | 0.14 | 0.17 |
| 9 | 90151362 | rs3028 | 1.91 | 0.653 | 0.788 | -0.065 | 0.016 | -0.026 | -0.026 | 0 | 0.37 | 0.25 | 0.29 | 0.41 | 0.32 | 0.4 |

| | | | | | | | | | | | | | | | | |
|---|---|---|---|---|---|---|---|---|---|---|---|---|---|---|---|---|
| 9 | 90155692 | rs4877365 | 1.929 | 0.503 | 0.79 | -0.097 | 0.021 | -0.028 | -0.028 | 0 | 0.36 | 0.25 | 0.28 | 0.41 | 0.31 | 0.39 |
| 9 | 90157451 | rs1014306 | 0.527 | 0.567 | 0.766 | -0.126 | -0.017 | 0.012 | 0.012 | 0 | 0.25 | 0.2 | 0.18 | 0.06 | 0.34 | 0.13 |
| 9 | 90160010 | rs7036598 | -0.753 | 0.477 | 0.78 | -0.14 | -0.003 | 0.001 | 0.001 | 0 | 0.42 | 0.54 | 0.41 | 0.19 | 0.47 | 0.3 |
| 9 | 90165926 | rs7866648 | 0.577 | 0.326 | 0.437 | -0.159 | 0.036 | 0.109 | 0.109 | 0.005 | 0.56 | 0.81 | 0.55 | 0.77 | 0.41 | 0.62 |
| 9 | 90168002 | rs1983973 | -0.585 | 0.297 | 0.62 | -0.121 | -0.026 | 0.229 | 0.229 | 0.563 | 0.13 | 0.43 | 0.22 | 0.34 | 0.09 | 0.21 |
| 9 | 90169540 | rs11141878 | -0.3 | 0.382 | 0.586 | -0.141 | 0.007 | 0.001 | 0.001 | 0 | 0.33 | 0.57 | 0.38 | 0.39 | 0.38 | 0.33 |
| 9 | 90170231 | rs6560001 | -1.269 | 0.433 | 0.286 | -0.182 | -0.032 | 0.024 | 0.024 | 0.44 | 0.89 | 0.98 | 0.9 | 0.95 | 0.85 | 0.91 |
| 9 | 90170885 | rs10512187 | 0.613 | 0.425 | 0.288 | -0.187 | 0.036 | 0.109 | 0.109 | 0.005 | 0.56 | 0.82 | 0.55 | 0.77 | 0.41 | 0.62 |
| 9 | 90170988 | rs11141879 | 0.44 | 0.413 | 0.247 | -0.164 | -0.074 | -0.027 | -0.027 | 0.16 | 0.76 | 0.96 | 0.71 | 0.82 | 0.71 | 0.74 |
| 9 | 90171408 | rs7852430 | 0.44 | 0.413 | 0.247 | -0.164 | -0.074 | -0.027 | -0.027 | 1.755 | 0.76 | 0.96 | 0.71 | 0.82 | 0.71 | 0.74 |
| 9 | 90172040 | rs17479142 | 0.257 | 0.422 | 0.268 | -0.152 | -0.042 | -0.047 | -0.047 | 0.037 | 0.13 | 0.01 | 0.19 | 0.13 | 0.13 | 0.17 |
| 9 | 90174169 | rs1861832 | 1.973 | 0.372 | 0.38 | -0.128 | 0.018 | -0.009 | -0.009 | 0 | 0.37 | 0.31 | 0.28 | 0.41 | 0.3 | 0.39 |
| 9 | 90177252 | rs1571515 | 0.906 | 0.489 | 0.062 | -0.137 | -0.072 | -0.142 | -0.142 | 1.649 | 0.68 | 0.93 | 0.52 | 0.47 | 0.63 | 0.53 |
| 9 | 90178124 | rs12001404 | 1.205 | 0.485 | 0.108 | -0.152 | -0.015 | 0.002 | 0.002 | 0 | 0.19 | 0.14 | 0.15 | 0.05 | 0.3 | 0.12 |
| 9 | 90179619 | rs888336 | 0.938 | 0.513 | 0.066 | -0.085 | -0.072 | -0.142 | -0.142 | 0.006 | 0.68 | 0.93 | 0.52 | 0.47 | 0.63 | 0.53 |
| 9 | 90183367 | rs9314763 | -1.147 | 0.633 | 0.242 | 0.004 | 0.008 | -0.029 | -0.029 | 0 | 0.29 | 0.22 | 0.44 | 0.49 | 0.24 | 0.4 |
| 9 | 90184453 | rs12340431 | 0.46 | 0.765 | 0.145 | 0.098 | -0.006 | 0.064 | 0.064 | 0 | 0.09 | 0.14 | 0.2 | 0.34 | 0.09 | 0.21 |
| 9 | 90186205 | rs3128519 | -1.081 | 0.752 | 0.165 | 0.028 | 0.011 | 0.001 | 0.001 | 0 | 0.47 | 0.39 | 0.57 | 0.54 | 0.54 | 0.53 |
| 9 | 90186976 | rs11141899 | -1.171 | 0.473 | 0.313 | 0.541 | -0.002 | 0.014 | 0.014 | 0 | 0.27 | 0.29 | 0.43 | 0.48 | 0.24 | 0.36 |
| 9 | 90192991 | rs4878104 | 0.43 | 0.563 | 0.124 | 0.063 | 0.013 | 0.02 | 0.02 | 0 | 0.45 | 0.42 | 0.48 | 0.75 | 0.39 | 0.59 |
| 9 | 90195672 | rs888333 | 0.436 | 0.666 | -0.174 | 0.113 | -0.005 | -0.009 | -0.009 | 0 | 0.46 | 0.52 | 0.48 | 0.75 | 0.4 | 0.59 |
| 9 | 90198251 | rs4877367 | -1.037 | 1.06 | -0.287 | 0.949 | -0.028 | -0.048 | -0.048 | 1.039 | 0.1 | 0.01 | 0.16 | 0.13 | 0.1 | 0.14 |
| 9 | 90209137 | rs3095748 | 0.786 | 0.899 | 0.577 | 1.494 | -0.003 | -0.029 | -0.029 | 0 | 0.92 | 0.98 | 0.91 | 0.76 | 0.93 | 0.87 |
| 9 | 90210508 | rs12551072 | -1.443 | 0.691 | 0.682 | 1.942 | 0.005 | 0.142 | 0.142 | 0 | 0.08 | 0.28 | 0.08 | 0.23 | 0.06 | 0.15 |
| 9 | 90211490 | rs3124236 | 2.127 | 0.58 | 0.985 | 1.884 | -0.028 | 0.216 | 0.216 | 0 | 0.82 | 0.56 | 0.82 | 0.53 | 0.87 | 0.72 |
| 9 | 90216094 | rs3739784 | -1.129 | 0.847 | 0.252 | 1.512 | -0.014 | -0.029 | -0.029 | 0.498 | 0.06 | 0 | 0.04 | 0.24 | 0.05 | 0.16 |
| 9 | 90218218 | rs3124237 | -1.547 | 0.873 | 1.029 | 0.751 | 0.006 | -0.098 | -0.098 | 1.005 | 0.14 | 0.37 | 0.1 | 0 | 0.11 | 0.12 |
| 9 | 90218347 | rs7027958 | 0.835 | 1.253 | 0.635 | 0.775 | 0.008 | -0.003 | -0.003 | 0 | 0.61 | 0.57 | 0.59 | 0.39 | 0.58 | 0.51 |
| 9 | 90218678 | rs3095747 | 1.004 | 1.68 | 0.517 | 0.918 | 0.006 | 0.004 | 0.004 | 0.009 | 0.67 | 0.65 | 0.64 | 0.42 | 0.64 | 0.62 |
| 9 | 90220370 | rs13288561 | -0.738 | 1.314 | 0.047 | 1.144 | 0.002 | -0.008 | -0.008 | 0 | 0.15 | 0.1 | 0.11 | 0.22 | 0.11 | 0.21 |
| 9 | 90224267 | rs2111554 | -1.687 | 1.72 | 0.32 | 1.37 | -0.003 | 0.007 | 0.007 | 0.611 | 0.33 | 0.6 | 0.38 | 0.63 | 0.31 | 0.43 |
| 9 | 90234714 | rs11141914 | -0.255 | 1.7 | 0.221 | 1.482 | -0.015 | -0.065 | -0.065 | 0.014 | 0.14 | 0.01 | 0.1 | 0.24 | 0.11 | 0.26 |
| 9 | 90235142 | rs13283404 | -1.074 | 1.566 | 0.289 | 1.237 | -0.004 | 0.119 | 0.119 | 0.032 | 0.21 | 0.6 | 0.29 | 0.41 | 0.2 | 0.18 |
| 9 | 90235794 | rs11141915 | -0.912 | 1.779 | 0.184 | 1.309 | 0.006 | 0.067 | 0.067 | 0 | 0.23 | 0.31 | 0.23 | 0.36 | 0.2 | 0.4 |
| 9 | 90240377 | rs4878112 | -0.587 | 1.82 | -0.277 | 1.43 | 0.004 | 0.024 | 0.024 | 0 | 0.16 | 0.21 | 0.27 | 0.41 | 0.25 | 0.22 |
| 9 | 90240813 | rs3095755 | 1.09 | 1.287 | -1.751 | 0.388 | -0.071 | 0.102 | 0.102 | 2.97 | 0.32 | 0.08 | 0.21 | 0.19 | 0.35 | 0.25 |
| 9 | 90246317 | rs3124238 | -0.315 | 1.31 | -1.722 | 0.967 | -0.089 | -0.121 | -0.121 | 0 | 0.2 | 0.04 | 0.31 | 0.39 | 0.36 | 0.18 |
| 9 | 90250053 | rs1861829 | 0.58 | 0.873 | -1.913 | 0.972 | -0.053 | -0.069 | -0.069 | 0.014 | 0.69 | 0.87 | 0.46 | 0.53 | 0.53 | 0.7 |
| 9 | 90258153 | rs2378753 | -0.362 | 0.6 | -1.113 | 0.742 | 0.006 | 0.06 | 0.06 | 0.503 | 0.22 | 0.42 | 0.2 | 0.3 | 0.18 | 0.31 |
| 9 | 90258633 | rs1927976 | -1.817 | -0.017 | -1.774 | 0.533 | -0.068 | 0.14 | 0.14 | 0 | 0.57 | 0.87 | 0.75 | 0.79 | 0.53 | 0.53 |
| 9 | 90261695 | rs3118867 | -0.522 | 0.077 | -1.595 | 0.74 | -0.031 | -0.075 | -0.075 | 0 | 0.33 | 0.19 | 0.61 | 0.51 | 0.48 | 0.31 |
| 9 | 90262425 | rs2274606 | 1.159 | -0.016 | -1.804 | 0.532 | 0.003 | 0.047 | 0.047 | 0 | 0.11 | 0.01 | 0.05 | 0.04 | 0.12 | 0.04 |
| 9 | 90264023 | rs3118870 | -1.844 | 0.052 | -1.043 | 0.572 | -0.004 | 0.018 | 0.018 | 0 | 0.57 | 0.61 | 0.77 | 0.78 | 0.61 | 0.54 |
| 9 | 90265015 | rs3818584 | 0.073 | -0.024 | -1.052 | 0.614 | 0.006 | -0.065 | -0.065 | 0.039 | 0.3 | 0.12 | 0.54 | 0.58 | 0.37 | 0.28 |
| 9 | 90265697 | rs10868642 | -0.623 | 0.377 | -0.229 | 0.869 | -0.019 | -0.026 | -0.026 | 0.4 | 0.25 | 0.47 | 0.19 | 0.21 | 0.16 | 0.25 |
| 9 | 90274869 | rs3128471 | -0.728 | 0.913 | -0.248 | 1.193 | 0.014 | 0.009 | 0.009 | 0 | 0.15 | 0.22 | 0.17 | 0.18 | 0.2 | 0.26 |
| 9 | 90276666 | rs11141934 | -1.166 | 0.72 | -0.339 | 1.218 | 0.004 | 0.002 | 0.002 | 0 | 0.09 | 0.06 | 0.05 | 0.16 | 0.08 | 0.09 |
| 9 | 90278786 | rs3118848 | -1.922 | 0.786 | -0.156 | 1.696 | 0.003 | 0.002 | 0.002 | 0.476 | 0.34 | 0.62 | 0.45 | 0.63 | 0.39 | 0.44 |
| 9 | 90280569 | rs12335796 | -1.128 | 0.707 | 0.575 | 2.193 | 0.036 | 0.222 | 0.222 | 0.984 | 0.1 | 0.39 | 0.22 | 0.24 | 0.12 | 0.14 |
| 9 | 90284962 | rs11141937 | -0.2 | 0.926 | 0.157 | 1.804 | -0.032 | -0.024 | -0.024 | 0 | 0.06 | 0.01 | 0.05 | 0.12 | 0.1 | 0.12 |
| 9 | 90285464 | rs4878117 | 1.906 | 0.862 | -0.451 | 1.48 | -0.041 | -0.001 | -0.001 | 0 | 0.51 | 0.1 | 0.57 | 0.51 | 0.49 | 0.6 |
| 9 | 90289936 | rs10868644 | -1.267 | 0.772 | 0.287 | 1.748 | 0.018 | 0.05 | 0.05 | 0 | 0.12 | 0.27 | 0.09 | 0.27 | 0.1 | 0.16 |
| 9 | 90290816 | rs3118850 | 1.342 | 0.674 | 0.092 | 1.559 | 0.006 | -0.001 | -0.001 | 0 | 0.69 | 0.31 | 0.81 | 0.64 | 0.78 | 0.81 |
| 9 | 90291121 | rs3118851 | -0.827 | 1.158 | 0.381 | 1.437 | 0.054 | 0.005 | 0.005 | 0 | 0.21 | 0.4 | 0.24 | 0.24 | 0.3 | 0.22 |
| 9 | 90297680 | rs3128479 | 0.872 | 0.494 | 0.012 | 1.292 | -0.001 | 0.052 | 0.052 | 0 | 0.53 | 0.33 | 0.61 | 0.71 | 0.5 | 0.62 |
| 9 | 90308000 | rs3128489 | -0.139 | 0.501 | 0.502 | 0.748 | -0.01 | 0.022 | 0.022 | 0 | 0.71 | 0.78 | 0.78 | 0.98 | 0.67 | 0.86 |
| 9 | 90312577 | rs3128494 | 0.532 | 0.511 | 0.696 | 0.714 | -0.01 | 0.015 | 0.015 | 0 | 0.65 | 0.69 | 0.75 | 0.98 | 0.64 | 0.84 |
| 9 | 90317668 | rs7025760 | -1.798 | 0.304 | 1.066 | 1.473 | -0.015 | 0.08 | 0.08 | 0 | 0.26 | 0.39 | 0.21 | 0.43 | 0.21 | 0.34 |
| 9 | 90318855 | rs3793647 | -0.796 | 0.335 | 0.876 | 1.615 | 0.006 | 0.074 | 0.074 | 0 | 0.09 | 0.09 | 0.17 | 0.44 | 0.16 | 0.28 |
| 9 | 90319906 | rs3118862 | 1.652 | 0.272 | 0.834 | 1.436 | -0.002 | 0.044 | 0.044 | 0 | 0.5 | 0.28 | 0.52 | 0.42 | 0.48 | 0.5 |
| 9 | 90321583 | rs3118863 | 1.849 | 0.27 | 0.692 | 1.441 | -0.012 | 0.02 | 0.02 | 0 | 0.46 | 0.12 | 0.4 | 0.41 | 0.46 | 0.49 |
| 9 | 90322023 | rs1056719 | 0.688 | 0.166 | 1.009 | 1.523 | 0 | -0.001 | -0.001 | 0 | 0.62 | 0.4 | 0.7 | 0.83 | 0.64 | 0.81 |
| 9 | 90324200 | rs3118866 | 1.76 | 0.222 | 0.786 | 1.516 | -0.019 | 0.044 | 0.044 | 0 | 0.47 | 0.13 | 0.52 | 0.39 | 0.48 | 0.48 |
| 9 | 90324869 | rs4877369 | -1.115 | 0.277 | 0.822 | 1.537 | -0.006 | -0.006 | -0.006 | 0 | 0.11 | 0.07 | 0.16 | 0.59 | 0.13 | 0.32 |
| 9 | 90327095 | rs3128499 | 1.761 | 0.242 | 0.786 | 1.513 | -0.019 | 0.044 | 0.044 | 0 | 0.47 | 0.13 | 0.53 | 0.39 | 0.48 | 0.48 |
| 9 | 90328478 | rs4877370 | 0.542 | 0.266 | 0.892 | 1.384 | -0.001 | -0.077 | -0.077 | 0 | 0.63 | 0.44 | 0.73 | 0.99 | 0.64 | 0.84 |
| 9 | 90334530 | rs3128506 | 0.756 | 0.15 | 0.584 | 1.526 | -0.041 | -0.032 | -0.032 | 0 | 0.59 | 0.8 | 0.64 | 0.53 | 0.53 | 0.56 |
| 9 | 90340863 | rs3118869 | 1.342 | 0.289 | 0.685 | 1.208 | -0.001 | -0.003 | -0.003 | 0 | 0.54 | 0.49 | 0.4 | 0.34 | 0.45 | 0.43 |
| 9 | 90343780 | rs2378757 | -0.277 | 0.272 | 0.861 | 0.878 | -0.023 | 0.021 | 0.021 | 0 | 0.77 | 0.83 | 0.63 | 0.98 | 0.64 | 0.83 |

| | | | | | | | | | | | | | | | | |
|---|---|---|---|---|---|---|---|---|---|---|---|---|---|---|---|---|
| 9 | 90344316 | rs2296958 | -1.514 | 0.038 | 0.874 | 1.301 | -0.027 | 0.044 | 0.044 | 0 | 0.11 | 0.22 | 0.09 | 0.15 | 0.05 | 0.07 |
| 9 | 90355120 | rs7870268 | -0.866 | 0.204 | 0.651 | 0.868 | -0.006 | -0.058 | -0.058 | 0 | 0.11 | 0.46 | 0.06 | 0 | 0.05 | 0.01 |
| 9 | 90362777 | rs3118839 | 0.314 | 0.116 | -0.23 | 1.03 | -0.011 | -0.002 | -0.002 | 0 | 0.57 | 0.71 | 0.49 | 0.46 | 0.53 | 0.49 |
| 9 | 90363086 | rs3118840 | 0.193 | -0.018 | -0.266 | 1.107 | -0.011 | -0.01 | -0.01 | 0 | 0.59 | 0.76 | 0.64 | 0.45 | 0.57 | 0.54 |
| 9 | 90364627 | rs1927982 | 0.443 | 0.314 | -0.055 | 1.035 | -0.031 | 0.077 | 0.077 | 0 | 0.63 | 0.22 | 0.71 | 0.84 | 0.59 | 0.73 |
| 9 | 90366947 | rs9410948 | 0.423 | 0.37 | 0 | 1.526 | -0.003 | -0.002 | -0.002 | 0.488 | 0.66 | 0.31 | 0.73 | 0.84 | 0.61 | 0.76 |
| 9 | 90370106 | rs2149383 | 0.911 | 0.313 | 0.001 | 1.506 | -0.059 | -0.041 | -0.041 | 0 | 0.58 | 0.14 | 0.59 | 0.56 | 0.57 | 0.68 |
| 9 | 90370511 | rs943856 | 0.532 | 0.35 | 0.127 | 1.056 | -0.003 | -0.002 | -0.002 | 0 | 0.66 | 0.3 | 0.73 | 0.85 | 0.61 | 0.76 |
| 9 | 90380485 | rs4598339 | -1.237 | 0.183 | 0.464 | 1.8 | 0.008 | 0.219 | 0.219 | 0.089 | 0.07 | 0.2 | 0.15 | 0.3 | 0.05 | 0.04 |
| 9 | 90383163 | rs4878122 | 0.104 | 0.172 | 0.602 | 1.235 | 0.005 | 0.004 | 0.004 | 0 | 0.67 | 0.59 | 0.65 | 0.69 | 0.63 | 0.74 |
| 9 | 90384418 | rs9410624 | 0.684 | -0.042 | 0.573 | 1.29 | -0.025 | -0.032 | -0.032 | 0 | 0.58 | 0.14 | 0.59 | 0.58 | 0.58 | 0.64 |
| 9 | 90387296 | rs7037401 | -0.402 | -0.038 | 1.028 | 1.329 | -0.035 | 0.04 | 0.04 | 0.221 | 0.71 | 0.8 | 0.77 | 0.83 | 0.64 | 0.66 |
| 9 | 90390559 | rs9285055 | -0.402 | -0.038 | 1.028 | 1.329 | -0.035 | 0.04 | 0.04 | 0.221 | 0.71 | 0.79 | 0.77 | 0.83 | 0.64 | 0.66 |
| 9 | 90392042 | rs4878130 | 0.06 | 0.062 | 1.583 | 1.766 | -0.003 | -0.005 | -0.005 | 0 | 0.67 | 0.68 | 0.65 | 0.59 | 0.63 | 0.62 |
| 9 | 90392406 | rs7848164 | 0.935 | -0.566 | 1.283 | 1.908 | 0.006 | -0.009 | -0.009 | 0 | 0.41 | 0.51 | 0.4 | 0.34 | 0.44 | 0.39 |
| 9 | 90395684 | rs10123732 | 0.188 | -0.57 | 1.266 | 1.879 | 0.024 | 0.006 | 0.006 | 0.255 | 0.25 | 0.16 | 0.26 | 0.25 | 0.19 | 0.23 |
| 9 | 90400909 | rs9410632 | 0.794 | -0.457 | 1.441 | 1.938 | 0.006 | -0.004 | -0.004 | 0 | 0.42 | 0.46 | 0.5 | 0.59 | 0.44 | 0.43 |
| 9 | 90403094 | rs4878132 | -0.778 | -0.434 | 1.188 | 1.892 | -0.011 | 0.015 | 0.015 | 0.099 | 0.71 | 0.75 | 0.79 | 0.85 | 0.64 | 0.69 |
| 9 | 90405210 | rs9410964 | 0.473 | 0.168 | 0.97 | 1.596 | -0.078 | -0.032 | -0.032 | 0.225 | 0.38 | 0.1 | 0.33 | 0.37 | 0.43 | 0.38 |
| 9 | 90410305 | rs7044255 | 0.302 | 0.148 | 0.906 | 1.135 | 0 | -0.004 | -0.004 | 0 | 0.41 | 0.41 | 0.33 | 0.28 | 0.44 | 0.37 |
| 9 | 90425335 | rs9410636 | -0.592 | 0.29 | -0.141 | 1.056 | -0.057 | 0.008 | 0.008 | 0.287 | 0.66 | 0.89 | 0.71 | 0.71 | 0.62 | 0.67 |
| 9 | 90428537 | rs6420260 | -0.368 | 0.208 | 0.446 | 0.997 | 0.002 | -0.008 | -0.008 | 0 | 0.61 | 0.61 | 0.69 | 0.7 | 0.61 | 0.58 |
| 9 | 90431177 | rs11141976 | 0.071 | -0.309 | 0.337 | 0.752 | 0.002 | 0.03 | 0.03 | 2.612 | 0.52 | 0.47 | 0.42 | 0.26 | 0.53 | 0.3 |
| 9 | 90432159 | rs11141977 | 0.26 | -0.376 | 0.412 | 0.723 | 0.013 | 0.11 | 0.11 | 2.966 | 0.5 | 0.33 | 0.4 | 0.26 | 0.52 | 0.28 |
| 9 | 90432515 | rs7867409 | 0.26 | -0.376 | 0.412 | 0.723 | 0.013 | 0.11 | 0.11 | 2.966 | 0.5 | 0.34 | 0.4 | 0.26 | 0.53 | 0.28 |
| 9 | 90440658 | rs11794730 | 0.248 | -0.514 | 0.34 | 0.563 | 0.003 | 0.053 | 0.053 | 2.681 | 0.48 | 0.35 | 0.4 | 0.01 | 0.51 | 0.26 |
| 9 | 90442020 | rs11141981 | -0.092 | -0.541 | 0.175 | 1.101 | 0.014 | 0.057 | 0.057 | 0.972 | 0.54 | 0.36 | 0.54 | 0.23 | 0.58 | 0.57 |
| 9 | 90448266 | rs7025119 | -0.881 | -0.595 | 0.422 | 1.568 | 0.013 | 0.206 | 0.206 | 1.472 | 0.1 | 0.44 | 0.09 | 0.21 | 0.07 | 0.06 |
| 9 | 90450644 | rs10116815 | -1.177 | -0.565 | 0.074 | 1.584 | -0.077 | -0.107 | -0.107 | 0.306 | 0.72 | 0.98 | 0.67 | 0.71 | 0.69 | 0.67 |
| 9 | 90455686 | rs7045334 | -1.228 | -0.514 | 0.514 | 1.649 | -0.006 | 0.097 | 0.097 | 0 | 0.1 | 0.16 | 0.16 | 0.48 | 0.09 | 0.35 |
| 9 | 90462253 | rs7018666 | -1.078 | -0.254 | 0.659 | 1.12 | 0.117 | 0.151 | 0.151 | 0 | 0.07 | 0.32 | 0.11 | 0.1 | 0.11 | 0.04 |
| 9 | 90466696 | rs11141992 | -0.015 | -0.442 | 0.349 | 1.074 | -0.002 | -0.01 | -0.01 | 0 | 0.59 | 0.49 | 0.46 | 0.23 | 0.61 | 0.32 |
| 9 | 90470780 | rs3935944 | -1.336 | -0.259 | 0.511 | 1.464 | -0.006 | 0.068 | 0.068 | 1.251 | 0.18 | 0.26 | 0.31 | 0.44 | 0.19 | 0.49 |
| 9 | 90481245 | rs11142008 | -1.34 | -0.177 | 0.475 | 1.496 | -0.009 | 0.148 | 0.148 | 2.033 | 0.19 | 0.39 | 0.34 | 0.44 | 0.19 | 0.49 |
| 9 | 90487391 | rs12001141 | 0.032 | 0.164 | -0.108 | 1.338 | -0.022 | 0.001 | 0.001 | 0 | 0.36 | 0.32 | 0.38 | 0.41 | 0.46 | 0.23 |
| 9 | 90487924 | rs4072799 | -0.537 | -0.284 | -0.091 | 0.96 | -0.017 | 0.044 | 0.044 | 0.374 | 0.2 | 0.35 | 0.29 | 0.29 | 0.16 | 0.38 |
| 9 | 90493423 | rs10868668 | -0.172 | -0.17 | -0.83 | 0.569 | -0.021 | 0.249 | 0.249 | 6.55 | 0.46 | 0.12 | 0.4 | 0.17 | 0.53 | 0.41 |
| 9 | 90494889 | rs10780878 | -0.498 | -0.255 | -0.797 | 0.503 | -0.006 | 0.175 | 0.175 | 0.185 | 0.42 | 0.17 | 0.47 | 0.17 | 0.56 | 0.48 |
| 9 | 90495370 | rs6560041 | -0.639 | 0.22 | -0.607 | 1.429 | -0.042 | -0.024 | -0.024 | 0.038 | 0.38 | 0.18 | 0.57 | 0.44 | 0.51 | 0.54 |
| 9 | 90503451 | rs11142017 | -0.837 | 0.281 | 0.397 | 1.587 | 0.004 | -0.007 | -0.007 | 0 | 0.4 | 0.52 | 0.4 | 0.34 | 0.52 | 0.49 |
| 9 | 90514482 | rs12552120 | -0.452 | 0.136 | -0.066 | 1.386 | -0.029 | 0.046 | 0.046 | 0.122 | 0.37 | 0.21 | 0.37 | 0.2 | 0.45 | 0.4 |
| 9 | 90515619 | rs7470248 | -0.383 | 0.163 | -0.272 | 1.398 | -0.06 | 0.072 | 0.072 | 0.491 | 0.36 | 0.14 | 0.36 | 0.2 | 0.45 | 0.4 |
| 9 | 90571740 | rs655500 | 0.551 | -0.228 | 0.325 | 2.028 | -0.025 | 0.044 | 0.044 | 0 | 0.83 | 0.74 | 0.71 | 0.33 | 0.92 | 0.84 |
| 9 | 90581425 | rs668108 | 0.673 | -0.237 | 0.439 | 1.975 | -0.029 | 0.044 | 0.044 | 0 | 0.82 | 0.7 | 0.71 | 0.33 | 0.92 | 0.84 |
| 9 | 90581746 | rs1167 | 0.656 | -0.236 | 0.435 | 1.975 | -0.025 | 0.039 | 0.039 | 0 | 0.82 | 0.72 | 0.71 | 0.33 | 0.92 | 0.84 |
| 9 | 90583521 | rs560019 | 1.125 | 0.111 | 0.427 | 1.911 | -0.002 | 0 | 0 | 1.533 | 0.62 | 0.51 | 0.46 | 0.38 | 0.62 | 0.72 |
| 9 | 90584478 | rs2271992 | -0.122 | -0.199 | 0.576 | 1.819 | -0.012 | 0.024 | 0.024 | 0 | 0.08 | 0.08 | 0.07 | 0.2 | 0.06 | 0.25 |
| 9 | 90586043 | rs622418 | 2.108 | -0.055 | 0.816 | 1.466 | 0.007 | 0.158 | 0.158 | 0.145 | 0.52 | 0.19 | 0.34 | 0.18 | 0.51 | 0.46 |
| 9 | 90586221 | rs9410653 | 1.392 | -0.195 | 0.786 | 1.494 | 0.034 | 0.034 | 0.034 | 0.006 | 0.06 | 0.01 | 0.01 | 0 | 0.01 | 0.01 |
| 9 | 90588903 | rs665983 | 0.768 | -0.184 | 1.1 | 1.615 | -0.036 | 0.01 | 0.01 | 0.134 | 0.75 | 0.64 | 0.69 | 0.31 | 0.9 | 0.83 |
| 9 | 90590650 | rs572992 | 0.563 | 0.117 | 1.101 | 1.614 | 0.006 | 0.002 | 0.002 | 2.118 | 0.65 | 0.77 | 0.49 | 0.32 | 0.65 | 0.73 |
| 9 | 90601003 | rs3898637 | 0.011 | 0.336 | 1.001 | 1.504 | 0.002 | -0.004 | -0.004 | 0 | 0.09 | 0.03 | 0.09 | 0.13 | 0.1 | 0.27 |
| 9 | 90603115 | rs473082 | 0.774 | 0.293 | 1.364 | 1.489 | -0.005 | -0.004 | -0.004 | 0 | 0.63 | 0.42 | 0.46 | 0.32 | 0.65 | 0.73 |
| 9 | 90611872 | rs10868677 | 1.926 | 0.38 | 1.353 | 1.458 | -0.001 | 0.137 | 0.137 | 0.032 | 0.52 | 0.25 | 0.33 | 0.19 | 0.5 | 0.46 |
| 9 | 90613915 | rs652529 | 0.696 | 0.424 | 1.446 | 1.43 | 0.003 | 0.002 | 0.002 | 2.153 | 0.65 | 0.67 | 0.48 | 0.32 | 0.65 | 0.73 |
| 9 | 90621460 | rs7043672 | 0.261 | 0.658 | 1.554 | 1.417 | 0.104 | 0.039 | 0.039 | 0 | 0.17 | 0.42 | 0.28 | 0.25 | 0.28 | 0.15 |
| 9 | 90627994 | rs13296013 | 0.527 | 0.721 | 1.445 | 1.362 | 0.073 | -0.09 | -0.09 | 0 | 0.14 | 0.31 | 0.19 | 0 | 0.27 | 0.09 |
| 9 | 90630239 | rs7864469 | 0.51 | 0.746 | 1.443 | 1.433 | 0.049 | 0.044 | 0.044 | 0 | 0.14 | 0.36 | 0.23 | 0.25 | 0.21 | 0.13 |
| 9 | 90641936 | rs10115989 | -1.332 | 0.631 | 1.046 | 1.584 | -0.01 | -0.007 | -0.007 | 0 | 0.39 | 0.83 | 0.55 | 0.67 | 0.37 | 0.27 |
| 9 | 90645727 | rs10780886 | -1.03 | 0.582 | 1.934 | 1.82 | 0.008 | 0.033 | 0.033 | 2.997 | 0.34 | 0.46 | 0.51 | 0.59 | 0.36 | 0.25 |
| 9 | 90646449 | rs11142050 | 0.798 | 0.605 | 2.063 | 1.753 | 0.048 | -0.071 | -0.071 | 0 | 0.14 | 0.28 | 0.19 | 0 | 0.27 | 0.08 |
| 9 | 90647962 | rs7038292 | -1.45 | 0.58 | 1.972 | 1.928 | -0.008 | -0.049 | -0.049 | 0.001 | 0.39 | 0.83 | 0.55 | 0.62 | 0.37 | 0.28 |
| 9 | 90648222 | rs1331220 | -1.042 | 0.583 | 2.195 | 1.936 | 0.001 | 0.009 | 0.009 | 2.208 | 0.36 | 0.6 | 0.52 | 0.53 | 0.36 | 0.26 |
| 9 | 90650132 | rs10780887 | -1.105 | 0.524 | 2.235 | 1.745 | -0.005 | 0.005 | 0.005 | 1.901 | 0.36 | 0.6 | 0.41 | 0.43 | 0.32 | 0.2 |
| 9 | 90650880 | rs10125370 | -0.222 | 0.311 | 2 | 1.876 | -0.019 | 0.022 | 0.022 | 0.333 | 0.2 | 0.24 | 0.25 | 0.34 | 0.09 | 0.12 |
| 9 | 90658749 | rs11142062 | 1.497 | 0.374 | 0.61 | 0.75 | -0.039 | 0.123 | 0.123 | 0 | 0.19 | 0.01 | 0.09 | 0.01 | 0.14 | 0.08 |
| 9 | 90665712 | rs7048705 | -0.048 | 0.489 | 0.853 | 1.175 | 0.018 | 0.007 | 0.007 | 0 | 0.24 | 0.66 | 0.26 | 0.33 | 0.32 | 0.16 |
| 9 | 90667427 | rs11142070 | 0.994 | 0.27 | 0.52 | 1.388 | -0.006 | -0.066 | -0.066 | 0 | 0.11 | 0.01 | 0.21 | 0.31 | 0.07 | 0.12 |
| 9 | 90668342 | rs9969798 | -0.024 | 0.449 | 0.948 | 1.102 | -0.003 | -0.006 | -0.006 | 0 | 0.23 | 0.76 | 0.39 | 0.45 | 0.28 | 0.16 |

| | | | | | | | | | | | | | | | | |
|---|---|---|---|---|---|---|---|---|---|---|---|---|---|---|---|---|
| 9 | 90669303 | rs10868691 | -0.05 | 0.44 | 0.943 | 1.075 | -0.003 | -0.007 | -0.007 | 0 | 0.24 | 0.76 | 0.39 | 0.45 | 0.28 | 0.16 |
| 9 | 90669897 | rs2026920 | 0.904 | 0.694 | 0.469 | 0.548 | 0.014 | -0.058 | -0.058 | 0.147 | 0.17 | 0.23 | 0.18 | 0 | 0.26 | 0.08 |
| 9 | 90674178 | rs538970 | -1.329 | 0.406 | 0.285 | 0.886 | -0.03 | 0.031 | 0.031 | 0 | 0.94 | 0.99 | 0.92 | 1 | 0.87 | 0.98 |
| 9 | 90688689 | rs10868700 | -1.88 | -0.507 | 1.309 | 1.736 | 0.028 | 0.522 | 0.522 | 0.523 | 0.05 | 0.33 | 0.19 | 0.47 | 0.01 | 0.09 |
| 9 | 90689210 | rs11142081 | -0.033 | -0.642 | 0.743 | 1.512 | -0.007 | -0.005 | -0.005 | 0.017 | 0.2 | 0.21 | 0.12 | 0.17 | 0.14 | 0.13 |
| 9 | 90697532 | rs2104827 | -1.137 | -0.405 | 1.182 | 1.801 | -0.026 | 0.249 | 0.249 | 3.161 | 0.17 | 0.55 | 0.26 | 0.52 | 0.11 | 0.22 |
| 9 | 90697669 | rs12001770 | -0.358 | -0.393 | 1.396 | 1.458 | 0.015 | 0.106 | 0.106 | 3.044 | 0.11 | 0.38 | 0.21 | 0.17 | 0.09 | 0.08 |
| 9 | 90703042 | rs4877394 | -1.179 | -0.374 | 1.085 | 1.456 | -0.018 | 0.056 | 0.056 | 3.118 | 0.16 | 0.55 | 0.23 | 0.19 | 0.11 | 0.16 |
| 9 | 90710965 | rs7020924 | -0.709 | -0.274 | 0.094 | 1.086 | 0.01 | 0.018 | 0.018 | 0 | 0.52 | 0.64 | 0.46 | 0.41 | 0.37 | 0.42 |
| 9 | 90720715 | rs11142112 | 0.107 | -0.174 | -0.538 | 0.506 | 0.028 | 0.125 | 0.125 | 0.017 | 0.38 | 0.2 | 0.27 | 0.1 | 0.27 | 0.29 |
| 9 | 90721446 | rs12237927 | 0.877 | -0.099 | -0.617 | 0.643 | -0.026 | -0.059 | -0.059 | 0.009 | 0.11 | 0.02 | 0.16 | 0.23 | 0.11 | 0.13 |
| 9 | 90722248 | rs11142115 | -0.302 | 0.134 | -0.099 | 0.355 | 0.032 | 0.131 | 0.131 | 0 | 0.39 | 0.2 | 0.27 | 0.1 | 0.27 | 0.29 |
| 9 | 90760805 | rs4877406 | -1.915 | 1.216 | 1.43 | 1.954 | 0.087 | 0.026 | 0.026 | 0.099 | 0.17 | 0.35 | 0.36 | 0.23 | 0.28 | 0.36 |
| 9 | 90762031 | rs10512193 | -1.145 | 1.311 | 1.318 | 1.984 | -0.056 | 0.006 | 0.006 | 0 | 0.11 | 0.05 | 0.12 | 0.11 | 0.2 | 0.2 |
| 9 | 90765354 | rs11142135 | -1.669 | 1.383 | 1.429 | 2.637 | 0.116 | 0.029 | 0.029 | 0.941 | 0.17 | 0.42 | 0.38 | 0.24 | 0.31 | 0.36 |
| 9 | 90780784 | rs7863242 | -2.49 | 0.938 | 1.056 | 2.429 | -0.084 | 0.04 | 0.04 | 1.593 | 0.35 | 0.89 | 0.47 | 0.26 | 0.39 | 0.45 |
| 9 | 90792721 | rs2814851 | -1.594 | 1.214 | 1.324 | 2.597 | 0.009 | 0.009 | 0.009 | 1.425 | 0.26 | 0.33 | 0.44 | 0.65 | 0.36 | 0.44 |
| 9 | 90792863 | rs2778019 | -1.527 | 1.3 | 1.366 | 2.639 | 0.012 | 0.012 | 0.012 | 0.6 | 0.26 | 0.33 | 0.43 | 0.65 | 0.36 | 0.44 |
| 9 | 90799534 | rs2578245 | 1.287 | 1.264 | 1.488 | 2.398 | 0.023 | 0.021 | 0.021 | 0.683 | 0.75 | 0.64 | 0.66 | 0.37 | 0.61 | 0.56 |
| 9 | 90801739 | rs7034024 | -2.16 | 1.083 | 2.106 | 2.482 | 0.004 | 0.049 | 0.049 | 0 | 0.11 | 0.24 | 0.14 | 0.16 | 0.11 | 0.1 |
| 9 | 90802614 | rs17080774 | -1.673 | 1.082 | 2.162 | 2.523 | -0.001 | 0.012 | 0.012 | 0 | 0.08 | 0.14 | 0.12 | 0.08 | 0.07 | 0.09 |
| 9 | 90802721 | rs7865433 | -1.497 | 1.158 | 2.861 | 2.773 | 0.025 | 0.031 | 0.031 | 0 | 0.16 | 0.26 | 0.28 | 0.22 | 0.23 | 0.29 |
| 9 | 90803472 | rs17053864 | -2.343 | 1.184 | 2.552 | 2.776 | 0.004 | 0.049 | 0.049 | 0 | 0.11 | 0.24 | 0.14 | 0.16 | 0.11 | 0.1 |
| 9 | 90808582 | rs4877127 | -2.495 | 1.177 | 2.58 | 2.757 | 0.016 | 0.04 | 0.04 | 0 | 0.26 | 0.3 | 0.39 | 0.37 | 0.33 | 0.39 |
| 9 | 90811182 | rs2814828 | 3.196 | 0.975 | 2.629 | 3.054 | 0.024 | 0.062 | 0.062 | 0 | 0.81 | 0.73 | 0.64 | 0.41 | 0.79 | 0.67 |
| 9 | 90816711 | rs7045211 | -1.654 | 1.009 | 2.66 | 3.002 | 0.002 | 0.019 | 0.019 | 0 | 0.08 | 0.16 | 0.12 | 0.08 | 0.07 | 0.08 |
| 9 | 90817188 | rs2814837 | 1.304 | 1.022 | 2.752 | 3.116 | 0.015 | 0.141 | 0.141 | 0 | 0.93 | 0.93 | 0.78 | 0.56 | 0.9 | 0.78 |
| 9 | 90819763 | rs2814839 | 1.304 | 1.022 | 2.752 | 3.116 | 0.015 | 0.141 | 0.141 | 0 | 0.93 | 0.93 | 0.78 | 0.56 | 0.9 | 0.78 |
| 9 | 90820013 | rs2492849 | 1.304 | 1.022 | 2.752 | 3.116 | 0.015 | 0.141 | 0.141 | 0 | 0.93 | 0.93 | 0.78 | 0.56 | 0.9 | 0.78 |
| 9 | 90821421 | rs2482488 | 3.205 | 0.973 | 3.271 | 3.037 | 0.058 | 0.209 | 0.209 | 0.013 | 0.81 | 0.57 | 0.63 | 0.48 | 0.8 | 0.69 |
| 9 | 90831332 | rs17053965 | -1.822 | 0.93 | 2.723 | 3.077 | -0.011 | 0.074 | 0.074 | 0 | 0.05 | 0.07 | 0.13 | 0.16 | 0.04 | 0.18 |
| 9 | 90832856 | rs2814845 | 2.596 | 0.958 | 2.836 | 3.133 | 0.04 | 0.225 | 0.225 | 0 | 0.87 | 0.7 | 0.67 | 0.49 | 0.85 | 0.72 |
| 9 | 90835589 | rs2578246 | 2.493 | 1.052 | 2.72 | 2.953 | 0.057 | 0.153 | 0.153 | 1.99 | 0.84 | 0.41 | 0.72 | 0.63 | 0.82 | 0.83 |
| 9 | 90839031 | rs17053978 | -0.666 | 0.958 | 1.988 | 2.731 | -0.021 | -0.009 | -0.009 | 0 | 0.11 | 0.07 | 0.2 | 0.12 | 0.18 | 0.15 |
| 9 | 90847356 | rs1550286 | 1.199 | 0.897 | 2.546 | 3.027 | 0.079 | 0.285 | 0.285 | 0.288 | 0.93 | 0.76 | 0.78 | 0.64 | 0.9 | 0.81 |
| 9 | 90852161 | rs17054006 | -2.445 | 0.879 | 2.49 | 2.868 | 0.033 | 0.074 | 0.074 | 0.434 | 0.13 | 0.56 | 0.15 | 0.17 | 0.14 | 0.14 |
| 9 | 90857957 | rs7028051 | 1.273 | 0.708 | 1.962 | 2.758 | 0 | -0.001 | -0.001 | 0 | 0.72 | 0.68 | 0.53 | 0.5 | 0.74 | 0.63 |
| 9 | 90865387 | rs17054025 | -1.087 | 0.705 | 2.071 | 2.763 | 0.002 | 0.001 | 0.001 | 0 | 0.07 | 0.06 | 0.06 | 0.08 | 0.12 | 0.06 |
| 9 | 90865441 | rs1438467 | -0.811 | 0.706 | 2.072 | 2.808 | 0.023 | -0.024 | -0.024 | 0.194 | 0.22 | 0.15 | 0.3 | 0.24 | 0.18 | 0.27 |
| 9 | 90866595 | rs11142202 | 1.879 | 0.705 | 2.081 | 2.805 | -0.001 | 0 | 0 | 0.236 | 0.65 | 0.63 | 0.47 | 0.41 | 0.62 | 0.58 |
| 9 | 90867612 | rs10780914 | 1.964 | 0.709 | 2.13 | 2.807 | 0.008 | 0 | 0 | 0 | 0.64 | 0.53 | 0.45 | 0.42 | 0.63 | 0.58 |
| 9 | 90868617 | rs6560070 | 0.665 | 0.683 | 1.282 | 2.536 | 0.045 | 0.144 | 0.144 | 0.42 | 0.93 | 0.83 | 0.83 | 0.74 | 0.92 | 0.91 |
| 9 | 90875879 | rs893494 | 1.138 | 0.576 | 0.626 | 2.248 | 0.017 | 0.046 | 0.046 | 0.704 | 0.86 | 0.85 | 0.77 | 0.65 | 0.8 | 0.82 |
| 9 | 90882154 | rs3937594 | 1.208 | 0.556 | -0.006 | 1.411 | -0.004 | 0.045 | 0.045 | 0 | 0.6 | 0.31 | 0.36 | 0.31 | 0.61 | 0.49 |
| 9 | 90891114 | rs4877434 | -0.781 | 0.542 | -0.543 | 1.426 | 0 | -0.017 | -0.017 | 0 | 0.36 | 0.19 | 0.37 | 0.33 | 0.35 | 0.51 |
| 9 | 90893431 | rs10780919 | 0.957 | 0.279 | 0.341 | 1.355 | 0.019 | 0.073 | 0.073 | 1.815 | 0.72 | 0.44 | 0.65 | 0.53 | 0.71 | 0.56 |
| 9 | 90895184 | rs4877132 | -1.567 | -0.529 | 0.801 | 0.692 | 0.006 | -0.022 | -0.022 | 0 | 0.11 | 0.28 | 0.13 | 0.05 | 0.08 | 0.14 |
| 9 | 90896018 | rs7851719 | -1.069 | -1.569 | 0.831 | 0.396 | -0.046 | -0.034 | -0.034 | 0 | 0.23 | 0.6 | 0.19 | 0.1 | 0.11 | 0.21 |
| 9 | 90896077 | rs10868746 | -0.835 | -1.94 | -0.101 | 0.418 | 0.013 | -0.012 | -0.012 | 0.035 | 0.14 | 0.06 | 0.08 | 0.2 | 0.09 | 0.13 |
| 9 | 90899627 | rs7867717 | -1.465 | -2.25 | -0.072 | 0.439 | 0.095 | 0.012 | 0.012 | 1.965 | 0.36 | 0.87 | 0.32 | 0.31 | 0.17 | 0.32 |
| 9 | 90905263 | rs4515651 | -1.193 | -2.402 | 0.902 | 0.372 | -0.084 | 0.018 | 0.018 | 0.445 | 0.25 | 0.53 | 0.22 | 0.28 | 0.1 | 0.25 |
| 9 | 90905665 | rs7870611 | -1.962 | -2.242 | 1.091 | 0.128 | -0.006 | -0.032 | -0.032 | 0 | 0.1 | 0.16 | 0.06 | 0.03 | 0.06 | 0.09 |
| 9 | 90908761 | rs6560071 | -2.327 | -2.504 | 1.848 | -0.006 | -0.012 | -0.025 | -0.025 | 0 | 0.3 | 0.65 | 0.27 | 0.13 | 0.19 | 0.32 |
| 9 | 90917530 | rs7865913 | -2.285 | -3.376 | 1.195 | -0.471 | 0.091 | 0.157 | 0.157 | 2.386 | 0.39 | 0.92 | 0.32 | 0.12 | 0.23 | 0.32 |
| 9 | 90917641 | rs7868935 | -2.24 | -3.337 | 2.035 | -0.502 | -0.024 | -0.078 | -0.078 | 0 | 0.12 | 0.63 | 0.13 | 0 | 0.06 | 0.08 |
| 9 | 90925255 | rs12377675 | -1.335 | -3.34 | 1.866 | -0.23 | 0.008 | 0.004 | 0.004 | 1.056 | 0.26 | 0.25 | 0.18 | 0.15 | 0.18 | 0.26 |
| 9 | 90925394 | rs11142238 | -1.325 | -3.351 | 1.86 | 0.09 | 0.009 | -0.006 | -0.006 | 0 | 0.26 | 0.24 | 0.19 | 0.26 | 0.18 | 0.26 |
| 9 | 90927030 | rs11142239 | -1.299 | -3.349 | 1.86 | -0.089 | 0.007 | 0.003 | 0.003 | 0.977 | 0.26 | 0.25 | 0.18 | 0.15 | 0.18 | 0.25 |
| 9 | 90927206 | rs10868754 | -2.215 | -3.373 | 2.623 | -0.103 | -0.008 | -0.01 | -0.01 | 2.266 | 0.34 | 0.61 | 0.27 | 0.15 | 0.23 | 0.33 |
| 9 | 90939033 | rs12005022 | -1.458 | -3.301 | 2.816 | -0.421 | -0.006 | -0.036 | -0.036 | 0.014 | 0.08 | 0.15 | 0.06 | 0 | 0.04 | 0.09 |
| 9 | 90945833 | rs10125017 | -2.296 | -3.376 | 3.579 | -0.456 | 0.002 | -0.052 | -0.052 | 0 | 0.1 | 0.37 | 0.08 | 0 | 0.04 | 0.08 |
| 9 | 90948749 | rs7046181 | -2.682 | -3.683 | 3.363 | -0.501 | 0.062 | 0.154 | 0.154 | 1.016 | 0.35 | 0.92 | 0.25 | 0.1 | 0.21 | 0.3 |
| 9 | 90955547 | rs12348836 | -2.258 | -3.787 | 3.613 | -0.498 | 0.014 | 0.015 | 0.015 | 0.348 | 0.29 | 0.75 | 0.22 | 0.1 | 0.16 | 0.29 |
| 9 | 90966143 | rs10121008 | -2.399 | -3.901 | 3.543 | -0.413 | 0.093 | 0.115 | 0.115 | 2.298 | 0.3 | 0.89 | 0.19 | 0.1 | 0.13 | 0.28 |
| 9 | 91002287 | rs3750493 | -2.365 | -3.915 | 3.383 | -0.42 | 0.062 | 0.072 | 0.072 | 1.876 | 0.29 | 0.8 | 0.2 | 0.1 | 0.13 | 0.27 |
| 9 | 91008303 | rs11142277 | -1.526 | -3.797 | 3.217 | -0.382 | -0.044 | -0.061 | -0.061 | 0 | 0.23 | 0.38 | 0.15 | 0.1 | 0.11 | 0.22 |
| 9 | 91024421 | rs10123897 | -1.537 | -3.797 | 3.217 | -0.378 | -0.044 | -0.058 | -0.058 | 0 | 0.23 | 0.38 | 0.15 | 0.1 | 0.11 | 0.22 |
| 9 | 91099778 | rs3010859 | -0.347 | -3.976 | 2.452 | -0.243 | -0.013 | -0.044 | -0.044 | 0.003 | 0.11 | 0.01 | 0.11 | 0.13 | 0.08 | 0.11 |
| 9 | 91104414 | rs12337803 | -1.198 | -3.986 | 2.68 | -0.634 | -0.028 | -0.037 | -0.037 | 0.169 | 0.22 | 0.33 | 0.14 | 0.1 | 0.11 | 0.22 |

| | | | | | | | | | | | | | | | | |
|---|---|---|---|---|---|---|---|---|---|---|---|---|---|---|---|---|
| 9 | 91109302 | rs11142322 | -1.794 | -4.553 | 2.406 | -0.285 | -0.085 | -0.009 | -0.009 | 0.403 | 0.3 | 0.52 | 0.29 | 0.76 | 0.1 | 0.43 |
| 9 | 91152492 | rs12685193 | -1.29 | -4.424 | 1.442 | -0.4 | 0 | 0.001 | 0.001 | 0 | 0.26 | 0.22 | 0.19 | 0.78 | 0.1 | 0.42 |
| 9 | 91154126 | rs10868791 | -2.106 | -4.405 | 0.656 | -0.443 | 0.156 | 0.063 | 0.063 | 0.113 | 0.36 | 0.94 | 0.27 | 0.8 | 0.12 | 0.43 |
| 9 | 91161674 | rs7860074 | -2.038 | -4.214 | 0.857 | -0.359 | 0.12 | 0.067 | 0.067 | 0.573 | 0.33 | 0.87 | 0.26 | 0.8 | 0.11 | 0.37 |
| 9 | 91175742 | rs13292086 | -0.851 | -3.976 | 0.737 | -0.287 | 0.066 | -0.037 | -0.037 | 0 | 0.25 | 0.02 | 0.18 | 0.79 | 0.1 | 0.32 |
| 9 | 91183368 | rs12346988 | -1.797 | -3.88 | 1.332 | -0.386 | 0.004 | 0.005 | 0.005 | 0 | 0.35 | 0.68 | 0.32 | 0.87 | 0.12 | 0.46 |
| 9 | 91186956 | rs6560097 | -1.735 | -3.878 | 1.297 | -0.405 | 0.005 | 0.005 | 0.005 | 0 | 0.34 | 0.68 | 0.31 | 0.87 | 0.13 | 0.44 |
| 9 | 91189922 | rs4263860 | -2.232 | -2.915 | 1.621 | -0.201 | -0.022 | -0.065 | -0.065 | 0.019 | 0.12 | 0.6 | 0.11 | 0 | 0.05 | 0.05 |
| 9 | 91190077 | rs4269628 | -1.559 | -2.84 | 1.413 | 0.198 | -0.083 | 0.052 | 0.052 | 0 | 0.22 | 0.5 | 0.25 | 0.71 | 0.08 | 0.26 |
| 9 | 91197917 | rs4877453 | -1.164 | -2.656 | 0.949 | 0.236 | -0.019 | 0.015 | 0.015 | 0.011 | 0.2 | 0.21 | 0.24 | 0.7 | 0.08 | 0.36 |
| 9 | 91203868 | rs4877136 | -0.725 | -1.394 | 0.236 | -0.115 | -0.024 | -0.018 | -0.018 | 0 | 0.3 | 0.41 | 0.27 | 0.2 | 0.22 | 0.26 |
| 9 | 91210293 | rs6560099 | -1.269 | -0.189 | -1.191 | -0.13 | -0.137 | -0.055 | -0.055 | 0 | 0.55 | 0.94 | 0.56 | 0.44 | 0.44 | 0.45 |
| 9 | 91211783 | rs4877459 | -0.601 | -0.077 | -1.128 | -0.739 | 0 | 0.032 | 0.032 | 0 | 0.33 | 0.79 | 0.3 | 0.24 | 0.26 | 0.24 |
| 9 | 91219303 | rs4242609 | -1.721 | -0.111 | -1.247 | -0.744 | -0.115 | 0.133 | 0.133 | 0 | 0.52 | 0.92 | 0.45 | 0.24 | 0.44 | 0.42 |
| 9 | 91229667 | rs4877464 | -1.285 | -0.205 | -0.678 | -1.037 | 0.033 | 0.11 | 0.11 | 0 | 0.41 | 0.24 | 0.36 | 0.18 | 0.34 | 0.31 |
| 9 | 91254321 | rs10512195 | -0.677 | -0.188 | -0.027 | -1.066 | -0.007 | -0.049 | -0.049 | 0 | 0.17 | 0.21 | 0.12 | 0 | 0.14 | 0.14 |
| 9 | 91255143 | rs17054259 | -0.633 | -0.238 | 0.236 | -1.179 | 0.001 | -0.07 | -0.07 | 0 | 0.16 | 0.69 | 0.11 | 0.01 | 0.1 | 0.12 |
| 9 | 91256579 | rs4075961 | -2.591 | -0.267 | -0.062 | -1.251 | -0.096 | 0.221 | 0.221 | 0 | 0.54 | 0.92 | 0.46 | 0.17 | 0.45 | 0.4 |
| 9 | 91259439 | rs4626701 | 0.291 | -0.115 | 0.904 | -1.155 | 0.038 | -0.065 | -0.065 | 0.042 | 0.11 | 0.25 | 0.06 | 0.01 | 0.1 | 0.11 |
| 9 | 91260562 | rs17054265 | -1.135 | -0.123 | 0.734 | -1.127 | 0.008 | -0.159 | -0.159 | 0 | 0.28 | 0.47 | 0.19 | 0.01 | 0.25 | 0.24 |
| 9 | 91260612 | rs7869919 | -1.925 | -0.135 | 1.264 | -1.144 | 0.015 | 0.173 | 0.173 | 0 | 0.49 | 0.76 | 0.42 | 0.17 | 0.41 | 0.37 |
| 9 | 91262215 | rs7048023 | 0.144 | -0.11 | 1.248 | -1.095 | -0.016 | -0.013 | -0.013 | 0 | 0.2 | 0.3 | 0.22 | 0.17 | 0.16 | 0.13 |
| 9 | 91262296 | rs4543636 | -1.474 | -0.025 | 0.704 | -0.906 | 0.012 | 0.178 | 0.178 | 0 | 0.49 | 0.77 | 0.42 | 0.18 | 0.41 | 0.37 |
| 9 | 91268541 | rs7855348 | -1.395 | 0.597 | 1.281 | -0.649 | 0.015 | -0.061 | -0.061 | 0 | 0.2 | 0.39 | 0.17 | 0.07 | 0.19 | 0.18 |
| 9 | 91281294 | rs9777071 | -1.616 | 0.648 | 1.004 | -0.487 | 0 | 0.013 | 0.013 | 0 | 0.49 | 0.58 | 0.62 | 0.69 | 0.46 | 0.47 |
| 9 | 91290319 | rs11516165 | -0.455 | 0.699 | 0.869 | -0.493 | -0.008 | -0.001 | -0.001 | 0 | 0.31 | 0.21 | 0.45 | 0.62 | 0.28 | 0.3 |
| 9 | 91293885 | rs7872811 | -0.26 | 0.708 | 0.834 | -0.432 | -0.022 | -0.023 | -0.023 | 0 | 0.29 | 0.16 | 0.45 | 0.62 | 0.28 | 0.31 |
| 9 | 91295698 | rs9410306 | 1.427 | 0.465 | 1.033 | -0.384 | 0.001 | 0.018 | 0.018 | 0 | 0.5 | 0.43 | 0.38 | 0.31 | 0.54 | 0.53 |
| 9 | 91296517 | rs9410313 | 1.296 | 0.488 | 0.439 | -0.365 | 0.001 | 0.112 | 0.112 | 0 | 0.49 | 0.26 | 0.38 | 0.32 | 0.54 | 0.53 |
| 9 | 91311747 | rs17054328 | -1.328 | 0.622 | 0.611 | -0.184 | -0.018 | -0.013 | -0.013 | 0.515 | 0.13 | 0.48 | 0.1 | 0.06 | 0.09 | 0.05 |
| 9 | 91313899 | rs4559317 | 0.379 | 0.903 | -0.375 | 0.174 | -0.065 | -0.018 | -0.018 | 0 | 0.69 | 0.86 | 0.54 | 0.37 | 0.66 | 0.61 |
| 9 | 91317827 | rs4495506 | 0.818 | 1.185 | -0.057 | 0.624 | -0.001 | -0.001 | -0.001 | 0 | 0.62 | 0.63 | 0.53 | 0.42 | 0.56 | 0.53 |
| 9 | 91318984 | rs9410282 | 0.588 | 1.45 | -0.364 | 0.501 | -0.042 | 0.04 | 0.04 | 0 | 0.64 | 0.78 | 0.46 | 0.27 | 0.57 | 0.43 |
| 9 | 91330562 | rs4877047 | 0.598 | 1.493 | -0.376 | 0.466 | -0.036 | 0.035 | 0.035 | 0 | 0.68 | 0.8 | 0.49 | 0.25 | 0.62 | 0.47 |
| 9 | 91337035 | rs7042278 | -0.183 | 1.438 | -0.5 | 1.147 | -0.031 | -0.046 | -0.046 | 0 | 0.74 | 0.86 | 0.53 | 0.41 | 0.66 | 0.61 |
| 9 | 91344293 | rs4877059 | -1.192 | 0.578 | 0.63 | 1.647 | -0.028 | 0.078 | 0.078 | 0 | 0.27 | 0.52 | 0.39 | 0.57 | 0.25 | 0.39 |
| 9 | 91344655 | rs4609263 | -1.517 | 0.527 | 1.423 | 1.919 | 0.021 | 0.181 | 0.181 | 0 | 0.17 | 0.32 | 0.36 | 0.55 | 0.23 | 0.28 |
| 9 | 91348970 | rs4877066 | -0.996 | 0.492 | 1.657 | 1.83 | 0.002 | 0.212 | 0.212 | 0 | 0.15 | 0.39 | 0.29 | 0.4 | 0.17 | 0.26 |
| 9 | 91359502 | rs1857490 | -1.995 | 0.454 | 0.845 | 1.461 | 0.016 | -0.016 | -0.016 | 0 | 0.35 | 0.74 | 0.44 | 0.56 | 0.29 | 0.46 |
| 9 | 91362843 | rs7869563 | -1.469 | 0.635 | 1.074 | 1.56 | 0.004 | -0.009 | -0.009 | 0.542 | 0.25 | 0.74 | 0.33 | 0.37 | 0.2 | 0.32 |
| 9 | 91365367 | rs4247466 | -0.715 | 0.963 | 1.662 | 1.753 | 0.008 | 0.106 | 0.106 | 0 | 0.15 | 0.27 | 0.35 | 0.37 | 0.18 | 0.23 |
| 9 | 91368043 | rs1857496 | -1.521 | 1.176 | 1.674 | 1.669 | 0.026 | 0.05 | 0.05 | 0 | 0.26 | 0.38 | 0.46 | 0.6 | 0.34 | 0.38 |
| 9 | 91370155 | rs17054421 | -1.171 | 1.184 | 1.744 | 1.632 | -0.001 | 0.058 | 0.058 | 0 | 0.15 | 0.19 | 0.19 | 0.39 | 0.15 | 0.21 |
| 9 | 91371086 | rs17054423 | -0.142 | 1.194 | 1.543 | 1.631 | 0 | 0.004 | 0.004 | 0 | 0.1 | 0.09 | 0.08 | 0.22 | 0.08 | 0.18 |
| 9 | 91377672 | rs7867153 | -0.986 | 1.391 | 1.636 | 1.571 | 0.036 | 0.112 | 0.112 | 0 | 0.17 | 0.31 | 0.38 | 0.4 | 0.26 | 0.29 |
| 9 | 91386221 | rs7048152 | -1.18 | 0.839 | 0.46 | 0.666 | 0.003 | 0.034 | 0.034 | 0 | 0.05 | 0.07 | 0.11 | 0.18 | 0.07 | 0.03 |
| 9 | 91392349 | rs7851181 | -0.571 | 0.511 | 0.672 | -0.268 | -0.008 | -0.142 | -0.142 | 0.405 | 0.22 | 0.44 | 0.14 | 0.01 | 0.18 | 0.1 |
| 9 | 91397359 | rs7038959 | -1.432 | 0.178 | 0.904 | -0.447 | -0.002 | -0.013 | -0.013 | 0 | 0.37 | 0.62 | 0.37 | 0.17 | 0.31 | 0.18 |
| 9 | 91399821 | rs9410340 | -1.375 | 0.232 | 1.257 | -0.404 | -0.01 | -0.041 | -0.041 | 0 | 0.37 | 0.53 | 0.35 | 0.18 | 0.31 | 0.17 |
| 9 | 91401893 | rs9410344 | -0.84 | 0.149 | 1.592 | -0.27 | -0.011 | 0.007 | 0.007 | 0 | 0.16 | 0.21 | 0.21 | 0.17 | 0.13 | 0.08 |
| 9 | 91415414 | rs7867530 | -0.998 | 0.119 | 1.99 | -0.294 | 0.081 | 0.184 | 0.184 | 0 | 0.06 | 0.24 | 0.16 | 0.17 | 0.09 | 0.03 |
| 9 | 91416645 | rs1288652 | 0.8 | 0.76 | 1.358 | -0.614 | -0.013 | 0.006 | 0.006 | 0 | 0.63 | 0.27 | 0.54 | 0.66 | 0.63 | 0.7 |
| 9 | 91423470 | rs2818251 | 0.791 | 0.756 | 1.355 | -0.616 | -0.014 | 0.005 | 0.005 | 0 | 0.63 | 0.28 | 0.55 | 0.67 | 0.63 | 0.7 |
| 9 | 91423611 | rs2653814 | -0.813 | 0.649 | 1.539 | -0.584 | 0.003 | -0.001 | -0.001 | 2.102 | 0.82 | 0.82 | 0.8 | 0.84 | 0.77 | 0.8 |
| 9 | 91425294 | rs7036143 | 1.177 | -0.086 | 0.527 | -0.881 | 0.08 | -0.032 | -0.032 | 0 | 0.34 | 0.17 | 0.3 | 0.61 | 0.22 | 0.41 |
| 9 | 91434617 | rs9410214 | -0.454 | 0.065 | 0.995 | -1.262 | -0.001 | -0.092 | -0.092 | 0 | 0.17 | 0.26 | 0.11 | 0 | 0.17 | 0.13 |
| 9 | 91438244 | rs879850 | 1.209 | 0.059 | 0.461 | -1.258 | 0.07 | 0.007 | 0.007 | 0 | 0.35 | 0.19 | 0.3 | 0.68 | 0.22 | 0.39 |
| 9 | 91444623 | rs9410375 | 1.372 | 0.309 | 0.496 | -1.267 | 0 | -0.007 | -0.007 | 0 | 0.05 | 0.05 | 0.04 | 0 | 0.11 | 0.07 |
| 9 | 91449366 | rs1779307 | -0.956 | 0.655 | 1.151 | -1.58 | 0.011 | -0.026 | -0.026 | 0 | 0.35 | 0.39 | 0.42 | 0.14 | 0.47 | 0.37 |
| 9 | 91456777 | rs2818249 | -0.982 | 0.423 | 1.324 | -1.563 | 0.004 | -0.014 | -0.014 | 0 | 0.7 | 0.66 | 0.72 | 0.81 | 0.71 | 0.76 |
| 9 | 91457113 | rs10780205 | 1.495 | 0.41 | 1.324 | -1.568 | 0.06 | 0.002 | 0.002 | 0 | 0.34 | 0.16 | 0.29 | 0.67 | 0.23 | 0.4 |
| 9 | 91460248 | rs2174057 | -1.002 | 0.419 | 1.421 | -1.565 | -0.003 | -0.003 | -0.003 | 2.787 | 0.35 | 0.35 | 0.42 | 0.13 | 0.47 | 0.36 |
| 9 | 91485533 | rs11137433 | 0.392 | 0.505 | 0.9 | -1.616 | -0.003 | -0.014 | -0.014 | 0.052 | 0.48 | 0.47 | 0.49 | 0.74 | 0.41 | 0.5 |
| 9 | 91485589 | rs7863591 | -0.152 | 0.564 | 0.881 | -1.593 | -0.029 | 0.057 | 0.057 | 0.809 | 0.78 | 0.88 | 0.77 | 0.93 | 0.7 | 0.73 |
| 9 | 91486658 | rs868073 | 0.258 | 0.539 | 0.881 | -1.578 | -0.022 | 0.002 | 0.002 | 0 | 0.11 | 0.04 | 0.17 | 0.06 | 0.18 | 0.1 |
| 9 | 91487768 | rs10867126 | 0.624 | 0.707 | 0.861 | -1.593 | 0.001 | -0.002 | -0.002 | 3.984 | 0.5 | 0.47 | 0.51 | 0.73 | 0.44 | 0.5 |
| 9 | 91488725 | rs987824 | 1.037 | 0.909 | 0.652 | -1.501 | -0.006 | 0.011 | 0.011 | 0 | 0.26 | 0.22 | 0.24 | 0.19 | 0.26 | 0.23 |
| 9 | 91489053 | rs6559300 | -0.122 | 0.91 | 0.601 | -1.522 | -0.013 | 0.022 | 0.022 | 0 | 0.61 | 0.65 | 0.58 | 0.73 | 0.51 | 0.56 |
| 9 | 91494640 | rs905363 | 0.313 | 0.638 | 0.166 | -1.41 | -0.021 | 0.055 | 0.055 | 0 | 0.31 | 0.17 | 0.29 | 0.19 | 0.32 | 0.26 |

| | | | | | | | | | | | | | | | | |
|---|---|---|---|---|---|---|---|---|---|---|---|---|---|---|---|---|
| 9 | 91502970 | rs1393570 | 0.612 | -0.339 | -0.207 | -1.731 | 0.002 | 0.089 | 0.089 | 0.41 | 0.42 | 0.29 | 0.28 | 0.2 | 0.37 | 0.32 |
| 9 | 91505370 | rs11137439 | 0.692 | -0.39 | -1.014 | -1.718 | -0.013 | -0.06 | -0.06 | 0.122 | 0.33 | 0.06 | 0.31 | 0.67 | 0.28 | 0.42 |
| 9 | 91511000 | rs10867131 | -0.362 | -0.179 | 0.096 | -1.506 | 0.065 | -0.058 | -0.058 | 0 | 0.84 | 0.56 | 0.79 | 0.93 | 0.77 | 0.79 |
| 9 | 91528751 | rs4128575 | -0.273 | -0.237 | -0.121 | -1.174 | 0.011 | 0.023 | 0.023 | 0 | 0.47 | 0.57 | 0.64 | 0.75 | 0.4 | 0.45 |
| 9 | 91530173 | rs1156619 | -1.066 | -0.131 | -0.187 | -1.21 | -0.087 | 0.072 | 0.072 | 0.504 | 0.62 | 0.91 | 0.8 | 0.77 | 0.61 | 0.64 |
| 9 | 91535231 | rs6559311 | -0.617 | -0.104 | 0 | -1.142 | -0.002 | 0.001 | 0.001 | 0 | 0.54 | 0.6 | 0.63 | 0.63 | 0.55 | 0.61 |
| 9 | 91535469 | rs17054591 | -0.298 | -0.114 | 0.075 | -1.105 | 0.085 | 0.196 | 0.196 | 0.647 | 0.08 | 0.37 | 0.18 | 0.14 | 0.06 | 0.03 |
| 9 | 91538158 | rs1502693 | 1.243 | 0.192 | -0.372 | -1.181 | -0.101 | 0.012 | 0.012 | 1.597 | 0.33 | 0.01 | 0.17 | 0.23 | 0.32 | 0.29 |
| 9 | 91539876 | rs7036574 | -0.182 | 0.317 | -0.151 | -1.063 | 0.02 | 0.038 | 0.038 | 0 | 0.46 | 0.62 | 0.58 | 0.71 | 0.4 | 0.44 |
| 9 | 91540059 | rs1875620 | 0.584 | 0.327 | -0.112 | -1.074 | 0.004 | 0.098 | 0.098 | 0 | 0.46 | 0.25 | 0.32 | 0.26 | 0.54 | 0.48 |
| 9 | 91541169 | rs7019570 | -0.232 | 0.314 | -0.139 | -1.028 | -0.015 | -0.009 | -0.009 | 0.227 | 0.95 | 0.99 | 0.92 | 0.97 | 0.94 | 0.92 |
| 9 | 91562311 | rs9410425 | 0.27 | 0.282 | -0.039 | -1.086 | 0.027 | -0.028 | -0.028 | 0 | 0.39 | 0.28 | 0.41 | 0.57 | 0.33 | 0.42 |
| 9 | 91562866 | rs9410427 | 0.035 | 0.436 | -0.301 | -1.071 | 0.01 | 0.059 | 0.059 | 0 | 0.48 | 0.36 | 0.34 | 0.27 | 0.54 | 0.48 |
| 9 | 91563034 | rs9410429 | 0.915 | 0.601 | -0.657 | -0.973 | -0.062 | 0.063 | 0.063 | 0.124 | 0.12 | 0.01 | 0.12 | 0.03 | 0.22 | 0.19 |
| 9 | 91569331 | rs6559318 | -0.601 | -0.037 | -0.282 | -0.722 | 0.01 | 0.086 | 0.086 | 0 | 0.52 | 0.71 | 0.68 | 0.73 | 0.46 | 0.52 |
| 9 | 91570795 | rs11137468 | -0.463 | 0.538 | 0.16 | -0.141 | -0.004 | -0.013 | -0.013 | 0.076 | 0.09 | 0.26 | 0.08 | 0.05 | 0.09 | 0.03 |
| 9 | 91575235 | rs12338537 | 0.046 | 0.951 | -0.203 | 0.07 | -0.021 | -0.004 | -0.004 | 0 | 0.71 | 0.74 | 0.47 | 0.29 | 0.57 | 0.52 |
| 9 | 91578102 | rs7854677 | -0.172 | 1.046 | -0.18 | 0.295 | -0.02 | -0.006 | -0.006 | 0 | 0.73 | 0.74 | 0.48 | 0.3 | 0.58 | 0.53 |
| 9 | 91581627 | rs6559328 | -0.36 | 1.015 | -0.358 | 0.312 | -0.039 | -0.067 | -0.067 | 0 | 0.77 | 0.85 | 0.7 | 0.44 | 0.63 | 0.57 |
| 9 | 91587228 | rs4570242 | -0.483 | 0.522 | -0.325 | -0.337 | -0.133 | -0.101 | -0.101 | 0.154 | 0.73 | 0.94 | 0.49 | 0.36 | 0.57 | 0.59 |
| 9 | 91591582 | rs7869712 | -0.28 | 0.514 | -0.219 | -0.498 | -0.066 | -0.029 | -0.029 | 0.22 | 0.71 | 0.83 | 0.48 | 0.33 | 0.55 | 0.51 |
| 9 | 91591680 | rs11137472 | -0.87 | 0.443 | 0.151 | -0.731 | 0.037 | -0.083 | -0.083 | 1.111 | 0.15 | 0.42 | 0.14 | 0.03 | 0.16 | 0.11 |
| 9 | 91599103 | rs4075163 | 0.957 | 0.392 | -0.115 | -0.735 | -0.002 | -0.007 | -0.007 | 0 | 0.57 | 0.52 | 0.36 | 0.31 | 0.42 | 0.44 |
| 9 | 91602796 | rs11791652 | 1.368 | 0.158 | -0.313 | -0.902 | -0.005 | -0.009 | -0.009 | 0.436 | 0.46 | 0.45 | 0.33 | 0.31 | 0.38 | 0.38 |
| 9 | 91604994 | rs7022664 | -0.692 | 0.042 | -0.875 | -0.749 | -0.128 | -0.085 | -0.085 | 0 | 0.73 | 0.95 | 0.5 | 0.34 | 0.58 | 0.54 |
| 9 | 91607869 | rs6559331 | 1.022 | 0.097 | -0.693 | -0.728 | 0.005 | 0.038 | 0.038 | 0 | 0.51 | 0.64 | 0.38 | 0.34 | 0.44 | 0.45 |
| 9 | 91608628 | rs10867149 | 0.226 | 0.457 | -0.759 | -0.303 | -0.025 | -0.002 | -0.002 | 1.359 | 0.66 | 0.77 | 0.47 | 0.35 | 0.54 | 0.51 |
| 9 | 91610543 | rs3934594 | 0.592 | 0.229 | -0.883 | -0.902 | 0.022 | 0.14 | 0.14 | 0 | 0.49 | 0.29 | 0.35 | 0.18 | 0.43 | 0.43 |
| 9 | 91610595 | rs11137481 | -0.024 | 0.492 | -0.897 | -1.52 | -0.008 | -0.004 | -0.004 | 0.426 | 0.06 | 0.09 | 0.03 | 0.05 | 0.05 | 0.05 |
| 9 | 91610929 | rs7045576 | -0.187 | 0.521 | -1.28 | -1.781 | -0.139 | 0.075 | 0.075 | 0.469 | 0.71 | 0.93 | 0.49 | 0.17 | 0.51 | 0.56 |
| 9 | 91613066 | rs7038457 | -0.403 | 0.185 | -0.479 | -1.946 | -0.011 | -0.021 | -0.021 | 0 | 0.15 | 0.34 | 0.09 | 0.09 | 0.1 | 0.12 |
| 9 | 91618102 | rs7853537 | -0.452 | 0.169 | -0.339 | -1.953 | -0.007 | -0.014 | -0.014 | 0 | 0.14 | 0.26 | 0.08 | 0.09 | 0.1 | 0.13 |
| 9 | 91620124 | rs9314668 | 0.809 | -0.433 | -0.818 | -2.872 | 0.033 | 0.084 | 0.084 | 0.078 | 0.29 | 0.13 | 0.24 | 0.03 | 0.24 | 0.17 |
| 9 | 91621197 | rs7865979 | 0.317 | 0.309 | -0.667 | -2.895 | -0.014 | 0.071 | 0.071 | 0.131 | 0.39 | 0.23 | 0.52 | 0.08 | 0.43 | 0.27 |
| 9 | 91622177 | rs3750396 | -0.755 | 0.126 | -0.37 | -2.982 | -0.045 | 0.142 | 0.142 | 0.06 | 0.57 | 0.84 | 0.63 | 0.17 | 0.53 | 0.4 |
| 9 | 91622691 | rs9410438 | 0.246 | 0.127 | -0.314 | -2.978 | -0.009 | -0.018 | -0.018 | 0 | 0.15 | 0.3 | 0.08 | 0.09 | 0.1 | 0.13 |
| 9 | 91623709 | rs2150611 | -0.727 | 0.133 | -0.381 | -2.965 | -0.042 | 0.138 | 0.138 | 0.042 | 0.57 | 0.83 | 0.63 | 0.17 | 0.53 | 0.39 |
| 9 | 91623811 | rs1125467 | 0.39 | 0.155 | -0.416 | -2.948 | -0.014 | 0.07 | 0.07 | 0.154 | 0.39 | 0.23 | 0.52 | 0.08 | 0.43 | 0.27 |
| 9 | 91625979 | rs944482 | -0.279 | 0.281 | -0.946 | -2.803 | -0.015 | -0.013 | -0.013 | 1.34 | 0.86 | 0.75 | 0.92 | 0.91 | 0.92 | 0.88 |
| 9 | 91630548 | rs1331180 | 0.791 | 0.178 | -1.721 | -2.963 | -0.02 | -0.008 | -0.008 | 0 | 0.24 | 0.08 | 0.27 | 0.79 | 0.29 | 0.4 |
| 9 | 91635785 | rs2316280 | 1.094 | 0.061 | -1.866 | -3.039 | -0.044 | 0.103 | 0.103 | 4.664 | 0.21 | 0.03 | 0.1 | 0.06 | 0.2 | 0.21 |
| 9 | 91637304 | rs10780221 | -0.046 | 0.119 | -1.369 | -2.698 | -0.007 | 0.064 | 0.064 | 0.063 | 0.45 | 0.58 | 0.57 | 0.15 | 0.47 | 0.34 |
| 9 | 91638212 | rs7029876 | -0.247 | 0.164 | -0.897 | -2.177 | -0.002 | -0.023 | -0.023 | 0 | 0.9 | 0.88 | 0.96 | 1 | 0.95 | 0.94 |
| 9 | 91641563 | rs12115280 | -0.639 | 0.371 | -1.16 | -2.232 | 0.009 | 0.014 | 0.014 | 0 | 0.15 | 0.13 | 0.07 | 0 | 0.11 | 0.09 |
| 9 | 91645705 | rs9410448 | 0.457 | 0.801 | -1.626 | -2.786 | 0.012 | 0.198 | 0.198 | 1.225 | 0.47 | 0.28 | 0.67 | 0.99 | 0.55 | 0.69 |
| 9 | 91653749 | rs4877041 | 0.64 | 1.425 | -1.705 | -2.829 | -0.029 | 0.267 | 0.267 | 1.753 | 0.44 | 0.16 | 0.49 | 0.98 | 0.45 | 0.63 |
| 9 | 91654117 | rs12337493 | -0.041 | 1.176 | -1.112 | -2.99 | 0.003 | -0.068 | -0.068 | 2.074 | 0.11 | 0.45 | 0.08 | 0 | 0.05 | 0.05 |
| 9 | 91656963 | rs3750399 | -0.295 | 1.217 | -1.23 | -3.093 | 0.016 | 0.098 | 0.098 | 1.021 | 0.62 | 0.8 | 0.77 | 0.99 | 0.63 | 0.76 |
| 9 | 91660158 | rs4877042 | -0.761 | 1.397 | -1.067 | -3.022 | 0.051 | -0.083 | -0.083 | 0.186 | 0.16 | 0.58 | 0.28 | 0 | 0.17 | 0.11 |
| 9 | 91662534 | rs6559335 | -1.375 | 1.385 | -1.322 | -3.136 | -0.03 | 0.031 | 0.031 | 0.243 | 0.3 | 0.69 | 0.4 | 0.01 | 0.34 | 0.16 |
| 9 | 91666487 | rs7855350 | 0.317 | 1.43 | -1.267 | -3.111 | 0.005 | -0.037 | -0.037 | 0 | 0.14 | 0.15 | 0.12 | 0.01 | 0.18 | 0.05 |
| 9 | 91666812 | rs7021743 | 0.252 | 1.397 | -1.658 | -3.18 | -0.017 | 0.072 | 0.072 | 0 | 0.65 | 0.72 | 0.65 | 1 | 0.67 | 0.71 |
| 9 | 91669362 | rs2297313 | 1.333 | 1.413 | -1.941 | -3.167 | -0.008 | 0.263 | 0.263 | 1.953 | 0.46 | 0.2 | 0.49 | 0.99 | 0.47 | 0.64 |
| 9 | 91672117 | rs9410307 | 1.405 | 1.103 | -2.007 | -3.147 | -0.029 | 0.291 | 0.291 | 2.563 | 0.45 | 0.15 | 0.49 | 0.98 | 0.47 | 0.63 |
| 9 | 91676380 | rs11789781 | 0.167 | 0.551 | -1.682 | -2.7 | -0.001 | -0.007 | -0.007 | 0 | 0.09 | 0.11 | 0.06 | 0 | 0.07 | 0.07 |
| 9 | 91677941 | rs9410453 | -0.579 | 0.382 | -1.747 | -2.53 | 0.026 | 0.138 | 0.138 | 0.169 | 0.77 | 0.96 | 0.9 | 1 | 0.83 | 0.82 |
| 9 | 91680600 | rs9410455 | 1.162 | 1.05 | -1.177 | -2.114 | -0.012 | 0.068 | 0.068 | 0.304 | 0.69 | 0.27 | 0.6 | 0.97 | 0.69 | 0.84 |
| 9 | 91710338 | rs7037454 | -1.03 | 1.049 | -0.409 | -2.108 | 0.001 | -0.083 | -0.083 | 0.052 | 0.23 | 0.36 | 0.27 | 0.01 | 0.19 | 0.14 |
| 9 | 91720619 | rs1331190 | 0.014 | 1.14 | -0.486 | -2.116 | 0.006 | -0.007 | -0.007 | 0 | 0.05 | 0.05 | 0.19 | 0 | 0.08 | 0.03 |
| 9 | 91726877 | rs4876968 | -1.008 | 1.132 | -0.479 | -2.142 | 0.004 | -0.011 | -0.011 | 0 | 0.08 | 0.09 | 0.2 | 0.01 | 0.09 | 0.06 |
| 9 | 91729926 | rs2183665 | -0.427 | 1.244 | -0.559 | -2.202 | -0.005 | 0.004 | 0.004 | 0 | 0.94 | 0.94 | 0.89 | 0.98 | 0.89 | 0.98 |
| 9 | 91768648 | rs4534195 | 2.076 | 1.607 | -0.639 | -2.425 | 0.002 | -0.051 | -0.051 | 0 | 0.68 | 0.6 | 0.6 | 0.97 | 0.66 | 0.78 |
| 9 | 91771148 | rs4876971 | -1.243 | 1.619 | -0.682 | -2.358 | 0.005 | -0.079 | -0.079 | 0.513 | 0.22 | 0.36 | 0.31 | 0 | 0.19 | 0.15 |
| 9 | 91771642 | rs10867158 | -1.595 | 1.66 | -0.657 | -2.366 | 0.005 | -0.085 | -0.085 | 0 | 0.26 | 0.4 | 0.32 | 0.01 | 0.21 | 0.19 |
| 9 | 91801585 | rs13285368 | -0.537 | 1.663 | -0.673 | -2.355 | 0.026 | -0.028 | -0.028 | 0 | 0.07 | 0.16 | 0.24 | 0 | 0.1 | 0.04 |
| 9 | 91805113 | rs11137533 | -0.097 | 1.649 | -0.68 | -2.345 | -0.001 | -0.03 | -0.03 | 0 | 0.14 | 0.14 | 0.07 | 0 | 0.1 | 0.1 |
| 9 | 91820998 | rs4877062 | -1.813 | 2.309 | -1.065 | -0.6 | 0.007 | 0.011 | 0.011 | 0.009 | 0.29 | 0.36 | 0.56 | 0.51 | 0.34 | 0.35 |
| 9 | 91838806 | rs17437738 | 0.355 | 2.352 | -1.252 | -0.603 | -0.036 | 0.025 | 0.025 | 0 | 0.06 | 0.01 | 0.06 | 0.01 | 0.13 | 0.07 |
| 9 | 91841972 | rs11137545 | 1.492 | 2.166 | -1.989 | -1.006 | 0.024 | -0.014 | -0.014 | 2.386 | 0.33 | 0.11 | 0.16 | 0.27 | 0.3 | 0.37 |

| | | | | | | | | | | | | | | | | |
|---|---|---|---|---|---|---|---|---|---|---|---|---|---|---|---|---|
| 9 | 91878702 | rs10441779 | -0.185 | 1.81 | -1.714 | -0.731 | -0.027 | 0.056 | 0.056 | 0 | 0.2 | 0.13 | 0.13 | 0.01 | 0.23 | 0.18 |
| 9 | 91900556 | rs1980888 | -0.37 | 1.896 | -1.6 | -0.274 | 0.029 | 0.188 | 0.188 | 0 | 0.09 | 0.19 | 0.43 | 0.46 | 0.11 | 0.17 |
| 9 | 91908783 | rs1556046 | -0.714 | 1.332 | -2.303 | -0.061 | -0.02 | 0.019 | 0.019 | 0.197 | 0.6 | 0.79 | 0.73 | 0.36 | 0.56 | 0.45 |
| 9 | 91920946 | rs11137559 | -0.259 | 0.913 | -2.385 | 0.376 | -0.008 | -0.087 | -0.087 | 0.143 | 0.23 | 0.07 | 0.21 | 0.39 | 0.28 | 0.36 |
| 9 | 91926088 | rs3211663 | 0.564 | 0.466 | -2.56 | -0.216 | -0.016 | 0.021 | 0.021 | 0 | 0.06 | 0.02 | 0.05 | 0.02 | 0.09 | 0.03 |
| 9 | 91932309 | rs3211703 | 0.152 | 0.946 | -2.525 | 0.08 | 0.004 | 0.008 | 0.008 | 0 | 0.09 | 0.06 | 0.38 | 0.24 | 0.15 | 0.16 |
| 9 | 91934787 | rs3763616 | -0.929 | 1.28 | -2.848 | -0.064 | -0.072 | -0.1 | -0.1 | 0.278 | 0.81 | 0.97 | 0.78 | 0.7 | 0.71 | 0.75 |
| 9 | 91953252 | rs7851988 | -0.97 | 0.862 | -2.069 | 0.29 | -0.002 | -0.004 | -0.004 | 0 | 0.28 | 0.31 | 0.46 | 0.41 | 0.29 | 0.44 |
| 9 | 91960826 | rs7470318 | 0.52 | 0.692 | -2.106 | 0.158 | 0.007 | 0.087 | 0.087 | 0 | 0.14 | 0 | 0.04 | 0 | 0.09 | 0.1 |
| 9 | 91976520 | rs4876978 | -1.053 | 0.828 | -2.058 | 0.274 | -0.003 | -0.006 | -0.006 | 0 | 0.28 | 0.31 | 0.47 | 0.41 | 0.3 | 0.44 |
| 9 | 91978159 | rs4876979 | -0.404 | 0.76 | -1.999 | 0.31 | 0.011 | 0.052 | 0.052 | 0 | 0.12 | 0.15 | 0.4 | 0.41 | 0.17 | 0.34 |
| 9 | 91979438 | rs13283164 | -0.528 | 0.842 | -1.872 | 0.361 | 0 | 0.002 | 0.002 | 0 | 0.28 | 0.29 | 0.46 | 0.41 | 0.26 | 0.43 |
| 9 | 91981877 | rs1134004 | 0.789 | 0.094 | -2.421 | 0.082 | -0.06 | -0.07 | -0.07 | 1.811 | 0.21 | 0.02 | 0.13 | 0.27 | 0.22 | 0.17 |
| 9 | 91984571 | rs7856532 | -0.435 | -0.697 | -1.525 | -0.444 | -0.004 | 0 | 0 | 0 | 0.42 | 0.4 | 0.33 | 0.41 | 0.24 | 0.49 |
| 9 | 91989539 | rs4877077 | -0.326 | -0.523 | -1.338 | -0.421 | -0.017 | 0.014 | 0.014 | 0 | 0.23 | 0.32 | 0.24 | 0.27 | 0.14 | 0.38 |
| 9 | 91990950 | rs34731447 | 0.339 | 0.31 | -1.48 | -0.703 | -0.05 | 0.016 | 0.016 | 0 | 0.08 | 0.01 | 0.25 | 0.03 | 0.14 | 0.12 |
| 9 | 91991069 | rs36050908 | -0.042 | -0.56 | -0.969 | -0.813 | 0.008 | -0.002 | -0.002 | 0.008 | 0.25 | 0.16 | 0.14 | 0 | 0.21 | 0.11 |
| 9 | 91997938 | rs7469438 | -0.657 | -0.719 | -0.844 | -0.762 | 0.008 | 0.006 | 0.006 | 2.375 | 0.34 | 0.3 | 0.18 | 0.09 | 0.27 | 0.22 |
| 9 | 92002060 | rs28402866 | -0.696 | -1.125 | -0.497 | -0.262 | -0.033 | 0.057 | 0.057 | 0 | 0.1 | 0.23 | 0.13 | 0.2 | 0.03 | 0.23 |
| 9 | 92007225 | rs6559349 | -0.608 | -1.375 | -0.59 | -0.491 | -0.057 | -0.025 | -0.025 | 0.01 | 0.31 | 0.38 | 0.22 | 0.2 | 0.16 | 0.29 |
| 9 | 92010796 | rs6559350 | 0.568 | -0.618 | -0.458 | -0.771 | 0.017 | -0.024 | -0.024 | 0 | 0.79 | 0.72 | 0.75 | 0.91 | 0.74 | 0.78 |
| 9 | 92016916 | rs7045789 | 0.384 | -1.705 | -0.577 | -0.563 | -0.051 | 0.038 | 0.038 | 2.84 | 0.29 | 0.5 | 0.3 | 0.59 | 0.16 | 0.25 |
| 9 | 92021256 | rs9410475 | 0.825 | -1.56 | -0.589 | -0.702 | -0.021 | 0.008 | 0.008 | 0 | 0.12 | 0.1 | 0.12 | 0 | 0.23 | 0.08 |
| 9 | 92030186 | rs9410321 | -0.103 | -1.75 | -0.675 | -0.792 | -0.019 | 0.031 | 0.031 | 0.017 | 0.23 | 0.16 | 0.18 | 0.09 | 0.3 | 0.21 |
| 9 | 92034938 | rs4074459 | -0.442 | -1.41 | -1.174 | -1.284 | 0.009 | 0.068 | 0.068 | 0.723 | 0.46 | 0.36 | 0.35 | 0.09 | 0.53 | 0.39 |
| 9 | 92036427 | rs4132699 | -0.614 | -1.671 | -0.96 | -0.528 | -0.003 | -0.001 | -0.001 | 0 | 0.41 | 0.51 | 0.25 | 0.39 | 0.39 | 0.31 |
| 9 | 92039416 | rs11265876 | -0.273 | -1.548 | -1.179 | -0.621 | -0.003 | 0.004 | 0.004 | 0 | 0.3 | 0.31 | 0.2 | 0.1 | 0.33 | 0.21 |
| 9 | 92040475 | rs10512197 | 1.679 | -1.327 | -1.356 | -0.416 | -0.029 | -0.021 | -0.021 | 0 | 0.17 | 0.11 | 0.22 | 0.23 | 0.27 | 0.22 |
| 9 | 92043184 | rs10797126 | 0.188 | -1.39 | -1.336 | -0.577 | -0.015 | 0.02 | 0.02 | 0 | 0.21 | 0.2 | 0.15 | 0.1 | 0.26 | 0.16 |
| 9 | 92047433 | rs4078338 | -0.814 | -1.573 | -1.019 | -0.485 | -0.01 | -0.014 | -0.014 | 0 | 0.7 | 0.8 | 0.64 | 0.38 | 0.67 | 0.59 |
| 9 | 92052258 | rs4073713 | 0.712 | -1.875 | -0.617 | -0.523 | -0.043 | -0.017 | -0.017 | 0 | 0.33 | 0.39 | 0.34 | 0.29 | 0.2 | 0.24 |
| 9 | 92053124 | rs7041468 | -0.1 | -1.796 | -0.668 | -0.548 | 0.007 | 0.028 | 0.028 | 0 | 0.57 | 0.69 | 0.51 | 0.38 | 0.43 | 0.51 |
| 9 | 92062038 | rs9330435 | -0.069 | -1.79 | -0.758 | -0.493 | 0.011 | 0.004 | 0.004 | 0 | 0.29 | 0.38 | 0.19 | 0.27 | 0.31 | 0.15 |
| 9 | 92067803 | rs7871577 | 0.547 | -1.7 | -1.118 | -0.685 | -0.044 | 0.007 | 0.007 | 3.466 | 0.41 | 0.48 | 0.37 | 0.57 | 0.26 | 0.31 |
| 9 | 92069671 | rs9410486 | 0.398 | -1.072 | -1.965 | -0.847 | -0.103 | 0.131 | 0.131 | 3.037 | 0.3 | 0.13 | 0.27 | 0 | 0.45 | 0.29 |
| 9 | 92070391 | rs11999884 | 1.211 | -0.688 | -1.772 | -0.289 | -0.047 | -0.045 | -0.045 | 0.311 | 0.17 | 0.06 | 0.22 | 0.25 | 0.27 | 0.22 |
| 9 | 92074429 | rs4360376 | 1.124 | -0.794 | -1.6 | -0.442 | -0.079 | 0.101 | 0.101 | 1.178 | 0.17 | 0.01 | 0.14 | 0 | 0.21 | 0.21 |
| 9 | 92076340 | rs4560885 | -0.025 | -2.015 | -1.397 | 0.078 | 0.007 | -0.017 | -0.017 | 0 | 0.73 | 0.75 | 0.6 | 0.54 | 0.86 | 0.69 |
| 9 | 92077132 | rs11265908 | -0.156 | -1.885 | -1.497 | -0.118 | 0.039 | -0.021 | -0.021 | 0 | 0.44 | 0.24 | 0.43 | 0.53 | 0.42 | 0.4 |
| 9 | 92081623 | rs17054916 | -0.807 | -1.757 | -0.677 | 0.132 | 0.057 | 0.136 | 0.136 | 0 | 0.9 | 0.67 | 0.85 | 0.82 | 0.92 | 0.8 |
| 9 | 92093576 | rs7859211 | 1.111 | -1.868 | -0.841 | 0.282 | -0.024 | 0.004 | 0.004 | 0.006 | 0.1 | 0.16 | 0.17 | 0.16 | 0.06 | 0.06 |
| 9 | 92100804 | rs4405007 | 0.645 | -1.837 | -1.181 | 0.169 | -0.014 | 0.018 | 0.018 | 0 | 0.23 | 0.24 | 0.16 | 0.11 | 0.28 | 0.13 |
| 9 | 92105483 | rs9695510 | 0.919 | -1.861 | -1.266 | 0.567 | -0.024 | 0.043 | 0.043 | 0.653 | 0.11 | 0.16 | 0.24 | 0.34 | 0.06 | 0.14 |
| 9 | 92111105 | rs13298567 | 1.136 | -1.144 | -1.66 | 0.28 | -0.003 | -0.003 | -0.003 | 0 | 0.15 | 0.14 | 0.17 | 0.33 | 0.21 | 0.19 |
| 9 | 92111838 | rs10908892 | 1.492 | -0.97 | -1.799 | -0.26 | 0.006 | 0.013 | 0.013 | 0 | 0.13 | 0.05 | 0.08 | 0 | 0.11 | 0.03 |
| 9 | 92116564 | rs11265780 | 0.55 | -0.842 | -2.003 | -0.452 | -0.032 | 0.206 | 0.206 | 1.402 | 0.37 | 0.09 | 0.25 | 0 | 0.35 | 0.22 |
| 9 | 92117102 | rs10908893 | 1.824 | -0.648 | -1.866 | 0.28 | 0.003 | -0.032 | -0.032 | 0.017 | 0.08 | 0.01 | 0.1 | 0.31 | 0.06 | 0.14 |
| 9 | 92117752 | rs1571534 | 0.18 | -0.521 | -1.473 | 0.199 | 0.003 | 0.034 | 0.034 | 0.556 | 0.52 | 0.4 | 0.44 | 0.27 | 0.55 | 0.45 |
| 9 | 92121724 | rs11265783 | 0.5 | 0.108 | -1.66 | 0.076 | -0.042 | -0.017 | -0.017 | 0.256 | 0.32 | 0.17 | 0.31 | 0.31 | 0.34 | 0.34 |
| 9 | 92125867 | rs2988071 | 0.607 | 0.334 | -1.435 | 0.248 | -0.017 | -0.078 | -0.078 | 0.213 | 0.25 | 0.07 | 0.26 | 0.35 | 0.28 | 0.27 |
| 9 | 92128645 | rs2149879 | 0.349 | 0.525 | -0.703 | 0.15 | -0.02 | -0.045 | -0.045 | 1.298 | 0.29 | 0.2 | 0.31 | 0.45 | 0.32 | 0.4 |
| 9 | 92143868 | rs13290596 | 0.779 | 0.61 | -0.699 | -0.644 | -0.016 | 0.106 | 0.106 | 0.559 | 0.23 | 0.08 | 0.17 | 0.01 | 0.21 | 0.18 |
| 9 | 92148993 | rs3002331 | -0.13 | 0.129 | -0.449 | -0.503 | 0.007 | 0.002 | 0.002 | 1.227 | 0.53 | 0.61 | 0.51 | 0.39 | 0.57 | 0.53 |
| 9 | 92154969 | rs1983976 | -1.008 | -0.42 | -0.693 | -0.615 | -0.025 | -0.027 | -0.027 | 4.125 | 0.7 | 0.78 | 0.68 | 0.6 | 0.63 | 0.63 |
| 9 | 92156146 | rs12000581 | 1.336 | -0.417 | -0.625 | -0.868 | -0.004 | 0.073 | 0.073 | 0.524 | 0.23 | 0.12 | 0.17 | 0.04 | 0.21 | 0.3 |
| 9 | 92158225 | rs2480490 | 0.195 | -0.441 | -0.135 | -0.758 | -0.001 | 0 | 0 | 0 | 0.34 | 0.4 | 0.33 | 0.4 | 0.34 | 0.28 |
| 9 | 92158829 | rs913807 | 1.208 | -0.63 | -0.637 | -0.68 | -0.027 | -0.143 | -0.143 | 1.339 | 0.24 | 0.04 | 0.25 | 0.4 | 0.25 | 0.24 |
| 9 | 92167606 | rs12345210 | 0.084 | -0.987 | 0.069 | -1.13 | 0.004 | -0.075 | -0.075 | 0 | 0.2 | 0.32 | 0.18 | 0.03 | 0.16 | 0.1 |
| 9 | 92170123 | rs10908894 | -0.14 | -1.072 | -0.098 | -1.12 | 0.013 | 0.03 | 0.03 | 0 | 0.35 | 0.32 | 0.34 | 0.21 | 0.28 | 0.36 |
| 9 | 92173910 | rs6559362 | -0.995 | -1.063 | 0.15 | -1.119 | 0.022 | 0.033 | 0.033 | 0 | 0.44 | 0.66 | 0.39 | 0.2 | 0.31 | 0.37 |
| 9 | 92186458 | rs7042552 | -0.585 | -0.501 | 0.039 | -1.202 | -0.006 | 0.011 | 0.011 | 0 | 0.62 | 0.65 | 0.51 | 0.21 | 0.55 | 0.45 |
| 9 | 92191467 | rs10797116 | -0.61 | -0.498 | 0.036 | -1.162 | -0.008 | 0.018 | 0.018 | 0 | 0.62 | 0.66 | 0.52 | 0.22 | 0.55 | 0.43 |
| 9 | 92196438 | rs12343854 | 0.869 | -0.575 | -0.44 | -1.479 | 0.005 | -0.022 | -0.022 | 0.377 | 0.34 | 0.31 | 0.22 | 0.02 | 0.28 | 0.27 |
| 9 | 92196724 | rs10908898 | 0.421 | 0.162 | -0.824 | -0.421 | -0.08 | -0.095 | -0.095 | 0.005 | 0.69 | 0.9 | 0.52 | 0.48 | 0.54 | 0.69 |
| 9 | 92204172 | rs2031970 | 0.126 | -0.217 | -0.303 | -0.818 | -0.021 | 0.035 | 0.035 | 1.968 | 0.19 | 0.07 | 0.24 | 0.14 | 0.24 | 0.09 |
| 9 | 92209587 | rs7871395 | 0.177 | 0.107 | 0.197 | -0.462 | 0.054 | 0.04 | 0.04 | 0 | 0.16 | 0.34 | 0.25 | 0.24 | 0.22 | 0.15 |
| 9 | 92212750 | rs1475537 | -0.235 | 0.031 | 0.261 | -0.508 | 0.032 | 0.01 | 0.01 | 0 | 0.19 | 0.3 | 0.26 | 0.24 | 0.24 | 0.16 |
| 9 | 92213923 | rs1055595 | -0.56 | -0.177 | -0.14 | -0.228 | 0 | 0.003 | 0.003 | 0 | 0.53 | 0.51 | 0.51 | 0.38 | 0.52 | 0.43 |
| 9 | 92213967 | rs1007966 | -0.561 | -0.223 | -0.137 | -0.867 | 0 | 0.005 | 0.005 | 0 | 0.53 | 0.51 | 0.47 | 0.26 | 0.52 | 0.43 |

| | | | | | | | | | | | | | | | | |
|---|---|---|---|---|---|---|---|---|---|---|---|---|---|---|---|---|
| 9 | 92215638 | rs1571536 | -0.5 | -0.249 | -0.118 | -0.811 | -0.002 | -0.016 | -0.016 | 4.84 | 0.52 | 0.42 | 0.35 | 0.26 | 0.53 | 0.4 |
| 9 | 92218862 | rs3138489 | -0.684 | -0.272 | -0.333 | -0.822 | 0 | 0.02 | 0.02 | 0 | 0.56 | 0.59 | 0.37 | 0.26 | 0.53 | 0.4 |
| 9 | 92222495 | rs3138513 | 1.399 | 0.021 | -0.534 | -0.396 | -0.033 | 0.027 | 0.027 | 0 | 0.06 | 0.01 | 0.12 | 0 | 0.14 | 0.09 |
| 9 | 92228559 | rs10908903 | -0.586 | -0.461 | -0.407 | -0.432 | 0.002 | 0.035 | 0.035 | 0 | 0.49 | 0.59 | 0.34 | 0.27 | 0.49 | 0.39 |
| 9 | 92231020 | rs870926 | 0.975 | -0.696 | -0.699 | -0.41 | 0.015 | 0.123 | 0.123 | 0.012 | 0.22 | 0.01 | 0.06 | 0.02 | 0.13 | 0.19 |
| 9 | 92231894 | rs10908904 | -0.095 | 0.04 | 0.383 | -0.025 | 0.127 | 0.019 | 0.019 | 0.21 | 0.14 | 0.47 | 0.13 | 0.17 | 0.22 | 0.07 |
| 9 | 92234041 | rs7851960 | -0.729 | -0.425 | -0.057 | 0.236 | 0 | 0 | 0 | 0 | 0.55 | 0.49 | 0.4 | 0.51 | 0.44 | 0.52 |
| 9 | 92237200 | rs7850947 | 0.012 | -0.662 | -0.142 | 0.187 | 0.001 | -0.001 | -0.001 | 0 | 0.15 | 0.12 | 0.2 | 0.33 | 0.09 | 0.3 |
| 9 | 92238244 | rs10908905 | 0.906 | -0.096 | -0.11 | 0.247 | -0.007 | -0.003 | -0.003 | 0.175 | 0.1 | 0.07 | 0.09 | 0.01 | 0.21 | 0.04 |
| 9 | 92239995 | rs7851166 | -0.396 | -0.098 | -0.15 | 0.365 | -0.002 | -0.002 | -0.002 | 0 | 0.53 | 0.46 | 0.39 | 0.49 | 0.44 | 0.52 |
| 9 | 92248027 | rs10124343 | 0.787 | 0.518 | -0.397 | 0.816 | 0.007 | -0.001 | -0.001 | 0 | 0.66 | 0.37 | 0.5 | 0.63 | 0.47 | 0.5 |
| 9 | 92249584 | rs10908907 | 0.051 | 0.249 | -0.611 | 0.954 | 0.086 | -0.007 | -0.007 | 0 | 0.53 | 0.68 | 0.54 | 0.51 | 0.28 | 0.64 |
| 9 | 92250365 | rs4877115 | -0.571 | 1.125 | -0.685 | 1.047 | -0.057 | -0.055 | -0.055 | 0.012 | 0.91 | 0.99 | 0.83 | 0.74 | 0.79 | 0.83 |
| 9 | 92254212 | rs4876989 | -0.372 | -0.335 | -0.103 | 1.113 | 0.032 | 0.171 | 0.171 | 1.337 | 0.09 | 0.37 | 0.2 | 0.23 | 0.07 | 0.25 |
| 9 | 92254248 | rs4877116 | -0.129 | -0.317 | -0.112 | 0.765 | 0.036 | 0.032 | 0.032 | 1.553 | 0.09 | 0.35 | 0.2 | 0.07 | 0.06 | 0.18 |
| 9 | 92261864 | rs1411156 | -0.36 | -0.043 | -0.588 | 0.797 | -0.078 | -0.021 | -0.021 | 0 | 0.81 | 0.9 | 0.73 | 0.76 | 0.56 | 0.78 |
| 9 | 92270655 | rs6559371 | -0.029 | -0.349 | -0.513 | 0.703 | -0.002 | 0.009 | 0.009 | 0 | 0.36 | 0.44 | 0.47 | 0.52 | 0.35 | 0.33 |
| 9 | 92273175 | rs10908909 | -0.2 | -0.568 | -0.56 | 0.415 | 0.005 | 0 | 0 | 8.193 | 0.66 | 0.64 | 0.57 | 0.63 | 0.51 | 0.58 |
| 9 | 92276116 | rs7848364 | 0.531 | -0.562 | -0.77 | 0.743 | 0.009 | 0.18 | 0.18 | 0 | 0.07 | 0.23 | 0.23 | 0.21 | 0.04 | 0.14 |
| 9 | 92277210 | rs1109998 | 0.96 | -0.514 | -0.87 | 0.505 | -0.013 | 0.005 | 0.005 | 0 | 0.22 | 0.15 | 0.2 | 0.19 | 0.28 | 0.13 |
| 9 | 92279448 | rs4877120 | 0.938 | -0.45 | -1.027 | 0.494 | -0.035 | 0.01 | 0.01 | 0 | 0.23 | 0.06 | 0.2 | 0.2 | 0.29 | 0.16 |
| 9 | 92281255 | rs11265860 | 0.187 | 0.393 | -0.213 | 0.727 | 0.008 | 0.158 | 0.158 | 0 | 0.07 | 0.24 | 0.24 | 0.19 | 0.04 | 0.15 |
| 9 | 92281403 | rs4876993 | 0.368 | 0.377 | -0.331 | 0.679 | 0.011 | -0.005 | -0.005 | 0 | 0.61 | 0.52 | 0.54 | 0.63 | 0.51 | 0.62 |
| 9 | 92285084 | rs10908911 | -0.708 | 0.57 | 0.094 | 0.468 | 0.006 | 0.176 | 0.176 | 0 | 0.09 | 0.33 | 0.24 | 0.19 | 0.04 | 0.15 |
| 9 | 92290874 | rs4876951 | 0.509 | 1.099 | -0.595 | 0.003 | 0.044 | 0.02 | 0.02 | 0 | 0.68 | 0.4 | 0.48 | 0.62 | 0.56 | 0.63 |
| 9 | 92301982 | rs4074312 | -0.567 | 0.613 | -1.062 | -0.933 | -0.003 | -0.031 | -0.031 | 0 | 0.11 | 0.11 | 0.08 | 0 | 0.09 | 0.01 |
| 9 | 92308660 | rs11265870 | -1.166 | 0.483 | -1.316 | -1.233 | 0.006 | -0.01 | -0.01 | 0 | 0.2 | 0.15 | 0.11 | 0.01 | 0.16 | 0.08 |
| 9 | 92309687 | rs10114707 | -0.117 | 0.736 | -2.04 | -1.504 | -0.076 | 0.178 | 0.178 | 0.235 | 0.71 | 0.97 | 0.88 | 0.98 | 0.73 | 0.85 |
| 9 | 92319527 | rs11265877 | -0.494 | 0.415 | -1.017 | -1.394 | 0.004 | 0.01 | 0.01 | 1.622 | 0.11 | 0.04 | 0.09 | 0 | 0.09 | 0.01 |
| 9 | 92328208 | rs11265882 | -0.701 | 0.329 | -0.785 | -1.436 | 0 | -0.023 | -0.023 | 0.318 | 0.12 | 0.11 | 0.09 | 0 | 0.09 | 0.01 |
| 9 | 92336018 | rs11790339 | -0.477 | 0.364 | -1.056 | -1.733 | -0.001 | -0.013 | -0.013 | 0 | 0.12 | 0.1 | 0.09 | 0.01 | 0.13 | 0.03 |
| 9 | 92444702 | rs12115250 | -0.057 | 0.072 | -0.594 | -1.786 | -0.007 | -0.035 | -0.035 | 0.133 | 0.07 | 0.14 | 0.05 | 0 | 0.05 | 0.03 |
| 9 | 92444823 | rs10797129 | -1.339 | 0.155 | -0.515 | -2.127 | -0.002 | -0.06 | -0.06 | 0.256 | 0.42 | 0.53 | 0.29 | 0.13 | 0.36 | 0.14 |
| 9 | 92448136 | rs12376091 | -0.17 | 0.226 | -0.511 | -2.228 | -0.012 | 0.017 | 0.017 | 0 | 0.23 | 0.16 | 0.19 | 0.09 | 0.25 | 0.06 |
| 9 | 92449055 | rs9722251 | -1.482 | 0.165 | -0.418 | -2.225 | -0.002 | -0.064 | -0.064 | 0 | 0.42 | 0.53 | 0.29 | 0.12 | 0.36 | 0.14 |
| 9 | 92454568 | rs9696601 | -0.702 | 0.145 | -0.473 | -2.134 | -0.018 | -0.064 | -0.064 | 0.998 | 0.15 | 0.23 | 0.08 | 0.03 | 0.11 | 0.08 |
| 9 | 92456953 | rs9722389 | -1.2 | 0.169 | -0.691 | -2.179 | -0.017 | 0.114 | 0.114 | 0 | 0.39 | 0.23 | 0.25 | 0.11 | 0.36 | 0.15 |
| 9 | 92457327 | rs11265920 | -0.997 | 0.288 | -0.623 | -2.121 | -0.001 | 0.006 | 0.006 | 0.019 | 0.34 | 0.32 | 0.23 | 0.11 | 0.3 | 0.12 |
| 9 | 92459463 | rs10908933 | 0.084 | -0.13 | -0.334 | -1.556 | -0.001 | 0.03 | 0.03 | 0 | 0.46 | 0.38 | 0.48 | 0.24 | 0.52 | 0.38 |
| 9 | 92464383 | rs9330440 | -0.991 | -0.263 | -0.487 | -1.205 | -0.003 | 0.003 | 0.003 | 0 | 0.65 | 0.64 | 0.67 | 0.43 | 0.68 | 0.47 |
| 9 | 92485878 | rs9777548 | 0.447 | 0.373 | -0.701 | -0.882 | -0.088 | -0.055 | -0.055 | 4.765 | 0.31 | 0.14 | 0.37 | 0.43 | 0.49 | 0.36 |
| 9 | 92486638 | rs10797131 | -0.905 | 0.202 | -0.308 | -1.012 | -0.01 | -0.002 | -0.002 | 0 | 0.55 | 0.48 | 0.51 | 0.43 | 0.68 | 0.43 |
| 9 | 92489679 | rs10116554 | -0.1 | 0.289 | -0.598 | -0.974 | -0.026 | -0.025 | -0.025 | 0 | 0.41 | 0.14 | 0.42 | 0.43 | 0.58 | 0.39 |
| 9 | 92492597 | rs10908940 | -1.068 | 0.105 | -0.445 | -0.918 | 0 | -0.001 | -0.001 | 0 | 0.57 | 0.6 | 0.55 | 0.46 | 0.68 | 0.47 |
| 9 | 92494316 | rs9330447 | -0.72 | 0.573 | -0.2 | -0.428 | -0.001 | -0.001 | -0.001 | 0 | 0.44 | 0.48 | 0.46 | 0.42 | 0.57 | 0.36 |
| 9 | 92494807 | rs9696730 | 0.379 | 0.914 | -0.865 | -0.349 | -0.07 | -0.057 | -0.057 | 0.015 | 0.29 | 0.13 | 0.34 | 0.42 | 0.42 | 0.33 |
| 9 | 92495076 | rs10908943 | -0.186 | 0.528 | -0.836 | 0.072 | 0.01 | 0.002 | 0.002 | 0 | 0.46 | 0.56 | 0.51 | 0.45 | 0.58 | 0.41 |
| 9 | 92498089 | rs11265957 | -0.478 | 0.769 | -1.138 | 0.713 | -0.005 | -0.033 | -0.033 | 0.377 | 0.33 | 0.22 | 0.37 | 0.48 | 0.32 | 0.42 |
| 9 | 92499064 | rs11265961 | 0.043 | 0.334 | -1.064 | 0.79 | 0.03 | -0.004 | -0.004 | 0 | 0.43 | 0.3 | 0.3 | 0.41 | 0.34 | 0.53 |
| 9 | 92499739 | rs12003053 | 0.22 | 0.928 | -1.241 | 0.569 | -0.021 | 0.012 | 0.012 | 0.298 | 0.23 | 0.12 | 0.3 | 0.16 | 0.27 | 0.08 |
| 9 | 92499771 | rs11265963 | 0.107 | 0.81 | -1.29 | 0.519 | -0.034 | 0.013 | 0.013 | 0 | 0.24 | 0.07 | 0.29 | 0.16 | 0.27 | 0.08 |
| 9 | 92500586 | rs9775766 | -0.335 | 1.045 | -0.65 | 0.645 | 0.036 | 0.003 | 0.003 | 0.006 | 0.72 | 0.53 | 0.64 | 0.68 | 0.65 | 0.65 |
| 9 | 92501369 | rs9330449 | 0.566 | 0.685 | -0.358 | 0.202 | -0.002 | -0.047 | -0.047 | 0.159 | 0.65 | 0.55 | 0.64 | 0.84 | 0.65 | 0.91 |
| 9 | 92501780 | rs9775506 | 0.164 | 0.313 | -0.24 | 0.422 | -0.006 | -0.013 | -0.013 | 0 | 0.23 | 0.38 | 0.29 | 0.16 | 0.2 | 0.07 |
| 9 | 92503272 | rs9330451 | 0.099 | 0.529 | -1.054 | 0.879 | -0.073 | -0.072 | -0.072 | 0 | 0.67 | 0.82 | 0.56 | 0.54 | 0.52 | 0.64 |
| 9 | 92509057 | rs12342215 | 0.796 | -0.299 | -1.224 | 0.239 | 0.039 | 0.078 | 0.078 | 0 | 0.19 | 0.05 | 0.1 | 0 | 0.14 | 0.22 |
| 9 | 92510619 | rs9777581 | 0.43 | 0.604 | -0.902 | 1.016 | -0.019 | -0.018 | -0.018 | 0.021 | 0.6 | 0.79 | 0.44 | 0.43 | 0.46 | 0.57 |
| 9 | 92515514 | rs9330454 | 0.522 | 0.465 | -0.632 | 0.737 | 0.004 | 0.004 | 0.004 | 0.023 | 0.57 | 0.58 | 0.39 | 0.35 | 0.43 | 0.52 |
| 9 | 92524438 | rs10122995 | -0.372 | 0.31 | -0.73 | 0.559 | 0.001 | 0.004 | 0.004 | 3.027 | 0.14 | 0.14 | 0.1 | 0.1 | 0.17 | 0.08 |
| 9 | 92526756 | rs10114488 | -0.945 | 0.105 | -0.48 | 0.527 | -0.018 | -0.02 | -0.02 | 0 | 0.3 | 0.45 | 0.31 | 0.25 | 0.28 | 0.29 |
| 9 | 92679450 | rs16905869 | 0.308 | 0.209 | -0.992 | 0.974 | -0.002 | -0.037 | -0.037 | 0.022 | 0.16 | 0.12 | 0.25 | 0.36 | 0.18 | 0.19 |
| 9 | 92681245 | rs7029990 | -0.183 | -0.231 | -1.459 | 0.321 | 0.013 | 0.04 | 0.04 | 0 | 0.19 | 0.11 | 0.16 | 0.05 | 0.15 | 0.2 |
| 9 | 92685454 | rs7021867 | -0.63 | -0.295 | -1.313 | 0.314 | 0.014 | -0.034 | -0.034 | 0 | 0.33 | 0.19 | 0.32 | 0.41 | 0.3 | 0.37 |
| 9 | 92686142 | rs4744475 | 0.102 | -0.381 | -1.285 | 0.222 | -0.017 | 0.035 | 0.035 | 0 | 0.14 | 0.01 | 0.07 | 0.05 | 0.12 | 0.18 |
| 9 | 92689363 | rs7863076 | -0.434 | -1.48 | -1.176 | -0.077 | 0.045 | 0.004 | 0.004 | 0 | 0.44 | 0.64 | 0.36 | 0.52 | 0.25 | 0.35 |
| 9 | 92691598 | rs10991825 | 0.667 | -1.321 | -1.344 | 0.38 | 0.05 | -0.038 | -0.038 | 1.132 | 0.17 | 0.03 | 0.1 | 0.18 | 0.11 | 0.25 |
| 9 | 92693362 | rs10820846 | -0.743 | -1.061 | -1.571 | 0.12 | 0.1 | 0.058 | 0.058 | 0.108 | 0.53 | 0.84 | 0.43 | 0.57 | 0.35 | 0.42 |
| 9 | 92693434 | rs10761114 | -0.743 | -1.061 | -1.571 | 0.12 | 0.1 | 0.058 | 0.058 | 0.108 | 0.53 | 0.84 | 0.43 | 0.57 | 0.35 | 0.42 |
| 9 | 92700500 | rs10992228 | 1.496 | -0.584 | -1.91 | 0.044 | -0.035 | 0.012 | 0.012 | 0 | 0.19 | 0.04 | 0.19 | 0.14 | 0.21 | 0.09 |

| | | | | | | | | | | | | | | | |
|---|---|---|---|---|---|---|---|---|---|---|---|---|---|---|---|
| 9 | 92704074 | rs10992338 | 1.018 | -0.975 | -0.579 | 0.218 | -0.026 | 0.056 | 0.056 | 0.493 | 0.26 | 0.38 | 0.18 | 0.35 | 0.2 | 0.1 |
| 9 | 92711789 | rs2196923 | 0.402 | -0.945 | -0.809 | 0.238 | -0.004 | 0.001 | 0.001 | 0.659 | 0.32 | 0.61 | 0.21 | 0.35 | 0.2 | 0.07 |
| 9 | 92713999 | rs7045961 | 1.007 | -0.526 | -0.906 | 0.089 | 0.003 | -0.026 | -0.026 | 0.029 | 0.12 | 0.19 | 0.13 | 0.07 | 0.14 | 0.29 |
| 9 | 92716596 | rs16905942 | 0.706 | -0.464 | -1.354 | 0.15 | 0.023 | -0.062 | -0.062 | 0 | 0.18 | 0.01 | 0.09 | 0.19 | 0.12 | 0.32 |
| 9 | 92722593 | rs7035196 | -0.902 | 0.077 | -1.104 | 0.14 | -0.042 | 0.012 | 0.012 | 0 | 0.61 | 0.83 | 0.51 | 0.61 | 0.47 | 0.69 |
| 9 | 92723125 | rs10993172 | 0.155 | 0.065 | -0.838 | 0.167 | -0.005 | -0.002 | -0.002 | 0.242 | 0.45 | 0.4 | 0.28 | 0.54 | 0.32 | 0.39 |
| 9 | 92725441 | rs7030989 | 1.059 | 0.181 | -0.63 | 0.108 | 0.056 | -0.037 | -0.037 | 0.11 | 0.13 | 0.41 | 0.16 | 0.07 | 0.15 | 0.27 |
| 9 | 92727048 | rs10993333 | 0.191 | 0.1 | -0.68 | 0.23 | 0.066 | -0.017 | -0.017 | 1.292 | 0.45 | 0.26 | 0.28 | 0.44 | 0.32 | 0.39 |
| 9 | 92727336 | rs10993342 | 0.269 | 0.103 | -0.679 | 0.232 | 0.065 | -0.015 | -0.015 | 0.755 | 0.45 | 0.27 | 0.34 | 0.44 | 0.32 | 0.4 |
| 9 | 92728873 | rs6479587 | 0.93 | 0.204 | -0.601 | 0.251 | 0.052 | -0.04 | -0.04 | 1.739 | 0.13 | 0.41 | 0.16 | 0.07 | 0.15 | 0.27 |
| 9 | 92739891 | rs10821475 | -0.744 | 0.202 | -0.931 | 0.37 | -0.003 | -0.006 | -0.006 | 0 | 0.59 | 0.68 | 0.5 | 0.51 | 0.47 | 0.67 |
| 9 | 92742478 | rs2890110 | 0.468 | -0.067 | -1.057 | 0.371 | 0.015 | 0.01 | 0.01 | 0.024 | 0.2 | 0.17 | 0.11 | 0.1 | 0.12 | 0.32 |
| 9 | 92748385 | rs12685019 | 0.671 | 1.047 | -1.183 | 0.435 | -0.024 | -0.026 | -0.026 | 0.023 | 0.06 | 0.01 | 0.21 | 0.22 | 0.21 | 0.15 |
| 9 | 92755133 | rs10821487 | -0.325 | 0.84 | -1.158 | 0.382 | -0.003 | -0.006 | -0.006 | 1.872 | 0.29 | 0.24 | 0.37 | 0.31 | 0.31 | 0.47 |
| 9 | 92763235 | rs7866392 | -0.699 | 1.043 | -1.228 | 0.376 | -0.009 | -0.004 | -0.004 | 0 | 0.6 | 0.64 | 0.51 | 0.51 | 0.52 | 0.67 |
| 9 | 92765694 | rs1036699 | 0.82 | 1.027 | -1.315 | 0.383 | -0.027 | 0.001 | 0.001 | 0.273 | 0.19 | 0.01 | 0.12 | 0.1 | 0.17 | 0.31 |
| 9 | 92769678 | rs10993701 | 1.423 | 0.755 | -1.226 | 0.132 | 0.017 | -0.031 | -0.031 | 0.733 | 0.23 | 0.17 | 0.17 | 0.34 | 0.18 | 0.07 |
| 9 | 92773125 | rs535599 | 1.1 | 0.949 | -1.341 | 0.141 | -0.007 | 0.021 | 0.021 | 0 | 0.33 | 0.21 | 0.28 | 0.28 | 0.33 | 0.18 |
| 9 | 92773905 | rs620311 | -1.375 | 0.789 | -1.773 | -0.015 | 0.003 | 0.105 | 0.105 | 0.14 | 0.83 | 1 | 0.97 | 1 | 0.9 | 0.83 |
| 9 | 92776497 | rs652454 | 0.533 | 0.915 | -1.717 | 0.512 | -0.011 | -0.011 | -0.011 | 0.008 | 0.41 | 0.34 | 0.49 | 0.6 | 0.47 | 0.31 |
| 9 | 92776614 | rs652913 | -1.003 | 0.919 | -1.771 | 0.554 | -0.027 | -0.047 | -0.047 | 2.819 | 0.79 | 0.99 | 0.83 | 0.85 | 0.82 | 0.68 |
| 9 | 92776695 | rs601472 | 0.933 | 1.137 | -1.408 | 0.855 | -0.002 | -0.005 | -0.005 | 0 | 0.32 | 0.26 | 0.29 | 0.4 | 0.32 | 0.17 |
| 9 | 92789486 | rs535414 | 1.502 | 1.357 | -1.598 | 0.253 | -0.018 | 0.04 | 0.04 | 0 | 0.15 | 0.09 | 0.13 | 0.04 | 0.18 | 0.09 |
| 9 | 92793780 | rs10991758 | 0.317 | 1.447 | -1.132 | 0.767 | -0.004 | -0.005 | -0.005 | 0 | 0.07 | 0.05 | 0.2 | 0.2 | 0.15 | 0.15 |
| 9 | 92794117 | rs10991761 | 1.375 | 1.27 | -1.226 | 0.741 | 0.004 | -0.059 | -0.059 | 0 | 0.19 | 0.04 | 0.18 | 0.2 | 0.17 | 0.06 |
| 9 | 92794619 | rs684476 | 0.493 | 1.316 | -1.139 | 0.758 | -0.017 | 0.007 | 0.007 | 0 | 0.41 | 0.31 | 0.5 | 0.46 | 0.48 | 0.33 |
| 9 | 92803759 | rs7049060 | -0.642 | 1.502 | -1.145 | 0.757 | -0.004 | 0.004 | 0.004 | 0 | 0.51 | 0.8 | 0.56 | 0.47 | 0.54 | 0.58 |
| 9 | 92804166 | rs10991799 | -0.507 | 1.632 | -1.276 | 0.87 | -0.006 | -0.006 | -0.006 | 0.808 | 0.08 | 0.06 | 0.2 | 0.21 | 0.15 | 0.15 |
| 9 | 92809471 | rs10122211 | -0.079 | 0.672 | -0.748 | 0.993 | -0.004 | -0.019 | -0.019 | 0.746 | 0.4 | 0.55 | 0.35 | 0.27 | 0.4 | 0.45 |
| 9 | 92814420 | rs997758 | -0.141 | 0.632 | -0.745 | 1.028 | 0.002 | -0.012 | -0.012 | 0 | 0.39 | 0.49 | 0.36 | 0.27 | 0.41 | 0.45 |
| 9 | 92814783 | rs997759 | -0.263 | 0.841 | -1.127 | 1.154 | 0 | -0.001 | -0.001 | 0.223 | 0.43 | 0.61 | 0.45 | 0.43 | 0.41 | 0.54 |
| 9 | 92816727 | rs10761113 | 0.902 | 0.815 | -1.267 | 1.149 | 0.005 | -0.014 | -0.014 | 0 | 0.18 | 0.16 | 0.19 | 0.2 | 0.16 | 0.07 |
| 9 | 92817191 | rs599721 | 0.674 | 0.908 | -1.624 | 0.894 | -0.002 | 0.007 | 0.007 | 0 | 0.43 | 0.34 | 0.34 | 0.31 | 0.45 | 0.2 |
| 9 | 92830743 | rs544950 | -0.568 | 1.008 | -2.121 | 0.577 | -0.036 | -0.079 | -0.079 | 0 | 0.87 | 0.99 | 0.86 | 0.75 | 0.86 | 0.76 |
| 9 | 92838827 | rs287860 | 0.356 | 0.877 | -1.533 | 0.068 | -0.003 | 0.051 | 0.051 | 0 | 0.4 | 0.26 | 0.31 | 0.19 | 0.44 | 0.2 |
| 9 | 92847174 | rs645706 | 0.026 | 0.971 | -1.165 | 0.537 | -0.003 | -0.007 | -0.007 | 0 | 0.58 | 0.33 | 0.56 | 0.6 | 0.61 | 0.37 |
| 9 | 92848755 | rs11792411 | -0.601 | 1.347 | -0.96 | 0.475 | 0.011 | -0.005 | -0.005 | 0 | 0.35 | 0.4 | 0.46 | 0.34 | 0.49 | 0.4 |
| 9 | 92850878 | rs4744088 | -0.195 | 0.598 | -0.852 | 0.725 | 0.005 | -0.007 | -0.007 | 0.348 | 0.23 | 0.2 | 0.25 | 0.26 | 0.22 | 0.11 |
| 9 | 92852199 | rs1329045 | -0.304 | 0.87 | -0.594 | 0.855 | 0.009 | 0.08 | 0.08 | 0.015 | 0.86 | 0.81 | 0.67 | 0.51 | 0.84 | 0.68 |
| 9 | 92852944 | rs7874990 | 1.066 | 0.874 | -0.342 | 0.433 | -0.007 | 0.052 | 0.052 | 0 | 0.45 | 0.34 | 0.27 | 0.2 | 0.45 | 0.48 |
| 9 | 92855420 | rs10123553 | 0.394 | 0.768 | -0.554 | 0.348 | 0.01 | 0.027 | 0.027 | 1.193 | 0.46 | 0.29 | 0.38 | 0.37 | 0.46 | 0.21 |
| 9 | 92855977 | rs10992049 | 1.056 | 0.804 | -0.588 | 0.234 | -0.006 | -0.063 | -0.063 | 0.053 | 0.17 | 0.25 | 0.15 | 0.05 | 0.16 | 0.09 |
| 9 | 92856946 | rs423239 | 1.098 | 0.794 | -0.83 | 0.232 | -0.003 | -0.042 | -0.042 | 0.043 | 0.17 | 0.21 | 0.15 | 0.05 | 0.17 | 0.09 |
| 9 | 92858197 | rs10761128 | 1.175 | 0.801 | -1.163 | 0.246 | 0.015 | -0.044 | -0.044 | 0 | 0.15 | 0.05 | 0.08 | 0.21 | 0.12 | 0.07 |
| 9 | 92859704 | rs16906191 | 0.992 | 0.917 | -1.07 | -0.026 | -0.006 | -0.076 | -0.076 | 0 | 0.18 | 0.28 | 0.15 | 0.05 | 0.16 | 0.09 |
| 9 | 92862486 | rs4744096 | -1.062 | 1.486 | -1.763 | 0.677 | -0.057 | -0.07 | -0.07 | 0.709 | 0.89 | 0.99 | 0.75 | 0.77 | 0.86 | 0.85 |
| 9 | 92863041 | rs10992083 | 0.516 | 0.83 | -1.322 | 0.254 | 0.002 | -0.025 | -0.025 | 0 | 0.22 | 0.22 | 0.13 | 0.09 | 0.22 | 0.38 |
| 9 | 92867363 | rs4743853 | -0.199 | 0.337 | -0.742 | 0.682 | -0.012 | -0.004 | -0.004 | 0 | 0.35 | 0.62 | 0.26 | 0.33 | 0.29 | 0.18 |
| 9 | 92869370 | rs16906203 | -0.892 | 0.852 | -0.684 | 1.029 | 0.006 | 0.053 | 0.053 | 0.11 | 0.07 | 0.11 | 0.28 | 0.27 | 0.12 | 0.17 |
| 9 | 92870113 | rs1854444 | 0.421 | 0.569 | -0.835 | 0.022 | -0.001 | -0.007 | -0.007 | 0.869 | 0.65 | 0.69 | 0.57 | 0.81 | 0.64 | 0.63 |
| 9 | 92870296 | rs1329047 | -0.383 | 0.247 | -1.204 | -0.807 | 0.018 | 0.067 | 0.067 | 0.22 | 0.4 | 0.33 | 0.26 | 0.11 | 0.36 | 0.42 |
| 9 | 92870425 | rs7852983 | 0.104 | 0.466 | -1.21 | -0.12 | -0.005 | 0.015 | 0.015 | 0 | 0.51 | 0.34 | 0.42 | 0.58 | 0.48 | 0.45 |
| 9 | 92872227 | rs7860989 | 0.107 | 0.102 | -1.258 | -0.566 | 0.021 | 0.048 | 0.048 | 0 | 0.67 | 0.79 | 0.81 | 0.85 | 0.76 | 0.66 |
| 9 | 92872306 | rs7851675 | -0.482 | -0.019 | -1.468 | -0.731 | -0.008 | 0.071 | 0.071 | 0.52 | 0.88 | 0.99 | 0.96 | 1 | 0.92 | 0.84 |
| 9 | 92872751 | rs17460947 | 1.162 | 0.651 | -1.98 | -1.11 | -0.08 | 0.086 | 0.086 | 0 | 0.19 | 0.04 | 0.16 | 0.03 | 0.26 | 0.1 |
| 9 | 92873665 | rs4744106 | 0.373 | 0.498 | -1.425 | -0.399 | -0.005 | 0 | 0 | 0 | 0.42 | 0.38 | 0.64 | 0.43 | 0.51 | 0.41 |
| 9 | 92875390 | rs7042866 | 1.507 | 0.237 | -1.176 | -0.158 | -0.003 | 0.046 | 0.046 | 0 | 0.18 | 0.21 | 0.12 | 0.32 | 0.17 | 0.18 |
| 9 | 92876060 | rs7848094 | -0.829 | 0.545 | -1.224 | 0.19 | -0.011 | 0.006 | 0.006 | 1.289 | 0.68 | 0.66 | 0.83 | 0.68 | 0.75 | 0.65 |
| 9 | 92876666 | rs4743859 | 0.2 | 0.273 | -1.169 | 0.809 | -0.002 | -0.028 | -0.028 | 0.02 | 0.23 | 0.2 | 0.47 | 0.46 | 0.26 | 0.32 |
| 9 | 92887130 | rs10992192 | 0.425 | 0.213 | -1.62 | 0.453 | -0.019 | -0.053 | -0.053 | 0 | 0.19 | 0.05 | 0.44 | 0.3 | 0.2 | 0.26 |
| 9 | 92887231 | rs2081670 | -0.16 | 0.159 | -0.929 | 0.089 | 0.002 | 0.003 | 0.003 | 0.096 | 0.22 | 0.22 | 0.15 | 0.16 | 0.17 | 0.18 |
| 9 | 92891077 | rs964912 | 0.108 | 0.59 | -1.006 | 0.237 | -0.004 | 0.021 | 0.021 | 0.024 | 0.44 | 0.27 | 0.57 | 0.64 | 0.49 | 0.47 |
| 9 | 92891865 | rs17445269 | 0.993 | 0.693 | -1.055 | 0.509 | -0.013 | -0.001 | -0.001 | 0 | 0.2 | 0.04 | 0.12 | 0.22 | 0.22 | 0.19 |
| 9 | 92892015 | rs10992211 | -0.151 | 0.458 | -0.734 | -0.25 | -0.012 | -0.039 | -0.039 | 0 | 0.27 | 0.4 | 0.14 | 0.21 | 0.24 | 0.13 |
| 9 | 92896723 | rs4744117 | 0.436 | 0.436 | -0.56 | -0.752 | -0.007 | -0.049 | -0.049 | 0 | 0.6 | 0.44 | 0.68 | 0.81 | 0.64 | 0.79 |
| 9 | 92897297 | rs7867301 | -0.295 | 0.338 | -0.518 | -0.789 | -0.007 | -0.003 | -0.003 | 0 | 0.12 | 0.35 | 0.09 | 0.08 | 0.08 | 0.06 |
| 9 | 92897338 | rs10481717 | -1.434 | 0.515 | -0.569 | -0.748 | -0.04 | 0.086 | 0.086 | 0 | 0.86 | 0.99 | 0.84 | 0.99 | 0.85 | 0.93 |
| 9 | 92898149 | rs7875016 | 0.32 | 0.455 | -0.084 | -0.41 | 0.038 | 0.031 | 0.031 | 0 | 0.07 | 0.33 | 0.06 | 0.06 | 0.06 | 0.03 |
| 9 | 92898177 | rs12352194 | -0.561 | 0.313 | -0.29 | -0.541 | -0.007 | -0.045 | -0.045 | 0.431 | 0.23 | 0.32 | 0.15 | 0.15 | 0.22 | 0.36 |

| | | | | | | | | | | | | | | | | |
|---|---|---|---|---|---|---|---|---|---|---|---|---|---|---|---|---|
| 9 | 92900609 | rs10992247 | 0.601 | 0.38 | -0.927 | -0.313 | -0.021 | -0.003 | -0.003 | 0.169 | 0.08 | 0 | 0.06 | 0.05 | 0.09 | 0.28 |
| 9 | 92903522 | rs7025302 | -1.803 | 0.632 | -1.104 | 0.498 | -0.016 | -0.011 | -0.011 | 0.008 | 0.44 | 0.83 | 0.37 | 0.57 | 0.41 | 0.51 |
| 9 | 92903671 | rs10820944 | -0.147 | 0.675 | -0.954 | 0.738 | 0.003 | 0.014 | 0.014 | 0 | 0.09 | 0.07 | 0.16 | 0.19 | 0.13 | 0.09 |
| 9 | 92906025 | rs12337287 | 1.535 | 0.649 | -0.892 | 0.769 | 0.002 | -0.003 | -0.003 | 0 | 0.05 | 0.06 | 0.01 | 0.1 | 0.04 | 0.02 |
| 9 | 92907061 | rs7860895 | -1.812 | 0.659 | -0.883 | 0.725 | -0.014 | -0.047 | -0.047 | 2.878 | 0.39 | 0.77 | 0.36 | 0.47 | 0.37 | 0.49 |
| 9 | 92907960 | rs10820953 | -0.121 | 0.666 | -0.919 | 0.73 | 0.003 | 0.015 | 0.015 | 0 | 0.08 | 0.07 | 0.16 | 0.19 | 0.13 | 0.07 |
| 9 | 92909654 | rs10820958 | -1.822 | 0.652 | -0.933 | 0.721 | -0.016 | -0.051 | -0.051 | 0.225 | 0.39 | 0.79 | 0.36 | 0.47 | 0.37 | 0.49 |
| 9 | 92910367 | rs10992278 | -0.21 | 0.73 | -0.938 | 0.603 | 0.001 | 0.011 | 0.011 | 0 | 0.09 | 0.07 | 0.16 | 0.19 | 0.13 | 0.07 |
| 9 | 92925179 | rs7468504 | -1.324 | 0.549 | -1.047 | 0.879 | -0.015 | -0.025 | -0.025 | 0 | 0.34 | 0.81 | 0.35 | 0.61 | 0.33 | 0.23 |
| 9 | 92927997 | rs10992326 | -1.604 | 0.564 | -0.957 | 0.979 | -0.001 | 0.033 | 0.033 | 0 | 0.43 | 0.82 | 0.4 | 0.66 | 0.4 | 0.53 |
| 9 | 92931861 | rs4744131 | 1.958 | 0.513 | -1.18 | 1.022 | -0.016 | 0.029 | 0.029 | 0.069 | 0.23 | 0.04 | 0.15 | 0.21 | 0.25 | 0.2 |
| 9 | 92941210 | rs7048754 | -1.343 | 0.307 | 0.276 | 1.414 | -0.011 | 0.041 | 0.041 | 0 | 0.28 | 0.61 | 0.32 | 0.55 | 0.22 | 0.46 |
| 9 | 92943286 | rs1854923 | -1.008 | 0.407 | 0.503 | 1.079 | 0.009 | -0.023 | -0.023 | 0 | 0.38 | 0.55 | 0.31 | 0.29 | 0.43 | 0.14 |
| 9 | 92946648 | rs10761166 | -0.885 | 0.076 | 0.79 | 0.993 | 0.002 | -0.041 | -0.041 | 0 | 0.18 | 0.31 | 0.14 | 0.13 | 0.22 | 0.2 |
| 9 | 92950272 | rs1539519 | -2.042 | -0.281 | 0.725 | 1.25 | -0.017 | -0.02 | -0.02 | 1.065 | 0.42 | 0.45 | 0.29 | 0.32 | 0.36 | 0.54 |
| 9 | 92960794 | rs1854924 | 0.514 | 0.014 | 0.604 | 1.428 | -0.036 | -0.009 | -0.009 | 0 | 0.12 | 0.02 | 0.14 | 0.07 | 0.17 | 0.03 |
| 9 | 92961558 | rs12347445 | 0.195 | 0.142 | 1.099 | 1.682 | 0.005 | -0.004 | -0.004 | 0 | 0.09 | 0.14 | 0.07 | 0.1 | 0.14 | 0.08 |
| 9 | 92962977 | rs6479430 | -0.711 | 0.103 | 1.163 | 1.988 | 0.059 | 0.34 | 0.34 | 0 | 0.06 | 0.31 | 0.18 | 0.36 | 0.07 | 0.14 |
| 9 | 92963339 | rs10992452 | -0.755 | 0.1 | 0.416 | 1.396 | 0.006 | -0.01 | -0.01 | 0.054 | 0.1 | 0.06 | 0.11 | 0.12 | 0.06 | 0.02 |
| 9 | 92967125 | rs7023271 | -0.688 | 0.062 | 0.532 | 1.036 | -0.02 | -0.037 | -0.037 | 0.951 | 0.09 | 0.11 | 0.02 | 0 | 0.03 | 0.03 |
| 9 | 92992878 | rs12345191 | -0.105 | 0.065 | 0.223 | 1.088 | -0.005 | -0.001 | -0.001 | 0 | 0.12 | 0 | 0.04 | 0.08 | 0.1 | 0.3 |
| 9 | 93005808 | rs7023466 | -0.324 | 0.223 | 0.353 | 1.28 | -0.031 | -0.019 | -0.019 | 0 | 0.17 | 0.11 | 0.21 | 0.22 | 0.26 | 0.17 |
| 9 | 93008041 | rs10821091 | -1.991 | 0.123 | 0.983 | 1.402 | 0.011 | -0.018 | -0.018 | 0.228 | 0.39 | 0.47 | 0.49 | 0.67 | 0.42 | 0.59 |
| 9 | 93010541 | rs7043530 | -2.376 | 0.084 | 1.04 | 1.385 | 0.002 | 0.06 | 0.06 | 0 | 0.49 | 0.61 | 0.51 | 0.67 | 0.45 | 0.62 |
| 9 | 93015989 | rs7865330 | -2.182 | 0.084 | 1.056 | 1.384 | -0.001 | 0.02 | 0.02 | 0 | 0.47 | 0.55 | 0.5 | 0.67 | 0.45 | 0.59 |
| 9 | 93025338 | rs1777018 | -2.803 | 0.011 | 0.898 | 1.308 | -0.014 | 0.087 | 0.087 | 0 | 0.58 | 0.76 | 0.62 | 0.76 | 0.59 | 0.7 |
| 9 | 93025656 | rs1777019 | -2.695 | 0.02 | 1.184 | 1.313 | 0 | -0.006 | -0.006 | 0 | 0.54 | 0.54 | 0.6 | 0.76 | 0.59 | 0.68 |
| 9 | 93029002 | rs2380291 | -1.851 | -0.255 | 1.167 | 1.464 | -0.064 | 0.062 | 0.062 | 0 | 0.31 | 0.43 | 0.3 | 0.45 | 0.19 | 0.44 |
| 9 | 93029282 | rs11792210 | -0.656 | -0.26 | 0.62 | 1.396 | 0.02 | -0.016 | -0.016 | 0 | 0.16 | 0.08 | 0.11 | 0.2 | 0.09 | 0.34 |
| 9 | 93031263 | rs12346947 | 0.391 | -0.252 | 0.522 | 1.335 | 0.029 | -0.009 | -0.009 | 0 | 0.11 | 0 | 0.03 | 0.08 | 0.07 | 0.29 |
| 9 | 93033040 | rs1410856 | -2.608 | -0.076 | 0.789 | 1.413 | -0.002 | -0.002 | -0.002 | 0.977 | 0.52 | 0.6 | 0.45 | 0.49 | 0.53 | 0.61 |
| 9 | 93036856 | rs10992761 | 0.15 | -0.09 | 0.378 | 1.35 | 0.033 | 0.013 | 0.013 | 0 | 0.12 | 0.03 | 0.04 | 0.08 | 0.07 | 0.32 |
| 9 | 93039262 | rs10821149 | -0.284 | 0.052 | 0.597 | 1.395 | 0.01 | 0.033 | 0.033 | 0.668 | 0.12 | 0.22 | 0.16 | 0.2 | 0.17 | 0.11 |
| 9 | 93039369 | rs2771091 | -0.64 | 0.096 | -0.122 | 1.426 | -0.034 | -0.04 | -0.04 | 0 | 0.21 | 0.14 | 0.28 | 0.33 | 0.28 | 0.2 |
| 9 | 93046322 | rs2380292 | -0.944 | -0.027 | -0.16 | 1.251 | 0.035 | 0.032 | 0.032 | 0.051 | 0.2 | 0.09 | 0.06 | 0.08 | 0.09 | 0.32 |
| 9 | 93049422 | rs1777003 | -0.661 | 0.369 | 0.093 | 1.439 | 0.004 | 0.001 | 0.001 | 0 | 0.19 | 0.21 | 0.23 | 0.23 | 0.27 | 0.18 |
| 9 | 93051826 | rs2771095 | -2.518 | 0.427 | -0.103 | 1.7 | -0.011 | 0.095 | 0.095 | 0.009 | 0.47 | 0.79 | 0.6 | 0.69 | 0.55 | 0.63 |
| 9 | 93058963 | rs2771098 | -2.276 | 0.459 | -0.141 | 1.721 | -0.027 | 0.014 | 0.014 | 0.261 | 0.45 | 0.76 | 0.55 | 0.58 | 0.55 | 0.62 |
| 9 | 93065137 | rs10217176 | -0.648 | 0.653 | -0.187 | 1.654 | -0.016 | 0.017 | 0.017 | 0 | 0.15 | 0.09 | 0.12 | 0.11 | 0.21 | 0.1 |
| 9 | 93067144 | rs2771102 | -2.096 | 0.647 | -0.072 | 1.745 | 0 | 0.002 | 0.002 | 0 | 0.32 | 0.6 | 0.48 | 0.59 | 0.35 | 0.54 |
| 9 | 93067790 | rs1754069 | -0.29 | 0.837 | -0.099 | 1.85 | 0.007 | 0.013 | 0.013 | 0 | 0.1 | 0.1 | 0.19 | 0.2 | 0.19 | 0.11 |
| 9 | 93071368 | rs729009 | -1.621 | 0.653 | -0.066 | 1.657 | 0.075 | 0.12 | 0.12 | 0.3 | 0.23 | 0.54 | 0.42 | 0.38 | 0.34 | 0.27 |
| 9 | 93071559 | rs1318387 | 0.363 | 0.52 | -0.56 | 1.512 | -0.02 | -0.008 | -0.008 | 1.465 | 0.1 | 0.02 | 0.07 | 0.1 | 0.12 | 0.09 |
| 9 | 93080054 | rs2771090 | -0.891 | 0.651 | -0.345 | 1.654 | 0.06 | 0.07 | 0.07 | 1.924 | 0.1 | 0.27 | 0.21 | 0.19 | 0.17 | 0.07 |
| 9 | 93080194 | rs2246522 | -2.228 | 0.3 | -0.745 | 1.587 | -0.041 | -0.099 | -0.099 | 0 | 0.43 | 0.95 | 0.46 | 0.42 | 0.42 | 0.53 |
| 9 | 93081851 | rs16906332 | 0.345 | 0.248 | -0.687 | 1.516 | 0.025 | 0.01 | 0.01 | 0 | 0.1 | 0.03 | 0.04 | 0.07 | 0.07 | 0.23 |
| 9 | 93083867 | rs16936969 | 0.345 | 0.248 | -0.687 | 1.516 | 0.025 | 0.01 | 0.01 | 0 | 0.1 | 0.03 | 0.04 | 0.07 | 0.07 | 0.24 |
| 9 | 93085382 | rs10993008 | -0.654 | 0.325 | -0.486 | 1.555 | -0.029 | -0.009 | -0.009 | 0.179 | 0.22 | 0.51 | 0.21 | 0.18 | 0.18 | 0.22 |
| 9 | 93090915 | rs2798025 | -1.709 | 0.356 | -0.674 | 1.553 | -0.013 | -0.033 | -0.033 | 0 | 0.32 | 0.82 | 0.42 | 0.37 | 0.35 | 0.3 |
| 9 | 93093378 | rs6479525 | -1.418 | 0.527 | -0.546 | 1.27 | 0.016 | -0.012 | -0.012 | 0 | 0.46 | 0.65 | 0.4 | 0.49 | 0.4 | 0.58 |
| 9 | 93095891 | rs1777045 | -1.382 | 0.44 | -1.045 | 0.92 | -0.058 | 0.03 | 0.03 | 1.89 | 0.43 | 0.88 | 0.58 | 0.61 | 0.5 | 0.42 |
| 9 | 93098295 | rs2798032 | 0.108 | 0.48 | -1.259 | 0.941 | 0.038 | 0.017 | 0.017 | 2.119 | 0.06 | 0.16 | 0.14 | 0.05 | 0.12 | 0.02 |
| 9 | 93099104 | rs1590811 | -1.768 | 0.148 | -1.828 | 0.687 | -0.103 | 0.064 | 0.064 | 1.062 | 0.55 | 0.96 | 0.62 | 0.68 | 0.58 | 0.67 |
| 9 | 93100550 | rs12348363 | -0.872 | -0.093 | -0.635 | 0.686 | 0.025 | 0.021 | 0.021 | 0 | 0.12 | 0.02 | 0.04 | 0.07 | 0.08 | 0.25 |
| 9 | 93102793 | rs1012161 | -1.185 | 0.422 | 0.072 | 1.014 | 0.041 | 0.061 | 0.061 | 0.058 | 0.2 | 0.57 | 0.28 | 0.32 | 0.26 | 0.25 |
| 9 | 93103388 | rs979464 | -0.395 | -0.384 | -0.398 | 0.753 | -0.009 | -0.01 | -0.01 | 0 | 0.15 | 0.02 | 0.08 | 0.15 | 0.13 | 0.11 |
| 9 | 93104329 | rs7048728 | -1.152 | -0.616 | 0.853 | 0.549 | 0.03 | 0.021 | 0.021 | 3.884 | 0.08 | 0.35 | 0.07 | 0.07 | 0.06 | 0.03 |
| 9 | 93105008 | rs1329092 | -1.879 | -0.714 | 0.93 | 0.595 | -0.013 | -0.004 | -0.004 | 0.045 | 0.12 | 0.25 | 0.08 | 0.11 | 0.1 | 0.17 |
| 9 | 93106543 | rs979814 | -2.869 | -1.025 | 1.637 | 0.656 | -0.035 | -0.006 | -0.006 | 0.885 | 0.19 | 0.64 | 0.14 | 0.16 | 0.12 | 0.27 |
| 9 | 93106801 | rs979816 | -2.472 | -1.024 | 1.734 | 0.656 | -0.038 | -0.017 | -0.017 | 0 | 0.17 | 0.58 | 0.11 | 0.14 | 0.1 | 0.25 |
| 9 | 93111241 | rs874977 | -1.159 | -1.647 | 1.251 | 0.554 | -0.032 | 0.001 | 0.001 | 0 | 0.37 | 0.47 | 0.3 | 0.32 | 0.29 | 0.29 |
| 9 | 93116944 | rs1777036 | -0.448 | -1.644 | 0.625 | 0.504 | 0.03 | 0.029 | 0.029 | 0 | 0.28 | 0.16 | 0.14 | 0.16 | 0.2 | 0.16 |
| 9 | 93129283 | rs1623337 | -0.469 | -1.628 | 0.616 | 0.512 | 0.032 | 0.031 | 0.031 | 0 | 0.29 | 0.15 | 0.14 | 0.17 | 0.2 | 0.19 |
| 9 | 93130383 | rs12004191 | -2.29 | -1.63 | 0.592 | 0.558 | 0.003 | -0.005 | -0.005 | 0 | 0.19 | 0.76 | 0.25 | 0.27 | 0.17 | 0.29 |
| 9 | 93130796 | rs1777052 | 0.062 | -1.548 | 0.489 | 0.516 | -0.002 | 0.007 | 0.007 | 0 | 0.19 | 0.04 | 0.09 | 0.15 | 0.16 | 0.14 |
| 9 | 93141541 | rs1648477 | -0.977 | -1.952 | 0.792 | 0.229 | 0.035 | 0.006 | 0.006 | 0 | 0.39 | 0.32 | 0.32 | 0.31 | 0.31 | 0.27 |
| 9 | 93148230 | rs1889204 | 0.845 | -1.904 | 1.206 | 0.242 | 0.006 | 0.005 | 0.005 | 0.019 | 0.65 | 0.65 | 0.79 | 0.75 | 0.75 | 0.8 |
| 9 | 93148579 | rs1426506 | 0.084 | -1.898 | 1.116 | 0.236 | 0.032 | 0.025 | 0.025 | 0 | 0.71 | 0.78 | 0.85 | 0.81 | 0.81 | 0.83 |
| 9 | 93152933 | rs10739979 | 0.993 | -1.954 | 1.447 | 0.226 | -0.01 | -0.006 | -0.006 | 0.118 | 0.6 | 0.45 | 0.76 | 0.71 | 0.72 | 0.72 |

| | | | | | | | | | | | | | | | | |
|---|---|---|---|---|---|---|---|---|---|---|---|---|---|---|---|---|
| 9 | 93154530 | rs2004205 | 1.492 | -1.959 | 1.06 | 0.235 | 0.108 | -0.041 | -0.041 | 0.078 | 0.5 | 0.1 | 0.58 | 0.56 | 0.61 | 0.51 |
| 9 | 93158434 | rs10993323 | 1.326 | -2.205 | 1.256 | -0.009 | 0.048 | -0.006 | -0.006 | 0 | 0.45 | 0.23 | 0.51 | 0.57 | 0.56 | 0.49 |
| 9 | 93162602 | rs1576290 | -0.161 | -2.275 | 1.616 | -0.095 | 0.024 | 0.028 | 0.028 | 0 | 0.64 | 0.7 | 0.81 | 0.82 | 0.77 | 0.76 |
| 9 | 93163805 | rs2162333 | 1.167 | -2.179 | 1.438 | -0.026 | 0 | 0.002 | 0.002 | 0 | 0.08 | 0.07 | 0.17 | 0.14 | 0.11 | 0.09 |
| 9 | 93168446 | rs10993355 | 1.026 | -2.18 | 1.572 | -0.026 | -0.007 | 0.039 | 0.039 | 0 | 0.08 | 0.17 | 0.18 | 0.14 | 0.11 | 0.09 |
| 9 | 93169441 | rs10821400 | 0.794 | -2.211 | 1.635 | -0.064 | 0.013 | 0.013 | 0.013 | 0.125 | 0.54 | 0.44 | 0.73 | 0.73 | 0.68 | 0.68 |
| 9 | 93183404 | rs4744418 | 1.337 | -2.208 | 1.349 | -0.037 | 0.05 | -0.001 | -0.001 | 0 | 0.45 | 0.21 | 0.55 | 0.59 | 0.57 | 0.58 |
| 9 | 93187329 | rs10993452 | -0.016 | -2.243 | 1.458 | -0.048 | -0.007 | 0.016 | 0.016 | 0 | 0.12 | 0.18 | 0.23 | 0.2 | 0.17 | 0.11 |
| 9 | 93187808 | rs10821428 | 1.017 | -2.217 | 1.305 | -0.044 | 0.098 | 0.026 | 0.026 | 0.305 | 0.48 | 0.11 | 0.56 | 0.6 | 0.6 | 0.52 |
| 9 | 93192521 | rs10821444 | 1.298 | -2.165 | 1.33 | -0.067 | 0.048 | 0.003 | 0.003 | 1.016 | 0.45 | 0.27 | 0.55 | 0.59 | 0.57 | 0.58 |
| 9 | 93193062 | rs6479594 | -0.609 | -2.244 | 1.011 | -0.113 | 0.017 | 0.027 | 0.027 | 0 | 0.76 | 0.87 | 0.88 | 0.84 | 0.82 | 0.8 |
| 9 | 93199571 | rs10821452 | 0.336 | -2.19 | 1.135 | -0.065 | 0.017 | 0.009 | 0.009 | 0.02 | 0.59 | 0.64 | 0.64 | 0.68 | 0.66 | 0.65 |
| 9 | 93205277 | rs17450197 | 1.514 | -1.539 | 0.2 | 0.268 | -0.138 | 0.01 | 0.01 | 0.85 | 0.34 | 0.07 | 0.36 | 0.27 | 0.46 | 0.22 |
| 9 | 93214896 | rs7858582 | 0.634 | -1.379 | 0.69 | -0.156 | 0.004 | 0.004 | 0.004 | 0.006 | 0.49 | 0.56 | 0.47 | 0.65 | 0.56 | 0.6 |
| 9 | 93219568 | rs4743992 | -0.528 | -1.502 | 0.671 | -0.229 | 0 | 0.015 | 0.015 | 0 | 0.69 | 0.8 | 0.71 | 0.81 | 0.73 | 0.71 |
| 9 | 93222465 | rs10761383 | 0.988 | -1.067 | 0.594 | -0.083 | -0.018 | -0.04 | -0.04 | 0 | 0.43 | 0.17 | 0.41 | 0.56 | 0.54 | 0.45 |
| 9 | 93225765 | rs1549508 | 0.974 | -1.304 | 0.597 | -0.08 | -0.001 | -0.039 | -0.039 | 0 | 0.43 | 0.16 | 0.42 | 0.56 | 0.57 | 0.45 |
| 9 | 93231161 | rs1426507 | 0.479 | -1.155 | 0.207 | -0.139 | 0.008 | 0.008 | 0.008 | 0.012 | 0.49 | 0.57 | 0.48 | 0.65 | 0.59 | 0.6 |
| 9 | 93240193 | rs1863559 | 0.123 | -0.948 | 0.152 | -0.012 | -0.01 | -0.003 | -0.003 | 0 | 0.62 | 0.53 | 0.51 | 0.61 | 0.64 | 0.63 |
| 9 | 93246553 | rs10761389 | 0.924 | -1.275 | -0.039 | -0.18 | 0.02 | 0.003 | 0.003 | 0.313 | 0.46 | 0.19 | 0.43 | 0.65 | 0.58 | 0.47 |
| 9 | 93248431 | rs12339486 | 0.526 | -1.443 | 0.137 | -0.248 | -0.004 | 0.001 | 0.001 | 0 | 0.1 | 0.14 | 0.18 | 0.09 | 0.11 | 0.07 |
| 9 | 93252163 | rs4743995 | -0.694 | -1.577 | 0.992 | -0.35 | 0.004 | -0.019 | -0.019 | 1.1 | 0.73 | 0.68 | 0.71 | 0.82 | 0.76 | 0.72 |
| 9 | 93256092 | rs7862334 | -0.02 | -1.621 | 1.064 | -0.374 | 0.009 | 0.026 | 0.026 | 0 | 0.63 | 0.67 | 0.66 | 0.81 | 0.72 | 0.71 |
| 9 | 93260040 | rs16906632 | -0.395 | -1.447 | 0.812 | -0.228 | -0.011 | 0.007 | 0.007 | 0 | 0.14 | 0.21 | 0.19 | 0.12 | 0.13 | 0.2 |
| 9 | 93265378 | rs12353086 | -0.375 | -1.445 | 0.787 | -0.215 | -0.009 | 0.006 | 0.006 | 0 | 0.14 | 0.19 | 0.19 | 0.12 | 0.13 | 0.2 |
| 9 | 93266508 | rs1346211 | 1.168 | -0.667 | -0.024 | -0.625 | -0.069 | 0.072 | 0.072 | 0 | 0.14 | 0.01 | 0.13 | 0.02 | 0.18 | 0.09 |
| 9 | 93272277 | rs10993564 | -0.474 | -1.091 | 0.725 | -0.352 | -0.073 | -0.013 | -0.013 | 0 | 0.26 | 0.46 | 0.27 | 0.14 | 0.18 | 0.24 |
| 9 | 93284804 | rs4744469 | 0.761 | -0.975 | 0.175 | -0.427 | 0.05 | -0.013 | -0.013 | 0.79 | 0.06 | 0.14 | 0.18 | 0.02 | 0.1 | 0.07 |
| 9 | 93285878 | rs10993570 | -1.137 | -1.018 | -0.143 | -0.417 | 0.007 | 0.004 | 0.004 | 0.715 | 0.34 | 0.68 | 0.45 | 0.2 | 0.28 | 0.32 |
| 9 | 93289728 | rs7870746 | 0.67 | -1.152 | -0.467 | -0.231 | 0.005 | 0.005 | 0.005 | 0 | 0.09 | 0.09 | 0.17 | 0.04 | 0.1 | 0.07 |
| 9 | 93309082 | rs690211 | -1.271 | -0.994 | -0.915 | -0.252 | -0.017 | -0.022 | -0.022 | 0 | 0.82 | 1 | 0.9 | 0.84 | 0.87 | 0.8 |
| 9 | 93316821 | rs12002017 | -0.515 | -1.032 | -0.236 | -0.221 | -0.055 | -0.008 | -0.008 | 2.012 | 0.28 | 0.59 | 0.27 | 0.18 | 0.18 | 0.25 |
| 9 | 93331003 | rs10993593 | 0.657 | 0.323 | -0.78 | -0.303 | -0.078 | 0.073 | 0.073 | 0 | 0.15 | 0.06 | 0.15 | 0.02 | 0.24 | 0.1 |
| 9 | 93332475 | rs4743997 | 0.949 | -0.54 | -0.927 | -0.499 | 0.049 | 0.031 | 0.031 | 0 | 0.29 | 0.12 | 0.13 | 0.18 | 0.16 | 0.22 |
| 9 | 93333075 | rs690140 | 0.948 | -0.484 | -1.184 | 0.099 | -0.022 | -0.033 | -0.033 | 0.839 | 0.18 | 0.07 | 0.24 | 0.35 | 0.29 | 0.19 |
| 9 | 93343832 | rs1331497 | -0.349 | -0.608 | -0.42 | 0.068 | 0.019 | -0.007 | -0.007 | 0 | 0.73 | 0.64 | 0.67 | 0.84 | 0.67 | 0.69 |
| 9 | 93344344 | rs690552 | 0.35 | -0.926 | -0.14 | 0.169 | -0.001 | -0.003 | -0.003 | 0 | 0.33 | 0.25 | 0.33 | 0.58 | 0.41 | 0.3 |
| 9 | 93348404 | rs16906689 | 0.511 | -0.775 | 0.112 | 0.068 | 0.01 | 0.011 | 0.011 | 0.093 | 0.08 | 0.28 | 0.2 | 0.08 | 0.09 | 0.13 |
| 9 | 93348786 | rs2613339 | -0.404 | -0.849 | -0.081 | -0.014 | 0.003 | 0.013 | 0.013 | 0 | 0.31 | 0.35 | 0.44 | 0.52 | 0.4 | 0.47 |
| 9 | 93350920 | rs525843 | 0.262 | -0.892 | -0.05 | -0.062 | -0.004 | 0 | 0 | 0 | 0.63 | 0.61 | 0.53 | 0.86 | 0.55 | 0.6 |
| 9 | 93351868 | rs555362 | 0.868 | -0.887 | -0.04 | -0.031 | 0.024 | 0.076 | 0.076 | 0 | 0.2 | 0.33 | 0.31 | 0.45 | 0.3 | 0.28 |
| 9 | 93353987 | rs690567 | 0.868 | -0.887 | -0.04 | -0.031 | 0.024 | 0.076 | 0.076 | 0 | 0.2 | 0.33 | 0.31 | 0.45 | 0.3 | 0.28 |
| 9 | 93357338 | rs7856157 | 1.081 | -0.841 | -0.09 | -0.072 | 0.04 | 0.006 | 0.006 | 1.227 | 0.43 | 0.27 | 0.21 | 0.41 | 0.25 | 0.33 |
| 9 | 93359512 | rs689917 | -0.735 | -0.408 | -0.166 | -0.062 | 0.018 | -0.039 | -0.039 | 1.185 | 0.75 | 0.67 | 0.66 | 0.93 | 0.66 | 0.8 |
| 9 | 93359638 | rs690150 | -0.742 | -0.397 | -0.165 | -0.057 | 0.009 | -0.027 | -0.027 | 0.162 | 0.75 | 0.7 | 0.66 | 0.93 | 0.66 | 0.8 |
| 9 | 93366369 | rs12004340 | 1.272 | -0.635 | -0.58 | 0.221 | 0.023 | -0.012 | -0.012 | 0 | 0.17 | 0.08 | 0.1 | 0.23 | 0.12 | 0.11 |
| 9 | 93369784 | rs7868194 | -0.294 | -0.418 | -0.549 | 0.121 | 0.001 | -0.007 | -0.007 | 0 | 0.43 | 0.49 | 0.45 | 0.34 | 0.48 | 0.39 |
| 9 | 93377290 | rs690111 | -1.157 | -0.456 | -0.783 | 0.378 | -0.02 | -0.002 | -0.002 | 0 | 0.63 | 0.49 | 0.68 | 0.67 | 0.77 | 0.59 |
| 9 | 93377901 | rs1017753 | -0.363 | -0.474 | -0.753 | 0.428 | -0.002 | 0.002 | 0.002 | 0 | 0.47 | 0.47 | 0.46 | 0.43 | 0.51 | 0.51 |
| 9 | 93379328 | rs16906717 | 0.009 | 0.218 | -1.07 | 0.552 | 0.006 | 0.055 | 0.055 | 1.439 | 0.05 | 0.22 | 0.09 | 0.1 | 0.03 | 0.02 |
| 9 | 93382438 | rs690232 | -0.176 | 0.344 | -1.045 | 0.602 | -0.008 | -0.014 | -0.014 | 0 | 0.25 | 0.25 | 0.32 | 0.43 | 0.33 | 0.28 |
| 9 | 93387516 | rs563726 | 0.446 | 0.486 | -1.85 | 0.476 | -0.055 | -0.061 | -0.061 | 0 | 0.2 | 0.05 | 0.23 | 0.32 | 0.29 | 0.2 |
| 9 | 93392332 | rs1331501 | 0.602 | 0.827 | -1.492 | 0.154 | -0.071 | 0.104 | 0.104 | 0 | 0.2 | 0.06 | 0.17 | 0.02 | 0.28 | 0.13 |
| 9 | 93402190 | rs4744473 | 0.496 | 0.333 | -1.127 | 0.774 | 0.021 | -0.017 | -0.017 | 0 | 0.58 | 0.26 | 0.26 | 0.62 | 0.45 | 0.38 | 0.56 |
| 9 | 93409161 | rs1412005 | 0.351 | 0.932 | -0.586 | 0.76 | 0.082 | 0.225 | 0.225 | 0.037 | 0.83 | 0.58 | 0.62 | 0.47 | 0.74 | 0.79 |
| 9 | 93410201 | rs689687 | 0.771 | 1.053 | -0.672 | 1.148 | -0.053 | -0.051 | -0.051 | 0 | 0.09 | 0.01 | 0.18 | 0.35 | 0.19 | 0.1 |
| 9 | 93412927 | rs1331507 | 1.405 | 0.629 | -1.007 | 0.482 | -0.016 | 0.066 | 0.066 | 0 | 0.11 | 0.01 | 0.05 | 0 | 0.08 | 0.09 |
| 9 | 93413258 | rs10993618 | 0.193 | 0.595 | -0.618 | 0.524 | -0.021 | -0.039 | -0.039 | 0.063 | 0.74 | 0.82 | 0.61 | 0.57 | 0.68 | 0.72 |
| 9 | 93418243 | rs4744000 | 0.707 | 0.165 | -0.564 | 0.46 | 0.096 | 0.022 | 0.022 | 0.018 | 0.42 | 0.19 | 0.18 | 0.37 | 0.27 | 0.35 |
| 9 | 93434093 | rs753593 | 0.69 | 0.253 | -1.418 | -0.013 | -0.001 | -0.001 | -0.001 | 0 | 0.14 | 0.16 | 0.1 | 0.13 | 0.12 | 0.18 |
| 9 | 93441674 | rs1173099 | -0.08 | 0.9 | -1.5 | -0.328 | -0.053 | 0.076 | 0.076 | 0 | 0.66 | 0.83 | 0.6 | 0.97 | 0.56 | 0.67 |
| 9 | 93443531 | rs1173100 | -0.111 | 0.994 | -1.396 | 0.811 | -0.014 | -0.05 | -0.05 | 0 | 0.22 | 0.13 | 0.3 | 0.52 | 0.24 | 0.25 |
| 9 | 93445063 | rs1173102 | -0.695 | 0.452 | -0.845 | 0.842 | 0 | -0.002 | -0.002 | 0.004 | 0.39 | 0.61 | 0.5 | 0.54 | 0.35 | 0.47 |
| 9 | 93445145 | rs10993628 | 0.868 | 0.412 | -1.754 | -0.035 | 0.013 | 0.169 | 0.169 | 0 | 0.33 | 0.05 | 0.19 | 0.04 | 0.28 | 0.13 |
| 9 | 93445335 | rs1173104 | -0.06 | 0.262 | -1.753 | -0.049 | 0.001 | -0.004 | -0.004 | 0 | 0.43 | 0.6 | 0.52 | 0.52 | 0.4 | 0.5 |
| 9 | 93455943 | rs16906767 | 0.941 | 0.859 | -1.649 | 0.133 | 0.008 | -0.002 | -0.002 | 0 | 0.23 | 0.29 | 0.18 | 0.24 | 0.24 | 0.25 |
| 9 | 93461649 | rs1173107 | 0.403 | 0.578 | -1.035 | 0.31 | -0.003 | -0.002 | -0.002 | 0.055 | 0.37 | 0.35 | 0.5 | 0.42 | 0.39 | 0.39 |
| 9 | 93466631 | rs10993639 | 1.731 | 0.163 | -1.488 | -0.058 | -0.021 | 0.061 | 0.061 | 0 | 0.19 | 0.02 | 0.11 | 0.03 | 0.19 | 0.09 |
| 9 | 93469911 | rs1172896 | -1.046 | 0.136 | -1.246 | -0.08 | -0.018 | -0.012 | -0.012 | 0 | 0.64 | 0.78 | 0.51 | 0.59 | 0.61 | 0.61 |

| | | | | | | | | | | | | | | | | |
|---|---|---|---|---|---|---|---|---|---|---|---|---|---|---|---|---|
| 9 | 93471627 | rs4744489 | 0.68 | 0.087 | -0.819 | -0.048 | -0.039 | 0.008 | 0.008 | 0 | 0.15 | 0.27 | 0.06 | 0.11 | 0.05 | 0.11 |
| 9 | 93476005 | rs2613319 | 1.021 | 0.098 | -0.621 | -0.024 | 0.018 | 0.005 | 0.005 | 0 | 0.25 | 0.31 | 0.21 | 0.29 | 0.32 | 0.29 |
| 9 | 93482553 | rs12336030 | 0.722 | 0.051 | -0.631 | 0.009 | 0.006 | 0.145 | 0.145 | 0.825 | 0.05 | 0.25 | 0.02 | 0.11 | 0.01 | 0.1 |
| 9 | 93486569 | rs2810919 | -0.298 | 0.09 | -0.725 | -0.022 | 0 | 0.001 | 0.001 | 0 | 0.4 | 0.56 | 0.28 | 0.41 | 0.37 | 0.4 |
| 9 | 93514742 | rs34993748 | 1.878 | -0.045 | -1.19 | -0.198 | 0.037 | -0.019 | -0.019 | 0.773 | 0.14 | 0.03 | 0.05 | 0.13 | 0.06 | 0.05 |
| 9 | 93526206 | rs1148851 | 0.551 | -0.017 | -1.073 | -0.316 | -0.01 | -0.014 | -0.014 | 0 | 0.36 | 0.2 | 0.48 | 0.4 | 0.36 | 0.39 |
| 9 | 93526329 | rs914926 | 0.528 | -0.024 | -1.076 | -0.32 | -0.009 | -0.013 | -0.013 | 0 | 0.37 | 0.2 | 0.48 | 0.4 | 0.36 | 0.39 |
| 9 | 93530889 | rs10821495 | 0.44 | -0.028 | -0.603 | -0.505 | -0.02 | -0.091 | -0.091 | 0 | 0.29 | 0.38 | 0.17 | 0.02 | 0.21 | 0.12 |
| 9 | 93530909 | rs1148853 | -0.159 | 0.115 | -0.532 | -0.446 | -0.01 | -0.032 | -0.032 | 0 | 0.64 | 0.56 | 0.78 | 0.87 | 0.69 | 0.74 |
| 9 | 93546691 | rs290275 | 0.582 | 0.879 | -0.771 | -0.59 | -0.008 | -0.021 | -0.021 | 3.909 | 0.17 | 0.13 | 0.19 | 0.41 | 0.21 | 0.25 |
| 9 | 93547691 | rs4744498 | 0.899 | 0.828 | -0.926 | -0.724 | -0.005 | 0.019 | 0.019 | 0.301 | 0.13 | 0.09 | 0.28 | 0.08 | 0.1 | 0.17 |
| 9 | 93550754 | rs10993688 | 1.226 | 0.802 | -0.776 | -0.829 | -0.031 | -0.042 | -0.042 | 0 | 0.3 | 0.17 | 0.21 | 0.32 | 0.33 | 0.32 |
| 9 | 93552550 | rs290267 | -0.346 | 0.926 | -0.964 | -0.807 | -0.024 | 0.036 | 0.036 | 0.031 | 0.83 | 0.91 | 0.68 | 0.92 | 0.81 | 0.79 |
| 9 | 93557698 | rs10993693 | 0.209 | 0.907 | -0.494 | -0.737 | 0 | 0.002 | 0.002 | 0 | 0.47 | 0.55 | 0.51 | 0.42 | 0.43 | 0.51 |
| 9 | 93561450 | rs290991 | -0.06 | 0.674 | -1.013 | -0.689 | 0 | 0.002 | 0.002 | 0.014 | 0.47 | 0.21 | 0.42 | 0.59 | 0.48 | 0.44 |
| 9 | 93561614 | rs10821507 | 1.362 | 0.711 | -0.851 | -0.791 | 0.041 | 0.115 | 0.115 | 0 | 0.27 | 0.07 | 0.12 | 0.02 | 0.2 | 0.11 |
| 9 | 93563536 | rs290986 | 0.428 | 0.94 | -0.599 | -0.435 | -0.013 | -0.026 | -0.026 | 0 | 0.17 | 0.13 | 0.2 | 0.42 | 0.22 | 0.25 |
| 9 | 93567572 | rs1333633 | -0.429 | 1.067 | -0.202 | -0.424 | -0.002 | 0 | 0 | 0 | 0.61 | 0.4 | 0.71 | 0.68 | 0.57 | 0.6 |
| 9 | 93570012 | rs7868795 | 0.748 | 0.853 | -0.439 | -0.546 | 0.004 | 0.011 | 0.011 | 0.18 | 0.14 | 0.11 | 0.28 | 0.08 | 0.09 | 0.15 |
| 9 | 93570505 | rs7036417 | 1.065 | 0.973 | -0.459 | -0.638 | -0.077 | -0.073 | -0.073 | 1.355 | 0.3 | 0.08 | 0.23 | 0.32 | 0.34 | 0.36 |
| 9 | 93571341 | rs3789889 | 0.14 | 0.444 | -0.229 | -0.563 | 0.044 | 0.065 | 0.065 | 0 | 0.48 | 0.35 | 0.48 | 0.26 | 0.33 | 0.37 |
| 9 | 93572543 | rs7020331 | 0.796 | 0.42 | 0.016 | -0.394 | 0.027 | 0.035 | 0.035 | 0 | 0.33 | 0.22 | 0.2 | 0.18 | 0.24 | 0.2 |
| 9 | 93573542 | rs290240 | -0.583 | 0.416 | 0.286 | -0.401 | -0.013 | -0.032 | -0.032 | 0 | 0.6 | 0.59 | 0.75 | 0.82 | 0.68 | 0.76 |
| 9 | 93573597 | rs12550873 | 1.358 | 0.361 | 0.266 | -0.488 | -0.015 | -0.072 | -0.072 | 2.44 | 0.24 | 0.24 | 0.08 | 0.01 | 0.11 | 0.13 |
| 9 | 93574650 | rs290237 | -0.96 | 0.391 | 0.078 | -0.463 | -0.02 | -0.117 | -0.117 | 1.597 | 0.71 | 0.65 | 0.87 | 0.99 | 0.81 | 0.84 |
| 9 | 93575966 | rs10993697 | 1.324 | 0.43 | 0.035 | -0.382 | 0.003 | -0.013 | -0.013 | 1.179 | 0.1 | 0.06 | 0.12 | 0.17 | 0.13 | 0.08 |
| 9 | 93577781 | rs10993698 | 1.234 | 0.428 | 0.034 | -0.383 | 0.004 | -0.014 | -0.014 | 0 | 0.11 | 0.06 | 0.12 | 0.17 | 0.13 | 0.08 |
| 9 | 93579997 | rs4744505 | -0.324 | 0.533 | 0.235 | -0.319 | -0.009 | 0.001 | 0.001 | 0 | 0.61 | 0.57 | 0.71 | 0.58 | 0.64 | 0.67 |
| 9 | 93581182 | rs1333630 | 0.234 | 0.603 | 0.265 | -0.268 | 0.006 | 0.006 | 0.006 | 0 | 0.5 | 0.5 | 0.59 | 0.41 | 0.51 | 0.59 |
| 9 | 93584793 | rs914925 | 0.487 | 0.597 | -0.189 | -0.278 | -0.021 | -0.01 | -0.01 | 0 | 0.47 | 0.14 | 0.56 | 0.41 | 0.5 | 0.58 |
| 9 | 93586759 | rs1081767 | 0.804 | 0.794 | -0.127 | -0.289 | -0.009 | 0.014 | 0.014 | 0 | 0.1 | 0.06 | 0.17 | 0.42 | 0.16 | 0.14 |
| 9 | 93587089 | rs187306 | -0.768 | 0.734 | 0.067 | -0.289 | 0.009 | 0.099 | 0.099 | 0 | 0.9 | 0.85 | 0.83 | 0.58 | 0.84 | 0.86 |
| 9 | 93590403 | rs290974 | 0.845 | 0.307 | -0.261 | -0.276 | -0.002 | 0.069 | 0.069 | 0 | 0.09 | 0.08 | 0.13 | 0.42 | 0.12 | 0.14 |
| 9 | 93595807 | rs290213 | -0.521 | 0.407 | -0.525 | -0.346 | -0.012 | 0.044 | 0.044 | 0 | 0.86 | 0.94 | 0.91 | 1 | 0.83 | 0.9 |
| 9 | 93596067 | rs290212 | 0.182 | -0.022 | -0.334 | -0.213 | 0.08 | -0.031 | -0.031 | 0 | 0.38 | 0.8 | 0.26 | 0.43 | 0.27 | 0.29 |
| 9 | 93597669 | rs1025325 | -0.769 | 0.635 | -0.095 | -0.267 | 0.017 | -0.003 | -0.003 | 0 | 0.63 | 0.76 | 0.68 | 0.4 | 0.66 | 0.62 |
| 9 | 93599890 | rs4744512 | 0.443 | 0.897 | 0.187 | -0.764 | -0.02 | -0.11 | -0.11 | 2.08 | 0.39 | 0.46 | 0.38 | 0.06 | 0.26 | 0.23 |
| 9 | 93601955 | rs1864202 | 0.483 | 1.429 | -0.022 | -0.495 | 0.026 | 0.002 | 0.002 | 0.49 | 0.15 | 0.17 | 0.23 | 0.16 | 0.28 | 0.32 |
| 9 | 93603138 | rs17489214 | 1.456 | 0.562 | -0.09 | -0.742 | 0.014 | 0.028 | 0.028 | 0 | 0.23 | 0.11 | 0.11 | 0.02 | 0.16 | 0.16 |
| 9 | 93605269 | rs10993707 | 0.486 | 0.96 | 0.066 | -0.57 | 0.051 | 0 | 0 | 0 | 0.17 | 0.29 | 0.23 | 0.16 | 0.32 | 0.36 |
| 9 | 93606103 | rs7855964 | 0.643 | 0.685 | 0.203 | -0.371 | -0.024 | -0.131 | -0.131 | 0.099 | 0.37 | 0.45 | 0.39 | 0.04 | 0.24 | 0.19 |
| 9 | 93608739 | rs2278278 | -1.34 | 0.607 | -0.402 | -0.099 | -0.002 | -0.001 | -0.001 | 0 | 0.88 | 0.94 | 0.84 | 0.63 | 0.84 | 0.88 |
| 9 | 93609419 | rs2035073 | 0.417 | 0.703 | -0.096 | -0.577 | 0.003 | 0.008 | 0.008 | 1.218 | 0.19 | 0.21 | 0.23 | 0.4 | 0.34 | 0.4 |
| 9 | 93610761 | rs4744514 | -0.507 | 0.595 | -0.566 | -0.451 | 0.021 | -0.007 | -0.007 | 0 | 0.56 | 0.73 | 0.63 | 0.44 | 0.57 | 0.6 |
| 9 | 93612959 | rs10993713 | -0.082 | 0.978 | -0.276 | -0.61 | -0.011 | -0.059 | -0.059 | 1.131 | 0.2 | 0.26 | 0.31 | 0.03 | 0.14 | 0.09 |
| 9 | 93616303 | rs7863711 | -0.572 | 0.952 | -1.098 | -0.378 | -0.046 | 0.069 | 0.069 | 0 | 0.38 | 0.23 | 0.53 | 0.18 | 0.46 | 0.46 |
| 9 | 93617445 | rs16905730 | -0.138 | 0.541 | -0.869 | -0.108 | 0.013 | 0.011 | 0.011 | 0 | 0.14 | 0.05 | 0.08 | 0.07 | 0.1 | 0.22 |
| 9 | 93618236 | rs10993725 | -1.186 | 0.53 | 0.28 | 0.357 | 0.026 | 0.127 | 0.127 | 0 | 0.07 | 0.32 | 0.32 | 0.19 | 0.07 | 0.06 |
| 9 | 93619736 | rs10124093 | -0.787 | 0.538 | 0.387 | 0.42 | 0.012 | 0.099 | 0.099 | 0 | 0.05 | 0.18 | 0.32 | 0.2 | 0.06 | 0.06 |
| 9 | 93622150 | rs12001366 | -1.239 | 0.535 | 0.419 | 0.419 | 0.02 | 0.124 | 0.124 | 0 | 0.06 | 0.24 | 0.32 | 0.2 | 0.07 | 0.06 |
| 9 | 93622835 | rs1760125 | 0.133 | 1.132 | 0.179 | -0.233 | -0.038 | 0.05 | 0.05 | 0 | 0.79 | 0.9 | 0.82 | 0.96 | 0.68 | 0.85 |
| 9 | 93623238 | rs10993728 | -0.712 | 1.099 | 0.174 | -0.233 | -0.01 | 0.014 | 0.014 | 0 | 0.15 | 0.28 | 0.36 | 0.2 | 0.13 | 0.08 |
| 9 | 93624896 | rs10821515 | 1.563 | 0.916 | 0.143 | -0.968 | 0.006 | -0.029 | -0.029 | 0 | 0.33 | 0.2 | 0.22 | 0 | 0.28 | 0.1 |
| 9 | 93626241 | rs4529574 | 0.322 | 0.934 | 0.367 | -0.956 | 0.003 | 0.029 | 0.029 | 0 | 0.13 | 0.31 | 0.37 | 0.21 | 0.14 | 0.08 |
| 9 | 93627484 | rs4744516 | 0.597 | 0.953 | 0.321 | -0.93 | 0.001 | -0.007 | -0.007 | 0 | 0.47 | 0.45 | 0.59 | 0.21 | 0.42 | 0.18 |
| 9 | 93628027 | rs2991216 | 0.311 | 1.181 | 0.248 | 0.106 | -0.008 | -0.006 | -0.006 | 0 | 0.14 | 0.12 | 0.14 | 0.27 | 0.26 | 0.24 |
| 9 | 93633166 | rs7855064 | 0.265 | 0.672 | 0.303 | 0.436 | 0.002 | 0.002 | 0.002 | 0 | 0.2 | 0.17 | 0.12 | 0.4 | 0.15 | 0.43 |
| 9 | 93633948 | rs7864625 | 1.118 | 0.638 | 0.293 | 0.37 | -0.004 | -0.001 | -0.001 | 0 | 0.09 | 0.1 | 0.05 | 0.07 | 0.06 | 0.2 |
| 9 | 93634413 | rs290229 | 0.053 | 0.659 | 0.231 | 0.426 | 0 | -0.018 | -0.018 | 0 | 0.24 | 0.16 | 0.15 | 0.65 | 0.22 | 0.4 |
| 9 | 93635975 | rs10512202 | 1.421 | 0.67 | 0.215 | 0.304 | 0 | -0.003 | -0.003 | 0 | 0.1 | 0.06 | 0.05 | 0.11 | 0.07 | 0.12 |
| 9 | 93636664 | rs290227 | -0.524 | 0.627 | 0.412 | 0.308 | -0.005 | 0.005 | 0.005 | 0 | 0.3 | 0.37 | 0.21 | 0.63 | 0.25 | 0.57 |
| 9 | 93637015 | rs2290890 | 0.335 | 0.593 | 0.466 | 0.279 | -0.005 | 0.016 | 0.016 | 0 | 0.14 | 0.21 | 0.37 | 0.23 | 0.14 | 0.08 |
| 9 | 93640009 | rs2290887 | 0.335 | 0.593 | 0.466 | 0.279 | -0.005 | 0.016 | 0.016 | 0 | 0.14 | 0.21 | 0.37 | 0.23 | 0.14 | 0.08 |
| 9 | 93641175 | rs2306041 | 0.318 | 0.596 | 0.456 | 0.281 | -0.005 | 0.017 | 0.017 | 0 | 0.14 | 0.22 | 0.37 | 0.23 | 0.14 | 0.08 |
| 9 | 93642897 | rs10761395 | 1.695 | 1.043 | 0.284 | 0.031 | -0.004 | -0.107 | -0.107 | 0 | 0.34 | 0.38 | 0.24 | 0 | 0.32 | 0.11 |
| 9 | 93644948 | rs1755938 | 0.917 | 1.461 | 0.011 | 0.596 | -0.001 | -0.003 | -0.003 | 0 | 0.12 | 0.11 | 0.09 | 0.31 | 0.14 | 0.16 |
| 9 | 93651615 | rs290256 | -0.746 | 1.557 | -0.228 | 0.516 | -0.072 | 0.056 | 0.056 | 0 | 0.88 | 0.99 | 0.87 | 0.97 | 0.78 | 0.88 |
| 9 | 93651657 | rs290255 | 0.894 | 1.145 | -0.123 | 0.638 | -0.003 | -0.02 | -0.02 | 0 | 0.12 | 0.07 | 0.08 | 0.31 | 0.13 | 0.16 |
| 9 | 93653237 | rs10993746 | 1.047 | 1.262 | -0.147 | 0.464 | 0 | -0.005 | -0.005 | 0 | 0.1 | 0.07 | 0.06 | 0.18 | 0.07 | 0.12 |

| | | | | | | | | | | | | | | | | |
|---|---|---|---|---|---|---|---|---|---|---|---|---|---|---|---|---|
| 9 | 93654090 | rs2613320 | -0.224 | 1.666 | 0.22 | 0.535 | -0.013 | 0.004 | 0.004 | 0 | 0.85 | 0.87 | 0.84 | 0.97 | 0.74 | 0.84 |
| 9 | 93654504 | rs1172665 | -0.721 | 1.407 | 0.424 | 0.797 | -0.012 | 0.016 | 0.016 | 0 | 0.37 | 0.57 | 0.34 | 0.49 | 0.34 | 0.47 |
| 9 | 93654697 | rs1172663 | -0.74 | 1.277 | 0.445 | 0.828 | -0.017 | 0.022 | 0.022 | 0 | 0.37 | 0.55 | 0.33 | 0.5 | 0.33 | 0.46 |
| 9 | 93655385 | rs726757 | 0.588 | 1.091 | 0.387 | 0.859 | -0.022 | 0.025 | 0.025 | 0 | 0.65 | 0.43 | 0.68 | 0.5 | 0.72 | 0.56 |
| 9 | 93658206 | rs1049164 | 0.321 | 1.215 | -0.218 | 0.855 | 0.011 | 0.013 | 0.013 | 0 | 0.21 | 0.12 | 0.19 | 0.12 | 0.19 | 0.28 |
| 9 | 93658854 | rs158688 | -0.42 | 2.123 | 0.225 | 1.446 | 0.043 | 0.023 | 0.023 | 0.27 | 0.2 | 0.28 | 0.4 | 0.27 | 0.35 | 0.27 |
| 9 | 93658952 | rs12344755 | 0.089 | 1.733 | 0.381 | 0.727 | -0.01 | 0.051 | 0.051 | 0.079 | 0.13 | 0.23 | 0.12 | 0.39 | 0.09 | 0.19 |
| 9 | 93658978 | rs158687 | -0.824 | 2.257 | 0.082 | 0 | 0.04 | -0.032 | -0.032 | 0.981 | 0.19 | 0.28 | 0.38 | 0.11 | 0.33 | 0.17 |
| 9 | 93661352 | rs1551747 | -0.504 | 1.226 | 0.128 | -0.059 | 0.009 | -0.048 | -0.048 | 2.76 | 0.17 | 0.25 | 0.11 | 0.04 | 0.19 | 0.12 |
| 9 | 93662205 | rs1760118 | 0.023 | 1.541 | -0.032 | -0.281 | 0 | -0.001 | -0.001 | 0 | 0.24 | 0.27 | 0.31 | 0.25 | 0.29 | 0.3 |
| 9 | 93662641 | rs10993751 | -0.509 | 1.042 | -0.14 | -1.078 | 0 | -0.011 | -0.011 | 0 | 0.46 | 0.46 | 0.47 | 0.73 | 0.54 | 0.53 |
| 9 | 93662762 | rs10993752 | 0.14 | 0.956 | -0.183 | -0.904 | 0 | 0.002 | 0.002 | 0 | 0.3 | 0.28 | 0.37 | 0.71 | 0.35 | 0.42 |
| 9 | 93663201 | rs4744520 | 0.3 | 0.699 | -0.437 | -0.219 | 0.008 | -0.07 | -0.07 | 0 | 0.23 | 0.15 | 0.33 | 0.56 | 0.23 | 0.24 |
| 9 | 93665389 | rs10993755 | -0.683 | 0.229 | 0.377 | -0.073 | -0.011 | 0.079 | 0.079 | 0 | 0.16 | 0.3 | 0.2 | 0.28 | 0.15 | 0.2 |
| 9 | 93665453 | rs2774241 | 0.342 | -0.464 | 0.616 | -0.306 | -0.02 | -0.004 | -0.004 | 0 | 0.91 | 0.79 | 0.96 | 0.88 | 0.96 | 0.84 |
| 9 | 93667010 | rs291765 | 1.615 | -0.8 | 0.242 | -0.677 | 0.013 | 0 | 0 | 0 | 0.66 | 0.32 | 0.75 | 0.66 | 0.81 | 0.62 |
| 9 | 93667457 | rs1585461 | 0.335 | -0.795 | 0.49 | -0.617 | 0.006 | 0.142 | 0.142 | 0 | 0.06 | 0.15 | 0.06 | 0.19 | 0.06 | 0.12 |
| 9 | 93667894 | rs10125876 | 0.1 | -0.85 | 0.619 | -0.663 | -0.007 | -0.083 | -0.083 | 1.704 | 0.14 | 0.41 | 0.07 | 0.01 | 0.08 | 0.09 |
| 9 | 93668008 | rs291768 | 1.074 | -0.596 | 0.256 | -0.803 | -0.021 | 0.015 | 0.015 | 0 | 0.2 | 0.14 | 0.18 | 0.15 | 0.29 | 0.19 |
| 9 | 93668751 | rs10993760 | 1.304 | -0.708 | 0.444 | -0.607 | 0.001 | 0 | 0 | 0 | 0.07 | 0.07 | 0.11 | 0.09 | 0.03 | 0.04 |
| 9 | 93668891 | rs10993761 | 0.458 | -0.625 | 0.382 | -0.558 | 0.008 | 0.142 | 0.142 | 0 | 0.06 | 0.15 | 0.06 | 0.19 | 0.06 | 0.11 |
| 9 | 93671163 | rs1675315 | 1.285 | -0.631 | 0.039 | -0.614 | 0.015 | -0.001 | -0.001 | 0 | 0.67 | 0.31 | 0.75 | 0.66 | 0.81 | 0.62 |
| 9 | 93681295 | rs291753 | 0.042 | -0.231 | 0.193 | -0.741 | 0.031 | -0.047 | -0.047 | 0 | 0.32 | 0.4 | 0.24 | 0.18 | 0.37 | 0.29 |
| 9 | 93685457 | rs10991692 | 1.054 | -0.356 | 0.001 | -0.3 | 0.01 | 0 | 0 | 0 | 0.07 | 0.06 | 0.03 | 0.09 | 0.03 | 0.04 |
| 9 | 93689819 | rs10991696 | 1.054 | -0.401 | 0.033 | -0.43 | 0.01 | 0 | 0 | 0 | 0.07 | 0.07 | 0.03 | 0.09 | 0.03 | 0.04 |
| 9 | 93692552 | rs169744 | 0.694 | -0.5 | 0.047 | -0.439 | -0.023 | -0.004 | -0.004 | 0.325 | 0.84 | 0.76 | 0.83 | 0.83 | 0.9 | 0.8 |
| 9 | 93692988 | rs10123208 | -0.124 | -0.416 | -0.179 | -0.413 | -0.015 | 0 | 0 | 0 | 0.12 | 0.21 | 0.08 | 0.13 | 0.09 | 0.1 |
| 9 | 93695725 | rs192650 | -0.085 | -0.808 | -0.238 | -0.954 | -0.003 | -0.007 | -0.007 | 0.34 | 0.37 | 0.34 | 0.27 | 0.1 | 0.39 | 0.31 |
| 9 | 93701052 | rs291748 | 0.557 | -0.692 | -0.502 | -0.7 | -0.045 | 0.009 | 0.009 | 0.014 | 0.84 | 0.7 | 0.82 | 0.75 | 0.9 | 0.81 |
| 9 | 93704904 | rs10991704 | 1.146 | -0.672 | -0.646 | -0.7 | 0.015 | -0.002 | -0.002 | 0.164 | 0.08 | 0.08 | 0.03 | 0.09 | 0.04 | 0.05 |
| 9 | 93704926 | rs10991705 | 1.064 | -0.672 | -0.637 | -0.7 | -0.013 | 0 | 0 | 1.862 | 0.08 | 0.12 | 0.04 | 0.09 | 0.04 | 0.05 |
| 9 | 93710061 | rs296663 | -0.431 | -0.985 | -1.273 | -0.547 | 0.021 | 0.015 | 0.015 | 0 | 0.58 | 0.77 | 0.39 | 0.38 | 0.58 | 0.48 |
| 9 | 93712104 | rs12341958 | 0.82 | -1.074 | -1.362 | -0.46 | -0.03 | -0.12 | -0.12 | 0 | 0.36 | 0.09 | 0.47 | 0.47 | 0.37 | 0.44 |
| 9 | 93712701 | rs17569206 | 0.942 | -0.786 | -1.37 | -0.631 | -0.029 | 0.029 | 0.029 | 0 | 0.14 | 0.04 | 0.09 | 0.06 | 0.16 | 0.23 |
| 9 | 93714080 | rs296650 | 0.437 | -0.789 | -1.245 | -0.634 | -0.009 | 0.008 | 0.008 | 0 | 0.38 | 0.62 | 0.32 | 0.27 | 0.43 | 0.19 |
| 9 | 93714388 | rs177040 | 0.304 | -1.173 | -0.964 | -0.373 | 0.036 | 0.059 | 0.059 | 0 | 0.51 | 0.63 | 0.37 | 0.29 | 0.54 | 0.38 |
| 9 | 93718266 | rs10991722 | -0.154 | -1.021 | -1.251 | -0.345 | 0.027 | 0.03 | 0.03 | 0 | 0.43 | 0.25 | 0.53 | 0.6 | 0.45 | 0.53 |
| 9 | 93719846 | rs10739911 | 0.452 | -0.778 | -1.588 | -0.197 | -0.042 | -0.101 | -0.101 | 0 | 0.35 | 0.1 | 0.5 | 0.51 | 0.41 | 0.47 |
| 9 | 93724431 | rs382847 | 0.201 | -1.296 | -1.188 | -0.202 | 0.05 | -0.036 | -0.036 | 0 | 0.63 | 0.23 | 0.68 | 0.5 | 0.72 | 0.69 |
| 9 | 93724516 | rs368754 | -0.236 | -1.488 | -0.744 | -0.193 | -0.016 | 0.006 | 0.006 | 0.332 | 0.85 | 0.8 | 0.81 | 0.69 | 0.89 | 0.8 |
| 9 | 93724845 | rs7032584 | 0.16 | -1.485 | -0.728 | -0.193 | 0.012 | -0.001 | -0.001 | 0 | 0.3 | 0.73 | 0.17 | 0.28 | 0.21 | 0.16 |
| 9 | 93727745 | rs169851 | 0.736 | -1.478 | -1.023 | -0.445 | -0.082 | 0.084 | 0.084 | 0.057 | 0.35 | 0.15 | 0.3 | 0.17 | 0.44 | 0.35 |
| 9 | 93728360 | rs296690 | 1.169 | -1.479 | -1.042 | -0.449 | -0.092 | 0.125 | 0.125 | 0.031 | 0.32 | 0.11 | 0.25 | 0.07 | 0.39 | 0.33 |
| 9 | 93730292 | rs192703 | 0.494 | -0.979 | -0.869 | 0.043 | 0.031 | -0.045 | -0.045 | 0 | 0.61 | 0.19 | 0.56 | 0.43 | 0.64 | 0.67 |
| 9 | 93731058 | rs12352804 | 1.429 | -1.236 | -0.755 | -0.146 | -0.033 | 0.002 | 0.002 | 0.589 | 0.08 | 0.2 | 0.05 | 0.09 | 0.04 | 0.06 |
| 9 | 93731198 | rs10820820 | 1.308 | -1.138 | -0.194 | -0.217 | -0.001 | -0.001 | -0.001 | 0.192 | 0.19 | 0.16 | 0.1 | 0.2 | 0.16 | 0.1 |
| 9 | 93736643 | rs7865547 | 0.998 | -1.117 | -0.186 | -0.224 | -0.001 | -0.001 | -0.001 | 0 | 0.2 | 0.2 | 0.11 | 0.2 | 0.16 | 0.1 |
| 9 | 93740708 | rs296713 | -0.682 | -1.254 | -0.163 | -0.199 | 0.004 | -0.006 | -0.006 | 0 | 0.77 | 0.79 | 0.81 | 0.66 | 0.83 | 0.78 |
| 9 | 93741233 | rs9696872 | 0.822 | -0.908 | -0.31 | -0.458 | -0.058 | -0.125 | -0.125 | 0 | 0.29 | 0.07 | 0.45 | 0.4 | 0.35 | 0.36 |
| 9 | 93745381 | rs296724 | -0.41 | -1.273 | -0.068 | -0.452 | 0.002 | -0.004 | -0.004 | 0 | 0.66 | 0.67 | 0.71 | 0.49 | 0.73 | 0.69 |
| 9 | 93752522 | rs7047670 | -0.71 | -0.991 | 0.6 | -0.535 | 0.057 | 0.157 | 0.157 | 0.823 | 0.05 | 0.38 | 0.09 | 0.12 | 0.01 | 0.11 |
| 9 | 93763731 | rs296610 | 0.309 | -0.823 | 0.064 | -0.662 | -0.062 | 0.096 | 0.096 | 0 | 0.33 | 0.15 | 0.25 | 0.06 | 0.39 | 0.3 |
| 9 | 93774385 | rs296646 | -0.163 | -0.995 | 0.517 | -0.644 | 0.01 | -0.04 | -0.04 | 0 | 0.38 | 0.55 | 0.34 | 0.18 | 0.41 | 0.4 |
| 9 | 93787482 | rs296630 | 0.273 | -0.91 | -0.042 | -0.496 | -0.042 | 0.045 | 0.045 | 0 | 0.33 | 0.23 | 0.26 | 0.06 | 0.41 | 0.28 |
| 9 | 93791410 | rs10121342 | -1.567 | -1.121 | 0.628 | 0.366 | 0.012 | 0.025 | 0.025 | 0.295 | 0.13 | 0.52 | 0.14 | 0.15 | 0.1 | 0.22 |
| 9 | 93793696 | rs12685800 | 0.122 | -0.67 | -0.253 | 0.493 | -0.011 | 0 | 0 | 0 | 0.06 | 0.04 | 0.05 | 0.04 | 0.09 | 0.1 |
| 9 | 93804208 | rs10761105 | -0.13 | -1.309 | -0.556 | 0.827 | -0.021 | -0.113 | -0.113 | 0 | 0.23 | 0.08 | 0.4 | 0.43 | 0.27 | 0.28 |
| 9 | 93824836 | rs7027785 | 0.026 | -1.382 | -0.446 | 0.939 | -0.038 | -0.078 | -0.078 | 0 | 0.18 | 0.08 | 0.39 | 0.44 | 0.28 | 0.27 |
| 9 | 93828321 | rs7025659 | -1.487 | -2.067 | 0.505 | 0.795 | 0.042 | 0.137 | 0.137 | 2.923 | 0.06 | 0.49 | 0.03 | 0.11 | 0.01 | 0.11 |
| 9 | 93829019 | rs296648 | -0.088 | -2.091 | -0.405 | 0.766 | -0.005 | -0.001 | -0.001 | 1.028 | 0.1 | 0.12 | 0.03 | 0.06 | 0.05 | 0.11 |
| 9 | 93834060 | rs10820828 | -0.241 | -2.088 | -0.358 | 1.011 | 0.008 | -0.029 | -0.029 | 0 | 0.23 | 0.15 | 0.25 | 0.46 | 0.2 | 0.35 |
| 9 | 93837417 | rs296614 | -1.172 | -2.94 | -0.419 | 0.843 | 0.03 | 0.015 | 0.015 | 0.009 | 0.46 | 0.37 | 0.34 | 0.66 | 0.29 | 0.56 |
| 9 | 93837497 | rs3853607 | -0.2 | -2.898 | -0.092 | 1.229 | -0.02 | 0.005 | 0.005 | 0.021 | 0.24 | 0.32 | 0.26 | 0.23 | 0.18 | 0.23 |
| 9 | 93840659 | rs296639 | -1.78 | -3.152 | -0.215 | 0.991 | -0.008 | -0.005 | -0.005 | 0.271 | 0.55 | 0.48 | 0.4 | 0.62 | 0.39 | 0.6 |
| 9 | 93840753 | rs7859968 | 0.431 | -3.153 | 0.017 | 1.145 | -0.002 | 0.002 | 0.002 | 0 | 0.2 | 0.16 | 0.24 | 0.22 | 0.16 | 0.2 |
| 9 | 93842021 | rs296638 | -0.655 | -2.952 | -0.125 | 1.137 | 0.044 | -0.032 | -0.032 | 0 | 0.27 | 0.15 | 0.12 | 0.37 | 0.17 | 0.25 |
| 9 | 93843942 | rs296636 | -0.983 | -2.941 | -0.121 | 1.163 | -0.022 | -0.002 | -0.002 | 0 | 0.28 | 0.32 | 0.09 | 0.23 | 0.12 | 0.32 |
| 9 | 93844314 | rs296634 | -0.418 | -2.768 | -0.807 | 1.117 | 0.063 | -0.052 | -0.052 | 0 | 0.23 | 0.11 | 0.08 | 0.36 | 0.12 | 0.21 |
| 9 | 93849856 | rs395612 | -1.705 | -2.501 | -0.619 | 1.015 | 0.005 | 0.002 | 0.002 | 0.205 | 0.54 | 0.46 | 0.42 | 0.59 | 0.39 | 0.61 |

| | | | | | | | | | | | | | | | | |
|---|---|---|---|---|---|---|---|---|---|---|---|---|---|---|---|---|
| 9 | 93851440 | rs393369 | -0.395 | -2.481 | -1.022 | 1.192 | 0.036 | -0.006 | -0.006 | 0 | 0.26 | 0.19 | 0.12 | 0.31 | 0.16 | 0.22 |
| 9 | 93852815 | rs430794 | -0.815 | -2.229 | -0.957 | 1.205 | -0.018 | 0.004 | 0.004 | 0 | 0.44 | 0.5 | 0.38 | 0.51 | 0.32 | 0.43 |
| 9 | 93853457 | rs582337 | -0.411 | -1.69 | -1.121 | 1.296 | 0.024 | -0.029 | -0.029 | 0 | 0.1 | 0.03 | 0.02 | 0.16 | 0.05 | 0.11 |
| 9 | 93858333 | rs365426 | 0.688 | -1.712 | -1.154 | 1.307 | 0.054 | -0.057 | -0.057 | 0 | 0.86 | 0.98 | 0.98 | 0.77 | 0.95 | 0.74 |
| 9 | 93861920 | rs296620 | -1.179 | -1.573 | -0.911 | 0.975 | -0.028 | -0.019 | -0.019 | 0 | 0.39 | 0.52 | 0.26 | 0.74 | 0.27 | 0.63 |
| 9 | 93862080 | rs296618 | -0.094 | -1.037 | -1.255 | 0.978 | 0.011 | -0.016 | -0.016 | 0 | 0.1 | 0 | 0.02 | 0.08 | 0.06 | 0.14 |
| 9 | 93874879 | rs773283 | 0.868 | -0.995 | -1.795 | 0.867 | -0.075 | -0.02 | -0.02 | 0.042 | 0.42 | 0.15 | 0.58 | 0.35 | 0.53 | 0.3 |
| 9 | 93881354 | rs2296751 | -1.383 | -1.306 | -0.744 | 1.265 | -0.01 | 0.008 | 0.008 | 0 | 0.2 | 0.26 | 0.12 | 0.3 | 0.12 | 0.31 |
| 9 | 93887045 | rs2274464 | -0.366 | -1.066 | -1.157 | 1.397 | 0.023 | -0.032 | -0.032 | 0 | 0.1 | 0.01 | 0.02 | 0.11 | 0.05 | 0.14 |
| 9 | 93893941 | rs7036424 | -0.569 | -0.665 | -0.702 | 1.231 | -0.001 | 0.028 | 0.028 | 0.158 | 0.07 | 0.11 | 0.09 | 0.19 | 0.05 | 0.17 |
| 9 | 93899625 | rs10820830 | -0.656 | -0.908 | -0.748 | 0.715 | 0 | 0 | 0 | 0 | 0.23 | 0.26 | 0.27 | 0.17 | 0.16 | 0.19 |
| 9 | 93901159 | rs7033430 | 1.157 | -1.003 | -1.193 | 1.009 | 0.017 | -0.004 | -0.004 | 0.031 | 0.67 | 0.73 | 0.62 | 0.73 | 0.77 | 0.68 |
| 9 | 93901785 | rs7034207 | 0.85 | -1.115 | -1.096 | 1.055 | 0.012 | -0.04 | -0.04 | 0.082 | 0.73 | 0.85 | 0.65 | 0.72 | 0.8 | 0.7 |
| 9 | 93904257 | rs4604577 | 1.463 | -1.324 | -0.74 | 0.312 | 0.024 | 0.085 | 0.085 | 0.475 | 0.51 | 0.25 | 0.5 | 0.65 | 0.58 | 0.57 |
| 9 | 93909589 | rs10991781 | 0.125 | -1.448 | -0.433 | 0.236 | -0.003 | -0.031 | -0.031 | 0 | 0.19 | 0.22 | 0.25 | 0.03 | 0.14 | 0.16 |
| 9 | 93920745 | rs943502 | 0.864 | -1.659 | 0.153 | 0.258 | -0.009 | 0.003 | 0.003 | 1.01 | 0.71 | 0.52 | 0.76 | 0.68 | 0.72 | 0.72 |
| 9 | 93923617 | rs16907199 | -0.779 | -1.587 | 0.437 | 0.014 | 0.007 | -0.08 | -0.08 | 0 | 0.2 | 0.5 | 0.15 | 0.07 | 0.19 | 0.13 |
| 9 | 93925207 | rs12351170 | 0.304 | -1.553 | 0.233 | 0.024 | 0.013 | 0.043 | 0.043 | 0 | 0.16 | 0.08 | 0.21 | 0.04 | 0.11 | 0.14 |
| 9 | 93926928 | rs1887369 | 0.386 | -1.665 | 0.795 | -0.031 | 0.022 | -0.047 | -0.047 | 0 | 0.86 | 0.68 | 0.89 | 0.94 | 0.85 | 0.89 |
| 9 | 93927913 | rs16907201 | 0.366 | -1.631 | 0.3 | -0.032 | 0.012 | 0.039 | 0.039 | 0 | 0.16 | 0.08 | 0.22 | 0.04 | 0.11 | 0.14 |
| 9 | 93930671 | rs10991797 | 1.321 | -1.634 | 0.445 | -0.024 | -0.011 | -0.058 | -0.058 | 2.55 | 0.77 | 0.45 | 0.84 | 0.89 | 0.81 | 0.8 |
| 9 | 93930913 | rs10820838 | 1.03 | -1.502 | 0.338 | -0.028 | -0.001 | -0.096 | -0.096 | 0 | 0.79 | 0.52 | 0.84 | 0.93 | 0.81 | 0.86 |
| 9 | 93934851 | rs4744038 | -0.237 | -1.324 | -0.671 | 0.463 | -0.029 | -0.008 | -0.008 | 0 | 0.21 | 0.01 | 0.29 | 0.22 | 0.16 | 0.29 |
| 9 | 93944244 | rs773519 | 1.316 | -1.456 | -0.27 | 1.022 | 0.004 | -0.007 | -0.007 | 0.072 | 0.79 | 0.18 | 0.8 | 0.72 | 0.83 | 0.79 |
| 9 | 93946379 | rs10991808 | -1.174 | -1.42 | 0.842 | 1.063 | 0.059 | 0.177 | 0.177 | 0 | 0.1 | 0.41 | 0.13 | 0.24 | 0.11 | 0.13 |
| 9 | 93950513 | rs10991816 | -0.544 | -2.212 | 0.165 | 0.713 | 0.024 | 0.019 | 0.019 | 0 | 0.37 | 0.25 | 0.26 | 0.64 | 0.28 | 0.36 |
| 9 | 93952940 | rs10991820 | -0.757 | -2.12 | 0.491 | 0.655 | 0.009 | 0.008 | 0.008 | 0 | 0.39 | 0.36 | 0.28 | 0.63 | 0.3 | 0.38 |
| 9 | 93961118 | rs10512206 | -0.264 | -2.324 | 0.443 | 0.553 | 0.005 | 0.001 | 0.001 | 0 | 0.33 | 0.28 | 0.2 | 0.64 | 0.21 | 0.32 |
| 9 | 93963953 | rs7025775 | -0.252 | -2.332 | 0.437 | 0.549 | 0.005 | 0.001 | 0.001 | 0 | 0.33 | 0.28 | 0.2 | 0.64 | 0.21 | 0.32 |
| 9 | 93969769 | rs7025567 | 0.033 | -2.473 | 0.368 | 1.03 | -0.001 | 0.045 | 0.045 | 0.003 | 0.13 | 0.08 | 0.08 | 0.4 | 0.11 | 0.2 |
| 9 | 93972658 | rs16907277 | 0.095 | -2.481 | 0.472 | 1.104 | 0.006 | -0.019 | -0.019 | 0 | 0.07 | 0.11 | 0.08 | 0.01 | 0.1 | 0.04 |
| 9 | 93974553 | rs773507 | 1.849 | -2.535 | 0.503 | 1.028 | 0.015 | -0.063 | -0.063 | 0 | 0.68 | 0.19 | 0.76 | 0.52 | 0.73 | 0.66 |
| 9 | 93975471 | rs773506 | 1.447 | -2.548 | 0.472 | 0.99 | 0.062 | 0.151 | 0.151 | 0 | 0.49 | 0.14 | 0.65 | 0.28 | 0.63 | 0.54 |
| 9 | 93982492 | rs9785225 | 0.59 | -2.537 | 0.505 | 1.045 | 0.033 | -0.035 | -0.035 | 0 | 0.23 | 0.1 | 0.12 | 0.32 | 0.12 | 0.13 |
| 9 | 93982932 | rs773515 | 1.378 | -2.506 | 0.465 | 0.932 | 0.063 | 0.127 | 0.127 | 0 | 0.49 | 0.16 | 0.65 | 0.34 | 0.63 | 0.57 |
| 9 | 93997764 | rs773518 | 0.182 | -2.406 | 0.617 | 0.586 | 0.013 | -0.044 | -0.044 | 0 | 0.88 | 0.53 | 0.93 | 0.94 | 0.94 | 0.89 |
| 9 | 94004897 | rs7857616 | -0.542 | -2.402 | 0.549 | 0.668 | 0.004 | 0.055 | 0.055 | 0 | 0.17 | 0.26 | 0.16 | 0.34 | 0.19 | 0.23 |
| 9 | 94019410 | rs16907311 | 0.911 | -2.4 | 0.52 | 0.672 | -0.019 | -0.052 | -0.052 | 0.529 | 0.09 | 0 | 0.07 | 0.27 | 0.09 | 0.15 |
| 9 | 94023575 | rs10991853 | -1.037 | -2.397 | 0.589 | 0.774 | 0.004 | 0.075 | 0.075 | 0.027 | 0.19 | 0.27 | 0.17 | 0.35 | 0.2 | 0.23 |
| 9 | 94025991 | rs7043552 | 0.034 | -2.386 | 0.576 | 0.77 | 0.039 | -0.028 | -0.028 | 0 | 0.06 | 0.19 | 0.08 | 0.01 | 0.1 | 0.03 |
| 9 | 94029256 | rs1615811 | -2.903 | -2.632 | -0.07 | 0.491 | -0.171 | 0.145 | 0.145 | 0.471 | 0.73 | 0.96 | 0.65 | 0.95 | 0.5 | 0.66 |
| 9 | 94034568 | rs12238387 | 1.003 | -2.603 | -0.037 | 0.503 | -0.005 | -0.037 | -0.037 | 0 | 0.09 | 0.01 | 0.07 | 0.27 | 0.09 | 0.15 |
| 9 | 94037842 | rs2482340 | -1.631 | -2.662 | -0.014 | 0.499 | 0.042 | 0.082 | 0.082 | 0 | 0.52 | 0.91 | 0.36 | 0.64 | 0.37 | 0.46 |
| 9 | 94037873 | rs2482338 | -1.534 | -2.639 | -0.03 | 0.51 | 0.063 | 0.095 | 0.095 | 0 | 0.51 | 0.84 | 0.36 | 0.64 | 0.37 | 0.46 |
| 9 | 94049879 | rs10991869 | 1.391 | -2.335 | 0.032 | 0.72 | -0.017 | 0.079 | 0.079 | 0 | 0.13 | 0.01 | 0.22 | 0 | 0.09 | 0.09 |
| 9 | 94062978 | rs298728 | -0.018 | -2.294 | 0.537 | 0.964 | 0.006 | -0.035 | -0.035 | 0 | 0.1 | 0.3 | 0.06 | 0.05 | 0.06 | 0.1 |
| 9 | 94076827 | rs10991880 | -1.106 | -2.087 | -0.15 | 1.103 | 0.002 | 0.113 | 0.113 | 0.179 | 0.19 | 0.33 | 0.16 | 0.4 | 0.18 | 0.27 |
| 9 | 94087622 | rs7874056 | -0.222 | -1.978 | -0.294 | 1.046 | 0.04 | -0.023 | -0.023 | 0 | 0.06 | 0.19 | 0.09 | 0.01 | 0.1 | 0.03 |
| 9 | 94108757 | rs7868771 | -0.164 | -2.149 | -0.927 | 1.156 | 0.002 | 0.011 | 0.011 | 0.062 | 0.13 | 0.08 | 0.08 | 0.34 | 0.1 | 0.19 |
| 9 | 94120749 | rs12552364 | 0.83 | -2.011 | -1.106 | 0.979 | 0.012 | -0.085 | -0.085 | 0 | 0.2 | 0.01 | 0.28 | 0.3 | 0.12 | 0.21 |
| 9 | 94141918 | rs774224 | -1.197 | -1.671 | -0.456 | 0.504 | -0.005 | 0.012 | 0.012 | 0 | 0.3 | 0.49 | 0.2 | 0.36 | 0.24 | 0.29 |
| 9 | 94153356 | rs2004308 | -0.735 | -1.667 | -0.476 | 0.552 | -0.001 | -0.038 | -0.038 | 0 | 0.25 | 0.46 | 0.14 | 0.19 | 0.18 | 0.22 |
| 9 | 94153503 | rs10991917 | -0.62 | -1.548 | -0.728 | 0.476 | 0.017 | -0.077 | -0.077 | 4.215 | 0.23 | 0.35 | 0.15 | 0.09 | 0.22 | 0.16 |
| 9 | 94154803 | rs16907447 | 0.29 | -1.29 | -0.954 | 0.558 | 0.01 | 0.099 | 0.099 | 0 | 0.12 | 0.23 | 0.1 | 0.25 | 0.13 | 0.15 |
| 9 | 94157846 | rs10991921 | 0.042 | -1.282 | -0.958 | 0.555 | 0.008 | 0.086 | 0.086 | 0 | 0.13 | 0.23 | 0.11 | 0.25 | 0.13 | 0.16 |
| 9 | 94161700 | rs894962 | -1.655 | -0.906 | -1.074 | -0.499 | -0.117 | 0.127 | 0.127 | 1.912 | 0.73 | 0.91 | 0.65 | 0.95 | 0.51 | 0.67 |
| 9 | 94162081 | rs883121 | -0.19 | -0.843 | -0.588 | 0.283 | 0.007 | -0.002 | -0.002 | 0 | 0.34 | 0.24 | 0.39 | 0.59 | 0.28 | 0.36 |
| 9 | 94164373 | rs7024443 | -0.954 | -0.537 | 0.024 | 0.168 | -0.011 | -0.048 | -0.048 | 1.928 | 0.19 | 0.48 | 0.13 | 0.15 | 0.12 | 0.17 |
| 9 | 94165889 | rs1758695 | 0.126 | -0.096 | -0.496 | 0.136 | -0.001 | -0.004 | -0.004 | 0 | 0.57 | 0.59 | 0.48 | 0.28 | 0.58 | 0.54 |
| 9 | 94176261 | rs7021746 | -0.273 | -1.079 | -0.951 | 0.736 | 0.074 | 0.03 | 0.03 | 0.647 | 0.44 | 0.77 | 0.33 | 0.3 | 0.26 | 0.35 |
| 9 | 94179978 | rs2482357 | -0.943 | -1.314 | -1.395 | 1.028 | -0.028 | -0.033 | -0.033 | 0.001 | 0.62 | 0.86 | 0.51 | 0.52 | 0.44 | 0.48 |
| 9 | 94181951 | rs2440592 | 0.162 | -1.371 | -1.41 | 0.93 | -0.002 | -0.011 | -0.011 | 2.233 | 0.16 | 0.02 | 0.17 | 0.14 | 0.16 | 0.13 |
| 9 | 94182363 | rs2482704 | -0.703 | -1.374 | -1.139 | 0.965 | -0.002 | -0.042 | -0.042 | 0 | 0.61 | 0.79 | 0.5 | 0.44 | 0.43 | 0.48 |
| 9 | 94188141 | rs13297268 | 0.101 | -1.49 | -0.665 | 1.364 | 0.029 | 0.015 | 0.015 | 0.773 | 0.11 | 0.02 | 0.03 | 0.06 | 0.07 | 0.05 |
| 9 | 94190241 | rs10820853 | -0.665 | -1.951 | -0.189 | 1.699 | -0.037 | 0.052 | 0.052 | 0.617 | 0.16 | 0.32 | 0.09 | 0.26 | 0.1 | 0.28 |
| 9 | 94190673 | rs7032571 | -1.366 | -2.119 | -0.111 | 1.72 | -0.026 | 0.011 | 0.011 | 0.12 | 0.28 | 0.34 | 0.12 | 0.34 | 0.17 | 0.33 |
| 9 | 94192804 | rs1467780 | -0.069 | -2.064 | -0.203 | 1.696 | -0.031 | 0.009 | 0.009 | 0.153 | 0.11 | 0.19 | 0.04 | 0.1 | 0.05 | 0.19 |
| 9 | 94193941 | rs774227 | 1.622 | -2.48 | -0.816 | 1.636 | 0.161 | -0.007 | -0.007 | 3.268 | 0.54 | 0.14 | 0.75 | 0.5 | 0.74 | 0.56 |
| 9 | 94199485 | rs809568 | 0.036 | -1.985 | -0.114 | 1.394 | -0.001 | -0.03 | -0.03 | 0.072 | 0.94 | 0.91 | 0.96 | 1 | 0.95 | 0.99 |

| | | | | | | | | | | | | | | | | |
|---|---|---|---|---|---|---|---|---|---|---|---|---|---|---|---|---|
| 9 | 94202927 | rs11535536 | -0.657 | -2.246 | -0.172 | 1.971 | 0.011 | -0.014 | -0.014 | 0 | 0.19 | 0.13 | 0.08 | 0.32 | 0.12 | 0.22 |
| 9 | 94214002 | rs1412470 | -2.26 | -2.695 | -0.445 | 2.053 | 0.155 | 0.013 | 0.013 | 0.443 | 0.43 | 0.88 | 0.26 | 0.63 | 0.28 | 0.42 |
| 9 | 94217301 | rs1981030 | -1.61 | -2.763 | -0.044 | 2.185 | -0.034 | 0.041 | 0.041 | 0.902 | 0.25 | 0.66 | 0.17 | 0.54 | 0.17 | 0.35 |
| 9 | 94221535 | rs7867022 | -1.294 | -2.81 | 0.39 | 2.329 | -0.028 | 0.028 | 0.028 | 0 | 0.25 | 0.29 | 0.11 | 0.47 | 0.13 | 0.26 |
| 9 | 94222921 | rs10820857 | -0.971 | -2.8 | 0.389 | 2.336 | -0.037 | 0.079 | 0.079 | 0 | 0.17 | 0.28 | 0.1 | 0.38 | 0.1 | 0.22 |
| 9 | 94229429 | rs12345170 | -0.694 | -2.752 | 0.455 | 2.29 | -0.032 | 0.012 | 0.012 | 0 | 0.16 | 0.21 | 0.04 | 0.25 | 0.05 | 0.09 |
| 9 | 94236664 | rs6479339 | -1.868 | -2.812 | 0.379 | 2.212 | -0.013 | -0.001 | -0.001 | 0 | 0.29 | 0.67 | 0.19 | 0.29 | 0.18 | 0.23 |
| 9 | 94240052 | rs12685188 | 1.069 | -2.655 | 0.265 | 2.189 | 0.033 | -0.018 | -0.018 | 0.25 | 0.08 | 0.01 | 0.01 | 0.09 | 0.03 | 0.04 |
| 9 | 94245669 | rs16907524 | -0.662 | -2.719 | 0.579 | 2.46 | -0.06 | 0.148 | 0.148 | 0.076 | 0.15 | 0.53 | 0.09 | 0.31 | 0.06 | 0.11 |
| 9 | 94246676 | rs4743836 | -0.747 | -2.757 | 0.444 | 2.416 | -0.002 | -0.001 | -0.001 | 3.551 | 0.21 | 0.2 | 0.16 | 0.13 | 0.18 | 0.24 |
| 9 | 94249082 | rs4744055 | -2.37 | -2.427 | 0.213 | 2.362 | 0.108 | 0.056 | 0.056 | 0 | 0.45 | 0.88 | 0.3 | 0.67 | 0.33 | 0.46 |
| 9 | 94258836 | rs1571892 | -2.17 | -2.276 | 0.286 | 2.413 | 0.041 | -0.127 | -0.127 | 0 | 0.34 | 0.87 | 0.26 | 0.48 | 0.26 | 0.37 |
| 9 | 94259474 | rs7032300 | -1.501 | -2.291 | 0.577 | 2.52 | -0.06 | 0.092 | 0.092 | 0 | 0.22 | 0.59 | 0.12 | 0.44 | 0.1 | 0.21 |
| 9 | 94260376 | rs7035655 | -0.956 | -2.281 | 0.671 | 2.469 | 0.066 | 0.111 | 0.111 | 3.632 | 0.07 | 0.47 | 0.05 | 0.07 | 0.03 | 0.07 |
| 9 | 94261753 | rs6479342 | -1.95 | -2.31 | 0.639 | 2.453 | -0.043 | 0.051 | 0.051 | 0 | 0.24 | 0.63 | 0.13 | 0.45 | 0.11 | 0.25 |
| 9 | 94264346 | rs12350612 | -1.351 | -2.299 | 0.631 | 2.439 | -0.028 | 0.037 | 0.037 | 0 | 0.19 | 0.23 | 0.08 | 0.38 | 0.08 | 0.18 |
| 9 | 94264616 | rs12350727 | -0.206 | -2.274 | 0.619 | 2.152 | 0.013 | -0.008 | -0.008 | 0 | 0.13 | 0.11 | 0.06 | 0.19 | 0.06 | 0.13 |
| 9 | 94266286 | rs768056 | -0.617 | -2.356 | 0.534 | 2.139 | -0.017 | 0.005 | 0.005 | 0 | 0.19 | 0.24 | 0.15 | 0.19 | 0.16 | 0.16 |
| 9 | 94267243 | rs16907551 | -0.398 | -2.356 | 0.475 | 2.129 | 0.012 | 0.021 | 0.021 | 0 | 0.18 | 0.1 | 0.12 | 0.04 | 0.16 | 0.16 |
| 9 | 94269967 | rs7868550 | -2.14 | -2.313 | 0.395 | 2.182 | 0.048 | -0.063 | -0.063 | 0 | 0.35 | 0.86 | 0.26 | 0.43 | 0.27 | 0.37 |
| 9 | 94270414 | rs7025323 | -0.178 | -2.309 | 0.424 | 2.188 | -0.016 | 0.056 | 0.056 | 0 | 0.06 | 0.12 | 0.02 | 0.14 | 0.02 | 0.05 |
| 9 | 94273334 | rs7038274 | -1.665 | -2.3 | 0.425 | 2.168 | 0.008 | 0 | 0 | 1.4 | 0.29 | 0.74 | 0.24 | 0.29 | 0.25 | 0.32 |
| 9 | 94279402 | rs10820865 | -2.58 | -1.875 | 0.207 | 2.128 | 0.06 | -0.042 | -0.042 | 0.095 | 0.45 | 0.89 | 0.31 | 0.59 | 0.34 | 0.46 |
| 9 | 94281499 | rs7856591 | -2.085 | -1.825 | 0.341 | 2.127 | 0.022 | -0.049 | -0.049 | 0 | 0.27 | 0.78 | 0.17 | 0.54 | 0.18 | 0.29 |
| 9 | 94291378 | rs10820870 | 0.694 | -1.761 | 0.296 | 1.983 | -0.003 | -0.041 | -0.041 | 0.887 | 0.1 | 0.01 | 0.04 | 0.16 | 0.08 | 0.09 |
| 9 | 94310766 | rs10481715 | 0.486 | -1.769 | 0.371 | 1.903 | 0 | -0.001 | -0.001 | 0.487 | 0.1 | 0.12 | 0.05 | 0.14 | 0.08 | 0.1 |
| 9 | 94312726 | rs9409602 | -1.823 | -1.889 | 0.573 | 1.882 | -0.028 | 0.016 | 0.016 | 0 | 0.37 | 0.44 | 0.23 | 0.45 | 0.28 | 0.31 |
| 9 | 94313839 | rs10991978 | -0.248 | -1.879 | 0.464 | 1.897 | -0.017 | 0.053 | 0.053 | 0 | 0.06 | 0.15 | 0.02 | 0.13 | 0.02 | 0.05 |
| 9 | 94314110 | rs6479358 | -2.421 | -1.85 | 0.43 | 1.896 | 0.069 | -0.041 | -0.041 | 0.326 | 0.47 | 0.91 | 0.31 | 0.57 | 0.36 | 0.48 |
| 9 | 94320054 | rs1546101 | -2.368 | -1.774 | 0.407 | 1.896 | 0.057 | -0.044 | -0.044 | 0.012 | 0.47 | 0.91 | 0.32 | 0.57 | 0.37 | 0.48 |
| 9 | 94331204 | rs10991990 | -0.467 | -1.722 | 0.392 | 1.854 | -0.001 | 0.001 | 0.001 | 0 | 0.17 | 0.16 | 0.09 | 0.26 | 0.15 | 0.15 |
| 9 | 94331504 | rs10991991 | -0.598 | -1.719 | 0.386 | 1.855 | -0.005 | 0.005 | 0.005 | 0 | 0.18 | 0.2 | 0.1 | 0.26 | 0.15 | 0.15 |
| 9 | 94333426 | rs1388966 | -2.494 | -1.723 | 0.203 | 1.864 | 0.061 | -0.017 | -0.017 | 0 | 0.49 | 0.91 | 0.34 | 0.57 | 0.41 | 0.48 |
| 9 | 94333599 | rs902923 | -0.614 | -1.223 | 0.6 | 1.577 | 0.009 | -0.001 | -0.001 | 0 | 0.2 | 0.16 | 0.15 | 0.19 | 0.18 | 0.17 |
| 9 | 94333889 | rs902921 | -1.463 | -0.973 | 0.746 | 1.456 | 0.073 | 0.063 | 0.063 | 2.359 | 0.09 | 0.42 | 0.08 | 0.12 | 0.08 | 0.16 |
| 9 | 94339510 | rs9409609 | -0.853 | -1.34 | 0.592 | 1.433 | -0.033 | -0.003 | -0.003 | 0 | 0.25 | 0.35 | 0.18 | 0.21 | 0.19 | 0.17 |
| 9 | 94341698 | rs9409610 | -0.889 | -1.367 | 0.579 | 1.42 | -0.033 | -0.004 | -0.004 | 0 | 0.26 | 0.35 | 0.18 | 0.21 | 0.19 | 0.17 |
| 9 | 94343125 | rs10991995 | -1.489 | -0.595 | 0.405 | 1.373 | 0.006 | -0.006 | -0.006 | 0 | 0.12 | 0.17 | 0.11 | 0.1 | 0.17 | 0.14 |
| 9 | 94363366 | rs7033293 | -1.218 | -0.995 | 0.535 | 1.252 | 0.005 | -0.006 | -0.006 | 3.093 | 0.2 | 0.2 | 0.21 | 0.19 | 0.27 | 0.21 |
| 9 | 94368561 | rs1027269 | -2.545 | -1.145 | 0.699 | 1.178 | 0.026 | 0.014 | 0.014 | 0.57 | 0.44 | 0.82 | 0.35 | 0.4 | 0.42 | 0.41 |
| 9 | 94369120 | rs4744070 | -1.186 | -1.148 | 0.708 | 1.178 | -0.05 | 0.003 | 0.003 | 1.112 | 0.24 | 0.62 | 0.14 | 0.21 | 0.15 | 0.2 |
| 9 | 94370675 | rs1319000 | -1.163 | -1.131 | 0.706 | 1.177 | 0.005 | -0.006 | -0.006 | 0.479 | 0.2 | 0.2 | 0.21 | 0.19 | 0.27 | 0.21 |
| 9 | 94372455 | rs7032779 | -1.018 | -1.133 | 0.782 | 1.164 | -0.006 | -0.025 | -0.025 | 0 | 0.18 | 0.23 | 0.13 | 0.1 | 0.15 | 0.18 |
| 9 | 94373024 | rs7850575 | -1.27 | -1.082 | 0.716 | 1.157 | -0.042 | -0.001 | -0.001 | 0.685 | 0.25 | 0.62 | 0.14 | 0.21 | 0.15 | 0.22 |
| 9 | 94374714 | rs10512215 | 2.006 | -0.3 | -0.215 | 0.911 | -0.052 | -0.046 | -0.046 | 0 | 0.28 | 0.04 | 0.13 | 0.22 | 0.27 | 0.15 |
| 9 | 94379035 | rs1021439 | 0.293 | -0.384 | 0.461 | 0.565 | -0.042 | -0.054 | -0.054 | 0.769 | 0.16 | 0.31 | 0.1 | 0.04 | 0.11 | 0.13 |
| 9 | 94379459 | rs1021441 | -1.536 | -0.362 | 0.463 | 0.567 | 0.003 | 0.021 | 0.021 | 0 | 0.37 | 0.64 | 0.32 | 0.23 | 0.39 | 0.36 |
| 9 | 94387597 | rs4744084 | -1.521 | -0.365 | 0.462 | 0.566 | 0.003 | 0.02 | 0.02 | 0 | 0.37 | 0.64 | 0.32 | 0.23 | 0.39 | 0.36 |
| 9 | 94391378 | rs12340837 | 0.114 | -0.365 | 0.529 | 0.559 | -0.053 | -0.078 | -0.078 | 0 | 0.17 | 0.44 | 0.11 | 0.04 | 0.12 | 0.14 |
| 9 | 94391746 | rs9695578 | -2.155 | -0.388 | 0.516 | 0.617 | 0.027 | 0.012 | 0.012 | 0.57 | 0.44 | 0.81 | 0.35 | 0.4 | 0.42 | 0.41 |
| 9 | 94394640 | rs12339155 | -0.863 | -0.104 | 0.482 | 0.706 | -0.004 | -0.065 | -0.065 | 0 | 0.24 | 0.45 | 0.2 | 0.14 | 0.26 | 0.23 |
| 9 | 94395522 | rs12340405 | -0.022 | -0.089 | 0.556 | 0.72 | -0.045 | -0.064 | -0.064 | 7.082 | 0.17 | 0.35 | 0.1 | 0.04 | 0.12 | 0.14 |
| 9 | 94396059 | rs1492682 | -0.136 | -0.086 | 0.344 | 0.722 | -0.057 | -0.082 | -0.082 | 0.524 | 0.18 | 0.44 | 0.11 | 0.04 | 0.12 | 0.14 |
| 9 | 94396676 | rs10992022 | 0.102 | 0.018 | 0.269 | 0.766 | -0.034 | -0.011 | -0.011 | 0 | 0.06 | 0.02 | 0.1 | 0.1 | 0.15 | 0.09 |
| 9 | 94403445 | rs2131304 | -0.246 | -0.168 | 0.382 | 0.322 | -0.002 | -0.02 | -0.02 | 0 | 0.14 | 0.18 | 0.12 | 0.09 | 0.13 | 0.13 |
| 9 | 94405559 | rs10992026 | -0.248 | -0.504 | 0.433 | 0.152 | -0.057 | -0.082 | -0.082 | 0 | 0.18 | 0.44 | 0.11 | 0.04 | 0.12 | 0.14 |
| 9 | 94406084 | rs1492685 | -0.495 | 0.081 | 0.01 | 0.624 | -0.002 | -0.019 | -0.019 | 0 | 0.14 | 0.18 | 0.11 | 0.1 | 0.13 | 0.13 |
| 9 | 94406606 | rs9409636 | -0.495 | 0.081 | 0.01 | 0.618 | -0.002 | -0.006 | -0.006 | 1.274 | 0.14 | 0.18 | 0.12 | 0.12 | 0.13 | 0.13 |
| 9 | 94408117 | rs10992029 | 0.378 | 0.114 | -0.34 | 0.639 | -0.001 | -0.001 | -0.001 | 0 | 0.43 | 0.41 | 0.26 | 0.44 | 0.41 | 0.4 |
| 9 | 94408473 | rs7851193 | -0.954 | 0.064 | -0.761 | 0.121 | -0.017 | -0.07 | -0.07 | 0 | 0.33 | 0.52 | 0.22 | 0.09 | 0.29 | 0.34 |
| 9 | 94412540 | rs9409449 | -0.29 | -0.034 | -0.089 | 0.041 | 0.014 | -0.048 | -0.048 | 0.293 | 0.15 | 0.29 | 0.13 | 0.04 | 0.15 | 0.13 |
| 9 | 94414784 | rs2076773 | 0.145 | -0.117 | -0.168 | 0.043 | -0.012 | -0.027 | -0.027 | 0 | 0.16 | 0.23 | 0.09 | 0.05 | 0.13 | 0.22 |
| 9 | 94417069 | rs9299395 | -1.151 | 0.472 | -0.953 | 0.002 | -0.063 | -0.065 | -0.065 | 0.269 | 0.68 | 0.92 | 0.6 | 0.57 | 0.66 | 0.69 |
| 9 | 94420140 | rs7032274 | 0.008 | 0.389 | -0.349 | -0.075 | -0.001 | 0.019 | 0.019 | 0 | 0.51 | 0.39 | 0.46 | 0.53 | 0.51 | 0.6 |
| 9 | 94422821 | rs17506543 | 0.403 | 0.598 | -0.384 | 0.012 | -0.01 | -0.005 | -0.005 | 0 | 0.05 | 0.02 | 0.22 | 0.11 | 0.12 | 0.12 |
| 9 | 94427820 | rs9299396 | 0.15 | 0.383 | -0.219 | -0.074 | 0.01 | -0.054 | -0.054 | 0 | 0.16 | 0.3 | 0.13 | 0.04 | 0.16 | 0.15 |
| 9 | 94428149 | rs10992039 | 1.706 | 0.486 | -0.296 | -0.036 | 0.028 | -0.076 | -0.076 | 8.682 | 0.24 | 0.08 | 0.09 | 0.3 | 0.19 | 0.17 |
| 9 | 94437294 | rs10046802 | 1.721 | 0.432 | -0.232 | -0.06 | -0.007 | 0.01 | 0.01 | 0.143 | 0.25 | 0.27 | 0.1 | 0.3 | 0.19 | 0.17 |
| 9 | 94441201 | rs1388967 | -1.169 | 0.459 | -0.2 | -0.108 | -0.006 | 0.005 | 0.005 | 0 | 0.6 | 0.65 | 0.58 | 0.57 | 0.58 | 0.65 |

| | | | | | | | | | | | | | | | | |
|---|---|---|---|---|---|---|---|---|---|---|---|---|---|---|---|---|
| 9 | 94441745 | rs10992043 | -0.341 | 0.517 | -0.227 | -0.076 | 0 | 0.004 | 0.004 | 0 | 0.46 | 0.57 | 0.5 | 0.53 | 0.45 | 0.4 |
| 9 | 94444830 | rs10820896 | 1.599 | 0.6 | -0.416 | 0.281 | 0.025 | -0.043 | -0.043 | 0 | 0.22 | 0.13 | 0.09 | 0.29 | 0.17 | 0.16 |
| 9 | 94445809 | rs2892376 | 0.297 | 0.665 | -0.476 | 0.252 | -0.002 | -0.004 | -0.004 | 0 | 0.33 | 0.29 | 0.42 | 0.51 | 0.36 | 0.32 |
| 9 | 94445943 | rs2387871 | -0.798 | 1.128 | -0.495 | 0.148 | -0.007 | -0.008 | -0.008 | 1.264 | 0.1 | 0.08 | 0.33 | 0.22 | 0.18 | 0.16 |
| 9 | 94448777 | rs12115281 | -0.109 | 0.482 | -0.53 | -0.265 | 0.008 | 0.019 | 0.019 | 0 | 0.17 | 0.12 | 0.07 | 0.04 | 0.13 | 0.25 |
| 9 | 94454217 | rs1201041 | 0.542 | 0.581 | -0.067 | -0.124 | 0.019 | -0.015 | -0.015 | 0 | 0.74 | 0.63 | 0.76 | 0.78 | 0.71 | 0.57 |
| 9 | 94455171 | rs10992053 | 0.884 | 0.155 | -0.969 | -0.158 | 0.017 | -0.022 | -0.022 | 0 | 0.29 | 0.24 | 0.15 | 0.33 | 0.2 | 0.22 |
| 9 | 94456714 | rs923769 | -0.347 | 0.153 | -0.699 | -0.188 | 0 | -0.008 | -0.008 | 0 | 0.8 | 0.77 | 0.87 | 0.92 | 0.79 | 0.67 |
| 9 | 94458669 | rs12348947 | 1.001 | 0.139 | -0.763 | -0.014 | 0.02 | -0.039 | -0.039 | 0 | 0.27 | 0.16 | 0.15 | 0.33 | 0.2 | 0.22 |
| 9 | 94482041 | rs10820899 | 0.939 | 0.309 | -0.806 | -0.226 | 0.008 | -0.001 | -0.001 | 0.019 | 0.37 | 0.5 | 0.54 | 0.42 | 0.42 | 0.28 |
| 9 | 94483198 | rs7466603 | -0.112 | 0.249 | -0.871 | -0.276 | -0.002 | 0.035 | 0.035 | 0 | 0.7 | 0.78 | 0.79 | 0.91 | 0.71 | 0.6 |
| 9 | 94483759 | rs10992058 | 1.012 | 0.28 | -0.833 | -0.241 | 0.007 | -0.002 | -0.002 | 0 | 0.37 | 0.5 | 0.56 | 0.43 | 0.42 | 0.28 |
| 9 | 94486321 | rs10761129 | 0.238 | 0.675 | -1.283 | -0.304 | -0.005 | 0.041 | 0.041 | 0 | 0.69 | 0.78 | 0.77 | 0.91 | 0.69 | 0.58 |
| 9 | 94487459 | rs10992065 | 0.47 | 0.499 | -1.719 | 0.059 | -0.019 | -0.031 | -0.031 | 0.569 | 0.31 | 0.11 | 0.36 | 0.38 | 0.27 | 0.27 |
| 9 | 94488415 | rs4073735 | -1.061 | -0.074 | -1.375 | -0.047 | -0.018 | 0.006 | 0.006 | 0.66 | 0.55 | 0.74 | 0.38 | 0.55 | 0.44 | 0.6 |
| 9 | 94489212 | rs10992066 | 1.059 | -0.345 | -1.028 | -0.323 | -0.003 | -0.036 | -0.036 | 0 | 0.12 | 0.11 | 0.1 | 0 | 0.11 | 0.03 |
| 9 | 94497680 | rs4237215 | 0.61 | -0.493 | -0.988 | -0.388 | 0.007 | -0.03 | -0.03 | 0 | 0.62 | 0.75 | 0.52 | 0.53 | 0.66 | 0.68 |
| 9 | 94498550 | rs9409651 | 1.879 | -0.502 | -0.864 | -0.435 | -0.036 | -0.018 | -0.018 | 0.064 | 0.41 | 0.27 | 0.41 | 0.46 | 0.51 | 0.38 |
| 9 | 94499797 | rs16907720 | 1.58 | -0.395 | -0.149 | -0.503 | 0.007 | -0.043 | -0.043 | 0 | 0.08 | 0.2 | 0.05 | 0 | 0.05 | 0.01 |
| 9 | 94501891 | rs10992075 | -0.507 | -0.455 | 0.098 | -0.306 | -0.004 | 0.012 | 0.012 | 0 | 0.29 | 0.41 | 0.35 | 0.31 | 0.26 | 0.24 |
| 9 | 94506153 | rs9409653 | -0.171 | -0.541 | -0.014 | -0.025 | -0.003 | -0.023 | -0.023 | 0 | 0.76 | 0.86 | 0.76 | 0.71 | 0.76 | 0.77 |
| 9 | 94508700 | rs3935601 | 1.327 | -0.724 | 0.148 | 0.082 | 0.006 | 0.005 | 0.005 | 0 | 0.49 | 0.52 | 0.48 | 0.44 | 0.57 | 0.43 |
| 9 | 94510145 | rs16907725 | -0.19 | -0.613 | -0.042 | 0.191 | -0.012 | 0.026 | 0.026 | 0 | 0.18 | 0.28 | 0.24 | 0.27 | 0.12 | 0.33 |
| 9 | 94514322 | rs10739919 | -0.113 | -0.292 | -0.601 | 0.287 | -0.006 | -0.028 | -0.028 | 0 | 0.77 | 0.87 | 0.76 | 0.71 | 0.76 | 0.77 |
| 9 | 94517715 | rs4467996 | 0.678 | -0.264 | -0.411 | 0.321 | 0.009 | 0.007 | 0.007 | 0.094 | 0.56 | 0.72 | 0.69 | 0.6 | 0.62 | 0.48 |
| 9 | 94525452 | rs7860482 | 0.123 | 0.048 | -0.256 | 0.265 | -0.001 | -0.078 | -0.078 | 0 | 0.15 | 0.21 | 0.08 | 0 | 0.11 | 0.04 |
| 9 | 94525991 | rs17585941 | -0.879 | -0.026 | -0.316 | 0.32 | -0.005 | 0.016 | 0.016 | 0 | 0.2 | 0.1 | 0.26 | 0.16 | 0.15 | 0.1 |
| 9 | 94528401 | rs10992085 | 0.029 | -0.435 | 0.227 | 0.588 | -0.006 | -0.011 | -0.011 | 4.312 | 0.22 | 0.28 | 0.1 | 0.16 | 0.13 | 0.29 |
| 9 | 94530989 | rs9409461 | 2.339 | -0.238 | -0.663 | 0.664 | -0.099 | 0.029 | 0.029 | 0.813 | 0.37 | 0.03 | 0.4 | 0.23 | 0.49 | 0.36 |
| 9 | 94532382 | rs7026175 | -0.94 | -0.752 | -0.145 | 0.683 | 0.004 | 0.027 | 0.027 | 0 | 0.22 | 0.33 | 0.35 | 0.26 | 0.18 | 0.11 |
| 9 | 94533121 | rs10992087 | -1.044 | -0.745 | -0.135 | 0.814 | 0.006 | 0.112 | 0.112 | 0 | 0.07 | 0.12 | 0.27 | 0.25 | 0.08 | 0.08 |
| 9 | 94539018 | rs4467997 | -1.446 | -0.892 | -0.202 | 0.72 | -0.002 | -0.002 | -0.002 | 0 | 0.4 | 0.33 | 0.4 | 0.33 | 0.31 | 0.36 |
| 9 | 94542019 | rs10992097 | -0.917 | -0.884 | -0.589 | 0.607 | 0.009 | 0.016 | 0.016 | 0.089 | 0.34 | 0.23 | 0.15 | 0.11 | 0.23 | 0.27 |
| 9 | 94542162 | rs7029673 | -0.806 | -0.862 | -0.605 | 0.602 | 0.007 | 0.015 | 0.015 | 0.153 | 0.33 | 0.22 | 0.14 | 0.07 | 0.23 | 0.27 |
| 9 | 94546564 | rs3935382 | -0.831 | -0.794 | -0.676 | 0.676 | 0.016 | 0.016 | 0.016 | 0 | 0.29 | 0.75 | 0.35 | 0.24 | 0.16 | 0.14 |
| 9 | 94547058 | rs4378021 | 1.322 | -0.575 | -0.695 | 1.379 | -0.009 | 0.022 | 0.022 | 0.421 | 0.43 | 0.14 | 0.48 | 0.3 | 0.62 | 0.4 |
| 9 | 94561730 | rs2103301 | -2.383 | -0.952 | 0.768 | 2.38 | 0.024 | 0.519 | 0.519 | 4.251 | 0.09 | 0.5 | 0.11 | 0.46 | 0.05 | 0.16 |
| 9 | 94562681 | rs7029814 | -1.582 | -0.907 | 0.162 | 2.068 | -0.015 | 0.03 | 0.03 | 2.078 | 0.47 | 0.94 | 0.56 | 0.69 | 0.31 | 0.42 |
| 9 | 94570125 | rs7867961 | -2.012 | -1.022 | 1.133 | 2.487 | 0.054 | 0.313 | 0.313 | 2.284 | 0.07 | 0.35 | 0.09 | 0.3 | 0.05 | 0.09 |
| 9 | 94573753 | rs10992121 | -2.587 | -1.019 | 1.045 | 2.067 | 0.016 | 0.374 | 0.374 | 7.515 | 0.1 | 0.57 | 0.07 | 0.35 | 0.05 | 0.17 |
| 9 | 94577828 | rs7041224 | -1.892 | -1.011 | 0.757 | 2.056 | 0.038 | 0.365 | 0.365 | 2.681 | 0.05 | 0.38 | 0.03 | 0.21 | 0.01 | 0.08 |
| 9 | 94581258 | rs4743855 | -1.711 | -1.275 | -0.654 | 1.352 | -0.023 | -0.004 | -0.004 | 0 | 0.52 | 0.95 | 0.57 | 0.62 | 0.38 | 0.44 |
| 9 | 94589135 | rs12551140 | -1.351 | -1.262 | -0.03 | 1.361 | 0.047 | -0.009 | -0.009 | 1.336 | 0.44 | 0.69 | 0.55 | 0.45 | 0.29 | 0.33 |
| 9 | 94597646 | rs7867707 | -0.207 | -1.451 | 0.086 | 1.349 | -0.018 | -0.044 | -0.044 | 5.386 | 0.22 | 0.14 | 0.48 | 0.34 | 0.25 | 0.25 |
| 9 | 94600207 | rs17587223 | -0.055 | -1.368 | -0.385 | 1.082 | -0.073 | 0.011 | 0.011 | 0 | 0.18 | 0.01 | 0.1 | 0.08 | 0.21 | 0.17 |
| 9 | 94601233 | rs12003273 | -0.009 | -1.886 | 0.869 | 1.009 | -0.09 | -0.023 | -0.023 | 0 | 0.14 | 0.38 | 0.05 | 0.11 | 0.03 | 0.08 |
| 9 | 94603970 | rs12002478 | -0.718 | -1.903 | 1.005 | 1.043 | -0.11 | -0.01 | -0.01 | 0 | 0.18 | 0.5 | 0.09 | 0.19 | 0.08 | 0.15 |
| 9 | 94609451 | rs2141368 | -0.028 | -1.791 | 0.947 | 1.138 | -0.003 | 0.002 | 0.002 | 0 | 0.2 | 0.14 | 0.3 | 0.18 | 0.22 | 0.15 |
| 9 | 94614626 | rs10992132 | -0.327 | -1.803 | 0.996 | 0.882 | 0.003 | 0.017 | 0.017 | 0 | 0.23 | 0.25 | 0.49 | 0.37 | 0.25 | 0.25 |
| 9 | 94625960 | rs10992142 | -1.047 | -1.854 | 1.028 | 0.801 | -0.009 | 0.009 | 0.009 | 0.696 | 0.35 | 0.55 | 0.54 | 0.42 | 0.28 | 0.33 |
| 9 | 94626543 | rs7850118 | -1.984 | -1.933 | 1.036 | 0.772 | 0.088 | -0.017 | -0.017 | 0.167 | 0.47 | 0.79 | 0.59 | 0.52 | 0.33 | 0.39 |
| 9 | 94629592 | rs12552858 | -0.001 | -1.918 | 1.065 | 0.782 | 0 | 0.004 | 0.004 | 0 | 0.22 | 0.19 | 0.49 | 0.34 | 0.25 | 0.24 |
| 9 | 94631630 | rs10992144 | -1.466 | -1.944 | 1.125 | 0.775 | -0.002 | 0.002 | 0.002 | 0 | 0.38 | 0.55 | 0.57 | 0.52 | 0.32 | 0.39 |
| 9 | 94631730 | rs11789730 | -1.036 | -1.772 | 0.886 | 0.839 | -0.002 | 0.014 | 0.014 | 0 | 0.32 | 0.37 | 0.53 | 0.44 | 0.3 | 0.3 |
| 9 | 94639795 | rs1528363 | -1.535 | -1.682 | 0.475 | 0.803 | 0.063 | 0.002 | 0.002 | 0.118 | 0.46 | 0.71 | 0.57 | 0.55 | 0.31 | 0.39 |
| 9 | 94640621 | rs1881389 | -2.646 | -1.506 | -0.228 | 0.783 | -0.184 | -0.073 | -0.073 | 1.524 | 0.64 | 0.96 | 0.7 | 0.63 | 0.5 | 0.56 |
| 9 | 94642450 | rs4744110 | -0.702 | -1.268 | 0.758 | 0.936 | 0.013 | 0.072 | 0.072 | 0 | 0.24 | 0.36 | 0.5 | 0.38 | 0.24 | 0.24 |
| 9 | 94646958 | rs10761133 | -1.705 | -1.408 | 0.114 | 0.851 | 0.069 | -0.056 | -0.056 | 0 | 0.43 | 0.81 | 0.57 | 0.44 | 0.29 | 0.37 |
| 9 | 94651415 | rs7858435 | -2.259 | -1.347 | -0.061 | 0.707 | -0.041 | -0.067 | -0.067 | 0.131 | 0.56 | 0.89 | 0.66 | 0.53 | 0.42 | 0.45 |
| 9 | 94656016 | rs10512219 | -0.897 | -1.3 | 0.744 | 0.583 | -0.021 | -0.04 | -0.04 | 1.523 | 0.27 | 0.32 | 0.11 | 0.11 | 0.16 | 0.09 |
| 9 | 94662474 | rs4430151 | -1.294 | -1.106 | 1.396 | 1.106 | 0 | 0.007 | 0.007 | 0 | 0.29 | 0.26 | 0.54 | 0.42 | 0.26 | 0.34 |
| 9 | 94668733 | rs3905385 | -0.669 | -1.057 | 1.418 | 1.213 | -0.02 | -0.051 | -0.051 | 2.245 | 0.22 | 0.11 | 0.5 | 0.35 | 0.22 | 0.28 |
| 9 | 94683050 | rs16907979 | -1.998 | -1.047 | 2.026 | 1.398 | 0.037 | 0.217 | 0.217 | 0 | 0.08 | 0.27 | 0.24 | 0.28 | 0.06 | 0.13 |
| 9 | 94685926 | rs2001917 | -1.861 | -1.381 | 1.884 | 1.291 | 0.044 | -0.032 | -0.032 | 0 | 0.26 | 0.8 | 0.31 | 0.41 | 0.11 | 0.2 |
| 9 | 94686736 | rs7045073 | -0.671 | -1.385 | 2.105 | 1.157 | -0.103 | -0.05 | -0.05 | 0 | 0.18 | 0.46 | 0.07 | 0.14 | 0.04 | 0.07 |
| 9 | 94687601 | rs1892263 | -0.382 | -1.174 | 1.742 | 1.218 | -0.029 | -0.003 | -0.003 | 0.17 | 0.2 | 0.06 | 0.29 | 0.15 | 0.2 | 0.2 |
| 9 | 94687927 | rs10992161 | -0.352 | -1.172 | 1.743 | 1.219 | -0.03 | -0.004 | -0.004 | 0.187 | 0.2 | 0.06 | 0.29 | 0.15 | 0.2 | 0.2 |
| 9 | 94696779 | rs7034006 | -1.115 | -0.869 | 1.974 | 1.016 | -0.017 | 0.03 | 0.03 | 0 | 0.23 | 0.14 | 0.3 | 0.11 | 0.26 | 0.21 |
| 9 | 94696954 | rs7037255 | -2.838 | -0.992 | 1.969 | 1.083 | 0.036 | -0.05 | -0.05 | 0 | 0.5 | 0.87 | 0.6 | 0.53 | 0.36 | 0.48 |

| | | | | | | | | | | | | | | | | |
|---|---|---|---|---|---|---|---|---|---|---|---|---|---|---|---|---|
| 9 | 94697108 | rs7038397 | -1.111 | -0.959 | 2.009 | 1.034 | -0.008 | 0.014 | 0.014 | 0 | 0.23 | 0.18 | 0.3 | 0.11 | 0.26 | 0.21 |
| 9 | 94697487 | rs7038823 | -1.695 | -0.955 | 2.069 | 1.04 | 0.004 | -0.016 | -0.016 | 2.027 | 0.28 | 0.28 | 0.33 | 0.18 | 0.29 | 0.32 |
| 9 | 94705717 | rs7863425 | -0.735 | -1.027 | 2.163 | 1.028 | -0.088 | -0.028 | -0.028 | 0 | 0.16 | 0.46 | 0.06 | 0.13 | 0.04 | 0.07 |
| 9 | 94708403 | rs7048699 | -3.07 | -1.054 | 2.07 | 1.063 | 0.038 | -0.042 | -0.042 | 0.113 | 0.5 | 0.87 | 0.6 | 0.54 | 0.36 | 0.47 |
| 9 | 94710847 | rs9774945 | -2.311 | -1.021 | 2.323 | 0.515 | 0.005 | 0.021 | 0.021 | 0.149 | 0.35 | 0.5 | 0.54 | 0.41 | 0.32 | 0.4 |
| 9 | 94728959 | rs10761137 | -1.522 | -1.043 | 2.265 | 0.467 | 0 | -0.007 | -0.007 | 0 | 0.28 | 0.23 | 0.46 | 0.33 | 0.23 | 0.28 |
| 9 | 94758241 | rs10992190 | 0.719 | -1.115 | 2.295 | 0.339 | 0.002 | 0.11 | 0.11 | 0 | 0.05 | 0.12 | 0.22 | 0.25 | 0.03 | 0.08 |
| 9 | 94758498 | rs6479395 | -1.819 | -1.154 | 2.446 | 0.277 | -0.027 | -0.059 | -0.059 | 0 | 0.18 | 0.44 | 0.12 | 0.11 | 0.13 | 0.19 |
| 9 | 94763136 | rs7867394 | -0.843 | -1.154 | 2.457 | 0.277 | -0.011 | -0.025 | -0.025 | 0 | 0.09 | 0.35 | 0.05 | 0.03 | 0.03 | 0.07 |
| 9 | 94764233 | rs7045602 | -0.632 | -1.159 | 2.704 | 0.275 | -0.015 | -0.027 | -0.027 | 0 | 0.08 | 0.28 | 0.04 | 0.03 | 0.03 | 0.07 |
| 9 | 94765564 | rs7018784 | 0.989 | -1.158 | 2.541 | 0.269 | 0.001 | -0.016 | -0.016 | 0 | 0.05 | 0.09 | 0.05 | 0 | 0.07 | 0 |
| 9 | 94769170 | rs6479397 | -1.354 | -1.17 | 2.569 | 0.277 | -0.021 | -0.084 | -0.084 | 0 | 0.15 | 0.54 | 0.11 | 0.04 | 0.1 | 0.07 |
| 9 | 94775387 | rs7039002 | -2.383 | -1.206 | 2.507 | 0.262 | 0.027 | 0.029 | 0.029 | 1.945 | 0.28 | 0.74 | 0.15 | 0.12 | 0.14 | 0.19 |
| 9 | 94794302 | rs7944 | 1.094 | -1.193 | 2.422 | 0.263 | -0.01 | 0.015 | 0.015 | 0 | 0.06 | 0.02 | 0.04 | 0 | 0.07 | 0 |
| 9 | 94796416 | rs12006507 | 0.837 | -1.152 | 2.402 | 0.265 | -0.029 | -0.037 | -0.037 | 0.053 | 0.07 | 0.11 | 0.01 | 0 | 0.01 | 0 |
| 9 | 94796491 | rs10820934 | -0.305 | -1.147 | 2.287 | 0.247 | -0.031 | 0.034 | 0.034 | 0 | 0.17 | 0.06 | 0.23 | 0.08 | 0.19 | 0.21 |
| 9 | 94800146 | rs3847308 | -3.333 | -1.155 | 2.272 | 0.286 | 0.011 | -0.052 | -0.052 | 0 | 0.51 | 0.92 | 0.61 | 0.45 | 0.36 | 0.47 |
| 9 | 94822146 | rs7858659 | -2.033 | -1.233 | 2.339 | 0.3 | 0.009 | -0.003 | -0.003 | 0 | 0.27 | 0.67 | 0.33 | 0.29 | 0.14 | 0.15 |
| 9 | 94841258 | rs10992224 | -0.464 | -1.09 | 2.072 | 0.363 | -0.019 | 0.033 | 0.033 | 0 | 0.17 | 0.07 | 0.23 | 0.08 | 0.18 | 0.21 |
| 9 | 94842715 | rs16908135 | 0.694 | -1.028 | 2.419 | 0.664 | 0.014 | 0.17 | 0.17 | 0 | 0.05 | 0.14 | 0.23 | 0.25 | 0.03 | 0.07 |
| 9 | 94864322 | rs7029539 | -2.358 | -1.034 | 2.167 | 0.631 | -0.013 | 0.017 | 0.017 | 0.012 | 0.25 | 0.65 | 0.36 | 0.37 | 0.16 | 0.26 |
| 9 | 94880110 | rs7858974 | -1.247 | -0.81 | 1.839 | 0.516 | 0.008 | -0.042 | -0.042 | 0 | 0.14 | 0.29 | 0.11 | 0.03 | 0.13 | 0.07 |
| 9 | 94882488 | rs7024547 | -1.87 | -1.008 | 1.558 | 0.435 | -0.029 | 0.012 | 0.012 | 0 | 0.36 | 0.44 | 0.5 | 0.38 | 0.25 | 0.35 |
| 9 | 94883880 | rs7048339 | -2.489 | -0.674 | 0.78 | 0.462 | -0.033 | -0.057 | -0.057 | 0 | 0.49 | 0.94 | 0.62 | 0.44 | 0.39 | 0.47 |
| 9 | 94902575 | rs10992250 | -0.925 | -0.762 | 0.905 | 0.567 | -0.002 | 0.005 | 0.005 | 0.002 | 0.15 | 0.02 | 0.2 | 0.11 | 0.15 | 0.23 |
| 9 | 94907008 | rs2274921 | -2.337 | -0.773 | 0.97 | 0.612 | 0.044 | -0.019 | -0.019 | 1.123 | 0.4 | 0.78 | 0.53 | 0.37 | 0.27 | 0.38 |
| 9 | 94909996 | rs7044976 | -1.416 | -0.22 | 0.994 | 0.843 | 0.006 | 0.019 | 0.019 | 0 | 0.14 | 0.31 | 0.19 | 0.13 | 0.12 | 0.23 |
| 9 | 94919766 | rs7874985 | -0.55 | -0.067 | 0.209 | 0.878 | -0.013 | -0.057 | -0.057 | 0.227 | 0.13 | 0.15 | 0.06 | 0 | 0.09 | 0.05 |
| 9 | 94920495 | rs10465156 | -0.276 | 0.06 | -0.017 | 1.241 | 0 | 0.047 | 0.047 | 0 | 0.07 | 0.06 | 0.25 | 0.26 | 0.05 | 0.13 |
| 9 | 94922307 | rs1970119 | -0.644 | 0.063 | -0.136 | 0.753 | -0.033 | -0.041 | -0.041 | 0 | 0.09 | 0 | 0.17 | 0.11 | 0.1 | 0.18 |
| 9 | 94927187 | rs6479409 | -0.811 | -0.265 | 0.455 | 0.417 | -0.035 | -0.049 | -0.049 | 0 | 0.16 | 0.23 | 0.09 | 0.05 | 0.08 | 0.13 |
| 9 | 94927564 | rs7035248 | -1.308 | 0.181 | 0.675 | 0.569 | -0.001 | 0.01 | 0.01 | 1.206 | 0.3 | 0.35 | 0.39 | 0.29 | 0.26 | 0.17 |
| 9 | 94927735 | rs7849530 | -1.409 | -0.113 | 1.056 | 0.357 | -0.005 | 0.026 | 0.026 | 0.111 | 0.14 | 0.37 | 0.17 | 0.13 | 0.09 | 0.23 |
| 9 | 94939615 | rs4744122 | -1.5 | -0.131 | -0.57 | -0.388 | -0.105 | 0.096 | 0.096 | 0.437 | 0.57 | 0.93 | 0.46 | 0.22 | 0.48 | 0.57 |
| 9 | 94944778 | rs4743863 | -0.679 | -0.186 | 0.015 | -0.59 | 0.001 | -0.056 | -0.056 | 0 | 0.33 | 0.46 | 0.3 | 0.09 | 0.31 | 0.35 |
| 9 | 94947595 | rs4743864 | 0.055 | -0.128 | 0.034 | -0.645 | -0.011 | -0.003 | -0.003 | 0.039 | 0.1 | 0.05 | 0.17 | 0.07 | 0.13 | 0.28 |
| 9 | 94949715 | rs11793122 | -0.164 | -0.192 | 0.306 | -0.684 | -0.005 | -0.042 | -0.042 | 0.431 | 0.18 | 0.26 | 0.11 | 0.09 | 0.16 | 0.13 |
| 9 | 94949905 | rs7023755 | -1.027 | -0.215 | 0.326 | -0.688 | 0 | -0.041 | -0.041 | 2.599 | 0.36 | 0.48 | 0.33 | 0.13 | 0.33 | 0.44 |
| 9 | 94951235 | rs11790258 | -0.286 | -0.288 | 0.417 | -0.69 | -0.008 | -0.042 | -0.042 | 0 | 0.19 | 0.28 | 0.11 | 0.09 | 0.15 | 0.13 |
| 9 | 94951819 | rs3858097 | -0.508 | -0.11 | 0.09 | -0.429 | -0.01 | -0.08 | -0.08 | 0 | 0.26 | 0.42 | 0.16 | 0.06 | 0.2 | 0.16 |
| 9 | 94962103 | rs10123080 | -0.21 | -0.113 | 0.025 | -0.449 | -0.008 | -0.088 | -0.088 | 0 | 0.22 | 0.36 | 0.11 | 0.02 | 0.18 | 0.06 |
| 9 | 94974188 | rs3780346 | -1.486 | -0.022 | -0.974 | -0.58 | -0.102 | 0.188 | 0.188 | 0.236 | 0.52 | 0.93 | 0.42 | 0.18 | 0.45 | 0.43 |
| 9 | 94985637 | rs556155 | -0.669 | -0.061 | -0.989 | -0.673 | -0.013 | -0.012 | -0.012 | 0 | 0.13 | 0.05 | 0.2 | 0.12 | 0.15 | 0.33 |
| 9 | 95007976 | rs895639 | -0.002 | -0.06 | -0.987 | -0.672 | -0.008 | -0.005 | -0.005 | 0 | 0.1 | 0.05 | 0.16 | 0.08 | 0.12 | 0.24 |
| 9 | 95041636 | rs10820964 | -1.127 | -0.381 | -1.167 | 0.394 | 0.007 | 0.235 | 0.235 | 0.056 | 0.46 | 0.9 | 0.3 | 0.15 | 0.35 | 0.28 |
| 9 | 95044229 | rs10119535 | -0.376 | -0.532 | -0.266 | 0.225 | -0.025 | -0.126 | -0.126 | 0 | 0.3 | 0.57 | 0.15 | 0.02 | 0.22 | 0.12 |
| 9 | 95052081 | rs10739927 | -1.435 | -0.525 | -0.283 | 0.232 | 0.007 | 0.236 | 0.236 | 0.048 | 0.46 | 0.9 | 0.3 | 0.15 | 0.35 | 0.28 |
| 9 | 95059804 | rs710163 | -2.438 | -0.429 | -0.504 | 0.214 | -0.129 | 0.067 | 0.067 | 0 | 0.62 | 0.97 | 0.74 | 0.69 | 0.52 | 0.75 |
| 9 | 95070095 | rs9409664 | -1.897 | -0.446 | -0.493 | 0.183 | -0.109 | 0.105 | 0.105 | 0.001 | 0.56 | 0.94 | 0.46 | 0.22 | 0.48 | 0.53 |
| 9 | 95087824 | rs10123342 | -0.943 | -0.511 | -0.072 | -0.001 | 0.015 | 0.091 | 0.091 | 0.266 | 0.41 | 0.74 | 0.24 | 0.1 | 0.33 | 0.19 |
| 9 | 95096811 | rs10992308 | 0.19 | -0.588 | -0.091 | -0.048 | -0.026 | -0.128 | -0.128 | 0.174 | 0.25 | 0.57 | 0.13 | 0.02 | 0.17 | 0.06 |
| 9 | 95105638 | rs10992312 | 0.182 | -0.606 | -0.099 | -0.057 | -0.027 | -0.128 | -0.128 | 0 | 0.25 | 0.57 | 0.13 | 0.02 | 0.17 | 0.06 |
| 9 | 95138564 | rs7026361 | -1.372 | -0.608 | -0.126 | -0.058 | 0.008 | 0.237 | 0.237 | 0.069 | 0.46 | 0.89 | 0.3 | 0.15 | 0.36 | 0.29 |
| 9 | 95140351 | rs1121978 | -0.6 | -0.613 | -0.1 | -0.061 | 0.004 | 0.029 | 0.029 | 0.794 | 0.36 | 0.71 | 0.22 | 0.1 | 0.29 | 0.13 |
| 9 | 95147434 | rs1380 | 0.53 | -0.632 | -0.088 | -0.05 | 0.005 | -0.007 | -0.007 | 1.22 | 0.11 | 0.14 | 0.09 | 0.08 | 0.11 | 0.07 |
| 9 | 95159196 | rs10992322 | -1.072 | -0.663 | -0.229 | 0.001 | 0.021 | 0.128 | 0.128 | 0 | 0.45 | 0.78 | 0.28 | 0.15 | 0.36 | 0.29 |
| 9 | 95160477 | rs2147060 | 2.001 | -0.664 | -0.417 | 0.039 | -0.148 | 0.036 | 0.036 | 0 | 0.4 | 0.03 | 0.28 | 0.35 | 0.5 | 0.4 |
| 9 | 95171852 | rs10761156 | 1.636 | -0.629 | -0.426 | -0.077 | -0.116 | 0.124 | 0.124 | 0.004 | 0.44 | 0.06 | 0.54 | 0.79 | 0.52 | 0.48 |
| 9 | 95193001 | rs10733744 | -0.355 | -0.571 | -0.341 | -0.034 | -0.037 | -0.012 | -0.012 | 0 | 0.9 | 1 | 0.83 | 0.94 | 0.88 | 0.76 |
| 9 | 95196273 | rs10125510 | -0.326 | -0.572 | -0.26 | -0.035 | -0.009 | 0.004 | 0.004 | 0 | 0.9 | 0.96 | 0.84 | 0.94 | 0.88 | 0.76 |
| 9 | 95218829 | rs8067 | -1.075 | -0.761 | -0.233 | -0.182 | 0.023 | 0.132 | 0.132 | 0 | 0.46 | 0.78 | 0.29 | 0.15 | 0.36 | 0.29 |
| 9 | 95219248 | rs3174352 | 1.654 | -0.783 | -0.202 | -0.193 | -0.078 | 0.141 | 0.141 | 0 | 0.44 | 0.08 | 0.54 | 0.79 | 0.52 | 0.47 |
| 9 | 95221475 | rs10992348 | 0.467 | -0.854 | 0.236 | -0.263 | -0.02 | -0.148 | -0.148 | 0.694 | 0.23 | 0.5 | 0.13 | 0.02 | 0.18 | 0.06 |
| 9 | 95229047 | rs13301537 | -0.569 | -0.859 | 0.188 | -0.268 | 0.004 | 0.028 | 0.028 | 0.18 | 0.35 | 0.71 | 0.23 | 0.1 | 0.29 | 0.13 |
| 9 | 95237222 | rs3739606 | -1.053 | -0.865 | 0.185 | -0.27 | 0.01 | 0.068 | 0.068 | 0.223 | 0.4 | 0.73 | 0.27 | 0.15 | 0.32 | 0.23 |
| 9 | 95240546 | rs10116241 | -1.46 | -0.868 | 0.186 | -0.271 | 0.014 | 0.247 | 0.247 | 0.023 | 0.47 | 0.9 | 0.31 | 0.15 | 0.36 | 0.29 |
| 9 | 95243549 | rs331377 | 2.502 | -0.817 | 0.147 | -0.135 | -0.144 | 0.055 | 0.055 | 0 | 0.37 | 0.02 | 0.25 | 0.31 | 0.48 | 0.25 |
| 9 | 95246833 | rs4744134 | -1.466 | -0.798 | 0.185 | -0.124 | 0.014 | 0.247 | 0.247 | 0.023 | 0.47 | 0.89 | 0.31 | 0.15 | 0.36 | 0.29 |
| 9 | 95256519 | rs331380 | 1.979 | -0.798 | 0.184 | -0.124 | -0.119 | 0.124 | 0.124 | 0 | 0.43 | 0.06 | 0.54 | 0.79 | 0.52 | 0.47 |

| | | | | | | | | | | | | | | | | |
|---|---|---|---|---|---|---|---|---|---|---|---|---|---|---|---|---|
| 9 | 95258119 | rs10429459 | -1.067 | -0.806 | 0.183 | -0.127 | 0.01 | 0.069 | 0.069 | 0.213 | 0.4 | 0.73 | 0.27 | 0.15 | 0.31 | 0.23 |
| 9 | 95271861 | rs2296665 | -0.497 | -0.819 | 0.346 | -0.125 | -0.003 | 0.002 | 0.002 | 0 | 0.9 | 0.93 | 0.84 | 0.94 | 0.88 | 0.76 |
| 9 | 95274338 | rs2296667 | -1.388 | -0.875 | 0.373 | -0.142 | 0.024 | 0.145 | 0.145 | 0 | 0.46 | 0.78 | 0.29 | 0.15 | 0.36 | 0.29 |
| 9 | 95279588 | rs2296668 | -1.34 | -1.189 | -0.934 | -1.05 | -0.105 | 0.027 | 0.027 | 0 | 0.74 | 0.94 | 0.66 | 0.81 | 0.54 | 0.64 |
| 9 | 95279985 | rs968040 | -0.059 | -1.242 | -0.95 | -1.079 | 0.007 | 0.071 | 0.071 | 0.374 | 0.36 | 0.76 | 0.24 | 0.09 | 0.29 | 0.13 |
| 9 | 95284212 | rs10491806 | 1.632 | -1.494 | -1.357 | -1.219 | 0.03 | -0.062 | -0.062 | 0 | 0.24 | 0.04 | 0.35 | 0.67 | 0.17 | 0.33 |
| 9 | 95284873 | rs9299405 | 0.058 | -1.479 | -1.347 | -1.178 | 0.008 | 0.072 | 0.072 | 0.333 | 0.37 | 0.76 | 0.24 | 0.09 | 0.29 | 0.13 |
| 9 | 95284982 | rs10120210 | -1.539 | -1.504 | -1.359 | -1.209 | -0.113 | 0.024 | 0.024 | 0 | 0.74 | 0.95 | 0.66 | 0.81 | 0.54 | 0.64 |
| 9 | 95293641 | rs10118939 | 0.038 | -1.466 | -1.324 | -1.19 | 0.008 | 0.069 | 0.069 | 0.387 | 0.37 | 0.75 | 0.24 | 0.09 | 0.29 | 0.13 |
| 9 | 95309243 | rs7036459 | 1.069 | -1.468 | -1.214 | -1.193 | -0.024 | -0.135 | -0.135 | 0 | 0.25 | 0.6 | 0.15 | 0 | 0.18 | 0.06 |
| 9 | 95323135 | rs10992367 | 1.546 | -1.656 | -1.246 | -1.125 | 0.03 | -0.062 | -0.062 | 0 | 0.24 | 0.04 | 0.35 | 0.67 | 0.17 | 0.33 |
| 9 | 95323669 | rs1536423 | -1.449 | -1.634 | -1.232 | -1.121 | -0.117 | 0.023 | 0.023 | 0 | 0.74 | 0.95 | 0.66 | 0.8 | 0.54 | 0.64 |
| 9 | 95325631 | rs7865019 | 0.17 | -1.64 | -1.211 | -1.124 | 0.008 | 0.063 | 0.063 | 0.408 | 0.37 | 0.74 | 0.24 | 0.09 | 0.29 | 0.13 |
| 9 | 95353785 | rs2131918 | -1.424 | -1.575 | -1.226 | -1.097 | -0.113 | 0.023 | 0.023 | 0 | 0.74 | 0.94 | 0.66 | 0.81 | 0.54 | 0.64 |
| 9 | 95360764 | rs7863882 | 0.916 | -1.686 | -1.19 | -1.074 | 0.046 | 0.003 | 0.003 | 0.36 | 0.31 | 0.14 | 0.38 | 0.68 | 0.22 | 0.39 |
| 9 | 95362665 | rs1874128 | 0.206 | -1.731 | -1.215 | -1.104 | 0.067 | 0.055 | 0.055 | 0.486 | 0.38 | 0.19 | 0.42 | 0.73 | 0.25 | 0.52 |
| 9 | 95372761 | rs10992384 | 1.58 | -1.768 | -1.233 | -1.125 | 0.036 | -0.016 | -0.016 | 0 | 0.26 | 0.12 | 0.35 | 0.67 | 0.17 | 0.33 |
| 9 | 95374368 | rs1980705 | 0.182 | -1.764 | -1.226 | -1.124 | 0.067 | 0.055 | 0.055 | 0.429 | 0.38 | 0.2 | 0.42 | 0.73 | 0.25 | 0.52 |
| 9 | 95383290 | rs724102 | 0.681 | -1.804 | -1.241 | -1.246 | 0.008 | -0.143 | -0.143 | 0 | 0.3 | 0.57 | 0.2 | 0 | 0.28 | 0.05 |
| 9 | 95384198 | rs1117670 | 0.987 | -1.805 | -1.315 | -1.236 | 0.047 | 0.004 | 0.004 | 0.266 | 0.31 | 0.15 | 0.38 | 0.67 | 0.22 | 0.39 |
| 9 | 95393049 | rs1323523 | 1.651 | -1.863 | -1.343 | -1.255 | 0.04 | -0.036 | -0.036 | 0.042 | 0.26 | 0.05 | 0.34 | 0.71 | 0.18 | 0.33 |
| 9 | 95419667 | rs10992392 | 1.654 | -1.838 | -1.318 | -1.424 | 0.036 | -0.003 | -0.003 | 0.2 | 0.26 | 0.12 | 0.35 | 0.73 | 0.18 | 0.33 |
| 9 | 95420983 | rs10992393 | 0.791 | -1.876 | -0.748 | -1.501 | 0.012 | -0.139 | -0.139 | 0 | 0.29 | 0.43 | 0.19 | 0 | 0.28 | 0.05 |
| 9 | 95425746 | rs7863890 | 0.273 | -1.838 | -1.034 | -1.471 | 0.068 | 0.077 | 0.077 | 0.493 | 0.39 | 0.2 | 0.41 | 0.78 | 0.26 | 0.52 |
| 9 | 95429120 | rs9969804 | -1.496 | -1.879 | -1.018 | -1.468 | -0.11 | 0.058 | 0.058 | 0 | 0.74 | 0.94 | 0.65 | 0.87 | 0.56 | 0.64 |
| 9 | 95450808 | rs10992420 | 1.612 | -1.685 | -0.77 | -1.26 | 0.027 | 0.003 | 0.003 | 0 | 0.25 | 0.12 | 0.34 | 0.72 | 0.18 | 0.31 |
| 9 | 95467111 | rs10992427 | 0.196 | -1.617 | -1.451 | -1.245 | 0.062 | 0.088 | 0.088 | 0.45 | 0.38 | 0.14 | 0.4 | 0.77 | 0.27 | 0.52 |
| 9 | 95472584 | rs12686771 | 2.074 | -1.66 | -1.478 | -0.897 | -0.013 | -0.081 | -0.081 | 0 | 0.21 | 0.01 | 0.09 | 0.3 | 0.16 | 0.25 |
| 9 | 95473719 | rs556 | 0.213 | -1.583 | -1.433 | -1.54 | 0.063 | 0.09 | 0.09 | 0.45 | 0.38 | 0.14 | 0.39 | 0.77 | 0.27 | 0.52 |
| 9 | 95475937 | rs3739605 | 1.697 | -1.588 | -1.432 | -1.515 | 0.031 | -0.006 | -0.006 | 1.413 | 0.25 | 0.08 | 0.33 | 0.72 | 0.18 | 0.31 |
| 9 | 95476958 | rs3203916 | 0.216 | -1.581 | -1.428 | -1.511 | 0.063 | 0.09 | 0.09 | 0.45 | 0.38 | 0.14 | 0.39 | 0.77 | 0.27 | 0.52 |
| 9 | 95488414 | rs10821009 | 0.228 | -1.582 | -1.427 | -1.508 | 0.063 | 0.089 | 0.089 | 0.522 | 0.38 | 0.13 | 0.39 | 0.77 | 0.27 | 0.52 |
| 9 | 95502571 | rs10821012 | 0.238 | -1.58 | -1.427 | -1.525 | 0.061 | 0.087 | 0.087 | 0.542 | 0.38 | 0.13 | 0.39 | 0.77 | 0.27 | 0.52 |
| 9 | 95527885 | rs7031163 | 1.696 | -1.571 | -1.419 | -1.509 | 0.031 | -0.007 | -0.007 | 0 | 0.25 | 0.07 | 0.33 | 0.72 | 0.18 | 0.31 |
| 9 | 95529408 | rs7043984 | 0.244 | -1.363 | -1.377 | -1.633 | 0.002 | 0.032 | 0.032 | 0.188 | 0.36 | 0.72 | 0.24 | 0.1 | 0.3 | 0.12 |
| 9 | 95551617 | rs10992469 | 0.786 | -1.335 | -1.489 | -1.351 | 0.046 | 0.017 | 0.017 | 0.028 | 0.32 | 0.09 | 0.36 | 0.7 | 0.22 | 0.45 |
| 9 | 95551830 | rs2181585 | 0.229 | -1.38 | -1.392 | -1.431 | -0.001 | -0.095 | -0.095 | 0 | 0.36 | 0.61 | 0.24 | 0.01 | 0.33 | 0.17 |
| 9 | 95555939 | rs7868651 | -0.297 | -1.246 | -1.533 | -1.29 | -0.01 | 0.011 | 0.011 | 0 | 0.63 | 0.72 | 0.56 | 0.71 | 0.5 | 0.56 |
| 9 | 95559505 | rs10992471 | 0.347 | -1.108 | -1.829 | -1.135 | -0.006 | 0.043 | 0.043 | 0 | 0.41 | 0.69 | 0.31 | 0.09 | 0.41 | 0.25 |
| 9 | 95559852 | rs1810696 | 0.401 | -0.742 | -2.289 | -0.885 | 0.039 | 0.01 | 0.01 | 0 | 0.31 | 0.07 | 0.35 | 0.71 | 0.21 | 0.44 |
| 9 | 95560534 | rs12554627 | 1.087 | -0.402 | -2.249 | -0.632 | -0.052 | 0.01 | 0.01 | 0 | 0.22 | 0.09 | 0.31 | 0.19 | 0.35 | 0.28 |
| 9 | 95561537 | rs7850075 | 0.448 | -0.633 | -1.709 | 0.199 | 0.025 | -0.052 | -0.052 | 0 | 0.24 | 0.05 | 0.32 | 0.65 | 0.18 | 0.29 |
| 9 | 95572545 | rs4141966 | 0.569 | -0.446 | -1.589 | 0.166 | 0.018 | -0.038 | -0.038 | 2.104 | 0.22 | 0.05 | 0.33 | 0.71 | 0.18 | 0.3 |
| 9 | 95580376 | rs10992480 | -0.444 | -0.039 | -1.508 | 0.543 | 0 | -0.025 | -0.025 | 0 | 0.37 | 0.24 | 0.24 | 0.43 | 0.35 | 0.35 |
| 9 | 95583472 | rs7038387 | -0.347 | -0.43 | -2.19 | -0.406 | 0.058 | 0.157 | 0.157 | 0 | 0.49 | 0.74 | 0.48 | 0.75 | 0.39 | 0.57 |
| 9 | 95586719 | rs1972214 | 1.461 | -0.84 | -1.798 | -0.494 | -0.035 | 0.007 | 0.007 | 0 | 0.26 | 0.57 | 0.38 | 0.32 | 0.2 | 0.29 |
| 9 | 95590093 | rs10821023 | 1.482 | -0.847 | -1.666 | -0.498 | -0.038 | 0.011 | 0.011 | 0 | 0.26 | 0.56 | 0.38 | 0.32 | 0.2 | 0.29 |
| 9 | 95599825 | rs7045305 | 1.76 | -1.067 | -1.167 | -0.491 | -0.022 | -0.04 | -0.04 | 0 | 0.21 | 0.41 | 0.34 | 0.12 | 0.16 | 0.13 |
| 9 | 95616149 | rs7859380 | -0.022 | -0.939 | -1.131 | 0.309 | 0.025 | 0.05 | 0.05 | 0 | 0.47 | 0.7 | 0.52 | 0.33 | 0.39 | 0.39 |
| 9 | 95618766 | rs10761174 | -1.796 | -0.88 | -1.243 | 0.31 | -0.059 | 0.032 | 0.032 | 0 | 0.7 | 0.86 | 0.63 | 0.75 | 0.58 | 0.67 |
| 9 | 95621504 | rs10739931 | -1.832 | -0.931 | -1.275 | 0.293 | -0.079 | 0.038 | 0.038 | 0 | 0.7 | 0.88 | 0.63 | 0.77 | 0.58 | 0.67 |
| 9 | 95624890 | rs10992498 | 0.749 | -0.892 | -1.21 | 0.351 | 0 | -0.034 | -0.034 | 0 | 0.2 | 0.11 | 0.1 | 0.42 | 0.19 | 0.28 |
| 9 | 95630363 | rs4744150 | 0.269 | -0.974 | -1.004 | 0.026 | 0.006 | 0 | 0 | 0 | 0.43 | 0.61 | 0.49 | 0.12 | 0.34 | 0.23 |
| 9 | 95652951 | rs10992512 | -0.747 | -0.846 | -1.184 | 0.039 | -0.022 | 0.042 | 0.042 | 0 | 0.54 | 0.76 | 0.56 | 0.62 | 0.48 | 0.44 |
| 9 | 95683366 | rs7035868 | 0.391 | -0.969 | -1.047 | -0.322 | 0.006 | 0.004 | 0.004 | 0 | 0.43 | 0.61 | 0.49 | 0.12 | 0.36 | 0.23 |
| 9 | 95688099 | rs10992535 | 0.136 | -1.213 | -0.803 | -0.231 | -0.004 | -0.004 | -0.004 | 0 | 0.29 | 0.32 | 0.17 | 0.23 | 0.23 | 0.41 |
| 9 | 95691037 | rs13294147 | 0.427 | -1.291 | -0.633 | -0.552 | 0.001 | -0.036 | -0.036 | 3.51 | 0.38 | 0.41 | 0.43 | 0.12 | 0.36 | 0.14 |
| 9 | 95694499 | rs10821041 | -0.22 | -1.231 | -0.751 | -0.151 | 0.003 | 0.01 | 0.01 | 0 | 0.47 | 0.57 | 0.46 | 0.38 | 0.42 | 0.33 |
| 9 | 95699292 | rs7856228 | -0.097 | -1.099 | -0.831 | -0.123 | -0.002 | -0.002 | -0.002 | 0 | 0.44 | 0.58 | 0.45 | 0.38 | 0.37 | 0.32 |
| 9 | 95708192 | rs10821046 | -1.67 | -1.097 | -1.29 | 0.136 | -0.075 | -0.104 | -0.104 | 0 | 0.74 | 0.92 | 0.64 | 0.6 | 0.63 | 0.69 |
| 9 | 95715717 | rs6479442 | -0.199 | -1.322 | -1.062 | -0.303 | -0.005 | 0.006 | 0.006 | 0.025 | 0.59 | 0.67 | 0.58 | 0.34 | 0.53 | 0.52 |
| 9 | 95720621 | rs7023321 | -1.861 | -1.357 | -1.133 | -0.24 | -0.096 | 0.117 | 0.117 | 0.004 | 0.77 | 0.96 | 0.66 | 0.92 | 0.63 | 0.72 |
| 9 | 95723163 | rs7035881 | 1.441 | -1.408 | -0.941 | -0.626 | -0.04 | -0.037 | -0.037 | 0 | 0.23 | 0.38 | 0.31 | 0.12 | 0.14 | 0.13 |
| 9 | 95727883 | rs4147166 | -0.306 | -0.866 | -0.645 | -0.157 | -0.026 | 0.015 | 0.015 | 0.001 | 0.1 | 0.18 | 0.03 | 0.14 | 0.05 | 0.16 |
| 9 | 95737649 | rs9942965 | -0.727 | -0.807 | -0.762 | -0.246 | 0.017 | 0.043 | 0.043 | 0.021 | 0.68 | 0.58 | 0.62 | 0.51 | 0.6 | 0.68 |
| 9 | 95746936 | rs10992564 | -0.711 | -0.806 | -0.762 | -0.245 | 0.016 | 0.041 | 0.041 | 0.039 | 0.68 | 0.58 | 0.62 | 0.51 | 0.61 | 0.68 |
| 9 | 95747389 | rs7850247 | 0.082 | -0.921 | -0.86 | -0.318 | 0.002 | -0.009 | -0.009 | 0.269 | 0.57 | 0.5 | 0.59 | 0.36 | 0.51 | 0.53 |
| 9 | 95753319 | rs10992568 | -0.436 | -1.084 | -0.823 | -0.284 | -0.021 | -0.004 | -0.004 | 0 | 0.16 | 0.25 | 0.07 | 0.15 | 0.12 | 0.13 |
| 9 | 95767060 | rs4744164 | -0.173 | -0.95 | -1.043 | -0.213 | -0.011 | -0.006 | -0.006 | 0.198 | 0.59 | 0.7 | 0.58 | 0.41 | 0.49 | 0.55 |

| | | | | | | | | | | | | | | | | |
|---|---|---|---|---|---|---|---|---|---|---|---|---|---|---|---|---|
| 9 | 95770161 | rs10761188 | 0.621 | -1.08 | -1.351 | -0.325 | 0.001 | -0.019 | -0.019 | 0 | 0.41 | 0.52 | 0.52 | 0.28 | 0.41 | 0.3 |
| 9 | 95772228 | rs10761189 | -1.298 | -0.622 | -2.06 | -0.064 | -0.101 | -0.115 | -0.115 | 0.76 | 0.75 | 0.94 | 0.65 | 0.56 | 0.6 | 0.69 |
| 9 | 95775561 | rs3934851 | 0.536 | -0.652 | -1.565 | -0.106 | -0.004 | -0.034 | -0.034 | 0 | 0.08 | 0 | 0.02 | 0.12 | 0.06 | 0.17 |
| 9 | 95781040 | rs10761191 | -1.017 | -0.513 | -1.783 | -0.23 | -0.085 | -0.091 | -0.091 | 0 | 0.63 | 0.92 | 0.61 | 0.46 | 0.56 | 0.44 |
| 9 | 95782667 | rs11790517 | 0.827 | -0.43 | -1.263 | -0.349 | -0.011 | -0.001 | -0.001 | 0.119 | 0.29 | 0.38 | 0.39 | 0.24 | 0.29 | 0.11 |
| 9 | 95785819 | rs7855476 | 0.866 | -0.091 | -1.333 | -0.687 | 0.005 | -0.01 | -0.01 | 0.077 | 0.13 | 0.19 | 0.15 | 0.07 | 0.17 | 0.16 |
| 9 | 95786065 | rs7854887 | 0.668 | -0.107 | -1.347 | -0.695 | 0.003 | -0.008 | -0.008 | 0.27 | 0.14 | 0.19 | 0.15 | 0.07 | 0.17 | 0.16 |
| 9 | 95788623 | rs11788432 | -0.359 | -0.051 | -1.424 | -0.685 | -0.026 | 0.017 | 0.017 | 0 | 0.23 | 0.17 | 0.17 | 0.21 | 0.26 | 0.29 |
| 9 | 95793816 | rs13290740 | 0.45 | -0.282 | -1.205 | -0.471 | -0.031 | -0.021 | -0.021 | 0 | 0.31 | 0.43 | 0.37 | 0.23 | 0.28 | 0.14 |
| 9 | 95797446 | rs7022396 | -0.143 | -0.179 | -1.243 | -0.861 | -0.013 | -0.004 | -0.004 | 0.655 | 0.41 | 0.46 | 0.4 | 0.37 | 0.36 | 0.27 |
| 9 | 95799446 | rs6479447 | -1.607 | 0.171 | -1.972 | -0.686 | -0.072 | -0.119 | -0.119 | 0.103 | 0.72 | 0.94 | 0.64 | 0.53 | 0.63 | 0.64 |
| 9 | 95802287 | rs11787849 | -0.288 | 0.024 | -1.268 | -0.413 | -0.001 | 0.001 | 0.001 | 0.79 | 0.16 | 0.17 | 0.08 | 0.12 | 0.14 | 0.12 |
| 9 | 95806365 | rs4573339 | -0.107 | 0.259 | -1.099 | -0.18 | -0.003 | 0.006 | 0.006 | 0 | 0.22 | 0.2 | 0.18 | 0.16 | 0.2 | 0.29 |
| 9 | 95806823 | rs10992608 | -0.751 | 0.229 | -1.525 | -0.101 | -0.083 | -0.071 | -0.071 | 0 | 0.58 | 0.89 | 0.58 | 0.5 | 0.5 | 0.51 |
| 9 | 95812707 | rs7027092 | -0.822 | -0.292 | -0.512 | -0.052 | 0.004 | -0.003 | -0.003 | 0.139 | 0.44 | 0.59 | 0.47 | 0.48 | 0.34 | 0.4 |
| 9 | 95818641 | rs4744167 | 0.002 | -0.26 | -0.126 | 0.044 | 0.002 | 0.097 | 0.097 | 0 | 0.17 | 0.36 | 0.32 | 0.24 | 0.14 | 0.12 |
| 9 | 95830197 | rs10992622 | -0.865 | -0.897 | 0.404 | -0.961 | -0.041 | -0.042 | -0.042 | 0 | 0.26 | 0.46 | 0.15 | 0.21 | 0.15 | 0.29 |
| 9 | 95839418 | rs10761192 | -1.574 | -0.993 | 0.47 | -1.137 | 0.009 | -0.069 | -0.069 | 0 | 0.14 | 0.41 | 0.11 | 0.07 | 0.09 | 0.13 |
| 9 | 95840256 | rs1131773 | -1.412 | -1.104 | 0.484 | -1.17 | -0.029 | -0.033 | -0.033 | 0 | 0.23 | 0.49 | 0.12 | 0.21 | 0.14 | 0.33 |
| 9 | 95845152 | rs9696357 | -1.418 | -1.077 | 0.37 | -1.244 | -0.03 | -0.05 | -0.05 | 0 | 0.25 | 0.41 | 0.12 | 0.2 | 0.15 | 0.32 |
| 9 | 95854444 | rs10992629 | -1.454 | -0.724 | -0.576 | -1.368 | 0.001 | -0.03 | -0.03 | 0 | 0.28 | 0.69 | 0.17 | 0.08 | 0.17 | 0.23 |
| 9 | 95855100 | rs10992630 | -1.025 | 0.168 | -0.786 | -1.039 | -0.006 | 0.005 | 0.005 | 0 | 0.16 | 0.15 | 0.13 | 0.23 | 0.16 | 0.21 |
| 9 | 95857333 | rs13288200 | -0.193 | 0.088 | -1.299 | -1.427 | -0.044 | 0.062 | 0.062 | 0.06 | 0.15 | 0.04 | 0.1 | 0 | 0.23 | 0.05 |
| 9 | 95860100 | rs4237219 | 0.595 | -0.866 | -1.219 | -1.349 | -0.027 | 0.014 | 0.014 | 1.508 | 0.77 | 0.39 | 0.85 | 0.61 | 0.88 | 0.74 |
| 9 | 95860415 | rs4744171 | 0.866 | -1.499 | -1.788 | -1.895 | 0.092 | 0.021 | 0.021 | 1.606 | 0.62 | 0.29 | 0.78 | 0.6 | 0.77 | 0.57 |
| 9 | 95869726 | rs4744175 | 1.665 | -1.61 | -1.917 | -2.029 | -0.061 | 0.02 | 0.02 | 0 | 0.43 | 0.06 | 0.67 | 0.6 | 0.54 | 0.52 |
| 9 | 95869943 | rs2275849 | -0.072 | -1.509 | -1.742 | -1.885 | -0.041 | 0.027 | 0.027 | 2.86 | 0.18 | 0.6 | 0.1 | 0.27 | 0.09 | 0.12 |
| 9 | 95872885 | rs17593496 | 0.599 | -0.964 | -1.66 | -1.989 | -0.013 | 0.051 | 0.051 | 0.014 | 0.14 | 0.04 | 0.06 | 0 | 0.16 | 0.05 |
| 9 | 95879087 | rs3802386 | 1.027 | -1.419 | -1.449 | -2.154 | 0.013 | 0.088 | 0.088 | 0.081 | 0.63 | 0.77 | 0.78 | 0.83 | 0.64 | 0.62 |
| 9 | 95881054 | rs13285107 | 0.185 | -1.413 | -1.065 | -2.099 | -0.001 | -0.033 | -0.033 | 0 | 0.13 | 0.2 | 0.06 | 0.01 | 0.15 | 0.04 |
| 9 | 95883363 | rs1954435 | -0.512 | -1.743 | -0.554 | -2.418 | -0.049 | -0.173 | -0.173 | 0 | 0.23 | 0.44 | 0.12 | 0 | 0.15 | 0.21 |
| 9 | 95887154 | rs13298119 | 0.248 | -1.627 | -0.35 | -2.12 | -0.003 | -0.045 | -0.045 | 0.147 | 0.13 | 0.2 | 0.06 | 0.01 | 0.16 | 0.04 |
| 9 | 95887320 | rs2275848 | 0.82 | -1.76 | -0.307 | -2.14 | -0.022 | -0.061 | -0.061 | 0 | 0.8 | 0.62 | 0.88 | 0.87 | 0.81 | 0.84 |
| 9 | 95888667 | rs12235416 | -0.489 | -1.599 | 0.132 | -1.756 | -0.006 | 0.019 | 0.019 | 0 | 0.19 | 0.35 | 0.1 | 0.25 | 0.19 | 0.14 |
| 9 | 95898531 | rs7025554 | -0.821 | -1.545 | 0.328 | -1.703 | -0.01 | -0.046 | -0.046 | 1.695 | 0.09 | 0.33 | 0.05 | 0.01 | 0.04 | 0.04 |
| 9 | 95899771 | rs10821076 | 1.459 | -1.296 | -0.823 | -2.209 | -0.04 | 0.07 | 0.07 | 0.18 | 0.42 | 0.19 | 0.65 | 0.63 | 0.49 | 0.55 |
| 9 | 95905552 | rs4743894 | 1.154 | -1.112 | -0.543 | -2.083 | 0.026 | 0.074 | 0.074 | 0.153 | 0.57 | 0.26 | 0.72 | 0.67 | 0.58 | 0.7 |
| 9 | 95910508 | rs6479457 | -0.687 | -1.207 | -0.452 | -2.001 | -0.008 | -0.036 | -0.036 | 0 | 0.29 | 0.64 | 0.15 | 0.17 | 0.27 | 0.22 |
| 9 | 95912016 | rs4335184 | 0.493 | -0.931 | -0.606 | -1.923 | -0.003 | -0.019 | -0.019 | 0.49 | 0.7 | 0.33 | 0.81 | 0.86 | 0.71 | 0.85 |
| 9 | 95917252 | rs10821077 | -0.061 | -1.382 | -1.143 | -1.935 | 0.013 | 0.015 | 0.015 | 0.342 | 0.21 | 0.14 | 0.11 | 0.11 | 0.14 | 0.23 |
| 9 | 95918643 | rs10992658 | 1.096 | -1.268 | -1.973 | -1.903 | -0.139 | 0.029 | 0.029 | 0.18 | 0.38 | 0.05 | 0.62 | 0.67 | 0.5 | 0.52 |
| 9 | 95919314 | rs12378014 | 0.092 | -0.841 | -0.934 | -1.579 | -0.008 | 0.016 | 0.016 | 0 | 0.16 | 0.1 | 0.11 | 0.07 | 0.2 | 0.06 |
| 9 | 95930334 | rs6479458 | -0.367 | -0.701 | -0.25 | -1.269 | 0.025 | -0.016 | -0.016 | 0 | 0.06 | 0.15 | 0.03 | 0.05 | 0.07 | 0.05 |
| 9 | 95930572 | rs10821080 | 0.914 | -0.904 | -1.075 | -0.716 | -0.118 | -0.105 | -0.105 | 0 | 0.27 | 0.09 | 0.5 | 0.51 | 0.49 | 0.3 |
| 9 | 95931685 | rs4743898 | 0.842 | -1.647 | -1.537 | -0.983 | -0.028 | -0.014 | -0.014 | 0.055 | 0.16 | 0.03 | 0.2 | 0.09 | 0.17 | 0.06 |
| 9 | 95939766 | rs2398821 | -1.22 | -2.239 | -1.936 | -1.329 | 0 | 0.029 | 0.029 | 0 | 0.88 | 0.99 | 0.93 | 0.91 | 0.9 | 0.9 |
| 9 | 95940368 | rs1106024 | 0.997 | -2.423 | -1.976 | -1.467 | -0.02 | -0.018 | -0.018 | 0 | 0.31 | 0.25 | 0.53 | 0.57 | 0.51 | 0.33 |
| 9 | 95951477 | rs10821083 | -0.04 | -2.447 | -1.926 | -1.479 | 0.023 | 0.015 | 0.015 | 0 | 0.56 | 0.72 | 0.4 | 0.34 | 0.38 | 0.56 |
| 9 | 95951769 | rs7041800 | -0.938 | -1.369 | -2.254 | -1.534 | -0.162 | 0.135 | 0.135 | 0 | 0.74 | 0.98 | 0.69 | 0.91 | 0.56 | 0.83 |
| 9 | 95952870 | rs10992671 | 0.144 | -2.671 | -1.557 | -1.669 | 0.012 | 0.038 | 0.038 | 0 | 0.34 | 0.27 | 0.32 | 0.17 | 0.32 | 0.27 |
| 9 | 95953341 | rs10512222 | -0.71 | -2.771 | -0.691 | -1.698 | -0.004 | -0.001 | -0.001 | 0.478 | 0.67 | 0.68 | 0.86 | 0.74 | 0.83 | 0.61 |
| 9 | 95955365 | rs7858813 | -1.677 | -2.743 | -0.803 | -1.689 | 0 | 0.033 | 0.033 | 0 | 0.88 | 0.99 | 0.93 | 0.91 | 0.9 | 0.9 |
| 9 | 95955986 | rs7031721 | 1.324 | -2.745 | -0.708 | -1.703 | 0.004 | 0 | 0 | 0 | 0.18 | 0.15 | 0.11 | 0.08 | 0.15 | 0.23 |
| 9 | 95958095 | rs6479466 | -0.753 | -2.67 | -0.512 | -1.683 | -0.025 | -0.016 | -0.016 | 0.44 | 0.67 | 0.62 | 0.86 | 0.74 | 0.83 | 0.63 |
| 9 | 95962890 | rs12336667 | 1.551 | -2.371 | -0.65 | -1.447 | -0.106 | -0.108 | -0.108 | 0 | 0.26 | 0.09 | 0.5 | 0.51 | 0.47 | 0.3 |
| 9 | 95963114 | rs10992674 | 0.281 | -2.295 | -0.458 | -1.39 | 0.009 | 0.005 | 0.005 | 0 | 0.47 | 0.4 | 0.63 | 0.59 | 0.62 | 0.52 |
| 9 | 95971876 | rs1539669 | -0.356 | -1.921 | -0.123 | -1.302 | -0.011 | -0.049 | -0.049 | 0 | 0.82 | 0.72 | 0.9 | 0.91 | 0.88 | 0.89 |
| 9 | 95983986 | rs7873162 | 0.819 | -2.074 | 0.137 | -1.281 | -0.003 | -0.004 | -0.004 | 0 | 0.19 | 0.21 | 0.12 | 0.13 | 0.15 | 0.31 |
| 9 | 95985281 | rs4744195 | 1.39 | -2.361 | 0.056 | -0.923 | -0.06 | -0.049 | -0.049 | 0.124 | 0.28 | 0.16 | 0.52 | 0.44 | 0.49 | 0.27 |
| 9 | 95988968 | rs4617217 | -0.985 | -2.662 | 0.293 | -1.015 | 0.027 | -0.002 | -0.002 | 0 | 0.68 | 0.78 | 0.88 | 0.65 | 0.84 | 0.67 |
| 9 | 95990501 | rs4744199 | 0.663 | -2.668 | 0.365 | -1.019 | 0.009 | -0.031 | -0.031 | 0.359 | 0.2 | 0.37 | 0.23 | 0.08 | 0.21 | 0.09 |
| 9 | 95991895 | rs7874101 | 1.052 | -2.647 | 0.286 | -1.007 | -0.008 | -0.011 | -0.011 | 0 | 0.2 | 0.24 | 0.13 | 0.13 | 0.15 | 0.31 |
| 9 | 95993170 | rs7048527 | 1.881 | -2.754 | 0.201 | -1.016 | -0.004 | 0.001 | 0.001 | 0.346 | 0.22 | 0.22 | 0.08 | 0.25 | 0.12 | 0.23 |
| 9 | 95993326 | rs11787888 | 1.53 | -2.732 | -0.031 | -1.011 | 0.01 | 0.015 | 0.015 | 0 | 0.18 | 0.1 | 0.11 | 0.08 | 0.15 | 0.3 |
| 9 | 95994155 | rs11793893 | 1.12 | -2.714 | -0.066 | -1.004 | -0.005 | -0.007 | -0.007 | 0.009 | 0.2 | 0.22 | 0.12 | 0.13 | 0.16 | 0.31 |
| 9 | 95994554 | rs7864826 | 1.464 | -2.616 | -0.179 | -0.971 | -0.015 | 0.002 | 0.002 | 0 | 0.15 | 0.06 | 0.2 | 0.08 | 0.17 | 0.06 |
| 9 | 95996573 | rs2398820 | 0.317 | -2.629 | -0.05 | -0.95 | 0.016 | 0.001 | 0.001 | 0 | 0.44 | 0.38 | 0.73 | 0.52 | 0.65 | 0.33 |
| 9 | 96000622 | rs1539670 | -1.584 | -2.48 | -0.443 | -0.834 | 0.023 | 0.004 | 0.004 | 3.05 | 0.89 | 0.98 | 0.96 | 0.91 | 0.96 | 0.9 |
| 9 | 96000921 | rs11788911 | 1.657 | -2.392 | -0.285 | -0.802 | 0.008 | -0.005 | -0.005 | 0.786 | 0.21 | 0.17 | 0.08 | 0.26 | 0.11 | 0.23 |

| | | | | | | | | | | | | | | | | |
|---|---|---|---|---|---|---|---|---|---|---|---|---|---|---|---|---|
| 9 | 96002983 | rs7022051 | 1.281 | -2.122 | -0.186 | -0.833 | -0.014 | -0.025 | -0.025 | 0 | 0.14 | 0.19 | 0.1 | 0.05 | 0.14 | 0.06 |
| 9 | 96003435 | rs12341303 | 1.344 | -1.983 | -0.467 | -0.751 | -0.094 | -0.077 | -0.077 | 0 | 0.27 | 0.1 | 0.52 | 0.44 | 0.47 | 0.27 |
| 9 | 96012145 | rs12554962 | 1.428 | -2.161 | -0.603 | -0.856 | -0.01 | -0.003 | -0.003 | 0 | 0.13 | 0.07 | 0.19 | 0.08 | 0.17 | 0.05 |
| 9 | 96015256 | rs4744201 | 1.415 | -2.103 | -0.596 | -0.789 | -0.125 | -0.098 | -0.098 | 0 | 0.26 | 0.06 | 0.51 | 0.43 | 0.47 | 0.24 |
| 9 | 96017751 | rs10992689 | 0.469 | -2.343 | -0.223 | -0.894 | 0.002 | -0.001 | -0.001 | 0 | 0.43 | 0.48 | 0.31 | 0.42 | 0.31 | 0.57 |
| 9 | 96019970 | rs10761202 | 1.598 | -2.292 | -0.391 | -0.881 | -0.117 | -0.094 | -0.094 | 0.081 | 0.25 | 0.06 | 0.51 | 0.43 | 0.46 | 0.24 |
| 9 | 96022116 | rs1110335 | 0.603 | -2.316 | -0.175 | -0.912 | 0.012 | -0.008 | -0.008 | 0 | 0.42 | 0.33 | 0.3 | 0.42 | 0.31 | 0.57 |
| 9 | 96038524 | rs4744213 | -0.432 | -2.388 | -0.017 | -0.976 | -0.038 | -0.134 | -0.134 | 0 | 0.79 | 0.46 | 0.85 | 1 | 0.82 | 0.9 |
| 9 | 96041713 | rs1931367 | -0.286 | -2.328 | 0.199 | -0.933 | 0.047 | -0.009 | -0.009 | 0 | 0.93 | 0.64 | 0.96 | 1 | 0.95 | 0.96 |
| 9 | 96053928 | rs1889339 | 0.53 | -2.461 | 0.067 | -0.867 | 0.009 | 0.007 | 0.007 | 0 | 0.67 | 0.31 | 0.8 | 0.85 | 0.78 | 0.81 |
| 9 | 96063456 | rs16936750 | -0.08 | -2.42 | 0.296 | -0.787 | -0.005 | 0 | 0 | 1.204 | 0.15 | 0.2 | 0.05 | 0.15 | 0.04 | 0.1 |
| 9 | 96068234 | rs10992701 | 1.465 | -2.52 | -0.08 | -0.82 | 0.022 | -0.014 | -0.014 | 0 | 0.28 | 0.17 | 0.1 | 0.36 | 0.15 | 0.52 |
| 9 | 96068328 | rs16909016 | 1.14 | -2.259 | 0.022 | -0.742 | 0.03 | -0.011 | -0.011 | 0.007 | 0.11 | 0 | 0.03 | 0.08 | 0.03 | 0.09 |
| 9 | 96077465 | rs10761209 | 0.55 | -2.251 | -0.173 | -0.646 | 0.017 | -0.016 | -0.016 | 0 | 0.42 | 0.25 | 0.29 | 0.44 | 0.31 | 0.58 |
| 9 | 96082752 | rs6479474 | 1.585 | -2.081 | 0.003 | -0.697 | -0.013 | -0.055 | -0.055 | 0.234 | 0.13 | 0.18 | 0.09 | 0 | 0.14 | 0.06 |
| 9 | 96082919 | rs13293217 | -1.105 | -2.039 | -0.041 | -0.602 | 0.026 | 0.015 | 0.015 | 0 | 0.7 | 0.71 | 0.45 | 0.58 | 0.5 | 0.74 |
| 9 | 96091688 | rs10821109 | 1.096 | -2.225 | -0.393 | -0.732 | 0.017 | -0.03 | -0.03 | 0.275 | 0.3 | 0.1 | 0.1 | 0.36 | 0.19 | 0.59 |
| 9 | 96093535 | rs3933798 | 1.261 | -2.231 | -0.395 | -0.734 | 0.006 | -0.046 | -0.046 | 0 | 0.29 | 0.07 | 0.1 | 0.36 | 0.19 | 0.59 |
| 9 | 96093865 | rs4633120 | 0.094 | -2.04 | -0.246 | -0.762 | -0.059 | -0.021 | -0.021 | 0 | 0.29 | 0.49 | 0.28 | 0.21 | 0.19 | 0.11 |
| 9 | 96093924 | rs4534181 | -0.634 | -2.039 | -0.417 | -0.818 | -0.017 | -0.024 | -0.024 | 0 | 0.4 | 0.51 | 0.35 | 0.25 | 0.34 | 0.17 |
| 9 | 96094499 | rs10992707 | 0.086 | -2.036 | -0.398 | -0.8 | -0.058 | -0.023 | -0.023 | 0 | 0.29 | 0.48 | 0.27 | 0.21 | 0.19 | 0.12 |
| 9 | 96094943 | rs2991377 | -0.37 | -2.063 | -0.385 | -0.842 | 0.001 | 0.003 | 0.003 | 0 | 0.55 | 0.62 | 0.79 | 0.61 | 0.63 | 0.32 |
| 9 | 96096243 | rs2927567 | 1.68 | -1.973 | -0.325 | -0.795 | -0.07 | -0.061 | -0.061 | 0 | 0.26 | 0.13 | 0.51 | 0.39 | 0.44 | 0.21 |
| 9 | 96099377 | rs2927572 | 0.291 | -1.999 | -0.267 | -0.818 | 0.001 | 0 | 0 | 0 | 0.46 | 0.52 | 0.74 | 0.49 | 0.59 | 0.27 |
| 9 | 96117149 | rs10761215 | -0.383 | -1.917 | -0.232 | -0.846 | 0.03 | 0.023 | 0.023 | 0 | 0.49 | 0.45 | 0.65 | 0.61 | 0.62 | 0.29 |
| 9 | 96117348 | rs10761216 | 1.227 | -1.921 | -0.394 | -0.827 | 0.013 | -0.013 | -0.013 | 0 | 0.31 | 0.11 | 0.11 | 0.35 | 0.22 | 0.62 |
| 9 | 96125819 | rs4744226 | -0.026 | -1.902 | -0.431 | -0.738 | 0.016 | 0.03 | 0.03 | 0.266 | 0.36 | 0.23 | 0.36 | 0.22 | 0.34 | 0.13 |
| 9 | 96126075 | rs4744227 | 1.315 | -1.831 | -0.319 | -0.71 | -0.035 | -0.039 | -0.039 | 2.751 | 0.28 | 0.23 | 0.48 | 0.43 | 0.41 | 0.21 |
| 9 | 96133597 | rs10992715 | 1.487 | -1.823 | -0.261 | -0.765 | -0.02 | -0.017 | -0.017 | 0 | 0.18 | 0.19 | 0.21 | 0.04 | 0.13 | 0.06 |
| 9 | 96138557 | rs1931371 | 0.761 | -1.895 | -0.504 | -0.798 | 0.094 | 0.105 | 0.105 | 0.838 | 0.53 | 0.18 | 0.56 | 0.73 | 0.63 | 0.84 |
| 9 | 96144349 | rs2086106 | -0.82 | -2.038 | -0.235 | -0.825 | 0.01 | -0.005 | -0.005 | 0 | 0.9 | 0.74 | 0.93 | 0.92 | 0.91 | 0.94 |
| 9 | 96145271 | rs10821119 | -0.386 | -1.926 | -0.372 | -0.767 | 0.031 | 0.028 | 0.028 | 0 | 0.72 | 0.23 | 0.73 | 0.9 | 0.84 | 0.9 |
| 9 | 96150272 | rs10739948 | 0.365 | -1.854 | -0.331 | -0.654 | -0.043 | -0.038 | -0.038 | 0 | 0.35 | 0.21 | 0.55 | 0.52 | 0.48 | 0.25 |
| 9 | 96154352 | rs11999013 | 0.64 | -1.433 | -0.188 | -0.791 | -0.004 | -0.042 | -0.042 | 0 | 0.25 | 0.26 | 0.15 | 0.02 | 0.21 | 0.07 |
| 9 | 96158277 | rs2266010 | -0.78 | -1.105 | 0.192 | -0.447 | -0.046 | -0.039 | -0.039 | 0 | 0.18 | 0.43 | 0.16 | 0.14 | 0.11 | 0.09 |
| 9 | 96163260 | rs7020852 | -0.713 | -1.127 | -0.157 | -0.321 | 0.005 | 0.009 | 0.009 | 0 | 0.46 | 0.66 | 0.66 | 0.56 | 0.58 | 0.29 |
| 9 | 96166579 | rs10761224 | 0.196 | -0.749 | 0.301 | -0.341 | 0.013 | 0.014 | 0.014 | 0.01 | 0.32 | 0.4 | 0.5 | 0.4 | 0.46 | 0.2 |
| 9 | 96169561 | rs10821123 | 1.14 | -0.71 | 0.241 | -0.281 | -0.07 | -0.071 | -0.071 | 1.246 | 0.24 | 0.11 | 0.44 | 0.35 | 0.37 | 0.16 |
| 9 | 96176203 | rs2184255 | -0.971 | -0.81 | 0.299 | -0.171 | 0.006 | 0.032 | 0.032 | 0 | 0.47 | 0.66 | 0.77 | 0.62 | 0.58 | 0.29 |
| 9 | 96178967 | rs13287066 | 0.578 | -0.701 | 0.215 | -0.131 | -0.005 | -0.005 | -0.005 | 0 | 0.29 | 0.31 | 0.49 | 0.41 | 0.43 | 0.17 |
| 9 | 96182487 | rs12379660 | 0.655 | -0.114 | 0.152 | -0.005 | 0.02 | 0.033 | 0.033 | 0.06 | 0.21 | 0.3 | 0.28 | 0.37 | 0.34 | 0.11 |
| 9 | 96191004 | rs944990 | 1.572 | -0.104 | 0.054 | -0.008 | -0.031 | -0.037 | -0.037 | 1.181 | 0.17 | 0.1 | 0.23 | 0.3 | 0.28 | 0.1 |
| 9 | 96192620 | rs2249747 | 1.402 | -0.434 | 0.002 | -0.48 | 0.009 | -0.009 | -0.009 | 0 | 0.2 | 0.16 | 0.1 | 0 | 0.13 | 0.06 |
| 9 | 96196166 | rs11789176 | -0.665 | -0.558 | 0.383 | -0.291 | 0.013 | -0.009 | -0.009 | 0 | 0.38 | 0.54 | 0.54 | 0.59 | 0.49 | 0.23 |
| 9 | 96200540 | rs7036984 | -0.106 | -0.509 | 0.508 | -0.318 | 0.015 | 0.08 | 0.08 | 0 | 0.14 | 0.43 | 0.23 | 0.16 | 0.13 | 0.11 |
| 9 | 96209943 | rs10821128 | 1.514 | -0.491 | 0.334 | -0.306 | -0.099 | -0.131 | -0.131 | 0 | 0.2 | 0.04 | 0.31 | 0.41 | 0.35 | 0.11 |
| 9 | 96214928 | rs1055710 | 1.289 | -0.212 | 0.437 | -0.32 | -0.002 | 0.002 | 0.002 | 0.198 | 0.21 | 0.22 | 0.33 | 0.42 | 0.35 | 0.15 |
| 9 | 96227100 | rs10512225 | 0.952 | -0.451 | 0.108 | -0.487 | -0.054 | 0.036 | 0.036 | 0 | 0.43 | 0.12 | 0.36 | 0.4 | 0.42 | 0.7 |
| 9 | 96228822 | rs10821131 | -0.563 | -0.315 | 0.225 | -0.436 | 0.016 | -0.004 | -0.004 | 0 | 0.37 | 0.51 | 0.54 | 0.56 | 0.48 | 0.2 |
| 9 | 96228952 | rs10821132 | 2.067 | -0.332 | 0.007 | -0.315 | -0.083 | -0.114 | -0.114 | 3.836 | 0.17 | 0.02 | 0.27 | 0.37 | 0.32 | 0.09 |
| 9 | 96229440 | rs10512224 | 0.674 | -0.867 | 0.216 | -0.325 | 0.058 | 0.146 | 0.146 | 0.048 | 0.05 | 0.37 | 0.15 | 0.06 | 0.03 | 0.07 |
| 9 | 96238218 | rs10821134 | 1.615 | -0.982 | -0.003 | -0.469 | -0.101 | -0.133 | -0.133 | 0.065 | 0.2 | 0.03 | 0.31 | 0.41 | 0.35 | 0.12 |
| 9 | 96242455 | rs4744244 | -2.068 | -1.327 | 0.055 | -0.541 | 0.049 | 0.076 | 0.076 | 0 | 0.85 | 0.98 | 0.93 | 1 | 0.92 | 0.94 |
| 9 | 96243745 | rs7025805 | -0.111 | -0.848 | 0.534 | -0.183 | 0.009 | 0.074 | 0.074 | 0 | 0.13 | 0.37 | 0.23 | 0.16 | 0.13 | 0.07 |
| 9 | 96265258 | rs10821144 | -0.764 | -0.766 | 0.158 | -0.231 | -0.015 | 0.008 | 0.008 | 0.014 | 0.39 | 0.67 | 0.56 | 0.59 | 0.51 | 0.24 |
| 9 | 96271752 | rs4744250 | 1.214 | -0.821 | 0.293 | -0.082 | -0.069 | -0.09 | -0.09 | 0.832 | 0.2 | 0.07 | 0.31 | 0.42 | 0.36 | 0.16 |
| 9 | 96291224 | rs7019001 | -0.881 | -0.878 | 0.303 | -0.184 | 0.007 | -0.001 | -0.001 | 0 | 0.38 | 0.58 | 0.56 | 0.58 | 0.51 | 0.24 |
| 9 | 96326127 | rs10821160 | 1.227 | -0.913 | -0.203 | -0.369 | -0.111 | -0.11 | -0.11 | 0 | 0.2 | 0.03 | 0.3 | 0.35 | 0.37 | 0.16 |
| 9 | 96336993 | rs7023577 | -0.735 | -0.918 | -0.022 | -0.435 | -0.032 | -0.012 | -0.012 | 0.009 | 0.38 | 0.72 | 0.56 | 0.53 | 0.52 | 0.24 |
| 9 | 96347718 | rs1000850 | 0.906 | -1.008 | 0.011 | -0.744 | 0.008 | -0.019 | -0.019 | 0 | 0.4 | 0.25 | 0.48 | 0.54 | 0.34 | 0.68 |
| 9 | 96351923 | rs10821166 | 0.186 | -1.143 | 0.133 | -0.745 | -0.064 | -0.025 | -0.025 | 0 | 0.3 | 0.16 | 0.32 | 0.34 | 0.47 | 0.21 |
| 9 | 96353119 | rs2150752 | -0.813 | -1.148 | 0.398 | -0.736 | 0.004 | 0.003 | 0.003 | 2.117 | 0.72 | 0.74 | 0.82 | 0.88 | 0.82 | 0.89 |
| 9 | 96353485 | rs10122014 | -0.077 | -1.125 | 0.965 | -0.289 | 0.082 | 0.177 | 0.177 | 0.002 | 0.06 | 0.46 | 0.23 | 0.06 | 0.03 | 0.03 |
| 9 | 96357739 | rs10116422 | 0.185 | -1.073 | 0.663 | -0.303 | 0.042 | 0.028 | 0.028 | 1.344 | 0.26 | 0.6 | 0.49 | 0.31 | 0.41 | 0.16 |
| 9 | 96359884 | rs10761239 | 0.204 | -1.377 | 0.512 | -0.309 | -0.051 | -0.016 | -0.016 | 0 | 0.29 | 0.19 | 0.33 | 0.34 | 0.48 | 0.21 |
| 9 | 96366415 | rs2254389 | 0.452 | -1.853 | 0.36 | -0.465 | -0.052 | -0.017 | -0.017 | 0 | 0.63 | 0.53 | 0.58 | 0.74 | 0.77 | 0.8 |
| 9 | 96366513 | rs2994369 | 1.342 | -1.845 | 0.361 | -0.463 | -0.023 | -0.04 | -0.04 | 0 | 0.4 | 0.14 | 0.29 | 0.49 | 0.37 | 0.66 |
| 9 | 96375217 | rs10123378 | 1.057 | -1.745 | 0.277 | -0.455 | 0.091 | 0.021 | 0.021 | 0 | 0.22 | 0.37 | 0.29 | 0.25 | 0.4 | 0.14 |
| 9 | 96379505 | rs9409478 | -1.048 | -1.929 | 0.042 | -0.437 | 0.013 | -0.009 | -0.009 | 0 | 0.8 | 0.86 | 0.92 | 1 | 0.91 | 0.95 |

| | | | | | | | | | | | | | | | | |
|---|---|---|---|---|---|---|---|---|---|---|---|---|---|---|---|---|
| 9 | 96379797 | rs12551314 | -0.874 | -1.434 | -0.173 | -0.1 | 0.001 | -0.002 | -0.002 | 0 | 0.12 | 0.12 | 0.13 | 0.2 | 0.11 | 0.11 |
| 9 | 96382100 | rs7033146 | 0.216 | -1.41 | -0.368 | -0.36 | -0.008 | -0.001 | -0.001 | 3.333 | 0.48 | 0.23 | 0.37 | 0.6 | 0.47 | 0.79 |
| 9 | 96382336 | rs7033439 | 0.193 | -1.538 | 0.086 | -0.262 | -0.014 | -0.011 | -0.011 | 0.111 | 0.65 | 0.61 | 0.65 | 0.85 | 0.76 | 0.84 |
| 9 | 96391990 | rs7849752 | 0.569 | -0.956 | 0.238 | -0.264 | -0.001 | -0.002 | -0.002 | 0 | 0.47 | 0.37 | 0.51 | 0.54 | 0.43 | 0.72 |
| 9 | 96401464 | rs11534279 | 0.226 | -0.279 | 0.724 | 0.081 | 0.047 | 0.049 | 0.049 | 0.064 | 0.29 | 0.54 | 0.36 | 0.39 | 0.4 | 0.15 |
| 9 | 96403367 | rs10761247 | -0.568 | -0.297 | 0.713 | -0.11 | -0.001 | 0.003 | 0.003 | 0 | 0.45 | 0.66 | 0.44 | 0.44 | 0.57 | 0.25 |
| 9 | 96407983 | rs9695734 | 1.385 | -0.279 | 0.375 | -0.104 | -0.001 | -0.017 | -0.017 | 1.69 | 0.16 | 0.12 | 0.08 | 0.05 | 0.16 | 0.1 |
| 9 | 96415899 | rs11793637 | -0.127 | -0.363 | 0.477 | -0.01 | 0.031 | 0.033 | 0.033 | 0 | 0.34 | 0.54 | 0.41 | 0.46 | 0.48 | 0.19 |
| 9 | 96425138 | rs3750358 | -0.701 | -0.291 | 0.099 | 0.035 | 0.019 | -0.003 | -0.003 | 1.558 | 0.47 | 0.66 | 0.48 | 0.49 | 0.61 | 0.27 |
| 9 | 96427505 | rs10992839 | 1.076 | 0.046 | -0.403 | 0.153 | 0.003 | 0 | 0 | 0 | 0.22 | 0.23 | 0.28 | 0.29 | 0.37 | 0.12 |
| 9 | 96440808 | rs1132004 | 1.072 | 0.056 | -0.398 | 0.146 | 0.003 | 0 | 0 | 0 | 0.22 | 0.24 | 0.28 | 0.29 | 0.37 | 0.12 |
| 9 | 96447178 | rs7023781 | 0.844 | -0.407 | -1.341 | 0.026 | 0.005 | -0.096 | -0.096 | 0.156 | 0.41 | 0.08 | 0.36 | 0.54 | 0.33 | 0.69 |
| 9 | 96451784 | rs4287000 | -0.349 | -0.415 | -0.926 | -0.163 | -0.007 | 0.007 | 0.007 | 0 | 0.64 | 0.33 | 0.64 | 0.76 | 0.56 | 0.84 |
| 9 | 96461013 | rs10992867 | 0.904 | -0.363 | -1.615 | -0.063 | -0.008 | -0.11 | -0.11 | 0 | 0.39 | 0.06 | 0.35 | 0.55 | 0.31 | 0.67 |
| 9 | 96463094 | rs12683791 | 0.888 | -0.136 | -1.564 | -0.002 | -0.027 | -0.105 | -0.105 | 1.117 | 0.39 | 0.07 | 0.37 | 0.55 | 0.32 | 0.67 |
| 9 | 96469075 | rs4744272 | 0.377 | -0.418 | -1.205 | -0.219 | 0.01 | 0.012 | 0.012 | 0 | 0.67 | 0.68 | 0.71 | 0.84 | 0.74 | 0.83 |
| 9 | 96484209 | rs11794188 | 0.315 | -0.464 | -1.327 | -0.051 | -0.013 | -0.031 | -0.031 | 0 | 0.1 | 0.13 | 0.04 | 0 | 0.03 | 0.02 |
| 9 | 96484220 | rs2398871 | 0.205 | -0.119 | -1.704 | 0.301 | -0.027 | -0.123 | -0.123 | 0.039 | 0.39 | 0.06 | 0.36 | 0.55 | 0.31 | 0.65 |
| 9 | 96488780 | rs10992886 | 0.427 | 0.38 | -1.272 | 0.615 | -0.042 | -0.041 | -0.041 | 0 | 0.08 | 0.01 | 0.22 | 0.16 | 0.17 | 0.13 |
| 9 | 96488923 | rs10821207 | 0.021 | 0.343 | -0.719 | 0.593 | 0.011 | 0.011 | 0.011 | 0 | 0.1 | 0.12 | 0.23 | 0.16 | 0.17 | 0.13 |
| 9 | 96493773 | rs2255840 | -0.296 | 0.249 | -0.201 | 0.929 | 0.035 | 0.1 | 0.1 | 0 | 0.11 | 0.26 | 0.24 | 0.29 | 0.17 | 0.16 |
| 9 | 96493970 | rs7049001 | -0.1 | -0.377 | -0.502 | 0.684 | -0.004 | 0.008 | 0.008 | 0.197 | 0.37 | 0.42 | 0.34 | 0.48 | 0.28 | 0.48 |
| 9 | 96522109 | rs2490997 | -1.227 | -1.043 | -0.602 | 0.523 | -0.008 | 0.008 | 0.008 | 1.178 | 0.61 | 0.64 | 0.52 | 0.73 | 0.5 | 0.67 |
| 9 | 96522775 | rs2490996 | 1.326 | -1.193 | -0.956 | 0.418 | -0.055 | -0.019 | -0.019 | 0 | 0.12 | 0.01 | 0.11 | 0.15 | 0.18 | 0.08 |
| 9 | 96524226 | rs10761266 | -0.236 | -1.279 | -0.708 | 0.97 | 0.069 | 0.042 | 0.042 | 0.392 | 0.44 | 0.23 | 0.36 | 0.28 | 0.28 | 0.48 |
| 9 | 96526835 | rs12351676 | -0.204 | -1.27 | -0.7 | 1.101 | 0.066 | 0.054 | 0.054 | 0.497 | 0.43 | 0.22 | 0.35 | 0.24 | 0.28 | 0.46 |
| 9 | 96531707 | rs2487013 | 1.347 | -1.267 | -0.723 | 1.066 | -0.055 | -0.035 | -0.035 | 0 | 0.12 | 0.01 | 0.11 | 0.19 | 0.18 | 0.08 |
| 9 | 96534118 | rs10119990 | -0.897 | -1.269 | -0.721 | 1.067 | 0.039 | 0.015 | 0.015 | 0.608 | 0.49 | 0.63 | 0.41 | 0.55 | 0.32 | 0.58 |
| 9 | 96534292 | rs2490992 | 0.633 | -1.263 | -0.819 | 0.974 | -0.074 | -0.017 | -0.017 | 0 | 0.17 | 0.01 | 0.15 | 0.19 | 0.26 | 0.09 |
| 9 | 96544917 | rs10821216 | 0.078 | -1.476 | -0.893 | 1.174 | 0.107 | -0.02 | -0.02 | 1.052 | 0.37 | 0.12 | 0.28 | 0.29 | 0.2 | 0.33 |
| 9 | 96559370 | rs7874836 | 1.317 | -1.151 | -0.752 | 1.239 | -0.006 | 0.008 | 0.008 | 1.264 | 0.38 | 0.37 | 0.48 | 0.18 | 0.49 | 0.36 |
| 9 | 96562563 | rs7027203 | 2.428 | -1.187 | -0.884 | 1.155 | 0.015 | -0.009 | -0.009 | 0 | 0.27 | 0.35 | 0.32 | 0.19 | 0.37 | 0.25 |
| 9 | 96563349 | rs10761275 | -0.506 | -1.195 | -0.788 | 1.161 | 0.027 | 0.018 | 0.018 | 0.617 | 0.49 | 0.61 | 0.41 | 0.58 | 0.32 | 0.56 |
| 9 | 96564575 | rs10821224 | -0.104 | -1.208 | -0.77 | 1.113 | 0.064 | -0.008 | -0.008 | 1.247 | 0.44 | 0.21 | 0.36 | 0.37 | 0.3 | 0.45 |
| 9 | 96569343 | rs10761280 | -0.516 | -1.188 | -0.813 | 1.088 | 0.027 | 0.036 | 0.036 | 0.476 | 0.5 | 0.61 | 0.41 | 0.67 | 0.33 | 0.56 |
| 9 | 96571211 | rs7854075 | 2.379 | -1.105 | -0.758 | 1.113 | 0.014 | -0.029 | -0.029 | 0 | 0.27 | 0.35 | 0.32 | 0.1 | 0.36 | 0.25 |
| 9 | 96577178 | rs4744283 | -1.503 | -1.954 | -0.701 | 0.783 | 0.001 | -0.003 | -0.003 | 0 | 0.46 | 0.57 | 0.39 | 0.59 | 0.42 | 0.47 |
| 9 | 96578169 | rs11999651 | -0.686 | -1.925 | -0.674 | 0.788 | -0.01 | 0.008 | 0.008 | 0 | 0.35 | 0.52 | 0.3 | 0.43 | 0.28 | 0.4 |
| 9 | 96582511 | rs11789114 | 0.673 | -1.805 | -0.566 | 0.817 | 0.003 | 0.011 | 0.011 | 0 | 0.12 | 0.1 | 0.16 | 0.08 | 0.12 | 0.11 |
| 9 | 96583746 | rs4744285 | -0.444 | -1.81 | -0.467 | 0.857 | 0.009 | 0.11 | 0.11 | 0 | 0.15 | 0.32 | 0.19 | 0.39 | 0.15 | 0.22 |
| 9 | 96591066 | rs3996253 | -0.937 | -1.877 | -0.509 | 0.831 | 0 | -0.008 | -0.008 | 1.498 | 0.57 | 0.49 | 0.49 | 0.63 | 0.43 | 0.6 |
| 9 | 96591546 | rs10992913 | -1.622 | -1.972 | -0.524 | 0.793 | 0 | 0 | 0 | 0 | 0.51 | 0.47 | 0.54 | 0.77 | 0.45 | 0.58 |
| 9 | 96600298 | rs4344139 | -2.689 | -1.92 | -0.798 | 0.788 | 0.018 | -0.013 | -0.013 | 0 | 0.71 | 0.62 | 0.66 | 0.82 | 0.59 | 0.75 |
| 9 | 96612770 | rs3957512 | -0.474 | -1.955 | -0.837 | 0.817 | -0.009 | 0 | 0 | 0 | 0.46 | 0.48 | 0.4 | 0.48 | 0.29 | 0.52 |
| 9 | 96614312 | rs10821237 | 0.464 | -2.027 | -0.937 | 0.869 | 0.043 | -0.042 | -0.042 | 0 | 0.25 | 0.08 | 0.27 | 0.23 | 0.15 | 0.27 |
| 9 | 96617463 | rs10821238 | 0.487 | -2.019 | -0.933 | 0.873 | 0.041 | -0.042 | -0.042 | 0 | 0.24 | 0.07 | 0.27 | 0.23 | 0.15 | 0.29 |
| 9 | 96617700 | rs10992927 | 0.727 | -1.335 | -0.034 | 1.035 | -0.04 | -0.023 | -0.023 | 0.118 | 0.11 | 0.01 | 0.09 | 0.15 | 0.14 | 0.08 |
| 9 | 96623453 | rs4428725 | -0.569 | -1.662 | 0.169 | 0.877 | 0.001 | 0.009 | 0.009 | 0.023 | 0.55 | 0.53 | 0.51 | 0.73 | 0.42 | 0.65 |
| 9 | 96624526 | rs4075732 | -0.667 | -1.658 | 0.22 | 0.83 | -0.001 | 0 | 0 | 0.06 | 0.54 | 0.43 | 0.5 | 0.54 | 0.42 | 0.57 |
| 9 | 96624645 | rs4075733 | -1.322 | -1.287 | -0.023 | 0.345 | -0.002 | 0 | 0 | 0 | 0.67 | 0.65 | 0.65 | 0.82 | 0.54 | 0.74 |
| 9 | 96630555 | rs10733750 | -1.427 | -1.845 | 0.439 | 0.631 | -0.034 | -0.007 | -0.007 | 1.552 | 0.34 | 0.45 | 0.29 | 0.32 | 0.34 | 0.32 |
| 9 | 96631287 | rs10115047 | -1.543 | -2.117 | 0.48 | 0.573 | -0.005 | -0.018 | -0.018 | 0 | 0.4 | 0.36 | 0.27 | 0.21 | 0.38 | 0.24 |
| 9 | 96631459 | rs10114476 | -0.593 | -2.248 | 0.497 | 0.608 | -0.017 | 0.148 | 0.148 | 7.261 | 0.08 | 0.28 | 0.06 | 0.17 | 0.02 | 0.17 |
| 9 | 96631796 | rs10117869 | -1.279 | -2.265 | 0.39 | 0.589 | -0.049 | 0.012 | 0.012 | 0.372 | 0.31 | 0.45 | 0.27 | 0.34 | 0.26 | 0.31 |
| 9 | 96635264 | rs9644971 | -0.949 | -2.305 | 0.333 | 0.575 | -0.021 | 0.026 | 0.026 | 0.578 | 0.22 | 0.36 | 0.19 | 0.26 | 0.22 | 0.27 |
| 9 | 96644841 | rs7855646 | -2.212 | -2.327 | 0.443 | 0.684 | 0.063 | 0.002 | 0.002 | 0.935 | 0.48 | 0.63 | 0.35 | 0.6 | 0.4 | 0.5 |
| 9 | 96646070 | rs11793716 | -1.041 | -2.139 | 0.175 | -0.303 | 0.043 | 0.045 | 0.045 | 0 | 0.24 | 0.16 | 0.15 | 0.12 | 0.18 | 0.14 |
| 9 | 96648679 | rs2185110 | -2.02 | -2.13 | 0.183 | -0.298 | -0.015 | -0.007 | -0.007 | 1.194 | 0.45 | 0.45 | 0.33 | 0.38 | 0.4 | 0.41 |
| 9 | 96654346 | rs12376257 | 0.269 | -2.055 | -0.081 | -0.326 | -0.043 | 0.021 | 0.021 | 0 | 0.13 | 0.01 | 0.11 | 0.1 | 0.19 | 0.11 |
| 9 | 96654597 | rs11794892 | -0.213 | -2.001 | -0.046 | -0.307 | 0.017 | 0.083 | 0.083 | 0 | 0.16 | 0.01 | 0.05 | 0.04 | 0.12 | 0.1 |
| 9 | 96658325 | rs10992966 | 0.095 | -1.706 | 0.161 | -0.185 | -0.043 | -0.003 | -0.003 | 0 | 0.11 | 0.29 | 0.12 | 0.08 | 0.07 | 0.05 |
| 9 | 96661535 | rs1890376 | -1.463 | -1.395 | -0.284 | -0.315 | 0.05 | 0.033 | 0.033 | 2.085 | 0.49 | 0.59 | 0.35 | 0.36 | 0.41 | 0.42 |
| 9 | 96663863 | rs7047570 | -1.268 | -1.332 | -0.197 | -0.249 | 0.01 | 0.007 | 0.007 | 1.025 | 0.47 | 0.41 | 0.33 | 0.36 | 0.4 | 0.42 |
| 9 | 96665767 | rs1890379 | -1.332 | -1.211 | -0.99 | -0.226 | 0.043 | 0.028 | 0.028 | 1.681 | 0.48 | 0.58 | 0.35 | 0.36 | 0.4 | 0.42 |
| 9 | 96670213 | rs10117505 | -0.492 | -0.072 | -1.505 | -0.44 | -0.005 | 0.033 | 0.033 | 0.01 | 0.15 | 0.09 | 0.11 | 0.09 | 0.19 | 0.12 |
| 9 | 96671646 | rs12343357 | -0.888 | -0.261 | -1.186 | -0.301 | 0 | -0.006 | -0.006 | 0 | 0.14 | 0.11 | 0.05 | 0.05 | 0.08 | 0.07 |
| 9 | 96671698 | rs7872495 | -0.854 | -0.026 | -1.269 | -0.153 | -0.004 | -0.004 | -0.004 | 1.499 | 0.14 | 0.16 | 0.13 | 0.12 | 0.11 | 0.08 |
| 9 | 96672124 | rs10119262 | -0.398 | 0.83 | -1.71 | 0.178 | -0.019 | 0.01 | 0.01 | 0 | 0.42 | 0.18 | 0.47 | 0.35 | 0.41 | 0.28 |
| 9 | 96679166 | rs13283140 | -0.916 | 1.862 | -1.383 | -0.261 | -0.014 | 0.008 | 0.008 | 0.007 | 0.19 | 0.12 | 0.29 | 0.03 | 0.32 | 0.19 |

| | | | | | | | | | | | | | | | | |
|---|---|---|---|---|---|---|---|---|---|---|---|---|---|---|---|---|
| 9 | 96679448 | rs10992979 | 0.082 | 1.298 | -2.163 | -0.392 | 0.012 | 0.009 | 0.009 | 0 | 0.26 | 0.12 | 0.25 | 0.2 | 0.22 | 0.18 |
| 9 | 96680040 | rs12353447 | -0.545 | 1.333 | -2.199 | 0.427 | -0.011 | -0.01 | -0.01 | 0.153 | 0.31 | 0.25 | 0.4 | 0.44 | 0.44 | 0.54 |
| 9 | 96680113 | rs12380877 | 1.146 | 0.447 | -2.993 | 0.022 | 0.002 | -0.015 | -0.015 | 0 | 0.13 | 0.01 | 0.12 | 0.14 | 0.11 | 0.14 |
| 9 | 96681388 | rs4743938 | -1.476 | -0.145 | -2.348 | -0.375 | -0.01 | 0.008 | 0.008 | 0 | 0.86 | 0.81 | 0.87 | 0.83 | 0.92 | 0.93 |
| 9 | 96682611 | rs10821266 | 0.338 | -0.589 | -1.469 | -0.606 | -0.008 | -0.024 | -0.024 | 0 | 0.26 | 0.27 | 0.2 | 0.18 | 0.25 | 0.19 |
| 9 | 96682813 | rs12156596 | 0.393 | -0.589 | -1.34 | -0.563 | -0.01 | -0.034 | -0.034 | 0 | 0.25 | 0.3 | 0.21 | 0.18 | 0.24 | 0.19 |
| 9 | 96685120 | rs10992984 | 1.079 | -0.727 | -1.495 | -0.695 | 0.013 | 0.005 | 0.005 | 0 | 0.17 | 0.01 | 0.12 | 0.14 | 0.13 | 0.14 |
| 9 | 96685308 | rs12376441 | 1.775 | -0.856 | -1.621 | -0.847 | 0.016 | -0.003 | -0.003 | 0 | 0.21 | 0.16 | 0.23 | 0.2 | 0.18 | 0.17 |
| 9 | 96691023 | rs7028921 | 0.2 | -0.733 | -1.509 | -0.808 | -0.017 | -0.041 | -0.041 | 0.923 | 0.29 | 0.31 | 0.21 | 0.18 | 0.26 | 0.2 |
| 9 | 96693709 | rs12341268 | -0.691 | -0.79 | -1.537 | -0.699 | 0.008 | 0.003 | 0.003 | 0 | 0.55 | 0.48 | 0.61 | 0.62 | 0.69 | 0.73 |
| 9 | 96697234 | rs1933678 | 1.282 | -0.527 | -0.657 | -0.621 | 0.006 | -0.027 | -0.027 | 0 | 0.11 | 0.18 | 0.09 | 0.04 | 0.13 | 0.06 |
| 9 | 96699676 | rs10992995 | 0.022 | 0.747 | -0.601 | -0.103 | 0.008 | -0.002 | -0.002 | 0 | 0.23 | 0.19 | 0.36 | 0.15 | 0.34 | 0.42 |
| 9 | 96709022 | rs1933679 | -0.14 | 0.136 | -1.117 | -0.139 | -0.085 | -0.015 | -0.015 | 0.546 | 0.61 | 0.88 | 0.75 | 0.63 | 0.54 | 0.42 |
| 9 | 96709483 | rs1933680 | 0.407 | -0.449 | -0.989 | -0.292 | 0 | -0.011 | -0.011 | 0 | 0.39 | 0.37 | 0.46 | 0.19 | 0.42 | 0.29 |
| 9 | 96709839 | rs10821272 | 2.077 | -0.589 | -0.8 | -0.031 | -0.001 | 0.009 | 0.009 | 0 | 0.11 | 0.12 | 0.1 | 0.23 | 0.1 | 0.07 |
| 9 | 96713690 | rs11793324 | 2.055 | -0.589 | -0.8 | -0.031 | -0.001 | 0.008 | 0.008 | 0 | 0.11 | 0.12 | 0.1 | 0.23 | 0.1 | 0.07 |
| 9 | 96714161 | rs1057713 | 0.397 | -0.636 | -0.685 | -0.064 | 0 | -0.034 | -0.034 | 0 | 0.39 | 0.45 | 0.46 | 0.21 | 0.41 | 0.3 |
| 9 | 96714981 | rs12684081 | 1.128 | -1.132 | -0.442 | -0.079 | -0.039 | 0.076 | 0.076 | 0 | 0.24 | 0.08 | 0.31 | 0.09 | 0.28 | 0.19 |
| 9 | 96716028 | rs11789015 | 1.027 | -1.136 | -0.427 | -0.092 | 0.014 | -0.065 | -0.065 | 0.078 | 0.26 | 0.35 | 0.34 | 0.09 | 0.29 | 0.19 |
| 9 | 96716158 | rs7872123 | -0.06 | -1.123 | -0.428 | -0.078 | 0.001 | -0.03 | -0.03 | 0 | 0.39 | 0.44 | 0.47 | 0.2 | 0.42 | 0.3 |
| 9 | 96740079 | rs10821278 | 1.054 | -1.221 | -0.193 | -0.002 | -0.035 | 0.08 | 0.08 | 0 | 0.2 | 0.3 | 0.27 | 0.44 | 0.12 | 0.11 |
| 9 | 96741641 | rs10821279 | 1.955 | -0.749 | -0.319 | -0.007 | -0.021 | -0.017 | -0.017 | 0 | 0.11 | 0.01 | 0.06 | 0.11 | 0.13 | 0.09 |
| 9 | 96757868 | rs987511 | -0.884 | -0.713 | -0.197 | -0.056 | -0.034 | 0.014 | 0.014 | 0 | 0.59 | 0.77 | 0.75 | 0.65 | 0.54 | 0.41 |
| 9 | 96763614 | rs1411861 | 1.205 | -0.709 | -0.302 | -0.067 | -0.041 | 0.08 | 0.08 | 0.033 | 0.23 | 0.07 | 0.32 | 0.09 | 0.28 | 0.19 |
| 9 | 96766248 | rs1331219 | -0.846 | -0.664 | -0.376 | -0.002 | -0.056 | 0.01 | 0.01 | 0 | 0.6 | 0.81 | 0.75 | 0.65 | 0.54 | 0.41 |
| 9 | 96769237 | rs10761312 | 0.615 | -0.631 | -0.366 | 0.105 | -0.023 | 0.032 | 0.032 | 0 | 0.24 | 0.29 | 0.35 | 0.43 | 0.15 | 0.2 |
| 9 | 96773760 | rs4743942 | 1.552 | -0.612 | -0.322 | 0.25 | -0.029 | 0.072 | 0.072 | 0 | 0.15 | 0.28 | 0.25 | 0.26 | 0.11 | 0.11 |
| 9 | 96776609 | rs2150624 | -1.391 | -0.57 | -0.75 | -0.006 | -0.075 | 0.036 | 0.036 | 0 | 0.78 | 0.98 | 0.88 | 0.89 | 0.77 | 0.6 |
| 9 | 96792054 | rs10116489 | 2.21 | -0.639 | -0.623 | -0.166 | -0.033 | 0.048 | 0.048 | 0 | 0.17 | 0.28 | 0.25 | 0.26 | 0.11 | 0.11 |
| 9 | 96792790 | rs7047684 | 0.471 | -0.618 | -0.515 | -0.227 | 0.002 | -0.022 | -0.022 | 0 | 0.32 | 0.35 | 0.44 | 0.26 | 0.33 | 0.28 |
| 9 | 96795797 | rs10116746 | 2.652 | -0.631 | -0.586 | -0.22 | -0.027 | 0.062 | 0.062 | 0 | 0.16 | 0.27 | 0.25 | 0.26 | 0.11 | 0.11 |
| 9 | 96805635 | rs10429556 | -0.625 | -0.967 | -0.553 | -0.355 | -0.018 | 0.002 | 0.002 | 0 | 0.62 | 0.65 | 0.74 | 0.65 | 0.55 | 0.47 |
| 9 | 96827424 | rs1539282 | 0.434 | -0.917 | -0.619 | -0.347 | 0 | 0.009 | 0.009 | 0 | 0.33 | 0.26 | 0.43 | 0.26 | 0.33 | 0.28 |
| 9 | 96830635 | rs6479526 | 1.582 | -1.003 | -0.556 | -0.31 | -0.009 | 0.012 | 0.012 | 0 | 0.27 | 0.32 | 0.31 | 0.39 | 0.22 | 0.2 |
| 9 | 96837996 | rs7020044 | 2.139 | -0.959 | -0.51 | -0.259 | -0.038 | 0.053 | 0.053 | 0 | 0.18 | 0.31 | 0.26 | 0.26 | 0.11 | 0.11 |
| 9 | 96846061 | rs4744313 | 0.526 | -0.886 | -0.855 | -0.298 | -0.003 | -0.002 | -0.002 | 0 | 0.41 | 0.39 | 0.57 | 0.35 | 0.39 | 0.3 |
| 9 | 96846500 | rs10993047 | 2.211 | -0.802 | -1.206 | -0.29 | -0.002 | -0.004 | -0.004 | 0.014 | 0.23 | 0.17 | 0.32 | 0.09 | 0.28 | 0.19 |
| 9 | 96858411 | rs6479527 | 0.039 | -0.525 | -1.684 | -0.104 | -0.019 | -0.033 | -0.033 | 0.149 | 0.54 | 0.79 | 0.66 | 0.48 | 0.5 | 0.38 |
| 9 | 96861798 | rs10993052 | 0.64 | -0.466 | -1.607 | -0.048 | -0.006 | -0.005 | -0.005 | 0 | 0.41 | 0.48 | 0.58 | 0.35 | 0.39 | 0.3 |
| 9 | 96864227 | rs10122334 | -0.52 | -0.174 | -1.814 | 0.074 | -0.175 | -0.076 | -0.076 | 0.118 | 0.64 | 0.96 | 0.77 | 0.65 | 0.56 | 0.47 |
| 9 | 96869147 | rs10821295 | 2.089 | 0.159 | -1.599 | -0.126 | -0.011 | 0.008 | 0.008 | 0 | 0.14 | 0.09 | 0.13 | 0.03 | 0.24 | 0.11 |
| 9 | 96873151 | rs10993058 | -0.228 | -0.352 | -1.918 | 0.116 | -0.088 | -0.087 | -0.087 | 0 | 0.57 | 0.87 | 0.68 | 0.48 | 0.54 | 0.45 |
| 9 | 96883271 | rs7875583 | 0.045 | -0.279 | -1.322 | 0.09 | 0.004 | 0.003 | 0.003 | 0 | 0.42 | 0.57 | 0.59 | 0.35 | 0.39 | 0.3 |
| 9 | 96889505 | rs3909256 | 1.158 | 0.21 | -1.796 | -0.228 | -0.017 | 0.054 | 0.054 | 0.016 | 0.23 | 0.12 | 0.31 | 0.09 | 0.25 | 0.2 |
| 9 | 96893369 | rs3858085 | 0.941 | -0.209 | -1.768 | 0.102 | 0.007 | -0.009 | -0.009 | 0 | 0.12 | 0.09 | 0.11 | 0.18 | 0.07 | 0.05 |
| 9 | 96893543 | rs7035599 | 0.464 | 0.198 | -1.941 | -0.217 | -0.03 | 0.037 | 0.037 | 0.472 | 0.34 | 0.23 | 0.43 | 0.15 | 0.47 | 0.34 |
| 9 | 96895108 | rs10821299 | 1.655 | 0.147 | -2.171 | 0.246 | -0.015 | -0.108 | -0.108 | 0 | 0.18 | 0.01 | 0.1 | 0.34 | 0.17 | 0.28 |
| 9 | 96895320 | rs10761316 | 0.719 | 0.572 | -2.413 | 0.018 | -0.025 | -0.024 | -0.024 | 0.007 | 0.26 | 0.08 | 0.28 | 0.27 | 0.32 | 0.18 |
| 9 | 96896292 | rs10739970 | -0.36 | 0.381 | -2.373 | -0.073 | 0.017 | -0.03 | -0.03 | 2.622 | 0.58 | 0.24 | 0.7 | 0.52 | 0.65 | 0.43 |
| 9 | 96897033 | rs2398901 | -1.787 | 0.665 | -2.126 | -0.165 | -0.01 | 0.006 | 0.006 | 0 | 0.87 | 0.91 | 0.95 | 0.91 | 0.85 | 0.81 |
| 9 | 96898436 | rs10761317 | 0.344 | 0.557 | -2.033 | -0.055 | -0.027 | -0.022 | -0.022 | 0 | 0.26 | 0.08 | 0.28 | 0.27 | 0.33 | 0.19 |
| 9 | 96902444 | rs959802 | 0.982 | 0.416 | -1.588 | 0.16 | 0.012 | -0.024 | -0.024 | 0 | 0.08 | 0.02 | 0.18 | 0.22 | 0.08 | 0.05 |
| 9 | 96906751 | rs10993068 | 0.19 | 0.576 | -1.112 | -0.548 | -0.002 | -0.01 | -0.01 | 0 | 0.39 | 0.31 | 0.36 | 0.11 | 0.4 | 0.25 |
| 9 | 96913764 | rs13439971 | 0.233 | 0.433 | -0.766 | -0.622 | -0.009 | -0.049 | -0.049 | 0 | 0.36 | 0.41 | 0.21 | 0.11 | 0.29 | 0.22 |
| 9 | 96916230 | rs10512230 | 1.823 | 0.676 | -1.478 | -0.29 | -0.112 | -0.145 | -0.145 | 0.852 | 0.23 | 0.01 | 0.28 | 0.41 | 0.33 | 0.34 |
| 9 | 96919182 | rs10114341 | 0.04 | 0.568 | -1.591 | -0.494 | -0.002 | -0.032 | -0.032 | 0 | 0.45 | 0.45 | 0.29 | 0.13 | 0.44 | 0.36 |
| 9 | 96930485 | rs1362820 | 0.99 | 0.396 | -1.083 | -0.372 | 0.001 | -0.001 | -0.001 | 0 | 0.12 | 0.08 | 0.19 | 0.22 | 0.08 | 0.09 |
| 9 | 96931807 | rs2026618 | -1.035 | 0.477 | -0.827 | -0.377 | 0.023 | 0.047 | 0.047 | 0 | 0.93 | 0.86 | 0.88 | 0.79 | 0.9 | 0.85 |
| 9 | 96939718 | rs10993081 | 1.087 | 0.405 | -0.922 | -0.532 | 0.009 | -0.063 | -0.063 | 0.27 | 0.11 | 0.01 | 0.19 | 0.22 | 0.08 | 0.09 |
| 9 | 96944581 | rs10821312 | 0.034 | 0.502 | -0.503 | -0.961 | 0.021 | -0.063 | -0.063 | 0 | 0.14 | 0.32 | 0.11 | 0.02 | 0.16 | 0.14 |
| 9 | 96945748 | rs10512231 | 1.208 | 0.485 | -0.507 | -0.987 | -0.006 | 0.12 | 0.12 | 0 | 0.32 | 0.1 | 0.18 | 0.11 | 0.29 | 0.22 |
| 9 | 96959415 | rs7038156 | -0.081 | 0.264 | -0.365 | -0.792 | 0.019 | -0.062 | -0.062 | 0 | 0.14 | 0.32 | 0.11 | 0.02 | 0.17 | 0.14 |
| 9 | 96970197 | rs7872931 | 1.188 | 0.367 | -1.073 | -0.904 | 0.009 | 0.113 | 0.113 | 0.016 | 0.31 | 0.09 | 0.17 | 0.11 | 0.26 | 0.2 |
| 9 | 96973995 | rs12379483 | -0.024 | 0.521 | -0.443 | -0.672 | 0.001 | -0.012 | -0.012 | 0 | 0.14 | 0.18 | 0.18 | 0.04 | 0.17 | 0.13 |
| 9 | 96990968 | rs9409706 | 2.151 | 0.521 | -0.848 | -0.503 | -0.108 | -0.146 | -0.146 | 0 | 0.22 | 0.01 | 0.2 | 0.44 | 0.32 | 0.36 |
| 9 | 97000398 | rs2675278 | -0.951 | 0.422 | -0.586 | -0.58 | 0.005 | -0.1 | -0.1 | 0.282 | 0.24 | 0.53 | 0.22 | 0.05 | 0.23 | 0.18 |
| 9 | 97000863 | rs7029157 | 0.846 | 0.455 | -0.469 | -0.602 | 0.026 | -0.061 | -0.061 | 0.029 | 0.11 | 0.38 | 0.05 | 0 | 0.07 | 0.05 |
| 9 | 97008212 | rs2675275 | 0.06 | 0.314 | -0.855 | -0.404 | 0.006 | -0.004 | -0.004 | 3.488 | 0.47 | 0.66 | 0.45 | 0.49 | 0.56 | 0.49 |
| 9 | 97015630 | rs10993117 | -0.745 | 0.124 | -0.902 | -0.535 | 0 | 0 | 0 | 1.856 | 0.23 | 0.19 | 0.37 | 0.25 | 0.25 | 0.21 |

| | | | | | | | | | | | | | | | | |
|---|---|---|---|---|---|---|---|---|---|---|---|---|---|---|---|---|
| 9 | 97016558 | rs9299416 | -0.445 | -0.349 | -1.522 | -1.133 | 0.042 | 0.101 | 0.101 | 0 | 0.63 | 0.83 | 0.78 | 0.83 | 0.69 | 0.6 |
| 9 | 97017858 | rs11789548 | 0.206 | -0.723 | -1.738 | -1.801 | 0.006 | 0.048 | 0.048 | 0 | 0.26 | 0.16 | 0.17 | 0.02 | 0.25 | 0.09 |
| 9 | 97018657 | rs10821325 | 2.318 | -0.069 | -2.061 | -1.568 | -0.103 | -0.151 | -0.151 | 0 | 0.22 | 0.01 | 0.31 | 0.51 | 0.32 | 0.34 |
| 9 | 97024028 | rs10993121 | -0.502 | -0.827 | -2.02 | -1.848 | 0.002 | 0.085 | 0.085 | 0 | 0.37 | 0.21 | 0.21 | 0.02 | 0.34 | 0.2 |
| 9 | 97025298 | rs3118766 | -0.131 | -0.754 | -1.597 | -1.768 | -0.004 | -0.002 | -0.002 | 0 | 0.31 | 0.29 | 0.46 | 0.4 | 0.39 | 0.29 |
| 9 | 97040683 | rs11793128 | 0.164 | -1.857 | -1.007 | -1.638 | 0.032 | 0.052 | 0.052 | 0 | 0.47 | 0.34 | 0.3 | 0.07 | 0.4 | 0.29 |
| 9 | 97042132 | rs10435926 | 0.13 | -1.655 | -0.799 | -1.28 | 0.002 | 0.112 | 0.112 | 0 | 0.06 | 0.08 | 0.26 | 0.41 | 0.07 | 0.19 |
| 9 | 97052841 | rs3118752 | 0.163 | -1.753 | -0.256 | -1.738 | -0.025 | -0.051 | -0.051 | 0.111 | 0.89 | 0.56 | 0.94 | 0.98 | 0.96 | 0.95 |
| 9 | 97053678 | rs3118749 | 0.031 | -1.737 | 0.045 | -1.73 | -0.016 | -0.039 | -0.039 | 0.037 | 0.91 | 0.68 | 0.95 | 0.99 | 0.96 | 0.95 |
| 9 | 97055310 | rs1536690 | 0.615 | -1.922 | -0.433 | -1.907 | 0.007 | -0.002 | -0.002 | 0 | 0.29 | 0.25 | 0.16 | 0.06 | 0.18 | 0.25 |
| 9 | 97056183 | rs3131910 | 2.12 | -1.483 | -0.911 | -1.73 | -0.135 | -0.12 | -0.12 | 0 | 0.22 | 0.05 | 0.33 | 0.56 | 0.39 | 0.33 |
| 9 | 97061544 | rs12343413 | 0.221 | -1.554 | -0.232 | -1.738 | -0.011 | -0.059 | -0.059 | 0 | 0.14 | 0.16 | 0.05 | 0 | 0.09 | 0.08 |
| 9 | 97061577 | rs9409511 | 0.77 | -1.443 | -0.755 | -1.717 | -0.149 | -0.098 | -0.098 | 0.051 | 0.34 | 0.05 | 0.36 | 0.56 | 0.47 | 0.46 |
| 9 | 97062981 | rs12236219 | 0.476 | -1.238 | -0.285 | -1.506 | 0.003 | 0.115 | 0.115 | 0 | 0.05 | 0.09 | 0.26 | 0.35 | 0.05 | 0.17 |
| 9 | 97064273 | rs7042481 | -0.324 | -1.364 | 0.016 | -1.534 | 0.001 | 0.002 | 0.002 | 0 | 0.74 | 0.71 | 0.58 | 0.65 | 0.68 | 0.74 |
| 9 | 97066584 | rs9299419 | -0.194 | -1.373 | 0.159 | -1.538 | 0.004 | 0.013 | 0.013 | 0 | 0.73 | 0.6 | 0.57 | 0.65 | 0.68 | 0.75 |
| 9 | 97068939 | rs10993161 | 0.371 | -1.143 | 0.135 | -1.379 | 0.022 | 0.213 | 0.213 | 0 | 0.05 | 0.16 | 0.26 | 0.35 | 0.05 | 0.17 |
| 9 | 97107088 | rs9409727 | -0.679 | -1.249 | -0.306 | -1.592 | 0 | -0.044 | -0.044 | 0 | 0.79 | 0.74 | 0.78 | 1 | 0.7 | 0.87 |
| 9 | 97123522 | rs9696076 | 0.753 | -1.338 | -0.375 | -1.51 | -0.02 | -0.021 | -0.021 | 0 | 0.05 | 0.14 | 0.03 | 0.01 | 0.02 | 0.02 |
| 9 | 97161890 | rs10993188 | 0.313 | -1.316 | -0.301 | -1.376 | 0.002 | 0.168 | 0.168 | 0 | 0.06 | 0.15 | 0.34 | 0.34 | 0.05 | 0.16 |
| 9 | 97177826 | rs1752784 | 0.229 | -1.349 | -0.36 | -1.459 | -0.002 | -0.006 | -0.006 | 0 | 0.25 | 0.26 | 0.5 | 0.34 | 0.32 | 0.28 |
| 9 | 97208231 | rs16910109 | 0.332 | -1.328 | -0.316 | -1.416 | 0.001 | 0.165 | 0.165 | 0 | 0.06 | 0.15 | 0.34 | 0.34 | 0.05 | 0.16 |
| 9 | 97216163 | rs7037298 | 0.694 | -1.732 | -0.358 | -1.711 | -0.026 | -0.052 | -0.052 | 0 | 0.31 | 0.26 | 0.13 | 0.05 | 0.15 | 0.23 |
| 9 | 97221434 | rs9409765 | -0.53 | -1.613 | -0.41 | -1.663 | 0.001 | 0.004 | 0.004 | 0 | 0.56 | 0.55 | 0.64 | 0.39 | 0.47 | 0.51 |
| 9 | 97228918 | rs9409552 | -0.671 | -1.775 | -0.202 | -1.51 | 0.001 | 0.004 | 0.004 | 0 | 0.56 | 0.55 | 0.64 | 0.39 | 0.47 | 0.51 |
| 9 | 97242304 | rs7020879 | -0.758 | -1.704 | -0.691 | -1.478 | 0.001 | 0.01 | 0.01 | 0 | 0.57 | 0.69 | 0.65 | 0.41 | 0.47 | 0.52 |
| 9 | 97246817 | rs9409562 | 1.113 | -1.669 | -0.782 | -1.492 | -0.046 | 0.104 | 0.104 | 0 | 0.19 | 0.03 | 0.16 | 0 | 0.27 | 0.11 |
| 9 | 97252505 | rs9409567 | -0.456 | -1.731 | -0.68 | -1.526 | 0.002 | 0.002 | 0.002 | 0 | 0.32 | 0.31 | 0.55 | 0.36 | 0.36 | 0.31 |
| 9 | 97253578 | rs7031088 | -1.116 | -1.996 | -0.074 | -1.088 | 0.102 | 0.047 | 0.047 | 0.202 | 0.49 | 0.7 | 0.55 | 0.43 | 0.28 | 0.5 |
| 9 | 97257314 | rs10821352 | 0.003 | -1.662 | 0.127 | -0.697 | 0.002 | 0.174 | 0.174 | 0 | 0.06 | 0.16 | 0.35 | 0.35 | 0.05 | 0.16 |
| 9 | 97257732 | rs6479549 | -0.137 | -1.546 | 0.017 | -1.266 | -0.064 | -0.104 | -0.104 | 1.186 | 0.31 | 0.4 | 0.15 | 0.07 | 0.15 | 0.25 |
| 9 | 97278389 | rs12343547 | -0.854 | -1.319 | -0.018 | -0.785 | -0.005 | -0.004 | -0.004 | 0.264 | 0.26 | 0.32 | 0.2 | 0.18 | 0.17 | 0.22 |
| 9 | 97279870 | rs12236456 | 1.2 | -1.372 | -0.302 | -0.909 | -0.001 | 0.027 | 0.027 | 0 | 0.3 | 0.17 | 0.49 | 0.21 | 0.35 | 0.18 |
| 9 | 97280338 | rs12342274 | 0.717 | -1.114 | -0.359 | -0.823 | -0.007 | 0.037 | 0.037 | 0 | 0.1 | 0.03 | 0.04 | 0 | 0.1 | 0.04 |
| 9 | 97283226 | rs9299425 | -0.468 | -0.975 | 0.14 | -0.722 | -0.022 | -0.046 | -0.046 | 0 | 0.78 | 0.75 | 0.92 | 0.94 | 0.87 | 0.79 |
| 9 | 97286964 | rs2772020 | -0.468 | -0.975 | 0.14 | -0.722 | -0.022 | -0.046 | -0.046 | 0.132 | 0.78 | 0.75 | 0.92 | 0.94 | 0.87 | 0.79 |
| 9 | 97290466 | rs605323 | 1.269 | -0.586 | 0.1 | -0.483 | -0.07 | -0.082 | -0.082 | 0 | 0.21 | 0.07 | 0.21 | 0.55 | 0.35 | 0.39 |
| 9 | 97310865 | rs543731 | 1.322 | -0.574 | -0.078 | -0.476 | -0.142 | -0.16 | -0.16 | 0 | 0.21 | 0.01 | 0.21 | 0.55 | 0.35 | 0.38 |
| 9 | 97312584 | rs10761333 | -0.802 | -0.731 | 0.321 | -0.522 | 0.006 | 0.008 | 0.008 | 0 | 0.73 | 0.74 | 0.76 | 0.45 | 0.61 | 0.57 |
| 9 | 97321127 | rs1048510 | 0.362 | -1.081 | 0.15 | -0.73 | 0.023 | 0.101 | 0.101 | 0 | 0.5 | 0.28 | 0.57 | 0.26 | 0.49 | 0.29 |
| 9 | 97321172 | rs1543185 | 0.054 | -1.131 | 0.569 | -0.669 | -0.013 | 0.056 | 0.056 | 0.038 | 0.07 | 0.24 | 0.11 | 0.12 | 0.03 | 0.1 |
| 9 | 97324198 | rs965474 | -0.83 | -1.126 | 0.609 | -0.711 | 0.079 | 0.085 | 0.085 | 0 | 0.71 | 0.56 | 0.75 | 0.46 | 0.6 | 0.56 |
| 9 | 97325998 | rs6479554 | -0.185 | -1.124 | 0.537 | -0.737 | -0.015 | 0.004 | 0.004 | 0 | 0.64 | 0.32 | 0.64 | 0.33 | 0.57 | 0.45 |
| 9 | 97328313 | rs10761339 | 0.497 | -1.175 | 0.474 | -0.768 | 0.026 | 0.097 | 0.097 | 0 | 0.49 | 0.3 | 0.57 | 0.26 | 0.48 | 0.29 |
| 9 | 97330693 | rs609901 | -0.096 | -1.18 | 0.5 | -0.751 | -0.01 | 0.006 | 0.006 | 0 | 0.42 | 0.66 | 0.33 | 0.61 | 0.47 | 0.55 |
| 9 | 97333275 | rs3852401 | 0.184 | -1.173 | 0.488 | -0.771 | -0.034 | -0.048 | -0.048 | 2.172 | 0.2 | 0.65 | 0.12 | 0.07 | 0.1 | 0.17 |
| 9 | 97337835 | rs1754432 | -0.019 | -1.109 | 0.61 | -0.726 | -0.005 | 0.002 | 0.002 | 0 | 0.42 | 0.64 | 0.32 | 0.61 | 0.47 | 0.55 |
| 9 | 97343620 | rs7870433 | 0.811 | -1.329 | 0.288 | -0.826 | 0.016 | 0.037 | 0.037 | 0 | 0.4 | 0.24 | 0.54 | 0.22 | 0.41 | 0.25 |
| 9 | 97343939 | rs663385 | -0.031 | -1.193 | 0.35 | -0.717 | -0.002 | 0.003 | 0.003 | 0 | 0.9 | 0.87 | 0.89 | 0.87 | 0.93 | 0.84 |
| 9 | 97345467 | rs802918 | 1.285 | -1.163 | 0.103 | -0.744 | -0.163 | -0.173 | -0.173 | 0.649 | 0.23 | 0.01 | 0.21 | 0.55 | 0.37 | 0.38 |
| 9 | 97346437 | rs10739976 | 0.6 | -1.415 | 0.053 | -0.988 | 0.041 | 0.151 | 0.151 | 0 | 0.5 | 0.23 | 0.51 | 0.2 | 0.48 | 0.37 |
| 9 | 97349520 | rs2280302 | 0.162 | -1.232 | 0.409 | -0.761 | -0.007 | 0.001 | 0.001 | 0 | 0.08 | 0.26 | 0.05 | 0.07 | 0.04 | 0.07 |
| 9 | 97349591 | rs2280301 | 0.147 | -1.233 | 0.408 | -0.762 | -0.008 | 0 | 0 | 0 | 0.08 | 0.26 | 0.05 | 0.07 | 0.04 | 0.07 |
| 9 | 97354528 | rs548348 | 0.896 | -0.742 | 0.576 | -0.328 | 0.003 | 0.008 | 0.008 | 0 | 0.24 | 0.28 | 0.2 | 0.55 | 0.35 | 0.37 |
| 9 | 97360148 | rs601389 | -0.362 | -0.926 | 0.283 | -0.476 | 0.014 | 0.086 | 0.086 | 0 | 0.79 | 0.95 | 0.92 | 0.95 | 0.85 | 0.8 |
| 9 | 97365642 | rs9695 | 0.327 | -0.946 | 0.509 | -0.479 | 0 | 0 | 0 | 3.164 | 0.53 | 0.47 | 0.7 | 0.4 | 0.5 | 0.44 |
| 9 | 97366400 | rs1754436 | -0.338 | -0.929 | 0.163 | -0.488 | 0.016 | 0.085 | 0.085 | 0 | 0.79 | 0.95 | 0.92 | 0.95 | 0.85 | 0.8 |
| 9 | 97380043 | rs8192689 | 0.821 | -0.703 | 0.473 | -0.271 | -0.006 | -0.007 | -0.007 | 1.084 | 0.26 | 0.28 | 0.21 | 0.56 | 0.35 | 0.38 |
| 9 | 97380597 | rs7867069 | 0.722 | -0.754 | 0.451 | -0.419 | 0.001 | 0.007 | 0.007 | 0 | 0.1 | 0.08 | 0.17 | 0.2 | 0.07 | 0.13 |
| 9 | 97388953 | rs10993269 | -0.559 | -1.025 | 0.485 | -0.713 | 0.002 | -0.038 | -0.038 | 0 | 0.44 | 0.57 | 0.47 | 0.8 | 0.5 | 0.55 |
| 9 | 97390098 | rs4129220 | 1.084 | -1.135 | 0.489 | -0.68 | -0.002 | -0.027 | -0.027 | 0 | 0.38 | 0.35 | 0.48 | 0.15 | 0.37 | 0.27 |
| 9 | 97390288 | rs4129219 | -0.237 | -1.083 | 0.416 | -0.507 | -0.005 | 0.002 | 0.002 | 0 | 0.26 | 0.3 | 0.24 | 0.27 | 0.17 | 0.27 |
| 9 | 97398792 | rs16910850 | 0.536 | -0.922 | -0.01 | -0.384 | -0.005 | 0.039 | 0.039 | 0 | 0.11 | 0.12 | 0.2 | 0.23 | 0.08 | 0.14 |
| 9 | 97403254 | rs7031924 | 0.54 | -1.033 | 0.14 | -0.575 | 0.009 | 0.034 | 0.034 | 0 | 0.44 | 0.37 | 0.49 | 0.15 | 0.43 | 0.32 |
| 9 | 97407889 | rs10761350 | 0.23 | -0.973 | -0.05 | -0.489 | 0.003 | 0.005 | 0.005 | 0 | 0.48 | 0.43 | 0.67 | 0.38 | 0.45 | 0.42 |
| 9 | 97409437 | rs7035325 | -0.066 | -0.431 | -0.303 | -0.007 | -0.005 | 0.073 | 0.073 | 0 | 0.12 | 0.22 | 0.21 | 0.26 | 0.08 | 0.14 |
| 9 | 97409771 | rs10993280 | 0.334 | -0.485 | -0.718 | -0.343 | 0 | 0.12 | 0.12 | 0 | 0.54 | 0.72 | 0.54 | 0.2 | 0.47 | 0.41 |
| 9 | 97412037 | rs10993282 | 0.276 | -0.173 | -0.677 | 0.317 | -0.001 | 0.001 | 0.001 | 0 | 0.08 | 0.1 | 0.04 | 0.03 | 0.08 | 0.05 |
| 9 | 97415529 | rs913185 | 0.112 | 0.33 | -0.646 | 0.868 | -0.02 | -0.039 | -0.039 | 0 | 0.8 | 0.85 | 0.82 | 0.4 | 0.76 | 0.76 |

| | | | | | | | | | | | | | | | | |
|---|---|---|---|---|---|---|---|---|---|---|---|---|---|---|---|---|
| 9 | 97419146 | rs12685159 | 0.279 | 0.21 | -0.615 | 1.548 | -0.01 | -0.041 | -0.041 | 2.552 | 0.11 | 0.02 | 0.15 | 0.37 | 0.09 | 0.08 |
| 9 | 97429160 | rs6479562 | -0.958 | 0.145 | -1.008 | 0.508 | -0.026 | 0.087 | 0.087 | 0 | 0.59 | 0.84 | 0.66 | 0.74 | 0.54 | 0.62 |
| 9 | 97430831 | rs10081702 | 0.167 | -0.027 | -0.001 | 0.716 | -0.002 | 0.062 | 0.062 | 0 | 0.13 | 0.33 | 0.14 | 0.2 | 0.09 | 0.08 |
| 9 | 97443341 | rs4460442 | -1.372 | -0.181 | -0.225 | 0.569 | -0.062 | 0.144 | 0.144 | 0.005 | 0.66 | 0.95 | 0.76 | 0.86 | 0.63 | 0.75 |
| 9 | 97445112 | rs12335609 | 0.5 | -0.333 | -0.217 | 0.366 | 0.013 | 0.075 | 0.075 | 0.346 | 0.15 | 0.03 | 0.06 | 0.02 | 0.11 | 0.11 |
| 9 | 97447430 | rs10124217 | 0.341 | -0.299 | 0.045 | 0.41 | -0.009 | 0.035 | 0.035 | 0 | 0.14 | 0.27 | 0.13 | 0.2 | 0.09 | 0.07 |
| 9 | 97463435 | rs12554486 | 0.211 | -0.088 | 0.568 | 0.514 | 0.001 | -0.007 | -0.007 | 0 | 0.37 | 0.37 | 0.47 | 0.56 | 0.35 | 0.46 |
| 9 | 97464381 | rs12349751 | 0.102 | 0.075 | 0.767 | 0.593 | 0.012 | 0.076 | 0.076 | 0.373 | 0.15 | 0.03 | 0.06 | 0.02 | 0.11 | 0.11 |
| 9 | 97466882 | rs12344604 | -0.41 | 0.073 | 1.121 | 0.933 | 0.122 | 0.252 | 0.252 | 0.376 | 0.08 | 0.42 | 0.12 | 0.18 | 0.08 | 0.07 |
| 9 | 97476866 | rs6479565 | -0.666 | -0.424 | 0.962 | 0.926 | -0.015 | 0.052 | 0.052 | 0.106 | 0.15 | 0.58 | 0.22 | 0.21 | 0.12 | 0.23 |
| 9 | 97487704 | rs396538 | -0.577 | -0.363 | 1.438 | 0.955 | 0.002 | 0.002 | 0.002 | 0 | 0.14 | 0.47 | 0.21 | 0.14 | 0.11 | 0.23 |
| 9 | 97503985 | rs7851726 | -1.322 | -0.555 | 0.782 | 0.898 | -0.021 | 0.041 | 0.041 | 0 | 0.61 | 0.9 | 0.66 | 0.76 | 0.6 | 0.78 |
| 9 | 97505608 | rs7866848 | -0.989 | -0.644 | 0.818 | 1.012 | 0.018 | 0.021 | 0.021 | 0 | 0.57 | 0.8 | 0.64 | 0.66 | 0.59 | 0.71 |
| 9 | 97522915 | rs12341814 | 0.587 | -0.594 | 1.115 | 0.923 | -0.026 | -0.023 | -0.023 | 0 | 0.11 | 0.13 | 0.1 | 0.07 | 0.06 | 0.12 |
| 9 | 97549812 | rs10512235 | 1.407 | -0.754 | 0.667 | 0.716 | 0.027 | 0.071 | 0.071 | 0 | 0.15 | 0.01 | 0.06 | 0.03 | 0.1 | 0.26 |
| 9 | 97586542 | rs10732405 | -0.386 | -0.608 | 0.986 | 0.891 | -0.005 | -0.018 | -0.018 | 0.209 | 0.88 | 0.83 | 0.81 | 0.89 | 0.9 | 0.81 |
| 9 | 97613428 | rs7847277 | -0.061 | -0.531 | 0.837 | 0.977 | -0.026 | -0.03 | -0.03 | 0.323 | 0.42 | 0.3 | 0.43 | 0.52 | 0.49 | 0.49 |
| 9 | 97639463 | rs10993381 | 0.833 | -0.558 | 0.889 | 0.577 | 0.017 | 0.05 | 0.05 | 0 | 0.1 | 0.01 | 0.02 | 0.03 | 0.06 | 0.11 |
| 9 | 97648587 | rs4385527 | 1.175 | -0.611 | 1.063 | 0.622 | -0.022 | 0.111 | 0.111 | 0 | 0.39 | 0.1 | 0.35 | 0.17 | 0.4 | 0.23 |
| 9 | 97662496 | rs10993387 | -0.092 | -0.559 | 1.171 | 0.65 | -0.019 | 0.014 | 0.014 | 0 | 0.42 | 0.36 | 0.51 | 0.63 | 0.49 | 0.49 |
| 9 | 97675840 | rs10993391 | 0.652 | -0.619 | 0.48 | 0.587 | 0.017 | 0.05 | 0.05 | 0 | 0.11 | 0.01 | 0.02 | 0.03 | 0.07 | 0.15 |
| 9 | 97677556 | rs12235316 | 0.993 | -0.626 | 0.47 | 0.58 | 0.008 | 0.079 | 0.079 | 0 | 0.43 | 0.13 | 0.35 | 0.27 | 0.4 | 0.29 |
| 9 | 97691104 | rs12344211 | -0.164 | -0.462 | 0.623 | 0.729 | -0.039 | 0.005 | 0.005 | 0 | 0.59 | 0.75 | 0.49 | 0.41 | 0.51 | 0.55 |
| 9 | 97723868 | rs2404295 | -1.095 | -0.89 | 0.741 | 1.103 | -0.021 | 0.013 | 0.013 | 0.149 | 0.22 | 0.68 | 0.15 | 0.28 | 0.12 | 0.31 |
| 9 | 97729136 | rs10993413 | 1.17 | -1.119 | 0.456 | 0.838 | 0.02 | 0.071 | 0.071 | 0 | 0.15 | 0.01 | 0.06 | 0.03 | 0.11 | 0.26 |
| 9 | 97734605 | rs16911887 | -0.253 | -0.76 | 1.194 | 1.22 | -0.02 | 0.054 | 0.054 | 0.004 | 0.06 | 0.28 | 0.03 | 0.12 | 0.01 | 0.06 |
| 9 | 97735639 | rs10993422 | -0.253 | -0.76 | 1.194 | 1.22 | -0.02 | 0.054 | 0.054 | 0.004 | 0.06 | 0.28 | 0.03 | 0.12 | 0.01 | 0.06 |
| 9 | 97738174 | rs10993428 | -0.253 | -0.76 | 1.194 | 1.22 | -0.02 | 0.054 | 0.054 | 0.004 | 0.06 | 0.28 | 0.03 | 0.12 | 0.01 | 0.06 |
| 9 | 97739465 | rs10993429 | -0.253 | -0.76 | 1.194 | 1.22 | -0.02 | 0.054 | 0.054 | 0.004 | 0.06 | 0.28 | 0.03 | 0.12 | 0.01 | 0.06 |
| 9 | 97749379 | rs7866661 | -1.137 | -0.775 | 1.661 | 1.229 | 0.015 | 0.172 | 0.172 | 0.727 | 0.08 | 0.48 | 0.04 | 0.15 | 0.02 | 0.09 |
| 9 | 97754447 | rs7042271 | -1.559 | -0.756 | 1.802 | 1.261 | -0.004 | 0.143 | 0.143 | 1.245 | 0.09 | 0.63 | 0.12 | 0.2 | 0.04 | 0.15 |
| 9 | 97759920 | rs10993446 | -0.02 | -0.756 | 1.773 | 1.251 | -0.018 | 0.05 | 0.05 | 0 | 0.05 | 0.23 | 0.02 | 0.12 | 0.01 | 0.06 |
| 9 | 97768286 | rs10761364 | 1.295 | -0.73 | 1.488 | 1.231 | -0.027 | 0.025 | 0.025 | 0.168 | 0.81 | 0.41 | 0.85 | 0.73 | 0.88 | 0.72 |
| 9 | 97771852 | rs3802456 | 1.277 | -0.725 | 1.527 | 1.233 | -0.029 | 0.032 | 0.032 | 0.099 | 0.81 | 0.43 | 0.85 | 0.73 | 0.88 | 0.72 |
| 9 | 97783630 | rs10993456 | -0.619 | -0.725 | 1.532 | 1.236 | -0.006 | -0.001 | -0.001 | 0.022 | 0.15 | 0.17 | 0.04 | 0.18 | 0.08 | 0.19 |
| 9 | 97785248 | rs7023719 | 0.262 | -0.741 | 1.53 | 1.234 | 0.003 | 0.078 | 0.078 | 0 | 0.94 | 0.82 | 0.91 | 0.84 | 0.96 | 0.88 |
| 9 | 97789745 | rs1020963 | 0.715 | -0.653 | 1.493 | 1.707 | -0.016 | 0.019 | 0.019 | 0 | 0.84 | 0.73 | 0.87 | 0.78 | 0.89 | 0.75 |
| 9 | 97831471 | rs4744435 | 1.354 | -0.715 | 1.297 | 1.734 | -0.027 | 0.072 | 0.072 | 0.015 | 0.81 | 0.49 | 0.85 | 0.69 | 0.88 | 0.7 |
| 9 | 97831581 | rs4744436 | 1.166 | -0.691 | 1.246 | 1.887 | -0.02 | 0.167 | 0.167 | 0.008 | 0.82 | 0.53 | 0.85 | 0.64 | 0.86 | 0.71 |
| 9 | 97834573 | rs10761374 | 0.057 | -1.061 | 0.604 | 0.96 | -0.036 | 0.012 | 0.012 | 0 | 0.66 | 0.54 | 0.7 | 0.6 | 0.78 | 0.63 |
| 9 | 97844513 | rs10491560 | 0.013 | -1.128 | 0.633 | 0.917 | 0.063 | 0.017 | 0.017 | 0 | 0.32 | 0.18 | 0.2 | 0.29 | 0.17 | 0.26 |
| 9 | 97845505 | rs10821447 | 0.039 | -1.138 | 0.682 | 0.919 | -0.035 | 0.008 | 0.008 | 0 | 0.64 | 0.53 | 0.69 | 0.6 | 0.77 | 0.61 |
| 9 | 97849090 | rs4657 | -0.055 | -1.111 | 0.689 | 0.928 | -0.009 | 0.001 | 0.001 | 0 | 0.65 | 0.6 | 0.7 | 0.61 | 0.78 | 0.61 |
| 9 | 97898220 | rs3780567 | 0.111 | -0.157 | 0.469 | 0.434 | -0.007 | 0.02 | 0.02 | 0.047 | 0.45 | 0.63 | 0.51 | 0.63 | 0.45 | 0.48 |
| 9 | 97913026 | rs640628 | -0.962 | -0.341 | 0.365 | 0.32 | -0.046 | 0.046 | 0.046 | 0 | 0.75 | 0.89 | 0.71 | 0.92 | 0.62 | 0.75 |
| 9 | 97923592 | rs565293 | 0.518 | -0.401 | 0.431 | 0.32 | 0.028 | 0.004 | 0.004 | 0.421 | 0.29 | 0.25 | 0.19 | 0.3 | 0.16 | 0.26 |
| 9 | 97928719 | rs657637 | 0.759 | -0.374 | 0.414 | 0.333 | 0.069 | 0.005 | 0.005 | 0.497 | 0.28 | 0.11 | 0.19 | 0.29 | 0.16 | 0.26 |
| 9 | 97940807 | rs2297597 | 0.393 | -0.332 | 0.536 | 0.268 | -0.025 | 0.001 | 0.001 | 0 | 0.42 | 0.23 | 0.48 | 0.59 | 0.45 | 0.44 |
| 9 | 97989207 | rs12380720 | 1.541 | 0.184 | 0.26 | 0.469 | -0.02 | -0.013 | -0.013 | 0.007 | 0.11 | 0.05 | 0.31 | 0.38 | 0.19 | 0.14 |
| 9 | 97992323 | rs16912035 | 1.667 | 0.171 | 0.04 | 0.311 | 0.037 | 0.126 | 0.126 | 0 | 0.21 | 0.01 | 0.07 | 0.02 | 0.13 | 0.25 |
| 9 | 97998601 | rs1326188 | 0.965 | 0.347 | 0.472 | 0.613 | -0.027 | -0.027 | -0.027 | 0 | 0.08 | 0.04 | 0.1 | 0.18 | 0.13 | 0.06 |
| 9 | 98004495 | rs4647432 | 1.029 | -0.183 | 0.726 | 0.253 | -0.101 | 0.141 | 0.141 | 0.031 | 0.24 | 0.01 | 0.21 | 0.02 | 0.33 | 0.17 |
| 9 | 98006368 | rs356659 | -1.116 | -0.157 | 0.773 | 0.273 | -0.048 | 0.05 | 0.05 | 0 | 0.75 | 0.88 | 0.71 | 0.93 | 0.62 | 0.74 |
| 9 | 98058759 | rs4647382 | 0.629 | -0.074 | 0.674 | 0.368 | -0.032 | 0 | 0 | 0 | 0.42 | 0.21 | 0.48 | 0.59 | 0.45 | 0.44 |
| 9 | 98060631 | rs356665 | -0.644 | -0.192 | 0.444 | 0.366 | 0.009 | 0.024 | 0.024 | 0 | 0.54 | 0.71 | 0.41 | 0.4 | 0.48 | 0.47 |
| 9 | 98094129 | rs10993512 | -0.704 | -0.155 | -0.25 | 0.277 | 0.033 | 0.045 | 0.045 | 0.912 | 0.74 | 0.87 | 0.84 | 0.82 | 0.83 | 0.78 |
| 9 | 98102983 | rs4743993 | 0.806 | 0.011 | -0.339 | 0.407 | 0.006 | 0.006 | 0.006 | 0 | 0.4 | 0.43 | 0.56 | 0.56 | 0.44 | 0.43 |
| 9 | 98110248 | rs6479604 | 0.089 | -0.05 | -1.11 | 0.443 | 0.044 | 0.008 | 0.008 | 0.316 | 0.69 | 0.86 | 0.82 | 0.68 | 0.8 | 0.74 |
| 9 | 98122445 | rs10761384 | -0.7 | 0.116 | -1.317 | 0.356 | 0.005 | 0.012 | 0.012 | 0.289 | 0.79 | 0.99 | 0.87 | 0.83 | 0.87 | 0.81 |
| 9 | 98131102 | rs2036516 | 0.048 | 0.674 | -0.454 | 0.612 | 0.004 | 0.005 | 0.005 | 0 | 0.41 | 0.41 | 0.59 | 0.5 | 0.46 | 0.5 |
| 9 | 98138938 | rs357547 | -0.097 | 1.228 | -0.184 | 0.234 | 0.004 | -0.007 | -0.007 | 1.077 | 0.73 | 0.74 | 0.74 | 0.86 | 0.69 | 0.72 |
| 9 | 98143642 | rs357542 | -0.112 | 1.263 | -0.214 | -0.046 | 0.015 | -0.017 | -0.017 | 0 | 0.33 | 0.42 | 0.28 | 0.23 | 0.41 | 0.2 |
| 9 | 98146478 | rs357539 | -0.776 | 1.202 | -0.241 | -0.111 | 0 | -0.002 | -0.002 | 0 | 0.75 | 0.81 | 0.75 | 0.87 | 0.71 | 0.72 |
| 9 | 98147804 | rs7860361 | 0.458 | 1.691 | -0.836 | -0.07 | -0.079 | 0.122 | 0.122 | 0.608 | 0.24 | 0.07 | 0.18 | 0.05 | 0.32 | 0.14 |
| 9 | 98171417 | rs7388704 | 1.11 | 1.458 | -0.689 | 0.108 | 0.079 | -0.057 | -0.057 | 0 | 0.39 | 0.15 | 0.45 | 0.55 | 0.26 | 0.51 |
| 9 | 98175928 | rs2026077 | 0.502 | 1.689 | -0.57 | 0.02 | -0.02 | 0.049 | 0.049 | 0.051 | 0.25 | 0.18 | 0.15 | 0.14 | 0.3 | 0.17 |
| 9 | 98177173 | rs357561 | 0.974 | 1.852 | -0.611 | 0.129 | -0.032 | -0.03 | -0.03 | 1.306 | 0.05 | 0 | 0.07 | 0.12 | 0.11 | 0.04 |
| 9 | 98177956 | rs16912133 | 0.003 | 1.445 | -0.396 | 0.082 | 0.024 | -0.005 | -0.005 | 0 | 0.66 | 0.48 | 0.61 | 0.75 | 0.57 | 0.69 |
| 9 | 98189458 | rs357522 | -1.072 | 1.003 | -1.059 | 0.191 | 0.087 | -0.044 | -0.044 | 0 | 0.58 | 0.85 | 0.52 | 0.44 | 0.7 | 0.44 |

| | | | | | | | | | | | | | | | | |
|---|---|---|---|---|---|---|---|---|---|---|---|---|---|---|---|---|
| 9 | 98204199 | rs357563 | -0.571 | 1.535 | -0.844 | 0.332 | -0.055 | -0.054 | -0.054 | 0 | 0.32 | 0.9 | 0.36 | 0.31 | 0.36 | 0.32 |
| 9 | 98204792 | rs16909859 | -0.825 | 1.404 | 0.47 | 0.642 | 0.096 | 0.165 | 0.165 | 0 | 0.07 | 0.31 | 0.07 | 0.16 | 0.08 | 0.07 |
| 9 | 98209594 | rs357564 | 0.506 | 1.153 | -0.187 | 0.343 | -0.002 | -0.026 | -0.026 | 0 | 0.33 | 0.23 | 0.41 | 0.59 | 0.35 | 0.47 |
| 9 | 98231008 | rs16909898 | -0.651 | 1.382 | 0.772 | 0.596 | 0.036 | 0.009 | 0.009 | 0 | 0.06 | 0.13 | 0.07 | 0.08 | 0.1 | 0.04 |
| 9 | 98238358 | rs2066836 | -0.663 | 1.464 | 0.349 | 0.348 | -0.038 | 0.061 | 0.061 | 0.064 | 0.19 | 0.01 | 0.1 | 0.08 | 0.19 | 0.1 |
| 9 | 98240120 | rs2297086 | -2.021 | 1.309 | 0.822 | 0.426 | -0.002 | 0.002 | 0.002 | 0.643 | 0.35 | 0.72 | 0.23 | 0.34 | 0.35 | 0.21 |
| 9 | 98242925 | rs2297087 | -0.945 | 1.411 | 1.084 | 0.345 | 0 | -0.061 | -0.061 | 0 | 0.23 | 0.26 | 0.12 | 0.08 | 0.22 | 0.11 |
| 9 | 98266370 | rs4448343 | -1.865 | 1.662 | 0.905 | 0.61 | 0.029 | 0.015 | 0.015 | 0 | 0.28 | 0.52 | 0.21 | 0.32 | 0.32 | 0.17 |
| 9 | 98273791 | rs28412122 | -1.039 | 1.683 | 0.381 | 0.658 | 0 | -0.004 | -0.004 | 0.065 | 0.2 | 0.17 | 0.12 | 0.21 | 0.21 | 0.14 |
| 9 | 98283093 | rs28536742 | -2.132 | 1.678 | -0.138 | 0.364 | 0.015 | -0.012 | -0.012 | 0.299 | 0.3 | 0.44 | 0.23 | 0.27 | 0.34 | 0.23 |
| 9 | 98293208 | rs16909955 | -1.735 | 1.949 | -0.421 | 0.425 | 0.006 | 0.08 | 0.08 | 1.707 | 0.05 | 0.3 | 0.04 | 0.07 | 0.01 | 0.05 |
| 9 | 98302535 | rs2897050 | -0.949 | 2.166 | -1.489 | 0.439 | -0.051 | 0.056 | 0.056 | 0.063 | 0.17 | 0.03 | 0.11 | 0.07 | 0.19 | 0.14 |
| 9 | 98307129 | rs16909975 | -1.359 | 1.589 | -1.071 | 0.756 | -0.009 | -0.009 | -0.009 | 0.339 | 0.09 | 0.11 | 0.1 | 0.15 | 0.11 | 0.07 |
| 9 | 98318926 | rs1329393 | -0.993 | 1.543 | -1.168 | 0.841 | -0.019 | -0.002 | -0.002 | 0.127 | 0.14 | 0.07 | 0.11 | 0.13 | 0.17 | 0.14 |
| 9 | 98319211 | rs576594 | -1.2 | 1.257 | -2.039 | 0.834 | -0.021 | 0.093 | 0.093 | 0.035 | 0.54 | 0.88 | 0.65 | 0.7 | 0.61 | 0.69 |
| 9 | 98326767 | rs7875436 | -0.608 | 0.833 | -1.061 | 1.449 | 0.018 | -0.013 | -0.013 | 0.525 | 0.4 | 0.67 | 0.47 | 0.43 | 0.34 | 0.54 |
| 9 | 98348885 | rs7040739 | -1.385 | 1.249 | -0.245 | 2.059 | 0.017 | 0.116 | 0.116 | 1.209 | 0.17 | 0.39 | 0.27 | 0.48 | 0.17 | 0.3 |
| 9 | 98349921 | rs1806706 | -0.827 | 1.22 | -0.154 | 2.222 | -0.001 | 0.062 | 0.062 | 0.343 | 0.12 | 0.25 | 0.09 | 0.34 | 0.09 | 0.1 |
| 9 | 98355001 | rs4590564 | -0.944 | 1.18 | -0.057 | 2.22 | -0.002 | 0.097 | 0.097 | 0.649 | 0.12 | 0.35 | 0.1 | 0.33 | 0.09 | 0.1 |
| 9 | 98355145 | rs10981328 | -0.939 | 1.152 | 0.846 | 2.151 | 0.077 | 0.149 | 0.149 | 0 | 0.05 | 0.17 | 0.19 | 0.14 | 0.08 | 0.21 |
| 9 | 98358166 | rs882130 | -1.506 | 0.845 | 0.488 | 2.115 | -0.015 | -0.002 | -0.002 | 0 | 0.29 | 0.58 | 0.23 | 0.28 | 0.24 | 0.22 |
| 9 | 98358683 | rs10115098 | -1.523 | 0.815 | 0.86 | 2.182 | -0.003 | 0.041 | 0.041 | 0.048 | 0.25 | 0.6 | 0.3 | 0.5 | 0.24 | 0.32 |
| 9 | 98359188 | rs1889616 | -2.04 | 0.822 | 0.823 | 2.167 | -0.009 | -0.028 | -0.028 | 2.51 | 0.36 | 0.86 | 0.38 | 0.63 | 0.32 | 0.43 |
| 9 | 98360213 | rs6478058 | -1.654 | 0.826 | 1.431 | 2.229 | 0.118 | 0.44 | 0.44 | 3.611 | 0.07 | 0.38 | 0.15 | 0.34 | 0.08 | 0.21 |
| 9 | 98362492 | rs928618 | -0.741 | 0.761 | 1.192 | 1.876 | -0.012 | -0.013 | -0.013 | 0 | 0.17 | 0.24 | 0.16 | 0.15 | 0.16 | 0.12 |
| 9 | 98362742 | rs928619 | -0.493 | 0.772 | 1.205 | 1.896 | -0.003 | 0.002 | 0.002 | 0 | 0.14 | 0.23 | 0.13 | 0.16 | 0.12 | 0.11 |
| 9 | 98364547 | rs10120219 | -0.525 | 0.403 | 0.679 | 1.859 | -0.019 | 0.068 | 0.068 | 0 | 0.27 | 0.44 | 0.27 | 0.53 | 0.23 | 0.4 |
| 9 | 98367717 | rs10818991 | -1.617 | 0.675 | 0.077 | 1.866 | 0.039 | 0.175 | 0.175 | 0 | 0.46 | 0.83 | 0.66 | 0.79 | 0.42 | 0.53 |
| 9 | 98368761 | rs1984119 | -0.503 | 1.581 | 0.597 | 2.009 | 0.019 | 0.024 | 0.024 | 0 | 0.15 | 0.2 | 0.38 | 0.26 | 0.27 | 0.13 |
| 9 | 98371526 | rs6478154 | -0.536 | 0.777 | 0.489 | 1.596 | -0.007 | 0.023 | 0.023 | 0 | 0.33 | 0.39 | 0.45 | 0.44 | 0.33 | 0.39 |
| 9 | 98376367 | rs4743090 | 0.53 | 0.738 | 0.333 | 1.656 | 0.012 | -0.007 | -0.007 | 0 | 0.15 | 0.09 | 0.08 | 0.15 | 0.13 | 0.19 |
| 9 | 98380588 | rs7036222 | 0.283 | 0.679 | 0.391 | 1.612 | -0.004 | -0.027 | -0.027 | 0 | 0.18 | 0.1 | 0.36 | 0.28 | 0.2 | 0.19 |
| 9 | 98381360 | rs7864457 | -0.594 | 0.651 | 0.678 | 1.814 | -0.029 | 0.025 | 0.025 | 0.01 | 0.23 | 0.4 | 0.14 | 0.27 | 0.19 | 0.22 |
| 9 | 98382950 | rs7854560 | -0.886 | 0.861 | 0.633 | 1.953 | 0 | 0.066 | 0.066 | 0 | 0.26 | 0.41 | 0.35 | 0.52 | 0.29 | 0.39 |
| 9 | 98383097 | rs7860540 | -2.068 | 0.748 | 0.265 | 1.624 | -0.083 | 0.165 | 0.165 | 0.087 | 0.58 | 0.92 | 0.82 | 0.8 | 0.59 | 0.61 |
| 9 | 98387529 | rs4742697 | -0.782 | 0.747 | 0.476 | 1.762 | -0.007 | 0.011 | 0.011 | 0 | 0.24 | 0.3 | 0.14 | 0.35 | 0.21 | 0.26 |
| 9 | 98391111 | rs930280 | -1.372 | 0.76 | 0.691 | 1.774 | -0.001 | -0.003 | -0.003 | 0.136 | 0.49 | 0.56 | 0.54 | 0.66 | 0.46 | 0.47 |
| 9 | 98392340 | rs6478302 | -0.635 | 0.866 | 0.656 | 1.928 | 0.002 | -0.001 | -0.001 | 0 | 0.33 | 0.37 | 0.62 | 0.63 | 0.37 | 0.44 |
| 9 | 98393411 | rs16910070 | 0.624 | 0.859 | 0.608 | 1.883 | -0.001 | 0 | 0 | 0 | 0.14 | 0.11 | 0.08 | 0.08 | 0.14 | 0.16 |
| 9 | 98393954 | rs16910073 | -0.51 | 0.87 | 0.212 | 1.867 | -0.001 | 0.002 | 0.002 | 0.158 | 0.31 | 0.33 | 0.43 | 0.38 | 0.3 | 0.28 |
| 9 | 98398956 | rs10819953 | -0.611 | 0.669 | 0.202 | 1.786 | -0.02 | 0.047 | 0.047 | 0 | 0.23 | 0.34 | 0.12 | 0.36 | 0.18 | 0.29 |
| 9 | 98402621 | rs10989747 | -0.202 | 1.016 | -0.493 | 1.691 | -0.01 | -0.027 | -0.027 | 0 | 0.27 | 0.14 | 0.42 | 0.31 | 0.3 | 0.22 |
| 9 | 98406181 | rs7027192 | -0.25 | 0.958 | -0.493 | 1.687 | -0.001 | -0.013 | -0.013 | 7.004 | 0.27 | 0.2 | 0.43 | 0.3 | 0.3 | 0.22 |
| 9 | 98407526 | rs9695781 | -0.208 | 0.88 | -0.257 | 1.65 | -0.004 | 0.002 | 0.002 | 0.5 | 0.15 | 0.17 | 0.11 | 0.18 | 0.14 | 0.16 |
| 9 | 98410405 | rs10990303 | -0.617 | 1.037 | -0.274 | 1.787 | 0.002 | 0.024 | 0.024 | 0.003 | 0.18 | 0.19 | 0.3 | 0.44 | 0.22 | 0.37 |
| 9 | 98414711 | rs10990614 | -0.032 | 0.091 | -0.813 | 1.027 | 0.033 | -0.004 | -0.004 | 0 | 0.18 | 0.1 | 0.07 | 0.17 | 0.12 | 0.14 |
| 9 | 98420881 | rs1582073 | -1.118 | 0.079 | -1.024 | 0.009 | -0.003 | -0.01 | -0.01 | 0 | 0.64 | 0.69 | 0.62 | 0.85 | 0.57 | 0.69 |
| 9 | 98426006 | rs7047000 | -1.072 | -0.062 | -1.537 | 0.288 | -0.011 | 0.016 | 0.016 | 0 | 0.57 | 0.82 | 0.39 | 0.62 | 0.5 | 0.47 |
| 9 | 98428619 | rs6478489 | -0.256 | -0.254 | -0.653 | 0.273 | -0.006 | 0.014 | 0.014 | 0 | 0.37 | 0.51 | 0.31 | 0.45 | 0.37 | 0.33 |
| 9 | 98430574 | rs4743148 | -0.753 | -0.424 | -0.849 | 0.132 | -0.003 | 0.008 | 0.008 | 0 | 0.39 | 0.36 | 0.49 | 0.68 | 0.4 | 0.54 |
| 9 | 98430835 | rs10991380 | 0.856 | -0.612 | -0.695 | 0.074 | 0.002 | -0.039 | -0.039 | 0 | 0.19 | 0.09 | 0.18 | 0.41 | 0.18 | 0.24 |
| 9 | 98431637 | rs12686467 | 1.307 | -0.549 | -0.852 | 0.134 | -0.024 | -0.047 | -0.047 | 0 | 0.09 | 0.01 | 0.23 | 0.27 | 0.09 | 0.25 |
| 9 | 98432200 | rs10820751 | 1.465 | -0.415 | -0.871 | -0.024 | 0.03 | -0.003 | -0.003 | 4.852 | 0.18 | 0.09 | 0.07 | 0.17 | 0.14 | 0.14 |
| 9 | 98435273 | rs700998 | 2.439 | 0.142 | -1.175 | -0.46 | -0.024 | 0.046 | 0.046 | 0.084 | 0.25 | 0.15 | 0.19 | 0.1 | 0.32 | 0.19 |
| 9 | 98435489 | rs17372214 | 1.733 | 0.157 | -1.29 | -0.228 | -0.03 | -0.019 | -0.019 | 0 | 0.06 | 0.01 | 0.04 | 0.09 | 0.09 | 0.14 |
| 9 | 98439240 | rs354276 | -0.163 | 0.071 | -1.1 | -0.166 | -0.003 | -0.005 | -0.005 | 0 | 0.81 | 0.8 | 0.58 | 0.72 | 0.81 | 0.66 |
| 9 | 98444303 | rs915228 | 1.566 | -0.099 | -1.242 | -0.191 | 0.016 | -0.081 | -0.081 | 0 | 0.13 | 0.02 | 0.14 | 0.32 | 0.09 | 0.1 |
| 9 | 98445060 | rs458477 | 1.043 | -0.048 | -0.98 | -0.345 | -0.002 | 0.009 | 0.009 | 0.046 | 0.41 | 0.42 | 0.29 | 0.2 | 0.53 | 0.38 |
| 9 | 98445610 | rs817182 | -1.087 | -0.011 | -0.959 | -0.271 | -0.019 | -0.025 | -0.025 | 0 | 0.88 | 0.96 | 0.81 | 0.85 | 0.89 | 0.89 |
| 9 | 98447172 | rs817184 | 2.587 | 0.069 | -0.934 | -0.389 | -0.067 | 0.096 | 0.096 | 0 | 0.25 | 0.05 | 0.18 | 0.12 | 0.32 | 0.19 |
| 9 | 98447883 | rs1335049 | -0.638 | 0.062 | -0.733 | -0.275 | 0.003 | 0.005 | 0.005 | 0 | 0.44 | 0.42 | 0.32 | 0.42 | 0.44 | 0.41 |
| 9 | 98451928 | rs10816456 | 0.852 | 0.231 | -1.16 | -0.349 | 0 | 0.042 | 0.042 | 0 | 0.25 | 0.19 | 0.13 | 0.02 | 0.26 | 0.17 |
| 9 | 98455545 | rs10978784 | 1.633 | -0.085 | -1.199 | -0.236 | 0.013 | 0.073 | 0.073 | 0.157 | 0.17 | 0.02 | 0.06 | 0.02 | 0.13 | 0.13 |
| 9 | 98466346 | rs2782951 | -0.207 | 0.73 | -0.977 | -0.034 | -0.022 | 0.003 | 0.003 | 0 | 0.43 | 0.34 | 0.34 | 0.42 | 0.49 | 0.4 |
| 9 | 98468885 | rs16910150 | 1.319 | 0.277 | -1.279 | 0.081 | -0.024 | -0.058 | -0.058 | 0.301 | 0.09 | 0.01 | 0.21 | 0.22 | 0.09 | 0.25 |
| 9 | 98473646 | rs700992 | 1.86 | 0.379 | -1.133 | -0.71 | -0.013 | 0.057 | 0.057 | 0 | 0.29 | 0.17 | 0.2 | 0.14 | 0.32 | 0.24 |
| 9 | 98475674 | rs1977620 | 0.886 | 0.298 | -1.193 | -0.761 | -0.007 | 0.055 | 0.055 | 0 | 0.28 | 0.19 | 0.17 | 0.04 | 0.32 | 0.18 |
| 9 | 98476064 | rs700990 | 0.931 | 0.515 | -0.891 | -0.443 | -0.001 | 0.037 | 0.037 | 0.2 | 0.36 | 0.29 | 0.24 | 0.15 | 0.36 | 0.26 |
| 9 | 98478118 | rs460175 | -0.24 | 0.671 | -1.324 | -0.148 | -0.036 | -0.033 | -0.033 | 0 | 0.86 | 0.92 | 0.77 | 0.76 | 0.81 | 0.77 |

| | | | | | | | | | | | | | | | | |
|---|---|---|---|---|---|---|---|---|---|---|---|---|---|---|---|---|
| 9 | 98479529 | rs1967908 | -0.048 | 0.219 | -0.898 | -0.15 | 0.008 | 0.028 | 0.028 | 0 | 0.46 | 0.36 | 0.51 | 0.63 | 0.44 | 0.52 |
| 9 | 98483660 | rs354271 | 1.271 | 0.452 | -1.108 | -0.164 | -0.022 | -0.022 | -0.022 | 0 | 0.47 | 0.13 | 0.48 | 0.53 | 0.49 | 0.58 |
| 9 | 98488882 | rs817166 | 0.498 | 0.137 | -0.394 | -0.195 | -0.009 | -0.025 | -0.025 | 0 | 0.83 | 0.8 | 0.92 | 0.96 | 0.9 | 0.94 |
| 9 | 98493505 | rs17373603 | -0.293 | 0.825 | -0.613 | 0.338 | -0.009 | -0.002 | -0.002 | 0.124 | 0.07 | 0.04 | 0.11 | 0.12 | 0.1 | 0.13 |
| 9 | 98496651 | rs700979 | 0.844 | 0.682 | -0.454 | 0.237 | 0.002 | 0.008 | 0.008 | 0 | 0.55 | 0.31 | 0.65 | 0.62 | 0.59 | 0.57 |
| 9 | 98496807 | rs700978 | 0.718 | 0.27 | 0.187 | 0.004 | -0.006 | -0.035 | -0.035 | 0 | 0.64 | 0.45 | 0.77 | 0.78 | 0.69 | 0.74 |
| 9 | 98497395 | rs700976 | 0.459 | 0.223 | 0.215 | -0.131 | 0.007 | 0.017 | 0.017 | 0 | 0.76 | 0.78 | 0.84 | 0.89 | 0.82 | 0.88 |
| 9 | 98497814 | rs700975 | 1.43 | 0.242 | 0.415 | 0.07 | 0.013 | 0.013 | 0.013 | 0 | 0.5 | 0.2 | 0.57 | 0.58 | 0.57 | 0.54 |
| 9 | 98501892 | rs16910200 | 0.068 | 0.419 | 0.554 | -0.012 | 0.016 | -0.016 | -0.016 | 0 | 0.08 | 0.17 | 0.13 | 0.07 | 0.08 | 0.04 |
| 9 | 98502921 | rs7035549 | 0.061 | 0.365 | 0.848 | -0.164 | 0 | -0.025 | -0.025 | 0 | 0.1 | 0.15 | 0.06 | 0 | 0.1 | 0.03 |
| 9 | 98504905 | rs414610 | 1.403 | 0.626 | 1.333 | -0.167 | -0.001 | -0.01 | -0.01 | 1.011 | 0.54 | 0.47 | 0.53 | 0.72 | 0.57 | 0.54 |
| 9 | 98507588 | rs700966 | 1.539 | 0.618 | 1.122 | -0.141 | 0.006 | 0.015 | 0.015 | 0.05 | 0.53 | 0.32 | 0.52 | 0.65 | 0.55 | 0.54 |
| 9 | 98508628 | rs4743271 | 0.044 | 0.457 | 0.939 | 0.055 | -0.002 | 0.01 | 0.01 | 0.612 | 0.12 | 0.15 | 0.16 | 0.19 | 0.12 | 0.21 |
| 9 | 98515578 | rs4742743 | -1.27 | 0.42 | 0.701 | -0.062 | 0.001 | 0.008 | 0.008 | 0 | 0.34 | 0.65 | 0.38 | 0.16 | 0.31 | 0.27 |
| 9 | 98516196 | rs933013 | -1.747 | 0.425 | 0.528 | -0.02 | -0.002 | -0.019 | -0.019 | 0 | 0.43 | 0.71 | 0.54 | 0.44 | 0.43 | 0.46 |
| 9 | 98516889 | rs10981787 | -0.183 | 0.468 | -0.032 | -0.121 | -0.023 | 0.094 | 0.094 | 0 | 0.22 | 0.06 | 0.2 | 0.05 | 0.22 | 0.21 |
| 9 | 98525304 | rs4743293 | -1.236 | 0.6 | 0.475 | 0.039 | -0.001 | -0.001 | -0.001 | 0 | 0.37 | 0.3 | 0.5 | 0.36 | 0.4 | 0.44 |
| 9 | 98533383 | rs7042092 | -0.391 | 0.606 | 0.006 | -0.28 | 0.016 | 0.041 | 0.041 | 0 | 0.24 | 0.16 | 0.2 | 0.12 | 0.17 | 0.2 |
| 9 | 98534372 | rs4743306 | -2.198 | 0.504 | -0.243 | -0.053 | -0.093 | -0.062 | -0.062 | 0.266 | 0.54 | 0.92 | 0.59 | 0.43 | 0.48 | 0.48 |
| 9 | 98534462 | rs10817721 | -1.832 | 0.649 | -1.009 | -0.275 | -0.062 | -0.123 | -0.123 | 0.254 | 0.82 | 0.96 | 0.75 | 0.51 | 0.73 | 0.73 |
| 9 | 98535990 | rs7035466 | -1.225 | 0.194 | -0.053 | -0.012 | 0.02 | 0.019 | 0.019 | 0 | 0.43 | 0.71 | 0.45 | 0.31 | 0.34 | 0.39 |
| 9 | 98536017 | rs7031550 | -0.861 | -0.324 | 0.364 | -0.25 | -0.033 | -0.036 | -0.036 | 0.242 | 0.34 | 0.5 | 0.28 | 0.2 | 0.21 | 0.26 |
| 9 | 98538954 | rs7854146 | -0.454 | -0.231 | -0.061 | -0.292 | -0.023 | -0.022 | -0.022 | 0.198 | 0.22 | 0.66 | 0.22 | 0.15 | 0.13 | 0.16 |
| 9 | 98541232 | rs7047415 | -0.008 | 0.087 | -0.573 | -0.194 | -0.034 | -0.057 | -0.057 | 0.028 | 0.17 | 0.5 | 0.11 | 0.04 | 0.09 | 0.08 |
| 9 | 98542044 | rs1836406 | -0.733 | 0.358 | -1.667 | 0.026 | -0.008 | 0.101 | 0.101 | 0.083 | 0.43 | 0.92 | 0.42 | 0.26 | 0.34 | 0.33 |
| 9 | 98546523 | rs10759835 | -0.014 | 0.647 | -0.888 | 0.67 | -0.001 | 0.028 | 0.028 | 0 | 0.27 | 0.35 | 0.33 | 0.52 | 0.29 | 0.22 |
| 9 | 98550264 | rs1836404 | -0.033 | 0.439 | -1.586 | 0.423 | -0.032 | 0.003 | 0.003 | 0.511 | 0.46 | 0.16 | 0.38 | 0.4 | 0.51 | 0.4 |
| 9 | 98550467 | rs1434729 | 0.899 | 0.731 | -1.622 | 0.603 | -0.002 | 0.007 | 0.007 | 0 | 0.05 | 0.03 | 0.14 | 0.16 | 0.1 | 0.08 |
| 9 | 98554957 | rs17393861 | 0.247 | 0.638 | -0.761 | 0.569 | 0.039 | 0.093 | 0.093 | 0 | 0.07 | 0.16 | 0.15 | 0.16 | 0.1 | 0.08 |
| 9 | 98555942 | rs10983622 | 0.209 | 0.532 | -0.749 | 0.456 | -0.047 | 0.078 | 0.078 | 0.689 | 0.41 | 0.17 | 0.25 | 0.24 | 0.41 | 0.33 |
| 9 | 98559246 | rs10983739 | 0.134 | 0.455 | -0.538 | 0.39 | -0.028 | 0.078 | 0.078 | 0.359 | 0.42 | 0.22 | 0.25 | 0.24 | 0.41 | 0.32 |
| 9 | 98568543 | rs816677 | -0.838 | -0.02 | -0.008 | 0.396 | 0.012 | 0.037 | 0.037 | 0 | 0.68 | 0.56 | 0.6 | 0.55 | 0.63 | 0.65 |
| 9 | 98573509 | rs17303045 | 1.236 | 0.013 | -0.108 | 0.423 | -0.01 | -0.012 | -0.012 | 0.182 | 0.05 | 0.02 | 0.15 | 0.16 | 0.1 | 0.08 |
| 9 | 98575014 | rs816671 | 1.453 | -0.321 | -0.474 | -0.046 | -0.101 | -0.035 | -0.035 | 1.358 | 0.26 | 0.04 | 0.19 | 0.24 | 0.33 | 0.25 |
| 9 | 98583426 | rs816685 | -1.162 | -0.123 | -0.463 | 0.303 | -0.054 | -0.088 | -0.088 | 0 | 0.7 | 0.85 | 0.57 | 0.5 | 0.59 | 0.64 |
| 9 | 98596745 | rs10985233 | 0.721 | -0.1 | -0.436 | 0.277 | -0.03 | -0.057 | -0.057 | 0 | 0.32 | 0.24 | 0.45 | 0.5 | 0.42 | 0.36 |
| 9 | 98603006 | rs10985499 | 1.35 | -0.083 | -0.326 | 0.296 | 0.002 | 0.007 | 0.007 | 0 | 0.27 | 0.32 | 0.31 | 0.34 | 0.32 | 0.28 |
| 9 | 98607989 | rs2149564 | 0.204 | -0.145 | -0.249 | 0.248 | 0.02 | -0.003 | -0.003 | 0.397 | 0.37 | 0.48 | 0.54 | 0.61 | 0.48 | 0.43 |
| 9 | 98623567 | rs10481647 | -0.529 | -0.415 | 0.151 | 0.165 | 0 | -0.005 | -0.005 | 0 | 0.55 | 0.57 | 0.72 | 0.76 | 0.55 | 0.66 |
| 9 | 98635675 | rs12004422 | -0.526 | -0.422 | 0.147 | 0.162 | 0 | -0.005 | -0.005 | 0 | 0.55 | 0.57 | 0.72 | 0.76 | 0.55 | 0.66 |
| 9 | 98637430 | rs12001 | 0.729 | -0.471 | 0.085 | 0.182 | -0.018 | -0.002 | -0.002 | 0.091 | 0.23 | 0.25 | 0.27 | 0.25 | 0.13 | 0.3 |
| 9 | 98640671 | rs589458 | 0.612 | -0.556 | 0.395 | 0.156 | 0.008 | -0.029 | -0.029 | 0.518 | 0.43 | 0.23 | 0.49 | 0.49 | 0.42 | 0.52 |
| 9 | 98645674 | rs4742781 | 1.099 | -0.621 | 0.428 | -0.02 | 0.021 | 0.02 | 0.02 | 0 | 0.2 | 0.12 | 0.2 | 0.15 | 0.11 | 0.24 |
| 9 | 98648085 | rs7868270 | 1.089 | -0.621 | 0.431 | -0.02 | -0.03 | -0.029 | -0.029 | 2.371 | 0.2 | 0.25 | 0.2 | 0.15 | 0.11 | 0.24 |
| 9 | 98663517 | rs667018 | -1.807 | -0.617 | 0.214 | 0.062 | -0.142 | 0.041 | 0.041 | 0.082 | 0.73 | 0.98 | 0.79 | 0.85 | 0.64 | 0.73 |
| 9 | 98668370 | rs12345187 | 1.73 | -0.555 | 0.284 | 0.157 | -0.044 | -0.085 | -0.085 | 0.184 | 0.24 | 0.12 | 0.29 | 0.38 | 0.31 | 0.27 |
| 9 | 98670456 | rs12684321 | -0.161 | -0.573 | 0.394 | 0.16 | -0.019 | 0.007 | 0.007 | 0 | 0.29 | 0.37 | 0.41 | 0.38 | 0.24 | 0.39 |
| 9 | 98691137 | rs2274654 | 0.756 | -0.575 | 0.62 | 0.163 | 0.002 | 0 | 0 | 0.177 | 0.22 | 0.19 | 0.26 | 0.26 | 0.13 | 0.3 |
| 9 | 98695299 | rs16910370 | 1.047 | -0.558 | 0.62 | 0.167 | 0.027 | 0.034 | 0.034 | 0.005 | 0.07 | 0.17 | 0.15 | 0.11 | 0.1 | 0.08 |
| 9 | 98719799 | rs10988792 | 0.753 | -0.436 | 0.409 | 0.156 | 0.001 | 0 | 0 | 1.125 | 0.22 | 0.19 | 0.26 | 0.27 | 0.13 | 0.3 |
| 9 | 98722130 | rs10819668 | 1.69 | -0.309 | 0.163 | 0.186 | -0.046 | -0.085 | -0.085 | 1.629 | 0.24 | 0.11 | 0.29 | 0.38 | 0.31 | 0.27 |
| 9 | 98733782 | rs16910396 | 0.245 | -0.212 | 0.183 | 0.233 | 0.018 | 0.059 | 0.059 | 0 | 0.09 | 0.16 | 0.21 | 0.22 | 0.13 | 0.14 |
| 9 | 98739498 | rs10739787 | -1.5 | -0.265 | 0.116 | 0.151 | -0.12 | -0.058 | -0.058 | 0.419 | 0.72 | 0.97 | 0.79 | 0.69 | 0.62 | 0.73 |
| 9 | 98745485 | rs4743443 | -0.123 | 0.005 | 0.381 | 0.403 | 0.018 | 0.044 | 0.044 | 0 | 0.1 | 0.2 | 0.22 | 0.16 | 0.12 | 0.14 |
| 9 | 98748034 | rs10512244 | 0.165 | 0.03 | 0.342 | 0.415 | 0.019 | 0.043 | 0.043 | 0 | 0.09 | 0.16 | 0.21 | 0.16 | 0.11 | 0.14 |
| 9 | 98750606 | rs690484 | -0.539 | -0.005 | 0.262 | 0.33 | -0.018 | 0.056 | 0.056 | 0.322 | 0.56 | 0.74 | 0.73 | 0.69 | 0.54 | 0.66 |
| 9 | 98765778 | rs657451 | 1.41 | 0.082 | 0.388 | 0.45 | 0.003 | 0.012 | 0.012 | 0 | 0.25 | 0.29 | 0.31 | 0.38 | 0.31 | 0.25 |
| 9 | 98770928 | rs913407 | 0.666 | 0.096 | 0.586 | 0.436 | 0.034 | 0.065 | 0.065 | 0 | 0.31 | 0.49 | 0.46 | 0.42 | 0.4 | 0.33 |
| 9 | 98781179 | rs2672813 | 0.344 | -0.015 | 0.422 | 0.266 | 0.021 | 0.009 | 0.009 | 0 | 0.35 | 0.54 | 0.52 | 0.53 | 0.42 | 0.41 |
| 9 | 98786276 | rs238 | -0.646 | -0.064 | 0.164 | 0.215 | -0.057 | 0.101 | 0.101 | 0.385 | 0.57 | 0.84 | 0.7 | 0.76 | 0.55 | 0.65 |
| 9 | 98794448 | rs7023522 | -0.499 | -0.192 | 0.406 | 0.241 | 0 | 0.017 | 0.017 | 0.002 | 0.54 | 0.57 | 0.62 | 0.72 | 0.56 | 0.59 |
| 9 | 98794679 | rs10760718 | 0.057 | -0.203 | 0.212 | 0.222 | 0.003 | 0.01 | 0.01 | 0 | 0.49 | 0.44 | 0.52 | 0.65 | 0.46 | 0.5 |
| 9 | 98800048 | rs690575 | -0.733 | -0.222 | -0.431 | -0.135 | -0.012 | 0.151 | 0.151 | 0.722 | 0.58 | 0.81 | 0.7 | 0.85 | 0.59 | 0.65 |
| 9 | 98805260 | rs12684445 | 0.354 | -0.308 | -0.389 | -0.118 | 0.015 | 0.021 | 0.021 | 0 | 0.23 | 0.14 | 0.22 | 0.21 | 0.16 | 0.26 |
| 9 | 98810506 | rs10819754 | -0.295 | -0.449 | -0.846 | -0.662 | 0.089 | 0.198 | 0.198 | 0 | 0.43 | 0.13 | 0.33 | 0.13 | 0.28 | 0.37 |
| 9 | 98817497 | rs4592147 | 0.567 | 0.035 | -0.606 | -0.138 | -0.001 | -0.003 | -0.003 | 0 | 0.38 | 0.33 | 0.26 | 0.43 | 0.39 | 0.33 |
| 9 | 98818646 | rs734632 | -0.157 | 0.045 | -0.564 | -0.124 | -0.009 | 0.022 | 0.022 | 0.032 | 0.62 | 0.71 | 0.6 | 0.78 | 0.6 | 0.65 |
| 9 | 98823407 | rs10989088 | 1.947 | -0.116 | -0.808 | -0.091 | -0.029 | -0.065 | -0.065 | 0 | 0.21 | 0.11 | 0.19 | 0.43 | 0.29 | 0.23 |
| 9 | 98823528 | rs7021333 | -0.127 | -0.163 | -0.362 | -0.165 | 0.006 | -0.009 | -0.009 | 0.143 | 0.63 | 0.63 | 0.68 | 0.78 | 0.57 | 0.62 |

| | | | | | | | | | | | | | | | | |
|---|---|---|---|---|---|---|---|---|---|---|---|---|---|---|---|---|
| 9 | 98824475 | rs4743501 | 0.135 | -0.472 | -0.242 | -0.039 | 0.012 | 0.008 | 0.008 | 0 | 0.1 | 0.16 | 0.12 | 0.1 | 0.14 | 0.1 |
| 9 | 98826252 | rs7041827 | 0.079 | -0.437 | -0.154 | 0.043 | -0.026 | -0.015 | -0.015 | 0.003 | 0.36 | 0.6 | 0.35 | 0.28 | 0.21 | 0.32 |
| 9 | 98827451 | rs4743505 | -1.182 | 0.117 | -0.838 | -0.107 | -0.097 | -0.064 | -0.064 | 0 | 0.82 | 0.99 | 0.72 | 0.78 | 0.73 | 0.75 |
| 9 | 98828510 | rs4743506 | 0.591 | 0.164 | 0.001 | 0.282 | 0.037 | 0.071 | 0.071 | 0 | 0.09 | 0.18 | 0.14 | 0.17 | 0.14 | 0.17 |
| 9 | 98828889 | rs10739797 | 0.04 | -0.231 | -0.45 | 0.098 | 0.008 | -0.014 | -0.014 | 0 | 0.42 | 0.26 | 0.3 | 0.51 | 0.4 | 0.34 |
| 9 | 98835067 | rs4075991 | -0.688 | -0.56 | 0.44 | -0.089 | -0.043 | -0.117 | -0.117 | 0 | 0.19 | 0.38 | 0.11 | 0 | 0.11 | 0.06 |
| 9 | 98839535 | rs4422885 | 0.219 | -0.607 | 0.684 | 0.433 | 0.002 | -0.019 | -0.019 | 0.008 | 0.13 | 0.09 | 0.08 | 0 | 0.07 | 0.05 |
| 9 | 98840177 | rs4237201 | -0.806 | -0.416 | 0.631 | 0.662 | 0.003 | 0.006 | 0.006 | 0 | 0.91 | 0.91 | 0.9 | 0.88 | 0.86 | 0.9 |
| 9 | 98840796 | rs4562454 | 0.019 | -0.193 | 0.658 | 0.517 | -0.006 | 0.003 | 0.003 | 0 | 0.42 | 0.6 | 0.35 | 0.68 | 0.48 | 0.45 |
| 9 | 98843334 | rs3925103 | 0.174 | -0.188 | 0.629 | 0.515 | -0.005 | -0.001 | -0.001 | 5.113 | 0.38 | 0.35 | 0.33 | 0.66 | 0.48 | 0.45 |
| 9 | 98843638 | rs4291387 | 1.288 | 0.269 | -0.086 | 0.688 | -0.117 | -0.139 | -0.139 | 0 | 0.23 | 0.02 | 0.21 | 0.43 | 0.34 | 0.31 |
| 9 | 98844752 | rs4743521 | 0.221 | 0.066 | 0.557 | 0.614 | 0.002 | 0.041 | 0.041 | 0 | 0.73 | 0.45 | 0.77 | 0.64 | 0.73 | 0.76 |
| 9 | 98849009 | rs4743525 | 0.922 | -0.302 | 0.322 | 0.538 | 0.007 | 0.002 | 0.002 | 0.812 | 0.3 | 0.36 | 0.28 | 0.41 | 0.44 | 0.37 |
| 9 | 98851230 | rs4443781 | 1.175 | -1.056 | 0.053 | 0.139 | 0.003 | 0.005 | 0.005 | 0 | 0.2 | 0.16 | 0.38 | 0.14 | 0.1 | 0.2 |
| 9 | 98851504 | rs10760747 | 1.591 | -1.045 | 0.192 | 0.134 | -0.066 | -0.035 | -0.035 | 2.836 | 0.27 | 0.16 | 0.27 | 0.41 | 0.44 | 0.37 |
| 9 | 98851777 | rs4620409 | 0.054 | -1.307 | -0.017 | 0.014 | 0.019 | -0.009 | -0.009 | 4.128 | 0.51 | 0.78 | 0.38 | 0.56 | 0.54 | 0.57 |
| 9 | 98856744 | rs7862829 | 1.504 | -1.197 | -0.137 | -0.063 | 0.026 | 0.011 | 0.011 | 0.815 | 0.3 | 0.38 | 0.28 | 0.43 | 0.44 | 0.37 |
| 9 | 98858735 | rs10760758 | -2.059 | -0.937 | -0.218 | 0.093 | -0.035 | -0.034 | -0.034 | 0 | 0.92 | 1 | 0.91 | 0.91 | 0.86 | 0.94 |
| 9 | 98864367 | rs16910480 | 0.218 | -1.727 | 0.193 | -0.053 | -0.052 | -0.115 | -0.115 | 0 | 0.19 | 0.36 | 0.09 | 0 | 0.09 | 0.06 |
| 9 | 98867813 | rs10819861 | 0.03 | -1.547 | -0.671 | 0.02 | 0.016 | 0.036 | 0.036 | 0 | 0.51 | 0.7 | 0.37 | 0.42 | 0.53 | 0.44 |
| 9 | 98874639 | rs10819882 | -0.038 | -1.421 | -0.48 | 0.126 | -0.041 | 0.034 | 0.034 | 0.041 | 0.24 | 0.4 | 0.43 | 0.33 | 0.17 | 0.38 |
| 9 | 98877206 | rs10733713 | -0.218 | -1.505 | -0.756 | 0.035 | 0.001 | -0.013 | -0.013 | 0 | 0.57 | 0.51 | 0.66 | 0.74 | 0.54 | 0.72 |
| 9 | 98879734 | rs12115436 | 0.577 | -1.461 | -0.678 | 0.102 | 0.002 | 0.002 | 0.002 | 0 | 0.2 | 0.14 | 0.37 | 0.15 | 0.16 | 0.25 |
| 9 | 98884945 | rs4743556 | 0.555 | -1.093 | -0.761 | -0.003 | -0.019 | 0.018 | 0.018 | 0 | 0.22 | 0.28 | 0.41 | 0.33 | 0.15 | 0.33 |
| 9 | 98885144 | rs4743558 | -0.478 | -0.54 | -0.597 | -0.189 | -0.074 | -0.052 | -0.052 | 0.947 | 0.72 | 0.93 | 0.75 | 0.74 | 0.64 | 0.72 |
| 9 | 98888628 | rs10760787 | 0.455 | -0.395 | -0.506 | -0.044 | 0.001 | -0.014 | -0.014 | 0 | 0.56 | 0.53 | 0.65 | 0.74 | 0.54 | 0.71 |
| 9 | 98891743 | rs7041700 | -0.585 | -1.042 | 0.202 | -0.21 | -0.034 | -0.121 | -0.121 | 0 | 0.17 | 0.39 | 0.11 | 0 | 0.11 | 0.06 |
| 9 | 98895004 | rs7871616 | -0.545 | -1.122 | -0.191 | -0.112 | -0.006 | -0.019 | -0.019 | 0 | 0.55 | 0.56 | 0.41 | 0.74 | 0.38 | 0.63 |
| 9 | 98904169 | rs12341515 | -0.708 | -1.29 | -0.223 | 0.099 | 0.004 | 0.04 | 0.04 | 0.208 | 0.11 | 0.37 | 0.09 | 0.14 | 0.08 | 0.09 |
| 9 | 98905141 | rs2010398 | -0.035 | -1.791 | -0.385 | 0.379 | 0.021 | -0.005 | -0.005 | 0 | 0.24 | 0.14 | 0.23 | 0.22 | 0.2 | 0.26 |
| 9 | 98907515 | rs3892812 | -1.085 | -1.496 | -0.507 | 0.355 | 0.037 | 0.026 | 0.026 | 0 | 0.54 | 0.66 | 0.38 | 0.6 | 0.38 | 0.52 |
| 9 | 98918219 | rs4743597 | -0.96 | -1.418 | -0.805 | -0.025 | 0.012 | 0.008 | 0.008 | 0 | 0.49 | 0.55 | 0.25 | 0.36 | 0.33 | 0.35 |
| 9 | 98918433 | rs4743598 | -0.353 | -0.732 | -1.22 | 0.797 | 0.043 | -0.045 | -0.045 | 0 | 0.27 | 0.11 | 0.12 | 0.35 | 0.16 | 0.26 |
| 9 | 98923066 | rs10989494 | -0.104 | -0.242 | -0.935 | 0.528 | -0.001 | -0.002 | -0.002 | 0 | 0.4 | 0.34 | 0.26 | 0.27 | 0.34 | 0.33 |
| 9 | 98923268 | rs10819930 | -0.214 | -0.565 | -0.205 | 1.068 | -0.001 | 0.002 | 0.002 | 0.507 | 0.39 | 0.55 | 0.46 | 0.53 | 0.35 | 0.43 |
| 9 | 98924140 | rs10989497 | 0.49 | 0.026 | -0.603 | 0.753 | -0.007 | 0.048 | 0.048 | 0.043 | 0.34 | 0.26 | 0.37 | 0.21 | 0.41 | 0.32 |
| 9 | 98925146 | rs10989503 | 0.421 | 0 | -0.455 | 0.766 | -0.003 | 0.029 | 0.029 | 0.053 | 0.35 | 0.29 | 0.37 | 0.22 | 0.41 | 0.33 |
| 9 | 98925677 | rs4742865 | -0.15 | 0.182 | -0.015 | 0.961 | -0.002 | 0.009 | 0.009 | 0 | 0.24 | 0.24 | 0.3 | 0.47 | 0.14 | 0.36 |
| 9 | 98928974 | rs12353207 | 0.317 | 0.515 | -0.026 | 0.773 | 0.002 | 0.037 | 0.037 | 0 | 0.37 | 0.33 | 0.38 | 0.22 | 0.42 | 0.33 |
| 9 | 98929986 | rs10819946 | -0.022 | 0.577 | 0.006 | 0.857 | -0.002 | 0.002 | 0.002 | 0 | 0.23 | 0.24 | 0.15 | 0.36 | 0.14 | 0.26 |
| 9 | 98932408 | rs4743611 | -1.188 | 0.221 | -0.077 | 0.816 | -0.01 | 0 | 0 | 0 | 0.62 | 0.74 | 0.69 | 0.69 | 0.58 | 0.69 |
| 9 | 98932749 | rs7862977 | -0.499 | 0.017 | -0.068 | 0.88 | -0.014 | 0.01 | 0.01 | 0 | 0.28 | 0.33 | 0.17 | 0.36 | 0.17 | 0.25 |
| 9 | 98935333 | rs10819950 | -0.554 | 0.568 | -0.299 | 0.769 | 0.002 | -0.018 | -0.018 | 0.471 | 0.29 | 0.18 | 0.18 | 0.36 | 0.24 | 0.22 |
| 9 | 98936604 | rs7857532 | -0.579 | 0.72 | -0.547 | 0.809 | 0.025 | -0.019 | -0.019 | 0 | 0.77 | 0.7 | 0.66 | 0.79 | 0.63 | 0.64 |
| 9 | 98936624 | rs7857544 | 0.649 | 0.567 | -0.584 | 1.019 | 0.008 | 0.001 | 0.001 | 0.039 | 0.5 | 0.55 | 0.47 | 0.46 | 0.43 | 0.43 |
| 9 | 98938479 | rs3892706 | -0.59 | 0.719 | -0.674 | 0.999 | 0.047 | -0.029 | -0.029 | 0 | 0.75 | 0.66 | 0.64 | 0.79 | 0.57 | 0.63 |
| 9 | 98940178 | rs714676 | 0.224 | 0.757 | -0.875 | 1.215 | -0.011 | -0.098 | -0.098 | 0.023 | 0.26 | 0.02 | 0.17 | 0.33 | 0.2 | 0.21 |
| 9 | 98952847 | rs10819981 | -0.442 | 0.543 | -0.885 | 1.013 | -0.106 | 0.026 | 0.026 | 0.183 | 0.77 | 0.89 | 0.64 | 0.79 | 0.57 | 0.64 |
| 9 | 98957977 | rs7048593 | 0.821 | 0.73 | -0.579 | 1.253 | 0.014 | 0.016 | 0.016 | 0.217 | 0.46 | 0.6 | 0.28 | 0.26 | 0.32 | 0.34 |
| 9 | 98958949 | rs912465 | 0.798 | 0.587 | -0.571 | 1.278 | 0.02 | 0.007 | 0.007 | 0.07 | 0.46 | 0.6 | 0.43 | 0.36 | 0.29 | 0.42 |
| 9 | 98962102 | rs4743638 | -0.649 | 0.541 | -0.907 | 1.358 | -0.103 | -0.041 | -0.041 | 0.003 | 0.79 | 0.92 | 0.7 | 0.74 | 0.62 | 0.67 |
| 9 | 98962994 | rs7857474 | 0.036 | 0.545 | -0.675 | 1.388 | 0.006 | -0.011 | -0.011 | 0 | 0.29 | 0.24 | 0.22 | 0.39 | 0.27 | 0.22 |
| 9 | 98963734 | rs10117895 | -0.32 | 0.557 | -0.657 | 1.378 | 0.005 | -0.018 | -0.018 | 0.234 | 0.32 | 0.3 | 0.4 | 0.48 | 0.3 | 0.3 |
| 9 | 98968142 | rs13300278 | 0.328 | 0.636 | -0.812 | 1.512 | -0.022 | -0.066 | -0.066 | 3.721 | 0.28 | 0.06 | 0.21 | 0.33 | 0.26 | 0.22 |
| 9 | 98968292 | rs13300603 | -0.007 | 0.63 | -0.794 | 1.443 | -0.025 | -0.103 | -0.103 | 1.296 | 0.3 | 0.06 | 0.39 | 0.41 | 0.29 | 0.29 |
| 9 | 98969746 | rs7869882 | -0.561 | 0.619 | -0.911 | 1.399 | -0.064 | -0.02 | -0.02 | 0 | 0.81 | 0.92 | 0.72 | 0.72 | 0.68 | 0.67 |
| 9 | 98972430 | rs11791270 | -0.142 | 0.494 | -0.752 | 1.427 | 0.002 | -0.013 | -0.013 | 0 | 0.31 | 0.31 | 0.41 | 0.41 | 0.29 | 0.29 |
| 9 | 98974141 | rs7020106 | 0.52 | 0.423 | -0.486 | 1.376 | 0.008 | 0.012 | 0.012 | 0.405 | 0.48 | 0.55 | 0.33 | 0.29 | 0.4 | 0.38 |
| 9 | 98978660 | rs10820020 | 0.038 | 0.396 | -1.067 | 1.37 | -0.018 | -0.115 | -0.115 | 0 | 0.2 | 0.07 | 0.21 | 0.48 | 0.19 | 0.34 |
| 9 | 98985321 | rs412147 | 1.162 | 0.009 | -0.772 | 0.876 | -0.007 | -0.048 | -0.048 | 0 | 0.31 | 0.32 | 0.16 | 0.05 | 0.27 | 0.13 |
| 9 | 98988429 | rs2479827 | 0.35 | 0.125 | -0.61 | 1.057 | 0 | 0 | 0 | 0 | 0.4 | 0.57 | 0.23 | 0.3 | 0.34 | 0.26 |
| 9 | 98991303 | rs2031321 | 0.292 | 0.17 | -0.595 | 1.149 | -0.011 | 0.002 | 0.002 | 0 | 0.43 | 0.32 | 0.43 | 0.47 | 0.5 | 0.43 |
| 9 | 98991519 | rs280661 | -1.241 | 0.175 | -0.667 | 1.14 | -0.018 | -0.027 | -0.027 | 0 | 0.86 | 0.96 | 0.83 | 0.84 | 0.86 | 0.77 |
| 9 | 98995852 | rs8190581 | -0.438 | 0.199 | -0.597 | 1.164 | -0.012 | 0.131 | 0.131 | 0 | 0.1 | 0.26 | 0.07 | 0.26 | 0.07 | 0.13 |
| 9 | 98996534 | rs7048229 | 0.175 | 0.213 | -0.716 | 1.099 | 0 | -0.012 | -0.012 | 0 | 0.45 | 0.36 | 0.61 | 0.53 | 0.52 | 0.51 |
| 9 | 99012277 | rs407179 | 0.944 | 0.379 | -0.852 | 0.658 | -0.021 | -0.122 | -0.122 | 0 | 0.31 | 0.4 | 0.16 | 0.03 | 0.27 | 0.13 |
| 9 | 99016095 | rs2026001 | -0.086 | 0.575 | -1.544 | 0.751 | 0.01 | 0.016 | 0.016 | 0 | 0.61 | 0.78 | 0.48 | 0.72 | 0.64 | 0.6 |
| 9 | 99016896 | rs370438 | -1.143 | 0.513 | -1.893 | 0.543 | -0.011 | -0.008 | -0.008 | 0.019 | 0.84 | 0.95 | 0.83 | 0.86 | 0.86 | 0.77 |
| 9 | 99018442 | rs2257157 | 0.318 | 0.411 | -1.986 | 0.439 | 0 | -0.002 | -0.002 | 0.885 | 0.5 | 0.5 | 0.36 | 0.34 | 0.46 | 0.44 |

| | | | | | | | | | | | | | | | | |
|---|---|---|---|---|---|---|---|---|---|---|---|---|---|---|---|---|
| 9 | 99021582 | rs7039978 | 0.317 | 0.484 | -2.22 | 0.759 | -0.046 | 0.019 | 0.019 | 0.195 | 0.56 | 0.83 | 0.38 | 0.34 | 0.52 | 0.29 |
| 9 | 99021661 | rs2476923 | 0.09 | 0.569 | -2.083 | 0.684 | -0.073 | -0.044 | -0.044 | 0 | 0.59 | 0.83 | 0.57 | 0.39 | 0.55 | 0.38 |
| 9 | 99029431 | rs280654 | 0.01 | -0.052 | -1.767 | 0.235 | 0.027 | 0.032 | 0.032 | 0 | 0.4 | 0.64 | 0.42 | 0.31 | 0.37 | 0.33 |
| 9 | 99031628 | rs11788785 | 0.108 | -0.055 | -2.232 | 0.071 | 0.022 | -0.066 | -0.066 | 2.01 | 0.24 | 0.13 | 0.2 | 0.56 | 0.16 | 0.3 |
| 9 | 99038030 | rs2476927 | 0.773 | -0.249 | -1.703 | -0.117 | -0.014 | -0.041 | -0.041 | 0 | 0.37 | 0.48 | 0.24 | 0.23 | 0.39 | 0.26 |
| 9 | 99040501 | rs1983828 | 0.769 | 0.541 | -1.745 | -0.066 | -0.027 | 0.02 | 0.02 | 0 | 0.23 | 0.15 | 0.2 | 0.06 | 0.34 | 0.11 |
| 9 | 99050100 | rs8190512 | -0.399 | 0.318 | -1.906 | -0.114 | -0.006 | 0.044 | 0.044 | 0 | 0.42 | 0.29 | 0.56 | 0.67 | 0.52 | 0.47 |
| 9 | 99050794 | rs2479828 | -0.857 | 0.277 | -1.951 | -0.133 | 0 | 0.021 | 0.021 | 0 | 0.88 | 0.97 | 0.83 | 0.92 | 0.9 | 0.76 |
| 9 | 99054604 | rs1119864 | 0.806 | 0.458 | -1.962 | 0.496 | -0.007 | -0.008 | -0.008 | 0 | 0.24 | 0.2 | 0.2 | 0.06 | 0.34 | 0.11 |
| 9 | 99056209 | rs11788083 | 0.361 | 0.033 | -1.727 | 0.679 | -0.002 | -0.002 | -0.002 | 0 | 0.15 | 0.15 | 0.34 | 0.32 | 0.15 | 0.25 |
| 9 | 99060612 | rs7029101 | 0.097 | -0.288 | -1.487 | 0.247 | 0.008 | 0.01 | 0.01 | 0 | 0.45 | 0.52 | 0.26 | 0.25 | 0.38 | 0.28 |
| 9 | 99069230 | rs11788621 | 0.32 | -0.128 | -1.368 | 1.091 | 0.004 | 0.007 | 0.007 | 0 | 0.25 | 0.19 | 0.16 | 0.19 | 0.2 | 0.14 |
| 9 | 99069944 | rs4743713 | -0.404 | 0.131 | -2.15 | 0.682 | -0.098 | -0.035 | -0.035 | 0 | 0.68 | 0.92 | 0.73 | 0.67 | 0.52 | 0.7 |
| 9 | 99073152 | rs4742910 | -0.578 | 0.116 | -2.119 | 0.683 | -0.113 | -0.013 | -0.013 | 0 | 0.69 | 0.94 | 0.73 | 0.71 | 0.52 | 0.77 |
| 9 | 99075707 | rs4743716 | 0.269 | -0.248 | -1.774 | 0.746 | -0.005 | -0.002 | -0.002 | 0 | 0.55 | 0.66 | 0.58 | 0.52 | 0.47 | 0.48 |
| 9 | 99077386 | rs7022202 | 0.269 | -0.248 | -1.774 | 0.746 | -0.005 | -0.002 | -0.002 | 0 | 0.55 | 0.66 | 0.58 | 0.52 | 0.47 | 0.48 |
| 9 | 99078150 | rs10990403 | -0.515 | -0.265 | -1.761 | 0.74 | -0.075 | -0.015 | -0.015 | 0.208 | 0.67 | 0.88 | 0.72 | 0.66 | 0.52 | 0.69 |
| 9 | 99079114 | rs12685240 | 0.147 | -0.103 | -1.712 | 0.6 | -0.005 | -0.002 | -0.002 | 0 | 0.55 | 0.66 | 0.57 | 0.52 | 0.47 | 0.48 |
| 9 | 99081512 | rs4743722 | -0.227 | 0.207 | -1.58 | 0.585 | -0.023 | -0.01 | -0.01 | 0 | 0.66 | 0.71 | 0.65 | 0.6 | 0.51 | 0.76 |
| 9 | 99085363 | rs10126039 | -0.46 | 0.283 | -0.679 | 1.003 | 0.019 | 0.117 | 0.117 | 0 | 0.12 | 0.22 | 0.29 | 0.3 | 0.13 | 0.28 |
| 9 | 99087446 | rs7858292 | -0.401 | 0.464 | -1.394 | 0.264 | -0.081 | 0.188 | 0.188 | 0 | 0.61 | 0.89 | 0.6 | 0.87 | 0.55 | 0.54 |
| 9 | 99087685 | rs10990503 | -0.002 | -0.139 | -0.889 | -0.112 | -0.01 | 0.018 | 0.018 | 0 | 0.34 | 0.39 | 0.35 | 0.32 | 0.28 | 0.36 |
| 9 | 99097527 | rs6479272 | -0.907 | -0.189 | -0.979 | -0.615 | -0.096 | 0.186 | 0.186 | 0.019 | 0.63 | 0.92 | 0.61 | 0.87 | 0.57 | 0.56 |
| 9 | 99108678 | rs10820410 | -0.046 | -0.49 | -0.206 | -0.385 | 0.001 | -0.042 | -0.042 | 0 | 0.3 | 0.62 | 0.12 | 0.04 | 0.18 | 0.13 |
| 9 | 99148152 | rs10820507 | 0.757 | -0.635 | 0.215 | -0.409 | 0.018 | 0.019 | 0.019 | 0 | 0.23 | 0.1 | 0.06 | 0.02 | 0.15 | 0.09 |
| 9 | 99171631 | rs10820564 | 0.807 | -0.538 | 0.349 | -0.782 | -0.033 | -0.118 | -0.118 | 1.382 | 0.23 | 0.31 | 0.08 | 0.02 | 0.15 | 0.09 |
| 9 | 99174951 | rs10760999 | 0.516 | -0.542 | 0.131 | -0.784 | -0.032 | -0.102 | -0.102 | 0 | 0.26 | 0.54 | 0.1 | 0.02 | 0.16 | 0.09 |
| 9 | 99188248 | rs1885427 | -0.006 | -0.465 | 0.178 | -0.748 | 0.004 | -0.036 | -0.036 | 1.835 | 0.3 | 0.64 | 0.13 | 0.03 | 0.19 | 0.12 |
| 9 | 99188347 | rs10820597 | 0.522 | -0.422 | 0.189 | -0.713 | -0.028 | -0.102 | -0.102 | 0 | 0.25 | 0.54 | 0.1 | 0.02 | 0.15 | 0.09 |
| 9 | 99189255 | rs10820598 | 0.522 | -0.422 | 0.189 | -0.713 | -0.028 | -0.102 | -0.102 | 4.934 | 0.25 | 0.54 | 0.1 | 0.02 | 0.15 | 0.09 |
| 9 | 99210584 | rs7870106 | -0.131 | -0.354 | 0.076 | -0.757 | 0.01 | -0.024 | -0.024 | 1.398 | 0.32 | 0.64 | 0.14 | 0.03 | 0.21 | 0.13 |
| 9 | 99247758 | rs10991346 | 0.651 | -0.379 | 0.489 | -0.729 | -0.04 | -0.147 | -0.147 | 0 | 0.26 | 0.45 | 0.11 | 0.02 | 0.17 | 0.09 |
| 9 | 99250463 | rs7030316 | 0.65 | -0.39 | 0.5 | -0.734 | -0.041 | -0.151 | -0.151 | 0.651 | 0.26 | 0.44 | 0.11 | 0.02 | 0.17 | 0.09 |
| 9 | 99252464 | rs1057687 | 0.523 | -0.386 | 0.441 | -0.732 | -0.004 | -0.048 | -0.048 | 0 | 0.27 | 0.64 | 0.12 | 0.02 | 0.17 | 0.09 |
| 9 | 99260928 | rs13302712 | -0.319 | -0.784 | 0.207 | -0.842 | 0.044 | 0.024 | 0.024 | 0.036 | 0.13 | 0.01 | 0.16 | 0.1 | 0.08 | 0.29 |
| 9 | 99262296 | rs7871404 | 0.59 | -0.377 | 0.67 | -0.6 | -0.026 | -0.104 | -0.104 | 0 | 0.27 | 0.55 | 0.11 | 0.02 | 0.17 | 0.1 |
| 9 | 99268429 | rs10118793 | 0.831 | -0.389 | 0.869 | -0.606 | 0.001 | -0.03 | -0.03 | 0 | 0.23 | 0.18 | 0.09 | 0.02 | 0.16 | 0.09 |
| 9 | 99288951 | rs16911097 | 0.699 | 0.204 | 1.584 | 0.5 | 0 | -0.047 | -0.047 | 0 | 0.25 | 0.19 | 0.31 | 0.56 | 0.26 | 0.16 |
| 9 | 99311177 | rs7869415 | -1.362 | 0.532 | 2.642 | 1.242 | -0.004 | -0.007 | -0.007 | 0 | 0.06 | 0.62 | 0.05 | 0 | 0 | 0.01 |
| 9 | 99346570 | rs7042473 | -1.049 | 0.959 | 1.877 | 1.359 | -0.009 | -0.014 | -0.014 | 0.969 | 0.14 | 0.1 | 0.27 | 0.29 | 0.19 | 0.34 |
| 9 | 99400209 | rs10978436 | -0.736 | 0.84 | 1.889 | 1.642 | -0.044 | -0.029 | -0.029 | 0.094 | 0.08 | 0.01 | 0.21 | 0.29 | 0.14 | 0.24 |
| 9 | 99448564 | rs2165890 | -0.936 | 0.689 | 1.25 | 1.717 | 0.039 | 0.086 | 0.086 | 0 | 0.09 | 0.22 | 0.23 | 0.22 | 0.13 | 0.17 |
| 9 | 99452245 | rs6477621 | -0.2 | 0.809 | 0.934 | 1.408 | -0.028 | 0.001 | 0.001 | 0 | 0.89 | 0.99 | 0.88 | 0.91 | 0.86 | 0.76 |
| 9 | 99461810 | rs2083825 | -1.223 | 0.744 | 1.247 | 2.178 | 0.028 | 0.181 | 0.181 | 0 | 0.1 | 0.24 | 0.25 | 0.44 | 0.14 | 0.21 |
| 9 | 99487607 | rs1596359 | -0.874 | 0.22 | 1.919 | 1.527 | 0.015 | 0.134 | 0.134 | 1.371 | 0.07 | 0.33 | 0.04 | 0.11 | 0.02 | 0.1 |
| 9 | 99488406 | rs16911482 | -0.874 | 0.22 | 1.959 | 1.527 | 0.012 | 0.128 | 0.128 | 1.371 | 0.07 | 0.31 | 0.04 | 0.11 | 0.02 | 0.1 |
| 9 | 99567384 | rs10124911 | -1.81 | 0.115 | 2.695 | 2.015 | 0.008 | 0.256 | 0.256 | 0.017 | 0.08 | 0.44 | 0.06 | 0.21 | 0.03 | 0.08 |
| 9 | 99581811 | rs4645656 | -2.317 | 0.057 | 3.174 | 2.001 | -0.035 | 0.205 | 0.205 | 1.772 | 0.1 | 0.63 | 0.07 | 0.32 | 0.03 | 0.13 |
| 9 | 99584880 | rs7859940 | -2.688 | 0.056 | 3.282 | 1.986 | -0.039 | 0.13 | 0.13 | 1.545 | 0.11 | 0.69 | 0.09 | 0.32 | 0.04 | 0.13 |
| 9 | 99589296 | rs7848710 | -2.191 | 0.072 | 3.407 | 1.995 | -0.02 | 0.285 | 0.285 | 1.418 | 0.1 | 0.56 | 0.06 | 0.32 | 0.03 | 0.13 |
| 9 | 99606529 | rs7037100 | -2.673 | 0.056 | 3.345 | 1.985 | -0.039 | 0.112 | 0.112 | 3.65 | 0.11 | 0.7 | 0.09 | 0.32 | 0.03 | 0.14 |
| 9 | 99619081 | rs2296812 | -0.161 | 0.287 | 3.208 | 2.151 | -0.042 | -0.026 | -0.026 | 0.056 | 0.07 | 0.01 | 0.21 | 0.1 | 0.13 | 0.14 |
| 9 | 99620277 | rs7040493 | -0.723 | -0.055 | 3.381 | 2.099 | 0.023 | 0.014 | 0.014 | 0.166 | 0.06 | 0.31 | 0.02 | 0 | 0.01 | 0.05 |
| 9 | 99625063 | rs6477733 | -2.527 | -0.08 | 3.391 | 2.088 | -0.037 | 0.167 | 0.167 | 2.001 | 0.1 | 0.66 | 0.08 | 0.32 | 0.03 | 0.14 |
| 9 | 99655358 | rs1046795 | -0.902 | -0.058 | 3.019 | 2.045 | 0.016 | 0.008 | 0.008 | 0.221 | 0.06 | 0.31 | 0.03 | 0 | 0.01 | 0.05 |
| 9 | 99792208 | rs10123754 | -1.101 | -0.148 | 2.402 | 1.784 | -0.024 | 0.193 | 0.193 | 0 | 0.2 | 0.52 | 0.19 | 0.43 | 0.15 | 0.28 |
| 9 | 99793583 | rs16919034 | -1.122 | -0.15 | 2.407 | 1.819 | -0.027 | 0.13 | 0.13 | 0 | 0.21 | 0.54 | 0.19 | 0.54 | 0.16 | 0.32 |
| 9 | 99795125 | rs15394 | -1.083 | -0.143 | 2.407 | 1.806 | -0.023 | 0.196 | 0.196 | 0 | 0.2 | 0.52 | 0.19 | 0.43 | 0.15 | 0.28 |
| 9 | 99798293 | rs10118886 | -1.099 | -0.138 | 2.407 | 1.806 | -0.027 | 0.195 | 0.195 | 0 | 0.2 | 0.52 | 0.19 | 0.43 | 0.15 | 0.28 |
| 9 | 99806169 | rs12236690 | -1.119 | -0.136 | 2.409 | 1.812 | -0.021 | 0.193 | 0.193 | 0 | 0.2 | 0.53 | 0.19 | 0.43 | 0.16 | 0.28 |
| 9 | 99811206 | rs10816962 | -0.204 | -0.132 | 2.468 | 1.851 | 0.001 | -0.05 | -0.05 | 2.912 | 0.15 | 0.07 | 0.15 | 0.42 | 0.15 | 0.24 |
| 9 | 99814841 | rs4510972 | -0.967 | -0.127 | 2.372 | 1.853 | -0.024 | 0.202 | 0.202 | 0 | 0.19 | 0.51 | 0.19 | 0.42 | 0.15 | 0.24 |
| 9 | 99815929 | rs10759417 | -0.967 | -0.127 | 2.372 | 1.853 | -0.02 | 0.202 | 0.202 | 0 | 0.19 | 0.51 | 0.19 | 0.42 | 0.15 | 0.24 |
| 9 | 99821611 | rs4743056 | 0.597 | -0.212 | 1.07 | 0.842 | 0.002 | 0.005 | 0.005 | 0 | 0.5 | 0.46 | 0.66 | 0.42 | 0.52 | 0.51 |
| 9 | 99824697 | rs4361859 | -0.336 | -0.009 | 0.923 | 0.22 | -0.072 | 0.207 | 0.207 | 0 | 0.69 | 0.97 | 0.85 | 0.98 | 0.67 | 0.8 |
| 9 | 99832156 | rs6477872 | -0.279 | -0.016 | 0.935 | 0.217 | -0.024 | 0.13 | 0.13 | 0 | 0.69 | 0.88 | 0.84 | 0.98 | 0.67 | 0.8 |
| 9 | 99856397 | rs3802470 | 0.047 | 0.177 | 0.858 | 0.381 | 0.008 | -0.064 | -0.064 | 0 | 0.27 | 0.47 | 0.15 | 0.14 | 0.28 | 0.2 |
| 9 | 99874486 | rs10980643 | -0.193 | -0.126 | 0.725 | 1.112 | 0.002 | -0.005 | -0.005 | 0 | 0.19 | 0.16 | 0.1 | 0.28 | 0.18 | 0.3 |
| 9 | 99883513 | rs13283681 | 0.646 | -0.286 | -0.23 | 0.995 | -0.055 | -0.024 | -0.024 | 0 | 0.37 | 0.17 | 0.53 | 0.39 | 0.52 | 0.39 |

| | | | | | | | | | | | | | | | | |
|---|---|---|---|---|---|---|---|---|---|---|---|---|---|---|---|---|
| 9 | 99890895 | rs4399037 | -0.763 | -0.625 | -0.394 | 0.744 | 0.011 | 0.022 | 0.022 | 0 | 0.64 | 0.84 | 0.68 | 0.67 | 0.72 | 0.74 |
| 9 | 99895662 | rs2001382 | 0.52 | -0.137 | 0.028 | 1.257 | -0.111 | 0.003 | 0.003 | 0 | 0.36 | 0.09 | 0.53 | 0.29 | 0.52 | 0.36 |
| 9 | 99899313 | rs4743059 | -0.114 | -0.033 | 0.379 | 1.368 | 0.003 | 0.077 | 0.077 | 0.223 | 0.09 | 0.14 | 0.13 | 0.31 | 0.1 | 0.05 |
| 9 | 99946041 | rs10980992 | -0.075 | 0.461 | -0.014 | 1.014 | 0.003 | -0.003 | -0.003 | 0 | 0.14 | 0.11 | 0.18 | 0.1 | 0.18 | 0.22 |
| 9 | 99973826 | rs4743061 | -0.684 | 0.252 | 0.846 | 1.292 | 0.023 | 0.024 | 0.024 | 0.222 | 0.19 | 0.56 | 0.34 | 0.21 | 0.18 | 0.26 |
| 9 | 100005128 | rs11791310 | 0.544 | 0.157 | 0.344 | 1.165 | -0.022 | -0.012 | -0.012 | 0 | 0.39 | 0.23 | 0.28 | 0.37 | 0.51 | 0.42 |
| 9 | 100016763 | rs7863530 | -1.207 | 0.141 | 0.252 | 1.089 | 0.015 | 0.194 | 0.194 | 0.017 | 0.66 | 0.96 | 0.8 | 0.94 | 0.8 | 0.75 |
| 9 | 100038067 | rs3994560 | 0.422 | 0.133 | 0.35 | 1.291 | -0.001 | 0.004 | 0.004 | 0 | 0.16 | 0.14 | 0.34 | 0.29 | 0.18 | 0.26 |
| 9 | 100043031 | rs7855042 | -0.085 | 0.131 | 0.795 | 1.284 | 0.054 | 0.069 | 0.069 | 0.261 | 0.09 | 0.39 | 0.16 | 0.16 | 0.1 | 0.03 |
| 9 | 100049972 | rs2487874 | 1.008 | 0.131 | 0.795 | 1.283 | 0.015 | 0.194 | 0.194 | 0.017 | 0.34 | 0.04 | 0.2 | 0.06 | 0.2 | 0.25 |
| 9 | 100051415 | rs2818242 | 1.008 | 0.131 | 0.795 | 1.283 | 0.015 | 0.194 | 0.194 | 0.017 | 0.34 | 0.04 | 0.2 | 0.06 | 0.2 | 0.25 |
| 9 | 100054429 | rs10817378 | -0.118 | 0.342 | 0.795 | 1.283 | 0.072 | 0.067 | 0.067 | 0.306 | 0.1 | 0.39 | 0.18 | 0.16 | 0.13 | 0.03 |
| 9 | 100061226 | rs1903635 | -0.932 | 0.166 | 0.807 | 1.297 | -0.02 | -0.014 | -0.014 | 0 | 0.61 | 0.77 | 0.72 | 0.62 | 0.5 | 0.58 |
| 9 | 100074446 | rs3747504 | 0.035 | 0.278 | 0.95 | 1.381 | 0.02 | 0.051 | 0.051 | 0 | 0.17 | 0.3 | 0.35 | 0.29 | 0.19 | 0.27 |
| 9 | 100076896 | rs3747502 | -0.11 | 0.276 | 0.707 | 1.373 | 0.056 | 0.071 | 0.071 | 0.337 | 0.09 | 0.39 | 0.1 | 0.16 | 0.1 | 0.03 |
| 9 | 100087220 | rs3747498 | 0.136 | 0.21 | 0.876 | 1.415 | 0.008 | 0.022 | 0.022 | 0 | 0.16 | 0.19 | 0.35 | 0.29 | 0.19 | 0.26 |
| 9 | 100088971 | rs10981625 | 0.645 | 0.171 | 0.736 | 1.288 | -0.02 | -0.05 | -0.05 | 0 | 0.34 | 0.41 | 0.3 | 0.18 | 0.2 | 0.26 |
| 9 | 100105735 | rs1947717 | -0.472 | 0.262 | 0.478 | 1.3 | 0.002 | 0.006 | 0.006 | 0.955 | 0.52 | 0.75 | 0.66 | 0.58 | 0.39 | 0.55 |
| 9 | 100107236 | rs10435865 | -0.472 | 0.262 | 0.478 | 1.3 | 0.002 | 0.006 | 0.006 | 0.955 | 0.52 | 0.75 | 0.66 | 0.58 | 0.39 | 0.55 |
| 9 | 100114057 | rs4313271 | -0.316 | 0.267 | 0.6 | 1.302 | 0.009 | 0.01 | 0.01 | 1.235 | 0.5 | 0.6 | 0.66 | 0.59 | 0.39 | 0.55 |
| 9 | 100116059 | rs4743082 | 0.198 | 0.259 | 0.643 | 1.315 | -0.002 | 0.01 | 0.01 | 0 | 0.16 | 0.14 | 0.35 | 0.4 | 0.19 | 0.29 |
| 9 | 100122291 | rs3747495 | 0.285 | 0.258 | 0.641 | 1.368 | -0.001 | 0.002 | 0.002 | 0 | 0.16 | 0.14 | 0.34 | 0.29 | 0.19 | 0.26 |
| 9 | 100137394 | rs2242187 | -0.485 | 0.246 | 0.635 | 1.365 | 0.008 | 0.007 | 0.007 | 1.029 | 0.51 | 0.72 | 0.66 | 0.58 | 0.39 | 0.55 |
| 9 | 100139396 | rs898870 | 0.087 | 0.43 | -0.057 | 1.289 | 0.009 | 0.046 | 0.046 | 0 | 0.16 | 0.18 | 0.35 | 0.4 | 0.19 | 0.29 |
| 9 | 100152684 | rs12115447 | -0.164 | 0.372 | -0.111 | 1.18 | 0.015 | 0.062 | 0.062 | 0 | 0.18 | 0.29 | 0.36 | 0.37 | 0.19 | 0.29 |
| 9 | 100153594 | rs10817513 | 0.911 | 0.362 | -0.645 | 0.986 | 0 | 0.131 | 0.131 | 0.358 | 0.31 | 0.03 | 0.2 | 0.08 | 0.19 | 0.25 |
| 9 | 100158052 | rs1020884 | 0.1 | 0.292 | -0.273 | 1.12 | -0.008 | 0.057 | 0.057 | 2.787 | 0.47 | 0.17 | 0.54 | 0.35 | 0.38 | 0.51 |
| 9 | 100161524 | rs10759658 | 0.072 | 0.216 | -0.077 | 1.219 | -0.011 | 0.01 | 0.01 | 0.426 | 0.73 | 0.4 | 0.73 | 0.66 | 0.72 | 0.7 |
| 9 | 100163998 | rs11791849 | 0.315 | 0.485 | 0.066 | 1.099 | -0.002 | -0.011 | -0.011 | 0 | 0.29 | 0.38 | 0.21 | 0.25 | 0.33 | 0.17 |
| 9 | 100167429 | rs1381533 | 0.387 | 0.446 | -0.004 | 1.166 | -0.001 | -0.004 | -0.004 | 0 | 0.28 | 0.34 | 0.25 | 0.29 | 0.34 | 0.24 |
| 9 | 100169852 | rs1462091 | 0.381 | 0.443 | -0.005 | 1.165 | -0.001 | -0.004 | -0.004 | 0 | 0.28 | 0.34 | 0.25 | 0.29 | 0.34 | 0.24 |
| 9 | 100172620 | rs12551443 | -0.007 | 0.977 | -0.494 | 0.953 | -0.055 | 0.027 | 0.027 | 0 | 0.1 | 0.02 | 0.05 | 0.03 | 0.13 | 0.04 |
| 9 | 100188129 | rs7853581 | -0.18 | 0.789 | -0.102 | 1.257 | 0.004 | 0.001 | 0.001 | 0 | 0.39 | 0.6 | 0.6 | 0.58 | 0.51 | 0.41 |
| 9 | 100192986 | rs13286028 | 1.279 | 0.698 | 0.005 | 1.326 | -0.004 | 0.004 | 0.004 | 1.546 | 0.23 | 0.19 | 0.23 | 0.18 | 0.3 | 0.17 |
| 9 | 100194406 | rs2045732 | -0.235 | 0.631 | -0.093 | 1.267 | 0.002 | 0.001 | 0.001 | 0 | 0.39 | 0.61 | 0.6 | 0.58 | 0.51 | 0.41 |
| 9 | 100203708 | rs17423994 | 1.803 | 0.203 | -0.46 | 0.82 | -0.028 | 0.121 | 0.121 | 0 | 0.29 | 0.01 | 0.17 | 0.07 | 0.19 | 0.22 |
| 9 | 100211869 | rs4557815 | -0.99 | 0.628 | 0.143 | 1.288 | -0.001 | -0.025 | -0.025 | 0 | 0.87 | 0.94 | 0.93 | 0.8 | 0.86 | 0.74 |
| 9 | 100220058 | rs7871839 | 0.869 | 0.974 | 0.432 | 1.317 | 0 | -0.004 | -0.004 | 0 | 0.23 | 0.23 | 0.23 | 0.18 | 0.3 | 0.17 |
| 9 | 100226631 | rs7020902 | -0.224 | 0.438 | 0.269 | 1.061 | 0.001 | 0 | 0 | 0.025 | 0.39 | 0.62 | 0.6 | 0.58 | 0.52 | 0.43 |
| 9 | 100233189 | rs4742693 | 0.087 | 0.693 | -0.082 | 0.955 | 0.021 | 0.025 | 0.025 | 0.24 | 0.34 | 0.42 | 0.57 | 0.47 | 0.49 | 0.36 |
| 9 | 100253819 | rs10817671 | 0.062 | 0.76 | -0.272 | 0.86 | 0.008 | 0.008 | 0.008 | 0 | 0.34 | 0.38 | 0.57 | 0.5 | 0.5 | 0.37 |
| 9 | 100269291 | rs2149981 | -0.774 | 0.675 | -0.427 | 0.419 | 0.001 | -0.016 | -0.016 | 0 | 0.16 | 0.19 | 0.08 | 0.05 | 0.17 | 0.09 |
| 9 | 100270198 | rs10982444 | -0.212 | 0.515 | -0.26 | 0.725 | 0.005 | -0.004 | -0.004 | 0.071 | 0.41 | 0.5 | 0.55 | 0.64 | 0.51 | 0.45 |
| 9 | 100270886 | rs11788529 | -0.448 | 0.888 | -0.526 | 1.056 | -0.01 | -0.011 | -0.011 | 0 | 0.33 | 0.24 | 0.45 | 0.39 | 0.41 | 0.43 |
| 9 | 100279009 | rs7862883 | 0.981 | 0.845 | -0.618 | 1.151 | -0.011 | -0.04 | -0.04 | 0 | 0.16 | 0.07 | 0.13 | 0.3 | 0.21 | 0.22 |
| 9 | 100282313 | rs953417 | -0.054 | 0.887 | -0.416 | 1.243 | -0.002 | -0.004 | -0.004 | 2.48 | 0.73 | 0.73 | 0.63 | 0.63 | 0.66 | 0.63 |
| 9 | 100285297 | rs10759754 | 0.572 | 0.476 | 0.017 | 1.142 | 0 | 0.021 | 0.021 | 0 | 0.65 | 0.45 | 0.49 | 0.54 | 0.64 | 0.51 |
| 9 | 100285653 | rs10982508 | 0.653 | 0.651 | -0.186 | 0.744 | 0 | 0 | 0 | 0 | 0.25 | 0.26 | 0.18 | 0.27 | 0.34 | 0.21 |
| 9 | 100287073 | rs7019830 | -0.15 | 0.565 | -0.428 | 0.532 | -0.003 | 0.004 | 0.004 | 0.46 | 0.73 | 0.76 | 0.64 | 0.79 | 0.67 | 0.63 |
| 9 | 100296219 | rs2149987 | -0.311 | 0.473 | -0.591 | 0.434 | 0.006 | 0.012 | 0.012 | 0 | 0.51 | 0.56 | 0.43 | 0.64 | 0.61 | 0.54 |
| 9 | 100297592 | rs913913 | -1.188 | 0.45 | -1.278 | 0.376 | -0.048 | -0.04 | -0.04 | 0.178 | 0.87 | 0.96 | 0.8 | 0.82 | 0.77 | 0.83 |
| 9 | 100297695 | rs1007070 | 0.953 | 0.111 | -1.124 | 0.043 | 0.075 | 0.113 | 0.113 | 0 | 0.32 | 0.13 | 0.22 | 0.05 | 0.15 | 0.2 |
| 9 | 100299775 | rs10982602 | 0.747 | 1.025 | -1.051 | 0.509 | -0.038 | -0.015 | -0.015 | 1.308 | 0.06 | 0 | 0.06 | 0.05 | 0.13 | 0.05 |
| 9 | 100305357 | rs1571567 | -0.079 | 0.967 | -0.773 | 0.408 | 0.006 | 0.008 | 0.008 | 0 | 0.47 | 0.39 | 0.36 | 0.61 | 0.44 | 0.41 |
| 9 | 100305698 | rs10817736 | 1.494 | 0.916 | -0.976 | 0.18 | 0.006 | 0.121 | 0.121 | 0 | 0.28 | 0.08 | 0.26 | 0.06 | 0.23 | 0.25 |
| 9 | 100306267 | rs10817737 | -0.036 | 1.17 | -1.043 | 0.026 | -0.001 | -0.001 | -0.001 | 0 | 0.48 | 0.43 | 0.34 | 0.61 | 0.41 | 0.4 |
| 9 | 100307789 | rs7045553 | 0.006 | 1.097 | -1.015 | 0.209 | 0 | 0.001 | 0.001 | 0 | 0.48 | 0.48 | 0.36 | 0.61 | 0.45 | 0.4 |
| 9 | 100308231 | rs10817739 | 1.413 | 1.213 | -1.762 | 0.096 | 0.007 | 0.109 | 0.109 | 0 | 0.28 | 0.09 | 0.26 | 0.05 | 0.23 | 0.25 |
| 9 | 100310534 | rs7867486 | -0.794 | 1.577 | -1.608 | 0.608 | -0.079 | -0.046 | -0.046 | 1.763 | 0.86 | 0.99 | 0.79 | 0.84 | 0.76 | 0.86 |
| 9 | 100310666 | rs10982661 | 0.903 | 0.891 | -1.333 | 1.234 | -0.011 | -0.088 | -0.088 | 0.409 | 0.16 | 0.03 | 0.08 | 0.31 | 0.14 | 0.07 |
| 9 | 100310711 | rs7855669 | -0.494 | 0.983 | -0.487 | 0.843 | -0.002 | -0.002 | -0.002 | 0 | 0.28 | 0.29 | 0.23 | 0.29 | 0.31 | 0.26 |
| 9 | 100312284 | rs4742698 | 0.508 | 1.405 | 0.093 | 1.265 | 0.014 | 0.134 | 0.134 | 0.063 | 0.85 | 0.69 | 0.58 | 0.62 | 0.82 | 0.87 |
| 9 | 100313725 | rs7875496 | 0.932 | 1.268 | -0.724 | 0.774 | 0.027 | 0 | 0 | 1.205 | 0.36 | 0.14 | 0.2 | 0.29 | 0.26 | 0.28 |
| 9 | 100313793 | rs4743106 | -0.146 | 1.335 | -0.518 | 1.083 | 0.011 | 0.066 | 0.066 | 0.039 | 0.86 | 0.73 | 0.71 | 0.74 | 0.81 | 0.85 |
| 9 | 100313903 | rs4743107 | 0.888 | 1.767 | -0.42 | 0.663 | -0.018 | 0.007 | 0.007 | 0 | 0.3 | 0.14 | 0.18 | 0.26 | 0.31 | 0.33 |
| 9 | 100316230 | rs1475545 | 0.026 | 1.699 | -0.377 | 0.602 | -0.003 | 0.002 | 0.002 | 0.165 | 0.57 | 0.59 | 0.41 | 0.36 | 0.51 | 0.53 |
| 9 | 100318458 | rs913911 | -0.558 | 2.21 | -0.306 | 0.655 | 0.026 | 0.035 | 0.035 | 0.01 | 0.86 | 0.83 | 0.7 | 0.74 | 0.75 | 0.85 |
| 9 | 100319031 | rs10817759 | 1.147 | 2.037 | -0.374 | 0.562 | 0.009 | -0.006 | -0.006 | 0 | 0.27 | 0.12 | 0.14 | 0.23 | 0.23 | 0.32 |
| 9 | 100319894 | rs7871185 | -0.213 | 1.908 | -0.143 | 0.361 | 0.015 | -0.054 | -0.054 | 0.352 | 0.23 | 0.44 | 0.33 | 0.2 | 0.26 | 0.25 |

| | | | | | | | | | | | | | | | | |
|---|---|---|---|---|---|---|---|---|---|---|---|---|---|---|---|---|
| 9 | 100324654 | rs10759781 | 0.006 | 1.829 | 0.337 | 0.518 | 0 | 0.001 | 0.001 | 0.199 | 0.57 | 0.51 | 0.53 | 0.51 | 0.53 | 0.55 |
| 9 | 100326824 | rs4743110 | -0.981 | 2.044 | 0.543 | 0.636 | 0.025 | 0.075 | 0.075 | 0.099 | 0.89 | 0.83 | 0.73 | 0.76 | 0.83 | 0.9 |
| 9 | 100327714 | rs4743114 | 0.99 | 1.82 | 0.451 | 0.509 | 0.046 | 0.009 | 0.009 | 0 | 0.29 | 0.11 | 0.14 | 0.25 | 0.21 | 0.44 |
| 9 | 100330140 | rs10817781 | 0.468 | 1.829 | 1.078 | 0.479 | 0.004 | -0.029 | -0.029 | 1.674 | 0.34 | 0.49 | 0.24 | 0.25 | 0.35 | 0.46 |
| 9 | 100333671 | rs1475543 | 1.245 | 1.46 | 1.015 | 0.294 | 0.021 | -0.004 | -0.004 | 0 | 0.33 | 0.25 | 0.15 | 0.32 | 0.17 | 0.22 |
| 9 | 100333772 | rs1536950 | 0.197 | 1.726 | 1.068 | 0.192 | -0.006 | -0.006 | -0.006 | 0 | 0.19 | 0.19 | 0.2 | 0.06 | 0.28 | 0.21 |
| 9 | 100334586 | rs2149986 | -0.257 | 1.752 | 1.081 | 0.297 | 0.001 | -0.001 | -0.001 | 0 | 0.54 | 0.47 | 0.48 | 0.52 | 0.45 | 0.44 |
| 9 | 100341038 | rs913912 | -0.247 | 1.267 | 1.257 | 0.38 | 0.019 | 0.004 | 0.004 | 1.927 | 0.5 | 0.63 | 0.52 | 0.51 | 0.44 | 0.4 |
| 9 | 100353801 | rs2805792 | -0.285 | 1.368 | 1.449 | 0.308 | 0.008 | -0.083 | -0.083 | 4.715 | 0.14 | 0.4 | 0.19 | 0.06 | 0.18 | 0.18 |
| 9 | 100357256 | rs2805785 | -0.433 | 0.842 | 1.321 | 0.362 | -0.002 | -0.013 | -0.013 | 1.849 | 0.16 | 0.45 | 0.32 | 0.21 | 0.2 | 0.19 |
| 9 | 100357658 | rs2773343 | -0.172 | 0.848 | 1.31 | 0.336 | 0 | -0.065 | -0.065 | 0.017 | 0.14 | 0.31 | 0.18 | 0.05 | 0.2 | 0.18 |
| 9 | 100361864 | rs958346 | -0.714 | 0.938 | 0.801 | 0.215 | -0.003 | -0.023 | -0.023 | 0 | 0.65 | 0.77 | 0.67 | 0.51 | 0.65 | 0.5 |
| 9 | 100367306 | rs952765 | 1.188 | 0.748 | 0.602 | -0.011 | -0.067 | 0.061 | 0.061 | 0 | 0.17 | 0.06 | 0.23 | 0 | 0.32 | 0.09 |
| 9 | 100381034 | rs2805777 | -0.22 | 0.612 | 0.693 | 0.119 | 0.007 | 0.003 | 0.003 | 0 | 0.6 | 0.36 | 0.53 | 0.37 | 0.67 | 0.48 |
| 9 | 100388119 | rs10817858 | 1.17 | 0.612 | 0.528 | 0.282 | -0.009 | -0.011 | -0.011 | 0.374 | 0.24 | 0.1 | 0.12 | 0.25 | 0.22 | 0.41 |
| 9 | 100388197 | rs2773347 | -0.582 | 0.598 | 0.784 | 0.342 | -0.001 | -0.023 | -0.023 | 0 | 0.65 | 0.77 | 0.7 | 0.52 | 0.67 | 0.5 |
| 9 | 100399132 | rs2773350 | 0.496 | 0.556 | 0.902 | 0.203 | -0.001 | -0.05 | -0.05 | 3.314 | 0.13 | 0.21 | 0.17 | 0.06 | 0.19 | 0.18 |
| 9 | 100420142 | rs715749 | -1.267 | 0.691 | 0.987 | 0.322 | 0.001 | 0.06 | 0.06 | 0.006 | 0.89 | 0.87 | 0.81 | 0.77 | 0.88 | 0.9 |
| 9 | 100428977 | rs10116536 | 1.241 | 0.732 | 0.936 | 0.227 | -0.067 | 0.062 | 0.062 | 0 | 0.17 | 0.06 | 0.22 | 0 | 0.31 | 0.09 |
| 9 | 100437058 | rs3176757 | 1.485 | 0.519 | 0.885 | 0.203 | -0.016 | -0.018 | -0.018 | 0 | 0.22 | 0.1 | 0.11 | 0.25 | 0.22 | 0.39 |
| 9 | 100459578 | rs1800975 | -0.641 | 0.661 | 0.803 | -0.011 | -0.004 | -0.039 | -0.039 | 0 | 0.66 | 0.81 | 0.7 | 0.51 | 0.67 | 0.5 |
| 9 | 100462409 | rs10817938 | 2.162 | 0.057 | 0.082 | -0.164 | 0.015 | -0.06 | -0.06 | 0.606 | 0.11 | 0 | 0.06 | 0.19 | 0.03 | 0.07 |
| 9 | 100467594 | rs2808672 | 0.286 | 0.28 | 0.502 | 0.062 | 0.052 | 0.03 | 0.03 | 3.4 | 0.46 | 0.32 | 0.3 | 0.34 | 0.35 | 0.37 |
| 9 | 100468218 | rs2808673 | -0.83 | 0.417 | 0.621 | 0.065 | 0.001 | -0.016 | -0.016 | 0 | 0.65 | 0.77 | 0.69 | 0.49 | 0.67 | 0.47 |
| 9 | 100480125 | rs7849509 | 1.05 | 0.57 | 0.555 | -0.026 | -0.035 | 0.022 | 0.022 | 0 | 0.16 | 0.13 | 0.22 | 0 | 0.32 | 0.09 |
| 9 | 100482503 | rs2668797 | 0.218 | 0.255 | 0.683 | -0.102 | 0.03 | -0.028 | -0.028 | 0 | 0.26 | 0.36 | 0.51 | 0.21 | 0.43 | 0.25 |
| 9 | 100482976 | rs953199 | 0.748 | 0.104 | 0.213 | -0.009 | 0.048 | 0.013 | 0.013 | 0.837 | 0.37 | 0.22 | 0.16 | 0.27 | 0.24 | 0.22 |
| 9 | 100498831 | rs2668802 | 0.088 | 0.517 | 0.439 | -0.166 | 0.037 | -0.034 | -0.034 | 0 | 0.26 | 0.39 | 0.5 | 0.21 | 0.43 | 0.26 |
| 9 | 100505830 | rs2808699 | 0.133 | 0.543 | 0.258 | -0.454 | 0 | -0.024 | -0.024 | 0.004 | 0.24 | 0.27 | 0.37 | 0.06 | 0.43 | 0.24 |
| 9 | 100507159 | rs2805838 | -0.289 | 0.127 | 0.086 | -0.206 | -0.008 | -0.015 | -0.015 | 0 | 0.45 | 0.5 | 0.54 | 0.79 | 0.4 | 0.63 |
| 9 | 100511331 | rs16924016 | 1.181 | -0.12 | 0.313 | 0.445 | -0.01 | 0.068 | 0.068 | 0 | 0.08 | 0.11 | 0.1 | 0.21 | 0.07 | 0.07 |
| 9 | 100512420 | rs10818021 | -0.049 | -0.118 | 0.174 | 0.255 | -0.019 | -0.003 | -0.003 | 0.128 | 0.34 | 0.46 | 0.39 | 0.7 | 0.27 | 0.51 |
| 9 | 100514919 | rs10818023 | 1.326 | -0.509 | -0.267 | 0.112 | 0.007 | -0.059 | -0.059 | 0 | 0.24 | 0.01 | 0.12 | 0.18 | 0.15 | 0.2 |
| 9 | 100520013 | rs1398230 | 0.099 | -0.572 | -0.116 | -0.023 | -0.011 | -0.006 | -0.006 | 0 | 0.34 | 0.42 | 0.38 | 0.72 | 0.26 | 0.52 |
| 9 | 100521585 | rs12377067 | 1.16 | -0.299 | -0.7 | 0.215 | 0.024 | 0.005 | 0.005 | 0.066 | 0.25 | 0.07 | 0.15 | 0.17 | 0.16 | 0.2 |
| 9 | 100527157 | rs7034310 | 0.14 | -0.288 | -0.414 | 0.164 | -0.001 | -0.001 | -0.001 | 0 | 0.44 | 0.39 | 0.35 | 0.43 | 0.36 | 0.32 |
| 9 | 100528320 | rs7357631 | -0.193 | 0.264 | -0.697 | -0.648 | -0.021 | 0.024 | 0.024 | 0 | 0.69 | 0.77 | 0.64 | 0.91 | 0.55 | 0.77 |
| 9 | 100543880 | rs1877432 | 0.367 | 0.109 | -0.855 | -0.16 | -0.001 | 0 | 0 | 0.01 | 0.36 | 0.34 | 0.29 | 0.33 | 0.36 | 0.28 |
| 9 | 100549013 | rs7850258 | -0.57 | 0.252 | -1.34 | -0.16 | -0.025 | 0.078 | 0.078 | 0 | 0.69 | 0.87 | 0.69 | 0.87 | 0.66 | 0.81 |
| 9 | 100550028 | rs1443438 | -0.56 | 0.239 | -1.346 | 0.066 | -0.025 | 0.079 | 0.079 | 0 | 0.69 | 0.87 | 0.69 | 0.87 | 0.66 | 0.81 |
| 9 | 100550227 | rs12347079 | 0.686 | 0.16 | -1.168 | 0.003 | -0.004 | 0.026 | 0.026 | 0 | 0.27 | 0.32 | 0.36 | 0.54 | 0.23 | 0.53 |
| 9 | 100550253 | rs10818071 | 0.788 | 0.237 | -1.34 | -0.213 | -0.009 | 0.025 | 0.025 | 0 | 0.09 | 0.07 | 0.22 | 0.42 | 0.1 | 0.26 |
| 9 | 100556109 | rs965513 | -0.537 | 0.267 | -1.309 | -0.332 | -0.027 | 0.084 | 0.084 | 0 | 0.69 | 0.89 | 0.7 | 0.87 | 0.66 | 0.81 |
| 9 | 100556972 | rs10759944 | -0.517 | 0.273 | -1.31 | -0.33 | -0.024 | 0.08 | 0.08 | 0 | 0.69 | 0.89 | 0.7 | 0.87 | 0.66 | 0.81 |
| 9 | 100557291 | rs16924274 | 0.478 | 0.321 | -1.084 | 0.199 | -0.003 | 0.029 | 0.029 | 0.011 | 0.27 | 0.31 | 0.32 | 0.49 | 0.23 | 0.51 |
| 9 | 100571442 | rs7871887 | -0.569 | 0.729 | -1.442 | -0.568 | -0.003 | -0.051 | -0.051 | 0.054 | 0.17 | 0.24 | 0.18 | 0.01 | 0.16 | 0.1 |
| 9 | 100574120 | rs7870795 | -0.419 | 0.888 | -1.8 | -0.795 | -0.014 | 0.034 | 0.034 | 4.166 | 0.74 | 0.88 | 0.8 | 0.89 | 0.72 | 0.76 |
| 9 | 100585818 | rs10983871 | 0.264 | 0.802 | -1.798 | -0.814 | -0.002 | 0.09 | 0.09 | 0 | 0.05 | 0.18 | 0.04 | 0.1 | 0.04 | 0.06 |
| 9 | 100588839 | rs7848973 | 0.538 | 0.748 | -1.993 | -1.112 | -0.026 | 0.143 | 0.143 | 0 | 0.6 | 0.8 | 0.65 | 0.98 | 0.59 | 0.7 |
| 9 | 100596165 | rs10983893 | -0.03 | 0.771 | -2.128 | -1.01 | 0.007 | -0.008 | -0.008 | 0 | 0.17 | 0.17 | 0.14 | 0.01 | 0.14 | 0.12 |
| 9 | 100608682 | rs12348691 | 0.879 | 0.799 | -2.107 | -1.015 | 0.002 | 0.058 | 0.058 | 0.029 | 0.57 | 0.67 | 0.65 | 0.88 | 0.58 | 0.64 |
| 9 | 100608745 | rs10759960 | -0.386 | 0.792 | -2.114 | -1.008 | -0.011 | 0.032 | 0.032 | 4.364 | 0.74 | 0.83 | 0.8 | 0.95 | 0.72 | 0.77 |
| 9 | 100617433 | rs7043516 | -0.574 | 0.798 | -2.117 | -1.008 | -0.019 | -0.053 | -0.053 | 0.007 | 0.21 | 0.3 | 0.17 | 0.05 | 0.18 | 0.16 |
| 9 | 100617583 | rs1443435 | 0.899 | 0.786 | -2.118 | -1.01 | 0.002 | 0.059 | 0.059 | 0 | 0.57 | 0.67 | 0.65 | 0.88 | 0.58 | 0.64 |
| 9 | 100618660 | rs10984009 | 0.068 | 0.746 | -2.137 | -0.971 | 0.014 | 0.029 | 0.029 | 0 | 0.17 | 0.09 | 0.14 | 0.01 | 0.14 | 0.12 |
| 9 | 100636398 | rs925487 | 0.376 | 0.811 | -2.057 | -0.92 | -0.013 | 0.11 | 0.11 | 0 | 0.62 | 0.78 | 0.67 | 0.98 | 0.63 | 0.7 |
| 9 | 100637339 | rs10119795 | 0.995 | 1.082 | -2.376 | -0.999 | 0.027 | 0.081 | 0.081 | 0 | 0.46 | 0.67 | 0.5 | 0.82 | 0.39 | 0.59 |
| 9 | 100650096 | rs7866436 | 0.216 | 0.923 | -2.416 | -1.056 | -0.026 | 0.17 | 0.17 | 0 | 0.61 | 0.87 | 0.67 | 0.93 | 0.63 | 0.68 |
| 9 | 100664194 | rs7040232 | 0.73 | 1.169 | -2.45 | -0.988 | 0.007 | -0.005 | -0.005 | 0.541 | 0.15 | 0.16 | 0.18 | 0.11 | 0.25 | 0.09 |
| 9 | 100666931 | rs6586 | 1.05 | 1.149 | -2.471 | -1.018 | 0.044 | 0.112 | 0.112 | 0 | 0.46 | 0.74 | 0.49 | 0.82 | 0.38 | 0.59 |
| 9 | 100667599 | rs1561961 | 0.2 | 1.144 | -2.485 | -1.013 | -0.041 | 0.192 | 0.192 | 0 | 0.61 | 0.91 | 0.67 | 0.93 | 0.63 | 0.68 |
| 9 | 100669799 | rs1561958 | 1.002 | 1.134 | -2.38 | -0.966 | 0.036 | 0.087 | 0.087 | 0 | 0.46 | 0.71 | 0.49 | 0.82 | 0.37 | 0.59 |
| 9 | 100675816 | rs1127703 | 0.172 | 1.301 | -2.111 | -0.306 | -0.002 | 0.001 | 0.001 | 0 | 0.16 | 0.17 | 0.19 | 0.15 | 0.27 | 0.07 |
| 9 | 100681490 | rs10984361 | -1.126 | 1.237 | -2.232 | -0.407 | -0.063 | 0.105 | 0.105 | 0.282 | 0.32 | 0.13 | 0.34 | 0.16 | 0.41 | 0.19 |
| 9 | 100684757 | rs3183927 | -1.19 | 1.265 | -2.213 | -0.364 | -0.035 | 0.048 | 0.048 | 0.393 | 0.32 | 0.2 | 0.34 | 0.21 | 0.41 | 0.21 |
| 9 | 100688511 | rs7030887 | -1.097 | 1.321 | -2.377 | -0.36 | -0.064 | 0.101 | 0.101 | 0.477 | 0.32 | 0.13 | 0.34 | 0.16 | 0.41 | 0.19 |
| 9 | 100689693 | rs1059003 | -0.162 | 1.329 | -2.384 | -0.382 | 0.003 | 0.093 | 0.093 | 0 | 0.17 | 0.02 | 0.14 | 0.01 | 0.15 | 0.12 |
| 9 | 100691397 | rs1475696 | -1.097 | 1.321 | -2.374 | -0.363 | -0.064 | 0.086 | 0.086 | 0.464 | 0.32 | 0.13 | 0.34 | 0.16 | 0.41 | 0.2 |
| 9 | 100721555 | rs17687621 | -0.533 | 0.605 | -2.006 | -0.096 | 0.004 | 0.041 | 0.041 | 0 | 0.17 | 0.03 | 0.22 | 0.07 | 0.15 | 0.12 |

| | | | | | | | | | | | | | | | | |
|---|---|---|---|---|---|---|---|---|---|---|---|---|---|---|---|---|
| 9 | 100726574 | rs4743149 | -0.567 | 0.581 | -1.984 | -0.364 | 0.017 | 0.048 | 0.048 | 0 | 0.18 | 0.07 | 0.15 | 0.06 | 0.15 | 0.13 |
| 9 | 100739888 | rs10116976 | -0.186 | 0.914 | -2.544 | -0.899 | -0.076 | 0.15 | 0.15 | 0.621 | 0.77 | 0.99 | 0.86 | 1 | 0.72 | 0.84 |
| 9 | 100741115 | rs1572025 | 0.265 | 1.247 | -2.716 | -0.25 | -0.083 | -0.017 | -0.017 | 0 | 0.67 | 0.89 | 0.49 | 0.32 | 0.59 | 0.66 |
| 9 | 100742253 | rs7855088 | 0.265 | 1.247 | -2.662 | -0.25 | -0.075 | -0.013 | -0.013 | 0 | 0.67 | 0.88 | 0.49 | 0.32 | 0.59 | 0.66 |
| 9 | 100750623 | rs10984650 | 0.796 | 1.529 | -2.536 | -0.909 | -0.001 | 0 | 0 | 0 | 0.17 | 0.17 | 0.19 | 0.16 | 0.3 | 0.07 |
| 9 | 100763358 | rs7041878 | 1.535 | 0.586 | -2.535 | -1.614 | 0.037 | 0.046 | 0.046 | 0 | 0.25 | 0.14 | 0.06 | 0.09 | 0.12 | 0.44 |
| 9 | 100767860 | rs1010777 | -0.181 | 0.661 | -1.386 | -1.565 | 0.004 | -0.013 | -0.013 | 0 | 0.59 | 0.55 | 0.41 | 0.27 | 0.52 | 0.63 |
| 9 | 100784050 | rs10984811 | 0.83 | 1.032 | -1.513 | -1.283 | 0.012 | -0.002 | -0.002 | 0.005 | 0.18 | 0.22 | 0.19 | 0.16 | 0.3 | 0.07 |
| 9 | 100818344 | rs6478537 | -0.472 | 1.257 | -1.69 | -0.564 | -0.073 | 0.03 | 0.03 | 0.134 | 0.51 | 0.97 | 0.61 | 0.33 | 0.43 | 0.64 |
| 9 | 100818720 | rs6478539 | 0.517 | 1.276 | -1.371 | -0.567 | 0.047 | 0.062 | 0.062 | 0 | 0.4 | 0.68 | 0.26 | 0.12 | 0.21 | 0.56 |
| 9 | 100819070 | rs3199064 | -0.561 | 1.276 | -1.452 | -0.556 | -0.077 | 0.026 | 0.026 | 0.149 | 0.5 | 0.97 | 0.61 | 0.33 | 0.43 | 0.63 |
| 9 | 100823084 | rs13049 | -0.927 | 1.228 | -1.479 | -0.572 | -0.107 | 0.01 | 0.01 | 0 | 0.55 | 0.93 | 0.66 | 0.33 | 0.48 | 0.71 |
| 9 | 100828866 | rs4412474 | -0.922 | 1.256 | -1.446 | -0.359 | -0.122 | -0.04 | -0.04 | 0 | 0.54 | 0.96 | 0.66 | 0.36 | 0.47 | 0.72 |
| 9 | 100830392 | rs9918951 | -0.952 | 1.256 | -1.447 | -0.359 | -0.15 | -0.064 | -0.064 | 0 | 0.54 | 0.97 | 0.67 | 0.36 | 0.47 | 0.73 |
| 9 | 100834294 | rs7021804 | -0.717 | 1.266 | -1.309 | -0.354 | -0.084 | -0.006 | -0.006 | 0 | 0.53 | 0.93 | 0.66 | 0.36 | 0.47 | 0.72 |
| 9 | 100835163 | rs16912590 | -0.09 | 1.306 | -1.229 | -0.34 | 0.107 | 0.101 | 0.101 | 0 | 0.08 | 0.21 | 0.27 | 0.2 | 0.22 | 0.06 |
| 9 | 100836889 | rs10818468 | -0.09 | 1.309 | -1.229 | -0.336 | 0.104 | 0.098 | 0.098 | 0 | 0.08 | 0.21 | 0.27 | 0.2 | 0.22 | 0.06 |
| 9 | 100842005 | rs6478556 | 0.444 | 1.182 | -1.22 | -0.395 | 0.007 | 0.002 | 0.002 | 0 | 0.39 | 0.58 | 0.24 | 0.13 | 0.2 | 0.57 |
| 9 | 100843417 | rs4237190 | -0.959 | 1.194 | -1.234 | -0.373 | -0.123 | -0.05 | -0.05 | 0.413 | 0.54 | 0.96 | 0.67 | 0.36 | 0.48 | 0.72 |
| 9 | 100843660 | rs16912622 | 0.205 | 0.522 | -0.924 | -0.578 | 0.009 | 0.004 | 0.004 | 0 | 0.39 | 0.58 | 0.24 | 0.12 | 0.2 | 0.57 |
| 9 | 100868189 | rs7855251 | -0.186 | 0.759 | -1.052 | -0.669 | 0.052 | 0.084 | 0.084 | 0.001 | 0.48 | 0.6 | 0.27 | 0.11 | 0.28 | 0.59 |
| 9 | 100868900 | rs10760120 | 0.315 | 0.957 | -0.86 | -0.451 | 0.003 | -0.01 | -0.01 | 0 | 0.57 | 0.45 | 0.4 | 0.68 | 0.54 | 0.44 |
| 9 | 100869920 | rs9792455 | 0.062 | 1.249 | -0.991 | -0.257 | 0.007 | -0.006 | -0.006 | 0 | 0.64 | 0.6 | 0.4 | 0.69 | 0.53 | 0.49 |
| 9 | 100871717 | rs2417731 | -0.084 | 0.614 | -0.973 | -0.867 | -0.006 | -0.011 | -0.011 | 0 | 0.67 | 0.71 | 0.62 | 0.91 | 0.68 | 0.61 |
| 9 | 100884896 | rs12238311 | 0.377 | 0.163 | -0.687 | 0.142 | 0.026 | -0.047 | -0.047 | 0.99 | 0.24 | 0.05 | 0.06 | 0.27 | 0.14 | 0.18 |
| 9 | 100889340 | rs942165 | -0.317 | 0.847 | -0.403 | -0.174 | 0 | -0.037 | -0.037 | 0 | 0.15 | 0.25 | 0.08 | 0.09 | 0.13 | 0.23 |
| 9 | 100890591 | rs11794723 | 0.351 | 0.99 | -0.946 | 0.213 | -0.03 | -0.005 | -0.005 | 1.856 | 0.06 | 0 | 0.08 | 0.04 | 0.11 | 0.1 |
| 9 | 100895443 | rs735111 | 0.365 | 0.9 | -0.947 | 0.21 | -0.028 | -0.002 | -0.002 | 0.333 | 0.06 | 0 | 0.09 | 0.03 | 0.1 | 0.1 |
| 9 | 100896048 | rs10985166 | 0.318 | 0.899 | -0.948 | 0.212 | -0.028 | -0.005 | -0.005 | 0 | 0.06 | 0 | 0.09 | 0.04 | 0.1 | 0.1 |
| 9 | 100904179 | rs2795487 | -0.392 | 0.654 | -0.845 | 0.353 | 0.001 | -0.005 | -0.005 | 0.307 | 0.68 | 0.72 | 0.73 | 0.56 | 0.74 | 0.69 |
| 9 | 100905658 | rs3780465 | 0.354 | 0.685 | -1.318 | 0.636 | -0.007 | -0.075 | -0.075 | 1.415 | 0.12 | 0.02 | 0.19 | 0.3 | 0.13 | 0.08 |
| 9 | 100909931 | rs2795489 | -0.808 | 0.535 | -0.847 | 0.693 | 0 | -0.002 | -0.002 | 0 | 0.73 | 0.75 | 0.74 | 0.56 | 0.76 | 0.73 |
| 9 | 100909996 | rs2795490 | -0.287 | 0.637 | -0.895 | 0.644 | -0.007 | -0.014 | -0.014 | 0.028 | 0.68 | 0.74 | 0.66 | 0.51 | 0.66 | 0.66 |
| 9 | 100911422 | rs2795491 | 0.235 | 0.886 | -1.107 | -0.232 | -0.002 | -0.012 | -0.012 | 0.165 | 0.31 | 0.33 | 0.22 | 0.17 | 0.32 | 0.3 |
| 9 | 100911671 | rs814027 | -0.169 | 0.979 | -1.089 | -0.21 | -0.001 | 0.001 | 0.001 | 0 | 0.37 | 0.35 | 0.31 | 0.21 | 0.42 | 0.37 |
| 9 | 100913376 | rs2795492 | -0.076 | 1.173 | -0.824 | 0.537 | -0.052 | 0.063 | 0.063 | 0 | 0.63 | 0.82 | 0.55 | 0.73 | 0.55 | 0.62 |
| 9 | 100919318 | rs2250494 | 0.17 | 0.652 | -0.738 | 0.512 | -0.003 | -0.002 | -0.002 | 0 | 0.56 | 0.46 | 0.44 | 0.57 | 0.42 | 0.48 |
| 9 | 100922407 | rs3780457 | -0.137 | 0.238 | -1.187 | 0.413 | 0.011 | 0.006 | 0.006 | 0 | 0.15 | 0.07 | 0.09 | 0.1 | 0.12 | 0.06 |
| 9 | 100925136 | rs7862309 | -0.094 | -0.462 | -1.008 | 0.749 | -0.021 | 0.042 | 0.042 | 0 | 0.25 | 0.55 | 0.19 | 0.41 | 0.15 | 0.26 |
| 9 | 100927255 | rs10985257 | -0.142 | -1.012 | -1.616 | -0.091 | 0.025 | 0.042 | 0.042 | 0 | 0.31 | 0.23 | 0.17 | 0.11 | 0.25 | 0.15 |
| 9 | 100927632 | rs10491723 | -0.915 | -0.393 | -2.225 | 0.335 | -0.039 | 0.108 | 0.108 | 0.016 | 0.5 | 0.15 | 0.52 | 0.68 | 0.5 | 0.5 |
| 9 | 100927851 | rs10491724 | 1.292 | -0.293 | -2.402 | 0.222 | 0.004 | 0.043 | 0.043 | 0 | 0.06 | 0.08 | 0.08 | 0.18 | 0.05 | 0.09 |
| 9 | 100927886 | rs13300050 | -0.842 | -0.721 | -1.688 | -0.283 | 0 | 0.004 | 0.004 | 0 | 0.43 | 0.49 | 0.57 | 0.45 | 0.42 | 0.45 |
| 9 | 100928982 | rs2417733 | 0.923 | -0.779 | -1.674 | -0.255 | 0.011 | -0.066 | -0.066 | 0 | 0.28 | 0.13 | 0.21 | 0.43 | 0.22 | 0.24 |
| 9 | 100930253 | rs4743174 | -0.148 | -0.635 | -1.309 | -0.111 | -0.001 | -0.003 | -0.003 | 0 | 0.45 | 0.52 | 0.46 | 0.36 | 0.47 | 0.39 |
| 9 | 100930947 | rs10985279 | 1.68 | -0.536 | -1.302 | -0.107 | -0.008 | -0.063 | -0.063 | 2.364 | 0.14 | 0.03 | 0.12 | 0.32 | 0.1 | 0.16 |
| 9 | 100934033 | rs4610873 | -0.498 | -0.652 | -1.27 | -0.563 | 0.028 | 0.039 | 0.039 | 0 | 0.51 | 0.62 | 0.44 | 0.3 | 0.39 | 0.31 |
| 9 | 100935050 | rs2275710 | -0.58 | -0.495 | -1.197 | -0.431 | -0.043 | 0.072 | 0.072 | 0.425 | 0.62 | 0.79 | 0.6 | 0.76 | 0.56 | 0.61 |
| 9 | 100941315 | rs16913573 | -0.881 | -0.529 | 0.352 | 0.317 | -0.012 | 0.032 | 0.032 | 0 | 0.15 | 0.33 | 0.22 | 0.2 | 0.1 | 0.11 |
| 9 | 100942499 | rs16913581 | 0.283 | -0.51 | 0.057 | 0.244 | -0.003 | -0.015 | -0.015 | 1.56 | 0.08 | 0.09 | 0.13 | 0.01 | 0.05 | 0.02 |
| 9 | 100943397 | rs12552786 | 0.048 | -0.681 | -0.232 | 0.355 | -0.01 | -0.026 | -0.026 | 0 | 0.06 | 0.02 | 0.08 | 0.2 | 0.04 | 0.09 |
| 9 | 100945146 | rs10818575 | -1.21 | -0.679 | 0.321 | 0.201 | -0.008 | -0.011 | -0.011 | 0.486 | 0.26 | 0.6 | 0.28 | 0.16 | 0.27 | 0.3 |
| 9 | 100947796 | rs16913644 | 0.292 | -0.525 | 0.075 | 0.318 | -0.03 | -0.023 | -0.023 | 0 | 0.05 | 0 | 0.17 | 0.16 | 0.13 | 0.19 |
| 9 | 100951838 | rs7870439 | -0.013 | -0.709 | 0.004 | 0.371 | -0.001 | -0.003 | -0.003 | 0 | 0.23 | 0.26 | 0.2 | 0.14 | 0.21 | 0.2 |
| 9 | 100955714 | rs4743184 | -0.397 | -0.611 | -0.091 | 0.394 | -0.05 | -0.049 | -0.049 | 0 | 0.12 | 0.02 | 0.21 | 0.16 | 0.22 | 0.25 |
| 9 | 100960607 | rs10985391 | 0.606 | -0.643 | 0.387 | 0.224 | 0.016 | -0.04 | -0.04 | 0.019 | 0.08 | 0.21 | 0.06 | 0 | 0.11 | 0.08 |
| 9 | 100963653 | rs3739668 | -1.429 | -1.017 | 0.389 | 0.311 | -0.003 | -0.002 | -0.002 | 0.046 | 0.39 | 0.48 | 0.44 | 0.35 | 0.37 | 0.44 |
| 9 | 100964651 | rs3780453 | -2.016 | -1.001 | 0.288 | 0.302 | 0.009 | 0.049 | 0.049 | 0 | 0.49 | 0.75 | 0.5 | 0.36 | 0.49 | 0.52 |
| 9 | 100965472 | rs3739667 | -2.626 | -1.089 | 0.049 | 0.378 | -0.098 | -0.097 | -0.097 | 0.107 | 0.58 | 0.91 | 0.58 | 0.49 | 0.57 | 0.51 |
| 9 | 100970289 | rs3780447 | 1.005 | -1.059 | 0.091 | 0.421 | -0.041 | -0.065 | -0.065 | 0 | 0.14 | 0.01 | 0.16 | 0.2 | 0.11 | 0.12 |
| 9 | 100975694 | rs7854249 | 0.559 | -1.075 | 0.23 | 0.392 | 0.001 | -0.018 | -0.018 | 0 | 0.06 | 0.11 | 0.04 | 0 | 0.06 | 0.07 |
| 9 | 100976086 | rs4743186 | -2.493 | -1.114 | 0.232 | 0.406 | -0.034 | -0.028 | -0.028 | 7.246 | 0.56 | 0.78 | 0.57 | 0.5 | 0.57 | 0.54 |
| 9 | 100981757 | rs7041802 | -1.263 | -1.026 | 0.387 | 0.367 | 0.002 | -0.002 | -0.002 | 0.036 | 0.37 | 0.46 | 0.34 | 0.35 | 0.37 | 0.29 |
| 9 | 100984946 | rs12351801 | -1.555 | -0.728 | 0.628 | 0.74 | 0.009 | 0.005 | 0.005 | 0 | 0.36 | 0.65 | 0.37 | 0.29 | 0.39 | 0.35 |
| 9 | 100985312 | rs10818621 | -0.697 | -0.917 | 0.463 | 0.69 | -0.015 | -0.027 | -0.027 | 0 | 0.26 | 0.33 | 0.28 | 0.16 | 0.3 | 0.33 |
| 9 | 100985388 | rs10985507 | -0.646 | -0.746 | -0.118 | 0.816 | 0.001 | -0.015 | -0.015 | 0.003 | 0.24 | 0.27 | 0.27 | 0.32 | 0.21 | 0.13 |
| 9 | 100987622 | rs7868451 | -1.255 | -0.702 | -0.391 | 0.364 | 0.031 | 0.129 | 0.129 | 0 | 0.39 | 0.82 | 0.34 | 0.17 | 0.34 | 0.26 |
| 9 | 100991430 | rs1573025 | 0.437 | -0.558 | -0.26 | 0.449 | -0.006 | 0.004 | 0.004 | 0 | 0.17 | 0.14 | 0.2 | 0.16 | 0.18 | 0.16 |
| 9 | 100995758 | rs879368 | -0.363 | -0.564 | -0.173 | 0.412 | -0.01 | 0.019 | 0.019 | 0 | 0.24 | 0.17 | 0.25 | 0.16 | 0.27 | 0.22 |

| | | | | | | | | | | | | | | | | |
|---|---|---|---|---|---|---|---|---|---|---|---|---|---|---|---|---|
| 9 | 101002868 | rs10985573 | 0.064 | -0.539 | -0.149 | 0.425 | -0.001 | -0.026 | -0.026 | 0 | 0.08 | 0.13 | 0.05 | 0 | 0.1 | 0.06 |
| 9 | 101008541 | rs4743193 | -2.337 | -0.579 | -0.36 | 0.562 | -0.088 | -0.063 | -0.063 | 0 | 0.52 | 0.93 | 0.57 | 0.45 | 0.53 | 0.39 |
| 9 | 101008885 | rs7853442 | -1.581 | -1.373 | 0.086 | 0.139 | 0.026 | 0.115 | 0.115 | 0 | 0.39 | 0.78 | 0.34 | 0.18 | 0.34 | 0.26 |
| 9 | 101010704 | rs12349672 | -0.061 | -1.082 | 0.162 | 0.325 | 0.003 | -0.008 | -0.008 | 0 | 0.07 | 0.07 | 0.07 | 0.18 | 0.08 | 0.02 |
| 9 | 101024887 | rs10115971 | 0.213 | -1.177 | 0.313 | 0.238 | -0.002 | 0.015 | 0.015 | 0.745 | 0.2 | 0.12 | 0.2 | 0.16 | 0.18 | 0.17 |
| 9 | 101026356 | rs10818693 | 0.244 | -1.187 | 0.269 | 0.241 | -0.014 | 0.013 | 0.013 | 1.055 | 0.19 | 0.05 | 0.2 | 0.16 | 0.18 | 0.17 |
| 9 | 101034477 | rs10985680 | 0.389 | -1.184 | 0.26 | 0.242 | -0.02 | 0.008 | 0.008 | 1.018 | 0.19 | 0.05 | 0.2 | 0.16 | 0.18 | 0.17 |
| 9 | 101034626 | rs10760233 | -0.365 | -1.215 | 0.518 | 0.251 | -0.003 | -0.016 | -0.016 | 0 | 0.25 | 0.12 | 0.28 | 0.33 | 0.26 | 0.19 |
| 9 | 101040915 | rs10739615 | -1.163 | -1.22 | 0.536 | 0.252 | -0.001 | -0.002 | -0.002 | 0 | 0.35 | 0.4 | 0.35 | 0.33 | 0.35 | 0.25 |
| 9 | 101046441 | rs1435259 | -0.746 | -1.124 | 0.768 | 0.407 | -0.043 | 0.001 | 0.001 | 0 | 0.17 | 0.32 | 0.12 | 0.18 | 0.12 | 0.05 |
| 9 | 101056009 | rs10818743 | -0.39 | -0.34 | 0.529 | 0.526 | -0.008 | -0.027 | -0.027 | 0 | 0.22 | 0.38 | 0.22 | 0.16 | 0.19 | 0.17 |
| 9 | 101056122 | rs10985765 | -0.39 | -0.34 | 0.529 | 0.526 | -0.008 | -0.027 | -0.027 | 0 | 0.22 | 0.38 | 0.22 | 0.16 | 0.19 | 0.17 |
| 9 | 101063070 | rs7850445 | -0.795 | -0.227 | 0.316 | 0.829 | -0.005 | 0.002 | 0.002 | 0.055 | 0.32 | 0.39 | 0.32 | 0.36 | 0.26 | 0.34 |
| 9 | 101065807 | rs10985795 | -0.985 | -1.222 | 0.39 | 0.731 | 0.04 | 0.006 | 0.006 | 0 | 0.37 | 0.27 | 0.3 | 0.37 | 0.25 | 0.41 |
| 9 | 101066578 | rs13290518 | -0.985 | -1.203 | 0.39 | 0.812 | 0.039 | -0.001 | -0.001 | 0 | 0.37 | 0.27 | 0.3 | 0.4 | 0.25 | 0.41 |
| 9 | 101068580 | rs2304389 | -0.057 | -1.265 | 0.232 | 0.487 | 0.035 | 0.029 | 0.029 | 0.034 | 0.26 | 0.18 | 0.21 | 0.19 | 0.17 | 0.23 |
| 9 | 101071090 | rs10985817 | -0.051 | -1.242 | 0.213 | 0.498 | 0.025 | 0.021 | 0.021 | 0.128 | 0.26 | 0.2 | 0.21 | 0.2 | 0.17 | 0.23 |
| 9 | 101071562 | rs3780446 | -0.759 | -0.729 | -0.085 | 0.708 | 0 | -0.001 | -0.001 | 0.788 | 0.44 | 0.58 | 0.38 | 0.54 | 0.32 | 0.43 |
| 9 | 101084042 | rs13440299 | -0.733 | -0.1 | 0.369 | 0.403 | 0.002 | -0.021 | -0.021 | 1.573 | 0.06 | 0.2 | 0.04 | 0 | 0.06 | 0.1 |
| 9 | 101084087 | rs7037275 | -0.426 | -0.104 | -0.971 | 0.687 | -0.014 | -0.086 | -0.086 | 1.734 | 0.32 | 0.15 | 0.3 | 0.55 | 0.29 | 0.41 |
| 9 | 101092018 | rs3780438 | -1.21 | -0.253 | -1.629 | 0.729 | -0.077 | -0.08 | -0.08 | 0.003 | 0.61 | 0.86 | 0.43 | 0.54 | 0.57 | 0.63 |
| 9 | 101092316 | rs16914931 | 0.41 | -0.461 | -1.113 | 0.464 | 0.005 | -0.006 | -0.006 | 0.451 | 0.14 | 0.09 | 0.04 | 0.01 | 0.11 | 0.11 |
| 9 | 101094353 | rs16914943 | 0.23 | -0.275 | -1.01 | 0.347 | 0 | -0.022 | -0.022 | 0 | 0.16 | 0.12 | 0.07 | 0 | 0.16 | 0.12 |
| 9 | 101094546 | rs879354 | -0.697 | -0.226 | -0.31 | 0.307 | 0 | -0.002 | -0.002 | 0.387 | 0.43 | 0.55 | 0.32 | 0.5 | 0.45 | 0.41 |
| 9 | 101097869 | rs7871923 | 0.473 | -0.165 | 0.052 | 0.25 | -0.008 | -0.02 | -0.02 | 0 | 0.08 | 0.18 | 0.04 | 0.02 | 0.05 | 0.08 |
| 9 | 101101888 | rs7874366 | -1.954 | -0.278 | -0.266 | 0.383 | -0.078 | 0.073 | 0.073 | 0.794 | 0.64 | 0.94 | 0.52 | 0.76 | 0.62 | 0.75 |
| 9 | 101102091 | rs12236069 | -0.894 | -0.336 | 0.342 | 0.401 | 0.003 | -0.02 | -0.02 | 0 | 0.42 | 0.47 | 0.41 | 0.77 | 0.45 | 0.62 |
| 9 | 101106398 | rs653873 | -0.393 | -0.4 | -0.29 | -0.014 | -0.03 | -0.047 | -0.047 | 0 | 0.69 | 0.83 | 0.67 | 0.47 | 0.63 | 0.61 |
| 9 | 101113241 | rs471722 | 0.246 | -0.693 | 0.425 | 0.201 | -0.025 | 0.01 | 0.01 | 0 | 0.61 | 0.46 | 0.77 | 0.57 | 0.71 | 0.58 |
| 9 | 101117385 | rs608900 | 0.166 | -0.725 | 0.71 | -0.183 | -0.017 | -0.019 | -0.019 | 0 | 0.61 | 0.5 | 0.78 | 0.79 | 0.71 | 0.61 |
| 9 | 101125148 | rs514993 | -0.478 | 0.01 | 0.857 | -0.728 | 0.013 | -0.04 | -0.04 | 0 | 0.15 | 0.16 | 0.09 | 0.04 | 0.19 | 0.24 |
| 9 | 101126504 | rs2572364 | -1.965 | -0.437 | 0.124 | -0.996 | 0.022 | 0.142 | 0.142 | 0.574 | 0.42 | 0.78 | 0.24 | 0.19 | 0.36 | 0.38 |
| 9 | 101127379 | rs660148 | -1.235 | -0.467 | 0.5 | -1.016 | 0.006 | -0.035 | -0.035 | 0.614 | 0.28 | 0.56 | 0.17 | 0.17 | 0.28 | 0.34 |
| 9 | 101128858 | rs3780429 | -1.324 | -0.263 | -0.002 | -0.985 | 0.007 | 0.015 | 0.015 | 3.488 | 0.29 | 0.71 | 0.19 | 0.16 | 0.22 | 0.18 |
| 9 | 101128916 | rs3780428 | -1.35 | -0.237 | 0.01 | -0.97 | 0 | 0.006 | 0.006 | 4.423 | 0.29 | 0.68 | 0.22 | 0.16 | 0.28 | 0.16 |
| 9 | 101132441 | rs585819 | -1.619 | -0.332 | 0.099 | -1.016 | 0.043 | 0.063 | 0.063 | 0 | 0.39 | 0.7 | 0.18 | 0.16 | 0.27 | 0.38 |
| 9 | 101132481 | rs487910 | -1.275 | -0.341 | 0.231 | -1.011 | -0.013 | -0.024 | -0.024 | 0.045 | 0.33 | 0.55 | 0.14 | 0.16 | 0.19 | 0.33 |
| 9 | 101132921 | rs583641 | -1.101 | 0.055 | 0.281 | -1.026 | -0.005 | -0.014 | -0.014 | 0 | 0.27 | 0.23 | 0.18 | 0.04 | 0.33 | 0.35 |
| 9 | 101133138 | rs571569 | -1.228 | -0.015 | 0.297 | -0.964 | 0 | -0.036 | -0.036 | 1.607 | 0.27 | 0.29 | 0.15 | 0.09 | 0.28 | 0.37 |
| 9 | 101133518 | rs567931 | -1.842 | -0.115 | 0.033 | -0.996 | 0.022 | 0.04 | 0.04 | 0 | 0.38 | 0.7 | 0.19 | 0.17 | 0.3 | 0.38 |
| 9 | 101135643 | rs10818832 | -0.056 | -0.195 | -0.309 | -0.632 | 0.002 | 0.002 | 0.002 | 0 | 0.12 | 0.21 | 0.08 | 0.15 | 0.11 | 0.13 |
| 9 | 101137452 | rs661105 | -0.596 | -0.273 | -0.327 | -0.628 | -0.003 | -0.001 | -0.001 | 0 | 0.14 | 0.2 | 0.06 | 0.15 | 0.11 | 0.27 |
| 9 | 101138682 | rs10818845 | -0.655 | -0.057 | -0.095 | -0.296 | -0.012 | -0.016 | -0.016 | 0.445 | 0.26 | 0.68 | 0.29 | 0.17 | 0.2 | 0.12 |
| 9 | 101138898 | rs10760292 | -1.381 | 0.303 | -0.249 | -0.292 | 0.015 | 0.03 | 0.03 | 0 | 0.38 | 0.78 | 0.25 | 0.31 | 0.31 | 0.25 |
| 9 | 101143028 | rs3780421 | -0.7 | 0.21 | 0.333 | -0.209 | 0.004 | 0.015 | 0.015 | 2.184 | 0.16 | 0.5 | 0.1 | 0.17 | 0.12 | 0.11 |
| 9 | 101147503 | rs10986084 | 0.243 | 0.224 | 0.335 | -0.188 | 0.017 | 0.025 | 0.025 | 0.309 | 0.11 | 0.23 | 0.08 | 0.15 | 0.1 | 0.09 |
| 9 | 101168577 | rs7020345 | 0.285 | 0.446 | -0.554 | -0.179 | -0.006 | -0.002 | -0.002 | 0.252 | 0.19 | 0.15 | 0.17 | 0.19 | 0.24 | 0.15 |
| 9 | 101171682 | rs12235930 | -0.456 | -0.047 | -0.292 | 0.119 | -0.019 | -0.003 | -0.003 | 0 | 0.14 | 0.42 | 0.09 | 0.11 | 0.1 | 0.12 |
| 9 | 101172234 | rs1537960 | 0.71 | -0.04 | -0.919 | 0.221 | -0.073 | -0.137 | -0.137 | 1.972 | 0.22 | 0.02 | 0.51 | 0.37 | 0.28 | 0.21 |
| 9 | 101172694 | rs1537959 | -0.8 | -0.051 | -0.808 | 0.261 | -0.001 | -0.001 | -0.001 | 0.785 | 0.43 | 0.6 | 0.62 | 0.48 | 0.4 | 0.34 |
| 9 | 101174188 | rs10818896 | -0.871 | -0.215 | 0.119 | 0.004 | -0.045 | -0.014 | -0.014 | 0.349 | 0.16 | 0.46 | 0.09 | 0.11 | 0.1 | 0.13 |
| 9 | 101176060 | rs7022828 | -0.581 | 0.281 | -0.669 | -0.183 | -0.007 | -0.019 | -0.019 | 0 | 0.36 | 0.37 | 0.2 | 0.24 | 0.3 | 0.24 |
| 9 | 101177852 | rs2900382 | -0.388 | 0.512 | -1.704 | -1.089 | -0.031 | 0.186 | 0.186 | 0 | 0.31 | 0.09 | 0.17 | 0.01 | 0.33 | 0.27 |
| 9 | 101178629 | rs4743221 | -0.804 | 0.078 | -1.446 | -1.56 | -0.007 | -0.013 | -0.013 | 0 | 0.72 | 0.72 | 0.83 | 0.88 | 0.8 | 0.78 |
| 9 | 101179298 | rs10986167 | 0.679 | 0.247 | -0.764 | -1.099 | -0.002 | -0.001 | -0.001 | 0.018 | 0.3 | 0.23 | 0.5 | 0.7 | 0.36 | 0.45 |
| 9 | 101179742 | rs8181080 | -0.7 | 0.144 | -0.229 | -1.02 | -0.047 | -0.056 | -0.056 | 0 | 0.69 | 0.49 | 0.8 | 0.85 | 0.78 | 0.78 |
| 9 | 101188123 | rs4743222 | -0.215 | 0.298 | 0.038 | -0.402 | 0.012 | 0.156 | 0.156 | 0.829 | 0.08 | 0.22 | 0.23 | 0.34 | 0.07 | 0.08 |
| 9 | 101192990 | rs1537957 | -0.047 | 0.194 | -0.146 | -0.455 | 0.057 | -0.028 | -0.028 | 0.373 | 0.59 | 0.18 | 0.57 | 0.54 | 0.7 | 0.69 |
| 9 | 101202803 | rs7466271 | -0.987 | 1.352 | -0.299 | -0.248 | -0.032 | -0.002 | -0.002 | 0 | 0.94 | 0.99 | 0.91 | 0.91 | 0.85 | 0.88 |
| 9 | 101207390 | rs1415116 | 0.585 | 1.038 | -0.405 | -0.488 | 0.003 | 0.011 | 0.011 | 0 | 0.13 | 0.12 | 0.09 | 0.03 | 0.07 | 0.15 |
| 9 | 101210609 | rs10986308 | -0.168 | 0.926 | -0.335 | -0.538 | 0 | 0.009 | 0.009 | 0 | 0.2 | 0.26 | 0.3 | 0.36 | 0.16 | 0.35 |
| 9 | 101211767 | rs10818941 | 0.241 | 1.149 | -0.19 | -0.494 | -0.019 | -0.026 | -0.026 | 0 | 0.17 | 0.13 | 0.31 | 0.51 | 0.26 | 0.28 |
| 9 | 101212597 | rs7042032 | -0.781 | 0.782 | 0.053 | -0.591 | 0.017 | 0.127 | 0.127 | 0.782 | 0.07 | 0.31 | 0.15 | 0.19 | 0.04 | 0.05 |
| 9 | 101218784 | rs10818955 | 0.554 | 0.957 | -0.597 | -0.865 | -0.036 | 0.087 | 0.087 | 0 | 0.31 | 0.1 | 0.29 | 0.18 | 0.39 | 0.22 |
| 9 | 101219548 | rs1930420 | -0.585 | 0.762 | -0.738 | -0.6 | 0.017 | 0.015 | 0.015 | 0 | 0.58 | 0.36 | 0.7 | 0.68 | 0.66 | 0.74 |
| 9 | 101219931 | rs2808542 | -1.523 | 0.485 | -0.383 | -0.982 | 0.019 | -0.051 | -0.051 | 0.027 | 0.78 | 0.59 | 0.85 | 0.9 | 0.76 | 0.8 |
| 9 | 101220651 | rs1930419 | -0.323 | 0.675 | -0.707 | -1.278 | -0.006 | 0.045 | 0.045 | 0 | 0.55 | 0.39 | 0.44 | 0.21 | 0.49 | 0.41 |
| 9 | 101221143 | rs2808543 | -0.653 | 0.701 | -0.73 | -1.077 | -0.003 | 0.011 | 0.011 | 0 | 0.59 | 0.44 | 0.52 | 0.38 | 0.55 | 0.52 |
| 9 | 101229396 | rs2808553 | -0.725 | 0.716 | -0.801 | -1.042 | -0.002 | 0.004 | 0.004 | 0 | 0.6 | 0.43 | 0.52 | 0.37 | 0.55 | 0.53 |

| | | | | | | | | | | | | | | | | |
|---|---|---|---|---|---|---|---|---|---|---|---|---|---|---|---|---|
| 9 | 101230221 | rs2151214 | -1.858 | 0.757 | -0.805 | -1.118 | -0.034 | -0.069 | -0.069 | 0 | 0.75 | 0.58 | 0.87 | 0.9 | 0.82 | 0.79 |
| 9 | 101231892 | rs2779531 | -1.185 | 0.671 | -0.901 | -1.194 | 0.015 | 0.02 | 0.02 | 0 | 0.68 | 0.3 | 0.78 | 0.75 | 0.77 | 0.64 |
| 9 | 101235046 | rs12554364 | 1.167 | 0.775 | -0.91 | -1.129 | -0.019 | -0.018 | -0.018 | 0 | 0.14 | 0.11 | 0.35 | 0.51 | 0.27 | 0.24 |
| 9 | 101238684 | rs1930416 | -0.114 | 0.728 | -0.938 | -1.163 | -0.037 | 0.197 | 0.197 | 0 | 0.53 | 0.18 | 0.42 | 0.23 | 0.5 | 0.42 |
| 9 | 101238974 | rs1930415 | 0.945 | 0.863 | -0.752 | -0.884 | -0.002 | 0.009 | 0.009 | 0 | 0.15 | 0.13 | 0.35 | 0.52 | 0.27 | 0.26 |
| 9 | 101243852 | rs2779548 | -1.387 | 0.706 | -1.058 | -1.618 | 0 | -0.025 | -0.025 | 0.016 | 0.52 | 0.57 | 0.71 | 0.9 | 0.54 | 0.72 |
| 9 | 101248083 | rs13295431 | 1.558 | 0.683 | -1.321 | -1.675 | -0.026 | 0.089 | 0.089 | 0 | 0.15 | 0.01 | 0.11 | 0.02 | 0.12 | 0.15 |
| 9 | 101249102 | rs872091 | 0.542 | 0.813 | -1.189 | -1.628 | -0.021 | -0.015 | -0.015 | 1.13 | 0.19 | 0.15 | 0.43 | 0.67 | 0.32 | 0.4 |
| 9 | 101249423 | rs874278 | -0.165 | 0.62 | -1.006 | -1.483 | -0.013 | -0.013 | -0.013 | 0 | 0.27 | 0.33 | 0.26 | 0.22 | 0.19 | 0.2 |
| 9 | 101253226 | rs2779551 | 0.406 | 0.828 | -1 | -1.309 | -0.009 | 0.013 | 0.013 | 0.716 | 0.06 | 0.09 | 0.12 | 0.16 | 0.05 | 0.04 |
| 9 | 101254476 | rs2779552 | -0.594 | 0.731 | -0.839 | -1.743 | -0.001 | -0.007 | -0.007 | 0 | 0.7 | 0.79 | 0.78 | 0.91 | 0.65 | 0.74 |
| 9 | 101258803 | rs16916507 | 0.234 | 1.025 | -1.077 | -1.903 | 0 | -0.052 | -0.052 | 0 | 0.29 | 0.37 | 0.14 | 0.04 | 0.27 | 0.27 |
| 9 | 101258881 | rs1000440 | -0.89 | 1.106 | -1.19 | -1.78 | -0.001 | -0.005 | -0.005 | 3.127 | 0.6 | 0.7 | 0.46 | 0.2 | 0.61 | 0.42 |
| 9 | 101266669 | rs12552067 | 1.142 | 1.829 | -1.026 | -0.602 | -0.033 | -0.066 | -0.066 | 0.715 | 0.11 | 0.02 | 0.25 | 0.5 | 0.14 | 0.22 |
| 9 | 101267927 | rs10986473 | 0.443 | 1.738 | -0.83 | -0.939 | -0.005 | -0.011 | -0.011 | 0 | 0.26 | 0.24 | 0.19 | 0.01 | 0.3 | 0.14 |
| 9 | 101269571 | rs2779558 | -0.058 | 1.876 | -0.648 | -0.864 | 0.001 | -0.018 | -0.018 | 3.071 | 0.29 | 0.3 | 0.29 | 0.12 | 0.33 | 0.26 |
| 9 | 101270230 | rs13295101 | 1.359 | 2.127 | -0.556 | -0.993 | 0.025 | 0.04 | 0.04 | 0 | 0.47 | 0.34 | 0.19 | 0.18 | 0.35 | 0.42 |
| 9 | 101277609 | rs7857375 | 0.004 | 2.106 | -0.57 | -0.941 | 0 | -0.006 | -0.006 | 1.967 | 0.31 | 0.32 | 0.31 | 0.17 | 0.33 | 0.16 |
| 9 | 101277681 | rs2779562 | -1.155 | 2.212 | -0.587 | -0.913 | -0.008 | 0.073 | 0.073 | 0 | 0.46 | 0.35 | 0.66 | 0.76 | 0.51 | 0.52 |
| 9 | 101279351 | rs944761 | 0 | 2.196 | -0.573 | -0.915 | 0 | -0.007 | -0.007 | 0 | 0.31 | 0.32 | 0.31 | 0.17 | 0.33 | 0.16 |
| 9 | 101281710 | rs2779572 | -1.11 | 2.196 | -0.57 | -0.905 | -0.011 | 0.077 | 0.077 | 0 | 0.46 | 0.34 | 0.66 | 0.76 | 0.51 | 0.51 |
| 9 | 101284111 | rs2095121 | 0.402 | 2.31 | -0.383 | -0.509 | 0.137 | 0.04 | 0.04 | 0.097 | 0.93 | 0.68 | 0.85 | 0.94 | 0.86 | 0.94 |
| 9 | 101289991 | rs10760389 | 1.066 | 2.268 | -0.487 | -0.508 | 0.149 | 0.072 | 0.072 | 0.468 | 0.9 | 0.63 | 0.74 | 0.82 | 0.81 | 0.81 |
| 9 | 101293382 | rs10986577 | 0.647 | 1.313 | -1.119 | -0.333 | -0.032 | -0.083 | -0.083 | 0.032 | 0.12 | 0.02 | 0.24 | 0.48 | 0.14 | 0.22 |
| 9 | 101297144 | rs4237192 | 1.129 | 1.031 | -0.983 | -0.469 | -0.033 | -0.026 | -0.026 | 1.832 | 0.1 | 0.01 | 0.16 | 0.14 | 0.09 | 0.17 |
| 9 | 101298707 | rs11787646 | 0.339 | 1.231 | -1.066 | -0.734 | -0.013 | 0.02 | 0.02 | 0 | 0.14 | 0.08 | 0.09 | 0 | 0.18 | 0.11 |
| 9 | 101300786 | rs2779600 | -1.13 | 1.301 | -1.004 | -0.819 | -0.004 | 0.011 | 0.011 | 0 | 0.46 | 0.62 | 0.55 | 0.65 | 0.53 | 0.42 |
| 9 | 101303077 | rs2779602 | -1.752 | 1.403 | -0.814 | -0.634 | 0.012 | 0.035 | 0.035 | 0 | 0.52 | 0.64 | 0.71 | 0.79 | 0.59 | 0.54 |
| 9 | 101303770 | rs2808521 | 0.989 | 1.507 | -0.808 | -0.525 | -0.077 | -0.116 | -0.116 | 0 | 0.16 | 0.05 | 0.3 | 0.49 | 0.25 | 0.22 |
| 9 | 101308111 | rs1999498 | 1.25 | 1.047 | -0.556 | -0.352 | 0.012 | 0.022 | 0.022 | 0 | 0.47 | 0.37 | 0.2 | 0.15 | 0.37 | 0.42 |
| 9 | 101313371 | rs16917019 | -0.793 | 1.038 | -0.459 | -0.265 | -0.028 | -0.083 | -0.083 | 0.138 | 0.24 | 0.53 | 0.25 | 0.15 | 0.24 | 0.12 |
| 9 | 101317401 | rs2259639 | 0.823 | 1.107 | -0.658 | -0.116 | -0.097 | -0.138 | -0.138 | 3.287 | 0.23 | 0.05 | 0.34 | 0.48 | 0.31 | 0.24 |
| 9 | 101317921 | rs2900512 | -0.397 | 1.286 | -0.238 | 0.13 | 0.111 | 0.052 | 0.052 | 0 | 0.08 | 0.28 | 0.19 | 0.15 | 0.19 | 0.18 |
| 9 | 101318075 | rs884886 | 0.321 | 1.124 | -0.377 | 0.059 | 0.154 | 0.1 | 0.1 | 0 | 0.89 | 0.6 | 0.7 | 0.79 | 0.77 | 0.77 |
| 9 | 101320582 | rs10986740 | 0.823 | 0.804 | -1.249 | -0.311 | 0.022 | 0.036 | 0.036 | 0 | 0.43 | 0.24 | 0.27 | 0.29 | 0.35 | 0.45 |
| 9 | 101328200 | rs1167767 | -0.41 | 1.102 | -1.735 | 0.157 | -0.019 | -0.014 | -0.014 | 0 | 0.38 | 0.19 | 0.52 | 0.62 | 0.39 | 0.27 |
| 9 | 101333453 | rs928425 | -0.316 | 0.738 | -1.879 | 0.069 | -0.034 | 0.039 | 0.039 | 0 | 0.43 | 0.13 | 0.57 | 0.64 | 0.61 | 0.54 |
| 9 | 101333901 | rs7044074 | -0.886 | 0.408 | -0.937 | -0.325 | -0.016 | -0.059 | -0.059 | 1.304 | 0.1 | 0.31 | 0.04 | 0 | 0.05 | 0 |
| 9 | 101334556 | rs1167769 | 0.266 | 0.768 | -1.467 | -0.3 | 0.001 | 0.004 | 0.004 | 0 | 0.55 | 0.52 | 0.35 | 0.32 | 0.45 | 0.43 |
| 9 | 101334945 | rs2779523 | 0.173 | 0.547 | -1.791 | -0.676 | 0.054 | 0.086 | 0.086 | 0 | 0.56 | 0.84 | 0.3 | 0.26 | 0.4 | 0.42 |
| 9 | 101336041 | rs1179740 | -0.428 | 0.516 | -1.605 | -1.256 | 0.01 | 0.046 | 0.046 | 0 | 0.48 | 0.62 | 0.4 | 0.31 | 0.45 | 0.52 |
| 9 | 101339690 | rs1758799 | -1.422 | 0.646 | -1.692 | -0.955 | -0.1 | -0.117 | -0.117 | 0.346 | 0.77 | 0.98 | 0.68 | 0.39 | 0.72 | 0.78 |
| 9 | 101339947 | rs2808534 | -0.568 | 0.501 | -1.686 | -1.19 | -0.031 | 0.057 | 0.057 | 0 | 0.63 | 0.79 | 0.57 | 0.27 | 0.57 | 0.55 |
| 9 | 101340316 | rs3750344 | 1.12 | 0.415 | -1.197 | -1.06 | 0 | -0.01 | -0.01 | 0 | 0.19 | 0.24 | 0.09 | 0.15 | 0.18 | 0.1 |
| 9 | 101341534 | rs10986816 | 1.05 | 0.453 | -1.346 | -1.041 | 0 | -0.015 | -0.015 | 0 | 0.19 | 0.33 | 0.1 | 0.15 | 0.18 | 0.1 |
| 9 | 101344340 | rs2779527 | 0.487 | 0.497 | -1.624 | -0.804 | -0.025 | -0.017 | -0.017 | 0 | 0.34 | 0.2 | 0.54 | 0.62 | 0.39 | 0.4 |
| 9 | 101345778 | rs2778908 | 0.779 | 0.274 | -1.829 | -0.711 | -0.044 | 0.011 | 0.011 | 0 | 0.11 | 0.01 | 0.3 | 0.09 | 0.13 | 0.21 |
| 9 | 101346168 | rs1167763 | -0.543 | -0.034 | -1.015 | -0.541 | -0.012 | 0.018 | 0.018 | 0 | 0.89 | 0.76 | 0.86 | 0.86 | 0.92 | 0.86 |
| 9 | 101346548 | rs10123407 | 0.124 | 0.342 | -1.448 | -0.598 | -0.008 | -0.008 | -0.008 | 0 | 0.13 | 0.03 | 0.1 | 0.12 | 0.13 | 0.2 |
| 9 | 101347219 | rs12337255 | 0.193 | 0.653 | -0.902 | -0.603 | -0.001 | 0.003 | 0.003 | 0 | 0.22 | 0.15 | 0.12 | 0 | 0.21 | 0.17 |
| 9 | 101350077 | rs2779529 | -0.114 | 0.685 | -0.319 | 0.096 | 0.007 | -0.009 | -0.009 | 0 | 0.34 | 0.49 | 0.51 | 0.66 | 0.36 | 0.31 |
| 9 | 101351051 | rs7860298 | 0.107 | 0.435 | 0.054 | -0.048 | -0.019 | -0.022 | -0.022 | 1.925 | 0.08 | 0.3 | 0.03 | 0 | 0.02 | 0.03 |
| 9 | 101351818 | rs10986852 | 0.943 | 0.579 | -0.049 | 0.382 | -0.03 | 0.035 | 0.035 | 0 | 0.22 | 0.14 | 0.26 | 0.14 | 0.29 | 0.19 |
| 9 | 101355624 | rs2808539 | -0.48 | 0.358 | 0.092 | 0.578 | 0 | -0.006 | -0.006 | 0 | 0.37 | 0.57 | 0.51 | 0.67 | 0.4 | 0.32 |
| 9 | 101356515 | rs1999500 | -0.911 | 0.242 | -0.234 | 0.046 | -0.001 | -0.001 | -0.001 | 0.75 | 0.72 | 0.7 | 0.71 | 0.79 | 0.69 | 0.69 |
| 9 | 101356663 | rs7035372 | 0.737 | 0.247 | -0.396 | 0.09 | 0.029 | 0.123 | 0.123 | 0 | 0.34 | 0.12 | 0.2 | 0.11 | 0.29 | 0.35 |
| 9 | 101361447 | rs10819071 | -0.141 | 0.645 | -0.337 | 0.232 | 0.008 | -0.003 | -0.003 | 0 | 0.67 | 0.62 | 0.5 | 0.3 | 0.65 | 0.6 |
| 9 | 101361584 | rs10819072 | -0.178 | 0.659 | -0.34 | 0.405 | 0.009 | 0.004 | 0.004 | 0 | 0.67 | 0.62 | 0.51 | 0.36 | 0.65 | 0.6 |
| 9 | 101363329 | rs914662 | 0.524 | 0.9 | -0.693 | 0.934 | -0.038 | 0.008 | 0.008 | 0 | 0.24 | 0.12 | 0.28 | 0.18 | 0.34 | 0.21 |
| 9 | 101366538 | rs944689 | -1.366 | 0.678 | 0.042 | 1.684 | 0 | 0.119 | 0.119 | 0.23 | 0.07 | 0.08 | 0.25 | 0.43 | 0.08 | 0.16 |
| 9 | 101379780 | rs2778913 | 1.25 | 1.114 | 0.664 | 1.494 | 0.095 | 0.024 | 0.024 | 0.209 | 0.9 | 0.67 | 0.82 | 0.85 | 0.87 | 0.83 |
| 9 | 101382838 | rs1167774 | 1.544 | 1.06 | 0.638 | 1.546 | 0.075 | 0.077 | 0.077 | 0.186 | 0.89 | 0.67 | 0.81 | 0.8 | 0.87 | 0.78 |
| 9 | 101384719 | rs6478785 | 1.125 | 1.101 | 0.496 | 1.584 | -0.016 | 0.035 | 0.035 | 0 | 0.93 | 0.96 | 0.65 | 0.41 | 0.9 | 0.69 |
| 9 | 101389408 | rs1930131 | 1.49 | 1.191 | 0.625 | 1.421 | 0.005 | 0.048 | 0.048 | 0.005 | 0.88 | 0.87 | 0.61 | 0.42 | 0.84 | 0.65 |
| 9 | 101406038 | rs7864144 | -1.846 | 1.113 | 0.869 | 1.393 | 0.084 | 0.042 | 0.042 | 0.289 | 0.12 | 0.39 | 0.11 | 0.09 | 0.14 | 0.09 |
| 9 | 101417785 | rs1967890 | -1.46 | 1.03 | 0.79 | 1.479 | 0.073 | 0.39 | 0.39 | 0 | 0.06 | 0.23 | 0.22 | 0.5 | 0.09 | 0.13 |
| 9 | 101427651 | rs12002737 | 0.5 | 0.666 | 0.235 | 0.556 | 0.028 | 0.03 | 0.03 | 0 | 0.28 | 0.09 | 0.14 | 0.19 | 0.22 | 0.27 |
| 9 | 101427796 | rs7865785 | -0.976 | 1.309 | 0.986 | 0.842 | 0.013 | 0.017 | 0.017 | 0 | 0.07 | 0.18 | 0.06 | 0.04 | 0.09 | 0.08 |
| 9 | 101433268 | rs1930136 | 1.446 | 1.345 | 0.961 | 0.839 | 0.017 | -0.014 | -0.014 | 0 | 0.89 | 0.81 | 0.81 | 0.89 | 0.84 | 0.75 |

| | | | | | | | | | | | | | | | | |
|---|---|---|---|---|---|---|---|---|---|---|---|---|---|---|---|---|
| 9 | 101436021 | rs10116255 | -0.842 | 1.266 | 1.075 | 0.839 | 0.093 | 0.015 | 0.015 | 1.88 | 0.05 | 0.26 | 0.05 | 0.04 | 0.06 | 0.01 |
| 9 | 101436463 | rs914667 | 0.576 | 0.993 | 0.804 | 0.886 | -0.016 | -0.02 | -0.02 | 0.034 | 0.94 | 0.98 | 0.87 | 0.86 | 0.93 | 0.79 |
| 9 | 101436894 | rs1105931 | 1.689 | 0.981 | 0.822 | 0.868 | 0.016 | 0.032 | 0.032 | 0 | 0.88 | 0.76 | 0.82 | 0.79 | 0.85 | 0.67 |
| 9 | 101437009 | rs1547271 | 0.76 | 1.003 | 0.151 | 1.043 | -0.019 | -0.024 | -0.024 | 0 | 0.94 | 0.99 | 0.87 | 0.83 | 0.93 | 0.71 |
| 9 | 101454321 | rs4517239 | 1.282 | 0.976 | 0.332 | 1.038 | 0.064 | 0.12 | 0.12 | 0.43 | 0.88 | 0.58 | 0.82 | 0.8 | 0.89 | 0.74 |
| 9 | 101491066 | rs380857 | -0.167 | 0.999 | -0.734 | 1.146 | -0.027 | -0.048 | -0.048 | 0 | 0.9 | 0.98 | 0.83 | 0.76 | 0.88 | 0.87 |
| 9 | 101507149 | rs16917884 | 0.044 | 1.284 | -0.856 | 1.921 | -0.024 | -0.025 | -0.025 | 0 | 0.08 | 0.02 | 0.05 | 0.13 | 0.12 | 0.14 |
| 9 | 101518442 | rs580809 | 0.106 | 1.054 | -0.758 | 1.812 | -0.024 | -0.054 | -0.054 | 0 | 0.81 | 0.91 | 0.74 | 0.56 | 0.77 | 0.73 |
| 9 | 101519554 | rs422531 | 0.085 | 1.053 | -0.758 | 1.508 | -0.024 | -0.04 | -0.04 | 1.859 | 0.81 | 0.91 | 0.78 | 0.64 | 0.77 | 0.73 |
| 9 | 101533973 | rs10819300 | 0.694 | 0.85 | -1.244 | 0.895 | -0.011 | 0.048 | 0.048 | 0 | 0.21 | 0.07 | 0.13 | 0.13 | 0.19 | 0.19 |
| 9 | 101534397 | rs12686258 | 1.192 | 0.946 | -1.757 | 0.532 | -0.077 | -0.05 | -0.05 | 0 | 0.17 | 0.02 | 0.26 | 0.22 | 0.24 | 0.16 |
| 9 | 101539601 | rs4742741 | 0.476 | 1.001 | -0.823 | 1.16 | -0.004 | -0.006 | -0.006 | 0 | 0.76 | 0.76 | 0.6 | 0.38 | 0.74 | 0.58 |
| 9 | 101550367 | rs7853904 | 0.042 | 0.918 | -0.777 | 1.183 | -0.041 | -0.036 | -0.036 | 0 | 0.71 | 0.86 | 0.54 | 0.4 | 0.66 | 0.55 |
| 9 | 101552955 | rs7027771 | 0.598 | 0.904 | -0.538 | 1.176 | 0.015 | -0.005 | -0.005 | 0.001 | 0.66 | 0.47 | 0.51 | 0.38 | 0.66 | 0.52 |
| 9 | 101553014 | rs7038640 | 0.254 | 0.978 | -1.229 | 1.216 | -0.015 | -0.014 | -0.014 | 0 | 0.69 | 0.75 | 0.53 | 0.4 | 0.66 | 0.55 |
| 9 | 101585992 | rs2295925 | -0.05 | 0.638 | -0.861 | 1.059 | -0.028 | -0.007 | -0.007 | 0.262 | 0.46 | 0.2 | 0.42 | 0.5 | 0.49 | 0.56 |
| 9 | 101590318 | rs911928 | 0.025 | 0.715 | -0.568 | 0.311 | 0.002 | 0.02 | 0.02 | 0 | 0.48 | 0.38 | 0.43 | 0.23 | 0.49 | 0.34 |
| 9 | 101593825 | rs2295926 | -0.488 | 1.044 | -0.262 | 0.184 | 0.022 | -0.039 | -0.039 | 0 | 0.33 | 0.52 | 0.34 | 0.2 | 0.39 | 0.28 |
| 9 | 101595443 | rs10987898 | -0.618 | 0.935 | 0.112 | 0.663 | -0.007 | -0.012 | -0.012 | 0 | 0.28 | 0.24 | 0.36 | 0.35 | 0.3 | 0.35 |
| 9 | 101598609 | rs1407503 | -0.411 | 0.739 | 0.16 | 0.58 | 0 | 0.003 | 0.003 | 0 | 0.3 | 0.36 | 0.21 | 0.36 | 0.3 | 0.37 |
| 9 | 101599758 | rs4743285 | -0.367 | 0.7 | 0.352 | 0.919 | 0.012 | 0.077 | 0.077 | 0 | 0.2 | 0.34 | 0.32 | 0.42 | 0.23 | 0.34 |
| 9 | 101601645 | rs999471 | -0.444 | 0.701 | 0.56 | 0.87 | 0.004 | 0.017 | 0.017 | 0 | 0.15 | 0.18 | 0.15 | 0.31 | 0.19 | 0.28 |
| 9 | 101603712 | rs10819377 | -0.23 | 0.247 | 0.546 | 0.389 | -0.021 | 0.007 | 0.007 | 0.853 | 0.42 | 0.3 | 0.44 | 0.39 | 0.44 | 0.4 |
| 9 | 101607247 | rs10987949 | -1.003 | -0.15 | 0.708 | 0.565 | -0.035 | 0.069 | 0.069 | 0 | 0.17 | 0.34 | 0.12 | 0.31 | 0.11 | 0.29 |
| 9 | 101608025 | rs10987952 | -0.115 | -0.126 | 0.581 | 0.626 | -0.017 | 0.053 | 0.053 | 0 | 0.11 | 0.17 | 0.1 | 0.29 | 0.08 | 0.25 |
| 9 | 101609706 | rs2295928 | -0.118 | -0.2 | 0.568 | 0.547 | -0.025 | 0.071 | 0.071 | 0 | 0.1 | 0.17 | 0.1 | 0.24 | 0.07 | 0.11 |
| 9 | 101609768 | rs2295929 | -0.826 | -0.195 | 0.645 | 0.546 | -0.033 | 0.062 | 0.062 | 0 | 0.15 | 0.26 | 0.11 | 0.24 | 0.1 | 0.12 |
| 9 | 101619191 | rs1322540 | -1.387 | -0.247 | 0.549 | 0.436 | -0.012 | -0.008 | -0.008 | 0 | 0.62 | 0.56 | 0.6 | 0.68 | 0.66 | 0.69 |
| 9 | 101619403 | rs7864581 | -1.401 | -0.252 | 0.62 | 0.433 | -0.014 | -0.01 | -0.01 | 0.029 | 0.62 | 0.55 | 0.6 | 0.68 | 0.66 | 0.69 |
| 9 | 101623189 | rs1157248 | -1.816 | -0.265 | 0.619 | 0.37 | 0.02 | 0.076 | 0.076 | 0.281 | 0.68 | 0.84 | 0.8 | 1 | 0.71 | 0.76 |
| 9 | 101625377 | rs10987993 | -0.174 | -0.06 | 0.64 | 0.36 | -0.005 | 0.039 | 0.039 | 0 | 0.13 | 0.17 | 0.13 | 0.24 | 0.14 | 0.11 |
| 9 | 101627022 | rs7875364 | -1.904 | 0.025 | 0.439 | 0.24 | 0.015 | 0.022 | 0.022 | 0.274 | 0.67 | 0.79 | 0.8 | 1 | 0.71 | 0.75 |
| 9 | 101629751 | rs10819389 | -0.161 | 0.018 | 0.08 | 0.272 | -0.038 | -0.006 | -0.006 | 0 | 0.39 | 0.3 | 0.45 | 0.44 | 0.47 | 0.52 |
| 9 | 101629892 | rs10739724 | -1.637 | -0.2 | 0.091 | 0.202 | 0.011 | 0.032 | 0.032 | 0.06 | 0.64 | 0.75 | 0.75 | 0.82 | 0.67 | 0.74 |
| 9 | 101634716 | rs2295932 | -0.652 | -0.279 | 0.197 | 0.832 | -0.052 | 0.071 | 0.071 | 0.031 | 0.31 | 0.46 | 0.38 | 0.56 | 0.25 | 0.49 |
| 9 | 101639369 | rs10819414 | -0.742 | 0.01 | -0.038 | 1.242 | -0.012 | 0.002 | 0.002 | 0 | 0.39 | 0.31 | 0.48 | 0.44 | 0.5 | 0.42 |
| 9 | 101647469 | rs16917989 | -0.406 | 0.131 | 0.318 | 0.411 | -0.013 | -0.082 | -0.082 | 0 | 0.16 | 0.25 | 0.05 | 0.01 | 0.1 | 0.03 |
| 9 | 101649752 | rs1007892 | 0.14 | 0.244 | 0.214 | 0.895 | -0.02 | -0.028 | -0.028 | 0.211 | 0.29 | 0.25 | 0.39 | 0.4 | 0.33 | 0.31 |
| 9 | 101651948 | rs1028653 | -1.089 | 0.126 | 0.34 | 0.979 | 0.001 | 0.018 | 0.018 | 0 | 0.27 | 0.5 | 0.22 | 0.36 | 0.29 | 0.24 |
| 9 | 101652106 | rs1028655 | -1.63 | 0.002 | -0.007 | 0.968 | 0.042 | 0.096 | 0.096 | 0 | 0.58 | 0.79 | 0.62 | 0.76 | 0.63 | 0.56 |
| 9 | 101655952 | rs2416541 | -0.108 | -0.074 | -0.307 | 1.039 | -0.005 | -0.006 | -0.006 | 0 | 0.35 | 0.35 | 0.42 | 0.4 | 0.38 | 0.33 |
| 9 | 101661037 | rs927468 | -1.467 | -0.308 | -1.375 | 0.688 | 0.02 | 0.112 | 0.112 | 0 | 0.58 | 0.87 | 0.61 | 0.79 | 0.61 | 0.56 |
| 9 | 101667612 | rs10819451 | -0.788 | -0.131 | -0.708 | 0.976 | -0.002 | 0.009 | 0.009 | 0 | 0.17 | 0.16 | 0.15 | 0.26 | 0.19 | 0.17 |
| 9 | 101678036 | rs7470221 | 0.098 | -0.136 | -1.796 | 0.52 | -0.139 | -0.112 | -0.112 | 0 | 0.37 | 0.04 | 0.37 | 0.55 | 0.5 | 0.4 |
| 9 | 101678096 | rs10123556 | -0.893 | -0.254 | -0.456 | 0.547 | -0.004 | -0.023 | -0.023 | 0 | 0.06 | 0.23 | 0.03 | 0 | 0.04 | 0.01 |
| 9 | 101678273 | rs716573 | -0.357 | -0.279 | -0.917 | 0.679 | 0.006 | -0.008 | -0.008 | 0.021 | 0.11 | 0.07 | 0.06 | 0.26 | 0.07 | 0.19 |
| 9 | 101685520 | rs10819460 | -0.889 | -0.326 | -0.256 | 0.455 | -0.028 | 0.016 | 0.016 | 0.244 | 0.16 | 0.33 | 0.12 | 0.25 | 0.12 | 0.19 |
| 9 | 101687361 | rs1555521 | 0.04 | 0.151 | -0.509 | 0.525 | -0.02 | -0.017 | -0.017 | 0 | 0.29 | 0.25 | 0.51 | 0.39 | 0.37 | 0.36 |
| 9 | 101688707 | rs10114761 | 0.12 | 0.056 | -0.515 | 0.585 | -0.037 | -0.045 | -0.045 | 0 | 0.29 | 0.15 | 0.52 | 0.43 | 0.37 | 0.38 |
| 9 | 101691988 | rs2065598 | -1.442 | -0.555 | -0.434 | -0.118 | 0.021 | 0.078 | 0.078 | 0 | 0.62 | 0.74 | 0.81 | 1 | 0.7 | 0.72 |
| 9 | 101694958 | rs10819472 | 0.665 | -0.439 | -0.317 | 0.256 | -0.089 | -0.039 | -0.039 | 0 | 0.26 | 0.13 | 0.31 | 0.28 | 0.37 | 0.3 |
| 9 | 101697289 | rs17783501 | 0.316 | -0.361 | -0.229 | 0.273 | -0.011 | -0.013 | -0.013 | 0 | 0.28 | 0.27 | 0.51 | 0.43 | 0.37 | 0.38 |
| 9 | 101707174 | rs911933 | 0.687 | -0.306 | -0.105 | 0.269 | -0.06 | -0.026 | -0.026 | 0 | 0.26 | 0.12 | 0.3 | 0.28 | 0.36 | 0.3 |
| 9 | 101708150 | rs6478963 | -0.206 | -0.349 | -0.166 | 0.292 | -0.012 | 0.04 | 0.04 | 0 | 0.31 | 0.38 | 0.3 | 0.57 | 0.32 | 0.34 |
| 9 | 101720921 | rs7874124 | -0.166 | 0.351 | -0.248 | 0.583 | 0.009 | 0.004 | 0.004 | 0.022 | 0.29 | 0.33 | 0.32 | 0.32 | 0.37 | 0.32 |
| 9 | 101727615 | rs2050257 | -0.014 | -0.07 | -0.339 | 0.4 | 0.006 | 0.011 | 0.011 | 0 | 0.42 | 0.33 | 0.22 | 0.32 | 0.37 | 0.41 |
| 9 | 101729103 | rs2151616 | 0.684 | -0.093 | -0.592 | 0.12 | -0.011 | 0.165 | 0.165 | 0.696 | 0.29 | 0.04 | 0.12 | 0 | 0.26 | 0.18 |
| 9 | 101734465 | rs10819537 | -1.345 | 0.651 | 0.939 | 1.543 | 0.041 | 0.035 | 0.035 | 0 | 0.22 | 0.51 | 0.44 | 0.29 | 0.3 | 0.34 |
| 9 | 101743336 | rs989393 | -1.844 | 0.371 | 1.068 | 2.015 | -0.005 | 0.052 | 0.052 | 0.023 | 0.29 | 0.48 | 0.36 | 0.49 | 0.31 | 0.46 |
| 9 | 101748234 | rs2075662 | -1.118 | 0.423 | 1.068 | 1.987 | -0.011 | 0.011 | 0.011 | 0 | 0.17 | 0.34 | 0.26 | 0.2 | 0.16 | 0.25 |
| 9 | 101758352 | rs10988510 | -0.51 | 0.379 | 0.699 | 2.127 | -0.028 | -0.047 | -0.047 | 0.021 | 0.1 | 0.03 | 0.09 | 0.3 | 0.14 | 0.2 |
| 9 | 101763633 | rs3780622 | -2.014 | 0.344 | 1.207 | 2.183 | 0.021 | 0.088 | 0.088 | 0 | 0.17 | 0.3 | 0.25 | 0.4 | 0.25 | 0.34 |
| 9 | 101779824 | rs4743309 | -1.405 | 0.146 | 1.059 | 2.647 | -0.01 | 0.052 | 0.052 | 0 | 0.12 | 0.15 | 0.25 | 0.42 | 0.08 | 0.33 |
| 9 | 101781301 | rs10819587 | 0.376 | 0.299 | 0.831 | 2.858 | -0.029 | -0.033 | -0.033 | 0 | 0.06 | 0.01 | 0.12 | 0.21 | 0.09 | 0.03 |
| 9 | 101781858 | rs1041631 | -1.943 | -0.08 | 0.995 | 2.775 | -0.005 | 0.039 | 0.039 | 0 | 0.17 | 0.23 | 0.28 | 0.42 | 0.16 | 0.37 |
| 9 | 101782535 | rs4743310 | -1.092 | -0.142 | 0.749 | 2.833 | 0.009 | -0.009 | -0.009 | 0 | 0.11 | 0.1 | 0.14 | 0.31 | 0.08 | 0.22 |
| 9 | 101784777 | rs3739799 | -0.22 | -0.139 | 0.684 | 2.84 | -0.039 | -0.049 | -0.049 | 0.173 | 0.08 | 0.01 | 0.09 | 0.29 | 0.14 | 0.19 |
| 9 | 101785207 | rs12552765 | -1.382 | -0.305 | 0.603 | 2.805 | 0.02 | -0.03 | -0.03 | 0 | 0.14 | 0.1 | 0.15 | 0.28 | 0.09 | 0.24 |
| 9 | 101785558 | rs7848837 | -0.768 | -0.197 | 1.104 | 2.668 | 0.027 | 0.134 | 0.134 | 0 | 0.1 | 0.24 | 0.12 | 0.29 | 0.14 | 0.19 |

| | | | | | | | | | | | | | | | | |
|---|---|---|---|---|---|---|---|---|---|---|---|---|---|---|---|---|
| 9 | 101785633 | rs7851787 | -0.768 | -0.197 | 1.104 | 2.666 | 0.027 | 0.147 | 0.147 | 0 | 0.1 | 0.24 | 0.16 | 0.32 | 0.14 | 0.19 |
| 9 | 101792716 | rs12684946 | -0.049 | -0.588 | 0.129 | 2.487 | -0.001 | -0.057 | -0.057 | 0 | 0.11 | 0.01 | 0.12 | 0.2 | 0.1 | 0.05 |
| 9 | 101797043 | rs976240 | -1.249 | -0.354 | 1 | 2.854 | -0.027 | 0.046 | 0.046 | 0 | 0.13 | 0.19 | 0.15 | 0.24 | 0.08 | 0.24 |
| 9 | 101804329 | rs7854112 | -0.762 | -0.372 | 0.845 | 2.525 | -0.002 | -0.011 | -0.011 | 0 | 0.15 | 0.08 | 0.14 | 0.17 | 0.15 | 0.23 |
| 9 | 101807463 | rs928522 | -1.296 | -0.395 | 1.082 | 2.554 | -0.012 | 0.036 | 0.036 | 0 | 0.18 | 0.29 | 0.22 | 0.3 | 0.15 | 0.32 |
| 9 | 101810339 | rs869776 | -2.716 | -0.472 | 1.131 | 2.567 | -0.043 | 0.007 | 0.007 | 0 | 0.31 | 0.5 | 0.38 | 0.66 | 0.23 | 0.57 |
| 9 | 101812072 | rs13296166 | -0.933 | -0.333 | 0.63 | 2.546 | 0.001 | -0.006 | -0.006 | 0 | 0.15 | 0.09 | 0.15 | 0.17 | 0.15 | 0.23 |
| 9 | 101815339 | rs16918167 | -2.762 | -0.418 | 0.563 | 2.476 | -0.035 | 0.001 | 0.001 | 0.494 | 0.32 | 0.42 | 0.37 | 0.65 | 0.21 | 0.6 |
| 9 | 101816399 | rs2067986 | -0.006 | -0.362 | 0.38 | 2.429 | -0.033 | -0.057 | -0.057 | 0.623 | 0.11 | 0.01 | 0.19 | 0.18 | 0.12 | 0.22 |
| 9 | 101816552 | rs3824511 | -1.629 | -0.388 | 0.406 | 2.448 | -0.024 | 0.058 | 0.058 | 0.31 | 0.16 | 0.2 | 0.15 | 0.35 | 0.09 | 0.21 |
| 9 | 101820545 | rs12001981 | -0.328 | -0.158 | 0.45 | 2.528 | -0.004 | -0.013 | -0.013 | 0.189 | 0.11 | 0.06 | 0.19 | 0.18 | 0.13 | 0.22 |
| 9 | 101821169 | rs1537506 | -1.157 | -0.172 | 0.44 | 2.521 | -0.001 | 0 | 0 | 4.804 | 0.15 | 0.16 | 0.2 | 0.18 | 0.13 | 0.23 |
| 9 | 101822302 | rs7031588 | -2.263 | -0.203 | 0.476 | 2.344 | -0.029 | 0.032 | 0.032 | 0.002 | 0.27 | 0.41 | 0.34 | 0.57 | 0.19 | 0.42 |
| 9 | 101823327 | rs1360686 | -1.21 | 0.161 | 0.308 | 2.19 | -0.086 | -0.075 | -0.075 | 0 | 0.15 | 0.01 | 0.22 | 0.23 | 0.24 | 0.23 |
| 9 | 101823373 | rs1413298 | -2.784 | 0.12 | 0.302 | 1.94 | -0.002 | 0.001 | 0.001 | 0 | 0.34 | 0.4 | 0.39 | 0.64 | 0.33 | 0.47 |
| 9 | 101832442 | rs1051105 | -2.506 | -0.458 | 0.568 | 2.117 | -0.014 | 0.023 | 0.023 | 0.047 | 0.29 | 0.4 | 0.32 | 0.63 | 0.22 | 0.46 |
| 9 | 101834340 | rs914991 | -2.512 | -0.401 | 0.31 | 2.151 | -0.016 | 0.006 | 0.006 | 0.361 | 0.3 | 0.45 | 0.33 | 0.68 | 0.23 | 0.61 |
| 9 | 101835165 | rs4743322 | -1.815 | 0.183 | -0.169 | 2.113 | 0 | -0.007 | -0.007 | 0 | 0.24 | 0.25 | 0.31 | 0.63 | 0.22 | 0.46 |
| 9 | 101844549 | rs2416666 | -0.731 | 0.232 | -0.333 | 2.177 | -0.053 | -0.08 | -0.08 | 0 | 0.12 | 0.01 | 0.19 | 0.29 | 0.15 | 0.25 |
| 9 | 101848586 | rs7874183 | -1.06 | 0.305 | -0.182 | 1.269 | -0.004 | 0.031 | 0.031 | 1.434 | 0.24 | 0.38 | 0.33 | 0.63 | 0.22 | 0.46 |
| 9 | 101858382 | rs7034716 | -0.523 | 0.465 | -0.688 | 0.106 | -0.001 | -0.025 | -0.025 | 0.014 | 0.25 | 0.26 | 0.19 | 0.04 | 0.28 | 0.14 |
| 9 | 101860076 | rs10819634 | -0.44 | 0.147 | -0.495 | 0.936 | -0.005 | 0.034 | 0.034 | 0 | 0.22 | 0.28 | 0.23 | 0.39 | 0.18 | 0.23 |
| 9 | 101861767 | rs12686783 | -1.438 | 0.111 | -0.46 | 0.967 | -0.002 | 0.068 | 0.068 | 0 | 0.1 | 0.11 | 0.17 | 0.4 | 0.1 | 0.17 |
| 9 | 101875789 | rs10739778 | -1.417 | -0.056 | -0.61 | 0.699 | -0.003 | 0.007 | 0.007 | 0 | 0.33 | 0.36 | 0.33 | 0.46 | 0.33 | 0.38 |
| 9 | 101882199 | rs10739779 | 0.593 | -0.086 | -0.621 | 0.661 | -0.006 | -0.009 | -0.009 | 2.688 | 0.16 | 0.11 | 0.12 | 0.01 | 0.19 | 0.1 |
| 9 | 101886071 | rs10512263 | 1.276 | 0.062 | -0.936 | 0.717 | -0.022 | -0.04 | -0.04 | 0.556 | 0.07 | 0.01 | 0.14 | 0.23 | 0.08 | 0.08 |
| 9 | 101905643 | rs10283455 | 0.235 | -0.041 | -0.825 | 0.569 | -0.003 | -0.016 | -0.016 | 0.167 | 0.18 | 0.16 | 0.13 | 0.01 | 0.19 | 0.1 |
| 9 | 101907424 | rs10733710 | 0.775 | 0.038 | -1.177 | -0.157 | -0.01 | 0.011 | 0.011 | 0.191 | 0.21 | 0.16 | 0.3 | 0.21 | 0.22 | 0.4 |
| 9 | 101912471 | rs334348 | -0.712 | 0.054 | -1.099 | -0.071 | -0.004 | 0.018 | 0.018 | 0 | 0.27 | 0.31 | 0.28 | 0.46 | 0.25 | 0.34 |
| 9 | 101914387 | rs334349 | -0.69 | 0.043 | -1.1 | -0.065 | 0 | 0 | 0 | 0 | 0.27 | 0.29 | 0.28 | 0.46 | 0.25 | 0.34 |
| 9 | 101916076 | rs7850895 | 0.133 | 0.115 | -1.066 | -0.057 | 0 | -0.005 | -0.005 | 0 | 0.06 | 0.05 | 0.05 | 0 | 0.08 | 0.04 |
| 9 | 101922412 | rs385057 | 0.313 | 0.084 | -1.045 | -0.049 | -0.001 | -0.033 | -0.033 | 2.997 | 0.18 | 0.18 | 0.13 | 0.01 | 0.19 | 0.1 |
| 9 | 101926594 | rs334362 | -0.017 | 0.175 | -0.993 | 0.022 | -0.003 | 0.029 | 0.029 | 0 | 0.22 | 0.27 | 0.24 | 0.43 | 0.19 | 0.24 |
| 9 | 101928702 | rs334363 | -0.58 | 0.295 | -0.959 | 0.13 | 0.003 | -0.011 | -0.011 | 0.008 | 0.31 | 0.3 | 0.32 | 0.43 | 0.32 | 0.36 |
| 9 | 101931740 | rs7035064 | -0.227 | -0.152 | -1.011 | 0.537 | 0 | -0.006 | -0.006 | 0 | 0.06 | 0.05 | 0.05 | 0 | 0.08 | 0.04 |
| 9 | 101933170 | rs7022989 | -1.026 | -0.21 | -0.619 | 0.929 | 0.028 | 0.522 | 0.522 | 0.592 | 0.05 | 0.29 | 0.13 | 0.43 | 0.02 | 0.06 |
| 9 | 101936231 | rs10819641 | -0.924 | -0.194 | -0.869 | 0.542 | -0.015 | 0.217 | 0.217 | 0 | 0.11 | 0.27 | 0.2 | 0.44 | 0.06 | 0.13 |
| 9 | 101952798 | rs10988744 | -0.907 | -0.132 | -1.002 | 0.008 | -0.011 | 0.21 | 0.21 | 0 | 0.11 | 0.29 | 0.21 | 0.42 | 0.06 | 0.13 |
| 9 | 101974824 | rs701376 | 0.347 | 0.041 | -2.326 | -1.799 | -0.06 | 0.147 | 0.147 | 0 | 0.28 | 0.01 | 0.17 | 0.05 | 0.26 | 0.13 |
| 9 | 101986627 | rs7036164 | -0.149 | 0.037 | -1.061 | -0.967 | 0.011 | 0.05 | 0.05 | 0 | 0.31 | 0.23 | 0.36 | 0.13 | 0.31 | 0.42 |
| 9 | 101987760 | rs3780616 | -1.667 | 0.434 | -0.533 | -0.074 | -0.001 | 0.032 | 0.032 | 0 | 0.09 | 0.03 | 0.08 | 0.01 | 0.07 | 0.05 |
| 9 | 101995174 | rs17725608 | -0.919 | 0.433 | -0.539 | -0.074 | -0.023 | 0.03 | 0.03 | 0 | 0.07 | 0 | 0.08 | 0.01 | 0.07 | 0.05 |
| 9 | 101999398 | rs17725644 | -1.383 | 0.446 | -0.5 | -0.068 | 0.002 | 0.006 | 0.006 | 0 | 0.1 | 0.07 | 0.09 | 0.01 | 0.07 | 0.07 |
| 9 | 102021226 | rs1746154 | 1.27 | 0.333 | 0.432 | -0.184 | 0.016 | -0.086 | -0.086 | 0.105 | 0.87 | 0.57 | 0.79 | 1 | 0.9 | 0.9 |
| 9 | 102029791 | rs228635 | 1.233 | 0.33 | 0.838 | -0.186 | 0.01 | -0.079 | -0.079 | 0.898 | 0.88 | 0.62 | 0.79 | 1 | 0.9 | 0.91 |
| 9 | 102033964 | rs162628 | -0.378 | 0.03 | -0.124 | -0.327 | -0.01 | -0.024 | -0.024 | 0 | 0.2 | 0.04 | 0.1 | 0.17 | 0.18 | 0.25 |
| 9 | 102036913 | rs182762 | 0.628 | 0.004 | 0.355 | -0.344 | 0.003 | -0.075 | -0.075 | 0 | 0.88 | 0.7 | 0.79 | 1 | 0.9 | 0.91 |
| 9 | 102039043 | rs189042 | 0.175 | -0.439 | -0.283 | -1.042 | 0.002 | -0.01 | -0.01 | 0 | 0.6 | 0.39 | 0.61 | 0.88 | 0.69 | 0.8 |
| 9 | 102049363 | rs692905 | -0.645 | -0.472 | 0.342 | -1.067 | -0.019 | -0.049 | -0.049 | 0 | 0.71 | 0.59 | 0.65 | 0.88 | 0.78 | 0.87 |
| 9 | 102054850 | rs12001129 | 1.242 | 0.313 | 0.053 | -0.568 | -0.051 | -0.08 | -0.08 | 0 | 0.21 | 0.08 | 0.31 | 0.47 | 0.29 | 0.19 |
| 9 | 102056001 | rs451008 | 0.335 | 0.233 | 0.475 | -0.708 | -0.054 | 0.013 | 0.013 | 0 | 0.41 | 0.17 | 0.4 | 0.64 | 0.47 | 0.44 |
| 9 | 102059951 | rs162624 | 0.329 | 0.297 | 0.539 | -0.717 | -0.059 | 0.014 | 0.014 | 0 | 0.41 | 0.17 | 0.4 | 0.64 | 0.48 | 0.45 |
| 9 | 102063150 | rs1622127 | 0.283 | 0.299 | 0.698 | -0.738 | 0 | 0.001 | 0.001 | 0.377 | 0.67 | 0.35 | 0.57 | 0.76 | 0.69 | 0.55 |
| 9 | 102066983 | rs7047384 | 0.795 | 0.435 | 0.483 | -0.533 | 0.001 | 0.001 | 0.001 | 0 | 0.19 | 0.2 | 0.2 | 0.23 | 0.21 | 0.35 |
| 9 | 102076659 | rs10988763 | 0.26 | 1.039 | 0.306 | -0.821 | -0.052 | 0.048 | 0.048 | 0 | 0.26 | 0.14 | 0.24 | 0.14 | 0.35 | 0.36 |
| 9 | 102079390 | rs170463 | 1.271 | 0.576 | 0.705 | -0.988 | -0.009 | -0.016 | -0.016 | 0 | 0.28 | 0.3 | 0.15 | 0.12 | 0.19 | 0.09 |
| 9 | 102079670 | rs379029 | 0.337 | 0.729 | 0.733 | -0.973 | 0.002 | -0.01 | -0.01 | 0 | 0.6 | 0.45 | 0.5 | 0.75 | 0.54 | 0.45 |
| 9 | 102079941 | rs10760678 | 0.351 | 0.714 | 0.834 | -1.002 | -0.047 | 0.051 | 0.051 | 0 | 0.29 | 0.19 | 0.25 | 0.12 | 0.39 | 0.38 |
| 9 | 102081002 | rs4268186 | -0.291 | 0.638 | 1.234 | -1.033 | -0.024 | 0.026 | 0.026 | 0 | 0.69 | 0.74 | 0.56 | 0.87 | 0.6 | 0.52 |
| 9 | 102084670 | rs10117464 | 0.345 | 0.655 | 1.396 | -1.038 | 0.026 | 0.066 | 0.066 | 0 | 0.07 | 0.24 | 0.06 | 0.12 | 0.07 | 0.07 |
| 9 | 102086148 | rs390312 | 0.49 | 0.631 | 1.372 | -1.077 | -0.003 | 0.01 | 0.01 | 0 | 0.58 | 0.4 | 0.48 | 0.76 | 0.49 | 0.41 |
| 9 | 102092282 | rs420258 | 1.377 | 0.739 | 1.288 | -1.184 | -0.003 | -0.011 | -0.011 | 0 | 0.3 | 0.3 | 0.16 | 0.12 | 0.23 | 0.09 |
| 9 | 102100078 | rs1323468 | 1.262 | 0.84 | 1.39 | -0.732 | -0.001 | 0.034 | 0.034 | 0.039 | 0.19 | 0.16 | 0.46 | 0.5 | 0.18 | 0.31 |
| 9 | 102101485 | rs7042978 | 0.338 | 0.854 | 1.433 | -0.741 | 0.021 | 0.061 | 0.061 | 0 | 0.07 | 0.24 | 0.06 | 0.12 | 0.07 | 0.08 |
| 9 | 102116594 | rs323729 | 0.539 | 0.993 | 1.452 | -0.757 | 0.016 | 0.012 | 0.012 | 0 | 0.89 | 0.7 | 0.92 | 0.88 | 0.88 | 0.88 |
| 9 | 102119090 | rs12551906 | 1.28 | 1.131 | 1.104 | -0.939 | -0.085 | 0.055 | 0.055 | 0.02 | 0.19 | 0.03 | 0.19 | 0.11 | 0.28 | 0.3 |
| 9 | 102123746 | rs323721 | -1.214 | 0.925 | 1.328 | -0.753 | -0.001 | 0.042 | 0.042 | 0 | 0.81 | 0.79 | 0.54 | 0.5 | 0.81 | 0.69 |
| 9 | 102127067 | rs1746153 | -0.095 | 0.884 | 1.115 | -0.803 | -0.005 | 0.028 | 0.028 | 0.331 | 0.7 | 0.47 | 0.46 | 0.38 | 0.7 | 0.57 |
| 9 | 102128509 | rs10819657 | 0.371 | 0.817 | 1.123 | -0.767 | -0.011 | 0.047 | 0.047 | 1.484 | 0.27 | 0.38 | 0.51 | 0.62 | 0.26 | 0.39 |

| | | | | | | | | | | | | | | | | |
|---|---|---|---|---|---|---|---|---|---|---|---|---|---|---|---|---|
| 9 | 102139440 | rs7024617 | 0.113 | 0.81 | 1.135 | -0.757 | 0.032 | 0.062 | 0.062 | 0 | 0.08 | 0.3 | 0.1 | 0.11 | 0.07 | 0.07 |
| 9 | 102139527 | rs10491654 | -0.176 | 0.821 | 1.173 | -0.739 | -0.005 | 0.024 | 0.024 | 0.089 | 0.71 | 0.44 | 0.47 | 0.38 | 0.71 | 0.6 |
| 9 | 102140053 | rs7871334 | -0.176 | 0.821 | 1.173 | -0.739 | -0.005 | 0.024 | 0.024 | 0.089 | 0.71 | 0.44 | 0.47 | 0.38 | 0.71 | 0.61 |
| 9 | 102143101 | rs894674 | 0.54 | 0.763 | 1.098 | -0.798 | -0.008 | -0.005 | -0.005 | 0 | 0.42 | 0.43 | 0.22 | 0.27 | 0.31 | 0.22 |
| 9 | 102146915 | rs9299331 | -0.085 | 0.757 | 1.152 | -0.796 | 0.003 | -0.016 | -0.016 | 0 | 0.37 | 0.37 | 0.35 | 0.22 | 0.48 | 0.46 |
| 9 | 102151801 | rs12555122 | 1.554 | 0.731 | 1.03 | -0.829 | -0.005 | -0.013 | -0.013 | 1.355 | 0.3 | 0.26 | 0.17 | 0.12 | 0.22 | 0.1 |
| 9 | 102152364 | rs10988794 | 1.315 | 0.795 | 0.755 | -0.761 | -0.007 | -0.074 | -0.074 | 0 | 0.19 | 0.08 | 0.41 | 0.51 | 0.19 | 0.32 |
| 9 | 102159610 | rs2416771 | 0.441 | 0.215 | 0.619 | -0.722 | 0.012 | -0.033 | -0.033 | 0 | 0.11 | 0.05 | 0.04 | 0.16 | 0.09 | 0.12 |
| 9 | 102168293 | rs880597 | 0.483 | 0.229 | 0.731 | -0.726 | 0.014 | 0.01 | 0.01 | 0.547 | 0.42 | 0.32 | 0.38 | 0.3 | 0.34 | 0.23 |
| 9 | 102177648 | rs4743350 | 1.154 | 0.102 | 0.701 | -0.782 | -0.011 | -0.02 | -0.02 | 0 | 0.32 | 0.29 | 0.19 | 0.14 | 0.25 | 0.13 |
| 9 | 102179061 | rs10760682 | 0.065 | 0.644 | 0.639 | -0.73 | 0.006 | 0.043 | 0.043 | 0 | 0.33 | 0.2 | 0.26 | 0.22 | 0.31 | 0.45 |
| 9 | 102182953 | rs10512264 | 1.208 | 0.528 | 0.441 | -0.78 | -0.018 | -0.03 | -0.03 | 0 | 0.32 | 0.31 | 0.19 | 0.14 | 0.25 | 0.12 |
| 9 | 102190129 | rs10512267 | 1.048 | 0.69 | 0.106 | -0.692 | 0.017 | 0.027 | 0.027 | 0.162 | 0.22 | 0.22 | 0.33 | 0.51 | 0.33 | 0.32 |
| 9 | 102214169 | rs608510 | -1.106 | 0.155 | -0.156 | -0.793 | -0.044 | -0.057 | -0.057 | 5.233 | 0.63 | 0.59 | 0.79 | 0.86 | 0.75 | 0.83 |
| 9 | 102233287 | rs634202 | 0.199 | 0.32 | -0.86 | -0.367 | 0 | -0.003 | -0.003 | 0 | 0.08 | 0.11 | 0.22 | 0.16 | 0.07 | 0.09 |
| 9 | 102234323 | rs649057 | 0.313 | 0.272 | -0.855 | -0.361 | 0.001 | -0.001 | -0.001 | 1.259 | 0.08 | 0.11 | 0.21 | 0.16 | 0.05 | 0.09 |
| 9 | 102247368 | rs10760688 | -0.485 | 0.173 | -1.87 | -0.509 | -0.019 | 0.007 | 0.007 | 0 | 0.35 | 0.17 | 0.47 | 0.31 | 0.37 | 0.52 |
| 9 | 102271150 | rs9299333 | -0.458 | 0.186 | -1.833 | -0.484 | -0.005 | 0.017 | 0.017 | 0 | 0.37 | 0.27 | 0.48 | 0.31 | 0.38 | 0.52 |
| 9 | 102278511 | rs7874483 | -0.429 | -0.33 | -1.742 | -0.024 | 0.013 | -0.035 | -0.035 | 0 | 0.87 | 0.99 | 0.96 | 0.86 | 0.92 | 0.95 |
| 9 | 102289651 | rs7044788 | -0.215 | -0.265 | -1.83 | -0.434 | 0.013 | 0.061 | 0.061 | 0 | 0.22 | 0.11 | 0.23 | 0.09 | 0.24 | 0.14 |
| 9 | 102304304 | rs13298309 | -0.114 | -0.381 | -1.834 | -0.384 | -0.009 | 0.082 | 0.082 | 0 | 0.2 | 0.07 | 0.17 | 0.07 | 0.23 | 0.14 |
| 9 | 102337331 | rs11788118 | -0.374 | -0.407 | -1.686 | -0.494 | 0.017 | 0.037 | 0.037 | 0 | 0.22 | 0.13 | 0.17 | 0.05 | 0.22 | 0.15 |
| 9 | 102342950 | rs10988865 | 0.993 | 1.422 | -2.386 | -1.074 | -0.082 | 0.067 | 0.067 | 0 | 0.11 | 0.01 | 0.13 | 0 | 0.27 | 0.03 |
| 9 | 102346029 | rs10819687 | 0.137 | 0.696 | -2.083 | -0.924 | -0.024 | 0.011 | 0.011 | 0 | 0.19 | 0.13 | 0.21 | 0.13 | 0.31 | 0.14 |
| 9 | 102374475 | rs4743361 | -1.225 | -0.27 | -2.521 | -1.248 | -0.077 | 0.014 | 0.014 | 0 | 0.77 | 0.98 | 0.81 | 0.88 | 0.77 | 0.86 |
| 9 | 102383474 | rs4743363 | 1.205 | -0.85 | -1.48 | -1.331 | 0.031 | -0.008 | -0.008 | 0 | 0.31 | 0.24 | 0.39 | 0.66 | 0.24 | 0.56 |
| 9 | 102400263 | rs10819689 | 0.936 | -0.793 | -0.911 | -1.317 | 0.002 | -0.007 | -0.007 | 0 | 0.22 | 0.21 | 0.17 | 0.06 | 0.22 | 0.13 |
| 9 | 102464577 | rs1578463 | 0.236 | -0.823 | -0.698 | -1.573 | -0.036 | 0.045 | 0.045 | 0.003 | 0.35 | 0.21 | 0.25 | 0.16 | 0.38 | 0.14 |
| 9 | 102473442 | rs1147312 | 0.807 | -0.895 | -0.844 | -1.599 | -0.057 | -0.017 | -0.017 | 0 | 0.1 | 0.01 | 0.23 | 0.1 | 0.16 | 0.25 |
| 9 | 102502572 | rs17798928 | 1.15 | -0.985 | -0.838 | -1.649 | 0.012 | 0.045 | 0.045 | 0 | 0.21 | 0.11 | 0.18 | 0.06 | 0.23 | 0.11 |
| 9 | 102540372 | rs10819694 | 2.39 | -1.21 | -1.073 | -1.818 | -0.004 | 0.015 | 0.015 | 0.498 | 0.1 | 0.01 | 0.1 | 0.06 | 0.13 | 0.04 |
| 9 | 102562216 | rs7026022 | 0.103 | -1.239 | -0.791 | -1.743 | 0.017 | -0.044 | -0.044 | 0.066 | 0.39 | 0.49 | 0.28 | 0.17 | 0.38 | 0.16 |
| 9 | 102575680 | rs13295822 | 0.951 | -1.229 | -0.852 | -1.736 | -0.057 | -0.02 | -0.02 | 0 | 0.1 | 0.01 | 0.23 | 0.11 | 0.16 | 0.25 |
| 9 | 102585545 | rs1405209 | 0.429 | -1.24 | -0.427 | -1.678 | -0.004 | 0.003 | 0.003 | 0 | 0.36 | 0.32 | 0.26 | 0.17 | 0.38 | 0.16 |
| 9 | 102595685 | rs10988904 | 0.131 | -1.166 | -0.548 | -1.674 | 0.004 | -0.018 | -0.018 | 0 | 0.4 | 0.42 | 0.27 | 0.17 | 0.39 | 0.16 |
| 9 | 102596321 | rs2900223 | -1.463 | -1.301 | -0.37 | -1.372 | -0.004 | 0 | 0 | 0 | 0.69 | 0.63 | 0.68 | 0.34 | 0.77 | 0.52 |
| 9 | 102622237 | rs7847479 | -0.2 | -1.176 | -1.023 | -1.254 | 0.035 | 0.044 | 0.044 | 2.094 | 0.49 | 0.26 | 0.41 | 0.34 | 0.59 | 0.23 |
| 9 | 102627302 | rs1131339 | -0.2 | -1.176 | -1.013 | -1.254 | 0.035 | 0.043 | 0.043 | 2.094 | 0.49 | 0.26 | 0.41 | 0.34 | 0.59 | 0.23 |
| 9 | 102635520 | rs10819700 | -0.379 | -1.023 | -1.031 | -1.254 | 0.056 | 0.031 | 0.031 | 0.134 | 0.53 | 0.26 | 0.6 | 0.34 | 0.67 | 0.3 |
| 9 | 102679173 | rs4742777 | -0.088 | -0.043 | -1.278 | -1.59 | 0.048 | 0.044 | 0.044 | 1.293 | 0.56 | 0.74 | 0.64 | 0.74 | 0.68 | 0.49 |
| 9 | 102686487 | rs4428721 | 0.816 | 0.668 | -1.575 | -2.137 | -0.06 | 0.079 | 0.079 | 1.229 | 0.29 | 0.15 | 0.21 | 0.02 | 0.36 | 0.14 |
| 9 | 102687703 | rs10124366 | -1.055 | 0.246 | -1.619 | -2.092 | 0.028 | 0.028 | 0.028 | 0.217 | 0.66 | 0.85 | 0.68 | 0.77 | 0.75 | 0.61 |
| 9 | 102714466 | rs10512268 | -0.497 | 0.529 | -1.638 | -1.973 | 0.048 | 0.044 | 0.044 | 1.172 | 0.56 | 0.76 | 0.64 | 0.74 | 0.68 | 0.49 |
| 9 | 102723692 | rs10760706 | -0.487 | 0.531 | -1.598 | -1.972 | 0.044 | 0.04 | 0.04 | 1.226 | 0.56 | 0.74 | 0.64 | 0.74 | 0.68 | 0.49 |
| 9 | 102732237 | rs9556 | -0.489 | 0.695 | -1.557 | -1.894 | 0.042 | 0.031 | 0.031 | 0.165 | 0.57 | 0.76 | 0.63 | 0.73 | 0.68 | 0.47 |
| 9 | 102735865 | rs2698999 | -0.311 | 0.907 | -1.502 | -1.348 | -0.002 | 0.075 | 0.075 | 0.634 | 0.43 | 0.71 | 0.33 | 0.14 | 0.55 | 0.25 |
| 9 | 102741475 | rs2795575 | -0.395 | 0.929 | -1.496 | -1.37 | 0 | 0.076 | 0.076 | 0.272 | 0.44 | 0.71 | 0.33 | 0.13 | 0.55 | 0.24 |
| 9 | 102748197 | rs10760708 | 0.3 | 1.008 | -1.134 | -1.42 | 0.079 | -0.023 | -0.023 | 0.321 | 0.09 | 0.48 | 0.08 | 0 | 0.1 | 0.03 |
| 9 | 102777978 | rs10819717 | -0.471 | 1.037 | -1.219 | -1.273 | -0.077 | 0.128 | 0.128 | 0 | 0.66 | 0.85 | 0.75 | 0.87 | 0.55 | 0.79 |
| 9 | 102816535 | rs7867803 | 0.391 | 1.063 | -1.204 | -1.297 | 0.066 | -0.026 | -0.026 | 0.424 | 0.09 | 0.55 | 0.08 | 0 | 0.1 | 0.03 |
| 9 | 102826067 | rs10739795 | -0.474 | 1.068 | -1.226 | -1.312 | -0.077 | 0.128 | 0.128 | 0 | 0.66 | 0.86 | 0.75 | 0.87 | 0.55 | 0.79 |
| 9 | 102849539 | rs10988967 | 0.382 | 1.062 | -1.205 | -1.291 | 0.065 | -0.027 | -0.027 | 0.389 | 0.09 | 0.55 | 0.08 | 0 | 0.1 | 0.03 |
| 9 | 102861613 | rs7024375 | 1.187 | 0.882 | -1.548 | -1.347 | 0.009 | -0.125 | -0.125 | 0 | 0.21 | 0.03 | 0.35 | 0.62 | 0.17 | 0.33 |
| 9 | 102868769 | rs12346672 | 0.687 | 2.241 | -0.782 | -0.979 | -0.052 | 0.086 | 0.086 | 0 | 0.25 | 0.09 | 0.18 | 0.01 | 0.3 | 0.13 |
| 9 | 102886948 | rs846761 | 0.028 | 2.011 | -0.638 | -0.85 | -0.063 | 0.121 | 0.121 | 0 | 0.65 | 0.85 | 0.75 | 0.87 | 0.57 | 0.78 |
| 9 | 102911706 | rs1415927 | -0.182 | 2.017 | -0.43 | -0.858 | 0.062 | -0.029 | -0.029 | 0 | 0.09 | 0.55 | 0.08 | 0.01 | 0.11 | 0.03 |
| 9 | 102939193 | rs10988996 | -0.184 | 2.035 | -0.443 | -0.849 | 0.084 | -0.019 | -0.019 | 0 | 0.09 | 0.48 | 0.08 | 0.01 | 0.11 | 0.03 |
| 9 | 102958736 | rs7020780 | -0.042 | 2.054 | -0.434 | -0.839 | 0.074 | -0.02 | -0.02 | 0 | 0.09 | 0.41 | 0.07 | 0.01 | 0.11 | 0.03 |
| 9 | 102973719 | rs1999877 | -0.891 | 1.986 | -0.701 | -0.857 | -0.004 | 0.028 | 0.028 | 0.547 | 0.44 | 0.64 | 0.33 | 0.13 | 0.54 | 0.24 |
| 9 | 103007414 | rs2787394 | -0.21 | 1.666 | -1.402 | -1.628 | -0.066 | 0.139 | 0.139 | 0 | 0.33 | 0.12 | 0.23 | 0.13 | 0.4 | 0.21 |
| 9 | 103013986 | rs2787390 | 0.757 | 1.56 | -1.285 | -1.283 | -0.021 | -0.026 | -0.026 | 0 | 0.05 | 0 | 0.05 | 0.13 | 0.11 | 0.05 |
| 9 | 103050407 | rs2787372 | -0.246 | 1.658 | -1.212 | -1.6 | -0.036 | 0.057 | 0.057 | 0.012 | 0.34 | 0.22 | 0.25 | 0.14 | 0.43 | 0.19 |
| 9 | 103054951 | rs2787374 | -1.216 | 1.437 | -0.735 | -1.929 | 0.003 | -0.003 | -0.003 | 0 | 0.47 | 0.62 | 0.35 | 0.16 | 0.57 | 0.25 |
| 9 | 103055997 | rs2806700 | -0.233 | 1.784 | -1.303 | -1.854 | -0.061 | 0.111 | 0.111 | 0.232 | 0.35 | 0.18 | 0.24 | 0.15 | 0.44 | 0.23 |
| 9 | 103063645 | rs10123866 | 1.158 | 1.685 | -1.269 | -2.028 | 0.055 | 0.082 | 0.082 | 0.831 | 0.51 | 0.62 | 0.61 | 0.85 | 0.33 | 0.72 |
| 9 | 103063826 | rs10123913 | 1.158 | 1.685 | -1.269 | -2.028 | 0.055 | 0.082 | 0.082 | 0.831 | 0.51 | 0.62 | 0.61 | 0.85 | 0.33 | 0.72 |
| 9 | 103064153 | rs10989049 | 0.155 | 1.69 | -1.324 | -2.06 | -0.01 | 0.114 | 0.114 | 0.197 | 0.6 | 0.79 | 0.7 | 0.86 | 0.47 | 0.78 |
| 9 | 103065884 | rs3923501 | 0.11 | 1.767 | -1.325 | -2.022 | -0.006 | 0.073 | 0.073 | 0.023 | 0.61 | 0.75 | 0.69 | 0.86 | 0.46 | 0.78 |
| 9 | 103085570 | rs10819758 | 0.55 | 1.738 | -1.337 | -1.968 | -0.026 | 0.054 | 0.054 | 0 | 0.1 | 0 | 0.08 | 0 | 0.14 | 0.05 |

| | | | | | | | | | | | | | | | | |
|---|---|---|---|---|---|---|---|---|---|---|---|---|---|---|---|---|
| 9 | 103091805 | rs6479013 | -0.404 | 2.04 | -1.395 | -2.014 | -0.048 | -0.022 | -0.022 | 0 | 0.92 | 0.99 | 0.89 | 0.87 | 0.78 | 0.93 |
| 9 | 103138400 | rs10989072 | 0.445 | 1.309 | -1.27 | -1.961 | -0.001 | -0.028 | -0.028 | 0 | 0.1 | 0.14 | 0.08 | 0 | 0.14 | 0.05 |
| 9 | 103140157 | rs2026999 | 0.814 | 1.141 | -1.281 | -1.915 | 0.044 | 0.058 | 0.058 | 0.337 | 0.49 | 0.6 | 0.61 | 0.86 | 0.33 | 0.72 |
| 9 | 103147531 | rs10819769 | 0.653 | 0.41 | -1.088 | -1.414 | -0.001 | -0.007 | -0.007 | 0.084 | 0.79 | 0.79 | 0.85 | 0.86 | 0.75 | 0.87 |
| 9 | 103149987 | rs13284428 | -0.512 | -0.227 | -1.528 | -1.412 | -0.008 | 0.037 | 0.037 | 0 | 0.54 | 0.37 | 0.31 | 0.26 | 0.48 | 0.43 |
| 9 | 103151524 | rs953381 | -0.841 | -0.092 | -0.726 | -0.897 | 0.001 | -0.002 | -0.002 | 0 | 0.94 | 0.95 | 0.95 | 0.87 | 0.91 | 0.93 |
| 9 | 103157515 | rs12000794 | -0.299 | -0.082 | -0.735 | -0.915 | -0.002 | -0.033 | -0.033 | 0.285 | 0.14 | 0.17 | 0.1 | 0.01 | 0.16 | 0.06 |
| 9 | 103174029 | rs7031884 | 0.066 | -0.609 | -0.796 | -0.973 | -0.008 | -0.014 | -0.014 | 0 | 0.31 | 0.37 | 0.18 | 0.21 | 0.23 | 0.39 |
| 9 | 103179614 | rs10819774 | -0.221 | -0.576 | -0.759 | -0.975 | -0.003 | -0.01 | -0.01 | 1.044 | 0.32 | 0.34 | 0.19 | 0.2 | 0.23 | 0.36 |
| 9 | 103181596 | rs2026998 | 0.638 | -0.603 | -0.795 | -1.083 | 0.006 | 0.042 | 0.042 | 0.213 | 0.22 | 0.09 | 0.13 | 0 | 0.17 | 0.08 |
| 9 | 103211325 | rs7046659 | -0.209 | -0.804 | -0.12 | -1.044 | -0.011 | -0.014 | -0.014 | 0.406 | 0.29 | 0.35 | 0.2 | 0.21 | 0.19 | 0.38 |
| 9 | 103211802 | rs10989097 | -0.698 | -0.804 | -0.121 | -1.049 | -0.011 | -0.024 | -0.024 | 0 | 0.33 | 0.39 | 0.22 | 0.21 | 0.23 | 0.38 |
| 9 | 103212528 | rs12683548 | 0.903 | -0.802 | -0.157 | -1.042 | -0.001 | 0.037 | 0.037 | 0 | 0.05 | 0.07 | 0.05 | 0.21 | 0.03 | 0.29 |
| 9 | 103213947 | rs1556487 | -0.303 | -1.444 | -0.345 | -1.177 | 0.017 | 0.006 | 0.006 | 0 | 0.79 | 0.83 | 0.87 | 0.82 | 0.76 | 0.75 |
| 9 | 103218013 | rs17813801 | 0.743 | -1.073 | -0.263 | -0.971 | 0.005 | 0.09 | 0.09 | 0.126 | 0.22 | 0.06 | 0.13 | 0 | 0.17 | 0.08 |
| 9 | 103227676 | rs7869933 | 0.068 | -0.922 | -0.069 | -0.794 | 0.006 | 0.061 | 0.061 | 0 | 0.09 | 0.15 | 0.07 | 0.21 | 0.06 | 0.28 |
| 9 | 103240259 | rs7041189 | 0.876 | -0.969 | -0.16 | -0.873 | 0.008 | 0.009 | 0.009 | 0.638 | 0.19 | 0.14 | 0.13 | 0 | 0.13 | 0.08 |
| 9 | 103252812 | rs10989112 | -0.328 | -0.973 | -0.081 | -0.854 | 0.013 | 0.092 | 0.092 | 0 | 0.09 | 0.22 | 0.08 | 0.21 | 0.06 | 0.29 |
| 9 | 103262228 | rs10819788 | 0.133 | -0.948 | -0.415 | -0.843 | 0.004 | 0.003 | 0.003 | 0.063 | 0.25 | 0.24 | 0.18 | 0.21 | 0.16 | 0.36 |
| 9 | 103299435 | rs10989146 | -0.294 | -0.955 | -0.353 | -0.833 | 0 | 0 | 0 | 0.746 | 0.28 | 0.3 | 0.2 | 0.22 | 0.19 | 0.38 |
| 9 | 103312290 | rs2274882 | 0.058 | -0.949 | -0.36 | -0.828 | 0.004 | 0.003 | 0.003 | 0.026 | 0.24 | 0.24 | 0.18 | 0.2 | 0.16 | 0.36 |
| 9 | 103319797 | rs10989153 | -0.387 | -1.017 | -0.231 | -0.899 | -0.01 | -0.013 | -0.013 | 0 | 0.3 | 0.38 | 0.21 | 0.23 | 0.19 | 0.36 |
| 9 | 103324710 | rs17741974 | 0.822 | -0.906 | -0.688 | -1.001 | 0.007 | 0.048 | 0.048 | 1 | 0.19 | 0.08 | 0.12 | 0 | 0.13 | 0.08 |
| 9 | 103331141 | rs12005789 | 0.528 | -0.958 | -0.728 | -0.824 | 0 | -0.001 | -0.001 | 6.577 | 0.07 | 0.08 | 0.04 | 0.08 | 0.08 | 0.18 |
| 9 | 103338012 | rs2800284 | 1.511 | -0.575 | -1.446 | -1.25 | -0.123 | 0.131 | 0.131 | 0 | 0.21 | 0.03 | 0.43 | 0 | 0.35 | 0.16 |
| 9 | 103338172 | rs13295029 | 0.865 | -0.738 | -1.551 | -1.475 | 0.01 | 0.099 | 0.099 | 3.27 | 0.23 | 0.07 | 0.13 | 0 | 0.16 | 0.09 |
| 9 | 103340309 | rs1226591 | 0.709 | -0.796 | -1.144 | -1.471 | -0.009 | 0.003 | 0.003 | 0 | 0.3 | 0.26 | 0.48 | 0.08 | 0.42 | 0.33 |
| 9 | 103341014 | rs1226588 | -0.583 | -0.847 | -1.041 | -1.188 | 0.004 | 0.004 | 0.004 | 0 | 0.58 | 0.41 | 0.66 | 0.3 | 0.6 | 0.67 |
| 9 | 103342983 | rs2151598 | -1.348 | -1.284 | -1.072 | -1.288 | -0.004 | 0.06 | 0.06 | 0 | 0.74 | 0.87 | 0.83 | 0.85 | 0.71 | 0.84 |
| 9 | 103343852 | rs2780958 | -0.355 | -0.758 | -1.042 | -1.132 | 0.021 | 0.034 | 0.034 | 0 | 0.53 | 0.37 | 0.64 | 0.3 | 0.57 | 0.66 |
| 9 | 103351014 | rs17742650 | 1.432 | -0.683 | -1.147 | -1.182 | 0.004 | 0.096 | 0.096 | 0.959 | 0.18 | 0.04 | 0.11 | 0 | 0.12 | 0.07 |
| 9 | 103353017 | rs1226593 | 1.405 | -0.503 | -0.828 | -1.085 | 0.012 | -0.053 | -0.053 | 0 | 0.22 | 0.22 | 0.44 | 0 | 0.35 | 0.16 |
| 9 | 103361114 | rs10989168 | -0.403 | -0.688 | -0.895 | -0.805 | 0.027 | 0.002 | 0.002 | 0.509 | 0.35 | 0.18 | 0.21 | 0.3 | 0.26 | 0.51 |
| 9 | 103364064 | rs10989173 | 0.159 | -0.623 | -0.54 | -0.732 | 0.005 | 0.004 | 0.004 | 1.761 | 0.28 | 0.23 | 0.18 | 0.22 | 0.18 | 0.33 |
| 9 | 103365585 | rs7022319 | 0.189 | -0.306 | -0.193 | 0.495 | -0.011 | 0.019 | 0.019 | 0 | 0.36 | 0.48 | 0.29 | 0.47 | 0.34 | 0.31 |
| 9 | 103367615 | rs4474081 | -0.23 | -0.231 | -0.742 | 0.119 | 0.015 | 0.031 | 0.031 | 0 | 0.42 | 0.7 | 0.35 | 0.67 | 0.37 | 0.57 |
| 9 | 103370505 | rs10118164 | -0.242 | 0.084 | -0.46 | 0.226 | 0 | 0 | 0 | 0 | 0.36 | 0.33 | 0.23 | 0.35 | 0.32 | 0.44 |
| 9 | 103370532 | rs10115383 | -0.946 | 0.457 | -0.322 | 0.279 | -0.005 | 0 | 0 | 0 | 0.64 | 0.61 | 0.73 | 0.63 | 0.69 | 0.62 |
| 9 | 103371976 | rs10760729 | -0.611 | 0.387 | -0.511 | 0.091 | 0.013 | 0.039 | 0.039 | 0 | 0.45 | 0.61 | 0.36 | 0.72 | 0.41 | 0.67 |
| 9 | 103375024 | rs4743404 | -1.448 | 0.286 | -0.35 | -0.015 | 0.006 | -0.002 | -0.002 | 0 | 0.72 | 0.75 | 0.85 | 1 | 0.78 | 0.83 |
| 9 | 103377514 | rs7869768 | -0.189 | 0.32 | -0.458 | 0.283 | 0.001 | -0.002 | -0.002 | 0 | 0.35 | 0.59 | 0.28 | 0.45 | 0.28 | 0.38 |
| 9 | 103383109 | rs10819800 | 0.108 | 0.66 | -0.469 | 0.617 | -0.037 | -0.053 | -0.053 | 0 | 0.1 | 0.02 | 0.09 | 0.27 | 0.13 | 0.28 |
| 9 | 103386660 | rs6479020 | 0.814 | 0.756 | -0.338 | 0.244 | -0.002 | -0.027 | -0.027 | 2.888 | 0.32 | 0.33 | 0.19 | 0.08 | 0.3 | 0.16 |
| 9 | 103388556 | rs7036906 | 1.295 | 1.005 | -0.526 | 0.34 | -0.011 | -0.013 | -0.013 | 0 | 0.21 | 0.18 | 0.44 | 0.27 | 0.27 | 0.16 |
| 9 | 103403174 | rs1106382 | -1.299 | 1.005 | -0.413 | 0.344 | -0.014 | 0.131 | 0.131 | 0.321 | 0.73 | 0.92 | 0.86 | 1 | 0.77 | 0.84 |
| 9 | 103405197 | rs1936641 | -0.738 | 0.992 | -0.197 | 0.407 | 0 | -0.001 | -0.001 | 0.051 | 0.68 | 0.67 | 0.76 | 0.73 | 0.73 | 0.72 |
| 9 | 103412206 | rs4330746 | 0.025 | 1.106 | -0.397 | 0.479 | -0.027 | -0.032 | -0.032 | 0.962 | 0.32 | 0.24 | 0.53 | 0.55 | 0.4 | 0.46 |
| 9 | 103427155 | rs12238759 | 0.51 | 1.039 | -0.565 | 0.672 | -0.007 | -0.033 | -0.033 | 0 | 0.07 | 0.01 | 0.06 | 0.24 | 0.05 | 0.3 |
| 9 | 103427659 | rs10114244 | -0.677 | 1.154 | -0.493 | 0.606 | -0.025 | 0.037 | 0.037 | 0 | 0.66 | 0.91 | 0.8 | 0.76 | 0.72 | 0.55 |
| 9 | 103427741 | rs10114272 | 1.251 | 1.505 | -0.452 | 0.604 | -0.009 | -0.014 | -0.014 | 0 | 0.2 | 0.17 | 0.44 | 0.28 | 0.25 | 0.14 |
| 9 | 103437984 | rs10819806 | 0.198 | 1.43 | -0.188 | 0.605 | -0.013 | -0.045 | -0.045 | 0 | 0.39 | 0.49 | 0.23 | 0.2 | 0.35 | 0.28 |
| 9 | 103439019 | rs4512437 | 0.198 | 1.43 | -0.188 | 0.605 | -0.013 | -0.045 | -0.045 | 0.661 | 0.39 | 0.49 | 0.24 | 0.2 | 0.35 | 0.28 |
| 9 | 103446253 | rs10819808 | 0.846 | 1.411 | -0.255 | 0.805 | -0.02 | -0.028 | -0.028 | 0 | 0.22 | 0.17 | 0.46 | 0.41 | 0.33 | 0.24 |
| 9 | 103452068 | rs10123021 | 0.266 | 1.112 | -0.385 | 0.348 | 0 | 0.002 | 0.002 | 0 | 0.33 | 0.3 | 0.22 | 0.3 | 0.34 | 0.18 |
| 9 | 103459867 | rs7875828 | 0.525 | 0.87 | -0.149 | 0.31 | -0.001 | -0.01 | -0.01 | 0 | 0.37 | 0.32 | 0.25 | 0.08 | 0.38 | 0.18 |
| 9 | 103474052 | rs1892128 | -0.117 | 0.921 | -0.088 | 0.428 | 0.003 | -0.002 | -0.002 | 0 | 0.57 | 0.53 | 0.69 | 0.38 | 0.57 | 0.3 |
| 9 | 103480387 | rs12346541 | 0.709 | 0.992 | -0.247 | 0.728 | 0.001 | -0.013 | -0.013 | 12.7 | 0.2 | 0.19 | 0.44 | 0.31 | 0.19 | 0.11 |
| 9 | 103485133 | rs6479028 | 0.182 | 0.972 | -0.143 | 0.761 | 0.001 | -0.019 | -0.019 | 0 | 0.43 | 0.61 | 0.33 | 0.47 | 0.43 | 0.45 |
| 9 | 103485819 | rs7853739 | 0.653 | 0.683 | 0.17 | 0.516 | -0.021 | 0.066 | 0.066 | 0 | 0.23 | 0.38 | 0.43 | 0.32 | 0.2 | 0.11 |
| 9 | 103486169 | rs10989224 | 1.04 | 0.74 | 0.134 | 0.386 | 0 | -0.02 | -0.02 | 0 | 0.34 | 0.31 | 0.23 | 0.05 | 0.35 | 0.3 |
| 9 | 103486336 | rs4415379 | -0.824 | 0.975 | -0.201 | 0.312 | -0.004 | 0.18 | 0.18 | 0 | 0.69 | 0.91 | 0.83 | 1 | 0.74 | 0.83 |
| 9 | 103494116 | rs4742792 | 0.087 | 0.913 | 0.364 | 0.942 | -0.003 | 0.068 | 0.068 | 0 | 0.27 | 0.4 | 0.5 | 0.47 | 0.28 | 0.24 |
| 9 | 103495396 | rs7854792 | -0.69 | 0.433 | 0.235 | 0.642 | -0.007 | -0.055 | -0.055 | 0 | 0.37 | 0.48 | 0.62 | 0.8 | 0.38 | 0.4 |
| 9 | 103495889 | rs10989231 | 0.2 | 0.555 | -0.125 | 1.033 | -0.001 | 0 | 0 | 0 | 0.23 | 0.26 | 0.4 | 0.29 | 0.21 | 0.11 |
| 9 | 103513029 | rs1578041 | 0.409 | 0.78 | 0.004 | 1.073 | 0.012 | -0.016 | -0.016 | 0 | 0.57 | 0.7 | 0.49 | 0.49 | 0.61 | 0.59 |
| 9 | 103514183 | rs10819824 | 1.24 | 0.928 | 0.154 | 0.887 | 0.003 | 0.12 | 0.12 | 0.801 | 0.39 | 0.17 | 0.27 | 0.06 | 0.4 | 0.29 |
| 9 | 103514405 | rs10739801 | 0.261 | 0.912 | 0.195 | 0.895 | 0.012 | -0.008 | -0.008 | 0 | 0.56 | 0.68 | 0.48 | 0.51 | 0.61 | 0.57 |
| 9 | 103517896 | rs10739802 | 1.203 | 0.896 | -0.014 | 1.181 | -0.002 | 0.084 | 0.084 | 0.716 | 0.39 | 0.14 | 0.27 | 0.19 | 0.4 | 0.3 |
| 9 | 103518490 | rs7019439 | 0.505 | 0.877 | 0.263 | 1.261 | 0.022 | 0.12 | 0.12 | 0 | 0.15 | 0.32 | 0.16 | 0.3 | 0.14 | 0.23 |

| | | | | | | | | | | | | | | | | |
|---|---|---|---|---|---|---|---|---|---|---|---|---|---|---|---|---|
| 9 | 103521788 | rs7024267 | -1.234 | 0.836 | -0.372 | 0.635 | -0.025 | 0.063 | 0.063 | 0.349 | 0.83 | 0.95 | 0.64 | 0.95 | 0.8 | 0.91 |
| 9 | 103524447 | rs10819829 | 1.167 | 0.924 | -0.345 | 0.65 | -0.008 | 0.164 | 0.164 | 1.302 | 0.4 | 0.13 | 0.27 | 0.09 | 0.41 | 0.34 |
| 9 | 103534708 | rs10760755 | 0.969 | 0.269 | -0.167 | 0.868 | -0.001 | 0.001 | 0.001 | 0.624 | 0.44 | 0.41 | 0.33 | 0.36 | 0.48 | 0.36 |
| 9 | 103536801 | rs1933794 | -0.949 | 0.205 | -0.715 | 0.654 | -0.033 | 0.068 | 0.068 | 0.713 | 0.81 | 0.98 | 0.64 | 0.94 | 0.8 | 0.91 |
| 9 | 103539701 | rs2211079 | -0.146 | 0.385 | -0.196 | 0.989 | 0 | -0.001 | -0.001 | 0.052 | 0.58 | 0.57 | 0.49 | 0.53 | 0.64 | 0.61 |
| 9 | 103545759 | rs4742795 | -0.647 | 0.414 | 0.602 | 1.599 | 0.01 | 0.141 | 0.141 | 0 | 0.17 | 0.37 | 0.19 | 0.36 | 0.17 | 0.3 |
| 9 | 103556016 | rs10113967 | -0.349 | 0.293 | 0.956 | 1.88 | 0.008 | 0.089 | 0.089 | 0.017 | 0.12 | 0.24 | 0.19 | 0.34 | 0.14 | 0.26 |
| 9 | 103558414 | rs10739807 | -0.51 | 0.343 | 0.831 | 1.487 | 0.001 | -0.003 | -0.003 | 0.153 | 0.69 | 0.71 | 0.72 | 0.67 | 0.72 | 0.83 |
| 9 | 103571388 | rs1338729 | -0.536 | 0.306 | 0.956 | 1.474 | 0.001 | -0.005 | -0.005 | 0 | 0.69 | 0.74 | 0.71 | 0.67 | 0.72 | 0.85 |
| 9 | 103573103 | rs1338736 | -0.064 | -0.065 | 0.823 | 1.464 | -0.028 | 0.019 | 0.019 | 0 | 0.58 | 0.76 | 0.5 | 0.4 | 0.53 | 0.4 |
| 9 | 103578202 | rs1538410 | 0.751 | -0.223 | 1.089 | 1.344 | -0.005 | 0.185 | 0.185 | 0.362 | 0.45 | 0.14 | 0.4 | 0.12 | 0.5 | 0.33 |
| 9 | 103582699 | rs7861739 | -1.109 | -0.231 | 1.039 | 1.444 | -0.106 | -0.022 | -0.022 | 0 | 0.71 | 0.95 | 0.7 | 0.77 | 0.68 | 0.66 |
| 9 | 103585379 | rs7034844 | -0.107 | -0.255 | 1.693 | 1.443 | -0.028 | 0.086 | 0.086 | 1.568 | 0.09 | 0.35 | 0.04 | 0.19 | 0.01 | 0.05 |
| 9 | 103591484 | rs10819849 | 0.77 | -0.251 | 1.545 | 1.444 | -0.012 | 0.139 | 0.139 | 0.299 | 0.46 | 0.14 | 0.42 | 0.2 | 0.52 | 0.35 |
| 9 | 103598789 | rs7868230 | -1.199 | -0.244 | 1.642 | 1.498 | -0.045 | 0.126 | 0.126 | 3.928 | 0.13 | 0.59 | 0.09 | 0.28 | 0.03 | 0.06 |
| 9 | 103603201 | rs10512274 | 0.809 | -0.216 | 1.4 | 1.469 | 0.001 | 0.025 | 0.025 | 0 | 0.15 | 0.18 | 0.23 | 0.37 | 0.16 | 0.25 |
| 9 | 103603365 | rs1953805 | -1.253 | -0.215 | 1.337 | 1.408 | -0.123 | -0.042 | -0.042 | 0 | 0.72 | 0.97 | 0.7 | 0.76 | 0.68 | 0.65 |
| 9 | 103603761 | rs1953807 | -0.053 | -0.595 | 1.415 | 1.274 | 0.01 | 0.075 | 0.075 | 0 | 0.54 | 0.69 | 0.44 | 0.3 | 0.5 | 0.38 |
| 9 | 103620015 | rs10819859 | 0.678 | -0.55 | 1.476 | 1.191 | 0.005 | 0.021 | 0.021 | 0 | 0.46 | 0.35 | 0.4 | 0.1 | 0.49 | 0.33 |
| 9 | 103624177 | rs16919567 | 0.012 | -0.47 | 1.623 | 1.283 | -0.024 | -0.014 | -0.014 | 0.052 | 0.08 | 0.29 | 0.02 | 0.04 | 0.01 | 0.04 |
| 9 | 103644950 | rs10760775 | 0.63 | -0.457 | 1.53 | 1.246 | -0.036 | 0.215 | 0.215 | 0 | 0.42 | 0.08 | 0.38 | 0.1 | 0.49 | 0.34 |
| 9 | 103651056 | rs10819865 | -0.136 | -0.386 | 1.747 | 1.399 | -0.009 | 0.051 | 0.051 | 0 | 0.22 | 0.31 | 0.26 | 0.56 | 0.17 | 0.48 |
| 9 | 103658148 | rs1338733 | 0.166 | -0.381 | 1.741 | 1.424 | 0.003 | -0.025 | -0.025 | 0 | 0.21 | 0.16 | 0.22 | 0.47 | 0.17 | 0.47 |
| 9 | 103660338 | rs10819866 | -0.563 | -0.358 | 1.762 | 1.386 | 0.011 | -0.008 | -0.008 | 0.505 | 0.65 | 0.51 | 0.64 | 0.66 | 0.66 | 0.82 |
| 9 | 103665363 | rs1547689 | -0.447 | -0.371 | 1.783 | 1.396 | -0.026 | 0.04 | 0.04 | 0 | 0.24 | 0.46 | 0.28 | 0.7 | 0.17 | 0.49 |
| 9 | 103675913 | rs12552974 | -0.606 | -0.314 | 1.696 | 1.354 | 0.007 | -0.004 | -0.004 | 0 | 0.66 | 0.55 | 0.64 | 0.64 | 0.66 | 0.81 |
| 9 | 103691860 | rs13285273 | 0.898 | -0.288 | 1.548 | 1.471 | -0.001 | -0.025 | -0.025 | 0 | 0.16 | 0.11 | 0.21 | 0.37 | 0.16 | 0.24 |
| 9 | 103701886 | rs10819885 | -0.642 | -0.321 | 1.544 | 1.355 | 0.016 | -0.01 | -0.01 | 0 | 0.66 | 0.47 | 0.65 | 0.66 | 0.67 | 0.82 |
| 9 | 103704579 | rs12551278 | -0.262 | -0.337 | 1.458 | 1.377 | -0.01 | 0.042 | 0.042 | 0 | 0.23 | 0.31 | 0.26 | 0.55 | 0.17 | 0.48 |
| 9 | 103717654 | rs607449 | 0.671 | -0.238 | 1.354 | 1.501 | -0.007 | 0.006 | 0.006 | 0 | 0.32 | 0.32 | 0.32 | 0.27 | 0.34 | 0.18 |
| 9 | 103728088 | rs2245719 | 0.582 | -0.318 | 1.338 | 1.08 | -0.001 | -0.002 | -0.002 | 0 | 0.33 | 0.36 | 0.36 | 0.2 | 0.35 | 0.17 |
| 9 | 103741404 | rs2250068 | 0.435 | -0.445 | 1.459 | 1.032 | 0.026 | -0.054 | -0.054 | 0 | 0.34 | 0.51 | 0.37 | 0.19 | 0.35 | 0.17 |
| 9 | 103749454 | rs2486370 | -0.922 | -0.415 | 0.982 | 0.726 | -0.071 | -0.003 | -0.003 | 0 | 0.88 | 0.99 | 0.82 | 0.92 | 0.77 | 0.86 |
| 9 | 103763948 | rs13297102 | 0.043 | -0.507 | 1.378 | 0.972 | 0.007 | -0.026 | -0.026 | 2.603 | 0.21 | 0.15 | 0.22 | 0.47 | 0.17 | 0.48 |
| 9 | 103769373 | rs1330193 | 0.605 | -0.475 | 1.381 | 0.894 | 0.023 | -0.047 | -0.047 | 0 | 0.33 | 0.48 | 0.37 | 0.19 | 0.35 | 0.16 |
| 9 | 103779613 | rs628371 | -0.681 | -0.529 | 1.305 | 0.864 | -0.006 | 0.157 | 0.157 | 0 | 0.57 | 0.83 | 0.64 | 0.9 | 0.52 | 0.64 |
| 9 | 103787416 | rs654414 | -0.484 | -0.562 | 1.28 | 0.922 | 0 | 0.032 | 0.032 | 0 | 0.53 | 0.68 | 0.62 | 0.65 | 0.5 | 0.44 |
| 9 | 103790352 | rs2486381 | 0.687 | -0.374 | 0.819 | 0.809 | -0.002 | 0 | 0 | 0 | 0.32 | 0.33 | 0.36 | 0.18 | 0.35 | 0.16 |
| 9 | 103793544 | rs2251885 | 0.87 | -0.38 | 0.813 | 0.822 | -0.035 | 0.043 | 0.043 | 0 | 0.31 | 0.19 | 0.36 | 0.19 | 0.36 | 0.18 |
| 9 | 103794247 | rs649414 | -0.639 | -0.493 | 0.784 | 0.788 | -0.006 | 0.149 | 0.149 | 0 | 0.57 | 0.82 | 0.65 | 0.9 | 0.53 | 0.65 |
| 9 | 103794389 | rs649892 | 0.794 | -0.46 | 0.832 | 0.81 | -0.009 | 0.013 | 0.013 | 0 | 0.32 | 0.31 | 0.37 | 0.19 | 0.36 | 0.18 |
| 9 | 103796067 | rs2252151 | -0.097 | -0.732 | 0.932 | 0.946 | 0.009 | 0.054 | 0.054 | 0 | 0.51 | 0.64 | 0.6 | 0.73 | 0.53 | 0.45 |
| 9 | 103797594 | rs2480736 | -0.624 | -0.544 | 0.655 | 1.381 | 0.005 | 0.045 | 0.045 | 0.116 | 0.59 | 0.72 | 0.65 | 0.73 | 0.59 | 0.71 |
| 9 | 103801806 | rs2012728 | 0.659 | -0.487 | 0.774 | 1.452 | -0.019 | 0.057 | 0.057 | 0 | 0.19 | 0.25 | 0.21 | 0.36 | 0.17 | 0.25 |
| 9 | 103805207 | rs2256081 | -0.133 | -0.328 | 0.78 | 1.466 | 0.009 | -0.002 | -0.002 | 0 | 0.54 | 0.58 | 0.62 | 0.54 | 0.57 | 0.44 |
| 9 | 103809117 | rs476599 | -0.384 | -0.31 | 0.717 | 1.406 | 0.009 | 0.065 | 0.065 | 0.089 | 0.58 | 0.7 | 0.67 | 0.79 | 0.59 | 0.48 |
| 9 | 103835968 | rs495837 | -0.245 | -0.316 | 0.543 | 1.395 | 0 | 0.001 | 0.001 | 0.253 | 0.57 | 0.57 | 0.65 | 0.62 | 0.59 | 0.48 |
| 9 | 103840814 | rs624169 | -0.07 | -0.34 | 0.562 | 1.467 | -0.002 | -0.001 | -0.001 | 0 | 0.53 | 0.5 | 0.62 | 0.54 | 0.57 | 0.46 |
| 9 | 103848006 | rs1339422 | -0.723 | -0.336 | 0.56 | 1.379 | 0 | 0.084 | 0.084 | 0.863 | 0.62 | 0.78 | 0.68 | 0.89 | 0.61 | 0.72 |
| 9 | 103848261 | rs1339424 | 0.502 | -0.788 | 0.688 | 1.529 | -0.01 | 0.027 | 0.027 | 0 | 0.2 | 0.25 | 0.22 | 0.35 | 0.18 | 0.27 |
| 9 | 103858565 | rs7848935 | -0.749 | -0.496 | -0.218 | 0.492 | 0.018 | 0.067 | 0.067 | 1.113 | 0.63 | 0.77 | 0.71 | 0.89 | 0.65 | 0.72 |
| 9 | 103859537 | rs10819899 | 0.501 | -0.403 | -0.174 | 0.569 | -0.018 | 0.008 | 0.008 | 0 | 0.37 | 0.35 | 0.45 | 0.28 | 0.45 | 0.2 |
| 9 | 103861352 | rs2993805 | 0.509 | -0.402 | -0.173 | 0.585 | -0.018 | 0.006 | 0.006 | 0 | 0.37 | 0.35 | 0.45 | 0.28 | 0.45 | 0.2 |
| 9 | 103864180 | rs1329082 | -0.622 | -0.351 | -0.387 | 0.198 | 0.021 | 0.071 | 0.071 | 1.232 | 0.62 | 0.75 | 0.71 | 0.89 | 0.65 | 0.72 |
| 9 | 103871680 | rs12236239 | -0.542 | -0.194 | -0.106 | 0.823 | -0.03 | 0.015 | 0.015 | 0.181 | 0.29 | 0.47 | 0.3 | 0.71 | 0.24 | 0.54 |
| 9 | 103872681 | rs7467823 | 0.288 | -0.045 | -0.273 | 0.634 | -0.021 | -0.002 | -0.002 | 0 | 0.68 | 0.59 | 0.65 | 0.7 | 0.72 | 0.57 |
| 9 | 103873057 | rs10819903 | -0.486 | -0.036 | -0.561 | 0.309 | -0.004 | -0.009 | -0.009 | 3.356 | 0.44 | 0.44 | 0.48 | 0.76 | 0.35 | 0.67 |
| 9 | 103873786 | rs1410854 | -0.592 | 0.724 | -0.543 | 0.95 | 0.001 | 0.018 | 0.018 | 0 | 0.1 | 0.13 | 0.26 | 0.31 | 0.14 | 0.14 |
| 9 | 103874937 | rs2993807 | 0.077 | 0.186 | -1.251 | 0.934 | 0.026 | -0.021 | -0.021 | 1.646 | 0.24 | 0.12 | 0.1 | 0.29 | 0.17 | 0.22 |
| 9 | 103875396 | rs10156478 | 0.385 | 0.397 | -1.268 | 0.848 | -0.05 | -0.052 | -0.052 | 0 | 0.08 | 0.01 | 0.25 | 0.27 | 0.14 | 0.14 |
| 9 | 103876273 | rs2993808 | 0.591 | 0.492 | -1.211 | 0.973 | -0.007 | -0.028 | -0.028 | 0 | 0.64 | 0.76 | 0.63 | 0.39 | 0.67 | 0.58 |
| 9 | 103877068 | rs3983746 | 0.531 | 0.419 | -0.586 | 0.46 | 0.011 | -0.042 | -0.042 | 0 | 0.21 | 0.37 | 0.26 | 0.13 | 0.22 | 0.11 |
| 9 | 103878207 | rs10760791 | 0.744 | 0.354 | -0.046 | 0.275 | -0.001 | 0.036 | 0.036 | 0 | 0.46 | 0.59 | 0.41 | 0.26 | 0.48 | 0.48 |
| 9 | 103894388 | rs10989415 | 0.97 | 0.119 | -0.015 | 0.16 | -0.005 | 0.045 | 0.045 | 0.456 | 0.47 | 0.36 | 0.4 | 0.25 | 0.54 | 0.48 |
| 9 | 103910778 | rs3118941 | -0.946 | -0.003 | 0.155 | 0.228 | 0.035 | -0.049 | -0.049 | 0 | 0.79 | 0.89 | 0.91 | 0.63 | 0.89 | 0.76 |
| 9 | 103912602 | rs1431473 | 0.046 | 0.011 | 0.149 | -0.036 | 0.008 | -0.028 | -0.028 | 0 | 0.68 | 0.79 | 0.65 | 0.35 | 0.76 | 0.62 |
| 9 | 103920578 | rs2771064 | -1.135 | 0.037 | -0.004 | -0.212 | -0.008 | 0.11 | 0.11 | 1.409 | 0.8 | 0.99 | 0.95 | 0.97 | 0.87 | 0.84 |
| 9 | 103921194 | rs2987746 | 1.073 | -0.022 | 0.294 | -0.232 | 0.005 | 0.046 | 0.046 | 0.047 | 0.47 | 0.62 | 0.41 | 0.27 | 0.5 | 0.47 |
| 9 | 103922830 | rs2771063 | 0.261 | -0.008 | 0.174 | -0.216 | -0.002 | 0 | 0 | 0.679 | 0.63 | 0.48 | 0.62 | 0.35 | 0.71 | 0.6 |

| | | | | | | | | | | | | | | | | |
|---|---|---|---|---|---|---|---|---|---|---|---|---|---|---|---|---|
| 9 | 103922841 | rs16919990 | 1.934 | -0.498 | -0.268 | -0.524 | -0.029 | -0.034 | -0.034 | 0.763 | 0.07 | 0.15 | 0.02 | 0 | 0.01 | 0 |
| 9 | 103923185 | rs438403 | 0.347 | 0.113 | -0.264 | -0.173 | 0.001 | 0.001 | 0.001 | 0.622 | 0.63 | 0.37 | 0.62 | 0.33 | 0.71 | 0.6 |
| 9 | 103923601 | rs382245 | 1.285 | 0.118 | -0.265 | -0.169 | -0.004 | 0.06 | 0.06 | 0.176 | 0.44 | 0.29 | 0.39 | 0.26 | 0.5 | 0.47 |
| 9 | 103927286 | rs1431463 | 1.292 | 0.112 | -0.268 | -0.171 | -0.004 | 0.059 | 0.059 | 0.186 | 0.44 | 0.29 | 0.39 | 0.26 | 0.5 | 0.47 |
| 9 | 103930908 | rs1367754 | 0.33 | 0.282 | -0.638 | -1.035 | -0.014 | 0.049 | 0.049 | 0 | 0.19 | 0.08 | 0.23 | 0.08 | 0.21 | 0.13 |
| 9 | 103934113 | rs10989419 | 0.409 | 0.287 | -0.46 | -1.003 | 0 | 0.021 | 0.021 | 0 | 0.18 | 0.1 | 0.21 | 0.07 | 0.18 | 0.12 |
| 9 | 103937432 | rs1346941 | 1.136 | 0.185 | -0.576 | -1.281 | -0.001 | -0.026 | -0.026 | 4.192 | 0.39 | 0.61 | 0.36 | 0.04 | 0.44 | 0.24 |
| 9 | 103939007 | rs9697078 | 0.127 | -0.031 | -0.538 | -0.757 | 0.025 | 0.001 | 0.001 | 0 | 0.67 | 0.82 | 0.83 | 0.37 | 0.75 | 0.5 |
| 9 | 103946357 | rs1864662 | -0.348 | -0.22 | -0.195 | -0.837 | -0.013 | -0.05 | -0.05 | 0.935 | 0.78 | 0.78 | 0.93 | 0.97 | 0.89 | 0.85 |
| 9 | 103952270 | rs7029916 | -0.002 | -0.008 | -0.246 | -0.772 | -0.002 | 0.053 | 0.053 | 0.022 | 0.13 | 0.21 | 0.27 | 0.28 | 0.1 | 0.18 |
| 9 | 103955515 | rs2987735 | 0.664 | 0.009 | -0.491 | -0.849 | -0.019 | -0.014 | -0.014 | 0 | 0.65 | 0.6 | 0.66 | 0.68 | 0.79 | 0.63 |
| 9 | 103955777 | rs12347855 | 0.179 | 0.238 | -1.388 | -0.586 | -0.004 | -0.047 | -0.047 | 0 | 0.12 | 0.07 | 0.26 | 0.3 | 0.1 | 0.21 |
| 9 | 103960732 | rs1346942 | 0.484 | 0.211 | -1.397 | -0.651 | -0.002 | 0.075 | 0.075 | 0 | 0.12 | 0.17 | 0.08 | 0.62 | 0.13 | 0.31 |
| 9 | 103964075 | rs2567310 | -0.514 | 0.23 | -1.734 | -0.464 | 0.01 | 0.007 | 0.007 | 3.543 | 0.81 | 0.85 | 0.94 | 0.96 | 0.9 | 0.85 |
| 9 | 103968906 | rs12351067 | -0.573 | 0.935 | -1.639 | 0.137 | -0.013 | -0.044 | -0.044 | 0.575 | 0.12 | 0.06 | 0.27 | 0.27 | 0.11 | 0.21 |
| 9 | 103969339 | rs17831028 | 0.135 | 1.04 | -1.502 | -0.389 | -0.036 | 0.042 | 0.042 | 0 | 0.11 | 0.04 | 0.23 | 0.02 | 0.18 | 0.06 |
| 9 | 103969740 | rs10125791 | -0.124 | 0.529 | -1.491 | -0.772 | 0.024 | 0.051 | 0.051 | 0 | 0.36 | 0.71 | 0.23 | 0.09 | 0.28 | 0.32 |
| 9 | 103969852 | rs7047561 | -0.836 | 1.544 | -1.107 | 0.302 | 0.007 | 0.018 | 0.018 | 0.002 | 0.1 | 0.15 | 0.45 | 0.31 | 0.2 | 0.26 |
| 9 | 103976442 | rs2987742 | -0.283 | 1.209 | -0.896 | -0.965 | -0.005 | -0.042 | -0.042 | 3.282 | 0.7 | 0.8 | 0.77 | 0.99 | 0.65 | 0.93 |
| 9 | 103993352 | rs1367756 | -0.189 | 0.107 | -1.234 | -0.591 | 0.05 | 0.06 | 0.06 | 2.237 | 0.51 | 0.84 | 0.46 | 0.38 | 0.4 | 0.58 |
| 9 | 104010913 | rs4743454 | 0.551 | -0.628 | -0.791 | -1.002 | -0.005 | 0.01 | 0.01 | 0 | 0.3 | 0.3 | 0.35 | 0.28 | 0.33 | 0.31 |
| 9 | 104012075 | rs10819912 | -0.199 | -0.933 | -0.43 | -1.041 | 0.005 | 0 | 0 | 0 | 0.41 | 0.67 | 0.43 | 0.4 | 0.37 | 0.52 |
| 9 | 104013824 | rs750662 | 0.326 | -0.807 | -0.346 | -1.097 | -0.001 | -0.002 | -0.002 | 0 | 0.33 | 0.38 | 0.37 | 0.29 | 0.33 | 0.32 |
| 9 | 104021756 | rs12237965 | 0.212 | -0.79 | -0.73 | -1.038 | 0.03 | 0.01 | 0.01 | 2.52 | 0.16 | 0.1 | 0.07 | 0.1 | 0.06 | 0.27 |
| 9 | 104030793 | rs2033394 | -0.528 | -0.436 | -0.066 | -1.287 | 0.003 | 0.005 | 0.005 | 3.977 | 0.5 | 0.59 | 0.43 | 0.31 | 0.39 | 0.4 |
| 9 | 104032801 | rs2567312 | -1.572 | 0.038 | -0.147 | -1.484 | -0.07 | 0.087 | 0.087 | 0.002 | 0.79 | 0.95 | 0.8 | 1 | 0.68 | 0.94 |
| 9 | 104033649 | rs7034058 | -0.461 | -0.246 | 0.388 | -1.176 | 0.013 | 0.041 | 0.041 | 2.736 | 0.46 | 0.39 | 0.39 | 0.29 | 0.39 | 0.37 |
| 9 | 104033919 | rs7035407 | -0.348 | -0.568 | 0.089 | -1.458 | -0.001 | -0.002 | -0.002 | 0 | 0.35 | 0.39 | 0.35 | 0.28 | 0.31 | 0.26 |
| 9 | 104034955 | rs1329077 | -0.778 | -0.461 | 0.215 | -1.396 | 0.008 | 0.022 | 0.022 | 2.859 | 0.46 | 0.42 | 0.39 | 0.29 | 0.39 | 0.36 |
| 9 | 104039202 | rs10124153 | 0.03 | -0.944 | -0.092 | -1.474 | 0.055 | 0.108 | 0.108 | 14.425 | 0.21 | 0.03 | 0.07 | 0.01 | 0.11 | 0.17 |
| 9 | 104047608 | rs10116399 | -0.172 | -1.02 | 0.178 | -1.489 | 0.004 | -0.028 | -0.028 | 0 | 0.22 | 0.23 | 0.06 | 0.01 | 0.09 | 0.15 |
| 9 | 104066880 | rs10512279 | -1.054 | -0.738 | 0.832 | -1.425 | 0.003 | -0.004 | -0.004 | 0 | 0.44 | 0.58 | 0.34 | 0.22 | 0.36 | 0.36 |
| 9 | 104068545 | rs6479048 | 0.08 | -0.779 | 1.074 | -1.505 | -0.011 | -0.061 | -0.061 | 0 | 0.2 | 0.19 | 0.07 | 0.01 | 0.11 | 0.18 |
| 9 | 104072889 | rs11790209 | 0.556 | -0.683 | 0.991 | -1.413 | 0 | 0 | 0 | 0 | 0.2 | 0.21 | 0.25 | 0.2 | 0.23 | 0.18 |
| 9 | 104075766 | rs7869625 | 0.01 | -0.728 | 1.377 | -1.432 | -0.011 | -0.023 | -0.023 | 0 | 0.24 | 0.48 | 0.27 | 0.22 | 0.23 | 0.18 |
| 9 | 104078061 | rs7848242 | 0.512 | -0.774 | 1.345 | -1.415 | -0.004 | -0.007 | -0.007 | 0 | 0.2 | 0.35 | 0.26 | 0.22 | 0.23 | 0.18 |
| 9 | 104090736 | rs11790501 | 0.359 | -0.709 | 1.447 | -1.361 | -0.006 | -0.006 | -0.006 | 0.108 | 0.21 | 0.35 | 0.26 | 0.22 | 0.23 | 0.19 |
| 9 | 104091467 | rs7040055 | -1.132 | -0.724 | 0.928 | -1.372 | 0.03 | 0.048 | 0.048 | 2.977 | 0.45 | 0.67 | 0.35 | 0.24 | 0.35 | 0.37 |
| 9 | 104093054 | rs13284177 | -0.663 | -0.293 | 0.768 | -0.987 | -0.009 | -0.025 | -0.025 | 0.616 | 0.28 | 0.4 | 0.26 | 0.24 | 0.23 | 0.31 |
| 9 | 104108650 | rs2417250 | -0.043 | -0.063 | 0.173 | -0.008 | 0 | 0.015 | 0.015 | 0.016 | 0.81 | 0.78 | 0.54 | 0.43 | 0.76 | 0.66 |
| 9 | 104109389 | rs16920222 | 0.086 | -0.42 | 0.331 | -0.445 | -0.014 | -0.059 | -0.059 | 0 | 0.24 | 0.29 | 0.16 | 0.05 | 0.16 | 0.19 |
| 9 | 104110677 | rs1417999 | 1.273 | -0.436 | 0.128 | -0.41 | 0.021 | 0.172 | 0.172 | 4.64 | 0.52 | 0.12 | 0.35 | 0.24 | 0.6 | 0.38 |
| 9 | 104123762 | rs7043029 | -0.165 | -0.19 | 0.372 | -0.051 | 0.019 | 0.06 | 0.06 | 0.814 | 0.86 | 0.75 | 0.77 | 0.67 | 0.85 | 0.62 |
| 9 | 104124664 | rs2229594 | -0.134 | -0.182 | 0.088 | -0.063 | 0.036 | 0.097 | 0.097 | 0.901 | 0.86 | 0.68 | 0.77 | 0.67 | 0.85 | 0.62 |
| 9 | 104126523 | rs4322060 | 1.258 | -0.282 | 0.03 | -0.155 | -0.011 | -0.03 | -0.03 | 0 | 0.11 | 0.17 | 0.05 | 0.03 | 0.07 | 0.06 |
| 9 | 104126546 | rs1341762 | 0.344 | -0.174 | 0.048 | -0.073 | -0.016 | -0.07 | -0.07 | 0 | 0.22 | 0.3 | 0.15 | 0.05 | 0.16 | 0.17 |
| 9 | 104133628 | rs1572983 | -0.042 | -0.149 | -0.612 | 0.2 | -0.003 | -0.003 | -0.003 | 0 | 0.63 | 0.45 | 0.63 | 0.63 | 0.69 | 0.47 |
| 9 | 104134777 | rs1891218 | 0.027 | -0.062 | -1.235 | 0.246 | 0.002 | 0.002 | 0.002 | 7.207 | 0.62 | 0.36 | 0.61 | 0.63 | 0.69 | 0.49 |
| 9 | 104142941 | rs11496612 | 1.356 | -0.307 | -1.753 | 0.071 | -0.114 | 0.1 | 0.1 | 0.48 | 0.28 | 0.08 | 0.23 | 0.08 | 0.41 | 0.15 |
| 9 | 104152155 | rs10118966 | -0.393 | -0.172 | -1.058 | 0.482 | 0.002 | 0.003 | 0.003 | 7.613 | 0.38 | 0.66 | 0.39 | 0.37 | 0.31 | 0.51 |
| 9 | 104161154 | rs2253258 | -0.221 | -0.206 | -1.033 | 0.72 | 0.019 | -0.042 | -0.042 | 0 | 0.87 | 0.96 | 0.81 | 0.77 | 0.88 | 0.94 |
| 9 | 104163789 | rs10819926 | 0.44 | 0.148 | -0.808 | -0.207 | -0.003 | 0.003 | 0.003 | 0 | 0.3 | 0.26 | 0.21 | 0.12 | 0.38 | 0.25 |
| 9 | 104164162 | rs2777336 | 0.134 | -0.276 | -0.589 | -0.404 | 0.008 | 0.017 | 0.017 | 0 | 0.39 | 0.51 | 0.49 | 0.51 | 0.39 | 0.61 |
| 9 | 104165360 | rs17185855 | 0.44 | -0.494 | -0.621 | -0.909 | 0.002 | -0.023 | -0.023 | 0 | 0.16 | 0.12 | 0.1 | 0 | 0.12 | 0.1 |
| 9 | 104167718 | rs2777345 | -0.455 | -0.683 | -0.362 | -1.025 | 0.002 | 0.012 | 0.012 | 0 | 0.49 | 0.45 | 0.62 | 0.41 | 0.49 | 0.72 |
| 9 | 104169089 | rs3739719 | 1.395 | -0.691 | -0.438 | -1.114 | -0.01 | 0.034 | 0.034 | 0 | 0.31 | 0.2 | 0.21 | 0.07 | 0.39 | 0.13 |
| 9 | 104172936 | rs546577 | -0.583 | -0.625 | -0.36 | -0.955 | 0.001 | 0.004 | 0.004 | 0.011 | 0.51 | 0.57 | 0.63 | 0.41 | 0.49 | 0.72 |
| 9 | 104184022 | rs4577 | 0.311 | -0.399 | -0.262 | -0.509 | 0.001 | 0 | 0 | 0.006 | 0.32 | 0.37 | 0.52 | 0.4 | 0.37 | 0.6 |
| 9 | 104191392 | rs550915 | 1.077 | 0.075 | -1.06 | -0.8 | -0.039 | 0.095 | 0.095 | 0 | 0.17 | 0.01 | 0.08 | 0.01 | 0.16 | 0.3 |
| 9 | 104194532 | rs533017 | 0.529 | 0.209 | -0.465 | -0.345 | -0.004 | 0.037 | 0.037 | 0 | 0.33 | 0.47 | 0.36 | 0.45 | 0.31 | 0.58 |
| 9 | 104202803 | rs479600 | 1.143 | 0.285 | -0.464 | 0.173 | 0.005 | -0.064 | -0.064 | 0.584 | 0.69 | 0.83 | 0.63 | 0.53 | 0.75 | 0.74 |
| 9 | 104204614 | rs573816 | 1.661 | 0.258 | -0.43 | 0.12 | 0 | -0.001 | -0.001 | 0 | 0.63 | 0.59 | 0.6 | 0.53 | 0.71 | 0.71 |
| 9 | 104210432 | rs10760796 | -0.112 | 0.327 | -0.763 | 0.169 | -0.029 | -0.011 | -0.011 | 0.563 | 0.92 | 0.99 | 0.72 | 0.53 | 0.9 | 0.84 |
| 9 | 104211810 | rs7035564 | -0.436 | 0.395 | -0.642 | 0.203 | 0.005 | -0.031 | -0.031 | 0 | 0.06 | 0.16 | 0.03 | 0 | 0.05 | 0.03 |
| 9 | 104217909 | rs2065395 | 0.153 | 0.443 | -1.012 | 0.333 | 0.005 | -0.025 | -0.025 | 8.606 | 0.79 | 0.83 | 0.68 | 0.54 | 0.84 | 0.81 |
| 9 | 104222989 | rs7872599 | -0.613 | 0.718 | -0.714 | -0.188 | 0 | -0.04 | -0.04 | 1.286 | 0.16 | 0.21 | 0.09 | 0 | 0.14 | 0.09 |
| 9 | 104226808 | rs4742814 | 0.04 | 0.738 | -0.991 | 0.323 | -0.007 | 0 | 0 | 0 | 0.53 | 0.69 | 0.51 | 0.59 | 0.5 | 0.58 |
| 9 | 104228493 | rs9409333 | 0.213 | 0.841 | -0.576 | 0.528 | 0.032 | 0.021 | 0.021 | 0 | 0.78 | 0.54 | 0.72 | 0.75 | 0.74 | 0.84 |
| 9 | 104236774 | rs1929494 | 0.969 | 0.404 | -0.497 | 0.444 | -0.015 | 0.002 | 0.002 | 0 | 0.42 | 0.24 | 0.37 | 0.47 | 0.45 | 0.55 |

| | | | | | | | | | | | | | | | | |
|---|---|---|---|---|---|---|---|---|---|---|---|---|---|---|---|---|
| 9 | 104237055 | rs13292501 | -0.104 | 0.191 | -0.786 | 0.089 | -0.03 | 0.102 | 0.102 | 0 | 0.69 | 0.9 | 0.74 | 0.88 | 0.61 | 0.83 |
| 9 | 104243757 | rs2297355 | 0.001 | 0.324 | -0.343 | 0.691 | 0.058 | 0.238 | 0.238 | 0 | 0.05 | 0.23 | 0.18 | 0.22 | 0.05 | 0.17 |
| 9 | 104246779 | rs1929482 | 0.44 | 0.254 | -0.776 | 0.479 | 0 | -0.011 | -0.011 | 0 | 0.34 | 0.36 | 0.37 | 0.57 | 0.36 | 0.5 |
| 9 | 104253681 | rs7867496 | 0.628 | 0.086 | -0.721 | 0.284 | 0.02 | 0.019 | 0.019 | 0 | 0.36 | 0.25 | 0.18 | 0.36 | 0.3 | 0.35 |
| 9 | 104262460 | rs7045287 | 1.046 | 0.015 | -0.474 | 0.226 | -0.015 | -0.055 | -0.055 | 1.007 | 0.14 | 0.16 | 0.09 | 0 | 0.1 | 0.04 |
| 9 | 104263799 | rs1929478 | -0.861 | 0.555 | -0.597 | 0.125 | -0.007 | 0.061 | 0.061 | 0.21 | 0.61 | 0.77 | 0.66 | 0.82 | 0.61 | 0.68 |
| 9 | 104266953 | rs1929496 | 0.082 | 0.695 | -0.529 | 0.281 | 0.026 | 0.116 | 0.116 | 0 | 0.1 | 0.19 | 0.2 | 0.26 | 0.13 | 0.15 |
| 9 | 104268342 | rs16920447 | 1.901 | 0.753 | -0.615 | 0.356 | -0.065 | -0.11 | -0.11 | 0 | 0.23 | 0.01 | 0.16 | 0.43 | 0.25 | 0.38 |
| 9 | 104269009 | rs13290720 | 0.087 | 0.762 | -0.541 | 0.388 | 0.026 | 0.116 | 0.116 | 0 | 0.1 | 0.19 | 0.2 | 0.26 | 0.13 | 0.15 |
| 9 | 104273901 | rs1407869 | 0.667 | 0.739 | -0.432 | 0.37 | 0.062 | 0.144 | 0.144 | 0 | 0.06 | 0.3 | 0.19 | 0.13 | 0.1 | 0.08 |
| 9 | 104274944 | rs1407868 | 1.601 | 0.706 | -0.625 | 0.307 | -0.027 | -0.08 | -0.08 | 3.958 | 0.26 | 0.03 | 0.17 | 0.43 | 0.24 | 0.38 |
| 9 | 104275622 | rs4743466 | -0.753 | 0.648 | -0.606 | 0.231 | -0.014 | 0.084 | 0.084 | 0 | 0.57 | 0.76 | 0.65 | 0.82 | 0.57 | 0.67 |
| 9 | 104277937 | rs12684621 | 1.96 | 0.606 | -0.742 | 0.224 | -0.013 | -0.064 | -0.064 | 0.022 | 0.22 | 0.03 | 0.09 | 0.37 | 0.2 | 0.32 |
| 9 | 104279275 | rs11790317 | 1.349 | 0.565 | -0.598 | 0.152 | 0.002 | -0.02 | -0.02 | 0.113 | 0.13 | 0.1 | 0.09 | 0 | 0.1 | 0.04 |
| 9 | 104291812 | rs7023854 | -0.177 | 0.773 | -0.432 | 0.261 | 0.007 | 0.036 | 0.036 | 0.443 | 0.5 | 0.38 | 0.45 | 0.68 | 0.47 | 0.59 |
| 9 | 104292666 | rs10989551 | 1.584 | 0.903 | -0.771 | -0.125 | -0.016 | 0.053 | 0.053 | 1.065 | 0.1 | 0.01 | 0.09 | 0 | 0.09 | 0.09 |
| 9 | 104295764 | rs3818337 | -0.511 | 1.073 | -0.507 | 0.04 | -0.001 | 0.026 | 0.026 | 0 | 0.57 | 0.66 | 0.66 | 0.82 | 0.58 | 0.66 |
| 9 | 104308365 | rs12343288 | -0.343 | 1.073 | -0.541 | 0.315 | 0.019 | 0.109 | 0.109 | 0 | 0.12 | 0.25 | 0.2 | 0.26 | 0.12 | 0.13 |
| 9 | 104309445 | rs2273976 | 0.717 | 1.055 | -0.639 | 0.516 | 0.013 | 0.004 | 0.004 | 0.571 | 0.17 | 0.08 | 0.15 | 0.11 | 0.15 | 0.08 |
| 9 | 104315255 | rs16920483 | 2.034 | 1.081 | -0.712 | 0.363 | -0.029 | -0.088 | -0.088 | 0 | 0.21 | 0 | 0.09 | 0.36 | 0.19 | 0.32 |
| 9 | 104315931 | rs10989553 | -0.544 | 1.053 | -0.67 | 0.345 | 0.005 | 0.07 | 0.07 | 0 | 0.14 | 0.24 | 0.21 | 0.26 | 0.12 | 0.13 |
| 9 | 104324545 | rs10521057 | 2.026 | 1.049 | -0.759 | 0.327 | -0.033 | -0.088 | -0.088 | 0.255 | 0.21 | 0 | 0.09 | 0.36 | 0.19 | 0.32 |
| 9 | 104334333 | rs10512282 | 1.608 | 1.028 | -0.677 | 0.224 | 0.011 | 0.009 | 0.009 | 0.408 | 0.13 | 0.07 | 0.08 | 0 | 0.11 | 0.04 |
| 9 | 104334777 | rs3824473 | -0.26 | 1.044 | -0.611 | 0.258 | -0.005 | 0.005 | 0.005 | 2.15 | 0.26 | 0.29 | 0.29 | 0.26 | 0.22 | 0.17 |
| 9 | 104335682 | rs3739722 | -0.524 | 1.046 | -0.606 | 0.27 | 0.003 | 0.049 | 0.049 | 0 | 0.14 | 0.22 | 0.2 | 0.26 | 0.12 | 0.13 |
| 9 | 104337744 | rs7859863 | 1.104 | 1.174 | -0.61 | 0.23 | -0.006 | 0.014 | 0.014 | 0.289 | 0.14 | 0.08 | 0.18 | 0.07 | 0.18 | 0.15 |
| 9 | 104338416 | rs11791337 | 1.592 | 1.002 | -0.617 | 0.117 | 0.01 | 0.009 | 0.009 | 0.437 | 0.13 | 0.07 | 0.08 | 0 | 0.11 | 0.04 |
| 9 | 104338456 | rs10081707 | -0.609 | 1.078 | -0.382 | 0.115 | 0.005 | 0.041 | 0.041 | 0 | 0.59 | 0.68 | 0.68 | 0.84 | 0.62 | 0.67 |
| 9 | 104343719 | rs10123961 | 0.562 | 0.478 | 0.055 | 0.723 | 0.001 | 0.001 | 0.001 | 0 | 0.35 | 0.33 | 0.31 | 0.28 | 0.32 | 0.25 |
| 9 | 104350424 | rs7873495 | 0.834 | 0.555 | 0.074 | 0.946 | 0.015 | 0.084 | 0.084 | 0 | 0.06 | 0.18 | 0.19 | 0.15 | 0.1 | 0.07 |
| 9 | 104352574 | rs4743473 | -0.21 | 0.376 | -0.081 | 0.77 | 0.011 | 0.004 | 0.004 | 0.665 | 0.62 | 0.54 | 0.52 | 0.39 | 0.57 | 0.5 |
| 9 | 104355455 | rs1407877 | 0.75 | 0.327 | -0.315 | 0.897 | 0.004 | 0.031 | 0.031 | 0.382 | 0.12 | 0.21 | 0.11 | 0.28 | 0.09 | 0.1 |
| 9 | 104356439 | rs3739724 | 0.093 | -0.03 | -0.356 | 0.794 | 0.014 | -0.008 | -0.008 | 0 | 0.23 | 0.19 | 0.21 | 0.26 | 0.13 | 0.22 |
| 9 | 104358072 | rs1323433 | -0.279 | -0.122 | -0.368 | 0.617 | 0.047 | 0.006 | 0.006 | 1.909 | 0.34 | 0.21 | 0.29 | 0.27 | 0.22 | 0.26 |
| 9 | 104359035 | rs7858998 | -0.32 | -0.123 | -0.366 | 0.617 | 0.042 | 0.007 | 0.007 | 2.091 | 0.34 | 0.23 | 0.29 | 0.27 | 0.23 | 0.26 |
| 9 | 104359261 | rs6479056 | -1.46 | 0.077 | -0.429 | 0.531 | -0.024 | 0.013 | 0.013 | 0 | 0.81 | 0.95 | 0.83 | 0.84 | 0.76 | 0.68 |
| 9 | 104362937 | rs1323432 | 2.042 | -0.407 | -0.349 | 0.595 | 0.011 | 0.032 | 0.032 | 0.014 | 0.12 | 0.03 | 0.08 | 0 | 0.11 | 0.04 |
| 9 | 104368397 | rs9969726 | -0.755 | -0.354 | -0.25 | 0.68 | -0.004 | -0.003 | -0.003 | 0 | 0.68 | 0.72 | 0.71 | 0.55 | 0.65 | 0.57 |
| 9 | 104371037 | rs1885979 | -0.178 | -1.011 | 0.163 | 0.619 | 0.012 | 0 | 0 | 0.008 | 0.48 | 0.61 | 0.49 | 0.43 | 0.44 | 0.39 |
| 9 | 104371736 | rs942141 | -0.26 | -0.981 | -0.054 | 0.576 | 0.022 | 0.003 | 0.003 | 0 | 0.48 | 0.65 | 0.43 | 0.43 | 0.41 | 0.45 |
| 9 | 104373787 | rs9299346 | 0.673 | -0.98 | 0.003 | 0.585 | -0.004 | -0.06 | -0.06 | 0.008 | 0.33 | 0.2 | 0.34 | 0.39 | 0.34 | 0.35 |
| 9 | 104374738 | rs7875712 | -0.768 | -0.867 | -0.008 | 0.658 | -0.022 | -0.026 | -0.026 | 0 | 0.61 | 0.76 | 0.52 | 0.43 | 0.51 | 0.48 |
| 9 | 104378003 | rs10121600 | 0.344 | -0.501 | -0.367 | 0.684 | -0.006 | -0.041 | -0.041 | 0.263 | 0.38 | 0.56 | 0.35 | 0.25 | 0.36 | 0.31 |
| 9 | 104378479 | rs10117679 | 0.51 | -0.235 | -0.185 | 0.878 | -0.045 | -0.09 | -0.09 | 0.471 | 0.15 | 0.39 | 0.1 | 0.04 | 0.06 | 0.12 |
| 9 | 104385711 | rs10989563 | 1.713 | -0.168 | -0.282 | 0.888 | -0.006 | 0.129 | 0.129 | 0 | 0.26 | 0.07 | 0.17 | 0.05 | 0.24 | 0.17 |
| 9 | 104385800 | rs10989565 | 0.343 | -0.234 | -0.261 | 0.987 | 0.036 | 0.055 | 0.055 | 0 | 0.43 | 0.2 | 0.33 | 0.29 | 0.38 | 0.32 |
| 9 | 104388863 | rs1570514 | -0.654 | -0.143 | -0.233 | 1.112 | 0.003 | 0.007 | 0.007 | 0 | 0.61 | 0.59 | 0.45 | 0.52 | 0.47 | 0.43 |
| 9 | 104390195 | rs2417290 | -0.694 | -0.172 | -0.218 | 1.138 | -0.036 | 0.024 | 0.024 | 1.151 | 0.35 | 0.44 | 0.28 | 0.48 | 0.23 | 0.26 |
| 9 | 104399139 | rs10989574 | -0.831 | -0.284 | -0.363 | 1.047 | 0.016 | 0.015 | 0.015 | 0 | 0.64 | 0.6 | 0.56 | 0.51 | 0.53 | 0.53 |
| 9 | 104404261 | rs7856074 | -0.716 | -0.355 | -0.384 | 0.878 | -0.024 | 0.017 | 0.017 | 0.534 | 0.37 | 0.53 | 0.39 | 0.45 | 0.3 | 0.36 |
| 9 | 104409985 | rs12342601 | 0.042 | -0.353 | -0.403 | 0.899 | -0.014 | 0.106 | 0.106 | 0 | 0.07 | 0.18 | 0.09 | 0.22 | 0.04 | 0.12 |
| 9 | 104413034 | rs13296742 | 1.626 | -0.324 | -0.535 | 0.767 | -0.008 | 0.133 | 0.133 | 0 | 0.26 | 0.07 | 0.17 | 0.05 | 0.24 | 0.17 |
| 9 | 104417812 | rs1924032 | -0.167 | -0.313 | -0.414 | 0.836 | -0.001 | -0.002 | -0.002 | 0 | 0.56 | 0.47 | 0.46 | 0.28 | 0.5 | 0.41 |
| 9 | 104420176 | rs12685672 | 0.272 | -0.304 | -0.367 | 0.942 | -0.012 | 0.078 | 0.078 | 0 | 0.07 | 0.12 | 0.09 | 0.22 | 0.04 | 0.11 |
| 9 | 104425362 | rs16920640 | 0.437 | -0.348 | -0.511 | 0.251 | -0.042 | -0.071 | -0.071 | 0.05 | 0.17 | 0.34 | 0.1 | 0.06 | 0.09 | 0.1 |
| 9 | 104425619 | rs1323416 | 0.548 | -0.252 | -0.676 | 0.301 | -0.026 | 0.105 | 0.105 | 0 | 0.39 | 0.13 | 0.36 | 0.22 | 0.41 | 0.3 |
| 9 | 104432873 | rs942142 | 0.564 | -0.24 | -0.702 | 0.308 | -0.029 | 0.103 | 0.103 | 0 | 0.39 | 0.12 | 0.36 | 0.22 | 0.41 | 0.3 |
| 9 | 104433034 | rs10512285 | 0.564 | -0.24 | -0.702 | 0.308 | -0.029 | 0.103 | 0.103 | 0 | 0.39 | 0.12 | 0.36 | 0.22 | 0.41 | 0.3 |
| 9 | 104433347 | rs13284303 | 1.276 | -0.14 | -0.533 | 0.605 | -0.046 | -0.054 | -0.054 | 0 | 0.13 | 0.01 | 0.19 | 0.17 | 0.17 | 0.13 |
| 9 | 104437373 | rs7469637 | 0.211 | -0.255 | -0.301 | 0.354 | -0.043 | -0.072 | -0.072 | 0 | 0.17 | 0.34 | 0.1 | 0.06 | 0.09 | 0.1 |
| 9 | 104439370 | rs1323423 | 0.033 | -0.07 | -0.4 | 0.828 | 0 | -0.016 | -0.016 | 0 | 0.2 | 0.21 | 0.3 | 0.58 | 0.21 | 0.25 |
| 9 | 104443319 | rs2506352 | 0.009 | -0.057 | -0.411 | 0.835 | -0.001 | -0.028 | -0.028 | 0.733 | 0.2 | 0.13 | 0.29 | 0.57 | 0.21 | 0.25 |
| 9 | 104445493 | rs2485531 | -0.096 | -0.059 | -0.412 | 0.834 | -0.001 | -0.029 | -0.029 | 0.549 | 0.8 | 0.87 | 0.71 | 0.43 | 0.79 | 0.74 |
| 9 | 104449063 | rs2506354 | -0.096 | -0.059 | -0.412 | 0.834 | -0.001 | -0.029 | -0.029 | 0 | 0.8 | 0.87 | 0.71 | 0.43 | 0.79 | 0.74 |
| 9 | 104449098 | rs10989591 | 0.429 | -0.142 | -0.448 | 0.483 | -0.007 | -0.067 | -0.067 | 0 | 0.44 | 0.53 | 0.28 | 0.11 | 0.33 | 0.27 |
| 9 | 104454025 | rs2485522 | 0.327 | 0.093 | -0.3 | 0.724 | 0.01 | 0 | 0 | 0 | 0.37 | 0.33 | 0.43 | 0.32 | 0.46 | 0.47 |
| 9 | 104459296 | rs1415645 | 1.034 | 0.523 | -0.613 | 0.753 | -0.009 | -0.002 | -0.002 | 0 | 0.13 | 0.05 | 0.13 | 0.17 | 0.16 | 0.29 |
| 9 | 104474060 | rs7043519 | 0.894 | -0.314 | -0.497 | 1.065 | -0.021 | -0.049 | -0.049 | 0 | 0.12 | 0.19 | 0.04 | 0.01 | 0.05 | 0.09 |
| 9 | 104475496 | rs945870 | 0.543 | -0.283 | -0.385 | 1.081 | 0.003 | 0.181 | 0.181 | 0 | 0.92 | 0.85 | 0.89 | 0.61 | 0.93 | 0.85 |

| | | | | | | | | | | | | | | | | |
|---|---|---|---|---|---|---|---|---|---|---|---|---|---|---|---|---|
| 9 | 104482845 | rs7863180 | -0.094 | -0.3 | -0.525 | 0.952 | 0.009 | 0.048 | 0.048 | 0 | 0.51 | 0.69 | 0.52 | 0.39 | 0.54 | 0.55 |
| 9 | 104489367 | rs2050641 | -1.17 | -0.285 | -0.619 | 0.784 | 0.017 | -0.058 | -0.058 | 0 | 0.68 | 0.92 | 0.72 | 0.57 | 0.74 | 0.71 |
| 9 | 104492591 | rs7041051 | 0.576 | -0.339 | -0.45 | 0.955 | -0.049 | 0.09 | 0.09 | 0 | 0.15 | 0.37 | 0.07 | 0.28 | 0.06 | 0.17 |
| 9 | 104493766 | rs1415644 | 0.521 | -0.347 | -0.492 | 1.017 | -0.007 | 0 | 0 | 0.014 | 0.17 | 0.21 | 0.23 | 0.17 | 0.21 | 0.13 |
| 9 | 104495190 | rs4278209 | 0.782 | -0.357 | -0.486 | 1.015 | 0.015 | 0.007 | 0.007 | 0 | 0.36 | 0.31 | 0.43 | 0.37 | 0.48 | 0.45 |
| 9 | 104497569 | rs7848096 | 0.676 | -0.266 | -0.417 | 1.034 | 0.009 | -0.003 | -0.003 | 0 | 0.39 | 0.38 | 0.45 | 0.32 | 0.49 | 0.46 |
| 9 | 104498664 | rs12340365 | -0.59 | -0.562 | -0.375 | 1.083 | 0.004 | 0.018 | 0.018 | 0 | 0.83 | 0.85 | 0.72 | 0.67 | 0.79 | 0.8 |
| 9 | 104504756 | rs2900317 | -1.042 | -0.587 | -0.394 | 1.072 | 0.019 | -0.095 | -0.095 | 0.699 | 0.68 | 0.92 | 0.72 | 0.5 | 0.74 | 0.71 |
| 9 | 104514018 | rs11788625 | 1.202 | -0.479 | -0.385 | 1.009 | -0.057 | -0.038 | -0.038 | 1.93 | 0.13 | 0.01 | 0.2 | 0.15 | 0.18 | 0.13 |
| 9 | 104515222 | rs10819985 | 0.254 | -0.471 | -0.369 | 1.011 | 0.024 | 0.004 | 0.004 | 0 | 0.42 | 0.37 | 0.25 | 0.1 | 0.27 | 0.29 |
| 9 | 104521883 | rs7864316 | 0.157 | -0.623 | -0.334 | 1.065 | 0.02 | 0.014 | 0.014 | 0 | 0.42 | 0.37 | 0.27 | 0.34 | 0.26 | 0.32 |
| 9 | 104525970 | rs1415652 | 0.362 | -0.534 | -0.409 | 1.049 | -0.005 | 0.022 | 0.022 | 0 | 0.7 | 0.64 | 0.66 | 0.41 | 0.74 | 0.64 |
| 9 | 104534755 | rs7874128 | 0.786 | -0.513 | -0.438 | 1.043 | 0.005 | -0.005 | -0.005 | 0.446 | 0.41 | 0.48 | 0.45 | 0.35 | 0.53 | 0.49 |
| 9 | 104535125 | rs10819990 | -0.859 | -0.411 | -0.977 | 0.741 | 0.01 | 0.083 | 0.083 | 0 | 0.55 | 0.38 | 0.46 | 0.24 | 0.43 | 0.4 |
| 9 | 104543022 | rs11791821 | 1.106 | -0.467 | -1.273 | 0.773 | -0.057 | -0.039 | -0.039 | 0.015 | 0.13 | 0.01 | 0.2 | 0.15 | 0.18 | 0.13 |
| 9 | 104550554 | rs1885387 | 1.073 | -0.512 | -1.251 | 0.757 | 0.016 | 0.163 | 0.163 | 0 | 0.27 | 0.04 | 0.19 | 0.06 | 0.21 | 0.16 |
| 9 | 104550927 | rs4743489 | -0.111 | 0.015 | 0 | 1.863 | -0.054 | 0.105 | 0.105 | 0 | 0.14 | 0.37 | 0.07 | 0.28 | 0.05 | 0.16 |
| 9 | 104552015 | rs4742825 | -0.407 | 0.244 | -0.135 | 1.844 | -0.017 | 0.052 | 0.052 | 0 | 0.28 | 0.52 | 0.3 | 0.43 | 0.23 | 0.29 |
| 9 | 104554756 | rs10819994 | -0.101 | 0.08 | -0.789 | 1.632 | 0.033 | -0.014 | -0.014 | 0 | 0.43 | 0.71 | 0.25 | 0.46 | 0.24 | 0.32 |
| 9 | 104555316 | rs1337706 | -0.38 | 0.579 | -1.38 | 0.968 | -0.064 | -0.075 | -0.075 | 0 | 0.14 | 0.01 | 0.17 | 0.22 | 0.21 | 0.14 |
| 9 | 104557532 | rs10989626 | 0.006 | 0.334 | -0.577 | 0.941 | -0.011 | 0.225 | 0.225 | 2.158 | 0.18 | 0.51 | 0.22 | 0.45 | 0.13 | 0.15 |
| 9 | 104563211 | rs10739815 | -0.168 | -0.122 | -0.926 | 0.277 | -0.008 | 0.002 | 0.002 | 1.07 | 0.37 | 0.46 | 0.28 | 0.56 | 0.23 | 0.36 |
| 9 | 104563334 | rs1184818 | 0.877 | 0.224 | -0.975 | -1.04 | -0.011 | 0.006 | 0.006 | 0 | 0.22 | 0.11 | 0.32 | 0.08 | 0.34 | 0.29 |
| 9 | 104567544 | rs7857093 | -0.668 | 0.01 | -0.875 | -1.086 | 0.001 | -0.02 | -0.02 | 0.089 | 0.85 | 0.83 | 0.83 | 0.96 | 0.83 | 0.87 |
| 9 | 104569036 | rs6479065 | -0.969 | -0.015 | -1.123 | -1.056 | -0.037 | 0.05 | 0.05 | 0 | 0.88 | 0.95 | 0.84 | 0.96 | 0.83 | 0.88 |
| 9 | 104569198 | rs7034450 | 0.737 | 0.021 | -1.179 | -1.134 | -0.015 | 0.023 | 0.023 | 0 | 0.23 | 0.11 | 0.33 | 0.14 | 0.32 | 0.3 |
| 9 | 104570604 | rs2417297 | -0.691 | -0.021 | -0.851 | -0.872 | -0.015 | 0.002 | 0.002 | 0 | 0.87 | 0.9 | 0.83 | 0.83 | 0.82 | 0.84 |
| 9 | 104576030 | rs2013563 | -0.336 | -0.596 | -0.234 | -1.029 | 0.009 | 0.036 | 0.036 | 0 | 0.42 | 0.3 | 0.51 | 0.24 | 0.58 | 0.39 |
| 9 | 104584492 | rs10989636 | 0.995 | -0.36 | 0.09 | -1.091 | -0.084 | -0.02 | -0.02 | 0 | 0.15 | 0.01 | 0.14 | 0.14 | 0.22 | 0.13 |
| 9 | 104588732 | rs2786715 | -1.421 | -0.648 | 0.238 | -1.258 | 0.025 | 0.085 | 0.085 | 0 | 0.68 | 0.83 | 0.78 | 0.89 | 0.72 | 0.67 |
| 9 | 104594799 | rs9409348 | 0.984 | -0.627 | 0.352 | -1.264 | 0.036 | -0.053 | -0.053 | 0 | 0.22 | 0.27 | 0.36 | 0.1 | 0.36 | 0.21 |
| 9 | 104595324 | rs1337687 | 1.181 | -0.64 | 0.268 | -1.27 | -0.055 | 0.055 | 0.055 | 0 | 0.21 | 0.06 | 0.35 | 0.1 | 0.35 | 0.21 |
| 9 | 104595776 | rs1022749 | 0.643 | -0.717 | 0.274 | -1.239 | -0.028 | 0.025 | 0.025 | 0 | 0.23 | 0.39 | 0.24 | 0.53 | 0.12 | 0.23 |
| 9 | 104596969 | rs2786717 | -1.514 | -0.73 | 0.265 | -1.524 | 0.027 | 0.109 | 0.109 | 0 | 0.68 | 0.86 | 0.78 | 0.89 | 0.72 | 0.67 |
| 9 | 104597349 | rs2181040 | -0.64 | -0.591 | 0.116 | -1.474 | 0.001 | 0 | 0 | 0 | 0.44 | 0.47 | 0.53 | 0.27 | 0.6 | 0.44 |
| 9 | 104613466 | rs10989657 | 1.117 | -0.47 | -0.072 | -1.366 | -0.082 | -0.017 | -0.017 | 0 | 0.15 | 0.01 | 0.15 | 0.14 | 0.23 | 0.13 |
| 9 | 104616470 | rs10739816 | -1.294 | -0.35 | 0.09 | -1.364 | 0.031 | 0.076 | 0.076 | 0 | 0.66 | 0.8 | 0.77 | 0.89 | 0.73 | 0.67 |
| 9 | 104619833 | rs10820002 | -0.332 | -0.239 | 0.204 | -1.322 | 0.003 | 0.004 | 0.004 | 0 | 0.42 | 0.34 | 0.51 | 0.15 | 0.59 | 0.38 |
| 9 | 104619905 | rs7868836 | -0.099 | -0.206 | 0.053 | -1.221 | 0.005 | -0.008 | -0.008 | 0 | 0.42 | 0.5 | 0.58 | 0.62 | 0.47 | 0.45 |
| 9 | 104622868 | rs10989661 | 0.738 | -0.116 | -0.444 | -1.171 | -0.005 | 0.006 | 0.006 | 0.486 | 0.21 | 0.17 | 0.16 | 0.14 | 0.25 | 0.16 |
| 9 | 104623359 | rs12552818 | 0.755 | -0.023 | -1.248 | -1.169 | 0.002 | -0.002 | -0.002 | 0 | 0.2 | 0.26 | 0.23 | 0.6 | 0.12 | 0.24 |
| 9 | 104623443 | rs2181038 | -0.523 | 0.473 | -1.615 | -1.599 | 0.029 | 0.063 | 0.063 | 0 | 0.66 | 0.78 | 0.77 | 0.89 | 0.73 | 0.67 |
| 9 | 104624002 | rs17792220 | -0.238 | 1.331 | -1.286 | -0.608 | -0.054 | 0.057 | 0.057 | 0 | 0.21 | 0.06 | 0.35 | 0.01 | 0.35 | 0.21 |
| 9 | 104625704 | rs16920898 | -0.646 | 0.289 | -0.828 | -0.213 | -0.008 | -0.003 | -0.003 | 0 | 0.22 | 0.27 | 0.29 | 0.18 | 0.18 | 0.2 |
| 9 | 104626607 | rs7047881 | -0.2 | -0.176 | -0.709 | -0.926 | -0.029 | -0.041 | -0.041 | 0 | 0.39 | 0.5 | 0.29 | 0.25 | 0.28 | 0.23 |
| 9 | 104626708 | rs1012203 | -0.069 | 0.316 | -0.707 | -0.465 | -0.002 | 0.002 | 0.002 | 0 | 0.08 | 0.07 | 0.13 | 0.15 | 0.09 | 0.13 |
| 9 | 104630136 | rs1002665 | -0.457 | 0.939 | -1.206 | -0.621 | -0.054 | -0.091 | -0.091 | 0.16 | 0.86 | 0.96 | 0.62 | 0.49 | 0.77 | 0.67 |
| 9 | 104637950 | rs10989672 | 0.178 | 0.192 | -0.816 | -1.197 | -0.014 | -0.092 | -0.092 | 0.615 | 0.14 | 0.3 | 0.09 | 0 | 0.08 | 0.13 |
| 9 | 104638067 | rs10989674 | 1.056 | 0.001 | -0.665 | -1.517 | -0.003 | -0.035 | -0.035 | 0.027 | 0.11 | 0.15 | 0.06 | 0.02 | 0.09 | 0.02 |
| 9 | 104640499 | rs12236471 | 1.766 | -0.14 | -1.329 | -1.895 | -0.025 | 0.096 | 0.096 | 0.003 | 0.23 | 0.06 | 0.13 | 0.06 | 0.26 | 0.21 |
| 9 | 104642966 | rs9299352 | 0.694 | -0.089 | -1.341 | -1.907 | -0.009 | -0.094 | -0.094 | 0 | 0.36 | 0.35 | 0.21 | 0.06 | 0.34 | 0.34 |
| 9 | 104650296 | rs1555461 | -0.421 | -0.274 | -0.857 | -1.646 | -0.003 | -0.023 | -0.023 | 0 | 0.26 | 0.54 | 0.22 | 0.2 | 0.24 | 0.2 |
| 9 | 104652159 | rs1743300 | -0.791 | -0.242 | -0.967 | -1.678 | -0.045 | -0.08 | -0.08 | 0 | 0.71 | 0.85 | 0.51 | 0.48 | 0.65 | 0.51 |
| 9 | 104655954 | rs11793523 | 1.425 | -0.356 | -0.952 | -1.691 | 0.001 | 0 | 0 | 0 | 0.07 | 0.03 | 0.03 | 0.02 | 0.05 | 0.08 |
| 9 | 104657274 | rs823922 | -0.975 | -0.322 | -0.997 | -1.692 | -0.056 | -0.105 | -0.105 | 0 | 0.72 | 0.91 | 0.53 | 0.5 | 0.66 | 0.52 |
| 9 | 104657657 | rs823921 | -0.546 | -0.503 | 0.313 | -0.898 | 0.001 | -0.026 | -0.026 | 0 | 0.31 | 0.16 | 0.31 | 0.4 | 0.3 | 0.28 |
| 9 | 104662585 | rs823920 | 0.876 | -0.526 | 0.219 | -0.876 | -0.034 | -0.054 | -0.054 | 0.279 | 0.17 | 0.01 | 0.17 | 0.23 | 0.16 | 0.11 |
| 9 | 104668084 | rs1165806 | -1.374 | -0.524 | 0.261 | -0.875 | 0.008 | 0.005 | 0.005 | 3.35 | 0.46 | 0.65 | 0.41 | 0.44 | 0.41 | 0.31 |
| 9 | 104674217 | rs10989691 | 0.385 | -0.361 | 0.962 | -0.854 | 0.035 | -0.064 | -0.064 | 0 | 0.12 | 0.35 | 0.07 | 0.02 | 0.11 | 0.02 |
| 9 | 104675628 | rs2795379 | -0.867 | -0.439 | 0.768 | -0.89 | 0.01 | -0.027 | -0.027 | 1.062 | 0.36 | 0.23 | 0.32 | 0.42 | 0.31 | 0.3 |
| 9 | 104676140 | rs2756919 | 0.753 | -0.445 | 0.533 | -0.901 | -0.037 | -0.054 | -0.054 | 0.082 | 0.17 | 0.01 | 0.16 | 0.23 | 0.16 | 0.1 |
| 9 | 104678175 | rs16920991 | -1.04 | -0.373 | 1.016 | -0.941 | -0.025 | -0.089 | -0.089 | 0 | 0.34 | 0.58 | 0.15 | 0.05 | 0.2 | 0.11 |
| 9 | 104680797 | rs11794511 | 0.492 | -0.348 | 0.458 | -0.918 | -0.01 | -0.025 | -0.025 | 0 | 0.08 | 0.11 | 0.03 | 0.02 | 0.05 | 0.08 |
| 9 | 104681509 | rs12343857 | 0.316 | -0.28 | -0.24 | -0.815 | -0.023 | -0.046 | -0.046 | 0 | 0.09 | 0.02 | 0.12 | 0.14 | 0.1 | 0.12 |
| 9 | 104682988 | rs10820007 | -0.708 | -0.279 | -0.152 | -0.849 | -0.033 | -0.109 | -0.109 | 0 | 0.33 | 0.55 | 0.14 | 0.05 | 0.2 | 0.11 |
| 9 | 104690092 | rs7867903 | -0.911 | -0.214 | -0.66 | -0.776 | -0.003 | -0.008 | -0.008 | 1.86 | 0.31 | 0.62 | 0.24 | 0.19 | 0.26 | 0.23 |
| 9 | 104705253 | rs1418236 | 0.476 | 0.245 | -0.671 | -0.497 | -0.023 | -0.028 | -0.028 | 0.188 | 0.86 | 0.92 | 0.75 | 0.61 | 0.79 | 0.79 |
| 9 | 104705331 | rs10820015 | 0.869 | 0.167 | -0.699 | -0.543 | 0.012 | 0.01 | 0.01 | 2.335 | 0.53 | 0.36 | 0.6 | 0.56 | 0.58 | 0.68 |
| 9 | 104706100 | rs1418232 | 1.026 | 0.15 | -0.697 | -0.551 | 0.023 | 0.02 | 0.02 | 4.433 | 0.51 | 0.32 | 0.6 | 0.56 | 0.58 | 0.68 |

| | | | | | | | | | | | | | | | | |
|---|---|---|---|---|---|---|---|---|---|---|---|---|---|---|---|---|
| 9 | 104707762 | rs3012580 | 1.154 | 0.173 | -0.694 | -0.54 | 0.013 | 0.008 | 0.008 | 3.752 | 0.5 | 0.1 | 0.59 | 0.56 | 0.58 | 0.68 |
| 9 | 104708430 | rs7041807 | -0.116 | 0.361 | -0.743 | -0.457 | -0.026 | -0.088 | -0.088 | 0 | 0.33 | 0.55 | 0.15 | 0.05 | 0.2 | 0.12 |
| 9 | 104708457 | rs1361665 | 1.357 | 0.943 | -0.915 | 0.572 | -0.013 | -0.017 | -0.017 | 0.057 | 0.84 | 0.88 | 0.74 | 0.61 | 0.78 | 0.79 |
| 9 | 104713471 | rs10989718 | -0.615 | 0.723 | -1.209 | 0.52 | -0.016 | -0.045 | -0.045 | 0.197 | 0.1 | 0.03 | 0.13 | 0.15 | 0.11 | 0.09 |
| 9 | 104728905 | rs1007468 | -0.382 | 0.611 | -1.116 | 0.655 | 0.005 | 0.01 | 0.01 | 4.636 | 0.3 | 0.19 | 0.24 | 0.29 | 0.28 | 0.14 |
| 9 | 104730823 | rs7850788 | -0.442 | 0.848 | -0.165 | 1.656 | 0.013 | 0.368 | 0.368 | 0 | 0.06 | 0.18 | 0.2 | 0.44 | 0.04 | 0.37 |
| 9 | 104759737 | rs4743514 | -0.412 | 1.508 | -0.387 | 0.819 | 0.003 | -0.003 | -0.003 | 0 | 0.36 | 0.35 | 0.52 | 0.3 | 0.53 | 0.33 |
| 9 | 104760894 | rs1418243 | 0.01 | 1.661 | -0.511 | 1.127 | -0.001 | 0.094 | 0.094 | 0 | 0.07 | 0.07 | 0.2 | 0.52 | 0.07 | 0.31 |
| 9 | 104765867 | rs1418247 | -0.73 | 2.71 | -0.355 | 1.057 | -0.011 | -0.001 | -0.001 | 1.265 | 0.38 | 0.24 | 0.57 | 0.63 | 0.41 | 0.56 |
| 9 | 104766075 | rs2995190 | 0.177 | 2.576 | 0.186 | 0.733 | 0.035 | 0.068 | 0.068 | 0 | 0.93 | 0.7 | 0.91 | 0.87 | 0.93 | 0.93 |
| 9 | 104768230 | rs10820027 | -0.229 | 2.706 | -0.135 | 0.605 | -0.001 | -0.037 | -0.037 | 0 | 0.32 | 0.32 | 0.37 | 0.11 | 0.34 | 0.25 |
| 9 | 104780164 | rs2995192 | -0.415 | 2.314 | -0.707 | 0.619 | -0.058 | -0.041 | -0.041 | 0 | 0.89 | 0.99 | 0.85 | 0.86 | 0.85 | 0.91 |
| 9 | 104781700 | rs1935376 | 0.249 | 2.361 | -0.762 | 0.353 | 0 | 0 | 0 | 0 | 0.22 | 0.2 | 0.24 | 0.19 | 0.31 | 0.28 |
| 9 | 104782096 | rs2995195 | 0.351 | 1.184 | -0.577 | -0.271 | 0 | 0.002 | 0.002 | 0 | 0.72 | 0.75 | 0.54 | 0.24 | 0.76 | 0.59 |
| 9 | 104789573 | rs1935383 | -0.531 | 0.497 | -0.741 | -0.97 | -0.002 | 0.025 | 0.025 | 0 | 0.74 | 0.8 | 0.78 | 0.86 | 0.78 | 0.89 |
| 9 | 104794273 | rs1342112 | 1.119 | 0.56 | -0.731 | -1.043 | -0.008 | 0.009 | 0.009 | 1.362 | 0.2 | 0.16 | 0.22 | 0.13 | 0.29 | 0.28 |
| 9 | 104795624 | rs7041890 | 0.816 | 0.17 | -0.46 | -1.281 | -0.001 | 0.047 | 0.047 | 0.566 | 0.38 | 0.21 | 0.42 | 0.68 | 0.38 | 0.53 |
| 9 | 104799619 | rs1342119 | -0.732 | 0.185 | -0.479 | -1.253 | -0.029 | 0.011 | 0.011 | 0.648 | 0.7 | 0.77 | 0.62 | 0.73 | 0.56 | 0.63 |
| 9 | 104801485 | rs10118278 | 0.493 | -0.023 | 0.016 | -1.272 | 0.009 | 0.044 | 0.044 | 0 | 0.44 | 0.3 | 0.29 | 0.12 | 0.41 | 0.3 |
| 9 | 104802828 | rs12551744 | 0.519 | -0.153 | 0.189 | -0.95 | -0.004 | 0.133 | 0.133 | 0 | 0.07 | 0.14 | 0.24 | 0.6 | 0.06 | 0.27 |
| 9 | 104808575 | rs1449655 | -0.32 | -0.21 | 0.054 | -0.994 | -0.074 | 0.093 | 0.093 | 0.049 | 0.21 | 0.5 | 0.35 | 0.64 | 0.1 | 0.35 |
| 9 | 104811726 | rs2839950 | -0.332 | -0.215 | 0.051 | -0.997 | -0.074 | 0.092 | 0.092 | 0.054 | 0.21 | 0.5 | 0.35 | 0.64 | 0.1 | 0.35 |
| 9 | 104819532 | rs16921199 | 1.044 | -0.246 | -0.253 | -1.121 | 0.038 | 0.038 | 0.038 | 0.272 | 0.12 | 0.05 | 0.08 | 0.04 | 0.03 | 0.08 |
| 9 | 104819955 | rs1935373 | 0.888 | 0.121 | -0.245 | -1.36 | 0.016 | -0.038 | -0.038 | 0.019 | 0.22 | 0.3 | 0.25 | 0.13 | 0.3 | 0.3 |
| 9 | 104830117 | rs320296 | -0.497 | 0.425 | -0.536 | -1.574 | -0.034 | 0.093 | 0.093 | 0.358 | 0.64 | 0.81 | 0.66 | 0.87 | 0.54 | 0.67 |
| 9 | 104830919 | rs320294 | -1.796 | 0.544 | -0.637 | -1.56 | -0.056 | 0.076 | 0.076 | 1.6 | 0.83 | 0.97 | 0.79 | 0.96 | 0.75 | 0.8 |
| 9 | 104834950 | rs1449653 | 1.593 | 0.572 | -0.689 | -1.67 | -0.04 | 0.117 | 0.117 | 0.08 | 0.31 | 0.12 | 0.18 | 0.08 | 0.35 | 0.25 |
| 9 | 104836267 | rs320291 | -1.612 | 0.359 | -0.491 | -1.407 | -0.074 | 0.058 | 0.058 | 1.508 | 0.82 | 0.95 | 0.77 | 0.92 | 0.68 | 0.73 |
| 9 | 104837536 | rs320290 | 1.609 | 0.261 | -0.492 | -1.246 | -0.058 | -0.045 | -0.045 | 0.049 | 0.1 | 0.01 | 0.14 | 0.15 | 0.15 | 0.09 |
| 9 | 104841979 | rs320286 | -1.106 | 0.343 | -0.361 | -1.321 | -0.012 | 0.009 | 0.009 | 0.702 | 0.79 | 0.83 | 0.71 | 0.87 | 0.64 | 0.7 |
| 9 | 104842043 | rs11790712 | 0.967 | -0.031 | -0.99 | -1.808 | 0.011 | 0.091 | 0.091 | 2.167 | 0.25 | 0.09 | 0.14 | 0.03 | 0.2 | 0.06 |
| 9 | 104845283 | rs16921254 | 1.153 | 0.128 | -0.19 | -1.469 | -0.039 | -0.059 | -0.059 | 0.013 | 0.13 | 0.21 | 0.1 | 0.03 | 0.05 | 0.04 |
| 9 | 104863013 | rs320261 | -0.923 | 0.339 | -0.231 | -1.362 | -0.001 | -0.001 | -0.001 | 0 | 0.77 | 0.76 | 0.7 | 0.84 | 0.64 | 0.65 |
| 9 | 104863180 | rs10760840 | 1.489 | 0.28 | -0.177 | -1.143 | 0.005 | 0.153 | 0.153 | 0 | 0.06 | 0.13 | 0.21 | 0.6 | 0.06 | 0.25 |
| 9 | 104879699 | rs10448296 | 0.332 | 0.195 | -0.309 | -1.326 | 0.024 | 0.095 | 0.095 | 0 | 0.51 | 0.24 | 0.44 | 0.66 | 0.44 | 0.53 |
| 9 | 104882097 | rs1962673 | 0.27 | 0.147 | -0.143 | -0.971 | 0.021 | 0.018 | 0.018 | 0 | 0.52 | 0.31 | 0.44 | 0.58 | 0.44 | 0.53 |
| 9 | 104904454 | rs10760846 | 1.627 | 0.235 | -0.007 | -0.799 | 0.003 | 0.214 | 0.214 | 0 | 0.05 | 0.13 | 0.21 | 0.53 | 0.05 | 0.25 |
| 9 | 104906690 | rs320245 | 0.15 | 0.172 | 0.026 | -1.032 | -0.01 | 0.012 | 0.012 | 0 | 0.26 | 0.38 | 0.26 | 0.27 | 0.19 | 0.17 |
| 9 | 104926878 | rs10118990 | 0.378 | 0.138 | -0.207 | -1.126 | 0.012 | -0.011 | -0.011 | 0 | 0.48 | 0.41 | 0.25 | 0.06 | 0.38 | 0.28 |
| 9 | 104940829 | rs10820084 | 1.281 | 0.256 | -0.408 | -0.579 | -0.001 | 0.06 | 0.06 | 0 | 0.07 | 0.06 | 0.21 | 0.54 | 0.07 | 0.3 |
| 9 | 104942186 | rs10117916 | -0.215 | -0.31 | -0.581 | -1.143 | -0.007 | 0.001 | 0.001 | 1.472 | 0.41 | 0.46 | 0.39 | 0.59 | 0.29 | 0.39 |
| 9 | 104943139 | rs320202 | -1.141 | -0.109 | -0.969 | -1.331 | -0.069 | 0.129 | 0.129 | 3.935 | 0.61 | 0.83 | 0.63 | 0.82 | 0.47 | 0.53 |
| 9 | 104943226 | rs320203 | 1.008 | -0.06 | -0.709 | -1.336 | 0.02 | 0.043 | 0.043 | 0.039 | 0.13 | 0.22 | 0.17 | 0.2 | 0.15 | 0.13 |
| 9 | 104943443 | rs320204 | -0.931 | -0.096 | -1.128 | -1.943 | -0.031 | 0.174 | 0.174 | 2.547 | 0.57 | 0.83 | 0.64 | 0.86 | 0.49 | 0.54 |
| 9 | 104943675 | rs320205 | -0.67 | 0.548 | -1.079 | -1.695 | -0.042 | 0.07 | 0.07 | 0 | 0.76 | 0.88 | 0.77 | 0.9 | 0.68 | 0.77 |
| 9 | 104944497 | rs7028042 | 0.338 | 0.11 | -0.914 | -1.757 | 0.025 | 0.054 | 0.054 | 0 | 0.52 | 0.58 | 0.29 | 0.19 | 0.43 | 0.3 |
| 9 | 104945543 | rs320213 | -0.678 | -0.003 | -0.681 | -1.764 | -0.003 | -0.027 | -0.027 | 0 | 0.38 | 0.52 | 0.41 | 0.25 | 0.33 | 0.28 |
| 9 | 104945929 | rs16921296 | 0.365 | 0.04 | -0.676 | -1.767 | -0.011 | -0.074 | -0.074 | 0 | 0.2 | 0.34 | 0.2 | 0.08 | 0.18 | 0.15 |
| 9 | 104946242 | rs320214 | 1.71 | 0.299 | -0.414 | -1.242 | -0.042 | -0.035 | -0.035 | 0 | 0.1 | 0.02 | 0.13 | 0.15 | 0.12 | 0.11 |
| 9 | 104947395 | rs9918994 | -0.17 | 0.194 | -0.048 | -0.884 | 0.007 | 0.132 | 0.132 | 0.334 | 0.17 | 0.36 | 0.44 | 0.54 | 0.16 | 0.35 |
| 9 | 104949975 | rs10820085 | 0.929 | 0.289 | -0.127 | -0.804 | 0 | 0.072 | 0.072 | 0 | 0.09 | 0.11 | 0.26 | 0.52 | 0.09 | 0.3 |
| 9 | 104950176 | rs10760848 | 0.006 | 0.066 | -0.106 | -0.866 | -0.017 | 0.08 | 0.08 | 0.988 | 0.2 | 0.34 | 0.35 | 0.54 | 0.14 | 0.32 |
| 9 | 104950962 | rs10739833 | -1.586 | 0.282 | -0.72 | -0.937 | -0.068 | 0.036 | 0.036 | 0.425 | 0.8 | 0.93 | 0.85 | 0.87 | 0.68 | 0.77 |
| 9 | 104951839 | rs10122572 | -1.037 | 0.252 | -0.73 | -0.947 | -0.065 | 0.015 | 0.015 | 0.837 | 0.77 | 0.9 | 0.73 | 0.81 | 0.64 | 0.66 |
| 9 | 104954524 | rs10760851 | -1.686 | 0.255 | -0.729 | -0.947 | -0.077 | 0.03 | 0.03 | 0.563 | 0.8 | 0.94 | 0.85 | 0.88 | 0.69 | 0.78 |
| 9 | 104955990 | rs7029138 | -0.864 | -0.098 | -0.603 | -1.048 | 0.011 | -0.01 | -0.01 | 0 | 0.31 | 0.7 | 0.39 | 0.61 | 0.17 | 0.37 |
| 9 | 104958057 | rs12336913 | 1.035 | -0.039 | -0.606 | -1.021 | -0.005 | 0.097 | 0.097 | 0 | 0.46 | 0.23 | 0.34 | 0.23 | 0.48 | 0.31 |
| 9 | 104959452 | rs10820088 | 0.801 | -0.594 | -0.161 | -0.565 | -0.001 | 0.058 | 0.058 | 0 | 0.1 | 0.11 | 0.26 | 0.54 | 0.1 | 0.32 |
| 9 | 104960302 | rs1932169 | -0.437 | -0.682 | 0.124 | -0.933 | -0.102 | -0.097 | -0.097 | 0 | 0.2 | 0.47 | 0.13 | 0.07 | 0.06 | 0.06 |
| 9 | 104963105 | rs16921360 | 1.656 | 0.113 | -0.406 | -0.455 | 0.021 | 0 | 0 | 0 | 0.05 | 0 | 0.02 | 0.06 | 0.02 | 0.03 |
| 9 | 104965221 | rs10119241 | -0.591 | 0.853 | -0.336 | -0.524 | -0.025 | 0.015 | 0.015 | 0.018 | 0.72 | 0.8 | 0.69 | 0.81 | 0.62 | 0.59 |
| 9 | 104965863 | rs7857876 | -0.261 | 0.979 | -0.369 | -0.391 | -0.032 | 0.002 | 0.002 | 0.042 | 0.69 | 0.77 | 0.6 | 0.73 | 0.56 | 0.56 |
| 9 | 104971300 | rs16921368 | -0.556 | 1.13 | -0.084 | -0.459 | 0.019 | 0.038 | 0.038 | 0 | 0.19 | 0.47 | 0.26 | 0.21 | 0.16 | 0.2 |
| 9 | 104971320 | rs7040955 | -1.081 | 1.615 | -0.589 | -0.213 | -0.103 | 0.019 | 0.019 | 0.558 | 0.81 | 0.98 | 0.83 | 0.88 | 0.7 | 0.74 |
| 9 | 104986223 | rs6479093 | -0.502 | 1.001 | 0.297 | -0.015 | 0.01 | 0.047 | 0.047 | 0.202 | 0.18 | 0.23 | 0.38 | 0.67 | 0.18 | 0.44 |
| 9 | 104990394 | rs10989904 | 0.562 | 0.393 | 0.079 | -0.509 | -0.011 | -0.022 | -0.022 | 2.049 | 0.16 | 0.18 | 0.1 | 0.02 | 0.1 | 0.01 |
| 9 | 104991804 | rs12336157 | 0.013 | 0.966 | 0.425 | -0.231 | 0.057 | 0.04 | 0.04 | 0 | 0.1 | 0.3 | 0.13 | 0.07 | 0.09 | 0.06 |
| 9 | 104992089 | rs9644962 | 0.064 | 1.029 | -0.25 | -0.155 | -0.035 | -0.054 | -0.054 | 0 | 0.11 | 0.04 | 0.35 | 0.67 | 0.14 | 0.4 |
| 9 | 104994909 | rs7861666 | -0.513 | 0.756 | -0.13 | -0.254 | -0.002 | -0.003 | -0.003 | 2.159 | 0.18 | 0.16 | 0.36 | 0.71 | 0.18 | 0.39 |

| | | | | | | | | | | | | | | | | |
|---|---|---|---|---|---|---|---|---|---|---|---|---|---|---|---|---|
| 9 | 105000062 | rs7869148 | -0.359 | 0.384 | 0.293 | -0.878 | 0.02 | -0.11 | -0.11 | 0.018 | 0.18 | 0.44 | 0.17 | 0.03 | 0.17 | 0.06 |
| 9 | 105009828 | rs10989911 | 0.435 | 0.431 | 0.178 | -0.357 | -0.002 | 0.015 | 0.015 | 0.058 | 0.09 | 0.09 | 0.14 | 0.71 | 0.08 | 0.36 |
| 9 | 105012816 | rs2067538 | 0.631 | 0.303 | 0.875 | -0.441 | -0.001 | 0.062 | 0.062 | 0 | 0.08 | 0.11 | 0.13 | 0.71 | 0.06 | 0.34 |
| 9 | 105016960 | rs912211 | -0.417 | 0.247 | 0.124 | -1.391 | -0.01 | 0 | 0 | 0.068 | 0.67 | 0.73 | 0.56 | 0.26 | 0.61 | 0.49 |
| 9 | 105021136 | rs1923983 | 1.288 | 0.361 | -0.152 | -1.281 | -0.057 | -0.053 | -0.053 | 0 | 0.11 | 0.02 | 0.07 | 0.11 | 0.18 | 0.22 |
| 9 | 105024768 | rs6479096 | -1.182 | -0.044 | -0.686 | -1.676 | -0.158 | 0.239 | 0.239 | 0.289 | 0.6 | 0.97 | 0.6 | 0.9 | 0.47 | 0.77 |
| 9 | 105025138 | rs12006153 | 1.182 | 0.108 | -0.27 | -1.453 | 0.015 | 0.014 | 0.014 | 0 | 0.12 | 0.18 | 0.08 | 0.11 | 0.18 | 0.19 |
| 9 | 105026764 | rs7031871 | 0.419 | -0.356 | 0.189 | -1.376 | 0.009 | -0.01 | -0.01 | 0 | 0.08 | 0.19 | 0.16 | 0.07 | 0.07 | 0.04 |
| 9 | 105029381 | rs10118114 | 0.534 | -0.491 | 0.036 | -1.421 | -0.01 | 0.084 | 0.084 | 0.099 | 0.13 | 0.26 | 0.15 | 0.71 | 0.08 | 0.39 |
| 9 | 105034238 | rs7036669 | 0.112 | -0.419 | -0.058 | -1.454 | 0.048 | 0.006 | 0.006 | 0.14 | 0.08 | 0.33 | 0.14 | 0.05 | 0.07 | 0.05 |
| 9 | 105035936 | rs1923987 | -0.568 | -0.502 | -1.162 | -1.4 | 0.063 | 0.065 | 0.065 | 0.212 | 0.46 | 0.78 | 0.52 | 0.74 | 0.3 | 0.56 |
| 9 | 105036478 | rs13296598 | 0.169 | 0.377 | -1.076 | -1.39 | -0.108 | 0.017 | 0.017 | 0 | 0.22 | 0.03 | 0.15 | 0.11 | 0.31 | 0.23 |
| 9 | 105037839 | rs7466994 | -0.424 | 0.302 | -0.824 | -1.551 | -0.05 | 0.07 | 0.07 | 0 | 0.33 | 0.16 | 0.28 | 0.82 | 0.38 | 0.6 |
| 9 | 105041206 | rs959065 | -1.122 | 0.092 | -0.674 | -1.709 | 0.02 | 0.113 | 0.113 | 0 | 0.55 | 0.7 | 0.63 | 0.9 | 0.57 | 0.71 |
| 9 | 105041263 | rs959066 | -0.378 | 0.168 | -0.16 | -1.104 | -0.005 | 0.039 | 0.039 | 0.148 | 0.29 | 0.49 | 0.32 | 0.6 | 0.27 | 0.4 |
| 9 | 105041375 | rs959068 | -0.511 | 0.131 | -0.221 | -1.517 | -0.004 | 0.044 | 0.044 | 0 | 0.47 | 0.36 | 0.45 | 0.73 | 0.48 | 0.62 |
| 9 | 105041756 | rs1040980 | -0.208 | 0.071 | -0.143 | -1.294 | -0.001 | -0.01 | -0.01 | 0 | 0.31 | 0.35 | 0.33 | 0.74 | 0.29 | 0.6 |
| 9 | 105042090 | rs10989950 | 0.052 | -0.355 | -0.237 | -1.36 | -0.002 | -0.01 | -0.01 | 0 | 0.22 | 0.23 | 0.22 | 0.08 | 0.16 | 0.09 |
| 9 | 105042198 | rs16921474 | 0.899 | -0.148 | -0.269 | -1.213 | 0.006 | 0.009 | 0.009 | 0 | 0.08 | 0.04 | 0.09 | 0.02 | 0.07 | 0.04 |
| 9 | 105043748 | rs10820120 | -0.889 | -0.213 | 0.259 | -1.322 | 0.005 | -0.021 | -0.021 | 1.324 | 0.18 | 0.31 | 0.3 | 0.14 | 0.19 | 0.12 |
| 9 | 105045726 | rs10820123 | -0.869 | -0.363 | 0.023 | -1.295 | -0.001 | -0.008 | -0.008 | 0 | 0.28 | 0.3 | 0.32 | 0.14 | 0.24 | 0.17 |
| 9 | 105046060 | rs7865273 | -0.338 | 0.089 | -0.191 | -0.956 | 0.013 | 0.005 | 0.005 | 0 | 0.08 | 0.14 | 0.2 | 0.12 | 0.12 | 0.08 |
| 9 | 105050664 | rs4376547 | 0.046 | 0.169 | -0.358 | -1.47 | 0.002 | 0.001 | 0.001 | 0 | 0.53 | 0.46 | 0.47 | 0.72 | 0.57 | 0.67 |
| 9 | 105055586 | rs1924004 | 1.092 | 1.092 | -0.54 | -1.625 | 0.018 | -0.022 | -0.022 | 0 | 0.1 | 0.13 | 0.13 | 0.02 | 0.21 | 0.06 |
| 9 | 105056347 | rs1924005 | 1.282 | 1.092 | -0.591 | -1.625 | -0.042 | 0.039 | 0.039 | 0 | 0.09 | 0.04 | 0.12 | 0.02 | 0.21 | 0.06 |
| 9 | 105058421 | rs2417335 | -0.496 | 0.757 | -0.612 | -2.209 | 0.001 | -0.022 | -0.022 | 0.011 | 0.41 | 0.57 | 0.53 | 0.86 | 0.46 | 0.6 |
| 9 | 105058753 | rs9409365 | -1.065 | 0.274 | -0.873 | -2.46 | -0.033 | 0.204 | 0.204 | 0.003 | 0.55 | 0.86 | 0.59 | 0.86 | 0.52 | 0.67 |
| 9 | 105058946 | rs10733718 | -1.554 | -0.054 | -0.986 | -2.548 | -0.004 | 0.121 | 0.121 | 0 | 0.7 | 0.95 | 0.75 | 0.88 | 0.7 | 0.73 |
| 9 | 105060163 | rs10989973 | 1.404 | -0.252 | -0.717 | -1.585 | 0.031 | -0.073 | -0.073 | 0 | 0.19 | 0.08 | 0.16 | 0.48 | 0.11 | 0.41 |
| 9 | 105074656 | rs1022838 | 1.235 | -0.014 | -0.777 | -1.933 | -0.075 | 0.073 | 0.073 | 0 | 0.24 | 0.11 | 0.27 | 0.03 | 0.4 | 0.12 |
| 9 | 105077400 | rs12005744 | 0.907 | -0.172 | -0.466 | -1.748 | -0.02 | 0.009 | 0.009 | 0 | 0.23 | 0.24 | 0.27 | 0.7 | 0.13 | 0.48 |
| 9 | 105077531 | rs4742843 | 0.467 | -0.202 | -0.248 | -1.778 | -0.042 | 0.009 | 0.009 | 0 | 0.27 | 0.58 | 0.3 | 0.7 | 0.14 | 0.49 |
| 9 | 105077543 | rs1543533 | 1.374 | -0.204 | -0.267 | -1.784 | -0.02 | -0.057 | -0.057 | 0.348 | 0.14 | 0.23 | 0.06 | 0 | 0.06 | 0.07 |
| 9 | 105086690 | rs10990004 | 1.964 | -0.104 | -0.376 | -1.75 | -0.033 | -0.03 | -0.03 | 1.269 | 0.05 | 0.01 | 0.12 | 0.14 | 0.11 | 0.05 |
| 9 | 105090815 | rs9409373 | 1.314 | -0.082 | -0.41 | -1.825 | -0.098 | 0.101 | 0.101 | 0 | 0.24 | 0.08 | 0.27 | 0.03 | 0.4 | 0.12 |
| 9 | 105093304 | rs10114491 | -0.329 | 0.129 | 0.007 | -1.434 | -0.021 | -0.057 | -0.057 | 0 | 0.31 | 0.5 | 0.41 | 0.84 | 0.24 | 0.53 |
| 9 | 105094335 | rs9409038 | 0.252 | 0.149 | -0.123 | -1.422 | 0.008 | 0.019 | 0.019 | 0 | 0.44 | 0.45 | 0.47 | 0.17 | 0.56 | 0.25 |
| 9 | 105096394 | rs1924024 | 0.628 | 0.146 | -0.098 | -1.421 | -0.019 | 0.009 | 0.009 | 0 | 0.23 | 0.25 | 0.27 | 0.71 | 0.13 | 0.49 |
| 9 | 105097068 | rs1924026 | -0.094 | 0.698 | -0.437 | -1.229 | -0.004 | -0.019 | -0.019 | 0 | 0.45 | 0.53 | 0.47 | 0.17 | 0.56 | 0.25 |
| 9 | 105108011 | rs1425279 | 0.855 | 0.803 | -0.824 | -1.139 | 0.007 | 0.012 | 0.012 | 0 | 0.07 | 0.1 | 0.14 | 0.13 | 0.11 | 0.05 |
| 9 | 105108224 | rs9409041 | 0.433 | 0.789 | -0.866 | -1.243 | -0.015 | -0.005 | -0.005 | 0 | 0.37 | 0.35 | 0.33 | 0.04 | 0.45 | 0.19 |
| 9 | 105111298 | rs1477530 | 0.692 | 0.786 | -0.699 | -0.338 | -0.037 | -0.035 | -0.035 | 0 | 0.06 | 0.01 | 0.12 | 0.18 | 0.11 | 0.06 |
| 9 | 105117488 | rs7038470 | -1.229 | 0.776 | -0.951 | -0.75 | 0.005 | 0.164 | 0.164 | 0.823 | 0.58 | 0.78 | 0.68 | 0.95 | 0.58 | 0.77 |
| 9 | 105120779 | rs1425280 | 0.063 | 0.688 | -0.772 | -0.471 | 0.013 | 0.043 | 0.043 | 0 | 0.14 | 0.18 | 0.21 | 0.62 | 0.16 | 0.49 |
| 9 | 105122350 | rs10441769 | 0.355 | 0.521 | -0.439 | -0.581 | 0.004 | -0.004 | -0.004 | 0 | 0.32 | 0.32 | 0.37 | 0.31 | 0.38 | 0.2 |
| 9 | 105135141 | rs2417361 | -0.935 | 0.553 | -0.6 | -0.579 | 0.005 | 0 | 0 | 0.569 | 0.51 | 0.59 | 0.59 | 0.51 | 0.52 | 0.34 |
| 9 | 105137537 | rs9299361 | 1.669 | 0.562 | -0.748 | -0.47 | -0.012 | 0.019 | 0.019 | 3.88 | 0.14 | 0.01 | 0.23 | 0.1 | 0.13 | 0.12 |
| 9 | 105143745 | rs1364820 | 1.297 | 0.525 | -0.346 | -0.249 | 0.014 | 0.229 | 0.229 | 0.118 | 0.07 | 0.17 | 0.09 | 0.44 | 0.06 | 0.42 |
| 9 | 105144688 | rs1923975 | -0.441 | 0.498 | -0.319 | -0.277 | -0.003 | -0.004 | -0.004 | 0 | 0.47 | 0.53 | 0.56 | 0.51 | 0.5 | 0.3 |
| 9 | 105146406 | rs12005061 | 1.888 | 0.338 | -0.416 | -0.503 | -0.045 | -0.047 | -0.047 | 0.265 | 0.08 | 0.21 | 0.03 | 0 | 0.01 | 0 |
| 9 | 105146568 | rs10990059 | 0.016 | 0.982 | -1.177 | -0.188 | -0.018 | -0.023 | -0.023 | 0.08 | 0.1 | 0.08 | 0.19 | 0.23 | 0.14 | 0.15 |
| 9 | 105146883 | rs10760887 | -0.157 | 0.878 | -1.087 | 0.024 | 0.008 | 0.097 | 0.097 | 1.247 | 0.48 | 0.69 | 0.49 | 0.73 | 0.45 | 0.62 |
| 9 | 105167688 | rs6479112 | -0.324 | 0.54 | -0.883 | 0.116 | 0.006 | -0.002 | -0.002 | 0 | 0.5 | 0.63 | 0.6 | 0.52 | 0.52 | 0.38 |
| 9 | 105172992 | rs10733724 | -0.741 | 0.352 | -0.137 | -0.027 | 0.002 | 0.004 | 0.004 | 0 | 0.44 | 0.56 | 0.38 | 0.48 | 0.47 | 0.31 |
| 9 | 105173098 | rs10760891 | -0.352 | 0.703 | 0.055 | 0.115 | -0.004 | 0.007 | 0.007 | 0.056 | 0.37 | 0.26 | 0.25 | 0.31 | 0.36 | 0.24 |
| 9 | 105175187 | rs10119950 | 2.611 | 0.439 | -0.07 | -0.009 | -0.006 | -0.007 | -0.007 | 0 | 0.05 | 0.04 | 0.01 | 0 | 0.01 | 0 |
| 9 | 105186697 | rs2031200 | -1.119 | 0.467 | 0.213 | -0.007 | 0.008 | -0.003 | -0.003 | 0.089 | 0.51 | 0.7 | 0.4 | 0.48 | 0.49 | 0.31 |
| 9 | 105202644 | rs16921686 | -0.196 | 0.459 | 0.438 | 0.023 | -0.019 | 0.012 | 0.012 | 0 | 0.27 | 0.3 | 0.14 | 0.34 | 0.2 | 0.15 |
| 9 | 105204612 | rs7847867 | -0.237 | 0.366 | 0.502 | 0.013 | -0.028 | 0.012 | 0.012 | 0 | 0.27 | 0.32 | 0.14 | 0.34 | 0.2 | 0.15 |
| 9 | 105207670 | rs4410962 | -0.659 | 0.631 | 0.696 | -0.043 | 0.076 | 0.018 | 0.018 | 0 | 0.16 | 0.38 | 0.23 | 0.2 | 0.25 | 0.22 |
| 9 | 105208847 | rs7026565 | 0.203 | 0.744 | 0.451 | -0.056 | -0.043 | 0.03 | 0.03 | 0 | 0.1 | 0.04 | 0.12 | 0.02 | 0.18 | 0.15 |
| 9 | 105214028 | rs10990103 | -1.112 | 0.394 | 0.666 | -0.031 | 0.004 | -0.007 | -0.007 | 0.019 | 0.38 | 0.64 | 0.28 | 0.51 | 0.29 | 0.23 |
| 9 | 105216600 | rs10990106 | -0.201 | 0.393 | 0.766 | 0.019 | -0.024 | 0.013 | 0.013 | 0 | 0.27 | 0.3 | 0.14 | 0.34 | 0.19 | 0.15 |
| 9 | 105219505 | rs10760894 | -0.974 | 0.282 | 0.405 | 0.032 | -0.002 | 0.003 | 0.003 | 0 | 0.37 | 0.59 | 0.28 | 0.51 | 0.29 | 0.23 |
| 9 | 105246175 | rs1946334 | -0.683 | 0.32 | 0.782 | -0.14 | 0.004 | -0.018 | -0.018 | 0 | 0.23 | 0.56 | 0.25 | 0.21 | 0.2 | 0.19 |
| 9 | 105259717 | rs7849848 | 0.59 | 0.142 | 0.614 | -0.107 | 0.001 | 0.001 | 0.001 | 0 | 0.07 | 0.07 | 0.07 | 0.03 | 0.03 | 0.08 |
| 9 | 105266273 | rs7868372 | 0.865 | -0.134 | 0.415 | 0.095 | -0.006 | 0.033 | 0.033 | 0 | 0.1 | 0.07 | 0.06 | 0.34 | 0.03 | 0.05 |
| 9 | 105279735 | rs7029589 | -1.444 | 0.112 | 0.416 | -0.068 | -0.017 | -0.029 | -0.029 | 2.334 | 0.39 | 0.78 | 0.37 | 0.57 | 0.38 | 0.31 |
| 9 | 105292037 | rs4514038 | -0.751 | 0.152 | 0.453 | -0.161 | 0.013 | -0.012 | -0.012 | 0 | 0.28 | 0.7 | 0.31 | 0.21 | 0.35 | 0.27 |

| | | | | | | | | | | | | | | | | |
|---|---|---|---|---|---|---|---|---|---|---|---|---|---|---|---|---|
| 9 | 105298240 | rs10115055 | -0.297 | 0.318 | 0.736 | -0.162 | -0.002 | 0.039 | 0.039 | 0 | 0.29 | 0.51 | 0.18 | 0.38 | 0.29 | 0.18 |
| 9 | 105306425 | rs16921790 | 0.729 | 0.534 | 0.319 | -0.32 | -0.003 | -0.007 | -0.007 | 0 | 0.1 | 0.06 | 0.1 | 0.39 | 0.09 | 0.3 |
| 9 | 105315604 | rs4521822 | -2.009 | 0.228 | -0.057 | -0.676 | -0.089 | 0.224 | 0.224 | 0 | 0.64 | 0.97 | 0.53 | 0.97 | 0.58 | 0.72 |
| 9 | 105323932 | rs1412799 | -0.989 | 0.491 | 0.526 | -0.382 | -0.01 | -0.01 | -0.01 | 0.012 | 0.47 | 0.79 | 0.34 | 0.55 | 0.44 | 0.32 |
| 9 | 105325292 | rs7846871 | -0.205 | 0.496 | 0.583 | -0.377 | -0.007 | 0.023 | 0.023 | 0 | 0.32 | 0.44 | 0.19 | 0.38 | 0.3 | 0.16 |
| 9 | 105342266 | rs10760899 | -0.684 | 0.545 | 0.1 | -0.22 | -0.01 | -0.012 | -0.012 | 0.189 | 0.45 | 0.71 | 0.34 | 0.54 | 0.44 | 0.32 |
| 9 | 105347447 | rs4743564 | 0.742 | 0.481 | 0.064 | -0.171 | 0.001 | 0 | 0 | 0 | 0.1 | 0.06 | 0.1 | 0.38 | 0.09 | 0.31 |
| 9 | 105350958 | rs1332984 | -0.869 | 0.535 | 0.274 | -0.122 | -0.006 | -0.01 | -0.01 | 0 | 0.44 | 0.65 | 0.32 | 0.54 | 0.42 | 0.32 |
| 9 | 105355578 | rs16921840 | -0.784 | 0.676 | 0.632 | 0.243 | 0.054 | 0.037 | 0.037 | 0 | 0.13 | 0.39 | 0.12 | 0.15 | 0.12 | 0.11 |
| 9 | 105371812 | rs7875594 | 0.184 | 0.643 | 0.565 | 0.206 | 0.014 | -0.028 | -0.028 | 0 | 0.08 | 0.29 | 0.04 | 0.02 | 0.04 | 0.02 |
| 9 | 105376813 | rs10739849 | -0.296 | 0.828 | -0.149 | 0.533 | 0.004 | -0.007 | -0.007 | 0 | 0.9 | 0.94 | 0.9 | 0.62 | 0.92 | 0.67 |
| 9 | 105390418 | rs1115634 | -0.444 | 1.169 | -0.024 | 0.335 | 0.009 | 0.03 | 0.03 | 0 | 0.21 | 0.24 | 0.2 | 0.51 | 0.27 | 0.24 |
| 9 | 105395352 | rs10820213 | -0.349 | 1.188 | -0.748 | -0.102 | -0.073 | -0.076 | -0.076 | 0.001 | 0.27 | 0.09 | 0.19 | 0.38 | 0.36 | 0.22 |
| 9 | 105399431 | rs7046513 | -0.884 | 1.035 | 0.4 | 0.258 | 0.063 | 0.11 | 0.11 | 0 | 0.26 | 0.49 | 0.25 | 0.51 | 0.33 | 0.27 |
| 9 | 105400552 | rs16926142 | 0.911 | 0.887 | 0.57 | 0.095 | -0.007 | -0.03 | -0.03 | 0 | 0.07 | 0.13 | 0.03 | 0.02 | 0.05 | 0.02 |
| 9 | 105408515 | rs1332999 | -0.731 | 0.884 | 0.416 | 0.087 | 0.037 | 0.065 | 0.065 | 0 | 0.26 | 0.42 | 0.25 | 0.51 | 0.33 | 0.28 |
| 9 | 105412545 | rs7865059 | -1.515 | 0.894 | -0.056 | 0.088 | 0.005 | 0.007 | 0.007 | 0 | 0.34 | 0.63 | 0.29 | 0.53 | 0.37 | 0.3 |
| 9 | 105414408 | rs4743569 | 0.992 | 0.812 | -0.635 | -0.118 | -0.011 | -0.054 | -0.054 | 0 | 0.15 | 0.18 | 0.08 | 0.01 | 0.12 | 0.1 |
| 9 | 105414477 | rs13283389 | -0.779 | 1.06 | -0.709 | 0.233 | -0.007 | -0.008 | -0.008 | 0 | 0.25 | 0.25 | 0.24 | 0.51 | 0.33 | 0.29 |
| 9 | 105420968 | rs12553605 | 0.252 | 1.569 | -0.88 | 0.246 | -0.035 | -0.033 | -0.033 | 0 | 0.06 | 0.01 | 0.13 | 0.14 | 0.12 | 0.08 |
| 9 | 105422136 | rs12553989 | -0.67 | 1.853 | -0.972 | 0.404 | -0.044 | -0.043 | -0.043 | 0 | 0.11 | 0.08 | 0.18 | 0.22 | 0.3 | 0.15 |
| 9 | 105422608 | rs12553388 | 0.414 | 1.161 | -0.96 | 0.519 | -0.042 | -0.038 | -0.038 | 0 | 0.07 | 0.01 | 0.13 | 0.18 | 0.14 | 0.1 |
| 9 | 105424988 | rs10760900 | -0.119 | 0.223 | -1.109 | -0.064 | -0.046 | -0.006 | -0.006 | 0 | 0.36 | 0.48 | 0.33 | 0.74 | 0.2 | 0.57 |
| 9 | 105425207 | rs1332988 | 0.559 | 0.701 | -1.956 | 0.082 | -0.075 | 0.175 | 0.175 | 0 | 0.39 | 0.13 | 0.46 | 0.07 | 0.55 | 0.27 |
| 9 | 105426977 | rs4294244 | 0.668 | 0.694 | -1.946 | 0.667 | -0.004 | -0.05 | -0.05 | 0 | 0.09 | 0.01 | 0.1 | 0.32 | 0.08 | 0.38 |
| 9 | 105430482 | rs12686229 | 0.4 | 0.712 | -1.567 | 0.643 | 0.019 | 0.035 | 0.035 | 0 | 0.07 | 0.09 | 0.13 | 0.18 | 0.14 | 0.1 |
| 9 | 105431576 | rs4313213 | 0.102 | 0.308 | -1.935 | -0.093 | 0.03 | 0.056 | 0.056 | 0 | 0.49 | 0.15 | 0.56 | 0.38 | 0.63 | 0.65 |
| 9 | 105442117 | rs2210533 | -0.4 | 0.036 | -1.786 | -0.907 | 0.015 | 0.014 | 0.014 | 0 | 0.74 | 0.85 | 0.87 | 1 | 0.77 | 0.93 |
| 9 | 105444722 | rs10990238 | 0.265 | -0.58 | -1.625 | -0.668 | -0.002 | 0.003 | 0.003 | 0 | 0.24 | 0.18 | 0.3 | 0.42 | 0.22 | 0.57 |
| 9 | 105445184 | rs7866103 | -1.101 | -0.594 | -1.632 | -0.676 | 0.012 | -0.006 | -0.006 | 0 | 0.77 | 0.85 | 0.89 | 1 | 0.83 | 0.94 |
| 9 | 105446609 | rs10820230 | 2.141 | 0.27 | -1.975 | -0.611 | -0.046 | -0.059 | -0.059 | 0.48 | 0.12 | 0.02 | 0.13 | 0.38 | 0.17 | 0.12 |
| 9 | 105454102 | rs1556503 | 0.539 | 0.008 | -2.279 | -0.887 | 0.006 | -0.007 | -0.007 | 0.394 | 0.27 | 0.17 | 0.3 | 0.42 | 0.21 | 0.59 |
| 9 | 105464554 | rs10820241 | 1.399 | 0.171 | -1.704 | -1.294 | 0.006 | 0.005 | 0.005 | 0 | 0.27 | 0.36 | 0.37 | 0.39 | 0.32 | 0.21 |
| 9 | 105465785 | rs10820244 | 1.615 | 0.15 | -1.645 | -1.269 | -0.007 | 0.015 | 0.015 | 0 | 0.21 | 0.17 | 0.27 | 0.42 | 0.17 | 0.57 |
| 9 | 105468208 | rs4743571 | -0.359 | -0.265 | -1.897 | -1.688 | 0.001 | 0.001 | 0.001 | 0 | 0.53 | 0.38 | 0.43 | 0.42 | 0.43 | 0.68 |
| 9 | 105474366 | rs7869982 | 2.279 | -0.312 | -1.946 | -1.714 | -0.01 | -0.042 | -0.042 | 0.288 | 0.16 | 0.15 | 0.07 | 0 | 0.11 | 0.05 |
| 9 | 105474384 | rs7870093 | 1.021 | -0.311 | -1.908 | -1.715 | 0.008 | -0.028 | -0.028 | 0 | 0.25 | 0.23 | 0.13 | 0 | 0.22 | 0.09 |
| 9 | 105474758 | rs1576949 | -1.484 | -0.186 | -1.788 | -1.644 | 0.008 | 0.004 | 0.004 | 0 | 0.79 | 0.73 | 0.79 | 0.82 | 0.75 | 0.89 |
| 9 | 105476561 | rs16922059 | 2.729 | -0.454 | -1.764 | -1.547 | 0.004 | -0.005 | -0.005 | 0 | 0.08 | 0.05 | 0.04 | 0.23 | 0.05 | 0.3 |
| 9 | 105482137 | rs10990259 | 2.669 | -0.441 | -1.549 | -1.523 | -0.005 | 0 | 0 | 0 | 0.07 | 0.08 | 0.07 | 0.07 | 0.13 | 0.08 |
| 9 | 105485375 | rs1602690 | -1.333 | -0.569 | -1.344 | -1.423 | -0.001 | 0 | 0 | 0 | 0.67 | 0.61 | 0.59 | 0.8 | 0.64 | 0.8 |
| 9 | 105489249 | rs10820255 | 2.022 | -0.74 | -1.16 | -1.202 | -0.064 | -0.073 | -0.073 | 0.295 | 0.24 | 0.14 | 0.33 | 0.4 | 0.28 | 0.21 |
| 9 | 105495207 | rs7870366 | 2.137 | -0.791 | -0.838 | -1.213 | -0.003 | -0.047 | -0.047 | 0 | 0.1 | 0.18 | 0.06 | 0 | 0.11 | 0.03 |
| 9 | 105495671 | rs1492594 | 1.421 | -0.881 | -0.892 | -1.253 | -0.006 | 0.014 | 0.014 | 0 | 0.21 | 0.17 | 0.27 | 0.42 | 0.17 | 0.57 |
| 9 | 105496890 | rs985453 | 1.237 | -0.84 | -0.905 | -1.209 | 0.009 | 0.004 | 0.004 | 0 | 0.17 | 0.14 | 0.1 | 0.23 | 0.16 | 0.34 |
| 9 | 105502498 | rs10990268 | 1.292 | -0.818 | -0.649 | -1.243 | -0.005 | 0.002 | 0.002 | 0 | 0.2 | 0.21 | 0.26 | 0.19 | 0.16 | 0.3 |
| 9 | 105502882 | rs1492595 | 0.359 | -0.823 | -0.704 | -1.31 | -0.003 | 0.009 | 0.009 | 0 | 0.45 | 0.47 | 0.53 | 0.74 | 0.48 | 0.55 |
| 9 | 105503137 | rs1492596 | -1.903 | -0.593 | -0.807 | -1.347 | -0.007 | 0.007 | 0.007 | 0 | 0.81 | 0.84 | 0.85 | 0.93 | 0.74 | 0.9 |
| 9 | 105503295 | rs10990270 | 1.558 | -0.39 | -0.8 | -0.947 | -0.02 | -0.023 | -0.023 | 0.039 | 0.25 | 0.22 | 0.33 | 0.39 | 0.28 | 0.21 |
| 9 | 105503706 | rs1388919 | 1.697 | -0.404 | -0.907 | -0.941 | -0.011 | -0.044 | -0.044 | 0.106 | 0.16 | 0.15 | 0.07 | 0 | 0.11 | 0.06 |
| 9 | 105507920 | rs4743577 | 0.298 | 0.359 | -1.029 | -0.289 | -0.015 | -0.01 | -0.01 | 0.114 | 0.33 | 0.31 | 0.4 | 0.4 | 0.36 | 0.24 |
| 9 | 105512975 | rs4354369 | 0.178 | 0.477 | -1.128 | -0.441 | -0.015 | 0.05 | 0.05 | 0.237 | 0.44 | 0.32 | 0.58 | 0.82 | 0.43 | 0.77 |
| 9 | 105518491 | rs10820273 | -0.561 | 0.42 | -1.406 | -0.428 | 0.004 | 0.064 | 0.064 | 0 | 0.55 | 0.74 | 0.61 | 0.64 | 0.58 | 0.54 |
| 9 | 105521808 | rs10990300 | 0.598 | 0.385 | -0.786 | -0.091 | -0.012 | 0.04 | 0.04 | 0 | 0.24 | 0.32 | 0.25 | 0.39 | 0.21 | 0.15 |
| 9 | 105523821 | rs1580650 | 1.353 | 0.466 | -0.376 | -0.239 | 0.014 | -0.059 | -0.059 | 0 | 0.1 | 0.22 | 0.08 | 0 | 0.13 | 0.04 |
| 9 | 105524913 | rs7469903 | 1.572 | 0.465 | -0.375 | -0.243 | -0.009 | -0.011 | -0.011 | 0 | 0.21 | 0.13 | 0.27 | 0.25 | 0.24 | 0.35 |
| 9 | 105526362 | rs7858600 | -0.842 | 0.053 | -0.465 | -0.433 | -0.017 | 0.137 | 0.137 | 0 | 0.54 | 0.73 | 0.63 | 0.76 | 0.49 | 0.52 |
| 9 | 105526510 | rs7047925 | 1.669 | 0.195 | -0.659 | -0.681 | -0.003 | 0.031 | 0.031 | 0 | 0.26 | 0.14 | 0.25 | 0.24 | 0.24 | 0.42 |
| 9 | 105527511 | rs10820274 | -1.091 | 0.083 | -0.618 | -0.607 | -0.009 | 0.058 | 0.058 | 0 | 0.62 | 0.7 | 0.68 | 0.76 | 0.62 | 0.57 |
| 9 | 105529745 | rs10990302 | 1.37 | 0.497 | -0.092 | -0.507 | -0.013 | 0.059 | 0.059 | 0 | 0.15 | 0.17 | 0.18 | 0.39 | 0.11 | 0.14 |
| 9 | 105534377 | rs4743578 | -0.047 | 0.525 | -0.026 | -0.419 | -0.026 | 0.079 | 0.079 | 0 | 0.25 | 0.43 | 0.27 | 0.39 | 0.21 | 0.16 |
| 9 | 105541245 | rs1388920 | 1.597 | 0.574 | -0.07 | -0.381 | -0.007 | -0.008 | -0.008 | 0 | 0.21 | 0.13 | 0.27 | 0.25 | 0.24 | 0.35 |
| 9 | 105541307 | rs12237372 | -0.379 | 0.542 | -0.07 | -0.175 | -0.008 | 0.031 | 0.031 | 0 | 0.32 | 0.48 | 0.33 | 0.39 | 0.33 | 0.2 |
| 9 | 105542534 | rs1492602 | -0.457 | 0.459 | -0.162 | -0.135 | 0.011 | 0.044 | 0.044 | 0 | 0.53 | 0.61 | 0.6 | 0.64 | 0.58 | 0.56 |
| 9 | 105545039 | rs4743582 | -1.042 | 0.593 | -0.098 | -0.195 | -0.004 | -0.048 | -0.048 | 0 | 0.87 | 0.8 | 0.95 | 1 | 0.89 | 0.99 |
| 9 | 105545483 | rs994864 | 0.253 | 0.655 | 0.098 | 0.128 | -0.003 | 0.02 | 0.02 | 0 | 0.71 | 0.56 | 0.78 | 0.7 | 0.72 | 0.85 |
| 9 | 105545528 | rs994865 | -1.161 | -0.265 | 0.195 | -0.099 | -0.005 | -0.051 | -0.051 | 1.058 | 0.88 | 0.85 | 0.96 | 1 | 0.89 | 0.98 |
| 9 | 105545596 | rs1388921 | 0.97 | -0.533 | -0.165 | -0.1 | 0.003 | 0.004 | 0.004 | 0 | 0.28 | 0.33 | 0.32 | 0.19 | 0.26 | 0.36 |
| 9 | 105545781 | rs10820282 | 0.587 | -0.301 | -0.251 | 0.31 | 0.018 | 0.007 | 0.007 | 0.137 | 0.37 | 0.31 | 0.48 | 0.66 | 0.3 | 0.34 |

| | | | | | | | | | | | | | | | | |
|---|---|---|---|---|---|---|---|---|---|---|---|---|---|---|---|---|
| 9 | 105546074 | rs7854657 | 0.783 | 0.167 | 0.088 | 0.199 | -0.012 | 0.009 | 0.009 | 0 | 0.17 | 0.09 | 0.12 | 0.05 | 0.25 | 0.14 |
| 9 | 105546493 | rs10990317 | 0.214 | 0.217 | 0.289 | 0.156 | 0.05 | -0.065 | -0.065 | 0 | 0.23 | 0.35 | 0.21 | 0.05 | 0.38 | 0.14 |
| 9 | 105547307 | rs12551330 | 1.429 | -0.115 | 0.274 | 0.129 | 0.004 | 0.001 | 0.001 | 0 | 0.18 | 0.09 | 0.19 | 0.18 | 0.15 | 0.27 |
| 9 | 105548952 | rs7869941 | 0.293 | 1.274 | 1.185 | 1.027 | 0.008 | -0.058 | -0.058 | 0 | 0.13 | 0.21 | 0.09 | 0 | 0.15 | 0.05 |
| 9 | 105563495 | rs12376672 | 1.152 | 1.298 | 1.179 | 1.088 | 0 | 0 | 0 | 0.032 | 0.07 | 0.08 | 0.07 | 0.08 | 0.07 | 0.08 |
| 9 | 105566691 | rs10217639 | 1.627 | 1.254 | 1.135 | 1.402 | 0.065 | -0.009 | -0.009 | 0.069 | 0.53 | 0.23 | 0.49 | 0.48 | 0.37 | 0.73 |
| 9 | 105573037 | rs7848024 | -1.081 | 1.237 | 1.225 | 1.488 | 0.004 | 0.004 | 0.004 | 0.129 | 0.25 | 0.69 | 0.29 | 0.27 | 0.27 | 0.16 |
| 9 | 105583926 | rs13294991 | 0.193 | 1.229 | 1.24 | 1.445 | 0.008 | -0.053 | -0.053 | 0.025 | 0.13 | 0.24 | 0.1 | 0 | 0.15 | 0.05 |
| 9 | 105585487 | rs1552766 | -1.412 | 1.303 | 1.007 | 1.474 | -0.02 | -0.027 | -0.027 | 3.797 | 0.35 | 0.75 | 0.46 | 0.52 | 0.5 | 0.24 |
| 9 | 105587136 | rs7033449 | 1.653 | 1.276 | 1.059 | 1.453 | 0.065 | -0.009 | -0.009 | 0.21 | 0.53 | 0.23 | 0.49 | 0.48 | 0.37 | 0.73 |
| 9 | 105591790 | rs2812594 | -1.429 | 1.379 | 1.123 | 1.557 | -0.018 | -0.024 | -0.024 | 0 | 0.35 | 0.73 | 0.46 | 0.52 | 0.5 | 0.24 |
| 9 | 105593380 | rs10990353 | 0.761 | 1.383 | 0.904 | 1.575 | -0.027 | -0.039 | -0.039 | 0 | 0.09 | 0.02 | 0.11 | 0.3 | 0.1 | 0.36 |
| 9 | 105595173 | rs1492598 | -0.146 | 1.492 | 1.32 | 1.385 | 0.041 | -0.012 | -0.012 | 0 | 0.18 | 0.22 | 0.18 | 0.13 | 0.3 | 0.17 |
| 9 | 105595627 | rs10512305 | 1.211 | 1.447 | 1.053 | 1.392 | -0.026 | 0.015 | 0.015 | 0 | 0.1 | 0.04 | 0.11 | 0.05 | 0.22 | 0.09 |
| 9 | 105603760 | rs843270 | -1.028 | 0.578 | 0.79 | 0.308 | 0.011 | -0.093 | -0.093 | 0 | 0.22 | 0.5 | 0.16 | 0.08 | 0.23 | 0.13 |
| 9 | 105608043 | rs10990359 | 0.303 | 1.03 | 0.051 | 0.83 | 0 | 0 | 0 | 0.167 | 0.07 | 0.06 | 0.06 | 0.08 | 0.07 | 0.09 |
| 9 | 105619674 | rs10990361 | 0.214 | 1.04 | 0.067 | 0.794 | 0.003 | 0.004 | 0.004 | 0.02 | 0.07 | 0.16 | 0.07 | 0.08 | 0.07 | 0.08 |
| 9 | 105620884 | rs2639345 | -0.615 | 1.116 | -0.542 | 0.749 | -0.015 | 0.047 | 0.047 | 0 | 0.9 | 0.98 | 0.95 | 1 | 0.86 | 0.97 |
| 9 | 105623942 | rs843283 | 1.061 | 0.872 | -0.312 | 0.917 | -0.005 | 0.001 | 0.001 | 0 | 0.05 | 0.02 | 0.05 | 0 | 0.12 | 0.01 |
| 9 | 105632182 | rs1352979 | 0.884 | 0.435 | -0.332 | 1.168 | 0.067 | 0.02 | 0.02 | 0 | 0.43 | 0.26 | 0.44 | 0.38 | 0.31 | 0.37 |
| 9 | 105633963 | rs2220706 | 0.985 | 0.418 | -0.248 | 1.244 | -0.001 | 0 | 0 | 0 | 0.06 | 0.05 | 0.04 | 0.08 | 0.05 | 0.1 |
| 9 | 105634309 | rs843275 | -0.314 | 0.431 | -0.168 | 1.403 | 0.007 | 0.04 | 0.04 | 0 | 0.27 | 0.61 | 0.34 | 0.43 | 0.28 | 0.41 |
| 9 | 105635106 | rs843272 | -0.519 | 0.43 | -0.173 | 1.406 | -0.057 | 0.034 | 0.034 | 0.019 | 0.7 | 0.86 | 0.79 | 0.82 | 0.58 | 0.77 |
| 9 | 105639770 | rs1492600 | 0.081 | 0.418 | -0.173 | 1.415 | -0.024 | 0.131 | 0.131 | 0 | 0.22 | 0.58 | 0.3 | 0.49 | 0.16 | 0.4 |
| 9 | 105675653 | rs11496631 | 0.28 | 0.449 | -0.197 | 1.246 | -0.057 | -0.031 | -0.031 | 0 | 0.31 | 0.14 | 0.26 | 0.36 | 0.42 | 0.24 |
| 9 | 105685265 | rs10990381 | 1.157 | 0.387 | 0.048 | 1.225 | 0.074 | 0.097 | 0.097 | 0 | 0.42 | 0.18 | 0.39 | 0.21 | 0.31 | 0.37 |
| 9 | 105695150 | rs10990384 | 0.154 | 0.405 | -0.053 | 1.193 | -0.057 | -0.029 | -0.029 | 0 | 0.32 | 0.15 | 0.27 | 0.36 | 0.42 | 0.24 |
| 9 | 105700390 | rs1935861 | 0.201 | 0.535 | 0.048 | 1.243 | 0.004 | 0.033 | 0.033 | 0 | 0.73 | 0.34 | 0.65 | 0.55 | 0.72 | 0.61 |
| 9 | 105704669 | rs10124970 | 0.205 | 0.54 | 0.021 | 1.24 | 0.019 | 0.085 | 0.085 | 0 | 0.75 | 0.47 | 0.67 | 0.57 | 0.73 | 0.62 |
| 9 | 105706469 | rs2067711 | 0.993 | 0.537 | 0.021 | 1.241 | 0.059 | 0.078 | 0.078 | 0.097 | 0.43 | 0.25 | 0.4 | 0.21 | 0.31 | 0.38 |
| 9 | 105707984 | rs10820319 | 0.953 | 0.584 | -0.017 | 1.488 | 0.003 | -0.011 | -0.011 | 0 | 0.16 | 0.07 | 0.09 | 0.22 | 0.15 | 0.36 |
| 9 | 105708946 | rs12005200 | 0.377 | 0.586 | -0.006 | 1.45 | 0.082 | 0.359 | 0.359 | 0 | 0.05 | 0.42 | 0.2 | 0.22 | 0.02 | 0.02 |
| 9 | 105711552 | rs10820320 | 1.21 | 0.589 | -0.122 | 1.407 | 0.08 | 0.108 | 0.108 | 0 | 0.41 | 0.1 | 0.38 | 0.22 | 0.31 | 0.37 |
| 9 | 105725193 | rs4409450 | 0.634 | 1.15 | 0.484 | 1.515 | 0.011 | 0.038 | 0.038 | 0 | 0.73 | 0.39 | 0.64 | 0.57 | 0.71 | 0.62 |
| 9 | 105734158 | rs10990402 | -0.007 | 0.922 | -0.921 | 0.923 | -0.001 | 0.002 | 0.002 | 0 | 0.16 | 0.13 | 0.09 | 0.23 | 0.15 | 0.35 |
| 9 | 105740221 | rs10990405 | -0.021 | 1.016 | -1.183 | 0.829 | 0.004 | -0.018 | -0.018 | 0 | 0.16 | 0.08 | 0.13 | 0.3 | 0.15 | 0.36 |
| 9 | 105746701 | rs1934995 | -0.372 | 1.694 | -1.27 | 0.156 | -0.058 | 0.052 | 0.052 | 0.386 | 0.25 | 0.13 | 0.18 | 0.09 | 0.37 | 0.12 |
| 9 | 105753694 | rs10124146 | -0.592 | 2.195 | -0.855 | 0.073 | 0.053 | -0.03 | -0.03 | 0.693 | 0.94 | 0.86 | 0.91 | 1 | 0.86 | 0.99 |
| 9 | 105755977 | rs1341204 | 0.658 | 2.45 | -1.514 | -0.488 | -0.043 | 0.047 | 0.047 | 0.351 | 0.76 | 0.85 | 0.82 | 0.92 | 0.69 | 0.9 |
| 9 | 105758477 | rs12156626 | 0.884 | 0.967 | -1.328 | -0.161 | 0.002 | -0.003 | -0.003 | 0 | 0.27 | 0.25 | 0.29 | 0.38 | 0.19 | 0.27 |
| 9 | 105763301 | rs10990416 | 0.009 | 1.009 | -0.868 | 0.053 | 0.033 | -0.009 | -0.009 | 0.012 | 0.74 | 0.57 | 0.72 | 0.79 | 0.63 | 0.83 |
| 9 | 105764116 | rs10760932 | 0.357 | 0.951 | -0.719 | -0.002 | -0.021 | 0.005 | 0.005 | 1.385 | 0.45 | 0.2 | 0.42 | 0.42 | 0.45 | 0.56 |
| 9 | 105766479 | rs10990423 | 0.355 | 0.949 | -0.719 | 0.039 | -0.021 | -0.02 | -0.02 | 0.035 | 0.46 | 0.2 | 0.51 | 0.47 | 0.45 | 0.56 |
| 9 | 105768503 | rs10119488 | 0.964 | 1.241 | -0.667 | 0.239 | 0.003 | 0.009 | 0.009 | 0 | 0.07 | 0.02 | 0.1 | 0.04 | 0.07 | 0.13 |
| 9 | 105770875 | rs10739862 | -0.193 | 1.307 | -0.37 | 0.297 | 0.035 | 0.011 | 0.011 | 0 | 0.83 | 0.74 | 0.84 | 0.79 | 0.71 | 0.96 |
| 9 | 105771136 | rs1107540 | -0.193 | 1.282 | -0.362 | 0.297 | 0.032 | 0.01 | 0.01 | 0 | 0.83 | 0.74 | 0.84 | 0.79 | 0.71 | 0.96 |
| 9 | 105772065 | rs2211279 | -0.193 | 1.282 | -0.366 | 0.297 | 0.032 | 0.01 | 0.01 | 0 | 0.83 | 0.74 | 0.84 | 0.79 | 0.71 | 0.96 |
| 9 | 105778790 | rs7856837 | -0.419 | 1.287 | -0.332 | 0.134 | 0.035 | -0.065 | -0.065 | 0 | 0.84 | 0.76 | 0.84 | 1 | 0.72 | 0.97 |
| 9 | 105780618 | rs13291989 | 0.531 | 1.297 | -0.328 | 0.206 | 0.02 | 0.002 | 0.002 | 0 | 0.76 | 0.69 | 0.73 | 0.75 | 0.63 | 0.83 |
| 9 | 105785533 | rs1341207 | 0.333 | 1.055 | -0.459 | 0.132 | -0.013 | -0.022 | -0.022 | 0 | 0.45 | 0.2 | 0.51 | 0.48 | 0.43 | 0.55 |
| 9 | 105788456 | rs10121525 | 0.039 | 1.013 | -0.254 | 0.148 | -0.012 | 0.059 | 0.059 | 0 | 0.1 | 0.19 | 0.12 | 0.26 | 0.08 | 0.13 |
| 9 | 105789727 | rs10512309 | 0.693 | 1.039 | -0.4 | 0.031 | -0.013 | 0.027 | 0.027 | 1.571 | 0.21 | 0.08 | 0.14 | 0.1 | 0.23 | 0.13 |
| 9 | 105790119 | rs947184 | 1.371 | 0.232 | -0.833 | -0.307 | -0.013 | 0.051 | 0.051 | 1.126 | 0.09 | 0 | 0.05 | 0 | 0.06 | 0.11 |
| 9 | 105790199 | rs11788777 | 1.238 | 0.246 | -0.679 | -0.266 | 0 | -0.011 | -0.011 | 0 | 0.25 | 0.19 | 0.38 | 0.37 | 0.21 | 0.42 |
| 9 | 105794662 | rs10990460 | 0.606 | 0.033 | -0.423 | -0.153 | 0.008 | 0.004 | 0.004 | 2.257 | 0.28 | 0.24 | 0.2 | 0.27 | 0.2 | 0.28 |
| 9 | 105797832 | rs10820351 | 0.066 | 0.341 | -0.202 | 0.159 | -0.006 | 0.05 | 0.05 | 0 | 0.1 | 0.23 | 0.12 | 0.18 | 0.08 | 0.12 |
| 9 | 105800243 | rs1341211 | 0.476 | 0.306 | -0.444 | 0.051 | 0 | -0.001 | -0.001 | 0 | 0.43 | 0.39 | 0.51 | 0.44 | 0.41 | 0.44 |
| 9 | 105803672 | rs7038696 | 0.269 | 0.228 | -0.471 | 0.108 | 0 | 0 | 0 | 0.561 | 0.46 | 0.43 | 0.54 | 0.55 | 0.44 | 0.49 |
| 9 | 105815502 | rs716450 | 0.95 | 0.12 | -0.575 | 0.038 | -0.013 | -0.008 | -0.008 | 0 | 0.34 | 0.23 | 0.48 | 0.4 | 0.36 | 0.43 |
| 9 | 105822396 | rs1099151 | 0.06 | -0.151 | -0.519 | -0.001 | 0 | -0.001 | -0.001 | 0.007 | 0.46 | 0.54 | 0.36 | 0.45 | 0.5 | 0.39 |
| 9 | 105827744 | rs12377905 | 0.237 | -0.14 | -0.505 | -0.029 | 0 | -0.001 | -0.001 | 0.075 | 0.46 | 0.44 | 0.54 | 0.52 | 0.44 | 0.47 |
| 9 | 105827815 | rs843192 | 1.775 | 0.203 | -0.477 | -0.097 | 0.015 | -0.02 | -0.02 | 0 | 0.07 | 0.1 | 0.08 | 0 | 0.14 | 0.01 |
| 9 | 105832538 | rs12347896 | 0.884 | 0.083 | -0.244 | -0.023 | -0.037 | -0.047 | -0.047 | 1.056 | 0.09 | 0.2 | 0.04 | 0.01 | 0.03 | 0.01 |
| 9 | 105837587 | rs843167 | 0.657 | 0.174 | -0.394 | 0.01 | -0.032 | 0.021 | 0.021 | 0 | 0.22 | 0.14 | 0.19 | 0.09 | 0.31 | 0.14 |
| 9 | 105839544 | rs10990512 | 0.22 | 0.306 | -0.394 | 0.154 | -0.04 | 0.007 | 0.007 | 0 | 0.13 | 0.19 | 0.06 | 0.1 | 0.05 | 0.04 |
| 9 | 105844979 | rs843162 | -0.001 | 0.656 | -0.712 | 0.442 | -0.004 | 0.001 | 0.001 | 0 | 0.43 | 0.41 | 0.33 | 0.39 | 0.48 | 0.36 |
| 9 | 105847351 | rs843159 | -0.206 | 0.506 | -0.777 | -0.005 | 0.004 | -0.048 | -0.048 | 0 | 0.43 | 0.45 | 0.29 | 0.14 | 0.43 | 0.39 |
| 9 | 105847555 | rs843158 | 0.192 | -0.008 | -0.478 | 0.403 | -0.002 | 0.009 | 0.009 | 0 | 0.83 | 0.78 | 0.83 | 0.75 | 0.84 | 0.64 |
| 9 | 105848723 | rs10990524 | 0.589 | -0.131 | -1.086 | -0.35 | -0.014 | -0.018 | -0.018 | 0.217 | 0.18 | 0.03 | 0.1 | 0.17 | 0.15 | 0.12 |

| | | | | | | | | | | | | | | | | |
|---|---|---|---|---|---|---|---|---|---|---|---|---|---|---|---|---|
| 9 | 105850270 | rs10123704 | 0.068 | -0.125 | -1.116 | -0.598 | -0.047 | -0.049 | -0.049 | 0.248 | 0.36 | 0.21 | 0.47 | 0.47 | 0.44 | 0.47 |
| 9 | 105853222 | rs1375209 | 1.152 | -0.103 | -1.19 | -0.577 | -0.013 | -0.05 | -0.05 | 0 | 0.15 | 0.07 | 0.21 | 0.38 | 0.15 | 0.26 |
| 9 | 105858882 | rs10990533 | 1.127 | -0.102 | -1.189 | -0.578 | -0.012 | -0.046 | -0.046 | 0.469 | 0.15 | 0.08 | 0.22 | 0.38 | 0.15 | 0.26 |
| 9 | 105864747 | rs1417702 | -0.519 | -0.404 | -0.629 | -0.816 | -0.028 | -0.038 | -0.038 | 0.048 | 0.93 | 0.81 | 0.99 | 0.99 | 0.99 | 0.9 |
| 9 | 105868652 | rs6479155 | -0.303 | -0.564 | -0.734 | -0.923 | -0.04 | -0.045 | -0.045 | 0 | 0.38 | 0.22 | 0.48 | 0.47 | 0.45 | 0.5 |
| 9 | 105879111 | rs7048164 | -1.44 | -0.835 | -0.394 | -1.023 | -0.022 | -0.004 | -0.004 | 0 | 0.7 | 0.59 | 0.84 | 0.71 | 0.75 | 0.75 |
| 9 | 105881438 | rs1597210 | 1.253 | -0.853 | -0.24 | -1.05 | 0 | 0.008 | 0.008 | 0 | 0.14 | 0.09 | 0.27 | 0.11 | 0.15 | 0.16 |
| 9 | 105882459 | rs10990553 | 1.316 | -0.81 | -0.266 | -1.072 | -0.028 | 0.028 | 0.028 | 0 | 0.2 | 0.13 | 0.23 | 0.03 | 0.29 | 0.21 |
| 9 | 105883836 | rs7873512 | -0.448 | -0.802 | -0.261 | -1.056 | -0.001 | -0.009 | -0.009 | 0 | 0.48 | 0.47 | 0.61 | 0.68 | 0.45 | 0.52 |
| 9 | 105884526 | rs843180 | 1.917 | -0.736 | -0.391 | -0.942 | 0.005 | -0.004 | -0.004 | 0 | 0.24 | 0.27 | 0.14 | 0.29 | 0.24 | 0.15 |
| 9 | 105893479 | rs10990559 | 1.007 | -0.498 | -0.554 | -0.898 | 0 | 0.006 | 0.006 | 0 | 0.14 | 0.1 | 0.27 | 0.11 | 0.15 | 0.16 |
| 9 | 105900043 | rs10120196 | 1.175 | -0.547 | -0.329 | -0.935 | 0.004 | 0.008 | 0.008 | 0 | 0.19 | 0.11 | 0.08 | 0.13 | 0.16 | 0.09 |
| 9 | 105900413 | rs4743624 | -0.126 | -0.548 | -0.221 | -0.899 | -0.001 | -0.001 | -0.001 | 0.455 | 0.33 | 0.29 | 0.36 | 0.29 | 0.3 | 0.25 |
| 9 | 105904199 | rs4743626 | 0.92 | -0.531 | -0.266 | -0.893 | 0.009 | 0.014 | 0.014 | 0 | 0.21 | 0.12 | 0.08 | 0.13 | 0.16 | 0.11 |
| 9 | 105920416 | rs2218687 | 0.367 | -0.478 | -0.038 | -0.59 | 0.007 | 0.063 | 0.063 | 0 | 0.18 | 0.36 | 0.31 | 0.29 | 0.17 | 0.19 |
| 9 | 105923585 | rs12378919 | 0.617 | 0.017 | -0.318 | -0.49 | -0.03 | 0.029 | 0.029 | 0 | 0.19 | 0.12 | 0.22 | 0.03 | 0.29 | 0.21 |
| 9 | 105937834 | rs1839166 | -0.198 | -0.101 | -0.35 | -0.324 | -0.039 | -0.046 | -0.046 | 0 | 0.68 | 0.79 | 0.53 | 0.6 | 0.52 | 0.64 |
| 9 | 105939715 | rs10990596 | 0.216 | -0.165 | -0.363 | -0.364 | 0.014 | -0.004 | -0.004 | 0 | 0.05 | 0.02 | 0.02 | 0.05 | 0.04 | 0.04 |
| 9 | 105940438 | rs1448579 | -0.266 | -0.031 | 0.051 | -0.418 | -0.001 | -0.002 | -0.002 | 0 | 0.61 | 0.62 | 0.62 | 0.77 | 0.55 | 0.73 |
| 9 | 105942389 | rs10990599 | 1.392 | -0.022 | -0.116 | -0.418 | 0.02 | 0.024 | 0.024 | 0 | 0.21 | 0.07 | 0.09 | 0.14 | 0.17 | 0.11 |
| 9 | 105942403 | rs4742875 | -1.556 | 0.083 | -0.35 | -0.393 | -0.031 | 0.026 | 0.026 | 0 | 0.86 | 0.91 | 0.75 | 0.97 | 0.75 | 0.86 |
| 9 | 105946455 | rs10820387 | 0.228 | -0.081 | -0.186 | -0.442 | -0.035 | -0.025 | -0.025 | 0 | 0.17 | 0.23 | 0.25 | 0.14 | 0.09 | 0.24 |
| 9 | 105947343 | rs10990603 | 1.157 | 0.023 | -0.235 | -0.373 | -0.013 | -0.024 | -0.024 | 0.358 | 0.17 | 0.12 | 0.23 | 0.35 | 0.23 | 0.21 |
| 9 | 105951481 | rs10739871 | -0.406 | -0.15 | -0.318 | -0.651 | 0.002 | 0.008 | 0.008 | 0 | 0.51 | 0.53 | 0.39 | 0.56 | 0.49 | 0.44 |
| 9 | 105951980 | rs7047105 | -0.352 | -0.049 | -0.81 | -0.645 | 0.008 | -0.031 | -0.031 | 0 | 0.77 | 0.76 | 0.73 | 0.96 | 0.71 | 0.77 |
| 9 | 105955511 | rs12339905 | 0.132 | -1.435 | -0.366 | -0.373 | -0.015 | -0.021 | -0.021 | 0 | 0.29 | 0.28 | 0.13 | 0.16 | 0.22 | 0.2 |
| 9 | 105964224 | rs10990617 | -0.462 | -0.929 | 0.187 | 0.043 | -0.026 | -0.001 | -0.001 | 0 | 0.36 | 0.55 | 0.38 | 0.4 | 0.27 | 0.42 |
| 9 | 105964658 | rs11789118 | -0.176 | -0.4 | -0.76 | 0.285 | -0.021 | -0.039 | -0.039 | 0 | 0.09 | 0.13 | 0.03 | 0.01 | 0.05 | 0.09 |
| 9 | 105965663 | rs10990619 | 0.065 | 0.202 | -1.114 | 0.762 | -0.01 | 0.006 | 0.006 | 0 | 0.09 | 0.11 | 0.22 | 0.13 | 0.03 | 0.15 |
| 9 | 105965683 | rs16922610 | -0.142 | 0.502 | -1.544 | 0.762 | 0.01 | -0.015 | -0.015 | 0 | 0.24 | 0.13 | 0.15 | 0.27 | 0.24 | 0.25 |
| 9 | 105969051 | rs10990625 | -0.451 | 1.249 | -1.321 | 1.111 | -0.016 | -0.021 | -0.021 | 0 | 0.21 | 0.16 | 0.37 | 0.63 | 0.26 | 0.39 |
| 9 | 105969101 | rs1823279 | -0.239 | 0.869 | -1.292 | 0.764 | -0.001 | -0.003 | -0.003 | 0 | 0.12 | 0.13 | 0.13 | 0.15 | 0.16 | 0.17 |
| 9 | 105969217 | rs1823281 | -0.803 | -0.767 | -1.677 | -0.415 | 0.053 | 0.039 | 0.039 | 0 | 0.76 | 0.89 | 0.83 | 0.99 | 0.88 | 0.85 |
| 9 | 105971199 | rs10990626 | -0.063 | -0.806 | -1.847 | -0.293 | 0.001 | -0.008 | -0.008 | 0 | 0.19 | 0.12 | 0.15 | 0.21 | 0.19 | 0.19 |
| 9 | 105971237 | rs7850677 | -0.548 | -1.395 | -1.227 | -0.744 | 0.003 | 0.004 | 0.004 | 0.137 | 0.65 | 0.64 | 0.73 | 0.86 | 0.8 | 0.76 |
| 9 | 105971829 | rs10820391 | 0.491 | -1.407 | -1.093 | -0.736 | 0.015 | 0.013 | 0.013 | 0 | 0.46 | 0.52 | 0.57 | 0.69 | 0.59 | 0.57 |
| 9 | 105977018 | rs4743634 | 0.209 | -1.272 | -1.01 | -0.714 | 0.001 | 0.001 | 0.001 | 0.184 | 0.19 | 0.12 | 0.15 | 0.17 | 0.19 | 0.19 |
| 9 | 105977028 | rs17775713 | 0.694 | -1.649 | -1.033 | -0.797 | 0.013 | 0.037 | 0.037 | 0 | 0.1 | 0.05 | 0.06 | 0 | 0.09 | 0.1 |
| 9 | 105983538 | rs7044631 | -0.096 | -1.63 | -1.037 | -0.893 | 0.005 | 0.016 | 0.016 | 0 | 0.27 | 0.21 | 0.2 | 0.16 | 0.24 | 0.31 |
| 9 | 105983780 | rs1448577 | 0.739 | -1.486 | -0.573 | -0.371 | 0.001 | 0 | 0 | 0 | 0.33 | 0.32 | 0.52 | 0.41 | 0.45 | 0.32 |
| 9 | 105983864 | rs10990632 | 0.754 | -1.558 | -0.424 | -0.288 | -0.008 | -0.029 | -0.029 | 0 | 0.14 | 0.06 | 0.11 | 0.28 | 0.19 | 0.22 |
| 9 | 105991203 | rs10739874 | -1.852 | -1.605 | -0.305 | -0.354 | 0.058 | 0.051 | 0.051 | 0 | 0.77 | 0.88 | 0.93 | 0.99 | 0.89 | 0.87 |
| 9 | 105993121 | rs10760966 | 0.579 | -1.597 | -0.194 | -0.316 | 0.002 | 0.042 | 0.042 | 0 | 0.16 | 0.17 | 0.2 | 0.41 | 0.2 | 0.23 |
| 9 | 105993267 | rs1176191 | -1.933 | -1.578 | -0.224 | -0.382 | 0.073 | 0.083 | 0.083 | 0.002 | 0.78 | 0.93 | 0.93 | 0.99 | 0.89 | 0.87 |
| 9 | 105999196 | rs7865219 | 0.61 | -1.465 | -0.106 | -0.301 | -0.001 | -0.001 | -0.001 | 0 | 0.57 | 0.57 | 0.6 | 0.43 | 0.56 | 0.47 |
| 9 | 106005044 | rs10990645 | 0.443 | -1.509 | -0.472 | -0.256 | 0.036 | -0.015 | -0.015 | 0 | 0.38 | 0.26 | 0.27 | 0.39 | 0.3 | 0.35 |
| 9 | 106006075 | rs10820403 | 1.045 | -1.37 | -0.46 | -0.171 | -0.033 | 0.029 | 0.029 | 0 | 0.3 | 0.2 | 0.22 | 0.08 | 0.41 | 0.28 |
| 9 | 106009535 | rs1456476 | -0.256 | -1.249 | -0.436 | 0.019 | 0.002 | -0.067 | -0.067 | 0.577 | 0.71 | 0.83 | 0.53 | 0.47 | 0.72 | 0.64 |
| 9 | 106013174 | rs7033431 | 0.315 | -1.201 | -0.388 | 0.052 | -0.002 | -0.021 | -0.021 | 0 | 0.16 | 0.11 | 0.12 | 0.24 | 0.19 | 0.24 |
| 9 | 106015010 | rs167334 | -0.508 | -1.411 | -0.103 | -0.068 | -0.001 | 0.048 | 0.048 | 0.046 | 0.72 | 0.53 | 0.75 | 0.62 | 0.7 | 0.59 |
| 9 | 106017371 | rs10990654 | 0.965 | -1.566 | -0.054 | -0.262 | -0.027 | 0.018 | 0.018 | 0 | 0.29 | 0.19 | 0.22 | 0.08 | 0.42 | 0.29 |
| 9 | 106021388 | rs303545 | 1.602 | -1.599 | -0.298 | -0.238 | 0.074 | 0.088 | 0.088 | 0.25 | 0.26 | 0.1 | 0.08 | 0.07 | 0.13 | 0.16 |
| 9 | 106024220 | rs16922670 | 0.409 | -1.358 | -0.186 | -0.026 | 0 | 0.03 | 0.03 | 0 | 0.11 | 0.13 | 0.34 | 0.33 | 0.14 | 0.13 |
| 9 | 106034122 | rs303585 | -0.838 | -1.371 | -0.348 | -0.181 | -0.003 | -0.019 | -0.019 | 1.378 | 0.73 | 0.79 | 0.78 | 0.62 | 0.7 | 0.59 |
| 9 | 106038803 | rs4295718 | 1.613 | -1.611 | -0.739 | -0.5 | 0.081 | 0.13 | 0.13 | 0.055 | 0.22 | 0.04 | 0.07 | 0.01 | 0.11 | 0.12 |
| 9 | 106046072 | rs1124647 | -0.399 | -1.502 | -0.269 | -0.375 | -0.021 | 0.007 | 0.007 | 0 | 0.15 | 0.39 | 0.19 | 0.14 | 0.09 | 0.11 |
| 9 | 106052433 | rs12337792 | 0.341 | -1.497 | -0.302 | -0.655 | -0.008 | -0.066 | -0.066 | 0.047 | 0.11 | 0.3 | 0.09 | 0 | 0.08 | 0.1 |
| 9 | 106053927 | rs12004858 | 0.879 | -0.974 | -1.281 | -0.816 | -0.085 | 0.091 | 0.091 | 0 | 0.24 | 0.08 | 0.26 | 0.05 | 0.36 | 0.2 |
| 9 | 106060322 | rs303615 | 1.564 | -1.252 | -1.199 | -0.75 | 0.081 | 0.115 | 0.115 | 0 | 0.22 | 0.07 | 0.08 | 0.01 | 0.11 | 0.12 |
| 9 | 106067533 | rs4743641 | 0.244 | -0.655 | -0.728 | -0.262 | -0.091 | 0.118 | 0.118 | 0 | 0.31 | 0.11 | 0.3 | 0.07 | 0.43 | 0.29 |
| 9 | 106068965 | rs1378974 | -0.07 | -0.819 | -0.668 | 0.004 | 0.006 | 0.005 | 0.005 | 0 | 0.36 | 0.39 | 0.61 | 0.45 | 0.51 | 0.38 |
| 9 | 106082463 | rs445722 | 1.179 | -1.212 | -1.06 | -0.293 | 0.053 | 0.096 | 0.096 | 0 | 0.2 | 0.07 | 0.08 | 0.01 | 0.12 | 0.11 |
| 9 | 106082492 | rs16922767 | 1.041 | -1.418 | -1.368 | -0.028 | -0.013 | -0.043 | -0.043 | 0 | 0.07 | 0.01 | 0.05 | 0.14 | 0.09 | 0.19 |
| 9 | 106083827 | rs370386 | -0.088 | -1.902 | -0.756 | 0.182 | 0.033 | 0.03 | 0.03 | 0 | 0.34 | 0.32 | 0.2 | 0.19 | 0.22 | 0.2 |
| 9 | 106087206 | rs423433 | 1.666 | -1.867 | -0.827 | 0.184 | 0.053 | 0.096 | 0.096 | 0 | 0.2 | 0.07 | 0.08 | 0.01 | 0.12 | 0.11 |
| 9 | 106087352 | rs373637 | -0.724 | -1.528 | -0.543 | 0.039 | -0.01 | -0.004 | -0.004 | 0 | 0.61 | 0.76 | 0.63 | 0.59 | 0.46 | 0.45 |
| 9 | 106091787 | rs379786 | 0.338 | -1.566 | -0.242 | 0.034 | -0.003 | -0.001 | -0.001 | 0.126 | 0.36 | 0.38 | 0.23 | 0.23 | 0.25 | 0.24 |
| 9 | 106093936 | rs450441 | 0.253 | -1.568 | -0.247 | 0.032 | 0.002 | 0.003 | 0.003 | 0 | 0.37 | 0.38 | 0.23 | 0.23 | 0.25 | 0.23 |
| 9 | 106098966 | rs414964 | 1.476 | -1.746 | 0.076 | 0.105 | 0.03 | 0.028 | 0.028 | 0 | 0.25 | 0.17 | 0.1 | 0.05 | 0.12 | 0.17 |

| | | | | | | | | | | | | | | | | |
|---|---|---|---|---|---|---|---|---|---|---|---|---|---|---|---|---|
| 9 | 106100469 | rs12002497 | 0.996 | -1.704 | 0.187 | 0.123 | -0.013 | 0.152 | 0.152 | 0.142 | 0.05 | 0.16 | 0.06 | 0.17 | 0.01 | 0.02 |
| 9 | 106101057 | rs16922813 | 1.206 | -1.672 | 0.054 | 0.031 | 0 | -0.013 | -0.013 | 0 | 0.09 | 0.13 | 0.07 | 0 | 0.09 | 0.07 |
| 9 | 106101691 | rs7037370 | -0.951 | -1.731 | 0.138 | 0.193 | -0.005 | -0.015 | -0.015 | 4.303 | 0.61 | 0.77 | 0.61 | 0.57 | 0.43 | 0.44 |
| 9 | 106104878 | rs303575 | 1.521 | -1.833 | 0.38 | 0.387 | 0.058 | 0.031 | 0.031 | 0 | 0.2 | 0.06 | 0.07 | 0.11 | 0.12 | 0.1 |
| 9 | 106106116 | rs11999327 | 0.155 | -1.852 | 0.522 | 0.483 | -0.03 | 0.077 | 0.077 | 0 | 0.08 | 0.17 | 0.07 | 0.21 | 0.05 | 0.03 |
| 9 | 106120670 | rs1904928 | 0.09 | -1.857 | 0.571 | 0.48 | -0.031 | 0.097 | 0.097 | 0 | 0.09 | 0.21 | 0.08 | 0.21 | 0.05 | 0.03 |
| 9 | 106122565 | rs303596 | -0.251 | -1.956 | 0.634 | 0.469 | -0.058 | 0.011 | 0.011 | 0 | 0.34 | 0.43 | 0.17 | 0.36 | 0.17 | 0.19 |
| 9 | 106123271 | rs303598 | 0.888 | -1.831 | 0.094 | 0.471 | 0.015 | 0.011 | 0.011 | 0 | 0.26 | 0.23 | 0.09 | 0.15 | 0.12 | 0.16 |
| 9 | 106124641 | rs303603 | 1.304 | -1.768 | 0.001 | 0.431 | 0.027 | 0.022 | 0.022 | 0 | 0.21 | 0.17 | 0.08 | 0.11 | 0.12 | 0.1 |
| 9 | 106126869 | rs10820413 | 0.771 | -0.563 | 0.045 | 0.514 | 0.077 | -0.023 | -0.023 | 0 | 0.07 | 0.1 | 0.19 | 0.03 | 0.21 | 0.09 |
| 9 | 106137424 | rs16922872 | -0.362 | -2.18 | -0.19 | 0.82 | 0 | 0 | 0 | 0 | 0.12 | 0.11 | 0.07 | 0.24 | 0.05 | 0.09 |
| 9 | 106148215 | rs303635 | -0.355 | -2.491 | -0.136 | 0.555 | -0.01 | -0.001 | -0.001 | 0 | 0.38 | 0.37 | 0.21 | 0.35 | 0.17 | 0.19 |
| 9 | 106153263 | rs16922886 | 0.009 | -2.005 | -0.616 | 0.692 | 0.017 | -0.02 | -0.02 | 0 | 0.08 | 0.02 | 0.09 | 0.2 | 0.03 | 0.07 |
| 9 | 106163708 | rs303620 | 1.085 | -2.225 | -0.771 | 0.258 | 0.091 | 0.13 | 0.13 | 0 | 0.22 | 0.03 | 0.07 | 0.03 | 0.1 | 0.09 |
| 9 | 106169268 | rs9695167 | -1.125 | -2.669 | 0.033 | 0.173 | -0.088 | 0.009 | 0.009 | 0 | 0.19 | 0.43 | 0.15 | 0.17 | 0.08 | 0.13 |
| 9 | 106173210 | rs303543 | -0.69 | -2.686 | -0.046 | 0.37 | -0.014 | 0.001 | 0.001 | 0 | 0.43 | 0.57 | 0.22 | 0.32 | 0.18 | 0.24 |
| 9 | 106173793 | rs303542 | 1.264 | -2.588 | -0.301 | 0.45 | 0.082 | 0.019 | 0.019 | 0 | 0.22 | 0.06 | 0.07 | 0.15 | 0.1 | 0.1 |
| 9 | 106177362 | rs6479168 | -0.39 | -2.36 | -0.011 | 0.329 | -0.017 | 0.006 | 0.006 | 0 | 0.12 | 0.14 | 0.1 | 0.17 | 0.04 | 0.1 |
| 9 | 106181050 | rs6479169 | -1.018 | -1.899 | -0.332 | 0.324 | 0.032 | 0.041 | 0.041 | 0 | 0.57 | 0.66 | 0.58 | 0.69 | 0.36 | 0.41 |
| 9 | 106205224 | rs10990701 | 0.975 | -1.951 | -0.795 | 0.115 | 0.074 | 0.038 | 0.038 | 0.094 | 0.24 | 0.07 | 0.1 | 0.15 | 0.13 | 0.11 |
| 9 | 106205827 | rs12682841 | -0.924 | -1.292 | -0.068 | 0.483 | -0.048 | 0.008 | 0.008 | 0 | 0.21 | 0.31 | 0.18 | 0.17 | 0.08 | 0.22 |
| 9 | 106207271 | rs12683176 | -0.605 | -0.786 | -0.548 | 0.87 | 0.006 | -0.005 | -0.005 | 0 | 0.09 | 0.06 | 0.13 | 0.17 | 0.02 | 0.11 |
| 9 | 106235615 | rs1936739 | -0.317 | 0.333 | -0.363 | 0.614 | 0.019 | 0.02 | 0.02 | 0 | 0.22 | 0.3 | 0.3 | 0.29 | 0.32 | 0.38 |
| 9 | 106236224 | rs1415686 | 0.587 | 0.538 | -0.514 | 0.686 | -0.021 | -0.017 | -0.017 | 0 | 0.07 | 0.02 | 0.18 | 0.21 | 0.1 | 0.14 |
| 9 | 106238925 | rs17790711 | 0.044 | -0.044 | -0.229 | 0.14 | -0.014 | -0.036 | -0.036 | 0 | 0.15 | 0.18 | 0.05 | 0.02 | 0.08 | 0.2 |
| 9 | 106240743 | rs16922936 | 0.664 | 0.536 | -1.024 | 0.475 | -0.063 | -0.113 | -0.113 | 0 | 0.15 | 0.02 | 0.32 | 0.37 | 0.19 | 0.09 |
| 9 | 106242831 | rs10820423 | 0.424 | 0.363 | -0.73 | -0.121 | 0 | 0 | 0 | 0.009 | 0.37 | 0.36 | 0.47 | 0.54 | 0.41 | 0.33 |
| 9 | 106243701 | rs12379035 | 1.948 | 0.072 | -0.928 | -0.395 | 0.06 | 0.054 | 0.054 | 0 | 0.23 | 0.1 | 0.09 | 0.14 | 0.12 | 0.11 |
| 9 | 106248670 | rs12351109 | -0.688 | 0.104 | -0.729 | -0.437 | 0.034 | 0.015 | 0.015 | 0 | 0.78 | 0.73 | 0.61 | 0.71 | 0.62 | 0.68 |
| 9 | 106261364 | rs10820441 | 1.876 | 0.072 | -0.473 | -0.478 | -0.001 | -0.004 | -0.004 | 2.413 | 0.21 | 0.19 | 0.16 | 0.18 | 0.23 | 0.22 |
| 9 | 106268589 | rs840250 | -1.229 | 0.72 | -0.506 | -0.473 | -0.014 | -0.01 | -0.01 | 0 | 0.9 | 0.92 | 0.73 | 0.76 | 0.83 | 0.77 |
| 9 | 106268919 | rs13285576 | 1.42 | -0.025 | -0.597 | -0.498 | -0.032 | -0.061 | -0.061 | 0.215 | 0.14 | 0.04 | 0.31 | 0.34 | 0.18 | 0.09 |
| 9 | 106270798 | rs10820445 | 1.178 | -0.512 | -0.423 | -0.791 | 0.062 | 0.055 | 0.055 | 0 | 0.29 | 0.17 | 0.11 | 0.17 | 0.14 | 0.21 |
| 9 | 106283105 | rs1703481 | 1.709 | -0.538 | -0.395 | -0.829 | 0.007 | -0.003 | -0.003 | 0.034 | 0.11 | 0.09 | 0.03 | 0 | 0.07 | 0.11 |
| 9 | 106287146 | rs10760969 | -0.657 | -0.12 | -0.189 | -0.514 | 0.035 | 0.016 | 0.016 | 0 | 0.78 | 0.73 | 0.61 | 0.7 | 0.63 | 0.7 |
| 9 | 106289427 | rs2202828 | 0.274 | -0.112 | -0.218 | -0.505 | -0.002 | 0.001 | 0.001 | 0 | 0.48 | 0.43 | 0.5 | 0.53 | 0.49 | 0.46 |
| 9 | 106295428 | rs10760970 | 1.59 | -0.29 | -0.463 | -0.601 | 0.056 | 0.046 | 0.046 | 0.079 | 0.21 | 0.08 | 0.08 | 0.14 | 0.12 | 0.09 |
| 9 | 106302762 | rs10990763 | 1.345 | 0.025 | -0.432 | -0.609 | 0.025 | 0.046 | 0.046 | 0.061 | 0.23 | 0.1 | 0.12 | 0.14 | 0.16 | 0.1 |
| 9 | 106306038 | rs840256 | -1.126 | 0.174 | -0.17 | -0.549 | 0.03 | 0.016 | 0.016 | 0 | 0.82 | 0.76 | 0.67 | 0.74 | 0.69 | 0.72 |
| 9 | 106306486 | rs840255 | -1.069 | -0.212 | -0.154 | -0.532 | 0.033 | 0.011 | 0.011 | 0 | 0.81 | 0.75 | 0.63 | 0.73 | 0.65 | 0.72 |
| 9 | 106311685 | rs840241 | -0.816 | -0.273 | -0.15 | -0.503 | 0.078 | 0.025 | 0.025 | 0 | 0.78 | 0.67 | 0.61 | 0.73 | 0.63 | 0.69 |
| 9 | 106326245 | rs2786798 | -0.097 | -0.38 | -0.276 | -0.549 | -0.001 | -0.001 | -0.001 | 0 | 0.53 | 0.5 | 0.52 | 0.6 | 0.5 | 0.57 |
| 9 | 106327061 | rs1995789 | 0.199 | -0.36 | -0.444 | -0.554 | -0.004 | 0.017 | 0.017 | 0 | 0.48 | 0.37 | 0.5 | 0.56 | 0.49 | 0.46 |
| 9 | 106331688 | rs10820459 | 1.153 | -1.536 | -0.765 | -0.874 | 0.044 | 0.038 | 0.038 | 0 | 0.22 | 0.11 | 0.08 | 0.14 | 0.12 | 0.09 |
| 9 | 106333748 | rs1509040 | 0.461 | -1.234 | -0.854 | -0.606 | 0.007 | -0.074 | -0.074 | 1.053 | 0.18 | 0.08 | 0.21 | 0.37 | 0.16 | 0.16 |
| 9 | 106335903 | rs1577902 | 0.107 | -1.444 | -0.504 | -0.96 | 0.007 | 0.005 | 0.005 | 0.225 | 0.44 | 0.38 | 0.39 | 0.57 | 0.42 | 0.48 |
| 9 | 106337200 | rs1577905 | -0.646 | -1.622 | -0.475 | -1.102 | 0.048 | -0.009 | -0.009 | 0.135 | 0.72 | 0.61 | 0.5 | 0.76 | 0.57 | 0.67 |
| 9 | 106344556 | rs4540458 | -0.313 | -1.703 | -0.331 | -1.133 | 0.004 | 0.006 | 0.006 | 0 | 0.44 | 0.6 | 0.71 | 0.6 | 0.56 | 0.44 |
| 9 | 106345031 | rs7871238 | 0.708 | -1.746 | -0.438 | -1.151 | 0.007 | 0.006 | 0.006 | 0 | 0.28 | 0.25 | 0.12 | 0.18 | 0.16 | 0.2 |
| 9 | 106348522 | rs7865138 | -1.277 | -1.686 | -0.35 | -1.078 | 0.009 | 0.026 | 0.026 | 0.001 | 0.7 | 0.81 | 0.82 | 0.78 | 0.7 | 0.62 |
| 9 | 106359311 | rs7048428 | 1.151 | -1.647 | -0.337 | -1.029 | -0.002 | 0.037 | 0.037 | 0 | 0.15 | 0.19 | 0.21 | 0.36 | 0.13 | 0.11 |
| 9 | 106375480 | rs10990824 | 0.413 | -1.705 | -0.241 | -1.113 | 0.009 | -0.001 | -0.001 | 0 | 0.28 | 0.28 | 0.5 | 0.27 | 0.43 | 0.37 |
| 9 | 106375915 | rs10990825 | 0.83 | -1.664 | -0.178 | -1.085 | 0 | 0.001 | 0.001 | 0 | 0.26 | 0.25 | 0.11 | 0.18 | 0.14 | 0.18 |
| 9 | 106375973 | rs10990826 | 0.83 | -1.664 | -0.178 | -1.085 | 0 | 0.001 | 0.001 | 0 | 0.26 | 0.25 | 0.11 | 0.18 | 0.14 | 0.18 |
| 9 | 106381192 | rs10990837 | 1.29 | -1.329 | -0.326 | -1.14 | 0.049 | -0.062 | -0.062 | 2.124 | 0.16 | 0.25 | 0.14 | 0.04 | 0.27 | 0.13 |
| 9 | 106385442 | rs10990840 | 1.051 | -2.483 | -0.537 | -0.902 | 0.03 | 0.018 | 0.018 | 0 | 0.2 | 0.05 | 0.08 | 0.16 | 0.13 | 0.11 |
| 9 | 106385994 | rs10990842 | 0.483 | -2.405 | -0.426 | -0.864 | 0.05 | 0.035 | 0.035 | 0 | 0.25 | 0.12 | 0.1 | 0.18 | 0.14 | 0.18 |
| 9 | 106386149 | rs10760978 | -0.296 | -2.365 | -0.407 | -0.852 | -0.013 | 0.002 | 0.002 | 0.05 | 0.41 | 0.46 | 0.33 | 0.53 | 0.27 | 0.3 |
| 9 | 106391797 | rs2221127 | -0.927 | -2.359 | -0.874 | -0.881 | 0.013 | 0.021 | 0.021 | 0 | 0.7 | 0.83 | 0.82 | 0.78 | 0.71 | 0.63 |
| 9 | 106396321 | rs10820503 | 1.006 | -2.059 | -0.568 | -0.664 | 0.005 | -0.001 | -0.001 | 0 | 0.31 | 0.33 | 0.52 | 0.25 | 0.43 | 0.34 |
| 9 | 106399281 | rs2055749 | -0.402 | -2.264 | -0.58 | -0.682 | -0.022 | 0.005 | 0.005 | 0 | 0.41 | 0.49 | 0.3 | 0.54 | 0.27 | 0.3 |
| 9 | 106400878 | rs11794668 | 1.13 | -2.326 | -0.614 | -0.501 | -0.002 | -0.01 | -0.01 | 0 | 0.13 | 0.09 | 0.18 | 0.35 | 0.13 | 0.12 |
| 9 | 106407456 | rs11788173 | -0.151 | -2.522 | -0.635 | -0.557 | 0.029 | 0.014 | 0.014 | 0 | 0.6 | 0.72 | 0.79 | 0.61 | 0.68 | 0.52 |
| 9 | 106409952 | rs7867943 | -0.796 | -2.559 | -0.625 | -0.564 | -0.021 | 0.004 | 0.004 | 0 | 0.42 | 0.49 | 0.33 | 0.53 | 0.27 | 0.3 |
| 9 | 106417844 | rs10990869 | 0.071 | -2.454 | -0.607 | -0.498 | -0.011 | 0.016 | 0.016 | 0 | 0.3 | 0.37 | 0.27 | 0.35 | 0.24 | 0.16 |
| 9 | 106422027 | rs11793044 | 1.163 | -2.196 | -0.634 | -0.331 | -0.002 | -0.011 | -0.011 | 0 | 0.14 | 0.1 | 0.18 | 0.35 | 0.13 | 0.13 |
| 9 | 106423464 | rs10760980 | 1.033 | -1.968 | -0.829 | -0.267 | 0.009 | -0.002 | -0.002 | 0.18 | 0.3 | 0.33 | 0.52 | 0.26 | 0.43 | 0.37 |
| 9 | 106424148 | rs10990877 | 1.11 | -1.651 | -0.898 | 0.098 | -0.003 | -0.001 | -0.001 | 0 | 0.07 | 0.08 | 0.02 | 0.16 | 0.01 | 0.11 |
| 9 | 106441872 | rs7860970 | -0.143 | -1.724 | -0.927 | 0.013 | -0.034 | 0.068 | 0.068 | 0.023 | 0.25 | 0.43 | 0.23 | 0.38 | 0.17 | 0.22 |

| | | | | | | | | | | | | | | | | |
|---|---|---|---|---|---|---|---|---|---|---|---|---|---|---|---|---|
| 9 | 106459537 | rs10820531 | 1.209 | -2.123 | -1.182 | -0.088 | 0.02 | -0.004 | -0.004 | 0 | 0.15 | 0.06 | 0.06 | 0.17 | 0.11 | 0.07 |
| 9 | 106459868 | rs368956 | 0.64 | -2.423 | -1.245 | -0.327 | 0.023 | 0.046 | 0.046 | 0 | 0.32 | 0.16 | 0.19 | 0.19 | 0.3 | 0.34 |
| 9 | 106460892 | rs412741 | -0.715 | -2.454 | -1.144 | -0.352 | 0.002 | 0.001 | 0.001 | 0 | 0.57 | 0.61 | 0.43 | 0.57 | 0.47 | 0.57 |
| 9 | 106464499 | rs970603 | 0.049 | -2.379 | -1.274 | -0.304 | -0.04 | 0.082 | 0.082 | 0 | 0.25 | 0.49 | 0.23 | 0.37 | 0.16 | 0.22 |
| 9 | 106470714 | rs449475 | 1.145 | -2.139 | -2.076 | -0.164 | 0.014 | 0.004 | 0.004 | 0 | 0.12 | 0.06 | 0.06 | 0 | 0.08 | 0.12 |
| 9 | 106477496 | rs403258 | -1.2 | -1.687 | -2.294 | 0.1 | 0.01 | -0.001 | -0.001 | 0.016 | 0.73 | 0.7 | 0.49 | 0.75 | 0.58 | 0.65 |
| 9 | 106481839 | rs10990914 | 0.876 | -1.308 | -2.858 | 0.395 | 0.041 | 0.004 | 0.004 | 0 | 0.08 | 0 | 0.02 | 0.05 | 0.01 | 0.07 |
| 9 | 106483432 | rs4743664 | 0.165 | -0.494 | -2.543 | 0.67 | -0.04 | -0.095 | -0.095 | 0.375 | 0.29 | 0.06 | 0.28 | 0.36 | 0.31 | 0.29 |
| 9 | 106487753 | rs4436177 | -0.468 | -0.34 | -1.305 | 0.6 | 0.015 | 0.004 | 0.004 | 0 | 0.15 | 0.09 | 0.07 | 0.17 | 0.11 | 0.08 |
| 9 | 106493088 | rs433341 | -0.487 | -0.348 | -1.394 | 0.89 | 0.021 | -0.02 | -0.02 | 0.155 | 0.25 | 0.13 | 0.16 | 0.28 | 0.18 | 0.21 |
| 9 | 106498091 | rs455178 | -0.575 | 0.101 | -2.137 | 0.229 | -0.059 | 0.075 | 0.075 | 0 | 0.67 | 0.84 | 0.59 | 0.8 | 0.55 | 0.7 |
| 9 | 106505693 | rs1659546 | 0.218 | -0.004 | -2.039 | 0.191 | 0.027 | -0.015 | -0.015 | 0 | 0.25 | 0.12 | 0.1 | 0.25 | 0.16 | 0.2 |
| 9 | 106512148 | rs428072 | -0.478 | -0.327 | -2.046 | 0.851 | -0.048 | -0.059 | -0.059 | 0.097 | 0.62 | 0.79 | 0.43 | 0.44 | 0.51 | 0.56 |
| 9 | 106530763 | rs10990941 | 0.802 | -0.51 | -1.856 | 0.433 | 0 | -0.028 | -0.028 | 0 | 0.28 | 0.56 | 0.26 | 0.16 | 0.27 | 0.28 |
| 9 | 106531629 | rs10739877 | -0.535 | -0.56 | -1.931 | 0.383 | -0.049 | -0.046 | -0.046 | 0.92 | 0.57 | 0.79 | 0.39 | 0.42 | 0.48 | 0.52 |
| 9 | 106535762 | rs10739878 | -1.807 | 0.225 | -2.423 | 0.46 | -0.035 | -0.051 | -0.051 | 0 | 0.88 | 0.97 | 0.69 | 0.61 | 0.85 | 0.87 |
| 9 | 106545276 | rs16923311 | 0.603 | 0.199 | -2.33 | 0.449 | 0.009 | -0.003 | -0.003 | 0.139 | 0.29 | 0.23 | 0.13 | 0.27 | 0.21 | 0.24 |
| 9 | 106546851 | rs16923314 | 1.075 | 0.183 | -1.636 | 0.066 | -0.002 | 0.021 | 0.021 | 0.107 | 0.11 | 0.19 | 0.16 | 0.15 | 0.12 | 0.08 |
| 9 | 106547099 | rs16923317 | 1.716 | -0.087 | -1.973 | 0.295 | 0.007 | -0.046 | -0.046 | 0 | 0.17 | 0.01 | 0.07 | 0.17 | 0.12 | 0.07 |
| 9 | 106554028 | rs7864801 | 0.663 | 0.078 | -1.077 | 0.35 | 0.008 | -0.035 | -0.035 | 0 | 0.27 | 0.41 | 0.25 | 0.15 | 0.28 | 0.28 |
| 9 | 106565689 | rs6479189 | -0.409 | 0.182 | -1.219 | 0.517 | -0.021 | -0.025 | -0.025 | 0 | 0.58 | 0.66 | 0.44 | 0.46 | 0.47 | 0.53 |
| 9 | 106571202 | rs4742892 | -0.1 | 0.13 | -1.249 | 0.418 | 0.009 | 0.026 | 0.026 | 0.18 | 0.53 | 0.66 | 0.4 | 0.37 | 0.43 | 0.48 |
| 9 | 106573399 | rs1439186 | -0.394 | 0.259 | -1.23 | 0.423 | -0.024 | -0.005 | -0.005 | 0 | 0.59 | 0.67 | 0.44 | 0.39 | 0.47 | 0.53 |
| 9 | 106581950 | rs1439190 | -0.307 | 0.264 | -0.983 | 0.203 | -0.005 | 0.003 | 0.003 | 0.045 | 0.28 | 0.3 | 0.27 | 0.18 | 0.38 | 0.33 |
| 9 | 106584972 | rs1371169 | -0.417 | -0.005 | -0.542 | 0.143 | -0.001 | -0.001 | -0.001 | 1.894 | 0.29 | 0.29 | 0.35 | 0.31 | 0.42 | 0.31 |
| 9 | 106585651 | rs1439189 | -1.625 | 0.137 | -0.581 | 0.012 | 0.006 | -0.017 | -0.017 | 2.827 | 0.85 | 0.89 | 0.66 | 0.51 | 0.89 | 0.82 |
| 9 | 106593860 | rs7037107 | 0.678 | 0.237 | -0.73 | -0.198 | 0.004 | 0.107 | 0.107 | 0 | 0.08 | 0.24 | 0.14 | 0.16 | 0.08 | 0.04 |
| 9 | 106600479 | rs17802452 | 1.749 | 0.106 | -1.162 | -0.546 | -0.08 | 0.095 | 0.095 | 0 | 0.16 | 0.01 | 0.1 | 0 | 0.21 | 0.2 |
| 9 | 106601859 | rs12555838 | 0.393 | 0.385 | -1.044 | -0.194 | -0.078 | -0.029 | -0.029 | 3.419 | 0.21 | 0.04 | 0.25 | 0.2 | 0.32 | 0.28 |
| 9 | 106604077 | rs10512319 | 0.339 | 0.418 | -0.824 | -0.388 | 0.025 | 0.08 | 0.08 | 0.03 | 0.44 | 0.31 | 0.24 | 0.14 | 0.38 | 0.33 |
| 9 | 106604233 | rs7046385 | -0.468 | 0.286 | -0.677 | -0.261 | 0.007 | 0.012 | 0.012 | 0.262 | 0.28 | 0.33 | 0.39 | 0.36 | 0.4 | 0.32 |
| 9 | 106610918 | rs13298450 | 0.297 | 0.297 | -0.716 | -0.252 | -0.078 | -0.032 | -0.032 | 0 | 0.21 | 0.04 | 0.25 | 0.21 | 0.32 | 0.28 |
| 9 | 106611898 | rs3930054 | 1.221 | 0.194 | -0.609 | -0.314 | 0.031 | 0.068 | 0.068 | 0.609 | 0.31 | 0.14 | 0.15 | 0.12 | 0.24 | 0.25 |
| 9 | 106631068 | rs7864255 | 1.111 | 0.309 | -1.144 | -0.23 | 0.003 | 0.083 | 0.083 | 0 | 0.06 | 0.15 | 0.09 | 0.15 | 0.07 | 0.04 |
| 9 | 106631091 | rs7036142 | 1.181 | 0.281 | -1.28 | -0.411 | 0.033 | 0.095 | 0.095 | 1.186 | 0.31 | 0.08 | 0.2 | 0.12 | 0.25 | 0.24 |
| 9 | 106637033 | rs10820559 | -0.675 | 0.443 | -1.068 | -0.277 | 0.007 | 0.03 | 0.03 | 0.04 | 0.66 | 0.47 | 0.58 | 0.5 | 0.68 | 0.61 |
| 9 | 106649139 | rs7037581 | -0.782 | 0.834 | -0.434 | 0.266 | 0.012 | 0.018 | 0.018 | 0.321 | 0.28 | 0.35 | 0.35 | 0.36 | 0.39 | 0.32 |
| 9 | 106659562 | rs920810 | -0.34 | 0.959 | -0.551 | -0.092 | -0.004 | 0.021 | 0.021 | 0 | 0.36 | 0.3 | 0.59 | 0.81 | 0.4 | 0.64 |
| 9 | 106659817 | rs920809 | -0.314 | 1.208 | -0.521 | -0.574 | -0.006 | -0.001 | -0.001 | 0 | 0.84 | 0.85 | 0.9 | 1 | 0.8 | 0.93 |
| 9 | 106659985 | rs9657690 | 1.325 | 0.77 | -0.065 | -0.826 | 0.003 | 0.023 | 0.023 | 0.323 | 0.45 | 0.37 | 0.3 | 0.17 | 0.42 | 0.3 |
| 9 | 106660496 | rs716533 | 1.325 | 0.724 | 0.065 | -0.862 | -0.004 | -0.036 | -0.036 | 0 | 0.45 | 0.48 | 0.31 | 0.17 | 0.42 | 0.28 |
| 9 | 106665541 | rs10990985 | 0.638 | 0.926 | 0.281 | -0.714 | 0.017 | 0.013 | 0.013 | 0 | 0.1 | 0.18 | 0.07 | 0.11 | 0.14 | 0.13 |
| 9 | 106667534 | rs1348451 | 1.513 | 0.889 | 0.2 | -0.74 | -0.004 | -0.035 | -0.035 | 0 | 0.45 | 0.48 | 0.31 | 0.17 | 0.42 | 0.28 |
| 9 | 106674242 | rs7025066 | -0.697 | 0.838 | 0.026 | -0.674 | -0.052 | 0.086 | 0.086 | 0 | 0.86 | 0.99 | 0.91 | 1 | 0.82 | 0.94 |
| 9 | 106674435 | rs7045335 | -0.114 | 0.83 | 0.24 | -0.594 | 0.003 | 0.021 | 0.021 | 0 | 0.61 | 0.73 | 0.44 | 0.3 | 0.62 | 0.46 |
| 9 | 106674850 | rs10990991 | 0.852 | 0.849 | 0.23 | -0.637 | -0.004 | -0.006 | -0.006 | 1.603 | 0.09 | 0.03 | 0.05 | 0.1 | 0.12 | 0.1 |
| 9 | 106675290 | rs718856 | 1.746 | 0.829 | 0.515 | -0.664 | -0.011 | 0.091 | 0.091 | 1.29 | 0.44 | 0.24 | 0.31 | 0.17 | 0.45 | 0.31 |
| 9 | 106681686 | rs10760997 | -0.021 | 0.803 | 0.484 | -0.618 | 0.006 | 0.021 | 0.021 | 0 | 0.59 | 0.67 | 0.44 | 0.29 | 0.62 | 0.45 |
| 9 | 106693503 | rs10217653 | -0.64 | 0.769 | 0.552 | 0.241 | 0.002 | -0.019 | -0.019 | 1.434 | 0.72 | 0.77 | 0.65 | 0.57 | 0.74 | 0.68 |
| 9 | 106694482 | rs10820566 | 0.126 | 0.748 | 0.584 | 0.208 | -0.002 | 0.004 | 0.004 | 1.932 | 0.64 | 0.36 | 0.57 | 0.55 | 0.68 | 0.63 |
| 9 | 106703809 | rs4743670 | 0.077 | 0.728 | 0.573 | 0.193 | -0.016 | 0.034 | 0.034 | 4.321 | 0.29 | 0.64 | 0.36 | 0.46 | 0.21 | 0.36 |
| 9 | 106707757 | rs2197295 | 0.531 | 0.727 | 0.602 | 0.222 | -0.027 | 0.09 | 0.09 | 1.12 | 0.24 | 0.57 | 0.31 | 0.44 | 0.17 | 0.34 |
| 9 | 106714767 | rs10820576 | 0.727 | 0.759 | 0.441 | 0.213 | -0.003 | -0.002 | -0.002 | 0 | 0.09 | 0.05 | 0.06 | 0.1 | 0.13 | 0.1 |
| 9 | 106715594 | rs7042666 | 0.723 | 0.791 | 0.603 | 0.307 | -0.006 | 0.016 | 0.016 | 0.684 | 0.22 | 0.26 | 0.28 | 0.44 | 0.16 | 0.31 |
| 9 | 106719165 | rs747029 | -0.247 | 0.872 | 0.687 | 0.317 | -0.001 | 0.002 | 0.002 | 0.096 | 0.7 | 0.68 | 0.65 | 0.56 | 0.73 | 0.68 |
| 9 | 106720681 | rs10820583 | -0.336 | 0.825 | 0.641 | 0.317 | 0.007 | -0.03 | -0.03 | 0.032 | 0.71 | 0.78 | 0.66 | 0.57 | 0.74 | 0.68 |
| 9 | 106721804 | rs10512321 | 0.96 | 1.14 | 0.448 | 0.125 | 0.058 | 0.045 | 0.045 | 0 | 0.11 | 0.36 | 0.14 | 0.13 | 0.12 | 0.09 |
| 9 | 106722204 | rs10820588 | 0.671 | 1.123 | 0.303 | 0.099 | -0.002 | 0.001 | 0.001 | 0.989 | 0.56 | 0.35 | 0.46 | 0.42 | 0.56 | 0.57 |
| 9 | 106723148 | rs2218384 | 0.171 | 1.053 | 0.214 | -0.178 | 0 | 0.004 | 0.004 | 0 | 0.32 | 0.36 | 0.4 | 0.62 | 0.29 | 0.45 |
| 9 | 106730859 | rs1348455 | -0.134 | 0.722 | 0.566 | -0.629 | 0 | -0.007 | -0.007 | 0 | 0.41 | 0.48 | 0.51 | 0.76 | 0.42 | 0.53 |
| 9 | 106736021 | rs16923598 | 1.667 | 0.711 | 0.507 | -0.696 | 0 | 0.002 | 0.002 | 0 | 0.08 | 0.08 | 0.1 | 0.13 | 0.1 | 0.09 |
| 9 | 106741410 | rs1443843 | -0.211 | 0.749 | 0.457 | -0.722 | -0.006 | -0.015 | -0.015 | 0 | 0.39 | 0.54 | 0.48 | 0.77 | 0.34 | 0.52 |
| 9 | 106741855 | rs1822539 | 0.6 | 0.724 | 0.402 | -0.691 | -0.025 | 0.011 | 0.011 | 0 | 0.31 | 0.52 | 0.42 | 0.7 | 0.21 | 0.46 |
| 9 | 106742792 | rs535966 | 0.672 | 0.771 | 0.28 | -0.754 | -0.029 | 0.182 | 0.182 | 0.636 | 0.47 | 0.18 | 0.39 | 0.1 | 0.52 | 0.36 |
| 9 | 106744641 | rs10991022 | 0.105 | 0.681 | 0.669 | -0.606 | 0.018 | -0.008 | -0.008 | 0 | 0.11 | 0.28 | 0.08 | 0.07 | 0.13 | 0.06 |
| 9 | 106745051 | rs10991023 | 0.214 | 0.682 | 0.683 | -0.605 | 0.011 | -0.01 | -0.01 | 0 | 0.1 | 0.22 | 0.08 | 0.07 | 0.13 | 0.06 |
| 9 | 106761243 | rs13301641 | 0.628 | 0.656 | 0.564 | -0.677 | -0.009 | -0.002 | -0.002 | 0 | 0.34 | 0.18 | 0.45 | 0.7 | 0.32 | 0.47 |
| 9 | 106765319 | rs7857804 | 0.589 | 0.684 | 0.495 | -0.672 | 0 | 0.003 | 0.003 | 0 | 0.36 | 0.32 | 0.46 | 0.7 | 0.32 | 0.47 |
| 9 | 106769158 | rs687586 | 0.726 | 0.761 | 0.054 | -0.658 | -0.037 | 0.136 | 0.136 | 0 | 0.43 | 0.18 | 0.43 | 0.13 | 0.53 | 0.4 |

| | | | | | | | | | | | | | | | | |
|---|---|---|---|---|---|---|---|---|---|---|---|---|---|---|---|---|
| 9 | 106772759 | rs1867347 | -0.327 | 0.688 | 0.129 | -0.646 | 0.002 | 0.031 | 0.031 | 0 | 0.47 | 0.61 | 0.5 | 0.78 | 0.45 | 0.55 |
| 9 | 106786496 | rs628345 | -0.643 | 0.725 | 0.099 | -0.528 | 0.005 | 0.006 | 0.006 | 0 | 0.86 | 0.76 | 0.87 | 0.85 | 0.86 | 0.89 |
| 9 | 106786864 | rs630103 | 0.267 | 0.699 | 0.324 | -0.668 | -0.001 | 0.012 | 0.012 | 0.034 | 0.51 | 0.55 | 0.46 | 0.16 | 0.55 | 0.42 |
| 9 | 106791364 | rs868619 | -1.074 | 0.706 | 0.531 | -0.718 | 0.003 | 0.026 | 0.026 | 0.028 | 0.58 | 0.69 | 0.61 | 0.91 | 0.57 | 0.64 |
| 9 | 106797388 | rs10991043 | 0.334 | 0.727 | 0.397 | -0.701 | -0.01 | 0.022 | 0.022 | 0.492 | 0.38 | 0.21 | 0.43 | 0.72 | 0.35 | 0.49 |
| 9 | 106798942 | rs12115654 | 1.283 | 0.717 | 0.409 | -0.626 | 0.007 | 0.021 | 0.021 | 0 | 0.1 | 0.23 | 0.11 | 0.13 | 0.1 | 0.09 |
| 9 | 106801187 | rs12345652 | 1.211 | 0.712 | 0.409 | -0.625 | -0.003 | 0.024 | 0.024 | 0 | 0.1 | 0.23 | 0.11 | 0.13 | 0.08 | 0.09 |
| 9 | 106813132 | rs10991055 | -0.5 | 0.785 | 0.472 | -0.538 | 0.019 | -0.035 | -0.035 | 0 | 0.16 | 0.33 | 0.11 | 0.08 | 0.19 | 0.08 |
| 9 | 106813465 | rs1411101 | 0.46 | 0.858 | -0.006 | -0.472 | -0.003 | 0.002 | 0.002 | 0 | 0.34 | 0.19 | 0.41 | 0.71 | 0.31 | 0.47 |
| 9 | 106822968 | rs4742899 | -1.278 | 1.043 | -0.316 | -0.568 | -0.032 | -0.029 | -0.029 | 0.27 | 0.92 | 0.99 | 0.82 | 0.91 | 0.91 | 0.89 |
| 9 | 106835052 | rs10991090 | -0.13 | 1.137 | -0.128 | -0.466 | 0.01 | -0.015 | -0.015 | 0 | 0.14 | 0.22 | 0.09 | 0.08 | 0.18 | 0.08 |
| 9 | 106850457 | rs10991099 | 0.561 | 1.057 | -0.089 | -0.411 | 0.023 | -0.001 | -0.001 | 0 | 0.1 | 0.22 | 0.07 | 0.08 | 0.15 | 0.06 |
| 9 | 106850787 | rs1411105 | -0.337 | 1.001 | -0.016 | -0.541 | 0.004 | -0.027 | -0.027 | 0 | 0.45 | 0.48 | 0.49 | 0.79 | 0.46 | 0.53 |
| 9 | 106854958 | rs1411103 | -0.764 | 1.101 | -0.017 | -0.505 | 0.005 | -0.002 | -0.002 | 0 | 0.89 | 0.81 | 0.84 | 0.94 | 0.87 | 0.89 |
| 9 | 106856043 | rs4742901 | 0.498 | 0.977 | 0.014 | -0.614 | 0 | 0 | 0 | 0 | 0.31 | 0.27 | 0.46 | 0.75 | 0.3 | 0.46 |
| 9 | 106856633 | rs3818626 | -0.019 | 0.765 | 0.223 | -0.742 | 0.004 | -0.022 | -0.022 | 3.242 | 0.57 | 0.5 | 0.38 | 0.19 | 0.57 | 0.42 |
| 9 | 106856910 | rs4743687 | -0.242 | 0.77 | 0.15 | -0.735 | -0.027 | 0.053 | 0.053 | 0 | 0.59 | 0.75 | 0.4 | 0.2 | 0.58 | 0.43 |
| 9 | 106857078 | rs4742906 | -0.224 | 0.771 | 0.127 | -0.734 | -0.017 | 0.035 | 0.035 | 0 | 0.59 | 0.72 | 0.39 | 0.2 | 0.58 | 0.43 |
| 9 | 106858851 | rs12236470 | 0.312 | 0.852 | 0.068 | -0.75 | -0.006 | 0.004 | 0.004 | 0 | 0.1 | 0.07 | 0.07 | 0.06 | 0.16 | 0.07 |
| 9 | 106859811 | rs7028408 | -0.129 | 0.81 | 0.236 | -0.886 | -0.008 | 0.023 | 0.023 | 0 | 0.57 | 0.58 | 0.39 | 0.19 | 0.57 | 0.43 |
| 9 | 106860969 | rs10991107 | 0.6 | 0.776 | 0.508 | -0.898 | 0.013 | 0.024 | 0.024 | 0 | 0.46 | 0.35 | 0.31 | 0.17 | 0.42 | 0.36 |
| 9 | 106865027 | rs10991109 | 0.659 | 0.409 | 0.267 | -1.529 | -0.001 | -0.001 | -0.001 | 0.712 | 0.2 | 0.18 | 0.16 | 0.17 | 0.17 | 0.25 |
| 9 | 106877939 | rs10820603 | -0.563 | 0.744 | 0.185 | -1.79 | -0.039 | 0.071 | 0.071 | 0 | 0.59 | 0.79 | 0.4 | 0.2 | 0.58 | 0.41 |
| 9 | 106896809 | rs7872034 | -0.57 | 0.726 | 0.177 | -1.79 | -0.041 | 0.071 | 0.071 | 0 | 0.59 | 0.79 | 0.41 | 0.2 | 0.57 | 0.43 |
| 9 | 106901039 | rs6479222 | 1.608 | 1.076 | -0.155 | -2.023 | -0.035 | 0.032 | 0.032 | 0 | 0.07 | 0 | 0.07 | 0 | 0.11 | 0.01 |
| 9 | 106901877 | rs15532 | 0.357 | 0.639 | 0.009 | -1.878 | -0.006 | 0.007 | 0.007 | 0 | 0.1 | 0.07 | 0.07 | 0.02 | 0.16 | 0.07 |
| 9 | 106921023 | rs6479223 | 0.162 | 0.554 | -0.314 | -2.384 | 0.013 | 0.017 | 0.017 | 0 | 0.48 | 0.52 | 0.32 | 0.16 | 0.42 | 0.34 |
| 9 | 106925008 | rs10991137 | 0.526 | 0.618 | -0.436 | -2.379 | -0.005 | 0.006 | 0.006 | 0 | 0.1 | 0.05 | 0.06 | 0.03 | 0.15 | 0.07 |
| 9 | 106926807 | rs12351427 | 0.577 | 0.615 | -0.409 | -2.381 | -0.002 | -0.002 | -0.002 | 0 | 0.1 | 0.11 | 0.07 | 0.03 | 0.15 | 0.07 |
| 9 | 106935585 | rs12351429 | 0.527 | 0.615 | -0.409 | -2.381 | -0.007 | 0.011 | 0.011 | 0 | 0.1 | 0.05 | 0.06 | 0.03 | 0.15 | 0.07 |
| 9 | 106944031 | rs966390 | -0.589 | 0.586 | -0.402 | -2.441 | -0.031 | 0.067 | 0.067 | 0 | 0.6 | 0.74 | 0.4 | 0.19 | 0.56 | 0.41 |
| 9 | 106956123 | rs1507506 | 0.106 | 0.554 | -0.379 | -2.393 | 0.034 | 0.078 | 0.078 | 0 | 0.49 | 0.62 | 0.33 | 0.16 | 0.42 | 0.34 |
| 9 | 106976151 | rs10991172 | -0.384 | 0.715 | -0.493 | -2.139 | -0.019 | 0.043 | 0.043 | 0 | 0.59 | 0.7 | 0.39 | 0.2 | 0.57 | 0.42 |
| 9 | 106989653 | rs1507515 | -0.339 | 0.694 | -0.403 | -2.148 | -0.008 | 0.014 | 0.014 | 0 | 0.59 | 0.65 | 0.39 | 0.2 | 0.57 | 0.41 |
| 9 | 106993898 | rs10124553 | -0.298 | 0.72 | -0.421 | -2.146 | -0.007 | 0.012 | 0.012 | 0 | 0.59 | 0.63 | 0.38 | 0.2 | 0.56 | 0.4 |
| 9 | 107009131 | rs2122582 | -0.047 | 1.067 | -0.651 | -1.989 | 0 | -0.007 | -0.007 | 0.724 | 0.63 | 0.42 | 0.42 | 0.29 | 0.63 | 0.39 |
| 9 | 107013530 | rs10120365 | 0.19 | 1.655 | -1.031 | -2.325 | 0.012 | -0.035 | -0.035 | 0 | 0.09 | 0.17 | 0.07 | 0.01 | 0.13 | 0.06 |
| 9 | 107013881 | rs4743702 | 0.52 | 1.51 | -1.26 | -2.682 | 0.003 | -0.063 | -0.063 | 0.154 | 0.59 | 0.43 | 0.56 | 0.94 | 0.53 | 0.66 |
| 9 | 107021689 | rs7034022 | -0.365 | 2.274 | -1.376 | -1.689 | 0.029 | -0.049 | -0.049 | 0 | 0.1 | 0.3 | 0.07 | 0.01 | 0.13 | 0.06 |
| 9 | 107026812 | rs1450679 | -0.464 | 2.316 | -1.474 | -1.695 | -0.04 | 0.016 | 0.016 | 0 | 0.88 | 0.96 | 0.76 | 0.96 | 0.85 | 0.82 |
| 9 | 107028394 | rs1597513 | -0.842 | 2.357 | -1.494 | -1.687 | -0.047 | -0.001 | -0.001 | 0.816 | 0.9 | 1 | 0.8 | 0.96 | 0.89 | 0.87 |
| 9 | 107029309 | rs1450675 | -0.014 | 2.263 | -1.409 | -1.634 | 0.003 | -0.017 | -0.017 | 0 | 0.09 | 0.09 | 0.06 | 0.01 | 0.13 | 0.06 |
| 9 | 107038190 | rs10991205 | 0.309 | 2.265 | -1.427 | -1.634 | -0.006 | 0.002 | 0.002 | 0 | 0.08 | 0.04 | 0.05 | 0.01 | 0.13 | 0.06 |
| 9 | 107041827 | rs4743703 | 0.884 | 2.328 | -1.654 | -1.786 | -0.019 | 0.081 | 0.081 | 0 | 0.13 | 0.01 | 0.09 | 0 | 0.14 | 0.14 |
| 9 | 107045058 | rs1507511 | 1.149 | 2.508 | -1.106 | -1.355 | 0.026 | -0.047 | -0.047 | 0.16 | 0.78 | 0.64 | 0.68 | 0.94 | 0.72 | 0.76 |
| 9 | 107047590 | rs1395832 | -0.725 | 1.984 | -0.782 | -0.937 | 0.018 | -0.039 | -0.039 | 1.274 | 0.1 | 0.18 | 0.07 | 0.01 | 0.13 | 0.06 |
| 9 | 107050093 | rs1354892 | -0.238 | 2.003 | -1.163 | -0.947 | 0.031 | -0.043 | -0.043 | 0 | 0.08 | 0.19 | 0.06 | 0.01 | 0.13 | 0.06 |
| 9 | 107054520 | rs10521067 | -0.537 | 2.293 | -1.671 | -0.738 | -0.063 | 0.008 | 0.008 | 0 | 0.17 | 0.03 | 0.11 | 0.1 | 0.22 | 0.06 |
| 9 | 107055840 | rs3012359 | 0.479 | 1.579 | -1.726 | -1.141 | -0.059 | 0.092 | 0.092 | 0 | 0.32 | 0.09 | 0.19 | 0.1 | 0.34 | 0.34 |
| 9 | 107056086 | rs2991282 | 0.183 | 0.315 | -1.2 | -0.78 | -0.008 | -0.04 | -0.04 | 0 | 0.46 | 0.44 | 0.33 | 0.11 | 0.58 | 0.38 |
| 9 | 107056847 | rs867977 | -0.059 | 0.73 | -0.851 | -0.427 | -0.009 | -0.026 | -0.026 | 0 | 0.53 | 0.45 | 0.35 | 0.17 | 0.62 | 0.41 |
| 9 | 107057177 | rs884258 | -0.073 | 0.518 | -1.139 | -0.217 | 0.015 | 0.018 | 0.018 | 0 | 0.56 | 0.38 | 0.44 | 0.17 | 0.69 | 0.44 |
| 9 | 107063817 | rs10820631 | -0.598 | 1.559 | -1.313 | 0.929 | -0.081 | -0.101 | -0.101 | 0 | 0.14 | 0.03 | 0.33 | 0.52 | 0.27 | 0.28 |
| 9 | 107073849 | rs7847263 | -0.284 | 1.173 | -0.615 | 0.472 | -0.005 | -0.001 | -0.001 | 0 | 0.21 | 0.38 | 0.22 | 0.16 | 0.18 | 0.24 |
| 9 | 107076147 | rs4743707 | -1.371 | 1.139 | -0.133 | 0.449 | 0 | 0 | 0 | 0 | 0.34 | 0.34 | 0.49 | 0.67 | 0.44 | 0.52 |
| 9 | 107083282 | rs13299767 | -0.466 | 0.944 | -0.054 | 0.244 | 0 | 0 | 0 | 0 | 0.19 | 0.22 | 0.22 | 0.16 | 0.18 | 0.24 |
| 9 | 107084182 | rs13291235 | -0.085 | 0.925 | -0.189 | 0.225 | -0.028 | -0.035 | -0.035 | 0 | 0.14 | 0.04 | 0.2 | 0.14 | 0.15 | 0.22 |
| 9 | 107097929 | rs1688666 | -1.161 | -0.14 | 0.052 | -0.812 | -0.009 | -0.1 | -0.1 | 0.309 | 0.63 | 0.53 | 0.69 | 0.97 | 0.66 | 0.88 |
| 9 | 107100866 | rs7857557 | -1.403 | -0.938 | 0.793 | -0.419 | -0.071 | 0.09 | 0.09 | 0.732 | 0.13 | 0.52 | 0.06 | 0.19 | 0.05 | 0.11 |
| 9 | 107100917 | rs6479233 | -0.071 | -1.182 | 0.394 | -0.593 | -0.009 | 0.045 | 0.045 | 0 | 0.27 | 0.08 | 0.27 | 0.16 | 0.24 | 0.34 |
| 9 | 107101626 | rs12238305 | -1.439 | -1.235 | 0.852 | -0.231 | -0.048 | 0.061 | 0.061 | 0 | 0.12 | 0.37 | 0.06 | 0.19 | 0.07 | 0.12 |
| 9 | 107104909 | rs10991225 | -1.152 | -1.256 | 0.889 | -0.192 | -0.052 | 0.074 | 0.074 | 0 | 0.11 | 0.37 | 0.05 | 0.18 | 0.05 | 0.11 |
| 9 | 107108324 | rs10991227 | 0.287 | -1.103 | 0.545 | -0.143 | -0.028 | 0.072 | 0.072 | 0 | 0.06 | 0.2 | 0.03 | 0.16 | 0.02 | 0.07 |
| 9 | 107113739 | rs2933323 | 0.817 | -1.716 | 0.577 | -0.141 | -0.07 | 0.001 | 0.001 | 0 | 0.73 | 0.45 | 0.88 | 0.74 | 0.85 | 0.78 |
| 9 | 107125805 | rs3012352 | 0.99 | -1.216 | 0.917 | -0.111 | -0.025 | -0.016 | -0.016 | 0 | 0.67 | 0.49 | 0.68 | 0.73 | 0.75 | 0.69 |
| 9 | 107128724 | rs3000671 | 0.235 | -1.332 | 1.138 | -0.163 | 0.004 | 0.001 | 0.001 | 0 | 0.73 | 0.78 | 0.71 | 0.75 | 0.76 | 0.72 |
| 9 | 107129400 | rs3012353 | 0.492 | -1.338 | 1.37 | -0.172 | -0.015 | -0.005 | -0.005 | 1.365 | 0.72 | 0.64 | 0.7 | 0.73 | 0.76 | 0.72 |
| 9 | 107130684 | rs7048700 | -0.769 | -1.362 | 1.225 | -0.206 | -0.009 | 0.016 | 0.016 | 0.441 | 0.93 | 1 | 0.95 | 0.94 | 0.93 | 0.97 |
| 9 | 107138679 | rs7029615 | 1.161 | -1.345 | 1.431 | 0.477 | -0.002 | 0.002 | 0.002 | 0 | 0.64 | 0.43 | 0.66 | 0.49 | 0.74 | 0.56 |

| | | | | | | | | | | | | | | | | |
|---|---|---|---|---|---|---|---|---|---|---|---|---|---|---|---|---|
| 9 | 107154109 | rs6479234 | 0.42 | -1.312 | 1.305 | 0.297 | 0.006 | 0.024 | 0.024 | 0 | 0.71 | 0.88 | 0.71 | 0.77 | 0.74 | 0.7 |
| 9 | 107154562 | rs1565603 | 0.519 | -1.317 | 1.321 | 0.295 | 0.008 | 0.018 | 0.018 | 0 | 0.7 | 0.82 | 0.71 | 0.77 | 0.74 | 0.7 |
| 9 | 107154760 | rs1565604 | 0.965 | -1.328 | 1.629 | 0.352 | -0.025 | 0.026 | 0.026 | 0 | 0.67 | 0.54 | 0.68 | 0.59 | 0.74 | 0.67 |
| 9 | 107180109 | rs10991262 | 1.75 | -1.05 | 0.951 | -0.293 | 0.001 | 0.002 | 0.002 | 0 | 0.37 | 0.35 | 0.31 | 0.07 | 0.36 | 0.21 |
| 9 | 107199088 | rs9299380 | 1.063 | -1.224 | 1.269 | 0.132 | -0.02 | 0.025 | 0.025 | 0 | 0.65 | 0.52 | 0.67 | 0.43 | 0.72 | 0.57 |
| 9 | 107227109 | rs902788 | 0.2 | -1.244 | 1.128 | 0.073 | 0.004 | -0.008 | -0.008 | 0 | 0.21 | 0.09 | 0.24 | 0.21 | 0.19 | 0.3 |
| 9 | 107227614 | rs1388752 | 0.161 | -1.25 | 1.282 | 0.065 | -0.027 | -0.052 | -0.052 | 0 | 0.86 | 0.75 | 0.92 | 0.94 | 0.91 | 0.94 |
| 9 | 107234158 | rs698459 | -1.184 | -1.187 | 1.006 | 0.102 | -0.02 | -0.017 | -0.017 | 0 | 0.35 | 0.4 | 0.33 | 0.28 | 0.28 | 0.36 |
| 9 | 107242297 | rs10820643 | -0.163 | -1.145 | 0.82 | 0.14 | 0.001 | -0.001 | -0.001 | 2.062 | 0.22 | 0.2 | 0.24 | 0.23 | 0.19 | 0.28 |
| 9 | 107245354 | rs10820644 | 0.036 | -1.193 | 0.792 | 0.117 | 0.004 | -0.01 | -0.01 | 0 | 0.21 | 0.12 | 0.24 | 0.21 | 0.19 | 0.29 |
| 9 | 107246760 | rs1916678 | -0.502 | -1.194 | 0.871 | 0.176 | -0.01 | 0.057 | 0.057 | 0 | 0.24 | 0.31 | 0.27 | 0.51 | 0.2 | 0.41 |
| 9 | 107257406 | rs10820650 | -1.281 | -1.076 | 0.414 | 0.14 | -0.017 | 0.026 | 0.026 | 0 | 0.35 | 0.42 | 0.35 | 0.55 | 0.3 | 0.47 |
| 9 | 107259865 | rs7849956 | -1.128 | -1.073 | 0.392 | 0.133 | -0.018 | 0.026 | 0.026 | 0 | 0.35 | 0.43 | 0.35 | 0.57 | 0.3 | 0.47 |
| 9 | 107263182 | rs2417531 | -0.641 | -1.345 | 0.339 | 0.067 | -0.052 | 0.058 | 0.058 | 0.034 | 0.21 | 0.35 | 0.17 | 0.35 | 0.12 | 0.17 |
| 9 | 107265452 | rs1464674 | -0.178 | -1.061 | 0.217 | -0.04 | -0.028 | -0.011 | -0.011 | 0 | 0.77 | 0.91 | 0.77 | 0.78 | 0.75 | 0.67 |
| 9 | 107265989 | rs10820661 | 1.433 | -1.056 | 0.218 | -0.037 | -0.006 | 0 | 0 | 0 | 0.56 | 0.56 | 0.59 | 0.43 | 0.62 | 0.49 |
| 9 | 107266844 | rs1403812 | -0.123 | -1.047 | 0.123 | -0.035 | -0.026 | -0.009 | -0.009 | 0.018 | 0.77 | 0.91 | 0.77 | 0.78 | 0.75 | 0.67 |
| 9 | 107266943 | rs1403811 | -0.109 | -1.048 | 0.123 | -0.036 | -0.026 | -0.009 | -0.009 | 0 | 0.77 | 0.91 | 0.77 | 0.78 | 0.75 | 0.67 |
| 9 | 107267304 | rs7030820 | 0.953 | -0.799 | 0.167 | -0.126 | -0.006 | 0 | 0 | 0.355 | 0.55 | 0.54 | 0.58 | 0.43 | 0.62 | 0.49 |
| 9 | 107273553 | rs7025484 | -0.367 | -0.12 | 0.298 | 0.145 | 0.042 | 0.04 | 0.04 | 0 | 0.28 | 0.44 | 0.35 | 0.41 | 0.36 | 0.42 |
| 9 | 107279558 | rs2900371 | 0.591 | -0.134 | 0.034 | 0.157 | -0.1 | -0.103 | -0.103 | 0.007 | 0.17 | 0.03 | 0.28 | 0.33 | 0.31 | 0.33 |
| 9 | 107280314 | rs7869860 | 0.485 | -1.029 | -0.696 | -0.108 | -0.041 | 0.018 | 0.018 | 0 | 0.2 | 0.34 | 0.13 | 0.24 | 0.11 | 0.23 |
| 9 | 107281379 | rs6479245 | -1.896 | -1.219 | -1.188 | -0.235 | -0.084 | 0.158 | 0.158 | 0.013 | 0.56 | 0.92 | 0.52 | 0.81 | 0.51 | 0.7 |
| 9 | 107292001 | rs10761033 | -0.944 | -1.046 | -0.518 | -0.022 | -0.008 | -0.021 | -0.021 | 0 | 0.38 | 0.47 | 0.33 | 0.73 | 0.33 | 0.56 |
| 9 | 107293126 | rs1523668 | -0.944 | -1.046 | -0.518 | -0.022 | -0.008 | -0.021 | -0.021 | 0 | 0.38 | 0.47 | 0.33 | 0.73 | 0.33 | 0.56 |
| 9 | 107293318 | rs1523667 | -0.637 | -1.112 | -0.506 | 0.215 | -0.006 | 0.003 | 0.003 | 0 | 0.34 | 0.43 | 0.26 | 0.37 | 0.3 | 0.43 |
| 9 | 107293507 | rs12349993 | -0.729 | -1.089 | -0.514 | 0.264 | -0.005 | 0.004 | 0.004 | 0 | 0.37 | 0.44 | 0.31 | 0.57 | 0.31 | 0.51 |
| 9 | 107293713 | rs7872276 | 0.157 | -1.235 | -0.23 | -0.149 | -0.012 | -0.028 | -0.028 | 0 | 0.12 | 0.14 | 0.07 | 0.07 | 0.08 | 0.11 |
| 9 | 107294021 | rs12003257 | 0.523 | -1.29 | -0.259 | -0.134 | -0.048 | 0.029 | 0.029 | 0 | 0.2 | 0.39 | 0.13 | 0.24 | 0.12 | 0.22 |
| 9 | 107294722 | rs10761034 | 0.756 | -1.236 | -0.037 | -0.09 | 0.017 | 0.097 | 0.097 | 0 | 0.14 | 0.24 | 0.18 | 0.49 | 0.2 | 0.34 |
| 9 | 107297337 | rs6479247 | -2.238 | -1.549 | -0.812 | -0.629 | -0.076 | 0.11 | 0.11 | 0 | 0.61 | 0.91 | 0.45 | 0.81 | 0.49 | 0.72 |
| 9 | 107300612 | rs6479250 | 0.014 | -1.29 | -0.375 | -0.382 | -0.023 | -0.042 | -0.042 | 0 | 0.14 | 0.21 | 0.07 | 0.08 | 0.08 | 0.12 |
| 9 | 107304806 | rs10991297 | 0.827 | -0.881 | -0.679 | -0.507 | 0.006 | -0.001 | -0.001 | 0 | 0.18 | 0.29 | 0.16 | 0.17 | 0.17 | 0.16 |
| 9 | 107312272 | rs6479251 | 1.255 | -1.387 | -0.56 | -0.665 | -0.058 | -0.021 | -0.021 | 0 | 0.18 | 0.38 | 0.1 | 0.12 | 0.07 | 0.18 |
| 9 | 107312475 | rs4742914 | -0.616 | -1.315 | -0.821 | -0.746 | -0.001 | -0.007 | -0.007 | 0 | 0.39 | 0.47 | 0.32 | 0.63 | 0.38 | 0.61 |
| 9 | 107313406 | rs10991298 | 0.821 | -1.176 | -1.046 | -0.613 | -0.042 | -0.057 | -0.057 | 0.032 | 0.23 | 0.07 | 0.37 | 0.35 | 0.25 | 0.23 |
| 9 | 107324200 | rs13287980 | 0.509 | -1.35 | -0.604 | -0.38 | -0.067 | 0.054 | 0.054 | 0.016 | 0.23 | 0.53 | 0.13 | 0.3 | 0.12 | 0.25 |
| 9 | 107324247 | rs13289787 | 0.392 | -1.279 | -0.583 | -0.348 | -0.05 | 0.019 | 0.019 | 0 | 0.28 | 0.53 | 0.17 | 0.3 | 0.19 | 0.29 |
| 9 | 107327794 | rs1403822 | -0.948 | -1.06 | -0.387 | -0.077 | 0.013 | -0.051 | -0.051 | 0.342 | 0.39 | 0.46 | 0.56 | 0.87 | 0.45 | 0.59 |
| 9 | 107341255 | rs10125033 | 0.293 | -0.578 | -0.263 | 0.584 | -0.005 | -0.006 | -0.006 | 0 | 0.19 | 0.2 | 0.37 | 0.38 | 0.25 | 0.19 |
| 9 | 107349640 | rs10761048 | 1.554 | -0.565 | -0.437 | 0.613 | -0.057 | -0.068 | -0.068 | 0 | 0.11 | 0.02 | 0.15 | 0.31 | 0.17 | 0.3 |
| 9 | 107370498 | rs10115731 | 0.537 | -0.571 | -0.215 | 0.569 | -0.019 | -0.032 | -0.032 | 0.294 | 0.17 | 0.14 | 0.37 | 0.38 | 0.24 | 0.19 |
| 9 | 107390277 | rs929963 | 0.529 | -0.556 | -0.21 | 0.575 | -0.019 | -0.033 | -0.033 | 0 | 0.18 | 0.13 | 0.37 | 0.38 | 0.24 | 0.19 |
| 9 | 107401344 | rs10739902 | -1.221 | -0.922 | -0.135 | 0.345 | -0.014 | 0.108 | 0.108 | 0 | 0.48 | 0.78 | 0.63 | 0.76 | 0.49 | 0.68 |
| 9 | 107415828 | rs2049347 | 0.17 | -0.914 | 0.125 | 0.351 | -0.004 | 0.007 | 0.007 | 0 | 0.28 | 0.37 | 0.25 | 0.35 | 0.21 | 0.47 |
| 9 | 107419241 | rs7872725 | -0.277 | -1.063 | 0.286 | 0.396 | 0.009 | -0.009 | -0.009 | 0 | 0.32 | 0.72 | 0.27 | 0.37 | 0.23 | 0.49 |
| 9 | 107422668 | rs13298262 | 0.868 | -0.994 | -0.16 | 0.321 | -0.046 | -0.07 | -0.07 | 0 | 0.16 | 0.05 | 0.36 | 0.39 | 0.25 | 0.19 |
| 9 | 107424198 | rs10820704 | 1.72 | -0.922 | -0.112 | 0.349 | -0.056 | -0.059 | -0.059 | 0.315 | 0.11 | 0.03 | 0.16 | 0.5 | 0.16 | 0.32 |
| 9 | 107429678 | rs10820705 | -1.881 | -1.291 | -0.261 | -0.474 | -0.139 | 0.204 | 0.204 | 0 | 0.71 | 0.98 | 0.75 | 1 | 0.63 | 0.84 |
| 9 | 107430373 | rs7848078 | 0.09 | -0.968 | 0.997 | 0.528 | -0.086 | -0.061 | -0.061 | 0 | 0.2 | 0.59 | 0.11 | 0.07 | 0.07 | 0.21 |
| 9 | 107436408 | rs7043099 | 0.09 | -0.968 | 0.997 | 0.528 | -0.086 | -0.061 | -0.061 | 0 | 0.2 | 0.59 | 0.11 | 0.07 | 0.07 | 0.21 |
| 9 | 107439675 | rs7025373 | 0.053 | -0.87 | 0.918 | 0.58 | -0.08 | -0.061 | -0.061 | 0.051 | 0.22 | 0.58 | 0.11 | 0.06 | 0.07 | 0.27 |
| 9 | 107440880 | rs1582708 | 0.064 | -0.872 | 0.917 | 0.579 | -0.08 | -0.061 | -0.061 | 0.599 | 0.21 | 0.57 | 0.11 | 0.06 | 0.07 | 0.27 |
| 9 | 107450409 | rs10991353 | 0.442 | -0.5 | 0.503 | 0.824 | -0.042 | -0.082 | -0.082 | 0.494 | 0.15 | 0.04 | 0.36 | 0.49 | 0.24 | 0.15 |
| 9 | 107457759 | rs1364946 | -0.464 | -0.552 | 0.818 | 0.616 | -0.002 | 0.006 | 0.006 | 0 | 0.31 | 0.29 | 0.43 | 0.45 | 0.31 | 0.29 |
| 9 | 107463151 | rs10761074 | 0.657 | -0.531 | 0.763 | 0.634 | -0.01 | 0.022 | 0.022 | 0 | 0.63 | 0.48 | 0.55 | 0.51 | 0.64 | 0.64 |
| 9 | 107475166 | rs10761076 | 0.896 | -0.365 | 0.048 | 0.417 | -0.027 | -0.049 | -0.049 | 4.195 | 0.14 | 0.08 | 0.16 | 0.52 | 0.17 | 0.33 |
| 9 | 107476188 | rs1425567 | -0.843 | -0.347 | -0.063 | 0.407 | -0.089 | -0.138 | -0.138 | 0 | 0.83 | 0.99 | 0.63 | 0.55 | 0.71 | 0.79 |
| 9 | 107477485 | rs10820714 | 1.186 | -0.171 | 0.129 | 0.308 | -0.047 | -0.063 | -0.063 | 0.114 | 0.12 | 0.04 | 0.14 | 0.3 | 0.17 | 0.3 |
| 9 | 107485577 | rs7027419 | 1.53 | -0.287 | 0.246 | -0.27 | -0.041 | 0.195 | 0.195 | 0 | 0.41 | 0.14 | 0.32 | 0.01 | 0.45 | 0.21 |
| 9 | 107505000 | rs4743756 | -0.272 | -0.307 | 0.389 | 0.434 | -0.009 | -0.012 | -0.012 | 0 | 0.72 | 0.69 | 0.8 | 0.81 | 0.72 | 0.7 |
| 9 | 107505262 | rs7034016 | -1.239 | -0.334 | 0.224 | 0.401 | -0.008 | -0.013 | -0.013 | 0 | 0.87 | 0.89 | 0.87 | 0.81 | 0.79 | 0.85 |
| 9 | 107506029 | rs10761081 | -1.116 | -0.336 | 0.267 | 0.4 | 0.003 | 0.009 | 0.009 | 0 | 0.87 | 0.85 | 0.86 | 0.81 | 0.79 | 0.85 |
| 9 | 107506122 | rs10739905 | 1.369 | -0.267 | -0.072 | 0.528 | -0.057 | -0.08 | -0.08 | 0 | 0.14 | 0.02 | 0.14 | 0.29 | 0.18 | 0.3 |
| 9 | 107506490 | rs1331926 | 0.658 | -0.32 | 0.039 | 0.585 | 0.014 | -0.065 | -0.065 | 0.355 | 0.62 | 0.17 | 0.5 | 0.55 | 0.64 | 0.6 |
| 9 | 107509404 | rs10820720 | 1.051 | -0.279 | -0.108 | 0.642 | -0.036 | -0.096 | -0.096 | 0.163 | 0.16 | 0.06 | 0.17 | 0.51 | 0.18 | 0.33 |
| 9 | 107509671 | rs2417563 | 0.328 | -0.298 | 0.154 | 0.658 | 0 | 0 | 0 | 0 | 0.64 | 0.41 | 0.55 | 0.56 | 0.65 | 0.67 |
| 9 | 107512888 | rs2417564 | 0.907 | 0.01 | -0.23 | 0.825 | -0.036 | -0.096 | -0.096 | 0 | 0.16 | 0.06 | 0.17 | 0.51 | 0.18 | 0.33 |
| 9 | 107516115 | rs4742921 | 0.534 | -0.087 | -0.245 | 0.747 | 0.017 | -0.012 | -0.012 | 0 | 0.06 | 0.04 | 0.04 | 0.22 | 0.02 | 0.05 |

| | | | | | | | | | | | | | | | | |
|---|---|---|---|---|---|---|---|---|---|---|---|---|---|---|---|---|
| 9 | 107516169 | rs10761082 | -0.248 | 0.048 | -0.214 | 0.766 | -0.03 | -0.059 | -0.059 | 0 | 0.73 | 0.83 | 0.72 | 0.52 | 0.62 | 0.55 |
| 9 | 107527830 | rs12552611 | 1.294 | 0.053 | -0.267 | 0.837 | 0.039 | 0.142 | 0.142 | 0 | 0.07 | 0.23 | 0.25 | 0.24 | 0.08 | 0.03 |
| 9 | 107529689 | rs10991376 | 1.176 | 0.244 | -1.153 | 0.359 | -0.059 | -0.081 | -0.081 | 0 | 0.14 | 0.03 | 0.15 | 0.29 | 0.18 | 0.3 |
| 9 | 107531956 | rs2472476 | -0.106 | 0.212 | -1.068 | 0.278 | -0.003 | -0.004 | -0.004 | 0 | 0.72 | 0.74 | 0.74 | 0.52 | 0.61 | 0.54 |
| 9 | 107536895 | rs2515606 | 1.487 | 0.045 | -2.12 | 0.037 | -0.089 | 0.231 | 0.231 | 0 | 0.41 | 0.08 | 0.34 | 0.08 | 0.45 | 0.23 |
| 9 | 107539352 | rs2482430 | -0.739 | 0.955 | -1.584 | 0.805 | -0.012 | -0.015 | -0.015 | 1.876 | 0.87 | 0.92 | 0.9 | 0.85 | 0.8 | 0.84 |
| 9 | 107544285 | rs4149341 | 0.784 | 1.024 | -1.652 | 0.804 | -0.059 | -0.083 | -0.083 | 0 | 0.14 | 0.03 | 0.14 | 0.29 | 0.18 | 0.29 |
| 9 | 107544700 | rs363717 | -0.703 | 0.906 | -1.527 | -0.154 | -0.016 | -0.029 | -0.029 | 0 | 0.87 | 0.94 | 0.89 | 0.8 | 0.79 | 0.84 |
| 9 | 107545156 | rs4149339 | -0.332 | 0.889 | -0.877 | 0.293 | 0.018 | 0.023 | 0.023 | 0.028 | 0.24 | 0.41 | 0.42 | 0.75 | 0.25 | 0.38 |
| 9 | 107545903 | rs4149338 | -0.338 | 0.989 | -0.816 | 0.352 | 0.005 | 0.006 | 0.006 | 0.009 | 0.28 | 0.41 | 0.43 | 0.75 | 0.27 | 0.38 |
| 9 | 107546901 | rs2482433 | 0.228 | 1.039 | -1.558 | -0.141 | -0.103 | 0.104 | 0.104 | 0 | 0.22 | 0.02 | 0.25 | 0.04 | 0.29 | 0.14 |
| 9 | 107547622 | rs1331924 | -0.763 | 0.763 | -0.12 | 0.724 | 0.103 | 0.143 | 0.143 | 0 | 0.12 | 0.4 | 0.31 | 0.26 | 0.16 | 0.06 |
| 9 | 107548375 | rs2777797 | 0.184 | -0.363 | -0.947 | 0.03 | 0.036 | 0.045 | 0.045 | 0 | 0.83 | 0.95 | 0.92 | 1 | 0.89 | 0.88 |
| 9 | 107548493 | rs2740485 | 0.156 | -0.358 | -0.945 | 0.032 | 0.035 | 0.044 | 0.044 | 0 | 0.83 | 0.95 | 0.92 | 1 | 0.89 | 0.88 |
| 9 | 107548566 | rs2066881 | -0.206 | -0.385 | -1.228 | 1.077 | -0.007 | -0.006 | -0.006 | 0 | 0.07 | 0.01 | 0.03 | 0.41 | 0.05 | 0.24 |
| 9 | 107548818 | rs4149337 | -0.206 | -0.385 | -1.228 | 1.077 | -0.007 | -0.006 | -0.006 | 0 | 0.07 | 0.01 | 0.03 | 0.41 | 0.05 | 0.24 |
| 9 | 107550639 | rs4149336 | -1.308 | -0.433 | -1.016 | 0.85 | -0.005 | -0.035 | -0.035 | 0 | 0.21 | 0.13 | 0.32 | 0.67 | 0.2 | 0.27 |
| 9 | 107551247 | rs2297405 | 0.106 | -0.501 | -0.973 | 1.296 | -0.004 | -0.027 | -0.027 | 0 | 0.07 | 0.01 | 0.03 | 0.28 | 0.05 | 0.1 |
| 9 | 107551538 | rs2297406 | 0.766 | -0.725 | -2.063 | 0.675 | -0.104 | -0.046 | -0.046 | 0.218 | 0.27 | 0.02 | 0.2 | 0.25 | 0.3 | 0.4 |
| 9 | 107552924 | rs4149333 | -1.33 | -1.901 | -0.326 | 0.998 | -0.057 | 0.084 | 0.084 | 2.076 | 0.23 | 0.54 | 0.15 | 0.43 | 0.1 | 0.27 |
| 9 | 107553185 | rs2020927 | -1.478 | -2.093 | -0.321 | 0.91 | -0.051 | 0.044 | 0.044 | 3.233 | 0.26 | 0.57 | 0.16 | 0.43 | 0.1 | 0.27 |
| 9 | 107553524 | rs10121901 | -1.478 | -2.093 | -0.321 | 0.91 | -0.051 | 0.044 | 0.044 | 3.233 | 0.26 | 0.57 | 0.15 | 0.43 | 0.1 | 0.27 |
| 9 | 107554069 | rs2066720 | -1.465 | -2.085 | -0.317 | 0.914 | -0.051 | 0.045 | 0.045 | 3.048 | 0.26 | 0.57 | 0.15 | 0.43 | 0.1 | 0.26 |
| 9 | 107554744 | rs4149330 | 0.104 | -2.104 | -0.011 | 0.986 | 0.004 | -0.009 | -0.009 | 2.836 | 0.11 | 0.08 | 0.04 | 0.25 | 0.06 | 0.1 |
| 9 | 107555306 | rs2297408 | 0.073 | -2.089 | 0.487 | 0.92 | -0.025 | 0.098 | 0.098 | 0.186 | 0.08 | 0.21 | 0.07 | 0.18 | 0.02 | 0.12 |
| 9 | 107556349 | rs4149326 | -0.943 | -2.129 | 0.807 | 0.9 | -0.064 | 0.105 | 0.105 | 0.766 | 0.13 | 0.41 | 0.11 | 0.18 | 0.03 | 0.12 |
| 9 | 107558983 | rs4149325 | 0.36 | -2.089 | 0.375 | 0.888 | -0.012 | -0.062 | -0.062 | 0 | 0.1 | 0 | 0.03 | 0.24 | 0.06 | 0.09 |
| 9 | 107560291 | rs4149323 | 0.287 | -2.093 | 0.418 | 0.928 | 0.003 | -0.042 | -0.042 | 0.086 | 0.1 | 0.04 | 0.03 | 0.23 | 0.06 | 0.09 |
| 9 | 107562804 | rs2230808 | 1.938 | -2.391 | 0.333 | 0.708 | 0.137 | -0.034 | -0.034 | 0 | 0.59 | 0.13 | 0.75 | 0.59 | 0.79 | 0.63 |
| 9 | 107563269 | rs2777800 | 1.066 | -2.442 | 0.856 | 0.682 | -0.05 | -0.027 | -0.027 | 0 | 0.71 | 0.55 | 0.86 | 0.77 | 0.83 | 0.76 |
| 9 | 107563437 | rs2740479 | 2.202 | -2.407 | 0.279 | 0.719 | 0.06 | 0.092 | 0.092 | 0 | 0.49 | 0.07 | 0.5 | 0.3 | 0.68 | 0.57 |
| 9 | 107564676 | rs1999431 | -0.293 | -2.425 | 0.389 | 0.732 | -0.036 | 0.024 | 0.024 | 0.891 | 0.1 | 0.18 | 0.09 | 0.18 | 0.03 | 0.12 |
| 9 | 107564839 | rs914546 | 0.25 | -2.403 | 0.349 | 0.743 | -0.026 | 0.037 | 0.037 | 0.013 | 0.08 | 0.16 | 0.08 | 0.18 | 0.02 | 0.12 |
| 9 | 107564846 | rs1883023 | 1.958 | -2.485 | -0.022 | 0.713 | 0.154 | -0.024 | -0.024 | 0 | 0.57 | 0.13 | 0.75 | 0.58 | 0.79 | 0.61 |
| 9 | 107564939 | rs4149321 | 0.445 | -2.384 | 0.04 | 0.771 | 0.006 | -0.029 | -0.029 | 0 | 0.1 | 0.05 | 0.04 | 0.23 | 0.06 | 0.1 |
| 9 | 107568705 | rs2066716 | 0.996 | -2.395 | -0.086 | 0.736 | -0.022 | -0.023 | -0.023 | 0 | 0.09 | 0.03 | 0.23 | 0.29 | 0.11 | 0.06 |
| 9 | 107569337 | rs2777802 | 1.618 | -2.432 | 0.073 | 0.718 | 0.047 | 0.001 | 0.001 | 0 | 0.6 | 0.31 | 0.76 | 0.59 | 0.79 | 0.62 |
| 9 | 107570714 | rs2740476 | -1.711 | -2.362 | -0.541 | 0.77 | -0.047 | 0.004 | 0.004 | 0 | 0.37 | 0.51 | 0.21 | 0.4 | 0.18 | 0.32 |
| 9 | 107571241 | rs2777804 | 1.735 | -2.121 | -1.514 | 0.867 | 0.037 | 0.06 | 0.06 | 0 | 0.48 | 0.04 | 0.51 | 0.3 | 0.68 | 0.56 |
| 9 | 107573757 | rs4149319 | -0.189 | -1.527 | -0.484 | 1.137 | -0.025 | 0.025 | 0.025 | 0 | 0.07 | 0.11 | 0.07 | 0.16 | 0.02 | 0.12 |
| 9 | 107576246 | rs2297409 | -1.536 | -1.358 | -0.607 | 1.18 | -0.001 | -0.001 | -0.001 | 0 | 0.29 | 0.3 | 0.19 | 0.4 | 0.16 | 0.33 |
| 9 | 107578103 | rs3780541 | -0.706 | -0.458 | -1.118 | 1.325 | -0.008 | -0.063 | -0.063 | 0.201 | 0.1 | 0 | 0.03 | 0.24 | 0.07 | 0.1 |
| 9 | 107580657 | rs3818688 | -1.09 | -0.428 | -1.323 | 1.547 | 0.013 | -0.068 | -0.068 | 0 | 0.1 | 0 | 0.04 | 0.31 | 0.06 | 0.12 |
| 9 | 107580842 | rs2020926 | -1.07 | -0.439 | -1.321 | 1.549 | 0.013 | -0.067 | -0.067 | 0 | 0.1 | 0 | 0.04 | 0.31 | 0.06 | 0.12 |
| 9 | 107581307 | rs4149316 | -1.498 | -0.402 | -1.349 | 1.782 | 0.008 | -0.057 | -0.057 | 1.5 | 0.09 | 0.01 | 0.08 | 0.33 | 0.07 | 0.14 |
| 9 | 107581749 | rs2254884 | -1.04 | 0.435 | -1.88 | 1.8 | -0.073 | -0.106 | -0.106 | 0.004 | 0.24 | 0.08 | 0.25 | 0.57 | 0.29 | 0.28 |
| 9 | 107582780 | rs3780542 | -0.258 | -0.65 | -1.583 | 1.905 | 0.007 | -0.003 | -0.003 | 0 | 0.11 | 0.1 | 0.14 | 0.24 | 0.06 | 0.09 |
| 9 | 107584309 | rs4149315 | 0.098 | -0.656 | -1.622 | 1.989 | 0.018 | -0.047 | -0.047 | 0 | 0.09 | 0.01 | 0.12 | 0.27 | 0.05 | 0.09 |
| 9 | 107584351 | rs4149314 | -0.009 | -0.872 | -1.927 | 1.628 | 0.003 | 0.021 | 0.021 | 0 | 0.11 | 0.05 | 0.05 | 0.06 | 0.1 | 0.09 |
| 9 | 107588328 | rs3824477 | 0.488 | -0.79 | -1.749 | 1.692 | -0.016 | -0.015 | -0.015 | 0 | 0.09 | 0.12 | 0.14 | 0.03 | 0.02 | 0.06 |
| 9 | 107589630 | rs4149308 | -0.772 | -1.832 | -0.758 | 1.213 | -0.039 | 0.048 | 0.048 | 0.044 | 0.21 | 0.46 | 0.21 | 0.68 | 0.12 | 0.14 |
| 9 | 107589744 | rs4149307 | -1.945 | -1.966 | -0.562 | 1.144 | 0.037 | 0.019 | 0.019 | 0 | 0.33 | 0.75 | 0.36 | 0.72 | 0.16 | 0.2 |
| 9 | 107594364 | rs2515629 | -0.436 | -2.21 | -0.719 | 0.638 | 0.001 | -0.005 | -0.005 | 0 | 0.19 | 0.18 | 0.1 | 0.07 | 0.17 | 0.16 |
| 9 | 107594515 | rs4149303 | -0.37 | -2.094 | 0.178 | 0.808 | -0.023 | 0.023 | 0.023 | 0 | 0.2 | 0.29 | 0.19 | 0.69 | 0.12 | 0.14 |
| 9 | 107598740 | rs2065412 | -2.321 | -2.287 | -0.843 | 0.003 | -0.171 | 0.103 | 0.103 | 0 | 0.76 | 0.98 | 0.61 | 0.95 | 0.57 | 0.84 |
| 9 | 107598884 | rs2297398 | 0.972 | -2.909 | 0.005 | 0.889 | 0.021 | -0.035 | -0.035 | 0 | 0.1 | 0.02 | 0.13 | 0.27 | 0.05 | 0.09 |
| 9 | 107599481 | rs2297400 | -0.673 | -2.952 | 0.404 | 0.482 | -0.026 | 0.011 | 0.011 | 0 | 0.22 | 0.36 | 0.21 | 0.69 | 0.13 | 0.18 |
| 9 | 107600193 | rs2515601 | -0.479 | -3.188 | 0.439 | 0.319 | -0.027 | -0.05 | -0.05 | 0.28 | 0.17 | 0.26 | 0.18 | 0.04 | 0.09 | 0.15 |
| 9 | 107600251 | rs4149299 | 0.757 | -3.217 | 0.576 | 0.277 | -0.001 | -0.006 | -0.006 | 0 | 0.11 | 0.1 | 0.05 | 0.07 | 0.1 | 0.08 |
| 9 | 107601512 | rs6479282 | -1.436 | -3.23 | 0.608 | 0.278 | -0.028 | -0.035 | -0.035 | 0 | 0.33 | 0.46 | 0.25 | 0.75 | 0.23 | 0.26 |
| 9 | 107602666 | rs2246841 | -0.476 | -3.196 | 0.411 | 0.264 | -0.029 | -0.053 | -0.053 | 0 | 0.17 | 0.27 | 0.19 | 0.04 | 0.09 | 0.15 |
| 9 | 107602678 | rs2274873 | 0.749 | -3.183 | 0.03 | 0.272 | -0.001 | -0.006 | -0.006 | 0 | 0.11 | 0.1 | 0.05 | 0.07 | 0.1 | 0.08 |
| 9 | 107603278 | rs2472385 | -0.242 | -3.176 | 0.073 | 0.275 | 0.014 | 0.015 | 0.015 | 0 | 0.16 | 0.13 | 0.18 | 0.04 | 0.09 | 0.15 |
| 9 | 107604104 | rs4149297 | 0.689 | -3.168 | 0.026 | 0.271 | -0.001 | -0.004 | -0.004 | 0 | 0.11 | 0.1 | 0.05 | 0.07 | 0.1 | 0.09 |
| 9 | 107604197 | rs2472449 | -2.453 | -3.197 | -0.068 | 0.231 | 0.102 | 0.203 | 0.203 | 0 | 0.51 | 0.83 | 0.45 | 0.79 | 0.33 | 0.41 |
| 9 | 107604507 | rs2472448 | -0.545 | -3.55 | 0.184 | 0.108 | 0.011 | 0.009 | 0.009 | 0 | 0.16 | 0.13 | 0.18 | 0.04 | 0.09 | 0.15 |
| 9 | 107607061 | rs4149295 | -1.309 | -3.447 | 0.549 | 0.141 | -0.01 | -0.011 | -0.011 | 0.191 | 0.22 | 0.72 | 0.12 | 0.07 | 0.12 | 0.12 |
| 9 | 107609065 | rs4149292 | 0.296 | -3.358 | 0.133 | 0.186 | 0.009 | 0.007 | 0.007 | 0 | 0.12 | 0.06 | 0.05 | 0.08 | 0.1 | 0.1 |
| 9 | 107611827 | rs4149290 | 0.294 | -3.355 | 0.147 | 0.188 | 0.009 | 0.006 | 0.006 | 0 | 0.12 | 0.06 | 0.05 | 0.08 | 0.1 | 0.1 |

| | | | | | | | | | | | | | | | | |
|---|---|---|---|---|---|---|---|---|---|---|---|---|---|---|---|---|
| 9 | 107617433 | rs2487039 | -0.883 | -3.574 | -0.076 | 0.238 | 0.055 | -0.111 | -0.111 | 0.822 | 0.24 | 0.07 | 0.24 | 0.34 | 0.13 | 0.25 |
| 9 | 107618681 | rs2472384 | -1.891 | -3.456 | 0.269 | 0.31 | 0.132 | -0.017 | -0.017 | 0 | 0.47 | 0.71 | 0.35 | 0.44 | 0.24 | 0.37 |
| 9 | 107619063 | rs2253304 | -2.084 | -3.462 | 0.274 | 0.248 | 0.195 | -0.028 | -0.028 | 0.069 | 0.49 | 0.85 | 0.36 | 0.42 | 0.25 | 0.37 |
| 9 | 107620299 | rs2253175 | -2.073 | -3.447 | 0.271 | 0.257 | 0.188 | -0.022 | -0.022 | 0 | 0.48 | 0.84 | 0.37 | 0.42 | 0.25 | 0.37 |
| 9 | 107620332 | rs2253174 | -2.032 | -3.432 | 0.238 | 0.265 | 0.161 | -0.01 | -0.01 | 0 | 0.48 | 0.79 | 0.36 | 0.42 | 0.25 | 0.37 |
| 9 | 107620387 | rs2253172 | -1.486 | -3.383 | 0.476 | 0.283 | 0.046 | -0.02 | -0.02 | 0.102 | 0.4 | 0.27 | 0.32 | 0.42 | 0.24 | 0.37 |
| 9 | 107620867 | rs2230806 | -1.701 | -3.191 | -0.165 | 0.388 | 0.124 | -0.013 | -0.013 | 0 | 0.46 | 0.71 | 0.35 | 0.41 | 0.24 | 0.37 |
| 9 | 107621140 | rs2243313 | -1.701 | -3.191 | -0.165 | 0.382 | 0.124 | -0.016 | -0.016 | 0.981 | 0.46 | 0.71 | 0.35 | 0.42 | 0.24 | 0.37 |
| 9 | 107621638 | rs2482420 | -1.695 | -3.227 | -0.165 | 0.383 | 0.125 | -0.016 | -0.016 | 0 | 0.46 | 0.71 | 0.35 | 0.42 | 0.24 | 0.37 |
| 9 | 107621672 | rs4149287 | 0.831 | -3.037 | -0.205 | 0.554 | 0.034 | -0.051 | -0.051 | 0 | 0.09 | 0 | 0.06 | 0.3 | 0.03 | 0.09 |
| 9 | 107623626 | rs2487058 | -1.514 | -3.091 | -0.056 | 0.415 | 0.123 | -0.017 | -0.017 | 0 | 0.45 | 0.71 | 0.34 | 0.42 | 0.23 | 0.37 |
| 9 | 107623750 | rs2487059 | -1.34 | -3.035 | 0.26 | 0.446 | 0.038 | -0.018 | -0.018 | 0 | 0.39 | 0.28 | 0.32 | 0.42 | 0.23 | 0.39 |
| 9 | 107624029 | rs2230805 | -1.326 | -2.995 | 0.354 | 0.341 | -0.046 | 0.005 | 0.005 | 0 | 0.4 | 0.49 | 0.32 | 0.4 | 0.23 | 0.37 |
| 9 | 107624242 | rs2249891 | -1.338 | -2.544 | -0.235 | 0.67 | 0.019 | -0.012 | -0.012 | 0 | 0.3 | 0.68 | 0.29 | 0.33 | 0.13 | 0.25 |
| 9 | 107625803 | rs4149286 | -0.237 | -2.078 | -0.517 | 0.92 | 0.052 | -0.065 | -0.065 | 0 | 0.13 | 0.05 | 0.06 | 0.29 | 0.03 | 0.11 |
| 9 | 107625932 | rs4149285 | -0.311 | -2.079 | -0.33 | 0.922 | 0.017 | -0.019 | -0.019 | 0 | 0.13 | 0.11 | 0.07 | 0.29 | 0.03 | 0.1 |
| 9 | 107626108 | rs4149284 | 0.001 | -2.116 | -0.358 | 0.93 | 0.054 | -0.067 | -0.067 | 0 | 0.12 | 0.03 | 0.06 | 0.29 | 0.02 | 0.1 |
| 9 | 107626189 | rs4149283 | -0.237 | -2.117 | -0.358 | 0.948 | 0.053 | -0.063 | -0.063 | 0 | 0.13 | 0.05 | 0.06 | 0.29 | 0.03 | 0.1 |
| 9 | 107626284 | rs4149282 | -0.384 | -2.127 | -0.372 | 0.934 | 0.027 | -0.033 | -0.033 | 0.122 | 0.14 | 0.09 | 0.06 | 0.29 | 0.03 | 0.1 |
| 9 | 107626389 | rs4149281 | -1.768 | -2.475 | 0.321 | 0.638 | -0.08 | 0.03 | 0.03 | 0 | 0.29 | 0.53 | 0.3 | 0.3 | 0.13 | 0.21 |
| 9 | 107626542 | rs4149280 | -0.238 | -2.179 | -0.055 | 0.898 | -0.02 | 0.027 | 0.027 | 0 | 0.13 | 0.13 | 0.06 | 0.28 | 0.02 | 0.1 |
| 9 | 107627259 | rs4149279 | -0.845 | -1.59 | -0.782 | 0.207 | 0.01 | -0.004 | -0.004 | 0 | 0.41 | 0.37 | 0.39 | 0.58 | 0.34 | 0.4 |
| 9 | 107629104 | rs4743764 | -1.074 | -1.596 | -1.115 | -0.345 | 0.103 | 0.073 | 0.073 | 0 | 0.54 | 0.86 | 0.38 | 0.37 | 0.4 | 0.53 |
| 9 | 107634640 | rs4742927 | -0.558 | -1.796 | -0.799 | -0.448 | 0.015 | 0.011 | 0.011 | 0 | 0.32 | 0.7 | 0.28 | 0.27 | 0.22 | 0.37 |
| 9 | 107635869 | rs2000069 | -1.431 | -1.717 | -1.481 | -0.514 | 0.031 | 0.051 | 0.051 | 0 | 0.55 | 0.93 | 0.4 | 0.35 | 0.41 | 0.59 |
| 9 | 107638507 | rs4149275 | -0.034 | -1.989 | -1.221 | -0.323 | 0.016 | -0.058 | -0.058 | 0.01 | 0.2 | 0.08 | 0.18 | 0.34 | 0.16 | 0.4 |
| 9 | 107639414 | rs4149274 | -0.963 | -2.372 | -0.768 | -0.909 | 0.06 | 0.056 | 0.056 | 0 | 0.41 | 0.26 | 0.27 | 0.32 | 0.29 | 0.54 |
| 9 | 107640786 | rs1999429 | -1.361 | -2.517 | 0.405 | -0.986 | -0.04 | -0.053 | -0.053 | 0.105 | 0.1 | 0.34 | 0.04 | 0 | 0.03 | 0.02 |
| 9 | 107642211 | rs4149273 | -1.641 | -2.563 | 0.948 | -1.041 | 0.111 | 0.121 | 0.121 | 0.011 | 0.5 | 0.75 | 0.32 | 0.32 | 0.35 | 0.55 |
| 9 | 107642287 | rs4149272 | -1.778 | -2.585 | 1.133 | -1.052 | 0.136 | 0.152 | 0.152 | 0.114 | 0.51 | 0.84 | 0.33 | 0.32 | 0.35 | 0.56 |
| 9 | 107644287 | rs4149271 | -0.001 | -2.585 | 1.234 | -1.018 | -0.001 | 0.001 | 0.001 | 0.226 | 0.19 | 0.15 | 0.2 | 0.26 | 0.2 | 0.4 |
| 9 | 107644830 | rs3858075 | 0.068 | -2.595 | 1.233 | -1.023 | -0.002 | 0 | 0 | 0 | 0.19 | 0.15 | 0.2 | 0.26 | 0.2 | 0.33 |
| 9 | 107645080 | rs2275542 | -0.904 | -2.671 | 1.264 | -1.063 | 0.037 | 0.035 | 0.035 | 0 | 0.37 | 0.21 | 0.26 | 0.32 | 0.3 | 0.46 |
| 9 | 107646260 | rs3904998 | 0.06 | -2.644 | 1.25 | -1.049 | -0.002 | 0 | 0 | 0 | 0.19 | 0.15 | 0.2 | 0.26 | 0.2 | 0.35 |
| 9 | 107647019 | rs11789603 | 0.092 | -2.749 | 1.164 | -1.111 | -0.001 | -0.013 | -0.013 | 0 | 0.08 | 0.15 | 0.07 | 0.07 | 0.09 | 0.18 |
| 9 | 107647121 | rs4149269 | -1.73 | -2.77 | 1.778 | -1.119 | 0.012 | 0.012 | 0.012 | 0 | 0.45 | 0.58 | 0.31 | 0.32 | 0.35 | 0.6 |
| 9 | 107647220 | rs4149268 | -1.924 | -2.778 | 1.785 | -1.124 | 0.096 | 0.101 | 0.101 | 0.02 | 0.49 | 0.74 | 0.32 | 0.32 | 0.35 | 0.6 |
| 9 | 107647655 | rs3890182 | 0.331 | -2.821 | 1.924 | -1.143 | 0.001 | 0 | 0 | 2.932 | 0.17 | 0.12 | 0.07 | 0.06 | 0.12 | 0.1 |
| 9 | 107648430 | rs3847300 | 0.331 | -2.821 | 1.924 | -1.143 | 0.001 | 0 | 0 | 2.932 | 0.17 | 0.12 | 0.07 | 0.06 | 0.12 | 0.1 |
| 9 | 107648652 | rs3847303 | 0.13 | -2.823 | 2.098 | -1.144 | -0.041 | -0.043 | -0.043 | 0.52 | 0.18 | 0.26 | 0.07 | 0.06 | 0.12 | 0.1 |
| 9 | 107650843 | rs10115928 | -2.035 | -2.841 | 1.686 | -1.154 | 0.121 | 0.131 | 0.131 | 0 | 0.49 | 0.8 | 0.33 | 0.32 | 0.35 | 0.6 |
| 9 | 107651174 | rs2275543 | 0.681 | -2.845 | 1.694 | -1.156 | 0.026 | 0.024 | 0.024 | 3.009 | 0.16 | 0.06 | 0.05 | 0.06 | 0.1 | 0.1 |
| 9 | 107651945 | rs4149267 | -2.012 | -2.815 | 1.646 | -1.14 | 0.122 | 0.132 | 0.132 | 0.034 | 0.49 | 0.8 | 0.33 | 0.32 | 0.35 | 0.6 |
| 9 | 107653426 | rs12686004 | 1.108 | -2.751 | 1.538 | -1.103 | -0.029 | -0.043 | -0.043 | 0 | 0.11 | 0.01 | 0.13 | 0.19 | 0.11 | 0.25 |
| 9 | 107655848 | rs3847304 | -0.289 | -2.76 | 1.578 | -1.1 | 0 | 0.003 | 0.003 | 0.776 | 0.2 | 0.16 | 0.21 | 0.26 | 0.2 | 0.45 |
| 9 | 107657070 | rs3905000 | -0.217 | -2.729 | 1.409 | -1.08 | -0.015 | -0.017 | -0.017 | 0.952 | 0.2 | 0.19 | 0.09 | 0.06 | 0.14 | 0.11 |
| 9 | 107658910 | rs12341993 | -0.2 | -2.755 | 1.012 | -1.092 | -0.054 | -0.054 | -0.054 | 0 | 0.2 | 0.31 | 0.08 | 0.06 | 0.12 | 0.1 |
| 9 | 107662494 | rs2575875 | -1.821 | -2.647 | 0.934 | -1.097 | 0.086 | 0.096 | 0.096 | 0.652 | 0.49 | 0.74 | 0.37 | 0.31 | 0.36 | 0.57 |
| 9 | 107662691 | rs2791950 | -1.892 | -2.645 | 0.795 | -1.096 | 0.12 | 0.135 | 0.135 | 0.493 | 0.5 | 0.79 | 0.38 | 0.32 | 0.36 | 0.57 |
| 9 | 107662701 | rs2777788 | -1.892 | -2.645 | 0.795 | -1.096 | 0.12 | 0.135 | 0.135 | 0.493 | 0.5 | 0.79 | 0.38 | 0.32 | 0.36 | 0.57 |
| 9 | 107662705 | rs4149266 | -1.892 | -2.645 | 0.795 | -1.096 | 0.12 | 0.135 | 0.135 | 0.493 | 0.5 | 0.79 | 0.38 | 0.32 | 0.36 | 0.57 |
| 9 | 107663094 | rs2777792 | -1.9 | -2.645 | 0.795 | -1.096 | 0.12 | 0.136 | 0.136 | 0.531 | 0.5 | 0.79 | 0.38 | 0.32 | 0.36 | 0.59 |
| 9 | 107664301 | rs1883025 | -0.792 | -2.5 | 0.835 | -1.028 | -0.005 | -0.005 | -0.005 | 0 | 0.33 | 0.35 | 0.27 | 0.24 | 0.24 | 0.41 |
| 9 | 107665511 | rs2777793 | -1.282 | -1.314 | 0.016 | -0.964 | 0.013 | 0.11 | 0.11 | 0 | 0.5 | 0.68 | 0.43 | 0.73 | 0.45 | 0.48 |
| 9 | 107665739 | rs2575876 | -0.183 | -1.309 | -0.355 | -1.02 | 0.017 | 0.018 | 0.018 | 0 | 0.3 | 0.2 | 0.27 | 0.24 | 0.24 | 0.4 |
| 9 | 107665751 | rs2482424 | 0.789 | -0.513 | -1.344 | -1.242 | -0.041 | -0.042 | -0.042 | 0.173 | 0.07 | 0.01 | 0.23 | 0.19 | 0.14 | 0.15 |
| 9 | 107666513 | rs2740486 | -1.081 | -1.244 | -1.081 | -1.004 | 0.003 | 0.136 | 0.136 | 0 | 0.51 | 0.76 | 0.44 | 0.73 | 0.46 | 0.51 |
| 9 | 107667079 | rs2575878 | -1.695 | -0.987 | -1.148 | -0.893 | -0.05 | 0.099 | 0.099 | 0.093 | 0.62 | 0.83 | 0.5 | 0.8 | 0.54 | 0.7 |
| 9 | 107668825 | rs2575879 | -1.711 | -0.478 | -0.888 | -0.57 | -0.045 | 0.139 | 0.139 | 0.042 | 0.58 | 0.81 | 0.51 | 0.81 | 0.55 | 0.68 |
| 9 | 107669073 | rs13284054 | -0.553 | -0.086 | -0.902 | 0.132 | -0.011 | -0.016 | -0.016 | 0.087 | 0.13 | 0.18 | 0.07 | 0.06 | 0.13 | 0.09 |
| 9 | 107669241 | rs4100654 | -0.002 | 0.075 | -1.197 | 0.298 | 0 | 0 | 0 | 0 | 0.1 | 0.08 | 0.05 | 0.06 | 0.1 | 0.08 |
| 9 | 107671659 | rs10820743 | -0.43 | 0.087 | -1.112 | 0.312 | 0 | 0.001 | 0.001 | 0 | 0.29 | 0.27 | 0.21 | 0.58 | 0.29 | 0.2 |
| 9 | 107672365 | rs2777795 | 0.323 | 0.263 | -1.921 | 0.385 | -0.024 | -0.04 | -0.04 | 0.124 | 0.11 | 0.01 | 0.14 | 0.17 | 0.1 | 0.24 |
| 9 | 107672498 | rs4149265 | -0.56 | 0.39 | -1.367 | 0.728 | -0.008 | -0.023 | -0.023 | 0 | 0.28 | 0.16 | 0.2 | 0.57 | 0.29 | 0.2 |
| 9 | 107676940 | rs12237054 | -0.676 | 0.442 | -1.357 | 1.523 | -0.014 | -0.037 | -0.037 | 0 | 0.08 | 0.01 | 0.11 | 0.33 | 0.07 | 0.07 |
| 9 | 107677289 | rs4149263 | -0.224 | -0.319 | -2.136 | 1.216 | 0.033 | 0.038 | 0.038 | 0 | 0.3 | 0.12 | 0.15 | 0.15 | 0.21 | 0.19 |
| 9 | 107678278 | rs4149261 | -0.769 | -0.687 | -2.188 | 0.977 | 0.009 | -0.045 | -0.045 | 0 | 0.13 | 0.07 | 0.11 | 0.33 | 0.09 | 0.09 |
| 9 | 107678797 | rs2515618 | 1.173 | -0.093 | -2.804 | 0.514 | -0.021 | -0.027 | -0.027 | 0 | 0.25 | 0.12 | 0.13 | 0.33 | 0.3 | 0.37 |
| 9 | 107678916 | rs10991408 | 1.015 | -0.03 | -2.714 | 0.525 | -0.008 | -0.025 | -0.025 | 0 | 0.09 | 0.02 | 0.11 | 0.33 | 0.07 | 0.07 |

| | | | | | | | | | | | | | | | | |
|---|---|---|---|---|---|---|---|---|---|---|---|---|---|---|---|---|
| 9 | 107679425 | rs13291117 | 1.124 | -0.004 | -1.849 | 0.204 | 0.008 | -0.024 | -0.024 | 0 | 0.06 | 0.15 | 0.06 | 0 | 0.07 | 0.02 |
| 9 | 107679500 | rs12336969 | 1.381 | 0.298 | -0.788 | -0.023 | 0.035 | -0.022 | -0.022 | 0 | 0.06 | 0.23 | 0.11 | 0 | 0.06 | 0.06 |
| 9 | 107680458 | rs2437811 | -1.46 | 0.516 | -0.364 | -0.171 | 0.013 | 0 | 0 | 1.15 | 0.87 | 0.84 | 0.76 | 0.84 | 0.79 | 0.8 |
| 9 | 107680730 | rs10512337 | 1.506 | 0.513 | -0.367 | -0.167 | 0.035 | -0.022 | -0.022 | 0 | 0.06 | 0.23 | 0.11 | 0 | 0.06 | 0.06 |
| 9 | 107680915 | rs2515617 | 0.627 | 0.505 | -0.315 | -0.18 | 0.005 | -0.016 | -0.016 | 2.632 | 0.69 | 0.34 | 0.43 | 0.83 | 0.6 | 0.64 |
| 9 | 107681995 | rs2515616 | -1.314 | 0.56 | -0.378 | -0.026 | 0.012 | 0 | 0 | 0 | 0.87 | 0.84 | 0.8 | 0.83 | 0.78 | 0.81 |
| 9 | 107682012 | rs7861459 | 0.966 | 0.462 | 0.045 | -0.057 | 0.023 | -0.017 | -0.017 | 0.147 | 0.05 | 0.23 | 0.1 | 0 | 0.04 | 0.05 |
| 9 | 107682517 | rs2791952 | -0.577 | 0.669 | -0.2 | -0.013 | 0.001 | -0.017 | -0.017 | 0 | 0.87 | 0.93 | 0.89 | 0.67 | 0.91 | 0.9 |
| 9 | 107684230 | rs2472509 | 1.032 | 0.321 | -0.388 | -0.22 | -0.029 | -0.027 | -0.027 | 0.646 | 0.28 | 0.12 | 0.34 | 0.35 | 0.36 | 0.46 |
| 9 | 107684253 | rs2472508 | 0.903 | 0.381 | -0.182 | -0.186 | 0.005 | -0.001 | -0.001 | 0.04 | 0.14 | 0.17 | 0.24 | 0.18 | 0.21 | 0.19 |
| 9 | 107684276 | rs10991411 | 0.192 | 0.372 | -0.084 | -0.196 | 0.042 | -0.022 | -0.022 | 0 | 0.07 | 0.37 | 0.13 | 0 | 0.07 | 0.06 |
| 9 | 107684286 | rs2487049 | 0.896 | 0.339 | -0.476 | -0.08 | -0.017 | 0 | 0 | 0 | 0.13 | 0.09 | 0.24 | 0.18 | 0.21 | 0.19 |
| 9 | 107686354 | rs10991415 | -0.051 | -1.154 | -0.533 | -0.476 | -0.004 | 0.001 | 0.001 | 0 | 0.2 | 0.16 | 0.09 | 0.13 | 0.12 | 0.11 |
| 9 | 107686405 | rs2487052 | 0.938 | -1.035 | -1.093 | -0.739 | 0.005 | -0.062 | -0.062 | 1.17 | 0.23 | 0.07 | 0.17 | 0.35 | 0.23 | 0.37 |
| 9 | 107690057 | rs2437817 | 0.23 | -1.153 | -0.925 | -1.007 | 0.015 | 0.011 | 0.011 | 0 | 0.37 | 0.26 | 0.44 | 0.35 | 0.37 | 0.46 |
| 9 | 107690450 | rs1800977 | 0.244 | -1.166 | -0.937 | -1.017 | 0.014 | 0.01 | 0.01 | 0 | 0.37 | 0.25 | 0.44 | 0.34 | 0.37 | 0.46 |
| 9 | 107690995 | rs2422493 | -0.228 | -1.063 | -0.991 | -1.008 | 0.007 | 0.001 | 0.001 | 0 | 0.52 | 0.45 | 0.53 | 0.46 | 0.49 | 0.53 |
| 9 | 107694245 | rs2043664 | -1.001 | -0.603 | -1.768 | -0.876 | -0.035 | 0.007 | 0.007 | 0 | 0.8 | 0.86 | 0.77 | 0.78 | 0.71 | 0.77 |
| 9 | 107695396 | rs2472495 | 1.422 | -0.874 | -1.944 | -0.692 | -0.014 | -0.096 | -0.096 | 0.765 | 0.22 | 0.03 | 0.2 | 0.34 | 0.24 | 0.38 |
| 9 | 107698322 | rs12553329 | 0.238 | 0.106 | -1.949 | -0.134 | -0.06 | -0.086 | -0.086 | 0 | 0.16 | 0.03 | 0.29 | 0.3 | 0.21 | 0.16 |
| 9 | 107698612 | rs3887137 | 1.205 | 0.193 | -1.885 | -0.071 | -0.035 | -0.049 | -0.049 | 0 | 0.1 | 0.03 | 0.1 | 0.3 | 0.11 | 0.1 |
| 9 | 107699196 | rs2437819 | -0.96 | 0.109 | -1.697 | -0.366 | -0.02 | 0.008 | 0.008 | 0 | 0.77 | 0.84 | 0.77 | 0.79 | 0.71 | 0.72 |
| 9 | 107701183 | rs7848232 | 1.361 | -0.078 | -1.162 | -0.418 | 0.037 | -0.01 | -0.01 | 2.888 | 0.05 | 0.21 | 0.05 | 0.01 | 0.04 | 0.01 |
| 9 | 107704228 | rs2487031 | -0.826 | -0.088 | -1.215 | -0.452 | 0.017 | -0.011 | -0.011 | 0 | 0.74 | 0.68 | 0.73 | 0.79 | 0.69 | 0.71 |
| 9 | 107707353 | rs2164560 | 1.102 | -0.758 | -1.316 | -0.485 | 0.047 | -0.033 | -0.033 | 0 | 0.36 | 0.08 | 0.27 | 0.36 | 0.32 | 0.42 |
| 9 | 107710562 | rs2472459 | 1.396 | -0.397 | -0.566 | -0.187 | -0.026 | 0.043 | 0.043 | 0 | 0.24 | 0.36 | 0.2 | 0.35 | 0.23 | 0.29 |
| 9 | 107715878 | rs2472519 | 1.574 | -0.161 | -0.991 | -0.017 | 0.007 | -0.017 | -0.017 | 0 | 0.24 | 0.22 | 0.21 | 0.37 | 0.24 | 0.29 |
| 9 | 107716396 | rs2472516 | 0.557 | -0.167 | -0.962 | -0.166 | -0.014 | 0.02 | 0.02 | 0 | 0.39 | 0.26 | 0.51 | 0.67 | 0.43 | 0.44 |
| 9 | 107722578 | rs4397467 | 0.782 | -0.305 | -0.506 | -0.12 | -0.015 | 0 | 0 | 0 | 0.36 | 0.53 | 0.43 | 0.35 | 0.34 | 0.32 |
| 9 | 107726070 | rs10991432 | 0.855 | -0.49 | -1.051 | -0.317 | -0.007 | -0.004 | -0.004 | 0 | 0.18 | 0.19 | 0.12 | 0.14 | 0.14 | 0.16 |
| 9 | 107726821 | rs9919066 | 1.275 | -0.698 | -1.36 | -0.489 | 0.046 | 0.058 | 0.058 | 0 | 0.16 | 0.05 | 0.11 | 0 | 0.09 | 0.12 |
| 9 | 107731568 | rs12380341 | 0.733 | 0.022 | -1.142 | -0.217 | -0.001 | -0.001 | -0.001 | 0 | 0.16 | 0.16 | 0.12 | 0.14 | 0.16 | 0.14 |
| 9 | 107731789 | rs7023634 | -1.008 | 0.469 | -1.179 | 0.027 | -0.037 | 0.015 | 0.015 | 0 | 0.77 | 0.89 | 0.77 | 0.8 | 0.72 | 0.68 |
| 9 | 107735920 | rs10991437 | 1.242 | 0.642 | -0.978 | 0.619 | -0.032 | -0.028 | -0.028 | 0.371 | 0.08 | 0.03 | 0.1 | 0.3 | 0.1 | 0.11 |
| 9 | 107736034 | rs1808789 | 0.219 | 0.312 | -0.607 | 0.404 | 0.005 | -0.008 | -0.008 | 0 | 0.39 | 0.52 | 0.34 | 0.63 | 0.39 | 0.4 |
| 9 | 107737393 | rs715119 | 1.387 | 0.243 | -0.364 | 0.303 | 0.005 | -0.016 | -0.016 | 0 | 0.28 | 0.22 | 0.21 | 0.33 | 0.26 | 0.29 |
| 9 | 107739312 | rs12379205 | 0.788 | 0.039 | -0.668 | -0.311 | -0.001 | -0.001 | -0.001 | 0 | 0.17 | 0.17 | 0.17 | 0.14 | 0.17 | 0.15 |
| 9 | 107740191 | rs10761089 | -0.419 | 0.075 | -0.69 | -0.29 | 0.004 | 0.045 | 0.045 | 0 | 0.56 | 0.69 | 0.52 | 0.78 | 0.56 | 0.56 |
| 9 | 107743081 | rs10991440 | 0.351 | 0.203 | -0.807 | -0.126 | -0.001 | -0.002 | -0.002 | 0 | 0.17 | 0.17 | 0.17 | 0.14 | 0.17 | 0.15 |
| 9 | 107744661 | rs4743784 | -0.379 | 0.358 | -1.361 | -0.429 | -0.058 | 0.102 | 0.102 | 0 | 0.63 | 0.83 | 0.57 | 0.82 | 0.57 | 0.59 |
| 9 | 107744958 | rs12342875 | 0.291 | 0.501 | -1.061 | 0.114 | 0.002 | 0.03 | 0.03 | 0 | 0.13 | 0.18 | 0.19 | 0.34 | 0.15 | 0.08 |
| 9 | 107748958 | rs1039406 | -0.28 | 0.054 | -0.969 | -0.412 | 0.002 | 0.05 | 0.05 | 0 | 0.47 | 0.58 | 0.47 | 0.85 | 0.47 | 0.47 |
| 9 | 107751571 | rs10991446 | 0 | -0.049 | -0.799 | -0.44 | 0.051 | 0.107 | 0.107 | 0 | 0.22 | 0.51 | 0.32 | 0.34 | 0.26 | 0.12 |
| 9 | 107751593 | rs7019610 | -1.052 | -0.19 | -1.384 | -0.813 | -0.023 | 0.015 | 0.015 | 0 | 0.48 | 0.81 | 0.64 | 0.36 | 0.48 | 0.21 |
| 9 | 107754330 | rs4743785 | -1.616 | -0.18 | -1.467 | -0.99 | -0.062 | 0.075 | 0.075 | 0 | 0.73 | 0.9 | 0.79 | 0.87 | 0.68 | 0.57 |
| 9 | 107760227 | rs10820754 | 0.507 | -0.133 | -1.273 | -1.082 | 0.012 | 0.086 | 0.086 | 0 | 0.39 | 0.21 | 0.33 | 0.86 | 0.35 | 0.43 |
| 9 | 107767036 | rs6479289 | -1.541 | -0.127 | -0.776 | -1.074 | -0.01 | 0.014 | 0.014 | 0 | 0.72 | 0.77 | 0.78 | 0.87 | 0.68 | 0.58 |
| 9 | 107770227 | rs13283838 | 1.987 | 0.14 | -0.599 | -0.677 | -0.024 | -0.034 | -0.034 | 0.408 | 0.11 | 0.05 | 0.16 | 0.31 | 0.13 | 0.07 |
| 9 | 107770256 | rs12000585 | 1.54 | 0.013 | -0.568 | -0.788 | 0.001 | -0.016 | -0.016 | 0.978 | 0.13 | 0.1 | 0.07 | 0 | 0.09 | 0.05 |
| 9 | 107778690 | rs12340129 | 1.11 | 0.125 | -0.504 | -0.662 | 0.008 | 0.045 | 0.045 | 0 | 0.14 | 0.18 | 0.19 | 0.32 | 0.15 | 0.07 |
| 9 | 107782100 | rs2417598 | -0.965 | 0.036 | -0.613 | -0.761 | -0.007 | 0.029 | 0.029 | 0 | 0.47 | 0.67 | 0.63 | 0.32 | 0.48 | 0.23 |
| 9 | 107782109 | rs4551427 | 1.574 | 0.222 | -0.163 | -0.292 | 0.023 | -0.045 | -0.045 | 0.011 | 0.09 | 0.27 | 0.14 | 0 | 0.11 | 0.07 |
| 9 | 107784172 | rs1827324 | -0.33 | 0.188 | -0.447 | -0.253 | 0.02 | 0.042 | 0.042 | 0 | 0.26 | 0.55 | 0.36 | 0.32 | 0.28 | 0.14 |
| 9 | 107786886 | rs10991473 | 1.583 | -0.266 | -1.216 | -0.011 | 0.011 | -0.093 | -0.093 | 0 | 0.2 | 0.03 | 0.15 | 0.39 | 0.16 | 0.25 |
| 9 | 107786913 | rs328875 | 0.086 | 0.066 | -1.187 | -0.535 | 0 | -0.002 | -0.002 | 0 | 0.43 | 0.44 | 0.42 | 0.31 | 0.47 | 0.3 |
| 9 | 107787214 | rs328876 | 0.023 | 0.124 | -0.985 | -0.499 | -0.008 | -0.005 | -0.005 | 1.738 | 0.59 | 0.33 | 0.54 | 0.61 | 0.54 | 0.57 |
| 9 | 107789252 | rs328879 | 0.165 | 0.098 | -0.72 | -0.422 | -0.001 | 0.003 | 0.003 | 0.056 | 0.65 | 0.52 | 0.57 | 0.7 | 0.64 | 0.55 |
| 9 | 107790065 | rs328880 | 0.048 | 0.152 | -0.811 | -0.665 | -0.011 | -0.007 | -0.007 | 0 | 0.6 | 0.63 | 0.48 | 0.6 | 0.53 | 0.6 |
| 9 | 107791702 | rs498275 | 0.893 | 0.014 | -0.858 | -1.093 | -0.001 | -0.036 | -0.036 | 0 | 0.31 | 0.35 | 0.26 | 0.06 | 0.32 | 0.24 |
| 9 | 107792622 | rs12346607 | -0.181 | 0.01 | -0.893 | -1.078 | 0.009 | 0.047 | 0.047 | 0 | 0.5 | 0.29 | 0.57 | 0.8 | 0.5 | 0.46 |
| 9 | 107793314 | rs7865004 | 1.814 | 0.05 | -0.989 | -0.694 | -0.026 | -0.036 | -0.036 | 0 | 0.12 | 0.03 | 0.15 | 0.24 | 0.15 | 0.06 |
| 9 | 107793365 | rs443757 | -1.325 | -0.204 | -1.175 | -1.043 | -0.002 | 0.003 | 0.003 | 0 | 0.76 | 0.77 | 0.81 | 0.86 | 0.74 | 0.69 |
| 9 | 107793488 | rs7865251 | 0.694 | -0.561 | -1.32 | -0.947 | 0.002 | -0.048 | -0.048 | 0 | 0.28 | 0.13 | 0.26 | 0.64 | 0.24 | 0.3 |
| 9 | 107795869 | rs375920 | 0.954 | -0.28 | -1.238 | -1.034 | -0.003 | -0.065 | -0.065 | 0.03 | 0.31 | 0.4 | 0.25 | 0.07 | 0.3 | 0.24 |
| 9 | 107796548 | rs419490 | -1.617 | -0.35 | -1.254 | -1.078 | 0.006 | -0.004 | -0.004 | 0.194 | 0.77 | 0.75 | 0.8 | 0.86 | 0.72 | 0.69 |
| 9 | 107797889 | rs7850052 | 0.472 | -0.405 | -1.288 | -1.024 | -0.007 | 0.004 | 0.004 | 0 | 0.17 | 0.15 | 0.28 | 0.17 | 0.18 | 0.15 |
| 9 | 107799042 | rs328881 | 1.971 | -0.467 | -1.42 | -1.068 | 0.002 | 0.072 | 0.072 | 0 | 0.22 | 0.08 | 0.1 | 0.06 | 0.19 | 0.18 |
| 9 | 107799223 | rs4743792 | 2.061 | -0.482 | -1.33 | -0.893 | 0.02 | -0.11 | -0.11 | 0 | 0.15 | 0.02 | 0.11 | 0.4 | 0.09 | 0.24 |
| 9 | 107802222 | rs7023868 | 1.819 | -0.361 | -0.874 | -1.613 | 0.033 | -0.046 | -0.046 | 0 | 0.09 | 0.32 | 0.15 | 0 | 0.12 | 0.06 |

| | | | | | | | | | | | | | | | | |
|---|---|---|---|---|---|---|---|---|---|---|---|---|---|---|---|---|
| 9 | 107804674 | rs7043894 | 0.056 | -0.392 | -0.86 | -1.615 | -0.004 | 0.039 | 0.039 | 0 | 0.17 | 0.42 | 0.2 | 0.17 | 0.16 | 0.17 |
| 9 | 107809019 | rs448085 | -0.191 | -0.328 | -1.367 | -1.6 | 0.005 | 0.007 | 0.007 | 0 | 0.68 | 0.51 | 0.73 | 0.7 | 0.67 | 0.59 |
| 9 | 107809131 | rs387394 | 0.569 | -0.413 | -1.31 | -1.571 | -0.004 | -0.081 | -0.081 | 0 | 0.31 | 0.41 | 0.25 | 0.07 | 0.3 | 0.24 |
| 9 | 107809777 | rs435986 | 0.376 | -0.39 | -1.431 | -1.557 | -0.009 | -0.11 | -0.11 | 0 | 0.33 | 0.47 | 0.26 | 0.07 | 0.3 | 0.24 |
| 9 | 107810223 | rs10820767 | 0.459 | -0.366 | -1.639 | -1.566 | -0.025 | 0.013 | 0.013 | 0.306 | 0.34 | 0.05 | 0.29 | 0.78 | 0.29 | 0.41 |
| 9 | 107810644 | rs446303 | -0.843 | -0.177 | -1.467 | -1.406 | 0.019 | -0.033 | -0.033 | 1.314 | 0.69 | 0.61 | 0.58 | 0.85 | 0.61 | 0.65 |
| 9 | 107814671 | rs1809921 | 0.042 | -0.397 | -1.289 | -1.648 | 0.003 | 0.008 | 0.008 | 0 | 0.59 | 0.39 | 0.69 | 0.78 | 0.65 | 0.68 |
| 9 | 107814816 | rs2417603 | -0.491 | 0.205 | -1.766 | -1.142 | -0.017 | 0.084 | 0.084 | 0.327 | 0.53 | 0.86 | 0.64 | 0.28 | 0.57 | 0.53 |
| 9 | 107816813 | rs1532677 | -0.089 | 0.514 | -1.388 | -1.238 | 0.003 | 0.038 | 0.038 | 0 | 0.62 | 0.74 | 0.72 | 0.85 | 0.64 | 0.71 |
| 9 | 107819715 | rs10991509 | -0.1 | 0.676 | -1.874 | -1.097 | -0.002 | 0.053 | 0.053 | 0 | 0.27 | 0.13 | 0.39 | 0.18 | 0.25 | 0.22 |
| 9 | 107819733 | rs10991510 | 1.811 | 0.988 | -1.247 | -0.815 | 0.003 | -0.017 | -0.017 | 0 | 0.06 | 0.08 | 0.04 | 0 | 0.09 | 0.03 |
| 9 | 107821021 | rs1501460 | 0.186 | 0.86 | -1.431 | -0.907 | -0.009 | 0.082 | 0.082 | 0 | 0.41 | 0.23 | 0.35 | 0.18 | 0.46 | 0.48 |
| 9 | 107826038 | rs28497830 | -0.675 | 1.132 | -0.67 | 0.12 | 0.033 | 0.043 | 0.043 | 0 | 0.3 | 0.55 | 0.35 | 0.46 | 0.39 | 0.21 |
| 9 | 107826647 | rs2937372 | -0.31 | 0.537 | -0.664 | -0.422 | -0.012 | 0.065 | 0.065 | 0 | 0.82 | 0.93 | 0.9 | 0.96 | 0.81 | 0.85 |
| 9 | 107827911 | rs882992 | 0.274 | 0.545 | -0.411 | -0.371 | 0.01 | -0.107 | -0.107 | 0 | 0.71 | 0.5 | 0.8 | 0.92 | 0.7 | 0.8 |
| 9 | 107830552 | rs11999215 | -1.078 | -0.029 | -0.604 | 0.26 | -0.006 | 0 | 0 | 0 | 0.12 | 0.11 | 0.07 | 0.21 | 0.07 | 0.1 |
| 9 | 107833459 | rs10991516 | -0.832 | 0.036 | -0.283 | 0.105 | -0.007 | -0.027 | -0.027 | 0 | 0.05 | 0.11 | 0.03 | 0 | 0.06 | 0.04 |
| 9 | 107833736 | rs3132117 | 0.742 | -0.228 | -0.581 | 0.054 | 0.005 | -0.001 | -0.001 | 0 | 0.61 | 0.65 | 0.77 | 0.58 | 0.67 | 0.46 |
| 9 | 107835785 | rs7047382 | 1.316 | -0.624 | -0.004 | -0.453 | -0.013 | -0.055 | -0.055 | 0.175 | 0.17 | 0.24 | 0.1 | 0.01 | 0.11 | 0.08 |
| 9 | 107839290 | rs3005724 | -0.945 | -0.764 | -0.421 | -0.477 | -0.006 | 0.034 | 0.034 | 0 | 0.83 | 0.93 | 0.94 | 0.96 | 0.85 | 0.86 |
| 9 | 107839841 | rs2937359 | 0.485 | 0.13 | -0.429 | 0.046 | -0.012 | 0.001 | 0.001 | 0 | 0.56 | 0.3 | 0.58 | 0.47 | 0.56 | 0.64 |
| 9 | 107841964 | rs2937352 | 0.344 | 0.126 | -0.421 | 0.052 | -0.013 | -0.013 | -0.013 | 0.053 | 0.57 | 0.32 | 0.59 | 0.57 | 0.58 | 0.69 |
| 9 | 107842490 | rs3005720 | -0.481 | 0.182 | -0.375 | -0.039 | 0 | 0.006 | 0.006 | 0 | 0.81 | 0.87 | 0.93 | 0.85 | 0.84 | 0.79 |
| 9 | 107851745 | rs10820780 | -0.368 | -0.095 | -0.011 | 0.437 | 0.028 | 0.001 | 0.001 | 0.294 | 0.39 | 0.69 | 0.39 | 0.64 | 0.27 | 0.51 |
| 9 | 107856028 | rs7855885 | 0.493 | 0.792 | 0.675 | 0.932 | 0.001 | -0.001 | -0.001 | 0 | 0.14 | 0.15 | 0.1 | 0.17 | 0.15 | 0.15 |
| 9 | 107856582 | rs1039407 | 0.013 | 0.623 | 0.727 | 0.885 | -0.063 | 0.164 | 0.164 | 0 | 0.22 | 0.49 | 0.26 | 0.5 | 0.11 | 0.34 |
| 9 | 107857255 | rs2173884 | -1.083 | 0.624 | 0.712 | 0.865 | 0.065 | -0.02 | -0.02 | 0 | 0.37 | 0.84 | 0.37 | 0.67 | 0.26 | 0.49 |
| 9 | 107858481 | rs3005716 | 0.764 | 0.934 | 0.708 | 0.805 | -0.02 | -0.027 | -0.027 | 0 | 0.65 | 0.75 | 0.72 | 0.59 | 0.6 | 0.65 |
| 9 | 107860817 | rs989125 | -0.064 | 1.163 | 0.887 | 0.946 | -0.033 | 0.016 | 0.016 | 0 | 0.23 | 0.46 | 0.19 | 0.3 | 0.16 | 0.3 |
| 9 | 107861083 | rs4609251 | -1.735 | 1.232 | 0.535 | 0.864 | 0.04 | -0.003 | -0.003 | 0 | 0.44 | 0.92 | 0.42 | 0.67 | 0.36 | 0.5 |
| 9 | 107863050 | rs7854755 | -0.349 | 1.125 | 0.655 | 0.901 | -0.02 | 0.063 | 0.063 | 0 | 0.28 | 0.54 | 0.29 | 0.5 | 0.22 | 0.37 |
| 9 | 107866405 | rs4742941 | -1.14 | 1.106 | 0.706 | 0.88 | 0.003 | 0 | 0 | 0 | 0.41 | 0.62 | 0.37 | 0.64 | 0.36 | 0.5 |
| 9 | 107869224 | rs2047701 | -0.496 | 1.061 | 0.738 | 0.929 | -0.011 | -0.012 | -0.012 | 0 | 0.2 | 0.42 | 0.15 | 0.18 | 0.19 | 0.16 |
| 9 | 107869834 | rs4743805 | -0.018 | 1.087 | 0.775 | 1.007 | -0.034 | 0.121 | 0.121 | 0.021 | 0.24 | 0.51 | 0.27 | 0.49 | 0.15 | 0.34 |
| 9 | 107870014 | rs4743806 | -0.831 | 0.384 | 0.485 | 0.892 | 0.04 | 0.004 | 0.004 | 0.839 | 0.4 | 0.77 | 0.37 | 0.66 | 0.31 | 0.51 |
| 9 | 107873231 | rs3124016 | 1.019 | 0.853 | 1.375 | 1.197 | -0.006 | 0.071 | 0.071 | 0 | 0.73 | 0.59 | 0.7 | 0.48 | 0.74 | 0.71 |
| 9 | 107873320 | rs10820784 | 0.666 | 0.402 | 1.413 | 0.267 | -0.002 | 0.101 | 0.101 | 0 | 0.49 | 0.34 | 0.52 | 0.21 | 0.53 | 0.42 |
| 9 | 107877624 | rs10991552 | -1.922 | 0.382 | 2.865 | 1.089 | 0.007 | 0.007 | 0.007 | 0 | 0.13 | 0.21 | 0.11 | 0.14 | 0.15 | 0.11 |
| 9 | 107878643 | rs12343892 | -0.76 | 0.321 | 2.724 | 1.005 | -0.004 | -0.017 | -0.017 | 0.317 | 0.06 | 0.05 | 0.02 | 0 | 0.06 | 0.03 |
| 9 | 107879948 | rs7861505 | -2.52 | 0.252 | 3.074 | 1.005 | -0.012 | -0.016 | -0.016 | 0 | 0.09 | 0.33 | 0.06 | 0.03 | 0.04 | 0.08 |
| 9 | 107881736 | rs7858499 | -2.134 | 0.339 | 2.586 | 1.218 | 0.004 | 0.069 | 0.069 | 0 | 0.2 | 0.5 | 0.17 | 0.3 | 0.23 | 0.17 |
| 9 | 107893381 | rs7022410 | -1.994 | -0.273 | 2.134 | 1.02 | 0 | 0.109 | 0.109 | 0 | 0.07 | 0.35 | 0.08 | 0.17 | 0.04 | 0.07 |
| 9 | 107910242 | rs35211598 | -2.108 | -0.157 | 1.462 | 0.973 | -0.019 | 0.093 | 0.093 | 0.115 | 0.15 | 0.54 | 0.12 | 0.21 | 0.09 | 0.16 |
| 9 | 107917632 | rs10991561 | -2.173 | -0.083 | 1.593 | 0.988 | -0.006 | 0.014 | 0.014 | 0.06 | 0.2 | 0.69 | 0.2 | 0.32 | 0.18 | 0.24 |
| 9 | 107920445 | rs7870426 | -2.364 | -0.07 | 2.036 | 0.98 | 0.01 | 0.076 | 0.076 | 0 | 0.18 | 0.56 | 0.2 | 0.32 | 0.18 | 0.24 |
| 9 | 107921428 | rs7856399 | -0.991 | -0.026 | 1.704 | 0.56 | 0.002 | -0.023 | -0.023 | 0 | 0.05 | 0.07 | 0.06 | 0 | 0.05 | 0.1 |
| 9 | 107923203 | rs765399 | -1.41 | -0.122 | 1.361 | 0.51 | -0.027 | -0.025 | -0.025 | 0 | 0.08 | 0.01 | 0.08 | 0.13 | 0.12 | 0.09 |
| 9 | 107929180 | rs3005738 | -2.732 | -0.268 | 1.37 | 0.416 | 0.031 | 0.057 | 0.057 | 0.046 | 0.25 | 0.87 | 0.21 | 0.15 | 0.2 | 0.22 |
| 9 | 107929206 | rs13295440 | -2.334 | -0.21 | 1.939 | 0.631 | -0.018 | 0.003 | 0.003 | 0 | 0.11 | 0.26 | 0.1 | 0.14 | 0.11 | 0.09 |
| 9 | 107931886 | rs12338452 | -2.406 | -0.218 | 2.284 | 0.637 | -0.016 | 0.033 | 0.033 | 0 | 0.11 | 0.41 | 0.08 | 0.14 | 0.08 | 0.09 |
| 9 | 107934729 | rs10991567 | 0.25 | -0.464 | 1.705 | 0.571 | -0.015 | 0.021 | 0.021 | 0 | 0.64 | 0.52 | 0.48 | 0.44 | 0.7 | 0.68 |
| 9 | 107954622 | rs10991580 | -1.367 | -0.497 | 1.668 | 0.37 | -0.005 | -0.006 | -0.006 | 0.141 | 0.05 | 0.13 | 0.03 | 0.05 | 0.04 | 0.03 |
| 9 | 107956549 | rs12335477 | -1.849 | -0.457 | 2.308 | 0.264 | 0.032 | -0.007 | -0.007 | 0 | 0.11 | 0.33 | 0.13 | 0.09 | 0.13 | 0.16 |
| 9 | 107958534 | rs327971 | -2.407 | -0.721 | 2.554 | 0.38 | -0.016 | -0.021 | -0.021 | 0.082 | 0.19 | 0.49 | 0.33 | 0.15 | 0.19 | 0.21 |
| 9 | 107959548 | rs10124147 | 0.881 | -0.674 | 2.094 | 0.351 | 0.003 | 0.002 | 0.002 | 0 | 0.43 | 0.4 | 0.37 | 0.34 | 0.58 | 0.46 |
| 9 | 107969815 | rs390253 | -3.304 | -0.191 | 2.907 | 0.145 | -0.014 | -0.036 | -0.036 | 0.055 | 0.19 | 0.62 | 0.2 | 0.11 | 0.19 | 0.2 |
| 9 | 107973277 | rs395992 | -3.205 | -0.115 | 2.989 | 0.096 | -0.007 | -0.023 | -0.023 | 0.106 | 0.2 | 0.45 | 0.29 | 0.13 | 0.19 | 0.21 |
| 9 | 107974737 | rs7854392 | -2.354 | -0.177 | 2.762 | 0.209 | 0.003 | 0.038 | 0.038 | 0 | 0.06 | 0.17 | 0.16 | 0.13 | 0.07 | 0.05 |
| 9 | 107975234 | rs412647 | -2.901 | -0.182 | 2.829 | 0.207 | -0.013 | 0.011 | 0.011 | 0.059 | 0.12 | 0.28 | 0.18 | 0.13 | 0.09 | 0.08 |
| 9 | 107986092 | rs384924 | -2.792 | -0.04 | 3.164 | 0.295 | 0 | 0.054 | 0.054 | 0 | 0.1 | 0.36 | 0.2 | 0.13 | 0.08 | 0.09 |
| 9 | 107988197 | rs446641 | -2.528 | -0.044 | 3.221 | 0.302 | 0.002 | 0.056 | 0.056 | 0 | 0.09 | 0.24 | 0.2 | 0.13 | 0.08 | 0.08 |
| 9 | 107989797 | rs370629 | -3.977 | -0.184 | 3.468 | 0.232 | -0.05 | -0.002 | -0.002 | 0.124 | 0.17 | 0.57 | 0.23 | 0.13 | 0.1 | 0.1 |
| 9 | 107990126 | rs438299 | -3.977 | -0.184 | 3.473 | 0.21 | -0.051 | -0.005 | -0.005 | 0.005 | 0.17 | 0.56 | 0.23 | 0.13 | 0.1 | 0.1 |
| 9 | 107990623 | rs366144 | -3.977 | -0.184 | 3.473 | 0.21 | -0.051 | -0.005 | -0.005 | 0 | 0.17 | 0.56 | 0.22 | 0.13 | 0.1 | 0.1 |
| 9 | 107990922 | rs422084 | -3.977 | -0.184 | 3.473 | 0.21 | -0.051 | -0.005 | -0.005 | 0.016 | 0.17 | 0.56 | 0.22 | 0.13 | 0.1 | 0.1 |
| 9 | 107991283 | rs384947 | -3.977 | -0.184 | 3.458 | 0.21 | -0.05 | -0.006 | -0.006 | 0 | 0.17 | 0.57 | 0.23 | 0.13 | 0.1 | 0.1 |
| 9 | 108008916 | rs327956 | -3.603 | -0.017 | 3.576 | 0.26 | -0.009 | 0.066 | 0.066 | 0 | 0.12 | 0.48 | 0.22 | 0.13 | 0.08 | 0.09 |
| 9 | 108042806 | rs193008 | -4.317 | 0.018 | 3.742 | 0.253 | -0.034 | 0.007 | 0.007 | 0.213 | 0.14 | 0.67 | 0.24 | 0.13 | 0.08 | 0.09 |
| 9 | 108043166 | rs193007 | -2.944 | 0.017 | 3.749 | 0.253 | 0.017 | 0.086 | 0.086 | 0 | 0.1 | 0.41 | 0.22 | 0.13 | 0.08 | 0.08 |

| | | | | | | | | | | | | | | | | |
|---|---|---|---|---|---|---|---|---|---|---|---|---|---|---|---|---|
| 9 | 108043636 | rs227887 | -4.082 | 0.019 | 3.475 | 0.253 | -0.034 | 0.007 | 0.007 | 0.213 | 0.14 | 0.67 | 0.24 | 0.13 | 0.08 | 0.09 |
| 9 | 108054966 | rs327984 | -2.966 | 0.113 | 2.821 | 0.297 | -0.005 | 0.039 | 0.039 | 0 | 0.11 | 0.32 | 0.21 | 0.13 | 0.09 | 0.08 |
| 9 | 108058559 | rs376926 | -3.259 | 0.113 | 2.854 | 0.315 | -0.002 | 0.059 | 0.059 | 0 | 0.11 | 0.41 | 0.22 | 0.13 | 0.08 | 0.08 |
| 9 | 108063472 | rs12378850 | -1.5 | 0.206 | 2.27 | 0.351 | -0.002 | -0.01 | -0.01 | 0 | 0.05 | 0.06 | 0.02 | 0 | 0.02 | 0.02 |
| 9 | 108084792 | rs10820798 | -2.592 | 1.384 | 2.285 | 0.194 | 0.12 | 0.005 | 0.005 | 0 | 0.12 | 0.59 | 0.15 | 0.06 | 0.19 | 0.08 |
| 9 | 108094256 | rs10991629 | -2.855 | 1.824 | 2.308 | -0.051 | 0.033 | -0.082 | -0.082 | 0.002 | 0.13 | 0.41 | 0.16 | 0 | 0.12 | 0.15 |
| 9 | 108111284 | rs10120853 | -2.626 | 1.825 | 2.438 | -0.059 | 0.049 | -0.031 | -0.031 | 0 | 0.09 | 0.39 | 0.07 | 0 | 0.06 | 0.02 |
| 9 | 108147763 | rs3199966 | -3.368 | 1.785 | 2.6 | -0.083 | 0.046 | -0.036 | -0.036 | 0.097 | 0.1 | 0.49 | 0.15 | 0 | 0.08 | 0.13 |
| 9 | 108150015 | rs12344130 | -2.963 | 1.801 | 2.611 | -0.071 | 0.066 | 0.004 | 0.004 | 0.186 | 0.08 | 0.47 | 0.13 | 0 | 0.06 | 0.11 |
| 9 | 108150827 | rs957903 | -0.73 | 2.531 | 2.164 | -0.312 | -0.064 | -0.03 | -0.03 | 0.364 | 0.1 | 0.01 | 0.19 | 0.09 | 0.29 | 0.09 |
| 9 | 108152199 | rs2771040 | 3.199 | 1.678 | 2.34 | -0.36 | 0.044 | -0.088 | -0.088 | 0 | 0.87 | 0.49 | 0.84 | 1 | 0.88 | 0.87 |
| 9 | 108166606 | rs2812314 | 3.117 | 1.573 | 2.25 | -0.416 | 0.034 | -0.113 | -0.113 | 0 | 0.85 | 0.51 | 0.83 | 1 | 0.88 | 0.9 |
| 9 | 108166638 | rs12340765 | -2.707 | 1.492 | 2.263 | -0.332 | 0.061 | -0.039 | -0.039 | 0 | 0.1 | 0.4 | 0.14 | 0 | 0.08 | 0.08 |
| 9 | 108207197 | rs2812297 | -0.504 | 1.014 | 0.833 | -0.657 | 0.019 | 0.043 | 0.043 | 0 | 0.49 | 0.47 | 0.66 | 0.7 | 0.56 | 0.47 |
| 9 | 108209066 | rs7037141 | -0.193 | 1.348 | 0.598 | 0.225 | -0.003 | 0.011 | 0.011 | 0 | 0.11 | 0.12 | 0.21 | 0.44 | 0.11 | 0.19 |
| 9 | 108211179 | rs7872832 | 0.135 | 0.495 | 0.658 | -0.645 | -0.031 | -0.099 | -0.099 | 2.39 | 0.17 | 0.26 | 0.07 | 0 | 0.11 | 0.11 |
| 9 | 108218852 | rs10991660 | 0.234 | 0.479 | 0.802 | -0.652 | -0.025 | -0.087 | -0.087 | 0 | 0.17 | 0.23 | 0.07 | 0 | 0.11 | 0.11 |
| 9 | 108223786 | rs10991661 | 1.336 | 0.288 | 0.189 | -0.895 | 0.024 | -0.048 | -0.048 | 0 | 0.16 | 0.01 | 0.24 | 0.17 | 0.11 | 0.16 |
| 9 | 108224424 | rs10991662 | 1.336 | 0.288 | 0.189 | -0.895 | 0.024 | -0.048 | -0.048 | 0.009 | 0.16 | 0.01 | 0.23 | 0.17 | 0.11 | 0.16 |
| 9 | 108240797 | rs2812310 | -1.885 | 0.25 | 0.177 | -0.886 | 0 | 0.033 | 0.033 | 0 | 0.59 | 0.74 | 0.7 | 0.66 | 0.56 | 0.53 |
| 9 | 108243493 | rs2771045 | 0.832 | 0.175 | 0.281 | -0.894 | -0.036 | 0.026 | 0.026 | 0 | 0.16 | 0.11 | 0.18 | 0.06 | 0.28 | 0.11 |
| 9 | 108244576 | rs16924672 | 0.299 | 0.152 | 0.422 | -0.901 | 0.033 | 0.008 | 0.008 | 0 | 0.22 | 0.11 | 0.25 | 0.17 | 0.16 | 0.18 |
| 9 | 108251400 | rs2812312 | 0.791 | 0.209 | 0.378 | -0.805 | -0.03 | 0.015 | 0.015 | 0 | 0.16 | 0.13 | 0.18 | 0.09 | 0.28 | 0.11 |
| 9 | 108276704 | rs10820813 | -2.406 | -0.009 | 0.396 | -0.849 | -0.011 | 0.014 | 0.014 | 0.074 | 0.66 | 0.93 | 0.74 | 0.69 | 0.63 | 0.57 |
| 9 | 108298183 | rs1327351 | 1.422 | 0.048 | 0.231 | -0.82 | 0.019 | -0.049 | -0.049 | 0 | 0.16 | 0.01 | 0.24 | 0.17 | 0.12 | 0.16 |
| 9 | 108324930 | rs12337780 | 0.609 | 0.06 | 0.265 | -0.779 | -0.004 | 0.009 | 0.009 | 0 | 0.11 | 0.12 | 0.21 | 0.44 | 0.11 | 0.19 |
| 9 | 108350181 | rs2768291 | -2.046 | -0.228 | 0.015 | -1.141 | 0.006 | 0.015 | 0.015 | 0 | 0.63 | 0.76 | 0.73 | 0.67 | 0.62 | 0.56 |
| 9 | 108371337 | rs2518129 | 1.668 | 0.075 | -0.665 | -0.921 | 0.01 | 0.072 | 0.072 | 0 | 0.12 | 0 | 0.08 | 0 | 0.08 | 0.08 |
| 9 | 108373184 | rs2417625 | 0.349 | 0.434 | -0.385 | -0.582 | -0.001 | 0.039 | 0.039 | 0 | 0.11 | 0.16 | 0.22 | 0.44 | 0.11 | 0.18 |
| 9 | 108377097 | rs4742955 | 0.047 | 0.433 | -0.362 | -0.468 | 0.059 | -0.023 | -0.023 | 0 | 0.19 | 0.3 | 0.24 | 0.09 | 0.28 | 0.12 |
| 9 | 108380004 | rs10816284 | -1.013 | 0.433 | -0.385 | -0.467 | 0.042 | 0.032 | 0.032 | 0 | 0.3 | 0.47 | 0.45 | 0.53 | 0.39 | 0.31 |
| 9 | 108380355 | rs17309806 | 0.433 | 0.584 | -0.725 | -0.384 | 0.03 | -0.007 | -0.007 | 0 | 0.16 | 0.21 | 0.22 | 0.09 | 0.27 | 0.1 |
| 9 | 108398678 | rs2768282 | 0.272 | 0.512 | -0.772 | -0.404 | 0.017 | -0.039 | -0.039 | 0.003 | 0.2 | 0.01 | 0.25 | 0.18 | 0.15 | 0.17 |
| 9 | 108399825 | rs1024366 | 0.314 | 0.527 | -0.765 | -0.397 | 0.016 | -0.04 | -0.04 | 0.01 | 0.19 | 0.01 | 0.25 | 0.18 | 0.15 | 0.17 |
| 9 | 108404763 | rs10978179 | 0.28 | 0.562 | -0.74 | -0.386 | -0.003 | 0.013 | 0.013 | 0 | 0.11 | 0.12 | 0.22 | 0.44 | 0.11 | 0.19 |
| 9 | 108405344 | rs1858743 | 0.408 | 0.53 | -0.738 | -0.519 | 0.01 | 0.003 | 0.003 | 0.157 | 0.17 | 0.01 | 0.19 | 0.09 | 0.13 | 0.14 |
| 9 | 108409400 | rs885954 | -0.949 | 0.557 | -0.736 | -0.553 | -0.042 | 0.015 | 0.015 | 0 | 0.36 | 0.22 | 0.47 | 0.27 | 0.42 | 0.26 |
| 9 | 108409614 | rs17251166 | -0.361 | 1.203 | -0.083 | -0.318 | 0.044 | -0.01 | -0.01 | 0 | 0.17 | 0.24 | 0.23 | 0.1 | 0.28 | 0.1 |
| 9 | 108413323 | rs605419 | -0.198 | 1.256 | -0.823 | -0.361 | -0.095 | 0.021 | 0.021 | 0 | 0.14 | 0.01 | 0.15 | 0.06 | 0.27 | 0.06 |
| 9 | 108415556 | rs635712 | -0.949 | 0.759 | -0.249 | 0.18 | 0.051 | 0.072 | 0.072 | 0 | 0.26 | 0.53 | 0.41 | 0.57 | 0.32 | 0.23 |
| 9 | 108417743 | rs567831 | -1.164 | 0.125 | -0.875 | -0.199 | 0.002 | -0.004 | -0.004 | 0 | 0.69 | 0.67 | 0.68 | 0.66 | 0.64 | 0.58 |
| 9 | 108418212 | rs583973 | -0.566 | 0.239 | -0.905 | -0.668 | -0.061 | 0.085 | 0.085 | 0 | 0.31 | 0.1 | 0.3 | 0.03 | 0.43 | 0.16 |
| 9 | 108423366 | rs10816292 | -1.237 | -0.559 | 0.134 | -0.305 | -0.014 | 0.129 | 0.129 | 0 | 0.26 | 0.49 | 0.31 | 0.47 | 0.22 | 0.2 |
| 9 | 108426149 | rs7874737 | -1.443 | -0.691 | 0.013 | -0.622 | 0.019 | 0.047 | 0.047 | 0 | 0.48 | 0.32 | 0.52 | 0.6 | 0.41 | 0.34 |
| 9 | 108426784 | rs1463983 | -1.514 | -0.678 | 0.073 | -0.616 | 0.02 | 0.044 | 0.044 | 0 | 0.49 | 0.37 | 0.52 | 0.6 | 0.41 | 0.34 |
| 9 | 108427062 | rs1463984 | -0.51 | -0.688 | 0.16 | -0.643 | -0.004 | 0.077 | 0.077 | 0 | 0.15 | 0.17 | 0.2 | 0.47 | 0.09 | 0.17 |
| 9 | 108428303 | rs1409922 | 0.483 | -0.611 | 0.036 | -0.639 | 0.014 | 0.039 | 0.039 | 0.025 | 0.26 | 0.08 | 0.23 | 0.12 | 0.19 | 0.16 |
| 9 | 108431783 | rs16924823 | 0.244 | 0.034 | 0.486 | -0.183 | 0.004 | -0.048 | -0.048 | 0 | 0.09 | 0.18 | 0.1 | 0.01 | 0.12 | 0.02 |
| 9 | 108434007 | rs9299108 | -1.325 | 0.018 | 0.417 | -0.088 | 0.002 | 0.119 | 0.119 | 0 | 0.25 | 0.56 | 0.32 | 0.49 | 0.24 | 0.19 |
| 9 | 108442205 | rs7047678 | -0.4 | 0.189 | 0.093 | 0.396 | 0.004 | 0.132 | 0.132 | 0 | 0.06 | 0.09 | 0.19 | 0.44 | 0.08 | 0.12 |
| 9 | 108443332 | rs1567368 | -0.363 | -0.694 | 0.087 | -0.242 | 0.012 | 0.011 | 0.011 | 0.885 | 0.38 | 0.63 | 0.26 | 0.13 | 0.22 | 0.25 |
| 9 | 108449986 | rs10816295 | -0.128 | -0.382 | -0.101 | 0.018 | 0.004 | 0.132 | 0.132 | 0 | 0.06 | 0.09 | 0.19 | 0.44 | 0.08 | 0.12 |
| 9 | 108457806 | rs7039618 | -1.253 | -1.057 | 0.308 | -0.433 | 0.042 | 0.03 | 0.03 | 0.318 | 0.52 | 0.62 | 0.47 | 0.57 | 0.32 | 0.39 |
| 9 | 108468129 | rs3817141 | -1.24 | -1.264 | 1.106 | -0.522 | -0.024 | 0.157 | 0.157 | 1.137 | 0.19 | 0.56 | 0.25 | 0.44 | 0.1 | 0.16 |
| 9 | 108471563 | rs10733555 | -1.305 | -1.362 | 1.107 | -0.583 | 0.067 | 0.049 | 0.049 | 0.466 | 0.5 | 0.67 | 0.47 | 0.57 | 0.32 | 0.38 |
| 9 | 108471885 | rs10978213 | -1.461 | -1.337 | 1.116 | -0.573 | 0.059 | 0.042 | 0.042 | 0.727 | 0.51 | 0.67 | 0.47 | 0.57 | 0.32 | 0.38 |
| 9 | 108474679 | rs10739211 | -1.563 | -1.349 | 1.089 | -0.58 | 0.079 | 0.051 | 0.051 | 1.215 | 0.52 | 0.77 | 0.48 | 0.57 | 0.32 | 0.38 |
| 9 | 108479126 | rs9408800 | -0.601 | -0.516 | 0.792 | -0.708 | -0.079 | 0.01 | 0.01 | 0 | 0.89 | 0.99 | 0.82 | 0.95 | 0.71 | 0.88 |
| 9 | 108504621 | rs10978230 | 0.062 | -0.476 | 0.985 | -0.664 | 0.009 | 0.15 | 0.15 | 0 | 0.07 | 0.11 | 0.2 | 0.44 | 0.08 | 0.13 |
| 9 | 108521693 | rs10759135 | -0.564 | -0.486 | 1.115 | -0.641 | 0.024 | 0.232 | 0.232 | 0 | 0.08 | 0.19 | 0.21 | 0.44 | 0.08 | 0.13 |
| 9 | 108523710 | rs10465120 | 0.356 | -0.56 | 1.061 | -0.706 | 0.027 | 0.03 | 0.03 | 0.298 | 0.32 | 0.21 | 0.23 | 0.13 | 0.21 | 0.22 |
| 9 | 108534505 | rs1979993 | -1.362 | -0.299 | 0.918 | -0.41 | -0.025 | 0.203 | 0.203 | 0.089 | 0.19 | 0.47 | 0.25 | 0.44 | 0.1 | 0.16 |
| 9 | 108535272 | rs10978240 | -0.319 | -0.204 | 0.726 | -0.355 | 0.018 | 0.194 | 0.194 | 0 | 0.07 | 0.14 | 0.2 | 0.44 | 0.08 | 0.13 |
| 9 | 108568402 | rs3010417 | -2.115 | -0.442 | 0.408 | -0.487 | -0.12 | -0.06 | -0.06 | 0.048 | 0.68 | 0.98 | 0.67 | 0.64 | 0.61 | 0.53 |
| 9 | 108583206 | rs10978293 | -2.31 | -0.138 | 1.641 | 0.165 | 0.037 | 0.391 | 0.391 | 0.509 | 0.12 | 0.47 | 0.23 | 0.45 | 0.08 | 0.15 |
| 9 | 108592589 | rs872246 | -2.218 | -0.145 | 1.622 | 0.165 | 0.027 | 0.394 | 0.394 | 0.443 | 0.12 | 0.48 | 0.23 | 0.46 | 0.08 | 0.14 |
| 9 | 108595364 | rs10512341 | -1.505 | -0.133 | 1.625 | 0.172 | 0.017 | 0.255 | 0.255 | 0 | 0.09 | 0.23 | 0.21 | 0.46 | 0.08 | 0.14 |
| 9 | 108596042 | rs7872125 | -1.814 | -0.133 | 1.592 | 0.197 | 0.035 | 0.361 | 0.361 | 0 | 0.1 | 0.33 | 0.22 | 0.46 | 0.08 | 0.14 |
| 9 | 108603039 | rs6477473 | -1.755 | -0.789 | 1.383 | 0.207 | -0.041 | 0.147 | 0.147 | 0.3 | 0.2 | 0.55 | 0.25 | 0.46 | 0.1 | 0.17 |

| | | | | | | | | | | | | | | | | |
|---|---|---|---|---|---|---|---|---|---|---|---|---|---|---|---|---|
| 9 | 108605956 | rs12346407 | -1.01 | -0.633 | 0.716 | 0.333 | 0.006 | 0.156 | 0.156 | 0 | 0.07 | 0.12 | 0.2 | 0.46 | 0.08 | 0.14 |
| 9 | 108608674 | rs3105991 | -1.661 | -0.819 | 0.79 | 0.191 | -0.03 | 0.144 | 0.144 | 0.183 | 0.21 | 0.51 | 0.27 | 0.48 | 0.12 | 0.17 |
| 9 | 108626502 | rs1806664 | -1.268 | -0.678 | 0.639 | 0.313 | -0.001 | 0.269 | 0.269 | 0 | 0.08 | 0.2 | 0.19 | 0.48 | 0.06 | 0.13 |
| 9 | 108626919 | rs2997726 | 0.774 | -0.727 | 0.558 | 0.201 | 0.032 | 0.231 | 0.231 | 0 | 0.83 | 0.64 | 0.76 | 0.52 | 0.86 | 0.88 |
| 9 | 108629422 | rs6477476 | -0.298 | -0.75 | -0.21 | -0.206 | -0.001 | 0.047 | 0.047 | 0.005 | 0.91 | 0.98 | 0.95 | 1 | 0.94 | 0.98 |
| 9 | 108633515 | rs7872318 | -1.476 | -1.282 | 0.158 | 0.155 | -0.04 | 0.049 | 0.049 | 0.561 | 0.29 | 0.55 | 0.32 | 0.54 | 0.14 | 0.27 |
| 9 | 108641808 | rs3010959 | -0.718 | -0.975 | -0.048 | -0.141 | -0.001 | -0.011 | -0.011 | 0 | 0.16 | 0.21 | 0.11 | 0.08 | 0.19 | 0.16 |
| 9 | 108642738 | rs6477480 | 1.708 | -1.724 | -0.189 | -0.592 | 0.025 | 0.024 | 0.024 | 2.285 | 0.63 | 0.32 | 0.78 | 0.92 | 0.71 | 0.8 |
| 9 | 108643176 | rs2417654 | 0.758 | -0.682 | 0.302 | 0.018 | 0.002 | -0.014 | -0.014 | 0.054 | 0.83 | 0.78 | 0.87 | 0.92 | 0.78 | 0.87 |
| 9 | 108648392 | rs13440292 | 0.27 | -0.023 | -0.335 | 0.075 | -0.053 | -0.041 | -0.041 | 0 | 0.1 | 0.01 | 0.17 | 0.14 | 0.2 | 0.14 |
| 9 | 108655096 | rs4742971 | -0.036 | -0.439 | -0.071 | 0.404 | -0.01 | -0.104 | -0.104 | 0 | 0.13 | 0.01 | 0.25 | 0.51 | 0.08 | 0.13 |
| 9 | 108656238 | rs4598317 | 2.398 | -0.459 | -0.068 | 0.33 | 0.028 | -0.057 | -0.057 | 0 | 0.59 | 0.11 | 0.54 | 0.4 | 0.67 | 0.7 |
| 9 | 108663489 | rs10115085 | 2.086 | -0.557 | 0.34 | 0.36 | -0.011 | 0.019 | 0.019 | 0.022 | 0.66 | 0.46 | 0.6 | 0.4 | 0.7 | 0.74 |
| 9 | 108672309 | rs7030289 | -0.332 | -0.493 | 0.429 | 0.363 | -0.03 | 0.335 | 0.335 | 0 | 0.13 | 0.35 | 0.26 | 0.53 | 0.05 | 0.11 |
| 9 | 108674392 | rs2417658 | 2.05 | -0.536 | 0.425 | 0.302 | -0.017 | 0.052 | 0.052 | 0 | 0.69 | 0.49 | 0.61 | 0.46 | 0.73 | 0.78 |
| 9 | 108678284 | rs4742974 | 2.488 | -0.548 | 0.483 | 0.286 | 0.009 | -0.023 | -0.023 | 0.073 | 0.66 | 0.34 | 0.58 | 0.46 | 0.69 | 0.76 |
| 9 | 108680292 | rs2417660 | 2.776 | -0.569 | 0.447 | 0.269 | 0.031 | -0.085 | -0.085 | 1.701 | 0.6 | 0.13 | 0.55 | 0.46 | 0.67 | 0.74 |
| 9 | 108681706 | rs2417662 | 0.625 | -0.454 | 0.491 | 0.026 | -0.014 | 0.016 | 0.016 | 0 | 0.86 | 0.9 | 0.89 | 0.99 | 0.79 | 0.91 |
| 9 | 108683134 | rs1987249 | -0.927 | -0.456 | 0.525 | 0.027 | 0.01 | -0.038 | -0.038 | 4.089 | 0.07 | 0.17 | 0.05 | 0 | 0.06 | 0.03 |
| 9 | 108683365 | rs1987248 | 1.327 | -0.461 | 0.518 | 0.024 | -0.002 | -0.014 | -0.014 | 1.985 | 0.83 | 0.86 | 0.87 | 0.99 | 0.78 | 0.9 |
| 9 | 108684675 | rs10759153 | 2.132 | -0.482 | 0.547 | 0.018 | -0.001 | -0.156 | -0.156 | 0.15 | 0.74 | 0.53 | 0.81 | 0.99 | 0.72 | 0.86 |
| 9 | 108688554 | rs10816354 | -1.217 | -0.051 | -0.139 | 0.067 | -0.025 | 0.028 | 0.028 | 0.363 | 0.23 | 0.47 | 0.39 | 0.72 | 0.19 | 0.26 |
| 9 | 108689648 | rs10978345 | 2.742 | -0.061 | -0.173 | 0.04 | -0.007 | 0.098 | 0.098 | 0.039 | 0.56 | 0.2 | 0.44 | 0.26 | 0.52 | 0.59 |
| 9 | 108694397 | rs4742659 | -2.269 | -0.13 | -0.146 | 0.045 | 0.004 | 0.004 | 0.004 | 0.219 | 0.32 | 0.67 | 0.45 | 0.73 | 0.26 | 0.32 |
| 9 | 108702986 | rs13293907 | -1.421 | -0.476 | -0.018 | 0.167 | 0.014 | -0.043 | -0.043 | 0 | 0.08 | 0.2 | 0.06 | 0 | 0.07 | 0.03 |
| 9 | 108717969 | rs10816356 | -2.107 | -0.474 | -0.374 | 0.173 | 0.013 | 0.007 | 0.007 | 0.388 | 0.3 | 0.7 | 0.43 | 0.73 | 0.24 | 0.31 |
| 9 | 108732469 | rs7866652 | 2.48 | -0.442 | -0.43 | 0.292 | 0 | 0.112 | 0.112 | 0.049 | 0.57 | 0.15 | 0.46 | 0.25 | 0.56 | 0.61 |
| 9 | 108733453 | rs10978362 | -1.27 | -0.374 | -0.254 | 0.469 | -0.043 | 0.018 | 0.018 | 0.51 | 0.22 | 0.5 | 0.37 | 0.73 | 0.16 | 0.25 |
| 9 | 108739271 | rs16925085 | -0.993 | -0.395 | -0.58 | 0.377 | -0.007 | -0.068 | -0.068 | 0 | 0.11 | 0.28 | 0.06 | 0 | 0.09 | 0.04 |
| 9 | 108740661 | rs4336667 | 2.536 | -0.794 | -0.637 | 0.598 | 0.014 | 0.149 | 0.149 | 0.581 | 0.52 | 0.12 | 0.45 | 0.27 | 0.54 | 0.58 |
| 9 | 108742160 | rs10978363 | 0.123 | -0.712 | 0.007 | 1.25 | 0.017 | -0.002 | -0.002 | 0.154 | 0.09 | 0.04 | 0.19 | 0.47 | 0.04 | 0.1 |
| 9 | 108742371 | rs7870421 | -2.202 | -0.311 | 0.279 | 0.435 | -0.01 | -0.082 | -0.082 | 0.48 | 0.15 | 0.33 | 0.09 | 0 | 0.12 | 0.06 |
| 9 | 108749726 | rs12006289 | -0.893 | -0.052 | -0.013 | 0.588 | 0.008 | -0.042 | -0.042 | 0 | 0.08 | 0.23 | 0.05 | 0 | 0.09 | 0.03 |
| 9 | 108759480 | rs7867860 | -1.399 | 0.318 | 0.13 | 1.334 | 0.027 | 0.068 | 0.068 | 0.126 | 0.12 | 0.24 | 0.19 | 0.22 | 0.16 | 0.06 |
| 9 | 108761012 | rs10759158 | -2.075 | 0.274 | 0.324 | 1.429 | 0.031 | 0.095 | 0.095 | 0.041 | 0.14 | 0.31 | 0.21 | 0.25 | 0.17 | 0.1 |
| 9 | 108779678 | rs2417682 | 1.833 | -0.408 | -0.259 | 1.297 | -0.001 | 0.06 | 0.06 | 1.443 | 0.53 | 0.17 | 0.51 | 0.38 | 0.57 | 0.63 |
| 9 | 108785618 | rs9886710 | -1.563 | -0.534 | 0.077 | 1.169 | 0.006 | 0.023 | 0.023 | 0 | 0.26 | 0.4 | 0.36 | 0.61 | 0.28 | 0.22 |
| 9 | 108791327 | rs10816368 | -0.748 | -0.547 | 0.105 | 1.178 | 0.066 | 0.155 | 0.155 | 0.194 | 0.07 | 0.2 | 0.17 | 0.22 | 0.14 | 0.06 |
| 9 | 108793429 | rs4255200 | 0.259 | -0.644 | 0.585 | 1.107 | -0.012 | -0.116 | -0.116 | 0 | 0.81 | 0.7 | 0.89 | 0.96 | 0.85 | 0.87 |
| 9 | 108798258 | rs7468493 | -2.265 | -0.556 | 0.636 | 1.473 | 0.008 | 0.043 | 0.043 | 0 | 0.29 | 0.52 | 0.39 | 0.59 | 0.3 | 0.24 |
| 9 | 108807861 | rs4742661 | -0.663 | -0.556 | 0.516 | 1.499 | 0.007 | -0.008 | -0.008 | 0.059 | 0.14 | 0.13 | 0.18 | 0.37 | 0.13 | 0.15 |
| 9 | 108808480 | rs7852935 | -0.612 | -0.595 | 0.338 | 1.541 | -0.038 | -0.034 | -0.034 | 0.003 | 0.06 | 0.01 | 0.15 | 0.17 | 0.12 | 0.06 |
| 9 | 108814037 | rs10816370 | -2.364 | -0.615 | 0.393 | 1.396 | 0.002 | 0.012 | 0.012 | 0 | 0.29 | 0.36 | 0.37 | 0.59 | 0.3 | 0.24 |
| 9 | 108817987 | rs4418395 | -2.352 | -0.575 | 0.387 | 1.402 | 0.002 | 0.03 | 0.03 | 0 | 0.29 | 0.45 | 0.39 | 0.59 | 0.3 | 0.24 |
| 9 | 108822093 | rs10116405 | -2.282 | -0.566 | 0.402 | 1.401 | 0.03 | 0.073 | 0.073 | 0.12 | 0.16 | 0.39 | 0.22 | 0.23 | 0.18 | 0.11 |
| 9 | 108823761 | rs4128998 | -1.222 | -0.448 | 0.232 | 1.416 | 0.031 | 0.059 | 0.059 | 0 | 0.11 | 0.23 | 0.2 | 0.2 | 0.15 | 0.06 |
| 9 | 108827402 | rs4587393 | -0.751 | -0.349 | -0.165 | 1.489 | 0.007 | -0.027 | -0.027 | 0 | 0.15 | 0.08 | 0.18 | 0.37 | 0.14 | 0.15 |
| 9 | 108838212 | rs13299446 | -0.315 | -0.618 | 0.685 | 0.48 | -0.041 | -0.096 | -0.096 | 0 | 0.14 | 0.32 | 0.06 | 0.02 | 0.06 | 0.09 |
| 9 | 108839164 | rs10512343 | -2.032 | -0.825 | 0.717 | 0.716 | 0.011 | -0.003 | -0.003 | 0.007 | 0.33 | 0.64 | 0.19 | 0.39 | 0.23 | 0.27 |
| 9 | 108846468 | rs971416 | -0.779 | -1.275 | 0.279 | 0.641 | -0.02 | 0.073 | 0.073 | 0 | 0.26 | 0.37 | 0.27 | 0.37 | 0.2 | 0.19 |
| 9 | 108854018 | rs10978413 | -0.295 | -1.352 | 0.063 | 0.567 | -0.012 | -0.079 | -0.079 | 0 | 0.13 | 0.01 | 0.09 | 0.34 | 0.13 | 0.15 |
| 9 | 108857533 | rs10978416 | 0.062 | -1.348 | 0.065 | 0.569 | -0.015 | -0.07 | -0.07 | 0 | 0.11 | 0.01 | 0.08 | 0.33 | 0.13 | 0.14 |
| 9 | 108862046 | rs9409077 | -2.3 | -1.389 | 0.192 | 0.501 | 0.043 | 0.068 | 0.068 | 0 | 0.44 | 0.66 | 0.39 | 0.71 | 0.33 | 0.34 |
| 9 | 108866460 | rs885578 | 0.172 | -1.359 | 0.467 | 0.485 | -0.035 | -0.089 | -0.089 | 0 | 0.14 | 0.29 | 0.06 | 0.04 | 0.06 | 0.1 |
| 9 | 108868577 | rs925837 | 2.492 | -1.617 | -0.001 | 0.164 | 0.009 | 0.179 | 0.179 | 0.035 | 0.48 | 0.13 | 0.56 | 0.26 | 0.63 | 0.6 |
| 9 | 108877645 | rs1516893 | 0.851 | -1.573 | 0.114 | 0.16 | 0.01 | -0.059 | -0.059 | 0.659 | 0.88 | 0.79 | 0.9 | 0.99 | 0.87 | 0.94 |
| 9 | 108878030 | rs9299118 | 0.728 | -1.586 | -0.051 | 0.25 | -0.033 | -0.071 | -0.071 | 0.014 | 0.11 | 0.01 | 0.2 | 0.33 | 0.11 | 0.07 |
| 9 | 108878507 | rs12236453 | 0.228 | -1.554 | -0.016 | 0.229 | 0.012 | -0.016 | -0.016 | 0 | 0.08 | 0.02 | 0.03 | 0.32 | 0.05 | 0.11 |
| 9 | 108882218 | rs4742987 | 2.189 | -1.623 | -0.054 | 0.031 | 0.036 | 0.176 | 0.176 | 0.832 | 0.53 | 0.13 | 0.58 | 0.26 | 0.65 | 0.61 |
| 9 | 108893840 | rs1120147 | 1.16 | -1.656 | 0.255 | 0.093 | 0.021 | -0.069 | -0.069 | 0 | 0.87 | 0.74 | 0.89 | 0.99 | 0.84 | 0.94 |
| 9 | 108893850 | rs1120146 | 2.28 | -1.65 | -0.428 | -0.128 | -0.082 | 0.06 | 0.06 | 3.475 | 0.38 | 0.12 | 0.48 | 0.27 | 0.46 | 0.57 |
| 9 | 108895689 | rs4742988 | 2.235 | -1.672 | -0.581 | -0.118 | -0.132 | 0.087 | 0.087 | 2.313 | 0.39 | 0.07 | 0.5 | 0.25 | 0.49 | 0.52 |
| 9 | 108898935 | rs1400438 | 0.966 | -1.377 | 0.066 | 0.225 | -0.008 | -0.044 | -0.044 | 0 | 0.76 | 0.42 | 0.85 | 0.87 | 0.8 | 0.8 |
| 9 | 108901049 | rs9409082 | -0.331 | -2.039 | 0.125 | 0.657 | -0.058 | -0.119 | -0.119 | 0.219 | 0.21 | 0.04 | 0.27 | 0.31 | 0.26 | 0.13 |
| 9 | 108901466 | rs9409083 | -0.048 | -2.074 | 0.123 | 0.541 | -0.049 | -0.13 | -0.13 | 3.115 | 0.18 | 0.02 | 0.25 | 0.33 | 0.21 | 0.12 |
| 9 | 108902045 | rs1516885 | 0.195 | -2.259 | 0.641 | 0.4 | -0.031 | 0.012 | 0.012 | 4.213 | 0.14 | 0.41 | 0.06 | 0.14 | 0.06 | 0.14 |
| 9 | 108912911 | rs1516882 | -1.018 | -2.293 | 0.651 | 0.398 | -0.001 | 0.001 | 0.001 | 0 | 0.31 | 0.32 | 0.34 | 0.42 | 0.32 | 0.16 |
| 9 | 108916895 | rs7038938 | -0.226 | -2.357 | 0.712 | 0.406 | -0.087 | 0.073 | 0.073 | 1.919 | 0.2 | 0.46 | 0.08 | 0.34 | 0.08 | 0.25 |
| 9 | 108923289 | rs12353538 | 0.726 | -2.182 | 0.707 | 0.416 | -0.011 | -0.024 | -0.024 | 0 | 0.05 | 0.1 | 0.04 | 0 | 0.03 | 0.01 |

| | | | | | | | | | | | | | | | | |
|---|---|---|---|---|---|---|---|---|---|---|---|---|---|---|---|---|
| 9 | 108923457 | rs2417687 | -2.539 | -2.314 | 0.473 | 0.351 | -0.097 | 0.126 | 0.126 | 2.207 | 0.6 | 0.93 | 0.52 | 0.78 | 0.48 | 0.48 |
| 9 | 108926464 | rs13300395 | -0.738 | -1.856 | 0.925 | 0.585 | 0.002 | 0.02 | 0.02 | 0 | 0.27 | 0.3 | 0.31 | 0.41 | 0.29 | 0.17 |
| 9 | 108927068 | rs7028916 | -0.748 | -1.91 | 0.969 | 0.585 | 0.003 | 0.033 | 0.033 | 0 | 0.27 | 0.33 | 0.31 | 0.41 | 0.29 | 0.17 |
| 9 | 108981298 | rs7862517 | -0.243 | -1.892 | 0.73 | 0.614 | -0.075 | -0.141 | -0.141 | 0 | 0.23 | 0.04 | 0.29 | 0.37 | 0.29 | 0.13 |
| 9 | 109004064 | rs12005566 | -1.054 | -2.549 | 1.165 | 0.161 | -0.028 | -0.004 | -0.004 | 1.125 | 0.1 | 0.28 | 0.05 | 0.05 | 0.04 | 0.02 |
| 9 | 109010799 | rs1484372 | 0.873 | -2.625 | 1.738 | 0.211 | -0.029 | 0.027 | 0.027 | 0 | 0.7 | 0.48 | 0.65 | 0.45 | 0.76 | 0.86 |
| 9 | 109014185 | rs2627741 | -0.177 | -2.604 | 1.406 | 0.213 | -0.012 | -0.101 | -0.101 | 0 | 0.79 | 0.96 | 0.71 | 0.51 | 0.79 | 0.88 |
| 9 | 109018525 | rs2417688 | 1.686 | -2.572 | 1.74 | 0.088 | 0.03 | 0.081 | 0.081 | 5.805 | 0.44 | 0.28 | 0.54 | 0.24 | 0.59 | 0.59 |
| 9 | 109043885 | rs16936272 | -0.876 | -1.597 | 1.666 | 0.453 | -0.016 | 0.018 | 0.018 | 0 | 0.08 | 0.3 | 0.05 | 0.06 | 0.03 | 0.02 |
| 9 | 109046007 | rs12344579 | 0.013 | -1.508 | 0.966 | 0.489 | -0.007 | -0.017 | -0.017 | 0 | 0.06 | 0.08 | 0.03 | 0 | 0.03 | 0.01 |
| 9 | 109065718 | rs7043293 | 0.892 | -1.554 | 0.51 | 0.307 | 0.002 | 0.018 | 0.018 | 0.015 | 0.7 | 0.66 | 0.64 | 0.46 | 0.68 | 0.85 |
| 9 | 109067561 | rs1484375 | 0.57 | -1.575 | 0.342 | 0.31 | 0 | -0.05 | -0.05 | 1.894 | 0.75 | 0.88 | 0.67 | 0.47 | 0.77 | 0.9 |
| 9 | 109088848 | rs12683342 | -0.672 | -1.604 | 0.281 | 0.282 | -0.021 | -0.067 | -0.067 | 0 | 0.24 | 0.09 | 0.31 | 0.53 | 0.28 | 0.13 |
| 9 | 109094026 | rs1876376 | -0.23 | -1.582 | 0.123 | 0.306 | 0 | -0.01 | -0.01 | 0 | 0.21 | 0.12 | 0.3 | 0.54 | 0.2 | 0.1 |
| 9 | 109104850 | rs12686474 | -0.664 | -1.559 | -0.421 | 0.313 | -0.027 | -0.079 | -0.079 | 0 | 0.24 | 0.09 | 0.31 | 0.53 | 0.28 | 0.13 |
| 9 | 109118797 | rs6477505 | 0.283 | -1.598 | -0.486 | 0.214 | -0.001 | 0.001 | 0.001 | 0 | 0.9 | 0.92 | 0.95 | 0.99 | 0.88 | 0.95 |
| 9 | 109138185 | rs13293802 | -1.084 | -2.074 | -0.305 | 0.146 | -0.03 | 0.041 | 0.041 | 0.407 | 0.31 | 0.45 | 0.37 | 0.54 | 0.27 | 0.14 |
| 9 | 109147362 | rs955062 | -1.211 | -1.91 | -0.221 | 0.171 | -0.024 | -0.03 | -0.03 | 0 | 0.36 | 0.43 | 0.34 | 0.78 | 0.22 | 0.33 |
| 9 | 109149614 | rs10759178 | 1.415 | -1.892 | -0.325 | 0.291 | 0 | -0.007 | -0.007 | 0 | 0.55 | 0.42 | 0.62 | 0.22 | 0.67 | 0.64 |
| 9 | 109150272 | rs7846958 | -0.935 | -1.847 | 0.344 | 0.708 | -0.108 | 0.068 | 0.068 | 4.432 | 0.16 | 0.45 | 0.06 | 0.25 | 0.02 | 0.22 |
| 9 | 109153027 | rs10759179 | -0.864 | -1.216 | -0.492 | 0.564 | -0.011 | -0.006 | -0.006 | 0 | 0.29 | 0.3 | 0.46 | 0.76 | 0.21 | 0.27 |
| 9 | 109154423 | rs10816387 | 0.779 | -1.334 | -2.011 | 0.775 | 0.029 | -0.11 | -0.11 | 0.651 | 0.45 | 0.05 | 0.46 | 0.57 | 0.66 | 0.53 |
| 9 | 109154569 | rs7857609 | -0.275 | -1.98 | -1.651 | 0.929 | -0.021 | 0.006 | 0.006 | 0 | 0.1 | 0.14 | 0.08 | 0.12 | 0.03 | 0.05 |
| 9 | 109155467 | rs6477511 | -0.504 | -2.367 | -1.575 | 0.826 | 0.005 | 0.002 | 0.002 | 0 | 0.88 | 0.86 | 0.75 | 0.92 | 0.95 | 0.92 |
| 9 | 109158155 | rs12352728 | -0.095 | -2.778 | -1.038 | 0.702 | -0.018 | -0.014 | -0.014 | 0 | 0.15 | 0.16 | 0.1 | 0.04 | 0.07 | 0.07 |
| 9 | 109160164 | rs7866564 | -0.577 | -2.667 | -1.13 | 0.751 | 0.004 | 0.002 | 0.002 | 0 | 0.88 | 0.85 | 0.75 | 0.91 | 0.95 | 0.91 |
| 9 | 109162816 | rs10759180 | 1.973 | -2.581 | -1.287 | 0.922 | 0.046 | -0.094 | -0.094 | 2.535 | 0.46 | 0.04 | 0.44 | 0.58 | 0.68 | 0.56 |
| 9 | 109164956 | rs10739222 | 1.928 | -2.593 | -1.282 | 0.915 | 0.057 | -0.085 | -0.085 | 0 | 0.46 | 0.05 | 0.45 | 0.59 | 0.69 | 0.61 |
| 9 | 109166421 | rs10759181 | 2.072 | -2.11 | -1.404 | 1.524 | -0.045 | 0.028 | 0.028 | 0.74 | 0.44 | 0.04 | 0.34 | 0.36 | 0.61 | 0.46 |
| 9 | 109178575 | rs4434671 | -1.86 | -1.526 | -0.44 | 1.491 | -0.003 | -0.003 | -0.003 | 0 | 0.43 | 0.56 | 0.37 | 0.58 | 0.38 | 0.53 |
| 9 | 109179510 | rs13293858 | 0.009 | -1.675 | -0.404 | 1.575 | 0.006 | -0.001 | -0.001 | 0.182 | 0.1 | 0.08 | 0.08 | 0.1 | 0.05 | 0.04 |
| 9 | 109179823 | rs11794472 | 1.046 | -1.713 | -0.519 | 1.627 | 0.021 | -0.012 | -0.012 | 0 | 0.07 | 0.01 | 0.04 | 0.03 | 0.04 | 0.02 |
| 9 | 109181134 | rs17291025 | -0.392 | -1.871 | -0.19 | 1.778 | -0.013 | -0.005 | -0.005 | 0 | 0.14 | 0.11 | 0.12 | 0.16 | 0.18 | 0.2 |
| 9 | 109181911 | rs902143 | -2.618 | -1.347 | -0.502 | 1.658 | 0.005 | 0.041 | 0.041 | 0 | 0.45 | 0.69 | 0.42 | 0.59 | 0.41 | 0.56 |
| 9 | 109182762 | rs7866482 | -0.557 | -1.367 | -0.54 | 1.642 | 0.001 | 0.003 | 0.003 | 0 | 0.9 | 1 | 0.77 | 0.92 | 0.95 | 0.91 |
| 9 | 109184595 | rs7041138 | -2.086 | -1.039 | -0.351 | 1.435 | 0.002 | 0.024 | 0.024 | 0 | 0.35 | 0.52 | 0.33 | 0.53 | 0.33 | 0.47 |
| 9 | 109185446 | rs10113989 | -0.499 | -1.06 | -0.164 | 1.403 | 0.002 | -0.029 | -0.029 | 0 | 0.1 | 0.19 | 0.1 | 0.04 | 0.08 | 0.08 |
| 9 | 109185593 | rs10114641 | -1.05 | -1.045 | -0.156 | 1.415 | -0.014 | -0.015 | -0.015 | 0.556 | 0.17 | 0.2 | 0.13 | 0.07 | 0.12 | 0.1 |
| 9 | 109189050 | rs7042670 | 0.38 | -0.964 | -0.201 | 1.428 | -0.057 | -0.049 | -0.049 | 0 | 0.11 | 0.01 | 0.1 | 0.14 | 0.14 | 0.14 |
| 9 | 109189121 | rs6477516 | -2.151 | -0.5 | -0.328 | 1.348 | -0.007 | 0 | 0 | 0.02 | 0.34 | 0.55 | 0.29 | 0.32 | 0.32 | 0.33 |
| 9 | 109190563 | rs10978480 | -2.055 | -0.555 | -0.321 | 1.322 | 0.003 | -0.004 | -0.004 | 0 | 0.37 | 0.5 | 0.35 | 0.36 | 0.38 | 0.37 |
| 9 | 109193717 | rs1352593 | -2.195 | -0.577 | -0.344 | 1.308 | 0.031 | 0.014 | 0.014 | 0 | 0.35 | 0.5 | 0.42 | 0.58 | 0.43 | 0.53 |
| 9 | 109196180 | rs13294274 | 0.251 | -0.544 | -0.343 | 1.297 | 0.013 | -0.001 | -0.001 | 0.143 | 0.1 | 0.07 | 0.08 | 0.1 | 0.05 | 0.04 |
| 9 | 109197082 | rs7024043 | -2.998 | -0.56 | -0.319 | 1.275 | -0.009 | 0.058 | 0.058 | 0 | 0.48 | 0.7 | 0.49 | 0.59 | 0.49 | 0.6 |
| 9 | 109197883 | rs10978485 | -1.568 | -0.491 | -0.285 | 1.312 | 0.039 | -0.002 | -0.002 | 0 | 0.28 | 0.44 | 0.27 | 0.25 | 0.35 | 0.3 |
| 9 | 109201514 | rs16925492 | -0.608 | -0.073 | -0.39 | 1.35 | -0.018 | -0.023 | -0.023 | 0 | 0.11 | 0.08 | 0.16 | 0.28 | 0.17 | 0.29 |
| 9 | 109205296 | rs4742996 | -2.109 | -0.171 | -0.4 | 0.98 | -0.012 | 0.052 | 0.052 | 0 | 0.47 | 0.69 | 0.48 | 0.59 | 0.47 | 0.59 |
| 9 | 109211561 | rs2417709 | -2.076 | 0.624 | -0.618 | 0.365 | 0.038 | 0.03 | 0.03 | 0.882 | 0.52 | 0.75 | 0.63 | 0.42 | 0.58 | 0.61 |
| 9 | 109211719 | rs2417710 | -1.092 | 1.108 | -0.078 | 0.25 | -0.002 | -0.002 | -0.002 | 0 | 0.32 | 0.29 | 0.4 | 0.19 | 0.36 | 0.34 |
| 9 | 109215469 | rs4742669 | -2.388 | 1.305 | -0.456 | 0.043 | -0.032 | -0.007 | -0.007 | 0 | 0.42 | 0.76 | 0.54 | 0.41 | 0.45 | 0.57 |
| 9 | 109216259 | rs12552379 | -0.661 | 1.397 | -0.447 | 0.209 | -0.041 | -0.034 | -0.034 | 0 | 0.07 | 0.02 | 0.16 | 0.21 | 0.14 | 0.26 |
| 9 | 109218799 | rs7855719 | -2.109 | 1.189 | -0.076 | -0.279 | 0.001 | 0.044 | 0.044 | 0.841 | 0.43 | 0.64 | 0.45 | 0.21 | 0.42 | 0.36 |
| 9 | 109222859 | rs17291806 | -1.257 | 1.371 | -0.171 | -0.152 | -0.003 | 0.004 | 0.004 | 0.441 | 0.33 | 0.31 | 0.39 | 0.21 | 0.36 | 0.3 |
| 9 | 109230037 | rs989980 | -1.981 | 2.124 | -0.474 | -0.038 | -0.021 | 0.031 | 0.031 | 0 | 0.43 | 0.76 | 0.55 | 0.31 | 0.47 | 0.48 |
| 9 | 109233625 | rs10512345 | -0.011 | 2.164 | -0.638 | 0.1 | -0.041 | -0.031 | -0.031 | 0 | 0.07 | 0.02 | 0.16 | 0.19 | 0.15 | 0.22 |
| 9 | 109233842 | rs13288836 | -0.385 | 2.124 | -0.269 | -0.142 | -0.004 | 0.01 | 0.01 | 0.243 | 0.25 | 0.24 | 0.35 | 0.18 | 0.3 | 0.21 |
| 9 | 109240561 | rs16925511 | 0.074 | 1.255 | -0.428 | -0.157 | -0.017 | -0.002 | -0.002 | 0 | 0.11 | 0.03 | 0.12 | 0.1 | 0.11 | 0.11 |
| 9 | 109252423 | rs1387595 | -1.288 | 1.541 | -0.285 | -0.075 | -0.002 | 0.001 | 0.001 | 0.548 | 0.46 | 0.75 | 0.61 | 0.41 | 0.56 | 0.53 |
| 9 | 109260333 | rs12338993 | -1.232 | 1.477 | 0.021 | 0.091 | -0.001 | 0 | 0 | 0.331 | 0.42 | 0.41 | 0.58 | 0.39 | 0.55 | 0.53 |
| 9 | 109265076 | rs6477523 | -1.247 | 1.257 | -0.53 | 0.615 | -0.013 | -0.002 | -0.002 | 0 | 0.4 | 0.66 | 0.57 | 0.39 | 0.48 | 0.49 |
| 9 | 109267590 | rs16925523 | -1.316 | 0.876 | -0.478 | 0.584 | -0.01 | 0.015 | 0.015 | 0 | 0.1 | 0.03 | 0.17 | 0.07 | 0.1 | 0.12 |
| 9 | 109270732 | rs1472029 | 0.604 | 0.581 | -0.393 | 0.651 | 0.026 | 0.031 | 0.031 | 0 | 0.59 | 0.29 | 0.56 | 0.67 | 0.69 | 0.62 |
| 9 | 109273253 | rs10978504 | -1.016 | 0.66 | -0.015 | 1.117 | -0.004 | 0.004 | 0.004 | 0 | 0.42 | 0.48 | 0.44 | 0.44 | 0.38 | 0.56 |
| 9 | 109274146 | rs9696881 | -0.345 | 0.671 | -0.489 | 1.198 | 0.02 | 0.105 | 0.105 | 0.127 | 0.11 | 0.24 | 0.19 | 0.26 | 0.15 | 0.25 |
| 9 | 109274388 | rs7873699 | 1.457 | 0.785 | -0.556 | 1.117 | -0.118 | -0.017 | -0.017 | 0 | 0.47 | 0.03 | 0.35 | 0.42 | 0.54 | 0.35 |
| 9 | 109275588 | rs6477524 | 0.265 | 0.779 | 0.665 | 1.469 | 0.02 | 0.055 | 0.055 | 0 | 0.92 | 0.73 | 0.83 | 0.87 | 0.94 | 0.92 |
| 9 | 109277184 | rs1602111 | -0.912 | 0.772 | 0.545 | 1.49 | 0.014 | 0.078 | 0.078 | 0.219 | 0.11 | 0.25 | 0.19 | 0.25 | 0.14 | 0.25 |
| 9 | 109290849 | rs7847257 | 0.216 | 0.737 | 0.575 | 1.417 | 0.005 | 0.023 | 0.023 | 0 | 0.92 | 0.84 | 0.83 | 0.87 | 0.93 | 0.92 |
| 9 | 109291197 | rs16925535 | -0.72 | 0.573 | 0.389 | 1.396 | 0 | 0.017 | 0.017 | 0.611 | 0.13 | 0.18 | 0.16 | 0.23 | 0.14 | 0.24 |

| | | | | | | | | | | | | | | | | |
|---|---|---|---|---|---|---|---|---|---|---|---|---|---|---|---|---|
| 9 | 109291610 | rs1532115 | 0.501 | 0.509 | 0.226 | 0.919 | 0.021 | 0.058 | 0.058 | 0 | 0.91 | 0.72 | 0.83 | 0.88 | 0.94 | 0.92 |
| 9 | 109292568 | rs931425 | 0.382 | 0.375 | -0.734 | 0.81 | 0.022 | 0.009 | 0.009 | 0 | 0.67 | 0.2 | 0.58 | 0.73 | 0.75 | 0.65 |
| 9 | 109296536 | rs7018699 | -0.702 | 0.731 | -0.628 | 0.58 | -0.001 | 0.042 | 0.042 | 0 | 0.52 | 0.68 | 0.62 | 0.64 | 0.51 | 0.63 |
| 9 | 109297588 | rs7042522 | -0.47 | 0.439 | -0.547 | 0.491 | 0.003 | -0.005 | -0.005 | 0 | 0.52 | 0.65 | 0.45 | 0.44 | 0.58 | 0.53 |
| 9 | 109298040 | rs10978516 | -1.086 | -0.327 | -0.046 | -0.016 | -0.022 | -0.04 | -0.04 | 0 | 0.17 | 0.28 | 0.07 | 0.03 | 0.11 | 0.34 |
| 9 | 109299254 | rs10978518 | -1.116 | -0.394 | 0.515 | -0.022 | -0.02 | -0.035 | -0.035 | 0 | 0.1 | 0.27 | 0.05 | 0.03 | 0.08 | 0.2 |
| 9 | 109308266 | rs10978525 | -0.845 | -0.365 | 0.577 | -0.028 | -0.007 | -0.026 | -0.026 | 0 | 0.08 | 0.12 | 0.04 | 0.01 | 0.07 | 0.19 |
| 9 | 109312922 | rs7865489 | -1.476 | -0.579 | 0.482 | -0.079 | 0.026 | 0.021 | 0.021 | 0.272 | 0.48 | 0.71 | 0.33 | 0.42 | 0.43 | 0.66 |
| 9 | 109317378 | rs10978534 | -0.385 | -0.588 | 0.668 | -0.085 | 0.004 | 0.013 | 0.013 | 0 | 0.3 | 0.32 | 0.26 | 0.37 | 0.31 | 0.28 |
| 9 | 109319711 | rs10978535 | -0.844 | -0.631 | 0.72 | -0.028 | 0.002 | 0.007 | 0.007 | 0 | 0.34 | 0.52 | 0.28 | 0.38 | 0.34 | 0.35 |
| 9 | 109321284 | rs10759187 | -0.596 | -0.561 | 0.193 | -0.571 | -0.003 | 0.002 | 0.002 | 0 | 0.35 | 0.32 | 0.29 | 0.32 | 0.38 | 0.25 |
| 9 | 109323247 | rs962395 | -0.19 | -0.61 | 0.363 | -0.367 | -0.003 | 0.019 | 0.019 | 0 | 0.27 | 0.35 | 0.23 | 0.39 | 0.26 | 0.25 |
| 9 | 109323457 | rs6477527 | -2.059 | -0.891 | 0.336 | -0.558 | 0.072 | 0.003 | 0.003 | 0.972 | 0.51 | 0.88 | 0.32 | 0.43 | 0.39 | 0.66 |
| 9 | 109325757 | rs10115173 | -0.726 | -1.041 | 0.389 | -0.538 | 0.008 | 0.006 | 0.006 | 0 | 0.23 | 0.21 | 0.15 | 0.16 | 0.15 | 0.29 |
| 9 | 109327591 | rs4743005 | 0.033 | -0.974 | 0.519 | -0.505 | 0.005 | 0.016 | 0.016 | 0 | 0.16 | 0.23 | 0.12 | 0.26 | 0.19 | 0.19 |
| 9 | 109339950 | rs7024815 | -1.382 | -1.009 | 0.438 | -0.517 | 0.053 | 0.104 | 0.104 | 1.395 | 0.07 | 0.43 | 0.04 | 0.07 | 0.03 | 0.16 |
| 9 | 109344847 | rs1489667 | -1.541 | -1.051 | 0.5 | -0.535 | 0.046 | 0.11 | 0.11 | 1.623 | 0.07 | 0.44 | 0.04 | 0.08 | 0.03 | 0.16 |
| 9 | 109345993 | rs10978550 | 0.034 | -1.043 | 0.414 | -0.543 | -0.064 | -0.046 | -0.046 | 0 | 0.15 | 0.01 | 0.12 | 0.17 | 0.22 | 0.13 |
| 9 | 109348980 | rs1907438 | 0.042 | -1.034 | 0.499 | -0.594 | 0.009 | 0.01 | 0.01 | 0.075 | 0.1 | 0.1 | 0.03 | 0.02 | 0.05 | 0.23 |
| 9 | 109350497 | rs10759188 | -1.426 | -1 | 0.476 | -0.585 | 0 | 0 | 0 | 0.031 | 0.39 | 0.42 | 0.26 | 0.34 | 0.34 | 0.46 |
| 9 | 109362226 | rs7871327 | -2.134 | -1.04 | 0.361 | -0.57 | 0.069 | 0.019 | 0.019 | 1.572 | 0.48 | 0.86 | 0.31 | 0.41 | 0.38 | 0.64 |
| 9 | 109364146 | rs10816399 | -0.145 | -1.204 | 0.433 | -0.445 | 0 | 0 | 0 | 0 | 0.25 | 0.28 | 0.21 | 0.3 | 0.26 | 0.18 |
| 9 | 109365223 | rs10816400 | -0.069 | -1.197 | 0.398 | -0.518 | 0.01 | 0.011 | 0.011 | 0.059 | 0.1 | 0.07 | 0.03 | 0 | 0.05 | 0.22 |
| 9 | 109373732 | rs12379601 | -0.392 | -1.233 | 0.374 | -0.535 | 0.005 | 0.002 | 0.002 | 0 | 0.11 | 0.11 | 0.03 | 0.04 | 0.05 | 0.27 |
| 9 | 109379687 | rs10978560 | -1.859 | -1.473 | 0.591 | -0.647 | -0.068 | -0.072 | -0.072 | 0 | 0.24 | 0.44 | 0.09 | 0.1 | 0.13 | 0.43 |
| 9 | 109386686 | rs12350093 | 0.468 | -1.27 | 0.335 | -0.7 | -0.006 | 0.008 | 0.008 | 0 | 0.1 | 0.07 | 0.06 | 0.05 | 0.11 | 0.07 |
| 9 | 109389147 | rs11792106 | -0.497 | -1.196 | 0.673 | -0.642 | -0.001 | 0.003 | 0.003 | 0 | 0.28 | 0.32 | 0.23 | 0.33 | 0.31 | 0.23 |
| 9 | 109390820 | rs984551 | -2.756 | -1.193 | -0.048 | -0.673 | -0.137 | -0.066 | -0.066 | 0.066 | 0.59 | 0.94 | 0.37 | 0.41 | 0.51 | 0.66 |
| 9 | 109391774 | rs2220623 | 0.058 | -1.203 | 0.108 | -0.676 | 0.001 | -0.005 | -0.005 | 0 | 0.1 | 0.11 | 0.03 | 0.02 | 0.05 | 0.21 |
| 9 | 109393881 | rs7865945 | 0.122 | -1.201 | 0.094 | -0.672 | -0.006 | -0.015 | -0.015 | 0.695 | 0.1 | 0.12 | 0.03 | 0.04 | 0.05 | 0.25 |
| 9 | 109393957 | rs12002108 | -0.231 | -1.176 | 0.145 | -0.65 | 0.005 | 0.03 | 0.03 | 0 | 0.24 | 0.39 | 0.21 | 0.3 | 0.26 | 0.15 |
| 9 | 109394613 | rs1491113 | -2.251 | -1.021 | -0.746 | -0.555 | -0.129 | -0.057 | -0.057 | 0.188 | 0.57 | 0.96 | 0.37 | 0.41 | 0.5 | 0.66 |
| 9 | 109403237 | rs7874140 | -0.934 | -0.874 | -0.051 | -0.552 | -0.027 | -0.048 | -0.048 | 0 | 0.08 | 0.17 | 0.03 | 0.01 | 0.04 | 0.16 |
| 9 | 109403758 | rs7044278 | -0.525 | -1.073 | 0.1 | -0.391 | -0.004 | 0.037 | 0.037 | 0.242 | 0.27 | 0.51 | 0.23 | 0.33 | 0.26 | 0.18 |
| 9 | 109404422 | rs12349610 | 0.447 | -0.985 | 0.057 | -0.461 | -0.009 | 0.023 | 0.023 | 0 | 0.1 | 0.04 | 0.06 | 0 | 0.1 | 0.05 |
| 9 | 109405667 | rs2417753 | -1.035 | -0.97 | 0.127 | -0.461 | -0.015 | -0.111 | -0.111 | 1.323 | 0.2 | 0.31 | 0.12 | 0.01 | 0.15 | 0.22 |
| 9 | 109410667 | rs7855642 | -0.567 | -0.692 | 0.226 | -0.054 | -0.002 | 0.058 | 0.058 | 0.437 | 0.24 | 0.49 | 0.21 | 0.3 | 0.25 | 0.16 |
| 9 | 109417359 | rs2417713 | -1.056 | -0.691 | -0.677 | -0.096 | 0.008 | 0.017 | 0.017 | 0.763 | 0.41 | 0.73 | 0.38 | 0.39 | 0.42 | 0.46 |
| 9 | 109423202 | rs10118378 | -1.118 | -0.715 | -0.528 | -0.097 | 0.004 | 0.011 | 0.011 | 0.721 | 0.4 | 0.67 | 0.37 | 0.39 | 0.42 | 0.46 |
| 9 | 109426449 | rs10978589 | 0.239 | -0.369 | -0.906 | -0.236 | 0.003 | -0.013 | -0.013 | 0.009 | 0.07 | 0.08 | 0.04 | 0 | 0.07 | 0.07 |
| 9 | 109427261 | rs7864733 | -2.009 | -0.529 | -1.01 | -0.118 | -0.127 | -0.118 | -0.118 | 0 | 0.52 | 0.96 | 0.48 | 0.47 | 0.51 | 0.58 |
| 9 | 109446277 | rs12377606 | -0.421 | -0.449 | -0.755 | 0.015 | -0.001 | -0.002 | -0.002 | 0 | 0.1 | 0.15 | 0.09 | 0.08 | 0.09 | 0.11 |
| 9 | 109459475 | rs10816421 | 1.281 | -0.338 | -0.898 | -0.712 | 0.007 | 0.005 | 0.005 | 0 | 0.42 | 0.47 | 0.51 | 0.55 | 0.47 | 0.6 |
| 9 | 109459699 | rs10759193 | -0.377 | -0.285 | -1.016 | -0.711 | 0.003 | -0.011 | -0.011 | 0.049 | 0.71 | 0.83 | 0.71 | 0.64 | 0.67 | 0.75 |
| 9 | 109461883 | rs12340308 | 0.098 | -0.296 | -0.991 | -0.746 | -0.001 | -0.03 | -0.03 | 0 | 0.3 | 0.16 | 0.3 | 0.41 | 0.35 | 0.28 |
| 9 | 109465397 | rs16925749 | 0.236 | -0.212 | -0.83 | -0.857 | 0.021 | 0.006 | 0.006 | 0 | 0.07 | 0.12 | 0.06 | 0.05 | 0.08 | 0.18 |
| 9 | 109468107 | rs1491103 | 2.378 | -0.02 | -1.203 | -0.985 | -0.09 | -0.159 | -0.159 | 0.007 | 0.31 | 0.04 | 0.45 | 0.51 | 0.35 | 0.34 |
| 9 | 109469564 | rs16925750 | -0.462 | -0.059 | -0.892 | -1.01 | 0.095 | 0.066 | 0.066 | 0.536 | 0.1 | 0.48 | 0.08 | 0.05 | 0.08 | 0.21 |
| 9 | 109470527 | rs10816422 | 0.229 | -0.219 | -0.968 | -1.086 | 0.018 | 0.01 | 0.01 | 0.2 | 0.54 | 0.84 | 0.62 | 0.6 | 0.54 | 0.67 |
| 9 | 109473227 | rs1491101 | 1.338 | -0.089 | -0.802 | -1.103 | 0.004 | 0.001 | 0.001 | 0 | 0.06 | 0.06 | 0.04 | 0.05 | 0.07 | 0.22 |
| 9 | 109473787 | rs10122952 | 0.461 | -0.098 | -0.817 | -1.105 | 0.025 | -0.037 | -0.037 | 0 | 0.09 | 0.19 | 0.06 | 0.03 | 0.09 | 0.09 |
| 9 | 109475636 | rs1388235 | -1.307 | -0.11 | -0.845 | -1.11 | 0.02 | -0.034 | -0.034 | 0 | 0.19 | 0.67 | 0.14 | 0.08 | 0.17 | 0.31 |
| 9 | 109478331 | rs12002898 | -1.074 | -0.129 | -1.053 | -0.972 | -0.002 | 0.002 | 0.002 | 0 | 0.45 | 0.47 | 0.37 | 0.49 | 0.46 | 0.39 |
| 9 | 109480510 | rs13291742 | -0.225 | -0.108 | -1.665 | -0.955 | 0.015 | -0.043 | -0.043 | 0 | 0.34 | 0.21 | 0.3 | 0.46 | 0.36 | 0.3 |
| 9 | 109482027 | rs3936578 | 0.874 | -0.068 | -1.689 | -0.997 | -0.013 | 0.008 | 0.008 | 0.342 | 0.47 | 0.73 | 0.57 | 0.49 | 0.45 | 0.43 |
| 9 | 109485116 | rs10978610 | 0.655 | 0.356 | -1.667 | -0.501 | 0.007 | -0.018 | -0.018 | 0.353 | 0.08 | 0.11 | 0.05 | 0.03 | 0.09 | 0.09 |
| 9 | 109492294 | rs2498861 | 1.304 | 0.254 | -2.171 | -0.418 | -0.115 | -0.068 | -0.068 | 0.699 | 0.46 | 0.04 | 0.55 | 0.47 | 0.48 | 0.39 |
| 9 | 109494624 | rs7041150 | -1.332 | -0.038 | -1.916 | -0.21 | -0.049 | -0.048 | -0.048 | 0 | 0.48 | 0.96 | 0.38 | 0.55 | 0.39 | 0.6 |
| 9 | 109497528 | rs7863929 | -0.604 | -0.076 | -1.856 | -0.218 | -0.001 | 0.017 | 0.017 | 1.863 | 0.24 | 0.83 | 0.26 | 0.09 | 0.18 | 0.34 |
| 9 | 109506245 | rs7036275 | 0.157 | -0.089 | -1.954 | -0.188 | 0.013 | -0.04 | -0.04 | 0.128 | 0.23 | 0.12 | 0.13 | 0.46 | 0.21 | 0.26 |
| 9 | 109507545 | rs10978620 | -0.374 | -0.078 | -1.952 | -0.268 | -0.001 | -0.001 | -0.001 | 0 | 0.23 | 0.77 | 0.25 | 0.09 | 0.18 | 0.34 |
| 9 | 109508320 | rs1491099 | -1.411 | 0.185 | -2.109 | -0.196 | -0.102 | -0.01 | -0.01 | 0 | 0.5 | 0.96 | 0.41 | 0.55 | 0.46 | 0.67 |
| 9 | 109513291 | rs2172111 | -0.534 | -0.132 | -1.196 | 0.393 | 0.013 | -0.035 | -0.035 | 0.019 | 0.25 | 0.15 | 0.13 | 0.46 | 0.21 | 0.28 |
| 9 | 109518208 | rs2451948 | 0.038 | -0.287 | -1.501 | -0.254 | -0.019 | 0.039 | 0.039 | 0 | 0.17 | 0.03 | 0.13 | 0.09 | 0.17 | 0.12 |
| 9 | 109519849 | rs725773 | -0.922 | -1.085 | -1.495 | 0.039 | 0.041 | -0.044 | -0.044 | 0 | 0.32 | 0.14 | 0.27 | 0.43 | 0.24 | 0.27 |
| 9 | 109520131 | rs716875 | -0.461 | -1.032 | -1.489 | 0.075 | 0.001 | 0.003 | 0.003 | 0 | 0.18 | 0.14 | 0.18 | 0.43 | 0.11 | 0.24 |
| 9 | 109520821 | rs2493636 | 1.24 | -0.912 | -1.307 | -0.767 | 0.007 | 0 | 0 | 0 | 0.55 | 0.31 | 0.62 | 0.56 | 0.57 | 0.49 |
| 9 | 109521257 | rs2493637 | -0.341 | -1.275 | -1.186 | -0.561 | 0.033 | -0.045 | -0.045 | 0 | 0.3 | 0.14 | 0.27 | 0.43 | 0.24 | 0.27 |
| 9 | 109522849 | rs17194561 | -0.003 | -1.873 | -0.537 | -0.691 | -0.011 | -0.003 | -0.003 | 0 | 0.13 | 0.14 | 0.07 | 0.05 | 0.05 | 0.16 |

| | | | | | | | | | | | | | | | | |
|---|---|---|---|---|---|---|---|---|---|---|---|---|---|---|---|---|
| 9 | 109529338 | rs10978628 | -0.259 | -1.911 | 0.09 | -0.679 | -0.035 | 0.012 | 0.012 | 0 | 0.21 | 0.27 | 0.2 | 0.3 | 0.14 | 0.29 |
| 9 | 109531515 | rs7022991 | 1.025 | -0.867 | 1.042 | -1.038 | -0.033 | -0.108 | -0.108 | 0.67 | 0.69 | 0.49 | 0.81 | 0.94 | 0.76 | 0.67 |
| 9 | 109531966 | rs10978631 | 1.583 | -0.707 | 1.01 | -1.084 | 0.002 | -0.005 | -0.005 | 0 | 0.61 | 0.34 | 0.75 | 0.93 | 0.66 | 0.59 |
| 9 | 109533269 | rs7873059 | -0.367 | -0.673 | 1.341 | -1.103 | 0.001 | -0.027 | -0.027 | 1.096 | 0.13 | 0.19 | 0.1 | 0.02 | 0.14 | 0.19 |
| 9 | 109538577 | rs7858184 | -0.069 | -0.604 | 1.17 | -1.039 | 0.001 | -0.008 | -0.008 | 8.199 | 0.07 | 0.08 | 0.05 | 0 | 0.09 | 0.08 |
| 9 | 109538962 | rs7048770 | -0.291 | -0.618 | 1.161 | -1.03 | 0.008 | 0.009 | 0.009 | 0 | 0.84 | 0.86 | 0.93 | 0.96 | 0.9 | 0.87 |
| 9 | 109542204 | rs4237185 | 0.523 | -0.545 | 1.042 | -0.965 | -0.004 | -0.058 | -0.058 | 0.116 | 0.76 | 0.66 | 0.87 | 0.95 | 0.8 | 0.79 |
| 9 | 109548485 | rs10978639 | 0.968 | -0.397 | 0.505 | -0.817 | 0.003 | 0.015 | 0.015 | 0.284 | 0.61 | 0.31 | 0.74 | 0.92 | 0.66 | 0.61 |
| 9 | 109557941 | rs4128264 | -1.582 | 0.297 | 1.555 | -0.028 | 0.03 | -0.033 | -0.033 | 0 | 0.22 | 0.55 | 0.17 | 0.18 | 0.26 | 0.27 |
| 9 | 109566506 | rs10978647 | -0.447 | 0.302 | 1.515 | 0.069 | 0 | 0 | 0 | 0 | 0.1 | 0.11 | 0.05 | 0.14 | 0.1 | 0.23 |
| 9 | 109579213 | rs10978651 | -1.022 | 0.357 | 1.975 | 0.039 | -0.001 | 0 | 0 | 0.038 | 0.17 | 0.15 | 0.13 | 0.17 | 0.21 | 0.25 |
| 9 | 109581132 | rs4742675 | -1.319 | 0.354 | 2.224 | 0.038 | 0.016 | -0.015 | -0.015 | 0 | 0.19 | 0.35 | 0.14 | 0.17 | 0.21 | 0.25 |
| 9 | 109592778 | rs4625093 | -3.01 | 0.207 | 1.707 | 0.151 | -0.072 | -0.006 | -0.006 | 0.361 | 0.42 | 0.93 | 0.29 | 0.41 | 0.43 | 0.52 |
| 9 | 109598171 | rs9792598 | -0.367 | 0.265 | 2.108 | 0.444 | 0.002 | -0.002 | -0.002 | 0 | 0.08 | 0.11 | 0.05 | 0.1 | 0.1 | 0.16 |
| 9 | 109614849 | rs10283476 | -0.345 | 0.249 | 2.158 | 0.473 | 0.003 | -0.002 | -0.002 | 0 | 0.08 | 0.15 | 0.06 | 0.1 | 0.09 | 0.16 |
| 9 | 109618474 | rs10978661 | -0.411 | 0.258 | 2.014 | 0.457 | -0.03 | 0.028 | 0.028 | 0 | 0.09 | 0.02 | 0.06 | 0.03 | 0.11 | 0.13 |
| 9 | 109624255 | rs7863663 | -2.473 | 0.186 | 2.409 | 0.452 | 0.02 | -0.022 | -0.022 | 0 | 0.31 | 0.54 | 0.22 | 0.28 | 0.36 | 0.43 |
| 9 | 109632353 | rs4743034 | -2.216 | -0.276 | 2.498 | 0.461 | -0.008 | -0.028 | -0.028 | 0 | 0.24 | 0.43 | 0.14 | 0.19 | 0.22 | 0.35 |
| 9 | 109638167 | rs9299129 | -0.483 | -0.247 | 2.291 | 0.478 | -0.003 | 0.012 | 0.012 | 0 | 0.09 | 0.12 | 0.06 | 0.15 | 0.11 | 0.16 |
| 9 | 109640836 | rs10978664 | -1.635 | -0.228 | 2.265 | 0.484 | -0.001 | -0.011 | -0.011 | 2.86 | 0.19 | 0.26 | 0.12 | 0.18 | 0.19 | 0.28 |
| 9 | 109644865 | rs6477547 | -2.637 | -0.351 | 1.4 | 0.453 | 0.013 | 0.012 | 0.012 | 2.95 | 0.34 | 0.8 | 0.24 | 0.31 | 0.3 | 0.44 |
| 9 | 109653221 | rs3814542 | -1.374 | 0.083 | 1.818 | 0.292 | -0.004 | -0.029 | -0.029 | 0 | 0.22 | 0.44 | 0.14 | 0.18 | 0.19 | 0.36 |
| 9 | 109658333 | rs907064 | -1.967 | 0.015 | 1.468 | 0.326 | -0.003 | 0.024 | 0.024 | 2.153 | 0.35 | 0.79 | 0.25 | 0.33 | 0.35 | 0.55 |
| 9 | 109661016 | rs7020870 | -2.066 | 0.035 | 1.521 | 0.337 | -0.002 | 0.01 | 0.01 | 1.41 | 0.34 | 0.74 | 0.25 | 0.33 | 0.35 | 0.55 |
| 9 | 109673244 | rs16925920 | -1.383 | -0.135 | 1.548 | 0.608 | 0.03 | 0.008 | 0.008 | 0.776 | 0.05 | 0.27 | 0.03 | 0 | 0.04 | 0.01 |
| 9 | 109675354 | rs1567451 | -1.387 | -0.023 | 1.245 | 0.768 | 0.002 | 0.053 | 0.053 | 0.551 | 0.13 | 0.39 | 0.08 | 0.16 | 0.11 | 0.22 |
| 9 | 109676893 | rs1858231 | -0.977 | -0.123 | 0.863 | 1.245 | -0.01 | 0.023 | 0.023 | 0.29 | 0.27 | 0.63 | 0.3 | 0.38 | 0.2 | 0.23 |
| 9 | 109677417 | rs11789013 | -1.471 | -0.048 | 1.353 | 1.285 | 0.001 | 0.005 | 0.005 | 1.319 | 0.28 | 0.48 | 0.29 | 0.32 | 0.26 | 0.33 |
| 9 | 109679767 | rs10816455 | -0.008 | 0.147 | 1.356 | 1.38 | -0.001 | 0.004 | 0.004 | 0.925 | 0.13 | 0.17 | 0.15 | 0.18 | 0.13 | 0.15 |
| 9 | 109682501 | rs10978672 | -0.367 | 0.153 | 1.379 | 1.363 | -0.012 | 0.021 | 0.021 | 0 | 0.14 | 0.08 | 0.1 | 0.08 | 0.17 | 0.2 |
| 9 | 109687288 | rs1000152 | -0.838 | 0.113 | 1.793 | 1.357 | 0.006 | 0.021 | 0.021 | 1.408 | 0.14 | 0.43 | 0.17 | 0.13 | 0.09 | 0.08 |
| 9 | 109687403 | rs17723637 | -0.362 | 0.125 | 1.504 | 1.321 | -0.012 | 0.021 | 0.021 | 0 | 0.14 | 0.07 | 0.1 | 0.08 | 0.17 | 0.2 |
| 9 | 109689859 | rs3814540 | -1.914 | 0.111 | 1.528 | 1.257 | 0.001 | -0.005 | -0.005 | 0 | 0.41 | 0.71 | 0.45 | 0.48 | 0.39 | 0.48 |
| 9 | 109691676 | rs3814538 | -2.096 | 0.028 | 1.23 | 1.212 | 0.004 | -0.025 | -0.025 | 1.404 | 0.44 | 0.86 | 0.46 | 0.48 | 0.39 | 0.48 |
| 9 | 109699547 | rs4743039 | -1.777 | 0.598 | 1.002 | 1.347 | -0.019 | -0.04 | -0.04 | 0 | 0.42 | 0.82 | 0.42 | 0.48 | 0.42 | 0.46 |
| 9 | 109728072 | rs6477551 | -1.262 | 0.487 | 1.796 | 1.215 | -0.009 | -0.009 | -0.009 | 1.445 | 0.2 | 0.68 | 0.16 | 0.16 | 0.13 | 0.07 |
| 9 | 109733743 | rs1510285 | -0.708 | 0.533 | 1.665 | 1.047 | 0.004 | -0.101 | -0.101 | 0 | 0.16 | 0.37 | 0.14 | 0 | 0.12 | 0.06 |
| 9 | 109740520 | rs10978689 | -0.616 | 0.541 | 1.727 | 1.051 | 0.008 | -0.1 | -0.1 | 0 | 0.16 | 0.32 | 0.13 | 0 | 0.12 | 0.06 |
| 9 | 109741309 | rs10978691 | -0.616 | 0.541 | 1.727 | 1.051 | 0.008 | -0.1 | -0.1 | 0 | 0.16 | 0.32 | 0.13 | 0 | 0.12 | 0.06 |
| 9 | 109757432 | rs10978695 | 1.046 | 0.565 | 1.075 | 1.035 | -0.002 | -0.004 | -0.004 | 0.983 | 0.08 | 0.06 | 0.04 | 0 | 0.08 | 0.02 |
| 9 | 109762044 | rs1876985 | -1.124 | 0.42 | 0.91 | 1.279 | -0.045 | -0.064 | -0.064 | 0 | 0.3 | 0.13 | 0.36 | 0.44 | 0.34 | 0.47 |
| 9 | 109764204 | rs10978698 | -0.6 | 0.416 | 1.014 | 1.313 | -0.027 | -0.048 | -0.048 | 0 | 0.23 | 0.13 | 0.26 | 0.36 | 0.29 | 0.4 |
| 9 | 109770755 | rs12343909 | -0.088 | 0.363 | 1.275 | 0.902 | -0.002 | -0.059 | -0.059 | 0 | 0.12 | 0.15 | 0.09 | 0 | 0.09 | 0.06 |
| 9 | 109787178 | rs7853813 | -0.589 | 0.35 | 1.3 | 0.914 | -0.013 | -0.082 | -0.082 | 0 | 0.14 | 0.31 | 0.11 | 0 | 0.08 | 0.06 |
| 9 | 109793425 | rs10816461 | -0.282 | 0.318 | 0.599 | 1.145 | -0.015 | -0.067 | -0.067 | 0 | 0.21 | 0.13 | 0.23 | 0.54 | 0.23 | 0.36 |
| 9 | 109796197 | rs7390631 | -1.341 | 0.333 | 0.702 | 1.225 | 0.006 | -0.017 | -0.017 | 0 | 0.39 | 0.69 | 0.36 | 0.54 | 0.32 | 0.42 |
| 9 | 109806199 | rs10759208 | -1.337 | 0.672 | 0.605 | 1.35 | -0.005 | -0.007 | -0.007 | 0.015 | 0.41 | 0.71 | 0.54 | 0.55 | 0.38 | 0.45 |
| 9 | 109809259 | rs6477555 | -1.911 | 0.344 | -0.04 | 1.079 | -0.035 | 0.109 | 0.109 | 0.035 | 0.53 | 0.87 | 0.53 | 0.69 | 0.49 | 0.52 |
| 9 | 109809366 | rs696562 | -1.788 | -0.343 | -0.912 | 0.385 | -0.075 | 0.016 | 0.016 | 1.444 | 0.66 | 0.94 | 0.6 | 0.72 | 0.59 | 0.56 |
| 9 | 109812226 | rs784790 | -1.803 | -0.366 | -0.869 | 0.36 | -0.074 | 0.021 | 0.021 | 2.708 | 0.65 | 0.94 | 0.61 | 0.73 | 0.58 | 0.56 |
| 9 | 109813447 | rs10739230 | -0.434 | -0.653 | -0.16 | 0.197 | 0.006 | -0.005 | -0.005 | 0 | 0.36 | 0.45 | 0.37 | 0.64 | 0.39 | 0.45 |
| 9 | 109815352 | rs7874831 | 0.106 | -0.697 | 0.171 | 0.152 | 0.001 | -0.051 | -0.051 | 1.898 | 0.14 | 0.32 | 0.14 | 0.07 | 0.09 | 0.07 |
| 9 | 109819184 | rs4492441 | 0.587 | -0.774 | -0.116 | 0.139 | 0.002 | 0.013 | 0.013 | 0 | 0.13 | 0.09 | 0.13 | 0.07 | 0.09 | 0.09 |
| 9 | 109819621 | rs12345400 | -1.345 | -0.708 | -0.291 | 0.198 | -0.003 | 0.11 | 0.11 | 0 | 0.51 | 0.77 | 0.51 | 0.7 | 0.48 | 0.54 |
| 9 | 109827660 | rs784649 | -1.724 | -0.537 | -0.383 | 0.41 | -0.082 | 0.012 | 0.012 | 2.241 | 0.66 | 0.94 | 0.6 | 0.73 | 0.58 | 0.59 |
| 9 | 109838977 | rs1519040 | -0.275 | -0.504 | -0.083 | 0.78 | -0.001 | 0 | 0 | 0.119 | 0.14 | 0.17 | 0.11 | 0.12 | 0.1 | 0.15 |
| 9 | 109839490 | rs10978713 | 1.627 | -0.438 | -0.103 | 1.117 | -0.003 | -0.008 | -0.008 | 0 | 0.08 | 0.07 | 0.14 | 0.2 | 0.09 | 0.11 |
| 9 | 109843602 | rs4443716 | 0.793 | -0.417 | -0.085 | 1.018 | -0.001 | 0 | 0 | 1.225 | 0.1 | 0.11 | 0.07 | 0.06 | 0.07 | 0.09 |
| 9 | 109848914 | rs7861530 | 0.751 | -0.407 | -0.086 | 1.023 | -0.002 | -0.001 | -0.001 | 0 | 0.1 | 0.11 | 0.08 | 0.07 | 0.07 | 0.09 |
| 9 | 109857771 | rs1714460 | -1.858 | -0.473 | -0.285 | 1.136 | -0.092 | -0.022 | -0.022 | 0 | 0.66 | 0.96 | 0.6 | 0.68 | 0.58 | 0.58 |
| 9 | 109862984 | rs16912206 | 0.791 | -0.459 | -0.271 | 0.97 | -0.001 | 0 | 0 | 0 | 0.1 | 0.11 | 0.07 | 0.06 | 0.07 | 0.09 |
| 9 | 109863693 | rs784661 | 0.516 | -0.582 | 0.162 | 0.946 | 0 | 0.015 | 0.015 | 0.414 | 0.34 | 0.36 | 0.31 | 0.48 | 0.32 | 0.33 |
| 9 | 109868858 | rs7029669 | -1.513 | -0.433 | 0.17 | 0.912 | -0.013 | -0.027 | -0.027 | 0 | 0.32 | 0.57 | 0.3 | 0.23 | 0.26 | 0.25 |
| 9 | 109869532 | rs7044846 | -1.342 | -0.471 | 0.235 | 0.913 | -0.012 | -0.027 | -0.027 | 0 | 0.31 | 0.48 | 0.28 | 0.23 | 0.25 | 0.25 |
| 9 | 109872374 | rs784664 | -2.24 | -0.544 | -0.23 | 0.848 | -0.121 | -0.062 | -0.062 | 0 | 0.68 | 0.98 | 0.62 | 0.7 | 0.61 | 0.58 |
| 9 | 109873993 | rs1878735 | -0.323 | -0.492 | -0.03 | 0.836 | 0.002 | 0.003 | 0.003 | 0 | 0.19 | 0.16 | 0.19 | 0.18 | 0.18 | 0.16 |
| 9 | 109880985 | rs784660 | -1.885 | -0.405 | -0.113 | 0.852 | -0.087 | -0.017 | -0.017 | 0 | 0.66 | 0.96 | 0.6 | 0.71 | 0.58 | 0.58 |
| 9 | 109881346 | rs939094 | -0.355 | -0.47 | 0.261 | 0.668 | 0.001 | 0.001 | 0.001 | 0 | 0.17 | 0.16 | 0.14 | 0.12 | 0.16 | 0.16 |
| 9 | 109882332 | rs1007061 | 0.621 | -0.391 | 0.247 | 0.609 | -0.005 | 0.018 | 0.018 | 0 | 0.11 | 0.05 | 0.13 | 0.09 | 0.1 | 0.1 |

| | | | | | | | | | | | | | | | | |
|---|---|---|---|---|---|---|---|---|---|---|---|---|---|---|---|---|
| 9 | 109890250 | rs12002945 | -0.213 | -0.409 | 0.37 | 0.608 | -0.003 | -0.039 | -0.039 | 0.047 | 0.15 | 0.34 | 0.15 | 0.09 | 0.1 | 0.1 |
| 9 | 109890759 | rs10759209 | 0.398 | -0.708 | 0.136 | 0.773 | -0.008 | 0.012 | 0.012 | 1.75 | 0.34 | 0.38 | 0.3 | 0.42 | 0.28 | 0.34 |
| 9 | 109892061 | rs7031681 | -1.594 | -0.652 | -0.102 | 0.851 | -0.111 | -0.02 | -0.02 | 0 | 0.62 | 0.94 | 0.57 | 0.63 | 0.49 | 0.56 |
| 9 | 109892770 | rs1105848 | 0.447 | -0.692 | 0.045 | 0.849 | -0.005 | 0.006 | 0.006 | 0.87 | 0.34 | 0.36 | 0.31 | 0.42 | 0.28 | 0.34 |
| 9 | 109894432 | rs10978723 | -1.425 | -0.366 | -0.399 | 0.387 | 0.006 | 0 | 0 | 0 | 0.65 | 0.62 | 0.6 | 0.63 | 0.56 | 0.58 |
| 9 | 109895391 | rs7864018 | 0.208 | -0.353 | -0.282 | 0.345 | 0.013 | -0.009 | -0.009 | 0.174 | 0.2 | 0.31 | 0.24 | 0.14 | 0.22 | 0.17 |
| 9 | 109897161 | rs7858186 | -0.985 | -0.407 | -0.304 | 0.304 | -0.012 | -0.009 | -0.009 | 0.021 | 0.59 | 0.81 | 0.57 | 0.57 | 0.51 | 0.51 |
| 9 | 109898877 | rs784683 | -1.075 | -0.099 | -0.383 | 0.195 | -0.008 | -0.012 | -0.012 | 0.238 | 0.64 | 0.8 | 0.57 | 0.59 | 0.56 | 0.59 |
| 9 | 109900756 | rs9696036 | 0.077 | -0.271 | -0.375 | 0.402 | 0.026 | 0.024 | 0.024 | 0.739 | 0.46 | 0.3 | 0.35 | 0.38 | 0.37 | 0.46 |
| 9 | 109903423 | rs12351052 | 1.274 | -0.099 | -0.4 | 0.376 | -0.001 | 0.003 | 0.003 | 0 | 0.12 | 0.11 | 0.16 | 0.13 | 0.14 | 0.11 |
| 9 | 109905613 | rs956578 | -0.198 | -0.149 | -0.361 | 0.229 | 0.005 | 0.014 | 0.014 | 0 | 0.19 | 0.28 | 0.13 | 0.21 | 0.13 | 0.17 |
| 9 | 109907509 | rs1714474 | 0.498 | -0.209 | -0.443 | 0.168 | 0.016 | 0.017 | 0.017 | 0.541 | 0.39 | 0.26 | 0.3 | 0.29 | 0.33 | 0.41 |
| 9 | 109910721 | rs1714473 | 0.881 | -0.23 | -0.558 | 0.311 | 0.01 | 0.035 | 0.035 | 0.416 | 0.35 | 0.26 | 0.29 | 0.19 | 0.3 | 0.27 |
| 9 | 109911084 | rs7849325 | -0.103 | -0.133 | -0.54 | 0.211 | 0.007 | 0.024 | 0.024 | 0 | 0.18 | 0.3 | 0.13 | 0.24 | 0.13 | 0.16 |
| 9 | 109912451 | rs784677 | 0.366 | -0.059 | -1.27 | 0.154 | 0.011 | 0.022 | 0.022 | 0.248 | 0.42 | 0.31 | 0.33 | 0.29 | 0.4 | 0.42 |
| 9 | 109912752 | rs784676 | -2.181 | -0.037 | -1.302 | 0.15 | -0.063 | -0.102 | -0.102 | 0 | 0.74 | 0.96 | 0.64 | 0.59 | 0.67 | 0.68 |
| 9 | 109915716 | rs784670 | 0.459 | -0.043 | -1.104 | 0.005 | 0.011 | 0.014 | 0.014 | 0.802 | 0.37 | 0.23 | 0.3 | 0.31 | 0.33 | 0.41 |
| 9 | 109921231 | rs871625 | 0.721 | 0.187 | -0.959 | 0.559 | -0.009 | 0.03 | 0.03 | 0 | 0.11 | 0.07 | 0.16 | 0.05 | 0.14 | 0.11 |
| 9 | 109923051 | rs10978733 | 0.262 | 0.258 | -0.709 | 0.718 | -0.007 | -0.046 | -0.046 | 0 | 0.15 | 0.06 | 0.15 | 0.27 | 0.14 | 0.31 |
| 9 | 109926563 | rs17195683 | 0.702 | 0.377 | -0.576 | 0.631 | 0.002 | 0.041 | 0.041 | 0.3 | 0.05 | 0.06 | 0.13 | 0.21 | 0.1 | 0.21 |
| 9 | 109926894 | rs10978734 | 0.494 | 0.349 | -0.354 | 0.284 | -0.001 | 0.023 | 0.023 | 0 | 0.18 | 0.18 | 0.22 | 0.43 | 0.19 | 0.33 |
| 9 | 109928973 | rs16912263 | 0.398 | 0.651 | -0.643 | 0.315 | 0.015 | -0.004 | -0.004 | 0 | 0.1 | 0.18 | 0.07 | 0.04 | 0.08 | 0.08 |
| 9 | 109931412 | rs6477562 | -1.259 | 0.51 | -1.139 | 0.057 | -0.082 | 0.007 | 0.007 | 0 | 0.55 | 0.92 | 0.55 | 0.65 | 0.53 | 0.6 |
| 9 | 109940271 | rs4443717 | -1.455 | 0.592 | -1.068 | 0.097 | -0.091 | -0.032 | -0.032 | 0 | 0.61 | 0.92 | 0.57 | 0.65 | 0.59 | 0.61 |
| 9 | 109942554 | rs10978740 | -1.473 | 0.554 | -0.7 | 0.079 | -0.033 | 0.008 | 0.008 | 0 | 0.6 | 0.81 | 0.56 | 0.65 | 0.59 | 0.61 |
| 9 | 109943165 | rs2417773 | -0.541 | 1.362 | 0.547 | -0.301 | 0.015 | 0.033 | 0.033 | 0.74 | 0.13 | 0.26 | 0.15 | 0.19 | 0.09 | 0.09 |
| 9 | 109949144 | rs7032927 | -0.165 | 1.253 | -0.21 | -0.615 | -0.004 | 0.037 | 0.037 | 0.007 | 0.15 | 0.07 | 0.11 | 0.01 | 0.16 | 0.1 |
| 9 | 109953472 | rs1464250 | -0.441 | 1.771 | 0.603 | -0.094 | 0.04 | -0.08 | -0.08 | 0 | 0.12 | 0.26 | 0.1 | 0 | 0.16 | 0.05 |
| 9 | 109960553 | rs12336793 | -0.275 | 1.977 | 0.735 | 0.285 | 0.022 | 0.113 | 0.113 | 0 | 0.1 | 0.22 | 0.16 | 0.36 | 0.18 | 0.21 |
| 9 | 109960591 | rs12336762 | -0.792 | 1.826 | 0.36 | 0.524 | 0.049 | -0.092 | -0.092 | 0 | 0.13 | 0.33 | 0.11 | 0 | 0.17 | 0.06 |
| 9 | 109971491 | rs2680198 | 0.301 | 1.691 | -0.658 | 0.736 | -0.002 | 0.006 | 0.006 | 0 | 0.84 | 0.84 | 0.61 | 0.41 | 0.87 | 0.59 |
| 9 | 109972066 | rs2680197 | 0.774 | 1.288 | -0.813 | 0.554 | 0.027 | 0.012 | 0.012 | 0.241 | 0.66 | 0.62 | 0.47 | 0.43 | 0.56 | 0.44 |
| 9 | 109973450 | rs2680195 | 0.387 | 0.68 | -0.743 | 0.724 | -0.017 | 0.054 | 0.054 | 0 | 0.89 | 0.79 | 0.85 | 0.76 | 0.95 | 0.84 |
| 9 | 109978917 | rs2680208 | -0.13 | 0.853 | -1.319 | -0.01 | -0.004 | -0.021 | -0.021 | 0.029 | 0.65 | 0.48 | 0.7 | 0.76 | 0.69 | 0.75 |
| 9 | 109981481 | rs10978757 | -0.426 | 0.503 | -2.333 | -0.285 | -0.042 | 0.127 | 0.127 | 0 | 0.32 | 0.01 | 0.17 | 0.1 | 0.32 | 0.22 |
| 9 | 109986449 | rs13292544 | -0.535 | 0.782 | -2.011 | 0.091 | -0.056 | 0.048 | 0.048 | 0 | 0.22 | 0.01 | 0.14 | 0.1 | 0.26 | 0.19 |
| 9 | 109986827 | rs10978761 | -0.564 | 0.698 | -1.417 | 1.099 | 0.001 | 0.055 | 0.055 | 1.286 | 0.09 | 0.12 | 0.19 | 0.37 | 0.14 | 0.21 |
| 9 | 109987079 | rs2808374 | -1.261 | 0.716 | -1.33 | 1.055 | -0.003 | 0.001 | 0.001 | 1.128 | 0.18 | 0.23 | 0.34 | 0.44 | 0.24 | 0.31 |
| 9 | 109989135 | rs7859324 | -0.223 | 0.732 | -0.816 | 0.404 | -0.005 | -0.033 | -0.033 | 0 | 0.07 | 0.13 | 0.09 | 0.03 | 0.05 | 0.05 |
| 9 | 109991942 | rs1519049 | -0.715 | 0.806 | -0.447 | 0.606 | 0.005 | 0.004 | 0.004 | 0 | 0.12 | 0.12 | 0.32 | 0.2 | 0.2 | 0.18 |
| 9 | 109995551 | rs2680216 | -1.775 | 0.449 | -0.52 | -0.009 | -0.005 | 0.03 | 0.03 | 0 | 0.4 | 0.31 | 0.42 | 0.21 | 0.42 | 0.45 |
| 9 | 109998563 | rs2808363 | -0.04 | 0.491 | -0.466 | 0.008 | -0.009 | -0.009 | -0.009 | 0 | 0.11 | 0.05 | 0.32 | 0.22 | 0.2 | 0.18 |
| 9 | 110000354 | rs17724552 | -0.011 | 0.404 | -0.527 | -0.075 | 0.005 | 0.03 | 0.03 | 0.129 | 0.13 | 0.05 | 0.08 | 0.01 | 0.13 | 0.19 |
| 9 | 110001533 | rs10978764 | -1.559 | 0.562 | -0.53 | -0.075 | 0 | -0.002 | -0.002 | 0 | 0.32 | 0.31 | 0.49 | 0.26 | 0.4 | 0.42 |
| 9 | 110002647 | rs4978764 | 2.016 | 0.285 | 0.523 | -0.286 | -0.001 | 0 | 0 | 0 | 0.57 | 0.43 | 0.42 | 0.71 | 0.52 | 0.51 |
| 9 | 110006333 | rs12004883 | -0.482 | 0.321 | 0.495 | -0.318 | 0 | -0.014 | -0.014 | 0 | 0.19 | 0.15 | 0.17 | 0.04 | 0.19 | 0.24 |
| 9 | 110008143 | rs10978769 | -1.079 | 0.253 | 0.462 | -0.351 | -0.024 | 0.007 | 0.007 | 0 | 0.22 | 0.11 | 0.37 | 0.21 | 0.28 | 0.28 |
| 9 | 110013980 | rs7867544 | -0.023 | 0.65 | 0.751 | -0.104 | 0.007 | 0.008 | 0.008 | 0.002 | 0.12 | 0.12 | 0.32 | 0.2 | 0.2 | 0.18 |
| 9 | 110022842 | rs10816477 | 0.165 | 0.614 | 0.716 | -0.088 | -0.016 | -0.016 | -0.016 | 1.426 | 0.11 | 0.04 | 0.31 | 0.21 | 0.18 | 0.16 |
| 9 | 110031745 | rs942291 | -1.132 | 0.524 | 0.938 | -0.138 | 0.012 | -0.005 | -0.005 | 0 | 0.21 | 0.27 | 0.38 | 0.21 | 0.25 | 0.26 |
| 9 | 110032498 | rs12345455 | -1.256 | 0.499 | 0.917 | -0.151 | 0.018 | -0.009 | -0.009 | 0.517 | 0.22 | 0.32 | 0.39 | 0.21 | 0.25 | 0.26 |
| 9 | 110042484 | rs10816481 | 1.385 | 0.054 | 1.15 | 0.031 | -0.034 | 0.014 | 0.014 | 0.107 | 0.67 | 0.46 | 0.72 | 0.39 | 0.71 | 0.48 |
| 9 | 110044572 | rs7041137 | -1.231 | -0.012 | 1.329 | -0.101 | 0.015 | -0.025 | -0.025 | 0 | 0.24 | 0.35 | 0.39 | 0.2 | 0.25 | 0.28 |
| 9 | 110052660 | rs12005522 | 1.166 | -0.021 | 1.359 | -0.093 | -0.001 | -0.024 | -0.024 | 0 | 0.06 | 0.11 | 0.08 | 0.03 | 0.05 | 0.04 |
| 9 | 110056739 | rs10759222 | 1.54 | 0.114 | 1.372 | -0.14 | -0.071 | 0.066 | 0.066 | 3.173 | 0.65 | 0.91 | 0.52 | 0.75 | 0.57 | 0.51 |
| 9 | 110063197 | rs11573662 | 0.429 | 0.103 | 1.255 | -0.149 | 0.001 | 0.008 | 0.008 | 0.115 | 0.13 | 0.11 | 0.08 | 0.01 | 0.14 | 0.17 |
| 9 | 110075437 | rs7023656 | 2.073 | 0.246 | 1.452 | -0.167 | 0.005 | -0.048 | -0.048 | 0 | 0.62 | 0.49 | 0.49 | 0.2 | 0.61 | 0.37 |
| 9 | 110076032 | rs942292 | 1.983 | 0.209 | 1.749 | -0.295 | 0.004 | -0.027 | -0.027 | 0 | 0.61 | 0.56 | 0.49 | 0.75 | 0.57 | 0.49 |
| 9 | 110078582 | rs7025563 | 0.286 | 0.232 | 1.732 | -0.275 | -0.025 | -0.021 | -0.021 | 0.753 | 0.85 | 0.96 | 0.65 | 0.76 | 0.79 | 0.78 |
| 9 | 110079453 | rs16912372 | 0.992 | 0.227 | 1.749 | -0.281 | -0.001 | -0.027 | -0.027 | 0 | 0.06 | 0.12 | 0.08 | 0.03 | 0.05 | 0.04 |
| 9 | 110083764 | rs2417784 | 0.223 | 0.24 | 1.47 | -0.276 | -0.016 | -0.013 | -0.013 | 0.063 | 0.12 | 0.04 | 0.3 | 0.2 | 0.18 | 0.16 |
| 9 | 110084261 | rs1805330 | -0.556 | 0.235 | 1.494 | -0.285 | 0 | -0.054 | -0.054 | 0 | 0.1 | 0.21 | 0.07 | 0 | 0.07 | 0.11 |
| 9 | 110084832 | rs1323809 | 2.511 | 0.132 | 1.41 | -0.343 | -0.033 | 0.168 | 0.168 | 2.218 | 0.53 | 0.18 | 0.39 | 0.72 | 0.52 | 0.45 |
| 9 | 110085255 | rs11573709 | -1.457 | 0.327 | 1.279 | -0.235 | -0.007 | -0.117 | -0.117 | 0 | 0.24 | 0.39 | 0.15 | 0.01 | 0.21 | 0.29 |
| 9 | 110086951 | rs11573721 | 0.263 | 0.204 | 1.241 | -0.191 | -0.004 | 0.039 | 0.039 | 0.167 | 0.12 | 0.05 | 0.07 | 0.01 | 0.14 | 0.16 |
| 9 | 110092856 | rs10868 | 0.375 | 0.212 | 1.151 | -0.188 | -0.027 | 0.064 | 0.064 | 0.13 | 0.12 | 0 | 0.07 | 0.01 | 0.14 | 0.17 |
| 9 | 110100745 | rs10816493 | 1.354 | 0.283 | 1.374 | -0.001 | 0.012 | -0.018 | -0.018 | 0.111 | 0.73 | 0.66 | 0.57 | 0.76 | 0.71 | 0.66 |
| 9 | 110101440 | rs817844 | -0.891 | 0.402 | 0.4 | 0.048 | -0.041 | 0.005 | 0.005 | 0 | 0.19 | 0.04 | 0.35 | 0.2 | 0.25 | 0.27 |
| 9 | 110106222 | rs817852 | 1.356 | -0.202 | 1.015 | 0.791 | -0.01 | 0.118 | 0.118 | 0.046 | 0.89 | 0.78 | 0.91 | 0.58 | 0.93 | 0.79 |

| | | | | | | | | | | | | | | | | |
|---|---|---|---|---|---|---|---|---|---|---|---|---|---|---|---|---|
| 9 | 110107862 | rs716004 | -0.077 | -0.163 | 0.744 | 0.496 | -0.041 | 0.077 | 0.077 | 0 | 0.14 | 0.01 | 0.09 | 0.01 | 0.17 | 0.17 |
| 9 | 110112190 | rs10816494 | -0.539 | -0.058 | 1.082 | 0.619 | -0.002 | 0.035 | 0.035 | 0 | 0.21 | 0.29 | 0.15 | 0.39 | 0.22 | 0.31 |
| 9 | 110113919 | rs817854 | 0.224 | -0.097 | 1.305 | 0.57 | 0.019 | 0.013 | 0.013 | 0 | 0.95 | 0.76 | 0.94 | 0.96 | 0.96 | 0.95 |
| 9 | 110114432 | rs10978803 | -0.611 | -0.071 | 0.785 | 0.591 | -0.002 | 0.018 | 0.018 | 0 | 0.22 | 0.3 | 0.15 | 0.39 | 0.22 | 0.32 |
| 9 | 110116369 | rs10816499 | -1.13 | -0.084 | 0.719 | 0.556 | -0.004 | 0.013 | 0.013 | 0 | 0.28 | 0.34 | 0.19 | 0.39 | 0.27 | 0.39 |
| 9 | 110127130 | rs10759229 | 1.725 | 0.027 | -0.011 | 0.318 | 0.011 | 0.035 | 0.035 | 0.111 | 0.52 | 0.15 | 0.5 | 0.37 | 0.58 | 0.41 |
| 9 | 110130919 | rs10739243 | 2.279 | -0.116 | 0.721 | 0.059 | -0.041 | 0.006 | 0.006 | 0 | 0.81 | 0.41 | 0.69 | 0.78 | 0.88 | 0.8 |
| 9 | 110131397 | rs1538985 | 1.787 | 0.149 | 0.241 | 0.427 | -0.023 | 0.067 | 0.067 | 0 | 0.87 | 0.58 | 0.75 | 0.78 | 0.92 | 0.87 |
| 9 | 110132198 | rs817861 | -0.724 | 0.505 | -1.047 | 0.283 | 0.003 | -0.026 | -0.026 | 0 | 0.05 | 0.01 | 0.18 | 0.14 | 0.04 | 0.07 |
| 9 | 110136208 | rs12375540 | -1.145 | 0.789 | -0.932 | -0.045 | -0.003 | -0.008 | -0.008 | 0.971 | 0.11 | 0.1 | 0.08 | 0 | 0.17 | 0.03 |
| 9 | 110141811 | rs1746837 | 0.005 | 0.464 | -1.401 | -0.678 | 0 | 0.127 | 0.127 | 0.006 | 0.47 | 0.29 | 0.71 | 0.85 | 0.56 | 0.48 |
| 9 | 110142277 | rs10122670 | -0.687 | -0.394 | -0.146 | -0.793 | 0.046 | 0.106 | 0.106 | 0.467 | 0.06 | 0.4 | 0.04 | 0.03 | 0.01 | 0.1 |
| 9 | 110144329 | rs11792269 | -0.908 | -0.405 | -0.624 | -1.121 | -0.007 | 0.018 | 0.018 | 0 | 0.15 | 0.06 | 0.16 | 0.1 | 0.16 | 0.26 |
| 9 | 110144688 | rs817870 | 1.339 | -0.872 | -1.134 | -1.258 | -0.007 | -0.006 | -0.006 | 0 | 0.6 | 0.46 | 0.68 | 0.67 | 0.72 | 0.45 |
| 9 | 110150168 | rs1771723 | 1.458 | -1.48 | -1.225 | -0.065 | 0.064 | 0.011 | 0.011 | 0 | 0.55 | 0.28 | 0.53 | 0.38 | 0.7 | 0.42 |
| 9 | 110150659 | rs1622267 | -0.192 | -1.53 | -0.916 | -0.244 | -0.045 | -0.033 | -0.033 | 0 | 0.79 | 0.59 | 0.85 | 0.8 | 0.89 | 0.81 |
| 9 | 110151512 | rs1771717 | -0.471 | -1.743 | -1.108 | -0.311 | -0.002 | 0.001 | 0.001 | 0 | 0.89 | 0.94 | 0.89 | 0.86 | 0.9 | 0.87 |
| 9 | 110152533 | rs7029300 | -0.538 | -2.099 | -0.84 | -0.484 | -0.048 | -0.053 | -0.053 | 0 | 0.21 | 0.36 | 0.11 | 0.09 | 0.11 | 0.31 |
| 9 | 110155358 | rs817829 | 0.24 | -2.576 | -0.82 | -0.308 | -0.025 | 0.008 | 0.008 | 0 | 0.8 | 0.6 | 0.85 | 0.76 | 0.89 | 0.82 |
| 9 | 110156279 | rs817827 | 1.357 | -2.349 | -1.845 | -0.614 | -0.093 | -0.089 | -0.089 | 1.005 | 0.32 | 0.08 | 0.5 | 0.51 | 0.54 | 0.35 |
| 9 | 110156300 | rs817826 | -0.797 | -2.619 | -1.736 | -0.841 | -0.011 | -0.054 | -0.054 | 0 | 0.8 | 0.65 | 0.89 | 0.88 | 0.86 | 0.79 |
| 9 | 110159566 | rs817824 | 1.364 | -3.291 | -2.01 | -0.479 | 0.064 | -0.07 | -0.07 | 0 | 0.42 | 0.1 | 0.55 | 0.5 | 0.68 | 0.42 |
| 9 | 110167103 | rs12004443 | -0.218 | -2.506 | -0.756 | -0.367 | -0.015 | -0.014 | -0.014 | 0 | 0.2 | 0.27 | 0.14 | 0.15 | 0.16 | 0.21 |
| 9 | 110167168 | rs9299131 | -0.641 | -2.703 | -0.814 | -0.312 | -0.016 | -0.006 | -0.006 | 0 | 0.28 | 0.31 | 0.14 | 0.24 | 0.17 | 0.38 |
| 9 | 110167768 | rs947125 | 0.434 | -2.622 | -1.1 | -0.367 | 0.007 | 0.014 | 0.014 | 0 | 0.1 | 0.07 | 0.1 | 0.06 | 0.08 | 0.07 |
| 9 | 110168438 | rs10978837 | 0.387 | -2.318 | -0.785 | -0.257 | 0.031 | -0.013 | -0.013 | 0 | 0.13 | 0.03 | 0.03 | 0.17 | 0.04 | 0.21 |
| 9 | 110168617 | rs16912495 | -0.892 | -1.787 | -0.339 | -0.318 | -0.012 | -0.016 | -0.016 | 0 | 0.24 | 0.36 | 0.16 | 0.22 | 0.18 | 0.24 |
| 9 | 110171107 | rs7861976 | 0.017 | -1.611 | -0.599 | -0.244 | -0.006 | -0.007 | -0.007 | 0.08 | 0.15 | 0.2 | 0.11 | 0.13 | 0.14 | 0.16 |
| 9 | 110183891 | rs4978954 | 0.464 | -1.642 | -1.182 | -0.1 | 0.018 | -0.013 | -0.013 | 0 | 0.09 | 0.05 | 0.03 | 0.14 | 0.07 | 0.02 |
| 9 | 110184048 | rs4978955 | 0.374 | -1.657 | -1.19 | 0.074 | 0.02 | -0.027 | -0.027 | 0 | 0.09 | 0.05 | 0.03 | 0.18 | 0.07 | 0.02 |
| 9 | 110184645 | rs1572905 | 1.195 | -1.189 | -1.089 | -0.669 | -0.008 | -0.014 | -0.014 | 0.207 | 0.62 | 0.44 | 0.72 | 0.74 | 0.69 | 0.56 |
| 9 | 110185695 | rs12336348 | 0.371 | -1.257 | -0.967 | -0.735 | -0.05 | -0.049 | -0.049 | 0 | 0.16 | 0.41 | 0.09 | 0.06 | 0.07 | 0.22 |
| 9 | 110185887 | rs13288073 | 1.037 | -1.2 | -0.927 | -0.711 | -0.007 | -0.012 | -0.012 | 0 | 0.62 | 0.44 | 0.71 | 0.74 | 0.7 | 0.56 |
| 9 | 110187151 | rs10978847 | 0.354 | -0.926 | -1.048 | -0.737 | -0.003 | -0.05 | -0.05 | 1.114 | 0.71 | 0.56 | 0.76 | 0.88 | 0.76 | 0.57 |
| 9 | 110193442 | rs17196438 | 0.536 | -1.039 | -1.216 | -0.479 | 0.036 | 0.012 | 0.012 | 0.59 | 0.18 | 0.1 | 0.07 | 0.17 | 0.11 | 0.18 |
| 9 | 110195120 | rs10816506 | 0.456 | -0.742 | -1.23 | -0.391 | -0.003 | -0.001 | -0.001 | 0 | 0.06 | 0.02 | 0.05 | 0.11 | 0.07 | 0.1 |
| 9 | 110195358 | rs947126 | -0.684 | -0.977 | -1.306 | -0.904 | 0.035 | 0.05 | 0.05 | 3.021 | 0.45 | 0.34 | 0.22 | 0.29 | 0.34 | 0.37 |
| 9 | 110195598 | rs12379012 | -0.124 | -0.574 | -1.07 | -0.437 | 0.02 | 0.025 | 0.025 | 0.128 | 0.15 | 0.06 | 0.07 | 0.11 | 0.08 | 0.1 |
| 9 | 110196197 | rs12342660 | -0.271 | -0.488 | -1.047 | -0.396 | -0.001 | -0.004 | -0.004 | 0 | 0.17 | 0.17 | 0.08 | 0.12 | 0.1 | 0.12 |
| 9 | 110197029 | rs2417791 | -0.698 | -0.32 | -1.122 | -0.353 | 0.007 | 0.014 | 0.014 | 0 | 0.27 | 0.23 | 0.17 | 0.15 | 0.25 | 0.23 |
| 9 | 110204166 | rs10759234 | 0.261 | -0.54 | -0.774 | 0.297 | -0.009 | 0 | 0 | 0.748 | 0.54 | 0.44 | 0.68 | 0.46 | 0.66 | 0.52 |
| 9 | 110205831 | rs10978858 | -0.141 | -0.146 | -0.77 | 0.332 | -0.009 | -0.006 | -0.006 | 0 | 0.93 | 0.94 | 0.91 | 0.89 | 0.87 | 0.9 |
| 9 | 110206622 | rs10816508 | 0.664 | -0.124 | -0.103 | 0.635 | 0.065 | 0.064 | 0.064 | 0 | 0.9 | 0.8 | 0.89 | 0.84 | 0.85 | 0.87 |
| 9 | 110210362 | rs10978861 | -0.574 | -0.653 | -0.278 | 1.154 | -0.001 | 0.044 | 0.044 | 0 | 0.11 | 0.06 | 0.3 | 0.39 | 0.07 | 0.25 |
| 9 | 110214347 | rs10759237 | 0.267 | -0.588 | -0.477 | 0.406 | -0.013 | -0.02 | -0.02 | 0 | 0.33 | 0.44 | 0.25 | 0.23 | 0.29 | 0.19 |
| 9 | 110214615 | rs12555837 | 1.453 | -0.264 | -1.203 | 0.389 | -0.034 | -0.035 | -0.035 | 0 | 0.1 | 0.03 | 0.18 | 0.2 | 0.19 | 0.08 |
| 9 | 110217097 | rs947122 | -0.595 | -0.169 | -0.735 | 0.393 | 0.025 | -0.009 | -0.009 | 0.982 | 0.94 | 0.88 | 0.89 | 0.97 | 0.87 | 0.94 |
| 9 | 110217742 | rs882942 | 0.84 | -0.807 | -1.21 | 0.167 | 0.01 | 0.001 | 0.001 | 0 | 0.58 | 0.18 | 0.49 | 0.41 | 0.68 | 0.49 |
| 9 | 110226138 | rs7035256 | 0.543 | -0.419 | -0.585 | 0.084 | -0.006 | -0.028 | -0.028 | 0.097 | 0.25 | 0.17 | 0.48 | 0.42 | 0.32 | 0.36 |
| 9 | 110229609 | rs1832881 | -1.314 | -0.514 | -1.158 | -0.049 | -0.053 | -0.02 | -0.02 | 0.558 | 0.59 | 0.86 | 0.65 | 0.63 | 0.56 | 0.75 |
| 9 | 110231170 | rs10759238 | 0.277 | -0.004 | -1.37 | -0.145 | -0.034 | -0.094 | -0.094 | 0 | 0.25 | 0.09 | 0.47 | 0.41 | 0.32 | 0.36 |
| 9 | 110231863 | rs1819730 | -0.497 | -0.175 | -0.475 | -0.503 | -0.016 | -0.099 | -0.099 | 0 | 0.22 | 0.27 | 0.12 | 0 | 0.16 | 0.12 |
| 9 | 110238361 | rs17725250 | -0.79 | 0.2 | -0.9 | 0.242 | -0.011 | 0.005 | 0.005 | 0 | 0.05 | 0.02 | 0.02 | 0.05 | 0.05 | 0.07 |
| 9 | 110241529 | rs1832741 | 0.188 | 0.284 | -0.67 | 0.382 | 0.009 | 0.011 | 0.011 | 0 | 0.6 | 0.54 | 0.37 | 0.51 | 0.48 | 0.46 |
| 9 | 110242157 | rs10759240 | 0.097 | 0.192 | -1.16 | 0.688 | -0.122 | -0.034 | -0.034 | 0.469 | 0.67 | 0.9 | 0.54 | 0.68 | 0.5 | 0.6 |
| 9 | 110243031 | rs10816518 | 0.769 | -0.609 | -1.142 | 0.287 | 0.003 | -0.015 | -0.015 | 0.547 | 0.46 | 0.31 | 0.35 | 0.52 | 0.46 | 0.46 |
| 9 | 110271554 | rs4448339 | -0.436 | 0.08 | -1.524 | 0.222 | -0.158 | 0.029 | 0.029 | 0 | 0.74 | 0.98 | 0.69 | 0.84 | 0.55 | 0.65 |
| 9 | 110274062 | rs10816523 | -0.355 | -0.612 | -0.773 | 0.078 | -0.012 | 0.004 | 0.004 | 0 | 0.31 | 0.37 | 0.35 | 0.66 | 0.25 | 0.37 |
| 9 | 110277340 | rs4979041 | 0.067 | -0.436 | -0.414 | 0.169 | -0.001 | 0.008 | 0.008 | 0 | 0.58 | 0.72 | 0.44 | 0.67 | 0.4 | 0.45 |
| 9 | 110280496 | rs7041758 | -0.703 | -0.992 | 0.13 | 0.588 | -0.045 | 0.08 | 0.08 | 0.202 | 0.17 | 0.49 | 0.27 | 0.24 | 0.09 | 0.17 |
| 9 | 110293247 | rs10816526 | -0.226 | -0.475 | -0.428 | 0.509 | -0.051 | 0.021 | 0.021 | 0 | 0.7 | 0.83 | 0.61 | 0.78 | 0.51 | 0.65 |
| 9 | 110296686 | rs10119193 | 0.285 | -0.694 | 0.186 | 0.869 | 0.003 | 0.004 | 0.004 | 0.353 | 0.63 | 0.63 | 0.32 | 0.42 | 0.47 | 0.38 |
| 9 | 110297442 | rs10978905 | -1.064 | -0.937 | 0.239 | 0.893 | -0.018 | 0.196 | 0.196 | 0 | 0.1 | 0.27 | 0.21 | 0.37 | 0.05 | 0.2 |
| 9 | 110300994 | rs4979065 | 0.129 | -1.016 | 0.083 | 0.695 | -0.039 | 0.03 | 0.03 | 0.004 | 0.67 | 0.8 | 0.51 | 0.78 | 0.5 | 0.63 |
| 9 | 110303463 | rs2185857 | 0.271 | -1.408 | 0.157 | 0.964 | 0.056 | 0.014 | 0.014 | 0 | 0.49 | 0.64 | 0.37 | 0.45 | 0.28 | 0.4 |
| 9 | 110311124 | rs10122326 | 0.201 | -1.452 | -0.01 | 0.615 | 0.009 | -0.022 | -0.022 | 0.302 | 0.06 | 0.24 | 0.03 | 0.02 | 0.05 | 0.11 |
| 9 | 110311539 | rs10512383 | 1.469 | -1.874 | -1.041 | 0.521 | 0.077 | 0.003 | 0.003 | 0.627 | 0.3 | 0.06 | 0.22 | 0.2 | 0.14 | 0.18 |
| 9 | 110318782 | rs2258083 | -0.838 | 0.063 | -1.246 | 0.342 | -0.021 | 0.005 | 0.005 | 0 | 0.75 | 0.86 | 0.79 | 0.78 | 0.67 | 0.78 |
| 9 | 110319927 | rs2258170 | -0.557 | 0.494 | -1.123 | 0.449 | -0.024 | 0.006 | 0.006 | 0.03 | 0.25 | 0.18 | 0.35 | 0.21 | 0.34 | 0.34 |

| | | | | | | | | | | | | | | | | |
|---|---|---|---|---|---|---|---|---|---|---|---|---|---|---|---|---|
| 9 | 110326874 | rs2258660 | 0.665 | 1.599 | -1.113 | 0.411 | -0.04 | 0.038 | 0.038 | 0 | 0.07 | 0.01 | 0.21 | 0 | 0.21 | 0.11 |
| 9 | 110328726 | rs1573048 | -0.368 | 0.998 | -1.044 | 0.531 | -0.026 | -0.011 | -0.011 | 0 | 0.2 | 0.13 | 0.35 | 0.26 | 0.34 | 0.32 |
| 9 | 110332569 | rs7045952 | -0.635 | -0.24 | -0.732 | 0.175 | 0.004 | 0.007 | 0.007 | 0 | 0.05 | 0.18 | 0.04 | 0.05 | 0.06 | 0.03 |
| 9 | 110346728 | rs10978931 | -0.178 | 0.014 | -2.193 | 0.048 | -0.159 | -0.069 | -0.069 | 0 | 0.69 | 0.98 | 0.5 | 0.35 | 0.45 | 0.41 |
| 9 | 110350727 | rs11792587 | 0.14 | -1.095 | -2.746 | -0.644 | -0.021 | 0.123 | 0.123 | 0 | 0.19 | 0.01 | 0.12 | 0 | 0.16 | 0.07 |
| 9 | 110354209 | rs1935973 | -0.657 | -0.992 | -2.676 | -0.559 | -0.008 | 0.058 | 0.058 | 0 | 0.9 | 1 | 0.97 | 1 | 0.94 | 0.97 |
| 9 | 110360883 | rs4979136 | -0.055 | -1.756 | -2.047 | -0.839 | 0.007 | 0.006 | 0.006 | 0 | 0.41 | 0.6 | 0.39 | 0.33 | 0.26 | 0.26 |
| 9 | 110363564 | rs7034018 | 0.179 | -1.643 | -1.423 | -0.782 | -0.015 | -0.094 | -0.094 | 0 | 0.26 | 0.38 | 0.11 | 0.03 | 0.18 | 0.14 |
| 9 | 110369095 | rs10816539 | -0.436 | -2.194 | -1.16 | 0.152 | -0.019 | 0.026 | 0.026 | 0 | 0.25 | 0.34 | 0.19 | 0.41 | 0.13 | 0.16 |
| 9 | 110370455 | rs10816540 | -0.389 | -2.852 | -1.476 | -0.169 | -0.008 | 0.006 | 0.006 | 0 | 0.58 | 0.58 | 0.38 | 0.51 | 0.37 | 0.38 |
| 9 | 110376654 | rs7869457 | -1.017 | -3.011 | -2.369 | -0.099 | -0.07 | -0.069 | -0.069 | 0.096 | 0.72 | 0.91 | 0.53 | 0.63 | 0.44 | 0.44 |
| 9 | 110377174 | rs10978945 | 1.036 | -3.532 | -2.772 | -0.343 | 0.016 | 0.058 | 0.058 | 1.343 | 0.23 | 0.01 | 0.11 | 0.07 | 0.16 | 0.1 |
| 9 | 110379567 | rs12682930 | 0.368 | -3.147 | -2.199 | 0.032 | -0.009 | 0.013 | 0.013 | 0.038 | 0.17 | 0.2 | 0.15 | 0.33 | 0.07 | 0.12 |
| 9 | 110382117 | rs2017419 | -1.266 | -2.973 | -2.812 | 0.076 | -0.125 | -0.115 | -0.115 | 0.387 | 0.76 | 0.96 | 0.49 | 0.63 | 0.46 | 0.51 |
| 9 | 110383280 | rs7022385 | 0.251 | -3.561 | -2.705 | 0.338 | 0.031 | -0.009 | -0.009 | 0 | 0.18 | 0.09 | 0.21 | 0.17 | 0.1 | 0.13 |
| 9 | 110383583 | rs1110403 | -1.051 | -3.121 | -3.012 | 0.358 | -0.075 | 0.012 | 0.012 | 1.109 | 0.67 | 0.93 | 0.61 | 0.76 | 0.4 | 0.46 |
| 9 | 110384379 | rs10759245 | -0.191 | -2.233 | -2.526 | 0.653 | 0.006 | -0.124 | -0.124 | 0.829 | 0.36 | 0.07 | 0.34 | 0.5 | 0.28 | 0.19 |
| 9 | 110386734 | rs10978947 | 0.467 | -1.998 | -2.381 | 0.845 | -0.014 | -0.046 | -0.046 | 0 | 0.1 | 0.01 | 0.22 | 0.48 | 0.07 | 0.1 |
| 9 | 110389815 | rs10978950 | 0.25 | -2.159 | -2.481 | -0.134 | -0.028 | 0.122 | 0.122 | 0.349 | 0.25 | 0.01 | 0.12 | 0.04 | 0.2 | 0.09 |
| 9 | 110396073 | rs11794132 | 0.343 | -2.377 | -2.509 | -0.244 | -0.015 | 0.101 | 0.101 | 0 | 0.19 | 0.01 | 0.09 | 0.03 | 0.16 | 0.08 |
| 9 | 110397107 | rs7864185 | -0.69 | -3.159 | -1.823 | -0.595 | -0.004 | -0.015 | -0.015 | 0.765 | 0.42 | 0.43 | 0.18 | 0.16 | 0.31 | 0.28 |
| 9 | 110407586 | rs12001098 | 0.189 | -3.081 | -1.498 | -0.463 | -0.002 | -0.001 | -0.001 | 1.255 | 0.19 | 0.19 | 0.1 | 0.06 | 0.13 | 0.15 |
| 9 | 110416006 | rs4978532 | -1.115 | -1.392 | -1.595 | -0.298 | -0.084 | -0.01 | -0.01 | 0 | 0.82 | 0.9 | 0.76 | 0.87 | 0.53 | 0.61 |
| 9 | 110419649 | rs11790667 | 0.282 | -1.852 | -1.005 | -0.535 | -0.011 | -0.022 | -0.022 | 0.177 | 0.2 | 0.28 | 0.13 | 0.13 | 0.17 | 0.15 |
| 9 | 110422723 | rs10816543 | 0.396 | -1.905 | -1.199 | -0.561 | -0.003 | -0.006 | -0.006 | 0.739 | 0.2 | 0.22 | 0.12 | 0.13 | 0.17 | 0.15 |
| 9 | 110422729 | rs10816544 | 0.396 | -1.905 | -1.199 | -0.561 | -0.003 | -0.006 | -0.006 | 0 | 0.2 | 0.22 | 0.12 | 0.13 | 0.17 | 0.15 |
| 9 | 110424020 | rs4979219 | 1.46 | -2.49 | -1.921 | 0.042 | -0.028 | -0.062 | -0.062 | 1.419 | 0.18 | 0.03 | 0.23 | 0.26 | 0.15 | 0.24 |
| 9 | 110424575 | rs7040614 | -0.281 | -2.719 | -1.17 | -0.161 | -0.014 | -0.019 | -0.019 | 0 | 0.38 | 0.52 | 0.2 | 0.17 | 0.29 | 0.25 |
| 9 | 110431994 | rs12351531 | 0.603 | -1.96 | -1.431 | 0.387 | -0.015 | -0.018 | -0.018 | 0.307 | 0.14 | 0.01 | 0.07 | 0.09 | 0.11 | 0.07 |
| 9 | 110432131 | rs10978988 | 1.025 | -2.289 | -1.606 | 0.567 | 0.005 | -0.038 | -0.038 | 0 | 0.15 | 0.01 | 0.28 | 0.22 | 0.07 | 0.19 |
| 9 | 110435868 | rs10816550 | -0.521 | -1.862 | -1.308 | 0.551 | 0.045 | -0.016 | -0.016 | 0 | 0.47 | 0.28 | 0.45 | 0.51 | 0.34 | 0.39 |
| 9 | 110436899 | rs940290 | -1.403 | -1.104 | -1.546 | 0.581 | -0.051 | -0.008 | -0.008 | 0 | 0.83 | 0.92 | 0.79 | 0.87 | 0.58 | 0.67 |
| 9 | 110436968 | rs940289 | -1.123 | -1.143 | -1.297 | 0.562 | 0.021 | 0 | 0 | 0 | 0.82 | 0.85 | 0.79 | 0.87 | 0.58 | 0.67 |
| 9 | 110436996 | rs940288 | -0.11 | -1.428 | -1.527 | 0.528 | -0.023 | 0.008 | 0.008 | 0 | 0.62 | 0.84 | 0.69 | 0.74 | 0.4 | 0.56 |
| 9 | 110438418 | rs10978995 | 0.784 | -1.43 | -1.832 | 0.43 | -0.019 | -0.023 | -0.023 | 0 | 0.2 | 0.01 | 0.1 | 0.13 | 0.18 | 0.11 |
| 9 | 110438441 | rs10978996 | -0.09 | -1.427 | -1.521 | 0.423 | 0.014 | -0.016 | -0.016 | 0 | 0.37 | 0.16 | 0.42 | 0.39 | 0.29 | 0.32 |
| 9 | 110438476 | rs10816552 | -0.09 | -1.427 | -1.533 | 0.423 | 0.014 | -0.017 | -0.017 | 0 | 0.37 | 0.15 | 0.42 | 0.39 | 0.29 | 0.35 |
| 9 | 110448724 | rs4979249 | -0.547 | -0.808 | -0.85 | 0.451 | 0.063 | -0.004 | -0.004 | 0.108 | 0.81 | 0.73 | 0.79 | 0.83 | 0.61 | 0.68 |
| 9 | 110451516 | rs4978541 | 1.496 | -1.518 | -1.507 | -0.077 | 0.017 | 0.067 | 0.067 | 0 | 0.26 | 0.11 | 0.13 | 0.02 | 0.19 | 0.19 |
| 9 | 110451579 | rs10816556 | 1.143 | -1.134 | -1.804 | 0.604 | -0.002 | -0.069 | -0.069 | 0 | 0.17 | 0 | 0.1 | 0.18 | 0.16 | 0.09 |
| 9 | 110451637 | rs10759254 | 1.143 | -1.134 | -1.804 | 0.604 | -0.002 | -0.069 | -0.069 | 0 | 0.17 | 0 | 0.1 | 0.18 | 0.16 | 0.09 |
| 9 | 110451936 | rs11789097 | 1.005 | -1.292 | -1.911 | 0.711 | -0.048 | -0.056 | -0.056 | 0.74 | 0.2 | 0.02 | 0.34 | 0.24 | 0.18 | 0.26 |
| 9 | 110452511 | rs1403856 | -1.377 | -1.149 | -1.797 | 0.7 | 0.027 | 0 | 0 | 0 | 0.85 | 0.85 | 0.8 | 0.83 | 0.61 | 0.73 |
| 9 | 110453475 | rs1523784 | 0.325 | -1.485 | -1.463 | 0.619 | 0.062 | 0.042 | 0.042 | 0.299 | 0.45 | 0.67 | 0.34 | 0.35 | 0.24 | 0.36 |
| 9 | 110454145 | rs2049366 | -0.213 | -0.738 | -0.916 | 0.557 | -0.013 | -0.015 | -0.015 | 0.211 | 0.65 | 0.69 | 0.68 | 0.59 | 0.42 | 0.62 |
| 9 | 110459321 | rs4557764 | -0.218 | -1.148 | -0.723 | 0.407 | -0.015 | 0 | 0 | 0 | 0.39 | 0.62 | 0.57 | 0.63 | 0.25 | 0.4 |
| 9 | 110460993 | rs10512382 | -0.349 | -0.9 | -0.719 | 0.31 | 0.132 | -0.012 | -0.012 | 0 | 0.78 | 0.61 | 0.78 | 0.82 | 0.59 | 0.69 |
| 9 | 110462320 | rs12685971 | 1.24 | -1.108 | -1.157 | 0.35 | 0.03 | -0.039 | -0.039 | 1.761 | 0.15 | 0.02 | 0.1 | 0.16 | 0.13 | 0.08 |
| 9 | 110462930 | rs2149998 | -1.283 | -0.84 | -0.97 | 0.204 | 0.027 | 0.001 | 0.001 | 0.222 | 0.85 | 0.84 | 0.81 | 0.83 | 0.62 | 0.74 |
| 9 | 110463371 | rs7032602 | 0.797 | -1.676 | -0.886 | 0.421 | 0.003 | 0.053 | 0.053 | 0.047 | 0.46 | 0.27 | 0.49 | 0.32 | 0.42 | 0.49 |
| 9 | 110466214 | rs4979264 | -1.337 | -1.09 | -1.084 | 0.463 | 0.031 | 0.001 | 0.001 | 0.286 | 0.85 | 0.84 | 0.81 | 0.83 | 0.62 | 0.74 |
| 9 | 110467351 | rs6477589 | -0.882 | -1.137 | -1.387 | 0.299 | -0.044 | 0.012 | 0.012 | 0.462 | 0.67 | 0.84 | 0.69 | 0.73 | 0.44 | 0.55 |
| 9 | 110467423 | rs6477590 | -1.526 | -0.853 | -1.249 | 0.443 | 0.012 | -0.002 | -0.002 | 0 | 0.87 | 0.88 | 0.83 | 0.82 | 0.65 | 0.76 |
| 9 | 110467498 | rs6477591 | 0.066 | -1.308 | -1.344 | 0.704 | -0.031 | -0.029 | -0.029 | 0 | 0.65 | 0.75 | 0.73 | 0.51 | 0.5 | 0.62 |
| 9 | 110469181 | rs10979018 | 0.684 | -1.203 | -1.397 | 0.778 | 0.022 | -0.058 | -0.058 | 0 | 0.2 | 0.07 | 0.11 | 0.31 | 0.15 | 0.14 |
| 9 | 110469925 | rs1880146 | -1.11 | -0.811 | -1.371 | 0.426 | -0.01 | -0.005 | -0.005 | 0.253 | 0.85 | 0.87 | 0.81 | 0.82 | 0.62 | 0.73 |
| 9 | 110472296 | rs1411355 | 1.2 | -1.304 | -0.952 | 0.099 | 0.007 | 0.024 | 0.024 | 0 | 0.27 | 0.24 | 0.17 | 0.1 | 0.24 | 0.23 |
| 9 | 110474940 | rs7874022 | -0.101 | -1.355 | -1.402 | 0.152 | 0.027 | 0.017 | 0.017 | 0 | 0.55 | 0.77 | 0.42 | 0.4 | 0.35 | 0.43 |
| 9 | 110477630 | rs11789693 | -0.706 | -1.063 | -0.916 | 0.227 | 0.009 | 0.004 | 0.004 | 0 | 0.52 | 0.55 | 0.42 | 0.56 | 0.41 | 0.42 |
| 9 | 110477887 | rs10979023 | 0.952 | -0.857 | -0.9 | 0.338 | -0.002 | 0.038 | 0.038 | 0 | 0.16 | 0.06 | 0.08 | 0.09 | 0.18 | 0.07 |
| 9 | 110480149 | rs7038436 | -0.789 | -0.669 | -0.745 | 0.479 | -0.001 | -0.002 | -0.002 | 10.599 | 0.58 | 0.54 | 0.44 | 0.56 | 0.44 | 0.49 |
| 9 | 110488648 | rs10217338 | -0.536 | -0.907 | -0.382 | 0.699 | -0.01 | 0.004 | 0.004 | 0 | 0.4 | 0.42 | 0.35 | 0.55 | 0.3 | 0.36 |
| 9 | 110489551 | rs12555298 | 0.761 | -0.819 | -1.08 | 0.981 | -0.011 | -0.097 | -0.097 | 0.485 | 0.18 | 0.03 | 0.31 | 0.33 | 0.18 | 0.2 |
| 9 | 110489771 | rs6477592 | 0.396 | 0.175 | -0.957 | 0.729 | -0.006 | -0.004 | -0.004 | 0.148 | 0.61 | 0.68 | 0.45 | 0.36 | 0.53 | 0.69 |
| 9 | 110494208 | rs6477593 | -0.702 | 0.586 | -1.254 | 0.446 | -0.079 | 0.023 | 0.023 | 0 | 0.84 | 0.99 | 0.89 | 0.9 | 0.75 | 0.92 |
| 9 | 110495044 | rs4979291 | -0.005 | 0.491 | -1.234 | 0.318 | -0.019 | -0.009 | -0.009 | 0 | 0.47 | 0.26 | 0.58 | 0.47 | 0.48 | 0.64 |
| 9 | 110497235 | rs7863292 | -0.426 | 0.151 | -0.602 | 0.525 | 0.025 | 0.067 | 0.067 | 0 | 0.05 | 0.22 | 0.02 | 0.1 | 0.03 | 0.04 |
| 9 | 110497951 | rs10759255 | -0.104 | -0.371 | -1.154 | 0.108 | 0.064 | -0.008 | -0.008 | 0.045 | 0.54 | 0.29 | 0.5 | 0.46 | 0.65 | 0.74 |
| 9 | 110498273 | rs2417804 | -1.04 | -0.213 | -1.517 | 0.005 | -0.039 | 0.018 | 0.018 | 0 | 0.86 | 0.99 | 0.92 | 0.91 | 0.82 | 0.92 |

| | | | | | | | | | | | | | | | | |
|---|---|---|---|---|---|---|---|---|---|---|---|---|---|---|---|---|
| 9 | 110501437 | rs1851741 | 0.053 | -0.166 | -1.608 | 0.341 | -0.041 | -0.081 | -0.081 | 0 | 0.36 | 0.16 | 0.47 | 0.54 | 0.54 | 0.6 |
| 9 | 110503765 | rs1880143 | -0.065 | 0.397 | -0.769 | 0.583 | 0.032 | 0.042 | 0.042 | 0 | 0.33 | 0.54 | 0.43 | 0.43 | 0.48 | 0.51 |
| 9 | 110504358 | rs944178 | 1.58 | -1.006 | -0.482 | 0.408 | -0.014 | -0.007 | -0.007 | 0 | 0.21 | 0.24 | 0.18 | 0.13 | 0.07 | 0.11 |
| 9 | 110504472 | rs2030641 | 0.681 | -0.412 | -0.333 | 0.666 | 0.08 | 0.067 | 0.067 | 1.307 | 0.28 | 0.53 | 0.44 | 0.38 | 0.48 | 0.34 |
| 9 | 110505424 | rs10979032 | 1.111 | -0.233 | -0.616 | 0.605 | -0.053 | -0.021 | -0.021 | 0.535 | 0.24 | 0.15 | 0.4 | 0.27 | 0.41 | 0.33 |
| 9 | 110509904 | rs7037931 | 0.332 | -0.405 | -0.43 | 0.361 | 0.005 | -0.021 | -0.021 | 0.036 | 0.72 | 0.66 | 0.81 | 0.82 | 0.68 | 0.67 |
| 9 | 110510862 | rs10733564 | -1.267 | -0.329 | -0.594 | 0.349 | -0.031 | -0.017 | -0.017 | 0 | 0.94 | 1 | 0.95 | 0.9 | 0.88 | 0.93 |
| 9 | 110511408 | rs7042939 | 1.343 | -0.443 | -0.383 | 0.454 | -0.005 | 0.001 | 0.001 | 0 | 0.67 | 0.34 | 0.78 | 0.71 | 0.6 | 0.61 |
| 9 | 110511936 | rs4978566 | 0.358 | -0.406 | -0.461 | 0.379 | -0.001 | -0.02 | -0.02 | 0 | 0.74 | 0.67 | 0.84 | 0.82 | 0.73 | 0.7 |
| 9 | 110512551 | rs7860653 | -0.936 | -0.31 | -0.778 | 0.377 | -0.047 | 0.025 | 0.025 | 0 | 0.85 | 0.99 | 0.9 | 0.9 | 0.8 | 0.9 |
| 9 | 110514284 | rs10979037 | 1.275 | -0.398 | -0.799 | 0.45 | -0.04 | -0.012 | -0.012 | 0 | 0.24 | 0.15 | 0.39 | 0.26 | 0.39 | 0.28 |
| 9 | 110517794 | rs944172 | -0.284 | -0.364 | -0.269 | 0.706 | 0.021 | 0.018 | 0.018 | 0 | 0.82 | 0.73 | 0.87 | 0.79 | 0.72 | 0.85 |
| 9 | 110519538 | rs10118459 | -0.909 | -0.311 | -0.391 | 0.713 | -0.051 | 0.028 | 0.028 | 0 | 0.84 | 0.99 | 0.89 | 0.9 | 0.78 | 0.9 |
| 9 | 110520428 | rs10512379 | 1.707 | -0.684 | -0.641 | 0.703 | -0.028 | -0.015 | -0.015 | 0 | 0.11 | 0 | 0.06 | 0.12 | 0.11 | 0.09 |
| 9 | 110520534 | rs12684481 | 0.566 | -0.501 | -0.415 | 0.824 | -0.014 | -0.025 | -0.025 | 4.14 | 0.06 | 0.02 | 0.02 | 0.08 | 0.05 | 0.19 |
| 9 | 110522741 | rs7040755 | 0.444 | -0.682 | -0.112 | 0.902 | 0.011 | 0.038 | 0.038 | 0 | 0.74 | 0.41 | 0.69 | 0.62 | 0.67 | 0.73 |
| 9 | 110531496 | rs10759257 | 1.139 | -0.675 | -0.475 | 0.886 | -0.007 | 0.058 | 0.058 | 0 | 0.68 | 0.53 | 0.72 | 0.53 | 0.65 | 0.58 |
| 9 | 110532522 | rs2140483 | 0.408 | -0.374 | -0.317 | 0.799 | 0.007 | -0.001 | -0.001 | 2.182 | 0.77 | 0.75 | 0.83 | 0.8 | 0.67 | 0.8 |
| 9 | 110534010 | rs7028227 | -0.802 | -0.344 | -0.587 | 0.797 | -0.048 | 0.025 | 0.025 | 0 | 0.84 | 0.99 | 0.89 | 0.9 | 0.79 | 0.9 |
| 9 | 110536932 | rs1467311 | -0.165 | -0.554 | -0.38 | 0.812 | -0.036 | -0.009 | -0.009 | 3.023 | 0.79 | 0.87 | 0.83 | 0.8 | 0.67 | 0.84 |
| 9 | 110542327 | rs1411357 | -0.258 | -0.529 | -0.381 | 0.838 | -0.04 | -0.014 | -0.014 | 0 | 0.79 | 0.88 | 0.83 | 0.8 | 0.67 | 0.84 |
| 9 | 110544407 | rs1330150 | 0.918 | -0.923 | -0.604 | 0.694 | -0.005 | 0.002 | 0.002 | 0.545 | 0.36 | 0.33 | 0.46 | 0.28 | 0.46 | 0.38 |
| 9 | 110544998 | rs4979374 | -0.2 | -1.512 | -1.03 | 0.543 | 0.035 | -0.013 | -0.013 | 0 | 0.42 | 0.35 | 0.36 | 0.52 | 0.21 | 0.43 |
| 9 | 110548764 | rs1330147 | 0.853 | -0.897 | -0.809 | 0.467 | -0.016 | -0.006 | -0.006 | 0 | 0.12 | 0.02 | 0.05 | 0.11 | 0.1 | 0.08 |
| 9 | 110551835 | rs953383 | 0.503 | -1.024 | -0.513 | -0.099 | -0.012 | -0.136 | -0.136 | 0.323 | 0.23 | 0.37 | 0.16 | 0.01 | 0.21 | 0.16 |
| 9 | 110552239 | rs1330146 | -1.348 | -1.241 | -0.778 | 0.22 | 0.006 | -0.019 | -0.019 | 0 | 0.57 | 0.71 | 0.44 | 0.55 | 0.47 | 0.57 |
| 9 | 110553667 | rs10816576 | -1.331 | -1.243 | -0.768 | 0.219 | 0.007 | -0.018 | -0.018 | 0 | 0.56 | 0.71 | 0.44 | 0.55 | 0.47 | 0.57 |
| 9 | 110556602 | rs1360257 | -0.532 | -1.32 | -0.638 | 0.291 | -0.007 | 0.002 | 0.002 | 0 | 0.29 | 0.3 | 0.26 | 0.43 | 0.21 | 0.3 |
| 9 | 110557377 | rs1411356 | -0.402 | -0.975 | -0.876 | 0.552 | 0.048 | 0.013 | 0.013 | 0.011 | 0.46 | 0.69 | 0.38 | 0.5 | 0.39 | 0.52 |
| 9 | 110570206 | rs12339041 | 0.711 | -0.538 | -1.311 | 0.499 | 0.007 | 0.013 | 0.013 | 0 | 0.11 | 0.07 | 0.06 | 0.07 | 0.09 | 0.05 |
| 9 | 110571327 | rs7028182 | -1.157 | -0.716 | -1.575 | 0.582 | -0.016 | 0.023 | 0.023 | 0 | 0.59 | 0.77 | 0.57 | 0.68 | 0.5 | 0.59 |
| 9 | 110571952 | rs9632903 | 0.631 | -0.45 | -1.431 | 0.676 | 0.008 | 0.014 | 0.014 | 0 | 0.11 | 0.07 | 0.06 | 0.07 | 0.09 | 0.05 |
| 9 | 110573303 | rs7848776 | 0.063 | -0.316 | -1.431 | 0.785 | 0.003 | -0.044 | -0.044 | 0 | 0.17 | 0.1 | 0.21 | 0.23 | 0.17 | 0.17 |
| 9 | 110581046 | rs10121193 | -0.555 | 0.162 | -1.275 | 0.515 | -0.048 | 0.015 | 0.015 | 0 | 0.62 | 0.77 | 0.59 | 0.67 | 0.56 | 0.61 |
| 9 | 110587368 | rs10733565 | -0.702 | 0.559 | -1.002 | 0.643 | -0.005 | -0.015 | -0.015 | 0 | 0.63 | 0.72 | 0.65 | 0.6 | 0.61 | 0.69 |
| 9 | 110587889 | rs7875395 | -1.276 | 0.513 | 0.093 | 1.402 | 0.005 | 0.12 | 0.12 | 1.1 | 0.14 | 0.37 | 0.23 | 0.24 | 0.15 | 0.22 |
| 9 | 110597519 | rs10979066 | -1.2 | 0.51 | -0.247 | 1.418 | -0.008 | 0.048 | 0.048 | 0 | 0.15 | 0.26 | 0.23 | 0.24 | 0.15 | 0.22 |
| 9 | 110612674 | rs7018799 | -1.229 | 0.485 | -0.381 | 1.571 | -0.005 | 0.067 | 0.067 | 0 | 0.17 | 0.25 | 0.32 | 0.59 | 0.16 | 0.31 |
| 9 | 110612675 | rs7018800 | -0.863 | 0.502 | -0.573 | 1.581 | 0 | 0.041 | 0.041 | 0 | 0.13 | 0.1 | 0.29 | 0.52 | 0.11 | 0.27 |
| 9 | 110615884 | rs4979494 | -0.884 | 0.497 | -0.563 | 1.588 | -0.001 | 0.047 | 0.047 | 0 | 0.13 | 0.12 | 0.29 | 0.53 | 0.11 | 0.27 |
| 9 | 110618805 | rs10979075 | -1.499 | 0.464 | -0.515 | 1.593 | -0.004 | 0.039 | 0.039 | 0 | 0.18 | 0.25 | 0.32 | 0.61 | 0.16 | 0.31 |
| 9 | 110628820 | rs4978629 | -0.961 | 0.451 | -0.89 | 1.338 | -0.001 | 0.003 | 0.003 | 0 | 0.29 | 0.26 | 0.36 | 0.48 | 0.26 | 0.47 |
| 9 | 110632874 | rs11794699 | -0.118 | 1.109 | -1.831 | 1.015 | -0.027 | -0.032 | -0.032 | 0 | 0.32 | 0.23 | 0.33 | 0.63 | 0.35 | 0.4 |
| 9 | 110635349 | rs10759261 | -0.239 | 1.145 | -2.019 | 0.947 | -0.017 | -0.002 | -0.002 | 0.866 | 0.49 | 0.23 | 0.46 | 0.59 | 0.55 | 0.55 |
| 9 | 110635883 | rs10816581 | -0.524 | 1.412 | -1.945 | 1.051 | -0.013 | 0.004 | 0.004 | 0 | 0.38 | 0.28 | 0.43 | 0.37 | 0.37 | 0.34 |
| 9 | 110637432 | rs7852099 | 0.831 | 1.21 | -1.714 | 1.497 | -0.053 | -0.04 | -0.04 | 0.259 | 0.92 | 0.98 | 0.7 | 0.66 | 0.84 | 0.82 |
| 9 | 110642150 | rs7045297 | 0.908 | 0.617 | -1.804 | 0.165 | -0.013 | 0.031 | 0.031 | 0 | 0.27 | 0.2 | 0.23 | 0.13 | 0.33 | 0.17 |
| 9 | 110645412 | rs7021395 | -0.054 | -0.09 | -1.322 | 0.089 | 0.003 | 0.003 | 0.003 | 0 | 0.71 | 0.72 | 0.6 | 0.42 | 0.67 | 0.55 |
| 9 | 110646522 | rs10979082 | 0.835 | -0.497 | -1.557 | 0.003 | 0.014 | -0.021 | -0.021 | 0.119 | 0.08 | 0 | 0.13 | 0.13 | 0.06 | 0.05 |
| 9 | 110649553 | rs10119909 | 0.529 | -0.937 | -1.371 | -0.323 | 0.001 | 0.042 | 0.042 | 0.382 | 0.25 | 0.12 | 0.21 | 0.16 | 0.24 | 0.3 |
| 9 | 110650284 | rs12335733 | 1.905 | -0.796 | -1.446 | -0.344 | -0.022 | 0.036 | 0.036 | 0.012 | 0.27 | 0.2 | 0.21 | 0.13 | 0.36 | 0.17 |
| 9 | 110657726 | rs944472 | -0.158 | -1.058 | -1.165 | -0.29 | 0.016 | 0.003 | 0.003 | 0 | 0.69 | 0.63 | 0.58 | 0.42 | 0.66 | 0.56 |
| 9 | 110658115 | rs1929273 | 1.431 | -0.783 | -1.05 | -0.362 | -0.022 | 0.043 | 0.043 | 0 | 0.3 | 0.18 | 0.21 | 0.13 | 0.38 | 0.17 |
| 9 | 110660983 | rs34048749 | 0.717 | -0.503 | -0.891 | -0.066 | 0.022 | -0.029 | -0.029 | 1.052 | 0.09 | 0 | 0.13 | 0.14 | 0.05 | 0.03 |
| 9 | 110663574 | rs10759262 | -0.838 | -0.669 | -0.59 | 0.118 | 0 | 0.002 | 0.002 | 0 | 0.57 | 0.64 | 0.6 | 0.65 | 0.45 | 0.68 |
| 9 | 110666409 | rs1929276 | -1.212 | -0.631 | -0.608 | 0.125 | -0.013 | 0.019 | 0.019 | 0 | 0.61 | 0.79 | 0.68 | 0.66 | 0.45 | 0.66 |
| 9 | 110668683 | rs1929278 | -0.376 | -0.807 | -0.363 | 0.326 | 0.01 | 0.009 | 0.009 | 0 | 0.44 | 0.45 | 0.46 | 0.38 | 0.37 | 0.43 |
| 9 | 110677097 | rs4978649 | 0.298 | -0.065 | -0.263 | -0.261 | 0.024 | 0 | 0 | 0 | 0.43 | 0.32 | 0.56 | 0.53 | 0.36 | 0.62 |
| 9 | 110683074 | rs12348520 | 1.02 | 0.476 | 0.152 | -0.386 | 0.001 | -0.009 | -0.009 | 0 | 0.07 | 0.08 | 0.03 | 0 | 0.04 | 0.01 |
| 9 | 110688674 | rs7020351 | -0.046 | 0.385 | 0.267 | -0.348 | 0.03 | -0.015 | -0.015 | 0 | 0.08 | 0.43 | 0.05 | 0 | 0.03 | 0.01 |
| 9 | 110688738 | rs1339764 | 0.138 | 0.861 | -0.543 | -0.133 | 0.025 | 0.045 | 0.045 | 0 | 0.55 | 0.74 | 0.64 | 0.67 | 0.41 | 0.68 |
| 9 | 110688855 | rs2039802 | 0.039 | 0.615 | -0.327 | -0.428 | -0.02 | 0.05 | 0.05 | 0 | 0.31 | 0.16 | 0.23 | 0.12 | 0.38 | 0.2 |
| 9 | 110694926 | rs1361369 | 0.246 | 0.74 | -0.334 | 0.716 | 0.003 | 0.002 | 0.002 | 0 | 0.62 | 0.64 | 0.4 | 0.55 | 0.54 | 0.44 |
| 9 | 110695062 | rs950143 | -0.861 | -0.381 | -0.799 | 0.923 | 0.041 | -0.031 | -0.031 | 0.021 | 0.15 | 0.06 | 0.05 | 0.21 | 0.08 | 0.24 |
| 9 | 110696206 | rs1361368 | 0.851 | -0.91 | 0.239 | 1.08 | -0.055 | 0.077 | 0.077 | 0 | 0.73 | 0.61 | 0.74 | 0.45 | 0.86 | 0.49 |
| 9 | 110700994 | rs10816591 | 0.12 | -0.928 | -0.339 | 0.276 | -0.022 | 0.06 | 0.06 | 0 | 0.38 | 0.31 | 0.35 | 0.13 | 0.52 | 0.32 |
| 9 | 110702554 | rs6477606 | 0.873 | -0.542 | -0.666 | -0.034 | -0.001 | -0.024 | -0.024 | 0 | 0.25 | 0.29 | 0.17 | 0.05 | 0.21 | 0.17 |
| 9 | 110703980 | rs12376694 | 0.65 | -0.651 | -0.618 | 0.059 | -0.09 | 0.019 | 0.019 | 0 | 0.12 | 0.01 | 0.18 | 0.08 | 0.31 | 0.13 |
| 9 | 110705646 | rs6477608 | -0.254 | -0.894 | -0.097 | 0.114 | -0.026 | 0.084 | 0.084 | 0 | 0.82 | 0.76 | 0.62 | 0.4 | 0.88 | 0.56 |

| | | | | | | | | | | | | | | | | |
|---|---|---|---|---|---|---|---|---|---|---|---|---|---|---|---|---|
| 9 | 110707050 | rs10512375 | 0.436 | -0.693 | -0.145 | -0.052 | -0.006 | -0.015 | -0.015 | 0 | 0.29 | 0.33 | 0.31 | 0.11 | 0.22 | 0.24 |
| 9 | 110707964 | rs7026539 | -0.384 | -0.608 | -0.122 | -0.232 | -0.016 | 0.048 | 0.048 | 0 | 0.39 | 0.35 | 0.36 | 0.13 | 0.52 | 0.31 |
| 9 | 110709587 | rs10521087 | -0.447 | -0.353 | 0.133 | 0.092 | 0.045 | -0.018 | -0.018 | 0 | 0.18 | 0.31 | 0.2 | 0.12 | 0.31 | 0.16 |
| 9 | 110710767 | rs10739251 | 0.546 | -0.898 | 0.488 | 0.355 | -0.057 | 0.163 | 0.163 | 0 | 0.78 | 0.54 | 0.72 | 0.44 | 0.85 | 0.54 |
| 9 | 110712028 | rs10979125 | -0.294 | -0.327 | 0.184 | 0.064 | -0.042 | 0.02 | 0.02 | 0 | 0.17 | 0.09 | 0.21 | 0.09 | 0.35 | 0.16 |
| 9 | 110712165 | rs10979126 | -0.381 | -0.398 | 0.249 | 0.198 | 0.005 | 0.064 | 0.064 | 0 | 0.07 | 0.23 | 0.05 | 0.09 | 0.04 | 0.15 |
| 9 | 110716274 | rs1888621 | -0.227 | -0.114 | -0.421 | 0.123 | -0.104 | 0.008 | 0.008 | 0 | 0.15 | 0.02 | 0.2 | 0.09 | 0.35 | 0.16 |
| 9 | 110724246 | rs9792589 | 0.982 | -1.696 | -0.853 | -0.092 | 0.011 | -0.053 | -0.053 | 0 | 0.15 | 0.03 | 0.11 | 0.35 | 0.08 | 0.1 |
| 9 | 110725754 | rs10979136 | 0.228 | -1.224 | -0.916 | -0.158 | -0.028 | 0.047 | 0.047 | 0 | 0.12 | 0.03 | 0.07 | 0.02 | 0.17 | 0.11 |
| 9 | 110729131 | rs2709418 | -1.012 | -1.502 | -1.209 | -0.121 | 0.033 | -0.07 | -0.07 | 1.123 | 0.85 | 0.97 | 0.8 | 0.77 | 0.9 | 0.77 |
| 9 | 110729237 | rs1554279 | 1.613 | -1.613 | -0.941 | -0.625 | -0.04 | -0.029 | -0.029 | 0.623 | 0.24 | 0.18 | 0.42 | 0.34 | 0.44 | 0.16 |
| 9 | 110731392 | rs10979145 | -0.095 | -1.629 | -0.435 | -0.732 | -0.002 | -0.046 | -0.046 | 0 | 0.15 | 0.25 | 0.09 | 0.02 | 0.18 | 0.12 |
| 9 | 110732326 | rs13288508 | 1.46 | -1.745 | -0.531 | -0.904 | 0.013 | 0.019 | 0.019 | 0 | 0.22 | 0.1 | 0.16 | 0.02 | 0.18 | 0.27 |
| 9 | 110732838 | rs498924 | 0.084 | -1.609 | -0.16 | -0.642 | 0.036 | -0.031 | -0.031 | 0 | 0.6 | 0.75 | 0.62 | 0.39 | 0.7 | 0.56 |
| 9 | 110734905 | rs560104 | -0.224 | -1.606 | -0.141 | -0.638 | 0.049 | -0.06 | -0.06 | 0 | 0.67 | 0.83 | 0.69 | 0.38 | 0.81 | 0.6 |
| 9 | 110735353 | rs1703007 | 0.015 | -1.723 | -0.038 | -0.597 | -0.004 | -0.004 | -0.004 | 0.093 | 0.75 | 0.75 | 0.75 | 0.82 | 0.78 | 0.72 |
| 9 | 110735536 | rs10120991 | 0.861 | -1.718 | 0.093 | -0.592 | 0.016 | -0.045 | -0.045 | 1.798 | 0.09 | 0.19 | 0.06 | 0 | 0.11 | 0.04 |
| 9 | 110735564 | rs10512374 | 1.113 | -1.749 | -0.009 | -0.589 | 0.022 | -0.066 | -0.066 | 0.219 | 0.18 | 0.09 | 0.12 | 0.44 | 0.08 | 0.16 |
| 9 | 110736719 | rs1703008 | -1.17 | -1.678 | 0.017 | -0.563 | 0.014 | -0.003 | -0.003 | 0.044 | 0.84 | 0.89 | 0.8 | 0.82 | 0.89 | 0.76 |
| 9 | 110736854 | rs7038277 | -0.579 | -1.735 | 0.248 | -0.639 | 0 | 0 | 0 | 0 | 0.58 | 0.52 | 0.37 | 0.51 | 0.45 | 0.59 |
| 9 | 110736914 | rs1769657 | 1.723 | -1.665 | 0.265 | -0.62 | 0.042 | 0.026 | 0.026 | 0 | 0.25 | 0.37 | 0.43 | 0.31 | 0.45 | 0.17 |
| 9 | 110738027 | rs10521086 | 0.976 | -1.841 | 0.268 | -0.634 | -0.005 | 0.029 | 0.029 | 0 | 0.05 | 0.14 | 0.02 | 0.06 | 0.02 | 0.1 |
| 9 | 110742521 | rs10979150 | 1.023 | -1.81 | -0.063 | -0.638 | 0.002 | -0.026 | -0.026 | 0.185 | 0.08 | 0.11 | 0.06 | 0 | 0.11 | 0.04 |
| 9 | 110743292 | rs10124568 | 0.986 | -1.803 | -0.232 | -0.635 | 0.013 | -0.044 | -0.044 | 0 | 0.09 | 0.18 | 0.06 | 0 | 0.11 | 0.04 |
| 9 | 110744722 | rs10119946 | -0.088 | -1.89 | -0.358 | -0.705 | -0.006 | -0.052 | -0.052 | 0 | 0.42 | 0.5 | 0.26 | 0.08 | 0.36 | 0.43 |
| 9 | 110751144 | rs7029391 | -0.084 | -1.818 | -0.501 | -0.615 | 0.05 | 0.023 | 0.023 | 0 | 0.49 | 0.32 | 0.31 | 0.5 | 0.33 | 0.55 |
| 9 | 110751734 | rs592069 | -1.672 | -1.704 | -0.421 | -0.488 | -0.046 | -0.033 | -0.033 | 0 | 0.76 | 0.83 | 0.57 | 0.69 | 0.55 | 0.83 |
| 9 | 110752106 | rs1009813 | 0.521 | -1.55 | -0.92 | -0.443 | 0.002 | -0.01 | -0.01 | 0 | 0.32 | 0.25 | 0.19 | 0.08 | 0.25 | 0.39 |
| 9 | 110754102 | rs1934419 | -0.366 | -0.372 | -0.879 | 0.204 | -0.032 | 0.021 | 0.021 | 0.004 | 0.34 | 0.51 | 0.21 | 0.49 | 0.21 | 0.31 |
| 9 | 110754778 | rs1769652 | 0.172 | -0.795 | -1.779 | 0.121 | 0.069 | 0.126 | 0.126 | 0.169 | 0.42 | 0.12 | 0.36 | 0.2 | 0.34 | 0.53 |
| 9 | 110757887 | rs10979157 | -0.293 | -0.743 | -1.238 | 0.87 | 0.023 | -0.028 | -0.028 | 0.858 | 0.21 | 0.14 | 0.19 | 0.46 | 0.09 | 0.25 |
| 9 | 110759922 | rs673985 | -0.233 | 0.028 | -1.588 | 0.778 | -0.049 | -0.052 | -0.052 | 0.686 | 0.71 | 0.88 | 0.8 | 0.68 | 0.65 | 0.87 |
| 9 | 110760598 | rs4979619 | -0.086 | -0.239 | -1.376 | 0.536 | -0.036 | -0.01 | -0.01 | 0.084 | 0.61 | 0.8 | 0.77 | 0.68 | 0.54 | 0.84 |
| 9 | 110761427 | rs12347482 | 0.675 | -0.673 | -1.198 | 0.331 | 0.022 | -0.02 | -0.02 | 0 | 0.45 | 0.23 | 0.63 | 0.57 | 0.45 | 0.58 |
| 9 | 110761513 | rs12236795 | 0.694 | -0.658 | -1.191 | 0.338 | 0.02 | -0.021 | -0.021 | 0 | 0.45 | 0.23 | 0.63 | 0.56 | 0.45 | 0.57 |
| 9 | 110762725 | rs7341756 | -0.281 | -0.63 | -0.431 | 0.316 | -0.01 | -0.007 | -0.007 | 0 | 0.15 | 0.47 | 0.13 | 0.11 | 0.08 | 0.26 |
| 9 | 110783036 | rs10512368 | -0.07 | -1.075 | -0.071 | 0.575 | 0.019 | 0.01 | 0.01 | 0 | 0.52 | 0.4 | 0.68 | 0.62 | 0.42 | 0.7 |
| 9 | 110787286 | rs7869206 | 0.213 | -1.106 | 0.081 | 0.584 | 0.043 | -0.01 | -0.01 | 0.013 | 0.47 | 0.21 | 0.63 | 0.52 | 0.41 | 0.56 |
| 9 | 110787730 | rs12551076 | 0.07 | -1.109 | 0.087 | 0.577 | 0.053 | 0.027 | 0.027 | 0 | 0.5 | 0.25 | 0.67 | 0.62 | 0.42 | 0.7 |
| 9 | 110801608 | rs10816618 | -0.074 | -1.186 | 0.033 | 0.59 | 0.028 | 0.007 | 0.007 | 0 | 0.51 | 0.37 | 0.67 | 0.6 | 0.42 | 0.7 |
| 9 | 110803324 | rs1914513 | 0.259 | -1.202 | 0.012 | 0.583 | 0.035 | -0.021 | -0.021 | 0 | 0.47 | 0.15 | 0.63 | 0.52 | 0.41 | 0.57 |
| 9 | 110808571 | rs10816621 | 0.783 | -1.861 | -0.472 | 0.431 | 0.022 | 0.003 | 0.003 | 0 | 0.11 | 0.01 | 0.1 | 0.1 | 0.07 | 0.12 |
| 9 | 110820132 | rs4978343 | -0.673 | -1.639 | 0.095 | 0.411 | -0.004 | -0.001 | -0.001 | 0 | 0.52 | 0.52 | 0.69 | 0.6 | 0.43 | 0.69 |
| 9 | 110821050 | rs10979183 | -0.562 | -1.598 | -0.286 | 0.422 | 0.026 | 0.008 | 0.008 | 0 | 0.51 | 0.33 | 0.67 | 0.6 | 0.43 | 0.69 |
| 9 | 110826460 | rs10512367 | 0.046 | -1.824 | -1.292 | 0.331 | 0.047 | -0.025 | -0.025 | 0 | 0.22 | 0.11 | 0.24 | 0.4 | 0.11 | 0.23 |
| 9 | 110829012 | rs13289116 | 0.638 | -1.138 | -1.137 | 0.398 | -0.057 | 0.02 | 0.02 | 0 | 0.25 | 0.08 | 0.39 | 0.12 | 0.31 | 0.32 |
| 9 | 110832584 | rs13291052 | 0.668 | -1.107 | -1.229 | 0.446 | -0.074 | -0.008 | -0.008 | 5.302 | 0.25 | 0.04 | 0.46 | 0.14 | 0.31 | 0.33 |
| 9 | 110837073 | rs10816625 | 1.082 | -1.051 | -0.951 | 1.191 | -0.026 | -0.022 | -0.022 | 0.076 | 0.08 | 0.01 | 0.19 | 0.38 | 0.09 | 0.08 |
| 9 | 110841593 | rs7042321 | -0.7 | -1.774 | -0.16 | -0.348 | -0.092 | -0.028 | -0.028 | 0 | 0.26 | 0.49 | 0.33 | 0.25 | 0.08 | 0.22 |
| 9 | 110843272 | rs10979187 | -1.4 | -1.693 | -0.274 | -0.616 | 0.002 | 0.002 | 0.002 | 1.571 | 0.33 | 0.64 | 0.4 | 0.28 | 0.18 | 0.32 |
| 9 | 110843426 | rs10816626 | -1.399 | -1.667 | -0.273 | -0.607 | 0.002 | 0.002 | 0.002 | 1.414 | 0.33 | 0.64 | 0.4 | 0.29 | 0.18 | 0.32 |
| 9 | 110843927 | rs10118883 | -0.092 | -1.592 | -0.429 | -0.683 | 0.012 | -0.005 | -0.005 | 0 | 0.08 | 0.15 | 0.07 | 0.03 | 0.1 | 0.1 |
| 9 | 110844942 | rs12006421 | -1.545 | -1.608 | -0.56 | -0.681 | 0.017 | 0.014 | 0.014 | 0.063 | 0.38 | 0.65 | 0.45 | 0.32 | 0.24 | 0.51 |
| 9 | 110850146 | rs10512366 | 0.096 | -1.458 | -0.667 | -0.666 | -0.006 | 0.002 | 0.002 | 0 | 0.07 | 0.06 | 0.06 | 0.03 | 0.1 | 0.1 |
| 9 | 110852176 | rs7875633 | -1.522 | -1.274 | -0.36 | -0.719 | 0.05 | 0.018 | 0.018 | 0.074 | 0.41 | 0.68 | 0.45 | 0.38 | 0.25 | 0.51 |
| 9 | 110857507 | rs10120248 | -1.506 | -1.293 | -0.445 | -0.782 | 0.053 | 0.035 | 0.035 | 0.243 | 0.42 | 0.73 | 0.45 | 0.33 | 0.25 | 0.5 |
| 9 | 110857960 | rs1434851 | -1.089 | -1.282 | -0.359 | -0.773 | -0.006 | -0.001 | -0.001 | 0 | 0.35 | 0.62 | 0.4 | 0.32 | 0.19 | 0.33 |
| 9 | 110864014 | rs1831590 | -1.072 | -1.377 | -0.213 | -0.758 | -0.016 | -0.007 | -0.007 | 0.203 | 0.35 | 0.59 | 0.39 | 0.29 | 0.19 | 0.34 |
| 9 | 110867734 | rs7024830 | -1.121 | -0.025 | -0.899 | -1.244 | -0.045 | 0.006 | 0.006 | 0 | 0.72 | 0.89 | 0.71 | 0.27 | 0.62 | 0.68 |
| 9 | 110868532 | rs10120623 | 1.21 | -0.235 | -0.476 | -1.445 | -0.036 | 0.127 | 0.127 | 0.624 | 0.36 | 0.21 | 0.29 | 0.08 | 0.39 | 0.35 |
| 9 | 110868550 | rs1412436 | 0.395 | -0.518 | 0.135 | -1.555 | 0 | -0.007 | -0.007 | 0 | 0.16 | 0.31 | 0.17 | 0.15 | 0.12 | 0.12 |
| 9 | 110871414 | rs10979214 | 1.415 | -0.442 | -0.432 | -1.553 | -0.049 | 0.151 | 0.151 | 0.744 | 0.36 | 0.18 | 0.29 | 0.08 | 0.39 | 0.35 |
| 9 | 110872527 | rs10124038 | -0.976 | -0.457 | -0.39 | -1.514 | -0.043 | 0.016 | 0.016 | 0 | 0.68 | 0.88 | 0.7 | 0.31 | 0.58 | 0.66 |
| 9 | 110874194 | rs10816631 | 1.384 | -0.178 | -0.335 | -1.576 | -0.02 | 0.042 | 0.042 | 0.864 | 0.3 | 0.27 | 0.26 | 0.03 | 0.34 | 0.18 |
| 9 | 110875895 | rs10979217 | -0.456 | -0.71 | -0.102 | -0.909 | -0.002 | 0 | 0 | 0 | 0.11 | 0.1 | 0.21 | 0.08 | 0.06 | 0.16 |
| 9 | 110881453 | rs837983 | -1.166 | 0.034 | 0.228 | -1.204 | 0.016 | -0.043 | -0.043 | 0.039 | 0.74 | 0.71 | 0.7 | 0.95 | 0.63 | 0.89 |
| 9 | 110882390 | rs667052 | 0.696 | -0.341 | 0.5 | -1.47 | -0.004 | 0.12 | 0.12 | 0 | 0.52 | 0.34 | 0.45 | 0.81 | 0.5 | 0.57 |
| 9 | 110885479 | rs630965 | -0.926 | -0.326 | 0.809 | -1.425 | 0.038 | -0.093 | -0.093 | 0.528 | 0.71 | 0.49 | 0.68 | 0.92 | 0.63 | 0.88 |
| 9 | 110887234 | rs1025572 | -0.437 | -0.295 | 0.247 | -1.392 | 0.005 | 0.006 | 0.006 | 0 | 0.11 | 0.06 | 0.19 | 0.1 | 0.07 | 0.16 |

| | | | | | | | | | | | | | | | | |
|---|---|---|---|---|---|---|---|---|---|---|---|---|---|---|---|---|
| 9 | 110887934 | rs580018 | 0.707 | -0.213 | -0.019 | -1.548 | -0.039 | 0.089 | 0.089 | 0 | 0.17 | 0.03 | 0.11 | 0.02 | 0.18 | 0.14 |
| 9 | 110891494 | rs7870781 | 0.72 | -0.337 | 0.37 | -1.508 | -0.012 | -0.081 | -0.081 | 0.096 | 0.41 | 0.46 | 0.27 | 0.07 | 0.32 | 0.36 |
| 9 | 110893829 | rs10979227 | 1.818 | -0.363 | -0.035 | -1.512 | -0.02 | 0.158 | 0.158 | 0.107 | 0.29 | 0.06 | 0.19 | 0.03 | 0.25 | 0.2 |
| 9 | 110896340 | rs516327 | -0.285 | -0.551 | 0.734 | -1.401 | 0 | 0.001 | 0.001 | 0.089 | 0.27 | 0.22 | 0.26 | 0.78 | 0.25 | 0.37 |
| 9 | 110896828 | rs595811 | -1.621 | -0.511 | 0.829 | -1.454 | 0.04 | 0.142 | 0.142 | 1.539 | 0.46 | 0.72 | 0.51 | 0.92 | 0.38 | 0.7 |
| 9 | 110901598 | rs7863999 | -0.858 | -0.466 | 1.196 | -1.272 | -0.008 | 0.04 | 0.04 | 0.66 | 0.1 | 0.31 | 0.09 | 0.12 | 0.06 | 0.14 |
| 9 | 110905570 | rs567032 | -1.165 | -0.535 | 0.841 | -0.417 | 0.026 | -0.022 | -0.022 | 0.029 | 0.39 | 0.74 | 0.32 | 0.48 | 0.32 | 0.56 |
| 9 | 110907467 | rs665556 | -0.002 | -0.52 | 1.1 | -0.398 | 0 | 0.002 | 0.002 | 0 | 0.26 | 0.22 | 0.25 | 0.45 | 0.25 | 0.32 |
| 9 | 110911316 | rs10512364 | 1.883 | -0.534 | 0.739 | -0.604 | 0.012 | 0.105 | 0.105 | 0.037 | 0.29 | 0.12 | 0.3 | 0.04 | 0.25 | 0.2 |
| 9 | 110911435 | rs566510 | -0.982 | -0.474 | 0.851 | -0.552 | -0.023 | -0.021 | -0.021 | 1.679 | 0.68 | 0.75 | 0.63 | 0.5 | 0.57 | 0.7 |
| 9 | 110917627 | rs614851 | -0.093 | -0.456 | 1.022 | -0.521 | -0.003 | 0.028 | 0.028 | 0 | 0.28 | 0.3 | 0.27 | 0.47 | 0.26 | 0.38 |
| 9 | 110923018 | rs4542003 | 0.049 | -0.409 | 0.784 | -0.517 | -0.01 | -0.032 | -0.032 | 0 | 0.09 | 0.15 | 0.05 | 0.01 | 0.06 | 0.11 |
| 9 | 110923883 | rs525803 | -0.848 | -0.19 | 0.879 | -0.297 | -0.037 | -0.031 | -0.031 | 0 | 0.69 | 0.79 | 0.63 | 0.52 | 0.58 | 0.73 |
| 9 | 110929030 | rs687992 | -0.317 | -0.224 | 0.995 | -0.193 | -0.007 | 0.063 | 0.063 | 0 | 0.28 | 0.4 | 0.32 | 0.48 | 0.26 | 0.38 |
| 9 | 110939597 | rs637635 | -0.367 | -0.196 | 0.874 | -0.261 | -0.008 | -0.009 | -0.009 | 0 | 0.65 | 0.69 | 0.57 | 0.52 | 0.56 | 0.73 |
| 9 | 110940013 | rs503238 | -0.326 | -0.182 | 0.885 | 0.071 | -0.007 | 0.089 | 0.089 | 0 | 0.27 | 0.43 | 0.27 | 0.45 | 0.25 | 0.33 |
| 9 | 110941390 | rs666303 | -0.388 | 0.21 | 0.553 | 0.384 | -0.002 | -0.002 | -0.002 | 0 | 0.71 | 0.71 | 0.69 | 0.46 | 0.68 | 0.8 |
| 9 | 110943815 | rs10979263 | -1.987 | -0.812 | 1.283 | 0.646 | -0.027 | 0.391 | 0.391 | 1.533 | 0.11 | 0.44 | 0.1 | 0.51 | 0.05 | 0.19 |
| 9 | 110946296 | rs10759273 | -1.605 | -0.93 | 1.234 | 0.536 | -0.033 | 0.297 | 0.297 | 1.353 | 0.12 | 0.37 | 0.16 | 0.51 | 0.07 | 0.16 |
| 9 | 110946380 | rs559836 | -0.202 | -0.376 | 0.457 | -0.313 | -0.018 | 0.025 | 0.025 | 0.268 | 0.17 | 0.09 | 0.18 | 0.04 | 0.24 | 0.16 |
| 9 | 110947155 | rs553333 | -0.617 | -0.397 | 0.487 | -0.324 | 0.018 | -0.096 | -0.096 | 0.817 | 0.22 | 0.37 | 0.21 | 0.05 | 0.24 | 0.21 |
| 9 | 110953051 | rs10512362 | -0.296 | -0.448 | -0.117 | 0.612 | 0.004 | 0.066 | 0.066 | 0 | 0.06 | 0.04 | 0.07 | 0.45 | 0.05 | 0.11 |
| 9 | 110954627 | rs13287243 | -0.296 | -0.448 | -0.117 | 0.612 | 0.004 | 0.066 | 0.066 | 0 | 0.06 | 0.03 | 0.07 | 0.47 | 0.05 | 0.1 |
| 9 | 110954663 | rs550891 | -1 | -0.34 | -0.56 | -0.342 | -0.042 | 0.058 | 0.058 | 0 | 0.73 | 0.85 | 0.68 | 0.92 | 0.65 | 0.85 |
| 9 | 110955288 | rs556399 | 0.148 | -0.795 | -1.202 | -0.733 | -0.024 | 0.306 | 0.306 | 0 | 0.48 | 0.16 | 0.49 | 0.92 | 0.5 | 0.65 |
| 9 | 110959588 | rs569859 | 0.022 | -0.8 | -0.947 | -0.814 | 0.014 | 0.247 | 0.247 | 0 | 0.5 | 0.22 | 0.5 | 0.92 | 0.48 | 0.71 |
| 9 | 110969777 | rs838012 | 0.614 | -0.815 | -1.187 | -0.562 | -0.049 | -0.092 | -0.092 | 0.049 | 0.4 | 0.06 | 0.42 | 0.43 | 0.43 | 0.57 |
| 9 | 110977335 | rs655655 | -0.385 | -0.887 | -0.213 | -0.66 | -0.004 | -0.005 | -0.005 | 0.004 | 0.6 | 0.39 | 0.58 | 0.42 | 0.6 | 0.67 |
| 9 | 110978841 | rs7026619 | -1.288 | -1.073 | 0.312 | -0.666 | -0.024 | 0.094 | 0.094 | 0 | 0.23 | 0.41 | 0.18 | 0.49 | 0.17 | 0.21 |
| 9 | 110978859 | rs672586 | 0.173 | -1.298 | -0.008 | -1.166 | 0.043 | 0.004 | 0.004 | 0 | 0.48 | 0.33 | 0.37 | 0.61 | 0.36 | 0.51 |
| 9 | 110981431 | rs2571504 | -1.878 | -1.36 | -0.337 | -1.048 | -0.096 | 0.091 | 0.091 | 0 | 0.77 | 0.95 | 0.71 | 0.95 | 0.67 | 0.89 |
| 9 | 110986403 | rs519144 | -0.921 | -1.525 | -0.298 | -0.989 | -0.065 | -0.064 | -0.064 | 0 | 0.69 | 0.84 | 0.67 | 0.46 | 0.62 | 0.75 |
| 9 | 110987236 | rs10979283 | -0.315 | -1.473 | -0.263 | -0.919 | 0.004 | 0.056 | 0.056 | 0 | 0.09 | 0.09 | 0.04 | 0.5 | 0.05 | 0.14 |
| 9 | 110989319 | rs837999 | 0.815 | -1.742 | -0.51 | -1.152 | -0.073 | -0.044 | -0.044 | 0 | 0.43 | 0.06 | 0.41 | 0.39 | 0.47 | 0.59 |
| 9 | 110989880 | rs868895 | -1.271 | -1.774 | -0.207 | -1.182 | -0.034 | -0.028 | -0.028 | 0 | 0.7 | 0.79 | 0.68 | 0.42 | 0.62 | 0.76 |
| 9 | 110990323 | rs1888913 | -1.111 | -1.8 | 0.146 | -1.067 | 0 | -0.037 | -0.037 | 0 | 0.26 | 0.73 | 0.26 | 0.03 | 0.15 | 0.18 |
| 9 | 110991069 | rs1819463 | -1.109 | -1.745 | 0.315 | -1.075 | -0.022 | -0.086 | -0.086 | 2.779 | 0.25 | 0.66 | 0.25 | 0.02 | 0.16 | 0.17 |
| 9 | 111000640 | rs10122058 | -1.734 | -1.429 | 1.018 | -0.641 | -0.008 | 0.358 | 0.358 | 0.195 | 0.12 | 0.55 | 0.12 | 0.55 | 0.06 | 0.16 |
| 9 | 111008699 | rs4644328 | -1.736 | -1.361 | 0.652 | -1.065 | -0.017 | 0.012 | 0.012 | 0 | 0.76 | 0.8 | 0.65 | 0.92 | 0.67 | 0.87 |
| 9 | 111014118 | rs13302091 | -0.258 | -1.469 | 0.719 | -0.938 | 0.004 | 0.049 | 0.049 | 0 | 0.09 | 0.09 | 0.04 | 0.54 | 0.07 | 0.14 |
| 9 | 111018343 | rs7043626 | -1.691 | -1.331 | 1.002 | -1.003 | 0.054 | -0.055 | -0.055 | 0 | 0.75 | 0.65 | 0.64 | 0.92 | 0.65 | 0.84 |
| 9 | 111022525 | rs7030526 | -0.77 | -1.365 | 1.526 | -1.017 | 0.032 | 0.013 | 0.013 | 0 | 0.67 | 0.49 | 0.59 | 0.38 | 0.6 | 0.74 |
| 9 | 111023899 | rs10739255 | 0.37 | -1.59 | 1.339 | -1.232 | 0.004 | 0.027 | 0.027 | 0 | 0.45 | 0.21 | 0.41 | 0.37 | 0.45 | 0.58 |
| 9 | 111027827 | rs3117888 | 0.032 | -1.582 | 1.41 | -1.273 | 0 | 0.054 | 0.054 | 0 | 0.47 | 0.6 | 0.53 | 0.09 | 0.49 | 0.29 |
| 9 | 111029103 | rs1330591 | 0.176 | -1.584 | 1.364 | -1.242 | -0.004 | 0.072 | 0.072 | 0 | 0.46 | 0.3 | 0.47 | 0.1 | 0.49 | 0.29 |
| 9 | 111033028 | rs16912823 | -0.575 | -1.517 | 1.285 | -1.119 | -0.002 | 0.279 | 0.279 | 0 | 0.09 | 0.28 | 0.06 | 0.55 | 0.07 | 0.17 |
| 9 | 111034788 | rs1323397 | -1.726 | -1.74 | 0.373 | -1.65 | -0.07 | 0.182 | 0.182 | 0 | 0.71 | 0.98 | 0.73 | 0.94 | 0.68 | 0.71 |
| 9 | 111037295 | rs1575450 | -1.562 | -1.839 | 0.365 | -1.65 | -0.08 | 0.194 | 0.194 | 0 | 0.7 | 0.96 | 0.72 | 0.93 | 0.65 | 0.73 |
| 9 | 111042233 | rs12004427 | -1.085 | -1.625 | 0.913 | -1.252 | -0.01 | -0.015 | -0.015 | 0 | 0.09 | 0.47 | 0.05 | 0.02 | 0.02 | 0.04 |
| 9 | 111043009 | rs1323377 | 1.897 | -1.543 | 0.503 | -1.303 | -0.013 | 0.008 | 0.008 | 1.455 | 0.26 | 0.21 | 0.34 | 0.08 | 0.33 | 0.14 |
| 9 | 111044933 | rs10979301 | 1.817 | -1.534 | 0.392 | -1.32 | -0.025 | 0.023 | 0.023 | 1.68 | 0.26 | 0.18 | 0.34 | 0.05 | 0.33 | 0.14 |
| 9 | 111044950 | rs7029971 | -1.19 | -1.52 | 0.407 | -1.359 | 0.001 | -0.027 | -0.027 | 0.268 | 0.1 | 0.44 | 0.07 | 0.02 | 0.05 | 0.04 |
| 9 | 111050366 | rs7021201 | -0.111 | -1.607 | 0.244 | -1.468 | -0.065 | 0.326 | 0.326 | 0 | 0.48 | 0.86 | 0.53 | 0.07 | 0.47 | 0.31 |
| 9 | 111050810 | rs12353336 | 1.7 | -1.633 | 0.345 | -1.468 | 0.028 | -0.096 | -0.096 | 0.255 | 0.28 | 0.35 | 0.33 | 0.05 | 0.31 | 0.15 |
| 9 | 111051418 | rs10117055 | -0.699 | -1.667 | 0.097 | -1.263 | -0.037 | 0.009 | 0.009 | 0 | 0.56 | 0.87 | 0.57 | 0.62 | 0.54 | 0.45 |
| 9 | 111054809 | rs1570504 | -0.835 | -1.602 | 0.405 | -1.122 | -0.038 | -0.133 | -0.133 | 0 | 0.2 | 0.46 | 0.18 | 0.02 | 0.14 | 0.15 |
| 9 | 111060921 | rs1323380 | -0.162 | -1.494 | 0.204 | -1.056 | -0.001 | -0.027 | -0.027 | 0.107 | 0.14 | 0.16 | 0.16 | 0.01 | 0.14 | 0.16 |
| 9 | 111062515 | rs10816656 | -1.288 | -1.572 | 0.145 | -0.995 | 0.013 | -0.033 | -0.033 | 0.256 | 0.24 | 0.18 | 0.2 | 0.56 | 0.2 | 0.3 |
| 9 | 111066833 | rs17543333 | -0.131 | -1.39 | 0.439 | -0.864 | -0.002 | -0.017 | -0.017 | 0 | 0.15 | 0.17 | 0.16 | 0.03 | 0.14 | 0.2 |
| 9 | 111068483 | rs1323384 | -1.221 | -1.493 | 0.661 | -0.798 | -0.038 | 0.046 | 0.046 | 0.173 | 0.27 | 0.44 | 0.21 | 0.58 | 0.19 | 0.31 |
| 9 | 111072429 | rs10979314 | 0.147 | -1.235 | -0.024 | -0.304 | -0.012 | -0.005 | -0.005 | 0 | 0.07 | 0.02 | 0.12 | 0.26 | 0.09 | 0.21 |
| 9 | 111072668 | rs12349823 | 0.513 | -1.183 | 0.074 | -0.838 | 0.002 | -0.039 | -0.039 | 1.332 | 0.43 | 0.51 | 0.58 | 0.09 | 0.45 | 0.3 |
| 9 | 111074108 | rs10816658 | 0.139 | -1.203 | -0.848 | -0.307 | -0.03 | 0.03 | 0.03 | 0 | 0.3 | 0.09 | 0.28 | 0.18 | 0.33 | 0.39 |
| 9 | 111074243 | rs10283642 | -1.085 | -0.66 | -0.649 | -0.567 | -0.054 | 0.151 | 0.151 | 0 | 0.6 | 0.84 | 0.58 | 0.92 | 0.56 | 0.77 |
| 9 | 111077062 | rs7033347 | -0.949 | -0.027 | 0.151 | 0.66 | 0.006 | -0.01 | -0.01 | 1.151 | 0.39 | 0.7 | 0.46 | 0.4 | 0.36 | 0.58 |
| 9 | 111082271 | rs4978345 | -1.047 | -0.06 | -0.032 | 0.437 | 0.013 | -0.021 | -0.021 | 0.797 | 0.41 | 0.87 | 0.37 | 0.35 | 0.37 | 0.59 |
| 9 | 111085981 | rs7860612 | -0.642 | -0.04 | 0.631 | 0.433 | -0.006 | 0.001 | 0.001 | 0 | 0.35 | 0.61 | 0.41 | 0.38 | 0.23 | 0.56 |
| 9 | 111098115 | rs7031855 | -0.058 | -0.137 | 1.112 | 0.349 | -0.032 | 0.066 | 0.066 | 0 | 0.26 | 0.41 | 0.38 | 0.38 | 0.19 | 0.46 |
| 9 | 111100388 | rs1323391 | -1.131 | -0.097 | 0.99 | 0.378 | 0.087 | -0.041 | -0.041 | 1.152 | 0.38 | 0.83 | 0.43 | 0.38 | 0.23 | 0.55 |

| | | | | | | | | | | | | | | | | |
|---|---|---|---|---|---|---|---|---|---|---|---|---|---|---|---|---|
| 9 | 111107191 | rs1323396 | -0.875 | 0.076 | 0.963 | 0.453 | -0.026 | 0.012 | 0.012 | 0 | 0.35 | 0.55 | 0.39 | 0.38 | 0.22 | 0.54 |
| 9 | 111109673 | rs7045550 | -0.946 | 0.075 | 0.966 | 0.463 | -0.022 | 0.009 | 0.009 | 0 | 0.36 | 0.57 | 0.39 | 0.38 | 0.22 | 0.54 |
| 9 | 111114728 | rs10979321 | -0.404 | 0.596 | 0.62 | 0.625 | -0.072 | -0.077 | -0.077 | 1.346 | 0.13 | 0.03 | 0.1 | 0.52 | 0.26 | 0.15 |
| 9 | 111118695 | rs12338469 | 0.062 | -0.435 | 1.056 | 0.632 | -0.038 | -0.011 | -0.011 | 0 | 0.11 | 0.22 | 0.14 | 0.05 | 0.04 | 0.11 |
| 9 | 111120353 | rs10979326 | -0.201 | -0.107 | 1.08 | 0.926 | -0.034 | 0.062 | 0.062 | 0 | 0.25 | 0.47 | 0.27 | 0.32 | 0.2 | 0.44 |
| 9 | 111120695 | rs10979327 | 0.129 | -0.074 | 0.808 | 0.95 | -0.044 | -0.055 | -0.055 | 0 | 0.1 | 0.01 | 0.05 | 0.5 | 0.12 | 0.11 |
| 9 | 111123697 | rs12379749 | -0.268 | -0.304 | 0.663 | 0.843 | -0.006 | 0.014 | 0.014 | 0 | 0.23 | 0.28 | 0.25 | 0.31 | 0.19 | 0.38 |
| 9 | 111140133 | rs1885968 | -0.664 | -0.095 | -0.138 | 0.303 | -0.014 | -0.001 | -0.001 | 0 | 0.36 | 0.54 | 0.31 | 0.36 | 0.27 | 0.53 |
| 9 | 111142265 | rs1323719 | -0.734 | -0.332 | 0.102 | -1.008 | 0 | -0.008 | -0.008 | 0 | 0.37 | 0.56 | 0.39 | 0.88 | 0.31 | 0.51 |
| 9 | 111143660 | rs10115131 | -1.654 | -0.732 | -0.746 | -1.704 | 0.036 | 0.312 | 0.312 | 0.13 | 0.54 | 0.95 | 0.56 | 0.93 | 0.35 | 0.7 |
| 9 | 111151700 | rs10816666 | -0.267 | -0.851 | -0.328 | -1.705 | -0.039 | -0.072 | -0.072 | 0 | 0.34 | 0.48 | 0.34 | 0.88 | 0.23 | 0.5 |
| 9 | 111152200 | rs10114425 | -0.519 | -0.749 | 0.083 | -1.61 | -0.011 | -0.002 | -0.002 | 0 | 0.09 | 0.24 | 0.17 | 0.05 | 0.08 | 0.11 |
| 9 | 111153825 | rs16913038 | -0.293 | -0.618 | -0.326 | -1.548 | 0.015 | 0.009 | 0.009 | 0 | 0.1 | 0.08 | 0.03 | 0 | 0.03 | 0.08 |
| 9 | 111154361 | rs16913043 | -0.382 | -0.619 | -0.295 | -1.549 | -0.004 | -0.023 | -0.023 | 0.452 | 0.1 | 0.13 | 0.04 | 0 | 0.03 | 0.08 |
| 9 | 111160260 | rs657749 | 0.412 | 0.102 | -0.733 | -1.833 | -0.04 | 0.057 | 0.057 | 0 | 0.11 | 0.02 | 0.08 | 0.02 | 0.17 | 0.03 |
| 9 | 111161855 | rs640278 | 0.277 | 0.221 | -0.795 | -2.607 | -0.003 | 0.082 | 0.082 | 0 | 0.55 | 0.33 | 0.38 | 0.11 | 0.56 | 0.4 |
| 9 | 111167197 | rs1998537 | 1.238 | 0.492 | -0.866 | -2.683 | 0.029 | 0.164 | 0.164 | 0 | 0.39 | 0.15 | 0.25 | 0.08 | 0.32 | 0.31 |
| 9 | 111175438 | rs10979346 | 0.388 | 0.452 | -0.018 | -2.682 | 0.011 | 0.003 | 0.003 | 0.036 | 0.2 | 0.29 | 0.28 | 0.76 | 0.21 | 0.28 |
| 9 | 111175475 | rs10979347 | -0.12 | 0.338 | 0.315 | -2.737 | 0.006 | -0.01 | -0.01 | 0.019 | 0.23 | 0.5 | 0.29 | 0.76 | 0.21 | 0.28 |
| 9 | 111176217 | rs518095 | -1.41 | 0.474 | 0.155 | -2.818 | 0 | 0.14 | 0.14 | 0 | 0.47 | 0.74 | 0.58 | 0.82 | 0.54 | 0.47 |
| 9 | 111182085 | rs2093679 | 0.159 | 0.254 | 0.296 | -3.023 | 0.029 | -0.018 | -0.018 | 0 | 0.25 | 0.37 | 0.32 | 0.76 | 0.28 | 0.31 |
| 9 | 111182146 | rs506707 | -0.934 | 0.199 | 0.244 | -3.11 | 0.01 | -0.05 | -0.05 | 0.855 | 0.4 | 0.5 | 0.44 | 0.79 | 0.51 | 0.4 |
| 9 | 111197736 | rs539864 | -1.3 | 0.102 | 0.279 | -3.157 | -0.01 | 0.072 | 0.072 | 0.106 | 0.44 | 0.68 | 0.56 | 0.84 | 0.52 | 0.43 |
| 9 | 111201554 | rs478208 | 0.333 | 0.293 | 0.138 | -3.103 | -0.051 | 0.054 | 0.054 | 0.021 | 0.16 | 0.07 | 0.15 | 0.02 | 0.29 | 0.05 |
| 9 | 111202210 | rs10979352 | 0.358 | 0.169 | 0.201 | -3.108 | 0.009 | -0.015 | -0.015 | 0.204 | 0.23 | 0.41 | 0.29 | 0.76 | 0.22 | 0.31 |
| 9 | 111204349 | rs488085 | 0.636 | 0.253 | 0.111 | -3.078 | -0.029 | 0.038 | 0.038 | 0 | 0.13 | 0.07 | 0.11 | 0.02 | 0.23 | 0.04 |
| 9 | 111230318 | rs9644923 | 0.457 | 0.176 | 0.092 | -3.018 | 0.01 | -0.01 | -0.01 | 0 | 0.22 | 0.33 | 0.28 | 0.78 | 0.22 | 0.3 |
| 9 | 111231987 | rs9644924 | -0.74 | -0.087 | -0.167 | -3.306 | -0.002 | 0.025 | 0.025 | 0 | 0.46 | 0.57 | 0.5 | 0.88 | 0.43 | 0.54 |
| 9 | 111234117 | rs7859673 | 1.496 | -0.122 | -0.296 | -3.371 | 0.011 | 0.044 | 0.044 | 0 | 0.4 | 0.33 | 0.42 | 0.1 | 0.4 | 0.38 |
| 9 | 111236780 | rs2417865 | 0.4 | 0.056 | -0.193 | -3.296 | -0.001 | -0.02 | -0.02 | 0.689 | 0.64 | 0.66 | 0.7 | 0.88 | 0.63 | 0.68 |
| 9 | 111236905 | rs7044687 | 0.374 | -0.01 | -0.185 | -3.291 | 0.009 | 0.016 | 0.016 | 0.001 | 0.21 | 0.19 | 0.18 | 0.1 | 0.14 | 0.23 |
| 9 | 111238914 | rs7039103 | -0.385 | 0.239 | -0.315 | -3.051 | 0.011 | -0.046 | -0.046 | 0 | 0.86 | 0.76 | 0.91 | 0.97 | 0.83 | 0.92 |
| 9 | 111250709 | rs1408066 | 1.843 | -0.002 | -0.801 | -3.228 | -0.012 | 0.142 | 0.142 | 0.009 | 0.29 | 0.07 | 0.35 | 0.03 | 0.31 | 0.31 |
| 9 | 111258189 | rs7029016 | 1.945 | 0.121 | -0.912 | -3.187 | -0.005 | 0.12 | 0.12 | 0 | 0.26 | 0.07 | 0.29 | 0.03 | 0.29 | 0.29 |
| 9 | 111259345 | rs10979372 | 0.236 | 0.097 | -0.462 | -3.283 | 0.001 | -0.022 | -0.022 | 0 | 0.44 | 0.46 | 0.46 | 0.09 | 0.45 | 0.41 |
| 9 | 111275801 | rs6477655 | -0.355 | 0.537 | -0.262 | -2.913 | 0.048 | -0.037 | -0.037 | 0.573 | 0.08 | 0.22 | 0.09 | 0.02 | 0.13 | 0.02 |
| 9 | 111277519 | rs9918937 | 0.308 | 0.351 | -0.009 | -3.192 | 0.016 | -0.106 | -0.106 | 0 | 0.62 | 0.54 | 0.63 | 0.9 | 0.6 | 0.72 |
| 9 | 111279713 | rs7042096 | 0.083 | 0.085 | 0.039 | -3.195 | 0.014 | -0.084 | -0.084 | 0 | 0.29 | 0.47 | 0.32 | 0.88 | 0.28 | 0.42 |
| 9 | 111281164 | rs7030851 | 0.152 | 0.062 | 0.025 | -3.004 | -0.017 | -0.071 | -0.071 | 0.646 | 0.24 | 0.34 | 0.25 | 0.09 | 0.17 | 0.23 |
| 9 | 111282789 | rs10979383 | 0.182 | -0.05 | -0.005 | -3.041 | -0.005 | -0.046 | -0.046 | 4.47 | 0.27 | 0.28 | 0.29 | 0.12 | 0.21 | 0.25 |
| 9 | 111283188 | rs10816679 | 1.079 | 0.258 | -0.314 | -3.34 | -0.017 | 0.103 | 0.103 | 0.018 | 0.55 | 0.65 | 0.49 | 0.88 | 0.52 | 0.69 |
| 9 | 111288037 | rs7865368 | 0.582 | 0.261 | -0.045 | -3.077 | -0.001 | -0.007 | -0.007 | 0 | 0.19 | 0.2 | 0.18 | 0.08 | 0.15 | 0.21 |
| 9 | 111291784 | rs7852410 | -1.034 | 0.391 | 0.089 | -3.054 | -0.009 | 0.014 | 0.014 | 0 | 0.88 | 0.88 | 0.9 | 0.98 | 0.84 | 0.91 |
| 9 | 111295630 | rs7018616 | 1.185 | 0.278 | 0.232 | -3.336 | -0.024 | -0.127 | -0.127 | 0 | 0.39 | 0.44 | 0.27 | 0.04 | 0.31 | 0.33 |
| 9 | 111302146 | rs7044959 | 1.134 | 0.282 | 0.474 | -3.328 | -0.008 | -0.06 | -0.06 | 0.228 | 0.4 | 0.41 | 0.27 | 0.06 | 0.31 | 0.34 |
| 9 | 111311096 | rs4978700 | -0.311 | 0.31 | 0.485 | -3.36 | 0 | -0.047 | -0.047 | 0.263 | 0.45 | 0.43 | 0.6 | 0.93 | 0.48 | 0.56 |
| 9 | 111311996 | rs10979395 | 0.903 | 0.32 | 0.377 | -3.42 | -0.02 | 0.036 | 0.036 | 0 | 0.28 | 0.17 | 0.42 | 0.84 | 0.34 | 0.36 |
| 9 | 111315303 | rs16913269 | 0.241 | 0.181 | 0.47 | -3.373 | 0 | -0.023 | -0.023 | 0.062 | 0.2 | 0.25 | 0.22 | 0.11 | 0.19 | 0.26 |
| 9 | 111315597 | rs10979398 | -0.051 | 0.161 | 0.45 | -3.334 | -0.022 | 0.047 | 0.047 | 0 | 0.6 | 0.68 | 0.49 | 0.16 | 0.5 | 0.6 |
| 9 | 111318495 | rs16913282 | 0.197 | 0.172 | 0.462 | -3.315 | 0.001 | -0.02 | -0.02 | 0.096 | 0.2 | 0.23 | 0.21 | 0.1 | 0.18 | 0.25 |
| 9 | 111319491 | rs16913284 | 0.325 | 0.1 | 0.445 | -3.35 | -0.028 | -0.067 | -0.067 | 0 | 0.2 | 0.45 | 0.19 | 0.11 | 0.14 | 0.2 |
| 9 | 111321154 | rs12349538 | -0.311 | 0.09 | 0.293 | -3.211 | -0.051 | 0.074 | 0.074 | 0 | 0.63 | 0.76 | 0.6 | 0.17 | 0.51 | 0.62 |
| 9 | 111322964 | rs7856330 | -0.483 | 0.047 | 0.364 | -2.934 | -0.015 | -0.035 | -0.035 | 0 | 0.25 | 0.48 | 0.25 | 0.16 | 0.19 | 0.27 |
| 9 | 111324602 | rs871424 | 1.87 | -0.072 | 0.125 | -2.989 | 0.001 | 0.182 | 0.182 | 0 | 0.33 | 0.07 | 0.23 | 0.01 | 0.31 | 0.31 |
| 9 | 111334514 | rs7858857 | -0.081 | 0.072 | 0.262 | -2.894 | -0.046 | 0.079 | 0.079 | 0 | 0.61 | 0.75 | 0.52 | 0.17 | 0.5 | 0.59 |
| 9 | 111335268 | rs10979405 | -0.036 | 0.067 | 0.288 | -2.896 | -0.028 | 0.06 | 0.06 | 0 | 0.6 | 0.68 | 0.51 | 0.17 | 0.5 | 0.59 |
| 9 | 111342144 | rs10217573 | -0.386 | 0.174 | 0.232 | -2.903 | -0.08 | 0.055 | 0.055 | 0 | 0.65 | 0.88 | 0.66 | 0.21 | 0.54 | 0.63 |
| 9 | 111342197 | rs10217224 | 0.003 | 0.125 | 0.281 | -2.906 | -0.032 | 0.061 | 0.061 | 0 | 0.6 | 0.7 | 0.52 | 0.18 | 0.5 | 0.6 |
| 9 | 111345751 | rs1930100 | -0.124 | 0.174 | 0.301 | -2.923 | 0.001 | -0.039 | -0.039 | 0.412 | 0.23 | 0.49 | 0.25 | 0.14 | 0.19 | 0.21 |
| 9 | 111345810 | rs4552986 | 1.24 | 0.214 | -0.019 | -2.903 | -0.03 | 0.006 | 0.006 | 0.018 | 0.25 | 0.12 | 0.27 | 0.79 | 0.3 | 0.36 |
| 9 | 111351436 | rs7848857 | 0.996 | 0.084 | 0.11 | -2.792 | 0.008 | 0.16 | 0.16 | 0 | 0.39 | 0.16 | 0.28 | 0.03 | 0.35 | 0.39 |
| 9 | 111359745 | rs4978715 | -0.695 | 0.158 | 0.003 | -2.803 | -0.084 | 0.079 | 0.079 | 0 | 0.64 | 0.87 | 0.54 | 0.19 | 0.51 | 0.62 |
| 9 | 111362601 | rs16913324 | 0.252 | 0.027 | 0.397 | -2.72 | -0.009 | -0.015 | -0.015 | 0 | 0.18 | 0.22 | 0.19 | 0.11 | 0.13 | 0.27 |
| 9 | 111367630 | rs1930108 | 1.481 | -0.068 | 0.054 | -2.821 | 0.017 | 0.181 | 0.181 | 0 | 0.35 | 0.08 | 0.24 | 0.01 | 0.31 | 0.32 |
| 9 | 111367795 | rs4623515 | 0.157 | 0.172 | 0.127 | -2.767 | -0.009 | -0.029 | -0.029 | 0.364 | 0.18 | 0.32 | 0.21 | 0.11 | 0.13 | 0.21 |
| 9 | 111368151 | rs959244 | 0.012 | -0.022 | -0.414 | -2.519 | -0.003 | 0.102 | 0.102 | 0 | 0.58 | 0.74 | 0.48 | 0.14 | 0.44 | 0.59 |
| 9 | 111369668 | rs4978717 | 0.686 | -0.083 | 0.183 | -2.045 | -0.012 | -0.017 | -0.017 | 0.037 | 0.71 | 0.36 | 0.72 | 0.77 | 0.77 | 0.74 |
| 9 | 111369823 | rs4978718 | 0.769 | 0.156 | 0.303 | -1.773 | -0.001 | -0.01 | -0.01 | 0.09 | 0.7 | 0.37 | 0.69 | 0.75 | 0.71 | 0.78 |
| 9 | 111370795 | rs7857549 | 0.343 | 0.414 | 0.448 | -1.998 | -0.001 | -0.012 | -0.012 | 0 | 0.77 | 0.78 | 0.75 | 0.94 | 0.71 | 0.78 |

| | | | | | | | | | | | | | | | | |
|---|---|---|---|---|---|---|---|---|---|---|---|---|---|---|---|---|
| 9 | 111372759 | rs10816689 | 0.558 | 0.382 | 0.168 | -2.045 | -0.007 | 0.003 | 0.003 | 0.356 | 0.3 | 0.24 | 0.43 | 0.7 | 0.35 | 0.33 |
| 9 | 111376151 | rs7851754 | 0.346 | 0.264 | 0.345 | -2.072 | 0.009 | -0.054 | -0.054 | 0.976 | 0.73 | 0.56 | 0.74 | 0.89 | 0.73 | 0.69 |
| 9 | 111376977 | rs7027855 | -0.152 | 0.289 | 0.333 | -2.016 | 0.017 | -0.042 | -0.042 | 0.379 | 0.79 | 0.71 | 0.76 | 0.93 | 0.74 | 0.78 |
| 9 | 111379252 | rs12340315 | 0.46 | 0.243 | 0.339 | -2.017 | 0.007 | 0.008 | 0.008 | 0.162 | 0.48 | 0.57 | 0.33 | 0.23 | 0.38 | 0.45 |
| 9 | 111379330 | rs1958943 | -0.116 | 0.282 | 0.206 | -1.991 | -0.002 | -0.009 | -0.009 | 0.047 | 0.78 | 0.8 | 0.77 | 0.94 | 0.71 | 0.78 |
| 9 | 111382775 | rs12336075 | 0.013 | 0.274 | 0.487 | -1.875 | 0.018 | -0.02 | -0.02 | 0.009 | 0.35 | 0.4 | 0.56 | 0.71 | 0.41 | 0.34 |
| 9 | 111387353 | rs10979414 | 0.629 | 0.385 | 0.212 | -1.717 | -0.006 | 0.008 | 0.008 | 0.136 | 0.33 | 0.2 | 0.33 | 0.69 | 0.34 | 0.36 |
| 9 | 111387462 | rs13289732 | 0.78 | 0.384 | 0.206 | -1.699 | -0.014 | -0.004 | -0.004 | 0.314 | 0.32 | 0.14 | 0.32 | 0.67 | 0.34 | 0.36 |
| 9 | 111388183 | rs4978359 | -0.112 | 0.402 | 0.127 | -1.785 | -0.017 | 0.049 | 0.049 | 0 | 0.57 | 0.68 | 0.48 | 0.26 | 0.51 | 0.54 |
| 9 | 111389865 | rs4978720 | -0.339 | 0.185 | 0.568 | -1.094 | -0.009 | 0.008 | 0.008 | 0 | 0.91 | 0.91 | 0.81 | 0.96 | 0.85 | 0.89 |
| 9 | 111396742 | rs10115769 | -1.201 | 0.033 | 0.927 | -1.178 | -0.008 | -0.002 | -0.002 | 0 | 0.11 | 0.31 | 0.08 | 0.13 | 0.1 | 0.05 |
| 9 | 111397766 | rs4978360 | 0.748 | 0.146 | 0.59 | -1.094 | 0.015 | 0.012 | 0.012 | 0 | 0.19 | 0.08 | 0.22 | 0.1 | 0.14 | 0.24 |
| 9 | 111400131 | rs13302880 | 0.844 | 0.085 | 0.784 | -1.178 | 0 | -0.006 | -0.006 | 0.107 | 0.72 | 0.63 | 0.82 | 0.87 | 0.69 | 0.68 |
| 9 | 111401766 | rs4978721 | 1.279 | 0.103 | 0.813 | -1.185 | 0 | 0.019 | 0.019 | 0 | 0.54 | 0.53 | 0.6 | 0.78 | 0.54 | 0.46 |
| 9 | 111402410 | rs4412431 | -0.641 | 0.082 | 0.853 | -1.188 | -0.004 | -0.013 | -0.013 | 0 | 0.21 | 0.28 | 0.13 | 0.12 | 0.2 | 0.25 |
| 9 | 111404834 | rs10816693 | 1.466 | 0.16 | 0.98 | -1.167 | 0.004 | 0.057 | 0.057 | 0 | 0.5 | 0.39 | 0.46 | 0.78 | 0.47 | 0.44 |
| 9 | 111405228 | rs10979424 | -0.129 | 0.082 | 0.54 | -1.048 | 0.002 | 0.009 | 0.009 | 0.031 | 0.25 | 0.21 | 0.29 | 0.13 | 0.24 | 0.26 |
| 9 | 111409524 | rs1873764 | -0.466 | -0.072 | 0.351 | -0.79 | 0.001 | 0.001 | 0.001 | 0 | 0.14 | 0.08 | 0.09 | 0.14 | 0.14 | 0.19 |
| 9 | 111413448 | rs7865339 | 0.984 | -0.109 | 0.334 | -0.73 | 0.013 | 0.011 | 0.011 | 0 | 0.18 | 0.09 | 0.22 | 0.1 | 0.14 | 0.21 |
| 9 | 111413719 | rs7018492 | -1.056 | -0.096 | 0.455 | -0.844 | 0.001 | 0.001 | 0.001 | 0 | 0.28 | 0.31 | 0.16 | 0.23 | 0.25 | 0.4 |
| 9 | 111414999 | rs10979429 | -0.576 | -0.188 | 0.337 | -1.001 | 0.011 | 0.01 | 0.01 | 0.167 | 0.32 | 0.23 | 0.33 | 0.26 | 0.3 | 0.4 |
| 9 | 111417991 | rs937160 | -0.588 | 0.105 | 0.488 | -1.219 | 0.017 | 0.02 | 0.02 | 0 | 0.34 | 0.29 | 0.18 | 0.22 | 0.25 | 0.41 |
| 9 | 111420155 | rs3983541 | -0.358 | 0.291 | 0.589 | -1.499 | 0.009 | 0.016 | 0.016 | 0 | 0.32 | 0.29 | 0.18 | 0.11 | 0.25 | 0.37 |
| 9 | 111423688 | rs1319976 | -0.916 | 0.41 | 0.914 | -1.564 | 0.024 | -0.035 | -0.035 | 0 | 0.92 | 0.88 | 0.95 | 0.99 | 0.89 | 0.92 |
| 9 | 111427754 | rs2417919 | 0.625 | 0.384 | 0.851 | -1.582 | 0.002 | 0.032 | 0.032 | 0 | 0.61 | 0.67 | 0.77 | 0.88 | 0.65 | 0.55 |
| 9 | 111429924 | rs7034647 | 0.034 | 0.378 | 1.142 | -1.597 | 0.007 | 0.007 | 0.007 | 0 | 0.11 | 0.16 | 0.18 | 0.14 | 0.12 | 0.16 |
| 9 | 111430589 | rs2417921 | -1.512 | 0.482 | 1.073 | -1.548 | 0.016 | 0.122 | 0.122 | 0 | 0.49 | 0.71 | 0.45 | 0.27 | 0.46 | 0.57 |
| 9 | 111446046 | rs4978725 | -1.38 | 0.542 | 1.129 | -1.444 | 0.073 | 0.004 | 0.004 | 0.104 | 0.15 | 0.39 | 0.24 | 0.15 | 0.19 | 0.17 |
| 9 | 111453191 | rs13286509 | 1.585 | 0.543 | 0.833 | -1.472 | -0.007 | 0.091 | 0.091 | 0 | 0.45 | 0.24 | 0.53 | 0.72 | 0.44 | 0.35 |
| 9 | 111455575 | rs10512385 | -0.032 | 0.488 | 0.857 | -1.569 | -0.014 | -0.018 | -0.018 | 0 | 0.17 | 0.25 | 0.08 | 0.06 | 0.11 | 0.19 |
| 9 | 111455892 | rs7874723 | -0.061 | 0.556 | 1.159 | -1.375 | 0.029 | 0.037 | 0.037 | 0 | 0.08 | 0.17 | 0.15 | 0.13 | 0.1 | 0.12 |
| 9 | 111460075 | rs10759298 | 0.854 | 0.567 | 1.08 | -1.424 | -0.008 | 0.012 | 0.012 | 1.045 | 0.61 | 0.44 | 0.6 | 0.8 | 0.55 | 0.55 |
| 9 | 111467834 | rs7031739 | -0.49 | 0.724 | 0.702 | -1.29 | -0.004 | 0.006 | 0.006 | 0 | 0.15 | 0.2 | 0.18 | 0.19 | 0.14 | 0.18 |
| 9 | 111468234 | rs1556950 | 1.61 | 0.634 | 0.688 | -1.336 | -0.001 | 0.036 | 0.036 | 0.371 | 0.47 | 0.37 | 0.53 | 0.75 | 0.46 | 0.36 |
| 9 | 111469933 | rs6477669 | -1.051 | 0.676 | 0.45 | -1.321 | 0.005 | -0.003 | -0.003 | 0.361 | 0.19 | 0.23 | 0.22 | 0.19 | 0.23 | 0.22 |
| 9 | 111481932 | rs13299079 | 0.697 | 0.28 | 0.11 | -1.119 | 0 | -0.054 | -0.054 | 0.013 | 0.61 | 0.4 | 0.69 | 0.92 | 0.6 | 0.54 |
| 9 | 111484899 | rs10979455 | -0.101 | 0.052 | -0.294 | -0.552 | 0.013 | -0.015 | -0.015 | 0 | 0.11 | 0.05 | 0.03 | 0.2 | 0.07 | 0.17 |
| 9 | 111485851 | rs10122805 | 0.385 | 0.554 | -0.091 | -0.714 | 0.007 | -0.059 | -0.059 | 0 | 0.87 | 0.74 | 0.9 | 1 | 0.85 | 0.85 |
| 9 | 111498383 | rs1934460 | 0.667 | 0.356 | -0.173 | -0.936 | -0.005 | -0.064 | -0.064 | 0.015 | 0.66 | 0.58 | 0.75 | 0.93 | 0.69 | 0.58 |
| 9 | 111504523 | rs10816707 | 0.008 | 0.362 | -0.253 | -1.213 | -0.002 | -0.012 | -0.012 | 0 | 0.44 | 0.45 | 0.54 | 0.73 | 0.45 | 0.55 |
| 9 | 111507261 | rs4978734 | 0.605 | 0.578 | -0.32 | -0.88 | -0.001 | -0.013 | -0.013 | 0 | 0.3 | 0.27 | 0.47 | 0.54 | 0.33 | 0.44 |
| 9 | 111511670 | rs2039810 | 0.508 | 0.434 | -0.469 | -1.615 | -0.001 | -0.034 | -0.034 | 0.377 | 0.23 | 0.3 | 0.17 | 0.07 | 0.21 | 0.21 |
| 9 | 111511992 | rs10816710 | -0.545 | 0.421 | -0.473 | -1.655 | -0.002 | -0.024 | -0.024 | 1.349 | 0.63 | 0.59 | 0.68 | 0.79 | 0.64 | 0.7 |
| 9 | 111518150 | rs10759310 | 0.359 | 0.457 | -0.888 | -1.473 | -0.008 | 0.05 | 0.05 | 0 | 0.39 | 0.24 | 0.51 | 0.72 | 0.42 | 0.49 |
| 9 | 111520844 | rs11789156 | 0.638 | 0.092 | -0.67 | -1.182 | -0.008 | -0.001 | -0.001 | 0 | 0.09 | 0.05 | 0.08 | 0 | 0.12 | 0.1 |
| 9 | 111523941 | rs1538658 | 0.227 | -0.303 | -0.69 | -0.718 | 0.013 | 0.014 | 0.014 | 0 | 0.22 | 0.12 | 0.24 | 0.2 | 0.19 | 0.22 |
| 9 | 111524704 | rs4978364 | 0.073 | -0.561 | -1.003 | -0.377 | 0.004 | -0.054 | -0.054 | 0 | 0.09 | 0 | 0.06 | 0.22 | 0.07 | 0.12 |
| 9 | 111528013 | rs9886734 | 1.037 | -0.633 | -0.463 | -0.689 | -0.007 | -0.003 | -0.003 | 0 | 0.44 | 0.36 | 0.5 | 0.37 | 0.51 | 0.47 |
| 9 | 111528780 | rs10118705 | -0.422 | -0.645 | -0.023 | -0.795 | -0.035 | -0.074 | -0.074 | 0 | 0.83 | 0.61 | 0.89 | 0.94 | 0.92 | 0.88 |
| 9 | 111538497 | rs10739269 | -1.252 | -0.588 | -0.186 | -0.773 | 0.007 | 0.04 | 0.04 | 0 | 0.88 | 0.98 | 0.93 | 0.97 | 0.93 | 0.91 |
| 9 | 111540653 | rs870913 | 0.459 | -0.616 | -0.034 | -0.67 | -0.037 | -0.013 | -0.013 | 0.125 | 0.26 | 0.11 | 0.31 | 0.3 | 0.29 | 0.28 |
| 9 | 111542647 | rs1416900 | 0.861 | -0.856 | 0.346 | -0.691 | -0.019 | -0.004 | -0.004 | 0 | 0.68 | 0.37 | 0.8 | 0.66 | 0.8 | 0.64 |
| 9 | 111542950 | rs10816720 | 1.169 | -0.719 | 0.286 | -0.719 | 0.012 | -0.006 | -0.006 | 0 | 0.61 | 0.31 | 0.71 | 0.5 | 0.68 | 0.51 |
| 9 | 111543572 | rs10448261 | -1.208 | -0.421 | 0.768 | -0.446 | -0.001 | -0.002 | -0.002 | 0.012 | 0.43 | 0.35 | 0.4 | 0.56 | 0.41 | 0.52 |
| 9 | 111547599 | rs7019881 | -1.309 | -0.418 | 0.857 | -0.615 | 0.043 | 0.04 | 0.04 | 0.118 | 0.18 | 0.57 | 0.16 | 0.23 | 0.2 | 0.28 |
| 9 | 111547669 | rs7034453 | -1.528 | -0.487 | 0.866 | -0.56 | 0.007 | -0.01 | -0.01 | 0.178 | 0.48 | 0.72 | 0.43 | 0.58 | 0.43 | 0.54 |
| 9 | 111547818 | rs7020124 | 0.807 | -0.248 | 1.026 | -0.506 | -0.003 | 0.024 | 0.024 | 0 | 0.05 | 0.13 | 0.02 | 0.1 | 0.04 | 0.06 |
| 9 | 111550543 | rs10118169 | 0.501 | -0.306 | 1.135 | -0.531 | -0.007 | -0.021 | -0.021 | 0 | 0.22 | 0.28 | 0.26 | 0.15 | 0.17 | 0.25 |
| 9 | 111551344 | rs10979489 | 1.069 | -0.324 | 1.11 | -0.53 | -0.016 | 0.025 | 0.025 | 0 | 0.18 | 0.03 | 0.24 | 0.15 | 0.16 | 0.16 |
| 9 | 111551724 | rs10512387 | -0.242 | -0.26 | 1.175 | -0.505 | 0.018 | 0.026 | 0.026 | 0 | 0.11 | 0.19 | 0.11 | 0.24 | 0.17 | 0.16 |
| 9 | 111552022 | rs1416898 | -0.776 | -0.263 | 1.156 | -0.507 | 0.006 | -0.001 | -0.001 | 0 | 0.33 | 0.45 | 0.37 | 0.38 | 0.33 | 0.41 |
| 9 | 111552269 | rs12377816 | 1.145 | -0.007 | 1.002 | -0.394 | -0.034 | -0.029 | -0.029 | 0 | 0.06 | 0.01 | 0.09 | 0.12 | 0.13 | 0.09 |
| 9 | 111555057 | rs10979496 | 0.701 | -0.041 | 1.031 | -0.401 | -0.004 | 0.026 | 0.026 | 0 | 0.05 | 0.13 | 0.02 | 0.11 | 0.04 | 0.06 |
| 9 | 111563327 | rs10979502 | -0.568 | -0.087 | 1.085 | -0.176 | 0.008 | 0.006 | 0.006 | 0 | 0.31 | 0.41 | 0.36 | 0.41 | 0.33 | 0.4 |
| 9 | 111563577 | rs10979504 | 0.603 | -0.232 | 1.087 | -0.307 | -0.001 | -0.003 | -0.003 | 0 | 0.2 | 0.21 | 0.25 | 0.17 | 0.16 | 0.24 |
| 9 | 111577140 | rs10979523 | 1.244 | -0.202 | 0.648 | -0.23 | -0.003 | -0.003 | -0.003 | 0.234 | 0.15 | 0.19 | 0.21 | 0.16 | 0.1 | 0.13 |
| 9 | 111584123 | rs10816732 | -1.027 | -0.084 | 0.802 | -0.229 | 0.046 | 0.024 | 0.024 | 0 | 0.2 | 0.42 | 0.18 | 0.25 | 0.26 | 0.29 |
| 9 | 111592442 | rs6477683 | -0.294 | -0.25 | 0.345 | -0.212 | -0.011 | -0.05 | -0.05 | 0.011 | 0.89 | 0.99 | 0.96 | 0.83 | 0.91 | 0.91 |
| 9 | 111594273 | rs10816733 | 0.001 | -0.218 | 0.736 | -0.246 | 0.003 | 0.004 | 0.004 | 0 | 0.26 | 0.3 | 0.32 | 0.41 | 0.27 | 0.31 |

| | | | | | | | | | | | | | | | | |
|---|---|---|---|---|---|---|---|---|---|---|---|---|---|---|---|---|
| 9 | 111595187 | rs10816735 | -0.493 | -0.164 | 0.785 | -0.232 | 0.019 | -0.003 | -0.003 | 0 | 0.13 | 0.17 | 0.16 | 0.14 | 0.22 | 0.26 |
| 9 | 111596085 | rs1417712 | 0.668 | -0.196 | 0.799 | -0.226 | -0.001 | 0 | 0 | 0 | 0.19 | 0.2 | 0.22 | 0.27 | 0.13 | 0.19 |
| 9 | 111598970 | rs2039923 | -0.152 | -0.466 | 0.548 | -0.861 | 0.022 | 0.042 | 0.042 | 0 | 0.66 | 0.81 | 0.73 | 0.78 | 0.72 | 0.74 |
| 9 | 111599066 | rs4354381 | 1.942 | -0.471 | 0.555 | -0.846 | -0.021 | -0.058 | -0.058 | 0 | 0.31 | 0.14 | 0.34 | 0.38 | 0.37 | 0.28 |
| 9 | 111603106 | rs16913584 | 0.706 | -0.438 | 0.599 | -0.824 | -0.001 | -0.032 | -0.032 | 0 | 0.08 | 0.15 | 0.06 | 0 | 0.07 | 0.14 |
| 9 | 111604933 | rs10979545 | 2.005 | -0.442 | 0.52 | -0.824 | 0.003 | 0.005 | 0.005 | 0 | 0.14 | 0.06 | 0.21 | 0.16 | 0.09 | 0.13 |
| 9 | 111605283 | rs12552409 | 1.143 | -0.386 | 0.782 | -0.726 | -0.009 | -0.005 | -0.005 | 0 | 0.38 | 0.39 | 0.45 | 0.51 | 0.52 | 0.4 |
| 9 | 111605612 | rs7034770 | 0.736 | -0.385 | 0.799 | -0.726 | -0.001 | 0.003 | 0.003 | 0 | 0.43 | 0.46 | 0.47 | 0.62 | 0.56 | 0.46 |
| 9 | 111606792 | rs7024768 | -1.112 | -0.467 | 0.628 | -0.775 | 0.027 | 0.029 | 0.029 | 0 | 0.45 | 0.67 | 0.33 | 0.33 | 0.34 | 0.46 |
| 9 | 111606875 | rs7039521 | -0.177 | -0.512 | 0.601 | -0.773 | -0.022 | -0.008 | -0.008 | 3.319 | 0.34 | 0.46 | 0.27 | 0.33 | 0.25 | 0.31 |
| 9 | 111608280 | rs1539028 | 1.131 | -0.457 | 0.252 | -0.758 | -0.005 | -0.009 | -0.009 | 0 | 0.38 | 0.34 | 0.4 | 0.38 | 0.43 | 0.46 |
| 9 | 111612990 | rs4978368 | -1.911 | -0.322 | -0.2 | -0.857 | -0.033 | -0.013 | -0.013 | 0.019 | 0.83 | 0.99 | 0.78 | 0.83 | 0.86 | 0.95 |
| 9 | 111613145 | rs10979554 | 0.539 | -0.404 | 0.062 | -0.846 | 0.005 | 0.005 | 0.005 | 0 | 0.25 | 0.19 | 0.24 | 0.21 | 0.21 | 0.25 |
| 9 | 111617341 | rs3750467 | 0.434 | -0.236 | -0.057 | -0.833 | -0.012 | -0.003 | -0.003 | 0 | 0.08 | 0.05 | 0.1 | 0.14 | 0.15 | 0.23 |
| 9 | 111619707 | rs6415805 | -0.351 | -0.289 | -0.045 | -0.809 | 0.001 | -0.002 | -0.002 | 0.259 | 0.32 | 0.36 | 0.34 | 0.32 | 0.36 | 0.38 |
| 9 | 111620728 | rs10816744 | -0.748 | -1.003 | 0.13 | -1.131 | -0.018 | -0.051 | -0.051 | 0 | 0.35 | 0.54 | 0.3 | 0.15 | 0.28 | 0.42 |
| 9 | 111621439 | rs10979564 | -0.257 | -0.667 | -0.368 | -0.97 | -0.007 | -0.037 | -0.037 | 0 | 0.29 | 0.38 | 0.27 | 0.15 | 0.27 | 0.35 |
| 9 | 111622975 | rs11794127 | 1.664 | -0.37 | -0.804 | -1.068 | -0.025 | 0.042 | 0.042 | 0.088 | 0.08 | 0 | 0.06 | 0 | 0.1 | 0.02 |
| 9 | 111623055 | rs4978746 | 0.861 | -0.537 | -0.506 | -0.779 | 0.006 | 0.039 | 0.039 | 0 | 0.49 | 0.59 | 0.5 | 0.68 | 0.55 | 0.55 |
| 9 | 111623941 | rs9299164 | 0.704 | -0.272 | -0.172 | -0.835 | 0.005 | 0.024 | 0.024 | 0 | 0.55 | 0.6 | 0.51 | 0.78 | 0.58 | 0.69 |
| 9 | 111623989 | rs10979570 | -0.258 | 0.081 | -0.092 | -0.603 | 0.003 | -0.07 | -0.07 | 0 | 0.23 | 0.29 | 0.25 | 0.05 | 0.24 | 0.19 |
| 9 | 111626090 | rs7872727 | -0.094 | 0.455 | -1.063 | 0.012 | -0.028 | 0.022 | 0.022 | 0 | 0.77 | 0.94 | 0.75 | 0.82 | 0.77 | 0.86 |
| 9 | 111628667 | rs12351317 | -0.866 | 0.668 | 0.325 | 0.446 | 0.127 | 0.156 | 0.156 | 0 | 0.08 | 0.42 | 0.2 | 0.09 | 0.13 | 0.09 |
| 9 | 111637304 | rs2027433 | -1.093 | 0.582 | 0.104 | 0.763 | -0.004 | -0.011 | -0.011 | 0 | 0.27 | 0.74 | 0.37 | 0.38 | 0.3 | 0.31 |
| 9 | 111637331 | rs1981268 | -0.099 | 0.33 | -0.416 | 0.733 | 0.012 | -0.009 | -0.009 | 0.644 | 0.88 | 0.94 | 0.91 | 0.86 | 0.91 | 0.66 |
| 9 | 111638684 | rs7874795 | -0.788 | 0.417 | -0.409 | 0.729 | -0.004 | -0.011 | -0.011 | 0.076 | 0.28 | 0.74 | 0.37 | 0.38 | 0.3 | 0.31 |
| 9 | 111640631 | rs10435834 | -0.733 | 0.402 | -0.164 | 0.244 | 0.007 | -0.022 | -0.022 | 0 | 0.35 | 0.57 | 0.28 | 0.32 | 0.38 | 0.3 |
| 9 | 111640667 | rs10435835 | 0.514 | 0.027 | -0.253 | -0.163 | -0.004 | 0.007 | 0.007 | 0 | 0.81 | 0.81 | 0.73 | 0.77 | 0.84 | 0.61 |
| 9 | 111641627 | rs2275496 | 1.994 | 0.117 | -0.421 | -0.102 | -0.106 | -0.094 | -0.094 | 0 | 0.46 | 0.06 | 0.46 | 0.48 | 0.45 | 0.27 |
| 9 | 111641825 | rs1538660 | 0.621 | 0.124 | -0.331 | -0.106 | 0.002 | 0.015 | 0.015 | 0 | 0.19 | 0.32 | 0.16 | 0.29 | 0.17 | 0.23 |
| 9 | 111642535 | rs7874280 | 0.974 | 0.114 | -0.078 | -0.119 | -0.004 | -0.061 | -0.061 | 0 | 0.14 | 0.19 | 0.09 | 0 | 0.16 | 0.05 |
| 9 | 111645198 | rs10117384 | 0.234 | 0.117 | -0.075 | -0.105 | 0.004 | 0.026 | 0.026 | 0 | 0.22 | 0.33 | 0.31 | 0.38 | 0.23 | 0.32 |
| 9 | 111646406 | rs6477685 | -1.043 | 0.11 | -0.106 | -0.117 | -0.007 | -0.038 | -0.038 | 0.372 | 0.92 | 0.99 | 0.93 | 0.86 | 0.94 | 0.73 |
| 9 | 111646983 | rs4978374 | 0.445 | 0.126 | -0.068 | -0.112 | -0.006 | -0.013 | -0.013 | 0 | 0.75 | 0.75 | 0.84 | 0.86 | 0.78 | 0.63 |
| 9 | 111647581 | rs10816756 | 0.073 | 0.185 | 0.037 | -0.044 | 0.103 | 0.136 | 0.136 | 0 | 0.08 | 0.37 | 0.2 | 0.09 | 0.13 | 0.08 |
| 9 | 111648732 | rs10816758 | -0.369 | 0.182 | -0.004 | -0.047 | 0.002 | 0.009 | 0.009 | 0.016 | 0.27 | 0.69 | 0.36 | 0.38 | 0.3 | 0.31 |
| 9 | 111650007 | rs10120358 | 0.549 | 0.17 | 0.02 | -0.053 | 0.004 | 0.015 | 0.015 | 0 | 0.19 | 0.32 | 0.16 | 0.29 | 0.17 | 0.23 |
| 9 | 111650048 | rs10117105 | 0.549 | 0.17 | 0.001 | -0.053 | 0.004 | 0.015 | 0.015 | 0 | 0.19 | 0.32 | 0.16 | 0.29 | 0.17 | 0.23 |
| 9 | 111650788 | rs11792254 | 1.182 | 0.119 | -0.022 | -0.144 | -0.005 | -0.051 | -0.051 | 0.021 | 0.13 | 0.17 | 0.08 | 0 | 0.14 | 0.05 |
| 9 | 111652477 | rs12344049 | 1.096 | 0.233 | -0.097 | -0.098 | -0.001 | -0.01 | -0.01 | 0 | 0.13 | 0.24 | 0.11 | 0.12 | 0.1 | 0.09 |
| 9 | 111654395 | rs3780512 | 0.476 | 0.222 | -0.225 | -0.056 | 0.013 | -0.01 | -0.01 | 0 | 0.11 | 0.06 | 0.09 | 0.14 | 0.08 | 0.34 |
| 9 | 111659021 | rs2275626 | -0.816 | 0.32 | -0.086 | 0.046 | 0.075 | 0.182 | 0.182 | 0.756 | 0.14 | 0.48 | 0.25 | 0.26 | 0.2 | 0.22 |
| 9 | 111659483 | rs2230793 | -0.816 | 0.323 | -0.087 | 0.041 | 0.075 | 0.187 | 0.187 | 0.756 | 0.14 | 0.48 | 0.25 | 0.26 | 0.2 | 0.22 |
| 9 | 111661150 | rs4978760 | -2.441 | 0.272 | -0.555 | -0.082 | -0.123 | -0.102 | -0.102 | 0 | 0.54 | 0.95 | 0.54 | 0.52 | 0.54 | 0.72 |
| 9 | 111663793 | rs10979605 | 0.369 | 0.267 | -0.099 | -0.075 | 0.024 | -0.029 | -0.029 | 0.238 | 0.06 | 0.18 | 0.03 | 0 | 0.09 | 0.05 |
| 9 | 111664178 | rs3818932 | 2.271 | 0.258 | -0.958 | -0.189 | -0.09 | -0.114 | -0.114 | 1.155 | 0.26 | 0.02 | 0.33 | 0.31 | 0.27 | 0.16 |
| 9 | 111670100 | rs838826 | 0.199 | 0.211 | -0.742 | -0.263 | 0 | 0.001 | 0.001 | 0 | 0.25 | 0.28 | 0.2 | 0.25 | 0.19 | 0.42 |
| 9 | 111678016 | rs1772043 | -1.891 | 0.382 | -0.873 | -0.135 | -0.027 | -0.023 | -0.023 | 0.131 | 0.7 | 0.84 | 0.64 | 0.69 | 0.66 | 0.82 |
| 9 | 111688767 | rs2275635 | 2.111 | 0.352 | -0.94 | -0.105 | -0.009 | 0.015 | 0.015 | 0.056 | 0.21 | 0.02 | 0.13 | 0.17 | 0.18 | 0.11 |
| 9 | 111697054 | rs1759842 | -1.188 | 0.488 | -0.536 | -0.017 | 0.019 | 0.038 | 0.038 | 0 | 0.64 | 0.57 | 0.46 | 0.59 | 0.58 | 0.72 |
| 9 | 111697427 | rs1616727 | -1.231 | 0.474 | -0.428 | -0.024 | 0.018 | 0.035 | 0.035 | 0 | 0.64 | 0.59 | 0.46 | 0.59 | 0.58 | 0.72 |
| 9 | 111703561 | rs13299328 | 1.459 | 0.33 | -0.419 | -0.121 | -0.015 | 0.012 | 0.012 | 0 | 0.12 | 0.03 | 0.1 | 0.1 | 0.1 | 0.09 |
| 9 | 111705501 | rs2289480 | 1.036 | 0.232 | -0.383 | -0.13 | 0.002 | 0.011 | 0.011 | 0 | 0.34 | 0.3 | 0.52 | 0.41 | 0.36 | 0.26 |
| 9 | 111744771 | rs10979638 | 1.218 | 0.295 | -0.144 | -0.078 | -0.001 | 0.01 | 0.01 | 0 | 0.05 | 0.05 | 0.05 | 0.17 | 0.07 | 0.16 |
| 9 | 111746097 | rs866311 | -0.663 | 0.252 | -0.154 | -0.177 | 0.006 | -0.006 | -0.006 | 0 | 0.59 | 0.53 | 0.41 | 0.42 | 0.51 | 0.56 |
| 9 | 111765737 | rs12555920 | -0.076 | 0.264 | 0.065 | 0.097 | 0.064 | 0.181 | 0.181 | 0 | 0.06 | 0.23 | 0.18 | 0.12 | 0.09 | 0.09 |
| 9 | 111774393 | rs10816766 | 0.391 | 0.294 | -0.08 | -0.109 | 0 | -0.015 | -0.015 | 0 | 0.72 | 0.58 | 0.76 | 0.76 | 0.75 | 0.6 |
| 9 | 111776224 | rs1333344 | 2.145 | 0.381 | -0.311 | -0.086 | -0.076 | -0.092 | -0.092 | 0.164 | 0.26 | 0.02 | 0.32 | 0.3 | 0.28 | 0.16 |
| 9 | 111782531 | rs7776 | 1.046 | 0.269 | -0.113 | -0.127 | 0.007 | -0.022 | -0.022 | 1.991 | 0.28 | 0.18 | 0.19 | 0.36 | 0.26 | 0.28 |
| 9 | 111782670 | rs10816767 | 0.635 | 0.265 | 0.032 | -0.119 | -0.003 | -0.002 | -0.002 | 0 | 0.46 | 0.51 | 0.59 | 0.4 | 0.5 | 0.35 |
| 9 | 111790432 | rs10979660 | 0.602 | 0.339 | 0.028 | -0.132 | -0.004 | -0.002 | -0.002 | 0 | 0.46 | 0.51 | 0.6 | 0.4 | 0.51 | 0.35 |
| 9 | 111795565 | rs7852550 | 1.887 | 0.272 | 0.197 | -0.172 | -0.009 | -0.065 | -0.065 | 0 | 0.14 | 0.21 | 0.08 | 0 | 0.14 | 0.09 |
| 9 | 111798751 | rs2256253 | -0.97 | 0.286 | 0.216 | -0.205 | -0.002 | -0.117 | -0.117 | 0 | 0.8 | 0.65 | 0.88 | 1 | 0.77 | 0.85 |
| 9 | 111799369 | rs489283 | 1.97 | 0.265 | 0.018 | -0.206 | -0.022 | 0 | 0 | 0.168 | 0.2 | 0.01 | 0.13 | 0.17 | 0.18 | 0.11 |
| 9 | 111800234 | rs534286 | -0.511 | 0.267 | 0.09 | -0.12 | -0.002 | -0.005 | -0.005 | 0 | 0.75 | 0.37 | 0.68 | 0.86 | 0.71 | 0.73 |
| 9 | 111801344 | rs3750453 | 1.24 | 0.283 | 0.032 | -0.102 | -0.009 | -0.032 | -0.032 | 0 | 0.08 | 0.01 | 0.08 | 0.13 | 0.07 | 0.19 |
| 9 | 111807394 | rs646234 | -0.599 | 0.333 | 0.261 | -0.117 | 0.009 | 0.003 | 0.003 | 0 | 0.5 | 0.43 | 0.38 | 0.6 | 0.43 | 0.62 |
| 9 | 111815340 | rs1003346 | 0.978 | 0.303 | 0.203 | -0.15 | 0 | -0.004 | -0.004 | 0 | 0.43 | 0.48 | 0.43 | 0.29 | 0.43 | 0.29 |
| 9 | 111831600 | rs11787869 | 0.95 | 0.313 | 0.075 | -0.168 | 0.01 | 0.038 | 0.038 | 0 | 0.34 | 0.23 | 0.21 | 0.17 | 0.33 | 0.22 |

| | | | | | | | | | | | | | | | | |
|---|---|---|---|---|---|---|---|---|---|---|---|---|---|---|---|---|
| 9 | 111834983 | rs7855980 | 1.895 | 0.391 | 0.193 | -0.004 | -0.008 | -0.064 | -0.064 | 0 | 0.14 | 0.2 | 0.09 | 0 | 0.14 | 0.09 |
| 9 | 111854246 | rs523340 | 0.943 | 0.244 | 0.036 | 0.112 | -0.006 | -0.028 | -0.028 | 0.06 | 0.09 | 0.03 | 0.08 | 0.13 | 0.09 | 0.25 |
| 9 | 111868885 | rs2271878 | 1.609 | 0.269 | -0.253 | 0.095 | -0.081 | -0.101 | -0.101 | 0.583 | 0.27 | 0.02 | 0.33 | 0.29 | 0.28 | 0.17 |
| 9 | 111880736 | rs2805888 | -0.189 | 0.074 | -0.218 | -0.047 | 0.007 | 0.013 | 0.013 | 0 | 0.74 | 0.81 | 0.79 | 0.76 | 0.79 | 0.65 |
| 9 | 111881927 | rs6825 | 1.07 | 0.116 | -0.136 | -0.154 | -0.029 | 0.036 | 0.036 | 0 | 0.43 | 0.2 | 0.42 | 0.29 | 0.47 | 0.32 |
| 9 | 111885869 | rs10816772 | 1.245 | 0.157 | 0.119 | -0.111 | -0.006 | 0.03 | 0.03 | 0 | 0.23 | 0.34 | 0.17 | 0.36 | 0.21 | 0.23 |
| 9 | 111886177 | rs17627977 | 2.153 | 0.176 | 0.085 | -0.132 | 0 | -0.055 | -0.055 | 0 | 0.15 | 0.19 | 0.09 | 0 | 0.17 | 0.12 |
| 9 | 111886575 | rs12379316 | 2.789 | 0.173 | 0.068 | -0.115 | -0.008 | -0.033 | -0.033 | 1.081 | 0.14 | 0.01 | 0.11 | 0.17 | 0.12 | 0.07 |
| 9 | 111887219 | rs4978770 | 1.064 | 0.164 | 0.074 | -0.149 | 0 | -0.001 | -0.001 | 0 | 0.16 | 0.16 | 0.12 | 0.1 | 0.12 | 0.1 |
| 9 | 111890197 | rs1416957 | 0.477 | 0.257 | 0.096 | -0.093 | -0.001 | -0.005 | -0.005 | 0 | 0.37 | 0.41 | 0.26 | 0.36 | 0.38 | 0.38 |
| 9 | 111890349 | rs2417976 | 1.311 | 0.257 | 0.095 | -0.092 | 0 | -0.001 | -0.001 | 0 | 0.22 | 0.23 | 0.17 | 0.36 | 0.21 | 0.23 |
| 9 | 111899198 | rs11794020 | 1.917 | 0.279 | 0.115 | -0.183 | 0 | -0.055 | -0.055 | 0.208 | 0.15 | 0.19 | 0.09 | 0 | 0.17 | 0.15 |
| 9 | 111899928 | rs10759329 | 0.718 | 0.349 | 0.021 | -0.016 | 0 | 0.002 | 0.002 | 0 | 0.66 | 0.55 | 0.59 | 0.66 | 0.67 | 0.55 |
| 9 | 111915474 | rs10979712 | 0.564 | 0.339 | -0.143 | -0.015 | 0 | -0.002 | -0.002 | 0 | 0.37 | 0.38 | 0.39 | 0.28 | 0.43 | 0.31 |
| 9 | 111918365 | rs6477693 | 0.498 | 0.216 | -0.26 | -0.026 | 0.007 | -0.006 | -0.006 | 0 | 0.69 | 0.69 | 0.62 | 0.65 | 0.74 | 0.55 |
| 9 | 111925315 | rs4978774 | 0.449 | 1.048 | -0.382 | 0.13 | 0 | -0.003 | -0.003 | 0 | 0.79 | 0.76 | 0.8 | 0.67 | 0.8 | 0.63 |
| 9 | 111928136 | rs7468610 | 0.755 | 0.96 | -0.308 | -0.094 | -0.002 | -0.161 | -0.161 | 0 | 0.3 | 0.51 | 0.16 | 0.01 | 0.3 | 0.15 |
| 9 | 111928593 | rs2139350 | 1.229 | 0.864 | -0.431 | -0.131 | -0.001 | -0.055 | -0.055 | 0 | 0.24 | 0.23 | 0.12 | 0 | 0.24 | 0.12 |
| 9 | 111934261 | rs10759332 | 2.191 | 0.928 | -0.673 | 0.218 | -0.008 | -0.038 | -0.038 | 0 | 0.14 | 0.01 | 0.1 | 0.18 | 0.13 | 0.07 |
| 9 | 111936788 | rs10979727 | 1.132 | 1.522 | 0.31 | 1.025 | -0.007 | -0.057 | -0.057 | 0 | 0.44 | 0.51 | 0.27 | 0.16 | 0.43 | 0.21 |
| 9 | 111938125 | rs10119402 | 0.568 | 1.749 | -0.143 | 1.213 | -0.006 | -0.011 | -0.011 | 2.541 | 0.86 | 0.87 | 0.85 | 0.75 | 0.83 | 0.58 |
| 9 | 111942070 | rs885506 | 0.312 | 2.06 | -0.239 | 1.171 | -0.066 | 0.001 | 0.001 | 0 | 0.63 | 0.89 | 0.44 | 0.38 | 0.57 | 0.38 |
| 9 | 111942504 | rs10816777 | 1.64 | 1.712 | -0.015 | 1.028 | 0.005 | 0.013 | 0.013 | 0.997 | 0.41 | 0.31 | 0.23 | 0.26 | 0.38 | 0.19 |
| 9 | 111945049 | rs3750450 | 1.069 | 1.785 | 0.047 | 1.077 | 0.014 | 0.013 | 0.013 | 2.244 | 0.81 | 0.74 | 0.83 | 0.75 | 0.77 | 0.56 |
| 9 | 111945992 | rs3983526 | 0.93 | 1.78 | 0.062 | 1.533 | 0.006 | 0.011 | 0.011 | 0 | 0.49 | 0.6 | 0.53 | 0.4 | 0.5 | 0.31 |
| 9 | 111946103 | rs10816779 | -0.644 | 1.741 | 0.138 | 1.017 | 0.013 | -0.044 | -0.044 | 0.256 | 0.08 | 0.28 | 0.05 | 0 | 0.07 | 0.04 |
| 9 | 111950088 | rs10116353 | -0.017 | 1.764 | 0.073 | 1.121 | 0 | -0.003 | -0.003 | 0 | 0.35 | 0.33 | 0.42 | 0.36 | 0.31 | 0.25 |
| 9 | 111955592 | rs12346595 | -0.79 | 1.706 | 0.034 | 0.906 | 0.023 | -0.035 | -0.035 | 0.923 | 0.09 | 0.45 | 0.05 | 0 | 0.06 | 0.05 |
| 9 | 111958345 | rs6415807 | 0.479 | 2.262 | -0.741 | 1.22 | -0.041 | -0.048 | -0.048 | 0.55 | 0.87 | 0.96 | 0.86 | 0.76 | 0.81 | 0.59 |
| 9 | 111961389 | rs7875754 | -0.601 | 2.273 | -0.713 | 1.219 | -0.008 | -0.01 | -0.01 | 0 | 0.4 | 0.34 | 0.48 | 0.5 | 0.41 | 0.39 |
| 9 | 111963923 | rs7021903 | -0.364 | 2.18 | -0.664 | 1.234 | 0.002 | 0.006 | 0.006 | 0 | 0.37 | 0.42 | 0.45 | 0.38 | 0.35 | 0.28 |
| 9 | 111967572 | rs4978780 | 0.744 | 1.977 | -0.95 | 1.082 | 0 | 0.033 | 0.033 | 0 | 0.54 | 0.67 | 0.43 | 0.36 | 0.49 | 0.36 |
| 9 | 111969201 | rs7044279 | 0.097 | 2.045 | -1.036 | 1.17 | -0.018 | -0.035 | -0.035 | 0 | 0.9 | 0.99 | 0.92 | 0.86 | 0.89 | 0.72 |
| 9 | 111972423 | rs10739274 | 0.787 | 1.809 | -0.682 | 0.845 | 0.003 | 0.004 | 0.004 | 0.772 | 0.48 | 0.49 | 0.39 | 0.36 | 0.42 | 0.3 |
| 9 | 111972497 | rs10979749 | -0.482 | 1.689 | -0.439 | 0.942 | 0.035 | -0.021 | -0.021 | 0.423 | 0.06 | 0.23 | 0.04 | 0 | 0.06 | 0.04 |
| 9 | 111973020 | rs7858370 | -0.877 | 1.707 | -0.725 | 0.955 | 0.06 | 0.046 | 0.046 | 0 | 0.1 | 0.32 | 0.2 | 0.09 | 0.13 | 0.14 |
| 9 | 111978049 | rs4978783 | 0.332 | 1.525 | -2.148 | 1.202 | -0.047 | -0.092 | -0.092 | 0 | 0.8 | 0.99 | 0.73 | 0.75 | 0.8 | 0.8 |
| 9 | 111979807 | rs7860076 | 0.788 | 2.078 | -1.95 | 0.619 | -0.11 | 0.01 | 0.01 | 0 | 0.81 | 0.97 | 0.75 | 0.88 | 0.71 | 0.89 |
| 9 | 111980234 | rs4483229 | -0.778 | 0.9 | -1.137 | 0.645 | -0.008 | -0.008 | -0.008 | 0.002 | 0.15 | 0.2 | 0.13 | 0.18 | 0.1 | 0.36 |
| 9 | 111984614 | rs10979760 | -0.716 | 0.944 | -1.072 | 0.84 | -0.001 | 0 | 0 | 0 | 0.25 | 0.22 | 0.29 | 0.33 | 0.29 | 0.16 |
| 9 | 111991700 | rs1879069 | -0.27 | 1.187 | -1.304 | 0.907 | -0.006 | -0.015 | -0.015 | 0 | 0.35 | 0.28 | 0.56 | 0.44 | 0.4 | 0.39 |
| 9 | 111991981 | rs2012370 | -0.231 | 0.687 | -0.921 | 0.586 | 0 | 0 | 0 | 0 | 0.43 | 0.44 | 0.6 | 0.5 | 0.44 | 0.51 |
| 9 | 111992509 | rs10816789 | 0.491 | 0.702 | -1.733 | 0.461 | -0.034 | -0.068 | -0.068 | 0 | 0.19 | 0.01 | 0.14 | 0.24 | 0.17 | 0.18 |
| 9 | 111994722 | rs3849111 | 0.19 | 0.73 | -1.31 | 0.329 | -0.011 | 0.008 | 0.008 | 0.02 | 0.64 | 0.76 | 0.69 | 0.65 | 0.61 | 0.69 |
| 9 | 111996188 | rs17200841 | -0.138 | 0.745 | -1.164 | 0.21 | -0.002 | 0.016 | 0.016 | 0 | 0.11 | 0.04 | 0.09 | 0.08 | 0.12 | 0.07 |
| 9 | 112020668 | rs745388 | -0.31 | 1.306 | -0.727 | 0.195 | -0.007 | 0.006 | 0.006 | 0 | 0.36 | 0.25 | 0.51 | 0.29 | 0.42 | 0.33 |
| 9 | 112021585 | rs750723 | 0.348 | 0.774 | -1.22 | -0.291 | 0.023 | 0.027 | 0.027 | 0 | 0.12 | 0.03 | 0.06 | 0.07 | 0.1 | 0.09 |
| 9 | 112021708 | rs893918 | 0.522 | 1.052 | -0.379 | 0.074 | -0.003 | 0.002 | 0.002 | 0 | 0.61 | 0.42 | 0.43 | 0.68 | 0.49 | 0.61 |
| 9 | 112021852 | rs2004094 | 0.244 | 1.081 | -0.224 | 0.107 | 0.12 | 0.147 | 0.147 | 0 | 0.95 | 0.66 | 0.92 | 0.87 | 0.89 | 0.93 |
| 9 | 112024242 | rs1837549 | 0.405 | 1.116 | -0.436 | -0.026 | 0.01 | -0.004 | -0.004 | 0 | 0.64 | 0.44 | 0.45 | 0.68 | 0.53 | 0.62 |
| 9 | 112024496 | rs4978787 | 0.405 | 1.116 | -0.436 | -0.026 | 0.01 | -0.004 | -0.004 | 0.102 | 0.64 | 0.44 | 0.45 | 0.68 | 0.53 | 0.63 |
| 9 | 112033697 | rs3793555 | 0.662 | 1.649 | -0.433 | 0.159 | 0.033 | -0.005 | -0.005 | 0 | 0.64 | 0.5 | 0.45 | 0.68 | 0.53 | 0.64 |
| 9 | 112039400 | rs3793552 | 0.54 | 1.607 | -0.512 | 0.086 | -0.06 | -0.004 | -0.004 | 0.288 | 0.16 | 0.24 | 0.18 | 0.14 | 0.07 | 0.14 |
| 9 | 112044763 | rs10979798 | 0.235 | 1.973 | -0.6 | 0.132 | 0.052 | 0.009 | 0.009 | 0 | 0.87 | 0.81 | 0.67 | 0.82 | 0.66 | 0.8 |
| 9 | 112056059 | rs10512392 | -0.546 | 1.995 | -0.706 | 0.12 | -0.092 | 0.05 | 0.05 | 0 | 0.15 | 0.03 | 0.3 | 0.04 | 0.3 | 0.15 |
| 9 | 112058964 | rs12352460 | -0.091 | 1.888 | -0.699 | 0.117 | -0.002 | -0.006 | -0.006 | 0 | 0.13 | 0.12 | 0.28 | 0.18 | 0.26 | 0.19 |
| 9 | 112062247 | rs3793549 | 0.649 | 1.803 | -0.686 | 0.05 | -0.046 | 0.021 | 0.021 | 0 | 0.82 | 0.88 | 0.64 | 0.92 | 0.58 | 0.78 |
| 9 | 112070162 | rs3763640 | 0.153 | 0.806 | -0.482 | 0.241 | -0.047 | -0.014 | -0.014 | 0.971 | 0.18 | 0.23 | 0.2 | 0.15 | 0.1 | 0.13 |
| 9 | 112072419 | rs7025125 | -0.7 | 1.001 | -0.605 | 0.355 | 0.009 | -0.004 | -0.004 | 0 | 0.34 | 0.49 | 0.49 | 0.33 | 0.35 | 0.32 |
| 9 | 112080128 | rs966697 | 0.15 | 0.913 | -1.947 | 0.796 | -0.08 | -0.125 | -0.125 | 0 | 0.7 | 0.96 | 0.74 | 0.62 | 0.63 | 0.79 |
| 9 | 112081994 | rs4978793 | 0.395 | 1.878 | -1.987 | 1.026 | -0.082 | -0.062 | -0.062 | 0.015 | 0.63 | 0.88 | 0.66 | 0.44 | 0.47 | 0.74 |
| 9 | 112089520 | rs4978795 | 0.292 | 0.788 | -2.147 | 0.366 | 0.049 | 0.026 | 0.026 | 0 | 0.42 | 0.7 | 0.32 | 0.34 | 0.27 | 0.57 |
| 9 | 112091175 | rs10979826 | 0.681 | 0.641 | -2.066 | 0.473 | 0.008 | 0.002 | 0.002 | 0.016 | 0.34 | 0.68 | 0.28 | 0.32 | 0.2 | 0.52 |
| 9 | 112097738 | rs12553743 | -0.379 | 0.324 | -1.106 | 0.56 | 0.008 | 0.096 | 0.096 | 0 | 0.08 | 0.34 | 0.18 | 0.13 | 0.07 | 0.09 |
| 9 | 112100958 | rs7857376 | -0.299 | 0.911 | -1.489 | 0.852 | -0.005 | -0.001 | -0.001 | 0 | 0.53 | 0.51 | 0.72 | 0.51 | 0.62 | 0.45 |
| 9 | 112103798 | rs7863010 | -1.356 | 0.908 | -1.628 | 0.851 | -0.057 | -0.083 | -0.083 | 0.725 | 0.87 | 0.99 | 0.87 | 0.73 | 0.82 | 0.93 |
| 9 | 112103828 | rs10979834 | 0.887 | 1.255 | -1.73 | 0.9 | -0.011 | 0.061 | 0.061 | 1.575 | 0.38 | 0.18 | 0.47 | 0.18 | 0.39 | 0.27 |
| 9 | 112104111 | rs7847955 | -0.752 | 1.222 | -1.642 | 0.909 | -0.014 | 0.009 | 0.009 | 0 | 0.6 | 0.5 | 0.76 | 0.53 | 0.7 | 0.48 |
| 9 | 112104315 | rs7848206 | -0.287 | 1.266 | -1.417 | 0.903 | 0.001 | 0 | 0 | 0 | 0.18 | 0.19 | 0.13 | 0.22 | 0.25 | 0.12 |

| | | | | | | | | | | | | | | | | |
|---|---|---|---|---|---|---|---|---|---|---|---|---|---|---|---|---|
| 9 | 112104544 | rs4292768 | 0.868 | 1.267 | -1.354 | 0.902 | 0.023 | 0.016 | 0.016 | 0.027 | 0.08 | 0.09 | 0.08 | 0.2 | 0.16 | 0.07 |
| 9 | 112105331 | rs7852279 | -0.221 | 1.257 | -1.363 | 0.897 | 0.001 | 0 | 0 | 0.397 | 0.17 | 0.18 | 0.13 | 0.22 | 0.25 | 0.12 |
| 9 | 112107196 | rs12685120 | -0.213 | 1.245 | -1.374 | 0.891 | 0.008 | 0.002 | 0.002 | 0 | 0.17 | 0.2 | 0.13 | 0.23 | 0.25 | 0.12 |
| 9 | 112108334 | rs12115792 | 1.716 | 1.044 | -1.16 | 0.815 | -0.04 | -0.031 | -0.031 | 0.719 | 0.26 | 0.37 | 0.1 | 0.2 | 0.12 | 0.45 |
| 9 | 112110035 | rs1371786 | 1.343 | 0.947 | -1.263 | 0.777 | -0.036 | 0.019 | 0.019 | 0 | 0.27 | 0.43 | 0.27 | 0.35 | 0.17 | 0.51 |
| 9 | 112112634 | rs10979838 | 0.995 | 1.047 | -1.325 | 0.813 | 0.006 | 0.006 | 0.006 | 0 | 0.1 | 0.05 | 0.05 | 0.07 | 0.08 | 0.05 |
| 9 | 112121089 | rs10759341 | 0.724 | 1.03 | -1.577 | 0.837 | -0.002 | 0.058 | 0.058 | 0.314 | 0.41 | 0.18 | 0.46 | 0.21 | 0.4 | 0.34 |
| 9 | 112131222 | rs7872998 | -0.692 | 1.354 | -1.372 | 1.224 | -0.021 | -0.023 | -0.023 | 2.169 | 0.86 | 0.89 | 0.88 | 0.8 | 0.81 | 0.93 |
| 9 | 112133083 | rs12551129 | 0.682 | 1.557 | -1.332 | 1.346 | 0.02 | 0.018 | 0.018 | 0.696 | 0.08 | 0.1 | 0.11 | 0.16 | 0.16 | 0.08 |
| 9 | 112137099 | rs3943672 | 1.463 | 1.366 | -1.255 | 1.323 | 0.017 | 0.022 | 0.022 | 0.264 | 0.66 | 0.51 | 0.59 | 0.44 | 0.54 | 0.79 |
| 9 | 112148314 | rs3793569 | 0.54 | 1.446 | -1.252 | 1.346 | 0.024 | 0.033 | 0.033 | 0.532 | 0.08 | 0.13 | 0.12 | 0.22 | 0.16 | 0.09 |
| 9 | 112149215 | rs3793567 | -0.054 | 1.384 | -1.192 | 1.253 | -0.001 | 0 | 0 | 0 | 0.2 | 0.18 | 0.14 | 0.21 | 0.21 | 0.13 |
| 9 | 112150516 | rs3793566 | 0.468 | 0.983 | -1.441 | 1.037 | -0.008 | -0.006 | -0.006 | 1.232 | 0.19 | 0.16 | 0.2 | 0.26 | 0.29 | 0.13 |
| 9 | 112151349 | rs1371787 | -0.555 | 1.088 | -1.537 | 0.981 | -0.058 | -0.049 | -0.049 | 0.061 | 0.84 | 0.97 | 0.85 | 0.75 | 0.75 | 0.88 |
| 9 | 112152419 | rs10816805 | 0.572 | 1.107 | -1.512 | 0.958 | -0.007 | 0.001 | 0.001 | 0 | 0.19 | 0.16 | 0.2 | 0.18 | 0.29 | 0.13 |
| 9 | 112153156 | rs10512393 | 1.134 | 0.902 | -1.461 | 0.979 | 0.004 | 0.001 | 0.001 | 0 | 0.56 | 0.46 | 0.51 | 0.42 | 0.43 | 0.72 |
| 9 | 112164045 | rs17201409 | 0.492 | 0.859 | -1.46 | 1.061 | -0.011 | -0.007 | -0.007 | 0 | 0.19 | 0.16 | 0.19 | 0.25 | 0.28 | 0.12 |
| 9 | 112165546 | rs10816807 | -0.472 | 0.899 | -1.59 | 1.071 | -0.033 | -0.034 | -0.034 | 0.185 | 0.82 | 0.97 | 0.87 | 0.82 | 0.8 | 0.9 |
| 9 | 112171184 | rs7847693 | 0.559 | 1.082 | -1.333 | 1.141 | 0.009 | 0.011 | 0.011 | 0 | 0.65 | 0.53 | 0.56 | 0.42 | 0.51 | 0.77 |
| 9 | 112172674 | rs17803052 | 0.786 | 1.468 | -1.434 | 1.097 | -0.038 | -0.021 | -0.021 | 0 | 0.08 | 0 | 0.05 | 0.06 | 0.07 | 0.09 |
| 9 | 112192338 | rs3793532 | 0.084 | 1.329 | -1.264 | 0.666 | -0.003 | -0.003 | -0.003 | 0.164 | 0.21 | 0.14 | 0.35 | 0.22 | 0.28 | 0.27 |
| 9 | 112193021 | rs7873088 | -0.084 | 1.069 | -0.991 | 0.473 | 0.002 | -0.003 | -0.003 | 0.192 | 0.19 | 0.25 | 0.2 | 0.19 | 0.25 | 0.12 |
| 9 | 112197634 | rs10979861 | 0.2 | 1.184 | -0.642 | 0.284 | 0.011 | -0.016 | -0.016 | 0 | 0.36 | 0.24 | 0.37 | 0.64 | 0.32 | 0.32 |
| 9 | 112199146 | rs4978809 | 1 | 1 | -0.602 | -0.198 | 0.005 | 0.012 | 0.012 | 0.168 | 0.25 | 0.16 | 0.35 | 0.07 | 0.24 | 0.17 |
| 9 | 112201603 | rs16914146 | 0.324 | 1.298 | -0.626 | -0.142 | 0.02 | -0.096 | -0.096 | 0 | 0.33 | 0.54 | 0.23 | 0.07 | 0.31 | 0.43 |
| 9 | 112207341 | rs1108496 | 0.315 | 1.313 | -0.627 | -0.109 | 0.019 | -0.116 | -0.116 | 0.321 | 0.33 | 0.54 | 0.23 | 0.05 | 0.31 | 0.43 |
| 9 | 112218035 | rs4978810 | -1.609 | 1.346 | -0.661 | -0.057 | -0.032 | -0.031 | -0.031 | 0 | 0.94 | 0.98 | 0.94 | 0.9 | 0.85 | 0.93 |
| 9 | 112218403 | rs7041037 | -0.008 | 1.091 | -0.457 | -0.2 | 0.012 | -0.003 | -0.003 | 0.046 | 0.36 | 0.53 | 0.3 | 0.32 | 0.34 | 0.45 |
| 9 | 112218841 | rs10816813 | -0.694 | 1.064 | -0.497 | -0.24 | -0.009 | 0.017 | 0.017 | 0 | 0.58 | 0.71 | 0.61 | 0.33 | 0.54 | 0.63 |
| 9 | 112220503 | rs7033652 | 0.748 | 0.966 | -0.046 | -0.931 | 0.016 | -0.037 | -0.037 | 0.66 | 0.07 | 0.23 | 0.05 | 0 | 0.07 | 0.2 |
| 9 | 112221052 | rs7850860 | 1.213 | 1.035 | -0.372 | -0.752 | 0.009 | 0.009 | 0.009 | 0.623 | 0.22 | 0.13 | 0.31 | 0.07 | 0.19 | 0.18 |
| 9 | 112236879 | rs7046039 | -0.627 | 0.67 | -0.279 | -0.754 | 0.005 | 0.014 | 0.014 | 0 | 0.58 | 0.56 | 0.59 | 0.27 | 0.59 | 0.61 |
| 9 | 112237679 | rs10979884 | 0.376 | 0.887 | -0.113 | -0.85 | -0.001 | 0.03 | 0.03 | 0 | 0.26 | 0.12 | 0.17 | 0.06 | 0.24 | 0.42 |
| 9 | 112243321 | rs10512395 | 0.193 | 0.638 | 0.198 | -0.85 | -0.004 | -0.08 | -0.08 | 0 | 0.28 | 0.26 | 0.18 | 0.04 | 0.24 | 0.43 |
| 9 | 112244438 | rs10759343 | -1.301 | 0.976 | -0.027 | -0.454 | -0.027 | -0.023 | -0.023 | 0 | 0.95 | 0.98 | 0.93 | 0.89 | 0.85 | 0.94 |
| 9 | 112251563 | rs4978813 | -0.106 | 0.743 | 0.271 | -0.925 | 0.019 | -0.048 | -0.048 | 0 | 0.65 | 0.48 | 0.63 | 0.9 | 0.53 | 0.6 |
| 9 | 112253555 | rs10979889 | 1.116 | 0.594 | 0.071 | -1.056 | 0.015 | 0.092 | 0.092 | 0 | 0.22 | 0.04 | 0.13 | 0.04 | 0.17 | 0.21 |
| 9 | 112266667 | rs4978394 | -0.611 | 0.677 | 0.315 | -1.015 | 0.025 | 0.029 | 0.029 | 0 | 0.92 | 0.82 | 0.91 | 0.91 | 0.85 | 0.94 |
| 9 | 112269480 | rs4978396 | 1.089 | 0.605 | 0.485 | -1.103 | 0.023 | -0.135 | -0.135 | 0 | 0.25 | 0.3 | 0.27 | 0.01 | 0.3 | 0.33 |
| 9 | 112271395 | rs4978816 | 0.874 | 0.509 | 0.384 | -0.854 | 0.001 | -0.004 | -0.004 | 0 | 0.37 | 0.39 | 0.47 | 0.63 | 0.31 | 0.33 |
| 9 | 112271805 | rs4978817 | 1.048 | 0.547 | 0.305 | -0.86 | 0.013 | -0.093 | -0.093 | 0 | 0.24 | 0.21 | 0.27 | 0.01 | 0.3 | 0.33 |
| 9 | 112277740 | rs10124300 | -0.543 | 0.525 | 0.305 | -0.808 | -0.002 | 0.008 | 0.008 | 1.285 | 0.54 | 0.32 | 0.44 | 0.28 | 0.54 | 0.61 |
| 9 | 112281863 | rs1109478 | -0.103 | 0.646 | 0.055 | -0.813 | 0.008 | 0.101 | 0.101 | 1.503 | 0.51 | 0.7 | 0.55 | 0.86 | 0.45 | 0.6 |
| 9 | 112282070 | rs881660 | 1.581 | 0.448 | 0.074 | -0.825 | -0.003 | 0.043 | 0.043 | 0.286 | 0.16 | 0.17 | 0.29 | 0.53 | 0.15 | 0.14 |
| 9 | 112284879 | rs10119758 | 1.034 | 0.543 | -0.008 | -1.018 | 0.018 | 0.077 | 0.077 | 0 | 0.22 | 0.06 | 0.13 | 0.04 | 0.17 | 0.21 |
| 9 | 112291281 | rs12554939 | 1.516 | 0.549 | -0.006 | -1.052 | -0.016 | 0.025 | 0.025 | 0.226 | 0.2 | 0.09 | 0.23 | 0.01 | 0.24 | 0.13 |
| 9 | 112291471 | rs9987721 | 1.027 | 0.53 | -0.008 | -1.038 | 0.018 | 0.077 | 0.077 | 0 | 0.22 | 0.06 | 0.13 | 0.04 | 0.17 | 0.21 |
| 9 | 112294766 | rs1999269 | 0.091 | 0.527 | 0.053 | -0.985 | 0.008 | 0.098 | 0.098 | 1.503 | 0.51 | 0.7 | 0.55 | 0.86 | 0.45 | 0.6 |
| 9 | 112302220 | rs7046565 | 0.289 | 0.537 | 0.037 | -0.981 | -0.003 | 0.111 | 0.111 | 1.029 | 0.41 | 0.18 | 0.36 | 0.05 | 0.4 | 0.33 |
| 9 | 112303530 | rs1541087 | 0.969 | 0.464 | 0.081 | -0.899 | -0.007 | 0.009 | 0.009 | 0.458 | 0.39 | 0.47 | 0.48 | 0.63 | 0.31 | 0.35 |
| 9 | 112303858 | rs17201835 | 0.547 | 0.499 | -0.319 | -0.751 | 0.002 | -0.049 | -0.049 | 0 | 0.08 | 0.01 | 0.04 | 0.22 | 0.08 | 0.07 |
| 9 | 112309305 | rs11788069 | 0.08 | 0.561 | -0.254 | -0.89 | -0.006 | 0.02 | 0.02 | 0 | 0.67 | 0.72 | 0.74 | 0.86 | 0.61 | 0.55 |
| 9 | 112315013 | rs10759349 | -0.212 | 0.947 | -0.09 | -0.571 | 0.026 | 0.01 | 0.01 | 0 | 0.89 | 0.81 | 0.88 | 0.91 | 0.79 | 0.74 |
| 9 | 112317462 | rs12554063 | 1.143 | 0.824 | -0.439 | -0.721 | -0.02 | 0.042 | 0.042 | 0.099 | 0.2 | 0.09 | 0.22 | 0.01 | 0.22 | 0.13 |
| 9 | 112325051 | rs7858161 | 0.159 | 0.802 | -0.344 | -0.654 | -0.001 | 0.001 | 0.001 | 0.081 | 0.29 | 0.23 | 0.26 | 0.23 | 0.29 | 0.2 |
| 9 | 112337709 | rs7032700 | 1.117 | 0.875 | -0.552 | -1.128 | -0.017 | 0.03 | 0.03 | 0.039 | 0.2 | 0.1 | 0.22 | 0.01 | 0.21 | 0.12 |
| 9 | 112339759 | rs13283237 | 0.931 | 0.82 | -0.345 | -1.155 | 0 | -0.052 | -0.052 | 0 | 0.22 | 0.21 | 0.23 | 0.01 | 0.21 | 0.12 |
| 9 | 112346022 | rs10512396 | 1.058 | 0.68 | -0.589 | -1.407 | 0.002 | 0.077 | 0.077 | 0.486 | 0.19 | 0.04 | 0.12 | 0.04 | 0.17 | 0.17 |
| 9 | 112346234 | rs10512397 | 1.058 | 0.68 | -0.589 | -1.407 | 0.002 | 0.077 | 0.077 | 0.486 | 0.19 | 0.04 | 0.12 | 0.04 | 0.17 | 0.17 |
| 9 | 112350261 | rs10979955 | 0.982 | 0.759 | -0.333 | -1.117 | 0.002 | 0.004 | 0.004 | 0.772 | 0.38 | 0.36 | 0.48 | 0.71 | 0.34 | 0.38 |
| 9 | 112353128 | rs7044539 | 0.388 | 1.002 | -0.411 | -0.49 | 0.001 | -0.073 | -0.073 | 1.237 | 0.23 | 0.24 | 0.23 | 0.01 | 0.21 | 0.13 |
| 9 | 112355217 | rs17794184 | -0.176 | 0.948 | -0.702 | -0.212 | 0.007 | -0.048 | -0.048 | 0 | 0.08 | 0.01 | 0.03 | 0.15 | 0.07 | 0.07 |
| 9 | 112358656 | rs1411683 | 0.014 | 1.333 | -0.244 | -0.53 | 0.031 | 0.019 | 0.019 | 0 | 0.93 | 0.85 | 0.92 | 0.93 | 0.86 | 0.94 |
| 9 | 112365670 | rs746376 | -0.042 | 0.812 | -0.592 | -0.844 | -0.003 | -0.038 | -0.038 | 0 | 0.26 | 0.22 | 0.24 | 0.02 | 0.25 | 0.13 |
| 9 | 112366444 | rs7852886 | -0.374 | 0.283 | -0.561 | -0.64 | -0.01 | -0.03 | -0.03 | 0 | 0.44 | 0.54 | 0.43 | 0.25 | 0.35 | 0.49 |
| 9 | 112366635 | rs4978839 | -0.384 | -0.098 | -0.293 | -0.164 | 0.003 | 0.005 | 0.005 | 3.694 | 0.44 | 0.41 | 0.39 | 0.4 | 0.42 | 0.52 |
| 9 | 112366665 | rs1169626 | 0.686 | -0.063 | -0.572 | -0.068 | 0.003 | 0.001 | 0.001 | 0 | 0.19 | 0.14 | 0.11 | 0.18 | 0.14 | 0.13 |
| 9 | 112367107 | rs10119049 | -0.051 | -0.651 | 0.07 | -0.262 | 0.011 | 0.036 | 0.036 | 2.77 | 0.41 | 0.25 | 0.23 | 0.19 | 0.35 | 0.48 |
| 9 | 112367400 | rs6477714 | -0.85 | -0.56 | 0.059 | -0.096 | 0.034 | 0.042 | 0.042 | 0.877 | 0.52 | 0.69 | 0.27 | 0.4 | 0.42 | 0.51 |

| | | | | | | | | | | | | | | | | |
|---|---|---|---|---|---|---|---|---|---|---|---|---|---|---|---|---|
| 9 | 112370353 | rs1576587 | -0.976 | -0.091 | -0.139 | -0.154 | -0.006 | 0.002 | 0.002 | 0.666 | 0.58 | 0.6 | 0.34 | 0.4 | 0.54 | 0.54 |
| 9 | 112377352 | rs4978842 | 0.36 | -0.292 | -0.101 | -0.081 | 0.002 | -0.028 | -0.028 | 0 | 0.06 | 0.05 | 0.02 | 0.18 | 0.06 | 0.04 |
| 9 | 112381955 | rs7049166 | -0.334 | -0.859 | -0.421 | -0.452 | -0.006 | -0.007 | -0.007 | 0 | 0.37 | 0.57 | 0.2 | 0.2 | 0.29 | 0.17 |
| 9 | 112382522 | rs10123567 | 0.737 | -0.868 | -0.119 | -0.679 | 0.013 | -0.009 | -0.009 | 0 | 0.26 | 0.15 | 0.14 | 0.01 | 0.21 | 0.13 |
| 9 | 112383429 | rs10979965 | 0.619 | -0.24 | -1.083 | -0.297 | -0.037 | -0.048 | -0.048 | 0 | 0.11 | 0 | 0.08 | 0.18 | 0.12 | 0.08 |
| 9 | 112390854 | rs6477716 | -0.951 | -0.191 | -0.786 | -0.716 | 0.001 | -0.013 | -0.013 | 0.014 | 0.5 | 0.5 | 0.31 | 0.15 | 0.47 | 0.44 |
| 9 | 112392697 | rs4344152 | 0.09 | -0.097 | -1.54 | -0.734 | -0.057 | 0.028 | 0.028 | 0.028 | 0.2 | 0.07 | 0.14 | 0.14 | 0.24 | 0.3 |
| 9 | 112393507 | rs7029517 | -1.302 | -0.11 | -1.358 | -0.413 | -0.018 | 0.032 | 0.032 | 0.157 | 0.59 | 0.73 | 0.35 | 0.32 | 0.56 | 0.49 |
| 9 | 112394924 | rs11791761 | 0.906 | 0.064 | -1.281 | -0.202 | -0.014 | 0.013 | 0.013 | 0.072 | 0.1 | 0.07 | 0.06 | 0.06 | 0.12 | 0.25 |
| 9 | 112396529 | rs1407393 | -1.362 | 0.139 | -1.942 | 0.08 | -0.078 | -0.123 | -0.123 | 0.39 | 0.67 | 0.94 | 0.4 | 0.5 | 0.62 | 0.64 |
| 9 | 112398477 | rs1407394 | -1.534 | 0.105 | -1.682 | -0.212 | -0.049 | -0.084 | -0.084 | 3.333 | 0.64 | 0.9 | 0.45 | 0.43 | 0.57 | 0.55 |
| 9 | 112398754 | rs1358917 | 0.84 | 0.163 | -1.281 | -0.456 | 0.003 | -0.007 | -0.007 | 0.232 | 0.11 | 0.09 | 0.07 | 0.04 | 0.11 | 0.23 |
| 9 | 112400897 | rs11790056 | 1.191 | -0.037 | -1.482 | -0.618 | -0.001 | -0.009 | -0.009 | 0.679 | 0.21 | 0.21 | 0.14 | 0.08 | 0.2 | 0.13 |
| 9 | 112401674 | rs10739280 | -1.995 | 0.169 | -1.769 | -0.34 | -0.041 | -0.102 | -0.102 | 0.018 | 0.69 | 0.9 | 0.53 | 0.46 | 0.62 | 0.59 |
| 9 | 112406744 | rs10979978 | -0.919 | -0.314 | -0.969 | -0.941 | -0.004 | -0.028 | -0.028 | 0 | 0.38 | 0.46 | 0.21 | 0.12 | 0.3 | 0.37 |
| 9 | 112411823 | rs7049116 | 0.039 | 0.763 | -1.284 | -0.086 | -0.012 | 0.011 | 0.011 | 1.11 | 0.11 | 0.05 | 0.07 | 0.04 | 0.11 | 0.23 |
| 9 | 112415744 | rs10512398 | 0.891 | 0.003 | -1.264 | -0.112 | 0.027 | 0.024 | 0.024 | 2.612 | 0.21 | 0.15 | 0.13 | 0.1 | 0.13 | 0.05 |
| 9 | 112418293 | rs16914315 | 0.464 | 0.326 | -1.186 | -0.138 | 0.027 | 0.047 | 0.047 | 0.536 | 0.35 | 0.27 | 0.22 | 0.16 | 0.26 | 0.12 |
| 9 | 112419244 | rs7026239 | 0.464 | 0.326 | -1.186 | -0.138 | 0.027 | 0.047 | 0.047 | 0.536 | 0.35 | 0.27 | 0.22 | 0.16 | 0.26 | 0.12 |
| 9 | 112429319 | rs10979990 | 0.1 | 0.791 | -1.365 | -0.554 | 0.024 | 0.074 | 0.074 | 9.126 | 0.49 | 0.38 | 0.29 | 0.2 | 0.37 | 0.21 |
| 9 | 112434587 | rs7875665 | 0.879 | 1.354 | -1.374 | -0.477 | -0.012 | 0.006 | 0.006 | 0 | 0.07 | 0.04 | 0.06 | 0.04 | 0.09 | 0.22 |
| 9 | 112438399 | rs10979994 | 0.619 | 1.429 | -1.125 | 0.115 | 0.012 | 0.082 | 0.082 | 0 | 0.06 | 0.08 | 0.09 | 0.23 | 0.09 | 0.09 |
| 9 | 112440123 | rs10979996 | 1.227 | 1.398 | -1.702 | -0.029 | -0.038 | -0.005 | -0.005 | 0 | 0.06 | 0.01 | 0.1 | 0.06 | 0.16 | 0.09 |
| 9 | 112443817 | rs4292771 | 1.744 | 1.176 | -1.665 | -0.011 | 0.03 | 0.032 | 0.032 | 2.296 | 0.3 | 0.26 | 0.18 | 0.13 | 0.19 | 0.06 |
| 9 | 112444460 | rs2146087 | -1.159 | 1.521 | -1.351 | 0.452 | -0.046 | -0.142 | -0.142 | 0.017 | 0.75 | 0.92 | 0.58 | 0.51 | 0.68 | 0.64 |
| 9 | 112451003 | rs10816855 | -0.719 | 1.054 | -1.048 | 0.479 | 0.019 | 0.068 | 0.068 | 0.674 | 0.19 | 0.48 | 0.22 | 0.27 | 0.2 | 0.21 |
| 9 | 112453013 | rs713145 | -0.24 | 1.246 | -1.697 | 0.813 | -0.076 | -0.113 | -0.113 | 0 | 0.69 | 0.88 | 0.52 | 0.47 | 0.59 | 0.42 |
| 9 | 112475587 | rs2031785 | -0.33 | 1.212 | -1.218 | 1.255 | -0.015 | -0.013 | -0.013 | 0 | 0.93 | 0.98 | 0.82 | 0.9 | 0.89 | 0.89 |
| 9 | 112477012 | rs10980011 | 0.098 | 1.078 | -0.714 | 1.068 | -0.002 | 0.001 | 0.001 | 0 | 0.42 | 0.46 | 0.43 | 0.41 | 0.38 | 0.54 |
| 9 | 112477726 | rs2208704 | 0.297 | 1.473 | -0.951 | 0.956 | -0.061 | -0.015 | -0.015 | 0 | 0.09 | 0.01 | 0.13 | 0.09 | 0.21 | 0.12 |
| 9 | 112479083 | rs239 | 0.982 | 1.204 | -0.691 | 0.872 | 0.044 | -0.005 | -0.005 | 0.319 | 0.37 | 0.29 | 0.22 | 0.38 | 0.26 | 0.18 |
| 9 | 112480192 | rs1322275 | 0.656 | 1.174 | -0.631 | 0.861 | -0.005 | -0.001 | -0.001 | 1.661 | 0.39 | 0.48 | 0.24 | 0.38 | 0.26 | 0.19 |
| 9 | 112483181 | rs1571267 | 0.612 | 1.179 | -0.627 | 0.862 | -0.003 | -0.001 | -0.001 | 0 | 0.41 | 0.48 | 0.27 | 0.41 | 0.31 | 0.23 |
| 9 | 112488155 | rs9409094 | -0.293 | 1.176 | -0.542 | 0.834 | 0.009 | 0 | 0 | 0.239 | 0.51 | 0.49 | 0.54 | 0.5 | 0.58 | 0.66 |
| 9 | 112492370 | rs1322272 | 0.868 | 1.063 | -0.522 | 0.701 | 0.033 | 0.002 | 0.002 | 0.053 | 0.37 | 0.34 | 0.2 | 0.36 | 0.25 | 0.15 |
| 9 | 112502765 | rs12343459 | 0.889 | 1.142 | -0.961 | 0.739 | 0.04 | 0.001 | 0.001 | 0.085 | 0.37 | 0.32 | 0.2 | 0.36 | 0.25 | 0.15 |
| 9 | 112508976 | rs7848345 | 1.019 | 1.125 | -0.964 | 0.731 | 0.043 | -0.047 | -0.047 | 0 | 0.34 | 0.1 | 0.18 | 0.36 | 0.25 | 0.15 |
| 9 | 112512382 | rs10512400 | -0.191 | 1.277 | -1.005 | 0.76 | -0.008 | -0.001 | -0.001 | 0 | 0.51 | 0.65 | 0.65 | 0.51 | 0.49 | 0.67 |
| 9 | 112513269 | rs4111120 | -0.257 | 1.476 | -1.058 | 0.627 | -0.066 | -0.021 | -0.021 | 0 | 0.9 | 0.99 | 0.87 | 0.89 | 0.79 | 0.88 |
| 9 | 112519888 | rs2149002 | 0.946 | 1.263 | -0.418 | 0.624 | 0.056 | -0.005 | -0.005 | 4.834 | 0.77 | 0.55 | 0.63 | 0.8 | 0.67 | 0.74 |
| 9 | 112521535 | rs2025878 | -0.88 | 0.75 | -0.748 | 0.564 | -0.053 | 0.086 | 0.086 | 2.731 | 0.58 | 0.87 | 0.7 | 0.74 | 0.57 | 0.78 |
| 9 | 112525489 | rs7028330 | 0.082 | 0.706 | -0.455 | 0.32 | 0.007 | -0.015 | -0.015 | 0 | 0.17 | 0.25 | 0.23 | 0.09 | 0.17 | 0.22 |
| 9 | 112525766 | rs11790000 | 0.64 | 0.142 | -0.43 | 0.598 | -0.001 | 0.001 | 0.001 | 0.482 | 0.26 | 0.27 | 0.22 | 0.35 | 0.25 | 0.38 |
| 9 | 112531466 | rs7023524 | -0.945 | 0.032 | -0.303 | 0.342 | -0.013 | -0.012 | -0.012 | 0.02 | 0.23 | 0.6 | 0.43 | 0.14 | 0.2 | 0.31 |
| 9 | 112531537 | rs7023411 | -0.849 | 0.783 | -0.501 | 0.658 | -0.042 | -0.032 | -0.032 | 0.681 | 0.93 | 0.99 | 0.86 | 0.69 | 0.81 | 0.9 |
| 9 | 112531693 | rs16914500 | -0.609 | 0.615 | -0.328 | 0.635 | -0.018 | -0.009 | -0.009 | 0.882 | 0.2 | 0.56 | 0.42 | 0.14 | 0.18 | 0.3 |
| 9 | 112532676 | rs2068439 | 0.144 | 0.482 | -0.686 | 0.638 | -0.008 | -0.026 | -0.026 | 0.123 | 0.05 | 0.01 | 0.02 | 0.2 | 0.05 | 0.04 |
| 9 | 112534856 | rs1327793 | 0.907 | 0.214 | -0.939 | 0.09 | -0.053 | -0.07 | -0.07 | 0 | 0.71 | 0.85 | 0.77 | 0.49 | 0.62 | 0.73 |
| 9 | 112534910 | rs1327794 | -0.71 | -0.237 | -1.208 | -0.146 | 0.048 | -0.089 | -0.089 | 0 | 0.54 | 0.89 | 0.62 | 0.45 | 0.39 | 0.67 |
| 9 | 112536455 | rs7022225 | -0.085 | -0.088 | -1.082 | -0.065 | 0.059 | -0.071 | -0.071 | 0 | 0.45 | 0.85 | 0.59 | 0.45 | 0.37 | 0.65 |
| 9 | 112539406 | rs10115708 | -0.058 | -0.299 | -0.838 | -0.207 | -0.025 | -0.055 | -0.055 | 0.003 | 0.13 | 0.52 | 0.23 | 0.03 | 0.07 | 0.17 |
| 9 | 112543228 | rs10816866 | 1.452 | -0.215 | -1.216 | -0.153 | 0.024 | -0.02 | -0.02 | 0 | 0.24 | 0.1 | 0.16 | 0.28 | 0.2 | 0.34 |
| 9 | 112546308 | rs2182815 | 1.466 | -0.224 | -1.218 | -0.163 | 0.024 | -0.026 | -0.026 | 0 | 0.24 | 0.1 | 0.16 | 0.28 | 0.2 | 0.35 |
| 9 | 112551867 | rs913359 | -1.882 | 0.054 | -1.198 | -0.346 | -0.019 | 0.134 | 0.134 | 0.206 | 0.75 | 0.93 | 0.83 | 0.96 | 0.76 | 0.91 |
| 9 | 112553003 | rs1887521 | -0.361 | 0.294 | -0.69 | -0.043 | 0.154 | 0.242 | 0.242 | 0.038 | 0.14 | 0.48 | 0.21 | 0.36 | 0.2 | 0.15 |
| 9 | 112554953 | rs13294989 | -0.424 | 0.233 | -0.74 | -0.061 | -0.001 | 0.013 | 0.013 | 0.08 | 0.16 | 0.17 | 0.31 | 0.3 | 0.17 | 0.27 |
| 9 | 112556848 | rs13286006 | -0.424 | 0.233 | -0.74 | -0.061 | -0.001 | 0.013 | 0.013 | 0.08 | 0.16 | 0.18 | 0.32 | 0.3 | 0.17 | 0.27 |
| 9 | 112556945 | rs10980041 | -0.387 | 0.374 | -0.81 | -0.001 | 0.154 | 0.242 | 0.242 | 0.038 | 0.14 | 0.48 | 0.21 | 0.36 | 0.2 | 0.15 |
| 9 | 112561012 | rs2209815 | -1.246 | 0.307 | -1.489 | -0.138 | -0.058 | -0.076 | -0.076 | 0 | 0.36 | 0.83 | 0.42 | 0.53 | 0.36 | 0.35 |
| 9 | 112569588 | rs10759367 | -1.838 | 0.33 | -1.576 | -0.158 | -0.036 | 0.093 | 0.093 | 0.004 | 0.47 | 0.89 | 0.6 | 0.69 | 0.49 | 0.53 |
| 9 | 112569923 | rs10759368 | 1.388 | 0.348 | -1.494 | -0.133 | 0.051 | 0.097 | 0.097 | 0 | 0.06 | 0.07 | 0.15 | 0.36 | 0.19 | 0.07 |
| 9 | 112574527 | rs730973 | -1.588 | 0.314 | -1.53 | -0.133 | 0.051 | 0.035 | 0.035 | 0 | 0.41 | 0.79 | 0.44 | 0.33 | 0.3 | 0.47 |
| 9 | 112576143 | rs12003677 | -1.579 | 0.338 | -1.609 | -0.122 | 0.053 | 0.036 | 0.036 | 0 | 0.41 | 0.8 | 0.44 | 0.32 | 0.3 | 0.47 |
| 9 | 112578906 | rs4295727 | -1.193 | 0.879 | -1.006 | 0.304 | -0.007 | 0.017 | 0.017 | 2.486 | 0.28 | 0.46 | 0.36 | 0.31 | 0.26 | 0.39 |
| 9 | 112579958 | rs2795058 | 0.706 | 1.348 | -1.425 | 0.722 | 0.005 | -0.03 | -0.03 | 2.07 | 0.28 | 0.07 | 0.23 | 0.27 | 0.27 | 0.37 |
| 9 | 112583729 | rs7032049 | -0.215 | 1.857 | -0.67 | 1.2 | -0.014 | 0.025 | 0.025 | 0 | 0.07 | 0.04 | 0.03 | 0.01 | 0.08 | 0.11 |
| 9 | 112584723 | rs10980070 | -0.75 | 1.779 | -0.342 | 1.613 | -0.008 | 0.05 | 0.05 | 0 | 0.1 | 0.28 | 0.08 | 0.14 | 0.08 | 0.11 |
| 9 | 112589201 | rs12001151 | -0.726 | 1.78 | -0.342 | 1.614 | -0.007 | 0.051 | 0.051 | 0 | 0.1 | 0.28 | 0.08 | 0.14 | 0.08 | 0.11 |
| 9 | 112589454 | rs2418050 | -1.577 | 1.818 | -0.345 | 1.641 | 0.077 | 0.154 | 0.154 | 0 | 0.17 | 0.33 | 0.23 | 0.5 | 0.26 | 0.17 |

| | | | | | | | | | | | | | | | | |
|---|---|---|---|---|---|---|---|---|---|---|---|---|---|---|---|---|
| 9 | 112591526 | rs10733573 | -1.988 | 1.812 | -0.389 | 1.632 | 0.071 | 0.114 | 0.114 | 0.45 | 0.23 | 0.45 | 0.4 | 0.54 | 0.32 | 0.31 |
| 9 | 112592855 | rs7867595 | -0.863 | 1.946 | -0.91 | 1.809 | -0.006 | 0.05 | 0.05 | 0 | 0.1 | 0.28 | 0.07 | 0.14 | 0.08 | 0.11 |
| 9 | 112593531 | rs10816891 | -0.518 | 2.332 | -1.629 | 1.723 | -0.026 | 0.004 | 0.004 | 0 | 0.06 | 0.01 | 0.15 | 0.36 | 0.19 | 0.07 |
| 9 | 112596911 | rs2025875 | 0.039 | 0.623 | -1.776 | 0.614 | -0.023 | 0.007 | 0.007 | 0 | 0.74 | 0.98 | 0.83 | 0.81 | 0.76 | 0.83 |
| 9 | 112604714 | rs2900498 | -0.63 | 0.518 | -0.534 | 0.741 | -0.004 | 0.011 | 0.011 | 0.039 | 0.1 | 0.27 | 0.07 | 0.09 | 0.07 | 0.13 |
| 9 | 112606655 | rs10980082 | -0.6 | 1.249 | -0.402 | 1.838 | 0.018 | 0.106 | 0.106 | 0 | 0.08 | 0.12 | 0.21 | 0.39 | 0.11 | 0.15 |
| 9 | 112608710 | rs2769151 | 0.21 | 0.385 | -0.325 | 1.373 | -0.005 | -0.003 | -0.003 | 0 | 0.55 | 0.42 | 0.77 | 0.68 | 0.68 | 0.73 |
| 9 | 112610830 | rs10816897 | -0.538 | 0.722 | -0.677 | 1.703 | -0.042 | -0.078 | -0.078 | 0 | 0.2 | 0.08 | 0.34 | 0.48 | 0.26 | 0.36 |
| 9 | 112611749 | rs2004426 | -1.107 | 0.07 | -0.446 | 1.396 | -0.027 | -0.08 | -0.08 | 0 | 0.71 | 0.59 | 0.87 | 0.92 | 0.77 | 0.89 |
| 9 | 112612553 | rs1077760 | 1.333 | 0.441 | -0.275 | 1.374 | -0.001 | 0.001 | 0.001 | 0 | 0.34 | 0.3 | 0.38 | 0.22 | 0.36 | 0.38 |
| 9 | 112613140 | rs17794822 | 0.584 | 0.44 | -0.626 | 1.558 | 0.007 | -0.014 | -0.014 | 0 | 0.09 | 0.04 | 0.09 | 0.2 | 0.08 | 0.2 |
| 9 | 112614108 | rs10980086 | 0.52 | 0.46 | -0.629 | 1.481 | 0.017 | -0.023 | -0.023 | 0 | 0.16 | 0.14 | 0.1 | 0.24 | 0.09 | 0.14 |
| 9 | 112618755 | rs10980089 | 0.142 | 0.374 | -0.621 | 1.44 | 0.007 | -0.018 | -0.018 | 0 | 0.18 | 0.15 | 0.14 | 0.24 | 0.15 | 0.15 |
| 9 | 112620371 | rs12338023 | 0.001 | 0.013 | -0.289 | 1.107 | -0.013 | 0.013 | 0.013 | 0 | 0.17 | 0.22 | 0.14 | 0.21 | 0.14 | 0.25 |
| 9 | 112621077 | rs2769143 | 1.756 | -0.044 | -0.369 | 1.016 | -0.101 | -0.015 | -0.015 | 0 | 0.25 | 0.05 | 0.36 | 0.23 | 0.36 | 0.32 |
| 9 | 112621234 | rs7875145 | -0.831 | 0.002 | -0.356 | 1.105 | -0.011 | 0.008 | 0.008 | 0 | 0.34 | 0.39 | 0.24 | 0.45 | 0.24 | 0.39 |
| 9 | 112621761 | rs2769145 | -1.885 | -0.01 | -0.255 | 0.977 | 0.014 | 0.047 | 0.047 | 0 | 0.74 | 0.85 | 0.88 | 0.92 | 0.77 | 0.89 |
| 9 | 112625522 | rs10980096 | -0.245 | 0.154 | -0.079 | 1.191 | -0.017 | 0.015 | 0.015 | 0 | 0.17 | 0.23 | 0.14 | 0.21 | 0.14 | 0.25 |
| 9 | 112628478 | rs11792458 | 0.416 | 0.539 | -0.401 | 1.151 | -0.053 | -0.052 | -0.052 | 0 | 0.08 | 0.01 | 0.25 | 0.26 | 0.16 | 0.13 |
| 9 | 112629397 | rs7044189 | -0.126 | 0.115 | -0.194 | 0.895 | 0.012 | 0.026 | 0.026 | 0 | 0.47 | 0.65 | 0.71 | 0.68 | 0.55 | 0.66 |
| 9 | 112630762 | rs10980100 | -0.748 | 0.655 | -0.367 | 0.947 | -0.006 | 0.003 | 0.003 | 0 | 0.28 | 0.29 | 0.35 | 0.27 | 0.33 | 0.23 |
| 9 | 112632327 | rs7042517 | 0.313 | -0.253 | -0.852 | 0.682 | 0.021 | -0.011 | -0.011 | 0 | 0.6 | 0.71 | 0.75 | 0.57 | 0.69 | 0.64 |
| 9 | 112632977 | rs2479314 | -0.84 | -0.575 | -1.064 | 0.585 | -0.001 | -0.046 | -0.046 | 0 | 0.89 | 0.94 | 0.9 | 0.72 | 0.88 | 0.88 |
| 9 | 112634210 | rs16914570 | -0.455 | -1.074 | -0.442 | 0.517 | -0.026 | -0.019 | -0.019 | 0 | 0.16 | 0.42 | 0.07 | 0.11 | 0.1 | 0.28 |
| 9 | 112634640 | rs10759382 | 2.089 | -1.438 | -0.493 | 0.747 | 0.099 | 0.088 | 0.088 | 0 | 0.48 | 0.23 | 0.69 | 0.38 | 0.65 | 0.38 |
| 9 | 112636653 | rs7040294 | 0.16 | -1.411 | -0.519 | 0.77 | 0.005 | -0.007 | -0.007 | 0 | 0.68 | 0.73 | 0.77 | 0.57 | 0.75 | 0.69 |
| 9 | 112638250 | rs1475433 | 1.629 | -1.205 | -0.415 | 0.894 | 0.042 | 0.007 | 0.007 | 0 | 0.53 | 0.37 | 0.7 | 0.47 | 0.65 | 0.41 |
| 9 | 112638465 | rs16914573 | 1.237 | -1.101 | -0.14 | 0.616 | 0.033 | 0.1 | 0.1 | 0 | 0.06 | 0.28 | 0.04 | 0.1 | 0.03 | 0.23 |
| 9 | 112640311 | rs9409096 | 2.256 | -0.565 | -0.705 | 0.292 | -0.047 | 0.103 | 0.103 | 0 | 0.36 | 0.2 | 0.4 | 0.15 | 0.41 | 0.24 |
| 9 | 112640616 | rs13283300 | 1.772 | -0.544 | -0.738 | 0.334 | -0.092 | -0.044 | -0.044 | 0.021 | 0.15 | 0.01 | 0.31 | 0.22 | 0.29 | 0.16 |
| 9 | 112641489 | rs16914575 | 1.455 | -0.537 | -0.413 | -0.254 | 0.041 | 0.097 | 0.097 | 0 | 0.06 | 0.26 | 0.04 | 0.1 | 0.04 | 0.23 |
| 9 | 112641903 | rs9408826 | 1.927 | -0.435 | -0.938 | -0.3 | -0.047 | 0.104 | 0.104 | 0 | 0.36 | 0.2 | 0.4 | 0.15 | 0.41 | 0.24 |
| 9 | 112642756 | rs10816904 | -0.64 | -0.85 | -0.036 | 0.025 | -0.061 | -0.02 | -0.02 | 0 | 0.31 | 0.44 | 0.17 | 0.26 | 0.18 | 0.46 |
| 9 | 112645868 | rs1341983 | -1.06 | -0.83 | -0.21 | 0.035 | -0.006 | -0.001 | -0.001 | 0 | 0.33 | 0.6 | 0.18 | 0.26 | 0.18 | 0.44 |
| 9 | 112648964 | rs10283686 | 0.797 | -0.696 | -0.378 | 0.086 | -0.041 | -0.023 | -0.023 | 0 | 0.65 | 0.52 | 0.8 | 0.74 | 0.78 | 0.53 |
| 9 | 112656046 | rs12343568 | 2.996 | -0.501 | -0.854 | -0.144 | 0.003 | 0.012 | 0.012 | 0.527 | 0.09 | 0.04 | 0.04 | 0 | 0.07 | 0.01 |
| 9 | 112660775 | rs6477730 | 0.188 | -1.103 | -0.686 | -0.559 | -0.02 | -0.055 | -0.055 | 1.128 | 0.7 | 0.66 | 0.81 | 0.84 | 0.77 | 0.76 |
| 9 | 112661908 | rs2095094 | 0.03 | -1.228 | -0.853 | -0.751 | -0.005 | 0.057 | 0.057 | 0.043 | 0.78 | 0.95 | 0.89 | 0.88 | 0.77 | 0.81 |
| 9 | 112663125 | rs1981035 | 1.517 | -0.582 | -1.186 | -0.169 | -0.053 | -0.051 | -0.051 | 0 | 0.08 | 0.01 | 0.25 | 0.27 | 0.18 | 0.15 |
| 9 | 112663525 | rs877640 | 2.024 | -0.805 | -1.553 | -0.292 | -0.094 | 0.143 | 0.143 | 0 | 0.31 | 0.02 | 0.37 | 0.09 | 0.38 | 0.22 |
| 9 | 112672877 | rs12115781 | -0.446 | -1.313 | -0.734 | 0.41 | -0.027 | 0.148 | 0.148 | 0 | 0.16 | 0.33 | 0.1 | 0.39 | 0.1 | 0.37 |
| 9 | 112684039 | rs7858354 | 0.041 | -1.273 | -0.441 | 0.456 | -0.013 | 0.236 | 0.236 | 0.569 | 0.07 | 0.26 | 0.07 | 0.24 | 0.03 | 0.15 |
| 9 | 112694486 | rs2277146 | -0.354 | -1.078 | -0.494 | 0.394 | 0.041 | 0.151 | 0.151 | 0 | 0.17 | 0.35 | 0.33 | 0.54 | 0.21 | 0.31 |
| 9 | 112702544 | rs13294193 | 2.05 | -0.85 | -1.061 | 0.476 | -0.042 | -0.042 | -0.042 | 0.353 | 0.07 | 0 | 0.12 | 0.19 | 0.13 | 0.1 |
| 9 | 112703215 | rs7029365 | 1.579 | -1.066 | -1.159 | 0.201 | -0.017 | 0.153 | 0.153 | 0.362 | 0.45 | 0.26 | 0.45 | 0.16 | 0.49 | 0.25 |
| 9 | 112706482 | rs7035935 | -0.176 | -1.202 | -0.978 | 0.414 | -0.013 | 0.061 | 0.061 | 0 | 0.12 | 0.2 | 0.09 | 0.26 | 0.09 | 0.19 |
| 9 | 112717505 | rs10120878 | 2.42 | -1.287 | -1.517 | -0.036 | 0.002 | 0.004 | 0.004 | 5.951 | 0.13 | 0.12 | 0.09 | 0.07 | 0.09 | 0.03 |
| 9 | 112722869 | rs12003083 | 0.544 | -1.319 | -1.169 | 0.205 | 0.001 | -0.003 | -0.003 | 0.239 | 0.09 | 0.07 | 0.03 | 0.16 | 0.07 | 0.21 |
| 9 | 112728014 | rs6477734 | 1.539 | -1.249 | -1.313 | 0.126 | -0.05 | 0.138 | 0.138 | 0.902 | 0.42 | 0.23 | 0.46 | 0.15 | 0.49 | 0.22 |
| 9 | 112731195 | rs7020474 | -1.127 | -0.969 | -1.335 | 0.335 | -0.043 | 0.001 | 0.001 | 0 | 0.87 | 0.99 | 0.92 | 0.92 | 0.84 | 0.92 |
| 9 | 112732288 | rs11794102 | 0.313 | -0.917 | -0.495 | 0.643 | -0.01 | -0.031 | -0.031 | 0 | 0.11 | 0.21 | 0.1 | 0.04 | 0.07 | 0.09 |
| 9 | 112734445 | rs4978864 | -0.456 | -0.505 | -0.971 | 0.91 | -0.004 | 0.011 | 0.011 | 0 | 0.13 | 0.12 | 0.25 | 0.44 | 0.16 | 0.34 |
| 9 | 112735377 | rs1930249 | -0.393 | -0.49 | -1.063 | 0.918 | -0.009 | -0.005 | -0.005 | 0 | 0.13 | 0.09 | 0.24 | 0.43 | 0.16 | 0.34 |
| 9 | 112735585 | rs4978866 | 0.487 | -0.24 | -1.282 | 0.682 | -0.05 | -0.056 | -0.056 | 0 | 0.09 | 0.01 | 0.23 | 0.27 | 0.15 | 0.17 |
| 9 | 112739344 | rs995294 | 1.222 | -0.418 | -1.447 | 0.023 | -0.083 | 0.12 | 0.12 | 1.118 | 0.29 | 0.12 | 0.4 | 0.11 | 0.43 | 0.2 |
| 9 | 112740024 | rs10739286 | -0.449 | -1.526 | -1.314 | 0.189 | 0.027 | 0.028 | 0.028 | 0 | 0.71 | 0.81 | 0.83 | 0.82 | 0.78 | 0.8 |
| 9 | 112741697 | rs7871446 | 2.045 | -1.529 | -1.307 | 0.342 | 0.001 | 0.003 | 0.003 | 5.413 | 0.12 | 0.08 | 0.07 | 0.08 | 0.1 | 0.04 |
| 9 | 112742786 | rs10980133 | 0.241 | -1.495 | -0.798 | 0.336 | -0.006 | 0.004 | 0.004 | 0.138 | 0.29 | 0.28 | 0.17 | 0.3 | 0.2 | 0.26 |
| 9 | 112750411 | rs4978411 | 0.377 | -1.525 | -1.076 | 0.169 | 0.039 | 0.049 | 0.049 | 0.255 | 0.57 | 0.36 | 0.75 | 0.78 | 0.7 | 0.68 |
| 9 | 112751482 | rs10217743 | -0.099 | -1.712 | -1.352 | 0.049 | 0.005 | 0.006 | 0.006 | 2.506 | 0.64 | 0.37 | 0.63 | 0.81 | 0.71 | 0.7 |
| 9 | 112751649 | rs10217567 | 1.727 | -1.543 | -1.539 | 0.059 | -0.128 | 0.117 | 0.117 | 1.956 | 0.28 | 0.04 | 0.24 | 0.12 | 0.41 | 0.2 |
| 9 | 112753883 | rs12000943 | -0.703 | -1.533 | -1.493 | 0.284 | 0.074 | 0.055 | 0.055 | 0.935 | 0.48 | 0.77 | 0.41 | 0.32 | 0.34 | 0.49 |
| 9 | 112756545 | rs10739287 | -1.466 | -1.316 | -1.431 | 0.313 | -0.041 | -0.044 | -0.044 | 0 | 0.93 | 0.99 | 0.77 | 0.72 | 0.88 | 0.81 |
| 9 | 112757373 | rs10816913 | -1.414 | -1.359 | -1.434 | 0.309 | -0.044 | -0.069 | -0.069 | 0.013 | 0.87 | 0.99 | 0.76 | 0.72 | 0.85 | 0.8 |
| 9 | 112757584 | rs7864336 | -1.4 | -0.964 | -0.998 | -0.211 | 0.002 | -0.049 | -0.049 | 0 | 0.71 | 0.83 | 0.63 | 0.44 | 0.73 | 0.67 |
| 9 | 112757604 | rs7864442 | 0.795 | -0.995 | -1.065 | -0.254 | -0.024 | -0.056 | -0.056 | 0 | 0.13 | 0.33 | 0.06 | 0.03 | 0.07 | 0.13 |
| 9 | 112758187 | rs10980136 | 0.722 | -0.871 | -1.229 | -0.182 | -0.118 | 0.054 | 0.054 | 1.996 | 0.35 | 0.07 | 0.25 | 0.23 | 0.44 | 0.27 |
| 9 | 112760293 | rs12555129 | 0.306 | -0.428 | -0.738 | -0.016 | 0 | 0 | 0 | 0 | 0.19 | 0.25 | 0.29 | 0.15 | 0.22 | 0.15 |
| 9 | 112769632 | rs6477736 | 0.328 | -1.122 | -0.974 | 0.012 | -0.071 | -0.112 | -0.112 | 2.908 | 0.26 | 0.58 | 0.1 | 0.04 | 0.09 | 0.18 |

| | | | | | | | | | | | | | | | | |
|---|---|---|---|---|---|---|---|---|---|---|---|---|---|---|---|---|
| 9 | 112775268 | rs10739288 | 0.55 | -0.706 | -2.235 | 0.355 | -0.04 | -0.079 | -0.079 | 0.219 | 0.43 | 0.05 | 0.53 | 0.4 | 0.64 | 0.36 |
| 9 | 112775738 | rs1571260 | -0.302 | -0.696 | -0.977 | 0.039 | -0.025 | -0.032 | -0.032 | 0.029 | 0.27 | 0.64 | 0.1 | 0.1 | 0.1 | 0.31 |
| 9 | 112778426 | rs723706 | -1.063 | 0.883 | -1.076 | 0.932 | -0.002 | 0.003 | 0.003 | 0 | 0.14 | 0.17 | 0.17 | 0.26 | 0.21 | 0.18 |
| 9 | 112778952 | rs7030564 | 0.607 | 0.93 | -1.674 | 0.307 | -0.046 | 0.018 | 0.018 | 0.063 | 0.79 | 0.95 | 0.87 | 0.89 | 0.81 | 0.88 |
| 9 | 112783824 | rs10512401 | -1.472 | 0.956 | -1.019 | 0.187 | 0.011 | -0.019 | -0.019 | 0 | 0.08 | 0.18 | 0.08 | 0.03 | 0.1 | 0.05 |
| 9 | 112787897 | rs10759387 | -0.152 | 1.381 | -1.756 | 0.427 | -0.068 | -0.07 | -0.07 | 0.076 | 0.15 | 0.01 | 0.33 | 0.24 | 0.22 | 0.2 |
| 9 | 112790036 | rs7024673 | -0.16 | 1.667 | -1.887 | 0.325 | -0.099 | -0.072 | -0.072 | 0.17 | 0.17 | 0.02 | 0.37 | 0.27 | 0.31 | 0.21 |
| 9 | 112791184 | rs260188 | -0.522 | 1.54 | -2.334 | 0.595 | 0 | 0 | 0 | 0 | 0.16 | 0.17 | 0.16 | 0.29 | 0.11 | 0.24 |
| 9 | 112796113 | rs260194 | -0.857 | 0.26 | -1.375 | 0.065 | -0.011 | 0.01 | 0.01 | 0 | 0.29 | 0.44 | 0.28 | 0.33 | 0.22 | 0.3 |
| 9 | 112802769 | rs260205 | -1.338 | -0.812 | -1.349 | -0.645 | -0.024 | -0.026 | -0.026 | 0 | 0.69 | 0.81 | 0.71 | 0.61 | 0.6 | 0.61 |
| 9 | 112807133 | rs10980159 | 1.564 | -0.926 | -1.222 | -0.815 | 0.033 | 0.044 | 0.044 | 0 | 0.2 | 0.07 | 0.09 | 0.03 | 0.09 | 0.1 |
| 9 | 112810330 | rs260215 | 0.447 | -0.149 | -1.209 | -0.649 | -0.048 | -0.016 | -0.016 | 0.062 | 0.14 | 0.03 | 0.17 | 0.13 | 0.19 | 0.09 |
| 9 | 112813375 | rs260218 | 0.424 | -0.151 | -0.504 | -0.657 | -0.014 | -0.002 | -0.002 | 0 | 0.14 | 0.08 | 0.21 | 0.12 | 0.19 | 0.09 |
| 9 | 112815997 | rs260223 | 0.424 | -0.151 | -0.504 | -0.676 | -0.014 | -0.001 | -0.001 | 0 | 0.14 | 0.08 | 0.21 | 0.12 | 0.19 | 0.09 |
| 9 | 112820620 | rs2017392 | 0.812 | -0.187 | -0.687 | -0.692 | -0.054 | -0.06 | -0.06 | 0 | 0.17 | 0.08 | 0.38 | 0.41 | 0.31 | 0.34 |
| 9 | 112823818 | rs541471 | 0.177 | -0.182 | -0.247 | -0.738 | 0.041 | -0.007 | -0.007 | 0 | 0.17 | 0.35 | 0.19 | 0.12 | 0.19 | 0.09 |
| 9 | 112825422 | rs552182 | 0.218 | -0.184 | -0.24 | -0.739 | 0.036 | -0.007 | -0.007 | 0 | 0.16 | 0.32 | 0.22 | 0.12 | 0.19 | 0.09 |
| 9 | 112826429 | rs521883 | -1.345 | -0.566 | -0.486 | -0.88 | -0.02 | -0.051 | -0.051 | 0.253 | 0.74 | 0.86 | 0.81 | 0.58 | 0.69 | 0.61 |
| 9 | 112832200 | rs12056992 | 1.189 | -1.129 | -0.44 | -0.821 | 0.03 | 0.035 | 0.035 | 0 | 0.22 | 0.12 | 0.1 | 0.03 | 0.1 | 0.1 |
| 9 | 112832326 | rs491533 | -1.315 | -1.119 | -0.344 | -0.868 | -0.015 | -0.02 | -0.02 | 0 | 0.73 | 0.78 | 0.75 | 0.58 | 0.61 | 0.65 |
| 9 | 112839238 | rs481661 | -0.625 | -1.102 | -0.334 | -0.855 | -0.006 | -0.007 | -0.007 | 0 | 0.51 | 0.72 | 0.66 | 0.55 | 0.51 | 0.54 |
| 9 | 112845614 | rs1752333 | 0.443 | -0.926 | -0.379 | -0.783 | 0.013 | 0.022 | 0.022 | 0.743 | 0.32 | 0.34 | 0.55 | 0.55 | 0.42 | 0.49 |
| 9 | 112847897 | rs564321 | -0.474 | -1.005 | -0.668 | -0.816 | -0.015 | -0.012 | -0.012 | 0 | 0.64 | 0.65 | 0.65 | 0.58 | 0.52 | 0.6 |
| 9 | 112850080 | rs1980874 | -0.906 | -1.613 | -0.432 | -1.169 | -0.048 | -0.11 | -0.11 | 0 | 0.43 | 0.54 | 0.21 | 0.03 | 0.2 | 0.15 |
| 9 | 112851492 | rs10512402 | -0.187 | -1.561 | -0.423 | -1.134 | -0.038 | -0.06 | -0.06 | 0.058 | 0.33 | 0.36 | 0.13 | 0.03 | 0.11 | 0.1 |
| 9 | 112853511 | rs7038399 | -0.571 | -0.516 | -0.403 | -0.617 | -0.004 | -0.035 | -0.035 | 0 | 0.1 | 0.18 | 0.06 | 0 | 0.08 | 0.04 |
| 9 | 112858901 | rs10980199 | -0.258 | -0.277 | -0.52 | -0.305 | -0.061 | -0.114 | -0.114 | 0.743 | 0.3 | 0.46 | 0.15 | 0.03 | 0.14 | 0.13 |
| 9 | 112864943 | rs494267 | -0.676 | 1.597 | -0.932 | 1.389 | -0.023 | 0.025 | 0.025 | 0 | 0.07 | 0.04 | 0.38 | 0.44 | 0.24 | 0.13 |
| 9 | 112868321 | rs7026192 | 0.09 | 1.159 | -0.903 | 0.653 | 0.003 | 0.001 | 0.001 | 0 | 0.43 | 0.4 | 0.28 | 0.43 | 0.33 | 0.39 |
| 9 | 112869277 | rs11788194 | 0.499 | 1.591 | -1.396 | 0.167 | -0.039 | 0.056 | 0.056 | 0 | 0.1 | 0.01 | 0.06 | 0.01 | 0.11 | 0.11 |
| 9 | 112873868 | rs11790833 | -0.168 | 1.486 | -1.345 | -0.21 | -0.009 | 0.046 | 0.046 | 0 | 0.2 | 0.1 | 0.11 | 0.02 | 0.18 | 0.16 |
| 9 | 112874767 | rs1327775 | 0.861 | 0.262 | -1.257 | -0.586 | -0.007 | -0.009 | -0.009 | 0 | 0.39 | 0.41 | 0.2 | 0.27 | 0.23 | 0.22 |
| 9 | 112879762 | rs525681 | -0.629 | 1.116 | -1.203 | 0.055 | -0.002 | -0.094 | -0.094 | 0 | 0.69 | 0.62 | 0.75 | 0.96 | 0.69 | 0.69 |
| 9 | 112879944 | rs496709 | -1.427 | 1.127 | -0.868 | 0.099 | -0.006 | 0.032 | 0.032 | 0 | 0.8 | 0.85 | 0.81 | 0.97 | 0.76 | 0.75 |
| 9 | 112884482 | rs4978870 | -0.639 | 0.671 | -1.017 | -0.316 | 0.011 | 0.055 | 0.055 | 0 | 0.57 | 0.66 | 0.69 | 0.75 | 0.6 | 0.69 |
| 9 | 112886940 | rs17806439 | 1.417 | 0.548 | -1.122 | -0.154 | -0.003 | -0.029 | -0.029 | 1.259 | 0.07 | 0.01 | 0.06 | 0.14 | 0.08 | 0.16 |
| 9 | 112889315 | rs914355 | 0.596 | 0.334 | -0.546 | -0.313 | -0.022 | -0.029 | -0.029 | 0.589 | 0.33 | 0.38 | 0.19 | 0.2 | 0.24 | 0.37 |
| 9 | 112891853 | rs2900505 | -0.447 | 0.435 | -0.606 | 0.191 | 0.001 | 0.044 | 0.044 | 0 | 0.51 | 0.6 | 0.36 | 0.31 | 0.46 | 0.5 |
| 9 | 112893850 | rs943616 | -1.773 | 0.335 | -0.645 | 0.035 | -0.001 | -0.022 | -0.022 | 0.285 | 0.77 | 0.76 | 0.78 | 0.97 | 0.73 | 0.76 |
| 9 | 112900199 | rs914358 | -0.325 | 0.406 | -0.676 | 0.135 | 0.005 | 0.043 | 0.043 | 0 | 0.51 | 0.59 | 0.36 | 0.31 | 0.45 | 0.5 |
| 9 | 112900785 | rs3739456 | 0.788 | 0.42 | -0.711 | 0.417 | -0.02 | -0.051 | -0.051 | 0 | 0.23 | 0.15 | 0.41 | 0.66 | 0.26 | 0.23 |
| 9 | 112908032 | rs16914814 | 0.978 | 0.202 | -0.084 | 0.289 | -0.005 | 0.015 | 0.015 | 0.805 | 0.17 | 0.41 | 0.1 | 0.21 | 0.14 | 0.09 |
| 9 | 112914082 | rs17203603 | 0.69 | -0.253 | -0.499 | 0.07 | 0.017 | 0.01 | 0.01 | 0 | 0.22 | 0.06 | 0.13 | 0.13 | 0.19 | 0.22 |
| 9 | 112916400 | rs4372077 | 0.985 | -0.232 | -0.635 | -0.026 | -0.051 | -0.002 | -0.002 | 0 | 0.15 | 0.04 | 0.15 | 0.11 | 0.21 | 0.16 |
| 9 | 112918360 | rs10816927 | -0.094 | -0.281 | -0.526 | -0.039 | 0.01 | -0.009 | -0.009 | 1.32 | 0.33 | 0.46 | 0.25 | 0.32 | 0.34 | 0.24 |
| 9 | 112918531 | rs10759392 | -1.356 | -0.276 | -0.349 | -0.297 | 0 | -0.051 | -0.051 | 0.554 | 0.61 | 0.63 | 0.71 | 0.96 | 0.66 | 0.73 |
| 9 | 112919288 | rs7847354 | -1.104 | -0.602 | -0.276 | -0.28 | -0.03 | 0.154 | 0.154 | 0 | 0.57 | 0.8 | 0.59 | 0.86 | 0.47 | 0.6 |
| 9 | 112920099 | rs7023507 | -1.001 | -0.557 | -0.26 | -0.279 | -0.007 | 0.172 | 0.172 | 0 | 0.53 | 0.78 | 0.58 | 0.86 | 0.46 | 0.57 |
| 9 | 112923119 | rs914359 | 1.305 | -0.615 | -0.304 | -0.428 | 0.014 | 0.035 | 0.035 | 0 | 0.18 | 0.11 | 0.08 | 0.06 | 0.12 | 0.2 |
| 9 | 112925906 | rs1571780 | -0.443 | -0.759 | -0.072 | -0.524 | 0 | 0.007 | 0.007 | 0 | 0.46 | 0.59 | 0.57 | 0.86 | 0.45 | 0.57 |
| 9 | 112925977 | rs10980229 | 0.837 | -0.719 | -0.07 | -0.38 | -0.019 | -0.071 | -0.071 | 0.345 | 0.22 | 0.12 | 0.38 | 0.51 | 0.23 | 0.35 |
| 9 | 112929130 | rs4978413 | -0.413 | -0.577 | 0.057 | -0.72 | 0.004 | 0.028 | 0.028 | 0 | 0.51 | 0.59 | 0.56 | 0.86 | 0.45 | 0.57 |
| 9 | 112931646 | rs1047469 | -0.693 | -0.346 | -0.343 | -0.642 | -0.033 | 0.144 | 0.144 | 0.033 | 0.58 | 0.8 | 0.59 | 0.86 | 0.47 | 0.59 |
| 9 | 112932182 | rs12340651 | 3.242 | -0.315 | -0.203 | -0.58 | -0.029 | -0.03 | -0.03 | 0 | 0.07 | 0.15 | 0.02 | 0.01 | 0.01 | 0 |
| 9 | 112933271 | rs4289885 | -0.549 | -0.133 | -0.582 | -0.539 | -0.024 | 0.134 | 0.134 | 0 | 0.57 | 0.76 | 0.58 | 0.86 | 0.47 | 0.6 |
| 9 | 112948620 | rs1041482 | -0.961 | -0.052 | -0.908 | -1.255 | 0.012 | -0.064 | -0.064 | 0 | 0.71 | 0.7 | 0.74 | 0.96 | 0.62 | 0.73 |
| 9 | 112949710 | rs7039011 | 0.582 | 0.342 | -1.393 | -1.175 | -0.09 | -0.079 | -0.079 | 0 | 0.18 | 0.02 | 0.2 | 0.22 | 0.3 | 0.16 |
| 9 | 112950305 | rs4978883 | -0.378 | -0.146 | -1.043 | -1.413 | 0 | -0.061 | -0.061 | 0 | 0.56 | 0.46 | 0.69 | 0.95 | 0.57 | 0.59 |
| 9 | 112952005 | rs10759396 | -0.162 | -0.108 | -1.058 | -1.176 | -0.001 | 0 | 0 | 0 | 0.51 | 0.44 | 0.38 | 0.47 | 0.53 | 0.38 |
| 9 | 112962809 | rs10512403 | 1.128 | -0.02 | -1.504 | -1.873 | -0.004 | 0.066 | 0.066 | 0 | 0.33 | 0.06 | 0.19 | 0.23 | 0.28 | 0.23 |
| 9 | 112962942 | rs17806752 | 1.857 | 0.231 | -1.525 | -1.866 | -0.039 | -0.016 | -0.016 | 0.001 | 0.08 | 0.01 | 0.06 | 0.08 | 0.16 | 0.09 |
| 9 | 112963702 | rs4978888 | 0.626 | 0.034 | -1.375 | -2.138 | -0.013 | -0.015 | -0.015 | 0 | 0.2 | 0.35 | 0.17 | 0.13 | 0.17 | 0.22 |
| 9 | 112966252 | rs7046745 | -2.129 | -0.226 | -0.881 | -1.666 | 0.008 | -0.036 | -0.036 | 0 | 0.76 | 0.77 | 0.77 | 0.97 | 0.65 | 0.75 |
| 9 | 112967140 | rs4978890 | 0.117 | -0.415 | -0.814 | -1.589 | 0.011 | 0.004 | 0.004 | 0 | 0.21 | 0.28 | 0.29 | 0.21 | 0.32 | 0.24 |
| 9 | 112970634 | rs4111656 | 1.163 | -0.337 | -0.86 | -1.522 | 0.038 | 0.047 | 0.047 | 0 | 0.3 | 0.16 | 0.17 | 0.23 | 0.22 | 0.2 |
| 9 | 112973648 | rs4314718 | -0.129 | -0.272 | -0.96 | -1.657 | 0.058 | 0.074 | 0.074 | 0.245 | 0.52 | 0.29 | 0.47 | 0.75 | 0.33 | 0.52 |
| 9 | 112974847 | rs6477748 | -0.167 | -0.211 | -0.631 | -1.501 | 0.047 | 0.124 | 0.124 | 6.57 | 0.11 | 0.49 | 0.22 | 0.13 | 0.12 | 0.13 |
| 9 | 112977595 | rs7866504 | -1.306 | -0.003 | -1.18 | -1.49 | -0.004 | 0.003 | 0.003 | 0.056 | 0.64 | 0.35 | 0.55 | 0.83 | 0.53 | 0.62 |
| 9 | 112978298 | rs7851800 | 0.041 | -0.074 | -1.308 | -1.529 | -0.069 | 0.178 | 0.178 | 0.043 | 0.47 | 0.1 | 0.49 | 0.83 | 0.48 | 0.54 |

| | | | | | | | | | | | | | | | | |
|---|---|---|---|---|---|---|---|---|---|---|---|---|---|---|---|---|
| 9 | 112983819 | rs10816943 | 0.62 | -0.471 | -1.823 | -1.271 | -0.026 | 0.038 | 0.038 | 0 | 0.06 | 0.01 | 0.27 | 0.5 | 0.06 | 0.22 |
| 9 | 112984666 | rs1411922 | -1.225 | -0.489 | -1.712 | -1.678 | -0.029 | 0.038 | 0.038 | 0.08 | 0.64 | 0.29 | 0.55 | 0.83 | 0.53 | 0.62 |
| 9 | 112984893 | rs1411921 | -0.104 | -0.176 | -2.193 | -1.356 | -0.069 | 0.178 | 0.178 | 0.045 | 0.47 | 0.1 | 0.49 | 0.83 | 0.48 | 0.54 |
| 9 | 112989951 | rs16914868 | 1.122 | -0.44 | -1.707 | -1.456 | 0.007 | 0.004 | 0.004 | 1.662 | 0.18 | 0.16 | 0.07 | 0.04 | 0.09 | 0.1 |
| 9 | 113000213 | rs10759402 | -0.006 | 0.012 | -2.31 | -1.609 | 0.011 | 0.098 | 0.098 | 0 | 0.39 | 0.23 | 0.61 | 0.86 | 0.37 | 0.5 |
| 9 | 113005260 | rs2026314 | -1.741 | 0.184 | -2.131 | -1.765 | 0.007 | -0.045 | -0.045 | 0.22 | 0.76 | 0.78 | 0.78 | 0.98 | 0.64 | 0.81 |
| 9 | 113005287 | rs2026313 | -0.97 | 0.845 | -2.678 | -0.748 | 0.007 | 0.056 | 0.056 | 0 | 0.47 | 0.25 | 0.39 | 0.39 | 0.46 | 0.45 |
| 9 | 113007499 | rs4135215 | 0.993 | 0.601 | -2.466 | -0.71 | 0.004 | -0.036 | -0.036 | 2.908 | 0.16 | 0.02 | 0.1 | 0.25 | 0.14 | 0.13 |
| 9 | 113009246 | rs4135208 | 0.124 | 0.795 | -2.22 | -0.662 | 0.013 | 0.01 | 0.01 | 0 | 0.33 | 0.21 | 0.33 | 0.32 | 0.3 | 0.29 |
| 9 | 113011039 | rs2031760 | -0.812 | 1.105 | -1.753 | 0.457 | -0.008 | -0.028 | -0.028 | 0 | 0.16 | 0.09 | 0.33 | 0.62 | 0.19 | 0.35 |
| 9 | 113013588 | rs4135192 | -0.428 | 1.404 | -1.995 | -0.374 | -0.092 | 0.091 | 0.091 | 0.675 | 0.24 | 0.02 | 0.14 | 0.08 | 0.32 | 0.2 |
| 9 | 113015602 | rs4135183 | -0.36 | 1.337 | -1.835 | -0.491 | -0.005 | 0.011 | 0.011 | 0 | 0.09 | 0.05 | 0.07 | 0.02 | 0.12 | 0.05 |
| 9 | 113016000 | rs4135179 | -0.243 | 1.685 | -2.084 | -0.678 | -0.072 | 0.024 | 0.024 | 0.015 | 0.12 | 0.01 | 0.09 | 0.05 | 0.19 | 0.11 |
| 9 | 113016399 | rs4135172 | 0.392 | 1.57 | -1.857 | -0.361 | -0.002 | 0.031 | 0.031 | 0 | 0.11 | 0.11 | 0.04 | 0.23 | 0.1 | 0.13 |
| 9 | 113016885 | rs4135168 | -1.188 | 1.346 | -1.194 | -0.591 | 0 | -0.042 | -0.042 | 0 | 0.2 | 0.24 | 0.23 | 0.09 | 0.24 | 0.22 |
| 9 | 113017352 | rs1410051 | 0.153 | 0.967 | -1.289 | -1.121 | 0.01 | 0.018 | 0.018 | 0 | 0.35 | 0.32 | 0.32 | 0.14 | 0.24 | 0.33 |
| 9 | 113017562 | rs4135165 | 1.459 | 0.818 | -1.323 | -1.212 | 0.041 | 0.087 | 0.087 | 0.364 | 0.23 | 0.09 | 0.06 | 0 | 0.09 | 0.15 |
| 9 | 113019508 | rs2301241 | -0.5 | 0.637 | -0.496 | -0.919 | -0.002 | -0.018 | -0.018 | 0 | 0.56 | 0.53 | 0.69 | 0.74 | 0.54 | 0.67 |
| 9 | 113020916 | rs3758192 | 1.015 | 0.626 | -0.708 | -0.881 | -0.001 | 0.037 | 0.037 | 0 | 0.11 | 0.11 | 0.05 | 0.23 | 0.1 | 0.13 |
| 9 | 113022794 | rs10733576 | -1.146 | 0.595 | -0.666 | -1.034 | -0.001 | -0.006 | -0.006 | 0 | 0.68 | 0.73 | 0.75 | 0.97 | 0.64 | 0.81 |
| 9 | 113043999 | rs1556047 | -0.855 | 0.184 | -0.353 | -0.761 | -0.01 | 0.017 | 0.017 | 0 | 0.64 | 0.65 | 0.67 | 0.84 | 0.53 | 0.77 |
| 9 | 113046742 | rs10816959 | 0.078 | 0.018 | -0.286 | -0.595 | 0.012 | 0.003 | 0.003 | 0 | 0.43 | 0.36 | 0.44 | 0.64 | 0.31 | 0.52 |
| 9 | 113050873 | rs10980301 | 0.166 | 0.179 | -0.652 | -0.711 | -0.037 | -0.107 | -0.107 | 0.074 | 0.14 | 0.01 | 0.35 | 0.64 | 0.13 | 0.37 |
| 9 | 113051296 | rs6477754 | 0.799 | 0.09 | -0.126 | -0.732 | -0.014 | -0.111 | -0.111 | 1.103 | 0.3 | 0.38 | 0.09 | 0 | 0.17 | 0.15 |
| 9 | 113058357 | rs10739293 | -1.065 | -0.141 | -0.009 | -0.014 | 0.018 | 0.021 | 0.021 | 0.023 | 0.42 | 0.59 | 0.37 | 0.73 | 0.29 | 0.51 |
| 9 | 113060439 | rs7869617 | -1.43 | 0.047 | -0.367 | 0.441 | 0.039 | 0.116 | 0.116 | 1.022 | 0.49 | 0.69 | 0.61 | 0.68 | 0.45 | 0.62 |
| 9 | 113063439 | rs10980309 | -0.334 | -0.116 | -0.671 | 0.738 | 0.003 | -0.136 | -0.136 | 0.15 | 0.24 | 0.04 | 0.25 | 0.44 | 0.21 | 0.34 |
| 9 | 113069090 | rs10816965 | -0.411 | 0.373 | 0.277 | 1.22 | -0.006 | 0.013 | 0.013 | 0 | 0.13 | 0.18 | 0.07 | 0.11 | 0.1 | 0.12 |
| 9 | 113080316 | rs17204340 | -0.89 | 0.751 | -0.15 | 1.775 | 0.001 | 0.092 | 0.092 | 0 | 0.06 | 0.06 | 0.16 | 0.42 | 0.04 | 0.21 |
| 9 | 113082105 | rs1327541 | -0.093 | 0.995 | -0.309 | 0.87 | 0.005 | -0.029 | -0.029 | 0 | 0.63 | 0.49 | 0.67 | 0.74 | 0.62 | 0.55 |
| 9 | 113083182 | rs16914942 | -0.287 | 1.002 | -0.286 | 1.074 | 0.007 | -0.059 | -0.059 | 2.132 | 0.12 | 0.24 | 0.06 | 0 | 0.1 | 0.05 |
| 9 | 113085232 | rs17807186 | -0.785 | 1.041 | -0.309 | 1.392 | 0.007 | 0.066 | 0.066 | 0 | 0.16 | 0.24 | 0.22 | 0.45 | 0.16 | 0.3 |
| 9 | 113089377 | rs10816972 | -0.294 | 1.1 | -0.552 | 0.749 | 0.001 | 0.001 | 0.001 | 0 | 0.1 | 0.08 | 0.17 | 0.07 | 0.12 | 0.12 |
| 9 | 113091523 | rs7041938 | -0.207 | 1.02 | -0.638 | 0.777 | -0.009 | -0.01 | -0.01 | 0.659 | 0.1 | 0.04 | 0.16 | 0.08 | 0.12 | 0.12 |
| 9 | 113099591 | rs7040502 | -1.211 | 0.639 | -0.333 | -0.126 | 0 | -0.012 | -0.012 | 0.528 | 0.33 | 0.67 | 0.33 | 0.07 | 0.32 | 0.25 |
| 9 | 113099863 | rs7040684 | -1.218 | 0.637 | -0.338 | -0.141 | 0 | -0.009 | -0.009 | 0 | 0.33 | 0.67 | 0.33 | 0.09 | 0.32 | 0.25 |
| 9 | 113103292 | rs6477755 | -1.304 | 0.455 | -1.351 | -0.901 | -0.01 | 0.248 | 0.248 | 0 | 0.5 | 0.85 | 0.52 | 0.13 | 0.49 | 0.37 |
| 9 | 113103597 | rs12686260 | 0.188 | 0.391 | -1.214 | -0.964 | -0.001 | 0.006 | 0.006 | 0.038 | 0.15 | 0.1 | 0.19 | 0.08 | 0.15 | 0.21 |
| 9 | 113109166 | rs7854346 | -1.666 | -0.218 | -2.084 | -0.541 | 0.01 | -0.124 | -0.124 | 0 | 0.84 | 0.99 | 0.84 | 0.6 | 0.9 | 0.75 |
| 9 | 113109179 | rs7854688 | -0.648 | -0.496 | -1.789 | -0.797 | 0.026 | 0.119 | 0.119 | 0.185 | 0.4 | 0.76 | 0.4 | 0.13 | 0.33 | 0.33 |
| 9 | 113110830 | rs6477758 | -0.958 | -0.435 | -1.741 | -0.702 | -0.012 | 0.249 | 0.249 | 0 | 0.51 | 0.84 | 0.52 | 0.13 | 0.49 | 0.37 |
| 9 | 113112572 | rs963143 | 1.266 | -0.222 | -1.675 | -0.611 | -0.133 | -0.107 | -0.107 | 0.012 | 0.33 | 0.05 | 0.32 | 0.47 | 0.41 | 0.38 |
| 9 | 113114946 | rs3949852 | -0.011 | -0.364 | -1.098 | -0.801 | -0.011 | -0.076 | -0.076 | 0 | 0.23 | 0.65 | 0.22 | 0.05 | 0.18 | 0.12 |
| 9 | 113124417 | rs13287615 | 1.744 | 0.211 | -1.691 | -1.171 | -0.015 | 0.024 | 0.024 | 0 | 0.11 | 0.03 | 0.08 | 0 | 0.12 | 0.08 |
| 9 | 113124562 | rs17733560 | 0.367 | -0.102 | -0.901 | -1.706 | 0.021 | -0.122 | -0.122 | 0 | 0.2 | 0.38 | 0.18 | 0 | 0.26 | 0.12 |
| 9 | 113125853 | rs10816993 | -0.156 | 0.201 | -1.282 | -1.775 | -0.003 | -0.048 | -0.048 | 0 | 0.46 | 0.46 | 0.35 | 0.15 | 0.53 | 0.38 |
| 9 | 113126314 | rs1327532 | -0.856 | -0.092 | -1.325 | -1.854 | 0.01 | 0.015 | 0.015 | 0.003 | 0.63 | 0.66 | 0.55 | 0.23 | 0.68 | 0.56 |
| 9 | 113126692 | rs12683685 | 0.994 | 0.021 | -1.377 | -1.764 | -0.003 | -0.007 | -0.007 | 0.042 | 0.17 | 0.21 | 0.2 | 0.07 | 0.15 | 0.19 |
| 9 | 113127924 | rs3739453 | 1.32 | 0.107 | -1.572 | -1.72 | -0.044 | 0.081 | 0.081 | 0 | 0.28 | 0.07 | 0.17 | 0.15 | 0.27 | 0.26 |
| 9 | 113128505 | rs2274483 | 1.316 | 0.011 | -1.448 | -1.035 | 0.008 | -0.015 | -0.015 | 0 | 0.1 | 0.07 | 0.21 | 0.38 | 0.08 | 0.15 |
| 9 | 113132221 | rs17204533 | 2.363 | -0.068 | -1.64 | -0.685 | -0.046 | -0.093 | -0.093 | 0 | 0.13 | 0.01 | 0.1 | 0.28 | 0.17 | 0.14 |
| 9 | 113133741 | rs10816998 | 0.134 | -0.097 | -1.46 | -0.888 | 0.004 | 0.01 | 0.01 | 0 | 0.3 | 0.41 | 0.39 | 0.4 | 0.34 | 0.28 |
| 9 | 113136177 | rs1327535 | -1.461 | 0.118 | -1.635 | -0.364 | -0.052 | -0.129 | -0.129 | 0.285 | 0.84 | 0.98 | 0.81 | 0.62 | 0.79 | 0.75 |
| 9 | 113147203 | rs12686142 | 0.083 | 0.19 | -1.401 | -1.062 | 0.007 | 0.024 | 0.024 | 0 | 0.46 | 0.6 | 0.36 | 0.23 | 0.38 | 0.31 |
| 9 | 113147855 | rs7855526 | -0.244 | -0.086 | -1.509 | -1.654 | -0.014 | -0.064 | -0.064 | 0 | 0.33 | 0.5 | 0.3 | 0.12 | 0.31 | 0.25 |
| 9 | 113149952 | rs2255007 | 2.196 | -0.052 | -1.564 | -1.079 | -0.005 | -0.017 | -0.017 | 1.841 | 0.08 | 0.01 | 0.22 | 0.43 | 0.09 | 0.17 |
| 9 | 113153159 | rs7847265 | -0.846 | -0.094 | -1.571 | -1.263 | -0.012 | 0.048 | 0.048 | 0 | 0.71 | 0.85 | 0.75 | 0.9 | 0.71 | 0.69 |
| 9 | 113155541 | rs10739300 | -0.939 | 0.053 | -1.62 | -1.083 | 0.005 | 0.02 | 0.02 | 0 | 0.72 | 0.65 | 0.69 | 0.65 | 0.65 | 0.66 |
| 9 | 113162199 | rs13294114 | 0.966 | 0.478 | -1.659 | -1.341 | 0.028 | -0.031 | -0.031 | 0 | 0.07 | 0.13 | 0.11 | 0.01 | 0.13 | 0.12 |
| 9 | 113162304 | rs1887457 | 1.196 | 0.259 | -2.035 | -1.182 | -0.017 | 0.067 | 0.067 | 0 | 0.25 | 0.13 | 0.16 | 0.12 | 0.24 | 0.15 |
| 9 | 113163239 | rs1410048 | 1.395 | 0.323 | -1.786 | -0.808 | -0.028 | -0.042 | -0.042 | 0 | 0.1 | 0.01 | 0.15 | 0.09 | 0.11 | 0.13 |
| 9 | 113167003 | rs12348838 | 1.382 | 0.4 | -1.781 | -0.842 | -0.02 | 0.043 | 0.043 | 0 | 0.16 | 0.04 | 0.11 | 0.03 | 0.16 | 0.25 |
| 9 | 113168073 | rs1327537 | -3.086 | -0.115 | -2.175 | -0.893 | 0.027 | 0 | 0 | 0.807 | 0.9 | 1 | 0.93 | 0.92 | 0.97 | 0.93 |
| 9 | 113169126 | rs2281937 | 0.582 | 0.026 | -2.241 | -2.086 | -0.021 | 0.114 | 0.114 | 0 | 0.41 | 0.21 | 0.34 | 0.09 | 0.43 | 0.28 |
| 9 | 113169631 | rs7030192 | 0.167 | -0.136 | -1.685 | -2.134 | -0.005 | -0.004 | -0.004 | 0 | 0.41 | 0.57 | 0.33 | 0.44 | 0.33 | 0.39 |
| 9 | 113171262 | rs16914997 | 1.343 | -0.018 | -1.474 | -2.143 | 0.003 | -0.021 | -0.021 | 0 | 0.21 | 0.23 | 0.15 | 0.08 | 0.21 | 0.28 |
| 9 | 113176113 | rs7027372 | -1.105 | 0.103 | -1.538 | -2.141 | -0.013 | -0.05 | -0.05 | 0 | 0.67 | 0.83 | 0.56 | 0.52 | 0.63 | 0.7 |
| 9 | 113177324 | rs4475572 | 2.3 | 0.087 | -1.546 | -2.137 | -0.022 | 0.045 | 0.045 | 0 | 0.16 | 0.02 | 0.11 | 0.03 | 0.16 | 0.26 |
| 9 | 113188572 | rs1410050 | -2.709 | 0.076 | -1.488 | -2.072 | -0.044 | -0.131 | -0.131 | 0.09 | 0.84 | 0.99 | 0.76 | 0.59 | 0.82 | 0.85 |

| | | | | | | | | | | | | | | | | |
|---|---|---|---|---|---|---|---|---|---|---|---|---|---|---|---|---|
| 9 | 113189213 | rs2254168 | -0.832 | 0.077 | -1.362 | -2.069 | -0.027 | -0.028 | -0.028 | 0 | 0.63 | 0.8 | 0.47 | 0.52 | 0.53 | 0.67 |
| 9 | 113192427 | rs3739450 | -0.088 | 0.138 | -1.318 | -2.101 | -0.002 | 0.046 | 0.046 | 0 | 0.25 | 0.44 | 0.29 | 0.43 | 0.25 | 0.32 |
| 9 | 113193574 | rs6477769 | 0.65 | 0.022 | -1.314 | -2.091 | -0.028 | 0.087 | 0.087 | 0 | 0.2 | 0.42 | 0.19 | 0.4 | 0.15 | 0.28 |
| 9 | 113196733 | rs7852962 | 1.067 | -0.023 | -0.982 | -2.091 | -0.01 | -0.051 | -0.051 | 0 | 0.17 | 0.31 | 0.18 | 0.09 | 0.14 | 0.12 |
| 9 | 113197263 | rs7042161 | -0.786 | 0.118 | -1.733 | -2.248 | -0.004 | -0.064 | -0.064 | 0 | 0.6 | 0.58 | 0.76 | 0.96 | 0.66 | 0.64 |
| 9 | 113202831 | rs7848767 | -0.381 | 0.14 | -1.749 | -2.235 | 0.006 | 0.006 | 0.006 | 0 | 0.43 | 0.52 | 0.5 | 0.51 | 0.48 | 0.47 |
| 9 | 113204681 | rs7864433 | 2.284 | 0.148 | -1.757 | -2.24 | -0.02 | 0.044 | 0.044 | 0 | 0.16 | 0.04 | 0.1 | 0.03 | 0.16 | 0.26 |
| 9 | 113208250 | rs7863519 | 0.242 | 0.126 | -1.758 | -2.254 | -0.01 | 0.075 | 0.075 | 0 | 0.24 | 0.46 | 0.22 | 0.45 | 0.19 | 0.3 |
| 9 | 113208581 | rs7851366 | -1.921 | 0.116 | -1.718 | -2.277 | 0.004 | -0.002 | -0.002 | 0.467 | 0.78 | 0.83 | 0.87 | 0.96 | 0.81 | 0.89 |
| 9 | 113209195 | rs1889323 | 2.116 | 0.167 | -1.728 | -2.139 | -0.019 | 0.045 | 0.045 | 0 | 0.16 | 0.04 | 0.1 | 0.03 | 0.16 | 0.26 |
| 9 | 113215792 | rs10081633 | 1.689 | 0.114 | -1.699 | -2.127 | 0 | -0.006 | -0.006 | 0 | 0.18 | 0.17 | 0.12 | 0.03 | 0.16 | 0.26 |
| 9 | 113217123 | rs7035805 | -0.592 | 0.21 | -1.68 | -2.072 | -0.002 | -0.002 | -0.002 | 0.038 | 0.44 | 0.58 | 0.51 | 0.55 | 0.48 | 0.49 |
| 9 | 113219779 | rs7875113 | -0.357 | -0.09 | -2.199 | -1.973 | 0 | -0.005 | -0.005 | 7.22 | 0.42 | 0.63 | 0.5 | 0.5 | 0.44 | 0.46 |
| 9 | 113227531 | rs7869377 | -0.286 | -0.051 | -2.357 | -1.944 | -0.001 | 0.001 | 0.001 | 0 | 0.41 | 0.59 | 0.49 | 0.5 | 0.44 | 0.46 |
| 9 | 113229411 | rs17806765 | 1.642 | 0.213 | -2.364 | -1.824 | -0.025 | 0.015 | 0.015 | 0 | 0.16 | 0.04 | 0.2 | 0.07 | 0.19 | 0.15 |
| 9 | 113234507 | rs10817025 | 0.963 | 0.037 | -2.168 | -1.689 | -0.021 | 0.032 | 0.032 | 0 | 0.24 | 0.14 | 0.21 | 0.08 | 0.29 | 0.31 |
| 9 | 113236797 | rs10512405 | 0.968 | 0.013 | -2.159 | -1.686 | -0.023 | 0.037 | 0.037 | 0 | 0.26 | 0.15 | 0.34 | 0.08 | 0.32 | 0.32 |
| 9 | 113239268 | rs10817026 | 0.931 | 0.037 | -2.117 | -1.686 | -0.012 | 0.016 | 0.016 | 0 | 0.26 | 0.19 | 0.35 | 0.08 | 0.32 | 0.32 |
| 9 | 113240481 | rs7028791 | 0.094 | 0.426 | -1.7 | -1.264 | 0.02 | -0.011 | -0.011 | 0 | 0.58 | 0.71 | 0.63 | 0.58 | 0.64 | 0.75 |
| 9 | 113246386 | rs1931304 | -0.476 | 0.584 | -2.475 | -1.204 | 0.003 | 0.066 | 0.066 | 0 | 0.79 | 0.89 | 0.89 | 0.99 | 0.82 | 0.9 |
| 9 | 113250699 | rs10739303 | -1.165 | 1.264 | -2.458 | -0.753 | -0.039 | 0.062 | 0.062 | 0 | 0.89 | 0.99 | 0.93 | 1 | 0.89 | 0.94 |
| 9 | 113254704 | rs4978934 | -0.208 | 0.901 | -2.067 | -0.1 | -0.03 | 0.01 | 0.01 | 1.098 | 0.46 | 0.16 | 0.58 | 0.52 | 0.57 | 0.42 |
| 9 | 113255402 | rs13283529 | -0.383 | 0.857 | -1.473 | -0.27 | -0.016 | 0.03 | 0.03 | 0 | 0.06 | 0.01 | 0.06 | 0.01 | 0.08 | 0.04 |
| 9 | 113255544 | rs1889325 | 0.128 | 0.79 | -1.184 | -0.417 | 0.001 | 0.057 | 0.057 | 0 | 0.52 | 0.39 | 0.51 | 0.17 | 0.52 | 0.5 |
| 9 | 113256301 | rs7021157 | -0.356 | 0.576 | -0.754 | -0.067 | -0.007 | -0.01 | -0.01 | 2.208 | 0.05 | 0.1 | 0.03 | 0.03 | 0.03 | 0.03 |
| 9 | 113259152 | rs10817033 | 0.156 | -0.028 | -0.766 | -0.773 | -0.002 | -0.043 | -0.043 | 0.532 | 0.22 | 0.19 | 0.08 | 0 | 0.12 | 0.04 |
| 9 | 113261483 | rs872665 | 0.834 | -0.188 | -0.62 | -1.187 | -0.003 | -0.009 | -0.009 | 0 | 0.33 | 0.35 | 0.26 | 0.18 | 0.25 | 0.26 |
| 9 | 113261645 | rs872666 | 0.272 | 0.782 | -0.696 | -0.164 | -0.001 | 0 | 0 | 0 | 0.43 | 0.44 | 0.47 | 0.54 | 0.44 | 0.47 |
| 9 | 113265860 | rs10980412 | -0.496 | 0.608 | -0.416 | -0.291 | -0.052 | -0.051 | -0.051 | 0 | 0.67 | 0.79 | 0.6 | 0.53 | 0.57 | 0.51 |
| 9 | 113267399 | rs7046184 | 1.37 | 0.529 | -0.283 | -0.281 | -0.017 | -0.064 | -0.064 | 0 | 0.19 | 0.21 | 0.06 | 0 | 0.09 | 0.03 |
| 9 | 113278199 | rs12346753 | -1.462 | 1.226 | -0.602 | 0.576 | -0.032 | -0.04 | -0.04 | 0 | 0.92 | 0.99 | 0.93 | 0.66 | 0.9 | 0.82 |
| 9 | 113279114 | rs4403484 | 0.025 | 1.301 | -0.47 | 0.326 | 0 | 0.005 | 0.005 | 0.492 | 0.84 | 0.83 | 0.72 | 0.31 | 0.73 | 0.61 |
| 9 | 113281255 | rs12553191 | 1.104 | 1.514 | -0.802 | 0.121 | -0.03 | -0.006 | -0.006 | 0 | 0.14 | 0.03 | 0.27 | 0.11 | 0.2 | 0.19 |
| 9 | 113282064 | rs10980424 | 0.732 | 0.711 | -0.728 | -0.313 | 0.015 | 0.003 | 0.003 | 0.012 | 0.28 | 0.18 | 0.11 | 0.01 | 0.17 | 0.1 |
| 9 | 113283373 | rs1889329 | 0.613 | 0.692 | -0.349 | 0.135 | -0.003 | -0.008 | -0.008 | 0 | 0.32 | 0.34 | 0.32 | 0.19 | 0.24 | 0.22 |
| 9 | 113287555 | rs10980428 | -0.117 | 1.165 | -0.437 | 0.766 | 0.005 | 0.059 | 0.059 | 0 | 0.1 | 0.08 | 0.14 | 0.36 | 0.16 | 0.23 |
| 9 | 113291642 | rs4521831 | 0.223 | 0.931 | -0.465 | 0.528 | 0.002 | -0.004 | -0.004 | 0.788 | 0.86 | 0.83 | 0.93 | 0.68 | 0.91 | 0.77 |
| 9 | 113300206 | rs10759440 | 0.797 | 0.893 | -0.256 | 0.319 | 0.014 | 0.032 | 0.032 | 0.057 | 0.73 | 0.57 | 0.73 | 0.29 | 0.71 | 0.51 |
| 9 | 113300835 | rs1889321 | 0.781 | 0.907 | -0.312 | 0.326 | 0.01 | 0.023 | 0.023 | 0.045 | 0.73 | 0.6 | 0.74 | 0.29 | 0.71 | 0.51 |
| 9 | 113307129 | rs4460449 | -0.198 | 0.558 | -0.607 | 0.353 | -0.001 | 0.016 | 0.016 | 0.057 | 0.79 | 0.79 | 0.69 | 0.49 | 0.83 | 0.71 |
| 9 | 113309263 | rs955833 | 0.628 | 0.651 | -1.874 | -0.049 | -0.027 | 0.082 | 0.082 | 0 | 0.36 | 0.24 | 0.4 | 0.14 | 0.42 | 0.3 |
| 9 | 113311710 | rs7035502 | -0.64 | 0.618 | -1.888 | 0.146 | -0.057 | -0.002 | -0.002 | 0 | 0.6 | 0.79 | 0.49 | 0.56 | 0.51 | 0.55 |
| 9 | 113313187 | rs10980438 | 1.134 | 0.874 | -1.41 | 0.543 | -0.03 | -0.018 | -0.018 | 0 | 0.22 | 0.38 | 0.26 | 0.18 | 0.16 | 0.2 |
| 9 | 113313503 | rs16915122 | 1.079 | 0.813 | -1.342 | 0.66 | 0 | -0.015 | -0.015 | 0.255 | 0.12 | 0.09 | 0.13 | 0.33 | 0.1 | 0.19 |
| 9 | 113316224 | rs10980439 | 1 | 0.954 | -1.436 | 0.585 | -0.015 | -0.028 | -0.028 | 3.097 | 0.24 | 0.4 | 0.28 | 0.18 | 0.2 | 0.25 |
| 9 | 113316793 | rs2991364 | -1.053 | 0.584 | -2.089 | 0.315 | -0.021 | 0.089 | 0.089 | 0 | 0.63 | 0.77 | 0.6 | 0.86 | 0.6 | 0.73 |
| 9 | 113319516 | rs3010808 | 0.58 | 0.728 | -1.903 | 0.592 | 0 | 0.006 | 0.006 | 0 | 0.13 | 0.12 | 0.13 | 0.33 | 0.09 | 0.19 |
| 9 | 113329058 | rs4978937 | -0.368 | 0.747 | -2.104 | -0.156 | -0.015 | 0.106 | 0.106 | 0 | 0.58 | 0.73 | 0.57 | 0.86 | 0.55 | 0.7 |
| 9 | 113331428 | rs11788554 | -0.081 | 0.859 | -1.571 | 0.654 | -0.009 | 0.019 | 0.019 | 0.515 | 0.13 | 0.06 | 0.08 | 0 | 0.15 | 0.03 |
| 9 | 113336184 | rs4075022 | -0.12 | 1.014 | -2.141 | 0.916 | -0.09 | -0.072 | -0.072 | 0 | 0.36 | 0.09 | 0.58 | 0.56 | 0.39 | 0.42 |
| 9 | 113343006 | rs7853273 | -0.333 | 1.579 | -1.504 | 0.881 | 0.056 | 0.04 | 0.04 | 0.181 | 0.06 | 0.15 | 0.07 | 0.08 | 0.1 | 0.05 |
| 9 | 113343555 | rs4978938 | -0.66 | 1.104 | -1.573 | -0.101 | 0.006 | 0.01 | 0.01 | 0 | 0.58 | 0.38 | 0.71 | 0.95 | 0.61 | 0.74 |
| 9 | 113345800 | rs16915134 | 0.643 | 0.904 | -1.705 | -0.044 | -0.008 | -0.063 | -0.063 | 3.531 | 0.18 | 0.02 | 0.07 | 0.26 | 0.17 | 0.24 |
| 9 | 113355719 | rs12340257 | 1.036 | 0.654 | -0.881 | -0.621 | 0.007 | -0.066 | -0.066 | 0.137 | 0.12 | 0.21 | 0.07 | 0.01 | 0.09 | 0.07 |
| 9 | 113360859 | rs10817044 | 0.967 | 0.674 | -1.098 | -0.49 | -0.07 | -0.071 | -0.071 | 0.021 | 0.35 | 0.09 | 0.57 | 0.55 | 0.38 | 0.42 |
| 9 | 113364308 | rs4978943 | -0.302 | 0.487 | -0.438 | -0.634 | 0.015 | 0.04 | 0.04 | 0 | 0.61 | 0.33 | 0.72 | 0.95 | 0.65 | 0.74 |
| 9 | 113364464 | rs12236915 | 0.757 | 0.457 | -0.471 | -0.594 | 0.016 | -0.024 | -0.024 | 0 | 0.19 | 0.11 | 0.08 | 0.31 | 0.17 | 0.22 |
| 9 | 113366508 | rs10817045 | 0.997 | 0.466 | -0.496 | -0.506 | -0.083 | -0.09 | -0.09 | 0 | 0.35 | 0.08 | 0.57 | 0.54 | 0.38 | 0.43 |
| 9 | 113366974 | rs10980480 | 0.753 | 0.434 | -0.457 | -0.465 | 0.018 | -0.024 | -0.024 | 0 | 0.19 | 0.11 | 0.08 | 0.31 | 0.17 | 0.23 |
| 9 | 113367300 | rs10980481 | 0.638 | 0.492 | -0.341 | -0.471 | 0.007 | 0.008 | 0.008 | 0 | 0.05 | 0.05 | 0.07 | 0.09 | 0.1 | 0.06 |
| 9 | 113368085 | rs7875689 | 0.182 | 0.336 | -0.302 | -0.547 | -0.012 | 0.123 | 0.123 | 0 | 0.57 | 0.16 | 0.66 | 0.82 | 0.57 | 0.68 |
| 9 | 113371069 | rs4978949 | 0.875 | 0.281 | -0.356 | -0.552 | -0.052 | -0.04 | -0.04 | 0.634 | 0.38 | 0.15 | 0.59 | 0.58 | 0.41 | 0.48 |
| 9 | 113372027 | rs10817051 | 0.829 | 0.217 | -0.285 | -0.807 | -0.011 | 0.017 | 0.017 | 0 | 0.12 | 0.06 | 0.11 | 0.02 | 0.17 | 0.09 |
| 9 | 113374865 | rs4302915 | -0.679 | 0.097 | 0.015 | -0.92 | 0 | 0 | 0 | 0 | 0.37 | 0.37 | 0.24 | 0.39 | 0.41 | 0.35 |
| 9 | 113374973 | rs4422837 | -0.069 | 0.089 | 0.015 | -0.927 | 0.001 | -0.002 | -0.002 | 2.095 | 0.69 | 0.3 | 0.77 | 0.89 | 0.74 | 0.74 |
| 9 | 113378789 | rs10817056 | 0.487 | 0.093 | 0.012 | -0.922 | 0.003 | -0.008 | -0.008 | 0 | 0.14 | 0.15 | 0.11 | 0.07 | 0.17 | 0.12 |
| 9 | 113380437 | rs12238269 | 1.716 | 0.089 | -0.02 | -0.962 | -0.052 | -0.07 | -0.07 | 0.003 | 0.33 | 0.13 | 0.57 | 0.58 | 0.36 | 0.47 |
| 9 | 113386179 | rs4509410 | 0.327 | -0.017 | -0.098 | -1.035 | 0.031 | 0.036 | 0.036 | 0.23 | 0.55 | 0.29 | 0.71 | 0.67 | 0.6 | 0.69 |
| 9 | 113388632 | rs10817061 | -0.123 | -0.02 | -0.096 | -1.033 | 0.004 | 0.006 | 0.006 | 0 | 0.32 | 0.2 | 0.19 | 0.31 | 0.34 | 0.26 |

| | | | | | | | | | | | | | | | | |
|---|---|---|---|---|---|---|---|---|---|---|---|---|---|---|---|---|
| 9 | 113390612 | rs13302218 | -0.01 | -0.038 | -0.045 | -1.01 | 0.036 | 0.075 | 0.075 | 0.129 | 0.06 | 0.23 | 0.06 | 0.09 | 0.07 | 0.08 |
| 9 | 113392182 | rs6477778 | -0.69 | -0.046 | 0.121 | -1.019 | -0.032 | -0.145 | -0.145 | 1.381 | 0.75 | 0.55 | 0.83 | 0.98 | 0.81 | 0.82 |
| 9 | 113393012 | rs6477780 | -1.228 | -0.087 | 0.668 | -1.026 | -0.017 | -0.125 | -0.125 | 0 | 0.81 | 0.63 | 0.85 | 1 | 0.84 | 0.92 |
| 9 | 113398561 | rs4978951 | -0.23 | -0.103 | 0.322 | -1.025 | -0.015 | -0.028 | -0.028 | 1.02 | 0.62 | 0.47 | 0.77 | 0.76 | 0.67 | 0.78 |
| 9 | 113403846 | rs3001115 | -2.209 | 0.988 | -0.047 | -1.301 | -0.016 | -0.112 | -0.112 | 0 | 0.72 | 0.95 | 0.5 | 0.43 | 0.77 | 0.57 |
| 9 | 113404293 | rs3001116 | -0.47 | 0.84 | 0.029 | -1.343 | 0.021 | 0.03 | 0.03 | 0.056 | 0.48 | 0.6 | 0.29 | 0.35 | 0.42 | 0.41 |
| 9 | 113409839 | rs16915239 | 0.343 | 0.834 | -0.011 | -1.322 | 0.006 | -0.027 | -0.027 | 0 | 0.27 | 0.14 | 0.14 | 0.34 | 0.25 | 0.33 |
| 9 | 113423432 | rs4269612 | -0.281 | 0.842 | -0.064 | -1.333 | 0.005 | -0.004 | -0.004 | 1.342 | 0.33 | 0.28 | 0.17 | 0.34 | 0.3 | 0.36 |
| 9 | 113431606 | rs3001125 | -1.237 | 1.125 | -0.377 | -1.128 | -0.085 | -0.033 | -0.033 | 0 | 0.63 | 0.91 | 0.41 | 0.36 | 0.59 | 0.52 |
| 9 | 113434178 | rs3010815 | -0.66 | 1.11 | -0.248 | -1.044 | 0.007 | 0.046 | 0.046 | 0.346 | 0.53 | 0.75 | 0.32 | 0.35 | 0.47 | 0.44 |
| 9 | 113454159 | rs7040569 | -0.661 | 1.002 | -0.298 | -0.931 | -0.021 | 0.031 | 0.031 | 0.682 | 0.61 | 0.69 | 0.38 | 0.28 | 0.57 | 0.45 |
| 9 | 113463246 | rs4498626 | -0.269 | 0.411 | -0.32 | -0.899 | 0.002 | -0.012 | -0.012 | 0.021 | 0.47 | 0.54 | 0.42 | 0.85 | 0.45 | 0.51 |
| 9 | 113466154 | rs4978952 | 1.505 | 0.306 | -0.712 | -0.208 | 0.027 | -0.099 | -0.099 | 0 | 0.14 | 0.01 | 0.21 | 0.43 | 0.09 | 0.17 |
| 9 | 113466941 | rs12347321 | 0.536 | 0.468 | -0.836 | -0.622 | -0.016 | 0.041 | 0.041 | 0 | 0.11 | 0.02 | 0.06 | 0.02 | 0.11 | 0.07 |
| 9 | 113467546 | rs7856889 | -0.298 | -0.065 | -0.393 | -0.582 | -0.001 | -0.009 | -0.009 | 0.027 | 0.46 | 0.59 | 0.42 | 0.76 | 0.42 | 0.52 |
| 9 | 113468600 | rs4623510 | 1.776 | 0.331 | -1.144 | -0.57 | -0.043 | 0.032 | 0.032 | 1.068 | 0.14 | 0.01 | 0.08 | 0.09 | 0.17 | 0.11 |
| 9 | 113468897 | rs4401940 | -0.12 | 0.237 | -0.735 | -0.549 | -0.006 | -0.012 | -0.012 | 0 | 0.54 | 0.52 | 0.39 | 0.32 | 0.61 | 0.36 |
| 9 | 113472810 | rs10980535 | 1.147 | 0.12 | -0.976 | -0.56 | 0.003 | -0.125 | -0.125 | 0.909 | 0.24 | 0.03 | 0.29 | 0.51 | 0.21 | 0.29 |
| 9 | 113475374 | rs10980536 | 0.681 | 0.184 | -0.751 | -0.744 | -0.059 | 0.031 | 0.031 | 0 | 0.34 | 0.18 | 0.27 | 0.22 | 0.43 | 0.19 |
| 9 | 113476105 | rs4297099 | 0.275 | 0.059 | -0.242 | -0.76 | -0.004 | 0.001 | 0.001 | 0 | 0.34 | 0.38 | 0.35 | 0.6 | 0.27 | 0.36 |
| 9 | 113480169 | rs4415391 | -1.049 | 0.186 | -0.17 | -0.875 | 0.005 | -0.025 | -0.025 | 0 | 0.86 | 0.78 | 0.71 | 0.92 | 0.88 | 0.69 |
| 9 | 113485961 | rs4336672 | -0.244 | 0.193 | -0.289 | -0.898 | 0.001 | -0.081 | -0.081 | 0 | 0.69 | 0.58 | 0.62 | 0.82 | 0.7 | 0.58 |
| 9 | 113486002 | rs4259484 | -1.317 | 0.33 | -0.716 | -0.723 | -0.014 | 0.016 | 0.016 | 0 | 0.88 | 0.96 | 0.72 | 0.92 | 0.88 | 0.72 |
| 9 | 113493735 | rs10759464 | 0.542 | 0.163 | -0.393 | -0.641 | 0.001 | -0.028 | -0.028 | 0 | 0.31 | 0.38 | 0.25 | 0.06 | 0.35 | 0.14 |
| 9 | 113501372 | rs4574919 | -0.131 | 0.058 | -0.312 | -0.578 | 0.006 | 0.042 | 0.042 | 0 | 0.48 | 0.63 | 0.33 | 0.11 | 0.43 | 0.27 |
| 9 | 113503673 | rs10980549 | 1.824 | -0.054 | -0.59 | 0.07 | -0.025 | -0.079 | -0.079 | 0 | 0.15 | 0.02 | 0.21 | 0.34 | 0.17 | 0.16 |
| 9 | 113503776 | rs4532674 | -0.82 | 0.207 | -0.535 | 0.035 | -0.003 | -0.004 | -0.004 | 2.464 | 0.63 | 0.66 | 0.54 | 0.45 | 0.6 | 0.39 |
| 9 | 113504910 | rs10980551 | -0.308 | -0.069 | -0.351 | -0.077 | 0.025 | 0.081 | 0.081 | 0 | 0.51 | 0.62 | 0.31 | 0.15 | 0.42 | 0.27 |
| 9 | 113506998 | rs4144418 | -0.311 | -0.047 | -0.342 | -0.066 | 0.024 | 0.074 | 0.074 | 0 | 0.51 | 0.61 | 0.31 | 0.15 | 0.42 | 0.27 |
| 9 | 113509778 | rs2846448 | -1.305 | -0.127 | -0.827 | 0.342 | 0 | -0.001 | -0.001 | 0 | 0.74 | 0.79 | 0.62 | 0.72 | 0.8 | 0.66 |
| 9 | 113512773 | rs2298492 | 0.946 | 0.134 | -1.593 | 0.435 | 0.002 | 0.014 | 0.014 | 0 | 0.1 | 0.11 | 0.08 | 0.26 | 0.19 | 0.26 |
| 9 | 113522917 | rs1940245 | 0.467 | -0.205 | -2.119 | 0.031 | 0.027 | -0.033 | -0.033 | 0 | 0.3 | 0.17 | 0.29 | 0.39 | 0.25 | 0.25 |
| 9 | 113523032 | rs1940247 | 1.876 | -0.37 | -2.081 | -0.239 | -0.033 | 0.086 | 0.086 | 0 | 0.3 | 0.18 | 0.24 | 0.05 | 0.36 | 0.17 |
| 9 | 113523448 | rs12001728 | 1.224 | -0.488 | -2.209 | -0.094 | -0.001 | -0.005 | -0.005 | 0 | 0.15 | 0.18 | 0.09 | 0.05 | 0.1 | 0.08 |
| 9 | 113526327 | rs1940251 | -0.994 | -0.202 | -1.65 | 0.467 | -0.015 | 0.016 | 0.016 | 0 | 0.73 | 0.46 | 0.6 | 0.68 | 0.82 | 0.7 |
| 9 | 113527226 | rs10817091 | 1.099 | -0.179 | -1.742 | 0.36 | -0.026 | 0.141 | 0.141 | 0 | 0.45 | 0.24 | 0.31 | 0.09 | 0.47 | 0.26 |
| 9 | 113531585 | rs2766998 | -1.333 | -0.059 | -1.751 | 0.503 | -0.006 | -0.056 | -0.056 | 0 | 0.83 | 0.93 | 0.77 | 0.64 | 0.79 | 0.81 |
| 9 | 113536274 | rs2148584 | -0.197 | 0.034 | -1.26 | 0.715 | -0.002 | 0.003 | 0.003 | 0 | 0.35 | 0.57 | 0.46 | 0.56 | 0.32 | 0.55 |
| 9 | 113538122 | rs2274419 | 1.697 | 0.309 | -1.236 | 0.805 | -0.032 | -0.026 | -0.026 | 0 | 0.05 | 0 | 0.04 | 0.19 | 0.13 | 0.16 |
| 9 | 113541125 | rs2766984 | -0.049 | 0.204 | -1.237 | 0.618 | 0 | 0.02 | 0.02 | 0 | 0.33 | 0.54 | 0.45 | 0.51 | 0.31 | 0.46 |
| 9 | 113547476 | rs1170327 | -0.639 | 0.157 | -1.164 | 0.656 | -0.008 | 0.008 | 0.008 | 0 | 0.18 | 0.69 | 0.11 | 0.25 | 0.13 | 0.21 |
| 9 | 113552952 | rs525344 | -1.102 | 0.186 | -1.647 | 0.697 | -0.035 | -0.093 | -0.093 | 0.05 | 0.85 | 0.98 | 0.78 | 0.64 | 0.8 | 0.83 |
| 9 | 113560071 | rs10759469 | 0.53 | 0.139 | -1.421 | 0.396 | 0.006 | 0.022 | 0.022 | 0 | 0.19 | 0.26 | 0.37 | 0.27 | 0.19 | 0.24 |
| 9 | 113560119 | rs2821143 | 0.787 | -0.023 | -1.511 | 0.316 | -0.011 | 0.02 | 0.02 | 0 | 0.65 | 0.5 | 0.68 | 0.38 | 0.65 | 0.5 |
| 9 | 113561251 | rs3780529 | 1.614 | -0.012 | -1.518 | 0.328 | -0.027 | -0.027 | -0.027 | 0 | 0.06 | 0.01 | 0.05 | 0.16 | 0.12 | 0.18 |
| 9 | 113563143 | rs578430 | 0.989 | -0.08 | -1.294 | 0.259 | -0.025 | -0.018 | -0.018 | 0 | 0.1 | 0.3 | 0.03 | 0.04 | 0.02 | 0.06 |
| 9 | 113565027 | rs521803 | -0.546 | -0.046 | -1.51 | 0.281 | -0.012 | -0.042 | -0.042 | 0 | 0.78 | 0.82 | 0.73 | 0.46 | 0.69 | 0.69 |
| 9 | 113566357 | rs2821142 | 1.438 | 0.006 | -1.494 | 0.226 | -0.027 | 0.165 | 0.165 | 0 | 0.46 | 0.23 | 0.31 | 0.11 | 0.46 | 0.26 |
| 9 | 113569096 | rs1409686 | 0.386 | 0.186 | -1.421 | 0.368 | -0.004 | 0.049 | 0.049 | 0 | 0.69 | 0.65 | 0.7 | 0.44 | 0.66 | 0.59 |
| 9 | 113570107 | rs2846444 | 0.554 | 0.123 | -1.558 | 0.351 | 0.01 | 0.011 | 0.011 | 0 | 0.56 | 0.41 | 0.38 | 0.33 | 0.58 | 0.53 |
| 9 | 113573073 | rs2846451 | 1.37 | 0.28 | -2.066 | -0.063 | -0.015 | 0.066 | 0.066 | 0 | 0.22 | 0.07 | 0.12 | 0.06 | 0.19 | 0.18 |
| 9 | 113579789 | rs16915435 | 0.495 | 0.499 | -1.4 | 0.384 | -0.002 | -0.06 | -0.06 | 0.947 | 0.14 | 0.03 | 0.17 | 0.34 | 0.16 | 0.14 |
| 9 | 113581359 | rs509276 | -0.577 | 0.608 | -0.823 | -0.345 | -0.005 | -0.11 | -0.11 | 0.087 | 0.25 | 0.46 | 0.15 | 0.04 | 0.22 | 0.17 |
| 9 | 113582050 | rs2821140 | 1.944 | 0.808 | -1.25 | -0.409 | 0.026 | 0.07 | 0.07 | 1.151 | 0.4 | 0.14 | 0.4 | 0.27 | 0.31 | 0.44 |
| 9 | 113582391 | rs540171 | 0.521 | 0.789 | -0.787 | -0.462 | -0.027 | -0.063 | -0.063 | 0.735 | 0.15 | 0.25 | 0.06 | 0.03 | 0.09 | 0.11 |
| 9 | 113582672 | rs12553421 | 1.424 | 0.846 | -0.848 | -0.097 | 0.01 | -0.03 | -0.03 | 0 | 0.11 | 0.04 | 0.12 | 0.38 | 0.09 | 0.11 |
| 9 | 113584012 | rs2767001 | 0.08 | 1.367 | -0.708 | -0.212 | 0.005 | -0.072 | -0.072 | 0 | 0.75 | 0.66 | 0.8 | 0.94 | 0.72 | 0.84 |
| 9 | 113585503 | rs10980575 | 1.533 | 1.282 | -0.816 | -0.029 | -0.041 | -0.037 | -0.037 | 0 | 0.1 | 0.02 | 0.15 | 0.17 | 0.18 | 0.15 |
| 9 | 113588332 | rs10980578 | 0.925 | 0.827 | -0.678 | -0.505 | -0.005 | 0.017 | 0.017 | 1.307 | 0.15 | 0.17 | 0.29 | 0.26 | 0.11 | 0.23 |
| 9 | 113590448 | rs2821134 | 0.593 | 0.864 | -0.626 | -0.656 | 0.017 | -0.078 | -0.078 | 0.375 | 0.66 | 0.52 | 0.71 | 0.93 | 0.58 | 0.78 |
| 9 | 113596088 | rs12684949 | 0.368 | 0.761 | -0.685 | -0.396 | 0.013 | 0.01 | 0.01 | 0.014 | 0.22 | 0.32 | 0.36 | 0.24 | 0.23 | 0.28 |
| 9 | 113597109 | rs2821153 | -0.331 | 0.692 | -0.201 | -0.616 | -0.006 | -0.012 | -0.012 | 0 | 0.82 | 0.81 | 0.86 | 0.91 | 0.84 | 0.9 |
| 9 | 113597449 | rs2846460 | 2.911 | 0.504 | -0.513 | -0.816 | 0.019 | 0.125 | 0.125 | 0.002 | 0.27 | 0.09 | 0.13 | 0.04 | 0.2 | 0.21 |
| 9 | 113597800 | rs4978958 | -0.616 | 0.771 | -0.035 | -0.334 | 0.006 | -0.022 | -0.022 | 0 | 0.3 | 0.44 | 0.4 | 0.24 | 0.29 | 0.36 |
| 9 | 113597938 | rs4978959 | -0.868 | 0.78 | -0.031 | -0.271 | -0.004 | 0.002 | 0.002 | 0.11 | 0.33 | 0.49 | 0.4 | 0.33 | 0.29 | 0.42 |
| 9 | 113599422 | rs943327 | -0.132 | 0.831 | -0.189 | -0.105 | 0.005 | 0.029 | 0.029 | 0 | 0.24 | 0.37 | 0.37 | 0.33 | 0.23 | 0.34 |
| 9 | 113602694 | rs3860159 | -0.32 | 1.023 | -0.161 | 0.003 | -0.004 | -0.009 | -0.009 | 0.088 | 0.23 | 0.19 | 0.28 | 0.65 | 0.27 | 0.34 |
| 9 | 113609826 | rs10980588 | -0.311 | 0.91 | -0.028 | -0.26 | -0.01 | -0.051 | -0.051 | 1.492 | 0.12 | 0.18 | 0.08 | 0 | 0.09 | 0.09 |
| 9 | 113619757 | rs10980596 | 0.409 | 1.043 | -0.334 | 0.108 | -0.046 | -0.048 | -0.048 | 0.327 | 0.14 | 0.05 | 0.18 | 0.25 | 0.24 | 0.27 |

| | | | | | | | | | | | | | | | | |
|---|---|---|---|---|---|---|---|---|---|---|---|---|---|---|---|---|
| 9 | 113622466 | rs474186 | 0.379 | 0.959 | -0.081 | -0.063 | -0.005 | 0.019 | 0.019 | 0 | 0.19 | 0.26 | 0.32 | 0.27 | 0.17 | 0.23 |
| 9 | 113625888 | rs576515 | -0.918 | 0.972 | -0.214 | -0.076 | 0.026 | 0.043 | 0.043 | 1.163 | 0.34 | 0.44 | 0.51 | 0.54 | 0.41 | 0.51 |
| 9 | 113629491 | rs472936 | 1.322 | 0.908 | -0.082 | -0.091 | -0.014 | -0.003 | -0.003 | 0 | 0.08 | 0.26 | 0.04 | 0.04 | 0.03 | 0.04 |
| 9 | 113631282 | rs512756 | 0.58 | 1.09 | -0.124 | -0.287 | 0.007 | -0.005 | -0.005 | 0 | 0.69 | 0.6 | 0.52 | 0.73 | 0.61 | 0.67 |
| 9 | 113631618 | rs516187 | -0.299 | 1.172 | -0.317 | -0.257 | 0.02 | 0.025 | 0.025 | 0.689 | 0.22 | 0.32 | 0.22 | 0.3 | 0.27 | 0.31 |
| 9 | 113632279 | rs693342 | -0.518 | 1.174 | -0.564 | -0.24 | 0.022 | 0.035 | 0.035 | 1.047 | 0.35 | 0.43 | 0.51 | 0.55 | 0.41 | 0.51 |
| 9 | 113633737 | rs12000617 | 2.337 | 1.03 | -0.976 | -0.455 | -0.01 | 0.178 | 0.178 | 0.036 | 0.36 | 0.11 | 0.18 | 0.03 | 0.31 | 0.26 |
| 9 | 113635180 | rs2766981 | 0.503 | 1.596 | -1.019 | -0.294 | -0.012 | 0.019 | 0.019 | 0.097 | 0.8 | 0.82 | 0.81 | 0.95 | 0.72 | 0.83 |
| 9 | 113635381 | rs505414 | -1.255 | 1.553 | -0.859 | -0.019 | 0.008 | 0.011 | 0.011 | 0.265 | 0.37 | 0.37 | 0.51 | 0.55 | 0.44 | 0.5 |
| 9 | 113635691 | rs6833 | -1.176 | 1.592 | -0.926 | 0.027 | 0.002 | 0.01 | 0.01 | 0 | 0.39 | 0.37 | 0.5 | 0.49 | 0.39 | 0.48 |
| 9 | 113635929 | rs1043128 | 1.3 | 1.728 | -1.069 | 0.065 | -0.05 | -0.046 | -0.046 | 0 | 0.09 | 0.01 | 0.17 | 0.19 | 0.2 | 0.15 |
| 9 | 113636353 | rs13094 | -1.275 | 1.704 | -1.046 | 0.068 | 0.002 | 0.019 | 0.019 | 0 | 0.39 | 0.41 | 0.5 | 0.49 | 0.38 | 0.48 |
| 9 | 113637662 | rs955888 | 0.167 | 1.948 | -1.138 | -0.011 | -0.047 | 0.06 | 0.06 | 0 | 0.14 | 0.08 | 0.17 | 0.01 | 0.27 | 0.12 |
| 9 | 113637854 | rs3739709 | 0.96 | 1.238 | -0.625 | 0.58 | -0.05 | -0.044 | -0.044 | 0 | 0.09 | 0.01 | 0.16 | 0.19 | 0.2 | 0.15 |
| 9 | 113640468 | rs544276 | -0.199 | 1.116 | -0.58 | 0.54 | -0.038 | 0.024 | 0.024 | 0 | 0.26 | 0.38 | 0.33 | 0.3 | 0.17 | 0.26 |
| 9 | 113642646 | rs576559 | -0.653 | 0.934 | -0.372 | 0.149 | 0.006 | 0.019 | 0.019 | 0 | 0.38 | 0.39 | 0.5 | 0.53 | 0.38 | 0.47 |
| 9 | 113648839 | rs1536435 | 0.623 | 0.624 | -0.611 | -1.321 | -0.026 | 0.181 | 0.181 | 0 | 0.54 | 0.78 | 0.42 | 0.06 | 0.56 | 0.36 |
| 9 | 113649462 | rs7039165 | 0.22 | 0.818 | -0.921 | -1.035 | -0.067 | -0.049 | -0.049 | 0 | 0.17 | 0.02 | 0.14 | 0.2 | 0.19 | 0.22 |
| 9 | 113654585 | rs7854710 | -0.728 | 0.629 | -0.527 | -1.438 | -0.002 | -0.053 | -0.053 | 0 | 0.18 | 0.66 | 0.15 | 0.04 | 0.14 | 0.2 |
| 9 | 113656170 | rs10817107 | 0.344 | 0.588 | -0.59 | -1.495 | -0.027 | -0.022 | -0.022 | 0.408 | 0.17 | 0.08 | 0.25 | 0.21 | 0.19 | 0.23 |
| 9 | 113656440 | rs1326893 | -0.486 | 0.417 | -0.778 | -1.361 | -0.021 | 0.01 | 0.01 | 0.065 | 0.77 | 0.98 | 0.75 | 0.83 | 0.78 | 0.74 |
| 9 | 113658671 | rs10817108 | 0.641 | 0.387 | -0.667 | -1.298 | -0.06 | -0.052 | -0.052 | 0 | 0.16 | 0.02 | 0.24 | 0.21 | 0.19 | 0.22 |
| 9 | 113660985 | rs10119345 | -0.587 | 0.379 | -0.633 | -1.307 | -0.002 | -0.055 | -0.055 | 2.499 | 0.18 | 0.66 | 0.15 | 0.04 | 0.15 | 0.19 |
| 9 | 113662681 | rs1007000 | 0.669 | 0.385 | -0.704 | -1.297 | -0.06 | -0.052 | -0.052 | 0 | 0.16 | 0.02 | 0.24 | 0.21 | 0.19 | 0.22 |
| 9 | 113665254 | rs1409684 | 0.426 | 0.382 | -0.701 | -1.299 | -0.036 | -0.03 | -0.03 | 0 | 0.17 | 0.06 | 0.25 | 0.21 | 0.19 | 0.23 |
| 9 | 113667277 | rs3780528 | -0.65 | 0.392 | -0.631 | -1.3 | -0.002 | -0.036 | -0.036 | 0.042 | 0.18 | 0.71 | 0.15 | 0.04 | 0.15 | 0.19 |
| 9 | 113674967 | rs1326899 | -1.151 | 0.384 | -0.678 | -1.294 | -0.023 | 0.035 | 0.035 | 0.106 | 0.35 | 0.77 | 0.4 | 0.26 | 0.34 | 0.43 |
| 9 | 113675287 | rs1326898 | -0.407 | 0.366 | -0.268 | -1.305 | -0.034 | 0 | 0 | 0 | 0.25 | 0.13 | 0.34 | 0.24 | 0.31 | 0.38 |
| 9 | 113680831 | rs2182577 | -0.384 | 0.237 | -0.148 | -1.401 | -0.002 | 0.033 | 0.033 | 0 | 0.75 | 0.89 | 0.74 | 0.83 | 0.76 | 0.7 |
| 9 | 113681255 | rs10980633 | -0.267 | -0.05 | -0.113 | -1.4 | -0.001 | 0.032 | 0.032 | 0.347 | 0.14 | 0.15 | 0.17 | 0.58 | 0.12 | 0.3 |
| 9 | 113683472 | rs2766993 | 1.917 | -0.058 | -0.191 | -1.47 | -0.029 | 0.144 | 0.144 | 0 | 0.38 | 0.17 | 0.27 | 0.02 | 0.44 | 0.17 |
| 9 | 113684143 | rs1547664 | 1.476 | -0.014 | -0.153 | -1.362 | -0.029 | -0.043 | -0.043 | 0.026 | 0.1 | 0.01 | 0.21 | 0.19 | 0.1 | 0.11 |
| 9 | 113686353 | rs1326890 | -0.28 | -0.027 | -0.165 | -1.393 | 0.018 | 0.016 | 0.016 | 0.183 | 0.77 | 0.89 | 0.78 | 0.83 | 0.85 | 0.72 |
| 9 | 113690028 | rs2254841 | 1.022 | -0.121 | 0.443 | -1.454 | -0.003 | -0.006 | -0.006 | 0.566 | 0.55 | 0.49 | 0.48 | 0.62 | 0.59 | 0.51 |
| 9 | 113694355 | rs4978968 | -1.107 | -0.122 | 0.674 | -1.454 | 0 | 0.002 | 0.002 | 0 | 0.44 | 0.41 | 0.51 | 0.38 | 0.41 | 0.49 |
| 9 | 113694568 | rs10817114 | 0.079 | -0.128 | 0.765 | -1.458 | 0.007 | -0.001 | -0.001 | 0.024 | 0.22 | 0.23 | 0.29 | 0.21 | 0.29 | 0.24 |
| 9 | 113696095 | rs2900519 | -0.157 | -0.127 | 0.891 | -1.487 | -0.009 | 0.094 | 0.094 | 0.011 | 0.13 | 0.21 | 0.17 | 0.59 | 0.1 | 0.29 |
| 9 | 113697520 | rs2900520 | 0.413 | -0.131 | 0.87 | -1.49 | 0.013 | 0.006 | 0.006 | 0.608 | 0.22 | 0.18 | 0.22 | 0.17 | 0.12 | 0.24 |
| 9 | 113697555 | rs4246884 | -2.174 | -0.114 | 0.754 | -1.48 | -0.059 | 0.16 | 0.16 | 0 | 0.61 | 0.82 | 0.7 | 0.98 | 0.51 | 0.79 |
| 9 | 113697686 | rs10980642 | -0.462 | -0.114 | 0.759 | -1.474 | -0.02 | 0.151 | 0.151 | 0 | 0.15 | 0.31 | 0.18 | 0.59 | 0.1 | 0.3 |
| 9 | 113698031 | rs4978970 | -2.113 | -0.056 | 0.283 | -1.486 | -0.058 | 0.154 | 0.154 | 0 | 0.61 | 0.83 | 0.7 | 0.98 | 0.5 | 0.79 |
| 9 | 113701264 | rs10817115 | -0.988 | -0.052 | 0.261 | -1.461 | -0.006 | 0.008 | 0.008 | 0.266 | 0.33 | 0.29 | 0.45 | 0.8 | 0.38 | 0.53 |
| 9 | 113703439 | rs12555560 | 0.125 | -0.068 | 0.126 | -1.486 | 0.003 | 0.003 | 0.003 | 0.86 | 0.25 | 0.2 | 0.24 | 0.18 | 0.15 | 0.28 |
| 9 | 113707225 | rs7866715 | -1.967 | 0.183 | -0.277 | -1.525 | -0.014 | 0.045 | 0.045 | 0 | 0.61 | 0.71 | 0.7 | 0.98 | 0.53 | 0.82 |
| 9 | 113709586 | rs10980653 | 2.22 | 0.04 | -0.658 | -1.749 | -0.079 | 0.122 | 0.122 | 0 | 0.3 | 0.08 | 0.26 | 0 | 0.44 | 0.16 |
| 9 | 113712132 | rs10817118 | 0.195 | 0.175 | -0.72 | -1.553 | -0.087 | -0.066 | -0.066 | 0 | 0.2 | 0.02 | 0.27 | 0.26 | 0.29 | 0.25 |
| 9 | 113722721 | rs12345986 | 1.445 | 0.15 | -0.634 | -1.627 | -0.014 | 0.049 | 0.049 | 0 | 0.39 | 0.31 | 0.3 | 0.02 | 0.47 | 0.19 |
| 9 | 113724346 | rs10817124 | -0.117 | -0.115 | -0.687 | -1.389 | -0.061 | -0.013 | -0.013 | 0 | 0.22 | 0.06 | 0.28 | 0.21 | 0.29 | 0.25 |
| 9 | 113732126 | rs10980665 | 0.665 | -0.083 | -0.693 | -1.537 | -0.037 | -0.035 | -0.035 | 0 | 0.13 | 0.01 | 0.21 | 0.17 | 0.14 | 0.12 |
| 9 | 113732167 | rs2418124 | 1.127 | -0.155 | -0.515 | -1.544 | 0.041 | 0.097 | 0.097 | 0 | 0.52 | 0.32 | 0.52 | 0.2 | 0.63 | 0.34 |
| 9 | 113734610 | rs2277179 | 0.716 | -0.315 | -0.7 | -1.606 | 0.006 | 0.015 | 0.015 | 0 | 0.15 | 0.11 | 0.11 | 0.03 | 0.11 | 0.15 |
| 9 | 113735263 | rs4978432 | 0.506 | -0.37 | -0.607 | -1.61 | -0.023 | 0.015 | 0.015 | 0 | 0.14 | 0.29 | 0.2 | 0.16 | 0.07 | 0.21 |
| 9 | 113737589 | rs10759478 | 0.151 | -0.229 | -1.176 | -1.526 | -0.001 | 0.017 | 0.017 | 0 | 0.82 | 0.79 | 0.83 | 0.38 | 0.82 | 0.68 |
| 9 | 113737831 | rs16915618 | -0.469 | 0.023 | -0.982 | -1.603 | -0.038 | -0.052 | -0.052 | 0 | 0.29 | 0.44 | 0.31 | 0.19 | 0.19 | 0.34 |
| 9 | 113747491 | rs17382569 | 0.679 | 0.179 | -1.315 | -1.641 | -0.008 | -0.031 | -0.031 | 0.156 | 0.11 | 0.18 | 0.08 | 0.02 | 0.06 | 0.14 |
| 9 | 113747902 | rs12552125 | 0.987 | 0.432 | -1.242 | -1.478 | -0.036 | -0.04 | -0.04 | 0 | 0.11 | 0.01 | 0.21 | 0.16 | 0.13 | 0.09 |
| 9 | 113751324 | rs12551377 | 0.19 | 0.491 | -1.204 | -1.537 | 0 | -0.031 | -0.031 | 0 | 0.25 | 0.26 | 0.28 | 0.12 | 0.25 | 0.3 |
| 9 | 113753524 | rs10980683 | -1.107 | 0.459 | -0.841 | -1.283 | -0.032 | 0.054 | 0.054 | 0 | 0.39 | 0.21 | 0.4 | 0.24 | 0.43 | 0.41 |
| 9 | 113766668 | rs7023788 | -0.262 | 0.602 | -1.496 | -1.117 | -0.106 | -0.07 | -0.07 | 0.779 | 0.22 | 0.02 | 0.29 | 0.26 | 0.32 | 0.21 |
| 9 | 113769925 | rs1030005 | 0.386 | 0.597 | -1.038 | -0.744 | -0.003 | -0.067 | -0.067 | 0 | 0.14 | 0.29 | 0.08 | 0.03 | 0.13 | 0.1 |
| 9 | 113779011 | rs1030006 | -0.542 | 0.622 | -0.999 | -0.747 | 0.001 | -0.004 | -0.004 | 0 | 0.45 | 0.56 | 0.43 | 0.33 | 0.45 | 0.43 |
| 9 | 113782852 | rs12353038 | 1.184 | 0.649 | -0.988 | -0.735 | -0.014 | 0.013 | 0.013 | 0 | 0.41 | 0.15 | 0.49 | 0.64 | 0.43 | 0.46 |
| 9 | 113783435 | rs723591 | 0.771 | 0.928 | -0.795 | -0.068 | 0.006 | -0.013 | -0.013 | 0 | 0.56 | 0.53 | 0.57 | 0.67 | 0.56 | 0.58 |
| 9 | 113784557 | rs10980694 | 0.258 | 1.051 | -0.971 | -0.043 | -0.028 | -0.042 | -0.042 | 2.802 | 0.09 | 0.01 | 0.2 | 0.19 | 0.1 | 0.06 |
| 9 | 113785983 | rs7039431 | 0.163 | 0.766 | -0.7 | -0.058 | 0.001 | -0.006 | -0.006 | 0 | 0.55 | 0.45 | 0.57 | 0.7 | 0.55 | 0.57 |
| 9 | 113791313 | rs2287005 | -0.972 | 0.651 | -0.665 | 0.036 | -0.003 | -0.017 | -0.017 | 0 | 0.06 | 0.1 | 0.05 | 0.03 | 0.03 | 0.1 |
| 9 | 113792706 | rs3739707 | 0.078 | 0.25 | -0.763 | -0.063 | 0 | 0 | 0 | 0 | 0.3 | 0.25 | 0.34 | 0.62 | 0.26 | 0.18 |
| 9 | 113794500 | rs1000115 | 0.385 | 0.736 | -0.907 | -1.049 | 0.008 | -0.069 | -0.069 | 0 | 0.28 | 0.39 | 0.32 | 0.13 | 0.3 | 0.3 |
| 9 | 113798729 | rs7041855 | -0.309 | 1.111 | -1.186 | -1.076 | 0 | -0.017 | -0.017 | 0 | 0.42 | 0.46 | 0.59 | 0.38 | 0.55 | 0.46 |

| | | | | | | | | | | | | | | | | |
|---|---|---|---|---|---|---|---|---|---|---|---|---|---|---|---|---|
| 9 | 113803185 | rs10980705 | 0.296 | 1.161 | -1.145 | -0.914 | -0.071 | -0.057 | -0.057 | 0 | 0.13 | 0.02 | 0.25 | 0.25 | 0.24 | 0.15 |
| 9 | 113811445 | rs4978434 | 0.981 | 1.511 | -1.191 | -0.819 | -0.032 | -0.031 | -0.031 | 0 | 0.05 | 0 | 0.05 | 0.15 | 0.11 | 0.07 |
| 9 | 113815164 | rs4576485 | -0.445 | 0.877 | -1.404 | -1.134 | 0.035 | 0.11 | 0.11 | 0 | 0.77 | 0.94 | 0.89 | 0.94 | 0.84 | 0.69 |
| 9 | 113815456 | rs7035625 | -0.94 | 1.183 | -0.628 | -0.64 | 0.102 | 0.052 | 0.052 | 0.003 | 0.18 | 0.41 | 0.29 | 0.29 | 0.27 | 0.19 |
| 9 | 113828759 | rs1008157 | 0.233 | 1.041 | -1.274 | -1.224 | -0.02 | 0.058 | 0.058 | 0 | 0.18 | 0.05 | 0.16 | 0 | 0.23 | 0.08 |
| 9 | 113831319 | rs4978979 | -1.132 | 0.697 | -1.066 | -1.884 | 0.004 | -0.013 | -0.013 | 3.585 | 0.6 | 0.61 | 0.77 | 0.89 | 0.71 | 0.69 |
| 9 | 113831644 | rs4246886 | -0.805 | 0.859 | -1.229 | -1.268 | 0 | 0.002 | 0.002 | 0 | 0.43 | 0.57 | 0.68 | 0.69 | 0.57 | 0.43 |
| 9 | 113831845 | rs10980720 | 0.541 | 1.29 | -1.403 | -0.524 | -0.082 | -0.077 | -0.077 | 0 | 0.13 | 0.01 | 0.26 | 0.29 | 0.23 | 0.24 |
| 9 | 113834438 | rs1978304 | 0.206 | 1.065 | -0.188 | -0.494 | 0.023 | 0.068 | 0.068 | 0 | 0.26 | 0.38 | 0.41 | 0.4 | 0.34 | 0.19 |
| 9 | 113837805 | rs10980729 | -0.156 | 1.213 | 0.367 | 0.313 | 0.001 | 0.013 | 0.013 | 1.079 | 0.24 | 0.26 | 0.36 | 0.58 | 0.25 | 0.37 |
| 9 | 113837908 | rs10759484 | -0.139 | 1.204 | 0.336 | 0.2 | -0.001 | -0.005 | -0.005 | 0 | 0.24 | 0.24 | 0.35 | 0.62 | 0.25 | 0.36 |
| 9 | 113838083 | rs10817138 | 1.771 | 0.894 | 0.316 | -0.441 | 0.042 | 0.096 | 0.096 | 0 | 0.4 | 0.24 | 0.22 | 0.13 | 0.32 | 0.36 |
| 9 | 113838967 | rs7873139 | 0.104 | 0.825 | 0.895 | -0.43 | 0.012 | -0.01 | -0.01 | 0 | 0.64 | 0.58 | 0.58 | 0.72 | 0.57 | 0.67 |
| 9 | 113844435 | rs2900522 | 1.113 | 1.022 | 0.617 | -0.499 | -0.081 | -0.077 | -0.077 | 0 | 0.13 | 0.01 | 0.26 | 0.27 | 0.22 | 0.24 |
| 9 | 113846467 | rs7851332 | 1.802 | 0.895 | 0.681 | -0.639 | 0.027 | 0.072 | 0.072 | 0 | 0.38 | 0.24 | 0.22 | 0.09 | 0.32 | 0.34 |
| 9 | 113848849 | rs7863097 | -1.012 | 0.84 | 0.685 | -0.634 | -0.002 | -0.088 | -0.088 | 0 | 0.79 | 0.71 | 0.85 | 1 | 0.79 | 0.92 |
| 9 | 113850877 | rs7857633 | 1.697 | 0.709 | 0.426 | -0.539 | 0.036 | 0.101 | 0.101 | 0 | 0.38 | 0.22 | 0.22 | 0.09 | 0.32 | 0.35 |
| 9 | 113852849 | rs10980755 | 0.143 | 0.794 | 0.524 | -0.043 | 0.003 | 0.011 | 0.011 | 0.343 | 0.24 | 0.24 | 0.35 | 0.63 | 0.26 | 0.33 |
| 9 | 113853766 | rs10980757 | 0.503 | 0.575 | 0.378 | -0.1 | -0.018 | -0.035 | -0.035 | 0 | 0.06 | 0.14 | 0.03 | 0 | 0.03 | 0.01 |
| 9 | 113857337 | rs7861175 | 0.659 | 0.956 | 0.381 | 0.035 | 0.005 | -0.05 | -0.05 | 0 | 0.17 | 0.22 | 0.14 | 0.02 | 0.19 | 0.08 |
| 9 | 113857380 | rs10491532 | 0.158 | 0.917 | 0.096 | 0.576 | -0.046 | -0.058 | -0.058 | 0 | 0.13 | 0.07 | 0.25 | 0.26 | 0.18 | 0.22 |
| 9 | 113858933 | rs1002607 | -0.959 | 0.491 | 0.467 | 0.71 | -0.007 | -0.002 | -0.002 | 0.066 | 0.53 | 0.49 | 0.61 | 0.55 | 0.59 | 0.51 |
| 9 | 113864119 | rs7852442 | -0.16 | 0.801 | 0.171 | 0.899 | -0.042 | -0.048 | -0.048 | 0.131 | 0.85 | 0.95 | 0.83 | 0.77 | 0.8 | 0.78 |
| 9 | 113864545 | rs9299191 | -0.549 | 0.513 | 0.416 | 0.8 | -0.006 | 0.02 | 0.02 | 0 | 0.65 | 0.77 | 0.63 | 0.72 | 0.63 | 0.74 |
| 9 | 113864749 | rs1476946 | -1.234 | 0.309 | 0.954 | 0.587 | -0.003 | 0.001 | 0.001 | 0 | 0.6 | 0.57 | 0.61 | 0.52 | 0.66 | 0.55 |
| 9 | 113870526 | rs11793292 | 0.212 | 0.382 | 0.892 | 0.487 | -0.018 | 0.058 | 0.058 | 0 | 0.38 | 0.24 | 0.42 | 0.11 | 0.46 | 0.3 |
| 9 | 113880073 | rs872069 | 0.08 | 0.608 | 0.845 | 1.123 | 0.001 | 0.006 | 0.006 | 0 | 0.15 | 0.15 | 0.17 | 0.29 | 0.19 | 0.23 |
| 9 | 113880116 | rs872068 | 0.409 | 0.599 | 0.839 | 1.146 | -0.075 | -0.011 | -0.011 | 0.206 | 0.13 | 0.01 | 0.15 | 0.12 | 0.19 | 0.22 |
| 9 | 113890977 | rs1888191 | 1.231 | 0.385 | 0.479 | 0.702 | -0.054 | 0.092 | 0.092 | 0 | 0.25 | 0.07 | 0.34 | 0.07 | 0.33 | 0.19 |
| 9 | 113892963 | rs1411424 | -0.715 | 0.261 | 1.218 | 0.963 | 0.004 | 0.002 | 0.002 | 0 | 0.49 | 0.48 | 0.56 | 0.43 | 0.52 | 0.46 |
| 9 | 113893460 | rs11789772 | -1.845 | 0.2 | 1.199 | 0.567 | 0.017 | -0.051 | -0.051 | 0 | 0.67 | 0.81 | 0.67 | 0.44 | 0.73 | 0.58 |
| 9 | 113894066 | rs10491882 | 0.33 | 0.193 | 1.157 | 0.51 | -0.01 | 0.001 | 0.001 | 0.003 | 0.36 | 0.49 | 0.41 | 0.32 | 0.33 | 0.24 |
| 9 | 113901309 | rs2418135 | -0.384 | -0.209 | 1.183 | 0.393 | 0.005 | 0.022 | 0.022 | 0 | 0.5 | 0.48 | 0.56 | 0.59 | 0.52 | 0.5 |
| 9 | 113902881 | rs12346499 | -0.382 | -0.223 | 1.176 | 0.385 | 0.005 | 0.027 | 0.027 | 0 | 0.5 | 0.48 | 0.56 | 0.63 | 0.52 | 0.5 |
| 9 | 113905451 | rs2026801 | -0.792 | -0.251 | 1.253 | 0.357 | -0.002 | -0.003 | -0.003 | 0 | 0.55 | 0.52 | 0.6 | 0.63 | 0.64 | 0.59 |
| 9 | 113911613 | rs7870066 | -1.75 | -0.203 | 1.224 | -0.394 | 0.014 | 0.008 | 0.008 | 0 | 0.67 | 0.79 | 0.51 | 0.24 | 0.72 | 0.55 |
| 9 | 113915905 | rs10980800 | -0.966 | -0.215 | 1.505 | -0.435 | 0.001 | 0.006 | 0.006 | 0 | 0.21 | 0.23 | 0.13 | 0.09 | 0.22 | 0.16 |
| 9 | 113919523 | rs10217157 | -1.363 | -0.238 | 1.815 | -0.413 | 0.003 | 0.003 | 0.003 | 0 | 0.62 | 0.35 | 0.48 | 0.24 | 0.71 | 0.54 |
| 9 | 113928554 | rs7848895 | 0.935 | 0.07 | 1.604 | -0.437 | -0.065 | 0.005 | 0.005 | 0 | 0.13 | 0.01 | 0.14 | 0.09 | 0.16 | 0.24 |
| 9 | 113932556 | rs17206709 | 1.137 | 0.216 | 1.566 | -0.478 | -0.04 | 0.031 | 0.031 | 0 | 0.09 | 0.01 | 0.12 | 0.03 | 0.11 | 0.17 |
| 9 | 113933206 | rs10980811 | 0.35 | 0.002 | 1.646 | -0.531 | 0.012 | 0.092 | 0.092 | 0 | 0.37 | 0.13 | 0.22 | 0.12 | 0.34 | 0.21 |
| 9 | 113940924 | rs16936387 | -0.582 | -0.062 | 1.789 | -0.587 | 0 | 0.001 | 0.001 | 0.044 | 0.5 | 0.51 | 0.35 | 0.14 | 0.55 | 0.31 |
| 9 | 113944432 | rs7871113 | -0.417 | -0.093 | 1.685 | -0.624 | -0.07 | 0.004 | 0.004 | 0.521 | 0.14 | 0.22 | 0.04 | 0.09 | 0.02 | 0.07 |
| 9 | 113949141 | rs7020211 | -2.351 | -0.076 | 1.709 | -0.647 | 0.003 | 0.036 | 0.036 | 0 | 0.42 | 0.7 | 0.32 | 0.21 | 0.39 | 0.41 |
| 9 | 113954413 | rs12335533 | -1.014 | 0.249 | 1.239 | -0.507 | -0.014 | 0.012 | 0.012 | 0 | 0.26 | 0.3 | 0.27 | 0.12 | 0.37 | 0.33 |
| 9 | 113957534 | rs7046956 | -1.414 | -0.049 | 1.379 | -0.274 | -0.007 | -0.005 | -0.005 | 0 | 0.66 | 0.65 | 0.5 | 0.23 | 0.71 | 0.55 |
| 9 | 113958351 | rs2039183 | -0.61 | 0.43 | 0.816 | -0.137 | 0.036 | -0.042 | -0.042 | 0 | 0.09 | 0.17 | 0.11 | 0 | 0.2 | 0.09 |
| 9 | 113973091 | rs824002 | -1.591 | -0.267 | 0.26 | -0.008 | 0.026 | -0.065 | -0.065 | 0 | 0.7 | 0.86 | 0.53 | 0.38 | 0.76 | 0.61 |
| 9 | 113973553 | rs823999 | -1.06 | -0.504 | 0.812 | -0.044 | 0.004 | 0.001 | 0.001 | 0.037 | 0.64 | 0.65 | 0.48 | 0.3 | 0.74 | 0.59 |
| 9 | 113973650 | rs10980825 | -0.791 | -0.332 | 0.661 | -0.263 | -0.046 | -0.03 | -0.03 | 0 | 0.19 | 0.07 | 0.21 | 0.21 | 0.26 | 0.3 |
| 9 | 113976743 | rs700131 | -0.791 | -0.332 | 0.661 | -0.258 | -0.046 | -0.028 | -0.028 | 0 | 0.19 | 0.08 | 0.2 | 0.21 | 0.26 | 0.3 |
| 9 | 113979182 | rs1433586 | -0.791 | -0.332 | 0.661 | -0.261 | -0.046 | -0.03 | -0.03 | 0 | 0.19 | 0.08 | 0.21 | 0.21 | 0.26 | 0.3 |
| 9 | 113983555 | rs2570380 | -0.791 | -0.332 | 0.661 | -0.261 | -0.046 | -0.03 | -0.03 | 0.196 | 0.19 | 0.08 | 0.2 | 0.21 | 0.26 | 0.3 |
| 9 | 113984805 | rs1330272 | -1.229 | -0.442 | 0.643 | -0.315 | 0.001 | 0 | 0 | 0 | 0.65 | 0.65 | 0.49 | 0.3 | 0.75 | 0.59 |
| 9 | 113989160 | rs7861100 | 0.533 | -0.292 | 1.078 | -0.352 | 0.025 | 0.002 | 0.002 | 0 | 0.07 | 0.16 | 0.04 | 0.08 | 0.1 | 0.08 |
| 9 | 113990853 | rs1836108 | -1.399 | -0.252 | 0.789 | -0.321 | 0.004 | 0.001 | 0.001 | 0.061 | 0.64 | 0.65 | 0.48 | 0.3 | 0.75 | 0.59 |
| 9 | 113991207 | rs2570387 | -1.829 | -0.178 | 0.497 | -0.291 | 0.012 | -0.02 | -0.02 | 0.044 | 0.69 | 0.75 | 0.51 | 0.38 | 0.76 | 0.61 |
| 9 | 113992598 | rs2570388 | -0.763 | -0.067 | -0.004 | -0.688 | -0.046 | -0.026 | -0.026 | 0 | 0.19 | 0.08 | 0.2 | 0.2 | 0.26 | 0.3 |
| 9 | 113993269 | rs10980833 | 1.383 | 0.022 | -0.072 | -0.679 | -0.034 | -0.023 | -0.023 | 0 | 0.07 | 0 | 0.1 | 0.06 | 0.09 | 0.12 |
| 9 | 113995017 | rs7036572 | 1.767 | -0.243 | -0.054 | -0.846 | -0.075 | 0.119 | 0.119 | 0 | 0.26 | 0.08 | 0.21 | 0 | 0.38 | 0.17 |
| 9 | 113997789 | rs700137 | -0.249 | -0.125 | -0.116 | -0.676 | -0.061 | -0.022 | -0.022 | 1.315 | 0.16 | 0.04 | 0.2 | 0.16 | 0.26 | 0.28 |
| 9 | 114002747 | rs824012 | -1.858 | -0.206 | -0.166 | -0.669 | 0.018 | -0.042 | -0.042 | 0 | 0.7 | 0.86 | 0.53 | 0.34 | 0.76 | 0.62 |
| 9 | 114003466 | rs17813724 | 0.166 | -0.123 | 0.052 | -0.784 | 0.001 | -0.033 | -0.033 | 0 | 0.11 | 0.13 | 0.08 | 0.01 | 0.12 | 0.1 |
| 9 | 114008711 | rs11788536 | 1.423 | 0.014 | -0.098 | -0.714 | -0.025 | 0.03 | 0.03 | 0 | 0.28 | 0.17 | 0.21 | 0 | 0.38 | 0.17 |
| 9 | 114021014 | rs700144 | -1.427 | -0.141 | -0.458 | -0.625 | 0.028 | -0.004 | -0.004 | 0 | 0.68 | 0.81 | 0.52 | 0.3 | 0.76 | 0.6 |
| 9 | 114027937 | rs12346444 | 1.485 | 0.257 | -0.676 | -0.594 | -0.088 | 0.149 | 0.149 | 0 | 0.26 | 0.06 | 0.2 | 0 | 0.38 | 0.17 |
| 9 | 114030915 | rs10980841 | 0.622 | 0.545 | -0.341 | -0.199 | -0.053 | -0.049 | -0.049 | 0 | 0.1 | 0.01 | 0.13 | 0.14 | 0.14 | 0.18 |
| 9 | 114038704 | rs534643 | -0.686 | 0.331 | 0.046 | -0.273 | -0.012 | -0.017 | -0.017 | 0.017 | 0.57 | 0.47 | 0.44 | 0.26 | 0.67 | 0.5 |
| 9 | 114040326 | rs691720 | -1.385 | 0.277 | -1.015 | -0.219 | 0.048 | -0.023 | -0.023 | 0.121 | 0.68 | 0.82 | 0.53 | 0.34 | 0.77 | 0.6 |

| | | | | | | | | | | | | | | | | |
|---|---|---|---|---|---|---|---|---|---|---|---|---|---|---|---|---|
| 9 | 114044933 | rs690722 | 0.775 | 1.082 | -0.95 | -0.289 | -0.084 | 0.164 | 0.164 | 0.061 | 0.28 | 0.07 | 0.21 | 0 | 0.37 | 0.15 |
| 9 | 114046614 | rs10980847 | 0.091 | 0.967 | -0.556 | 0.373 | -0.044 | -0.049 | -0.049 | 0 | 0.08 | 0.01 | 0.08 | 0.17 | 0.12 | 0.18 |
| 9 | 114050357 | rs10980854 | 0.061 | 0.549 | -0.123 | 0.935 | -0.016 | 0.179 | 0.179 | 8.25 | 0.1 | 0.19 | 0.32 | 0.48 | 0.05 | 0.13 |
| 9 | 114050929 | rs10980855 | -1.104 | 0.549 | -0.076 | 0.96 | 0 | 0.048 | 0.048 | 0.737 | 0.18 | 0.28 | 0.43 | 0.72 | 0.15 | 0.3 |
| 9 | 114052520 | rs700148 | 0.684 | 0.368 | -0.752 | 0.413 | -0.124 | 0.214 | 0.214 | 0.317 | 0.34 | 0.08 | 0.25 | 0.01 | 0.46 | 0.19 |
| 9 | 114056169 | rs695207 | -0.811 | 0.407 | -0.103 | 0.68 | 0.025 | 0.04 | 0.04 | 0 | 0.59 | 0.67 | 0.72 | 0.75 | 0.64 | 0.59 |
| 9 | 114059341 | rs10980857 | -0.84 | 0.539 | 0.315 | 0.961 | 0.06 | 0.044 | 0.044 | 0.024 | 0.08 | 0.4 | 0.06 | 0.03 | 0.04 | 0.11 |
| 9 | 114069273 | rs4978437 | -0.116 | 0.87 | -0.879 | 1.084 | -0.035 | -0.035 | -0.035 | 0 | 0.07 | 0.01 | 0.11 | 0.24 | 0.1 | 0.16 |
| 9 | 114078716 | rs2095108 | 0.142 | 0.541 | -0.886 | 0.663 | 0.035 | 0.03 | 0.03 | 0 | 0.54 | 0.65 | 0.45 | 0.36 | 0.36 | 0.52 |
| 9 | 114087402 | rs12347972 | -0.563 | 0.266 | -0.347 | 0.689 | 0.049 | 0.001 | 0.001 | 0.005 | 0.51 | 0.32 | 0.57 | 0.5 | 0.63 | 0.5 |
| 9 | 114087696 | rs866624 | -1.091 | -0.193 | -0.391 | 0.69 | 0.026 | 0.004 | 0.004 | 0 | 0.6 | 0.87 | 0.64 | 0.66 | 0.64 | 0.52 |
| 9 | 114119984 | rs2418157 | -0.396 | 0.243 | 0.378 | 0.516 | -0.001 | 0.024 | 0.024 | 0 | 0.48 | 0.44 | 0.33 | 0.35 | 0.5 | 0.4 |
| 9 | 114125835 | rs2297524 | -1.704 | 0.26 | 0.719 | 0.612 | 0.014 | 0.011 | 0.011 | 0 | 0.63 | 0.82 | 0.63 | 0.68 | 0.65 | 0.56 |
| 9 | 114129847 | rs4978992 | -0.592 | 0.485 | 0.629 | 0.528 | -0.026 | 0.103 | 0.103 | 0.132 | 0.14 | 0.32 | 0.11 | 0.35 | 0.08 | 0.15 |
| 9 | 114134198 | rs10980884 | 0.504 | 0.523 | 0.492 | 0.497 | -0.114 | 0.235 | 0.235 | 0.448 | 0.33 | 0.05 | 0.21 | 0 | 0.41 | 0.25 |
| 9 | 114138702 | rs2274898 | 1.129 | 0.523 | 0.501 | 0.518 | -0.02 | -0.039 | -0.039 | 1.153 | 0.07 | 0 | 0.08 | 0.19 | 0.07 | 0.11 |
| 9 | 114161657 | rs2418159 | -0.361 | 0.508 | 0.546 | 0.524 | -0.016 | 0.058 | 0.058 | 0 | 0.46 | 0.3 | 0.31 | 0.35 | 0.49 | 0.4 |
| 9 | 114173689 | rs3818719 | -0.024 | 0.509 | 0.781 | 0.567 | 0.029 | 0.101 | 0.101 | 0 | 0.13 | 0.18 | 0.28 | 0.33 | 0.16 | 0.15 |
| 9 | 114184108 | rs7038720 | -0.118 | 0.505 | 0.69 | 0.557 | -0.013 | 0.066 | 0.066 | 0.488 | 0.12 | 0.22 | 0.1 | 0.36 | 0.08 | 0.15 |
| 9 | 114185794 | rs1041576 | 0.36 | 0.722 | 0.257 | 0.25 | -0.141 | 0.236 | 0.236 | 0.286 | 0.36 | 0.09 | 0.27 | 0 | 0.5 | 0.26 |
| 9 | 114190337 | rs16916100 | 1.033 | 0.498 | 0.343 | 0.454 | -0.017 | -0.039 | -0.039 | 0 | 0.07 | 0 | 0.08 | 0.18 | 0.07 | 0.11 |
| 9 | 114191098 | rs12345793 | 1.034 | 1.06 | -0.012 | 0.107 | -0.073 | 0.125 | 0.125 | 0.45 | 0.19 | 0.01 | 0.14 | 0 | 0.26 | 0.15 |
| 9 | 114195587 | rs2297530 | 1.03 | 0.656 | 0.312 | 0.362 | -0.017 | -0.04 | -0.04 | 0 | 0.07 | 0 | 0.08 | 0.2 | 0.07 | 0.11 |
| 9 | 114220603 | rs10759497 | -1.14 | 0.699 | 0.398 | 0.258 | 0.014 | -0.002 | -0.002 | 0 | 0.53 | 0.64 | 0.54 | 0.5 | 0.59 | 0.45 |
| 9 | 114279497 | rs12003641 | 0.491 | 0.722 | 0.378 | 0.448 | 0 | 0.115 | 0.115 | 0 | 0.11 | 0.26 | 0.26 | 0.25 | 0.08 | 0.13 |
| 9 | 114287275 | rs9299199 | -0.626 | 0.539 | -0.225 | 0.223 | 0.001 | -0.02 | -0.02 | 0 | 0.47 | 0.58 | 0.33 | 0.26 | 0.51 | 0.37 |
| 9 | 114287818 | rs7851745 | -2.234 | 0.331 | -0.543 | -0.114 | 0.011 | -0.021 | -0.021 | 0 | 0.69 | 0.91 | 0.68 | 0.68 | 0.71 | 0.63 |
| 9 | 114301585 | rs10441737 | -1.044 | 0.333 | -0.503 | -0.106 | -0.007 | -0.002 | -0.002 | 0 | 0.57 | 0.56 | 0.42 | 0.42 | 0.63 | 0.5 |
| 9 | 114305576 | rs10759501 | -2.05 | 0.439 | -1.367 | -0.477 | 0.012 | -0.021 | -0.021 | 0 | 0.69 | 0.9 | 0.68 | 0.68 | 0.71 | 0.63 |
| 9 | 114316077 | rs12341420 | -0.26 | 0.862 | -1.248 | -0.868 | -0.03 | 0.076 | 0.076 | 0 | 0.35 | 0.25 | 0.24 | 0 | 0.43 | 0.26 |
| 9 | 114318911 | rs7027778 | -0.221 | 0.901 | -1.501 | -0.849 | -0.049 | 0.129 | 0.129 | 0 | 0.34 | 0.19 | 0.23 | 0 | 0.43 | 0.26 |
| 9 | 114335885 | rs10122842 | -0.079 | 1.025 | -1.597 | -0.731 | -0.093 | 0.191 | 0.191 | 0 | 0.32 | 0.08 | 0.21 | 0.02 | 0.4 | 0.24 |
| 9 | 114341033 | rs17320021 | -0.078 | 1.137 | -1.032 | -0.277 | 0.01 | -0.024 | -0.024 | 0.907 | 0.05 | 0.12 | 0.06 | 0 | 0.05 | 0.03 |
| 9 | 114341376 | rs17320055 | -0.292 | 1.392 | -1.591 | -0.143 | -0.049 | 0.008 | 0.008 | 0 | 0.28 | 0.13 | 0.22 | 0.27 | 0.34 | 0.29 |
| 9 | 114342146 | rs10491726 | 1.065 | 1.136 | -1.701 | -0.359 | -0.024 | 0.043 | 0.043 | 0 | 0.08 | 0.01 | 0.05 | 0 | 0.13 | 0.02 |
| 9 | 114342378 | rs10817189 | -0.654 | 1.149 | -0.744 | -0.107 | 0.013 | -0.034 | -0.034 | 0 | 0.32 | 0.5 | 0.35 | 0.2 | 0.34 | 0.33 |
| 9 | 114343376 | rs12555931 | 0.117 | 1.189 | -0.24 | -0.13 | 0.002 | -0.005 | -0.005 | 0 | 0.23 | 0.25 | 0.26 | 0.18 | 0.24 | 0.24 |
| 9 | 114348513 | rs2273786 | -0.827 | 1.154 | 0.06 | -0.176 | 0.005 | -0.021 | -0.021 | 0 | 0.33 | 0.55 | 0.34 | 0.18 | 0.33 | 0.33 |
| 9 | 114348617 | rs2273788 | 0.044 | 1.141 | 0.066 | -0.163 | 0.007 | -0.011 | -0.011 | 0 | 0.24 | 0.32 | 0.26 | 0.18 | 0.24 | 0.24 |
| 9 | 114352627 | rs6477828 | -1.102 | 0.945 | -0.487 | 0.111 | -0.001 | 0 | 0 | 0.012 | 0.42 | 0.57 | 0.45 | 0.41 | 0.4 | 0.45 |
| 9 | 114356367 | rs3739704 | -1.682 | 0.489 | -0.649 | -0.301 | 0.003 | 0.004 | 0.004 | 0 | 0.58 | 0.58 | 0.59 | 0.66 | 0.57 | 0.64 |
| 9 | 114363693 | rs10122597 | -0.622 | 0.269 | 0.031 | 0.259 | 0.033 | -0.058 | -0.058 | 0 | 0.1 | 0.38 | 0.09 | 0 | 0.09 | 0.1 |
| 9 | 114365046 | rs1475106 | -1.803 | 0.203 | -0.206 | 0.202 | -0.011 | -0.007 | -0.007 | 0 | 0.4 | 0.83 | 0.51 | 0.64 | 0.35 | 0.44 |
| 9 | 114366248 | rs966628 | -1.45 | 0.396 | -0.012 | 0.231 | -0.021 | -0.05 | -0.05 | 0 | 0.27 | 0.83 | 0.53 | 0.43 | 0.22 | 0.25 |
| 9 | 114371624 | rs12342716 | -1.105 | 0.423 | 0.149 | 0.298 | 0.036 | 0.09 | 0.09 | 0 | 0.24 | 0.44 | 0.5 | 0.45 | 0.23 | 0.24 |
| 9 | 114374729 | rs16916245 | -0.025 | 0.121 | -0.156 | 0.389 | -0.007 | -0.025 | -0.025 | 0.055 | 0.21 | 0.18 | 0.35 | 0.44 | 0.19 | 0.19 |
| 9 | 114382444 | rs10817197 | -1.641 | 0.014 | -0.177 | 0.404 | 0 | 0 | 0 | 0.233 | 0.47 | 0.56 | 0.66 | 0.61 | 0.4 | 0.44 |
| 9 | 114384386 | rs7036413 | -0.064 | 0.285 | -0.805 | 0.526 | -0.036 | -0.046 | -0.046 | 0.044 | 0.24 | 0.12 | 0.39 | 0.43 | 0.26 | 0.21 |
| 9 | 114385055 | rs10759503 | -0.853 | -0.027 | -0.463 | 0.38 | -0.016 | 0 | 0 | 0 | 0.14 | 0.18 | 0.12 | 0.17 | 0.07 | 0.16 |
| 9 | 114386599 | rs10121085 | -0.837 | -0.16 | -0.461 | 0.39 | -0.021 | 0.001 | 0.001 | 0 | 0.14 | 0.17 | 0.12 | 0.17 | 0.07 | 0.16 |
| 9 | 114393235 | rs2065099 | -1.188 | -0.164 | -0.769 | 0.39 | 0.021 | -0.031 | -0.031 | 0.78 | 0.34 | 0.75 | 0.47 | 0.49 | 0.28 | 0.29 |
| 9 | 114400998 | rs1570303 | 0.645 | -0.175 | -0.448 | 0.661 | 0.003 | -0.022 | -0.022 | 0 | 0.16 | 0.12 | 0.22 | 0.39 | 0.16 | 0.06 |
| 9 | 114406649 | rs2146079 | -1.391 | -0.184 | -0.567 | 0.56 | 0.028 | -0.025 | -0.025 | 0.8 | 0.36 | 0.76 | 0.49 | 0.52 | 0.29 | 0.34 |
| 9 | 114409550 | rs4978455 | -0.313 | -0.217 | 0.285 | 0.555 | -0.016 | 0.072 | 0.072 | 0 | 0.23 | 0.35 | 0.43 | 0.41 | 0.23 | 0.16 |
| 9 | 114416298 | rs1051423 | 0.926 | -0.068 | 0.241 | 0.655 | -0.031 | -0.11 | -0.11 | 1.056 | 0.15 | 0.01 | 0.22 | 0.39 | 0.16 | 0.06 |
| 9 | 114416802 | rs12553893 | -0.313 | 0.165 | -0.041 | 0.362 | -0.004 | -0.007 | -0.007 | 0 | 0.3 | 0.26 | 0.46 | 0.45 | 0.33 | 0.37 |
| 9 | 114417082 | rs16916271 | -0.296 | 0.145 | -0.146 | 0.252 | -0.009 | 0.027 | 0.027 | 0 | 0.13 | 0.41 | 0.06 | 0.11 | 0.07 | 0.19 |
| 9 | 114421608 | rs4979028 | -0.99 | 0.166 | -0.53 | 0.178 | 0.001 | 0.01 | 0.01 | 0 | 0.39 | 0.47 | 0.48 | 0.59 | 0.32 | 0.51 |
| 9 | 114421679 | rs4979030 | -1.829 | 0.528 | -0.582 | -0.011 | 0.017 | 0.091 | 0.091 | 0 | 0.6 | 0.75 | 0.78 | 0.92 | 0.63 | 0.65 |
| 9 | 114423433 | rs4979031 | -0.513 | 0.5 | -0.55 | 0.131 | -0.01 | 0.041 | 0.041 | 0 | 0.33 | 0.54 | 0.47 | 0.45 | 0.29 | 0.31 |
| 9 | 114426645 | rs12376497 | -0.402 | 0.741 | -0.181 | 0.502 | 0.007 | 0.044 | 0.044 | 0 | 0.29 | 0.55 | 0.45 | 0.47 | 0.27 | 0.31 |
| 9 | 114435668 | rs4979034 | -0.748 | 0.545 | -0.937 | 0.522 | 0.007 | -0.015 | -0.015 | 0 | 0.33 | 0.7 | 0.42 | 0.46 | 0.31 | 0.35 |
| 9 | 114444565 | rs7863832 | -0.791 | 0.493 | -1.269 | 0.171 | 0.003 | 0.143 | 0.143 | 0 | 0.49 | 0.82 | 0.64 | 0.79 | 0.45 | 0.43 |
| 9 | 114447531 | rs16916293 | 0.647 | 0.292 | -1.737 | -0.154 | -0.029 | -0.05 | -0.05 | 1.734 | 0.11 | 0 | 0.05 | 0.12 | 0.1 | 0.25 |
| 9 | 114449970 | rs6477842 | 0.057 | 0.059 | -0.458 | 0.391 | -0.001 | 0.003 | 0.003 | 0 | 0.42 | 0.46 | 0.59 | 0.61 | 0.41 | 0.3 |
| 9 | 114453006 | rs10981012 | 1.181 | -0.213 | -0.712 | 0.527 | 0.018 | -0.057 | -0.057 | 0 | 0.14 | 0.02 | 0.24 | 0.28 | 0.1 | 0.17 |
| 9 | 114458600 | rs10981014 | -0.47 | -0.265 | 0.656 | 0.426 | 0.006 | 0.123 | 0.123 | 0 | 0.17 | 0.41 | 0.08 | 0.29 | 0.14 | 0.28 |
| 9 | 114462322 | rs6477845 | -2.247 | -0.307 | 0.165 | -0.399 | -0.001 | -0.013 | -0.013 | 1.421 | 0.79 | 0.8 | 0.87 | 0.93 | 0.8 | 0.79 |
| 9 | 114464487 | rs1407390 | -2.214 | -0.324 | 0.525 | -0.407 | 0.002 | 0.025 | 0.025 | 1.825 | 0.79 | 0.86 | 0.87 | 0.93 | 0.8 | 0.79 |

| | | | | | | | | | | | | | | | | |
|---|---|---|---|---|---|---|---|---|---|---|---|---|---|---|---|---|
| 9 | 114464733 | rs10981022 | 0.498 | -0.367 | 0.634 | -0.449 | -0.04 | -0.049 | -0.049 | 0 | 0.39 | 0.13 | 0.56 | 0.61 | 0.4 | 0.3 |
| 9 | 114468966 | rs7036568 | 0.873 | -0.439 | 0.7 | -0.45 | 0.002 | -0.038 | -0.038 | 0.478 | 0.22 | 0.33 | 0.22 | 0.05 | 0.25 | 0.21 |
| 9 | 114481631 | rs9657640 | 1.298 | -0.404 | 0.779 | -0.614 | -0.007 | 0.035 | 0.035 | 0 | 0.19 | 0.15 | 0.2 | 0.03 | 0.23 | 0.14 |
| 9 | 114482980 | rs12683870 | -0.473 | -0.426 | 0.829 | -0.637 | 0.009 | 0.127 | 0.127 | 0 | 0.54 | 0.25 | 0.63 | 0.88 | 0.55 | 0.58 |
| 9 | 114484682 | rs1337581 | -2.257 | -0.365 | 0.618 | -0.599 | 0 | -0.018 | -0.018 | 0 | 0.79 | 0.79 | 0.87 | 0.93 | 0.79 | 0.79 |
| 9 | 114484783 | rs12352352 | 1.301 | -0.362 | 0.573 | -0.598 | -0.007 | 0.035 | 0.035 | 0 | 0.19 | 0.14 | 0.2 | 0.03 | 0.23 | 0.14 |
| 9 | 114489887 | rs953536 | -0.68 | -0.351 | 0.566 | -0.579 | 0 | -0.018 | -0.018 | 0.074 | 0.57 | 0.45 | 0.65 | 0.88 | 0.55 | 0.58 |
| 9 | 114542451 | rs823639 | -1.016 | -0.211 | 0.694 | -0.621 | -0.005 | 0.041 | 0.041 | 0 | 0.6 | 0.67 | 0.73 | 0.9 | 0.56 | 0.65 |
| 9 | 114542761 | rs823638 | -2.185 | -0.169 | 0.729 | -0.551 | 0 | -0.004 | -0.004 | 0 | 0.79 | 0.81 | 0.86 | 0.93 | 0.79 | 0.8 |
| 9 | 114545878 | rs559248 | -2.199 | -0.168 | 0.643 | -0.55 | -0.004 | 0.036 | 0.036 | 1.393 | 0.79 | 0.87 | 0.87 | 0.93 | 0.79 | 0.8 |
| 9 | 114557475 | rs823645 | 1.899 | -0.262 | 0.343 | -0.434 | 0.006 | -0.024 | -0.024 | 0 | 0.08 | 0 | 0.03 | 0.08 | 0.04 | 0.19 |
| 9 | 114561679 | rs10981078 | 1.184 | -0.154 | 0.37 | -0.588 | 0.003 | 0.068 | 0.068 | 0 | 0.2 | 0.11 | 0.14 | 0.03 | 0.21 | 0.16 |
| 9 | 114562289 | rs446465 | -0.744 | -0.191 | 0.58 | -0.682 | -0.001 | 0.026 | 0.026 | 0 | 0.57 | 0.62 | 0.71 | 0.9 | 0.56 | 0.64 |
| 9 | 114563069 | rs10981079 | 1.235 | -0.164 | 0.391 | -0.665 | 0.004 | 0.059 | 0.059 | 0 | 0.2 | 0.12 | 0.14 | 0.03 | 0.21 | 0.16 |
| 9 | 114567760 | rs667463 | -0.509 | -0.2 | 0.61 | -0.649 | 0.01 | 0.099 | 0.099 | 0 | 0.54 | 0.3 | 0.68 | 0.88 | 0.55 | 0.57 |
| 9 | 114572011 | rs7875230 | 1.129 | -0.107 | 0.483 | -0.653 | 0.003 | 0.02 | 0.02 | 0 | 0.21 | 0.15 | 0.14 | 0.02 | 0.23 | 0.16 |
| 9 | 114572203 | rs10981083 | 1.328 | -0.223 | 0.349 | -0.419 | -0.031 | -0.03 | -0.03 | 0.703 | 0.23 | 0.14 | 0.32 | 0.33 | 0.29 | 0.11 |
| 9 | 114577770 | rs7045809 | 0.225 | -0.372 | 0.08 | -0.265 | -0.002 | 0.001 | 0.001 | 0 | 0.22 | 0.25 | 0.27 | 0.43 | 0.15 | 0.23 |
| 9 | 114582649 | rs700049 | -2.119 | -0.219 | 0.009 | -0.363 | -0.013 | 0.034 | 0.034 | 1.141 | 0.79 | 0.88 | 0.86 | 0.93 | 0.78 | 0.8 |
| 9 | 114591865 | rs7036787 | 0.981 | -0.056 | 0.169 | 0 | 0 | 0.029 | 0.029 | 0 | 0.12 | 0.13 | 0.3 | 0.43 | 0.11 | 0.19 |
| 9 | 114593311 | rs10981100 | 1.136 | 0.29 | -0.36 | -0.304 | -0.088 | -0.037 | -0.037 | 0 | 0.15 | 0.01 | 0.12 | 0.12 | 0.24 | 0.12 |
| 9 | 114593488 | rs7042137 | -1.232 | -0.09 | -0.359 | -0.664 | 0.012 | 0.189 | 0.189 | 0 | 0.48 | 0.74 | 0.7 | 0.88 | 0.51 | 0.48 |
| 9 | 114599048 | rs4978466 | -1.996 | -0.227 | -0.514 | -0.769 | 0.029 | 0.159 | 0.159 | 0.859 | 0.64 | 0.88 | 0.81 | 0.91 | 0.71 | 0.61 |
| 9 | 114606411 | rs16916411 | 1.258 | -0.251 | -0.321 | -0.727 | 0.004 | 0.02 | 0.02 | 0.004 | 0.15 | 0.13 | 0.09 | 0.02 | 0.16 | 0.13 |
| 9 | 114607816 | rs4979054 | -0.259 | -0.346 | 0.204 | -0.664 | -0.014 | 0.067 | 0.067 | 0 | 0.2 | 0.31 | 0.34 | 0.45 | 0.14 | 0.27 |
| 9 | 114612458 | rs12376643 | 0.061 | -0.214 | -0.139 | -0.651 | -0.065 | 0.045 | 0.045 | 0 | 0.3 | 0.14 | 0.21 | 0.15 | 0.39 | 0.25 |
| 9 | 114615980 | rs10817227 | -1.468 | -0.246 | -0.139 | -0.645 | 0.016 | 0.006 | 0.006 | 1.238 | 0.52 | 0.79 | 0.5 | 0.48 | 0.55 | 0.41 |
| 9 | 114621854 | rs10981126 | 0.636 | 0.184 | 0.342 | -0.342 | 0.031 | 0.033 | 0.033 | 0 | 0.17 | 0.22 | 0.13 | 0.27 | 0.23 | 0.14 |
| 9 | 114627170 | rs315712 | -0.259 | -0.104 | 0.421 | -0.324 | -0.007 | 0.005 | 0.005 | 0 | 0.81 | 0.8 | 0.67 | 0.69 | 0.87 | 0.75 |
| 9 | 114629162 | rs12379047 | -0.071 | 0.275 | 0.695 | -0.138 | 0.07 | 0.222 | 0.222 | 0 | 0.16 | 0.36 | 0.25 | 0.46 | 0.2 | 0.14 |
| 9 | 114636015 | rs12001972 | -0.103 | 0.228 | 0.507 | -0.424 | 0.012 | -0.016 | -0.016 | 0 | 0.22 | 0.5 | 0.15 | 0.18 | 0.23 | 0.19 |
| 9 | 114638922 | rs10121681 | 0.731 | 0.218 | 0.525 | -0.447 | -0.006 | -0.034 | -0.034 | 0 | 0.17 | 0.21 | 0.1 | 0.02 | 0.16 | 0.13 |
| 9 | 114639407 | rs13295839 | 0.459 | 0.168 | 0.253 | -0.5 | -0.014 | -0.044 | -0.044 | 0 | 0.26 | 0.16 | 0.41 | 0.41 | 0.27 | 0.31 |
| 9 | 114639652 | rs7023375 | -2.835 | -0.048 | 0.159 | -0.68 | 0.004 | 0.126 | 0.126 | 0.13 | 0.68 | 0.9 | 0.82 | 0.92 | 0.7 | 0.66 |
| 9 | 114641291 | rs7043162 | -1.081 | 0.051 | 0.359 | -0.573 | 0 | -0.005 | -0.005 | 0 | 0.44 | 0.56 | 0.67 | 0.74 | 0.44 | 0.44 |
| 9 | 114643893 | rs2418192 | -1.58 | 0.048 | 0.675 | -0.519 | 0.001 | 0.006 | 0.006 | 0.225 | 0.58 | 0.64 | 0.76 | 0.76 | 0.6 | 0.58 |
| 9 | 114645077 | rs10981136 | 1.294 | 0.016 | 0.039 | -0.363 | -0.005 | 0.007 | 0.007 | 0 | 0.13 | 0.15 | 0.3 | 0.29 | 0.07 | 0.21 |
| 9 | 114654601 | rs9299201 | 0.193 | 0.223 | 0.381 | -0.201 | 0.005 | 0.023 | 0.023 | 0.547 | 0.25 | 0.31 | 0.31 | 0.34 | 0.3 | 0.21 |
| 9 | 114658407 | rs1413218 | -1.754 | 0.209 | -0.118 | -0.395 | 0.024 | 0.151 | 0.151 | 0.546 | 0.63 | 0.87 | 0.8 | 0.92 | 0.68 | 0.64 |
| 9 | 114661611 | rs7042346 | 0.482 | 0.593 | -0.041 | -0.004 | -0.001 | -0.004 | -0.004 | 0 | 0.27 | 0.25 | 0.42 | 0.39 | 0.27 | 0.3 |
| 9 | 114668639 | rs12554037 | 1.667 | 0.619 | -0.294 | 0.143 | -0.04 | -0.039 | -0.039 | 0 | 0.08 | 0.02 | 0.2 | 0.32 | 0.14 | 0.07 |
| 9 | 114672325 | rs10759516 | -0.576 | 0.498 | 0.08 | -0.062 | -0.006 | -0.006 | -0.006 | 0 | 0.38 | 0.66 | 0.5 | 0.62 | 0.39 | 0.39 |
| 9 | 114672865 | rs16916456 | -0.298 | 0.465 | 0.422 | -0.065 | 0.027 | 0.103 | 0.103 | 0.03 | 0.1 | 0.32 | 0.07 | 0.2 | 0.08 | 0.09 |
| 9 | 114688103 | rs1813202 | -2.145 | 0.217 | 0.024 | -0.234 | -0.001 | 0.189 | 0.189 | 0.343 | 0.63 | 0.94 | 0.8 | 0.93 | 0.67 | 0.67 |
| 9 | 114690765 | rs3824464 | 0.156 | 0.669 | 0.446 | 0.12 | 0.006 | -0.016 | -0.016 | 0 | 0.3 | 0.43 | 0.44 | 0.31 | 0.3 | 0.27 |
| 9 | 114694486 | rs7850023 | 0.438 | 0.615 | 0.484 | 0.106 | 0.005 | -0.011 | -0.011 | 0 | 0.27 | 0.38 | 0.43 | 0.3 | 0.28 | 0.29 |
| 9 | 114699654 | rs7873908 | -0.431 | 0.729 | 0.629 | 0.011 | -0.004 | 0.007 | 0.007 | 0 | 0.37 | 0.39 | 0.5 | 0.28 | 0.39 | 0.41 |
| 9 | 114699757 | rs7860208 | -2.071 | 0.789 | 0.736 | -0.057 | 0.016 | 0.065 | 0.065 | 0.098 | 0.57 | 0.76 | 0.76 | 0.94 | 0.6 | 0.68 |
| 9 | 114699872 | rs7860334 | 0.59 | 0.883 | 0.505 | 0.102 | -0.001 | 0.001 | 0.001 | 0 | 0.07 | 0.08 | 0.07 | 0.01 | 0.09 | 0.14 |
| 9 | 114700584 | rs10981165 | 0.208 | 0.928 | 0.633 | 0.297 | 0.118 | 0.252 | 0.252 | 2.494 | 0.05 | 0.31 | 0.04 | 0.17 | 0.05 | 0.03 |
| 9 | 114701630 | rs7041171 | -0.081 | 0.791 | 0.453 | 0.23 | -0.004 | 0.016 | 0.016 | 0.183 | 0.22 | 0.25 | 0.25 | 0.37 | 0.2 | 0.16 |
| 9 | 114702514 | rs7854170 | -0.01 | 0.813 | 0.263 | 0.154 | -0.01 | -0.016 | -0.016 | 0 | 0.34 | 0.28 | 0.58 | 0.6 | 0.4 | 0.37 |
| 9 | 114703029 | rs10125079 | -1.309 | 0.882 | 0.301 | 0.289 | 0.009 | 0.162 | 0.162 | 0 | 0.55 | 0.75 | 0.72 | 0.91 | 0.57 | 0.56 |
| 9 | 114703106 | rs10817241 | -0.419 | 0.893 | 0.34 | 0.355 | -0.004 | 0 | 0 | 0 | 0.44 | 0.38 | 0.63 | 0.6 | 0.47 | 0.4 |
| 9 | 114703215 | rs10125129 | 0.036 | 0.9 | 0.364 | 0.38 | 0.103 | 0.279 | 0.279 | 2.17 | 0.06 | 0.32 | 0.04 | 0.21 | 0.05 | 0.05 |
| 9 | 114703347 | rs10981168 | -0.45 | 0.861 | 0.031 | 0.314 | -0.004 | 0 | 0 | 0 | 0.44 | 0.38 | 0.63 | 0.6 | 0.47 | 0.4 |
| 9 | 114705137 | rs7031902 | -2.049 | 0.986 | -0.204 | 0.415 | -0.06 | -0.101 | -0.101 | 0 | 0.6 | 0.93 | 0.65 | 0.56 | 0.58 | 0.62 |
| 9 | 114705265 | rs7046626 | -0.845 | 0.869 | 0.224 | 0.277 | 0.001 | 0.002 | 0.002 | 0 | 0.44 | 0.47 | 0.52 | 0.32 | 0.39 | 0.4 |
| 9 | 114706396 | rs16916492 | -0.874 | 1.121 | 0.466 | 0.853 | 0.052 | 0.155 | 0.155 | 0.09 | 0.08 | 0.29 | 0.08 | 0.22 | 0.1 | 0.09 |
| 9 | 114706412 | rs7848697 | -0.599 | 0.728 | 0.039 | 0.187 | 0.003 | 0.004 | 0.004 | 0.14 | 0.2 | 0.23 | 0.12 | 0.13 | 0.17 | 0.24 |
| 9 | 114707212 | rs10817246 | -0.852 | 0.926 | 0.089 | 0.248 | 0.011 | 0.012 | 0.012 | 0 | 0.41 | 0.35 | 0.42 | 0.35 | 0.36 | 0.49 |
| 9 | 114711743 | rs419097 | 1.16 | 1.131 | -0.131 | 0.214 | -0.001 | -0.003 | -0.003 | 0.617 | 0.49 | 0.49 | 0.5 | 0.62 | 0.46 | 0.41 |
| 9 | 114714853 | rs170282 | 1.493 | 1.169 | 0.342 | 0.525 | 0.018 | -0.029 | -0.029 | 0 | 0.79 | 0.67 | 0.84 | 0.9 | 0.74 | 0.73 |
| 9 | 114721455 | rs13299128 | -0.69 | 0.797 | 0.167 | 0.622 | -0.002 | -0.009 | -0.009 | 1.746 | 0.1 | 0.1 | 0.13 | 0 | 0.16 | 0.03 |
| 9 | 114721483 | rs2900535 | 1.582 | 0.667 | 0.036 | 1.843 | -0.007 | -0.026 | -0.026 | 0.577 | 0.82 | 0.85 | 0.84 | 0.59 | 0.79 | 0.8 |
| 9 | 114721683 | rs1889224 | 2.358 | 0.374 | 0.966 | 2.029 | -0.007 | 0.043 | 0.043 | 0 | 0.73 | 0.58 | 0.79 | 0.41 | 0.75 | 0.69 |
| 9 | 114721737 | rs1889223 | -0.537 | 0.384 | 1.321 | 2.166 | 0.024 | 0.059 | 0.059 | 0 | 0.07 | 0.16 | 0.3 | 0.14 | 0.1 | 0.07 |
| 9 | 114724494 | rs4979067 | 3.127 | 0.134 | 1.224 | 1.814 | -0.005 | 0.11 | 0.11 | 0 | 0.58 | 0.16 | 0.45 | 0.27 | 0.62 | 0.51 |
| 9 | 114727220 | rs4978473 | 2.572 | 0.219 | 1.753 | 1.803 | -0.017 | 0.026 | 0.026 | 0 | 0.72 | 0.45 | 0.63 | 0.44 | 0.8 | 0.57 |

| | | | | | | | | | | | | | | | |
|---|---|---|---|---|---|---|---|---|---|---|---|---|---|---|---|
| 9 | 114731867 | rs10733581 | 2.485 | 0.085 | 1.809 | 2.54 | 0.029 | 0.064 | 0.064 | 0.061 | 0.73 | 0.53 | 0.52 | 0.65 | 0.67 | 0.71 |
| 9 | 114732385 | rs12005421 | -1.434 | -0.262 | 1.723 | 2.246 | 0.006 | 0.003 | 0.003 | 0 | 0.11 | 0.23 | 0.32 | 0.09 | 0.12 | 0.13 |
| 9 | 114733502 | rs6477867 | -2.547 | -0.28 | 1.534 | 2.177 | 0.003 | -0.062 | -0.062 | 0.265 | 0.28 | 0.6 | 0.49 | 0.09 | 0.31 | 0.29 |
| 9 | 114736410 | rs2418194 | 2.785 | -0.187 | 1.285 | 2.368 | -0.005 | -0.008 | -0.008 | 0 | 0.59 | 0.1 | 0.47 | 0.44 | 0.63 | 0.48 |
| 9 | 114747481 | rs12000633 | -2.294 | -0.076 | 1.651 | 1.928 | -0.004 | 0.013 | 0.013 | 0 | 0.28 | 0.68 | 0.47 | 0.13 | 0.31 | 0.18 |
| 9 | 114749320 | rs2805481 | -1.984 | 0.234 | 2.02 | 2.553 | 0.001 | -0.003 | -0.003 | 0 | 0.29 | 0.65 | 0.19 | 0.43 | 0.19 | 0.32 |
| 9 | 114753347 | rs12376779 | -0.672 | 0.222 | 1.914 | 2.062 | 0.004 | 0.049 | 0.049 | 2.497 | 0.17 | 0.29 | 0.15 | 0.23 | 0.14 | 0.15 |
| 9 | 114758073 | rs4979071 | -2.31 | 0.407 | 1.833 | 1.648 | -0.006 | 0.031 | 0.031 | 1.942 | 0.37 | 0.71 | 0.5 | 0.27 | 0.37 | 0.38 |
| 9 | 114759313 | rs4979072 | -2.64 | 0.557 | 1.648 | 1.642 | 0.025 | -0.021 | -0.021 | 0 | 0.19 | 0.4 | 0.37 | 0.12 | 0.24 | 0.22 |
| 9 | 114764995 | rs12004156 | -1.417 | 0.453 | 1.63 | 1.6 | 0.002 | -0.036 | -0.036 | 0.535 | 0.11 | 0.34 | 0.06 | 0.04 | 0.09 | 0.16 |
| 9 | 114765400 | rs13299968 | -0.978 | 0.641 | 1.175 | 1.848 | -0.043 | -0.033 | -0.033 | 0.001 | 0.07 | 0.01 | 0.31 | 0.09 | 0.16 | 0.07 |
| 9 | 114768887 | rs7859911 | -0.59 | 0.437 | 1.508 | 1.65 | 0.008 | 0.001 | 0.001 | 6.023 | 0.16 | 0.29 | 0.15 | 0.16 | 0.14 | 0.17 |
| 9 | 114777275 | rs10981208 | -0.694 | -0.052 | 1.478 | 2.504 | -0.004 | 0.01 | 0.01 | 0 | 0.15 | 0.12 | 0.06 | 0.36 | 0.08 | 0.19 |
| 9 | 114780290 | rs10981210 | -0.839 | 0.032 | 1.142 | 2.551 | -0.039 | -0.051 | -0.051 | 0 | 0.08 | 0.01 | 0.06 | 0.22 | 0.13 | 0.13 |
| 9 | 114791018 | rs2281787 | -1.652 | -0.077 | 1.785 | 2.356 | -0.009 | 0.095 | 0.095 | 0.035 | 0.07 | 0.28 | 0.17 | 0.13 | 0.02 | 0.23 |
| 9 | 114791150 | rs1324210 | -1.034 | -0.092 | 1.94 | 2.47 | -0.022 | 0.065 | 0.065 | 0 | 0.25 | 0.38 | 0.14 | 0.61 | 0.22 | 0.31 |
| 9 | 114798037 | rs10125054 | -0.561 | 0.142 | 1.549 | 2.327 | 0.002 | -0.004 | -0.004 | 0 | 0.31 | 0.29 | 0.3 | 0.42 | 0.26 | 0.35 |
| 9 | 114803286 | rs2281788 | -0.721 | 0.193 | 1.916 | 2.584 | -0.017 | -0.036 | -0.036 | 0.314 | 0.07 | 0.01 | 0.04 | 0.27 | 0.08 | 0.19 |
| 9 | 114803631 | rs2281790 | -2.653 | 0.235 | 2.486 | 2.679 | 0.018 | 0.047 | 0.047 | 0.01 | 0.14 | 0.28 | 0.31 | 0.23 | 0.2 | 0.36 |
| 9 | 114807641 | rs7854368 | -2.988 | 0.097 | 2.645 | 2.633 | 0.002 | -0.004 | -0.004 | 0 | 0.25 | 0.75 | 0.2 | 0.41 | 0.19 | 0.36 |
| 9 | 114810102 | rs10981228 | -0.63 | 0.127 | 2.529 | 2.54 | -0.019 | -0.039 | -0.039 | 0 | 0.07 | 0.01 | 0.04 | 0.27 | 0.09 | 0.2 |
| 9 | 114814828 | rs7034341 | -2.77 | 0.148 | 2.824 | 2.427 | 0.045 | 0.001 | 0.001 | 4.8 | 0.12 | 0.56 | 0.26 | 0.1 | 0.16 | 0.25 |
| 9 | 114821568 | rs16916623 | -0.624 | 0.155 | 1.816 | 2.52 | -0.019 | -0.043 | -0.043 | 0 | 0.07 | 0.01 | 0.04 | 0.23 | 0.09 | 0.19 |
| 9 | 114823121 | rs4979078 | -2.907 | 0.176 | 2.026 | 2.085 | -0.005 | 0.091 | 0.091 | 1.432 | 0.17 | 0.57 | 0.32 | 0.24 | 0.15 | 0.31 |
| 9 | 114823298 | rs4979079 | -1.534 | 0.241 | 1.188 | 1.945 | -0.005 | -0.009 | -0.009 | 0 | 0.07 | 0.06 | 0.19 | 0.17 | 0.1 | 0.25 |
| 9 | 114823991 | rs10981242 | -1.673 | -0.54 | 1.325 | 1.777 | -0.036 | 0.046 | 0.046 | 0.001 | 0.09 | 0.37 | 0.04 | 0.08 | 0.01 | 0.06 |
| 9 | 114828332 | rs10817260 | -1.239 | 0.024 | 0.292 | 1.801 | -0.017 | 0.045 | 0.045 | 1.716 | 0.22 | 0.65 | 0.31 | 0.34 | 0.2 | 0.47 |
| 9 | 114832050 | rs2762495 | -1.209 | 0.17 | -0.155 | 2.004 | 0.001 | 0.002 | 0.002 | 0 | 0.13 | 0.16 | 0.1 | 0.26 | 0.1 | 0.17 |
| 9 | 114832919 | rs2762496 | -1.413 | -0.043 | -0.009 | 2.221 | 0.001 | 0.002 | 0.002 | 0 | 0.13 | 0.16 | 0.19 | 0.24 | 0.09 | 0.17 |
| 9 | 114837321 | rs2043057 | 0.187 | -0.881 | -1.134 | 2.054 | 0.022 | -0.033 | -0.033 | 0 | 0.25 | 0.04 | 0.18 | 0.28 | 0.25 | 0.23 |
| 9 | 114837337 | rs10116017 | -1.589 | 0.078 | 0.013 | 1.649 | 0.024 | 0.035 | 0.035 | 0 | 0.2 | 0.43 | 0.2 | 0.23 | 0.22 | 0.2 |
| 9 | 114839679 | rs7862743 | -0.696 | 0.121 | 0.064 | 1.577 | 0.113 | -0.011 | -0.011 | 0.009 | 0.06 | 0.27 | 0.06 | 0 | 0.12 | 0.05 |
| 9 | 114843415 | rs2782942 | -1.598 | -0.277 | 0.09 | 1.323 | -0.055 | 0.116 | 0.116 | 0 | 0.6 | 0.81 | 0.69 | 0.81 | 0.56 | 0.63 |
| 9 | 114844254 | rs3849119 | -0.286 | -0.125 | 0.274 | 1.391 | 0.131 | -0.006 | -0.006 | 0.178 | 0.06 | 0.27 | 0.06 | 0 | 0.12 | 0.05 |
| 9 | 114844883 | rs9408834 | -1.006 | 0.313 | 0.145 | 1.335 | -0.035 | -0.017 | -0.017 | 0 | 0.21 | 0.15 | 0.24 | 0.24 | 0.32 | 0.2 |
| 9 | 114846751 | rs10981252 | 0.043 | 0.303 | 0.135 | 1.349 | -0.038 | -0.08 | -0.08 | 0 | 0.13 | 0.01 | 0.17 | 0.25 | 0.17 | 0.16 |
| 9 | 114850190 | rs2782931 | -0.981 | 0.294 | 0.195 | 1.316 | -0.02 | -0.009 | -0.009 | 0 | 0.21 | 0.19 | 0.25 | 0.23 | 0.32 | 0.19 |
| 9 | 114856721 | rs7032451 | -1.752 | 0.011 | 0.087 | 1.164 | -0.046 | 0.095 | 0.095 | 0.116 | 0.6 | 0.82 | 0.7 | 0.79 | 0.58 | 0.65 |
| 9 | 114858776 | rs3849120 | -0.621 | -0.009 | 0.282 | 1.205 | -0.006 | 0.002 | 0.002 | 0 | 0.43 | 0.45 | 0.5 | 0.56 | 0.4 | 0.48 |
| 9 | 114868037 | rs12686374 | -0.124 | -0.011 | 0.219 | 1.525 | -0.032 | -0.025 | -0.025 | 0.18 | 0.06 | 0.02 | 0.19 | 0.06 | 0.16 | 0.05 |
| 9 | 114870301 | rs12338324 | -0.154 | 0.052 | 0.413 | 1.581 | -0.001 | -0.001 | -0.001 | 0 | 0.4 | 0.54 | 0.34 | 0.45 | 0.28 | 0.43 |
| 9 | 114875617 | rs2476934 | -1.822 | -0.012 | 0.286 | 1.531 | -0.015 | 0.047 | 0.047 | 0 | 0.62 | 0.71 | 0.7 | 0.79 | 0.62 | 0.65 |
| 9 | 114877392 | rs2762472 | -0.962 | 0.116 | 0.363 | 1.701 | -0.019 | -0.021 | -0.021 | 0 | 0.22 | 0.17 | 0.3 | 0.35 | 0.34 | 0.22 |
| 9 | 114879755 | rs9408838 | -0.707 | -0.332 | 0.33 | 1.077 | 0.046 | 0.055 | 0.055 | 0 | 0.5 | 0.6 | 0.56 | 0.72 | 0.38 | 0.65 |
| 9 | 114888516 | rs2762475 | -0.712 | -0.326 | 0.85 | 1.112 | -0.002 | -0.002 | -0.002 | 0 | 0.47 | 0.5 | 0.5 | 0.67 | 0.39 | 0.65 |
| 9 | 114888773 | rs2177667 | -1.472 | -0.341 | 0.141 | 1.125 | -0.024 | 0.063 | 0.063 | 0 | 0.6 | 0.8 | 0.68 | 0.72 | 0.6 | 0.71 |
| 9 | 114902932 | rs7870176 | -1.601 | 0.14 | 0.702 | 1.626 | -0.004 | -0.005 | -0.005 | 0.362 | 0.35 | 0.61 | 0.45 | 0.47 | 0.42 | 0.26 |
| 9 | 114907131 | rs700098 | -0.359 | 0.299 | -0.088 | 1.737 | -0.041 | -0.12 | -0.12 | 0.673 | 0.14 | 0.01 | 0.17 | 0.42 | 0.19 | 0.15 |
| 9 | 114912752 | rs10817270 | -0.909 | 0.392 | 0.453 | 0.762 | 0.069 | -0.077 | -0.077 | 0 | 0.19 | 0.42 | 0.27 | 0.04 | 0.23 | 0.08 |
| 9 | 114917202 | rs2900537 | -1.255 | 0.042 | 0.019 | 1.262 | -0.008 | -0.011 | -0.011 | 0 | 0.36 | 0.63 | 0.41 | 0.38 | 0.41 | 0.24 |
| 9 | 114919707 | rs700105 | 0.422 | -2.02 | -0.05 | 0.635 | -0.01 | -0.011 | -0.011 | 0 | 0.77 | 0.79 | 0.7 | 0.9 | 0.85 | 0.82 |
| 9 | 114922067 | rs9886804 | 1.165 | -2.532 | -0.865 | 0.552 | -0.001 | 0 | 0 | 0.62 | 0.42 | 0.63 | 0.44 | 0.36 | 0.42 | 0.48 |
| 9 | 114924720 | rs10981291 | 0.542 | -2.737 | -1.384 | 0.95 | -0.032 | -0.065 | -0.065 | 0 | 0.14 | 0.01 | 0.09 | 0.24 | 0.2 | 0.17 |
| 9 | 114934114 | rs10739335 | -0.524 | -2.696 | -1.26 | 1.405 | -0.014 | -0.09 | -0.09 | 0.004 | 0.82 | 0.98 | 0.84 | 0.68 | 0.77 | 0.85 |
| 9 | 114934532 | rs10817277 | -0.6 | -2.921 | -1.326 | 1.197 | 0.032 | 0.032 | 0.032 | 0.119 | 0.34 | 0.12 | 0.33 | 0.27 | 0.34 | 0.33 |
| 9 | 114938931 | rs10759529 | 1.012 | -2.869 | -0.442 | 1.232 | 0.02 | 0.091 | 0.091 | 0.117 | 0.47 | 0.79 | 0.4 | 0.33 | 0.45 | 0.51 |
| 9 | 114942200 | rs4979087 | 0.874 | -2.822 | -0.493 | 1.241 | 0.019 | 0.083 | 0.083 | 0.274 | 0.48 | 0.87 | 0.44 | 0.38 | 0.45 | 0.52 |
| 9 | 114960138 | rs10981305 | 1.068 | -2.861 | -0.311 | 1.387 | 0.008 | 0.04 | 0.04 | 0 | 0.45 | 0.7 | 0.4 | 0.34 | 0.47 | 0.5 |
| 9 | 114963346 | rs12554866 | 1.626 | -3.06 | 0.096 | 1.352 | -0.054 | 0.113 | 0.113 | 0 | 0.36 | 0.17 | 0.34 | 0.15 | 0.45 | 0.4 |
| 9 | 114973383 | rs7874580 | -0.114 | -3.109 | -0.232 | 1.692 | 0.03 | -0.038 | -0.038 | 0 | 0.33 | 0.06 | 0.4 | 0.35 | 0.32 | 0.4 |
| 9 | 114976485 | rs10817291 | -0.126 | -3.109 | -0.231 | 1.693 | 0.034 | -0.037 | -0.037 | 0.135 | 0.33 | 0.06 | 0.4 | 0.35 | 0.32 | 0.4 |
| 9 | 114980967 | rs6575 | 0.778 | -3.138 | -0.086 | 1.761 | 0.025 | 0.055 | 0.055 | 0 | 0.49 | 0.77 | 0.45 | 0.39 | 0.47 | 0.52 |
| 9 | 114982463 | rs7855745 | 0.891 | -3.141 | -0.109 | 1.76 | 0.022 | 0.07 | 0.07 | 0 | 0.5 | 0.87 | 0.45 | 0.39 | 0.47 | 0.52 |
| 9 | 114982549 | rs7869523 | 0.906 | -3.153 | -0.115 | 1.754 | 0.023 | 0.071 | 0.071 | 0 | 0.5 | 0.87 | 0.45 | 0.39 | 0.47 | 0.52 |
| 9 | 114985497 | rs1534069 | -0.063 | -3.124 | -0.132 | 1.765 | -0.005 | -0.084 | -0.084 | 0 | 0.33 | 0.02 | 0.39 | 0.35 | 0.32 | 0.4 |
| 9 | 114987364 | rs7041185 | 0.72 | -3.187 | -0.123 | 1.77 | 0.03 | 0.066 | 0.066 | 0 | 0.5 | 0.8 | 0.45 | 0.39 | 0.47 | 0.52 |
| 9 | 114989616 | rs6477900 | -1.237 | -3.153 | -0.23 | 1.775 | -0.051 | -0.088 | -0.088 | 0.408 | 0.84 | 0.99 | 0.85 | 0.74 | 0.79 | 0.92 |
| 9 | 115011317 | rs1359878 | 0.956 | -3.15 | -0.155 | 1.759 | -0.017 | 0.036 | 0.036 | 0.087 | 0.47 | 0.9 | 0.46 | 0.39 | 0.47 | 0.52 |
| 9 | 115015710 | rs10817301 | -0.211 | -3.139 | -0.143 | 1.762 | 0.035 | -0.04 | -0.04 | 0 | 0.34 | 0.07 | 0.39 | 0.35 | 0.32 | 0.4 |

| | | | | | | | | | | | | | | | |
|---|---|---|---|---|---|---|---|---|---|---|---|---|---|---|---|
| 9 | 115023853 | rs7874660 | 1.395 | -3.025 | 0.089 | 1.717 | -0.003 | 0.008 | 0.008 | 0.043 | 0.42 | 0.64 | 0.39 | 0.31 | 0.45 | 0.48 |
| 9 | 115030570 | rs7870866 | 1.059 | -3.389 | 0.048 | 1.742 | 0.002 | 0.04 | 0.04 | 0.093 | 0.46 | 0.85 | 0.46 | 0.39 | 0.46 | 0.51 |
| 9 | 115035134 | rs10981337 | 1.183 | -3.624 | -0.126 | 1.755 | 0.033 | 0.009 | 0.009 | 0 | 0.19 | 0.02 | 0.3 | 0.09 | 0.13 | 0.21 |
| 9 | 115036021 | rs10981338 | 1.063 | -3.486 | -0.035 | 1.741 | -0.002 | 0.04 | 0.04 | 0.064 | 0.46 | 0.89 | 0.45 | 0.39 | 0.46 | 0.53 |
| 9 | 115039845 | rs7875779 | -1.098 | -3.476 | -0.096 | 1.851 | -0.019 | -0.055 | -0.055 | 0 | 0.83 | 0.95 | 0.85 | 0.75 | 0.79 | 0.93 |
| 9 | 115085206 | rs4310281 | -0.23 | -3.524 | 0.285 | 1.763 | 0.024 | -0.052 | -0.052 | 0 | 0.35 | 0.03 | 0.39 | 0.35 | 0.33 | 0.4 |
| 9 | 115088669 | rs4542000 | -1.219 | -3.324 | 0.365 | 1.809 | -0.049 | -0.093 | -0.093 | 0 | 0.84 | 0.99 | 0.85 | 0.74 | 0.79 | 0.93 |
| 9 | 115095743 | rs7040447 | -1.048 | -3.325 | 0.37 | 1.831 | -0.019 | -0.057 | -0.057 | 0.143 | 0.83 | 0.95 | 0.85 | 0.73 | 0.79 | 0.93 |
| 9 | 115103266 | rs10817319 | -0.768 | -3.067 | 0.702 | 1.656 | 0.011 | 0.004 | 0.004 | 0 | 0.4 | 0.33 | 0.42 | 0.39 | 0.34 | 0.43 |
| 9 | 115120143 | rs12685878 | -1.193 | -2.388 | 1.557 | 2.247 | -0.028 | 0.05 | 0.05 | 0 | 0.21 | 0.32 | 0.19 | 0.35 | 0.21 | 0.25 |
| 9 | 115137476 | rs7862690 | 1.667 | -2.423 | 1.649 | 2.508 | -0.026 | 0.066 | 0.066 | 0 | 0.73 | 0.46 | 0.76 | 0.45 | 0.72 | 0.71 |
| 9 | 115140133 | rs4979108 | -0.066 | -2.313 | 1.802 | 2.625 | -0.019 | -0.013 | -0.013 | 0 | 0.43 | 0.47 | 0.56 | 0.24 | 0.29 | 0.35 |
| 9 | 115140735 | rs10817330 | 0.029 | -2.221 | 1.597 | 2.855 | 0.003 | -0.001 | -0.001 | 0 | 0.56 | 0.57 | 0.41 | 0.57 | 0.64 | 0.63 |
| 9 | 115155264 | rs6477925 | -0.932 | -2.317 | 1.756 | 2.846 | 0.012 | -0.002 | -0.002 | 0 | 0.53 | 0.54 | 0.62 | 0.5 | 0.43 | 0.47 |
| 9 | 115156389 | rs7852741 | -0.368 | -2.28 | 1.728 | 2.916 | -0.004 | -0.019 | -0.019 | 3.905 | 0.12 | 0.06 | 0.1 | 0.21 | 0.18 | 0.11 |
| 9 | 115167146 | rs4978489 | 0.22 | -2.318 | 1.949 | 2.803 | 0.033 | 0.027 | 0.027 | 0 | 0.39 | 0.3 | 0.47 | 0.21 | 0.25 | 0.35 |
| 9 | 115170313 | rs1571639 | 0.27 | -2.325 | 1.946 | 2.725 | 0.026 | 0.024 | 0.024 | 0 | 0.38 | 0.3 | 0.47 | 0.19 | 0.25 | 0.35 |
| 9 | 115171986 | rs1537055 | 0.913 | -2.154 | 1.755 | 2.705 | 0.018 | 0.037 | 0.037 | 0 | 0.95 | 0.93 | 0.92 | 0.88 | 0.89 | 0.96 |
| 9 | 115178029 | rs10739349 | 0.921 | -2.136 | 1.713 | 2.722 | 0.001 | -0.001 | -0.001 | 0.36 | 0.44 | 0.47 | 0.35 | 0.49 | 0.46 | 0.51 |
| 9 | 115182612 | rs4979113 | -0.543 | -2.598 | 1.664 | 2.731 | 0.045 | 0.023 | 0.023 | 0.134 | 0.5 | 0.36 | 0.56 | 0.39 | 0.43 | 0.45 |
| 9 | 115193763 | rs7857000 | 0.816 | -2.404 | 1.243 | 2.346 | -0.051 | -0.073 | -0.073 | 0.514 | 0.86 | 0.98 | 0.86 | 0.76 | 0.77 | 0.87 |
| 9 | 115196114 | rs7038491 | 0.428 | -2.534 | 1.368 | 2.302 | -0.021 | -0.006 | -0.006 | 1.691 | 0.36 | 0.33 | 0.41 | 0.22 | 0.24 | 0.26 |
| 9 | 115196558 | rs10817341 | 0.334 | -2.53 | 1.387 | 2.325 | 0.036 | 0.028 | 0.028 | 0.031 | 0.5 | 0.67 | 0.45 | 0.47 | 0.52 | 0.61 |
| 9 | 115202068 | rs10759563 | 0.684 | -2.533 | 1.212 | 2.294 | 0.019 | 0.015 | 0.015 | 0.118 | 0.47 | 0.49 | 0.44 | 0.47 | 0.52 | 0.61 |
| 9 | 115205042 | rs10817344 | -0.378 | -2.531 | 1.197 | 2.315 | 0.011 | -0.005 | -0.005 | 0 | 0.61 | 0.69 | 0.55 | 0.67 | 0.7 | 0.71 |
| 9 | 115219619 | rs7862037 | 0.623 | -2.486 | 1.075 | 2.447 | -0.022 | -0.051 | -0.051 | 1.375 | 0.86 | 0.94 | 0.86 | 0.69 | 0.77 | 0.86 |
| 9 | 115234611 | rs7096 | 0.672 | -2.546 | 1.118 | 2.307 | 0.021 | 0.015 | 0.015 | 0.241 | 0.47 | 0.48 | 0.46 | 0.47 | 0.52 | 0.59 |
| 9 | 115265330 | rs4979125 | 0.421 | -2.536 | 1.081 | 2.211 | 0.004 | 0.006 | 0.006 | 0.327 | 0.36 | 0.28 | 0.38 | 0.19 | 0.25 | 0.25 |
| 9 | 115270889 | rs7018648 | 0.449 | -2.426 | 1.011 | 2.044 | 0.011 | 0.008 | 0.008 | 0.082 | 0.48 | 0.51 | 0.47 | 0.48 | 0.53 | 0.61 |
| 9 | 115289605 | rs4979128 | 0.269 | -2.285 | 1.089 | 2.099 | 0.042 | 0.027 | 0.027 | 0.039 | 0.38 | 0.24 | 0.34 | 0.2 | 0.26 | 0.31 |
| 9 | 115302544 | rs10759577 | 1.246 | -2.418 | 0.691 | 2.114 | -0.002 | 0.148 | 0.148 | 0 | 0.37 | 0.16 | 0.33 | 0.16 | 0.51 | 0.28 |
| 9 | 115316399 | rs10817357 | 0.595 | -1.891 | 0.868 | 2.196 | 0.006 | 0.003 | 0.003 | 0 | 0.48 | 0.58 | 0.49 | 0.5 | 0.54 | 0.55 |
| 9 | 115326187 | rs10981445 | 0.595 | -1.891 | 0.861 | 2.195 | 0.008 | 0.003 | 0.003 | 0 | 0.48 | 0.59 | 0.49 | 0.5 | 0.54 | 0.55 |
| 9 | 115335386 | rs2806320 | 1.377 | -1.89 | 0.608 | 2.173 | 0.016 | 0.157 | 0.157 | 0 | 0.83 | 0.54 | 0.8 | 0.7 | 0.79 | 0.86 |
| 9 | 115343222 | rs7040835 | 0.631 | -1.667 | 0.194 | 2.243 | 0.04 | 0.051 | 0.051 | 0 | 0.46 | 0.41 | 0.47 | 0.38 | 0.54 | 0.54 |
| 9 | 115347088 | rs2593684 | 0.154 | -2.013 | -0.264 | 1.59 | -0.041 | -0.022 | -0.022 | 0 | 0.4 | 0.38 | 0.36 | 0.19 | 0.26 | 0.3 |
| 9 | 115367078 | rs10739367 | 0.545 | -1.602 | -0.873 | 1.762 | 0.018 | 0.004 | 0.004 | 0 | 0.46 | 0.46 | 0.47 | 0.51 | 0.56 | 0.55 |
| 9 | 115371397 | rs2676629 | 1.253 | -0.019 | -1.12 | 1.94 | 0 | 0.001 | 0.001 | 0 | 0.89 | 0.93 | 0.85 | 0.7 | 0.84 | 0.87 |
| 9 | 115373695 | rs1517079 | -0.644 | -0.063 | -1.481 | 1.959 | 0 | -0.042 | -0.042 | 0.411 | 0.08 | 0.01 | 0.06 | 0.18 | 0.09 | 0.11 |
| 9 | 115399064 | rs2798317 | -0.782 | -0.187 | -1.676 | 0.96 | -0.003 | -0.008 | -0.008 | 0 | 0.2 | 0.01 | 0.09 | 0.17 | 0.17 | 0.25 |
| 9 | 115401368 | rs11788361 | -0.148 | 0.639 | -2.03 | 0.852 | -0.045 | -0.014 | -0.014 | 0.002 | 0.08 | 0.01 | 0.06 | 0.1 | 0.11 | 0.1 |
| 9 | 115403585 | rs2995806 | 0.607 | -0.488 | -1.096 | 1.413 | -0.042 | 0.109 | 0.109 | 0 | 0.76 | 0.58 | 0.75 | 0.48 | 0.83 | 0.6 |
| 9 | 115404746 | rs10739369 | -1.387 | -1.127 | -0.877 | 0.724 | 0.003 | 0 | 0 | 0 | 0.41 | 0.62 | 0.36 | 0.63 | 0.32 | 0.52 |
| 9 | 115416257 | rs786990 | -1.944 | -1.071 | -2.146 | 0.097 | -0.172 | -0.053 | -0.053 | 2.605 | 0.74 | 0.98 | 0.52 | 0.8 | 0.57 | 0.83 |
| 9 | 115416612 | rs1568125 | 0.636 | -1.378 | -1.606 | 0.823 | -0.02 | 0.093 | 0.093 | 0 | 0.16 | 0.29 | 0.24 | 0.37 | 0.14 | 0.11 |
| 9 | 115417171 | rs4979141 | -1.422 | -1.92 | -1.474 | 0.573 | 0.002 | -0.001 | -0.001 | 0 | 0.41 | 0.61 | 0.36 | 0.62 | 0.32 | 0.41 |
| 9 | 115421424 | rs4979142 | 0.702 | -1.998 | -1.462 | 0.466 | -0.033 | -0.004 | -0.004 | 0 | 0.1 | 0.23 | 0.03 | 0.1 | 0.03 | 0.12 |
| 9 | 115421788 | rs16917185 | 0.728 | -1.998 | -1.403 | 0.465 | -0.032 | -0.005 | -0.005 | 0.785 | 0.1 | 0.2 | 0.03 | 0.1 | 0.03 | 0.12 |
| 9 | 115422854 | rs7028901 | -1.669 | -1.972 | -1.412 | 0.609 | 0 | -0.001 | -0.001 | 0 | 0.43 | 0.59 | 0.35 | 0.68 | 0.32 | 0.59 |
| 9 | 115426603 | rs2040062 | -0.035 | -1.941 | -1.262 | 0.5 | -0.012 | 0.016 | 0.016 | 0.145 | 0.22 | 0.28 | 0.26 | 0.36 | 0.18 | 0.13 |
| 9 | 115429998 | rs12002812 | 0.482 | -1.746 | -1.123 | 0.629 | -0.033 | -0.001 | -0.001 | 0.047 | 0.1 | 0.2 | 0.03 | 0.1 | 0.03 | 0.12 |
| 9 | 115430157 | rs11998945 | 0.679 | -1.727 | -1.277 | 0.638 | 0.009 | 0.001 | 0.001 | 0 | 0.09 | 0.06 | 0.01 | 0.1 | 0.03 | 0.12 |
| 9 | 115430670 | rs7040287 | -0.818 | -1.57 | -1.264 | 0.523 | -0.009 | 0.014 | 0.014 | 0 | 0.37 | 0.59 | 0.35 | 0.53 | 0.3 | 0.33 |
| 9 | 115440165 | rs10759581 | -0.777 | -1.631 | -1.027 | 0.486 | -0.008 | 0.011 | 0.011 | 0 | 0.37 | 0.59 | 0.36 | 0.54 | 0.3 | 0.34 |
| 9 | 115444828 | rs786962 | -0.484 | -1.062 | -1.232 | 0.105 | 0.01 | 0 | 0 | 0 | 0.58 | 0.69 | 0.59 | 0.59 | 0.6 | 0.51 |
| 9 | 115446426 | rs2173196 | 1.257 | -1.172 | -1.377 | -0.094 | -0.009 | 0 | 0 | 0 | 0.18 | 0.07 | 0.28 | 0.16 | 0.17 | 0.13 |
| 9 | 115447935 | rs786963 | -0.148 | -0.947 | -1.018 | -0.3 | 0.002 | -0.054 | -0.054 | 0.283 | 0.22 | 0.36 | 0.16 | 0.08 | 0.22 | 0.21 |
| 9 | 115448341 | rs786966 | 0.609 | -0.284 | -0.906 | -0.363 | -0.006 | -0.045 | -0.045 | 0.364 | 0.12 | 0.18 | 0.06 | 0 | 0.09 | 0.06 |
| 9 | 115449110 | rs786968 | 1 | -0.099 | -0.744 | -0.273 | 0.008 | -0.054 | -0.054 | 1.097 | 0.09 | 0.26 | 0.07 | 0 | 0.09 | 0.06 |
| 9 | 115454504 | rs2676622 | -0.645 | -0.169 | -0.822 | 0.101 | 0.008 | -0.003 | -0.003 | 0 | 0.19 | 0.36 | 0.22 | 0.25 | 0.23 | 0.22 |
| 9 | 115460239 | rs786978 | 0.103 | -0.034 | -0.634 | 0.246 | -0.004 | 0.013 | 0.013 | 0 | 0.12 | 0.37 | 0.14 | 0.17 | 0.11 | 0.09 |
| 9 | 115462552 | rs2798316 | 0.582 | -0.053 | -0.645 | 0.251 | -0.001 | 0.013 | 0.013 | 0 | 0.11 | 0.25 | 0.13 | 0.17 | 0.11 | 0.09 |
| 9 | 115472230 | rs805050 | -0.342 | -0.17 | -0.819 | 0.195 | 0.013 | 0.004 | 0.004 | 0 | 0.32 | 0.46 | 0.49 | 0.41 | 0.39 | 0.35 |
| 9 | 115509156 | rs2173193 | 1.036 | -0.071 | -0.65 | 0.072 | 0.003 | -0.001 | -0.001 | 0 | 0.21 | 0.18 | 0.38 | 0.22 | 0.29 | 0.18 |
| 9 | 115517264 | rs2645986 | 0.342 | 0.071 | -0.998 | 0.07 | -0.013 | -0.019 | -0.019 | 0.836 | 0.46 | 0.48 | 0.23 | 0.22 | 0.33 | 0.51 |
| 9 | 115519675 | rs2796033 | -0.146 | 0.171 | -0.906 | 0.108 | 0.022 | -0.006 | -0.006 | 0 | 0.67 | 0.52 | 0.5 | 0.63 | 0.62 | 0.74 |
| 9 | 115519904 | rs2796032 | -0.836 | 0.109 | -0.933 | 0.103 | -0.003 | -0.004 | -0.004 | 1.207 | 0.72 | 0.65 | 0.6 | 0.7 | 0.72 | 0.79 |
| 9 | 115525259 | rs10981499 | 0.942 | -0.149 | -1.164 | -0.121 | -0.007 | 0.07 | 0.07 | 0 | 0.19 | 0.02 | 0.12 | 0.08 | 0.18 | 0.16 |
| 9 | 115527183 | rs1891403 | -1.032 | 0.803 | -0.411 | 0.561 | -0.005 | 0.001 | 0.001 | 0 | 0.89 | 0.78 | 0.89 | 0.86 | 0.9 | 0.92 |

| | | | | | | | | | | | | | | | | |
|---|---|---|---|---|---|---|---|---|---|---|---|---|---|---|---|---|
| 9 | 115531238 | rs7039744 | 0.795 | 0.911 | -0.429 | 0.553 | 0.013 | -0.003 | -0.003 | 0 | 0.22 | 0.26 | 0.39 | 0.22 | 0.29 | 0.18 |
| 9 | 115546442 | rs7044734 | 0.937 | 0.74 | -0.178 | 0.458 | 0.026 | 0.045 | 0.045 | 0 | 0.42 | 0.29 | 0.19 | 0.17 | 0.32 | 0.49 |
| 9 | 115549463 | rs787271 | -0.674 | 1.07 | -0.715 | 0.737 | 0.003 | 0.014 | 0.014 | 0 | 0.88 | 0.63 | 0.87 | 0.86 | 0.9 | 0.91 |
| 9 | 115550363 | rs787274 | -0.825 | 1.046 | -0.735 | 0.724 | 0.005 | 0.021 | 0.021 | 0 | 0.91 | 0.76 | 0.89 | 0.86 | 0.91 | 0.92 |
| 9 | 115558246 | rs10739371 | -0.007 | 1.096 | -0.722 | 0.753 | 0.029 | 0.009 | 0.009 | 0 | 0.86 | 0.75 | 0.81 | 0.79 | 0.79 | 0.87 |
| 9 | 115559659 | rs2153839 | 0.806 | 1.075 | -0.771 | 0.692 | 0.036 | -0.02 | -0.02 | 0.436 | 0.79 | 0.6 | 0.75 | 0.79 | 0.72 | 0.84 |
| 9 | 115585109 | rs12555600 | 1.72 | 1.254 | -0.935 | 0.862 | -0.005 | -0.009 | -0.009 | 0.029 | 0.16 | 0.06 | 0.25 | 0.17 | 0.15 | 0.09 |
| 9 | 115589251 | rs10120988 | 0.59 | 1.173 | -0.953 | 0.952 | 0.104 | 0.073 | 0.073 | 0 | 0.51 | 0.81 | 0.35 | 0.37 | 0.36 | 0.56 |
| 9 | 115593729 | rs1539339 | 1.063 | 1.66 | -0.762 | 1.087 | -0.108 | -0.146 | -0.146 | 0 | 0.2 | 0.01 | 0.28 | 0.39 | 0.29 | 0.24 |
| 9 | 115609941 | rs10817396 | 0.719 | 1.75 | -0.243 | 0.629 | 0.056 | 0.033 | 0.033 | 0 | 0.85 | 0.59 | 0.79 | 0.79 | 0.79 | 0.87 |
| 9 | 115616110 | rs1539341 | 1.366 | 1.811 | -0.255 | 0.546 | 0.011 | -0.013 | -0.013 | 0.557 | 0.77 | 0.71 | 0.75 | 0.79 | 0.73 | 0.85 |
| 9 | 115617291 | rs10481671 | 2.084 | 1.763 | 0.196 | 1.068 | 0.013 | 0.048 | 0.048 | 0 | 0.78 | 0.53 | 0.69 | 0.71 | 0.79 | 0.65 |
| 9 | 115617822 | rs10115576 | 0.486 | 1.723 | 0.127 | 0.938 | -0.015 | -0.02 | -0.02 | 0 | 0.87 | 0.76 | 0.87 | 0.89 | 0.91 | 0.92 |
| 9 | 115625699 | rs2182046 | 0.181 | 1.793 | 0.029 | 0.891 | 0.001 | 0.002 | 0.002 | 0 | 0.73 | 0.76 | 0.75 | 0.73 | 0.76 | 0.78 |
| 9 | 115627012 | rs4978508 | 1.107 | 1.854 | -0.182 | 0.95 | -0.103 | -0.143 | -0.143 | 0 | 0.17 | 0.01 | 0.31 | 0.49 | 0.3 | 0.22 |
| 9 | 115632898 | rs16917417 | 0.659 | 2.227 | 0.727 | 1.493 | -0.018 | -0.001 | -0.001 | 0 | 0.15 | 0.06 | 0.13 | 0.15 | 0.15 | 0.11 |
| 9 | 115632969 | rs2025544 | -0.231 | 2.193 | 0.774 | 1.477 | 0 | -0.01 | -0.01 | 0.212 | 0.92 | 0.95 | 0.97 | 1 | 0.93 | 0.94 |
| 9 | 115633479 | rs10116897 | 1.797 | 2.173 | 0.884 | 1.487 | -0.026 | 0.034 | 0.034 | 0 | 0.85 | 0.66 | 0.78 | 0.79 | 0.9 | 0.67 |
| 9 | 115634282 | rs1324931 | 0.33 | 2.217 | 0.809 | 1.384 | -0.002 | -0.025 | -0.025 | 0 | 0.91 | 0.93 | 0.96 | 1 | 0.92 | 0.94 |
| 9 | 115639748 | rs10759604 | 0.688 | 2.241 | 0.823 | 1.157 | -0.011 | -0.057 | -0.057 | 0 | 0.89 | 0.81 | 0.95 | 1 | 0.91 | 0.94 |
| 9 | 115641569 | rs16917440 | -0.567 | 2.305 | 0.677 | 1.196 | 0.034 | -0.007 | -0.007 | 0 | 0.06 | 0.1 | 0.07 | 0.06 | 0.11 | 0.02 |
| 9 | 115645184 | rs10156558 | 1.226 | 2.235 | 0.607 | 1.225 | 0.006 | -0.017 | -0.017 | 0 | 0.63 | 0.49 | 0.63 | 0.69 | 0.64 | 0.56 |
| 9 | 115654098 | rs4142852 | -1.093 | 2.455 | 0.566 | 1.283 | 0.017 | -0.004 | -0.004 | 0 | 0.31 | 0.4 | 0.52 | 0.75 | 0.34 | 0.47 |
| 9 | 115654246 | rs4978511 | -0.189 | 2.379 | 0.139 | 1.044 | 0 | 0.002 | 0.002 | 0 | 0.93 | 0.91 | 0.97 | 0.98 | 0.94 | 0.92 |
| 9 | 115654683 | rs10124939 | -0.561 | 2.489 | -0.041 | 1.105 | -0.009 | 0.021 | 0.021 | 0 | 0.94 | 0.98 | 0.98 | 0.98 | 0.94 | 0.92 |
| 9 | 115660368 | rs4978512 | 0.759 | 2.582 | 0.003 | 1.159 | -0.049 | 0.034 | 0.034 | 0 | 0.71 | 0.82 | 0.78 | 0.77 | 0.63 | 0.77 |
| 9 | 115666561 | rs10981540 | 0.613 | 2.601 | -0.044 | 1.306 | -0.009 | 0.071 | 0.071 | 0 | 0.86 | 0.8 | 0.66 | 0.41 | 0.9 | 0.63 |
| 9 | 115669613 | rs17751300 | 0.1 | 2.618 | -0.069 | 1.271 | -0.013 | 0.012 | 0.012 | 0 | 0.19 | 0.09 | 0.12 | 0.14 | 0.2 | 0.13 |
| 9 | 115669711 | rs912664 | 1.082 | 2.226 | -0.041 | 1.103 | -0.013 | 0.02 | 0.02 | 0 | 0.62 | 0.7 | 0.45 | 0.19 | 0.58 | 0.45 |
| 9 | 115673030 | rs912670 | 0.328 | 2.211 | -0.224 | 1.231 | -0.072 | 0.006 | 0.006 | 0 | 0.79 | 0.9 | 0.75 | 0.78 | 0.57 | 0.82 |
| 9 | 115674281 | rs1952004 | -1.273 | 0.432 | 0.382 | 1.584 | -0.033 | 0.273 | 0.273 | 0.524 | 0.1 | 0.35 | 0.21 | 0.44 | 0.04 | 0.26 |
| 9 | 115675673 | rs12377544 | -0.261 | 0.842 | -0.187 | 0.438 | 0 | 0.003 | 0.003 | 0 | 0.2 | 0.17 | 0.12 | 0.11 | 0.17 | 0.18 |
| 9 | 115677671 | rs11791620 | -0.213 | 2.034 | -0.252 | 0.959 | -0.049 | -0.049 | -0.049 | 0 | 0.08 | 0.02 | 0.15 | 0.19 | 0.26 | 0.06 |
| 9 | 115678033 | rs1925081 | -0.366 | 1.349 | -0.182 | 0.179 | -0.002 | -0.007 | -0.007 | 0.249 | 0.48 | 0.48 | 0.4 | 0.29 | 0.62 | 0.39 |
| 9 | 115678692 | rs10817404 | 0.461 | 1.251 | -0.944 | 0.13 | -0.02 | 0.144 | 0.144 | 0 | 0.32 | 0.09 | 0.18 | 0.09 | 0.31 | 0.32 |
| 9 | 115681924 | rs16917501 | 0.227 | 1.348 | -0.312 | 1.642 | 0.004 | 0.002 | 0.002 | 0 | 0.1 | 0.07 | 0.19 | 0.34 | 0.08 | 0.28 |
| 9 | 115682269 | rs11790986 | 0.219 | 1.595 | -0.307 | 1.123 | -0.042 | -0.044 | -0.044 | 1.504 | 0.07 | 0.01 | 0.13 | 0.2 | 0.22 | 0.06 |
| 9 | 115683429 | rs10759611 | -1.351 | 1.4 | -0.014 | 0.907 | 0.026 | 0.009 | 0.009 | 0 | 0.23 | 0.27 | 0.23 | 0.29 | 0.35 | 0.19 |
| 9 | 115684234 | rs10817407 | -1.374 | 1.39 | -0.059 | 0.9 | 0.053 | 0.021 | 0.021 | 0 | 0.23 | 0.32 | 0.23 | 0.29 | 0.35 | 0.19 |
| 9 | 115687192 | rs10759614 | -1.037 | 1.572 | -0.444 | 1.085 | 0.031 | 0.013 | 0.013 | 0 | 0.21 | 0.22 | 0.23 | 0.29 | 0.35 | 0.19 |
| 9 | 115687735 | rs10817408 | -0.53 | 0.24 | -1.09 | 1.29 | -0.015 | 0.003 | 0.003 | 0 | 0.11 | 0.03 | 0.08 | 0.09 | 0.13 | 0.14 |
| 9 | 115688494 | rs4571802 | -0.76 | -0.214 | -0.88 | 1.594 | 0.009 | -0.03 | -0.03 | 0 | 0.17 | 0.12 | 0.41 | 0.52 | 0.14 | 0.48 |
| 9 | 115694415 | rs1324952 | 0.358 | -0.675 | -0.718 | 1.32 | -0.004 | 0.004 | 0.004 | 0 | 0.75 | 0.75 | 0.85 | 0.58 | 0.82 | 0.8 |
| 9 | 115694490 | rs10981550 | 0.04 | -0.317 | -1.09 | 1.063 | -0.021 | 0.013 | 0.013 | 0 | 0.14 | 0.09 | 0.21 | 0.09 | 0.18 | 0.14 |
| 9 | 115694935 | rs6477967 | 1.002 | -0.272 | -0.921 | 1.051 | 0.004 | -0.006 | -0.006 | 0 | 0.61 | 0.61 | 0.59 | 0.45 | 0.64 | 0.64 |
| 9 | 115696187 | rs10817411 | -0.759 | -0.385 | -1.27 | 1.109 | -0.013 | -0.059 | -0.059 | 0 | 0.27 | 0.07 | 0.27 | 0.38 | 0.25 | 0.27 |
| 9 | 115698104 | rs747065 | 0.906 | -0.47 | -1.18 | 1.04 | -0.014 | 0.023 | 0.023 | 0 | 0.76 | 0.8 | 0.72 | 0.9 | 0.69 | 0.81 |
| 9 | 115698346 | rs747066 | 1.192 | -0.436 | -1.158 | 1.074 | 0.004 | -0.031 | -0.031 | 0 | 0.63 | 0.8 | 0.66 | 0.55 | 0.63 | 0.66 |
| 9 | 115702065 | rs11792260 | -0.046 | -0.285 | -1.034 | 0.935 | -0.022 | 0.033 | 0.033 | 0 | 0.14 | 0.07 | 0.21 | 0.03 | 0.18 | 0.12 |
| 9 | 115707599 | rs10817414 | 0.699 | -2.26 | -0.525 | 0.66 | 0.014 | 0.006 | 0.006 | 0.432 | 0.56 | 0.4 | 0.79 | 0.58 | 0.8 | 0.7 |
| 9 | 115707661 | rs7043761 | 0.392 | -2.312 | -0.164 | 0.617 | -0.02 | 0.044 | 0.044 | 0 | 0.92 | 0.8 | 0.97 | 0.85 | 0.97 | 0.94 |
| 9 | 115708164 | rs6477968 | 0.629 | -2.904 | -0.554 | 0.401 | -0.02 | 0 | 0 | 0 | 0.68 | 0.65 | 0.76 | 0.64 | 0.82 | 0.74 |
| 9 | 115709698 | rs10733601 | 0.666 | -3.026 | -0.342 | 0.492 | -0.018 | 0.005 | 0.005 | 0.352 | 0.62 | 0.6 | 0.69 | 0.39 | 0.75 | 0.66 |
| 9 | 115710102 | rs4532675 | 0.539 | -3.041 | -0.273 | 0.488 | 0.032 | -0.02 | -0.02 | 0 | 0.64 | 0.71 | 0.71 | 0.4 | 0.75 | 0.66 |
| 9 | 115712068 | rs7858808 | 0.099 | -3.345 | -0.239 | 0.374 | -0.076 | -0.095 | -0.095 | 0 | 0.2 | 0.39 | 0.09 | 0.04 | 0.07 | 0.13 |
| 9 | 115712252 | rs7874374 | 0.383 | -3.442 | -0.289 | 0.4 | -0.034 | 0 | 0 | 0 | 0.67 | 0.63 | 0.86 | 0.63 | 0.86 | 0.71 |
| 9 | 115714804 | rs7020864 | 0.704 | -3.409 | -0.292 | 0.416 | -0.024 | 0.007 | 0.007 | 0.362 | 0.63 | 0.59 | 0.71 | 0.4 | 0.77 | 0.67 |
| 9 | 115716155 | rs995366 | -1.106 | -3.394 | -0.39 | 0.426 | 0.032 | -0.097 | -0.097 | 0 | 0.85 | 1 | 0.95 | 0.65 | 0.93 | 0.84 |
| 9 | 115716647 | rs12115363 | 0.046 | -3.373 | -0.294 | 0.378 | -0.01 | 0.001 | 0.001 | 0 | 0.19 | 0.21 | 0.22 | 0.22 | 0.15 | 0.12 |
| 9 | 115720888 | rs979596 | 1.253 | -3.424 | -0.199 | 0.527 | 0.048 | -0.002 | -0.002 | 0 | 0.46 | 0.35 | 0.64 | 0.4 | 0.7 | 0.58 |
| 9 | 115722489 | rs10817416 | -0.193 | -2.759 | -0.074 | 0.715 | -0.036 | -0.018 | -0.018 | 0 | 0.2 | 0.31 | 0.23 | 0.17 | 0.15 | 0.12 |
| 9 | 115723429 | rs1408696 | -0.439 | -2.81 | -0.103 | 0.728 | 0.031 | -0.052 | -0.052 | 0 | 0.84 | 0.91 | 0.94 | 0.57 | 0.92 | 0.79 |
| 9 | 115724471 | rs10981559 | 1.003 | -2.797 | -0.211 | 0.81 | 0.033 | -0.063 | -0.063 | 0 | 0.13 | 0 | 0.05 | 0.25 | 0.06 | 0.11 |
| 9 | 115724490 | rs3903908 | -0.208 | -2.734 | -0.156 | 0.45 | -0.014 | -0.006 | -0.006 | 0 | 0.19 | 0.23 | 0.21 | 0.17 | 0.15 | 0.12 |
| 9 | 115724549 | rs3903907 | 0.169 | -2.418 | -0.237 | 0.214 | -0.004 | -0.001 | -0.001 | 0 | 0.64 | 0.66 | 0.71 | 0.44 | 0.77 | 0.72 |
| 9 | 115733682 | rs9886741 | 0.368 | -0.898 | -0.313 | 0.431 | 0.025 | 0.009 | 0.009 | 0 | 0.32 | 0.37 | 0.42 | 0.32 | 0.45 | 0.46 |
| 9 | 115737459 | rs7040359 | -0.356 | -0.5 | -0.842 | 0.354 | 0.001 | 0.002 | 0.002 | 0 | 0.33 | 0.66 | 0.28 | 0.27 | 0.28 | 0.34 |
| 9 | 115740086 | rs4618774 | -0.177 | -0.504 | -1.037 | 0.452 | -0.007 | 0.008 | 0.008 | 0 | 0.31 | 0.62 | 0.36 | 0.37 | 0.25 | 0.4 |
| 9 | 115745205 | rs1408691 | -1.264 | -0.319 | -1.584 | 0.326 | -0.042 | 0.011 | 0.011 | 0 | 0.54 | 0.83 | 0.53 | 0.54 | 0.55 | 0.6 |

| | | | | | | | | | | | | | | | | |
|---|---|---|---|---|---|---|---|---|---|---|---|---|---|---|---|---|
| 9 | 115745240 | rs2104391 | 0.46 | -0.373 | -1.165 | 0.282 | -0.027 | 0.053 | 0.053 | 0 | 0.29 | 0.46 | 0.33 | 0.37 | 0.23 | 0.4 |
| 9 | 115745656 | rs10981576 | 0.804 | -0.286 | -1.252 | 0.242 | -0.028 | 0.025 | 0.025 | 0 | 0.1 | 0.03 | 0.07 | 0.03 | 0.16 | 0.12 |
| 9 | 115747393 | rs3945761 | -0.571 | -0.258 | -1.338 | 0.261 | 0.011 | -0.013 | -0.013 | 0 | 0.42 | 0.68 | 0.43 | 0.4 | 0.4 | 0.5 |
| 9 | 115757581 | rs4979204 | -0.784 | -0.591 | -1.853 | 0.213 | 0.047 | 0.101 | 0.101 | 0 | 0.55 | 0.7 | 0.48 | 0.82 | 0.38 | 0.57 |
| 9 | 115759519 | rs10981589 | 1.075 | -0.703 | -1.918 | 0.173 | -0.013 | 0.029 | 0.029 | 0 | 0.07 | 0 | 0.02 | 0.01 | 0.04 | 0.06 |
| 9 | 115764076 | rs1324935 | -0.897 | -0.353 | -1.637 | 0.392 | 0.048 | 0.104 | 0.104 | 0 | 0.55 | 0.7 | 0.48 | 0.82 | 0.37 | 0.57 |
| 9 | 115764347 | rs10981592 | 0.817 | -0.128 | -1.415 | 0.464 | -0.028 | 0.038 | 0.038 | 0 | 0.3 | 0.47 | 0.34 | 0.35 | 0.23 | 0.4 |
| 9 | 115770285 | rs6477978 | -1.401 | 0.026 | -1.51 | 0.227 | -0.068 | 0.11 | 0.11 | 0 | 0.68 | 0.85 | 0.6 | 0.91 | 0.62 | 0.71 |
| 9 | 115775168 | rs1570756 | -1.112 | 0.043 | -1.515 | 0.235 | -0.025 | 0.059 | 0.059 | 0 | 0.64 | 0.74 | 0.55 | 0.85 | 0.53 | 0.68 |
| 9 | 115777556 | rs4246894 | -0.471 | 0.023 | -1.486 | 0.259 | 0.013 | 0.091 | 0.091 | 0 | 0.56 | 0.72 | 0.5 | 0.76 | 0.46 | 0.62 |
| 9 | 115781111 | rs2418246 | 0.372 | 0.04 | -1.414 | 0.267 | -0.029 | 0.036 | 0.036 | 0 | 0.33 | 0.5 | 0.36 | 0.39 | 0.25 | 0.44 |
| 9 | 115783589 | rs4979208 | -0.15 | 0.869 | -1.709 | -0.161 | -0.078 | 0.122 | 0.122 | 0 | 0.68 | 0.87 | 0.61 | 0.91 | 0.62 | 0.72 |
| 9 | 115788707 | rs1411600 | -0.516 | 0.255 | -0.928 | 0.821 | -0.01 | 0.029 | 0.029 | 0 | 0.17 | 0.2 | 0.19 | 0.29 | 0.1 | 0.18 |
| 9 | 115789373 | rs9299213 | 0.469 | -0.689 | -1.49 | 0.25 | 0.026 | 0.036 | 0.036 | 0 | 0.54 | 0.59 | 0.49 | 0.19 | 0.68 | 0.37 |
| 9 | 115790227 | rs16917698 | 0.361 | -0.939 | -1.561 | 0.774 | 0.002 | 0.074 | 0.074 | 0 | 0.06 | 0.04 | 0.11 | 0.39 | 0.04 | 0.23 |
| 9 | 115794341 | rs7863800 | 0.943 | -0.053 | -1.402 | -0.223 | 0.062 | -0.08 | -0.08 | 0 | 0.23 | 0.36 | 0.22 | 0.09 | 0.37 | 0.15 |
| 9 | 115794588 | rs10739386 | -0.499 | -0.127 | -1.454 | -0.202 | 0.008 | -0.002 | -0.002 | 0.068 | 0.6 | 0.6 | 0.61 | 0.58 | 0.72 | 0.59 |
| 9 | 115794614 | rs10817431 | -0.031 | -0.109 | -1.44 | -0.316 | 0.029 | 0.044 | 0.044 | 0 | 0.53 | 0.6 | 0.5 | 0.18 | 0.68 | 0.37 |
| 9 | 115794712 | rs16917707 | 1.356 | -0.065 | -0.927 | -0.451 | -0.007 | -0.031 | -0.031 | 0 | 0.09 | 0.2 | 0.06 | 0.05 | 0.05 | 0.04 |
| 9 | 115796553 | rs10981613 | 1.337 | -0.602 | -1.563 | -0.859 | 0.036 | 0.022 | 0.022 | 0 | 0.15 | 0.07 | 0.05 | 0.08 | 0.06 | 0.08 |
| 9 | 115802287 | rs10981618 | 0.007 | -0.542 | -1.412 | -0.897 | -0.009 | 0.053 | 0.053 | 0.703 | 0.45 | 0.6 | 0.53 | 0.78 | 0.51 | 0.55 |
| 9 | 115808300 | rs1411605 | -1.661 | -0.495 | -1.667 | -0.806 | 0.016 | 0.024 | 0.024 | 1.386 | 0.78 | 0.8 | 0.85 | 0.94 | 0.84 | 0.8 |
| 9 | 115811102 | rs12003447 | -0.637 | -0.395 | -1.424 | -0.477 | -0.029 | -0.016 | -0.016 | 0 | 0.56 | 0.74 | 0.48 | 0.46 | 0.53 | 0.44 |
| 9 | 115815398 | rs10981621 | -0.616 | -0.03 | -1.408 | 0.287 | 0.009 | 0.011 | 0.011 | 0 | 0.49 | 0.38 | 0.55 | 0.52 | 0.58 | 0.49 |
| 9 | 115817156 | rs10981622 | 0.682 | -0.126 | -1.513 | 0.248 | 0.044 | 0.02 | 0.02 | 0 | 0.14 | 0.03 | 0.05 | 0.08 | 0.07 | 0.1 |
| 9 | 115819250 | rs11791740 | 0.964 | -0.084 | -1.344 | 0.42 | -0.063 | 0.059 | 0.059 | 0.674 | 0.14 | 0.04 | 0.18 | 0 | 0.33 | 0.07 |
| 9 | 115821485 | rs10759623 | -0.13 | -0.771 | -0.919 | 1.037 | -0.003 | 0.078 | 0.078 | 0 | 0.21 | 0.29 | 0.32 | 0.49 | 0.18 | 0.35 |
| 9 | 115852715 | rs7022165 | -0.335 | -0.731 | -0.839 | 0.917 | -0.009 | 0.029 | 0.029 | 0.044 | 0.27 | 0.3 | 0.35 | 0.53 | 0.21 | 0.42 |
| 9 | 115864465 | rs7034705 | 0.201 | -0.841 | -1.057 | 0.973 | -0.003 | 0.003 | 0.003 | 0 | 0.21 | 0.28 | 0.12 | 0.35 | 0.15 | 0.25 |
| 9 | 115864525 | rs13294922 | 0.578 | -0.543 | -1.535 | 0.242 | -0.057 | 0.143 | 0.143 | 2.101 | 0.4 | 0.16 | 0.25 | 0.14 | 0.47 | 0.24 |
| 9 | 115865501 | rs17828145 | 0.855 | -0.89 | -1.575 | 0.09 | 0.034 | 0.029 | 0.029 | 0.055 | 0.19 | 0.08 | 0.06 | 0.13 | 0.11 | 0.17 |
| 9 | 115866017 | rs6477983 | 1.255 | -0.966 | -1.13 | 0.176 | 0.022 | 0.073 | 0.073 | 0 | 0.1 | 0.22 | 0.06 | 0.18 | 0.09 | 0.11 |
| 9 | 115866273 | rs10981630 | 1.339 | -0.998 | -1.1 | -0.093 | 0.005 | -0.075 | -0.075 | 0 | 0.16 | 0.27 | 0.24 | 0.01 | 0.15 | 0.11 |
| 9 | 115867276 | rs2009977 | 1.143 | -0.843 | -1.503 | 0.12 | -0.005 | -0.012 | -0.012 | 0 | 0.11 | 0.08 | 0.17 | 0.14 | 0.12 | 0.15 |
| 9 | 115874261 | rs2039221 | 1.595 | -0.865 | -1.445 | 0.12 | -0.044 | -0.041 | -0.041 | 0.374 | 0.28 | 0.13 | 0.37 | 0.37 | 0.4 | 0.28 |
| 9 | 115885060 | rs10817441 | -0.338 | -0.9 | -1.321 | 0.012 | -0.016 | -0.063 | -0.063 | 0 | 0.66 | 0.48 | 0.64 | 0.82 | 0.7 | 0.71 |
| 9 | 115893229 | rs4598325 | 0.674 | -0.754 | -1.211 | 0.562 | 0.011 | 0.187 | 0.187 | 0 | 0.11 | 0.26 | 0.18 | 0.53 | 0.1 | 0.28 |
| 9 | 115897653 | rs10981639 | 1.116 | -0.782 | -1.495 | 0.234 | -0.014 | -0.029 | -0.029 | 0 | 0.06 | 0.01 | 0.02 | 0.1 | 0.05 | 0.11 |
| 9 | 115897803 | rs10981640 | 1.342 | -0.748 | -1.466 | -0.467 | 0.005 | 0.008 | 0.008 | 0 | 0.21 | 0.22 | 0.16 | 0.16 | 0.19 | 0.2 |
| 9 | 115899317 | rs10513183 | 1.083 | -0.444 | -1.641 | 0.104 | -0.017 | 0.102 | 0.102 | 0 | 0.1 | 0.28 | 0.03 | 0.17 | 0.04 | 0.14 |
| 9 | 115900533 | rs9792498 | -0.932 | -0.478 | -1.649 | 0.024 | -0.009 | -0.029 | -0.029 | 0 | 0.42 | 0.45 | 0.29 | 0.8 | 0.3 | 0.67 |
| 9 | 115901241 | rs12336914 | 0.638 | -0.072 | -1.884 | 0.459 | -0.022 | 0.006 | 0.006 | 0.014 | 0.2 | 0.02 | 0.08 | 0.13 | 0.16 | 0.19 |
| 9 | 115905480 | rs4979216 | -0.132 | 0.574 | -0.834 | 0.67 | -0.069 | 0.091 | 0.091 | 0.27 | 0.76 | 0.92 | 0.69 | 0.97 | 0.64 | 0.93 |
| 9 | 115905837 | rs10981645 | -0.704 | -0.194 | -0.031 | 1.712 | -0.018 | 0.185 | 0.185 | 0 | 0.1 | 0.28 | 0.03 | 0.3 | 0.04 | 0.15 |
| 9 | 115905850 | rs12115536 | -0.649 | -0.173 | -0.021 | 1.092 | -0.016 | 0.104 | 0.104 | 0 | 0.09 | 0.28 | 0.03 | 0.17 | 0.04 | 0.14 |
| 9 | 115908452 | rs10981648 | -0.47 | -0.095 | -0.716 | 0.666 | 0.014 | 0.012 | 0.012 | 0 | 0.08 | 0.04 | 0.02 | 0.06 | 0.04 | 0.14 |
| 9 | 115910991 | rs10981650 | -0.835 | -0.215 | -0.446 | 0.6 | -0.005 | -0.012 | -0.012 | 0.115 | 0.08 | 0.1 | 0.03 | 0.04 | 0.04 | 0.14 |
| 9 | 115911904 | rs10817446 | -0.866 | -0.225 | -0.451 | 0.595 | -0.005 | -0.012 | -0.012 | 0 | 0.08 | 0.1 | 0.03 | 0.04 | 0.04 | 0.14 |
| 9 | 115915194 | rs11794048 | 0.321 | -0.118 | -1.343 | -0.174 | -0.052 | 0.115 | 0.115 | 0 | 0.43 | 0.15 | 0.34 | 0.2 | 0.49 | 0.33 |
| 9 | 115915624 | rs7027743 | -0.438 | -0.466 | -0.508 | 0.617 | 0.008 | 0.017 | 0.017 | 0 | 0.61 | 0.54 | 0.44 | 0.45 | 0.58 | 0.52 |
| 9 | 115917783 | rs4263842 | -0.631 | -0.806 | -0.675 | 1.161 | 0.04 | 0.002 | 0.002 | 0 | 0.72 | 0.57 | 0.56 | 0.67 | 0.66 | 0.73 |
| 9 | 115918552 | rs10121164 | -0.414 | -0.809 | -0.128 | 0.959 | -0.001 | -0.001 | -0.001 | 0.179 | 0.76 | 0.73 | 0.72 | 0.68 | 0.69 | 0.72 |
| 9 | 115919782 | rs3934969 | 1.371 | -0.331 | -0.546 | 0.349 | 0.015 | 0.05 | 0.05 | 0 | 0.39 | 0.27 | 0.29 | 0.16 | 0.35 | 0.26 |
| 9 | 115920099 | rs4344146 | 0.04 | -0.013 | -0.575 | 0.579 | -0.016 | -0.012 | -0.012 | 0 | 0.64 | 0.67 | 0.61 | 0.6 | 0.56 | 0.55 |
| 9 | 115921843 | rs4483236 | -0.183 | -0.066 | -0.427 | 0.575 | 0.001 | -0.005 | -0.005 | 0.295 | 0.25 | 0.21 | 0.4 | 0.61 | 0.25 | 0.37 |
| 9 | 115922694 | rs7849150 | 1.679 | -0.094 | -0.382 | 0.59 | 0.009 | 0.003 | 0.003 | 0 | 0.4 | 0.34 | 0.3 | 0.37 | 0.35 | 0.27 |
| 9 | 115924388 | rs4073133 | -0.273 | -0.099 | -0.385 | 0.587 | 0 | -0.004 | -0.004 | 0 | 0.25 | 0.21 | 0.39 | 0.41 | 0.25 | 0.36 |
| 9 | 115924811 | rs3827672 | 1.673 | -0.1 | -0.394 | 0.573 | -0.045 | 0.012 | 0.012 | 0 | 0.11 | 0.02 | 0.09 | 0.05 | 0.18 | 0.12 |
| 9 | 115925104 | rs3810906 | -0.265 | -0.104 | -0.39 | 0.573 | 0 | -0.004 | -0.004 | 0 | 0.25 | 0.21 | 0.39 | 0.41 | 0.25 | 0.36 |
| 9 | 115925396 | rs1128774 | -1.254 | -0.093 | -0.377 | 0.553 | 0.027 | -0.012 | -0.012 | 0 | 0.85 | 0.68 | 0.8 | 0.87 | 0.82 | 0.9 |
| 9 | 115926317 | rs2571 | 1.615 | -0.089 | -0.417 | 0.544 | -0.03 | -0.047 | -0.047 | 0 | 0.08 | 0.01 | 0.09 | 0.17 | 0.1 | 0.1 |
| 9 | 115927857 | rs1128150 | 1.187 | -0.088 | -0.352 | 0.542 | -0.012 | -0.021 | -0.021 | 0 | 0.09 | 0.11 | 0.03 | 0.04 | 0.04 | 0.15 |
| 9 | 115928275 | rs3810908 | -1.521 | -0.089 | -0.307 | 0.545 | 0.017 | -0.004 | -0.004 | 0 | 0.87 | 0.79 | 0.82 | 0.87 | 0.82 | 0.9 |
| 9 | 115934852 | rs10739387 | -1.241 | -0.096 | -0.292 | 0.546 | 0 | -0.004 | -0.004 | 0 | 0.47 | 0.46 | 0.53 | 0.71 | 0.47 | 0.64 |
| 9 | 115938318 | rs10981669 | -0.334 | -0.532 | -0.974 | -0.089 | 0.004 | 0.024 | 0.024 | 0 | 0.38 | 0.29 | 0.48 | 0.75 | 0.39 | 0.56 |
| 9 | 115938750 | rs17762224 | 0.76 | -0.518 | -0.975 | -0.317 | -0.011 | 0.02 | 0.02 | 0 | 0.2 | 0.13 | 0.12 | 0.09 | 0.23 | 0.27 |
| 9 | 115941144 | rs10118914 | 1.914 | -0.567 | -0.949 | -0.308 | 0 | -0.035 | -0.035 | 2.095 | 0.28 | 0.25 | 0.2 | 0.02 | 0.21 | 0.14 |
| 9 | 115946170 | rs10981674 | 1.562 | -0.537 | -0.953 | -0.269 | -0.009 | -0.019 | -0.019 | 0.049 | 0.09 | 0.1 | 0.04 | 0.04 | 0.05 | 0.15 |
| 9 | 115950156 | rs10465129 | 1.956 | -0.536 | -1.097 | -0.218 | 0.016 | -0.064 | -0.064 | 0.022 | 0.1 | 0.01 | 0.11 | 0.22 | 0.07 | 0.23 |

| | | | | | | | | | | | | | | | | |
|---|---|---|---|---|---|---|---|---|---|---|---|---|---|---|---|---|
| 9 | 115952895 | rs12684417 | 0.936 | -0.388 | -0.904 | -0.746 | 0.006 | 0.011 | 0.011 | 0 | 0.39 | 0.32 | 0.29 | 0.16 | 0.35 | 0.26 |
| 9 | 115960547 | rs10981683 | 1.601 | -0.38 | -0.879 | -0.684 | 0.013 | 0.021 | 0.021 | 0 | 0.08 | 0.04 | 0.03 | 0.04 | 0.05 | 0.15 |
| 9 | 115962551 | rs7852613 | 1.33 | -0.254 | -0.804 | -0.631 | -0.028 | -0.113 | -0.113 | 0 | 0.29 | 0.39 | 0.22 | 0.02 | 0.21 | 0.14 |
| 9 | 115963507 | rs7029139 | 1.319 | -0.259 | -0.806 | -0.634 | -0.029 | -0.112 | -0.112 | 0 | 0.3 | 0.39 | 0.22 | 0.02 | 0.21 | 0.14 |
| 9 | 115963802 | rs4979220 | -0.658 | -0.224 | -0.848 | -0.584 | 0.006 | -0.021 | -0.021 | 0.033 | 0.68 | 0.75 | 0.67 | 0.41 | 0.69 | 0.51 |
| 9 | 115964664 | rs12380325 | 0.371 | -0.277 | -0.796 | -0.521 | 0.003 | 0.002 | 0.002 | 0 | 0.27 | 0.27 | 0.36 | 0.33 | 0.31 | 0.26 |
| 9 | 115968797 | rs1133618 | 0.589 | -0.355 | -0.687 | -0.699 | -0.013 | -0.072 | -0.072 | 0.497 | 0.29 | 0.33 | 0.22 | 0.02 | 0.21 | 0.14 |
| 9 | 115981113 | rs6477997 | -0.293 | -0.086 | -0.453 | -0.751 | 0.008 | -0.066 | -0.066 | 3.97 | 0.3 | 0.4 | 0.3 | 0.15 | 0.3 | 0.35 |
| 9 | 115983216 | rs2233913 | 1.595 | 0.131 | -0.569 | -0.827 | -0.04 | -0.008 | -0.008 | 0 | 0.06 | 0.01 | 0.07 | 0.04 | 0.15 | 0.12 |
| 9 | 115983540 | rs2233914 | 0.554 | -0.352 | -0.819 | -0.24 | -0.001 | 0.023 | 0.023 | 0 | 0.12 | 0.16 | 0.1 | 0.32 | 0.13 | 0.29 |
| 9 | 115990221 | rs10513189 | -1.013 | -0.767 | -0.847 | -0.489 | 0.008 | 0.002 | 0.002 | 0.095 | 0.48 | 0.6 | 0.42 | 0.47 | 0.46 | 0.65 |
| 9 | 115995901 | rs10981699 | 1.889 | -0.989 | -1.405 | -0.102 | -0.012 | -0.068 | -0.068 | 1.228 | 0.11 | 0.01 | 0.16 | 0.24 | 0.12 | 0.1 |
| 9 | 115997249 | rs6477998 | -0.587 | -0.819 | -1.092 | -0.121 | -0.001 | 0.002 | 0.002 | 0 | 0.43 | 0.56 | 0.4 | 0.47 | 0.43 | 0.64 |
| 9 | 116002565 | rs7851623 | 0.973 | -0.935 | -0.66 | -0.26 | -0.002 | -0.004 | -0.004 | 0 | 0.2 | 0.2 | 0.26 | 0.08 | 0.25 | 0.2 |
| 9 | 116005783 | rs10817465 | -0.033 | -0.185 | -0.963 | -0.4 | -0.001 | 0 | 0 | 1.014 | 0.62 | 0.64 | 0.59 | 0.71 | 0.59 | 0.61 |
| 9 | 116011458 | rs16930959 | 1.821 | -0.263 | -0.957 | -0.433 | 0.005 | -0.021 | -0.021 | 0.024 | 0.07 | 0.01 | 0.04 | 0.32 | 0.07 | 0.22 |
| 9 | 116025024 | rs10759637 | -0.546 | -0.203 | -0.818 | -0.456 | 0 | 0.001 | 0.001 | 0 | 0.42 | 0.57 | 0.4 | 0.47 | 0.43 | 0.64 |
| 9 | 116029174 | rs3087548 | 0.634 | -0.376 | -0.312 | -0.538 | 0.003 | -0.056 | -0.056 | 0 | 0.22 | 0.33 | 0.14 | 0.06 | 0.21 | 0.2 |
| 9 | 116037894 | rs3810919 | 1.199 | -0.494 | -0.364 | 0.018 | -0.043 | -0.052 | -0.052 | 0 | 0.16 | 0.05 | 0.1 | 0.26 | 0.2 | 0.2 |
| 9 | 116041061 | rs12342549 | 1.493 | -0.768 | -0.028 | -0.144 | -0.015 | -0.026 | -0.026 | 1.135 | 0.09 | 0.22 | 0.04 | 0.05 | 0.04 | 0.17 |
| 9 | 116042263 | rs10733604 | -1.518 | -0.568 | -0.279 | -0.262 | 0.002 | -0.011 | -0.011 | 0 | 0.93 | 0.91 | 0.81 | 1 | 0.86 | 0.93 |
| 9 | 116042492 | rs10759638 | -0.737 | -0.995 | -0.084 | 0.035 | -0.017 | 0.012 | 0.012 | 0 | 0.38 | 0.46 | 0.21 | 0.43 | 0.32 | 0.57 |
| 9 | 116045074 | rs12352784 | 2.271 | -0.973 | -0.233 | 0.054 | -0.008 | -0.015 | -0.015 | 0 | 0.07 | 0.11 | 0.03 | 0.05 | 0.04 | 0.09 |
| 9 | 116045272 | rs16932083 | 1.49 | -0.914 | -0.354 | 0.097 | 0.018 | -0.046 | -0.046 | 0.535 | 0.09 | 0.01 | 0.05 | 0.12 | 0.08 | 0.2 |
| 9 | 116048874 | rs4979232 | 0.238 | -0.919 | -0.356 | 0.041 | -0.04 | -0.109 | -0.109 | 0.745 | 0.25 | 0.06 | 0.15 | 0.37 | 0.28 | 0.41 |
| 9 | 116048977 | rs10981714 | 1.472 | -0.918 | -0.328 | 0.075 | 0.018 | -0.048 | -0.048 | 0.04 | 0.09 | 0.01 | 0.05 | 0.32 | 0.08 | 0.2 |
| 9 | 116054040 | rs2418249 | 1.367 | -0.918 | -0.325 | 0.081 | 0.023 | -0.045 | -0.045 | 0 | 0.09 | 0.03 | 0.17 | 0.34 | 0.08 | 0.21 |
| 9 | 116059054 | rs10817475 | 0.257 | -0.908 | -0.344 | 0.014 | -0.041 | -0.109 | -0.109 | 0 | 0.24 | 0.06 | 0.15 | 0.37 | 0.27 | 0.41 |
| 9 | 116060124 | rs3750534 | -0.683 | -0.972 | -0.05 | -0.029 | -0.014 | 0.01 | 0.01 | 0 | 0.38 | 0.45 | 0.21 | 0.42 | 0.32 | 0.57 |
| 9 | 116060221 | rs3750533 | -0.482 | -0.981 | -0.055 | -0.034 | -0.012 | 0.017 | 0.017 | 0 | 0.36 | 0.45 | 0.2 | 0.42 | 0.3 | 0.57 |
| 9 | 116060487 | rs3750532 | -0.628 | -0.92 | -0.012 | -0.007 | -0.014 | 0.011 | 0.011 | 0 | 0.38 | 0.45 | 0.21 | 0.42 | 0.32 | 0.58 |
| 9 | 116065087 | rs7860625 | -0.287 | -0.961 | 0.059 | 0.129 | -0.08 | 0.087 | 0.087 | 0.21 | 0.23 | 0.37 | 0.11 | 0.37 | 0.12 | 0.38 |
| 9 | 116066958 | rs10817476 | 0.813 | -0.603 | -0.535 | -0.179 | -0.015 | -0.048 | -0.048 | 3.903 | 0.22 | 0.12 | 0.14 | 0.32 | 0.25 | 0.28 |
| 9 | 116073892 | rs10124515 | 0.082 | -0.875 | -0.411 | -0.411 | -0.029 | 0.016 | 0.016 | 0.203 | 0.28 | 0.54 | 0.16 | 0.37 | 0.17 | 0.4 |
| 9 | 116077694 | rs12684385 | 2.096 | -0.809 | -0.459 | -0.363 | 0.017 | -0.048 | -0.048 | 0 | 0.09 | 0.01 | 0.05 | 0.13 | 0.08 | 0.19 |
| 9 | 116078468 | rs17764563 | 1.675 | -0.824 | -0.265 | -0.497 | -0.001 | -0.014 | -0.014 | 0.525 | 0.12 | 0.12 | 0.07 | 0.03 | 0.08 | 0.2 |
| 9 | 116081449 | rs7038589 | 1.972 | -0.774 | -0.394 | -0.506 | -0.017 | 0.014 | 0.014 | 0.491 | 0.13 | 0.09 | 0.09 | 0 | 0.17 | 0.07 |
| 9 | 116084691 | rs16932649 | 1.424 | -1.042 | -0.156 | -0.358 | -0.013 | -0.039 | -0.039 | 0 | 0.13 | 0.19 | 0.07 | 0.05 | 0.09 | 0.2 |
| 9 | 116085423 | rs10817479 | 2.031 | -1.14 | -0.964 | -0.229 | 0 | -0.046 | -0.046 | 0 | 0.08 | 0 | 0.07 | 0.15 | 0.05 | 0.06 |
| 9 | 116091342 | rs3810921 | -0.078 | -0.877 | -0.75 | -0.535 | -0.009 | 0.139 | 0.139 | 0 | 0.47 | 0.64 | 0.29 | 0.08 | 0.45 | 0.48 |
| 9 | 116097493 | rs4636275 | 0.975 | -0.873 | -0.594 | -0.517 | -0.049 | -0.11 | -0.11 | 0 | 0.18 | 0.36 | 0.1 | 0.03 | 0.09 | 0.2 |
| 9 | 116102463 | rs10481674 | 0.442 | -0.878 | -0.676 | -0.603 | -0.037 | -0.119 | -0.119 | 0 | 0.22 | 0.48 | 0.13 | 0.04 | 0.12 | 0.2 |
| 9 | 116107087 | rs2093691 | -0.57 | -0.805 | -0.627 | -0.204 | -0.003 | -0.007 | -0.007 | 0 | 0.56 | 0.65 | 0.34 | 0.42 | 0.52 | 0.68 |
| 9 | 116111407 | rs1998503 | -0.095 | -0.366 | -1.107 | -0.505 | 0.001 | 0.029 | 0.029 | 0 | 0.48 | 0.57 | 0.29 | 0.08 | 0.46 | 0.49 |
| 9 | 116118952 | rs4979237 | 0.078 | 0.048 | -1.864 | -0.038 | -0.109 | 0.069 | 0.069 | 0 | 0.38 | 0.12 | 0.23 | 0.21 | 0.44 | 0.47 |
| 9 | 116121520 | rs4978539 | -0.241 | -0.894 | -1.738 | 0.193 | 0.017 | 0.024 | 0.024 | 0 | 0.49 | 0.55 | 0.43 | 0.35 | 0.39 | 0.48 |
| 9 | 116122326 | rs1077636 | -1.267 | -0.801 | -1.749 | 0.541 | -0.017 | -0.001 | -0.001 | 1 | 0.69 | 0.71 | 0.55 | 0.66 | 0.55 | 0.73 |
| 9 | 116122954 | rs752757 | 0.343 | -0.754 | -1.626 | 0.456 | -0.025 | 0.007 | 0.007 | 0.051 | 0.33 | 0.54 | 0.33 | 0.39 | 0.2 | 0.29 |
| 9 | 116124325 | rs7038037 | -0.768 | -0.753 | -1.672 | 0.399 | -0.025 | 0.01 | 0.01 | 0 | 0.6 | 0.67 | 0.5 | 0.35 | 0.47 | 0.53 |
| 9 | 116125789 | rs10981769 | -0.549 | -0.864 | -1.326 | 0.36 | 0.056 | 0.075 | 0.075 | 0.034 | 0.45 | 0.67 | 0.42 | 0.21 | 0.29 | 0.33 |
| 9 | 116128568 | rs12342334 | 0.131 | -0.243 | -1.058 | 0.462 | -0.008 | -0.006 | -0.006 | 0.753 | 0.23 | 0.32 | 0.15 | 0.22 | 0.17 | 0.17 |
| 9 | 116128884 | rs4979243 | 1.183 | -0.023 | -1.342 | 0.302 | 0.002 | 0.007 | 0.007 | 0 | 0.08 | 0.06 | 0.21 | 0.29 | 0.08 | 0.08 |
| 9 | 116130924 | rs7857932 | 1.292 | 0.029 | -1.196 | -0.079 | -0.001 | -0.033 | -0.033 | 0 | 0.11 | 0.15 | 0.07 | 0 | 0.08 | 0.04 |
| 9 | 116131453 | rs818710 | -0.708 | 0.512 | -1.214 | -0.294 | 0.036 | -0.073 | -0.073 | 0.112 | 0.81 | 0.7 | 0.73 | 0.98 | 0.72 | 0.86 |
| 9 | 116131695 | rs10121150 | 0.721 | 0.695 | -1.611 | 0.334 | 0 | 0.004 | 0.004 | 0 | 0.22 | 0.22 | 0.18 | 0.16 | 0.22 | 0.14 |
| 9 | 116131873 | rs3827661 | 0.359 | 0.784 | -1.652 | 0.816 | -0.004 | -0.016 | -0.016 | 0.062 | 0.11 | 0.07 | 0.23 | 0.28 | 0.13 | 0.1 |
| 9 | 116132275 | rs3750529 | 0.447 | 0.657 | -1.416 | 0.093 | -0.004 | -0.024 | -0.024 | 0 | 0.11 | 0.13 | 0.04 | 0.03 | 0.08 | 0.13 |
| 9 | 116132334 | rs3088235 | 0.808 | 0.911 | -1.668 | 0.442 | -0.065 | -0.04 | -0.04 | 0 | 0.11 | 0.01 | 0.09 | 0.14 | 0.18 | 0.24 |
| 9 | 116133095 | rs10817488 | -0.176 | -0.005 | -1.375 | 0.243 | 0.009 | 0.007 | 0.007 | 0 | 0.44 | 0.56 | 0.32 | 0.29 | 0.34 | 0.35 |
| 9 | 116134127 | rs818713 | 0.075 | 0.303 | -1.735 | 0.021 | -0.033 | 0.102 | 0.102 | 2.781 | 0.29 | 0.09 | 0.21 | 0.11 | 0.35 | 0.18 |
| 9 | 116136198 | rs1043836 | 0.739 | -0.259 | -0.777 | 0.895 | -0.002 | -0.001 | -0.001 | 0 | 0.52 | 0.49 | 0.7 | 0.47 | 0.62 | 0.72 |
| 9 | 116137673 | rs3747815 | -0.753 | -0.501 | -0.293 | 1.363 | -0.029 | 0.018 | 0.018 | 0.098 | 0.36 | 0.59 | 0.32 | 0.54 | 0.24 | 0.24 |
| 9 | 116138013 | rs3747816 | -0.411 | -0.666 | -0.252 | 1.559 | -0.009 | 0.005 | 0.005 | 0 | 0.25 | 0.29 | 0.2 | 0.34 | 0.18 | 0.17 |
| 9 | 116139835 | rs16936474 | -0.58 | -0.322 | -0.644 | 1.496 | -0.027 | 0.037 | 0.037 | 0 | 0.21 | 0.33 | 0.15 | 0.29 | 0.13 | 0.14 |
| 9 | 116144973 | rs818702 | 1.145 | 0.72 | -1.185 | 1.865 | -0.005 | -0.008 | -0.008 | 0 | 0.82 | 0.83 | 0.66 | 0.53 | 0.8 | 0.67 |
| 9 | 116148630 | rs818706 | 0.93 | 0.727 | -1.193 | 1.623 | -0.007 | -0.019 | -0.019 | 1.173 | 0.85 | 0.87 | 0.68 | 0.67 | 0.83 | 0.8 |
| 9 | 116150109 | rs818708 | -0.013 | 0.196 | -2.093 | 0.837 | 0.055 | 0.031 | 0.031 | 0 | 0.52 | 0.87 | 0.45 | 0.62 | 0.45 | 0.47 |
| 9 | 116151191 | rs1805313 | -0.184 | -0.135 | -1.379 | 0.576 | -0.011 | -0.003 | -0.003 | 0.136 | 0.42 | 0.51 | 0.56 | 0.44 | 0.31 | 0.46 |
| 9 | 116151527 | rs8177812 | -0.818 | 0.021 | -1.481 | 0.564 | 0.004 | -0.023 | -0.023 | 0 | 0.15 | 0.1 | 0.2 | 0.24 | 0.12 | 0.3 |

| | | | | | | | | | | | | | | | | |
|---|---|---|---|---|---|---|---|---|---|---|---|---|---|---|---|---|
| 9 | 116152940 | rs2228083 | -0.262 | 0.295 | -1.248 | 0.56 | 0.009 | 0.005 | 0.005 | 0.369 | 0.06 | 0.08 | 0.06 | 0.11 | 0.08 | 0.06 |
| 9 | 116153352 | rs2761016 | 0.198 | 0.713 | -1.383 | 0.222 | -0.01 | 0.023 | 0.023 | 0 | 0.52 | 0.67 | 0.72 | 0.61 | 0.53 | 0.44 |
| 9 | 116155428 | rs8177796 | -0.264 | 1.164 | -0.698 | 0.075 | 0.013 | -0.028 | -0.028 | 0.422 | 0.06 | 0.1 | 0.05 | 0 | 0.09 | 0.02 |
| 9 | 116159423 | rs818684 | -1.044 | 1.224 | -0.151 | -0.033 | 0.044 | -0.05 | -0.05 | 0.009 | 0.13 | 0.27 | 0.14 | 0.06 | 0.17 | 0.09 |
| 9 | 116162760 | rs818689 | -1.003 | 1.123 | -0.018 | -0.048 | 0.027 | -0.034 | -0.034 | 0 | 0.13 | 0.23 | 0.14 | 0.07 | 0.17 | 0.09 |
| 9 | 116171275 | rs3827674 | -0.448 | 0.717 | -0.281 | -0.163 | 0.019 | 0.028 | 0.028 | 0 | 0.08 | 0.02 | 0.02 | 0.01 | 0.05 | 0.11 |
| 9 | 116173206 | rs3810925 | -0.46 | 0.713 | -0.312 | -0.165 | 0.018 | 0.037 | 0.037 | 0 | 0.08 | 0.01 | 0.02 | 0.01 | 0.05 | 0.11 |
| 9 | 116183301 | rs818714 | -1.9 | 0.791 | 0.256 | -0.085 | 0.008 | -0.038 | -0.038 | 0 | 0.28 | 0.44 | 0.24 | 0.18 | 0.3 | 0.33 |
| 9 | 116184307 | rs10116799 | -0.435 | 0.924 | 0.227 | -0.349 | -0.002 | -0.008 | -0.008 | 0.206 | 0.08 | 0.08 | 0.06 | 0 | 0.1 | 0.04 |
| 9 | 116190841 | rs7021688 | -0.144 | 0.979 | 0.281 | -0.322 | 0.004 | -0.018 | -0.018 | 0.662 | 0.06 | 0.09 | 0.05 | 0 | 0.09 | 0.04 |
| 9 | 116194436 | rs1578592 | 0.478 | 0.975 | -0.098 | 0.116 | 0.009 | -0.04 | -0.04 | 0 | 0.24 | 0.12 | 0.39 | 0.34 | 0.19 | 0.16 |
| 9 | 116214058 | rs12338788 | -1.027 | 0.794 | 0.282 | -0.004 | -0.002 | -0.04 | -0.04 | 0.537 | 0.14 | 0.19 | 0.09 | 0.01 | 0.15 | 0.15 |
| 9 | 116266704 | rs3761820 | 0.299 | 0.937 | 0.601 | 0.286 | -0.001 | -0.001 | -0.001 | 0 | 0.07 | 0.05 | 0.05 | 0.11 | 0.09 | 0.07 |
| 9 | 116282114 | rs12342699 | -0.179 | 0.942 | 0.497 | 0.09 | 0.021 | -0.038 | -0.038 | 0 | 0.07 | 0.17 | 0.07 | 0 | 0.1 | 0.04 |
| 9 | 116288537 | rs7855434 | -1.25 | 0.964 | 0.466 | 0.069 | 0 | -0.03 | -0.03 | 0 | 0.16 | 0.19 | 0.13 | 0.01 | 0.18 | 0.15 |
| 9 | 116289417 | rs7046427 | -0.211 | 0.975 | 0.473 | 0.075 | 0.026 | -0.039 | -0.039 | 0 | 0.07 | 0.18 | 0.07 | 0 | 0.1 | 0.04 |
| 9 | 116304165 | rs7851718 | -0.211 | 0.975 | 0.469 | 0.075 | 0.025 | -0.039 | -0.039 | 0 | 0.07 | 0.18 | 0.07 | 0 | 0.1 | 0.04 |
| 9 | 116304304 | rs7851937 | -1.404 | 0.965 | 0.463 | 0.091 | 0 | -0.024 | -0.024 | 0 | 0.16 | 0.2 | 0.13 | 0.02 | 0.18 | 0.15 |
| 9 | 116334777 | rs4979257 | 0.656 | 0.951 | 0.309 | 0.161 | 0.016 | -0.047 | -0.047 | 0.537 | 0.87 | 0.81 | 0.86 | 1 | 0.81 | 0.95 |
| 9 | 116343044 | rs12339493 | -0.208 | 0.912 | 0.34 | 0.192 | 0.027 | -0.039 | -0.039 | 0 | 0.07 | 0.18 | 0.07 | 0 | 0.1 | 0.04 |
| 9 | 116346287 | rs3810927 | 0.708 | 0.969 | 0.133 | 0.418 | 0.015 | 0.011 | 0.011 | 0 | 0.23 | 0.11 | 0.24 | 0.21 | 0.19 | 0.14 |
| 9 | 116349298 | rs10981835 | -1.127 | 1.595 | 0.852 | 0.837 | -0.003 | 0.032 | 0.032 | 0 | 0.13 | 0.06 | 0.07 | 0.01 | 0.14 | 0.14 |
| 9 | 116355415 | rs4979261 | 2.666 | 1.584 | 0.97 | 0.878 | 0.008 | -0.048 | -0.048 | 0 | 0.65 | 0.42 | 0.69 | 0.81 | 0.61 | 0.64 |
| 9 | 116362738 | rs17187621 | -1.558 | 1.484 | 0.838 | 0.82 | -0.002 | -0.006 | -0.006 | 0 | 0.16 | 0.23 | 0.12 | 0.12 | 0.17 | 0.27 |
| 9 | 116363146 | rs518636 | 1.973 | 1.524 | 0.791 | 0.783 | -0.007 | 0.019 | 0.019 | 0 | 0.77 | 0.76 | 0.78 | 0.88 | 0.71 | 0.71 |
| 9 | 116365451 | rs10817494 | -1.176 | 1.601 | 1.108 | 1.039 | 0.002 | 0.14 | 0.14 | 0 | 0.13 | 0.19 | 0.16 | 0.45 | 0.16 | 0.2 |
| 9 | 116372579 | rs514309 | 1.212 | 2.241 | 1.101 | 0.733 | -0.045 | 0.074 | 0.074 | 0 | 0.85 | 0.98 | 0.86 | 0.99 | 0.8 | 0.87 |
| 9 | 116373700 | rs482318 | 2.245 | 2.25 | 1.423 | 0.702 | 0.016 | -0.078 | -0.078 | 0 | 0.78 | 0.69 | 0.77 | 0.99 | 0.69 | 0.83 |
| 9 | 116374874 | rs590963 | 0.078 | 2.137 | 1.342 | 0.812 | -0.014 | 0.013 | 0.013 | 0 | 0.93 | 0.98 | 0.89 | 1 | 0.88 | 0.98 |
| 9 | 116378169 | rs586265 | 1.564 | 2.138 | 1.369 | 0.786 | 0.017 | -0.051 | -0.051 | 0 | 0.85 | 0.8 | 0.82 | 1 | 0.77 | 0.94 |
| 9 | 116384750 | rs611335 | 1.665 | 2.059 | 1.52 | 0.763 | 0.022 | -0.068 | -0.068 | 0.283 | 0.84 | 0.76 | 0.82 | 1 | 0.76 | 0.94 |
| 9 | 116386438 | rs569679 | 2.248 | 2.096 | 2.115 | 0.642 | 0.029 | -0.154 | -0.154 | 0 | 0.73 | 0.5 | 0.75 | 0.99 | 0.67 | 0.81 |
| 9 | 116387254 | rs540599 | 2.904 | 2.08 | 1.719 | 1.005 | -0.049 | -0.062 | -0.062 | 0 | 0.66 | 0.15 | 0.61 | 0.61 | 0.58 | 0.72 |
| 9 | 116388816 | rs670719 | 1.811 | 1.951 | 1.702 | 0.035 | 0.046 | -0.104 | -0.104 | 0.827 | 0.83 | 0.63 | 0.81 | 1 | 0.76 | 0.94 |
| 9 | 116391069 | rs2127744 | -1.923 | 1.77 | 0.986 | 0.157 | 0.016 | -0.098 | -0.098 | 0 | 0.21 | 0.31 | 0.13 | 0.03 | 0.22 | 0.2 |
| 9 | 116391162 | rs2127745 | -1.297 | 1.602 | 0.863 | 0.516 | 0.021 | -0.05 | -0.05 | 0 | 0.09 | 0.2 | 0.05 | 0 | 0.09 | 0.07 |
| 9 | 116399607 | rs658580 | 1.34 | 1.689 | 0.459 | 0.896 | 0 | 0 | 0 | 0 | 0.72 | 0.76 | 0.79 | 0.76 | 0.7 | 0.81 |
| 9 | 116400152 | rs7862785 | -1.447 | 2.256 | 0.58 | 0.854 | 0.015 | 0.008 | 0.008 | 0 | 0.26 | 0.34 | 0.24 | 0.22 | 0.29 | 0.25 |
| 9 | 116400936 | rs4978549 | 1.105 | 2.097 | 0.247 | 0.669 | -0.004 | 0.002 | 0.002 | 0.107 | 0.72 | 0.31 | 0.73 | 0.75 | 0.67 | 0.81 |
| 9 | 116403013 | rs10981854 | -0.133 | 1.816 | -0.221 | 0.567 | -0.013 | -0.04 | -0.04 | 0 | 0.3 | 0.16 | 0.38 | 0.51 | 0.29 | 0.42 |
| 9 | 116404703 | rs584190 | -0.143 | 2.109 | -0.154 | 0.265 | -0.001 | -0.001 | -0.001 | 0.722 | 0.14 | 0.13 | 0.13 | 0.21 | 0.18 | 0.15 |
| 9 | 116405376 | rs7873198 | 1.328 | 1.981 | 0.43 | 0.11 | 0.023 | 0.058 | 0.058 | 0 | 0.46 | 0.34 | 0.38 | 0.22 | 0.42 | 0.44 |
| 9 | 116405475 | rs13297834 | 0.313 | 2.003 | 0.098 | 0.199 | -0.018 | 0.018 | 0.018 | 0 | 0.05 | 0.02 | 0.08 | 0 | 0.09 | 0.01 |
| 9 | 116405568 | rs2812401 | 0.093 | 1.736 | 0.461 | 0.316 | 0.002 | 0.019 | 0.019 | 0 | 0.68 | 0.51 | 0.6 | 0.45 | 0.71 | 0.61 |
| 9 | 116406907 | rs7862521 | 0.417 | 1.658 | 0.334 | 0.447 | -0.001 | 0.017 | 0.017 | 0 | 0.56 | 0.68 | 0.49 | 0.42 | 0.49 | 0.52 |
| 9 | 116407044 | rs7862352 | 0.649 | 1.772 | 0.422 | 0.272 | 0.003 | 0.012 | 0.012 | 0 | 0.51 | 0.48 | 0.42 | 0.23 | 0.51 | 0.46 |
| 9 | 116407227 | rs7035632 | -0.746 | 1.623 | 0.539 | 1.406 | 0.038 | 0.224 | 0.224 | 0.454 | 0.06 | 0.17 | 0.2 | 0.31 | 0.08 | 0.11 |
| 9 | 116409528 | rs602569 | -0.288 | 1.182 | 0.009 | 0.194 | -0.003 | -0.013 | -0.013 | 0 | 0.46 | 0.5 | 0.59 | 0.69 | 0.58 | 0.42 |
| 9 | 116410383 | rs6478027 | 0.277 | 0.381 | -0.147 | -0.17 | -0.022 | -0.029 | -0.029 | 0 | 0.18 | 0.3 | 0.14 | 0.18 | 0.07 | 0.17 |
| 9 | 116411416 | rs681218 | 1.027 | 0.421 | -1.042 | -0.126 | -0.033 | 0.012 | 0.012 | 0 | 0.22 | 0.02 | 0.18 | 0.15 | 0.2 | 0.16 |
| 9 | 116411753 | rs13291990 | 0.833 | 0.303 | -1.058 | 0.13 | 0.007 | -0.004 | -0.004 | 0 | 0.35 | 0.1 | 0.26 | 0.28 | 0.36 | 0.4 |
| 9 | 116412663 | rs7864770 | -0.153 | 0.56 | -0.093 | 0.255 | 0.034 | -0.082 | -0.082 | 0.022 | 0.3 | 0.53 | 0.28 | 0.17 | 0.31 | 0.25 |
| 9 | 116413354 | rs10981859 | -0.131 | -0.031 | 0.244 | 0.948 | 0 | 0.058 | 0.058 | 0 | 0.08 | 0.07 | 0.2 | 0.37 | 0.07 | 0.2 |
| 9 | 116413751 | rs10817497 | -0.774 | -0.058 | 0.352 | 0.956 | -0.005 | 0.091 | 0.091 | 0 | 0.1 | 0.16 | 0.29 | 0.37 | 0.07 | 0.21 |
| 9 | 116416163 | rs618624 | 0.41 | -0.354 | 0.322 | 0.358 | -0.038 | 0.035 | 0.035 | 0 | 0.71 | 0.54 | 0.6 | 0.45 | 0.81 | 0.68 |
| 9 | 116418881 | rs1077072 | -0.436 | -0.553 | 0.539 | -0.314 | -0.025 | -0.026 | -0.026 | 0 | 0.27 | 0.4 | 0.18 | 0.15 | 0.14 | 0.22 |
| 9 | 116420246 | rs2622266 | 0.815 | -0.577 | 0.833 | -0.425 | -0.006 | -0.011 | -0.011 | 0 | 0.69 | 0.29 | 0.8 | 0.87 | 0.84 | 0.81 |
| 9 | 116421786 | rs2101069 | -0.652 | 0.281 | -0.078 | -0.479 | -0.003 | 0.023 | 0.023 | 0 | 0.36 | 0.33 | 0.23 | 0.12 | 0.33 | 0.26 |
| 9 | 116432208 | rs586397 | -0.949 | 0.36 | -0.126 | -0.034 | 0.02 | 0.032 | 0.032 | 0 | 0.44 | 0.63 | 0.52 | 0.72 | 0.32 | 0.35 |
| 9 | 116432952 | rs536984 | -1.505 | 0.449 | -0.317 | -0.293 | -0.016 | 0.063 | 0.063 | 0 | 0.56 | 0.7 | 0.64 | 0.72 | 0.51 | 0.45 |
| 9 | 116433501 | rs10117193 | 0.135 | -0.166 | -0.054 | -0.212 | 0.018 | 0.037 | 0.037 | 0 | 0.23 | 0.12 | 0.11 | 0.1 | 0.15 | 0.16 |
| 9 | 116434346 | rs784935 | -0.247 | -0.034 | 0.484 | 0.605 | 0.001 | 0.178 | 0.178 | 0 | 0.15 | 0.31 | 0.37 | 0.4 | 0.12 | 0.1 |
| 9 | 116435724 | rs784932 | -1.123 | 0.225 | 0.211 | 0.193 | 0.032 | 0.081 | 0.081 | 0 | 0.47 | 0.7 | 0.54 | 0.73 | 0.37 | 0.38 |
| 9 | 116436877 | rs3860173 | -0.572 | 0.264 | 0.369 | 0.379 | -0.006 | 0.003 | 0.003 | 0 | 0.32 | 0.35 | 0.38 | 0.61 | 0.25 | 0.3 |
| 9 | 116438047 | rs554920 | -1.353 | 0.432 | 0.761 | 0.533 | 0.003 | -0.046 | -0.046 | 0 | 0.18 | 0.57 | 0.18 | 0.12 | 0.19 | 0.14 |
| 9 | 116439749 | rs7852392 | -1.421 | 0.531 | -0.171 | 1.074 | -0.001 | -0.057 | -0.057 | 0.726 | 0.14 | 0.03 | 0.15 | 0.29 | 0.13 | 0.1 |
| 9 | 116441603 | rs7860671 | -0.71 | 0.887 | 0.213 | 1.509 | 0.004 | 0.084 | 0.084 | 0 | 0.1 | 0.17 | 0.42 | 0.48 | 0.14 | 0.09 |
| 9 | 116443721 | rs9644931 | -1.859 | 0.734 | 0.548 | 1.432 | 0.019 | 0.156 | 0.156 | 0.011 | 0.17 | 0.45 | 0.48 | 0.56 | 0.19 | 0.14 |
| 9 | 116444269 | rs4979274 | -0.296 | 0.764 | -0.267 | 1.531 | -0.03 | -0.031 | -0.031 | 0 | 0.09 | 0.01 | 0.39 | 0.42 | 0.1 | 0.07 |

| | | | | | | | | | | | | | | | | |
|---|---|---|---|---|---|---|---|---|---|---|---|---|---|---|---|---|
| 9 | 116445988 | rs16907988 | -1.004 | 0.708 | -0.058 | 0.736 | 0.004 | -0.031 | -0.031 | 3.78 | 0.07 | 0.14 | 0.04 | 0.01 | 0.05 | 0.07 |
| 9 | 116446881 | rs3904317 | -1 | 0.83 | -0.517 | 1.219 | -0.018 | -0.058 | -0.058 | 0 | 0.14 | 0.06 | 0.41 | 0.49 | 0.17 | 0.16 |
| 9 | 116447723 | rs586348 | -0.803 | 0.552 | -1.332 | 0.106 | -0.085 | 0.054 | 0.054 | 0 | 0.55 | 0.92 | 0.72 | 0.64 | 0.5 | 0.43 |
| 9 | 116453001 | rs7855534 | -0.41 | 0.12 | -0.678 | -0.583 | -0.001 | -0.053 | -0.053 | 0 | 0.36 | 0.57 | 0.26 | 0.06 | 0.32 | 0.24 |
| 9 | 116460731 | rs11999448 | -1.135 | 0.206 | 0.139 | -0.448 | 0.02 | -0.028 | -0.028 | 1.58 | 0.07 | 0.25 | 0.07 | 0 | 0.05 | 0.02 |
| 9 | 116461477 | rs10981892 | -0.936 | -0.151 | -0.117 | -0.28 | 0.01 | 0.005 | 0.005 | 1.492 | 0.5 | 0.42 | 0.59 | 0.54 | 0.42 | 0.47 |
| 9 | 116462317 | rs10817507 | 0.445 | -0.228 | 0.079 | -0.586 | 0.002 | -0.026 | -0.026 | 0 | 0.28 | 0.22 | 0.15 | 0.04 | 0.24 | 0.18 |
| 9 | 116464160 | rs2170226 | -0.951 | -0.218 | 0.103 | -0.319 | -0.012 | 0.045 | 0.045 | 0 | 0.24 | 0.3 | 0.48 | 0.5 | 0.18 | 0.26 |
| 9 | 116468123 | rs16908163 | 0.426 | -0.445 | 0.195 | -0.578 | -0.01 | -0.087 | -0.087 | 0 | 0.29 | 0.29 | 0.15 | 0.03 | 0.23 | 0.18 |
| 9 | 116468685 | rs10123403 | -1.305 | -0.438 | 0.284 | -0.507 | 0.002 | -0.001 | -0.001 | 1.297 | 0.52 | 0.52 | 0.63 | 0.51 | 0.42 | 0.44 |
| 9 | 116475151 | rs10817510 | -1.663 | -0.303 | 0.356 | -0.448 | -0.021 | -0.027 | -0.027 | 0 | 0.57 | 0.78 | 0.67 | 0.53 | 0.48 | 0.49 |
| 9 | 116477070 | rs7852684 | -0.248 | -0.328 | 0.407 | -0.484 | -0.016 | -0.127 | -0.127 | 0.018 | 0.34 | 0.51 | 0.22 | 0.05 | 0.29 | 0.2 |
| 9 | 116478405 | rs10981899 | 0.19 | -0.311 | 0.331 | -0.442 | 0.009 | -0.105 | -0.105 | 0.504 | 0.17 | 0.05 | 0.41 | 0.48 | 0.14 | 0.25 |
| 9 | 116486584 | rs10123658 | -0.728 | -0.239 | 0.439 | -0.397 | 0.012 | -0.003 | -0.003 | 0 | 0.47 | 0.42 | 0.57 | 0.51 | 0.37 | 0.42 |
| 9 | 116493215 | rs1326328 | -1.635 | -0.174 | 0.216 | -0.411 | -0.022 | -0.021 | -0.021 | 0 | 0.58 | 0.72 | 0.67 | 0.53 | 0.49 | 0.49 |
| 9 | 116495444 | rs7866229 | -0.494 | -0.072 | 0.074 | -0.071 | 0.028 | -0.016 | -0.016 | 0 | 0.06 | 0.2 | 0.04 | 0.01 | 0.06 | 0.05 |
| 9 | 116498284 | rs10817516 | -0.312 | -0.107 | -0.211 | 0.04 | 0.011 | -0.03 | -0.03 | 0 | 0.19 | 0.12 | 0.42 | 0.48 | 0.14 | 0.25 |
| 9 | 116499027 | rs10817517 | 0.199 | 0.09 | -0.151 | -0.226 | 0.002 | 0.017 | 0.017 | 0 | 0.3 | 0.22 | 0.17 | 0.11 | 0.3 | 0.19 |
| 9 | 116501277 | rs1871692 | -1.211 | -0.346 | -0.077 | -0.055 | -0.022 | -0.007 | -0.007 | 0 | 0.37 | 0.56 | 0.55 | 0.64 | 0.29 | 0.35 |
| 9 | 116506979 | rs9792536 | 1.399 | 0.442 | -0.612 | -0.233 | -0.033 | 0.03 | 0.03 | 0.172 | 0.35 | 0.13 | 0.28 | 0.34 | 0.39 | 0.47 |
| 9 | 116507666 | rs7045057 | -0.536 | 0.219 | -0.39 | -0.281 | -0.006 | -0.021 | -0.021 | 0.214 | 0.24 | 0.34 | 0.14 | 0.1 | 0.21 | 0.15 |
| 9 | 116508048 | rs12003007 | -0.471 | 0.219 | -0.39 | -0.281 | -0.006 | -0.022 | -0.022 | 0 | 0.24 | 0.34 | 0.14 | 0.1 | 0.21 | 0.15 |
| 9 | 116508649 | rs7847036 | 0.227 | 0.552 | -0.446 | -0.296 | -0.002 | 0.014 | 0.014 | 0 | 0.14 | 0.14 | 0.39 | 0.5 | 0.14 | 0.19 |
| 9 | 116509853 | rs7866886 | 0.159 | 0.548 | -0.438 | -0.298 | -0.002 | 0.015 | 0.015 | 0 | 0.14 | 0.15 | 0.39 | 0.5 | 0.14 | 0.19 |
| 9 | 116512100 | rs7032818 | -1.435 | 0.549 | -0.386 | -0.331 | 0 | 0.003 | 0.003 | 0 | 0.37 | 0.42 | 0.52 | 0.63 | 0.35 | 0.34 |
| 9 | 116514497 | rs16908855 | -0.291 | 0.54 | -0.386 | -0.334 | 0.001 | 0.001 | 0.001 | 0 | 0.23 | 0.25 | 0.13 | 0.08 | 0.21 | 0.15 |
| 9 | 116517302 | rs10981912 | 0.389 | 0.672 | -0.194 | -0.213 | -0.001 | 0 | 0 | 0 | 0.59 | 0.45 | 0.41 | 0.46 | 0.6 | 0.62 |
| 9 | 116521430 | rs10981914 | 0.313 | 0.774 | -0.477 | -0.048 | -0.015 | -0.033 | -0.033 | 0.119 | 0.13 | 0.06 | 0.38 | 0.5 | 0.14 | 0.19 |
| 9 | 116528183 | rs7868736 | -1.107 | 0.741 | -0.223 | -0.534 | 0.003 | -0.111 | -0.111 | 0 | 0.75 | 0.71 | 0.81 | 0.97 | 0.74 | 0.82 |
| 9 | 116530593 | rs1994002 | 1.096 | 0.81 | -0.869 | 0.041 | -0.01 | 0.084 | 0.084 | 2.378 | 0.44 | 0.15 | 0.54 | 0.77 | 0.49 | 0.55 |
| 9 | 116535326 | rs16909256 | -0.394 | 0.784 | -0.914 | -0.091 | -0.006 | 0.023 | 0.023 | 0 | 0.24 | 0.23 | 0.17 | 0.1 | 0.26 | 0.14 |
| 9 | 116535410 | rs1359637 | 0.562 | 0.825 | -0.907 | 0.074 | -0.018 | 0.015 | 0.015 | 0 | 0.53 | 0.33 | 0.6 | 0.67 | 0.52 | 0.67 |
| 9 | 116545758 | rs1022949 | -0.917 | 0.938 | -0.931 | 0.175 | -0.017 | -0.027 | -0.027 | 0 | 0.89 | 0.96 | 0.85 | 0.82 | 0.88 | 0.86 |
| 9 | 116546918 | rs987073 | 1.307 | 0.548 | -0.818 | 0.555 | 0.009 | 0.018 | 0.018 | 0.042 | 0.38 | 0.22 | 0.3 | 0.41 | 0.38 | 0.53 |
| 9 | 116553683 | rs7030473 | 1.006 | 0.71 | -0.415 | 0.63 | 0.011 | 0.019 | 0.019 | 0 | 0.27 | 0.33 | 0.42 | 0.29 | 0.3 | 0.19 |
| 9 | 116555563 | rs913080 | 0.946 | 0.643 | -0.422 | 0.843 | 0.005 | -0.004 | -0.004 | 0.095 | 0.42 | 0.45 | 0.29 | 0.57 | 0.36 | 0.6 |
| 9 | 116555661 | rs10817520 | 1.421 | 0.621 | -0.658 | 0.919 | -0.035 | -0.031 | -0.031 | 0 | 0.36 | 0.07 | 0.25 | 0.41 | 0.34 | 0.52 |
| 9 | 116560635 | rs7037774 | -0.299 | 0.894 | -0.271 | 1.071 | -0.036 | 0.005 | 0.005 | 0.027 | 0.83 | 0.95 | 0.8 | 0.87 | 0.75 | 0.91 |
| 9 | 116564763 | rs988015 | -1.933 | 0.345 | 0.258 | 1.081 | -0.004 | 0.207 | 0.207 | 2.568 | 0.08 | 0.43 | 0.05 | 0.16 | 0.01 | 0.07 |
| 9 | 116565746 | rs10981934 | -1.191 | 0.355 | 0.317 | 1.089 | 0 | 0.197 | 0.197 | 0.587 | 0.06 | 0.3 | 0.04 | 0.16 | 0.01 | 0.07 |
| 9 | 116573081 | rs10981940 | -1.572 | 0.422 | 0.353 | 1.196 | 0.012 | 0.568 | 0.568 | 0.683 | 0.07 | 0.43 | 0.25 | 0.42 | 0.02 | 0.11 |
| 9 | 116629769 | rs1330165 | -1.6 | 0.511 | -0.059 | 1.105 | 0.024 | 0.433 | 0.433 | 0.755 | 0.08 | 0.43 | 0.3 | 0.4 | 0.05 | 0.11 |
| 9 | 116635490 | rs4978557 | -0.709 | 0.425 | -1.174 | 0.575 | 0.007 | -0.048 | -0.048 | 0 | 0.5 | 0.88 | 0.56 | 0.44 | 0.41 | 0.42 |
| 9 | 116652328 | rs1999203 | -0.581 | -0.015 | -0.736 | 1.224 | -0.004 | 0.052 | 0.052 | 0 | 0.1 | 0.1 | 0.23 | 0.51 | 0.05 | 0.16 |
| 9 | 116660999 | rs17200294 | 1.02 | -0.009 | -1.239 | 0.528 | -0.007 | 0.028 | 0.028 | 0 | 0.21 | 0.16 | 0.25 | 0.03 | 0.25 | 0.15 |
| 9 | 116668012 | rs10982000 | -0.308 | 0.039 | -0.954 | 0.351 | -0.001 | 0 | 0 | 0 | 0.25 | 0.22 | 0.15 | 0.36 | 0.27 | 0.26 |
| 9 | 116668681 | rs2150014 | -0.92 | 0.063 | -0.756 | 0.29 | -0.001 | 0 | 0 | 0 | 0.3 | 0.37 | 0.4 | 0.65 | 0.31 | 0.32 |
| 9 | 116670142 | rs7038501 | -1.389 | 0.218 | -0.304 | 0.321 | -0.002 | 0.028 | 0.028 | 0 | 0.54 | 0.67 | 0.66 | 0.67 | 0.58 | 0.47 |
| 9 | 116672801 | rs7034929 | 0.459 | -0.1 | -0.367 | 0.191 | 0.004 | -0.001 | -0.001 | 0 | 0.17 | 0.15 | 0.13 | 0.15 | 0.25 | 0.15 |
| 9 | 116680454 | rs7852787 | -0.034 | -0.254 | -0.257 | 0.286 | -0.03 | 0.086 | 0.086 | 0.462 | 0.39 | 0.18 | 0.36 | 0.15 | 0.49 | 0.31 |
| 9 | 116683645 | rs882980 | -0.56 | -0.305 | -0.184 | 0.481 | -0.002 | 0.012 | 0.012 | 0 | 0.48 | 0.45 | 0.44 | 0.35 | 0.54 | 0.41 |
| 9 | 116684240 | rs4979298 | -1.181 | -0.465 | -0.29 | 0.419 | -0.001 | 0.008 | 0.008 | 0 | 0.54 | 0.63 | 0.67 | 0.64 | 0.58 | 0.48 |
| 9 | 116697000 | rs10513223 | -0.096 | -0.394 | -0.674 | 0.395 | -0.001 | 0.029 | 0.029 | 0 | 0.4 | 0.28 | 0.32 | 0.26 | 0.41 | 0.39 |
| 9 | 116702866 | rs10448275 | 0.09 | -0.445 | -0.765 | 0.147 | 0.008 | 0.052 | 0.052 | 0 | 0.5 | 0.27 | 0.39 | 0.36 | 0.55 | 0.4 |
| 9 | 116704466 | rs11791817 | 1.007 | -0.458 | -0.85 | 0.143 | -0.009 | -0.024 | -0.024 | 0 | 0.06 | 0.11 | 0.03 | 0 | 0.03 | 0.01 |
| 9 | 116711241 | rs7853673 | 0.044 | -0.766 | -0.664 | 0.196 | 0 | 0.015 | 0.015 | 0.922 | 0.47 | 0.37 | 0.38 | 0.16 | 0.49 | 0.39 |
| 9 | 116715090 | rs1571578 | 1.619 | -0.362 | -1.081 | 0.765 | 0.02 | -0.03 | -0.03 | 0.651 | 0.13 | 0.02 | 0.05 | 0.21 | 0.13 | 0.11 |
| 9 | 116715706 | rs7046776 | 0.183 | 0.264 | -0.677 | 0.941 | 0.007 | 0.036 | 0.036 | 0 | 0.5 | 0.34 | 0.4 | 0.36 | 0.56 | 0.42 |
| 9 | 116722693 | rs10982017 | 0.099 | 0.369 | -0.952 | 0.564 | 0.002 | 0.024 | 0.024 | 0 | 0.51 | 0.4 | 0.61 | 0.69 | 0.56 | 0.48 |
| 9 | 116723095 | rs814694 | 0.868 | 0.423 | -1.144 | 0.702 | -0.012 | -0.018 | -0.018 | 0.022 | 0.05 | 0.13 | 0.03 | 0.01 | 0.03 | 0.01 |
| 9 | 116723488 | rs1544102 | -0.403 | 0.869 | -1.382 | 0.935 | 0 | -0.009 | -0.009 | 0.109 | 0.57 | 0.59 | 0.67 | 0.69 | 0.61 | 0.49 |
| 9 | 116724402 | rs16911006 | -0.365 | 0.982 | -1.463 | 1.394 | 0.001 | 0.006 | 0.006 | 0 | 0.24 | 0.33 | 0.39 | 0.3 | 0.25 | 0.21 |
| 9 | 116725481 | rs10817542 | 0.092 | 1.288 | -2.039 | 1.154 | -0.079 | -0.043 | -0.043 | 0 | 0.16 | 0.02 | 0.23 | 0.24 | 0.24 | 0.36 |
| 9 | 116725719 | rs16911010 | 0.001 | 0.35 | -1.405 | 1.3 | -0.013 | -0.011 | -0.011 | 0.228 | 0.19 | 0.09 | 0.14 | 0.17 | 0.2 | 0.18 |
| 9 | 116726904 | rs4979309 | -0.813 | 0.072 | -0.704 | 1.412 | -0.004 | 0.016 | 0.016 | 1.216 | 0.18 | 0.17 | 0.35 | 0.34 | 0.2 | 0.22 |
| 9 | 116733183 | rs1544065 | -1.081 | 0.107 | -0.703 | 1.035 | 0.002 | -0.002 | -0.002 | 0.029 | 0.18 | 0.2 | 0.14 | 0.19 | 0.13 | 0.23 |
| 9 | 116735001 | rs7021972 | -0.945 | 0.526 | -0.857 | 0.791 | -0.037 | -0.071 | -0.071 | 0 | 0.33 | 0.15 | 0.29 | 0.39 | 0.36 | 0.39 |
| 9 | 116736791 | rs10982023 | 0.992 | 0.225 | 0.174 | 0.009 | 0.005 | -0.053 | -0.053 | 0 | 0.91 | 0.77 | 0.91 | 1 | 0.93 | 0.95 |
| 9 | 116739981 | rs12351979 | 0.301 | -0.639 | -1.034 | -1.051 | 0.031 | 0.181 | 0.181 | 0 | 0.29 | 0.05 | 0.2 | 0.01 | 0.21 | 0.23 |

| | | | | | | | | | | | | | | | | |
|---|---|---|---|---|---|---|---|---|---|---|---|---|---|---|---|---|
| 9 | 116745655 | rs1006541 | 1.079 | 0.676 | -0.201 | -0.217 | 0.067 | 0.065 | 0.065 | 0.004 | 0.77 | 0.58 | 0.66 | 0.36 | 0.62 | 0.65 |
| 9 | 116749297 | rs4979310 | -0.357 | 0.705 | -0.171 | -0.21 | 0.015 | 0.032 | 0.032 | 0 | 0.14 | 0.19 | 0.27 | 0.6 | 0.31 | 0.3 |
| 9 | 116751850 | rs10817547 | 1.301 | 0.695 | 0.408 | -0.342 | 0.06 | 0.085 | 0.085 | 0 | 0.75 | 0.52 | 0.66 | 0.4 | 0.65 | 0.65 |
| 9 | 116752771 | rs11791155 | -0.099 | 0.795 | 0.122 | -0.275 | -0.064 | -0.063 | -0.063 | 0 | 0.11 | 0.03 | 0.25 | 0.6 | 0.27 | 0.29 |
| 9 | 116763905 | rs700125 | 0.239 | 0.387 | 0.298 | -0.387 | -0.007 | -0.036 | -0.036 | 0 | 0.92 | 0.93 | 0.96 | 1 | 0.95 | 0.97 |
| 9 | 116764283 | rs4978561 | -0.558 | 1.003 | 0.213 | -0.388 | -0.03 | -0.028 | -0.028 | 0.616 | 0.14 | 0.13 | 0.29 | 0.6 | 0.35 | 0.33 |
| 9 | 116766112 | rs942288 | 0.616 | 0.922 | 0.699 | -0.434 | -0.025 | -0.069 | -0.069 | 1.319 | 0.88 | 0.77 | 0.94 | 1 | 0.94 | 0.97 |
| 9 | 116766695 | rs7858120 | -0.594 | 0.999 | 0.698 | -0.348 | 0.02 | 0.029 | 0.029 | 0 | 0.14 | 0.18 | 0.28 | 0.6 | 0.35 | 0.33 |
| 9 | 116768637 | rs4978562 | -0.568 | 1.02 | 0.659 | -0.327 | -0.015 | -0.012 | -0.012 | 0 | 0.14 | 0.14 | 0.28 | 0.61 | 0.35 | 0.33 |
| 9 | 116770186 | rs2025391 | 0.635 | 0.402 | 0.523 | -1.016 | -0.005 | -0.021 | -0.021 | 0 | 0.35 | 0.39 | 0.36 | 0.16 | 0.3 | 0.26 |
| 9 | 116776289 | rs10982031 | -0.157 | 0.503 | 0.554 | -0.936 | 0.043 | 0.045 | 0.045 | 0 | 0.14 | 0.14 | 0.26 | 0.65 | 0.34 | 0.32 |
| 9 | 116778644 | rs1853546 | -0.034 | 0.518 | 0.511 | -1.154 | 0.051 | 0.058 | 0.058 | 0 | 0.53 | 0.36 | 0.61 | 0.78 | 0.68 | 0.64 |
| 9 | 116779813 | rs10817557 | 0.127 | 0.334 | 0.613 | -1.127 | 0.059 | 0.098 | 0.098 | 0 | 0.5 | 0.29 | 0.59 | 0.77 | 0.64 | 0.62 |
| 9 | 116780314 | rs4979325 | 0.138 | 0.328 | 0.557 | -1.129 | 0.029 | 0.047 | 0.047 | 0 | 0.51 | 0.4 | 0.6 | 0.77 | 0.64 | 0.62 |
| 9 | 116780465 | rs4979326 | 0.301 | 0.287 | 0.53 | -1.148 | -0.026 | -0.023 | -0.023 | 0 | 0.13 | 0.06 | 0.27 | 0.59 | 0.34 | 0.34 |
| 9 | 116780559 | rs13297916 | 0.882 | 0.295 | 0.541 | -1.13 | 0.014 | 0.024 | 0.024 | 0 | 0.38 | 0.35 | 0.33 | 0.17 | 0.3 | 0.28 |
| 9 | 116780992 | rs7023951 | 0.851 | 0.107 | 0.192 | -1.353 | -0.01 | -0.087 | -0.087 | 1.418 | 0.31 | 0.42 | 0.29 | 0.06 | 0.28 | 0.26 |
| 9 | 116782921 | rs10982035 | 0.942 | -0.577 | 0.095 | -1.302 | 0.029 | 0.138 | 0.138 | 0 | 0.37 | 0.1 | 0.3 | 0.12 | 0.29 | 0.28 |
| 9 | 116789571 | rs10982039 | -0.908 | -0.921 | 0.342 | -1.398 | -0.064 | -0.107 | -0.107 | 0.067 | 0.22 | 0.53 | 0.1 | 0.04 | 0.09 | 0.05 |
| 9 | 116789690 | rs10125470 | -0.136 | -0.687 | 0.35 | -1.285 | 0.012 | 0.047 | 0.047 | 0.311 | 0.57 | 0.63 | 0.4 | 0.17 | 0.38 | 0.31 |
| 9 | 116790352 | rs7040743 | -0.054 | -0.653 | 0.393 | -1.332 | 0.014 | 0.045 | 0.045 | 0.139 | 0.56 | 0.61 | 0.4 | 0.13 | 0.38 | 0.31 |
| 9 | 116790511 | rs10759679 | 0.79 | -0.583 | -0.022 | -1.242 | 0.001 | -0.01 | -0.01 | 0 | 0.43 | 0.6 | 0.36 | 0.13 | 0.3 | 0.27 |
| 9 | 116800606 | rs10121424 | -0.976 | -0.048 | -0.642 | -0.939 | 0.058 | 0.076 | 0.076 | 0 | 0.57 | 0.85 | 0.64 | 0.7 | 0.7 | 0.63 |
| 9 | 116800860 | rs10124293 | -0.929 | -0.036 | -0.521 | -0.931 | 0.056 | 0.068 | 0.068 | 0 | 0.57 | 0.8 | 0.64 | 0.69 | 0.7 | 0.63 |
| 9 | 116805634 | rs12555049 | -0.871 | 0.406 | -0.088 | -0.405 | -0.003 | -0.032 | -0.032 | 0 | 0.14 | 0.12 | 0.06 | 0 | 0.09 | 0.04 |
| 9 | 116810204 | rs12378906 | -0.884 | 0.406 | -0.088 | -0.404 | -0.003 | -0.031 | -0.031 | 0.187 | 0.14 | 0.12 | 0.06 | 0 | 0.09 | 0.04 |
| 9 | 116812084 | rs7856141 | -0.773 | 0.46 | -0.053 | -0.378 | -0.006 | -0.038 | -0.038 | 0 | 0.13 | 0.13 | 0.06 | 0 | 0.09 | 0.04 |
| 9 | 116814126 | rs7037095 | 0.871 | 1.445 | 0.132 | 0.483 | 0.064 | 0.139 | 0.139 | 0 | 0.86 | 0.74 | 0.68 | 0.42 | 0.78 | 0.69 |
| 9 | 116814661 | rs12683020 | -0.298 | 1.548 | -0.485 | 0.546 | -0.058 | -0.064 | -0.064 | 4.781 | 0.11 | 0.02 | 0.3 | 0.57 | 0.22 | 0.29 |
| 9 | 116815777 | rs3810928 | -1.111 | 1.378 | -0.881 | -0.05 | -0.133 | 0.029 | 0.029 | 0.005 | 0.29 | 0.07 | 0.43 | 0.75 | 0.43 | 0.4 |
| 9 | 116821526 | rs3860175 | -0.035 | 1.139 | -0.028 | 0.388 | 0.015 | 0.005 | 0.005 | 0 | 0.21 | 0.27 | 0.29 | 0.23 | 0.28 | 0.18 |
| 9 | 116825675 | rs7034903 | 0.222 | 0.837 | -0.799 | -0.231 | 0.003 | 0.054 | 0.054 | 0 | 0.27 | 0.17 | 0.13 | 0.11 | 0.21 | 0.23 |
| 9 | 116825745 | rs10759680 | 0.13 | 0.664 | -0.782 | 0.469 | -0.009 | -0.037 | -0.037 | 0 | 0.24 | 0.14 | 0.27 | 0.38 | 0.26 | 0.29 |
| 9 | 116826842 | rs10817563 | 0.125 | 0.647 | -0.778 | 0.463 | -0.01 | -0.033 | -0.033 | 0 | 0.24 | 0.16 | 0.27 | 0.38 | 0.26 | 0.29 |
| 9 | 116837745 | rs12344124 | 0.23 | 0.405 | -0.358 | 0.188 | 0.001 | -0.049 | -0.049 | 0 | 0.1 | 0.13 | 0.07 | 0 | 0.11 | 0.03 |
| 9 | 116837945 | rs2567698 | -0.128 | 0.595 | -0.307 | 0.536 | 0.004 | 0.042 | 0.042 | 0 | 0.15 | 0.12 | 0.33 | 0.44 | 0.22 | 0.23 |
| 9 | 116838743 | rs2567700 | -0.586 | 0.405 | -0.467 | 0.636 | -0.003 | -0.052 | -0.052 | 1.71 | 0.21 | 0.14 | 0.36 | 0.44 | 0.26 | 0.23 |
| 9 | 116840204 | rs2251680 | -0.066 | -0.03 | -1.008 | -0.231 | -0.016 | -0.007 | -0.007 | 0 | 0.55 | 0.19 | 0.49 | 0.51 | 0.46 | 0.57 |
| 9 | 116841180 | rs2285055 | -0.998 | -0.004 | -0.648 | -0.514 | -0.001 | 0 | 0 | 0 | 0.72 | 0.74 | 0.62 | 0.5 | 0.61 | 0.65 |
| 9 | 116844869 | rs13283242 | 1.497 | -0.209 | -0.941 | -0.758 | 0.063 | 0.132 | 0.132 | 0 | 0.31 | 0.1 | 0.1 | 0.06 | 0.14 | 0.28 |
| 9 | 116847201 | rs891725 | 1.122 | -0.167 | -0.615 | -0.665 | -0.029 | -0.059 | -0.059 | 0 | 0.1 | 0.01 | 0.27 | 0.4 | 0.13 | 0.25 |
| 9 | 116847365 | rs891724 | 0.537 | -0.105 | -0.53 | -1.341 | 0 | -0.08 | -0.08 | 0 | 0.19 | 0.08 | 0.35 | 0.64 | 0.2 | 0.39 |
| 9 | 116849069 | rs7862938 | 1.237 | -0.047 | -0.277 | -1.342 | 0 | -0.028 | -0.028 | 0 | 0.11 | 0.12 | 0.08 | 0 | 0.14 | 0.03 |
| 9 | 116849447 | rs728071 | 0.478 | -0.06 | -0.288 | -1.35 | -0.002 | -0.078 | -0.078 | 0 | 0.14 | 0.35 | 0.11 | 0 | 0.14 | 0.03 |
| 9 | 116850345 | rs10817565 | 1.755 | -0.139 | -0.86 | -1.551 | 0.066 | 0.135 | 0.135 | 0.211 | 0.29 | 0.1 | 0.08 | 0.04 | 0.11 | 0.23 |
| 9 | 116851076 | rs2093785 | -1.469 | 0.243 | -0.852 | -1.414 | 0 | -0.001 | -0.001 | 1.993 | 0.79 | 0.85 | 0.68 | 0.71 | 0.67 | 0.79 |
| 9 | 116853797 | rs7854030 | -0.174 | 0.069 | -0.627 | -1.429 | 0.014 | -0.016 | -0.016 | 0.635 | 0.62 | 0.55 | 0.56 | 0.71 | 0.5 | 0.7 |
| 9 | 116861763 | rs1886128 | -0.955 | 0.344 | -1.041 | -0.876 | 0.011 | 0.001 | 0.001 | 0.436 | 0.78 | 0.81 | 0.66 | 0.69 | 0.63 | 0.78 |
| 9 | 116865784 | rs2418265 | 0.083 | 0.274 | -1.053 | -0.917 | 0.002 | -0.007 | -0.007 | 0 | 0.61 | 0.52 | 0.54 | 0.67 | 0.46 | 0.69 |
| 9 | 116866605 | rs7856692 | 0.868 | 0.286 | -0.983 | -0.818 | -0.017 | -0.089 | -0.089 | 0 | 0.43 | 0.44 | 0.19 | 0.04 | 0.26 | 0.3 |
| 9 | 116867054 | rs7029198 | -0.105 | 1.12 | -0.626 | 0.008 | 0.026 | 0.003 | 0.003 | 0 | 0.73 | 0.65 | 0.63 | 0.67 | 0.6 | 0.72 |
| 9 | 116870232 | rs2900573 | -0.543 | 0.589 | -0.858 | -0.142 | -0.002 | 0.042 | 0.042 | 0.145 | 0.58 | 0.66 | 0.56 | 0.73 | 0.62 | 0.68 |
| 9 | 116870327 | rs2900574 | -0.618 | 1.419 | -0.395 | 0.83 | 0.022 | 0.041 | 0.041 | 2.099 | 0.21 | 0.18 | 0.33 | 0.45 | 0.37 | 0.43 |
| 9 | 116872524 | rs7047941 | 0.031 | 1.239 | -0.469 | 0.492 | 0.017 | 0.012 | 0.012 | 0 | 0.68 | 0.79 | 0.82 | 0.73 | 0.71 | 0.7 |
| 9 | 116873183 | rs10759682 | 0.225 | 0.542 | -0.505 | 0.191 | 0.001 | -0.006 | -0.006 | 0.977 | 0.57 | 0.6 | 0.56 | 0.32 | 0.44 | 0.43 |
| 9 | 116880005 | rs4978567 | 0.31 | 0.333 | -0.639 | 0.076 | 0.004 | 0.004 | 0.004 | 0 | 0.47 | 0.45 | 0.49 | 0.41 | 0.38 | 0.43 |
| 9 | 116880031 | rs10982068 | -0.774 | -0.056 | -0.682 | -0.139 | 0.002 | -0.004 | -0.004 | 0 | 0.13 | 0.15 | 0.07 | 0.17 | 0.1 | 0.21 |
| 9 | 116887405 | rs7856967 | -0.826 | -0.221 | -0.487 | -0.548 | -0.018 | -0.042 | -0.042 | 0.842 | 0.08 | 0.18 | 0.03 | 0 | 0.04 | 0.04 |
| 9 | 116887845 | rs7857688 | -0.672 | -0.565 | -1.044 | -1.096 | 0.017 | 0.006 | 0.006 | 0 | 0.22 | 0.19 | 0.11 | 0 | 0.19 | 0.08 |
| 9 | 116890596 | rs2418267 | 0.458 | -1.074 | -0.846 | -1.403 | 0.002 | -0.035 | -0.035 | 1.834 | 0.74 | 0.75 | 0.87 | 1 | 0.8 | 0.89 |
| 9 | 116895387 | rs13300798 | 0.499 | -1.022 | -1.123 | -1.374 | 0.009 | 0.08 | 0.08 | 0.008 | 0.14 | 0.02 | 0.08 | 0 | 0.16 | 0.03 |
| 9 | 116896966 | rs891721 | -0.7 | -0.721 | -0.858 | -1.767 | -0.017 | -0.094 | -0.094 | 0 | 0.39 | 0.44 | 0.28 | 0.08 | 0.33 | 0.34 |
| 9 | 116897213 | rs891720 | 0.891 | 0.318 | -1.546 | -1.381 | -0.084 | -0.103 | -0.103 | 0.017 | 0.16 | 0.02 | 0.34 | 0.6 | 0.26 | 0.34 |
| 9 | 116901926 | rs1017360 | -0.59 | 0.141 | -1.323 | -1.674 | -0.013 | 0.021 | 0.021 | 1.869 | 0.37 | 0.29 | 0.6 | 0.8 | 0.49 | 0.7 |
| 9 | 116902782 | rs7868176 | 1.446 | 0.086 | -1.649 | -1.626 | 0.029 | 0.054 | 0.054 | 0 | 0.2 | 0.09 | 0.12 | 0.03 | 0.14 | 0.15 |
| 9 | 116903410 | rs737174 | 0.63 | 0.282 | -1.383 | -1.394 | 0.014 | 0.081 | 0.081 | 0 | 0.33 | 0.17 | 0.27 | 0.07 | 0.26 | 0.37 |
| 9 | 116907448 | rs10817573 | 0.027 | 0.828 | -0.885 | -0.419 | 0.016 | 0.03 | 0.03 | 0 | 0.07 | 0.13 | 0.11 | 0.15 | 0.09 | 0.14 |
| 9 | 116908453 | rs2002284 | 0.017 | 0.573 | -1.267 | -0.562 | -0.024 | -0.005 | -0.005 | 0 | 0.84 | 0.87 | 0.87 | 0.84 | 0.81 | 0.96 |
| 9 | 116913145 | rs10739403 | 0.244 | -0.126 | -1.289 | -0.95 | 0.02 | 0.048 | 0.048 | 0 | 0.61 | 0.69 | 0.77 | 0.77 | 0.68 | 0.73 |

| | | | | | | | | | | | | | | | | |
|---|---|---|---|---|---|---|---|---|---|---|---|---|---|---|---|---|
| 9 | 116913717 | rs919641 | -0.387 | -1.353 | -0.853 | -1.979 | -0.057 | -0.16 | -0.16 | 0 | 0.33 | 0.55 | 0.17 | 0 | 0.18 | 0.24 |
| 9 | 116913794 | rs2808772 | -0.559 | -0.304 | -0.817 | -0.978 | -0.001 | 0 | 0 | 0 | 0.49 | 0.55 | 0.6 | 0.55 | 0.45 | 0.63 |
| 9 | 116916904 | rs1469599 | 1.122 | -0.571 | -0.967 | -1.124 | 0.012 | 0.017 | 0.017 | 0.058 | 0.12 | 0.09 | 0.06 | 0 | 0.09 | 0.05 |
| 9 | 116916931 | rs1469598 | 0.48 | -0.818 | -0.798 | -0.716 | -0.026 | -0.028 | -0.028 | 0.225 | 0.31 | 0.21 | 0.52 | 0.42 | 0.36 | 0.54 |
| 9 | 116920024 | rs2808781 | -0.485 | -1.055 | -0.812 | -0.855 | 0.001 | 0 | 0 | 0 | 0.49 | 0.36 | 0.61 | 0.51 | 0.48 | 0.61 |
| 9 | 116921849 | rs2808785 | 0.018 | -0.883 | -0.964 | -0.899 | 0.011 | 0.035 | 0.035 | 0 | 0.25 | 0.17 | 0.16 | 0.11 | 0.22 | 0.19 |
| 9 | 116929327 | rs737142 | 1.408 | -0.807 | -0.297 | -0.634 | 0.041 | 0.047 | 0.047 | 0 | 0.19 | 0.26 | 0.46 | 0.38 | 0.28 | 0.41 |
| 9 | 116930100 | rs2567707 | 1.178 | -0.917 | -0.744 | -1.288 | 0.003 | 0.054 | 0.054 | 0.35 | 0.27 | 0.16 | 0.14 | 0 | 0.27 | 0.09 |
| 9 | 116930194 | rs2567706 | 0.69 | -1.003 | -0.289 | -1.292 | -0.003 | -0.019 | -0.019 | 0 | 0.31 | 0.3 | 0.17 | 0.13 | 0.29 | 0.12 |
| 9 | 116931099 | rs2241671 | -0.133 | -1.234 | -0.33 | -1.323 | 0.02 | 0.007 | 0.007 | 0 | 0.45 | 0.3 | 0.32 | 0.51 | 0.43 | 0.32 |
| 9 | 116931445 | rs2808770 | 0.809 | -1.101 | -0.315 | -1.513 | -0.002 | -0.017 | -0.017 | 0 | 0.31 | 0.3 | 0.18 | 0.13 | 0.3 | 0.13 |
| 9 | 116931737 | rs2808771 | 0.82 | -1.1 | -0.316 | -1.512 | -0.002 | -0.015 | -0.015 | 0 | 0.31 | 0.3 | 0.18 | 0.13 | 0.3 | 0.13 |
| 9 | 116932906 | rs2567704 | -1.478 | -1.217 | -0.113 | -1.571 | 0.003 | -0.028 | -0.028 | 0 | 0.87 | 0.76 | 0.9 | 0.96 | 0.85 | 0.93 |
| 9 | 116934281 | rs2808777 | 0.749 | -1.174 | -0.202 | -1.454 | -0.005 | -0.021 | -0.021 | 0.392 | 0.3 | 0.32 | 0.17 | 0.16 | 0.27 | 0.1 |
| 9 | 116936905 | rs10982093 | 1.118 | -1.102 | -0.052 | -1.36 | 0.007 | -0.017 | -0.017 | 0.079 | 0.2 | 0.18 | 0.11 | 0 | 0.16 | 0.12 |
| 9 | 116937667 | rs10759684 | -0.752 | -1.026 | 0.054 | -1.321 | -0.001 | -0.011 | -0.011 | 0.301 | 0.62 | 0.63 | 0.77 | 0.8 | 0.61 | 0.82 |
| 9 | 116939620 | rs10759685 | -0.154 | -1.096 | 0.027 | -1.029 | -0.01 | 0.004 | 0.004 | 0.462 | 0.37 | 0.4 | 0.27 | 0.36 | 0.31 | 0.36 |
| 9 | 116941019 | rs3827676 | 0.766 | -1.182 | -0.267 | -1.068 | -0.013 | -0.017 | -0.017 | 0.19 | 0.31 | 0.25 | 0.54 | 0.6 | 0.35 | 0.55 |
| 9 | 116942796 | rs12001466 | -0.458 | -1.899 | 0.468 | -0.642 | -0.02 | 0.008 | 0.008 | 0.107 | 0.35 | 0.57 | 0.26 | 0.36 | 0.27 | 0.33 |
| 9 | 116943362 | rs1431818 | -0.761 | -1.95 | 0.39 | -0.577 | -0.017 | 0.249 | 0.249 | 0.546 | 0.16 | 0.47 | 0.16 | 0.36 | 0.12 | 0.2 |
| 9 | 116944333 | rs10759686 | 0.901 | -1.766 | 0.125 | -0.907 | 0.001 | 0.012 | 0.012 | 1.752 | 0.43 | 0.39 | 0.66 | 0.73 | 0.57 | 0.57 |
| 9 | 116944847 | rs10759687 | 0.353 | -1.754 | -0.363 | -0.879 | 0.03 | 0.053 | 0.053 | 1.098 | 0.52 | 0.59 | 0.73 | 0.77 | 0.62 | 0.66 |
| 9 | 116945454 | rs10759688 | -0.477 | -1.59 | -0.208 | -0.819 | -0.008 | -0.044 | -0.044 | 0 | 0.76 | 0.71 | 0.88 | 0.96 | 0.8 | 0.91 |
| 9 | 116947947 | rs10982094 | -0.514 | -1.242 | -0.099 | -0.389 | 0.028 | 0.044 | 0.044 | 0.32 | 0.32 | 0.19 | 0.2 | 0.19 | 0.25 | 0.3 |
| 9 | 116949449 | rs891718 | -1.239 | -1.193 | 0.952 | 0.013 | -0.01 | -0.01 | -0.01 | 0 | 0.17 | 0.33 | 0.14 | 0.12 | 0.15 | 0.21 |
| 9 | 116951459 | rs10817578 | -0.064 | -1.142 | 0.656 | 0.043 | 0.02 | -0.06 | -0.06 | 1.796 | 0.76 | 0.84 | 0.81 | 0.58 | 0.84 | 0.83 |
| 9 | 116958646 | rs2808788 | 0.289 | -0.865 | 1.163 | -0.182 | -0.007 | 0.052 | 0.052 | 0.106 | 0.71 | 0.53 | 0.76 | 0.51 | 0.77 | 0.82 |
| 9 | 116961156 | rs2567712 | 0.483 | -1.035 | 1.169 | -0.19 | -0.028 | 0.035 | 0.035 | 0 | 0.69 | 0.54 | 0.77 | 0.49 | 0.8 | 0.79 |
| 9 | 116963876 | rs2567713 | 0.298 | -0.888 | 1.173 | -0.232 | -0.015 | 0.008 | 0.008 | 0.778 | 0.63 | 0.58 | 0.74 | 0.46 | 0.75 | 0.78 |
| 9 | 116964061 | rs2567714 | 0.495 | -0.851 | 0.889 | -0.16 | 0.001 | 0 | 0 | 0.603 | 0.59 | 0.44 | 0.7 | 0.35 | 0.74 | 0.77 |
| 9 | 116965542 | rs2567716 | 0.482 | -0.88 | 0.171 | -0.317 | 0.011 | -0.027 | -0.027 | 0 | 0.61 | 0.74 | 0.73 | 0.48 | 0.67 | 0.8 |
| 9 | 116967263 | rs2567719 | 1.027 | -1.809 | 0.11 | -0.312 | 0.017 | 0.002 | 0.002 | 0 | 0.51 | 0.53 | 0.72 | 0.47 | 0.69 | 0.58 |
| 9 | 116967405 | rs13290696 | -0.245 | -1.751 | -0.658 | -0.279 | 0.021 | 0.055 | 0.055 | 0 | 0.14 | 0.04 | 0.03 | 0.02 | 0.08 | 0.23 |
| 9 | 116967952 | rs2567720 | 0.408 | -1.16 | -0.181 | 0.219 | 0.024 | -0.037 | -0.037 | 0 | 0.68 | 0.74 | 0.77 | 0.49 | 0.79 | 0.83 |
| 9 | 116971384 | rs12001716 | -1.333 | -0.967 | 0.583 | -0.1 | -0.028 | -0.105 | -0.105 | 0 | 0.17 | 0.48 | 0.1 | 0.03 | 0.12 | 0.12 |
| 9 | 116972955 | rs10982107 | -1.206 | -1.2 | 0.546 | 0.255 | -0.028 | 0.031 | 0.031 | 0 | 0.31 | 0.48 | 0.25 | 0.61 | 0.23 | 0.17 |
| 9 | 116973755 | rs4979354 | -0.269 | -1.421 | 0.697 | 0.227 | -0.013 | 0.039 | 0.039 | 0.241 | 0.22 | 0.32 | 0.18 | 0.56 | 0.14 | 0.1 |
| 9 | 116974295 | rs13300252 | -1.086 | -1.307 | 0.531 | 0.233 | -0.031 | 0.022 | 0.022 | 0 | 0.32 | 0.48 | 0.25 | 0.62 | 0.23 | 0.17 |
| 9 | 116979172 | rs7870474 | -0.451 | -1.935 | 0.681 | -0.539 | -0.054 | -0.089 | -0.089 | 0 | 0.28 | 0.41 | 0.09 | 0.1 | 0.13 | 0.31 |
| 9 | 116980052 | rs10982110 | 0.501 | -2.071 | 0.247 | -0.811 | 0.047 | 0.042 | 0.042 | 0.022 | 0.59 | 0.3 | 0.71 | 0.26 | 0.69 | 0.82 |
| 9 | 116980821 | rs10513237 | -0.285 | -2.205 | 0.581 | -0.657 | -0.004 | 0.002 | 0.002 | 0 | 0.09 | 0.12 | 0.06 | 0.14 | 0.08 | 0.05 |
| 9 | 116985526 | rs7027825 | -0.46 | -2.183 | 0.781 | -0.333 | -0.021 | 0.049 | 0.049 | 0.26 | 0.73 | 0.59 | 0.78 | 0.41 | 0.78 | 0.9 |
| 9 | 116986741 | rs16927476 | -0.639 | -2.219 | 0.828 | -0.486 | 0.013 | -0.001 | -0.001 | 0 | 0.11 | 0.07 | 0.05 | 0.15 | 0.09 | 0.08 |
| 9 | 116987587 | rs7855833 | -0.448 | -2.179 | 0.932 | -0.418 | -0.019 | 0.03 | 0.03 | 0.255 | 0.73 | 0.61 | 0.78 | 0.39 | 0.78 | 0.9 |
| 9 | 116989375 | rs871878 | 0.023 | -2.343 | 1.184 | -0.298 | -0.018 | 0.019 | 0.019 | 0.283 | 0.7 | 0.37 | 0.78 | 0.44 | 0.78 | 0.87 |
| 9 | 116989776 | rs902496 | -0.475 | -2.317 | 1.512 | -0.3 | -0.015 | 0.05 | 0.05 | 0.068 | 0.77 | 0.65 | 0.81 | 0.45 | 0.8 | 0.9 |
| 9 | 116989900 | rs11794759 | 1.5 | -2.461 | 1.018 | -0.501 | -0.041 | 0.05 | 0.05 | 0.199 | 0.37 | 0.14 | 0.64 | 0.21 | 0.57 | 0.57 |
| 9 | 116990995 | rs4979357 | -0.122 | -2.063 | 1.538 | -0.224 | -0.014 | 0.027 | 0.027 | 0.386 | 0.73 | 0.39 | 0.79 | 0.44 | 0.8 | 0.9 |
| 9 | 116991071 | rs7868992 | 0.683 | -1.807 | 1.537 | -0.305 | -0.013 | -0.003 | -0.003 | 0 | 0.64 | 0.45 | 0.68 | 0.33 | 0.7 | 0.84 |
| 9 | 116995483 | rs7875159 | -0.253 | -1.698 | 1.649 | -0.114 | -0.015 | 0.07 | 0.07 | 0 | 0.75 | 0.58 | 0.76 | 0.47 | 0.78 | 0.87 |
| 9 | 116996533 | rs7021841 | -1.391 | -1.795 | 1.38 | -0.249 | 0.025 | 0.004 | 0.004 | 0.942 | 0.08 | 0.39 | 0.03 | 0 | 0.01 | 0.02 |
| 9 | 116997048 | rs10982118 | 0.527 | -1.682 | 0.516 | -0.207 | 0.039 | -0.055 | -0.055 | 0.362 | 0.66 | 0.2 | 0.7 | 0.47 | 0.77 | 0.84 |
| 9 | 117000687 | rs1249715 | 0.358 | -1.41 | -0.401 | -0.804 | 0.045 | 0.071 | 0.071 | 0 | 0.42 | 0.21 | 0.22 | 0.29 | 0.36 | 0.34 |
| 9 | 117000801 | rs1982850 | 0.677 | -1.133 | -0.644 | -0.679 | -0.043 | 0 | 0 | 0.006 | 0.25 | 0.15 | 0.48 | 0.19 | 0.39 | 0.49 |
| 9 | 117004955 | rs7875437 | 0.359 | -1.608 | -1.224 | -0.919 | 0.031 | 0.049 | 0.049 | 0 | 0.37 | 0.1 | 0.19 | 0.29 | 0.34 | 0.28 |
| 9 | 117005624 | rs7850171 | 0.091 | -1.257 | -0.877 | -1.049 | 0.001 | 0.002 | 0.002 | 0 | 0.41 | 0.41 | 0.4 | 0.19 | 0.58 | 0.55 |
| 9 | 117009115 | rs1048469 | -0.507 | -0.303 | -0.708 | -0.537 | -0.001 | -0.008 | -0.008 | 0.014 | 0.48 | 0.46 | 0.33 | 0.36 | 0.45 | 0.4 |
| 9 | 117009953 | rs2029522 | 0.491 | -0.031 | -0.495 | 0.248 | -0.007 | 0.051 | 0.051 | 0 | 0.18 | 0.24 | 0.13 | 0.47 | 0.17 | 0.18 |
| 9 | 117010405 | rs1249710 | -0.259 | 0.511 | -0.559 | 0.637 | -0.004 | 0.011 | 0.011 | 0 | 0.35 | 0.3 | 0.25 | 0.35 | 0.36 | 0.31 |
| 9 | 117011595 | rs999599 | -0.386 | 0.553 | -0.798 | 0.646 | -0.006 | 0.015 | 0.015 | 0 | 0.35 | 0.25 | 0.25 | 0.35 | 0.36 | 0.31 |
| 9 | 117015956 | rs10982124 | -0.602 | 0.491 | -1.188 | 0.339 | 0.005 | 0.021 | 0.021 | 1.393 | 0.41 | 0.6 | 0.33 | 0.77 | 0.38 | 0.41 |
| 9 | 117019167 | rs4979361 | -0.612 | 0.447 | -1.249 | 1.23 | 0 | -0.029 | -0.029 | 0 | 0.09 | 0.02 | 0.11 | 0.38 | 0.07 | 0.16 |
| 9 | 117019801 | rs923601 | -0.338 | 0.446 | -1.48 | 0.085 | 0.007 | 0.094 | 0.094 | 0 | 0.54 | 0.74 | 0.51 | 0.22 | 0.39 | 0.43 |
| 9 | 117021282 | rs7869221 | 0.084 | 0.202 | -0.899 | -0.639 | -0.001 | -0.056 | -0.056 | 0 | 0.23 | 0.29 | 0.2 | 0 | 0.23 | 0.13 |
| 9 | 117022495 | rs10732377 | -0.019 | -0.017 | -1.045 | -0.549 | 0.002 | -0.003 | -0.003 | 0.224 | 0.55 | 0.55 | 0.59 | 0.61 | 0.51 | 0.51 |
| 9 | 117022609 | rs923602 | 0.005 | -0.088 | -1.057 | -0.657 | 0.035 | -0.131 | -0.131 | 0 | 0.22 | 0.44 | 0.2 | 0.03 | 0.24 | 0.17 |
| 9 | 117023606 | rs2418268 | 1.288 | 0.073 | -1.437 | -0.517 | -0.043 | -0.038 | -0.038 | 0 | 0.25 | 0.12 | 0.44 | 0.4 | 0.34 | 0.25 |
| 9 | 117024515 | rs2636856 | -1.666 | -0.08 | -0.937 | -0.492 | -0.043 | -0.143 | -0.143 | 0 | 0.77 | 0.98 | 0.8 | 0.63 | 0.76 | 0.69 |
| 9 | 117024627 | rs2418269 | 0.356 | -0.112 | -0.556 | -0.517 | 0.005 | -0.002 | -0.002 | 2.905 | 0.36 | 0.46 | 0.48 | 0.4 | 0.37 | 0.34 |

| | | | | | | | | | | | | | | | | |
|---|---|---|---|---|---|---|---|---|---|---|---|---|---|---|---|---|
| 9 | 117027068 | rs1623718 | -0.117 | -0.271 | -0.316 | -0.486 | -0.001 | -0.002 | -0.002 | 0 | 0.55 | 0.6 | 0.55 | 0.6 | 0.43 | 0.57 |
| 9 | 117030261 | rs7027370 | 1.212 | 0.077 | -0.303 | -0.239 | -0.037 | -0.03 | -0.03 | 0.145 | 0.24 | 0.11 | 0.44 | 0.35 | 0.34 | 0.24 |
| 9 | 117030582 | rs2636860 | -0.503 | 0.055 | -0.333 | -0.52 | 0.013 | 0.03 | 0.03 | 0 | 0.52 | 0.63 | 0.35 | 0.23 | 0.38 | 0.41 |
| 9 | 117032111 | rs2491426 | -1.878 | 0.064 | -0.628 | -0.496 | -0.014 | -0.087 | -0.087 | 0 | 0.8 | 0.95 | 0.82 | 0.62 | 0.77 | 0.7 |
| 9 | 117032189 | rs5003737 | -0.812 | 0.118 | -0.554 | -0.668 | -0.02 | 0.104 | 0.104 | 0 | 0.56 | 0.84 | 0.38 | 0.23 | 0.44 | 0.42 |
| 9 | 117032261 | rs1490740 | 0.536 | 0.163 | -0.272 | -0.709 | 0.083 | -0.063 | -0.063 | 0 | 0.1 | 0.3 | 0.11 | 0 | 0.19 | 0.07 |
| 9 | 117032340 | rs1490741 | -0.33 | 0.021 | -0.357 | -0.626 | -0.025 | -0.026 | -0.026 | 0 | 0.45 | 0.52 | 0.26 | 0.23 | 0.25 | 0.35 |
| 9 | 117033022 | rs4143245 | -0.171 | 0.69 | -0.441 | -0.837 | 0.009 | -0.009 | -0.009 | 0 | 0.35 | 0.42 | 0.49 | 0.32 | 0.42 | 0.32 |
| 9 | 117033379 | rs10982127 | 0.188 | 1.053 | -0.514 | -0.477 | 0.047 | -0.057 | -0.057 | 0 | 0.11 | 0.24 | 0.11 | 0 | 0.18 | 0.07 |
| 9 | 117034510 | rs10982130 | 0.214 | 1.047 | -0.587 | -0.477 | 0.044 | -0.052 | -0.052 | 0.492 | 0.11 | 0.22 | 0.11 | 0 | 0.19 | 0.07 |
| 9 | 117034674 | rs10982133 | 0.156 | 1.071 | -0.552 | -0.477 | 0.077 | -0.064 | -0.064 | 0 | 0.11 | 0.27 | 0.11 | 0 | 0.19 | 0.07 |
| 9 | 117034749 | rs1490742 | -0.14 | 0.691 | -1.282 | -0.208 | -0.058 | 0.019 | 0.019 | 0 | 0.34 | 0.13 | 0.53 | 0.32 | 0.45 | 0.29 |
| 9 | 117037820 | rs10513241 | -0.032 | 0.484 | -0.922 | 0.301 | 0 | 0 | 0 | 0.733 | 0.35 | 0.23 | 0.36 | 0.33 | 0.37 | 0.24 |
| 9 | 117039111 | rs1766059 | -0.452 | 0.917 | -0.928 | 0.092 | 0 | 0 | 0 | 0 | 0.27 | 0.29 | 0.27 | 0.29 | 0.34 | 0.32 |
| 9 | 117039222 | rs10817580 | 0.412 | 0.794 | -1.258 | 0.134 | -0.063 | -0.071 | -0.071 | 0 | 0.18 | 0.02 | 0.15 | 0.29 | 0.24 | 0.25 |
| 9 | 117042977 | rs1249745 | -0.54 | 0.397 | -0.604 | -0.828 | 0.024 | -0.163 | -0.163 | 0 | 0.78 | 0.53 | 0.81 | 1 | 0.76 | 0.91 |
| 9 | 117043352 | rs1249744 | 0.663 | -0.745 | -0.803 | -0.262 | 0.032 | -0.037 | -0.037 | 0 | 0.3 | 0.1 | 0.24 | 0.35 | 0.28 | 0.17 |
| 9 | 117044458 | rs1628968 | 0.423 | -0.752 | -0.674 | -0.303 | 0.026 | -0.008 | -0.008 | 0 | 0.32 | 0.19 | 0.25 | 0.35 | 0.3 | 0.18 |
| 9 | 117047502 | rs4381012 | 1.56 | -1.022 | -1.011 | -0.755 | 0.001 | 0.064 | 0.064 | 0 | 0.12 | 0.01 | 0.07 | 0 | 0.15 | 0.04 |
| 9 | 117047888 | rs2130722 | 0.896 | -0.926 | -1.097 | -0.41 | -0.011 | -0.046 | -0.046 | 0 | 0.11 | 0.01 | 0.24 | 0.16 | 0.07 | 0.17 |
| 9 | 117048609 | rs4978577 | 0.708 | -0.625 | -0.831 | -0.55 | 0.008 | -0.018 | -0.018 | 0 | 0.18 | 0.1 | 0.18 | 0.28 | 0.14 | 0.14 |
| 9 | 117049891 | rs946053 | -1.035 | -0.724 | -0.639 | -0.742 | -0.054 | 0.075 | 0.075 | 0 | 0.68 | 0.81 | 0.75 | 0.93 | 0.49 | 0.78 |
| 9 | 117050998 | rs10982134 | 0.509 | -0.902 | -0.194 | -0.714 | 0.054 | 0.041 | 0.041 | 0.404 | 0.4 | 0.24 | 0.45 | 0.63 | 0.29 | 0.57 |
| 9 | 117051020 | rs1687410 | 0.375 | -0.818 | -0.208 | -0.547 | -0.019 | 0.013 | 0.013 | 0 | 0.2 | 0.36 | 0.2 | 0.28 | 0.14 | 0.14 |
| 9 | 117053215 | rs1249738 | 0.186 | -0.578 | -0.654 | -0.325 | 0.006 | 0.005 | 0.005 | 0.005 | 0.42 | 0.62 | 0.47 | 0.63 | 0.29 | 0.58 |
| 9 | 117054195 | rs10429546 | -0.365 | 0.008 | -0.677 | 0.221 | 0.002 | 0.005 | 0.005 | 0 | 0.25 | 0.22 | 0.26 | 0.25 | 0.2 | 0.18 |
| 9 | 117060746 | rs4979365 | -0.938 | -0.122 | -0.385 | 0.815 | -0.015 | 0.043 | 0.043 | 0.072 | 0.09 | 0.17 | 0.04 | 0.26 | 0.03 | 0.24 |
| 9 | 117061237 | rs1687401 | 0.327 | 0.084 | -0.537 | 0.59 | 0.007 | 0.011 | 0.011 | 0 | 0.49 | 0.39 | 0.47 | 0.41 | 0.4 | 0.44 |
| 9 | 117062677 | rs1626295 | -0.297 | 0.466 | -0.95 | 0.312 | -0.08 | 0.162 | 0.162 | 0 | 0.63 | 0.85 | 0.6 | 0.92 | 0.46 | 0.76 |
| 9 | 117064306 | rs1546049 | 0.183 | 0.548 | -0.347 | 0.636 | -0.002 | -0.006 | -0.006 | 1.239 | 0.57 | 0.53 | 0.48 | 0.69 | 0.43 | 0.66 |
| 9 | 117066040 | rs10817588 | 0.455 | 1.342 | -0.901 | 0.256 | -0.024 | 0.047 | 0.047 | 0 | 0.25 | 0.13 | 0.29 | 0.1 | 0.38 | 0.22 |
| 9 | 117066467 | rs10759690 | -0.063 | 0.985 | -1.502 | 0.274 | -0.059 | 0.021 | 0.021 | 0 | 0.4 | 0.14 | 0.41 | 0.34 | 0.49 | 0.39 |
| 9 | 117066823 | rs10817590 | -0.001 | 1.099 | -1.513 | 0.277 | -0.045 | 0.03 | 0.03 | 0 | 0.39 | 0.13 | 0.27 | 0.31 | 0.46 | 0.35 |
| 9 | 117073588 | rs11537824 | -0.093 | 0.024 | -0.359 | 0.397 | -0.074 | 0.152 | 0.152 | 0 | 0.2 | 0.43 | 0.19 | 0.51 | 0.06 | 0.35 |
| 9 | 117075811 | rs10982144 | 1.18 | 1.017 | -0.956 | 0.25 | -0.071 | 0.074 | 0.074 | 0.39 | 0.12 | 0.01 | 0.17 | 0 | 0.15 | 0.03 |
| 9 | 117076684 | rs1631637 | -0.482 | 0.476 | -0.348 | 0.382 | -0.002 | -0.009 | -0.009 | 0.623 | 0.57 | 0.59 | 0.68 | 0.62 | 0.59 | 0.6 |
| 9 | 117078286 | rs7851482 | -0.131 | 0.561 | -0.688 | 0.171 | 0.024 | 0.131 | 0.131 | 0 | 0.5 | 0.28 | 0.48 | 0.26 | 0.59 | 0.46 |
| 9 | 117078838 | rs2787334 | -0.247 | 0.365 | -0.457 | 0.214 | 0 | 0.009 | 0.009 | 0 | 0.54 | 0.56 | 0.6 | 0.32 | 0.58 | 0.48 |
| 9 | 117079287 | rs2636885 | -0.645 | 0.647 | -1.464 | -0.376 | -0.102 | 0.077 | 0.077 | 0 | 0.73 | 0.95 | 0.72 | 0.9 | 0.66 | 0.8 |
| 9 | 117080388 | rs2636881 | 0.189 | 0.188 | -0.662 | 0.148 | -0.007 | 0.01 | 0.01 | 0 | 0.31 | 0.44 | 0.25 | 0.35 | 0.26 | 0.26 |
| 9 | 117084228 | rs2787337 | -0.015 | 0.203 | -0.851 | -0.634 | -0.001 | -0.032 | -0.032 | 0 | 0.36 | 0.49 | 0.39 | 0.81 | 0.32 | 0.59 |
| 9 | 117086271 | rs10982151 | 1.823 | 1.056 | -1.186 | -0.541 | -0.038 | 0.06 | 0.06 | 1.043 | 0.1 | 0 | 0.16 | 0 | 0.13 | 0.03 |
| 9 | 117093029 | rs17230081 | 0.818 | 0.992 | -1.085 | -0.193 | -0.031 | -0.01 | -0.01 | 0.92 | 0.18 | 0.08 | 0.17 | 0.19 | 0.19 | 0.27 |
| 9 | 117097755 | rs10739407 | -0.858 | 0.875 | -0.99 | -0.441 | -0.056 | 0.028 | 0.028 | 0 | 0.79 | 0.95 | 0.87 | 0.86 | 0.74 | 0.76 |
| 9 | 117099995 | rs10982171 | -0.019 | 0.705 | -0.456 | -0.128 | 0.047 | -0.042 | -0.042 | 0 | 0.4 | 0.12 | 0.29 | 0.57 | 0.39 | 0.4 |
| 9 | 117102046 | rs2636870 | -0.576 | 0.694 | -0.194 | -0.158 | 0.031 | 0.001 | 0.001 | 0 | 0.47 | 0.75 | 0.51 | 0.56 | 0.48 | 0.38 |
| 9 | 117103973 | rs2250242 | -0.801 | 0.702 | -0.17 | -0.131 | 0.029 | 0.048 | 0.048 | 0.059 | 0.5 | 0.88 | 0.54 | 0.61 | 0.48 | 0.42 |
| 9 | 117110046 | rs3748178 | 0.661 | 0.429 | -0.234 | -0.314 | 0.001 | 0.112 | 0.112 | 0.052 | 0.24 | 0.01 | 0.14 | 0.07 | 0.26 | 0.09 |
| 9 | 117110112 | rs1265891 | -0.783 | 0.536 | 0.085 | -0.14 | -0.035 | 0.02 | 0.02 | 8.581 | 0.08 | 0.69 | 0.06 | 0.05 | 0 | 0.04 |
| 9 | 117111707 | rs2787346 | -0.499 | 0.562 | 0.12 | -0.127 | 0.024 | 0.098 | 0.098 | 6.445 | 0.06 | 0.59 | 0.05 | 0.05 | 0 | 0.04 |
| 9 | 117111951 | rs10817593 | -0.141 | 0.628 | -0.575 | -0.15 | -0.014 | -0.03 | -0.03 | 0 | 0.44 | 0.14 | 0.42 | 0.57 | 0.48 | 0.43 |
| 9 | 117112814 | rs1249725 | -0.813 | 0.753 | -0.607 | -0.261 | 0.046 | 0.06 | 0.06 | 0.13 | 0.48 | 0.81 | 0.52 | 0.63 | 0.43 | 0.5 |
| 9 | 117112886 | rs1766069 | -0.44 | 0.49 | -0.281 | -0.156 | -0.008 | 0.046 | 0.046 | 5.53 | 0.06 | 0.67 | 0.06 | 0.05 | 0 | 0.04 |
| 9 | 117114168 | rs10982177 | 1.614 | 0.541 | -0.689 | -0.302 | 0.012 | -0.066 | -0.066 | 0 | 0.16 | 0.1 | 0.23 | 0.51 | 0.12 | 0.25 |
| 9 | 117120680 | rs7866925 | 1.7 | 0.573 | -1.078 | -0.619 | 0 | 0.052 | 0.052 | 0 | 0.3 | 0.05 | 0.15 | 0.15 | 0.23 | 0.25 |
| 9 | 117121840 | rs2787348 | -1.157 | 0.924 | -1.161 | -0.691 | -0.056 | 0.076 | 0.076 | 0 | 0.81 | 0.98 | 0.89 | 0.94 | 0.75 | 0.87 |
| 9 | 117122615 | rs2636897 | -0.475 | 0.749 | -0.579 | -0.674 | -0.029 | -0.015 | -0.015 | 0.455 | 0.63 | 0.71 | 0.53 | 0.42 | 0.53 | 0.6 |
| 9 | 117123494 | rs10817594 | 1.002 | 0.953 | -0.769 | -0.586 | -0.04 | -0.013 | -0.013 | 0 | 0.21 | 0.05 | 0.31 | 0.23 | 0.24 | 0.25 |
| 9 | 117125236 | rs12337816 | 1.445 | 1.082 | -0.738 | -0.565 | -0.031 | 0.025 | 0.025 | 0 | 0.08 | 0.01 | 0.13 | 0.01 | 0.13 | 0.02 |
| 9 | 117126307 | rs10125969 | 0.028 | 0.935 | -0.566 | -0.428 | -0.05 | 0.009 | 0.009 | 0 | 0.55 | 0.11 | 0.48 | 0.39 | 0.51 | 0.55 |
| 9 | 117148327 | rs10739409 | 0.94 | 0.976 | -0.439 | -0.269 | -0.032 | -0.041 | -0.041 | 0 | 0.23 | 0.05 | 0.35 | 0.3 | 0.25 | 0.29 |
| 9 | 117148430 | rs10733608 | 0.177 | 0.964 | -0.257 | -0.285 | -0.015 | 0.001 | 0.001 | 0 | 0.54 | 0.27 | 0.51 | 0.46 | 0.47 | 0.52 |
| 9 | 117149218 | rs2787328 | -1.456 | 0.977 | 0.047 | -0.685 | 0.017 | -0.019 | -0.019 | 1.16 | 0.84 | 0.74 | 0.87 | 0.88 | 0.77 | 0.88 |
| 9 | 117150070 | rs1324632 | 0.749 | 1.033 | 0.304 | -0.406 | 0 | 0.111 | 0.111 | 0 | 0.21 | 0.41 | 0.24 | 0.4 | 0.21 | 0.3 |
| 9 | 117150951 | rs2763196 | -0.435 | 1.026 | 0.179 | -0.558 | -0.001 | -0.001 | -0.001 | 0 | 0.44 | 0.47 | 0.67 | 0.8 | 0.43 | 0.6 |
| 9 | 117153206 | rs12351392 | 1.209 | 0.865 | -0.495 | -0.581 | 0.009 | 0.082 | 0.082 | 0 | 0.3 | 0.1 | 0.16 | 0.12 | 0.25 | 0.23 |
| 9 | 117156596 | rs4979375 | 0.961 | 0.934 | -0.529 | -0.494 | -0.031 | -0.076 | -0.076 | 0.398 | 0.22 | 0.05 | 0.32 | 0.36 | 0.22 | 0.26 |
| 9 | 117157334 | rs17231346 | 1.145 | 0.923 | -0.487 | -0.568 | 0.014 | 0.068 | 0.068 | 0 | 0.31 | 0.13 | 0.16 | 0.13 | 0.25 | 0.23 |
| 9 | 117161055 | rs10759692 | -0.011 | 0.995 | -0.426 | -0.507 | -0.031 | -0.002 | -0.002 | 0 | 0.54 | 0.19 | 0.49 | 0.47 | 0.47 | 0.5 |

| | | | | | | | | | | | | | | | | |
|---|---|---|---|---|---|---|---|---|---|---|---|---|---|---|---|---|
| 9 | 117162253 | rs10759693 | -0.15 | 0.991 | -0.406 | -0.509 | -0.016 | -0.025 | -0.025 | 0 | 0.57 | 0.21 | 0.51 | 0.5 | 0.51 | 0.56 |
| 9 | 117164006 | rs10759694 | 0.838 | 0.954 | -0.473 | -0.481 | -0.031 | -0.072 | -0.072 | 0.365 | 0.22 | 0.06 | 0.32 | 0.35 | 0.22 | 0.26 |
| 9 | 117166206 | rs2274158 | 0.907 | 0.957 | -0.471 | -0.48 | -0.032 | -0.072 | -0.072 | 0 | 0.22 | 0.05 | 0.32 | 0.35 | 0.22 | 0.26 |
| 9 | 117166246 | rs2274159 | 0.039 | 0.923 | -0.433 | -0.507 | -0.032 | -0.003 | -0.003 | 0.426 | 0.54 | 0.19 | 0.49 | 0.47 | 0.47 | 0.5 |
| 9 | 117169300 | rs2236388 | -1.686 | 0.666 | -0.175 | -0.402 | 0.005 | 0.003 | 0.003 | 0 | 0.84 | 0.87 | 0.81 | 0.58 | 0.88 | 0.73 |
| 9 | 117181430 | rs7849013 | 0.277 | 0.97 | 0.175 | -0.429 | -0.013 | -0.061 | -0.061 | 0.083 | 0.23 | 0.07 | 0.37 | 0.37 | 0.22 | 0.3 |
| 9 | 117183149 | rs10739411 | -0.176 | 1.063 | 0.462 | -0.454 | 0 | 0 | 0 | 0 | 0.27 | 0.29 | 0.52 | 0.37 | 0.3 | 0.36 |
| 9 | 117185395 | rs2181928 | -0.265 | 0.974 | -0.343 | -0.513 | 0.002 | -0.025 | -0.025 | 1.078 | 0.73 | 0.28 | 0.7 | 0.94 | 0.64 | 0.8 |
| 9 | 117186712 | rs4978584 | 0.17 | 0.389 | -0.734 | -0.1 | -0.026 | -0.065 | -0.065 | 0.207 | 0.22 | 0.06 | 0.37 | 0.34 | 0.21 | 0.27 |
| 9 | 117194505 | rs7851968 | 0.584 | 0.414 | -0.59 | -0.557 | -0.023 | 0.078 | 0.078 | 0 | 0.21 | 0.06 | 0.13 | 0.02 | 0.26 | 0.17 |
| 9 | 117197493 | rs7044547 | -0.58 | 0.086 | -0.599 | -0.565 | 0.001 | -0.005 | -0.005 | 0 | 0.36 | 0.73 | 0.56 | 0.39 | 0.33 | 0.33 |
| 9 | 117200079 | rs4979391 | 0.929 | 0.136 | -0.659 | -0.519 | -0.068 | -0.048 | -0.048 | 0.218 | 0.43 | 0.07 | 0.36 | 0.56 | 0.42 | 0.47 |
| 9 | 117200140 | rs10982218 | -0.647 | 0.1 | 0.459 | -0.489 | 0.042 | 0.058 | 0.058 | 0.1 | 0.06 | 0.35 | 0.05 | 0.05 | 0.05 | 0.03 |
| 9 | 117208985 | rs10982222 | 0.117 | -0.118 | 0.629 | -0.464 | -0.001 | -0.001 | -0.001 | 0 | 0.26 | 0.27 | 0.49 | 0.33 | 0.25 | 0.29 |
| 9 | 117214909 | rs942520 | -1.244 | -0.203 | 0.663 | -0.537 | -0.01 | 0.017 | 0.017 | 0 | 0.54 | 0.73 | 0.6 | 0.42 | 0.52 | 0.46 |
| 9 | 117218901 | rs10739418 | -0.751 | -0.236 | 0.558 | -0.599 | -0.008 | -0.1 | -0.1 | 0 | 0.75 | 0.7 | 0.85 | 0.96 | 0.74 | 0.84 |
| 9 | 117219888 | rs4979397 | -0.791 | -0.279 | 0.57 | -0.583 | -0.002 | -0.1 | -0.1 | 0 | 0.75 | 0.7 | 0.84 | 0.96 | 0.73 | 0.83 |
| 9 | 117222287 | rs10759707 | -0.446 | -0.283 | 0.563 | -0.556 | -0.01 | 0.023 | 0.023 | 0.056 | 0.3 | 0.63 | 0.45 | 0.38 | 0.25 | 0.31 |
| 9 | 117222675 | rs4979402 | -0.759 | -0.275 | 0.563 | -0.561 | -0.003 | -0.1 | -0.1 | 0 | 0.75 | 0.7 | 0.84 | 0.96 | 0.73 | 0.83 |
| 9 | 117228115 | rs4979407 | 0.627 | -0.218 | 0.561 | -0.57 | 0.001 | -0.036 | -0.036 | 0.246 | 0.18 | 0.13 | 0.32 | 0.31 | 0.14 | 0.28 |
| 9 | 117229400 | rs1535971 | -0.232 | -0.472 | 0.458 | -0.649 | -0.004 | -0.037 | -0.037 | 0 | 0.69 | 0.37 | 0.81 | 0.93 | 0.72 | 0.87 |
| 9 | 117231413 | rs10817618 | 0.559 | -0.102 | 0.386 | 0.029 | -0.005 | 0.008 | 0.008 | 0.122 | 0.71 | 0.32 | 0.56 | 0.65 | 0.75 | 0.67 |
| 9 | 117236557 | rs1123056 | -0.166 | -0.393 | -0.027 | 0.019 | -0.005 | 0.009 | 0.009 | 0 | 0.55 | 0.31 | 0.61 | 0.62 | 0.52 | 0.59 |
| 9 | 117237754 | rs1000709 | 0.237 | -0.565 | -0.085 | -0.072 | -0.026 | 0.005 | 0.005 | 0 | 0.46 | 0.21 | 0.4 | 0.6 | 0.49 | 0.56 |
| 9 | 117245132 | rs16929545 | 1.038 | -0.516 | -0.225 | 0.116 | -0.056 | -0.011 | -0.011 | 0 | 0.17 | 0.01 | 0.1 | 0.13 | 0.17 | 0.22 |
| 9 | 117249874 | rs725340 | -0.394 | -0.771 | 0.555 | 0.335 | -0.016 | 0.059 | 0.059 | 0 | 0.14 | 0.16 | 0.11 | 0.38 | 0.06 | 0.19 |
| 9 | 117251352 | rs1535960 | 0.594 | -0.813 | 0.458 | 0.225 | -0.046 | -0.021 | -0.021 | 0.053 | 0.49 | 0.11 | 0.4 | 0.59 | 0.5 | 0.58 |
| 9 | 117256896 | rs12551529 | -0.29 | -0.852 | 0.31 | 0.206 | -0.016 | -0.008 | -0.008 | 0 | 0.43 | 0.33 | 0.44 | 0.41 | 0.49 | 0.42 |
| 9 | 117259062 | rs1535962 | 0.937 | -0.773 | -0.268 | 0.264 | -0.123 | -0.128 | -0.128 | 0 | 0.46 | 0.04 | 0.38 | 0.57 | 0.5 | 0.47 |
| 9 | 117264083 | rs4979418 | 1.097 | -0.781 | -0.325 | 0.27 | -0.115 | -0.147 | -0.147 | 1.382 | 0.43 | 0.04 | 0.33 | 0.57 | 0.48 | 0.49 |
| 9 | 117265406 | rs2296262 | 0.917 | -0.7 | -0.129 | 0.53 | -0.025 | 0.025 | 0.025 | 0.133 | 0.28 | 0.22 | 0.33 | 0.14 | 0.4 | 0.22 |
| 9 | 117272858 | rs942518 | 1.027 | -0.761 | 0.032 | 0.252 | -0.019 | 0.007 | 0.007 | 0.015 | 0.76 | 0.35 | 0.67 | 0.71 | 0.91 | 0.76 |
| 9 | 117275012 | rs998547 | 0.525 | -0.621 | -0.24 | 0.349 | -0.028 | 0.082 | 0.082 | 0 | 0.49 | 0.18 | 0.39 | 0.69 | 0.52 | 0.54 |
| 9 | 117278844 | rs10513248 | -0.445 | -0.237 | 0.583 | 0.084 | 0.008 | -0.09 | -0.09 | 0 | 0.67 | 0.58 | 0.74 | 0.97 | 0.6 | 0.78 |
| 9 | 117282390 | rs2770154 | 1.137 | -0.545 | 0.254 | 0.85 | 0.038 | -0.055 | -0.055 | 0 | 0.73 | 0.19 | 0.65 | 0.47 | 0.87 | 0.7 |
| 9 | 117283232 | rs1272624 | 0.589 | -0.513 | 0.7 | 1.04 | -0.031 | 0.026 | 0.026 | 1.728 | 0.85 | 0.33 | 0.86 | 0.83 | 0.94 | 0.91 |
| 9 | 117287928 | rs10759716 | -0.878 | -0.471 | 0.149 | 1.105 | -0.017 | 0.015 | 0.015 | 0.992 | 0.17 | 0.68 | 0.15 | 0.21 | 0.09 | 0.13 |
| 9 | 117288169 | rs10817635 | 0.679 | -0.283 | -0.409 | 1.268 | -0.134 | -0.024 | -0.024 | 0 | 0.44 | 0.04 | 0.4 | 0.41 | 0.49 | 0.48 |
| 9 | 117289895 | rs4978594 | -0.687 | -0.128 | 0.839 | 2.228 | -0.006 | 0.098 | 0.098 | 0 | 0.08 | 0.09 | 0.19 | 0.35 | 0.04 | 0.2 |
| 9 | 117309052 | rs4979424 | -1.499 | 0.052 | 1.115 | 1.401 | 0.021 | 0.101 | 0.101 | 0.43 | 0.09 | 0.31 | 0.15 | 0.16 | 0.04 | 0.05 |
| 9 | 117312703 | rs10982285 | -1.018 | 0.173 | 0.565 | 1.468 | -0.005 | 0.007 | 0.007 | 0.027 | 0.31 | 0.59 | 0.39 | 0.58 | 0.24 | 0.37 |
| 9 | 117314774 | rs10982289 | -1.05 | -0.131 | 0.299 | 1.496 | 0.028 | 0.009 | 0.009 | 0 | 0.24 | 0.13 | 0.13 | 0.23 | 0.16 | 0.18 |
| 9 | 117315880 | rs10114562 | -0.748 | 0.143 | -0.085 | 1.216 | -0.018 | 0.005 | 0.005 | 0 | 0.63 | 0.71 | 0.72 | 0.69 | 0.53 | 0.53 |
| 9 | 117322542 | rs10817638 | 0.744 | 0.285 | 0.388 | 1.261 | 0 | 0 | 0 | 0 | 0.38 | 0.37 | 0.51 | 0.42 | 0.36 | 0.35 |
| 9 | 117323426 | rs4979427 | -1.028 | 0.523 | 0.057 | 1.002 | -0.009 | 0.015 | 0.015 | 0 | 0.34 | 0.39 | 0.34 | 0.53 | 0.29 | 0.33 |
| 9 | 117324845 | rs4979428 | -0.433 | 0.014 | -0.044 | 1.011 | 0.014 | -0.035 | -0.035 | 0 | 0.26 | 0.21 | 0.22 | 0.47 | 0.2 | 0.33 |
| 9 | 117325336 | rs7851259 | 0.912 | 0.161 | -0.223 | 0.687 | 0 | 0 | 0 | 0 | 0.49 | 0.51 | 0.48 | 0.43 | 0.46 | 0.47 |
| 9 | 117329675 | rs12343284 | 0.298 | 0.029 | -0.609 | 0.637 | 0.016 | -0.018 | -0.018 | 0.555 | 0.16 | 0.1 | 0.05 | 0.24 | 0.08 | 0.17 |
| 9 | 117339877 | rs10982300 | 1.56 | 0.127 | -0.531 | 0.381 | 0 | -0.039 | -0.039 | 0 | 0.11 | 0.15 | 0.05 | 0.01 | 0.1 | 0.13 |
| 9 | 117348466 | rs12238437 | 0.331 | 0.163 | -0.895 | 0.906 | 0.013 | -0.037 | -0.037 | 0 | 0.15 | 0.11 | 0.08 | 0.37 | 0.08 | 0.16 |
| 9 | 117349653 | rs10982317 | -0.514 | 0.182 | -1.03 | 0.749 | 0.019 | -0.045 | -0.045 | 0 | 0.24 | 0.13 | 0.15 | 0.37 | 0.17 | 0.18 |
| 9 | 117353464 | rs10124511 | -0.077 | 0.783 | -0.808 | 0.387 | -0.049 | 0.063 | 0.063 | 0 | 0.76 | 0.89 | 0.85 | 0.99 | 0.69 | 0.9 |
| 9 | 117360729 | rs7868842 | -0.149 | 0.641 | -0.922 | 0.572 | -0.042 | -0.018 | -0.018 | 0 | 0.75 | 0.86 | 0.72 | 0.76 | 0.66 | 0.76 |
| 9 | 117370538 | rs4978600 | -0.646 | 0.314 | -0.937 | 0.286 | 0.009 | 0.009 | 0.009 | 0 | 0.88 | 0.9 | 0.93 | 1 | 0.9 | 0.93 |
| 9 | 117375909 | rs4979444 | 0.745 | 0.315 | -0.787 | 0.619 | -0.001 | 0.007 | 0.007 | 0.255 | 0.41 | 0.5 | 0.55 | 0.53 | 0.46 | 0.4 |
| 9 | 117383919 | rs7027695 | 0.21 | -0.354 | -0.98 | 0.564 | -0.019 | -0.005 | -0.005 | 0 | 0.57 | 0.53 | 0.65 | 0.67 | 0.71 | 0.51 |
| 9 | 117385992 | rs7036626 | 0.888 | -0.11 | -0.927 | 0.468 | -0.009 | -0.018 | -0.018 | 2.12 | 0.31 | 0.21 | 0.45 | 0.42 | 0.38 | 0.26 |
| 9 | 117391325 | rs1536738 | 0.232 | -0.775 | -0.869 | 0.257 | 0.011 | 0.009 | 0.009 | 0 | 0.78 | 0.77 | 0.85 | 0.87 | 0.88 | 0.8 |
| 9 | 117393736 | rs4978602 | 0.509 | -0.79 | -0.796 | 0.32 | 0.022 | 0.015 | 0.015 | 0 | 0.59 | 0.65 | 0.66 | 0.74 | 0.75 | 0.55 |
| 9 | 117393934 | rs2418310 | 0.509 | -0.791 | -0.788 | 0.32 | 0.018 | 0.014 | 0.014 | 0 | 0.59 | 0.65 | 0.66 | 0.74 | 0.73 | 0.55 |
| 9 | 117394595 | rs12379659 | -0.612 | -0.942 | -0.629 | 0.449 | -0.005 | -0.001 | -0.001 | 0 | 0.34 | 0.34 | 0.31 | 0.26 | 0.21 | 0.43 |
| 9 | 117394754 | rs10982340 | -0.612 | -0.942 | -0.629 | 0.449 | -0.005 | -0.001 | -0.001 | 0 | 0.34 | 0.34 | 0.31 | 0.26 | 0.21 | 0.43 |
| 9 | 117396237 | rs3810935 | -0.171 | -1.028 | -0.602 | 0.334 | 0 | 0.001 | 0.001 | 0 | 0.29 | 0.29 | 0.12 | 0.14 | 0.18 | 0.19 |
| 9 | 117405189 | rs7027937 | -0.244 | -0.999 | -1.072 | 0.358 | 0.063 | 0.05 | 0.05 | 0 | 0.29 | 0.12 | 0.22 | 0.15 | 0.17 | 0.28 |
| 9 | 117406601 | rs1058278 | -0.244 | -0.999 | -1.072 | 0.358 | 0.063 | 0.05 | 0.05 | 0 | 0.29 | 0.12 | 0.22 | 0.15 | 0.17 | 0.28 |
| 9 | 117406965 | rs10759728 | -0.244 | -0.999 | -1.072 | 0.358 | 0.063 | 0.05 | 0.05 | 0 | 0.29 | 0.12 | 0.22 | 0.15 | 0.17 | 0.28 |
| 9 | 117407923 | rs1058280 | -0.252 | -0.977 | -1.072 | 0.42 | 0.065 | 0.037 | 0.037 | 0 | 0.29 | 0.12 | 0.22 | 0.16 | 0.17 | 0.28 |
| 9 | 117429227 | rs2282003 | 0.619 | -0.295 | -0.545 | 0.875 | 0.012 | 0 | 0 | 1.776 | 0.32 | 0.47 | 0.52 | 0.37 | 0.4 | 0.28 |
| 9 | 117430524 | rs2296947 | 0.534 | -0.344 | -0.214 | 1.033 | 0.006 | -0.029 | -0.029 | 0 | 0.33 | 0.43 | 0.46 | 0.27 | 0.4 | 0.28 |

| | | | | | | | | | | | | | | | | |
|---|---|---|---|---|---|---|---|---|---|---|---|---|---|---|---|---|
| 9 | 117432041 | rs11793719 | 0.749 | -0.709 | -0.418 | 0.779 | 0.001 | -0.009 | -0.009 | 0 | 0.11 | 0.13 | 0.04 | 0 | 0.04 | 0.04 |
| 9 | 117432944 | rs7846979 | -0.678 | -0.624 | -0.453 | 1.029 | -0.004 | 0.005 | 0.005 | 0 | 0.54 | 0.65 | 0.72 | 0.56 | 0.53 | 0.51 |
| 9 | 117433685 | rs2296948 | -0.567 | -0.62 | -0.431 | 1.036 | -0.014 | 0.002 | 0.002 | 0 | 0.55 | 0.67 | 0.74 | 0.56 | 0.57 | 0.51 |
| 9 | 117440143 | rs16930798 | 0.021 | -0.669 | -0.356 | 1.154 | -0.017 | 0.011 | 0.011 | 0 | 0.14 | 0.19 | 0.23 | 0.2 | 0.07 | 0.07 |
| 9 | 117442123 | rs6478100 | -0.594 | -0.762 | -0.433 | 1.145 | -0.007 | -0.014 | -0.014 | 0.507 | 0.57 | 0.74 | 0.6 | 0.47 | 0.51 | 0.62 |
| 9 | 117444227 | rs13286744 | -0.452 | -0.798 | -0.398 | 1.144 | 0.001 | 0.034 | 0.034 | 0 | 0.53 | 0.65 | 0.54 | 0.35 | 0.53 | 0.5 |
| 9 | 117445168 | rs10982370 | 0.975 | -0.875 | -0.323 | 0.978 | -0.003 | -0.024 | -0.024 | 0 | 0.34 | 0.42 | 0.43 | 0.29 | 0.37 | 0.39 |
| 9 | 117448713 | rs7853853 | -0.905 | -0.936 | -0.304 | 0.992 | -0.003 | 0.066 | 0.066 | 0.105 | 0.16 | 0.21 | 0.34 | 0.38 | 0.18 | 0.23 |
| 9 | 117450244 | rs10739424 | -1.169 | -0.902 | -0.412 | 0.982 | -0.074 | 0.015 | 0.015 | 0 | 0.61 | 0.85 | 0.82 | 0.67 | 0.59 | 0.66 |
| 9 | 117451913 | rs7039257 | 0.973 | -0.877 | -0.418 | 0.999 | -0.001 | -0.022 | -0.022 | 5.359 | 0.33 | 0.41 | 0.43 | 0.29 | 0.37 | 0.39 |
| 9 | 117453668 | rs16930907 | -0.393 | -0.898 | -0.537 | 1.132 | -0.018 | 0.145 | 0.145 | 0.058 | 0.08 | 0.14 | 0.27 | 0.31 | 0.05 | 0.17 |
| 9 | 117456045 | rs16930926 | 0.728 | -0.63 | -0.977 | 0.613 | -0.011 | -0.015 | -0.015 | 0 | 0.06 | 0.01 | 0.04 | 0.07 | 0.08 | 0.06 |
| 9 | 117464374 | rs17816047 | -0.225 | -0.706 | -0.945 | 0.69 | -0.001 | 0 | 0 | 0.153 | 0.14 | 0.17 | 0.07 | 0.07 | 0.08 | 0.1 |
| 9 | 117478697 | rs4979458 | -0.609 | -0.462 | -1.68 | 0.911 | -0.138 | -0.021 | -0.021 | 0 | 0.6 | 0.96 | 0.83 | 0.67 | 0.59 | 0.6 |
| 9 | 117481666 | rs4979459 | -0.084 | -0.936 | -1.548 | 0.617 | 0.012 | 0.004 | 0.004 | 0 | 0.54 | 0.42 | 0.4 | 0.59 | 0.53 | 0.72 |
| 9 | 117483015 | rs10817669 | -0.264 | -1.046 | -1.64 | 0.551 | -0.02 | -0.047 | -0.047 | 0 | 0.68 | 0.45 | 0.7 | 0.85 | 0.72 | 0.83 |
| 9 | 117483943 | rs7026324 | 1.092 | -0.731 | -1.863 | 0.884 | -0.117 | -0.102 | -0.102 | 0 | 0.39 | 0.06 | 0.25 | 0.48 | 0.42 | 0.49 |
| 9 | 117491859 | rs11790825 | 0.306 | -0.287 | -1.638 | 0.617 | 0.075 | -0.009 | -0.009 | 0 | 0.1 | 0.32 | 0.14 | 0.04 | 0.16 | 0.14 |
| 9 | 117502242 | rs2185935 | 1.257 | -0.534 | -2.077 | 0.629 | -0.09 | -0.099 | -0.099 | 0.984 | 0.36 | 0.05 | 0.22 | 0.47 | 0.32 | 0.36 |
| 9 | 117516418 | rs7038398 | -0.994 | -0.98 | -1.415 | -0.005 | 0.006 | -0.006 | -0.006 | 0 | 0.81 | 0.72 | 0.77 | 0.85 | 0.77 | 0.84 |
| 9 | 117517897 | rs10982395 | 1.25 | -0.977 | -1.613 | 0.062 | 0.018 | -0.032 | -0.032 | 0.41 | 0.06 | 0.01 | 0.01 | 0.13 | 0.01 | 0.15 |
| 9 | 117523018 | rs12683136 | 2.453 | -0.607 | -1.582 | -0.397 | -0.007 | -0.049 | -0.049 | 0 | 0.09 | 0.01 | 0.05 | 0.16 | 0.08 | 0.05 |
| 9 | 117523192 | rs10759736 | -1.051 | 0.12 | -0.977 | -0.889 | 0.017 | -0.047 | -0.047 | 0.703 | 0.92 | 0.8 | 0.86 | 1 | 0.89 | 0.98 |
| 9 | 117530133 | rs10513258 | -0.046 | -0.229 | -1.077 | -0.65 | -0.008 | -0.069 | -0.069 | 0 | 0.27 | 0.33 | 0.21 | 0 | 0.25 | 0.11 |
| 9 | 117530319 | rs4978609 | 1.021 | 0.092 | -1.469 | 0.004 | -0.018 | -0.041 | -0.041 | 0 | 0.09 | 0.02 | 0.09 | 0.15 | 0.12 | 0.14 |
| 9 | 117533062 | rs7032238 | -0.53 | 0.043 | -1.017 | -0.885 | -0.012 | -0.118 | -0.118 | 0 | 0.33 | 0.47 | 0.29 | 0.05 | 0.34 | 0.19 |
| 9 | 117533289 | rs7048659 | -0.731 | 0.042 | -0.887 | -0.981 | -0.006 | -0.103 | -0.103 | 0 | 0.38 | 0.49 | 0.32 | 0.05 | 0.41 | 0.23 |
| 9 | 117535054 | rs12238270 | -0.25 | 0.322 | -1.072 | -0.382 | -0.019 | 0.016 | 0.016 | 0 | 0.18 | 0.04 | 0.19 | 0.16 | 0.19 | 0.23 |
| 9 | 117540910 | rs7856856 | 0.393 | -0.081 | -0.891 | -0.037 | -0.001 | 0.004 | 0.004 | 0 | 0.22 | 0.2 | 0.36 | 0.31 | 0.28 | 0.33 |
| 9 | 117541450 | rs7860414 | 1.377 | -0.021 | -1.387 | -0.624 | 0.044 | 0.146 | 0.146 | 5.249 | 0.33 | 0.12 | 0.14 | 0.04 | 0.23 | 0.15 |
| 9 | 117545666 | rs6478106 | 0.431 | 0.036 | -0.786 | -0.557 | -0.003 | 0.059 | 0.059 | 0.416 | 0.24 | 0.43 | 0.39 | 0.31 | 0.29 | 0.33 |
| 9 | 117547772 | rs10114470 | -0.353 | 0.239 | -1.068 | -0.506 | -0.086 | -0.12 | -0.12 | 0.153 | 0.76 | 0.91 | 0.7 | 0.51 | 0.68 | 0.7 |
| 9 | 117552885 | rs3810936 | -0.364 | 0.248 | -1.202 | -0.536 | -0.086 | -0.119 | -0.119 | 0 | 0.76 | 0.91 | 0.7 | 0.51 | 0.68 | 0.69 |
| 9 | 117557872 | rs7867918 | 1.717 | -0.006 | -1.315 | -1.004 | 0.034 | 0.133 | 0.133 | 5.175 | 0.25 | 0.06 | 0.11 | 0 | 0.17 | 0.05 |
| 9 | 117558703 | rs6478108 | -0.161 | 0.457 | -0.835 | -0.275 | -0.063 | -0.082 | -0.082 | 0 | 0.77 | 0.89 | 0.72 | 0.51 | 0.66 | 0.74 |
| 9 | 117560265 | rs7847158 | 0.789 | 0.333 | -0.684 | -0.351 | 0.002 | -0.001 | -0.001 | 0 | 0.22 | 0.18 | 0.22 | 0.04 | 0.26 | 0.27 |
| 9 | 117566440 | rs4263839 | -0.38 | 0.541 | -0.59 | -0.111 | -0.078 | -0.1 | -0.1 | 2.549 | 0.78 | 0.91 | 0.73 | 0.51 | 0.68 | 0.76 |
| 9 | 117568766 | rs6478109 | -0.369 | 0.597 | -0.482 | -0.08 | -0.077 | -0.1 | -0.1 | 0 | 0.79 | 0.91 | 0.73 | 0.51 | 0.68 | 0.75 |
| 9 | 117569046 | rs7848647 | -0.37 | 0.598 | -0.482 | -0.08 | -0.077 | -0.1 | -0.1 | 0 | 0.79 | 0.91 | 0.73 | 0.51 | 0.67 | 0.76 |
| 9 | 117570856 | rs10982412 | 1.312 | 0.096 | -0.335 | -0.192 | -0.015 | 0.03 | 0.03 | 0 | 0.1 | 0.02 | 0.08 | 0.03 | 0.12 | 0.07 |
| 9 | 117584082 | rs7868351 | -0.248 | 0.149 | -0.347 | -0.121 | -0.007 | -0.018 | -0.018 | 0 | 0.2 | 0.28 | 0.2 | 0.18 | 0.16 | 0.27 |
| 9 | 117585482 | rs1322063 | -1.343 | 0.599 | -0.535 | -0.09 | -0.019 | 0.022 | 0.022 | 0 | 0.92 | 0.97 | 0.91 | 1 | 0.86 | 0.98 |
| 9 | 117597885 | rs34187507 | 1.63 | 0.182 | -0.72 | -0.297 | -0.008 | 0.162 | 0.162 | 5.318 | 0.29 | 0.05 | 0.13 | 0 | 0.24 | 0.13 |
| 9 | 117605070 | rs11554257 | 0.175 | 0.195 | -0.38 | -0.08 | 0.005 | -0.002 | -0.002 | 0 | 0.16 | 0.12 | 0.16 | 0.22 | 0.13 | 0.25 |
| 9 | 117608270 | rs12350737 | 0.003 | 0.274 | -0.131 | -0.072 | 0.007 | -0.025 | -0.025 | 0.061 | 0.08 | 0.33 | 0.04 | 0 | 0.02 | 0.02 |
| 9 | 117609919 | rs16931895 | 0.423 | 0.283 | -0.357 | -0.055 | 0.002 | 0 | 0 | 0 | 0.14 | 0.1 | 0.16 | 0.18 | 0.12 | 0.25 |
| 9 | 117611886 | rs12336469 | -0.331 | 0.287 | -0.164 | -0.032 | 0.01 | 0.022 | 0.022 | 0.177 | 0.08 | 0.36 | 0.04 | 0.05 | 0.02 | 0.04 |
| 9 | 117616813 | rs7037755 | -0.333 | 0.29 | -0.189 | 0.007 | 0.016 | 0.043 | 0.043 | 0.221 | 0.08 | 0.36 | 0.05 | 0.06 | 0.02 | 0.04 |
| 9 | 117618373 | rs10982432 | 0.535 | 0.318 | -0.499 | -0.016 | 0.007 | -0.022 | -0.022 | 0 | 0.06 | 0.22 | 0.03 | 0 | 0.02 | 0.02 |
| 9 | 117627569 | rs7866342 | -0.474 | 0.676 | -1.82 | 0.059 | -0.095 | -0.12 | -0.12 | 4.575 | 0.8 | 0.96 | 0.81 | 0.56 | 0.7 | 0.74 |
| 9 | 117630043 | rs4979467 | 0.694 | 0.672 | -1.597 | -0.411 | -0.007 | 0.009 | 0.009 | 0.008 | 0.57 | 0.4 | 0.6 | 0.31 | 0.55 | 0.46 |
| 9 | 117636879 | rs7874896 | 0.912 | 0.584 | -1.668 | -0.57 | 0.003 | 0.005 | 0.005 | 0 | 0.24 | 0.18 | 0.25 | 0.04 | 0.27 | 0.27 |
| 9 | 117643362 | rs13300483 | 1.331 | 0.524 | -1.736 | -0.46 | 0.01 | -0.015 | -0.015 | 0 | 0.31 | 0.07 | 0.27 | 0.25 | 0.25 | 0.19 |
| 9 | 117645015 | rs7028891 | 0.671 | 0.596 | -1.588 | -0.38 | -0.001 | 0.024 | 0.024 | 0.095 | 0.54 | 0.39 | 0.62 | 0.26 | 0.53 | 0.45 |
| 9 | 117651780 | rs1407309 | -0.934 | 0.396 | -0.866 | 0.234 | 0.029 | 0.047 | 0.047 | 0 | 0.46 | 0.8 | 0.7 | 0.31 | 0.44 | 0.43 |
| 9 | 117660933 | rs1322067 | -0.648 | 0.362 | -0.845 | 0.238 | 0.002 | 0.006 | 0.006 | 0.053 | 0.41 | 0.36 | 0.66 | 0.32 | 0.44 | 0.43 |
| 9 | 117664211 | rs2295800 | -0.907 | 0.444 | -0.982 | 0.278 | 0.028 | 0.046 | 0.046 | 0 | 0.46 | 0.8 | 0.7 | 0.32 | 0.44 | 0.43 |
| 9 | 117665435 | rs3181372 | -0.95 | 0.461 | -1.4 | 0.246 | 0.017 | -0.005 | -0.005 | 0.157 | 0.46 | 0.85 | 0.71 | 0.39 | 0.45 | 0.46 |
| 9 | 117665866 | rs3181369 | -0.393 | 0.446 | -1.39 | 0.301 | -0.005 | 0.023 | 0.023 | 0 | 0.35 | 0.2 | 0.5 | 0.25 | 0.4 | 0.4 |
| 9 | 117667443 | rs3181362 | -0.053 | 0.429 | -1.362 | 0.31 | -0.037 | 0.035 | 0.035 | 2.468 | 0.12 | 0.65 | 0.21 | 0.14 | 0.04 | 0.05 |
| 9 | 117668142 | rs3181195 | -0.393 | 0.449 | -1.512 | 0.304 | -0.005 | 0.023 | 0.023 | 0 | 0.35 | 0.2 | 0.5 | 0.25 | 0.4 | 0.4 |
| 9 | 117669585 | rs1322055 | -1.023 | 0.63 | -1.561 | 0.542 | -0.03 | -0.025 | -0.025 | 0 | 0.9 | 0.97 | 0.9 | 0.84 | 0.84 | 0.89 |
| 9 | 117674320 | rs1555457 | 0.383 | 0.007 | -1.004 | 0.283 | -0.017 | -0.034 | -0.034 | 5.012 | 0.12 | 0.06 | 0.23 | 0.2 | 0.14 | 0.13 |
| 9 | 117679474 | rs2418326 | 0.285 | 0.002 | -1.013 | 0.253 | 0.002 | 0.035 | 0.035 | 0 | 0.23 | 0.13 | 0.26 | 0.04 | 0.25 | 0.24 |
| 9 | 117685398 | rs4979472 | -0.883 | 0.049 | -0.819 | 0.446 | 0.011 | -0.006 | -0.006 | 0 | 0.46 | 0.86 | 0.71 | 0.39 | 0.45 | 0.43 |
| 9 | 117695094 | rs3181192 | -0.322 | -0.062 | -0.751 | -0.312 | -0.027 | -0.063 | -0.063 | 0 | 0.11 | 0.52 | 0.07 | 0 | 0.03 | 0.03 |
| 9 | 117696336 | rs726657 | -0.309 | 0.51 | -1.432 | 0.048 | 0.029 | 0.079 | 0.079 | 0 | 0.45 | 0.72 | 0.57 | 0.24 | 0.42 | 0.4 |
| 9 | 117699850 | rs10817686 | -0.001 | 0.705 | -1.145 | 0.094 | 0.002 | 0.035 | 0.035 | 0 | 0.53 | 0.4 | 0.4 | 0.27 | 0.44 | 0.38 |
| 9 | 117700710 | rs4979476 | 0.415 | 0.664 | -1.452 | 1.223 | -0.012 | -0.054 | -0.054 | 3.94 | 0.91 | 1 | 0.81 | 0.71 | 0.92 | 0.83 |

| | | | | | | | | | | | | | | | |
|---|---|---|---|---|---|---|---|---|---|---|---|---|---|---|---|
| 9 | 117701752 | rs10759743 | 0.259 | 0.094 | -2.032 | 1.162 | 0.024 | -0.023 | -0.023 | 0 | 0.21 | 0.06 | 0.23 | 0.27 | 0.15 | 0.22 |
| 9 | 117704155 | rs11791315 | 0.303 | 0.742 | -2.421 | 0.467 | -0.026 | 0.039 | 0.039 | 0 | 0.16 | 0.09 | 0.11 | 0.03 | 0.23 | 0.17 |
| 9 | 117708099 | rs10817689 | -0.581 | 0.953 | -1.897 | 0.741 | -0.013 | -0.02 | -0.02 | 0 | 0.4 | 0.28 | 0.39 | 0.46 | 0.47 | 0.52 |
| 9 | 117720262 | rs10817691 | -0.824 | 0.758 | -1.172 | 0.589 | 0.002 | 0.003 | 0.003 | 0 | 0.44 | 0.51 | 0.4 | 0.46 | 0.49 | 0.54 |
| 9 | 117722283 | rs12236266 | -0.901 | 0.749 | -1.348 | 0.301 | 0.001 | 0.001 | 0.001 | 0 | 0.5 | 0.55 | 0.46 | 0.46 | 0.55 | 0.53 |
| 9 | 117722690 | rs1265918 | -0.814 | 0.61 | -1.558 | 0.618 | -0.012 | -0.048 | -0.048 | 0 | 0.91 | 1 | 0.81 | 0.71 | 0.92 | 0.81 |
| 9 | 117723333 | rs7853354 | 0.504 | 0.662 | -1.484 | 0.47 | 0.015 | -0.042 | -0.042 | 1.18 | 0.24 | 0.13 | 0.18 | 0.43 | 0.22 | 0.36 |
| 9 | 117728895 | rs10123153 | -0.07 | 0.785 | -1.095 | 0.084 | -0.032 | -0.034 | -0.034 | 3.345 | 0.38 | 0.76 | 0.5 | 0.49 | 0.5 | 0.46 |
| 9 | 117739612 | rs10982474 | 0.671 | 0.769 | -1.233 | 0.261 | -0.001 | 0 | 0 | 0 | 0.17 | 0.15 | 0.15 | 0.16 | 0.14 | 0.21 |
| 9 | 117739703 | rs10982475 | 0.999 | 1.183 | -1.007 | 0.288 | 0.02 | 0.076 | 0.076 | 0 | 0.16 | 0.01 | 0.06 | 0.07 | 0.08 | 0.04 |
| 9 | 117740248 | rs12342713 | 0.13 | 2.443 | 0.311 | 1.201 | -0.006 | -0.029 | -0.029 | 0 | 0.23 | 0.34 | 0.13 | 0.2 | 0.2 | 0.22 |
| 9 | 117741304 | rs10982476 | 0.173 | 2.446 | 0.136 | 1.202 | -0.004 | -0.019 | -0.019 | 0 | 0.23 | 0.29 | 0.12 | 0.2 | 0.2 | 0.22 |
| 9 | 117743178 | rs1012823 | 0.602 | 0.992 | -0.473 | -0.084 | 0.029 | -0.03 | -0.03 | 0 | 0.08 | 0.15 | 0.09 | 0.01 | 0.15 | 0.1 |
| 9 | 117744430 | rs1330367 | -0.364 | 0.68 | -0.983 | 0.049 | 0 | -0.005 | -0.005 | 0 | 0.33 | 0.66 | 0.22 | 0.21 | 0.36 | 0.33 |
| 9 | 117750277 | rs4979482 | -0.06 | -0.212 | -1.272 | 0.467 | 0.095 | -0.001 | -0.001 | 0 | 0.48 | 0.67 | 0.58 | 0.54 | 0.34 | 0.39 |
| 9 | 117753004 | rs10817694 | 0.038 | -0.359 | -0.864 | 0.872 | 0.041 | 0.016 | 0.016 | 0.028 | 0.46 | 0.58 | 0.46 | 0.37 | 0.33 | 0.35 |
| 9 | 117754106 | rs16932027 | 0.69 | -0.409 | -1.225 | 0.679 | 0.004 | 0.017 | 0.017 | 0.115 | 0.09 | 0.04 | 0.05 | 0.09 | 0.08 | 0.16 |
| 9 | 117757075 | rs17817825 | 0.264 | 0.272 | -1.732 | 0.158 | -0.011 | 0.161 | 0.161 | 0 | 0.3 | 0.04 | 0.17 | 0.09 | 0.25 | 0.27 |
| 9 | 117758092 | rs17239323 | 1.239 | 0.332 | -1.735 | 0.165 | -0.024 | 0.132 | 0.132 | 0 | 0.21 | 0.01 | 0.12 | 0.01 | 0.16 | 0.11 |
| 9 | 117766101 | rs4979488 | 1.449 | -0.574 | -1.566 | 0.032 | 0.049 | 0.01 | 0.01 | 0.229 | 0.3 | 0.12 | 0.26 | 0.22 | 0.23 | 0.27 |
| 9 | 117768021 | rs10759751 | 0.82 | 1.308 | -1.391 | 0.588 | -0.027 | -0.026 | -0.026 | 0 | 0.09 | 0.06 | 0.09 | 0.23 | 0.16 | 0.17 |
| 9 | 117768680 | rs4979489 | 0.565 | 1.204 | -1.297 | 0.475 | 0.043 | 0.008 | 0.008 | 0 | 0.14 | 0.19 | 0.17 | 0.14 | 0.26 | 0.17 |
| 9 | 117768771 | rs10982490 | 1.563 | 1.057 | -0.532 | 0.124 | -0.012 | -0.034 | -0.034 | 0 | 0.06 | 0.13 | 0.05 | 0 | 0.05 | 0.01 |
| 9 | 117771310 | rs12006094 | 1.22 | 1.816 | -0.693 | 0.277 | 0.071 | -0.012 | -0.012 | 0 | 0.08 | 0.15 | 0.11 | 0.02 | 0.2 | 0.09 |
| 9 | 117771475 | rs16932055 | -0.065 | 1.143 | -0.416 | 0.461 | 0.009 | 0.034 | 0.034 | 0 | 0.12 | 0.16 | 0.22 | 0.41 | 0.19 | 0.2 |
| 9 | 117776052 | rs16932062 | 0.171 | 1.056 | -0.366 | 0.088 | 0.006 | 0.001 | 0.001 | 0 | 0.17 | 0.17 | 0.13 | 0.22 | 0.22 | 0.24 |
| 9 | 117778369 | rs10117421 | 1.211 | 0.68 | -0.002 | -0.39 | -0.006 | -0.051 | -0.051 | 0 | 0.22 | 0.19 | 0.11 | 0.01 | 0.14 | 0.12 |
| 9 | 117779668 | rs1330362 | 0.388 | 0.215 | 0.061 | -0.481 | -0.057 | -0.084 | -0.084 | 1.305 | 0.15 | 0.31 | 0.07 | 0.03 | 0.07 | 0.06 |
| 9 | 117779933 | rs1107373 | 1.176 | 0.563 | -0.397 | -0.274 | -0.002 | -0.023 | -0.023 | 0 | 0.1 | 0.01 | 0.04 | 0.12 | 0.09 | 0.16 |
| 9 | 117781656 | rs11787664 | 0.098 | 0.354 | -0.581 | -0.385 | -0.043 | -0.006 | -0.006 | 0.649 | 0.35 | 0.22 | 0.28 | 0.35 | 0.51 | 0.41 |
| 9 | 117795288 | rs3789875 | 1.284 | 0.016 | -0.578 | -0.212 | -0.042 | -0.009 | -0.009 | 0.129 | 0.18 | 0.07 | 0.15 | 0.13 | 0.28 | 0.15 |
| 9 | 117797597 | rs12347433 | 1.251 | 0.016 | -0.578 | -0.21 | -0.044 | -0.013 | -0.013 | 0 | 0.18 | 0.07 | 0.15 | 0.13 | 0.28 | 0.17 |
| 9 | 117803720 | rs10759752 | -0.629 | -0.105 | -0.1 | -0.307 | 0.016 | 0.037 | 0.037 | 0 | 0.51 | 0.64 | 0.38 | 0.37 | 0.59 | 0.44 |
| 9 | 117804027 | rs12343737 | 1.948 | -0.037 | -0.193 | -0.35 | 0.042 | -0.024 | -0.024 | 0 | 0.15 | 0.06 | 0.11 | 0.13 | 0.12 | 0.11 |
| 9 | 117804544 | rs1061495 | -0.091 | -0.05 | 0.278 | -0.416 | 0.037 | -0.064 | -0.064 | 0.154 | 0.21 | 0.53 | 0.28 | 0.15 | 0.26 | 0.12 |
| 9 | 117805719 | rs953288 | -1.076 | -0.07 | 0.183 | -0.407 | 0.02 | -0.002 | -0.002 | 0 | 0.54 | 0.77 | 0.54 | 0.48 | 0.62 | 0.44 |
| 9 | 117807508 | rs10122770 | 1.694 | -0.133 | 0.399 | -0.434 | -0.018 | -0.052 | -0.052 | 0 | 0.09 | 0.19 | 0.06 | 0 | 0.06 | 0.01 |
| 9 | 117810036 | rs2297181 | 1.931 | -0.145 | 0.185 | -0.421 | 0.036 | -0.012 | -0.012 | 0 | 0.16 | 0.08 | 0.11 | 0.13 | 0.12 | 0.14 |
| 9 | 117811205 | rs11789268 | 1.07 | -0.062 | 0.135 | -0.419 | -0.012 | 0.011 | 0.011 | 0 | 0.07 | 0.04 | 0.05 | 0.03 | 0.17 | 0.08 |
| 9 | 117812400 | rs1547691 | -1.43 | -0.159 | -0.025 | -0.579 | -0.022 | -0.076 | -0.076 | 0 | 0.77 | 0.89 | 0.63 | 0.5 | 0.75 | 0.59 |
| 9 | 117812791 | rs11789200 | 1.144 | -0.099 | -0.508 | -0.581 | -0.013 | 0.012 | 0.012 | 0 | 0.07 | 0.04 | 0.05 | 0.03 | 0.17 | 0.08 |
| 9 | 117812891 | rs2150101 | 1.653 | -0.166 | -0.521 | -0.59 | 0.016 | 0.004 | 0.004 | 0 | 0.22 | 0.13 | 0.09 | 0.01 | 0.13 | 0.11 |
| 9 | 117813335 | rs7875852 | -1.498 | 0.084 | 0.034 | 0.053 | 0.017 | 0.003 | 0.003 | 0 | 0.55 | 0.74 | 0.53 | 0.49 | 0.62 | 0.48 |
| 9 | 117815273 | rs11794797 | 0.483 | 0.197 | 0.115 | 0.266 | -0.003 | 0 | 0 | 0 | 0.18 | 0.17 | 0.14 | 0.21 | 0.24 | 0.21 |
| 9 | 117817683 | rs1330365 | 0.406 | 0.2 | 0.131 | 0.267 | 0.023 | -0.003 | -0.003 | 0.648 | 0.18 | 0.23 | 0.15 | 0.21 | 0.24 | 0.21 |
| 9 | 117821026 | rs1330368 | -1.069 | 0.189 | -0.086 | 0.253 | -0.012 | 0.021 | 0.021 | 0.052 | 0.39 | 0.69 | 0.42 | 0.36 | 0.5 | 0.33 |
| 9 | 117829987 | rs16932160 | -0.553 | 0.146 | -0.129 | 0.188 | 0.012 | -0.021 | -0.021 | 0 | 0.32 | 0.53 | 0.37 | 0.33 | 0.34 | 0.26 |
| 9 | 117830812 | rs7847271 | 0.16 | -0.438 | 0.21 | 0.336 | -0.016 | -0.026 | -0.026 | 0.064 | 0.13 | 0.31 | 0.22 | 0.11 | 0.1 | 0.05 |
| 9 | 117832416 | rs7043308 | -0.392 | -0.519 | 0.162 | 0.338 | -0.04 | -0.008 | -0.008 | 0 | 0.29 | 0.37 | 0.34 | 0.24 | 0.23 | 0.19 |
| 9 | 117834493 | rs11791562 | 0.057 | -0.499 | -0.008 | 0.335 | 0.032 | -0.036 | -0.036 | 0 | 0.08 | 0.16 | 0.07 | 0 | 0.17 | 0.08 |
| 9 | 117834840 | rs10982513 | -0.65 | -1.034 | -0.054 | 0.753 | -0.049 | -0.01 | -0.01 | 0.139 | 0.29 | 0.41 | 0.34 | 0.24 | 0.22 | 0.19 |
| 9 | 117835254 | rs3789871 | 0.469 | -0.513 | -0.988 | 0.934 | 0.041 | -0.019 | -0.019 | 0 | 0.15 | 0.06 | 0.12 | 0.13 | 0.12 | 0.1 |
| 9 | 117840922 | rs1330351 | 0.161 | 0.074 | -0.707 | 0.472 | -0.019 | 0.027 | 0.027 | 0 | 0.37 | 0.29 | 0.42 | 0.27 | 0.48 | 0.5 |
| 9 | 117841416 | rs12340324 | 1.016 | -0.486 | -1.026 | 0.196 | 0.023 | 0.009 | 0.009 | 0 | 0.14 | 0.05 | 0.09 | 0.08 | 0.12 | 0.05 |
| 9 | 117844239 | rs2989519 | 0.021 | -0.346 | -0.351 | 0.708 | 0.015 | 0.1 | 0.1 | 0 | 0.49 | 0.71 | 0.53 | 0.28 | 0.44 | 0.35 |
| 9 | 117845333 | rs1757106 | -0.67 | -0.346 | -0.245 | 0.854 | -0.005 | 0.001 | 0.001 | 0 | 0.63 | 0.65 | 0.65 | 0.35 | 0.56 | 0.45 |
| 9 | 117845347 | rs10982515 | 1.134 | -0.735 | -0.382 | 0.515 | 0.016 | 0.022 | 0.022 | 0.056 | 0.09 | 0.05 | 0.05 | 0 | 0.06 | 0.02 |
| 9 | 117846570 | rs2274836 | 0.662 | -0.792 | 0.474 | 0.55 | 0.013 | 0.005 | 0.005 | 0 | 0.37 | 0.39 | 0.36 | 0.59 | 0.44 | 0.53 |
| 9 | 117848394 | rs1757095 | -0.938 | -0.835 | 0.298 | 0.507 | 0.011 | 0.001 | 0.001 | 0.21 | 0.9 | 0.89 | 0.94 | 1 | 0.94 | 0.98 |
| 9 | 117853022 | rs944510 | -0.191 | -0.797 | 0.3 | 0.537 | 0.006 | 0.042 | 0.042 | 0 | 0.48 | 0.6 | 0.52 | 0.28 | 0.45 | 0.35 |
| 9 | 117856114 | rs1511 | -0.747 | -0.411 | 0.511 | 0.838 | -0.014 | -0.007 | -0.007 | 0 | 0.63 | 0.67 | 0.65 | 0.4 | 0.57 | 0.48 |
| 9 | 117856877 | rs1330360 | 0.145 | -0.47 | 0.489 | 0.791 | 0.013 | 0.034 | 0.034 | 0 | 0.51 | 0.55 | 0.5 | 0.25 | 0.44 | 0.33 |
| 9 | 117860951 | rs1250027 | -0.815 | -0.481 | 0.444 | 0.791 | -0.051 | -0.023 | -0.023 | 0 | 0.63 | 0.79 | 0.62 | 0.4 | 0.57 | 0.47 |
| 9 | 117865376 | rs2482078 | 0.177 | -0.424 | 0.53 | 0.825 | 0.004 | 0.006 | 0.006 | 0 | 0.51 | 0.5 | 0.5 | 0.24 | 0.45 | 0.35 |
| 9 | 117869417 | rs10982536 | 0.061 | -0.407 | 0.49 | 0.865 | -0.001 | 0.024 | 0.024 | 0 | 0.13 | 0.23 | 0.12 | 0.16 | 0.13 | 0.15 |
| 9 | 117869930 | rs1250023 | 0.039 | -0.319 | 0.574 | 0.596 | 0.014 | 0.044 | 0.044 | 0 | 0.5 | 0.4 | 0.49 | 0.24 | 0.45 | 0.35 |
| 9 | 117879414 | rs10118040 | 0.048 | -0.101 | 0.396 | 0.495 | 0.001 | -0.007 | -0.007 | 0.272 | 0.38 | 0.39 | 0.32 | 0.71 | 0.38 | 0.39 |
| 9 | 117880792 | rs2071520 | 0.681 | -0.091 | 0.578 | 0.955 | -0.012 | -0.006 | -0.006 | 0 | 0.28 | 0.22 | 0.25 | 0.32 | 0.32 | 0.32 |
| 9 | 117885635 | rs12380363 | 0.294 | -0.54 | 0.208 | 0.318 | 0.003 | 0.052 | 0.052 | 0 | 0.31 | 0.22 | 0.26 | 0.1 | 0.28 | 0.28 |

| | | | | | | | | | | | | | | | | |
|---|---|---|---|---|---|---|---|---|---|---|---|---|---|---|---|---|
| 9 | 117889206 | rs10817713 | 1.041 | -0.529 | 0.55 | 0.41 | 0.001 | -0.002 | -0.002 | 0 | 0.28 | 0.32 | 0.26 | 0.27 | 0.32 | 0.32 |
| 9 | 117896185 | rs10817715 | 0.108 | -0.412 | 0.893 | 0.598 | 0.003 | -0.05 | -0.05 | 0 | 0.31 | 0.2 | 0.26 | 0.5 | 0.29 | 0.33 |
| 9 | 117896349 | rs1537722 | -0.88 | -0.855 | 0.872 | 0.304 | -0.014 | 0.002 | 0.002 | 0 | 0.7 | 0.77 | 0.56 | 0.77 | 0.63 | 0.76 |
| 9 | 117897993 | rs7021057 | 0.266 | -0.869 | 0.897 | 0.3 | 0.001 | -0.006 | -0.006 | 0 | 0.32 | 0.31 | 0.28 | 0.5 | 0.29 | 0.33 |
| 9 | 117903975 | rs2012566 | 0.266 | -0.869 | 0.897 | 0.3 | 0.001 | -0.006 | -0.006 | 1.239 | 0.32 | 0.31 | 0.28 | 0.5 | 0.29 | 0.33 |
| 9 | 117905403 | rs7865971 | 0.388 | -0.851 | 0.804 | 0.28 | 0.002 | 0.038 | 0.038 | 0 | 0.31 | 0.24 | 0.26 | 0.1 | 0.28 | 0.28 |
| 9 | 117905653 | rs1414137 | 0.619 | -0.851 | 0.8 | 0.297 | -0.003 | -0.018 | -0.018 | 0 | 0.38 | 0.48 | 0.29 | 0.27 | 0.34 | 0.42 |
| 9 | 117909793 | rs1335334 | 1.103 | -0.598 | 0.501 | 0.399 | 0.042 | -0.012 | -0.012 | 0 | 0.19 | 0.06 | 0.34 | 0.14 | 0.16 | 0.09 |
| 9 | 117912654 | rs6478135 | -0.697 | -0.668 | 0.765 | 0.521 | 0.027 | 0.575 | 0.575 | 0 | 0.06 | 0.25 | 0.03 | 0.47 | 0.02 | 0.15 |
| 9 | 117912890 | rs1537728 | -0.639 | -0.608 | 0.765 | 0.475 | -0.025 | 0.033 | 0.033 | 0 | 0.72 | 0.79 | 0.56 | 0.85 | 0.64 | 0.85 |
| 9 | 117913870 | rs12683723 | -1.414 | -0.617 | 0.789 | 0.712 | 0.063 | 0.43 | 0.43 | 0 | 0.08 | 0.25 | 0.11 | 0.53 | 0.09 | 0.19 |
| 9 | 117919576 | rs10982575 | 0.804 | -0.954 | 0.526 | 0.197 | -0.046 | -0.025 | -0.025 | 0 | 0.18 | 0.23 | 0.1 | 0.09 | 0.12 | 0.13 |
| 9 | 117926650 | rs7851086 | 0.827 | -0.96 | 0.582 | 0.194 | -0.042 | -0.023 | -0.023 | 0 | 0.18 | 0.25 | 0.11 | 0.09 | 0.12 | 0.13 |
| 9 | 117927227 | rs10982590 | 1.464 | -0.945 | 0.363 | 0.2 | 0.038 | 0.012 | 0.012 | 0.57 | 0.15 | 0.05 | 0.09 | 0.08 | 0.12 | 0.12 |
| 9 | 117928419 | rs7866181 | 1.13 | -1 | 0.022 | 0.032 | 0.003 | 0.044 | 0.044 | 0 | 0.32 | 0.2 | 0.18 | 0.25 | 0.27 | 0.39 |
| 9 | 117928483 | rs7024589 | 0.375 | -0.934 | 0.417 | -0.025 | 0.015 | -0.058 | -0.058 | 0 | 0.32 | 0.46 | 0.35 | 0.15 | 0.35 | 0.29 |
| 9 | 117932692 | rs10982592 | 1.338 | -0.772 | 0.445 | 0.443 | -0.026 | 0.063 | 0.063 | 0.018 | 0.17 | 0.2 | 0.11 | 0.48 | 0.12 | 0.16 |
| 9 | 117932816 | rs4979508 | -0.95 | -0.727 | 0.01 | 0.233 | -0.026 | -0.006 | -0.006 | 0 | 0.84 | 0.86 | 0.65 | 0.86 | 0.74 | 0.86 |
| 9 | 117934415 | rs10817719 | 1.617 | -0.75 | -0.009 | -0.067 | 0.038 | 0.007 | 0.007 | 0.48 | 0.15 | 0.05 | 0.09 | 0.11 | 0.11 | 0.11 |
| 9 | 117950332 | rs7040243 | 1.305 | -0.686 | -0.063 | -0.209 | -0.028 | 0.008 | 0.008 | 0 | 0.14 | 0.07 | 0.22 | 0.11 | 0.19 | 0.19 |
| 9 | 117951073 | rs7027882 | -0.211 | -0.838 | 0.593 | -0.253 | 0.048 | 0.146 | 0.146 | 0 | 0.78 | 0.6 | 0.61 | 0.48 | 0.72 | 0.77 |
| 9 | 117956575 | rs1335344 | -0.75 | -0.866 | 0.578 | -0.255 | -0.011 | -0.013 | -0.013 | 0.767 | 0.82 | 0.81 | 0.64 | 0.48 | 0.73 | 0.78 |
| 9 | 117956828 | rs7851185 | 0.368 | -1.115 | 0.475 | -0.482 | -0.012 | 0.055 | 0.055 | 0.866 | 0.28 | 0.18 | 0.28 | 0.11 | 0.28 | 0.25 |
| 9 | 117956937 | rs16932349 | 1.985 | -1.202 | 0.348 | -0.506 | 0.04 | 0.009 | 0.009 | 0.24 | 0.15 | 0.05 | 0.1 | 0.1 | 0.11 | 0.12 |
| 9 | 117962526 | rs4978627 | 0.2 | -1.251 | 0.506 | -0.57 | -0.002 | 0.001 | 0.001 | 0.485 | 0.3 | 0.35 | 0.33 | 0.13 | 0.33 | 0.28 |
| 9 | 117964656 | rs4978628 | 1.658 | -1.256 | 0.48 | -0.559 | -0.053 | 0.02 | 0.02 | 0 | 0.31 | 0.04 | 0.18 | 0.25 | 0.29 | 0.37 |
| 9 | 117966294 | rs10759761 | -0.094 | -1.263 | 0.508 | -0.552 | 0.01 | 0.054 | 0.054 | 0 | 0.45 | 0.4 | 0.42 | 0.23 | 0.44 | 0.4 |
| 9 | 117967117 | rs7044001 | -0.823 | -1.084 | 0.333 | -0.217 | -0.016 | -0.023 | -0.023 | 0 | 0.81 | 0.83 | 0.64 | 0.52 | 0.73 | 0.75 |
| 9 | 117970568 | rs1889651 | 0.044 | -1.244 | 0.508 | -0.402 | -0.018 | 0.012 | 0.012 | 0 | 0.64 | 0.69 | 0.41 | 0.28 | 0.53 | 0.52 |
| 9 | 117970824 | rs4335203 | -0.282 | -1.167 | 0.76 | -0.395 | -0.001 | 0 | 0 | 0 | 0.64 | 0.63 | 0.48 | 0.36 | 0.56 | 0.63 |
| 9 | 117972155 | rs1335357 | -0.012 | -1.19 | 0.523 | -0.473 | 0.001 | -0.002 | -0.002 | 0 | 0.64 | 0.6 | 0.4 | 0.28 | 0.53 | 0.52 |
| 9 | 117972516 | rs4147052 | 1.529 | -1.157 | -0.05 | -0.361 | -0.024 | 0.085 | 0.085 | 0 | 0.31 | 0.07 | 0.18 | 0.16 | 0.28 | 0.34 |
| 9 | 117975864 | rs10982611 | 0.74 | -1.071 | 0.042 | -0.304 | 0.048 | 0.121 | 0.121 | 0.438 | 0.47 | 0.15 | 0.28 | 0.26 | 0.41 | 0.45 |
| 9 | 117979360 | rs13440344 | 0.475 | -1.027 | 0.511 | -0.134 | 0.027 | -0.003 | -0.003 | 0.079 | 0.2 | 0.33 | 0.28 | 0.22 | 0.24 | 0.26 |
| 9 | 117979410 | rs10817727 | 0.877 | -1.279 | 0.44 | -0.176 | -0.042 | -0.012 | -0.012 | 0 | 0.17 | 0.21 | 0.11 | 0.12 | 0.12 | 0.11 |
| 9 | 117981328 | rs10982615 | 1.244 | -1.121 | -0.127 | -0.149 | 0.003 | 0.001 | 0.001 | 0.341 | 0.14 | 0.08 | 0.09 | 0.11 | 0.12 | 0.09 |
| 9 | 117983998 | rs10759762 | -0.018 | -0.899 | -0.735 | -0.263 | -0.018 | -0.024 | -0.024 | 0 | 0.69 | 0.73 | 0.57 | 0.45 | 0.64 | 0.72 |
| 9 | 117985994 | rs10759763 | 0.557 | -0.799 | -0.723 | -0.845 | 0.009 | 0.034 | 0.034 | 0 | 0.35 | 0.27 | 0.21 | 0.19 | 0.29 | 0.42 |
| 9 | 117987615 | rs10982618 | 0.558 | -0.754 | -0.587 | -0.865 | 0.006 | 0.02 | 0.02 | 0.004 | 0.35 | 0.28 | 0.22 | 0.19 | 0.3 | 0.43 |
| 9 | 117992846 | rs17834626 | 1.406 | -0.294 | -0.616 | -1.117 | -0.009 | -0.001 | -0.001 | 0 | 0.09 | 0.06 | 0.06 | 0 | 0.17 | 0.05 |
| 9 | 117992860 | rs16932390 | 0.688 | -0.441 | -0.547 | -1.452 | -0.004 | -0.001 | -0.001 | 9.546 | 0.2 | 0.17 | 0.24 | 0.03 | 0.22 | 0.13 |
| 9 | 117994565 | rs2992140 | 0.865 | -0.543 | -0.373 | -0.869 | -0.002 | 0.008 | 0.008 | 0 | 0.21 | 0.19 | 0.37 | 0.33 | 0.17 | 0.18 |
| 9 | 117996625 | rs2992142 | 0.334 | -0.763 | -0.41 | -0.733 | 0.037 | 0.01 | 0.01 | 4.526 | 0.65 | 0.54 | 0.67 | 0.56 | 0.58 | 0.71 |
| 9 | 118001034 | rs10120911 | 1.502 | -1.926 | -0.339 | -0.49 | -0.047 | 0.001 | 0.001 | 0 | 0.16 | 0.24 | 0.06 | 0.12 | 0.05 | 0.19 |
| 9 | 118001670 | rs2989509 | -0.299 | -2.027 | -0.455 | -0.641 | -0.008 | -0.002 | -0.002 | 0 | 0.45 | 0.47 | 0.5 | 0.55 | 0.32 | 0.47 |
| 9 | 118007135 | rs7869917 | 2.378 | -1.993 | -0.586 | -0.592 | -0.009 | -0.026 | -0.026 | 1.081 | 0.1 | 0 | 0.09 | 0.1 | 0.13 | 0.11 |
| 9 | 118012841 | rs10125253 | -0.402 | -2.143 | -0.518 | -0.81 | -0.003 | -0.011 | -0.011 | 0.091 | 0.42 | 0.5 | 0.62 | 0.77 | 0.37 | 0.42 |
| 9 | 118013289 | rs883402 | 1.602 | -2.154 | -0.536 | -0.783 | 0 | 0.004 | 0.004 | 0 | 0.18 | 0.16 | 0.36 | 0.34 | 0.14 | 0.17 |
| 9 | 118029484 | rs4147051 | 0.87 | -2.023 | -1.711 | -0.78 | -0.015 | 0.024 | 0.024 | 6.816 | 0.16 | 0.18 | 0.07 | 0.4 | 0.07 | 0.12 |
| 9 | 118029729 | rs10817738 | 1.804 | -1.911 | -1.718 | -0.793 | -0.142 | 0.167 | 0.167 | 0 | 0.29 | 0.06 | 0.23 | 0.02 | 0.44 | 0.29 |
| 9 | 118039472 | rs10982650 | 0.276 | -1.332 | -0.821 | 0.279 | 0.01 | -0.001 | -0.001 | 3.918 | 0.27 | 0.23 | 0.14 | 0.21 | 0.18 | 0.29 |
| 9 | 118042353 | rs1414153 | 0.409 | -0.887 | -0.105 | 0.556 | -0.001 | -0.001 | -0.001 | 0 | 0.23 | 0.25 | 0.24 | 0.14 | 0.21 | 0.31 |
| 9 | 118043489 | rs7846857 | -0.669 | -0.727 | -0.274 | 0.91 | -0.014 | 0.027 | 0.027 | 4.696 | 0.16 | 0.17 | 0.07 | 0.4 | 0.07 | 0.13 |
| 9 | 118057870 | rs11998787 | -0.661 | -0.433 | -0.476 | -0.079 | -0.015 | -0.035 | -0.035 | 0.025 | 0.13 | 0.18 | 0.06 | 0.04 | 0.07 | 0.03 |
| 9 | 118068722 | rs1157836 | 0.008 | 0.006 | -1.584 | -0.117 | -0.04 | 0.138 | 0.138 | 0.091 | 0.53 | 0.24 | 0.62 | 0.76 | 0.56 | 0.58 |
| 9 | 118070259 | rs10982664 | 0.34 | -0.225 | -1.66 | 0.078 | 0.005 | -0.023 | -0.023 | 0 | 0.12 | 0 | 0.13 | 0.14 | 0.12 | 0.09 |
| 9 | 118073084 | rs1335335 | -0.128 | -0.37 | -2.235 | -0.176 | 0.07 | 0.099 | 0.099 | 0 | 0.56 | 0.18 | 0.47 | 0.23 | 0.69 | 0.48 |
| 9 | 118081458 | rs2027155 | -0.185 | -0.474 | -1.897 | 0.763 | 0.04 | 0.01 | 0.01 | 0 | 0.57 | 0.28 | 0.55 | 0.64 | 0.67 | 0.55 |
| 9 | 118087739 | rs11787534 | 0.65 | 0.235 | -2.089 | 0.623 | -0.005 | 0.009 | 0.009 | 0 | 0.06 | 0.05 | 0.06 | 0.01 | 0.15 | 0.05 |
| 9 | 118091712 | rs10817746 | 1.223 | 0.479 | -2.586 | 0.276 | -0.045 | -0.124 | -0.124 | 0 | 0.25 | 0.07 | 0.25 | 0.46 | 0.24 | 0.25 |
| 9 | 118092924 | rs7034237 | -0.775 | -0.112 | -1.478 | 0.073 | -0.006 | 0.004 | 0.004 | 0 | 0.67 | 0.52 | 0.57 | 0.69 | 0.68 | 0.6 |
| 9 | 118093007 | rs3814957 | 0.627 | -0.13 | -1.039 | 0.052 | -0.001 | 0.001 | 0.001 | 0 | 0.32 | 0.34 | 0.42 | 0.31 | 0.32 | 0.4 |
| 9 | 118094992 | rs7028273 | 1.163 | -0.219 | -1.107 | 0.062 | -0.002 | -0.001 | -0.001 | 0 | 0.23 | 0.2 | 0.15 | 0.23 | 0.24 | 0.33 |
| 9 | 118104142 | rs10982691 | 1.325 | -0.451 | -1.532 | -0.061 | -0.035 | 0.072 | 0.072 | 1.419 | 0.15 | 0.02 | 0.18 | 0.03 | 0.18 | 0.19 |
| 9 | 118107224 | rs1981499 | -0.69 | -0.311 | -1.307 | -0.03 | -0.005 | -0.001 | -0.001 | 0 | 0.66 | 0.52 | 0.63 | 0.7 | 0.67 | 0.6 |
| 9 | 118107547 | rs10817753 | 1.553 | -0.319 | -1.96 | 0.028 | -0.054 | 0.06 | 0.06 | 0 | 0.23 | 0.1 | 0.17 | 0.06 | 0.26 | 0.17 |
| 9 | 118113846 | rs13300423 | 1.689 | -0.355 | -1.854 | -0.014 | 0.02 | -0.024 | -0.024 | 0 | 0.1 | 0 | 0.11 | 0.14 | 0.1 | 0.08 |
| 9 | 118119122 | rs10982704 | 0.438 | -0.098 | -1.715 | -0.175 | -0.099 | 0.163 | 0.163 | 0 | 0.39 | 0.12 | 0.35 | 0.09 | 0.44 | 0.36 |
| 9 | 118121096 | rs10429586 | 0.456 | -0.079 | -1.592 | -0.171 | -0.067 | 0.142 | 0.142 | 0 | 0.39 | 0.16 | 0.35 | 0.09 | 0.44 | 0.36 |

| | | | | | | | | | | | | | | | | |
|---|---|---|---|---|---|---|---|---|---|---|---|---|---|---|---|---|
| 9 | 118121597 | rs10982710 | 0.573 | 0.182 | -1.748 | -0.321 | -0.065 | 0.13 | 0.13 | 0 | 0.37 | 0.14 | 0.32 | 0.1 | 0.39 | 0.34 |
| 9 | 118124242 | rs4978633 | -0.425 | -0.453 | -1.63 | -0.228 | 0.013 | 0.057 | 0.057 | 1.437 | 0.54 | 0.29 | 0.6 | 0.67 | 0.56 | 0.51 |
| 9 | 118129621 | rs9409144 | -0.511 | -0.365 | -1.909 | 0.003 | -0.052 | 0.039 | 0.039 | 0.831 | 0.59 | 0.9 | 0.54 | 0.78 | 0.55 | 0.63 |
| 9 | 118137835 | rs7850103 | -0.944 | -0.401 | -1.88 | -0.025 | -0.067 | 0.043 | 0.043 | 0 | 0.68 | 0.92 | 0.65 | 0.87 | 0.65 | 0.74 |
| 9 | 118148224 | rs10733614 | -0.812 | -0.391 | -1.616 | 0.009 | 0 | 0 | 0 | 0 | 0.91 | 0.9 | 0.8 | 0.95 | 0.91 | 0.93 |
| 9 | 118148690 | rs7859920 | -0.203 | -0.554 | -1.08 | 0.091 | -0.008 | 0.015 | 0.015 | 0 | 0.55 | 0.72 | 0.51 | 0.44 | 0.54 | 0.61 |
| 9 | 118152551 | rs10124685 | 0.005 | -0.351 | -0.871 | 0.353 | 0.007 | 0.087 | 0.087 | 0 | 0.14 | 0.24 | 0.14 | 0.44 | 0.13 | 0.14 |
| 9 | 118156112 | rs3893062 | 0.361 | -0.531 | -0.799 | 0.266 | 0.012 | 0.021 | 0.021 | 0 | 0.29 | 0.63 | 0.31 | 0.62 | 0.37 | 0.35 |
| 9 | 118157729 | rs10982724 | -0.005 | -0.493 | -0.328 | 0.314 | 0.137 | 0.311 | 0.311 | 0 | 0.06 | 0.31 | 0.14 | 0.35 | 0.11 | 0.1 |
| 9 | 118161597 | rs10817762 | -0.764 | -0.717 | -0.546 | -0.228 | -0.011 | 0.092 | 0.092 | 0 | 0.48 | 0.72 | 0.46 | 0.79 | 0.5 | 0.5 |
| 9 | 118163563 | rs2269700 | -1.474 | -0.669 | -0.691 | -0.237 | -0.055 | 0.051 | 0.051 | 0 | 0.66 | 0.91 | 0.6 | 0.85 | 0.65 | 0.71 |
| 9 | 118170889 | rs7862745 | -0.191 | -0.41 | -0.401 | -0.02 | 0.034 | 0.189 | 0.189 | 1.152 | 0.17 | 0.37 | 0.27 | 0.52 | 0.23 | 0.22 |
| 9 | 118173811 | rs10124376 | 1.092 | -0.484 | -0.394 | -0.188 | 0.01 | 0.001 | 0.001 | 0 | 0.23 | 0.31 | 0.18 | 0.27 | 0.26 | 0.25 |
| 9 | 118174830 | rs7037974 | -0.326 | -0.641 | -0.246 | -0.23 | 0.011 | 0.026 | 0.026 | 0.137 | 0.51 | 0.59 | 0.34 | 0.33 | 0.41 | 0.52 |
| 9 | 118179485 | rs721227 | 0.283 | -0.678 | -0.327 | -0.279 | 0 | 0 | 0 | 0 | 0.39 | 0.36 | 0.36 | 0.32 | 0.4 | 0.46 |
| 9 | 118184602 | rs6478163 | -0.04 | -0.699 | -0.485 | -0.32 | 0.032 | 0.081 | 0.081 | 3.212 | 0.79 | 0.95 | 0.9 | 0.94 | 0.85 | 0.82 |
| 9 | 118186082 | rs10124142 | 1.228 | -0.616 | -0.359 | -0.146 | -0.009 | 0.018 | 0.018 | 0 | 0.34 | 0.29 | 0.39 | 0.18 | 0.34 | 0.28 |
| 9 | 118189520 | rs4979527 | -0.125 | -0.614 | -0.342 | -0.152 | -0.021 | 0.045 | 0.045 | 1.986 | 0.45 | 0.69 | 0.5 | 0.76 | 0.51 | 0.55 |
| 9 | 118194134 | rs10982742 | 0.321 | -0.545 | -0.07 | -0.127 | 0.001 | 0.019 | 0.019 | 1.695 | 0.39 | 0.32 | 0.35 | 0.33 | 0.4 | 0.46 |
| 9 | 118196088 | rs10759780 | -1.6 | -0.546 | -0.1 | -0.127 | -0.016 | 0.012 | 0.012 | 3.092 | 0.89 | 1 | 0.91 | 0.94 | 0.86 | 0.89 |
| 9 | 118199668 | rs10982746 | 0.462 | -0.595 | -0.056 | -0.137 | -0.004 | 0.102 | 0.102 | 0.786 | 0.73 | 0.55 | 0.75 | 0.5 | 0.74 | 0.74 |
| 9 | 118202187 | rs7042718 | -1.263 | -0.432 | -0.031 | -0.053 | 0.001 | 0.021 | 0.021 | 3.603 | 0.88 | 0.95 | 0.91 | 0.94 | 0.86 | 0.89 |
| 9 | 118209542 | rs2157664 | -0.516 | -0.48 | -0.485 | -0.017 | -0.028 | 0.05 | 0.05 | 0.115 | 0.56 | 0.81 | 0.51 | 0.75 | 0.52 | 0.61 |
| 9 | 118213502 | rs2157663 | 1.458 | -0.412 | -0.943 | -0.243 | -0.024 | -0.075 | -0.075 | 0 | 0.12 | 0.01 | 0.13 | 0.27 | 0.11 | 0.15 |
| 9 | 118217828 | rs1632835 | -0.66 | -0.395 | -0.835 | -0.297 | -0.001 | 0.026 | 0.026 | 0.08 | 0.53 | 0.6 | 0.5 | 0.75 | 0.52 | 0.61 |
| 9 | 118223430 | rs1632812 | 1.521 | -0.403 | -1.028 | -0.126 | -0.021 | 0.001 | 0.001 | 0 | 0.32 | 0.17 | 0.42 | 0.21 | 0.34 | 0.28 |
| 9 | 118243754 | rs12380029 | 1.014 | -0.439 | -0.463 | 0.185 | 0.001 | -0.014 | -0.014 | 1.383 | 0.13 | 0.08 | 0.07 | 0.23 | 0.11 | 0.14 |
| 9 | 118244107 | rs12341727 | 0.371 | -0.644 | -0.297 | 0.069 | -0.011 | 0.005 | 0.005 | 0 | 0.08 | 0.15 | 0.1 | 0.09 | 0.07 | 0.05 |
| 9 | 118248522 | rs7852274 | -0.012 | -0.689 | -0.195 | 0.057 | 0.043 | 0.206 | 0.206 | 0.847 | 0.83 | 0.5 | 0.76 | 0.51 | 0.78 | 0.8 |
| 9 | 118250623 | rs7874499 | -0.04 | -0.709 | -0.27 | 0.033 | -0.014 | 0.056 | 0.056 | 0 | 0.09 | 0.26 | 0.18 | 0.15 | 0.08 | 0.06 |
| 9 | 118252779 | rs6478174 | -1.522 | -0.645 | -0.435 | 0.077 | -0.007 | 0 | 0 | 1.061 | 0.89 | 1 | 0.91 | 0.93 | 0.88 | 0.89 |
| 9 | 118255620 | rs12238100 | 0.146 | -0.555 | -0.073 | 0.213 | 0.223 | 0.569 | 0.569 | 0.106 | 0.05 | 0.35 | 0.14 | 0.42 | 0.1 | 0.08 |
| 9 | 118258940 | rs6478176 | -0.05 | -0.767 | -0.364 | -0.381 | -0.018 | -0.028 | -0.028 | 0.014 | 0.25 | 0.53 | 0.34 | 0.19 | 0.23 | 0.21 |
| 9 | 118259578 | rs1989767 | 1.148 | -0.943 | -0.579 | -0.549 | -0.037 | 0.017 | 0.017 | 0 | 0.59 | 0.16 | 0.44 | 0.32 | 0.56 | 0.56 |
| 9 | 118260090 | rs12551402 | 0.248 | -0.96 | -0.357 | -0.554 | 0.007 | 0.043 | 0.043 | 0 | 0.09 | 0.33 | 0.18 | 0.12 | 0.08 | 0.05 |
| 9 | 118261303 | rs7038462 | 0.461 | -0.989 | -0.349 | -0.524 | 0.006 | 0.026 | 0.026 | 0 | 0.68 | 0.49 | 0.62 | 0.43 | 0.64 | 0.61 |
| 9 | 118262641 | rs9299225 | 0.057 | -0.938 | -0.282 | -0.477 | 0.151 | 0.448 | 0.448 | 0.396 | 0.07 | 0.32 | 0.15 | 0.42 | 0.11 | 0.11 |
| 9 | 118266112 | rs9409167 | -0.446 | -0.891 | -0.433 | -0.468 | 0.056 | 0.194 | 0.194 | 0 | 0.8 | 0.58 | 0.71 | 0.52 | 0.76 | 0.74 |
| 9 | 118276864 | rs2418353 | -0.913 | -1.35 | -1.014 | -0.76 | 0.081 | 0.083 | 0.083 | 0.014 | 0.48 | 0.21 | 0.3 | 0.37 | 0.35 | 0.48 |
| 9 | 118277315 | rs9408872 | -0.3 | -1.329 | -0.956 | -0.719 | 0.058 | 0.028 | 0.028 | 0.269 | 0.35 | 0.2 | 0.17 | 0.31 | 0.22 | 0.32 |
| 9 | 118279910 | rs10982776 | -0.14 | -1.388 | -0.947 | -0.709 | 0.042 | 0.01 | 0.01 | 0 | 0.31 | 0.19 | 0.12 | 0.31 | 0.16 | 0.31 |
| 9 | 118286802 | rs10124673 | 0.229 | -1.324 | -0.915 | -0.775 | 0.012 | -0.005 | -0.005 | 0 | 0.63 | 0.35 | 0.7 | 0.33 | 0.73 | 0.7 |
| 9 | 118287508 | rs7033617 | -0.224 | -1.379 | -1.022 | -0.84 | -0.004 | -0.004 | -0.004 | 0 | 0.29 | 0.33 | 0.34 | 0.25 | 0.26 | 0.32 |
| 9 | 118298964 | rs2040480 | 2.033 | -1.185 | -1.167 | -0.78 | -0.137 | 0.107 | 0.107 | 0 | 0.35 | 0.07 | 0.37 | 0.08 | 0.47 | 0.38 |
| 9 | 118305438 | rs10119624 | 0.947 | -1.164 | -0.668 | -0.691 | 0.017 | 0.011 | 0.011 | 0.082 | 0.56 | 0.38 | 0.62 | 0.27 | 0.66 | 0.66 |
| 9 | 118311164 | rs1018845 | 1.351 | -1.188 | -0.73 | -0.721 | 0.037 | 0.097 | 0.097 | 0.486 | 0.48 | 0.18 | 0.56 | 0.21 | 0.59 | 0.54 |
| 9 | 118311318 | rs10817792 | -0.161 | -1.43 | -0.737 | -0.827 | -0.005 | 0.065 | 0.065 | 0.007 | 0.73 | 0.49 | 0.72 | 0.35 | 0.74 | 0.77 |
| 9 | 118311498 | rs7046356 | -0.161 | -1.43 | -0.737 | -0.827 | -0.005 | 0.065 | 0.065 | 0.007 | 0.73 | 0.49 | 0.73 | 0.35 | 0.74 | 0.77 |
| 9 | 118311709 | rs10982793 | 0.666 | -1.412 | -0.991 | -0.825 | 0.03 | 0.036 | 0.036 | 0 | 0.17 | 0.09 | 0.1 | 0.08 | 0.08 | 0.11 |
| 9 | 118312655 | rs10982794 | 2.93 | -1.189 | -1.199 | -0.952 | -0.108 | 0.01 | 0.01 | 0 | 0.24 | 0.02 | 0.29 | 0.07 | 0.28 | 0.22 |
| 9 | 118313925 | rs10982796 | 2.03 | -1.699 | -0.403 | -0.484 | -0.032 | 0.064 | 0.064 | 0 | 0.42 | 0.16 | 0.49 | 0.2 | 0.55 | 0.53 |
| 9 | 118314095 | rs10982797 | 1.955 | -1.699 | -0.354 | -0.48 | -0.007 | 0.04 | 0.04 | 0 | 0.43 | 0.3 | 0.5 | 0.21 | 0.56 | 0.54 |
| 9 | 118314283 | rs4979552 | 1.238 | -1.748 | -0.397 | -0.506 | 0.005 | -0.006 | -0.006 | 0 | 0.66 | 0.28 | 0.6 | 0.29 | 0.68 | 0.65 |
| 9 | 118314667 | rs2900601 | -0.76 | -1.733 | -0.33 | -0.4 | 0.016 | -0.005 | -0.005 | 0.218 | 0.84 | 0.85 | 0.93 | 0.76 | 0.91 | 0.86 |
| 9 | 118332688 | rs2157666 | 2.292 | -1.589 | -0.566 | -0.441 | -0.073 | 0.046 | 0.046 | 0 | 0.37 | 0.17 | 0.45 | 0.16 | 0.5 | 0.42 |
| 9 | 118335952 | rs7027632 | 2.494 | -1.584 | -0.642 | -0.45 | -0.144 | 0.069 | 0.069 | 0 | 0.36 | 0.07 | 0.37 | 0.12 | 0.49 | 0.41 |
| 9 | 118339551 | rs6478191 | 0.406 | -1.576 | -0.467 | -0.383 | -0.005 | 0.034 | 0.034 | 0 | 0.73 | 0.42 | 0.72 | 0.34 | 0.74 | 0.76 |
| 9 | 118339739 | rs2157668 | 0.805 | -1.639 | -0.527 | -0.439 | 0.001 | 0 | 0 | 0 | 0.61 | 0.39 | 0.6 | 0.29 | 0.62 | 0.64 |
| 9 | 118348955 | rs10982832 | -0.487 | -1.679 | -0.675 | -0.499 | 0.042 | 0.085 | 0.085 | 0 | 0.29 | 0.11 | 0.22 | 0.13 | 0.18 | 0.23 |
| 9 | 118352164 | rs1981500 | 2.022 | -1.648 | -0.642 | -0.468 | -0.018 | 0.089 | 0.089 | 0 | 0.43 | 0.23 | 0.42 | 0.16 | 0.54 | 0.53 |
| 9 | 118357126 | rs875069 | -0.666 | -1.618 | -0.688 | -0.468 | 0.034 | 0.048 | 0.048 | 0 | 0.23 | 0.13 | 0.14 | 0.12 | 0.12 | 0.22 |
| 9 | 118367195 | rs7859870 | -1.74 | -1.54 | -0.125 | -0.253 | 0.068 | 0.197 | 0.197 | 0 | 0.92 | 0.83 | 0.81 | 0.59 | 0.83 | 0.9 |
| 9 | 118368761 | rs12115665 | 0.995 | -1.485 | -0.555 | -0.458 | -0.011 | 0.01 | 0.01 | 0 | 0.07 | 0.01 | 0.04 | 0.04 | 0.06 | 0.12 |
| 9 | 118380578 | rs4593632 | 2.34 | -1.543 | -0.578 | -0.502 | -0.106 | 0.077 | 0.077 | 0 | 0.36 | 0.11 | 0.38 | 0.12 | 0.49 | 0.41 |
| 9 | 118392208 | rs4979570 | 1.447 | -1.651 | -0.181 | -0.316 | 0.017 | 0.004 | 0.004 | 0.204 | 0.56 | 0.41 | 0.68 | 0.32 | 0.67 | 0.58 |
| 9 | 118397913 | rs7850229 | -1.115 | -1.519 | 0.201 | -0.044 | -0.032 | -0.026 | -0.026 | 0.18 | 0.19 | 0.26 | 0.14 | 0.09 | 0.08 | 0.22 |
| 9 | 118405173 | rs10982853 | -0.79 | -0.961 | -0.246 | 0.16 | 0.002 | 0.002 | 0.002 | 0 | 0.13 | 0.06 | 0.12 | 0.14 | 0.11 | 0.18 |
| 9 | 118408803 | rs956325 | -1.014 | -1.354 | 0.058 | 0.126 | -0.015 | 0.004 | 0.004 | 0 | 0.21 | 0.28 | 0.18 | 0.3 | 0.13 | 0.29 |
| 9 | 118409944 | rs10982854 | -1.282 | -1.294 | 0.451 | -0.292 | 0.023 | 0.029 | 0.029 | 0 | 0.29 | 0.45 | 0.45 | 0.66 | 0.34 | 0.36 |

| | | | | | | | | | | | | | | | | |
|---|---|---|---|---|---|---|---|---|---|---|---|---|---|---|---|---|
| 9 | 118411240 | rs12378054 | -0.055 | -1.415 | 0.376 | -0.302 | 0.023 | -0.022 | -0.022 | 0 | 0.14 | 0.06 | 0.05 | 0.22 | 0.09 | 0.13 |
| 9 | 118413485 | rs9650715 | -0.382 | -1.11 | 0.261 | -0.14 | -0.002 | 0.052 | 0.052 | 0 | 0.1 | 0.08 | 0.2 | 0.52 | 0.17 | 0.12 |
| 9 | 118415208 | rs10982856 | 0.181 | -1.309 | 0.148 | -0.191 | -0.012 | -0.028 | -0.028 | 1.369 | 0.12 | 0.06 | 0.13 | 0.41 | 0.11 | 0.17 |
| 9 | 118433442 | rs10817809 | -0.309 | -1.334 | 0.205 | -0.33 | -0.012 | -0.015 | -0.015 | 0 | 0.21 | 0.17 | 0.33 | 0.58 | 0.32 | 0.25 |
| 9 | 118440540 | rs2900611 | -0.743 | -1.6 | 0.338 | -0.546 | -0.027 | -0.008 | -0.008 | 0 | 0.29 | 0.32 | 0.11 | 0.23 | 0.13 | 0.26 |
| 9 | 118448142 | rs10982873 | -2.541 | -1.583 | 0.348 | -0.552 | 0 | 0.011 | 0.011 | 0 | 0.53 | 0.61 | 0.44 | 0.81 | 0.46 | 0.51 |
| 9 | 118448207 | rs10982874 | -2.288 | -1.577 | 0.353 | -0.548 | 0.004 | 0.022 | 0.022 | 0 | 0.51 | 0.61 | 0.44 | 0.81 | 0.45 | 0.51 |
| 9 | 118451241 | rs10817817 | -0.719 | -1.685 | 0.441 | -0.405 | 0.001 | 0.028 | 0.028 | 0 | 0.24 | 0.31 | 0.23 | 0.58 | 0.22 | 0.3 |
| 9 | 118455827 | rs7863684 | 0.236 | -1.654 | 0.438 | -0.39 | -0.031 | 0.019 | 0.019 | 0 | 0.15 | 0.27 | 0.09 | 0.21 | 0.1 | 0.14 |
| 9 | 118456851 | rs2188073 | -3.101 | -1.678 | 0.282 | -0.454 | -0.034 | 0.097 | 0.097 | 0 | 0.6 | 0.82 | 0.51 | 0.88 | 0.48 | 0.6 |
| 9 | 118461146 | rs4978647 | -0.185 | -1.689 | 0.226 | -0.429 | -0.045 | 0.017 | 0.017 | 0 | 0.2 | 0.34 | 0.17 | 0.25 | 0.13 | 0.17 |
| 9 | 118462312 | rs7040032 | 0.822 | -1.588 | 0.326 | -0.414 | -0.006 | -0.029 | -0.029 | 0.825 | 0.08 | 0.13 | 0.1 | 0.01 | 0.06 | 0.02 |
| 9 | 118475932 | rs12380012 | -0.248 | -1.473 | -0.069 | -0.334 | 0.007 | 0.044 | 0.044 | 0 | 0.15 | 0.16 | 0.19 | 0.58 | 0.23 | 0.22 |
| 9 | 118484694 | rs7469580 | 1.367 | -1.341 | -0.265 | -0.335 | 0.001 | 0.001 | 0.001 | 0 | 0.57 | 0.41 | 0.56 | 0.33 | 0.62 | 0.53 |
| 9 | 118484799 | rs10118264 | -0.244 | -1.063 | -0.058 | 0.088 | -0.036 | 0.014 | 0.014 | 0 | 0.17 | 0.29 | 0.08 | 0.22 | 0.1 | 0.13 |
| 9 | 118485764 | rs7045079 | 1.539 | -0.222 | -0.557 | 0.307 | -0.001 | -0.001 | -0.001 | 0 | 0.05 | 0.05 | 0.09 | 0.03 | 0.04 | 0.04 |
| 9 | 118499474 | rs10982901 | -0.581 | -0.234 | -0.539 | 0.45 | -0.004 | 0.043 | 0.043 | 0 | 0.2 | 0.24 | 0.23 | 0.45 | 0.16 | 0.23 |
| 9 | 118503865 | rs2213931 | 0.816 | -0.165 | -0.646 | 0.358 | -0.008 | 0.004 | 0.004 | 0 | 0.06 | 0.03 | 0.02 | 0.08 | 0.07 | 0.12 |
| 9 | 118508283 | rs10982903 | -0.24 | -0.08 | -0.678 | 0.329 | -0.009 | -0.006 | -0.006 | 0 | 0.12 | 0.07 | 0.08 | 0.21 | 0.14 | 0.16 |
| 9 | 118512043 | rs7855735 | -0.355 | -0.499 | -0.362 | 0.159 | -0.012 | -0.03 | -0.03 | 0 | 0.14 | 0.24 | 0.13 | 0.05 | 0.09 | 0.11 |
| 9 | 118512297 | rs7856085 | -1.099 | -0.44 | -0.628 | 0.161 | -0.019 | 0.047 | 0.047 | 0.361 | 0.33 | 0.48 | 0.36 | 0.5 | 0.25 | 0.33 |
| 9 | 118518250 | rs7874534 | -1.505 | -0.067 | -1.313 | -0.056 | -0.022 | 0.016 | 0.016 | 0 | 0.68 | 0.84 | 0.45 | 0.72 | 0.61 | 0.6 |
| 9 | 118518733 | rs10429616 | -0.264 | -0.341 | -1.142 | -0.109 | 0.001 | 0 | 0 | 0 | 0.43 | 0.48 | 0.29 | 0.54 | 0.43 | 0.39 |
| 9 | 118526471 | rs689505 | -0.556 | -0.359 | -1.224 | -0.351 | -0.003 | -0.015 | -0.015 | 0 | 0.42 | 0.68 | 0.39 | 0.58 | 0.37 | 0.39 |
| 9 | 118526984 | rs10982908 | 1.259 | -0.193 | -1.425 | -0.205 | -0.012 | -0.02 | -0.02 | 0.186 | 0.09 | 0.04 | 0.06 | 0.08 | 0.09 | 0.04 |
| 9 | 118535406 | rs2188078 | 2.092 | -0.306 | -1.453 | -0.303 | -0.072 | 0.008 | 0.008 | 0.207 | 0.31 | 0.05 | 0.41 | 0.24 | 0.39 | 0.4 |
| 9 | 118540831 | rs2418410 | 0.506 | -0.364 | -1.392 | -0.166 | -0.02 | 0.022 | 0.022 | 0 | 0.27 | 0.19 | 0.18 | 0.17 | 0.27 | 0.25 |
| 9 | 118540913 | rs6478204 | -1.891 | -0.45 | -1.015 | 0.169 | -0.023 | 0.032 | 0.032 | 0 | 0.67 | 0.84 | 0.56 | 0.75 | 0.59 | 0.59 |
| 9 | 118541068 | rs2418412 | 0.057 | -0.468 | -0.929 | 0.179 | 0.004 | 0.003 | 0.003 | 0 | 0.42 | 0.51 | 0.41 | 0.57 | 0.42 | 0.38 |
| 9 | 118542030 | rs7855710 | -1.327 | -0.45 | -0.988 | 0.173 | -0.02 | 0.047 | 0.047 | 0 | 0.62 | 0.81 | 0.51 | 0.74 | 0.55 | 0.59 |
| 9 | 118550806 | rs7043919 | -0.533 | -0.442 | -1.153 | 0.094 | -0.009 | 0.014 | 0.014 | 0 | 0.83 | 0.69 | 0.89 | 0.83 | 0.88 | 0.88 |
| 9 | 118552995 | rs7865455 | -1.368 | -0.16 | -0.928 | 0.212 | -0.004 | 0.011 | 0.011 | 0 | 0.53 | 0.65 | 0.48 | 0.58 | 0.49 | 0.49 |
| 9 | 118559109 | rs2157673 | 0.528 | -0.013 | -0.761 | 0.465 | -0.001 | -0.002 | -0.002 | 0 | 0.27 | 0.29 | 0.32 | 0.41 | 0.31 | 0.27 |
| 9 | 118559403 | rs10982920 | 0.536 | 0.467 | -1.411 | 0.475 | -0.009 | -0.017 | -0.017 | 0.029 | 0.09 | 0.03 | 0.06 | 0.08 | 0.09 | 0.04 |
| 9 | 118567019 | rs9696301 | -0.403 | 0.45 | -1.403 | 0.468 | 0 | 0 | 0 | 0 | 0.46 | 0.59 | 0.42 | 0.57 | 0.44 | 0.48 |
| 9 | 118568823 | rs10982926 | -0.303 | 0.504 | -0.994 | 0.644 | -0.012 | -0.017 | -0.017 | 4.44 | 0.09 | 0.03 | 0.06 | 0.08 | 0.09 | 0.04 |
| 9 | 118570426 | rs7850583 | -0.7 | 0.436 | -0.704 | 0.559 | -0.001 | 0.002 | 0.002 | 0 | 0.16 | 0.19 | 0.1 | 0.17 | 0.11 | 0.12 |
| 9 | 118571428 | rs4979583 | -1.074 | 0.389 | -0.524 | 0.267 | 0 | -0.005 | -0.005 | 0 | 0.36 | 0.41 | 0.27 | 0.27 | 0.34 | 0.41 |
| 9 | 118573812 | rs7863556 | -0.531 | -0.175 | 0.436 | 0.27 | -0.023 | 0.015 | 0.015 | 0 | 0.1 | 0.34 | 0.09 | 0.12 | 0.05 | 0.1 |
| 9 | 118585450 | rs2106056 | -1.317 | -0.404 | 0.493 | 0.115 | -0.018 | -0.015 | -0.015 | 0.481 | 0.24 | 0.43 | 0.23 | 0.23 | 0.19 | 0.22 |
| 9 | 118592791 | rs6478207 | -2.273 | -0.483 | 0.853 | 0.264 | -0.027 | -0.005 | -0.005 | 0.573 | 0.33 | 0.55 | 0.32 | 0.35 | 0.26 | 0.38 |
| 9 | 118594766 | rs10982943 | 0.075 | -0.167 | 0.791 | 0.379 | 0.002 | -0.005 | -0.005 | 0.564 | 0.08 | 0.07 | 0.11 | 0.12 | 0.07 | 0.16 |
| 9 | 118597837 | rs10817839 | -1.884 | -0.273 | 0.977 | 0.145 | 0.032 | -0.008 | -0.008 | 0 | 0.19 | 0.44 | 0.23 | 0.16 | 0.2 | 0.29 |
| 9 | 118599720 | rs10982946 | -0.505 | -0.285 | 1.048 | 0.1 | 0.002 | 0 | 0 | 0.14 | 0.15 | 0.14 | 0.11 | 0.19 | 0.1 | 0.11 |
| 9 | 118607894 | rs966955 | 0.311 | -0.189 | 0.044 | 0.17 | -0.004 | 0.005 | 0.005 | 0 | 0.32 | 0.21 | 0.26 | 0.26 | 0.3 | 0.22 |
| 9 | 118615701 | rs10118806 | -0.299 | -0.196 | 0.04 | 0.168 | -0.005 | -0.001 | -0.001 | 0 | 0.39 | 0.21 | 0.34 | 0.33 | 0.38 | 0.29 |
| 9 | 118617531 | rs10982957 | 1.16 | -0.135 | -0.037 | 0.211 | -0.011 | -0.02 | -0.02 | 2.578 | 0.07 | 0.01 | 0.08 | 0.07 | 0.08 | 0.07 |
| 9 | 118624384 | rs10982970 | 0.832 | -0.124 | -0.002 | 0.187 | -0.002 | 0.002 | 0.002 | 0 | 0.08 | 0.07 | 0.11 | 0.11 | 0.08 | 0.16 |
| 9 | 118628595 | rs12379210 | 0.479 | -0.12 | 0.045 | 0.172 | 0.014 | 0.003 | 0.003 | 0 | 0.1 | 0.16 | 0.12 | 0.11 | 0.13 | 0.11 |
| 9 | 118630270 | rs2040483 | 2.089 | -0.187 | -0.626 | 0.192 | -0.052 | -0.058 | -0.058 | 0 | 0.33 | 0.11 | 0.4 | 0.39 | 0.4 | 0.39 |
| 9 | 118642639 | rs7858008 | 0.812 | -0.015 | -0.394 | 0.154 | -0.002 | 0.009 | 0.009 | 0.007 | 0.21 | 0.12 | 0.23 | 0.15 | 0.2 | 0.12 |
| 9 | 118644810 | rs995804 | -1.085 | 0.065 | -0.32 | 0.107 | -0.01 | 0.005 | 0.005 | 0 | 0.42 | 0.27 | 0.46 | 0.38 | 0.43 | 0.4 |
| 9 | 118649073 | rs11787606 | 0.224 | 0.089 | -0.574 | 0.101 | 0 | -0.012 | -0.012 | 0.013 | 0.05 | 0.05 | 0.04 | 0 | 0.06 | 0.03 |
| 9 | 118650653 | rs1243 | 0.416 | 0.077 | -0.588 | 0.121 | 0.011 | -0.016 | -0.016 | 0 | 0.08 | 0.03 | 0.09 | 0.1 | 0.07 | 0.16 |
| 9 | 118650689 | rs1989775 | -1.176 | 0.043 | -0.542 | 0.106 | 0.001 | -0.001 | -0.001 | 0 | 0.89 | 0.88 | 0.93 | 0.86 | 0.87 | 0.93 |
| 9 | 118650724 | rs2040485 | 0.447 | -0.167 | -1.068 | 0.156 | 0.007 | -0.031 | -0.031 | 0 | 0.08 | 0.02 | 0.09 | 0.1 | 0.07 | 0.16 |
| 9 | 118651401 | rs10982982 | 0.447 | -0.167 | -1.09 | 0.156 | 0.003 | -0.035 | -0.035 | 0 | 0.08 | 0.01 | 0.09 | 0.1 | 0.07 | 0.16 |
| 9 | 118653960 | rs10982987 | 0.035 | -0.152 | -1.081 | 0.147 | -0.023 | -0.005 | -0.005 | 0 | 0.28 | 0.15 | 0.33 | 0.28 | 0.3 | 0.22 |
| 9 | 118657072 | rs10817846 | 0.441 | -0.085 | -1.215 | 0.119 | -0.018 | -0.015 | -0.015 | 0 | 0.07 | 0.03 | 0.11 | 0.12 | 0.09 | 0.1 |
| 9 | 118658716 | rs10982990 | 0.45 | -0.068 | -1.206 | 0.127 | -0.018 | -0.015 | -0.015 | 0.604 | 0.07 | 0.03 | 0.11 | 0.12 | 0.09 | 0.11 |
| 9 | 118660341 | rs4979591 | 0.45 | -0.068 | -1.206 | 0.127 | -0.018 | -0.015 | -0.015 | 0 | 0.07 | 0.03 | 0.11 | 0.12 | 0.09 | 0.11 |
| 9 | 118662045 | rs10982996 | 0.084 | -0.343 | -0.884 | 0.178 | 0.003 | -0.035 | -0.035 | 0 | 0.08 | 0.01 | 0.09 | 0.1 | 0.07 | 0.16 |
| 9 | 118662804 | rs10982997 | -0.681 | -0.328 | -0.846 | 0.194 | 0.003 | 0.003 | 0.003 | 0 | 0.14 | 0.09 | 0.13 | 0.1 | 0.13 | 0.19 |
| 9 | 118665465 | rs9409176 | 0.311 | -0.298 | -0.72 | 0.304 | -0.018 | -0.046 | -0.046 | 0 | 0.43 | 0.22 | 0.42 | 0.47 | 0.44 | 0.49 |
| 9 | 118666158 | rs9408883 | 0.385 | -0.236 | -0.773 | 0.335 | -0.026 | -0.055 | -0.055 | 0.042 | 0.42 | 0.18 | 0.41 | 0.47 | 0.43 | 0.49 |
| 9 | 118677606 | rs6478213 | -1.335 | -0.583 | 0.345 | 0.458 | -0.038 | 0.005 | 0.005 | 1.294 | 0.17 | 0.53 | 0.11 | 0.13 | 0.1 | 0.12 |
| 9 | 118685872 | rs12683184 | -0.018 | -0.469 | 0.751 | 0.707 | 0.007 | 0.009 | 0.009 | 0 | 0.07 | 0.12 | 0.12 | 0.09 | 0.08 | 0.13 |
| 9 | 118694535 | rs7027312 | 0.725 | -0.49 | 0.686 | 0.984 | 0.015 | 0.133 | 0.133 | 0 | 0.92 | 0.79 | 0.95 | 0.78 | 0.93 | 0.91 |
| 9 | 118698901 | rs6478215 | -0.869 | -0.55 | 0.678 | 0.954 | 0.017 | 0.128 | 0.128 | 2.751 | 0.1 | 0.44 | 0.05 | 0.12 | 0.04 | 0.07 |

| | | | | | | | | | | | | | | | | |
|---|---|---|---|---|---|---|---|---|---|---|---|---|---|---|---|---|
| 9 | 118699375 | rs9409178 | 2.606 | -0.882 | 0.33 | 0.86 | 0.032 | -0.081 | -0.081 | 0 | 0.64 | 0.14 | 0.71 | 0.51 | 0.71 | 0.56 |
| 9 | 118711935 | rs7868474 | -0.773 | -0.873 | 0.323 | 0.833 | 0.017 | 0.013 | 0.013 | 2.736 | 0.1 | 0.46 | 0.06 | 0.05 | 0.04 | 0.07 |
| 9 | 118712965 | rs4979597 | 2.366 | -0.642 | 0.571 | 1.069 | -0.021 | 0.082 | 0.082 | 0 | 0.72 | 0.53 | 0.76 | 0.57 | 0.75 | 0.63 |
| 9 | 118713227 | rs9408887 | 2.525 | -0.645 | 0.838 | 1.067 | -0.022 | 0.061 | 0.061 | 0.006 | 0.7 | 0.46 | 0.75 | 0.57 | 0.75 | 0.63 |
| 9 | 118716486 | rs2106063 | 0.428 | -0.636 | 0.775 | 1.074 | 0.023 | 0.078 | 0.078 | 0.218 | 0.94 | 0.74 | 0.96 | 0.9 | 0.97 | 0.94 |
| 9 | 118719627 | rs7468745 | 0.943 | -0.641 | 0.761 | 1.071 | 0.024 | 0.082 | 0.082 | 0.631 | 0.93 | 0.66 | 0.96 | 0.9 | 0.97 | 0.94 |
| 9 | 118719754 | rs1993013 | 1.03 | -0.61 | 0.188 | 0.722 | -0.111 | -0.144 | -0.144 | 3.1 | 0.28 | 0.02 | 0.26 | 0.44 | 0.34 | 0.36 |
| 9 | 118720050 | rs12115454 | 0.187 | -0.456 | 0.895 | 1.192 | 0.006 | 0.006 | 0.006 | 0 | 0.07 | 0.12 | 0.13 | 0.09 | 0.08 | 0.13 |
| 9 | 118720333 | rs7468919 | 2.46 | -0.546 | 0.901 | 1.015 | -0.022 | 0.078 | 0.078 | 0 | 0.71 | 0.53 | 0.76 | 0.57 | 0.75 | 0.63 |
| 9 | 118725422 | rs10983034 | -1.773 | -0.516 | 0.542 | 1.031 | -0.006 | -0.02 | -0.02 | 4.719 | 0.17 | 0.13 | 0.14 | 0.22 | 0.18 | 0.21 |
| 9 | 118736263 | rs10123549 | -0.605 | -0.821 | -0.003 | -0.022 | -0.014 | -0.048 | -0.048 | 0.469 | 0.09 | 0.17 | 0.02 | 0 | 0.05 | 0.05 |
| 9 | 118738038 | rs4979603 | 0.744 | -0.633 | 0.177 | 0.11 | 0.11 | 0.16 | 0.16 | 0.419 | 0.94 | 0.51 | 0.93 | 0.88 | 0.92 | 0.86 |
| 9 | 118738595 | rs7469847 | 1.486 | -0.696 | 0.112 | 0.139 | -0.068 | 0.017 | 0.017 | 1.114 | 0.78 | 0.49 | 0.88 | 0.79 | 0.89 | 0.85 |
| 9 | 118741214 | rs12551620 | -0.394 | -0.541 | -0.09 | -0.275 | 0.013 | -0.008 | -0.008 | 0 | 0.67 | 0.63 | 0.61 | 0.24 | 0.6 | 0.53 |
| 9 | 118741718 | rs1349323 | 0.315 | -0.164 | -0.854 | -0.129 | -0.002 | 0.098 | 0.098 | 0 | 0.49 | 0.39 | 0.5 | 0.1 | 0.49 | 0.35 |
| 9 | 118744545 | rs1901069 | 0.115 | 0.282 | -0.521 | 0.475 | -0.005 | 0.015 | 0.015 | 0 | 0.57 | 0.45 | 0.57 | 0.23 | 0.51 | 0.48 |
| 9 | 118764806 | rs204190 | -1.197 | -0.132 | 0.32 | 1.287 | -0.02 | 0.147 | 0.147 | 0.799 | 0.13 | 0.53 | 0.26 | 0.3 | 0.11 | 0.22 |
| 9 | 118781688 | rs6478223 | 0.278 | -0.3 | -0.335 | 1.277 | 0 | 0.006 | 0.006 | 0.33 | 0.8 | 0.87 | 0.81 | 0.82 | 0.82 | 0.86 |
| 9 | 118790317 | rs204169 | 1.728 | -0.242 | -0.654 | 1.738 | -0.001 | 0.056 | 0.056 | 0.099 | 0.57 | 0.11 | 0.51 | 0.26 | 0.62 | 0.5 |
| 9 | 118790936 | rs204168 | 1.332 | -0.239 | -0.515 | 1.676 | 0.032 | -0.031 | -0.031 | 0 | 0.62 | 0.26 | 0.54 | 0.52 | 0.69 | 0.61 |
| 9 | 118817903 | rs1314016 | -0.805 | -0.265 | -0.092 | 1.658 | -0.003 | 0.127 | 0.127 | 1.317 | 0.15 | 0.49 | 0.27 | 0.3 | 0.14 | 0.28 |
| 9 | 118841680 | rs2418432 | 1.154 | -0.318 | 0.261 | 1.506 | 0.005 | 0.164 | 0.164 | 0.27 | 0.78 | 0.47 | 0.7 | 0.43 | 0.83 | 0.6 |
| 9 | 118843677 | rs7032067 | 1.39 | -0.564 | -0.217 | 1.009 | 0.025 | 0.026 | 0.026 | 0 | 0.59 | 0.28 | 0.58 | 0.27 | 0.69 | 0.56 |
| 9 | 118845404 | rs2213941 | -1.291 | -0.377 | -0.28 | 1.277 | -0.021 | 0.068 | 0.068 | 2.083 | 0.25 | 0.57 | 0.23 | 0.58 | 0.15 | 0.33 |
| 9 | 118871741 | rs6478230 | -0.861 | -0.009 | -1.49 | 0.158 | -0.008 | 0.023 | 0.023 | 0 | 0.58 | 0.63 | 0.67 | 0.88 | 0.55 | 0.69 |
| 9 | 118898718 | rs204514 | -0.355 | 0.136 | -2.214 | 0.895 | -0.019 | -0.105 | -0.105 | 0.115 | 0.28 | 0.05 | 0.18 | 0.45 | 0.24 | 0.4 |
| 9 | 118919075 | rs10983070 | -0.173 | 0.028 | -2.745 | 0.002 | 0.032 | 0.201 | 0.201 | 0 | 0.47 | 0.09 | 0.35 | 0.21 | 0.61 | 0.4 |
| 9 | 118927725 | rs10983073 | -0.309 | -0.262 | -1.785 | 0.328 | 0.058 | 0.09 | 0.09 | 0 | 0.49 | 0.32 | 0.37 | 0.32 | 0.62 | 0.39 |
| 9 | 118935491 | rs717963 | 0.147 | 1.037 | -2.389 | -0.244 | -0.076 | 0.134 | 0.134 | 0 | 0.32 | 0.05 | 0.19 | 0.13 | 0.35 | 0.29 |
| 9 | 118949998 | rs12375498 | 0.298 | 1.046 | -2.457 | -0.295 | -0.089 | -0.009 | -0.009 | 0 | 0.14 | 0.02 | 0.15 | 0.09 | 0.26 | 0.11 |
| 9 | 118953457 | rs3789281 | -0.953 | 0.394 | -1.148 | -0.37 | 0.007 | 0.009 | 0.009 | 0 | 0.51 | 0.54 | 0.38 | 0.33 | 0.62 | 0.42 |
| 9 | 118954624 | rs1405 | -0.518 | 0.417 | -1.242 | -0.372 | 0.034 | 0.177 | 0.177 | 0 | 0.47 | 0.12 | 0.35 | 0.23 | 0.62 | 0.41 |
| 9 | 118957985 | rs446783 | 0.24 | 0.01 | -1.003 | -0.61 | 0.019 | 0.11 | 0.11 | 0 | 0.53 | 0.79 | 0.68 | 0.75 | 0.42 | 0.61 |
| 9 | 118959731 | rs3789283 | -0.631 | -0.149 | -0.499 | -0.621 | 0.015 | 0.038 | 0.038 | 0 | 0.5 | 0.41 | 0.33 | 0.34 | 0.58 | 0.4 |
| 9 | 118966153 | rs452312 | 0.153 | -0.539 | -0.657 | -0.798 | 0.024 | 0.128 | 0.128 | 0.02 | 0.48 | 0.83 | 0.68 | 0.75 | 0.42 | 0.6 |
| 9 | 118980520 | rs433687 | 0.198 | -0.743 | -0.485 | -0.98 | 0.003 | -0.025 | -0.025 | 0 | 0.57 | 0.5 | 0.67 | 0.81 | 0.49 | 0.66 |
| 9 | 118981886 | rs412644 | 0.185 | -0.728 | -0.516 | -0.985 | 0.003 | -0.015 | -0.015 | 0 | 0.57 | 0.5 | 0.66 | 0.72 | 0.48 | 0.63 |
| 9 | 118984002 | rs398293 | 0.176 | -0.923 | -0.453 | -0.789 | 0.022 | 0.042 | 0.042 | 0 | 0.48 | 0.67 | 0.63 | 0.71 | 0.4 | 0.55 |
| 9 | 118988913 | rs390623 | 0.778 | -1.184 | -0.728 | -0.045 | -0.006 | 0.012 | 0.012 | 0 | 0.16 | 0.17 | 0.28 | 0.29 | 0.13 | 0.25 |
| 9 | 118998517 | rs10983084 | -0.137 | -0.938 | -0.964 | 0.336 | -0.006 | -0.003 | -0.003 | 0 | 0.54 | 0.47 | 0.37 | 0.44 | 0.6 | 0.45 |
| 9 | 119011391 | rs10983088 | 0.163 | -0.652 | -1.863 | 0.155 | -0.037 | -0.071 | -0.071 | 0 | 0.23 | 0.01 | 0.31 | 0.24 | 0.23 | 0.23 |
| 9 | 119022802 | rs13298102 | -0.291 | -0.861 | -1.73 | -0.696 | 0.031 | 0.068 | 0.068 | 0.091 | 0.28 | 0.15 | 0.13 | 0.11 | 0.22 | 0.24 |
| 9 | 119023867 | rs7847346 | -0.803 | -0.117 | -1.359 | -1.142 | 0.011 | 0.017 | 0.017 | 0.435 | 0.43 | 0.4 | 0.31 | 0.15 | 0.38 | 0.39 |
| 9 | 119024899 | rs17260413 | 0.001 | -0.048 | -1.598 | -0.961 | 0.021 | 0.092 | 0.092 | 0.153 | 0.27 | 0.08 | 0.14 | 0.09 | 0.22 | 0.21 |
| 9 | 119026907 | rs1411820 | -0.036 | 0.529 | -0.778 | -0.803 | 0 | -0.006 | -0.006 | 0 | 0.55 | 0.5 | 0.74 | 0.7 | 0.53 | 0.48 |
| 9 | 119027718 | rs1331140 | 0.609 | 0.749 | -0.713 | -0.96 | 0.002 | 0.003 | 0.003 | 2.598 | 0.43 | 0.37 | 0.4 | 0.68 | 0.35 | 0.38 |
| 9 | 119031497 | rs1999351 | -0.401 | 0.87 | -0.474 | -0.905 | -0.005 | 0.047 | 0.047 | 0 | 0.61 | 0.67 | 0.77 | 0.91 | 0.57 | 0.63 |
| 9 | 119034475 | rs4837294 | 0.23 | 1.029 | -0.083 | -0.851 | 0.025 | 0.03 | 0.03 | 0 | 0.15 | 0.35 | 0.37 | 0.22 | 0.2 | 0.21 |
| 9 | 119038034 | rs10435873 | 0.051 | 0.868 | -0.382 | -0.452 | 0.006 | 0.032 | 0.032 | 0 | 0.47 | 0.37 | 0.38 | 0.61 | 0.41 | 0.44 |
| 9 | 119041018 | rs7874262 | -1.368 | 1.779 | 0.49 | -0.291 | 0.16 | -0.017 | -0.017 | 0 | 0.07 | 0.33 | 0.09 | 0 | 0.17 | 0.02 |
| 9 | 119044580 | rs4836727 | 0.221 | 1.507 | 0.118 | -0.121 | 0.005 | -0.007 | -0.007 | 0 | 0.61 | 0.52 | 0.63 | 0.71 | 0.58 | 0.64 |
| 9 | 119065358 | rs7857793 | -1.156 | 1.15 | 0.179 | 0.288 | 0 | -0.023 | -0.023 | 0 | 0.09 | 0.07 | 0.18 | 0.15 | 0.07 | 0.16 |
| 9 | 119074532 | rs11794542 | -1.158 | 1.072 | 0.855 | 0.25 | 0 | -0.009 | -0.009 | 0 | 0.09 | 0.09 | 0.18 | 0.15 | 0.07 | 0.16 |
| 9 | 119078356 | rs1885983 | 2.316 | 1.555 | 1.813 | -0.144 | -0.001 | 0 | 0 | 0 | 0.77 | 0.24 | 0.68 | 0.82 | 0.68 | 0.75 |
| 9 | 119079076 | rs1570518 | 0.387 | 1.813 | 0.91 | 0.114 | -0.105 | -0.071 | -0.071 | 0.132 | 0.28 | 0.02 | 0.33 | 0.32 | 0.3 | 0.34 |
| 9 | 119085047 | rs10817871 | 2.454 | 1.761 | 1.861 | 1.368 | 0.014 | 0.042 | 0.042 | 0 | 0.73 | 0.34 | 0.63 | 0.51 | 0.67 | 0.67 |
| 9 | 119090744 | rs17301196 | -0.488 | 1.974 | 2.154 | 0.911 | 0.101 | -0.028 | -0.028 | 0 | 0.05 | 0.11 | 0.09 | 0 | 0.23 | 0.03 |
| 9 | 119093923 | rs2273977 | -1.787 | 1.795 | 1.961 | 0.977 | 0.02 | -0.013 | -0.013 | 0 | 0.17 | 0.2 | 0.25 | 0.11 | 0.32 | 0.22 |
| 9 | 119099873 | rs13294988 | 1.765 | 1.785 | 1.637 | 0.997 | 0.017 | 0.028 | 0.028 | 0 | 0.74 | 0.33 | 0.66 | 0.57 | 0.67 | 0.7 |
| 9 | 119106881 | rs7020782 | 1.906 | 1.902 | 1.223 | 0.619 | -0.009 | -0.001 | -0.001 | 0.133 | 0.75 | 0.26 | 0.64 | 0.7 | 0.69 | 0.69 |
| 9 | 119107708 | rs912213 | 2.215 | 1.918 | 1.188 | 0.638 | 0.001 | 0 | 0 | 0 | 0.75 | 0.28 | 0.65 | 0.7 | 0.7 | 0.69 |
| 9 | 119111270 | rs1475215 | 1.969 | 2.154 | 1.236 | 0.678 | 0.048 | 0.014 | 0.014 | 0.42 | 0.79 | 0.37 | 0.63 | 0.71 | 0.65 | 0.75 |
| 9 | 119112366 | rs2224768 | 0.874 | 1.502 | 0.238 | 0.713 | -0.006 | -0.06 | -0.06 | 0 | 0.46 | 0.05 | 0.36 | 0.34 | 0.39 | 0.43 |
| 9 | 119115449 | rs10983112 | -1.215 | 1.682 | 0.631 | 0.64 | 0.014 | -0.001 | -0.001 | 0.073 | 0.08 | 0.08 | 0.16 | 0.11 | 0.2 | 0.13 |
| 9 | 119117464 | rs7865534 | -0.317 | 1.699 | 0.602 | 0.62 | 0.042 | 0.003 | 0.003 | 0 | 0.05 | 0.06 | 0.16 | 0.08 | 0.19 | 0.04 |
| 9 | 119122342 | rs751543 | 1.827 | 1.657 | 0.719 | 0.556 | 0.054 | 0.022 | 0.022 | 0.271 | 0.77 | 0.47 | 0.71 | 0.7 | 0.72 | 0.73 |
| 9 | 119128575 | rs7390972 | 0.203 | 2.055 | 0.165 | 0.293 | -0.001 | 0.009 | 0.009 | 0 | 0.36 | 0.28 | 0.34 | 0.35 | 0.34 | 0.31 |
| 9 | 119130627 | rs12353147 | -0.057 | 1.634 | -1.645 | 0.326 | 0.01 | 0.012 | 0.012 | 0 | 0.53 | 0.17 | 0.48 | 0.55 | 0.47 | 0.58 |
| 9 | 119134443 | rs16933460 | -1.096 | 1.863 | -0.73 | 0.294 | -0.001 | 0 | 0 | 0 | 0.05 | 0.05 | 0.06 | 0.01 | 0.07 | 0.05 |

| | | | | | | | | | | | | | | | | |
|---|---|---|---|---|---|---|---|---|---|---|---|---|---|---|---|---|
| 9 | 119136171 | rs12339056 | -1.332 | 2.43 | -0.704 | 0.31 | 0.024 | -0.015 | -0.015 | 0 | 0.08 | 0.1 | 0.09 | 0 | 0.23 | 0.07 |
| 9 | 119138292 | rs9697050 | -0.892 | 1.501 | -1.103 | 0.578 | 0.004 | -0.018 | -0.018 | 0 | 0.17 | 0.08 | 0.17 | 0.2 | 0.13 | 0.2 |
| 9 | 119140488 | rs10817881 | 1.036 | 0.996 | -0.766 | 0.654 | 0.006 | 0.013 | 0.013 | 0 | 0.63 | 0.7 | 0.62 | 0.27 | 0.69 | 0.36 |
| 9 | 119142351 | rs4837261 | -0.37 | 0.875 | -0.643 | 1.609 | -0.001 | 0.017 | 0.017 | 0 | 0.84 | 0.85 | 0.79 | 0.64 | 0.83 | 0.76 |
| 9 | 119147754 | rs932705 | 0.42 | 0.671 | -0.792 | 1.513 | 0.005 | -0.024 | -0.024 | 0 | 0.75 | 0.85 | 0.72 | 0.62 | 0.77 | 0.69 |
| 9 | 119148669 | rs12236532 | -0.892 | 0.632 | -0.581 | 1.539 | 0.009 | -0.009 | -0.009 | 0 | 0.14 | 0.07 | 0.07 | 0.13 | 0.09 | 0.17 |
| 9 | 119150675 | rs3789304 | 0.979 | 0.439 | -0.559 | 0.384 | -0.014 | 0.023 | 0.023 | 0 | 0.38 | 0.36 | 0.35 | 0.13 | 0.45 | 0.23 |
| 9 | 119154200 | rs16933484 | -0.009 | 0.579 | -0.68 | 0.436 | -0.003 | -0.013 | -0.013 | 0 | 0.06 | 0.01 | 0.03 | 0.08 | 0.04 | 0.04 |
| 9 | 119155568 | rs10120005 | 1.175 | 0.499 | -0.14 | 0.37 | -0.004 | -0.022 | -0.022 | 2.026 | 0.25 | 0.24 | 0.17 | 0.1 | 0.24 | 0.11 |
| 9 | 119160150 | rs6478235 | -0.399 | 0.46 | 0.346 | 0.896 | 0.006 | 0.115 | 0.115 | 0 | 0.95 | 0.86 | 0.92 | 0.81 | 0.97 | 0.92 |
| 9 | 119160791 | rs3810939 | 0.191 | 0.346 | 0.154 | 0.452 | 0.01 | -0.018 | -0.018 | 0 | 0.31 | 0.16 | 0.43 | 0.36 | 0.27 | 0.33 |
| 9 | 119161253 | rs3810940 | 0.22 | 0.332 | 0.147 | 0.445 | 0.009 | -0.019 | -0.019 | 0.138 | 0.31 | 0.16 | 0.46 | 0.37 | 0.27 | 0.34 |
| 9 | 119163067 | rs10817885 | 1.592 | 0.061 | -0.001 | 0.264 | -0.017 | 0.08 | 0.08 | 0 | 0.22 | 0.03 | 0.14 | 0.09 | 0.23 | 0.11 |
| 9 | 119163509 | rs1054402 | -0.958 | 0.044 | 0.078 | 0.245 | -0.015 | -0.007 | -0.007 | 0.143 | 0.71 | 0.58 | 0.76 | 0.72 | 0.73 | 0.81 |
| 9 | 119163576 | rs16933502 | 0.573 | 0.049 | -0.004 | 0.217 | -0.008 | -0.015 | -0.015 | 0.536 | 0.05 | 0.01 | 0.03 | 0.08 | 0.04 | 0.04 |
| 9 | 119172047 | rs17302884 | 0.921 | 0.084 | 0.31 | 0.132 | -0.003 | -0.005 | -0.005 | 0 | 0.33 | 0.38 | 0.27 | 0.12 | 0.42 | 0.21 |
| 9 | 119174267 | rs10983142 | 1.558 | 0.077 | 0.234 | 0.141 | -0.015 | 0.068 | 0.068 | 0 | 0.22 | 0.03 | 0.14 | 0.1 | 0.23 | 0.11 |
| 9 | 119179876 | rs7866443 | 0.838 | 0.038 | 0.204 | 0.156 | 0.003 | -0.018 | -0.018 | 0 | 0.24 | 0.17 | 0.4 | 0.36 | 0.22 | 0.29 |
| 9 | 119181214 | rs4837514 | -0.137 | -0.055 | 0.295 | 0.051 | -0.003 | -0.003 | -0.003 | 0.093 | 0.59 | 0.63 | 0.68 | 0.64 | 0.65 | 0.71 |
| 9 | 119181794 | rs17303101 | 1.213 | 0.85 | 0.264 | -0.041 | -0.003 | 0.025 | 0.025 | 0 | 0.28 | 0.27 | 0.22 | 0.12 | 0.31 | 0.19 |
| 9 | 119186551 | rs10759837 | -0.488 | 0.528 | 0.331 | 0.029 | -0.013 | 0.013 | 0.013 | 0 | 0.87 | 0.82 | 0.88 | 0.82 | 0.94 | 0.89 |
| 9 | 119187220 | rs9299239 | 1.457 | 0.393 | 0.237 | -0.077 | 0.001 | -0.001 | -0.001 | 0 | 0.24 | 0.18 | 0.16 | 0.1 | 0.24 | 0.11 |
| 9 | 119187459 | rs1507909 | 0.728 | 0.395 | 0.231 | 0.043 | 0.001 | -0.004 | -0.004 | 0 | 0.25 | 0.24 | 0.4 | 0.52 | 0.22 | 0.51 |
| 9 | 119198324 | rs1396042 | -0.944 | 0.46 | 0.237 | 0.01 | -0.011 | -0.025 | -0.025 | 0 | 0.9 | 0.87 | 0.92 | 0.97 | 0.95 | 0.94 |
| 9 | 119200847 | rs7850952 | 0.473 | 0.245 | 0.009 | -0.038 | 0.003 | -0.018 | -0.018 | 0 | 0.29 | 0.26 | 0.46 | 0.5 | 0.25 | 0.36 |
| 9 | 119205302 | rs4837530 | -0.077 | 0.191 | 0.027 | -0.058 | 0.011 | -0.036 | -0.036 | 0 | 0.36 | 0.26 | 0.49 | 0.5 | 0.29 | 0.41 |
| 9 | 119230710 | rs13298648 | 1.108 | 0.146 | -0.043 | -0.136 | -0.051 | 0.022 | 0.022 | 0 | 0.27 | 0.02 | 0.17 | 0.18 | 0.28 | 0.16 |
| 9 | 119236373 | rs10759842 | 0.475 | 0.041 | 0.305 | -0.22 | -0.026 | -0.043 | -0.043 | 0 | 0.82 | 0.49 | 0.89 | 0.89 | 0.9 | 0.9 |
| 9 | 119238756 | rs12005639 | 1.581 | 0.042 | 0.18 | -0.26 | -0.009 | 0.064 | 0.064 | 0 | 0.22 | 0.05 | 0.15 | 0.1 | 0.24 | 0.1 |
| 9 | 119244942 | rs6478238 | 1.228 | 0.574 | 0.205 | 0.005 | -0.002 | 0 | 0 | 0 | 0.75 | 0.42 | 0.8 | 0.72 | 0.82 | 0.81 |
| 9 | 119247845 | rs10983190 | 0.208 | 0.61 | 0.233 | 0.161 | -0.009 | -0.016 | -0.016 | 0 | 0.06 | 0.02 | 0.07 | 0.19 | 0.05 | 0.08 |
| 9 | 119248059 | rs17220352 | 1.249 | 0.953 | 0.008 | -0.104 | -0.016 | 0.035 | 0.035 | 0 | 0.21 | 0.19 | 0.2 | 0.11 | 0.25 | 0.14 |
| 9 | 119252629 | rs6478241 | 0.286 | 0.872 | -0.209 | -0.108 | -0.011 | 0.003 | 0.003 | 0.581 | 0.62 | 0.27 | 0.64 | 0.62 | 0.62 | 0.71 |
| 9 | 119255010 | rs10983194 | 0.237 | 1.163 | -0.158 | -0.052 | -0.021 | -0.018 | -0.018 | 0 | 0.06 | 0.03 | 0.07 | 0.17 | 0.08 | 0.08 |
| 9 | 119258187 | rs767432 | 1.003 | 0.903 | -0.292 | -0.015 | -0.009 | -0.015 | -0.015 | 0 | 0.28 | 0.18 | 0.45 | 0.32 | 0.31 | 0.25 |
| 9 | 119258919 | rs4837566 | 0.143 | 0.792 | 0.012 | -0.152 | -0.001 | -0.022 | -0.022 | 0 | 0.23 | 0.64 | 0.22 | 0.17 | 0.2 | 0.14 |
| 9 | 119261166 | rs1475057 | 0.782 | 0.863 | -0.572 | -0.253 | -0.031 | -0.007 | -0.007 | 0 | 0.37 | 0.07 | 0.5 | 0.31 | 0.35 | 0.28 |
| 9 | 119266468 | rs4836731 | 0.835 | 0.91 | -0.101 | -0.176 | -0.006 | -0.006 | -0.006 | 0 | 0.61 | 0.3 | 0.66 | 0.42 | 0.62 | 0.36 |
| 9 | 119266695 | rs4836732 | -0.016 | 0.582 | 0.069 | -0.157 | -0.018 | 0.018 | 0.018 | 1.255 | 0.54 | 0.3 | 0.61 | 0.56 | 0.5 | 0.67 |
| 9 | 119277811 | rs4837584 | -0.255 | 0.706 | 0.273 | -0.119 | 0.016 | -0.002 | -0.002 | 0 | 0.93 | 0.86 | 0.95 | 0.89 | 0.92 | 0.92 |
| 9 | 119285144 | rs4837587 | 1.01 | 0.708 | -0.005 | -0.128 | -0.059 | -0.001 | -0.001 | 0.557 | 0.37 | 0.08 | 0.5 | 0.31 | 0.38 | 0.29 |
| 9 | 119287555 | rs12237388 | -0.409 | 0.76 | 0.374 | -0.092 | 0.084 | 0.132 | 0.132 | 0.033 | 0.1 | 0.25 | 0.16 | 0.21 | 0.16 | 0.16 |
| 9 | 119300776 | rs12379134 | -0.674 | 0.674 | 0.297 | -0.17 | -0.012 | 0.017 | 0.017 | 0.466 | 0.57 | 0.7 | 0.45 | 0.6 | 0.57 | 0.66 |
| 9 | 119305421 | rs7042951 | 0.754 | 0.59 | 0.329 | -0.241 | -0.008 | -0.072 | -0.072 | 1.437 | 0.19 | 0.31 | 0.13 | 0.01 | 0.15 | 0.08 |
| 9 | 119307773 | rs10983229 | 1.729 | 0.608 | 0.236 | -0.119 | 0.002 | 0.008 | 0.008 | 0 | 0.24 | 0.17 | 0.16 | 0.11 | 0.26 | 0.08 |
| 9 | 119308184 | rs7029693 | 0.503 | 0.602 | 0.243 | -0.065 | -0.003 | 0.01 | 0.01 | 0 | 0.35 | 0.37 | 0.31 | 0.56 | 0.39 | 0.56 |
| 9 | 119309260 | rs10983230 | -0.134 | 0.942 | 0.234 | 0.095 | -0.002 | 0.001 | 0.001 | 0.009 | 0.08 | 0.07 | 0.09 | 0.14 | 0.11 | 0.1 |
| 9 | 119314217 | rs13290997 | -0.572 | 0.568 | 0.311 | 0.139 | -0.001 | 0.019 | 0.019 | 0.086 | 0.53 | 0.69 | 0.46 | 0.55 | 0.58 | 0.66 |
| 9 | 119314851 | rs1407243 | 0.379 | 0.605 | 0.44 | 0.181 | 0.019 | 0.029 | 0.029 | 0 | 0.33 | 0.4 | 0.33 | 0.54 | 0.43 | 0.59 |
| 9 | 119323187 | rs10759848 | -0.856 | 0.957 | 0.306 | 0.272 | 0.009 | 0.001 | 0.001 | 0 | 0.92 | 0.9 | 0.95 | 0.87 | 0.88 | 0.91 |
| 9 | 119331057 | rs1467554 | 1.287 | 0.565 | -0.087 | 0.064 | -0.033 | -0.024 | -0.024 | 0 | 0.31 | 0.08 | 0.46 | 0.31 | 0.3 | 0.23 |
| 9 | 119339463 | rs16933690 | 0.495 | 0.97 | 0.55 | 0.103 | -0.027 | -0.147 | -0.147 | 0 | 0.21 | 0.43 | 0.14 | 0.01 | 0.15 | 0.07 |
| 9 | 119341037 | rs11787627 | 0.981 | 0.984 | 0.541 | 0.181 | 0.001 | 0 | 0 | 0.011 | 0.29 | 0.28 | 0.32 | 0.3 | 0.41 | 0.25 |
| 9 | 119348005 | rs12376892 | 2.282 | 1.055 | 0.377 | 0.163 | -0.059 | 0.026 | 0.026 | 0 | 0.19 | 0.01 | 0.14 | 0.11 | 0.26 | 0.07 |
| 9 | 119357964 | rs7866525 | 0.413 | 1.048 | 0.548 | 0.183 | -0.001 | 0.095 | 0.095 | 0 | 0.5 | 0.28 | 0.54 | 0.68 | 0.44 | 0.69 |
| 9 | 119367308 | rs10983262 | 1.759 | 1.017 | 0.5 | 0.135 | -0.007 | 0.051 | 0.051 | 0 | 0.22 | 0.1 | 0.15 | 0.12 | 0.26 | 0.08 |
| 9 | 119367476 | rs7855974 | 0.229 | 1.143 | 0.541 | 0.212 | 0 | -0.003 | -0.003 | 0.393 | 0.52 | 0.46 | 0.55 | 0.69 | 0.44 | 0.69 |
| 9 | 119382930 | rs4837682 | 0.043 | 1.157 | 0.489 | 0.198 | 0.003 | 0.007 | 0.007 | 0 | 0.91 | 0.89 | 0.83 | 0.8 | 0.84 | 0.82 |
| 9 | 119383888 | rs10491578 | 1.989 | 1.007 | 0.407 | -0.037 | -0.034 | 0.055 | 0.055 | 0 | 0.19 | 0.04 | 0.11 | 0.1 | 0.26 | 0.08 |
| 9 | 119397553 | rs9697091 | -0.117 | 0.966 | 0.47 | -0.036 | 0 | -0.005 | -0.005 | 0 | 0.56 | 0.56 | 0.65 | 0.86 | 0.51 | 0.79 |
| 9 | 119404723 | rs811690 | 0.67 | 1.017 | 0.483 | 0.063 | 0.01 | 0.007 | 0.007 | 0 | 0.09 | 0.15 | 0.08 | 0.15 | 0.12 | 0.08 |
| 9 | 119405271 | rs811458 | -1.411 | 1.039 | 0.488 | -0.021 | 0.002 | -0.011 | -0.011 | 0 | 0.76 | 0.75 | 0.78 | 0.87 | 0.66 | 0.86 |
| 9 | 119419325 | rs1742821 | -0.167 | 1.024 | 0.496 | 0.051 | 0 | -0.041 | -0.041 | 0 | 0.56 | 0.53 | 0.65 | 0.87 | 0.5 | 0.79 |
| 9 | 119453455 | rs803939 | -0.201 | 1.016 | 0.254 | 0.047 | 0.001 | 0.083 | 0.083 | 0.072 | 0.55 | 0.35 | 0.63 | 0.87 | 0.5 | 0.78 |
| 9 | 119458020 | rs803932 | -1.364 | 1.052 | 0.344 | 0.087 | 0.023 | -0.086 | -0.086 | 0 | 0.74 | 0.58 | 0.76 | 0.87 | 0.65 | 0.84 |
| 9 | 119461275 | rs1661300 | 0.84 | 1.04 | 0.404 | 0.272 | -0.001 | -0.002 | -0.002 | 0.399 | 0.08 | 0.08 | 0.07 | 0.15 | 0.12 | 0.09 |
| 9 | 119466436 | rs803919 | 0.012 | 1.017 | 0.417 | 0.087 | 0.12 | 0.043 | 0.043 | 0 | 0.93 | 0.56 | 0.87 | 0.97 | 0.91 | 0.94 |
| 9 | 119471881 | rs4141711 | -0.752 | 0.993 | 0.222 | 0.136 | 0 | -0.001 | -0.001 | 0 | 0.9 | 0.88 | 0.85 | 0.84 | 0.82 | 0.87 |
| 9 | 119474700 | rs1441748 | 0.842 | 0.71 | 0.006 | -0.301 | 0.007 | 0 | 0 | 0 | 0.18 | 0.18 | 0.12 | 0 | 0.15 | 0.06 |

| | | | | | | | | | | | | | | | | |
|---|---|---|---|---|---|---|---|---|---|---|---|---|---|---|---|---|
| 9 | 119476270 | rs719535 | 0.589 | 0.933 | 0.349 | 0.37 | 0.006 | 0.025 | 0.025 | 1.566 | 0.8 | 0.74 | 0.77 | 0.45 | 0.76 | 0.42 |
| 9 | 119483436 | rs10817926 | 0.286 | 0.663 | -0.963 | 0.291 | -0.009 | 0.043 | 0.043 | 1.937 | 0.55 | 0.33 | 0.6 | 0.32 | 0.45 | 0.3 |
| 9 | 119484528 | rs1372328 | 0.185 | 0.681 | -1.297 | 0.3 | -0.014 | 0.061 | 0.061 | 2.37 | 0.55 | 0.27 | 0.59 | 0.32 | 0.45 | 0.3 |
| 9 | 119487445 | rs10983326 | 0.626 | 0.318 | -1.438 | 0.43 | -0.003 | 0.001 | 0.001 | 0 | 0.26 | 0.13 | 0.39 | 0.23 | 0.28 | 0.19 |
| 9 | 119487811 | rs1441747 | 0.054 | 0.109 | -1.687 | 0.274 | 0.023 | 0.059 | 0.059 | 1.285 | 0.55 | 0.2 | 0.48 | 0.68 | 0.59 | 0.73 |
| 9 | 119489348 | rs9657672 | 0.447 | 0.474 | -1.692 | 0.483 | 0.043 | -0.025 | -0.025 | 0 | 0.45 | 0.24 | 0.34 | 0.58 | 0.36 | 0.43 |
| 9 | 119489471 | rs10125636 | -0.002 | 0.206 | -1.223 | 0.053 | -0.001 | 0.001 | 0.001 | 0 | 0.48 | 0.49 | 0.42 | 0.3 | 0.56 | 0.26 |
| 9 | 119490572 | rs7028544 | 0.404 | 0.029 | -1.508 | 0.524 | -0.034 | -0.027 | -0.027 | 4.329 | 0.36 | 0.14 | 0.25 | 0.5 | 0.39 | 0.44 |
| 9 | 119493194 | rs1441744 | -0.962 | -0.047 | -1.056 | 0.151 | -0.005 | 0.006 | 0.006 | 0 | 0.73 | 0.78 | 0.82 | 0.86 | 0.65 | 0.87 |
| 9 | 119495575 | rs2274414 | 0.73 | -0.578 | -0.944 | 0.356 | 0.003 | -0.004 | -0.004 | 2.143 | 0.16 | 0.16 | 0.09 | 0.2 | 0.06 | 0.06 |
| 9 | 119508795 | rs2416570 | 0.538 | -0.772 | -0.604 | 0.094 | 0.04 | 0.063 | 0.063 | 0 | 0.5 | 0.32 | 0.71 | 0.65 | 0.58 | 0.61 |
| 9 | 119510166 | rs894207 | -0.423 | -0.852 | -0.393 | 0.1 | 0 | -0.023 | -0.023 | 1.641 | 0.62 | 0.57 | 0.75 | 0.77 | 0.6 | 0.84 |
| 9 | 119510525 | rs894208 | 0.5 | -0.707 | -0.833 | 0.156 | 0.035 | 0.053 | 0.053 | 0 | 0.5 | 0.35 | 0.71 | 0.65 | 0.57 | 0.61 |
| 9 | 119512923 | rs16933879 | 1.247 | -0.509 | -1.028 | 0.168 | -0.002 | 0.01 | 0.01 | 0 | 0.12 | 0.09 | 0.09 | 0.09 | 0.1 | 0.07 |
| 9 | 119513452 | rs10732384 | -0.225 | -0.461 | -1.293 | 0.158 | 0.015 | 0.041 | 0.041 | 0 | 0.59 | 0.25 | 0.74 | 0.71 | 0.6 | 0.85 |
| 9 | 119516944 | rs961797 | 0.378 | 0.046 | -1.24 | 0.498 | -0.04 | -0.04 | -0.04 | 0.094 | 0.43 | 0.2 | 0.55 | 0.53 | 0.49 | 0.57 |
| 9 | 119518570 | rs10983338 | 0.899 | -0.091 | -1.088 | 0.499 | -0.018 | -0.005 | -0.005 | 0 | 0.23 | 0.31 | 0.18 | 0.21 | 0.14 | 0.09 |
| 9 | 119519588 | rs1372330 | 0.952 | 0.14 | -1.202 | 0.656 | 0.011 | 0.05 | 0.05 | 0.367 | 0.06 | 0.07 | 0.12 | 0.25 | 0.13 | 0.11 |
| 9 | 119522256 | rs10983339 | 0.738 | -0.643 | -1.099 | 0.748 | -0.069 | 0.091 | 0.091 | 0 | 0.41 | 0.14 | 0.47 | 0.28 | 0.48 | 0.5 |
| 9 | 119525648 | rs13283406 | 1.285 | 0.276 | -0.827 | 0.419 | -0.06 | 0.024 | 0.024 | 0 | 0.19 | 0.01 | 0.18 | 0.09 | 0.24 | 0.07 |
| 9 | 119525840 | rs930466 | -0.65 | -0.006 | -0.419 | 0.458 | 0.019 | -0.002 | -0.002 | 0 | 0.69 | 0.54 | 0.73 | 0.75 | 0.64 | 0.86 |
| 9 | 119526375 | rs4836849 | 0.504 | -0.033 | -0.372 | 0.395 | 0.004 | 0.01 | 0.01 | 0 | 0.36 | 0.42 | 0.33 | 0.52 | 0.41 | 0.66 |
| 9 | 119527755 | rs12338231 | 0.261 | 0.149 | -1.25 | 0.51 | -0.022 | -0.022 | -0.022 | 0.523 | 0.36 | 0.58 | 0.43 | 0.23 | 0.23 | 0.21 |
| 9 | 119528478 | rs10759855 | -0.011 | 0.868 | -1.021 | 0.693 | -0.011 | -0.001 | -0.001 | 0 | 0.71 | 0.81 | 0.75 | 0.75 | 0.64 | 0.86 |
| 9 | 119530849 | rs10983343 | -0.101 | 0.592 | -0.917 | 0.62 | -0.003 | 0.014 | 0.014 | 0 | 0.55 | 0.41 | 0.5 | 0.42 | 0.52 | 0.59 |
| 9 | 119531310 | rs7847485 | -0.291 | 0.065 | -0.623 | 0.616 | 0 | -0.003 | -0.003 | 0 | 0.48 | 0.52 | 0.47 | 0.76 | 0.4 | 0.52 |
| 9 | 119533737 | rs1441750 | 0.5 | -0.166 | -0.346 | 0.393 | -0.016 | 0.085 | 0.085 | 0 | 0.44 | 0.21 | 0.47 | 0.24 | 0.53 | 0.37 |
| 9 | 119538249 | rs4836850 | 0.56 | -0.249 | -0.346 | 0.438 | -0.045 | 0.169 | 0.169 | 0 | 0.43 | 0.13 | 0.47 | 0.15 | 0.52 | 0.36 |
| 9 | 119538823 | rs7858285 | 1.244 | -0.537 | 0.222 | 1.578 | -0.062 | 0.203 | 0.203 | 0 | 0.83 | 0.59 | 0.81 | 0.52 | 0.9 | 0.78 |
| 9 | 119539410 | rs4837849 | 0.138 | -0.535 | 0.219 | 1.553 | 0.006 | -0.027 | -0.027 | 0.271 | 0.91 | 0.69 | 0.96 | 1 | 0.96 | 0.94 |
| 9 | 119539531 | rs4836851 | 0.504 | -0.467 | 0.243 | 1.588 | -0.048 | 0.103 | 0.103 | 0 | 0.84 | 0.68 | 0.81 | 0.61 | 0.9 | 0.78 |
| 9 | 119539774 | rs4836852 | 0.595 | -0.485 | 0.215 | 1.578 | -0.057 | 0.11 | 0.11 | 0 | 0.83 | 0.62 | 0.81 | 0.61 | 0.9 | 0.78 |
| 9 | 119542324 | rs4837857 | 0.761 | -0.473 | -0.085 | 1.645 | -0.07 | 0.216 | 0.216 | 0 | 0.82 | 0.53 | 0.81 | 0.49 | 0.89 | 0.77 |
| 9 | 119543476 | rs10759859 | 0.992 | -0.32 | -0.461 | 1.046 | 0 | 0 | 0 | 1.401 | 0.66 | 0.4 | 0.62 | 0.28 | 0.73 | 0.52 |
| 9 | 119544164 | rs7021928 | -0.704 | -0.561 | -0.14 | 1.049 | -0.004 | 0.033 | 0.033 | 0 | 0.31 | 0.56 | 0.28 | 0.42 | 0.27 | 0.4 |
| 9 | 119544321 | rs10817933 | 0.523 | 0.29 | 0.33 | 0.65 | 0.057 | 0.026 | 0.026 | 0 | 0.73 | 0.62 | 0.53 | 0.37 | 0.58 | 0.66 |
| 9 | 119544433 | rs4837864 | 0.092 | -0.493 | 0.272 | 0.107 | -0.016 | -0.001 | -0.001 | 0.096 | 0.84 | 0.77 | 0.86 | 0.79 | 0.89 | 0.74 |
| 9 | 119544455 | rs10759860 | -0.887 | -0.622 | 0.348 | -0.683 | -0.005 | -0.042 | -0.042 | 0 | 0.69 | 0.65 | 0.73 | 0.92 | 0.73 | 0.75 |
| 9 | 119546021 | rs7030327 | 0.728 | -0.726 | 0.062 | -0.756 | 0.001 | -0.007 | -0.007 | 0 | 0.59 | 0.56 | 0.53 | 0.2 | 0.55 | 0.48 |
| 9 | 119550051 | rs10817934 | 1.743 | -0.661 | -0.473 | -0.885 | -0.042 | 0.073 | 0.073 | 0 | 0.26 | 0.04 | 0.26 | 0.1 | 0.3 | 0.24 |
| 9 | 119550176 | rs7852805 | -1.015 | -0.372 | -0.127 | -0.919 | 0.013 | -0.036 | -0.036 | 0 | 0.89 | 0.85 | 0.87 | 0.99 | 0.81 | 0.86 |
| 9 | 119554510 | rs2028495 | -1.043 | -0.37 | -0.204 | -0.918 | 0.003 | -0.021 | -0.021 | 0 | 0.89 | 0.88 | 0.87 | 0.99 | 0.81 | 0.86 |
| 9 | 119561084 | rs8181035 | 0.899 | -0.538 | 0.091 | -0.867 | -0.004 | -0.031 | -0.031 | 0 | 0.3 | 0.37 | 0.26 | 0.08 | 0.26 | 0.23 |
| 9 | 119564014 | rs36105701 | 1.243 | -0.414 | -0.36 | -0.78 | -0.05 | 0.091 | 0.091 | 0 | 0.28 | 0.04 | 0.26 | 0.08 | 0.26 | 0.26 |
| 9 | 119566533 | rs10983355 | 1.67 | -0.229 | -0.484 | -0.532 | -0.019 | 0.013 | 0.013 | 0 | 0.24 | 0.07 | 0.25 | 0.17 | 0.28 | 0.21 |
| 9 | 119572334 | rs1155556 | 0.525 | -0.355 | -0.075 | -0.412 | -0.001 | 0.001 | 0.001 | 0 | 0.41 | 0.39 | 0.47 | 0.67 | 0.48 | 0.46 |
| 9 | 119573867 | rs7021435 | 2.572 | -0.488 | 0.22 | -0.453 | -0.003 | -0.006 | -0.006 | 0.458 | 0.06 | 0.3 | 0.03 | 0 | 0 | 0 |
| 9 | 119575226 | rs1556480 | 1.073 | -0.178 | 0.213 | -0.385 | -0.003 | -0.022 | -0.022 | 0 | 0.34 | 0.38 | 0.29 | 0.08 | 0.26 | 0.26 |
| 9 | 119576473 | rs6478254 | -1.734 | -0.072 | 0.066 | -0.414 | 0.012 | -0.02 | -0.02 | 0.046 | 0.93 | 0.89 | 0.9 | 0.99 | 0.83 | 0.88 |
| 9 | 119576802 | rs7034397 | -0.087 | -0.201 | 0.083 | -0.35 | -0.003 | 0.031 | 0.031 | 0 | 0.51 | 0.69 | 0.51 | 0.58 | 0.45 | 0.51 |
| 9 | 119578333 | rs1156155 | 0.38 | -0.112 | 0.191 | 0.106 | 0.015 | 0.122 | 0.122 | 1.892 | 0.17 | 0.29 | 0.22 | 0.5 | 0.2 | 0.25 |
| 9 | 119580735 | rs4836864 | 0.684 | -0.083 | -0.22 | 0.073 | 0.001 | 0.009 | 0.009 | 0 | 0.76 | 0.8 | 0.83 | 0.78 | 0.76 | 0.7 |
| 9 | 119582571 | rs10491575 | 0.624 | 0.071 | -0.175 | 0.373 | 0.004 | 0.055 | 0.055 | 0.275 | 0.14 | 0.19 | 0.24 | 0.48 | 0.17 | 0.24 |
| 9 | 119582761 | rs3765536 | 0.624 | 0.068 | -0.154 | 0.341 | 0.006 | 0.068 | 0.068 | 0.318 | 0.14 | 0.2 | 0.25 | 0.49 | 0.17 | 0.24 |
| 9 | 119582872 | rs3765537 | 0.558 | -0.063 | -0.319 | 0.33 | -0.012 | -0.026 | -0.026 | 0 | 0.76 | 0.83 | 0.83 | 0.71 | 0.73 | 0.62 |
| 9 | 119585852 | rs1326807 | -0.106 | 0.221 | 0.147 | 0.371 | 0.028 | -0.016 | -0.016 | 0 | 0.87 | 0.72 | 0.89 | 0.91 | 0.8 | 0.79 |
| 9 | 119591344 | rs1590605 | 0.103 | 0.204 | 0.125 | 0.364 | 0.021 | -0.018 | -0.018 | 0 | 0.86 | 0.72 | 0.89 | 0.91 | 0.8 | 0.79 |
| 9 | 119598413 | rs7866077 | -0.019 | 0.212 | 0.243 | 0.519 | 0.013 | 0.154 | 0.154 | 0 | 0.17 | 0.38 | 0.26 | 0.52 | 0.17 | 0.2 |
| 9 | 119598710 | rs10125438 | 0.432 | 0.294 | 0.082 | 0.527 | -0.004 | 0.069 | 0.069 | 0 | 0.15 | 0.25 | 0.23 | 0.51 | 0.14 | 0.18 |
| 9 | 119599613 | rs943308 | -0.977 | 0.339 | -0.076 | 0.198 | -0.002 | 0.001 | 0.001 | 0 | 0.9 | 0.89 | 0.91 | 0.95 | 0.81 | 0.75 |
| 9 | 119612028 | rs1326814 | 0.571 | 0.192 | 0.128 | 0.41 | 0.043 | 0.08 | 0.08 | 0.36 | 0.72 | 0.46 | 0.56 | 0.52 | 0.64 | 0.55 |
| 9 | 119613865 | rs2296673 | 1.721 | -0.012 | 0.034 | 0.085 | -0.001 | -0.001 | -0.001 | 0 | 0.22 | 0.16 | 0.25 | 0.21 | 0.27 | 0.13 |
| 9 | 119616042 | rs1536428 | 0.459 | -0.545 | 0.172 | 0.468 | 0.04 | 0.082 | 0.082 | 1.193 | 0.73 | 0.46 | 0.56 | 0.53 | 0.65 | 0.57 |
| 9 | 119616074 | rs10124883 | 0.461 | -0.568 | -0.407 | 0.737 | 0.001 | 0.036 | 0.036 | 0.5 | 0.14 | 0.18 | 0.24 | 0.43 | 0.16 | 0.18 |
| 9 | 119617397 | rs10983378 | 0.147 | -0.947 | -0.797 | 0.336 | 0.038 | -0.071 | -0.071 | 0 | 0.15 | 0.01 | 0.1 | 0.22 | 0.07 | 0.18 |
| 9 | 119618830 | rs1359702 | -0.504 | -1.125 | -0.334 | 0.124 | -0.011 | 0 | 0 | 0 | 0.28 | 0.36 | 0.28 | 0.27 | 0.25 | 0.41 |
| 9 | 119619937 | rs4837952 | 1.195 | -1.273 | -0.724 | -0.211 | -0.073 | 0.06 | 0.06 | 0.113 | 0.35 | 0.16 | 0.3 | 0.21 | 0.48 | 0.17 |
| 9 | 119622438 | rs10491574 | 1.18 | -1.179 | -0.49 | -0.048 | -0.001 | 0.082 | 0.082 | 1.078 | 0.12 | 0.21 | 0.28 | 0.47 | 0.12 | 0.17 |
| 9 | 119624675 | rs943306 | -0.886 | -1.323 | -0.39 | -0.193 | -0.005 | -0.012 | -0.012 | 0 | 0.41 | 0.61 | 0.57 | 0.72 | 0.39 | 0.59 |

| | | | | | | | | | | | | | | | | |
|---|---|---|---|---|---|---|---|---|---|---|---|---|---|---|---|---|
| 9 | 119626289 | rs2026587 | -1.411 | -1.505 | -0.574 | -0.395 | 0.033 | 0.042 | 0.042 | 0 | 0.76 | 0.88 | 0.79 | 0.95 | 0.88 | 0.69 |
| 9 | 119626482 | rs4837955 | 1.216 | -1.154 | -0.425 | -0.059 | -0.075 | 0.02 | 0.02 | 0 | 0.33 | 0.15 | 0.4 | 0.28 | 0.48 | 0.17 |
| 9 | 119627005 | rs10817944 | 1.106 | -1.163 | -0.371 | -0.083 | 0.013 | -0.057 | -0.057 | 0 | 0.15 | 0.33 | 0.11 | 0.04 | 0.18 | 0.16 |
| 9 | 119627429 | rs7034422 | -0.488 | -1.222 | -0.674 | -0.079 | 0.03 | -0.028 | -0.028 | 0 | 0.85 | 0.94 | 0.91 | 0.78 | 0.92 | 0.8 |
| 9 | 119627987 | rs7862589 | 1.201 | -1.09 | -0.628 | -0.018 | -0.038 | 0.011 | 0.011 | 0 | 0.32 | 0.23 | 0.39 | 0.28 | 0.46 | 0.14 |
| 9 | 119629567 | rs4837957 | -0.972 | -1.013 | -0.914 | 0.131 | 0.019 | -0.026 | -0.026 | 0 | 0.89 | 0.94 | 0.93 | 0.76 | 0.93 | 0.83 |
| 9 | 119630148 | rs9696212 | 0.298 | -0.74 | -0.842 | 0.163 | 0.009 | -0.011 | -0.011 | 0 | 0.37 | 0.31 | 0.31 | 0.46 | 0.27 | 0.51 |
| 9 | 119633968 | rs2225067 | 0.275 | -0.375 | -0.891 | 0.177 | -0.014 | 0.016 | 0.016 | 0 | 0.35 | 0.31 | 0.42 | 0.26 | 0.5 | 0.19 |
| 9 | 119641176 | rs4837967 | -0.558 | -0.52 | -1.035 | 0.744 | -0.038 | -0.084 | -0.084 | 0 | 0.33 | 0.15 | 0.33 | 0.5 | 0.4 | 0.34 |
| 9 | 119641530 | rs7868701 | 0.636 | -1.017 | -0.941 | 0.812 | 0 | 0.017 | 0.017 | 0.517 | 0.12 | 0.14 | 0.19 | 0.4 | 0.07 | 0.19 |
| 9 | 119641763 | rs7868991 | -0.342 | -0.946 | -0.998 | 0.791 | -0.038 | -0.082 | -0.082 | 0 | 0.33 | 0.15 | 0.33 | 0.5 | 0.4 | 0.34 |
| 9 | 119645481 | rs12336175 | 0.075 | -1.208 | -0.757 | 1.038 | -0.021 | 0.046 | 0.046 | 0 | 0.15 | 0.2 | 0.21 | 0.38 | 0.09 | 0.15 |
| 9 | 119646682 | rs1998898 | 1.289 | -0.97 | -1.07 | 1.144 | -0.062 | -0.04 | -0.04 | 0.184 | 0.16 | 0.05 | 0.23 | 0.24 | 0.3 | 0.1 |
| 9 | 119649615 | rs4837980 | 1.532 | -1.816 | -1.412 | 0.499 | -0.11 | 0.179 | 0.179 | 0 | 0.4 | 0.09 | 0.4 | 0.08 | 0.51 | 0.5 |
| 9 | 119652622 | rs3843825 | 0.244 | -1.756 | -1.343 | 0.819 | -0.069 | -0.117 | -0.117 | 0.773 | 0.25 | 0.05 | 0.29 | 0.46 | 0.34 | 0.25 |
| 9 | 119654521 | rs4091697 | 0.296 | -2.043 | -0.19 | 0.981 | -0.006 | 0 | 0 | 1.811 | 0.73 | 0.31 | 0.79 | 0.68 | 0.9 | 0.83 |
| 9 | 119655679 | rs10124561 | 0.282 | -2.056 | 0.224 | 0.975 | -0.048 | 0.101 | 0.101 | 1.825 | 0.17 | 0.36 | 0.18 | 0.37 | 0.08 | 0.14 |
| 9 | 119657927 | rs16934033 | 1.511 | -1.86 | 0.377 | 1.02 | -0.039 | -0.049 | -0.049 | 0 | 0.09 | 0.15 | 0.03 | 0 | 0.02 | 0.03 |
| 9 | 119659517 | rs10983398 | -0.496 | -1.706 | -0.067 | 1.097 | -0.037 | -0.083 | -0.083 | 5.17 | 0.29 | 0.1 | 0.32 | 0.55 | 0.36 | 0.29 |
| 9 | 119665210 | rs10983400 | -0.76 | -1.838 | 0.119 | 1.056 | -0.003 | 0.004 | 0.004 | 0 | 0.16 | 0.1 | 0.1 | 0.31 | 0.08 | 0.21 |
| 9 | 119666575 | rs7042929 | -0.737 | -1.882 | 0.272 | 1.02 | -0.048 | 0.084 | 0.084 | 0 | 0.13 | 0.2 | 0.08 | 0.28 | 0.05 | 0.17 |
| 9 | 119667151 | rs7029188 | 1.328 | -1.967 | -0.471 | 0.343 | -0.061 | 0.226 | 0.226 | 0.136 | 0.45 | 0.05 | 0.47 | 0.09 | 0.55 | 0.52 |
| 9 | 119672835 | rs4838015 | 0.008 | -1.582 | 0.535 | 0.942 | -0.023 | 0.053 | 0.053 | 0.006 | 0.83 | 0.77 | 0.79 | 0.35 | 0.91 | 0.73 |
| 9 | 119673408 | rs10817958 | 1.268 | -1.449 | 0.479 | 0.813 | -0.044 | -0.04 | -0.04 | 0 | 0.14 | 0.06 | 0.23 | 0.24 | 0.28 | 0.1 |
| 9 | 119674722 | rs944323 | 1.183 | -1.608 | 1.068 | 0.678 | -0.002 | -0.046 | -0.046 | 0.101 | 0.59 | 0.47 | 0.51 | 0.1 | 0.57 | 0.54 |
| 9 | 119676324 | rs7029462 | 0.332 | -1.618 | 0.86 | 0.746 | -0.017 | 0.031 | 0.031 | 0 | 0.81 | 0.77 | 0.78 | 0.35 | 0.89 | 0.67 |
| 9 | 119683394 | rs2094807 | 1.685 | -1.588 | 0.698 | 0.667 | 0.008 | 0.286 | 0.286 | 0.067 | 0.52 | 0.09 | 0.47 | 0.11 | 0.58 | 0.57 |
| 9 | 119685232 | rs4838020 | 0.53 | -1.654 | 1.14 | 0.73 | -0.062 | 0.08 | 0.08 | 0.108 | 0.78 | 0.47 | 0.75 | 0.35 | 0.89 | 0.8 |
| 9 | 119685882 | rs10513279 | 1.607 | -1.503 | 0.72 | 0.525 | 0.008 | 0.286 | 0.286 | 0.067 | 0.52 | 0.09 | 0.47 | 0.11 | 0.58 | 0.57 |
| 9 | 119685920 | rs10513280 | 1.181 | -1.514 | 0.727 | 0.519 | -0.015 | -0.034 | -0.034 | 0 | 0.09 | 0.14 | 0.04 | 0 | 0.03 | 0.04 |
| 9 | 119688194 | rs1929010 | -0.684 | -1.403 | 0.408 | 0.893 | -0.022 | 0.098 | 0.098 | 0.902 | 0.87 | 0.78 | 0.8 | 0.59 | 0.93 | 0.89 |
| 9 | 119688600 | rs1929009 | 1.154 | -1.193 | -0.188 | 0.561 | 0.039 | 0.253 | 0.253 | 0.047 | 0.52 | 0.17 | 0.48 | 0.1 | 0.58 | 0.57 |
| 9 | 119691640 | rs12551186 | 1.048 | -1.119 | -0.589 | 0.564 | -0.011 | 0.221 | 0.221 | 0.01 | 0.49 | 0.08 | 0.46 | 0.16 | 0.56 | 0.56 |
| 9 | 119693164 | rs16934069 | -0.22 | -0.311 | 0.346 | 0.869 | 0.023 | 0.084 | 0.084 | 0 | 0.22 | 0.33 | 0.33 | 0.37 | 0.2 | 0.25 |
| 9 | 119693948 | rs10983411 | 0.396 | -0.365 | -0.04 | 0.692 | -0.002 | -0.006 | -0.006 | 0 | 0.2 | 0.18 | 0.25 | 0.22 | 0.25 | 0.1 |
| 9 | 119696088 | rs10983413 | 1.275 | -0.36 | -0.323 | 0.695 | -0.014 | -0.021 | -0.021 | 0 | 0.13 | 0.08 | 0.2 | 0.22 | 0.2 | 0.08 |
| 9 | 119696591 | rs10759875 | 1.13 | -0.546 | -0.466 | 0.303 | -0.02 | 0.244 | 0.244 | 0 | 0.48 | 0.18 | 0.38 | 0.1 | 0.47 | 0.44 |
| 9 | 119702496 | rs2150198 | 1.117 | -0.032 | -0.299 | 0.677 | -0.021 | -0.082 | -0.082 | 0 | 0.59 | 0.14 | 0.5 | 0.44 | 0.53 | 0.47 |
| 9 | 119707772 | rs10116120 | 0.201 | 0.021 | 0.638 | 0.747 | 0.022 | 0.03 | 0.03 | 0 | 0.18 | 0.4 | 0.34 | 0.23 | 0.24 | 0.2 |
| 9 | 119708335 | rs3849142 | 1.182 | 0.213 | 0.69 | 0.842 | 0.004 | 0.007 | 0.007 | 0 | 0.12 | 0.12 | 0.31 | 0.23 | 0.24 | 0.19 |
| 9 | 119710771 | rs10759876 | 0.777 | 0.038 | 1.035 | 0.847 | -0.015 | 0.035 | 0.035 | 4.214 | 0.78 | 0.64 | 0.87 | 0.67 | 0.8 | 0.66 |
| 9 | 119712333 | rs3849144 | 0.847 | 0.032 | 0.874 | 0.844 | -0.019 | 0.042 | 0.042 | 3.458 | 0.78 | 0.54 | 0.86 | 0.66 | 0.8 | 0.66 |
| 9 | 119712401 | rs10817967 | 1.047 | 0.285 | 0.597 | 0.8 | 0.005 | 0.01 | 0.01 | 0 | 0.12 | 0.13 | 0.31 | 0.23 | 0.23 | 0.18 |
| 9 | 119719068 | rs10983433 | 1.25 | 0.347 | 0.093 | 0.8 | -0.025 | -0.027 | -0.027 | 0.402 | 0.11 | 0.06 | 0.35 | 0.3 | 0.22 | 0.13 |
| 9 | 119719162 | rs10983434 | 0.145 | 0.227 | 0.293 | 0.655 | 0.01 | -0.029 | -0.029 | 0 | 0.13 | 0.28 | 0.1 | 0.05 | 0.14 | 0.23 |
| 9 | 119720895 | rs10983435 | 1.531 | 0.314 | -0.279 | 0.756 | -0.024 | -0.021 | -0.021 | 0 | 0.1 | 0.06 | 0.35 | 0.3 | 0.22 | 0.13 |
| 9 | 119720910 | rs7026444 | 1.175 | -0.017 | -0.347 | 0.65 | 0.022 | 0.026 | 0.026 | 3.825 | 0.72 | 0.24 | 0.83 | 0.75 | 0.78 | 0.66 |
| 9 | 119721232 | rs13283397 | 1.243 | -0.186 | -1.025 | 0.527 | -0.041 | -0.129 | -0.129 | 0 | 0.59 | 0.08 | 0.46 | 0.45 | 0.59 | 0.52 |
| 9 | 119722707 | rs16934119 | -0.032 | 0.093 | -0.912 | 0.481 | -0.011 | -0.018 | -0.018 | 5.686 | 0.1 | 0.01 | 0.04 | 0.16 | 0.1 | 0.13 |
| 9 | 119724111 | rs4838041 | 1.605 | -0.408 | -1.473 | 0.17 | -0.043 | -0.074 | -0.074 | 1.159 | 0.51 | 0.08 | 0.43 | 0.46 | 0.52 | 0.52 |
| 9 | 119730709 | rs10759878 | 1.57 | -0.365 | -1.119 | 0.155 | -0.062 | -0.077 | -0.077 | 0.897 | 0.13 | 0.02 | 0.38 | 0.28 | 0.27 | 0.12 |
| 9 | 119732360 | rs10759879 | 0.258 | -0.5 | -0.925 | 0.081 | -0.003 | -0.009 | -0.009 | 4.233 | 0.23 | 0.22 | 0.43 | 0.29 | 0.29 | 0.2 |
| 9 | 119735989 | rs11787916 | 2.209 | -0.849 | -2 | -0.202 | -0.068 | 0.029 | 0.029 | 0.018 | 0.3 | 0.01 | 0.26 | 0.13 | 0.3 | 0.29 |
| 9 | 119738890 | rs2274762 | 1.697 | -0.303 | -1.401 | -0.127 | -0.075 | -0.085 | -0.085 | 0 | 0.14 | 0.03 | 0.39 | 0.29 | 0.3 | 0.13 |
| 9 | 119745284 | rs13284239 | 0.82 | -0.365 | -1.455 | -0.199 | -0.069 | -0.069 | -0.069 | 0.114 | 0.6 | 0.09 | 0.45 | 0.63 | 0.57 | 0.63 |
| 9 | 119746836 | rs12337139 | 2.036 | -0.625 | -0.913 | -0.698 | -0.036 | -0.02 | -0.02 | 0 | 0.07 | 0.01 | 0.05 | 0.07 | 0.11 | 0.08 |
| 9 | 119753154 | rs7848841 | 0.203 | -0.653 | -0.215 | -0.726 | -0.049 | -0.083 | -0.083 | 0.295 | 0.13 | 0.41 | 0.07 | 0.01 | 0.03 | 0.16 |
| 9 | 119762707 | rs4144636 | 1.484 | -0.759 | -0.771 | -0.68 | 0.086 | 0.04 | 0.04 | 2.23 | 0.65 | 0.15 | 0.82 | 0.7 | 0.81 | 0.66 |
| 9 | 119766969 | rs7042036 | 1.421 | -0.433 | -1.71 | -0.76 | 0.005 | 0.1 | 0.1 | 1.011 | 0.58 | 0.14 | 0.74 | 0.7 | 0.69 | 0.61 |
| 9 | 119767269 | rs12686071 | -0.11 | -0.668 | -1.138 | -0.185 | 0.018 | -0.013 | -0.013 | 0 | 0.1 | 0.06 | 0.03 | 0.23 | 0.05 | 0.12 |
| 9 | 119768572 | rs10817972 | 0.176 | -0.394 | -1.068 | -0.442 | -0.007 | -0.006 | -0.006 | 0 | 0.83 | 0.27 | 0.88 | 0.93 | 0.88 | 0.86 |
| 9 | 119770480 | rs3761845 | 0.112 | -0.07 | -1.951 | -0.891 | -0.023 | 0.164 | 0.164 | 0 | 0.43 | 0.14 | 0.71 | 0.75 | 0.57 | 0.47 |
| 9 | 119771893 | rs1537528 | -0.168 | 0.574 | -1.284 | -0.603 | -0.012 | 0.015 | 0.015 | 0 | 0.12 | 0.09 | 0.07 | 0.14 | 0.19 | 0.12 |
| 9 | 119777844 | rs10739481 | 0.317 | 0.69 | -0.835 | -0.269 | 0.005 | 0.057 | 0.057 | 0 | 0.71 | 0.45 | 0.77 | 0.52 | 0.7 | 0.64 |
| 9 | 119779810 | rs2418446 | 0.256 | 0.63 | -0.748 | -0.269 | 0.004 | 0.092 | 0.092 | 0 | 0.74 | 0.45 | 0.8 | 0.52 | 0.72 | 0.64 |
| 9 | 119780936 | rs10817976 | 1.321 | 0.61 | -0.721 | -0.653 | -0.001 | -0.001 | -0.001 | 0 | 0.18 | 0.19 | 0.3 | 0.27 | 0.29 | 0.12 |
| 9 | 119783586 | rs7874684 | 0.224 | -0.144 | -1.347 | -1.186 | 0.002 | -0.01 | -0.01 | 0 | 0.27 | 0.22 | 0.26 | 0.12 | 0.18 | 0.35 |
| 9 | 119784980 | rs10817980 | 1.746 | -0.244 | -1.115 | -1.171 | -0.065 | -0.063 | -0.063 | 0.411 | 0.17 | 0.06 | 0.29 | 0.27 | 0.29 | 0.12 |
| 9 | 119785506 | rs12341741 | 1.695 | -0.316 | -1.003 | -1.233 | 0.003 | 0.091 | 0.091 | 0 | 0.31 | 0.11 | 0.25 | 0.14 | 0.25 | 0.2 |

| | | | | | | | | | | | | | | | | |
|---|---|---|---|---|---|---|---|---|---|---|---|---|---|---|---|---|
| 9 | 119786753 | rs949870 | -1.115 | -0.491 | -0.7 | -1.192 | 0 | 0 | 0 | 0 | 0.43 | 0.47 | 0.32 | 0.58 | 0.41 | 0.53 |
| 9 | 119789130 | rs7047287 | 0.412 | -0.217 | -0.986 | -0.932 | -0.039 | -0.103 | -0.103 | 0 | 0.19 | 0.05 | 0.16 | 0.47 | 0.26 | 0.21 |
| 9 | 119792047 | rs7857238 | 0.379 | -0.036 | -0.805 | -0.986 | -0.04 | 0.005 | 0.005 | 0 | 0.31 | 0.27 | 0.38 | 0.29 | 0.41 | 0.28 |
| 9 | 119799584 | rs9697192 | 0.645 | -0.042 | -0.284 | -0.648 | 0.009 | -0.058 | -0.058 | 0 | 0.11 | 0.26 | 0.09 | 0.01 | 0.11 | 0.17 |
| 9 | 119800808 | rs1334078 | -0.799 | -0.421 | -0.331 | -0.508 | -0.001 | -0.003 | -0.003 | 0.776 | 0.4 | 0.43 | 0.31 | 0.55 | 0.41 | 0.52 |
| 9 | 119802694 | rs1334077 | 0.837 | -0.558 | -0.221 | -0.327 | 0.002 | -0.005 | -0.005 | 0 | 0.57 | 0.27 | 0.57 | 0.42 | 0.55 | 0.36 |
| 9 | 119803284 | rs12351322 | 0.861 | -0.551 | -0.08 | -0.785 | 0.003 | 0.047 | 0.047 | 0 | 0.33 | 0.22 | 0.28 | 0.13 | 0.29 | 0.18 |
| 9 | 119803528 | rs7855056 | -0.01 | -0.632 | 0.297 | -0.691 | -0.007 | 0.13 | 0.13 | 0 | 0.49 | 0.78 | 0.55 | 0.24 | 0.48 | 0.27 |
| 9 | 119803933 | rs10481683 | 0.241 | -0.597 | 0.484 | -0.671 | 0 | -0.013 | -0.013 | 0.052 | 0.46 | 0.54 | 0.53 | 0.23 | 0.47 | 0.22 |
| 9 | 119806789 | rs10983472 | 0.833 | -0.548 | -0.102 | -0.56 | 0.004 | 0.01 | 0.01 | 0 | 0.07 | 0.06 | 0.17 | 0.02 | 0.07 | 0.15 |
| 9 | 119808697 | rs10817983 | 0.719 | -0.662 | -0.062 | -0.472 | -0.002 | 0.009 | 0.009 | 0 | 0.32 | 0.27 | 0.2 | 0.24 | 0.31 | 0.24 |
| 9 | 119808900 | rs4836966 | 0.229 | -0.516 | -0.149 | -0.218 | 0.007 | -0.036 | -0.036 | 0.117 | 0.13 | 0.07 | 0.1 | 0.49 | 0.15 | 0.35 |
| 9 | 119811790 | rs4838117 | -1.165 | -0.447 | 0.112 | -0.374 | 0 | -0.002 | -0.002 | 0 | 0.67 | 0.75 | 0.72 | 0.87 | 0.72 | 0.82 |
| 9 | 119811891 | rs13298897 | 0.939 | -0.17 | 0.278 | -0.291 | 0.023 | -0.046 | -0.046 | 0 | 0.11 | 0.19 | 0.12 | 0 | 0.16 | 0.04 |
| 9 | 119812580 | rs10817987 | 0.085 | -0.351 | 0.228 | -0.275 | -0.002 | -0.006 | -0.006 | 0 | 0.47 | 0.54 | 0.35 | 0.28 | 0.56 | 0.38 |
| 9 | 119815778 | rs1334070 | -0.612 | -0.392 | 0.244 | -0.198 | -0.011 | -0.02 | -0.02 | 0 | 0.6 | 0.61 | 0.46 | 0.71 | 0.7 | 0.71 |
| 9 | 119816239 | rs10983479 | 2.122 | -0.442 | 0.02 | -0.26 | -0.082 | 0.004 | 0.004 | 0.335 | 0.22 | 0.01 | 0.16 | 0.13 | 0.25 | 0.17 |
| 9 | 119816619 | rs10119031 | 0.95 | -0.236 | 0.688 | -0.281 | 0.048 | -0.063 | -0.063 | 0.054 | 0.11 | 0.27 | 0.12 | 0 | 0.17 | 0.05 |
| 9 | 119817038 | rs944976 | -1.49 | -0.277 | 0.357 | -0.292 | 0.005 | 0.014 | 0.014 | 0 | 0.72 | 0.82 | 0.74 | 0.87 | 0.79 | 0.88 |
| 9 | 119818979 | rs6478276 | 0.16 | -0.122 | 0.069 | -0.15 | -0.01 | 0.097 | 0.097 | 0 | 0.47 | 0.31 | 0.34 | 0.72 | 0.54 | 0.66 |
| 9 | 119821287 | rs6478277 | -0.135 | -0.605 | 0.273 | -0.036 | 0.014 | 0.019 | 0.019 | 0 | 0.56 | 0.43 | 0.43 | 0.72 | 0.7 | 0.68 |
| 9 | 119821314 | rs6478278 | 0.064 | -0.14 | 0.667 | 0.966 | -0.001 | 0.004 | 0.004 | 1.336 | 0.14 | 0.17 | 0.07 | 0.25 | 0.15 | 0.21 |
| 9 | 119822186 | rs7872915 | -1.128 | -0.314 | 0.745 | 0.798 | -0.006 | -0.01 | -0.01 | 0 | 0.79 | 0.8 | 0.77 | 0.87 | 0.84 | 0.88 |
| 9 | 119823868 | rs4087490 | 0.412 | -0.475 | 0.597 | 0.754 | 0.03 | 0.017 | 0.017 | 0 | 0.14 | 0.06 | 0.27 | 0.14 | 0.08 | 0.17 |
| 9 | 119824051 | rs10983487 | 0.986 | -0.476 | 0.902 | 0.806 | 0.046 | 0.001 | 0.001 | 0 | 0.5 | 0.34 | 0.41 | 0.46 | 0.61 | 0.5 |
| 9 | 119825237 | rs10121565 | -0.835 | -0.497 | 1.012 | 0.777 | -0.016 | 0.013 | 0.013 | 0.1 | 0.28 | 0.4 | 0.35 | 0.39 | 0.23 | 0.38 |
| 9 | 119826169 | rs11789058 | 0.706 | 0.201 | 1.274 | 1.089 | 0.005 | 0.021 | 0.021 | 0.064 | 0.53 | 0.63 | 0.57 | 0.32 | 0.46 | 0.42 |
| 9 | 119826602 | rs7848256 | -1.142 | -0.121 | 1.523 | 1.123 | -0.008 | 0.08 | 0.08 | 1.525 | 0.18 | 0.51 | 0.1 | 0.25 | 0.15 | 0.23 |
| 9 | 119827197 | rs7848689 | 0.119 | -0.072 | 1.153 | 1.014 | -0.002 | -0.006 | -0.006 | 0 | 0.12 | 0.1 | 0.06 | 0.21 | 0.13 | 0.16 |
| 9 | 119835353 | rs10983505 | -0.797 | -0.047 | 1.095 | 0.992 | -0.001 | 0.059 | 0.059 | 1.746 | 0.17 | 0.44 | 0.22 | 0.22 | 0.14 | 0.18 |
| 9 | 119836771 | rs10118188 | -0.341 | 0.018 | 0.698 | 0.852 | -0.027 | -0.019 | -0.019 | 0 | 0.79 | 0.69 | 0.89 | 0.86 | 0.87 | 0.88 |
| 9 | 119838444 | rs12344337 | -0.528 | 0.104 | 0.447 | 1.088 | 0.004 | -0.025 | -0.025 | 0 | 0.32 | 0.22 | 0.28 | 0.38 | 0.27 | 0.28 |
| 9 | 119839328 | rs16934284 | -0.55 | 0.297 | 0.976 | 1.611 | 0.063 | 0.249 | 0.249 | 0 | 0.07 | 0.3 | 0.09 | 0.3 | 0.13 | 0.18 |
| 9 | 119840344 | rs10759896 | -0.365 | -0.064 | 0.49 | 1.144 | 0.006 | 0.005 | 0.005 | 0.124 | 0.88 | 0.95 | 0.83 | 0.97 | 0.91 | 0.82 |
| 9 | 119843385 | rs1105890 | -0.504 | -0.226 | 0.68 | 1.492 | -0.017 | 0.091 | 0.091 | 0 | 0.13 | 0.23 | 0.27 | 0.41 | 0.07 | 0.23 |
| 9 | 119854198 | rs12339142 | -0.691 | 0.434 | 0.628 | 1.215 | -0.002 | 0 | 0 | 0 | 0.39 | 0.64 | 0.41 | 0.65 | 0.44 | 0.47 |
| 9 | 119856725 | rs10983516 | -0.052 | 0.431 | 0.048 | 1.45 | -0.023 | -0.029 | -0.029 | 1.295 | 0.17 | 0.11 | 0.23 | 0.33 | 0.23 | 0.16 |
| 9 | 119857460 | rs11791192 | 0.358 | 0.596 | -0.001 | 1.522 | 0.047 | 0.076 | 0.076 | 0 | 0.06 | 0.15 | 0.09 | 0.19 | 0.15 | 0.16 |
| 9 | 119858116 | rs10983517 | -0.606 | 0.4 | -0.489 | 1.479 | -0.003 | -0.01 | -0.01 | 0 | 0.23 | 0.2 | 0.36 | 0.52 | 0.23 | 0.28 |
| 9 | 119863851 | rs10983521 | 2.225 | 0.328 | -0.785 | 1.213 | -0.035 | 0.118 | 0.118 | 0.362 | 0.33 | 0.02 | 0.23 | 0.09 | 0.31 | 0.17 |
| 9 | 119864616 | rs7031714 | 0.592 | 0.255 | -0.332 | 1.332 | -0.047 | -0.008 | -0.008 | 0 | 0.79 | 0.71 | 0.7 | 0.78 | 0.9 | 0.68 |
| 9 | 119865441 | rs10983522 | -0.594 | 0.514 | -0.653 | 1.369 | -0.09 | -0.091 | -0.091 | 0 | 0.35 | 0.12 | 0.35 | 0.49 | 0.44 | 0.34 |
| 9 | 119865765 | rs13297784 | 1.043 | 0.583 | -0.597 | 1.243 | -0.05 | -0.018 | -0.018 | 0 | 0.09 | 0.01 | 0.08 | 0.11 | 0.16 | 0.07 |
| 9 | 119870336 | rs16934320 | 0.081 | 0.492 | -0.573 | 1.269 | 0.002 | -0.003 | -0.003 | 0 | 0.11 | 0.11 | 0.16 | 0.23 | 0.08 | 0.13 |
| 9 | 119872587 | rs13285598 | 0.952 | 0.669 | -0.571 | 1.276 | -0.051 | -0.018 | -0.018 | 0 | 0.09 | 0.01 | 0.08 | 0.11 | 0.16 | 0.07 |
| 9 | 119876954 | rs1889320 | 0.601 | 0.439 | -0.316 | 1.209 | 0.009 | 0.003 | 0.003 | 0 | 0.53 | 0.69 | 0.48 | 0.55 | 0.49 | 0.5 |
| 9 | 119878362 | rs11789158 | 0.513 | 0.455 | -0.708 | 1.248 | 0.002 | -0.003 | -0.003 | 0 | 0.54 | 0.76 | 0.49 | 0.55 | 0.5 | 0.5 |
| 9 | 119878770 | rs11789213 | -0.296 | 0.455 | -0.497 | 1.347 | -0.027 | 0.017 | 0.017 | 0 | 0.08 | 0.15 | 0.13 | 0.19 | 0.02 | 0.13 |
| 9 | 119880993 | rs3923774 | -0.345 | 1.064 | -0.858 | 0.97 | -0.029 | 0.026 | 0.026 | 0 | 0.2 | 0.07 | 0.12 | 0.08 | 0.22 | 0.16 |
| 9 | 119885715 | rs10983537 | -1.195 | 0.962 | -0.463 | 1.31 | 0 | 0.018 | 0.018 | 0 | 0.09 | 0.12 | 0.07 | 0.19 | 0.1 | 0.16 |
| 9 | 119887577 | rs1572155 | -0.813 | 1.394 | -0.433 | 1.341 | -0.002 | -0.002 | -0.002 | 0 | 0.19 | 0.17 | 0.31 | 0.25 | 0.23 | 0.3 |
| 9 | 119896151 | rs10817997 | 0.049 | 1.558 | -0.831 | 1.308 | -0.034 | -0.007 | -0.007 | 0 | 0.13 | 0.04 | 0.08 | 0.09 | 0.17 | 0.11 |
| 9 | 119899060 | rs1413413 | -0.605 | 0.976 | -0.627 | 1.17 | 0 | 0.001 | 0.001 | 0 | 0.2 | 0.21 | 0.11 | 0.16 | 0.19 | 0.22 |
| 9 | 119901378 | rs4090221 | 1.694 | 0.623 | -1.966 | 0.637 | -0.003 | 0.061 | 0.061 | 0 | 0.29 | 0.04 | 0.2 | 0.17 | 0.26 | 0.18 |
| 9 | 119903506 | rs10817999 | 0.146 | -0.026 | -1.281 | 0.35 | 0.001 | 0.024 | 0.024 | 0 | 0.42 | 0.31 | 0.65 | 0.67 | 0.54 | 0.6 |
| 9 | 119908494 | rs16934346 | 1.246 | 0.623 | -1.211 | 0.866 | -0.021 | -0.033 | -0.033 | 0 | 0.07 | 0.12 | 0.02 | 0 | 0.03 | 0.07 |
| 9 | 119913608 | rs10983551 | 0.548 | 0.951 | -1.331 | 0.984 | -0.002 | -0.002 | -0.002 | 0 | 0.18 | 0.16 | 0.28 | 0.24 | 0.22 | 0.3 |
| 9 | 119920031 | rs7019490 | 2.376 | 1.086 | -0.773 | 0.764 | 0.011 | 0.06 | 0.06 | 3.079 | 0.15 | 0.03 | 0.08 | 0 | 0.17 | 0.1 |
| 9 | 119921723 | rs1334081 | -0.189 | 1.146 | -0.692 | 0.916 | 0.001 | 0.031 | 0.031 | 0 | 0.42 | 0.29 | 0.64 | 0.67 | 0.54 | 0.61 |
| 9 | 119922519 | rs17309941 | 0.345 | 0.713 | -0.691 | 0.78 | -0.005 | 0.021 | 0.021 | 0.019 | 0.34 | 0.23 | 0.37 | 0.25 | 0.4 | 0.42 |
| 9 | 119923266 | rs1334082 | 0.705 | 0.919 | -0.715 | 0.593 | -0.001 | 0 | 0 | 0 | 0.16 | 0.18 | 0.09 | 0.08 | 0.17 | 0.11 |
| 9 | 119925095 | rs6478282 | -0.5 | 0.992 | -0.484 | 0.836 | 0.017 | 0.017 | 0.017 | 0 | 0.25 | 0.37 | 0.28 | 0.27 | 0.27 | 0.28 |
| 9 | 119932558 | rs10983556 | 0.298 | 0.975 | -0.562 | 0.846 | 0.001 | 0.007 | 0.007 | 0.35 | 0.09 | 0.13 | 0.18 | 0.19 | 0.08 | 0.17 |
| 9 | 119939389 | rs1413409 | 1.878 | 0.757 | -0.994 | 0.61 | -0.058 | 0.084 | 0.084 | 0 | 0.31 | 0.05 | 0.23 | 0.16 | 0.32 | 0.2 |
| 9 | 119942649 | rs977937 | 0.341 | 1.33 | -0.415 | 0.96 | 0.004 | -0.008 | -0.008 | 0.003 | 0.58 | 0.5 | 0.61 | 0.66 | 0.55 | 0.58 |
| 9 | 119945170 | rs10983562 | 1.748 | 1.247 | -0.656 | 1.255 | 0.014 | 0.049 | 0.049 | 3.592 | 0.16 | 0.05 | 0.09 | 0 | 0.17 | 0.09 |
| 9 | 119946330 | rs10983563 | -0.335 | 1.416 | -0.432 | 1.71 | 0.024 | 0.06 | 0.06 | 0.517 | 0.09 | 0.2 | 0.19 | 0.2 | 0.08 | 0.18 |
| 9 | 119946647 | rs10983564 | 0.581 | 1.006 | -0.761 | 1.602 | -0.052 | -0.029 | -0.029 | 0 | 0.08 | 0.01 | 0.19 | 0.43 | 0.15 | 0.13 |
| 9 | 119956288 | rs7858836 | -0.529 | 0.033 | -0.089 | 1.046 | -0.006 | 0.012 | 0.012 | 0 | 0.64 | 0.61 | 0.62 | 0.48 | 0.64 | 0.67 |

| | | | | | | | | | | | | | | | | |
|---|---|---|---|---|---|---|---|---|---|---|---|---|---|---|---|---|
| 9 | 119957909 | rs10818012 | -0.464 | -0.002 | 0.152 | 1.029 | -0.01 | 0.016 | 0.016 | 0 | 0.63 | 0.48 | 0.61 | 0.49 | 0.64 | 0.68 |
| 9 | 119958853 | rs958804 | 0.625 | -0.12 | 0.236 | 0.981 | -0.001 | -0.012 | -0.012 | 0 | 0.31 | 0.26 | 0.37 | 0.43 | 0.34 | 0.3 |
| 9 | 119967963 | rs915281 | 0.353 | -0.2 | -0.098 | 0.974 | 0.007 | 0.107 | 0.107 | 0 | 0.48 | 0.2 | 0.33 | 0.27 | 0.51 | 0.38 |
| 9 | 119968145 | rs6478290 | -0.16 | -0.13 | 0.033 | 0.695 | -0.004 | 0.032 | 0.032 | 0.343 | 0.15 | 0.23 | 0.28 | 0.22 | 0.12 | 0.29 |
| 9 | 119986987 | rs2184543 | 0.338 | -0.392 | -0.864 | -0.438 | -0.1 | 0.145 | 0.145 | 0 | 0.34 | 0.07 | 0.18 | 0.08 | 0.4 | 0.21 |
| 9 | 119989912 | rs4838299 | -0.503 | -0.397 | -0.791 | -0.436 | -0.063 | 0.108 | 0.108 | 0 | 0.66 | 0.9 | 0.81 | 0.92 | 0.6 | 0.76 |
| 9 | 119990164 | rs10121207 | 0.648 | -0.862 | -0.469 | -0.466 | 0.001 | -0.008 | -0.008 | 0 | 0.44 | 0.56 | 0.7 | 0.68 | 0.46 | 0.44 |
| 9 | 119990556 | rs10818014 | -0.016 | -0.883 | -0.433 | -0.411 | -0.032 | 0.018 | 0.018 | 0 | 0.16 | 0.29 | 0.09 | 0.16 | 0.11 | 0.25 |
| 9 | 119992025 | rs11787579 | 0.14 | -0.838 | -0.171 | -0.033 | -0.037 | 0.12 | 0.12 | 0.083 | 0.13 | 0.29 | 0.14 | 0.32 | 0.06 | 0.24 |
| 9 | 119992795 | rs6478297 | -1.175 | -0.907 | -0.105 | -0.232 | -0.001 | 0 | 0 | 0 | 0.6 | 0.41 | 0.3 | 0.33 | 0.52 | 0.63 |
| 9 | 119993022 | rs1335405 | 0.503 | -0.962 | -0.119 | -0.248 | -0.037 | 0.066 | 0.066 | 0 | 0.11 | 0.26 | 0.06 | 0.17 | 0.05 | 0.21 |
| 9 | 119993084 | rs4838301 | 1.06 | -1.115 | -0.325 | -0.331 | -0.085 | -0.005 | -0.005 | 0.429 | 0.4 | 0.12 | 0.68 | 0.67 | 0.51 | 0.37 |
| 9 | 119993094 | rs1160246 | -0.022 | -1.087 | -0.343 | -0.329 | -0.067 | 0.187 | 0.187 | 0.18 | 0.44 | 0.12 | 0.22 | 0.15 | 0.44 | 0.39 |
| 9 | 119994691 | rs6478298 | -0.938 | -1.042 | -0.293 | -0.295 | -0.018 | 0.016 | 0.016 | 0.525 | 0.58 | 0.7 | 0.31 | 0.33 | 0.49 | 0.63 |
| 9 | 120000826 | rs10513289 | 0.158 | -0.768 | -0.649 | -0.691 | -0.038 | 0.136 | 0.136 | 0.03 | 0.34 | 0.07 | 0.17 | 0.09 | 0.32 | 0.36 |
| 9 | 120001836 | rs16934422 | 0.974 | -0.462 | -0.44 | -0.433 | -0.035 | -0.036 | -0.036 | 0 | 0.08 | 0.01 | 0.11 | 0.14 | 0.08 | 0.02 |
| 9 | 120007625 | rs10465132 | 0.669 | -0.579 | -0.478 | -0.474 | 0.075 | 0.122 | 0.122 | 0.208 | 0.52 | 0.12 | 0.77 | 0.78 | 0.66 | 0.48 |
| 9 | 120011056 | rs10818015 | -0.634 | -0.571 | -0.162 | -0.492 | -0.005 | 0.014 | 0.014 | 0 | 0.21 | 0.27 | 0.18 | 0.28 | 0.17 | 0.23 |
| 9 | 120012421 | rs1335419 | 0.703 | -0.595 | -0.805 | -0.525 | 0.07 | 0.113 | 0.113 | 0.162 | 0.53 | 0.14 | 0.77 | 0.77 | 0.66 | 0.51 |
| 9 | 120017634 | rs10983584 | 0.553 | -0.455 | -1.299 | -0.549 | -0.039 | 0.007 | 0.007 | 0 | 0.28 | 0.07 | 0.2 | 0.21 | 0.28 | 0.28 |
| 9 | 120023135 | rs11794931 | -0.536 | -0.488 | -0.934 | -0.412 | -0.013 | -0.008 | -0.008 | 0 | 0.2 | 0.3 | 0.11 | 0.16 | 0.15 | 0.22 |
| 9 | 120023774 | rs10983589 | 0.284 | -0.635 | -0.849 | -0.25 | -0.046 | 0.046 | 0.046 | 0 | 0.12 | 0.31 | 0.06 | 0.16 | 0.06 | 0.2 |
| 9 | 120028948 | rs3900788 | 0.423 | -0.427 | -2.152 | -0.241 | -0.045 | -0.003 | -0.003 | 0 | 0.28 | 0.08 | 0.2 | 0.24 | 0.31 | 0.22 |
| 9 | 120029861 | rs1007834 | 0.335 | 0.233 | -1.979 | -0.069 | 0.05 | 0.006 | 0.006 | 0.597 | 0.5 | 0.11 | 0.69 | 0.62 | 0.62 | 0.47 |
| 9 | 120031254 | rs7043216 | -0.983 | 0.737 | -1.254 | -0.115 | -0.02 | 0.029 | 0.029 | 0.967 | 0.17 | 0.09 | 0.14 | 0.09 | 0.2 | 0.15 |
| 9 | 120032451 | rs7019790 | -1.195 | 0.733 | -1.192 | -0.146 | -0.102 | 0.027 | 0.027 | 0.252 | 0.25 | 0.05 | 0.23 | 0.16 | 0.36 | 0.33 |
| 9 | 120035769 | rs6478300 | -0.236 | 0.23 | -1.178 | -0.245 | 0.038 | 0.006 | 0.006 | 0 | 0.42 | 0.65 | 0.24 | 0.39 | 0.26 | 0.47 |
| 9 | 120038801 | rs1889655 | -0.742 | 0.626 | -1.123 | -0.176 | -0.027 | 0.01 | 0.01 | 0 | 0.17 | 0.08 | 0.19 | 0.14 | 0.26 | 0.3 |
| 9 | 120043171 | rs7026789 | -0.619 | 0.47 | -1.193 | -0.197 | -0.109 | 0.04 | 0.04 | 0.049 | 0.21 | 0.03 | 0.22 | 0.11 | 0.32 | 0.31 |
| 9 | 120050463 | rs10513297 | 1.082 | 0.196 | -0.682 | -0.101 | -0.004 | -0.008 | -0.008 | 0.046 | 0.38 | 0.39 | 0.43 | 0.55 | 0.45 | 0.22 |
| 9 | 120052529 | rs10759913 | 1.25 | 0.188 | -0.659 | -0.095 | -0.061 | -0.078 | -0.078 | 2.284 | 0.36 | 0.14 | 0.4 | 0.56 | 0.45 | 0.22 |
| 9 | 120054021 | rs7862566 | 0.623 | -0.182 | -0.694 | -0.076 | 0.008 | -0.005 | -0.005 | 0 | 0.26 | 0.2 | 0.3 | 0.32 | 0.17 | 0.32 |
| 9 | 120055202 | rs3893339 | 0.813 | 0.312 | -1.112 | 0.251 | -0.091 | -0.083 | -0.083 | 0 | 0.4 | 0.11 | 0.43 | 0.54 | 0.52 | 0.24 |
| 9 | 120065787 | rs10983613 | 0.852 | 0.717 | -1.574 | 0.36 | -0.066 | -0.109 | -0.109 | 0 | 0.37 | 0.07 | 0.45 | 0.47 | 0.35 | 0.21 |
| 9 | 120068520 | rs7022764 | 0.011 | 0.411 | -1.239 | 0.171 | -0.019 | -0.002 | -0.002 | 0 | 0.67 | 0.56 | 0.72 | 0.64 | 0.71 | 0.59 |
| 9 | 120071548 | rs1572306 | 0.708 | 0.389 | -1.56 | 0.042 | -0.041 | -0.044 | -0.044 | 0 | 0.18 | 0.02 | 0.22 | 0.2 | 0.22 | 0.24 |
| 9 | 120071949 | rs1572307 | 0.181 | 0.374 | -1.368 | 0.028 | 0 | 0.008 | 0.008 | 0.694 | 0.22 | 0.1 | 0.24 | 0.19 | 0.24 | 0.25 |
| 9 | 120078941 | rs11790950 | 1.159 | 0.467 | -1.087 | 0.292 | -0.057 | -0.12 | -0.12 | 0.481 | 0.36 | 0.09 | 0.43 | 0.47 | 0.36 | 0.21 |
| 9 | 120085704 | rs7021801 | 0.346 | 0.468 | -0.979 | 0.194 | -0.001 | -0.001 | -0.001 | 1.323 | 0.47 | 0.49 | 0.5 | 0.47 | 0.48 | 0.23 |
| 9 | 120086176 | rs10983619 | 1.004 | 0.575 | -1.478 | 0.25 | -0.052 | -0.083 | -0.083 | 0 | 0.37 | 0.16 | 0.42 | 0.47 | 0.38 | 0.21 |
| 9 | 120095530 | rs10983624 | -0.172 | 0.265 | -1.681 | 0.096 | -0.002 | -0.041 | -0.041 | 0.074 | 0.1 | 0 | 0.19 | 0.23 | 0.07 | 0.14 |
| 9 | 120099047 | rs1339921 | 0.402 | 0.9 | -1.444 | 0.142 | 0.001 | 0.018 | 0.018 | 2.22 | 0.45 | 0.34 | 0.35 | 0.55 | 0.47 | 0.31 |
| 9 | 120107961 | rs1934485 | 0.951 | 0.552 | -2.011 | 0.193 | 0.019 | -0.004 | -0.004 | 0 | 0.35 | 0.05 | 0.23 | 0.69 | 0.29 | 0.26 |
| 9 | 120108044 | rs10818024 | 0.826 | 0.537 | -2.081 | 0.234 | -0.004 | 0.104 | 0.104 | 1.443 | 0.37 | 0.04 | 0.3 | 0.84 | 0.32 | 0.29 |
| 9 | 120108170 | rs1934484 | 0.739 | 0.122 | -2.402 | 0.029 | 0.025 | 0.071 | 0.071 | 4.675 | 0.41 | 0.04 | 0.28 | 0.76 | 0.33 | 0.39 |
| 9 | 120109509 | rs4240415 | -0.188 | 0.704 | -1.974 | -0.416 | 0.008 | -0.037 | -0.037 | 0 | 0.89 | 0.84 | 0.92 | 0.99 | 0.85 | 0.97 |
| 9 | 120110190 | rs12552082 | 1.023 | 0.438 | -2.242 | 0.224 | 0.042 | -0.052 | -0.052 | 0.21 | 0.34 | 0.08 | 0.23 | 0.39 | 0.28 | 0.26 |
| 9 | 120114228 | rs10818029 | 0.27 | 1.058 | -1.37 | -0.614 | -0.01 | 0.062 | 0.062 | 0.839 | 0.57 | 0.32 | 0.64 | 0.94 | 0.49 | 0.75 |
| 9 | 120122802 | rs7029526 | 0.334 | 1.026 | -1.006 | -0.323 | 0.017 | -0.091 | -0.091 | 0 | 0.31 | 0.37 | 0.26 | 0.05 | 0.36 | 0.23 |
| 9 | 120126470 | rs10818030 | 1.091 | 0.803 | -1.152 | 0.424 | -0.003 | -0.07 | -0.07 | 0 | 0.26 | 0.1 | 0.19 | 0.47 | 0.25 | 0.15 |
| 9 | 120130206 | rs10513300 | -0.134 | 0.904 | -1.354 | 0.719 | -0.015 | -0.029 | -0.029 | 0 | 0.06 | 0 | 0.3 | 0.26 | 0.06 | 0.15 |
| 9 | 120133394 | rs1416940 | 0.094 | 1.311 | -0.951 | 0.11 | 0.022 | -0.07 | -0.07 | 0 | 0.79 | 0.72 | 0.82 | 0.97 | 0.7 | 0.73 |
| 9 | 120137252 | rs1339929 | 0.016 | 1.298 | -0.836 | 0.104 | -0.002 | -0.025 | -0.025 | 0.127 | 0.79 | 0.79 | 0.84 | 0.97 | 0.71 | 0.73 |
| 9 | 120137758 | rs2226006 | -0.013 | 1.232 | -0.865 | 0.155 | 0.005 | 0.002 | 0.002 | 4.315 | 0.53 | 0.61 | 0.63 | 0.5 | 0.47 | 0.58 |
| 9 | 120141746 | rs1339930 | 0.099 | 0.96 | -0.636 | -0.122 | -0.009 | -0.06 | -0.06 | 0 | 0.42 | 0.46 | 0.52 | 0.91 | 0.32 | 0.48 |
| 9 | 120141805 | rs1934487 | 0.116 | 1.302 | -0.387 | -0.126 | 0.005 | 0.025 | 0.025 | 0.88 | 0.48 | 0.44 | 0.67 | 0.31 | 0.56 | 0.41 |
| 9 | 120145848 | rs10818033 | 0.989 | 0.904 | -0.101 | 0.095 | 0.013 | -0.006 | -0.006 | 0 | 0.71 | 0.61 | 0.5 | 0.65 | 0.64 | 0.59 |
| 9 | 120146772 | rs16905575 | -0.189 | 1.006 | -0.329 | 0.188 | -0.032 | -0.04 | -0.04 | 0 | 0.07 | 0.01 | 0.34 | 0.32 | 0.12 | 0.15 |
| 9 | 120148425 | rs7042295 | -0.593 | 0.999 | -0.297 | 0.239 | -0.002 | -0.002 | -0.002 | 0 | 0.08 | 0.1 | 0.2 | 0.17 | 0.12 | 0.15 |
| 9 | 120150050 | rs1416943 | 0.218 | 0.874 | -0.078 | 0.327 | 0.014 | -0.019 | -0.019 | 0 | 0.79 | 0.7 | 0.71 | 0.82 | 0.76 | 0.73 |
| 9 | 120151034 | rs4419882 | -0.394 | 0.865 | -0.126 | 0.36 | -0.002 | -0.003 | -0.003 | 0 | 0.07 | 0.08 | 0.2 | 0.17 | 0.11 | 0.14 |
| 9 | 120153306 | rs12551446 | 1.692 | 0.612 | -0.301 | 0.155 | 0.016 | -0.06 | -0.06 | 0 | 0.24 | 0.1 | 0.16 | 0.42 | 0.21 | 0.14 |
| 9 | 120154325 | rs4837163 | 0.115 | 0.73 | -0.254 | 0.004 | 0.001 | -0.019 | -0.019 | 0 | 0.78 | 0.7 | 0.72 | 0.82 | 0.8 | 0.73 |
| 9 | 120154849 | rs1416942 | 0.046 | 0.52 | -0.169 | -0.085 | 0.002 | 0.004 | 0.004 | 0 | 0.42 | 0.42 | 0.4 | 0.76 | 0.38 | 0.46 |
| 9 | 120161777 | rs10513301 | 1.801 | 0.584 | -0.599 | -0.008 | -0.029 | -0.117 | -0.117 | 1.411 | 0.25 | 0.04 | 0.17 | 0.38 | 0.23 | 0.19 |
| 9 | 120162325 | rs7872945 | 1.535 | 0.622 | -0.475 | -0.213 | -0.004 | 0.023 | 0.023 | 0.128 | 0.32 | 0.23 | 0.28 | 0.06 | 0.35 | 0.22 |
| 9 | 120162380 | rs7875944 | 0.808 | 0.55 | -0.471 | -0.124 | -0.027 | 0.058 | 0.058 | 0.428 | 0.42 | 0.27 | 0.5 | 0.27 | 0.53 | 0.36 |
| 9 | 120163466 | rs4240422 | 0.366 | 0.598 | -0.699 | -0.196 | 0.001 | -0.005 | -0.005 | 0.139 | 0.45 | 0.44 | 0.67 | 0.42 | 0.55 | 0.37 |
| 9 | 120165049 | rs1934483 | -0.052 | 0.906 | -1.249 | -0.094 | 0.002 | -0.049 | -0.049 | 0 | 0.79 | 0.74 | 0.87 | 0.97 | 0.8 | 0.74 |

| | | | | | | | | | | | | | | | | |
|---|---|---|---|---|---|---|---|---|---|---|---|---|---|---|---|---|
| 9 | 120165611 | rs7467917 | -0.432 | 1.149 | -1.362 | 0.485 | -0.034 | -0.044 | -0.044 | 0.828 | 0.1 | 0.04 | 0.36 | 0.36 | 0.18 | 0.15 |
| 9 | 120169552 | rs10739493 | -0.232 | 0.91 | -1.217 | 0.178 | -0.026 | 0.003 | 0.003 | 1.761 | 0.58 | 0.79 | 0.7 | 0.59 | 0.58 | 0.6 |
| 9 | 120179586 | rs7023773 | -0.827 | 0.728 | -1.107 | 0.37 | 0.024 | 0.015 | 0.015 | 0.08 | 0.48 | 0.75 | 0.54 | 0.54 | 0.38 | 0.54 |
| 9 | 120185992 | rs955302 | 0.02 | 0.922 | -1.282 | 0.427 | -0.047 | -0.048 | -0.048 | 1.604 | 0.26 | 0.14 | 0.27 | 0.42 | 0.34 | 0.25 |
| 9 | 120189474 | rs4836590 | -0.76 | 0.733 | -1.459 | 0.215 | -0.062 | -0.039 | -0.039 | 0 | 0.33 | 0.14 | 0.31 | 0.41 | 0.39 | 0.45 |
| 9 | 120192435 | rs17419570 | -0.214 | 0.903 | -1.509 | 0.146 | -0.042 | 0.063 | 0.063 | 0 | 0.11 | 0.02 | 0.1 | 0.01 | 0.14 | 0.2 |
| 9 | 120194838 | rs4146493 | -1.657 | 0.73 | -0.698 | 0.447 | 0.001 | 0.01 | 0.01 | 4.254 | 0.44 | 0.41 | 0.64 | 0.76 | 0.55 | 0.6 |
| 9 | 120196215 | rs17419694 | -0.778 | 0.715 | -0.97 | 0.455 | -0.059 | -0.051 | -0.051 | 0 | 0.34 | 0.14 | 0.31 | 0.44 | 0.39 | 0.45 |
| 9 | 120199140 | rs16905625 | 0.852 | 0.737 | -1.183 | 0.515 | -0.077 | -0.117 | -0.117 | 0 | 0.2 | 0.05 | 0.21 | 0.41 | 0.26 | 0.25 |
| 9 | 120209679 | rs12685568 | 1.119 | 0.176 | -0.788 | 0.45 | 0.012 | -0.002 | -0.002 | 0 | 0.22 | 0.2 | 0.09 | 0.18 | 0.09 | 0.19 |
| 9 | 120210027 | rs9987535 | -1.38 | 0.282 | -0.64 | 0.374 | -0.041 | -0.001 | -0.001 | 0 | 0.6 | 0.8 | 0.44 | 0.59 | 0.48 | 0.65 |
| 9 | 120210307 | rs10513307 | 1.098 | 0.053 | -0.587 | 0.331 | -0.042 | 0.001 | 0.001 | 2.778 | 0.22 | 0.29 | 0.1 | 0.18 | 0.09 | 0.19 |
| 9 | 120211956 | rs10983659 | 0.008 | 0.602 | -1.017 | 0.523 | -0.037 | -0.035 | -0.035 | 0 | 0.08 | 0.02 | 0.31 | 0.31 | 0.15 | 0.14 |
| 9 | 120212123 | rs10818038 | -1.731 | 0.258 | -0.804 | -0.122 | -0.013 | 0.077 | 0.077 | 1.313 | 0.69 | 0.85 | 0.76 | 0.93 | 0.65 | 0.78 |
| 9 | 120216660 | rs4837254 | -0.953 | 0.157 | -0.816 | 0.181 | 0.02 | 0.049 | 0.049 | 0.074 | 0.56 | 0.8 | 0.74 | 0.73 | 0.6 | 0.62 |
| 9 | 120217280 | rs1572523 | 0.752 | 0.577 | -1.19 | 0.591 | -0.09 | -0.121 | -0.121 | 0 | 0.21 | 0.05 | 0.4 | 0.42 | 0.3 | 0.29 |
| 9 | 120225010 | rs13285674 | -0.114 | 0.311 | -0.816 | 0.588 | 0.018 | 0.003 | 0.003 | 0.13 | 0.31 | 0.25 | 0.19 | 0.27 | 0.21 | 0.39 |
| 9 | 120225715 | rs7025237 | 0.086 | 0.319 | -0.411 | 0.544 | -0.03 | -0.006 | -0.006 | 0 | 0.67 | 0.76 | 0.46 | 0.33 | 0.55 | 0.59 |
| 9 | 120226415 | rs1577808 | 0.164 | 0.397 | -0.275 | 0.37 | 0.02 | 0.092 | 0.092 | 0 | 0.07 | 0.14 | 0.13 | 0.25 | 0.12 | 0.12 |
| 9 | 120229555 | rs10983664 | 1.3 | 0.245 | -0.246 | -0.104 | -0.003 | -0.052 | -0.052 | 0 | 0.36 | 0.36 | 0.24 | 0.06 | 0.34 | 0.2 |
| 9 | 120239806 | rs12115318 | 0.549 | 0.097 | -0.375 | 0.213 | 0.007 | 0.002 | 0.002 | 0.054 | 0.2 | 0.18 | 0.09 | 0.13 | 0.11 | 0.22 |
| 9 | 120240163 | rs10983670 | 0.324 | 0.591 | -0.609 | 0.486 | -0.059 | -0.08 | -0.08 | 8.779 | 0.25 | 0.1 | 0.42 | 0.44 | 0.34 | 0.29 |
| 9 | 120248751 | rs1337206 | 0.494 | 0.612 | -0.492 | 0.454 | -0.051 | -0.073 | -0.073 | 0 | 0.23 | 0.11 | 0.42 | 0.44 | 0.34 | 0.27 |
| 9 | 120249360 | rs4633129 | 0.649 | 0.521 | -0.405 | 0.307 | 0.002 | 0.002 | 0.002 | 0.445 | 0.21 | 0.22 | 0.14 | 0.12 | 0.18 | 0.2 |
| 9 | 120249464 | rs10983673 | -0.747 | 0.461 | -0.306 | -0.052 | 0 | 0.044 | 0.044 | 0.022 | 0.17 | 0.22 | 0.18 | 0.39 | 0.16 | 0.32 |
| 9 | 120250392 | rs1316439 | -1.35 | 0.391 | -0.255 | -0.329 | 0.016 | 0.063 | 0.063 | 0.2 | 0.62 | 0.77 | 0.76 | 0.94 | 0.66 | 0.75 |
| 9 | 120254911 | rs1337215 | 0.713 | 0.619 | -0.121 | -0.113 | 0.009 | 0.014 | 0.014 | 0 | 0.46 | 0.37 | 0.32 | 0.21 | 0.37 | 0.38 |
| 9 | 120255966 | rs2786130 | 0.089 | 0.683 | -0.105 | 0.114 | 0.002 | -0.011 | -0.011 | 0 | 0.55 | 0.49 | 0.43 | 0.38 | 0.49 | 0.53 |
| 9 | 120263905 | rs12348211 | -0.089 | 1.218 | -0.087 | 0.131 | -0.044 | 0.044 | 0.044 | 0 | 0.09 | 0.02 | 0.09 | 0.01 | 0.18 | 0.1 |
| 9 | 120270109 | rs540657 | 1.107 | 0.966 | -0.162 | -0.569 | -0.001 | -0.074 | -0.074 | 0 | 0.37 | 0.35 | 0.28 | 0 | 0.4 | 0.24 |
| 9 | 120275146 | rs11794261 | 0.975 | 1.071 | -0.446 | -0.18 | -0.073 | 0.096 | 0.096 | 1.925 | 0.4 | 0.17 | 0.32 | 0.22 | 0.5 | 0.43 |
| 9 | 120275159 | rs12337381 | 0.245 | 1.245 | -0.351 | -0.127 | -0.016 | 0.016 | 0.016 | 0 | 0.09 | 0.05 | 0.07 | 0.01 | 0.16 | 0.1 |
| 9 | 120277454 | rs545966 | 0.122 | 1.059 | 0.051 | -0.08 | 0.015 | 0.032 | 0.032 | 4.282 | 0.47 | 0.41 | 0.4 | 0.23 | 0.66 | 0.53 |
| 9 | 120285351 | rs4510935 | 0.725 | 0.915 | -0.228 | 0.43 | -0.009 | -0.036 | -0.036 | 0 | 0.16 | 0.1 | 0.37 | 0.44 | 0.15 | 0.2 |
| 9 | 120292334 | rs1985317 | 0.635 | 0.984 | -0.179 | -0.285 | -0.004 | 0.093 | 0.093 | 3.624 | 0.43 | 0.22 | 0.36 | 0.23 | 0.62 | 0.48 |
| 9 | 120293454 | rs10983688 | 0.571 | 1.15 | -0.219 | -0.279 | -0.018 | 0.016 | 0.016 | 0 | 0.08 | 0.03 | 0.11 | 0.01 | 0.21 | 0.1 |
| 9 | 120299808 | rs495062 | -0.176 | 0.992 | 0.015 | -0.212 | 0.006 | 0.097 | 0.097 | 0.002 | 0.23 | 0.48 | 0.42 | 0.67 | 0.19 | 0.3 |
| 9 | 120302842 | rs559135 | -0.536 | 0.497 | -0.271 | -0.138 | 0.003 | 0 | 0 | 0 | 0.66 | 0.69 | 0.78 | 0.91 | 0.81 | 0.79 |
| 9 | 120303051 | rs11795161 | 1.301 | 0.752 | -1.322 | -0.271 | -0.022 | 0.071 | 0.071 | 1.322 | 0.33 | 0.17 | 0.24 | 0 | 0.39 | 0.23 |
| 9 | 120304076 | rs10117151 | -0.427 | 1.402 | -1.128 | 0.61 | -0.053 | -0.043 | -0.043 | 0 | 0.27 | 0.14 | 0.48 | 0.69 | 0.37 | 0.43 |
| 9 | 120309723 | rs540178 | 0.056 | 0.691 | -0.431 | 0.47 | -0.004 | 0.032 | 0.032 | 0.424 | 0.7 | 0.49 | 0.76 | 0.65 | 0.75 | 0.78 |
| 9 | 120315637 | rs12339717 | -0.469 | 0.645 | -0.361 | 0.446 | -0.004 | 0.003 | 0.003 | 0.045 | 0.1 | 0.15 | 0.23 | 0.14 | 0.11 | 0.23 |
| 9 | 120316896 | rs495083 | 0.921 | 0.514 | 0.231 | 0.145 | 0 | 0.008 | 0.008 | 0.002 | 0.58 | 0.51 | 0.53 | 0.56 | 0.6 | 0.54 |
| 9 | 120327559 | rs476999 | 0.195 | 0.557 | 0.299 | 0.14 | 0.004 | 0.007 | 0.007 | 0.164 | 0.68 | 0.68 | 0.75 | 0.7 | 0.7 | 0.77 |
| 9 | 120339304 | rs565055 | 0.717 | 0.619 | -0.037 | -0.119 | 0.02 | 0.061 | 0.061 | 0 | 0.5 | 0.37 | 0.42 | 0.38 | 0.47 | 0.47 |
| 9 | 120341514 | rs7858743 | 0.315 | 0.346 | -0.245 | -0.679 | -0.001 | -0.046 | -0.046 | 5.34 | 0.25 | 0.25 | 0.19 | 0.01 | 0.21 | 0.26 |
| 9 | 120353505 | rs7857205 | 0.801 | 0.144 | -0.465 | -0.764 | 0.002 | 0.005 | 0.005 | 0 | 0.44 | 0.43 | 0.4 | 0.37 | 0.39 | 0.34 |
| 9 | 120353610 | rs7857333 | 1.35 | 0.235 | -0.616 | -0.745 | 0.015 | 0.062 | 0.062 | 0 | 0.18 | 0.06 | 0.14 | 0 | 0.12 | 0.13 |
| 9 | 120358616 | rs10983707 | 1.907 | 0.404 | -0.397 | -0.507 | -0.048 | -0.079 | -0.079 | 0 | 0.23 | 0.06 | 0.23 | 0.38 | 0.26 | 0.22 |
| 9 | 120360094 | rs10983708 | 0.905 | 0.359 | -0.273 | -0.52 | 0.003 | 0.01 | 0.01 | 0 | 0.16 | 0.22 | 0.15 | 0.19 | 0.19 | 0.2 |
| 9 | 120368736 | rs10983712 | 0.974 | 0.464 | -0.498 | -0.777 | 0 | 0.001 | 0.001 | 0 | 0.39 | 0.37 | 0.34 | 0.39 | 0.36 | 0.26 |
| 9 | 120369028 | rs7875849 | 1.096 | 0.503 | -0.551 | -0.756 | 0.003 | 0.009 | 0.009 | 0 | 0.16 | 0.22 | 0.15 | 0.19 | 0.19 | 0.22 |
| 9 | 120372882 | rs1329055 | 0.422 | 0.613 | -0.801 | -0.808 | 0.013 | 0.063 | 0.063 | 0 | 0.12 | 0.14 | 0.1 | 0.28 | 0.16 | 0.17 |
| 9 | 120374480 | rs10491854 | 2.047 | 0.616 | -0.726 | -0.66 | -0.019 | -0.056 | -0.056 | 0.627 | 0.24 | 0.09 | 0.22 | 0.38 | 0.25 | 0.22 |
| 9 | 120378483 | rs1928295 | 0.553 | 0.582 | -0.7 | -0.651 | 0.001 | 0.007 | 0.007 | 0 | 0.46 | 0.45 | 0.39 | 0.39 | 0.44 | 0.39 |
| 9 | 120389465 | rs1155928 | 2.016 | 0.561 | -0.693 | -0.623 | -0.02 | -0.059 | -0.059 | 0 | 0.24 | 0.09 | 0.22 | 0.38 | 0.25 | 0.22 |
| 9 | 120390282 | rs1887904 | -0.585 | 0.607 | -0.661 | -0.549 | -0.002 | -0.003 | -0.003 | 1.857 | 0.65 | 0.77 | 0.71 | 0.72 | 0.68 | 0.69 |
| 9 | 120398385 | rs7026557 | 0.419 | 0.404 | -0.182 | -0.132 | 0.007 | 0.029 | 0.029 | 0 | 0.17 | 0.31 | 0.15 | 0.2 | 0.2 | 0.21 |
| 9 | 120410219 | rs4837462 | -1.159 | 0.429 | -0.668 | -0.52 | -0.013 | 0.079 | 0.079 | 0 | 0.48 | 0.78 | 0.52 | 0.61 | 0.51 | 0.56 |
| 9 | 120413192 | rs7034845 | -0.643 | 0.431 | -0.172 | -0.431 | 0.001 | 0.004 | 0.004 | 0 | 0.38 | 0.56 | 0.32 | 0.45 | 0.4 | 0.34 |
| 9 | 120416248 | rs10983736 | -0.476 | 0.573 | 0.377 | -0.221 | 0.001 | 0.017 | 0.017 | 0 | 0.14 | 0.24 | 0.13 | 0.19 | 0.16 | 0.14 |
| 9 | 120416402 | rs10983737 | 0.664 | 0.524 | 0.227 | -0.376 | -0.004 | 0.037 | 0.037 | 0 | 0.07 | 0.01 | 0.03 | 0 | 0.07 | 0.13 |
| 9 | 120418118 | rs913615 | 1.829 | 0.614 | 0.108 | -0.468 | -0.05 | 0.028 | 0.028 | 0 | 0.53 | 0.13 | 0.47 | 0.43 | 0.51 | 0.49 |
| 9 | 120427484 | rs1854448 | 1.023 | 0.6 | 0.22 | -0.462 | 0.004 | 0.008 | 0.008 | 0.066 | 0.64 | 0.31 | 0.75 | 0.75 | 0.67 | 0.68 |
| 9 | 120431545 | rs7039756 | -0.834 | 0.412 | 0.657 | -0.406 | 0.026 | 0.063 | 0.063 | 0 | 0.19 | 0.35 | 0.32 | 0.33 | 0.22 | 0.24 |
| 9 | 120433401 | rs10818066 | -1.181 | 0.434 | 0.521 | -0.402 | 0.003 | 0.024 | 0.024 | 0 | 0.23 | 0.34 | 0.34 | 0.33 | 0.24 | 0.26 |
| 9 | 120441946 | rs1329057 | 0.435 | -0.015 | 0.697 | -0.952 | 0.008 | 0.054 | 0.054 | 0.066 | 0.78 | 0.63 | 0.83 | 0.76 | 0.8 | 0.84 |
| 9 | 120443779 | rs960312 | -0.669 | -0.002 | 0.765 | -0.959 | 0.006 | 0.066 | 0.066 | 0.558 | 0.84 | 0.75 | 0.86 | 0.76 | 0.85 | 0.89 |
| 9 | 120448498 | rs1252038 | -0.044 | -0.049 | 0.436 | -1.118 | -0.006 | -0.039 | -0.039 | 0.003 | 0.08 | 0.16 | 0.04 | 0 | 0.07 | 0.13 |

| | | | | | | | | | | | | | | | | |
|---|---|---|---|---|---|---|---|---|---|---|---|---|---|---|---|---|
| 9 | 120449466 | rs1360094 | 1.441 | -0.058 | 0.377 | -1.143 | 0.011 | -0.035 | -0.035 | 0 | 0.61 | 0.16 | 0.6 | 0.62 | 0.67 | 0.61 |
| 9 | 120450798 | rs4837496 | 0.837 | -0.032 | 0.439 | -1.106 | 0.001 | -0.004 | -0.004 | 0 | 0.69 | 0.34 | 0.65 | 0.63 | 0.74 | 0.74 |
| 9 | 120456495 | rs10818070 | 0.008 | -0.025 | 0.418 | -1.143 | -0.006 | -0.038 | -0.038 | 0.016 | 0.08 | 0.14 | 0.04 | 0 | 0.07 | 0.13 |
| 9 | 120461621 | rs10759930 | 1.734 | -0.211 | -0.14 | -1.6 | -0.111 | -0.107 | -0.107 | 0.364 | 0.31 | 0.06 | 0.39 | 0.63 | 0.42 | 0.38 |
| 9 | 120462715 | rs2737191 | 0.811 | -0.321 | -0.071 | -1.581 | 0.017 | 0.109 | 0.109 | 0 | 0.3 | 0.11 | 0.23 | 0 | 0.25 | 0.23 |
| 9 | 120470054 | rs1927911 | 1.046 | -0.332 | 0.711 | -0.803 | 0 | -0.001 | -0.001 | 0 | 0.69 | 0.32 | 0.66 | 0.63 | 0.74 | 0.75 |
| 9 | 120473834 | rs5030717 | 0.446 | -0.258 | 0.24 | -0.803 | -0.002 | -0.063 | -0.063 | 0 | 0.14 | 0.19 | 0.11 | 0.01 | 0.11 | 0.03 |
| 9 | 120474199 | rs2149356 | 1.229 | -0.194 | -0.563 | -0.716 | 0.021 | -0.031 | -0.031 | 0 | 0.61 | 0.18 | 0.63 | 0.63 | 0.69 | 0.63 |
| 9 | 120480010 | rs11536897 | 0.076 | 0.059 | -0.11 | -1.227 | -0.001 | -0.02 | -0.02 | 0.086 | 0.06 | 0.07 | 0.03 | 0 | 0.05 | 0.05 |
| 9 | 120480115 | rs1927906 | -1.158 | -0.103 | 0.805 | -1.278 | 0.011 | -0.013 | -0.013 | 0 | 0.12 | 0.51 | 0.1 | 0.04 | 0.1 | 0.19 |
| 9 | 120480812 | rs1554973 | -2.457 | -0.154 | 0.669 | -1.363 | 0.006 | 0.036 | 0.036 | 1.729 | 0.27 | 0.79 | 0.19 | 0.14 | 0.23 | 0.3 |
| 9 | 120481856 | rs7856729 | -1.346 | -0.151 | 1.26 | -1.358 | 0.015 | -0.038 | -0.038 | 0 | 0.16 | 0.33 | 0.1 | 0.1 | 0.13 | 0.11 |
| 9 | 120482940 | rs7846989 | -0.831 | -0.16 | 1.287 | -1.348 | 0.021 | -0.002 | -0.002 | 0 | 0.11 | 0.4 | 0.09 | 0.04 | 0.1 | 0.19 |
| 9 | 120483639 | rs7037225 | -1.69 | -0.146 | 1.236 | -1.191 | -0.004 | -0.003 | -0.003 | 3.201 | 0.2 | 0.73 | 0.25 | 0.12 | 0.16 | 0.25 |
| 9 | 120485215 | rs2183016 | -1.382 | -0.254 | 1.19 | -1.251 | 0.017 | -0.037 | -0.037 | 0 | 0.15 | 0.34 | 0.1 | 0.1 | 0.13 | 0.11 |
| 9 | 120485795 | rs7045953 | -1.397 | -0.254 | 1.19 | -1.251 | 0.016 | -0.038 | -0.038 | 0.911 | 0.16 | 0.34 | 0.1 | 0.1 | 0.13 | 0.12 |
| 9 | 120496387 | rs7025144 | -0.118 | -0.966 | -0.008 | -2.263 | 0.043 | 0.081 | 0.081 | 0 | 0.67 | 0.87 | 0.74 | 0.99 | 0.74 | 0.78 |
| 9 | 120497958 | rs10465133 | 0.745 | -0.646 | 0.367 | -2.376 | 0.048 | 0.149 | 0.149 | 0.31 | 0.54 | 0.24 | 0.6 | 0.96 | 0.59 | 0.63 |
| 9 | 120502265 | rs7866830 | -0.944 | -0.801 | 0.439 | -2.432 | -0.019 | -0.075 | -0.075 | 0 | 0.23 | 0.64 | 0.17 | 0.03 | 0.2 | 0.18 |
| 9 | 120503274 | rs7870814 | 1.209 | -0.934 | 0.113 | -2.532 | 0.027 | 0.1 | 0.1 | 0 | 0.5 | 0.34 | 0.47 | 0.89 | 0.54 | 0.61 |
| 9 | 120510364 | rs10818081 | 1.788 | -1.162 | -0.104 | -2.733 | -0.061 | 0.156 | 0.156 | 2.108 | 0.4 | 0.13 | 0.43 | 0.89 | 0.49 | 0.58 |
| 9 | 120510899 | rs17330490 | 0.217 | -1.168 | -0.072 | -2.731 | 0 | 0 | 0 | 0 | 0.1 | 0.05 | 0.06 | 0 | 0.09 | 0.04 |
| 9 | 120512266 | rs756136 | 1.696 | -1.22 | -0.052 | -2.751 | -0.016 | 0.052 | 0.052 | 1.529 | 0.4 | 0.3 | 0.43 | 0.89 | 0.48 | 0.58 |
| 9 | 120514683 | rs10739501 | 0.663 | -1.105 | 0.076 | -2.706 | 0.027 | 0.05 | 0.05 | 1.616 | 0.58 | 0.34 | 0.53 | 0.9 | 0.67 | 0.67 |
| 9 | 120514770 | rs2225207 | 0.661 | -1.107 | 0.076 | -2.707 | 0.027 | 0.05 | 0.05 | 1.586 | 0.58 | 0.33 | 0.53 | 0.9 | 0.67 | 0.67 |
| 9 | 120518162 | rs10123941 | 0.289 | -1.187 | 0.118 | -2.58 | -0.004 | -0.065 | -0.065 | 0 | 0.66 | 0.68 | 0.72 | 0.98 | 0.73 | 0.77 |
| 9 | 120518859 | rs10818087 | 0.247 | -1.136 | 0.126 | -2.569 | -0.02 | 0.025 | 0.025 | 1.409 | 0.12 | 0.05 | 0.09 | 0 | 0.18 | 0.1 |
| 9 | 120519878 | rs1927896 | 0.258 | -1.158 | 0.183 | -2.516 | -0.001 | -0.054 | -0.054 | 0 | 0.67 | 0.68 | 0.72 | 0.98 | 0.74 | 0.78 |
| 9 | 120520277 | rs7026522 | -0.754 | -1.155 | 0.262 | -2.456 | 0.016 | 0.023 | 0.023 | 0.404 | 0.77 | 0.86 | 0.77 | 0.99 | 0.81 | 0.84 |
| 9 | 120522873 | rs10759943 | -0.754 | -1.155 | 0.262 | -2.456 | 0.016 | 0.023 | 0.023 | 0.404 | 0.77 | 0.86 | 0.77 | 0.99 | 0.81 | 0.84 |
| 9 | 120524178 | rs7027711 | -0.569 | -1.135 | -0.019 | -2.444 | -0.004 | -0.052 | -0.052 | 0 | 0.75 | 0.76 | 0.76 | 0.99 | 0.8 | 0.84 |
| 9 | 120526112 | rs10983778 | -0.219 | -1.037 | -1.025 | -2.39 | 0.026 | 0.033 | 0.033 | 3.78 | 0.7 | 0.25 | 0.62 | 0.9 | 0.79 | 0.73 |
| 9 | 120533422 | rs1475546 | 0.783 | -0.836 | -1.113 | -2.203 | 0.038 | 0.264 | 0.264 | 2.793 | 0.5 | 0.17 | 0.5 | 0.89 | 0.54 | 0.59 |
| 9 | 120536514 | rs10739502 | 0.259 | -0.525 | -0.887 | -2.377 | 0.009 | 0.005 | 0.005 | 0 | 0.65 | 0.74 | 0.78 | 0.99 | 0.71 | 0.78 |
| 9 | 120540290 | rs1927934 | -0.31 | -0.591 | -0.912 | -2.41 | 0.004 | 0.141 | 0.141 | 0 | 0.34 | 0.15 | 0.22 | 0.01 | 0.28 | 0.23 |
| 9 | 120543270 | rs7018934 | 0.623 | -0.588 | -0.762 | -2.581 | 0.009 | -0.077 | -0.077 | 0 | 0.67 | 0.6 | 0.78 | 0.91 | 0.65 | 0.75 |
| 9 | 120546133 | rs7046116 | 0.389 | -0.525 | -1.059 | -2.512 | -0.007 | 0.012 | 0.012 | 0.806 | 0.18 | 0.11 | 0.1 | 0.09 | 0.21 | 0.12 |
| 9 | 120547282 | rs12352822 | 1.06 | -0.406 | -1.328 | -2.446 | -0.02 | 0.054 | 0.054 | 0 | 0.13 | 0.03 | 0.1 | 0.01 | 0.14 | 0.13 |
| 9 | 120548934 | rs4836703 | 0.789 | -0.43 | -1.238 | -2.355 | 0.017 | 0.057 | 0.057 | 0 | 0.21 | 0.11 | 0.12 | 0 | 0.14 | 0.09 |
| 9 | 120556606 | rs2225208 | 0.933 | -0.261 | -1.089 | -2.238 | 0.041 | 0.107 | 0.107 | 1.199 | 0.26 | 0.09 | 0.12 | 0 | 0.15 | 0.09 |
| 9 | 120560559 | rs2149361 | 0.465 | 0.349 | -1.755 | -1.822 | -0.009 | 0.06 | 0.06 | 0 | 0.17 | 0.04 | 0.1 | 0.01 | 0.16 | 0.15 |
| 9 | 120571308 | rs2149358 | -0.471 | 0.385 | -1.695 | -1.756 | 0.038 | -0.013 | -0.013 | 0.027 | 0.88 | 0.8 | 0.9 | 0.94 | 0.79 | 0.92 |
| 9 | 120576387 | rs12342517 | 0.435 | 0.398 | -1.812 | -1.741 | -0.003 | 0.05 | 0.05 | 0 | 0.17 | 0.06 | 0.1 | 0.01 | 0.16 | 0.16 |
| 9 | 120576634 | rs10759948 | 1.018 | 0.375 | -1.903 | -0.988 | -0.072 | -0.129 | -0.129 | 0 | 0.29 | 0.03 | 0.38 | 0.61 | 0.33 | 0.45 |
| 9 | 120591239 | rs4837510 | 0.302 | 0.946 | -0.807 | -0.964 | -0.008 | 0.006 | 0.006 | 0 | 0.7 | 0.76 | 0.8 | 0.93 | 0.63 | 0.77 |
| 9 | 120613258 | rs694428 | -0.675 | 0.257 | 0.578 | 0.441 | 0.025 | 0.297 | 0.297 | 1.058 | 0.08 | 0.35 | 0.27 | 0.33 | 0.06 | 0.12 |
| 9 | 120613902 | rs1709578 | -0.259 | 0.579 | -0.186 | 0.186 | -0.058 | -0.083 | -0.083 | 0.178 | 0.71 | 0.89 | 0.64 | 0.57 | 0.63 | 0.44 |
| 9 | 120616052 | rs524074 | -0.923 | 0.223 | 0.265 | 0.01 | -0.054 | 0.148 | 0.148 | 2.503 | 0.17 | 0.54 | 0.33 | 0.34 | 0.11 | 0.16 |
| 9 | 120619093 | rs6478330 | 0.989 | 0.587 | -0.115 | -0.227 | -0.009 | 0.018 | 0.018 | 0.773 | 0.1 | 0.04 | 0.07 | 0 | 0.12 | 0.02 |
| 9 | 120626011 | rs10818096 | -0.259 | 0.674 | 0.06 | -0.215 | -0.016 | -0.026 | -0.026 | 0 | 0.75 | 0.81 | 0.68 | 0.56 | 0.69 | 0.51 |
| 9 | 120626599 | rs10818097 | -0.272 | 0.674 | 0.06 | -0.215 | -0.016 | -0.026 | -0.026 | 0 | 0.75 | 0.81 | 0.68 | 0.56 | 0.69 | 0.51 |
| 9 | 120629491 | rs963466 | -0.276 | 0.7 | 0.269 | -0.219 | -0.009 | -0.015 | -0.015 | 0.039 | 0.75 | 0.81 | 0.69 | 0.57 | 0.69 | 0.51 |
| 9 | 120630599 | rs11790155 | 0.478 | 0.672 | -0.035 | -0.415 | -0.029 | -0.027 | -0.027 | 6.211 | 0.17 | 0.04 | 0.14 | 0.2 | 0.23 | 0.15 |
| 9 | 120632054 | rs2296943 | 0.921 | 0.485 | 0.696 | -0.421 | 0.002 | 0.013 | 0.013 | 3.444 | 0.53 | 0.46 | 0.51 | 0.36 | 0.44 | 0.29 |
| 9 | 120636801 | rs908198 | 0.266 | 0.488 | 0.829 | -0.423 | 0 | 0 | 0 | 0.385 | 0.18 | 0.17 | 0.16 | 0.2 | 0.23 | 0.15 |
| 9 | 120637262 | rs2048741 | 1.036 | 0.427 | 0.688 | -0.416 | 0.01 | 0.046 | 0.046 | 4.207 | 0.51 | 0.41 | 0.51 | 0.32 | 0.44 | 0.29 |
| 9 | 120659844 | rs10818102 | 0.923 | 0.373 | 0.777 | -0.345 | 0.007 | 0.025 | 0.025 | 3.269 | 0.52 | 0.44 | 0.53 | 0.32 | 0.43 | 0.3 |
| 9 | 120660568 | rs1399428 | -0.821 | 0.395 | 0.806 | -0.328 | -0.025 | -0.034 | -0.034 | 2.204 | 0.77 | 0.84 | 0.7 | 0.56 | 0.68 | 0.53 |
| 9 | 120665221 | rs10435879 | 0.946 | 0.208 | 0.901 | -0.3 | 0.013 | 0.045 | 0.045 | 3.21 | 0.51 | 0.39 | 0.52 | 0.33 | 0.43 | 0.3 |
| 9 | 120669106 | rs7872095 | -0.114 | 0.167 | 0.965 | -0.261 | -0.011 | 0.012 | 0.012 | 0 | 0.2 | 0.32 | 0.12 | 0.23 | 0.19 | 0.19 |
| 9 | 120669183 | rs1399436 | 0.629 | 0.076 | 1.043 | -0.283 | -0.011 | 0.162 | 0.162 | 0.849 | 0.07 | 0.25 | 0.24 | 0.3 | 0.04 | 0.13 |
| 9 | 120678965 | rs7043830 | -0.409 | 0.279 | 0.753 | -0.141 | 0.002 | 0.004 | 0.004 | 3.085 | 0.72 | 0.72 | 0.63 | 0.59 | 0.61 | 0.49 |
| 9 | 120693160 | rs520466 | 0.852 | 0.303 | 0.596 | -0.258 | 0.014 | 0.036 | 0.036 | 3.171 | 0.51 | 0.39 | 0.53 | 0.36 | 0.43 | 0.3 |
| 9 | 120695091 | rs1514775 | 0.351 | 0.529 | 0.704 | -0.186 | 0.001 | 0.005 | 0.005 | 0 | 0.17 | 0.27 | 0.11 | 0.2 | 0.2 | 0.18 |
| 9 | 120704358 | rs1996215 | -0.559 | 0.559 | 0.544 | -0.144 | 0.002 | 0.004 | 0.004 | 4.786 | 0.72 | 0.72 | 0.65 | 0.56 | 0.63 | 0.52 |
| 9 | 120713296 | rs626584 | -0.48 | 0.505 | 0.379 | -0.161 | 0.006 | 0.011 | 0.011 | 3.295 | 0.71 | 0.69 | 0.64 | 0.56 | 0.61 | 0.51 |
| 9 | 120717586 | rs7028883 | -0.367 | 0.232 | 0.615 | 0.362 | -0.002 | 0.006 | 0.006 | 0.22 | 0.18 | 0.25 | 0.11 | 0.21 | 0.2 | 0.18 |
| 9 | 120728632 | rs10115978 | -0.148 | 0.321 | -0.181 | 0.405 | 0 | -0.003 | -0.003 | 0 | 0.17 | 0.12 | 0.1 | 0.21 | 0.2 | 0.18 |
| 9 | 120732125 | rs2722824 | 0.673 | 0.322 | -0.09 | 0.29 | -0.008 | 0.052 | 0.052 | 0.009 | 0.71 | 0.57 | 0.63 | 0.49 | 0.72 | 0.62 |

| | | | | | | | | | | | | | | | | |
|---|---|---|---|---|---|---|---|---|---|---|---|---|---|---|---|---|
| 9 | 120733712 | rs1930713 | -0.157 | 0.07 | -0.571 | -0.727 | 0.006 | 0.006 | 0.006 | 0 | 0.29 | 0.22 | 0.22 | 0.04 | 0.26 | 0.17 |
| 9 | 120737136 | rs2809398 | -0.733 | 0.219 | -0.825 | -0.787 | -0.002 | 0.012 | 0.012 | 0 | 0.63 | 0.48 | 0.6 | 0.51 | 0.66 | 0.6 |
| 9 | 120737349 | rs2416603 | 0.458 | 0.073 | -0.841 | -0.784 | -0.056 | -0.062 | -0.062 | 0 | 0.32 | 0.13 | 0.36 | 0.44 | 0.39 | 0.42 |
| 9 | 120739065 | rs955117 | 0.431 | 0.044 | -0.502 | -0.798 | -0.025 | -0.028 | -0.028 | 0 | 0.32 | 0.21 | 0.37 | 0.43 | 0.39 | 0.43 |
| 9 | 120745836 | rs2225600 | 0.608 | 0.05 | -0.422 | -0.815 | 0.001 | -0.011 | -0.011 | 0.383 | 0.28 | 0.27 | 0.22 | 0.07 | 0.25 | 0.17 |
| 9 | 120751155 | rs9408905 | -0.978 | -0.189 | -0.188 | -0.124 | 0.043 | 0.372 | 0.372 | 0.246 | 0.93 | 0.64 | 0.76 | 0.67 | 0.96 | 0.86 |
| 9 | 120756290 | rs17345248 | 1.231 | -0.526 | -0.597 | -0.459 | -0.091 | -0.002 | -0.002 | 0 | 0.26 | 0.04 | 0.17 | 0.17 | 0.32 | 0.23 |
| 9 | 120757370 | rs2245960 | 1.132 | -0.637 | -0.493 | -0.498 | 0.008 | 0.03 | 0.03 | 0 | 0.25 | 0.17 | 0.18 | 0.04 | 0.23 | 0.14 |
| 9 | 120763748 | rs1930697 | -0.509 | -0.449 | -0.372 | -0.227 | 0.009 | 0.032 | 0.032 | 0 | 0.68 | 0.59 | 0.61 | 0.49 | 0.69 | 0.63 |
| 9 | 120767777 | rs2806085 | -1.304 | -0.463 | 0.515 | -0.115 | 0.009 | 0.247 | 0.247 | 0 | 0.93 | 0.76 | 0.77 | 0.68 | 0.96 | 0.86 |
| 9 | 120769844 | rs2780246 | -0.742 | -0.464 | 0.462 | -0.115 | 0.048 | 0.393 | 0.393 | 0.485 | 0.92 | 0.58 | 0.76 | 0.68 | 0.96 | 0.86 |
| 9 | 120773585 | rs2806081 | -0.08 | -0.837 | -0.057 | -0.242 | -0.071 | 0.137 | 0.137 | 0.164 | 0.84 | 0.6 | 0.75 | 0.67 | 0.95 | 0.78 |
| 9 | 120786036 | rs2722841 | 0.52 | -0.713 | -0.693 | -0.724 | -0.012 | 0.134 | 0.134 | 0.341 | 0.52 | 0.26 | 0.38 | 0.21 | 0.53 | 0.37 |
| 9 | 120787025 | rs2809420 | 1.117 | -0.681 | -0.811 | -0.875 | -0.001 | -0.031 | -0.031 | 0.542 | 0.26 | 0.29 | 0.23 | 0.05 | 0.25 | 0.13 |
| 9 | 120795990 | rs2416604 | 0.637 | -0.535 | -0.741 | -0.706 | 0.037 | 0.037 | 0.037 | 0 | 0.53 | 0.26 | 0.49 | 0.62 | 0.64 | 0.61 |
| 9 | 120814952 | rs2809369 | -0.447 | -0.609 | -0.637 | -0.667 | -0.072 | 0.124 | 0.124 | 0.39 | 0.83 | 0.47 | 0.72 | 0.69 | 0.95 | 0.77 |
| 9 | 120831882 | rs2900142 | 1.095 | -0.373 | -0.794 | -0.558 | -0.021 | -0.018 | -0.018 | 0 | 0.3 | 0.23 | 0.37 | 0.43 | 0.41 | 0.38 |
| 9 | 120839644 | rs2722866 | 1.272 | -0.188 | -0.43 | -0.521 | 0.004 | 0.124 | 0.124 | 0 | 0.26 | 0.06 | 0.16 | 0.03 | 0.24 | 0.14 |
| 9 | 120841269 | rs2722865 | 0.623 | -0.182 | -0.398 | -0.488 | -0.011 | 0.042 | 0.042 | 0 | 0.3 | 0.23 | 0.23 | 0.07 | 0.31 | 0.16 |
| 9 | 120842790 | rs2252957 | 0.192 | -0.17 | -0.417 | -0.473 | 0.005 | -0.051 | -0.051 | 0 | 0.32 | 0.36 | 0.27 | 0.07 | 0.34 | 0.17 |
| 9 | 120851579 | rs2809380 | -0.192 | 0.095 | 0.033 | 0.21 | 0.103 | 0.334 | 0.334 | 1.41 | 0.89 | 0.51 | 0.68 | 0.67 | 0.89 | 0.85 |
| 9 | 120854939 | rs2722806 | -0.97 | 0.046 | -0.033 | 0.21 | 0.035 | 0.365 | 0.365 | 1.938 | 0.92 | 0.6 | 0.75 | 0.71 | 0.96 | 0.86 |
| 9 | 120862912 | rs2722814 | 0.491 | -0.075 | -0.569 | 0.188 | -0.002 | 0.055 | 0.055 | 2.522 | 0.81 | 0.33 | 0.65 | 0.7 | 0.87 | 0.83 |
| 9 | 120863737 | rs12349683 | 1.273 | 0.241 | -0.959 | 0.189 | -0.004 | -0.016 | -0.016 | 0 | 0.11 | 0.05 | 0.2 | 0.42 | 0.13 | 0.21 |
| 9 | 120869313 | rs2806110 | 0.385 | 0.261 | -0.777 | 0.241 | -0.002 | 0.057 | 0.057 | 2.522 | 0.81 | 0.33 | 0.65 | 0.7 | 0.87 | 0.83 |
| 9 | 120874982 | rs13299605 | 1.303 | 0.321 | -1.638 | -0.117 | -0.088 | -0.073 | -0.073 | 0 | 0.26 | 0.04 | 0.2 | 0.26 | 0.29 | 0.27 |
| 9 | 120889966 | rs2769164 | 1.249 | 0.017 | -2.192 | -0.719 | -0.016 | 0.067 | 0.067 | 1.231 | 0.18 | 0 | 0.07 | 0.08 | 0.15 | 0.23 |
| 9 | 120899084 | rs2769183 | 0.244 | -0.011 | -1.88 | -0.801 | -0.033 | 0.259 | 0.259 | 0.739 | 0.48 | 0.17 | 0.3 | 0.11 | 0.53 | 0.43 |
| 9 | 120913104 | rs2806117 | -0.867 | -0.042 | -0.643 | -0.119 | 0.028 | 0.225 | 0.225 | 0.668 | 0.9 | 0.59 | 0.72 | 0.75 | 0.92 | 0.82 |
| 9 | 120916741 | rs2025881 | 1.806 | -0.091 | -1.549 | 0.255 | -0.025 | -0.031 | -0.031 | 3.708 | 0.2 | 0.01 | 0.11 | 0.17 | 0.17 | 0.19 |
| 9 | 120922128 | rs10983899 | 1.138 | 0.167 | -1.451 | 0.062 | -0.017 | -0.005 | -0.005 | 0 | 0.25 | 0.17 | 0.18 | 0.21 | 0.25 | 0.23 |
| 9 | 120925960 | rs10818118 | 0.367 | 0.07 | -0.976 | -0.071 | -0.008 | -0.008 | -0.008 | 3.205 | 0.34 | 0.26 | 0.39 | 0.58 | 0.37 | 0.35 |
| 9 | 120929609 | rs2769186 | 0.795 | 0.169 | -1.718 | -0.5 | 0.012 | 0.067 | 0.067 | 2.314 | 0.26 | 0.07 | 0.14 | 0.16 | 0.24 | 0.36 |
| 9 | 120945445 | rs2769192 | 0.024 | 0.296 | -1.398 | -0.227 | -0.012 | 0.157 | 0.157 | 0.646 | 0.48 | 0.25 | 0.31 | 0.16 | 0.53 | 0.46 |
| 9 | 120953087 | rs913367 | 0.115 | 0.276 | -1.48 | -0.222 | -0.04 | 0.225 | 0.225 | 1.136 | 0.47 | 0.13 | 0.3 | 0.16 | 0.53 | 0.44 |
| 9 | 120962237 | rs12552190 | -0.109 | -0.002 | -1.352 | 0.109 | 0.023 | 0.067 | 0.067 | 0.399 | 0.52 | 0.32 | 0.35 | 0.35 | 0.57 | 0.49 |
| 9 | 120962282 | rs12551731 | 0.587 | 0.42 | -1.167 | 0.865 | -0.041 | -0.048 | -0.048 | 0 | 0.09 | 0.01 | 0.19 | 0.4 | 0.13 | 0.15 |
| 9 | 120967585 | rs1327797 | -0.066 | 0.259 | -1.123 | 0.257 | -0.007 | -0.005 | -0.005 | 0 | 0.3 | 0.19 | 0.21 | 0.35 | 0.3 | 0.4 |
| 9 | 120969775 | rs10818121 | -0.777 | -0.54 | -0.614 | 0.351 | -0.051 | 0.065 | 0.065 | 0.32 | 0.79 | 0.45 | 0.86 | 0.75 | 0.89 | 0.79 |
| 9 | 120970254 | rs1327798 | -1.023 | -0.537 | -0.552 | 0.323 | -0.034 | -0.084 | -0.084 | 0.729 | 0.81 | 0.6 | 0.88 | 0.97 | 0.89 | 0.82 |
| 9 | 120975586 | rs10983917 | 1.11 | -0.519 | -1.483 | 0.108 | -0.064 | 0.153 | 0.153 | 0 | 0.36 | 0.12 | 0.25 | 0 | 0.42 | 0.15 |
| 9 | 120978387 | rs10513326 | -0.017 | -0.568 | -1.458 | 0.383 | -0.056 | -0.151 | -0.151 | 2.079 | 0.27 | 0.04 | 0.48 | 0.55 | 0.3 | 0.47 |
| 9 | 120981380 | rs1571272 | 0.607 | -0.421 | -1.335 | -0.586 | 0.036 | 0.139 | 0.139 | 0 | 0.51 | 0.25 | 0.36 | 0.2 | 0.59 | 0.32 |
| 9 | 120985358 | rs10983923 | 0.842 | 0.222 | -1.331 | 0.059 | -0.03 | -0.056 | -0.056 | 0 | 0.1 | 0.01 | 0.4 | 0.44 | 0.13 | 0.1 |
| 9 | 120988194 | rs927953 | -0.424 | 0.011 | -1.121 | -1.193 | -0.021 | -0.028 | -0.028 | 0 | 0.65 | 0.51 | 0.77 | 0.76 | 0.72 | 0.64 |
| 9 | 120992769 | rs1410201 | 1.103 | 0.106 | -1.506 | -1.103 | -0.042 | -0.07 | -0.07 | 0 | 0.12 | 0.02 | 0.41 | 0.6 | 0.17 | 0.15 |
| 9 | 120994586 | rs10123927 | 0.629 | 0.114 | -1.147 | -1.401 | -0.044 | -0.001 | -0.001 | 0.941 | 0.33 | 0.04 | 0.49 | 0.74 | 0.3 | 0.48 |
| 9 | 121006917 | rs10983934 | 0.071 | 0.931 | -1.064 | -1.412 | -0.059 | 0.091 | 0.091 | 0 | 0.3 | 0.13 | 0.24 | 0.02 | 0.42 | 0.15 |
| 9 | 121008432 | rs10983936 | 0.08 | 0.942 | -1.29 | -1.398 | -0.085 | 0.138 | 0.138 | 0 | 0.3 | 0.09 | 0.24 | 0.02 | 0.42 | 0.15 |
| 9 | 121009165 | rs7867162 | -0.036 | 0.707 | -0.846 | -0.943 | -0.001 | -0.063 | -0.063 | 0 | 0.51 | 0.51 | 0.63 | 0.95 | 0.47 | 0.66 |
| 9 | 121010969 | rs10983944 | -0.032 | 0.607 | -0.893 | -0.268 | -0.003 | 0.018 | 0.018 | 0 | 0.42 | 0.29 | 0.44 | 0.34 | 0.42 | 0.45 |
| 9 | 121010987 | rs4836716 | -0.512 | 0.289 | -0.296 | 0.355 | -0.01 | 0.114 | 0.114 | 0 | 0.87 | 0.73 | 0.84 | 0.66 | 0.91 | 0.87 |
| 9 | 121016602 | rs10983945 | 0.372 | 0.22 | -0.142 | 0.308 | -0.001 | 0.003 | 0.003 | 0 | 0.43 | 0.36 | 0.45 | 0.34 | 0.42 | 0.45 |
| 9 | 121036075 | rs10983950 | 0.729 | -0.018 | -0.513 | 0.073 | 0.048 | 0.052 | 0.052 | 0 | 0.24 | 0.04 | 0.1 | 0.14 | 0.16 | 0.22 |
| 9 | 121037997 | rs1927320 | -0.768 | -0.388 | 0.055 | -0.06 | -0.016 | 0.121 | 0.121 | 0 | 0.85 | 0.69 | 0.82 | 0.67 | 0.89 | 0.87 |
| 9 | 121041753 | rs954070 | 1.201 | -0.395 | 0.081 | -0.104 | -0.022 | 0.036 | 0.036 | 0.078 | 0.41 | 0.28 | 0.36 | 0.32 | 0.47 | 0.41 |
| 9 | 121045042 | rs12236645 | 1.203 | -0.629 | -0.114 | -0.187 | 0.048 | 0.039 | 0.039 | 0 | 0.19 | 0.01 | 0.05 | 0.11 | 0.1 | 0.14 |
| 9 | 121051077 | rs10759964 | 1.561 | -0.66 | -0.343 | -0.745 | 0.01 | 0.003 | 0.003 | 0 | 0.29 | 0.38 | 0.31 | 0.31 | 0.35 | 0.36 |
| 9 | 121069222 | rs10983957 | 1.784 | -0.707 | -0.521 | -0.814 | -0.072 | -0.026 | -0.026 | 0 | 0.17 | 0.07 | 0.33 | 0.21 | 0.29 | 0.24 |
| 9 | 121071032 | rs7874874 | 0.543 | -0.96 | -0.359 | -0.833 | -0.071 | -0.094 | -0.094 | 0.053 | 0.3 | 0.45 | 0.13 | 0.11 | 0.15 | 0.24 |
| 9 | 121072539 | rs7025261 | -0.44 | -0.969 | -0.356 | -0.829 | 0.007 | -0.016 | -0.016 | 0 | 0.75 | 0.78 | 0.76 | 0.64 | 0.79 | 0.84 |
| 9 | 121074020 | rs10513328 | 0.57 | -0.619 | -0.431 | -0.679 | 0.052 | -0.016 | -0.016 | 0.019 | 0.45 | 0.31 | 0.63 | 0.53 | 0.63 | 0.61 |
| 9 | 121075273 | rs7862653 | 0.108 | -0.314 | -0.4 | -0.278 | 0.012 | 0.01 | 0.01 | 0 | 0.3 | 0.35 | 0.37 | 0.38 | 0.35 | 0.29 |
| 9 | 121076460 | rs12552250 | 1.528 | -0.347 | -0.518 | -0.491 | -0.108 | -0.052 | -0.052 | 0 | 0.19 | 0.03 | 0.34 | 0.22 | 0.29 | 0.24 |
| 9 | 121080374 | rs7873210 | 1.75 | -0.57 | -0.386 | -0.486 | -0.014 | -0.002 | -0.002 | 0 | 0.23 | 0.2 | 0.28 | 0.29 | 0.3 | 0.28 |
| 9 | 121080656 | rs12345715 | 0.487 | -0.679 | -0.275 | -0.562 | 0.057 | 0.023 | 0.023 | 0 | 0.07 | 0.3 | 0.04 | 0.04 | 0.07 | 0.07 |
| 9 | 121081100 | rs17364704 | 0.79 | -0.966 | -0.044 | -0.672 | 0.005 | -0.009 | -0.009 | 0.1 | 0.3 | 0.25 | 0.12 | 0.08 | 0.17 | 0.26 |
| 9 | 121081314 | rs7047454 | -0.64 | -0.874 | -0.278 | -0.565 | 0.001 | 0.004 | 0.004 | 0 | 0.8 | 0.82 | 0.78 | 0.63 | 0.83 | 0.84 |
| 9 | 121090791 | rs1418495 | 0.492 | -0.638 | -0.359 | -0.55 | 0.023 | -0.009 | -0.009 | 0 | 0.44 | 0.28 | 0.62 | 0.51 | 0.6 | 0.52 |

| | | | | | | | | | | | | | | | | |
|---|---|---|---|---|---|---|---|---|---|---|---|---|---|---|---|---|
| 9 | 121094973 | rs7866799 | 0.61 | -0.625 | -0.483 | -0.543 | 0.014 | -0.023 | -0.023 | 0 | 0.43 | 0.24 | 0.62 | 0.51 | 0.59 | 0.52 |
| 9 | 121110670 | rs10983993 | 1.874 | -0.621 | -0.649 | -0.634 | -0.094 | -0.045 | -0.045 | 0 | 0.22 | 0.02 | 0.26 | 0.25 | 0.31 | 0.27 |
| 9 | 121111790 | rs7036676 | -0.424 | -0.795 | -0.54 | -0.665 | -0.042 | 0.011 | 0.011 | 0 | 0.62 | 0.81 | 0.45 | 0.39 | 0.55 | 0.6 |
| 9 | 121125496 | rs10124472 | 0.09 | -0.866 | -0.115 | -0.718 | 0.064 | 0.02 | 0.02 | 0.005 | 0.08 | 0.35 | 0.04 | 0.04 | 0.07 | 0.07 |
| 9 | 121127354 | rs3893408 | 0.149 | -0.864 | -0.114 | -0.717 | 0.068 | 0.024 | 0.024 | 0 | 0.08 | 0.35 | 0.05 | 0.04 | 0.07 | 0.07 |
| 9 | 121133696 | rs10983999 | 0.69 | -1.015 | 0.044 | -0.803 | 0.057 | 0.072 | 0.072 | 0 | 0.29 | 0.14 | 0.1 | 0.1 | 0.16 | 0.26 |
| 9 | 121136695 | rs7864888 | -0.377 | -1.009 | 0.11 | -0.809 | 0.003 | -0.003 | -0.003 | 0 | 0.61 | 0.61 | 0.46 | 0.39 | 0.54 | 0.6 |
| 9 | 121137235 | rs17366047 | 0.865 | -1.11 | 0.081 | -0.869 | 0.072 | 0.12 | 0.12 | 0 | 0.28 | 0.06 | 0.1 | 0.1 | 0.16 | 0.25 |
| 9 | 121139437 | rs4593622 | -0.462 | -1.036 | 0.157 | -0.862 | -0.013 | 0.006 | 0.006 | 0 | 0.61 | 0.68 | 0.46 | 0.39 | 0.54 | 0.59 |
| 9 | 121140638 | rs10513331 | 0.471 | -0.694 | 0.229 | -0.618 | 0.023 | -0.013 | -0.013 | 0 | 0.32 | 0.45 | 0.35 | 0.29 | 0.38 | 0.34 |
| 9 | 121149060 | rs7855921 | -0.534 | -1.11 | -0.04 | -0.592 | -0.001 | 0.001 | 0.001 | 0 | 0.71 | 0.66 | 0.57 | 0.37 | 0.68 | 0.6 |
| 9 | 121151094 | rs12350323 | 1.482 | -1.137 | -0.052 | -0.526 | -0.017 | -0.028 | -0.028 | 0.294 | 0.17 | 0.15 | 0.29 | 0.26 | 0.24 | 0.32 |
| 9 | 121157043 | rs1342500 | 0.305 | -1.541 | 0.351 | -1.313 | -0.031 | 0.076 | 0.076 | 0 | 0.73 | 0.51 | 0.78 | 0.58 | 0.8 | 0.71 |
| 9 | 121158641 | rs1418514 | -1.397 | -1.535 | 0.289 | -1.706 | -0.017 | 0.112 | 0.112 | 0 | 0.88 | 0.83 | 0.87 | 0.61 | 0.92 | 0.92 |
| 9 | 121159542 | rs12350098 | -0.183 | -1.541 | 0.305 | -1.718 | 0.001 | -0.076 | -0.076 | 1.16 | 0.15 | 0.33 | 0.08 | 0.05 | 0.12 | 0.2 |
| 9 | 121162755 | rs10818145 | 0.87 | -1.693 | -0.76 | -1.963 | -0.044 | 0.046 | 0.046 | 0.934 | 0.36 | 0.18 | 0.43 | 0.24 | 0.43 | 0.25 |
| 9 | 121163842 | rs16907082 | 1.834 | -1.695 | -0.807 | -1.893 | -0.027 | -0.034 | -0.034 | 0 | 0.16 | 0.12 | 0.29 | 0.24 | 0.25 | 0.33 |
| 9 | 121168521 | rs10818147 | 0.34 | -1.69 | -0.998 | -1.925 | 0.003 | 0.028 | 0.028 | 0.035 | 0.57 | 0.36 | 0.49 | 0.32 | 0.54 | 0.38 |
| 9 | 121171878 | rs4836723 | 0.127 | -1.693 | -0.977 | -1.903 | -0.026 | 0.061 | 0.061 | 0 | 0.73 | 0.49 | 0.78 | 0.56 | 0.8 | 0.71 |
| 9 | 121173877 | rs1935624 | 1.621 | -1.503 | -1.185 | -2.138 | -0.098 | -0.009 | -0.009 | 0.037 | 0.26 | 0.05 | 0.36 | 0.24 | 0.34 | 0.21 |
| 9 | 121177842 | rs1935621 | 0.167 | -1.373 | -0.922 | -2.031 | 0.004 | 0.028 | 0.028 | 0.037 | 0.57 | 0.36 | 0.49 | 0.32 | 0.54 | 0.38 |
| 9 | 121179496 | rs4401938 | 0.469 | -1.215 | -0.682 | -1.863 | 0.02 | 0.056 | 0.056 | 2.215 | 0.25 | 0.14 | 0.08 | 0.09 | 0.18 | 0.22 |
| 9 | 121182554 | rs10818151 | 0.543 | -1.273 | -0.882 | -1.925 | -0.029 | 0.028 | 0.028 | 0.266 | 0.41 | 0.22 | 0.55 | 0.67 | 0.46 | 0.33 |
| 9 | 121183669 | rs10984022 | 1.848 | -0.634 | -1.129 | -2.067 | -0.042 | -0.056 | -0.056 | 0.05 | 0.09 | 0 | 0.15 | 0.23 | 0.13 | 0.15 |
| 9 | 121190095 | rs17367380 | 0.466 | -0.998 | -0.783 | -2.071 | 0.021 | 0.094 | 0.094 | 2.621 | 0.25 | 0.06 | 0.1 | 0.08 | 0.17 | 0.21 |
| 9 | 121191140 | rs10818153 | -0.095 | -1.071 | -0.376 | -2.14 | 0.003 | -0.021 | -0.021 | 0 | 0.32 | 0.44 | 0.32 | 0.26 | 0.4 | 0.48 |
| 9 | 121195316 | rs10435881 | 1.134 | -1.001 | -0.445 | -2.102 | -0.089 | 0.005 | 0.005 | 0.326 | 0.37 | 0.1 | 0.54 | 0.64 | 0.42 | 0.29 |
| 9 | 121201702 | rs1891442 | 0.285 | -0.335 | 0.329 | -1.454 | 0 | -0.021 | -0.021 | 0 | 0.14 | 0.11 | 0.11 | 0 | 0.15 | 0.07 |
| 9 | 121213149 | rs7022505 | 0.513 | -0.397 | 0.266 | -1.448 | 0.014 | 0.018 | 0.018 | 0.915 | 0.15 | 0.09 | 0.05 | 0.09 | 0.1 | 0.19 |
| 9 | 121213469 | rs1984268 | 1.376 | -0.327 | 0.296 | -1.437 | -0.022 | 0.069 | 0.069 | 1.02 | 0.47 | 0.12 | 0.6 | 0.64 | 0.49 | 0.31 |
| 9 | 121218719 | rs1418501 | -1.399 | -0.091 | 0.034 | -1.328 | -0.001 | -0.002 | -0.002 | 0 | 0.5 | 0.55 | 0.38 | 0.35 | 0.51 | 0.69 |
| 9 | 121241085 | rs7030074 | -1.422 | -0.348 | -0.109 | -1.269 | -0.001 | -0.003 | -0.003 | 0 | 0.49 | 0.57 | 0.38 | 0.32 | 0.5 | 0.67 |
| 9 | 121243073 | rs7022160 | -1.19 | -0.736 | -0.105 | -1.312 | 0.006 | 0.001 | 0.001 | 0.388 | 0.33 | 0.35 | 0.17 | 0.1 | 0.26 | 0.26 |
| 9 | 121244869 | rs7858444 | 0.097 | -0.725 | -0.107 | -1.315 | -0.003 | -0.044 | -0.044 | 0 | 0.16 | 0.21 | 0.12 | 0.01 | 0.17 | 0.07 |
| 9 | 121253842 | rs10818161 | 0.003 | -0.724 | -0.311 | -1.292 | 0.018 | 0.019 | 0.019 | 0.795 | 0.17 | 0.15 | 0.05 | 0.1 | 0.09 | 0.21 |
| 9 | 121254222 | rs12342183 | -1.176 | -0.65 | -0.279 | -1.24 | 0 | -0.015 | -0.015 | 0 | 0.34 | 0.38 | 0.18 | 0.1 | 0.26 | 0.28 |
| 9 | 121263967 | rs10984054 | 0.636 | -0.35 | -0.386 | -0.714 | 0.012 | -0.003 | -0.003 | 0 | 0.1 | 0.02 | 0.17 | 0.41 | 0.08 | 0.08 |
| 9 | 121277542 | rs7029465 | 0.412 | -0.192 | -0.221 | -0.82 | -0.019 | -0.008 | -0.008 | 6.585 | 0.12 | 0.2 | 0.04 | 0.09 | 0.05 | 0.16 |
| 9 | 121282616 | rs7864314 | -0.922 | -0.15 | -0.188 | -1.079 | 0 | -0.007 | -0.007 | 0 | 0.45 | 0.59 | 0.38 | 0.3 | 0.48 | 0.64 |
| 9 | 121284915 | rs10984077 | 0.763 | -0.134 | -0.259 | -1.045 | -0.012 | -0.005 | -0.005 | 0 | 0.12 | 0.15 | 0.03 | 0.09 | 0.05 | 0.16 |
| 9 | 121288997 | rs10984078 | 0.949 | 0.009 | -0.448 | -1.052 | -0.044 | -0.011 | -0.011 | 0 | 0.18 | 0.1 | 0.23 | 0.2 | 0.28 | 0.4 |
| 9 | 121301720 | rs10818184 | 0.589 | 0.011 | -0.413 | -1.052 | -0.026 | 0.003 | 0.003 | 0 | 0.21 | 0.18 | 0.24 | 0.2 | 0.28 | 0.43 |
| 9 | 121312200 | rs12378763 | -1.12 | 0.102 | -0.338 | -1.039 | 0.002 | 0.042 | 0.042 | 0.515 | 0.48 | 0.67 | 0.39 | 0.29 | 0.5 | 0.63 |
| 9 | 121312632 | rs7848936 | 0.872 | 0.109 | -0.785 | -1.07 | -0.021 | -0.009 | -0.009 | 0 | 0.12 | 0.19 | 0.04 | 0.09 | 0.06 | 0.16 |
| 9 | 121312824 | rs7848982 | -1.492 | 0.646 | -1.574 | -1.197 | -0.031 | 0.052 | 0.052 | 0.155 | 0.64 | 0.81 | 0.58 | 0.74 | 0.62 | 0.74 |
| 9 | 121314808 | rs7857107 | -1.377 | 1.487 | -1.446 | -0.966 | -0.002 | 0.003 | 0.003 | 0.015 | 0.35 | 0.39 | 0.34 | 0.2 | 0.44 | 0.48 |
| 9 | 121317025 | rs9409210 | -1.425 | 0.896 | -1.359 | -0.537 | -0.006 | 0.012 | 0.012 | 0 | 0.33 | 0.35 | 0.33 | 0.2 | 0.44 | 0.48 |
| 9 | 121318471 | rs10116305 | -0.631 | 0.348 | -1.592 | -0.297 | -0.001 | -0.029 | -0.029 | 0 | 0.13 | 0.14 | 0.1 | 0 | 0.16 | 0.08 |
| 9 | 121320294 | rs1360810 | 0.359 | 0.582 | -1.681 | 0.281 | -0.019 | -0.016 | -0.016 | 0 | 0.83 | 0.89 | 0.8 | 0.78 | 0.78 | 0.67 |
| 9 | 121323568 | rs11792957 | -0.571 | -0.488 | -2.128 | 0.016 | 0.061 | 0.011 | 0.011 | 0 | 0.24 | 0.04 | 0.32 | 0.22 | 0.14 | 0.17 |
| 9 | 121325521 | rs10818187 | 0.259 | 0.018 | -2.039 | -1.21 | -0.014 | 0.01 | 0.01 | 0 | 0.42 | 0.27 | 0.29 | 0.03 | 0.5 | 0.24 |
| 9 | 121326409 | rs7855145 | 0.153 | -0.19 | -1.389 | -1.455 | 0.002 | 0.02 | 0.02 | 1.494 | 0.49 | 0.38 | 0.32 | 0.15 | 0.51 | 0.29 |
| 9 | 121327323 | rs1335258 | -0.356 | -0.949 | -1.389 | -1.056 | 0.035 | 0.009 | 0.009 | 0 | 0.73 | 0.55 | 0.47 | 0.38 | 0.65 | 0.56 |
| 9 | 121327603 | rs7469341 | -0.288 | -1.081 | -1.122 | -1.106 | -0.01 | -0.002 | -0.002 | 0 | 0.26 | 0.28 | 0.15 | 0.22 | 0.15 | 0.27 |
| 9 | 121328526 | rs10759982 | -0.577 | -0.812 | -0.782 | -0.326 | 0.03 | 0.111 | 0.111 | 0 | 0.8 | 0.66 | 0.67 | 0.54 | 0.72 | 0.6 |
| 9 | 121334810 | rs10984091 | 1.389 | -0.805 | -0.851 | -0.364 | -0.004 | 0.085 | 0.085 | 0.781 | 0.47 | 0.27 | 0.35 | 0.17 | 0.51 | 0.27 |
| 9 | 121342907 | rs11790829 | 1.324 | -0.84 | -0.868 | -0.349 | -0.002 | 0.091 | 0.091 | 0.629 | 0.47 | 0.27 | 0.35 | 0.17 | 0.51 | 0.27 |
| 9 | 121344767 | rs1335255 | 0.181 | -1.137 | -0.974 | -0.473 | 0 | -0.014 | -0.014 | 0 | 0.1 | 0.11 | 0.07 | 0.03 | 0.1 | 0.02 |
| 9 | 121351852 | rs7864368 | 0.14 | -1.15 | -1.018 | -0.429 | 0.008 | -0.004 | -0.004 | 0 | 0.15 | 0.12 | 0.09 | 0.19 | 0.05 | 0.26 |
| 9 | 121351909 | rs7037161 | 0.351 | -0.934 | -0.584 | -0.45 | 0.027 | 0.086 | 0.086 | 0 | 0.08 | 0.19 | 0.2 | 0.17 | 0.07 | 0.06 |
| 9 | 121359692 | rs1414097 | 0.868 | -1.002 | -0.633 | -0.445 | 0.002 | 0.003 | 0.003 | 1.212 | 0.55 | 0.4 | 0.55 | 0.34 | 0.58 | 0.33 |
| 9 | 121364552 | rs10984116 | -1.008 | -1.442 | -0.811 | 0.471 | 0.031 | -0.001 | -0.001 | 0.244 | 0.17 | 0.11 | 0.06 | 0.14 | 0.1 | 0.1 |
| 9 | 121369402 | rs10124018 | 0.284 | -0.36 | -0.092 | 0.91 | 0.055 | 0.043 | 0.043 | 0 | 0.76 | 0.59 | 0.66 | 0.71 | 0.69 | 0.85 |
| 9 | 121370958 | rs1335246 | 1.261 | -0.784 | 0.144 | 1.167 | -0.02 | -0.079 | -0.079 | 0.015 | 0.65 | 0.16 | 0.54 | 0.52 | 0.67 | 0.7 |
| 9 | 121372582 | rs2416662 | -0.605 | -0.574 | 0.395 | 1.203 | 0.001 | 0 | 0 | 0 | 0.84 | 0.8 | 0.71 | 0.78 | 0.76 | 0.93 |
| 9 | 121375928 | rs16907415 | 0.106 | -0.75 | 0.375 | 1.063 | 0.021 | 0.038 | 0.038 | 0 | 0.09 | 0.23 | 0.05 | 0.08 | 0.08 | 0.08 |
| 9 | 121385397 | rs4301511 | 1.783 | -0.259 | -0.091 | 1.366 | 0.016 | 0.166 | 0.166 | 0.201 | 0.52 | 0.16 | 0.4 | 0.25 | 0.44 | 0.48 |
| 9 | 121387146 | rs7850923 | -0.476 | -0.036 | 0.142 | 1.565 | 0.015 | 0.051 | 0.051 | 0 | 0.22 | 0.27 | 0.19 | 0.37 | 0.29 | 0.27 |
| 9 | 121406435 | rs4837567 | -1.262 | -0.048 | 0.304 | 1.646 | 0.009 | 0.05 | 0.05 | 0.019 | 0.27 | 0.34 | 0.29 | 0.55 | 0.32 | 0.44 |

| | | | | | | | | | | | | | | | | |
|---|---|---|---|---|---|---|---|---|---|---|---|---|---|---|---|---|
| 9 | 121406910 | rs1508377 | -0.906 | -0.046 | 0.279 | 1.635 | -0.002 | -0.001 | -0.001 | 0 | 0.83 | 0.81 | 0.71 | 0.81 | 0.75 | 0.92 |
| 9 | 121407046 | rs1508378 | -0.941 | -0.045 | 0.252 | 1.649 | -0.006 | -0.003 | -0.003 | 0.528 | 0.83 | 0.81 | 0.71 | 0.81 | 0.75 | 0.92 |
| 9 | 121409332 | rs10118699 | 0.449 | -0.099 | 0.205 | 1.681 | -0.001 | 0.028 | 0.028 | 0 | 0.05 | 0.06 | 0.1 | 0.19 | 0.02 | 0.17 |
| 9 | 121417973 | rs4837570 | -1.198 | -0.092 | 0.371 | 1.571 | 0.008 | 0.055 | 0.055 | 0.028 | 0.25 | 0.53 | 0.2 | 0.37 | 0.29 | 0.27 |
| 9 | 121418763 | rs10759994 | 0.4 | 0.013 | -0.05 | 1.718 | -0.035 | -0.055 | -0.055 | 0.044 | 0.16 | 0.03 | 0.11 | 0.26 | 0.23 | 0.21 |
| 9 | 121424346 | rs1335259 | 0.401 | 0.298 | 0.436 | 1.86 | 0.005 | 0.012 | 0.012 | 0 | 0.73 | 0.33 | 0.57 | 0.62 | 0.72 | 0.74 |
| 9 | 121427985 | rs10984153 | 0.119 | 0.287 | 0.522 | 1.895 | -0.005 | 0.035 | 0.035 | 0 | 0.06 | 0.08 | 0.11 | 0.21 | 0.02 | 0.16 |
| 9 | 121428389 | rs1414096 | 0.939 | 0.071 | 0.447 | 1.647 | 0.001 | -0.006 | -0.006 | 0 | 0.54 | 0.47 | 0.46 | 0.36 | 0.49 | 0.52 |
| 9 | 121442207 | rs10759998 | -0.053 | 0.071 | -0.317 | 1.739 | 0.006 | -0.055 | -0.055 | 0 | 0.27 | 0.13 | 0.22 | 0.45 | 0.25 | 0.38 |
| 9 | 121450688 | rs2416680 | -0.463 | -0.012 | -0.17 | 1.816 | -0.013 | 0.02 | 0.02 | 0 | 0.3 | 0.38 | 0.25 | 0.57 | 0.25 | 0.38 |
| 9 | 121452706 | rs10984174 | 0.699 | -0.023 | -0.149 | 1.655 | 0.001 | -0.018 | -0.018 | 0 | 0.21 | 0.08 | 0.12 | 0.24 | 0.23 | 0.21 |
| 9 | 121452775 | rs10116611 | -1.073 | 0.088 | -0.227 | 1.276 | 0.05 | -0.073 | -0.073 | 0.11 | 0.82 | 0.68 | 0.69 | 0.93 | 0.75 | 0.91 |
| 9 | 121456562 | rs4836729 | 0.839 | 0.136 | -1.296 | 1.41 | -0.019 | 0.015 | 0.015 | 0 | 0.53 | 0.26 | 0.46 | 0.39 | 0.52 | 0.53 |
| 9 | 121458762 | rs10818219 | 0.003 | 0.124 | -1.375 | 1.394 | 0.012 | -0.043 | -0.043 | 0 | 0.28 | 0.15 | 0.22 | 0.45 | 0.24 | 0.38 |
| 9 | 121463210 | rs13288491 | 0.515 | 0.386 | -0.949 | 1.407 | -0.009 | -0.017 | -0.017 | 1.342 | 0.07 | 0.03 | 0.07 | 0.11 | 0.09 | 0.1 |
| 9 | 121463496 | rs1355034 | -0.235 | 0.373 | -0.822 | 0.999 | 0.05 | -0.018 | -0.018 | 0 | 0.79 | 0.48 | 0.68 | 0.84 | 0.76 | 0.9 |
| 9 | 121467481 | rs10513341 | 0.356 | -0.607 | -1.869 | 1.173 | 0.046 | -0.013 | -0.013 | 0.006 | 0.3 | 0.1 | 0.2 | 0.27 | 0.22 | 0.29 |
| 9 | 121467568 | rs10984184 | 0.481 | -0.649 | -2.094 | 1.231 | -0.003 | -0.058 | -0.058 | 0 | 0.24 | 0.09 | 0.22 | 0.32 | 0.24 | 0.28 |
| 9 | 121468282 | rs3750473 | -0.04 | -0.903 | -1.331 | 0.782 | 0.048 | -0.011 | -0.011 | 0 | 0.3 | 0.1 | 0.2 | 0.27 | 0.22 | 0.29 |
| 9 | 121469510 | rs16907545 | -1.118 | -0.096 | -0.166 | 0.841 | -0.013 | -0.008 | -0.008 | 3.674 | 0.13 | 0.22 | 0.05 | 0.12 | 0.08 | 0.08 |
| 9 | 121475729 | rs10984189 | -0.697 | -0.23 | -0.179 | 1.113 | -0.023 | 0.039 | 0.039 | 0 | 0.33 | 0.45 | 0.23 | 0.56 | 0.23 | 0.41 |
| 9 | 121482333 | rs10513342 | -0.31 | -0.277 | 0.108 | 1.531 | -0.049 | -0.109 | -0.109 | 0 | 0.28 | 0.03 | 0.21 | 0.41 | 0.26 | 0.41 |
| 9 | 121482477 | rs10818225 | -0.454 | -0.263 | 0.144 | 1.53 | -0.032 | -0.098 | -0.098 | 0 | 0.29 | 0.05 | 0.21 | 0.41 | 0.26 | 0.42 |
| 9 | 121494898 | rs1508389 | -0.617 | -0.326 | 0.242 | 1.537 | 0.01 | -0.037 | -0.037 | 0 | 0.3 | 0.21 | 0.21 | 0.56 | 0.23 | 0.41 |
| 9 | 121501251 | rs965853 | -1.426 | -0.264 | -0.234 | 1.174 | -0.072 | 0.148 | 0.148 | 0.602 | 0.57 | 0.89 | 0.53 | 0.79 | 0.5 | 0.71 |
| 9 | 121502691 | rs4433214 | -0.256 | -0.249 | -0.131 | 1.164 | -0.001 | -0.001 | -0.001 | 0 | 0.28 | 0.7 | 0.32 | 0.22 | 0.26 | 0.3 |
| 9 | 121506034 | rs7868904 | -0.116 | -0.267 | -0.148 | 1.247 | 0 | 0.088 | 0.088 | 0.367 | 0.26 | 0.57 | 0.31 | 0.39 | 0.24 | 0.31 |
| 9 | 121510009 | rs7021488 | 0.038 | -0.261 | -0.216 | 1.104 | 0.022 | 0.011 | 0.011 | 0 | 0.26 | 0.51 | 0.32 | 0.25 | 0.28 | 0.3 |
| 9 | 121522885 | rs9632909 | -0.156 | -0.274 | -0.137 | 1.144 | 0.01 | 0.049 | 0.049 | 0.123 | 0.27 | 0.59 | 0.33 | 0.38 | 0.28 | 0.36 |
| 9 | 121524247 | rs10984217 | 0.187 | -0.327 | -0.287 | 1.197 | -0.02 | -0.088 | -0.088 | 3.343 | 0.29 | 0.05 | 0.18 | 0.37 | 0.23 | 0.34 |
| 9 | 121524285 | rs7029069 | 0.168 | -0.305 | -0.262 | 1.131 | 0.022 | 0.006 | 0.006 | 0 | 0.26 | 0.48 | 0.32 | 0.23 | 0.28 | 0.35 |
| 9 | 121525411 | rs2136062 | -1.145 | -0.335 | -0.246 | 1.114 | -0.001 | 0.021 | 0.021 | 0 | 0.56 | 0.64 | 0.51 | 0.75 | 0.51 | 0.7 |
| 9 | 121529204 | rs2416686 | 0.148 | -0.232 | -0.754 | 1.017 | 0.001 | 0.005 | 0.005 | 1 | 0.71 | 0.22 | 0.64 | 0.77 | 0.73 | 0.65 |
| 9 | 121535396 | rs2025756 | -0.977 | -0.693 | 0.059 | 1.341 | 0.002 | -0.003 | -0.003 | 0 | 0.34 | 0.32 | 0.2 | 0.41 | 0.26 | 0.34 |
| 9 | 121536509 | rs1326628 | -0.765 | -0.62 | -0.501 | 1.381 | 0.013 | -0.025 | -0.025 | 0 | 0.33 | 0.25 | 0.19 | 0.41 | 0.26 | 0.34 |
| 9 | 121539063 | rs10984224 | -0.384 | -0.298 | -0.821 | 1.558 | -0.016 | -0.072 | -0.072 | 0 | 0.28 | 0.1 | 0.17 | 0.41 | 0.25 | 0.33 |
| 9 | 121545222 | rs2900156 | -0.307 | -0.361 | -0.673 | 1.863 | 0.004 | -0.032 | -0.032 | 0.41 | 0.26 | 0.19 | 0.17 | 0.4 | 0.22 | 0.33 |
| 9 | 121545457 | rs1326624 | -0.393 | -0.372 | -0.933 | 1.646 | -0.005 | -0.057 | -0.057 | 0 | 0.28 | 0.1 | 0.18 | 0.64 | 0.24 | 0.33 |
| 9 | 121546954 | rs1571091 | 0.503 | -0.906 | 0.14 | 1.603 | -0.037 | 0.003 | 0.003 | 2.578 | 0.72 | 0.55 | 0.72 | 0.72 | 0.82 | 0.61 |
| 9 | 121547660 | rs943259 | 1.199 | -0.967 | -0.38 | 1.796 | -0.075 | 0.095 | 0.095 | 0 | 0.42 | 0.13 | 0.5 | 0.22 | 0.56 | 0.29 |
| 9 | 121559353 | rs10984229 | 0.011 | -0.991 | -0.076 | 1.741 | -0.048 | -0.118 | -0.118 | 0 | 0.29 | 0.05 | 0.22 | 0.47 | 0.28 | 0.32 |
| 9 | 121559990 | rs1409550 | 0.037 | -1.065 | 0.209 | 2.033 | -0.038 | 0.016 | 0.016 | 2.551 | 0.72 | 0.55 | 0.72 | 0.69 | 0.82 | 0.61 |
| 9 | 121577481 | rs7025578 | -0.239 | -1.224 | 0.798 | 2.24 | 0.004 | 0.05 | 0.05 | 0 | 0.28 | 0.36 | 0.22 | 0.47 | 0.26 | 0.3 |
| 9 | 121588193 | rs10984245 | 0.109 | -1.273 | 0.881 | 2.278 | -0.005 | -0.028 | -0.028 | 0.01 | 0.25 | 0.15 | 0.19 | 0.46 | 0.26 | 0.29 |
| 9 | 121593535 | rs10818242 | -0.219 | -1.463 | 1.649 | 2.274 | 0.005 | -0.003 | -0.003 | 0.039 | 0.2 | 0.16 | 0.17 | 0.22 | 0.12 | 0.29 |
| 9 | 121597614 | rs1359679 | -0.603 | -1.446 | 1.536 | 2.304 | -0.011 | 0.048 | 0.048 | 0 | 0.32 | 0.47 | 0.26 | 0.5 | 0.27 | 0.31 |
| 9 | 121624958 | rs10818249 | -1.076 | -1.46 | 1.492 | 2.255 | 0.018 | 0.06 | 0.06 | 0 | 0.46 | 0.18 | 0.37 | 0.7 | 0.38 | 0.57 |
| 9 | 121641891 | rs1326631 | -0.159 | -1.058 | 1.214 | 1.97 | -0.007 | -0.05 | -0.05 | 1.84 | 0.28 | 0.12 | 0.19 | 0.48 | 0.27 | 0.31 |
| 9 | 121645267 | rs7853689 | -0.27 | -1.055 | 1.216 | 1.972 | -0.001 | -0.023 | -0.023 | 2.023 | 0.29 | 0.21 | 0.2 | 0.48 | 0.27 | 0.31 |
| 9 | 121667815 | rs4837588 | -0.495 | -1.479 | 1.402 | 1.131 | -0.012 | -0.079 | -0.079 | 0 | 0.33 | 0.47 | 0.13 | 0.15 | 0.24 | 0.29 |
| 9 | 121670536 | rs10984298 | -0.013 | -1.2 | 0.759 | 1.226 | -0.001 | -0.003 | -0.003 | 0 | 0.29 | 0.25 | 0.11 | 0.13 | 0.24 | 0.29 |
| 9 | 121698714 | rs12237612 | -0.385 | -1.163 | 0.314 | 1.585 | 0.019 | -0.099 | -0.099 | 0 | 0.16 | 0 | 0.11 | 0.28 | 0.11 | 0.22 |
| 9 | 121711420 | rs10984333 | 0.046 | -1.019 | 0.693 | 0.96 | -0.008 | 0.043 | 0.043 | 0.302 | 0.28 | 0.14 | 0.11 | 0.13 | 0.24 | 0.3 |
| 9 | 121719114 | rs10984339 | -0.646 | -1.397 | 0.882 | 1.399 | -0.01 | 0.024 | 0.024 | 0 | 0.16 | 0.15 | 0.1 | 0.33 | 0.08 | 0.17 |
| 9 | 121724665 | rs12376255 | 0.324 | -1.354 | 0.714 | 0.517 | -0.003 | 0.048 | 0.048 | 0 | 0.26 | 0.08 | 0.1 | 0.16 | 0.21 | 0.29 |
| 9 | 121724759 | rs2818326 | -1.733 | -1.54 | 1.361 | 0.509 | 0.068 | 0.055 | 0.055 | 0 | 0.5 | 0.69 | 0.27 | 0.38 | 0.33 | 0.5 |
| 9 | 121727404 | rs10818262 | -1.383 | -1.549 | 1.571 | 0.475 | 0.039 | 0.029 | 0.029 | 0 | 0.44 | 0.19 | 0.22 | 0.38 | 0.33 | 0.48 |
| 9 | 121731238 | rs7048230 | -0.967 | -1.544 | 1.532 | 0.482 | -0.054 | 0.023 | 0.023 | 0 | 0.2 | 0.4 | 0.14 | 0.22 | 0.12 | 0.2 |
| 9 | 121738737 | rs2818311 | 0.182 | -1.339 | 1.946 | 0.648 | -0.018 | -0.067 | -0.067 | 0 | 0.71 | 0.58 | 0.87 | 0.84 | 0.79 | 0.72 |
| 9 | 121751926 | rs2807714 | 0.788 | -1.339 | 1.495 | 0.692 | 0.014 | 0.028 | 0.028 | 0.079 | 0.67 | 0.31 | 0.84 | 0.81 | 0.77 | 0.71 |
| 9 | 121753075 | rs2807715 | 0.387 | -1.313 | 1.39 | 0.694 | -0.005 | -0.019 | -0.019 | 0 | 0.69 | 0.7 | 0.87 | 0.84 | 0.77 | 0.69 |
| 9 | 121753744 | rs2818299 | 1.433 | -1.371 | 1.27 | 0.722 | 0.054 | 0.011 | 0.011 | 0.008 | 0.55 | 0.33 | 0.76 | 0.56 | 0.71 | 0.54 |
| 9 | 121756868 | rs7850538 | -0.485 | -1.431 | 1.326 | 0.76 | -0.037 | 0.078 | 0.078 | 0 | 0.12 | 0.24 | 0.09 | 0.27 | 0.06 | 0.15 |
| 9 | 121759567 | rs2818294 | 1.056 | -1.396 | 0.92 | 0.752 | 0.021 | 0.018 | 0.018 | 3.299 | 0.94 | 0.69 | 0.95 | 0.97 | 0.98 | 0.99 |
| 9 | 121765918 | rs2818325 | 1.062 | -1.338 | -0.287 | 0.761 | -0.056 | 0.033 | 0.033 | 3.007 | 0.81 | 0.36 | 0.85 | 0.73 | 0.92 | 0.84 |
| 9 | 121774806 | rs10739528 | 1.042 | -0.918 | -0.65 | 0.816 | 0.023 | 0.007 | 0.007 | 0 | 0.57 | 0.35 | 0.67 | 0.58 | 0.71 | 0.53 |
| 9 | 121778640 | rs2416691 | -0.331 | 0.049 | -1.368 | 0.359 | -0.055 | 0.047 | 0.047 | 0 | 0.27 | 0.01 | 0.2 | 0.15 | 0.23 | 0.35 |
| 9 | 121798140 | rs2109619 | 0.039 | 0.3 | -0.711 | 0.813 | -0.022 | -0.046 | -0.046 | 1.038 | 0.31 | 0.2 | 0.36 | 0.45 | 0.33 | 0.2 |
| 9 | 121803609 | rs2191622 | 0.692 | 0.273 | -1.45 | 0.527 | -0.039 | 0.006 | 0.006 | 0 | 0.18 | 0.01 | 0.18 | 0.1 | 0.18 | 0.19 |

| | | | | | | | | | | | | | | | | |
|---|---|---|---|---|---|---|---|---|---|---|---|---|---|---|---|---|
| 9 | 121805021 | rs10760015 | 0.052 | 0.2 | -0.286 | 0.329 | 0.004 | 0.009 | 0.009 | 0 | 0.5 | 0.53 | 0.43 | 0.47 | 0.5 | 0.61 |
| 9 | 121808496 | rs10122991 | 0.78 | 0.048 | -0.555 | 0.036 | -0.033 | 0.075 | 0.075 | 0 | 0.28 | 0.12 | 0.2 | 0.12 | 0.32 | 0.35 |
| 9 | 121809618 | rs1158809 | 0.572 | -0.062 | -0.682 | 0.079 | 0.02 | -0.071 | -0.071 | 0 | 0.18 | 0.06 | 0.19 | 0.35 | 0.18 | 0.26 |
| 9 | 121836674 | rs6478394 | 0.449 | -0.1 | -0.47 | 0.029 | 0.005 | -0.048 | -0.048 | 0.13 | 0.32 | 0.35 | 0.25 | 0.12 | 0.34 | 0.35 |
| 9 | 121836984 | rs10984399 | 0.574 | 0.009 | -1.151 | 0.115 | 0.016 | -0.069 | -0.069 | 0 | 0.17 | 0.06 | 0.18 | 0.34 | 0.16 | 0.26 |
| 9 | 121839928 | rs10514837 | 0.412 | 0.403 | -0.795 | -0.104 | -0.005 | 0.016 | 0.016 | 0.372 | 0.33 | 0.26 | 0.35 | 0.17 | 0.3 | 0.17 |
| 9 | 121840391 | rs737640 | -0.133 | 0.67 | -0.801 | 0.071 | -0.016 | 0.036 | 0.036 | 0 | 0.72 | 0.85 | 0.77 | 0.91 | 0.67 | 0.71 |
| 9 | 121842266 | rs7867690 | -0.382 | 0.627 | -0.098 | 1.129 | 0.034 | 0.235 | 0.235 | 2.087 | 0.1 | 0.41 | 0.1 | 0.28 | 0.06 | 0.04 |
| 9 | 121844326 | rs7018768 | -0.412 | 0.876 | -0.781 | 0.475 | 0.006 | -0.011 | -0.011 | 0.664 | 0.39 | 0.49 | 0.29 | 0.29 | 0.41 | 0.25 |
| 9 | 121858094 | rs17292757 | -0.721 | 0.725 | -1.157 | 0.312 | 0.001 | 0.015 | 0.015 | 0 | 0.12 | 0.05 | 0.17 | 0.08 | 0.09 | 0.17 |
| 9 | 121861960 | rs2159639 | -1.307 | 1.737 | -1.528 | -0.008 | -0.063 | 0.043 | 0.043 | 0.576 | 0.18 | 0.04 | 0.25 | 0.08 | 0.25 | 0.21 |
| 9 | 121862016 | rs4836745 | 0.157 | 1.374 | -1.54 | -0.039 | -0.002 | 0 | 0 | 1.207 | 0.31 | 0.31 | 0.37 | 0.36 | 0.28 | 0.42 |
| 9 | 121878377 | rs10760019 | 0.873 | 0.615 | -0.686 | -0.341 | -0.008 | -0.037 | -0.037 | 1.747 | 0.44 | 0.18 | 0.31 | 0.52 | 0.43 | 0.3 |
| 9 | 121881005 | rs10984414 | 0.948 | 0.273 | -1.11 | -0.558 | -0.029 | -0.069 | -0.069 | 0 | 0.15 | 0.03 | 0.19 | 0.32 | 0.14 | 0.15 |
| 9 | 121883165 | rs115795 | 0.289 | 0.386 | -0.851 | -0.604 | 0.031 | -0.081 | -0.081 | 0.06 | 0.78 | 0.49 | 0.7 | 0.92 | 0.74 | 0.76 |
| 9 | 121911162 | rs17292799 | 0.551 | 0.194 | -1.024 | -0.723 | 0.015 | 0.028 | 0.028 | 0.162 | 0.1 | 0.01 | 0.11 | 0.04 | 0.06 | 0.2 |
| 9 | 121911404 | rs230123 | -0.757 | 0.313 | -0.907 | -0.756 | 0.027 | -0.084 | -0.084 | 0 | 0.28 | 0.55 | 0.36 | 0.1 | 0.34 | 0.29 |
| 9 | 121914488 | rs57440 | -1.733 | -0.404 | -1.318 | -0.98 | -0.126 | -0.085 | -0.085 | 0 | 0.58 | 0.97 | 0.68 | 0.41 | 0.54 | 0.63 |
| 9 | 121916627 | rs10984425 | -0.172 | -0.43 | -1.29 | -0.991 | -0.032 | 0.013 | 0.013 | 1.059 | 0.31 | 0.44 | 0.33 | 0.34 | 0.21 | 0.34 |
| 9 | 121918081 | rs110797 | -0.864 | -0.355 | -1.294 | -0.984 | 0.015 | -0.081 | -0.081 | 1.265 | 0.31 | 0.49 | 0.37 | 0.11 | 0.36 | 0.29 |
| 9 | 121927511 | rs10984426 | 1.564 | -0.414 | -1.274 | -0.867 | -0.121 | -0.164 | -0.164 | 0.068 | 0.37 | 0.03 | 0.29 | 0.54 | 0.42 | 0.35 |
| 9 | 121927666 | rs1571904 | 0.67 | -0.546 | -1.063 | -1.038 | -0.006 | 0.015 | 0.015 | 0 | 0.05 | 0.03 | 0.04 | 0 | 0.07 | 0.06 |
| 9 | 121929575 | rs28453 | -0.34 | -0.55 | -0.827 | -1.044 | -0.014 | -0.086 | -0.086 | 0.053 | 0.13 | 0.51 | 0.11 | 0 | 0.09 | 0.08 |
| 9 | 121951619 | rs230080 | -0.72 | 0.281 | -2.464 | -1.306 | -0.082 | -0.124 | -0.124 | 0.086 | 0.65 | 0.98 | 0.75 | 0.39 | 0.64 | 0.73 |
| 9 | 121956076 | rs230089 | 0.067 | -0.044 | -1.396 | -1.524 | -0.017 | -0.01 | -0.01 | 0.006 | 0.29 | 0.63 | 0.19 | 0.21 | 0.17 | 0.18 |
| 9 | 121961896 | rs7018536 | -0.521 | -0.152 | -0.293 | -1.723 | 0.027 | 0.003 | 0.003 | 0.92 | 0.07 | 0.3 | 0.04 | 0.01 | 0.04 | 0.04 |
| 9 | 121965240 | rs10984435 | -0.516 | -0.17 | -0.144 | -1.732 | 0.014 | -0.009 | -0.009 | 0.021 | 0.07 | 0.25 | 0.04 | 0.01 | 0.04 | 0.05 |
| 9 | 121967454 | rs1332452 | -0.753 | -0.043 | -0.314 | -1.671 | -0.005 | -0.005 | -0.005 | 0 | 0.86 | 0.88 | 0.87 | 0.8 | 0.82 | 0.83 |
| 9 | 121967597 | rs1332453 | 0.21 | 0.33 | -0.875 | -1.557 | -0.007 | 0.036 | 0.036 | 2.494 | 0.19 | 0.06 | 0.31 | 0.12 | 0.22 | 0.24 |
| 9 | 121967957 | rs10760025 | 1.427 | 0.071 | -0.712 | -1.669 | -0.006 | 0.073 | 0.073 | 5.89 | 0.56 | 0.18 | 0.48 | 0.67 | 0.56 | 0.55 |
| 9 | 121970419 | rs1332455 | -0.304 | 0.483 | -0.869 | -1.687 | -0.006 | 0 | 0 | 0 | 0.71 | 0.69 | 0.62 | 0.7 | 0.6 | 0.63 |
| 9 | 121973842 | rs9696959 | 1.733 | 0.405 | -0.7 | -1.694 | 0.045 | 0.024 | 0.024 | 0 | 0.46 | 0.24 | 0.26 | 0.58 | 0.37 | 0.39 |
| 9 | 121974448 | rs2183964 | -1.676 | 0.767 | -0.308 | -1.471 | 0.016 | 0.017 | 0.017 | 0 | 0.85 | 0.8 | 0.86 | 0.79 | 0.83 | 0.84 |
| 9 | 121976044 | rs7853697 | 0.973 | 0.8 | -0.315 | -1.429 | 0.005 | -0.015 | -0.015 | 0 | 0.06 | 0.2 | 0.03 | 0.01 | 0.04 | 0.05 |
| 9 | 121979181 | rs10984443 | 1.485 | 0.89 | -0.543 | -1.414 | -0.041 | 0.021 | 0.021 | 0 | 0.15 | 0.06 | 0.23 | 0.1 | 0.22 | 0.21 |
| 9 | 121982370 | rs875166 | 1.458 | 0.775 | -0.643 | -1.478 | -0.032 | -0.009 | -0.009 | 0 | 0.14 | 0.01 | 0.26 | 0.11 | 0.17 | 0.2 |
| 9 | 121984553 | rs10984447 | 1.47 | 1.036 | -0.271 | -1.358 | -0.044 | 0.023 | 0.023 | 0 | 0.15 | 0.07 | 0.24 | 0.11 | 0.23 | 0.21 |
| 9 | 121984673 | rs10760028 | -1.481 | 1.027 | -0.243 | -1.35 | 0.018 | 0.014 | 0.014 | 0 | 0.86 | 0.8 | 0.86 | 0.8 | 0.82 | 0.84 |
| 9 | 121988899 | rs10818288 | 0.935 | 0.861 | -0.227 | -1.36 | -0.059 | 0.056 | 0.056 | 0 | 0.6 | 0.17 | 0.51 | 0.69 | 0.51 | 0.56 |
| 9 | 121993508 | rs10818289 | 0.848 | 0.678 | -0.18 | -1.557 | 0.005 | 0.042 | 0.042 | 0 | 0.57 | 0.34 | 0.51 | 0.69 | 0.47 | 0.55 |
| 9 | 121996466 | rs989094 | -0.29 | 0.759 | -0.577 | -1.555 | -0.007 | 0.003 | 0.003 | 0.144 | 0.75 | 0.79 | 0.79 | 0.78 | 0.69 | 0.76 |
| 9 | 122001000 | rs2274157 | 1.03 | 0.683 | -0.677 | -1.591 | 0.021 | 0.099 | 0.099 | 0 | 0.54 | 0.17 | 0.47 | 0.67 | 0.45 | 0.54 |
| 9 | 122001714 | rs12378647 | -0.116 | 0.869 | -0.631 | -1.33 | -0.009 | 0.006 | 0.006 | 0.198 | 0.75 | 0.79 | 0.78 | 0.78 | 0.69 | 0.76 |
| 9 | 122008060 | rs1571903 | 0.82 | 0.64 | -0.489 | -1.352 | -0.006 | 0.024 | 0.024 | 0 | 0.57 | 0.69 | 0.55 | 0.67 | 0.5 | 0.54 |
| 9 | 122012643 | rs7027123 | 0.972 | 0.311 | -0.533 | -2.145 | -0.018 | 0.05 | 0.05 | 1.495 | 0.59 | 0.32 | 0.39 | 0.8 | 0.53 | 0.53 |
| 9 | 122014027 | rs539054 | 0.381 | 0.351 | -0.026 | -2.107 | -0.002 | -0.061 | -0.061 | 0 | 0.76 | 0.68 | 0.67 | 0.9 | 0.76 | 0.72 |
| 9 | 122014339 | rs600066 | 0.369 | 0.34 | 0.061 | -2.122 | -0.006 | -0.055 | -0.055 | 0.043 | 0.74 | 0.47 | 0.64 | 0.89 | 0.76 | 0.72 |
| 9 | 122016593 | rs1576836 | 0.566 | 0.407 | -0.405 | -2.047 | -0.002 | 0.006 | 0.006 | 0.31 | 0.17 | 0.1 | 0.28 | 0.09 | 0.17 | 0.2 |
| 9 | 122016656 | rs12238422 | 0.762 | 0.597 | -0.801 | -1.813 | -0.052 | -0.052 | -0.052 | 0 | 0.13 | 0.03 | 0.15 | 0.24 | 0.2 | 0.12 |
| 9 | 122023574 | rs2031380 | 0.103 | 0.225 | -0.26 | -1.71 | 0.006 | -0.027 | -0.027 | 0 | 0.93 | 0.76 | 0.95 | 0.99 | 0.94 | 0.92 |
| 9 | 122030076 | rs1221136 | 0.652 | 0.359 | -0.483 | -1.774 | -0.027 | 0.024 | 0.024 | 0.942 | 0.16 | 0.11 | 0.25 | 0.1 | 0.23 | 0.18 |
| 9 | 122030983 | rs10760030 | 0.083 | 0.353 | -0.435 | -1.748 | 0.012 | -0.059 | -0.059 | 0 | 0.74 | 0.59 | 0.68 | 0.89 | 0.71 | 0.72 |
| 9 | 122035992 | rs10818292 | 1.272 | -0.118 | -0.808 | -1.446 | 0.008 | -0.021 | -0.021 | 0.824 | 0.26 | 0.21 | 0.19 | 0.56 | 0.24 | 0.35 |
| 9 | 122036135 | rs487240 | -0.177 | -0.05 | -1.267 | -1.844 | -0.053 | 0.053 | 0.053 | 1.304 | 0.46 | 0.07 | 0.42 | 0.34 | 0.51 | 0.43 |
| 9 | 122036993 | rs914651 | 0.49 | 0.097 | -1.526 | -2.103 | -0.016 | 0.147 | 0.147 | 4.987 | 0.3 | 0.07 | 0.2 | 0.08 | 0.25 | 0.27 |
| 9 | 122039576 | rs680003 | -0.882 | -0.048 | -0.792 | -2.175 | 0.006 | 0.081 | 0.081 | 1.418 | 0.51 | 0.71 | 0.48 | 0.31 | 0.5 | 0.4 |
| 9 | 122039704 | rs7856245 | 0.279 | 0.439 | 0.049 | -1.858 | 0.034 | 0.014 | 0.014 | 0 | 0.17 | 0.25 | 0.23 | 0.22 | 0.25 | 0.14 |
| 9 | 122043820 | rs792933 | -0.487 | 0.127 | -0.745 | -1.854 | 0.009 | -0.014 | -0.014 | 0 | 0.41 | 0.53 | 0.42 | 0.32 | 0.46 | 0.36 |
| 9 | 122044643 | rs10217394 | 1.766 | 0.128 | -1.558 | -1.687 | -0.074 | -0.164 | -0.164 | 0.568 | 0.23 | 0.02 | 0.19 | 0.52 | 0.27 | 0.3 |
| 9 | 122047381 | rs560056 | 0.178 | 0.985 | -0.994 | -1.684 | -0.119 | -0.06 | -0.06 | 0.061 | 0.26 | 0.03 | 0.3 | 0.26 | 0.33 | 0.29 |
| 9 | 122059709 | rs4837628 | -1.057 | 0.813 | -0.479 | -1.727 | 0.014 | -0.009 | -0.009 | 0.872 | 0.39 | 0.64 | 0.43 | 0.32 | 0.45 | 0.33 |
| 9 | 122060357 | rs12683279 | 0.958 | 0.872 | -0.528 | -1.575 | -0.039 | -0.035 | -0.035 | 0.88 | 0.07 | 0 | 0.13 | 0.2 | 0.12 | 0.1 |
| 9 | 122068881 | rs16908223 | 0.727 | 0.691 | -0.172 | -1.646 | 0.005 | -0.016 | -0.016 | 0.054 | 0.09 | 0.2 | 0.07 | 0.05 | 0.1 | 0.02 |
| 9 | 122070829 | rs10818297 | 1.316 | 0.703 | -0.23 | -1.64 | 0.006 | -0.01 | -0.01 | 0 | 0.07 | 0.15 | 0.06 | 0.05 | 0.09 | 0.01 |
| 9 | 122071105 | rs10739534 | -0.659 | 0.301 | -0.288 | -1.675 | 0.012 | -0.012 | -0.012 | 0 | 0.4 | 0.6 | 0.43 | 0.32 | 0.46 | 0.32 |
| 9 | 122081818 | rs10118663 | 0.884 | 0.51 | -1.003 | -1.571 | 0.023 | -0.008 | -0.008 | 0 | 0.08 | 0.22 | 0.11 | 0.06 | 0.12 | 0.04 |
| 9 | 122091015 | rs17634284 | 2.03 | 0.522 | -1.227 | -1.59 | -0.036 | -0.01 | -0.01 | 0 | 0.06 | 0 | 0.07 | 0.05 | 0.11 | 0.04 |
| 9 | 122095833 | rs10984490 | 0.508 | 0.539 | -1.292 | -1.631 | -0.076 | -0.043 | -0.043 | 0.297 | 0.28 | 0.1 | 0.3 | 0.31 | 0.35 | 0.28 |
| 9 | 122096532 | rs10984491 | 0.658 | 0.857 | -1.48 | -1.494 | -0.026 | -0.073 | -0.073 | 0 | 0.4 | 0.02 | 0.33 | 0.62 | 0.34 | 0.4 |

| | | | | | | | | | | | | | | | | |
|---|---|---|---|---|---|---|---|---|---|---|---|---|---|---|---|---|
| 9 | 122098205 | rs10984492 | 0.026 | 1.465 | 0.248 | -1.311 | 0.037 | -0.058 | -0.058 | 0 | 0.12 | 0.17 | 0.19 | 0.01 | 0.16 | 0.13 |
| 9 | 122112563 | rs16908302 | 0.09 | 1.167 | -0.444 | -0.794 | 0.019 | -0.009 | -0.009 | 0 | 0.08 | 0.22 | 0.11 | 0.06 | 0.11 | 0.03 |
| 9 | 122120197 | rs1324623 | -0.293 | 0.961 | -0.706 | -1.682 | -0.002 | -0.051 | -0.051 | 0 | 0.62 | 0.68 | 0.72 | 1 | 0.62 | 0.81 |
| 9 | 122120555 | rs4540472 | 0.111 | 1.028 | -0.723 | -2.184 | 0 | -0.089 | -0.089 | 0 | 0.49 | 0.49 | 0.58 | 0.99 | 0.45 | 0.72 |
| 9 | 122132184 | rs1324627 | -0.568 | 0.753 | -0.802 | -2.294 | -0.002 | -0.002 | -0.002 | 0 | 0.58 | 0.68 | 0.69 | 1 | 0.59 | 0.75 |
| 9 | 122133105 | rs7020462 | -0.022 | 0.998 | -0.792 | -2.068 | 0.122 | 0.083 | 0.083 | 0 | 0.09 | 0.37 | 0.17 | 0.11 | 0.16 | 0.06 |
| 9 | 122140786 | rs2025501 | -0.252 | 0.599 | -1.384 | -2.022 | -0.016 | 0.183 | 0.183 | 0.262 | 0.43 | 0.15 | 0.48 | 0.86 | 0.4 | 0.63 |
| 9 | 122146403 | rs2210379 | -0.227 | 0.412 | -0.88 | -2.269 | -0.009 | 0.053 | 0.053 | 0.32 | 0.65 | 0.81 | 0.56 | 0.13 | 0.65 | 0.42 |
| 9 | 122150396 | rs7848896 | 0.378 | -0.13 | -0.556 | -2.797 | 0.004 | 0.122 | 0.122 | 0 | 0.55 | 0.77 | 0.49 | 0.13 | 0.51 | 0.43 |
| 9 | 122154649 | rs4455935 | -0.54 | -0.151 | -0.587 | -2.832 | 0.003 | 0.04 | 0.04 | 0 | 0.7 | 0.84 | 0.59 | 0.13 | 0.69 | 0.48 |
| 9 | 122156770 | rs17635586 | 0.986 | -0.132 | -0.626 | -2.809 | -0.012 | 0.045 | 0.045 | 0 | 0.14 | 0.07 | 0.1 | 0 | 0.18 | 0.06 |
| 9 | 122158444 | rs2150753 | 0.318 | -0.049 | -0.319 | -2.821 | 0.002 | 0.075 | 0.075 | 0 | 0.54 | 0.72 | 0.48 | 0.13 | 0.5 | 0.42 |
| 9 | 122158693 | rs2183761 | 0.318 | -0.049 | -0.319 | -2.821 | 0.002 | 0.075 | 0.075 | 0 | 0.54 | 0.72 | 0.48 | 0.13 | 0.5 | 0.42 |
| 9 | 122162502 | rs7847786 | 0.673 | -0.032 | -0.212 | -2.811 | 0.01 | 0.064 | 0.064 | 0 | 0.51 | 0.37 | 0.44 | 0.13 | 0.5 | 0.42 |
| 9 | 122166761 | rs17635892 | 0.717 | -0.101 | -0.184 | -2.514 | -0.005 | 0.012 | 0.012 | 0 | 0.14 | 0.08 | 0.1 | 0 | 0.18 | 0.06 |
| 9 | 122171470 | rs10984519 | 0.874 | -0.664 | -0.813 | -1.658 | 0.004 | -0.132 | -0.132 | 0.378 | 0.19 | 0.01 | 0.15 | 0.41 | 0.12 | 0.32 |
| 9 | 122174362 | rs1331621 | 0.951 | -0.53 | -0.445 | -1.648 | -0.003 | 0.031 | 0.031 | 0 | 0.67 | 0.43 | 0.61 | 0.53 | 0.63 | 0.72 |
| 9 | 122176793 | rs10491527 | 0.801 | -0.722 | -0.5 | -1.92 | -0.007 | 0.031 | 0.031 | 0 | 0.14 | 0.08 | 0.09 | 0 | 0.16 | 0.06 |
| 9 | 122189165 | rs9299259 | 0.275 | -0.696 | -0.321 | -1.832 | 0.018 | -0.028 | -0.028 | 0.361 | 0.06 | 0.24 | 0.06 | 0 | 0.05 | 0.02 |
| 9 | 122193951 | rs10491526 | 0.672 | -0.747 | -0.517 | -1.602 | -0.029 | 0.129 | 0.129 | 0 | 0.79 | 0.42 | 0.71 | 0.53 | 0.85 | 0.76 |
| 9 | 122198190 | rs10818313 | 0.996 | -0.685 | -0.508 | -1.636 | -0.006 | 0.02 | 0.02 | 0 | 0.13 | 0.08 | 0.09 | 0 | 0.14 | 0.05 |
| 9 | 122202000 | rs10120508 | 0.902 | -0.75 | -0.531 | -1.614 | -0.001 | 0.005 | 0.005 | 0 | 0.69 | 0.34 | 0.63 | 0.53 | 0.72 | 0.72 |
| 9 | 122204804 | rs1331624 | 0.806 | -0.502 | -0.53 | -1.643 | 0.022 | -0.008 | -0.008 | 0 | 0.57 | 0.32 | 0.64 | 0.54 | 0.69 | 0.53 |
| 9 | 122215618 | rs10818318 | 0.702 | -0.218 | -0.593 | -1.553 | -0.002 | -0.015 | -0.015 | 0 | 0.12 | 0.15 | 0.09 | 0 | 0.15 | 0.05 |
| 9 | 122222718 | rs16908409 | 0.672 | -0.129 | -0.538 | -0.962 | -0.003 | 0.076 | 0.076 | 0 | 0.09 | 0.11 | 0.17 | 0.46 | 0.06 | 0.22 |
| 9 | 122222874 | rs9299260 | 0.642 | -0.038 | -0.448 | -1.365 | 0.015 | 0.065 | 0.065 | 0.21 | 0.5 | 0.33 | 0.41 | 0 | 0.5 | 0.37 |
| 9 | 122222907 | rs9299261 | 0.14 | 0.002 | -0.441 | -1.326 | -0.001 | 0.001 | 0.001 | 0.255 | 0.59 | 0.45 | 0.58 | 0.46 | 0.56 | 0.59 |
| 9 | 122226426 | rs1331591 | 1.212 | 0.042 | -0.283 | -1.322 | 0.006 | -0.051 | -0.051 | 3.503 | 0.12 | 0.19 | 0.1 | 0 | 0.15 | 0.06 |
| 9 | 122233951 | rs10818319 | 0.261 | 0.101 | -0.489 | -1.275 | 0.036 | -0.031 | -0.031 | 0 | 0.17 | 0.4 | 0.15 | 0.09 | 0.21 | 0.09 |
| 9 | 122244046 | rs1990422 | -1.127 | -0.02 | -0.557 | -1.303 | 0.012 | -0.028 | -0.028 | 0.008 | 0.75 | 0.83 | 0.73 | 0.55 | 0.78 | 0.68 |
| 9 | 122247156 | rs7868280 | -0.257 | 0.073 | -0.918 | -1.316 | -0.013 | 0.022 | 0.022 | 0.11 | 0.65 | 0.56 | 0.63 | 0.54 | 0.65 | 0.64 |
| 9 | 122247734 | rs10984556 | 1.323 | 0.214 | -1.146 | -1.285 | 0.007 | 0.014 | 0.014 | 0 | 0.05 | 0.09 | 0.04 | 0.08 | 0.06 | 0.04 |
| 9 | 122249965 | rs1412055 | -0.014 | 0.18 | -1.158 | -1.321 | -0.003 | 0.076 | 0.076 | 0 | 0.14 | 0.2 | 0.21 | 0.53 | 0.12 | 0.26 |
| 9 | 122254248 | rs12555385 | 0.101 | 0.625 | -1.651 | -2.441 | 0.023 | 0.102 | 0.102 | 0.213 | 0.49 | 0.3 | 0.39 | 0 | 0.51 | 0.38 |
| 9 | 122257254 | rs10491524 | -0.494 | 0.608 | -2.127 | -1.931 | -0.007 | -0.029 | -0.029 | 0.385 | 0.67 | 0.67 | 0.54 | 0.09 | 0.71 | 0.47 |
| 9 | 122257627 | rs10984560 | -0.256 | 1.347 | -2.618 | -1.295 | -0.071 | 0.148 | 0.148 | 0.989 | 0.32 | 0.06 | 0.3 | 0 | 0.38 | 0.22 |
| 9 | 122259593 | rs10984564 | -0.535 | 1.136 | -2.043 | -0.594 | -0.043 | 0.076 | 0.076 | 0.031 | 0.32 | 0.11 | 0.28 | 0.14 | 0.36 | 0.28 |
| 9 | 122260749 | rs10984568 | 0.197 | 0.463 | -2.098 | -0.186 | 0.044 | -0.029 | -0.029 | 0.218 | 0.46 | 0.18 | 0.29 | 0.52 | 0.39 | 0.41 |
| 9 | 122260800 | rs1412056 | -0.108 | 0.805 | -1.853 | -0.231 | -0.011 | 0.008 | 0.008 | 1.486 | 0.4 | 0.66 | 0.59 | 0.29 | 0.44 | 0.46 |
| 9 | 122261157 | rs11790549 | 0.75 | 0.008 | -2.103 | -0.327 | 0.032 | -0.069 | -0.069 | 0 | 0.29 | 0.15 | 0.22 | 0.53 | 0.23 | 0.32 |
| 9 | 122261319 | rs2416707 | -0.292 | -0.48 | -1.483 | -0.684 | 0.072 | -0.05 | -0.05 | 0 | 0.61 | 0.21 | 0.61 | 0.48 | 0.73 | 0.65 |
| 9 | 122263352 | rs758875 | 0.232 | -0.128 | -1.413 | -0.595 | -0.049 | -0.029 | -0.029 | 3.712 | 0.12 | 0.01 | 0.09 | 0.19 | 0.15 | 0.1 |
| 9 | 122264713 | rs10491522 | -0.66 | -0.372 | -0.695 | -0.77 | 0.01 | -0.003 | -0.003 | 0 | 0.58 | 0.62 | 0.6 | 0.45 | 0.65 | 0.61 |
| 9 | 122272453 | rs17638213 | 0.328 | -0.256 | -0.99 | -0.742 | -0.044 | -0.028 | -0.028 | 0 | 0.11 | 0.01 | 0.1 | 0.18 | 0.14 | 0.11 |
| 9 | 122277651 | rs12345648 | -0.68 | -0.457 | -0.043 | -0.865 | 0.016 | -0.036 | -0.036 | 0.377 | 0.1 | 0.44 | 0.16 | 0 | 0.06 | 0.05 |
| 9 | 122279421 | rs10818328 | -0.942 | -0.39 | -0.082 | -0.702 | 0.001 | -0.001 | -0.001 | 0 | 0.59 | 0.59 | 0.67 | 0.43 | 0.67 | 0.66 |
| 9 | 122288239 | rs7849695 | -0.339 | -0.244 | -0.103 | -0.409 | 0.123 | -0.005 | -0.005 | 0 | 0.49 | 0.16 | 0.55 | 0.53 | 0.64 | 0.64 |
| 9 | 122288413 | rs1014162 | -0.363 | -0.239 | -0.1 | -0.413 | 0.126 | -0.007 | -0.007 | 0 | 0.49 | 0.16 | 0.55 | 0.52 | 0.64 | 0.64 |
| 9 | 122288939 | rs10760049 | 0.428 | 0.135 | 0.284 | -0.275 | 0.018 | -0.001 | -0.001 | 0 | 0.54 | 0.23 | 0.38 | 0.61 | 0.46 | 0.49 |
| 9 | 122289187 | rs1009470 | -0.541 | 0.134 | 0.27 | -0.276 | 0.002 | -0.002 | -0.002 | 0 | 0.44 | 0.57 | 0.6 | 0.4 | 0.54 | 0.51 |
| 9 | 122300146 | rs1331608 | -0.261 | -0.087 | 0.241 | -0.215 | -0.003 | 0.003 | 0.003 | 0 | 0.42 | 0.48 | 0.55 | 0.4 | 0.55 | 0.49 |
| 9 | 122301156 | rs2109664 | -0.421 | 0.003 | 0.246 | -0.1 | 0.002 | -0.003 | -0.003 | 0 | 0.45 | 0.44 | 0.63 | 0.41 | 0.55 | 0.51 |
| 9 | 122303614 | rs4644327 | -0.145 | -0.002 | 0.189 | -0.07 | -0.003 | -0.027 | -0.027 | 0 | 0.42 | 0.27 | 0.59 | 0.4 | 0.55 | 0.5 |
| 9 | 122312068 | rs9644909 | 0.301 | 0.268 | -1.09 | 0.096 | 0.044 | -0.014 | -0.014 | 0 | 0.5 | 0.18 | 0.33 | 0.59 | 0.43 | 0.48 |
| 9 | 122319889 | rs990622 | 0.371 | 0.74 | -0.646 | 0.711 | 0.001 | 0.012 | 0.012 | 0 | 0.8 | 0.77 | 0.76 | 0.6 | 0.76 | 0.69 |
| 9 | 122333861 | rs10818334 | 0.221 | 0.355 | -1.699 | -0.176 | -0.072 | 0.198 | 0.198 | 4.434 | 0.42 | 0.07 | 0.3 | 0.18 | 0.53 | 0.27 |
| 9 | 122337936 | rs12350573 | -0.154 | -0.031 | -0.091 | -0.382 | 0.007 | 0.027 | 0.027 | 2.024 | 0.48 | 0.57 | 0.37 | 0.18 | 0.55 | 0.3 |
| 9 | 122341917 | rs4412427 | 1.245 | -0.358 | -0.498 | 0.213 | -0.03 | -0.07 | -0.07 | 1.581 | 0.15 | 0.02 | 0.27 | 0.33 | 0.17 | 0.28 |
| 9 | 122343811 | rs10984605 | 0.239 | -0.154 | -0.237 | -0.261 | -0.001 | -0.016 | -0.016 | 0.147 | 0.46 | 0.45 | 0.36 | 0.18 | 0.53 | 0.29 |
| 9 | 122352612 | rs10818336 | -0.287 | -0.397 | -0.293 | -0.375 | 0 | 0.008 | 0.008 | 6.56 | 0.53 | 0.6 | 0.37 | 0.17 | 0.53 | 0.31 |
| 9 | 122360136 | rs12349509 | -0.171 | -0.404 | 0.113 | -0.34 | -0.026 | -0.03 | -0.03 | 1.576 | 0.09 | 0.34 | 0.04 | 0 | 0.01 | 0.02 |
| 9 | 122372872 | rs7021853 | -1.406 | -0.359 | 0.307 | -0.022 | 0.073 | 0.291 | 0.291 | 1.591 | 0.05 | 0.4 | 0.06 | 0.13 | 0.02 | 0.08 |
| 9 | 122375077 | rs11794047 | 0.769 | -0.321 | 0.036 | -0.06 | -0.145 | 0.187 | 0.187 | 3.329 | 0.41 | 0.04 | 0.29 | 0.16 | 0.51 | 0.27 |
| 9 | 122393306 | rs10984619 | 0.203 | -0.26 | 0.354 | 0.051 | -0.001 | -0.009 | -0.009 | 0.01 | 0.46 | 0.48 | 0.36 | 0.29 | 0.53 | 0.33 |
| 9 | 122394119 | rs17580614 | 1.178 | 0.065 | -0.345 | -0.351 | -0.058 | 0.066 | 0.066 | 0.916 | 0.25 | 0.02 | 0.14 | 0.13 | 0.26 | 0.2 |
| 9 | 122417221 | rs6478430 | -0.469 | 0.317 | 0.076 | -0.041 | 0.001 | 0.017 | 0.017 | 6.058 | 0.53 | 0.57 | 0.39 | 0.29 | 0.53 | 0.34 |
| 9 | 122422035 | rs1860665 | -0.448 | 0.345 | -0.071 | -0.111 | 0 | 0.006 | 0.006 | 6.202 | 0.53 | 0.5 | 0.48 | 0.3 | 0.53 | 0.34 |
| 9 | 122426409 | rs1013282 | -0.246 | 0.535 | -0.81 | -0.698 | 0.007 | 0.113 | 0.113 | 6.75 | 0.49 | 0.33 | 0.35 | 0.14 | 0.53 | 0.29 |
| 9 | 122431172 | rs10818361 | -0.271 | 0.538 | -0.817 | -0.687 | 0.007 | 0.116 | 0.116 | 6.927 | 0.49 | 0.33 | 0.35 | 0.15 | 0.53 | 0.28 |

| | | | | | | | | | | | | | | | | |
|---|---|---|---|---|---|---|---|---|---|---|---|---|---|---|---|---|
| 9 | 122434053 | rs4837656 | -0.167 | 1.104 | -0.724 | -0.472 | 0.001 | -0.023 | -0.023 | 0 | 0.21 | 0.23 | 0.21 | 0.03 | 0.31 | 0.09 |
| 9 | 122436623 | rs10125439 | 0.134 | 1.706 | -1.054 | -0.26 | -0.013 | 0.005 | 0.005 | 0 | 0.12 | 0.1 | 0.17 | 0.01 | 0.25 | 0.07 |
| 9 | 122439147 | rs10984635 | -0.31 | 1.033 | -1.016 | 0.31 | 0.005 | 0.004 | 0.004 | 3.835 | 0.51 | 0.48 | 0.66 | 0.56 | 0.57 | 0.67 |
| 9 | 122439531 | rs17648616 | -0.168 | 1.258 | -1.407 | 0.44 | -0.027 | 0.003 | 0.003 | 1.7 | 0.14 | 0.02 | 0.23 | 0.13 | 0.17 | 0.26 |
| 9 | 122442505 | rs6415829 | 0.279 | 1.424 | -1.282 | 0.574 | -0.02 | -0.009 | -0.009 | 0.487 | 0.59 | 0.72 | 0.49 | 0.44 | 0.52 | 0.47 |
| 9 | 122446737 | rs4466463 | -0.775 | 2.124 | -1.28 | 0.453 | -0.034 | -0.011 | -0.011 | 0 | 0.23 | 0.13 | 0.29 | 0.22 | 0.32 | 0.25 |
| 9 | 122451259 | rs12551982 | 1.39 | 1.837 | -0.766 | -0.336 | 0.002 | -0.015 | -0.015 | 0 | 0.57 | 0.45 | 0.58 | 0.75 | 0.51 | 0.43 |
| 9 | 122456873 | rs12376944 | -0.032 | 1.765 | -0.435 | -0.474 | -0.001 | -0.043 | -0.043 | 0.043 | 0.15 | 0.17 | 0.07 | 0 | 0.13 | 0.2 |
| 9 | 122463062 | rs4437722 | 0.161 | 1.851 | -0.35 | -0.346 | 0.014 | -0.015 | -0.015 | 0.459 | 0.93 | 0.84 | 0.9 | 0.97 | 0.93 | 0.88 |
| 9 | 122473586 | rs10984654 | -0.194 | 1.996 | -0.417 | -0.355 | -0.037 | -0.017 | -0.017 | 0 | 0.2 | 0.11 | 0.26 | 0.22 | 0.31 | 0.24 |
| 9 | 122474896 | rs4407966 | 0.222 | 1.649 | -0.343 | -0.47 | 0.012 | -0.02 | -0.02 | 1.35 | 0.93 | 0.86 | 0.9 | 0.97 | 0.92 | 0.88 |
| 9 | 122477415 | rs7042510 | 1.197 | 1.399 | -0.362 | -0.652 | -0.013 | -0.11 | -0.11 | 0.046 | 0.74 | 0.58 | 0.83 | 0.97 | 0.8 | 0.66 |
| 9 | 122479120 | rs4520239 | 0.71 | 1.644 | 0.079 | -0.365 | 0.061 | -0.013 | -0.013 | 0.225 | 0.92 | 0.73 | 0.89 | 0.97 | 0.92 | 0.89 |
| 9 | 122481907 | rs4642724 | 0.408 | 1.623 | -0.035 | -0.361 | 0.01 | -0.023 | -0.023 | 0.193 | 0.93 | 0.85 | 0.9 | 0.97 | 0.93 | 0.89 |
| 9 | 122486128 | rs12343952 | -0.475 | 2.111 | -0.276 | -0.351 | -0.051 | -0.026 | -0.026 | 1.932 | 0.19 | 0.08 | 0.27 | 0.22 | 0.31 | 0.26 |
| 9 | 122487783 | rs4255207 | -0.353 | 2.092 | -0.271 | -0.361 | -0.041 | -0.022 | -0.022 | 0 | 0.19 | 0.1 | 0.26 | 0.21 | 0.31 | 0.26 |
| 9 | 122487902 | rs4240461 | 1.094 | 2.104 | -0.026 | -0.303 | 0.052 | -0.013 | -0.013 | 0.862 | 0.9 | 0.69 | 0.88 | 0.95 | 0.91 | 0.89 |
| 9 | 122491590 | rs4240462 | 1.152 | 2.1 | 0.248 | -0.338 | 0.05 | 0.004 | 0.004 | 0 | 0.71 | 0.47 | 0.62 | 0.73 | 0.63 | 0.63 |
| 9 | 122500393 | rs17478596 | -0.809 | 2.129 | 0.379 | -0.341 | 0 | -0.001 | -0.001 | 0 | 0.06 | 0.06 | 0.09 | 0.03 | 0.09 | 0.1 |
| 9 | 122507529 | rs16908934 | -0.014 | 1.55 | 0.368 | -0.752 | 0.014 | 0.002 | 0.002 | 0 | 0.14 | 0.1 | 0.08 | 0 | 0.09 | 0.13 |
| 9 | 122510438 | rs4836779 | 1.699 | 1.425 | 0.292 | -0.921 | -0.002 | 0.007 | 0.007 | 0 | 0.56 | 0.45 | 0.51 | 0.76 | 0.53 | 0.51 |
| 9 | 122514849 | rs10760059 | -0.755 | 1.594 | 0.406 | -0.904 | 0.007 | 0 | 0 | 0 | 0.28 | 0.3 | 0.4 | 0.25 | 0.38 | 0.35 |
| 9 | 122522400 | rs3934932 | -0.082 | 1.456 | 0.446 | -0.842 | 0.009 | 0.007 | 0.007 | 0 | 0.22 | 0.25 | 0.35 | 0.21 | 0.29 | 0.26 |
| 9 | 122525203 | rs11795124 | 0.116 | 1.805 | 0.275 | -0.952 | 0.013 | 0.021 | 0.021 | 0 | 0.15 | 0.09 | 0.1 | 0 | 0.13 | 0.15 |
| 9 | 122525788 | rs10760062 | -0.44 | 1.91 | 0.35 | -0.858 | 0.004 | 0.001 | 0.001 | 0 | 0.26 | 0.27 | 0.39 | 0.24 | 0.34 | 0.33 |
| 9 | 122526774 | rs10513351 | 0.714 | 1.865 | 0.332 | -0.883 | 0.034 | -0.001 | -0.001 | 0 | 0.71 | 0.56 | 0.6 | 0.76 | 0.65 | 0.67 |
| 9 | 122528499 | rs7854124 | 0.665 | 1.88 | 0.33 | -0.876 | 0.034 | 0 | 0 | 1.844 | 0.71 | 0.57 | 0.6 | 0.77 | 0.65 | 0.67 |
| 9 | 122533070 | rs2781116 | 1.221 | 2.042 | 0.152 | 0.059 | 0.029 | 0.012 | 0.012 | 0 | 0.67 | 0.56 | 0.44 | 0.43 | 0.59 | 0.6 |
| 9 | 122533488 | rs2781115 | 1.069 | 1.998 | 0.092 | 0.053 | -0.006 | 0.019 | 0.019 | 0 | 0.73 | 0.66 | 0.68 | 0.61 | 0.74 | 0.7 |
| 9 | 122533710 | rs2809898 | 1.748 | 2.052 | 0.005 | 0.12 | 0.001 | 0.008 | 0.008 | 0 | 0.51 | 0.47 | 0.34 | 0.42 | 0.46 | 0.44 |
| 9 | 122535333 | rs10818382 | 0.34 | 2.052 | -0.061 | 0.082 | -0.001 | 0 | 0 | 0 | 0.06 | 0.06 | 0.04 | 0.04 | 0.06 | 0.08 |
| 9 | 122538345 | rs1360689 | 1.559 | 1.895 | 0.234 | 0.009 | -0.003 | 0.033 | 0.033 | 0 | 0.53 | 0.38 | 0.49 | 0.76 | 0.51 | 0.5 |
| 9 | 122538992 | rs1333891 | 0.435 | 2.338 | -0.054 | -0.078 | -0.051 | 0.046 | 0.046 | 0.008 | 0.08 | 0.01 | 0.07 | 0 | 0.15 | 0.05 |
| 9 | 122541037 | rs1413317 | 1.327 | 1.52 | -0.056 | -0.177 | -0.003 | 0.01 | 0.01 | 0 | 0.29 | 0.12 | 0.2 | 0.28 | 0.27 | 0.35 |
| 9 | 122544250 | rs12376610 | 0.031 | 2.009 | 1.024 | 0.279 | -0.002 | 0.001 | 0.001 | 0 | 0.06 | 0.06 | 0.04 | 0.04 | 0.06 | 0.08 |
| 9 | 122545704 | rs2809903 | 1.288 | 1.851 | 1.294 | 0.082 | -0.004 | 0.003 | 0.003 | 0 | 0.55 | 0.4 | 0.35 | 0.46 | 0.51 | 0.51 |
| 9 | 122545827 | rs10984698 | -0.879 | 1.511 | 0.898 | -0.282 | 0.006 | 0.039 | 0.039 | 0 | 0.37 | 0.27 | 0.42 | 0.17 | 0.35 | 0.41 |
| 9 | 122549068 | rs4837674 | 1.584 | 1.642 | 0.948 | -0.113 | -0.002 | 0.005 | 0.005 | 0 | 0.41 | 0.11 | 0.31 | 0.37 | 0.39 | 0.37 |
| 9 | 122549086 | rs1360695 | 0.569 | 1.694 | 1.312 | -0.18 | 0 | 0.001 | 0.001 | 0 | 0.64 | 0.33 | 0.65 | 0.6 | 0.64 | 0.63 |
| 9 | 122552939 | rs1333913 | -0.198 | 1.576 | 1.269 | -0.259 | -0.006 | 0.003 | 0.003 | 0 | 0.25 | 0.16 | 0.34 | 0.23 | 0.25 | 0.25 |
| 9 | 122556173 | rs13297781 | 1.472 | 1.624 | 1.209 | -0.272 | -0.059 | 0.139 | 0.139 | 0 | 0.44 | 0.12 | 0.43 | 0.77 | 0.48 | 0.44 |
| 9 | 122560262 | rs1333903 | -1.075 | 1.522 | 0.89 | -0.168 | -0.013 | 0.097 | 0.097 | 0 | 0.45 | 0.24 | 0.45 | 0.21 | 0.46 | 0.51 |
| 9 | 122562214 | rs10760064 | -0.188 | 1.488 | 0.841 | -0.155 | 0.008 | -0.007 | -0.007 | 0 | 0.24 | 0.29 | 0.34 | 0.21 | 0.25 | 0.25 |
| 9 | 122566359 | rs2151641 | 0.183 | 1.302 | 0.518 | -0.175 | 0 | -0.039 | -0.039 | 0.419 | 0.22 | 0.2 | 0.13 | 0 | 0.21 | 0.27 |
| 9 | 122568511 | rs2900162 | 1.09 | 1.298 | -0.234 | 0.071 | -0.034 | 0.033 | 0.033 | 0 | 0.47 | 0.15 | 0.35 | 0.4 | 0.49 | 0.44 |
| 9 | 122570847 | rs2809909 | -0.174 | 0.946 | -0.234 | 0.166 | -0.081 | -0.129 | -0.129 | 0 | 0.25 | 0.03 | 0.25 | 0.39 | 0.31 | 0.34 |
| 9 | 122571328 | rs1889274 | -0.643 | 0.771 | 0.935 | 0.15 | -0.008 | 0.034 | 0.034 | 0 | 0.29 | 0.39 | 0.25 | 0.41 | 0.27 | 0.32 |
| 9 | 122572896 | rs944961 | 1.456 | 1.014 | 0.515 | -0.079 | -0.055 | 0.036 | 0.036 | 0.084 | 0.42 | 0.12 | 0.31 | 0.36 | 0.43 | 0.38 |
| 9 | 122574527 | rs4378028 | -0.244 | 0.846 | 1.167 | -0.123 | -0.003 | -0.055 | -0.055 | 0 | 0.77 | 0.77 | 0.87 | 0.97 | 0.77 | 0.77 |
| 9 | 122577120 | rs4469535 | -0.162 | 0.808 | 1.493 | -0.118 | 0.002 | 0.005 | 0.005 | 0 | 0.76 | 0.79 | 0.85 | 0.97 | 0.75 | 0.77 |
| 9 | 122578632 | rs7045823 | 0.199 | 0.867 | 1.421 | -0.073 | 0.009 | 0.037 | 0.037 | 0 | 0.72 | 0.64 | 0.66 | 0.6 | 0.67 | 0.68 |
| 9 | 122578906 | rs7859254 | 0.003 | 0.881 | 1.263 | -0.17 | -0.018 | 0.005 | 0.005 | 0 | 0.28 | 0.19 | 0.35 | 0.24 | 0.31 | 0.33 |
| 9 | 122582415 | rs953841 | 0.05 | 0.966 | 1.032 | -0.14 | -0.002 | -0.003 | -0.003 | 0 | 0.73 | 0.77 | 0.68 | 0.58 | 0.67 | 0.67 |
| 9 | 122583234 | rs1333914 | 1.446 | 0.95 | 0.351 | -0.102 | -0.007 | 0.03 | 0.03 | 0.331 | 0.4 | 0.13 | 0.3 | 0.36 | 0.36 | 0.34 |
| 9 | 122589792 | rs7039987 | 0.019 | 1.145 | 0.021 | -0.195 | -0.076 | -0.122 | -0.122 | 0 | 0.76 | 0.96 | 0.69 | 0.59 | 0.67 | 0.7 |
| 9 | 122591297 | rs12551283 | 0.021 | 0.936 | 0.406 | -1.378 | 0.004 | -0.074 | -0.074 | 0.74 | 0.25 | 0.31 | 0.21 | 0 | 0.27 | 0.19 |
| 9 | 122595692 | rs13283350 | 0.456 | 0.552 | 0.208 | -0.584 | -0.004 | -0.006 | -0.006 | 0 | 0.72 | 0.56 | 0.68 | 0.77 | 0.69 | 0.6 |
| 9 | 122596036 | rs1333916 | 0.466 | 0.669 | -0.517 | -0.556 | -0.002 | -0.003 | -0.003 | 0 | 0.72 | 0.69 | 0.68 | 0.77 | 0.7 | 0.58 |
| 9 | 122607145 | rs7046054 | 0.185 | 1.217 | -1.491 | -0.754 | -0.104 | 0.15 | 0.15 | 0.201 | 0.76 | 0.99 | 0.84 | 0.97 | 0.68 | 0.79 |
| 9 | 122608692 | rs2225659 | 0.979 | 1.169 | -1.384 | -0.919 | 0.003 | 0.04 | 0.04 | 0 | 0.5 | 0.3 | 0.58 | 0.74 | 0.5 | 0.48 |
| 9 | 122609065 | rs4837681 | -0.121 | 1.282 | -1.455 | -1.583 | -0.102 | 0.158 | 0.158 | 0.393 | 0.75 | 0.99 | 0.84 | 0.98 | 0.67 | 0.79 |
| 9 | 122621100 | rs7857834 | 0.272 | 1.024 | -0.879 | -1.349 | 0 | -0.054 | -0.054 | 0 | 0.54 | 0.36 | 0.71 | 0.97 | 0.56 | 0.68 |
| 9 | 122623813 | rs10984725 | -0.519 | 1.587 | -1.094 | -1.252 | -0.018 | 0.013 | 0.013 | 0.176 | 0.28 | 0.14 | 0.23 | 0.06 | 0.38 | 0.28 |
| 9 | 122624757 | rs1360690 | -0.161 | 0.842 | -0.956 | -0.592 | -0.003 | 0.006 | 0.006 | 0.364 | 0.27 | 0.31 | 0.38 | 0.6 | 0.25 | 0.31 |
| 9 | 122625226 | rs10984729 | -1.051 | 1.336 | -0.678 | -1.061 | -0.002 | -0.008 | -0.008 | 0 | 0.08 | 0.09 | 0.06 | 0 | 0.11 | 0.08 |
| 9 | 122631415 | rs7864914 | -1.324 | 1.329 | 0.268 | -1.343 | 0.01 | -0.063 | -0.063 | 0 | 0.1 | 0.74 | 0.15 | 0 | 0.1 | 0.02 |
| 9 | 122656864 | rs10818398 | -0.229 | 0.349 | 0.828 | -1.487 | 0.001 | 0 | 0 | 0 | 0.5 | 0.55 | 0.61 | 0.29 | 0.61 | 0.51 |
| 9 | 122657312 | rs10984766 | 0.095 | 0.421 | 0.509 | -1.736 | 0.075 | -0.081 | -0.081 | 0 | 0.26 | 0.54 | 0.37 | 0.11 | 0.38 | 0.23 |
| 9 | 122664468 | rs10818400 | -0.181 | -0.086 | 1.082 | -1.254 | 0.02 | 0.044 | 0.044 | 0 | 0.52 | 0.68 | 0.62 | 0.3 | 0.61 | 0.52 |

| | | | | | | | | | | | | | | | | |
|---|---|---|---|---|---|---|---|---|---|---|---|---|---|---|---|---|
| 9 | 122673299 | rs1333908 | 0.054 | 0.074 | 0.369 | -1.23 | 0.01 | 0.019 | 0.019 | 0 | 0.5 | 0.57 | 0.63 | 0.28 | 0.61 | 0.5 |
| 9 | 122674412 | rs1333904 | -0.171 | 0.166 | -0.362 | -1.178 | 0.036 | 0.128 | 0.128 | 0 | 0.52 | 0.81 | 0.63 | 0.27 | 0.61 | 0.48 |
| 9 | 122676273 | rs7873013 | -0.144 | 0.224 | -0.336 | -0.82 | 0.024 | 0.061 | 0.061 | 0 | 0.55 | 0.72 | 0.62 | 0.27 | 0.63 | 0.48 |
| 9 | 122676538 | rs1537510 | 0.136 | 0.059 | -0.289 | -0.24 | -0.005 | 0.025 | 0.025 | 3.089 | 0.72 | 0.59 | 0.62 | 0.35 | 0.72 | 0.52 |
| 9 | 122677087 | rs7019090 | -0.547 | 0.372 | -0.633 | 0.082 | 0.001 | -0.011 | -0.011 | 0 | 0.41 | 0.55 | 0.42 | 0.68 | 0.42 | 0.47 |
| 9 | 122681580 | rs6478444 | -0.77 | -0.184 | -1.323 | 0.027 | 0.033 | 0.023 | 0.023 | 1.428 | 0.45 | 0.89 | 0.42 | 0.59 | 0.35 | 0.39 |
| 9 | 122688434 | rs12001524 | -0.819 | -0.389 | 0.001 | -0.203 | -0.027 | -0.037 | -0.037 | 0 | 0.1 | 0.46 | 0.07 | 0.02 | 0.02 | 0.02 |
| 9 | 122689304 | rs7020446 | -0.512 | -0.375 | 0.336 | -0.197 | -0.031 | -0.038 | -0.038 | 0 | 0.08 | 0.27 | 0.04 | 0.02 | 0.02 | 0.02 |
| 9 | 122710349 | rs11999958 | -0.273 | -0.425 | 0.507 | -0.244 | -0.027 | -0.032 | -0.032 | 0.125 | 0.07 | 0.15 | 0.03 | 0.02 | 0.02 | 0.03 |
| 9 | 122720191 | rs2416724 | -0.719 | -0.476 | 0.468 | 0.061 | 0.001 | 0.034 | 0.034 | 2 | 0.27 | 0.36 | 0.37 | 0.53 | 0.25 | 0.36 |
| 9 | 122720528 | rs2900163 | -0.973 | -0.509 | 0.161 | 0.046 | -0.003 | 0.052 | 0.052 | 2.527 | 0.29 | 0.55 | 0.4 | 0.54 | 0.25 | 0.36 |
| 9 | 122734446 | rs10435885 | 0.086 | -0.301 | -0.658 | 0.134 | -0.069 | -0.104 | -0.104 | 0.035 | 0.18 | 0.01 | 0.19 | 0.37 | 0.22 | 0.36 |
| 9 | 122739135 | rs6478449 | -0.435 | 0.206 | -1.397 | -0.668 | -0.04 | 0.058 | 0.058 | 0.224 | 0.78 | 0.9 | 0.78 | 0.95 | 0.7 | 0.65 |
| 9 | 122740885 | rs10984782 | 1.508 | -0.6 | -2.068 | -1.05 | -0.069 | -0.119 | -0.119 | 3.946 | 0.3 | 0.03 | 0.42 | 0.54 | 0.37 | 0.22 |
| 9 | 122762260 | rs10984786 | 0.618 | -0.423 | -2.073 | -1.039 | -0.057 | -0.105 | -0.105 | 0 | 0.21 | 0.02 | 0.19 | 0.35 | 0.23 | 0.37 |
| 9 | 122763224 | rs2416735 | -0.927 | -0.423 | -2.16 | -1.133 | -0.116 | 0.105 | 0.105 | 0.728 | 0.8 | 0.98 | 0.78 | 0.96 | 0.7 | 0.66 |
| 9 | 122763603 | rs7872205 | 1.486 | -1.004 | -1.618 | -1.448 | 0.017 | 0.001 | 0.001 | 4.616 | 0.41 | 0.27 | 0.43 | 0.58 | 0.35 | 0.24 |
| 9 | 122763797 | rs7872465 | -0.385 | -0.533 | -1.369 | -1.251 | -0.07 | 0.103 | 0.103 | 0.312 | 0.78 | 0.92 | 0.77 | 0.96 | 0.7 | 0.66 |
| 9 | 122767607 | rs7866747 | -0.643 | -0.652 | -1.236 | -1.477 | -0.035 | 0.049 | 0.049 | 1.188 | 0.74 | 0.8 | 0.73 | 0.95 | 0.65 | 0.66 |
| 9 | 122771568 | rs10818415 | 1.136 | -0.997 | -1.737 | -1.411 | 0.089 | -0.041 | -0.041 | 2.573 | 0.32 | 0.06 | 0.11 | 0.32 | 0.17 | 0.34 |
| 9 | 122773855 | rs2900167 | -0.421 | -0.852 | -1.017 | -1.469 | 0.036 | 0.021 | 0.021 | 0.147 | 0.51 | 0.61 | 0.32 | 0.4 | 0.32 | 0.43 |
| 9 | 122775371 | rs10984796 | 0.163 | -0.798 | -1.067 | -1.477 | -0.001 | -0.027 | -0.027 | 0 | 0.14 | 0.4 | 0.15 | 0.09 | 0.1 | 0.08 |
| 9 | 122776605 | rs2002821 | 0.716 | -0.841 | -0.744 | -1.486 | 0.012 | 0.008 | 0.008 | 0.206 | 0.11 | 0.34 | 0.12 | 0.09 | 0.06 | 0.06 |
| 9 | 122777336 | rs2416740 | 1.173 | -0.881 | -0.931 | -1.503 | 0.102 | -0.017 | -0.017 | 0 | 0.33 | 0.1 | 0.12 | 0.31 | 0.17 | 0.34 |
| 9 | 122777758 | rs2416742 | -0.356 | -0.862 | -0.882 | -1.511 | 0.019 | 0.011 | 0.011 | 0.183 | 0.5 | 0.53 | 0.32 | 0.4 | 0.32 | 0.43 |
| 9 | 122777961 | rs2416743 | -0.797 | -0.145 | -0.419 | -0.854 | 0.022 | -0.047 | -0.047 | 3.304 | 0.16 | 0.43 | 0.19 | 0.09 | 0.16 | 0.09 |
| 9 | 122779915 | rs16909325 | 0.154 | -0.088 | -0.733 | -0.583 | 0.013 | 0.006 | 0.006 | 0.061 | 0.1 | 0.26 | 0.12 | 0.09 | 0.07 | 0.06 |
| 9 | 122781549 | rs10984806 | 0.002 | -0.064 | -0.78 | -0.798 | 0.018 | 0.005 | 0.005 | 0.955 | 0.32 | 0.34 | 0.51 | 0.64 | 0.39 | 0.27 |
| 9 | 122786574 | rs7873896 | -0.2 | -0.747 | -0.794 | -0.864 | -0.002 | -0.012 | -0.012 | 0 | 0.18 | 0.22 | 0.2 | 0.1 | 0.13 | 0.2 |
| 9 | 122786815 | rs6478451 | -0.74 | -0.817 | -1.419 | -1.047 | -0.03 | 0.023 | 0.023 | 0.583 | 0.81 | 0.99 | 0.85 | 0.91 | 0.78 | 0.76 |
| 9 | 122791136 | rs12344392 | 0.14 | -0.767 | -0.998 | -0.997 | -0.011 | -0.04 | -0.04 | 2.446 | 0.18 | 0.27 | 0.2 | 0.1 | 0.13 | 0.2 |
| 9 | 122793550 | rs12350202 | 0.466 | -0.735 | -0.995 | -0.998 | 0 | -0.003 | -0.003 | 0 | 0.17 | 0.18 | 0.19 | 0.1 | 0.13 | 0.2 |
| 9 | 122795581 | rs6478453 | 0.48 | -0.705 | -0.914 | -0.971 | 0.009 | 0.023 | 0.023 | 0 | 0.62 | 0.69 | 0.65 | 0.81 | 0.65 | 0.55 |
| 9 | 122799150 | rs7851216 | 0.487 | -0.714 | -0.891 | -0.994 | 0.009 | 0.027 | 0.027 | 0 | 0.62 | 0.67 | 0.65 | 0.81 | 0.65 | 0.56 |
| 9 | 122810876 | rs4836798 | 0.332 | -1.849 | -0.641 | -0.341 | 0.11 | 0.04 | 0.04 | 0 | 0.49 | 0.25 | 0.58 | 0.49 | 0.64 | 0.49 |
| 9 | 122814965 | rs7865756 | 0.369 | -2.119 | -0.044 | -0.422 | -0.02 | -0.055 | -0.055 | 0.113 | 0.19 | 0.35 | 0.16 | 0.09 | 0.17 | 0.24 |
| 9 | 122817066 | rs10760075 | 0.4 | -2.381 | -0.697 | -0.128 | 0.13 | 0.088 | 0.088 | 0.019 | 0.53 | 0.26 | 0.6 | 0.74 | 0.69 | 0.68 |
| 9 | 122829607 | rs13302206 | 1.137 | -2.001 | -0.645 | -0.497 | 0.05 | 0.014 | 0.014 | 0 | 0.45 | 0.23 | 0.53 | 0.48 | 0.59 | 0.49 |
| 9 | 122829654 | rs7854483 | 1.066 | -2.039 | -0.457 | -0.399 | 0.068 | -0.115 | -0.115 | 4.88 | 0.28 | 0.1 | 0.23 | 0.41 | 0.15 | 0.24 |
| 9 | 122833468 | rs4837698 | 0.245 | -2.024 | -0.377 | -0.398 | -0.015 | -0.048 | -0.048 | 0 | 0.76 | 0.39 | 0.8 | 0.91 | 0.8 | 0.76 |
| 9 | 122839274 | rs7870808 | 0.768 | -1.76 | 0.032 | 0.041 | 0.087 | -0.099 | -0.099 | 0 | 0.25 | 0.1 | 0.2 | 0.42 | 0.1 | 0.19 |
| 9 | 122842260 | rs945249 | -0.092 | -1.683 | 0.16 | 0.064 | -0.042 | 0.038 | 0.038 | 0.536 | 0.67 | 0.44 | 0.71 | 0.57 | 0.82 | 0.76 |
| 9 | 122846710 | rs4837700 | 0.338 | -0.652 | -0.048 | 0.16 | 0.013 | -0.013 | -0.013 | 0 | 0.45 | 0.25 | 0.59 | 0.57 | 0.54 | 0.67 |
| 9 | 122853291 | rs6478456 | 0.083 | -0.817 | -0.05 | 0.06 | -0.013 | -0.034 | -0.034 | 0 | 0.06 | 0.17 | 0.03 | 0 | 0.03 | 0.01 |
| 9 | 122855414 | rs12003361 | -0.012 | -0.893 | 0.104 | 0.008 | 0.068 | 0.038 | 0.038 | 2 | 0.05 | 0.35 | 0.03 | 0 | 0.03 | 0.02 |
| 9 | 122856204 | rs1981122 | 0.861 | -0.897 | -0.927 | 0.176 | 0.021 | -0.061 | -0.061 | 0.197 | 0.38 | 0.06 | 0.29 | 0.34 | 0.34 | 0.27 |
| 9 | 122857993 | rs12554672 | 0.103 | -0.959 | -0.879 | -0.076 | -0.011 | -0.028 | -0.028 | 0 | 0.07 | 0.12 | 0.03 | 0 | 0.03 | 0.01 |
| 9 | 122859248 | rs7022811 | 0.237 | -0.714 | -0.735 | 0.171 | 0.045 | 0.002 | 0.002 | 1.738 | 0.51 | 0.32 | 0.61 | 0.57 | 0.56 | 0.67 |
| 9 | 122865267 | rs16909453 | 0.408 | -0.575 | -1.197 | 0.23 | -0.03 | -0.124 | -0.124 | 0 | 0.38 | 0.06 | 0.31 | 0.42 | 0.36 | 0.3 |
| 9 | 122868699 | rs5020365 | -0.045 | -1.059 | -0.78 | 0.288 | 0.044 | -0.017 | -0.017 | 0 | 0.43 | 0.66 | 0.33 | 0.43 | 0.33 | 0.28 |
| 9 | 122868788 | rs10984840 | 0.335 | -0.858 | -1.296 | 0.226 | 0.023 | -0.14 | -0.14 | 0 | 0.38 | 0.04 | 0.29 | 0.44 | 0.34 | 0.26 |
| 9 | 122873248 | rs2416750 | -1.17 | -0.12 | 0.575 | 0.036 | 0.037 | 0.034 | 0.034 | 1.178 | 0.06 | 0.38 | 0.04 | 0 | 0.01 | 0 |
| 9 | 122874156 | rs945246 | 0.15 | 0.323 | 0.253 | 0.147 | 0.015 | 0.001 | 0.001 | 0.02 | 0.63 | 0.59 | 0.58 | 0.6 | 0.56 | 0.52 |
| 9 | 122887345 | rs4837708 | 0.381 | -0.07 | 0.438 | 0.557 | -0.035 | -0.104 | -0.104 | 0 | 0.18 | 0.03 | 0.2 | 0.37 | 0.23 | 0.25 |
| 9 | 122894522 | rs1414063 | 1.161 | 0.341 | 1.612 | -0.011 | 0.021 | -0.019 | -0.019 | 3.566 | 0.75 | 0.53 | 0.7 | 0.78 | 0.76 | 0.68 |
| 9 | 122894800 | rs10760078 | 1.912 | 0.193 | 1.095 | -0.045 | -0.012 | -0.037 | -0.037 | 1.206 | 0.67 | 0.09 | 0.64 | 0.65 | 0.73 | 0.58 |
| 9 | 122908894 | rs4836804 | -1.264 | 0.014 | 1.731 | -0.311 | 0.004 | 0.008 | 0.008 | 0 | 0.24 | 0.37 | 0.3 | 0.27 | 0.26 | 0.43 |
| 9 | 122918266 | rs4837719 | -1.805 | 0.002 | 2.019 | -0.316 | -0.001 | -0.001 | -0.001 | 0 | 0.29 | 0.69 | 0.35 | 0.26 | 0.27 | 0.43 |
| 9 | 122937625 | rs1335210 | -1.543 | 0.016 | 2.195 | -0.309 | -0.003 | -0.001 | -0.001 | 0 | 0.27 | 0.53 | 0.32 | 0.26 | 0.26 | 0.43 |
| 9 | 122937656 | rs1335211 | -1.258 | 0.095 | 1.975 | -0.271 | 0.009 | 0.011 | 0.011 | 0.28 | 0.24 | 0.37 | 0.3 | 0.26 | 0.26 | 0.42 |
| 9 | 122954405 | rs4837728 | 1.525 | 0.087 | 0.602 | -0.352 | -0.006 | -0.008 | -0.008 | 0 | 0.72 | 0.11 | 0.65 | 0.74 | 0.76 | 0.64 |
| 9 | 122958532 | rs10984865 | -1.498 | 0.238 | 0.887 | -0.268 | 0.004 | 0.02 | 0.02 | 0 | 0.07 | 0.09 | 0.19 | 0.17 | 0.11 | 0.06 |
| 9 | 122959229 | rs10984866 | -2.046 | 0.08 | 1.115 | -0.319 | 0 | 0.033 | 0.033 | 0 | 0.09 | 0.16 | 0.19 | 0.17 | 0.11 | 0.06 |
| 9 | 122960228 | rs1860990 | -1.219 | -0.059 | 0.942 | -0.672 | -0.021 | 0.055 | 0.055 | 0 | 0.26 | 0.1 | 0.16 | 0.16 | 0.22 | 0.36 |
| 9 | 122965218 | rs4837732 | 1.796 | -0.308 | 1.483 | -0.718 | -0.029 | 0.043 | 0.043 | 1.581 | 0.86 | 0.45 | 0.78 | 0.84 | 0.89 | 0.92 |
| 9 | 122968584 | rs13292795 | -0.186 | -0.305 | 1.244 | -0.74 | -0.046 | -0.022 | -0.022 | 0.027 | 0.13 | 0.01 | 0.09 | 0.16 | 0.14 | 0.35 |
| 9 | 122972617 | rs4836809 | 2.196 | -0.27 | 1.406 | -0.703 | 0.007 | 0.014 | 0.014 | 0 | 0.59 | 0.35 | 0.62 | 0.68 | 0.67 | 0.54 |
| 9 | 122975509 | rs7040919 | 1.791 | -0.185 | 1.261 | -0.657 | -0.022 | 0.053 | 0.053 | 1.15 | 0.87 | 0.47 | 0.78 | 0.84 | 0.9 | 0.92 |
| 9 | 123005508 | rs4997435 | 1.531 | 0.228 | -0.616 | -1.144 | -0.021 | 0.184 | 0.184 | 0.409 | 0.6 | 0.14 | 0.68 | 0.91 | 0.62 | 0.59 |

| | | | | | | | | | | | | | | | | |
|---|---|---|---|---|---|---|---|---|---|---|---|---|---|---|---|---|
| 9 | 123011212 | rs10739558 | 1.587 | -0.365 | 0.159 | -0.447 | 0.017 | 0.021 | 0.021 | 4.582 | 0.76 | 0.23 | 0.83 | 0.92 | 0.87 | 0.88 |
| 9 | 123011594 | rs4837737 | 1.418 | -0.327 | 0.237 | -0.429 | -0.004 | -0.014 | -0.014 | 0 | 0.77 | 0.29 | 0.84 | 0.92 | 0.87 | 0.88 |
| 9 | 123025230 | rs2416755 | 1.672 | -0.806 | 0.627 | -0.337 | -0.074 | -0.049 | -0.049 | 0.669 | 0.8 | 0.31 | 0.9 | 0.93 | 0.95 | 0.89 |
| 9 | 123041996 | rs759125 | 1.627 | -0.603 | 1.381 | -0.392 | -0.024 | -0.02 | -0.02 | 0 | 0.87 | 0.66 | 0.92 | 0.92 | 0.91 | 0.86 |
| 9 | 123050797 | rs887807 | 2.374 | -0.584 | 1.982 | 0.277 | -0.016 | 0.14 | 0.14 | 0.951 | 0.87 | 0.53 | 0.7 | 0.72 | 0.93 | 0.85 |
| 9 | 123056637 | rs4837752 | -2.184 | -0.454 | 1.882 | 0.461 | 0.007 | 0.127 | 0.127 | 0 | 0.08 | 0.19 | 0.25 | 0.28 | 0.07 | 0.17 |
| 9 | 123074244 | rs917873 | -0.676 | -0.469 | 0.845 | -0.229 | -0.007 | -0.007 | -0.007 | 0.523 | 0.22 | 0.1 | 0.3 | 0.26 | 0.22 | 0.2 |
| 9 | 123075135 | rs2159994 | -0.544 | -0.457 | 0.882 | -0.213 | -0.003 | -0.015 | -0.015 | 0 | 0.18 | 0.12 | 0.27 | 0.26 | 0.18 | 0.19 |
| 9 | 123085369 | rs200551 | -2.476 | -0.542 | 2.222 | 0.05 | 0.007 | 0.127 | 0.127 | 2.732 | 0.1 | 0.51 | 0.07 | 0.13 | 0.03 | 0.09 |
| 9 | 123088244 | rs10984890 | -1.817 | -0.514 | 2.328 | 0.084 | 0.002 | 0.043 | 0.043 | 0 | 0.05 | 0.13 | 0.03 | 0.12 | 0.03 | 0.1 |
| 9 | 123093411 | rs10984892 | -2.262 | -0.517 | 2.424 | 0.262 | -0.004 | 0.05 | 0.05 | 0.013 | 0.06 | 0.2 | 0.04 | 0.14 | 0.03 | 0.1 |
| 9 | 123105974 | rs846826 | -3.194 | -0.608 | 3.592 | 0.231 | -0.027 | 0.036 | 0.036 | 3.356 | 0.11 | 0.66 | 0.08 | 0.13 | 0.04 | 0.1 |
| 9 | 123113475 | rs1332247 | -2.292 | -0.786 | 3.757 | 0.511 | 0.038 | 0.256 | 0.256 | 0.436 | 0.07 | 0.38 | 0.05 | 0.18 | 0.01 | 0.11 |
| 9 | 123114119 | rs10818441 | -1.033 | -0.408 | 3.077 | 1.097 | -0.005 | 0.02 | 0.02 | 1.028 | 0.27 | 0.67 | 0.36 | 0.5 | 0.2 | 0.42 |
| 9 | 123114345 | rs12376794 | -1.045 | -0.409 | 3.077 | 1.096 | -0.005 | 0.02 | 0.02 | 1.115 | 0.27 | 0.67 | 0.36 | 0.5 | 0.2 | 0.41 |
| 9 | 123115902 | rs10818442 | -0.193 | -0.106 | 2.948 | 1.1 | 0 | 0.003 | 0.003 | 0.027 | 0.2 | 0.27 | 0.32 | 0.33 | 0.2 | 0.3 |
| 9 | 123118155 | rs7024514 | -1.368 | -0.111 | 3.084 | 1.096 | 0.046 | 0.251 | 0.251 | 0.099 | 0.05 | 0.31 | 0.05 | 0.17 | 0.01 | 0.11 |
| 9 | 123119421 | rs1332245 | 0.825 | 0.269 | 2.503 | 0.953 | 0.014 | 0.061 | 0.061 | 0 | 0.72 | 0.45 | 0.62 | 0.5 | 0.74 | 0.58 |
| 9 | 123122187 | rs1112863 | -0.741 | 0.351 | 1.387 | 0.5 | -0.006 | 0.004 | 0.004 | 0 | 0.4 | 0.38 | 0.47 | 0.66 | 0.46 | 0.5 |
| 9 | 123124504 | rs1537282 | -0.814 | 0.813 | 0.515 | 0.595 | 0.003 | -0.001 | -0.001 | 0.073 | 0.35 | 0.61 | 0.47 | 0.66 | 0.4 | 0.48 |
| 9 | 123133385 | rs7852998 | -0.045 | 1.073 | -0.465 | 0.846 | -0.155 | -0.135 | -0.135 | 0.909 | 0.72 | 0.94 | 0.59 | 0.56 | 0.58 | 0.62 |
| 9 | 123133808 | rs10760096 | 0.319 | 0.248 | -0.55 | 0.539 | 0.006 | 0.008 | 0.008 | 1.092 | 0.25 | 0.21 | 0.32 | 0.16 | 0.21 | 0.1 |
| 9 | 123134187 | rs7040576 | 0.153 | 0.817 | -1.239 | 0.771 | -0.078 | -0.105 | -0.105 | 0.272 | 0.23 | 0.08 | 0.37 | 0.46 | 0.34 | 0.42 |
| 9 | 123137705 | rs10818449 | -0.034 | 0.094 | -0.905 | -0.522 | -0.009 | -0.068 | -0.068 | 10.228 | 0.41 | 0.41 | 0.24 | 0.11 | 0.35 | 0.15 |
| 9 | 123142848 | rs10818451 | -0.844 | -1.141 | -0.346 | -0.549 | -0.049 | -0.022 | -0.022 | 0.016 | 0.32 | 0.52 | 0.26 | 0.27 | 0.18 | 0.16 |
| 9 | 123144243 | rs7030519 | -0.917 | -1.234 | -0.692 | -0.684 | -0.003 | -0.014 | -0.014 | 0.009 | 0.7 | 0.64 | 0.77 | 0.79 | 0.69 | 0.65 |
| 9 | 123146316 | rs2183906 | -1.516 | -1.06 | -0.708 | -0.727 | -0.001 | -0.004 | -0.004 | 0 | 0.75 | 0.75 | 0.8 | 0.86 | 0.76 | 0.67 |
| 9 | 123147885 | rs7852100 | -0.799 | -1.228 | -0.453 | -0.786 | -0.008 | -0.034 | -0.034 | 1.729 | 0.69 | 0.59 | 0.76 | 0.82 | 0.7 | 0.66 |
| 9 | 123156591 | rs7859743 | -0.671 | -1.275 | -0.511 | -0.78 | -0.011 | -0.036 | -0.036 | 1.072 | 0.68 | 0.59 | 0.75 | 0.82 | 0.71 | 0.66 |
| 9 | 123161735 | rs10125592 | 0.2 | -1.301 | -0.436 | -0.721 | -0.005 | -0.001 | -0.001 | 0 | 0.27 | 0.3 | 0.21 | 0.26 | 0.14 | 0.15 |
| 9 | 123162666 | rs12379034 | 0.223 | -1.297 | -0.433 | -0.718 | -0.009 | -0.001 | -0.001 | 0 | 0.27 | 0.3 | 0.21 | 0.27 | 0.14 | 0.15 |
| 9 | 123170167 | rs914592 | -1.665 | -1.162 | -0.681 | -0.861 | 0.001 | -0.005 | -0.005 | 0 | 0.82 | 0.84 | 0.88 | 0.9 | 0.8 | 0.83 |
| 9 | 123171755 | rs2297454 | -0.937 | -1.427 | -0.599 | -0.834 | -0.024 | -0.008 | -0.008 | 0 | 0.42 | 0.6 | 0.35 | 0.35 | 0.25 | 0.32 |
| 9 | 123182202 | rs6478475 | 0.533 | -1.065 | -0.489 | -0.703 | -0.007 | 0.021 | 0.021 | 0 | 0.06 | 0.25 | 0.04 | 0.04 | 0.03 | 0 |
| 9 | 123185724 | rs2900169 | 1.613 | -0.867 | -0.898 | -0.768 | -0.094 | -0.138 | -0.138 | 0 | 0.26 | 0.04 | 0.42 | 0.46 | 0.37 | 0.49 |
| 9 | 123207702 | rs4837771 | 0.726 | -1.103 | -0.809 | -0.792 | 0.046 | 0.016 | 0.016 | 0 | 0.5 | 0.26 | 0.58 | 0.53 | 0.59 | 0.56 |
| 9 | 123211786 | rs10984917 | 1.062 | -1.077 | -1.051 | -0.716 | 0.012 | 0.062 | 0.062 | 1.704 | 0.14 | 0.03 | 0.07 | 0 | 0.11 | 0.04 |
| 9 | 123217710 | rs10984920 | 1.214 | -1.076 | -1.011 | -0.629 | 0.007 | -0.025 | -0.025 | 0 | 0.08 | 0.02 | 0.1 | 0.28 | 0.06 | 0.15 |
| 9 | 123220829 | rs2501727 | -0.356 | -1.167 | -0.779 | -0.671 | -0.042 | 0.093 | 0.093 | 0 | 0.16 | 0.36 | 0.16 | 0.32 | 0.1 | 0.16 |
| 9 | 123225926 | rs7869290 | -0.166 | -0.92 | -1.001 | -0.968 | -0.001 | 0.036 | 0.036 | 0.157 | 0.35 | 0.32 | 0.23 | 0.13 | 0.38 | 0.11 |
| 9 | 123234422 | rs10984922 | 0.512 | -1.033 | -0.695 | -0.568 | 0.019 | 0.05 | 0.05 | 1.473 | 0.16 | 0.06 | 0.09 | 0.04 | 0.15 | 0.04 |
| 9 | 123296964 | rs10984944 | 0.225 | -0.916 | 0.084 | -0.101 | -0.027 | 0.14 | 0.14 | 0 | 0.88 | 0.64 | 0.86 | 0.71 | 0.94 | 0.87 |
| 9 | 123298319 | rs7868597 | 0.091 | -0.89 | 0.035 | -0.086 | -0.015 | 0.109 | 0.109 | 0 | 0.1 | 0.25 | 0.12 | 0.29 | 0.06 | 0.13 |
| 9 | 123301346 | rs3750494 | 0.824 | -0.88 | -0.073 | -0.082 | 0.006 | -0.021 | -0.021 | 0 | 0.08 | 0.02 | 0.1 | 0.29 | 0.06 | 0.13 |
| 9 | 123305220 | rs7854663 | 0.004 | -0.871 | -0.001 | -0.054 | -0.014 | 0.15 | 0.15 | 0 | 0.89 | 0.66 | 0.87 | 0.71 | 0.95 | 0.88 |
| 9 | 123322700 | rs1475663 | 1.25 | -1.449 | 0.102 | -0.484 | 0.016 | 0.004 | 0.004 | 0.601 | 0.6 | 0.36 | 0.65 | 0.62 | 0.77 | 0.67 |
| 9 | 123324845 | rs10984956 | 0.313 | -0.616 | 0.176 | 0.102 | -0.008 | 0.092 | 0.092 | 0 | 0.09 | 0.16 | 0.11 | 0.29 | 0.06 | 0.13 |
| 9 | 123327435 | rs12348950 | 0.798 | -0.285 | -0.39 | 0.025 | -0.095 | -0.121 | -0.121 | 0.044 | 0.35 | 0.07 | 0.47 | 0.45 | 0.43 | 0.52 |
| 9 | 123330813 | rs4837782 | 0.421 | 0.063 | 0.908 | 0.709 | -0.02 | 0.102 | 0.102 | 0 | 0.89 | 0.75 | 0.87 | 0.71 | 0.95 | 0.88 |
| 9 | 123351121 | rs7861226 | -1.283 | 0.291 | 1.048 | 0.603 | -0.005 | 0.044 | 0.044 | 1.44 | 0.31 | 0.45 | 0.32 | 0.43 | 0.27 | 0.24 |
| 9 | 123370985 | rs1359329 | 1.385 | 1.323 | 2.008 | 0.872 | 0.063 | 0.153 | 0.153 | 0 | 0.84 | 0.62 | 0.9 | 0.65 | 0.84 | 0.85 |
| 9 | 123374639 | rs2416760 | 2.504 | 1.167 | 2.091 | 0.776 | -0.009 | -0.026 | -0.026 | 0.781 | 0.67 | 0.21 | 0.73 | 0.58 | 0.69 | 0.74 |
| 9 | 123404610 | rs1324473 | 2.437 | 1.148 | 2.17 | 0.797 | -0.003 | -0.015 | -0.015 | 0 | 0.68 | 0.25 | 0.73 | 0.58 | 0.7 | 0.74 |
| 9 | 123421879 | rs2274144 | -0.88 | 1.15 | 2.577 | 0.772 | -0.012 | -0.012 | -0.012 | 0.236 | 0.11 | 0.18 | 0.07 | 0.07 | 0.07 | 0.11 |
| 9 | 123453281 | rs966396 | 1.987 | 1.157 | 2.627 | 0.781 | 0.012 | 0.035 | 0.035 | 0.429 | 0.72 | 0.61 | 0.76 | 0.58 | 0.71 | 0.73 |
| 9 | 123480696 | rs4617229 | 2.545 | 1.055 | 2.029 | 0.726 | -0.001 | -0.018 | -0.018 | 0 | 0.66 | 0.25 | 0.74 | 0.58 | 0.7 | 0.74 |
| 9 | 123480866 | rs2416799 | 2.329 | 1.074 | 2 | 0.735 | 0.003 | 0.024 | 0.024 | 0.926 | 0.68 | 0.38 | 0.75 | 0.58 | 0.7 | 0.74 |
| 9 | 123497595 | rs10985011 | -1.231 | 1.091 | 1.744 | 0.592 | -0.004 | -0.032 | -0.032 | 0 | 0.13 | 0.17 | 0.15 | 0.02 | 0.14 | 0.12 |
| 9 | 123497645 | rs2900177 | 2.736 | 1.086 | 1.985 | 0.753 | -0.003 | -0.036 | -0.036 | 0 | 0.66 | 0.22 | 0.71 | 0.56 | 0.7 | 0.72 |
| 9 | 123498290 | rs10985014 | -1.286 | 1.042 | 1.954 | 0.58 | -0.005 | -0.045 | -0.045 | 0 | 0.13 | 0.2 | 0.15 | 0.02 | 0.14 | 0.12 |
| 9 | 123498335 | rs10985015 | -1.573 | 1.041 | 2.078 | 0.579 | -0.008 | -0.069 | -0.069 | 0 | 0.14 | 0.31 | 0.16 | 0.02 | 0.14 | 0.12 |
| 9 | 123514048 | rs10985021 | -0.891 | 1.18 | 1.749 | 0.593 | -0.008 | 0.02 | 0.02 | 3.166 | 0.12 | 0.06 | 0.14 | 0.02 | 0.16 | 0.08 |
| 9 | 123520577 | rs16910125 | -1.062 | 1.175 | 1.91 | 0.604 | 0.003 | -0.025 | -0.025 | 0 | 0.11 | 0.15 | 0.14 | 0.02 | 0.15 | 0.08 |
| 9 | 123522115 | rs3814536 | -0.97 | 1.176 | 1.793 | 0.605 | -0.003 | 0.001 | 0.001 | 3.034 | 0.11 | 0.1 | 0.14 | 0.02 | 0.15 | 0.08 |
| 9 | 123525175 | rs10985030 | -0.861 | 1.124 | 1.83 | 0.569 | -0.007 | 0.01 | 0.01 | 3.536 | 0.11 | 0.06 | 0.13 | 0.02 | 0.15 | 0.08 |
| 9 | 123539791 | rs16910134 | -0.141 | 1.131 | 1.734 | 0.903 | -0.012 | 0.028 | 0.028 | 0 | 0.05 | 0 | 0.01 | 0.33 | 0.03 | 0.12 |
| 9 | 123544003 | rs2416800 | 2.826 | 2.369 | 1.533 | 0.858 | -0.146 | -0.149 | -0.149 | 0 | 0.64 | 0.08 | 0.64 | 0.47 | 0.56 | 0.7 |
| 9 | 123548885 | rs10985036 | -0.956 | 2.158 | 1.58 | 0.918 | -0.003 | 0.003 | 0.003 | 2.421 | 0.12 | 0.1 | 0.13 | 0.02 | 0.15 | 0.08 |
| 9 | 123550027 | rs7026635 | 1.728 | 2.293 | 1.923 | 0.948 | 0.001 | 0.014 | 0.014 | 0 | 0.84 | 0.85 | 0.85 | 0.54 | 0.74 | 0.8 |

| | | | | | | | | | | | | | | | | |
|---|---|---|---|---|---|---|---|---|---|---|---|---|---|---|---|---|
| 9 | 123560833 | rs13285804 | -0.7 | 2.332 | 1.837 | 0.917 | -0.008 | 0.016 | 0.016 | 3.101 | 0.11 | 0.06 | 0.13 | 0.02 | 0.15 | 0.08 |
| 9 | 123563510 | rs10985044 | -0.7 | 2.332 | 1.837 | 0.917 | -0.008 | 0.016 | 0.016 | 3.101 | 0.11 | 0.06 | 0.13 | 0.02 | 0.15 | 0.08 |
| 9 | 123578964 | rs6602 | -0.7 | 2.332 | 1.825 | 0.917 | -0.009 | 0.019 | 0.019 | 3.101 | 0.11 | 0.05 | 0.13 | 0.02 | 0.15 | 0.08 |
| 9 | 123583192 | rs1060817 | 3.052 | 2.332 | 1.94 | 0.977 | -0.058 | -0.078 | -0.078 | 0 | 0.66 | 0.16 | 0.66 | 0.47 | 0.59 | 0.73 |
| 9 | 123586737 | rs10760117 | 3.052 | 2.332 | 1.94 | 0.956 | -0.058 | -0.08 | -0.08 | 0 | 0.66 | 0.16 | 0.65 | 0.47 | 0.59 | 0.73 |
| 9 | 123606101 | rs10739575 | 1.636 | 2.413 | 1.786 | 0.964 | 0.025 | 0.084 | 0.084 | 0 | 0.88 | 0.85 | 0.88 | 0.54 | 0.84 | 0.82 |
| 9 | 123607829 | rs933003 | -0.187 | 2.33 | 1.492 | 1.001 | -0.012 | 0.023 | 0.023 | 0 | 0.05 | 0 | 0.01 | 0.33 | 0.03 | 0.12 |
| 9 | 123607880 | rs10985051 | -0.742 | 2.289 | 1.525 | 0.876 | -0.009 | 0.019 | 0.019 | 3.121 | 0.11 | 0.05 | 0.13 | 0.02 | 0.16 | 0.08 |
| 9 | 123610288 | rs4837796 | 2.656 | 2.372 | 1.44 | 0.829 | -0.057 | -0.08 | -0.08 | 0.441 | 0.66 | 0.16 | 0.65 | 0.47 | 0.58 | 0.73 |
| 9 | 123616858 | rs12379604 | -0.888 | 2.719 | 1.144 | 0.849 | -0.015 | 0.021 | 0.021 | 2.94 | 0.11 | 0.05 | 0.13 | 0.02 | 0.17 | 0.08 |
| 9 | 123618229 | rs1837 | 1.587 | 2.588 | 1.35 | 0.867 | 0.018 | 0.04 | 0.04 | 0 | 0.81 | 0.75 | 0.84 | 0.55 | 0.73 | 0.79 |
| 9 | 123618971 | rs914842 | 1.274 | 2.468 | 1.407 | 0.923 | 0.012 | 0.06 | 0.06 | 0 | 0.83 | 0.75 | 0.85 | 0.55 | 0.77 | 0.82 |
| 9 | 123626533 | rs12377227 | -0.754 | 2.207 | 0.952 | 0.814 | -0.003 | 0.001 | 0.001 | 4.699 | 0.09 | 0.06 | 0.11 | 0.02 | 0.12 | 0.07 |
| 9 | 123632045 | rs1056567 | 1.611 | 2.348 | 1.272 | 1.054 | 0.077 | 0.171 | 0.171 | 0 | 0.77 | 0.53 | 0.78 | 0.5 | 0.68 | 0.78 |
| 9 | 123633948 | rs3933326 | 1.609 | 2.357 | 1.27 | 1.052 | 0.08 | 0.171 | 0.171 | 0 | 0.77 | 0.53 | 0.78 | 0.5 | 0.68 | 0.77 |
| 9 | 123640500 | rs1953126 | 1.64 | 2.498 | 1.517 | 1.174 | 0.007 | 0.005 | 0.005 | 0.007 | 0.77 | 0.78 | 0.68 | 0.67 | 0.64 | 0.78 |
| 9 | 123642351 | rs1609810 | 1.64 | 2.498 | 1.517 | 1.237 | 0.007 | 0.003 | 0.003 | 0.007 | 0.77 | 0.78 | 0.69 | 0.72 | 0.64 | 0.78 |
| 9 | 123643855 | rs10985073 | 2.603 | 2.25 | 1.241 | 1.181 | -0.138 | -0.152 | -0.152 | 0 | 0.63 | 0.07 | 0.57 | 0.5 | 0.56 | 0.65 |
| 9 | 123652898 | rs881375 | 1.427 | 2.259 | 1.248 | 1.179 | 0.007 | 0.003 | 0.003 | 0.009 | 0.77 | 0.78 | 0.69 | 0.72 | 0.64 | 0.78 |
| 9 | 123655329 | rs6478486 | 1.419 | 2.262 | 1.689 | 1.18 | 0.008 | 0.004 | 0.004 | 0.022 | 0.78 | 0.78 | 0.69 | 0.72 | 0.64 | 0.8 |
| 9 | 123665019 | rs9886724 | 2.522 | 2.26 | 1.89 | 1.182 | -0.102 | -0.118 | -0.118 | 0 | 0.64 | 0.11 | 0.56 | 0.5 | 0.56 | 0.67 |
| 9 | 123668199 | rs10435844 | 1.431 | 2.268 | 1.887 | 1.177 | 0.008 | 0.003 | 0.003 | 0.02 | 0.78 | 0.78 | 0.69 | 0.72 | 0.64 | 0.8 |
| 9 | 123671520 | rs2239657 | 1.431 | 2.268 | 1.887 | 1.177 | 0.008 | 0.003 | 0.003 | 0.02 | 0.78 | 0.78 | 0.69 | 0.72 | 0.64 | 0.8 |
| 9 | 123673890 | rs7021880 | -1.199 | 2.262 | 1.941 | 1.149 | 0.012 | 0.007 | 0.007 | 0 | 0.17 | 0.16 | 0.28 | 0.28 | 0.34 | 0.19 |
| 9 | 123677827 | rs2109895 | 1.461 | 2.277 | 1.888 | 1.173 | 0.008 | 0.003 | 0.003 | 0.02 | 0.78 | 0.78 | 0.69 | 0.72 | 0.64 | 0.8 |
| 9 | 123681275 | rs7038991 | -1.148 | 2.232 | 2.456 | 1.144 | 0.043 | -0.025 | -0.025 | 0 | 0.07 | 0.23 | 0.07 | 0 | 0.06 | 0.08 |
| 9 | 123683982 | rs7021049 | 2.684 | 2.244 | 2.403 | 1.17 | -0.105 | -0.115 | -0.115 | 0 | 0.64 | 0.12 | 0.56 | 0.5 | 0.56 | 0.67 |
| 9 | 123685092 | rs1014530 | 2.917 | 2.25 | 2.437 | 1.18 | -0.003 | -0.003 | -0.003 | 0 | 0.65 | 0.34 | 0.6 | 0.5 | 0.57 | 0.67 |
| 9 | 123690239 | rs3761847 | 2.886 | 2.25 | 2.613 | 1.184 | 0.016 | 0.017 | 0.017 | 0 | 0.66 | 0.39 | 0.6 | 0.52 | 0.57 | 0.68 |
| 9 | 123699083 | rs10985095 | -1.474 | 2.255 | 2.716 | 1.189 | -0.004 | 0.157 | 0.157 | 2.598 | 0.06 | 0.28 | 0.04 | 0.22 | 0.01 | 0.05 |
| 9 | 123700779 | rs10733648 | 1.644 | 2.249 | 2.716 | 1.197 | 0.058 | 0.032 | 0.032 | 0 | 0.77 | 0.65 | 0.68 | 0.73 | 0.64 | 0.79 |
| 9 | 123703894 | rs10985097 | -1.217 | 2.247 | 2.775 | 1.196 | -0.01 | 0.118 | 0.118 | 2.621 | 0.05 | 0.22 | 0.04 | 0.2 | 0.01 | 0.04 |
| 9 | 123705087 | rs10818488 | 2.953 | 2.252 | 2.702 | 1.194 | 0.005 | 0.007 | 0.007 | 0 | 0.66 | 0.36 | 0.6 | 0.5 | 0.57 | 0.67 |
| 9 | 123705945 | rs7039505 | 1.67 | 2.266 | 2.528 | 1.147 | 0.057 | 0.033 | 0.033 | 0 | 0.77 | 0.65 | 0.68 | 0.73 | 0.64 | 0.8 |
| 9 | 123716103 | rs41258306 | -1.127 | 1.886 | 2.524 | 0.671 | -0.001 | -0.009 | -0.009 | 5.133 | 0.06 | 0.07 | 0.04 | 0 | 0.06 | 0.08 |
| 9 | 123716681 | rs12004487 | -2.811 | 1.857 | 2.591 | 0.706 | 0.008 | 0.077 | 0.077 | 0.052 | 0.12 | 0.53 | 0.11 | 0.22 | 0.08 | 0.13 |
| 9 | 123716840 | rs2300929 | -2.811 | 1.857 | 2.591 | 0.706 | 0.008 | 0.077 | 0.077 | 0.052 | 0.12 | 0.53 | 0.11 | 0.22 | 0.08 | 0.13 |
| 9 | 123717840 | rs7045519 | -3.131 | 1.842 | 2.653 | 0.698 | -0.018 | 0.029 | 0.029 | 0.143 | 0.13 | 0.64 | 0.12 | 0.22 | 0.08 | 0.13 |
| 9 | 123727994 | rs7035682 | -2.523 | 1.877 | 2.472 | 0.716 | 0.007 | 0.076 | 0.076 | 0.047 | 0.12 | 0.53 | 0.11 | 0.22 | 0.08 | 0.13 |
| 9 | 123731408 | rs10985112 | -2.381 | 1.878 | 2.497 | 0.615 | 0.025 | -0.016 | -0.016 | 6.192 | 0.11 | 0.53 | 0.1 | 0.03 | 0.07 | 0.09 |
| 9 | 123733133 | rs7026551 | -2.795 | 2.362 | 2.746 | 0.665 | 0.064 | 0.024 | 0.024 | 0.698 | 0.15 | 0.59 | 0.13 | 0.25 | 0.19 | 0.15 |
| 9 | 123737018 | rs2269066 | -2.099 | 2.381 | 2.916 | 0.67 | 0.081 | 0.094 | 0.094 | 2.33 | 0.09 | 0.37 | 0.08 | 0.21 | 0.1 | 0.04 |
| 9 | 123739993 | rs10760132 | 1.612 | 2.228 | 1.803 | -0.057 | 0.043 | 0.046 | 0.046 | 0 | 0.49 | 0.24 | 0.5 | 0.57 | 0.42 | 0.6 |
| 9 | 123746255 | rs4837805 | 0.603 | 2.06 | 1.627 | -0.285 | -0.002 | 0.01 | 0.01 | 0 | 0.63 | 0.68 | 0.62 | 0.79 | 0.62 | 0.75 |
| 9 | 123756983 | rs16910255 | -0.741 | 2.035 | 1.699 | -0.312 | 0.033 | -0.016 | -0.016 | 0 | 0.06 | 0.13 | 0.04 | 0.04 | 0.09 | 0.11 |
| 9 | 123763006 | rs10760135 | 1.386 | 2.044 | 1.473 | -0.278 | -0.061 | 0.026 | 0.026 | 0 | 0.55 | 0.1 | 0.54 | 0.61 | 0.55 | 0.73 |
| 9 | 123769200 | rs17611 | 1.727 | 2.146 | 1.43 | -0.283 | -0.039 | -0.019 | -0.019 | 0 | 0.49 | 0.03 | 0.5 | 0.57 | 0.46 | 0.62 |
| 9 | 123770436 | rs2300932 | 0.67 | 2.229 | 1.565 | -0.31 | -0.001 | -0.016 | -0.016 | 1.168 | 0.62 | 0.63 | 0.6 | 0.82 | 0.61 | 0.76 |
| 9 | 123771799 | rs7027797 | -0.726 | 2.283 | 1.602 | -0.259 | 0.039 | -0.015 | -0.015 | 0 | 0.06 | 0.13 | 0.05 | 0.04 | 0.1 | 0.11 |
| 9 | 123783934 | rs10985126 | -2.388 | 2.286 | 1.681 | -0.253 | 0.131 | 0.084 | 0.084 | 1.016 | 0.13 | 0.44 | 0.1 | 0.24 | 0.17 | 0.14 |
| 9 | 123791936 | rs7031128 | 0.649 | 2.445 | 1.546 | -0.46 | 0.04 | -0.118 | -0.118 | 0 | 0.82 | 0.64 | 0.81 | 0.99 | 0.78 | 0.94 |
| 9 | 123800218 | rs17216529 | -1.371 | 2.473 | 1.531 | -0.453 | 0.078 | 0.111 | 0.111 | 3.993 | 0.07 | 0.31 | 0.05 | 0.18 | 0.07 | 0.03 |
| 9 | 123809890 | rs1468673 | 2.371 | 2.453 | 1.388 | -0.474 | -0.052 | 0.044 | 0.044 | 0 | 0.68 | 0.18 | 0.71 | 0.75 | 0.61 | 0.79 |
| 9 | 123817613 | rs7029523 | 0.657 | 2.456 | 1.506 | -0.441 | 0.039 | -0.098 | -0.098 | 0.129 | 0.82 | 0.65 | 0.81 | 0.97 | 0.78 | 0.94 |
| 9 | 123822837 | rs10985140 | 2.275 | 2.477 | 1.463 | -0.453 | -0.06 | 0.047 | 0.047 | 0 | 0.68 | 0.17 | 0.7 | 0.75 | 0.61 | 0.8 |
| 9 | 123823972 | rs1179766 | 0.172 | 2.433 | 1.617 | -0.393 | 0.117 | 0.014 | 0.014 | 0.059 | 0.95 | 0.75 | 0.94 | 0.99 | 0.94 | 0.98 |
| 9 | 123824021 | rs2416815 | -1.273 | 2.347 | 1.314 | -0.356 | 0.073 | 0.109 | 0.109 | 4.171 | 0.07 | 0.3 | 0.05 | 0.18 | 0.07 | 0.03 |
| 9 | 123835199 | rs16910300 | 0.159 | 2.239 | 0.85 | -0.38 | 0.018 | -0.005 | -0.005 | 0.317 | 0.19 | 0.09 | 0.2 | 0.17 | 0.15 | 0.16 |
| 9 | 123848837 | rs10985148 | -0.462 | 2.239 | 0.832 | -0.424 | -0.006 | 0.011 | 0.011 | 0 | 0.9 | 0.92 | 0.88 | 0.98 | 0.85 | 0.97 |
| 9 | 123850770 | rs10818503 | 1.338 | 2.398 | 0.902 | -0.415 | -0.016 | 0.015 | 0.015 | 0 | 0.74 | 0.26 | 0.74 | 0.78 | 0.7 | 0.88 |
| 9 | 123857085 | rs10760146 | 2.201 | 2.465 | 0.806 | -0.455 | 0.023 | 0.03 | 0.03 | 0 | 0.49 | 0.12 | 0.5 | 0.57 | 0.45 | 0.64 |
| 9 | 123858194 | rs10733651 | 1.274 | 2.472 | 0.811 | -0.452 | -0.014 | 0.013 | 0.013 | 0 | 0.74 | 0.26 | 0.74 | 0.78 | 0.7 | 0.91 |
| 9 | 123860689 | rs10818504 | 2.193 | 2.471 | 0.804 | -0.449 | 0.024 | 0.035 | 0.035 | 0 | 0.49 | 0.12 | 0.5 | 0.57 | 0.45 | 0.64 |
| 9 | 123867782 | rs10156413 | 1.043 | 2.513 | 0.839 | -0.464 | 0.086 | -0.138 | -0.138 | 0.016 | 0.78 | 0.54 | 0.78 | 0.97 | 0.71 | 0.93 |
| 9 | 123876305 | rs2146838 | 2.171 | 2.471 | 0.723 | -0.464 | -0.068 | 0.048 | 0.048 | 0 | 0.68 | 0.21 | 0.69 | 0.74 | 0.6 | 0.8 |
| 9 | 123914306 | rs3747843 | 2.318 | 2.233 | 0.409 | -0.649 | -0.023 | 0.137 | 0.137 | 0 | 0.58 | 0.17 | 0.65 | 0.74 | 0.51 | 0.78 |
| 9 | 123914592 | rs4836840 | 1.233 | 2.228 | 0.547 | -0.648 | 0.024 | -0.106 | -0.106 | 0 | 0.68 | 0.61 | 0.74 | 0.97 | 0.62 | 0.91 |
| 9 | 123930296 | rs2146836 | 1.319 | 2.003 | 0.517 | -0.569 | -0.016 | 0.014 | 0.014 | 0 | 0.73 | 0.26 | 0.74 | 0.78 | 0.7 | 0.92 |
| 9 | 123938998 | rs3789311 | -0.046 | 1.928 | 0.478 | -0.554 | 0.027 | -0.014 | -0.014 | 1.11 | 0.18 | 0.09 | 0.2 | 0.17 | 0.14 | 0.16 |

| | | | | | | | | | | | | | | | | |
|---|---|---|---|---|---|---|---|---|---|---|---|---|---|---|---|---|
| 9 | 123941122 | rs2057470 | 0.894 | 2.013 | 0.426 | -0.586 | 0.071 | -0.11 | -0.11 | 0 | 0.78 | 0.62 | 0.78 | 0.97 | 0.71 | 0.93 |
| 9 | 123943884 | rs2296078 | 2.337 | 1.585 | 0.049 | -0.545 | 0.007 | 0.015 | 0.015 | 0 | 0.49 | 0.09 | 0.49 | 0.57 | 0.45 | 0.64 |
| 9 | 123951685 | rs942152 | 1.319 | 1.606 | 0.114 | -0.565 | 0.071 | -0.114 | -0.114 | 2.15 | 0.7 | 0.49 | 0.73 | 0.93 | 0.59 | 0.81 |
| 9 | 123955756 | rs10818516 | 1.689 | 1.487 | -0.59 | -0.281 | -0.038 | -0.017 | -0.017 | 0 | 0.5 | 0.03 | 0.49 | 0.57 | 0.46 | 0.64 |
| 9 | 123961405 | rs2209076 | 1.869 | 1.516 | -0.556 | -0.268 | 0.011 | 0.027 | 0.027 | 0 | 0.51 | 0.12 | 0.5 | 0.57 | 0.46 | 0.64 |
| 9 | 123971612 | rs10739592 | 1.523 | 1.183 | -0.716 | -0.445 | 0.019 | 0.073 | 0.073 | 0 | 0.49 | 0.15 | 0.52 | 0.62 | 0.46 | 0.61 |
| 9 | 123985348 | rs1590345 | -1.1 | 0.851 | -0.044 | -0.224 | -0.011 | 0.054 | 0.054 | 0 | 0.08 | 0.22 | 0.24 | 0.12 | 0.05 | 0.11 |
| 9 | 123998342 | rs10760159 | 1.624 | 0.834 | -0.14 | -0.194 | -0.018 | 0.044 | 0.044 | 0 | 0.49 | 0.08 | 0.43 | 0.62 | 0.46 | 0.62 |
| 9 | 124003326 | rs10760160 | 1.723 | 0.811 | -0.161 | -0.199 | -0.071 | 0.004 | 0.004 | 0 | 0.48 | 0.03 | 0.43 | 0.62 | 0.46 | 0.62 |
| 9 | 124016909 | rs4556152 | 1.436 | 0.759 | -0.093 | -0.234 | 0.012 | 0.121 | 0.121 | 0 | 0.52 | 0.3 | 0.63 | 0.75 | 0.47 | 0.72 |
| 9 | 124021254 | rs10760165 | 1.436 | 0.759 | -0.094 | -0.234 | 0.012 | 0.118 | 0.118 | 0 | 0.52 | 0.3 | 0.64 | 0.75 | 0.47 | 0.72 |
| 9 | 124029155 | rs2239696 | 1.465 | 0.759 | -0.094 | -0.196 | 0.013 | 0.117 | 0.117 | 0 | 0.52 | 0.29 | 0.64 | 0.73 | 0.47 | 0.72 |
| 9 | 124030179 | rs10818524 | 1.747 | 0.792 | -0.131 | -0.189 | -0.07 | -0.001 | -0.001 | 0 | 0.47 | 0.03 | 0.42 | 0.61 | 0.45 | 0.62 |
| 9 | 124033044 | rs10985196 | -1.278 | 1.019 | 0.52 | -0.344 | 0.063 | 0.027 | 0.027 | 0 | 0.19 | 0.5 | 0.33 | 0.2 | 0.21 | 0.22 |
| 9 | 124043867 | rs10985200 | 2.156 | 0.966 | 0.443 | -0.388 | 0.021 | 0.085 | 0.085 | 0 | 0.5 | 0.19 | 0.43 | 0.63 | 0.46 | 0.62 |
| 9 | 124044995 | rs3810942 | 2.076 | 0.975 | 0.335 | -0.384 | -0.072 | 0.013 | 0.013 | 0.081 | 0.48 | 0.03 | 0.42 | 0.63 | 0.46 | 0.62 |
| 9 | 124047237 | rs7046030 | -0.592 | 1.006 | 0.545 | -0.428 | 0.066 | -0.066 | -0.066 | 0 | 0.15 | 0.38 | 0.13 | 0.06 | 0.19 | 0.11 |
| 9 | 124047246 | rs913767 | 0.654 | 0.99 | 0.178 | -0.391 | -0.001 | 0.004 | 0.004 | 0 | 0.69 | 0.73 | 0.77 | 0.81 | 0.66 | 0.83 |
| 9 | 124051400 | rs1078305 | 0.816 | 0.998 | 0.165 | -0.378 | -0.001 | -0.015 | -0.015 | 0 | 0.68 | 0.69 | 0.76 | 0.82 | 0.66 | 0.83 |
| 9 | 124060879 | rs3827677 | 0.482 | 0.975 | 0.077 | -0.389 | -0.004 | 0.011 | 0.011 | 0 | 0.71 | 0.78 | 0.77 | 0.81 | 0.66 | 0.83 |
| 9 | 124067789 | rs913763 | 1.793 | 0.962 | -0.45 | -0.146 | -0.074 | -0.034 | -0.034 | 0 | 0.47 | 0.03 | 0.42 | 0.6 | 0.45 | 0.62 |
| 9 | 124072652 | rs306784 | -1.434 | 0.687 | 0.022 | -0.393 | -0.001 | -0.011 | -0.011 | 0 | 0.34 | 0.66 | 0.26 | 0.12 | 0.39 | 0.22 |
| 9 | 124074420 | rs2039144 | 0.306 | 0.922 | -0.096 | -0.553 | -0.002 | -0.026 | -0.026 | 0 | 0.82 | 0.84 | 0.85 | 0.99 | 0.8 | 0.89 |
| 9 | 124075544 | rs306761 | -1.283 | 0.788 | 0.171 | -0.341 | 0.004 | -0.016 | -0.016 | 0 | 0.33 | 0.61 | 0.45 | 0.24 | 0.36 | 0.27 |
| 9 | 124083614 | rs2304393 | 0.604 | 0.391 | -0.05 | -0.275 | -0.01 | 0.004 | 0.004 | 0 | 0.1 | 0.08 | 0.04 | 0.18 | 0.05 | 0.07 |
| 9 | 124090784 | rs306770 | 0.491 | 0.471 | -0.017 | -0.426 | -0.005 | 0.007 | 0.007 | 0 | 0.8 | 0.85 | 0.85 | 0.99 | 0.81 | 0.89 |
| 9 | 124092355 | rs306772 | -0.189 | 0.525 | 0.342 | -0.5 | 0.087 | -0.031 | -0.031 | 0 | 0.16 | 0.43 | 0.12 | 0.11 | 0.2 | 0.11 |
| 9 | 124093940 | rs446038 | -0.272 | 0.46 | 0.261 | -0.5 | -0.021 | 0.044 | 0.044 | 0.001 | 0.85 | 0.92 | 0.88 | 0.99 | 0.83 | 0.93 |
| 9 | 124094014 | rs364814 | -0.742 | 0.461 | 0.262 | -0.5 | -0.022 | 0.031 | 0.031 | 0 | 0.87 | 0.92 | 0.88 | 0.99 | 0.84 | 0.93 |
| 9 | 124095271 | rs398720 | 0.952 | 0.427 | 0.334 | -0.463 | 0 | -0.016 | -0.016 | 0.781 | 0.78 | 0.67 | 0.61 | 0.87 | 0.79 | 0.79 |
| 9 | 124098336 | rs767769 | 2.594 | 0.341 | 0.14 | -0.472 | -0.045 | -0.06 | -0.06 | 0 | 0.39 | 0.08 | 0.36 | 0.58 | 0.39 | 0.55 |
| 9 | 124103759 | rs7853284 | -0.448 | 0.424 | 0.608 | -0.223 | -0.059 | 0.021 | 0.021 | 0.589 | 0.14 | 0.25 | 0.06 | 0.17 | 0.05 | 0.07 |
| 9 | 124109022 | rs306793 | -0.575 | 0.479 | 0.575 | -0.246 | -0.007 | -0.006 | -0.006 | 0.938 | 0.87 | 0.89 | 0.88 | 0.99 | 0.84 | 0.93 |
| 9 | 124118776 | rs3747846 | -1.014 | 0.442 | 0.598 | -0.233 | -0.044 | 0.002 | 0.002 | 0.134 | 0.18 | 0.28 | 0.08 | 0.17 | 0.08 | 0.11 |
| 9 | 124125520 | rs17086 | -2.26 | 0.455 | 0.637 | -0.261 | -0.01 | -0.013 | -0.013 | 0 | 0.45 | 0.5 | 0.51 | 0.31 | 0.39 | 0.34 |
| 9 | 124126618 | rs16910566 | -0.415 | 0.636 | 0.554 | -0.318 | 0.013 | 0.012 | 0.012 | 0 | 0.18 | 0.1 | 0.11 | 0.01 | 0.17 | 0.1 |
| 9 | 124127335 | rs306796 | -1.684 | 0.636 | 0.561 | -0.314 | 0.001 | 0.013 | 0.013 | 0 | 0.34 | 0.3 | 0.45 | 0.14 | 0.34 | 0.27 |
| 9 | 124128354 | rs167396 | -0.205 | 0.599 | 0.412 | -0.227 | -0.003 | -0.001 | -0.001 | 0.007 | 0.15 | 0.17 | 0.34 | 0.13 | 0.18 | 0.16 |
| 9 | 124131512 | rs367395 | 0.458 | -0.003 | 0.786 | 0.138 | -0.028 | 0.003 | 0.003 | 0 | 0.81 | 0.78 | 0.68 | 0.83 | 0.85 | 0.89 |
| 9 | 124135334 | rs3761853 | 0.376 | -0.257 | 0.686 | 0.122 | -0.015 | 0.003 | 0.003 | 0.205 | 0.1 | 0.08 | 0.04 | 0.12 | 0.05 | 0.08 |
| 9 | 124160616 | rs424829 | 0.183 | 0.452 | 0.533 | 0.052 | -0.052 | 0.045 | 0.045 | 0 | 0.17 | 0.09 | 0.16 | 0 | 0.29 | 0.12 |
| 9 | 124170490 | rs2196311 | 1.02 | 0.162 | 0.979 | 0.378 | -0.089 | 0.057 | 0.057 | 1.264 | 0.72 | 0.46 | 0.64 | 0.61 | 0.86 | 0.76 |
| 9 | 124189101 | rs12682698 | -0.05 | 0.552 | 0.799 | 0.384 | -0.028 | -0.021 | -0.021 | 1.336 | 0.07 | 0.03 | 0.06 | 0.12 | 0.14 | 0.08 |
| 9 | 124192482 | rs306777 | 0.014 | 0.318 | 1.289 | 0.312 | -0.048 | 0.074 | 0.074 | 0 | 0.85 | 0.67 | 0.7 | 0.73 | 0.91 | 0.84 |
| 9 | 124200417 | rs177698 | -1.046 | 0.394 | 1.303 | 0.229 | -0.001 | -0.005 | -0.005 | 0 | 0.31 | 0.3 | 0.22 | 0.12 | 0.34 | 0.2 |
| 9 | 124213962 | rs4837853 | 1.752 | 0.504 | 1.237 | 0.198 | -0.003 | 0.004 | 0.004 | 0 | 0.54 | 0.38 | 0.47 | 0.61 | 0.52 | 0.68 |
| 9 | 124238924 | rs10818548 | -0.993 | 0.15 | 1.226 | 0.297 | -0.007 | 0.015 | 0.015 | 0 | 0.3 | 0.23 | 0.21 | 0.12 | 0.34 | 0.2 |
| 9 | 124252503 | rs10985272 | -0.707 | 0.217 | 1.274 | 0.356 | -0.04 | 0.053 | 0.053 | 0 | 0.27 | 0.14 | 0.2 | 0.12 | 0.34 | 0.2 |
| 9 | 124253751 | rs12551532 | 0.457 | 0.191 | 1.248 | 0.282 | 0.011 | 0.005 | 0.005 | 0 | 0.13 | 0.06 | 0.07 | 0.01 | 0.08 | 0.02 |
| 9 | 124259751 | rs10985274 | 1.602 | 0.225 | 1.365 | 0.327 | 0.005 | 0.003 | 0.003 | 0 | 0.54 | 0.4 | 0.48 | 0.6 | 0.56 | 0.63 |
| 9 | 124280991 | rs4836853 | 1.453 | 0.471 | 1.512 | -0.033 | 0 | -0.004 | -0.004 | 0.36 | 0.55 | 0.46 | 0.68 | 0.67 | 0.51 | 0.72 |
| 9 | 124282782 | rs10760179 | -0.95 | 0.44 | 1.573 | -0.094 | -0.044 | 0.015 | 0.015 | 0 | 0.28 | 0.4 | 0.16 | 0.33 | 0.2 | 0.14 |
| 9 | 124283089 | rs12551209 | -0.222 | 0.493 | 1.672 | -0.1 | -0.006 | -0.001 | -0.001 | 0.134 | 0.18 | 0.21 | 0.11 | 0.21 | 0.15 | 0.06 |
| 9 | 124288948 | rs7854285 | 0.124 | 0.525 | 1.384 | -0.124 | 0.003 | -0.033 | -0.033 | 2.553 | 0.82 | 0.79 | 0.83 | 0.98 | 0.72 | 0.81 |
| 9 | 124295569 | rs4837876 | 1.702 | 0.426 | 1.331 | -0.085 | 0.001 | 0.018 | 0.018 | 0 | 0.53 | 0.43 | 0.67 | 0.65 | 0.51 | 0.67 |
| 9 | 124303632 | rs10985311 | 0.509 | 0.39 | 0.681 | -0.088 | -0.008 | -0.012 | -0.012 | 0 | 0.06 | 0.06 | 0.03 | 0.02 | 0.03 | 0.05 |
| 9 | 124303655 | rs478352 | -0.481 | 0.637 | 0.617 | -0.075 | -0.027 | 0.019 | 0.019 | 0 | 0.22 | 0.14 | 0.18 | 0.12 | 0.32 | 0.2 |
| 9 | 124308077 | rs7873850 | -0.612 | 0.495 | 0.614 | -0.133 | 0.002 | 0.002 | 0.002 | 0 | 0.23 | 0.23 | 0.15 | 0.24 | 0.17 | 0.11 |
| 9 | 124320762 | rs475972 | -0.115 | 1.235 | 0.247 | -0.268 | -0.065 | 0.062 | 0.062 | 0 | 0.17 | 0.08 | 0.17 | 0 | 0.3 | 0.14 |
| 9 | 124325541 | rs1107811 | 0.186 | 1.256 | 0.393 | -0.286 | -0.02 | -0.003 | -0.003 | 0 | 0.81 | 0.86 | 0.8 | 1 | 0.69 | 0.85 |
| 9 | 124326028 | rs7867606 | 0.075 | 0.761 | 0.77 | 0.09 | -0.018 | 0.027 | 0.027 | 0.203 | 0.07 | 0.1 | 0.02 | 0.1 | 0.02 | 0.05 |
| 9 | 124328194 | rs7042758 | -0.17 | 0.74 | 0.906 | 0.079 | -0.02 | 0.027 | 0.027 | 0.389 | 0.07 | 0.16 | 0.02 | 0.1 | 0.02 | 0.05 |
| 9 | 124330141 | rs657239 | 0.103 | 0.603 | 0.882 | 0.157 | -0.004 | -0.001 | -0.001 | 0 | 0.82 | 0.82 | 0.89 | 0.81 | 0.83 | 0.91 |
| 9 | 124330970 | rs10985320 | -0.862 | 0.806 | 1.168 | 0.641 | -0.001 | 0.058 | 0.058 | 0.427 | 0.19 | 0.35 | 0.32 | 0.27 | 0.15 | 0.26 |
| 9 | 124336665 | rs556331 | -0.076 | 1.136 | 0.754 | 0.362 | 0.028 | -0.09 | -0.09 | 0.246 | 0.2 | 0.32 | 0.19 | 0 | 0.28 | 0.11 |
| 9 | 124337517 | rs7045915 | -0.514 | 1.017 | 0.605 | 0.755 | -0.002 | -0.005 | -0.005 | 0 | 0.16 | 0.13 | 0.29 | 0.27 | 0.15 | 0.25 |
| 9 | 124343306 | rs914388 | 0.112 | 0.659 | 0.174 | 0.111 | -0.004 | 0.035 | 0.035 | 0 | 0.52 | 0.55 | 0.65 | 0.75 | 0.48 | 0.75 |
| 9 | 124343700 | rs603315 | 0.62 | 0.228 | 0.109 | 0.537 | -0.011 | 0.068 | 0.068 | 0.336 | 0.78 | 0.43 | 0.65 | 0.67 | 0.84 | 0.7 |
| 9 | 124344438 | rs883095 | 1.147 | 0.032 | -0.056 | 0.364 | -0.015 | -0.05 | -0.05 | 0.011 | 0.32 | 0.11 | 0.33 | 0.43 | 0.33 | 0.46 |

| | | | | | | | | | | | | | | | | |
|---|---|---|---|---|---|---|---|---|---|---|---|---|---|---|---|---|
| 9 | 124344832 | rs587936 | -0.33 | -0.154 | 0.364 | 0.243 | 0.035 | -0.009 | -0.009 | 0.021 | 0.52 | 0.36 | 0.57 | 0.46 | 0.61 | 0.38 |
| 9 | 124354826 | rs10513376 | 0.349 | 0.301 | 0.282 | -0.345 | 0.012 | -0.073 | -0.073 | 1.097 | 0.21 | 0.27 | 0.19 | 0 | 0.32 | 0.11 |
| 9 | 124362502 | rs2777319 | -0.606 | 0.065 | 0.107 | -0.516 | 0.002 | 0.007 | 0.007 | 0 | 0.55 | 0.63 | 0.56 | 0.42 | 0.62 | 0.56 |
| 9 | 124364349 | rs7021714 | 0.882 | -0.066 | 0.19 | -0.315 | -0.001 | 0.001 | 0.001 | 0 | 0.29 | 0.3 | 0.33 | 0.35 | 0.29 | 0.4 |
| 9 | 124364572 | rs10818569 | 0.417 | 0.016 | -0.097 | -0.339 | -0.007 | 0.007 | 0.007 | 0 | 0.33 | 0.43 | 0.38 | 0.42 | 0.31 | 0.45 |
| 9 | 124368674 | rs2777317 | -0.273 | 0.179 | -0.075 | -0.849 | 0.014 | -0.126 | -0.126 | 0.074 | 0.28 | 0.54 | 0.26 | 0.04 | 0.34 | 0.15 |
| 9 | 124372152 | rs10985332 | 0.435 | -0.024 | -0.38 | -0.174 | -0.003 | 0.032 | 0.032 | 0 | 0.25 | 0.27 | 0.42 | 0.56 | 0.19 | 0.31 |
| 9 | 124372312 | rs10985334 | 0.364 | -0.089 | -0.599 | -0.537 | 0.002 | 0.007 | 0.007 | 0 | 0.07 | 0 | 0.04 | 0.02 | 0.03 | 0.08 |
| 9 | 124376589 | rs605576 | 0.376 | -0.125 | -0.902 | -1.108 | -0.054 | 0.169 | 0.169 | 0 | 0.38 | 0.17 | 0.33 | 0.03 | 0.48 | 0.29 |
| 9 | 124379788 | rs1984035 | 0.092 | -0.188 | -0.661 | -1.239 | 0.016 | 0.019 | 0.019 | 0.178 | 0.51 | 0.56 | 0.58 | 0.97 | 0.39 | 0.6 |
| 9 | 124383244 | rs2789876 | 0.51 | -0.166 | -1.252 | -1.224 | -0.108 | 0.261 | 0.261 | 0 | 0.4 | 0.08 | 0.33 | 0.06 | 0.5 | 0.32 |
| 9 | 124384647 | rs2789875 | 0.549 | 0.099 | -0.336 | -0.595 | 0.015 | -0.08 | -0.08 | 0.536 | 0.88 | 0.42 | 0.89 | 1 | 0.9 | 0.91 |
| 9 | 124388306 | rs7863224 | 0.492 | 0.049 | -0.675 | -0.302 | -0.004 | 0.028 | 0.028 | 0 | 0.28 | 0.34 | 0.45 | 0.59 | 0.23 | 0.32 |
| 9 | 124390320 | rs520198 | -0.175 | 0.153 | -1.643 | -0.259 | 0.017 | -0.09 | -0.09 | 0 | 0.68 | 0.16 | 0.52 | 0.41 | 0.77 | 0.68 |
| 9 | 124394575 | rs871214 | -0.435 | 0.098 | -1.581 | -0.165 | -0.026 | -0.04 | -0.04 | 0.751 | 0.43 | 0.88 | 0.56 | 0.62 | 0.36 | 0.46 |
| 9 | 124396753 | rs2797349 | -0.11 | 0.4 | -0.834 | 0.339 | 0.023 | 0.031 | 0.031 | 0 | 0.74 | 0.65 | 0.58 | 0.69 | 0.63 | 0.8 |
| 9 | 124397290 | rs584985 | -0.135 | -0.229 | -0.586 | 0.461 | -0.011 | 0 | 0 | 0 | 0.36 | 0.42 | 0.24 | 0.32 | 0.25 | 0.4 |
| 9 | 124399756 | rs2777310 | -0.344 | -0.136 | -1.295 | 0.48 | 0.012 | -0.015 | -0.015 | 0.003 | 0.41 | 0.62 | 0.43 | 0.45 | 0.3 | 0.48 |
| 9 | 124407976 | rs12553641 | -0.088 | -0.139 | -1.947 | 0.301 | -0.017 | -0.026 | -0.026 | 0.235 | 0.32 | 0.03 | 0.2 | 0.26 | 0.25 | 0.43 |
| 9 | 124410605 | rs12554667 | -0.303 | 0.397 | -1.203 | 0.522 | 0.013 | 0.027 | 0.027 | 0.568 | 0.32 | 0.54 | 0.42 | 0.36 | 0.36 | 0.27 |
| 9 | 124412961 | rs10760182 | -0.727 | 0.593 | -1.184 | 0.42 | 0.005 | -0.007 | -0.007 | 0 | 0.47 | 0.63 | 0.53 | 0.53 | 0.49 | 0.64 |
| 9 | 124425243 | rs10985349 | -0.31 | 1.332 | -0.64 | 0.864 | -0.002 | -0.004 | -0.004 | 2.985 | 0.15 | 0.15 | 0.16 | 0.28 | 0.15 | 0.24 |
| 9 | 124427373 | rs1571801 | -0.55 | 1.698 | -0.314 | 0.378 | 0.009 | -0.004 | -0.004 | 0 | 0.16 | 0.15 | 0.19 | 0.06 | 0.26 | 0.13 |
| 9 | 124436035 | rs7029095 | 0.231 | 0.923 | 0.207 | 0.253 | -0.009 | -0.042 | -0.042 | 0 | 0.08 | 0.23 | 0.04 | 0 | 0.04 | 0.08 |
| 9 | 124436050 | rs7028742 | -0.533 | 0.898 | 0.837 | 0.146 | 0.008 | -0.04 | -0.04 | 0 | 0.22 | 0.31 | 0.21 | 0.11 | 0.24 | 0.18 |
| 9 | 124438732 | rs7875955 | -1.267 | 0.743 | 0.766 | 0.102 | 0 | -0.038 | -0.038 | 0.002 | 0.36 | 0.6 | 0.3 | 0.12 | 0.36 | 0.34 |
| 9 | 124441404 | rs7047340 | -0.514 | 0.519 | 1.227 | 0.844 | -0.006 | 0.139 | 0.139 | 0.978 | 0.12 | 0.5 | 0.21 | 0.2 | 0.05 | 0.14 |
| 9 | 124441657 | rs12350358 | -0.68 | 0.523 | 1.274 | 0.79 | -0.015 | 0.083 | 0.083 | 1.893 | 0.13 | 0.46 | 0.21 | 0.18 | 0.06 | 0.14 |
| 9 | 124444109 | rs870556 | -1.016 | 0.623 | 1.18 | 0.839 | 0.001 | -0.01 | -0.01 | 0 | 0.36 | 0.61 | 0.43 | 0.29 | 0.36 | 0.38 |
| 9 | 124445115 | rs2051053 | -1.854 | 0.429 | 0.519 | 0.969 | -0.063 | 0.031 | 0.031 | 0.17 | 0.53 | 0.88 | 0.6 | 0.66 | 0.48 | 0.56 |
| 9 | 124449142 | rs2900195 | -0.844 | 0.689 | 1.24 | 0.756 | 0 | -0.035 | -0.035 | 0 | 0.36 | 0.62 | 0.32 | 0.11 | 0.36 | 0.34 |
| 9 | 124449774 | rs16910935 | -0.472 | 0.73 | 1.153 | 0.777 | -0.006 | 0.022 | 0.022 | 1.7 | 0.31 | 0.22 | 0.28 | 0.11 | 0.35 | 0.32 |
| 9 | 124487495 | rs13290547 | 0.574 | 0.492 | 0.821 | 1.4 | -0.015 | 0.031 | 0.031 | 0 | 0.09 | 0 | 0.02 | 0.03 | 0.07 | 0.02 |
| 9 | 124516590 | rs12685856 | -1.494 | 0.566 | 1.316 | 2.11 | 0.064 | 0.283 | 0.283 | 0.683 | 0.08 | 0.51 | 0.07 | 0.23 | 0.04 | 0.04 |
| 9 | 124546143 | rs10985372 | 0.14 | 0.958 | -0.135 | 1.016 | -0.01 | 0.025 | 0.025 | 0 | 0.08 | 0.04 | 0.08 | 0 | 0.13 | 0.03 |
| 9 | 124550237 | rs10985376 | -1.018 | 0.773 | 0.089 | 1.07 | 0.002 | 0.083 | 0.083 | 2.968 | 0.53 | 0.62 | 0.43 | 0.87 | 0.49 | 0.66 |
| 9 | 124550994 | rs12238806 | -0.278 | 1.355 | -0.344 | 1.944 | -0.054 | -0.046 | -0.046 | 0.359 | 0.09 | 0.01 | 0.17 | 0.35 | 0.15 | 0.26 |
| 9 | 124560853 | rs10818593 | -0.77 | 1.074 | -0.541 | 1.705 | -0.012 | -0.019 | -0.019 | 0 | 0.16 | 0.14 | 0.21 | 0.35 | 0.22 | 0.29 |
| 9 | 124567667 | rs12347373 | -0.843 | 0.685 | -0.575 | 0.882 | 0.011 | 0.053 | 0.053 | 5.487 | 0.57 | 0.35 | 0.51 | 0.84 | 0.55 | 0.71 |
| 9 | 124571484 | rs4641136 | -0.426 | 0.655 | -0.792 | 1.303 | -0.003 | -0.001 | -0.001 | 0 | 0.32 | 0.26 | 0.22 | 0.37 | 0.29 | 0.45 |
| 9 | 124572031 | rs7858071 | -0.296 | 0.517 | -0.247 | 1.29 | -0.003 | 0.038 | 0.038 | 0.016 | 0.29 | 0.41 | 0.32 | 0.47 | 0.27 | 0.27 |
| 9 | 124573625 | rs10116392 | -0.475 | 0.57 | -0.972 | 1.346 | -0.015 | -0.029 | -0.029 | 0.022 | 0.25 | 0.24 | 0.33 | 0.5 | 0.3 | 0.35 |
| 9 | 124578338 | rs1129678 | 0.074 | 0.581 | -1.209 | 1.264 | 0.019 | -0.029 | -0.029 | 0 | 0.31 | 0.08 | 0.22 | 0.32 | 0.26 | 0.32 |
| 9 | 124583411 | rs3802488 | -0.468 | 1.055 | -0.796 | 1.601 | -0.053 | -0.065 | -0.065 | 0 | 0.21 | 0.13 | 0.24 | 0.41 | 0.3 | 0.33 |
| 9 | 124589809 | rs10818600 | -0.794 | 0.829 | 0.026 | 1.586 | 0.046 | 0.095 | 0.095 | 0.169 | 0.26 | 0.5 | 0.31 | 0.5 | 0.34 | 0.35 |
| 9 | 124600375 | rs7862029 | -0.598 | 0.367 | 0.096 | 1.091 | 0.009 | 0.046 | 0.046 | 3.687 | 0.56 | 0.31 | 0.39 | 0.73 | 0.6 | 0.64 |
| 9 | 124602102 | rs7039561 | -1.282 | 0.035 | 0.133 | 0.821 | -0.037 | 0.066 | 0.066 | 3.595 | 0.67 | 0.88 | 0.5 | 0.8 | 0.64 | 0.72 |
| 9 | 124606788 | rs10818602 | -1.204 | 0.058 | 0.661 | 0.906 | -0.005 | 0.033 | 0.033 | 2.894 | 0.66 | 0.77 | 0.49 | 0.79 | 0.64 | 0.72 |
| 9 | 124613105 | rs4308811 | -0.012 | 0.012 | 0.811 | 1.448 | 0.011 | 0 | 0 | 0.179 | 0.39 | 0.32 | 0.16 | 0.35 | 0.27 | 0.38 |
| 9 | 124614496 | rs10818605 | -0.786 | 0.195 | 0.918 | 1.342 | -0.001 | 0.008 | 0.008 | 1.586 | 0.59 | 0.4 | 0.31 | 0.73 | 0.59 | 0.66 |
| 9 | 124617259 | rs7864184 | -0.786 | 0.195 | 0.857 | 1.347 | 0 | 0.001 | 0.001 | 1.586 | 0.59 | 0.43 | 0.32 | 0.73 | 0.59 | 0.63 |
| 9 | 124619225 | rs13295845 | 0.775 | 0.415 | 0.64 | 1.107 | 0.021 | -0.028 | -0.028 | 0 | 0.05 | 0.09 | 0.03 | 0 | 0.08 | 0.04 |
| 9 | 124620787 | rs4836866 | -1.016 | 0.036 | 0.05 | 0.574 | -0.01 | 0.001 | 0.001 | 0.304 | 0.77 | 0.79 | 0.56 | 0.81 | 0.71 | 0.74 |
| 9 | 124623873 | rs10739600 | -1.045 | 0.08 | 0.038 | 0.245 | -0.001 | -0.009 | -0.009 | 0 | 0.75 | 0.74 | 0.58 | 0.9 | 0.69 | 0.7 |
| 9 | 124625023 | rs4538956 | 0.907 | -0.044 | -0.453 | 0.072 | -0.097 | 0.057 | 0.057 | 0 | 0.18 | 0.02 | 0.16 | 0.08 | 0.3 | 0.22 |
| 9 | 124627927 | rs10513383 | 0.228 | -0.437 | -0.271 | -0.195 | -0.001 | 0.001 | 0.001 | 0 | 0.13 | 0.12 | 0.17 | 0.03 | 0.12 | 0.09 |
| 9 | 124628199 | rs16911057 | 1.831 | -1.012 | -0.748 | -0.103 | -0.026 | -0.081 | -0.081 | 0 | 0.14 | 0.01 | 0.07 | 0.23 | 0.13 | 0.19 |
| 9 | 124630333 | rs7047365 | 0.445 | -0.719 | 0.149 | -0.685 | 0.001 | -0.035 | -0.035 | 0 | 0.55 | 0.47 | 0.37 | 0.87 | 0.51 | 0.57 |
| 9 | 124640841 | rs4503172 | -0.571 | -0.629 | 0.171 | -0.63 | 0.014 | -0.073 | -0.073 | 0 | 0.69 | 0.6 | 0.54 | 0.9 | 0.64 | 0.68 |
| 9 | 124643095 | rs7033700 | -0.183 | -0.455 | -0.39 | -0.417 | -0.017 | 0.032 | 0.032 | 0 | 0.64 | 0.68 | 0.43 | 0.87 | 0.57 | 0.61 |
| 9 | 124643367 | rs6478545 | -1.151 | -0.218 | -0.673 | -0.22 | -0.029 | 0.045 | 0.045 | 0 | 0.77 | 0.84 | 0.6 | 0.91 | 0.69 | 0.72 |
| 9 | 124650611 | rs7871736 | 0.036 | -0.194 | -0.576 | 0.201 | 0.016 | 0.006 | 0.006 | 0.371 | 0.61 | 0.44 | 0.6 | 0.54 | 0.52 | 0.58 |
| 9 | 124651790 | rs10985431 | 0.087 | -0.036 | -0.705 | -0.032 | 0 | -0.008 | -0.008 | 0 | 0.11 | 0.13 | 0.25 | 0.05 | 0.12 | 0.09 |
| 9 | 124661100 | rs4271039 | 0.177 | -0.036 | -0.66 | -0.014 | 0.001 | -0.013 | -0.013 | 0 | 0.11 | 0.15 | 0.25 | 0.05 | 0.12 | 0.08 |
| 9 | 124668198 | rs4518734 | 0.326 | -0.067 | -0.635 | -0.118 | 0.006 | -0.028 | -0.028 | 0 | 0.16 | 0.21 | 0.27 | 0.07 | 0.17 | 0.15 |
| 9 | 124675677 | rs9408931 | 1.115 | -0.262 | -1.94 | -0.024 | 0.021 | 0.016 | 0.016 | 0.118 | 0.57 | 0.08 | 0.51 | 0.65 | 0.67 | 0.6 |
| 9 | 124676496 | rs7019301 | 1.141 | -0.13 | -1.952 | 0.6 | -0.075 | -0.104 | -0.104 | 0 | 0.34 | 0.04 | 0.29 | 0.45 | 0.33 | 0.38 |
| 9 | 124678508 | rs11793905 | 0.28 | -0.301 | -1.135 | 0.612 | -0.004 | 0.009 | 0.009 | 0.64 | 0.13 | 0.16 | 0.09 | 0.28 | 0.06 | 0.2 |
| 9 | 124679214 | rs2416835 | 0.878 | -0.09 | -1.039 | 0.494 | -0.003 | -0.006 | -0.006 | 0 | 0.31 | 0.24 | 0.26 | 0.36 | 0.27 | 0.33 |

| | | | | | | | | | | | | | | | | |
|---|---|---|---|---|---|---|---|---|---|---|---|---|---|---|---|---|
| 9 | 124680367 | rs2416837 | -1.447 | -0.645 | 0.435 | 0.168 | -0.067 | 0.079 | 0.079 | 4.8 | 0.19 | 0.45 | 0.15 | 0.29 | 0.07 | 0.21 |
| 9 | 124685499 | rs13295747 | -0.081 | -0.675 | 0.002 | -0.508 | -0.069 | 0.156 | 0.156 | 0 | 0.4 | 0.17 | 0.47 | 0.11 | 0.56 | 0.34 |
| 9 | 124687446 | rs6478548 | 0.851 | -0.789 | 1.007 | -0.002 | -0.062 | 0.134 | 0.134 | 0.488 | 0.84 | 0.6 | 0.78 | 0.71 | 0.95 | 0.8 |
| 9 | 124693697 | rs4344151 | -0.063 | -0.765 | 1.285 | 0.006 | -0.017 | 0.152 | 0.152 | 0 | 0.92 | 0.79 | 0.86 | 0.74 | 0.98 | 0.82 |
| 9 | 124694838 | rs9409243 | 1.583 | -0.94 | 1.289 | -0.251 | -0.035 | -0.04 | -0.04 | 0 | 0.4 | 0.16 | 0.29 | 0.59 | 0.39 | 0.43 |
| 9 | 124697074 | rs11999315 | -0.758 | -0.769 | 1.252 | -0.546 | -0.001 | -0.008 | -0.008 | 0.676 | 0.32 | 0.35 | 0.29 | 0.09 | 0.36 | 0.22 |
| 9 | 124697877 | rs13295532 | -0.24 | -0.777 | 1.411 | -0.556 | -0.018 | -0.069 | -0.069 | 0 | 0.26 | 0.41 | 0.27 | 0.08 | 0.25 | 0.19 |
| 9 | 124701103 | rs2416841 | 0.365 | -0.678 | 1.27 | -0.581 | -0.055 | 0.065 | 0.065 | 0 | 0.2 | 0.07 | 0.18 | 0.06 | 0.28 | 0.14 |
| 9 | 124703332 | rs10985459 | 0.362 | -0.645 | 1.375 | -0.559 | -0.001 | 0.003 | 0.003 | 0 | 0.22 | 0.19 | 0.23 | 0.05 | 0.27 | 0.21 |
| 9 | 124704304 | rs10739601 | 1.919 | -0.659 | 1.409 | -0.567 | -0.014 | -0.017 | -0.017 | 0 | 0.4 | 0.26 | 0.29 | 0.59 | 0.39 | 0.44 |
| 9 | 124711549 | rs2416848 | 0.495 | -0.638 | 1.567 | -0.497 | -0.02 | 0.016 | 0.016 | 0 | 0.69 | 0.5 | 0.63 | 0.69 | 0.71 | 0.61 |
| 9 | 124712136 | rs7028210 | 1.998 | -0.637 | 1.428 | -0.514 | -0.023 | -0.024 | -0.024 | 0 | 0.42 | 0.18 | 0.31 | 0.6 | 0.41 | 0.49 |
| 9 | 124716151 | rs12379136 | 0.466 | -0.635 | 1.441 | -1.076 | -0.003 | 0.023 | 0.023 | 0 | 0.21 | 0.15 | 0.22 | 0.05 | 0.27 | 0.19 |
| 9 | 124718909 | rs9409246 | -0.324 | -0.657 | 1.493 | -1.019 | -0.019 | 0.084 | 0.084 | 0 | 0.92 | 0.78 | 0.86 | 0.83 | 0.98 | 0.8 |
| 9 | 124720937 | rs4837923 | 0.253 | -0.642 | 1.401 | -1.033 | -0.017 | 0.03 | 0.03 | 0.017 | 0.21 | 0.18 | 0.25 | 0.05 | 0.27 | 0.1 |
| 9 | 124723489 | rs10985469 | 0.473 | -0.627 | 1.364 | -1.019 | -0.003 | 0.022 | 0.022 | 0 | 0.21 | 0.15 | 0.22 | 0.05 | 0.27 | 0.19 |
| 9 | 124723960 | rs2416843 | -0.708 | -0.478 | 1.061 | -0.939 | 0.002 | -0.036 | -0.036 | 1.082 | 0.28 | 0.36 | 0.33 | 0.09 | 0.3 | 0.12 |
| 9 | 124736081 | rs4534187 | 1.609 | -0.745 | 1.015 | -0.806 | -0.031 | 0.026 | 0.026 | 0.019 | 0.42 | 0.17 | 0.31 | 0.69 | 0.41 | 0.48 |
| 9 | 124738711 | rs16911180 | -1.056 | -0.329 | 1.498 | -0.628 | -0.021 | -0.05 | -0.05 | 0.613 | 0.24 | 0.44 | 0.26 | 0.11 | 0.16 | 0.1 |
| 9 | 124740606 | rs6478551 | -0.291 | -0.183 | 1.727 | -0.074 | -0.012 | 0.022 | 0.022 | 0.313 | 0.06 | 0.22 | 0.07 | 0.05 | 0.01 | 0.02 |
| 9 | 124747323 | rs7019095 | -0.786 | 0.078 | 1.672 | -0.331 | 0.042 | -0.069 | -0.069 | 0.007 | 0.22 | 0.36 | 0.19 | 0.06 | 0.36 | 0.2 |
| 9 | 124747765 | rs17177082 | 0.924 | -0.096 | 1.423 | -0.539 | -0.001 | 0.023 | 0.023 | 0 | 0.08 | 0.02 | 0.03 | 0 | 0.06 | 0.04 |
| 9 | 124754815 | rs10739604 | 0.357 | 0.067 | 2.39 | 0.192 | -0.006 | 0.029 | 0.029 | 0 | 0.89 | 0.8 | 0.8 | 0.75 | 0.94 | 0.74 |
| 9 | 124756264 | rs4836872 | -1.985 | 0.736 | 2.713 | -0.428 | -0.017 | 0.042 | 0.042 | 0 | 0.3 | 0.73 | 0.33 | 0.08 | 0.38 | 0.22 |
| 9 | 124759431 | rs10985478 | -0.324 | 0.794 | 2.882 | -0.461 | -0.007 | 0.003 | 0.003 | 0 | 0.14 | 0.13 | 0.13 | 0.03 | 0.27 | 0.13 |
| 9 | 124778409 | rs16911266 | -2.62 | 0.766 | 3.139 | -0.455 | 0.088 | 0.021 | 0.021 | 0.041 | 0.09 | 0.42 | 0.22 | 0.04 | 0.06 | 0.02 |
| 9 | 124779322 | rs13289318 | -2.143 | 0.758 | 3.05 | -0.482 | -0.002 | -0.009 | -0.009 | 0 | 0.29 | 0.63 | 0.39 | 0.08 | 0.38 | 0.22 |
| 9 | 124786156 | rs11506930 | -1.117 | 0.599 | 3.091 | -0.491 | 0.002 | -0.011 | -0.011 | 0 | 0.05 | 0.08 | 0.03 | 0.02 | 0.04 | 0.01 |
| 9 | 124790623 | rs10125937 | -1.631 | 0.638 | 3.103 | -0.479 | 0.025 | -0.056 | -0.056 | 2.618 | 0.24 | 0.59 | 0.35 | 0.07 | 0.33 | 0.15 |
| 9 | 124810592 | rs4837936 | -2.279 | 0.583 | 2.769 | -0.403 | -0.014 | 0.038 | 0.038 | 0 | 0.29 | 0.73 | 0.41 | 0.1 | 0.37 | 0.21 |
| 9 | 124818049 | rs7045782 | -0.561 | 0.606 | 2.925 | -0.416 | 0.046 | -0.07 | -0.07 | 3.106 | 0.16 | 0.38 | 0.16 | 0.03 | 0.27 | 0.14 |
| 9 | 124821226 | rs10513389 | -0.428 | 0.617 | 2.842 | -0.415 | 0.01 | -0.027 | -0.027 | 0 | 0.15 | 0.2 | 0.15 | 0.03 | 0.26 | 0.13 |
| 9 | 124831273 | rs16911368 | -1.491 | 0.622 | 2.823 | -0.41 | 0.098 | 0.039 | 0.039 | 0 | 0.05 | 0.21 | 0.2 | 0.04 | 0.06 | 0.01 |
| 9 | 124835368 | rs10818623 | -0.165 | 0.622 | 1.934 | -0.404 | -0.007 | 0.005 | 0.005 | 0 | 0.14 | 0.12 | 0.13 | 0.03 | 0.27 | 0.14 |
| 9 | 124852286 | rs7860331 | -2.179 | 0.603 | 1.893 | -0.407 | -0.014 | 0.041 | 0.041 | 0 | 0.3 | 0.73 | 0.41 | 0.11 | 0.37 | 0.22 |
| 9 | 124863020 | rs10985526 | -2.077 | 0.427 | 1.817 | -0.337 | -0.021 | 0.036 | 0.036 | 0 | 0.31 | 0.71 | 0.41 | 0.11 | 0.41 | 0.29 |
| 9 | 124864385 | rs12551572 | -0.999 | 0.616 | 1.573 | -0.316 | -0.027 | 0.029 | 0.029 | 0 | 0.22 | 0.14 | 0.18 | 0.06 | 0.36 | 0.26 |
| 9 | 124869062 | rs10985529 | -0.208 | 0.526 | 1.485 | -0.286 | 0.01 | -0.023 | -0.023 | 0.014 | 0.15 | 0.16 | 0.13 | 0.03 | 0.27 | 0.17 |
| 9 | 124870192 | rs7034156 | -1.73 | 0.517 | 1.281 | -0.035 | 0.034 | -0.001 | -0.001 | 0 | 0.14 | 0.35 | 0.25 | 0.13 | 0.15 | 0.14 |
| 9 | 124870596 | rs7021304 | -1.444 | 0.31 | 0.638 | 0.139 | 0.002 | -0.01 | -0.01 | 0 | 0.27 | 0.64 | 0.37 | 0.14 | 0.34 | 0.24 |
| 9 | 124870612 | rs13287142 | -0.751 | 0.374 | -0.138 | 0.03 | -0.03 | 0.056 | 0.056 | 0 | 0.2 | 0.1 | 0.15 | 0.04 | 0.3 | 0.21 |
| 9 | 124871697 | rs2247680 | -1.513 | -0.451 | 0.878 | 0.971 | -0.002 | 0.124 | 0.124 | 0 | 0.14 | 0.46 | 0.13 | 0.23 | 0.11 | 0.1 |
| 9 | 124876341 | rs10985532 | -1.731 | -1.474 | -0.013 | 1.119 | 0.113 | 0.051 | 0.051 | 1.071 | 0.39 | 0.83 | 0.51 | 0.76 | 0.21 | 0.42 |
| 9 | 124877059 | rs7861047 | -1.475 | -1.476 | 0.526 | 1.125 | -0.028 | -0.011 | -0.011 | 0 | 0.34 | 0.6 | 0.48 | 0.76 | 0.2 | 0.42 |
| 9 | 124879007 | rs11793614 | -1.591 | -1.238 | 0.636 | 1.069 | -0.034 | 0.044 | 0.044 | 0 | 0.14 | 0.61 | 0.13 | 0.24 | 0.08 | 0.11 |
| 9 | 124884007 | rs10985538 | -1.356 | -1.286 | 0.98 | 0.596 | -0.002 | -0.027 | -0.027 | 0.056 | 0.09 | 0.51 | 0.08 | 0 | 0.03 | 0.02 |
| 9 | 124885254 | rs7038781 | 0.959 | -1.692 | 0.54 | 1.467 | -0.02 | 0.019 | 0.019 | 2.685 | 0.74 | 0.37 | 0.58 | 0.37 | 0.87 | 0.72 |
| 9 | 124885596 | rs7042153 | 1.481 | -1.812 | 0.353 | 1.333 | 0.059 | -0.003 | -0.003 | 0 | 0.64 | 0.28 | 0.52 | 0.31 | 0.8 | 0.64 |
| 9 | 124893355 | rs1411451 | -1.44 | -1.514 | 0.754 | 0.729 | -0.011 | -0.056 | -0.056 | 0.526 | 0.1 | 0.18 | 0.1 | 0 | 0.08 | 0.1 |
| 9 | 124895008 | rs1983985 | 0.386 | -0.903 | 0.117 | 1.085 | -0.009 | -0.066 | -0.066 | 0.023 | 0.77 | 0.87 | 0.7 | 0.46 | 0.78 | 0.69 |
| 9 | 124900274 | rs1124729 | 1.636 | -1.067 | 0.652 | 1.152 | 0.032 | 0.031 | 0.031 | 0 | 0.58 | 0.3 | 0.42 | 0.31 | 0.65 | 0.51 |
| 9 | 124900788 | rs9775695 | 1.821 | -1.089 | 0.639 | 1.138 | 0.031 | 0.044 | 0.044 | 0 | 0.56 | 0.28 | 0.4 | 0.3 | 0.63 | 0.51 |
| 9 | 124901941 | rs1536958 | 0.226 | -1.071 | 0.575 | 1.139 | -0.008 | -0.054 | -0.054 | 0 | 0.78 | 0.86 | 0.7 | 0.46 | 0.78 | 0.69 |
| 9 | 124905629 | rs4147660 | -1.755 | -1.194 | 0.698 | 0.784 | 0 | 0 | 0 | 0 | 0.23 | 0.73 | 0.34 | 0.18 | 0.15 | 0.17 |
| 9 | 124906456 | rs6822 | -1.432 | -1.205 | 1.095 | 0.778 | -0.045 | 0.028 | 0.028 | 0 | 0.19 | 0.44 | 0.31 | 0.18 | 0.14 | 0.17 |
| 9 | 124906723 | rs3818638 | 0.2 | -1.082 | 0.275 | 0.723 | -0.007 | -0.041 | -0.041 | 0.401 | 0.75 | 0.84 | 0.71 | 0.38 | 0.76 | 0.68 |
| 9 | 124910582 | rs3793616 | 1.966 | -1.007 | 0.053 | 0.547 | 0.017 | 0.228 | 0.228 | 0 | 0.51 | 0.11 | 0.36 | 0.2 | 0.58 | 0.51 |
| 9 | 124913308 | rs12686688 | 1.845 | -0.8 | -0.041 | 0.48 | -0.071 | 0.228 | 0.228 | 0 | 0.48 | 0.04 | 0.33 | 0.18 | 0.53 | 0.46 |
| 9 | 124918150 | rs10818639 | 1.943 | -0.842 | -0.006 | 0.502 | 0.016 | 0.237 | 0.237 | 0 | 0.5 | 0.1 | 0.36 | 0.2 | 0.58 | 0.49 |
| 9 | 124918545 | rs7875039 | -0.557 | -0.777 | -0.05 | 0.81 | -0.032 | -0.09 | -0.09 | 0.56 | 0.81 | 0.97 | 0.79 | 0.64 | 0.8 | 0.82 |
| 9 | 124918682 | rs1411445 | 1.598 | -0.624 | 0.067 | -0.268 | 0.023 | 0.207 | 0.207 | 0 | 0.51 | 0.11 | 0.36 | 0.22 | 0.59 | 0.51 |
| 9 | 124924543 | rs10985554 | 1.454 | -0.73 | -0.3 | -0.793 | -0.054 | 0.238 | 0.238 | 0 | 0.43 | 0.13 | 0.29 | 0.06 | 0.47 | 0.3 |
| 9 | 124925605 | rs950019 | 0.877 | -0.895 | 0.003 | -0.884 | 0.007 | 0.163 | 0.163 | 0 | 0.52 | 0.71 | 0.37 | 0.06 | 0.49 | 0.32 |
| 9 | 124929161 | rs7466389 | -0.029 | -1.018 | -0.135 | -0.802 | -0.016 | -0.028 | -0.028 | 0 | 0.05 | 0.1 | 0.06 | 0 | 0.02 | 0.05 |
| 9 | 124930771 | rs10985555 | 1.657 | -0.978 | -0.273 | -0.798 | -0.083 | 0.247 | 0.247 | 0 | 0.42 | 0.11 | 0.28 | 0.08 | 0.49 | 0.31 |
| 9 | 124935044 | rs10818642 | 1.762 | -0.977 | -0.36 | -0.798 | -0.106 | 0.247 | 0.247 | 0 | 0.41 | 0.07 | 0.28 | 0.08 | 0.49 | 0.29 |
| 9 | 124941854 | rs1010804 | 0.965 | -0.966 | 0.054 | -0.783 | 0.01 | 0.029 | 0.029 | 0 | 0.51 | 0.57 | 0.37 | 0.09 | 0.53 | 0.36 |
| 9 | 124945604 | rs10760211 | 0.766 | -0.421 | -0.211 | -0.854 | -0.002 | 0.035 | 0.035 | 0 | 0.52 | 0.57 | 0.44 | 0.26 | 0.51 | 0.48 |
| 9 | 124946021 | rs7869304 | -0.292 | -0.988 | 0.16 | -0.321 | -0.004 | 0.033 | 0.033 | 0 | 0.74 | 0.66 | 0.6 | 0.37 | 0.76 | 0.73 |

| | | | | | | | | | | | | | | | | |
|---|---|---|---|---|---|---|---|---|---|---|---|---|---|---|---|---|
| 9 | 124949046 | rs2039197 | 1.136 | -1.067 | -0.252 | -0.618 | 0.002 | 0 | 0 | 0 | 0.7 | 0.69 | 0.71 | 0.72 | 0.75 | 0.58 |
| 9 | 124949538 | rs913933 | 1.134 | -0.998 | -0.327 | -0.622 | 0.004 | 0.006 | 0.006 | 0 | 0.54 | 0.48 | 0.49 | 0.25 | 0.59 | 0.39 |
| 9 | 124949601 | rs2039196 | 0.149 | -0.968 | -0.304 | 0.125 | -0.016 | 0.038 | 0.038 | 0 | 0.21 | 0.26 | 0.34 | 0.65 | 0.19 | 0.21 |
| 9 | 124955028 | rs7855478 | -0.904 | -0.951 | -0.845 | -0.084 | -0.032 | -0.064 | -0.064 | 0 | 0.17 | 0.43 | 0.13 | 0.1 | 0.13 | 0.16 |
| 9 | 124957525 | rs1571586 | -0.88 | -0.673 | -2.267 | -0.081 | 0.085 | 0.09 | 0.09 | 0.07 | 0.45 | 0.86 | 0.53 | 0.73 | 0.36 | 0.42 |
| 9 | 124958572 | rs12683922 | -0.63 | -0.499 | -2.221 | -0.074 | 0.059 | 0.085 | 0.085 | 0 | 0.41 | 0.87 | 0.52 | 0.77 | 0.36 | 0.42 |
| 9 | 124966261 | rs3750486 | -0.57 | -0.308 | -2.102 | 0.247 | -0.006 | -0.04 | -0.04 | 0 | 0.14 | 0 | 0.06 | 0.13 | 0.1 | 0.24 |
| 9 | 124966449 | rs3793617 | -0.029 | 0.428 | -2.544 | 0.816 | -0.036 | -0.059 | -0.059 | 0 | 0.17 | 0.07 | 0.37 | 0.33 | 0.24 | 0.1 |
| 9 | 124967776 | rs10818647 | -0.057 | 0.412 | -2.08 | 0.808 | -0.004 | -0.005 | -0.005 | 0 | 0.18 | 0.18 | 0.38 | 0.33 | 0.23 | 0.1 |
| 9 | 124969315 | rs10818649 | 1.021 | -0.032 | -1.709 | 0.527 | 0.013 | 0.009 | 0.009 | 0 | 0.51 | 0.35 | 0.47 | 0.45 | 0.57 | 0.55 |
| 9 | 124974419 | rs10818651 | -0.164 | 0.628 | -2.708 | -0.031 | -0.107 | -0.044 | -0.044 | 0 | 0.32 | 0.03 | 0.22 | 0.33 | 0.36 | 0.12 |
| 9 | 124980026 | rs10985568 | 0.47 | 0.905 | -2.714 | -0.095 | -0.053 | 0.019 | 0.019 | 6.003 | 0.16 | 0.03 | 0.31 | 0.12 | 0.23 | 0.28 |
| 9 | 124984125 | rs3793621 | -0.204 | 0.553 | -1.976 | -0.204 | -0.003 | -0.006 | -0.006 | 0 | 0.32 | 0.63 | 0.36 | 0.28 | 0.28 | 0.12 |
| 9 | 124985250 | rs12345155 | -0.495 | -0.117 | -1.895 | -0.462 | -0.042 | -0.048 | -0.048 | 0 | 0.68 | 0.57 | 0.83 | 0.84 | 0.82 | 0.72 |
| 9 | 124985785 | rs10818652 | 0.463 | 0.999 | -2.611 | -0.02 | -0.168 | -0.179 | -0.179 | 0.395 | 0.3 | 0.03 | 0.4 | 0.48 | 0.46 | 0.47 |
| 9 | 124992764 | rs3808900 | -0.107 | 0.975 | -2.142 | 0.122 | -0.01 | -0.018 | -0.018 | 0.011 | 0.16 | 0.15 | 0.3 | 0.33 | 0.2 | 0.21 |
| 9 | 124993505 | rs7849627 | -0.26 | 0.728 | -2.396 | -0.871 | -0.039 | 0.076 | 0.076 | 0 | 0.35 | 0.14 | 0.27 | 0.19 | 0.44 | 0.32 |
| 9 | 124995282 | rs4837951 | 0.378 | 0.999 | -2.082 | -0.957 | -0.135 | 0.069 | 0.069 | 0 | 0.29 | 0.02 | 0.34 | 0.12 | 0.38 | 0.32 |
| 9 | 124998225 | rs1571585 | 0.297 | 0.903 | -1.031 | -0.71 | -0.072 | 0.133 | 0.133 | 0 | 0.39 | 0.11 | 0.35 | 0.16 | 0.43 | 0.47 |
| 9 | 125002275 | rs10818655 | 0.109 | 0.669 | 0.824 | -0.883 | -0.004 | 0.036 | 0.036 | 0 | 0.45 | 0.4 | 0.36 | 0.16 | 0.46 | 0.45 |
| 9 | 125003679 | rs1060586 | 0.104 | 0.612 | 0.351 | -0.807 | -0.003 | 0.13 | 0.13 | 0 | 0.48 | 0.27 | 0.59 | 0.84 | 0.5 | 0.52 |
| 9 | 125003751 | rs10739606 | 0.256 | 0.708 | -0.312 | -0.562 | -0.076 | 0.137 | 0.137 | 0 | 0.4 | 0.1 | 0.34 | 0.16 | 0.42 | 0.46 |
| 9 | 125024291 | rs12555156 | -0.169 | -0.558 | -0.169 | -0.032 | -0.037 | -0.081 | -0.081 | 0.429 | 0.13 | 0.03 | 0.32 | 0.32 | 0.17 | 0.12 |
| 9 | 125026664 | rs7034992 | 0.06 | -0.507 | 0.397 | -0.935 | -0.039 | 0.133 | 0.133 | 0 | 0.38 | 0.1 | 0.29 | 0.16 | 0.38 | 0.45 |
| 9 | 125043555 | rs9695213 | -2.555 | -0.465 | 2.498 | -0.009 | -0.021 | 0.045 | 0.045 | 0 | 0.15 | 0.36 | 0.16 | 0.22 | 0.1 | 0.2 |
| 9 | 125044448 | rs10818673 | -3.13 | -0.575 | 2.791 | -0.108 | -0.033 | 0.019 | 0.019 | 0.079 | 0.24 | 0.58 | 0.28 | 0.34 | 0.13 | 0.33 |
| 9 | 125061280 | rs10985588 | -2.447 | -0.18 | 1.959 | -0.015 | -0.01 | 0.039 | 0.039 | 0 | 0.13 | 0.23 | 0.15 | 0.23 | 0.1 | 0.13 |
| 9 | 125062104 | rs10985589 | -1.585 | -0.167 | 1.961 | -0.009 | -0.005 | -0.02 | -0.02 | 0 | 0.09 | 0.05 | 0.12 | 0.23 | 0.1 | 0.13 |
| 9 | 125063820 | rs2026152 | -1.551 | -0.167 | 1.959 | -0.009 | -0.004 | -0.016 | -0.016 | 0 | 0.09 | 0.06 | 0.12 | 0.23 | 0.1 | 0.13 |
| 9 | 125065423 | rs7019478 | -2.611 | -0.167 | 1.963 | -0.008 | -0.009 | 0.044 | 0.044 | 0 | 0.13 | 0.23 | 0.15 | 0.23 | 0.1 | 0.13 |
| 9 | 125075782 | rs3818711 | -2.731 | -0.201 | 2.692 | 0.764 | -0.018 | 0.051 | 0.051 | 0 | 0.12 | 0.22 | 0.12 | 0.22 | 0.08 | 0.12 |
| 9 | 125081147 | rs10985594 | -1.707 | -0.008 | 1.691 | 0.86 | 0.001 | -0.012 | -0.012 | 0.415 | 0.08 | 0.06 | 0.09 | 0.23 | 0.08 | 0.12 |
| 9 | 125102160 | rs7049102 | -2.538 | 0.044 | 1.864 | 0.167 | -0.028 | 0.013 | 0.013 | 0 | 0.18 | 0.27 | 0.11 | 0.25 | 0.1 | 0.2 |
| 9 | 125105790 | rs13283472 | -1.548 | 0.471 | 1.416 | 0.291 | -0.002 | 0.005 | 0.005 | 0.18 | 0.06 | 0.07 | 0.08 | 0.18 | 0.1 | 0.19 |
| 9 | 125111953 | rs7022025 | -2.653 | 0.178 | 2.236 | -0.324 | 0.029 | -0.007 | -0.007 | 0 | 0.1 | 0.39 | 0.08 | 0.05 | 0.09 | 0.11 |
| 9 | 125117174 | rs7029883 | -2.679 | 0.111 | 2.226 | -0.39 | -0.004 | -0.051 | -0.051 | 0 | 0.14 | 0.49 | 0.06 | 0.05 | 0.08 | 0.03 |
| 9 | 125123971 | rs4837960 | -2.718 | 0.239 | 2 | -0.473 | 0.007 | -0.111 | -0.111 | 0 | 0.16 | 0.42 | 0.11 | 0.04 | 0.13 | 0.2 |
| 9 | 125131480 | rs10306108 | -1.565 | 0.409 | 1.299 | -0.293 | 0.014 | -0.028 | -0.028 | 0 | 0.05 | 0.13 | 0.03 | 0 | 0.07 | 0.01 |
| 9 | 125131631 | rs10306109 | -1.565 | 0.409 | 1.299 | -0.293 | 0.014 | -0.028 | -0.028 | 0 | 0.05 | 0.13 | 0.03 | 0 | 0.07 | 0.01 |
| 9 | 125136884 | rs1213266 | 2.266 | -0.254 | 1.779 | -0.362 | -0.066 | -0.082 | -0.082 | 0.437 | 0.83 | 0.41 | 0.87 | 0.94 | 0.93 | 0.85 |
| 9 | 125140206 | rs3842788 | -2.08 | -0.205 | 2.178 | -0.335 | -0.042 | -0.023 | -0.023 | 0 | 0.12 | 0.32 | 0.03 | 0.07 | 0.04 | 0.04 |
| 9 | 125141786 | rs10306143 | -1.193 | 0.258 | 2.505 | -0.718 | 0.024 | -0.056 | -0.056 | 0 | 0.1 | 0.16 | 0.12 | 0 | 0.14 | 0.09 |
| 9 | 125143792 | rs5788 | -2.744 | -0.066 | 3.009 | -0.597 | -0.014 | -0.034 | -0.034 | 0.236 | 0.2 | 0.69 | 0.24 | 0.04 | 0.14 | 0.1 |
| 9 | 125146701 | rs10306156 | -1.359 | -0.075 | 3.411 | -0.607 | -0.028 | -0.03 | -0.03 | 0 | 0.07 | 0.25 | 0.02 | 0 | 0 | 0 |
| 9 | 125146895 | rs4836886 | -2.565 | -0.104 | 3.38 | -0.657 | -0.015 | -0.036 | -0.036 | 0 | 0.21 | 0.69 | 0.25 | 0.04 | 0.14 | 0.1 |
| 9 | 125147871 | rs9299281 | -2.565 | -0.104 | 3.38 | -0.657 | -0.015 | -0.036 | -0.036 | 0.647 | 0.21 | 0.69 | 0.25 | 0.04 | 0.14 | 0.1 |
| 9 | 125148632 | rs6478565 | -2.877 | -0.056 | 3.182 | -0.654 | -0.005 | -0.019 | -0.019 | 0.951 | 0.25 | 0.69 | 0.28 | 0.07 | 0.17 | 0.19 |
| 9 | 125149958 | rs10306162 | -2.647 | -0.05 | 3.042 | -0.771 | 0.013 | -0.051 | -0.051 | 1.073 | 0.19 | 0.41 | 0.26 | 0.07 | 0.17 | 0.19 |
| 9 | 125149970 | rs10306163 | -3.479 | -0.054 | 2.985 | -0.77 | 0.022 | 0.04 | 0.04 | 2.454 | 0.31 | 0.74 | 0.3 | 0.11 | 0.2 | 0.23 |
| 9 | 125150888 | rs12551233 | -3.252 | -0.053 | 3.023 | -0.718 | -0.04 | -0.059 | -0.059 | 0.051 | 0.21 | 0.65 | 0.11 | 0.07 | 0.07 | 0.14 |
| 9 | 125153352 | rs7874531 | -1.682 | 0.05 | 2.487 | -0.675 | -0.024 | -0.032 | -0.032 | 0 | 0.08 | 0.31 | 0.03 | 0 | 0.01 | 0 |
| 9 | 125155408 | rs8046 | -3.083 | 0.043 | 2.208 | -0.669 | -0.057 | -0.088 | -0.088 | 0 | 0.18 | 0.51 | 0.1 | 0.09 | 0.07 | 0.14 |
| 9 | 125155685 | rs10306188 | -1.941 | 0.043 | 2.204 | -0.667 | -0.037 | -0.047 | -0.047 | 0 | 0.1 | 0.5 | 0.04 | 0 | 0.01 | 0.01 |
| 9 | 125157198 | rs10306194 | 0.198 | 0.681 | 0.364 | -0.958 | -0.051 | 0.045 | 0.045 | 0 | 0.12 | 0.02 | 0.11 | 0.04 | 0.15 | 0.14 |
| 9 | 125157316 | rs10306196 | -1.58 | 0.497 | 0.667 | -0.642 | -0.014 | 0.004 | 0.004 | 0.048 | 0.08 | 0.01 | 0.05 | 0.09 | 0.06 | 0.13 |
| 9 | 125157718 | rs9233 | -1.58 | 0.497 | 0.667 | -0.642 | -0.014 | 0.004 | 0.004 | 0.048 | 0.08 | 0.01 | 0.05 | 0.09 | 0.06 | 0.14 |
| 9 | 125164936 | rs10513402 | -1.343 | 0.501 | 0.823 | -0.826 | 0.001 | -0.023 | -0.023 | 1.139 | 0.1 | 0.11 | 0.16 | 0 | 0.12 | 0.08 |
| 9 | 125165225 | rs950104 | -0.498 | -0.337 | 1.196 | -0.622 | -0.026 | 0.004 | 0.004 | 0 | 0.22 | 0.34 | 0.14 | 0.24 | 0.15 | 0.3 |
| 9 | 125169930 | rs10818684 | 0.963 | -0.478 | 1.162 | -0.611 | 0.004 | 0.001 | 0.001 | 1.537 | 0.64 | 0.36 | 0.6 | 0.64 | 0.74 | 0.58 |
| 9 | 125177491 | rs1758701 | -1.232 | -0.425 | 0.986 | -0.579 | -0.027 | -0.027 | -0.027 | 0.179 | 0.95 | 0.99 | 0.77 | 0.89 | 0.89 | 0.88 |
| 9 | 125186011 | rs1234909 | -0.966 | -0.947 | 1.097 | -0.265 | -0.061 | 0.015 | 0.015 | 0.565 | 0.31 | 0.53 | 0.17 | 0.33 | 0.17 | 0.3 |
| 9 | 125192159 | rs1332408 | -0.444 | -0.959 | 1.34 | -0.299 | -0.001 | 0 | 0 | 0 | 0.26 | 0.25 | 0.27 | 0.25 | 0.25 | 0.34 |
| 9 | 125209061 | rs2778624 | 1.349 | -0.984 | 1.188 | -0.317 | -0.024 | -0.007 | -0.007 | 0 | 0.61 | 0.5 | 0.69 | 0.66 | 0.72 | 0.62 |
| 9 | 125229799 | rs2778635 | 1.206 | -0.317 | 0.682 | 0.025 | -0.002 | 0.088 | 0.088 | 0 | 0.71 | 0.53 | 0.59 | 0.5 | 0.68 | 0.67 |
| 9 | 125230685 | rs1332406 | -0.284 | -0.323 | 0.921 | 0.15 | 0.009 | 0.074 | 0.074 | 0 | 0.14 | 0.19 | 0.15 | 0.38 | 0.18 | 0.19 |
| 9 | 125231092 | rs2778636 | 1.073 | -0.303 | 0.693 | 0.157 | -0.004 | 0.088 | 0.088 | 0 | 0.71 | 0.53 | 0.59 | 0.5 | 0.69 | 0.67 |
| 9 | 125233864 | rs7865902 | 2.075 | -0.307 | 0.594 | 0.178 | -0.065 | 0.199 | 0.199 | 0.641 | 0.42 | 0.12 | 0.31 | 0.13 | 0.42 | 0.3 |
| 9 | 125236148 | rs1888941 | -0.62 | -0.105 | 0.804 | 0.859 | 0.017 | 0.176 | 0.176 | 0 | 0.12 | 0.32 | 0.12 | 0.33 | 0.11 | 0.11 |
| 9 | 125238040 | rs1228138 | 0.135 | -0.024 | 0.312 | 0.816 | -0.017 | 0.063 | 0.063 | 0 | 0.17 | 0.29 | 0.14 | 0.33 | 0.13 | 0.26 |

| | | | | | | | | | | | | | | | | |
|---|---|---|---|---|---|---|---|---|---|---|---|---|---|---|---|---|
| 9 | 125246793 | rs1547929 | -0.466 | -0.001 | -0.165 | 0.597 | 0.007 | -0.001 | -0.001 | 0 | 0.2 | 0.37 | 0.23 | 0.2 | 0.23 | 0.23 |
| 9 | 125252977 | rs2416863 | 0.014 | -0.025 | -0.23 | 0.857 | -0.004 | 0.039 | 0.039 | 0.057 | 0.08 | 0.17 | 0.21 | 0.23 | 0.09 | 0.14 |
| 9 | 125253682 | rs7048727 | 0.14 | 0.463 | -1.238 | 0.404 | -0.015 | 0.097 | 0.097 | 0 | 0.19 | 0.05 | 0.09 | 0.02 | 0.13 | 0.18 |
| 9 | 125254955 | rs7022194 | -2.077 | 0.756 | -1.508 | 0.243 | -0.076 | -0.101 | -0.101 | 0 | 0.59 | 0.92 | 0.61 | 0.49 | 0.53 | 0.66 |
| 9 | 125255361 | rs10985655 | -1.046 | 0.744 | -1.379 | 0.21 | -0.001 | -0.004 | -0.004 | 0 | 0.47 | 0.55 | 0.53 | 0.46 | 0.46 | 0.54 |
| 9 | 125255623 | rs1592031 | -1.729 | 0.897 | -1.47 | 0.502 | -0.076 | -0.097 | -0.097 | 0.041 | 0.59 | 0.92 | 0.61 | 0.49 | 0.53 | 0.66 |
| 9 | 125257838 | rs11496595 | -0.306 | 0.823 | -1.179 | 0.707 | 0.003 | 0.046 | 0.046 | 0.079 | 0.27 | 0.46 | 0.44 | 0.44 | 0.32 | 0.36 |
| 9 | 125261468 | rs2096064 | -0.36 | 0.803 | -1.185 | 0.65 | 0.003 | 0.061 | 0.061 | 0.222 | 0.27 | 0.49 | 0.44 | 0.45 | 0.33 | 0.36 |
| 9 | 125265317 | rs10739609 | -1.213 | 0.812 | -2.226 | 0.78 | -0.031 | -0.101 | -0.101 | 0.044 | 0.59 | 0.91 | 0.62 | 0.53 | 0.55 | 0.65 |
| 9 | 125265496 | rs10985658 | 0.408 | 0.71 | -1.782 | 0.967 | 0.007 | -0.014 | -0.014 | 0.029 | 0.14 | 0.08 | 0.1 | 0.18 | 0.13 | 0.16 |
| 9 | 125266798 | rs12380552 | 0.57 | 0.802 | -2.403 | 1.139 | -0.036 | -0.112 | -0.112 | 0.926 | 0.34 | 0.06 | 0.34 | 0.44 | 0.39 | 0.28 |
| 9 | 125268884 | rs1556190 | 0.397 | 0.954 | -2.257 | 1.011 | -0.029 | 0.002 | 0.002 | 0.517 | 0.43 | 0.23 | 0.32 | 0.55 | 0.46 | 0.46 |
| 9 | 125271022 | rs7873246 | -0.586 | 1.04 | -2.077 | 1.05 | -0.021 | -0.029 | -0.029 | 0 | 0.89 | 0.92 | 0.69 | 0.78 | 0.8 | 0.78 |
| 9 | 125271839 | rs7019219 | 0.52 | 0.525 | -1.397 | 0.761 | 0.006 | 0.002 | 0.002 | 0 | 0.47 | 0.58 | 0.32 | 0.42 | 0.43 | 0.3 |
| 9 | 125272062 | rs10985662 | 0.297 | -0.223 | -1.344 | 0.669 | 0.001 | -0.041 | -0.041 | 0 | 0.12 | 0 | 0.05 | 0.17 | 0.06 | 0.11 |
| 9 | 125273574 | rs4836891 | 0.762 | -0.242 | -1.377 | 0.678 | 0.005 | -0.068 | -0.068 | 0 | 0.14 | 0 | 0.1 | 0.24 | 0.07 | 0.18 |
| 9 | 125274730 | rs1329954 | 0.552 | -0.098 | -0.681 | 0.918 | 0.013 | 0.068 | 0.068 | 0 | 0.12 | 0.26 | 0.25 | 0.25 | 0.17 | 0.16 |
| 9 | 125276105 | rs10985663 | -1.375 | -0.806 | -0.872 | 0.18 | -0.01 | 0.126 | 0.126 | 0 | 0.59 | 0.77 | 0.59 | 0.87 | 0.48 | 0.71 |
| 9 | 125276867 | rs12349388 | 0.646 | -1.024 | -0.583 | 0.178 | 0.002 | 0.002 | 0.002 | 0 | 0.1 | 0.09 | 0.13 | 0.09 | 0.06 | 0.22 |
| 9 | 125282970 | rs10985667 | 0.912 | -0.952 | 0.294 | 0.225 | 0.012 | 0.067 | 0.067 | 0 | 0.13 | 0.23 | 0.25 | 0.26 | 0.17 | 0.17 |
| 9 | 125289083 | rs16911867 | -0.237 | -1.013 | 0.636 | -0.198 | -0.001 | 0.003 | 0.003 | 0 | 0.36 | 0.37 | 0.39 | 0.51 | 0.32 | 0.38 |
| 9 | 125289521 | rs10818708 | -1.679 | -0.922 | 0.371 | -0.311 | -0.018 | 0.156 | 0.156 | 0 | 0.58 | 0.82 | 0.59 | 0.87 | 0.48 | 0.71 |
| 9 | 125290397 | rs10760228 | -1.708 | -0.935 | 0.394 | -0.305 | -0.021 | 0.149 | 0.149 | 0 | 0.59 | 0.82 | 0.59 | 0.87 | 0.48 | 0.71 |
| 9 | 125298845 | rs2104751 | -1.367 | -0.694 | 0.21 | -0.638 | -0.003 | -0.019 | -0.019 | 0.026 | 0.84 | 0.86 | 0.78 | 1 | 0.71 | 0.82 |
| 9 | 125303760 | rs7037327 | -0.329 | -0.906 | 0.324 | -0.57 | 0.007 | -0.015 | -0.015 | 0 | 0.19 | 0.14 | 0.13 | 0.33 | 0.14 | 0.34 |
| 9 | 125304750 | rs1041356 | -0.781 | -0.956 | 0.527 | -0.656 | -0.005 | -0.04 | -0.04 | 0.076 | 0.46 | 0.54 | 0.52 | 0.87 | 0.38 | 0.68 |
| 9 | 125307248 | rs10760235 | -0.768 | -0.937 | 0.495 | -0.646 | -0.004 | -0.037 | -0.037 | 0 | 0.46 | 0.52 | 0.52 | 0.87 | 0.38 | 0.68 |
| 9 | 125308131 | rs978477 | -0.762 | -0.94 | 0.494 | -0.648 | -0.004 | -0.038 | -0.038 | 5.026 | 0.46 | 0.52 | 0.52 | 0.87 | 0.37 | 0.68 |
| 9 | 125314024 | rs3983868 | 0.762 | -0.896 | 0.46 | -0.539 | -0.012 | 0.047 | 0.047 | 0.044 | 0.28 | 0.4 | 0.39 | 0.54 | 0.24 | 0.34 |
| 9 | 125314400 | rs2416875 | 0.815 | -0.933 | 0.608 | -0.644 | -0.002 | 0.003 | 0.003 | 0 | 0.56 | 0.6 | 0.59 | 0.67 | 0.48 | 0.46 |
| 9 | 125314885 | rs7859232 | 0.557 | -0.96 | 0.632 | -0.683 | 0.007 | 0.001 | 0.001 | 0 | 0.5 | 0.43 | 0.37 | 0.45 | 0.41 | 0.44 |
| 9 | 125315107 | rs7031561 | 0.834 | -0.941 | 0.626 | -0.659 | -0.002 | 0.003 | 0.003 | 0 | 0.56 | 0.59 | 0.59 | 0.67 | 0.48 | 0.46 |
| 9 | 125315393 | rs12684786 | 2.208 | -0.948 | 0.616 | -0.665 | 0.006 | 0.026 | 0.026 | 0.091 | 0.28 | 0.2 | 0.2 | 0.13 | 0.24 | 0.12 |
| 9 | 125315542 | rs1831369 | 0.85 | -0.934 | 0.64 | -0.655 | -0.012 | 0.046 | 0.046 | 0.048 | 0.28 | 0.4 | 0.39 | 0.54 | 0.24 | 0.34 |
| 9 | 125315557 | rs1831370 | 0.906 | -0.801 | 0.32 | -0.639 | -0.002 | 0.003 | 0.003 | 0 | 0.56 | 0.59 | 0.59 | 0.66 | 0.48 | 0.46 |
| 9 | 125316042 | rs1341043 | 0.915 | -0.801 | 0.32 | -0.639 | -0.002 | 0.003 | 0.003 | 0 | 0.56 | 0.6 | 0.59 | 0.66 | 0.48 | 0.46 |
| 9 | 125316179 | rs1341044 | -1.304 | -0.639 | 0.219 | -0.628 | -0.005 | -0.018 | -0.018 | 0 | 0.84 | 0.86 | 0.78 | 1 | 0.71 | 0.81 |
| 9 | 125316350 | rs1411272 | 0.089 | -0.773 | 0.327 | -0.427 | 0.002 | -0.001 | -0.001 | 0.049 | 0.62 | 0.64 | 0.6 | 0.67 | 0.55 | 0.5 |
| 9 | 125316850 | rs10985690 | 0.362 | -0.635 | 0.403 | -0.941 | -0.002 | -0.001 | -0.001 | 0 | 0.3 | 0.33 | 0.38 | 0.77 | 0.24 | 0.35 |
| 9 | 125317485 | rs10760239 | -0.325 | -0.623 | 0.464 | -1.166 | 0.007 | -0.08 | -0.08 | 2.152 | 0.7 | 0.7 | 0.64 | 0.98 | 0.61 | 0.62 |
| 9 | 125318521 | rs2183328 | 1.073 | -0.64 | 0.421 | -1.228 | 0.01 | -0.045 | -0.045 | 0 | 0.21 | 0.23 | 0.25 | 0 | 0.34 | 0.17 |
| 9 | 125318782 | rs2183326 | -0.067 | -0.298 | 0.305 | -0.993 | 0.02 | -0.134 | -0.134 | 0.013 | 0.63 | 0.57 | 0.62 | 0.98 | 0.55 | 0.58 |
| 9 | 125325182 | rs10985700 | 1.611 | -0.293 | 0.163 | -0.981 | -0.004 | 0.061 | 0.061 | 0 | 0.3 | 0.12 | 0.22 | 0.18 | 0.29 | 0.23 |
| 9 | 125328122 | rs4836894 | 1.571 | -0.264 | 0.132 | -0.967 | 0.003 | 0.054 | 0.054 | 0 | 0.31 | 0.15 | 0.22 | 0.18 | 0.29 | 0.23 |
| 9 | 125328865 | rs10513416 | 1.452 | -0.269 | 0.211 | -0.969 | 0.003 | 0.013 | 0.013 | 0 | 0.32 | 0.27 | 0.23 | 0.18 | 0.29 | 0.23 |
| 9 | 125330184 | rs10985702 | -0.273 | -0.266 | 0.251 | -1.005 | -0.01 | -0.035 | -0.035 | 0 | 0.38 | 0.45 | 0.41 | 0.81 | 0.32 | 0.4 |
| 9 | 125330678 | rs10985704 | -0.054 | 0.084 | 0.263 | -0.617 | 0.008 | -0.092 | -0.092 | 0 | 0.63 | 0.6 | 0.62 | 0.98 | 0.55 | 0.58 |
| 9 | 125330739 | rs1999182 | 0.066 | 0.09 | 0.24 | -0.614 | 0.013 | -0.117 | -0.117 | 0 | 0.62 | 0.48 | 0.61 | 0.98 | 0.55 | 0.58 |
| 9 | 125331205 | rs1411265 | -0.03 | 0.174 | -0.174 | -0.573 | 0.009 | -0.095 | -0.095 | 0 | 0.63 | 0.6 | 0.62 | 0.98 | 0.55 | 0.58 |
| 9 | 125332930 | rs7860446 | 0.038 | 0.116 | -0.101 | 0.701 | -0.021 | 0.057 | 0.057 | 0 | 0.2 | 0.25 | 0.17 | 0.55 | 0.09 | 0.2 |
| 9 | 125335216 | rs12376146 | -0.057 | 1.225 | 0.398 | 0.163 | 0.181 | -0.084 | -0.084 | 0 | 0.13 | 0.33 | 0.21 | 0 | 0.24 | 0.15 |
| 9 | 125339689 | rs10119609 | 1.061 | 1.284 | -0.196 | 0.558 | -0.036 | -0.04 | -0.04 | 0 | 0.07 | 0.03 | 0.21 | 0.25 | 0.13 | 0.12 |
| 9 | 125340264 | rs7867785 | -0.952 | 0.369 | -0.211 | 0.617 | -0.012 | 0.008 | 0.008 | 0 | 0.39 | 0.62 | 0.38 | 0.26 | 0.44 | 0.43 |
| 9 | 125342209 | rs2050898 | 0.535 | -0.612 | -0.37 | 0.5 | 0.015 | 0.047 | 0.047 | 0.277 | 0.19 | 0.12 | 0.09 | 0.01 | 0.18 | 0.21 |
| 9 | 125343746 | rs959550 | 0.925 | -0.784 | -0.557 | 0.189 | 0.001 | 0.008 | 0.008 | 0 | 0.43 | 0.28 | 0.45 | 0.4 | 0.53 | 0.37 |
| 9 | 125349081 | rs10818721 | -0.094 | -0.879 | -0.763 | 0.496 | 0.006 | -0.042 | -0.042 | 0 | 0.29 | 0.19 | 0.46 | 0.6 | 0.26 | 0.32 |
| 9 | 125364208 | rs10985729 | -0.019 | -1.016 | -0.486 | 0.755 | 0.007 | -0.007 | -0.007 | 2.902 | 0.3 | 0.21 | 0.38 | 0.36 | 0.23 | 0.35 |
| 9 | 125371916 | rs10818729 | 0.037 | -1.357 | -0.011 | 0.589 | -0.013 | -0.004 | -0.004 | 0 | 0.64 | 0.56 | 0.61 | 0.63 | 0.69 | 0.58 |
| 9 | 125372878 | rs1000510 | 0.571 | -1.409 | -0.109 | 0.716 | -0.001 | 0.038 | 0.038 | 0 | 0.1 | 0.08 | 0.18 | 0.3 | 0.1 | 0.22 |
| 9 | 125375428 | rs4836901 | 0.317 | -1.485 | -0.059 | 0.76 | -0.048 | 0.163 | 0.163 | 0.08 | 0.81 | 0.56 | 0.74 | 0.61 | 0.83 | 0.74 |
| 9 | 125376842 | rs972926 | -1.075 | -1.436 | -0.049 | 0.716 | -0.021 | -0.029 | -0.029 | 0 | 0.89 | 0.64 | 0.92 | 0.95 | 0.91 | 0.96 |
| 9 | 125377087 | rs972925 | -0.35 | -1.515 | -0.116 | 0.647 | -0.042 | -0.087 | -0.087 | 0 | 0.78 | 0.63 | 0.86 | 0.94 | 0.81 | 0.95 |
| 9 | 125384197 | rs10124314 | 0.773 | -1.12 | -0.448 | 0.723 | -0.022 | -0.041 | -0.041 | 0 | 0.24 | 0.06 | 0.3 | 0.3 | 0.21 | 0.37 |
| 9 | 125384788 | rs10739616 | 0.037 | -1.178 | -0.266 | 0.658 | -0.03 | -0.042 | -0.042 | 6.288 | 0.81 | 0.66 | 0.9 | 0.92 | 0.84 | 0.79 |
| 9 | 125385408 | rs10760244 | -0.061 | -1.378 | -0.468 | 0.467 | 0.028 | 0.041 | 0.041 | 0 | 0.41 | 0.27 | 0.45 | 0.31 | 0.34 | 0.4 |
| 9 | 125386182 | rs12000807 | 0.506 | -1.357 | -0.483 | 0.509 | -0.003 | 0.007 | 0.007 | 0 | 0.22 | 0.22 | 0.32 | 0.33 | 0.19 | 0.1 |
| 9 | 125387735 | rs11792470 | 0.239 | -1.094 | -0.636 | 0.622 | -0.025 | -0.083 | -0.083 | 0.171 | 0.3 | 0.09 | 0.48 | 0.61 | 0.29 | 0.45 |
| 9 | 125387853 | rs10491515 | 1.353 | -0.978 | -0.497 | 0.64 | -0.005 | 0 | 0 | 0 | 0.35 | 0.35 | 0.27 | 0.3 | 0.42 | 0.32 |
| 9 | 125390874 | rs1556189 | -0.915 | -1.379 | -0.205 | 0.793 | -0.036 | -0.094 | -0.094 | 0 | 0.77 | 0.64 | 0.85 | 0.95 | 0.8 | 0.94 |

| | | | | | | | | | | | | | | | |
|---|---|---|---|---|---|---|---|---|---|---|---|---|---|---|---|
| 9 | 125391026 | rs1476858 | -0.656 | -1.291 | -0.38 | 0.836 | -0.049 | -0.098 | -0.098 | 0 | 0.72 | 0.44 | 0.77 | 0.94 | 0.79 | 0.93 |
| 9 | 125391127 | rs1476859 | -0.059 | -1.381 | -0.504 | 0.96 | -0.038 | 0.001 | 0.001 | 0.419 | 0.29 | 0.06 | 0.49 | 0.79 | 0.28 | 0.47 |
| 9 | 125391241 | rs1476860 | 0.09 | -1.329 | -0.478 | 0.864 | -0.043 | -0.127 | -0.127 | 0.008 | 0.28 | 0.05 | 0.47 | 0.6 | 0.28 | 0.43 |
| 9 | 125391369 | rs1536929 | -0.286 | -1.301 | -0.476 | 0.831 | -0.024 | 0.003 | 0.003 | 0 | 0.68 | 0.44 | 0.6 | 0.64 | 0.72 | 0.87 |
| 9 | 125391409 | rs1536928 | 0.145 | -0.695 | 0.369 | 0.413 | -0.001 | 0.004 | 0.004 | 0 | 0.44 | 0.39 | 0.3 | 0.34 | 0.5 | 0.49 |
| 9 | 125398582 | rs2041474 | -1.31 | -0.26 | 1.532 | 1.038 | -0.033 | 0.043 | 0.043 | 2.125 | 0.28 | 0.49 | 0.47 | 0.66 | 0.21 | 0.4 |
| 9 | 125415365 | rs1928624 | -1.085 | -0.026 | 1.473 | 1.195 | -0.026 | 0.067 | 0.067 | 1.736 | 0.26 | 0.46 | 0.34 | 0.65 | 0.2 | 0.37 |
| 9 | 125418870 | rs2215532 | -1.058 | -0.029 | 1.475 | 1.221 | -0.029 | 0.127 | 0.127 | 1.654 | 0.24 | 0.46 | 0.32 | 0.6 | 0.18 | 0.36 |
| 9 | 125422424 | rs2416891 | -0.692 | 0.191 | 0.966 | 0.681 | 0 | -0.041 | -0.041 | 0 | 0.21 | 0.13 | 0.36 | 0.46 | 0.21 | 0.36 |
| 9 | 125422865 | rs237617 | 0.388 | 0.631 | 1.508 | 0.036 | 0.004 | -0.071 | -0.071 | 0 | 0.28 | 0.27 | 0.23 | 0.01 | 0.31 | 0.22 |
| 9 | 125426554 | rs7027867 | -1.425 | 0.523 | 2.489 | 1.72 | 0.026 | 0.173 | 0.173 | 0.016 | 0.08 | 0.17 | 0.19 | 0.47 | 0.13 | 0.1 |
| 9 | 125433467 | rs7035512 | -2.318 | 0.351 | 2.994 | 1.66 | 0.02 | 0.041 | 0.041 | 0.978 | 0.1 | 0.47 | 0.06 | 0.07 | 0.06 | 0.05 |
| 9 | 125436147 | rs7044945 | -2.346 | 0.339 | 3.168 | 1.654 | 0.019 | 0.041 | 0.041 | 0.844 | 0.1 | 0.48 | 0.06 | 0.08 | 0.06 | 0.05 |
| 9 | 125444045 | rs10985748 | 0.572 | 0.417 | 2.725 | 1.604 | -0.019 | 0.025 | 0.025 | 0 | 0.05 | 0 | 0.05 | 0 | 0.06 | 0.05 |
| 9 | 125453469 | rs12238140 | -0.869 | 0.478 | 2.818 | 1.757 | 0.007 | 0.105 | 0.105 | 0.173 | 0.05 | 0.06 | 0.17 | 0.37 | 0.11 | 0.07 |
| 9 | 125464010 | rs764373 | -0.455 | -0.424 | 2.677 | 1.28 | -0.006 | -0.163 | -0.163 | 2.767 | 0.21 | 0.02 | 0.36 | 0.53 | 0.18 | 0.3 |
| 9 | 125465183 | rs12002657 | -1.749 | 0.261 | 3.742 | 0.593 | 0.014 | -0.025 | -0.025 | 0 | 0.06 | 0.2 | 0.04 | 0 | 0.06 | 0.03 |
| 9 | 125478541 | rs4571792 | 1.929 | 0.266 | 3.799 | 0.595 | 0.016 | -0.026 | -0.026 | 1.49 | 0.93 | 0.78 | 0.96 | 1 | 0.94 | 0.97 |
| 9 | 125491426 | rs10429618 | -2.055 | -0.021 | 3.863 | 0.736 | 0.005 | -0.033 | -0.033 | 1.713 | 0.07 | 0.18 | 0.04 | 0 | 0.06 | 0.03 |
| 9 | 125502869 | rs7046355 | 2.881 | -0.026 | 3.845 | 0.67 | 0.023 | 0.045 | 0.045 | 0.31 | 0.9 | 0.55 | 0.95 | 0.93 | 0.94 | 0.96 |
| 9 | 125511661 | rs10120579 | 2.185 | 0.012 | 3.121 | 0.343 | 0.009 | -0.034 | -0.034 | 0 | 0.92 | 0.78 | 0.96 | 1 | 0.94 | 0.97 |
| 9 | 125517223 | rs2416893 | 2.764 | -0.005 | 2.551 | 0.335 | -0.03 | -0.074 | -0.074 | 0 | 0.88 | 0.35 | 0.92 | 1 | 0.95 | 0.97 |
| 9 | 125542513 | rs2050871 | -1.8 | 0.3 | 1.272 | 0.527 | -0.002 | -0.031 | -0.031 | 0.08 | 0.07 | 0.17 | 0.02 | 0 | 0.04 | 0.02 |
| 9 | 125554766 | rs2535739 | 2.108 | 0.066 | 2.006 | 0.882 | -0.006 | 0.097 | 0.097 | 3.716 | 0.91 | 0.49 | 0.91 | 0.85 | 0.98 | 0.75 |
| 9 | 125559860 | rs2535742 | 1.414 | -0.132 | 2.204 | 0.281 | -0.012 | -0.044 | -0.044 | 0.037 | 0.82 | 0.76 | 0.86 | 0.95 | 0.87 | 0.88 |
| 9 | 125573235 | rs11790713 | -1.334 | -0.097 | 2.626 | 0.263 | -0.018 | -0.093 | -0.093 | 0 | 0.17 | 0.27 | 0.11 | 0 | 0.13 | 0.07 |
| 9 | 125582173 | rs4997232 | -2.114 | 0.618 | 2.778 | 1.061 | 0.001 | 0.014 | 0.014 | 0.071 | 0.26 | 0.67 | 0.41 | 0.56 | 0.25 | 0.27 |
| 9 | 125585909 | rs10818752 | -1.721 | 0.724 | 2.449 | 1.056 | 0.033 | 0.099 | 0.099 | 0.029 | 0.21 | 0.45 | 0.37 | 0.54 | 0.24 | 0.15 |
| 9 | 125588724 | rs7037776 | -1.496 | 0.808 | 2.186 | 1.193 | 0.081 | 0.071 | 0.071 | 2.422 | 0.06 | 0.37 | 0.08 | 0.06 | 0.03 | 0.05 |
| 9 | 125590647 | rs3802487 | -1.831 | 1.09 | 1.584 | 1.349 | 0.094 | 0.267 | 0.267 | 0 | 0.1 | 0.27 | 0.29 | 0.47 | 0.18 | 0.06 |
| 9 | 125601801 | rs10818753 | -1.458 | 0.641 | 1.163 | 1.16 | 0.004 | 0.16 | 0.16 | 0 | 0.15 | 0.35 | 0.36 | 0.61 | 0.13 | 0.1 |
| 9 | 125605708 | rs10513414 | -0.778 | 0.625 | 1.197 | 1.382 | 0.001 | -0.002 | -0.002 | 0 | 0.08 | 0.02 | 0.28 | 0.6 | 0.06 | 0.09 |
| 9 | 125606191 | rs10985793 | -2.603 | 0.131 | 1.368 | 0.955 | -0.023 | 0.024 | 0.024 | 0 | 0.23 | 0.6 | 0.4 | 0.7 | 0.16 | 0.17 |
| 9 | 125608627 | rs10818754 | -0.114 | -0.513 | 0.072 | -0.199 | -0.018 | 0.124 | 0.124 | 0 | 0.2 | 0.01 | 0.1 | 0.01 | 0.21 | 0.2 |
| 9 | 125617037 | rs1375862 | -1.461 | -0.551 | 0.482 | -0.027 | 0.014 | 0.16 | 0.16 | 0 | 0.14 | 0.35 | 0.35 | 0.64 | 0.12 | 0.09 |
| 9 | 125652757 | rs10818759 | -1.582 | -0.57 | 0.646 | -0.05 | 0.017 | 0.173 | 0.173 | 0 | 0.14 | 0.39 | 0.36 | 0.64 | 0.12 | 0.09 |
| 9 | 125675559 | rs2239539 | -1.595 | -0.707 | 0.657 | -0.121 | 0.014 | 0.171 | 0.171 | 0 | 0.14 | 0.39 | 0.35 | 0.64 | 0.12 | 0.09 |
| 9 | 125709323 | rs10818768 | -0.037 | -0.743 | -0.909 | -1.23 | -0.126 | 0.098 | 0.098 | 0.089 | 0.54 | 0.03 | 0.4 | 0.29 | 0.52 | 0.62 |
| 9 | 125766114 | rs10985854 | 0.544 | -0.794 | -0.698 | -1.049 | 0.003 | 0.021 | 0.021 | 0 | 0.07 | 0.02 | 0.26 | 0.6 | 0.05 | 0.08 |
| 9 | 125799927 | rs10985869 | 0.009 | -0.81 | -0.718 | -1.063 | -0.002 | -0.014 | -0.014 | 0 | 0.08 | 0.03 | 0.27 | 0.6 | 0.06 | 0.09 |
| 9 | 125870466 | rs700085 | 1.324 | -1.002 | -0.82 | -1.324 | 0.015 | 0.027 | 0.027 | 1.451 | 0.84 | 0.14 | 0.86 | 0.91 | 0.91 | 0.92 |
| 9 | 125880828 | rs700091 | 1.856 | -1.398 | -1.11 | -1.542 | 0.063 | 0.019 | 0.019 | 0 | 0.65 | 0.16 | 0.55 | 0.32 | 0.8 | 0.81 |
| 9 | 125880929 | rs700092 | 1.582 | -1.397 | -1.136 | -1.542 | 0.045 | -0.02 | -0.02 | 0 | 0.65 | 0.06 | 0.54 | 0.32 | 0.8 | 0.81 |
| 9 | 125917138 | rs7032325 | -0.995 | -0.816 | -0.24 | -1.176 | -0.026 | -0.005 | -0.005 | 0.372 | 0.11 | 0.52 | 0.09 | 0.07 | 0.02 | 0.06 |
| 9 | 125963450 | rs803730 | 1.511 | -0.756 | -0.671 | -1.477 | -0.127 | 0.138 | 0.138 | 0.038 | 0.23 | 0.03 | 0.18 | 0.02 | 0.33 | 0.26 |
| 9 | 125975397 | rs7026688 | -1.6 | -0.824 | -0.484 | -1.309 | -0.014 | 0.037 | 0.037 | 1.066 | 0.21 | 0.63 | 0.39 | 0.62 | 0.15 | 0.09 |
| 9 | 125984527 | rs3860155 | -0.183 | -0.837 | -0.464 | -1.236 | 0.001 | 0.002 | 0.002 | 0.039 | 0.08 | 0.02 | 0.28 | 0.59 | 0.06 | 0.08 |
| 9 | 126017354 | rs6478601 | 0.442 | -0.785 | -0.377 | -1.179 | 0.065 | 0.09 | 0.09 | 6.196 | 0.05 | 0.53 | 0.06 | 0.03 | 0.01 | 0 |
| 9 | 126031850 | rs3739839 | 0.108 | -0.649 | -0.62 | -1.182 | -0.001 | 0.024 | 0.024 | 0 | 0.07 | 0.04 | 0.27 | 0.61 | 0.05 | 0.08 |
| 9 | 126037523 | rs10818801 | 0.045 | -0.389 | -0.886 | -1.274 | -0.046 | 0.084 | 0.084 | 0.862 | 0.59 | 0.12 | 0.45 | 0.28 | 0.59 | 0.67 |
| 9 | 126044010 | rs10760271 | -0.498 | -0.071 | -2.165 | -1.114 | -0.002 | -0.008 | -0.008 | 0 | 0.15 | 0.12 | 0.31 | 0.6 | 0.13 | 0.08 |
| 9 | 126048349 | rs999655 | -0.544 | -0.163 | -2.301 | -1.034 | -0.006 | 0.065 | 0.065 | 0.833 | 0.64 | 0.27 | 0.72 | 0.89 | 0.61 | 0.66 |
| 9 | 126049975 | rs868693 | 0.255 | -0.283 | -2.621 | -1.075 | -0.066 | 0.15 | 0.15 | 0.005 | 0.48 | 0.08 | 0.4 | 0.29 | 0.47 | 0.56 |
| 9 | 126050038 | rs10985938 | -0.135 | -0.355 | -1.363 | -0.569 | -0.014 | 0.048 | 0.048 | 0 | 0.09 | 0 | 0.04 | 0 | 0.11 | 0.09 |
| 9 | 126054988 | rs12004778 | -0.372 | -0.944 | -1.241 | -0.702 | 0.027 | 0.027 | 0.027 | 0.035 | 0.28 | 0.18 | 0.23 | 0.19 | 0.19 | 0.29 |
| 9 | 126056464 | rs10818803 | 0.385 | -0.505 | -1.414 | -1.14 | -0.043 | 0.041 | 0.041 | 0 | 0.28 | 0.2 | 0.28 | 0.1 | 0.43 | 0.4 |
| 9 | 126060193 | rs7854591 | -0.586 | -1.363 | -1.327 | -1.567 | 0.067 | 0.074 | 0.074 | 1.594 | 0.45 | 0.68 | 0.53 | 0.84 | 0.23 | 0.4 |
| 9 | 126063065 | rs10985950 | -0.24 | -1.493 | -1.251 | -0.835 | 0.02 | 0.011 | 0.011 | 0 | 0.53 | 0.32 | 0.54 | 0.45 | 0.57 | 0.7 |
| 9 | 126063746 | rs10985952 | -1.003 | -1.303 | -0.879 | -0.62 | -0.005 | -0.022 | -0.022 | 0 | 0.68 | 0.77 | 0.63 | 0.46 | 0.63 | 0.73 |
| 9 | 126064590 | rs7859203 | -0.233 | -1.235 | -0.577 | 0.074 | -0.027 | 0.115 | 0.115 | 0.738 | 0.07 | 0.25 | 0.07 | 0.17 | 0 | 0.06 |
| 9 | 126073680 | rs10739625 | -0.349 | -0.683 | -1.476 | -0.356 | 0.001 | -0.016 | -0.016 | 0 | 0.15 | 0.11 | 0.29 | 0.71 | 0.15 | 0.22 |
| 9 | 126073830 | rs10739626 | -0.667 | -0.762 | -1.645 | -0.749 | 0.043 | 0.077 | 0.077 | 0 | 0.4 | 0.24 | 0.48 | 0.84 | 0.28 | 0.49 |
| 9 | 126081738 | rs10985963 | 0.176 | -0.213 | -2.364 | -0.441 | -0.07 | -0.052 | -0.052 | 0.075 | 0.32 | 0.05 | 0.46 | 0.67 | 0.36 | 0.4 |
| 9 | 126082343 | rs7024188 | -0.536 | -0.584 | -1.89 | -1.021 | 0.006 | -0.036 | -0.036 | 0.043 | 0.62 | 0.54 | 0.69 | 0.8 | 0.66 | 0.69 |
| 9 | 126083799 | rs2416901 | -0.007 | -0.529 | -2.692 | -0.884 | -0.003 | 0.081 | 0.081 | 0 | 0.8 | 0.98 | 0.89 | 0.9 | 0.82 | 0.86 |
| 9 | 126087359 | rs3927757 | 0.141 | -0.21 | -2.56 | 0.249 | -0.026 | -0.117 | -0.117 | 0.608 | 0.16 | 0.03 | 0.34 | 0.55 | 0.19 | 0.26 |
| 9 | 126091085 | rs7341862 | 0.399 | 0.098 | -2.852 | -0.219 | -0.085 | 0.024 | 0.024 | 0 | 0.25 | 0.07 | 0.25 | 0.22 | 0.29 | 0.25 |
| 9 | 126094458 | rs10760278 | -0.713 | -0.293 | -2.672 | -0.254 | 0.001 | 0 | 0 | 0 | 0.88 | 0.93 | 0.94 | 0.94 | 0.89 | 0.95 |
| 9 | 126096522 | rs10818810 | 0.954 | -0.179 | -2.844 | 0.104 | -0.077 | -0.055 | -0.055 | 1.767 | 0.27 | 0.1 | 0.44 | 0.44 | 0.39 | 0.28 |

| | | | | | | | | | | | | | | | | |
|---|---|---|---|---|---|---|---|---|---|---|---|---|---|---|---|---|
| 9 | 126098374 | rs3932701 | 0.406 | -0.641 | -2.144 | 0.021 | -0.003 | -0.02 | -0.02 | 0 | 0.3 | 0.3 | 0.32 | 0.48 | 0.33 | 0.43 |
| 9 | 126099604 | rs1891636 | 0.38 | -0.684 | -1.106 | -0.637 | 0 | -0.009 | -0.009 | 0 | 0.37 | 0.33 | 0.34 | 0.24 | 0.42 | 0.35 |
| 9 | 126101008 | rs10114139 | 0.021 | -0.659 | -0.773 | -0.564 | 0 | 0.022 | 0.022 | 0 | 0.5 | 0.52 | 0.42 | 0.28 | 0.52 | 0.4 |
| 9 | 126102968 | rs7022169 | 0.366 | -0.466 | -0.698 | -0.502 | -0.016 | -0.062 | -0.062 | 0.009 | 0.86 | 0.75 | 0.92 | 1 | 0.9 | 0.94 |
| 9 | 126106114 | rs1971603 | 0.013 | -0.436 | -0.692 | -0.477 | -0.006 | -0.034 | -0.034 | 0 | 0.89 | 0.86 | 0.94 | 1 | 0.91 | 0.94 |
| 9 | 126106142 | rs7035562 | 0.256 | -0.379 | -0.745 | -0.404 | -0.003 | 0.006 | 0.006 | 0 | 0.66 | 0.63 | 0.72 | 0.68 | 0.63 | 0.7 |
| 9 | 126111351 | rs1970585 | -0.329 | -0.522 | -0.724 | -0.264 | 0.002 | -0.005 | -0.005 | 0 | 0.93 | 0.95 | 0.96 | 1 | 0.94 | 0.98 |
| 9 | 126113594 | rs10985976 | 0.214 | -0.611 | -0.54 | 0.032 | 0.001 | -0.001 | -0.001 | 0 | 0.4 | 0.34 | 0.49 | 0.5 | 0.4 | 0.51 |
| 9 | 126116865 | rs4838051 | 0.446 | -0.84 | -0.088 | -0.174 | -0.016 | -0.027 | -0.027 | 0 | 0.15 | 0.22 | 0.12 | 0.02 | 0.1 | 0.1 |
| 9 | 126117558 | rs1891632 | 0.458 | -0.588 | 0.233 | 0.58 | 0.019 | 0.11 | 0.11 | 0 | 0.09 | 0.15 | 0.29 | 0.26 | 0.16 | 0.2 |
| 9 | 126120159 | rs2808416 | -0.511 | -0.918 | -0.321 | 0.299 | 0.007 | 0.001 | 0.001 | 0 | 0.66 | 0.71 | 0.78 | 0.72 | 0.72 | 0.69 |
| 9 | 126123009 | rs2491353 | -1.064 | -0.374 | -0.424 | 0.27 | -0.055 | -0.002 | -0.002 | 0 | 0.68 | 0.87 | 0.64 | 0.59 | 0.57 | 0.66 |
| 9 | 126127998 | rs1105224 | 0.47 | -0.12 | -0.112 | 0.186 | 0 | -0.001 | -0.001 | 0 | 0.36 | 0.38 | 0.35 | 0.42 | 0.38 | 0.35 |
| 9 | 126142233 | rs1048616 | 0.262 | -0.335 | -0.077 | 0.855 | -0.005 | -0.049 | -0.049 | 0 | 0.14 | 0.04 | 0.09 | 0.29 | 0.12 | 0.18 |
| 9 | 126145002 | rs4836919 | 0.178 | -0.092 | 0.495 | 1.112 | 0.021 | 0.104 | 0.104 | 0 | 0.08 | 0.15 | 0.29 | 0.26 | 0.14 | 0.2 |
| 9 | 126150830 | rs2797945 | 1.76 | -0.66 | 0.745 | 0.618 | 0.009 | 0.024 | 0.024 | 0 | 0.57 | 0.33 | 0.45 | 0.39 | 0.62 | 0.44 |
| 9 | 126151949 | rs10985997 | 0.499 | -0.63 | 0.621 | 0.688 | -0.012 | -0.039 | -0.039 | 0.077 | 0.12 | 0.04 | 0.09 | 0.28 | 0.12 | 0.17 |
| 9 | 126152911 | rs1891640 | 0.846 | -0.702 | 0.869 | 0.651 | -0.022 | 0.043 | 0.043 | 0 | 0.7 | 0.49 | 0.55 | 0.67 | 0.75 | 0.62 |
| 9 | 126153636 | rs1891641 | 0.077 | -0.724 | 1.108 | 0.89 | 0.081 | 0.028 | 0.028 | 0.035 | 0.08 | 0.24 | 0.2 | 0.09 | 0.15 | 0.09 |
| 9 | 126154776 | rs1891642 | 1.944 | -0.745 | 1.036 | 0.87 | -0.001 | -0.004 | -0.004 | 0.85 | 0.58 | 0.44 | 0.45 | 0.38 | 0.63 | 0.44 |
| 9 | 126157360 | rs4838057 | -0.386 | -0.7 | 1 | 0.872 | 0.003 | -0.006 | -0.006 | 0.456 | 0.22 | 0.19 | 0.24 | 0.23 | 0.1 | 0.29 |
| 9 | 126162221 | rs4466467 | 0.335 | -0.57 | 0.903 | 0.679 | 0.009 | 0.011 | 0.011 | 0 | 0.16 | 0.13 | 0.13 | 0.04 | 0.08 | 0.1 |
| 9 | 126166739 | rs7865976 | -0.588 | -0.323 | 0.969 | 0.801 | 0.09 | 0.035 | 0.035 | 0.049 | 0.09 | 0.31 | 0.21 | 0.09 | 0.15 | 0.13 |
| 9 | 126173552 | rs16926471 | -0.802 | -0.295 | 0.738 | 0.949 | 0.031 | 0.028 | 0.028 | 0 | 0.11 | 0.2 | 0.21 | 0.18 | 0.16 | 0.17 |
| 9 | 126177860 | rs7048861 | 1.679 | -0.632 | 0.523 | 0.843 | 0.017 | 0.039 | 0.039 | 0 | 0.55 | 0.33 | 0.46 | 0.39 | 0.63 | 0.5 |
| 9 | 126178928 | rs12347955 | -1.009 | -0.491 | 0.751 | 1.033 | 0.001 | 0.006 | 0.006 | 0 | 0.18 | 0.19 | 0.23 | 0.24 | 0.18 | 0.24 |
| 9 | 126179204 | rs7851052 | -1.481 | -0.517 | 0.76 | 1.093 | -0.009 | 0.06 | 0.06 | 0 | 0.24 | 0.33 | 0.28 | 0.4 | 0.21 | 0.28 |
| 9 | 126179799 | rs7026808 | -0.001 | -0.379 | -0.051 | 0.43 | -0.005 | -0.058 | -0.058 | 0 | 0.14 | 0.23 | 0.05 | 0.03 | 0.09 | 0.1 |
| 9 | 126181097 | rs10124048 | -0.715 | -0.047 | -0.543 | 0.855 | -0.001 | -0.002 | -0.002 | 0 | 0.15 | 0.13 | 0.22 | 0.23 | 0.18 | 0.23 |
| 9 | 126181588 | rs10760286 | 1.403 | -0.243 | -1.081 | 0.365 | -0.122 | 0.058 | 0.058 | 0 | 0.44 | 0.05 | 0.39 | 0.39 | 0.55 | 0.4 |
| 9 | 126187698 | rs3924101 | 1.106 | 0.445 | 0.32 | 0.659 | 0.013 | 0.009 | 0.009 | 0.21 | 0.63 | 0.54 | 0.53 | 0.59 | 0.56 | 0.47 |
| 9 | 126196204 | rs10818820 | -1.859 | -0.796 | 1.175 | -0.133 | -0.006 | -0.015 | -0.015 | 0.494 | 0.37 | 0.54 | 0.38 | 0.3 | 0.37 | 0.35 |
| 9 | 126198147 | rs7044960 | 1.846 | -0.958 | 1.376 | -0.057 | 0.011 | 0.006 | 0.006 | 0 | 0.63 | 0.3 | 0.64 | 0.66 | 0.69 | 0.69 |
| 9 | 126202551 | rs10739633 | 1.796 | -0.917 | 1.259 | -0.028 | 0.014 | 0.003 | 0.003 | 0 | 0.63 | 0.3 | 0.63 | 0.65 | 0.68 | 0.68 |
| 9 | 126205754 | rs7859831 | -0.618 | -0.841 | 1.282 | -0.046 | 0.022 | 0.015 | 0.015 | 0 | 0.07 | 0.11 | 0.12 | 0.14 | 0.15 | 0.12 |
| 9 | 126227862 | rs10760289 | 2.28 | -1.014 | 1.432 | -0.163 | 0.039 | 0.023 | 0.023 | 0 | 0.6 | 0.23 | 0.61 | 0.66 | 0.68 | 0.68 |
| 9 | 126231899 | rs11999243 | -1.35 | -1.085 | 1.749 | -0.375 | -0.024 | -0.043 | -0.043 | 0.742 | 0.19 | 0.3 | 0.1 | 0.08 | 0.12 | 0.13 |
| 9 | 126234414 | rs10760290 | 2.325 | -1.067 | 1.593 | -0.347 | 0.038 | 0.04 | 0.04 | 0 | 0.6 | 0.23 | 0.61 | 0.68 | 0.68 | 0.67 |
| 9 | 126253077 | rs12350892 | -0.386 | -1.084 | 1.956 | -0.582 | -0.014 | -0.017 | -0.017 | 0 | 0.15 | 0.23 | 0.16 | 0.08 | 0.1 | 0.13 |
| 9 | 126259435 | rs2808397 | 0.721 | -1.107 | 1.639 | -0.592 | 0.003 | 0.002 | 0.002 | 0 | 0.85 | 0.86 | 0.92 | 0.88 | 0.88 | 0.87 |
| 9 | 126262932 | rs2772208 | 2.182 | -1.15 | 1.129 | -0.585 | 0.052 | 0.071 | 0.071 | 0 | 0.57 | 0.24 | 0.62 | 0.72 | 0.67 | 0.64 |
| 9 | 126282999 | rs2772215 | 1.386 | -0.989 | 0.64 | -0.549 | 0.008 | 0.008 | 0.008 | 0 | 0.7 | 0.73 | 0.79 | 0.87 | 0.83 | 0.8 |
| 9 | 126286695 | rs10818827 | -1.481 | -0.616 | 0.364 | -0.398 | -0.008 | -0.001 | -0.001 | 0 | 0.23 | 0.13 | 0.33 | 0.2 | 0.29 | 0.24 |
| 9 | 126287416 | rs2772213 | 1.342 | -0.53 | 0.957 | -0.407 | 0.06 | 0.019 | 0.019 | 0 | 0.92 | 0.83 | 0.84 | 0.88 | 0.84 | 0.87 |
| 9 | 126292586 | rs2772212 | 1.14 | -0.776 | 0.967 | -0.601 | -0.007 | -0.03 | -0.03 | 0 | 0.84 | 0.8 | 0.84 | 0.94 | 0.84 | 0.86 |
| 9 | 126297895 | rs7026921 | -2.701 | -1.064 | 1.607 | -0.721 | 0.001 | 0.002 | 0.002 | 0 | 0.38 | 0.57 | 0.38 | 0.27 | 0.3 | 0.3 |
| 9 | 126304695 | rs748994 | -1.066 | -1.068 | 1.861 | -0.719 | -0.012 | -0.019 | -0.019 | 0.022 | 0.13 | 0.23 | 0.16 | 0.09 | 0.06 | 0.11 |
| 9 | 126309872 | rs4836931 | -1.261 | -1.226 | 1.674 | -0.799 | -0.013 | -0.036 | -0.036 | 3.71 | 0.18 | 0.21 | 0.16 | 0.07 | 0.16 | 0.14 |
| 9 | 126312791 | rs10818831 | -1.943 | -1.245 | 1.726 | -0.809 | -0.034 | -0.065 | -0.065 | 0 | 0.28 | 0.38 | 0.29 | 0.12 | 0.19 | 0.22 |
| 9 | 126315123 | rs6478623 | -2.94 | -1.264 | 1.682 | -0.795 | -0.003 | -0.007 | -0.007 | 0.139 | 0.4 | 0.54 | 0.37 | 0.25 | 0.29 | 0.31 |
| 9 | 126319341 | rs10986051 | -1.219 | -1.166 | 1.691 | -0.744 | -0.005 | -0.019 | -0.019 | 0.87 | 0.17 | 0.18 | 0.16 | 0.06 | 0.16 | 0.15 |
| 9 | 126321003 | rs10986053 | -0.266 | -1.177 | 1.737 | -0.748 | 0.006 | 0.006 | 0.006 | 0 | 0.1 | 0.08 | 0.13 | 0.06 | 0.04 | 0.07 |
| 9 | 126321363 | rs4838070 | -1.258 | -1.152 | 1.817 | -0.733 | -0.016 | -0.041 | -0.041 | 0.328 | 0.18 | 0.23 | 0.17 | 0.07 | 0.16 | 0.15 |
| 9 | 126324050 | rs10119667 | -2.264 | -1.129 | 1.593 | -0.714 | -0.011 | -0.036 | -0.036 | 0.835 | 0.27 | 0.47 | 0.25 | 0.19 | 0.24 | 0.22 |
| 9 | 126326884 | rs10760291 | -2.399 | -1.359 | 1.371 | -0.906 | -0.01 | -0.024 | -0.024 | 0.453 | 0.39 | 0.53 | 0.37 | 0.21 | 0.31 | 0.26 |
| 9 | 126333104 | rs7048972 | -0.461 | -1.209 | 1.224 | -0.825 | -0.006 | -0.002 | -0.002 | 0 | 0.06 | 0.04 | 0.06 | 0.07 | 0.08 | 0.07 |
| 9 | 126338482 | rs7020626 | -2.997 | -1.492 | 1.155 | -0.891 | 0.019 | 0.135 | 0.135 | 0.277 | 0.49 | 0.92 | 0.46 | 0.29 | 0.38 | 0.32 |
| 9 | 126341310 | rs7857782 | -2.378 | -1.432 | 1.444 | -0.942 | -0.002 | -0.046 | -0.046 | 0.11 | 0.33 | 0.51 | 0.3 | 0.21 | 0.35 | 0.25 |
| 9 | 126343077 | rs17286621 | 0.079 | -1.363 | 1.225 | -1.11 | -0.012 | 0.001 | 0.001 | 0 | 0.05 | 0.01 | 0.04 | 0.05 | 0.04 | 0.05 |
| 9 | 126346105 | rs10818846 | -1.618 | -1.379 | 1.304 | -1.067 | -0.001 | 0 | 0 | 0 | 0.17 | 0.15 | 0.14 | 0.12 | 0.11 | 0.14 |
| 9 | 126351423 | rs10986064 | -0.341 | -1.341 | 1.222 | -1.053 | -0.029 | 0.023 | 0.023 | 2.234 | 0.11 | 0.04 | 0.18 | 0.07 | 0.21 | 0.1 |
| 9 | 126366529 | rs10986070 | -2.66 | -1.492 | 1.772 | -1.104 | 0.019 | 0.058 | 0.058 | 0.172 | 0.46 | 0.63 | 0.39 | 0.23 | 0.39 | 0.32 |
| 9 | 126369663 | rs7868205 | -1.216 | -1.427 | 1.719 | -1.059 | -0.001 | -0.011 | -0.011 | 0 | 0.07 | 0.08 | 0.04 | 0.04 | 0.04 | 0.05 |
| 9 | 126383362 | rs7037809 | -1.849 | -1.427 | 1.747 | -1.058 | -0.008 | -0.011 | -0.011 | 0 | 0.09 | 0.15 | 0.05 | 0.07 | 0.04 | 0.05 |
| 9 | 126396606 | rs1475527 | -2.035 | -1.472 | 1.786 | -1.23 | -0.013 | -0.061 | -0.061 | 0.038 | 0.36 | 0.45 | 0.33 | 0.17 | 0.34 | 0.26 |
| 9 | 126412353 | rs10760294 | -0.184 | -1.419 | 1.623 | -1.246 | 0.005 | 0.006 | 0.006 | 0 | 0.1 | 0.08 | 0.07 | 0.05 | 0.05 | 0.07 |
| 9 | 126415616 | rs1366810 | -1.562 | -1.407 | 1.664 | -1.236 | 0.003 | 0.041 | 0.041 | 1.433 | 0.25 | 0.14 | 0.25 | 0.09 | 0.29 | 0.17 |
| 9 | 126423340 | rs7027523 | -2.4 | -1.452 | 2.035 | -1.258 | -0.001 | -0.014 | -0.014 | 0 | 0.31 | 0.64 | 0.31 | 0.11 | 0.32 | 0.19 |
| 9 | 126435369 | rs7028815 | -1.68 | -1.429 | 1.804 | -1.246 | 0.003 | 0.04 | 0.04 | 1.407 | 0.25 | 0.15 | 0.24 | 0.09 | 0.29 | 0.17 |

| | | | | | | | | | | | | | | | | |
|---|---|---|---|---|---|---|---|---|---|---|---|---|---|---|---|---|
| 9 | 126439100 | rs12377595 | -1.304 | -1.42 | 1.735 | -1.24 | -0.004 | 0.053 | 0.053 | 0 | 0.22 | 0.11 | 0.22 | 0.1 | 0.27 | 0.17 |
| 9 | 126442550 | rs6478629 | -2.964 | -1.464 | 1.723 | -1.255 | 0.044 | 0.174 | 0.174 | 0.087 | 0.47 | 0.81 | 0.41 | 0.21 | 0.39 | 0.33 |
| 9 | 126448659 | rs12003386 | -0.85 | -1.715 | 2.127 | -0.997 | 0.012 | 0.013 | 0.013 | 0 | 0.11 | 0.08 | 0.04 | 0.02 | 0.07 | 0.06 |
| 9 | 126458789 | rs10760297 | -1.989 | -1.79 | 2.199 | -1.025 | -0.005 | -0.037 | -0.037 | 0 | 0.24 | 0.5 | 0.22 | 0.17 | 0.21 | 0.22 |
| 9 | 126459380 | rs10818859 | -1.515 | -1.81 | 2.219 | -1.034 | -0.031 | -0.058 | -0.058 | 0.055 | 0.18 | 0.43 | 0.11 | 0.08 | 0.07 | 0.14 |
| 9 | 126462707 | rs7860055 | -2.498 | -1.75 | 2.287 | -1.054 | -0.015 | -0.058 | -0.058 | 0 | 0.25 | 0.52 | 0.18 | 0.15 | 0.24 | 0.21 |
| 9 | 126470140 | rs7867502 | -1.091 | -1.647 | 2.306 | -1.002 | -0.034 | -0.056 | -0.056 | 0.702 | 0.13 | 0.24 | 0.05 | 0.03 | 0.07 | 0.07 |
| 9 | 126472839 | rs7864135 | -3.197 | -1.694 | 2.021 | -1.035 | 0.065 | 0.144 | 0.144 | 0 | 0.42 | 0.78 | 0.29 | 0.19 | 0.29 | 0.28 |
| 9 | 126476963 | rs7851425 | -2.218 | -1.765 | 2.109 | -1.071 | -0.027 | -0.051 | -0.051 | 2.633 | 0.29 | 0.62 | 0.16 | 0.06 | 0.11 | 0.15 |
| 9 | 126477127 | rs4838077 | -2.368 | -1.632 | 2.148 | -1.003 | 0.027 | -0.003 | -0.003 | 0 | 0.14 | 0.32 | 0.15 | 0.12 | 0.18 | 0.15 |
| 9 | 126509806 | rs7850520 | -3.065 | -1.627 | 1.788 | -0.997 | 0.056 | 0.134 | 0.134 | 0 | 0.41 | 0.83 | 0.3 | 0.18 | 0.3 | 0.3 |
| 9 | 126511085 | rs12339991 | -1.913 | -1.59 | 1.827 | -1.066 | -0.007 | -0.01 | -0.01 | 0 | 0.21 | 0.24 | 0.15 | 0.14 | 0.19 | 0.2 |
| 9 | 126520068 | rs9785285 | -2.828 | -1.672 | 1.887 | -1.052 | 0.044 | 0.066 | 0.066 | 0 | 0.36 | 0.74 | 0.22 | 0.11 | 0.19 | 0.23 |
| 9 | 126520497 | rs10818868 | -0.189 | -1.662 | 2.027 | -1.045 | -0.011 | -0.011 | -0.011 | 0 | 0.1 | 0.15 | 0.08 | 0.05 | 0.03 | 0.06 |
| 9 | 126525212 | rs2479106 | -3.25 | -1.668 | 1.943 | -1.062 | 0.057 | 0.133 | 0.133 | 0 | 0.4 | 0.82 | 0.3 | 0.18 | 0.3 | 0.3 |
| 9 | 126530988 | rs2799463 | -2.699 | -1.716 | 1.994 | -1.051 | 0.029 | 0.043 | 0.043 | 0 | 0.33 | 0.74 | 0.19 | 0.11 | 0.17 | 0.23 |
| 9 | 126537952 | rs1752168 | -1.906 | -1.727 | 2.32 | -1.056 | -0.066 | -0.114 | -0.114 | 1.625 | 0.24 | 0.45 | 0.13 | 0.06 | 0.11 | 0.15 |
| 9 | 126544609 | rs2799465 | -2.531 | -1.729 | 2.297 | -1.058 | -0.002 | -0.014 | -0.014 | 0 | 0.31 | 0.68 | 0.16 | 0.09 | 0.14 | 0.22 |
| 9 | 126548090 | rs2799466 | -1.351 | -1.69 | 2.567 | -1.085 | -0.027 | -0.073 | -0.073 | 0.003 | 0.13 | 0.25 | 0.05 | 0.01 | 0.08 | 0.08 |
| 9 | 126551037 | rs1752162 | -2.23 | -1.691 | 2.429 | -1.086 | -0.022 | -0.062 | -0.062 | 0 | 0.22 | 0.57 | 0.12 | 0.06 | 0.13 | 0.16 |
| 9 | 126558843 | rs1752166 | -2.642 | -1.708 | 2.297 | -1.086 | 0.025 | 0.033 | 0.033 | 0.222 | 0.31 | 0.72 | 0.18 | 0.08 | 0.16 | 0.19 |
| 9 | 126582578 | rs12000707 | -1.354 | -1.765 | 2.532 | -1.05 | -0.024 | -0.044 | -0.044 | 0 | 0.16 | 0.37 | 0.08 | 0.04 | 0.06 | 0.14 |
| 9 | 126593370 | rs2768819 | -3.38 | -1.779 | 2.36 | -1.056 | 0.06 | 0.126 | 0.126 | 0 | 0.39 | 0.81 | 0.26 | 0.15 | 0.28 | 0.3 |
| 9 | 126598435 | rs2458490 | -3.378 | -1.764 | 2.368 | -1.049 | 0.059 | 0.125 | 0.125 | 0 | 0.39 | 0.81 | 0.26 | 0.15 | 0.28 | 0.3 |
| 9 | 126601256 | rs7863739 | -0.732 | -1.738 | 2.385 | -1.037 | 0 | -0.002 | -0.002 | 0 | 0.09 | 0.09 | 0.1 | 0.07 | 0.13 | 0.11 |
| 9 | 126613028 | rs549779 | 3.026 | -1.737 | 2.203 | -0.772 | 0.057 | 0.078 | 0.078 | 0 | 0.61 | 0.18 | 0.74 | 0.73 | 0.73 | 0.7 |
| 9 | 126619233 | rs9696009 | -1.636 | -1.7 | 2.319 | -0.962 | -0.037 | -0.066 | -0.066 | 0 | 0.18 | 0.46 | 0.1 | 0.04 | 0.06 | 0.14 |
| 9 | 126664663 | rs7021780 | -0.962 | -1.597 | 2.375 | -0.904 | 0.013 | -0.001 | -0.001 | 0 | 0.07 | 0.24 | 0.03 | 0.02 | 0.02 | 0.08 |
| 9 | 126680050 | rs12001870 | -2.359 | -1.63 | 2.456 | -0.899 | -0.039 | -0.068 | -0.068 | 0.263 | 0.2 | 0.52 | 0.11 | 0.06 | 0.11 | 0.12 |
| 9 | 126685609 | rs12115297 | -1.486 | -1.552 | 2.135 | -0.888 | -0.003 | -0.005 | -0.005 | 0 | 0.07 | 0.22 | 0.05 | 0.02 | 0.05 | 0.01 |
| 9 | 126690157 | rs3739827 | -2.129 | -1.581 | 2.151 | -0.919 | -0.025 | -0.071 | -0.071 | 1.789 | 0.25 | 0.55 | 0.16 | 0.11 | 0.2 | 0.21 |
| 9 | 126695770 | rs10818894 | -1.824 | -1.577 | 1.688 | -0.934 | -0.008 | -0.02 | -0.02 | 0 | 0.28 | 0.68 | 0.19 | 0.11 | 0.2 | 0.18 |
| 9 | 126710886 | rs10760301 | -2.101 | -1.669 | 0.761 | -0.895 | 0.073 | 0.138 | 0.138 | 0 | 0.4 | 0.86 | 0.29 | 0.2 | 0.32 | 0.31 |
| 9 | 126715275 | rs4838084 | -1.379 | -1.313 | 1.111 | -0.638 | -0.019 | -0.043 | -0.043 | 0 | 0.25 | 0.44 | 0.19 | 0.16 | 0.23 | 0.25 |
| 9 | 126722647 | rs10760302 | 0.307 | -1.384 | 0.576 | -0.86 | -0.042 | -0.08 | -0.08 | 0 | 0.71 | 0.47 | 0.8 | 0.88 | 0.74 | 0.79 |
| 9 | 126722993 | rs10986159 | -0.072 | -0.282 | -0.318 | 0.317 | -0.03 | 0.052 | 0.052 | 0.209 | 0.05 | 0.01 | 0.27 | 0.4 | 0.14 | 0.33 |
| 9 | 126729111 | rs4523363 | 0.477 | -1.711 | -0.873 | -1.043 | 0.06 | 0.104 | 0.104 | 0.254 | 0.22 | 0.04 | 0.07 | 0.02 | 0.1 | 0.04 |
| 9 | 126732528 | rs3983857 | -0.315 | -1.254 | -0.447 | -0.999 | -0.008 | -0.046 | -0.046 | 0.197 | 0.7 | 0.59 | 0.81 | 0.85 | 0.69 | 0.73 |
| 9 | 126742752 | rs10986162 | -0.114 | -0.441 | -0.439 | -0.468 | -0.019 | -0.018 | -0.018 | 0.449 | 0.1 | 0.27 | 0.07 | 0.03 | 0.03 | 0.08 |
| 9 | 126749631 | rs4836950 | 0.704 | -0.339 | -0.988 | -0.466 | -0.13 | 0.032 | 0.032 | 0 | 0.34 | 0.05 | 0.63 | 0.76 | 0.53 | 0.58 |
| 9 | 126753312 | rs10986172 | -0.176 | -0.413 | -0.491 | -0.38 | -0.021 | -0.02 | -0.02 | 0 | 0.1 | 0.35 | 0.08 | 0.03 | 0.03 | 0.08 |
| 9 | 126756640 | rs1973 | 0.442 | 0.031 | -1.609 | -0.204 | -0.124 | 0.031 | 0.031 | 0 | 0.34 | 0.05 | 0.64 | 0.78 | 0.53 | 0.55 |
| 9 | 126758163 | rs7849890 | 0.147 | -0.073 | -1.469 | -0.033 | -0.023 | 0.047 | 0.047 | 0 | 0.38 | 0.16 | 0.66 | 0.72 | 0.56 | 0.6 |
| 9 | 126763128 | rs4314746 | -0.364 | 0.218 | -1.664 | 0.472 | -0.077 | -0.058 | -0.058 | 0 | 0.72 | 0.93 | 0.42 | 0.3 | 0.5 | 0.57 |
| 9 | 126767232 | rs7046300 | -0.793 | -1.145 | -1.471 | 0.322 | 0.022 | 0.033 | 0.033 | 0 | 0.59 | 0.77 | 0.74 | 0.72 | 0.6 | 0.61 |
| 9 | 126768048 | rs1327291 | -0.132 | -1.036 | -0.772 | 0.314 | -0.025 | 0.002 | 0.002 | 0 | 0.4 | 0.24 | 0.52 | 0.43 | 0.5 | 0.41 |
| 9 | 126768582 | rs969585 | -1.029 | -0.851 | -0.278 | 0.234 | 0.005 | 0.002 | 0.002 | 0 | 0.53 | 0.58 | 0.68 | 0.55 | 0.55 | 0.52 |
| 9 | 126771321 | rs7867942 | 0.452 | -0.701 | -0.273 | 0.12 | -0.005 | -0.014 | -0.014 | 0.035 | 0.09 | 0.15 | 0.08 | 0.02 | 0.05 | 0.09 |
| 9 | 126779494 | rs3780680 | -0.376 | -0.602 | -0.217 | 0.31 | 0.004 | 0.004 | 0.004 | 0 | 0.42 | 0.48 | 0.55 | 0.46 | 0.5 | 0.41 |
| 9 | 126800143 | rs4836959 | -0.534 | -0.277 | 0.014 | 0.378 | 0.011 | 0.019 | 0.019 | 0 | 0.49 | 0.45 | 0.58 | 0.7 | 0.53 | 0.52 |
| 9 | 126807459 | rs12555707 | 0.059 | 0.282 | -0.156 | 0.591 | -0.008 | 0.013 | 0.013 | 0 | 0.14 | 0.1 | 0.13 | 0.02 | 0.17 | 0.13 |
| 9 | 126811384 | rs12004441 | -0.541 | -0.058 | 0.431 | 0.907 | -0.041 | 0.031 | 0.031 | 0 | 0.21 | 0.3 | 0.23 | 0.26 | 0.13 | 0.16 |
| 9 | 126813100 | rs12002230 | -0.889 | -0.16 | 0.524 | 0.429 | -0.05 | 0.058 | 0.058 | 0 | 0.25 | 0.57 | 0.28 | 0.37 | 0.13 | 0.17 |
| 9 | 126817862 | rs7857883 | -0.921 | -0.261 | 0.602 | 0.386 | -0.032 | 0.084 | 0.084 | 0 | 0.22 | 0.41 | 0.29 | 0.37 | 0.15 | 0.22 |
| 9 | 126818818 | rs10986190 | 0.756 | -0.277 | 0.46 | 0.213 | -0.029 | 0.047 | 0.047 | 0 | 0.24 | 0.14 | 0.14 | 0.11 | 0.3 | 0.22 |
| 9 | 126820948 | rs1887342 | -2.166 | -0.153 | 0.443 | 0.338 | 0.025 | 0.044 | 0.044 | 0 | 0.64 | 0.85 | 0.71 | 0.75 | 0.64 | 0.65 |
| 9 | 126821992 | rs12682984 | -0.435 | -0.263 | 0.59 | 0.33 | -0.043 | 0.127 | 0.127 | 0.264 | 0.12 | 0.25 | 0.19 | 0.35 | 0.05 | 0.09 |
| 9 | 126823490 | rs10121243 | -1.718 | -0.171 | 0.529 | 0.314 | -0.023 | -0.035 | -0.035 | 0 | 0.59 | 0.79 | 0.51 | 0.51 | 0.54 | 0.52 |
| 9 | 126823752 | rs10118027 | 0.661 | -0.074 | 0.577 | 0.263 | 0.001 | -0.02 | -0.02 | 0 | 0.1 | 0.17 | 0.1 | 0.02 | 0.09 | 0.13 |
| 9 | 126824521 | rs2025822 | -1.146 | -0.154 | 0.632 | 0.339 | 0.036 | 0.031 | 0.031 | 0 | 0.52 | 0.6 | 0.52 | 0.39 | 0.59 | 0.55 |
| 9 | 126828640 | rs7867375 | -0.453 | -0.4 | 0.674 | 0.237 | -0.025 | -0.075 | -0.075 | 0 | 0.81 | 0.7 | 0.89 | 0.97 | 0.87 | 0.92 |
| 9 | 126829274 | rs7868236 | 1.134 | -0.474 | 0.55 | 0.248 | 0.051 | 0.033 | 0.033 | 0 | 0.45 | 0.3 | 0.44 | 0.61 | 0.37 | 0.44 |
| 9 | 126831118 | rs10760312 | -0.88 | 0.199 | 0.847 | 0.966 | -0.002 | 0.001 | 0.001 | 0 | 0.36 | 0.38 | 0.45 | 0.37 | 0.5 | 0.47 |
| 9 | 126835484 | rs3861868 | -1.497 | 0.138 | 0.583 | 1.009 | 0.003 | 0.01 | 0.01 | 0 | 0.47 | 0.57 | 0.63 | 0.7 | 0.53 | 0.58 |
| 9 | 126835798 | rs4836962 | -0.431 | 0.116 | 0.122 | 1.157 | -0.025 | 0.106 | 0.106 | 0 | 0.1 | 0.17 | 0.16 | 0.38 | 0.06 | 0.09 |
| 9 | 126848072 | rs7046686 | 0.137 | 0.361 | -0.046 | 1.02 | -0.002 | -0.009 | -0.009 | 0.018 | 0.11 | 0.13 | 0.09 | 0.03 | 0.09 | 0.14 |
| 9 | 126848249 | rs7858267 | -0.375 | 0.429 | -0.454 | 1.074 | -0.016 | -0.084 | -0.084 | 0 | 0.3 | 0.14 | 0.28 | 0.47 | 0.31 | 0.31 |
| 9 | 126850913 | rs10115776 | -0.961 | 0.796 | 0.456 | 0.97 | 0.072 | 0.043 | 0.043 | 0 | 0.18 | 0.44 | 0.33 | 0.25 | 0.21 | 0.26 |
| 9 | 126855342 | rs7858299 | -0.239 | 0.524 | -0.092 | 0.498 | 0.007 | 0.028 | 0.028 | 0 | 0.24 | 0.19 | 0.14 | 0.12 | 0.21 | 0.22 |

| | | | | | | | | | | | | | | | | |
|---|---|---|---|---|---|---|---|---|---|---|---|---|---|---|---|---|
| 9 | 126859995 | rs4838113 | -0.922 | 0.326 | 0.847 | 1.329 | 0.055 | 0.164 | 0.164 | 0.029 | 0.2 | 0.44 | 0.52 | 0.44 | 0.22 | 0.27 |
| 9 | 126860775 | rs7860363 | 0.018 | 0.288 | 0.638 | 1.243 | 0.007 | 0.043 | 0.043 | 0 | 0.24 | 0.16 | 0.14 | 0.12 | 0.22 | 0.22 |
| 9 | 126861389 | rs7863879 | -1.619 | 0.047 | 0.81 | 1.457 | -0.008 | 0.041 | 0.041 | 0.53 | 0.47 | 0.78 | 0.68 | 0.65 | 0.44 | 0.49 |
| 9 | 126862478 | rs7868364 | -0.209 | 0.081 | 0.729 | 1.584 | 0.011 | 0.048 | 0.048 | 0 | 0.24 | 0.15 | 0.13 | 0.12 | 0.21 | 0.22 |
| 9 | 126862735 | rs7868648 | -0.458 | 0.115 | 1.124 | 2.014 | -0.022 | 0.005 | 0.005 | 0.207 | 0.28 | 0.39 | 0.23 | 0.3 | 0.22 | 0.22 |
| 9 | 126862951 | rs6478644 | -1.171 | 0.246 | 1.084 | 1.992 | 0.002 | 0.03 | 0.03 | 0 | 0.37 | 0.47 | 0.59 | 0.53 | 0.37 | 0.35 |
| 9 | 126863443 | rs7030878 | -0.806 | -0.303 | 0.74 | 1.61 | -0.017 | 0.02 | 0.02 | 0.501 | 0.32 | 0.42 | 0.26 | 0.43 | 0.26 | 0.28 |
| 9 | 126871299 | rs2182660 | -0.495 | -0.231 | 0.454 | 1.813 | -0.006 | 0.005 | 0.005 | 0 | 0.31 | 0.38 | 0.27 | 0.35 | 0.29 | 0.32 |
| 9 | 126875893 | rs4838114 | -2.243 | -0.404 | 1.074 | 2.472 | -0.008 | 0.151 | 0.151 | 0 | 0.09 | 0.25 | 0.16 | 0.28 | 0.07 | 0.13 |
| 9 | 126879182 | rs2807610 | -0.074 | -0.818 | 1.277 | 1.884 | -0.02 | -0.017 | -0.017 | 0.437 | 0.19 | 0.28 | 0.21 | 0.15 | 0.16 | 0.08 |
| 9 | 126881448 | rs1463729 | -1.343 | -0.027 | 0.683 | 1.778 | 0.012 | -0.015 | -0.015 | 0 | 0.55 | 0.77 | 0.56 | 0.48 | 0.6 | 0.44 |
| 9 | 126885146 | rs1576011 | -1.531 | -0.108 | 0.862 | 1.877 | -0.021 | 0.011 | 0.011 | 0 | 0.58 | 0.8 | 0.68 | 0.65 | 0.62 | 0.56 |
| 9 | 126888086 | rs7029319 | 0.077 | -0.077 | 1.179 | 1.784 | -0.003 | 0.001 | 0.001 | 0 | 0.3 | 0.28 | 0.31 | 0.26 | 0.4 | 0.33 |
| 9 | 126888457 | rs2807602 | 0.061 | -0.168 | 1.328 | 1.649 | -0.017 | -0.004 | -0.004 | 0 | 0.21 | 0.3 | 0.31 | 0.21 | 0.18 | 0.17 |
| 9 | 126889267 | rs10118855 | 0.012 | -0.055 | 1.175 | 1.525 | 0.004 | -0.005 | -0.005 | 0 | 0.3 | 0.34 | 0.29 | 0.24 | 0.39 | 0.31 |
| 9 | 126890847 | rs2767755 | -1.713 | 0.049 | 0.878 | 1.601 | -0.002 | -0.015 | -0.015 | 0.238 | 0.62 | 0.71 | 0.72 | 0.59 | 0.64 | 0.62 |
| 9 | 126891306 | rs2638377 | -0.801 | 0.008 | 1.043 | 1.605 | -0.042 | 0.009 | 0.009 | 0.016 | 0.32 | 0.48 | 0.41 | 0.33 | 0.24 | 0.28 |
| 9 | 126891383 | rs2807597 | 0.31 | 0.109 | 0.93 | 1.638 | 0.012 | 0.003 | 0.003 | 0 | 0.21 | 0.16 | 0.28 | 0.16 | 0.17 | 0.16 |
| 9 | 126891946 | rs7873448 | 0.103 | 0.452 | 0.796 | 1.729 | -0.028 | 0.014 | 0.014 | 0 | 0.29 | 0.2 | 0.31 | 0.27 | 0.41 | 0.34 |
| 9 | 126893167 | rs2807593 | -0.119 | 1.007 | 0.443 | 1.228 | 0.019 | -0.029 | -0.029 | 0 | 0.24 | 0.31 | 0.47 | 0.13 | 0.32 | 0.22 |
| 9 | 126893436 | rs2807592 | -1.936 | 0.892 | -0.322 | 1.205 | -0.044 | -0.127 | -0.127 | 0 | 0.63 | 0.97 | 0.7 | 0.6 | 0.68 | 0.64 |
| 9 | 126894348 | rs2638381 | -1.717 | 0.213 | 0.191 | 1.536 | -0.021 | 0.03 | 0.03 | 0 | 0.59 | 0.75 | 0.54 | 0.67 | 0.58 | 0.64 |
| 9 | 126897258 | rs2767762 | -0.006 | -0.227 | 0.38 | 1.392 | -0.005 | -0.046 | -0.046 | 0 | 0.22 | 0.36 | 0.26 | 0.11 | 0.22 | 0.2 |
| 9 | 126905188 | rs2478894 | -0.539 | -0.129 | -0.645 | 1.427 | 0.016 | 0.009 | 0.009 | 3.391 | 0.28 | 0.74 | 0.31 | 0.2 | 0.23 | 0.23 |
| 9 | 126905298 | rs2452174 | -0.529 | -0.123 | -0.642 | 1.468 | 0.015 | -0.006 | -0.006 | 0.137 | 0.28 | 0.74 | 0.31 | 0.24 | 0.23 | 0.22 |
| 9 | 126913862 | rs10818924 | -0.607 | 0.397 | -0.573 | 1.096 | 0.002 | 0.003 | 0.003 | 0 | 0.36 | 0.45 | 0.38 | 0.59 | 0.38 | 0.42 |
| 9 | 126914307 | rs2767770 | -1.621 | 0.3 | -0.609 | 1.09 | -0.053 | 0.035 | 0.035 | 0.595 | 0.6 | 0.9 | 0.62 | 0.71 | 0.57 | 0.6 |
| 9 | 126918033 | rs2807584 | -0.456 | 0.058 | 0.262 | 1.033 | -0.043 | -0.057 | -0.057 | 2.732 | 0.24 | 0.45 | 0.24 | 0.13 | 0.18 | 0.17 |
| 9 | 126920280 | rs7865381 | -1.571 | 0.857 | -0.001 | 1.722 | 0.015 | 0.056 | 0.056 | 0 | 0.1 | 0.17 | 0.11 | 0.22 | 0.09 | 0.19 |
| 9 | 126935255 | rs4838118 | -1.447 | 0.523 | -0.161 | 1.324 | -0.001 | 0.003 | 0.003 | 0.133 | 0.07 | 0.08 | 0.11 | 0.07 | 0.04 | 0.12 |
| 9 | 126935364 | rs6478649 | -1.63 | -0.324 | -0.428 | 1.718 | 0.048 | -0.053 | -0.053 | 1.434 | 0.32 | 0.81 | 0.33 | 0.42 | 0.24 | 0.36 |
| 9 | 126935882 | rs1536510 | -0.885 | -0.311 | 0.934 | 1.797 | -0.02 | 0.07 | 0.07 | 0 | 0.18 | 0.26 | 0.15 | 0.32 | 0.17 | 0.13 |
| 9 | 126946822 | rs7860461 | -1.743 | -0.382 | 1.639 | 1.414 | 0.018 | -0.013 | -0.013 | 0 | 0.11 | 0.47 | 0.15 | 0.03 | 0.07 | 0.15 |
| 9 | 126949213 | rs4311746 | 0.005 | 0.162 | 0.773 | 1.768 | -0.068 | -0.078 | -0.078 | 1.748 | 0.13 | 0.01 | 0.12 | 0.24 | 0.17 | 0.13 |
| 9 | 126951284 | rs7042608 | -0.071 | -0.19 | 0.859 | 1.641 | -0.007 | 0 | 0 | 0.035 | 0.15 | 0.18 | 0.13 | 0.17 | 0.15 | 0.06 |
| 9 | 126957987 | rs10986266 | -1.634 | -0.331 | 0.988 | 1.852 | -0.042 | 0.068 | 0.068 | 0.843 | 0.22 | 0.64 | 0.19 | 0.33 | 0.11 | 0.21 |
| 9 | 126959681 | rs12003839 | -1.377 | -0.172 | 0.915 | 1.931 | -0.031 | 0.145 | 0.145 | 0.028 | 0.17 | 0.36 | 0.16 | 0.33 | 0.11 | 0.21 |
| 9 | 126965119 | rs7034837 | -1.794 | 0.295 | -0.034 | 0.99 | 0.015 | 0.181 | 0.181 | 0 | 0.52 | 0.82 | 0.56 | 0.77 | 0.58 | 0.51 |
| 9 | 126965396 | rs1106489 | -0.707 | 0.048 | 0.313 | 0.972 | -0.002 | -0.006 | -0.006 | 0.048 | 0.08 | 0.09 | 0.1 | 0.04 | 0.08 | 0.14 |
| 9 | 126967977 | rs10986270 | -0.965 | -0.161 | 0.609 | 1.282 | -0.037 | 0.072 | 0.072 | 0 | 0.28 | 0.39 | 0.27 | 0.46 | 0.2 | 0.36 |
| 9 | 126969243 | rs7043914 | -0.315 | 0.424 | 0.757 | 0.754 | -0.005 | -0.085 | -0.085 | 0 | 0.13 | 0.3 | 0.09 | 0 | 0.12 | 0.01 |
| 9 | 126969598 | rs10986273 | -0.365 | 0.85 | 0.084 | 1.276 | -0.057 | 0.013 | 0.013 | 0.068 | 0.1 | 0.03 | 0.11 | 0.07 | 0.16 | 0.15 |
| 9 | 126970903 | rs7021109 | 1.302 | -0.25 | 0.809 | 1.649 | -0.086 | 0.009 | 0.009 | 0.544 | 0.78 | 0.56 | 0.85 | 0.8 | 0.92 | 0.75 |
| 9 | 126971170 | rs4838130 | -1.71 | 0.94 | 0.605 | 2.05 | 0.016 | 0.051 | 0.051 | 0.024 | 0.11 | 0.2 | 0.16 | 0.26 | 0.14 | 0.14 |
| 9 | 126974767 | rs10760331 | -1.474 | 1.031 | 0.634 | 2.122 | 0.025 | 0.098 | 0.098 | 0 | 0.17 | 0.26 | 0.4 | 0.46 | 0.2 | 0.27 |
| 9 | 126975022 | rs10760332 | -1.814 | 0.772 | 1.211 | 1.984 | 0.018 | 0.166 | 0.166 | 1.879 | 0.21 | 0.53 | 0.43 | 0.53 | 0.21 | 0.27 |
| 9 | 126983801 | rs12339663 | -1.102 | 0.396 | 0.857 | 2.273 | 0.007 | 0.023 | 0.023 | 0 | 0.08 | 0.08 | 0.17 | 0.36 | 0.04 | 0.21 |
| 9 | 126983984 | rs16927289 | -1.883 | 0.111 | 1.712 | 2.261 | 0.054 | 0.091 | 0.091 | 2.202 | 0.1 | 0.43 | 0.21 | 0.15 | 0.09 | 0.13 |
| 9 | 126984382 | rs953470 | -1.915 | 0.641 | 1.323 | 2.033 | 0.011 | 0.063 | 0.063 | 0 | 0.24 | 0.37 | 0.3 | 0.5 | 0.24 | 0.31 |
| 9 | 126985334 | rs16927294 | -2.19 | 0.619 | 1.37 | 1.8 | 0.055 | 0.018 | 0.018 | 4.386 | 0.23 | 0.55 | 0.32 | 0.31 | 0.28 | 0.23 |
| 9 | 126987143 | rs6478652 | -2.208 | 0.74 | 1.441 | 1.782 | 0.103 | 0.032 | 0.032 | 4.456 | 0.17 | 0.51 | 0.3 | 0.25 | 0.25 | 0.2 |
| 9 | 126991185 | rs10818930 | 2.05 | -0.588 | 1.105 | 1.62 | 0.01 | -0.003 | -0.003 | 0 | 0.67 | 0.32 | 0.53 | 0.39 | 0.8 | 0.58 |
| 9 | 126997079 | rs7849959 | 1.341 | -0.384 | 1.176 | 1.626 | -0.015 | 0.013 | 0.013 | 0 | 0.74 | 0.7 | 0.74 | 0.47 | 0.84 | 0.7 |
| 9 | 127001460 | rs7861786 | 0.557 | -0.325 | 0.966 | 0.656 | 0.003 | -0.022 | -0.022 | 0 | 0.3 | 0.34 | 0.35 | 0.12 | 0.35 | 0.28 |
| 9 | 127005041 | rs10986285 | 0.74 | -0.476 | 1.443 | 0.752 | 0.001 | 0.039 | 0.039 | 0 | 0.46 | 0.37 | 0.39 | 0.35 | 0.49 | 0.43 |
| 9 | 127010114 | rs4838140 | -0.467 | -0.362 | 1.387 | 0.653 | 0.014 | 0 | 0 | 1.043 | 0.13 | 0.17 | 0.12 | 0.16 | 0.17 | 0.1 |
| 9 | 127020828 | rs2065221 | -0.14 | -0.465 | 1.422 | 0.767 | 0.001 | 0.008 | 0.008 | 0.531 | 0.58 | 0.63 | 0.56 | 0.79 | 0.56 | 0.63 |
| 9 | 127025040 | rs2282085 | 1.475 | -0.524 | 1.53 | 0.909 | 0.007 | 0.012 | 0.012 | 0 | 0.36 | 0.27 | 0.29 | 0.28 | 0.33 | 0.34 |
| 9 | 127026282 | rs10760344 | 1.578 | -0.527 | 1.517 | 0.91 | 0.011 | 0.036 | 0.036 | 0 | 0.34 | 0.22 | 0.28 | 0.19 | 0.31 | 0.33 |
| 9 | 127029736 | rs12379417 | 1.822 | -0.601 | 1.327 | 0.861 | -0.034 | 0.059 | 0.059 | 0.319 | 0.33 | 0.05 | 0.28 | 0.19 | 0.33 | 0.35 |
| 9 | 127031054 | rs16927321 | 0.8 | -0.603 | 1.371 | 0.876 | 0.002 | -0.005 | -0.005 | 0 | 0.07 | 0.04 | 0.05 | 0.09 | 0.03 | 0.08 |
| 9 | 127035761 | rs10986296 | 0.17 | -0.618 | 1.201 | 0.919 | -0.009 | -0.046 | -0.046 | 0 | 0.1 | 0.02 | 0.15 | 0.38 | 0.08 | 0.21 |
| 9 | 127039017 | rs7042263 | 0.001 | -0.624 | 1.357 | 0.713 | 0.001 | 0.007 | 0.007 | 0.422 | 0.58 | 0.63 | 0.56 | 0.79 | 0.57 | 0.63 |
| 9 | 127052272 | rs7872321 | 1.231 | -0.65 | 1.397 | 0.673 | -0.058 | 0.075 | 0.075 | 0 | 0.27 | 0.1 | 0.35 | 0.12 | 0.37 | 0.25 |
| 9 | 127054184 | rs4838153 | -0.002 | -0.328 | 1.442 | 0.907 | 0.002 | -0.001 | -0.001 | 0 | 0.1 | 0.11 | 0.11 | 0.13 | 0.16 | 0.07 |
| 9 | 127060431 | rs944340 | 0.588 | -0.382 | 1.676 | 1.03 | 0.018 | 0.039 | 0.039 | 0 | 0.45 | 0.27 | 0.44 | 0.65 | 0.4 | 0.6 |
| 9 | 127062488 | rs2026191 | -0.286 | -0.33 | 1.518 | 0.926 | 0.004 | 0.017 | 0.017 | 0.706 | 0.54 | 0.62 | 0.57 | 0.79 | 0.58 | 0.69 |
| 9 | 127065836 | rs17222478 | 0.541 | -0.168 | 1.131 | 1.089 | 0.027 | 0.011 | 0.011 | 0 | 0.06 | 0.15 | 0.09 | 0.13 | 0.13 | 0.06 |
| 9 | 127069629 | rs4838159 | -1.185 | -0.308 | 1.108 | 1.333 | -0.004 | 0.003 | 0.003 | 0 | 0.18 | 0.19 | 0.23 | 0.52 | 0.22 | 0.35 |

| | | | | | | | | | | | | | | | | |
|---|---|---|---|---|---|---|---|---|---|---|---|---|---|---|---|---|
| 9 | 127082375 | rs2150237 | 0.505 | -0.556 | 1.593 | 1.229 | 0.001 | -0.002 | -0.002 | 1.433 | 0.41 | 0.37 | 0.44 | 0.56 | 0.39 | 0.59 |
| 9 | 127083933 | rs2274781 | -0.622 | -0.552 | 1.552 | 1.247 | -0.014 | 0.101 | 0.101 | 0 | 0.12 | 0.23 | 0.16 | 0.38 | 0.07 | 0.28 |
| 9 | 127088029 | rs10986315 | 0.04 | -0.488 | 1.369 | 0.799 | 0.003 | 0.005 | 0.005 | 0 | 0.09 | 0.04 | 0.16 | 0.47 | 0.07 | 0.27 |
| 9 | 127089775 | rs748741 | 0.923 | -0.505 | 1.598 | 0.661 | -0.001 | -0.003 | -0.003 | 0.942 | 0.34 | 0.33 | 0.31 | 0.19 | 0.34 | 0.35 |
| 9 | 127090966 | rs912355 | -1.019 | -0.559 | 1.851 | 0.643 | -0.008 | -0.034 | -0.034 | 0 | 0.65 | 0.57 | 0.7 | 0.81 | 0.68 | 0.66 |
| 9 | 127112027 | rs10818945 | 0.077 | -0.695 | 1.519 | 0.839 | 0.031 | -0.023 | -0.023 | 0 | 0.1 | 0.03 | 0.14 | 0.46 | 0.04 | 0.26 |
| 9 | 127113660 | rs2416 | 0.062 | -0.777 | 1.708 | 0.686 | 0.034 | 0.073 | 0.073 | 0 | 0.5 | 0.31 | 0.49 | 0.33 | 0.58 | 0.36 |
| 9 | 127113732 | rs12555646 | 0.181 | -0.765 | 1.644 | 0.698 | 0.031 | -0.02 | -0.02 | 0.147 | 0.1 | 0.03 | 0.14 | 0.47 | 0.04 | 0.26 |
| 9 | 127114489 | rs13467 | -0.024 | -0.766 | 1.642 | 0.697 | 0.037 | -0.038 | -0.038 | 1.829 | 0.11 | 0.03 | 0.14 | 0.47 | 0.04 | 0.26 |
| 9 | 127117346 | rs1330811 | -0.032 | -0.691 | 1.854 | 0.712 | 0.033 | 0.082 | 0.082 | 0 | 0.51 | 0.32 | 0.48 | 0.32 | 0.57 | 0.35 |
| 9 | 127118386 | rs7873147 | -0.142 | -0.645 | 1.9 | 0.75 | 0 | 0.024 | 0.024 | 0 | 0.87 | 0.91 | 0.85 | 0.54 | 0.96 | 0.74 |
| 9 | 127130137 | rs10818950 | 0.933 | -0.571 | 1.891 | 0.568 | 0.001 | -0.005 | -0.005 | 0 | 0.34 | 0.31 | 0.33 | 0.21 | 0.37 | 0.38 |
| 9 | 127131062 | rs10818951 | 1.159 | -0.565 | 1.672 | 0.592 | -0.015 | 0.022 | 0.022 | 0 | 0.32 | 0.2 | 0.32 | 0.21 | 0.37 | 0.37 |
| 9 | 127162888 | rs7864941 | 1.165 | -0.516 | 1.292 | 0.631 | -0.021 | 0.028 | 0.028 | 0 | 0.32 | 0.2 | 0.31 | 0.22 | 0.38 | 0.39 |
| 9 | 127163623 | rs12344916 | 1.018 | -0.401 | 0.339 | 0.599 | -0.1 | 0.108 | 0.108 | 0.071 | 0.33 | 0.04 | 0.35 | 0.15 | 0.38 | 0.22 |
| 9 | 127173209 | rs7027515 | 0.334 | -0.699 | 0.989 | 1.056 | 0.022 | 0.065 | 0.065 | 0 | 0.5 | 0.34 | 0.48 | 0.31 | 0.56 | 0.34 |
| 9 | 127177161 | rs4574 | 0.404 | -0.588 | 0.876 | 1.073 | 0.001 | -0.017 | -0.017 | 3.257 | 0.37 | 0.61 | 0.38 | 0.23 | 0.4 | 0.39 |
| 9 | 127184067 | rs12343206 | -0.063 | -0.124 | 0.097 | 0.828 | 0.022 | 0.031 | 0.031 | 0 | 0.53 | 0.38 | 0.5 | 0.32 | 0.62 | 0.38 |
| 9 | 127190340 | rs10818964 | -0.011 | 0.343 | -0.069 | 1.404 | 0.017 | 0.006 | 0.006 | 0.86 | 0.61 | 0.26 | 0.56 | 0.36 | 0.67 | 0.44 |
| 9 | 127194892 | rs4836979 | -1.722 | 0.478 | 1.396 | 2.412 | 0.076 | 0.558 | 0.558 | 5.999 | 0.06 | 0.43 | 0.13 | 0.4 | 0.02 | 0.11 |
| 9 | 127195100 | rs4838174 | -1.42 | 0.51 | 1.18 | 2.322 | -0.024 | 0.187 | 0.187 | 0 | 0.2 | 0.45 | 0.33 | 0.48 | 0.15 | 0.21 |
| 9 | 127195873 | rs10986339 | -1.4 | 0.672 | 0.792 | 1.893 | 0.009 | 0.01 | 0.01 | 0 | 0.19 | 0.25 | 0.26 | 0.27 | 0.18 | 0.12 |
| 9 | 127204712 | rs11794849 | -0.736 | 0.254 | 1.009 | 2.214 | 0 | -0.001 | -0.001 | 0 | 0.22 | 0.24 | 0.1 | 0.2 | 0.15 | 0.1 |
| 9 | 127210541 | rs12003217 | -1.2 | 0.199 | 0.495 | 1.851 | -0.017 | 0.03 | 0.03 | 0.031 | 0.3 | 0.59 | 0.27 | 0.59 | 0.2 | 0.25 |
| 9 | 127215344 | rs7861556 | 0.551 | 1.47 | 0.299 | 1.675 | -0.012 | 0.01 | 0.01 | 1.891 | 0.8 | 0.81 | 0.78 | 0.86 | 0.73 | 0.67 |
| 9 | 127218258 | rs4838177 | 1.517 | 2.13 | 0.943 | 2.261 | 0.05 | 0.044 | 0.044 | 2.792 | 0.7 | 0.41 | 0.59 | 0.45 | 0.57 | 0.54 |
| 9 | 127220133 | rs10733669 | 1.093 | 2.239 | 1.3 | 2.263 | 0.051 | 0.085 | 0.085 | 2.806 | 0.73 | 0.53 | 0.63 | 0.46 | 0.64 | 0.58 |
| 9 | 127223127 | rs10760362 | 1.17 | 1.99 | 1.268 | 2.666 | -0.087 | -0.054 | -0.054 | 0 | 0.78 | 0.84 | 0.67 | 0.58 | 0.62 | 0.71 |
| 9 | 127223212 | rs4838181 | 1.704 | 1.94 | 1.345 | 2.799 | 0.076 | 0.078 | 0.078 | 1.707 | 0.72 | 0.4 | 0.62 | 0.44 | 0.6 | 0.61 |
| 9 | 127224865 | rs7851555 | 0.907 | 2.049 | 0.76 | 2.367 | -0.114 | -0.032 | -0.032 | 0 | 0.8 | 0.9 | 0.78 | 0.72 | 0.61 | 0.75 |
| 9 | 127230891 | rs10760365 | 1.095 | 0.859 | 0.645 | 1.874 | 0.003 | 0.01 | 0.01 | 0.018 | 0.53 | 0.54 | 0.53 | 0.62 | 0.42 | 0.5 |
| 9 | 127233823 | rs7852843 | 0.691 | 0.905 | 0.265 | 1.645 | -0.075 | 0.116 | 0.116 | 0.033 | 0.6 | 0.91 | 0.69 | 0.76 | 0.45 | 0.63 |
| 9 | 127244955 | rs915034 | 0.927 | 0.651 | 0.497 | 1.569 | -0.002 | 0.088 | 0.088 | 0 | 0.55 | 0.76 | 0.67 | 0.75 | 0.44 | 0.62 |
| 9 | 127247931 | rs7851737 | -0.965 | 0.814 | 0.452 | 1.786 | -0.017 | 0.049 | 0.049 | 0 | 0.42 | 0.18 | 0.31 | 0.29 | 0.53 | 0.33 |
| 9 | 127256223 | rs10114542 | -1.628 | 0.542 | 0.866 | 2.126 | -0.043 | 0.196 | 0.196 | 1.623 | 0.1 | 0.42 | 0.04 | 0.24 | 0.01 | 0.09 |
| 9 | 127267689 | rs3814134 | -2.394 | 0.434 | 0.552 | 2.327 | 0.033 | -0.036 | -0.036 | 0 | 0.15 | 0.83 | 0.09 | 0.31 | 0.02 | 0.12 |
| 9 | 127294185 | rs10986358 | -0.367 | 0.674 | 0.776 | 2.312 | -0.15 | -0.011 | -0.011 | 0 | 0.22 | 0.03 | 0.16 | 0.19 | 0.35 | 0.26 |
| 9 | 127320204 | rs12341407 | -1.497 | 0.456 | 1.022 | 2.401 | -0.044 | 0.213 | 0.213 | 1.823 | 0.1 | 0.41 | 0.06 | 0.24 | 0.01 | 0.09 |
| 9 | 127335754 | rs4590537 | 1.207 | 0.33 | 0.755 | 2.465 | -0.011 | -0.015 | -0.015 | 0 | 0.55 | 0.2 | 0.66 | 0.53 | 0.47 | 0.57 |
| 9 | 127338173 | rs7032937 | 0.93 | 0.377 | 0.793 | 2.538 | -0.018 | -0.019 | -0.019 | 0.825 | 0.58 | 0.75 | 0.69 | 0.47 | 0.47 | 0.58 |
| 9 | 127341110 | rs4240486 | 0.449 | 0.378 | 0.783 | 2.486 | -0.09 | -0.044 | -0.044 | 0 | 0.63 | 0.85 | 0.72 | 0.62 | 0.48 | 0.62 |
| 9 | 127406961 | rs10120039 | 1.438 | 0.404 | 0.777 | 2.304 | -0.08 | -0.005 | -0.005 | 0 | 0.49 | 0.04 | 0.63 | 0.39 | 0.47 | 0.53 |
| 9 | 127458468 | rs6478680 | 0.857 | 0.454 | 0.63 | 2.343 | -0.053 | -0.065 | -0.065 | 0.97 | 0.58 | 0.83 | 0.69 | 0.49 | 0.48 | 0.58 |
| 9 | 127495508 | rs10760374 | 1.174 | 0.512 | 0.681 | 2.374 | 0 | 0 | 0 | 0 | 0.53 | 0.48 | 0.66 | 0.39 | 0.47 | 0.53 |
| 9 | 127535992 | rs721862 | 1.273 | 0.567 | -0.236 | 2.18 | 0.002 | 0.012 | 0.012 | 0.005 | 0.52 | 0.43 | 0.66 | 0.39 | 0.47 | 0.52 |
| 9 | 127554070 | rs10760376 | 0.848 | 0.504 | -0.319 | 2.286 | 0.005 | 0 | 0 | 0 | 0.58 | 0.53 | 0.69 | 0.56 | 0.48 | 0.54 |
| 9 | 127562973 | rs913232 | 0.55 | 1.098 | -0.692 | 2.152 | -0.008 | 0.004 | 0.004 | 0.523 | 0.64 | 0.46 | 0.77 | 0.64 | 0.67 | 0.79 |
| 9 | 127564587 | rs10818989 | 1.057 | 0.855 | -1.974 | 1.501 | -0.08 | 0.074 | 0.074 | 0.636 | 0.48 | 0.04 | 0.62 | 0.31 | 0.47 | 0.52 |
| 9 | 127566378 | rs7874348 | 0.364 | 1.583 | -1.205 | 1.338 | -0.074 | 0.047 | 0.047 | 0 | 0.75 | 0.88 | 0.8 | 0.9 | 0.62 | 0.89 |
| 9 | 127570640 | rs10986432 | -0.524 | 0.704 | -1.28 | 1.638 | -0.02 | -0.018 | -0.018 | 0 | 0.19 | 0.04 | 0.28 | 0.22 | 0.17 | 0.16 |
| 9 | 127573167 | rs2416952 | -1.557 | 0.766 | -0.115 | 2.895 | 0.007 | 0.401 | 0.401 | 0 | 0.05 | 0.22 | 0.05 | 0.38 | 0.01 | 0.14 |
| 9 | 127573750 | rs4838215 | -1.618 | 0.757 | -0.121 | 2.891 | 0.001 | 0.379 | 0.379 | 0 | 0.06 | 0.21 | 0.05 | 0.38 | 0.01 | 0.14 |
| 9 | 127594371 | rs1887384 | 1.663 | 0.94 | 0.301 | 2.17 | 0.046 | 0.182 | 0.182 | 0 | 0.93 | 0.71 | 0.91 | 0.84 | 0.95 | 0.88 |
| 9 | 127602518 | rs1467676 | 0.222 | 0.803 | -0.032 | 1.744 | 0 | 0.004 | 0.004 | 0 | 0.42 | 0.37 | 0.44 | 0.35 | 0.41 | 0.33 |
| 9 | 127612399 | rs12552010 | 0.079 | -0.206 | 0.418 | 1.658 | -0.021 | -0.008 | -0.008 | 0 | 0.38 | 0.22 | 0.32 | 0.35 | 0.42 | 0.28 |
| 9 | 127615995 | rs3814131 | -0.349 | -0.174 | 1.123 | 1.723 | -0.001 | -0.025 | -0.025 | 0 | 0.46 | 0.45 | 0.51 | 0.76 | 0.45 | 0.5 |
| 9 | 127616537 | rs12375547 | -0.129 | -0.274 | 1.759 | 1.779 | 0 | -0.001 | -0.001 | 0.023 | 0.43 | 0.48 | 0.36 | 0.58 | 0.43 | 0.37 |
| 9 | 127620141 | rs876663 | 0.313 | -0.18 | 1.668 | 1.736 | -0.134 | -0.064 | -0.064 | 0 | 0.37 | 0.07 | 0.32 | 0.35 | 0.46 | 0.32 |
| 9 | 127621374 | rs2274969 | 0.222 | -0.706 | 1.748 | 0.979 | -0.014 | 0.005 | 0.005 | 0.551 | 0.24 | 0.33 | 0.26 | 0.23 | 0.15 | 0.21 |
| 9 | 127621644 | rs3780591 | 0.999 | -0.796 | 1.762 | 0.931 | -0.007 | 0.058 | 0.058 | 2.51 | 0.18 | 0.04 | 0.21 | 0.09 | 0.14 | 0.1 |
| 9 | 127634429 | rs1999589 | -0.942 | -0.81 | 1.883 | 0.935 | 0.002 | -0.004 | -0.004 | 0.8 | 0.58 | 0.51 | 0.51 | 0.39 | 0.57 | 0.62 |
| 9 | 127643168 | rs623096 | 0.478 | -0.876 | 1.56 | 0.636 | -0.006 | -0.006 | -0.006 | 0 | 0.55 | 0.47 | 0.48 | 0.68 | 0.7 | 0.58 |
| 9 | 127643456 | rs687434 | 1.809 | -1.075 | 1.526 | 0.438 | -0.019 | -0.007 | -0.007 | 0 | 0.34 | 0.21 | 0.32 | 0.34 | 0.4 | 0.26 |
| 9 | 127643946 | rs2304957 | 0.895 | -0.983 | 1.563 | 0.439 | -0.019 | 0.011 | 0.011 | 0 | 0.14 | 0.09 | 0.12 | 0.11 | 0.28 | 0.21 |
| 9 | 127645241 | rs17328115 | 1.726 | -1.102 | 1.467 | 0.373 | -0.007 | 0.096 | 0.096 | 0 | 0.16 | 0.01 | 0.1 | 0.01 | 0.13 | 0.16 |
| 9 | 127648182 | rs13283523 | 0.308 | -1.102 | 1.478 | 0.372 | -0.067 | 0.116 | 0.116 | 0 | 0.3 | 0.07 | 0.21 | 0.12 | 0.39 | 0.36 |
| 9 | 127661645 | rs634710 | 1.232 | -1.142 | 1.544 | 0.43 | -0.006 | -0.007 | -0.007 | 0 | 0.4 | 0.3 | 0.35 | 0.57 | 0.43 | 0.37 |
| 9 | 127662425 | rs4837003 | -0.633 | -1.14 | 1.352 | 0.651 | 0.016 | 0.02 | 0.02 | 0 | 0.44 | 0.38 | 0.37 | 0.28 | 0.31 | 0.4 |
| 9 | 127688614 | rs3780598 | 1.387 | -1.031 | 1.341 | 0.658 | -0.001 | 0.026 | 0.026 | 1.823 | 0.17 | 0.11 | 0.21 | 0.1 | 0.13 | 0.1 |

| | | | | | | | | | | | | | | | | |
|---|---|---|---|---|---|---|---|---|---|---|---|---|---|---|---|---|
| 9 | 127720397 | rs10986490 | -1.128 | -0.624 | 1.339 | 0.645 | 0.002 | 0.004 | 0.004 | 0 | 0.56 | 0.41 | 0.47 | 0.38 | 0.57 | 0.62 |
| 9 | 127733926 | rs2289631 | -1.667 | -0.588 | 1.37 | 0.672 | -0.002 | 0 | 0 | 0.011 | 0.62 | 0.67 | 0.66 | 0.64 | 0.57 | 0.71 |
| 9 | 127736933 | rs7045489 | -0.767 | -0.596 | 1.395 | 0.655 | 0.006 | -0.001 | -0.001 | 0 | 0.48 | 0.58 | 0.54 | 0.53 | 0.31 | 0.5 |
| 9 | 127764305 | rs764304 | 0.875 | -0.582 | 1.231 | 0.624 | -0.018 | 0.011 | 0.011 | 0 | 0.14 | 0.09 | 0.11 | 0.11 | 0.26 | 0.2 |
| 9 | 127765978 | rs4837005 | -0.727 | -0.661 | 1.344 | 0.771 | -0.001 | 0.002 | 0.002 | 0 | 0.39 | 0.42 | 0.36 | 0.33 | 0.41 | 0.44 |
| 9 | 127781259 | rs1017530 | 0.299 | -0.769 | 1.29 | 0.793 | -0.01 | -0.004 | -0.004 | 2.527 | 0.25 | 0.33 | 0.25 | 0.22 | 0.15 | 0.23 |
| 9 | 127784990 | rs12684101 | 0.828 | -0.774 | 1.299 | 0.776 | 0.006 | -0.007 | -0.007 | 0.019 | 0.21 | 0.1 | 0.14 | 0.24 | 0.17 | 0.25 |
| 9 | 127785819 | rs360198 | 0.642 | -0.772 | 1.348 | 0.776 | -0.008 | 0.002 | 0.002 | 0 | 0.22 | 0.28 | 0.16 | 0.24 | 0.17 | 0.25 |
| 9 | 127785957 | rs7857443 | 0.671 | -0.772 | 1.333 | 0.776 | 0 | 0 | 0 | 0 | 0.22 | 0.18 | 0.15 | 0.24 | 0.17 | 0.25 |
| 9 | 127792230 | rs4838242 | -0.651 | -0.188 | 1.096 | 0.491 | -0.011 | 0.008 | 0.008 | 0 | 0.42 | 0.4 | 0.54 | 0.41 | 0.47 | 0.49 |
| 9 | 127794843 | rs2052601 | 0.87 | -0.177 | 0.938 | 0.451 | -0.063 | 0.031 | 0.031 | 0 | 0.16 | 0.07 | 0.16 | 0.11 | 0.32 | 0.24 |
| 9 | 127801065 | rs394750 | -0.584 | -0.216 | 1.004 | 0.594 | -0.001 | 0 | 0 | 0 | 0.42 | 0.5 | 0.59 | 0.43 | 0.47 | 0.49 |
| 9 | 127809723 | rs608895 | 1.149 | -0.272 | 0.802 | 0.634 | 0 | -0.001 | -0.001 | 0.078 | 0.36 | 0.31 | 0.26 | 0.32 | 0.36 | 0.26 |
| 9 | 127818276 | rs589292 | 1.359 | -0.339 | 0.149 | 0.52 | -0.014 | -0.039 | -0.039 | 0 | 0.28 | 0.13 | 0.21 | 0.31 | 0.31 | 0.22 |
| 9 | 127820479 | rs603799 | -1.37 | -0.447 | 0.444 | 0.356 | 0 | 0.006 | 0.006 | 0.821 | 0.78 | 0.81 | 0.73 | 0.89 | 0.83 | 0.89 |
| 9 | 127821778 | rs662621 | 0.221 | -0.706 | 0.474 | 0.189 | 0.001 | -0.001 | -0.001 | 0 | 0.53 | 0.42 | 0.59 | 0.5 | 0.69 | 0.55 |
| 9 | 127825903 | rs11789451 | -0.338 | -0.796 | 0.51 | 0.274 | -0.014 | 0.016 | 0.016 | 0.378 | 0.4 | 0.34 | 0.33 | 0.35 | 0.51 | 0.54 |
| 9 | 127833905 | rs687654 | 1.164 | -0.777 | 0.602 | 0.304 | -0.007 | 0.023 | 0.023 | 0 | 0.33 | 0.38 | 0.26 | 0.53 | 0.32 | 0.3 |
| 9 | 127834277 | rs656322 | 1.675 | -0.524 | 0.453 | 0.127 | -0.064 | -0.057 | -0.057 | 0.486 | 0.31 | 0.07 | 0.23 | 0.31 | 0.39 | 0.24 |
| 9 | 127834878 | rs643228 | 0.521 | -0.668 | 0.555 | 0.119 | 0.001 | 0.001 | 0.001 | 0 | 0.45 | 0.38 | 0.33 | 0.54 | 0.44 | 0.43 |
| 9 | 127842701 | rs636033 | 0.171 | -0.702 | 0.718 | 0.132 | 0.012 | 0.039 | 0.039 | 0 | 0.5 | 0.59 | 0.5 | 0.65 | 0.44 | 0.47 |
| 9 | 127848785 | rs663153 | 1.964 | -0.694 | 0.715 | 0.154 | -0.024 | -0.055 | -0.055 | 0.147 | 0.28 | 0.1 | 0.21 | 0.31 | 0.31 | 0.22 |
| 9 | 127872298 | rs10760392 | 0.363 | -0.69 | 0.724 | 0.153 | 0.003 | 0.012 | 0.012 | 0 | 0.47 | 0.55 | 0.48 | 0.62 | 0.44 | 0.45 |
| 9 | 127910498 | rs1048251 | 0.499 | -0.686 | 0.495 | 0.15 | -0.001 | -0.001 | -0.001 | 0 | 0.46 | 0.47 | 0.35 | 0.54 | 0.45 | 0.43 |
| 9 | 127913837 | rs7019470 | 0.499 | -0.686 | 0.495 | 0.15 | -0.001 | -0.001 | -0.001 | 0 | 0.46 | 0.47 | 0.35 | 0.54 | 0.45 | 0.43 |
| 9 | 127942240 | rs10986614 | 1.885 | -0.694 | -0.078 | 0.076 | -0.052 | 0.04 | 0.04 | 2.345 | 0.2 | 0.01 | 0.26 | 0.1 | 0.17 | 0.12 |
| 9 | 127948321 | rs10986626 | 1.184 | -0.635 | 0.234 | 0.309 | -0.001 | -0.017 | -0.017 | 0.168 | 0.15 | 0.11 | 0.11 | 0.23 | 0.14 | 0.21 |
| 9 | 127958854 | rs17337635 | 1.602 | -0.627 | 0.186 | 0.289 | -0.011 | 0.03 | 0.03 | 0 | 0.13 | 0.08 | 0.09 | 0.01 | 0.14 | 0.13 |
| 9 | 127961771 | rs828095 | 1.799 | -0.619 | -0.156 | 0.965 | -0.074 | -0.039 | -0.039 | 0.531 | 0.27 | 0.03 | 0.19 | 0.23 | 0.31 | 0.22 |
| 9 | 127985775 | rs393721 | -1.673 | -0.565 | -0.328 | 0.686 | -0.018 | -0.049 | -0.049 | 0.805 | 0.71 | 0.89 | 0.79 | 0.69 | 0.69 | 0.78 |
| 9 | 127994103 | rs10986660 | 1.144 | -0.367 | 0.003 | 0.417 | -0.001 | -0.018 | -0.018 | 0 | 0.07 | 0.03 | 0.04 | 0.14 | 0.08 | 0.12 |
| 9 | 127996046 | rs599063 | -0.048 | -0.482 | 0.415 | 0.361 | -0.009 | 0.038 | 0.038 | 0.981 | 0.1 | 0.23 | 0.05 | 0.14 | 0.08 | 0.12 |
| 9 | 127997303 | rs12009 | 0.131 | -0.358 | 0.083 | 0.544 | 0.004 | -0.003 | -0.003 | 0.003 | 0.52 | 0.45 | 0.66 | 0.57 | 0.54 | 0.54 |
| 9 | 128004024 | rs391957 | -0.931 | -0.251 | -1.028 | 0.561 | -0.045 | -0.021 | -0.021 | 1.368 | 0.64 | 0.92 | 0.75 | 0.7 | 0.62 | 0.66 |
| 9 | 128021166 | rs662637 | -0.51 | -0.36 | -1.255 | 0.53 | -0.101 | -0.109 | -0.109 | 0.98 | 0.62 | 0.92 | 0.54 | 0.57 | 0.48 | 0.54 |
| 9 | 128024049 | rs372225 | -0.549 | -0.376 | -1.257 | 0.506 | -0.105 | -0.116 | -0.116 | 0.023 | 0.62 | 0.92 | 0.54 | 0.57 | 0.48 | 0.54 |
| 9 | 128035341 | rs359594 | -0.552 | -0.368 | -1.258 | 0.51 | -0.108 | -0.119 | -0.119 | 0.322 | 0.62 | 0.92 | 0.54 | 0.57 | 0.48 | 0.54 |
| 9 | 128068388 | rs13294021 | 0.791 | -0.811 | -0.89 | 0.607 | 0 | -0.016 | -0.016 | 0 | 0.46 | 0.52 | 0.31 | 0.23 | 0.45 | 0.35 |
| 9 | 128102865 | rs10114684 | 0.079 | -0.433 | -0.796 | 0.589 | -0.005 | -0.009 | -0.009 | 1.513 | 0.55 | 0.59 | 0.5 | 0.57 | 0.47 | 0.54 |
| 9 | 128104105 | rs828290 | -0.891 | -0.415 | -0.814 | 0.589 | -0.108 | -0.119 | -0.119 | 0 | 0.62 | 0.92 | 0.55 | 0.57 | 0.48 | 0.54 |
| 9 | 128116817 | rs937482 | -0.057 | -0.332 | -1.332 | 0.612 | -0.002 | 0.003 | 0.003 | 0 | 0.55 | 0.54 | 0.36 | 0.46 | 0.48 | 0.52 |
| 9 | 128122320 | rs2841334 | -2.968 | -0.151 | -1.479 | 0.824 | -0.065 | -0.066 | -0.066 | 0 | 0.88 | 0.98 | 0.81 | 0.76 | 0.79 | 0.85 |
| 9 | 128127412 | rs10819047 | 0.332 | -0.316 | -1.399 | 0.801 | -0.003 | -0.008 | -0.008 | 0 | 0.55 | 0.59 | 0.5 | 0.55 | 0.47 | 0.54 |
| 9 | 128133207 | rs10760397 | 0.365 | -0.443 | -1.389 | 0.776 | 0.003 | 0.001 | 0.001 | 0 | 0.55 | 0.59 | 0.49 | 0.6 | 0.45 | 0.54 |
| 9 | 128135215 | rs1463837 | 1.392 | -0.433 | -1.672 | 0.448 | -0.057 | 0.069 | 0.069 | 0.583 | 0.24 | 0.05 | 0.24 | 0.15 | 0.32 | 0.28 |
| 9 | 128137615 | rs10986727 | -0.368 | -0.09 | -0.785 | 0.737 | -0.013 | 0 | 0 | 0 | 0.68 | 0.7 | 0.57 | 0.71 | 0.58 | 0.68 |
| 9 | 128141348 | rs6478703 | 0.74 | -0.163 | -0.813 | 0.736 | 0.001 | 0.001 | 0.001 | 0 | 0.56 | 0.58 | 0.49 | 0.62 | 0.44 | 0.52 |
| 9 | 128148391 | rs10986735 | -0.168 | -0.547 | -0.767 | 0.728 | -0.05 | 0.075 | 0.075 | 0.019 | 0.1 | 0.23 | 0.18 | 0.17 | 0.04 | 0.08 |
| 9 | 128149249 | rs4838261 | 1.21 | -0.438 | -0.894 | 0.694 | 0.002 | 0.004 | 0.004 | 0 | 0.51 | 0.51 | 0.34 | 0.47 | 0.43 | 0.45 |
| 9 | 128150784 | rs2648315 | 0.344 | -0.118 | -1.162 | 0.631 | -0.065 | 0.037 | 0.037 | 0.185 | 0.27 | 0.09 | 0.28 | 0.13 | 0.32 | 0.32 |
| 9 | 128153831 | rs10986738 | -1.185 | -0.5 | -0.392 | 0.858 | -0.014 | 0.105 | 0.105 | 0.004 | 0.1 | 0.29 | 0.19 | 0.16 | 0.07 | 0.11 |
| 9 | 128156648 | rs2819625 | 0.012 | -0.516 | -1.297 | 0.569 | -0.081 | -0.029 | -0.029 | 0 | 0.58 | 0.93 | 0.52 | 0.65 | 0.46 | 0.51 |
| 9 | 128161614 | rs10986739 | 0.546 | -0.763 | -0.72 | 0.678 | -0.006 | -0.02 | -0.02 | 0 | 0.13 | 0.01 | 0.07 | 0.11 | 0.12 | 0.11 |
| 9 | 128175143 | rs12002063 | -0.233 | -0.712 | 0.169 | 0.69 | -0.03 | -0.038 | -0.038 | 0 | 0.08 | 0.34 | 0.05 | 0 | 0.03 | 0.02 |
| 9 | 128185330 | rs3122931 | -1.115 | -1.016 | 0.04 | 1.321 | -0.112 | 0.215 | 0.215 | 0 | 0.17 | 0.45 | 0.21 | 0.37 | 0.05 | 0.12 |
| 9 | 128193456 | rs1250505 | 0.062 | -0.659 | -0.653 | 0.955 | 0.025 | 0.034 | 0.034 | 0 | 0.53 | 0.71 | 0.49 | 0.4 | 0.43 | 0.39 |
| 9 | 128198422 | rs6478704 | 0.936 | -0.558 | -0.684 | 0.872 | 0.012 | -0.013 | -0.013 | 0 | 0.61 | 0.27 | 0.46 | 0.48 | 0.63 | 0.54 |
| 9 | 128207717 | rs907500 | -1.257 | -0.5 | -0.529 | 0.677 | 0.026 | -0.008 | -0.008 | 0 | 0.88 | 0.87 | 0.83 | 0.94 | 0.81 | 0.83 |
| 9 | 128208234 | rs10115550 | 0.475 | -0.774 | -0.198 | 0.61 | -0.04 | -0.046 | -0.046 | 0 | 0.08 | 0.16 | 0.01 | 0 | 0.01 | 0 |
| 9 | 128208927 | rs10760398 | -0.875 | -0.727 | -0.483 | 0.749 | -0.013 | -0.001 | -0.001 | 0 | 0.4 | 0.44 | 0.41 | 0.59 | 0.34 | 0.38 |
| 9 | 128224030 | rs7857424 | 0.308 | -0.707 | -0.279 | 0.717 | -0.043 | -0.052 | -0.052 | 0.119 | 0.1 | 0.36 | 0.04 | 0 | 0.01 | 0 |
| 9 | 128233506 | rs7018948 | 0.053 | -0.712 | -0.688 | 0.686 | 0.011 | -0.005 | -0.005 | 0 | 0.25 | 0.3 | 0.32 | 0.18 | 0.32 | 0.33 |
| 9 | 128239920 | rs7038668 | 0.075 | -0.895 | -0.702 | 0.536 | -0.005 | -0.008 | -0.008 | 0 | 0.58 | 0.69 | 0.5 | 0.57 | 0.47 | 0.49 |
| 9 | 128240922 | rs13291556 | 2.365 | -0.826 | -0.768 | 0.317 | -0.06 | 0.076 | 0.076 | 0 | 0.33 | 0.09 | 0.24 | 0.23 | 0.34 | 0.29 |
| 9 | 128268263 | rs7044663 | 0.49 | -0.817 | -0.634 | 0.444 | 0.018 | 0.031 | 0.031 | 0 | 0.53 | 0.67 | 0.47 | 0.38 | 0.43 | 0.41 |
| 9 | 128277742 | rs10986783 | 2.021 | -0.914 | -0.741 | 0.43 | 0.024 | -0.041 | -0.041 | 0.041 | 0.09 | 0.01 | 0.04 | 0.12 | 0.05 | 0.12 |
| 9 | 128282914 | rs7861903 | -0.711 | -0.907 | -0.252 | 0.224 | -0.058 | -0.099 | -0.099 | 0 | 0.14 | 0.45 | 0.07 | 0 | 0.06 | 0.07 |
| 9 | 128291664 | rs700115 | 0.2 | -0.861 | -0.181 | 0.249 | 0.006 | -0.028 | -0.028 | 0 | 0.92 | 0.66 | 0.95 | 1 | 0.94 | 0.93 |
| 9 | 128299605 | rs522894 | 1.665 | -0.602 | -0.516 | 0.233 | -0.028 | 0.026 | 0.026 | 0 | 0.09 | 0.02 | 0.11 | 0 | 0.18 | 0.13 |

| | | | | | | | | | | | | | | | | |
|---|---|---|---|---|---|---|---|---|---|---|---|---|---|---|---|---|
| 9 | 128303394 | rs17257143 | 2.76 | -1.052 | -1.119 | -0.076 | -0.022 | 0.059 | 0.059 | 3.702 | 0.18 | 0.01 | 0.08 | 0.07 | 0.14 | 0.1 |
| 9 | 128303425 | rs10114882 | 2.067 | -0.769 | -0.66 | 0.358 | -0.074 | 0.089 | 0.089 | 0 | 0.34 | 0.08 | 0.27 | 0.23 | 0.38 | 0.25 |
| 9 | 128316645 | rs829179 | 1.254 | -0.871 | -0.445 | 0.429 | -0.088 | 0.004 | 0.004 | 0.202 | 0.79 | 0.44 | 0.76 | 0.85 | 0.91 | 0.87 |
| 9 | 128329179 | rs512445 | -0.388 | -0.819 | -0.387 | 0.555 | 0.008 | -0.014 | -0.014 | 0 | 0.44 | 0.17 | 0.51 | 0.44 | 0.55 | 0.55 |
| 9 | 128341278 | rs545900 | -0.961 | -0.647 | -0.402 | 0.359 | -0.001 | -0.018 | -0.018 | 0.15 | 0.52 | 0.5 | 0.65 | 0.78 | 0.55 | 0.68 |
| 9 | 128373469 | rs7864945 | 2.385 | -0.52 | -0.778 | 0.437 | -0.094 | 0.059 | 0.059 | 0 | 0.31 | 0.05 | 0.24 | 0.22 | 0.35 | 0.25 |
| 9 | 128403509 | rs4838278 | 1.953 | -0.572 | -0.283 | 0.391 | -0.045 | 0.092 | 0.092 | 0 | 0.35 | 0.13 | 0.27 | 0.22 | 0.38 | 0.25 |
| 9 | 128406040 | rs1250733 | 0.155 | -0.628 | -0.231 | 0.303 | -0.007 | -0.044 | -0.044 | 0.494 | 0.91 | 0.67 | 0.94 | 1 | 0.94 | 0.93 |
| 9 | 128412676 | rs631287 | -0.726 | -0.702 | -0.17 | 0.71 | 0.003 | 0.001 | 0.001 | 0 | 0.48 | 0.46 | 0.51 | 0.43 | 0.54 | 0.58 |
| 9 | 128420381 | rs871391 | 0.697 | -0.41 | -0.284 | 0.765 | -0.012 | -0.009 | -0.009 | 0 | 0.23 | 0.15 | 0.33 | 0.29 | 0.29 | 0.31 |
| 9 | 128440014 | rs2416967 | 1.43 | -0.437 | -0.121 | 0.513 | -0.04 | 0.104 | 0.104 | 0 | 0.36 | 0.13 | 0.28 | 0.22 | 0.39 | 0.31 |
| 9 | 128449085 | rs7863341 | 0.264 | 0.138 | 0.129 | 0.808 | 0.02 | 0.034 | 0.034 | 0 | 0.53 | 0.69 | 0.48 | 0.37 | 0.44 | 0.36 |
| 9 | 128476464 | rs10118570 | -0.551 | 0.51 | -0.049 | 0.632 | -0.033 | 0.028 | 0.028 | 0 | 0.7 | 0.88 | 0.75 | 0.77 | 0.65 | 0.71 |
| 9 | 128488794 | rs10986857 | -0.11 | 0.678 | 0.189 | 0.935 | 0.01 | 0.06 | 0.06 | 0 | 0.2 | 0.27 | 0.32 | 0.39 | 0.21 | 0.36 |
| 9 | 128546359 | rs12341976 | 0.402 | 0.511 | 0.132 | 0.885 | 0.006 | -0.002 | -0.002 | 0 | 0.57 | 0.26 | 0.54 | 0.61 | 0.58 | 0.65 |
| 9 | 128569777 | rs10986936 | 0.424 | 0.515 | 0.134 | 0.889 | 0.007 | -0.001 | -0.001 | 0 | 0.56 | 0.25 | 0.54 | 0.61 | 0.58 | 0.65 |
| 9 | 128610269 | rs13295318 | 1.484 | 0.508 | 0.124 | 0.871 | -0.048 | 0.101 | 0.101 | 0.313 | 0.37 | 0.08 | 0.23 | 0.23 | 0.37 | 0.3 |
| 9 | 128616376 | rs11789020 | 0.323 | 0.512 | 0.39 | 0.86 | 0.003 | -0.002 | -0.002 | 0 | 0.58 | 0.32 | 0.54 | 0.61 | 0.58 | 0.66 |
| 9 | 128643415 | rs1411353 | 0.713 | 0.348 | 0.476 | 1.279 | 0.008 | 0.136 | 0.136 | 0 | 0.06 | 0.11 | 0.24 | 0.26 | 0.07 | 0.19 |
| 9 | 128664410 | rs1411351 | 1.554 | 0.256 | 0.311 | 0.918 | -0.079 | 0.151 | 0.151 | 0.6 | 0.4 | 0.08 | 0.26 | 0.21 | 0.42 | 0.32 |
| 9 | 128680148 | rs2026134 | -0.244 | 0.864 | 1.031 | 1.294 | 0.01 | -0.028 | -0.028 | 0 | 0.71 | 0.56 | 0.74 | 0.76 | 0.69 | 0.74 |
| 9 | 128680409 | rs2026133 | -0.098 | 0.889 | 0.688 | 1.349 | 0 | 0.013 | 0.013 | 0 | 0.18 | 0.13 | 0.3 | 0.39 | 0.21 | 0.36 |
| 9 | 128695180 | rs10819087 | -0.572 | 0.86 | 0.742 | 1.048 | -0.025 | 0.011 | 0.011 | 0 | 0.76 | 0.85 | 0.77 | 0.76 | 0.7 | 0.74 |
| 9 | 128719849 | rs10513456 | -0.055 | 0.94 | 0.486 | 1.092 | 0.002 | -0.02 | -0.02 | 0 | 0.69 | 0.38 | 0.73 | 0.76 | 0.69 | 0.74 |
| 9 | 128728459 | rs11792585 | -0.51 | 1.363 | 0.297 | 1.478 | -0.001 | 0.012 | 0.012 | 0 | 0.18 | 0.13 | 0.3 | 0.48 | 0.21 | 0.36 |
| 9 | 128740896 | rs10987069 | -0.922 | 1.128 | 0.867 | 1.774 | 0.032 | 0.293 | 0.293 | 0.361 | 0.09 | 0.37 | 0.25 | 0.35 | 0.08 | 0.19 |
| 9 | 128746044 | rs7868608 | -0.014 | 1.502 | 0.364 | 0.974 | 0.006 | -0.003 | -0.003 | 0.333 | 0.77 | 0.76 | 0.77 | 0.75 | 0.71 | 0.74 |
| 9 | 128754234 | rs2149990 | -0.307 | 0.583 | 0.173 | 1.098 | 0.037 | -0.004 | -0.004 | 0 | 0.41 | 0.71 | 0.58 | 0.52 | 0.32 | 0.3 |
| 9 | 128755979 | rs4240495 | -0.051 | 0.592 | 0.949 | 1.286 | -0.036 | 0.025 | 0.025 | 0 | 0.37 | 0.46 | 0.38 | 0.46 | 0.24 | 0.27 |
| 9 | 128756484 | rs4837027 | -0.051 | 0.521 | 0.847 | 1.266 | -0.036 | 0.025 | 0.025 | 0 | 0.37 | 0.46 | 0.38 | 0.46 | 0.24 | 0.27 |
| 9 | 128765264 | rs10987095 | 0.333 | 0.84 | 0.771 | 1.394 | 0.008 | 0.015 | 0.015 | 0 | 0.35 | 0.21 | 0.28 | 0.31 | 0.34 | 0.29 |
| 9 | 128765362 | rs7874953 | -0.433 | 0.83 | 0.348 | 1.463 | 0.009 | -0.018 | -0.018 | 0 | 0.45 | 0.8 | 0.62 | 0.51 | 0.43 | 0.34 |
| 9 | 128768663 | rs10819101 | 0.028 | 0.923 | 0.659 | 0.999 | 0.002 | 0.006 | 0.006 | 0 | 0.4 | 0.57 | 0.46 | 0.37 | 0.42 | 0.33 |
| 9 | 128773388 | rs13291454 | 0.186 | 0.918 | 0.4 | 0.818 | -0.004 | -0.007 | -0.007 | 0 | 0.43 | 0.4 | 0.35 | 0.31 | 0.4 | 0.32 |
| 9 | 128780293 | rs13290614 | 0.941 | 1.511 | -0.193 | 0.576 | 0.02 | -0.029 | -0.029 | 0 | 0.07 | 0.08 | 0.04 | 0 | 0.09 | 0.08 |
| 9 | 128780313 | rs13290624 | 0.941 | 1.511 | -0.193 | 0.576 | 0.02 | -0.029 | -0.029 | 0 | 0.07 | 0.08 | 0.04 | 0 | 0.09 | 0.08 |
| 9 | 128783188 | rs10987107 | -0.019 | 1.068 | -0.413 | 0.711 | -0.039 | -0.017 | -0.017 | 0 | 0.44 | 0.2 | 0.45 | 0.62 | 0.46 | 0.58 |
| 9 | 128788342 | rs4838311 | 0.157 | 0.99 | -0.805 | 0.887 | -0.023 | -0.037 | -0.037 | 0 | 0.62 | 0.73 | 0.66 | 0.58 | 0.5 | 0.61 |
| 9 | 128788957 | rs9299306 | 0.13 | 1.485 | -0.458 | 0.769 | 0.005 | 0.021 | 0.021 | 0 | 0.7 | 0.64 | 0.76 | 0.58 | 0.68 | 0.64 |
| 9 | 128791490 | rs10819102 | 1.099 | 0.575 | -0.713 | 0.772 | 0.023 | -0.004 | -0.004 | 0.011 | 0.32 | 0.21 | 0.21 | 0.28 | 0.26 | 0.33 |
| 9 | 128791503 | rs10819103 | 1.669 | 0.493 | -0.816 | 0.689 | -0.008 | -0.034 | -0.034 | 0 | 0.23 | 0.01 | 0.16 | 0.19 | 0.21 | 0.23 |
| 9 | 128792334 | rs7029545 | -0.114 | 1.037 | -0.001 | 1.103 | -0.004 | 0.007 | 0.007 | 0 | 0.32 | 0.3 | 0.52 | 0.24 | 0.37 | 0.25 |
| 9 | 128792958 | rs4838313 | 1.097 | 0.77 | -0.089 | 0.929 | 0.029 | 0.027 | 0.027 | 0 | 0.4 | 0.26 | 0.24 | 0.25 | 0.32 | 0.42 |
| 9 | 128792979 | rs10987111 | -0.811 | 1.071 | -0.146 | 1.002 | -0.012 | 0.022 | 0.022 | 0.861 | 0.45 | 0.31 | 0.45 | 0.39 | 0.46 | 0.43 |
| 9 | 128797551 | rs2416993 | 1.12 | 1.223 | -0.208 | 0.975 | -0.005 | 0 | 0 | 0 | 0.13 | 0.1 | 0.14 | 0.1 | 0.16 | 0.17 |
| 9 | 128798584 | rs2019199 | 0.603 | 1.144 | -0.175 | 1.069 | 0.041 | 0.068 | 0.068 | 0 | 0.71 | 0.45 | 0.43 | 0.48 | 0.62 | 0.69 |
| 9 | 128800424 | rs1558836 | -0.586 | 0.725 | -0.465 | 0.801 | -0.008 | 0.021 | 0.021 | 1.157 | 0.37 | 0.2 | 0.26 | 0.34 | 0.36 | 0.35 |
| 9 | 128805768 | rs4837031 | -0.022 | 0.669 | -0.456 | 0.951 | -0.008 | -0.046 | -0.046 | 0 | 0.29 | 0.13 | 0.37 | 0.41 | 0.3 | 0.26 |
| 9 | 128806385 | rs10819104 | -0.022 | 0.669 | -0.428 | 0.922 | -0.005 | -0.02 | -0.02 | 0 | 0.29 | 0.15 | 0.35 | 0.36 | 0.3 | 0.25 |
| 9 | 128807593 | rs2098440 | 0.232 | 0.859 | -0.257 | 1.025 | 0.045 | 0.146 | 0.146 | 0.747 | 0.86 | 0.66 | 0.78 | 0.6 | 0.83 | 0.83 |
| 9 | 128808825 | rs16928613 | 0.521 | 0.715 | -0.581 | 0.943 | -0.043 | -0.072 | -0.072 | 0 | 0.25 | 0.03 | 0.19 | 0.34 | 0.27 | 0.23 |
| 9 | 128810469 | rs7861741 | 0.787 | 0.697 | -0.387 | 0.554 | 0.008 | 0.015 | 0.015 | 0 | 0.49 | 0.52 | 0.37 | 0.24 | 0.42 | 0.5 |
| 9 | 128810846 | rs10217721 | 0.897 | 0.751 | -0.308 | 0.429 | -0.003 | -0.013 | -0.013 | 0 | 0.48 | 0.46 | 0.34 | 0.19 | 0.43 | 0.48 |
| 9 | 128812194 | rs10987119 | -0.916 | 1.151 | 0.274 | 1.038 | 0.044 | 0.151 | 0.151 | 0.544 | 0.19 | 0.41 | 0.38 | 0.41 | 0.21 | 0.22 |
| 9 | 128816587 | rs929673 | -0.184 | 0.787 | 0.071 | 0.85 | 0.042 | 0.059 | 0.059 | 0 | 0.78 | 0.63 | 0.6 | 0.57 | 0.71 | 0.74 |
| 9 | 128817516 | rs888236 | -0.333 | 0.93 | -0.588 | 0.757 | -0.002 | -0.002 | -0.002 | 0.342 | 0.29 | 0.26 | 0.22 | 0.38 | 0.31 | 0.23 |
| 9 | 128822026 | rs4838318 | 0.178 | 0.619 | -0.464 | 0.559 | 0 | -0.009 | -0.009 | 0 | 0.62 | 0.44 | 0.66 | 0.63 | 0.64 | 0.69 |
| 9 | 128826791 | rs888231 | -0.005 | 0.572 | -0.978 | 0.143 | 0.008 | 0.072 | 0.072 | 0 | 0.55 | 0.66 | 0.65 | 0.8 | 0.61 | 0.55 |
| 9 | 128827305 | rs7020350 | 0.675 | 0.345 | -0.883 | 0.54 | -0.008 | 0.018 | 0.018 | 0 | 0.76 | 0.6 | 0.81 | 0.73 | 0.8 | 0.73 |
| 9 | 128828716 | rs10987122 | 0.415 | 0.108 | -0.654 | 0.667 | 0.001 | 0 | 0 | 0 | 0.47 | 0.39 | 0.41 | 0.4 | 0.57 | 0.55 |
| 9 | 128831097 | rs4838320 | -0.652 | 0.069 | -0.451 | 0.669 | -0.005 | 0.004 | 0.004 | 0 | 0.89 | 0.8 | 0.96 | 0.9 | 0.92 | 0.86 |
| 9 | 128834294 | rs10987129 | 0.36 | 0.126 | -0.57 | 0.785 | 0.01 | 0.029 | 0.029 | 0 | 0.15 | 0.24 | 0.37 | 0.22 | 0.17 | 0.14 |
| 9 | 128835802 | rs999566 | 0.5 | -0.182 | -0.469 | 0.568 | -0.003 | 0.003 | 0.003 | 0 | 0.33 | 0.44 | 0.33 | 0.4 | 0.28 | 0.3 |
| 9 | 128836559 | rs4838322 | -0.114 | 0.145 | -0.849 | 0.814 | 0.043 | -0.005 | -0.005 | 0.009 | 0.51 | 0.27 | 0.63 | 0.5 | 0.63 | 0.56 |
| 9 | 128842857 | rs4837036 | -0.876 | 0.031 | -0.741 | 0.747 | 0.028 | 0.055 | 0.055 | 0 | 0.72 | 0.83 | 0.88 | 0.89 | 0.78 | 0.74 |
| 9 | 128843485 | rs10819112 | -0.33 | 0.869 | -0.229 | 1.32 | 0.05 | 0.057 | 0.057 | 0 | 0.21 | 0.3 | 0.41 | 0.39 | 0.32 | 0.24 |
| 9 | 128844657 | rs13295982 | 0.905 | 0.128 | -0.784 | 1.165 | -0.052 | 0.019 | 0.019 | 0 | 0.33 | 0.14 | 0.37 | 0.26 | 0.39 | 0.28 |
| 9 | 128844997 | rs7468639 | 0.834 | 0.093 | -0.411 | 1.45 | -0.022 | -0.003 | -0.003 | 0 | 0.34 | 0.21 | 0.39 | 0.35 | 0.39 | 0.33 |
| 9 | 128845089 | rs16928653 | 1.058 | -0.408 | -0.495 | 1.582 | 0.021 | -0.002 | -0.002 | 0 | 0.27 | 0.17 | 0.17 | 0.27 | 0.22 | 0.28 |

| | | | | | | | | | | | | | | | | |
|---|---|---|---|---|---|---|---|---|---|---|---|---|---|---|---|---|
| 9 | 128845092 | rs10819113 | 0.622 | -0.55 | -0.597 | 1.668 | -0.008 | 0.003 | 0.003 | 0 | 0.11 | 0.15 | 0.03 | 0.19 | 0.05 | 0.07 |
| 9 | 128846395 | rs17269441 | 1.355 | -0.519 | -0.624 | 1.652 | -0.052 | 0.019 | 0.019 | 0 | 0.33 | 0.14 | 0.37 | 0.26 | 0.39 | 0.28 |
| 9 | 128847015 | rs10987141 | 0.156 | -0.569 | -0.636 | 1.766 | 0.05 | 0.026 | 0.026 | 0 | 0.36 | 0.23 | 0.16 | 0.26 | 0.26 | 0.34 |
| 9 | 128847170 | rs13289050 | 1.065 | -0.532 | -0.909 | 1.253 | -0.043 | 0.027 | 0.027 | 0 | 0.35 | 0.17 | 0.38 | 0.26 | 0.39 | 0.28 |
| 9 | 128849488 | rs1420771 | -0.078 | -0.494 | -0.804 | 1.034 | -0.032 | 0.06 | 0.06 | 0 | 0.65 | 0.89 | 0.6 | 0.83 | 0.62 | 0.6 |
| 9 | 128858530 | rs10819120 | -0.989 | -0.665 | -0.529 | 1.299 | -0.066 | 0.138 | 0.138 | 0.516 | 0.15 | 0.39 | 0.09 | 0.43 | 0.06 | 0.12 |
| 9 | 128858795 | rs12552226 | 0.581 | -0.321 | -1.232 | 0.593 | 0.023 | 0.067 | 0.067 | 0 | 0.27 | 0.1 | 0.14 | 0.07 | 0.21 | 0.27 |
| 9 | 128866262 | rs2041944 | -0.789 | 0.131 | -1.096 | 0.58 | -0.014 | 0.045 | 0.045 | 0 | 0.54 | 0.67 | 0.59 | 0.3 | 0.51 | 0.56 |
| 9 | 128866390 | rs2111362 | 0.076 | -0.575 | -1.244 | 0.812 | 0.037 | 0.103 | 0.103 | 0 | 0.31 | 0.11 | 0.15 | 0.12 | 0.23 | 0.3 |
| 9 | 128869210 | rs10819122 | 0.669 | 0.038 | -0.454 | 1.717 | -0.002 | 0.098 | 0.098 | 0 | 0.91 | 0.87 | 0.96 | 0.56 | 0.91 | 0.84 |
| 9 | 128875526 | rs16928677 | -0.248 | -0.119 | 0.088 | 0.917 | -0.009 | -0.019 | -0.019 | 0 | 0.09 | 0.2 | 0.17 | 0.06 | 0.07 | 0.1 |
| 9 | 128876516 | rs6478724 | -1.217 | 0.312 | -0.21 | 1.487 | 0.007 | 0.043 | 0.043 | 0 | 0.22 | 0.25 | 0.27 | 0.35 | 0.26 | 0.31 |
| 9 | 128877531 | rs10987154 | 0.476 | -0.405 | -1.135 | 0.966 | 0.012 | 0.043 | 0.043 | 0 | 0.21 | 0.03 | 0.11 | 0.11 | 0.14 | 0.22 |
| 9 | 128879579 | rs2417001 | 0.561 | -0.751 | -1.137 | 0.72 | 0 | 0 | 0 | 0 | 0.19 | 0.22 | 0.09 | 0.24 | 0.12 | 0.17 |
| 9 | 128879934 | rs4595229 | 0.894 | -0.7 | -1.207 | 0.783 | -0.01 | -0.012 | -0.012 | 0 | 0.09 | 0.2 | 0.17 | 0.06 | 0.07 | 0.11 |
| 9 | 128880042 | rs888234 | 1.016 | -0.966 | -1.803 | 0.583 | -0.012 | -0.027 | -0.027 | 0 | 0.52 | 0.11 | 0.6 | 0.39 | 0.61 | 0.49 |
| 9 | 128880225 | rs4838340 | 0.111 | -0.732 | -1.088 | 0.738 | -0.01 | -0.008 | -0.008 | 0.202 | 0.55 | 0.48 | 0.76 | 0.65 | 0.72 | 0.6 |
| 9 | 128880277 | rs7871394 | 0.388 | -0.289 | -1.026 | 0.983 | -0.028 | -0.046 | -0.046 | 0 | 0.13 | 0.05 | 0.1 | 0.28 | 0.19 | 0.2 |
| 9 | 128883849 | rs7858074 | 1.375 | 0.128 | -1.108 | 0.721 | -0.072 | -0.053 | -0.053 | 0 | 0.26 | 0.08 | 0.42 | 0.29 | 0.34 | 0.17 |
| 9 | 128885548 | rs1861716 | 0.309 | 0.35 | -0.43 | 0.637 | -0.008 | -0.004 | -0.004 | 0 | 0.76 | 0.56 | 0.87 | 0.76 | 0.84 | 0.79 |
| 9 | 128893045 | rs7046411 | -0.895 | 0.187 | -0.229 | 0.934 | -0.002 | -0.002 | -0.002 | 0 | 0.4 | 0.61 | 0.44 | 0.57 | 0.46 | 0.48 |
| 9 | 128893372 | rs10217645 | 0.284 | 0.246 | -0.367 | 1.199 | -0.016 | -0.021 | -0.021 | 1.692 | 0.11 | 0.06 | 0.18 | 0.16 | 0.12 | 0.14 |
| 9 | 128900528 | rs1156715 | -1.001 | -0.246 | -0.595 | 0.931 | -0.001 | -0.007 | -0.007 | 0 | 0.87 | 0.9 | 0.92 | 0.85 | 0.9 | 0.85 |
| 9 | 128905129 | rs10760416 | -0.524 | -0.335 | -0.074 | 1.058 | -0.018 | -0.024 | -0.024 | 0 | 0.3 | 0.23 | 0.39 | 0.48 | 0.43 | 0.47 |
| 9 | 128910419 | rs1055352 | 1.635 | -0.562 | -0.262 | 1.019 | -0.027 | 0.061 | 0.061 | 0 | 0.44 | 0.12 | 0.46 | 0.25 | 0.44 | 0.38 |
| 9 | 128911346 | rs882061 | -0.397 | -0.478 | 0.274 | 1.193 | 0.024 | 0.068 | 0.068 | 0 | 0.27 | 0.57 | 0.25 | 0.42 | 0.32 | 0.32 |
| 9 | 128920470 | rs4838353 | 0.34 | -0.388 | 0.064 | 1.209 | -0.082 | -0.095 | -0.095 | 0 | 0.2 | 0.05 | 0.18 | 0.34 | 0.33 | 0.27 |
| 9 | 128921921 | rs10125891 | -0.591 | -0.718 | 0.234 | 0.918 | -0.005 | -0.009 | -0.009 | 0.009 | 0.31 | 0.35 | 0.3 | 0.32 | 0.24 | 0.3 |
| 9 | 128922682 | rs4837043 | -1.524 | -0.456 | -0.589 | 1.02 | 0.011 | 0.025 | 0.025 | 0 | 0.55 | 0.64 | 0.55 | 0.67 | 0.59 | 0.62 |
| 9 | 128923809 | rs7019762 | 1.701 | -0.468 | -1.007 | 0.977 | -0.038 | -0.047 | -0.047 | 0 | 0.38 | 0.12 | 0.45 | 0.36 | 0.37 | 0.35 |
| 9 | 128926091 | rs888225 | 0.362 | -0.296 | -0.745 | 0.906 | 0.001 | 0.018 | 0.018 | 0 | 0.09 | 0.12 | 0.19 | 0.18 | 0.1 | 0.12 |
| 9 | 128926239 | rs888224 | 0.127 | -0.256 | -0.949 | 0.512 | -0.039 | -0.018 | -0.018 | 3.16 | 0.24 | 0.14 | 0.19 | 0.24 | 0.35 | 0.3 |
| 9 | 128926989 | rs7018610 | -0.672 | -0.201 | -0.831 | 0.615 | 0.006 | 0.046 | 0.046 | 0 | 0.49 | 0.69 | 0.34 | 0.39 | 0.51 | 0.52 |
| 9 | 128928087 | rs10987177 | -0.363 | 0.06 | -0.79 | 0.032 | 0.023 | 0.022 | 0.022 | 0.302 | 0.5 | 0.48 | 0.5 | 0.4 | 0.59 | 0.54 |
| 9 | 128928438 | rs13290757 | 1.226 | -0.027 | -0.95 | -0.393 | -0.039 | 0.061 | 0.061 | 1.656 | 0.1 | 0.01 | 0.19 | 0 | 0.14 | 0.15 |
| 9 | 128937675 | rs1536960 | -0.186 | 0.037 | -0.493 | -0.56 | -0.002 | -0.007 | -0.007 | 6.894 | 0.67 | 0.63 | 0.73 | 0.72 | 0.69 | 0.64 |
| 9 | 128947475 | rs2809443 | -0.127 | -0.479 | -0.976 | -0.756 | -0.008 | 0.019 | 0.019 | 0 | 0.47 | 0.65 | 0.53 | 0.63 | 0.48 | 0.47 |
| 9 | 128952700 | rs11790107 | 1.17 | -0.696 | -1.581 | -0.732 | -0.023 | -0.073 | -0.073 | 7.048 | 0.29 | 0.07 | 0.22 | 0.37 | 0.3 | 0.37 |
| 9 | 128953978 | rs2773379 | 0.399 | -0.52 | -1.405 | -0.405 | -0.061 | -0.056 | -0.056 | 0 | 0.24 | 0.05 | 0.19 | 0.28 | 0.28 | 0.34 |
| 9 | 128954507 | rs7039848 | 0.593 | -0.171 | -0.283 | -0.304 | 0.001 | 0.007 | 0.007 | 0 | 0.22 | 0.23 | 0.3 | 0.34 | 0.23 | 0.15 |
| 9 | 128965556 | rs7034019 | -0.104 | -0.217 | 0.484 | -0.401 | -0.052 | -0.085 | -0.085 | 0 | 0.12 | 0.51 | 0.09 | 0 | 0.05 | 0.01 |
| 9 | 128966109 | rs10819136 | 0.526 | -0.505 | -0.037 | -0.462 | 0.008 | -0.006 | -0.006 | 0 | 0.36 | 0.22 | 0.24 | 0.38 | 0.32 | 0.45 |
| 9 | 128968531 | rs10987198 | 0.604 | -0.493 | -0.085 | -0.511 | 0.008 | -0.012 | -0.012 | 0 | 0.33 | 0.19 | 0.22 | 0.38 | 0.29 | 0.44 |
| 9 | 128968987 | rs10987199 | 0.459 | -0.479 | 0.089 | -0.499 | 0.006 | 0.007 | 0.007 | 0 | 0.13 | 0.13 | 0.09 | 0.11 | 0.09 | 0.06 |
| 9 | 128970219 | rs2417008 | -0.445 | -0.5 | 0.254 | -0.51 | -0.063 | -0.072 | -0.072 | 0 | 0.18 | 0.51 | 0.13 | 0.11 | 0.09 | 0.06 |
| 9 | 128970806 | rs10987202 | 0.955 | -0.496 | 0.237 | -0.508 | -0.003 | -0.001 | -0.001 | 0 | 0.07 | 0.11 | 0.03 | 0.11 | 0.04 | 0.05 |
| 9 | 128972210 | rs7021461 | -0.301 | -0.506 | 0.476 | -0.513 | -0.055 | -0.064 | -0.064 | 0 | 0.17 | 0.49 | 0.12 | 0.11 | 0.09 | 0.06 |
| 9 | 128975311 | rs2094763 | 0.102 | -0.474 | 0.591 | -0.497 | -0.026 | -0.03 | -0.03 | 0 | 0.15 | 0.22 | 0.09 | 0.11 | 0.09 | 0.06 |
| 9 | 128976978 | rs10987204 | 0.767 | -0.208 | 0.029 | -0.321 | -0.004 | -0.001 | -0.001 | 0.965 | 0.08 | 0.09 | 0.03 | 0.11 | 0.04 | 0.05 |
| 9 | 128978468 | rs2065183 | -0.891 | 0.262 | -0.323 | -0.26 | -0.009 | 0.011 | 0.011 | 0 | 0.65 | 0.71 | 0.51 | 0.67 | 0.62 | 0.69 |
| 9 | 128982650 | rs2150010 | 0.842 | 0.49 | -0.468 | 0.163 | -0.053 | -0.055 | -0.055 | 0.913 | 0.11 | 0.03 | 0.15 | 0.19 | 0.21 | 0.2 |
| 9 | 128983878 | rs2809423 | -1.061 | 0.539 | -0.533 | 0.062 | -0.006 | -0.019 | -0.019 | 0 | 0.76 | 0.81 | 0.66 | 0.67 | 0.76 | 0.86 |
| 9 | 128986041 | rs2809426 | 0.341 | 0.552 | -0.523 | 0.068 | 0.018 | 0.023 | 0.023 | 0 | 0.14 | 0.25 | 0.17 | 0.18 | 0.2 | 0.2 |
| 9 | 128987079 | rs1106477 | -0.42 | 0.333 | -0.632 | -0.156 | -0.009 | -0.012 | -0.012 | 0 | 0.62 | 0.59 | 0.49 | 0.48 | 0.56 | 0.66 |
| 9 | 128987905 | rs10987215 | 0.57 | 0.674 | -1.101 | 0.178 | -0.039 | -0.039 | -0.039 | 0.099 | 0.09 | 0.01 | 0.07 | 0.16 | 0.11 | 0.12 |
| 9 | 128990287 | rs12005670 | -0.285 | 0.352 | -0.713 | -0.042 | 0.002 | 0.013 | 0.013 | 0.002 | 0.51 | 0.37 | 0.42 | 0.38 | 0.51 | 0.58 |
| 9 | 128990760 | rs2773390 | -0.257 | 0.409 | -0.672 | -0.161 | -0.003 | -0.011 | -0.011 | 0 | 0.18 | 0.16 | 0.16 | 0.21 | 0.23 | 0.25 |
| 9 | 128991622 | rs16928802 | 0.295 | 0.231 | -0.528 | -0.658 | 0.008 | 0.041 | 0.041 | 0 | 0.08 | 0.03 | 0.06 | 0 | 0.07 | 0.03 |
| 9 | 128993470 | rs1537010 | 0.44 | 0.124 | 0.104 | -0.702 | 0.005 | -0.015 | -0.015 | 0.064 | 0.95 | 0.77 | 0.97 | 1 | 0.98 | 0.99 |
| 9 | 128993952 | rs16928804 | 0.236 | 0.37 | -0.121 | -0.9 | 0.007 | -0.042 | -0.042 | 0 | 0.1 | 0.15 | 0.14 | 0 | 0.12 | 0.12 |
| 9 | 128994668 | rs2039200 | -0.792 | 0.244 | 0.184 | -0.854 | 0.002 | -0.128 | -0.128 | 1.494 | 0.17 | 0.46 | 0.11 | 0.01 | 0.13 | 0.04 |
| 9 | 129001029 | rs2809428 | -0.145 | 0.262 | -1.276 | -0.703 | -0.045 | -0.015 | -0.015 | 1.989 | 0.32 | 0.1 | 0.25 | 0.31 | 0.35 | 0.46 |
| 9 | 129001487 | rs17272567 | 0.713 | -0.022 | -1.39 | -0.845 | -0.001 | 0.014 | 0.014 | 0 | 0.09 | 0.06 | 0.03 | 0.05 | 0.09 | 0.07 |
| 9 | 129018625 | rs10987225 | 0.195 | -0.444 | -1.003 | -1.012 | -0.031 | -0.013 | -0.013 | 0 | 0.1 | 0.16 | 0.04 | 0.1 | 0.04 | 0.11 |
| 9 | 129019248 | rs10760423 | -1.185 | -0.787 | -1.04 | -1.597 | 0.054 | 0.111 | 0.111 | 0.147 | 0.36 | 0.79 | 0.23 | 0.13 | 0.26 | 0.3 |
| 9 | 129029139 | rs2491650 | -1.051 | -0.62 | -0.974 | -1.448 | -0.008 | -0.019 | -0.019 | 0 | 0.63 | 0.72 | 0.56 | 0.42 | 0.58 | 0.7 |
| 9 | 129030818 | rs16928843 | 0.2 | -0.661 | -0.849 | -1.466 | -0.003 | -0.101 | -0.101 | 0.452 | 0.15 | 0.47 | 0.16 | 0.01 | 0.13 | 0.11 |
| 9 | 129031092 | rs2773371 | 0.299 | -0.655 | -1.431 | -1.391 | -0.11 | -0.022 | -0.022 | 0 | 0.38 | 0.07 | 0.36 | 0.32 | 0.4 | 0.47 |
| 9 | 129042345 | rs12686700 | 0.896 | -0.257 | -1.035 | -1.174 | 0.005 | -0.003 | -0.003 | 0.58 | 0.11 | 0.07 | 0.13 | 0.12 | 0.07 | 0.13 |

| | | | | | | | | | | | | | | | | |
|---|---|---|---|---|---|---|---|---|---|---|---|---|---|---|---|---|
| 9 | 129042699 | rs951214 | 1.544 | -0.387 | -1.143 | -1.101 | -0.039 | -0.128 | -0.128 | 0.404 | 0.27 | 0.07 | 0.36 | 0.51 | 0.3 | 0.22 |
| 9 | 129046512 | rs4836535 | 0.947 | -0.464 | -0.948 | -1.024 | 0.009 | -0.003 | -0.003 | 0.35 | 0.1 | 0.07 | 0.13 | 0.11 | 0.05 | 0.11 |
| 9 | 129046730 | rs1556218 | 0.646 | -0.43 | -0.357 | -1.096 | -0.007 | -0.001 | -0.001 | 0 | 0.39 | 0.3 | 0.47 | 0.6 | 0.45 | 0.31 |
| 9 | 129049692 | rs2026808 | -0.604 | -0.503 | -0.161 | -0.975 | -0.01 | 0.03 | 0.03 | 0 | 0.63 | 0.76 | 0.74 | 0.73 | 0.64 | 0.52 |
| 9 | 129053531 | rs1536959 | -0.458 | -0.496 | 0.492 | -0.682 | -0.034 | -0.019 | -0.019 | 0 | 0.15 | 0.43 | 0.15 | 0.12 | 0.05 | 0.11 |
| 9 | 129056597 | rs1317309 | 0.551 | -0.143 | -0.119 | -1.052 | -0.019 | -0.098 | -0.098 | 3.27 | 0.68 | 0.56 | 0.81 | 0.92 | 0.75 | 0.8 |
| 9 | 129057800 | rs4837051 | 0.098 | 0.901 | -0.533 | -0.844 | 0.001 | -0.001 | -0.001 | 0 | 0.21 | 0.23 | 0.23 | 0.18 | 0.27 | 0.23 |
| 9 | 129058255 | rs10987235 | 0.466 | 0.354 | -1.099 | -0.496 | 0.013 | 0.014 | 0.014 | 0 | 0.25 | 0.13 | 0.15 | 0.21 | 0.18 | 0.35 |
| 9 | 129058924 | rs12004434 | 0.739 | 0.358 | -0.857 | 0.687 | 0.013 | -0.068 | -0.068 | 0 | 0.26 | 0.13 | 0.36 | 0.37 | 0.25 | 0.18 |
| 9 | 129059870 | rs7466244 | -1.256 | 0.604 | 0.179 | 0.742 | -0.014 | 0.043 | 0.043 | 0.266 | 0.12 | 0.34 | 0.1 | 0.2 | 0.14 | 0.16 |
| 9 | 129060075 | rs944212 | -0.52 | 1.204 | -0.036 | 1.105 | 0.004 | 0.044 | 0.044 | 0 | 0.24 | 0.3 | 0.36 | 0.54 | 0.3 | 0.3 |
| 9 | 129061512 | rs539215 | -0.77 | 2.038 | -0.068 | 0.627 | 0.105 | 0.025 | 0.025 | 0 | 0.12 | 0.26 | 0.2 | 0.14 | 0.22 | 0.09 |
| 9 | 129061773 | rs10116388 | -0.215 | 1.332 | 0.009 | 1.087 | 0.033 | 0.227 | 0.227 | 0 | 0.16 | 0.37 | 0.31 | 0.47 | 0.2 | 0.15 |
| 9 | 129074984 | rs504184 | -0.363 | 1.177 | -1.346 | 0.269 | 0.005 | 0.048 | 0.048 | 0.342 | 0.34 | 0.2 | 0.26 | 0.24 | 0.32 | 0.31 |
| 9 | 129075914 | rs561769 | -0.184 | 1.177 | -1.346 | 0.201 | 0.023 | 0.076 | 0.076 | 0.975 | 0.32 | 0.2 | 0.2 | 0.11 | 0.25 | 0.26 |
| 9 | 129079854 | rs10987246 | -0.476 | 1.669 | -1.191 | 1.216 | -0.02 | -0.041 | -0.041 | 0.375 | 0.17 | 0.11 | 0.33 | 0.48 | 0.23 | 0.14 |
| 9 | 129087114 | rs562800 | 0.116 | 1.79 | -1.008 | 1.283 | -0.015 | -0.005 | -0.005 | 0 | 0.88 | 0.92 | 0.69 | 0.56 | 0.8 | 0.88 |
| 9 | 129088202 | rs10739669 | 0.537 | 1.49 | -1.204 | 0.347 | -0.013 | 0.007 | 0.007 | 0.13 | 0.59 | 0.7 | 0.49 | 0.38 | 0.56 | 0.65 |
| 9 | 129089985 | rs568240 | 0.369 | 1.4 | -1.186 | 0.335 | 0.003 | 0.037 | 0.037 | 2.525 | 0.28 | 0.17 | 0.15 | 0.11 | 0.23 | 0.23 |
| 9 | 129096474 | rs10819147 | 0.687 | 0.962 | -0.863 | -0.318 | 0.003 | 0.003 | 0.003 | 0.014 | 0.24 | 0.29 | 0.29 | 0.29 | 0.27 | 0.28 |
| 9 | 129102381 | rs554317 | 1.128 | 0.916 | -1.161 | -0.626 | 0.007 | 0.039 | 0.039 | 1.919 | 0.27 | 0.16 | 0.13 | 0.1 | 0.21 | 0.24 |
| 9 | 129102840 | rs1888156 | 0.503 | 0.703 | -1.001 | -0.773 | -0.005 | -0.008 | -0.008 | 0.692 | 0.29 | 0.39 | 0.3 | 0.29 | 0.29 | 0.28 |
| 9 | 129105723 | rs12554306 | 1.034 | 0.627 | -1.065 | -0.964 | -0.001 | -0.008 | -0.008 | 0.213 | 0.24 | 0.26 | 0.14 | 0.1 | 0.21 | 0.37 |
| 9 | 129115381 | rs568565 | -1.272 | 0.206 | -1.009 | -0.666 | -0.012 | 0.004 | 0.004 | 0 | 0.88 | 0.92 | 0.7 | 0.56 | 0.81 | 0.91 |
| 9 | 129115717 | rs1360289 | -0.728 | -0.022 | -1.147 | -0.897 | -0.003 | -0.015 | -0.015 | 0 | 0.69 | 0.76 | 0.58 | 0.38 | 0.69 | 0.74 |
| 9 | 129117662 | rs505442 | 0.203 | 0.15 | -1.259 | -1.121 | -0.004 | -0.019 | -0.019 | 0 | 0.56 | 0.5 | 0.43 | 0.27 | 0.53 | 0.66 |
| 9 | 129120430 | rs10987253 | 0.506 | 0.051 | -1.297 | -1.184 | 0.01 | -0.041 | -0.041 | 0.114 | 0.35 | 0.39 | 0.31 | 0.09 | 0.41 | 0.49 |
| 9 | 129121098 | rs548403 | 0.908 | 0.079 | -1.308 | -1.086 | 0.004 | 0.016 | 0.016 | 1.722 | 0.28 | 0.2 | 0.2 | 0.18 | 0.23 | 0.24 |
| 9 | 129139361 | rs10987261 | 1.419 | -0.007 | -1.28 | -1.149 | 0.003 | 0.004 | 0.004 | 0 | 0.07 | 0.05 | 0.04 | 0 | 0.07 | 0.02 |
| 9 | 129141450 | rs10124379 | -0.31 | 0.072 | -1.287 | -1.118 | -0.015 | 0.008 | 0.008 | 0 | 0.6 | 0.72 | 0.5 | 0.39 | 0.57 | 0.66 |
| 9 | 129143435 | rs2286889 | 2.355 | -0.185 | -1.616 | -1.255 | -0.013 | -0.069 | -0.069 | 0 | 0.11 | 0.01 | 0.14 | 0.26 | 0.11 | 0.17 |
| 9 | 129146344 | rs917776 | -0.369 | -0.198 | -1.385 | -1.252 | -0.001 | 0.001 | 0.001 | 0.537 | 0.58 | 0.54 | 0.39 | 0.39 | 0.47 | 0.49 |
| 9 | 129157541 | rs10491513 | 1.243 | -0.157 | -1.479 | -1.231 | 0.003 | 0.053 | 0.053 | 2.006 | 0.26 | 0.13 | 0.13 | 0.1 | 0.22 | 0.24 |
| 9 | 129158371 | rs11793111 | 1.526 | 0.13 | -1.427 | -1.067 | -0.003 | -0.012 | -0.012 | 0.132 | 0.09 | 0.06 | 0.07 | 0 | 0.11 | 0.08 |
| 9 | 129159558 | rs2241011 | 1.356 | -0.069 | -1.262 | -1.074 | -0.016 | 0.018 | 0.018 | 0.66 | 0.23 | 0.19 | 0.23 | 0.09 | 0.3 | 0.41 |
| 9 | 129159883 | rs2052156 | -1.358 | -0.14 | -1.311 | -0.847 | -0.025 | -0.024 | -0.024 | 0.438 | 0.88 | 0.94 | 0.69 | 0.56 | 0.79 | 0.91 |
| 9 | 129165182 | rs4837076 | 0.298 | -0.241 | -1.195 | -0.932 | 0.039 | -0.048 | -0.048 | 0.432 | 0.41 | 0.16 | 0.4 | 0.54 | 0.35 | 0.28 |
| 9 | 129166088 | rs10760430 | -0.676 | 0.018 | -0.602 | -0.695 | -0.008 | 0.018 | 0.018 | 0.015 | 0.78 | 0.68 | 0.76 | 0.7 | 0.79 | 0.78 |
| 9 | 129184309 | rs887659 | 0.355 | -0.241 | -0.714 | -0.045 | -0.002 | -0.012 | -0.012 | 0 | 0.47 | 0.51 | 0.43 | 0.25 | 0.44 | 0.53 |
| 9 | 129194773 | rs758970 | -0.257 | -0.292 | -0.719 | 0.09 | 0.006 | -0.01 | -0.01 | 3.778 | 0.62 | 0.48 | 0.65 | 0.66 | 0.59 | 0.62 |
| 9 | 129197579 | rs2041543 | 1.09 | -0.614 | -0.725 | -0.156 | -0.044 | 0.002 | 0.002 | 1.406 | 0.3 | 0.43 | 0.25 | 0.23 | 0.18 | 0.28 |
| 9 | 129204547 | rs10987278 | 1.228 | -0.457 | -0.715 | -0.149 | 0 | 0 | 0 | 0.299 | 0.23 | 0.24 | 0.26 | 0.23 | 0.29 | 0.27 |
| 9 | 129216848 | rs13287948 | 1.553 | -0.393 | -0.938 | -0.204 | -0.045 | 0.053 | 0.053 | 2.496 | 0.17 | 0.07 | 0.17 | 0.03 | 0.26 | 0.24 |
| 9 | 129221756 | rs10760437 | 1.656 | -0.522 | -1.025 | -0.042 | -0.043 | -0.097 | -0.097 | 0 | 0.13 | 0.01 | 0.23 | 0.47 | 0.16 | 0.09 |
| 9 | 129229878 | rs12683116 | 1.23 | -0.648 | -0.632 | -0.195 | -0.006 | -0.001 | -0.001 | 0 | 0.29 | 0.28 | 0.25 | 0.22 | 0.18 | 0.29 |
| 9 | 129230541 | rs12376387 | 1.166 | -0.843 | -0.613 | -0.074 | -0.017 | 0.014 | 0.014 | 0 | 0.1 | 0.18 | 0.18 | 0.08 | 0.07 | 0.14 |
| 9 | 129231957 | rs1468675 | 1.291 | -0.535 | -0.974 | 0.041 | -0.029 | -0.012 | -0.012 | 1.231 | 0.22 | 0.11 | 0.24 | 0.23 | 0.28 | 0.28 |
| 9 | 129234311 | rs887653 | -1.175 | -0.377 | -0.659 | -0.319 | 0.001 | -0.058 | -0.058 | 0 | 0.88 | 0.79 | 0.92 | 1 | 0.9 | 0.93 |
| 9 | 129236624 | rs7047946 | -0.199 | -0.382 | -0.658 | -0.403 | 0 | -0.052 | -0.052 | 0.004 | 0.7 | 0.68 | 0.75 | 0.97 | 0.64 | 0.7 |
| 9 | 129237931 | rs4837090 | -0.597 | -0.568 | -0.659 | -0.156 | 0.007 | 0.008 | 0.008 | 0.613 | 0.58 | 0.4 | 0.66 | 0.78 | 0.71 | 0.64 |
| 9 | 129242209 | rs10733677 | -0.022 | -0.452 | -0.886 | -0.161 | -0.011 | 0.022 | 0.022 | 0 | 0.4 | 0.32 | 0.49 | 0.75 | 0.46 | 0.4 |
| 9 | 129242770 | rs13297792 | 1.36 | -0.33 | -0.889 | -0.215 | -0.048 | 0.073 | 0.073 | 2.686 | 0.17 | 0.05 | 0.17 | 0.03 | 0.24 | 0.23 |
| 9 | 129244060 | rs11787581 | 1.379 | -0.336 | -0.891 | -0.477 | -0.048 | 0.072 | 0.072 | 2.525 | 0.16 | 0.05 | 0.17 | 0.03 | 0.24 | 0.23 |
| 9 | 129246177 | rs2286886 | 0.124 | -0.472 | -0.83 | -0.516 | -0.025 | 0.037 | 0.037 | 0 | 0.39 | 0.26 | 0.49 | 0.76 | 0.45 | 0.41 |
| 9 | 129249570 | rs1890008 | -0.079 | -0.593 | -0.654 | -0.621 | 0.001 | -0.027 | -0.027 | 0 | 0.69 | 0.67 | 0.65 | 0.9 | 0.66 | 0.58 |
| 9 | 129252874 | rs10819170 | 1.065 | -0.808 | -0.44 | -0.166 | -0.022 | 0.027 | 0.027 | 0 | 0.22 | 0.16 | 0.24 | 0.14 | 0.29 | 0.29 |
| 9 | 129253461 | rs2417022 | -1.143 | -0.952 | -0.391 | -0.143 | -0.011 | -0.005 | -0.005 | 0.028 | 0.87 | 0.82 | 0.82 | 0.92 | 0.91 | 0.82 |
| 9 | 129258837 | rs1548788 | 0.844 | -0.83 | -0.326 | -0.162 | -0.04 | -0.052 | -0.052 | 0 | 0.31 | 0.49 | 0.16 | 0.15 | 0.2 | 0.18 |
| 9 | 129260257 | rs10987305 | 0.003 | -0.737 | -0.764 | -0.085 | -0.02 | 0.015 | 0.015 | 0 | 0.56 | 0.72 | 0.43 | 0.34 | 0.48 | 0.5 |
| 9 | 129265742 | rs3814127 | 0.831 | -0.718 | -0.615 | -0.056 | 0.001 | -0.004 | -0.004 | 0 | 0.26 | 0.33 | 0.27 | 0.19 | 0.28 | 0.32 |
| 9 | 129268065 | rs10819174 | 0.799 | -0.718 | -0.616 | -0.059 | 0.002 | -0.013 | -0.013 | 0 | 0.25 | 0.33 | 0.27 | 0.15 | 0.27 | 0.28 |
| 9 | 129268204 | rs10739677 | -0.214 | -0.749 | -0.736 | -0.061 | -0.048 | 0 | 0 | 0 | 0.58 | 0.81 | 0.43 | 0.35 | 0.49 | 0.5 |
| 9 | 129269846 | rs16929088 | 0.524 | -0.645 | -0.407 | 0.049 | 0 | 0.03 | 0.03 | 0 | 0.37 | 0.3 | 0.44 | 0.23 | 0.36 | 0.47 |
| 9 | 129270114 | rs10987313 | 0.799 | -0.607 | -0.428 | 0.081 | -0.026 | -0.032 | -0.032 | 0 | 0.31 | 0.37 | 0.15 | 0.16 | 0.2 | 0.18 |
| 9 | 129270986 | rs7042444 | 0.457 | -0.317 | -1.135 | 0.183 | -0.004 | 0.036 | 0.036 | 0 | 0.36 | 0.25 | 0.43 | 0.23 | 0.35 | 0.47 |
| 9 | 129276052 | rs4837096 | -0.341 | 0.334 | -0.956 | 0.576 | -0.052 | -0.065 | -0.065 | 0 | 0.32 | 0.16 | 0.41 | 0.53 | 0.44 | 0.33 |
| 9 | 129284289 | rs13295594 | -0.532 | -0.135 | -0.423 | 0.433 | -0.014 | 0.02 | 0.02 | 0.083 | 0.36 | 0.43 | 0.27 | 0.54 | 0.34 | 0.29 |
| 9 | 129284495 | rs12001841 | 0.118 | 0.115 | -0.565 | 0.249 | 0 | -0.026 | -0.026 | 2.346 | 0.23 | 0.2 | 0.14 | 0.01 | 0.26 | 0.2 |
| 9 | 129286085 | rs10987324 | 0.343 | 0.122 | -0.777 | 0.865 | -0.008 | 0.023 | 0.023 | 0 | 0.07 | 0.04 | 0.11 | 0.45 | 0.07 | 0.08 |

| | | | | | | | | | | | | | | | | |
|---|---|---|---|---|---|---|---|---|---|---|---|---|---|---|---|---|
| 9 | 129294471 | rs10739679 | -0.608 | -0.022 | -0.801 | 0.148 | 0.008 | 0.011 | 0.011 | 0 | 0.58 | 0.63 | 0.5 | 0.67 | 0.65 | 0.62 |
| 9 | 129300073 | rs7860558 | -0.584 | -0.09 | -0.918 | 0.079 | 0.006 | 0.004 | 0.004 | 0.002 | 0.6 | 0.63 | 0.54 | 0.65 | 0.7 | 0.65 |
| 9 | 129302884 | rs10819180 | 0.017 | 0.051 | -0.651 | -0.066 | 0.003 | -0.006 | -0.006 | 0 | 0.31 | 0.34 | 0.27 | 0.23 | 0.37 | 0.37 |
| 9 | 129303487 | rs10987336 | 0.277 | 0.054 | -0.673 | -0.254 | -0.01 | 0.008 | 0.008 | 0.129 | 0.29 | 0.25 | 0.27 | 0.07 | 0.36 | 0.36 |
| 9 | 129310354 | rs7864931 | 0.133 | -0.765 | -0.268 | 0.168 | -0.031 | -0.037 | -0.037 | 0.006 | 0.07 | 0.21 | 0.01 | 0 | 0.02 | 0 |
| 9 | 129310775 | rs4557799 | 0.396 | -0.61 | -1.097 | 0.02 | -0.04 | 0.154 | 0.154 | 1.4 | 0.38 | 0.16 | 0.31 | 0.09 | 0.55 | 0.53 |
| 9 | 129311651 | rs945683 | 0.377 | -0.613 | -1.089 | 0.206 | -0.029 | 0.126 | 0.126 | 1.516 | 0.39 | 0.16 | 0.31 | 0.1 | 0.55 | 0.57 |
| 9 | 129312129 | rs2417025 | 0.803 | -0.05 | -0.987 | 1.078 | -0.052 | -0.081 | -0.081 | 0 | 0.12 | 0 | 0.12 | 0.4 | 0.13 | 0.09 |
| 9 | 129320566 | rs10819182 | -0.047 | -1.144 | -0.221 | 0.666 | -0.029 | 0.004 | 0.004 | 0.066 | 0.29 | 0.42 | 0.45 | 0.31 | 0.23 | 0.27 |
| 9 | 129323601 | rs3861869 | -0.69 | -1.321 | 0.111 | 0.944 | -0.031 | 0.066 | 0.066 | 0 | 0.23 | 0.37 | 0.16 | 0.39 | 0.16 | 0.18 |
| 9 | 129324021 | rs3861870 | -0.679 | -1.32 | 0.124 | 0.945 | -0.031 | 0.069 | 0.069 | 0 | 0.23 | 0.37 | 0.16 | 0.39 | 0.16 | 0.18 |
| 9 | 129324974 | rs13301795 | 0.159 | -1.303 | -0.154 | 1.005 | 0.019 | -0.052 | -0.052 | 0 | 0.13 | 0.04 | 0.09 | 0.29 | 0.09 | 0.07 |
| 9 | 129325376 | rs7021240 | -0.49 | -1.491 | 0.089 | 0.773 | -0.004 | 0.002 | 0.002 | 0 | 0.21 | 0.19 | 0.14 | 0.29 | 0.16 | 0.12 |
| 9 | 129325783 | rs13287122 | -1.17 | -1.507 | 0.018 | 0.698 | -0.031 | 0.017 | 0.017 | 0 | 0.32 | 0.53 | 0.21 | 0.4 | 0.23 | 0.23 |
| 9 | 129326937 | rs10987360 | -0.1 | -1.459 | -0.019 | 0.157 | -0.02 | 0 | 0 | 0 | 0.27 | 0.63 | 0.28 | 0.28 | 0.17 | 0.24 |
| 9 | 129332556 | rs7856616 | 1.254 | -0.34 | -0.744 | -0.836 | -0.091 | 0.103 | 0.103 | 0 | 0.17 | 0.01 | 0.13 | 0.01 | 0.27 | 0.2 |
| 9 | 129334700 | rs3904195 | 1.287 | -0.483 | -1.262 | -1.108 | -0.114 | 0.055 | 0.055 | 0 | 0.24 | 0.02 | 0.19 | 0.1 | 0.33 | 0.24 |
| 9 | 129340692 | rs7868155 | 0.058 | -0.973 | -0.376 | -0.77 | -0.002 | 0 | 0 | 0 | 0.4 | 0.53 | 0.37 | 0.44 | 0.4 | 0.32 |
| 9 | 129344763 | rs7855272 | -0.618 | -0.761 | -0.449 | -0.732 | 0.004 | -0.016 | -0.016 | 0.034 | 0.71 | 0.79 | 0.61 | 0.55 | 0.76 | 0.57 |
| 9 | 129345624 | rs10115026 | 0.414 | -0.519 | 0.184 | -0.159 | 0.03 | 0.052 | 0.052 | 0 | 0.22 | 0.31 | 0.31 | 0.42 | 0.31 | 0.27 |
| 9 | 129361764 | rs7040946 | 0.693 | -1.057 | 0.322 | 0.1 | -0.039 | -0.073 | -0.073 | 0 | 0.19 | 0.33 | 0.08 | 0.03 | 0.09 | 0.07 |
| 9 | 129361955 | rs10733680 | -0.25 | -1.109 | 0.056 | 0.205 | -0.027 | 0.016 | 0.016 | 0.471 | 0.74 | 0.33 | 0.73 | 0.57 | 0.87 | 0.88 |
| 9 | 129377235 | rs2235056 | -0.321 | -0.299 | -0.021 | 0.295 | 0.001 | 0.063 | 0.063 | 0 | 0.08 | 0.07 | 0.28 | 0.44 | 0.07 | 0.23 |
| 9 | 129383199 | rs4455975 | -0.111 | -0.109 | -0.422 | -0.363 | 0.008 | 0.039 | 0.039 | 0.72 | 0.45 | 0.18 | 0.52 | 0.57 | 0.45 | 0.55 |
| 9 | 129390800 | rs3829849 | 0.406 | -0.139 | -0.367 | -0.468 | 0.003 | 0.138 | 0.138 | 0.549 | 0.38 | 0.11 | 0.25 | 0.13 | 0.37 | 0.33 |
| 9 | 129396434 | rs10760444 | 0.194 | -0.161 | 0.202 | -0.456 | 0.008 | 0.015 | 0.015 | 0.631 | 0.49 | 0.43 | 0.45 | 0.43 | 0.55 | 0.45 |
| 9 | 129397014 | rs10448285 | 0.189 | -0.16 | 0.253 | -0.649 | 0.013 | 0.036 | 0.036 | 0.099 | 0.4 | 0.29 | 0.26 | 0.13 | 0.36 | 0.29 |
| 9 | 129397755 | rs12377679 | 0.824 | 0.666 | 0.032 | -1.051 | 0 | -0.052 | -0.052 | 0 | 0.34 | 0.35 | 0.18 | 0.02 | 0.27 | 0.19 |
| 9 | 129399870 | rs12336217 | 0.157 | 0.743 | 0.226 | -0.747 | 0.001 | 0.246 | 0.246 | 0.119 | 0.12 | 0.32 | 0.3 | 0.41 | 0.09 | 0.26 |
| 9 | 129406644 | rs7858338 | 0.723 | 0.9 | -0.217 | -0.85 | 0.047 | -0.056 | -0.056 | 0 | 0.15 | 0.21 | 0.16 | 0.01 | 0.25 | 0.21 |
| 9 | 129407543 | rs11793373 | 1.063 | 0.508 | -0.775 | -0.894 | 0.006 | -0.014 | -0.014 | 0.003 | 0.31 | 0.28 | 0.16 | 0.02 | 0.25 | 0.16 |
| 9 | 129408513 | rs10819190 | -0.085 | 0.879 | -1.238 | -0.208 | 0.039 | -0.058 | -0.058 | 0 | 0.42 | 0.14 | 0.37 | 0.56 | 0.39 | 0.35 |
| 9 | 129416317 | rs10987386 | -0.009 | 1.603 | -0.286 | -0.37 | -0.001 | -0.022 | -0.022 | 0 | 0.16 | 0.16 | 0.11 | 0.01 | 0.2 | 0.24 |
| 9 | 129418360 | rs7874916 | -1.041 | 1.909 | 0.208 | -0.652 | 0.04 | -0.142 | -0.142 | 0 | 0.27 | 0.53 | 0.23 | 0.01 | 0.35 | 0.33 |
| 9 | 129422783 | rs10819195 | -0.764 | 1.982 | 0.022 | -0.407 | 0 | -0.011 | -0.011 | 0 | 0.33 | 0.35 | 0.26 | 0.16 | 0.35 | 0.22 |
| 9 | 129428677 | rs4322101 | -1.24 | 1.666 | 0.369 | -0.137 | 0.002 | 0.006 | 0.006 | 0 | 0.43 | 0.5 | 0.39 | 0.54 | 0.44 | 0.42 |
| 9 | 129438872 | rs7030919 | -0.508 | 1.629 | 0.285 | -0.142 | -0.028 | -0.134 | -0.134 | 0 | 0.19 | 0.52 | 0.11 | 0 | 0.13 | 0.19 |
| 9 | 129453229 | rs2277158 | -0.933 | 1.57 | 0.358 | 0.109 | 0 | 0 | 0 | 0 | 0.44 | 0.55 | 0.41 | 0.56 | 0.44 | 0.4 |
| 9 | 129459478 | rs10987414 | -0.262 | 1.865 | -0.762 | -0.123 | -0.016 | 0.085 | 0.085 | 0 | 0.35 | 0.12 | 0.23 | 0.2 | 0.34 | 0.32 |
| 9 | 129459628 | rs10760450 | -0.54 | 1.892 | -0.683 | -0.017 | -0.034 | 0.034 | 0.034 | 0.066 | 0.06 | 0.02 | 0.09 | 0 | 0.18 | 0.1 |
| 9 | 129460914 | rs10733682 | 0.564 | 0.874 | -0.431 | 0.302 | -0.011 | 0.09 | 0.09 | 0.273 | 0.53 | 0.75 | 0.42 | 0.25 | 0.49 | 0.47 |
| 9 | 129461912 | rs3861878 | -0.283 | 1.606 | -0.105 | 0.978 | -0.001 | 0 | 0 | 0 | 0.31 | 0.32 | 0.2 | 0.29 | 0.29 | 0.38 |
| 9 | 129462611 | rs10987421 | -0.872 | 1.271 | -0.416 | 0.746 | -0.009 | -0.053 | -0.053 | 0.045 | 0.1 | 0.03 | 0.23 | 0.22 | 0.09 | 0.17 |
| 9 | 129463613 | rs3906146 | -0.844 | 1.504 | 0 | -0.435 | -0.003 | 0.011 | 0.011 | 0 | 0.44 | 0.35 | 0.55 | 0.86 | 0.46 | 0.59 |
| 9 | 129466456 | rs13287548 | -0.723 | 1.596 | -0.377 | -0.395 | -0.04 | 0.165 | 0.165 | 0.231 | 0.43 | 0.19 | 0.53 | 0.86 | 0.46 | 0.59 |
| 9 | 129471897 | rs7870976 | -0.401 | 1.539 | -0.307 | -0.122 | 0.003 | 0.014 | 0.014 | 0 | 0.37 | 0.28 | 0.51 | 0.78 | 0.39 | 0.47 |
| 9 | 129477410 | rs7041217 | -0.385 | 1.229 | -0.805 | -0.469 | -0.006 | 0.014 | 0.014 | 0 | 0.46 | 0.54 | 0.6 | 0.9 | 0.48 | 0.56 |
| 9 | 129481232 | rs9792682 | -0.614 | 1.283 | -1.201 | 0.153 | -0.043 | -0.003 | -0.003 | 0 | 0.14 | 0.01 | 0.1 | 0.11 | 0.19 | 0.25 |
| 9 | 129490258 | rs2417034 | -0.33 | 1.043 | -0.827 | 0.892 | -0.002 | 0.01 | 0.01 | 0 | 0.16 | 0.18 | 0.22 | 0.31 | 0.12 | 0.27 |
| 9 | 129506906 | rs10125011 | -0.418 | 1.111 | -0.302 | 0.975 | -0.004 | 0.053 | 0.053 | 0.791 | 0.25 | 0.62 | 0.15 | 0.47 | 0.25 | 0.22 |
| 9 | 129510823 | rs10819218 | -0.841 | 1.058 | -0.284 | 1.668 | -0.001 | 0.03 | 0.03 | 0 | 0.19 | 0.27 | 0.23 | 0.59 | 0.2 | 0.3 |
| 9 | 129512251 | rs10114029 | -0.841 | 1.058 | -0.259 | 1.668 | -0.001 | 0.032 | 0.032 | 0 | 0.19 | 0.28 | 0.23 | 0.59 | 0.2 | 0.3 |
| 9 | 129512633 | rs913716 | -1.363 | 1.094 | 0.104 | 1.519 | 0.007 | 0.005 | 0.005 | 0.27 | 0.29 | 0.49 | 0.29 | 0.73 | 0.32 | 0.38 |
| 9 | 129513586 | rs1005918 | -0.598 | 0.901 | -0.323 | 1.602 | -0.019 | -0.03 | -0.03 | 0.826 | 0.18 | 0.07 | 0.21 | 0.24 | 0.21 | 0.23 |
| 9 | 129523907 | rs7026747 | 0.855 | 1.237 | 0.643 | 1.177 | 0.012 | -0.003 | -0.003 | 0.439 | 0.85 | 0.72 | 0.62 | 0.91 | 0.82 | 0.83 |
| 9 | 129524753 | rs7858544 | 1.085 | 1.178 | 0.774 | 1.092 | 0.018 | -0.01 | -0.01 | 0 | 0.84 | 0.64 | 0.62 | 0.9 | 0.82 | 0.83 |
| 9 | 129540970 | rs12352423 | -0.272 | 1.71 | -0.15 | 0.415 | -0.083 | 0.071 | 0.071 | 0.276 | 0.18 | 0.03 | 0.15 | 0.03 | 0.32 | 0.13 |
| 9 | 129544443 | rs6478755 | 1.213 | 1.156 | 0.884 | 0.367 | 0.016 | -0.054 | -0.054 | 0 | 0.78 | 0.39 | 0.57 | 0.92 | 0.79 | 0.75 |
| 9 | 129551466 | rs2417038 | 0.55 | 1.068 | 0.888 | 1.141 | 0.014 | -0.013 | -0.013 | 0 | 0.86 | 0.77 | 0.61 | 0.95 | 0.81 | 0.84 |
| 9 | 129552881 | rs7036823 | -0.657 | 0.728 | 1.287 | 1.5 | 0.013 | 0.504 | 0.504 | 0.129 | 0.08 | 0.41 | 0.07 | 0.39 | 0.01 | 0.19 |
| 9 | 129567615 | rs873836 | -0.222 | 0.706 | 1.227 | 1.331 | 0.041 | 0.043 | 0.043 | 0 | 0.07 | 0.36 | 0.03 | 0.01 | 0.01 | 0.07 |
| 9 | 129567832 | rs873837 | -0.633 | 0.684 | 1.292 | 1.49 | -0.001 | 0.48 | 0.48 | 0.372 | 0.08 | 0.46 | 0.07 | 0.4 | 0.01 | 0.2 |
| 9 | 129571638 | rs1329407 | 0.415 | 0.78 | -0.315 | 1.171 | -0.052 | -0.085 | -0.085 | 0 | 0.2 | 0.04 | 0.16 | 0.41 | 0.23 | 0.23 |
| 9 | 129577902 | rs2900252 | 0.223 | 0.467 | -0.772 | 0.737 | -0.037 | 0.061 | 0.061 | 0.377 | 0.22 | 0.01 | 0.21 | 0.05 | 0.18 | 0.29 |
| 9 | 129595895 | rs3118994 | 0.123 | 0.84 | -0.106 | 0.986 | -0.014 | 0.012 | 0.012 | 0 | 0.85 | 0.85 | 0.66 | 0.97 | 0.78 | 0.89 |
| 9 | 129601835 | rs10760457 | -0.23 | 0.893 | -0.456 | 0.911 | -0.011 | 0.009 | 0.009 | 0 | 0.14 | 0.15 | 0.34 | 0.03 | 0.22 | 0.11 |
| 9 | 129609815 | rs7032601 | -0.7 | 0.102 | -1.242 | 0.24 | 0.017 | 0 | 0 | 0 | 0.88 | 0.99 | 0.96 | 0.93 | 0.92 | 0.91 |
| 9 | 129613696 | rs7035535 | -0.044 | 0.403 | -1.033 | 0.212 | 0 | 0.002 | 0.002 | 0 | 0.57 | 0.52 | 0.7 | 0.7 | 0.6 | 0.73 |
| 9 | 129616938 | rs10733686 | 0.035 | 0.278 | -1.196 | -0.118 | 0.007 | -0.005 | -0.005 | 5.672 | 0.57 | 0.35 | 0.42 | 0.48 | 0.37 | 0.66 |

| | | | | | | | | | | | | | | | | |
|---|---|---|---|---|---|---|---|---|---|---|---|---|---|---|---|---|
| 9 | 129617253 | rs1463810 | 0.422 | 0.139 | -0.995 | -0.084 | -0.006 | 0 | 0 | 0 | 0.44 | 0.43 | 0.39 | 0.47 | 0.29 | 0.52 |
| 9 | 129619875 | rs10987505 | 0.593 | 0.251 | -1.17 | -0.22 | 0.024 | -0.003 | -0.003 | 0 | 0.46 | 0.35 | 0.44 | 0.48 | 0.34 | 0.52 |
| 9 | 129625226 | rs7030157 | 0.003 | -0.265 | -1.323 | -0.745 | -0.008 | -0.023 | -0.023 | 0 | 0.37 | 0.6 | 0.43 | 0.19 | 0.27 | 0.21 |
| 9 | 129630126 | rs10819242 | 0.801 | -0.541 | -1.615 | -0.496 | -0.011 | 0.013 | 0.013 | 0 | 0.08 | 0.07 | 0.22 | 0.4 | 0.08 | 0.24 |
| 9 | 129633328 | rs10819243 | 0.335 | -1.174 | -1.049 | -0.744 | 0.011 | 0.054 | 0.054 | 0 | 0.53 | 0.3 | 0.45 | 0.33 | 0.62 | 0.47 |
| 9 | 129638142 | rs4291364 | 1.413 | -0.973 | -1.167 | -0.636 | -0.01 | 0.043 | 0.043 | 0 | 0.07 | 0.07 | 0.12 | 0.38 | 0.08 | 0.24 |
| 9 | 129643296 | rs10819246 | 1.281 | -0.935 | -1.154 | -0.622 | -0.015 | 0.002 | 0.002 | 0 | 0.08 | 0.07 | 0.11 | 0.38 | 0.08 | 0.24 |
| 9 | 129646775 | rs12178 | 0.97 | -0.942 | -1.114 | -0.662 | -0.005 | 0.037 | 0.037 | 0 | 0.09 | 0.09 | 0.11 | 0.49 | 0.08 | 0.31 |
| 9 | 129647028 | rs571606 | 1.728 | -1.245 | -1.171 | -0.974 | 0.003 | 0.01 | 0.01 | 0 | 0.13 | 0.09 | 0.13 | 0.01 | 0.12 | 0.02 |
| 9 | 129648531 | rs12235895 | 0.31 | 0.275 | -1.168 | -0.801 | 0.102 | 0.235 | 0.235 | 0 | 0.11 | 0.29 | 0.17 | 0.48 | 0.24 | 0.33 |
| 9 | 129649356 | rs3120029 | -0.31 | -0.17 | -1.111 | -0.991 | -0.018 | -0.032 | -0.032 | 0 | 0.68 | 0.41 | 0.63 | 0.83 | 0.78 | 0.82 |
| 9 | 129659918 | rs671261 | -0.188 | -0.048 | -1.366 | -0.884 | 0.032 | 0.061 | 0.061 | 0.041 | 0.61 | 0.2 | 0.56 | 0.75 | 0.73 | 0.72 |
| 9 | 129670028 | rs4836555 | 0.915 | 0.692 | -1.804 | -1.087 | -0.07 | -0.001 | -0.001 | 0 | 0.1 | 0.01 | 0.26 | 0.04 | 0.29 | 0.1 |
| 9 | 129676891 | rs3118831 | -0.47 | 0.355 | -1.722 | -1.017 | 0.033 | 0.078 | 0.078 | 0.046 | 0.62 | 0.14 | 0.56 | 0.82 | 0.73 | 0.79 |
| 9 | 129686561 | rs581444 | 0.668 | 0.401 | -1.939 | -0.862 | -0.034 | -0.132 | -0.132 | 0.006 | 0.29 | 0.03 | 0.21 | 0.42 | 0.26 | 0.41 |
| 9 | 129715736 | rs518116 | -0.871 | 0.389 | -1.927 | -0.883 | 0.033 | 0.076 | 0.076 | 0.154 | 0.62 | 0.14 | 0.56 | 0.82 | 0.73 | 0.79 |
| 9 | 129755509 | rs2031852 | -1.242 | 0.295 | -1.365 | -0.934 | 0.002 | 0.006 | 0.006 | 0 | 0.67 | 0.73 | 0.59 | 0.82 | 0.74 | 0.79 |
| 9 | 129762597 | rs11789307 | 0.573 | 0.448 | -1.396 | -0.81 | -0.072 | 0.01 | 0.01 | 0.179 | 0.33 | 0.1 | 0.35 | 0.31 | 0.48 | 0.3 |
| 9 | 129772495 | rs10739688 | 1.342 | 0.376 | -1.453 | -0.846 | -0.065 | -0.149 | -0.149 | 0 | 0.21 | 0.01 | 0.19 | 0.43 | 0.21 | 0.4 |
| 9 | 129789615 | rs7019291 | 0.056 | 0.44 | -1.211 | -0.796 | -0.009 | 0.002 | 0.002 | 0 | 0.38 | 0.7 | 0.38 | 0.4 | 0.48 | 0.36 |
| 9 | 129791911 | rs11793604 | 1.57 | 0.955 | -1.377 | -0.697 | -0.043 | -0.012 | -0.012 | 0 | 0.07 | 0.01 | 0.22 | 0.05 | 0.21 | 0.09 |
| 9 | 129807559 | rs7864425 | 0.103 | 0.593 | -0.885 | -0.851 | -0.001 | -0.001 | -0.001 | 0 | 0.43 | 0.35 | 0.37 | 0.4 | 0.52 | 0.38 |
| 9 | 129828036 | rs2789513 | 2.769 | 0.324 | -1.074 | -0.845 | 0.021 | 0.045 | 0.045 | 2.1 | 0.08 | 0 | 0.02 | 0 | 0.04 | 0.02 |
| 9 | 129833029 | rs2789514 | -2.672 | 0.223 | -1.305 | -1.148 | -0.007 | 0.036 | 0.036 | 0 | 0.88 | 0.96 | 0.89 | 1 | 0.88 | 0.98 |
| 9 | 129848136 | rs4837129 | 1.267 | 0.242 | -1.768 | -1.055 | -0.067 | -0.149 | -0.149 | 0.048 | 0.21 | 0.01 | 0.19 | 0.44 | 0.22 | 0.43 |
| 9 | 129849880 | rs8838 | 1.817 | 0.133 | -1.395 | -0.84 | -0.089 | -0.01 | -0.01 | 0 | 0.25 | 0.07 | 0.31 | 0.25 | 0.42 | 0.22 |
| 9 | 129854199 | rs2297866 | 0.714 | 0.296 | -1.573 | -0.558 | -0.065 | 0.016 | 0.016 | 0.098 | 0.33 | 0.11 | 0.34 | 0.3 | 0.48 | 0.3 |
| 9 | 129862996 | rs13298677 | 2.287 | 0.294 | -1.567 | -0.47 | -0.009 | -0.028 | -0.028 | 0 | 0.18 | 0.06 | 0.28 | 0.16 | 0.15 | 0.19 |
| 9 | 129877979 | rs1768374 | 2.742 | 0.411 | -1.427 | -0.756 | 0.021 | 0.045 | 0.045 | 2.1 | 0.08 | 0 | 0.02 | 0 | 0.04 | 0.02 |
| 9 | 129883806 | rs7045790 | 3.218 | 0.893 | -1.121 | -0.459 | -0.014 | 0.057 | 0.057 | 0 | 0.11 | 0.03 | 0.08 | 0 | 0.1 | 0.03 |
| 9 | 129906656 | rs12551115 | 0.33 | 1.211 | -0.882 | 0.027 | -0.081 | 0.023 | 0.023 | 0.102 | 0.35 | 0.1 | 0.35 | 0.3 | 0.49 | 0.29 |
| 9 | 129921983 | rs11789486 | 0.483 | 1.524 | -1.591 | 0.057 | -0.08 | 0.012 | 0.012 | 0.077 | 0.33 | 0.1 | 0.34 | 0.3 | 0.48 | 0.29 |
| 9 | 129954334 | rs10819272 | 0.045 | 1.388 | -1.787 | 0.192 | -0.109 | -0.105 | -0.105 | 0 | 0.35 | 0.09 | 0.32 | 0.48 | 0.43 | 0.47 |
| 9 | 129956147 | rs2779711 | -1.465 | 0.388 | -1.512 | 0.167 | 0.001 | 0.001 | 0.001 | 0 | 0.92 | 0.95 | 0.97 | 0.96 | 0.94 | 0.95 |
| 9 | 129968138 | rs10987574 | 0.984 | 0.551 | -1.16 | 0.273 | -0.058 | -0.131 | -0.131 | 0 | 0.18 | 0.01 | 0.2 | 0.41 | 0.19 | 0.39 |
| 9 | 129975605 | rs2779710 | 0.745 | 0.113 | -1.177 | -0.03 | -0.031 | 0.098 | 0.098 | 0 | 0.44 | 0.2 | 0.6 | 0.21 | 0.51 | 0.31 |
| 9 | 129979725 | rs2779708 | -1.243 | -0.289 | -0.984 | 0.022 | 0.012 | -0.013 | -0.013 | 0 | 0.72 | 0.92 | 0.85 | 0.73 | 0.74 | 0.78 |
| 9 | 129982879 | rs7040276 | 0.849 | -0.308 | -0.237 | -0.002 | 0.042 | 0.05 | 0.05 | 5.767 | 0.06 | 0.44 | 0.03 | 0.01 | 0 | 0 |
| 9 | 129984107 | rs3739553 | 0.365 | -0.322 | 0.065 | 0.012 | -0.004 | 0.081 | 0.081 | 4.394 | 0.07 | 0.65 | 0.04 | 0.09 | 0 | 0.07 |
| 9 | 129985108 | rs1058266 | 1.058 | -0.292 | -0.033 | 0.006 | 0.01 | -0.007 | -0.007 | 0.795 | 0.28 | 0.08 | 0.16 | 0.27 | 0.27 | 0.23 |
| 9 | 129992084 | rs2812289 | -1.102 | -0.32 | -0.105 | 0.008 | 0.017 | 0.007 | 0.007 | 0 | 0.71 | 0.88 | 0.84 | 0.73 | 0.74 | 0.77 |
| 9 | 130022540 | rs2779732 | -0.595 | -0.209 | -0.182 | 0.067 | 0.047 | 0.003 | 0.003 | 0 | 0.64 | 0.22 | 0.8 | 0.64 | 0.73 | 0.7 |
| 9 | 130046379 | rs10987587 | 0.979 | 0.045 | -0.838 | 0.406 | -0.046 | -0.124 | -0.124 | 0.062 | 0.16 | 0.01 | 0.16 | 0.42 | 0.17 | 0.36 |
| 9 | 130052614 | rs16929752 | 1.637 | -0.17 | -0.853 | 0.335 | -0.001 | 0.003 | 0.003 | 0 | 0.19 | 0.2 | 0.12 | 0.33 | 0.17 | 0.24 |
| 9 | 130053105 | rs2779727 | 0.81 | -0.144 | -1.378 | 0.213 | 0.019 | 0.119 | 0.119 | 0 | 0.47 | 0.22 | 0.65 | 0.21 | 0.56 | 0.32 |
| 9 | 130055785 | rs2812273 | -1.021 | -0.302 | -1.006 | 0.313 | -0.012 | -0.004 | -0.004 | 0 | 0.73 | 0.97 | 0.85 | 0.76 | 0.76 | 0.78 |
| 9 | 130064972 | rs2812283 | -0.117 | -0.088 | -0.914 | 0.697 | 0.048 | -0.05 | -0.05 | 1.065 | 0.64 | 0.23 | 0.8 | 0.49 | 0.73 | 0.68 |
| 9 | 130094526 | rs10987596 | 0.01 | 0.819 | -0.415 | 1.265 | -0.048 | -0.085 | -0.085 | 0 | 0.2 | 0.03 | 0.17 | 0.32 | 0.19 | 0.38 |
| 9 | 130100305 | rs3802370 | 1.965 | 0.708 | -1.034 | 0.868 | -0.05 | 0.098 | 0.098 | 0.27 | 0.34 | 0.05 | 0.19 | 0.21 | 0.32 | 0.21 |
| 9 | 130105618 | rs10987600 | 0.33 | 0.769 | -1.026 | 1.041 | -0.06 | -0.128 | -0.128 | 0 | 0.19 | 0.01 | 0.15 | 0.39 | 0.18 | 0.39 |
| 9 | 130106778 | rs7863063 | 1.414 | 0.656 | -0.124 | 1.108 | -0.008 | -0.009 | -0.009 | 1.974 | 0.39 | 0.65 | 0.21 | 0.36 | 0.31 | 0.21 |
| 9 | 130107964 | rs4130590 | 0.761 | 0.845 | -0.237 | 0.853 | 0.015 | 0.005 | 0.005 | 0 | 0.52 | 0.71 | 0.67 | 0.55 | 0.59 | 0.48 |
| 9 | 130110411 | rs7029536 | 0.048 | 0.842 | 0.106 | 0.858 | 0.071 | 0.144 | 0.144 | 0.137 | 0.16 | 0.39 | 0.45 | 0.34 | 0.26 | 0.27 |
| 9 | 130112231 | rs6478770 | 1.027 | 0.807 | -0.119 | 0.969 | 0.016 | 0.006 | 0.006 | 0 | 0.52 | 0.72 | 0.67 | 0.55 | 0.59 | 0.47 |
| 9 | 130114364 | rs4837142 | 0.304 | 1.06 | 0.645 | 1.104 | -0.008 | -0.034 | -0.034 | 0 | 0.71 | 0.77 | 0.81 | 0.94 | 0.77 | 0.86 |
| 9 | 130116358 | rs3934537 | -0.435 | 1.385 | 0.455 | 1.556 | -0.004 | -0.001 | -0.001 | 0.485 | 0.14 | 0.11 | 0.16 | 0.32 | 0.16 | 0.37 |
| 9 | 130123425 | rs4454390 | 0.916 | 1.572 | 0.811 | 1.531 | 0.002 | -0.009 | -0.009 | 0 | 0.59 | 0.54 | 0.4 | 0.74 | 0.54 | 0.71 |
| 9 | 130128056 | rs10819279 | -0.256 | 1.554 | 0.598 | 1.669 | -0.03 | -0.054 | -0.054 | 0 | 0.13 | 0.06 | 0.16 | 0.31 | 0.16 | 0.35 |
| 9 | 130132392 | rs7048256 | 1.917 | 1.475 | 0.847 | 1.655 | 0.016 | 0.179 | 0.179 | 1.451 | 0.45 | 0.16 | 0.24 | 0.23 | 0.39 | 0.34 |
| 9 | 130143113 | rs10819283 | -0.641 | 1.19 | 0.872 | 1.591 | 0.006 | -0.003 | -0.003 | 0 | 0.88 | 0.88 | 0.98 | 0.86 | 0.95 | 0.97 |
| 9 | 130145624 | rs3824415 | -0.28 | 1.386 | 0.82 | 1.748 | 0.01 | 0.073 | 0.073 | 0 | 0.13 | 0.18 | 0.14 | 0.38 | 0.15 | 0.35 |
| 9 | 130147979 | rs10819284 | 0.062 | 1.384 | 0.661 | 1.755 | -0.053 | -0.088 | -0.088 | 0 | 0.12 | 0.01 | 0.13 | 0.38 | 0.15 | 0.34 |
| 9 | 130152522 | rs10987630 | 0.55 | 1.097 | 0.71 | 1.382 | -0.053 | -0.032 | -0.032 | 0 | 0.69 | 0.53 | 0.81 | 0.8 | 0.8 | 0.85 |
| 9 | 130159333 | rs3808835 | 1.858 | 1.221 | 0.121 | 0.985 | 0.024 | 0.196 | 0.196 | 1.454 | 0.44 | 0.17 | 0.22 | 0.17 | 0.38 | 0.33 |
| 9 | 130165995 | rs1138740 | -0.027 | 1.947 | 0.151 | 1.427 | -0.062 | 0.035 | 0.035 | 0.232 | 0.1 | 0.01 | 0.21 | 0.01 | 0.23 | 0.12 |
| 9 | 130166857 | rs3824414 | 1.137 | 1.888 | 0.538 | 1.672 | 0 | 0.002 | 0.002 | 0 | 0.62 | 0.61 | 0.65 | 0.45 | 0.64 | 0.53 |
| 9 | 130166957 | rs3802364 | 1.137 | 1.887 | 0.545 | 1.65 | 0 | 0.002 | 0.002 | 0 | 0.62 | 0.61 | 0.65 | 0.47 | 0.64 | 0.53 |
| 9 | 130172485 | rs10987639 | -0.023 | 1.976 | 0.644 | 1.535 | 0.016 | 0.027 | 0.027 | 0.124 | 0.11 | 0.1 | 0.39 | 0.21 | 0.25 | 0.16 |
| 9 | 130181886 | rs17464948 | 1.738 | 0.981 | 0.586 | 1.602 | 0.009 | 0.098 | 0.098 | 0.485 | 0.55 | 0.28 | 0.39 | 0.2 | 0.58 | 0.39 |

| | | | | | | | | | | | | | | | |
|---|---|---|---|---|---|---|---|---|---|---|---|---|---|---|---|
| 9 | 130186700 | rs10819291 | 0.463 | 1.161 | 0.487 | 1.489 | -0.018 | -0.046 | -0.046 | 0.162 | 0.09 | 0.02 | 0.35 | 0.2 | 0.1 | 0.15 |
| 9 | 130191186 | rs13292096 | 1.244 | 1.149 | 0.531 | 1.489 | 0 | 0 | 0 | 0 | 0.61 | 0.61 | 0.65 | 0.41 | 0.63 | 0.5 |
| 9 | 130196674 | rs10819293 | 0.488 | 1.148 | 0.505 | 1.502 | -0.018 | -0.045 | -0.045 | 0 | 0.09 | 0.02 | 0.35 | 0.2 | 0.1 | 0.15 |
| 9 | 130197414 | rs4504745 | 1.234 | 1.152 | 0.531 | 1.493 | 0 | 0 | 0 | 0.008 | 0.62 | 0.62 | 0.65 | 0.41 | 0.63 | 0.5 |
| 9 | 130201678 | rs7869023 | 1.234 | 1.152 | 0.531 | 1.493 | 0 | 0 | 0 | 0 | 0.62 | 0.62 | 0.65 | 0.41 | 0.63 | 0.5 |
| 9 | 130215249 | rs2491101 | 2.166 | 1.147 | 0.592 | 1.428 | 0.002 | 0.105 | 0.105 | 0 | 0.51 | 0.36 | 0.28 | 0.21 | 0.53 | 0.35 |
| 9 | 130219669 | rs2243906 | 1.261 | 1.152 | 0.531 | 1.453 | 0 | 0 | 0 | 0 | 0.61 | 0.61 | 0.65 | 0.41 | 0.63 | 0.5 |
| 9 | 130221388 | rs2243767 | 1.265 | 1.138 | 0.524 | 1.448 | 0 | 0 | 0 | 0 | 0.61 | 0.61 | 0.65 | 0.41 | 0.63 | 0.5 |
| 9 | 130235053 | rs10819295 | -0.642 | 1.195 | 0.528 | 1.428 | 0 | 0.04 | 0.04 | 0.057 | 0.15 | 0.22 | 0.39 | 0.2 | 0.15 | 0.18 |
| 9 | 130241822 | rs1539569 | 0.925 | 1.358 | -0.111 | 1.971 | -0.023 | 0.007 | 0.007 | 0.513 | 0.72 | 0.63 | 0.81 | 0.69 | 0.8 | 0.77 |
| 9 | 130242166 | rs1539567 | 0.919 | 1.369 | -0.105 | 1.985 | -0.024 | 0.005 | 0.005 | 0.815 | 0.72 | 0.63 | 0.81 | 0.69 | 0.8 | 0.77 |
| 9 | 130246564 | rs2246270 | 0.44 | 1.341 | -0.03 | 1.002 | -0.001 | -0.047 | -0.047 | 0 | 0.8 | 0.85 | 0.83 | 0.46 | 0.83 | 0.67 |
| 9 | 130266566 | rs2245389 | 1.344 | 1.407 | -0.01 | 0.91 | 0 | 0.004 | 0.004 | 0.039 | 0.65 | 0.62 | 0.69 | 0.4 | 0.68 | 0.54 |
| 9 | 130280589 | rs2254868 | 1.029 | 1.445 | -0.003 | 0.896 | 0.009 | 0.028 | 0.028 | 0 | 0.74 | 0.63 | 0.77 | 0.35 | 0.77 | 0.55 |
| 9 | 130282839 | rs2254622 | 0.449 | 1.591 | 0.019 | 1.04 | -0.005 | -0.023 | -0.023 | 0 | 0.82 | 0.83 | 0.84 | 0.46 | 0.84 | 0.67 |
| 9 | 130286428 | rs1891732 | 0.818 | 1.563 | -0.064 | 0.905 | 0.001 | 0.002 | 0.002 | 0 | 0.66 | 0.59 | 0.69 | 0.37 | 0.68 | 0.55 |
| 9 | 130289951 | rs2298181 | -0.266 | 1.731 | 0.022 | 1.56 | -0.02 | -0.038 | -0.038 | 0 | 0.1 | 0.03 | 0.36 | 0.2 | 0.12 | 0.16 |
| 9 | 130297246 | rs2247374 | 0.899 | 1.765 | 0.293 | 1.604 | 0.04 | 0.137 | 0.137 | 0 | 0.84 | 0.62 | 0.83 | 0.4 | 0.82 | 0.64 |
| 9 | 130309024 | rs1891729 | -0.819 | 1.609 | 0.088 | 1.565 | 0.024 | 0.157 | 0.157 | 0 | 0.16 | 0.32 | 0.13 | 0.53 | 0.16 | 0.37 |
| 9 | 130310834 | rs7854003 | -0.782 | 1.406 | -0.421 | 1.477 | 0.006 | 0.041 | 0.041 | 0.084 | 0.13 | 0.2 | 0.37 | 0.25 | 0.14 | 0.16 |
| 9 | 130311404 | rs2257055 | -0.364 | 1.108 | -1.214 | 0.728 | -0.026 | 0.022 | 0.022 | 0.036 | 0.49 | 0.15 | 0.48 | 0.4 | 0.49 | 0.38 |
| 9 | 130313173 | rs7035401 | -0.606 | 1.08 | -0.203 | 0.891 | 0.028 | 0.046 | 0.046 | 0 | 0.12 | 0.25 | 0.25 | 0.17 | 0.14 | 0.13 |
| 9 | 130316561 | rs4378065 | -0.748 | 1 | -0.319 | 0.659 | 0.006 | -0.005 | -0.005 | 0 | 0.15 | 0.22 | 0.26 | 0.11 | 0.14 | 0.16 |
| 9 | 130320119 | rs2253112 | -0.189 | 0.534 | -0.148 | 0.17 | -0.008 | -0.028 | -0.028 | 0.821 | 0.29 | 0.37 | 0.17 | 0.22 | 0.26 | 0.3 |
| 9 | 130321117 | rs2252968 | -0.344 | 0.52 | 0.497 | 0.185 | -0.003 | 0.006 | 0.006 | 0 | 0.3 | 0.32 | 0.23 | 0.38 | 0.29 | 0.38 |
| 9 | 130323485 | rs2252730 | -1.112 | 0.706 | 0.739 | -0.19 | -0.006 | -0.003 | -0.003 | 0.005 | 0.42 | 0.63 | 0.52 | 0.48 | 0.45 | 0.59 |
| 9 | 130330131 | rs10987687 | -0.721 | 0.738 | 1 | -0.183 | 0.024 | -0.061 | -0.061 | 0 | 0.22 | 0.47 | 0.34 | 0.12 | 0.23 | 0.33 |
| 9 | 130334178 | rs4240418 | -0.147 | 0.704 | 0.964 | -0.174 | 0 | 0.013 | 0.013 | 0 | 0.74 | 0.74 | 0.79 | 0.63 | 0.75 | 0.67 |
| 9 | 130335418 | rs10760490 | 0.364 | 0.763 | 0.871 | -0.27 | -0.027 | 0.033 | 0.033 | 0 | 0.11 | 0.1 | 0.2 | 0.03 | 0.16 | 0.13 |
| 9 | 130342992 | rs1124602 | -0.896 | 0.676 | 1.229 | -0.284 | 0.02 | -0.041 | -0.041 | 0 | 0.21 | 0.39 | 0.33 | 0.12 | 0.23 | 0.31 |
| 9 | 130349014 | rs2417053 | -0.942 | 0.86 | 1.225 | -0.083 | -0.03 | -0.059 | -0.059 | 0 | 0.83 | 0.98 | 0.86 | 0.72 | 0.81 | 0.77 |
| 9 | 130358236 | rs12341585 | -1.677 | 0.691 | 1.729 | -0.598 | 0.01 | 0.02 | 0.02 | 0.432 | 0.16 | 0.61 | 0.2 | 0.16 | 0.13 | 0.28 |
| 9 | 130363183 | rs10987696 | -1.554 | 0.738 | 1.938 | -0.584 | 0.07 | 0.041 | 0.041 | 0 | 0.09 | 0.41 | 0.17 | 0.1 | 0.1 | 0.24 |
| 9 | 130372487 | rs1573178 | -0.104 | 0.534 | 1.942 | -0.35 | -0.037 | -0.018 | -0.018 | 0 | 0.62 | 0.78 | 0.6 | 0.4 | 0.57 | 0.56 |
| 9 | 130373596 | rs7852204 | -0.216 | 0.537 | 1.948 | -0.35 | -0.029 | -0.032 | -0.032 | 0 | 0.63 | 0.82 | 0.64 | 0.41 | 0.61 | 0.6 |
| 9 | 130376248 | rs6478788 | 0.557 | 0.486 | 2.086 | -0.378 | 0 | -0.005 | -0.005 | 0 | 0.54 | 0.48 | 0.48 | 0.31 | 0.51 | 0.36 |
| 9 | 130376287 | rs10819302 | -1.21 | 0.52 | 2.058 | -0.394 | 0.016 | 0.022 | 0.022 | 0 | 0.09 | 0.38 | 0.13 | 0.09 | 0.06 | 0.21 |
| 9 | 130402180 | rs36081179 | 0.943 | 0.172 | 1.461 | -1.073 | 0.009 | 0.09 | 0.09 | 0.013 | 0.17 | 0.04 | 0.07 | 0 | 0.12 | 0.03 |
| 9 | 130420813 | rs2287116 | -1.392 | 0.163 | 1.773 | -0.827 | -0.01 | 0.032 | 0.032 | 0.145 | 0.11 | 0.62 | 0.11 | 0.11 | 0.06 | 0.22 |
| 9 | 130430116 | rs2241167 | 0.302 | 0.132 | 1.748 | -0.837 | -0.015 | 0.069 | 0.069 | 0 | 0.55 | 0.28 | 0.47 | 0.25 | 0.52 | 0.39 |
| 9 | 130439542 | rs4837175 | -1.626 | 0.101 | 1.82 | -0.899 | 0.019 | 0.019 | 0.019 | 0.309 | 0.12 | 0.63 | 0.15 | 0.12 | 0.1 | 0.26 |
| 9 | 130460971 | rs10123531 | -1.6 | 0.104 | 1.58 | -0.898 | 0.02 | 0.02 | 0.02 | 0.321 | 0.11 | 0.63 | 0.15 | 0.11 | 0.1 | 0.26 |
| 9 | 130475011 | rs522328 | -0.602 | 0.128 | 1.394 | -0.89 | -0.067 | -0.058 | -0.058 | 0.803 | 0.64 | 0.9 | 0.61 | 0.37 | 0.61 | 0.61 |
| 9 | 130477160 | rs550045 | -0.322 | 0.156 | 1.594 | -0.895 | 0.007 | -0.005 | -0.005 | 0 | 0.62 | 0.64 | 0.54 | 0.29 | 0.55 | 0.58 |
| 9 | 130477768 | rs504434 | -0.322 | 0.096 | 1.594 | -0.895 | 0.007 | -0.005 | -0.005 | 0 | 0.62 | 0.64 | 0.54 | 0.29 | 0.56 | 0.58 |
| 9 | 130478653 | rs3802357 | 0.548 | 0.065 | 1.466 | -0.913 | 0.005 | 0.099 | 0.099 | 0 | 0.52 | 0.27 | 0.46 | 0.24 | 0.49 | 0.35 |
| 9 | 130486915 | rs2256139 | -0.59 | 0.097 | 1.498 | -0.891 | -0.014 | -0.015 | -0.015 | 0 | 0.66 | 0.78 | 0.6 | 0.37 | 0.62 | 0.67 |
| 9 | 130489558 | rs472144 | -0.603 | 0.116 | 1.503 | -0.894 | -0.015 | -0.015 | -0.015 | 1.136 | 0.66 | 0.78 | 0.6 | 0.37 | 0.62 | 0.67 |
| 9 | 130489743 | rs13298768 | 0.632 | 0.091 | 1.789 | -0.928 | 0.002 | 0.12 | 0.12 | 0.029 | 0.52 | 0.22 | 0.46 | 0.25 | 0.5 | 0.35 |
| 9 | 130492039 | rs557749 | -0.619 | 0.14 | 1.612 | -0.859 | 0.011 | -0.006 | -0.006 | 0.889 | 0.65 | 0.62 | 0.59 | 0.3 | 0.62 | 0.66 |
| 9 | 130493513 | rs482095 | -0.559 | 0.147 | 1.454 | -0.863 | -0.015 | -0.015 | -0.015 | 0.048 | 0.66 | 0.79 | 0.61 | 0.37 | 0.62 | 0.67 |
| 9 | 130494667 | rs534652 | -0.642 | 0.143 | 1.437 | -0.892 | -0.01 | -0.001 | -0.001 | 0.947 | 0.67 | 0.76 | 0.61 | 0.31 | 0.63 | 0.67 |
| 9 | 130495890 | rs3739812 | -0.664 | 0.143 | 1.308 | -0.896 | -0.018 | -0.023 | -0.023 | 0 | 0.67 | 0.81 | 0.61 | 0.37 | 0.63 | 0.67 |
| 9 | 130498120 | rs1999224 | -0.768 | -0.801 | 1.296 | -1.158 | -0.023 | -0.02 | -0.02 | 0 | 0.14 | 0.29 | 0.08 | 0.09 | 0.09 | 0.3 |
| 9 | 130502238 | rs7852090 | -1.393 | -0.838 | 1.14 | -1.13 | 0.044 | 0.078 | 0.078 | 0 | 0.07 | 0.34 | 0.04 | 0.06 | 0.04 | 0.05 |
| 9 | 130504070 | rs514024 | -0.322 | -0.672 | 0.942 | -1.144 | 0.002 | -0.018 | -0.018 | 0 | 0.63 | 0.57 | 0.58 | 0.28 | 0.61 | 0.62 |
| 9 | 130527786 | rs10739696 | -0.963 | -0.607 | 0.275 | -1.136 | -0.009 | 0.006 | 0.006 | 0 | 0.61 | 0.7 | 0.56 | 0.96 | 0.54 | 0.72 |
| 9 | 130528548 | rs4836579 | -0.486 | -0.514 | -0.503 | -1.035 | -0.011 | 0.108 | 0.108 | 0 | 0.55 | 0.29 | 0.51 | 0.89 | 0.52 | 0.64 |
| 9 | 130536717 | rs10760500 | -0.594 | -0.487 | -0.609 | -1.046 | -0.011 | 0.143 | 0.143 | 0 | 0.54 | 0.21 | 0.51 | 0.9 | 0.52 | 0.64 |
| 9 | 130538720 | rs913990 | 0.807 | -0.505 | -1.274 | -0.613 | -0.067 | -0.129 | -0.129 | 0.209 | 0.26 | 0.02 | 0.36 | 0.65 | 0.31 | 0.3 |
| 9 | 130545950 | rs1002095 | 0.453 | -0.427 | -0.853 | -0.496 | -0.017 | -0.085 | -0.085 | 0 | 0.4 | 0.07 | 0.42 | 0.57 | 0.38 | 0.35 |
| 9 | 130561688 | rs12379987 | 0.286 | -0.413 | -0.135 | -0.436 | 0.009 | 0.01 | 0.01 | 0 | 0.43 | 0.41 | 0.46 | 0.69 | 0.38 | 0.35 |
| 9 | 130562725 | rs1544105 | 0.194 | -0.477 | 0.104 | -0.48 | 0.002 | 0.002 | 0.002 | 0 | 0.47 | 0.62 | 0.48 | 0.69 | 0.4 | 0.4 |
| 9 | 130576075 | rs10106 | 0.256 | -0.159 | 0.146 | -0.406 | 0.013 | 0.018 | 0.018 | 0 | 0.47 | 0.5 | 0.47 | 0.69 | 0.39 | 0.4 |
| 9 | 130576597 | rs4451422 | 0.428 | -0.172 | 0.174 | -0.413 | 0.016 | 0.026 | 0.026 | 0 | 0.44 | 0.4 | 0.46 | 0.69 | 0.39 | 0.4 |
| 9 | 130578404 | rs10760503 | 0.326 | 0.072 | -0.027 | -0.254 | 0.022 | 0.05 | 0.05 | 0 | 0.51 | 0.43 | 0.49 | 0.69 | 0.44 | 0.4 |
| 9 | 130580093 | rs10987746 | 0.057 | -0.001 | -0.132 | -0.637 | 0.001 | 0.007 | 0.007 | 0 | 0.54 | 0.54 | 0.5 | 0.69 | 0.45 | 0.41 |
| 9 | 130586688 | rs3739817 | -0.723 | -0.136 | -0.515 | -0.506 | 0.004 | 0.01 | 0.01 | 0.001 | 0.09 | 0.04 | 0.06 | 0.02 | 0.07 | 0.23 |
| 9 | 130602124 | rs10819311 | -0.975 | -0.317 | 0.023 | -1.213 | -0.01 | -0.101 | -0.101 | 0 | 0.28 | 0.55 | 0.22 | 0.05 | 0.22 | 0.13 |

| | | | | | | | | | | | | | | | | |
|---|---|---|---|---|---|---|---|---|---|---|---|---|---|---|---|---|
| 9 | 130602408 | rs10121110 | 0.563 | -0.869 | -0.687 | -1.811 | 0.036 | 0.126 | 0.126 | 0.143 | 0.51 | 0.34 | 0.64 | 0.91 | 0.6 | 0.76 |
| 9 | 130607518 | rs11789185 | 0.914 | -1.13 | 0.036 | -1.354 | -0.024 | -0.031 | -0.031 | 0.29 | 0.8 | 0.37 | 0.87 | 0.93 | 0.9 | 0.9 |
| 9 | 130608296 | rs10819312 | -0.654 | -1.063 | 0.994 | -1.331 | -0.01 | -0.025 | -0.025 | 0 | 0.16 | 0.27 | 0.12 | 0.06 | 0.12 | 0.08 |
| 9 | 130612218 | rs12379490 | -0.563 | -1.119 | 1.029 | -1.351 | -0.011 | -0.063 | -0.063 | 0 | 0.14 | 0.22 | 0.07 | 0 | 0.09 | 0.02 |
| 9 | 130614912 | rs4837192 | 1.267 | -1.301 | 1.667 | -1.172 | -0.038 | -0.019 | -0.019 | 0.094 | 0.79 | 0.45 | 0.8 | 0.82 | 0.88 | 0.91 |
| 9 | 130615751 | rs10115613 | 1.18 | -1.251 | 1.688 | -1.32 | -0.033 | -0.116 | -0.116 | 0 | 0.79 | 0.4 | 0.88 | 1 | 0.89 | 0.97 |
| 9 | 130619637 | rs7865146 | 1.142 | -1.481 | 0.685 | -0.839 | 0.047 | 0.023 | 0.023 | 0 | 0.52 | 0.33 | 0.68 | 0.67 | 0.63 | 0.35 |
| 9 | 130622946 | rs4837197 | 1.267 | -1.081 | 1.072 | -0.943 | -0.024 | -0.048 | -0.048 | 0 | 0.73 | 0.39 | 0.82 | 0.91 | 0.82 | 0.78 |
| 9 | 130624723 | rs1537034 | -0.183 | -0.944 | 0.582 | -0.839 | -0.007 | -0.053 | -0.053 | 0 | 0.12 | 0.19 | 0.07 | 0 | 0.08 | 0.03 |
| 9 | 130629139 | rs913987 | 1.148 | -1.137 | 0.503 | -0.877 | -0.035 | -0.053 | -0.053 | 0 | 0.72 | 0.37 | 0.81 | 0.91 | 0.82 | 0.78 |
| 9 | 130631519 | rs3780663 | -0.339 | -0.484 | 0.204 | -1.354 | 0.002 | -0.083 | -0.083 | 1.974 | 0.31 | 0.41 | 0.18 | 0.05 | 0.32 | 0.08 |
| 9 | 130633010 | rs2094332 | 0.558 | -1.301 | 0.179 | -0.751 | 0.001 | -0.002 | -0.002 | 0 | 0.63 | 0.64 | 0.59 | 0.5 | 0.65 | 0.47 |
| 9 | 130633433 | rs1556246 | -1.012 | -1.48 | 1.09 | -0.432 | 0.021 | 0.043 | 0.043 | 0 | 0.42 | 0.35 | 0.24 | 0.13 | 0.34 | 0.44 |
| 9 | 130633904 | rs743539 | -1.191 | -1.382 | 1.266 | -0.324 | -0.014 | -0.047 | -0.047 | 0 | 0.44 | 0.48 | 0.25 | 0.14 | 0.35 | 0.48 |
| 9 | 130634944 | rs743537 | -1.195 | -1.379 | 1.163 | -0.313 | -0.007 | -0.028 | -0.028 | 0 | 0.44 | 0.51 | 0.26 | 0.14 | 0.35 | 0.49 |
| 9 | 130635937 | rs968463 | 1.045 | -1.364 | 0.851 | -0.305 | 0.033 | 0.065 | 0.065 | 0 | 0.54 | 0.35 | 0.74 | 0.86 | 0.65 | 0.51 |
| 9 | 130645262 | rs10819315 | 0.537 | -0.889 | 1.108 | 0.433 | 0.008 | 0.04 | 0.04 | 0 | 0.94 | 0.76 | 0.95 | 0.94 | 0.96 | 0.98 |
| 9 | 130657833 | rs3780670 | -1.262 | -1.174 | 0.758 | 0.271 | 0.005 | 0.004 | 0.004 | 0 | 0.23 | 0.2 | 0.14 | 0.07 | 0.17 | 0.28 |
| 9 | 130659406 | rs7027357 | 1.48 | -1.203 | 1.145 | 1.213 | 0 | 0.086 | 0.086 | 2.729 | 0.85 | 0.35 | 0.87 | 0.74 | 0.88 | 0.87 |
| 9 | 130660232 | rs3758330 | -0.834 | -1.629 | 1.18 | 1.516 | 0.014 | -0.014 | -0.014 | 0.019 | 0.12 | 0.04 | 0.08 | 0.15 | 0.08 | 0.23 |
| 9 | 130660810 | rs1571142 | 1.504 | -1.391 | 1.449 | 1.827 | -0.035 | 0.154 | 0.154 | 3.163 | 0.86 | 0.39 | 0.88 | 0.69 | 0.91 | 0.87 |
| 9 | 130668004 | rs4837204 | 1.449 | -1.467 | 1.652 | 2.03 | 0.022 | 0.201 | 0.201 | 1.951 | 0.93 | 0.5 | 0.95 | 0.86 | 0.99 | 0.94 |
| 9 | 130668957 | rs10819317 | -0.598 | -1.685 | 0.707 | 2.204 | 0.008 | -0.112 | -0.112 | 0.144 | 0.22 | 0.05 | 0.17 | 0.44 | 0.17 | 0.41 |
| 9 | 130670532 | rs3818509 | 1.711 | -1.482 | 1.651 | 2.219 | -0.041 | 0.09 | 0.09 | 0.874 | 0.83 | 0.43 | 0.91 | 0.72 | 0.9 | 0.83 |
| 9 | 130673419 | rs3780668 | 1.947 | -1.354 | 1.571 | 2.001 | -0.041 | -0.014 | -0.014 | 0 | 0.67 | 0.47 | 0.67 | 0.29 | 0.77 | 0.59 |
| 9 | 130696710 | rs4836593 | 1.044 | -1.789 | 2.015 | 2.633 | -0.058 | 0.061 | 0.061 | 0 | 0.85 | 0.79 | 0.92 | 0.65 | 0.96 | 0.75 |
| 9 | 130697328 | rs10819321 | 2.096 | -1.933 | 2.582 | 2.467 | -0.085 | 0.022 | 0.022 | 0 | 0.72 | 0.52 | 0.85 | 0.39 | 0.87 | 0.66 |
| 9 | 130702477 | rs3739821 | 3.035 | -2.067 | 2.771 | 2.293 | 0.04 | 0.034 | 0.034 | 0 | 0.59 | 0.26 | 0.75 | 0.26 | 0.76 | 0.44 |
| 9 | 130702642 | rs3824527 | -1.366 | -2.068 | 2.818 | 2.294 | -0.018 | -0.015 | -0.015 | 0 | 0.13 | 0.28 | 0.1 | 0.11 | 0.11 | 0.22 |
| 9 | 130703565 | rs7859 | 1.893 | -2.001 | 2.54 | 2.35 | -0.029 | 0.01 | 0.01 | 0 | 0.73 | 0.65 | 0.85 | 0.45 | 0.87 | 0.66 |
| 9 | 130705120 | rs1043060 | 1.965 | -2.007 | 2.562 | 2.339 | -0.047 | 0.007 | 0.007 | 0 | 0.72 | 0.59 | 0.85 | 0.37 | 0.87 | 0.66 |
| 9 | 130713308 | rs3802355 | 0.763 | -1.992 | 2.486 | 2.362 | -0.071 | 0.109 | 0.109 | 0 | 0.86 | 0.72 | 0.92 | 0.66 | 0.97 | 0.75 |
| 9 | 130716761 | rs6415846 | 1.037 | -1.986 | 2.507 | 2.366 | -0.074 | 0.087 | 0.087 | 0 | 0.84 | 0.69 | 0.92 | 0.65 | 0.96 | 0.75 |
| 9 | 130727677 | rs3780666 | 1.743 | -1.909 | 2.503 | 2.336 | -0.068 | 0.026 | 0.026 | 0 | 0.74 | 0.55 | 0.86 | 0.38 | 0.88 | 0.66 |
| 9 | 130735937 | rs7865476 | -0.703 | -1.867 | 2.418 | 2.219 | 0.011 | 0.02 | 0.02 | 0 | 0.06 | 0.19 | 0.02 | 0.08 | 0.05 | 0.08 |
| 9 | 130737250 | rs1078697 | 2.238 | -1.841 | 2.255 | 2.229 | -0.001 | -0.002 | -0.002 | 0 | 0.69 | 0.67 | 0.82 | 0.54 | 0.78 | 0.49 |
| 9 | 130755881 | rs7032368 | 1.393 | -1.824 | 2.245 | 2.166 | -0.078 | 0.124 | 0.124 | 0 | 0.8 | 0.56 | 0.89 | 0.63 | 0.92 | 0.76 |
| 9 | 130769800 | rs7873657 | -1.922 | -1.462 | 2.03 | 1.761 | -0.014 | 0.014 | 0.014 | 0 | 0.25 | 0.34 | 0.18 | 0.36 | 0.21 | 0.3 |
| 9 | 130770276 | rs7874551 | -0.955 | -1.41 | 2.133 | 1.928 | -0.004 | 0.118 | 0.118 | 0.026 | 0.07 | 0.22 | 0.06 | 0.26 | 0.04 | 0.02 |
| 9 | 130774369 | rs947626 | 1.54 | -1.304 | 2.355 | 2.074 | -0.059 | 0.073 | 0.073 | 0 | 0.73 | 0.55 | 0.82 | 0.48 | 0.83 | 0.53 |
| 9 | 130776407 | rs4837223 | 1.566 | -1.299 | 2.256 | 1.868 | -0.06 | -0.002 | -0.002 | 0 | 0.72 | 0.54 | 0.79 | 0.31 | 0.83 | 0.52 |
| 9 | 130776897 | rs10760523 | 1.76 | -1.3 | 1.513 | 1.99 | 0.028 | -0.019 | -0.019 | 1.127 | 0.7 | 0.2 | 0.76 | 0.42 | 0.84 | 0.52 |
| 9 | 130778514 | rs7026544 | 1.009 | -1.043 | 1.775 | 1.588 | -0.092 | 0.274 | 0.274 | 0 | 0.85 | 0.52 | 0.9 | 0.59 | 0.96 | 0.76 |
| 9 | 130782031 | rs7852168 | -1.284 | 0.561 | 0.966 | 0.556 | -0.027 | -0.002 | -0.002 | 0 | 0.19 | 0.14 | 0.21 | 0.19 | 0.3 | 0.23 |
| 9 | 130793484 | rs10987842 | 0.184 | 0.004 | 0.775 | 0.255 | -0.024 | -0.018 | -0.018 | 0 | 0.1 | 0.02 | 0.17 | 0.11 | 0.13 | 0.05 |
| 9 | 130808444 | rs10819341 | 1.219 | -0.319 | 1.922 | 0.004 | -0.005 | -0.003 | -0.003 | 0.014 | 0.72 | 0.34 | 0.68 | 0.77 | 0.81 | 0.63 |
| 9 | 130814366 | rs882895 | 1.224 | -0.294 | 1.935 | 0.062 | -0.005 | -0.003 | -0.003 | 0 | 0.72 | 0.34 | 0.68 | 0.75 | 0.81 | 0.61 |
| 9 | 130816609 | rs10987850 | -0.959 | -0.267 | 1.786 | -0.107 | 0.015 | 0.048 | 0.048 | 0 | 0.19 | 0.07 | 0.14 | 0.06 | 0.15 | 0.22 |
| 9 | 130819574 | rs10987852 | -1.278 | -0.154 | 2.212 | -0.265 | -0.006 | -0.016 | -0.016 | 0 | 0.2 | 0.24 | 0.2 | 0.12 | 0.15 | 0.26 |
| 9 | 130829564 | rs7852659 | 1.469 | -0.226 | 2.45 | -0.319 | -0.012 | -0.013 | -0.013 | 0.548 | 0.71 | 0.32 | 0.72 | 0.82 | 0.83 | 0.67 |
| 9 | 130835780 | rs4240425 | 1.369 | -0.263 | 2.408 | -0.345 | -0.018 | -0.015 | -0.015 | 0 | 0.71 | 0.32 | 0.75 | 0.82 | 0.86 | 0.72 |
| 9 | 130847209 | rs10739708 | 1.13 | -0.228 | 2.517 | -0.346 | -0.054 | -0.043 | -0.043 | 0 | 0.73 | 0.47 | 0.76 | 0.82 | 0.87 | 0.73 |
| 9 | 130851573 | rs10987871 | -0.9 | -0.228 | 2.578 | -0.346 | 0.019 | 0.016 | 0.016 | 0 | 0.22 | 0.15 | 0.2 | 0.16 | 0.13 | 0.26 |
| 9 | 130854968 | rs7042793 | -1.118 | -0.214 | 2.43 | -0.341 | -0.03 | -0.028 | -0.028 | 0 | 0.26 | 0.38 | 0.23 | 0.16 | 0.14 | 0.26 |
| 9 | 130870023 | rs7872702 | 1.21 | -0.218 | 1.109 | -0.411 | -0.028 | -0.033 | -0.033 | 0 | 0.72 | 0.36 | 0.73 | 0.83 | 0.84 | 0.69 |
| 9 | 130873428 | rs4837240 | 1.213 | -0.219 | 1.048 | -0.428 | -0.032 | -0.039 | -0.039 | 0 | 0.71 | 0.38 | 0.73 | 0.83 | 0.84 | 0.69 |
| 9 | 130891060 | rs12377832 | -0.848 | -0.398 | -0.415 | -0.282 | 0.009 | 0.004 | 0.004 | 0 | 0.2 | 0.13 | 0.16 | 0.06 | 0.11 | 0.16 |
| 9 | 130906346 | rs10819368 | 0.771 | -0.284 | -1.333 | 0.102 | 0.028 | -0.008 | -0.008 | 0.118 | 0.52 | 0.12 | 0.73 | 0.5 | 0.61 | 0.64 |
| 9 | 130909329 | rs10987895 | -0.679 | -0.488 | -0.185 | -0.033 | 0.007 | -0.017 | -0.017 | 0 | 0.38 | 0.3 | 0.2 | 0.03 | 0.33 | 0.19 |
| 9 | 130918370 | rs10987899 | 1.105 | -0.632 | -0.292 | 0.133 | 0.047 | -0.01 | -0.01 | 0.022 | 0.54 | 0.12 | 0.73 | 0.5 | 0.63 | 0.63 |
| 9 | 130931586 | rs10987905 | -0.659 | -1.096 | -0.194 | -0.269 | 0.019 | 0.155 | 0.155 | 0.817 | 0.35 | 0.13 | 0.18 | 0.04 | 0.3 | 0.18 |
| 9 | 130935956 | rs7847286 | -1.215 | -1.098 | 1.004 | -0.118 | -0.004 | 0.003 | 0.003 | 0 | 0.4 | 0.47 | 0.24 | 0.49 | 0.36 | 0.31 |
| 9 | 130936003 | rs7847709 | -1.274 | -1.253 | 1.542 | 0.199 | 0.039 | 0.063 | 0.063 | 0.515 | 0.4 | 0.69 | 0.22 | 0.17 | 0.29 | 0.25 |
| 9 | 130936530 | rs10987908 | 1.098 | -1.461 | 2.029 | -0.183 | 0 | -0.002 | -0.002 | 0 | 0.58 | 0.44 | 0.72 | 0.54 | 0.61 | 0.65 |
| 9 | 130937738 | rs870451 | 0.736 | -1.644 | 2.087 | -0.537 | 0.004 | -0.034 | -0.034 | 0 | 0.64 | 0.69 | 0.78 | 0.99 | 0.65 | 0.83 |
| 9 | 130940822 | rs3892074 | 1.274 | -1.577 | 2.224 | -0.472 | 0.007 | -0.035 | -0.035 | 0.983 | 0.57 | 0.42 | 0.76 | 0.94 | 0.65 | 0.77 |
| 9 | 130945298 | rs10987911 | -0.718 | -1.252 | 2.235 | -0.306 | -0.004 | -0.034 | -0.034 | 1.861 | 0.25 | 0.23 | 0.2 | 0.01 | 0.33 | 0.13 |
| 9 | 130967288 | rs3003569 | 1.853 | -1.234 | 2.271 | 0.272 | 0.046 | 0.474 | 0.474 | 0.342 | 0.91 | 0.6 | 0.91 | 0.62 | 0.94 | 0.82 |
| 9 | 130971586 | rs2502817 | 1.537 | -0.872 | 1.314 | -0.534 | -0.001 | -0.158 | -0.158 | 0 | 0.65 | 0.52 | 0.75 | 0.99 | 0.65 | 0.78 |

| | | | | | | | | | | | | | | | | |
|---|---|---|---|---|---|---|---|---|---|---|---|---|---|---|---|---|
| 9 | 130979181 | rs3003599 | 0.592 | -0.901 | 0.793 | -0.34 | -0.001 | -0.033 | -0.033 | 0 | 0.83 | 0.81 | 0.85 | 1 | 0.8 | 0.93 |
| 9 | 130984755 | rs3003609 | 1.569 | -1.324 | -0.386 | -0.112 | -0.063 | -0.052 | -0.052 | 0 | 0.48 | 0.06 | 0.63 | 0.61 | 0.56 | 0.6 |
| 9 | 130994179 | rs3003578 | 1.509 | -1.415 | -0.262 | -0.044 | -0.003 | 0.005 | 0.005 | 0 | 0.49 | 0.09 | 0.64 | 0.62 | 0.57 | 0.54 |
| 9 | 130997868 | rs13285411 | -0.955 | -1.465 | 0.755 | 0.362 | -0.007 | 0.095 | 0.095 | 0.006 | 0.15 | 0.23 | 0.11 | 0.3 | 0.12 | 0.29 |
| 9 | 130998850 | rs9644916 | -1.661 | -1.519 | 0.706 | 0.311 | -0.002 | 0.004 | 0.004 | 0 | 0.26 | 0.26 | 0.17 | 0.33 | 0.2 | 0.38 |
| 9 | 131003677 | rs1004464 | -1.515 | -1.554 | 0.75 | 0.505 | -0.009 | -0.011 | -0.011 | 0 | 0.22 | 0.23 | 0.12 | 0.12 | 0.15 | 0.27 |
| 9 | 131017962 | rs7023913 | -2.538 | -1.528 | 1.1 | 1.029 | 0.018 | -0.053 | -0.053 | 0 | 0.34 | 0.77 | 0.32 | 0.51 | 0.27 | 0.47 |
| 9 | 131018639 | rs4734 | -1.453 | -1.511 | 1.397 | 1.079 | -0.001 | -0.009 | -0.009 | 0 | 0.19 | 0.14 | 0.13 | 0.21 | 0.17 | 0.27 |
| 9 | 131039250 | rs2240960 | -1.812 | -1.645 | 1.901 | 1.192 | -0.013 | 0.065 | 0.065 | 0 | 0.21 | 0.51 | 0.15 | 0.31 | 0.15 | 0.27 |
| 9 | 131042734 | rs2270204 | -2.709 | -1.76 | 1.315 | 1.089 | 0.039 | -0.03 | -0.03 | 0 | 0.38 | 0.77 | 0.33 | 0.57 | 0.28 | 0.42 |
| 9 | 131055525 | rs10114273 | 0.305 | -2.048 | 1.43 | 0.465 | 0.014 | 0.016 | 0.016 | 0 | 0.06 | 0.03 | 0.03 | 0 | 0.05 | 0.02 |
| 9 | 131087837 | rs16930351 | 0.116 | -2.058 | 1.93 | 0.458 | -0.004 | -0.017 | -0.017 | 0.009 | 0.07 | 0.06 | 0.02 | 0 | 0.04 | 0.03 |
| 9 | 131088464 | rs3003600 | -0.386 | -2.043 | 1.759 | 0.337 | -0.01 | 0.009 | 0.009 | 0 | 0.84 | 0.85 | 0.88 | 1 | 0.77 | 0.95 |
| 9 | 131095112 | rs7850697 | -2.886 | -2.244 | 2.362 | 0.98 | 0.052 | 0.007 | 0.007 | 0 | 0.34 | 0.75 | 0.2 | 0.3 | 0.2 | 0.26 |
| 9 | 131101919 | rs7030121 | -2.862 | -2.23 | 2.397 | 0.987 | 0.048 | 0.006 | 0.006 | 0 | 0.33 | 0.75 | 0.2 | 0.3 | 0.2 | 0.26 |
| 9 | 131110443 | rs10987965 | -1.745 | -2.231 | 2.427 | 0.951 | -0.053 | -0.102 | -0.102 | 0.415 | 0.15 | 0.48 | 0.07 | 0 | 0.06 | 0.06 |
| 9 | 131118222 | rs2003560 | -1.248 | -2.279 | 2.233 | 0.835 | 0.009 | 0.09 | 0.09 | 0 | 0.15 | 0.01 | 0.06 | 0.01 | 0.09 | 0.06 |
| 9 | 131120452 | rs6478827 | -1.795 | -2.203 | 2.375 | 0.903 | -0.054 | -0.103 | -0.103 | 0.231 | 0.15 | 0.48 | 0.07 | 0 | 0.06 | 0.06 |
| 9 | 131123542 | rs7048106 | -2.661 | -2.18 | 2.007 | 0.976 | 0.033 | 0.003 | 0.003 | 0 | 0.34 | 0.76 | 0.22 | 0.32 | 0.24 | 0.35 |
| 9 | 131129207 | rs10124204 | -1.355 | -2.168 | 2.609 | 1.161 | 0.026 | 0.005 | 0.005 | 1.026 | 0.08 | 0.43 | 0.05 | 0.01 | 0.02 | 0.06 |
| 9 | 131132758 | rs7019382 | -1.442 | -2.135 | 2.218 | 1.09 | 0.022 | 0.103 | 0.103 | 0.024 | 0.2 | 0.05 | 0.07 | 0.01 | 0.15 | 0.06 |
| 9 | 131133983 | rs2286814 | -2.19 | -2.001 | 2.193 | 1.201 | 0.026 | 0.005 | 0.005 | 0 | 0.32 | 0.73 | 0.19 | 0.3 | 0.22 | 0.26 |
| 9 | 131169528 | rs12351561 | -1.679 | -1.983 | 1.782 | 1.314 | -0.056 | 0.007 | 0.007 | 0 | 0.23 | 0.46 | 0.14 | 0.18 | 0.14 | 0.19 |
| 9 | 131185961 | rs2072400 | -1.684 | -1.954 | 2.551 | 1.728 | -0.031 | 0.026 | 0.026 | 1.207 | 0.27 | 0.51 | 0.2 | 0.37 | 0.2 | 0.25 |
| 9 | 131191403 | rs2270202 | -1.949 | -2.013 | 2.802 | 1.688 | 0.016 | -0.01 | -0.01 | 0 | 0.29 | 0.71 | 0.22 | 0.38 | 0.21 | 0.26 |
| 9 | 131196695 | rs7258 | -1.652 | -1.978 | 3.08 | 1.871 | -0.021 | 0.041 | 0.041 | 1.655 | 0.25 | 0.37 | 0.18 | 0.41 | 0.21 | 0.25 |
| 9 | 131199576 | rs6918 | -1.847 | -1.821 | 3.218 | 1.948 | -0.026 | 0.038 | 0.038 | 0 | 0.24 | 0.45 | 0.2 | 0.35 | 0.22 | 0.26 |
| 9 | 131201895 | rs4837264 | -1.937 | -1.799 | 3.197 | 1.929 | -0.023 | 0.032 | 0.032 | 0 | 0.25 | 0.45 | 0.2 | 0.34 | 0.24 | 0.26 |
| 9 | 131210643 | rs7870167 | -1.872 | -2.217 | 3.028 | 1.767 | -0.046 | 0.023 | 0.023 | 0 | 0.29 | 0.45 | 0.21 | 0.38 | 0.24 | 0.34 |
| 9 | 131212372 | rs7849186 | 2.342 | -2.277 | 3.29 | 1.736 | 0.008 | 0 | 0 | 0.018 | 0.66 | 0.38 | 0.78 | 0.62 | 0.74 | 0.66 |
| 9 | 131224189 | rs7032246 | -1.968 | -2.294 | 3.311 | 1.935 | -0.047 | 0.043 | 0.043 | 0 | 0.29 | 0.47 | 0.25 | 0.48 | 0.24 | 0.36 |
| 9 | 131236201 | rs12347774 | 0.167 | -2.538 | 3.3 | 2.593 | -0.021 | 0.065 | 0.065 | 0.833 | 0.05 | 0.1 | 0.04 | 0.25 | 0 | 0.05 |
| 9 | 131285955 | rs2275260 | -1.726 | -2.094 | 3.398 | 2.196 | -0.035 | 0.04 | 0.04 | 0 | 0.28 | 0.38 | 0.24 | 0.48 | 0.25 | 0.35 |
| 9 | 131287352 | rs10988014 | 2.107 | -2.099 | 3.387 | 2.193 | 0.029 | -0.007 | -0.007 | 0.153 | 0.65 | 0.3 | 0.72 | 0.52 | 0.74 | 0.64 |
| 9 | 131299228 | rs7871140 | -2.359 | -2.087 | 3.243 | 2.2 | -0.016 | 0.004 | 0.004 | 0 | 0.34 | 0.55 | 0.26 | 0.48 | 0.26 | 0.35 |
| 9 | 131303522 | rs10760563 | -1.509 | -1.668 | 3.032 | 2.332 | 0.016 | -0.017 | -0.017 | 0 | 0.26 | 0.19 | 0.18 | 0.38 | 0.24 | 0.34 |
| 9 | 131311128 | rs10513497 | -1.701 | -1.71 | 3.056 | 2.412 | -0.03 | 0.024 | 0.024 | 0 | 0.27 | 0.38 | 0.2 | 0.39 | 0.24 | 0.34 |
| 9 | 131311333 | rs11999274 | -1.759 | -1.69 | 3.055 | 2.402 | -0.032 | 0.016 | 0.016 | 0 | 0.28 | 0.4 | 0.2 | 0.38 | 0.24 | 0.34 |
| 9 | 131327982 | rs10819410 | -1.458 | -1.26 | 3.235 | 3.124 | -0.011 | 0.048 | 0.048 | 0 | 0.16 | 0.25 | 0.17 | 0.42 | 0.11 | 0.24 |
| 9 | 131331320 | rs2152833 | 2.112 | -1.313 | 3.488 | 3.161 | -0.033 | 0.071 | 0.071 | 0 | 0.79 | 0.34 | 0.78 | 0.51 | 0.88 | 0.72 |
| 9 | 131334744 | rs10819411 | -1.712 | -1.262 | 3.525 | 3.186 | -0.029 | 0.101 | 0.101 | 0 | 0.18 | 0.35 | 0.18 | 0.43 | 0.12 | 0.24 |
| 9 | 131380225 | rs7864187 | -2.006 | -1.282 | 3.419 | 2.901 | -0.043 | 0.149 | 0.149 | 0 | 0.19 | 0.47 | 0.2 | 0.43 | 0.12 | 0.24 |
| 9 | 131397479 | rs4837291 | 2.11 | -1.26 | 2.422 | 2.944 | -0.039 | 0.126 | 0.126 | 0 | 0.8 | 0.37 | 0.79 | 0.49 | 0.87 | 0.72 |
| 9 | 131398636 | rs10760570 | 2.032 | -1.226 | 1.699 | 2.848 | -0.019 | 0.04 | 0.04 | 0 | 0.79 | 0.28 | 0.78 | 0.51 | 0.87 | 0.72 |
| 9 | 131403096 | rs2298045 | 1.554 | -0.901 | 0.573 | 2.866 | -0.018 | 0.064 | 0.064 | 0 | 0.83 | 0.69 | 0.82 | 0.61 | 0.87 | 0.76 |
| 9 | 131406739 | rs4397511 | 1.241 | -0.906 | 0.577 | 2.962 | -0.035 | 0.053 | 0.053 | 0 | 0.86 | 0.71 | 0.85 | 0.7 | 0.93 | 0.82 |
| 9 | 131406896 | rs16936868 | -0.487 | -0.613 | -0.526 | 2.684 | 0.023 | -0.066 | -0.066 | 1.613 | 0.11 | 0.01 | 0.12 | 0.2 | 0.06 | 0.11 |
| 9 | 131468740 | rs9697210 | 0.018 | 0.145 | -0.643 | 1.758 | -0.005 | 0.012 | 0.012 | 0 | 0.13 | 0.06 | 0.11 | 0.08 | 0.16 | 0.13 |
| 9 | 131482467 | rs3750325 | 0.599 | -0.689 | 0.495 | 1.624 | -0.043 | 0.07 | 0.07 | 1.753 | 0.89 | 0.54 | 0.94 | 0.92 | 0.96 | 0.93 |
| 9 | 131484526 | rs10988105 | 0.851 | -1.333 | -0.897 | 1.191 | -0.125 | 0.088 | 0.088 | 0 | 0.35 | 0.05 | 0.52 | 0.11 | 0.5 | 0.32 |
| 9 | 131493140 | rs1129169 | 0.641 | -1.394 | -0.321 | 1.467 | -0.07 | -0.056 | -0.056 | 1.234 | 0.37 | 0.06 | 0.53 | 0.27 | 0.55 | 0.39 |
| 9 | 131515547 | rs8507 | 0.566 | -1.442 | -0.256 | 1.524 | -0.006 | 0.021 | 0.021 | 0 | 0.38 | 0.17 | 0.54 | 0.24 | 0.55 | 0.39 |
| 9 | 131516838 | rs4836625 | 0.767 | -1.445 | -0.473 | 1.478 | -0.125 | 0.092 | 0.092 | 0 | 0.35 | 0.06 | 0.53 | 0.1 | 0.5 | 0.33 |
| 9 | 131527676 | rs10739734 | -0.098 | -1.535 | -0.38 | 1.416 | 0.015 | -0.002 | -0.002 | 0 | 0.52 | 0.55 | 0.41 | 0.63 | 0.41 | 0.54 |
| 9 | 131555306 | rs4836628 | 0.098 | -1.337 | -0.055 | 1.328 | -0.038 | 0.066 | 0.066 | 2.369 | 0.89 | 0.62 | 0.94 | 0.91 | 0.96 | 0.92 |
| 9 | 131562457 | rs2417126 | -0.248 | -2.259 | -0.988 | 1.139 | 0.064 | 0.063 | 0.063 | 0.251 | 0.33 | 0.81 | 0.22 | 0.26 | 0.25 | 0.41 |
| 9 | 131565554 | rs1572912 | -0.703 | -2.329 | -1.067 | 1.197 | 0.1 | -0.015 | -0.015 | 0 | 0.41 | 0.92 | 0.26 | 0.36 | 0.29 | 0.48 |
| 9 | 131571568 | rs11999937 | 0.423 | -2.327 | -0.719 | 1.189 | 0.01 | -0.012 | -0.012 | 0 | 0.08 | 0.07 | 0.04 | 0.1 | 0.04 | 0.08 |
| 9 | 131572027 | rs15676 | 0.515 | -2.388 | -0.778 | 1.195 | 0.107 | 0.019 | 0.019 | 0 | 0.59 | 0.17 | 0.75 | 0.64 | 0.71 | 0.52 |
| 9 | 131572215 | rs3829901 | 0.472 | -2.361 | -0.773 | 1.192 | 0.01 | -0.012 | -0.012 | 0 | 0.08 | 0.07 | 0.04 | 0.1 | 0.04 | 0.08 |
| 9 | 131580744 | rs2997922 | 0.558 | -2.424 | -0.803 | 1.197 | 0.105 | 0.023 | 0.023 | 0.114 | 0.59 | 0.17 | 0.74 | 0.64 | 0.71 | 0.52 |
| 9 | 131582104 | rs10819449 | -0.587 | -2.46 | -0.666 | 1.428 | -0.011 | 0.078 | 0.078 | 0.056 | 0.12 | 0.24 | 0.06 | 0.26 | 0.1 | 0.16 |
| 9 | 131582625 | rs2977996 | -0.02 | -2.468 | -0.631 | 1.214 | 0.026 | 0.023 | 0.023 | 0.234 | 0.67 | 0.3 | 0.79 | 0.73 | 0.75 | 0.6 |
| 9 | 131584631 | rs2280844 | 0.347 | -2.473 | -0.633 | 1.213 | -0.015 | 0.006 | 0.006 | 0.378 | 0.08 | 0.13 | 0.04 | 0.09 | 0.04 | 0.08 |
| 9 | 131585069 | rs2280843 | -0.012 | -2.477 | -0.634 | 1.205 | 0.026 | 0.023 | 0.023 | 0.234 | 0.67 | 0.31 | 0.79 | 0.73 | 0.75 | 0.59 |
| 9 | 131585873 | rs7029309 | 0.421 | -2.471 | -0.64 | 1.224 | -0.016 | -0.003 | -0.003 | 1.454 | 0.08 | 0.13 | 0.04 | 0.05 | 0.04 | 0.03 |
| 9 | 131586374 | rs7033070 | 0.033 | -2.48 | -0.171 | 1.212 | -0.026 | -0.022 | -0.022 | 0.787 | 0.31 | 0.56 | 0.2 | 0.27 | 0.25 | 0.41 |
| 9 | 131592150 | rs2293964 | 0.232 | -2.45 | -0.338 | 1.235 | -0.015 | 0.006 | 0.006 | 0.378 | 0.08 | 0.13 | 0.04 | 0.09 | 0.04 | 0.08 |
| 9 | 131595407 | rs10988134 | -0.115 | -2.468 | -0.347 | 1.218 | 0.011 | 0.01 | 0.01 | 0.078 | 0.32 | 0.64 | 0.21 | 0.27 | 0.25 | 0.41 |

| | | | | | | | | | | | | | | | | |
|---|---|---|---|---|---|---|---|---|---|---|---|---|---|---|---|---|
| 9 | 131609974 | rs3763597 | 0.28 | -2.186 | -0.29 | 1.241 | 0.003 | -0.005 | -0.005 | 1.026 | 0.08 | 0.09 | 0.04 | 0.09 | 0.04 | 0.08 |
| 9 | 131644307 | rs1923619 | 0.359 | -0.9 | -0.443 | 1.081 | -0.001 | 0.001 | 0.001 | 1.278 | 0.07 | 0.08 | 0.03 | 0.05 | 0.04 | 0.03 |
| 9 | 131670457 | rs3750320 | -0.509 | -2.018 | -0.133 | 0.362 | -0.039 | -0.021 | -0.021 | 0 | 0.29 | 0.43 | 0.15 | 0.17 | 0.2 | 0.28 |
| 9 | 131670919 | rs3750319 | 0.112 | -1.204 | -0.659 | 0.654 | 0.011 | -0.006 | -0.006 | 2.162 | 0.7 | 0.23 | 0.74 | 0.67 | 0.72 | 0.66 |
| 9 | 131672558 | rs10988151 | 0.299 | -1.024 | -0.852 | 0.817 | -0.004 | -0.088 | -0.088 | 1.018 | 0.69 | 0.08 | 0.71 | 0.62 | 0.72 | 0.61 |
| 9 | 131675360 | rs11789878 | 0.244 | -1.025 | -0.85 | 0.801 | 0.005 | -0.046 | -0.046 | 0 | 0.69 | 0.12 | 0.71 | 0.64 | 0.72 | 0.61 |
| 9 | 131676550 | rs16930764 | 2.09 | -0.346 | -1.261 | 0.411 | -0.03 | 0.029 | 0.029 | 0 | 0.06 | 0 | 0.02 | 0 | 0.06 | 0.06 |
| 9 | 131679892 | rs9031 | 0.048 | -0.366 | -1.274 | 0.456 | -0.003 | -0.069 | -0.069 | 0 | 0.69 | 0.08 | 0.71 | 0.64 | 0.72 | 0.61 |
| 9 | 131680177 | rs11542344 | 1.461 | 0.249 | -0.36 | 0.886 | -0.03 | 0.029 | 0.029 | 0 | 0.06 | 0 | 0.02 | 0 | 0.06 | 0.06 |
| 9 | 131687338 | rs7048932 | 1.375 | 0.291 | 0.668 | 1.614 | 0 | -0.023 | -0.023 | 0 | 0.45 | 0.42 | 0.44 | 0.24 | 0.41 | 0.35 |
| 9 | 131699571 | rs28365521 | 1.192 | 0.249 | 0.622 | 1.71 | -0.005 | -0.008 | -0.008 | 0.022 | 0.05 | 0.01 | 0.02 | 0.07 | 0.07 | 0.15 |
| 9 | 131702891 | rs2273866 | 1.523 | 0.263 | 0.852 | 1.627 | 0 | 0.025 | 0.025 | 0 | 0.45 | 0.36 | 0.43 | 0.24 | 0.41 | 0.35 |
| 9 | 131705763 | rs17432839 | 1.803 | 0.237 | 0.79 | 1.927 | -0.066 | 0.18 | 0.18 | 0.07 | 0.42 | 0.1 | 0.4 | 0.2 | 0.41 | 0.31 |
| 9 | 131737339 | rs10760583 | -0.053 | 0.317 | 1.049 | 2.44 | -0.053 | 0.06 | 0.06 | 0 | 0.69 | 0.94 | 0.77 | 0.88 | 0.68 | 0.81 |
| 9 | 131745548 | rs719703 | 2.033 | 0.308 | 1.112 | 2.588 | -0.076 | 0.13 | 0.13 | 0.051 | 0.42 | 0.09 | 0.4 | 0.25 | 0.41 | 0.31 |
| 9 | 131761899 | rs2302811 | 1.992 | 0.375 | 0.956 | 2.34 | -0.074 | 0.138 | 0.138 | 0.088 | 0.42 | 0.09 | 0.4 | 0.22 | 0.43 | 0.36 |
| 9 | 131767668 | rs2287363 | 1.449 | 0.658 | 1.067 | 2.647 | 0 | 0.005 | 0.005 | 0 | 0.49 | 0.47 | 0.5 | 0.41 | 0.47 | 0.42 |
| 9 | 131768191 | rs17485667 | 0.259 | 0.633 | 1.004 | 2.553 | -0.004 | 0.016 | 0.016 | 0 | 0.08 | 0.2 | 0.04 | 0.1 | 0.07 | 0.15 |
| 9 | 131772389 | rs3750340 | 0.034 | 0.828 | 1.05 | 2.348 | -0.043 | -0.03 | -0.03 | 0 | 0.93 | 0.99 | 0.79 | 0.66 | 0.85 | 0.84 |
| 9 | 131776041 | rs10819457 | 1.279 | 0.761 | 1.265 | 1.365 | -0.001 | -0.009 | -0.009 | 0 | 0.48 | 0.45 | 0.49 | 0.37 | 0.44 | 0.42 |
| 9 | 131788926 | rs7874044 | 1.092 | 0.207 | 1.177 | 1.343 | -0.006 | 0.032 | 0.032 | 0 | 0.57 | 0.62 | 0.57 | 0.36 | 0.54 | 0.44 |
| 9 | 131796961 | rs10760585 | -0.702 | 1.06 | 1.239 | 1.415 | -0.073 | 0.143 | 0.143 | 0 | 0.25 | 0.07 | 0.23 | 0 | 0.32 | 0.21 |
| 9 | 131799261 | rs11564101 | -2.979 | 0.896 | 2.292 | 2.147 | 0.024 | 0.129 | 0.129 | 0.982 | 0.08 | 0.37 | 0.27 | 0.15 | 0.05 | 0.17 |
| 9 | 131834811 | rs11564100 | -0.633 | 1.117 | 1.691 | 1.734 | 0.044 | -0.022 | -0.022 | 0 | 0.05 | 0.19 | 0.07 | 0 | 0.1 | 0.02 |
| 9 | 131838821 | rs4295766 | 1.427 | 0.642 | 1.641 | 2.54 | -0.002 | 0.006 | 0.006 | 0 | 0.89 | 0.89 | 0.76 | 0.76 | 0.83 | 0.76 |
| 9 | 131841887 | rs10819461 | 1.438 | 0.469 | 1.452 | 2.376 | -0.006 | -0.036 | -0.036 | 0.534 | 0.76 | 0.88 | 0.7 | 0.75 | 0.75 | 0.74 |
| 9 | 131846125 | rs10988198 | 2.223 | 0.241 | 2.222 | 2.544 | 0.027 | -0.014 | -0.014 | 0.76 | 0.65 | 0.29 | 0.63 | 0.63 | 0.73 | 0.66 |
| 9 | 131873222 | rs3814492 | 2.084 | 0.821 | 2.684 | 2.318 | 0.001 | -0.002 | -0.002 | 0 | 0.7 | 0.34 | 0.63 | 0.62 | 0.73 | 0.67 |
| 9 | 131873256 | rs3118625 | -1.507 | 0.882 | 2.121 | 2.281 | -0.018 | -0.048 | -0.048 | 0.426 | 0.25 | 0.1 | 0.31 | 0.27 | 0.27 | 0.27 |
| 9 | 131884802 | rs2541161 | -1.761 | 0.895 | 2.286 | 2.262 | 0.001 | -0.014 | -0.014 | 0 | 0.27 | 0.25 | 0.34 | 0.39 | 0.27 | 0.33 |
| 9 | 131923949 | rs4837345 | -1.584 | 1.003 | 2.408 | 1.828 | 0.004 | -0.018 | -0.018 | 5.968 | 0.25 | 0.34 | 0.21 | 0.15 | 0.28 | 0.25 |
| 9 | 131932940 | rs10739743 | 2.025 | 0.946 | 2.732 | 2.062 | 0.002 | -0.013 | -0.013 | 0 | 0.7 | 0.28 | 0.65 | 0.63 | 0.71 | 0.64 |
| 9 | 131938913 | rs1075650 | -1.393 | 0.883 | 2.686 | 1.92 | -0.014 | 0.013 | 0.013 | 0 | 0.24 | 0.14 | 0.19 | 0.15 | 0.28 | 0.25 |
| 9 | 131941309 | rs7034195 | 1.816 | 0.908 | 2.714 | 1.925 | -0.003 | 0.024 | 0.024 | 0 | 0.71 | 0.44 | 0.67 | 0.66 | 0.71 | 0.62 |
| 9 | 131944138 | rs913264 | -1.541 | 0.779 | 2.601 | 1.876 | 0.001 | 0.004 | 0.004 | 0 | 0.26 | 0.38 | 0.2 | 0.28 | 0.27 | 0.28 |
| 9 | 131948373 | rs7871824 | 1.796 | 0.781 | 2.003 | 1.857 | 0.002 | -0.005 | -0.005 | 1.906 | 0.71 | 0.33 | 0.67 | 0.65 | 0.72 | 0.62 |
| 9 | 131953598 | rs10760593 | -1.295 | 0.679 | 1.454 | 1.678 | 0.009 | 0.009 | 0.009 | 0 | 0.23 | 0.19 | 0.12 | 0.14 | 0.17 | 0.22 |
| 9 | 131956834 | rs6478869 | -1.611 | 1.022 | 1.744 | 1.538 | 0.075 | -0.057 | -0.057 | 0 | 0.17 | 0.48 | 0.27 | 0.07 | 0.29 | 0.16 |
| 9 | 131960107 | rs422134 | -1.529 | 0.98 | 1.632 | 1.493 | -0.002 | -0.003 | -0.003 | 0.003 | 0.29 | 0.28 | 0.26 | 0.21 | 0.26 | 0.33 |
| 9 | 131960131 | rs4836643 | 2.396 | 0.97 | 1.334 | 1.718 | -0.021 | 0.011 | 0.011 | 0.564 | 0.57 | 0.08 | 0.48 | 0.69 | 0.54 | 0.56 |
| 9 | 131963745 | rs10988248 | 2.475 | 1.016 | 1.16 | 1.8 | -0.048 | -0.072 | -0.072 | 0 | 0.56 | 0.06 | 0.46 | 0.58 | 0.54 | 0.54 |
| 9 | 131964881 | rs10760594 | -1.493 | 1.085 | 1.35 | 1.805 | 0.006 | 0.013 | 0.013 | 0 | 0.29 | 0.23 | 0.25 | 0.21 | 0.26 | 0.32 |
| 9 | 131969556 | rs7856501 | 2.341 | 1.081 | 1.188 | 1.823 | -0.02 | -0.05 | -0.05 | 0.022 | 0.56 | 0.11 | 0.46 | 0.57 | 0.54 | 0.52 |
| 9 | 131980567 | rs10988252 | -1.441 | 1.44 | 1.548 | 1.885 | 0.001 | -0.004 | -0.004 | 0 | 0.26 | 0.22 | 0.24 | 0.24 | 0.19 | 0.24 |
| 9 | 131982125 | rs2479030 | 1.938 | 1.44 | 1.402 | 1.677 | -0.046 | -0.075 | -0.075 | 0 | 0.62 | 0.1 | 0.5 | 0.6 | 0.62 | 0.6 |
| 9 | 131984217 | rs7873242 | -0.961 | 1.455 | 1.464 | 1.54 | 0.005 | 0.019 | 0.019 | 0 | 0.16 | 0.17 | 0.33 | 0.34 | 0.22 | 0.32 |
| 9 | 131984706 | rs4836644 | -0.875 | 1.163 | 1.208 | 1.526 | -0.029 | -0.118 | -0.118 | 0.143 | 0.18 | 0.39 | 0.14 | 0 | 0.16 | 0.04 |
| 9 | 131986809 | rs2479028 | 1.202 | 1.318 | 0.033 | 1.758 | -0.078 | -0.033 | -0.033 | 0 | 0.59 | 0.1 | 0.47 | 0.63 | 0.57 | 0.56 |
| 9 | 131992861 | rs716548 | 1.072 | 0.954 | 1.436 | 2.396 | 0.056 | 0.216 | 0.216 | 0 | 0.95 | 0.79 | 0.96 | 0.7 | 0.96 | 0.89 |
| 9 | 131993390 | rs1411080 | -1.146 | 0.65 | 1.867 | 2.604 | -0.011 | -0.006 | -0.006 | 0 | 0.23 | 0.3 | 0.27 | 0.17 | 0.23 | 0.18 |
| 9 | 131995919 | rs12352047 | -1.284 | 0.583 | 1.872 | 3.038 | -0.003 | 0.002 | 0.002 | 0 | 0.23 | 0.26 | 0.29 | 0.29 | 0.23 | 0.21 |
| 9 | 131996071 | rs395119 | 1.903 | 0.546 | 2.415 | 2.412 | 0.008 | -0.016 | -0.016 | 0 | 0.69 | 0.32 | 0.67 | 0.53 | 0.73 | 0.69 |
| 9 | 131996947 | rs7048616 | -1.066 | 0.559 | 2.137 | 2.35 | -0.004 | -0.007 | -0.007 | 0 | 0.22 | 0.25 | 0.26 | 0.13 | 0.23 | 0.12 |
| 9 | 131996962 | rs9409314 | 1.771 | 0.75 | 1.51 | 2.414 | 0.005 | -0.071 | -0.071 | 0 | 0.64 | 0.15 | 0.53 | 0.41 | 0.66 | 0.58 |
| 9 | 131998296 | rs11790192 | -1.773 | 0.904 | 1.782 | 2.293 | 0.007 | 0.036 | 0.036 | 0 | 0.31 | 0.49 | 0.34 | 0.48 | 0.34 | 0.38 |
| 9 | 131998330 | rs17456493 | -0.987 | 0.751 | 1.337 | 2.103 | -0.004 | -0.003 | -0.003 | 0 | 0.05 | 0.07 | 0.13 | 0.12 | 0.09 | 0.11 |
| 9 | 131999768 | rs11794903 | -1.382 | 0.8 | 2.312 | 1.883 | -0.01 | -0.064 | -0.064 | 0.165 | 0.22 | 0.43 | 0.23 | 0.1 | 0.19 | 0.15 |
| 9 | 131999876 | rs913775 | -2.438 | 0.692 | 2.527 | 2.461 | 0.003 | -0.041 | -0.041 | 0 | 0.33 | 0.81 | 0.43 | 0.53 | 0.31 | 0.36 |
| 9 | 132002616 | rs12237856 | -1.781 | 0.689 | 2.633 | 2.627 | 0.003 | 0.072 | 0.072 | 0.122 | 0.09 | 0.14 | 0.18 | 0.45 | 0.12 | 0.21 |
| 9 | 132008809 | rs4240433 | 1.505 | 0.75 | 3.213 | 2.796 | -0.005 | -0.008 | -0.008 | 0 | 0.76 | 0.36 | 0.74 | 0.82 | 0.79 | 0.78 |
| 9 | 132009096 | rs927926 | 1.537 | 0.742 | 3.236 | 2.792 | -0.008 | -0.013 | -0.013 | 0 | 0.76 | 0.42 | 0.75 | 0.82 | 0.79 | 0.78 |
| 9 | 132009235 | rs10116285 | -1.83 | 0.756 | 2.858 | 2.646 | 0.002 | 0.062 | 0.062 | 0.107 | 0.09 | 0.14 | 0.17 | 0.42 | 0.11 | 0.19 |
| 9 | 132010706 | rs12236289 | -1.234 | 0.615 | 2.884 | 2.556 | -0.005 | 0.157 | 0.157 | 0.02 | 0.06 | 0.25 | 0.05 | 0.2 | 0.02 | 0.08 |
| 9 | 132015425 | rs10988273 | 1.555 | 0.882 | 2.789 | 2.372 | 0.016 | 0.007 | 0.007 | 0 | 0.74 | 0.38 | 0.63 | 0.67 | 0.72 | 0.73 |
| 9 | 132020431 | rs7849837 | 1.502 | 0.769 | 2.851 | 2.4 | 0.01 | 0.012 | 0.012 | 0 | 0.74 | 0.41 | 0.7 | 0.66 | 0.74 | 0.75 |
| 9 | 132032637 | rs10819484 | 1.976 | 0.881 | 3.287 | 2.717 | 0.062 | 0.195 | 0.195 | 0 | 0.82 | 0.49 | 0.68 | 0.51 | 0.77 | 0.77 |
| 9 | 132041447 | rs10988296 | -2.444 | 0.651 | 3.44 | 2.794 | -0.026 | 0.341 | 0.341 | 0.072 | 0.1 | 0.54 | 0.17 | 0.43 | 0.02 | 0.15 |
| 9 | 132042502 | rs10988298 | -1.913 | 1.276 | 3.216 | 2.02 | 0.052 | -0.027 | -0.027 | 1.131 | 0.14 | 0.29 | 0.18 | 0.11 | 0.24 | 0.13 |
| 9 | 132044938 | rs11794563 | -1.801 | 1.104 | 2.816 | 1.975 | 0.014 | -0.012 | -0.012 | 1.476 | 0.14 | 0.17 | 0.17 | 0.11 | 0.23 | 0.14 |

| | | | | | | | | | | | | | | | | |
|---|---|---|---|---|---|---|---|---|---|---|---|---|---|---|---|---|
| 9 | 132046491 | rs17456938 | -2.237 | 1.062 | 3.186 | 1.942 | 0.07 | -0.022 | -0.022 | 0.406 | 0.18 | 0.57 | 0.22 | 0.14 | 0.24 | 0.18 |
| 9 | 132046625 | rs17456945 | -1.856 | 1.142 | 2.163 | 1.827 | 0.006 | -0.011 | -0.011 | 0 | 0.15 | 0.18 | 0.17 | 0.08 | 0.23 | 0.14 |
| 9 | 132048953 | rs7029664 | 0.66 | 1.181 | 1.506 | 1.912 | -0.008 | -0.003 | -0.003 | 0 | 0.63 | 0.35 | 0.54 | 0.58 | 0.52 | 0.55 |
| 9 | 132049842 | rs4366175 | -1.48 | 1.277 | 1.63 | 2.068 | 0.056 | 0.036 | 0.036 | 0.012 | 0.15 | 0.61 | 0.31 | 0.17 | 0.22 | 0.25 |
| 9 | 132053313 | rs10988304 | -0.862 | 1.211 | 1.6 | 1.742 | 0.067 | -0.049 | -0.049 | 0 | 0.23 | 0.43 | 0.36 | 0.11 | 0.37 | 0.13 |
| 9 | 132056318 | rs10819490 | 0.74 | 0.832 | 1.283 | 1.966 | 0.047 | 0.108 | 0.108 | 1.054 | 0.55 | 0.76 | 0.56 | 0.73 | 0.36 | 0.78 |
| 9 | 132063801 | rs4599906 | -0.685 | 1.154 | 0.728 | 1.921 | -0.085 | 0.04 | 0.04 | 6.242 | 0.75 | 0.96 | 0.81 | 0.8 | 0.67 | 0.86 |
| 9 | 132064209 | rs4523365 | 0.981 | 0.18 | 0.623 | 2.018 | 0.109 | 0.147 | 0.147 | 0.342 | 0.41 | 0.14 | 0.24 | 0.15 | 0.24 | 0.36 |
| 9 | 132071242 | rs10988311 | 0.214 | 0.143 | 1.062 | 2.122 | -0.001 | 0 | 0 | 0 | 0.44 | 0.47 | 0.31 | 0.6 | 0.31 | 0.56 |
| 9 | 132072646 | rs7867597 | 0.239 | 0.145 | 0.809 | 2.115 | -0.014 | -0.004 | -0.004 | 0 | 0.44 | 0.49 | 0.32 | 0.6 | 0.32 | 0.52 |
| 9 | 132073191 | rs10819494 | 0.231 | 0.144 | 0.809 | 2.102 | -0.014 | -0.004 | -0.004 | 0 | 0.44 | 0.49 | 0.32 | 0.61 | 0.32 | 0.53 |
| 9 | 132078047 | rs10988319 | 0.115 | 0.253 | 0.413 | 1.819 | -0.004 | -0.002 | -0.002 | 0.065 | 0.43 | 0.57 | 0.35 | 0.57 | 0.33 | 0.55 |
| 9 | 132078777 | rs10760607 | 0.009 | 0.233 | 0.19 | 1.654 | 0.025 | 0.014 | 0.014 | 0 | 0.44 | 0.71 | 0.38 | 0.57 | 0.34 | 0.55 |
| 9 | 132081623 | rs10819500 | 0.034 | 0.182 | -0.135 | 1.608 | 0.057 | 0.024 | 0.024 | 0 | 0.45 | 0.8 | 0.38 | 0.6 | 0.33 | 0.56 |
| 9 | 132082564 | rs11788098 | -0.04 | 0.168 | -0.127 | 1.602 | 0.059 | 0.015 | 0.015 | 0 | 0.45 | 0.8 | 0.36 | 0.59 | 0.32 | 0.55 |
| 9 | 132084621 | rs913770 | -0.71 | 0.155 | 0.433 | 1.523 | 0.032 | 0.143 | 0.143 | 0 | 0.1 | 0.42 | 0.12 | 0.15 | 0.05 | 0.1 |
| 9 | 132085237 | rs6478890 | -0.332 | 0.474 | -1.174 | 0.958 | -0.014 | 0.037 | 0.037 | 0 | 0.59 | 0.75 | 0.67 | 0.67 | 0.57 | 0.54 |
| 9 | 132085293 | rs7034728 | 0.069 | -0.201 | -1.544 | 1.188 | 0.059 | -0.033 | -0.033 | 0.297 | 0.42 | 0.83 | 0.4 | 0.54 | 0.31 | 0.47 |
| 9 | 132085618 | rs7850090 | -0.459 | -0.263 | -1.052 | 0.976 | -0.019 | 0.013 | 0.013 | 0 | 0.15 | 0.2 | 0.12 | 0.18 | 0.15 | 0.2 |
| 9 | 132085635 | rs913772 | 0.2 | 0.198 | -2.132 | 0.528 | 0.046 | 0.124 | 0.124 | 0 | 0.5 | 0.16 | 0.5 | 0.26 | 0.57 | 0.31 |
| 9 | 132088190 | rs4837357 | -0.712 | 0.091 | -1.755 | 0.707 | 0.032 | 0.007 | 0.007 | 0 | 0.39 | 0.75 | 0.43 | 0.55 | 0.32 | 0.45 |
| 9 | 132090371 | rs2282179 | -0.272 | 0.144 | -1.14 | 1.098 | 0.007 | 0.027 | 0.027 | 0 | 0.1 | 0.15 | 0.09 | 0.12 | 0.13 | 0.1 |
| 9 | 132098060 | rs17457362 | -0.143 | 0.015 | -0.37 | 1.37 | 0.004 | 0.105 | 0.105 | 0 | 0.12 | 0.34 | 0.13 | 0.18 | 0.1 | 0.1 |
| 9 | 132098528 | rs7859961 | -1.601 | 0.496 | -1.474 | 0.892 | -0.006 | 0.116 | 0.116 | 0 | 0.59 | 0.92 | 0.62 | 0.76 | 0.61 | 0.78 |
| 9 | 132105143 | rs10760611 | -2.301 | 0.351 | -1.937 | 0.441 | -0.011 | 0.02 | 0.02 | 0 | 0.75 | 0.93 | 0.71 | 0.83 | 0.72 | 0.82 |
| 9 | 132106034 | rs17441302 | -0.093 | 0.876 | -1.425 | 0.747 | 0.009 | 0.056 | 0.056 | 0 | 0.15 | 0.22 | 0.21 | 0.34 | 0.23 | 0.15 |
| 9 | 132109628 | rs4837360 | 0.143 | 0.819 | -1.713 | 0.636 | 0 | 0 | 0 | 0 | 0.36 | 0.34 | 0.4 | 0.48 | 0.34 | 0.57 |
| 9 | 132111298 | rs10988337 | 0.641 | 0.619 | -2.089 | 0.752 | 0.009 | 0.005 | 0.005 | 0.149 | 0.21 | 0.06 | 0.2 | 0.17 | 0.18 | 0.22 |
| 9 | 132111630 | rs17441351 | 0.042 | 0.564 | -1.837 | 0.68 | -0.01 | -0.015 | -0.015 | 0 | 0.11 | 0.08 | 0.11 | 0.18 | 0.18 | 0.12 |
| 9 | 132114326 | rs10819506 | -1.091 | 0.32 | -1.931 | -0.112 | -0.004 | 0.061 | 0.061 | 0 | 0.69 | 0.93 | 0.76 | 0.83 | 0.7 | 0.78 |
| 9 | 132115179 | rs10988341 | 0.377 | 0.638 | -1.245 | 0.63 | -0.023 | -0.025 | -0.025 | 0 | 0.05 | 0 | 0.02 | 0.13 | 0.04 | 0.08 |
| 9 | 132116060 | rs2011978 | -0.235 | 0.457 | -0.899 | 0.454 | -0.01 | -0.016 | -0.016 | 0 | 0.33 | 0.26 | 0.46 | 0.6 | 0.42 | 0.52 |
| 9 | 132127719 | rs4507869 | -1.289 | -0.347 | -0.06 | 0.294 | -0.056 | -0.002 | -0.002 | 0 | 0.15 | 0.67 | 0.06 | 0.1 | 0.04 | 0.12 |
| 9 | 132130971 | rs7868621 | -0.7 | -0.349 | 0.896 | 0.296 | -0.011 | 0.077 | 0.077 | 0.475 | 0.08 | 0.32 | 0.04 | 0.1 | 0.03 | 0.06 |
| 9 | 132131098 | rs10988346 | 0.827 | -0.276 | 0.065 | -0.367 | -0.044 | 0.107 | 0.107 | 0 | 0.21 | 0.02 | 0.29 | 0.06 | 0.25 | 0.08 |
| 9 | 132142411 | rs28361625 | -1.804 | -0.885 | 1.056 | 0.304 | -0.057 | 0.044 | 0.044 | 1.593 | 0.14 | 0.57 | 0.06 | 0.13 | 0.04 | 0.12 |
| 9 | 132145524 | rs10819507 | -0.144 | -0.565 | 1.171 | 0.391 | -0.006 | -0.009 | -0.009 | 0 | 0.61 | 0.47 | 0.66 | 0.73 | 0.65 | 0.61 |
| 9 | 132147812 | rs3853486 | 0.76 | -0.724 | 1.936 | 1.004 | 0.013 | -0.023 | -0.023 | 0 | 0.56 | 0.19 | 0.54 | 0.5 | 0.61 | 0.41 |
| 9 | 132147913 | rs10819508 | -0.123 | -0.768 | 1.946 | 1.017 | 0.035 | 0.076 | 0.076 | 0 | 0.07 | 0.22 | 0.14 | 0.2 | 0.09 | 0.21 |
| 9 | 132150169 | rs7047362 | 0.046 | -0.816 | 2.094 | 0.997 | -0.01 | 0 | 0 | 0 | 0.63 | 0.63 | 0.7 | 0.7 | 0.7 | 0.62 |
| 9 | 132150442 | rs4090173 | 0.203 | -0.455 | 2.1 | 1.173 | -0.012 | 0.004 | 0.004 | 0 | 0.65 | 0.49 | 0.69 | 0.7 | 0.7 | 0.61 |
| 9 | 132150995 | rs11789821 | -0.391 | -0.428 | 1.583 | 0.353 | -0.003 | -0.007 | -0.007 | 0.946 | 0.7 | 0.73 | 0.74 | 0.82 | 0.74 | 0.68 |
| 9 | 132155664 | rs10819510 | 0.847 | -0.438 | 1.093 | 0.368 | 0.013 | 0.019 | 0.019 | 0 | 0.55 | 0.2 | 0.55 | 0.61 | 0.62 | 0.44 |
| 9 | 132157537 | rs12684332 | 0.746 | -0.423 | 1.008 | 0.375 | 0.014 | 0.024 | 0.024 | 0 | 0.55 | 0.18 | 0.55 | 0.61 | 0.62 | 0.42 |
| 9 | 132159786 | rs12345504 | -1.429 | 0.275 | 1.244 | 0.523 | 0.043 | 0.064 | 0.064 | 0.015 | 0.14 | 0.53 | 0.19 | 0.21 | 0.12 | 0.24 |
| 9 | 132162419 | rs10118507 | -0.22 | 0.629 | 0.408 | 0.837 | 0.041 | 0.082 | 0.082 | 0 | 0.06 | 0.18 | 0.11 | 0.19 | 0.09 | 0.2 |
| 9 | 132162470 | rs10118527 | -0.316 | 0.618 | 0.358 | 0.837 | 0.036 | 0.07 | 0.07 | 0 | 0.06 | 0.18 | 0.11 | 0.19 | 0.09 | 0.2 |
| 9 | 132163254 | rs4837362 | -0.084 | -0.015 | -0.185 | 0.449 | 0.002 | 0.007 | 0.007 | 0.044 | 0.75 | 0.82 | 0.77 | 0.83 | 0.75 | 0.71 |
| 9 | 132167567 | rs10988356 | -1.194 | 0.393 | -0.119 | 0.542 | -0.057 | 0.023 | 0.023 | 0 | 0.13 | 0.02 | 0.12 | 0.05 | 0.18 | 0.14 |
| 9 | 132167952 | rs9918961 | -1.842 | -0.198 | 0.735 | 0.972 | -0.03 | 0.017 | 0.017 | 0.629 | 0.24 | 0.65 | 0.3 | 0.3 | 0.18 | 0.35 |
| 9 | 132168169 | rs7048833 | -1.199 | -0.315 | 1.14 | 0.574 | 0.001 | -0.054 | -0.054 | 0 | 0.17 | 0.38 | 0.15 | 0.05 | 0.17 | 0.3 |
| 9 | 132170673 | rs7046787 | -2.088 | -0.608 | -0.117 | 0.817 | 0.034 | -0.11 | -0.11 | 0 | 0.38 | 0.89 | 0.39 | 0.51 | 0.32 | 0.55 |
| 9 | 132176751 | rs3928291 | -0.832 | -0.579 | -0.807 | 0.932 | -0.031 | 0.008 | 0.008 | 0 | 0.19 | 0.01 | 0.13 | 0.19 | 0.18 | 0.36 |
| 9 | 132177003 | rs17441968 | -0.399 | -0.435 | -0.514 | 1.053 | -0.002 | 0.105 | 0.105 | 0 | 0.07 | 0.11 | 0.14 | 0.27 | 0.1 | 0.13 |
| 9 | 132177381 | rs1105551 | -1.241 | -0.477 | -0.717 | 1.009 | 0.032 | 0.055 | 0.055 | 2.219 | 0.31 | 0.79 | 0.26 | 0.23 | 0.22 | 0.42 |
| 9 | 132178213 | rs1810914 | -1.199 | -0.147 | -0.503 | 1.247 | -0.008 | 0.115 | 0.115 | 0 | 0.1 | 0.2 | 0.18 | 0.31 | 0.12 | 0.14 |
| 9 | 132180229 | rs4837365 | -0.759 | 0.185 | -1.449 | 0.841 | -0.013 | -0.064 | -0.064 | 0.045 | 0.42 | 0.92 | 0.38 | 0.43 | 0.36 | 0.52 |
| 9 | 132188431 | rs10819516 | -0.632 | 0.205 | -0.921 | 0.887 | 0.02 | -0.009 | -0.009 | 0 | 0.41 | 0.75 | 0.39 | 0.43 | 0.37 | 0.53 |
| 9 | 132191452 | rs9657682 | 0.155 | 0.39 | -0.744 | 0.933 | 0.002 | 0.007 | 0.007 | 1.681 | 0.59 | 0.51 | 0.49 | 0.37 | 0.56 | 0.61 |
| 9 | 132192986 | rs4837366 | -0.561 | 0.165 | -1.17 | 1.158 | -0.01 | -0.074 | -0.074 | 0 | 0.18 | 0.07 | 0.38 | 0.46 | 0.2 | 0.21 |
| 9 | 132194565 | rs10988373 | 0.762 | 0.342 | -1.753 | 0.629 | -0.002 | 0.034 | 0.034 | 0 | 0.28 | 0.18 | 0.3 | 0.12 | 0.24 | 0.33 |
| 9 | 132196531 | rs914708 | -0.206 | 0.242 | -1.303 | 0.611 | 0.001 | 0.049 | 0.049 | 0 | 0.49 | 0.67 | 0.54 | 0.7 | 0.52 | 0.47 |
| 9 | 132198135 | rs1147331 | 0.966 | 0.123 | -0.344 | 0.674 | 0.006 | -0.013 | -0.013 | 0 | 0.29 | 0.36 | 0.18 | 0.17 | 0.32 | 0.21 |
| 9 | 132198514 | rs1147332 | -0.704 | 0.209 | -0.672 | 0.721 | -0.058 | 0.019 | 0.019 | 0.383 | 0.8 | 0.96 | 0.86 | 0.87 | 0.77 | 0.85 |
| 9 | 132205249 | rs1147338 | -1.017 | 0.163 | 0.214 | 1.486 | 0.014 | 0.249 | 0.249 | 0 | 0.17 | 0.52 | 0.24 | 0.49 | 0.15 | 0.22 |
| 9 | 132218356 | rs932988 | -0.591 | -0.326 | -1.091 | 0.586 | -0.099 | 0.069 | 0.069 | 0 | 0.58 | 0.91 | 0.59 | 0.72 | 0.55 | 0.61 |
| 9 | 132232943 | rs10988396 | 0.888 | -0.056 | -1.009 | 0.244 | 0.009 | 0.073 | 0.073 | 0.038 | 0.28 | 0.16 | 0.16 | 0.09 | 0.25 | 0.14 |
| 9 | 132236708 | rs4837371 | -0.91 | -0.038 | -0.559 | 0.475 | -0.004 | 0.007 | 0.007 | 0 | 0.57 | 0.56 | 0.5 | 0.71 | 0.52 | 0.48 |
| 9 | 132239272 | rs7848479 | -0.954 | 0.038 | -0.579 | 0.523 | -0.004 | 0.002 | 0.002 | 0 | 0.57 | 0.57 | 0.54 | 0.68 | 0.54 | 0.57 |

| | | | | | | | | | | | | | | | | |
|---|---|---|---|---|---|---|---|---|---|---|---|---|---|---|---|---|
| 9 | 132241861 | rs968477 | 0.458 | 0.001 | -1.055 | 0.615 | 0.001 | -0.016 | -0.016 | 0 | 0.25 | 0.23 | 0.34 | 0.55 | 0.27 | 0.33 |
| 9 | 132253919 | rs1220795 | 0.909 | 0.155 | -1.868 | 0.362 | -0.138 | -0.092 | -0.092 | 0.157 | 0.41 | 0.05 | 0.41 | 0.35 | 0.45 | 0.42 |
| 9 | 132254665 | rs1220793 | -0.183 | 0.065 | -1.258 | 0.477 | 0 | 0.001 | 0.001 | 0 | 0.71 | 0.36 | 0.59 | 0.44 | 0.71 | 0.62 |
| 9 | 132257783 | rs1220789 | 0.533 | 0.366 | -1.819 | 0.518 | -0.123 | -0.088 | -0.088 | 0 | 0.42 | 0.05 | 0.42 | 0.35 | 0.44 | 0.42 |
| 9 | 132260350 | rs10819528 | -0.076 | 0.645 | -1.318 | 0.522 | -0.006 | -0.046 | -0.046 | 0 | 0.27 | 0.18 | 0.29 | 0.53 | 0.24 | 0.24 |
| 9 | 132268125 | rs1780456 | 0.284 | 0.278 | -0.715 | -0.081 | 0 | 0.01 | 0.01 | 0.459 | 0.68 | 0.68 | 0.65 | 0.51 | 0.69 | 0.74 |
| 9 | 132270020 | rs1721643 | -0.276 | 0.063 | -0.824 | -0.698 | 0.007 | 0.009 | 0.009 | 0.06 | 0.56 | 0.43 | 0.75 | 0.87 | 0.62 | 0.79 |
| 9 | 132271882 | rs1721642 | 1.056 | 0.423 | -0.432 | -0.63 | -0.002 | -0.025 | -0.025 | 0 | 0.19 | 0.13 | 0.33 | 0.37 | 0.21 | 0.49 |
| 9 | 132273421 | rs1780461 | 0.916 | 0.418 | -0.185 | -0.626 | -0.002 | 0.025 | 0.025 | 0 | 0.21 | 0.27 | 0.34 | 0.37 | 0.21 | 0.5 |
| 9 | 132281601 | rs10988407 | 0.692 | 0.48 | -0.202 | -0.828 | -0.015 | 0.011 | 0.011 | 0 | 0.12 | 0.09 | 0.12 | 0.01 | 0.17 | 0.04 |
| 9 | 132281720 | rs10429519 | -0.137 | 0.031 | 0.058 | -0.695 | 0.005 | 0.016 | 0.016 | 0.086 | 0.64 | 0.61 | 0.42 | 0.48 | 0.6 | 0.44 |
| 9 | 132287630 | rs4524899 | 0.58 | -0.037 | -0.134 | -0.727 | -0.111 | -0.073 | -0.073 | 3.715 | 0.35 | 0.12 | 0.3 | 0.41 | 0.42 | 0.31 |
| 9 | 132287716 | rs1533967 | 0.953 | 0.098 | 0.417 | -0.266 | -0.001 | 0.015 | 0.015 | 0 | 0.21 | 0.23 | 0.33 | 0.39 | 0.21 | 0.49 |
| 9 | 132292954 | rs3121625 | 0.102 | 0.037 | 0.874 | -0.34 | 0.013 | 0.114 | 0.114 | 0 | 0.92 | 0.73 | 0.8 | 0.86 | 0.94 | 0.97 |
| 9 | 132293940 | rs1780482 | 0.751 | 0.046 | 0.669 | -0.347 | -0.006 | 0.074 | 0.074 | 0 | 0.23 | 0.44 | 0.36 | 0.38 | 0.22 | 0.49 |
| 9 | 132298733 | rs1616208 | 0.178 | 0.025 | 0.604 | -0.64 | -0.012 | 0.061 | 0.061 | 0 | 0.29 | 0.55 | 0.54 | 0.52 | 0.28 | 0.52 |
| 9 | 132302923 | rs2498905 | -0.304 | 0.122 | 0.288 | -0.69 | 0.001 | 0.015 | 0.015 | 0.065 | 0.38 | 0.52 | 0.61 | 0.52 | 0.41 | 0.56 |
| 9 | 132303601 | rs2542250 | -0.409 | 0.272 | -0.204 | -0.646 | -0.05 | -0.051 | -0.051 | 0 | 0.65 | 0.84 | 0.74 | 0.58 | 0.61 | 0.71 |
| 9 | 132305053 | rs2542249 | 0.538 | 0.772 | 0.676 | -1.227 | 0.031 | -0.075 | -0.075 | 0 | 0.72 | 0.62 | 0.78 | 0.92 | 0.67 | 0.9 |
| 9 | 132308843 | rs10988420 | -0.032 | 0.243 | 0.332 | -1.366 | -0.004 | -0.045 | -0.045 | 0 | 0.3 | 0.36 | 0.18 | 0.07 | 0.27 | 0.12 |
| 9 | 132314273 | rs7868056 | 0.291 | 0.356 | -0.372 | -1.199 | 0.006 | 0.011 | 0.011 | 0 | 0.46 | 0.45 | 0.41 | 0.69 | 0.39 | 0.35 |
| 9 | 132315998 | rs17469896 | 0.428 | 0.716 | -1.031 | -1.054 | -0.001 | -0.016 | -0.016 | 0 | 0.11 | 0.13 | 0.08 | 0 | 0.12 | 0.05 |
| 9 | 132319215 | rs10819541 | 0.378 | 0.703 | -1.203 | -0.673 | 0.004 | -0.032 | -0.032 | 0 | 0.28 | 0.25 | 0.31 | 0.62 | 0.23 | 0.27 |
| 9 | 132321716 | rs1220688 | 0.149 | 0.933 | -1.018 | -0.281 | -0.019 | -0.007 | -0.007 | 0.009 | 0.25 | 0.13 | 0.41 | 0.26 | 0.34 | 0.52 |
| 9 | 132324366 | rs1220687 | -0.211 | 0.701 | -0.731 | -0.279 | -0.001 | -0.001 | -0.001 | 1.204 | 0.52 | 0.52 | 0.55 | 0.39 | 0.58 | 0.61 |
| 9 | 132336394 | rs1220680 | 0.149 | 0.731 | -1.337 | -0.498 | -0.082 | -0.051 | -0.051 | 0 | 0.32 | 0.14 | 0.47 | 0.66 | 0.44 | 0.37 |
| 9 | 132339872 | rs10819543 | 0.485 | 0.338 | -0.818 | -1.192 | 0.021 | 0.031 | 0.031 | 0 | 0.42 | 0.31 | 0.29 | 0.06 | 0.28 | 0.28 |
| 9 | 132340037 | rs4837389 | 0.695 | 0.384 | -0.439 | -1.221 | 0.006 | 0.027 | 0.027 | 0 | 0.13 | 0.18 | 0.16 | 0.22 | 0.14 | 0.14 |
| 9 | 132342393 | rs1220672 | 0.794 | 0.439 | -0.63 | -1.146 | -0.096 | -0.055 | -0.055 | 0 | 0.31 | 0.1 | 0.47 | 0.66 | 0.44 | 0.37 |
| 9 | 132344026 | rs17415618 | 0.776 | -0.157 | -0.463 | -1.485 | 0.01 | 0.083 | 0.083 | 0 | 0.29 | 0.13 | 0.13 | 0.04 | 0.22 | 0.2 |
| 9 | 132344256 | rs10988435 | 0.689 | -0.182 | -0.353 | -1.498 | 0.012 | 0.063 | 0.063 | 0 | 0.3 | 0.16 | 0.13 | 0.04 | 0.22 | 0.2 |
| 9 | 132349265 | rs10988437 | -0.698 | -0.005 | 0.08 | -1.435 | 0.004 | -0.018 | -0.018 | 0.012 | 0.57 | 0.47 | 0.45 | 0.28 | 0.48 | 0.6 |
| 9 | 132351047 | rs1220666 | 0.748 | -0.067 | -0.226 | -1.368 | -0.022 | 0.018 | 0.018 | 0.055 | 0.38 | 0.37 | 0.52 | 0.75 | 0.47 | 0.38 |
| 9 | 132352178 | rs1228439 | 0.465 | -0.22 | -0.377 | -1.347 | 0.001 | 0.006 | 0.006 | 0 | 0.43 | 0.45 | 0.65 | 0.76 | 0.57 | 0.43 |
| 9 | 132352219 | rs10988439 | 0.086 | -0.28 | -0.739 | -1.215 | 0.039 | 0.085 | 0.085 | 0 | 0.31 | 0.13 | 0.12 | 0.05 | 0.19 | 0.22 |
| 9 | 132357836 | rs1220651 | 0.837 | 0.265 | 0.116 | -0.591 | 0.017 | -0.007 | -0.007 | 0 | 0.88 | 0.7 | 0.85 | 0.92 | 0.87 | 0.85 |
| 9 | 132358827 | rs1220652 | 1.147 | -0.184 | -0.566 | -0.622 | 0.028 | 0.024 | 0.024 | 3.643 | 0.79 | 0.15 | 0.8 | 0.89 | 0.84 | 0.75 |
| 9 | 132363481 | rs10819548 | -0.374 | 0.045 | -0.551 | -1.032 | -0.003 | -0.029 | -0.029 | 0 | 0.36 | 0.38 | 0.37 | 0.15 | 0.39 | 0.4 |
| 9 | 132363874 | rs1220658 | -0.973 | 0.133 | -1.093 | -0.525 | -0.04 | -0.02 | -0.02 | 9.03 | 0.58 | 0.85 | 0.71 | 0.45 | 0.58 | 0.63 |
| 9 | 132366836 | rs2542242 | -0.687 | 0.571 | -1.471 | -0.44 | -0.021 | 0.017 | 0.017 | 0 | 0.48 | 0.8 | 0.62 | 0.42 | 0.5 | 0.32 |
| 9 | 132370360 | rs10988449 | 0.313 | -0.012 | -0.187 | -0.566 | -0.019 | -0.056 | -0.056 | 0 | 0.18 | 0.35 | 0.11 | 0.05 | 0.13 | 0.03 |
| 9 | 132374013 | rs1220701 | 0.073 | 0.296 | 0.152 | -0.544 | 0.009 | 0.032 | 0.032 | 0.1 | 0.14 | 0.15 | 0.37 | 0.33 | 0.21 | 0.2 |
| 9 | 132376859 | rs10988452 | 0.262 | 0.097 | 0.037 | -0.469 | -0.031 | -0.066 | -0.066 | 0.052 | 0.21 | 0.35 | 0.12 | 0.05 | 0.15 | 0.15 |
| 9 | 132380301 | rs7850102 | 1.101 | -0.37 | -0.153 | -0.725 | 0.021 | -0.001 | -0.001 | 0 | 0.52 | 0.35 | 0.5 | 0.57 | 0.56 | 0.5 |
| 9 | 132387665 | rs4837393 | 1.177 | -0.421 | -0.148 | -0.729 | 0.031 | 0.006 | 0.006 | 0 | 0.51 | 0.29 | 0.49 | 0.51 | 0.56 | 0.48 |
| 9 | 132394270 | rs2900277 | 0.267 | -0.501 | 0.238 | -0.712 | -0.001 | 0.025 | 0.025 | 0.197 | 0.72 | 0.65 | 0.61 | 0.55 | 0.71 | 0.62 |
| 9 | 132395316 | rs12335819 | 0.373 | -0.784 | 0.269 | -0.899 | -0.048 | -0.057 | -0.057 | 0 | 0.7 | 0.53 | 0.79 | 0.85 | 0.81 | 0.77 |
| 9 | 132397166 | rs9691 | 1.202 | -0.464 | -0.069 | -0.78 | 0.03 | -0.002 | -0.002 | 0 | 0.51 | 0.28 | 0.49 | 0.51 | 0.56 | 0.48 |
| 9 | 132397239 | rs3088095 | 0.387 | -0.522 | -0.05 | -0.788 | -0.024 | -0.051 | -0.051 | 0 | 0.21 | 0.3 | 0.13 | 0.05 | 0.15 | 0.15 |
| 9 | 132400480 | rs3739851 | 0.649 | -0.268 | -0.684 | -0.62 | 0.026 | -0.002 | -0.002 | 2.103 | 0.51 | 0.32 | 0.49 | 0.51 | 0.56 | 0.48 |
| 9 | 132412706 | rs4836665 | 0.181 | -0.345 | -1.074 | -0.866 | -0.001 | -0.008 | -0.008 | 0 | 0.78 | 0.78 | 0.74 | 0.92 | 0.79 | 0.8 |
| 9 | 132414185 | rs7866311 | -1.1 | -0.553 | -1.704 | -0.314 | 0.015 | 0 | 0 | 1.208 | 0.54 | 0.74 | 0.56 | 0.43 | 0.41 | 0.51 |
| 9 | 132416836 | rs7020887 | -1.891 | -0.079 | -2.158 | 0.073 | -0.047 | -0.081 | -0.081 | 0 | 0.82 | 0.95 | 0.8 | 0.64 | 0.77 | 0.79 |
| 9 | 132416914 | rs11793700 | 1.435 | -0.606 | -2.294 | 0.219 | 0.019 | 0.023 | 0.023 | 0 | 0.16 | 0.01 | 0.09 | 0.07 | 0.1 | 0.15 |
| 9 | 132417166 | rs16931378 | 1.102 | -0.305 | -1.968 | 0.533 | 0 | -0.061 | -0.061 | 0.43 | 0.09 | 0.01 | 0.16 | 0.24 | 0.05 | 0.12 |
| 9 | 132417991 | rs10124196 | -0.49 | 0.091 | -1.222 | -0.022 | 0.016 | -0.019 | -0.019 | 0 | 0.4 | 0.53 | 0.44 | 0.3 | 0.47 | 0.39 |
| 9 | 132425233 | rs10760632 | -0.17 | -0.034 | -0.86 | -0.033 | 0.003 | -0.015 | -0.015 | 0 | 0.16 | 0.2 | 0.17 | 0.06 | 0.19 | 0.23 |
| 9 | 132428927 | rs10988467 | -0.642 | -0.168 | -1.012 | 0.074 | -0.02 | -0.038 | -0.038 | 0 | 0.68 | 0.77 | 0.63 | 0.44 | 0.61 | 0.58 |
| 9 | 132430178 | rs7032080 | 0.4 | 0.018 | -0.955 | 0.068 | 0.007 | 0.009 | 0.009 | 0 | 0.48 | 0.57 | 0.46 | 0.35 | 0.41 | 0.3 |
| 9 | 132431625 | rs12343788 | -0.371 | 0.016 | -0.439 | 0.216 | -0.024 | 0.013 | 0.013 | 2.529 | 0.13 | 0.04 | 0.15 | 0.06 | 0.18 | 0.22 |
| 9 | 132444099 | rs7866070 | 0.158 | -0.443 | 0.359 | 0.511 | 0.003 | -0.004 | -0.004 | 0 | 0.4 | 0.58 | 0.35 | 0.49 | 0.36 | 0.39 |
| 9 | 132448019 | rs10125991 | -0.735 | -0.851 | 0.589 | 0.478 | -0.013 | -0.008 | -0.008 | 0 | 0.65 | 0.79 | 0.66 | 0.63 | 0.64 | 0.6 |
| 9 | 132449756 | rs10819557 | -0.177 | -0.758 | 1.207 | 0.446 | -0.012 | 0.019 | 0.019 | 0.052 | 0.54 | 0.34 | 0.58 | 0.62 | 0.56 | 0.57 |
| 9 | 132452336 | rs7848375 | -2.125 | -1.406 | 2.54 | 1.592 | 0.052 | 0.391 | 0.391 | 2.762 | 0.11 | 0.51 | 0.13 | 0.4 | 0.1 | 0.16 |
| 9 | 132458113 | rs894417 | -1.663 | -1.28 | 1.868 | 1.56 | 0.005 | 0.073 | 0.073 | 0 | 0.12 | 0.3 | 0.2 | 0.19 | 0.08 | 0.13 |
| 9 | 132463618 | rs11794459 | -1.755 | -1.352 | 1.774 | 1.525 | -0.014 | 0.004 | 0.004 | 0.022 | 0.35 | 0.5 | 0.36 | 0.6 | 0.27 | 0.36 |
| 9 | 132465237 | rs3844046 | -1.505 | -1.354 | 1.835 | 1.558 | 0.012 | 0.176 | 0.176 | 2.515 | 0.09 | 0.25 | 0.12 | 0.4 | 0.09 | 0.16 |
| 9 | 132465304 | rs3861882 | -1.784 | -1.364 | 1.722 | 1.562 | -0.02 | 0.007 | 0.007 | 0.022 | 0.35 | 0.52 | 0.37 | 0.61 | 0.27 | 0.36 |
| 9 | 132465556 | rs4836667 | -0.625 | -1.165 | 0.996 | 1.473 | 0.039 | -0.004 | -0.004 | 0 | 0.22 | 0.1 | 0.22 | 0.2 | 0.14 | 0.2 |

| | | | | | | | | | | | | | | | | |
|---|---|---|---|---|---|---|---|---|---|---|---|---|---|---|---|---|
| 9 | 132474185 | rs12685582 | -1.8 | -0.455 | 1.892 | 1.663 | 0.014 | 0.136 | 0.136 | 0 | 0.12 | 0.24 | 0.16 | 0.4 | 0.1 | 0.2 |
| 9 | 132478092 | rs11791419 | -1.692 | -0.524 | 1.721 | 1.409 | -0.017 | 0.164 | 0.164 | 0.666 | 0.18 | 0.52 | 0.17 | 0.36 | 0.11 | 0.23 |
| 9 | 132486736 | rs16931414 | -1.386 | -0.075 | 1.186 | 1.686 | -0.01 | 0.05 | 0.05 | 0.328 | 0.2 | 0.66 | 0.21 | 0.38 | 0.15 | 0.36 |
| 9 | 132497550 | rs16931419 | -0.997 | 0.127 | 1.013 | 1.136 | 0.014 | 0.083 | 0.083 | 0 | 0.15 | 0.47 | 0.11 | 0.21 | 0.13 | 0.19 |
| 9 | 132505659 | rs4837404 | -0.058 | 0.612 | -0.643 | 0.794 | -0.115 | -0.091 | -0.091 | 2.547 | 0.72 | 0.98 | 0.71 | 0.64 | 0.63 | 0.71 |
| 9 | 132507910 | rs10739757 | -1.215 | 0.193 | 0.582 | 1.141 | 0.011 | 0.096 | 0.096 | 0 | 0.13 | 0.32 | 0.13 | 0.21 | 0.1 | 0.2 |
| 9 | 132510366 | rs10448290 | -1.139 | 0.238 | 0.599 | 1.137 | -0.006 | 0.035 | 0.035 | 0 | 0.1 | 0.13 | 0.11 | 0.19 | 0.08 | 0.15 |
| 9 | 132511956 | rs10988493 | -1.034 | 0.222 | 0.59 | 1.14 | -0.009 | 0.034 | 0.034 | 0 | 0.1 | 0.13 | 0.1 | 0.19 | 0.06 | 0.15 |
| 9 | 132515081 | rs2241271 | -1.923 | 0.238 | 0.888 | 0.731 | 0 | -0.017 | -0.017 | 0 | 0.22 | 0.53 | 0.15 | 0.17 | 0.2 | 0.19 |
| 9 | 132515247 | rs11792431 | -1.216 | 0.301 | 1.372 | 0.846 | 0.018 | -0.018 | -0.018 | 0 | 0.07 | 0.21 | 0.05 | 0.02 | 0.06 | 0.05 |
| 9 | 132515730 | rs7872802 | -2.28 | 0.291 | 0.919 | 0.895 | 0.019 | 0.025 | 0.025 | 0 | 0.15 | 0.53 | 0.11 | 0.15 | 0.1 | 0.17 |
| 9 | 132522269 | rs3844048 | 0.505 | 0.091 | -0.178 | 0.298 | 0.026 | -0.044 | -0.044 | 0.003 | 0.6 | 0.25 | 0.63 | 0.55 | 0.66 | 0.67 |
| 9 | 132523860 | rs10118377 | -0.115 | 0.216 | -0.246 | 0.267 | -0.001 | -0.035 | -0.035 | 0 | 0.76 | 0.78 | 0.84 | 1 | 0.71 | 0.81 |
| 9 | 132529504 | rs7857431 | -1.173 | 0.037 | 0.201 | 1.044 | -0.029 | 0.138 | 0.138 | 1.13 | 0.18 | 0.67 | 0.21 | 0.47 | 0.09 | 0.21 |
| 9 | 132537865 | rs4837412 | -1.073 | 0.747 | -0.022 | 0.77 | 0.007 | 0.031 | 0.031 | 0 | 0.32 | 0.47 | 0.33 | 0.53 | 0.34 | 0.28 |
| 9 | 132539425 | rs4837413 | -0.843 | 0.766 | 0.239 | 0.865 | 0.014 | 0.1 | 0.1 | 0.2 | 0.16 | 0.28 | 0.21 | 0.53 | 0.21 | 0.23 |
| 9 | 132539989 | rs11999731 | -1.7 | 0.857 | 0.345 | 0.319 | 0.002 | -0.085 | -0.085 | 0.689 | 0.3 | 0.43 | 0.21 | 0.04 | 0.29 | 0.17 |
| 9 | 132541344 | rs11998910 | -0.633 | 0.531 | 0.268 | 0.295 | -0.007 | -0.057 | -0.057 | 0 | 0.16 | 0.22 | 0.13 | 0.01 | 0.13 | 0.07 |
| 9 | 132546316 | rs7873555 | -0.797 | 1.547 | -0.465 | 0.441 | -0.018 | 0 | 0 | 0 | 0.27 | 0.19 | 0.27 | 0.27 | 0.35 | 0.27 |
| 9 | 132549493 | rs10115114 | -1.139 | 0.574 | 0.344 | -0.199 | 0.025 | -0.021 | -0.021 | 1.846 | 0.27 | 0.51 | 0.25 | 0.22 | 0.3 | 0.33 |
| 9 | 132549740 | rs7867490 | 0.537 | 0.657 | 0.201 | -0.203 | -0.065 | -0.052 | -0.052 | 0 | 0.12 | 0.01 | 0.15 | 0.19 | 0.22 | 0.18 |
| 9 | 132552206 | rs11793587 | -2.094 | 0.402 | -0.529 | -1.338 | -0.007 | 0.17 | 0.17 | 0 | 0.66 | 0.89 | 0.71 | 0.97 | 0.68 | 0.76 |
| 9 | 132556974 | rs7040096 | -0.144 | 0.399 | -0.323 | -1.304 | -0.03 | 0.011 | 0.011 | 0 | 0.25 | 0.11 | 0.21 | 0.21 | 0.29 | 0.27 |
| 9 | 132557210 | rs7025992 | -1.092 | 0.337 | -0.307 | -1.401 | 0 | 0.006 | 0.006 | 0 | 0.41 | 0.6 | 0.31 | 0.23 | 0.37 | 0.4 |
| 9 | 132566666 | rs7029757 | 1.309 | 0.716 | -0.506 | -1.494 | -0.036 | -0.039 | -0.039 | 0 | 0.09 | 0 | 0.07 | 0.13 | 0.11 | 0.03 |
| 9 | 132568500 | rs10116248 | -0.485 | 0.565 | 0.151 | -1.478 | -0.006 | 0.002 | 0.002 | 0 | 0.26 | 0.22 | 0.21 | 0.22 | 0.29 | 0.34 |
| 9 | 132572793 | rs2287367 | -0.539 | 0.573 | 0 | -1.483 | 0.001 | -0.002 | -0.002 | 0 | 0.26 | 0.24 | 0.22 | 0.21 | 0.28 | 0.27 |
| 9 | 132573536 | rs12634 | -0.365 | 0.125 | -0.271 | -1.744 | 0.033 | 0.011 | 0.011 | 0 | 0.18 | 0.24 | 0.21 | 0.21 | 0.27 | 0.21 |
| 9 | 132574862 | rs13283584 | -0.066 | 0.128 | -0.3 | -1.742 | -0.056 | -0.031 | -0.031 | 0.016 | 0.17 | 0.06 | 0.19 | 0.21 | 0.27 | 0.21 |
| 9 | 132585058 | rs2296793 | -0.162 | 0.137 | -0.302 | -1.738 | 0.026 | 0.009 | 0.009 | 0 | 0.17 | 0.22 | 0.21 | 0.21 | 0.28 | 0.21 |
| 9 | 132589606 | rs8113 | -0.593 | 0.085 | -0.328 | -1.763 | -0.002 | 0 | 0 | 0 | 0.25 | 0.22 | 0.21 | 0.21 | 0.27 | 0.27 |
| 9 | 132591509 | rs3818553 | -0.593 | 0.085 | -0.331 | -1.763 | -0.001 | 0 | 0 | 0 | 0.25 | 0.22 | 0.21 | 0.21 | 0.27 | 0.27 |
| 9 | 132597021 | rs3814547 | -0.569 | 0.087 | -0.339 | -1.763 | -0.002 | 0 | 0 | 0 | 0.25 | 0.2 | 0.21 | 0.21 | 0.28 | 0.27 |
| 9 | 132601229 | rs7019254 | -0.496 | 0.096 | -0.567 | -1.758 | -0.078 | -0.024 | -0.024 | 0 | 0.24 | 0.03 | 0.2 | 0.21 | 0.27 | 0.27 |
| 9 | 132601903 | rs11790814 | -0.442 | 0.098 | -0.567 | -1.765 | -0.076 | -0.016 | -0.016 | 0.536 | 0.24 | 0.03 | 0.19 | 0.2 | 0.27 | 0.27 |
| 9 | 132602868 | rs7042087 | 1.638 | -0.175 | -0.65 | -2.151 | -0.024 | 0.225 | 0.225 | 0 | 0.37 | 0.11 | 0.33 | 0.03 | 0.38 | 0.24 |
| 9 | 132606981 | rs7045022 | 0.405 | -0.319 | -0.241 | -1.953 | -0.046 | -0.104 | -0.104 | 0 | 0.24 | 0.31 | 0.1 | 0 | 0.09 | 0.15 |
| 9 | 132614661 | rs11999367 | 0.38 | -0.322 | -0.196 | -1.955 | -0.051 | -0.113 | -0.113 | 0 | 0.24 | 0.32 | 0.1 | 0 | 0.09 | 0.15 |
| 9 | 132615077 | rs2296786 | 1.13 | -0.743 | -0.135 | -1.919 | 0.031 | 0.048 | 0.048 | 0.108 | 0.51 | 0.4 | 0.66 | 0.7 | 0.59 | 0.6 |
| 9 | 132616003 | rs4288464 | -0.819 | -0.422 | -0.089 | -2.144 | -0.006 | -0.11 | -0.11 | 1.035 | 0.8 | 0.65 | 0.87 | 0.98 | 0.82 | 0.87 |
| 9 | 132616209 | rs10819567 | 0.118 | 0.055 | -0.74 | -2.353 | -0.001 | -0.005 | -0.005 | 0 | 0.61 | 0.67 | 0.69 | 0.81 | 0.56 | 0.71 |
| 9 | 132616564 | rs7048514 | -1.947 | -0.128 | -0.651 | -2.322 | -0.039 | -0.092 | -0.092 | 0 | 0.84 | 0.73 | 0.92 | 1 | 0.92 | 0.91 |
| 9 | 132618578 | rs4836674 | -1.99 | -0.13 | -0.838 | -2.323 | -0.038 | -0.091 | -0.091 | 0 | 0.84 | 0.73 | 0.92 | 1 | 0.92 | 0.91 |
| 9 | 132618807 | rs7862315 | -1.6 | -0.142 | -0.839 | -2.332 | -0.034 | -0.08 | -0.08 | 0.565 | 0.83 | 0.73 | 0.91 | 0.98 | 0.91 | 0.89 |
| 9 | 132619089 | rs11788613 | 0.34 | -0.112 | -0.896 | -2.339 | -0.13 | -0.001 | -0.001 | 0 | 0.26 | 0.04 | 0.25 | 0.19 | 0.41 | 0.22 |
| 9 | 132620224 | rs2417150 | -2.03 | -0.139 | -0.862 | -2.34 | -0.037 | -0.09 | -0.09 | 0.16 | 0.84 | 0.73 | 0.92 | 1 | 0.92 | 0.91 |
| 9 | 132620364 | rs10819568 | -2.03 | -0.139 | -0.862 | -2.34 | -0.037 | -0.09 | -0.09 | 0 | 0.84 | 0.73 | 0.92 | 1 | 0.92 | 0.91 |
| 9 | 132623293 | rs1503375 | -1.614 | -0.139 | -0.862 | -2.34 | -0.034 | -0.08 | -0.08 | 0.17 | 0.83 | 0.73 | 0.9 | 0.98 | 0.91 | 0.89 |
| 9 | 132623868 | rs10115249 | -2.03 | -0.139 | -0.862 | -2.34 | -0.037 | -0.09 | -0.09 | 0 | 0.84 | 0.73 | 0.92 | 1 | 0.92 | 0.91 |
| 9 | 132625352 | rs4836675 | -0.729 | -0.247 | -0.789 | -2.399 | -0.082 | -0.147 | -0.147 | 0.163 | 0.72 | 0.55 | 0.88 | 0.98 | 0.89 | 0.83 |
| 9 | 132625818 | rs913316 | -2.059 | -0.23 | -0.834 | -2.387 | -0.037 | -0.09 | -0.09 | 0 | 0.84 | 0.73 | 0.92 | 1 | 0.92 | 0.91 |
| 9 | 132629029 | rs1108535 | -1.107 | -0.245 | -0.841 | -2.395 | -0.047 | -0.098 | -0.098 | 0.513 | 0.76 | 0.73 | 0.9 | 1 | 0.91 | 0.85 |
| 9 | 132629832 | rs880242 | 0.115 | -0.979 | -1.203 | -2.979 | -0.011 | -0.034 | -0.034 | 0 | 0.56 | 0.54 | 0.71 | 0.81 | 0.64 | 0.76 |
| 9 | 132631680 | rs6478920 | -2.169 | -0.79 | -0.823 | -2.684 | -0.036 | -0.088 | -0.088 | 0 | 0.85 | 0.75 | 0.92 | 1 | 0.92 | 0.91 |
| 9 | 132637712 | rs735115 | 0.572 | -0.724 | -0.981 | -2.689 | -0.112 | 0.012 | 0.012 | 0 | 0.26 | 0.05 | 0.25 | 0.19 | 0.41 | 0.2 |
| 9 | 132644138 | rs7849898 | -1.6 | -1.102 | -0.613 | -2.516 | -0.035 | -0.083 | -0.083 | 0.144 | 0.83 | 0.72 | 0.9 | 0.98 | 0.91 | 0.89 |
| 9 | 132649662 | rs1503372 | -1.514 | -0.955 | -0.558 | -2.337 | -0.028 | -0.055 | -0.055 | 0 | 0.89 | 0.6 | 0.96 | 1 | 0.98 | 0.94 |
| 9 | 132649857 | rs1057888 | -0.66 | -0.952 | -0.572 | -2.339 | -0.04 | -0.065 | -0.065 | 0.045 | 0.87 | 0.57 | 0.94 | 0.99 | 0.97 | 0.92 |
| 9 | 132655864 | rs13295392 | 1.065 | -0.866 | -1.128 | -2.386 | 0.027 | 0.13 | 0.13 | 0 | 0.22 | 0.04 | 0.15 | 0.03 | 0.18 | 0.1 |
| 9 | 132657519 | rs10760644 | 0.434 | -0.271 | -0.288 | -2.313 | 0.001 | -0.01 | -0.01 | 0 | 0.57 | 0.58 | 0.65 | 0.79 | 0.51 | 0.66 |
| 9 | 132662786 | rs1023000 | 0.454 | -0.264 | -0.284 | -2.31 | 0 | -0.009 | -0.009 | 0 | 0.57 | 0.58 | 0.65 | 0.79 | 0.51 | 0.67 |
| 9 | 132666628 | rs7044285 | 0.25 | -0.217 | -0.265 | -2.309 | 0.002 | -0.015 | -0.015 | 0.349 | 0.59 | 0.59 | 0.67 | 0.81 | 0.52 | 0.69 |
| 9 | 132673694 | rs10739766 | 0.134 | -0.866 | 0.284 | -1.663 | -0.024 | -0.045 | -0.045 | 0.34 | 0.74 | 0.64 | 0.83 | 0.89 | 0.82 | 0.69 |
| 9 | 132681784 | rs2295952 | 1.371 | -0.582 | 0.374 | -1.465 | -0.011 | -0.04 | -0.04 | 0.756 | 0.84 | 0.69 | 0.87 | 0.91 | 0.84 | 0.73 |
| 9 | 132698515 | rs7022657 | -0.568 | -0.574 | 0.402 | -1.464 | -0.008 | -0.007 | -0.007 | 0 | 0.92 | 0.84 | 0.93 | 0.93 | 0.93 | 0.89 |
| 9 | 132707164 | rs7870884 | 1.464 | -0.565 | 0.363 | -1.474 | -0.011 | -0.049 | -0.049 | 0 | 0.83 | 0.66 | 0.87 | 0.91 | 0.84 | 0.74 |
| 9 | 132720770 | rs17519205 | 0.061 | -0.543 | 0.013 | -1.273 | 0.002 | 0.008 | 0.008 | 0 | 0.09 | 0.05 | 0.15 | 0.08 | 0.08 | 0.12 |
| 9 | 132753534 | rs2102209 | 0.031 | -0.987 | 0.236 | -1.416 | -0.021 | -0.023 | -0.023 | 0 | 0.83 | 0.77 | 0.81 | 0.9 | 0.91 | 0.87 |
| 9 | 132758347 | rs2417202 | 1.571 | -1.008 | 0.684 | -1.067 | 0.023 | 0.259 | 0.259 | 0 | 0.06 | 0.17 | 0.25 | 0.64 | 0.06 | 0.16 |

| | | | | | | | | | | | | | | | | |
|---|---|---|---|---|---|---|---|---|---|---|---|---|---|---|---|---|
| 9 | 132758467 | rs1393904 | 0.088 | -0.975 | 0.184 | -1.339 | -0.02 | -0.021 | -0.021 | 0.933 | 0.83 | 0.77 | 0.81 | 0.9 | 0.91 | 0.87 |
| 9 | 132759808 | rs1322671 | 0.632 | -1.3 | 0.186 | -1.618 | -0.04 | -0.066 | -0.066 | 0 | 0.69 | 0.59 | 0.73 | 0.9 | 0.83 | 0.78 |
| 9 | 132771351 | rs2417192 | 0.026 | -1.17 | 0.229 | -1.437 | -0.036 | -0.006 | -0.006 | 0.01 | 0.83 | 0.71 | 0.81 | 0.84 | 0.91 | 0.87 |
| 9 | 132777024 | rs10760653 | 0.755 | -0.883 | 0.206 | -1.605 | -0.029 | -0.053 | -0.053 | 0 | 0.62 | 0.51 | 0.69 | 0.83 | 0.73 | 0.63 |
| 9 | 132786313 | rs1923664 | 0.371 | -0.651 | 0.512 | -1.013 | -0.021 | -0.048 | -0.048 | 0 | 0.11 | 0.2 | 0.05 | 0.01 | 0.04 | 0.07 |
| 9 | 132787528 | rs806861 | 0.901 | -0.718 | 0.395 | -1.045 | -0.043 | -0.02 | -0.02 | 0 | 0.8 | 0.63 | 0.8 | 0.84 | 0.88 | 0.83 |
| 9 | 132792272 | rs12353358 | -0.183 | -0.565 | 0.102 | -1.635 | -0.009 | 0.032 | 0.032 | 0 | 0.3 | 0.22 | 0.23 | 0.04 | 0.32 | 0.16 |
| 9 | 132803206 | rs1923662 | -0.002 | -0.358 | 0.509 | -1.125 | -0.04 | -0.056 | -0.056 | 0.651 | 0.88 | 0.7 | 0.94 | 0.97 | 0.97 | 0.95 |
| 9 | 132829354 | rs10819596 | -0.053 | -0.289 | 0.169 | -1.091 | -0.039 | -0.053 | -0.053 | 0 | 0.89 | 0.74 | 0.95 | 0.97 | 0.96 | 0.94 |
| 9 | 132830460 | rs2065159 | -1.11 | -0.28 | -0.073 | -1.084 | 0.008 | 0.003 | 0.003 | 0 | 0.92 | 0.97 | 0.98 | 1 | 0.99 | 0.96 |
| 9 | 132837130 | rs12380103 | -0.429 | -0.257 | -0.15 | -0.846 | -0.004 | -0.04 | -0.04 | 0 | 0.1 | 0.03 | 0.17 | 0.26 | 0.12 | 0.11 |
| 9 | 132837275 | rs10739772 | 0.787 | -0.133 | -0.118 | -1.424 | -0.002 | -0.091 | -0.091 | 0.78 | 0.72 | 0.6 | 0.78 | 0.95 | 0.74 | 0.78 |
| 9 | 132849126 | rs2159640 | 1.591 | -0.233 | -0.278 | -1.312 | -0.006 | -0.031 | -0.031 | 0.063 | 0.58 | 0.53 | 0.62 | 0.8 | 0.61 | 0.67 |
| 9 | 132850277 | rs12346450 | -0.964 | -0.09 | -0.906 | -1.204 | -0.009 | 0.022 | 0.022 | 0 | 0.3 | 0.23 | 0.33 | 0.17 | 0.35 | 0.27 |
| 9 | 132861672 | rs2286792 | 0.414 | -0.09 | -0.91 | -1.195 | -0.01 | -0.031 | -0.031 | 0 | 0.08 | 0.02 | 0.15 | 0.14 | 0.09 | 0.08 |
| 9 | 132869344 | rs10988593 | -0.118 | -0.033 | -0.812 | -1.138 | -0.004 | 0.002 | 0.002 | 0 | 0.22 | 0.2 | 0.18 | 0.04 | 0.26 | 0.19 |
| 9 | 132872686 | rs4837460 | -0.823 | -0.034 | -0.733 | -1.016 | -0.027 | -0.035 | -0.035 | 0 | 0.92 | 0.89 | 0.98 | 1 | 0.99 | 0.96 |
| 9 | 132882000 | rs7851378 | -0.861 | -0.029 | -0.791 | -1.013 | -0.018 | -0.026 | -0.026 | 4.915 | 0.92 | 0.93 | 0.98 | 1 | 0.99 | 0.96 |
| 9 | 132882141 | rs7851530 | 0.369 | 0.368 | -0.807 | -0.79 | 0.003 | 0.014 | 0.014 | 0 | 0.61 | 0.67 | 0.65 | 0.83 | 0.64 | 0.69 |
| 9 | 132895950 | rs1003415 | 0.448 | 0.539 | -0.801 | -0.829 | -0.012 | 0.139 | 0.139 | 0 | 0.48 | 0.33 | 0.41 | 0.13 | 0.53 | 0.29 |
| 9 | 132897036 | rs12344508 | -0.66 | 0.346 | 0.151 | -0.776 | 0.007 | -0.01 | -0.01 | 0.149 | 0.09 | 0.28 | 0.16 | 0.07 | 0.08 | 0.15 |
| 9 | 132897231 | rs1003416 | 0.685 | 0.367 | 0.525 | 0.08 | -0.009 | 0.057 | 0.057 | 0 | 0.69 | 0.55 | 0.6 | 0.47 | 0.72 | 0.57 |
| 9 | 132900076 | rs1306 | 1.745 | 0.272 | 0.518 | -0.309 | -0.051 | 0.116 | 0.116 | 0 | 0.38 | 0.18 | 0.41 | 0.15 | 0.43 | 0.26 |
| 9 | 132901403 | rs10819601 | -1.211 | 0.328 | 1.455 | 0.026 | 0.063 | 0.297 | 0.297 | 1.088 | 0.09 | 0.33 | 0.07 | 0.31 | 0.11 | 0.27 |
| 9 | 132901507 | rs10988603 | -0.96 | 0.443 | 1.591 | 0.071 | 0.12 | 0.324 | 0.324 | 0.128 | 0.1 | 0.34 | 0.2 | 0.37 | 0.18 | 0.16 |
| 9 | 132902487 | rs6478949 | -0.442 | 0.353 | 1.002 | 0.011 | 0.002 | -0.013 | -0.013 | 0.738 | 0.79 | 0.83 | 0.76 | 0.54 | 0.82 | 0.72 |
| 9 | 132906486 | rs17221008 | 1.354 | 0.689 | 1.131 | -0.703 | -0.001 | -0.01 | -0.01 | 0.396 | 0.56 | 0.43 | 0.4 | 0.17 | 0.51 | 0.41 |
| 9 | 132911947 | rs7851165 | -0.565 | 0.642 | 0.923 | -1.29 | 0.005 | -0.014 | -0.014 | 0 | 0.17 | 0.13 | 0.1 | 0 | 0.15 | 0.07 |
| 9 | 132912105 | rs7866942 | 0.184 | 0.924 | 1.52 | -1.187 | -0.004 | -0.02 | -0.02 | 0 | 0.67 | 0.61 | 0.68 | 0.71 | 0.7 | 0.68 |
| 9 | 132912150 | rs12347960 | -1.073 | 1.116 | 1.15 | -0.922 | 0.06 | 0.107 | 0.107 | 0.058 | 0.06 | 0.32 | 0.13 | 0.1 | 0.06 | 0.12 |
| 9 | 132914712 | rs4836691 | -0.202 | 1.174 | 0.29 | -1.01 | 0.01 | 0.034 | 0.034 | 0 | 0.71 | 0.89 | 0.8 | 0.72 | 0.76 | 0.73 |
| 9 | 132917235 | rs10819605 | -1.324 | 1.445 | 0.814 | -0.681 | 0.023 | 0.038 | 0.038 | 0 | 0.13 | 0.18 | 0.34 | 0.6 | 0.2 | 0.17 |
| 9 | 132920697 | rs12352544 | -0.623 | 1.401 | 0.567 | -0.703 | 0 | 0 | 0 | 0.038 | 0.09 | 0.07 | 0.13 | 0.07 | 0.09 | 0.04 |
| 9 | 132923772 | rs10760659 | -0.263 | 1.216 | 0.118 | -0.969 | 0.006 | 0.098 | 0.098 | 0.099 | 0.7 | 0.96 | 0.81 | 0.76 | 0.77 | 0.63 |
| 9 | 132926691 | rs11789529 | 1.032 | 1.348 | 0.594 | -1.107 | 0.002 | 0.036 | 0.036 | 0.261 | 0.46 | 0.34 | 0.32 | 0.12 | 0.46 | 0.28 |
| 9 | 132930343 | rs12378342 | 0.188 | 1.312 | 0.046 | -1.018 | -0.022 | 0.02 | 0.02 | 0 | 0.07 | 0.03 | 0.12 | 0.04 | 0.1 | 0.06 |
| 9 | 132935951 | rs3780708 | 0.318 | 1.252 | 0.349 | -0.965 | 0.026 | 0.061 | 0.061 | 0.295 | 0.62 | 0.79 | 0.74 | 0.72 | 0.71 | 0.53 |
| 9 | 132944545 | rs7849345 | 0.707 | 1.436 | 0.767 | -1.285 | 0.003 | -0.052 | -0.052 | 0 | 0.56 | 0.56 | 0.46 | 0.14 | 0.53 | 0.32 |
| 9 | 132946615 | rs10819610 | -1.418 | 1.892 | 1.904 | -1.026 | 0.018 | -0.062 | -0.062 | 2.007 | 0.27 | 0.5 | 0.41 | 0.14 | 0.26 | 0.16 |
| 9 | 132948550 | rs3824544 | 1.27 | 1.943 | 2.144 | -1.227 | -0.015 | -0.082 | -0.082 | 2.408 | 0.73 | 0.49 | 0.68 | 0.9 | 0.79 | 0.84 |
| 9 | 132955172 | rs740918 | -1.074 | 1.752 | 2 | -1.147 | 0.011 | 0.024 | 0.024 | 0 | 0.23 | 0.19 | 0.31 | 0.18 | 0.2 | 0.11 |
| 9 | 132955453 | rs10819611 | 1.965 | 1.928 | 2.033 | -1.486 | 0.007 | 0.053 | 0.053 | 0 | 0.64 | 0.29 | 0.52 | 0.87 | 0.71 | 0.78 |
| 9 | 132956229 | rs1009502 | -1.071 | 2.243 | 1.852 | -1.5 | -0.05 | 0.079 | 0.079 | 0 | 0.25 | 0.1 | 0.35 | 0.1 | 0.28 | 0.15 |
| 9 | 132958305 | rs1342043 | 2.698 | 2.5 | 2.171 | -1.416 | 0.013 | -0.051 | -0.051 | 0 | 0.84 | 0.33 | 0.78 | 0.96 | 0.85 | 0.86 |
| 9 | 132963129 | rs887535 | 1.22 | 2.517 | 1.728 | -1.384 | -0.005 | -0.031 | -0.031 | 0.585 | 0.92 | 0.88 | 0.95 | 1 | 0.96 | 0.94 |
| 9 | 132967196 | rs7024495 | 2.196 | 2.521 | 2.033 | -1.529 | -0.012 | 0.025 | 0.025 | 0 | 0.58 | 0.39 | 0.4 | 0.88 | 0.54 | 0.72 |
| 9 | 132967288 | rs7024934 | 1.783 | 1.788 | 2.209 | -1.465 | -0.012 | -0.055 | -0.055 | 0 | 0.89 | 0.7 | 0.93 | 0.98 | 0.95 | 0.88 |
| 9 | 132971820 | rs868964 | -1.203 | 1.854 | 1.956 | -1.444 | 0.095 | 0.011 | 0.011 | 0 | 0.05 | 0.15 | 0.12 | 0.03 | 0.09 | 0.03 |
| 9 | 132977438 | rs7848440 | 0.895 | 1.259 | 1.859 | -1.352 | 0 | -0.015 | -0.015 | 0.456 | 0.93 | 0.92 | 0.95 | 1 | 0.95 | 0.94 |
| 9 | 132978477 | rs4240447 | -1.202 | 1.199 | 1.962 | -1.366 | -0.004 | -0.041 | -0.041 | 0.513 | 0.28 | 0.33 | 0.39 | 0.1 | 0.27 | 0.18 |
| 9 | 132979088 | rs1001742 | 0.959 | 1.227 | 1.553 | -1.277 | 0 | -0.002 | -0.002 | 0 | 0.93 | 0.93 | 0.95 | 1 | 0.95 | 0.94 |
| 9 | 132985111 | rs3829905 | 1.686 | 1.357 | 1.609 | -1.256 | 0.001 | -0.027 | -0.027 | 0 | 0.61 | 0.45 | 0.48 | 0.86 | 0.63 | 0.76 |
| 9 | 132988709 | rs2277200 | -1.358 | 1.403 | 1.18 | -1.232 | -0.003 | -0.064 | -0.064 | 0.125 | 0.32 | 0.47 | 0.47 | 0.14 | 0.31 | 0.18 |
| 9 | 132993162 | rs7873936 | 2.067 | 1.465 | 0.779 | -0.426 | 0.037 | -0.013 | -0.013 | 0.479 | 0.84 | 0.66 | 0.72 | 0.89 | 0.84 | 0.86 |
| 9 | 132995160 | rs7852859 | -1.383 | 1.294 | 0.145 | -0.302 | 0.012 | 0 | 0 | 5.228 | 0.09 | 0.17 | 0.21 | 0.08 | 0.1 | 0.04 |
| 9 | 132998359 | rs709611 | 0.714 | 0.602 | -0.113 | -0.73 | -0.016 | -0.042 | -0.042 | 0 | 0.6 | 0.48 | 0.79 | 0.79 | 0.71 | 0.69 |
| 9 | 132999182 | rs7847694 | 0.706 | 0.239 | 0.128 | -0.776 | -0.01 | -0.019 | -0.019 | 0 | 0.63 | 0.44 | 0.77 | 0.74 | 0.7 | 0.62 |
| 9 | 133000282 | rs7867364 | 0.422 | 0.756 | 0.15 | -1.143 | -0.005 | 0.007 | 0.007 | 0 | 0.8 | 0.86 | 0.9 | 0.98 | 0.82 | 0.86 |
| 9 | 133002805 | rs7863731 | -0.379 | 0.853 | -0.145 | -1.501 | 0.001 | -0.012 | -0.012 | 0 | 0.35 | 0.53 | 0.56 | 0.72 | 0.41 | 0.44 |
| 9 | 133003804 | rs7867693 | -0.388 | 1.094 | -0.336 | -1.66 | -0.012 | 0.031 | 0.031 | 0 | 0.88 | 0.95 | 0.91 | 1 | 0.87 | 0.94 |
| 9 | 133006474 | rs10988653 | 1.178 | 0.956 | -0.395 | -1.844 | -0.01 | 0.18 | 0.18 | 0.634 | 0.45 | 0.21 | 0.29 | 0.06 | 0.37 | 0.44 |
| 9 | 133009150 | rs7866994 | 0.241 | 1.015 | 0.068 | -2.02 | 0.014 | -0.013 | -0.013 | 0 | 0.64 | 0.57 | 0.49 | 0.28 | 0.61 | 0.57 |
| 9 | 133011232 | rs4553010 | -0.762 | 1.013 | 0.039 | -2.046 | -0.002 | -0.015 | -0.015 | 0 | 0.9 | 0.93 | 0.95 | 1 | 0.9 | 0.94 |
| 9 | 133013233 | rs7026795 | 0.488 | 1.161 | 0.065 | -1.982 | 0.009 | 0.03 | 0.03 | 0 | 0.23 | 0.34 | 0.41 | 0.71 | 0.24 | 0.36 |
| 9 | 133014637 | rs13302591 | -0.08 | 1.046 | 0.105 | -1.8 | -0.013 | -0.044 | -0.044 | 0 | 0.11 | 0.19 | 0.12 | 0.01 | 0.13 | 0.07 |
| 9 | 133014733 | rs10988659 | -0.679 | 1.001 | 0.114 | -1.799 | -0.002 | -0.056 | -0.056 | 1.644 | 0.14 | 0.24 | 0.19 | 0.01 | 0.21 | 0.08 |
| 9 | 133019198 | rs7045878 | 0.354 | 1.066 | -0.114 | -1.836 | -0.006 | -0.014 | -0.014 | 0 | 0.75 | 0.82 | 0.76 | 0.99 | 0.67 | 0.86 |
| 9 | 133019752 | rs10115217 | -0.392 | 0.675 | -0.396 | -1.314 | -0.004 | 0.034 | 0.034 | 0 | 0.08 | 0.11 | 0.06 | 0.23 | 0.07 | 0.06 |
| 9 | 133021887 | rs10119818 | -0.871 | 0.644 | -0.39 | -1.306 | -0.014 | 0.059 | 0.059 | 0 | 0.1 | 0.23 | 0.08 | 0.23 | 0.08 | 0.07 |

| | | | | | | | | | | | | | | | | |
|---|---|---|---|---|---|---|---|---|---|---|---|---|---|---|---|---|
| 9 | 133025756 | rs10819626 | 0.592 | 0.647 | -1.007 | -1.518 | -0.001 | 0.039 | 0.039 | 0 | 0.45 | 0.3 | 0.29 | 0.3 | 0.39 | 0.48 |
| 9 | 133039956 | rs10988672 | -0.094 | 0.465 | -1.664 | -1.791 | -0.032 | 0.15 | 0.15 | 0 | 0.7 | 0.94 | 0.8 | 0.9 | 0.73 | 0.81 |
| 9 | 133042138 | rs10988675 | 1.746 | 0.535 | -1.994 | -1.983 | -0.034 | 0.114 | 0.114 | 0 | 0.23 | 0.02 | 0.17 | 0.03 | 0.21 | 0.2 |
| 9 | 133045648 | rs7866146 | 0.209 | 0.471 | -1.706 | -1.621 | -0.004 | 0.071 | 0.071 | 0.494 | 0.63 | 0.8 | 0.74 | 0.76 | 0.64 | 0.68 |
| 9 | 133050233 | rs1105621 | 0.18 | 0.677 | -1.41 | -0.426 | 0.039 | 0.193 | 0.193 | 0.025 | 0.2 | 0.39 | 0.29 | 0.51 | 0.27 | 0.26 |
| 9 | 133051110 | rs947510 | -0.148 | 0.779 | -2.498 | -0.508 | -0.064 | 0.118 | 0.118 | 1.269 | 0.71 | 0.97 | 0.8 | 0.89 | 0.73 | 0.8 |
| 9 | 133054868 | rs1935344 | 0.1 | 0.931 | -1.693 | 0.153 | -0.003 | 0.002 | 0.002 | 0 | 0.29 | 0.68 | 0.36 | 0.33 | 0.25 | 0.35 |
| 9 | 133055949 | rs4260979 | 0.593 | 1.045 | -1.845 | 0.105 | -0.028 | -0.046 | -0.046 | 0.076 | 0.15 | 0.06 | 0.25 | 0.48 | 0.28 | 0.27 |
| 9 | 133057805 | rs947507 | -1.499 | 0.953 | -1.933 | -0.165 | 0.01 | 0.085 | 0.085 | 0 | 0.62 | 0.78 | 0.76 | 0.96 | 0.68 | 0.72 |
| 9 | 133059626 | rs12555056 | -0.183 | 0.739 | -1.234 | 0.678 | -0.006 | 0.008 | 0.008 | 0 | 0.29 | 0.66 | 0.36 | 0.37 | 0.23 | 0.34 |
| 9 | 133070628 | rs10819633 | -0.52 | 1.035 | -1.22 | 1.374 | -0.046 | -0.071 | -0.071 | 0.021 | 0.14 | 0.05 | 0.28 | 0.53 | 0.27 | 0.26 |
| 9 | 133226582 | rs914977 | 0.793 | -0.142 | -0.515 | 1.006 | -0.027 | 0.039 | 0.039 | 1.828 | 0.68 | 0.41 | 0.76 | 0.51 | 0.76 | 0.67 |
| 9 | 133229311 | rs914976 | 0.5 | -0.177 | 0.339 | 0.68 | -0.008 | 0.014 | 0.014 | 0 | 0.9 | 0.78 | 0.93 | 0.88 | 0.92 | 0.9 |
| 9 | 133239496 | rs7047329 | 1.03 | -0.297 | 1.686 | 0.415 | -0.009 | -0.039 | -0.039 | 0 | 0.86 | 0.49 | 0.86 | 0.94 | 0.89 | 0.87 |
| 9 | 133241327 | rs4579578 | 0.52 | -0.664 | 0.917 | -0.017 | 0.003 | 0.009 | 0.009 | 0 | 0.43 | 0.28 | 0.58 | 0.56 | 0.44 | 0.52 |
| 9 | 133242154 | rs7874077 | -0.531 | -0.62 | 0.396 | 0.001 | 0.008 | 0.014 | 0.014 | 0 | 0.53 | 0.42 | 0.4 | 0.43 | 0.56 | 0.4 |
| 9 | 133245991 | rs914981 | 0.606 | -1.563 | 0.528 | 0.293 | -0.04 | 0.005 | 0.005 | 0 | 0.83 | 0.38 | 0.86 | 0.77 | 0.94 | 0.79 |
| 9 | 133246083 | rs1316391 | 0.588 | -1.562 | 0.531 | 0.294 | -0.042 | 0.007 | 0.007 | 0 | 0.83 | 0.39 | 0.86 | 0.77 | 0.94 | 0.79 |
| 9 | 133246381 | rs11243875 | -0.383 | -1.477 | 0.025 | 0.227 | 0.001 | -0.061 | -0.061 | 0 | 0.11 | 0.01 | 0.09 | 0.18 | 0.07 | 0.07 |
| 9 | 133250275 | rs7031579 | 0.725 | -1.703 | 0.329 | -0.34 | -0.052 | -0.022 | -0.022 | 0 | 0.77 | 0.42 | 0.85 | 0.78 | 0.91 | 0.78 |
| 9 | 133251986 | rs11244043 | 0.979 | -1.444 | 0.279 | -0.836 | 0.022 | 0.053 | 0.053 | 0 | 0.62 | 0.3 | 0.77 | 0.87 | 0.73 | 0.72 |
| 9 | 133252969 | rs7020021 | 1.091 | -1.429 | 0.246 | -0.844 | 0.036 | 0.099 | 0.099 | 0 | 0.61 | 0.24 | 0.77 | 0.87 | 0.73 | 0.71 |
| 9 | 133262821 | rs7855331 | -0.469 | -1.218 | 0.682 | -0.816 | 0.003 | -0.064 | -0.064 | 0 | 0.11 | 0.29 | 0.1 | 0 | 0.12 | 0.13 |
| 9 | 133264096 | rs11244127 | -0.931 | -1.289 | 0.388 | -0.831 | -0.006 | -0.041 | -0.041 | 0 | 0.27 | 0.36 | 0.17 | 0.09 | 0.22 | 0.19 |
| 9 | 133271928 | rs4372050 | -1.577 | -1.148 | 0.178 | -0.855 | 0.009 | 0.084 | 0.084 | 0.156 | 0.34 | 0.79 | 0.18 | 0.05 | 0.24 | 0.15 |
| 9 | 133274265 | rs4573325 | -0.465 | -1.821 | 0.317 | -0.374 | -0.006 | 0.005 | 0.005 | 0 | 0.27 | 0.34 | 0.42 | 0.37 | 0.21 | 0.36 |
| 9 | 133274806 | rs11244165 | 0.089 | -1.63 | 0.671 | -0.593 | 0.001 | -0.035 | -0.035 | 0.045 | 0.18 | 0.23 | 0.08 | 0.03 | 0.12 | 0.07 |
| 9 | 133276075 | rs885531 | -1.367 | -1.813 | 1.133 | -0.315 | -0.016 | -0.09 | -0.09 | 0 | 0.13 | 0.45 | 0.07 | 0.01 | 0.06 | 0.12 |
| 9 | 133277641 | rs914978 | -0.375 | -1.752 | 0.849 | 0.045 | 0.001 | -0.002 | -0.002 | 0.022 | 0.22 | 0.24 | 0.43 | 0.36 | 0.19 | 0.26 |
| 9 | 133279395 | rs10901300 | 1.011 | -2.405 | 1.15 | 0.077 | -0.054 | 0.052 | 0.052 | 0 | 0.7 | 0.59 | 0.63 | 0.53 | 0.8 | 0.61 |
| 9 | 133283255 | rs10736862 | 1.046 | -2.298 | 0.987 | 0.047 | -0.025 | 0.023 | 0.023 | 0 | 0.7 | 0.6 | 0.58 | 0.52 | 0.81 | 0.67 |
| 9 | 133283580 | rs6597658 | -0.176 | -2.158 | 0.839 | 0.029 | -0.007 | -0.024 | -0.024 | 0.457 | 0.9 | 0.85 | 0.96 | 1 | 0.96 | 0.94 |
| 9 | 133284310 | rs10793975 | 1.37 | -2.269 | 0.817 | -0.129 | -0.022 | 0.007 | 0.007 | 0.185 | 0.58 | 0.41 | 0.53 | 0.52 | 0.74 | 0.64 |
| 9 | 133285049 | rs10901311 | -0.549 | -1.838 | 1.017 | 0.36 | -0.031 | 0.155 | 0.155 | 1.274 | 0.18 | 0.41 | 0.22 | 0.39 | 0.11 | 0.19 |
| 9 | 133287097 | rs4740391 | 1.092 | -1.513 | 1.441 | -0.063 | -0.022 | -0.09 | -0.09 | 1.87 | 0.85 | 0.71 | 0.92 | 0.99 | 0.9 | 0.91 |
| 9 | 133288738 | rs11244241 | -0.79 | -1.587 | 1.656 | 0.65 | -0.007 | 0.16 | 0.16 | 0 | 0.11 | 0.22 | 0.21 | 0.4 | 0.1 | 0.25 |
| 9 | 133289174 | rs7044220 | 1.035 | -1.104 | 0.896 | -0.037 | -0.004 | 0.007 | 0.007 | 0.057 | 0.51 | 0.25 | 0.34 | 0.43 | 0.48 | 0.5 |
| 9 | 133290501 | rs7049027 | -0.515 | -1.022 | 1.305 | -0.079 | -0.012 | 0.016 | 0.016 | 0 | 0.7 | 0.73 | 0.6 | 0.84 | 0.61 | 0.78 |
| 9 | 133292993 | rs10901326 | -0.146 | -0.951 | 1.197 | 0.029 | 0 | 0.2 | 0.2 | 0.103 | 0.11 | 0.25 | 0.21 | 0.41 | 0.11 | 0.24 |
| 9 | 133294244 | rs13294886 | 0.631 | -0.999 | 0.414 | -0.228 | -0.01 | 0.073 | 0.073 | 0 | 0.5 | 0.15 | 0.34 | 0.36 | 0.48 | 0.48 |
| 9 | 133307984 | rs2282217 | -0.505 | -0.684 | -0.075 | -0.021 | 0.012 | -0.052 | -0.052 | 0 | 0.23 | 0.15 | 0.32 | 0.42 | 0.16 | 0.27 |
| 9 | 133308063 | rs2282218 | 1.128 | -0.212 | 0.059 | -0.498 | 0.002 | 0.013 | 0.013 | 1.155 | 0.46 | 0.34 | 0.28 | 0.34 | 0.44 | 0.41 |
| 9 | 133310815 | rs12342208 | 0.024 | -0.259 | 0.408 | -0.256 | -0.003 | -0.014 | -0.014 | 0 | 0.15 | 0.17 | 0.08 | 0.07 | 0.12 | 0.1 |
| 9 | 133313115 | rs4237122 | -0.883 | -0.18 | 0.857 | 0.148 | -0.022 | 0.176 | 0.176 | 0 | 0.14 | 0.28 | 0.32 | 0.5 | 0.08 | 0.2 |
| 9 | 133318751 | rs928518 | -0.809 | 0.178 | 0.808 | -0.273 | 0.001 | -0.009 | -0.009 | 0 | 0.34 | 0.33 | 0.44 | 0.48 | 0.27 | 0.28 |
| 9 | 133323675 | rs10901047 | 1.496 | 1.077 | 1.056 | -0.578 | 0.021 | 0.057 | 0.057 | 0 | 0.48 | 0.22 | 0.38 | 0.34 | 0.4 | 0.35 |
| 9 | 133324400 | rs11243372 | -1.087 | 1.036 | 1.354 | -0.613 | 0.014 | 0.007 | 0.007 | 0 | 0.33 | 0.43 | 0.49 | 0.56 | 0.47 | 0.39 |
| 9 | 133325472 | rs4740158 | 0.352 | 1.203 | 1.029 | -0.735 | 0.003 | 0.015 | 0.015 | 0 | 0.8 | 0.85 | 0.73 | 0.9 | 0.78 | 0.7 |
| 9 | 133326213 | rs914983 | 0.49 | 1.209 | 1.043 | -0.737 | -0.022 | 0.035 | 0.035 | 0 | 0.76 | 0.84 | 0.7 | 0.9 | 0.7 | 0.67 |
| 9 | 133328969 | rs1653332 | -1.366 | 0.703 | 1.076 | -0.667 | 0.015 | 0.051 | 0.051 | 0 | 0.46 | 0.26 | 0.69 | 0.67 | 0.6 | 0.66 |
| 9 | 133331226 | rs7860909 | 1.634 | 0.673 | 1.104 | -0.582 | 0.025 | 0.025 | 0.025 | 0 | 0.47 | 0.34 | 0.23 | 0.34 | 0.33 | 0.3 |
| 9 | 133335415 | rs1215985 | 1.492 | 0.098 | 1.308 | -0.409 | 0.01 | -0.003 | -0.003 | 0 | 0.7 | 0.25 | 0.8 | 0.96 | 0.7 | 0.8 |
| 9 | 133338393 | rs914971 | 1.51 | 0.191 | 1.178 | -0.15 | 0.026 | 0.059 | 0.059 | 0.001 | 0.73 | 0.16 | 0.82 | 0.96 | 0.74 | 0.79 |
| 9 | 133343659 | rs590086 | 1.486 | 0.166 | 1.055 | -0.079 | -0.011 | -0.049 | -0.049 | 0 | 0.84 | 0.22 | 0.89 | 0.99 | 0.9 | 0.92 |
| 9 | 133343744 | rs12551145 | -0.601 | 0.301 | 1.039 | 0.017 | -0.005 | -0.034 | -0.034 | 0 | 0.1 | 0.11 | 0.05 | 0 | 0.09 | 0.07 |
| 9 | 133346226 | rs10901072 | -0.16 | 0.287 | 0.922 | 0.078 | -0.004 | -0.022 | -0.022 | 0 | 0.11 | 0.11 | 0.09 | 0.04 | 0.15 | 0.14 |
| 9 | 133350608 | rs1215972 | 1.654 | 0.06 | 1.623 | 0.776 | -0.039 | 0.093 | 0.093 | 0 | 0.81 | 0.34 | 0.77 | 0.61 | 0.87 | 0.8 |
| 9 | 133351106 | rs476067 | 1.134 | 0.137 | 1.605 | 0.882 | 0.058 | 0.619 | 0.619 | 0.017 | 0.94 | 0.58 | 0.84 | 0.62 | 0.99 | 0.87 |
| 9 | 133355371 | rs540140 | -0.416 | 0.595 | 0.736 | 0.381 | -0.087 | 0.004 | 0.004 | 0 | 0.17 | 0.04 | 0.36 | 0.14 | 0.28 | 0.33 |
| 9 | 133360892 | rs474330 | -1.111 | 0.464 | 0.85 | 0.459 | 0.005 | 0.01 | 0.01 | 0 | 0.24 | 0.29 | 0.51 | 0.56 | 0.29 | 0.48 |
| 9 | 133361010 | rs17147023 | -0.746 | 0.119 | 0.605 | 0.036 | -0.019 | -0.102 | -0.102 | 0 | 0.25 | 0.32 | 0.14 | 0.03 | 0.28 | 0.22 |
| 9 | 133362055 | rs666971 | -1.605 | 0.345 | -0.396 | 0.158 | 0.014 | -0.001 | -0.001 | 0 | 0.34 | 0.52 | 0.5 | 0.61 | 0.43 | 0.56 |
| 9 | 133365094 | rs12375699 | 1.884 | 0.393 | -0.838 | 0.11 | -0.11 | 0.059 | 0.059 | 0.029 | 0.46 | 0.04 | 0.37 | 0.37 | 0.4 | 0.29 |
| 9 | 133367368 | rs634432 | 0.261 | 0.559 | -0.414 | 0.327 | -0.03 | -0.116 | -0.116 | 0 | 0.8 | 0.67 | 0.87 | 0.98 | 0.83 | 0.84 |
| 9 | 133367704 | rs7018779 | -0.489 | 1.185 | -1.058 | 1.229 | -0.002 | -0.037 | -0.037 | 0.123 | 0.08 | 0.11 | 0.05 | 0.01 | 0.12 | 0.06 |
| 9 | 133372383 | rs544701 | -1.174 | 1.388 | -1.417 | 1.861 | -0.034 | -0.007 | -0.007 | 0 | 0.08 | 0.01 | 0.25 | 0.46 | 0.12 | 0.16 |
| 9 | 133376153 | rs2282219 | -1.163 | 1.533 | -1.455 | 1.319 | -0.047 | -0.033 | -0.033 | 0 | 0.14 | 0.04 | 0.16 | 0.14 | 0.17 | 0.27 |
| 9 | 133387709 | rs2991274 | 1.424 | 1.727 | -0.934 | 1.858 | 0.024 | 0.224 | 0.224 | 0 | 0.92 | 0.78 | 0.63 | 0.48 | 0.9 | 0.81 |
| 9 | 133392659 | rs612380 | -1.39 | 1.515 | -1.23 | 0.839 | -0.005 | -0.002 | -0.002 | 0 | 0.09 | 0.09 | 0.15 | 0.1 | 0.15 | 0.22 |

| | | | | | | | | | | | | | | | | |
|---|---|---|---|---|---|---|---|---|---|---|---|---|---|---|---|---|
| 9 | 133400958 | rs4740339 | -0.2 | 1.416 | -2.11 | 0.356 | -0.098 | 0.019 | 0.019 | 0.415 | 0.37 | 0.07 | 0.51 | 0.72 | 0.41 | 0.4 |
| 9 | 133405176 | rs7874812 | -0.582 | 1.491 | -1.319 | 0.431 | 0.006 | 0.005 | 0.005 | 0 | 0.1 | 0.1 | 0.07 | 0.09 | 0.16 | 0.22 |
| 9 | 133410844 | rs7849694 | 0.035 | 1.323 | -1.125 | -0.507 | -0.001 | -0.003 | -0.003 | 0 | 0.42 | 0.62 | 0.36 | 0.17 | 0.42 | 0.37 |
| 9 | 133415084 | rs944605 | -0.645 | 1.215 | -1.113 | -0.756 | 0.001 | 0.109 | 0.109 | 0 | 0.44 | 0.75 | 0.61 | 0.81 | 0.41 | 0.43 |
| 9 | 133421313 | rs4740193 | -0.639 | 1.168 | -0.091 | -0.617 | 0.002 | -0.026 | -0.026 | 0.016 | 0.42 | 0.45 | 0.62 | 0.67 | 0.42 | 0.44 |
| 9 | 133422828 | rs4740194 | -0.861 | 1.18 | -0.026 | -0.662 | -0.008 | 0.041 | 0.041 | 0.053 | 0.44 | 0.63 | 0.68 | 0.77 | 0.46 | 0.51 |
| 9 | 133424020 | rs944604 | -0.72 | 1.126 | -0.046 | -0.711 | 0.007 | 0.067 | 0.067 | 0.005 | 0.48 | 0.62 | 0.72 | 0.77 | 0.56 | 0.52 |
| 9 | 133427274 | rs1560532 | -1.346 | 0.822 | -0.03 | -0.85 | -0.018 | 0.246 | 0.246 | 0.226 | 0.55 | 0.84 | 0.78 | 0.91 | 0.58 | 0.58 |
| 9 | 133427524 | rs11243759 | -0.371 | 0.751 | 0.533 | 0.244 | -0.001 | -0.01 | -0.01 | 0 | 0.38 | 0.53 | 0.46 | 0.55 | 0.42 | 0.27 |
| 9 | 133429006 | rs2313200 | -0.13 | 0.775 | 0.513 | 0.234 | 0.012 | 0.025 | 0.025 | 0 | 0.34 | 0.42 | 0.44 | 0.41 | 0.42 | 0.22 |
| 9 | 133436730 | rs10793932 | -1.074 | 0.744 | 0.151 | 0.092 | -0.023 | 0.109 | 0.109 | 0.412 | 0.49 | 0.82 | 0.72 | 0.73 | 0.53 | 0.56 |
| 9 | 133439577 | rs4740347 | -2.334 | 1.166 | 0.888 | 0.332 | 0.003 | 0.046 | 0.046 | 0 | 0.24 | 0.68 | 0.37 | 0.4 | 0.23 | 0.38 |
| 9 | 133440620 | rs7030440 | 0.574 | 1.028 | 0.816 | 0.279 | -0.007 | -0.014 | -0.014 | 0 | 0.27 | 0.18 | 0.42 | 0.5 | 0.37 | 0.27 |
| 9 | 133442778 | rs10793939 | -0.623 | 1.025 | 1.228 | 0.282 | 0.025 | -0.006 | -0.006 | 0.119 | 0.39 | 0.48 | 0.68 | 0.67 | 0.51 | 0.5 |
| 9 | 133453327 | rs4740196 | -1.24 | 0.669 | 1.903 | 1.184 | 0.003 | 0.031 | 0.031 | 0 | 0.1 | 0.2 | 0.23 | 0.18 | 0.1 | 0.22 |
| 9 | 133483217 | rs10901226 | -2.592 | 0.465 | 1.539 | 0.882 | 0.02 | 0.19 | 0.19 | 0.282 | 0.13 | 0.53 | 0.12 | 0.22 | 0.1 | 0.15 |
| 9 | 133488494 | rs11243976 | -1.132 | 0.545 | 1.148 | 1.028 | 0.018 | 0.244 | 0.244 | 0 | 0.05 | 0.24 | 0.06 | 0.16 | 0.01 | 0.07 |
| 9 | 133490496 | rs7040341 | -1.964 | 0.538 | 1.069 | 0.922 | 0.019 | 0.054 | 0.054 | 0 | 0.19 | 0.58 | 0.3 | 0.25 | 0.19 | 0.37 |
| 9 | 133516789 | rs10123316 | -1.39 | 0.205 | 0.535 | 0.693 | 0.011 | 0.205 | 0.205 | 0 | 0.06 | 0.24 | 0.06 | 0.17 | 0.01 | 0.06 |
| 9 | 133526315 | rs11244092 | -0.554 | 0.058 | -0.474 | -0.267 | 0.023 | 0.05 | 0.05 | 0 | 0.16 | 0.1 | 0.08 | 0 | 0.09 | 0.13 |
| 9 | 133532160 | rs7028896 | -1.73 | 0.295 | -0.154 | -0.109 | 0.002 | 0.003 | 0.003 | 0 | 0.07 | 0.22 | 0.18 | 0.07 | 0.07 | 0.2 |
| 9 | 133537672 | rs1890599 | -0.673 | 0.277 | -0.159 | -0.544 | 0.007 | 0.013 | 0.013 | 0 | 0.49 | 0.4 | 0.55 | 0.71 | 0.46 | 0.54 |
| 9 | 133539126 | rs2313583 | -0.487 | 0.285 | -0.143 | -0.571 | 0.006 | 0.008 | 0.008 | 0 | 0.47 | 0.38 | 0.49 | 0.69 | 0.42 | 0.41 |
| 9 | 133539661 | rs3808825 | -0.276 | 0.667 | -0.218 | -1.035 | 0.016 | 0.015 | 0.015 | 0 | 0.19 | 0.33 | 0.31 | 0.27 | 0.23 | 0.29 |
| 9 | 133543980 | rs2277186 | 0.489 | 0.075 | -0.592 | -1.247 | 0.046 | 0.031 | 0.031 | 0 | 0.18 | 0.01 | 0.09 | 0.09 | 0.08 | 0.2 |
| 9 | 133545578 | rs10901270 | -1.355 | 0.58 | 0.074 | -1.333 | -0.03 | 0.034 | 0.034 | 0 | 0.68 | 0.78 | 0.82 | 0.99 | 0.63 | 0.83 |
| 9 | 133548589 | rs10120700 | -1.191 | 0.515 | 0.142 | -1.379 | 0.002 | -0.058 | -0.058 | 0 | 0.65 | 0.59 | 0.79 | 0.99 | 0.61 | 0.82 |
| 9 | 133550133 | rs1545229 | -0.697 | 0.459 | -0.005 | -1.369 | 0.007 | -0.053 | -0.053 | 1.721 | 0.61 | 0.41 | 0.71 | 0.93 | 0.54 | 0.7 |
| 9 | 133552145 | rs10751506 | -1.551 | 0.371 | -0.492 | -1.389 | -0.075 | 0.196 | 0.196 | 0.094 | 0.7 | 0.98 | 0.85 | 0.99 | 0.7 | 0.83 |
| 9 | 133553187 | rs894243 | -1.286 | 0.416 | 0.411 | -1.066 | 0.085 | 0.044 | 0.044 | 0 | 0.07 | 0.32 | 0.11 | 0.07 | 0.08 | 0.12 |
| 9 | 133553993 | rs7021384 | 0.241 | 0.011 | 0.134 | -0.614 | -0.006 | 0.026 | 0.026 | 0 | 0.32 | 0.33 | 0.45 | 0.47 | 0.32 | 0.35 |
| 9 | 133554189 | rs7024667 | -0.314 | 0.217 | 0.027 | -0.769 | -0.011 | 0.138 | 0.138 | 0 | 0.14 | 0.25 | 0.34 | 0.56 | 0.15 | 0.33 |
| 9 | 133566997 | rs2855159 | 0.508 | -0.178 | -0.996 | -0.984 | -0.036 | 0.132 | 0.132 | 0 | 0.34 | 0.02 | 0.18 | 0.09 | 0.26 | 0.22 |
| 9 | 133593939 | rs2583839 | -1.003 | 0.016 | -1.325 | -0.879 | 0.022 | 0.06 | 0.06 | 0 | 0.55 | 0.8 | 0.58 | 0.65 | 0.42 | 0.57 |
| 9 | 133621773 | rs2253084 | 0.455 | 0.165 | -1.977 | -0.762 | 0.022 | 0.112 | 0.112 | 0 | 0.35 | 0.16 | 0.2 | 0.09 | 0.27 | 0.21 |
| 9 | 133629184 | rs4740363 | 0.15 | 0.169 | -1.847 | -0.073 | -0.011 | 0.005 | 0.005 | 0 | 0.07 | 0.01 | 0.25 | 0.48 | 0.08 | 0.22 |
| 9 | 133638943 | rs2855171 | -0.905 | 0.516 | -1.692 | -1.077 | -0.055 | 0.159 | 0.159 | 2.182 | 0.72 | 0.98 | 0.86 | 0.94 | 0.75 | 0.84 |
| 9 | 133646592 | rs4990084 | -0.207 | 0.624 | -1.015 | -0.987 | -0.001 | 0.006 | 0.006 | 0 | 0.39 | 0.36 | 0.31 | 0.19 | 0.41 | 0.27 |
| 9 | 133651648 | rs2855172 | -0.363 | 0.441 | -0.64 | -1.13 | 0 | -0.031 | -0.031 | 0 | 0.42 | 0.48 | 0.35 | 0.2 | 0.45 | 0.3 |
| 9 | 133660799 | rs11244139 | -0.016 | -0.012 | 0.281 | -1.549 | -0.025 | -0.112 | -0.112 | 0 | 0.16 | 0.42 | 0.1 | 0 | 0.08 | 0.02 |
| 9 | 133663058 | rs11244140 | -1.037 | -0.086 | 0.261 | -1.583 | -0.032 | -0.107 | -0.107 | 0 | 0.22 | 0.47 | 0.2 | 0.09 | 0.15 | 0.13 |
| 9 | 133667899 | rs7026988 | -0.78 | -0.141 | 0.446 | -1.603 | -0.033 | -0.099 | -0.099 | 2.016 | 0.2 | 0.44 | 0.19 | 0.08 | 0.13 | 0.13 |
| 9 | 133670874 | rs10901285 | 1.141 | 0.113 | -0.142 | -1.5 | -0.093 | -0.017 | -0.017 | 0 | 0.23 | 0.02 | 0.22 | 0.17 | 0.31 | 0.13 |
| 9 | 133675739 | rs4740372 | -0.161 | -0.206 | 1.036 | -1.351 | -0.008 | -0.029 | -0.029 | 0 | 0.16 | 0.16 | 0.16 | 0.08 | 0.11 | 0.13 |
| 9 | 133677168 | rs11244147 | -1.551 | -0.233 | 1.234 | -1.371 | -0.032 | -0.106 | -0.106 | 0 | 0.23 | 0.53 | 0.21 | 0.09 | 0.15 | 0.13 |
| 9 | 133707362 | rs2314342 | -0.99 | -0.166 | 1.339 | -1.335 | -0.02 | -0.087 | -0.087 | 0 | 0.21 | 0.35 | 0.19 | 0.09 | 0.15 | 0.11 |
| 9 | 133725767 | rs6597642 | -0.537 | -0.166 | 1.169 | -1.303 | -0.009 | -0.04 | -0.04 | 0 | 0.18 | 0.22 | 0.17 | 0.08 | 0.14 | 0.1 |
| 9 | 133732752 | rs10901294 | 0.695 | -0.166 | 1.178 | -1.299 | -0.01 | -0.048 | -0.048 | 1.226 | 0.12 | 0.15 | 0.07 | 0 | 0.07 | 0.03 |
| 9 | 133734075 | rs3824400 | -0.901 | -0.192 | 1.278 | -1.31 | -0.03 | -0.097 | -0.097 | 0.365 | 0.2 | 0.49 | 0.2 | 0.07 | 0.13 | 0.1 |
| 9 | 133735644 | rs2855194 | -1.509 | 0.075 | 0.177 | -1.575 | -0.031 | 0.073 | 0.073 | 0 | 0.63 | 0.82 | 0.67 | 0.76 | 0.6 | 0.63 |
| 9 | 133736116 | rs11244164 | -1.346 | -0.675 | 0.674 | -1.614 | 0.028 | 0.045 | 0.045 | 1.122 | 0.29 | 0.78 | 0.24 | 0.07 | 0.16 | 0.2 |
| 9 | 133743654 | rs3780282 | -1.376 | -0.621 | 0.68 | -1.604 | 0.021 | 0.031 | 0.031 | 1.52 | 0.29 | 0.77 | 0.25 | 0.08 | 0.16 | 0.14 |
| 9 | 133744672 | rs7848941 | -1.05 | -0.633 | 0.996 | -1.614 | -0.014 | -0.052 | -0.052 | 3.588 | 0.27 | 0.67 | 0.22 | 0.08 | 0.14 | 0.14 |
| 9 | 133749848 | rs7851994 | -1.323 | -0.62 | 0.797 | -1.606 | 0.017 | 0.022 | 0.022 | 1.527 | 0.29 | 0.77 | 0.25 | 0.08 | 0.16 | 0.14 |
| 9 | 133755528 | rs2227985 | 1.007 | -0.566 | 1.257 | -1.579 | -0.004 | -0.058 | -0.058 | 0 | 0.12 | 0.21 | 0.08 | 0 | 0.07 | 0.03 |
| 9 | 133755754 | rs12005009 | -1.702 | -0.518 | 0.869 | -1.68 | 0.025 | 0.135 | 0.135 | 3.464 | 0.32 | 0.84 | 0.28 | 0.08 | 0.21 | 0.18 |
| 9 | 133757785 | rs3808814 | 0.191 | -0.426 | 1.218 | -1.633 | 0.055 | 0.017 | 0.017 | 0 | 0.06 | 0.19 | 0.13 | 0.08 | 0.09 | 0.1 |
| 9 | 133761001 | rs1056171 | -1.858 | -0.539 | 0.47 | -1.248 | -0.094 | -0.027 | -0.027 | 0 | 0.58 | 0.88 | 0.7 | 0.6 | 0.49 | 0.55 |
| 9 | 133762558 | rs11991 | -1.068 | -0.505 | 1.458 | -1.27 | -0.037 | -0.137 | -0.137 | 0 | 0.24 | 0.49 | 0.14 | 0 | 0.12 | 0.09 |
| 9 | 133762826 | rs7457 | 0.439 | -0.48 | 1.311 | -1.255 | 0.002 | -0.074 | -0.074 | 0 | 0.12 | 0.29 | 0.09 | 0 | 0.07 | 0.03 |
| 9 | 133766008 | rs11244178 | -0.617 | -0.39 | 0.365 | -1.197 | -0.026 | -0.1 | -0.1 | 0.174 | 0.25 | 0.57 | 0.15 | 0 | 0.12 | 0.09 |
| 9 | 133769169 | rs3847193 | -1.096 | -0.121 | -0.21 | -1.279 | 0.01 | 0.044 | 0.044 | 2.241 | 0.35 | 0.74 | 0.29 | 0.07 | 0.25 | 0.28 |
| 9 | 133769786 | rs7034278 | 0.925 | -0.286 | -0.171 | -1.245 | 0.01 | 0.019 | 0.019 | 3.423 | 0.11 | 0.06 | 0.05 | 0.04 | 0.06 | 0.08 |
| 9 | 133770017 | rs7040543 | -1.396 | -0.507 | 0.197 | -0.773 | -0.017 | -0.001 | -0.001 | 0 | 0.28 | 0.67 | 0.45 | 0.64 | 0.19 | 0.31 |
| 9 | 133770222 | rs7865017 | -1.227 | -0.732 | 0.392 | -0.76 | -0.056 | -0.089 | -0.089 | 1.381 | 0.2 | 0.54 | 0.1 | 0.06 | 0.06 | 0.13 |
| 9 | 133770270 | rs7853654 | 0.266 | -0.427 | 0.76 | -0.163 | 0.014 | 0.213 | 0.213 | 0 | 0.08 | 0.19 | 0.34 | 0.44 | 0.13 | 0.17 |
| 9 | 133772910 | rs7026476 | -1.482 | -0.618 | 0.673 | -1.05 | -0.003 | 0.026 | 0.026 | 0 | 0.48 | 0.59 | 0.67 | 0.63 | 0.48 | 0.49 |
| 9 | 133775857 | rs2297091 | 0.151 | -0.287 | 0.535 | -0.653 | 0.041 | -0.027 | -0.027 | 0 | 0.08 | 0.15 | 0.15 | 0.03 | 0.13 | 0.12 |
| 9 | 133777993 | rs11789517 | 1.43 | -0.285 | 0.127 | -0.643 | 0.012 | 0.022 | 0.022 | 3.836 | 0.11 | 0.06 | 0.05 | 0 | 0.06 | 0.01 |

| | | | | | | | | | | | | | | | | |
|---|---|---|---|---|---|---|---|---|---|---|---|---|---|---|---|---|
| 9 | 133778117 | rs11791437 | 1.43 | -0.285 | 0.127 | -0.643 | 0.012 | 0.022 | 0.022 | 3.836 | 0.11 | 0.06 | 0.05 | 0 | 0.06 | 0.01 |
| 9 | 133779020 | rs3802347 | 0.068 | -0.151 | 0.128 | -0.565 | -0.011 | 0.014 | 0.014 | 0 | 0.11 | 0.09 | 0.36 | 0.49 | 0.19 | 0.21 |
| 9 | 133779448 | rs11792889 | -0.839 | -0.213 | 0.225 | -0.791 | 0.001 | -0.023 | -0.023 | 0 | 0.27 | 0.29 | 0.21 | 0.19 | 0.22 | 0.29 |
| 9 | 133787225 | rs6597650 | -1.956 | 0.115 | -0.016 | -0.328 | -0.05 | 0.007 | 0.007 | 0 | 0.4 | 0.84 | 0.6 | 0.61 | 0.4 | 0.47 |
| 9 | 133789420 | rs10901304 | -0.56 | 0.176 | 0.074 | -0.298 | -0.002 | -0.001 | -0.001 | 2.008 | 0.21 | 0.2 | 0.38 | 0.58 | 0.23 | 0.31 |
| 9 | 133790521 | rs3847195 | -1.468 | 0.064 | -0.587 | -0.261 | -0.031 | 0.01 | 0.01 | 0 | 0.4 | 0.77 | 0.59 | 0.61 | 0.4 | 0.47 |
| 9 | 133795610 | rs13293102 | -0.761 | -0.256 | -0.386 | -0.311 | 0 | -0.004 | -0.004 | 0.321 | 0.1 | 0.12 | 0.2 | 0.08 | 0.12 | 0.2 |
| 9 | 133796694 | rs6597655 | -0.395 | -0.481 | -0.695 | -0.519 | 0.002 | 0.003 | 0.003 | 0 | 0.84 | 0.88 | 0.92 | 0.84 | 0.86 | 0.71 |
| 9 | 133797557 | rs7035023 | -0.722 | -0.698 | -0.724 | -0.419 | -0.017 | 0.028 | 0.028 | 0 | 0.28 | 0.36 | 0.41 | 0.53 | 0.18 | 0.27 |
| 9 | 133808080 | rs7864465 | 1.013 | 0.024 | -1.361 | -0.233 | -0.051 | 0.105 | 0.105 | 0 | 0.41 | 0.21 | 0.33 | 0.23 | 0.55 | 0.28 |
| 9 | 133809775 | rs11244202 | -0.376 | -0.159 | -0.919 | -0.715 | 0.004 | -0.14 | -0.14 | 0.151 | 0.2 | 0.48 | 0.25 | 0.03 | 0.17 | 0.19 |
| 9 | 133810848 | rs10793974 | 0.205 | 0.393 | -1.213 | -0.134 | 0.031 | 0.07 | 0.07 | 0 | 0.51 | 0.39 | 0.58 | 0.75 | 0.66 | 0.53 |
| 9 | 133823473 | rs10901309 | -0.313 | 0.832 | -1.87 | 0.549 | -0.132 | -0.053 | -0.053 | 0 | 0.46 | 0.95 | 0.62 | 0.57 | 0.52 | 0.44 |
| 9 | 133828475 | rs7026302 | -1.47 | 1.1 | -0.158 | 0.419 | 0.089 | -0.025 | -0.025 | 0.003 | 0.07 | 0.27 | 0.07 | 0 | 0.1 | 0.03 |
| 9 | 133828515 | rs4568653 | -1.395 | 1.113 | -0.008 | 1.126 | 0.003 | 0.112 | 0.112 | 0 | 0.05 | 0.12 | 0.06 | 0.32 | 0.06 | 0.06 |
| 9 | 133848437 | rs12346096 | -1.616 | 0.773 | 0.611 | 0.326 | 0.001 | -0.079 | -0.079 | 0.304 | 0.16 | 0.59 | 0.12 | 0.03 | 0.15 | 0.07 |
| 9 | 133849498 | rs9886861 | -1.967 | 0.485 | 0.637 | 0.332 | 0.001 | 0.008 | 0.008 | 2.983 | 0.28 | 0.73 | 0.42 | 0.28 | 0.25 | 0.13 |
| 9 | 133849547 | rs9886826 | -1.148 | 0.789 | 0.883 | 0.512 | 0.019 | 0.013 | 0.013 | 0.228 | 0.13 | 0.15 | 0.16 | 0.25 | 0.2 | 0.06 |
| 9 | 133852742 | rs10901314 | -0.942 | 0.879 | 0.804 | 0.332 | 0.007 | 0.006 | 0.006 | 0.053 | 0.35 | 0.55 | 0.52 | 0.39 | 0.4 | 0.16 |
| 9 | 133856814 | rs12344376 | -1.095 | 0.539 | 1.306 | -0.195 | -0.03 | -0.024 | -0.024 | 0 | 0.12 | 0.47 | 0.07 | 0.08 | 0.03 | 0.06 |
| 9 | 133856962 | rs7048958 | -1.238 | 0.521 | 1.448 | -0.204 | -0.041 | -0.035 | -0.035 | 0.331 | 0.12 | 0.54 | 0.07 | 0.08 | 0.03 | 0.06 |
| 9 | 133857758 | rs17147386 | -1.794 | 0.38 | 1.455 | -0.176 | -0.007 | -0.004 | -0.004 | 0 | 0.17 | 0.71 | 0.12 | 0.15 | 0.11 | 0.07 |
| 9 | 133858122 | rs2178107 | -0.479 | 0.759 | 0.926 | -0.216 | -0.015 | -0.1 | -0.1 | 0 | 0.26 | 0.38 | 0.34 | 0.1 | 0.21 | 0.32 |
| 9 | 133860275 | rs10114591 | -0.838 | 0.739 | 0.728 | -0.235 | 0.002 | -0.009 | -0.009 | 0 | 0.29 | 0.63 | 0.4 | 0.17 | 0.28 | 0.34 |
| 9 | 133861632 | rs6597668 | -0.772 | 0.736 | 0.461 | -0.211 | -0.005 | 0.006 | 0.006 | 0.692 | 0.28 | 0.68 | 0.41 | 0.25 | 0.28 | 0.34 |
| 9 | 133862285 | rs7862053 | 0.039 | 1.141 | -0.021 | -0.218 | -0.041 | 0.011 | 0.011 | 0.019 | 0.85 | 0.97 | 0.91 | 0.9 | 0.8 | 0.8 |
| 9 | 133862958 | rs10901316 | 0.986 | 1.262 | -0.619 | -0.386 | -0.007 | 0.006 | 0.006 | 0.495 | 0.48 | 0.04 | 0.45 | 0.63 | 0.45 | 0.44 |
| 9 | 133863770 | rs12005819 | -0.36 | 1.299 | -0.16 | -0.166 | -0.002 | 0.002 | 0.002 | 0.319 | 0.36 | 0.6 | 0.44 | 0.22 | 0.36 | 0.35 |
| 9 | 133865384 | rs7852743 | -0.295 | 1.32 | -0.35 | -0.138 | 0.001 | 0.006 | 0.006 | 0.156 | 0.34 | 0.6 | 0.41 | 0.26 | 0.29 | 0.35 |
| 9 | 133865861 | rs11795033 | -0.241 | 0.965 | -1.101 | -0.143 | 0.01 | -0.02 | -0.02 | 0 | 0.35 | 0.55 | 0.43 | 0.23 | 0.36 | 0.35 |
| 9 | 133868398 | rs12683313 | 0.017 | 0.186 | -2.23 | 0.098 | 0.029 | 0.012 | 0.012 | 0.062 | 0.44 | 0.09 | 0.37 | 0.61 | 0.41 | 0.46 |
| 9 | 133872871 | rs11244233 | 0.006 | 0.14 | -2.143 | 0.101 | 0.039 | 0.034 | 0.034 | 1.305 | 0.41 | 0.11 | 0.47 | 0.64 | 0.36 | 0.52 |
| 9 | 133873991 | rs11244234 | -0.53 | 0.111 | -1.562 | 0.695 | -0.002 | 0.01 | 0.01 | 0 | 0.27 | 0.26 | 0.25 | 0.39 | 0.25 | 0.26 |
| 9 | 133874462 | rs10118337 | -0.252 | 0.39 | -1.382 | 0.478 | -0.005 | -0.006 | -0.006 | 0.021 | 0.24 | 0.25 | 0.24 | 0.62 | 0.39 | 0.36 |
| 9 | 133874568 | rs10116370 | -0.918 | -0.433 | -1.378 | -0.396 | -0.018 | 0.076 | 0.076 | 0 | 0.55 | 0.74 | 0.56 | 0.75 | 0.53 | 0.71 |
| 9 | 133874685 | rs11998813 | -0.549 | -0.921 | -0.826 | -0.448 | -0.003 | -0.026 | -0.026 | 0.36 | 0.15 | 0.14 | 0.08 | 0.02 | 0.11 | 0.12 |
| 9 | 133876113 | rs3857985 | -1.36 | -1.136 | -0.939 | -0.637 | 0.006 | 0.006 | 0.006 | 0 | 0.33 | 0.67 | 0.34 | 0.15 | 0.22 | 0.31 |
| 9 | 133876911 | rs10901317 | -0.947 | -0.922 | -0.656 | -0.467 | -0.02 | -0.025 | -0.025 | 0.138 | 0.23 | 0.66 | 0.29 | 0.12 | 0.15 | 0.23 |
| 9 | 133886042 | rs11244245 | -0.775 | -0.831 | -0.263 | -0.511 | -0.034 | -0.112 | -0.112 | 0.355 | 0.15 | 0.54 | 0.1 | 0.02 | 0.1 | 0.1 |
| 9 | 133887901 | rs869936 | -0.338 | -0.598 | -1.181 | -0.271 | -0.017 | -0.054 | -0.054 | 0.017 | 0.23 | 0.11 | 0.25 | 0.64 | 0.32 | 0.4 |
| 9 | 133888771 | rs915401 | -1.049 | -1.004 | -1.315 | -0.908 | 0.03 | 0.049 | 0.049 | 0 | 0.51 | 0.68 | 0.36 | 0.68 | 0.46 | 0.49 |
| 9 | 133891847 | rs881676 | -0.29 | -1.099 | -1.125 | -1.15 | -0.04 | -0.081 | -0.081 | 0.313 | 0.19 | 0.6 | 0.11 | 0.03 | 0.13 | 0.07 |
| 9 | 133897119 | rs4588908 | -0.598 | -1.038 | -1.52 | -1.147 | -0.011 | -0.013 | -0.013 | 0 | 0.29 | 0.67 | 0.31 | 0.14 | 0.2 | 0.29 |
| 9 | 133906060 | rs10793982 | -1.108 | -0.198 | -1.886 | -1.309 | -0.02 | 0.011 | 0.011 | 0 | 0.69 | 0.75 | 0.63 | 0.95 | 0.57 | 0.77 |
| 9 | 133906250 | rs3780280 | -0.422 | -0.413 | -2.478 | -1.159 | -0.006 | 0.093 | 0.093 | 0.038 | 0.39 | 0.11 | 0.33 | 0.78 | 0.42 | 0.44 |
| 9 | 133907229 | rs2275134 | -0.266 | -0.284 | -2.415 | -0.621 | -0.031 | -0.039 | -0.039 | 1.428 | 0.32 | 0.09 | 0.26 | 0.67 | 0.35 | 0.34 |
| 9 | 133908753 | rs3808810 | 0.187 | -0.676 | -1.557 | -0.839 | -0.015 | -0.033 | -0.033 | 0.714 | 0.19 | 0.2 | 0.08 | 0.04 | 0.08 | 0.1 |
| 9 | 133908961 | rs3780279 | 0.042 | 0.2 | -2.109 | -0.318 | -0.028 | -0.028 | -0.028 | 0 | 0.82 | 0.96 | 0.84 | 0.81 | 0.8 | 0.71 |
| 9 | 133911706 | rs2275131 | 0.271 | 0.504 | -2.006 | -0.196 | -0.072 | 0.026 | 0.026 | 0 | 0.4 | 0.16 | 0.49 | 0.66 | 0.48 | 0.29 |
| 9 | 133913434 | rs3780275 | 0.359 | 0.234 | -1.048 | -0.036 | -0.01 | -0.009 | -0.009 | 0.06 | 0.75 | 0.67 | 0.7 | 0.81 | 0.79 | 0.69 |
| 9 | 133914570 | rs2293518 | 0.3 | 0.561 | -1.612 | -0.049 | -0.004 | 0.033 | 0.033 | 0.09 | 0.22 | 0.11 | 0.21 | 0.14 | 0.24 | 0.1 |
| 9 | 133917277 | rs2293520 | 1.094 | 0.239 | -1.735 | 0.07 | 0.003 | -0.012 | -0.012 | 12.191 | 0.06 | 0.01 | 0.04 | 0.07 | 0.03 | 0.03 |
| 9 | 133917807 | rs4740396 | 0.092 | -0.028 | -2.472 | -0.285 | -0.005 | 0.037 | 0.037 | 0.114 | 0.27 | 0.02 | 0.25 | 0.15 | 0.24 | 0.23 |
| 9 | 133922072 | rs1414402 | 0.421 | 0.153 | -1.873 | 0.123 | 0 | -0.001 | -0.001 | 0 | 0.44 | 0.54 | 0.32 | 0.44 | 0.49 | 0.47 |
| 9 | 133924451 | rs869457 | 1.029 | 0.471 | -1.624 | 0.091 | -0.013 | 0.006 | 0.006 | 0 | 0.28 | 0.24 | 0.21 | 0.25 | 0.32 | 0.26 |
| 9 | 133925813 | rs11793535 | 0.288 | 0.125 | -1.092 | 0.616 | 0.017 | 0.065 | 0.065 | 0.065 | 0.08 | 0.18 | 0.05 | 0.15 | 0.08 | 0.12 |
| 9 | 133927286 | rs10793985 | -0.443 | -0.059 | -1.414 | 0.457 | -0.051 | -0.127 | -0.127 | 0 | 0.79 | 0.97 | 0.64 | 0.64 | 0.78 | 0.8 |
| 9 | 133927878 | rs10901333 | 0.188 | 0.126 | -1.034 | 0.42 | 0 | -0.001 | -0.001 | 0 | 0.44 | 0.55 | 0.33 | 0.43 | 0.5 | 0.48 |
| 9 | 133935704 | rs10901336 | 0.641 | 0.395 | -0.204 | 0.657 | 0.008 | 0.053 | 0.053 | 0 | 0.72 | 0.38 | 0.57 | 0.48 | 0.71 | 0.64 |
| 9 | 133937944 | rs6597674 | 0.29 | 0.347 | 0.428 | 0.697 | 0.017 | 0.06 | 0.06 | 0 | 0.75 | 0.66 | 0.6 | 0.5 | 0.72 | 0.64 |
| 9 | 133938973 | rs4740402 | 0.361 | 0.138 | -0.063 | 0.362 | 0.017 | 0.046 | 0.046 | 0 | 0.43 | 0.34 | 0.33 | 0.21 | 0.39 | 0.4 |
| 9 | 133938981 | rs4740403 | 1.127 | -0.235 | -0.146 | 0.386 | 0.068 | 0.144 | 0.144 | 0 | 0.33 | 0.09 | 0.25 | 0.08 | 0.26 | 0.26 |
| 9 | 133942844 | rs12553829 | -0.1 | -0.229 | 0.325 | 0.597 | -0.007 | 0.13 | 0.13 | 0.001 | 0.82 | 0.54 | 0.66 | 0.63 | 0.85 | 0.8 |
| 9 | 133946909 | rs10901344 | 0.546 | 0.172 | -0.202 | 0.331 | 0.019 | 0.145 | 0.145 | 0 | 0.44 | 0.16 | 0.33 | 0.19 | 0.41 | 0.42 |
| 9 | 133947345 | rs7858204 | 0.129 | 0.353 | 0.156 | 0.582 | 0.005 | 0.006 | 0.006 | 0.244 | 0.09 | 0.11 | 0.06 | 0.13 | 0.12 | 0.15 |
| 9 | 133948049 | rs2275140 | -0.22 | 0.382 | 0.556 | 0.69 | 0.005 | 0.143 | 0.143 | 0.12 | 0.83 | 0.56 | 0.66 | 0.64 | 0.86 | 0.81 |
| 9 | 133951180 | rs13301105 | -0.285 | 0.667 | 0.678 | 0.789 | 0.001 | 0.038 | 0.038 | 1.013 | 0.12 | 0.39 | 0.09 | 0.15 | 0.07 | 0.13 |
| 9 | 133951230 | rs3818800 | -0.375 | 0.656 | 0.642 | 0.783 | -0.007 | 0.026 | 0.026 | 1.115 | 0.12 | 0.38 | 0.1 | 0.16 | 0.07 | 0.12 |
| 9 | 133957759 | rs7860487 | -0.536 | 0.714 | 0.062 | 0.694 | -0.004 | -0.001 | -0.001 | 0 | 0.44 | 0.68 | 0.35 | 0.4 | 0.44 | 0.37 |

| | | | | | | | | | | | | | | | | |
|---|---|---|---|---|---|---|---|---|---|---|---|---|---|---|---|---|
| 9 | 133958148 | rs7873372 | -0.176 | 0.614 | -0.158 | 0.446 | -0.026 | 0.022 | 0.022 | 1.148 | 0.15 | 0.48 | 0.12 | 0.16 | 0.09 | 0.16 |
| 9 | 133958298 | rs1889749 | -0.969 | 0.753 | -1.321 | 0.247 | 0.024 | 0.026 | 0.026 | 0.019 | 0.51 | 0.87 | 0.43 | 0.55 | 0.57 | 0.54 |
| 9 | 133962930 | rs7024108 | 0.577 | 0.953 | -1.057 | 0.39 | -0.013 | 0 | 0 | 0 | 0.25 | 0.21 | 0.21 | 0.19 | 0.34 | 0.16 |
| 9 | 133963008 | rs4740412 | 1.5 | 0.476 | -1.068 | 0.008 | -0.013 | 0.159 | 0.159 | 0 | 0.3 | 0.06 | 0.24 | 0.08 | 0.27 | 0.21 |
| 9 | 133967240 | rs818056 | -0.359 | 0.628 | -0.884 | 0.157 | -0.006 | 0.013 | 0.013 | 1.701 | 0.27 | 0.64 | 0.22 | 0.36 | 0.22 | 0.37 |
| 9 | 133967510 | rs818055 | 0.127 | 0.584 | -0.971 | 0.155 | -0.004 | 0.008 | 0.008 | 1.705 | 0.73 | 0.35 | 0.78 | 0.64 | 0.78 | 0.63 |
| 9 | 133967828 | rs710075 | 0.142 | 0.604 | -0.967 | 0.159 | -0.004 | 0.008 | 0.008 | 1.686 | 0.73 | 0.35 | 0.78 | 0.64 | 0.78 | 0.63 |
| 9 | 133968546 | rs752929 | 1.505 | 0.548 | -1.181 | 0.061 | 0.01 | 0.139 | 0.139 | 0 | 0.31 | 0.07 | 0.24 | 0.08 | 0.28 | 0.21 |
| 9 | 133970178 | rs818051 | -0.333 | 1.054 | -0.829 | 0.594 | -0.045 | -0.014 | -0.014 | 0 | 0.41 | 0.24 | 0.53 | 0.55 | 0.5 | 0.4 |
| 9 | 133979352 | rs748697 | -0.17 | 1.116 | -0.298 | 0.364 | 0.01 | -0.011 | -0.011 | 0 | 0.09 | 0.22 | 0.07 | 0.03 | 0.1 | 0.05 |
| 9 | 133981491 | rs2872823 | -0.354 | 0.849 | 0.35 | 0.65 | -0.004 | 0.079 | 0.079 | 0 | 0.18 | 0.36 | 0.35 | 0.33 | 0.16 | 0.17 |
| 9 | 133981629 | rs2315075 | -0.294 | 0.859 | 0.357 | 0.655 | -0.006 | 0.046 | 0.046 | 0 | 0.17 | 0.3 | 0.34 | 0.33 | 0.16 | 0.17 |
| 9 | 133995353 | rs353514 | 0.422 | 1.117 | 0.643 | 0.109 | 0.011 | 0.051 | 0.051 | 0.154 | 0.74 | 0.42 | 0.8 | 0.62 | 0.75 | 0.73 |
| 9 | 133995959 | rs353513 | 0.429 | 1.102 | 0.634 | 0.091 | 0.01 | 0.046 | 0.046 | 0.169 | 0.74 | 0.42 | 0.8 | 0.63 | 0.75 | 0.7 |
| 9 | 133996184 | rs353512 | 0.494 | 1.231 | 0.628 | 0.04 | 0.024 | 0.055 | 0.055 | 0 | 0.74 | 0.46 | 0.78 | 0.62 | 0.73 | 0.72 |
| 9 | 134004912 | rs3736957 | -0.528 | 1.255 | 0.5 | 0.046 | 0 | 0.013 | 0.013 | 0.021 | 0.24 | 0.27 | 0.18 | 0.42 | 0.23 | 0.34 |
| 9 | 134012922 | rs7861678 | -0.502 | 1.258 | 0.508 | 0.047 | 0 | 0.009 | 0.009 | 0.028 | 0.24 | 0.25 | 0.17 | 0.42 | 0.24 | 0.34 |
| 9 | 134014000 | rs928675 | -0.664 | 1.254 | 0.534 | 0.045 | 0.003 | 0.041 | 0.041 | 0.103 | 0.25 | 0.34 | 0.18 | 0.42 | 0.24 | 0.34 |
| 9 | 134015072 | rs102993 | 1.385 | 1.145 | 0.026 | -0.527 | 0.005 | 0.135 | 0.135 | 0 | 0.33 | 0.12 | 0.29 | 0.03 | 0.3 | 0.18 |
| 9 | 134016121 | rs2281901 | 0.482 | 1.158 | 0.491 | 0.608 | -0.061 | -0.074 | -0.074 | 1.705 | 0.12 | 0.02 | 0.11 | 0.22 | 0.17 | 0.17 |
| 9 | 134020092 | rs103612 | 1.307 | 1.091 | 0.642 | 0.606 | -0.002 | -0.001 | -0.001 | 0 | 0.67 | 0.66 | 0.75 | 0.63 | 0.68 | 0.51 |
| 9 | 134025174 | rs7859782 | -1.403 | 1.28 | 0.82 | 0.945 | -0.002 | -0.017 | -0.017 | 0.324 | 0.3 | 0.27 | 0.21 | 0.42 | 0.29 | 0.45 |
| 9 | 134025595 | rs11244305 | -0.021 | 1.345 | 0.813 | 0.974 | -0.079 | -0.084 | -0.084 | 0 | 0.17 | 0.02 | 0.15 | 0.23 | 0.23 | 0.23 |
| 9 | 134064345 | rs2274427 | 1.072 | 1.286 | 0.987 | 0.975 | 0.007 | 0.037 | 0.037 | 0 | 0.68 | 0.57 | 0.76 | 0.57 | 0.68 | 0.59 |
| 9 | 134067474 | rs10901356 | 1.28 | 1.294 | 0.86 | 0.974 | 0.003 | 0.024 | 0.024 | 0.012 | 0.67 | 0.45 | 0.75 | 0.57 | 0.67 | 0.59 |
| 9 | 134096015 | rs4740420 | -0.881 | 1.428 | -0.737 | 1.052 | -0.017 | -0.069 | -0.069 | 0.01 | 0.29 | 0.17 | 0.2 | 0.42 | 0.3 | 0.39 |
| 9 | 134118740 | rs11244336 | 0.768 | 1.108 | -0.448 | 0.822 | 0.003 | 0.025 | 0.025 | 0.286 | 0.88 | 0.76 | 0.93 | 0.77 | 0.9 | 0.82 |
| 9 | 134136248 | rs9966 | -0.554 | 1.08 | -0.872 | 0.28 | -0.031 | 0.058 | 0.058 | 0 | 0.12 | 0.02 | 0.1 | 0.01 | 0.19 | 0.15 |
| 9 | 134139458 | rs3739508 | -0.371 | 1.594 | -0.236 | 0.768 | -0.004 | -0.003 | -0.003 | 0 | 0.26 | 0.21 | 0.29 | 0.67 | 0.36 | 0.47 |
| 9 | 134142444 | rs7041339 | -0.435 | 1.789 | 0.568 | 0.636 | -0.009 | -0.002 | -0.002 | 0 | 0.26 | 0.21 | 0.31 | 0.68 | 0.39 | 0.47 |
| 9 | 134149338 | rs10793992 | -0.567 | 1.766 | 0.556 | 1.472 | -0.005 | -0.007 | -0.007 | 0.013 | 0.28 | 0.2 | 0.31 | 0.5 | 0.39 | 0.46 |
| 9 | 134149767 | rs10901359 | -0.556 | 1.825 | 0.485 | 1.473 | -0.016 | -0.029 | -0.029 | 0 | 0.28 | 0.16 | 0.31 | 0.5 | 0.39 | 0.46 |
| 9 | 134156043 | rs10901363 | -0.547 | 1.904 | 0.498 | 1.488 | -0.031 | -0.057 | -0.057 | 1.226 | 0.28 | 0.14 | 0.31 | 0.49 | 0.4 | 0.48 |
| 9 | 134162558 | rs11244354 | 0.878 | 1.493 | -0.427 | 0.723 | -0.038 | -0.031 | -0.031 | 0.439 | 0.2 | 0.02 | 0.3 | 0.19 | 0.18 | 0.15 |
| 9 | 134164454 | rs11793913 | -0.155 | 1.578 | -0.086 | 0.887 | -0.03 | -0.056 | -0.056 | 0 | 0.28 | 0.14 | 0.31 | 0.51 | 0.39 | 0.46 |
| 9 | 134165248 | rs7041276 | -0.169 | 1.579 | 0.129 | 0.887 | -0.01 | -0.018 | -0.018 | 0 | 0.28 | 0.19 | 0.31 | 0.51 | 0.39 | 0.46 |
| 9 | 134169176 | rs2986608 | 0.769 | 1.446 | 0.353 | 1.361 | 0.009 | 0.159 | 0.159 | 0.051 | 0.88 | 0.7 | 0.81 | 0.65 | 0.86 | 0.8 |
| 9 | 134169616 | rs2483469 | 0.272 | 1.363 | -0.134 | 0.694 | -0.001 | -0.001 | -0.001 | 0 | 0.52 | 0.53 | 0.53 | 0.5 | 0.48 | 0.44 |
| 9 | 134183379 | rs2966332 | 0.464 | 1.642 | -0.321 | 0.635 | 0.029 | 0.015 | 0.015 | 2.215 | 0.76 | 0.4 | 0.65 | 0.73 | 0.74 | 0.67 |
| 9 | 134183503 | rs2986606 | 0.432 | 1.636 | -0.322 | 0.637 | 0.034 | 0.016 | 0.016 | 2.066 | 0.76 | 0.44 | 0.65 | 0.73 | 0.74 | 0.67 |
| 9 | 134183913 | rs2483476 | 0.446 | 1.649 | -0.315 | 0.644 | 0.033 | 0.015 | 0.015 | 2.183 | 0.76 | 0.43 | 0.65 | 0.73 | 0.74 | 0.67 |
| 9 | 134184191 | rs2483477 | 0.451 | 1.645 | -0.368 | 0.641 | 0.03 | 0.014 | 0.014 | 2.264 | 0.76 | 0.42 | 0.65 | 0.73 | 0.74 | 0.67 |
| 9 | 134195966 | rs913218 | 0.575 | 1.196 | -1.497 | 0.582 | -0.047 | -0.115 | -0.115 | 0 | 0.32 | 0.03 | 0.38 | 0.36 | 0.37 | 0.34 |
| 9 | 134198900 | rs2986599 | 0.077 | 1.13 | -0.986 | 0.49 | 0 | 0.011 | 0.011 | 0 | 0.36 | 0.58 | 0.38 | 0.55 | 0.32 | 0.38 |
| 9 | 134209707 | rs10751521 | 0.428 | 1.025 | -1.253 | 0.487 | 0 | -0.045 | -0.045 | 0 | 0.4 | 0.21 | 0.46 | 0.47 | 0.49 | 0.36 |
| 9 | 134212002 | rs4740438 | 0.351 | 0.711 | -0.424 | 0.411 | -0.004 | 0.009 | 0.009 | 0.857 | 0.7 | 0.42 | 0.73 | 0.73 | 0.73 | 0.57 |
| 9 | 134214383 | rs522464 | 0.289 | 0.399 | -0.114 | -0.025 | -0.013 | -0.062 | -0.062 | 1.475 | 0.72 | 0.54 | 0.78 | 0.93 | 0.77 | 0.67 |
| 9 | 134222085 | rs2966373 | 0.321 | 0.613 | -0.273 | 0.571 | -0.001 | 0 | 0 | 0 | 0.37 | 0.38 | 0.47 | 0.45 | 0.36 | 0.31 |
| 9 | 134223191 | rs11243356 | 0.843 | 0.867 | -0.591 | 0.439 | -0.017 | -0.016 | -0.016 | 1.258 | 0.1 | 0.06 | 0.17 | 0.15 | 0.12 | 0.12 |
| 9 | 134225572 | rs2050243 | -0.65 | 0.698 | -0.132 | 0.333 | -0.001 | 0.006 | 0.006 | 0 | 0.48 | 0.5 | 0.42 | 0.4 | 0.5 | 0.45 |
| 9 | 134228471 | rs11243362 | -0.005 | 0.852 | 0.032 | 0.181 | -0.006 | -0.045 | -0.045 | 0 | 0.36 | 0.42 | 0.24 | 0.13 | 0.35 | 0.32 |
| 9 | 134233268 | rs7852711 | -1.24 | 0.289 | 0.139 | 0.446 | 0.008 | 0.015 | 0.015 | 0 | 0.36 | 0.52 | 0.51 | 0.57 | 0.35 | 0.34 |
| 9 | 134233569 | rs4740227 | 0.83 | 0.225 | -0.03 | 0.409 | -0.001 | -0.001 | -0.001 | 0 | 0.08 | 0.04 | 0.14 | 0.09 | 0.09 | 0.06 |
| 9 | 134237148 | rs10901052 | -0.226 | -0.142 | 0.145 | 0.306 | -0.001 | 0.031 | 0.031 | 0 | 0.45 | 0.37 | 0.33 | 0.27 | 0.44 | 0.42 |
| 9 | 134238611 | rs11243375 | -0.379 | 0.11 | -0.247 | 0.41 | 0.002 | 0.154 | 0.154 | 0.029 | 0.06 | 0.22 | 0.13 | 0.15 | 0.03 | 0.08 |
| 9 | 134240052 | rs10901054 | -0.76 | 0.275 | -0.407 | -0.41 | -0.002 | 0.003 | 0.003 | 0 | 0.54 | 0.59 | 0.49 | 0.43 | 0.52 | 0.48 |
| 9 | 134247571 | rs6597494 | -1.019 | 0.055 | -0.3 | -0.795 | -0.013 | -0.051 | -0.051 | 0.415 | 0.18 | 0.19 | 0.07 | 0.02 | 0.09 | 0.07 |
| 9 | 134254451 | rs500446 | -1.018 | 0.355 | -0.542 | -0.89 | 0.001 | 0 | 0 | 0 | 0.3 | 0.34 | 0.28 | 0.14 | 0.26 | 0.33 |
| 9 | 134256675 | rs2483468 | 0.697 | 0.878 | -0.958 | -1.211 | 0.005 | 0.152 | 0.152 | 0 | 0.55 | 0.23 | 0.57 | 0.83 | 0.6 | 0.86 |
| 9 | 134258129 | rs11243383 | 0.722 | 1.618 | -0.813 | -0.794 | 0.009 | 0 | 0 | 0.113 | 0.08 | 0.1 | 0.07 | 0.08 | 0.14 | 0.11 |
| 9 | 134258779 | rs542305 | 0.558 | 0.918 | -0.496 | -1.04 | -0.001 | 0.055 | 0.055 | 0 | 0.58 | 0.32 | 0.6 | 0.83 | 0.65 | 0.84 |
| 9 | 134272423 | rs517961 | 1.078 | 1.299 | 0.578 | -0.604 | 0.092 | -0.052 | -0.052 | 0 | 0.85 | 0.47 | 0.76 | 0.92 | 0.79 | 0.94 |
| 9 | 134290898 | rs2994040 | 1.039 | 1.305 | 0.615 | -0.646 | 0.102 | -0.066 | -0.066 | 0.423 | 0.85 | 0.49 | 0.75 | 0.94 | 0.78 | 0.94 |
| 9 | 134313625 | rs7049110 | -0.524 | 1.392 | 1.506 | -0.417 | 0.038 | -0.035 | -0.035 | 0.774 | 0.07 | 0.18 | 0.07 | 0 | 0.15 | 0.03 |
| 9 | 134314882 | rs2966344 | 1.781 | 1.384 | 1.839 | -0.436 | 0.099 | -0.059 | -0.059 | 0 | 0.86 | 0.55 | 0.76 | 0.94 | 0.78 | 0.94 |
| 9 | 134316004 | rs2994060 | 0.457 | 1.372 | 1.821 | -0.428 | 0.044 | 0.007 | 0.007 | 0 | 0.92 | 0.74 | 0.82 | 0.94 | 0.93 | 0.97 |
| 9 | 134326950 | rs2994066 | -0.456 | 1.537 | 1.441 | -1.013 | -0.02 | 0.031 | 0.031 | 0 | 0.37 | 0.29 | 0.25 | 0.12 | 0.4 | 0.26 |
| 9 | 134342938 | rs10793870 | 0.66 | 1.82 | 1.264 | -1.108 | -0.092 | 0.02 | 0.02 | 0 | 0.14 | 0.04 | 0.15 | 0.11 | 0.29 | 0.18 |
| 9 | 134345401 | rs10793871 | 1.483 | 1.434 | 1.651 | -0.794 | 0.101 | -0.054 | -0.054 | 0 | 0.86 | 0.52 | 0.75 | 0.93 | 0.78 | 0.94 |

| | | | | | | | | | | | | | | | | |
|---|---|---|---|---|---|---|---|---|---|---|---|---|---|---|---|---|
| 9 | 134351343 | rs11243403 | -0.605 | 1.364 | 1.666 | -0.804 | 0.018 | 0.042 | 0.042 | 0 | 0.43 | 0.37 | 0.32 | 0.12 | 0.56 | 0.29 |
| 9 | 134353724 | rs4076428 | 0.985 | 1.382 | 0.737 | -0.742 | 0.143 | 0.243 | 0.243 | 0 | 0.48 | 0.12 | 0.48 | 0.81 | 0.37 | 0.68 |
| 9 | 134355421 | rs7854462 | 1.088 | 1.722 | 1.165 | -0.181 | 0.046 | 0.007 | 0.007 | 0 | 0.92 | 0.66 | 0.82 | 0.94 | 0.93 | 0.98 |
| 9 | 134358108 | rs2275498 | -0.503 | 1.722 | 1.198 | -0.182 | 0.012 | -0.006 | -0.006 | 0 | 0.06 | 0.12 | 0.16 | 0.06 | 0.07 | 0.02 |
| 9 | 134361627 | rs10901062 | 1.251 | 1.722 | 1.179 | -0.182 | 0.048 | 0.008 | 0.008 | 0 | 0.92 | 0.65 | 0.82 | 0.94 | 0.93 | 0.98 |
| 9 | 134362878 | rs3802349 | 1.215 | 1.722 | 1.173 | -0.181 | 0.046 | 0.007 | 0.007 | 0 | 0.92 | 0.66 | 0.82 | 0.94 | 0.93 | 0.98 |
| 9 | 134385436 | rs2296949 | 0.77 | 1.791 | 0.96 | -0.075 | 0.042 | 0.039 | 0.039 | 0 | 0.93 | 0.72 | 0.83 | 0.92 | 0.94 | 0.97 |
| 9 | 134387488 | rs3739494 | 0.73 | 1.799 | 0.964 | -0.07 | 0.044 | 0.041 | 0.041 | 0 | 0.93 | 0.72 | 0.83 | 0.92 | 0.94 | 0.97 |
| 9 | 134397624 | rs4740165 | 2.009 | 1.459 | 1.04 | -0.047 | 0.042 | 0.025 | 0.025 | 0.101 | 0.9 | 0.6 | 0.82 | 0.92 | 0.92 | 0.97 |
| 9 | 134398534 | rs3739495 | 0.804 | 1.573 | 0.754 | 0.035 | 0.045 | 0.038 | 0.038 | 0 | 0.93 | 0.72 | 0.83 | 0.93 | 0.94 | 0.97 |
| 9 | 134398719 | rs3824395 | 0.804 | 1.573 | 0.754 | 0.035 | 0.045 | 0.038 | 0.038 | 0 | 0.93 | 0.72 | 0.83 | 0.93 | 0.94 | 0.97 |
| 9 | 134398771 | rs10793885 | 0.804 | 1.573 | 0.754 | 0.035 | 0.045 | 0.034 | 0.034 | 0 | 0.93 | 0.72 | 0.83 | 0.93 | 0.94 | 0.97 |
| 9 | 134398841 | rs7857419 | 1.082 | 1.573 | 0.754 | 0.035 | 0.043 | 0.032 | 0.032 | 0 | 0.93 | 0.72 | 0.83 | 0.92 | 0.93 | 0.97 |
| 9 | 134402338 | rs6597502 | -1.231 | 1.45 | 0.185 | -0.072 | 0.031 | -0.035 | -0.035 | 0.237 | 0.11 | 0.25 | 0.09 | 0.04 | 0.16 | 0.07 |
| 9 | 134405598 | rs2282010 | -0.349 | 1.558 | 0.035 | 0.202 | 0.023 | 0.016 | 0.016 | 0 | 0.05 | 0.12 | 0.15 | 0.07 | 0.06 | 0.02 |
| 9 | 134408434 | rs3808830 | 1.233 | 1.657 | 0.068 | 0.143 | 0.126 | -0.006 | -0.006 | 0 | 0.88 | 0.64 | 0.75 | 0.93 | 0.77 | 0.94 |
| 9 | 134411018 | rs7874376 | -0.891 | 0.476 | -1.472 | 0.579 | 0.104 | -0.021 | -0.021 | 0 | 0.55 | 0.92 | 0.69 | 0.57 | 0.63 | 0.57 |
| 9 | 134413298 | rs4740263 | -0.849 | 0.478 | -1.469 | 0.643 | 0.099 | -0.014 | -0.014 | 0 | 0.54 | 0.92 | 0.68 | 0.57 | 0.62 | 0.57 |
| 9 | 134420585 | rs4740268 | -0.126 | 0.157 | -1.259 | 0.73 | 0.017 | 0.02 | 0.02 | 0 | 0.06 | 0.12 | 0.17 | 0.08 | 0.09 | 0.05 |
| 9 | 134428539 | rs10117759 | -0.748 | -0.056 | -1.713 | 0.734 | 0.074 | -0.04 | -0.04 | 0 | 0.56 | 0.96 | 0.68 | 0.6 | 0.62 | 0.58 |
| 9 | 134434362 | rs11243426 | -0.082 | -0.702 | -1.954 | 1.035 | 0.046 | -0.089 | -0.089 | 0.05 | 0.35 | 0.09 | 0.4 | 0.49 | 0.27 | 0.26 |
| 9 | 134437086 | rs4740280 | -0.715 | -0.821 | -0.623 | 1.322 | -0.019 | -0.014 | -0.014 | 0 | 0.77 | 0.75 | 0.82 | 0.8 | 0.83 | 0.81 |
| 9 | 134440917 | rs4740282 | 0.146 | -0.828 | 0.411 | 1.509 | 0.054 | 0.186 | 0.186 | 0 | 0.07 | 0.21 | 0.23 | 0.21 | 0.09 | 0.06 |
| 9 | 134442561 | rs12002074 | 0.083 | -0.566 | 0.079 | 0.969 | 0.007 | 0.069 | 0.069 | 0 | 0.33 | 0.22 | 0.21 | 0.09 | 0.29 | 0.22 |
| 9 | 134443228 | rs12378905 | 1.188 | -0.758 | 0.309 | 1.026 | -0.04 | 0.015 | 0.015 | 0.091 | 0.11 | 0.06 | 0.16 | 0.1 | 0.26 | 0.13 |
| 9 | 134445326 | rs6597513 | -0.589 | -0.88 | 0.597 | 1.014 | 0.056 | 0.023 | 0.023 | 0.183 | 0.22 | 0.59 | 0.41 | 0.31 | 0.35 | 0.2 |
| 9 | 134448296 | rs4740283 | -1.602 | -0.926 | 0.439 | 0.928 | 0.038 | -0.001 | -0.001 | 0 | 0.79 | 0.95 | 0.84 | 0.76 | 0.84 | 0.8 |
| 9 | 134449275 | rs17538157 | 1.543 | -0.973 | 0.409 | 0.866 | -0.058 | 0.03 | 0.03 | 0 | 0.26 | 0.08 | 0.21 | 0.15 | 0.36 | 0.33 |
| 9 | 134449467 | rs11243436 | 1.375 | -0.951 | 0.736 | 0.894 | 0.02 | -0.001 | -0.001 | 0 | 0.11 | 0.1 | 0.16 | 0.11 | 0.25 | 0.14 |
| 9 | 134451843 | rs7030401 | 0.514 | -1.003 | 1.171 | 0.906 | -0.013 | -0.019 | -0.019 | 1.897 | 0.31 | 0.34 | 0.12 | 0.09 | 0.12 | 0.17 |
| 9 | 134452681 | rs943851 | -0.909 | -1.002 | 1.342 | 1.036 | -0.001 | 0.001 | 0.001 | 0.861 | 0.72 | 0.76 | 0.62 | 0.55 | 0.74 | 0.74 |
| 9 | 134453976 | rs3739497 | 1.471 | -0.996 | 1.123 | 1.007 | -0.059 | 0.034 | 0.034 | 0 | 0.27 | 0.07 | 0.21 | 0.15 | 0.36 | 0.33 |
| 9 | 134458109 | rs3765545 | 0.714 | -0.995 | 1.21 | 0.977 | 0.029 | 0.032 | 0.032 | 0 | 0.29 | 0.23 | 0.11 | 0.09 | 0.12 | 0.17 |
| 9 | 134458378 | rs3765542 | -0.707 | -0.731 | 1.205 | 1.127 | -0.052 | 0.027 | 0.027 | 0 | 0.69 | 0.85 | 0.68 | 0.66 | 0.59 | 0.66 |
| 9 | 134460326 | rs744313 | 0.333 | -0.802 | 1.434 | 1.03 | 0.031 | 0.143 | 0.143 | 0 | 0.07 | 0.18 | 0.22 | 0.22 | 0.1 | 0.11 |
| 9 | 134467153 | rs10736852 | -1.232 | -1.023 | 1.317 | 0.985 | -0.008 | -0.003 | -0.003 | 1.102 | 0.77 | 0.77 | 0.83 | 0.77 | 0.84 | 0.81 |
| 9 | 134467314 | rs2296952 | -1.178 | -1.03 | 1.296 | 0.981 | -0.023 | -0.011 | -0.011 | 0.388 | 0.77 | 0.72 | 0.82 | 0.77 | 0.84 | 0.81 |
| 9 | 134470351 | rs3780263 | -1.519 | -0.977 | 1.186 | 0.978 | 0.033 | 0.004 | 0.004 | 0.81 | 0.79 | 0.95 | 0.84 | 0.77 | 0.83 | 0.81 |
| 9 | 134471153 | rs3808803 | -0.709 | -0.843 | 1.189 | 0.96 | 0.011 | 0.041 | 0.041 | 0 | 0.5 | 0.65 | 0.58 | 0.61 | 0.46 | 0.53 |
| 9 | 134473517 | rs2296950 | -0.222 | -0.417 | 1.056 | 1.084 | 0.119 | 0.105 | 0.105 | 0 | 0.19 | 0.39 | 0.38 | 0.37 | 0.34 | 0.24 |
| 9 | 134473655 | rs11243446 | 0.179 | -1.073 | 0.561 | 0.58 | -0.008 | 0.044 | 0.044 | 0 | 0.06 | 0.19 | 0.04 | 0.09 | 0.02 | 0.08 |
| 9 | 134483805 | rs1410562 | -1.134 | -1.2 | 0.121 | 0.493 | 0.048 | 0.032 | 0.032 | 0.411 | 0.78 | 0.94 | 0.79 | 0.83 | 0.86 | 0.8 |
| 9 | 134489763 | rs1887787 | -0.329 | -1.304 | 0.338 | 0.489 | 0.02 | 0.005 | 0.005 | 0.02 | 0.7 | 0.72 | 0.73 | 0.73 | 0.81 | 0.7 |
| 9 | 134492235 | rs2282006 | -0.853 | -1.357 | 0.517 | 0.631 | 0.031 | 0 | 0 | 0.014 | 0.76 | 0.83 | 0.72 | 0.73 | 0.85 | 0.7 |
| 9 | 134495007 | rs10901075 | -0.834 | -1.362 | 0.515 | 0.659 | 0.032 | 0.012 | 0.012 | 0.413 | 0.76 | 0.84 | 0.72 | 0.76 | 0.85 | 0.7 |
| 9 | 134497703 | rs10121026 | 1.017 | -1.239 | 0.664 | 0.755 | 0.007 | 0.074 | 0.074 | 0 | 0.06 | 0.06 | 0.1 | 0.24 | 0.12 | 0.1 |
| 9 | 134499896 | rs2031878 | -1.045 | -1.265 | 0.313 | 0.563 | -0.02 | 0.062 | 0.062 | 0 | 0.56 | 0.9 | 0.63 | 0.67 | 0.55 | 0.56 |
| 9 | 134501124 | rs2026006 | 0.018 | -1.241 | 0.383 | 0.578 | -0.005 | 0.029 | 0.029 | 0 | 0.08 | 0.21 | 0.07 | 0.1 | 0.05 | 0.09 |
| 9 | 134503810 | rs17148121 | -0.874 | -1.229 | 0.658 | 0.511 | 0.022 | 0.217 | 0.217 | 0 | 0.16 | 0.49 | 0.2 | 0.33 | 0.18 | 0.2 |
| 9 | 134508442 | rs7036709 | 0.435 | -1.172 | 0.563 | 0.452 | -0.032 | 0.056 | 0.056 | 0 | 0.36 | 0.2 | 0.34 | 0.24 | 0.36 | 0.25 |
| 9 | 134530248 | rs1887786 | 1.713 | -1.18 | 0.455 | 0.423 | -0.085 | -0.011 | -0.011 | 0 | 0.23 | 0.03 | 0.17 | 0.16 | 0.31 | 0.24 |
| 9 | 134536621 | rs17148166 | 1.105 | -1.228 | 0.431 | 0.293 | 0.056 | 0.071 | 0.071 | 0.015 | 0.24 | 0.09 | 0.08 | 0 | 0.1 | 0.1 |
| 9 | 134540980 | rs6597517 | 1.429 | -1.189 | 0.521 | 0.341 | -0.002 | -0.001 | -0.001 | 0 | 0.27 | 0.26 | 0.26 | 0.26 | 0.32 | 0.34 |
| 9 | 134555186 | rs12001816 | 0.499 | -1.319 | 0.492 | 0.302 | 0.03 | 0.067 | 0.067 | 0 | 0.51 | 0.34 | 0.33 | 0.26 | 0.41 | 0.44 |
| 9 | 134555407 | rs17148186 | 1.067 | -1.255 | 0.504 | 0.32 | 0.027 | 0.03 | 0.03 | 0 | 0.25 | 0.17 | 0.14 | 0.1 | 0.1 | 0.2 |
| 9 | 134582408 | rs11243479 | -0.456 | -1.009 | 0.684 | 0.504 | -0.005 | 0.096 | 0.096 | 0 | 0.1 | 0.29 | 0.15 | 0.2 | 0.05 | 0.2 |
| 9 | 134585181 | rs11243480 | 1.215 | -0.94 | 0.584 | 0.441 | 0.005 | -0.005 | -0.005 | 0 | 0.12 | 0.11 | 0.27 | 0.24 | 0.27 | 0.15 |
| 9 | 134591644 | rs1952422 | -0.706 | -1.241 | 0.607 | 0.442 | -0.064 | -0.018 | -0.018 | 0 | 0.69 | 0.85 | 0.53 | 0.6 | 0.59 | 0.65 |
| 9 | 134593583 | rs7020704 | 1.008 | -1.093 | 0.694 | 0.518 | -0.002 | 0.001 | 0.001 | 0 | 0.29 | 0.26 | 0.28 | 0.26 | 0.34 | 0.38 |
| 9 | 134596993 | rs11791414 | 0.559 | -1.419 | 0.488 | 0.273 | 0.002 | -0.003 | -0.003 | 0.004 | 0.41 | 0.38 | 0.57 | 0.5 | 0.61 | 0.49 |
| 9 | 134600009 | rs4740304 | -0.458 | -1.396 | 0.558 | 0.258 | 0.028 | 0.027 | 0.027 | 1.737 | 0.56 | 0.64 | 0.76 | 0.83 | 0.78 | 0.72 |
| 9 | 134604548 | rs11243490 | 0.516 | -1.202 | 0.303 | 0.328 | 0.001 | 0.008 | 0.008 | 0 | 0.34 | 0.45 | 0.33 | 0.35 | 0.36 | 0.45 |
| 9 | 134611544 | rs11243494 | 0.652 | -1.17 | 0.222 | 0.407 | 0.005 | 0.006 | 0.006 | 1.637 | 0.4 | 0.39 | 0.52 | 0.38 | 0.6 | 0.41 |
| 9 | 134621332 | rs4740309 | 0.979 | -0.645 | 0.397 | 0.806 | -0.003 | -0.004 | -0.004 | 0 | 0.16 | 0.15 | 0.33 | 0.25 | 0.29 | 0.17 |
| 9 | 134621809 | rs1123551 | -0.751 | -0.805 | 0.412 | 0.89 | -0.018 | 0.151 | 0.151 | 0.006 | 0.12 | 0.36 | 0.14 | 0.21 | 0.06 | 0.21 |
| 9 | 134630976 | rs2987406 | -1.345 | -0.755 | 0.173 | 0.8 | 0.019 | -0.01 | -0.01 | 0 | 0.85 | 0.98 | 0.85 | 0.83 | 0.89 | 0.82 |
| 9 | 134631095 | rs3012771 | 0.32 | -0.718 | 0.315 | 0.849 | -0.015 | 0.042 | 0.042 | 0.004 | 0.7 | 0.64 | 0.68 | 0.51 | 0.72 | 0.61 |
| 9 | 134633824 | rs12340210 | 0.349 | -0.41 | -0.041 | 0.379 | 0.007 | 0.127 | 0.127 | 0 | 0.08 | 0.16 | 0.18 | 0.36 | 0.13 | 0.21 |
| 9 | 134634208 | rs2987409 | 0.706 | -0.6 | -0.338 | -0.328 | -0.003 | -0.005 | -0.005 | 1.695 | 0.39 | 0.34 | 0.37 | 0.5 | 0.46 | 0.52 |

| | | | | | | | | | | | | | | | | |
|---|---|---|---|---|---|---|---|---|---|---|---|---|---|---|---|---|
| 9 | 134634566 | rs2987410 | -0.644 | -0.696 | -0.385 | -0.398 | -0.027 | -0.081 | -0.081 | 0 | 0.77 | 0.77 | 0.92 | 1 | 0.92 | 0.94 |
| 9 | 134635034 | rs10901086 | -0.154 | 0.179 | -0.48 | 0.422 | -0.003 | 0.008 | 0.008 | 0 | 0.08 | 0.12 | 0.06 | 0.08 | 0.07 | 0.1 |
| 9 | 134635088 | rs1887778 | -0.133 | 0.324 | -0.608 | 0.076 | 0 | -0.002 | -0.002 | 2.015 | 0.48 | 0.48 | 0.41 | 0.59 | 0.54 | 0.62 |
| 9 | 134642577 | rs869382 | 0.399 | 0.663 | -0.644 | -0.262 | -0.006 | -0.042 | -0.042 | 0 | 0.6 | 0.5 | 0.37 | 0.15 | 0.65 | 0.53 |
| 9 | 134643466 | rs10901089 | 0.321 | 0.872 | -0.526 | 0.832 | -0.018 | 0.054 | 0.054 | 0 | 0.68 | 0.55 | 0.62 | 0.51 | 0.74 | 0.63 |
| 9 | 134645218 | rs2031872 | -0.059 | 0.766 | -0.624 | 0.907 | 0.022 | -0.028 | -0.028 | 0.151 | 0.85 | 0.56 | 0.88 | 0.85 | 0.86 | 0.83 |
| 9 | 134645623 | rs3012723 | 0.325 | 1.308 | -0.301 | 0.903 | 0.034 | -0.017 | -0.017 | 0 | 0.83 | 0.43 | 0.83 | 0.84 | 0.8 | 0.8 |
| 9 | 134647656 | rs2987423 | -0.009 | 0.909 | -1.046 | 0.241 | 0.056 | -0.066 | -0.066 | 0 | 0.39 | 0.14 | 0.38 | 0.5 | 0.3 | 0.44 |
| 9 | 134649676 | rs10751488 | 1.163 | 1.817 | -0.515 | -0.094 | -0.004 | -0.04 | -0.04 | 0.012 | 0.33 | 0.39 | 0.21 | 0.09 | 0.3 | 0.2 |
| 9 | 134651561 | rs3012735 | -0.017 | 1.844 | -0.803 | -0.054 | -0.011 | -0.032 | -0.032 | 0 | 0.45 | 0.21 | 0.47 | 0.51 | 0.45 | 0.47 |
| 9 | 134652749 | rs3012738 | -0.712 | 1.813 | -0.727 | -0.076 | 0.008 | 0.078 | 0.078 | 0 | 0.54 | 0.32 | 0.66 | 0.76 | 0.55 | 0.63 |
| 9 | 134655176 | rs3012740 | 0.224 | 1.793 | -1.041 | -0.225 | 0.068 | -0.054 | -0.054 | 0.316 | 0.43 | 0.14 | 0.39 | 0.5 | 0.35 | 0.48 |
| 9 | 134656555 | rs2987430 | -0.162 | 1.887 | -1.019 | -0.245 | 0.109 | -0.071 | -0.071 | 0 | 0.81 | 0.51 | 0.77 | 0.9 | 0.72 | 0.81 |
| 9 | 134657367 | rs10793897 | 1.089 | 1.513 | -1.043 | -0.153 | -0.003 | 0.076 | 0.076 | 0 | 0.32 | 0.16 | 0.19 | 0.15 | 0.32 | 0.2 |
| 9 | 134660273 | rs3012748 | -0.621 | 1.559 | -1.17 | 0.048 | 0.001 | -0.028 | -0.028 | 0 | 0.56 | 0.54 | 0.67 | 0.82 | 0.57 | 0.65 |
| 9 | 134663526 | rs3012750 | 0.382 | 1.274 | -1.567 | 0.657 | 0.026 | -0.08 | -0.08 | 0 | 0.15 | 0.03 | 0.2 | 0.33 | 0.1 | 0.19 |
| 9 | 134663576 | rs4740313 | 0.093 | 1.671 | -1.117 | 0.605 | -0.035 | -0.048 | -0.048 | 0 | 0.08 | 0 | 0.18 | 0.23 | 0.11 | 0.16 |
| 9 | 134667753 | rs10901096 | 0.036 | 1.387 | -1.149 | -0.345 | 0 | 0.059 | 0.059 | 0 | 0.37 | 0.29 | 0.28 | 0.1 | 0.35 | 0.26 |
| 9 | 134674287 | rs10125655 | -0.319 | 1.348 | -1.248 | 0.391 | 0.01 | 0.001 | 0.001 | 0 | 0.5 | 0.22 | 0.38 | 0.53 | 0.47 | 0.38 |
| 9 | 134675670 | rs11243536 | -0.651 | 1.503 | -0.633 | 0.809 | -0.005 | 0.046 | 0.046 | 0 | 0.2 | 0.27 | 0.21 | 0.43 | 0.21 | 0.15 |
| 9 | 134676400 | rs10901101 | -1.071 | 1.93 | -0.118 | 0.716 | 0.024 | 0.084 | 0.084 | 0 | 0.23 | 0.36 | 0.31 | 0.43 | 0.29 | 0.18 |
| 9 | 134676845 | rs12683140 | -0.867 | 1.279 | -0.237 | 0.704 | -0.005 | 0.024 | 0.024 | 0 | 0.22 | 0.28 | 0.28 | 0.43 | 0.22 | 0.19 |
| 9 | 134677332 | rs4740186 | -1.078 | 1.652 | 0.201 | 0.96 | 0.047 | 0.168 | 0.168 | 0 | 0.05 | 0.21 | 0.35 | 0.27 | 0.15 | 0.13 |
| 9 | 134678778 | rs2987364 | -0.543 | 0.535 | -0.331 | -0.17 | 0.027 | 0.042 | 0.042 | 0 | 0.62 | 0.77 | 0.81 | 0.92 | 0.75 | 0.83 |
| 9 | 134680532 | rs3012755 | 0.44 | 0.729 | -0.336 | 0.288 | -0.005 | 0.011 | 0.011 | 0.022 | 0.31 | 0.35 | 0.22 | 0.49 | 0.25 | 0.27 |
| 9 | 134682384 | rs3012757 | 0.477 | 0.684 | -0.396 | 0.29 | 0.003 | -0.007 | -0.007 | 0 | 0.31 | 0.31 | 0.21 | 0.49 | 0.25 | 0.27 |
| 9 | 134683985 | rs2987367 | -0.012 | 0.792 | -0.47 | 0.306 | 0.003 | -0.009 | -0.009 | 0.231 | 0.35 | 0.34 | 0.24 | 0.48 | 0.31 | 0.38 |
| 9 | 134686240 | rs11243539 | -0.194 | 0.412 | -0.455 | -0.686 | -0.001 | -0.014 | -0.014 | 0 | 0.45 | 0.54 | 0.29 | 0.19 | 0.39 | 0.43 |
| 9 | 134687735 | rs3012761 | 0.366 | 0.095 | -0.168 | -1.076 | -0.003 | -0.021 | -0.021 | 0 | 0.36 | 0.37 | 0.24 | 0.19 | 0.38 | 0.22 |
| 9 | 134693169 | rs10901103 | 0.436 | 0.381 | -0.985 | -1.001 | -0.045 | -0.028 | -0.028 | 0 | 0.27 | 0.12 | 0.5 | 0.7 | 0.34 | 0.32 |
| 9 | 134696200 | rs2031874 | 0.168 | 0.636 | -0.736 | -0.958 | -0.001 | 0.083 | 0.083 | 0.245 | 0.31 | 0.15 | 0.2 | 0.1 | 0.25 | 0.42 |
| 9 | 134698342 | rs4740188 | 0.114 | 0.816 | -0.102 | -0.63 | 0.02 | -0.01 | -0.01 | 0 | 0.32 | 0.46 | 0.55 | 0.7 | 0.4 | 0.33 |
| 9 | 134708706 | rs943837 | -0.045 | 0.592 | -0.383 | -0.805 | -0.004 | -0.041 | -0.041 | 0 | 0.6 | 0.49 | 0.65 | 0.86 | 0.67 | 0.53 |
| 9 | 134710059 | rs2987389 | 1.013 | 0.228 | -1.113 | -0.661 | -0.034 | 0.002 | 0.002 | 0.084 | 0.23 | 0.06 | 0.15 | 0.18 | 0.27 | 0.19 |
| 9 | 134711461 | rs2987392 | -0.188 | -0.503 | -0.905 | -1.17 | -0.026 | -0.009 | -0.009 | 0 | 0.76 | 0.31 | 0.89 | 0.9 | 0.93 | 0.82 |
| 9 | 134715999 | rs7850959 | 0.005 | 1.145 | -0.949 | -0.533 | -0.018 | 0.006 | 0.006 | 0 | 0.17 | 0.12 | 0.26 | 0.14 | 0.23 | 0.32 |
| 9 | 134717658 | rs913506 | 0.256 | 0.793 | -0.536 | -0.531 | 0.001 | -0.07 | -0.07 | 0 | 0.27 | 0.41 | 0.21 | 0.09 | 0.31 | 0.17 |
| 9 | 134719002 | rs2987401 | -0.083 | 0.83 | -0.642 | -0.46 | -0.001 | -0.082 | -0.082 | 0 | 0.33 | 0.47 | 0.25 | 0.1 | 0.35 | 0.21 |
| 9 | 134721070 | rs10122844 | 0.128 | 1.098 | -1.464 | 0.461 | -0.074 | -0.002 | -0.002 | 0 | 0.62 | 0.89 | 0.64 | 0.7 | 0.49 | 0.56 |
| 9 | 134722109 | rs11243551 | -0.773 | 0.749 | -0.428 | 1.641 | 0.007 | 0.225 | 0.225 | 0 | 0.07 | 0.19 | 0.26 | 0.32 | 0.05 | 0.2 |
| 9 | 134722733 | rs1547209 | 0.482 | 0.881 | -0.703 | 1.026 | -0.009 | 0 | 0 | 0.354 | 0.78 | 0.81 | 0.85 | 0.79 | 0.79 | 0.8 |
| 9 | 134723854 | rs3012780 | -0.201 | 0.826 | -0.803 | 0.858 | 0 | 0.004 | 0.004 | 1.9 | 0.14 | 0.18 | 0.14 | 0.21 | 0.12 | 0.16 |
| 9 | 134724256 | rs3012782 | -0.273 | 0.797 | -0.647 | 1.183 | 0.001 | 0.03 | 0.03 | 1.789 | 0.15 | 0.2 | 0.14 | 0.32 | 0.12 | 0.17 |
| 9 | 134725562 | rs7350156 | 1.723 | 0.042 | -0.051 | 1.037 | 0.008 | 0.032 | 0.032 | 0 | 0.57 | 0.3 | 0.52 | 0.29 | 0.68 | 0.46 |
| 9 | 134732553 | rs4740321 | 0.166 | 0.189 | 0.293 | 1.293 | -0.005 | 0.205 | 0.205 | 0 | 0.05 | 0.13 | 0.22 | 0.38 | 0.03 | 0.2 |
| 9 | 134734108 | rs7036013 | -1.143 | 0.113 | 0.717 | 1.135 | 0.003 | 0.062 | 0.062 | 0 | 0.22 | 0.33 | 0.32 | 0.42 | 0.19 | 0.37 |
| 9 | 134755291 | rs7848165 | -0.017 | 0.321 | 0.549 | 0.999 | -0.009 | 0.005 | 0.005 | 0 | 0.12 | 0.12 | 0.09 | 0.02 | 0.15 | 0.13 |
| 9 | 134767080 | rs4579577 | 1.459 | 0.038 | 0.777 | 0.941 | 0.005 | 0.016 | 0.016 | 0 | 0.61 | 0.28 | 0.52 | 0.31 | 0.69 | 0.47 |
| 9 | 134772966 | rs3924215 | -0.217 | 0.165 | 0.783 | 1.167 | 0 | 0.005 | 0.005 | 1.881 | 0.86 | 0.83 | 0.86 | 0.72 | 0.87 | 0.83 |
| 9 | 134781550 | rs9411321 | 0.767 | 0.662 | -0.083 | 0.511 | -0.058 | -0.06 | -0.06 | 3.602 | 0.13 | 0.01 | 0.15 | 0.19 | 0.15 | 0.15 |
| 9 | 134788993 | rs9411419 | 0.975 | 0.415 | 0.095 | 0.226 | -0.011 | -0.001 | -0.001 | 0 | 0.32 | 0.17 | 0.23 | 0.3 | 0.38 | 0.22 |
| 9 | 134789096 | rs10901112 | 0.014 | 0.394 | 0.59 | 0.151 | 0.015 | 0.035 | 0.035 | 0 | 0.34 | 0.46 | 0.41 | 0.41 | 0.33 | 0.42 |
| 9 | 134791779 | rs6597536 | -1.077 | -0.017 | 0.126 | 0.117 | -0.008 | -0.023 | -0.023 | 0 | 0.87 | 0.91 | 0.8 | 0.73 | 0.87 | 0.85 |
| 9 | 134799366 | rs7047055 | 0.554 | 0.041 | 0.153 | -0.043 | -0.004 | 0.083 | 0.083 | 0 | 0.51 | 0.22 | 0.35 | 0.32 | 0.54 | 0.42 |
| 9 | 134813428 | rs4579576 | -0.269 | -0.119 | 0.252 | 0.004 | 0.002 | -0.017 | -0.017 | 0 | 0.82 | 0.88 | 0.76 | 0.69 | 0.84 | 0.8 |
| 9 | 134818418 | rs4128957 | 0.821 | -0.11 | 0.515 | 0.016 | -0.02 | -0.036 | -0.036 | 0 | 0.26 | 0.16 | 0.35 | 0.35 | 0.29 | 0.36 |
| 9 | 134818509 | rs4128956 | -1.963 | -0.213 | 0.409 | -0.111 | -0.029 | -0.003 | -0.003 | 0 | 0.69 | 0.84 | 0.78 | 0.69 | 0.62 | 0.78 |
| 9 | 134824764 | rs11243574 | 1.142 | -0.185 | 0.361 | 0.037 | 0.018 | -0.01 | -0.01 | 0 | 0.12 | 0.06 | 0.17 | 0.18 | 0.08 | 0.06 |
| 9 | 134829149 | rs10121309 | 1.897 | -0.257 | 0.718 | -0.065 | -0.017 | -0.029 | -0.029 | 0 | 0.05 | 0.13 | 0.03 | 0 | 0.01 | 0.01 |
| 9 | 134836197 | rs7032759 | 0.252 | -0.104 | 1.039 | 0.019 | 0.01 | 0.036 | 0.036 | 0 | 0.29 | 0.44 | 0.37 | 0.37 | 0.3 | 0.39 |
| 9 | 134844839 | rs11243585 | 0.825 | -0.147 | 0.86 | 0.038 | -0.006 | -0.059 | -0.059 | 0.11 | 0.23 | 0.09 | 0.23 | 0.34 | 0.21 | 0.34 |
| 9 | 134866354 | rs7043386 | 0.863 | 0.149 | 1.868 | 0.691 | -0.008 | -0.002 | -0.002 | 2.098 | 0.6 | 0.5 | 0.6 | 0.59 | 0.68 | 0.6 |
| 9 | 134881443 | rs7025089 | 0.991 | 0.149 | 1.763 | 0.694 | 0.007 | 0.002 | 0.002 | 0.016 | 0.59 | 0.4 | 0.59 | 0.59 | 0.68 | 0.6 |
| 9 | 134890430 | rs4363310 | 1.279 | 0.141 | 1.346 | 0.671 | 0.062 | 0.013 | 0.013 | 1.413 | 0.56 | 0.19 | 0.57 | 0.58 | 0.69 | 0.59 |
| 9 | 134899743 | rs11243598 | 1.123 | 0.142 | 1.208 | 0.717 | -0.009 | -0.014 | -0.014 | 0 | 0.11 | 0.07 | 0.18 | 0.26 | 0.11 | 0.13 |
| 9 | 134912614 | rs7467576 | 0.199 | 0.117 | 1.284 | 0.689 | -0.002 | -0.008 | -0.008 | 0 | 0.16 | 0.13 | 0.22 | 0.32 | 0.13 | 0.2 |
| 9 | 134917623 | rs4592148 | 1.26 | 0.01 | 1.339 | 0.275 | 0.047 | 0.013 | 0.013 | 0.611 | 0.57 | 0.25 | 0.57 | 0.56 | 0.68 | 0.57 |
| 9 | 134934986 | rs4526483 | 0.124 | 0.045 | 0.922 | 0.228 | 0.007 | 0.007 | 0.007 | 0 | 0.21 | 0.15 | 0.13 | 0.09 | 0.15 | 0.18 |
| 9 | 134944428 | rs7851262 | -0.872 | 0.416 | 1.2 | 0.445 | 0.008 | 0 | 0 | 0 | 0.37 | 0.29 | 0.37 | 0.4 | 0.31 | 0.4 |

| | | | | | | | | | | | | | | | |
|---|---|---|---|---|---|---|---|---|---|---|---|---|---|---|---|
| 9 | 134961834 | rs4962112 | 1.114 | 0.334 | 0.344 | 0.601 | -0.033 | 0.04 | 0.04 | 0.481 | 0.52 | 0.16 | 0.46 | 0.58 | 0.54 | 0.49 |
| 9 | 134965643 | rs7038996 | 0.709 | 0.221 | 1.056 | 0.435 | -0.001 | 0 | 0 | 0.72 | 0.68 | 0.33 | 0.69 | 0.85 | 0.67 | 0.64 |
| 9 | 134987645 | rs9411345 | -1.269 | 0.539 | 0.644 | 1.008 | -0.013 | -0.02 | -0.02 | 0.275 | 0.39 | 0.62 | 0.41 | 0.45 | 0.46 | 0.58 |
| 9 | 134993563 | rs1331630 | -0.484 | 0.611 | 0.497 | 1.026 | -0.006 | 0.01 | 0.01 | 0 | 0.27 | 0.21 | 0.29 | 0.13 | 0.31 | 0.35 |
| 9 | 134995268 | rs7865653 | -0.511 | 0.821 | 1.009 | 1.38 | 0.019 | 0.144 | 0.144 | 0 | 0.12 | 0.28 | 0.15 | 0.33 | 0.14 | 0.24 |
| 9 | 134997981 | rs1571832 | -0.744 | 0.789 | 0.347 | 0.994 | 0.009 | 0.005 | 0.005 | 0 | 0.21 | 0.25 | 0.25 | 0.21 | 0.25 | 0.34 |
| 9 | 135001703 | rs11243634 | -0.908 | 0.124 | 0.52 | 0.864 | -0.026 | 0.011 | 0.011 | 0.073 | 0.19 | 0.37 | 0.21 | 0.24 | 0.14 | 0.19 |
| 9 | 135003565 | rs1018691 | -0.098 | 0.209 | -0.024 | 0.682 | -0.001 | -0.004 | -0.004 | 0.099 | 0.67 | 0.7 | 0.66 | 0.63 | 0.65 | 0.69 |
| 9 | 135009024 | rs7858174 | 0.549 | -0.056 | -0.241 | 0.291 | 0.039 | -0.006 | -0.006 | 0 | 0.46 | 0.33 | 0.36 | 0.5 | 0.33 | 0.3 |
| 9 | 135015927 | rs9411446 | -0.441 | 0.349 | -0.517 | 0.247 | -0.001 | -0.003 | -0.003 | 0 | 0.63 | 0.66 | 0.59 | 0.56 | 0.6 | 0.65 |
| 9 | 135018146 | rs12684726 | 0.128 | -1.292 | -1.08 | 0.121 | 0.051 | 0.019 | 0.019 | 0 | 0.47 | 0.27 | 0.36 | 0.42 | 0.34 | 0.32 |
| 9 | 135020849 | rs10901130 | 1.102 | -2.269 | -1.708 | -0.891 | 0.008 | 0.115 | 0.115 | 0 | 0.22 | 0.01 | 0.1 | 0.03 | 0.17 | 0.09 |
| 9 | 135021608 | rs11243638 | 0.07 | -2.046 | -1.686 | -0.697 | 0.009 | 0.056 | 0.056 | 0 | 0.25 | 0.01 | 0.2 | 0.11 | 0.2 | 0.13 |
| 9 | 135030039 | rs7862560 | -0.083 | -2.657 | -0.949 | -0.725 | 0.04 | 0.054 | 0.054 | 0 | 0.52 | 0.62 | 0.31 | 0.16 | 0.37 | 0.25 |
| 9 | 135032890 | rs1412076 | -0.669 | -2.564 | -0.209 | -0.064 | 0.02 | -0.02 | -0.02 | 0 | 0.61 | 0.72 | 0.37 | 0.47 | 0.38 | 0.36 |
| 9 | 135034618 | rs11243648 | -0.113 | -3.054 | 0.103 | -0.155 | -0.029 | -0.025 | -0.025 | 0 | 0.38 | 0.52 | 0.22 | 0.21 | 0.23 | 0.25 |
| 9 | 135039556 | rs1105685 | 0.192 | -2.793 | 0.052 | -0.51 | 0.014 | 0.003 | 0.003 | 0 | 0.32 | 0.23 | 0.16 | 0.03 | 0.28 | 0.21 |
| 9 | 135048889 | rs6597547 | -0.825 | -2.487 | -0.169 | -0.315 | 0.017 | -0.013 | -0.013 | 0 | 0.58 | 0.68 | 0.33 | 0.39 | 0.43 | 0.38 |
| 9 | 135048978 | rs11788925 | 0.82 | -1.929 | -0.485 | -0.223 | -0.034 | 0.041 | 0.041 | 0 | 0.1 | 0.01 | 0.07 | 0 | 0.16 | 0.07 |
| 9 | 135053756 | rs7039329 | -0.026 | -1.907 | 0.008 | 0.132 | -0.021 | 0.016 | 0.016 | 0 | 0.25 | 0.31 | 0.16 | 0.27 | 0.17 | 0.13 |
| 9 | 135064788 | rs11243657 | -0.332 | -1.279 | -0.005 | 0.365 | -0.002 | 0.004 | 0.004 | 0 | 0.57 | 0.44 | 0.35 | 0.51 | 0.45 | 0.46 |
| 9 | 135070104 | rs7870760 | -0.857 | -1.486 | -0.559 | 0.252 | -0.1 | -0.159 | -0.159 | 4.092 | 0.64 | 0.94 | 0.43 | 0.51 | 0.46 | 0.46 |
| 9 | 135072009 | rs1041504 | -0.844 | -1.502 | -0.434 | 0.295 | -0.057 | -0.109 | -0.109 | 0 | 0.65 | 0.87 | 0.41 | 0.5 | 0.45 | 0.46 |
| 9 | 135073877 | rs3824574 | 0.632 | -1.96 | -0.571 | 0.098 | 0.013 | -0.025 | -0.025 | 1.261 | 0.33 | 0.19 | 0.25 | 0.38 | 0.32 | 0.33 |
| 9 | 135073967 | rs3739930 | 0.496 | -1.95 | -1.235 | -0.028 | 0.038 | 0.062 | 0.062 | 0 | 0.2 | 0.05 | 0.08 | 0.12 | 0.12 | 0.12 |
| 9 | 135077296 | rs871462 | -0.219 | -0.847 | -0.58 | 0.713 | -0.058 | -0.056 | -0.056 | 0.058 | 0.62 | 0.82 | 0.45 | 0.44 | 0.49 | 0.49 |
| 9 | 135077361 | rs871461 | -0.966 | -0.75 | -0.351 | 0.814 | -0.061 | 0.028 | 0.028 | 5.864 | 0.11 | 0.55 | 0.07 | 0.11 | 0.01 | 0.07 |
| 9 | 135081514 | rs7865565 | -0.522 | -0.297 | -0.64 | 1.032 | -0.034 | 0.153 | 0.153 | 3.453 | 0.09 | 0.54 | 0.05 | 0.19 | 0.01 | 0.07 |
| 9 | 135082979 | rs7026497 | 0.583 | -0.168 | -0.838 | 0.952 | -0.006 | 0.017 | 0.017 | 0.002 | 0.56 | 0.41 | 0.42 | 0.31 | 0.49 | 0.49 |
| 9 | 135087892 | rs7025107 | 0.31 | 0.854 | -1.781 | 0.995 | -0.136 | -0.142 | -0.142 | 0 | 0.65 | 0.95 | 0.49 | 0.52 | 0.51 | 0.59 |
| 9 | 135089172 | rs7018926 | 0.311 | 0.831 | -1.865 | 1 | -0.158 | -0.16 | -0.16 | 0 | 0.65 | 0.95 | 0.49 | 0.52 | 0.51 | 0.59 |
| 9 | 135102254 | rs2149171 | 0.621 | 1.657 | -0.83 | 1.295 | 0.048 | 0.046 | 0.046 | 0 | 0.72 | 0.58 | 0.59 | 0.43 | 0.62 | 0.66 |
| 9 | 135104786 | rs914420 | -0.326 | 2.679 | -0.855 | 2.11 | -0.023 | -0.016 | -0.016 | 0.597 | 0.11 | 0.06 | 0.28 | 0.41 | 0.19 | 0.23 |
| 9 | 135105059 | rs914419 | 0.631 | 2.514 | -0.571 | 1.88 | 0.045 | -0.022 | -0.022 | 2.094 | 0.84 | 0.66 | 0.87 | 0.86 | 0.82 | 0.91 |
| 9 | 135108510 | rs2854926 | 0.187 | 2.938 | -1.102 | 1.883 | -0.038 | -0.025 | -0.025 | 0 | 0.07 | 0 | 0.12 | 0.06 | 0.11 | 0.12 |
| 9 | 135108551 | rs2854925 | 0.234 | 1.491 | -1.466 | 0.699 | 0.022 | 0.021 | 0.021 | 0.046 | 0.47 | 0.24 | 0.44 | 0.39 | 0.43 | 0.54 |
| 9 | 135109833 | rs17148812 | 0.739 | 0.376 | -0.998 | -0.08 | -0.002 | -0.022 | -0.022 | 0 | 0.26 | 0.22 | 0.11 | 0.04 | 0.11 | 0.06 |
| 9 | 135110034 | rs10901140 | -0.486 | 0.015 | -1.124 | 0.448 | 0.011 | -0.038 | -0.038 | 0 | 0.22 | 0.14 | 0.35 | 0.38 | 0.16 | 0.39 |
| 9 | 135110456 | rs1810887 | -1.052 | -0.198 | -0.819 | -0.172 | -0.003 | 0.015 | 0.015 | 0 | 0.49 | 0.63 | 0.57 | 0.56 | 0.47 | 0.53 |
| 9 | 135110611 | rs10901141 | -0.55 | -0.424 | -0.311 | 0.188 | -0.004 | 0.005 | 0.005 | 0 | 0.32 | 0.35 | 0.43 | 0.43 | 0.23 | 0.45 |
| 9 | 135120155 | rs914418 | -0.815 | -0.31 | -0.893 | 0.287 | 0.044 | -0.003 | -0.003 | 1.606 | 0.45 | 0.79 | 0.49 | 0.48 | 0.32 | 0.45 |
| 9 | 135126458 | rs4962060 | -0.733 | -0.329 | -0.485 | 0.563 | -0.014 | -0.006 | -0.006 | 0 | 0.58 | 0.68 | 0.59 | 0.53 | 0.47 | 0.5 |
| 9 | 135132081 | rs4962061 | -0.233 | -0.15 | -0.649 | 0.979 | 0.005 | -0.01 | -0.01 | 0 | 0.27 | 0.21 | 0.41 | 0.38 | 0.23 | 0.33 |
| 9 | 135132930 | rs4962062 | -0.526 | -0.149 | -0.48 | 0.797 | 0.002 | -0.001 | -0.001 | 0 | 0.57 | 0.58 | 0.57 | 0.57 | 0.46 | 0.52 |
| 9 | 135133039 | rs7031969 | -0.477 | -0.306 | -0.35 | 1.067 | 0.001 | 0.001 | 0.001 | 0.435 | 0.35 | 0.39 | 0.28 | 0.32 | 0.32 | 0.25 |
| 9 | 135136162 | rs1571336 | -0.591 | -1.806 | -0.011 | 1.292 | -0.052 | 0.023 | 0.023 | 0 | 0.27 | 0.33 | 0.15 | 0.41 | 0.1 | 0.28 |
| 9 | 135138261 | rs7875801 | -0.244 | -2.005 | 0.154 | 1.37 | -0.02 | 0.102 | 0.102 | 0 | 0.07 | 0.13 | 0.06 | 0.36 | 0.03 | 0.15 |
| 9 | 135139064 | rs11787894 | -0.168 | -2.335 | 0.014 | 1.494 | 0 | -0.019 | -0.019 | 0 | 0.18 | 0.15 | 0.16 | 0.28 | 0.16 | 0.15 |
| 9 | 135139826 | rs3739927 | 0.067 | -2.335 | 0.144 | 1.495 | -0.024 | 0.164 | 0.164 | 0 | 0.08 | 0.18 | 0.07 | 0.36 | 0.03 | 0.15 |
| 9 | 135139901 | rs1056899 | -2.134 | -2.906 | 0.649 | 1.115 | 0.117 | 0.088 | 0.088 | 0 | 0.52 | 0.81 | 0.35 | 0.68 | 0.27 | 0.44 |
| 9 | 135150461 | rs17148869 | -0.424 | -2.853 | 0.691 | 1.153 | -0.022 | 0.006 | 0.006 | 0 | 0.27 | 0.29 | 0.14 | 0.4 | 0.1 | 0.28 |
| 9 | 135150907 | rs2296867 | -0.003 | -2.798 | 0.532 | 1.07 | 0 | -0.023 | -0.023 | 4.911 | 0.18 | 0.16 | 0.17 | 0.28 | 0.16 | 0.15 |
| 9 | 135152439 | rs17148873 | 0.195 | -2.674 | 0.445 | 0.928 | 0.019 | 0.019 | 0.019 | 0 | 0.19 | 0.14 | 0.08 | 0.04 | 0.07 | 0.13 |
| 9 | 135159200 | rs11243707 | 1.858 | -2.745 | -0.083 | 0.709 | -0.037 | 0.109 | 0.109 | 0 | 0.37 | 0.09 | 0.39 | 0.18 | 0.59 | 0.53 |
| 9 | 135185355 | rs744133 | 0.969 | -2.924 | 0.388 | 1.159 | -0.017 | -0.023 | -0.023 | 0 | 0.07 | 0.01 | 0.22 | 0.07 | 0.1 | 0.03 |
| 9 | 135188935 | rs10793918 | -1.082 | -3.199 | 0.716 | 1.27 | 0.042 | -0.002 | -0.002 | 0 | 0.37 | 0.69 | 0.23 | 0.64 | 0.14 | 0.37 |
| 9 | 135191473 | rs3758350 | -0.768 | -3.29 | 1.191 | 1.606 | -0.037 | 0.169 | 0.169 | 0 | 0.14 | 0.31 | 0.11 | 0.42 | 0.07 | 0.19 |
| 9 | 135192063 | rs12343347 | 0.891 | -3.294 | 1.198 | 1.608 | -0.021 | 0.035 | 0.035 | 0 | 0.06 | 0.14 | 0.02 | 0.18 | 0.03 | 0.1 |
| 9 | 135193946 | rs17148925 | 0.554 | -3.342 | 1.262 | 1.493 | 0.064 | -0.025 | -0.025 | 0 | 0.19 | 0.04 | 0.09 | 0.21 | 0.06 | 0.19 |
| 9 | 135200861 | rs10901164 | -0.297 | -3.335 | 1.289 | 1.499 | -0.024 | 0.172 | 0.172 | 0 | 0.11 | 0.24 | 0.09 | 0.41 | 0.06 | 0.17 |
| 9 | 135202829 | rs543573 | 1.148 | -3.359 | 1.115 | 1.467 | 0.044 | -0.002 | -0.002 | 0 | 0.63 | 0.31 | 0.77 | 0.36 | 0.86 | 0.63 |
| 9 | 135203231 | rs1183768 | 1.119 | -3.354 | 1.118 | 1.469 | 0.042 | -0.002 | -0.002 | 0 | 0.63 | 0.31 | 0.77 | 0.36 | 0.86 | 0.63 |
| 9 | 135203409 | rs1185193 | 0.528 | -3.328 | 1.155 | 1.485 | -0.06 | 0.021 | 0.021 | 0 | 0.69 | 0.58 | 0.81 | 0.37 | 0.88 | 0.64 |
| 9 | 135203838 | rs3739921 | 0.561 | -3.326 | 1.154 | 1.503 | 0.06 | -0.021 | -0.021 | 0 | 0.19 | 0.04 | 0.09 | 0.21 | 0.06 | 0.19 |
| 9 | 135206460 | rs9411449 | 0.885 | -3.231 | 1.133 | 1.285 | -0.062 | 0.009 | 0.009 | 0 | 0.66 | 0.55 | 0.79 | 0.37 | 0.86 | 0.63 |
| 9 | 135216186 | rs501377 | 0.95 | -3.212 | 1.117 | 1.274 | -0.061 | 0.007 | 0.007 | 0 | 0.66 | 0.55 | 0.78 | 0.36 | 0.85 | 0.6 |
| 9 | 135218343 | rs514279 | 1.957 | -2.725 | 0.562 | 1.159 | -0.035 | 0.103 | 0.103 | 0 | 0.39 | 0.09 | 0.4 | 0.19 | 0.58 | 0.51 |
| 9 | 135226280 | rs508322 | 1.08 | -3.106 | 1.292 | 1.351 | -0.035 | 0.017 | 0.017 | 0 | 0.6 | 0.5 | 0.71 | 0.5 | 0.76 | 0.69 |
| 9 | 135227241 | rs488568 | 1.027 | -3.198 | 1.495 | 1.492 | 0.011 | -0.002 | -0.002 | 1.305 | 0.64 | 0.36 | 0.78 | 0.36 | 0.84 | 0.62 |

| | | | | | | | | | | | | | | | | |
|---|---|---|---|---|---|---|---|---|---|---|---|---|---|---|---|---|
| 9 | 135227581 | rs492082 | 0.068 | -3.152 | 1.28 | 1.562 | -0.053 | 0.083 | 0.083 | 0 | 0.74 | 0.63 | 0.81 | 0.55 | 0.87 | 0.75 |
| 9 | 135230834 | rs544674 | 0.063 | -3.189 | 1.257 | 1.544 | -0.035 | 0.049 | 0.049 | 0 | 0.74 | 0.68 | 0.81 | 0.56 | 0.87 | 0.75 |
| 9 | 135234019 | rs477012 | 0.128 | -3.25 | 1.554 | 1.762 | -0.01 | -0.002 | -0.002 | 0 | 0.73 | 0.72 | 0.88 | 0.63 | 0.91 | 0.71 |
| 9 | 135235457 | rs10217641 | -1.171 | -2.651 | 1.767 | 1.805 | 0.01 | 0.194 | 0.194 | 0 | 0.07 | 0.23 | 0.1 | 0.26 | 0.05 | 0.08 |
| 9 | 135241588 | rs484932 | 0.863 | -2.662 | 1.772 | 1.822 | 0.013 | 0.188 | 0.188 | 0 | 0.94 | 0.8 | 0.91 | 0.74 | 0.95 | 0.94 |
| 9 | 135245867 | rs1185995 | 1.635 | -2.898 | 1.719 | 1.647 | -0.017 | 0.082 | 0.082 | 0 | 0.88 | 0.77 | 0.88 | 0.73 | 0.93 | 0.89 |
| 9 | 135250473 | rs7030233 | -1.489 | -2.847 | 1.745 | 1.629 | 0.007 | -0.012 | -0.012 | 0 | 0.24 | 0.22 | 0.19 | 0.33 | 0.16 | 0.16 |
| 9 | 135251616 | rs17149015 | 0.615 | -3.035 | 1.683 | 1.689 | 0.034 | -0.042 | -0.042 | 0 | 0.15 | 0.04 | 0.09 | 0.25 | 0.08 | 0.16 |
| 9 | 135257703 | rs11243755 | -1.426 | -2.929 | 1.683 | 1.67 | 0.007 | -0.013 | -0.013 | 0.347 | 0.24 | 0.22 | 0.19 | 0.35 | 0.16 | 0.16 |
| 9 | 135271536 | rs554118 | -1.703 | -2.885 | 1.267 | 1.584 | -0.045 | 0.032 | 0.032 | 0 | 0.35 | 0.5 | 0.26 | 0.55 | 0.2 | 0.31 |
| 9 | 135271692 | rs1166319 | -1.177 | -2.887 | 1.15 | 1.663 | -0.054 | 0.001 | 0.001 | 0 | 0.3 | 0.38 | 0.17 | 0.29 | 0.15 | 0.24 |
| 9 | 135277007 | rs3739916 | 0.071 | -2.242 | 0.439 | 1.61 | -0.008 | 0.032 | 0.032 | 0 | 0.08 | 0.01 | 0.06 | 0.03 | 0.05 | 0.05 |
| 9 | 135285010 | rs10901173 | 1.342 | -2.198 | 1.034 | 1.234 | 0.002 | -0.005 | -0.005 | 0 | 0.92 | 0.64 | 0.96 | 0.97 | 0.97 | 0.95 |
| 9 | 135286130 | rs11795094 | 2.145 | -2.743 | 0.506 | 1.427 | 0.067 | 0.023 | 0.023 | 1.312 | 0.67 | 0.11 | 0.7 | 0.28 | 0.78 | 0.75 |
| 9 | 135288807 | rs2885495 | 2.655 | -2.809 | 0.263 | 1.094 | 0.076 | 0.259 | 0.259 | 0.04 | 0.51 | 0.1 | 0.59 | 0.19 | 0.66 | 0.64 |
| 9 | 135289797 | rs10125557 | -1.647 | -2.36 | 0.849 | 1.524 | -0.011 | -0.003 | -0.003 | 1.138 | 0.25 | 0.59 | 0.28 | 0.74 | 0.2 | 0.2 |
| 9 | 135292355 | rs10156646 | -1.422 | -2.358 | 0.714 | 1.997 | -0.036 | 0.199 | 0.199 | 0.038 | 0.19 | 0.42 | 0.11 | 0.41 | 0.13 | 0.11 |
| 9 | 135297764 | rs944614 | -0.908 | -2.11 | 0.082 | 2.14 | -0.001 | 0.046 | 0.046 | 0 | 0.1 | 0.12 | 0.17 | 0.37 | 0.08 | 0.14 |
| 9 | 135298061 | rs944616 | -1.27 | -2.133 | 0.37 | 2.043 | -0.013 | 0.033 | 0.033 | 0 | 0.22 | 0.41 | 0.34 | 0.29 | 0.15 | 0.11 |
| 9 | 135298195 | rs11795012 | 2.171 | -2.431 | -1.916 | 0.822 | -0.092 | 0.132 | 0.132 | 0 | 0.35 | 0.1 | 0.32 | 0.12 | 0.55 | 0.43 |
| 9 | 135298232 | rs2362365 | -1.367 | -1.993 | -1.594 | 1.383 | 0.008 | -0.004 | -0.004 | 0 | 0.42 | 0.36 | 0.4 | 0.53 | 0.29 | 0.46 |
| 9 | 135298917 | rs1412272 | -0.663 | -1.381 | -2.205 | 1.086 | -0.027 | -0.001 | -0.001 | 2.631 | 0.16 | 0.02 | 0.29 | 0.09 | 0.14 | 0.17 |
| 9 | 135300917 | rs1001606 | 1.029 | -1.607 | -2.192 | 1.185 | 0.059 | 0.007 | 0.007 | 0.035 | 0.64 | 0.18 | 0.53 | 0.67 | 0.77 | 0.73 |
| 9 | 135301389 | rs999483 | -0.879 | -0.03 | -2.137 | 0.976 | -0.014 | 0.002 | 0.002 | 0 | 0.26 | 0.13 | 0.15 | 0.18 | 0.22 | 0.3 |
| 9 | 135302624 | rs557495 | -1.148 | 0.238 | -2.153 | 1.196 | -0.004 | 0.058 | 0.058 | 0 | 0.54 | 0.82 | 0.47 | 0.63 | 0.49 | 0.53 |
| 9 | 135308773 | rs4551482 | -0.301 | -0.021 | -2.007 | 1.382 | 0.001 | -0.031 | -0.031 | 0 | 0.11 | 0.06 | 0.26 | 0.26 | 0.07 | 0.23 |
| 9 | 135310415 | rs563521 | -0.88 | 0.2 | -1.839 | 0.982 | 0.042 | 0.046 | 0.046 | 0.817 | 0.51 | 0.85 | 0.56 | 0.6 | 0.4 | 0.63 |
| 9 | 135322596 | rs507371 | -0.823 | -0.241 | -1.371 | 1.246 | 0.018 | -0.017 | -0.017 | 0 | 0.22 | 0.19 | 0.13 | 0.31 | 0.11 | 0.27 |
| 9 | 135326806 | rs10429632 | -0.523 | 0.093 | -0.657 | 0.893 | 0.021 | -0.031 | -0.031 | 1.438 | 0.06 | 0.16 | 0.09 | 0 | 0.09 | 0.05 |
| 9 | 135329771 | rs558356 | -1.214 | -0.074 | -0.452 | 0.657 | -0.005 | -0.029 | -0.029 | 0 | 0.1 | 0.13 | 0.08 | 0.04 | 0.08 | 0.11 |
| 9 | 135330157 | rs562007 | -1.782 | 1.137 | 0.646 | 1.237 | 0.048 | 0.044 | 0.044 | 0 | 0.23 | 0.53 | 0.42 | 0.36 | 0.28 | 0.28 |
| 9 | 135331958 | rs529317 | -2.07 | 1 | 0.66 | 1.197 | -0.004 | 0.002 | 0.002 | 0 | 0.26 | 0.25 | 0.45 | 0.24 | 0.36 | 0.31 |
| 9 | 135336094 | rs787887 | -2.238 | 1.346 | 1.145 | 1.665 | -0.002 | 0.055 | 0.055 | 0.884 | 0.28 | 0.58 | 0.27 | 0.44 | 0.28 | 0.28 |
| 9 | 135338596 | rs541142 | -2.637 | 1.442 | 0.938 | 1.794 | 0.008 | -0.004 | -0.004 | 0 | 0.37 | 0.43 | 0.54 | 0.63 | 0.46 | 0.45 |
| 9 | 135338786 | rs498495 | -1.681 | 1.187 | 1.063 | 1.816 | -0.003 | 0.125 | 0.125 | 0.299 | 0.21 | 0.39 | 0.19 | 0.44 | 0.19 | 0.23 |
| 9 | 135343839 | rs540407 | 0.618 | 0.608 | 0.106 | 0.509 | -0.013 | 0.073 | 0.073 | 0 | 0.46 | 0.31 | 0.5 | 0.72 | 0.52 | 0.47 |
| 9 | 135344407 | rs569434 | 0.45 | 0.424 | 0.306 | 0.083 | 0.058 | 0.166 | 0.166 | 0 | 0.51 | 0.29 | 0.7 | 0.87 | 0.6 | 0.6 |
| 9 | 135350866 | rs472207 | 0.526 | 0.292 | 0.444 | 0.225 | -0.005 | 0.058 | 0.058 | 0 | 0.46 | 0.68 | 0.65 | 0.77 | 0.48 | 0.51 |
| 9 | 135351031 | rs482275 | -0.264 | 0.104 | 0.286 | 0.068 | -0.028 | 0.119 | 0.119 | 0.271 | 0.6 | 0.85 | 0.74 | 0.84 | 0.58 | 0.58 |
| 9 | 135351678 | rs565325 | 0.653 | 0.103 | 0.459 | 0.549 | -0.008 | 0.028 | 0.028 | 0 | 0.46 | 0.72 | 0.58 | 0.39 | 0.47 | 0.5 |
| 9 | 135365028 | rs530582 | -0.209 | 0.613 | 1.207 | 0.855 | 0.15 | 0.144 | 0.144 | 0.546 | 0.06 | 0.4 | 0.2 | 0.13 | 0.14 | 0.06 |
| 9 | 135365135 | rs10122574 | -1.325 | 0.437 | 1.174 | 0.769 | -0.012 | 0.031 | 0.031 | 0 | 0.27 | 0.3 | 0.25 | 0.52 | 0.2 | 0.2 |
| 9 | 135365590 | rs559127 | -0.693 | 0.044 | 1.121 | 0.548 | -0.001 | 0.012 | 0.012 | 0.736 | 0.18 | 0.59 | 0.27 | 0.18 | 0.17 | 0.17 |
| 9 | 135371169 | rs12002251 | 0.182 | -0.017 | 0.631 | 0.491 | -0.041 | -0.029 | -0.029 | 0 | 0.13 | 0.22 | 0.07 | 0.05 | 0.03 | 0.11 |
| 9 | 135372489 | rs561974 | -2.014 | 0.075 | 0.379 | 0.521 | 0.032 | 0.081 | 0.081 | 0.076 | 0.46 | 0.93 | 0.52 | 0.7 | 0.37 | 0.38 |
| 9 | 135372852 | rs10120389 | -0.604 | 0.077 | 0.854 | 0.551 | -0.008 | 0.073 | 0.073 | 0 | 0.16 | 0.21 | 0.16 | 0.45 | 0.1 | 0.13 |
| 9 | 135373497 | rs528460 | -0.02 | -0.087 | 1.153 | 0.372 | 0.156 | 0.222 | 0.222 | 0.207 | 0.06 | 0.44 | 0.2 | 0.14 | 0.12 | 0.06 |
| 9 | 135374445 | rs557874 | -0.81 | -0.149 | 0.922 | 0.089 | -0.005 | -0.005 | -0.005 | 0 | 0.22 | 0.29 | 0.15 | 0.16 | 0.17 | 0.27 |
| 9 | 135374764 | rs7047726 | -0.334 | -0.171 | 0.647 | 0.133 | 0.014 | -0.023 | -0.023 | 2.088 | 0.13 | 0.1 | 0.14 | 0.48 | 0.07 | 0.15 |
| 9 | 135374898 | rs562350 | -2.495 | -0.121 | 0.64 | -0.111 | 0.005 | 0.191 | 0.191 | 1.509 | 0.52 | 0.85 | 0.57 | 0.83 | 0.48 | 0.57 |
| 9 | 135377218 | rs4962067 | -0.243 | -0.138 | 0.627 | -0.039 | 0.013 | 0.017 | 0.017 | 0 | 0.13 | 0.1 | 0.05 | 0.08 | 0.06 | 0.12 |
| 9 | 135377764 | rs515721 | 0.126 | -0.071 | 0.755 | 0.014 | -0.026 | -0.04 | -0.04 | 0 | 0.14 | 0.22 | 0.1 | 0.04 | 0.07 | 0.12 |
| 9 | 135379520 | rs787904 | -1.179 | 0.062 | 0.812 | 0.098 | 0.009 | 0.056 | 0.056 | 0.412 | 0.27 | 0.53 | 0.34 | 0.35 | 0.32 | 0.32 |
| 9 | 135381734 | rs475027 | -1.553 | -0.035 | 0.726 | 0.058 | -0.012 | 0.027 | 0.027 | 0.538 | 0.32 | 0.54 | 0.34 | 0.35 | 0.32 | 0.32 |
| 9 | 135382198 | rs7021144 | -0.399 | 0.076 | 0.534 | -0.147 | -0.004 | -0.03 | -0.03 | 0 | 0.18 | 0.27 | 0.14 | 0.09 | 0.16 | 0.17 |
| 9 | 135384783 | rs17149204 | 0.155 | 0.391 | 0.273 | -0.348 | -0.007 | -0.023 | -0.023 | 0 | 0.14 | 0.2 | 0.09 | 0.07 | 0.1 | 0.14 |
| 9 | 135386515 | rs518457 | -0.506 | 0.366 | 0.407 | -0.344 | -0.039 | -0.072 | -0.072 | 0.093 | 0.2 | 0.37 | 0.11 | 0.08 | 0.11 | 0.21 |
| 9 | 135387021 | rs4962183 | 1.941 | 0.659 | -0.101 | -0.311 | -0.107 | 0.031 | 0.031 | 0 | 0.27 | 0.02 | 0.3 | 0.15 | 0.33 | 0.28 |
| 9 | 135389030 | rs12005531 | -0.271 | 1.12 | 1.051 | 0.692 | 0.002 | -0.003 | -0.003 | 0 | 0.16 | 0.21 | 0.11 | 0.14 | 0.15 | 0.27 |
| 9 | 135391963 | rs10793935 | -0.972 | 1.172 | 0.657 | 0.74 | 0.018 | 0.049 | 0.049 | 0 | 0.25 | 0.45 | 0.42 | 0.64 | 0.31 | 0.3 |
| 9 | 135394526 | rs2363020 | -0.953 | 1.115 | -0.186 | 0.202 | 0.015 | -0.029 | -0.029 | 0.032 | 0.32 | 0.49 | 0.43 | 0.78 | 0.35 | 0.52 |
| 9 | 135406613 | rs11243819 | -0.91 | 1.12 | -0.554 | 0.191 | -0.013 | -0.002 | -0.002 | 0 | 0.07 | 0.04 | 0.06 | 0.07 | 0.11 | 0.14 |
| 9 | 135408571 | rs944812 | -0.679 | 0.829 | -0.347 | -0.094 | -0.006 | -0.012 | -0.012 | 0 | 0.41 | 0.53 | 0.33 | 0.33 | 0.37 | 0.38 |
| 9 | 135414981 | rs7023572 | -0.292 | 0.597 | -0.207 | -0.281 | 0.003 | 0 | 0 | 0.039 | 0.28 | 0.39 | 0.28 | 0.29 | 0.31 | 0.24 |
| 9 | 135424607 | rs870596 | -0.752 | 0.909 | -0.471 | -0.843 | -0.024 | 0.024 | 0.024 | 1.328 | 0.79 | 0.88 | 0.69 | 0.94 | 0.65 | 0.8 |
| 9 | 135431295 | rs869890 | 1.253 | 0.633 | -0.406 | -0.833 | 0.01 | -0.003 | -0.003 | 0.014 | 0.4 | 0.31 | 0.38 | 0.58 | 0.3 | 0.44 |
| 9 | 135435742 | rs11243826 | -0.726 | 0.364 | -0.035 | -0.701 | -0.044 | -0.074 | -0.074 | 0.361 | 0.18 | 0.57 | 0.13 | 0.04 | 0.12 | 0.14 |
| 9 | 135435808 | rs17149282 | 0.019 | 0.675 | -0.55 | -0.74 | -0.075 | -0.08 | -0.08 | 0 | 0.21 | 0.03 | 0.19 | 0.31 | 0.27 | 0.25 |
| 9 | 135442556 | rs10122218 | -1.067 | 0.542 | -0.113 | -0.601 | -0.006 | -0.01 | -0.01 | 0 | 0.37 | 0.5 | 0.34 | 0.36 | 0.39 | 0.36 |

| | | | | | | | | | | | | | | | | |
|---|---|---|---|---|---|---|---|---|---|---|---|---|---|---|---|---|
| 9 | 135448320 | rs433311 | 0.071 | 0.86 | -0.259 | -0.829 | 0.016 | -0.033 | -0.033 | 0 | 0.67 | 0.59 | 0.69 | 0.91 | 0.56 | 0.79 |
| 9 | 135450156 | rs811504 | -0.502 | 0.845 | -0.757 | -0.775 | -0.065 | 0.065 | 0.065 | 0.017 | 0.73 | 0.91 | 0.77 | 0.91 | 0.65 | 0.84 |
| 9 | 135450636 | rs17149310 | -0.486 | 0.564 | -0.228 | -0.025 | -0.002 | -0.004 | -0.004 | 0 | 0.22 | 0.18 | 0.38 | 0.38 | 0.22 | 0.24 |
| 9 | 135462633 | rs880182 | -1.096 | 0.077 | -0.056 | -0.037 | -0.034 | 0.004 | 0.004 | 0 | 0.15 | 0.25 | 0.15 | 0.17 | 0.08 | 0.15 |
| 9 | 135467359 | rs1328403 | 0.678 | 0.061 | 0.058 | 0.266 | -0.022 | -0.023 | -0.023 | 0.553 | 0.63 | 0.44 | 0.77 | 0.7 | 0.72 | 0.84 |
| 9 | 135470414 | rs306547 | 0.625 | -0.017 | -0.1 | 0.331 | 0.026 | 0 | 0 | 2.594 | 0.67 | 0.94 | 0.84 | 0.72 | 0.77 | 0.83 |
| 9 | 135473647 | rs306542 | 0.542 | 0.258 | -0.173 | 0.431 | -0.009 | -0.013 | -0.013 | 0 | 0.79 | 0.95 | 0.84 | 0.78 | 0.8 | 0.88 |
| 9 | 135479687 | rs7037273 | -0.363 | 0.792 | 0.589 | 0.468 | 0.07 | 0.025 | 0.025 | 1.762 | 0.07 | 0.34 | 0.2 | 0.04 | 0.1 | 0.05 |
| 9 | 135483721 | rs445990 | 1.027 | 0.73 | -0.099 | 0.568 | -0.101 | 0.087 | 0.087 | 1.386 | 0.48 | 0.93 | 0.67 | 0.25 | 0.49 | 0.54 |
| 9 | 135483846 | rs10901193 | 0.322 | 0.724 | 0.059 | 0.566 | 0.035 | 0.008 | 0.008 | 0.689 | 0.06 | 0.2 | 0.2 | 0.04 | 0.1 | 0.06 |
| 9 | 135484385 | rs10901194 | 0.889 | 0.759 | -0.174 | 0.884 | -0.036 | -0.029 | -0.029 | 0.019 | 0.06 | 0.01 | 0.07 | 0.1 | 0.13 | 0.18 |
| 9 | 135489660 | rs509762 | 0.127 | 0.562 | 0.061 | 0.521 | -0.01 | -0.015 | -0.015 | 1.806 | 0.67 | 0.95 | 0.83 | 0.69 | 0.74 | 0.81 |
| 9 | 135490787 | rs10736856 | 0.211 | 0.547 | 0.102 | 0.52 | 0.022 | -0.026 | -0.026 | 0 | 0.63 | 0.81 | 0.71 | 0.57 | 0.69 | 0.74 |
| 9 | 135493386 | rs1028892 | 2.046 | 0.453 | 0.408 | 0.478 | -0.002 | 0.004 | 0.004 | 0 | 0.35 | 0.25 | 0.33 | 0.1 | 0.35 | 0.39 |
| 9 | 135494669 | rs306568 | -0.304 | 0.456 | 0.387 | 0.585 | 0 | 0.015 | 0.015 | 0.17 | 0.24 | 0.26 | 0.29 | 0.55 | 0.29 | 0.35 |
| 9 | 135521679 | rs3739905 | 0.432 | 0.509 | -0.112 | 0.612 | -0.067 | -0.128 | -0.128 | 0 | 0.18 | 0.02 | 0.17 | 0.42 | 0.26 | 0.28 |
| 9 | 135521917 | rs306538 | 0.142 | 0.418 | -0.089 | 0.514 | 0.033 | 0.017 | 0.017 | 2.424 | 0.66 | 0.86 | 0.83 | 0.68 | 0.75 | 0.82 |
| 9 | 135535929 | rs11243869 | 0.315 | 0.503 | -0.406 | 0.569 | -0.066 | -0.128 | -0.128 | 0 | 0.19 | 0.02 | 0.17 | 0.41 | 0.26 | 0.27 |
| 9 | 135550773 | rs10491904 | -0.2 | 0.392 | -0.314 | 0.61 | -0.051 | -0.078 | -0.078 | 0 | 0.24 | 0.11 | 0.35 | 0.44 | 0.36 | 0.33 |
| 9 | 135555341 | rs424416 | -1.638 | 0.32 | -0.296 | 0.432 | 0.002 | 0.105 | 0.105 | 0 | 0.5 | 0.72 | 0.61 | 0.87 | 0.54 | 0.49 |
| 9 | 135561944 | rs3780813 | 0.493 | 0.29 | -0.248 | 0.45 | 0.005 | -0.042 | -0.042 | 1.591 | 0.17 | 0.07 | 0.11 | 0.3 | 0.14 | 0.1 |
| 9 | 135562800 | rs456396 | -0.805 | 0.32 | -0.241 | 0.476 | 0.005 | 0.009 | 0.009 | 0 | 0.33 | 0.65 | 0.5 | 0.57 | 0.4 | 0.39 |
| 9 | 135568632 | rs302917 | -1.182 | 0.699 | -0.374 | -0.152 | -0.002 | 0.02 | 0.02 | 1.05 | 0.86 | 0.98 | 0.94 | 0.94 | 0.89 | 0.94 |
| 9 | 135573396 | rs302925 | -1.011 | 0.579 | -0.284 | -0.097 | 0.001 | 0.115 | 0.115 | 0 | 0.51 | 0.74 | 0.61 | 0.87 | 0.54 | 0.49 |
| 9 | 135579101 | rs11243880 | -0.234 | 0.984 | -0.333 | 0.295 | 0.051 | 0.064 | 0.064 | 0 | 0.08 | 0.16 | 0.32 | 0.16 | 0.17 | 0.09 |
| 9 | 135587540 | rs2772002 | -0.197 | 0.091 | -0.727 | 0.114 | 0.035 | 0.052 | 0.052 | 2.353 | 0.82 | 0.92 | 0.94 | 0.95 | 0.9 | 0.95 |
| 9 | 135588382 | rs7466085 | 0.497 | 0.749 | -0.299 | 0.104 | -0.009 | 0.023 | 0.023 | 0.296 | 0.42 | 0.33 | 0.61 | 0.21 | 0.49 | 0.53 |
| 9 | 135590998 | rs2772000 | -0.599 | 0.906 | 0.239 | 1.001 | 0.064 | 0.209 | 0.209 | 0.507 | 0.14 | 0.5 | 0.32 | 0.35 | 0.15 | 0.2 |
| 9 | 135598603 | rs306530 | -0.19 | 0.718 | -0.681 | 0.039 | 0.008 | 0.102 | 0.102 | 0.486 | 0.51 | 0.67 | 0.61 | 0.77 | 0.54 | 0.49 |
| 9 | 135600320 | rs306525 | 0.172 | 0.221 | -0.696 | 0.166 | -0.007 | 0.024 | 0.024 | 1.635 | 0.68 | 0.59 | 0.81 | 0.51 | 0.72 | 0.65 |
| 9 | 135602456 | rs10901205 | 0.552 | -0.288 | -0.842 | -0.374 | 0.001 | 0 | 0 | 0 | 0.51 | 0.48 | 0.72 | 0.5 | 0.58 | 0.56 |
| 9 | 135602727 | rs1999121 | -0.521 | -0.666 | -0.855 | -0.193 | -0.002 | -0.041 | -0.041 | 0 | 0.78 | 0.58 | 0.87 | 0.85 | 0.81 | 0.83 |
| 9 | 135603540 | rs10901206 | 0.194 | -0.663 | -0.948 | -0.151 | 0.054 | -0.104 | -0.104 | 4.486 | 0.28 | 0.05 | 0.15 | 0.36 | 0.23 | 0.27 |
| 9 | 135605439 | rs12550888 | 1.924 | -0.025 | -1.325 | -0.431 | -0.045 | -0.058 | -0.058 | 0 | 0.18 | 0.04 | 0.36 | 0.33 | 0.28 | 0.23 |
| 9 | 135605490 | rs693692 | -1.185 | -0.408 | -0.898 | -0.451 | -0.004 | 0.009 | 0.009 | 1.391 | 0.83 | 0.88 | 0.9 | 0.85 | 0.82 | 0.86 |
| 9 | 135608113 | rs386472 | 0.472 | -0.568 | -1.087 | -0.414 | 0.033 | 0.024 | 0.024 | 0.151 | 0.51 | 0.58 | 0.73 | 0.43 | 0.58 | 0.56 |
| 9 | 135615309 | rs574544 | -0.947 | -0.467 | -1.047 | -0.656 | -0.009 | 0.018 | 0.018 | 1.005 | 0.84 | 0.91 | 0.91 | 0.86 | 0.83 | 0.86 |
| 9 | 135616246 | rs4962210 | 0.064 | -0.45 | -1.148 | -0.837 | -0.022 | 0.033 | 0.033 | 0 | 0.42 | 0.73 | 0.56 | 0.38 | 0.42 | 0.56 |
| 9 | 135616700 | rs913705 | -0.445 | -1.073 | -0.585 | -1.089 | 0.012 | 0.006 | 0.006 | 0 | 0.49 | 0.6 | 0.27 | 0.56 | 0.31 | 0.55 |
| 9 | 135618325 | rs10901207 | 0.486 | -1.06 | -0.429 | -1.202 | -0.018 | -0.057 | -0.057 | 0.317 | 0.39 | 0.14 | 0.29 | 0.47 | 0.4 | 0.47 |
| 9 | 135624197 | rs11243900 | 0.443 | -0.893 | -0.096 | -1.147 | 0.004 | -0.004 | -0.004 | 0 | 0.37 | 0.37 | 0.58 | 0.36 | 0.41 | 0.33 |
| 9 | 135626396 | rs10901209 | 1.52 | -1.15 | 0.793 | -1.128 | 0.09 | 0.22 | 0.22 | 1.621 | 0.51 | 0.15 | 0.63 | 0.75 | 0.65 | 0.72 |
| 9 | 135632056 | rs954390 | -0.263 | -1.018 | 1.52 | -0.856 | -0.017 | -0.043 | -0.043 | 2.851 | 0.22 | 0.26 | 0.22 | 0.07 | 0.16 | 0.08 |
| 9 | 135632480 | rs1887964 | -0.615 | -1.003 | 0.973 | -0.848 | -0.047 | -0.093 | -0.093 | 0 | 0.25 | 0.57 | 0.26 | 0.07 | 0.17 | 0.09 |
| 9 | 135635852 | rs4962078 | -0.788 | -0.978 | 0.623 | -0.836 | -0.04 | -0.078 | -0.078 | 0 | 0.27 | 0.6 | 0.25 | 0.07 | 0.17 | 0.1 |
| 9 | 135647021 | rs10901212 | -0.011 | -0.391 | 0.373 | -0.688 | 0.003 | -0.011 | -0.011 | 0 | 0.82 | 0.79 | 0.88 | 0.88 | 0.83 | 0.89 |
| 9 | 135648905 | rs932887 | 1.474 | -0.881 | 0.162 | -0.969 | 0.054 | 0.204 | 0.204 | 1.65 | 0.55 | 0.15 | 0.62 | 0.82 | 0.66 | 0.81 |
| 9 | 135651456 | rs4962214 | -0.326 | -0.776 | 0.55 | -0.927 | -0.058 | -0.093 | -0.093 | 0 | 0.25 | 0.38 | 0.24 | 0.06 | 0.17 | 0.08 |
| 9 | 135654476 | rs2809247 | 1.121 | -0.689 | -0.33 | -0.876 | 0.029 | 0.084 | 0.084 | 0.289 | 0.58 | 0.69 | 0.66 | 0.82 | 0.67 | 0.81 |
| 9 | 135669228 | rs1410969 | -0.602 | -0.865 | -0.453 | -0.535 | 0.051 | 0.071 | 0.071 | 0.535 | 0.23 | 0.12 | 0.22 | 0.06 | 0.16 | 0.09 |
| 9 | 135672580 | rs215187 | -1.016 | -0.308 | 0.061 | 0.349 | 0.028 | 0.147 | 0.147 | 0 | 0.24 | 0.47 | 0.26 | 0.47 | 0.28 | 0.27 |
| 9 | 135675674 | rs10901214 | 1.206 | -0.609 | -0.96 | 0.15 | 0.023 | 0.057 | 0.057 | 0 | 0.52 | 0.08 | 0.49 | 0.41 | 0.55 | 0.64 |
| 9 | 135682019 | rs369511 | -1.2 | -0.736 | -0.357 | 0.043 | 0.008 | 0.052 | 0.052 | 0 | 0.23 | 0.29 | 0.24 | 0.48 | 0.28 | 0.27 |
| 9 | 135692783 | rs2809253 | -0.77 | -1.256 | -0.738 | -0.227 | 0.006 | -0.036 | -0.036 | 0 | 0.84 | 0.99 | 0.83 | 0.86 | 0.9 | 0.9 |
| 9 | 135701424 | rs2809255 | 0.866 | -1.179 | -0.505 | -0.075 | 0.053 | 0.004 | 0.004 | 0 | 0.06 | 0.14 | 0.06 | 0.05 | 0.09 | 0.08 |
| 9 | 135713036 | rs4962220 | 2.367 | -1.159 | -0.617 | -0.183 | -0.155 | -0.019 | -0.019 | 0.157 | 0.37 | 0.03 | 0.43 | 0.38 | 0.48 | 0.58 |
| 9 | 135719758 | rs17149822 | -0.535 | -1.185 | -0.415 | -0.16 | -0.003 | -0.02 | -0.02 | 0 | 0.09 | 0.19 | 0.1 | 0.04 | 0.06 | 0.05 |
| 9 | 135722652 | rs2771997 | 0.069 | -1.23 | 0.43 | -0.207 | 0.001 | 0.001 | 0.001 | 0.351 | 0.67 | 0.68 | 0.73 | 0.86 | 0.8 | 0.87 |
| 9 | 135724901 | rs2771994 | 1.255 | -1.261 | 0.406 | -0.207 | 0.082 | 0.118 | 0.118 | 0.047 | 0.53 | 0.21 | 0.57 | 0.82 | 0.69 | 0.79 |
| 9 | 135737088 | rs10491535 | -0.122 | -1.224 | 0.428 | -0.183 | -0.001 | -0.003 | -0.003 | 0 | 0.08 | 0.11 | 0.1 | 0.04 | 0.06 | 0.05 |
| 9 | 135740858 | rs10901217 | -0.672 | -1.209 | 0.395 | -0.153 | 0.01 | 0.08 | 0.08 | 0 | 0.16 | 0.17 | 0.14 | 0.45 | 0.21 | 0.31 |
| 9 | 135742518 | rs2526009 | 0.688 | -1.246 | 0.45 | -0.206 | -0.004 | -0.007 | -0.007 | 0 | 0.59 | 0.57 | 0.63 | 0.82 | 0.73 | 0.82 |
| 9 | 135753629 | rs2231400 | -1.271 | -1.153 | 0.472 | -0.198 | 0.035 | 0.001 | 0.001 | 0 | 0.06 | 0.36 | 0.06 | 0 | 0.04 | 0.03 |
| 9 | 135754010 | rs2231403 | 2.187 | -1.173 | 0.082 | -0.181 | -0.151 | -0.023 | -0.023 | 0.411 | 0.37 | 0.04 | 0.43 | 0.38 | 0.48 | 0.57 |
| 9 | 135763816 | rs2073869 | -0.688 | -1.137 | 0.515 | 0.424 | 0.012 | 0.047 | 0.047 | 0 | 0.15 | 0.2 | 0.13 | 0.27 | 0.2 | 0.19 |
| 9 | 135764081 | rs1062218 | 1.281 | -1.223 | 0.651 | 0.1 | 0.027 | 0.039 | 0.039 | 0 | 0.51 | 0.37 | 0.58 | 0.41 | 0.58 | 0.66 |
| 9 | 135765261 | rs2231410 | 0.781 | -1.378 | 0.933 | -0.53 | 0.007 | -0.008 | -0.008 | 2.197 | 0.14 | 0.13 | 0.08 | 0 | 0.09 | 0.03 |
| 9 | 135767185 | rs11553763 | -0.884 | -1.253 | 1.038 | -0.461 | -0.005 | -0.026 | -0.026 | 9.103 | 0.1 | 0.21 | 0.12 | 0.04 | 0.07 | 0.04 |
| 9 | 135767943 | rs1050700 | 0.4 | -1.289 | 1.211 | -0.412 | -0.01 | -0.012 | -0.012 | 0 | 0.63 | 0.59 | 0.65 | 0.83 | 0.75 | 0.83 |

| | | | | | | | | | | | | | | | | |
|---|---|---|---|---|---|---|---|---|---|---|---|---|---|---|---|---|
| 9 | 135769688 | rs10491534 | -0.87 | -1.062 | 1.388 | -0.378 | 0.078 | 0.012 | 0.012 | 0 | 0.07 | 0.24 | 0.07 | 0.07 | 0.1 | 0.1 |
| 9 | 135770115 | rs739441 | 0.537 | -1.498 | 1.646 | -0.496 | -0.006 | -0.003 | -0.003 | 0 | 0.68 | 0.66 | 0.75 | 0.82 | 0.83 | 0.83 |
| 9 | 135770134 | rs739442 | 1.814 | -1.443 | 0.948 | -0.338 | -0.029 | -0.069 | -0.069 | 0 | 0.44 | 0.06 | 0.5 | 0.58 | 0.61 | 0.65 |
| 9 | 135770300 | rs2809243 | 1.246 | -1.472 | 1.26 | -0.36 | 0.04 | 0.015 | 0.015 | 0 | 0.52 | 0.35 | 0.6 | 0.6 | 0.63 | 0.67 |
| 9 | 135770347 | rs2809244 | 0.095 | -1.391 | 1.234 | -0.316 | 0.006 | -0.004 | -0.004 | 0 | 0.68 | 0.78 | 0.74 | 0.68 | 0.75 | 0.78 |
| 9 | 135772717 | rs4962081 | 0.464 | -1.043 | 0.486 | -0.212 | 0 | -0.01 | -0.01 | 0 | 0.08 | 0.1 | 0.1 | 0.02 | 0.06 | 0.03 |
| 9 | 135775203 | rs2250057 | 0.926 | -1.337 | 0.532 | -0.34 | 0.009 | 0.007 | 0.007 | 0 | 0.5 | 0.51 | 0.58 | 0.63 | 0.66 | 0.72 |
| 9 | 135778827 | rs10901219 | 0.397 | -1.213 | 0.586 | -0.299 | 0.047 | 0.022 | 0.022 | 0 | 0.05 | 0.11 | 0.06 | 0.07 | 0.09 | 0.07 |
| 9 | 135786112 | rs6597586 | -0.911 | -1.221 | 0.599 | -0.32 | 0.036 | -0.011 | -0.011 | 0.497 | 0.1 | 0.23 | 0.09 | 0.07 | 0.13 | 0.1 |
| 9 | 135786904 | rs1073123 | -0.911 | -1.221 | 0.599 | -0.32 | 0.037 | -0.01 | -0.01 | 0.052 | 0.1 | 0.23 | 0.09 | 0.07 | 0.13 | 0.1 |
| 9 | 135799776 | rs13295430 | 0.179 | -1.179 | 0.542 | -0.339 | 0 | -0.017 | -0.017 | 0 | 0.09 | 0.13 | 0.12 | 0.02 | 0.1 | 0.04 |
| 9 | 135804735 | rs3761840 | 1.811 | -1.297 | 0.534 | -0.371 | -0.049 | -0.031 | -0.031 | 0 | 0.4 | 0.18 | 0.46 | 0.56 | 0.49 | 0.62 |
| 9 | 135813001 | rs7874234 | -1.111 | -1.386 | 0.312 | -0.367 | 0.017 | -0.044 | -0.044 | 1.128 | 0.19 | 0.33 | 0.2 | 0.09 | 0.21 | 0.13 |
| 9 | 135815269 | rs4419933 | 0.307 | -1.498 | 0.155 | -0.343 | 0.012 | 0.002 | 0.002 | 0 | 0.6 | 0.64 | 0.69 | 0.64 | 0.73 | 0.75 |
| 9 | 135820608 | rs4962083 | -1.21 | -1.228 | -0.003 | -0.305 | 0.007 | -0.026 | -0.026 | 0 | 0.18 | 0.24 | 0.2 | 0.1 | 0.21 | 0.14 |
| 9 | 135826950 | rs7040593 | 0.013 | -1.175 | -0.008 | -0.28 | 0 | -0.004 | -0.004 | 1.028 | 0.08 | 0.06 | 0.07 | 0.01 | 0.09 | 0.03 |
| 9 | 135831893 | rs3011289 | 0.182 | -1.106 | -0.339 | -0.398 | 0.011 | 0.004 | 0.004 | 0.009 | 0.59 | 0.63 | 0.65 | 0.66 | 0.69 | 0.76 |
| 9 | 135835334 | rs2905082 | 0.832 | -0.986 | -0.43 | -0.232 | 0.06 | 0.039 | 0.039 | 0 | 0.48 | 0.23 | 0.55 | 0.63 | 0.6 | 0.74 |
| 9 | 135840535 | rs2905078 | 0.028 | -0.738 | -1.762 | -0.765 | 0.002 | -0.007 | -0.007 | 0 | 0.43 | 0.5 | 0.45 | 0.4 | 0.45 | 0.41 |
| 9 | 135840945 | rs11243956 | 0.697 | -0.584 | -1.894 | -0.654 | -0.027 | 0.001 | 0.001 | 0 | 0.22 | 0.39 | 0.17 | 0.19 | 0.18 | 0.11 |
| 9 | 135841420 | rs11998995 | 0.652 | 0.021 | -2.523 | -0.243 | -0.037 | -0.048 | -0.048 | 0 | 0.29 | 0.12 | 0.27 | 0.31 | 0.34 | 0.36 |
| 9 | 135841437 | rs2905077 | 1.054 | 0.628 | -1.824 | -0.056 | -0.092 | -0.016 | -0.016 | 0 | 0.17 | 0.02 | 0.26 | 0.2 | 0.26 | 0.29 |
| 9 | 135842952 | rs2905076 | 1.084 | 0.68 | -1.787 | -0.026 | -0.093 | -0.018 | -0.018 | 0 | 0.17 | 0.02 | 0.27 | 0.2 | 0.26 | 0.29 |
| 9 | 135845035 | rs2905072 | 0.051 | 0.072 | -1.105 | 0.379 | 0 | -0.003 | -0.003 | 0 | 0.79 | 0.84 | 0.78 | 0.7 | 0.82 | 0.78 |
| 9 | 135849248 | rs6597589 | -0.012 | -0.064 | -1.223 | 0.151 | 0.012 | -0.002 | -0.002 | 0 | 0.43 | 0.31 | 0.3 | 0.49 | 0.42 | 0.52 |
| 9 | 135850574 | rs7036106 | 0.42 | -0.15 | -1.71 | 0.5 | -0.001 | -0.08 | -0.08 | 0 | 0.24 | 0.06 | 0.17 | 0.3 | 0.26 | 0.4 |
| 9 | 135852953 | rs3011260 | -0.033 | 0.152 | -0.889 | 0.373 | 0.014 | 0.085 | 0.085 | 0 | 0.72 | 0.58 | 0.61 | 0.48 | 0.7 | 0.78 |
| 9 | 135855420 | rs1964196 | 0.129 | -0.288 | -0.762 | 0.075 | 0.001 | 0 | 0 | 0.223 | 0.5 | 0.52 | 0.65 | 0.53 | 0.46 | 0.6 |
| 9 | 135855768 | rs685783 | -0.409 | -0.745 | -1.219 | -0.281 | 0.014 | -0.01 | -0.01 | 0 | 0.2 | 0.19 | 0.08 | 0.01 | 0.1 | 0.08 |
| 9 | 135859519 | rs2773818 | -0.236 | -0.322 | -1.185 | -0.072 | -0.012 | 0 | 0 | 0 | 0.45 | 0.32 | 0.37 | 0.41 | 0.5 | 0.4 |
| 9 | 135860839 | rs2905069 | -0.027 | 0.131 | -1.297 | 0.246 | -0.015 | -0.009 | -0.009 | 0.178 | 0.75 | 0.82 | 0.77 | 0.7 | 0.67 | 0.75 |
| 9 | 135861033 | rs2773822 | -0.555 | 0.074 | -1.389 | 0.26 | -0.021 | 0.006 | 0.006 | 0 | 0.11 | 0.02 | 0.19 | 0.09 | 0.11 | 0.15 |
| 9 | 135861241 | rs649651 | -0.612 | -0.123 | -1.295 | 0.221 | 0.017 | 0.026 | 0.026 | 0 | 0.15 | 0.06 | 0.19 | 0.1 | 0.11 | 0.22 |
| 9 | 135861998 | rs2073577 | 0.973 | 0.066 | -1.652 | 0.401 | -0.13 | -0.051 | -0.051 | 0.038 | 0.28 | 0.04 | 0.38 | 0.28 | 0.43 | 0.19 |
| 9 | 135862479 | rs606141 | -0.032 | -0.131 | -1.555 | 0.302 | 0.004 | 0.024 | 0.024 | 0 | 0.19 | 0.12 | 0.22 | 0.12 | 0.17 | 0.37 |
| 9 | 135862632 | rs633153 | -0.272 | -0.117 | -1.194 | 0.365 | 0.025 | 0.035 | 0.035 | 0 | 0.48 | 0.31 | 0.61 | 0.39 | 0.59 | 0.62 |
| 9 | 135864190 | rs2073819 | 0.498 | -0.288 | -0.361 | 0.914 | -0.012 | 0.015 | 0.015 | 0 | 0.73 | 0.35 | 0.79 | 0.7 | 0.78 | 0.57 |
| 9 | 135865496 | rs8193003 | 0.885 | -0.213 | -0.675 | 0.826 | 0.026 | 0.01 | 0.01 | 0.55 | 0.55 | 0.22 | 0.65 | 0.41 | 0.63 | 0.37 |
| 9 | 135866693 | rs667805 | -0.429 | -0.157 | 0.079 | 0.764 | -0.011 | 0.035 | 0.035 | 0.011 | 0.19 | 0.32 | 0.17 | 0.31 | 0.13 | 0.23 |
| 9 | 135867687 | rs678946 | -0.757 | -0.447 | 0.103 | 0.203 | -0.008 | -0.041 | -0.041 | 0 | 0.17 | 0.26 | 0.11 | 0.06 | 0.16 | 0.24 |
| 9 | 135867989 | rs7019702 | -0.38 | -0.738 | -0.053 | 0.434 | -0.025 | 0.027 | 0.027 | 0 | 0.28 | 0.58 | 0.34 | 0.4 | 0.18 | 0.29 |
| 9 | 135872063 | rs507586 | -0.962 | -0.448 | -0.449 | 0.521 | 0.035 | 0.003 | 0.003 | 0 | 0.42 | 0.77 | 0.45 | 0.38 | 0.35 | 0.43 |
| 9 | 135877287 | rs628094 | -0.223 | -0.704 | -0.439 | 0.292 | -0.016 | -0.001 | -0.001 | 1.77 | 0.65 | 0.43 | 0.62 | 0.65 | 0.73 | 0.76 |
| 9 | 135877904 | rs11243977 | 0.883 | -0.214 | -0.796 | 0.016 | -0.064 | 0.029 | 0.029 | 0 | 0.28 | 0.13 | 0.37 | 0.19 | 0.44 | 0.32 |
| 9 | 135879020 | rs9942933 | 1.097 | -0.311 | -0.835 | -0.007 | -0.09 | 0.027 | 0.027 | 0 | 0.27 | 0.13 | 0.38 | 0.18 | 0.45 | 0.22 |
| 9 | 135881044 | rs603328 | -0.692 | -0.57 | -0.428 | -0.051 | -0.006 | -0.03 | -0.03 | 0 | 0.09 | 0.11 | 0.16 | 0.01 | 0.06 | 0.19 |
| 9 | 135881193 | rs3011267 | -0.949 | -1.218 | -0.401 | 0.123 | 0.021 | -0.021 | -0.021 | 0 | 0.77 | 0.84 | 0.83 | 0.65 | 0.83 | 0.78 |
| 9 | 135883620 | rs3011271 | -0.934 | -1.347 | -0.786 | -0.16 | 0 | 0.031 | 0.031 | 0 | 0.2 | 0.12 | 0.24 | 0.05 | 0.21 | 0.31 |
| 9 | 135888407 | rs520812 | 0.06 | -1.332 | -0.84 | -0.164 | -0.031 | 0.048 | 0.048 | 0.065 | 0.12 | 0.03 | 0.07 | 0.04 | 0.16 | 0.12 |
| 9 | 135888516 | rs3011274 | -0.034 | -1.872 | -0.418 | -0.165 | 0.012 | 0.009 | 0.009 | 0 | 0.55 | 0.43 | 0.69 | 0.29 | 0.72 | 0.53 |
| 9 | 135889579 | rs7021235 | 0.96 | -1.635 | -0.464 | -0.008 | -0.008 | 0.012 | 0.012 | 0 | 0.37 | 0.33 | 0.45 | 0.24 | 0.52 | 0.23 |
| 9 | 135897389 | rs12685748 | 0.214 | -1.506 | 0.119 | 0.087 | -0.027 | -0.008 | -0.008 | 0 | 0.6 | 0.47 | 0.6 | 0.61 | 0.65 | 0.6 |
| 9 | 135899888 | rs2988566 | -0.158 | -1.181 | -0.605 | 0.283 | 0.013 | -0.043 | -0.043 | 0 | 0.33 | 0.15 | 0.24 | 0.41 | 0.33 | 0.4 |
| 9 | 135902962 | rs608604 | 0.43 | -0.712 | -0.196 | 0.238 | -0.003 | 0.006 | 0.006 | 0 | 0.68 | 0.36 | 0.74 | 0.63 | 0.7 | 0.57 |
| 9 | 135903335 | rs7035838 | -0.402 | -0.481 | -0.591 | 0.252 | 0.001 | 0 | 0 | 0 | 0.37 | 0.61 | 0.47 | 0.65 | 0.34 | 0.43 |
| 9 | 135903997 | rs623489 | 0.628 | 0.037 | -0.672 | 0.052 | 0.003 | -0.018 | -0.018 | 0 | 0.31 | 0.42 | 0.32 | 0.21 | 0.37 | 0.18 |
| 9 | 135904566 | rs2988571 | 1.176 | 0.099 | -1.037 | 0.145 | -0.025 | 0.056 | 0.056 | 0 | 0.28 | 0.1 | 0.17 | 0.21 | 0.25 | 0.42 |
| 9 | 135905701 | rs649168 | 1.123 | 0.264 | -0.525 | 0.58 | -0.026 | 0.008 | 0.008 | 2.644 | 0.24 | 0.13 | 0.29 | 0.17 | 0.32 | 0.11 |
| 9 | 135913483 | rs675167 | 0.896 | 0.271 | -0.216 | 0.597 | 0.021 | -0.02 | -0.02 | 3.916 | 0.27 | 0.42 | 0.32 | 0.16 | 0.34 | 0.1 |
| 9 | 135926017 | rs1541331 | -1.174 | 0.159 | -0.107 | 0.708 | 0.03 | 0.064 | 0.064 | 0 | 0.07 | 0.34 | 0.03 | 0.09 | 0.04 | 0.08 |
| 9 | 135929200 | rs7863758 | -0.741 | 0.163 | -0.027 | 0.676 | 0.022 | 0.042 | 0.042 | 0 | 0.06 | 0.26 | 0.03 | 0.08 | 0.04 | 0.08 |
| 9 | 135933122 | rs2073821 | 1.317 | -0.019 | -0.691 | 0.948 | -0.004 | -0.072 | -0.072 | 0.144 | 0.11 | 0.01 | 0.23 | 0.3 | 0.09 | 0.21 |
| 9 | 135933761 | rs7324 | 1.028 | 0.106 | -0.453 | 0.493 | -0.043 | 0.014 | 0.014 | 2.831 | 0.25 | 0.13 | 0.3 | 0.16 | 0.33 | 0.1 |
| 9 | 135934431 | rs586977 | 0.257 | 0.109 | -0.463 | 0.507 | 0.009 | -0.017 | -0.017 | 5.922 | 0.32 | 0.45 | 0.33 | 0.24 | 0.37 | 0.18 |
| 9 | 135951231 | rs2283134 | 1.214 | 0.312 | -1.133 | 0.769 | -0.067 | -0.043 | -0.043 | 0 | 0.2 | 0.07 | 0.31 | 0.2 | 0.32 | 0.07 |
| 9 | 135954310 | rs509064 | 0.032 | 0.073 | -0.609 | 0.615 | 0.007 | -0.017 | -0.017 | 0 | 0.41 | 0.58 | 0.47 | 0.28 | 0.47 | 0.22 |
| 9 | 135955672 | rs563864 | 0.604 | 0.014 | -1.303 | 0.57 | -0.089 | 0.017 | 0.017 | 0 | 0.3 | 0.09 | 0.42 | 0.21 | 0.42 | 0.1 |
| 9 | 135957091 | rs3761829 | 0.947 | -0.257 | -1.417 | 1.032 | 0.012 | -0.013 | -0.013 | 0.031 | 0.11 | 0.08 | 0.18 | 0.19 | 0.07 | 0.19 |
| 9 | 135957452 | rs694388 | 0.566 | 0.15 | -1.248 | 0.671 | -0.089 | 0.033 | 0.033 | 0 | 0.3 | 0.09 | 0.31 | 0.2 | 0.42 | 0.1 |

| | | | | | | | | | | | | | | | | |
|---|---|---|---|---|---|---|---|---|---|---|---|---|---|---|---|---|
| 9 | 135966362 | rs591790 | 0.162 | 0.222 | -0.502 | 0.743 | 0.009 | 0.028 | 0.028 | 0.397 | 0.51 | 0.66 | 0.47 | 0.63 | 0.48 | 0.65 |
| 9 | 135972407 | rs7023365 | -0.178 | -0.095 | -0.405 | 0.904 | -0.002 | 0.002 | 0.002 | 0 | 0.84 | 0.87 | 0.85 | 0.79 | 0.82 | 0.81 |
| 9 | 135976679 | rs2519090 | -0.237 | 0.417 | -0.39 | 1.172 | -0.011 | -0.032 | -0.032 | 0.738 | 0.19 | 0.09 | 0.28 | 0.37 | 0.22 | 0.24 |
| 9 | 135982088 | rs2073822 | 0.668 | 0.69 | 0.074 | 0.416 | 0.002 | 0.077 | 0.077 | 4.446 | 0.32 | 0.14 | 0.32 | 0.16 | 0.3 | 0.11 |
| 9 | 135983504 | rs2285375 | 0.39 | 0.703 | 0.694 | 0.39 | -0.014 | -0.053 | -0.053 | 0.012 | 0.35 | 0.48 | 0.35 | 0.16 | 0.3 | 0.11 |
| 9 | 135985796 | rs3761824 | 0.884 | 0.602 | 0.095 | 0.461 | -0.024 | -0.007 | -0.007 | 0 | 0.27 | 0.13 | 0.22 | 0.32 | 0.27 | 0.34 |
| 9 | 135995166 | rs10901235 | 0.84 | 0.624 | 0.09 | 0.486 | -0.025 | -0.029 | -0.029 | 0 | 0.27 | 0.13 | 0.24 | 0.36 | 0.27 | 0.34 |
| 9 | 136000445 | rs602060 | -0.081 | 0.838 | 0.642 | 0.274 | 0 | -0.01 | -0.01 | 2.389 | 0.57 | 0.57 | 0.5 | 0.69 | 0.5 | 0.7 |
| 9 | 136001966 | rs671050 | -0.868 | 1.174 | 0.756 | -0.739 | -0.012 | -0.077 | -0.077 | 3.377 | 0.67 | 0.61 | 0.72 | 0.88 | 0.71 | 0.68 |
| 9 | 136007358 | rs482670 | -0.925 | 0.698 | 0.539 | -0.552 | 0.007 | -0.011 | -0.011 | 0.866 | 0.62 | 0.62 | 0.67 | 0.63 | 0.6 | 0.61 |
| 9 | 136007988 | rs626713 | -0.433 | 1.129 | 0.764 | 0.301 | -0.003 | -0.025 | -0.025 | 0.221 | 0.28 | 0.21 | 0.32 | 0.51 | 0.29 | 0.5 |
| 9 | 136009191 | rs658853 | -0.626 | 1.074 | 0.686 | 0.242 | 0 | -0.001 | -0.001 | 0.391 | 0.33 | 0.38 | 0.32 | 0.48 | 0.34 | 0.51 |
| 9 | 136023053 | rs7022583 | -1.387 | 0.811 | 1.141 | -0.211 | 0.044 | -0.095 | -0.095 | 0 | 0.14 | 0.41 | 0.1 | 0 | 0.11 | 0.05 |
| 9 | 136029645 | rs35898523 | -0.973 | 0.687 | 1.077 | -0.196 | -0.007 | 0.012 | 0.012 | 0 | 0.1 | 0.03 | 0.04 | 0 | 0.08 | 0.06 |
| 9 | 136031346 | rs2073927 | -0.419 | 0.531 | -0.011 | -0.114 | -0.024 | 0.014 | 0.014 | 0 | 0.54 | 0.8 | 0.53 | 0.54 | 0.51 | 0.42 |
| 9 | 136037259 | rs2073925 | -0.046 | 0.052 | 0.165 | 0.55 | -0.008 | 0.111 | 0.111 | 0.168 | 0.08 | 0.11 | 0.19 | 0.49 | 0.07 | 0.25 |
| 9 | 136037742 | rs2073924 | -0.765 | -0.1 | -0.063 | 0.221 | -0.003 | 0.015 | 0.015 | 0 | 0.22 | 0.26 | 0.3 | 0.53 | 0.18 | 0.33 |
| 9 | 136040134 | rs10901239 | -0.738 | 0.509 | -0.894 | -1.05 | -0.026 | 0.1 | 0.1 | 0 | 0.34 | 0.22 | 0.29 | 0.04 | 0.39 | 0.19 |
| 9 | 136040899 | rs7044834 | -0.864 | -0.355 | -0.345 | -1 | -0.047 | -0.05 | -0.05 | 0.022 | 0.28 | 0.47 | 0.28 | 0.14 | 0.15 | 0.15 |
| 9 | 136042694 | rs11244017 | 0.547 | -0.292 | -1.851 | -1.438 | -0.078 | 0.11 | 0.11 | 0 | 0.34 | 0.08 | 0.33 | 0.17 | 0.34 | 0.29 |
| 9 | 136043270 | rs7022409 | -0.243 | -0.213 | -1.725 | -1.936 | 0.032 | 0.066 | 0.066 | 0 | 0.6 | 0.33 | 0.67 | 0.18 | 0.67 | 0.47 |
| 9 | 136047139 | rs4284147 | -0.836 | -0.396 | -1.587 | -0.962 | -0.001 | -0.013 | -0.013 | 0.561 | 0.78 | 0.81 | 0.82 | 0.7 | 0.77 | 0.83 |
| 9 | 136047393 | rs2039184 | -0.971 | -0.528 | -1.439 | -0.521 | 0.012 | 0.023 | 0.023 | 0.069 | 0.45 | 0.68 | 0.49 | 0.37 | 0.45 | 0.53 |
| 9 | 136049780 | rs6597604 | -0.011 | -0.206 | -1.103 | -0.517 | -0.013 | 0.003 | 0.003 | 0 | 0.25 | 0.17 | 0.33 | 0.01 | 0.32 | 0.16 |
| 9 | 136057106 | rs10751500 | 0.683 | -0.406 | -1.084 | -0.458 | 0.004 | -0.003 | -0.003 | 0 | 0.52 | 0.61 | 0.39 | 0.52 | 0.45 | 0.67 |
| 9 | 136061648 | rs7028259 | -0.891 | -0.359 | -0.751 | -0.357 | 0 | -0.005 | -0.005 | 0 | 0.8 | 0.8 | 0.83 | 0.71 | 0.81 | 0.85 |
| 9 | 136075679 | rs13295207 | 0.18 | 0.028 | -0.664 | -0.75 | -0.02 | 0.002 | 0.002 | 0 | 0.21 | 0.14 | 0.35 | 0.01 | 0.34 | 0.15 |
| 9 | 136079463 | rs10901244 | 1.127 | -0.238 | -0.406 | -0.703 | -0.009 | 0.025 | 0.025 | 0 | 0.46 | 0.3 | 0.43 | 0.56 | 0.43 | 0.37 |
| 9 | 136080272 | rs11244034 | -0.156 | -0.646 | 0.489 | 0.062 | -0.019 | 0.042 | 0.042 | 0 | 0.8 | 0.79 | 0.85 | 0.64 | 0.83 | 0.58 |
| 9 | 136081319 | rs11244035 | 1.567 | -0.261 | -0.009 | -0.024 | -0.017 | 0.018 | 0.018 | 0 | 0.05 | 0.01 | 0.07 | 0 | 0.11 | 0.03 |
| 9 | 136084941 | rs4489379 | 0.721 | -1.075 | 0.297 | 0.184 | 0.02 | 0.084 | 0.084 | 0 | 0.45 | 0.27 | 0.4 | 0.23 | 0.57 | 0.39 |
| 9 | 136085783 | rs3761821 | 0.484 | -0.461 | 1.251 | 0.812 | 0.005 | -0.034 | -0.034 | 0 | 0.8 | 0.93 | 0.71 | 0.63 | 0.81 | 0.58 |
| 9 | 136086916 | rs7864821 | 0.503 | -0.461 | 1.251 | 0.812 | 0.006 | -0.034 | -0.034 | 0.911 | 0.8 | 0.93 | 0.71 | 0.63 | 0.81 | 0.58 |
| 9 | 136101999 | rs886082 | 0.912 | -0.909 | 1.75 | 0.714 | -0.034 | 0.044 | 0.044 | 0 | 0.74 | 0.49 | 0.67 | 0.57 | 0.8 | 0.45 |
| 9 | 136112413 | rs4246170 | -0.425 | -0.887 | 1.521 | 0.646 | -0.02 | 0.002 | 0.002 | 0.01 | 0.21 | 0.19 | 0.05 | 0.23 | 0.1 | 0.28 |
| 9 | 136115440 | rs13301148 | -1.452 | -0.673 | 1.443 | 0.359 | -0.006 | -0.038 | -0.038 | 2.135 | 0.11 | 0.24 | 0.11 | 0.07 | 0.12 | 0.1 |
| 9 | 136116759 | rs7036324 | -1.83 | -0.797 | 1.758 | 0.271 | -0.026 | -0.095 | -0.095 | 0 | 0.16 | 0.46 | 0.13 | 0.07 | 0.14 | 0.1 |
| 9 | 136118513 | rs9411367 | -0.944 | -0.663 | 1.36 | 0.605 | -0.034 | 0.009 | 0.009 | 0 | 0.27 | 0.6 | 0.18 | 0.34 | 0.22 | 0.38 |
| 9 | 136119527 | rs9919007 | -0.984 | -0.665 | 0.867 | 0.582 | -0.044 | 0.003 | 0.003 | 0 | 0.27 | 0.47 | 0.14 | 0.31 | 0.22 | 0.39 |
| 9 | 136119888 | rs9411468 | -0.984 | -0.63 | 0.867 | 0.582 | -0.042 | 0.003 | 0.003 | 0 | 0.27 | 0.47 | 0.14 | 0.31 | 0.22 | 0.39 |
| 9 | 136120091 | rs9411471 | -0.984 | -0.664 | 0.867 | 0.582 | -0.044 | 0.003 | 0.003 | 0 | 0.27 | 0.47 | 0.14 | 0.31 | 0.22 | 0.39 |
| 9 | 136121441 | rs7046863 | -0.906 | -0.629 | 0.885 | 0.599 | -0.043 | 0.005 | 0.005 | 0 | 0.27 | 0.47 | 0.14 | 0.3 | 0.22 | 0.39 |
| 9 | 136127268 | rs9411475 | -1.076 | -0.064 | 0.74 | 0.982 | 0.004 | 0.045 | 0.045 | 0.463 | 0.11 | 0.21 | 0.07 | 0.26 | 0.11 | 0.13 |
| 9 | 136127641 | rs4962113 | -0.322 | 0.263 | 0.679 | 0.886 | 0.014 | -0.007 | -0.007 | 0 | 0.39 | 0.54 | 0.57 | 0.4 | 0.42 | 0.3 |
| 9 | 136128546 | rs7857390 | -1.164 | 0.179 | 0.459 | 0.997 | 0.007 | -0.001 | -0.001 | 0.008 | 0.63 | 0.81 | 0.68 | 0.67 | 0.62 | 0.63 |
| 9 | 136131188 | rs8176749 | 1.094 | -0.042 | 0.992 | 0.99 | -0.006 | 0.001 | 0.001 | 0 | 0.17 | 0.17 | 0.05 | 0.19 | 0.08 | 0.23 |
| 9 | 136131322 | rs8176746 | 1.094 | -0.036 | 0.984 | 0.982 | -0.006 | 0.001 | 0.001 | 0 | 0.17 | 0.17 | 0.05 | 0.19 | 0.08 | 0.23 |
| 9 | 136131415 | rs8176743 | 1.094 | -0.036 | 0.984 | 0.982 | -0.006 | 0.001 | 0.001 | 0 | 0.17 | 0.17 | 0.05 | 0.19 | 0.08 | 0.23 |
| 9 | 136131651 | rs1053878 | -0.066 | 0.436 | 1.51 | 1.005 | 0.004 | 0.026 | 0.026 | 2.798 | 0.11 | 0.26 | 0.07 | 0.15 | 0.1 | 0.03 |
| 9 | 136131905 | rs7873522 | 0.021 | 0.755 | 1.387 | 0.868 | -0.006 | 0.003 | 0.003 | 0 | 0.42 | 0.49 | 0.5 | 0.48 | 0.34 | 0.47 |
| 9 | 136132303 | rs8176732 | 1.167 | 0.761 | 1.359 | 0.865 | -0.005 | 0.001 | 0.001 | 0 | 0.24 | 0.24 | 0.43 | 0.29 | 0.22 | 0.22 |
| 9 | 136132633 | rs2073824 | 0.066 | 0.762 | 1.31 | 0.954 | 0.002 | -0.002 | -0.002 | 0 | 0.42 | 0.47 | 0.5 | 0.55 | 0.34 | 0.47 |
| 9 | 136132754 | rs8176722 | 1.172 | 0.733 | 1.285 | 0.762 | 0.015 | -0.011 | -0.011 | 0 | 0.17 | 0.14 | 0.06 | 0.19 | 0.11 | 0.24 |
| 9 | 136132852 | rs8176721 | 2.161 | 0.739 | 1.4 | 0.722 | -0.017 | -0.023 | -0.023 | 0 | 0.05 | 0.16 | 0.02 | 0 | 0.01 | 0 |
| 9 | 136133034 | rs8176717 | 0.996 | 0.79 | 1.27 | 0.744 | 0 | 0 | 0 | 0.02 | 0.25 | 0.27 | 0.43 | 0.29 | 0.22 | 0.22 |
| 9 | 136133506 | rs512770 | -1.231 | 1.075 | 1.245 | 0.709 | -0.001 | 0 | 0 | 0 | 0.75 | 0.72 | 0.58 | 0.71 | 0.81 | 0.77 |
| 9 | 136133699 | rs641959 | -1.076 | 0.786 | 1.309 | 0.705 | 0.005 | -0.007 | -0.007 | 0 | 0.74 | 0.63 | 0.55 | 0.7 | 0.75 | 0.77 |
| 9 | 136133743 | rs514708 | -1.1 | 0.79 | 1.285 | 0.707 | 0.005 | -0.006 | -0.006 | 0 | 0.74 | 0.63 | 0.55 | 0.7 | 0.75 | 0.77 |
| 9 | 136135238 | rs549446 | -1.052 | 0.792 | 1.282 | 0.709 | 0.005 | -0.003 | -0.003 | 0 | 0.75 | 0.67 | 0.57 | 0.71 | 0.78 | 0.77 |
| 9 | 136135659 | rs574347 | -1.052 | 0.842 | 1.329 | 0.771 | 0.005 | -0.003 | -0.003 | 0 | 0.75 | 0.67 | 0.57 | 0.71 | 0.78 | 0.78 |
| 9 | 136136770 | rs688976 | -1.302 | 0.787 | 1.373 | 0.739 | 0.004 | 0 | 0 | 0 | 0.77 | 0.69 | 0.58 | 0.71 | 0.79 | 0.78 |
| 9 | 136137106 | rs687289 | -0.653 | 0.892 | 1.292 | 0.738 | 0 | 0.002 | 0.002 | 0 | 0.37 | 0.44 | 0.23 | 0.38 | 0.37 | 0.39 |
| 9 | 136137646 | rs8176694 | 1.776 | 0.784 | 1.108 | 0.701 | -0.039 | 0.071 | 0.071 | 0 | 0.18 | 0.08 | 0.12 | 0.06 | 0.17 | 0.17 |
| 9 | 136138125 | rs672316 | -1.234 | 0.837 | 1.1 | 0.733 | 0.003 | -0.004 | -0.004 | 0 | 0.74 | 0.67 | 0.55 | 0.7 | 0.75 | 0.77 |
| 9 | 136139265 | rs657152 | -0.802 | 0.851 | 1.167 | 0.709 | 0 | 0 | 0 | 0 | 0.38 | 0.46 | 0.25 | 0.37 | 0.4 | 0.4 |
| 9 | 136139297 | rs8176682 | 1.172 | 0.851 | 1.096 | 0.74 | 0.007 | 0 | 0 | 0 | 0.37 | 0.17 | 0.32 | 0.32 | 0.38 | 0.37 |
| 9 | 136139617 | rs474279 | -1.041 | 1.148 | 0.923 | 0.819 | 0.004 | 0 | 0 | 0 | 0.76 | 0.7 | 0.57 | 0.7 | 0.78 | 0.77 |
| 9 | 136149399 | rs507666 | 0.372 | 1.155 | 0.872 | 0.808 | -0.009 | -0.008 | -0.008 | 0.019 | 0.11 | 0.09 | 0.12 | 0.19 | 0.19 | 0.13 |
| 9 | 136149722 | rs630014 | -0.448 | 0.899 | 0.459 | 0.446 | -0.001 | 0.008 | 0.008 | 0 | 0.49 | 0.57 | 0.42 | 0.56 | 0.54 | 0.63 |

| | | | | | | | | | | | | | | | | |
|---|---|---|---|---|---|---|---|---|---|---|---|---|---|---|---|---|
| 9 | 136151445 | rs568203 | -1.3 | 0.934 | 0.1 | 0.503 | -0.003 | -0.001 | -0.001 | 0 | 0.76 | 0.75 | 0.57 | 0.69 | 0.78 | 0.77 |
| 9 | 136152722 | rs8176630 | 1.048 | 0.825 | -0.039 | 0.433 | 0.007 | 0.043 | 0.043 | 0 | 0.25 | 0.08 | 0.14 | 0.12 | 0.21 | 0.13 |
| 9 | 136153481 | rs552148 | -1.273 | 0.957 | 0.109 | 0.504 | -0.004 | -0.001 | -0.001 | 0 | 0.75 | 0.75 | 0.57 | 0.69 | 0.77 | 0.77 |
| 9 | 136154168 | rs579459 | 0.287 | 1.12 | 0.124 | 0.591 | 0.006 | 0.003 | 0.003 | 0 | 0.12 | 0.13 | 0.14 | 0.19 | 0.21 | 0.14 |
| 9 | 136155359 | rs7030248 | 0.347 | 0.98 | 0.206 | 0.607 | 0.018 | 0.025 | 0.025 | 1.025 | 0.46 | 0.33 | 0.23 | 0.3 | 0.38 | 0.38 |
| 9 | 136155444 | rs633862 | -0.193 | 0.66 | 0.318 | 0.471 | 0 | -0.003 | -0.003 | 0.414 | 0.57 | 0.42 | 0.35 | 0.49 | 0.57 | 0.52 |
| 9 | 136156067 | rs9411381 | 1.023 | 0.695 | -0.11 | 0.442 | -0.006 | 0.008 | 0.008 | 0.114 | 0.16 | 0.16 | 0.2 | 0.19 | 0.17 | 0.25 |
| 9 | 136157133 | rs558240 | 0.175 | 0.919 | -0.088 | 0.045 | -0.021 | 0.062 | 0.062 | 0 | 0.36 | 0.17 | 0.25 | 0.15 | 0.39 | 0.24 |
| 9 | 136160228 | rs1752337 | 0.342 | 0.979 | -0.015 | 0.299 | 0.004 | 0.001 | 0.001 | 0 | 0.27 | 0.21 | 0.45 | 0.31 | 0.25 | 0.25 |
| 9 | 136161851 | rs9411493 | 0.145 | 1.304 | 0.362 | 0.242 | 0.013 | -0.017 | -0.017 | 0.333 | 0.67 | 0.54 | 0.5 | 0.67 | 0.66 | 0.73 |
| 9 | 136175051 | rs624960 | -0.465 | 0.855 | 0.117 | 0.311 | -0.016 | -0.002 | -0.002 | 0 | 0.73 | 0.69 | 0.56 | 0.69 | 0.78 | 0.74 |
| 9 | 136179407 | rs7357661 | 0.451 | 1.09 | -0.424 | 0.326 | 0.007 | 0.001 | 0.001 | 0 | 0.26 | 0.2 | 0.43 | 0.31 | 0.23 | 0.26 |
| 9 | 136184598 | rs17150319 | 1.288 | 1.045 | -1.235 | 0.4 | -0.017 | -0.046 | -0.046 | 1.618 | 0.11 | 0.02 | 0.09 | 0.16 | 0.12 | 0.11 |
| 9 | 136187163 | rs493014 | 0.413 | 0.927 | -1.257 | -0.254 | 0.01 | -0.043 | -0.043 | 0.473 | 0.7 | 0.4 | 0.82 | 0.9 | 0.66 | 0.77 |
| 9 | 136187652 | rs9411396 | 1.473 | 0.409 | -1.74 | 0.407 | 0.009 | -0.077 | -0.077 | 0 | 0.15 | 0.01 | 0.41 | 0.28 | 0.11 | 0.21 |
| 9 | 136194595 | rs17150482 | 0.712 | 0.855 | -2.075 | 0.293 | -0.057 | -0.064 | -0.064 | 0 | 0.2 | 0.04 | 0.32 | 0.3 | 0.27 | 0.44 |
| 9 | 136197834 | rs12335 | 0.368 | 0.456 | -1.374 | -0.391 | 0 | -0.003 | -0.003 | 0.345 | 0.55 | 0.41 | 0.52 | 0.68 | 0.46 | 0.5 |
| 9 | 136199503 | rs886090 | 0.887 | 0.742 | -1.592 | -0.414 | -0.097 | -0.052 | -0.052 | 0 | 0.25 | 0.04 | 0.35 | 0.31 | 0.36 | 0.46 |
| 9 | 136207843 | rs11655 | 2.173 | 0.737 | -0.897 | -0.542 | -0.042 | 0.017 | 0.017 | 0.032 | 0.07 | 0.01 | 0.05 | 0.02 | 0.11 | 0.02 |
| 9 | 136208044 | rs621907 | 2.74 | 0.509 | -1.315 | -0.941 | -0.013 | 0.037 | 0.037 | 0 | 0.07 | 0.01 | 0.05 | 0 | 0.09 | 0.01 |
| 9 | 136209319 | rs7870707 | 1.336 | 0.211 | -1.382 | -0.92 | 0.004 | -0.008 | -0.008 | 0 | 0.25 | 0.17 | 0.36 | 0.33 | 0.23 | 0.39 |
| 9 | 136210046 | rs513822 | -1.143 | 0.114 | -0.326 | -1.371 | 0.031 | 0.018 | 0.018 | 0.047 | 0.95 | 0.68 | 0.98 | 1 | 0.99 | 1 |
| 9 | 136214955 | rs553877 | -1.44 | 0.318 | -0.393 | -1.4 | -0.043 | 0.002 | 0.002 | 0.102 | 0.73 | 0.83 | 0.63 | 0.7 | 0.61 | 0.53 |
| 9 | 136217958 | rs28693986 | 2.627 | 0.308 | -0.337 | -1.396 | -0.023 | 0.021 | 0.021 | 0.075 | 0.07 | 0 | 0.04 | 0.02 | 0.08 | 0.02 |
| 9 | 136222789 | rs2051680 | 0.98 | 0.303 | -0.198 | -1.398 | -0.006 | -0.052 | -0.052 | 0 | 0.11 | 0.31 | 0.06 | 0.02 | 0.09 | 0.02 |
| 9 | 136227017 | rs2296809 | 2.953 | 0.244 | -0.676 | -1.301 | -0.025 | -0.032 | -0.032 | 0 | 0.1 | 0 | 0.07 | 0.1 | 0.11 | 0.06 |
| 9 | 136227260 | rs12763 | -0.844 | 0.75 | -0.143 | -1.699 | -0.045 | 0 | 0 | 0.13 | 0.73 | 0.84 | 0.63 | 0.7 | 0.6 | 0.53 |
| 9 | 136229950 | rs11543366 | 1.826 | 0.475 | -0.05 | -1.867 | -0.023 | 0.021 | 0.021 | 0.075 | 0.07 | 0 | 0.04 | 0.02 | 0.08 | 0.02 |
| 9 | 136246666 | rs28602591 | 0.573 | 0.362 | -0.117 | -1.946 | -0.006 | 0.025 | 0.025 | 0 | 0.09 | 0.04 | 0.07 | 0.02 | 0.1 | 0.1 |
| 9 | 136259809 | rs36091105 | 2.007 | 0.466 | -0.02 | -1.82 | -0.017 | 0.033 | 0.033 | 0.233 | 0.06 | 0 | 0.03 | 0 | 0.06 | 0.02 |
| 9 | 136263584 | rs7040821 | 0.357 | 0.428 | 0.171 | -1.831 | 0 | -0.042 | -0.042 | 0.231 | 0.1 | 0.24 | 0.05 | 0.02 | 0.09 | 0.03 |
| 9 | 136269143 | rs17150554 | 1.685 | 0.445 | 0.045 | -1.825 | -0.025 | 0.021 | 0.021 | 0.075 | 0.07 | 0 | 0.04 | 0.02 | 0.08 | 0.03 |
| 9 | 136270996 | rs3124782 | 0.732 | 0.435 | 0.437 | -1.888 | -0.001 | 0.004 | 0.004 | 0 | 0.59 | 0.36 | 0.5 | 0.73 | 0.47 | 0.51 |
| 9 | 136275313 | rs3124781 | -0.864 | 0.452 | 0.28 | -1.922 | -0.044 | 0.011 | 0.011 | 0.064 | 0.74 | 0.83 | 0.63 | 0.75 | 0.61 | 0.55 |
| 9 | 136277482 | rs2285487 | 1.76 | 0.471 | 0.15 | -1.889 | -0.017 | 0.024 | 0.024 | 0.075 | 0.07 | 0.01 | 0.04 | 0.02 | 0.08 | 0.03 |
| 9 | 136281966 | rs3124779 | 0.224 | 0.493 | 0.252 | -1.871 | 0.041 | -0.039 | -0.039 | 0 | 0.66 | 0.45 | 0.57 | 0.75 | 0.55 | 0.54 |
| 9 | 136287582 | rs34024143 | 0.305 | 0.397 | -0.136 | -1.771 | -0.014 | 0.031 | 0.031 | 0.005 | 0.1 | 0.04 | 0.07 | 0.01 | 0.13 | 0.03 |
| 9 | 136290672 | rs28571612 | 1.86 | 0.437 | -0.159 | -1.853 | -0.028 | 0.032 | 0.032 | 0.319 | 0.06 | 0 | 0.04 | 0 | 0.07 | 0.02 |
| 9 | 136291063 | rs3118667 | 0.306 | 0.408 | 0.157 | -2.079 | 0.016 | -0.051 | -0.051 | 0 | 0.63 | 0.49 | 0.53 | 0.83 | 0.51 | 0.56 |
| 9 | 136291361 | rs34054981 | 1.37 | 0.411 | -0.017 | -2.047 | -0.026 | 0.041 | 0.041 | 0.146 | 0.08 | 0.01 | 0.04 | 0 | 0.08 | 0.03 |
| 9 | 136299394 | rs681092 | 0.873 | 0.529 | 0.249 | -2.009 | 0 | -0.049 | -0.049 | 0 | 0.09 | 0.16 | 0.08 | 0 | 0.1 | 0.02 |
| 9 | 136303253 | rs2028002 | 0.323 | 0.507 | 0.068 | -2.057 | -0.144 | 0.024 | 0.024 | 0 | 0.28 | 0.04 | 0.39 | 0.19 | 0.43 | 0.43 |
| 9 | 136304497 | rs3124768 | 1.315 | 0.403 | 0.353 | -2.12 | 0.003 | 0.032 | 0.032 | 0 | 0.53 | 0.37 | 0.48 | 0.81 | 0.42 | 0.53 |
| 9 | 136305275 | rs2073933 | 0.915 | 0.518 | 0.077 | -2.058 | -0.103 | -0.005 | -0.005 | 0 | 0.22 | 0.03 | 0.35 | 0.17 | 0.34 | 0.4 |
| 9 | 136305530 | rs28647808 | 1.675 | 0.413 | 0.267 | -2.169 | -0.027 | 0.025 | 0.025 | 0.055 | 0.07 | 0.01 | 0.04 | 0.02 | 0.08 | 0.03 |
| 9 | 136307089 | rs603551 | 0.737 | 0.529 | 0.358 | -1.984 | -0.002 | -0.048 | -0.048 | 0 | 0.09 | 0.15 | 0.08 | 0 | 0.1 | 0.02 |
| 9 | 136307408 | rs602184 | 0.737 | 0.529 | 0.358 | -1.984 | -0.002 | -0.048 | -0.048 | 0.005 | 0.09 | 0.15 | 0.08 | 0 | 0.1 | 0.02 |
| 9 | 136309841 | rs592514 | 1.157 | 0.681 | 0.093 | -1.94 | -0.007 | -0.002 | -0.002 | 0 | 0.07 | 0.04 | 0.06 | 0 | 0.13 | 0.03 |
| 9 | 136311017 | rs652600 | -1.705 | -0.166 | -0.041 | -2.109 | -0.009 | -0.023 | -0.023 | 0 | 0.59 | 0.58 | 0.6 | 0.2 | 0.72 | 0.58 |
| 9 | 136311159 | rs34063534 | 1.936 | 0.191 | -0.366 | -1.864 | 0.006 | 0.004 | 0.004 | 0.052 | 0.06 | 0.03 | 0.03 | 0 | 0.03 | 0.02 |
| 9 | 136311391 | rs4962150 | 0.224 | 0.777 | -0.676 | -1.522 | -0.145 | -0.058 | -0.058 | 0 | 0.26 | 0.04 | 0.37 | 0.28 | 0.41 | 0.43 |
| 9 | 136319068 | rs3094375 | -0.476 | 1.168 | -0.091 | -1.468 | 0 | -0.001 | -0.001 | 0.291 | 0.29 | 0.31 | 0.44 | 0.2 | 0.43 | 0.41 |
| 9 | 136323754 | rs4962153 | -0.542 | 1.326 | -0.392 | -1.387 | 0.015 | -0.049 | -0.049 | 2.266 | 0.83 | 0.66 | 0.89 | 0.94 | 0.84 | 0.9 |
| 9 | 136324239 | rs1055432 | 0.19 | 1.763 | -1.144 | -0.982 | -0.113 | -0.048 | -0.048 | 0 | 0.18 | 0.03 | 0.33 | 0.19 | 0.33 | 0.37 |
| 9 | 136335824 | rs1054379 | -0.719 | 0.914 | -0.16 | -0.77 | -0.004 | -0.035 | -0.035 | 2.973 | 0.1 | 0.12 | 0.08 | 0 | 0.1 | 0.02 |
| 9 | 136337168 | rs3094378 | 0.066 | 0.917 | -0.173 | -0.768 | -0.009 | 0.03 | 0.03 | 0 | 0.07 | 0.01 | 0.06 | 0 | 0.09 | 0.02 |
| 9 | 136339959 | rs3124760 | -1.315 | 0.908 | -0.171 | -0.783 | -0.001 | -0.031 | -0.031 | 0.332 | 0.13 | 0.14 | 0.11 | 0 | 0.15 | 0.05 |
| 9 | 136344613 | rs968471 | -0.451 | 1.009 | -0.139 | -0.718 | 0.012 | -0.037 | -0.037 | 1.254 | 0.08 | 0.14 | 0.09 | 0 | 0.11 | 0.02 |
| 9 | 136344853 | rs3124758 | -0.74 | 0.889 | -0.198 | -0.7 | 0 | -0.029 | -0.029 | 0 | 0.14 | 0.16 | 0.14 | 0 | 0.16 | 0.06 |
| 9 | 136359085 | rs736417 | 0.171 | 0.201 | -0.494 | -0.172 | 0 | 0.007 | 0.007 | 0 | 0.38 | 0.34 | 0.42 | 0.38 | 0.37 | 0.53 |
| 9 | 136365075 | rs17810852 | 0.527 | 1.979 | -0.339 | -0.201 | 0.042 | -0.061 | -0.061 | 0 | 0.67 | 0.48 | 0.64 | 0.88 | 0.49 | 0.8 |
| 9 | 136395973 | rs3763650 | -0.609 | 2.099 | -0.744 | 0.539 | -0.016 | -0.009 | -0.009 | 0.908 | 0.16 | 0.1 | 0.14 | 0.24 | 0.25 | 0.31 |
| 9 | 136397834 | rs1008250 | -1.061 | 2.647 | 0.373 | 0.519 | 0.072 | 0.027 | 0.027 | 0 | 0.19 | 0.34 | 0.22 | 0.26 | 0.31 | 0.31 |
| 9 | 136399075 | rs9696672 | -0.957 | 2.359 | 0.096 | 0.623 | 0.006 | 0.014 | 0.014 | 0 | 0.3 | 0.3 | 0.2 | 0.44 | 0.35 | 0.49 |
| 9 | 136402067 | rs28569945 | -1.016 | 2.363 | 0.532 | -0.102 | 0.093 | -0.061 | -0.061 | 0.745 | 0.12 | 0.34 | 0.19 | 0.02 | 0.24 | 0.06 |
| 9 | 136407208 | rs28520600 | 0.002 | 1.572 | -0.049 | -0.722 | -0.004 | -0.084 | -0.084 | 0 | 0.26 | 0.35 | 0.17 | 0.02 | 0.26 | 0.15 |
| 9 | 136408841 | rs11537116 | 0.375 | 1.527 | -0.521 | -0.289 | -0.017 | -0.026 | -0.026 | 0 | 0.36 | 0.23 | 0.3 | 0.45 | 0.38 | 0.42 |
| 9 | 136408876 | rs11516159 | -0.396 | 1.194 | -0.605 | -0.503 | -0.009 | 0.064 | 0.064 | 0.035 | 0.8 | 0.75 | 0.6 | 0.44 | 0.85 | 0.63 |
| 9 | 136409854 | rs9330463 | -0.916 | 0.867 | -0.503 | -0.224 | -0.025 | 0.193 | 0.193 | 0 | 0.87 | 0.75 | 0.69 | 0.51 | 0.91 | 0.72 |

| | | | | | | | | | | | | | | | | |
|---|---|---|---|---|---|---|---|---|---|---|---|---|---|---|---|---|
| 9 | 136410525 | rs10119879 | 0.504 | 0.958 | -0.595 | -0.313 | -0.015 | -0.01 | -0.01 | 0 | 0.36 | 0.25 | 0.27 | 0.42 | 0.38 | 0.42 |
| 9 | 136411785 | rs11507718 | 0.637 | 0.731 | -0.888 | -0.443 | 0.007 | -0.008 | -0.008 | 0 | 0.24 | 0.17 | 0.16 | 0 | 0.24 | 0.06 |
| 9 | 136413041 | rs10120207 | -0.642 | 0.686 | -0.571 | 0.117 | -0.025 | 0.198 | 0.198 | 0 | 0.87 | 0.76 | 0.69 | 0.52 | 0.91 | 0.72 |
| 9 | 136414531 | rs9696616 | -0.65 | 0.69 | -0.569 | 0.129 | -0.025 | 0.195 | 0.195 | 0 | 0.87 | 0.76 | 0.68 | 0.52 | 0.91 | 0.72 |
| 9 | 136414937 | rs11535670 | 0.555 | 1.336 | -0.909 | 0.098 | -0.01 | -0.018 | -0.018 | 0 | 0.21 | 0.14 | 0.15 | 0.3 | 0.23 | 0.31 |
| 9 | 136415608 | rs9777051 | -0.209 | 1.058 | -0.975 | 0.233 | 0 | 0.001 | 0.001 | 0 | 0.5 | 0.58 | 0.43 | 0.38 | 0.51 | 0.56 |
| 9 | 136454105 | rs28666751 | 0.2 | 0.808 | -1.748 | 0.673 | -0.074 | -0.155 | -0.155 | 0.101 | 0.2 | 0.03 | 0.18 | 0.43 | 0.25 | 0.34 |
| 9 | 136454700 | rs10993774 | -0.739 | -0.217 | -1.348 | -0.311 | 0.009 | 0.02 | 0.02 | 0.073 | 0.45 | 0.37 | 0.38 | 0.9 | 0.42 | 0.64 |
| 9 | 136456607 | rs7040825 | -0.841 | -0.43 | -1.141 | -0.017 | -0.005 | -0.036 | -0.036 | 0 | 0.78 | 0.88 | 0.81 | 0.59 | 0.74 | 0.74 |
| 9 | 136457146 | rs7856830 | 0.02 | -0.819 | -0.802 | -0.953 | -0.026 | -0.173 | -0.173 | 0.094 | 0.23 | 0.52 | 0.15 | 0 | 0.19 | 0.02 |
| 9 | 136458311 | rs9409872 | -0.571 | -0.873 | -0.532 | -0.952 | -0.001 | -0.003 | -0.003 | 0 | 0.64 | 0.66 | 0.72 | 0.59 | 0.59 | 0.7 |
| 9 | 136458404 | rs3025299 | 0.984 | -1.019 | -0.872 | -1.042 | -0.003 | 0 | 0 | 0 | 0.35 | 0.32 | 0.53 | 0.26 | 0.41 | 0.37 |
| 9 | 136460220 | rs10821577 | -0.162 | -0.659 | -1.309 | -1.259 | -0.013 | -0.001 | -0.001 | 0.393 | 0.29 | 0.35 | 0.19 | 0.32 | 0.18 | 0.34 |
| 9 | 136462863 | rs1079783 | -1.992 | -0.434 | -1.324 | -1.097 | -0.031 | -0.098 | -0.098 | 0 | 0.79 | 0.95 | 0.82 | 0.59 | 0.74 | 0.72 |
| 9 | 136466648 | rs3025324 | -1.069 | -0.195 | -1.028 | -1.148 | -0.004 | 0.023 | 0.023 | 0 | 0.5 | 0.78 | 0.47 | 0.47 | 0.48 | 0.63 |
| 9 | 136466832 | rs3025325 | 0.612 | -0.114 | -0.545 | -1.063 | -0.009 | -0.098 | -0.098 | 0 | 0.18 | 0.62 | 0.14 | 0 | 0.15 | 0.02 |
| 9 | 136469322 | rs3025336 | 1.156 | 0.047 | -0.462 | -0.587 | -0.007 | 0 | 0 | 0 | 0.12 | 0.13 | 0.21 | 0.11 | 0.05 | 0.06 |
| 9 | 136471193 | rs7025648 | -0.324 | 0.108 | -0.707 | -0.589 | 0.019 | 0.005 | 0.005 | 0 | 0.11 | 0.09 | 0.04 | 0.11 | 0.07 | 0.07 |
| 9 | 136478442 | rs1029372 | -1.675 | 0.745 | -0.425 | -0.76 | 0.005 | 0.001 | 0.001 | 0 | 0.52 | 0.59 | 0.65 | 0.45 | 0.52 | 0.67 |
| 9 | 136479571 | rs3025351 | 0.503 | 1.246 | 0.644 | -0.802 | -0.003 | 0.005 | 0.005 | 0 | 0.06 | 0.02 | 0.03 | 0 | 0.07 | 0.03 |
| 9 | 136482068 | rs3025363 | -1.681 | 1.244 | 0.64 | -0.767 | 0.016 | -0.022 | -0.022 | 0 | 0.29 | 0.59 | 0.33 | 0.16 | 0.36 | 0.37 |
| 9 | 136482476 | rs3025365 | -0.613 | 1.345 | 0.585 | -0.662 | 0.104 | 0.115 | 0.115 | 0 | 0.1 | 0.36 | 0.17 | 0.16 | 0.14 | 0.31 |
| 9 | 136484292 | rs7872903 | -0.966 | 0.434 | 0.015 | -1.014 | 0.003 | -0.122 | -0.122 | 0 | 0.2 | 0.53 | 0.18 | 0.01 | 0.21 | 0.06 |
| 9 | 136487721 | rs916658 | 0.271 | 0.681 | -0.215 | -1.295 | 0.005 | -0.117 | -0.117 | 0.016 | 0.68 | 0.56 | 0.73 | 0.98 | 0.65 | 0.86 |
| 9 | 136488144 | rs11998869 | -0.23 | 0.223 | -0.386 | -0.876 | -0.001 | 0.005 | 0.005 | 0 | 0.1 | 0.11 | 0.05 | 0.18 | 0.06 | 0.12 |
| 9 | 136489539 | rs10993874 | 0.551 | -0.506 | -0.995 | -0.602 | 0.03 | -0.071 | -0.071 | 0 | 0.2 | 0.05 | 0.19 | 0.53 | 0.14 | 0.33 |
| 9 | 136494425 | rs2519765 | -1.186 | -0.706 | -0.64 | -0.973 | -0.005 | -0.021 | -0.021 | 1.289 | 0.79 | 0.8 | 0.79 | 1 | 0.71 | 0.9 |
| 9 | 136494566 | rs2519148 | 0.606 | -0.922 | 0.022 | -1.06 | -0.005 | -0.009 | -0.009 | 0 | 0.45 | 0.37 | 0.52 | 0.44 | 0.5 | 0.53 |
| 9 | 136494594 | rs3025369 | 0.253 | -0.706 | 0.159 | -1.18 | -0.007 | 0.031 | 0.031 | 0 | 0.09 | 0.15 | 0.04 | 0.18 | 0.05 | 0.11 |
| 9 | 136496708 | rs3025373 | -0.574 | -0.22 | -0.142 | -1.418 | -0.018 | -0.005 | -0.005 | 0 | 0.16 | 0.04 | 0.06 | 0.12 | 0.13 | 0.18 |
| 9 | 136498143 | rs1076153 | 0.597 | -0.136 | -0.263 | -1.598 | 0.017 | -0.029 | -0.029 | 0.029 | 0.29 | 0.2 | 0.23 | 0.66 | 0.21 | 0.49 |
| 9 | 136498761 | rs1076150 | 0.005 | -0.231 | 0.209 | -1.856 | 0.029 | -0.067 | -0.067 | 1.844 | 0.37 | 0.46 | 0.36 | 0.09 | 0.5 | 0.26 |
| 9 | 136500203 | rs1611114 | 0.575 | -0.298 | 0.289 | -1.878 | 0.005 | -0.008 | -0.008 | 0 | 0.34 | 0.31 | 0.3 | 0.67 | 0.27 | 0.5 |
| 9 | 136500515 | rs1611115 | -1.114 | -0.32 | 0.345 | -1.783 | 0.017 | 0.016 | 0.016 | 0 | 0.73 | 0.81 | 0.68 | 0.81 | 0.79 | 0.8 |
| 9 | 136502321 | rs3025382 | -0.891 | -0.144 | 0.669 | -1.795 | 0.1 | 0.013 | 0.013 | 0.192 | 0.08 | 0.29 | 0.08 | 0.03 | 0.12 | 0.08 |
| 9 | 136502328 | rs2519143 | -1.119 | -0.059 | 0.421 | -1.854 | 0.042 | 0.023 | 0.023 | 0 | 0.74 | 0.88 | 0.7 | 0.82 | 0.81 | 0.8 |
| 9 | 136503819 | rs2007153 | 0.24 | 0.023 | 0.546 | -1.858 | -0.006 | -0.004 | -0.004 | 0.158 | 0.58 | 0.53 | 0.59 | 0.68 | 0.63 | 0.65 |
| 9 | 136504580 | rs2797851 | 0.74 | 0.583 | 0.998 | -2.014 | -0.005 | -0.03 | -0.03 | 0 | 0.23 | 0.25 | 0.26 | 0 | 0.33 | 0.12 |
| 9 | 136505114 | rs1108580 | -0.966 | 0.594 | 1.104 | -2.064 | -0.01 | 0.047 | 0.047 | 0 | 0.42 | 0.68 | 0.4 | 0.12 | 0.54 | 0.32 |
| 9 | 136505241 | rs1108581 | -1.368 | 0.6 | 1.145 | -2.066 | 0.02 | 0.004 | 0.004 | 0 | 0.2 | 0.48 | 0.14 | 0.14 | 0.2 | 0.17 |
| 9 | 136505644 | rs2873804 | -0.966 | 0.502 | 0.912 | -2.108 | -0.01 | 0.033 | 0.033 | 0 | 0.42 | 0.68 | 0.39 | 0.14 | 0.53 | 0.29 |
| 9 | 136507473 | rs5320 | -0.148 | 0.648 | 1.26 | -1.919 | -0.011 | 0.04 | 0.04 | 0 | 0.08 | 0.17 | 0.04 | 0.11 | 0.06 | 0.08 |
| 9 | 136509083 | rs1611123 | -0.551 | 0.697 | 1.035 | -1.917 | 0.018 | -0.042 | -0.042 | 0 | 0.34 | 0.43 | 0.33 | 0.12 | 0.46 | 0.25 |
| 9 | 136509370 | rs4531 | -0.809 | 0.159 | 0.531 | -1.788 | 0.004 | -0.026 | -0.026 | 0 | 0.06 | 0.13 | 0.03 | 0 | 0.07 | 0.06 |
| 9 | 136511516 | rs1541332 | -0.09 | -0.351 | -0.047 | -2.054 | 0.001 | -0.091 | -0.091 | 0.31 | 0.58 | 0.47 | 0.6 | 0.97 | 0.47 | 0.74 |
| 9 | 136512275 | rs2519154 | 0.085 | -0.194 | -0.47 | -1.868 | 0.031 | 0.166 | 0.166 | 0 | 0.51 | 0.28 | 0.61 | 0.92 | 0.46 | 0.68 |
| 9 | 136514539 | rs7876027 | -1.127 | -0.332 | 0.731 | -1.353 | 0.027 | 0.008 | 0.008 | 0 | 0.07 | 0.33 | 0.04 | 0 | 0.03 | 0.05 |
| 9 | 136515297 | rs2283123 | -0.467 | -0.195 | 0.238 | -0.745 | -0.005 | -0.011 | -0.011 | 0 | 0.11 | 0.06 | 0.05 | 0.13 | 0.11 | 0.21 |
| 9 | 136515330 | rs2283124 | -0.805 | -0.203 | 0.527 | -0.748 | -0.002 | 0.003 | 0.003 | 0 | 0.13 | 0.19 | 0.07 | 0.13 | 0.11 | 0.24 |
| 9 | 136518097 | rs77905 | -0.378 | -0.53 | 0.11 | -1.505 | -0.001 | -0.035 | -0.035 | 0 | 0.58 | 0.57 | 0.59 | 0.9 | 0.52 | 0.52 |
| 9 | 136519702 | rs2097628 | 0.076 | -0.716 | 0.215 | -0.59 | 0.001 | -0.001 | -0.001 | 0 | 0.42 | 0.43 | 0.39 | 0.54 | 0.41 | 0.3 |
| 9 | 136522426 | rs3025422 | -0.116 | -0.949 | 0.1 | -0.3 | -0.022 | 0.01 | 0.01 | 0 | 0.05 | 0.07 | 0.08 | 0.1 | 0.02 | 0.2 |
| 9 | 136523669 | rs129882 | 0.315 | -0.797 | 0.018 | -0.635 | 0.009 | -0.019 | -0.019 | 0 | 0.27 | 0.21 | 0.22 | 0.36 | 0.17 | 0.32 |
| 9 | 136527226 | rs129885 | 0.163 | -0.979 | 0.164 | -0.428 | -0.006 | 0.01 | 0.01 | 0 | 0.28 | 0.33 | 0.23 | 0.38 | 0.18 | 0.32 |
| 9 | 136528721 | rs129886 | 0.261 | -1.036 | 0.109 | -0.425 | -0.01 | 0.015 | 0.015 | 0.006 | 0.28 | 0.34 | 0.24 | 0.38 | 0.17 | 0.32 |
| 9 | 136532183 | rs2073835 | 0.408 | -0.672 | -0.68 | -0.658 | 0.033 | -0.039 | -0.039 | 0.181 | 0.37 | 0.13 | 0.36 | 0.42 | 0.35 | 0.22 |
| 9 | 136536633 | rs2797840 | -0.311 | -0.31 | -0.399 | -0.655 | -0.035 | 0.049 | 0.049 | 0.638 | 0.63 | 0.69 | 0.6 | 0.8 | 0.48 | 0.52 |
| 9 | 136542868 | rs129889 | 0.506 | -0.189 | -0.12 | 0.005 | -0.008 | -0.032 | -0.032 | 0 | 0.27 | 0.13 | 0.32 | 0.41 | 0.31 | 0.08 |
| 9 | 136544943 | rs129956 | -0.863 | -0.585 | 0.645 | -0.344 | -0.025 | -0.076 | -0.076 | 0.078 | 0.13 | 0.54 | 0.07 | 0.02 | 0.06 | 0.18 |
| 9 | 136545363 | rs129895 | -0.681 | -0.518 | 0.309 | -0.399 | 0.002 | -0.025 | -0.025 | 0 | 0.46 | 0.81 | 0.45 | 0.44 | 0.41 | 0.4 |
| 9 | 136545423 | rs129940 | -0.349 | -0.551 | 1.094 | -0.412 | -0.01 | -0.048 | -0.048 | 0.713 | 0.1 | 0.27 | 0.05 | 0.02 | 0.06 | 0.18 |
| 9 | 136550817 | rs2519072 | -0.522 | -0.523 | 0.933 | -0.366 | 0.007 | -0.022 | -0.022 | 0 | 0.45 | 0.81 | 0.45 | 0.44 | 0.41 | 0.4 |
| 9 | 136552263 | rs129901 | -0.319 | -0.529 | 1.193 | -0.362 | 0.004 | -0.015 | -0.015 | 0 | 0.42 | 0.63 | 0.43 | 0.44 | 0.4 | 0.3 |
| 9 | 136552362 | rs4744533 | -0.001 | -0.58 | 0.915 | -0.375 | -0.041 | -0.075 | -0.075 | 0 | 0.37 | 0.1 | 0.39 | 0.44 | 0.4 | 0.28 |
| 9 | 136554167 | rs129904 | -0.257 | -0.583 | 0.91 | -0.346 | 0.002 | -0.006 | -0.006 | 0 | 0.43 | 0.29 | 0.5 | 0.44 | 0.43 | 0.31 |
| 9 | 136555629 | rs886016 | -0.73 | -0.536 | 0.257 | -0.343 | 0.019 | -0.012 | -0.012 | 0 | 0.5 | 0.84 | 0.49 | 0.44 | 0.44 | 0.44 |
| 9 | 136559460 | rs2073817 | -0.277 | -0.406 | 0.774 | -0.277 | 0.002 | -0.003 | -0.003 | 0 | 0.43 | 0.31 | 0.42 | 0.42 | 0.43 | 0.3 |
| 9 | 136566959 | rs2427995 | 0.126 | -0.339 | 1.096 | -0.482 | -0.022 | -0.076 | -0.076 | 0 | 0.87 | 0.7 | 0.89 | 1 | 0.92 | 0.8 |
| 9 | 136569061 | rs2427989 | 0.01 | -0.387 | 1.208 | -0.536 | -0.023 | -0.074 | -0.074 | 0.058 | 0.87 | 0.68 | 0.89 | 1 | 0.93 | 0.82 |

| | | | | | | | | | | | | | | | | |
|---|---|---|---|---|---|---|---|---|---|---|---|---|---|---|---|---|
| 9 | 136570740 | rs694821 | 0.157 | -0.192 | 0.912 | -0.537 | -0.001 | -0.001 | -0.001 | 0.758 | 0.5 | 0.46 | 0.47 | 0.55 | 0.5 | 0.66 |
| 9 | 136571089 | rs2510245 | 0.101 | -0.486 | 0.717 | -0.648 | 0.001 | 0 | 0 | 0.006 | 0.53 | 0.41 | 0.5 | 0.5 | 0.56 | 0.53 |
| 9 | 136571886 | rs4979633 | 1.38 | -0.442 | 0.47 | -0.46 | 0.008 | -0.091 | -0.091 | 1.839 | 0.27 | 0.08 | 0.31 | 0.38 | 0.28 | 0.09 |
| 9 | 136572750 | rs4979631 | 0.387 | -0.332 | 0.816 | -0.44 | 0.005 | -0.01 | -0.01 | 0.242 | 0.37 | 0.3 | 0.41 | 0.48 | 0.37 | 0.37 |
| 9 | 136573412 | rs2073815 | -0.125 | 0.187 | 0.869 | -0.124 | 0.034 | -0.007 | -0.007 | 0 | 0.47 | 0.23 | 0.46 | 0.49 | 0.56 | 0.46 |
| 9 | 136577094 | rs511293 | 0.195 | 0.586 | 0.527 | -0.29 | -0.001 | 0.009 | 0.009 | 0 | 0.23 | 0.16 | 0.24 | 0.14 | 0.22 | 0.21 |
| 9 | 136579589 | rs522676 | -0.626 | 0.679 | 0.621 | -0.147 | -0.006 | -0.002 | -0.002 | 0.01 | 0.73 | 0.76 | 0.69 | 0.68 | 0.66 | 0.74 |
| 9 | 136581419 | rs2427981 | -0.983 | 0.608 | 0.605 | 0.056 | -0.004 | -0.001 | -0.001 | 0 | 0.53 | 0.47 | 0.54 | 0.56 | 0.56 | 0.48 |
| 9 | 136584196 | rs2027963 | -1.027 | 0.667 | 0.588 | 0.04 | -0.003 | -0.001 | -0.001 | 0 | 0.54 | 0.49 | 0.6 | 0.56 | 0.56 | 0.47 |
| 9 | 136586367 | rs7040170 | 0.138 | 0.754 | 0.057 | -0.212 | -0.002 | -0.004 | -0.004 | 0 | 0.22 | 0.17 | 0.23 | 0.09 | 0.22 | 0.19 |
| 9 | 136590369 | rs2510234 | 0.375 | 0.747 | 0.002 | -0.155 | -0.013 | -0.002 | -0.002 | 0 | 0.3 | 0.22 | 0.31 | 0.34 | 0.35 | 0.28 |
| 9 | 136592014 | rs2251217 | -0.419 | 0.544 | -0.249 | 0.05 | -0.002 | 0.022 | 0.022 | 0 | 0.89 | 0.73 | 0.95 | 0.86 | 0.9 | 0.75 |
| 9 | 136592342 | rs2283125 | 1.096 | 0.536 | -0.597 | 0.202 | 0.021 | -0.034 | -0.034 | 0 | 0.34 | 0.13 | 0.36 | 0.36 | 0.34 | 0.27 |
| 9 | 136598128 | rs3780804 | -0.637 | 1.113 | 0.012 | 0.473 | 0.005 | 0.001 | 0.001 | 0 | 0.16 | 0.16 | 0.2 | 0.16 | 0.17 | 0.19 |
| 9 | 136599146 | rs573904 | 0.299 | 1.286 | -0.213 | 0.009 | -0.05 | -0.038 | -0.038 | 0 | 0.21 | 0.08 | 0.24 | 0.31 | 0.3 | 0.22 |
| 9 | 136600201 | rs493901 | -0.874 | 1.211 | -0.203 | -0.062 | 0.013 | -0.006 | -0.006 | 0 | 0.38 | 0.55 | 0.49 | 0.39 | 0.46 | 0.4 |
| 9 | 136606230 | rs10465165 | -0.746 | 1.113 | -0.194 | -0.297 | 0.034 | -0.02 | -0.02 | 0 | 0.11 | 0.17 | 0.17 | 0.09 | 0.16 | 0.18 |
| 9 | 136608669 | rs2510257 | 0.791 | 0.884 | -0.907 | -0.323 | -0.003 | 0.029 | 0.029 | 2.368 | 0.74 | 0.13 | 0.78 | 0.8 | 0.78 | 0.47 |
| 9 | 136612426 | rs646708 | 0.849 | 0.948 | -0.685 | -0.511 | 0 | -0.001 | -0.001 | 0 | 0.67 | 0.43 | 0.74 | 0.72 | 0.68 | 0.45 |
| 9 | 136617292 | rs491220 | 0.551 | 1.295 | -1.36 | -0.948 | -0.017 | 0.134 | 0.134 | 0.193 | 0.54 | 0.19 | 0.56 | 0.73 | 0.46 | 0.48 |
| 9 | 136620568 | rs667420 | 0.599 | 1.43 | -1.1 | -0.999 | -0.008 | 0.113 | 0.113 | 0.289 | 0.54 | 0.21 | 0.55 | 0.73 | 0.44 | 0.49 |
| 9 | 136622634 | rs10993787 | 0.307 | 1.661 | -0.789 | -1.141 | -0.01 | 0.034 | 0.034 | 0 | 0.6 | 0.33 | 0.61 | 0.88 | 0.53 | 0.53 |
| 9 | 136626906 | rs642193 | 0.413 | 1.448 | -0.673 | -1.047 | -0.004 | 0.032 | 0.032 | 0 | 0.6 | 0.31 | 0.6 | 0.78 | 0.56 | 0.52 |
| 9 | 136629298 | rs2797825 | -0.184 | 1.04 | -0.473 | -1.511 | -0.012 | 0.002 | 0.002 | 0 | 0.41 | 0.32 | 0.41 | 0.07 | 0.45 | 0.36 |
| 9 | 136632662 | rs622035 | 0.004 | 0.948 | -0.67 | -0.251 | -0.033 | -0.103 | -0.103 | 0 | 0.15 | 0.01 | 0.14 | 0.53 | 0.14 | 0.2 |
| 9 | 136643994 | rs602990 | -0.355 | 1.207 | -0.974 | -1.984 | -0.022 | 0.328 | 0.328 | 0 | 0.5 | 0.84 | 0.48 | 0.06 | 0.52 | 0.46 |
| 9 | 136647466 | rs9409864 | -0.288 | 1.12 | -0.31 | -2.331 | 0 | -0.057 | -0.057 | 1.239 | 0.2 | 0.25 | 0.18 | 0.02 | 0.19 | 0.16 |
| 9 | 136647624 | rs525060 | 0.675 | 1.029 | 0.516 | -2.545 | 0 | -0.073 | -0.073 | 0 | 0.57 | 0.5 | 0.65 | 0.96 | 0.52 | 0.55 |
| 9 | 136647817 | rs650830 | 0.559 | 0.978 | 0.487 | -2.461 | 0.008 | -0.066 | -0.066 | 0.208 | 0.22 | 0.18 | 0.16 | 0.01 | 0.29 | 0.29 |
| 9 | 136648535 | rs2488563 | -0.071 | 0.849 | 0.74 | -2.328 | 0.01 | -0.058 | -0.058 | 0 | 0.88 | 0.82 | 0.92 | 1 | 0.87 | 0.94 |
| 9 | 136650740 | rs2023665 | -0.685 | -0.131 | 0.42 | -2.052 | 0.002 | 0.002 | 0.002 | 0 | 0.55 | 0.53 | 0.5 | 0.56 | 0.61 | 0.55 |
| 9 | 136654038 | rs2520003 | 1.099 | 0.108 | 0.023 | -2.045 | -0.131 | 0.152 | 0.152 | 0 | 0.27 | 0.05 | 0.24 | 0.01 | 0.45 | 0.22 |
| 9 | 136657960 | rs3824561 | -0.627 | -0.372 | 0.665 | -1.465 | -0.001 | -0.001 | -0.001 | 0 | 0.52 | 0.54 | 0.56 | 0.44 | 0.44 | 0.53 |
| 9 | 136658987 | rs10993803 | -0.785 | -0.343 | 0.548 | -1.382 | 0.004 | 0.004 | 0.004 | 0 | 0.56 | 0.68 | 0.62 | 0.48 | 0.45 | 0.54 |
| 9 | 136661934 | rs2428099 | -1.017 | -0.355 | 0.539 | -1.388 | -0.004 | -0.003 | -0.003 | 3.777 | 0.58 | 0.74 | 0.62 | 0.48 | 0.45 | 0.54 |
| 9 | 136662638 | rs10821525 | 0.655 | -0.482 | 0.575 | -1.537 | 0.037 | -0.01 | -0.01 | 0 | 0.35 | 0.16 | 0.32 | 0.38 | 0.28 | 0.3 |
| 9 | 136662928 | rs7865299 | -1.894 | 0.039 | 1.748 | -1.151 | -0.001 | -0.104 | -0.104 | 0.126 | 0.21 | 0.51 | 0.25 | 0.09 | 0.17 | 0.2 |
| 9 | 136666483 | rs2510262 | -0.706 | -0.201 | 1.479 | -1.08 | 0.026 | -0.002 | -0.002 | 0.01 | 0.22 | 0.16 | 0.1 | 0.16 | 0.14 | 0.23 |
| 9 | 136666560 | rs10123394 | -0.791 | -0.188 | 1.83 | -1.102 | 0.048 | -0.044 | -0.044 | 0 | 0.1 | 0.18 | 0.17 | 0.03 | 0.12 | 0.14 |
| 9 | 136669004 | rs2488552 | -2.049 | -0.194 | 1.789 | -1.104 | 0.008 | 0.015 | 0.015 | 0.146 | 0.37 | 0.69 | 0.37 | 0.25 | 0.28 | 0.43 |
| 9 | 136676014 | rs7021663 | -2.361 | -0.029 | 2.095 | -1.196 | -0.005 | -0.089 | -0.089 | 0.468 | 0.21 | 0.47 | 0.3 | 0.1 | 0.18 | 0.21 |
| 9 | 136681696 | rs12337328 | -2.179 | 0.182 | 2.197 | -0.954 | 0.05 | -0.042 | -0.042 | 0 | 0.14 | 0.42 | 0.27 | 0.1 | 0.14 | 0.2 |
| 9 | 136682408 | rs7026263 | -1.823 | 0.197 | 2.093 | -1.031 | 0.027 | -0.056 | -0.056 | 1.791 | 0.12 | 0.23 | 0.22 | 0.03 | 0.13 | 0.17 |
| 9 | 136682482 | rs7026295 | -1.474 | 0.315 | 2.133 | -0.973 | 0.042 | -0.047 | -0.047 | 1.897 | 0.11 | 0.24 | 0.22 | 0.03 | 0.13 | 0.17 |
| 9 | 136686242 | rs3858101 | -2.006 | 0.288 | 2.043 | -0.998 | 0.054 | -0.073 | -0.073 | 0 | 0.13 | 0.37 | 0.23 | 0.02 | 0.12 | 0.18 |
| 9 | 136689278 | rs2486335 | -1.728 | 0.341 | 1.835 | -0.963 | 0.016 | -0.058 | -0.058 | 0 | 0.1 | 0.28 | 0.15 | 0.02 | 0.09 | 0.13 |
| 9 | 136693906 | rs3893493 | 1.891 | 0.316 | 1.803 | -1.163 | 0.004 | -0.065 | -0.065 | 0 | 0.71 | 0.36 | 0.69 | 0.9 | 0.74 | 0.63 |
| 9 | 136705956 | rs2260095 | -1.959 | -0.074 | 1.839 | -0.965 | -0.016 | -0.074 | -0.074 | 0.13 | 0.21 | 0.48 | 0.14 | 0.12 | 0.14 | 0.17 |
| 9 | 136710514 | rs12000549 | -0.964 | -0.024 | 1.485 | -0.909 | -0.007 | 0.009 | 0.009 | 0 | 0.12 | 0.1 | 0.07 | 0.12 | 0.11 | 0.14 |
| 9 | 136719045 | rs10993826 | -1.55 | -0.061 | 1.406 | -0.94 | -0.028 | 0.031 | 0.031 | 0 | 0.15 | 0.04 | 0.08 | 0.12 | 0.13 | 0.15 |
| 9 | 136721078 | rs3819502 | -2.806 | -0.137 | 2.071 | -0.964 | -0.018 | -0.053 | -0.053 | 0 | 0.23 | 0.7 | 0.17 | 0.15 | 0.15 | 0.25 |
| 9 | 136723520 | rs12344583 | -2.821 | 0.126 | 2.253 | -0.899 | -0.003 | -0.073 | -0.073 | 0.606 | 0.24 | 0.57 | 0.18 | 0.17 | 0.2 | 0.26 |
| 9 | 136733975 | rs10821527 | -2.46 | 0.216 | 2.111 | -0.875 | -0.004 | -0.064 | -0.064 | 0 | 0.2 | 0.61 | 0.15 | 0.11 | 0.13 | 0.19 |
| 9 | 136740971 | rs12341506 | -2.76 | 0.246 | 1.869 | -0.85 | 0.036 | 0.038 | 0.038 | 0.725 | 0.06 | 0.41 | 0.03 | 0.03 | 0.02 | 0.09 |
| 9 | 136747090 | rs7869522 | -2.151 | 0.164 | 1.568 | -0.915 | -0.007 | -0.071 | -0.071 | 0 | 0.2 | 0.43 | 0.1 | 0.09 | 0.14 | 0.31 |
| 9 | 136753246 | rs2789853 | 2.298 | 0.337 | 1.155 | -0.31 | 0.016 | 0 | 0 | 0 | 0.79 | 0.35 | 0.73 | 0.79 | 0.75 | 0.63 |
| 9 | 136754076 | rs2073883 | 0.229 | -0.038 | 0.587 | 0.035 | 0.022 | -0.036 | -0.036 | 0 | 0.29 | 0.06 | 0.22 | 0.34 | 0.22 | 0.26 |
| 9 | 136755590 | rs2519766 | 1.914 | -0.01 | 0.61 | -0.09 | 0.015 | 0.02 | 0.02 | 0.394 | 0.85 | 0.35 | 0.81 | 0.86 | 0.84 | 0.73 |
| 9 | 136758586 | rs2428082 | 0.361 | -0.023 | 0.813 | -0.066 | -0.008 | 0.004 | 0.004 | 2.23 | 0.66 | 0.58 | 0.6 | 0.62 | 0.66 | 0.66 |
| 9 | 136759508 | rs2519777 | 1.517 | -0.213 | 0.407 | -0.096 | 0.028 | 0.018 | 0.018 | 0.372 | 0.85 | 0.36 | 0.81 | 0.87 | 0.83 | 0.76 |
| 9 | 136760639 | rs2810472 | 1.889 | -0.254 | 0.371 | -0.118 | 0.012 | -0.009 | -0.009 | 0 | 0.78 | 0.33 | 0.76 | 0.82 | 0.76 | 0.66 |
| 9 | 136760719 | rs13287070 | 0.557 | -0.273 | 0.055 | -0.055 | 0.016 | -0.051 | -0.051 | 0 | 0.26 | 0.02 | 0.25 | 0.28 | 0.2 | 0.25 |
| 9 | 136762458 | rs3780751 | -1.785 | 0.211 | 0.579 | 0.099 | 0.021 | -0.002 | -0.002 | 0 | 0.2 | 0.36 | 0.23 | 0.18 | 0.25 | 0.32 |
| 9 | 136762800 | rs10993845 | 0.413 | -0.341 | 0.054 | 0.028 | -0.012 | 0.005 | 0.005 | 0 | 0.05 | 0.01 | 0.04 | 0.05 | 0.06 | 0.1 |
| 9 | 136762812 | rs13291383 | -0.278 | -0.607 | -0.179 | -0.167 | 0.064 | 0.032 | 0.032 | 0.111 | 0.4 | 0.11 | 0.29 | 0.3 | 0.3 | 0.29 |
| 9 | 136767301 | rs2157834 | 0.953 | 0.19 | 0.88 | -0.088 | 0.044 | -0.029 | -0.029 | 0 | 0.9 | 0.69 | 0.94 | 0.96 | 0.9 | 0.87 |
| 9 | 136769348 | rs2519996 | 1.18 | -0.178 | 0.41 | 0.027 | -0.018 | -0.009 | -0.009 | 0 | 0.88 | 0.66 | 0.93 | 0.94 | 0.92 | 0.84 |
| 9 | 136771319 | rs10114577 | -0.332 | -0.18 | -0.126 | -0.555 | -0.001 | 0.009 | 0.009 | 0 | 0.19 | 0.14 | 0.09 | 0.03 | 0.17 | 0.16 |
| 9 | 136771432 | rs679923 | 0.238 | -0.547 | -0.319 | -0.414 | 0.017 | 0.062 | 0.062 | 0 | 0.43 | 0.26 | 0.46 | 0.65 | 0.43 | 0.45 |

| | | | | | | | | | | | | | | | | |
|---|---|---|---|---|---|---|---|---|---|---|---|---|---|---|---|---|
| 9 | 136774217 | rs2789825 | 0.141 | -0.46 | -0.241 | -0.559 | -0.011 | -0.025 | -0.025 | 0 | 0.94 | 0.88 | 0.94 | 1 | 0.98 | 0.96 |
| 9 | 136779153 | rs608402 | 0.355 | -0.511 | -0.354 | -0.341 | 0.013 | 0.05 | 0.05 | 0 | 0.41 | 0.3 | 0.43 | 0.68 | 0.38 | 0.41 |
| 9 | 136785840 | rs608968 | 0.067 | -0.496 | -0.849 | 0.17 | 0.063 | 0.025 | 0.025 | 0 | 0.45 | 0.72 | 0.56 | 0.57 | 0.36 | 0.5 |
| 9 | 136790282 | rs2265520 | 0.854 | -0.535 | -0.549 | 0.109 | 0.011 | 0.005 | 0.005 | 0.311 | 0.68 | 0.31 | 0.56 | 0.61 | 0.77 | 0.54 |
| 9 | 136798370 | rs2519098 | 1.274 | -0.84 | 0.69 | -0.07 | -0.033 | -0.05 | -0.05 | 1.583 | 0.83 | 0.57 | 0.78 | 0.92 | 0.89 | 0.84 |
| 9 | 136803161 | rs654387 | 0.826 | -0.777 | -0.124 | -0.334 | -0.053 | 0.069 | 0.069 | 0 | 0.46 | 0.15 | 0.35 | 0.34 | 0.43 | 0.39 |
| 9 | 136803249 | rs2428003 | 0.515 | -0.858 | 0.475 | -0.391 | 0.009 | -0.007 | -0.007 | 0 | 0.93 | 0.72 | 0.83 | 0.97 | 0.94 | 0.92 |
| 9 | 136804211 | rs2073886 | 0.463 | -0.694 | 0.932 | -0.344 | 0.023 | 0.018 | 0.018 | 0.391 | 0.82 | 0.59 | 0.71 | 0.76 | 0.77 | 0.73 |
| 9 | 136805443 | rs2810531 | -0.398 | -0.833 | 0.852 | -0.504 | -0.003 | -0.033 | -0.033 | 0 | 0.73 | 0.69 | 0.71 | 0.8 | 0.69 | 0.76 |
| 9 | 136805789 | rs2519803 | -0.229 | -0.843 | 0.888 | -0.508 | -0.009 | -0.058 | -0.058 | 0 | 0.71 | 0.57 | 0.7 | 0.8 | 0.69 | 0.76 |
| 9 | 136807238 | rs520308 | 0.251 | -0.826 | 0.38 | -0.155 | 0.007 | -0.015 | -0.015 | 0 | 0.32 | 0.26 | 0.31 | 0.37 | 0.29 | 0.34 |
| 9 | 136810857 | rs2519799 | -0.564 | -0.68 | -0.163 | -0.084 | 0 | 0.014 | 0.014 | 0.475 | 0.73 | 0.78 | 0.71 | 0.79 | 0.7 | 0.76 |
| 9 | 136813341 | rs481964 | 0.558 | -0.164 | -0.448 | 0.178 | -0.06 | -0.023 | -0.023 | 2.445 | 0.38 | 0.13 | 0.37 | 0.39 | 0.4 | 0.37 |
| 9 | 136822827 | rs648353 | 0.543 | -0.256 | -0.472 | 0.325 | -0.07 | -0.045 | -0.045 | 2.497 | 0.34 | 0.12 | 0.37 | 0.39 | 0.4 | 0.36 |
| 9 | 136824023 | rs664026 | 0.493 | -0.409 | -0.196 | 0.271 | -0.02 | 0.073 | 0.073 | 0 | 0.27 | 0.5 | 0.28 | 0.4 | 0.23 | 0.33 |
| 9 | 136826457 | rs2073829 | -0.089 | -0.329 | -0.383 | -0.128 | 0.006 | 0.012 | 0.012 | 0 | 0.6 | 0.62 | 0.65 | 0.76 | 0.63 | 0.65 |
| 9 | 136829071 | rs10993877 | 0.066 | 0.093 | -0.365 | -0.098 | -0.005 | -0.042 | -0.042 | 0.176 | 0.36 | 0.48 | 0.36 | 0.25 | 0.31 | 0.36 |
| 9 | 136832528 | rs452999 | 0.12 | 0.452 | -1.04 | 0.182 | -0.002 | -0.028 | -0.028 | 0.056 | 0.23 | 0.19 | 0.24 | 0.44 | 0.27 | 0.39 |
| 9 | 136832577 | rs452943 | 0.009 | 0.143 | -0.65 | -0.415 | 0 | -0.004 | -0.004 | 0 | 0.43 | 0.51 | 0.39 | 0.37 | 0.43 | 0.23 |
| 9 | 136835343 | rs3780792 | -0.287 | 0.511 | -1.195 | -1.039 | -0.011 | 0.068 | 0.068 | 0 | 0.35 | 0.19 | 0.21 | 0.15 | 0.32 | 0.46 |
| 9 | 136835918 | rs2506686 | 0.126 | 0.006 | -0.542 | -1.274 | -0.012 | -0.091 | -0.091 | 1.369 | 0.67 | 0.61 | 0.77 | 0.92 | 0.67 | 0.86 |
| 9 | 136836196 | rs618439 | -0.512 | 0.401 | -0.426 | -0.415 | 0.005 | 0.014 | 0.014 | 0 | 0.16 | 0.31 | 0.16 | 0.16 | 0.15 | 0.13 |
| 9 | 136839840 | rs679106 | -0.404 | 0.29 | -0.95 | -0.673 | 0.002 | 0.009 | 0.009 | 0 | 0.16 | 0.12 | 0.1 | 0.13 | 0.16 | 0.07 |
| 9 | 136843974 | rs425850 | -0.199 | 0.292 | -1.373 | -1.12 | -0.01 | 0.075 | 0.075 | 0 | 0.3 | 0.14 | 0.2 | 0.13 | 0.36 | 0.18 |
| 9 | 136844973 | rs435586 | -0.783 | -0.117 | -0.565 | -1.493 | 0 | -0.009 | -0.009 | 0 | 0.4 | 0.39 | 0.26 | 0.13 | 0.41 | 0.18 |
| 9 | 136848821 | rs7018770 | -0.888 | -0.227 | -0.796 | -1.965 | -0.015 | -0.089 | -0.089 | 0 | 0.23 | 0.32 | 0.13 | 0 | 0.16 | 0.12 |
| 9 | 136851763 | rs420897 | -0.233 | -0.23 | -0.278 | -1.958 | -0.002 | -0.025 | -0.025 | 0 | 0.38 | 0.4 | 0.25 | 0.13 | 0.41 | 0.16 |
| 9 | 136852891 | rs2789864 | 0.047 | -0.139 | -0.308 | -1.904 | 0.008 | 0.079 | 0.079 | 0 | 0.54 | 0.65 | 0.7 | 0.84 | 0.5 | 0.71 |
| 9 | 136865879 | rs460126 | 0.337 | 0.294 | -0.503 | -2.033 | -0.001 | 0.026 | 0.026 | 1.747 | 0.52 | 0.51 | 0.41 | 0.12 | 0.45 | 0.33 |
| 9 | 136870787 | rs419218 | -0.1 | 0.589 | -0.331 | -1.169 | -0.002 | 0.003 | 0.003 | 0 | 0.16 | 0.18 | 0.34 | 0.6 | 0.17 | 0.45 |
| 9 | 136874066 | rs17664384 | -0.28 | 1.006 | -0.173 | -1.325 | 0.027 | -0.05 | -0.05 | 0 | 0.09 | 0.19 | 0.07 | 0.01 | 0.08 | 0.07 |
| 9 | 136874158 | rs464930 | 0.18 | 0.603 | -0.592 | -1.175 | -0.021 | -0.046 | -0.046 | 0 | 0.14 | 0.09 | 0.33 | 0.6 | 0.17 | 0.44 |
| 9 | 136876021 | rs441233 | 0.436 | 0.376 | -1.049 | -1.566 | -0.064 | -0.081 | -0.081 | 0.07 | 0.33 | 0.02 | 0.37 | 0.61 | 0.31 | 0.48 |
| 9 | 136892523 | rs408307 | 0.204 | 0.26 | -0.722 | -1.637 | 0.046 | 0.054 | 0.054 | 0 | 0.39 | 0.13 | 0.28 | 0.65 | 0.3 | 0.48 |
| 9 | 136898236 | rs1051467 | 1.785 | 0.54 | -0.334 | -1.775 | 0.025 | -0.032 | -0.032 | 0 | 0.07 | 0.14 | 0.04 | 0.01 | 0.08 | 0.08 |
| 9 | 136898922 | rs443876 | -0.711 | 0.829 | 0.194 | -1.96 | -0.006 | -0.004 | -0.004 | 1.207 | 0.8 | 0.85 | 0.55 | 0.68 | 0.68 | 0.75 |
| 9 | 136906752 | rs3847320 | 1.232 | 0.588 | 0.831 | -2.112 | -0.007 | -0.107 | -0.107 | 0.04 | 0.3 | 0.39 | 0.21 | 0.03 | 0.29 | 0.16 |
| 9 | 136909261 | rs457420 | -0.066 | 0.722 | 1.057 | -2.078 | 0.025 | 0.069 | 0.069 | 0 | 0.49 | 0.37 | 0.35 | 0.67 | 0.42 | 0.52 |
| 9 | 136909309 | rs460154 | 0.825 | 0.671 | 1.044 | -2.032 | 0.036 | 0.041 | 0.041 | 0 | 0.41 | 0.32 | 0.29 | 0.67 | 0.3 | 0.48 |
| 9 | 136913355 | rs464826 | -1.046 | 1.028 | 0.726 | -1.885 | -0.007 | -0.005 | -0.005 | 4.272 | 0.81 | 0.87 | 0.54 | 0.68 | 0.68 | 0.75 |
| 9 | 136915508 | rs2301576 | 2.078 | 1.263 | 0.45 | -1.906 | -0.026 | -0.01 | -0.01 | 4.976 | 0.07 | 0.01 | 0.14 | 0.13 | 0.08 | 0.13 |
| 9 | 136918662 | rs2073818 | 1.116 | 1.099 | 0.429 | -1.966 | 0.025 | 0.041 | 0.041 | 0.057 | 0.36 | 0.08 | 0.26 | 0.66 | 0.29 | 0.49 |
| 9 | 136921015 | rs17566226 | 1.019 | 1.476 | 1.043 | -1.444 | -0.007 | 0.031 | 0.031 | 0 | 0.29 | 0.15 | 0.19 | 0.02 | 0.32 | 0.17 |
| 9 | 136923972 | rs2810492 | 1.722 | 1.147 | 0.785 | -1.236 | 0.013 | -0.031 | -0.031 | 0 | 0.29 | 0.18 | 0.17 | 0.52 | 0.2 | 0.46 |
| 9 | 136925451 | rs1124317 | -0.031 | 1.126 | 0.96 | -1.029 | 0.041 | 0.02 | 0.02 | 0 | 0.81 | 0.72 | 0.54 | 0.68 | 0.68 | 0.75 |
| 9 | 136927657 | rs1076148 | 1.272 | 1.15 | 0.571 | -1.045 | 0.004 | -0.001 | -0.001 | 0 | 0.37 | 0.33 | 0.24 | 0.54 | 0.31 | 0.5 |
| 9 | 136928276 | rs1075654 | 1.492 | 0.969 | 0.4 | -1.069 | -0.026 | -0.023 | -0.023 | 0.934 | 0.07 | 0.01 | 0.2 | 0.15 | 0.08 | 0.14 |
| 9 | 136928644 | rs10993907 | -0.302 | 0.488 | 0.693 | -1.112 | 0.003 | -0.02 | -0.02 | 0 | 0.44 | 0.44 | 0.24 | 0.12 | 0.37 | 0.24 |
| 9 | 136938961 | rs2078778 | -0.323 | 0.502 | 0.685 | -1.084 | -0.001 | -0.027 | -0.027 | 0.022 | 0.44 | 0.47 | 0.24 | 0.12 | 0.37 | 0.24 |
| 9 | 136943067 | rs10993924 | 0.51 | 0.618 | 0.881 | -1.065 | -0.029 | -0.121 | -0.121 | 0.528 | 0.25 | 0.41 | 0.15 | 0.02 | 0.18 | 0.13 |
| 9 | 136950303 | rs2427951 | 0.323 | 0.741 | 0.596 | -1.003 | -0.006 | 0.028 | 0.028 | 0 | 0.56 | 0.41 | 0.34 | 0.64 | 0.51 | 0.62 |
| 9 | 136950691 | rs2506696 | -0.391 | 0.91 | 0.298 | -0.735 | 0.002 | 0.001 | 0.001 | 0 | 0.81 | 0.83 | 0.48 | 0.66 | 0.69 | 0.76 |
| 9 | 136961298 | rs10761407 | 0.386 | 0.206 | 0.461 | -0.235 | -0.001 | -0.004 | -0.004 | 0 | 0.34 | 0.37 | 0.24 | 0.46 | 0.33 | 0.47 |
| 9 | 136961808 | rs2427971 | 0.427 | 0.303 | -0.03 | -0.201 | -0.017 | 0.006 | 0.006 | 0 | 0.4 | 0.17 | 0.62 | 0.41 | 0.46 | 0.34 |
| 9 | 136965553 | rs12344192 | -0.252 | 0.728 | 0.396 | 0.543 | 0.002 | 0.014 | 0.014 | 0 | 0.37 | 0.49 | 0.26 | 0.5 | 0.35 | 0.5 |
| 9 | 136986911 | rs4292757 | 0.504 | 0.395 | 0.751 | 0.366 | -0.012 | -0.088 | -0.088 | 0 | 0.87 | 0.68 | 0.93 | 1 | 0.91 | 0.97 |
| 9 | 136990836 | rs10993944 | -0.446 | 0.729 | 0.052 | 1.122 | -0.01 | -0.014 | -0.014 | 0 | 0.13 | 0.1 | 0.24 | 0.35 | 0.21 | 0.36 |
| 9 | 136991034 | rs10993945 | -0.042 | 0.84 | -0.045 | 1.425 | -0.005 | -0.014 | -0.014 | 0 | 0.1 | 0.07 | 0.05 | 0.18 | 0.11 | 0.26 |
| 9 | 136992903 | rs4358859 | 0.752 | 0.588 | -0.504 | 0.977 | -0.08 | 0.052 | 0.052 | 0 | 0.39 | 0.15 | 0.37 | 0.28 | 0.45 | 0.28 |
| 9 | 136993590 | rs4744560 | 0.681 | 0.582 | -0.224 | 0.947 | -0.025 | 0.055 | 0.055 | 0 | 0.41 | 0.31 | 0.39 | 0.26 | 0.45 | 0.26 |
| 9 | 136994129 | rs28577297 | 0.127 | 0.716 | -0.801 | 0.77 | -0.026 | -0.045 | -0.045 | 0.948 | 0.11 | 0.03 | 0.12 | 0.22 | 0.14 | 0.24 |
| 9 | 136999786 | rs28464304 | 0.545 | 0.616 | -0.656 | 0.497 | -0.054 | -0.004 | -0.004 | 0.516 | 0.38 | 0.19 | 0.44 | 0.35 | 0.42 | 0.26 |
| 9 | 137012032 | rs34856382 | -0.299 | 0.373 | 0.222 | 0.363 | 0.004 | -0.022 | -0.022 | 0 | 0.07 | 0.25 | 0.04 | 0 | 0.03 | 0.01 |
| 9 | 137015216 | rs34775966 | -0.937 | 0.223 | 0.32 | 0.145 | -0.031 | -0.104 | -0.104 | 0 | 0.15 | 0.32 | 0.08 | 0 | 0.09 | 0.03 |
| 9 | 137018130 | rs28604373 | 0.401 | 0.435 | -0.04 | 0.335 | -0.001 | 0.002 | 0.002 | 0 | 0.46 | 0.42 | 0.47 | 0.36 | 0.45 | 0.27 |
| 9 | 137019776 | rs28694247 | 0.013 | 0.568 | -0.508 | 0.394 | -0.008 | -0.018 | -0.018 | 2.054 | 0.12 | 0.06 | 0.15 | 0.19 | 0.16 | 0.23 |
| 9 | 137026934 | rs28516666 | -0.607 | 0.51 | -0.389 | 0.343 | 0 | 0 | 0 | 0 | 0.2 | 0.15 | 0.19 | 0.19 | 0.22 | 0.25 |
| 9 | 137031454 | rs28498266 | 0.811 | 0.884 | -0.795 | 0.533 | 0.005 | -0.087 | -0.087 | 0.596 | 0.41 | 0.14 | 0.42 | 0.5 | 0.37 | 0.44 |
| 9 | 137037262 | rs28619507 | -0.187 | 1.215 | -0.486 | 0.334 | 0.044 | 0.075 | 0.075 | 0 | 0.62 | 0.82 | 0.75 | 0.78 | 0.72 | 0.73 |

| | | | | | | | | | | | | | | | | |
|---|---|---|---|---|---|---|---|---|---|---|---|---|---|---|---|---|
| 9 | 137037501 | rs28379755 | -0.211 | 1.341 | -0.334 | 0.504 | -0.011 | 0.024 | 0.024 | 0 | 0.65 | 0.77 | 0.65 | 0.71 | 0.61 | 0.68 |
| 9 | 137038257 | rs28404378 | -0.438 | 1.552 | -0.302 | 0.69 | -0.022 | -0.014 | -0.014 | 4.467 | 0.15 | 0.1 | 0.17 | 0.2 | 0.21 | 0.26 |
| 9 | 137092388 | rs12376985 | -0.598 | 1.084 | -0.565 | 0.343 | -0.028 | 0.02 | 0.02 | 0 | 0.07 | 0.01 | 0.09 | 0.02 | 0.12 | 0.04 |
| 9 | 137098042 | rs7872110 | 1.006 | 2.57 | -0.201 | 0.554 | 0.005 | -0.001 | -0.001 | 0 | 0.75 | 0.71 | 0.66 | 0.77 | 0.51 | 0.71 |
| 9 | 137100317 | rs4917376 | 0.034 | 1.371 | -0.302 | 0.913 | -0.003 | 0.026 | 0.026 | 0 | 0.19 | 0.2 | 0.23 | 0.46 | 0.15 | 0.21 |
| 9 | 137101104 | rs11185726 | 1.255 | -0.568 | -1.444 | 0.112 | 0.056 | 0.009 | 0.009 | 0 | 0.24 | 0.07 | 0.18 | 0.14 | 0.16 | 0.27 |
| 9 | 137102473 | rs7874315 | 1.211 | -0.695 | -0.297 | 0.011 | 0.005 | 0.005 | 0.005 | 0.34 | 0.3 | 0.29 | 0.26 | 0.18 | 0.2 | 0.23 |
| 9 | 137103126 | rs7027150 | -0.339 | -0.735 | -0.099 | 0.001 | 0.013 | 0.029 | 0.029 | 0 | 0.53 | 0.36 | 0.54 | 0.66 | 0.44 | 0.56 |
| 9 | 137104155 | rs7031566 | -1.092 | -0.731 | 0.291 | 0.141 | 0.083 | -0.001 | -0.001 | 0 | 0.83 | 0.76 | 0.77 | 0.87 | 0.63 | 0.77 |
| 9 | 137106879 | rs11791747 | 0.209 | -0.984 | 0.662 | 0.489 | 0.014 | 0.002 | 0.002 | 0.019 | 0.49 | 0.55 | 0.45 | 0.54 | 0.33 | 0.4 |
| 9 | 137106964 | rs17567190 | 1.445 | -1.146 | 0.676 | 0.436 | 0.027 | 0.029 | 0.029 | 0.017 | 0.29 | 0.23 | 0.24 | 0.13 | 0.19 | 0.21 |
| 9 | 137109245 | rs7033108 | 1.124 | -1.182 | 0.792 | 0.512 | -0.008 | 0.016 | 0.016 | 0.152 | 0.2 | 0.23 | 0.21 | 0.4 | 0.14 | 0.2 |
| 9 | 137110257 | rs11185664 | 1.849 | -1.22 | 0.579 | 0.478 | -0.023 | -0.008 | -0.008 | 0 | 0.07 | 0.01 | 0.1 | 0.04 | 0.12 | 0.04 |
| 9 | 137110907 | rs13302475 | 1.141 | -1.284 | 0.579 | 0.398 | 0.029 | 0.036 | 0.036 | 0 | 0.31 | 0.22 | 0.23 | 0.14 | 0.2 | 0.22 |
| 9 | 137111146 | rs11185665 | 1.821 | -1.204 | 0.53 | 0.411 | -0.021 | -0.007 | -0.007 | 0 | 0.07 | 0.01 | 0.1 | 0.04 | 0.12 | 0.04 |
| 9 | 137112421 | rs10881585 | 0.088 | -1.345 | 0.677 | 0.361 | -0.026 | 0.063 | 0.063 | 0.338 | 0.29 | 0.53 | 0.35 | 0.49 | 0.26 | 0.27 |
| 9 | 137112655 | rs7873963 | -1.267 | -0.81 | 0.439 | 0.475 | 0.111 | 0.021 | 0.021 | 0.607 | 0.83 | 0.74 | 0.76 | 0.83 | 0.64 | 0.77 |
| 9 | 137113356 | rs7041347 | 1.152 | -0.892 | 0.046 | 0.475 | 0.017 | -0.007 | -0.007 | 0 | 0.29 | 0.07 | 0.28 | 0.19 | 0.3 | 0.22 |
| 9 | 137113880 | rs11185668 | -1.253 | -0.703 | 0.408 | 0.512 | 0.007 | -0.003 | -0.003 | 0 | 0.83 | 0.86 | 0.77 | 0.83 | 0.64 | 0.76 |
| 9 | 137115785 | rs11185670 | 1.553 | -1.059 | 0.812 | 0.397 | -0.014 | -0.004 | -0.004 | 0 | 0.24 | 0.23 | 0.19 | 0.16 | 0.18 | 0.21 |
| 9 | 137117194 | rs4994332 | 0.852 | -1.156 | 1.23 | 0.264 | 0.028 | 0.015 | 0.015 | 0.057 | 0.51 | 0.44 | 0.49 | 0.34 | 0.43 | 0.36 |
| 9 | 137117496 | rs7028191 | -0.28 | -1.228 | 1.215 | 0.256 | 0 | 0.008 | 0.008 | 0.566 | 0.62 | 0.58 | 0.56 | 0.45 | 0.49 | 0.53 |
| 9 | 137117770 | rs4917357 | -1.268 | -1.155 | 1.359 | 0.343 | 0.211 | 0.056 | 0.056 | 0.082 | 0.85 | 0.66 | 0.76 | 0.82 | 0.64 | 0.79 |
| 9 | 137119281 | rs17093638 | 1.637 | 0.129 | 1.216 | 0.23 | 0.069 | 0.04 | 0.04 | 0.973 | 0.43 | 0.21 | 0.38 | 0.31 | 0.32 | 0.33 |
| 9 | 137120661 | rs7848481 | -0.126 | 0.059 | 1.235 | 0.213 | -0.077 | 0.01 | 0.01 | 0 | 0.76 | 0.9 | 0.74 | 0.8 | 0.59 | 0.65 |
| 9 | 137121862 | rs7848058 | 1.028 | -0.17 | 1.62 | 0.262 | -0.014 | 0.024 | 0.024 | 0 | 0.21 | 0.25 | 0.19 | 0.4 | 0.14 | 0.24 |
| 9 | 137122827 | rs10881587 | 0.795 | 0.077 | 1.845 | 0.375 | 0.011 | -0.016 | -0.016 | 0 | 0.64 | 0.49 | 0.57 | 0.71 | 0.45 | 0.58 |
| 9 | 137125501 | rs10785877 | 0.412 | 0.738 | 0.175 | 0.65 | 0.067 | -0.001 | -0.001 | 0 | 0.78 | 0.72 | 0.66 | 0.79 | 0.61 | 0.68 |
| 9 | 137126551 | rs11185681 | 0.188 | -0.205 | -1.853 | 0.694 | 0.054 | -0.021 | -0.021 | 0 | 0.39 | 0.21 | 0.37 | 0.39 | 0.32 | 0.51 |
| 9 | 137131611 | rs1889157 | 0.761 | 0.387 | -1.759 | 0.95 | -0.028 | 0.002 | 0.002 | 0 | 0.56 | 0.22 | 0.45 | 0.42 | 0.52 | 0.49 |
| 9 | 137137245 | rs4917360 | -0.1 | 0.648 | -1.621 | 0.972 | -0.011 | -0.01 | -0.01 | 0 | 0.8 | 0.84 | 0.82 | 0.72 | 0.7 | 0.79 |
| 9 | 137138000 | rs4917362 | 1.002 | 0.631 | -1.541 | 0.954 | -0.005 | -0.001 | -0.001 | 0 | 0.35 | 0.27 | 0.42 | 0.35 | 0.37 | 0.36 |
| 9 | 137138414 | rs10785879 | -0.126 | -0.169 | -1.244 | 0.376 | -0.053 | -0.011 | -0.011 | 0 | 0.17 | 0.49 | 0.16 | 0.14 | 0.06 | 0.09 |
| 9 | 137142353 | rs11185689 | -0.146 | 0.889 | -1.916 | 0.471 | -0.035 | -0.035 | -0.035 | 1.313 | 0.29 | 0.03 | 0.24 | 0.23 | 0.27 | 0.34 |
| 9 | 137147081 | rs4917339 | 0.506 | 1.621 | -0.582 | 0.529 | 0.025 | 0.037 | 0.037 | 0.316 | 0.7 | 0.54 | 0.56 | 0.55 | 0.62 | 0.53 |
| 9 | 137153870 | rs10748520 | 0.769 | 1.597 | -0.406 | -0.214 | 0 | -0.006 | -0.006 | 0 | 0.89 | 0.86 | 0.87 | 0.96 | 0.87 | 0.93 |
| 9 | 137153882 | rs10785882 | 1.396 | 1.63 | -0.513 | -0.993 | -0.002 | 0.022 | 0.022 | 0.061 | 0.66 | 0.74 | 0.74 | 0.94 | 0.7 | 0.66 |
| 9 | 137155945 | rs11789899 | 1.871 | 1.938 | -0.75 | -1.112 | 0.002 | 0.14 | 0.14 | 2.761 | 0.45 | 0.13 | 0.41 | 0.75 | 0.45 | 0.41 |
| 9 | 137159154 | rs7867061 | 1.633 | 1.652 | 0.357 | -0.925 | -0.002 | -0.061 | -0.061 | 0.022 | 0.61 | 0.47 | 0.58 | 0.85 | 0.61 | 0.56 |
| 9 | 137160637 | rs1889151 | -0.305 | 1.847 | 0.675 | -0.776 | 0.02 | 0.014 | 0.014 | 0 | 0.08 | 0.16 | 0.14 | 0.09 | 0.12 | 0.13 |
| 9 | 137160810 | rs1889152 | 1.513 | 1.854 | 0.783 | -0.823 | -0.017 | -0.071 | -0.071 | 0 | 0.67 | 0.53 | 0.64 | 0.85 | 0.7 | 0.61 |
| 9 | 137163255 | rs7848672 | -0.537 | 1.857 | 0.951 | -0.731 | 0.049 | -0.057 | -0.057 | 0.02 | 0.12 | 0.36 | 0.13 | 0.05 | 0.16 | 0.11 |
| 9 | 137163718 | rs1930788 | 0.652 | 1.821 | 0.784 | -0.57 | -0.005 | 0 | 0 | 0 | 0.83 | 0.86 | 0.68 | 0.85 | 0.76 | 0.81 |
| 9 | 137163978 | rs10785883 | 0.804 | 1.191 | 0.454 | -0.323 | -0.007 | 0.026 | 0.026 | 2.406 | 0.55 | 0.39 | 0.46 | 0.7 | 0.5 | 0.44 |
| 9 | 137165349 | rs12346747 | -1.72 | 0.886 | 1.165 | 0.037 | 0.126 | 0.055 | 0.055 | 0.834 | 0.05 | 0.37 | 0.06 | 0.01 | 0.06 | 0.02 |
| 9 | 137165429 | rs17093685 | -1.545 | 1.121 | 0.247 | 0.271 | 0.036 | 0.023 | 0.023 | 0 | 0.14 | 0.33 | 0.27 | 0.22 | 0.2 | 0.18 |
| 9 | 137166136 | rs7858041 | -1.084 | 0.421 | -0.495 | 0.324 | -0.006 | -0.021 | -0.021 | 0.522 | 0.1 | 0.06 | 0.23 | 0.15 | 0.13 | 0.09 |
| 9 | 137169720 | rs7874131 | -1.022 | 0.207 | -0.55 | 0.028 | 0.002 | 0.005 | 0.005 | 0 | 0.1 | 0.08 | 0.09 | 0.09 | 0.09 | 0.21 |
| 9 | 137170414 | rs12350120 | -1.286 | -1.379 | -0.378 | -0.76 | -0.047 | -0.056 | -0.056 | 0 | 0.34 | 0.59 | 0.17 | 0.13 | 0.16 | 0.32 |
| 9 | 137174079 | rs11185709 | -0.21 | -1.079 | -0.754 | -0.747 | -0.024 | 0.034 | 0.034 | 0.005 | 0.07 | 0 | 0.11 | 0.01 | 0.08 | 0.17 |
| 9 | 137174180 | rs875464 | -0.84 | -2.044 | -0.735 | -0.68 | 0.043 | 0.027 | 0.027 | 2.163 | 0.34 | 0.22 | 0.17 | 0.19 | 0.17 | 0.16 |
| 9 | 137182099 | rs11185717 | -1.093 | -1.748 | -0.703 | -0.483 | 0.002 | -0.004 | -0.004 | 0 | 0.3 | 0.31 | 0.23 | 0.35 | 0.25 | 0.35 |
| 9 | 137185046 | rs7032984 | -0.867 | -1.883 | -0.406 | -0.596 | -0.044 | -0.028 | -0.028 | 0 | 0.27 | 0.46 | 0.16 | 0.18 | 0.18 | 0.28 |
| 9 | 137186036 | rs3750548 | -0.81 | -1.112 | -0.791 | -0.312 | -0.006 | -0.009 | -0.009 | 0 | 0.13 | 0.2 | 0.07 | 0.12 | 0.09 | 0.27 |
| 9 | 137186524 | rs7871088 | -0.581 | -0.73 | -1.321 | -0.117 | -0.03 | -0.05 | -0.05 | 1.194 | 0.6 | 0.86 | 0.48 | 0.44 | 0.45 | 0.53 |
| 9 | 137195632 | rs12006542 | -0.934 | -1.096 | -0.823 | -0.586 | 0.027 | 0.045 | 0.045 | 0.813 | 0.37 | 0.75 | 0.35 | 0.22 | 0.31 | 0.31 |
| 9 | 137196129 | rs11794220 | -0.618 | -1.241 | -0.528 | -0.692 | 0 | 0.001 | 0.001 | 0 | 0.08 | 0.12 | 0.17 | 0.08 | 0.09 | 0.02 |
| 9 | 137196155 | rs11792663 | -1.126 | -1.289 | 0.706 | -0.716 | 0.026 | 0.035 | 0.035 | 0.407 | 0.09 | 0.31 | 0.2 | 0.08 | 0.09 | 0.02 |
| 9 | 137205188 | rs12339163 | -0.786 | -1.488 | 1.356 | -0.849 | -0.002 | -0.019 | -0.019 | 0 | 0.16 | 0.34 | 0.1 | 0.14 | 0.12 | 0.24 |
| 9 | 137205222 | rs10881574 | -0.56 | -1.499 | 1.346 | -0.852 | 0 | 0.002 | 0.002 | 0 | 0.08 | 0.12 | 0.17 | 0.08 | 0.09 | 0.02 |
| 9 | 137210863 | rs11185644 | -1.086 | -1.545 | 1.716 | -0.879 | -0.039 | -0.07 | -0.07 | 0 | 0.22 | 0.56 | 0.27 | 0.08 | 0.19 | 0.06 |
| 9 | 137215828 | rs11185647 | -1.683 | -1.426 | 1.755 | -0.82 | 0.033 | 0.05 | 0.05 | 1.055 | 0.36 | 0.77 | 0.35 | 0.21 | 0.29 | 0.29 |
| 9 | 137222137 | rs872298 | -1.612 | -1.504 | 2.031 | -0.842 | -0.013 | -0.021 | -0.021 | 0 | 0.34 | 0.38 | 0.31 | 0.21 | 0.29 | 0.29 |
| 9 | 137223240 | rs10881576 | -1.608 | -1.502 | 2.032 | -0.841 | -0.013 | -0.021 | -0.021 | 0 | 0.33 | 0.38 | 0.31 | 0.21 | 0.29 | 0.29 |
| 9 | 137226094 | rs3818740 | -1.675 | -1.495 | 2.102 | -0.838 | 0.003 | 0.003 | 0.003 | 0.431 | 0.36 | 0.64 | 0.34 | 0.21 | 0.3 | 0.3 |
| 9 | 137232535 | rs10881578 | -1.678 | -1.489 | 2.119 | -0.835 | -0.011 | -0.017 | -0.017 | 0 | 0.34 | 0.4 | 0.31 | 0.21 | 0.29 | 0.3 |
| 9 | 137234671 | rs10881580 | -1.718 | -1.507 | 2.094 | -0.844 | -0.017 | -0.032 | -0.032 | 0 | 0.34 | 0.44 | 0.32 | 0.21 | 0.3 | 0.31 |
| 9 | 137237661 | rs4917354 | -1.788 | -1.516 | 2.307 | -0.848 | -0.026 | -0.048 | -0.048 | 0.019 | 0.35 | 0.49 | 0.32 | 0.21 | 0.3 | 0.31 |
| 9 | 137239163 | rs914853 | -1.758 | -1.506 | 2.307 | -0.829 | -0.025 | -0.045 | -0.045 | 0.03 | 0.35 | 0.49 | 0.32 | 0.21 | 0.3 | 0.32 |

| | | | | | | | | | | | | | | | | |
|---|---|---|---|---|---|---|---|---|---|---|---|---|---|---|---|---|
| 9 | 137243383 | rs11185659 | -0.883 | -1.638 | 2.223 | -0.889 | -0.028 | -0.067 | -0.067 | 0 | 0.25 | 0.3 | 0.13 | 0.08 | 0.19 | 0.2 |
| 9 | 137248674 | rs7041449 | -1.54 | -1.441 | 2.03 | -0.875 | 0.016 | 0.023 | 0.023 | 0.102 | 0.3 | 0.75 | 0.19 | 0.13 | 0.23 | 0.29 |
| 9 | 137252760 | rs7853934 | -0.6 | -1.42 | 2.26 | -0.835 | -0.017 | -0.041 | -0.041 | 0 | 0.22 | 0.28 | 0.13 | 0.11 | 0.2 | 0.18 |
| 9 | 137253447 | rs7048602 | -2.425 | -1.311 | 2.338 | -0.676 | 0.044 | 0.083 | 0.083 | 0.633 | 0.07 | 0.47 | 0.05 | 0.05 | 0.01 | 0.1 |
| 9 | 137256078 | rs10881582 | -1.264 | -1.797 | 1.772 | -0.852 | 0.03 | 0.033 | 0.033 | 0.063 | 0.36 | 0.72 | 0.42 | 0.18 | 0.24 | 0.22 |
| 9 | 137259992 | rs11185660 | -1.244 | -1.75 | 1.661 | -0.829 | -0.047 | -0.055 | -0.055 | 0 | 0.34 | 0.4 | 0.39 | 0.19 | 0.24 | 0.23 |
| 9 | 137266704 | rs7039190 | -0.492 | -1.171 | 1.405 | -0.553 | 0.018 | 0.009 | 0.009 | 0 | 0.06 | 0.15 | 0.12 | 0.05 | 0.06 | 0.01 |
| 9 | 137269456 | rs34312136 | 0.085 | -0.285 | 0.45 | -0.205 | 0.039 | -0.009 | -0.009 | 0 | 0.31 | 0.53 | 0.39 | 0.23 | 0.41 | 0.16 |
| 9 | 137280939 | rs35079168 | 1.006 | -0.661 | -0.034 | -0.144 | 0.064 | 0.062 | 0.062 | 0 | 0.57 | 0.22 | 0.55 | 0.73 | 0.65 | 0.44 |
| 9 | 137285503 | rs11102986 | 0.078 | -0.994 | -0.141 | -0.308 | 0.046 | 0.038 | 0.038 | 0 | 0.76 | 0.97 | 0.78 | 0.86 | 0.82 | 0.71 |
| 9 | 137290725 | rs12004589 | -0.787 | -0.973 | 0.715 | -0.315 | 0.013 | -0.012 | -0.012 | 0 | 0.09 | 0.18 | 0.09 | 0.06 | 0.1 | 0.19 |
| 9 | 137292505 | rs6537998 | -0.951 | -1.103 | 0.918 | -0.359 | 0.067 | 0.063 | 0.063 | 2.907 | 0.06 | 0.52 | 0.05 | 0 | 0 | 0.01 |
| 9 | 137294030 | rs3132301 | 0.048 | -1.026 | 0.172 | -0.249 | 0.051 | 0.031 | 0.031 | 0 | 0.74 | 0.98 | 0.77 | 0.85 | 0.82 | 0.66 |
| 9 | 137295304 | rs875444 | 1.644 | -0.752 | 0.101 | 0.086 | 0.054 | -0.001 | -0.001 | 0 | 0.53 | 0.08 | 0.52 | 0.6 | 0.61 | 0.41 |
| 9 | 137299949 | rs1805352 | 1.273 | -0.831 | 1.736 | -0.061 | 0.039 | 0.032 | 0.032 | 0.395 | 0.6 | 0.27 | 0.62 | 0.75 | 0.7 | 0.42 |
| 9 | 137302631 | rs3132296 | 1.285 | -0.823 | 1.9 | -0.039 | 0.016 | 0.013 | 0.013 | 0.379 | 0.61 | 0.32 | 0.62 | 0.74 | 0.7 | 0.48 |
| 9 | 137304915 | rs3118529 | 1.287 | -0.915 | 1.861 | -0.032 | 0.065 | 0.046 | 0.046 | 0.525 | 0.6 | 0.18 | 0.62 | 0.74 | 0.7 | 0.48 |
| 9 | 137308462 | rs3118536 | 0.012 | -0.931 | 1.347 | 0.049 | 0.02 | -0.059 | -0.059 | 0 | 0.77 | 0.99 | 0.8 | 0.8 | 0.84 | 0.7 |
| 9 | 137311400 | rs4240705 | 1.45 | -0.591 | 1.702 | -0.171 | -0.004 | -0.009 | -0.009 | 0 | 0.6 | 0.55 | 0.6 | 0.7 | 0.66 | 0.49 |
| 9 | 137312828 | rs6537944 | -1.303 | -0.727 | 1.416 | 0.04 | 0.004 | 0.027 | 0.027 | 0.156 | 0.09 | 0.27 | 0.1 | 0.13 | 0.07 | 0.17 |
| 9 | 137318178 | rs3118571 | 1.523 | -0.584 | -0.181 | 0.009 | 0.043 | 0.051 | 0.051 | 0.654 | 0.59 | 0.13 | 0.56 | 0.7 | 0.66 | 0.45 |
| 9 | 137321156 | rs1536475 | 0.166 | -0.262 | -0.045 | 0.131 | 0.021 | -0.03 | -0.03 | 0.266 | 0.75 | 0.98 | 0.79 | 0.76 | 0.82 | 0.67 |
| 9 | 137324120 | rs3132294 | 0.199 | -0.302 | -0.126 | 0.109 | -0.033 | -0.02 | -0.02 | 0 | 0.74 | 0.98 | 0.76 | 0.81 | 0.77 | 0.67 |
| 9 | 137326408 | rs877954 | 1.035 | -0.369 | -0.02 | 0.147 | 0.024 | 0.015 | 0.015 | 0 | 0.6 | 0.24 | 0.58 | 0.66 | 0.67 | 0.46 |
| 9 | 137328286 | rs1805343 | 1.139 | -0.313 | -0.318 | 0.146 | 0.036 | 0.031 | 0.031 | 0 | 0.58 | 0.19 | 0.57 | 0.67 | 0.66 | 0.44 |
| 9 | 137337357 | rs3118526 | 0.704 | -0.311 | -0.667 | 0.293 | -0.01 | -0.054 | -0.054 | 0.919 | 0.8 | 0.99 | 0.82 | 0.82 | 0.89 | 0.69 |
| 9 | 137340513 | rs9409929 | -0.759 | 0.294 | -0.377 | 0.277 | -0.014 | 0.061 | 0.061 | 0 | 0.33 | 0.18 | 0.25 | 0.22 | 0.36 | 0.35 |
| 9 | 137341600 | rs11103603 | -1.342 | 0.025 | 0.382 | 0.078 | 0.018 | 0.061 | 0.061 | 0 | 0.43 | 0.7 | 0.31 | 0.22 | 0.38 | 0.42 |
| 9 | 137359635 | rs10776936 | -2.019 | 0.558 | 2.179 | 1.36 | 0.001 | 0.015 | 0.015 | 0.275 | 0.24 | 0.61 | 0.18 | 0.26 | 0.24 | 0.39 |
| 9 | 137359948 | rs11103714 | -0.844 | 0.604 | 2.134 | 1.235 | -0.014 | -0.015 | -0.015 | 0.616 | 0.13 | 0.1 | 0.14 | 0.18 | 0.18 | 0.3 |
| 9 | 137360936 | rs4842229 | 2.492 | 0.546 | 2.338 | 1.632 | -0.006 | 0.018 | 0.018 | 1.471 | 0.81 | 0.3 | 0.76 | 0.72 | 0.88 | 0.75 |
| 9 | 137366540 | rs943430 | 2.046 | 0.51 | 2.7 | 1.458 | -0.001 | 0.013 | 0.013 | 0.166 | 0.7 | 0.38 | 0.65 | 0.57 | 0.75 | 0.49 |
| 9 | 137369200 | rs7033773 | -1.735 | 0.288 | 2.782 | 1.415 | 0.028 | 0.007 | 0.007 | 0 | 0.07 | 0.28 | 0.03 | 0.03 | 0.05 | 0.08 |
| 9 | 137373598 | rs7033608 | -2.221 | 0.476 | 2.722 | 1.418 | 0 | 0.002 | 0.002 | 0.213 | 0.33 | 0.67 | 0.38 | 0.45 | 0.32 | 0.59 |
| 9 | 137374584 | rs3818518 | -1.598 | 0.574 | 2.682 | 1.563 | 0.004 | 0.022 | 0.022 | 0.577 | 0.27 | 0.3 | 0.33 | 0.44 | 0.26 | 0.53 |
| 9 | 137377445 | rs1555949 | -1.103 | 0.607 | 2.701 | 1.677 | 0.009 | 0.099 | 0.099 | 0.525 | 0.11 | 0.26 | 0.17 | 0.26 | 0.08 | 0.17 |
| 9 | 137379635 | rs7871031 | -1.231 | 0.574 | 2.586 | 1.417 | 0.009 | -0.004 | -0.004 | 0 | 0.23 | 0.35 | 0.19 | 0.21 | 0.23 | 0.44 |
| 9 | 137380516 | rs13295631 | -0.957 | 0.547 | 2.57 | 1.401 | -0.001 | -0.01 | -0.01 | 0.471 | 0.22 | 0.34 | 0.18 | 0.18 | 0.19 | 0.36 |
| 9 | 137382629 | rs10776949 | -2.1 | 0.601 | 2.359 | 1.583 | -0.008 | -0.057 | -0.057 | 0 | 0.37 | 0.82 | 0.37 | 0.52 | 0.33 | 0.62 |
| 9 | 137384140 | rs12336911 | -1.925 | 0.715 | 2.289 | 1.617 | 0.018 | 0.197 | 0.197 | 1.728 | 0.14 | 0.46 | 0.19 | 0.35 | 0.13 | 0.26 |
| 9 | 137385159 | rs943434 | -1.171 | 0.684 | 1.746 | 1.594 | 0.001 | 0.131 | 0.131 | 0.911 | 0.11 | 0.29 | 0.16 | 0.31 | 0.09 | 0.17 |
| 9 | 137392705 | rs7863795 | -1.294 | 0.66 | 1.39 | 0.959 | -0.003 | -0.019 | -0.019 | 0.061 | 0.25 | 0.37 | 0.19 | 0.18 | 0.21 | 0.35 |
| 9 | 137406529 | rs9409898 | -2.076 | 0.705 | 1.878 | 1.547 | 0.055 | 0.483 | 0.483 | 0.631 | 0.1 | 0.37 | 0.16 | 0.51 | 0.08 | 0.08 |
| 9 | 137406638 | rs9409899 | -1.586 | 0.695 | 2.263 | 1.543 | 0.048 | 0.376 | 0.376 | 0.028 | 0.07 | 0.19 | 0.15 | 0.51 | 0.08 | 0.07 |
| 9 | 137406713 | rs9409970 | -2.753 | 0.448 | 2.153 | 1.314 | -0.014 | 0.093 | 0.093 | 3.016 | 0.16 | 0.66 | 0.22 | 0.51 | 0.12 | 0.09 |
| 9 | 137406985 | rs11103096 | -1.317 | 0.59 | 1.811 | 0.48 | -0.005 | -0.024 | -0.024 | 2.185 | 0.05 | 0.18 | 0.04 | 0 | 0.03 | 0.02 |
| 9 | 137408432 | rs943427 | -1.048 | 1.535 | 1.227 | -0.167 | 0.054 | -0.082 | -0.082 | 0 | 0.3 | 0.42 | 0.23 | 0.08 | 0.43 | 0.47 |
| 9 | 137408717 | rs3132330 | -0.764 | 1.622 | 0.52 | -0.79 | 0.009 | -0.079 | -0.079 | 0.086 | 0.34 | 0.37 | 0.25 | 0.06 | 0.43 | 0.34 |
| 9 | 137413400 | rs3132329 | 1.137 | 1.982 | 0.21 | -0.314 | -0.001 | 0 | 0 | 0 | 0.75 | 0.71 | 0.75 | 0.82 | 0.64 | 0.78 |
| 9 | 137415319 | rs4842031 | 1.203 | 1.341 | 0.204 | -0.106 | 0.011 | 0 | 0 | 0 | 0.46 | 0.3 | 0.42 | 0.45 | 0.38 | 0.3 |
| 9 | 137415342 | rs3118580 | -0.036 | 1.603 | 0.092 | -0.126 | -0.078 | 0.017 | 0.017 | 0 | 0.85 | 0.97 | 0.85 | 0.88 | 0.71 | 0.82 |
| 9 | 137417305 | rs7037930 | 0.686 | 1.602 | 0.201 | -0.146 | -0.004 | -0.001 | -0.001 | 5.553 | 0.81 | 0.78 | 0.79 | 0.81 | 0.69 | 0.8 |
| 9 | 137417308 | rs10776839 | 1.015 | 1.59 | 0.189 | -0.15 | 0 | 0.001 | 0.001 | 0 | 0.53 | 0.37 | 0.46 | 0.45 | 0.42 | 0.34 |
| 9 | 137420521 | rs3118586 | 0.75 | 1.613 | 0.2 | -0.138 | -0.045 | 0.044 | 0.044 | 0 | 0.79 | 0.88 | 0.8 | 0.88 | 0.66 | 0.78 |
| 9 | 137422484 | rs3118589 | 0.902 | 1.589 | 0.299 | -0.141 | -0.026 | 0.006 | 0.006 | 0 | 0.78 | 0.84 | 0.77 | 0.82 | 0.66 | 0.78 |
| 9 | 137424233 | rs4841898 | 1.855 | 1.307 | 0.021 | -0.43 | 0.004 | -0.009 | -0.009 | 0 | 0.39 | 0.28 | 0.41 | 0.44 | 0.32 | 0.23 |
| 9 | 137425441 | rs3132321 | -0.544 | 0.981 | 0.268 | -0.474 | -0.029 | -0.01 | -0.01 | 0.09 | 0.38 | 0.53 | 0.37 | 0.36 | 0.31 | 0.5 |
| 9 | 137426119 | rs3132317 | 0.953 | 0.662 | -0.094 | -0.423 | -0.016 | -0.005 | -0.005 | 0 | 0.12 | 0.01 | 0.12 | 0.05 | 0.13 | 0.28 |
| 9 | 137426334 | rs3118593 | -0.538 | 0.614 | 0.309 | -0.242 | 0.004 | 0.002 | 0.002 | 0 | 0.4 | 0.67 | 0.38 | 0.36 | 0.3 | 0.5 |
| 9 | 137426935 | rs1006418 | 1.987 | 0.757 | 0.622 | -0.226 | 0.009 | -0.022 | -0.022 | 0 | 0.64 | 0.42 | 0.75 | 0.79 | 0.6 | 0.67 |
| 9 | 137427442 | rs3132315 | -0.085 | 0.855 | 0.424 | -0.195 | -0.036 | -0.02 | -0.02 | 0 | 0.85 | 0.9 | 0.84 | 0.8 | 0.72 | 0.81 |
| 9 | 137427767 | rs3118595 | 1.012 | 1.032 | 0.32 | -0.105 | 0.011 | -0.009 | -0.009 | 0.066 | 0.8 | 0.72 | 0.81 | 0.82 | 0.71 | 0.77 |
| 9 | 137428425 | rs1409832 | 0.676 | 0.904 | -0.204 | -0.051 | 0.02 | -0.018 | -0.018 | 0.334 | 0.28 | 0.2 | 0.31 | 0.35 | 0.25 | 0.49 |
| 9 | 137429719 | rs4240698 | 1.488 | 1.015 | -0.099 | 0.109 | 0 | 0 | 0 | 0 | 0.39 | 0.36 | 0.44 | 0.4 | 0.36 | 0.2 |
| 9 | 137431373 | rs3132310 | -0.523 | 1.259 | 0.267 | 0.24 | -0.062 | -0.037 | -0.037 | 0 | 0.91 | 0.98 | 0.87 | 0.83 | 0.77 | 0.81 |
| 9 | 137431424 | rs1316404 | 1.127 | 0.842 | 0.389 | 0.288 | 0 | 0.001 | 0.001 | 0 | 0.45 | 0.4 | 0.48 | 0.36 | 0.41 | 0.24 |
| 9 | 137436314 | rs3118515 | -0.37 | 0.302 | 0.048 | 0.266 | 0.034 | -0.015 | -0.015 | 0 | 0.35 | 0.24 | 0.34 | 0.39 | 0.3 | 0.56 |
| 9 | 137437183 | rs943423 | 0.093 | 0.126 | 0.01 | -0.107 | 0.028 | 0.03 | 0.03 | 0 | 0.64 | 0.71 | 0.71 | 0.74 | 0.69 | 0.43 |
| 9 | 137441595 | rs3118520 | -0.807 | 0.289 | -0.15 | -0.266 | 0.072 | 0.164 | 0.164 | 0 | 0.48 | 0.8 | 0.37 | 0.22 | 0.39 | 0.57 |

| | | | | | | | | | | | | | | | | |
|---|---|---|---|---|---|---|---|---|---|---|---|---|---|---|---|---|
| 9 | 137443896 | rs3132304 | 0.547 | 0.447 | 1.212 | 0.12 | 0.001 | 0.019 | 0.019 | 0 | 0.68 | 0.58 | 0.51 | 0.62 | 0.68 | 0.78 |
| 9 | 137447293 | rs11103276 | -2.162 | 0.391 | 1.784 | -0.277 | -0.038 | -0.101 | -0.101 | 0.118 | 0.2 | 0.6 | 0.11 | 0.02 | 0.12 | 0.24 |
| 9 | 137469004 | rs7025493 | -0.318 | 0.463 | 1.203 | -0.125 | -0.001 | -0.019 | -0.019 | 0 | 0.06 | 0.1 | 0.04 | 0 | 0.06 | 0.06 |
| 9 | 137470639 | rs7855550 | -1.757 | 0.439 | 1.483 | -0.137 | -0.001 | -0.031 | -0.031 | 0 | 0.09 | 0.28 | 0.06 | 0.01 | 0.07 | 0.1 |
| 9 | 137476593 | rs11103404 | -0.323 | 0.59 | 0.834 | -0.077 | 0.003 | -0.019 | -0.019 | 0 | 0.05 | 0.1 | 0.05 | 0 | 0.08 | 0.06 |
| 9 | 137477402 | rs11103407 | -1.792 | 0.833 | 0.395 | -0.025 | 0.013 | -0.022 | -0.022 | 0 | 0.22 | 0.58 | 0.16 | 0.13 | 0.23 | 0.18 |
| 9 | 137485476 | rs10858254 | 0.103 | -0.018 | 0.028 | -0.237 | 0.001 | -0.004 | -0.004 | 5.861 | 0.29 | 0.29 | 0.45 | 0.33 | 0.27 | 0.16 |
| 9 | 137485536 | rs10735245 | -1.162 | 0.009 | 0.027 | -0.353 | 0.024 | -0.016 | -0.016 | 0.457 | 0.43 | 0.8 | 0.55 | 0.36 | 0.38 | 0.31 |
| 9 | 137485876 | rs7862997 | -1.17 | 0.002 | 0.03 | -0.346 | 0.016 | -0.057 | -0.057 | 0 | 0.12 | 0.38 | 0.08 | 0.01 | 0.11 | 0.14 |
| 9 | 137489483 | rs4842136 | -0.748 | 0.161 | -1.081 | -0.234 | 0.024 | -0.008 | -0.008 | 0 | 0.43 | 0.76 | 0.54 | 0.34 | 0.37 | 0.3 |
| 9 | 137490048 | rs12348251 | -1.013 | 0.243 | -0.712 | -0.483 | -0.006 | -0.021 | -0.021 | 1.179 | 0.07 | 0.09 | 0.03 | 0 | 0.04 | 0.06 |
| 9 | 137494787 | rs11103417 | -0.373 | 0.445 | -1.134 | -0.8 | -0.008 | 0.044 | 0.044 | 0.354 | 0.24 | 0.16 | 0.15 | 0.08 | 0.25 | 0.28 |
| 9 | 137494819 | rs7020813 | -0.254 | 0.289 | -1.641 | -1.298 | -0.091 | 0.183 | 0.183 | 0 | 0.62 | 0.83 | 0.71 | 0.95 | 0.51 | 0.76 |
| 9 | 137503695 | rs7856818 | -0.974 | -0.626 | 0.114 | -0.798 | -0.025 | -0.022 | -0.022 | 0 | 0.14 | 0.7 | 0.1 | 0.09 | 0.08 | 0.08 |
| 9 | 137506578 | rs11103429 | -0.387 | -0.658 | 0.406 | -0.784 | -0.002 | -0.008 | -0.008 | 0 | 0.07 | 0.19 | 0.05 | 0.05 | 0.06 | 0.07 |
| 9 | 137511278 | rs6537934 | -0.931 | -0.918 | -1.144 | -0.454 | -0.014 | -0.094 | -0.094 | 0 | 0.55 | 0.94 | 0.45 | 0.6 | 0.37 | 0.61 |
| 9 | 137512483 | rs12344933 | -1.058 | -0.918 | -0.042 | -0.378 | -0.053 | -0.053 | -0.053 | 0.021 | 0.21 | 0.58 | 0.1 | 0.1 | 0.12 | 0.13 |
| 9 | 137515158 | rs4424343 | -1.006 | -0.503 | -0.239 | 0.154 | -0.004 | 0.018 | 0.018 | 0.364 | 0.27 | 0.69 | 0.34 | 0.43 | 0.25 | 0.41 |
| 9 | 137516213 | rs4841920 | -0.949 | -0.85 | -0.401 | 0.063 | -0.005 | 0.003 | 0.003 | 0 | 0.22 | 0.25 | 0.18 | 0.27 | 0.18 | 0.38 |
| 9 | 137516625 | rs7341858 | -1.726 | -0.806 | -1.044 | 0.08 | 0.052 | -0.02 | -0.02 | 0.674 | 0.39 | 0.92 | 0.4 | 0.37 | 0.31 | 0.45 |
| 9 | 137517022 | rs4076173 | -1.323 | -0.248 | -0.875 | -0.16 | 0.006 | 0.013 | 0.013 | 0.668 | 0.39 | 0.7 | 0.32 | 0.34 | 0.38 | 0.46 |
| 9 | 137518733 | rs7865626 | -1.045 | -0.867 | -0.411 | -0.699 | -0.007 | -0.067 | -0.067 | 0 | 0.2 | 0.23 | 0.1 | 0 | 0.18 | 0.1 |
| 9 | 137520936 | rs4304399 | -0.979 | -1.628 | -0.424 | -0.899 | 0.009 | 0.004 | 0.004 | 0.97 | 0.34 | 0.36 | 0.29 | 0.26 | 0.2 | 0.36 |
| 9 | 137530346 | rs7021140 | -0.93 | -1.568 | -0.125 | -1.601 | -0.009 | -0.046 | -0.046 | 0 | 0.32 | 0.59 | 0.27 | 0.09 | 0.28 | 0.36 |
| 9 | 137538057 | rs11103455 | 1.336 | -0.763 | -0.81 | -1.027 | -0.127 | -0.115 | -0.115 | 0.045 | 0.29 | 0.03 | 0.41 | 0.39 | 0.38 | 0.38 |
| 9 | 137543874 | rs10858265 | -0.881 | -0.685 | -0.253 | -0.906 | 0.002 | 0.039 | 0.039 | 0.099 | 0.39 | 0.69 | 0.31 | 0.1 | 0.39 | 0.41 |
| 9 | 137552398 | rs4335205 | 1.419 | -0.707 | 1.067 | 0.268 | 0 | -0.002 | -0.002 | 0 | 0.7 | 0.33 | 0.56 | 0.46 | 0.74 | 0.61 |
| 9 | 137554581 | rs4842138 | -0.045 | -0.552 | 1.24 | 0.33 | 0.009 | 0.236 | 0.236 | 0.01 | 0.86 | 0.7 | 0.74 | 0.54 | 0.89 | 0.91 |
| 9 | 137559556 | rs6537942 | -0.991 | -0.548 | 0.821 | 0.318 | 0.005 | -0.004 | -0.004 | 0 | 0.17 | 0.44 | 0.19 | 0.13 | 0.15 | 0.23 |
| 9 | 137563692 | rs4842139 | -0.069 | -0.402 | 0.741 | 0.232 | 0.006 | 0.214 | 0.214 | 0 | 0.87 | 0.7 | 0.74 | 0.54 | 0.89 | 0.9 |
| 9 | 137568051 | rs7044529 | -1.225 | -0.245 | 0.486 | -0.493 | 0.008 | -0.029 | -0.029 | 1.069 | 0.16 | 0.36 | 0.18 | 0.11 | 0.15 | 0.23 |
| 9 | 137573609 | rs12554098 | -0.493 | -0.123 | -0.755 | -0.299 | -0.014 | 0 | 0 | 0 | 0.11 | 0.03 | 0.15 | 0.08 | 0.13 | 0.14 |
| 9 | 137586995 | rs3922982 | -0.702 | -0.26 | -0.139 | -0.329 | -0.04 | -0.088 | -0.088 | 0.526 | 0.21 | 0.36 | 0.09 | 0.05 | 0.12 | 0.12 |
| 9 | 137592961 | rs4341231 | -0.145 | 0.548 | -0.842 | 0.09 | -0.024 | 0.009 | 0.009 | 0 | 0.6 | 0.7 | 0.39 | 0.73 | 0.54 | 0.66 |
| 9 | 137594829 | rs4529530 | -0.386 | 1.13 | -0.375 | -0.476 | 0.021 | -0.043 | -0.043 | 0 | 0.25 | 0.31 | 0.16 | 0.11 | 0.29 | 0.14 |
| 9 | 137595859 | rs4240702 | 0.713 | 0.807 | -0.194 | 0.056 | -0.001 | -0.001 | -0.001 | 0 | 0.6 | 0.64 | 0.72 | 0.48 | 0.52 | 0.56 |
| 9 | 137595928 | rs4240703 | -0.169 | 0.914 | 0.139 | 0.07 | -0.017 | 0.084 | 0.084 | 0 | 0.84 | 0.77 | 0.88 | 0.58 | 0.85 | 0.72 |
| 9 | 137596271 | rs4319175 | 0.906 | 0.19 | 0.651 | -0.056 | 0.009 | -0.002 | -0.002 | 0 | 0.6 | 0.46 | 0.75 | 0.58 | 0.52 | 0.6 |
| 9 | 137596949 | rs4842145 | 1.158 | 0.121 | 0.968 | -0.071 | 0.006 | 0.001 | 0.001 | 0 | 0.57 | 0.53 | 0.69 | 0.47 | 0.5 | 0.44 |
| 9 | 137598731 | rs3922914 | 0.797 | -0.225 | 0.875 | -0.234 | -0.045 | 0.071 | 0.071 | 0 | 0.8 | 0.5 | 0.89 | 0.7 | 0.85 | 0.71 |
| 9 | 137599444 | rs4548258 | 0.013 | -0.183 | 0.866 | -0.272 | -0.019 | 0.021 | 0.021 | 0 | 0.25 | 0.19 | 0.17 | 0.12 | 0.33 | 0.19 |
| 9 | 137599992 | rs3109687 | 1.594 | -0.084 | 0.621 | -0.399 | 0.027 | -0.002 | -0.002 | 1.742 | 0.47 | 0.21 | 0.63 | 0.46 | 0.42 | 0.4 |
| 9 | 137601285 | rs4841926 | 0.537 | 0.269 | 0.842 | -0.069 | -0.007 | -0.027 | -0.027 | 0 | 0.8 | 0.88 | 0.85 | 0.59 | 0.77 | 0.73 |
| 9 | 137602119 | rs3128591 | 1.017 | 0.28 | 0.837 | -0.048 | -0.002 | 0.015 | 0.015 | 0 | 0.75 | 0.69 | 0.82 | 0.58 | 0.76 | 0.57 |
| 9 | 137610423 | rs11103479 | -1.309 | 0.548 | 0.758 | -0.099 | 0.017 | 0.016 | 0.016 | 0 | 0.29 | 0.45 | 0.21 | 0.41 | 0.27 | 0.46 |
| 9 | 137612284 | rs12004951 | -1.503 | 0.499 | 0.639 | -0.102 | 0.001 | -0.006 | -0.006 | 0 | 0.3 | 0.76 | 0.23 | 0.4 | 0.25 | 0.48 |
| 9 | 137613772 | rs7875140 | -1.109 | 0.74 | 0.649 | 0.005 | -0.001 | -0.004 | -0.004 | 0 | 0.26 | 0.69 | 0.2 | 0.42 | 0.23 | 0.46 |
| 9 | 137619862 | rs3124300 | 1.728 | 0.763 | 0.289 | -0.104 | 0.002 | 0.004 | 0.004 | 0 | 0.54 | 0.18 | 0.63 | 0.48 | 0.47 | 0.44 |
| 9 | 137620847 | rs3128597 | 0.524 | 0.812 | 0.48 | -0.118 | 0.041 | -0.07 | -0.07 | 0 | 0.79 | 0.59 | 0.84 | 0.9 | 0.73 | 0.83 |
| 9 | 137630568 | rs3109677 | -1.627 | 1.115 | -0.183 | 0.305 | -0.038 | -0.008 | -0.008 | 0.339 | 0.43 | 0.84 | 0.32 | 0.51 | 0.5 | 0.5 |
| 9 | 137631198 | rs3109676 | -0.949 | 1.258 | 0.099 | 0.723 | -0.009 | 0.019 | 0.019 | 0 | 0.27 | 0.54 | 0.26 | 0.29 | 0.25 | 0.37 |
| 9 | 137641411 | rs3128615 | -2.065 | 1.286 | -0.38 | 0.63 | 0.024 | 0.109 | 0.109 | 0 | 0.51 | 0.95 | 0.49 | 0.77 | 0.55 | 0.59 |
| 9 | 137642577 | rs12686426 | -0.897 | 1.063 | 0.402 | 0.953 | 0 | 0.001 | 0.001 | 0 | 0.22 | 0.16 | 0.18 | 0.33 | 0.23 | 0.16 |
| 9 | 137646046 | rs11103501 | -0.537 | 1.272 | 0.241 | 1.07 | -0.016 | -0.022 | -0.022 | 0 | 0.18 | 0.09 | 0.16 | 0.35 | 0.22 | 0.16 |
| 9 | 137650824 | rs7849193 | -1.053 | 0.57 | 1.399 | 1.11 | -0.021 | 0.153 | 0.153 | 0.669 | 0.16 | 0.47 | 0.24 | 0.3 | 0.09 | 0.14 |
| 9 | 137654103 | rs11103509 | -0.658 | 0.621 | 1.538 | 1.581 | -0.002 | 0.246 | 0.246 | 0.169 | 0.1 | 0.35 | 0.22 | 0.28 | 0.04 | 0.12 |
| 9 | 137658608 | rs4842157 | -1.938 | 0.965 | 0.987 | 1.522 | 0.008 | 0.007 | 0.007 | 0 | 0.32 | 0.41 | 0.38 | 0.69 | 0.32 | 0.43 |
| 9 | 137662523 | rs10745384 | -2.155 | 0.954 | 0.142 | 0.808 | -0.069 | 0.328 | 0.328 | 0 | 0.49 | 0.9 | 0.48 | 0.93 | 0.43 | 0.52 |
| 9 | 137665445 | rs4842164 | -1.807 | 1.171 | 0.638 | 1.282 | 0.008 | 0.023 | 0.023 | 0 | 0.32 | 0.42 | 0.38 | 0.65 | 0.32 | 0.43 |
| 9 | 137666414 | rs4842165 | -0.621 | 1.269 | 0.823 | 0.936 | -0.002 | -0.007 | -0.007 | 0 | 0.19 | 0.18 | 0.14 | 0.35 | 0.19 | 0.18 |
| 9 | 137670002 | rs4563961 | 1.16 | 1.202 | 0.765 | 0.975 | 0.029 | 0.074 | 0.074 | 0 | 0.74 | 0.58 | 0.8 | 0.51 | 0.71 | 0.67 |
| 9 | 137672012 | rs3124932 | 1.379 | 0.835 | 0.186 | -0.003 | -0.027 | 0.284 | 0.284 | 0 | 0.47 | 0.09 | 0.47 | 0.07 | 0.49 | 0.44 |
| 9 | 137672801 | rs7849777 | -0.671 | 0.743 | 0.692 | 0.183 | 0.028 | 0.049 | 0.049 | 0.792 | 0.09 | 0.3 | 0.06 | 0.13 | 0.07 | 0.15 |
| 9 | 137673861 | rs6537949 | -1.673 | 0.771 | 0.699 | 0.173 | 0.038 | 0.019 | 0.019 | 0 | 0.12 | 0.36 | 0.09 | 0.13 | 0.13 | 0.17 |
| 9 | 137677090 | rs12005196 | -1.044 | 0.831 | 0.493 | 0.202 | 0.027 | 0.014 | 0.014 | 0 | 0.11 | 0.28 | 0.08 | 0.13 | 0.12 | 0.08 |
| 9 | 137677489 | rs4072077 | 0.138 | 0.776 | 0.392 | 0.183 | 0.003 | 0.03 | 0.03 | 0.756 | 0.07 | 0.18 | 0.02 | 0.11 | 0.04 | 0.04 |
| 9 | 137678306 | rs3128621 | 1.343 | 0.801 | -0.121 | 0.089 | -0.045 | 0.243 | 0.243 | 0 | 0.48 | 0.07 | 0.48 | 0.14 | 0.51 | 0.46 |
| 9 | 137679703 | rs11998946 | -1.474 | 0.806 | 0.46 | 0.093 | 0.044 | 0.025 | 0.025 | 0.913 | 0.13 | 0.46 | 0.08 | 0.12 | 0.11 | 0.06 |
| 9 | 137681458 | rs9409992 | 0.799 | 0.579 | 0.16 | -0.038 | -0.002 | 0.017 | 0.017 | 0 | 0.56 | 0.41 | 0.51 | 0.23 | 0.58 | 0.47 |

| | | | | | | | | | | | | | | | | |
|---|---|---|---|---|---|---|---|---|---|---|---|---|---|---|---|---|
| 9 | 137687314 | rs4842167 | -1.383 | 0.295 | -0.571 | -0.285 | -0.028 | 0.128 | 0.128 | 0.874 | 0.6 | 0.87 | 0.55 | 0.83 | 0.55 | 0.6 |
| 9 | 137688657 | rs3811159 | -0.896 | 0.366 | -0.146 | -0.269 | -0.001 | 0.035 | 0.035 | 0 | 0.52 | 0.55 | 0.44 | 0.75 | 0.41 | 0.46 |
| 9 | 137694369 | rs3827851 | -1.166 | 0.073 | -0.454 | -0.371 | -0.017 | 0.119 | 0.119 | 0 | 0.57 | 0.78 | 0.54 | 0.83 | 0.5 | 0.56 |
| 9 | 137699002 | rs4842169 | -0.064 | -0.558 | -0.478 | 0.454 | -0.019 | -0.02 | -0.02 | 2.535 | 0.37 | 0.18 | 0.31 | 0.43 | 0.35 | 0.28 |
| 9 | 137700749 | rs11103534 | -1.161 | -0.569 | 0.088 | 0.575 | -0.023 | 0.13 | 0.13 | 0.001 | 0.18 | 0.51 | 0.18 | 0.31 | 0.13 | 0.21 |
| 9 | 137700999 | rs11103535 | -0.942 | -0.53 | -0.472 | 0.089 | 0.002 | 0.001 | 0.001 | 0.005 | 0.09 | 0.11 | 0.07 | 0.12 | 0.08 | 0.14 |
| 9 | 137703923 | rs10776908 | -0.614 | -0.805 | -0.334 | 0.285 | 0 | 0 | 0 | 0.25 | 0.45 | 0.53 | 0.42 | 0.46 | 0.37 | 0.32 |
| 9 | 137704782 | rs7874142 | -1.274 | -0.718 | -1.206 | 0.415 | -0.042 | -0.015 | -0.015 | 0 | 0.58 | 0.85 | 0.5 | 0.49 | 0.51 | 0.46 |
| 9 | 137707834 | rs3827848 | 1.251 | -0.579 | -1.724 | 0.349 | 0.003 | 0 | 0 | 0 | 0.12 | 0.1 | 0.1 | 0.17 | 0.14 | 0.14 |
| 9 | 137707942 | rs4841934 | -1.397 | -0.672 | -1.979 | 0.485 | -0.058 | -0.034 | -0.034 | 0 | 0.59 | 0.9 | 0.52 | 0.49 | 0.52 | 0.46 |
| 9 | 137713649 | rs10858284 | -0.047 | -0.749 | -1.609 | 0.644 | -0.019 | -0.025 | -0.025 | 0.322 | 0.16 | 0.02 | 0.14 | 0.17 | 0.13 | 0.07 |
| 9 | 137718113 | rs4842173 | -0.469 | -0.369 | -1.289 | 0.809 | -0.009 | -0.002 | -0.002 | 0 | 0.75 | 0.8 | 0.7 | 0.75 | 0.69 | 0.82 |
| 9 | 137722170 | rs3128578 | 1.014 | -1.799 | -1.008 | 1.268 | 0.084 | -0.004 | -0.004 | 0.019 | 0.51 | 0.3 | 0.61 | 0.46 | 0.67 | 0.53 |
| 9 | 137722311 | rs4842174 | -1.154 | -1.995 | -0.54 | 1.981 | -0.035 | 0.026 | 0.026 | 0 | 0.3 | 0.34 | 0.28 | 0.48 | 0.16 | 0.41 |
| 9 | 137722644 | rs11103541 | -0.668 | -1.223 | -1.144 | 2.159 | 0.033 | -0.064 | -0.064 | 0 | 0.13 | 0.04 | 0.17 | 0.36 | 0.05 | 0.11 |
| 9 | 137724371 | rs7864699 | -1.05 | -0.707 | -0.698 | 1.609 | 0.05 | -0.062 | -0.062 | 0 | 0.21 | 0.87 | 0.25 | 0.34 | 0.09 | 0.28 |
| 9 | 137730384 | rs10858286 | 0.598 | -0.276 | 0 | 1.199 | 0 | 0 | 0 | 0 | 0.59 | 0.41 | 0.45 | 0.39 | 0.62 | 0.58 |
| 9 | 137730956 | rs9410002 | 0.886 | -0.288 | -0.565 | 1.073 | -0.016 | 0.035 | 0.035 | 0.545 | 0.53 | 0.07 | 0.4 | 0.36 | 0.59 | 0.49 |
| 9 | 137734232 | rs13946 | 0.919 | -0.216 | -0.312 | 1.424 | -0.018 | -0.05 | -0.05 | 0 | 0.8 | 0.85 | 0.62 | 0.63 | 0.78 | 0.81 |
| 9 | 137735043 | rs11103544 | -1.092 | -0.529 | 0.483 | 1.307 | -0.033 | 0.007 | 0.007 | 0 | 0.2 | 0.38 | 0.12 | 0.23 | 0.15 | 0.29 |
| 9 | 137736544 | rs4504708 | 0.649 | -0.256 | 0.515 | 1.442 | -0.031 | -0.053 | -0.053 | 0 | 0.82 | 0.87 | 0.62 | 0.64 | 0.77 | 0.81 |
| 9 | 137736650 | rs3128575 | 1.555 | -0.386 | 1.41 | 1.584 | 0.009 | 0.046 | 0.046 | 1.471 | 0.7 | 0.49 | 0.56 | 0.46 | 0.73 | 0.66 |
| 9 | 137736729 | rs4428739 | -1.027 | -0.388 | 1.091 | 1.576 | -0.023 | 0.057 | 0.057 | 0 | 0.12 | 0.27 | 0.04 | 0.19 | 0.04 | 0.15 |
| 9 | 137736913 | rs3128574 | 0.617 | -0.146 | 0.372 | 1.675 | -0.029 | -0.05 | -0.05 | 0 | 0.82 | 0.87 | 0.62 | 0.65 | 0.78 | 0.81 |
| 9 | 137737438 | rs12551466 | -0.42 | -0.148 | 0.414 | 1.372 | 0 | 0.003 | 0.003 | 0 | 0.09 | 0.09 | 0.07 | 0.19 | 0.12 | 0.12 |
| 9 | 137739980 | rs11103545 | 1.412 | -0.261 | 0.41 | 1.191 | 0.009 | 0.011 | 0.011 | 0 | 0.75 | 0.64 | 0.58 | 0.31 | 0.77 | 0.66 |
| 9 | 137743309 | rs3128570 | 0.805 | -0.234 | -0.148 | 1.034 | -0.009 | -0.013 | -0.013 | 0.716 | 0.79 | 0.83 | 0.59 | 0.67 | 0.74 | 0.76 |
| 9 | 137747707 | rs9410004 | 1.433 | -0.323 | -0.696 | 0.067 | -0.057 | 0.211 | 0.211 | 1.08 | 0.49 | 0.12 | 0.43 | 0.2 | 0.56 | 0.35 |
| 9 | 137749015 | rs7045674 | -0.497 | -0.702 | -0.442 | -0.639 | -0.004 | -0.062 | -0.062 | 0 | 0.47 | 0.51 | 0.55 | 0.93 | 0.37 | 0.57 |
| 9 | 137754467 | rs11103552 | -0.457 | -1.387 | -1.108 | -0.475 | 0.028 | 0.046 | 0.046 | 0 | 0.28 | 0.16 | 0.39 | 0.13 | 0.17 | 0.21 |
| 9 | 137754738 | rs13289166 | -0.052 | -1.454 | -1.199 | -0.595 | 0.038 | 0.039 | 0.039 | 0.098 | 0.2 | 0.09 | 0.17 | 0.01 | 0.09 | 0.14 |
| 9 | 137755163 | rs4842176 | -0.022 | -1.438 | -1.202 | -0.552 | 0.027 | 0.049 | 0.049 | 0.007 | 0.21 | 0.09 | 0.28 | 0.06 | 0.12 | 0.18 |
| 9 | 137757212 | rs4841940 | -0.407 | -1.866 | -0.808 | -0.805 | 0.014 | 0.013 | 0.013 | 1.472 | 0.26 | 0.22 | 0.29 | 0.06 | 0.15 | 0.15 |
| 9 | 137757545 | rs11103559 | -1.344 | -2.005 | 0.136 | -0.028 | -0.024 | 0.064 | 0.064 | 0 | 0.14 | 0.27 | 0.14 | 0.27 | 0.07 | 0.22 |
| 9 | 137761104 | rs4073505 | -1.762 | -2.623 | -0.427 | -0.083 | 0.104 | -0.007 | -0.007 | 0.273 | 0.54 | 0.76 | 0.49 | 0.42 | 0.28 | 0.43 |
| 9 | 137762857 | rs4436197 | -1.301 | -2.073 | 0.323 | 0.231 | -0.046 | 0.125 | 0.125 | 0 | 0.19 | 0.39 | 0.15 | 0.36 | 0.1 | 0.18 |
| 9 | 137764457 | rs6537952 | -0.256 | -0.772 | -0.372 | -0.155 | -0.008 | -0.001 | -0.001 | 0.075 | 0.75 | 0.79 | 0.73 | 0.73 | 0.67 | 0.83 |
| 9 | 137764945 | rs4347056 | -0.605 | -0.827 | -0.944 | -0.58 | -0.04 | 0.118 | 0.118 | 0.09 | 0.8 | 0.96 | 0.8 | 0.98 | 0.8 | 0.89 |
| 9 | 137767197 | rs3128632 | 0.707 | -0.659 | -0.52 | -0.117 | 0.001 | -0.002 | -0.002 | 0 | 0.42 | 0.35 | 0.31 | 0.42 | 0.43 | 0.51 |
| 9 | 137773556 | rs3128626 | 1.692 | -0.747 | -0.971 | 0.151 | -0.056 | -0.098 | -0.098 | 0 | 0.23 | 0.03 | 0.23 | 0.32 | 0.29 | 0.42 |
| 9 | 137775084 | rs7024491 | -0.202 | -0.411 | -0.381 | 0.003 | 0.002 | 0.009 | 0.009 | 0 | 0.4 | 0.53 | 0.47 | 0.56 | 0.39 | 0.39 |
| 9 | 137775139 | rs3128624 | 0.918 | -0.227 | -0.348 | 0.229 | 0.003 | -0.008 | -0.008 | 0.34 | 0.36 | 0.32 | 0.3 | 0.42 | 0.37 | 0.47 |
| 9 | 137775155 | rs4520243 | 0.91 | -0.23 | -0.349 | 0.23 | 0.003 | -0.008 | -0.008 | 0 | 0.36 | 0.32 | 0.3 | 0.41 | 0.37 | 0.47 |
| 9 | 137777504 | rs12684512 | 0.734 | 0.025 | -0.334 | 0.174 | 0.028 | 0.063 | 0.063 | 0 | 0.08 | 0.19 | 0.12 | 0.19 | 0.1 | 0.13 |
| 9 | 137779091 | rs7851696 | 0.667 | 0.022 | -0.448 | 0.161 | 0.023 | 0.053 | 0.053 | 0 | 0.08 | 0.21 | 0.12 | 0.19 | 0.1 | 0.13 |
| 9 | 137779875 | rs11103564 | 0.065 | -0.38 | -0.486 | 0.057 | -0.005 | -0.12 | -0.12 | 1.928 | 0.33 | 0.46 | 0.26 | 0.08 | 0.29 | 0.23 |
| 9 | 137780823 | rs6537958 | 0.32 | -0.373 | -0.438 | 0.426 | -0.015 | -0.056 | -0.056 | 0 | 0.3 | 0.43 | 0.21 | 0.18 | 0.24 | 0.26 |
| 9 | 137782277 | rs11103567 | 0.133 | -0.503 | -0.753 | 0.871 | 0.008 | -0.042 | -0.042 | 0 | 0.35 | 0.21 | 0.3 | 0.46 | 0.38 | 0.49 |
| 9 | 137784268 | rs10858290 | 0.025 | -0.317 | -0.545 | 0.654 | 0.004 | -0.003 | -0.003 | 0.005 | 0.35 | 0.32 | 0.35 | 0.41 | 0.31 | 0.37 |
| 9 | 137787248 | rs11103569 | -0.178 | -0.397 | 0.005 | -0.08 | -0.024 | -0.073 | -0.073 | 0 | 0.23 | 0.44 | 0.26 | 0.14 | 0.18 | 0.27 |
| 9 | 137788134 | rs4842188 | 0.813 | -0.475 | -1.014 | -0.227 | -0.119 | -0.129 | -0.129 | 0 | 0.4 | 0.04 | 0.36 | 0.43 | 0.41 | 0.51 |
| 9 | 137789793 | rs10776911 | -0.321 | -0.091 | -1.151 | -0.138 | 0.005 | -0.033 | -0.033 | 0 | 0.67 | 0.48 | 0.79 | 0.87 | 0.63 | 0.89 |
| 9 | 137790376 | rs4841944 | -0.945 | -0.288 | -0.635 | -0.157 | 0.009 | -0.025 | -0.025 | 0 | 0.73 | 0.64 | 0.82 | 0.87 | 0.7 | 0.89 |
| 9 | 137790916 | rs4384059 | 0.003 | -0.502 | -0.608 | 0.175 | 0.003 | -0.028 | -0.028 | 0 | 0.24 | 0.19 | 0.39 | 0.44 | 0.21 | 0.38 |
| 9 | 137800247 | rs1537189 | -0.129 | -0.814 | -0.989 | -0.068 | -0.067 | -0.106 | -0.106 | 0.016 | 0.66 | 0.91 | 0.47 | 0.47 | 0.62 | 0.74 |
| 9 | 137800731 | rs3012787 | -0.129 | -0.814 | -0.989 | -0.081 | -0.067 | -0.109 | -0.109 | 0.543 | 0.66 | 0.91 | 0.47 | 0.47 | 0.62 | 0.74 |
| 9 | 137801416 | rs2989735 | 0.199 | -0.961 | -0.536 | -0.145 | 0 | 0.005 | 0.005 | 0 | 0.6 | 0.56 | 0.43 | 0.43 | 0.62 | 0.55 |
| 9 | 137801800 | rs1071583 | -0.389 | -0.933 | -0.917 | -0.149 | -0.094 | -0.142 | -0.142 | 0 | 0.68 | 0.95 | 0.48 | 0.47 | 0.63 | 0.74 |
| 9 | 137809374 | rs2989726 | 1.225 | -0.547 | -1.451 | -0.313 | -0.009 | -0.024 | -0.024 | 2.673 | 0.27 | 0.18 | 0.2 | 0.3 | 0.28 | 0.4 |
| 9 | 137810259 | rs10120023 | -0.16 | -0.939 | -1.328 | -0.698 | -0.006 | 0.075 | 0.075 | 1.065 | 0.32 | 0.15 | 0.22 | 0.13 | 0.34 | 0.17 |
| 9 | 137816459 | rs2094962 | -0.581 | -1.729 | -0.722 | -0.56 | -0.027 | -0.004 | -0.004 | 0.37 | 0.3 | 0.66 | 0.11 | 0.15 | 0.11 | 0.13 |
| 9 | 137824613 | rs10858294 | -0.082 | -0.322 | -0.77 | -0.728 | -0.019 | 0.045 | 0.045 | 1.641 | 0.27 | 0.15 | 0.2 | 0.01 | 0.33 | 0.15 |
| 9 | 137829465 | rs2274838 | 0.283 | 0.173 | -1.15 | -0.082 | -0.074 | 0.028 | 0.028 | 0 | 0.57 | 0.9 | 0.37 | 0.6 | 0.48 | 0.52 |
| 9 | 137831204 | rs7849296 | 0.81 | -0.223 | -0.156 | -0.055 | 0.004 | 0.012 | 0.012 | 0 | 0.47 | 0.39 | 0.31 | 0.5 | 0.42 | 0.46 |
| 9 | 137834280 | rs17570546 | 0.718 | -0.209 | -0.569 | -0.549 | -0.047 | 0.133 | 0.133 | 0.405 | 0.22 | 0.02 | 0.14 | 0.01 | 0.24 | 0.13 |
| 9 | 137834828 | rs2274844 | 0.84 | -0.271 | 0.387 | -0.024 | -0.001 | -0.003 | -0.003 | 1.146 | 0.45 | 0.51 | 0.3 | 0.37 | 0.41 | 0.44 |
| 9 | 137835248 | rs1331429 | 0.936 | -0.377 | 0.599 | 0.009 | 0.001 | -0.005 | -0.005 | 0 | 0.42 | 0.45 | 0.27 | 0.37 | 0.42 | 0.46 |
| 9 | 137837317 | rs10119380 | 0.556 | -0.574 | 0.53 | 0.165 | -0.006 | 0.01 | 0.01 | 3.239 | 0.68 | 0.41 | 0.45 | 0.38 | 0.69 | 0.65 |
| 9 | 137840809 | rs10776914 | 0.568 | -0.58 | 0.677 | 0.211 | -0.005 | -0.001 | -0.001 | 0 | 0.63 | 0.37 | 0.44 | 0.39 | 0.67 | 0.64 |

| | | | | | | | | | | | | | | | | |
|---|---|---|---|---|---|---|---|---|---|---|---|---|---|---|---|---|
| 9 | 137842500 | rs10858300 | 0.367 | -0.642 | 0.391 | 0.708 | -0.043 | -0.061 | -0.061 | 0.851 | 0.24 | 0.07 | 0.5 | 0.38 | 0.29 | 0.26 |
| 9 | 137846865 | rs10858302 | 0.846 | -0.581 | 0.16 | 0.424 | -0.088 | 0.14 | 0.14 | 1.626 | 0.24 | 0.05 | 0.18 | 0.01 | 0.31 | 0.16 |
| 9 | 137848020 | rs12554353 | 0.41 | -0.578 | 0.197 | 0.464 | -0.023 | -0.031 | -0.031 | 0 | 0.25 | 0.13 | 0.51 | 0.37 | 0.29 | 0.26 |
| 9 | 137850460 | rs10776916 | 1.395 | -0.522 | 0.476 | 0.532 | 0.001 | 0.004 | 0.004 | 0.13 | 0.38 | 0.28 | 0.24 | 0.34 | 0.35 | 0.46 |
| 9 | 137852506 | rs7851502 | -1.157 | -0.421 | 0.572 | 0.596 | 0.006 | 0.004 | 0.004 | 0 | 0.54 | 0.45 | 0.72 | 0.47 | 0.63 | 0.5 |
| 9 | 137856025 | rs1041498 | 0.359 | -0.304 | 0.037 | 0.78 | 0 | -0.003 | -0.003 | 0.047 | 0.69 | 0.65 | 0.45 | 0.4 | 0.7 | 0.67 |
| 9 | 137856280 | rs4841950 | 0.103 | -0.147 | 0.301 | 0.641 | -0.019 | 0.156 | 0.156 | 0 | 0.93 | 0.75 | 0.96 | 0.81 | 0.98 | 0.96 |
| 9 | 137858418 | rs10858305 | 0.487 | 0.032 | -0.046 | 0.564 | -0.001 | 0.004 | 0.004 | 3.54 | 0.67 | 0.64 | 0.79 | 0.77 | 0.66 | 0.76 |
| 9 | 137864163 | rs7025292 | 0.328 | 0.21 | -0.2 | 1.29 | -0.022 | 0.168 | 0.168 | 0 | 0.93 | 0.78 | 0.97 | 0.77 | 0.98 | 0.96 |
| 9 | 137873099 | rs899833 | 0.413 | 0.787 | -0.939 | 0.721 | -0.048 | -0.053 | -0.053 | 0 | 0.81 | 0.92 | 0.54 | 0.76 | 0.74 | 0.7 |
| 9 | 137880493 | rs10858311 | -0.342 | 0.892 | -0.576 | -0.586 | 0 | -0.022 | -0.022 | 1.065 | 0.62 | 0.46 | 0.61 | 0.23 | 0.64 | 0.55 |
| 9 | 137883639 | rs10124926 | -1.103 | 1.064 | -0.297 | -0.794 | -0.007 | -0.026 | -0.026 | 0.067 | 0.73 | 0.54 | 0.69 | 0.24 | 0.76 | 0.63 |
| 9 | 137885561 | rs2279264 | -1.121 | 0.893 | -0.084 | -0.962 | -0.001 | 0.003 | 0.003 | 0 | 0.72 | 0.76 | 0.67 | 0.25 | 0.71 | 0.66 |
| 9 | 137886960 | rs13290953 | 0.511 | 0.683 | -0.062 | -1.347 | -0.021 | 0.144 | 0.144 | 1.705 | 0.44 | 0.2 | 0.47 | 0.21 | 0.45 | 0.45 |
| 9 | 137894409 | rs914392 | -0.473 | 0.931 | 0.902 | -0.993 | 0.034 | -0.035 | -0.035 | 0.492 | 0.07 | 0.25 | 0.06 | 0 | 0.07 | 0.07 |
| 9 | 137895354 | rs7046791 | -1.233 | 1.118 | 0.389 | -1.233 | -0.073 | 0.118 | 0.118 | 0.953 | 0.6 | 0.87 | 0.62 | 0.26 | 0.55 | 0.55 |
| 9 | 137899436 | rs11103625 | 0.885 | 1.538 | 0.872 | -0.477 | -0.011 | -0.012 | -0.012 | 0 | 0.41 | 0.27 | 0.34 | 0.54 | 0.38 | 0.53 |
| 9 | 137900695 | rs10448312 | -1.258 | 1.664 | 1.897 | -0.729 | 0 | -0.004 | -0.004 | 0 | 0.33 | 0.67 | 0.47 | 0.24 | 0.36 | 0.38 |
| 9 | 137901678 | rs7035792 | -2.368 | 1.028 | 2.127 | -0.489 | -0.005 | -0.081 | -0.081 | 0 | 0.15 | 0.35 | 0.08 | 0.02 | 0.1 | 0.14 |
| 9 | 137904004 | rs4841952 | -1.413 | 1.023 | 2.061 | -0.262 | -0.011 | -0.001 | -0.001 | 0.112 | 0.32 | 0.49 | 0.49 | 0.36 | 0.34 | 0.48 |
| 9 | 137906694 | rs11103631 | -1.783 | 1.048 | 1.843 | -0.231 | 0.008 | 0.009 | 0.009 | 1.56 | 0.38 | 0.68 | 0.51 | 0.37 | 0.38 | 0.52 |
| 9 | 137907348 | rs10776925 | -1.923 | 1.398 | 1.82 | -0.218 | -0.002 | 0.001 | 0.001 | 0 | 0.4 | 0.58 | 0.51 | 0.37 | 0.47 | 0.46 |
| 9 | 137907399 | rs11788133 | -0.537 | 0.594 | 1.343 | -0.054 | -0.002 | -0.003 | -0.003 | 2.117 | 0.08 | 0.08 | 0.15 | 0.11 | 0.1 | 0.09 |
| 9 | 137907690 | rs4842206 | -0.726 | 0.426 | 1.492 | -0.004 | 0 | 0 | 0 | 0 | 0.18 | 0.16 | 0.21 | 0.23 | 0.2 | 0.3 |
| 9 | 137909610 | rs6537976 | -1.486 | 0.823 | 1.53 | -0.46 | 0.057 | 0.009 | 0.009 | 0 | 0.14 | 0.32 | 0.21 | 0.11 | 0.18 | 0.12 |
| 9 | 137919491 | rs4641145 | -1.641 | 0.295 | 0.982 | -0.419 | -0.013 | -0.045 | -0.045 | 1.78 | 0.27 | 0.62 | 0.31 | 0.1 | 0.26 | 0.27 |
| 9 | 137919909 | rs7025267 | -1.193 | 0.296 | 0.288 | -0.142 | -0.02 | -0.048 | -0.048 | 0 | 0.21 | 0.35 | 0.24 | 0.1 | 0.23 | 0.25 |
| 9 | 137925300 | rs4842212 | -0.989 | 0.426 | -0.781 | 0.058 | 0 | 0.02 | 0.02 | 0.645 | 0.15 | 0.03 | 0.19 | 0.09 | 0.17 | 0.23 |
| 9 | 137932668 | rs607223 | -0.711 | 0.524 | -0.56 | -0.098 | -0.004 | 0.021 | 0.021 | 1.716 | 0.45 | 0.63 | 0.58 | 0.74 | 0.49 | 0.61 |
| 9 | 137933642 | rs17493408 | -0.121 | 0.899 | 0.124 | 0.318 | 0.026 | 0.01 | 0.01 | 0 | 0.14 | 0.25 | 0.13 | 0.2 | 0.17 | 0.2 |
| 9 | 137934787 | rs637087 | -0.52 | 0.786 | 0.455 | 0.055 | -0.01 | -0.01 | -0.01 | 0.531 | 0.3 | 0.24 | 0.45 | 0.54 | 0.37 | 0.42 |
| 9 | 137937541 | rs1009305 | -1.611 | 0.337 | 0.66 | 0.898 | 0.003 | 0.001 | 0.001 | 0 | 0.45 | 0.39 | 0.48 | 0.49 | 0.5 | 0.52 |
| 9 | 137937988 | rs664866 | -2.159 | -0.06 | 0.716 | 0.771 | -0.002 | -0.012 | -0.012 | 0 | 0.53 | 0.57 | 0.58 | 0.76 | 0.53 | 0.63 |
| 9 | 137938621 | rs7048977 | -1.041 | -0.14 | 0.401 | 0.757 | -0.039 | -0.033 | -0.033 | 0 | 0.38 | 0.21 | 0.45 | 0.49 | 0.48 | 0.5 |
| 9 | 137941500 | rs10776927 | -0.185 | -0.039 | 0.221 | 0.788 | -0.093 | -0.04 | -0.04 | 0 | 0.23 | 0.06 | 0.34 | 0.28 | 0.33 | 0.21 |
| 9 | 137941854 | rs616656 | -1.12 | -0.093 | 0.532 | 0.833 | -0.043 | 0.13 | 0.13 | 0.065 | 0.12 | 0.27 | 0.12 | 0.27 | 0.05 | 0.12 |
| 9 | 137942956 | rs17039656 | -0.711 | -0.027 | 0.978 | 0.828 | -0.001 | -0.005 | -0.005 | 0 | 0.19 | 0.38 | 0.13 | 0.2 | 0.14 | 0.21 |
| 9 | 137946707 | rs12554348 | -0.319 | 0.127 | 0.454 | 0.988 | -0.093 | -0.039 | -0.039 | 0 | 0.23 | 0.06 | 0.34 | 0.27 | 0.33 | 0.21 |
| 9 | 137963187 | rs521942 | -1.383 | -0.085 | 0.735 | 0.844 | -0.002 | -0.001 | -0.001 | 0.435 | 0.39 | 0.59 | 0.5 | 0.54 | 0.37 | 0.36 |
| 9 | 137965410 | rs9308281 | 0.792 | -0.179 | 0.383 | 0.257 | 0.01 | 0.03 | 0.03 | 0.086 | 0.47 | 0.56 | 0.37 | 0.3 | 0.45 | 0.34 |
| 9 | 137970302 | rs546682 | -0.529 | -0.433 | 0.02 | 0.267 | 0.013 | -0.009 | -0.009 | 2.843 | 0.29 | 0.21 | 0.23 | 0.29 | 0.2 | 0.24 |
| 9 | 137971918 | rs472048 | -0.236 | -0.027 | -0.211 | 0.554 | 0 | 0.007 | 0.007 | 0 | 0.13 | 0.13 | 0.3 | 0.34 | 0.13 | 0.26 |
| 9 | 137975041 | rs563961 | 1.011 | 0.19 | -0.615 | 0.23 | -0.017 | -0.036 | -0.036 | 0 | 0.44 | 0.25 | 0.36 | 0.47 | 0.5 | 0.39 |
| 9 | 137982127 | rs500461 | -0.338 | 0.42 | -0.498 | 0.643 | 0 | 0.009 | 0.009 | 0 | 0.89 | 0.88 | 0.63 | 0.68 | 0.88 | 0.76 |
| 9 | 137982621 | rs660701 | -0.769 | 0.699 | -0.942 | 0.288 | 0 | 0 | 0 | 1.621 | 0.37 | 0.38 | 0.31 | 0.52 | 0.4 | 0.43 |
| 9 | 137995557 | rs4842220 | -0.109 | 0.696 | -1.105 | 0.203 | 0.006 | -0.03 | -0.03 | 0 | 0.81 | 0.85 | 0.83 | 0.48 | 0.79 | 0.76 |
| 9 | 137995811 | rs899842 | 1.116 | 0.806 | -1.495 | -0.406 | 0.014 | 0.015 | 0.015 | 1.506 | 0.53 | 0.24 | 0.53 | 0.41 | 0.55 | 0.49 |
| 9 | 137996712 | rs11103672 | 0.087 | 1.435 | -1.667 | -0.579 | -0.014 | 0.004 | 0.004 | 0 | 0.13 | 0.06 | 0.13 | 0.09 | 0.16 | 0.14 |
| 9 | 137999847 | rs7853782 | -0.077 | 1.092 | -1.653 | -0.573 | -0.006 | 0.002 | 0.002 | 0 | 0.18 | 0.16 | 0.24 | 0.05 | 0.23 | 0.25 |
| 9 | 138001526 | rs12001599 | 0.694 | 1.104 | -1.566 | -0.547 | 0.008 | -0.044 | -0.044 | 0 | 0.11 | 0.19 | 0.07 | 0.02 | 0.14 | 0.13 |
| 9 | 138002096 | rs1536661 | -0.415 | 0.66 | -1.784 | -0.056 | -0.003 | -0.126 | -0.126 | 0.048 | 0.81 | 0.95 | 0.85 | 0.54 | 0.81 | 0.77 |
| 9 | 138002714 | rs1571332 | -0.796 | 0.596 | -1.085 | -0.487 | -0.024 | -0.087 | -0.087 | 0.336 | 0.19 | 0.52 | 0.23 | 0.05 | 0.13 | 0.17 |
| 9 | 138002918 | rs4842221 | 0.012 | 1.441 | -1.698 | -0.211 | -0.003 | 0.003 | 0.003 | 0.057 | 0.1 | 0.09 | 0.23 | 0.05 | 0.2 | 0.15 |
| 9 | 138002994 | rs4262379 | 0.186 | 0.675 | -1.996 | -0.011 | -0.017 | -0.037 | -0.037 | 0 | 0.39 | 0.16 | 0.39 | 0.42 | 0.38 | 0.38 |
| 9 | 138004530 | rs943733 | -0.06 | 0.329 | -1.589 | -0.304 | 0.009 | 0.024 | 0.024 | 0 | 0.25 | 0.22 | 0.18 | 0.13 | 0.2 | 0.13 |
| 9 | 138005260 | rs11788357 | 0.071 | 0.293 | -1.322 | -0.321 | -0.002 | -0.007 | -0.007 | 0 | 0.25 | 0.28 | 0.19 | 0.13 | 0.2 | 0.13 |
| 9 | 138005478 | rs11789656 | 0.071 | 0.293 | -1.305 | -0.324 | -0.002 | -0.009 | -0.009 | 0.437 | 0.25 | 0.28 | 0.16 | 0.12 | 0.2 | 0.13 |
| 9 | 138006871 | rs10858338 | -0.394 | 0.207 | -1.826 | -0.789 | 0.002 | 0.049 | 0.049 | 1.131 | 0.39 | 0.66 | 0.33 | 0.17 | 0.34 | 0.26 |
| 9 | 138006947 | rs11103681 | 0.715 | 0.128 | -1.973 | -0.911 | -0.044 | 0.016 | 0.016 | 0 | 0.18 | 0.03 | 0.21 | 0.12 | 0.22 | 0.18 |
| 9 | 138008243 | rs11790027 | 0.65 | -0.221 | -1.883 | -0.94 | -0.012 | -0.023 | -0.023 | 3.727 | 0.16 | 0.25 | 0.12 | 0.07 | 0.13 | 0.11 |
| 9 | 138009779 | rs11103683 | 1.157 | -0.2 | -2.053 | -0.986 | -0.006 | 0.062 | 0.062 | 0 | 0.14 | 0.05 | 0.07 | 0 | 0.1 | 0.06 |
| 9 | 138012070 | rs11103684 | 1.111 | -0.238 | -2.076 | -1.024 | 0.001 | 0.05 | 0.05 | 0 | 0.15 | 0.06 | 0.08 | 0 | 0.11 | 0.06 |
| 9 | 138012647 | rs2298249 | 0.694 | -0.383 | -2.113 | -1.058 | 0.004 | -0.002 | -0.002 | 0 | 0.17 | 0.15 | 0.2 | 0.19 | 0.14 | 0.19 |
| 9 | 138013365 | rs7035473 | 1.057 | -0.348 | -1.974 | -1.084 | 0.005 | 0.005 | 0.005 | 0 | 0.16 | 0.11 | 0.09 | 0 | 0.11 | 0.06 |
| 9 | 138013614 | rs11103685 | -0.118 | -0.349 | -2.007 | -1.161 | -0.031 | 0.108 | 0.108 | 0 | 0.33 | 0.15 | 0.21 | 0 | 0.34 | 0.16 |
| 9 | 138014017 | rs7039212 | -1.416 | -0.286 | -2.101 | -1.01 | -0.099 | 0.062 | 0.062 | 1.653 | 0.56 | 0.91 | 0.46 | 0.28 | 0.48 | 0.37 |
| 9 | 138015713 | rs647128 | -0.477 | 0.05 | -1.975 | -0.519 | 0.004 | 0.036 | 0.036 | 0 | 0.51 | 0.61 | 0.44 | 0.37 | 0.5 | 0.43 |
| 9 | 138016657 | rs11103693 | 0.578 | 0.204 | -2.174 | -0.55 | -0.036 | -0.06 | -0.06 | 0 | 0.28 | 0.07 | 0.28 | 0.37 | 0.25 | 0.33 |
| 9 | 138016676 | rs11103694 | -0.144 | -0.029 | -1.699 | -0.659 | -0.018 | -0.005 | -0.005 | 0 | 0.34 | 0.45 | 0.3 | 0.37 | 0.25 | 0.33 |

| | | | | | | | | | | | | | | | | |
|---|---|---|---|---|---|---|---|---|---|---|---|---|---|---|---|---|
| 9 | 138016698 | rs875230 | -1.586 | -0.186 | -1.841 | -0.646 | -0.013 | -0.033 | -0.033 | 0.058 | 0.68 | 0.79 | 0.65 | 0.55 | 0.63 | 0.64 |
| 9 | 138017196 | rs11103699 | -0.199 | -0.369 | -1.272 | -0.594 | -0.062 | -0.017 | -0.017 | 0 | 0.23 | 0.54 | 0.23 | 0.18 | 0.13 | 0.19 |
| 9 | 138019726 | rs12380578 | -0.261 | -0.031 | -1.651 | -0.576 | 0.003 | 0.039 | 0.039 | 0 | 0.36 | 0.26 | 0.36 | 0.2 | 0.38 | 0.31 |
| 9 | 138022634 | rs745757 | -0.804 | 0.589 | -1.917 | -0.548 | -0.007 | -0.041 | -0.041 | 0 | 0.68 | 0.82 | 0.66 | 0.55 | 0.67 | 0.67 |
| 9 | 138026999 | rs2031825 | -0.876 | 0.546 | -2.173 | -0.633 | 0.003 | -0.065 | -0.065 | 0 | 0.69 | 0.89 | 0.66 | 0.47 | 0.73 | 0.68 |
| 9 | 138027657 | rs3860249 | 0.197 | 0.499 | -2.164 | -0.678 | -0.002 | -0.003 | -0.003 | 3.185 | 0.44 | 0.23 | 0.44 | 0.46 | 0.42 | 0.52 |
| 9 | 138032691 | rs1778993 | -0.654 | 0.395 | -2.108 | -0.546 | 0.005 | -0.093 | -0.093 | 0 | 0.7 | 0.92 | 0.63 | 0.47 | 0.73 | 0.7 |
| 9 | 138033333 | rs11103711 | 0.179 | 0.612 | -1.994 | -0.585 | -0.007 | 0.032 | 0.032 | 0 | 0.47 | 0.27 | 0.42 | 0.45 | 0.45 | 0.54 |
| 9 | 138036671 | rs1778992 | 0.11 | 0.568 | -1.96 | -0.672 | 0.002 | 0.059 | 0.059 | 0.076 | 0.5 | 0.72 | 0.41 | 0.38 | 0.44 | 0.45 |
| 9 | 138037129 | rs1752664 | -0.674 | 1.364 | -1.408 | -1.485 | -0.002 | -0.037 | -0.037 | 0.856 | 0.39 | 0.36 | 0.32 | 0.05 | 0.49 | 0.23 |
| 9 | 138037372 | rs11103712 | -0.499 | 1.337 | -1.389 | -1.434 | -0.001 | -0.041 | -0.041 | 0.298 | 0.37 | 0.36 | 0.3 | 0.05 | 0.46 | 0.21 |
| 9 | 138038093 | rs17495030 | 0.41 | 1.27 | -1.394 | -1.418 | 0.022 | -0.051 | -0.051 | 0 | 0.15 | 0.17 | 0.16 | 0.01 | 0.27 | 0.1 |
| 9 | 138039129 | rs10858353 | 0.387 | 0.696 | -1.415 | -0.899 | -0.008 | 0.028 | 0.028 | 0.025 | 0.2 | 0.14 | 0.12 | 0.02 | 0.17 | 0.08 |
| 9 | 138045318 | rs1778990 | 0.181 | 0.368 | -1.567 | -0.739 | 0.019 | 0.034 | 0.034 | 0.021 | 0.19 | 0.05 | 0.11 | 0.13 | 0.16 | 0.15 |
| 9 | 138050248 | rs10858356 | -0.187 | 0.903 | -0.63 | -0.854 | 0.021 | -0.094 | -0.094 | 1.172 | 0.28 | 0.43 | 0.21 | 0.08 | 0.28 | 0.12 |
| 9 | 138053217 | rs6537988 | -0.126 | 0.473 | -0.278 | -0.112 | 0 | 0.047 | 0.047 | 0.306 | 0.77 | 0.69 | 0.55 | 0.42 | 0.73 | 0.71 |
| 9 | 138054178 | rs6537989 | -0.256 | -0.039 | -0.727 | -0.469 | 0.054 | 0.029 | 0.029 | 1.345 | 0.57 | 0.84 | 0.6 | 0.39 | 0.6 | 0.54 |
| 9 | 138056027 | rs1892001 | -0.21 | 0.299 | 0.656 | -0.562 | -0.002 | 0.026 | 0.026 | 0.307 | 0.26 | 0.58 | 0.36 | 0.36 | 0.22 | 0.35 |
| 9 | 138058487 | rs12379748 | 0.29 | 0.438 | 0.95 | -0.503 | -0.001 | -0.001 | -0.001 | 0.014 | 0.2 | 0.19 | 0.31 | 0.24 | 0.18 | 0.32 |
| 9 | 138059746 | rs7025467 | 0.726 | 0.388 | 1.115 | -0.645 | -0.001 | -0.01 | -0.01 | 0 | 0.09 | 0.08 | 0.06 | 0 | 0.07 | 0.02 |
| 9 | 138059937 | rs10858359 | -0.16 | 0.14 | 1.411 | -0.818 | -0.035 | -0.017 | -0.017 | 0 | 0.27 | 0.51 | 0.34 | 0.24 | 0.16 | 0.33 |
| 9 | 138061259 | rs7049140 | -0.148 | 0.352 | 1.244 | -0.717 | 0.027 | 0.007 | 0.007 | 0 | 0.13 | 0.34 | 0.1 | 0.11 | 0.11 | 0.06 |
| 9 | 138063959 | rs12236583 | 0.242 | 0.266 | 0.875 | -0.723 | -0.007 | 0.001 | 0.001 | 0.325 | 0.21 | 0.21 | 0.33 | 0.26 | 0.15 | 0.31 |
| 9 | 138065012 | rs7034486 | -0.975 | 0.262 | 0.872 | -0.731 | -0.002 | 0 | 0 | 0 | 0.35 | 0.61 | 0.44 | 0.36 | 0.27 | 0.38 |
| 9 | 138066092 | rs12555792 | -0.295 | 0.729 | 0.27 | -0.726 | 0.012 | 0.005 | 0.005 | 1.084 | 0.25 | 0.26 | 0.38 | 0.3 | 0.25 | 0.37 |
| 9 | 138067653 | rs2382864 | -0.265 | 0.265 | -0.438 | -0.927 | -0.003 | 0.036 | 0.036 | 0 | 0.38 | 0.27 | 0.4 | 0.25 | 0.37 | 0.41 |
| 9 | 138068124 | rs7047526 | -0.761 | 0.391 | -1.151 | -0.627 | -0.017 | 0.054 | 0.054 | 0 | 0.54 | 0.67 | 0.52 | 0.35 | 0.46 | 0.51 |
| 9 | 138072689 | rs10858363 | -0.219 | 0.137 | -1.057 | -0.77 | 0.003 | -0.025 | -0.025 | 0 | 0.17 | 0.15 | 0.08 | 0.02 | 0.14 | 0.18 |
| 9 | 138074372 | rs7849389 | 0.729 | 0.036 | -1.978 | -1.12 | 0.018 | 0.081 | 0.081 | 0 | 0.28 | 0.12 | 0.2 | 0.08 | 0.25 | 0.19 |
| 9 | 138074576 | rs1891988 | -0.334 | 0.971 | -1.973 | -0.032 | -0.021 | -0.036 | -0.036 | 0.07 | 0.76 | 0.84 | 0.69 | 0.55 | 0.67 | 0.74 |
| 9 | 138074968 | rs1891987 | 0.09 | 0.465 | -2.092 | 0.274 | -0.062 | -0.148 | -0.148 | 0 | 0.21 | 0.02 | 0.33 | 0.45 | 0.23 | 0.32 |
| 9 | 138076081 | rs10858364 | -0.453 | 0.281 | -1.532 | 0.281 | -0.011 | 0.011 | 0.011 | 0 | 0.34 | 0.4 | 0.29 | 0.39 | 0.28 | 0.26 |
| 9 | 138076167 | rs10858365 | 0.088 | 0.867 | -1.901 | 0.575 | -0.014 | -0.028 | -0.028 | 0 | 0.78 | 0.83 | 0.7 | 0.68 | 0.75 | 0.77 |
| 9 | 138077427 | rs10858366 | 0.438 | 0.217 | -1.81 | -0.357 | 0 | -0.001 | -0.001 | 0 | 0.1 | 0.09 | 0.12 | 0.11 | 0.07 | 0.17 |
| 9 | 138078679 | rs4842226 | 0.725 | 0.26 | -2.675 | -1.223 | -0.071 | 0.099 | 0.099 | 0 | 0.51 | 0.09 | 0.44 | 0.34 | 0.51 | 0.38 |
| 9 | 138081976 | rs17495922 | 0.448 | 0.159 | -2.011 | -1.746 | 0.011 | 0.037 | 0.037 | 0.008 | 0.48 | 0.39 | 0.41 | 0.24 | 0.41 | 0.28 |
| 9 | 138082201 | rs11103734 | 1.07 | 0.028 | -1.805 | -1.507 | 0.011 | 0.026 | 0.026 | 2.487 | 0.11 | 0.05 | 0.12 | 0.03 | 0.06 | 0.15 |
| 9 | 138082759 | rs11103735 | 1.171 | 0.235 | -1.898 | -1.473 | -0.031 | 0.019 | 0.019 | 0 | 0.13 | 0.03 | 0.1 | 0.05 | 0.18 | 0.12 |
| 9 | 138083351 | rs7028628 | -0.85 | -0.04 | -1.446 | -1.599 | 0.012 | 0.033 | 0.033 | 0.032 | 0.41 | 0.68 | 0.33 | 0.12 | 0.34 | 0.36 |
| 9 | 138086589 | rs1891999 | 0.955 | 0.192 | -1.761 | -1.611 | -0.022 | 0.081 | 0.081 | 0 | 0.32 | 0.05 | 0.33 | 0.24 | 0.31 | 0.29 |
| 9 | 138087879 | rs11793247 | 0.514 | 0.251 | -1 | -1.675 | 0 | -0.002 | -0.002 | 0 | 0.17 | 0.16 | 0.09 | 0.06 | 0.16 | 0.07 |
| 9 | 138091594 | rs11794621 | 0.02 | 0.595 | -0.34 | -2.071 | 0.01 | -0.081 | -0.081 | 0 | 0.22 | 0.32 | 0.11 | 0.04 | 0.22 | 0.13 |
| 9 | 138093089 | rs7023348 | 1.75 | 0.519 | -0.676 | -2.331 | -0.052 | 0.087 | 0.087 | 0 | 0.33 | 0.06 | 0.2 | 0.23 | 0.34 | 0.3 |
| 9 | 138093430 | rs6538001 | -0.517 | 0.554 | -0.226 | -2.484 | 0.006 | 0.005 | 0.005 | 0 | 0.78 | 0.74 | 0.52 | 0.33 | 0.76 | 0.62 |
| 9 | 138093775 | rs1891996 | 0.557 | 0.585 | -0.048 | -2.444 | -0.002 | -0.021 | -0.021 | 0 | 0.55 | 0.45 | 0.32 | 0.27 | 0.56 | 0.43 |
| 9 | 138093796 | rs1891995 | 1.896 | 0.612 | -0.236 | -2.393 | -0.015 | 0.084 | 0.084 | 0 | 0.33 | 0.14 | 0.21 | 0.23 | 0.34 | 0.3 |
| 9 | 138097443 | rs11103745 | -0.467 | 0.495 | 0.981 | -2.481 | 0.002 | -0.044 | -0.044 | 0 | 0.24 | 0.27 | 0.12 | 0.04 | 0.24 | 0.13 |
| 9 | 138100852 | rs12006325 | -0.024 | 0.394 | 1.113 | -2.519 | -0.013 | -0.034 | -0.034 | 0 | 0.22 | 0.25 | 0.18 | 0.06 | 0.2 | 0.19 |
| 9 | 138101987 | rs7866572 | -0.277 | 0.452 | 1.311 | -2.528 | -0.009 | -0.035 | -0.035 | 9.065 | 0.24 | 0.31 | 0.22 | 0.07 | 0.23 | 0.18 |
| 9 | 138102304 | rs11103755 | -1.345 | 0.527 | 1.237 | -2.543 | 0.02 | 0.066 | 0.066 | 0 | 0.47 | 0.66 | 0.33 | 0.11 | 0.42 | 0.35 |
| 9 | 138104368 | rs7042583 | -0.564 | 0.558 | 0.534 | -2.503 | -0.02 | -0.099 | -0.099 | 2.303 | 0.17 | 0.59 | 0.11 | 0.02 | 0.09 | 0.08 |
| 9 | 138104497 | rs1891981 | -1.433 | 0.808 | 0.058 | -2.253 | -0.05 | 0.224 | 0.224 | 0 | 0.56 | 0.91 | 0.37 | 0.11 | 0.48 | 0.37 |
| 9 | 138105075 | rs1891980 | -0.982 | 0.877 | 0.686 | -1.647 | 0.007 | 0.008 | 0.008 | 0.062 | 0.35 | 0.67 | 0.26 | 0.1 | 0.28 | 0.21 |
| 9 | 138114573 | rs7022353 | 0.652 | 2.228 | 0.413 | -0.897 | -0.016 | -0.005 | -0.005 | 0 | 0.81 | 0.87 | 0.61 | 0.76 | 0.74 | 0.77 |
| 9 | 138114837 | rs7043651 | -0.577 | 2.102 | 1.289 | -1.095 | 0.097 | 0.086 | 0.086 | 1.87 | 0.07 | 0.39 | 0.04 | 0.03 | 0.06 | 0.07 |
| 9 | 138120574 | rs7854386 | 1.577 | 2.376 | 1.368 | -1.193 | 0.004 | -0.001 | -0.001 | 0 | 0.7 | 0.34 | 0.51 | 0.71 | 0.61 | 0.61 |
| 9 | 138123236 | rs7875079 | 2.003 | 2.221 | 1.024 | -1.618 | 0.105 | 0.089 | 0.089 | 0.896 | 0.51 | 0.25 | 0.38 | 0.69 | 0.34 | 0.55 |
| 9 | 138125763 | rs7043655 | -1.115 | 2.545 | 1.134 | -1.636 | 0.042 | -0.077 | -0.077 | 0 | 0.3 | 0.57 | 0.26 | 0.05 | 0.44 | 0.15 |
| 9 | 138126646 | rs12380243 | -0.026 | 2.578 | 1.148 | -1.634 | -0.03 | 0.062 | 0.062 | 0 | 0.2 | 0.09 | 0.14 | 0.01 | 0.28 | 0.07 |
| 9 | 138127492 | rs10858372 | -0.596 | 2.574 | 1.433 | -1.638 | 0.001 | -0.034 | -0.034 | 0.006 | 0.24 | 0.26 | 0.21 | 0.01 | 0.38 | 0.09 |
| 9 | 138128536 | rs7865457 | 0.518 | 2.422 | 1.338 | -1.595 | 0.004 | 0.002 | 0.002 | 0.064 | 0.8 | 0.81 | 0.64 | 0.74 | 0.77 | 0.69 |
| 9 | 138128603 | rs7038247 | 1.988 | 2.384 | 1.312 | -1.6 | 0.107 | 0.09 | 0.09 | 0.909 | 0.51 | 0.25 | 0.38 | 0.69 | 0.33 | 0.54 |
| 9 | 138129028 | rs4842231 | 1.095 | 2.418 | 1.279 | -1.591 | 0.047 | 0 | 0 | 0 | 0.76 | 0.64 | 0.57 | 0.74 | 0.67 | 0.67 |
| 9 | 138129933 | rs11103764 | -0.38 | 2.566 | 1.211 | -1.367 | 0.014 | -0.021 | -0.021 | 0.405 | 0.24 | 0.31 | 0.18 | 0.06 | 0.33 | 0.13 |
| 9 | 138134313 | rs1891983 | -0.435 | 2.517 | 0.574 | -1.479 | 0.078 | -0.024 | -0.024 | 0 | 0.06 | 0.22 | 0.08 | 0 | 0.11 | 0.03 |
| 9 | 138135660 | rs1106259 | -0.102 | 1.943 | 0.042 | -1.444 | -0.001 | -0.001 | -0.001 | 0 | 0.87 | 0.89 | 0.82 | 0.76 | 0.85 | 0.81 |
| 9 | 138136568 | rs1106261 | 1.429 | 1.675 | -0.377 | -1.551 | 0.041 | 0.049 | 0.049 | 0.959 | 0.5 | 0.38 | 0.54 | 0.69 | 0.36 | 0.51 |
| 9 | 138137715 | rs2382859 | 1.261 | 1.728 | -0.514 | -1.35 | 0.008 | 0.02 | 0.02 | 0.212 | 0.55 | 0.39 | 0.57 | 0.7 | 0.41 | 0.53 |
| 9 | 138139506 | rs10858374 | -0.859 | 0.56 | -1.2 | -1.009 | 0.066 | 0.139 | 0.139 | 0.09 | 0.38 | 0.82 | 0.18 | 0.09 | 0.23 | 0.3 |

| | | | | | | | | | | | | | | | | |
|---|---|---|---|---|---|---|---|---|---|---|---|---|---|---|---|---|
| 9 | 138139987 | rs4842232 | -0.032 | 1.106 | -1.003 | -0.191 | -0.034 | -0.044 | -0.044 | 0 | 0.93 | 1 | 0.84 | 0.77 | 0.89 | 0.85 |
| 9 | 138143145 | rs7033668 | 0.423 | 0.695 | -0.702 | -0.415 | -0.015 | -0.013 | -0.013 | 3.539 | 0.64 | 0.53 | 0.7 | 0.7 | 0.67 | 0.66 |
| 9 | 138146809 | rs11792631 | -0.921 | 0.707 | -1.128 | -0.796 | 0.005 | 0 | 0 | 1.003 | 0.16 | 0.09 | 0.07 | 0 | 0.11 | 0.08 |
| 9 | 138149166 | rs7858253 | 0.612 | 0.753 | -1.754 | -0.69 | 0.026 | 0.054 | 0.054 | 0.135 | 0.46 | 0.27 | 0.53 | 0.67 | 0.39 | 0.53 |
| 9 | 138149317 | rs7858399 | 0.534 | 1.311 | -1.182 | -0.608 | 0.007 | -0.01 | -0.01 | 0.004 | 0.67 | 0.44 | 0.73 | 0.79 | 0.61 | 0.72 |
| 9 | 138151500 | rs12347149 | 0.471 | 1.847 | -1.45 | -0.219 | -0.054 | -0.01 | -0.01 | 0.348 | 0.8 | 0.86 | 0.74 | 0.81 | 0.64 | 0.74 |
| 9 | 138155396 | rs11103782 | 0.31 | 0.638 | -1.343 | -0.109 | 0.005 | 0.007 | 0.007 | 3.408 | 0.68 | 0.7 | 0.69 | 0.27 | 0.73 | 0.55 |
| 9 | 138156321 | rs7038150 | 0.448 | 0.504 | -1.724 | 0.502 | -0.003 | -0.031 | -0.031 | 0.14 | 0.63 | 0.91 | 0.64 | 0.57 | 0.65 | 0.55 |
| 9 | 138159376 | rs10858377 | -0.007 | 0.662 | -0.926 | 0.337 | 0.017 | 0.016 | 0.016 | 0 | 0.74 | 0.66 | 0.53 | 0.65 | 0.67 | 0.62 |
| 9 | 138160077 | rs2989741 | 0.945 | 0.268 | -0.801 | -0.207 | 0.002 | 0.015 | 0.015 | 0 | 0.46 | 0.59 | 0.43 | 0.64 | 0.44 | 0.47 |
| 9 | 138164038 | rs1778982 | -0.41 | 0.265 | -0.944 | -0.27 | -0.061 | -0.034 | -0.034 | 1.138 | 0.76 | 0.87 | 0.54 | 0.66 | 0.64 | 0.65 |
| 9 | 138165193 | rs1778981 | 1.186 | -0.637 | -0.308 | -0.116 | 0.001 | -0.002 | -0.002 | 0 | 0.37 | 0.42 | 0.3 | 0.41 | 0.31 | 0.25 |
| 9 | 138165503 | rs1757350 | 0.818 | -0.215 | -0.144 | 0.228 | -0.022 | -0.022 | -0.022 | 0 | 0.57 | 0.25 | 0.4 | 0.44 | 0.52 | 0.4 |
| 9 | 138166444 | rs11103786 | -1.298 | -0.535 | 0.827 | 0.181 | 0.06 | 0.019 | 0.019 | 0.038 | 0.07 | 0.33 | 0.03 | 0.01 | 0.03 | 0.02 |
| 9 | 138170334 | rs11103787 | -0.575 | -0.454 | 0.305 | 0.406 | -0.025 | 0.062 | 0.062 | 0 | 0.07 | 0.13 | 0.07 | 0.2 | 0.04 | 0.21 |
| 9 | 138171694 | rs914405 | -1.243 | -0.921 | -0.03 | -0.303 | 0.014 | 0.005 | 0.005 | 0 | 0.38 | 0.64 | 0.28 | 0.35 | 0.24 | 0.45 |
| 9 | 138172097 | rs1106182 | -0.104 | -0.965 | 0.643 | -0.36 | 0.012 | -0.02 | -0.02 | 0 | 0.06 | 0.26 | 0.06 | 0 | 0.05 | 0.07 |
| 9 | 138172789 | rs7026842 | -1.569 | -1.154 | 0.573 | -0.742 | -0.014 | -0.012 | -0.012 | 0.064 | 0.41 | 0.48 | 0.3 | 0.34 | 0.32 | 0.43 |
| 9 | 138173238 | rs11103789 | -1.562 | -1.17 | 0.504 | -0.735 | 0 | 0 | 0 | 0 | 0.39 | 0.34 | 0.26 | 0.35 | 0.3 | 0.53 |
| 9 | 138173409 | rs11103792 | -0.513 | -1.024 | 0.538 | -0.603 | -0.025 | 0.037 | 0.037 | 0 | 0.19 | 0.25 | 0.19 | 0.32 | 0.13 | 0.25 |
| 9 | 138174163 | rs7041675 | -0.544 | -0.96 | 0.51 | -0.656 | -0.018 | 0.029 | 0.029 | 0 | 0.15 | 0.19 | 0.11 | 0.26 | 0.13 | 0.12 |
| 9 | 138174978 | rs1806807 | -1.651 | -1 | 0.604 | -0.676 | -0.008 | -0.005 | -0.005 | 0 | 0.35 | 0.62 | 0.18 | 0.27 | 0.24 | 0.25 |
| 9 | 138175847 | rs7874626 | -0.425 | -0.937 | 0.537 | -0.583 | -0.02 | 0.044 | 0.044 | 0 | 0.14 | 0.22 | 0.08 | 0.24 | 0.07 | 0.07 |
| 9 | 138176724 | rs914407 | -1.591 | -0.94 | 0.07 | -0.652 | 0.016 | 0.014 | 0.014 | 0 | 0.39 | 0.68 | 0.19 | 0.29 | 0.27 | 0.37 |
| 9 | 138180621 | rs4842244 | 0.99 | -0.378 | -0.401 | -0.918 | 0.012 | 0.031 | 0.031 | 0 | 0.76 | 0.8 | 0.77 | 0.85 | 0.74 | 0.77 |
| 9 | 138181228 | rs11103796 | -0.493 | 0.175 | -0.419 | -1.073 | -0.003 | 0.007 | 0.007 | 0 | 0.24 | 0.15 | 0.24 | 0.11 | 0.21 | 0.26 |
| 9 | 138181270 | rs10776941 | 0.804 | 0.2 | -0.559 | -1.037 | 0.01 | 0.031 | 0.031 | 0 | 0.76 | 0.8 | 0.77 | 0.85 | 0.74 | 0.76 |
| 9 | 138182186 | rs10745401 | 1.347 | 0.334 | -0.372 | -0.883 | -0.001 | -0.009 | -0.009 | 0 | 0.49 | 0.5 | 0.53 | 0.72 | 0.52 | 0.47 |
| 9 | 138182939 | rs7026673 | 0.318 | 0.014 | -1.265 | -0.698 | 0.005 | 0 | 0 | 0 | 0.87 | 0.91 | 0.81 | 0.85 | 0.84 | 0.82 |
| 9 | 138184453 | rs7867539 | -0.979 | 0.516 | -1.282 | -0.763 | 0.01 | 0.102 | 0.102 | 0 | 0.36 | 0.17 | 0.25 | 0.1 | 0.31 | 0.32 |
| 9 | 138188447 | rs7853412 | -0.845 | 1.27 | -0.54 | -0.019 | 0.019 | -0.015 | -0.015 | 0 | 0.22 | 0.28 | 0.2 | 0.14 | 0.24 | 0.18 |
| 9 | 138189561 | rs7869911 | -0.298 | 1.036 | -1.617 | -0.051 | -0.063 | -0.014 | -0.014 | 0 | 0.27 | 0.06 | 0.31 | 0.23 | 0.33 | 0.27 |
| 9 | 138189578 | rs1778986 | -0.305 | 1.072 | -0.689 | -0.451 | 0.013 | -0.058 | -0.058 | 0.25 | 0.1 | 0.26 | 0.09 | 0 | 0.1 | 0.05 |
| 9 | 138191389 | rs10776946 | -0.121 | 1.317 | -1.066 | -0.092 | -0.001 | -0.002 | -0.002 | 0 | 0.59 | 0.6 | 0.64 | 0.61 | 0.61 | 0.54 |
| 9 | 138193283 | rs10858383 | 0.176 | 1.23 | -1.089 | -0.18 | 0 | 0 | 0 | 0.022 | 0.55 | 0.58 | 0.52 | 0.61 | 0.52 | 0.5 |
| 9 | 138195203 | rs4841962 | -0.072 | 1.244 | -1.214 | -0.107 | -0.022 | 0.02 | 0.02 | 1.911 | 0.6 | 0.75 | 0.57 | 0.66 | 0.57 | 0.62 |
| 9 | 138195529 | rs10858385 | -0.06 | 1.281 | -1.216 | -0.089 | -0.016 | 0.022 | 0.022 | 2.254 | 0.59 | 0.72 | 0.57 | 0.66 | 0.57 | 0.62 |
| 9 | 138197209 | rs3884535 | 0.067 | 0.886 | -0.781 | 0.29 | 0.007 | -0.002 | -0.002 | 0 | 0.63 | 0.75 | 0.5 | 0.65 | 0.63 | 0.69 |
| 9 | 138199317 | rs4842247 | 0.461 | 0.469 | -0.394 | 0.312 | 0.017 | 0.02 | 0.02 | 0 | 0.42 | 0.26 | 0.25 | 0.37 | 0.34 | 0.48 |
| 9 | 138202418 | rs7032559 | 0.753 | -0.041 | -0.108 | 0.097 | 0.012 | -0.031 | -0.031 | 0 | 0.4 | 0.16 | 0.32 | 0.49 | 0.35 | 0.51 |
| 9 | 138210649 | rs7047604 | 0.447 | -0.086 | 0.222 | -0.058 | -0.042 | -0.007 | -0.007 | 0.011 | 0.31 | 0.18 | 0.48 | 0.32 | 0.39 | 0.3 |
| 9 | 138211368 | rs4841964 | -0.639 | 0.167 | 0.414 | 0.037 | 0 | -0.001 | -0.001 | 0 | 0.84 | 0.84 | 0.85 | 0.81 | 0.78 | 0.84 |
| 9 | 138211783 | rs4842249 | -0.283 | 0.095 | 0.565 | 0.065 | -0.001 | 0.005 | 0.005 | 0 | 0.41 | 0.68 | 0.52 | 0.32 | 0.4 | 0.3 |
| 9 | 138219743 | rs4842251 | -0.048 | 0.22 | 0.091 | 0.121 | -0.034 | 0.011 | 0.011 | 0 | 0.36 | 0.23 | 0.48 | 0.32 | 0.4 | 0.3 |
| 9 | 138219780 | rs4841967 | -0.138 | 0.198 | 0.489 | 0.111 | -0.006 | 0.003 | 0.003 | 0 | 0.37 | 0.33 | 0.48 | 0.32 | 0.4 | 0.3 |
| 9 | 138221647 | rs11103820 | -0.114 | 0.195 | 0.862 | 0.13 | 0.012 | -0.008 | -0.008 | 0 | 0.36 | 0.46 | 0.51 | 0.32 | 0.4 | 0.28 |
| 9 | 138222860 | rs12555689 | -0.054 | 0.218 | 0.475 | 0.142 | -0.036 | 0.003 | 0.003 | 0.003 | 0.35 | 0.23 | 0.48 | 0.32 | 0.4 | 0.28 |
| 9 | 138229678 | rs4842253 | 0.224 | -0.054 | 0.834 | 0.275 | -0.002 | -0.003 | -0.003 | 0 | 0.81 | 0.67 | 0.84 | 0.77 | 0.77 | 0.73 |
| 9 | 138232307 | rs6538017 | 0.118 | -0.018 | 0.889 | 0.285 | 0.004 | 0.001 | 0.001 | 0.28 | 0.81 | 0.67 | 0.83 | 0.77 | 0.77 | 0.73 |
| 9 | 138236305 | rs4842254 | 0.593 | -0.372 | 0.324 | 0.078 | 0.016 | -0.011 | -0.011 | 0 | 0.39 | 0.36 | 0.29 | 0.48 | 0.29 | 0.4 |
| 9 | 138237120 | rs11103828 | -0.021 | -0.106 | 0.024 | 0.125 | -0.029 | 0.007 | 0.007 | 0 | 0.39 | 0.23 | 0.53 | 0.34 | 0.51 | 0.38 |
| 9 | 138239887 | rs6538018 | 0.413 | -0.262 | 0.237 | 0.099 | 0.054 | 0.01 | 0.01 | 0 | 0.46 | 0.65 | 0.34 | 0.39 | 0.3 | 0.4 |
| 9 | 138241815 | rs7039860 | 0.678 | -0.056 | -0.006 | 0.094 | 0.062 | -0.036 | -0.036 | 0 | 0.44 | 0.16 | 0.34 | 0.39 | 0.36 | 0.42 |
| 9 | 138243149 | rs7036094 | 0.411 | 0.031 | 1.129 | -0.035 | 0.006 | 0.013 | 0.013 | 0.448 | 0.81 | 0.67 | 0.85 | 0.75 | 0.78 | 0.75 |
| 9 | 138244411 | rs9409933 | 1.466 | 0.24 | 0.976 | -0.056 | 0.003 | 0.002 | 0.002 | 0 | 0.62 | 0.4 | 0.55 | 0.57 | 0.58 | 0.63 |
| 9 | 138245050 | rs11789038 | -0.408 | 0.398 | 0.362 | -0.097 | 0.005 | -0.03 | -0.03 | 0 | 0.12 | 0.14 | 0.22 | 0 | 0.15 | 0.09 |
| 9 | 138246910 | rs7467652 | -0.095 | 0.302 | 0.042 | 0.057 | -0.051 | -0.058 | -0.058 | 0.15 | 0.84 | 0.96 | 0.87 | 0.77 | 0.78 | 0.76 |
| 9 | 138252877 | rs7853278 | 1.591 | 0.129 | 0.505 | -0.087 | -0.001 | -0.001 | -0.001 | 0 | 0.59 | 0.39 | 0.54 | 0.58 | 0.58 | 0.62 |
| 9 | 138258205 | rs1571823 | 0.946 | 0.127 | 0.642 | -0.078 | 0 | 0.099 | 0.099 | 0.032 | 0.75 | 0.53 | 0.77 | 0.6 | 0.73 | 0.72 |
| 9 | 138263731 | rs7040000 | -1.368 | -0.033 | 1.023 | -0.416 | -0.014 | -0.044 | -0.044 | 0.033 | 0.25 | 0.5 | 0.34 | 0.16 | 0.2 | 0.13 |
| 9 | 138270241 | rs11103850 | -1.196 | -0.043 | 0.94 | -0.33 | 0.021 | 0.006 | 0.006 | 0.008 | 0.19 | 0.5 | 0.33 | 0.16 | 0.18 | 0.14 |
| 9 | 138275572 | rs11789484 | -1.374 | -0.039 | 0.644 | -0.636 | -0.004 | -0.029 | -0.029 | 0 | 0.3 | 0.67 | 0.36 | 0.13 | 0.24 | 0.18 |
| 9 | 138288052 | rs9409884 | -0.727 | -0.37 | 0.831 | -0.678 | -0.067 | -0.069 | -0.069 | 0 | 0.12 | 0.23 | 0.02 | 0 | 0.01 | 0.01 |
| 9 | 138292538 | rs13294986 | -1.151 | 0.278 | 0.432 | -0.752 | -0.001 | -0.022 | -0.022 | 0 | 0.43 | 0.52 | 0.42 | 0.22 | 0.42 | 0.43 |
| 9 | 138298164 | rs9409948 | -1.23 | 0.125 | 0.361 | -0.746 | -0.001 | -0.004 | -0.004 | 1.333 | 0.44 | 0.58 | 0.4 | 0.22 | 0.4 | 0.44 |
| 9 | 138306060 | rs6537893 | -1.153 | 0.526 | 0.738 | -0.793 | 0.016 | -0.078 | -0.078 | 0.555 | 0.12 | 0.39 | 0.07 | 0 | 0.12 | 0.03 |
| 9 | 138312393 | rs7019162 | -0.965 | 0.593 | 0.171 | -0.852 | -0.002 | -0.103 | -0.103 | 0 | 0.25 | 0.61 | 0.29 | 0.01 | 0.2 | 0.24 |
| 9 | 138318759 | rs12377350 | -0.365 | 0.386 | -0.829 | -0.07 | 0.031 | 0.036 | 0.036 | 3.991 | 0.41 | 0.28 | 0.65 | 0.31 | 0.66 | 0.47 |
| 9 | 138324679 | rs2384792 | -0.466 | 0.878 | -0.614 | 0.696 | 0.014 | 0.008 | 0.008 | 0 | 0.37 | 0.46 | 0.5 | 0.45 | 0.44 | 0.42 |

| | | | | | | | | | | | | | | | | |
|---|---|---|---|---|---|---|---|---|---|---|---|---|---|---|---|---|
| 9 | 138325937 | rs4841983 | -0.316 | 1.014 | -0.383 | 1.086 | 0.005 | 0.006 | 0.006 | 0 | 0.3 | 0.43 | 0.43 | 0.27 | 0.34 | 0.27 |
| 9 | 138327242 | rs10776810 | -0.613 | 0.846 | -0.505 | 0.857 | 0 | 0 | 0 | 0 | 0.38 | 0.48 | 0.46 | 0.27 | 0.39 | 0.35 |
| 9 | 138332424 | rs10858106 | -0.304 | 0.91 | -1.229 | 1.46 | -0.035 | -0.085 | -0.085 | 0 | 0.83 | 0.95 | 0.82 | 0.59 | 0.79 | 0.7 |
| 9 | 138337411 | rs10858108 | -0.297 | 0.927 | -1.218 | 1.796 | -0.037 | -0.084 | -0.084 | 0 | 0.83 | 0.95 | 0.82 | 0.58 | 0.79 | 0.72 |
| 9 | 138340774 | rs10776811 | -0.147 | 0.801 | -0.669 | 1.158 | 0 | 0.002 | 0.002 | 0 | 0.34 | 0.32 | 0.37 | 0.13 | 0.36 | 0.24 |
| 9 | 138343253 | rs4841986 | -0.084 | 0.558 | -0.259 | 1.606 | -0.012 | 0.017 | 0.017 | 0 | 0.74 | 0.56 | 0.68 | 0.39 | 0.79 | 0.65 |
| 9 | 138345912 | rs895191 | -0.015 | 0.543 | -0.825 | 1.339 | -0.009 | 0.002 | 0.002 | 0.176 | 0.65 | 0.45 | 0.73 | 0.6 | 0.66 | 0.66 |
| 9 | 138349162 | rs10776815 | 0.156 | 0.432 | -0.788 | 0.585 | -0.001 | 0.002 | 0.002 | 0.019 | 0.54 | 0.41 | 0.66 | 0.36 | 0.53 | 0.51 |
| 9 | 138355370 | rs4841875 | -0.052 | 0.588 | -1.459 | -0.301 | 0 | 0.146 | 0.146 | 1.165 | 0.46 | 0.25 | 0.38 | 0.12 | 0.51 | 0.47 |
| 9 | 138355410 | rs4841876 | 0.116 | 0.578 | -1.414 | -0.196 | -0.036 | 0.112 | 0.112 | 0.385 | 0.41 | 0.24 | 0.33 | 0.11 | 0.44 | 0.47 |
| 9 | 138357705 | rs10745363 | 0.155 | 0.317 | -1.308 | 0.137 | -0.01 | 0.007 | 0.007 | 1.028 | 0.61 | 0.61 | 0.71 | 0.57 | 0.64 | 0.64 |
| 9 | 138359002 | rs17040232 | -0.493 | 0.469 | -1.64 | -0.164 | -0.005 | -0.006 | -0.006 | 0.184 | 0.41 | 0.48 | 0.37 | 0.23 | 0.32 | 0.39 |
| 9 | 138361170 | rs10858115 | -0.377 | 0.829 | -2.746 | -0.355 | 0.004 | 0.096 | 0.096 | 0 | 0.36 | 0.13 | 0.34 | 0.12 | 0.31 | 0.33 |
| 9 | 138363933 | rs3748191 | 0.055 | 0.7 | -3.1 | -0.31 | -0.012 | 0.047 | 0.047 | 1.175 | 0.29 | 0.04 | 0.31 | 0.13 | 0.23 | 0.32 |
| 9 | 138365495 | rs2590504 | 0.251 | 0.803 | -3.521 | -0.155 | -0.113 | -0.02 | -0.02 | 0 | 0.47 | 0.95 | 0.59 | 0.6 | 0.5 | 0.48 |
| 9 | 138366254 | rs895192 | -0.494 | 0.799 | -3.041 | -0.342 | -0.037 | -0.049 | -0.049 | 0 | 0.81 | 0.99 | 0.93 | 0.78 | 0.8 | 0.78 |
| 9 | 138370613 | rs3762066 | 0.47 | 1.4 | -3.055 | -0.698 | -0.088 | 0.081 | 0.081 | 0 | 0.18 | 0.02 | 0.17 | 0.07 | 0.26 | 0.12 |
| 9 | 138372737 | rs3789911 | -0.562 | 0.267 | -3.162 | -0.497 | -0.017 | -0.019 | -0.019 | 0.276 | 0.67 | 0.98 | 0.88 | 0.73 | 0.74 | 0.78 |
| 9 | 138373864 | rs1412512 | -0.331 | -0.009 | -1.678 | 0.2 | -0.006 | 0.003 | 0.003 | 0 | 0.26 | 0.29 | 0.49 | 0.65 | 0.21 | 0.51 |
| 9 | 138375861 | rs2254813 | -0.448 | 0.023 | -1.557 | 0.144 | -0.023 | 0.014 | 0.014 | 0 | 0.27 | 0.37 | 0.52 | 0.64 | 0.21 | 0.49 |
| 9 | 138383213 | rs7026686 | -0.3 | 0.598 | -2.07 | -0.198 | -0.004 | 0.037 | 0.037 | 0 | 0.43 | 0.13 | 0.39 | 0.66 | 0.37 | 0.56 |
| 9 | 138386597 | rs4841991 | -0.337 | 0.797 | -0.71 | -0.752 | 0.001 | -0.001 | -0.001 | 0 | 0.48 | 0.49 | 0.51 | 0.84 | 0.43 | 0.48 |
| 9 | 138389740 | rs3762067 | -0.746 | 0.873 | -0.322 | -0.92 | -0.008 | 0.12 | 0.12 | 0.042 | 0.54 | 0.7 | 0.59 | 0.9 | 0.49 | 0.59 |
| 9 | 138391517 | rs2274651 | -0.166 | 0.855 | 0.157 | -0.906 | 0.003 | 0.031 | 0.031 | 0 | 0.48 | 0.37 | 0.5 | 0.83 | 0.43 | 0.48 |
| 9 | 138395969 | rs3748199 | -0.566 | 0.964 | 0.636 | -0.798 | 0.084 | 0.064 | 0.064 | 0 | 0.07 | 0.36 | 0.08 | 0.07 | 0.06 | 0.12 |
| 9 | 138396198 | rs3748201 | -0.571 | 0.958 | 0.663 | -0.801 | 0.064 | 0.046 | 0.046 | 0 | 0.07 | 0.31 | 0.08 | 0.07 | 0.06 | 0.12 |
| 9 | 138396251 | rs535 | -0.79 | 0.809 | -0.22 | -1.127 | -0.078 | 0.275 | 0.275 | 0.145 | 0.54 | 0.91 | 0.6 | 0.9 | 0.49 | 0.64 |
| 9 | 138404555 | rs2188210 | -1.392 | 0.856 | 0.607 | -0.922 | 0.015 | -0.035 | -0.035 | 0.284 | 0.16 | 0.64 | 0.26 | 0.09 | 0.14 | 0.25 |
| 9 | 138406444 | rs739395 | -1.372 | 0.956 | 0.494 | -0.897 | 0.057 | -0.027 | -0.027 | 0 | 0.14 | 0.49 | 0.27 | 0.09 | 0.14 | 0.26 |
| 9 | 138406460 | rs756684 | -1.326 | 0.97 | 0.35 | -0.89 | 0.055 | -0.025 | -0.025 | 0 | 0.14 | 0.47 | 0.26 | 0.09 | 0.14 | 0.26 |
| 9 | 138406565 | rs13290613 | -0.605 | 1.249 | -1.011 | -0.424 | -0.041 | -0.044 | -0.044 | 0 | 0.14 | 0.01 | 0.16 | 0.13 | 0.15 | 0.09 |
| 9 | 138417830 | rs13284515 | -0.765 | 1.143 | -1.058 | -0.425 | -0.035 | -0.024 | -0.024 | 0.051 | 0.15 | 0.01 | 0.15 | 0.12 | 0.14 | 0.09 |
| 9 | 138425308 | rs13295025 | -0.952 | 1.265 | -0.723 | -0.594 | -0.006 | 0.001 | 0.001 | 0 | 0.13 | 0.14 | 0.12 | 0.15 | 0.16 | 0.21 |
| 9 | 138428645 | rs210081 | 0.406 | 1.423 | -1.247 | -1.115 | -0.07 | 0.27 | 0.27 | 0 | 0.4 | 0.07 | 0.41 | 0.02 | 0.44 | 0.28 |
| 9 | 138437163 | rs462537 | 0.115 | 1.469 | -0.672 | -0.464 | -0.004 | 0.067 | 0.067 | 0 | 0.46 | 0.26 | 0.6 | 0.67 | 0.46 | 0.57 |
| 9 | 138437736 | rs7045084 | -0.453 | 1.37 | -0.102 | -0.408 | 0.008 | 0.113 | 0.113 | 0 | 0.07 | 0.27 | 0.13 | 0.13 | 0.06 | 0.09 |
| 9 | 138440045 | rs2590498 | 0.036 | 1.749 | -1.112 | -0.384 | -0.004 | 0.052 | 0.052 | 0 | 0.43 | 0.24 | 0.57 | 0.69 | 0.42 | 0.58 |
| 9 | 138445655 | rs10858121 | 0.001 | 1.773 | -1.171 | -0.267 | -0.004 | 0.05 | 0.05 | 0 | 0.43 | 0.22 | 0.58 | 0.66 | 0.44 | 0.56 |
| 9 | 138447855 | rs7023309 | 0.593 | 1.86 | -0.725 | 0.102 | 0.055 | -0.011 | -0.011 | 0.983 | 0.7 | 0.53 | 0.75 | 0.72 | 0.55 | 0.71 |
| 9 | 138450193 | rs11103062 | -0.549 | 0.694 | -1.41 | -0.259 | -0.01 | 0.062 | 0.062 | 0 | 0.18 | 0.01 | 0.12 | 0.09 | 0.16 | 0.12 |
| 9 | 138452026 | rs10858123 | -0.648 | 0.742 | -0.417 | 0.792 | 0.034 | 0.429 | 0.429 | 0 | 0.07 | 0.28 | 0.26 | 0.49 | 0.07 | 0.21 |
| 9 | 138454301 | rs3748206 | -0.771 | -0.166 | -1.214 | -0.443 | 0.052 | 0.206 | 0.206 | 0.01 | 0.42 | 0.14 | 0.23 | 0.11 | 0.31 | 0.32 |
| 9 | 138455746 | rs783776 | 0.036 | -0.258 | -0.835 | -0.108 | 0.022 | 0.468 | 0.468 | 0 | 0.93 | 0.7 | 0.75 | 0.49 | 0.95 | 0.8 |
| 9 | 138456727 | rs11103063 | 0.658 | 0.249 | -1.615 | -0.679 | -0.072 | 0 | 0 | 0.264 | 0.15 | 0.01 | 0.13 | 0.07 | 0.17 | 0.27 |
| 9 | 138460445 | rs783771 | 0.073 | -0.577 | -1.286 | -0.695 | 0.027 | 0.065 | 0.065 | 0 | 0.27 | 0.16 | 0.12 | 0.04 | 0.18 | 0.25 |
| 9 | 138467959 | rs697449 | 0.234 | -0.984 | -0.434 | -0.661 | 0 | -0.022 | -0.022 | 0 | 0.13 | 0.16 | 0.1 | 0.04 | 0.14 | 0.15 |
| 9 | 138468394 | rs705669 | 0.697 | -1.288 | -0.502 | -0.802 | -0.005 | 0 | 0 | 0 | 0.62 | 0.59 | 0.59 | 0.37 | 0.72 | 0.62 |
| 9 | 138469460 | rs3811133 | 0.404 | -1.194 | -0.483 | -0.816 | -0.008 | -0.042 | -0.042 | 0 | 0.67 | 0.56 | 0.81 | 0.87 | 0.7 | 0.7 |
| 9 | 138473753 | rs10776827 | 1.502 | -1.014 | -0.613 | -0.752 | 0.031 | 0.058 | 0.058 | 0 | 0.5 | 0.36 | 0.5 | 0.26 | 0.58 | 0.45 |
| 9 | 138474785 | rs4842007 | 0.2 | -0.747 | -0.571 | -0.831 | -0.003 | 0.053 | 0.053 | 0 | 0.71 | 0.83 | 0.83 | 0.96 | 0.71 | 0.72 |
| 9 | 138476823 | rs7029507 | 1.048 | -0.31 | -0.49 | -0.63 | -0.002 | 0.066 | 0.066 | 0 | 0.45 | 0.31 | 0.47 | 0.22 | 0.53 | 0.45 |
| 9 | 138480486 | rs3849218 | 0.871 | -0.358 | -0.264 | -0.568 | 0.014 | 0.035 | 0.035 | 0 | 0.49 | 0.36 | 0.46 | 0.28 | 0.56 | 0.47 |
| 9 | 138482453 | rs2151376 | -0.654 | -0.383 | -0.262 | -0.159 | 0.004 | 0.015 | 0.015 | 0 | 0.26 | 0.19 | 0.16 | 0.12 | 0.22 | 0.22 |
| 9 | 138482709 | rs4842008 | 0.59 | -0.214 | -0.179 | 0.095 | -0.014 | -0.02 | -0.02 | 0.097 | 0.71 | 0.6 | 0.78 | 0.75 | 0.76 | 0.69 |
| 9 | 138490710 | rs7873626 | 0.109 | -0.163 | -0.594 | 0.7 | 0.033 | 0.014 | 0.014 | 0 | 0.59 | 0.31 | 0.7 | 0.64 | 0.69 | 0.72 |
| 9 | 138496035 | rs11103084 | -0.562 | 1.397 | 0.687 | 1.606 | 0.031 | 0.025 | 0.025 | 0 | 0.12 | 0.19 | 0.11 | 0.2 | 0.17 | 0.1 |
| 9 | 138499206 | rs4842012 | 0.379 | 1.26 | 0.412 | 1.593 | -0.002 | 0.015 | 0.015 | 0 | 0.72 | 0.39 | 0.67 | 0.67 | 0.71 | 0.8 |
| 9 | 138503416 | rs872005 | -0.352 | 1.199 | 0.566 | 1.662 | 0.007 | 0.029 | 0.029 | 0 | 0.07 | 0.1 | 0.16 | 0.2 | 0.09 | 0.03 |
| 9 | 138505985 | rs11792145 | 0.964 | 0.923 | 0.429 | 1.656 | 0.004 | 0.126 | 0.126 | 1.676 | 0.76 | 0.54 | 0.73 | 0.56 | 0.75 | 0.67 |
| 9 | 138510121 | rs4841889 | -1.367 | 0.967 | 0.049 | 1.45 | 0.018 | 0.021 | 0.021 | 0.189 | 0.49 | 0.63 | 0.4 | 0.58 | 0.55 | 0.51 |
| 9 | 138513956 | rs11103105 | -0.478 | 0.591 | 0.039 | 1.26 | -0.005 | 0.026 | 0.026 | 0.054 | 0.06 | 0.1 | 0.09 | 0.08 | 0.07 | 0.1 |
| 9 | 138515929 | rs12336965 | -1.384 | 0.496 | -0.049 | 1.06 | -0.008 | 0.001 | 0.001 | 0 | 0.21 | 0.26 | 0.24 | 0.22 | 0.2 | 0.18 |
| 9 | 138516119 | rs17040344 | -0.358 | 0.58 | -0.287 | 0.843 | -0.01 | 0.053 | 0.053 | 0 | 0.09 | 0.02 | 0.04 | 0 | 0.07 | 0.06 |
| 9 | 138518114 | rs11103106 | -0.457 | 0.555 | -0.25 | 0.337 | 0 | -0.001 | -0.001 | 1.445 | 0.46 | 0.43 | 0.53 | 0.6 | 0.48 | 0.61 |
| 9 | 138518827 | rs10858139 | -0.081 | 0.707 | -0.382 | 0.194 | 0.007 | -0.003 | -0.003 | 0 | 0.37 | 0.43 | 0.49 | 0.6 | 0.41 | 0.56 |
| 9 | 138519360 | rs7037835 | 1.04 | 0.421 | -0.878 | -0.459 | 0.005 | 0.09 | 0.09 | 0 | 0.34 | 0.13 | 0.28 | 0.21 | 0.31 | 0.21 |
| 9 | 138523332 | rs1333234 | -1.488 | 0.335 | -0.714 | -0.563 | -0.001 | 0.018 | 0.018 | 0 | 0.68 | 0.78 | 0.81 | 0.94 | 0.72 | 0.8 |
| 9 | 138525778 | rs10858143 | 0.318 | 0.151 | -0.761 | -0.399 | 0.022 | 0.002 | 0.002 | 0.02 | 0.35 | 0.51 | 0.47 | 0.6 | 0.39 | 0.47 |
| 9 | 138527271 | rs11103115 | -0.347 | 0.05 | -1.417 | -0.866 | 0.013 | 0.028 | 0.028 | 0.335 | 0.27 | 0.07 | 0.23 | 0.18 | 0.24 | 0.31 |

| | | | | | | | | | | | | | | | | |
|---|---|---|---|---|---|---|---|---|---|---|---|---|---|---|---|---|
| 9 | 138527656 | rs11103116 | 1.764 | 0.017 | -1.521 | -0.926 | -0.014 | 0.127 | 0.127 | 0 | 0.33 | 0.13 | 0.19 | 0.05 | 0.29 | 0.2 |
| 9 | 138529219 | rs7855164 | -0.031 | 0.067 | -1.349 | -0.738 | 0.016 | -0.038 | -0.038 | 0.04 | 0.36 | 0.53 | 0.54 | 0.76 | 0.43 | 0.5 |
| 9 | 138529777 | rs1537414 | 0.65 | -0.095 | -1.422 | -0.132 | -0.001 | 0.004 | 0.004 | 0 | 0.68 | 0.36 | 0.66 | 0.43 | 0.68 | 0.62 |
| 9 | 138536400 | rs13286163 | -0.664 | 0.344 | -1.199 | -0.376 | 0 | -0.003 | -0.003 | 1.03 | 0.08 | 0.09 | 0.07 | 0.02 | 0.08 | 0.09 |
| 9 | 138537416 | rs11103123 | 0.606 | 0.332 | -1.479 | -0.451 | -0.032 | -0.03 | -0.03 | 0 | 0.15 | 0.02 | 0.14 | 0.17 | 0.16 | 0.08 |
| 9 | 138540992 | rs13293756 | -0.965 | 0.614 | -1.066 | -0.234 | 0.035 | 0.046 | 0.046 | 0 | 0.22 | 0.15 | 0.14 | 0.06 | 0.15 | 0.22 |
| 9 | 138544017 | rs488948 | 1.038 | 1.176 | -0.542 | -0.011 | -0.005 | 0.095 | 0.095 | 0.76 | 0.53 | 0.3 | 0.38 | 0.23 | 0.49 | 0.33 |
| 9 | 138545090 | rs7020804 | -0.029 | 1.422 | -0.778 | 0.153 | -0.001 | 0 | 0 | 0 | 0.05 | 0.06 | 0.09 | 0.02 | 0.08 | 0.08 |
| 9 | 138552234 | rs541131 | 0.456 | 1.586 | -0.571 | 0.467 | 0.033 | 0.011 | 0.011 | 0 | 0.66 | 0.55 | 0.57 | 0.39 | 0.59 | 0.43 |
| 9 | 138555161 | rs10776838 | -0.257 | 1.757 | -0.654 | 0.703 | -0.058 | -0.035 | -0.035 | 0.004 | 0.13 | 0.02 | 0.15 | 0.17 | 0.19 | 0.11 |
| 9 | 138560311 | rs514157 | 1.381 | 1.17 | 0.296 | 0.537 | -0.025 | -0.044 | -0.044 | 0 | 0.84 | 0.68 | 0.86 | 0.92 | 0.92 | 0.92 |
| 9 | 138560560 | rs510386 | 1.354 | 1.164 | 0.544 | 0.534 | -0.021 | -0.039 | -0.039 | 0 | 0.85 | 0.71 | 0.87 | 0.92 | 0.92 | 0.92 |
| 9 | 138567059 | rs480554 | 1.712 | 1.142 | 1.11 | 0.467 | -0.023 | -0.062 | -0.062 | 0.8 | 0.83 | 0.53 | 0.82 | 0.91 | 0.91 | 0.91 |
| 9 | 138576462 | rs480224 | 1.527 | 1.29 | 1.074 | 0.458 | 0.006 | -0.064 | -0.064 | 0.092 | 0.88 | 0.68 | 0.92 | 0.98 | 0.93 | 0.93 |
| 9 | 138580775 | rs480422 | -0.348 | 2.303 | 1.471 | 1.382 | 0.073 | 0.147 | 0.147 | 0 | 0.1 | 0.27 | 0.26 | 0.28 | 0.19 | 0.23 |
| 9 | 138580847 | rs528472 | 1.467 | 2.344 | 1.002 | 1.294 | 0.001 | 0.009 | 0.009 | 4.015 | 0.75 | 0.73 | 0.63 | 0.44 | 0.69 | 0.63 |
| 9 | 138581671 | rs13284789 | -0.501 | 2.229 | 0.918 | 1.181 | -0.039 | -0.034 | -0.034 | 0 | 0.07 | 0 | 0.1 | 0.12 | 0.16 | 0.17 |
| 9 | 138584199 | rs485779 | 1.261 | 1.955 | 1.226 | 1.105 | 0.018 | -0.044 | -0.044 | 0.138 | 0.91 | 0.75 | 0.93 | 0.97 | 0.95 | 0.96 |
| 9 | 138586309 | rs484524 | 1.691 | 1.927 | 1.39 | 1.071 | 0.032 | -0.06 | -0.06 | 0 | 0.89 | 0.63 | 0.92 | 0.97 | 0.94 | 0.95 |
| 9 | 138591230 | rs558113 | 2.155 | 2.013 | 1 | 1.4 | 0.004 | 0.021 | 0.021 | 0 | 0.85 | 0.77 | 0.83 | 0.66 | 0.91 | 0.89 |
| 9 | 138592785 | rs523140 | 1.077 | 1.715 | 1.016 | 0.895 | 0.003 | -0.031 | -0.031 | 0 | 0.91 | 0.85 | 0.94 | 0.97 | 0.95 | 0.96 |
| 9 | 138595441 | rs571842 | 1.027 | 1.562 | 0.959 | 0.828 | -0.002 | -0.037 | -0.037 | 0 | 0.89 | 0.83 | 0.94 | 0.97 | 0.95 | 0.95 |
| 9 | 138596974 | rs496503 | 1.832 | 1.492 | 0.943 | 1.118 | -0.022 | 0.008 | 0.008 | 0.013 | 0.84 | 0.72 | 0.84 | 0.84 | 0.94 | 0.92 |
| 9 | 138608747 | rs10858154 | -1.388 | 1.923 | 0.559 | 0.91 | -0.015 | 0.01 | 0.01 | 1.445 | 0.16 | 0.15 | 0.16 | 0.12 | 0.23 | 0.29 |
| 9 | 138610028 | rs492361 | -0.356 | 1.731 | 0.555 | 0.745 | 0.008 | 0.044 | 0.044 | 0 | 0.27 | 0.36 | 0.4 | 0.58 | 0.35 | 0.43 |
| 9 | 138615966 | rs498618 | -1.17 | 1.363 | 0.327 | 0.638 | 0.007 | 0.02 | 0.02 | 0 | 0.48 | 0.63 | 0.62 | 0.76 | 0.62 | 0.73 |
| 9 | 138617303 | rs1537416 | 0.564 | 1.746 | -0.018 | 0.808 | 0.009 | 0.001 | 0.001 | 0 | 0.6 | 0.44 | 0.49 | 0.36 | 0.53 | 0.47 |
| 9 | 138620891 | rs11103147 | -1.036 | 1.393 | -0.172 | 0.747 | 0.007 | 0.008 | 0.008 | 0 | 0.07 | 0.14 | 0.09 | 0.11 | 0.12 | 0.18 |
| 9 | 138628816 | rs10121357 | 1.188 | 1.319 | -0.251 | 0.654 | -0.005 | 0.004 | 0.004 | 0 | 0.84 | 0.78 | 0.84 | 0.88 | 0.8 | 0.78 |
| 9 | 138640273 | rs7020956 | -1.512 | 1.315 | 0.116 | 1.243 | 0.056 | 0.211 | 0.211 | 0.085 | 0.09 | 0.25 | 0.19 | 0.43 | 0.14 | 0.33 |
| 9 | 138641780 | rs11103161 | -1.05 | 1.283 | 0.141 | 1.13 | 0.081 | 0.059 | 0.059 | 0 | 0.07 | 0.31 | 0.12 | 0.11 | 0.13 | 0.2 |
| 9 | 138643920 | rs7037014 | -0.138 | 1.082 | -0.267 | 0.42 | -0.001 | -0.046 | -0.046 | 0.073 | 0.36 | 0.34 | 0.21 | 0.07 | 0.31 | 0.33 |
| 9 | 138644203 | rs10735239 | -0.492 | 0.872 | -0.308 | 0.748 | -0.001 | -0.004 | -0.004 | 0 | 0.46 | 0.55 | 0.42 | 0.51 | 0.46 | 0.65 |
| 9 | 138662309 | rs10776844 | -0.253 | 0.401 | -1.031 | 0.537 | 0.008 | 0.017 | 0.017 | 0 | 0.66 | 0.78 | 0.65 | 0.71 | 0.71 | 0.83 |
| 9 | 138668073 | rs1318383 | -0.25 | -0.067 | -0.935 | 0.349 | 0.006 | 0.008 | 0.008 | 0 | 0.69 | 0.72 | 0.69 | 0.78 | 0.74 | 0.84 |
| 9 | 138669261 | rs914428 | -0.128 | -0.343 | -0.539 | 0.627 | 0.017 | 0.005 | 0.005 | 1.388 | 0.52 | 0.55 | 0.69 | 0.57 | 0.68 | 0.82 |
| 9 | 138669976 | rs10776845 | -0.937 | 0.041 | 0.13 | 0.704 | -0.009 | -0.004 | -0.004 | 0.229 | 0.24 | 0.41 | 0.16 | 0.23 | 0.21 | 0.21 |
| 9 | 138672652 | rs1556417 | -0.923 | -0.071 | 0.26 | 0.757 | -0.02 | 0.009 | 0.009 | 0 | 0.26 | 0.42 | 0.22 | 0.27 | 0.21 | 0.28 |
| 9 | 138695235 | rs7852055 | -1.266 | 0.098 | 0.953 | 0.932 | -0.003 | 0.002 | 0.002 | 0 | 0.3 | 0.63 | 0.21 | 0.35 | 0.25 | 0.37 |
| 9 | 138701523 | rs7838 | -0.193 | 0.104 | 1.629 | 0.851 | -0.001 | -0.025 | -0.025 | 0.198 | 0.13 | 0.14 | 0.08 | 0 | 0.13 | 0.15 |
| 9 | 138706844 | rs3739465 | -2.125 | -0.178 | 1.625 | 1.005 | -0.001 | 0 | 0 | 0 | 0.33 | 0.64 | 0.22 | 0.44 | 0.27 | 0.4 |
| 9 | 138707074 | rs3739464 | -0.316 | -0.141 | 1.852 | 0.962 | 0.02 | 0.134 | 0.134 | 0 | 0.1 | 0.22 | 0.1 | 0.33 | 0.11 | 0.18 |
| 9 | 138711942 | rs12554173 | -0.161 | -0.141 | 2.131 | 0.945 | 0.001 | 0.036 | 0.036 | 0 | 0.09 | 0.08 | 0.1 | 0.33 | 0.11 | 0.18 |
| 9 | 138718981 | rs10114302 | -0.442 | -0.14 | 2.148 | 0.947 | 0.027 | 0.168 | 0.168 | 0 | 0.11 | 0.29 | 0.11 | 0.33 | 0.11 | 0.18 |
| 9 | 138756168 | rs13299821 | 0.368 | 1.046 | 1.537 | 0.697 | 0.027 | -0.001 | -0.001 | 0.178 | 0.12 | 0.14 | 0.22 | 0.1 | 0.23 | 0.24 |
| 9 | 138761762 | rs11792453 | 0.334 | 1.08 | 1.532 | 0.665 | 0.029 | -0.002 | -0.002 | 1.284 | 0.12 | 0.14 | 0.22 | 0.09 | 0.23 | 0.24 |
| 9 | 138772579 | rs11793168 | 0.542 | 1.064 | 1.543 | 0.668 | -0.003 | -0.001 | -0.001 | 3.81 | 0.11 | 0.08 | 0.22 | 0.09 | 0.23 | 0.24 |
| 9 | 138775419 | rs7861677 | -0.325 | 0.997 | 1.537 | 0.618 | 0.009 | -0.087 | -0.087 | 0.042 | 0.15 | 0.31 | 0.1 | 0 | 0.13 | 0.14 |
| 9 | 138777312 | rs1009473 | -1.401 | 0.951 | 1.212 | 0.652 | 0.02 | -0.044 | -0.044 | 0.027 | 0.34 | 0.59 | 0.38 | 0.14 | 0.4 | 0.47 |
| 9 | 138781516 | rs3119680 | -1.848 | 1.208 | 0.449 | 0.657 | 0.035 | -0.039 | -0.039 | 0 | 0.47 | 0.85 | 0.53 | 0.52 | 0.54 | 0.66 |
| 9 | 138803846 | rs4842042 | 0.744 | 0.834 | -0.729 | 0.313 | -0.027 | -0.013 | -0.013 | 0 | 0.19 | 0.01 | 0.22 | 0.14 | 0.22 | 0.16 |
| 9 | 138813713 | rs11103218 | -1.409 | 1.264 | 0.242 | 0.642 | 0.011 | 0.009 | 0.009 | 0 | 0.28 | 0.54 | 0.24 | 0.34 | 0.28 | 0.34 |
| 9 | 138816252 | rs10114289 | -0.846 | 1.248 | 0.168 | 0.293 | 0.025 | -0.114 | -0.114 | 0.322 | 0.15 | 0.45 | 0.1 | 0 | 0.13 | 0.08 |
| 9 | 138816796 | rs4842047 | 1.014 | 1.315 | -0.469 | 0.677 | 0.003 | 0.007 | 0.007 | 0 | 0.71 | 0.42 | 0.75 | 0.62 | 0.74 | 0.66 |
| 9 | 138817142 | rs11103226 | 0.649 | 1.355 | -1.169 | 0.439 | -0.01 | -0.023 | -0.023 | 0 | 0.53 | 0.24 | 0.54 | 0.48 | 0.56 | 0.47 |
| 9 | 138817178 | rs10776854 | 0.711 | 1.577 | -0.282 | 0.261 | 0.048 | -0.058 | -0.058 | 0 | 0.74 | 0.53 | 0.66 | 0.86 | 0.68 | 0.69 |
| 9 | 138818500 | rs7030382 | 0.483 | 0.949 | -0.554 | 0.579 | 0 | -0.002 | -0.002 | 0 | 0.53 | 0.48 | 0.57 | 0.53 | 0.55 | 0.49 |
| 9 | 138822770 | rs10858181 | 0.087 | 0.746 | -0.578 | 0.581 | -0.013 | 0.02 | 0.02 | 0.062 | 0.73 | 0.81 | 0.76 | 0.78 | 0.73 | 0.77 |
| 9 | 138824310 | rs10745376 | 0.095 | 0.648 | -0.495 | 0.538 | -0.001 | 0 | 0 | 0 | 0.74 | 0.67 | 0.76 | 0.78 | 0.75 | 0.79 |
| 9 | 138825117 | rs6569 | 1.166 | 0.683 | -0.407 | 0.817 | 0.001 | -0.002 | -0.002 | 0 | 0.53 | 0.48 | 0.57 | 0.5 | 0.55 | 0.49 |
| 9 | 138837113 | rs10776856 | 1.156 | 0.683 | -0.416 | 0.802 | 0.001 | -0.001 | -0.001 | 0 | 0.53 | 0.48 | 0.57 | 0.5 | 0.55 | 0.49 |
| 9 | 138838305 | rs2297690 | 1.94 | 0.781 | -0.538 | 0.52 | 0.003 | 0.111 | 0.111 | 0.163 | 0.35 | 0.13 | 0.24 | 0.12 | 0.32 | 0.21 |
| 9 | 138850610 | rs7868264 | 0.229 | 0.989 | -0.337 | 0.775 | -0.001 | 0.003 | 0.003 | 0 | 0.72 | 0.65 | 0.74 | 0.78 | 0.73 | 0.77 |
| 9 | 138862766 | rs4842059 | 1.447 | 0.975 | 0.047 | 0.853 | -0.002 | -0.007 | -0.007 | 0 | 0.42 | 0.39 | 0.29 | 0.26 | 0.36 | 0.39 |
| 9 | 138865612 | rs17040586 | -0.027 | 1.19 | 0.178 | 0.983 | 0.051 | 0.076 | 0.076 | 0.112 | 0.11 | 0.22 | 0.25 | 0.22 | 0.17 | 0.14 |
| 9 | 138866836 | rs11789650 | 0.049 | 1.231 | -0.344 | 0.887 | -0.08 | -0.065 | -0.065 | 1.948 | 0.14 | 0.01 | 0.15 | 0.24 | 0.18 | 0.14 |
| 9 | 138867842 | rs4841903 | 2.045 | 1.094 | -0.188 | 0.591 | 0.014 | 0.081 | 0.081 | 0.243 | 0.35 | 0.17 | 0.25 | 0.16 | 0.31 | 0.26 |
| 9 | 138875850 | rs2152174 | -0.78 | 1.165 | -0.186 | 0.518 | -0.043 | 0.008 | 0.008 | 0 | 0.42 | 0.25 | 0.45 | 0.64 | 0.48 | 0.5 |
| 9 | 138876604 | rs4842064 | 1.909 | 0.961 | 0.228 | 0.308 | 0.015 | 0.053 | 0.053 | 1.02 | 0.37 | 0.25 | 0.27 | 0.11 | 0.32 | 0.26 |

| | | | | | | | | | | | | | | | | |
|---|---|---|---|---|---|---|---|---|---|---|---|---|---|---|---|---|
| 9 | 138877266 | rs1475784 | -0.411 | 1.101 | -0.062 | 0.468 | -0.001 | -0.001 | -0.001 | 1.269 | 0.73 | 0.69 | 0.7 | 0.74 | 0.72 | 0.68 |
| 9 | 138877724 | rs7875089 | -1.525 | 0.962 | -0.628 | 0.315 | -0.003 | -0.035 | -0.035 | 2.769 | 0.56 | 0.57 | 0.72 | 0.88 | 0.66 | 0.7 |
| 9 | 138878884 | rs1475783 | 0.092 | 1.091 | -0.615 | 0.812 | -0.019 | -0.014 | -0.014 | 0 | 0.72 | 0.88 | 0.71 | 0.74 | 0.71 | 0.64 |
| 9 | 138878949 | rs10858194 | -0.639 | 1.205 | 0.324 | 0.859 | -0.017 | -0.052 | -0.052 | 0 | 0.1 | 0.26 | 0.05 | 0.01 | 0.04 | 0.05 |
| 9 | 138880743 | rs870164 | -1.106 | 1.865 | -0.59 | 1.376 | -0.069 | -0.068 | -0.068 | 0 | 0.1 | 0.01 | 0.14 | 0.35 | 0.21 | 0.12 |
| 9 | 138884700 | rs1932090 | 1.027 | 1.033 | 0.127 | 1.082 | -0.005 | 0.046 | 0.046 | 0 | 0.78 | 0.56 | 0.79 | 0.65 | 0.76 | 0.75 |
| 9 | 138892195 | rs10858197 | 0.448 | 1.055 | -0.586 | 0.658 | 0.009 | 0.009 | 0.009 | 0 | 0.55 | 0.38 | 0.42 | 0.41 | 0.48 | 0.4 |
| 9 | 138893874 | rs7869795 | 0.954 | 1.212 | -0.129 | 0.883 | 0.044 | 0.041 | 0.041 | 0 | 0.66 | 0.51 | 0.54 | 0.41 | 0.5 | 0.53 |
| 9 | 138894400 | rs7867262 | 0.83 | 0.438 | 0.536 | 0.474 | 0.051 | 0.042 | 0.042 | 0.038 | 0.92 | 0.65 | 0.87 | 0.94 | 0.92 | 0.83 |
| 9 | 138896566 | rs11103259 | 1.041 | 0.205 | 0.462 | 0.403 | -0.059 | -0.034 | -0.034 | 0 | 0.82 | 0.54 | 0.78 | 0.88 | 0.89 | 0.7 |
| 9 | 138900839 | rs10858200 | -0.518 | 0.269 | 0.081 | 0.329 | 0.007 | 0.008 | 0.008 | 0.564 | 0.15 | 0.12 | 0.13 | 0.06 | 0.08 | 0.07 |
| 9 | 138905136 | rs2280486 | -1.291 | 1.071 | 0.575 | 0.647 | 0.006 | 0.002 | 0.002 | 0 | 0.37 | 0.55 | 0.47 | 0.45 | 0.43 | 0.4 |
| 9 | 138908250 | rs2275107 | 0.027 | 1.052 | 0.736 | 0.546 | -0.041 | -0.041 | -0.041 | 0 | 0.17 | 0.33 | 0.22 | 0.09 | 0.12 | 0.18 |
| 9 | 138908957 | rs11103274 | 1.976 | 0.814 | -0.481 | 0.241 | 0.01 | 0.127 | 0.127 | 0.196 | 0.33 | 0.08 | 0.19 | 0.12 | 0.29 | 0.27 |
| 9 | 138911431 | rs10776870 | -0.405 | 1.484 | 0.472 | 0.617 | 0.007 | 0.004 | 0.004 | 0 | 0.17 | 0.14 | 0.24 | 0.36 | 0.3 | 0.2 |
| 9 | 138913314 | rs4842073 | -1.688 | 1.529 | -0.022 | 0.603 | -0.02 | -0.009 | -0.009 | 0.722 | 0.38 | 0.67 | 0.47 | 0.45 | 0.44 | 0.4 |
| 9 | 138913404 | rs4842074 | -0.655 | 1.086 | 0.483 | 0.004 | -0.036 | -0.049 | -0.049 | 0 | 0.18 | 0.52 | 0.23 | 0.09 | 0.14 | 0.18 |
| 9 | 138923780 | rs4409473 | 0.682 | 2.304 | 0.144 | 0.453 | 0.06 | -0.004 | -0.004 | 0 | 0.71 | 0.51 | 0.65 | 0.73 | 0.64 | 0.71 |
| 9 | 138947167 | rs4842091 | -0.919 | 2.427 | -0.425 | 0.687 | -0.034 | -0.051 | -0.051 | 1.533 | 0.19 | 0.09 | 0.33 | 0.42 | 0.26 | 0.24 |
| 9 | 138955201 | rs2385188 | 1.016 | 1.667 | -0.586 | -0.094 | 0.047 | -0.015 | -0.015 | 0 | 0.4 | 0.21 | 0.34 | 0.53 | 0.3 | 0.59 |
| 9 | 138955246 | rs871095 | 0.653 | 1.876 | 0.064 | -0.156 | -0.005 | 0.066 | 0.066 | 0 | 0.75 | 0.53 | 0.8 | 0.56 | 0.81 | 0.8 |
| 9 | 138958552 | rs11103302 | -0.48 | 2.346 | 0.26 | -0.535 | 0.002 | -0.044 | -0.044 | 0 | 0.34 | 0.36 | 0.43 | 0.08 | 0.49 | 0.19 |
| 9 | 138964892 | rs10776875 | 1.029 | 2.57 | 1.133 | 0.147 | 0.011 | 0.008 | 0.008 | 0 | 0.63 | 0.57 | 0.39 | 0.52 | 0.51 | 0.48 |
| 9 | 138965868 | rs10776877 | 1.441 | 3.139 | 1.718 | 0.182 | 0.127 | -0.013 | -0.013 | 0 | 0.88 | 0.44 | 0.75 | 0.94 | 0.78 | 0.75 |
| 9 | 138972104 | rs4842108 | -0.855 | 1.829 | 1.422 | 0.149 | 0.005 | -0.02 | -0.02 | 0 | 0.3 | 0.19 | 0.37 | 0.45 | 0.26 | 0.26 |
| 9 | 138974889 | rs7468351 | 1.319 | 2.088 | 1.544 | 0.177 | -0.024 | -0.02 | -0.02 | 0 | 0.61 | 0.27 | 0.48 | 0.55 | 0.6 | 0.7 |
| 9 | 138985434 | rs1717453 | -1.518 | 1.998 | 1.248 | 0.358 | 0.07 | 0.029 | 0.029 | 0.995 | 0.07 | 0.57 | 0.12 | 0.02 | 0.07 | 0.02 |
| 9 | 138988568 | rs7019243 | -1.106 | 2.099 | 0.058 | 0.443 | 0.004 | 0.006 | 0.006 | 0 | 0.31 | 0.37 | 0.4 | 0.43 | 0.34 | 0.26 |
| 9 | 138992466 | rs7030214 | -0.717 | 1.737 | -0.304 | 0.627 | -0.004 | -0.027 | -0.027 | 0 | 0.25 | 0.18 | 0.34 | 0.42 | 0.26 | 0.24 |
| 9 | 138999710 | rs1164893 | -0.443 | 1.739 | -0.016 | 0.32 | 0.045 | -0.008 | -0.008 | 0 | 0.28 | 0.53 | 0.26 | 0.27 | 0.33 | 0.25 |
| 9 | 139000752 | rs1620864 | -0.904 | 1.246 | -1.133 | 0.403 | -0.07 | -0.059 | -0.059 | 0 | 0.75 | 0.94 | 0.77 | 0.7 | 0.68 | 0.81 |
| 9 | 139015411 | rs1747839 | 0.779 | 0.179 | -1.537 | 0.286 | -0.001 | -0.058 | -0.058 | 0.115 | 0.17 | 0.07 | 0.24 | 0.31 | 0.17 | 0.34 |
| 9 | 139016326 | rs10858225 | 0.863 | -0.084 | -1.538 | -0.021 | 0.041 | 0.007 | 0.007 | 2.701 | 0.18 | 0.25 | 0.21 | 0.2 | 0.29 | 0.22 |
| 9 | 139018847 | rs1164884 | -1.584 | -0.283 | -1.263 | -0.127 | -0.057 | -0.034 | -0.034 | 0 | 0.71 | 0.84 | 0.68 | 0.66 | 0.63 | 0.75 |
| 9 | 139019445 | rs1164883 | -0.609 | -0.421 | -0.695 | -0.184 | 0 | 0 | 0 | 0 | 0.44 | 0.46 | 0.47 | 0.55 | 0.47 | 0.57 |
| 9 | 139020613 | rs1164882 | -0.895 | -0.397 | -0.918 | -0.198 | -0.012 | 0.008 | 0.008 | 0.17 | 0.49 | 0.79 | 0.51 | 0.55 | 0.49 | 0.62 |
| 9 | 139025615 | rs10858226 | -1.645 | -0.321 | -0.728 | -0.031 | -0.033 | -0.028 | -0.028 | 0.824 | 0.73 | 0.8 | 0.77 | 0.66 | 0.68 | 0.8 |
| 9 | 139029017 | rs1747846 | 0.149 | -0.102 | -0.387 | 0.056 | 0.016 | 0.055 | 0.055 | 0.028 | 0.45 | 0.25 | 0.39 | 0.3 | 0.44 | 0.36 |
| 9 | 139035609 | rs11103350 | 0.298 | -0.035 | -0.222 | 0.061 | 0.001 | 0.01 | 0.01 | 0.016 | 0.41 | 0.56 | 0.42 | 0.28 | 0.43 | 0.4 |
| 9 | 139048452 | rs11792632 | 0.282 | 0.138 | 0.018 | 0.293 | 0 | 0.11 | 0.11 | 0.16 | 0.19 | 0.34 | 0.32 | 0.37 | 0.18 | 0.38 |
| 9 | 139048974 | rs17038777 | -0.199 | 0.094 | -0.034 | 0.259 | -0.001 | 0.002 | 0.002 | 0.186 | 0.23 | 0.2 | 0.35 | 0.37 | 0.23 | 0.42 |
| 9 | 139049366 | rs7858442 | 0.956 | 0.214 | 0.03 | 0.324 | 0.014 | 0.01 | 0.01 | 0.132 | 0.1 | 0.05 | 0.01 | 0.06 | 0.04 | 0.03 |
| 9 | 139049385 | rs9409914 | 0.824 | 0.314 | 0.096 | 0.232 | 0 | -0.003 | -0.003 | 0.01 | 0.32 | 0.32 | 0.39 | 0.23 | 0.42 | 0.33 |
| 9 | 139050961 | rs912496 | -0.297 | 0.312 | 0.104 | 0.22 | 0.001 | 0.004 | 0.004 | 0.171 | 0.49 | 0.43 | 0.62 | 0.6 | 0.58 | 0.66 |
| 9 | 139051266 | rs748750 | -0.161 | 0.54 | -0.071 | 0.324 | -0.001 | 0.001 | 0.001 | 0.18 | 0.42 | 0.4 | 0.46 | 0.34 | 0.45 | 0.4 |
| 9 | 139061474 | rs4842123 | -0.277 | 1.296 | -1.095 | 0.925 | -0.003 | 0.026 | 0.026 | 0 | 0.53 | 0.73 | 0.7 | 0.6 | 0.58 | 0.7 |
| 9 | 139069713 | rs11103366 | 0.075 | 1.295 | -1.533 | 0.88 | -0.029 | 0.08 | 0.08 | 0.053 | 0.44 | 0.07 | 0.45 | 0.32 | 0.44 | 0.57 |
| 9 | 139077343 | rs4842126 | -0.917 | 1.876 | 0.225 | 1.592 | 0.142 | 0.247 | 0.247 | 0.069 | 0.14 | 0.44 | 0.28 | 0.32 | 0.21 | 0.15 |
| 9 | 139082796 | rs10858244 | -0.528 | 1.474 | 0.426 | 1.466 | -0.002 | 0.032 | 0.032 | 0 | 0.33 | 0.53 | 0.42 | 0.56 | 0.31 | 0.27 |
| 9 | 139083019 | rs11103372 | 0.112 | 1.439 | -0.02 | 1.036 | -0.006 | -0.047 | -0.047 | 0 | 0.24 | 0.03 | 0.13 | 0.17 | 0.16 | 0.17 |
| 9 | 139087521 | rs11103374 | 0.363 | 1.384 | 0.053 | 1.1 | -0.01 | -0.057 | -0.057 | 0.274 | 0.21 | 0.02 | 0.12 | 0.17 | 0.15 | 0.15 |
| 9 | 139089679 | rs7860634 | -0.945 | 1.454 | 0.408 | 0.567 | -0.001 | 0.002 | 0.002 | 0 | 0.56 | 0.59 | 0.41 | 0.33 | 0.55 | 0.44 |
| 9 | 139097645 | rs3758199 | -0.809 | 1.151 | 0.611 | 0.499 | 0.002 | 0.01 | 0.01 | 0 | 0.51 | 0.4 | 0.38 | 0.39 | 0.48 | 0.5 |
| 9 | 139099073 | rs3780190 | -1.099 | 1.175 | 0.599 | 0.506 | 0.001 | -0.001 | -0.001 | 0 | 0.57 | 0.42 | 0.39 | 0.41 | 0.5 | 0.53 |
| 9 | 139100413 | rs7024579 | 0.908 | 1.006 | 0.14 | 0.514 | -0.082 | 0.001 | 0.001 | 0.201 | 0.19 | 0.02 | 0.21 | 0.12 | 0.29 | 0.21 |
| 9 | 139105229 | rs3824359 | 0.27 | 0.831 | 0.299 | 0.51 | 0.041 | 0.001 | 0.001 | 0.293 | 0.25 | 0.11 | 0.11 | 0.22 | 0.14 | 0.16 |
| 9 | 139107879 | rs11103387 | -0.706 | 0.494 | 0.664 | 0.519 | 0.002 | 0.018 | 0.018 | 0.143 | 0.33 | 0.29 | 0.27 | 0.19 | 0.34 | 0.41 |
| 9 | 139109861 | rs10122824 | -1.363 | 0.841 | 0.465 | 0.74 | 0.008 | -0.03 | -0.03 | 0 | 0.62 | 0.77 | 0.68 | 0.47 | 0.63 | 0.55 |
| 9 | 139110654 | rs12684650 | 0.678 | 1.147 | 0.248 | 0.895 | -0.087 | -0.022 | -0.022 | 0.765 | 0.19 | 0.01 | 0.21 | 0.12 | 0.29 | 0.2 |
| 9 | 139119215 | rs10858250 | -0.021 | 0.856 | 1.129 | 0.995 | -0.009 | 0 | 0 | 0.046 | 0.28 | 0.2 | 0.38 | 0.25 | 0.29 | 0.16 |
| 9 | 139124856 | rs7032944 | -1.228 | 0.863 | 0.396 | 1.034 | -0.028 | -0.018 | -0.018 | 0 | 0.1 | 0.23 | 0.04 | 0.08 | 0.04 | 0.18 |
| 9 | 139134981 | rs7038554 | -0.509 | 1.299 | -0.323 | 1.08 | -0.004 | 0.017 | 0.017 | 0.461 | 0.37 | 0.72 | 0.45 | 0.28 | 0.34 | 0.2 |
| 9 | 139220876 | rs28661870 | -0.773 | 1.626 | 0.206 | 1.651 | 0.01 | 0.058 | 0.058 | 0 | 0.22 | 0.31 | 0.21 | 0.62 | 0.26 | 0.19 |
| 9 | 139234512 | rs28536668 | -1.02 | 1.657 | 0.682 | 1.096 | 0.044 | 0.046 | 0.046 | 1.092 | 0.15 | 0.35 | 0.13 | 0.26 | 0.16 | 0.2 |
| 9 | 139240630 | rs28539249 | 1.545 | 2.446 | 0.425 | 0.732 | 0.023 | -0.007 | -0.007 | 0.276 | 0.66 | 0.48 | 0.55 | 0.63 | 0.57 | 0.59 |
| 9 | 139243790 | rs28439345 | -1.101 | 2.818 | 0.54 | 0.958 | 0.052 | 0.096 | 0.096 | 0 | 0.06 | 0.13 | 0.12 | 0.31 | 0.15 | 0.2 |
| 9 | 139245289 | rs28642252 | 1.1 | 2.787 | 0.815 | 0.601 | 0.049 | -0.054 | -0.054 | 0 | 0.81 | 0.72 | 0.72 | 0.95 | 0.73 | 0.79 |
| 9 | 139252495 | rs3812547 | 1.179 | 3.543 | 0.506 | 0.365 | 0.027 | -0.033 | -0.033 | 1.414 | 0.8 | 0.79 | 0.53 | 0.95 | 0.67 | 0.73 |
| 9 | 139253839 | rs1128905 | -1.859 | 2.935 | 0.448 | 0.231 | 0.001 | -0.002 | -0.002 | 0.424 | 0.35 | 0.35 | 0.62 | 0.35 | 0.49 | 0.43 |

| | | | | | | | | | | | | | | | | |
|---|---|---|---|---|---|---|---|---|---|---|---|---|---|---|---|---|
| 9 | 139256141 | rs10747047 | -1.226 | 4.296 | -0.056 | -0.14 | -0.068 | 0.039 | 0.039 | 1.963 | 0.17 | 0.07 | 0.41 | 0.12 | 0.33 | 0.32 |
| 9 | 139257411 | rs10870202 | -2.057 | 4.17 | 0.701 | -0.253 | 0.033 | -0.006 | -0.006 | 0.571 | 0.31 | 0.43 | 0.64 | 0.33 | 0.5 | 0.43 |
| 9 | 139259249 | rs11145750 | -1.83 | 4.107 | 0.566 | -0.367 | 0.067 | 0.026 | 0.026 | 0.857 | 0.23 | 0.34 | 0.59 | 0.34 | 0.43 | 0.44 |
| 9 | 139261086 | rs10781496 | -1.898 | 3.929 | 0.644 | -0.687 | 0.013 | 0.001 | 0.001 | 0.91 | 0.24 | 0.25 | 0.56 | 0.33 | 0.42 | 0.44 |
| 9 | 139261991 | rs3812556 | -1.016 | 3.962 | 0.697 | -0.681 | 0.003 | 0.011 | 0.011 | 0.005 | 0.15 | 0.2 | 0.17 | 0.28 | 0.17 | 0.21 |
| 9 | 139266405 | rs10781499 | -1.687 | 4.051 | 0.666 | -0.717 | 0.045 | 0.009 | 0.009 | 0.455 | 0.21 | 0.25 | 0.57 | 0.32 | 0.4 | 0.39 |
| 9 | 139266496 | rs4077515 | -1.674 | 4.045 | 0.666 | -0.718 | 0.046 | 0.011 | 0.011 | 0.414 | 0.21 | 0.25 | 0.57 | 0.32 | 0.4 | 0.39 |
| 9 | 139269338 | rs10781500 | -1.685 | 4.054 | 0.666 | -0.718 | 0.045 | 0.01 | 0.01 | 0.455 | 0.21 | 0.25 | 0.57 | 0.32 | 0.39 | 0.4 |
| 9 | 139269905 | rs3812560 | -2.287 | 4.048 | 0.683 | -0.721 | 0.024 | -0.001 | -0.001 | 0 | 0.29 | 0.36 | 0.61 | 0.32 | 0.42 | 0.4 |
| 9 | 139272058 | rs10781507 | -1.687 | 4.062 | 0.716 | -0.71 | 0.048 | 0.011 | 0.011 | 0.455 | 0.21 | 0.26 | 0.57 | 0.32 | 0.4 | 0.4 |
| 9 | 139273288 | rs3829112 | -1.294 | 4.205 | 0.686 | -0.718 | 0.014 | 0.026 | 0.026 | 0 | 0.08 | 0.09 | 0.26 | 0.27 | 0.16 | 0.21 |
| 9 | 139274523 | rs11145793 | -0.746 | 4.208 | 0.759 | -0.723 | -0.002 | 0.003 | 0.003 | 0 | 0.13 | 0.11 | 0.31 | 0.04 | 0.23 | 0.19 |
| 9 | 139275204 | rs3812570 | -2.423 | 4.153 | 0.76 | -0.737 | 0.038 | -0.002 | -0.002 | 0 | 0.3 | 0.39 | 0.61 | 0.32 | 0.42 | 0.4 |
| 9 | 139279173 | rs10781510 | -1.839 | 4.208 | 0.776 | -0.694 | 0.044 | 0.007 | 0.007 | 0.374 | 0.21 | 0.26 | 0.57 | 0.32 | 0.39 | 0.4 |
| 9 | 139289825 | rs4266763 | -1.925 | 4.191 | 0.771 | -0.669 | 0.091 | 0.025 | 0.025 | 0.35 | 0.22 | 0.36 | 0.58 | 0.32 | 0.39 | 0.4 |
| 9 | 139296485 | rs563 | -0.882 | 4.217 | 0.71 | -0.7 | 0.07 | -0.034 | -0.034 | 0.65 | 0.13 | 0.23 | 0.32 | 0.05 | 0.23 | 0.19 |
| 9 | 139299637 | rs3812577 | -0.726 | 4.053 | 0.591 | -0.674 | -0.037 | -0.036 | -0.036 | 0 | 0.06 | 0.01 | 0.23 | 0.14 | 0.11 | 0.13 |
| 9 | 139302200 | rs11145917 | -0.653 | 4.23 | 0.059 | -1.194 | -0.042 | 0.032 | 0.032 | 0.083 | 0.2 | 0.12 | 0.35 | 0.05 | 0.34 | 0.2 |
| 9 | 139304016 | rs11145930 | -1.298 | 4.155 | 0.685 | -1.16 | 0.058 | -0.051 | -0.051 | 0 | 0.24 | 0.42 | 0.37 | 0.16 | 0.3 | 0.26 |
| 9 | 139306467 | rs10870144 | -0.62 | 4.238 | 0.15 | -1.323 | -0.042 | -0.023 | -0.023 | 0 | 0.1 | 0.03 | 0.27 | 0.14 | 0.16 | 0.17 |
| 9 | 139316744 | rs11145974 | -0.31 | 4.26 | 0.148 | -1.333 | -0.078 | 0.037 | 0.037 | 0 | 0.13 | 0.03 | 0.3 | 0.05 | 0.24 | 0.18 |
| 9 | 139317823 | rs9548 | -0.614 | 4.248 | 0.146 | -1.332 | -0.042 | -0.024 | -0.024 | 0.012 | 0.1 | 0.03 | 0.27 | 0.14 | 0.16 | 0.17 |
| 9 | 139317869 | rs1130635 | -0.539 | 4.195 | 0.167 | -1.349 | -0.069 | -0.009 | -0.009 | 0 | 0.14 | 0.06 | 0.31 | 0.17 | 0.27 | 0.26 |
| 9 | 139323201 | rs1128877 | -0.412 | 3.946 | 0.134 | -1.351 | -0.033 | -0.028 | -0.028 | 0 | 0.09 | 0.04 | 0.24 | 0.15 | 0.12 | 0.16 |
| 9 | 139323311 | rs8413 | -1.669 | 4.027 | 0.347 | -1.376 | 0.05 | -0.012 | -0.012 | 0 | 0.25 | 0.38 | 0.59 | 0.19 | 0.39 | 0.35 |
| 9 | 139327034 | rs10870194 | -1.883 | 4.084 | 0.342 | -1.397 | 0.01 | -0.011 | -0.011 | 0 | 0.33 | 0.39 | 0.62 | 0.19 | 0.41 | 0.35 |
| 9 | 139327439 | rs10781542 | -1.892 | 3.997 | 0.329 | -1.405 | 0.007 | -0.01 | -0.01 | 0 | 0.33 | 0.39 | 0.62 | 0.19 | 0.41 | 0.35 |
| 9 | 139335206 | rs1127162 | -0.354 | 4.055 | 0.233 | -1.361 | -0.035 | -0.029 | -0.029 | 0 | 0.09 | 0.04 | 0.3 | 0.16 | 0.13 | 0.16 |
| 9 | 139335599 | rs1127152 | -1.725 | 3.899 | 0.173 | -1.419 | -0.073 | 0.042 | 0.042 | 1.31 | 0.32 | 0.2 | 0.58 | 0.21 | 0.4 | 0.3 |
| 9 | 139335879 | rs3812589 | -0.172 | 3.9 | 0.155 | -1.393 | -0.033 | -0.028 | -0.028 | 0 | 0.09 | 0.04 | 0.3 | 0.16 | 0.12 | 0.16 |
| 9 | 139368953 | rs3812594 | -0.106 | 3.911 | 0.311 | -1.421 | -0.071 | 0.037 | 0.037 | 0 | 0.13 | 0.04 | 0.26 | 0.05 | 0.25 | 0.14 |
| 9 | 139369062 | rs3812595 | -0.185 | 3.916 | 0.338 | -1.419 | -0.035 | 0.026 | 0.026 | 0 | 0.14 | 0.1 | 0.26 | 0.06 | 0.25 | 0.14 |
| 9 | 139369212 | rs17567909 | 1.009 | 2.405 | -0.851 | -2.398 | 0.025 | 0.115 | 0.115 | 0 | 0.18 | 0 | 0.06 | 0 | 0.12 | 0.29 |
| 9 | 139371234 | rs10781497 | 0.406 | 2.471 | -0.812 | -2.36 | -0.033 | -0.028 | -0.028 | 0 | 0.09 | 0.04 | 0.3 | 0.16 | 0.12 | 0.16 |
| 9 | 139374952 | rs10870070 | -0.232 | 2.451 | -0.761 | -2.361 | -0.001 | 0 | 0 | 0 | 0.17 | 0.18 | 0.32 | 0.16 | 0.14 | 0.16 |
| 9 | 139375962 | rs4378078 | -1.205 | 2.414 | -0.644 | -2.342 | 0.025 | -0.021 | -0.021 | 0 | 0.33 | 0.42 | 0.61 | 0.22 | 0.41 | 0.3 |
| 9 | 139380320 | rs1139057 | 0.085 | 1.607 | -0.718 | -2.43 | 0.021 | 0.053 | 0.053 | 0 | 0.24 | 0.11 | 0.12 | 0.04 | 0.21 | 0.36 |
| 9 | 139389184 | rs6563 | -1.105 | 1.949 | -0.563 | -2.104 | 0.012 | -0.01 | -0.01 | 0 | 0.34 | 0.6 | 0.47 | 0.69 | 0.48 | 0.24 |
| 9 | 139391636 | rs2229974 | -1.058 | 0.934 | -0.734 | -1.118 | 0.001 | 0.025 | 0.025 | 4.578 | 0.54 | 0.63 | 0.65 | 0.95 | 0.58 | 0.67 |
| 9 | 139401504 | rs3124596 | -1.077 | 0.809 | -0.691 | -0.578 | -0.011 | 0.079 | 0.079 | 2.156 | 0.57 | 0.72 | 0.6 | 0.91 | 0.53 | 0.63 |
| 9 | 139403102 | rs3812605 | -1.246 | 0.705 | -0.96 | -0.556 | -0.133 | 0.207 | 0.207 | 3.896 | 0.6 | 0.94 | 0.64 | 0.91 | 0.58 | 0.64 |
| 9 | 139404895 | rs11145766 | -0.698 | -0.007 | -0.462 | 0.623 | -0.019 | -0.055 | -0.055 | 1.028 | 0.13 | 0.2 | 0.07 | 0.01 | 0.09 | 0.18 |
| 9 | 139408892 | rs3812609 | -0.836 | -0.119 | -1.127 | -0.123 | 0.043 | 0.088 | 0.088 | 0 | 0.24 | 0.08 | 0.11 | 0.1 | 0.16 | 0.28 |
| 9 | 139425634 | rs13290979 | -0.963 | 0.418 | -0.461 | 0.562 | -0.002 | -0.002 | -0.002 | 0.022 | 0.4 | 0.58 | 0.47 | 0.37 | 0.35 | 0.63 |
| 9 | 139431057 | rs3013300 | -0.405 | 0.298 | -0.259 | 0.061 | 0 | 0.013 | 0.013 | 0 | 0.36 | 0.24 | 0.45 | 0.72 | 0.34 | 0.6 |
| 9 | 139431874 | rs3013302 | -0.787 | 0.443 | -0.48 | 0.18 | 0.022 | 0.036 | 0.036 | 0 | 0.45 | 0.77 | 0.5 | 0.74 | 0.38 | 0.7 |
| 9 | 139443253 | rs3013309 | 0.901 | 0.789 | -0.412 | 0.074 | 0.001 | 0.015 | 0.015 | 0.817 | 0.7 | 0.23 | 0.7 | 0.72 | 0.72 | 0.67 |
| 9 | 139443956 | rs710411 | 1.011 | 0.883 | 0.056 | -0.254 | 0.005 | -0.009 | -0.009 | 0 | 0.6 | 0.47 | 0.52 | 0.66 | 0.63 | 0.55 |
| 9 | 139443989 | rs885446 | 0.197 | 0.915 | 0.054 | -0.299 | 0.002 | -0.007 | -0.007 | 0 | 0.92 | 0.82 | 0.95 | 0.95 | 0.93 | 0.96 |
| 9 | 139444102 | rs1058233 | -0.661 | 0.814 | 0.176 | -0.165 | -0.028 | 0.01 | 0.01 | 0 | 0.29 | 0.16 | 0.4 | 0.26 | 0.28 | 0.41 |
| 9 | 139456212 | rs7034799 | 0.583 | -0.112 | 0.423 | -0.477 | 0.001 | 0.047 | 0.047 | 0 | 0.93 | 0.75 | 0.96 | 0.91 | 0.95 | 0.97 |
| 9 | 139457506 | rs7849014 | 1.358 | -0.158 | 0.385 | -0.515 | -0.008 | 0.022 | 0.022 | 0.063 | 0.9 | 0.7 | 0.94 | 0.91 | 0.93 | 0.88 |
| 9 | 139468185 | rs10870085 | 0.565 | -0.178 | 0.172 | -0.465 | -0.028 | -0.033 | -0.033 | 0.841 | 0.68 | 0.48 | 0.81 | 0.81 | 0.79 | 0.65 |
| 9 | 139477521 | rs10870088 | 0.18 | -0.1 | 0.105 | -0.403 | -0.009 | -0.011 | -0.011 | 0.073 | 0.71 | 0.62 | 0.84 | 0.81 | 0.79 | 0.71 |
| 9 | 139483954 | rs944758 | 0.732 | -0.322 | -0.825 | -0.902 | 0.001 | -0.014 | -0.014 | 0 | 0.51 | 0.5 | 0.72 | 0.83 | 0.66 | 0.55 |
| 9 | 139487092 | rs4880055 | 0.363 | -0.045 | -0.471 | -0.307 | -0.017 | -0.028 | -0.028 | 0 | 0.64 | 0.4 | 0.77 | 0.84 | 0.77 | 0.63 |
| 9 | 139500617 | rs11145791 | -0.539 | 0.541 | -0.641 | -0.066 | -0.001 | 0.006 | 0.006 | 0 | 0.43 | 0.57 | 0.52 | 0.32 | 0.4 | 0.58 |
| 9 | 139500886 | rs10870094 | -0.322 | 0.164 | -0.737 | -1.326 | -0.001 | -0.043 | -0.043 | 0 | 0.44 | 0.47 | 0.3 | 0.09 | 0.41 | 0.51 |
| 9 | 139501085 | rs7041089 | 0.611 | -0.409 | -0.315 | -0.952 | -0.024 | -0.019 | -0.019 | 0.247 | 0.77 | 0.36 | 0.85 | 0.81 | 0.85 | 0.7 |
| 9 | 139511247 | rs7852910 | 0.523 | -0.322 | -1.089 | -0.947 | 0.05 | 0.158 | 0.158 | 0 | 0.53 | 0.26 | 0.56 | 0.14 | 0.66 | 0.35 |
| 9 | 139513538 | rs12378485 | 0.641 | -0.441 | -0.916 | 0.191 | 0.079 | 0.235 | 0.235 | 0 | 0.52 | 0.16 | 0.55 | 0.14 | 0.66 | 0.36 |
| 9 | 139518718 | rs12376605 | 0.029 | -0.497 | -1.176 | -0.134 | 0.006 | 0.044 | 0.044 | 0.355 | 0.25 | 0.15 | 0.23 | 0.14 | 0.21 | 0.3 |
| 9 | 139518965 | rs7863705 | 0.702 | -0.422 | -1.269 | -0.251 | 0.002 | 0.212 | 0.212 | 0 | 0.44 | 0.15 | 0.4 | 0.17 | 0.58 | 0.38 |
| 9 | 139522101 | rs914737 | -0.831 | 0.291 | -0.172 | 0.871 | 0.006 | 0.127 | 0.127 | 0 | 0.18 | 0.35 | 0.37 | 0.53 | 0.19 | 0.34 |
| 9 | 139526086 | rs10870101 | 0.271 | 0.341 | -0.182 | -0.051 | -0.003 | 0.009 | 0.009 | 0 | 0.69 | 0.45 | 0.59 | 0.42 | 0.66 | 0.49 |
| 9 | 139534703 | rs10747029 | 0.068 | 0.194 | -0.358 | -0.624 | -0.005 | 0.017 | 0.017 | 0 | 0.72 | 0.77 | 0.76 | 0.78 | 0.69 | 0.87 |
| 9 | 139535714 | rs10735657 | 0.881 | -0.034 | -0.128 | -0.662 | 0.012 | 0.037 | 0.037 | 0 | 0.51 | 0.35 | 0.57 | 0.64 | 0.52 | 0.69 |
| 9 | 139540385 | rs7854618 | 0.063 | -0.449 | 0.2 | -0.827 | 0.001 | 0.001 | 0.001 | 0 | 0.19 | 0.18 | 0.17 | 0.15 | 0.15 | 0.15 |
| 9 | 139544437 | rs7871194 | 0.771 | -0.471 | 0.094 | -0.818 | 0.031 | 0.044 | 0.044 | 0 | 0.51 | 0.35 | 0.58 | 0.64 | 0.54 | 0.68 |

| | | | | | | | | | | | | | | | | |
|---|---|---|---|---|---|---|---|---|---|---|---|---|---|---|---|---|
| 9 | 139556771 | rs1332793 | 0.737 | -0.347 | 0.151 | -0.789 | 0.024 | 0.026 | 0.026 | 0 | 0.53 | 0.38 | 0.62 | 0.67 | 0.6 | 0.68 |
| 9 | 139564668 | rs2297538 | -0.322 | -0.105 | 0.527 | -0.892 | -0.004 | -0.012 | -0.012 | 0 | 0.22 | 0.21 | 0.19 | 0.13 | 0.21 | 0.09 |
| 9 | 139566889 | rs11558192 | -0.531 | 0.291 | 0.239 | -0.968 | 0.009 | 0.008 | 0.008 | 0.229 | 0.1 | 0.06 | 0.15 | 0 | 0.11 | 0.05 |
| 9 | 139568047 | rs4880119 | 1.47 | 0.385 | 0.632 | -0.79 | -0.008 | -0.076 | -0.076 | 0 | 0.81 | 0.58 | 0.82 | 0.95 | 0.82 | 0.91 |
| 9 | 139570617 | rs10217363 | -0.772 | 0.547 | -0.165 | -0.721 | 0 | -0.003 | -0.003 | 0 | 0.14 | 0.15 | 0.12 | 0.05 | 0.13 | 0.03 |
| 9 | 139575487 | rs9411216 | -0.086 | 1.233 | -0.08 | -0.788 | 0.008 | -0.035 | -0.035 | 1.313 | 0.11 | 0.18 | 0.07 | 0.02 | 0.12 | 0.19 |
| 9 | 139578880 | rs1889047 | 0.564 | 0.768 | 0.241 | -0.819 | -0.002 | -0.053 | -0.053 | 0 | 0.75 | 0.45 | 0.65 | 0.88 | 0.78 | 0.81 |
| 9 | 139590934 | rs4880127 | 0.698 | 0.721 | 0.255 | -0.883 | -0.001 | -0.002 | -0.002 | 0 | 0.77 | 0.73 | 0.67 | 0.88 | 0.8 | 0.81 |
| 9 | 139596420 | rs2385891 | -1.296 | 1.237 | 0.122 | -0.745 | -0.022 | 0.05 | 0.05 | 0 | 0.45 | 0.74 | 0.46 | 0.27 | 0.42 | 0.54 |
| 9 | 139598108 | rs10870112 | -1.186 | 0.786 | -0.008 | -0.994 | -0.004 | 0.053 | 0.053 | 0 | 0.49 | 0.65 | 0.46 | 0.27 | 0.42 | 0.55 |
| 9 | 139607966 | rs11793385 | -1.501 | 1.684 | 0.839 | -0.788 | 0.145 | 0.036 | 0.036 | 0 | 0.12 | 0.46 | 0.15 | 0.09 | 0.19 | 0.16 |
| 9 | 139615549 | rs7029239 | -1.105 | 2.086 | 0.801 | -0.956 | 0.038 | 0.009 | 0.009 | 0.229 | 0.16 | 0.31 | 0.31 | 0.18 | 0.22 | 0.24 |
| 9 | 139617379 | rs11793962 | -1.754 | 2.068 | 0.602 | -1.232 | 0.027 | -0.014 | -0.014 | 0.421 | 0.24 | 0.36 | 0.38 | 0.18 | 0.34 | 0.33 |
| 9 | 139617836 | rs6874 | -0.268 | 1.803 | 0.231 | -1.11 | -0.019 | 0.019 | 0.019 | 0 | 0.08 | 0.04 | 0.07 | 0 | 0.12 | 0.09 |
| 9 | 139623701 | rs11999246 | -0.348 | 1.772 | 0.216 | -1.124 | -0.019 | 0.021 | 0.021 | 0 | 0.09 | 0.04 | 0.07 | 0 | 0.12 | 0.09 |
| 9 | 139633733 | rs10118736 | -0.4 | 1.001 | -0.126 | -1.39 | 0.008 | -0.013 | -0.013 | 0 | 0.2 | 0.17 | 0.24 | 0.27 | 0.15 | 0.17 |
| 9 | 139634495 | rs9886752 | -0.262 | 0.995 | -0.129 | -1.393 | 0.009 | -0.013 | -0.013 | 0 | 0.2 | 0.18 | 0.24 | 0.27 | 0.15 | 0.17 |
| 9 | 139638740 | rs2811724 | 0.098 | 0.683 | 0.176 | -1.864 | -0.014 | -0.118 | -0.118 | 0.186 | 0.78 | 0.42 | 0.78 | 0.98 | 0.82 | 0.69 |
| 9 | 139642961 | rs2811728 | 1.66 | 0.505 | -0.111 | -2.062 | -0.02 | 0.048 | 0.048 | 0.04 | 0.43 | 0.22 | 0.41 | 0.71 | 0.5 | 0.45 |
| 9 | 139643023 | rs1360879 | -0.077 | 0.37 | 0.547 | -1.695 | -0.017 | -0.006 | -0.006 | 1.146 | 0.19 | 0.27 | 0.24 | 0.15 | 0.13 | 0.11 |
| 9 | 139648721 | rs1108607 | 1.198 | 0.488 | 0.241 | -1.876 | 0.027 | 0.041 | 0.041 | 0 | 0.55 | 0.3 | 0.54 | 0.71 | 0.65 | 0.54 |
| 9 | 139649273 | rs2248077 | 0.326 | 0.599 | 0.818 | -1.846 | -0.02 | -0.048 | -0.048 | 1.53 | 0.77 | 0.56 | 0.79 | 0.87 | 0.82 | 0.65 |
| 9 | 139656706 | rs2297723 | 0.278 | 0.845 | 0.684 | -1.884 | -0.004 | -0.031 | -0.031 | 0 | 0.76 | 0.66 | 0.78 | 0.85 | 0.78 | 0.63 |
| 9 | 139660402 | rs4880142 | 0.42 | 0.846 | 0.768 | -1.89 | -0.006 | -0.042 | -0.042 | 3.32 | 0.75 | 0.57 | 0.78 | 0.85 | 0.78 | 0.62 |
| 9 | 139679717 | rs2784054 | -0.52 | 0.903 | 0.499 | -1.94 | 0 | -0.05 | -0.05 | 0 | 0.29 | 0.43 | 0.28 | 0.15 | 0.28 | 0.4 |
| 9 | 139685136 | rs2811787 | -1.424 | 0.858 | 0.119 | -2.212 | 0.004 | 0.04 | 0.04 | 0 | 0.52 | 0.65 | 0.57 | 0.27 | 0.46 | 0.52 |
| 9 | 139686247 | rs3739950 | 0.227 | 0.745 | -0.25 | -2.136 | -0.004 | 0.05 | 0.05 | 0.498 | 0.22 | 0.03 | 0.28 | 0.12 | 0.19 | 0.16 |
| 9 | 139690840 | rs1055995 | 0.507 | 0.915 | -0.463 | -2.254 | -0.024 | 0.097 | 0.097 | 0 | 0.6 | 0.22 | 0.65 | 0.8 | 0.62 | 0.52 |
| 9 | 139693596 | rs945386 | -0.336 | 0.306 | -0.045 | -1.947 | -0.004 | -0.002 | -0.002 | 0 | 0.28 | 0.34 | 0.3 | 0.24 | 0.21 | 0.26 |
| 9 | 139693992 | rs945387 | 1.367 | 0.503 | -0.098 | -2.082 | 0.013 | 0.024 | 0.024 | 0 | 0.36 | 0.25 | 0.3 | 0.74 | 0.3 | 0.32 |
| 9 | 139694521 | rs4546744 | -0.614 | 0.392 | 0.116 | -1.101 | -0.001 | 0 | 0 | 0 | 0.28 | 0.3 | 0.3 | 0.24 | 0.22 | 0.27 |
| 9 | 139694569 | rs7859194 | -0.725 | 0.358 | 0.093 | -1.038 | -0.003 | -0.002 | -0.002 | 0 | 0.29 | 0.34 | 0.3 | 0.24 | 0.22 | 0.27 |
| 9 | 139700605 | rs2811795 | 0.205 | 0.757 | 0.032 | -0.819 | -0.04 | -0.017 | -0.017 | 4.298 | 0.82 | 0.98 | 0.72 | 0.86 | 0.8 | 0.82 |
| 9 | 139701108 | rs2254143 | 0.433 | 0.736 | 0.251 | -0.829 | 0.001 | 0.005 | 0.005 | 0 | 0.81 | 0.82 | 0.7 | 0.86 | 0.8 | 0.81 |
| 9 | 139709976 | rs2484661 | 1.186 | 0.547 | 0.813 | -0.63 | -0.027 | -0.005 | -0.005 | 0 | 0.76 | 0.54 | 0.66 | 0.75 | 0.8 | 0.67 |
| 9 | 139711726 | rs7856666 | -1.337 | 0.51 | 0.736 | -0.603 | 0.005 | 0.074 | 0.074 | 3.349 | 0.05 | 0.25 | 0.04 | 0.11 | 0 | 0.14 |
| 9 | 139719671 | rs2811754 | 0.242 | 0.647 | 0.226 | -0.607 | -0.001 | 0.012 | 0.012 | 0 | 0.81 | 0.91 | 0.71 | 0.86 | 0.8 | 0.81 |
| 9 | 139721513 | rs2811751 | 1.243 | 1.219 | 0.761 | -0.768 | 0.106 | 0.015 | 0.015 | 0.051 | 0.87 | 0.55 | 0.77 | 0.82 | 0.82 | 0.83 |
| 9 | 139725227 | rs2811749 | 0.706 | 0.712 | 0.312 | -0.432 | -0.009 | -0.012 | -0.012 | 0 | 0.78 | 0.69 | 0.69 | 0.82 | 0.79 | 0.72 |
| 9 | 139726239 | rs2811748 | 0.772 | 0.269 | -0.096 | -0.525 | 0.037 | -0.005 | -0.005 | 0.276 | 0.79 | 0.87 | 0.73 | 0.76 | 0.89 | 0.79 |
| 9 | 139730129 | rs2811744 | 0.77 | 0.695 | 0.013 | -0.601 | -0.009 | -0.019 | -0.019 | 0 | 0.79 | 0.73 | 0.71 | 0.87 | 0.82 | 0.79 |
| 9 | 139736198 | rs2077581 | 0.228 | 0.746 | -0.657 | -0.633 | -0.006 | 0.029 | 0.029 | 0.368 | 0.78 | 0.97 | 0.7 | 0.87 | 0.81 | 0.78 |
| 9 | 139743497 | rs12555238 | 0.585 | 1.595 | -0.571 | -0.768 | -0.019 | 0.034 | 0.034 | 0.047 | 0.71 | 0.82 | 0.59 | 0.87 | 0.66 | 0.79 |
| 9 | 139746992 | rs10123375 | 0.359 | 1.181 | -0.671 | -0.79 | -0.004 | 0.048 | 0.048 | 0.336 | 0.73 | 0.84 | 0.55 | 0.88 | 0.75 | 0.76 |
| 9 | 139754110 | rs2236544 | 0.325 | 1.18 | -0.696 | -0.806 | -0.007 | 0.055 | 0.055 | 0.476 | 0.73 | 0.86 | 0.55 | 0.88 | 0.75 | 0.78 |
| 9 | 139779462 | rs10781520 | 0.131 | 1.19 | -0.645 | -0.778 | -0.017 | 0.052 | 0.052 | 1.389 | 0.74 | 0.88 | 0.55 | 0.88 | 0.74 | 0.78 |
| 9 | 139792926 | rs2784075 | 0.306 | 1.244 | -0.626 | -0.79 | -0.006 | 0.039 | 0.039 | 0.715 | 0.76 | 0.88 | 0.58 | 0.88 | 0.78 | 0.82 |
| 9 | 139796419 | rs10870140 | 0.261 | 1.227 | -0.547 | -0.793 | -0.004 | 0.031 | 0.031 | 0.364 | 0.73 | 0.82 | 0.55 | 0.88 | 0.74 | 0.81 |
| 9 | 139801932 | rs10119096 | 0.107 | 1.231 | -0.562 | -0.791 | -0.012 | 0.044 | 0.044 | 0.551 | 0.74 | 0.86 | 0.56 | 0.88 | 0.74 | 0.81 |
| 9 | 139804679 | rs4880158 | 0.323 | 1.232 | -0.577 | -0.79 | -0.024 | 0.051 | 0.051 | 1.135 | 0.75 | 0.91 | 0.57 | 0.88 | 0.75 | 0.81 |
| 9 | 139815053 | rs10781522 | 0.995 | 1.535 | -0.066 | -0.823 | 0.004 | -0.031 | -0.031 | 0 | 0.64 | 0.57 | 0.44 | 0.72 | 0.63 | 0.74 |
| 9 | 139817353 | rs4880073 | -0.729 | 1.461 | -0.562 | -0.928 | -0.008 | 0.008 | 0.008 | 0.735 | 0.34 | 0.69 | 0.42 | 0.37 | 0.4 | 0.5 |
| 9 | 139820421 | rs7852970 | 0.832 | 1.162 | -0.22 | -0.902 | 0 | -0.032 | -0.032 | 0 | 0.62 | 0.53 | 0.43 | 0.71 | 0.62 | 0.65 |
| 9 | 139821068 | rs3750512 | 0.603 | 1.35 | -0.16 | -1.003 | 0.005 | -0.03 | -0.03 | 0 | 0.67 | 0.66 | 0.47 | 0.86 | 0.64 | 0.73 |
| 9 | 139823670 | rs7019752 | 0.156 | 1.434 | -0.014 | -0.87 | 0.066 | -0.005 | -0.005 | 0 | 0.07 | 0.2 | 0.09 | 0.02 | 0.11 | 0.04 |
| 9 | 139830787 | rs10115595 | 0.348 | 1.056 | -0.635 | -1.058 | 0 | 0.064 | 0.064 | 0 | 0.53 | 0.64 | 0.44 | 0.84 | 0.53 | 0.62 |
| 9 | 139830940 | rs6560652 | 0.245 | 1.056 | -0.64 | -1.057 | -0.003 | 0.06 | 0.06 | 0 | 0.54 | 0.7 | 0.45 | 0.84 | 0.53 | 0.67 |
| 9 | 139834195 | rs4880169 | 0.163 | 0.778 | -0.576 | -1.007 | 0.02 | -0.023 | -0.023 | 0 | 0.09 | 0.17 | 0.1 | 0.03 | 0.13 | 0.08 |
| 9 | 139836362 | rs7040705 | 0.465 | 0.776 | -0.531 | -1.008 | 0 | 0.004 | 0.004 | 0 | 0.52 | 0.56 | 0.41 | 0.7 | 0.54 | 0.62 |
| 9 | 139841558 | rs884112 | 0.572 | 1.038 | -0.551 | -1.112 | 0.059 | -0.054 | -0.054 | 0 | 0.87 | 0.71 | 0.86 | 0.97 | 0.81 | 0.9 |
| 9 | 139842262 | rs1879011 | 0.109 | 1.019 | -0.617 | -1.058 | 0.022 | -0.023 | -0.023 | 0 | 0.09 | 0.17 | 0.11 | 0.03 | 0.13 | 0.08 |
| 9 | 139844151 | rs4880172 | 0.089 | 0.748 | -0.859 | -1.268 | 0.015 | 0.082 | 0.082 | 0 | 0.57 | 0.74 | 0.5 | 0.86 | 0.6 | 0.65 |
| 9 | 139847570 | rs884218 | -0.157 | 0.452 | -0.996 | -1.065 | 0.024 | 0.093 | 0.093 | 0 | 0.63 | 0.86 | 0.58 | 0.86 | 0.69 | 0.67 |
| 9 | 139850119 | rs868102 | 0.714 | 0.615 | -0.952 | -1.209 | 0.002 | 0.048 | 0.048 | 0 | 0.48 | 0.64 | 0.39 | 0.84 | 0.44 | 0.56 |
| 9 | 139854099 | rs10781528 | -0.384 | 0.599 | -1.22 | -1.2 | 0.024 | 0.085 | 0.085 | 0 | 0.64 | 0.86 | 0.58 | 0.86 | 0.71 | 0.66 |
| 9 | 139860264 | rs11145951 | -0.829 | 0.54 | -1.312 | -1.111 | -0.015 | 0.156 | 0.156 | 0 | 0.49 | 0.21 | 0.56 | 0.17 | 0.52 | 0.41 |
| 9 | 139860631 | rs11793787 | -0.589 | 1.028 | -0.857 | -0.966 | -0.005 | -0.002 | -0.002 | 0 | 0.16 | 0.12 | 0.17 | 0.03 | 0.26 | 0.1 |
| 9 | 139869600 | rs10114487 | 0.181 | 0.256 | -0.672 | -0.37 | 0.009 | -0.028 | -0.028 | 0.073 | 0.28 | 0.13 | 0.4 | 0.33 | 0.26 | 0.21 |
| 9 | 139869914 | rs3814499 | 0.187 | 0.218 | -0.11 | -0.752 | -0.006 | -0.023 | -0.023 | 0.879 | 0.67 | 0.61 | 0.55 | 0.81 | 0.69 | 0.77 |

| | | | | | | | | | | | | | | | | |
|---|---|---|---|---|---|---|---|---|---|---|---|---|---|---|---|---|
| 9 | 139876141 | rs6926 | -0.139 | 0.241 | -0.126 | -0.822 | 0 | 0.017 | 0.017 | 0.022 | 0.7 | 0.76 | 0.57 | 0.83 | 0.69 | 0.77 |
| 9 | 139877129 | rs2049043 | 0.053 | 0.24 | -0.102 | -0.825 | -0.001 | -0.011 | -0.011 | 0 | 0.69 | 0.65 | 0.55 | 0.83 | 0.69 | 0.77 |
| 9 | 139878317 | rs7029565 | 0.066 | 0.25 | -0.098 | -0.821 | -0.002 | -0.012 | -0.012 | 0 | 0.68 | 0.64 | 0.55 | 0.83 | 0.69 | 0.77 |
| 9 | 139878828 | rs4880082 | 0.054 | 0.25 | -0.098 | -0.82 | -0.002 | -0.013 | -0.013 | 0 | 0.69 | 0.64 | 0.55 | 0.83 | 0.69 | 0.77 |
| 9 | 139879170 | rs17578859 | 0.175 | 0.248 | -0.148 | -0.816 | 0.007 | 0.026 | 0.026 | 0 | 0.28 | 0.18 | 0.39 | 0.17 | 0.26 | 0.21 |
| 9 | 139885820 | rs10870158 | 0.165 | 0.285 | -0.044 | -0.834 | -0.008 | -0.058 | -0.058 | 0 | 0.66 | 0.57 | 0.55 | 0.82 | 0.68 | 0.76 |
| 9 | 139885948 | rs10781530 | 0.165 | 0.285 | -0.044 | -0.834 | -0.008 | -0.058 | -0.058 | 1.08 | 0.66 | 0.57 | 0.55 | 0.82 | 0.68 | 0.76 |
| 9 | 139911983 | rs12348881 | -0.215 | 0.703 | -0.802 | -0.723 | -0.003 | -0.037 | -0.037 | 0 | 0.15 | 0.19 | 0.1 | 0.03 | 0.11 | 0.07 |
| 9 | 139923937 | rs4880191 | 0.315 | 0.619 | -1.141 | -0.775 | -0.002 | 0.017 | 0.017 | 0 | 0.64 | 0.67 | 0.6 | 0.77 | 0.62 | 0.72 |
| 9 | 139924637 | rs13301354 | 0.295 | 0.616 | -1.165 | -0.786 | -0.002 | 0.016 | 0.016 | 0 | 0.65 | 0.67 | 0.59 | 0.77 | 0.61 | 0.72 |
| 9 | 139926960 | rs10732706 | 0.451 | 0.666 | -1.14 | -0.755 | -0.004 | 0.02 | 0.02 | 0 | 0.66 | 0.69 | 0.6 | 0.77 | 0.62 | 0.74 |
| 9 | 139929435 | rs12337910 | -1.417 | 0.654 | -1.122 | -0.747 | 0.046 | -0.007 | -0.007 | 0 | 0.07 | 0.16 | 0.09 | 0.03 | 0.1 | 0.03 |
| 9 | 139930945 | rs7047770 | 0.519 | 0.53 | -1.212 | -0.712 | -0.001 | 0.008 | 0.008 | 0 | 0.65 | 0.66 | 0.61 | 0.76 | 0.64 | 0.74 |
| 9 | 139932211 | rs1138628 | 0.482 | 0.589 | -1.326 | -0.677 | -0.03 | -0.006 | -0.006 | 0.003 | 0.25 | 0.01 | 0.29 | 0.21 | 0.24 | 0.21 |
| 9 | 139933244 | rs7038280 | -0.16 | 0.594 | -1.29 | -0.679 | 0.004 | 0.039 | 0.039 | 0 | 0.71 | 0.84 | 0.68 | 0.79 | 0.73 | 0.77 |
| 9 | 139934020 | rs6605 | 0.365 | 0.581 | -1.346 | -0.636 | -0.038 | -0.015 | -0.015 | 0 | 0.25 | 0.01 | 0.29 | 0.21 | 0.24 | 0.21 |
| 9 | 139934107 | rs6604 | 0.073 | 0.626 | -1.316 | -0.607 | -0.009 | 0.017 | 0.017 | 0 | 0.28 | 0.11 | 0.35 | 0.25 | 0.3 | 0.23 |
| 9 | 139936141 | rs13289619 | -0.248 | 0.294 | -1.293 | -0.696 | -0.018 | 0.023 | 0.023 | 0 | 0.05 | 0.02 | 0.07 | 0 | 0.09 | 0.02 |
| 9 | 139938681 | rs3829107 | 0.426 | 0.435 | -1.407 | -0.558 | 0.026 | 0.047 | 0.047 | 0 | 0.3 | 0.56 | 0.4 | 0.53 | 0.33 | 0.29 |
| 9 | 139947473 | rs7861020 | 0.135 | -0.085 | -1.122 | -0.125 | -0.004 | 0.007 | 0.007 | 0 | 0.63 | 0.61 | 0.59 | 0.54 | 0.66 | 0.68 |
| 9 | 139958296 | rs3750517 | 0.097 | -0.344 | -1.72 | -0.106 | 0.045 | -0.042 | -0.042 | 0 | 0.32 | 0.04 | 0.31 | 0.34 | 0.23 | 0.19 |
| 9 | 139959853 | rs4880195 | 0.105 | -0.347 | -1.751 | -0.11 | 0.036 | -0.058 | -0.058 | 8.219 | 0.31 | 0.01 | 0.31 | 0.34 | 0.22 | 0.19 |
| 9 | 139973820 | rs7037849 | -0.019 | 0.025 | -0.892 | -0.725 | 0.004 | -0.012 | -0.012 | 2.849 | 0.37 | 0.48 | 0.45 | 0.65 | 0.4 | 0.45 |
| 9 | 139980385 | rs1018330 | -0.113 | 0.071 | -0.754 | -0.657 | 0.006 | -0.013 | -0.013 | 0 | 0.65 | 0.26 | 0.59 | 0.42 | 0.67 | 0.68 |
| 9 | 139987350 | rs10870177 | 0.825 | 0.176 | -1.28 | -0.818 | -0.11 | 0.163 | 0.163 | 0.149 | 0.29 | 0.03 | 0.24 | 0.03 | 0.37 | 0.32 |
| 9 | 139990530 | rs7871579 | 1.065 | 0.16 | -1.131 | -0.534 | 0.011 | -0.057 | -0.057 | 0 | 0.2 | 0.11 | 0.33 | 0.43 | 0.21 | 0.15 |
| 9 | 139994102 | rs4880200 | -0.9 | 0.075 | -1.423 | -1.077 | 0.051 | 0.016 | 0.016 | 0 | 0.67 | 0.87 | 0.68 | 0.68 | 0.74 | 0.77 |
| 9 | 139995065 | rs4880201 | 0.995 | 0.191 | -1.687 | -1.071 | -0.078 | 0.111 | 0.111 | 0.004 | 0.27 | 0.07 | 0.22 | 0.01 | 0.35 | 0.31 |
| 9 | 140002989 | rs4880091 | -1.156 | 0.122 | -1.848 | -1.033 | 0.012 | -0.032 | -0.032 | 0 | 0.69 | 0.99 | 0.75 | 0.7 | 0.76 | 0.78 |
| 9 | 140003427 | rs4567 | -0.339 | 0.377 | -1.842 | -1.212 | 0.001 | -0.017 | -0.017 | 0 | 0.45 | 0.55 | 0.53 | 0.7 | 0.46 | 0.51 |
| 9 | 140003640 | rs7853336 | 0.613 | 0.133 | -1.762 | -1.044 | -0.004 | 0.034 | 0.034 | 0.013 | 0.3 | 0.46 | 0.44 | 0.57 | 0.3 | 0.25 |
| 9 | 140004341 | rs11145996 | -0.89 | -0.026 | -1.752 | -1.086 | 0.026 | -0.017 | -0.017 | 0 | 0.69 | 0.98 | 0.75 | 0.7 | 0.75 | 0.78 |
| 9 | 140006202 | rs13631 | 0.084 | 0.138 | -1.715 | -1.067 | -0.002 | -0.024 | -0.024 | 0 | 0.39 | 0.53 | 0.5 | 0.68 | 0.39 | 0.46 |
| 9 | 140006491 | rs10870180 | 0.461 | 0.73 | -2.533 | -1.069 | -0.114 | 0.158 | 0.158 | 0.017 | 0.27 | 0.04 | 0.22 | 0 | 0.35 | 0.3 |
| 9 | 140018138 | rs4880094 | -0.563 | 0.065 | -2.43 | -0.477 | -0.002 | -0.033 | -0.033 | 1.352 | 0.74 | 0.99 | 0.77 | 0.8 | 0.74 | 0.78 |
| 9 | 140018892 | rs9411311 | -0.229 | 0.335 | -2.041 | -0.293 | -0.002 | -0.003 | -0.003 | 3.125 | 0.06 | 0.07 | 0.06 | 0.09 | 0.07 | 0.09 |
| 9 | 140034431 | rs2301363 | -0.629 | -0.245 | -1.987 | -0.492 | -0.001 | -0.001 | -0.001 | 0 | 0.16 | 0.17 | 0.25 | 0.22 | 0.12 | 0.33 |
| 9 | 140037029 | rs4880215 | 0.481 | 0.184 | -2.725 | -1.569 | -0.096 | 0.226 | 0.226 | 0.2 | 0.36 | 0.07 | 0.26 | 0 | 0.41 | 0.19 |
| 9 | 140042320 | rs10870198 | 0.046 | 0.24 | -2.677 | -1.533 | -0.025 | 0.133 | 0.133 | 0 | 0.59 | 0.8 | 0.72 | 0.92 | 0.62 | 0.58 |
| 9 | 140054136 | rs11794201 | 0.492 | 0.338 | -2.691 | -1.639 | -0.065 | 0.135 | 0.135 | 0 | 0.29 | 0.09 | 0.2 | 0 | 0.34 | 0.19 |
| 9 | 140055876 | rs10747050 | -0.611 | 0.192 | -2.414 | -1.32 | 0.009 | 0.045 | 0.045 | 0 | 0.89 | 0.98 | 0.94 | 0.98 | 0.94 | 0.8 |
| 9 | 140068342 | rs4074426 | -0.173 | 0.11 | -1.595 | -1.485 | -0.003 | -0.043 | -0.043 | 0 | 0.34 | 0.37 | 0.21 | 0.08 | 0.27 | 0.4 |
| 9 | 140071683 | rs28544087 | -0.202 | 0.172 | -1.297 | -1.549 | -0.007 | -0.06 | -0.06 | 1.167 | 0.34 | 0.39 | 0.16 | 0.07 | 0.25 | 0.37 |
| 9 | 140073120 | rs28560209 | -0.023 | 0.169 | -1.244 | -1.566 | 0 | -0.02 | -0.02 | 0 | 0.32 | 0.34 | 0.15 | 0.07 | 0.24 | 0.35 |
| 9 | 140075719 | rs28451678 | -0.281 | 0.159 | -1.249 | -1.571 | -0.004 | -0.04 | -0.04 | 0 | 0.34 | 0.37 | 0.16 | 0.08 | 0.25 | 0.37 |
| 9 | 140076439 | rs28665875 | 0.05 | 0.17 | -1.28 | -1.582 | 0.004 | 0.009 | 0.009 | 0 | 0.31 | 0.3 | 0.14 | 0.04 | 0.24 | 0.34 |
| 9 | 140085991 | rs13292648 | 0.199 | 0.158 | -1.43 | -1.847 | 0.001 | 0.084 | 0.084 | 0.015 | 0.61 | 0.75 | 0.81 | 0.96 | 0.7 | 0.65 |
| 9 | 140101687 | rs6606564 | 0.074 | 0.39 | -1.147 | -2.05 | -0.021 | 0.023 | 0.023 | 0 | 0.88 | 0.93 | 0.94 | 0.91 | 0.84 | 0.88 |
| 9 | 140106538 | rs34383296 | 0.795 | 1.227 | -0.513 | -1.768 | 0.001 | -0.001 | -0.001 | 0 | 0.46 | 0.41 | 0.57 | 0.55 | 0.39 | 0.46 |
| 9 | 140110274 | rs7390663 | -0.824 | 1.226 | -0.514 | -1.768 | -0.021 | 0.024 | 0.024 | 0 | 0.88 | 0.92 | 0.94 | 0.91 | 0.84 | 0.88 |
| 9 | 140128085 | rs28407527 | 0.691 | 1.672 | -0.271 | -1.223 | 0.007 | 0.002 | 0.002 | 0 | 0.48 | 0.4 | 0.58 | 0.47 | 0.4 | 0.48 |
| 9 | 140133600 | rs9722600 | 0.339 | 2.133 | -0.52 | -0.869 | 0.036 | -0.061 | -0.061 | 0 | 0.24 | 0.07 | 0.16 | 0.31 | 0.16 | 0.27 |
| 9 | 140133719 | rs9722604 | 0.958 | 2.162 | -0.529 | -0.873 | 0.027 | -0.025 | -0.025 | 0 | 0.19 | 0.05 | 0.08 | 0.16 | 0.13 | 0.18 |
| 9 | 140141794 | rs11497277 | 2.081 | 1.592 | 0.24 | -0.427 | -0.003 | 0.058 | 0.058 | 0.842 | 0.51 | 0.33 | 0.46 | 0.34 | 0.48 | 0.43 |
| 9 | 140143801 | rs11535662 | 1.484 | 1.506 | 0.361 | -0.19 | 0.003 | 0.001 | 0.001 | 0 | 0.09 | 0.02 | 0.07 | 0.31 | 0.08 | 0.17 |
| 9 | 140144215 | rs1142749 | 1.065 | 2.253 | 0.685 | -0.293 | 0.054 | -0.056 | -0.056 | 0 | 0.9 | 0.64 | 0.93 | 1 | 0.85 | 0.94 |
| 9 | 140144649 | rs9802576 | 2.684 | 2.154 | 0.7 | -0.199 | -0.013 | 0.05 | 0.05 | 0 | 0.71 | 0.52 | 0.76 | 0.48 | 0.72 | 0.68 |
| 9 | 140148051 | rs28631372 | 2.64 | 2.117 | 0.682 | -0.18 | -0.007 | 0.068 | 0.068 | 0.059 | 0.73 | 0.54 | 0.76 | 0.49 | 0.72 | 0.7 |
| 9 | 140148189 | rs28455716 | 2.912 | 2.117 | 0.682 | -0.18 | -0.012 | 0.046 | 0.046 | 0 | 0.71 | 0.54 | 0.76 | 0.48 | 0.72 | 0.68 |
| 9 | 140167877 | rs8281 | 2.73 | 2.187 | 0.802 | -0.166 | -0.011 | 0.061 | 0.061 | 0.008 | 0.72 | 0.54 | 0.76 | 0.49 | 0.73 | 0.7 |
| 9 | 140188225 | rs7850758 | -2.491 | 1.399 | 1.367 | -0.411 | -0.026 | -0.026 | -0.026 | 0 | 0.14 | 0.62 | 0.21 | 0.05 | 0.09 | 0.09 |
| 9 | 140291708 | rs7469429 | 1.838 | -0.261 | -0.688 | -0.731 | -0.033 | 0.041 | 0.041 | 0.233 | 0.27 | 0.13 | 0.21 | 0.19 | 0.31 | 0.18 |
| 9 | 140301447 | rs11516126 | 2.6 | -1.202 | -0.01 | -0.081 | -0.071 | -0.037 | -0.037 | 0 | 0.23 | 0.09 | 0.14 | 0.27 | 0.29 | 0.3 |
| 9 | 140311092 | rs11516128 | -1.503 | -1.915 | 0.065 | -0.302 | -0.005 | -0.016 | -0.016 | 0 | 0.7 | 0.76 | 0.7 | 0.45 | 0.68 | 0.62 |
| 9 | 140313025 | rs6422889 | -1.299 | -1.895 | 0.073 | -0.293 | 0.008 | 0.043 | 0.043 | 0 | 0.68 | 0.61 | 0.7 | 0.45 | 0.67 | 0.61 |
| 9 | 140316819 | rs6606588 | 1.395 | -1.835 | -0.374 | -0.782 | -0.002 | -0.011 | -0.011 | 0.423 | 0.3 | 0.24 | 0.35 | 0.61 | 0.3 | 0.37 |
| 9 | 140317084 | rs13288824 | 2.577 | -1.816 | -0.393 | -0.773 | -0.087 | -0.05 | -0.05 | 0 | 0.23 | 0.09 | 0.13 | 0.25 | 0.32 | 0.31 |
| 9 | 140321666 | rs11534413 | 1.253 | -1.78 | -0.467 | -0.756 | 0 | 0.017 | 0.017 | 0 | 0.31 | 0.3 | 0.35 | 0.61 | 0.3 | 0.33 |
| 9 | 140325962 | rs11534415 | 1.287 | -1.787 | -0.57 | -0.791 | 0 | 0.011 | 0.011 | 0 | 0.3 | 0.29 | 0.35 | 0.62 | 0.29 | 0.33 |

| | | | | | | | | | | | | | | | | |
|---|---|---|---|---|---|---|---|---|---|---|---|---|---|---|---|---|
| 9 | 140327663 | rs34155071 | 2.601 | -1.649 | -1.111 | -0.764 | -0.085 | -0.046 | -0.046 | 0 | 0.23 | 0.08 | 0.13 | 0.25 | 0.32 | 0.31 |
| 9 | 140329141 | rs28649497 | 0.929 | -0.983 | -0.826 | -0.406 | -0.001 | -0.002 | -0.002 | 0 | 0.31 | 0.25 | 0.35 | 0.7 | 0.3 | 0.34 |
| 9 | 140331833 | rs9414731 | -1.271 | -0.975 | -1.05 | -0.733 | -0.021 | 0.033 | 0.033 | 0 | 0.89 | 0.74 | 0.95 | 0.89 | 0.93 | 0.88 |
| 9 | 140333859 | rs9722391 | -0.343 | -0.033 | -1.815 | -0.812 | -0.002 | 0.046 | 0.046 | 0 | 0.29 | 0.16 | 0.45 | 0.01 | 0.31 | 0.22 |
| 9 | 140344328 | rs28699612 | 0.757 | -0.105 | -1.732 | -0.611 | -0.053 | -0.139 | -0.139 | 0.177 | 0.22 | 0.01 | 0.24 | 0.46 | 0.22 | 0.28 |
| 9 | 140348401 | rs9410194 | -0.947 | -0.023 | -0.647 | -1.576 | 0.009 | 0.018 | 0.018 | 0 | 0.32 | 0.22 | 0.41 | 0.01 | 0.31 | 0.24 |
| 9 | 140358581 | rs4962237 | -0.979 | 0.066 | 0.136 | -0.494 | 0.002 | -0.005 | -0.005 | 0.069 | 0.1 | 0.07 | 0.03 | 0 | 0.08 | 0.04 |
| 9 | 140382705 | rs11137287 | -0.054 | 0.16 | 0.268 | -0.557 | 0.001 | -0.018 | -0.018 | 0.174 | 0.56 | 0.52 | 0.53 | 0.67 | 0.55 | 0.49 |
| 9 | 140389613 | rs13283256 | -0.199 | 0.185 | -0.369 | -0.323 | 0 | 0.009 | 0.009 | 0 | 0.11 | 0.04 | 0.05 | 0.05 | 0.11 | 0.04 |
| 9 | 140392494 | rs4962232 | 0.309 | 0.031 | -0.048 | -0.137 | 0.002 | -0.008 | -0.008 | 0.102 | 0.44 | 0.48 | 0.47 | 0.66 | 0.48 | 0.43 |
| 9 | 140396212 | rs1891629 | 0.357 | 0.247 | -0.199 | -0.198 | 0.017 | 0 | 0 | 0 | 0.16 | 0.18 | 0.29 | 0.16 | 0.18 | 0.32 |
| 9 | 140399983 | rs7037200 | -0.303 | 0.128 | -0.293 | -0.127 | 0.003 | 0.054 | 0.054 | 0 | 0.61 | 0.76 | 0.59 | 0.74 | 0.65 | 0.58 |
| 9 | 140408189 | rs17065442 | -0.068 | -0.014 | 0.434 | 0.334 | -0.002 | 0.005 | 0.005 | 0 | 0.11 | 0.13 | 0.07 | 0.21 | 0.09 | 0.07 |
| 9 | 140410881 | rs2480105 | -0.967 | -0.209 | 1.161 | 0.298 | -0.028 | 0.097 | 0.097 | 0 | 0.15 | 0.46 | 0.09 | 0.23 | 0.09 | 0.13 |
| 9 | 140414410 | rs11137410 | -0.234 | -0.116 | 1.054 | 0.227 | -0.011 | 0.032 | 0.032 | 0 | 0.12 | 0.18 | 0.07 | 0.21 | 0.09 | 0.07 |
| 9 | 140418677 | rs821304 | -0.76 | 0.214 | 0.836 | 0.07 | -0.007 | 0.003 | 0.003 | 0.006 | 0.35 | 0.55 | 0.22 | 0.63 | 0.3 | 0.45 |
| 9 | 140422452 | rs28875616 | -0.204 | 0.234 | 0.418 | -0.139 | -0.001 | 0.006 | 0.006 | 0.246 | 0.25 | 0.26 | 0.18 | 0.69 | 0.21 | 0.34 |
| 9 | 140432318 | rs10867123 | -0.181 | 0.251 | 0.411 | -0.127 | -0.001 | 0.006 | 0.006 | 0.424 | 0.24 | 0.26 | 0.18 | 0.69 | 0.2 | 0.34 |
| 9 | 140436548 | rs1099298 | -0.479 | 0.375 | 0.038 | -0.146 | -0.009 | -0.004 | -0.004 | 0 | 0.32 | 0.55 | 0.21 | 0.71 | 0.26 | 0.43 |
| 9 | 140449582 | rs10110 | -0.763 | 0.633 | 0.236 | -0.941 | 0 | -0.016 | -0.016 | 1.476 | 0.18 | 0.21 | 0.41 | 0.09 | 0.19 | 0.19 |
| 9 | 140450159 | rs1105221 | -0.763 | 0.633 | 0.236 | -0.941 | 0 | -0.016 | -0.016 | 0.743 | 0.18 | 0.21 | 0.41 | 0.09 | 0.19 | 0.19 |
| 9 | 140466816 | rs950078 | -0.724 | 0.922 | 0.433 | -0.989 | 0 | -0.008 | -0.008 | 0.053 | 0.17 | 0.16 | 0.4 | 0.09 | 0.19 | 0.19 |
| 9 | 140483076 | rs2259657 | -1.256 | 1.289 | 0.165 | -1.889 | 0.006 | 0.252 | 0.252 | 3.32 | 0.49 | 0.85 | 0.63 | 0.16 | 0.51 | 0.48 |
| 9 | 140507688 | rs4962227 | 0.013 | 1.172 | 0.543 | -1.767 | -0.001 | 0.005 | 0.005 | 0 | 0.11 | 0.1 | 0.38 | 0.09 | 0.13 | 0.15 |
| 9 | 140512555 | rs1045777 | -1.372 | 1.106 | 1.597 | -1.851 | -0.005 | -0.062 | -0.062 | 0.547 | 0.2 | 0.6 | 0.44 | 0.1 | 0.18 | 0.38 |
| 9 | 140515875 | rs10867031 | 0.127 | 1.016 | 1.175 | -1.826 | -0.032 | 0.005 | 0.005 | 0 | 0.1 | 0.01 | 0.37 | 0.08 | 0.13 | 0.14 |
| 9 | 140526439 | rs2987621 | -0.425 | 0.99 | 1.205 | -1.812 | 0 | -0.017 | -0.017 | 0 | 0.12 | 0.26 | 0.39 | 0.09 | 0.13 | 0.15 |
| 9 | 140542378 | rs11137167 | 0.003 | 1.136 | 0.958 | -1.802 | -0.009 | 0.005 | 0.005 | 0.022 | 0.09 | 0.08 | 0.38 | 0.08 | 0.13 | 0.17 |
| 9 | 140566936 | rs17496354 | -0.695 | 1.117 | 1.454 | -1.874 | -0.004 | 0.004 | 0.004 | 0 | 0.13 | 0.14 | 0.39 | 0.1 | 0.15 | 0.24 |
| 9 | 140569260 | rs2501557 | -0.57 | 1.234 | 1.959 | -2.291 | -0.048 | -0.18 | -0.18 | 0 | 0.27 | 0.44 | 0.11 | 0 | 0.16 | 0.24 |
| 9 | 140581700 | rs10732688 | -1.097 | 1.801 | 1.831 | -2.271 | 0.055 | -0.034 | -0.034 | 0.491 | 0.17 | 0.3 | 0.46 | 0.1 | 0.28 | 0.24 |
| 9 | 140587156 | rs975978 | -1.403 | 1.14 | 1.945 | -2.283 | 0.009 | 0.003 | 0.003 | 0.029 | 0.36 | 0.69 | 0.49 | 0.11 | 0.26 | 0.31 |
| 9 | 140601645 | rs4634736 | 0.286 | 1.042 | 1.916 | -2.203 | 0.011 | -0.008 | -0.008 | 1.02 | 0.18 | 0.13 | 0.07 | 0 | 0.1 | 0.07 |
| 9 | 140602887 | rs3125785 | 0.303 | 1.063 | 1.972 | -2.202 | 0.008 | -0.015 | -0.015 | 0 | 0.18 | 0.14 | 0.07 | 0 | 0.1 | 0.07 |
| 9 | 140605334 | rs7039441 | 0.292 | 0.361 | 1.362 | -2.908 | 0.01 | 0.012 | 0.012 | 0 | 0.49 | 0.45 | 0.3 | 0.83 | 0.37 | 0.69 |
| 9 | 140610042 | rs7037783 | -1.323 | 0.998 | 1.412 | -2.383 | 0.049 | 0.179 | 0.179 | 0.542 | 0.49 | 0.79 | 0.58 | 0.17 | 0.37 | 0.35 |
| 9 | 140611436 | rs3812497 | 0.1 | 0.869 | 1.237 | -2.48 | -0.006 | 0.076 | 0.076 | 1.074 | 0.28 | 0.09 | 0.45 | 0.14 | 0.26 | 0.31 |
| 9 | 140613053 | rs9314636 | -0.093 | 0.986 | 1.586 | -2.532 | -0.038 | -0.109 | -0.109 | 0 | 0.15 | 0.43 | 0.08 | 0 | 0.06 | 0.01 |
| 9 | 140623027 | rs3125782 | -0.129 | 0.98 | 1.607 | -2.517 | -0.004 | -0.056 | -0.056 | 0 | 0.1 | 0.44 | 0.07 | 0 | 0.04 | 0 |
| 9 | 140624798 | rs10867056 | 0.1 | 1.009 | 1.507 | -2.556 | -0.022 | 0.096 | 0.096 | 1.144 | 0.29 | 0.09 | 0.46 | 0.12 | 0.28 | 0.31 |
| 9 | 140629940 | rs9410127 | -0.16 | 1.026 | 1.615 | -2.455 | -0.007 | -0.054 | -0.054 | 0 | 0.1 | 0.37 | 0.06 | 0 | 0.04 | 0 |
| 9 | 140636333 | rs3123510 | -1.533 | 1.095 | 1.204 | -2.542 | -0.039 | 0.314 | 0.314 | 3.059 | 0.5 | 0.91 | 0.6 | 0.12 | 0.4 | 0.52 |
| 9 | 140642331 | rs6559221 | 0.231 | 0.73 | 1.316 | -2.273 | -0.005 | 0.088 | 0.088 | 0.551 | 0.67 | 0.84 | 0.52 | 0.88 | 0.69 | 0.65 |
| 9 | 140661826 | rs7034879 | -0.482 | 0.746 | 1.52 | -2.16 | -0.001 | -0.05 | -0.05 | 1.62 | 0.34 | 0.39 | 0.49 | 0.12 | 0.3 | 0.35 |
| 9 | 140677678 | rs12004293 | -0.531 | 0.662 | 1.587 | -2.165 | -0.007 | -0.079 | -0.079 | 0.64 | 0.36 | 0.46 | 0.5 | 0.12 | 0.29 | 0.35 |
| 9 | 140682841 | rs4487855 | 0.343 | 0.731 | 1.268 | -1.951 | 0 | -0.031 | -0.031 | 0 | 0.55 | 0.52 | 0.46 | 0.87 | 0.65 | 0.65 |
| 9 | 140698856 | rs7852475 | 0.127 | 0.844 | 0.589 | -1.586 | -0.026 | 0.159 | 0.159 | 0.13 | 0.58 | 0.85 | 0.46 | 0.81 | 0.64 | 0.64 |
| 9 | 140700868 | rs7028850 | -0.26 | 0.67 | 1.016 | -1.713 | 0.021 | 0.018 | 0.018 | 1.004 | 0.06 | 0.6 | 0.05 | 0.01 | 0 | 0 |
| 9 | 140706127 | rs11789684 | 0.743 | 0.648 | 0.904 | -1.706 | 0.043 | 0.252 | 0.252 | 1.181 | 0.5 | 0.18 | 0.41 | 0.89 | 0.65 | 0.6 |
| 9 | 140720117 | rs12344858 | -0.775 | 0.303 | 0.57 | -1.116 | 0.002 | -0.001 | -0.001 | 0.51 | 0.06 | 0.62 | 0.05 | 0.01 | 0 | 0 |
| 9 | 140734161 | rs4526432 | 0.152 | 0.865 | -0.548 | -0.725 | -0.002 | 0.005 | 0.005 | 0 | 0.61 | 0.63 | 0.34 | 0.76 | 0.47 | 0.59 |
| 9 | 140738306 | rs7390989 | 0.585 | 0.666 | -1.621 | -0.602 | 0.034 | 0.121 | 0.121 | 0.78 | 0.46 | 0.08 | 0.24 | 0.7 | 0.38 | 0.4 |
| 9 | 140751501 | rs10867083 | -0.014 | 1.252 | -0.544 | 0.256 | 0.057 | 0.078 | 0.078 | 0 | 0.77 | 0.58 | 0.61 | 0.65 | 0.67 | 0.64 |
| 9 | 140755119 | rs7389298 | -0.109 | 0.754 | -1.064 | 0.332 | 0.002 | 0.003 | 0.003 | 0 | 0.28 | 0.27 | 0.23 | 0.27 | 0.22 | 0.26 |
| 9 | 140756825 | rs7862101 | -0.134 | 1.095 | -0.943 | 0.501 | -0.001 | 0.001 | 0.001 | 0 | 0.13 | 0.17 | 0.19 | 0.24 | 0.14 | 0.15 |
| 10 | 426966 | rs4242758 | -0.069 | -0.839 | -0.745 | -0.472 | -0.002 | 0.008 | 0.008 | 0 | 0.38 | 0.42 | 0.51 | 0.51 | 0.36 | 0.5 |
| 10 | 431161 | rs10904086 | 0.029 | -0.85 | -0.715 | -0.488 | -0.002 | 0.004 | 0.004 | 0 | 0.32 | 0.66 | 0.49 | 0.51 | 0.29 | 0.46 |
| 10 | 468599 | rs12415961 | 0.367 | -1.056 | -0.356 | 0.089 | 0.002 | -0.004 | -0.004 | 0 | 0.41 | 0.63 | 0.52 | 0.52 | 0.37 | 0.49 |
| 10 | 544349 | rs11252791 | -0.38 | -0.591 | 0.617 | -0.201 | 0.003 | 0.078 | 0.078 | 0.671 | 0.08 | 0.41 | 0.05 | 0.1 | 0.04 | 0.07 |
| 10 | 547567 | rs816599 | 0.655 | -0.675 | 0.671 | -0.256 | -0.029 | 0.094 | 0.094 | 0.38 | 0.9 | 0.46 | 0.93 | 0.87 | 0.94 | 0.93 |
| 10 | 554186 | rs12251997 | -0.3 | -0.639 | 0.678 | -0.227 | 0.004 | 0.079 | 0.079 | 0.666 | 0.08 | 0.39 | 0.05 | 0.1 | 0.05 | 0.06 |
| 10 | 570172 | rs1672503 | -1.634 | -1.132 | 0.099 | -0.99 | -0.008 | 0.041 | 0.041 | 0 | 0.59 | 0.83 | 0.49 | 0.34 | 0.61 | 0.37 |
| 10 | 588126 | rs11253082 | -1.205 | -0.603 | 0.558 | -0.776 | -0.022 | 0.027 | 0.027 | 0.148 | 0.14 | 0.53 | 0.09 | 0.13 | 0.1 | 0.11 |
| 10 | 588406 | rs10904450 | -1.736 | -0.637 | 0.467 | -0.792 | -0.043 | 0.02 | 0.02 | 0.64 | 0.21 | 0.66 | 0.16 | 0.27 | 0.14 | 0.13 |
| 10 | 592283 | rs816646 | -1.653 | -0.573 | 0.242 | -0.881 | -0.023 | 0.038 | 0.038 | 0 | 0.59 | 0.77 | 0.47 | 0.33 | 0.57 | 0.34 |
| 10 | 595504 | rs865289 | -1.861 | -0.254 | 0.994 | -0.214 | 0.029 | 0.098 | 0.098 | 0 | 0.48 | 0.78 | 0.42 | 0.34 | 0.45 | 0.29 |
| 10 | 601089 | rs816650 | -1.124 | -0.206 | 1.709 | 0.309 | 0.042 | 0.07 | 0.07 | 0.408 | 0.09 | 0.45 | 0.08 | 0.1 | 0.08 | 0.06 |
| 10 | 605525 | rs2794954 | -2.334 | -0.237 | 1.278 | 0.29 | 0.021 | -0.007 | -0.007 | 0 | 0.23 | 0.81 | 0.25 | 0.28 | 0.16 | 0.1 |
| 10 | 609945 | rs816623 | -1.599 | -0.263 | 1.767 | 0.365 | -0.066 | 0.074 | 0.074 | 0 | 0.13 | 0.43 | 0.1 | 0.18 | 0.04 | 0.05 |

| | | | | | | | | | | | | | | | | |
|---|---|---|---|---|---|---|---|---|---|---|---|---|---|---|---|---|
| 10 | 615782 | rs17221309 | -0.339 | -0.014 | 1.408 | 0.065 | -0.032 | 0.044 | 0.044 | 0 | 0.3 | 0.22 | 0.31 | 0.07 | 0.37 | 0.22 |
| 10 | 623410 | rs11253256 | 1.393 | -0.291 | 1.133 | 0.251 | -0.096 | 0.036 | 0.036 | 0 | 0.48 | 0.03 | 0.43 | 0.63 | 0.47 | 0.7 |
| 10 | 626782 | rs7095189 | 1.442 | -0.869 | 1.713 | 0.409 | -0.031 | 0.043 | 0.043 | 0.522 | 0.85 | 0.43 | 0.83 | 0.83 | 0.9 | 0.92 |
| 10 | 627211 | rs1672535 | 1.599 | -0.885 | 1.674 | 0.402 | -0.059 | 0.044 | 0.044 | 0.795 | 0.79 | 0.37 | 0.78 | 0.7 | 0.88 | 0.89 |
| 10 | 645916 | rs816620 | 1.245 | -0.792 | 1.167 | 0.269 | -0.009 | 0.085 | 0.085 | 0 | 0.49 | 0.13 | 0.47 | 0.63 | 0.54 | 0.66 |
| 10 | 662193 | rs2386709 | -0.241 | -0.953 | 1.418 | 0.016 | 0.003 | 0.032 | 0.032 | 0 | 0.08 | 0.25 | 0.13 | 0.1 | 0.06 | 0.05 |
| 10 | 673037 | rs816563 | 0.842 | -1.101 | 1.56 | 0.007 | -0.034 | 0.121 | 0.121 | 0 | 0.89 | 0.61 | 0.92 | 0.82 | 0.96 | 0.97 |
| 10 | 678000 | rs7077311 | -0.144 | -1.088 | 1.678 | 0.014 | -0.015 | 0.074 | 0.074 | 0 | 0.07 | 0.2 | 0.06 | 0.14 | 0.02 | 0.02 |
| 10 | 680428 | rs2124585 | 0.851 | -1.099 | 1.557 | -0.067 | 0.001 | 0.072 | 0.072 | 0 | 0.58 | 0.28 | 0.59 | 0.75 | 0.61 | 0.75 |
| 10 | 680475 | rs816570 | -0.123 | -1.052 | 1.524 | -0.061 | -0.048 | 0.085 | 0.085 | 0.034 | 0.28 | 0.13 | 0.3 | 0.07 | 0.35 | 0.2 |
| 10 | 700623 | rs883728 | 1.233 | -0.326 | 1.731 | 0.189 | -0.044 | 0.073 | 0.073 | 1.129 | 0.88 | 0.52 | 0.91 | 0.85 | 0.97 | 0.96 |
| 10 | 712617 | rs17136372 | 0.295 | -0.247 | 1.682 | 0.226 | -0.011 | 0.091 | 0.091 | 0.072 | 0.92 | 0.73 | 0.94 | 0.86 | 0.98 | 0.98 |
| 10 | 719803 | rs4880737 | 0.572 | -0.233 | 0.599 | 0.222 | -0.015 | 0.115 | 0.115 | 0.308 | 0.9 | 0.53 | 0.93 | 0.85 | 0.98 | 0.96 |
| 10 | 738630 | rs1750792 | -1.37 | -0.201 | -0.17 | 0.26 | -0.012 | 0.002 | 0.002 | 3.036 | 0.2 | 0.74 | 0.2 | 0.26 | 0.08 | 0.08 |
| 10 | 743836 | rs11253324 | -0.472 | 0.063 | -0.153 | 0.256 | 0.008 | 0.137 | 0.137 | 0 | 0.08 | 0.35 | 0.07 | 0.15 | 0.02 | 0.05 |
| 10 | 744591 | rs12763675 | -0.23 | 0.434 | -1.675 | 0.003 | -0.067 | 0.094 | 0.094 | 1.969 | 0.27 | 0.14 | 0.31 | 0.09 | 0.36 | 0.19 |
| 10 | 749238 | rs553326 | -0.593 | 0.626 | -1.714 | 0.408 | -0.019 | 0.074 | 0.074 | 0 | 0.48 | 0.88 | 0.54 | 0.36 | 0.46 | 0.27 |
| 10 | 752193 | rs1581843 | -0.661 | 1.117 | -0.99 | 0.551 | 0.019 | 0.013 | 0.013 | 0 | 0.32 | 0.4 | 0.34 | 0.42 | 0.39 | 0.26 |
| 10 | 754033 | rs2265090 | -0.107 | 0.735 | -1.253 | 0.086 | 0.01 | 0.019 | 0.019 | 0 | 0.35 | 0.22 | 0.32 | 0.74 | 0.3 | 0.49 |
| 10 | 757222 | rs10904546 | 0.587 | 0.996 | -1.414 | -0.534 | -0.059 | 0.072 | 0.072 | 0 | 0.12 | 0.01 | 0.09 | 0 | 0.17 | 0.05 |
| 10 | 759268 | rs7916968 | -0.334 | 0.746 | -1.068 | -0.746 | -0.001 | 0.002 | 0.002 | 0 | 0.09 | 0.19 | 0.19 | 0.1 | 0.1 | 0.12 |
| 10 | 761907 | rs2486581 | 0.425 | 0.258 | -0.935 | -0.622 | -0.04 | 0.004 | 0.004 | 2.779 | 0.09 | 0.14 | 0.01 | 0.08 | 0.03 | 0.02 |
| 10 | 762872 | rs816314 | -1.079 | 0.178 | -0.726 | -1.163 | -0.01 | -0.045 | -0.045 | 0 | 0.79 | 0.71 | 0.85 | 0.92 | 0.82 | 0.83 |
| 10 | 766105 | rs7906313 | 0.431 | 0.112 | -0.837 | -1.332 | -0.023 | 0.081 | 0.081 | 0 | 0.45 | 0.28 | 0.43 | 0.74 | 0.48 | 0.55 |
| 10 | 767395 | rs816318 | 0.046 | 0.494 | -0.499 | -1.303 | 0.005 | -0.013 | -0.013 | 0 | 0.2 | 0.2 | 0.31 | 0.07 | 0.31 | 0.14 |
| 10 | 776027 | rs11253377 | -0.589 | 0.096 | -0.363 | -1.364 | -0.005 | 0.081 | 0.081 | 0 | 0.45 | 0.32 | 0.46 | 0.17 | 0.55 | 0.29 |
| 10 | 776394 | rs11253380 | -0.589 | 0.055 | -0.332 | -1.369 | -0.007 | 0.11 | 0.11 | 0 | 0.46 | 0.3 | 0.45 | 0.17 | 0.56 | 0.29 |
| 10 | 776544 | rs943204 | -0.979 | -0.168 | 0.005 | -1.453 | -0.022 | -0.083 | -0.083 | 0 | 0.78 | 0.65 | 0.74 | 0.93 | 0.84 | 0.79 |
| 10 | 778318 | rs754034 | -0.536 | 0.053 | -0.135 | -1.356 | -0.012 | 0.124 | 0.124 | 0 | 0.45 | 0.24 | 0.45 | 0.17 | 0.55 | 0.29 |
| 10 | 787896 | rs1769242 | 0.581 | -0.016 | -0.096 | -1.255 | -0.007 | 0.012 | 0.012 | 0 | 0.18 | 0.08 | 0.19 | 0 | 0.21 | 0.06 |
| 10 | 800213 | rs2254501 | 0.142 | -0.31 | -0.236 | -1.393 | -0.016 | 0.02 | 0.02 | 0.658 | 0.37 | 0.29 | 0.44 | 0.23 | 0.52 | 0.32 |
| 10 | 802272 | rs2256711 | -0.001 | -0.415 | -0.164 | -0.619 | -0.018 | -0.033 | -0.033 | 0 | 0.73 | 0.57 | 0.68 | 0.78 | 0.77 | 0.81 |
| 10 | 802445 | rs1769215 | 0.864 | -0.14 | 0.054 | -0.283 | 0.018 | -0.024 | -0.024 | 0 | 0.36 | 0.22 | 0.25 | 0.55 | 0.28 | 0.5 |
| 10 | 803721 | rs2790381 | 0.086 | -0.156 | 0.174 | -0.184 | 0.016 | 0.143 | 0.143 | 0 | 0.9 | 0.69 | 0.88 | 0.78 | 0.94 | 0.86 |
| 10 | 804315 | rs497338 | 0.759 | -0.096 | -0.647 | -0.222 | 0.029 | -0.035 | -0.035 | 0 | 0.36 | 0.22 | 0.25 | 0.54 | 0.27 | 0.52 |
| 10 | 811876 | rs12414585 | 0.146 | 0.11 | -0.425 | -0.397 | 0.001 | 0.006 | 0.006 | 0 | 0.34 | 0.3 | 0.28 | 0.16 | 0.31 | 0.27 |
| 10 | 813426 | rs11253424 | 1.084 | 0.34 | -0.748 | 0.03 | -0.049 | 0.06 | 0.06 | 0 | 0.1 | 0.01 | 0.17 | 0 | 0.19 | 0.04 |
| 10 | 815149 | rs7084027 | -1.018 | 0.292 | -0.759 | 0.307 | 0.003 | 0.018 | 0.018 | 0.128 | 0.76 | 0.77 | 0.72 | 0.59 | 0.75 | 0.57 |
| 10 | 819904 | rs17159886 | 0.44 | -0.16 | -0.929 | 0.804 | 0.003 | -0.005 | -0.005 | 0 | 0.13 | 0.06 | 0.12 | 0.17 | 0.12 | 0.04 |
| 10 | 820868 | rs10752019 | 0.167 | -0.207 | -0.898 | 0.813 | -0.002 | 0.009 | 0.009 | 0.404 | 0.46 | 0.44 | 0.42 | 0.41 | 0.45 | 0.38 |
| 10 | 824693 | rs11815869 | -0.573 | -0.794 | -1 | 1.04 | -0.015 | 0.016 | 0.016 | 0 | 0.2 | 0.21 | 0.13 | 0.34 | 0.13 | 0.11 |
| 10 | 831747 | rs4881527 | -0.15 | -0.47 | -1.299 | 0.479 | 0.029 | 0.022 | 0.022 | 0.836 | 0.19 | 0.09 | 0.17 | 0.11 | 0.12 | 0.08 |
| 10 | 834736 | rs4881530 | -0.035 | 0.234 | -1.723 | 0.543 | -0.02 | -0.021 | -0.021 | 0.003 | 0.34 | 0.14 | 0.36 | 0.37 | 0.38 | 0.53 |
| 10 | 836115 | rs11253444 | 0.025 | 1 | -0.903 | 0.049 | 0.061 | -0.062 | -0.062 | 0 | 0.11 | 0.28 | 0.14 | 0 | 0.19 | 0.09 |
| 10 | 837079 | rs12772869 | 0.212 | 1.338 | -1.181 | 0.346 | -0.01 | -0.016 | -0.016 | 0 | 0.36 | 0.23 | 0.35 | 0.43 | 0.42 | 0.58 |
| 10 | 837450 | rs4881531 | 0.532 | 1.278 | -1.431 | 0.084 | -0.005 | -0.01 | -0.01 | 0 | 0.2 | 0.16 | 0.24 | 0.46 | 0.21 | 0.44 |
| 10 | 838179 | rs1536337 | 0.373 | 0.091 | -2.167 | -0.001 | 0.043 | -0.038 | -0.038 | 0.542 | 0.29 | 0.02 | 0.27 | 0.31 | 0.18 | 0.13 |
| 10 | 853106 | rs1057304 | -0.229 | 0.321 | -1.728 | -0.159 | -0.025 | -0.012 | -0.012 | 0.083 | 0.33 | 0.27 | 0.41 | 0.38 | 0.47 | 0.5 |
| 10 | 854011 | rs7071801 | -0.044 | 0.326 | -1.748 | -0.184 | -0.076 | -0.042 | -0.042 | 0 | 0.32 | 0.16 | 0.31 | 0.37 | 0.44 | 0.49 |
| 10 | 858022 | rs9124 | 0.64 | 0.405 | -1.807 | -0.259 | -0.035 | -0.08 | -0.08 | 0.005 | 0.23 | 0.07 | 0.2 | 0.35 | 0.25 | 0.44 |
| 10 | 858394 | rs7703 | -0.226 | 0.712 | -1.916 | -0.389 | -0.067 | -0.037 | -0.037 | 0.846 | 0.32 | 0.18 | 0.41 | 0.37 | 0.48 | 0.5 |
| 10 | 862889 | rs748112 | -0.075 | 0.695 | -1.918 | -0.369 | -0.082 | -0.055 | -0.055 | 0 | 0.31 | 0.13 | 0.31 | 0.37 | 0.43 | 0.47 |
| 10 | 863964 | rs2296625 | 1.097 | 0.301 | -0.929 | -0.894 | 0.003 | -0.091 | -0.091 | 0.715 | 0.18 | 0.3 | 0.11 | 0.02 | 0.17 | 0.27 |
| 10 | 888073 | rs1341741 | 0.355 | 0.247 | -0.453 | -0.749 | 0.087 | -0.05 | -0.05 | 0 | 0.09 | 0.23 | 0.22 | 0 | 0.21 | 0.06 |
| 10 | 894838 | rs12249828 | 0.314 | 0.214 | -0.37 | -0.764 | 0.056 | -0.046 | -0.046 | 0.303 | 0.1 | 0.18 | 0.21 | 0 | 0.21 | 0.05 |
| 10 | 903950 | rs11253471 | -0.335 | -0.234 | -0.153 | -0.746 | 0.005 | 0.002 | 0.002 | 0.948 | 0.48 | 0.46 | 0.51 | 0.41 | 0.63 | 0.7 |
| 10 | 927331 | rs11253489 | 0.366 | -0.318 | -0.274 | -0.439 | 0.091 | -0.015 | -0.015 | 0 | 0.45 | 0.14 | 0.39 | 0.49 | 0.3 | 0.25 |
| 10 | 936103 | rs17221435 | 1.588 | -0.486 | -0.554 | -0.396 | -0.022 | -0.07 | -0.07 | 0 | 0.14 | 0 | 0.11 | 0.19 | 0.13 | 0.08 |
| 10 | 944496 | rs7077992 | 0.145 | -0.074 | -0.073 | -0.398 | 0.003 | 0.001 | 0.001 | 0.911 | 0.4 | 0.43 | 0.3 | 0.41 | 0.42 | 0.66 |
| 10 | 964306 | rs11253509 | -0.253 | 0.231 | -0.119 | -0.662 | -0.009 | -0.006 | -0.006 | 0 | 0.13 | 0.15 | 0.22 | 0 | 0.24 | 0.05 |
| 10 | 969377 | rs12259338 | -0.253 | 0.231 | -0.111 | -0.662 | -0.001 | -0.018 | -0.018 | 0.132 | 0.13 | 0.17 | 0.23 | 0 | 0.24 | 0.05 |
| 10 | 991737 | rs10904588 | -1.02 | 0.537 | -0.002 | -0.776 | 0.031 | -0.088 | -0.088 | 1.801 | 0.35 | 0.52 | 0.39 | 0.1 | 0.44 | 0.35 |
| 10 | 1013535 | rs6560863 | 0.611 | 0.876 | -0.107 | -0.665 | -0.017 | 0.026 | 0.026 | 0 | 0.8 | 0.82 | 0.8 | 0.9 | 0.72 | 0.84 |
| 10 | 1017325 | rs11253543 | 0.519 | 0.1 | 0.483 | -0.577 | 0.001 | -0.001 | -0.001 | 0.262 | 0.23 | 0.22 | 0.24 | 0.26 | 0.19 | 0.22 |
| 10 | 1018585 | rs4880745 | 1.009 | 0.1 | 0.545 | -0.58 | -0.003 | 0.012 | 0.012 | 0.09 | 0.56 | 0.59 | 0.56 | 0.64 | 0.52 | 0.62 |
| 10 | 1032568 | rs11253552 | 0.841 | 0.075 | 0.454 | -0.643 | 0.004 | 0.008 | 0.008 | 0 | 0.1 | 0.04 | 0.07 | 0.1 | 0.09 | 0.12 |
| 10 | 1034412 | rs12772979 | 1.139 | 0.107 | 0.446 | -0.62 | -0.024 | 0.021 | 0.021 | 0 | 0.09 | 0.03 | 0.12 | 0 | 0.19 | 0.04 |
| 10 | 1037091 | rs7094916 | 0.727 | 0.033 | 0.496 | -0.489 | 0.008 | -0.051 | -0.051 | 0 | 0.22 | 0.07 | 0.23 | 0.29 | 0.19 | 0.22 |
| 10 | 1040798 | rs7095853 | 0.251 | 0.047 | 0.477 | -0.505 | -0.021 | 0.031 | 0.031 | 0 | 0.8 | 0.87 | 0.8 | 0.9 | 0.71 | 0.84 |

| | | | | | | | | | | | | | | | | |
|---|---|---|---|---|---|---|---|---|---|---|---|---|---|---|---|---|
| 10 | 1043469 | rs2242271 | 1.516 | -0.364 | 1.038 | -0.663 | 0.014 | 0.041 | 0.041 | 0 | 0.49 | 0.22 | 0.52 | 0.58 | 0.51 | 0.55 |
| 10 | 1043798 | rs10752022 | 0.104 | -0.194 | 0.775 | -0.61 | -0.018 | 0.024 | 0.024 | 0 | 0.8 | 0.86 | 0.8 | 0.9 | 0.71 | 0.84 |
| 10 | 1046712 | rs2306409 | 1.149 | -0.209 | 0.859 | -0.61 | 0 | 0.002 | 0.002 | 0 | 0.54 | 0.41 | 0.54 | 0.61 | 0.52 | 0.61 |
| 10 | 1050754 | rs12415759 | 1.191 | -0.199 | 0.562 | -0.606 | 0.001 | 0.008 | 0.008 | 0 | 0.54 | 0.39 | 0.54 | 0.61 | 0.52 | 0.61 |
| 10 | 1055271 | rs10904597 | 0.82 | -0.009 | -0.294 | -0.552 | 0.006 | 0.005 | 0.005 | 0 | 0.1 | 0.05 | 0.06 | 0.1 | 0.08 | 0.12 |
| 10 | 1056252 | rs17293608 | 0.973 | 0.027 | -0.312 | -0.561 | -0.06 | 0.054 | 0.054 | 0 | 0.1 | 0.01 | 0.14 | 0 | 0.21 | 0.04 |
| 10 | 1058837 | rs2306404 | 0.728 | -0.064 | -0.224 | -0.54 | 0.009 | -0.013 | -0.013 | 0 | 0.22 | 0.15 | 0.24 | 0.29 | 0.19 | 0.22 |
| 10 | 1060218 | rs3207775 | 0.593 | 0.279 | -0.316 | -0.515 | 0.006 | 0.006 | 0.006 | 0 | 0.1 | 0.05 | 0.06 | 0.1 | 0.08 | 0.12 |
| 10 | 1065191 | rs17221491 | 0.686 | 0.524 | -0.325 | -0.455 | -0.06 | 0.054 | 0.054 | 0 | 0.1 | 0.01 | 0.14 | 0 | 0.21 | 0.04 |
| 10 | 1065710 | rs1044261 | 0.826 | 0.407 | -0.362 | -0.553 | -0.003 | 0.012 | 0.012 | 0.348 | 0.07 | 0.04 | 0.04 | 0 | 0.07 | 0.01 |
| 10 | 1067284 | rs3812656 | -1.526 | 0.665 | -0.427 | -0.449 | -0.031 | 0.028 | 0.028 | 0.85 | 0.34 | 0.79 | 0.38 | 0.25 | 0.43 | 0.27 |
| 10 | 1068639 | rs13377037 | 0.034 | 0.799 | 0.023 | -0.352 | 0.003 | 0.014 | 0.014 | 0 | 0.08 | 0.15 | 0.12 | 0.11 | 0.08 | 0.03 |
| 10 | 1068802 | rs3750729 | -1.916 | 0.518 | 0.219 | -0.528 | -0.005 | 0.003 | 0.003 | 0 | 0.59 | 0.68 | 0.54 | 0.43 | 0.57 | 0.46 |
| 10 | 1073684 | rs10751789 | 0.078 | 0.377 | -0.024 | -0.52 | 0.026 | 0.041 | 0.041 | 2.857 | 0.33 | 0.09 | 0.23 | 0.28 | 0.26 | 0.34 |
| 10 | 1074710 | rs11250227 | 0.078 | 0.377 | -0.041 | -0.499 | 0.026 | 0.038 | 0.038 | 2.857 | 0.33 | 0.09 | 0.23 | 0.28 | 0.26 | 0.34 |
| 10 | 1081293 | rs3793781 | 0.213 | 0.452 | 0.217 | -0.479 | -0.004 | 0.013 | 0.013 | 0 | 0.08 | 0.15 | 0.11 | 0.11 | 0.07 | 0.05 |
| 10 | 1081882 | rs7075695 | 0.654 | 0.33 | 0.552 | -0.553 | 0.003 | 0.005 | 0.005 | 0.631 | 0.23 | 0.24 | 0.19 | 0.18 | 0.18 | 0.17 |
| 10 | 1084360 | rs10508204 | -1.269 | 0.495 | 0.667 | -0.647 | -0.001 | -0.001 | -0.001 | 0 | 0.45 | 0.61 | 0.48 | 0.34 | 0.5 | 0.32 |
| 10 | 1086116 | rs1285 | 1.073 | 0.483 | 0.592 | -0.669 | -0.006 | 0.023 | 0.023 | 1.187 | 0.21 | 0.05 | 0.17 | 0.18 | 0.18 | 0.17 |
| 10 | 1087551 | rs12768746 | 1.81 | 0.401 | 0.85 | -0.63 | -0.013 | 0.015 | 0.015 | 0 | 0.43 | 0.23 | 0.45 | 0.56 | 0.43 | 0.55 |
| 10 | 1089616 | rs11250238 | -0.748 | 0.268 | 0.655 | -0.541 | -0.004 | -0.002 | -0.002 | 0.112 | 0.88 | 0.88 | 0.93 | 0.88 | 0.92 | 0.87 |
| 10 | 1096538 | rs12359250 | -0.171 | 0.764 | 0.716 | -0.469 | 0.006 | -0.007 | -0.007 | 0 | 0.15 | 0.19 | 0.24 | 0.09 | 0.23 | 0.08 |
| 10 | 1096932 | rs7100196 | 0.436 | 0.509 | 0.902 | -0.343 | -0.008 | 0.005 | 0.005 | 0.336 | 0.05 | 0.17 | 0.02 | 0.05 | 0.02 | 0.06 |
| 10 | 1100064 | rs11250242 | 0.294 | 0.623 | 0.644 | -0.334 | 0.02 | 0.047 | 0.047 | 0 | 0.05 | 0.15 | 0.11 | 0.09 | 0.07 | 0.05 |
| 10 | 1101535 | rs10903357 | 0.816 | 0.639 | 0.23 | -0.377 | -0.014 | 0.022 | 0.022 | 1.315 | 0.21 | 0.04 | 0.17 | 0.17 | 0.18 | 0.17 |
| 10 | 1116671 | rs2387217 | 0.936 | 0.548 | 0.264 | -0.34 | -0.006 | 0.02 | 0.02 | 0.725 | 0.2 | 0.03 | 0.17 | 0.17 | 0.15 | 0.16 |
| 10 | 1119203 | rs10508203 | -1.208 | 0.463 | 0.734 | -0.387 | -0.028 | 0.016 | 0.016 | 0.028 | 0.22 | 0.42 | 0.26 | 0.27 | 0.17 | 0.25 |
| 10 | 1126293 | rs7087381 | -1.281 | 0.44 | 0.745 | -0.4 | -0.013 | -0.007 | -0.007 | 0 | 0.42 | 0.49 | 0.43 | 0.44 | 0.32 | 0.41 |
| 10 | 1128216 | rs7074721 | 0.985 | 0.426 | 0.903 | -0.406 | 0.008 | 0.005 | 0.005 | 2.238 | 0.57 | 0.4 | 0.56 | 0.56 | 0.68 | 0.59 |
| 10 | 1128812 | rs11819230 | 0.236 | 0.453 | 0.947 | -0.399 | 0.004 | 0.02 | 0.02 | 0 | 0.06 | 0.13 | 0.12 | 0.09 | 0.06 | 0.05 |
| 10 | 1129814 | rs7080500 | -0.68 | 0.424 | 0.987 | -0.413 | 0 | -0.001 | -0.001 | 0.669 | 0.16 | 0.18 | 0.23 | 0.26 | 0.14 | 0.19 |
| 10 | 1132852 | rs2279256 | -0.032 | 0.523 | 1.041 | -0.364 | 0.002 | 0.01 | 0.01 | 0 | 0.06 | 0.14 | 0.12 | 0.07 | 0.06 | 0.05 |
| 10 | 1135221 | rs4880473 | -0.564 | 0.486 | 1.109 | -0.407 | -0.028 | -0.009 | -0.009 | 0.628 | 0.16 | 0.29 | 0.14 | 0.18 | 0.11 | 0.2 |
| 10 | 1139183 | rs10159718 | -1.104 | 0.5 | 1.065 | -0.308 | -0.018 | -0.008 | -0.008 | 0 | 0.43 | 0.47 | 0.43 | 0.44 | 0.32 | 0.41 |
| 10 | 1141749 | rs3816735 | -1.822 | 1.798 | 1.376 | -0.071 | -0.013 | 0.05 | 0.05 | 0.005 | 0.54 | 0.72 | 0.51 | 0.36 | 0.53 | 0.44 |
| 10 | 1142424 | rs10794717 | -1.111 | 0.693 | 0.924 | -0.575 | -0.004 | 0 | 0 | 0 | 0.23 | 0.23 | 0.28 | 0.27 | 0.2 | 0.26 |
| 10 | 1143827 | rs12784270 | 1.385 | 1.012 | 1.151 | -0.054 | -0.055 | -0.004 | -0.004 | 0 | 0.45 | 0.12 | 0.36 | 0.53 | 0.48 | 0.54 |
| 10 | 1144049 | rs7067536 | -0.124 | 1.227 | 1.051 | 0.049 | 0.025 | 0.012 | 0.012 | 0.433 | 0.06 | 0.28 | 0.03 | 0.01 | 0.03 | 0.07 |
| 10 | 1145182 | rs17155886 | 0.418 | 1.46 | 0.633 | 0.169 | 0.004 | 0.018 | 0.018 | 0 | 0.05 | 0.08 | 0.12 | 0.09 | 0.06 | 0.05 |
| 10 | 1151723 | rs11250264 | 0.364 | 1.468 | 0.637 | 0.172 | 0.004 | 0.018 | 0.018 | 0 | 0.05 | 0.09 | 0.12 | 0.1 | 0.06 | 0.05 |
| 10 | 1156487 | rs11250267 | 0.712 | 1.626 | 0.624 | 0.078 | 0.027 | -0.04 | -0.04 | 0.185 | 0.11 | 0.18 | 0.17 | 0 | 0.23 | 0.04 |
| 10 | 1157410 | rs7088740 | -0.582 | 0.641 | 0.55 | 0.689 | -0.014 | 0 | 0 | 0.866 | 0.37 | 0.39 | 0.35 | 0.45 | 0.26 | 0.37 |
| 10 | 1159566 | rs11250268 | 0.036 | 0.211 | 0.153 | 0.718 | -0.011 | 0.052 | 0.052 | 0 | 0.1 | 0.15 | 0.16 | 0.26 | 0.07 | 0.13 |
| 10 | 1166400 | rs1871626 | 0.31 | 0.386 | -0.3 | 0.491 | 0.018 | 0.001 | 0.001 | 1.279 | 0.57 | 0.4 | 0.53 | 0.54 | 0.71 | 0.59 |
| 10 | 1168267 | rs7899518 | 0.124 | 1.09 | -1.162 | 0.67 | -0.003 | -0.028 | -0.028 | 0 | 0.65 | 0.25 | 0.6 | 0.4 | 0.68 | 0.6 |
| 10 | 1171354 | rs11250274 | -0.59 | 1.385 | -0.682 | 0.475 | 0.019 | -0.027 | -0.027 | 0 | 0.05 | 0.14 | 0.05 | 0.01 | 0.07 | 0.04 |
| 10 | 1171734 | rs10903372 | -0.013 | 0.916 | -1.222 | 0.468 | 0.019 | 0.027 | 0.027 | 4.843 | 0.22 | 0.04 | 0.15 | 0.19 | 0.15 | 0.17 |
| 10 | 1179519 | rs7924182 | -0.539 | 1.621 | -0.921 | 0.748 | -0.016 | 0.001 | 0.001 | 0.675 | 0.41 | 0.34 | 0.43 | 0.44 | 0.46 | 0.27 |
| 10 | 1181623 | rs6560712 | -0.539 | 1.552 | -0.655 | 0.847 | -0.025 | -0.016 | -0.016 | 0 | 0.41 | 0.31 | 0.43 | 0.48 | 0.46 | 0.27 |
| 10 | 1183077 | rs12762464 | 0.555 | 1.604 | -0.093 | 0.727 | -0.003 | 0.026 | 0.026 | 0.246 | 0.53 | 0.2 | 0.5 | 0.57 | 0.5 | 0.71 |
| 10 | 1183713 | rs7918800 | 0.425 | 1.561 | -0.202 | 0.371 | 0.022 | -0.033 | -0.033 | 0 | 0.33 | 0.51 | 0.23 | 0.26 | 0.38 | 0.37 |
| 10 | 1183739 | rs11250283 | 0.703 | 2.564 | 0.142 | 0.199 | 0.055 | -0.139 | -0.139 | 1.75 | 0.75 | 0.45 | 0.79 | 0.96 | 0.69 | 0.86 |
| 10 | 1185306 | rs6560713 | -0.2 | 2.122 | -0.081 | 0.81 | 0.008 | -0.015 | -0.015 | 1.616 | 0.35 | 0.31 | 0.47 | 0.52 | 0.3 | 0.39 |
| 10 | 1187512 | rs4880476 | -1.351 | 2.218 | 0.351 | 0.756 | -0.002 | 0.001 | 0.001 | 0.28 | 0.38 | 0.64 | 0.61 | 0.3 | 0.44 | 0.28 |
| 10 | 1188555 | rs4880479 | 0.421 | 2.254 | 0.537 | 0.705 | -0.039 | 0.049 | 0.049 | 0.056 | 0.19 | 0.1 | 0.24 | 0.1 | 0.24 | 0.07 |
| 10 | 1195959 | rs4310544 | 1.27 | 2.248 | 0.865 | 0.728 | -0.018 | 0.021 | 0.021 | 0.297 | 0.62 | 0.28 | 0.39 | 0.7 | 0.58 | 0.73 |
| 10 | 1199313 | rs10903382 | -0.289 | 2.208 | 0.882 | 0.717 | 0.014 | 0.019 | 0.019 | 0 | 0.17 | 0.31 | 0.34 | 0.18 | 0.18 | 0.19 |
| 10 | 1200445 | rs1876899 | -0.594 | 3.41 | 1.043 | 0.83 | -0.004 | 0.005 | 0.005 | 0.119 | 0.81 | 0.84 | 0.78 | 0.9 | 0.78 | 0.94 |
| 10 | 1204296 | rs11250295 | -0.098 | 2.847 | 0.866 | 0.816 | 0.008 | 0.03 | 0.03 | 0 | 0.16 | 0.31 | 0.25 | 0.18 | 0.15 | 0.17 |
| 10 | 1205850 | rs12570526 | 0.989 | 2.674 | 0.599 | 0.898 | 0.012 | -0.07 | -0.07 | 0 | 0.23 | 0.06 | 0.17 | 0.34 | 0.18 | 0.21 |
| 10 | 1207809 | rs10903387 | 0.164 | 2.728 | 0.743 | 0.907 | -0.001 | 0 | 0 | 0.912 | 0.55 | 0.42 | 0.56 | 0.55 | 0.57 | 0.71 |
| 10 | 1211558 | rs11250304 | 0.953 | 2.704 | 0.643 | 0.949 | 0 | 0.005 | 0.005 | 0 | 0.1 | 0.08 | 0.29 | 0.18 | 0.11 | 0.18 |
| 10 | 1216861 | rs10903395 | -0.28 | 2.715 | 0.588 | 0.927 | -0.012 | -0.011 | -0.011 | 0 | 0.59 | 0.76 | 0.59 | 0.57 | 0.58 | 0.74 |
| 10 | 1218030 | rs10903396 | 0.015 | 3.32 | 0.798 | 1.607 | -0.005 | -0.001 | -0.001 | 0 | 0.57 | 0.62 | 0.58 | 0.57 | 0.56 | 0.74 |
| 10 | 1219013 | rs12771555 | 1.218 | 2.161 | 0.178 | 0.916 | 0 | 0.014 | 0.014 | 0 | 0.38 | 0.1 | 0.2 | 0.35 | 0.37 | 0.5 |
| 10 | 1223742 | rs4880783 | -0.325 | 1.529 | 0.565 | 0.726 | -0.037 | 0.045 | 0.045 | 1.12 | 0.71 | 0.5 | 0.75 | 0.64 | 0.79 | 0.76 |
| 10 | 1224336 | rs6560716 | -0.964 | 1.641 | 0.664 | 0.646 | -0.005 | 0.012 | 0.012 | 5.675 | 0.78 | 0.75 | 0.82 | 0.7 | 0.82 | 0.81 |
| 10 | 1224459 | rs1500966 | -0.32 | 1.417 | 0.582 | 0.535 | -0.001 | 0 | 0 | 0 | 0.71 | 0.71 | 0.78 | 0.7 | 0.75 | 0.78 |
| 10 | 1224599 | rs7901513 | -0.049 | 1.047 | 0.626 | 0.229 | 0.001 | -0.001 | -0.001 | 0 | 0.29 | 0.3 | 0.22 | 0.3 | 0.24 | 0.22 |

| | | | | | | | | | | | | | | | | |
|---|---|---|---|---|---|---|---|---|---|---|---|---|---|---|---|---|
| 10 | 1226783 | rs17293692 | 1.426 | 1.079 | 0.523 | 0.164 | -0.045 | -0.003 | -0.003 | 0 | 0.35 | 0.08 | 0.27 | 0.37 | 0.37 | 0.47 |
| 10 | 1226944 | rs4880484 | -0.685 | 0.633 | 1.26 | 0.235 | 0 | 0 | 0 | 0 | 0.29 | 0.34 | 0.45 | 0.29 | 0.32 | 0.29 |
| 10 | 1228206 | rs1046914 | -0.676 | 0.647 | 1.264 | 0.241 | 0 | 0 | 0 | 0 | 0.28 | 0.34 | 0.45 | 0.29 | 0.32 | 0.29 |
| 10 | 1228296 | rs904957 | 0.446 | 0.64 | 1.298 | 0.24 | 0.001 | 0.002 | 0.002 | 0.49 | 0.69 | 0.34 | 0.53 | 0.67 | 0.68 | 0.7 |
| 10 | 1228905 | rs11250317 | 0.923 | 0.698 | 0.989 | 0.175 | -0.044 | 0.068 | 0.068 | 0 | 0.15 | 0.06 | 0.13 | 0 | 0.21 | 0.04 |
| 10 | 1229009 | rs904960 | 0.548 | 0.698 | 1.023 | 0.184 | 0.014 | 0.009 | 0.009 | 0.052 | 0.72 | 0.59 | 0.54 | 0.71 | 0.68 | 0.71 |
| 10 | 1229663 | rs882312 | 0.542 | 0.644 | 0.5 | 0.17 | 0.007 | -0.033 | -0.033 | 0 | 0.11 | 0.05 | 0.25 | 0.25 | 0.09 | 0.25 |
| 10 | 1230968 | rs2271275 | 0.775 | 0.607 | 0.524 | 0.16 | -0.001 | -0.004 | -0.004 | 0 | 0.67 | 0.25 | 0.51 | 0.67 | 0.68 | 0.69 |
| 10 | 1232500 | rs7924253 | 0.122 | 0.538 | 0.458 | 0.289 | -0.008 | 0.053 | 0.053 | 0 | 0.14 | 0.26 | 0.32 | 0.29 | 0.11 | 0.25 |
| 10 | 1240519 | rs7081523 | -0.16 | 0.54 | 0.533 | 0.292 | -0.012 | 0.135 | 0.135 | 0 | 0.15 | 0.39 | 0.34 | 0.34 | 0.11 | 0.27 |
| 10 | 1240966 | rs7099511 | -0.137 | 0.769 | 0.6 | -0.116 | 0.03 | -0.108 | -0.108 | 0.565 | 0.82 | 0.68 | 0.85 | 1 | 0.78 | 0.96 |
| 10 | 1244414 | rs1392827 | 0.081 | 0.527 | 0.377 | 0.184 | -0.013 | 0.01 | 0.01 | 0.017 | 0.69 | 0.45 | 0.72 | 0.7 | 0.72 | 0.76 |
| 10 | 1246883 | rs4880487 | 0.653 | 0.827 | 0.224 | -0.207 | 0.05 | -0.114 | -0.114 | 0 | 0.19 | 0.37 | 0.16 | 0 | 0.26 | 0.05 |
| 10 | 1250184 | rs883248 | 0.702 | 0.715 | -0.101 | -0.124 | -0.045 | 0.033 | 0.033 | 0 | 0.63 | 0.13 | 0.47 | 0.73 | 0.59 | 0.67 |
| 10 | 1252224 | rs10751794 | -0.814 | 0.172 | -0.28 | -0.2 | 0.061 | -0.013 | -0.013 | 0 | 0.42 | 0.24 | 0.53 | 0.55 | 0.34 | 0.48 |
| 10 | 1256632 | rs3817043 | 1.89 | 0.218 | -0.721 | -0.296 | -0.116 | -0.115 | -0.115 | 0.34 | 0.33 | 0.03 | 0.27 | 0.43 | 0.37 | 0.47 |
| 10 | 1259485 | rs2387308 | -0.151 | 0.013 | -0.505 | -0.277 | -0.007 | 0 | 0 | 0 | 0.71 | 0.74 | 0.8 | 0.71 | 0.78 | 0.8 |
| 10 | 1274538 | rs11250349 | 0.342 | 0.096 | -0.638 | -0.238 | 0.008 | -0.025 | -0.025 | 0.597 | 0.14 | 0.12 | 0.31 | 0.28 | 0.12 | 0.28 |
| 10 | 1275157 | rs4880490 | 0.378 | 0.106 | -0.552 | -0.228 | 0.047 | -0.003 | -0.003 | 0 | 0.58 | 0.77 | 0.5 | 0.44 | 0.65 | 0.53 |
| 10 | 1275314 | rs4880491 | 0.766 | 0.187 | -0.465 | -0.257 | 0.063 | -0.145 | -0.145 | 0 | 0.21 | 0.53 | 0.18 | 0.01 | 0.26 | 0.05 |
| 10 | 1277716 | rs1995852 | -0.518 | -0.479 | -1.062 | -0.155 | 0.002 | -0.002 | -0.002 | 0 | 0.84 | 0.89 | 0.85 | 0.71 | 0.84 | 0.83 |
| 10 | 1280409 | rs4880802 | 0.42 | -0.337 | -0.916 | -0.168 | 0.045 | -0.004 | -0.004 | 0 | 0.59 | 0.77 | 0.51 | 0.44 | 0.66 | 0.53 |
| 10 | 1282828 | rs3793735 | 1.286 | 0.364 | -0.996 | -0.104 | -0.098 | -0.102 | -0.102 | 0 | 0.38 | 0.02 | 0.29 | 0.44 | 0.36 | 0.54 |
| 10 | 1286740 | rs7900341 | 0.131 | 0.632 | -0.599 | -0.133 | -0.001 | 0.003 | 0.003 | 0.147 | 0.13 | 0.12 | 0.31 | 0.22 | 0.14 | 0.24 |
| 10 | 1293100 | rs7100066 | 1.058 | 0.686 | -0.544 | -0.393 | -0.01 | 0.002 | 0.002 | 0 | 0.21 | 0.16 | 0.19 | 0.02 | 0.31 | 0.06 |
| 10 | 1300253 | rs6560721 | -0.449 | 0.812 | -1.39 | -0.151 | 0 | -0.009 | -0.009 | 0 | 0.58 | 0.59 | 0.7 | 0.66 | 0.54 | 0.8 |
| 10 | 1306745 | rs4880807 | 1.482 | 0.607 | -1.521 | -0.15 | -0.092 | -0.061 | -0.061 | 0 | 0.36 | 0.02 | 0.28 | 0.41 | 0.33 | 0.5 |
| 10 | 1307456 | rs11250372 | 0.208 | 0.642 | -1.457 | -0.124 | -0.006 | -0.007 | -0.007 | 0 | 0.13 | 0.12 | 0.35 | 0.24 | 0.17 | 0.25 |
| 10 | 1310657 | rs4880494 | 0.743 | 0.515 | -1.408 | -0.153 | 0.009 | 0.012 | 0.012 | 0.015 | 0.43 | 0.41 | 0.35 | 0.42 | 0.35 | 0.55 |
| 10 | 1322617 | rs17156146 | 1.174 | 0.896 | -1.913 | -0.029 | -0.067 | -0.042 | -0.042 | 0 | 0.4 | 0.01 | 0.28 | 0.42 | 0.33 | 0.55 |
| 10 | 1325411 | rs2804102 | -0.133 | 1.63 | -1.496 | 0.34 | 0.001 | -0.012 | -0.012 | 0 | 0.59 | 0.53 | 0.69 | 0.67 | 0.54 | 0.8 |
| 10 | 1328249 | rs12573544 | -0.264 | 1.835 | -2.097 | 0.141 | 0.014 | 0.047 | 0.047 | 0.015 | 0.11 | 0.16 | 0.35 | 0.24 | 0.18 | 0.22 |
| 10 | 1329985 | rs7096088 | 0.028 | 1.074 | -2.299 | -0.63 | -0.039 | 0.106 | 0.106 | 0 | 0.21 | 0.03 | 0.16 | 0 | 0.27 | 0.04 |
| 10 | 1332768 | rs4880811 | -0.217 | 0.706 | -2.527 | -0.176 | -0.018 | -0.026 | -0.026 | 0.04 | 0.39 | 0.03 | 0.25 | 0.66 | 0.31 | 0.52 |
| 10 | 1333590 | rs2387639 | -0.467 | 0.802 | -1.621 | -0.975 | 0.001 | -0.066 | -0.066 | 4.81 | 0.64 | 0.45 | 0.72 | 0.95 | 0.6 | 0.79 |
| 10 | 1335849 | rs1027604 | -0.135 | 0.323 | -1.978 | -1.392 | -0.008 | 0.193 | 0.193 | 1.341 | 0.51 | 0.74 | 0.67 | 0.87 | 0.47 | 0.65 |
| 10 | 1337709 | rs1542874 | -0.266 | 0.665 | -0.743 | -1.412 | 0.036 | -0.045 | -0.045 | 0 | 0.75 | 0.64 | 0.67 | 0.81 | 0.7 | 0.61 |
| 10 | 1338168 | rs4880813 | 0.762 | 0.948 | -0.791 | -1.331 | -0.03 | 0.115 | 0.115 | 0 | 0.26 | 0.05 | 0.19 | 0 | 0.31 | 0.09 |
| 10 | 1339022 | rs11250389 | 1.583 | 0.79 | -0.424 | -0.453 | -0.032 | -0.035 | -0.035 | 0 | 0.06 | 0 | 0.03 | 0.1 | 0.11 | 0.04 |
| 10 | 1340620 | rs3849975 | 0.238 | 0.596 | -0.233 | -0.29 | 0.024 | 0.068 | 0.068 | 1.613 | 0.51 | 0.18 | 0.55 | 0.69 | 0.46 | 0.72 |
| 10 | 1341691 | rs4471349 | 0.119 | 0.651 | -0.152 | -0.213 | 0.025 | 0.071 | 0.071 | 1.893 | 0.49 | 0.18 | 0.55 | 0.69 | 0.46 | 0.7 |
| 10 | 1343017 | rs11599315 | -0.546 | 0.664 | 0.112 | -0.468 | -0.001 | -0.007 | -0.007 | 0.036 | 0.61 | 0.32 | 0.72 | 0.93 | 0.63 | 0.82 |
| 10 | 1343198 | rs11594721 | -0.008 | 0.596 | 0.164 | -0.398 | 0.028 | 0.124 | 0.124 | 2.207 | 0.5 | 0.24 | 0.64 | 0.8 | 0.47 | 0.71 |
| 10 | 1359774 | rs2242423 | -0.224 | 0.541 | 0.351 | 0.328 | -0.021 | 0.072 | 0.072 | 0 | 0.48 | 0.8 | 0.35 | 0.33 | 0.47 | 0.32 |
| 10 | 1371610 | rs884949 | 0.807 | 0.606 | 0.015 | 0.42 | -0.06 | 0.19 | 0.19 | 0 | 0.33 | 0.05 | 0.21 | 0.01 | 0.35 | 0.14 |
| 10 | 1372501 | rs904962 | -0.547 | 0.708 | 0.134 | 0.576 | 0.012 | 0.162 | 0.162 | 0.104 | 0.08 | 0.25 | 0.07 | 0.31 | 0.07 | 0.07 |
| 10 | 1374007 | rs7073579 | -0.191 | 0.569 | 0.25 | 0.538 | -0.001 | 0.006 | 0.006 | 0 | 0.48 | 0.61 | 0.34 | 0.32 | 0.47 | 0.32 |
| 10 | 1376968 | rs7084512 | -0.054 | 0.723 | -0.99 | 0.607 | -0.002 | 0.093 | 0.093 | 0.287 | 0.06 | 0.24 | 0.05 | 0.19 | 0.04 | 0.05 |
| 10 | 1387508 | rs11250434 | -0.084 | 0.742 | -2.225 | -0.096 | -0.033 | 0.037 | 0.037 | 0 | 0.46 | 0.08 | 0.5 | 0.69 | 0.46 | 0.64 |
| 10 | 1396155 | rs2892333 | 0.017 | 0.837 | -1.539 | -0.637 | -0.023 | 0.169 | 0.169 | 3.232 | 0.45 | 0.2 | 0.37 | 0.11 | 0.5 | 0.37 |
| 10 | 1404314 | rs11250460 | -0.738 | 1.006 | -1.188 | 0.759 | -0.038 | -0.097 | -0.097 | 0.276 | 0.17 | 0.05 | 0.34 | 0.48 | 0.2 | 0.33 |
| 10 | 1415366 | rs12766316 | -1.055 | 0.888 | 0.615 | 1.116 | 0.003 | 0.017 | 0.017 | 0 | 0.1 | 0.12 | 0.1 | 0.21 | 0.11 | 0.24 |
| 10 | 1421516 | rs10794748 | -0.555 | 0.584 | 0.362 | 0.203 | 0.005 | 0.022 | 0.022 | 0.016 | 0.59 | 0.29 | 0.59 | 0.66 | 0.6 | 0.64 |
| 10 | 1424216 | rs11596049 | -0.164 | 0.615 | 0.125 | -0.019 | -0.019 | -0.095 | -0.095 | 0 | 0.2 | 0.05 | 0.32 | 0.58 | 0.2 | 0.3 |
| 10 | 1426013 | rs17293789 | 0.52 | 0.494 | 0.2 | -0.063 | -0.042 | 0.039 | 0.039 | 0.541 | 0.32 | 0.05 | 0.35 | 0.2 | 0.34 | 0.28 |
| 10 | 1426303 | rs4880842 | 0.294 | 0.442 | 0.192 | -0.128 | -0.003 | 0.063 | 0.063 | 0 | 0.5 | 0.31 | 0.33 | 0.34 | 0.5 | 0.51 |
| 10 | 1430761 | rs2387667 | 0.619 | 0.344 | 0.092 | -0.216 | 0 | 0.107 | 0.107 | 0 | 0.47 | 0.27 | 0.3 | 0.21 | 0.45 | 0.44 |
| 10 | 1436204 | rs1986608 | 0.19 | 0.066 | -0.221 | -0.572 | 0.003 | 0.155 | 0.155 | 0 | 0.49 | 0.22 | 0.29 | 0.21 | 0.42 | 0.47 |
| 10 | 1436858 | rs4880503 | 1.254 | 0.158 | 0.137 | -0.658 | 0.016 | 0.082 | 0.082 | 0 | 0.35 | 0.2 | 0.19 | 0.01 | 0.28 | 0.21 |
| 10 | 1443016 | rs2820590 | -1.459 | 0.202 | 0.503 | -0.664 | 0.003 | -0.042 | -0.042 | 0.535 | 0.64 | 0.65 | 0.8 | 1 | 0.72 | 0.79 |
| 10 | 1445849 | rs7067912 | 0.74 | 0.217 | 0.762 | -0.557 | 0.032 | 0.143 | 0.143 | 0 | 0.84 | 0.48 | 0.85 | 0.67 | 0.85 | 0.71 |
| 10 | 1447951 | rs2001197 | -0.22 | -0.063 | 0.421 | -0.274 | -0.028 | -0.127 | -0.127 | 0 | 0.22 | 0.03 | 0.32 | 0.44 | 0.22 | 0.22 |
| 10 | 1452786 | rs17293817 | 0.464 | -0.175 | -0.242 | -0.234 | -0.029 | 0.013 | 0.013 | 2.034 | 0.27 | 0.09 | 0.31 | 0.18 | 0.35 | 0.3 |
| 10 | 1454864 | rs10903443 | -0.487 | -0.032 | -0.229 | -0.046 | 0.008 | -0.073 | -0.073 | 0 | 0.23 | 0.11 | 0.34 | 0.45 | 0.2 | 0.22 |
| 10 | 1455543 | rs2255595 | -0.322 | 0.227 | 0.229 | 0.119 | 0.004 | 0.001 | 0.001 | 0.14 | 0.35 | 0.59 | 0.43 | 0.38 | 0.41 | 0.36 |
| 10 | 1460627 | rs4447096 | -1.279 | 0.158 | 0.881 | 1.626 | -0.003 | 0.077 | 0.077 | 0 | 0.16 | 0.3 | 0.45 | 0.6 | 0.15 | 0.18 |
| 10 | 1460688 | rs2892336 | -1.335 | 0.199 | 0.899 | 1.644 | 0.002 | 0.092 | 0.092 | 0 | 0.15 | 0.3 | 0.45 | 0.6 | 0.15 | 0.18 |
| 10 | 1463208 | rs7079958 | -0.515 | 0.195 | 0.999 | 1.42 | 0.007 | 0.024 | 0.024 | 0 | 0.05 | 0.13 | 0.05 | 0.1 | 0.06 | 0.03 |
| 10 | 1468196 | rs4448619 | -0.357 | 0.2 | 1.099 | 1.691 | -0.001 | 0.02 | 0.02 | 0.165 | 0.09 | 0.09 | 0.05 | 0.17 | 0.08 | 0.22 |

| | | | | | | | | | | | | | | | | |
|---|---|---|---|---|---|---|---|---|---|---|---|---|---|---|---|---|
| 10 | 1468978 | rs17156303 | -0.768 | 0.202 | 1.118 | 2.112 | -0.003 | 0.001 | 0.001 | 0 | 0.13 | 0.14 | 0.2 | 0.17 | 0.08 | 0.13 |
| 10 | 1471765 | rs2820651 | -0.39 | 0.17 | 1.381 | 2.086 | 0.019 | 0.085 | 0.085 | 0.169 | 0.1 | 0.26 | 0.06 | 0.17 | 0.08 | 0.22 |
| 10 | 1475286 | rs12415114 | 0.171 | 0.076 | 1.11 | 2.203 | 0.008 | 0.068 | 0.068 | 0 | 0.05 | 0.06 | 0.16 | 0.25 | 0.1 | 0.05 |
| 10 | 1475811 | rs7923036 | -1.851 | -0.01 | 1.113 | 2.157 | 0 | 0.013 | 0.013 | 0 | 0.18 | 0.2 | 0.36 | 0.42 | 0.18 | 0.18 |
| 10 | 1477677 | rs17156334 | -1.346 | -0.038 | 1.175 | 2.48 | -0.007 | 0.1 | 0.1 | 0.118 | 0.16 | 0.28 | 0.17 | 0.36 | 0.16 | 0.27 |
| 10 | 1478107 | rs4880508 | -0.871 | 0.019 | 0.999 | 2.466 | 0.007 | 0.047 | 0.047 | 0 | 0.1 | 0.13 | 0.17 | 0.32 | 0.15 | 0.13 |
| 10 | 1478171 | rs12267070 | -2.466 | -0.055 | 1.253 | 2.424 | -0.004 | 0.009 | 0.009 | 1.527 | 0.31 | 0.44 | 0.43 | 0.68 | 0.3 | 0.49 |
| 10 | 1479767 | rs2387675 | -1.75 | 0.029 | 1.206 | 2.378 | 0.024 | 0.207 | 0.207 | 0 | 0.2 | 0.47 | 0.23 | 0.48 | 0.22 | 0.37 |
| 10 | 1491872 | rs2820623 | -1.112 | -0.069 | 0.547 | 2.096 | -0.001 | -0.001 | -0.001 | 0 | 0.16 | 0.15 | 0.17 | 0.14 | 0.1 | 0.16 |
| 10 | 1492512 | rs10903450 | 0.721 | 1.286 | -0.081 | 1.767 | -0.051 | -0.056 | -0.056 | 0.027 | 0.19 | 0.04 | 0.3 | 0.31 | 0.3 | 0.26 |
| 10 | 1492807 | rs2813427 | 0.948 | -0.185 | -0.237 | 0.989 | 0.014 | -0.022 | -0.022 | 3.488 | 0.63 | 0.26 | 0.5 | 0.47 | 0.68 | 0.54 |
| 10 | 1495331 | rs2820629 | -0.307 | 0.506 | 0.374 | 1.001 | 0.073 | 0.13 | 0.13 | 0 | 0.94 | 0.7 | 0.83 | 0.85 | 0.93 | 0.88 |
| 10 | 1500525 | rs2820636 | 0.692 | 0.09 | 0.087 | 1.038 | -0.026 | 0.045 | 0.045 | 3.468 | 0.67 | 0.5 | 0.67 | 0.46 | 0.75 | 0.51 |
| 10 | 1503564 | rs10903458 | 0.607 | 0.796 | -0.296 | 0.441 | 0.004 | -0.005 | -0.005 | 0.609 | 0.48 | 0.58 | 0.36 | 0.16 | 0.4 | 0.27 |
| 10 | 1503838 | rs11250534 | 0.607 | 0.796 | -0.296 | 0.441 | 0.004 | -0.005 | -0.005 | 0 | 0.48 | 0.58 | 0.35 | 0.16 | 0.4 | 0.27 |
| 10 | 1505491 | rs10903459 | -0.995 | 0.314 | -0.316 | 0.826 | -0.036 | 0.008 | 0.008 | 0 | 0.32 | 0.48 | 0.22 | 0.41 | 0.18 | 0.54 |
| 10 | 1505573 | rs17156353 | 0.006 | 0.585 | -0.913 | 0.989 | 0.019 | -0.034 | -0.034 | 0 | 0.1 | 0.05 | 0.09 | 0.23 | 0.07 | 0.13 |
| 10 | 1507960 | rs11250544 | -0.452 | 0.839 | -0.995 | 1.098 | -0.003 | -0.064 | -0.064 | 0 | 0.14 | 0.04 | 0.1 | 0.33 | 0.12 | 0.17 |
| 10 | 1509658 | rs12782764 | 0.863 | -0.055 | -1.568 | 0.175 | 0.052 | 0.058 | 0.058 | 2.923 | 0.23 | 0.13 | 0.09 | 0 | 0.11 | 0.06 |
| 10 | 1510208 | rs12778089 | 0.826 | 0.004 | -1.576 | 0.167 | 0.047 | 0.061 | 0.061 | 2.644 | 0.23 | 0.13 | 0.1 | 0 | 0.12 | 0.06 |
| 10 | 1511786 | rs1533486 | 0.518 | 0.481 | -1.362 | -0.075 | 0.01 | 0.009 | 0.009 | 0 | 0.39 | 0.52 | 0.6 | 0.51 | 0.46 | 0.42 |
| 10 | 1512546 | rs5024862 | -0.174 | 0.351 | -1.229 | 0.001 | -0.025 | -0.067 | -0.067 | 1.69 | 0.36 | 0.16 | 0.28 | 0.49 | 0.41 | 0.52 |
| 10 | 1515126 | rs2676199 | -0.2 | 1.175 | -1.134 | 0.424 | -0.003 | -0.018 | -0.018 | 0 | 0.58 | 0.64 | 0.72 | 0.51 | 0.61 | 0.69 |
| 10 | 1517243 | rs2676753 | -0.502 | 0.51 | -0.728 | 0.286 | -0.005 | 0.017 | 0.017 | 0 | 0.47 | 0.39 | 0.38 | 0.37 | 0.51 | 0.73 |
| 10 | 1518682 | rs7906088 | 0.419 | 0.187 | -0.727 | 0.354 | 0.046 | -0.021 | -0.021 | 0 | 0.38 | 0.29 | 0.22 | 0.45 | 0.21 | 0.34 |
| 10 | 1519151 | rs3849976 | 0.324 | 0.781 | -1.147 | 0.37 | 0.009 | 0.006 | 0.006 | 0 | 0.36 | 0.34 | 0.56 | 0.42 | 0.45 | 0.28 |
| 10 | 1521858 | rs2813404 | -0.49 | 1.394 | -1.416 | 0.207 | -0.025 | 0.022 | 0.022 | 0 | 0.24 | 0.18 | 0.35 | 0.15 | 0.35 | 0.38 |
| 10 | 1524409 | rs2176377 | -0.189 | 0.262 | -1.308 | 0.485 | 0 | -0.001 | -0.001 | 0 | 0.38 | 0.61 | 0.2 | 0.32 | 0.39 | 0.23 |
| 10 | 1524715 | rs2138690 | 0.186 | 0.119 | -1.6 | 0.473 | -0.051 | -0.027 | -0.027 | 0.183 | 0.28 | 0.03 | 0.16 | 0.22 | 0.36 | 0.2 |
| 10 | 1525186 | rs11250563 | 0.053 | 0.805 | -1.637 | 0.437 | -0.013 | -0.007 | -0.007 | 0.01 | 0.19 | 0.1 | 0.22 | 0.23 | 0.24 | 0.22 |
| 10 | 1527241 | rs12784235 | -0.236 | 0.138 | -1.649 | 0.431 | 0.01 | 0.027 | 0.027 | 0 | 0.54 | 0.67 | 0.61 | 0.64 | 0.47 | 0.48 |
| 10 | 1530884 | rs4880511 | 0.102 | 0.13 | -1.703 | 0.914 | 0.002 | -0.062 | -0.062 | 0 | 0.15 | 0.03 | 0.39 | 0.33 | 0.09 | 0.2 |
| 10 | 1531776 | rs4353199 | -0.067 | 0.008 | -1.036 | 0.277 | 0.05 | -0.034 | -0.034 | 0 | 0.33 | 0.45 | 0.32 | 0.25 | 0.4 | 0.39 |
| 10 | 1535202 | rs11817394 | 0.063 | -0.295 | -0.641 | 0.473 | -0.011 | 0.02 | 0.02 | 0 | 0.17 | 0.19 | 0.29 | 0.31 | 0.08 | 0.22 |
| 10 | 1535739 | rs10903467 | 0.564 | -0.266 | -0.817 | 0.081 | -0.019 | -0.001 | -0.001 | 1.645 | 0.35 | 0.23 | 0.31 | 0.33 | 0.41 | 0.3 |
| 10 | 1543614 | rs2063941 | 0.002 | -0.604 | -0.88 | -0.171 | 0.006 | -0.001 | -0.001 | 0.543 | 0.38 | 0.38 | 0.42 | 0.35 | 0.3 | 0.38 |
| 10 | 1546782 | rs2813433 | 1.393 | -0.601 | -0.903 | -0.19 | -0.001 | 0.052 | 0.052 | 0.564 | 0.19 | 0.08 | 0.13 | 0.05 | 0.22 | 0.16 |
| 10 | 1549881 | rs2676719 | -0.498 | -0.582 | -0.796 | -0.152 | 0.007 | -0.005 | -0.005 | 0.039 | 0.72 | 0.61 | 0.73 | 0.68 | 0.71 | 0.68 |
| 10 | 1553772 | rs2676774 | 0.365 | -0.702 | -0.436 | 0.02 | -0.034 | 0.055 | 0.055 | 0 | 0.19 | 0.36 | 0.3 | 0.31 | 0.09 | 0.24 |
| 10 | 1560743 | rs11250578 | 0.368 | -0.586 | -0.066 | -0.01 | 0.014 | -0.004 | -0.004 | 0 | 0.37 | 0.5 | 0.31 | 0.32 | 0.4 | 0.33 |
| 10 | 1566851 | rs11250582 | 1.579 | -0.762 | -0.384 | -0.075 | -0.032 | -0.043 | -0.043 | 1.816 | 0.13 | 0.02 | 0.08 | 0.15 | 0.18 | 0.13 |
| 10 | 1567107 | rs7905434 | 0.496 | -0.846 | -0.198 | 0.005 | -0.042 | 0.052 | 0.052 | 0 | 0.33 | 0.15 | 0.2 | 0.17 | 0.4 | 0.3 |
| 10 | 1567138 | rs2813451 | -0.067 | -0.635 | 0.123 | -0.059 | -0.001 | 0.007 | 0.007 | 0 | 0.6 | 0.35 | 0.64 | 0.7 | 0.57 | 0.66 |
| 10 | 1567300 | rs2813452 | -0.397 | -0.463 | -0.288 | 0.019 | 0 | -0.001 | -0.001 | 0 | 0.63 | 0.62 | 0.65 | 0.7 | 0.57 | 0.66 |
| 10 | 1574057 | rs11250593 | -0.003 | -0.711 | -0.627 | -0.335 | 0.043 | 0.08 | 0.08 | 0 | 0.43 | 0.12 | 0.43 | 0.74 | 0.37 | 0.48 |
| 10 | 1574443 | rs2813458 | -0.397 | -0.509 | -0.093 | -0.656 | 0.01 | 0.034 | 0.034 | 0 | 0.81 | 0.45 | 0.81 | 0.77 | 0.83 | 0.78 |
| 10 | 1576292 | rs10903470 | 0.449 | -0.602 | -0.149 | -0.455 | 0.032 | -0.016 | -0.016 | 0 | 0.42 | 0.11 | 0.42 | 0.58 | 0.37 | 0.48 |
| 10 | 1581728 | rs2138692 | 0.418 | -0.909 | 0.278 | -0.585 | -0.001 | -0.001 | -0.001 | 0 | 0.37 | 0.39 | 0.43 | 0.74 | 0.35 | 0.48 |
| 10 | 1581875 | rs2813462 | -0.247 | -0.892 | 0.26 | -0.522 | -0.009 | -0.005 | -0.005 | 0 | 0.77 | 0.72 | 0.81 | 0.79 | 0.81 | 0.78 |
| 10 | 1589508 | rs2012196 | -1.933 | -0.978 | 0.605 | -0.506 | -0.045 | -0.056 | -0.056 | 0 | 0.9 | 0.74 | 0.97 | 1 | 0.98 | 0.99 |
| 10 | 1590256 | rs2813352 | -2.088 | -0.976 | 0.567 | -0.505 | -0.042 | -0.052 | -0.052 | 0 | 0.91 | 0.79 | 0.97 | 1 | 0.98 | 0.99 |
| 10 | 1590440 | rs11250609 | 0.542 | -0.767 | 0.096 | -0.713 | -0.044 | 0.036 | 0.036 | 0 | 0.33 | 0.12 | 0.31 | 0.24 | 0.42 | 0.35 |
| 10 | 1591562 | rs11591776 | 1.232 | -0.625 | 0.168 | -0.654 | -0.016 | 0.028 | 0.028 | 0.36 | 0.19 | 0.14 | 0.21 | 0.03 | 0.21 | 0.15 |
| 10 | 1593558 | rs4880867 | -0.104 | -0.673 | 0.217 | -0.57 | -0.005 | -0.029 | -0.029 | 0 | 0.56 | 0.53 | 0.65 | 0.77 | 0.57 | 0.6 |
| 10 | 1595155 | rs4880516 | 0.119 | -0.308 | -0.236 | -0.081 | -0.007 | -0.011 | -0.011 | 0 | 0.39 | 0.21 | 0.31 | 0.58 | 0.43 | 0.53 |
| 10 | 1595481 | rs7087167 | -1.268 | -0.42 | 0.588 | -0.19 | -0.047 | 0.053 | 0.053 | 0.075 | 0.88 | 0.58 | 0.94 | 0.83 | 0.96 | 0.99 |
| 10 | 1601316 | rs10903486 | 1.127 | -0.098 | 0.464 | -0.285 | 0.011 | -0.049 | -0.049 | 0 | 0.2 | 0.31 | 0.21 | 0.03 | 0.2 | 0.12 |
| 10 | 1601900 | rs4440958 | 0.704 | -0.205 | 0.165 | -0.266 | -0.015 | 0.037 | 0.037 | 0 | 0.36 | 0.2 | 0.32 | 0.23 | 0.42 | 0.42 |
| 10 | 1601990 | rs2813363 | -2.116 | -0.19 | -0.34 | -0.224 | -0.034 | -0.052 | -0.052 | 0 | 0.9 | 0.78 | 0.96 | 1 | 0.96 | 0.99 |
| 10 | 1602635 | rs7918979 | 0.383 | 0.332 | -0.583 | 0.072 | -0.004 | -0.035 | -0.035 | 0 | 0.34 | 0.16 | 0.43 | 0.59 | 0.34 | 0.43 |
| 10 | 1605715 | rs12268030 | 0.382 | 0.632 | -0.276 | 0.385 | -0.001 | -0.012 | -0.012 | 0.475 | 0.19 | 0.23 | 0.2 | 0.03 | 0.2 | 0.12 |
| 10 | 1610839 | rs11250633 | -0.096 | 0.517 | -1.178 | 0.764 | -0.002 | -0.054 | -0.054 | 0 | 0.4 | 0.14 | 0.44 | 0.46 | 0.34 | 0.53 |
| 10 | 1611165 | rs10508213 | 0.998 | 0.362 | -1.156 | 0.316 | -0.022 | -0.024 | -0.024 | 0.005 | 0.12 | 0.01 | 0.08 | 0.12 | 0.15 | 0.11 |
| 10 | 1612405 | rs17293949 | 0.92 | 0.451 | -1.629 | 0.01 | -0.068 | 0.019 | 0.019 | 0.016 | 0.2 | 0.01 | 0.21 | 0.1 | 0.21 | 0.12 |
| 10 | 1612952 | rs2676760 | -0.485 | 0.082 | -0.806 | -0.22 | -0.016 | -0.052 | -0.052 | 0 | 0.34 | 0.56 | 0.25 | 0.21 | 0.3 | 0.33 |
| 10 | 1616438 | rs17293956 | 0.85 | -0.138 | -0.971 | 0.045 | 0.005 | 0.005 | 0.005 | 0 | 0.18 | 0.1 | 0.14 | 0.11 | 0.15 | 0.21 |
| 10 | 1618717 | rs1574269 | 1.167 | 0.447 | -0.767 | 0.13 | -0.024 | 0.018 | 0.018 | 0 | 0.22 | 0.12 | 0.19 | 0.15 | 0.34 | 0.21 |
| 10 | 1618731 | rs1320077 | -0.048 | 0.431 | -0.642 | 0.123 | -0.004 | -0.009 | -0.009 | 0 | 0.34 | 0.4 | 0.25 | 0.25 | 0.31 | 0.33 |
| 10 | 1620356 | rs6560740 | -0.551 | 0.261 | -0.84 | 0.331 | 0.023 | 0.006 | 0.006 | 0.361 | 0.57 | 0.69 | 0.44 | 0.4 | 0.65 | 0.54 |

| | | | | | | | | | | | | | | | | |
|---|---|---|---|---|---|---|---|---|---|---|---|---|---|---|---|---|
| 10 | 1621219 | rs4880875 | 0.464 | 0.453 | -0.826 | 0.339 | -0.007 | -0.003 | -0.003 | 0 | 0.1 | 0.1 | 0.1 | 0.15 | 0.16 | 0.09 |
| 10 | 1621412 | rs4880876 | -0.405 | -0.009 | -0.937 | 0.244 | -0.004 | -0.006 | -0.006 | 0 | 0.26 | 0.15 | 0.24 | 0.28 | 0.32 | 0.34 |
| 10 | 1621911 | rs7894920 | 0.828 | 0.184 | -0.806 | 0.256 | 0.003 | 0.004 | 0.004 | 2.191 | 0.16 | 0.09 | 0.15 | 0.09 | 0.17 | 0.21 |
| 10 | 1622189 | rs17156544 | 0.887 | -0.122 | -1.088 | 0.35 | 0.002 | -0.081 | -0.081 | 0 | 0.26 | 0.11 | 0.32 | 0.36 | 0.2 | 0.35 |
| 10 | 1623406 | rs10903495 | 0.871 | -0.162 | -1.042 | 0.28 | 0.006 | 0.007 | 0.007 | 1.091 | 0.18 | 0.11 | 0.15 | 0.09 | 0.18 | 0.21 |
| 10 | 1623734 | rs10903496 | -0.359 | -0.269 | -0.64 | 0.101 | 0.011 | -0.003 | -0.003 | 0 | 0.61 | 0.43 | 0.65 | 0.63 | 0.52 | 0.55 |
| 10 | 1624284 | rs4880878 | -1.042 | -0.198 | -0.401 | 0.191 | -0.02 | 0.004 | 0.004 | 0.555 | 0.81 | 0.61 | 0.88 | 0.86 | 0.82 | 0.78 |
| 10 | 1624900 | rs10903497 | 0.775 | -0.045 | -1.094 | 0.391 | -0.014 | -0.11 | -0.11 | 0.059 | 0.26 | 0.07 | 0.29 | 0.38 | 0.21 | 0.34 |
| 10 | 1625294 | rs10794765 | 0.636 | -0.029 | -0.951 | 0.212 | -0.008 | 0.018 | 0.018 | 0 | 0.19 | 0.1 | 0.15 | 0.18 | 0.21 | 0.12 |
| 10 | 1628905 | rs2813392 | -0.603 | 0.046 | -0.575 | 0.317 | 0.011 | -0.008 | -0.008 | 0 | 0.6 | 0.49 | 0.61 | 0.67 | 0.5 | 0.52 |
| 10 | 1630612 | rs4880882 | 1.051 | 0.222 | -0.81 | 1.049 | -0.017 | -0.067 | -0.067 | 0.204 | 0.11 | 0.01 | 0.2 | 0.33 | 0.05 | 0.16 |
| 10 | 1630821 | rs2999399 | -1.076 | -0.28 | -0.635 | 0.565 | -0.038 | -0.042 | -0.042 | 0.033 | 0.67 | 0.6 | 0.76 | 0.89 | 0.78 | 0.62 |
| 10 | 1637209 | rs2676187 | 0.299 | 0.61 | -0.977 | 0.588 | -0.029 | -0.005 | -0.005 | 0 | 0.3 | 0.19 | 0.45 | 0.41 | 0.47 | 0.31 |
| 10 | 1637345 | rs2676186 | 0.827 | 0.287 | -1.017 | 0.634 | 0.007 | 0.002 | 0.002 | 0 | 0.12 | 0.05 | 0.16 | 0.18 | 0.12 | 0.07 |
| 10 | 1639362 | rs7903852 | 1.235 | 0.461 | -1.062 | 0.528 | -0.047 | 0.011 | 0.011 | 0 | 0.17 | 0.07 | 0.14 | 0.1 | 0.23 | 0.12 |
| 10 | 1641948 | rs2813397 | -0.681 | 0.16 | -0.619 | 0.77 | -0.002 | -0.007 | -0.007 | 0 | 0.45 | 0.57 | 0.54 | 0.68 | 0.41 | 0.35 |
| 10 | 1642428 | rs4880885 | 0.404 | -0.249 | -0.723 | 1.213 | 0.015 | -0.026 | -0.026 | 0 | 0.14 | 0.07 | 0.14 | 0.25 | 0.04 | 0.15 |
| 10 | 1642998 | rs2813400 | 0.827 | -0.134 | -0.25 | 1.058 | -0.023 | 0.044 | 0.044 | 0 | 0.11 | 0.2 | 0.22 | 0.28 | 0.07 | 0.06 |
| 10 | 1644363 | rs1533482 | 0.448 | 0.386 | -0.648 | 0.643 | 0.02 | -0.034 | -0.034 | 0 | 0.36 | 0.52 | 0.24 | 0.21 | 0.41 | 0.37 |
| 10 | 1644771 | rs12778518 | 0.458 | 0.535 | -0.678 | 0.978 | 0 | -0.001 | -0.001 | 0 | 0.21 | 0.2 | 0.2 | 0.19 | 0.26 | 0.18 |
| 10 | 1646899 | rs1112787 | 0.318 | 0.499 | -0.82 | 0.978 | 0 | -0.001 | -0.001 | 0 | 0.22 | 0.21 | 0.2 | 0.19 | 0.27 | 0.18 |
| 10 | 1646959 | rs1112786 | -0.569 | 0.173 | -0.905 | 1.064 | 0.001 | -0.002 | -0.002 | 0 | 0.9 | 0.92 | 0.79 | 0.8 | 0.93 | 0.94 |
| 10 | 1647114 | rs10437529 | -0.569 | 0.173 | -0.928 | 1.046 | 0.003 | 0.002 | 0.002 | 0 | 0.9 | 0.93 | 0.86 | 0.91 | 0.93 | 0.94 |
| 10 | 1648296 | rs11597244 | 0.582 | 0.735 | -1.026 | 0.86 | -0.033 | -0.007 | -0.007 | 0 | 0.14 | 0.07 | 0.11 | 0.1 | 0.16 | 0.09 |
| 10 | 1649762 | rs10903510 | -0.579 | 0.372 | -0.508 | 1.34 | -0.02 | 0.079 | 0.079 | 0 | 0.2 | 0.35 | 0.27 | 0.4 | 0.14 | 0.19 |
| 10 | 1650460 | rs17156607 | 0.82 | 0.505 | -1.007 | 1.037 | -0.02 | -0.008 | -0.008 | 0 | 0.11 | 0.02 | 0.12 | 0.12 | 0.13 | 0.14 |
| 10 | 1653960 | rs11250664 | -0.254 | 0.112 | -0.236 | 1.252 | -0.02 | 0.094 | 0.094 | 0 | 0.14 | 0.37 | 0.13 | 0.26 | 0.1 | 0.13 |
| 10 | 1655849 | rs6560744 | -1.121 | -0.073 | -0.79 | 0.936 | 0.008 | -0.006 | -0.006 | 0 | 0.73 | 0.82 | 0.65 | 0.71 | 0.75 | 0.69 |
| 10 | 1655969 | rs7900711 | -0.777 | -0.313 | -0.904 | 0.913 | 0.017 | -0.007 | -0.007 | 0.345 | 0.7 | 0.82 | 0.63 | 0.68 | 0.74 | 0.67 |
| 10 | 1657184 | rs1874995 | -0.777 | -0.313 | -0.94 | 0.912 | 0.018 | -0.019 | -0.019 | 0.931 | 0.7 | 0.82 | 0.63 | 0.63 | 0.74 | 0.67 |
| 10 | 1661213 | rs4880887 | 0.902 | 0.713 | -0.972 | 0.604 | 0.021 | -0.009 | -0.009 | 0 | 0.09 | 0.13 | 0.12 | 0.08 | 0.2 | 0.12 |
| 10 | 1661398 | rs1875005 | -0.671 | 1.006 | -1.153 | 0.226 | 0 | 0 | 0 | 0.843 | 0.6 | 0.42 | 0.57 | 0.67 | 0.57 | 0.64 |
| 10 | 1663908 | rs7077892 | -0.301 | 0.949 | -1.162 | 0.162 | 0.08 | -0.097 | -0.097 | 0 | 0.14 | 0.4 | 0.08 | 0 | 0.17 | 0.05 |
| 10 | 1664242 | rs1909437 | 0.346 | 0.607 | -1.272 | 0.02 | -0.022 | 0.041 | 0.041 | 1.148 | 0.27 | 0.49 | 0.29 | 0.33 | 0.22 | 0.21 |
| 10 | 1665029 | rs10794773 | -2.056 | 0.623 | -1.318 | 0.037 | 0.009 | -0.016 | -0.016 | 0.241 | 0.73 | 0.82 | 0.65 | 0.67 | 0.75 | 0.68 |
| 10 | 1665211 | rs10082419 | -1.132 | 0.385 | -1.229 | 0.157 | -0.016 | -0.004 | -0.004 | 0 | 0.63 | 0.78 | 0.58 | 0.63 | 0.59 | 0.64 |
| 10 | 1665466 | rs11250672 | 1.23 | 0.526 | -1.332 | 0.204 | 0.021 | -0.009 | -0.009 | 0 | 0.09 | 0.13 | 0.12 | 0.08 | 0.2 | 0.12 |
| 10 | 1665821 | rs7086346 | 1.162 | 0.515 | -1.332 | 0.199 | 0.039 | -0.016 | -0.016 | 0 | 0.1 | 0.18 | 0.12 | 0.08 | 0.2 | 0.12 |
| 10 | 1667365 | rs2387952 | 0.996 | 0.478 | -1.814 | 0.311 | 0.039 | 0.002 | 0.002 | 0.267 | 0.26 | 0.1 | 0.17 | 0.21 | 0.17 | 0.31 |
| 10 | 1669110 | rs12572078 | 0.369 | 0.547 | -1.739 | 0.144 | 0.016 | -0.054 | -0.054 | 0 | 0.3 | 0.13 | 0.37 | 0.53 | 0.28 | 0.32 |
| 10 | 1669347 | rs17294019 | 1.875 | 0.452 | -1.492 | -0.116 | 0.024 | 0.072 | 0.072 | 0.152 | 0.17 | 0.05 | 0.06 | 0.02 | 0.09 | 0.19 |
| 10 | 1670434 | rs6560748 | -0.421 | 0.923 | -1.17 | 0.226 | 0 | -0.003 | -0.003 | 1.075 | 0.58 | 0.63 | 0.45 | 0.52 | 0.56 | 0.62 |
| 10 | 1672408 | rs7078498 | -0.208 | 0.404 | -1.367 | -0.376 | 0.02 | 0.088 | 0.088 | 0 | 0.57 | 0.22 | 0.73 | 0.83 | 0.63 | 0.63 |
| 10 | 1674132 | rs1909440 | 0.446 | 0.681 | -1.469 | 0.039 | -0.041 | -0.051 | -0.051 | 0.527 | 0.3 | 0.12 | 0.38 | 0.57 | 0.37 | 0.27 |
| 10 | 1674607 | rs7092046 | -0.91 | 0.307 | -1.312 | -0.113 | 0.01 | 0.002 | 0.002 | 0 | 0.69 | 0.78 | 0.63 | 0.73 | 0.71 | 0.66 |
| 10 | 1675548 | rs12267329 | 0.188 | 0.105 | -1.045 | -0.342 | 0.017 | 0.027 | 0.027 | 0 | 0.5 | 0.57 | 0.26 | 0.18 | 0.37 | 0.53 |
| 10 | 1676185 | rs7903381 | 0.521 | 0.132 | -1.226 | -0.02 | -0.001 | -0.001 | -0.001 | 0.203 | 0.13 | 0.11 | 0.23 | 0.25 | 0.18 | 0.09 |
| 10 | 1676417 | rs7920777 | 1.868 | -0.538 | -1.072 | -0.094 | -0.045 | -0.098 | -0.098 | 0 | 0.16 | 0.41 | 0.05 | 0.02 | 0.06 | 0.24 |
| 10 | 1677579 | rs11250686 | 2.298 | -0.253 | -1.482 | 0.016 | -0.093 | -0.061 | -0.061 | 0 | 0.15 | 0.01 | 0.2 | 0.21 | 0.26 | 0.19 |
| 10 | 1678572 | rs10794774 | 1.294 | -0.745 | -1.232 | -0.433 | -0.091 | -0.11 | -0.11 | 0 | 0.24 | 0.06 | 0.32 | 0.45 | 0.37 | 0.29 |
| 10 | 1679189 | rs7917694 | -0.852 | -0.827 | -1.228 | -0.5 | -0.022 | -0.014 | -0.014 | 0 | 0.74 | 0.32 | 0.79 | 0.85 | 0.91 | 0.71 |
| 10 | 1680786 | rs10903516 | 1.555 | -0.76 | -1.277 | -0.467 | -0.11 | -0.131 | -0.131 | 0 | 0.23 | 0.04 | 0.31 | 0.45 | 0.37 | 0.29 |
| 10 | 1681656 | rs4880890 | 0.678 | -0.688 | -1.28 | -0.477 | -0.026 | -0.025 | -0.025 | 0 | 0.17 | 0.11 | 0.27 | 0.25 | 0.25 | 0.16 |
| 10 | 1684797 | rs4880897 | 0.336 | -0.794 | -1.245 | -0.522 | 0.003 | 0.042 | 0.042 | 0.211 | 0.41 | 0.14 | 0.59 | 0.7 | 0.62 | 0.44 |
| 10 | 1685180 | rs7097535 | 0.729 | -0.637 | -1.352 | -0.44 | 0.018 | 0.069 | 0.069 | 0 | 0.32 | 0.13 | 0.2 | 0.16 | 0.29 | 0.27 |
| 10 | 1687799 | rs2173524 | 0.011 | -0.538 | -1.179 | -0.351 | 0.008 | 0.006 | 0.006 | 0.381 | 0.61 | 0.23 | 0.53 | 0.65 | 0.67 | 0.57 |
| 10 | 1688405 | rs11592943 | 0.459 | -0.844 | -1.491 | -0.455 | 0.062 | 0.036 | 0.036 | 0.036 | 0.51 | 0.15 | 0.51 | 0.61 | 0.65 | 0.55 |
| 10 | 1692147 | rs17221916 | 0.616 | -0.792 | -1.013 | -0.477 | 0.008 | 0.008 | 0.008 | 0 | 0.18 | 0.2 | 0.29 | 0.25 | 0.24 | 0.16 |
| 10 | 1692663 | rs10903534 | 0.355 | -0.818 | -1.165 | -0.551 | -0.001 | 0 | 0 | 0 | 0.48 | 0.52 | 0.38 | 0.51 | 0.45 | 0.55 |
| 10 | 1692796 | rs7072699 | 0.68 | -0.825 | -0.934 | -0.6 | -0.003 | 0.046 | 0.046 | 0 | 0.28 | 0.12 | 0.2 | 0.16 | 0.29 | 0.26 |
| 10 | 1694750 | rs11814112 | 1.583 | -0.788 | -0.781 | -0.591 | -0.05 | -0.101 | -0.101 | 0 | 0.17 | 0.39 | 0.05 | 0.02 | 0.07 | 0.24 |
| 10 | 1696510 | rs4303163 | -0.461 | -0.702 | -1.012 | -0.495 | 0.003 | 0.007 | 0.007 | 0 | 0.67 | 0.71 | 0.67 | 0.76 | 0.7 | 0.71 |
| 10 | 1701872 | rs11250713 | 1.106 | -0.719 | -1.206 | -0.498 | 0.012 | 0.011 | 0.011 | 0 | 0.22 | 0.1 | 0.26 | 0.17 | 0.22 | 0.27 |
| 10 | 1704555 | rs7070710 | 0.554 | -0.984 | -1.064 | -0.543 | 0.012 | 0.034 | 0.034 | 0 | 0.42 | 0.28 | 0.61 | 0.69 | 0.61 | 0.45 |
| 10 | 1704740 | rs12265257 | 0.31 | -1.065 | -1.064 | -0.584 | 0.024 | 0.056 | 0.056 | 0 | 0.35 | 0.13 | 0.31 | 0.22 | 0.29 | 0.3 |
| 10 | 1705247 | rs1909428 | -0.339 | -1.072 | -1.01 | -0.589 | 0.029 | 0.045 | 0.045 | 0 | 0.5 | 0.58 | 0.36 | 0.26 | 0.38 | 0.54 |
| 10 | 1706151 | rs4880907 | -2.638 | -0.857 | -0.924 | -0.565 | -0.027 | -0.015 | -0.015 | 0 | 0.92 | 0.87 | 0.97 | 0.95 | 0.99 | 0.98 |
| 10 | 1706725 | rs11250717 | 1.265 | -0.413 | -0.666 | -0.431 | -0.055 | -0.103 | -0.103 | 0 | 0.18 | 0.38 | 0.06 | 0.02 | 0.08 | 0.24 |
| 10 | 1710677 | rs7073160 | 0.08 | 0.553 | -0.622 | 0.249 | -0.006 | 0.031 | 0.031 | 0 | 0.27 | 0.11 | 0.29 | 0.17 | 0.28 | 0.28 |

| | | | | | | | | | | | | | | | | |
|---|---|---|---|---|---|---|---|---|---|---|---|---|---|---|---|---|
| 10 | 1712209 | rs3750685 | -0.649 | 0.67 | -0.806 | 0.773 | -0.003 | -0.002 | -0.002 | 0 | 0.19 | 0.18 | 0.29 | 0.33 | 0.24 | 0.16 |
| 10 | 1715867 | rs10219144 | -0.091 | 0.048 | -0.721 | 0.192 | -0.001 | 0 | 0 | 0 | 0.18 | 0.18 | 0.21 | 0.16 | 0.13 | 0.21 |
| 10 | 1718040 | rs7076664 | -0.16 | -0.02 | -0.3 | 0.098 | -0.013 | -0.009 | -0.009 | 0.19 | 0.86 | 0.83 | 0.91 | 0.88 | 0.9 | 0.94 |
| 10 | 1724276 | rs12766674 | -0.916 | 0.352 | -0.103 | -0.19 | -0.002 | 0.006 | 0.006 | 0 | 0.21 | 0.18 | 0.16 | 0.13 | 0.23 | 0.16 |
| 10 | 1726719 | rs11250727 | -0.028 | 0.69 | -0.51 | 0.1 | -0.025 | -0.032 | -0.032 | 0 | 0.08 | 0.02 | 0.11 | 0.21 | 0.1 | 0.1 |
| 10 | 1727152 | rs11250728 | -0.178 | 0.673 | -0.06 | -0.033 | 0.027 | -0.038 | -0.038 | 0 | 0.08 | 0.17 | 0.1 | 0.02 | 0.14 | 0.11 |
| 10 | 1727178 | rs4880531 | 1.034 | 0.398 | -0.218 | 0.071 | 0.014 | 0.044 | 0.044 | 2.545 | 0.11 | 0 | 0.05 | 0.04 | 0.07 | 0.03 |
| 10 | 1729026 | rs7896371 | 0.776 | 0.2 | -0.143 | -0.039 | -0.003 | -0.008 | -0.008 | 0 | 0.41 | 0.32 | 0.45 | 0.46 | 0.44 | 0.34 |
| 10 | 1734473 | rs7915178 | 0.405 | 0.121 | -0.07 | -0.18 | -0.001 | 0 | 0 | 0 | 0.53 | 0.47 | 0.49 | 0.57 | 0.51 | 0.54 |
| 10 | 1735559 | rs2173523 | -0.498 | 0.177 | 0.06 | -0.597 | -0.008 | -0.045 | -0.045 | 0 | 0.22 | 0.4 | 0.2 | 0.13 | 0.18 | 0.26 |
| 10 | 1736632 | rs7924155 | 0.333 | 0.085 | 0.019 | -0.94 | 0.022 | 0.052 | 0.052 | 0 | 0.24 | 0.08 | 0.21 | 0.06 | 0.18 | 0.27 |
| 10 | 1736783 | rs11250736 | -0.123 | 0.307 | 0.663 | -0.208 | 0.043 | -0.022 | -0.022 | 0 | 0.16 | 0.06 | 0.05 | 0.19 | 0.08 | 0.09 |
| 10 | 1737244 | rs11250739 | 1.183 | 0.689 | 1.314 | -0.257 | 0.086 | -0.074 | -0.074 | 0 | 0.87 | 0.6 | 0.86 | 0.97 | 0.81 | 0.91 |
| 10 | 1737850 | rs4440960 | 1.381 | 0.674 | 1.417 | -0.263 | 0.091 | -0.084 | -0.084 | 0 | 0.86 | 0.45 | 0.85 | 0.97 | 0.81 | 0.91 |
| 10 | 1739576 | rs11250741 | 0.17 | 0.619 | 0.935 | -0.232 | 0.008 | -0.001 | -0.001 | 0.606 | 0.06 | 0.03 | 0.01 | 0.15 | 0.03 | 0.06 |
| 10 | 1739627 | rs12779596 | -0.151 | 1.199 | 1.051 | -0.066 | 0.025 | 0.018 | 0.018 | 0 | 0.07 | 0.12 | 0.1 | 0.12 | 0.14 | 0.09 |
| 10 | 1740278 | rs2892394 | 1.193 | 1.179 | 1.071 | -0.084 | 0.085 | -0.075 | -0.075 | 0 | 0.87 | 0.6 | 0.86 | 0.97 | 0.81 | 0.91 |
| 10 | 1744971 | rs11250748 | -0.682 | 1.247 | 0.458 | 0.041 | 0.01 | -0.009 | -0.009 | 0 | 0.06 | 0.07 | 0.07 | 0.02 | 0.13 | 0.05 |
| 10 | 1745254 | rs6560762 | 1.107 | 1.176 | 0.232 | 0.083 | -0.014 | 0.007 | 0.007 | 0 | 0.6 | 0.23 | 0.7 | 0.66 | 0.57 | 0.72 |
| 10 | 1745303 | rs6560763 | 0.629 | 0.212 | 0.368 | -0.15 | -0.009 | -0.015 | -0.015 | 0 | 0.68 | 0.35 | 0.8 | 0.78 | 0.71 | 0.81 |
| 10 | 1746219 | rs4880914 | 1.174 | 1.113 | 0.748 | -0.055 | 0.13 | -0.035 | -0.035 | 0 | 0.89 | 0.52 | 0.88 | 0.95 | 0.81 | 0.95 |
| 10 | 1747614 | rs1909434 | 0.782 | 0.313 | 0.573 | 0.023 | -0.016 | -0.015 | -0.015 | 0 | 0.72 | 0.64 | 0.59 | 0.67 | 0.75 | 0.82 |
| 10 | 1747923 | rs1909433 | 0.541 | -0.087 | 0.792 | -0.16 | -0.039 | -0.022 | -0.022 | 0 | 0.86 | 0.61 | 0.76 | 0.88 | 0.9 | 0.93 |
| 10 | 1750471 | rs11250752 | 1.264 | -0.234 | 1.075 | -0.231 | -0.027 | -0.082 | -0.082 | 0 | 0.69 | 0.51 | 0.63 | 0.82 | 0.69 | 0.84 |
| 10 | 1750763 | rs17156773 | 0.332 | -0.157 | 0.878 | 0.61 | -0.001 | 0.086 | 0.086 | 0.477 | 0.07 | 0.06 | 0.18 | 0.37 | 0.08 | 0.25 |
| 10 | 1751710 | rs17221972 | 1.575 | -0.39 | 0.554 | 0.102 | -0.035 | 0.105 | 0.105 | 0 | 0.41 | 0.21 | 0.21 | 0.15 | 0.43 | 0.35 |
| 10 | 1753115 | rs12572345 | 0.359 | -0.473 | 0.43 | 0.086 | -0.032 | -0.095 | -0.095 | 0 | 0.24 | 0.08 | 0.31 | 0.37 | 0.24 | 0.45 |
| 10 | 1753194 | rs9702737 | 0.984 | -0.838 | 1.231 | 0.407 | -0.028 | -0.03 | -0.03 | 0 | 0.69 | 0.55 | 0.55 | 0.67 | 0.69 | 0.82 |
| 10 | 1755177 | rs4880535 | -1.717 | -0.892 | 0.868 | 0.381 | 0.007 | -0.003 | -0.003 | 0 | 0.37 | 0.67 | 0.49 | 0.48 | 0.35 | 0.2 |
| 10 | 1756440 | rs11250757 | 0.354 | -0.944 | 0.622 | 0.292 | -0.056 | 0.038 | 0.038 | 0 | 0.11 | 0.01 | 0.1 | 0.05 | 0.19 | 0.08 |
| 10 | 1758915 | rs866987 | -0.429 | -0.823 | 1.042 | 0.396 | 0.024 | 0.16 | 0.16 | 0 | 0.05 | 0.24 | 0.05 | 0.15 | 0.04 | 0.01 |
| 10 | 1760992 | rs7918362 | -1.204 | -0.739 | 0.39 | 0.369 | 0.016 | 0.015 | 0.015 | 0 | 0.36 | 0.71 | 0.41 | 0.32 | 0.36 | 0.17 |
| 10 | 1763798 | rs10903552 | 0.643 | -0.36 | 0.313 | 0.56 | -0.011 | 0.013 | 0.013 | 0.244 | 0.68 | 0.36 | 0.55 | 0.52 | 0.66 | 0.81 |
| 10 | 1765025 | rs7907903 | 0.651 | -0.249 | 0.249 | 0.316 | -0.004 | -0.005 | -0.005 | 0 | 0.68 | 0.36 | 0.62 | 0.67 | 0.65 | 0.83 |
| 10 | 1774564 | rs1316273 | 0.645 | -0.49 | 0.169 | 0.083 | -0.006 | -0.027 | -0.027 | 0 | 0.56 | 0.49 | 0.46 | 0.17 | 0.58 | 0.53 |
| 10 | 1775906 | rs884863 | 0.748 | -0.532 | 0.549 | 0.002 | 0.017 | 0.047 | 0.047 | 0 | 0.55 | 0.32 | 0.45 | 0.12 | 0.58 | 0.51 |
| 10 | 1779498 | rs3750683 | -0.809 | -0.673 | 0.372 | 1.173 | 0 | -0.051 | -0.051 | 1.602 | 0.27 | 0.12 | 0.27 | 0.47 | 0.3 | 0.34 |
| 10 | 1784653 | rs6560767 | 0.796 | -0.482 | 0.639 | 1.233 | -0.001 | -0.007 | -0.007 | 0 | 0.7 | 0.76 | 0.66 | 0.54 | 0.65 | 0.62 |
| 10 | 1785731 | rs11250769 | -0.312 | -0.564 | 0.422 | 1.377 | -0.063 | -0.094 | -0.094 | 0 | 0.19 | 0.03 | 0.27 | 0.34 | 0.29 | 0.25 |
| 10 | 1795194 | rs997934 | 0.858 | -0.49 | 0.664 | 1.307 | -0.002 | -0.021 | -0.021 | 0.718 | 0.71 | 0.77 | 0.67 | 0.54 | 0.65 | 0.61 |
| 10 | 1799598 | rs6560768 | 1.226 | -0.832 | 0.591 | 1.015 | 0.006 | 0.014 | 0.014 | 0 | 0.54 | 0.37 | 0.45 | 0.11 | 0.55 | 0.48 |
| 10 | 1805990 | rs7095591 | 0.53 | -0.916 | 0.333 | 1.463 | -0.005 | -0.019 | -0.019 | 0 | 0.67 | 0.76 | 0.64 | 0.54 | 0.61 | 0.59 |
| 10 | 1813242 | rs11250778 | 1.31 | -1.052 | 0.359 | 1.428 | 0.017 | 0.063 | 0.063 | 0 | 0.53 | 0.35 | 0.49 | 0.25 | 0.58 | 0.48 |
| 10 | 1816298 | rs12570664 | 0.216 | -0.547 | 0.441 | 1.616 | 0.036 | 0.148 | 0.148 | 0 | 0.13 | 0.36 | 0.18 | 0.29 | 0.13 | 0.14 |
| 10 | 1820854 | rs11250783 | 0.556 | -0.551 | -0.203 | 1.608 | 0.001 | -0.003 | -0.003 | 0 | 0.71 | 0.77 | 0.68 | 0.57 | 0.72 | 0.68 |
| 10 | 1830144 | rs7923476 | 0.577 | -0.829 | 0.21 | 1.703 | -0.025 | -0.002 | -0.002 | 0 | 0.92 | 0.83 | 0.89 | 0.91 | 0.96 | 0.89 |
| 10 | 1831930 | rs4457675 | 0.32 | -0.696 | -0.18 | 2.051 | -0.03 | -0.046 | -0.046 | 0.164 | 0.08 | 0.01 | 0.08 | 0.19 | 0.1 | 0.07 |
| 10 | 1831994 | rs11250787 | 1.594 | -1.539 | -0.542 | 1.243 | 0.009 | 0.179 | 0.179 | 0.067 | 0.46 | 0.1 | 0.48 | 0.15 | 0.57 | 0.41 |
| 10 | 1836223 | rs17156958 | -0.215 | -0.717 | 0.317 | 1.854 | 0.059 | 0.088 | 0.088 | 0 | 0.09 | 0.18 | 0.14 | 0.21 | 0.15 | 0.16 |
| 10 | 1836595 | rs9633666 | -0.438 | -0.917 | 0.65 | 1.756 | -0.001 | 0.035 | 0.035 | 0 | 0.16 | 0.24 | 0.19 | 0.55 | 0.17 | 0.29 |
| 10 | 1840995 | rs7475202 | 1.862 | -1.158 | 0.81 | 1.538 | 0.014 | 0.063 | 0.063 | 0 | 0.57 | 0.25 | 0.52 | 0.15 | 0.61 | 0.43 |
| 10 | 1844321 | rs4345892 | -1.179 | -1.058 | 0.819 | 1.524 | -0.007 | -0.031 | -0.031 | 1.305 | 0.9 | 0.85 | 0.95 | 1 | 0.94 | 0.99 |
| 10 | 1847006 | rs7070304 | -1.02 | -1.068 | 0.844 | 1.613 | 0.005 | -0.01 | -0.01 | 0 | 0.19 | 0.24 | 0.13 | 0.14 | 0.22 | 0.21 |
| 10 | 1848263 | rs4880925 | 0.138 | -1.089 | 0.958 | 1.68 | -0.024 | 0.214 | 0.214 | 0 | 0.85 | 0.65 | 0.82 | 0.52 | 0.87 | 0.77 |
| 10 | 1849532 | rs4880537 | 2.104 | -0.772 | 0.725 | 1.604 | 0.01 | 0.095 | 0.095 | 0 | 0.54 | 0.24 | 0.52 | 0.15 | 0.57 | 0.38 |
| 10 | 1854595 | rs10751816 | 0.752 | -0.327 | 0.593 | 2.057 | 0.038 | 0.008 | 0.008 | 0 | 0.85 | 0.75 | 0.85 | 0.8 | 0.81 | 0.75 |
| 10 | 1855203 | rs11250801 | -0.818 | -0.19 | 0.437 | 1.771 | 0.013 | 0.002 | 0.002 | 0 | 0.14 | 0.17 | 0.15 | 0.2 | 0.19 | 0.25 |
| 10 | 1863720 | rs10903564 | 0.815 | -0.659 | -0.2 | 1.427 | -0.005 | 0.127 | 0.127 | 0 | 0.83 | 0.68 | 0.81 | 0.52 | 0.84 | 0.7 |
| 10 | 1868349 | rs7905930 | 0.754 | -0.858 | -0.079 | 1.525 | -0.027 | 0.134 | 0.134 | 0 | 0.83 | 0.64 | 0.8 | 0.49 | 0.86 | 0.7 |
| 10 | 1872726 | rs6560772 | -1.301 | -0.597 | -0.51 | 0.733 | -0.026 | 0.01 | 0.01 | 0.264 | 0.32 | 0.57 | 0.32 | 0.41 | 0.26 | 0.29 |
| 10 | 1874354 | rs7912438 | 0.851 | 0.027 | -1.212 | 0.581 | 0.011 | -0.002 | -0.002 | 2.165 | 0.59 | 0.3 | 0.61 | 0.55 | 0.6 | 0.56 |
| 10 | 1875945 | rs11250815 | -0.563 | 0.492 | -1.391 | 0.936 | -0.01 | -0.063 | -0.063 | 0 | 0.1 | 0.01 | 0.17 | 0.29 | 0.11 | 0.18 |
| 10 | 1882338 | rs7904909 | -0.662 | 0.758 | -0.9 | 0.54 | 0.009 | -0.023 | -0.023 | 0 | 0.07 | 0.15 | 0.07 | 0.03 | 0.08 | 0.1 |
| 10 | 1891174 | rs6560779 | -0.588 | 2.049 | -1.52 | 0.765 | -0.108 | -0.045 | -0.045 | 0.03 | 0.16 | 0.02 | 0.3 | 0.17 | 0.33 | 0.19 |
| 10 | 1891815 | rs4880539 | -1.071 | 1.753 | -2.062 | 0.009 | -0.028 | 0.099 | 0.099 | 0.44 | 0.4 | 0.11 | 0.47 | 0.21 | 0.56 | 0.37 |
| 10 | 1895607 | rs10903567 | -0.769 | 1.157 | -2.187 | 0.419 | 0.02 | 0.001 | 0.001 | 0 | 0.38 | 0.17 | 0.22 | 0.39 | 0.32 | 0.37 |
| 10 | 1895851 | rs7893262 | -1.619 | 0.997 | -2.205 | 0.443 | 0.088 | -0.034 | -0.034 | 0 | 0.54 | 0.17 | 0.53 | 0.55 | 0.65 | 0.56 |
| 10 | 1898716 | rs11250832 | -0.195 | 0.84 | -2.032 | 0.588 | -0.006 | 0.016 | 0.016 | 0 | 0.15 | 0.21 | 0.06 | 0.35 | 0.11 | 0.17 |
| 10 | 1899620 | rs10508219 | 0.238 | 1.185 | -1.945 | 0.632 | -0.043 | -0.017 | -0.017 | 0 | 0.16 | 0.1 | 0.29 | 0.17 | 0.31 | 0.19 |

| | | | | | | | | | | | | | | | | |
|---|---|---|---|---|---|---|---|---|---|---|---|---|---|---|---|---|
| 10 | 1901330 | rs4076720 | -0.824 | 1.089 | -1.959 | 0.568 | 0.019 | -0.002 | -0.002 | 0.181 | 0.4 | 0.26 | 0.26 | 0.4 | 0.36 | 0.36 |
| 10 | 1902429 | rs4077784 | -0.894 | 1.081 | -1.962 | 0.536 | 0.013 | 0.018 | 0.018 | 0 | 0.39 | 0.26 | 0.26 | 0.36 | 0.34 | 0.35 |
| 10 | 1904598 | rs10903574 | 0.004 | 1.184 | -1.918 | 0.487 | -0.012 | 0.061 | 0.061 | 0 | 0.27 | 0.21 | 0.21 | 0.05 | 0.25 | 0.24 |
| 10 | 1906265 | rs10430757 | -0.994 | 1.331 | -1.992 | 0.554 | -0.11 | -0.14 | -0.14 | 0 | 0.28 | 0.05 | 0.33 | 0.48 | 0.4 | 0.3 |
| 10 | 1907280 | rs7080557 | 0.049 | 1.058 | -1.847 | 0.62 | 0 | 0.008 | 0.008 | 0 | 0.13 | 0.15 | 0.05 | 0.31 | 0.09 | 0.11 |
| 10 | 1912306 | rs4545463 | -1.592 | 0.877 | -1.814 | 0.625 | 0.06 | -0.005 | -0.005 | 0.213 | 0.55 | 0.37 | 0.55 | 0.53 | 0.65 | 0.54 |
| 10 | 1914990 | rs11250867 | -0.849 | 1.301 | -1.639 | 0.657 | -0.042 | 0.026 | 0.026 | 1.283 | 0.4 | 0.23 | 0.48 | 0.3 | 0.52 | 0.43 |
| 10 | 1918528 | rs7096499 | -0.282 | 1.458 | -1.49 | 0.688 | -0.034 | -0.022 | -0.022 | 0.008 | 0.84 | 0.9 | 0.72 | 0.8 | 0.7 | 0.81 |
| 10 | 1921133 | rs7084945 | -1.754 | 0.797 | -0.927 | 0.61 | 0.058 | 0.028 | 0.028 | 0.246 | 0.54 | 0.75 | 0.57 | 0.62 | 0.66 | 0.58 |
| 10 | 1921146 | rs17222168 | 0.925 | 0.643 | -1.003 | 0.523 | 0.012 | 0.012 | 0.012 | 0 | 0.06 | 0.05 | 0.04 | 0 | 0.04 | 0.02 |
| 10 | 1921530 | rs7098283 | -1.299 | 0.634 | -0.992 | 0.528 | 0.036 | 0.007 | 0.007 | 1.034 | 0.46 | 0.66 | 0.53 | 0.57 | 0.6 | 0.55 |
| 10 | 1922296 | rs2492866 | 1.59 | 0.642 | -0.995 | 0.531 | 0.07 | 0.029 | 0.029 | 0.334 | 0.46 | 0.22 | 0.43 | 0.38 | 0.34 | 0.42 |
| 10 | 1923792 | rs7073281 | 0.526 | 0.634 | -0.685 | 0.511 | -0.003 | -0.087 | -0.087 | 5.516 | 0.19 | 0.4 | 0.17 | 0.1 | 0.16 | 0.24 |
| 10 | 1923821 | rs11250877 | 0.154 | 0.433 | -0.582 | 0.612 | -0.002 | 0.012 | 0.012 | 0 | 0.12 | 0.11 | 0.07 | 0.32 | 0.17 | 0.13 |
| 10 | 1925816 | rs10903587 | 0.36 | 0.452 | -0.581 | 0.611 | 0.005 | 0.022 | 0.022 | 0 | 0.1 | 0.12 | 0.07 | 0.27 | 0.14 | 0.11 |
| 10 | 1926697 | rs11250880 | 0.604 | 0.41 | -0.548 | 0.602 | -0.009 | -0.023 | -0.023 | 0.014 | 0.07 | 0.14 | 0.05 | 0 | 0.04 | 0.02 |
| 10 | 1926738 | rs7088760 | -0.467 | 0.399 | -0.541 | 0.595 | 0.033 | -0.016 | -0.016 | 0 | 0.36 | 0.47 | 0.45 | 0.3 | 0.46 | 0.44 |
| 10 | 1932182 | rs2387965 | -1.017 | 0.438 | -0.688 | 0.618 | -0.004 | 0.01 | 0.01 | 0.015 | 0.43 | 0.63 | 0.49 | 0.3 | 0.49 | 0.46 |
| 10 | 1934068 | rs7923512 | -0.483 | 0.684 | -0.598 | 0.831 | 0.023 | 0.001 | 0.001 | 1.588 | 0.36 | 0.55 | 0.46 | 0.37 | 0.48 | 0.45 |
| 10 | 1934824 | rs12354635 | -0.292 | 0.477 | -0.881 | 0.589 | 0 | 0.002 | 0.002 | 0 | 0.17 | 0.19 | 0.12 | 0.27 | 0.22 | 0.12 |
| 10 | 1935402 | rs10128131 | -1.144 | 0.13 | -0.98 | 0.445 | 0.085 | 0.045 | 0.045 | 0.51 | 0.52 | 0.74 | 0.58 | 0.64 | 0.7 | 0.58 |
| 10 | 1945103 | rs10751822 | 0.553 | 0.679 | -0.724 | 0.675 | -0.001 | -0.002 | -0.002 | 0 | 0.71 | 0.62 | 0.75 | 0.73 | 0.7 | 0.67 |
| 10 | 1948064 | rs10794801 | 1.226 | 0.169 | -0.77 | 0.954 | -0.004 | 0.001 | 0.001 | 0.138 | 0.56 | 0.46 | 0.62 | 0.53 | 0.6 | 0.55 |
| 10 | 1949101 | rs11250907 | 0.625 | 0.5 | -0.622 | 1.25 | 0.026 | -0.046 | -0.046 | 3.179 | 0.08 | 0.02 | 0.01 | 0.17 | 0.03 | 0.07 |
| 10 | 1953525 | rs2039568 | 0.586 | 0.57 | -0.429 | 1.147 | -0.008 | 0.017 | 0.017 | 0.433 | 0.75 | 0.47 | 0.79 | 0.73 | 0.78 | 0.68 |
| 10 | 1953826 | rs2039566 | 1.022 | 0.578 | -0.437 | 1.144 | 0.006 | 0.013 | 0.013 | 0 | 0.72 | 0.46 | 0.7 | 0.7 | 0.71 | 0.62 |
| 10 | 1956003 | rs17157295 | 0.701 | 0.905 | -0.127 | 1.251 | 0.001 | -0.021 | -0.021 | 0 | 0.06 | 0.16 | 0.04 | 0.02 | 0.05 | 0.07 |
| 10 | 1959231 | rs10903595 | 0.701 | 1.513 | -0.21 | 1.466 | -0.021 | -0.02 | -0.02 | 7.969 | 0.07 | 0.03 | 0.14 | 0.21 | 0.18 | 0.08 |
| 10 | 1959367 | rs10903596 | 1.384 | 1.485 | -0.14 | 1.45 | 0.059 | 0.047 | 0.047 | 0 | 0.68 | 0.43 | 0.55 | 0.48 | 0.53 | 0.54 |
| 10 | 1963068 | rs11592109 | 1.726 | 1.037 | 0.208 | 1.098 | 0.027 | 0.048 | 0.048 | 1.624 | 0.4 | 0.2 | 0.36 | 0.24 | 0.32 | 0.26 |
| 10 | 1965937 | rs1029182 | 1.044 | 0.86 | 0.469 | 1.307 | 0.026 | 0.095 | 0.095 | 0 | 0.72 | 0.44 | 0.66 | 0.52 | 0.67 | 0.61 |
| 10 | 1969555 | rs7909259 | -0.096 | 0.818 | 0.582 | 1.121 | 0.011 | 0.049 | 0.049 | 0 | 0.84 | 0.66 | 0.77 | 0.73 | 0.83 | 0.77 |
| 10 | 1972320 | rs10751825 | 0.698 | 0.811 | 0.647 | 1.116 | 0.006 | 0.042 | 0.042 | 0 | 0.78 | 0.35 | 0.71 | 0.71 | 0.78 | 0.71 |
| 10 | 1972657 | rs2387960 | 0.625 | 0.828 | 0.519 | 1.124 | 0.012 | 0.055 | 0.055 | 0 | 0.78 | 0.47 | 0.72 | 0.71 | 0.78 | 0.7 |
| 10 | 1973370 | rs10160165 | -0.147 | 0.958 | 0.045 | 1.209 | 0.002 | 0.013 | 0.013 | 0 | 0.85 | 0.78 | 0.78 | 0.73 | 0.83 | 0.77 |
| 10 | 1975035 | rs12220734 | -0.422 | 1.019 | -0.027 | 1.182 | 0 | 0.002 | 0.002 | 3.455 | 0.35 | 0.33 | 0.37 | 0.46 | 0.38 | 0.35 |
| 10 | 1975142 | rs10903599 | 0.287 | 1.108 | 0.092 | 1.28 | 0.023 | 0.065 | 0.065 | 0 | 0.81 | 0.62 | 0.73 | 0.69 | 0.77 | 0.66 |
| 10 | 1975754 | rs10903601 | -0.472 | -0.033 | -0.542 | 0.941 | -0.028 | -0.088 | -0.088 | 0 | 0.34 | 0.08 | 0.27 | 0.44 | 0.35 | 0.31 |
| 10 | 1976399 | rs915265 | -0.168 | -0.005 | -0.169 | 1.004 | -0.039 | 0.061 | 0.061 | 0.832 | 0.87 | 0.47 | 0.84 | 0.86 | 0.93 | 0.8 |
| 10 | 1976876 | rs915267 | 1.177 | 0.546 | -0.568 | 0.946 | -0.023 | 0.033 | 0.033 | 0.746 | 0.43 | 0.2 | 0.43 | 0.36 | 0.45 | 0.37 |
| 10 | 1977041 | rs11250933 | 0.264 | 1.303 | -0.452 | 1.19 | -0.036 | -0.033 | -0.033 | 0 | 0.07 | 0.01 | 0.07 | 0.12 | 0.13 | 0.09 |
| 10 | 1981965 | rs10903603 | -0.651 | 0.883 | -0.188 | 1.219 | -0.009 | 0.031 | 0.031 | 0 | 0.12 | 0.19 | 0.05 | 0.2 | 0.08 | 0.05 |
| 10 | 1982392 | rs7096375 | -1.724 | 0.865 | -0.109 | 1.171 | -0.001 | 0.03 | 0.03 | 0 | 0.19 | 0.33 | 0.18 | 0.27 | 0.18 | 0.15 |
| 10 | 1984431 | rs11250938 | -1.724 | 0.865 | -0.109 | 1.171 | -0.001 | 0.03 | 0.03 | 0 | 0.19 | 0.33 | 0.18 | 0.27 | 0.18 | 0.15 |
| 10 | 1986322 | rs11250945 | -0.643 | 0.64 | -0.193 | 1.148 | -0.005 | 0.018 | 0.018 | 0 | 0.12 | 0.16 | 0.05 | 0.2 | 0.08 | 0.05 |
| 10 | 1988598 | rs7073023 | 0.583 | 0.623 | -0.408 | 0.945 | -0.001 | -0.015 | -0.015 | 0.631 | 0.05 | 0.11 | 0.04 | 0.02 | 0.05 | 0.06 |
| 10 | 1991225 | rs10751826 | -1.065 | 0.961 | -0.48 | 1.092 | 0.009 | 0.01 | 0.01 | 0 | 0.33 | 0.34 | 0.39 | 0.51 | 0.47 | 0.36 |
| 10 | 1993586 | rs11250954 | -0.134 | 0.495 | -0.65 | 1.241 | 0.016 | -0.053 | -0.053 | 0 | 0.08 | 0 | 0.02 | 0.24 | 0.04 | 0.04 |
| 10 | 1994126 | rs12414918 | -0.246 | 0.911 | -0.416 | 1.042 | 0.008 | -0.027 | -0.027 | 0 | 0.16 | 0.2 | 0.18 | 0.08 | 0.2 | 0.16 |
| 10 | 1994789 | rs11250955 | 0.865 | 0.058 | -0.762 | 0.942 | 0.008 | 0.005 | 0.005 | 1.089 | 0.46 | 0.25 | 0.52 | 0.44 | 0.55 | 0.56 |
| 10 | 1995891 | rs12415378 | -0.215 | -0.638 | -1.18 | 0.812 | 0.065 | -0.036 | -0.036 | 0 | 0.33 | 0.06 | 0.2 | 0.34 | 0.22 | 0.18 |
| 10 | 1997959 | rs17294368 | 0.268 | -0.756 | -1.065 | 0.705 | 0.054 | 0.096 | 0.096 | 5.765 | 0.28 | 0.09 | 0.19 | 0.09 | 0.19 | 0.13 |
| 10 | 2000068 | rs4362074 | -0.272 | -0.975 | -0.206 | 0.844 | -0.055 | 0.038 | 0.038 | 3.652 | 0.84 | 0.47 | 0.84 | 0.83 | 0.92 | 0.84 |
| 10 | 2003368 | rs2892403 | 0.882 | -0.98 | -0.479 | 0.84 | -0.005 | 0.007 | 0.007 | 3.116 | 0.75 | 0.28 | 0.79 | 0.53 | 0.84 | 0.73 |
| 10 | 2005604 | rs7912369 | 0.699 | -1.005 | -0.668 | 0.808 | 0.144 | -0.024 | -0.024 | 0.194 | 0.51 | 0.21 | 0.63 | 0.49 | 0.69 | 0.66 |
| 10 | 2005963 | rs7895688 | 0.307 | -0.208 | -0.944 | 1.079 | -0.042 | -0.039 | -0.039 | 0 | 0.11 | 0.04 | 0.2 | 0.18 | 0.22 | 0.13 |
| 10 | 2006209 | rs7919602 | 1.154 | -0.09 | -1.032 | 0.881 | -0.043 | 0.013 | 0.013 | 0.173 | 0.4 | 0.16 | 0.46 | 0.35 | 0.5 | 0.53 |
| 10 | 2007658 | rs7893323 | 0.829 | -0.423 | -0.595 | 0.844 | -0.023 | 0.032 | 0.032 | 0.034 | 0.71 | 0.43 | 0.67 | 0.38 | 0.72 | 0.74 |
| 10 | 2008013 | rs12218258 | 0.241 | -0.433 | -0.645 | 1.003 | 0.011 | -0.024 | -0.024 | 0.069 | 0.07 | 0.04 | 0.03 | 0.26 | 0.05 | 0.06 |
| 10 | 2008811 | rs11250965 | 0.202 | -0.434 | -0.646 | 1.003 | 0.012 | -0.025 | -0.025 | 0 | 0.07 | 0.04 | 0.03 | 0.26 | 0.05 | 0.06 |
| 10 | 2011233 | rs6560783 | -0.399 | -0.478 | -0.454 | 0.817 | -0.012 | 0.139 | 0.139 | 0.019 | 0.82 | 0.5 | 0.72 | 0.69 | 0.81 | 0.86 |
| 10 | 2014089 | rs11250969 | 0.407 | -0.469 | -0.623 | 0.981 | -0.013 | 0.05 | 0.05 | 0.217 | 0.73 | 0.39 | 0.67 | 0.46 | 0.73 | 0.79 |
| 10 | 2015854 | rs3763685 | 1.467 | -0.587 | -0.932 | 0.924 | -0.077 | -0.049 | -0.049 | 0 | 0.34 | 0.12 | 0.41 | 0.35 | 0.41 | 0.48 |
| 10 | 2021054 | rs1335314 | -0.424 | -0.473 | -0.686 | 1.253 | 0.002 | 0.004 | 0.004 | 0.2 | 0.09 | 0.07 | 0.09 | 0.42 | 0.09 | 0.13 |
| 10 | 2022323 | rs11250972 | 0.279 | -0.396 | -0.603 | 1.087 | -0.001 | 0 | 0 | 0.016 | 0.68 | 0.34 | 0.63 | 0.42 | 0.66 | 0.7 |
| 10 | 2027751 | rs7915592 | 0.395 | -0.587 | -0.642 | 1.019 | 0.006 | 0 | 0 | 0 | 0.57 | 0.32 | 0.61 | 0.38 | 0.58 | 0.61 |
| 10 | 2042174 | rs1414129 | 0.02 | -0.565 | -0.385 | 1.124 | 0.007 | 0.014 | 0.014 | 0.009 | 0.62 | 0.34 | 0.68 | 0.76 | 0.66 | 0.72 |
| 10 | 2047386 | rs11591721 | -0.076 | -0.558 | -0.412 | 1.344 | -0.029 | 0.021 | 0.021 | 0 | 0.05 | 0.01 | 0.08 | 0.28 | 0.08 | 0.12 |
| 10 | 2049668 | rs11250988 | 0.83 | -0.463 | -0.547 | 0.878 | 0.005 | 0.105 | 0.105 | 0.494 | 0.28 | 0.07 | 0.22 | 0.07 | 0.23 | 0.16 |

| | | | | | | | | | | | | | | | | |
|---|---|---|---|---|---|---|---|---|---|---|---|---|---|---|---|---|
| 10 | 2051048 | rs6560785 | -0.419 | -0.425 | -0.364 | 0.879 | -0.028 | -0.072 | -0.072 | 0 | 0.65 | 0.53 | 0.72 | 0.8 | 0.69 | 0.76 |
| 10 | 2055591 | rs1414125 | 0.743 | -0.266 | -0.414 | 1.105 | -0.03 | 0.029 | 0.029 | 0.442 | 0.36 | 0.26 | 0.47 | 0.69 | 0.43 | 0.6 |
| 10 | 2057844 | rs11250992 | 0.172 | -0.207 | -0.193 | 1.132 | 0.005 | -0.015 | -0.015 | 0 | 0.41 | 0.47 | 0.45 | 0.69 | 0.46 | 0.65 |
| 10 | 2061812 | rs10794826 | 0.745 | -0.111 | -0.245 | 1.713 | -0.017 | -0.017 | -0.017 | 0 | 0.36 | 0.28 | 0.36 | 0.4 | 0.4 | 0.48 |
| 10 | 2064234 | rs7095469 | 0.263 | -0.139 | -0.311 | 1.554 | 0 | -0.003 | -0.003 | 0 | 0.39 | 0.39 | 0.43 | 0.68 | 0.45 | 0.61 |
| 10 | 2065094 | rs12220122 | -0.267 | 0.073 | -0.532 | 1.722 | -0.029 | 0.007 | 0.007 | 0 | 0.05 | 0.01 | 0.03 | 0.23 | 0.1 | 0.11 |
| 10 | 2067049 | rs7911388 | -0.253 | -0.239 | -0.117 | 1.434 | 0.005 | 0.005 | 0.005 | 0 | 0.66 | 0.63 | 0.71 | 0.69 | 0.72 | 0.75 |
| 10 | 2067523 | rs11250995 | 0.684 | -0.319 | -0.253 | 1.274 | -0.009 | 0.12 | 0.12 | 0.116 | 0.28 | 0.06 | 0.29 | 0.07 | 0.26 | 0.17 |
| 10 | 2069021 | rs17234214 | 1.713 | -0.289 | -0.194 | 1.442 | -0.065 | 0.057 | 0.057 | 0.02 | 0.28 | 0.08 | 0.3 | 0.16 | 0.32 | 0.37 |
| 10 | 2072663 | rs17808220 | 0.543 | -0.324 | -0.108 | 1.554 | 0.004 | 0.029 | 0.029 | 0.011 | 0.29 | 0.11 | 0.34 | 0.19 | 0.28 | 0.26 |
| 10 | 2073711 | rs10430746 | 0.835 | -0.324 | 0.391 | 1.511 | -0.016 | -0.015 | -0.015 | 9.22 | 0.35 | 0.27 | 0.34 | 0.4 | 0.41 | 0.47 |
| 10 | 2073927 | rs10430747 | -0.219 | -0.463 | 0.461 | 1.406 | -0.012 | -0.013 | -0.013 | 0 | 0.65 | 0.39 | 0.7 | 0.69 | 0.72 | 0.75 |
| 10 | 2074022 | rs10430765 | -0.366 | -0.461 | 0.497 | 1.407 | -0.006 | -0.006 | -0.006 | 0 | 0.66 | 0.57 | 0.7 | 0.69 | 0.72 | 0.75 |
| 10 | 2078596 | rs4880959 | 0.435 | -0.378 | -0.104 | 1.602 | -0.002 | 0.042 | 0.042 | 0 | 0.32 | 0.16 | 0.31 | 0.19 | 0.33 | 0.22 |
| 10 | 2080204 | rs7083890 | 0.528 | 0.126 | 0.14 | 1.742 | -0.009 | 0.004 | 0.004 | 0.086 | 0.32 | 0.27 | 0.34 | 0.23 | 0.34 | 0.41 |
| 10 | 2080271 | rs7099247 | 0.258 | 0.078 | 0.034 | 1.42 | 0.002 | 0.002 | 0.002 | 0.398 | 0.35 | 0.36 | 0.4 | 0.5 | 0.38 | 0.52 |
| 10 | 2082974 | rs10903639 | -0.102 | 0.079 | -0.159 | 0.769 | -0.009 | 0.019 | 0.019 | 0 | 0.46 | 0.58 | 0.31 | 0.37 | 0.46 | 0.59 |
| 10 | 2084135 | rs1335310 | -0.48 | 0.032 | -0.117 | 0.713 | -0.002 | -0.004 | -0.004 | 0 | 0.56 | 0.51 | 0.56 | 0.56 | 0.54 | 0.67 |
| 10 | 2085671 | rs870041 | 0.394 | 0.234 | 0.071 | 0.443 | -0.002 | -0.004 | -0.004 | 0.454 | 0.39 | 0.35 | 0.47 | 0.43 | 0.4 | 0.56 |
| 10 | 2086923 | rs10794839 | -1.138 | 0.073 | -0.236 | 0.307 | 0.004 | -0.008 | -0.008 | 0 | 0.7 | 0.72 | 0.76 | 0.61 | 0.73 | 0.78 |
| 10 | 2087246 | rs11251015 | 1.519 | 0.454 | -0.474 | 0.328 | 0 | -0.001 | -0.001 | 0 | 0.06 | 0.05 | 0.08 | 0.15 | 0.08 | 0.1 |
| 10 | 2090551 | rs7913242 | 1.648 | 0.518 | -0.473 | 0.234 | -0.007 | 0.018 | 0.018 | 0.385 | 0.23 | 0.15 | 0.13 | 0.07 | 0.23 | 0.33 |
| 10 | 2091021 | rs17157564 | 1.055 | 0.455 | -0.617 | 0.277 | -0.01 | 0.03 | 0.03 | 0 | 0.07 | 0.01 | 0.02 | 0.04 | 0.05 | 0.05 |
| 10 | 2091393 | rs17235357 | 0.314 | 0.458 | -0.594 | 0.284 | 0.006 | 0.006 | 0.006 | 0 | 0.41 | 0.48 | 0.5 | 0.44 | 0.43 | 0.61 |
| 10 | 2091485 | rs11594876 | 0.988 | 0.466 | -0.541 | 0.357 | -0.021 | 0.023 | 0.023 | 0 | 0.2 | 0.08 | 0.24 | 0.13 | 0.25 | 0.12 |
| 10 | 2092920 | rs17157580 | -0.759 | 0.711 | -0.304 | 0.635 | -0.001 | 0.001 | 0.001 | 0 | 0.48 | 0.54 | 0.52 | 0.48 | 0.48 | 0.66 |
| 10 | 2093110 | rs12266113 | -0.262 | 0.721 | -0.293 | 0.613 | 0.002 | -0.009 | -0.009 | 0 | 0.42 | 0.5 | 0.44 | 0.33 | 0.4 | 0.56 |
| 10 | 2093997 | rs7895736 | 0.058 | 0.755 | -0.07 | 0.879 | -0.002 | 0.042 | 0.042 | 0 | 0.07 | 0.12 | 0.09 | 0.15 | 0.08 | 0.11 |
| 10 | 2094235 | rs11251021 | -0.907 | 0.795 | -0.44 | 0.834 | 0 | 0.006 | 0.006 | 0 | 0.18 | 0.24 | 0.14 | 0.21 | 0.19 | 0.23 |
| 10 | 2095216 | rs945254 | 0.876 | 1.168 | -0.682 | 0.847 | -0.005 | -0.007 | -0.007 | 0.657 | 0.63 | 0.27 | 0.64 | 0.52 | 0.6 | 0.65 |
| 10 | 2095580 | rs17235747 | -0.783 | 1.553 | -1.039 | 0.872 | -0.025 | 0.001 | 0.001 | 0 | 0.24 | 0.24 | 0.28 | 0.16 | 0.33 | 0.15 |
| 10 | 2095726 | rs11251024 | -0.783 | 1.553 | -1.039 | 0.872 | -0.025 | 0.001 | 0.001 | 0 | 0.24 | 0.24 | 0.28 | 0.16 | 0.33 | 0.15 |
| 10 | 2098466 | rs7902045 | -0.636 | 1.697 | -1.02 | 0.624 | -0.002 | 0.002 | 0.002 | 0 | 0.18 | 0.23 | 0.27 | 0.09 | 0.3 | 0.1 |
| 10 | 2101136 | rs11251034 | 0.111 | 1.818 | -1.048 | 0.598 | -0.058 | 0.053 | 0.053 | 0 | 0.11 | 0.04 | 0.2 | 0.01 | 0.23 | 0.09 |
| 10 | 2101561 | rs4880970 | -1.445 | 1.447 | -1.117 | 0.552 | -0.023 | 0.019 | 0.019 | 0 | 0.33 | 0.33 | 0.54 | 0.22 | 0.45 | 0.27 |
| 10 | 2101863 | rs4880550 | -0.55 | 1.452 | -1.118 | 0.561 | 0 | 0.017 | 0.017 | 0 | 0.16 | 0.1 | 0.27 | 0.14 | 0.15 | 0.17 |
| 10 | 2104199 | rs4342949 | -0.351 | 1.435 | -1.202 | 0.568 | -0.006 | 0.004 | 0.004 | 0 | 0.22 | 0.28 | 0.29 | 0.09 | 0.33 | 0.12 |
| 10 | 2104415 | rs2388121 | -0.469 | 0.825 | -1.472 | 0.605 | 0.038 | 0.033 | 0.033 | 0 | 0.51 | 0.76 | 0.65 | 0.58 | 0.55 | 0.55 |
| 10 | 2107812 | rs6560794 | 0.132 | 1.034 | -1.632 | 0.569 | -0.032 | 0.007 | 0.007 | 1.262 | 0.59 | 0.17 | 0.35 | 0.63 | 0.51 | 0.66 |
| 10 | 2108579 | rs7922880 | 0.299 | 0.418 | -0.467 | 0.009 | -0.034 | -0.046 | -0.046 | 0 | 0.91 | 0.75 | 0.97 | 1 | 0.97 | 0.93 |
| 10 | 2113191 | rs7921086 | -0.329 | 0.99 | 0.346 | 0.056 | 0.073 | 0.021 | 0.021 | 15.802 | 0.07 | 0.29 | 0.19 | 0.05 | 0.13 | 0.03 |
| 10 | 2113429 | rs12415258 | -0.375 | 1.256 | -0.364 | -0.196 | -0.068 | 0.009 | 0.009 | 0 | 0.13 | 0.04 | 0.3 | 0.11 | 0.25 | 0.08 |
| 10 | 2113955 | rs17155596 | 0.211 | 0.687 | -0.584 | -0.351 | 0.027 | 0.013 | 0.013 | 0 | 0.11 | 0.03 | 0.04 | 0.07 | 0.08 | 0.15 |
| 10 | 2120407 | rs11251048 | -0.725 | 0.844 | -0.628 | -0.405 | -0.025 | 0.005 | 0.005 | 0 | 0.19 | 0.17 | 0.41 | 0.19 | 0.29 | 0.19 |
| 10 | 2125211 | rs11251050 | 2.088 | 0.195 | -1.262 | -0.868 | -0.068 | 0.225 | 0.225 | 0 | 0.39 | 0.06 | 0.25 | 0.1 | 0.35 | 0.43 |
| 10 | 2126066 | rs10903668 | 0.903 | -0.166 | -0.261 | -1.172 | -0.065 | -0.05 | -0.05 | 0.044 | 0.66 | 0.55 | 0.8 | 0.84 | 0.83 | 0.71 |
| 10 | 2126324 | rs7924123 | 0.023 | -0.256 | 0.012 | -1.215 | -0.038 | -0.086 | -0.086 | 0.568 | 0.82 | 0.72 | 0.95 | 1 | 0.92 | 0.93 |
| 10 | 2128355 | rs11251055 | 0.8 | -0.277 | -0.003 | -1.21 | 0.019 | 0.011 | 0.011 | 0.128 | 0.12 | 0.04 | 0.04 | 0.07 | 0.08 | 0.15 |
| 10 | 2128951 | rs10794850 | 1.566 | -0.142 | 0.331 | -1.163 | 0.021 | 0.078 | 0.078 | 0 | 0.49 | 0.32 | 0.48 | 0.73 | 0.54 | 0.58 |
| 10 | 2131912 | rs1333203 | -0.571 | -0.248 | 0.434 | -1.119 | -0.043 | -0.059 | -0.059 | 0.066 | 0.89 | 0.73 | 0.97 | 1 | 0.97 | 0.93 |
| 10 | 2134606 | rs7096214 | -0.841 | -0.375 | 0.11 | -1.201 | 0.017 | 0.013 | 0.013 | 0.909 | 0.22 | 0.15 | 0.16 | 0.15 | 0.13 | 0.26 |
| 10 | 2135810 | rs7101173 | -0.634 | 0.264 | -0.087 | -1.425 | -0.001 | 0.011 | 0.011 | 0.118 | 0.15 | 0.12 | 0.14 | 0.08 | 0.12 | 0.07 |
| 10 | 2136063 | rs11251056 | 1.6 | 1.24 | -0.16 | -1.486 | 0.025 | 0.101 | 0.101 | 1.126 | 0.48 | 0.16 | 0.52 | 0.64 | 0.47 | 0.44 |
| 10 | 2143308 | rs11251057 | -0.992 | 1.477 | -0.102 | -1.448 | 0.005 | -0.002 | -0.002 | 0.169 | 0.19 | 0.25 | 0.3 | 0.16 | 0.28 | 0.16 |
| 10 | 2148077 | rs7087819 | 0.936 | 1.361 | 0.17 | -1.491 | 0.001 | -0.023 | -0.023 | 0 | 0.58 | 0.46 | 0.68 | 0.77 | 0.59 | 0.62 |
| 10 | 2148432 | rs4880555 | 1.285 | 1.387 | 0.013 | -1.476 | 0 | 0.061 | 0.061 | 0.595 | 0.55 | 0.33 | 0.59 | 0.72 | 0.59 | 0.56 |
| 10 | 2148732 | rs7071344 | 0.98 | 1.222 | 0.064 | -1.45 | 0.017 | 0.011 | 0.011 | 0.453 | 0.13 | 0.09 | 0.06 | 0.09 | 0.1 | 0.21 |
| 10 | 2148963 | rs10903673 | -0.781 | 1.533 | 0.735 | -1.481 | -0.039 | -0.049 | -0.049 | 0 | 0.9 | 0.79 | 0.96 | 1 | 0.97 | 0.93 |
| 10 | 2151354 | rs10751841 | 1.296 | 1.745 | 0.254 | -1.516 | -0.002 | -0.025 | -0.025 | 0 | 0.67 | 0.42 | 0.74 | 0.84 | 0.74 | 0.74 |
| 10 | 2158398 | rs7080520 | 0.57 | 1.564 | -0.033 | -1.705 | -0.002 | -0.067 | -0.067 | 0.05 | 0.74 | 0.39 | 0.57 | 0.89 | 0.77 | 0.77 |
| 10 | 2159967 | rs11251067 | 1.049 | 0.856 | -1.027 | -2.06 | -0.004 | 0.02 | 0.02 | 0.195 | 0.16 | 0.01 | 0.08 | 0.06 | 0.14 | 0.07 |
| 10 | 2165018 | rs10903679 | 0.323 | 0.915 | -0.996 | -2.07 | -0.033 | 0.033 | 0.033 | 0 | 0.22 | 0.05 | 0.15 | 0.13 | 0.26 | 0.19 |
| 10 | 2165774 | rs1412914 | -0.832 | 1.073 | -1.187 | -2.263 | -0.117 | 0.094 | 0.094 | 0.037 | 0.35 | 0.07 | 0.45 | 0.19 | 0.49 | 0.27 |
| 10 | 2165880 | rs4357617 | 0.297 | 0.984 | -0.928 | -2.344 | 0.001 | -0.093 | -0.093 | 0.302 | 0.75 | 0.46 | 0.57 | 0.89 | 0.77 | 0.77 |
| 10 | 2167727 | rs7071544 | 0.394 | 0.981 | -1.248 | -2.345 | -0.002 | -0.04 | -0.04 | 0 | 0.74 | 0.33 | 0.57 | 0.89 | 0.77 | 0.77 |
| 10 | 2167778 | rs7086965 | 1.812 | 0.984 | -1.366 | -2.332 | 0.019 | 0.088 | 0.088 | 0 | 0.46 | 0.3 | 0.39 | 0.76 | 0.42 | 0.57 |
| 10 | 2168400 | rs12413439 | 0.964 | 0.986 | -2.028 | -2.251 | -0.044 | 0.006 | 0.006 | 0 | 0.07 | 0.01 | 0.27 | 0.06 | 0.15 | 0.07 |
| 10 | 2172881 | rs1999531 | 1.798 | 0.944 | -1.982 | -2.247 | 0.014 | 0.141 | 0.141 | 0 | 0.43 | 0.13 | 0.34 | 0.76 | 0.39 | 0.58 |
| 10 | 2173156 | rs10903680 | -0.726 | 0.759 | -1.673 | -1.792 | -0.084 | 0.084 | 0.084 | 0 | 0.29 | 0.06 | 0.21 | 0.13 | 0.38 | 0.2 |

| | | | | | | | | | | | | | | | | |
|---|---|---|---|---|---|---|---|---|---|---|---|---|---|---|---|---|
| 10 | 2178468 | rs12261574 | -0.256 | 0.288 | -1.477 | -1.539 | -0.046 | 0.045 | 0.045 | 0 | 0.24 | 0.05 | 0.18 | 0.13 | 0.3 | 0.19 |
| 10 | 2184223 | rs1041595 | -0.303 | 0.169 | -0.496 | -1.409 | -0.037 | -0.052 | -0.052 | 0.152 | 0.9 | 0.73 | 0.96 | 1 | 0.97 | 0.93 |
| 10 | 2185839 | rs7088792 | 0.318 | 0.303 | -1.566 | -1.336 | -0.002 | 0.05 | 0.05 | 0 | 0.33 | 0.09 | 0.29 | 0.76 | 0.3 | 0.53 |
| 10 | 2186466 | rs4277032 | -0.574 | 0.602 | -1.009 | -1.495 | 0.02 | -0.023 | -0.023 | 0 | 0.91 | 0.83 | 0.76 | 0.95 | 0.87 | 0.95 |
| 10 | 2187043 | rs7085369 | 0.15 | 0.36 | -1.145 | -1.778 | 0.001 | 0.003 | 0.003 | 0 | 0.79 | 0.79 | 0.71 | 0.93 | 0.8 | 0.87 |
| 10 | 2187737 | rs7086382 | 0.402 | 0.556 | -1.583 | -1.951 | -0.055 | 0.033 | 0.033 | 1.673 | 0.33 | 0.19 | 0.32 | 0.24 | 0.45 | 0.37 |
| 10 | 2188486 | rs10159898 | 0.815 | 0.487 | -1.372 | -2.187 | -0.001 | -0.017 | -0.017 | 0 | 0.2 | 0.13 | 0.08 | 0 | 0.19 | 0.04 |
| 10 | 2188538 | rs7068690 | 0.385 | 0.572 | -1.272 | -1.894 | -0.04 | 0.031 | 0.031 | 1.729 | 0.32 | 0.19 | 0.3 | 0.24 | 0.43 | 0.35 |
| 10 | 2189155 | rs12780291 | 0.01 | 0.621 | -1.321 | -1.92 | -0.055 | 0.044 | 0.044 | 0.498 | 0.34 | 0.2 | 0.32 | 0.24 | 0.46 | 0.35 |
| 10 | 2189259 | rs17157759 | 0.192 | 0.11 | -0.841 | -1.726 | 0.001 | -0.015 | -0.015 | 0 | 0.14 | 0.18 | 0.11 | 0.04 | 0.16 | 0.12 |
| 10 | 2189971 | rs7074107 | -0.59 | -0.606 | -0.358 | -1.915 | -0.024 | -0.108 | -0.108 | 0 | 0.24 | 0.44 | 0.16 | 0.07 | 0.2 | 0.21 |
| 10 | 2190850 | rs10903684 | 1.191 | -0.82 | -1.015 | -2.223 | -0.133 | 0.052 | 0.052 | 0 | 0.32 | 0.11 | 0.3 | 0.2 | 0.45 | 0.24 |
| 10 | 2190873 | rs1333200 | -0.85 | -1.056 | -0.696 | -2.338 | 0.018 | -0.054 | -0.054 | 0 | 0.87 | 0.7 | 0.74 | 0.95 | 0.85 | 0.94 |
| 10 | 2191007 | rs1930656 | -1.385 | -0.917 | -0.875 | -2.277 | 0.007 | -0.023 | -0.023 | 0 | 0.89 | 0.84 | 0.75 | 0.95 | 0.85 | 0.94 |
| 10 | 2192149 | rs17817879 | 1.476 | -0.98 | -1.139 | -2.257 | -0.026 | 0.053 | 0.053 | 0 | 0.09 | 0 | 0.03 | 0 | 0.11 | 0.04 |
| 10 | 2197103 | rs2095159 | 1.414 | -1.16 | -1.008 | -2.284 | -0.007 | 0 | 0 | 0 | 0.2 | 0.15 | 0.17 | 0.68 | 0.08 | 0.45 |
| 10 | 2200068 | rs10508223 | 1.3 | -1.045 | -0.774 | -2.315 | -0.013 | 0.014 | 0.014 | 0 | 0.36 | 0.31 | 0.32 | 0.2 | 0.45 | 0.24 |
| 10 | 2200592 | rs10903687 | 0.26 | -1.114 | -0.826 | -1.963 | 0.012 | 0.001 | 0.001 | 0 | 0.35 | 0.25 | 0.28 | 0.63 | 0.23 | 0.58 |
| 10 | 2200634 | rs10508224 | 0.472 | -0.473 | -1.04 | -1.801 | 0.01 | 0.04 | 0.04 | 0.022 | 0.19 | 0.11 | 0.12 | 0.04 | 0.16 | 0.15 |
| 10 | 2204321 | rs7918782 | 0.48 | 0.083 | -0.651 | -0.793 | 0.002 | 0.062 | 0.062 | 0 | 0.78 | 0.58 | 0.53 | 0.37 | 0.77 | 0.72 |
| 10 | 2208519 | rs1333201 | 0.35 | 0.12 | -0.688 | -1.503 | -0.001 | -0.065 | -0.065 | 0 | 0.71 | 0.64 | 0.61 | 0.93 | 0.7 | 0.8 |
| 10 | 2212037 | rs1984070 | -0.109 | 0.147 | -0.941 | -1.057 | 0.007 | 0.017 | 0.017 | 0.008 | 0.56 | 0.62 | 0.6 | 0.81 | 0.62 | 0.48 |
| 10 | 2215512 | rs11596966 | 0.332 | -0.692 | -1.278 | -1.07 | 0.03 | 0.097 | 0.097 | 0 | 0.22 | 0.06 | 0.13 | 0.01 | 0.15 | 0.15 |
| 10 | 2215931 | rs10444197 | 0.699 | -0.38 | -1.009 | -1.215 | 0.04 | 0.104 | 0.104 | 0 | 0.51 | 0.28 | 0.47 | 0.74 | 0.37 | 0.56 |
| 10 | 2219930 | rs12781711 | -0.361 | 0.041 | -0.862 | -1.365 | 0.013 | 0.133 | 0.133 | 0 | 0.34 | 0.13 | 0.3 | 0.09 | 0.31 | 0.36 |
| 10 | 2220505 | rs17157831 | -0.603 | 0.009 | -0.419 | -1.391 | 0.145 | -0.007 | -0.007 | 0 | 0.85 | 0.38 | 0.79 | 0.83 | 0.67 | 0.91 |
| 10 | 2222546 | rs12413933 | -0.183 | -0.207 | -0.592 | -0.649 | -0.003 | -0.027 | -0.027 | 0 | 0.17 | 0.1 | 0.34 | 0.58 | 0.19 | 0.3 |
| 10 | 2224013 | rs10751855 | -0.919 | -0.143 | 0.152 | -0.621 | 0.156 | 0.006 | 0.006 | 0 | 0.87 | 0.62 | 0.82 | 0.83 | 0.67 | 0.91 |
| 10 | 2226082 | rs11251084 | 1.039 | -0.318 | -0.33 | -0.631 | 0.048 | 0.114 | 0.114 | 0.221 | 0.46 | 0.16 | 0.3 | 0.25 | 0.33 | 0.46 |
| 10 | 2241511 | rs2151358 | 0.945 | -0.494 | -0.343 | -0.596 | 0.053 | 0.097 | 0.097 | 0.382 | 0.45 | 0.19 | 0.3 | 0.22 | 0.3 | 0.42 |
| 10 | 2256399 | rs11597710 | 0.831 | -0.482 | -0.292 | -0.644 | 0.028 | 0.101 | 0.101 | 0 | 0.23 | 0.07 | 0.14 | 0.01 | 0.18 | 0.16 |
| 10 | 2256731 | rs1000416 | 1.239 | -0.639 | -0.532 | -0.767 | 0.021 | 0.061 | 0.061 | 1.205 | 0.37 | 0.2 | 0.17 | 0.11 | 0.26 | 0.35 |
| 10 | 2260774 | rs914817 | -0.097 | -0.564 | -0.434 | -0.673 | 0.105 | 0 | 0 | 0.493 | 0.77 | 0.44 | 0.76 | 0.74 | 0.59 | 0.73 |
| 10 | 2261571 | rs7091338 | 0.105 | -0.763 | -0.636 | -0.388 | 0.047 | 0.129 | 0.129 | 0.153 | 0.34 | 0.12 | 0.3 | 0.11 | 0.24 | 0.23 |
| 10 | 2265880 | rs17157889 | 0.48 | -0.151 | -0.37 | 0.136 | 0.002 | 0.111 | 0.111 | 0 | 0.07 | 0.1 | 0.29 | 0.52 | 0.09 | 0.15 |
| 10 | 2268245 | rs10903703 | -0.142 | -0.025 | -0.102 | -0.246 | 0.029 | 0.006 | 0.006 | 0.941 | 0.2 | 0.26 | 0.3 | 0.26 | 0.34 | 0.14 |
| 10 | 2268247 | rs10794877 | -1.318 | -0.307 | -0.218 | 0.359 | -0.017 | -0.041 | -0.041 | 0 | 0.92 | 0.97 | 0.87 | 0.78 | 0.92 | 0.86 |
| 10 | 2268356 | rs10794878 | -0.206 | -0.741 | -0.475 | 0.023 | -0.011 | 0.028 | 0.028 | 0.456 | 0.41 | 0.31 | 0.5 | 0.27 | 0.56 | 0.3 |
| 10 | 2270504 | rs12573685 | -0.233 | -0.755 | -0.097 | 0.007 | 0.016 | 0.01 | 0.01 | 0.491 | 0.6 | 0.55 | 0.56 | 0.51 | 0.48 | 0.7 |
| 10 | 2279963 | rs10903712 | -0.314 | -0.74 | -0.108 | 0.02 | 0.006 | -0.016 | -0.016 | 0 | 0.82 | 0.88 | 0.73 | 0.65 | 0.88 | 0.78 |
| 10 | 2281519 | rs2014892 | 1.266 | -1.099 | -0.227 | 0.083 | 0.018 | 0.066 | 0.066 | 0 | 0.35 | 0.16 | 0.15 | 0.18 | 0.28 | 0.37 |
| 10 | 2286078 | rs7915509 | 0.389 | -1.096 | 0.006 | 0.074 | 0.022 | 0.035 | 0.035 | 0 | 0.48 | 0.36 | 0.36 | 0.29 | 0.36 | 0.47 |
| 10 | 2286495 | rs7916063 | -1.202 | -0.728 | 0.368 | 0.415 | 0.01 | 0.01 | 0.01 | 0 | 0.56 | 0.63 | 0.7 | 0.63 | 0.63 | 0.49 |
| 10 | 2287457 | rs17241211 | 1.104 | -0.746 | 0.122 | 0.207 | 0.012 | 0.074 | 0.074 | 0 | 0.41 | 0.24 | 0.27 | 0.16 | 0.34 | 0.47 |
| 10 | 2290244 | rs17819557 | 1.149 | -0.736 | 0.053 | 0.212 | 0.011 | 0.091 | 0.091 | 0.047 | 0.41 | 0.21 | 0.27 | 0.16 | 0.34 | 0.47 |
| 10 | 2290566 | rs7910363 | 0.649 | -0.429 | 0.092 | 0.131 | 0.016 | -0.015 | -0.015 | 0 | 0.13 | 0.03 | 0.16 | 0.15 | 0.08 | 0.08 |
| 10 | 2291522 | rs11817443 | 1.635 | -0.233 | -0.087 | 0.008 | -0.006 | 0.009 | 0.009 | 0 | 0.14 | 0.09 | 0.08 | 0.08 | 0.15 | 0.16 |
| 10 | 2292180 | rs10903713 | 0.777 | 0.185 | 0.372 | 0.315 | 0.131 | 0.024 | 0.024 | 0 | 0.09 | 0.23 | 0.16 | 0.13 | 0.29 | 0.07 |
| 10 | 2292684 | rs10903714 | -0.789 | -0.315 | -0.204 | 0.109 | 0.008 | 0.008 | 0.008 | 0.361 | 0.4 | 0.37 | 0.38 | 0.17 | 0.31 | 0.25 |
| 10 | 2292862 | rs11251116 | 0.857 | -0.28 | -0.996 | 0.197 | -0.031 | 0.094 | 0.094 | 0 | 0.19 | 0.01 | 0.13 | 0.01 | 0.18 | 0.13 |
| 10 | 2294754 | rs10903715 | 0.133 | 0.189 | -0.622 | 0.494 | 0.04 | 0.074 | 0.074 | 0.405 | 0.36 | 0.14 | 0.3 | 0.17 | 0.27 | 0.22 |
| 10 | 2297365 | rs7909838 | 0 | -0.14 | -0.445 | 0.744 | 0.014 | -0.006 | -0.006 | 0 | 0.48 | 0.27 | 0.46 | 0.5 | 0.38 | 0.57 |
| 10 | 2298593 | rs7090855 | -0.047 | -0.134 | 0.084 | 1.21 | 0.023 | 0.183 | 0.183 | 0 | 0.94 | 0.8 | 0.95 | 0.75 | 0.95 | 0.85 |
| 10 | 2301039 | rs7919805 | 0.149 | -0.759 | 0.477 | 1.176 | 0.009 | 0.024 | 0.024 | 0 | 0.51 | 0.43 | 0.39 | 0.19 | 0.37 | 0.37 |
| 10 | 2305056 | rs10903719 | 0.514 | -0.891 | 0.925 | 1.293 | -0.103 | 0.001 | 0.001 | 0 | 0.36 | 0.13 | 0.34 | 0.29 | 0.53 | 0.39 |
| 10 | 2317755 | rs1926039 | 0.727 | -1.027 | 1.129 | 0.989 | -0.013 | -0.014 | -0.014 | 0 | 0.07 | 0.08 | 0.11 | 0.05 | 0.03 | 0 |
| 10 | 2321173 | rs10751862 | 0.261 | -1.07 | 1.066 | 1.021 | -0.023 | -0.01 | -0.01 | 0 | 0.38 | 0.31 | 0.48 | 0.52 | 0.54 | 0.44 |
| 10 | 2322896 | rs10751864 | 0.279 | -0.954 | 1.259 | 0.992 | 0.001 | 0.005 | 0.005 | 0 | 0.92 | 0.81 | 0.89 | 0.91 | 0.93 | 0.88 |
| 10 | 2323505 | rs2148201 | -0.225 | -1.19 | 0.612 | 0.668 | 0.002 | -0.014 | -0.014 | 0 | 0.57 | 0.47 | 0.45 | 0.26 | 0.47 | 0.44 |
| 10 | 2328469 | rs11251139 | 0.626 | -1.19 | 0.347 | 1.045 | 0.013 | 0.122 | 0.122 | 0.162 | 0.48 | 0.25 | 0.29 | 0.15 | 0.38 | 0.34 |
| 10 | 2331425 | rs2025685 | 0.491 | -1.226 | 0.209 | 1.044 | -0.12 | -0.118 | -0.118 | 0.003 | 0.33 | 0.1 | 0.34 | 0.46 | 0.53 | 0.46 |
| 10 | 2332465 | rs10903735 | 0.004 | -1.238 | 0.287 | 1.039 | -0.043 | -0.03 | -0.03 | 0 | 0.4 | 0.22 | 0.45 | 0.56 | 0.55 | 0.47 |
| 10 | 2336035 | rs12256419 | 0.925 | -1.197 | 0.328 | 0.951 | -0.019 | -0.011 | -0.011 | 0 | 0.07 | 0.11 | 0.11 | 0.1 | 0.03 | 0.01 |
| 10 | 2338795 | rs7904173 | 0.597 | -1.256 | 0.373 | 1.061 | 0.014 | 0.044 | 0.044 | 0.093 | 0.48 | 0.29 | 0.29 | 0.3 | 0.38 | 0.39 |
| 10 | 2340038 | rs7919956 | -0.388 | -1.252 | 0.459 | 1.067 | -0.02 | -0.01 | -0.01 | 0.26 | 0.63 | 0.63 | 0.49 | 0.45 | 0.46 | 0.53 |
| 10 | 2341947 | rs11251149 | -1.078 | -0.998 | 1.253 | 1.603 | -0.061 | 0.105 | 0.105 | 2.459 | 0.16 | 0.46 | 0.2 | 0.31 | 0.08 | 0.19 |
| 10 | 2351551 | rs10794901 | 0.555 | -0.757 | -0.023 | 1.229 | -0.064 | 0.108 | 0.108 | 2.158 | 0.84 | 0.38 | 0.69 | 0.61 | 0.95 | 0.87 |
| 10 | 2351833 | rs768628 | 0.525 | 0.133 | -0.886 | 0.613 | -0.058 | -0.013 | -0.013 | 1.55 | 0.14 | 0.01 | 0.08 | 0.11 | 0.16 | 0.11 |
| 10 | 2355384 | rs7093387 | -0.166 | -0.442 | -1.092 | 0.78 | 0.007 | -0.073 | -0.073 | 0 | 0.81 | 0.93 | 0.78 | 0.64 | 0.84 | 0.85 |

| | | | | | | | | | | | | | | | | |
|---|---|---|---|---|---|---|---|---|---|---|---|---|---|---|---|---|
| 10 | 2356008 | rs1570951 | 0.431 | -1.501 | -0.229 | 0.386 | -0.044 | 0.018 | 0.018 | 0 | 0.69 | 0.61 | 0.71 | 0.6 | 0.81 | 0.76 |
| 10 | 2357702 | rs12412942 | 0.589 | -1.153 | -0.526 | 0.069 | 0.015 | -0.006 | -0.006 | 0 | 0.47 | 0.36 | 0.52 | 0.51 | 0.59 | 0.55 |
| 10 | 2358911 | rs17243233 | 1.719 | -0.61 | -0.701 | 0.07 | -0.016 | 0 | 0 | 0 | 0.12 | 0.05 | 0.07 | 0.12 | 0.15 | 0.11 |
| 10 | 2360003 | rs12570649 | 1.015 | -0.757 | -0.128 | 0.055 | -0.029 | -0.02 | -0.02 | 0 | 0.1 | 0.14 | 0.06 | 0.04 | 0.04 | 0.08 |
| 10 | 2363832 | rs7099701 | 0.726 | -0.767 | 0.051 | 0.112 | -0.05 | 0.051 | 0.051 | 0 | 0.11 | 0.27 | 0.07 | 0.12 | 0.04 | 0.16 |
| 10 | 2367939 | rs11251166 | 1.011 | -0.341 | -0.406 | -0.045 | 0 | 0.001 | 0.001 | 0 | 0.25 | 0.29 | 0.29 | 0.28 | 0.32 | 0.33 |
| 10 | 2372763 | rs7100859 | -0.124 | -0.13 | -0.608 | -0.308 | -0.002 | -0.002 | -0.002 | 0.147 | 0.38 | 0.37 | 0.24 | 0.36 | 0.36 | 0.47 |
| 10 | 2373295 | rs7087615 | 0.573 | -0.086 | -0.551 | -0.279 | -0.027 | 0.002 | 0.002 | 0 | 0.25 | 0.32 | 0.14 | 0.24 | 0.19 | 0.32 |
| 10 | 2374315 | rs17832702 | 0.999 | -0.037 | -0.976 | -0.469 | -0.029 | 0.068 | 0.068 | 0.328 | 0.21 | 0.11 | 0.17 | 0.01 | 0.23 | 0.14 |
| 10 | 2375069 | rs11251170 | -0.562 | -0.066 | -0.878 | -0.408 | -0.002 | 0 | 0 | 0 | 0.6 | 0.42 | 0.39 | 0.38 | 0.56 | 0.59 |
| 10 | 2375730 | rs12784984 | 0.536 | -0.223 | -1.245 | -0.036 | -0.004 | -0.006 | -0.006 | 0.943 | 0.32 | 0.18 | 0.14 | 0.33 | 0.29 | 0.37 |
| 10 | 2378501 | rs7095513 | -0.253 | -0.28 | -1.7 | -0.007 | 0.011 | 0.079 | 0.079 | 0 | 0.52 | 0.84 | 0.65 | 0.29 | 0.57 | 0.51 |
| 10 | 2380746 | rs10903747 | -1.115 | -0.227 | -1.388 | 0.242 | 0.013 | -0.066 | -0.066 | 0.04 | 0.84 | 0.94 | 0.78 | 0.62 | 0.85 | 0.88 |
| 10 | 2383461 | rs10508227 | 1.748 | 0.066 | -1.441 | 0.162 | -0.024 | 0.025 | 0.025 | 0 | 0.05 | 0 | 0.14 | 0 | 0.08 | 0.1 |
| 10 | 2383782 | rs10444195 | 0.325 | -0.359 | -1.134 | 0.274 | 0.009 | -0.018 | -0.018 | 0.11 | 0.19 | 0.12 | 0.22 | 0.38 | 0.16 | 0.22 |
| 10 | 2385405 | rs7086099 | 0.43 | -0.279 | -1.312 | -0.194 | -0.007 | 0.03 | 0.03 | 0 | 0.28 | 0.18 | 0.19 | 0.13 | 0.3 | 0.29 |
| 10 | 2385486 | rs11251178 | -0.078 | -0.793 | -1.017 | -0.033 | 0.014 | -0.006 | -0.006 | 0.565 | 0.38 | 0.3 | 0.29 | 0.64 | 0.27 | 0.35 |
| 10 | 2386983 | rs11251185 | 1.489 | -0.652 | -1.46 | 0.031 | -0.021 | 0.039 | 0.039 | 0 | 0.13 | 0 | 0.06 | 0.04 | 0.15 | 0.11 |
| 10 | 2387114 | rs4415681 | 0.898 | -0.895 | -1.388 | 0.177 | 0.03 | -0.073 | -0.073 | 3.413 | 0.2 | 0.11 | 0.21 | 0.57 | 0.12 | 0.21 |
| 10 | 2387282 | rs12779267 | -0.315 | -1.072 | -0.973 | -0.064 | -0.004 | -0.044 | -0.044 | 0.032 | 0.08 | 0.21 | 0.04 | 0 | 0.05 | 0.09 |
| 10 | 2387307 | rs4506560 | 1.644 | -1.152 | -1.016 | 0.188 | 0.019 | -0.028 | -0.028 | 0.042 | 0.17 | 0.07 | 0.06 | 0.23 | 0.11 | 0.13 |
| 10 | 2390581 | rs12764167 | 0.048 | -1.049 | -0.955 | -0.051 | -0.003 | 0.003 | 0.003 | 0 | 0.27 | 0.32 | 0.24 | 0.57 | 0.15 | 0.28 |
| 10 | 2393152 | rs10508230 | 0.01 | -1.059 | -0.96 | -0.055 | -0.002 | 0.002 | 0.002 | 0 | 0.28 | 0.32 | 0.24 | 0.57 | 0.15 | 0.29 |
| 10 | 2394016 | rs12411602 | 0.216 | -1.282 | -1.454 | -0.2 | 0.028 | 0.062 | 0.062 | 0 | 0.44 | 0.13 | 0.37 | 0.72 | 0.38 | 0.42 |
| 10 | 2394389 | rs4611129 | 0.942 | -1.131 | -0.474 | 0.055 | -0.008 | 0.021 | 0.021 | 0 | 0.22 | 0.25 | 0.22 | 0.57 | 0.12 | 0.2 |
| 10 | 2394444 | rs12257283 | -0.798 | -0.991 | -0.397 | 0.053 | -0.001 | -0.037 | -0.037 | 0 | 0.08 | 0.35 | 0.04 | 0 | 0.03 | 0.08 |
| 10 | 2405229 | rs7084211 | -0.209 | -1.056 | -0.187 | -0.003 | 0.026 | -0.023 | -0.023 | 0 | 0.63 | 0.5 | 0.48 | 0.76 | 0.55 | 0.57 |
| 10 | 2408852 | rs826474 | -1.038 | -1.101 | -0.247 | -0.117 | 0.004 | 0.096 | 0.096 | 0.58 | 0.62 | 0.88 | 0.77 | 0.81 | 0.59 | 0.67 |
| 10 | 2415439 | rs862282 | -0.115 | -0.793 | -0.081 | -0.127 | 0.041 | -0.034 | -0.034 | 0.017 | 0.33 | 0.46 | 0.51 | 0.16 | 0.43 | 0.39 |
| 10 | 2422803 | rs12761358 | 0.384 | -0.064 | -0.317 | -0.486 | 0.006 | -0.05 | -0.05 | 0 | 0.15 | 0.23 | 0.2 | 0 | 0.16 | 0.17 |
| 10 | 2424556 | rs12571964 | -0.577 | -0.128 | -0.24 | -0.512 | 0 | -0.006 | -0.006 | 0.14 | 0.2 | 0.17 | 0.2 | 0.08 | 0.18 | 0.23 |
| 10 | 2424806 | rs2638458 | -0.611 | -0.52 | -0.261 | -0.316 | 0.01 | 0.009 | 0.009 | 0 | 0.54 | 0.63 | 0.74 | 0.63 | 0.6 | 0.57 |
| 10 | 2429307 | rs826489 | -0.64 | -0.246 | -0.458 | -0.203 | 0.001 | 0.001 | 0.001 | 0 | 0.54 | 0.54 | 0.72 | 0.63 | 0.6 | 0.58 |
| 10 | 2432266 | rs7895038 | -0.595 | 0.077 | -0.523 | -0.014 | 0.009 | 0.001 | 0.001 | 0 | 0.62 | 0.59 | 0.46 | 0.38 | 0.56 | 0.61 |
| 10 | 2436043 | rs10903756 | 0.189 | -0.251 | -0.936 | 0.011 | -0.047 | -0.057 | -0.057 | 1.029 | 0.31 | 0.14 | 0.38 | 0.45 | 0.32 | 0.24 |
| 10 | 2437962 | rs17158243 | 1.146 | -0.114 | -1.141 | 0.053 | -0.015 | -0.063 | -0.063 | 1.037 | 0.1 | 0.01 | 0.16 | 0.37 | 0.06 | 0.11 |
| 10 | 2439846 | rs826479 | 0.448 | -0.519 | -0.872 | -0.168 | -0.018 | -0.009 | -0.009 | 0.001 | 0.1 | 0.13 | 0.02 | 0.07 | 0.06 | 0.17 |
| 10 | 2441043 | rs10903757 | 1.134 | -0.483 | -1.02 | -0.168 | -0.088 | 0.048 | 0.048 | 0 | 0.21 | 0.04 | 0.2 | 0.08 | 0.26 | 0.12 |
| 10 | 2441593 | rs826482 | 0.508 | -0.69 | -1.053 | -0.421 | 0.001 | -0.002 | -0.002 | 0 | 0.2 | 0.16 | 0.2 | 0.07 | 0.2 | 0.29 |
| 10 | 2443043 | rs861155 | 1.608 | -0.654 | -1.018 | -0.35 | -0.017 | 0.011 | 0.011 | 0 | 0.06 | 0.13 | 0.02 | 0.07 | 0.04 | 0.12 |
| 10 | 2445563 | rs7069544 | 1.76 | -0.947 | -0.83 | -0.393 | -0.006 | -0.014 | -0.014 | 2.52 | 0.37 | 0.28 | 0.37 | 0.41 | 0.45 | 0.36 |
| 10 | 2447002 | rs2638450 | 1.614 | -0.794 | -0.923 | -0.313 | -0.017 | 0.028 | 0.028 | 0 | 0.06 | 0.15 | 0.03 | 0.08 | 0.04 | 0.12 |
| 10 | 2447411 | rs2638449 | -0.062 | -0.756 | -0.758 | -0.315 | -0.017 | -0.036 | -0.036 | 0 | 0.26 | 0.32 | 0.22 | 0.13 | 0.21 | 0.35 |
| 10 | 2448609 | rs11251228 | 1.223 | -0.493 | -1.169 | -0.125 | -0.098 | 0.038 | 0.038 | 0 | 0.21 | 0.03 | 0.2 | 0.09 | 0.25 | 0.13 |
| 10 | 2454195 | rs11251231 | 0.187 | -0.544 | -1.13 | -0.115 | -0.126 | -0.145 | -0.145 | 0.036 | 0.3 | 0.05 | 0.37 | 0.46 | 0.32 | 0.24 |
| 10 | 2463464 | rs12262142 | 0.744 | -0.649 | -0.51 | -0.074 | -0.02 | -0.001 | -0.001 | 0 | 0.09 | 0.16 | 0.04 | 0.06 | 0.06 | 0.04 |
| 10 | 2463585 | rs4880562 | 1.302 | -0.516 | -0.604 | -0.205 | 0.031 | 0.006 | 0.006 | 0.049 | 0.51 | 0.34 | 0.49 | 0.47 | 0.59 | 0.41 |
| 10 | 2466713 | rs10508236 | 1.21 | -0.542 | -0.248 | -0.217 | 0.01 | 0.001 | 0.001 | 0.035 | 0.52 | 0.47 | 0.51 | 0.47 | 0.59 | 0.41 |
| 10 | 2467249 | rs7906049 | 2.213 | -0.409 | -0.625 | -0.265 | -0.039 | -0.01 | -0.01 | 0.074 | 0.25 | 0.18 | 0.33 | 0.26 | 0.38 | 0.28 |
| 10 | 2468358 | rs11251234 | 2.16 | -0.472 | -0.897 | -0.361 | -0.051 | 0.048 | 0.048 | 0 | 0.11 | 0.02 | 0.09 | 0.04 | 0.14 | 0.12 |
| 10 | 2468408 | rs10903763 | -1.166 | -0.637 | -0.61 | -0.301 | -0.032 | -0.009 | -0.009 | 0.132 | 0.88 | 0.8 | 0.96 | 0.94 | 0.94 | 0.92 |
| 10 | 2471961 | rs10903766 | 0.906 | -0.656 | -0.945 | -0.313 | -0.047 | -0.142 | -0.142 | 0.855 | 0.21 | 0.03 | 0.25 | 0.47 | 0.19 | 0.35 |
| 10 | 2472145 | rs11251237 | 1.423 | -0.626 | -0.95 | -0.251 | -0.063 | -0.114 | -0.114 | 0 | 0.19 | 0.03 | 0.25 | 0.4 | 0.19 | 0.23 |
| 10 | 2476773 | rs11251240 | 1.385 | -0.422 | -0.726 | -0.378 | -0.007 | 0.056 | 0.056 | 0 | 0.24 | 0.08 | 0.15 | 0.1 | 0.21 | 0.1 |
| 10 | 2480445 | rs12780201 | 1.501 | -0.375 | -0.09 | -0.186 | -0.001 | -0.022 | -0.022 | 0 | 0.13 | 0.12 | 0.17 | 0 | 0.14 | 0.16 |
| 10 | 2480963 | rs2029602 | 2.13 | -0.419 | -0.303 | -0.164 | -0.043 | -0.039 | -0.039 | 0 | 0.08 | 0.01 | 0.24 | 0.13 | 0.26 | 0.12 |
| 10 | 2483585 | rs7900071 | 1.531 | -0.791 | 0.164 | -0.262 | 0.001 | -0.077 | -0.077 | 0 | 0.14 | 0.25 | 0.19 | 0 | 0.15 | 0.16 |
| 10 | 2484526 | rs7923870 | -1.259 | -0.784 | -0.117 | -0.2 | -0.02 | -0.004 | -0.004 | 0 | 0.9 | 0.86 | 0.96 | 0.95 | 0.95 | 0.93 |
| 10 | 2488754 | rs7910547 | 0.08 | -0.796 | -0.176 | -0.233 | -0.004 | -0.017 | -0.017 | 0 | 0.25 | 0.26 | 0.22 | 0.05 | 0.2 | 0.21 |
| 10 | 2505690 | rs993651 | -0.385 | -0.808 | -0.271 | -0.239 | 0.018 | 0.036 | 0.036 | 0 | 0.76 | 0.84 | 0.79 | 0.95 | 0.8 | 0.79 |
| 10 | 2509159 | rs17158371 | 2.39 | -0.681 | -0.087 | -0.019 | 0.004 | -0.002 | -0.002 | 0.764 | 0.14 | 0.09 | 0.06 | 0.19 | 0.1 | 0.09 |
| 10 | 2511754 | rs1388723 | 0.751 | -0.688 | -0.079 | 0.074 | -0.016 | -0.009 | -0.009 | 0 | 0.39 | 0.32 | 0.39 | 0.46 | 0.48 | 0.36 |
| 10 | 2512919 | rs2452213 | -0.142 | -0.392 | 0.569 | 0.329 | -0.027 | 0.007 | 0.007 | 0 | 0.89 | 0.7 | 0.95 | 0.95 | 0.94 | 0.96 |
| 10 | 2524045 | rs9423540 | 0.459 | -0.11 | -0.058 | 0.324 | 0.003 | 0.008 | 0.008 | 0 | 0.5 | 0.45 | 0.48 | 0.63 | 0.44 | 0.45 |
| 10 | 2529326 | rs9419445 | 0.633 | -0.051 | -0.227 | 0.233 | -0.038 | 0.092 | 0.092 | 0 | 0.52 | 0.15 | 0.47 | 0.68 | 0.44 | 0.46 |
| 10 | 2530583 | rs2458713 | 0.144 | 0.035 | 0.186 | 0.152 | 0.022 | -0.008 | -0.008 | 0.054 | 0.27 | 0.36 | 0.32 | 0.24 | 0.4 | 0.22 |
| 10 | 2532247 | rs9419446 | 0.124 | -0.076 | -0.024 | 0.052 | 0.018 | 0.011 | 0.011 | 0 | 0.69 | 0.31 | 0.64 | 0.71 | 0.73 | 0.6 |
| 10 | 2536538 | rs2452200 | -0.345 | 0.127 | 0.883 | 0.719 | 0.01 | -0.009 | -0.009 | 0 | 0.39 | 0.54 | 0.47 | 0.7 | 0.44 | 0.47 |
| 10 | 2537042 | rs2452199 | -0.345 | 0.133 | 0.886 | 0.722 | 0.01 | -0.009 | -0.009 | 0.113 | 0.39 | 0.54 | 0.47 | 0.7 | 0.44 | 0.47 |

| | | | | | | | | | | | | | | | | |
|---|---|---|---|---|---|---|---|---|---|---|---|---|---|---|---|---|
| 10 | 2537313 | rs2452198 | -0.426 | 0.064 | 1.113 | 0.708 | -0.002 | 0.004 | 0.004 | 0 | 0.4 | 0.62 | 0.49 | 0.7 | 0.44 | 0.47 |
| 10 | 2541377 | rs2458705 | 0.872 | -0.248 | 1.755 | 1.288 | -0.024 | -0.004 | -0.004 | 0 | 0.69 | 0.43 | 0.66 | 0.31 | 0.74 | 0.7 |
| 10 | 2541943 | rs10903784 | 1.9 | -0.197 | 1.55 | 1.08 | 0.072 | 0.158 | 0.158 | 0 | 0.55 | 0.22 | 0.5 | 0.08 | 0.66 | 0.46 |
| 10 | 2544480 | rs9663489 | -1.127 | -0.147 | 1.522 | 1.121 | 0.024 | -0.025 | -0.025 | 0 | 0.08 | 0.14 | 0.06 | 0.02 | 0.12 | 0.07 |
| 10 | 2547022 | rs9423666 | -0.191 | -0.218 | 1.763 | 1.494 | -0.012 | 0.209 | 0.209 | 0 | 0.06 | 0.11 | 0.16 | 0.47 | 0.06 | 0.12 |
| 10 | 2548452 | rs2458698 | 0.891 | -0.315 | 1.957 | 1.482 | -0.046 | 0.017 | 0.017 | 0 | 0.68 | 0.49 | 0.79 | 0.7 | 0.76 | 0.64 |
| 10 | 2549436 | rs1492184 | -1.789 | -0.224 | 1.828 | 1.391 | 0.016 | 0.004 | 0.004 | 0 | 0.13 | 0.22 | 0.19 | 0.18 | 0.15 | 0.13 |
| 10 | 2553240 | rs1155447 | 2.475 | -0.187 | 1.678 | 1.344 | 0.015 | 0.301 | 0.301 | 0 | 0.48 | 0.13 | 0.43 | 0.05 | 0.55 | 0.4 |
| 10 | 2555773 | rs9665557 | -0.01 | -0.231 | 2.086 | 1.412 | 0 | 0.039 | 0.039 | 0 | 0.07 | 0.2 | 0.03 | 0.06 | 0.04 | 0.16 |
| 10 | 2557076 | rs2458688 | 0.985 | -0.276 | 2.31 | 1.392 | -0.01 | 0.043 | 0.043 | 0.06 | 0.89 | 0.56 | 0.93 | 0.93 | 0.95 | 0.83 |
| 10 | 2557691 | rs11814052 | 0.463 | -0.269 | 2.323 | 1.622 | -0.004 | 0.021 | 0.021 | 0 | 0.06 | 0.12 | 0.03 | 0.06 | 0.04 | 0.16 |
| 10 | 2558675 | rs1155134 | -1.71 | -0.258 | 2.237 | 1.629 | 0.017 | 0.006 | 0.006 | 0 | 0.13 | 0.2 | 0.19 | 0.18 | 0.15 | 0.13 |
| 10 | 2559558 | rs9665581 | -0.591 | -0.274 | 1.754 | 1.594 | 0.014 | -0.024 | -0.024 | 0 | 0.23 | 0.16 | 0.23 | 0.34 | 0.22 | 0.23 |
| 10 | 2561216 | rs11251290 | 2.052 | -0.142 | 1.297 | 1.499 | 0.043 | 0.258 | 0.258 | 0 | 0.49 | 0.15 | 0.43 | 0.11 | 0.56 | 0.4 |
| 10 | 2562328 | rs11251294 | -1.47 | -0.115 | 1.606 | 1.638 | -0.027 | 0.007 | 0.007 | 0 | 0.34 | 0.47 | 0.24 | 0.29 | 0.28 | 0.38 |
| 10 | 2562642 | rs1472135 | -0.849 | -0.114 | 1.215 | 1.637 | 0.002 | 0.002 | 0.002 | 0 | 0.08 | 0.08 | 0.05 | 0.05 | 0.04 | 0.02 |
| 10 | 2562836 | rs1472138 | -1.096 | -0.057 | 1.184 | 1.689 | -0.007 | 0.113 | 0.113 | 0 | 0.1 | 0.12 | 0.21 | 0.53 | 0.06 | 0.13 |
| 10 | 2566085 | rs2388097 | 1.702 | -0.047 | 1.173 | 1.579 | 0.057 | 0.014 | 0.014 | 0.159 | 0.56 | 0.24 | 0.53 | 0.44 | 0.66 | 0.49 |
| 10 | 2571838 | rs10903794 | 0.9 | -0.194 | 1.448 | 1.56 | -0.028 | 0.011 | 0.011 | 0 | 0.7 | 0.42 | 0.68 | 0.41 | 0.79 | 0.55 |
| 10 | 2573937 | rs11251322 | -0.78 | -0.173 | 1.417 | 1.612 | -0.001 | 0.018 | 0.018 | 0 | 0.06 | 0.06 | 0.14 | 0.16 | 0.05 | 0.05 |
| 10 | 2574759 | rs17158469 | -1.586 | -0.178 | 1.419 | 1.599 | -0.001 | 0.01 | 0.01 | 4.136 | 0.08 | 0.08 | 0.15 | 0.16 | 0.06 | 0.05 |
| 10 | 2577639 | rs17158495 | -1.058 | -0.098 | 1.398 | 1.542 | -0.003 | 0.003 | 0.003 | 0 | 0.1 | 0.1 | 0.09 | 0.06 | 0.14 | 0.09 |
| 10 | 2580216 | rs11251328 | -1.563 | -0.085 | 1.254 | 1.54 | -0.001 | 0.01 | 0.01 | 4.136 | 0.08 | 0.08 | 0.15 | 0.16 | 0.06 | 0.05 |
| 10 | 2582917 | rs11251339 | -1.865 | -0.075 | 1.098 | 1.533 | -0.041 | -0.047 | -0.047 | 5.697 | 0.15 | 0.05 | 0.21 | 0.23 | 0.18 | 0.14 |
| 10 | 2584073 | rs2388257 | 2.379 | -0.173 | 1.183 | 1.493 | 0.048 | 0.151 | 0.151 | 0 | 0.51 | 0.3 | 0.45 | 0.18 | 0.6 | 0.4 |
| 10 | 2584375 | rs2388258 | 1.097 | -0.817 | 1.447 | 1.593 | -0.045 | 0.023 | 0.023 | 0 | 0.7 | 0.46 | 0.68 | 0.41 | 0.79 | 0.54 |
| 10 | 2589776 | rs10508237 | -1.084 | -0.623 | 1.776 | 1.842 | -0.021 | 0.033 | 0.033 | 0 | 0.26 | 0.38 | 0.26 | 0.3 | 0.2 | 0.36 |
| 10 | 2590305 | rs2388260 | 2.038 | -0.558 | 1.745 | 1.848 | -0.009 | 0.07 | 0.07 | 0 | 0.81 | 0.72 | 0.71 | 0.48 | 0.83 | 0.77 |
| 10 | 2592123 | rs9919456 | 0.417 | -0.552 | 1.523 | 1.857 | -0.004 | 0.054 | 0.054 | 0 | 0.91 | 0.86 | 0.89 | 0.7 | 0.91 | 0.84 |
| 10 | 2593077 | rs7093049 | -0.353 | -0.585 | 1.516 | 1.855 | -0.013 | 0.013 | 0.013 | 0 | 0.19 | 0.24 | 0.22 | 0.25 | 0.14 | 0.21 |
| 10 | 2595331 | rs2388268 | 1.961 | -0.652 | 1.201 | 1.856 | -0.02 | 0.142 | 0.142 | 0 | 0.79 | 0.58 | 0.69 | 0.48 | 0.82 | 0.77 |
| 10 | 2602415 | rs2388273 | 1.993 | -0.345 | 1.068 | 1.869 | 0.03 | 0.173 | 0.173 | 0.022 | 0.8 | 0.49 | 0.68 | 0.51 | 0.8 | 0.77 |
| 10 | 2605786 | rs17179961 | 0.504 | -0.968 | 0.91 | 1.817 | -0.022 | -0.027 | -0.027 | 0.57 | 0.06 | 0.01 | 0.04 | 0.05 | 0.06 | 0.16 |
| 10 | 2606133 | rs4433480 | 1.956 | -0.958 | 0.978 | 1.626 | 0.066 | 0.311 | 0.311 | 0 | 0.5 | 0.15 | 0.43 | 0.06 | 0.61 | 0.37 |
| 10 | 2606565 | rs2388279 | 1.137 | -1.27 | 1.295 | 1.737 | -0.008 | 0.213 | 0.213 | 0.483 | 0.89 | 0.53 | 0.88 | 0.71 | 0.95 | 0.9 |
| 10 | 2606931 | rs2388282 | 0.841 | -1.222 | 1.048 | 1.723 | 0.008 | 0.199 | 0.199 | 0.466 | 0.89 | 0.6 | 0.87 | 0.73 | 0.94 | 0.9 |
| 10 | 2607487 | rs11595982 | -1.052 | -1.271 | 0.998 | 1.762 | -0.012 | 0.01 | 0.01 | 0 | 0.31 | 0.35 | 0.3 | 0.36 | 0.24 | 0.41 |
| 10 | 2610176 | rs17158651 | -0.418 | -1.238 | 0.836 | 1.684 | 0 | 0.008 | 0.008 | 0 | 0.07 | 0.08 | 0.16 | 0.19 | 0.08 | 0.09 |
| 10 | 2611192 | rs12783377 | 2.009 | -1.272 | 0.765 | 1.626 | 0.063 | 0.304 | 0.304 | 0 | 0.49 | 0.15 | 0.42 | 0.02 | 0.61 | 0.36 |
| 10 | 2616253 | rs7918919 | 0.715 | -0.956 | 0.921 | 2.221 | -0.001 | 0.139 | 0.139 | 0.208 | 0.91 | 0.78 | 0.9 | 0.73 | 0.95 | 0.88 |
| 10 | 2619314 | rs2388287 | 2.085 | -0.971 | 0.967 | 2.239 | 0 | 0.203 | 0.203 | 0 | 0.82 | 0.51 | 0.72 | 0.53 | 0.85 | 0.78 |
| 10 | 2619695 | rs7920309 | 0.795 | -1.046 | 0.747 | 2.246 | 0.002 | 0.154 | 0.154 | 0.136 | 0.9 | 0.76 | 0.9 | 0.73 | 0.95 | 0.88 |
| 10 | 2620063 | rs11595442 | -0.232 | -1.03 | 0.317 | 2.357 | 0.021 | -0.061 | -0.061 | 0 | 0.16 | 0.03 | 0.11 | 0.22 | 0.1 | 0.26 |
| 10 | 2621756 | rs11251374 | -1.932 | -0.8 | 1.049 | 2.79 | 0.002 | -0.004 | -0.004 | 0 | 0.12 | 0.08 | 0.25 | 0.35 | 0.11 | 0.14 |
| 10 | 2627161 | rs2388288 | 1.554 | -0.78 | 1.304 | 2.873 | -0.017 | 0.319 | 0.319 | 0.454 | 0.86 | 0.57 | 0.78 | 0.5 | 0.88 | 0.77 |
| 10 | 2634007 | rs7896692 | 1.557 | -0.737 | 0.742 | 2.543 | -0.041 | 0.101 | 0.101 | 0 | 0.8 | 0.49 | 0.67 | 0.34 | 0.85 | 0.72 |
| 10 | 2634831 | rs17158676 | -1.236 | -0.195 | -0.119 | 2.12 | -0.013 | 0.108 | 0.108 | 2.418 | 0.09 | 0.15 | 0.13 | 0.34 | 0.08 | 0.17 |
| 10 | 2635105 | rs11251391 | -0.821 | -0.19 | -0.161 | 2.123 | -0.019 | -0.03 | -0.03 | 0 | 0.05 | 0 | 0.14 | 0.18 | 0.07 | 0.08 |
| 10 | 2636063 | rs17155640 | -1.375 | -0.284 | 0.13 | 1.595 | -0.013 | 0.105 | 0.105 | 2.418 | 0.09 | 0.15 | 0.13 | 0.34 | 0.08 | 0.17 |
| 10 | 2642800 | rs11812571 | -0.653 | -0.319 | -0.054 | 0.819 | -0.007 | -0.021 | -0.021 | 0.001 | 0.13 | 0.18 | 0.08 | 0.04 | 0.1 | 0.1 |
| 10 | 2646621 | rs2892499 | 0.294 | -0.615 | -0.374 | 0.728 | -0.029 | -0.023 | -0.023 | 0 | 0.39 | 0.26 | 0.44 | 0.52 | 0.51 | 0.51 |
| 10 | 2648164 | rs10903830 | 0.087 | -1.416 | -0.551 | 0.402 | -0.001 | 0 | 0 | 0.01 | 0.66 | 0.38 | 0.7 | 0.75 | 0.79 | 0.81 |
| 10 | 2653147 | rs10794929 | 1.015 | -1.549 | -0.475 | 0.878 | -0.041 | -0.012 | -0.012 | 0 | 0.38 | 0.22 | 0.44 | 0.33 | 0.51 | 0.47 |
| 10 | 2658796 | rs7923950 | 0.175 | -1.632 | -0.493 | 0.84 | 0.019 | 0.01 | 0.01 | 0.198 | 0.65 | 0.29 | 0.69 | 0.74 | 0.78 | 0.77 |
| 10 | 2661196 | rs6601797 | -0.347 | -1.676 | -0.326 | 0.842 | 0.014 | 0.054 | 0.054 | 0.073 | 0.39 | 0.23 | 0.34 | 0.25 | 0.37 | 0.37 |
| 10 | 2662611 | rs11598665 | 0.929 | -1.652 | 0.583 | 0.847 | -0.008 | 0.008 | 0.008 | 0 | 0.18 | 0.24 | 0.18 | 0.2 | 0.1 | 0.09 |
| 10 | 2666834 | rs7080305 | 0.743 | -2.068 | 0.378 | 0.601 | 0.008 | 0.035 | 0.035 | 0 | 0.47 | 0.23 | 0.47 | 0.34 | 0.55 | 0.49 |
| 10 | 2668786 | rs2388390 | 0.247 | -2.107 | 0.475 | 0.596 | 0 | 0 | 0 | 0 | 0.58 | 0.42 | 0.54 | 0.38 | 0.64 | 0.55 |
| 10 | 2670553 | rs4278423 | 0.978 | -2.122 | 0.533 | 0.617 | -0.021 | 0.018 | 0.018 | 0 | 0.19 | 0.31 | 0.19 | 0.2 | 0.11 | 0.1 |
| 10 | 2672202 | rs17158714 | 1.02 | -2.098 | 0.322 | 0.545 | -0.018 | -0.051 | -0.051 | 0.942 | 0.12 | 0.01 | 0.07 | 0.23 | 0.1 | 0.2 |
| 10 | 2674458 | rs6601818 | 1.104 | -2.096 | 0.474 | 0.506 | -0.009 | 0.03 | 0.03 | 0 | 0.46 | 0.13 | 0.47 | 0.34 | 0.55 | 0.47 |
| 10 | 2676299 | rs10903834 | 0.212 | -2.46 | 0.146 | 0.343 | -0.002 | 0 | 0 | 0 | 0.55 | 0.47 | 0.54 | 0.58 | 0.66 | 0.63 |
| 10 | 2678825 | rs7915957 | -0.549 | -2.449 | 0.166 | 0.286 | -0.007 | -0.003 | -0.003 | 0 | 0.48 | 0.49 | 0.45 | 0.42 | 0.36 | 0.38 |
| 10 | 2679240 | rs7900654 | -0.482 | -2.271 | 0.114 | 0.265 | 0.006 | 0.003 | 0.003 | 0.193 | 0.47 | 0.45 | 0.46 | 0.41 | 0.37 | 0.38 |
| 10 | 2691576 | rs7074375 | 0.472 | -2.255 | -0.369 | 0.289 | 0.02 | 0.019 | 0.019 | 0 | 0.25 | 0.04 | 0.24 | 0.2 | 0.24 | 0.24 |
| 10 | 2692656 | rs7079384 | 1.433 | -2.261 | -0.047 | 0.27 | 0.003 | -0.003 | -0.003 | 0 | 0.18 | 0.17 | 0.18 | 0.2 | 0.11 | 0.1 |
| 10 | 2695158 | rs7899257 | 1.277 | -2.077 | 0.031 | 0.174 | 0.034 | 0.01 | 0.01 | 1.119 | 0.3 | 0.37 | 0.41 | 0.31 | 0.44 | 0.38 |
| 10 | 2697071 | rs2068455 | 0.544 | -2.118 | 0.027 | 0.166 | -0.001 | -0.001 | -0.001 | 0 | 0.42 | 0.52 | 0.48 | 0.35 | 0.54 | 0.47 |
| 10 | 2701181 | rs1413759 | 1.059 | -1.537 | -0.233 | 0.19 | -0.079 | 0.06 | 0.06 | 0.162 | 0.23 | 0.09 | 0.26 | 0.08 | 0.33 | 0.22 |

| | | | | | | | | | | | | | | | | |
|---|---|---|---|---|---|---|---|---|---|---|---|---|---|---|---|---|
| 10 | 2703572 | rs17158770 | 1.4 | -1.643 | -0.195 | 0.313 | 0.001 | -0.009 | -0.009 | 1.581 | 0.11 | 0.05 | 0.07 | 0.2 | 0.11 | 0.15 |
| 10 | 2704059 | rs7072489 | 0.078 | -1.954 | -0.079 | 0.261 | -0.026 | 0.022 | 0.022 | 0 | 0.32 | 0.56 | 0.36 | 0.51 | 0.23 | 0.33 |
| 10 | 2706717 | rs7072084 | 0.449 | -1.54 | -0.115 | 0.142 | 0.023 | 0.05 | 0.05 | 1.796 | 0.34 | 0.21 | 0.3 | 0.21 | 0.33 | 0.26 |
| 10 | 2707468 | rs10903840 | 1.377 | -1.479 | -0.168 | 0.145 | -0.077 | 0.047 | 0.047 | 1.482 | 0.21 | 0.07 | 0.25 | 0.08 | 0.33 | 0.26 |
| 10 | 2711041 | rs7893477 | -0.271 | -1.626 | -0.093 | 0.261 | -0.005 | 0 | 0 | 0.482 | 0.43 | 0.53 | 0.45 | 0.45 | 0.42 | 0.4 |
| 10 | 2713713 | rs10794932 | -1.267 | -1.614 | -0.132 | 0.256 | -0.001 | 0.002 | 0.002 | 0 | 0.7 | 0.69 | 0.73 | 0.56 | 0.79 | 0.67 |
| 10 | 2714278 | rs17158795 | 0.858 | -1.26 | -0.529 | 0.397 | -0.002 | -0.001 | -0.001 | 0.047 | 0.06 | 0.04 | 0.16 | 0.16 | 0.13 | 0.1 |
| 10 | 2714749 | rs884289 | 0.742 | -1.542 | -0.32 | 0.39 | 0.017 | 0.196 | 0.196 | 0 | 0.08 | 0.28 | 0.15 | 0.25 | 0.08 | 0.15 |
| 10 | 2715418 | rs11597478 | 0.255 | -1.124 | -1.298 | -0.244 | 0.035 | 0.08 | 0.08 | 0.536 | 0.28 | 0.15 | 0.14 | 0.04 | 0.2 | 0.16 |
| 10 | 2720453 | rs12764543 | 0.164 | -1.049 | -1.597 | -0.193 | 0.021 | 0.026 | 0.026 | 0.616 | 0.3 | 0.24 | 0.14 | 0.06 | 0.2 | 0.16 |
| 10 | 2721782 | rs12359239 | -0.309 | 0.694 | -1.457 | 0.635 | -0.141 | -0.009 | -0.009 | 0.098 | 0.28 | 0.06 | 0.43 | 0.25 | 0.5 | 0.36 |
| 10 | 2731235 | rs10794936 | -0.13 | 0.416 | -0.618 | -0.003 | -0.008 | -0.101 | -0.101 | 0 | 0.69 | 0.42 | 0.79 | 0.97 | 0.73 | 0.78 |
| 10 | 2733495 | rs11817995 | -0.112 | 0.969 | -0.549 | 0.017 | -0.007 | -0.108 | -0.108 | 0 | 0.3 | 0.42 | 0.2 | 0.03 | 0.27 | 0.19 |
| 10 | 2737238 | rs1008859 | -0.367 | 1.65 | -1.233 | 0.992 | -0.056 | -0.075 | -0.075 | 0 | 0.25 | 0.1 | 0.29 | 0.52 | 0.34 | 0.3 |
| 10 | 2738665 | rs11251436 | -0.199 | 0.906 | -1.24 | 0.447 | 0.008 | 0.054 | 0.054 | 0 | 0.31 | 0.09 | 0.33 | 0.15 | 0.23 | 0.36 |
| 10 | 2739606 | rs6601863 | -0.291 | 0.837 | -0.961 | 0.371 | 0.015 | 0.047 | 0.047 | 0 | 0.33 | 0.2 | 0.34 | 0.13 | 0.23 | 0.36 |
| 10 | 2740608 | rs7920763 | -0.372 | 0.837 | -0.371 | 0.725 | 0.001 | -0.004 | -0.004 | 0.172 | 0.58 | 0.61 | 0.54 | 0.38 | 0.54 | 0.53 |
| 10 | 2743333 | rs7906405 | 0.334 | 0.143 | -0.66 | 0.662 | 0.013 | 0.004 | 0.004 | 6.736 | 0.63 | 0.32 | 0.62 | 0.33 | 0.71 | 0.53 |
| 10 | 2743897 | rs2050343 | -0.717 | -0.269 | -0.928 | 1.03 | 0.02 | -0.031 | -0.031 | 1.767 | 0.23 | 0.17 | 0.31 | 0.41 | 0.17 | 0.32 |
| 10 | 2744038 | rs11592580 | -0.792 | -0.553 | -1.017 | 0.435 | 0.03 | 0.042 | 0.042 | 0 | 0.26 | 0.15 | 0.16 | 0.11 | 0.16 | 0.28 |
| 10 | 2744342 | rs4881012 | 0.376 | -0.254 | -1.295 | -0.219 | -0.006 | 0.053 | 0.053 | 3.055 | 0.3 | 0.11 | 0.19 | 0.12 | 0.29 | 0.17 |
| 10 | 2746308 | rs17158867 | -0.013 | -0.185 | -0.894 | -0.37 | 0.001 | 0.002 | 0.002 | 1.04 | 0.22 | 0.14 | 0.13 | 0.15 | 0.17 | 0.15 |
| 10 | 2747283 | rs2065682 | 0.455 | -0.213 | -0.936 | -0.382 | -0.017 | -0.025 | -0.025 | 1.228 | 0.19 | 0.01 | 0.12 | 0.15 | 0.14 | 0.14 |
| 10 | 2749234 | rs7909413 | -1.328 | -0.083 | -0.729 | -0.322 | 0.018 | 0.025 | 0.025 | 0 | 0.52 | 0.35 | 0.32 | 0.35 | 0.37 | 0.55 |
| 10 | 2749491 | rs11251440 | -0.378 | -0.145 | -0.746 | -0.341 | 0.016 | 0.018 | 0.018 | 0 | 0.29 | 0.22 | 0.18 | 0.19 | 0.19 | 0.34 |
| 10 | 2751834 | rs7919069 | -0.266 | 0.478 | -0.741 | -0.385 | 0.032 | 0.053 | 0.053 | 0 | 0.28 | 0.16 | 0.17 | 0.11 | 0.18 | 0.31 |
| 10 | 2754463 | rs7904477 | -0.023 | 0.51 | -0.761 | -0.372 | 0.001 | 0.001 | 0.001 | 0.748 | 0.23 | 0.16 | 0.15 | 0.16 | 0.18 | 0.22 |
| 10 | 2754671 | rs1004641 | -0.272 | 0.329 | -0.825 | -0.548 | 0.025 | 0.045 | 0.045 | 0 | 0.28 | 0.19 | 0.18 | 0.13 | 0.18 | 0.31 |
| 10 | 2755927 | rs945142 | -1.654 | 0.725 | -0.107 | -0.682 | -0.004 | 0.032 | 0.032 | 0 | 0.62 | 0.62 | 0.81 | 0.89 | 0.64 | 0.71 |
| 10 | 2757555 | rs11251448 | -0.766 | 0.79 | 0.243 | -0.316 | 0.015 | -0.024 | -0.024 | 0 | 0.33 | 0.35 | 0.63 | 0.76 | 0.46 | 0.4 |
| 10 | 2762427 | rs1889507 | -0.3 | 0.318 | 0.634 | 0.128 | 0.001 | 0.15 | 0.15 | 0 | 0.07 | 0.14 | 0.36 | 0.55 | 0.11 | 0.12 |
| 10 | 2770276 | rs12764463 | 0.126 | -0.009 | 0.834 | -0.318 | 0.023 | 0.037 | 0.037 | 0 | 0.2 | 0.05 | 0.1 | 0.13 | 0.14 | 0.3 |
| 10 | 2779797 | rs11251480 | -0.742 | 0.505 | 2.06 | 0.092 | 0 | 0.031 | 0.031 | 0 | 0.25 | 0.19 | 0.27 | 0.13 | 0.28 | 0.28 |
| 10 | 2780603 | rs7904050 | -1.226 | 0.507 | 2.611 | 0.041 | 0 | -0.022 | -0.022 | 0 | 0.31 | 0.39 | 0.3 | 0.18 | 0.34 | 0.3 |
| 10 | 2782166 | rs7918571 | -0.942 | 0.792 | 2.419 | 0.082 | 0.034 | 0.032 | 0.032 | 0 | 0.06 | 0.23 | 0.03 | 0.05 | 0.06 | 0.02 |
| 10 | 2782509 | rs7922741 | -0.942 | 0.797 | 2.419 | 0.155 | 0.038 | 0.082 | 0.082 | 0 | 0.06 | 0.23 | 0.05 | 0.11 | 0.07 | 0.03 |
| 10 | 2782803 | rs1537617 | 1.481 | 0.85 | 2.56 | 0.152 | 0.1 | 0.098 | 0.098 | 0.032 | 0.9 | 0.55 | 0.83 | 0.84 | 0.81 | 0.91 |
| 10 | 2783225 | rs7093433 | -1.565 | 0.804 | 2.097 | -0.103 | 0.023 | 0.032 | 0.032 | 0.189 | 0.07 | 0.25 | 0.03 | 0.09 | 0.06 | 0.07 |
| 10 | 2804109 | rs2152001 | -0.922 | 0.223 | 1.602 | -0.263 | 0.001 | 0.003 | 0.003 | 0 | 0.34 | 0.3 | 0.63 | 0.75 | 0.39 | 0.43 |
| 10 | 2804997 | rs2184413 | -0.72 | 0.314 | 1.444 | -0.219 | 0 | 0.022 | 0.022 | 0 | 0.32 | 0.22 | 0.62 | 0.71 | 0.39 | 0.39 |
| 10 | 2809279 | rs7095226 | -0.75 | 0.257 | 1.89 | -0.125 | 0.008 | -0.041 | -0.041 | 3.619 | 0.08 | 0.2 | 0.04 | 0 | 0.06 | 0.01 |
| 10 | 2815824 | rs7912465 | 0.677 | 0.391 | 1.149 | -0.062 | 0.039 | -0.025 | -0.025 | 0.001 | 0.9 | 0.65 | 0.93 | 0.93 | 0.92 | 0.94 |
| 10 | 2823407 | rs7915938 | -1.207 | 0.04 | 0.59 | 0.123 | -0.02 | -0.02 | -0.02 | 0 | 0.13 | 0.22 | 0.08 | 0.04 | 0.07 | 0.11 |
| 10 | 2826548 | rs9943312 | -1.369 | -0.04 | 0.756 | 0.106 | -0.026 | -0.042 | -0.042 | 0 | 0.17 | 0.33 | 0.17 | 0.07 | 0.14 | 0.22 |
| 10 | 2831743 | rs10736960 | 1.027 | -0.154 | 0.937 | 0.193 | -0.027 | -0.014 | -0.014 | 0 | 0.81 | 0.53 | 0.87 | 0.81 | 0.87 | 0.82 |
| 10 | 2835293 | rs945141 | -0.944 | 0.344 | 1.103 | 1.001 | 0 | 0.071 | 0.071 | 0 | 0.09 | 0.08 | 0.34 | 0.5 | 0.12 | 0.14 |
| 10 | 2835603 | rs1889504 | 0.834 | 0.036 | 1.895 | 0.626 | -0.01 | -0.075 | -0.075 | 0 | 0.87 | 0.7 | 0.92 | 0.99 | 0.92 | 0.95 |
| 10 | 2839591 | rs1577255 | 0.92 | 0.032 | 2.12 | 0.596 | -0.001 | -0.085 | -0.085 | 0.059 | 0.87 | 0.57 | 0.92 | 0.99 | 0.92 | 0.95 |
| 10 | 2840150 | rs1591313 | 0.992 | 0.02 | 2.069 | 0.591 | 0.008 | -0.093 | -0.093 | 0 | 0.87 | 0.49 | 0.92 | 0.99 | 0.92 | 0.95 |
| 10 | 2842782 | rs1999647 | 0.568 | 0.505 | 2.158 | 0.577 | -0.006 | -0.059 | -0.059 | 0 | 0.89 | 0.79 | 0.93 | 0.99 | 0.92 | 0.95 |
| 10 | 2845542 | rs4132076 | 1.09 | 0.502 | 2.119 | 0.576 | 0.008 | -0.093 | -0.093 | 0 | 0.87 | 0.5 | 0.92 | 0.99 | 0.92 | 0.95 |
| 10 | 2854697 | rs10794947 | 0.794 | 0.299 | 0.906 | -0.225 | -0.002 | -0.048 | -0.048 | 0.834 | 0.35 | 0.42 | 0.21 | 0.11 | 0.28 | 0.24 |
| 10 | 2864447 | rs7080882 | -0.245 | 0.197 | 0.725 | -0.39 | 0.008 | 0.04 | 0.04 | 0 | 0.51 | 0.61 | 0.73 | 0.89 | 0.62 | 0.71 |
| 10 | 2867879 | rs1875164 | -0.392 | 0.197 | 0.575 | -0.369 | 0.004 | -0.01 | -0.01 | 0 | 0.84 | 0.86 | 0.92 | 1 | 0.89 | 0.95 |
| 10 | 2877311 | rs11251532 | 1.101 | 0.288 | 0.582 | -0.279 | -0.003 | 0.061 | 0.061 | 0 | 0.32 | 0.18 | 0.18 | 0.1 | 0.26 | 0.23 |
| 10 | 2877343 | rs2065681 | -0.151 | 0.679 | 0.34 | -0.357 | 0.007 | 0.064 | 0.064 | 0 | 0.51 | 0.61 | 0.68 | 0.88 | 0.53 | 0.69 |
| 10 | 2880121 | rs12220754 | 0.379 | 0.764 | -0.121 | 0.154 | -0.009 | 0.044 | 0.044 | 0 | 0.07 | 0.04 | 0.36 | 0.52 | 0.14 | 0.16 |
| 10 | 2881331 | rs10903880 | 0.135 | 0.76 | 0.302 | 0.07 | 0.019 | 0.085 | 0.085 | 0 | 0.11 | 0.23 | 0.13 | 0.28 | 0.13 | 0.31 |
| 10 | 2882338 | rs7911151 | -1.511 | 0.784 | -0.267 | -0.449 | -0.003 | -0.039 | -0.039 | 0 | 0.39 | 0.52 | 0.58 | 0.79 | 0.4 | 0.57 |
| 10 | 2882816 | rs7899347 | -1.073 | 0.767 | -0.487 | -0.771 | -0.001 | -0.043 | -0.043 | 0 | 0.7 | 0.68 | 0.76 | 0.89 | 0.67 | 0.78 |
| 10 | 2883078 | rs17280444 | -0.122 | 0.972 | -1.216 | -0.468 | -0.02 | 0.097 | 0.097 | 0 | 0.17 | 0.02 | 0.08 | 0 | 0.13 | 0.07 |
| 10 | 2887947 | rs4436465 | 0.132 | 0.845 | -1.062 | -0.089 | -0.017 | -0.016 | -0.016 | 0.907 | 0.62 | 0.15 | 0.63 | 0.58 | 0.58 | 0.47 |
| 10 | 2891279 | rs1909692 | 0.098 | 1.35 | -0.778 | -0.61 | -0.008 | -0.006 | -0.006 | 0.005 | 0.16 | 0.11 | 0.11 | 0 | 0.17 | 0.11 |
| 10 | 2895361 | rs10903886 | -0.329 | 1.585 | -1.152 | -0.397 | -0.015 | -0.033 | -0.033 | 0 | 0.21 | 0.16 | 0.5 | 0.67 | 0.27 | 0.35 |
| 10 | 2901165 | rs6601988 | -0.478 | 1.592 | -0.619 | -0.487 | 0.008 | -0.054 | -0.054 | 0.218 | 0.83 | 0.82 | 0.88 | 1 | 0.83 | 0.89 |
| 10 | 2902609 | rs11251540 | 0.401 | 1.584 | -0.877 | -0.811 | 0.001 | 0 | 0 | 0 | 0.18 | 0.21 | 0.21 | 0.19 | 0.3 | 0.35 |
| 10 | 2909884 | rs10903888 | -0.316 | 1.553 | -0.85 | -0.559 | -0.004 | -0.012 | -0.012 | 0.316 | 0.2 | 0.2 | 0.5 | 0.68 | 0.25 | 0.35 |
| 10 | 2910624 | rs1909690 | -0.088 | 1.619 | -0.939 | -0.72 | -0.023 | 0.034 | 0.034 | 0 | 0.19 | 0.08 | 0.13 | 0 | 0.2 | 0.13 |
| 10 | 2932234 | rs10751896 | 0.351 | 1.331 | -0.482 | -0.576 | -0.002 | 0.001 | 0.001 | 0 | 0.64 | 0.66 | 0.38 | 0.25 | 0.54 | 0.54 |

| | | | | | | | | | | | | | | | | |
|---|---|---|---|---|---|---|---|---|---|---|---|---|---|---|---|---|
| 10 | 2933230 | rs10794964 | -1.104 | -0.058 | -0.74 | -1.243 | 0.005 | -0.006 | -0.006 | 7.548 | 0.67 | 0.34 | 0.86 | 0.93 | 0.81 | 0.87 |
| 10 | 2933969 | rs4242741 | -0.305 | 0.759 | -0.38 | -0.996 | -0.024 | 0.03 | 0.03 | 0.38 | 0.35 | 0.22 | 0.66 | 0.76 | 0.53 | 0.46 |
| 10 | 2953012 | rs11251563 | -0.218 | 0.69 | 0.103 | -1.072 | -0.033 | 0.063 | 0.063 | 0.4 | 0.34 | 0.18 | 0.64 | 0.81 | 0.51 | 0.47 |
| 10 | 2960256 | rs4526685 | 0.087 | 0.331 | 0.186 | -1.326 | -0.002 | -0.012 | -0.012 | 0 | 0.11 | 0.16 | 0.11 | 0.03 | 0.11 | 0.2 |
| 10 | 2960317 | rs10751898 | 0.064 | 0.53 | 0.454 | -0.632 | 0.006 | 0.003 | 0.003 | 0.31 | 0.33 | 0.34 | 0.58 | 0.55 | 0.42 | 0.43 |
| 10 | 2961101 | rs11251567 | 0.946 | 1.015 | 0.21 | -0.264 | -0.046 | -0.001 | -0.001 | 0.645 | 0.07 | 0.01 | 0.3 | 0.41 | 0.16 | 0.21 |
| 10 | 2964727 | rs7904324 | -1.018 | 0.247 | 0.548 | -0.62 | -0.031 | -0.005 | -0.005 | 0 | 0.29 | 0.56 | 0.23 | 0.28 | 0.29 | 0.27 |
| 10 | 2965692 | rs7084464 | -1.306 | 0.093 | 0.505 | -0.801 | 0.021 | 0.055 | 0.055 | 4.662 | 0.52 | 0.68 | 0.77 | 0.81 | 0.64 | 0.7 |
| 10 | 2966418 | rs1909684 | -0.83 | 0.067 | 0.478 | -0.761 | -0.01 | 0.022 | 0.022 | 1.268 | 0.19 | 0.31 | 0.19 | 0.29 | 0.24 | 0.26 |
| 10 | 2967402 | rs1574317 | 0.327 | 0.118 | 0.387 | -0.72 | -0.05 | -0.033 | -0.033 | 0.222 | 0.32 | 0.19 | 0.57 | 0.57 | 0.4 | 0.42 |
| 10 | 2967536 | rs11251570 | 0.95 | 0.126 | 0.303 | -0.691 | -0.028 | -0.024 | -0.024 | 0.022 | 0.06 | 0 | 0.06 | 0.25 | 0.12 | 0.06 |
| 10 | 2967596 | rs1574318 | 0.533 | 0.092 | 0.299 | -0.599 | -0.075 | -0.055 | -0.055 | 0 | 0.3 | 0.12 | 0.56 | 0.57 | 0.39 | 0.4 |
| 10 | 2967660 | rs11813223 | 1.83 | -0.108 | 0.514 | -1.143 | 0.033 | 0.063 | 0.063 | 0.75 | 0.31 | 0.12 | 0.11 | 0.07 | 0.18 | 0.14 |
| 10 | 2971227 | rs1875172 | 0.793 | -0.093 | 0.602 | -1.04 | -0.004 | -0.018 | -0.018 | 0 | 0.07 | 0.12 | 0.08 | 0.03 | 0.09 | 0.12 |
| 10 | 2972261 | rs2936896 | -0.626 | -0.293 | 0.721 | -1.139 | -0.019 | -0.099 | -0.099 | 3.557 | 0.78 | 0.57 | 0.86 | 0.92 | 0.81 | 0.84 |
| 10 | 2973425 | rs3003514 | -0.626 | -0.293 | 0.721 | -1.139 | -0.019 | -0.099 | -0.099 | 3.266 | 0.78 | 0.57 | 0.86 | 0.92 | 0.81 | 0.84 |
| 10 | 2977047 | rs2502068 | -0.695 | -0.339 | 0.674 | -1.199 | 0.011 | 0.012 | 0.012 | 2.595 | 0.49 | 0.49 | 0.75 | 0.85 | 0.62 | 0.69 |
| 10 | 2977918 | rs2454827 | 1.737 | -0.322 | 0.644 | -1.083 | 0.018 | 0.036 | 0.036 | 0.848 | 0.31 | 0.16 | 0.11 | 0.07 | 0.19 | 0.14 |
| 10 | 2978321 | rs7073091 | -0.938 | -0.01 | 0.834 | -1.114 | 0.038 | 0.119 | 0.119 | 0 | 0.58 | 0.25 | 0.75 | 0.9 | 0.68 | 0.75 |
| 10 | 2979537 | rs11251583 | 0.511 | -0.361 | 0.939 | -1.111 | 0.001 | -0.049 | -0.049 | 0 | 0.11 | 0.25 | 0.1 | 0.03 | 0.12 | 0.12 |
| 10 | 2979826 | rs2454824 | 0.094 | -0.208 | 0.589 | -1.094 | -0.035 | -0.062 | -0.062 | 0 | 0.18 | 0.05 | 0.36 | 0.63 | 0.26 | 0.31 |
| 10 | 2981236 | rs17281778 | 1.994 | -0.177 | 0.694 | -1.151 | 0.007 | 0.049 | 0.049 | 0.291 | 0.28 | 0.14 | 0.13 | 0.06 | 0.21 | 0.14 |
| 10 | 2981486 | rs7087799 | -0.408 | -0.166 | 0.832 | -1.228 | -0.025 | 0.075 | 0.075 | 0 | 0.6 | 0.73 | 0.38 | 0.19 | 0.53 | 0.48 |
| 10 | 2983047 | rs7923826 | -0.71 | -0.171 | 0.916 | -1.228 | -0.005 | -0.078 | -0.078 | 0 | 0.3 | 0.38 | 0.23 | 0.12 | 0.31 | 0.34 |
| 10 | 2983081 | rs2454823 | -0.038 | -0.133 | 0.683 | -1.16 | -0.059 | 0.144 | 0.144 | 0 | 0.4 | 0.16 | 0.62 | 0.81 | 0.48 | 0.51 |
| 10 | 2986450 | rs11251588 | -0.355 | 0.094 | 0.384 | -0.66 | -0.009 | -0.03 | -0.03 | 0 | 0.21 | 0.19 | 0.39 | 0.39 | 0.26 | 0.34 |
| 10 | 2987086 | rs17281966 | 1.745 | 0.354 | 0.543 | -0.586 | 0.006 | 0.048 | 0.048 | 0.333 | 0.28 | 0.14 | 0.13 | 0.06 | 0.21 | 0.14 |
| 10 | 2988202 | rs10794975 | -0.572 | 0.308 | 0.458 | -0.502 | -0.002 | 0.054 | 0.054 | 0.196 | 0.67 | 0.49 | 0.65 | 0.51 | 0.66 | 0.7 |
| 10 | 2989572 | rs12220191 | -0.073 | 0.365 | 0.201 | -0.292 | -0.013 | -0.037 | -0.037 | 0.179 | 0.29 | 0.17 | 0.43 | 0.41 | 0.34 | 0.44 |
| 10 | 2990190 | rs11251594 | 0.393 | 0.39 | 0.687 | 0.298 | -0.016 | -0.016 | -0.016 | 0 | 0.06 | 0 | 0.06 | 0.3 | 0.11 | 0.08 |
| 10 | 2990261 | rs10903900 | 0.21 | 0.288 | 0.989 | -0.295 | 0.001 | -0.005 | -0.005 | 0.148 | 0.57 | 0.35 | 0.56 | 0.48 | 0.54 | 0.58 |
| 10 | 2991774 | rs2454837 | -0.375 | 0.244 | 1.612 | 0.685 | 0.01 | 0.057 | 0.057 | 0 | 0.07 | 0.1 | 0.31 | 0.32 | 0.11 | 0.24 |
| 10 | 2992399 | rs2936891 | 1.031 | 0.276 | 1.811 | 0.295 | -0.002 | -0.056 | -0.056 | 1.093 | 0.89 | 0.65 | 0.9 | 0.96 | 0.91 | 0.87 |
| 10 | 2992761 | rs2454838 | -0.197 | 0.315 | 1.65 | 0.343 | 0.016 | 0.102 | 0.102 | 0 | 0.07 | 0.11 | 0.31 | 0.34 | 0.11 | 0.25 |
| 10 | 2993191 | rs11251596 | -1.354 | -0.628 | 1.592 | -0.874 | -0.01 | -0.051 | -0.051 | 0.021 | 0.11 | 0.27 | 0.09 | 0.03 | 0.09 | 0.12 |
| 10 | 3003449 | rs2454842 | 1.724 | -1.051 | 1.951 | -0.915 | -0.018 | -0.09 | -0.09 | 0 | 0.86 | 0.44 | 0.92 | 0.97 | 0.93 | 0.86 |
| 10 | 3013719 | rs12357680 | -1.54 | -1.031 | 2.487 | -0.906 | -0.006 | -0.065 | -0.065 | 0 | 0.12 | 0.26 | 0.06 | 0.03 | 0.07 | 0.14 |
| 10 | 3013810 | rs12357696 | -1.189 | -1.031 | 2.554 | -0.907 | -0.025 | -0.05 | -0.05 | 0.381 | 0.1 | 0.22 | 0.03 | 0.03 | 0.03 | 0.12 |
| 10 | 3020453 | rs10794978 | -2.279 | -1.549 | 2.401 | -1.109 | -0.052 | -0.099 | -0.099 | 0 | 0.22 | 0.45 | 0.11 | 0.07 | 0.11 | 0.19 |
| 10 | 3022195 | rs7896008 | -2.158 | -1.492 | 2.087 | -1.084 | -0.013 | -0.032 | -0.032 | 0 | 0.2 | 0.24 | 0.1 | 0.07 | 0.1 | 0.16 |
| 10 | 3031753 | rs4353181 | 2.389 | -1.637 | 1.922 | -1.144 | 0.045 | 0.059 | 0.059 | 0 | 0.75 | 0.17 | 0.85 | 0.93 | 0.89 | 0.8 |
| 10 | 3040889 | rs7073415 | -1.401 | -1.71 | 2.049 | -1.175 | -0.018 | -0.071 | -0.071 | 0.132 | 0.15 | 0.27 | 0.06 | 0.02 | 0.06 | 0.15 |
| 10 | 3041788 | rs10903912 | -2.321 | -1.762 | 2.159 | -1.195 | -0.002 | -0.016 | -0.016 | 0 | 0.25 | 0.71 | 0.14 | 0.06 | 0.1 | 0.2 |
| 10 | 3049757 | rs7914724 | -1.251 | -1.841 | 1.917 | -1.286 | -0.035 | -0.115 | -0.115 | 4.324 | 0.19 | 0.38 | 0.1 | 0.02 | 0.09 | 0.14 |
| 10 | 3054104 | rs12260533 | -2.315 | -1.977 | 1.901 | -1.326 | 0.023 | 0.018 | 0.018 | 0.144 | 0.29 | 0.7 | 0.16 | 0.06 | 0.13 | 0.19 |
| 10 | 3055401 | rs10903927 | -2.309 | -1.975 | 1.797 | -1.325 | 0.03 | 0.028 | 0.028 | 0.145 | 0.29 | 0.71 | 0.16 | 0.06 | 0.13 | 0.19 |
| 10 | 3056078 | rs12098591 | -1.085 | -2.05 | 1.689 | -1.351 | -0.04 | -0.024 | -0.024 | 0 | 0.1 | 0.18 | 0.05 | 0.04 | 0.04 | 0.14 |
| 10 | 3058348 | rs10903929 | 1.32 | -1.837 | 0.535 | -1.26 | 0.103 | 0.162 | 0.162 | 1.216 | 0.69 | 0.11 | 0.82 | 0.94 | 0.87 | 0.79 |
| 10 | 3058941 | rs7095229 | -1.659 | -1.719 | 0.592 | -1.208 | -0.017 | -0.042 | -0.042 | 0.042 | 0.29 | 0.63 | 0.16 | 0.06 | 0.13 | 0.19 |
| 10 | 3060901 | rs7070854 | -1.662 | -1.723 | 0.589 | -1.211 | -0.016 | -0.042 | -0.042 | 0 | 0.29 | 0.63 | 0.16 | 0.06 | 0.13 | 0.19 |
| 10 | 3067312 | rs10794989 | 0.263 | -0.486 | -0.554 | -0.938 | -0.013 | 0.114 | 0.114 | 1.633 | 0.44 | 0.21 | 0.67 | 0.81 | 0.6 | 0.63 |
| 10 | 3068685 | rs7087749 | 1.187 | -0.385 | 0.82 | -0.418 | 0.025 | 0.146 | 0.146 | 0.332 | 0.9 | 0.55 | 0.93 | 0.85 | 0.94 | 0.88 |
| 10 | 3071566 | rs6422398 | 0.853 | -0.693 | 0.666 | -0.793 | -0.034 | -0.087 | -0.087 | 0 | 0.79 | 0.43 | 0.78 | 0.93 | 0.88 | 0.78 |
| 10 | 3074091 | rs4880585 | 0.999 | -0.767 | 0.627 | -0.825 | -0.038 | -0.089 | -0.089 | 0.072 | 0.78 | 0.43 | 0.78 | 0.93 | 0.88 | 0.78 |
| 10 | 3075076 | rs17135074 | -1.374 | -0.52 | 0.748 | -0.298 | -0.003 | 0.096 | 0.096 | 0 | 0.08 | 0.22 | 0.05 | 0.15 | 0.05 | 0.05 |
| 10 | 3075870 | rs7096174 | 0.333 | -0.356 | 0.144 | -1.132 | 0.009 | 0.065 | 0.065 | 0 | 0.53 | 0.37 | 0.36 | 0.21 | 0.43 | 0.39 |
| 10 | 3076294 | rs10903937 | -0.047 | 0.874 | -0.101 | -1.061 | -0.008 | -0.012 | -0.012 | 0 | 0.15 | 0.08 | 0.12 | 0 | 0.19 | 0.08 |
| 10 | 3077299 | rs10903938 | -0.184 | 0.883 | -0.071 | -1.057 | -0.002 | -0.025 | -0.025 | 0.461 | 0.15 | 0.09 | 0.12 | 0 | 0.19 | 0.08 |
| 10 | 3083649 | rs7089499 | -0.744 | 1.286 | -0.108 | -0.689 | -0.005 | -0.003 | -0.003 | 1.387 | 0.09 | 0.08 | 0.1 | 0.1 | 0.13 | 0.12 |
| 10 | 3095046 | rs3814582 | -0.566 | 0.794 | -0.105 | -1.077 | 0.009 | 0.019 | 0.019 | 0 | 0.28 | 0.21 | 0.27 | 0.05 | 0.27 | 0.14 |
| 10 | 3095185 | rs3814585 | -0.61 | 1.302 | 0.971 | 0.254 | 0.012 | 0.123 | 0.123 | 0 | 0.11 | 0.38 | 0.23 | 0.19 | 0.12 | 0.12 |
| 10 | 3095281 | rs3829910 | 0.672 | 2.142 | 0.859 | 1.185 | 0.13 | 0.21 | 0.21 | 0 | 0.88 | 0.66 | 0.8 | 0.6 | 0.78 | 0.75 |
| 10 | 3096702 | rs7087613 | -0.747 | 1.853 | 0.294 | 1.086 | 0.008 | 0.034 | 0.034 | 0 | 0.2 | 0.28 | 0.33 | 0.5 | 0.27 | 0.4 |
| 10 | 3097212 | rs10732921 | 0.383 | 1.474 | 0.11 | 0.723 | -0.001 | -0.001 | -0.001 | 0 | 0.59 | 0.47 | 0.5 | 0.44 | 0.55 | 0.49 |
| 10 | 3097745 | rs10903949 | 0.488 | 0.926 | -0.575 | -0.181 | 0.012 | 0.091 | 0.091 | 0 | 0.47 | 0.37 | 0.45 | 0.13 | 0.46 | 0.39 |
| 10 | 3098711 | rs11813585 | -0.564 | 0.638 | -0.835 | 0.032 | -0.006 | 0.013 | 0.013 | 0 | 0.36 | 0.22 | 0.36 | 0.27 | 0.37 | 0.3 |
| 10 | 3099351 | rs7091953 | -0.175 | -0.671 | -0.832 | -0.427 | -0.023 | -0.046 | -0.046 | 0 | 0.69 | 0.4 | 0.84 | 0.92 | 0.86 | 0.81 |
| 10 | 3106857 | rs7923030 | 0.181 | -0.826 | -2.016 | 0.018 | 0.038 | -0.073 | -0.073 | 0 | 0.47 | 0.08 | 0.55 | 0.51 | 0.67 | 0.54 |
| 10 | 3114645 | rs11251694 | -0.721 | -0.632 | -1.327 | 0.363 | 0.003 | -0.003 | -0.003 | 0 | 0.25 | 0.26 | 0.29 | 0.28 | 0.17 | 0.31 |

| | | | | | | | | | | | | | | | | |
|---|---|---|---|---|---|---|---|---|---|---|---|---|---|---|---|---|
| 10 | 3114994 | rs10795000 | 0.06 | -0.502 | -1.444 | 0.47 | 0.019 | -0.029 | -0.029 | 0 | 0.18 | 0.12 | 0.27 | 0.28 | 0.12 | 0.26 |
| 10 | 3118871 | rs7909171 | 0.343 | -0.109 | -1.319 | 0.266 | -0.028 | -0.046 | -0.046 | 0 | 0.38 | 0.24 | 0.51 | 0.45 | 0.57 | 0.41 |
| 10 | 3119753 | rs7080058 | 0.297 | 0.129 | -0.34 | 0.852 | -0.02 | 0.037 | 0.037 | 0 | 0.79 | 0.66 | 0.73 | 0.7 | 0.85 | 0.71 |
| 10 | 3123433 | rs882849 | -1.155 | 1.294 | -0.832 | 1.481 | -0.032 | -0.032 | -0.032 | 0 | 0.24 | 0.17 | 0.38 | 0.36 | 0.36 | 0.36 |
| 10 | 3124319 | rs11251702 | -0.615 | -0.028 | -0.41 | 1.549 | -0.001 | 0.022 | 0.022 | 0.224 | 0.11 | 0.08 | 0.19 | 0.3 | 0.08 | 0.27 |
| 10 | 3124516 | rs4881074 | 0.781 | 0.022 | -0.487 | 1.417 | -0.025 | 0.046 | 0.046 | 0.149 | 0.74 | 0.6 | 0.69 | 0.59 | 0.78 | 0.54 |
| 10 | 3131553 | rs10795002 | 0.923 | 0.152 | -0.769 | 1.412 | -0.03 | 0.076 | 0.076 | 0.144 | 0.71 | 0.43 | 0.51 | 0.49 | 0.74 | 0.5 |
| 10 | 3134083 | rs11251711 | -0.761 | 0.131 | -1.549 | 1.636 | -0.005 | -0.064 | -0.064 | 0 | 0.1 | 0.01 | 0.18 | 0.28 | 0.09 | 0.3 |
| 10 | 3134681 | rs7090199 | 0.328 | 0.316 | -2.205 | 0.737 | -0.092 | 0.089 | 0.089 | 0 | 0.44 | 0.08 | 0.33 | 0.28 | 0.46 | 0.3 |
| 10 | 3137472 | rs7076989 | 0.645 | 0.446 | -1.99 | 0.558 | -0.078 | 0.125 | 0.125 | 0 | 0.32 | 0.07 | 0.22 | 0.1 | 0.35 | 0.25 |
| 10 | 3138009 | rs2388595 | 0.614 | 0.24 | -1.367 | 0.854 | 0.004 | -0.004 | -0.004 | 5.284 | 0.38 | 0.45 | 0.29 | 0.33 | 0.38 | 0.35 |
| 10 | 3138265 | rs7074733 | 1.169 | -0.037 | -0.581 | 1.697 | -0.016 | 0.044 | 0.044 | 0 | 0.73 | 0.62 | 0.54 | 0.43 | 0.75 | 0.61 |
| 10 | 3139701 | rs10903963 | 0.466 | 0.264 | -0.791 | 1.65 | -0.008 | 0.053 | 0.053 | 0 | 0.91 | 0.88 | 0.76 | 0.76 | 0.94 | 0.82 |
| 10 | 3145541 | rs2892550 | 0.759 | 0.759 | -0.007 | 1.291 | 0.028 | 0.086 | 0.086 | 0.027 | 0.82 | 0.54 | 0.72 | 0.67 | 0.75 | 0.82 |
| 10 | 3145906 | rs11251719 | 0.504 | 0.405 | 0.261 | 1.483 | -0.012 | 0.054 | 0.054 | 0 | 0.73 | 0.62 | 0.52 | 0.55 | 0.75 | 0.67 |
| 10 | 3147192 | rs4256893 | 0.245 | 0.105 | -0.125 | 1.11 | 0.004 | -0.017 | -0.017 | 0.026 | 0.28 | 0.27 | 0.32 | 0.54 | 0.26 | 0.36 |
| 10 | 3148116 | rs2388569 | 0.186 | 0.057 | 0.221 | 1.016 | -0.002 | 0.005 | 0.005 | 0 | 0.32 | 0.34 | 0.34 | 0.5 | 0.28 | 0.36 |
| 10 | 3149402 | rs10903967 | -0.031 | 0.275 | -0.441 | 0.397 | 0.001 | 0.001 | 0.001 | 0 | 0.61 | 0.39 | 0.53 | 0.81 | 0.66 | 0.63 |
| 10 | 3150371 | rs4881084 | 0.646 | -0.277 | -0.024 | 0.173 | -0.025 | -0.01 | -0.01 | 0.195 | 0.15 | 0.19 | 0.13 | 0.1 | 0.06 | 0.24 |
| 10 | 3150898 | rs11251721 | 0.54 | -0.117 | 0.148 | -0.103 | -0.001 | -0.005 | -0.005 | 0 | 0.2 | 0.15 | 0.21 | 0.05 | 0.2 | 0.09 |
| 10 | 3150973 | rs1132173 | 1.277 | -0.389 | -0.018 | -0.209 | 0.003 | -0.03 | -0.03 | 0 | 0.33 | 0.28 | 0.4 | 0.51 | 0.33 | 0.37 |
| 10 | 3151325 | rs2306297 | -0.077 | -0.326 | 0.178 | -0.078 | -0.004 | -0.003 | -0.003 | 1.26 | 0.52 | 0.54 | 0.54 | 0.6 | 0.42 | 0.63 |
| 10 | 3153888 | rs10508245 | 1.49 | -0.301 | 0.014 | -0.101 | -0.133 | -0.087 | -0.087 | 0 | 0.25 | 0.02 | 0.21 | 0.31 | 0.37 | 0.26 |
| 10 | 3154339 | rs2306301 | 1.202 | -0.247 | 0.158 | -0.104 | -0.036 | -0.024 | -0.024 | 1.163 | 0.27 | 0.14 | 0.22 | 0.33 | 0.37 | 0.27 |
| 10 | 3154461 | rs2306302 | 0.745 | -0.328 | 0.19 | -0.116 | -0.024 | -0.011 | -0.011 | 0 | 0.14 | 0.18 | 0.11 | 0.09 | 0.06 | 0.24 |
| 10 | 3154534 | rs2306304 | -0.221 | -0.315 | 0.302 | -0.124 | 0.004 | 0.003 | 0.003 | 0 | 0.47 | 0.44 | 0.46 | 0.4 | 0.56 | 0.37 |
| 10 | 3155217 | rs6602023 | -0.701 | -0.099 | 0.514 | -0.21 | 0.026 | -0.029 | -0.029 | 0 | 0.92 | 0.76 | 0.88 | 0.98 | 0.9 | 0.94 |
| 10 | 3155237 | rs6602024 | -0.835 | -0.1 | 0.514 | -0.211 | 0.03 | -0.026 | -0.026 | 0 | 0.92 | 0.76 | 0.88 | 0.98 | 0.9 | 0.94 |
| 10 | 3155353 | rs3816699 | 0.994 | -0.083 | 0.433 | -0.218 | 0.003 | -0.014 | -0.014 | 0.314 | 0.16 | 0.15 | 0.19 | 0.05 | 0.16 | 0.07 |
| 10 | 3155428 | rs3816701 | 0.874 | -0.063 | 0.486 | -0.213 | 0.015 | 0.072 | 0.072 | 0.051 | 0.59 | 0.29 | 0.56 | 0.81 | 0.66 | 0.61 |
| 10 | 3155698 | rs4881086 | -0.645 | 0.045 | 0.276 | -0.262 | -0.004 | -0.016 | -0.016 | 0 | 0.32 | 0.32 | 0.31 | 0.13 | 0.25 | 0.32 |
| 10 | 3155770 | rs4881088 | 0.46 | -0.1 | 0.22 | -0.265 | -0.011 | -0.009 | -0.009 | 0 | 0.16 | 0.18 | 0.12 | 0.08 | 0.09 | 0.26 |
| 10 | 3155790 | rs4881089 | -0.519 | 0.045 | 0.149 | -0.214 | -0.003 | -0.025 | -0.025 | 0 | 0.32 | 0.33 | 0.34 | 0.14 | 0.29 | 0.33 |
| 10 | 3156214 | rs2388566 | -0.24 | 0.241 | 0.204 | 0.21 | 0.006 | -0.027 | -0.027 | 0 | 0.92 | 0.73 | 0.91 | 0.98 | 0.95 | 0.95 |
| 10 | 3158646 | rs7916889 | -0.102 | 0.168 | -0.233 | 0.388 | 0.006 | 0 | 0 | 0.068 | 0.53 | 0.43 | 0.53 | 0.44 | 0.63 | 0.41 |
| 10 | 3159234 | rs2306305 | 0.636 | 1.291 | -0.263 | 0.105 | 0.02 | -0.031 | -0.031 | 0.035 | 0.81 | 0.73 | 0.77 | 0.91 | 0.76 | 0.76 |
| 10 | 3160127 | rs12257705 | 0.056 | 1.095 | -0.254 | 0.325 | -0.004 | 0.001 | 0.001 | 0.332 | 0.1 | 0.12 | 0.17 | 0.14 | 0.08 | 0.05 |
| 10 | 3160726 | rs9423462 | -0.179 | 0.619 | -0.33 | 0.236 | -0.007 | 0.006 | 0.006 | 0 | 0.39 | 0.44 | 0.4 | 0.43 | 0.36 | 0.27 |
| 10 | 3160847 | rs4881098 | -1.759 | 1.213 | -0.657 | -0.156 | 0.009 | -0.011 | -0.011 | 0.399 | 0.71 | 0.75 | 0.74 | 0.63 | 0.78 | 0.58 |
| 10 | 3161131 | rs3829914 | -1.223 | 0.924 | -0.736 | -0.426 | -0.001 | -0.011 | -0.011 | 0 | 0.57 | 0.58 | 0.64 | 0.84 | 0.68 | 0.72 |
| 10 | 3161253 | rs9423464 | 1.276 | 0.889 | -0.769 | -0.357 | 0.004 | 0.003 | 0.003 | 0 | 0.17 | 0.12 | 0.16 | 0.03 | 0.13 | 0.12 |
| 10 | 3162423 | rs6602026 | -2.891 | 0.818 | -0.437 | -0.344 | 0.005 | 0.004 | 0.004 | 0 | 0.76 | 0.78 | 0.86 | 0.87 | 0.85 | 0.85 |
| 10 | 3162525 | rs3814590 | 1.492 | 0.891 | -0.619 | -0.16 | -0.016 | -0.078 | -0.078 | 0.728 | 0.16 | 0.06 | 0.13 | 0.39 | 0.19 | 0.12 |
| 10 | 3163068 | rs7086721 | -1.402 | 1.24 | -0.506 | -0.359 | 0.009 | 0.015 | 0.015 | 0 | 0.61 | 0.65 | 0.65 | 0.71 | 0.68 | 0.76 |
| 10 | 3163175 | rs7075876 | 0.386 | 1.436 | -0.29 | -0.417 | 0 | -0.006 | -0.006 | 0.075 | 0.21 | 0.21 | 0.28 | 0.26 | 0.24 | 0.3 |
| 10 | 3163452 | rs11251728 | -0.371 | 1.411 | 0.032 | -0.431 | -0.009 | -0.005 | -0.005 | 2.014 | 0.39 | 0.32 | 0.28 | 0.41 | 0.44 | 0.44 |
| 10 | 3163585 | rs9423467 | 1.662 | 1.428 | -0.013 | -0.413 | 0.014 | 0.027 | 0.027 | 0 | 0.17 | 0.09 | 0.11 | 0.02 | 0.13 | 0.12 |
| 10 | 3163973 | rs9423685 | 0.27 | 1.429 | -0.022 | -0.387 | -0.001 | 0 | 0 | 0.098 | 0.22 | 0.24 | 0.28 | 0.26 | 0.24 | 0.3 |
| 10 | 3164719 | rs6602028 | -1.465 | 1.547 | 0.087 | -0.231 | -0.024 | -0.018 | -0.018 | 0 | 0.59 | 0.53 | 0.65 | 0.66 | 0.72 | 0.77 |
| 10 | 3165316 | rs12268267 | 1.455 | 1.492 | 0.097 | -0.409 | 0.001 | 0 | 0 | 0.288 | 0.11 | 0.12 | 0.2 | 0.16 | 0.12 | 0.07 |
| 10 | 3165972 | rs9423690 | -1.536 | 1.671 | -0.174 | -0.327 | -0.025 | -0.03 | -0.03 | 0.633 | 0.59 | 0.53 | 0.65 | 0.7 | 0.72 | 0.77 |
| 10 | 3166109 | rs9423474 | -0.739 | 1.878 | -0.097 | -0.478 | 0 | -0.008 | -0.008 | 0.717 | 0.44 | 0.49 | 0.52 | 0.37 | 0.53 | 0.65 |
| 10 | 3166345 | rs9423479 | -0.343 | 2.098 | 0.021 | -0.499 | 0.007 | -0.004 | -0.004 | 0 | 0.38 | 0.44 | 0.47 | 0.35 | 0.43 | 0.62 |
| 10 | 3166624 | rs11591441 | 1.031 | 1.752 | -0.124 | -0.903 | 0.008 | -0.021 | -0.021 | 0 | 0.18 | 0.18 | 0.14 | 0.07 | 0.22 | 0.26 |
| 10 | 3167463 | rs2306309 | -0.198 | 0.859 | 0.013 | -0.651 | -0.001 | 0.019 | 0.019 | 0 | 0.23 | 0.3 | 0.3 | 0.32 | 0.26 | 0.3 |
| 10 | 3167902 | rs7091322 | -0.1 | 0.87 | 0.011 | -0.646 | 0 | 0.009 | 0.009 | 0 | 0.22 | 0.27 | 0.3 | 0.32 | 0.26 | 0.3 |
| 10 | 3169771 | rs6602032 | -1.292 | 0.698 | -0.052 | -0.763 | 0.016 | 0.027 | 0.027 | 0 | 0.6 | 0.66 | 0.64 | 0.74 | 0.68 | 0.74 |
| 10 | 3169803 | rs7080240 | 0.363 | 0.614 | -0.144 | -0.641 | -0.045 | 0.005 | 0.005 | 0 | 0.33 | 0.19 | 0.28 | 0.27 | 0.43 | 0.43 |
| 10 | 3171043 | rs2279208 | 0.253 | 0.803 | -0.006 | -0.627 | -0.013 | 0.004 | 0.004 | 0 | 0.08 | 0.08 | 0.17 | 0.3 | 0.17 | 0.23 |
| 10 | 3172121 | rs1052333 | -0.488 | 0.948 | -0.078 | -0.752 | 0.02 | 0.014 | 0.014 | 0 | 0.34 | 0.47 | 0.38 | 0.37 | 0.39 | 0.42 |
| 10 | 3173581 | rs4880592 | -0.567 | 0.943 | -0.076 | -0.757 | 0.006 | 0.019 | 0.019 | 0 | 0.36 | 0.53 | 0.45 | 0.51 | 0.4 | 0.45 |
| 10 | 3176363 | rs2306310 | 1.56 | 0.974 | -0.133 | -0.749 | 0.026 | -0.04 | -0.04 | 1.621 | 0.12 | 0.19 | 0.08 | 0.05 | 0.13 | 0.12 |
| 10 | 3176389 | rs2306311 | 0.79 | 0.988 | -0.212 | -0.754 | -0.037 | -0.067 | -0.067 | 0.761 | 0.22 | 0.12 | 0.27 | 0.36 | 0.29 | 0.25 |
| 10 | 3176456 | rs2306312 | -1.728 | 0.895 | -0.205 | -0.832 | 0.004 | 0.006 | 0.006 | 0 | 0.66 | 0.67 | 0.66 | 0.74 | 0.75 | 0.7 |
| 10 | 3177320 | rs9423490 | -1.589 | 0.853 | -0.183 | -0.753 | 0.011 | 0.024 | 0.024 | 0 | 0.63 | 0.68 | 0.64 | 0.74 | 0.71 | 0.68 |
| 10 | 3178881 | rs1053000 | 1.263 | 0.865 | -0.284 | -0.731 | -0.109 | -0.088 | -0.088 | 0.746 | 0.18 | 0.01 | 0.18 | 0.22 | 0.28 | 0.27 |
| 10 | 3179197 | rs12772817 | 1.174 | 0.769 | -0.235 | -0.712 | 0.026 | 0.068 | 0.068 | 0.104 | 0.17 | 0.04 | 0.11 | 0.02 | 0.14 | 0.08 |
| 10 | 3180227 | rs6901 | -1.543 | 0.769 | -0.202 | -0.708 | 0.013 | 0.024 | 0.024 | 0 | 0.63 | 0.69 | 0.65 | 0.74 | 0.72 | 0.72 |
| 10 | 3180298 | rs10175 | -0.326 | 0.774 | -0.345 | -0.722 | 0.01 | 0.058 | 0.058 | 0.022 | 0.34 | 0.2 | 0.35 | 0.16 | 0.31 | 0.2 |

| | | | | | | | | | | | | | | | | |
|---|---|---|---|---|---|---|---|---|---|---|---|---|---|---|---|---|
| 10 | 3180689 | rs2388558 | -0.955 | 1.38 | 0.253 | -0.086 | -0.006 | 0.012 | 0.012 | 0 | 0.29 | 0.38 | 0.31 | 0.32 | 0.32 | 0.32 |
| 10 | 3180759 | rs9423495 | 0.356 | 1.382 | 0.13 | -0.09 | 0.017 | 0.021 | 0.021 | 0 | 0.19 | 0.11 | 0.12 | 0.02 | 0.15 | 0.08 |
| 10 | 3180951 | rs2279216 | 1.235 | 1.411 | 0.107 | -0.07 | -0.091 | -0.087 | -0.087 | 0.118 | 0.16 | 0.02 | 0.17 | 0.21 | 0.26 | 0.21 |
| 10 | 3184507 | rs7898695 | -1.48 | 1.351 | 0.142 | -0.109 | 0 | -0.007 | -0.007 | 0.182 | 0.5 | 0.53 | 0.59 | 0.68 | 0.61 | 0.59 |
| 10 | 3185237 | rs4881109 | -0.719 | 0.755 | 0.215 | -0.054 | -0.001 | 0.001 | 0.001 | 0 | 0.28 | 0.3 | 0.31 | 0.32 | 0.31 | 0.31 |
| 10 | 3185393 | rs4881110 | -1.075 | 0.768 | 0.192 | -0.049 | -0.014 | 0.005 | 0.005 | 0 | 0.32 | 0.42 | 0.32 | 0.33 | 0.31 | 0.33 |
| 10 | 3189115 | rs7924157 | 1.021 | 0.829 | 0.237 | -0.061 | 0.035 | -0.049 | -0.049 | 0 | 0.14 | 0.23 | 0.1 | 0.05 | 0.16 | 0.13 |
| 10 | 3189380 | rs3740607 | -1.302 | 0.849 | 0.236 | -0.05 | -0.003 | 0.025 | 0.025 | 0 | 0.48 | 0.45 | 0.54 | 0.69 | 0.52 | 0.55 |
| 10 | 3190265 | rs4881111 | -1.114 | 0.819 | 0.213 | -0.031 | -0.009 | 0.033 | 0.033 | 0 | 0.47 | 0.33 | 0.47 | 0.55 | 0.55 | 0.55 |
| 10 | 3191823 | rs2388556 | 1.172 | 0.829 | 0.152 | -0.007 | 0.012 | -0.025 | -0.025 | 0 | 0.13 | 0.13 | 0.09 | 0.05 | 0.16 | 0.13 |
| 10 | 3200292 | rs3182535 | -1.009 | -0.537 | 0.507 | 0.321 | -0.005 | -0.003 | -0.003 | 0 | 0.6 | 0.57 | 0.57 | 0.6 | 0.71 | 0.68 |
| 10 | 3201097 | rs3814594 | -1.108 | -0.536 | 0.512 | 0.322 | -0.021 | -0.013 | -0.013 | 0 | 0.6 | 0.5 | 0.57 | 0.6 | 0.71 | 0.68 |
| 10 | 3202065 | rs4242746 | -0.826 | -0.497 | 0.235 | 0.333 | 0.011 | 0.008 | 0.008 | 0.225 | 0.56 | 0.59 | 0.56 | 0.6 | 0.67 | 0.67 |
| 10 | 3202682 | rs9423709 | -0.569 | -0.542 | 0.292 | 0.401 | 0.001 | 0.014 | 0.014 | 0 | 0.21 | 0.25 | 0.29 | 0.32 | 0.27 | 0.31 |
| 10 | 3204682 | rs5017749 | -1.252 | -1.024 | -0.205 | 0.613 | -0.016 | 0.003 | 0.003 | 0 | 0.66 | 0.57 | 0.59 | 0.58 | 0.75 | 0.72 |
| 10 | 3208557 | rs12359035 | -1.315 | -0.234 | 0.106 | 0.942 | -0.005 | -0.003 | -0.003 | 0 | 0.62 | 0.59 | 0.58 | 0.6 | 0.7 | 0.72 |
| 10 | 3218457 | rs9423506 | -1.318 | -0.167 | 0.25 | 0.463 | 0.009 | 0.021 | 0.021 | 0.362 | 0.57 | 0.63 | 0.62 | 0.74 | 0.63 | 0.7 |
| 10 | 3219171 | rs9423508 | -0.202 | 0.296 | 0.129 | -0.115 | 0.013 | 0.017 | 0.017 | 0 | 0.18 | 0.1 | 0.11 | 0.02 | 0.13 | 0.12 |
| 10 | 3219445 | rs7078068 | -0.351 | 0.292 | 0.194 | 0.36 | -0.001 | -0.002 | -0.002 | 0 | 0.48 | 0.48 | 0.43 | 0.29 | 0.57 | 0.47 |
| 10 | 3230052 | rs7093944 | -1.411 | 0.453 | 0.673 | 0.146 | 0.027 | -0.043 | -0.043 | 0 | 0.15 | 0.66 | 0.17 | 0.08 | 0.17 | 0.03 |
| 10 | 3230337 | rs17131865 | -1.214 | -0.393 | 0.388 | 0.119 | -0.079 | -0.075 | -0.075 | 0 | 0.21 | 0.53 | 0.1 | 0.09 | 0.11 | 0.09 |
| 10 | 3245862 | rs10751916 | -0.567 | 0.077 | -0.458 | 0.338 | 0.009 | 0.007 | 0.007 | 0 | 0.35 | 0.25 | 0.56 | 0.66 | 0.36 | 0.49 |
| 10 | 3260462 | rs2477765 | 0.63 | 0.398 | -0.251 | -0.192 | -0.017 | 0.035 | 0.035 | 0 | 0.44 | 0.32 | 0.35 | 0.32 | 0.49 | 0.36 |
| 10 | 3264590 | rs7099179 | -0.964 | 0.389 | 0.261 | 0.055 | -0.008 | 0.012 | 0.012 | 0 | 0.55 | 0.6 | 0.64 | 0.64 | 0.49 | 0.64 |
| 10 | 3264809 | rs7088720 | -0.582 | 0.32 | 0.668 | 0.003 | 0.001 | 0.006 | 0.006 | 0.006 | 0.5 | 0.51 | 0.62 | 0.64 | 0.48 | 0.61 |
| 10 | 3270102 | rs2477764 | 0.362 | 0.257 | 0.362 | 0.09 | -0.004 | 0.02 | 0.02 | 0 | 0.49 | 0.33 | 0.46 | 0.38 | 0.49 | 0.38 |
| 10 | 3271675 | rs12269271 | -0.869 | 0.314 | 0.257 | 0.119 | -0.002 | 0.008 | 0.008 | 0 | 0.08 | 0.07 | 0.16 | 0.02 | 0.09 | 0.09 |
| 10 | 3273550 | rs7084176 | -0.812 | 0.526 | -0.131 | 0.332 | 0 | 0.021 | 0.021 | 0 | 0.5 | 0.63 | 0.54 | 0.44 | 0.52 | 0.62 |
| 10 | 3274756 | rs10795014 | -0.861 | 0.465 | -0.337 | 0.32 | -0.011 | 0.021 | 0.021 | 0 | 0.51 | 0.74 | 0.55 | 0.44 | 0.52 | 0.63 |
| 10 | 3282738 | rs11251793 | 0.001 | 0.287 | -0.423 | -0.143 | -0.001 | -0.004 | -0.004 | 0.105 | 0.29 | 0.27 | 0.18 | 0.19 | 0.26 | 0.21 |
| 10 | 3284061 | rs2814925 | 0.02 | 0.09 | -0.665 | 0.22 | -0.018 | 0.025 | 0.025 | 0 | 0.47 | 0.26 | 0.6 | 0.57 | 0.5 | 0.56 |
| 10 | 3286815 | rs2764986 | 0.286 | 0.431 | -0.583 | 0.451 | -0.009 | -0.012 | -0.012 | 0 | 0.21 | 0.18 | 0.32 | 0.26 | 0.24 | 0.21 |
| 10 | 3287305 | rs11251799 | 0.883 | 0.371 | -1.259 | 0.139 | -0.015 | 0.022 | 0.022 | 0 | 0.1 | 0.01 | 0.08 | 0.05 | 0.1 | 0.05 |
| 10 | 3288594 | rs11251800 | 0.738 | 0.651 | -0.885 | 0.704 | -0.017 | -0.025 | -0.025 | 0 | 0.21 | 0.09 | 0.31 | 0.29 | 0.28 | 0.35 |
| 10 | 3289771 | rs2814930 | -0.282 | 0.57 | -0.177 | 0.481 | 0.054 | 0.046 | 0.046 | 0 | 0.07 | 0.2 | 0.04 | 0.1 | 0.08 | 0.04 |
| 10 | 3290142 | rs10508250 | 0.766 | 0.612 | -0.463 | 0.598 | 0 | 0.013 | 0.013 | 0 | 0.11 | 0.09 | 0.08 | 0.22 | 0.08 | 0.1 |
| 10 | 3290827 | rs11251803 | 0.463 | 0.579 | -0.449 | 0.564 | 0 | 0.003 | 0.003 | 0 | 0.13 | 0.11 | 0.1 | 0.22 | 0.1 | 0.16 |
| 10 | 3295673 | rs4881131 | 0.711 | 0.967 | -0.701 | 0.31 | -0.009 | -0.002 | -0.002 | 0 | 0.19 | 0.14 | 0.28 | 0.22 | 0.23 | 0.32 |
| 10 | 3303486 | rs912826 | -0.864 | 0.92 | -0.531 | 0.149 | -0.004 | -0.037 | -0.037 | 0 | 0.91 | 0.81 | 0.94 | 1 | 0.95 | 0.98 |
| 10 | 3304955 | rs4489652 | -0.036 | 0.957 | -0.303 | 0.16 | 0.002 | -0.018 | -0.018 | 0 | 0.25 | 0.34 | 0.3 | 0.23 | 0.25 | 0.39 |
| 10 | 3307294 | rs12220117 | 0.522 | 0.773 | 0.098 | 0.516 | -0.007 | 0.068 | 0.068 | 0 | 0.16 | 0.29 | 0.13 | 0.21 | 0.12 | 0.11 |
| 10 | 3308716 | rs11251814 | 0.877 | 0.778 | 0.107 | 0.464 | -0.002 | -0.022 | -0.022 | 0.118 | 0.08 | 0.13 | 0.06 | 0 | 0.05 | 0.03 |
| 10 | 3308846 | rs963355 | 0.208 | 0.723 | 0.134 | 0.505 | 0.007 | 0.113 | 0.113 | 0 | 0.8 | 0.56 | 0.83 | 0.68 | 0.82 | 0.86 |
| 10 | 3309647 | rs2296465 | 0.968 | 0.836 | -0.21 | 0.438 | -0.001 | -0.001 | -0.001 | 0 | 0.2 | 0.17 | 0.33 | 0.26 | 0.25 | 0.43 |
| 10 | 3310378 | rs12784859 | -0.055 | 0.731 | -0.498 | 0.193 | 0.002 | 0.016 | 0.016 | 0.586 | 0.51 | 0.52 | 0.51 | 0.3 | 0.43 | 0.4 |
| 10 | 3310566 | rs12766168 | 1.452 | 0.684 | -0.718 | 0.181 | -0.01 | 0.082 | 0.082 | 0 | 0.28 | 0.11 | 0.34 | 0.09 | 0.26 | 0.26 |
| 10 | 3313456 | rs9804298 | 0.695 | 0.673 | -0.467 | 0.447 | 0.045 | 0.069 | 0.069 | 0 | 0.05 | 0.22 | 0.04 | 0.11 | 0.06 | 0.03 |
| 10 | 3314267 | rs7082002 | 0.71 | 0.636 | -0.849 | 0.172 | -0.005 | 0.041 | 0.041 | 0 | 0.16 | 0.25 | 0.13 | 0.21 | 0.13 | 0.12 |
| 10 | 3316388 | rs12776746 | 1.166 | 0.758 | -1.343 | 0.079 | -0.008 | 0.037 | 0.037 | 0.549 | 0.07 | 0.01 | 0.03 | 0 | 0.04 | 0.03 |
| 10 | 3319474 | rs1009543 | 0.419 | 0.773 | -1.34 | 0.081 | -0.003 | 0.012 | 0.012 | 1.605 | 0.37 | 0.55 | 0.42 | 0.45 | 0.39 | 0.56 |
| 10 | 3319978 | rs1570856 | 0.306 | 0.731 | -0.935 | 0.872 | 0 | -0.016 | -0.016 | 0 | 0.68 | 0.72 | 0.78 | 0.54 | 0.64 | 0.8 |
| 10 | 3320113 | rs912837 | 0.245 | 0.801 | -1.298 | 0.937 | -0.036 | -0.006 | -0.006 | 0 | 0.11 | 0.01 | 0.08 | 0.07 | 0.15 | 0.05 |
| 10 | 3322307 | rs2147939 | 0.455 | 0.903 | -1.038 | 0.888 | -0.003 | 0.009 | 0.009 | 0 | 0.59 | 0.52 | 0.74 | 0.54 | 0.58 | 0.78 |
| 10 | 3323315 | rs11251827 | 0.49 | 1.075 | -1.308 | 0.767 | -0.007 | 0.028 | 0.028 | 0.175 | 0.07 | 0.04 | 0.03 | 0 | 0.05 | 0.03 |
| 10 | 3323773 | rs1325586 | -0.043 | 1.093 | -1.127 | 0.976 | -0.001 | -0.001 | -0.001 | 0 | 0.6 | 0.58 | 0.74 | 0.5 | 0.59 | 0.78 |
| 10 | 3327813 | rs10904017 | -0.361 | 1.216 | -1.423 | 1.056 | -0.047 | -0.088 | -0.088 | 0 | 0.17 | 0.02 | 0.12 | 0.29 | 0.22 | 0.19 |
| 10 | 3327931 | rs17135001 | -0.899 | 1.05 | -0.79 | 0.768 | 0.007 | -0.012 | -0.012 | 0 | 0.08 | 0.15 | 0.05 | 0.03 | 0.06 | 0.06 |
| 10 | 3330360 | rs10904018 | 0.212 | 0.824 | -0.934 | 0.48 | 0 | 0 | 0 | 0.064 | 0.57 | 0.58 | 0.58 | 0.48 | 0.64 | 0.48 |
| 10 | 3330587 | rs7070200 | 0.176 | 0.419 | -0.978 | 0.564 | 0.016 | -0.037 | -0.037 | 0.239 | 0.77 | 0.83 | 0.78 | 0.62 | 0.85 | 0.71 |
| 10 | 3331092 | rs728939 | -0.018 | -0.319 | -1.039 | 0.067 | 0.018 | 0.021 | 0.021 | 0 | 0.62 | 0.33 | 0.68 | 0.32 | 0.72 | 0.61 |
| 10 | 3332360 | rs12765323 | 0.989 | -0.44 | -0.947 | -0.036 | 0.014 | -0.035 | -0.035 | 2.577 | 0.24 | 0.11 | 0.12 | 0.29 | 0.17 | 0.19 |
| 10 | 3333530 | rs17135017 | -0.245 | -0.453 | -0.962 | -0.095 | 0.013 | -0.085 | -0.085 | 0 | 0.18 | 0.43 | 0.16 | 0.04 | 0.17 | 0.11 |
| 10 | 3335061 | rs12766986 | 0.954 | -0.313 | -1.022 | 0.01 | 0.014 | -0.035 | -0.035 | 0 | 0.24 | 0.11 | 0.12 | 0.29 | 0.17 | 0.19 |
| 10 | 3335447 | rs11251836 | 0.748 | -0.16 | -0.632 | 0.112 | 0.058 | 0.018 | 0.018 | 0 | 0.07 | 0.19 | 0.08 | 0.04 | 0.09 | 0.07 |
| 10 | 3342774 | rs4881147 | -0.184 | -0.353 | -0.953 | 0.24 | 0.048 | 0.009 | 0.009 | 0 | 0.55 | 0.68 | 0.68 | 0.42 | 0.71 | 0.56 |
| 10 | 3344048 | rs4881149 | -0.716 | -0.564 | -1.047 | 0.312 | -0.005 | 0.009 | 0.009 | 0 | 0.88 | 0.92 | 0.88 | 0.66 | 0.96 | 0.85 |
| 10 | 3347548 | rs502397 | -0.158 | -0.528 | -1.138 | 0.31 | -0.018 | 0.012 | 0.012 | 0 | 0.17 | 0.18 | 0.15 | 0.35 | 0.08 | 0.19 |
| 10 | 3348124 | rs549676 | 0.031 | 0.842 | -1.283 | -0.046 | -0.011 | 0.029 | 0.029 | 0 | 0.56 | 0.62 | 0.46 | 0.66 | 0.46 | 0.56 |
| 10 | 3350027 | rs17291583 | 0.319 | 1.904 | -1.764 | -0.104 | -0.04 | -0.001 | -0.001 | 0 | 0.16 | 0.09 | 0.21 | 0.18 | 0.3 | 0.22 |

| | | | | | | | | | | | | | | | | |
|---|---|---|---|---|---|---|---|---|---|---|---|---|---|---|---|---|
| 10 | 3350441 | rs17219336 | -0.638 | 1.194 | -1.556 | -0.188 | -0.02 | 0.038 | 0.038 | 0 | 0.15 | 0.05 | 0.14 | 0.08 | 0.16 | 0.13 |
| 10 | 3355303 | rs1337382 | -0.766 | 2.284 | -1.78 | -0.114 | -0.091 | 0.02 | 0.02 | 0 | 0.22 | 0.06 | 0.38 | 0.15 | 0.34 | 0.25 |
| 10 | 3363137 | rs737533 | 1.21 | 1.327 | -1.462 | -0.923 | -0.027 | 0.072 | 0.072 | 0 | 0.59 | 0.31 | 0.45 | 0.82 | 0.49 | 0.68 |
| 10 | 3363208 | rs1807437 | -0.167 | 1.371 | -1.402 | -0.867 | -0.08 | 0.023 | 0.023 | 0 | 0.22 | 0.07 | 0.38 | 0.15 | 0.34 | 0.25 |
| 10 | 3368555 | rs11594429 | -0.217 | 1.36 | -1.413 | -0.821 | -0.082 | 0.024 | 0.024 | 0 | 0.22 | 0.07 | 0.38 | 0.15 | 0.35 | 0.26 |
| 10 | 3368595 | rs4881155 | 1.047 | 1.189 | -0.264 | -0.917 | 0.015 | -0.036 | -0.036 | 0.082 | 0.66 | 0.61 | 0.55 | 0.82 | 0.6 | 0.71 |
| 10 | 3381474 | rs10751920 | 0.215 | 0.785 | 0.034 | -0.814 | -0.038 | -0.055 | -0.055 | 0 | 0.88 | 0.69 | 0.94 | 0.97 | 0.95 | 0.96 |
| 10 | 3387978 | rs7903644 | 1.601 | 1.131 | 0.193 | -0.976 | -0.014 | -0.11 | -0.11 | 0.15 | 0.78 | 0.41 | 0.82 | 0.98 | 0.76 | 0.89 |
| 10 | 3389934 | rs722828 | 0.045 | 1.204 | 0.806 | -0.913 | 0.037 | -0.03 | -0.03 | 0 | 0.94 | 0.86 | 0.91 | 1 | 0.88 | 0.98 |
| 10 | 3390320 | rs1337381 | 0.889 | 0.426 | 0.15 | -0.734 | 0.03 | 0.019 | 0.019 | 0 | 0.49 | 0.31 | 0.48 | 0.56 | 0.53 | 0.55 |
| 10 | 3390575 | rs7079506 | -1.048 | 1.361 | 0.26 | -0.903 | 0.007 | -0.007 | -0.007 | 0 | 0.24 | 0.27 | 0.28 | 0.16 | 0.37 | 0.27 |
| 10 | 3391621 | rs6602045 | 0.749 | -0.116 | 0.104 | -1.017 | 0.001 | 0.004 | 0.004 | 0.09 | 0.68 | 0.64 | 0.62 | 0.63 | 0.6 | 0.67 |
| 10 | 3395216 | rs2182195 | -0.122 | -0.189 | 0.035 | -1.05 | -0.006 | -0.027 | -0.027 | 0 | 0.21 | 0.16 | 0.28 | 0.35 | 0.23 | 0.28 |
| 10 | 3397417 | rs1408985 | 1.66 | -0.34 | 0.124 | -1.159 | 0.035 | 0.03 | 0.03 | 0 | 0.53 | 0.19 | 0.47 | 0.58 | 0.54 | 0.62 |
| 10 | 3397728 | rs11251861 | 1.66 | -0.34 | 0.124 | -1.159 | 0.035 | 0.03 | 0.03 | 0 | 0.53 | 0.19 | 0.47 | 0.58 | 0.54 | 0.62 |
| 10 | 3399877 | rs7901930 | -0.281 | -0.311 | 0.309 | -1.089 | -0.049 | -0.056 | -0.056 | 0 | 0.14 | 0.3 | 0.13 | 0.05 | 0.06 | 0.04 |
| 10 | 3400516 | rs4271299 | -0.824 | -0.044 | -0.069 | -0.85 | -0.006 | -0.036 | -0.036 | 0 | 0.24 | 0.17 | 0.29 | 0.37 | 0.26 | 0.31 |
| 10 | 3416737 | rs7090790 | 0.174 | -0.028 | -0.194 | -0.721 | -0.014 | -0.051 | -0.051 | 0.409 | 0.16 | 0.06 | 0.26 | 0.37 | 0.17 | 0.26 |
| 10 | 3418124 | rs1570854 | 1.387 | -0.392 | -0.225 | -0.867 | 0.016 | -0.016 | -0.016 | 0 | 0.63 | 0.19 | 0.56 | 0.61 | 0.63 | 0.64 |
| 10 | 3421414 | rs7080295 | -0.547 | -0.515 | 0.275 | -0.905 | 0.063 | -0.022 | -0.022 | 3.135 | 0.93 | 0.75 | 0.91 | 1 | 0.88 | 0.98 |
| 10 | 3429328 | rs7898173 | 0.677 | -0.867 | -0.518 | -0.9 | -0.052 | 0.018 | 0.018 | 0 | 0.13 | 0 | 0.09 | 0.05 | 0.16 | 0.11 |
| 10 | 3431825 | rs7894960 | -0.529 | -0.903 | -0.079 | -0.8 | -0.037 | -0.05 | -0.05 | 0.176 | 0.1 | 0.35 | 0.08 | 0 | 0.04 | 0.04 |
| 10 | 3442533 | rs17135209 | 1.336 | -1.061 | -1.224 | -0.595 | 0.064 | -0.088 | -0.088 | 0 | 0.31 | 0.09 | 0.11 | 0.39 | 0.18 | 0.26 |
| 10 | 3444489 | rs10508254 | 0.017 | -0.93 | -0.894 | -0.647 | 0.005 | -0.006 | -0.006 | 0 | 0.55 | 0.4 | 0.27 | 0.44 | 0.41 | 0.42 |
| 10 | 3444851 | rs10508255 | 0.907 | -1.061 | -1.281 | -0.525 | 0.087 | -0.086 | -0.086 | 0 | 0.35 | 0.12 | 0.11 | 0.4 | 0.2 | 0.27 |
| 10 | 3445668 | rs7079024 | -0.207 | -1.071 | -1.306 | -0.647 | 0.108 | -0.018 | -0.018 | 0.193 | 0.49 | 0.19 | 0.71 | 0.57 | 0.64 | 0.64 |
| 10 | 3447072 | rs10795033 | -0.217 | -1.061 | -1.387 | -0.643 | 0.091 | -0.005 | -0.005 | 0.041 | 0.49 | 0.25 | 0.71 | 0.57 | 0.63 | 0.64 |
| 10 | 3448187 | rs11591499 | 1.165 | -0.925 | -1.44 | -0.602 | 0.059 | 0.137 | 0.137 | 5.753 | 0.32 | 0.03 | 0.1 | 0.12 | 0.18 | 0.22 |
| 10 | 3457288 | rs7901480 | 0.626 | -0.018 | -0.465 | -0.064 | -0.007 | -0.037 | -0.037 | 0 | 0.83 | 0.27 | 0.82 | 0.98 | 0.83 | 0.9 |
| 10 | 3466699 | rs11599597 | 0.256 | -0.023 | -0.715 | -0.075 | -0.022 | -0.092 | -0.092 | 0 | 0.81 | 0.4 | 0.83 | 1 | 0.82 | 0.89 |
| 10 | 3468208 | rs10904036 | 0.185 | -0.005 | -0.706 | -0.067 | -0.021 | -0.092 | -0.092 | 0.333 | 0.81 | 0.4 | 0.84 | 1 | 0.82 | 0.88 |
| 10 | 3475318 | rs10904040 | 0.262 | 1.013 | -1.386 | 0.725 | -0.052 | -0.078 | -0.078 | 1.139 | 0.11 | 0.01 | 0.22 | 0.35 | 0.17 | 0.23 |
| 10 | 3482498 | rs7084941 | 0.421 | 1.254 | -1.308 | 0.166 | -0.082 | 0.008 | 0.008 | 0 | 0.19 | 0.05 | 0.35 | 0.17 | 0.28 | 0.28 |
| 10 | 3484881 | rs7920115 | -0.031 | 0.157 | -0.66 | 0.17 | 0.003 | 0.005 | 0.005 | 0.247 | 0.54 | 0.59 | 0.38 | 0.61 | 0.49 | 0.45 |
| 10 | 3490932 | rs7894316 | -1.165 | -0.499 | 0.029 | 0.241 | -0.007 | 0.018 | 0.018 | 0 | 0.29 | 0.67 | 0.31 | 0.5 | 0.29 | 0.21 |
| 10 | 3493718 | rs10795049 | -0.664 | -0.516 | 0.281 | 0.258 | -0.027 | 0.112 | 0.112 | 0 | 0.25 | 0.47 | 0.28 | 0.5 | 0.26 | 0.18 |
| 10 | 3494886 | rs11251898 | -0.562 | -0.628 | -0.017 | 0.287 | 0.005 | -0.023 | -0.023 | 0 | 0.16 | 0.12 | 0.27 | 0.69 | 0.13 | 0.3 |
| 10 | 3495382 | rs1890073 | -0.087 | -0.554 | 0.008 | 0.317 | -0.001 | 0.013 | 0.013 | 0 | 0.08 | 0.09 | 0.15 | 0.2 | 0.05 | 0.19 |
| 10 | 3496426 | rs17135269 | -0.623 | -0.519 | 0.153 | 0.199 | -0.001 | -0.001 | -0.001 | 0 | 0.1 | 0.1 | 0.14 | 0.13 | 0.07 | 0.21 |
| 10 | 3502282 | rs4881173 | 0.006 | -0.883 | 0.825 | 0.211 | -0.031 | 0.152 | 0.152 | 0 | 0.16 | 0.32 | 0.17 | 0.48 | 0.12 | 0.16 |
| 10 | 3505030 | rs1964428 | -1.384 | -1.021 | 1.054 | 0.139 | -0.031 | 0.076 | 0.076 | 0 | 0.27 | 0.56 | 0.29 | 0.5 | 0.27 | 0.19 |
| 10 | 3506070 | rs1036747 | -0.018 | -1.014 | 0.938 | 0.21 | -0.032 | 0.149 | 0.149 | 0 | 0.16 | 0.33 | 0.17 | 0.48 | 0.12 | 0.16 |
| 10 | 3509300 | rs2053889 | -1.663 | -1.253 | 0.626 | 0.027 | 0.027 | 0.005 | 0.005 | 0 | 0.39 | 0.7 | 0.47 | 0.62 | 0.33 | 0.4 |
| 10 | 3518880 | rs11251910 | -0.602 | -1.204 | 0.62 | 0.104 | -0.02 | 0.03 | 0.03 | 0 | 0.23 | 0.31 | 0.22 | 0.46 | 0.14 | 0.17 |
| 10 | 3520151 | rs483881 | 0.191 | -1.007 | 0.627 | 0.05 | 0.012 | -0.009 | -0.009 | 5.54 | 0.88 | 0.8 | 0.79 | 0.86 | 0.83 | 0.8 |
| 10 | 3522302 | rs622898 | 1.044 | -1.097 | 0.332 | -0.026 | 0.006 | 0 | 0 | 0 | 0.63 | 0.33 | 0.54 | 0.38 | 0.68 | 0.62 |
| 10 | 3523872 | rs10904048 | 0.215 | -0.257 | 0.242 | -0.32 | 0.001 | -0.012 | -0.012 | 0 | 0.73 | 0.76 | 0.6 | 0.45 | 0.79 | 0.56 |
| 10 | 3525645 | rs10904049 | 1.437 | -0.327 | -0.268 | -1.127 | -0.024 | -0.033 | -0.033 | 0 | 0.28 | 0.14 | 0.28 | 0.33 | 0.29 | 0.15 |
| 10 | 3526096 | rs17135292 | -0.103 | -0.78 | 0.349 | -1.124 | 0.001 | -0.005 | -0.005 | 0 | 0.07 | 0.18 | 0.06 | 0.02 | 0.08 | 0.03 |
| 10 | 3527900 | rs7099538 | -0.229 | -0.92 | 0.553 | -1.284 | -0.011 | -0.031 | -0.031 | 0 | 0.07 | 0.16 | 0.04 | 0.01 | 0.03 | 0.1 |
| 10 | 3529103 | rs1660980 | 0.675 | -0.929 | 0.536 | -1.326 | -0.003 | 0 | 0 | 0 | 0.57 | 0.49 | 0.43 | 0.41 | 0.55 | 0.52 |
| 10 | 3530108 | rs7095342 | 0.914 | -1.089 | 0.088 | -1.398 | 0.019 | 0.009 | 0.009 | 0 | 0.53 | 0.31 | 0.42 | 0.4 | 0.55 | 0.51 |
| 10 | 3535233 | rs964162 | 0.168 | -0.972 | 0.153 | -1.185 | -0.04 | 0.029 | 0.029 | 0 | 0.69 | 0.56 | 0.56 | 0.4 | 0.76 | 0.62 |
| 10 | 3536296 | rs959509 | -0.129 | -0.95 | -0.096 | -1.396 | 0.021 | -0.019 | -0.019 | 0 | 0.66 | 0.7 | 0.6 | 0.41 | 0.77 | 0.61 |
| 10 | 3545431 | rs10751929 | -0.335 | -1.065 | 0.248 | -1.296 | -0.018 | 0.025 | 0.025 | 0 | 0.75 | 0.67 | 0.62 | 0.4 | 0.84 | 0.66 |
| 10 | 3545917 | rs7907091 | 1.232 | -0.967 | -0.112 | -1.557 | 0.018 | 0.02 | 0.02 | 0.518 | 0.54 | 0.16 | 0.39 | 0.37 | 0.53 | 0.42 |
| 10 | 3549366 | rs4880616 | 1.687 | -1.122 | -0.281 | -1.215 | 0.026 | -0.081 | -0.081 | 0 | 0.12 | 0.02 | 0.31 | 0.48 | 0.09 | 0.19 |
| 10 | 3549609 | rs7910111 | 1.892 | -0.401 | -0.54 | -1.589 | 0.054 | -0.087 | -0.087 | 0 | 0.25 | 0.04 | 0.19 | 0.32 | 0.12 | 0.1 |
| 10 | 3555578 | rs17135332 | 0.164 | -0.105 | 0.285 | -1.141 | 0.05 | -0.008 | -0.008 | 1.921 | 0.05 | 0.16 | 0.05 | 0.01 | 0.09 | 0.04 |
| 10 | 3558799 | rs684240 | -0.286 | -0.284 | 0.134 | -1.201 | 0.032 | -0.052 | -0.052 | 0 | 0.75 | 0.83 | 0.63 | 0.41 | 0.83 | 0.65 |
| 10 | 3558971 | rs476079 | -0.416 | 0.049 | 0.602 | -1.228 | 0.095 | -0.074 | -0.074 | 0.037 | 0.13 | 0.46 | 0.19 | 0.01 | 0.25 | 0.06 |
| 10 | 3560223 | rs563341 | 0.184 | -0.403 | -0.279 | -1.244 | 0.048 | -0.044 | -0.044 | 0.091 | 0.66 | 0.81 | 0.6 | 0.4 | 0.79 | 0.5 |
| 10 | 3564591 | rs11598081 | 0.311 | 0.075 | 0.039 | -1.215 | 0.015 | -0.032 | -0.032 | 0 | 0.09 | 0.18 | 0.15 | 0.01 | 0.23 | 0.05 |
| 10 | 3580253 | rs7914011 | 1.226 | 0.107 | 0.203 | -1.428 | -0.005 | 0.033 | 0.033 | 0.514 | 0.51 | 0.21 | 0.37 | 0.37 | 0.44 | 0.42 |
| 10 | 3580801 | rs11251934 | 1.016 | 0.064 | 0.112 | -1.51 | -0.028 | 0.05 | 0.05 | 0 | 0.13 | 0.03 | 0.08 | 0.01 | 0.16 | 0.19 |
| 10 | 3584829 | rs24401 | -0.549 | -0.026 | 0.418 | -1.542 | 0.019 | -0.031 | -0.031 | 0.178 | 0.76 | 0.82 | 0.59 | 0.39 | 0.84 | 0.67 |
| 10 | 3592082 | rs10795059 | 1.441 | 0.001 | 0.351 | -1.533 | -0.005 | 0.035 | 0.035 | 0.751 | 0.5 | 0.2 | 0.33 | 0.37 | 0.44 | 0.42 |
| 10 | 3596526 | rs7086377 | 0.58 | -0.088 | 0.521 | -1.531 | 0.001 | -0.002 | -0.002 | 0 | 0.64 | 0.38 | 0.42 | 0.38 | 0.6 | 0.61 |
| 10 | 3597405 | rs7914587 | 1.406 | 0.013 | 0.019 | -1.895 | 0 | -0.015 | -0.015 | 0 | 0.19 | 0.17 | 0.09 | 0.06 | 0.16 | 0.28 |

| | | | | | | | | | | | | | | | | |
|---|---|---|---|---|---|---|---|---|---|---|---|---|---|---|---|---|
| 10 | 3597542 | rs551307 | -0.903 | 0.014 | 0.059 | -1.875 | 0.031 | -0.06 | -0.06 | 0 | 0.77 | 0.91 | 0.6 | 0.39 | 0.84 | 0.67 |
| 10 | 3600741 | rs7068219 | 0.385 | 0.386 | 0.693 | -1.879 | 0.043 | -0.055 | -0.055 | 0 | 0.1 | 0.26 | 0.16 | 0.01 | 0.23 | 0.05 |
| 10 | 3606072 | rs1441265 | -1.25 | 0.406 | 0.098 | -1.865 | -0.019 | 0.123 | 0.123 | 0 | 0.45 | 0.79 | 0.36 | 0.08 | 0.55 | 0.52 |
| 10 | 3620969 | rs11812097 | 0.789 | 0.505 | 0.101 | -1.618 | -0.002 | -0.034 | -0.034 | 0 | 0.2 | 0.19 | 0.1 | 0.06 | 0.16 | 0.28 |
| 10 | 3625624 | rs12772528 | -0.145 | 0.586 | 0.083 | -1.631 | 0.019 | -0.118 | -0.118 | 0 | 0.3 | 0.46 | 0.26 | 0.07 | 0.39 | 0.33 |
| 10 | 3627637 | rs17224806 | 1.444 | 0.008 | -0.447 | -1.425 | -0.022 | 0.059 | 0.059 | 0 | 0.1 | 0.02 | 0.07 | 0.01 | 0.12 | 0.09 |
| 10 | 3628517 | rs11251954 | 1.367 | -0.257 | -0.545 | -1.34 | -0.006 | -0.005 | -0.005 | 0 | 0.29 | 0.23 | 0.29 | 0.29 | 0.29 | 0.13 |
| 10 | 3635369 | rs17135436 | 0.635 | -0.599 | -0.295 | -1.052 | 0.001 | -0.024 | -0.024 | 0 | 0.17 | 0.23 | 0.19 | 0.11 | 0.14 | 0.35 |
| 10 | 3635518 | rs17135438 | 0.489 | -0.683 | -0.475 | -1.045 | 0.004 | -0.012 | -0.012 | 0 | 0.09 | 0 | 0.04 | 0.13 | 0.05 | 0.14 |
| 10 | 3640752 | rs1668538 | 0.317 | -0.677 | -0.145 | -1.225 | 0 | 0.001 | 0.001 | 1.367 | 0.58 | 0.4 | 0.44 | 0.43 | 0.53 | 0.38 |
| 10 | 3642555 | rs7901181 | 0.28 | -0.659 | 0.071 | -1.272 | 0.044 | -0.085 | -0.085 | 0.002 | 0.19 | 0.48 | 0.13 | 0.05 | 0.18 | 0.36 |
| 10 | 3644879 | rs619778 | -1.329 | -0.702 | -0.082 | -1.273 | 0.007 | -0.061 | -0.061 | 0 | 0.8 | 0.91 | 0.58 | 0.48 | 0.75 | 0.76 |
| 10 | 3652078 | rs11816894 | 1.017 | -0.675 | 0.044 | -1.299 | 0.016 | -0.037 | -0.037 | 0 | 0.15 | 0.23 | 0.09 | 0.05 | 0.14 | 0.35 |
| 10 | 3657243 | rs11251965 | 0.561 | -0.68 | -0.122 | -1.277 | -0.003 | 0 | 0 | 0.042 | 0.09 | 0 | 0.04 | 0.12 | 0.06 | 0.14 |
| 10 | 3657375 | rs11251967 | 0.621 | -0.7 | -0.098 | -1.293 | 0.003 | 0.001 | 0.001 | 0 | 0.07 | 0.08 | 0.04 | 0.12 | 0.05 | 0.12 |
| 10 | 3658434 | rs705468 | -0.791 | -0.713 | -0.074 | -1.285 | 0.009 | -0.033 | -0.033 | 0 | 0.77 | 0.8 | 0.56 | 0.48 | 0.71 | 0.74 |
| 10 | 3661064 | rs11251971 | 0.073 | -0.638 | -0.12 | -1.231 | 0.007 | 0.007 | 0.007 | 0 | 0.09 | 0.08 | 0.05 | 0.12 | 0.06 | 0.14 |
| 10 | 3664150 | rs11251973 | 0.049 | -0.638 | -0.119 | -1.23 | 0.003 | 0.003 | 0.003 | 0 | 0.1 | 0.11 | 0.05 | 0.12 | 0.06 | 0.14 |
| 10 | 3667726 | rs705471 | 1.53 | -0.419 | -0.208 | -1.209 | -0.004 | 0.008 | 0.008 | 1.267 | 0.39 | 0.3 | 0.32 | 0.3 | 0.36 | 0.17 |
| 10 | 3669970 | rs7919625 | 1.072 | -0.704 | -0.295 | -1.309 | -0.007 | 0.027 | 0.027 | 0.438 | 0.45 | 0.32 | 0.39 | 0.31 | 0.47 | 0.24 |
| 10 | 3676325 | rs10904062 | 1.445 | -0.013 | -0.191 | -0.961 | -0.021 | 0.055 | 0.055 | 0 | 0.1 | 0.02 | 0.07 | 0.01 | 0.13 | 0.08 |
| 10 | 3682002 | rs7098140 | -0.307 | -0.021 | 0.158 | -0.762 | -0.005 | -0.039 | -0.039 | 0.776 | 0.24 | 0.34 | 0.14 | 0.17 | 0.18 | 0.47 |
| 10 | 3683786 | rs7909512 | 1.065 | 0.066 | 0.023 | -0.438 | 0.016 | 0.054 | 0.054 | 4.127 | 0.51 | 0.26 | 0.39 | 0.33 | 0.51 | 0.35 |
| 10 | 3684990 | rs12262514 | 1.226 | -0.056 | 0.393 | -0.446 | -0.007 | -0.091 | -0.091 | 1.141 | 0.16 | 0.28 | 0.11 | 0 | 0.19 | 0.07 |
| 10 | 3686435 | rs4881184 | -0.539 | -0.093 | 0.537 | -0.29 | -0.024 | 0.146 | 0.146 | 1.759 | 0.87 | 0.76 | 0.65 | 0.51 | 0.91 | 0.77 |
| 10 | 3686485 | rs4880619 | 0.127 | -0.07 | 0.303 | -0.367 | 0.001 | -0.05 | -0.05 | 0 | 0.3 | 0.31 | 0.21 | 0.05 | 0.32 | 0.3 |
| 10 | 3689257 | rs10904063 | 1.858 | 0.123 | 0.166 | -0.476 | -0.002 | -0.01 | -0.01 | 0 | 0.1 | 0.09 | 0.08 | 0.01 | 0.13 | 0.06 |
| 10 | 3689466 | rs11251988 | 0.606 | 0.13 | 0.231 | -0.446 | 0 | 0.002 | 0.002 | 0 | 0.05 | 0.09 | 0.04 | 0.13 | 0.07 | 0.13 |
| 10 | 3690181 | rs12573672 | 1.57 | 0.169 | 0.249 | -0.455 | -0.007 | -0.013 | -0.013 | 0.133 | 0.44 | 0.21 | 0.34 | 0.45 | 0.42 | 0.41 |
| 10 | 3690408 | rs11251992 | 1.382 | 0.17 | 0.228 | -0.465 | -0.022 | 0.069 | 0.069 | 4.481 | 0.49 | 0.17 | 0.38 | 0.32 | 0.48 | 0.33 |
| 10 | 3691304 | rs2397327 | -0.574 | -0.099 | 0.317 | -0.533 | 0.005 | -0.097 | -0.097 | 0.003 | 0.41 | 0.47 | 0.3 | 0.06 | 0.45 | 0.35 |
| 10 | 3692191 | rs2129441 | -0.556 | -0.132 | 0.214 | -0.545 | 0.006 | -0.101 | -0.101 | 0.053 | 0.41 | 0.43 | 0.3 | 0.06 | 0.45 | 0.35 |
| 10 | 3695386 | rs11251995 | 0.31 | 0.013 | -0.121 | -0.292 | -0.003 | -0.007 | -0.007 | 0 | 0.05 | 0.08 | 0.04 | 0.13 | 0.07 | 0.13 |
| 10 | 3697104 | rs11251998 | 1.215 | -0.186 | -0.065 | -0.388 | -0.004 | -0.064 | -0.064 | 0 | 0.15 | 0.19 | 0.11 | 0 | 0.19 | 0.07 |
| 10 | 3699620 | rs10904068 | -0.816 | -0.232 | -0.061 | 0.147 | -0.011 | 0.036 | 0.036 | 3.346 | 0.3 | 0.35 | 0.33 | 0.55 | 0.33 | 0.4 |
| 10 | 3699729 | rs10508257 | 0.761 | -0.233 | -0.051 | 0.227 | 0.005 | 0.041 | 0.041 | 0 | 0.05 | 0.11 | 0.03 | 0.13 | 0.05 | 0.13 |
| 10 | 3700290 | rs12765657 | -0.093 | -0.172 | -0.578 | 0.309 | -0.001 | 0.002 | 0.002 | 3.789 | 0.24 | 0.24 | 0.3 | 0.41 | 0.27 | 0.27 |
| 10 | 3701654 | rs12412149 | 1.078 | -0.144 | -0.614 | 0.26 | -0.007 | -0.007 | -0.007 | 0 | 0.16 | 0.1 | 0.23 | 0.17 | 0.14 | 0.26 |
| 10 | 3703206 | rs6601742 | -1.066 | -0.179 | -0.572 | 0.209 | 0.001 | -0.002 | -0.002 | 0 | 0.4 | 0.34 | 0.53 | 0.58 | 0.41 | 0.53 |
| 10 | 3706894 | rs7095969 | 0.707 | -0.137 | -0.232 | 0.446 | 0.003 | 0.014 | 0.014 | 0.285 | 0.57 | 0.35 | 0.46 | 0.28 | 0.61 | 0.36 |
| 10 | 3707650 | rs7909212 | 0.044 | -0.153 | -0.179 | 0.549 | -0.022 | 0.091 | 0.091 | 0.276 | 0.72 | 0.5 | 0.68 | 0.48 | 0.75 | 0.61 |
| 10 | 3708015 | rs4279919 | 0.211 | -0.105 | -0.388 | 0.196 | 0.001 | 0.006 | 0.006 | 0.302 | 0.57 | 0.37 | 0.46 | 0.28 | 0.61 | 0.36 |
| 10 | 3711122 | rs10508258 | -0.277 | 0.377 | -0.668 | 0.339 | 0 | -0.007 | -0.007 | 0 | 0.54 | 0.47 | 0.58 | 0.79 | 0.43 | 0.69 |
| 10 | 3715765 | rs11252010 | -0.329 | 0.649 | -0.648 | 1.426 | -0.006 | 0.001 | 0.001 | 0.336 | 0.07 | 0.01 | 0.15 | 0.38 | 0.07 | 0.18 |
| 10 | 3719164 | rs6601745 | 0.87 | 1.533 | -0.127 | 0.945 | 0.046 | 0.045 | 0.045 | 0.051 | 0.9 | 0.77 | 0.9 | 0.81 | 0.85 | 0.71 |
| 10 | 3723698 | rs11252021 | -0.516 | 1.24 | -0.05 | 0.793 | -0.005 | 0.018 | 0.018 | 0.307 | 0.23 | 0.23 | 0.25 | 0.42 | 0.21 | 0.19 |
| 10 | 3724611 | rs12412703 | 0.808 | 1.208 | -0.48 | 0.393 | -0.025 | 0.009 | 0.009 | 0 | 0.23 | 0.13 | 0.17 | 0.16 | 0.26 | 0.19 |
| 10 | 3724752 | rs873207 | 0.842 | 1.194 | -0.486 | 0.387 | -0.054 | 0.003 | 0.003 | 0 | 0.23 | 0.08 | 0.17 | 0.16 | 0.26 | 0.19 |
| 10 | 3727947 | rs7100211 | 0.38 | 1.169 | -0.114 | 0.337 | -0.024 | 0.132 | 0.132 | 0.152 | 0.18 | 0.38 | 0.24 | 0.41 | 0.17 | 0.13 |
| 10 | 3728834 | rs11252026 | 0.765 | 1.162 | -0.075 | 0.31 | -0.001 | -0.003 | -0.003 | 1.109 | 0.06 | 0.05 | 0.03 | 0.02 | 0.05 | 0.06 |
| 10 | 3729202 | rs10795063 | -0.289 | 1.19 | -0.127 | 0.272 | -0.043 | -0.006 | -0.006 | 3.609 | 0.37 | 0.2 | 0.3 | 0.36 | 0.45 | 0.54 |
| 10 | 3733386 | rs2129446 | -0.097 | 1.195 | -0.133 | 0.264 | -0.018 | -0.012 | -0.012 | 1.136 | 0.35 | 0.18 | 0.25 | 0.36 | 0.35 | 0.45 |
| 10 | 3733731 | rs10508261 | -1.469 | 0.932 | -0.192 | 0.126 | -0.005 | -0.017 | -0.017 | 0.022 | 0.59 | 0.59 | 0.55 | 0.77 | 0.63 | 0.68 |
| 10 | 3734857 | rs4881194 | 0.644 | 0.951 | -0.275 | 0.138 | -0.029 | 0.033 | 0.033 | 0.42 | 0.27 | 0.14 | 0.19 | 0.16 | 0.27 | 0.19 |
| 10 | 3737388 | rs11252040 | 0.216 | 1.007 | -0.124 | 0.285 | -0.011 | 0.011 | 0.011 | 0.406 | 0.07 | 0.11 | 0.05 | 0.16 | 0.04 | 0.2 |
| 10 | 3738103 | rs7895630 | -0.16 | 0.998 | -0.079 | 0.283 | -0.018 | 0.065 | 0.065 | 0.661 | 0.23 | 0.33 | 0.27 | 0.43 | 0.21 | 0.19 |
| 10 | 3739510 | rs4881197 | -0.995 | 1.026 | -0.082 | 0.278 | 0.009 | 0.033 | 0.033 | 5.05 | 0.32 | 0.44 | 0.36 | 0.6 | 0.35 | 0.48 |
| 10 | 3743084 | rs12218626 | 0.181 | 0.752 | -0.102 | 0.295 | -0.011 | 0.01 | 0.01 | 0.354 | 0.07 | 0.11 | 0.05 | 0.16 | 0.04 | 0.2 |
| 10 | 3746775 | rs2129445 | -0.758 | 0.771 | -0.197 | 0.279 | -0.037 | 0.044 | 0.044 | 4.105 | 0.3 | 0.44 | 0.32 | 0.6 | 0.25 | 0.39 |
| 10 | 3754781 | rs11597829 | -0.551 | 0.957 | -0.174 | 0.173 | 0.013 | 0.02 | 0.02 | 0 | 0.47 | 0.37 | 0.46 | 0.61 | 0.56 | 0.46 |
| 10 | 3761111 | rs17325819 | 0.773 | 0.99 | 0.023 | -0.029 | 0.009 | -0.007 | -0.007 | 0.064 | 0.32 | 0.25 | 0.15 | 0.05 | 0.22 | 0.18 |
| 10 | 3764605 | rs17228197 | 0.682 | 1.076 | -0.222 | 0.021 | -0.085 | 0.034 | 0.034 | 0.043 | 0.26 | 0.11 | 0.21 | 0.14 | 0.36 | 0.28 |
| 10 | 3765346 | rs4488095 | -0.886 | 1.16 | 0.04 | 0.069 | -0.029 | -0.1 | -0.1 | 0.141 | 0.67 | 0.58 | 0.84 | 0.95 | 0.78 | 0.81 |
| 10 | 3766472 | rs10751938 | -0.289 | 1.198 | 0.034 | 0.094 | -0.005 | -0.017 | -0.017 | 0.312 | 0.58 | 0.43 | 0.75 | 0.78 | 0.7 | 0.6 |
| 10 | 3766644 | rs2388861 | -0.442 | 1.417 | 0.014 | 0.075 | -0.001 | -0.005 | -0.005 | 0 | 0.6 | 0.4 | 0.73 | 0.78 | 0.63 | 0.55 |
| 10 | 3766959 | rs11252055 | 0.662 | 1.422 | 0.027 | 0.15 | 0.01 | -0.011 | -0.011 | 0 | 0.32 | 0.18 | 0.53 | 0.65 | 0.34 | 0.32 |
| 10 | 3769134 | rs11252058 | -0.398 | 1.445 | 0.835 | 0.739 | -0.004 | 0.011 | 0.011 | 1.114 | 0.09 | 0.15 | 0.09 | 0.17 | 0.08 | 0.22 |
| 10 | 3769727 | rs7094467 | 1.316 | 1.466 | 0.852 | 0.713 | 0.041 | -0.005 | -0.005 | 0 | 0.83 | 0.63 | 0.82 | 0.81 | 0.78 | 0.67 |
| 10 | 3770014 | rs7094797 | 0.593 | 1.51 | 0.614 | 0.633 | 0.021 | 0.033 | 0.033 | 0 | 0.55 | 0.37 | 0.74 | 0.75 | 0.68 | 0.58 |

| | | | | | | | | | | | | | | | | |
|---|---|---|---|---|---|---|---|---|---|---|---|---|---|---|---|---|
| 10 | 3771084 | rs7069135 | 1.167 | 1.771 | 0.524 | 0.373 | 0.005 | 0.024 | 0.024 | 0 | 0.46 | 0.24 | 0.53 | 0.36 | 0.45 | 0.45 |
| 10 | 3771779 | rs11592787 | -1.111 | 1.583 | 0.523 | 0.07 | -0.003 | -0.005 | -0.005 | 0 | 0.48 | 0.5 | 0.38 | 0.65 | 0.4 | 0.47 |
| 10 | 3772249 | rs11599366 | 0.39 | 1.2 | 0.487 | 0.107 | -0.013 | -0.086 | -0.086 | 0 | 0.12 | 0.01 | 0.2 | 0.57 | 0.13 | 0.27 |
| 10 | 3774779 | rs11252061 | 0.271 | 1.166 | 0.612 | 0.133 | -0.004 | -0.005 | -0.005 | 0 | 0.07 | 0.15 | 0.05 | 0.05 | 0.05 | 0.06 |
| 10 | 3778825 | rs7094917 | 0.59 | 1.176 | 0.155 | 0.02 | 0.003 | 0.023 | 0.023 | 0 | 0.49 | 0.4 | 0.57 | 0.34 | 0.49 | 0.47 |
| 10 | 3782419 | rs10795068 | -0.297 | 1.431 | -0.548 | -0.124 | 0.011 | -0.089 | -0.089 | 0 | 0.71 | 0.66 | 0.82 | 0.97 | 0.7 | 0.83 |
| 10 | 3785290 | rs12254418 | 0.598 | 0.787 | -0.684 | 0.047 | -0.004 | 0.085 | 0.085 | 0 | 0.46 | 0.18 | 0.54 | 0.28 | 0.47 | 0.46 |
| 10 | 3785605 | rs10508263 | -0.604 | 0.69 | -0.737 | -0.343 | -0.016 | -0.079 | -0.079 | 0 | 0.77 | 0.7 | 0.91 | 0.98 | 0.85 | 0.9 |
| 10 | 3786476 | rs10904088 | -0.13 | 0.9 | -1.465 | 0.289 | -0.008 | 0.097 | 0.097 | 0 | 0.46 | 0.17 | 0.52 | 0.28 | 0.48 | 0.47 |
| 10 | 3789223 | rs10795070 | -0.624 | 0.656 | -0.928 | 0.562 | -0.007 | 0.002 | 0.002 | 0.498 | 0.33 | 0.56 | 0.4 | 0.26 | 0.29 | 0.24 |
| 10 | 3791224 | rs10904089 | -0.582 | 1.117 | -1.475 | 0.4 | -0.051 | 0.023 | 0.023 | 0.494 | 0.4 | 0.8 | 0.57 | 0.35 | 0.49 | 0.45 |
| 10 | 3792608 | rs2279415 | -0.59 | 1.157 | -1.541 | 0.406 | -0.059 | 0.031 | 0.031 | 0.574 | 0.41 | 0.81 | 0.58 | 0.35 | 0.5 | 0.46 |
| 10 | 3798495 | rs7896849 | -0.493 | 1.787 | -1.65 | 0.46 | -0.006 | 0.055 | 0.055 | 0.307 | 0.38 | 0.81 | 0.45 | 0.29 | 0.36 | 0.39 |
| 10 | 3806376 | rs7078292 | 0.461 | 1.622 | -1.69 | 0.375 | -0.038 | 0.071 | 0.071 | 2.842 | 0.67 | 0.11 | 0.64 | 0.8 | 0.7 | 0.69 |
| 10 | 3807852 | rs4259743 | 0.042 | 1.539 | -1.697 | 0.392 | 0.001 | -0.001 | -0.001 | 0 | 0.07 | 0.06 | 0.1 | 0.07 | 0.08 | 0.1 |
| 10 | 3809243 | rs11252077 | -0.271 | 1.536 | -0.703 | 0.218 | 0.049 | 0.009 | 0.009 | 0 | 0.17 | 0.31 | 0.29 | 0.2 | 0.22 | 0.24 |
| 10 | 3812240 | rs7076536 | -1.865 | 1.497 | -0.751 | 0.268 | -0.026 | 0.022 | 0.022 | 1.014 | 0.39 | 0.79 | 0.49 | 0.36 | 0.44 | 0.51 |
| 10 | 3815292 | rs2171302 | -0.169 | 1.682 | -0.452 | 0.331 | 0.026 | -0.017 | -0.017 | 2.75 | 0.09 | 0.17 | 0.15 | 0.05 | 0.17 | 0.09 |
| 10 | 3818058 | rs2279414 | -1.322 | 1.709 | -0.573 | 0.465 | 0.006 | 0.002 | 0.002 | 0 | 0.23 | 0.24 | 0.4 | 0.29 | 0.31 | 0.34 |
| 10 | 3818497 | rs2129439 | -0.378 | 1.79 | -1.12 | 0.478 | -0.016 | -0.004 | -0.004 | 0 | 0.15 | 0.11 | 0.28 | 0.2 | 0.23 | 0.22 |
| 10 | 3820787 | rs1043003 | 0.849 | 1.201 | -2.078 | 0.178 | -0.12 | -0.162 | -0.162 | 0.055 | 0.35 | 0.03 | 0.28 | 0.49 | 0.35 | 0.3 |
| 10 | 3824433 | rs3750861 | -0.298 | 1.776 | -1.151 | 0.298 | 0.008 | 0.006 | 0.006 | 0 | 0.08 | 0.02 | 0.1 | 0.04 | 0.08 | 0.12 |
| 10 | 3825343 | rs10795076 | -0.502 | 2.286 | -0.886 | 0.405 | -0.008 | -0.005 | -0.005 | 0 | 0.13 | 0.08 | 0.25 | 0.18 | 0.16 | 0.23 |
| 10 | 3833232 | rs17249262 | -0.93 | 2.554 | -0.794 | 0.498 | 0.002 | -0.001 | -0.001 | 0 | 0.17 | 0.15 | 0.35 | 0.19 | 0.31 | 0.24 |
| 10 | 3838993 | rs12773202 | -0.267 | 2.349 | -1.422 | 0.71 | -0.038 | -0.005 | -0.005 | 0 | 0.25 | 0.15 | 0.21 | 0.29 | 0.31 | 0.17 |
| 10 | 3839024 | rs10508265 | 0.784 | 0.937 | -1.155 | 0.388 | -0.034 | -0.016 | -0.016 | 0 | 0.24 | 0.28 | 0.22 | 0.16 | 0.12 | 0.28 |
| 10 | 3839642 | rs7919841 | 0.201 | 0.734 | -1.186 | 0.43 | 0.031 | 0.013 | 0.013 | 0.11 | 0.55 | 0.18 | 0.38 | 0.37 | 0.58 | 0.43 |
| 10 | 3840014 | rs10508266 | 1.365 | 0.553 | -1.439 | 0.311 | -0.056 | -0.113 | -0.113 | 0 | 0.28 | 0.01 | 0.21 | 0.31 | 0.25 | 0.29 |
| 10 | 3842321 | rs11252108 | -0.392 | 0.61 | -0.667 | 0.409 | -0.013 | -0.004 | -0.004 | 0.006 | 0.63 | 0.47 | 0.57 | 0.65 | 0.7 | 0.52 |
| 10 | 3844038 | rs7898258 | 0.349 | 0.828 | -0.518 | 0.367 | 0.028 | 0.004 | 0.004 | 0 | 0.79 | 0.67 | 0.65 | 0.76 | 0.66 | 0.75 |
| 10 | 3846327 | rs7906552 | 0.692 | 0.81 | -0.745 | 0.248 | 0.032 | 0 | 0 | 0 | 0.52 | 0.34 | 0.42 | 0.47 | 0.35 | 0.57 |
| 10 | 3848340 | rs11252110 | 0.731 | 0.757 | -0.775 | 0.287 | -0.017 | -0.029 | -0.029 | 0 | 0.33 | 0.22 | 0.32 | 0.36 | 0.36 | 0.36 |
| 10 | 3848431 | rs10508268 | 1.236 | 0.906 | -0.701 | 0.209 | -0.063 | 0.089 | 0.089 | 3.406 | 0.22 | 0.1 | 0.15 | 0.06 | 0.29 | 0.09 |
| 10 | 3853031 | rs4881220 | 0.192 | 0.847 | -0.545 | 0.47 | 0.032 | -0.026 | -0.026 | 0 | 0.34 | 0.19 | 0.36 | 0.42 | 0.25 | 0.47 |
| 10 | 3856293 | rs12245875 | 1.149 | 1.333 | -0.766 | 0.457 | 0 | -0.006 | -0.006 | 0 | 0.21 | 0.21 | 0.16 | 0.11 | 0.28 | 0.08 |
| 10 | 3857746 | rs7077239 | 0.454 | 1.354 | -0.842 | 0.462 | 0.014 | 0.002 | 0.002 | 0 | 0.06 | 0.09 | 0.13 | 0.09 | 0.14 | 0.05 |
| 10 | 3857816 | rs7076208 | 0.534 | 0.454 | -0.947 | 0.597 | -0.04 | -0.064 | -0.064 | 1.302 | 0.11 | 0.02 | 0.14 | 0.19 | 0.13 | 0.18 |
| 10 | 3858778 | rs11252116 | 0.424 | 0.778 | -0.467 | 0.255 | 0.027 | -0.021 | -0.021 | 4.956 | 0.27 | 0.33 | 0.3 | 0.19 | 0.42 | 0.13 |
| 10 | 3860167 | rs11252118 | 0.017 | 0.358 | -1.298 | 0.678 | -0.001 | -0.035 | -0.035 | 1.003 | 0.31 | 0.11 | 0.21 | 0.27 | 0.26 | 0.32 |
| 10 | 3861214 | rs7082849 | -0.064 | -0.128 | -0.405 | 0.64 | -0.045 | -0.011 | -0.011 | 0 | 0.76 | 0.57 | 0.72 | 0.81 | 0.85 | 0.78 |
| 10 | 3861344 | rs11252119 | -0.471 | 0.223 | -0.708 | 0.263 | -0.001 | -0.006 | -0.006 | 0.041 | 0.34 | 0.33 | 0.26 | 0.22 | 0.31 | 0.33 |
| 10 | 3861861 | rs7904671 | -0.863 | 0.796 | -0.812 | 0.436 | 0.024 | 0.001 | 0.001 | 0 | 0.21 | 0.24 | 0.17 | 0.2 | 0.34 | 0.33 |
| 10 | 3862320 | rs17842279 | 0.195 | -0.066 | -1.47 | 0.575 | 0.026 | -0.071 | -0.071 | 0 | 0.15 | 0.01 | 0.24 | 0.24 | 0.11 | 0.17 |
| 10 | 3862514 | rs11252121 | -0.689 | 0.558 | -0.767 | 0.485 | 0.064 | 0.029 | 0.029 | 0.288 | 0.07 | 0.17 | 0.08 | 0.07 | 0.13 | 0.09 |
| 10 | 3862542 | rs6601764 | 0.03 | 0.079 | -1.496 | 0.043 | -0.007 | 0.151 | 0.151 | 0 | 0.48 | 0.23 | 0.45 | 0.22 | 0.53 | 0.31 |
| 10 | 3862940 | rs12266938 | 0.03 | -0.475 | -1.265 | -0.175 | -0.078 | 0.019 | 0.019 | 0 | 0.3 | 0.44 | 0.33 | 0.39 | 0.15 | 0.35 |
| 10 | 3864942 | rs11252124 | 0.083 | 0.764 | -2.143 | -0.992 | 0.024 | 0.034 | 0.034 | 0.153 | 0.39 | 0.14 | 0.43 | 0.67 | 0.36 | 0.56 |
| 10 | 3865519 | rs10904101 | 0.568 | 0.589 | -2.188 | -1.196 | -0.083 | 0.133 | 0.133 | 0.095 | 0.36 | 0.13 | 0.32 | 0.11 | 0.42 | 0.16 |
| 10 | 3866284 | rs4880634 | -0.051 | 0.478 | -1.819 | -0.551 | 0.024 | 0.022 | 0.022 | 0 | 0.36 | 0.14 | 0.41 | 0.68 | 0.31 | 0.58 |
| 10 | 3868056 | rs4880635 | -0.087 | 0.72 | -1.273 | -0.798 | 0.006 | -0.022 | -0.022 | 0 | 0.38 | 0.4 | 0.37 | 0.33 | 0.44 | 0.44 |
| 10 | 3868226 | rs3829203 | 0.306 | 0.568 | -1.469 | -0.858 | 0.003 | 0.009 | 0.009 | 0 | 0.66 | 0.7 | 0.71 | 0.75 | 0.7 | 0.78 |
| 10 | 3868392 | rs7914924 | -0.274 | 0.742 | -1.762 | -0.714 | -0.022 | 0.159 | 0.159 | 0 | 0.4 | 0.13 | 0.33 | 0.1 | 0.42 | 0.3 |
| 10 | 3872876 | rs10904103 | -0.729 | 0.395 | -0.837 | -0.359 | -0.017 | -0.04 | -0.04 | 2.657 | 0.3 | 0.47 | 0.33 | 0.26 | 0.3 | 0.18 |
| 10 | 3876063 | rs12245903 | -0.357 | 0.274 | -1.186 | -0.679 | 0.028 | 0.1 | 0.1 | 0 | 0.52 | 0.68 | 0.45 | 0.27 | 0.57 | 0.35 |
| 10 | 3877968 | rs11597773 | -0.005 | 0.195 | -0.764 | -0.144 | 0.014 | 0.024 | 0.024 | 0 | 0.31 | 0.55 | 0.41 | 0.4 | 0.34 | 0.33 |
| 10 | 3887162 | rs12411753 | -0.098 | -0.349 | -1.019 | 0.155 | 0.018 | -0.01 | -0.01 | 0 | 0.08 | 0.04 | 0.22 | 0.24 | 0.04 | 0.16 |
| 10 | 3887605 | rs7901658 | -1.184 | -0.441 | -1.134 | -0.245 | 0.029 | 0.039 | 0.039 | 0.72 | 0.41 | 0.73 | 0.44 | 0.3 | 0.38 | 0.39 |
| 10 | 3887689 | rs10904104 | 0.985 | -0.413 | -1.247 | -0.251 | -0.087 | -0.05 | -0.05 | 0 | 0.18 | 0.01 | 0.23 | 0.22 | 0.23 | 0.12 |
| 10 | 3888845 | rs7906433 | 0.309 | -0.444 | -1.165 | -0.309 | -0.001 | -0.011 | -0.011 | 0.808 | 0.22 | 0.17 | 0.14 | 0.02 | 0.24 | 0.09 |
| 10 | 3892083 | rs7907906 | 0.571 | -0.416 | -0.356 | -0.192 | -0.012 | -0.003 | -0.003 | 0 | 0.2 | 0.13 | 0.24 | 0.22 | 0.23 | 0.13 |
| 10 | 3892180 | rs7923118 | -0.418 | -0.553 | -0.041 | -0.335 | 0.036 | -0.02 | -0.02 | 1.577 | 0.28 | 0.36 | 0.32 | 0.22 | 0.38 | 0.16 |
| 10 | 3893980 | rs7897383 | 0.443 | -0.387 | 0.496 | -0.304 | 0.079 | -0.047 | -0.047 | 0 | 0.09 | 0.24 | 0.08 | 0 | 0.15 | 0.03 |
| 10 | 3896229 | rs7902434 | -1.85 | -0.544 | 0.454 | -0.179 | -0.002 | -0.017 | -0.017 | 0.367 | 0.6 | 0.71 | 0.68 | 0.54 | 0.61 | 0.51 |
| 10 | 3899157 | rs17135906 | -0.11 | -0.557 | 0.553 | 0.113 | -0.032 | 0.031 | 0.031 | 0 | 0.08 | 0.17 | 0.13 | 0.12 | 0.02 | 0.07 |
| 10 | 3900125 | rs7100255 | -0.391 | -0.423 | 0.236 | -0.024 | 0.001 | 0.012 | 0.012 | 0 | 0.25 | 0.29 | 0.36 | 0.38 | 0.25 | 0.22 |
| 10 | 3906911 | rs7902252 | 1.009 | -0.998 | -0.262 | -0.141 | -0.035 | -0.016 | -0.016 | 0 | 0.4 | 0.24 | 0.4 | 0.59 | 0.45 | 0.63 |
| 10 | 3910198 | rs4881226 | 0.664 | -1.122 | -0.208 | 0.045 | -0.024 | -0.022 | -0.022 | 0.498 | 0.43 | 0.29 | 0.44 | 0.54 | 0.46 | 0.63 |
| 10 | 3913700 | rs6601770 | -0.007 | -1.424 | 0.039 | -0.303 | -0.002 | -0.027 | -0.027 | 0 | 0.78 | 0.56 | 0.79 | 0.86 | 0.8 | 0.88 |
| 10 | 3915501 | rs6601771 | 0.302 | -1.434 | 0.218 | -0.318 | 0.004 | 0.001 | 0.001 | 0 | 0.62 | 0.67 | 0.6 | 0.65 | 0.66 | 0.75 |

| | | | | | | | | | | | | | | | | |
|---|---|---|---|---|---|---|---|---|---|---|---|---|---|---|---|---|
| 10 | 3917237 | rs11252148 | 1.854 | -1.455 | 0.081 | -0.329 | -0.039 | -0.038 | -0.038 | 0.864 | 0.4 | 0.24 | 0.4 | 0.51 | 0.45 | 0.64 |
| 10 | 3918640 | rs943029 | -0.044 | -1.573 | 0.158 | -0.383 | 0.007 | 0.002 | 0.002 | 0 | 0.29 | 0.21 | 0.26 | 0.27 | 0.3 | 0.23 |
| 10 | 3921082 | rs943034 | -0.39 | -1.583 | 0.108 | -0.388 | 0.023 | 0.004 | 0.004 | 0 | 0.34 | 0.2 | 0.3 | 0.31 | 0.3 | 0.23 |
| 10 | 3925238 | rs10904120 | 1.805 | -1.535 | -0.022 | -0.359 | -0.04 | -0.036 | -0.036 | 0.554 | 0.4 | 0.24 | 0.39 | 0.5 | 0.45 | 0.65 |
| 10 | 3925728 | rs3932533 | 0.824 | -1.359 | 0.236 | -0.293 | -0.003 | 0 | 0 | 1.389 | 0.68 | 0.37 | 0.63 | 0.7 | 0.71 | 0.8 |
| 10 | 3928995 | rs12240700 | -0.096 | -0.895 | 0.47 | -0.372 | 0.001 | -0.079 | -0.079 | 0 | 0.2 | 0.31 | 0.12 | 0.02 | 0.21 | 0.09 |
| 10 | 3929526 | rs10795086 | -0.895 | -0.997 | 0.017 | -0.157 | -0.017 | 0.01 | 0.01 | 0 | 0.69 | 0.57 | 0.64 | 0.33 | 0.76 | 0.45 |
| 10 | 3930424 | rs10795088 | -1.011 | -1.066 | 0.019 | -0.139 | -0.013 | 0.011 | 0.011 | 0 | 0.7 | 0.63 | 0.66 | 0.41 | 0.8 | 0.5 |
| 10 | 3932870 | rs12412574 | -0.118 | -1.052 | -0.117 | -0.327 | -0.004 | 0.028 | 0.028 | 0 | 0.37 | 0.24 | 0.39 | 0.24 | 0.44 | 0.23 |
| 10 | 3934581 | rs10795092 | -1.223 | -1.185 | -0.24 | -0.249 | 0.018 | -0.025 | -0.025 | 0 | 0.72 | 0.72 | 0.76 | 0.47 | 0.83 | 0.53 |
| 10 | 3935402 | rs11252168 | 0.423 | -1.215 | 0.166 | -0.3 | 0.014 | 0.014 | 0.014 | 0 | 0.28 | 0.17 | 0.24 | 0.2 | 0.27 | 0.16 |
| 10 | 3935589 | rs11252169 | 1.824 | -1.019 | 0.799 | -0.063 | -0.065 | 0.001 | 0.001 | 0.492 | 0.19 | 0.09 | 0.36 | 0.17 | 0.3 | 0.21 |
| 10 | 3935964 | rs943026 | -0.823 | -1.45 | 0.968 | 0.173 | -0.001 | -0.002 | -0.002 | 0 | 0.72 | 0.69 | 0.76 | 0.47 | 0.83 | 0.54 |
| 10 | 3939728 | rs10795093 | -0.597 | -0.673 | 1.535 | 0.596 | 0 | 0.246 | 0.246 | 0 | 0.13 | 0.46 | 0.33 | 0.34 | 0.11 | 0.14 |
| 10 | 3945400 | rs4881232 | -0.41 | -0.513 | 0.896 | 0.357 | 0.003 | -0.008 | -0.008 | 0 | 0.45 | 0.47 | 0.57 | 0.39 | 0.44 | 0.38 |
| 10 | 3946610 | rs10732922 | -0.953 | -0.698 | 0.339 | 0.327 | 0.051 | -0.036 | -0.036 | 0 | 0.55 | 0.76 | 0.74 | 0.5 | 0.61 | 0.43 |
| 10 | 3948464 | rs7899935 | -1.031 | -0.692 | 0.318 | 0.378 | 0.056 | 0.018 | 0.018 | 0 | 0.55 | 0.77 | 0.75 | 0.58 | 0.61 | 0.47 |
| 10 | 3948928 | rs10904134 | -0.376 | -0.828 | 0.648 | 0.163 | -0.012 | -0.003 | -0.003 | 0 | 0.45 | 0.63 | 0.48 | 0.38 | 0.46 | 0.42 |
| 10 | 3949016 | rs4881234 | -1.194 | -0.873 | 0.463 | 0.18 | 0.059 | -0.029 | -0.029 | 0 | 0.57 | 0.79 | 0.76 | 0.51 | 0.64 | 0.47 |
| 10 | 3949085 | rs11252180 | 1.504 | -0.443 | 0.481 | -0.144 | -0.111 | -0.022 | -0.022 | 0 | 0.19 | 0.05 | 0.28 | 0.14 | 0.3 | 0.19 |
| 10 | 3949166 | rs9651513 | -0.225 | -1.08 | 1.219 | -0.057 | 0.003 | -0.001 | -0.001 | 0 | 0.42 | 0.62 | 0.45 | 0.37 | 0.56 | 0.35 |
| 10 | 3950093 | rs4457649 | 0.889 | -1.381 | 2.211 | 0.02 | 0.027 | 0.024 | 0.024 | 2.264 | 0.84 | 0.37 | 0.87 | 0.85 | 0.81 | 0.91 |
| 10 | 3950427 | rs10795100 | 1.485 | -1.723 | 1.775 | -0.599 | 0.017 | 0.029 | 0.029 | 0.093 | 0.71 | 0.2 | 0.6 | 0.79 | 0.76 | 0.83 |
| 10 | 3950446 | rs4364955 | 0.946 | -0.954 | 2.683 | -0.03 | 0.026 | -0.012 | -0.012 | 0 | 0.87 | 0.73 | 0.91 | 0.92 | 0.83 | 0.94 |
| 10 | 3954953 | rs12217962 | -0.172 | -0.87 | 1.546 | -0.822 | 0.037 | 0.048 | 0.048 | 0 | 0.16 | 0.04 | 0.09 | 0.07 | 0.14 | 0.12 |
| 10 | 3956841 | rs17136022 | -0.878 | -0.033 | 2.316 | -0.419 | 0.014 | -0.051 | -0.051 | 0 | 0.13 | 0.21 | 0.09 | 0 | 0.16 | 0.02 |
| 10 | 3962232 | rs4881235 | 1.328 | -0.095 | 2.609 | -0.44 | 0.02 | -0.102 | -0.102 | 0.013 | 0.83 | 0.42 | 0.8 | 0.98 | 0.84 | 0.93 |
| 10 | 3982319 | rs4880646 | 0.862 | 0.063 | 1.127 | -0.304 | 0.026 | -0.078 | -0.078 | 2.209 | 0.85 | 0.68 | 0.81 | 0.98 | 0.84 | 0.95 |
| 10 | 3986323 | rs4881251 | 0.678 | -0.124 | 0.946 | -0.228 | -0.001 | -0.022 | -0.022 | 0.33 | 0.82 | 0.82 | 0.77 | 0.91 | 0.81 | 0.89 |
| 10 | 3986376 | rs7070280 | -1.459 | -0.27 | 1.571 | -0.093 | 0.013 | 0.047 | 0.047 | 0 | 0.07 | 0.29 | 0.03 | 0.11 | 0.05 | 0.04 |
| 10 | 3988146 | rs10904140 | 1.475 | -0.324 | 1.539 | -0.113 | -0.007 | -0.017 | -0.017 | 1.101 | 0.76 | 0.45 | 0.74 | 0.8 | 0.77 | 0.85 |
| 10 | 3989427 | rs4630187 | 0.501 | -0.358 | 0.756 | -0.204 | 0.006 | 0.032 | 0.032 | 0 | 0.83 | 0.89 | 0.77 | 0.95 | 0.82 | 0.9 |
| 10 | 3991102 | rs6601785 | -1.554 | -0.78 | 0.431 | -0.236 | 0.017 | 0.033 | 0.033 | 0.128 | 0.37 | 0.72 | 0.29 | 0.29 | 0.33 | 0.31 |
| 10 | 3992163 | rs4880650 | 0.084 | -1.387 | -0.136 | -0.7 | 0.053 | 0.068 | 0.068 | 0 | 0.52 | 0.32 | 0.7 | 0.69 | 0.57 | 0.6 |
| 10 | 3997672 | rs10795105 | 1.301 | -1.315 | -0.498 | -0.631 | -0.11 | -0.068 | -0.068 | 0 | 0.32 | 0.1 | 0.52 | 0.6 | 0.38 | 0.43 |
| 10 | 3999496 | rs7898425 | -0.031 | -2.157 | 0.089 | -1.064 | -0.002 | -0.062 | -0.062 | 0 | 0.27 | 0.41 | 0.2 | 0.12 | 0.32 | 0.3 |
| 10 | 4000140 | rs7098821 | 0.956 | -2.368 | 0.256 | -1.153 | 0.001 | 0.14 | 0.14 | 0 | 0.51 | 0.19 | 0.59 | 0.72 | 0.49 | 0.56 |
| 10 | 4000708 | rs4881260 | -0.013 | -2.376 | 0.348 | -1.171 | -0.004 | -0.023 | -0.023 | 0 | 0.27 | 0.48 | 0.21 | 0.23 | 0.32 | 0.31 |
| 10 | 4001897 | rs2031573 | 0.558 | -2.383 | 0.188 | -1.175 | -0.041 | 0.08 | 0.08 | 0 | 0.18 | 0.01 | 0.16 | 0.04 | 0.17 | 0.13 |
| 10 | 4002033 | rs7895964 | 0.575 | -2.601 | 0.111 | -1.105 | -0.007 | -0.014 | -0.014 | 0.199 | 0.67 | 0.62 | 0.73 | 0.78 | 0.74 | 0.77 |
| 10 | 4003340 | rs12217980 | 0.673 | -2.461 | 0.047 | -0.88 | 0.014 | -0.041 | -0.041 | 0 | 0.1 | 0.03 | 0.07 | 0.18 | 0.07 | 0.1 |
| 10 | 4006579 | rs7079191 | -0.251 | -2.604 | 0.064 | -1.008 | -0.04 | 0.065 | 0.065 | 0 | 0.38 | 0.14 | 0.32 | 0.26 | 0.47 | 0.41 |
| 10 | 4020133 | rs11252232 | -0.191 | -2.671 | -0.102 | -1.041 | -0.076 | 0.143 | 0.143 | 0.064 | 0.38 | 0.08 | 0.32 | 0.15 | 0.47 | 0.41 |
| 10 | 4021078 | rs10751951 | -1.115 | -2.717 | 0.261 | -1.031 | 0 | -0.001 | -0.001 | 0 | 0.53 | 0.45 | 0.43 | 0.43 | 0.55 | 0.47 |
| 10 | 4023054 | rs11252236 | 0.766 | -2.738 | 0.232 | -1.041 | -0.006 | -0.048 | -0.048 | 3.273 | 0.1 | 0.01 | 0.07 | 0.17 | 0.07 | 0.06 |
| 10 | 4023855 | rs10795115 | -0.346 | -2.726 | 0.351 | -1.049 | 0.001 | 0.026 | 0.026 | 0 | 0.41 | 0.32 | 0.35 | 0.15 | 0.48 | 0.41 |
| 10 | 4026125 | rs11252237 | 0.34 | -2.565 | 0.436 | -0.984 | -0.008 | 0.031 | 0.031 | 0 | 0.12 | 0.14 | 0.08 | 0.29 | 0.08 | 0.07 |
| 10 | 4029755 | rs4242768 | -0.325 | -2.533 | 0.599 | -1.041 | 0.028 | -0.007 | -0.007 | 0.173 | 0.94 | 0.73 | 0.96 | 1 | 0.97 | 0.97 |
| 10 | 4030590 | rs10795118 | -0.447 | -2.773 | 0.225 | -1.052 | 0.003 | 0.031 | 0.031 | 0 | 0.43 | 0.33 | 0.34 | 0.16 | 0.48 | 0.43 |
| 10 | 4037207 | rs11252249 | 0.046 | -2.784 | 0.226 | -0.804 | -0.027 | 0.039 | 0.039 | 0 | 0.17 | 0.05 | 0.14 | 0.09 | 0.22 | 0.23 |
| 10 | 4038347 | rs9664331 | -1.006 | -2.552 | 0.428 | -0.631 | -0.031 | 0.016 | 0.016 | 0 | 0.18 | 0.41 | 0.09 | 0.2 | 0.11 | 0.09 |
| 10 | 4044541 | rs1540992 | 1.25 | -2.553 | -0.674 | -0.641 | 0.091 | 0.016 | 0.016 | 0 | 0.53 | 0.09 | 0.72 | 0.6 | 0.64 | 0.63 |
| 10 | 4050003 | rs2031577 | 1.119 | -2.528 | -0.629 | -0.632 | 0.089 | 0.053 | 0.053 | 0 | 0.54 | 0.24 | 0.73 | 0.62 | 0.64 | 0.64 |
| 10 | 4050058 | rs2031575 | -0.06 | -2.483 | -0.93 | -0.63 | -0.011 | 0.032 | 0.032 | 0 | 0.88 | 0.85 | 0.95 | 0.71 | 0.92 | 0.93 |
| 10 | 4061039 | rs10904169 | -0.874 | -2.506 | -1.048 | -0.846 | 0.068 | 0.115 | 0.115 | 0.023 | 0.36 | 0.81 | 0.22 | 0.1 | 0.29 | 0.28 |
| 10 | 4062327 | rs7919822 | -0.83 | -2.38 | -1.006 | -0.816 | 0.031 | 0.044 | 0.044 | 0 | 0.34 | 0.72 | 0.22 | 0.08 | 0.28 | 0.28 |
| 10 | 4065704 | rs11252277 | -0.25 | -1.76 | -0.902 | -0.214 | -0.001 | -0.042 | -0.042 | 0 | 0.11 | 0.03 | 0.04 | 0.21 | 0.07 | 0.07 |
| 10 | 4069271 | rs2765654 | 0.161 | -1.932 | -0.829 | -0.111 | 0.003 | -0.006 | -0.006 | 0.482 | 0.1 | 0.08 | 0.02 | 0.18 | 0.04 | 0.05 |
| 10 | 4070444 | rs2291788 | -0.065 | -1.694 | -0.496 | -0.125 | 0.009 | -0.011 | -0.011 | 0 | 0.15 | 0.2 | 0.14 | 0.08 | 0.23 | 0.12 |
| 10 | 4075656 | rs1886946 | -1.602 | -2.405 | -0.39 | -0.424 | 0.076 | 0.052 | 0.052 | 2.102 | 0.51 | 0.78 | 0.28 | 0.35 | 0.39 | 0.3 |
| 10 | 4076159 | rs2765648 | -0.671 | -2.408 | -0.641 | -0.442 | -0.033 | -0.01 | -0.01 | 0 | 0.36 | 0.57 | 0.14 | 0.26 | 0.16 | 0.15 |
| 10 | 4076637 | rs11252287 | 0.362 | -2.276 | -1.582 | -0.392 | -0.037 | 0.03 | 0.03 | 0 | 0.14 | 0.03 | 0.12 | 0.05 | 0.23 | 0.1 |
| 10 | 4079520 | rs7093608 | 0.891 | -1.962 | -1.407 | -0.185 | 0.02 | -0.012 | -0.012 | 0 | 0.48 | 0.05 | 0.71 | 0.65 | 0.61 | 0.7 |
| 10 | 4091774 | rs10904174 | 0.551 | -2.026 | -0.813 | -0.095 | 0.092 | 0.056 | 0.056 | 0 | 0.47 | 0.22 | 0.7 | 0.71 | 0.67 | 0.56 |
| 10 | 4092494 | rs1325998 | 0.506 | -2.053 | -0.905 | -0.146 | 0.121 | 0.171 | 0.171 | 0 | 0.51 | 0.23 | 0.77 | 0.81 | 0.69 | 0.61 |
| 10 | 4094234 | rs2645211 | -0.079 | -2.818 | -0.817 | -0.456 | -0.037 | -0.007 | -0.007 | 0 | 0.74 | 0.47 | 0.79 | 0.79 | 0.83 | 0.75 |
| 10 | 4095177 | rs4880661 | 0.76 | -3.123 | -0.967 | -0.593 | 0.04 | 0.049 | 0.049 | 0 | 0.66 | 0.29 | 0.76 | 0.89 | 0.78 | 0.7 |
| 10 | 4097118 | rs4147111 | 0.225 | -3.519 | -1.405 | -0.889 | 0.015 | -0.005 | -0.005 | 0 | 0.16 | 0.01 | 0.06 | 0.08 | 0.07 | 0.11 |
| 10 | 4098453 | rs2645210 | -0.259 | -3.509 | -1.194 | -0.904 | -0.038 | -0.057 | -0.057 | 0 | 0.75 | 0.48 | 0.8 | 0.89 | 0.82 | 0.76 |

| | | | | | | | | | | | | | | | | |
|---|---|---|---|---|---|---|---|---|---|---|---|---|---|---|---|---|
| 10 | 4101955 | rs2935284 | -1.189 | -3.518 | -0.179 | -0.912 | -0.007 | -0.017 | -0.017 | 0 | 0.82 | 0.8 | 0.85 | 0.91 | 0.84 | 0.78 |
| 10 | 4103418 | rs7078122 | 1.634 | -3.48 | -0.235 | -0.495 | 0.073 | -0.013 | -0.013 | 0 | 0.44 | 0.21 | 0.68 | 0.62 | 0.67 | 0.58 |
| 10 | 4107596 | rs4397729 | 0.361 | -2.684 | 0.293 | -0.81 | -0.021 | -0.041 | -0.041 | 0.542 | 0.07 | 0.14 | 0.03 | 0 | 0.04 | 0.05 |
| 10 | 4115502 | rs11252318 | 1.666 | -2.83 | -0.271 | -0.702 | -0.048 | -0.049 | -0.049 | 0 | 0.29 | 0.21 | 0.56 | 0.55 | 0.46 | 0.47 |
| 10 | 4115602 | rs10508274 | -1.796 | -2.875 | -0.138 | -0.74 | 0.007 | 0.002 | 0.002 | 0 | 0.82 | 0.91 | 0.83 | 0.83 | 0.81 | 0.87 |
| 10 | 4121705 | rs7341996 | 0.101 | -3.041 | 0.068 | -0.824 | -0.001 | -0.001 | -0.001 | 0 | 0.37 | 0.36 | 0.27 | 0.3 | 0.35 | 0.31 |
| 10 | 4128024 | rs2069178 | 0.76 | -3.162 | 0.172 | -0.873 | 0.024 | 0.024 | 0.024 | 0 | 0.45 | 0.37 | 0.65 | 0.62 | 0.54 | 0.56 |
| 10 | 4133306 | rs11252333 | 1.195 | -1.49 | 0.731 | -1.109 | -0.014 | -0.072 | -0.072 | 0 | 0.66 | 0.49 | 0.82 | 0.91 | 0.76 | 0.82 |
| 10 | 4134507 | rs4881281 | 0.198 | -1.504 | 0.835 | -1.144 | 0 | -0.013 | -0.013 | 0 | 0.82 | 0.81 | 0.88 | 0.93 | 0.83 | 0.86 |
| 10 | 4134653 | rs17132053 | 0.709 | -1.522 | 0.796 | -1.152 | 0.014 | 0.014 | 0.014 | 0 | 0.07 | 0.03 | 0.02 | 0 | 0.04 | 0.05 |
| 10 | 4135785 | rs4881282 | 1.207 | -1.51 | 0.796 | -1.123 | -0.01 | -0.04 | -0.04 | 0 | 0.67 | 0.59 | 0.8 | 0.86 | 0.76 | 0.78 |
| 10 | 4138683 | rs10904182 | 1.606 | -1.574 | 0.582 | -1.206 | 0.013 | 0.147 | 0.147 | 0 | 0.47 | 0.25 | 0.7 | 0.82 | 0.59 | 0.6 |
| 10 | 4138794 | rs7087387 | 1.064 | -1.508 | 0.568 | -1.157 | -0.001 | -0.018 | -0.018 | 0.665 | 0.18 | 0.14 | 0.08 | 0.04 | 0.16 | 0.17 |
| 10 | 4140086 | rs6601810 | -0.339 | -1.564 | 0.535 | -1.191 | -0.032 | -0.034 | -0.034 | 0 | 0.18 | 0.3 | 0.09 | 0.07 | 0.08 | 0.08 |
| 10 | 4140580 | rs11252339 | 0.36 | -1.459 | 0.369 | -1.164 | -0.019 | 0.014 | 0.014 | 0 | 0.08 | 0.03 | 0.06 | 0.02 | 0.11 | 0.06 |
| 10 | 4146692 | rs7079146 | -0.829 | -1.56 | 0.417 | -1.187 | -0.04 | -0.044 | -0.044 | 0 | 0.27 | 0.4 | 0.14 | 0.12 | 0.14 | 0.16 |
| 10 | 4150876 | rs10795128 | -1.283 | -1.486 | 0.195 | -1.151 | -0.014 | -0.038 | -0.038 | 0 | 0.36 | 0.44 | 0.2 | 0.14 | 0.24 | 0.23 |
| 10 | 4151051 | rs4881291 | 1.397 | -1.339 | -0.041 | -1.102 | 0.006 | 0.134 | 0.134 | 0 | 0.46 | 0.22 | 0.7 | 0.8 | 0.59 | 0.58 |
| 10 | 4157839 | rs10795132 | 1.391 | -1.329 | -0.037 | -1.098 | 0.007 | 0.134 | 0.134 | 0 | 0.46 | 0.22 | 0.7 | 0.8 | 0.59 | 0.6 |
| 10 | 4163987 | rs10904194 | -1.064 | -1.237 | 0.124 | -0.77 | -0.022 | -0.025 | -0.025 | 0 | 0.27 | 0.37 | 0.13 | 0.11 | 0.13 | 0.15 |
| 10 | 4164331 | rs7101224 | 0.259 | -1.288 | 0.12 | -0.798 | -0.039 | -0.082 | -0.082 | 0 | 0.13 | 0.34 | 0.04 | 0 | 0.05 | 0.03 |
| 10 | 4164366 | rs1885477 | -0.078 | -0.606 | 0.136 | -0.972 | 0.05 | -0.013 | -0.013 | 6.439 | 0.88 | 0.83 | 0.87 | 0.92 | 0.81 | 0.88 |
| 10 | 4164550 | rs10904195 | 1.725 | -0.564 | 0.114 | -0.972 | 0.002 | -0.034 | -0.034 | 0 | 0.61 | 0.46 | 0.76 | 0.83 | 0.67 | 0.77 |
| 10 | 4168660 | rs11252364 | -0.051 | -0.744 | 0.018 | -0.896 | 0.009 | 0.009 | 0.009 | 0 | 0.13 | 0.03 | 0.07 | 0.07 | 0.08 | 0.08 |
| 10 | 4169040 | rs10904196 | -0.28 | -0.381 | 0.316 | -0.721 | -0.04 | -0.087 | -0.087 | 0.07 | 0.87 | 0.62 | 0.95 | 1 | 0.95 | 0.97 |
| 10 | 4171967 | rs10904197 | -0.255 | -0.37 | 0.258 | -0.713 | -0.04 | -0.052 | -0.052 | 0.049 | 0.87 | 0.57 | 0.93 | 0.95 | 0.95 | 0.93 |
| 10 | 4178976 | rs12247651 | -0.095 | -0.274 | -0.008 | -0.605 | -0.007 | 0.007 | 0.007 | 0 | 0.11 | 0.2 | 0.05 | 0.08 | 0.05 | 0.06 |
| 10 | 4179366 | rs12244138 | -0.321 | -0.309 | 0.008 | -0.621 | -0.011 | 0.005 | 0.005 | 0.074 | 0.11 | 0.23 | 0.06 | 0.08 | 0.05 | 0.06 |
| 10 | 4184635 | rs4881298 | -1.182 | -0.452 | 0.186 | -0.5 | -0.053 | -0.012 | -0.012 | 0.012 | 0.23 | 0.48 | 0.1 | 0.15 | 0.11 | 0.11 |
| 10 | 4186228 | rs11252374 | 0.895 | -0.434 | 0.066 | -0.497 | 0.007 | 0.001 | 0.001 | 0.37 | 0.1 | 0.07 | 0.02 | 0.07 | 0.05 | 0.05 |
| 10 | 4187316 | rs4881299 | -1.79 | 0.129 | -0.308 | -0.473 | -0.001 | 0.017 | 0.017 | 0 | 0.71 | 0.65 | 0.49 | 0.5 | 0.72 | 0.56 |
| 10 | 4190058 | rs6601817 | -1.691 | 0.345 | -0.289 | -0.556 | -0.001 | -0.001 | -0.001 | 0 | 0.35 | 0.59 | 0.22 | 0.21 | 0.31 | 0.24 |
| 10 | 4203580 | rs7913465 | -0.318 | 0.071 | -0.544 | -0.53 | -0.009 | -0.014 | -0.014 | 5.289 | 0.18 | 0.23 | 0.08 | 0.08 | 0.1 | 0.07 |
| 10 | 4206194 | rs7897041 | 1.137 | 0.172 | -0.601 | -0.574 | 0.002 | -0.01 | -0.01 | 0 | 0.06 | 0.09 | 0.05 | 0.01 | 0.05 | 0.01 |
| 10 | 4207398 | rs7916036 | 1.106 | 0.113 | -0.639 | -0.558 | -0.003 | -0.015 | -0.015 | 0 | 0.11 | 0.1 | 0.02 | 0.02 | 0.05 | 0.04 |
| 10 | 4207411 | rs11252381 | 1.709 | 0.131 | -0.99 | -0.605 | -0.013 | -0.049 | -0.049 | 1.689 | 0.2 | 0.02 | 0.17 | 0.22 | 0.23 | 0.14 |
| 10 | 4208924 | rs10508273 | 1.343 | 0.319 | -0.915 | -0.718 | -0.049 | 0.04 | 0.04 | 0.52 | 0.15 | 0.01 | 0.08 | 0.1 | 0.18 | 0.17 |
| 10 | 4209856 | rs7069837 | -1.271 | 1.457 | -0.156 | -0.33 | 0.001 | -0.099 | -0.099 | 0 | 0.29 | 0.48 | 0.17 | 0.08 | 0.24 | 0.13 |
| 10 | 4212277 | rs7907648 | -0.17 | 1.452 | -0.098 | -0.325 | -0.004 | -0.022 | -0.022 | 0 | 0.2 | 0.23 | 0.11 | 0.06 | 0.15 | 0.06 |
| 10 | 4216511 | rs17132251 | -0.446 | 1.472 | -0.037 | -0.312 | 0.073 | -0.024 | -0.024 | 0 | 0.09 | 0.25 | 0.07 | 0.02 | 0.1 | 0.08 |
| 10 | 4217047 | rs11252385 | -1.227 | 1.494 | -0.112 | -0.329 | -0.001 | -0.103 | -0.103 | 0 | 0.31 | 0.49 | 0.18 | 0.08 | 0.25 | 0.15 |
| 10 | 4219333 | rs4275521 | 0.575 | 1.808 | -0.179 | 0.082 | -0.02 | 0.042 | 0.042 | 0 | 0.33 | 0.47 | 0.42 | 0.48 | 0.26 | 0.49 |
| 10 | 4222387 | rs7072145 | -0.989 | 1.836 | -0.403 | 0.164 | -0.042 | 0.032 | 0.032 | 0 | 0.6 | 0.86 | 0.64 | 0.67 | 0.54 | 0.66 |
| 10 | 4222609 | rs1923405 | -0.71 | 1.921 | -0.235 | 0.199 | -0.004 | 0.004 | 0.004 | 0 | 0.56 | 0.6 | 0.58 | 0.57 | 0.52 | 0.59 |
| 10 | 4232538 | rs1885476 | -0.486 | 2.141 | -0.235 | 0.56 | 0.076 | -0.009 | -0.009 | 0 | 0.2 | 0.28 | 0.23 | 0.15 | 0.32 | 0.13 |
| 10 | 4232937 | rs1108079 | -0.482 | 1.48 | -0.791 | 0.65 | 0.008 | 0.022 | 0.022 | 0 | 0.49 | 0.57 | 0.65 | 0.6 | 0.51 | 0.59 |
| 10 | 4233037 | rs7922038 | 0.531 | 1.549 | -0.798 | 0.401 | 0.011 | 0.038 | 0.038 | 0 | 0.34 | 0.4 | 0.51 | 0.53 | 0.37 | 0.5 |
| 10 | 4233048 | rs1108080 | 0.079 | 1.325 | -1.191 | 0.113 | 0.012 | 0.048 | 0.048 | 0 | 0.49 | 0.59 | 0.71 | 0.66 | 0.52 | 0.59 |
| 10 | 4233932 | rs2397411 | -0.214 | 1.064 | -1.577 | 0.282 | -0.03 | -0.053 | -0.053 | 0 | 0.27 | 0.06 | 0.32 | 0.29 | 0.3 | 0.11 |
| 10 | 4237510 | rs9664754 | -0.868 | 1.733 | -0.417 | -0.499 | -0.005 | 0.005 | 0.005 | 0 | 0.13 | 0.1 | 0.16 | 0.08 | 0.16 | 0.08 |
| 10 | 4238622 | rs879913 | -1.056 | 1.703 | -0.794 | -0.804 | -0.022 | 0.044 | 0.044 | 0 | 0.23 | 0.13 | 0.25 | 0.09 | 0.28 | 0.14 |
| 10 | 4238934 | rs1810649 | -0.978 | 1.24 | -0.061 | -1.093 | 0.017 | -0.055 | -0.055 | 0 | 0.21 | 0.33 | 0.26 | 0.08 | 0.26 | 0.13 |
| 10 | 4240572 | rs2181124 | -1.522 | 0.701 | 0.147 | -1.362 | 0 | -0.038 | -0.038 | 0 | 0.36 | 0.68 | 0.36 | 0.08 | 0.38 | 0.2 |
| 10 | 4241547 | rs7096293 | -1.241 | 0.721 | 0.125 | -1.354 | 0.016 | -0.096 | -0.096 | 0.037 | 0.31 | 0.6 | 0.34 | 0.08 | 0.37 | 0.17 |
| 10 | 4241572 | rs11252395 | 0.7 | 0.855 | -0.893 | -1.657 | -0.104 | -0.059 | -0.059 | 0 | 0.22 | 0.02 | 0.2 | 0.74 | 0.27 | 0.54 |
| 10 | 4249881 | rs7900699 | -0.851 | 0.725 | -0.335 | -1.587 | -0.006 | 0.069 | 0.069 | 0 | 0.31 | 0.19 | 0.28 | 0.08 | 0.31 | 0.17 |
| 10 | 4252171 | rs6601822 | -0.75 | 0.676 | -0.11 | -1.677 | 0.032 | -0.137 | -0.137 | 0 | 0.29 | 0.6 | 0.34 | 0.02 | 0.37 | 0.15 |
| 10 | 4256975 | rs1475097 | -1.463 | 0.197 | -0.465 | -1.568 | -0.025 | -0.019 | -0.019 | 0 | 0.09 | 0.25 | 0.06 | 0.06 | 0.03 | 0.05 |
| 10 | 4257317 | rs1475098 | -1.168 | 0.525 | -0.228 | -1.758 | 0.009 | -0.119 | -0.119 | 0.871 | 0.34 | 0.47 | 0.33 | 0.08 | 0.35 | 0.17 |
| 10 | 4259123 | rs10508277 | -2.09 | -0.016 | 0.139 | -1.066 | 0.023 | -0.036 | -0.036 | 0 | 0.11 | 0.3 | 0.11 | 0.07 | 0.13 | 0.07 |
| 10 | 4259467 | rs17132334 | -1.193 | -0.967 | -0.586 | -0.996 | 0.032 | 0.006 | 0.006 | 2.971 | 0.17 | 0.06 | 0.12 | 0.13 | 0.09 | 0.11 |
| 10 | 4264793 | rs7475965 | 1.333 | -1.582 | -0.377 | -0.991 | 0.07 | 0.032 | 0.032 | 5.284 | 0.68 | 0.21 | 0.75 | 0.77 | 0.88 | 0.85 |
| 10 | 4265108 | rs1998328 | -1.533 | -1.61 | 0.122 | -0.986 | -0.047 | -0.001 | -0.001 | 0 | 0.2 | 0.3 | 0.15 | 0.19 | 0.08 | 0.11 |
| 10 | 4271121 | rs17132378 | -1.484 | -1.632 | -0.004 | -1.047 | -0.033 | -0.004 | -0.004 | 0.157 | 0.19 | 0.27 | 0.1 | 0.16 | 0.07 | 0.11 |
| 10 | 4274736 | rs12262827 | -1.457 | -1.687 | 0.055 | -1.078 | 0.014 | 0.014 | 0.014 | 0.937 | 0.11 | 0.12 | 0.05 | 0.06 | 0.03 | 0.05 |
| 10 | 4275733 | rs6601823 | -0.475 | -1.613 | 0.131 | -1.067 | -0.024 | 0 | 0 | 0 | 0.11 | 0.17 | 0.07 | 0.11 | 0.05 | 0.06 |
| 10 | 4280637 | rs12264229 | -1.049 | -1.584 | 0.092 | -1.151 | -0.03 | -0.022 | -0.022 | 0.108 | 0.15 | 0.28 | 0.11 | 0.08 | 0.07 | 0.08 |
| 10 | 4286897 | rs10904207 | 1.834 | -1.744 | -0.269 | -1.233 | 0.09 | 0.072 | 0.072 | 5.737 | 0.69 | 0.17 | 0.8 | 0.84 | 0.89 | 0.88 |
| 10 | 4287879 | rs4881303 | 1.435 | -1.703 | -0.043 | -1.217 | -0.006 | -0.007 | -0.007 | 0.549 | 0.78 | 0.34 | 0.82 | 0.89 | 0.9 | 0.91 |

| | | | | | | | | | | | | | | | | |
|---|---|---|---|---|---|---|---|---|---|---|---|---|---|---|---|---|
| 10 | 4288645 | rs10795146 | 2.127 | -1.708 | -0.047 | -1.219 | 0.052 | 0.044 | 0.044 | 4.899 | 0.7 | 0.26 | 0.81 | 0.84 | 0.89 | 0.88 |
| 10 | 4289738 | rs6601824 | 1.468 | -1.549 | 0.017 | -1.175 | -0.018 | -0.021 | -0.021 | 1.277 | 0.78 | 0.36 | 0.81 | 0.89 | 0.89 | 0.91 |
| 10 | 4294071 | rs1535447 | -0.677 | -1.428 | -0.307 | -1.084 | 0.014 | 0.011 | 0.011 | 0 | 0.13 | 0.08 | 0.08 | 0.07 | 0.06 | 0.08 |
| 10 | 4300320 | rs10904211 | 0.427 | -1.386 | -1.104 | -1.392 | -0.012 | -0.01 | -0.01 | 0 | 0.19 | 0.04 | 0.17 | 0.8 | 0.16 | 0.58 |
| 10 | 4301143 | rs2065089 | 1.07 | -1.594 | -0.2 | -1.295 | -0.028 | -0.022 | -0.022 | 2.173 | 0.71 | 0.37 | 0.82 | 0.84 | 0.88 | 0.89 |
| 10 | 4306421 | rs10904218 | -0.74 | -0.539 | -0.016 | -0.257 | 0.004 | 0.027 | 0.027 | 0 | 0.14 | 0.19 | 0.32 | 0.64 | 0.14 | 0.43 |
| 10 | 4321300 | rs1923396 | 0.711 | -1.242 | 0.093 | -1.461 | 0.009 | -0.003 | -0.003 | 0 | 0.56 | 0.38 | 0.52 | 0.06 | 0.66 | 0.4 |
| 10 | 4322833 | rs1923397 | 0.591 | -1.365 | 0.385 | -1.452 | 0.003 | 0.011 | 0.011 | 0 | 0.63 | 0.62 | 0.58 | 0.13 | 0.68 | 0.45 |
| 10 | 4324728 | rs2065092 | 1.012 | -0.486 | -0.633 | -1.717 | -0.095 | 0.098 | 0.098 | 1.114 | 0.18 | 0.03 | 0.22 | 0.02 | 0.28 | 0.25 |
| 10 | 4330575 | rs11252445 | -0.481 | -1.166 | 0.149 | -1.514 | -0.009 | -0.026 | -0.026 | 0 | 0.39 | 0.43 | 0.43 | 0.77 | 0.32 | 0.52 |
| 10 | 4332824 | rs7917410 | 0.228 | -1.501 | 0.167 | -1.863 | -0.009 | -0.032 | -0.032 | 0 | 0.66 | 0.42 | 0.56 | 0.19 | 0.74 | 0.46 |
| 10 | 4334578 | rs9423526 | 0.745 | -1.576 | 0.259 | -1.935 | 0.005 | 0.006 | 0.006 | 1.107 | 0.55 | 0.42 | 0.51 | 0.23 | 0.67 | 0.45 |
| 10 | 4335760 | rs9423365 | -0.44 | -1.711 | 0.119 | -1.843 | 0.012 | -0.024 | -0.024 | 0 | 0.94 | 0.97 | 0.96 | 0.83 | 0.96 | 0.89 |
| 10 | 4338361 | rs7096543 | -0.529 | -1.695 | 0.425 | -1.757 | 0.008 | 0.008 | 0.008 | 0 | 0.37 | 0.63 | 0.46 | 0.73 | 0.27 | 0.57 |
| 10 | 4341279 | rs7901676 | -0.536 | -1.752 | 0.536 | -1.774 | -0.019 | -0.016 | -0.016 | 0.02 | 0.35 | 0.52 | 0.45 | 0.72 | 0.25 | 0.55 |
| 10 | 4341874 | rs11252451 | 1.042 | -2.009 | 0.397 | -1.894 | 0.079 | 0.194 | 0.194 | 0.739 | 0.49 | 0.26 | 0.45 | 0.09 | 0.65 | 0.33 |
| 10 | 4343285 | rs11813559 | 0.316 | -1.978 | 0.321 | -1.862 | 0.025 | -0.014 | -0.014 | 0 | 0.15 | 0.12 | 0.08 | 0.19 | 0.1 | 0.13 |
| 10 | 4343318 | rs17353967 | 1.141 | -1.662 | 0.024 | -2.046 | -0.101 | 0.094 | 0.094 | 0.201 | 0.18 | 0.02 | 0.23 | 0.02 | 0.29 | 0.2 |
| 10 | 4351582 | rs4475832 | -0.739 | -2.525 | 1.138 | -1.515 | -0.019 | -0.014 | -0.014 | 0 | 0.34 | 0.44 | 0.44 | 0.72 | 0.25 | 0.55 |
| 10 | 4352698 | rs10904225 | 0.858 | -2.636 | 1.119 | -1.518 | 0.011 | 0.007 | 0.007 | 0.48 | 0.52 | 0.48 | 0.47 | 0.13 | 0.65 | 0.34 |
| 10 | 4355798 | rs12777785 | 1.007 | -2.6 | 1.055 | -1.546 | 0.046 | 0.176 | 0.176 | 0.013 | 0.47 | 0.3 | 0.45 | 0.13 | 0.62 | 0.34 |
| 10 | 4359834 | rs12261707 | 0.035 | -2.565 | 0.788 | -1.376 | -0.027 | -0.002 | -0.002 | 0 | 0.21 | 0.23 | 0.11 | 0.19 | 0.11 | 0.15 |
| 10 | 4361265 | rs10904228 | 0.188 | -2.626 | 0.526 | -1.37 | 0.041 | -0.004 | -0.004 | 0 | 0.2 | 0.07 | 0.09 | 0.19 | 0.1 | 0.15 |
| 10 | 4362720 | rs4881316 | 1.121 | -2.67 | 0.481 | -1.465 | 0.048 | 0.188 | 0.188 | 0.047 | 0.47 | 0.23 | 0.44 | 0.13 | 0.62 | 0.34 |
| 10 | 4363142 | rs1926787 | -0.292 | -2.388 | 0.639 | -1.286 | -0.005 | -0.011 | -0.011 | 0 | 0.24 | 0.21 | 0.41 | 0.66 | 0.25 | 0.48 |
| 10 | 4365399 | rs10904233 | 0.428 | -2.017 | 0.977 | -1.331 | 0 | -0.023 | -0.023 | 0 | 0.54 | 0.43 | 0.54 | 0.13 | 0.67 | 0.37 |
| 10 | 4374838 | rs10904234 | 0.54 | -1.747 | 1.07 | -1.251 | 0.013 | 0.002 | 0.002 | 0.002 | 0.51 | 0.45 | 0.53 | 0.12 | 0.67 | 0.4 |
| 10 | 4379270 | rs9988693 | 0.387 | -1.723 | 0.882 | -1.241 | 0.034 | 0.059 | 0.059 | 0 | 0.53 | 0.68 | 0.55 | 0.12 | 0.67 | 0.4 |
| 10 | 4380623 | rs4261192 | 0.237 | -1.132 | 1.072 | -0.642 | -0.001 | 0.012 | 0.012 | 0 | 0.77 | 0.72 | 0.64 | 0.32 | 0.77 | 0.56 |
| 10 | 4382152 | rs10795158 | 0.392 | -1.137 | 1.069 | -0.642 | -0.003 | 0.025 | 0.025 | 0 | 0.76 | 0.65 | 0.63 | 0.33 | 0.77 | 0.56 |
| 10 | 4384056 | rs9633757 | -0.131 | -0.942 | 0.566 | -0.501 | -0.017 | -0.053 | -0.053 | 0 | 0.2 | 0.11 | 0.35 | 0.67 | 0.22 | 0.43 |
| 10 | 4390097 | rs11252477 | -0.123 | -0.915 | 1.045 | -0.45 | -0.002 | 0.017 | 0.017 | 0.023 | 0.19 | 0.21 | 0.34 | 0.68 | 0.2 | 0.4 |
| 10 | 4395682 | rs7082366 | -0.086 | -0.845 | 1.3 | -0.644 | 0.025 | -0.007 | -0.007 | 0 | 0.93 | 0.79 | 0.82 | 0.95 | 0.94 | 0.95 |
| 10 | 4398913 | rs1926810 | -0.049 | -0.85 | 1.307 | -0.646 | 0.025 | -0.007 | -0.007 | 0.374 | 0.93 | 0.79 | 0.82 | 0.95 | 0.94 | 0.95 |
| 10 | 4408326 | rs7095119 | 0.222 | -1.088 | 1.416 | -0.549 | -0.004 | -0.037 | -0.037 | 0 | 0.18 | 0.13 | 0.34 | 0.66 | 0.19 | 0.39 |
| 10 | 4409895 | rs4881319 | 0.055 | -1.117 | 1.337 | -0.555 | -0.008 | -0.057 | -0.057 | 1.565 | 0.19 | 0.11 | 0.34 | 0.66 | 0.19 | 0.4 |
| 10 | 4415882 | rs7894528 | -0.424 | -0.669 | 0.831 | -0.963 | 0.028 | -0.072 | -0.072 | 0 | 0.7 | 0.53 | 0.67 | 0.88 | 0.68 | 0.77 |
| 10 | 4424952 | rs11814820 | -0.283 | -0.942 | 0.179 | -1.044 | -0.002 | -0.026 | -0.026 | 0.007 | 0.56 | 0.57 | 0.51 | 0.83 | 0.37 | 0.67 |
| 10 | 4428563 | rs1679446 | 0.695 | -0.856 | -0.042 | -1.033 | -0.004 | 0.003 | 0.003 | 0.083 | 0.27 | 0.24 | 0.3 | 0.17 | 0.31 | 0.21 |
| 10 | 4428799 | rs7072047 | 0.738 | -1.019 | -0.774 | -0.971 | 0 | 0.008 | 0.008 | 0.028 | 0.2 | 0.22 | 0.38 | 0.65 | 0.21 | 0.42 |
| 10 | 4429530 | rs12267660 | 0.812 | -0.977 | -0.779 | -0.974 | 0.105 | 0.068 | 0.068 | 1.291 | 0.32 | 0.08 | 0.1 | 0.15 | 0.14 | 0.24 |
| 10 | 4431428 | rs1751312 | 0.913 | -0.776 | -1.231 | -0.915 | -0.061 | 0.036 | 0.036 | 0.213 | 0.25 | 0.1 | 0.28 | 0.18 | 0.31 | 0.22 |
| 10 | 4431610 | rs1952231 | 0.887 | 0.423 | -0.99 | -0.398 | -0.013 | -0.009 | -0.009 | 0 | 0.19 | 0.13 | 0.2 | 0 | 0.33 | 0.12 |
| 10 | 4432617 | rs12770204 | 0.804 | 0.434 | -1.031 | -0.393 | -0.001 | -0.021 | -0.021 | 0 | 0.19 | 0.18 | 0.2 | 0.02 | 0.33 | 0.12 |
| 10 | 4433008 | rs10904247 | -0.499 | 1.048 | -1.111 | 0.597 | -0.001 | 0.015 | 0.015 | 0.072 | 0.22 | 0.23 | 0.38 | 0.59 | 0.22 | 0.43 |
| 10 | 4435485 | rs12784701 | -0.403 | 1.312 | -1.571 | 0.326 | -0.083 | 0.065 | 0.065 | 0 | 0.24 | 0.03 | 0.24 | 0.09 | 0.27 | 0.29 |
| 10 | 4437754 | rs17132814 | -1.079 | 1.089 | -1.157 | 1.422 | -0.034 | -0.065 | -0.065 | 0 | 0.14 | 0.07 | 0.29 | 0.56 | 0.18 | 0.37 |
| 10 | 4444195 | rs10751968 | -1.056 | -0.003 | -0.629 | 1.257 | -0.017 | 0.061 | 0.061 | 0 | 0.2 | 0.35 | 0.19 | 0.49 | 0.12 | 0.26 |
| 10 | 4444521 | rs1751308 | -0.747 | -0.318 | -0.726 | 1.159 | 0.003 | -0.008 | -0.008 | 0 | 0.14 | 0.1 | 0.13 | 0.3 | 0.1 | 0.3 |
| 10 | 4445549 | rs10904249 | -0.67 | -0.417 | -0.717 | 1.24 | 0.015 | 0.03 | 0.03 | 0 | 0.08 | 0.05 | 0.13 | 0.44 | 0.04 | 0.2 |
| 10 | 4447552 | rs10508282 | -1.166 | -0.498 | -0.48 | 1.216 | -0.014 | 0.11 | 0.11 | 0 | 0.09 | 0.13 | 0.13 | 0.44 | 0.04 | 0.2 |
| 10 | 4450357 | rs993175 | -1.236 | -0.315 | -1.211 | 0.454 | 0.013 | -0.057 | -0.057 | 0 | 0.3 | 0.18 | 0.36 | 0.5 | 0.25 | 0.34 |
| 10 | 4451300 | rs1033992 | 0.304 | -0.328 | -1.194 | 0.457 | -0.001 | 0.009 | 0.009 | 1.011 | 0.12 | 0.11 | 0.11 | 0.3 | 0.08 | 0.28 |
| 10 | 4452786 | rs10795163 | -0.685 | -0.34 | -1.218 | 0.445 | 0.002 | -0.087 | -0.087 | 0 | 0.24 | 0.12 | 0.33 | 0.46 | 0.21 | 0.3 |
| 10 | 4453312 | rs1751307 | -2.134 | -0.417 | -0.371 | 0.341 | 0.005 | 0.013 | 0.013 | 0.227 | 0.48 | 0.58 | 0.52 | 0.81 | 0.38 | 0.64 |
| 10 | 4458818 | rs10795164 | -0.871 | -0.748 | 0.014 | 0.673 | -0.02 | 0.084 | 0.084 | 0 | 0.16 | 0.25 | 0.17 | 0.49 | 0.08 | 0.25 |
| 10 | 4458882 | rs1679409 | -2.186 | -0.75 | 0.033 | 0.64 | -0.025 | -0.032 | -0.032 | 0 | 0.32 | 0.55 | 0.33 | 0.79 | 0.21 | 0.54 |
| 10 | 4464252 | rs1751284 | -0.817 | -0.722 | 0.033 | 0.65 | -0.002 | 0.053 | 0.053 | 0 | 0.17 | 0.31 | 0.16 | 0.3 | 0.12 | 0.3 |
| 10 | 4466940 | rs1751282 | -0.062 | -0.732 | 0.414 | 0.665 | -0.006 | 0.033 | 0.033 | 0.843 | 0.13 | 0.14 | 0.12 | 0.29 | 0.09 | 0.28 |
| 10 | 4469292 | rs1679434 | -0.276 | -0.733 | 0.15 | 0.685 | -0.009 | 0.047 | 0.047 | 0.535 | 0.13 | 0.19 | 0.13 | 0.3 | 0.09 | 0.28 |
| 10 | 4469878 | rs1751280 | -1.889 | -0.754 | 0.108 | 0.594 | -0.04 | -0.031 | -0.031 | 0 | 0.3 | 0.47 | 0.31 | 0.78 | 0.18 | 0.53 |
| 10 | 4480571 | rs10904254 | -0.523 | -0.738 | -0.021 | 0.711 | 0.021 | 0 | 0 | 0 | 0.09 | 0.04 | 0.13 | 0.4 | 0.04 | 0.21 |
| 10 | 4480902 | rs1751275 | -2.383 | -0.863 | 0.038 | 0.549 | -0.023 | -0.04 | -0.04 | 2.909 | 0.35 | 0.54 | 0.34 | 0.78 | 0.22 | 0.56 |
| 10 | 4482137 | rs1679442 | -2.035 | -0.854 | -0.06 | 0.554 | -0.026 | -0.029 | -0.029 | 0 | 0.32 | 0.44 | 0.32 | 0.78 | 0.18 | 0.54 |
| 10 | 4484135 | rs1679444 | -1.499 | -0.942 | -0.732 | -0.284 | 0.005 | -0.044 | -0.044 | 0 | 0.58 | 0.5 | 0.52 | 0.78 | 0.49 | 0.59 |
| 10 | 4492612 | rs1751291 | -0.655 | -1.343 | -0.441 | 0.166 | 0.003 | -0.008 | -0.008 | 0 | 0.19 | 0.16 | 0.17 | 0.33 | 0.15 | 0.27 |
| 10 | 4495777 | rs11815537 | 1.183 | -1.349 | -0.384 | 0.16 | 0.006 | -0.005 | -0.005 | 0 | 0.07 | 0.15 | 0.04 | 0.05 | 0.05 | 0.03 |
| 10 | 4496292 | rs1751295 | -0.109 | -1.385 | -0.431 | 0.153 | 0.02 | -0.038 | -0.038 | 0 | 0.16 | 0.09 | 0.14 | 0.33 | 0.11 | 0.26 |
| 10 | 4496690 | rs1625110 | -2.151 | -1.545 | -0.457 | 0.071 | -0.026 | -0.044 | -0.044 | 0.012 | 0.42 | 0.5 | 0.37 | 0.79 | 0.26 | 0.52 |

| | | | | | | | | | | | | | | | | |
|---|---|---|---|---|---|---|---|---|---|---|---|---|---|---|---|---|
| 10 | 4499085 | rs1751300 | -1.127 | -1.445 | -0.426 | -0.023 | -0.004 | 0.01 | 0.01 | 0.691 | 0.26 | 0.3 | 0.2 | 0.38 | 0.2 | 0.3 |
| 10 | 4501924 | rs1615752 | -2.104 | -1.481 | -0.338 | 0.129 | -0.027 | -0.062 | -0.062 | 0 | 0.4 | 0.5 | 0.4 | 0.83 | 0.27 | 0.5 |
| 10 | 4502809 | rs1926797 | -0.62 | -1.497 | -0.411 | 0.184 | -0.02 | 0.08 | 0.08 | 0 | 0.12 | 0.12 | 0.14 | 0.44 | 0.04 | 0.19 |
| 10 | 4505344 | rs1679414 | -0.011 | -1.499 | -0.445 | 0.177 | 0.016 | -0.033 | -0.033 | 0 | 0.15 | 0.08 | 0.18 | 0.33 | 0.12 | 0.25 |
| 10 | 4505667 | rs2127598 | 1.216 | -1.452 | -0.403 | 0.222 | 0.001 | -0.004 | -0.004 | 0 | 0.07 | 0.12 | 0.04 | 0.05 | 0.05 | 0.02 |
| 10 | 4507840 | rs10904261 | -0.738 | -1.421 | -0.633 | 0.339 | -0.02 | 0.079 | 0.079 | 0 | 0.12 | 0.12 | 0.14 | 0.44 | 0.04 | 0.19 |
| 10 | 4508341 | rs17133009 | -0.623 | -1.349 | -0.644 | 0.354 | 0.006 | -0.015 | -0.015 | 0 | 0.2 | 0.15 | 0.21 | 0.33 | 0.18 | 0.28 |
| 10 | 4511571 | rs2077668 | -1.709 | -1.337 | -0.656 | 0.252 | -0.016 | -0.016 | -0.016 | 0 | 0.32 | 0.36 | 0.36 | 0.77 | 0.22 | 0.47 |
| 10 | 4516515 | rs17133040 | -0.799 | -0.763 | -0.948 | 0.605 | 0.005 | -0.015 | -0.015 | 0.476 | 0.2 | 0.15 | 0.22 | 0.32 | 0.18 | 0.28 |
| 10 | 4517381 | rs9663977 | -0.738 | -0.368 | -0.97 | 0.567 | -0.001 | -0.018 | -0.018 | 0.535 | 0.22 | 0.16 | 0.24 | 0.32 | 0.24 | 0.25 |
| 10 | 4518806 | rs11593634 | 0.031 | -0.318 | -1.122 | 0.479 | -0.001 | 0.001 | 0.001 | 0 | 0.06 | 0 | 0.12 | 0.06 | 0.03 | 0.03 |
| 10 | 4519338 | rs17358959 | -1.025 | -0.235 | -0.649 | 0.57 | 0 | -0.003 | -0.003 | 0 | 0.3 | 0.29 | 0.28 | 0.38 | 0.29 | 0.32 |
| 10 | 4522299 | rs11252558 | -0.921 | -0.358 | -0.013 | 1.154 | -0.019 | 0.136 | 0.136 | 1.445 | 0.16 | 0.39 | 0.2 | 0.34 | 0.1 | 0.09 |
| 10 | 4526742 | rs12415117 | 0.023 | 0.105 | -0.734 | 1.298 | -0.028 | -0.037 | -0.037 | 0.96 | 0.08 | 0.01 | 0.15 | 0.41 | 0.06 | 0.19 |
| 10 | 4527027 | rs10751969 | 0.245 | 0.618 | -1.004 | 0.304 | -0.054 | 0.058 | 0.058 | 0.055 | 0.36 | 0.13 | 0.42 | 0.2 | 0.39 | 0.39 |
| 10 | 4529263 | rs1079110 | 0.184 | 0.59 | -0.913 | 0.295 | -0.044 | 0.056 | 0.056 | 0.09 | 0.36 | 0.14 | 0.42 | 0.2 | 0.39 | 0.39 |
| 10 | 4538492 | rs4551658 | -1.014 | 0.152 | -0.743 | -0.066 | 0.008 | 0.022 | 0.022 | 0.727 | 0.74 | 0.79 | 0.85 | 0.91 | 0.78 | 0.64 |
| 10 | 4543335 | rs4313471 | -1.196 | 0.162 | -0.798 | -0.061 | 0.006 | 0.063 | 0.063 | 0.734 | 0.75 | 0.9 | 0.86 | 0.91 | 0.78 | 0.68 |
| 10 | 4547174 | rs11252568 | 0.304 | 0.29 | -0.529 | 0.204 | -0.002 | 0.012 | 0.012 | 0 | 0.2 | 0.22 | 0.2 | 0.4 | 0.1 | 0.24 |
| 10 | 4554674 | rs11252571 | 0.305 | 0.284 | -0.525 | 0.192 | -0.002 | 0.011 | 0.011 | 0 | 0.2 | 0.22 | 0.2 | 0.4 | 0.1 | 0.25 |
| 10 | 4555068 | rs17133097 | 0.695 | 0.461 | -0.997 | 0.229 | 0.022 | 0.059 | 0.059 | 0.635 | 0.29 | 0.18 | 0.15 | 0.09 | 0.22 | 0.32 |
| 10 | 4555301 | rs17133099 | 0.593 | 0.542 | -1.081 | 0.264 | 0.011 | 0.019 | 0.019 | 0.728 | 0.3 | 0.26 | 0.16 | 0.09 | 0.23 | 0.37 |
| 10 | 4559656 | rs11252576 | 0.021 | 1.065 | -0.913 | 0.6 | -0.045 | 0.055 | 0.055 | 0 | 0.16 | 0.06 | 0.24 | 0.02 | 0.27 | 0.1 |
| 10 | 4564553 | rs1904667 | -0.438 | 0.708 | -0.594 | 0.848 | 0.005 | 0.001 | 0.001 | 0.027 | 0.42 | 0.49 | 0.48 | 0.43 | 0.44 | 0.37 |
| 10 | 4576042 | rs313479 | 0.714 | 0.302 | -0.399 | 1.241 | -0.005 | 0.072 | 0.072 | 0.194 | 0.86 | 0.87 | 0.82 | 0.48 | 0.88 | 0.76 |
| 10 | 4576240 | rs313480 | 0.215 | -0.629 | -0.818 | 0.494 | 0.042 | 0.032 | 0.032 | 0 | 0.34 | 0.18 | 0.17 | 0.25 | 0.17 | 0.2 |
| 10 | 4577247 | rs313483 | 0.215 | -0.629 | -0.812 | 0.494 | 0.046 | 0.035 | 0.035 | 0 | 0.34 | 0.17 | 0.17 | 0.25 | 0.17 | 0.2 |
| 10 | 4579540 | rs313484 | 0.251 | -0.651 | -0.847 | 0.476 | 0.059 | 0.044 | 0.044 | 0 | 0.33 | 0.13 | 0.17 | 0.24 | 0.17 | 0.2 |
| 10 | 4582015 | rs7079674 | 0.283 | -0.57 | -0.095 | 0.111 | 0.025 | 0.042 | 0.042 | 0 | 0.47 | 0.42 | 0.56 | 0.2 | 0.66 | 0.38 |
| 10 | 4585838 | rs313474 | -0.206 | -1.141 | -0.314 | -0.053 | 0.113 | 0.127 | 0.127 | 0 | 0.5 | 0.19 | 0.35 | 0.78 | 0.3 | 0.46 |
| 10 | 4586244 | rs313475 | -0.48 | 0.179 | 0.104 | -0.434 | 0.037 | -0.05 | -0.05 | 0 | 0.75 | 0.68 | 0.63 | 0.94 | 0.56 | 0.79 |
| 10 | 4588261 | rs11814411 | 0.113 | 0.398 | 0.282 | -0.545 | 0.004 | -0.03 | -0.03 | 0 | 0.19 | 0.24 | 0.33 | 0.06 | 0.25 | 0.2 |
| 10 | 4594032 | rs387673 | -0.277 | 0.191 | 0.169 | -0.685 | 0.003 | 0.006 | 0.006 | 0 | 0.8 | 0.72 | 0.85 | 0.83 | 0.76 | 0.85 |
| 10 | 4596608 | rs416990 | 0.96 | -0.24 | 0.13 | -0.904 | 0.007 | 0.055 | 0.055 | 0 | 0.52 | 0.34 | 0.49 | 0.76 | 0.49 | 0.46 |
| 10 | 4598433 | rs372744 | 1.282 | -0.04 | 0.043 | -0.568 | 0.022 | 0.036 | 0.036 | 0 | 0.37 | 0.26 | 0.19 | 0.19 | 0.23 | 0.22 |
| 10 | 4598903 | rs715987 | 0.998 | 0.257 | -0.224 | -0.471 | -0.037 | -0.043 | -0.043 | 0.318 | 0.13 | 0.04 | 0.12 | 0.15 | 0.22 | 0.14 |
| 10 | 4604549 | rs374802 | 1.15 | 0.153 | -0.104 | -0.12 | 0.054 | 0.054 | 0.054 | 0 | 0.39 | 0.19 | 0.23 | 0.27 | 0.24 | 0.34 |
| 10 | 4604986 | rs745782 | -1.038 | 0.482 | 0.248 | 0.084 | 0 | 0 | 0 | 0 | 0.42 | 0.57 | 0.62 | 0.59 | 0.51 | 0.52 |
| 10 | 4606853 | rs7906409 | -0.947 | 0.608 | -0.185 | 0.054 | 0.009 | 0.011 | 0.011 | 0 | 0.41 | 0.45 | 0.61 | 0.56 | 0.51 | 0.52 |
| 10 | 4609790 | rs10795187 | -0.202 | 1.078 | -0.87 | 0.19 | -0.002 | 0 | 0 | 0 | 0.21 | 0.2 | 0.27 | 0.45 | 0.27 | 0.31 |
| 10 | 4610644 | rs12413638 | 0.134 | 0.548 | -1.01 | -0.461 | -0.033 | 0.048 | 0.048 | 0 | 0.12 | 0.03 | 0.19 | 0 | 0.18 | 0.05 |
| 10 | 4611536 | rs7089195 | -0.598 | -0.919 | -1.079 | -0.707 | -0.007 | -0.007 | -0.007 | 0 | 0.32 | 0.38 | 0.23 | 0.13 | 0.16 | 0.36 |
| 10 | 4612690 | rs313500 | 0.906 | -1.54 | -0.709 | -0.271 | -0.027 | 0.033 | 0.033 | 0 | 0.74 | 0.34 | 0.83 | 0.4 | 0.85 | 0.56 |
| 10 | 4616862 | rs10904290 | 0.379 | -1.607 | -0.205 | -0.02 | -0.015 | 0.231 | 0.231 | 0 | 0.06 | 0.17 | 0.05 | 0.55 | 0.02 | 0.27 |
| 10 | 4619601 | rs2028690 | -0.634 | -1.186 | -0.714 | -1.135 | 0.012 | -0.077 | -0.077 | 0.026 | 0.34 | 0.34 | 0.31 | 0.11 | 0.34 | 0.39 |
| 10 | 4621751 | rs381443 | -0.277 | -1.228 | -0.773 | -1.181 | -0.003 | -0.038 | -0.038 | 0 | 0.29 | 0.2 | 0.29 | 0.04 | 0.33 | 0.28 |
| 10 | 4625573 | rs1008042 | 1.257 | -1.181 | -1.192 | -1.325 | 0 | 0.139 | 0.139 | 0 | 0.49 | 0.13 | 0.33 | 0.29 | 0.46 | 0.3 |
| 10 | 4627791 | rs1498271 | 1.356 | -1.199 | -1.002 | -1.333 | 0.028 | 0.132 | 0.132 | 0 | 0.5 | 0.25 | 0.34 | 0.29 | 0.46 | 0.3 |
| 10 | 4628581 | rs11252602 | 1.016 | -1.212 | -1.012 | -1.339 | -0.002 | 0.085 | 0.085 | 0 | 0.54 | 0.22 | 0.34 | 0.31 | 0.47 | 0.4 |
| 10 | 4628616 | rs1498274 | -0.104 | -1.133 | -1.134 | -1.289 | -0.024 | 0.026 | 0.026 | 0 | 0.73 | 0.59 | 0.63 | 0.4 | 0.78 | 0.62 |
| 10 | 4632979 | rs569744 | -0.549 | -0.984 | -1.093 | -1.423 | 0.074 | -0.087 | -0.087 | 0.09 | 0.24 | 0.45 | 0.3 | 0.11 | 0.32 | 0.32 |
| 10 | 4635292 | rs17295905 | 1.26 | -1.083 | -1.838 | -1.498 | 0.015 | 0.05 | 0.05 | 3.199 | 0.1 | 0.03 | 0.03 | 0 | 0.06 | 0.01 |
| 10 | 4636054 | rs2895006 | 0.758 | -0.927 | -1.661 | -1.327 | -0.003 | 0.01 | 0.01 | 0 | 0.18 | 0.09 | 0.3 | 0.08 | 0.19 | 0.11 |
| 10 | 4638643 | rs11252607 | 1.202 | -0.753 | -1.822 | -1.442 | 0.04 | -0.006 | -0.006 | 0 | 0.31 | 0.11 | 0.26 | 0.32 | 0.22 | 0.3 |
| 10 | 4641211 | rs1443852 | 0.409 | -0.623 | -1.683 | -1.632 | -0.023 | 0.017 | 0.017 | 0 | 0.56 | 0.35 | 0.63 | 0.35 | 0.57 | 0.42 |
| 10 | 4642156 | rs11252615 | -0.623 | -0.688 | -1.324 | -1.732 | 0.004 | 0.006 | 0.006 | 0 | 0.52 | 0.51 | 0.45 | 0.65 | 0.48 | 0.55 |
| 10 | 4646167 | rs10904299 | 1.569 | -0.429 | -1.421 | -1.598 | 0.026 | -0.007 | -0.007 | 0.239 | 0.29 | 0.11 | 0.25 | 0.28 | 0.23 | 0.26 |
| 10 | 4647269 | rs12775823 | 0.227 | -0.582 | -1.25 | -1.696 | 0.021 | 0.006 | 0.006 | 0 | 0.62 | 0.52 | 0.63 | 0.38 | 0.59 | 0.47 |
| 10 | 4647506 | rs1838238 | 0.452 | -0.611 | -1.26 | -1.711 | -0.007 | 0.098 | 0.098 | 0 | 0.29 | 0.15 | 0.36 | 0.06 | 0.35 | 0.2 |
| 10 | 4648430 | rs7096965 | 0.924 | -0.636 | -1.275 | -1.723 | -0.035 | 0.087 | 0.087 | 0 | 0.25 | 0.09 | 0.35 | 0.05 | 0.34 | 0.16 |
| 10 | 4648434 | rs10904300 | 1.522 | -0.447 | -1.138 | -1.602 | 0.024 | -0.011 | -0.011 | 0.059 | 0.3 | 0.17 | 0.25 | 0.33 | 0.23 | 0.26 |
| 10 | 4650036 | rs11252622 | 1.603 | -0.308 | -1.412 | -1.558 | -0.005 | -0.001 | -0.001 | 0 | 0.3 | 0.11 | 0.27 | 0.28 | 0.3 | 0.27 |
| 10 | 4653578 | rs1027504 | 0.236 | 0.546 | -1.029 | -1.473 | 0.019 | 0.093 | 0.093 | 0 | 0.28 | 0.15 | 0.34 | 0.06 | 0.29 | 0.19 |
| 10 | 4655565 | rs10458787 | -0.415 | 1.645 | -0.239 | -0.97 | 0.148 | -0.095 | -0.095 | 1.195 | 0.86 | 0.68 | 0.76 | 1 | 0.76 | 0.86 |
| 10 | 4659197 | rs1039197 | 0.158 | 1.369 | -0.399 | -0.936 | 0.163 | -0.081 | -0.081 | 0.714 | 0.82 | 0.6 | 0.72 | 0.94 | 0.72 | 0.82 |
| 10 | 4672693 | rs6601853 | -0.675 | 0.465 | -0.01 | -0.632 | -0.05 | -0.143 | -0.143 | 0 | 0.21 | 0.5 | 0.26 | 0.01 | 0.15 | 0.22 |
| 10 | 4674335 | rs313462 | -0.89 | 0.025 | -0.856 | -1.503 | 0.127 | 0.265 | 0.265 | 3.511 | 0.5 | 0.84 | 0.49 | 0.06 | 0.38 | 0.31 |
| 10 | 4676196 | rs7086888 | -0.234 | -0.198 | -0.175 | -1.409 | 0.015 | -0.059 | -0.059 | 0.781 | 0.1 | 0.32 | 0.05 | 0.01 | 0.07 | 0.14 |
| 10 | 4680910 | rs313445 | -0.012 | -0.132 | -0.128 | -1.391 | 0.003 | -0.004 | -0.004 | 0 | 0.32 | 0.29 | 0.25 | 0.07 | 0.29 | 0.21 |

| | | | | | | | | | | | | | | | | |
|---|---|---|---|---|---|---|---|---|---|---|---|---|---|---|---|---|
| 10 | 4683858 | rs313438 | 0.361 | -0.304 | -0.396 | -1.326 | 0.036 | 0.077 | 0.077 | 0 | 0.27 | 0.13 | 0.21 | 0.05 | 0.23 | 0.08 |
| 10 | 4684916 | rs17298157 | 0.46 | -0.281 | -0.544 | -1.354 | 0.045 | 0.113 | 0.113 | 0 | 0.26 | 0.05 | 0.2 | 0.06 | 0.21 | 0.13 |
| 10 | 4685231 | rs313435 | 0.177 | 0.455 | -0.486 | -1.716 | 0.018 | -0.026 | -0.026 | 0.415 | 0.63 | 0.52 | 0.54 | 0.26 | 0.55 | 0.57 |
| 10 | 4687532 | rs2088500 | -0.506 | -0.317 | -0.749 | -2.228 | 0.004 | 0.042 | 0.042 | 0.094 | 0.51 | 0.42 | 0.7 | 0.87 | 0.54 | 0.59 |
| 10 | 4689138 | rs12255709 | 1.67 | -0.399 | -0.664 | -2.384 | 0.007 | 0.03 | 0.03 | 0 | 0.19 | 0.12 | 0.12 | 0.07 | 0.2 | 0.1 |
| 10 | 4689842 | rs313452 | -1.119 | -0.48 | -0.548 | -2.512 | 0.001 | -0.042 | -0.042 | 0.399 | 0.75 | 0.74 | 0.84 | 0.95 | 0.76 | 0.76 |
| 10 | 4694013 | rs313455 | 0.817 | -0.207 | -0.383 | -2.205 | -0.002 | -0.022 | -0.022 | 0 | 0.24 | 0.27 | 0.14 | 0.08 | 0.23 | 0.15 |
| 10 | 4698360 | rs12249207 | 1.685 | -0.283 | -0.597 | -2.235 | 0.005 | 0.028 | 0.028 | 0 | 0.18 | 0.12 | 0.11 | 0.08 | 0.19 | 0.1 |
| 10 | 4699392 | rs11598678 | 1.528 | -0.358 | -0.776 | -1.997 | 0.002 | -0.057 | -0.057 | 0 | 0.09 | 0 | 0.18 | 0.24 | 0.08 | 0.13 |
| 10 | 4700968 | rs10508287 | 1.074 | 0.214 | -0.637 | -2.52 | -0.016 | 0.003 | 0.003 | 0 | 0.17 | 0.1 | 0.23 | 0.01 | 0.31 | 0.11 |
| 10 | 4711764 | rs313430 | 1.684 | -0.277 | -0.115 | -1.911 | 0.015 | -0.019 | -0.019 | 0 | 0.05 | 0.18 | 0.03 | 0 | 0.03 | 0.05 |
| 10 | 4716018 | rs1838239 | 0.601 | -0.365 | -1.073 | -1.943 | -0.022 | -0.084 | -0.084 | 0.384 | 0.16 | 0.31 | 0.14 | 0.02 | 0.12 | 0.2 |
| 10 | 4716542 | rs2165953 | -0.416 | 0.008 | -0.805 | -1.548 | -0.021 | -0.066 | -0.066 | 3.008 | 0.72 | 0.44 | 0.82 | 0.89 | 0.76 | 0.81 |
| 10 | 4719796 | rs1391511 | 0.594 | 0.842 | -1.276 | -0.183 | -0.031 | -0.086 | -0.086 | 0.152 | 0.58 | 0.1 | 0.53 | 0.49 | 0.6 | 0.53 |
| 10 | 4721471 | rs7916170 | 0.968 | 0.671 | -0.616 | -0.302 | -0.017 | -0.049 | -0.049 | 0.068 | 0.09 | 0.18 | 0.02 | 0 | 0.04 | 0.08 |
| 10 | 4725340 | rs6601855 | -0.443 | 0.965 | -0.34 | -0.312 | -0.006 | -0.09 | -0.09 | 0 | 0.77 | 0.62 | 0.74 | 0.95 | 0.8 | 0.83 |
| 10 | 4728091 | rs1498273 | -0.054 | 1.048 | -0.916 | 0.224 | 0.006 | 0.103 | 0.103 | 0 | 0.07 | 0.11 | 0.18 | 0.47 | 0.15 | 0.19 |
| 10 | 4731081 | rs7080649 | 0.371 | 0.873 | -1.315 | -0.064 | 0.013 | -0.055 | -0.055 | 0.132 | 0.48 | 0.09 | 0.47 | 0.48 | 0.46 | 0.39 |
| 10 | 4734128 | rs433430 | 0.578 | 0.852 | -0.083 | -0.069 | 0.033 | -0.021 | -0.021 | 0 | 0.93 | 0.7 | 0.95 | 1 | 0.93 | 0.93 |
| 10 | 4741842 | rs10904319 | 0.126 | 0.645 | -0.642 | -0.348 | 0.004 | 0.126 | 0.126 | 0 | 0.45 | 0.21 | 0.4 | 0.18 | 0.53 | 0.42 |
| 10 | 4746345 | rs10904321 | 0.029 | 0.646 | -0.269 | -0.344 | 0.031 | 0.094 | 0.094 | 0 | 0.46 | 0.28 | 0.43 | 0.19 | 0.57 | 0.46 |
| 10 | 4746670 | rs7068675 | -0.102 | 0.328 | -0.045 | 0.087 | 0 | 0.015 | 0.015 | 0 | 0.15 | 0.13 | 0.23 | 0.4 | 0.17 | 0.26 |
| 10 | 4748240 | rs10904323 | 0.98 | 0.029 | -0.064 | -0.01 | 0.048 | -0.1 | -0.1 | 2.545 | 0.31 | 0.03 | 0.29 | 0.42 | 0.24 | 0.21 |
| 10 | 4752471 | rs11252672 | 0.558 | 0.154 | 0.317 | 0.079 | 0.02 | -0.027 | -0.027 | 0 | 0.1 | 0.03 | 0.19 | 0.4 | 0.07 | 0.21 |
| 10 | 4753422 | rs11252674 | -0.027 | 0.299 | 0.436 | -0.013 | 0.007 | 0.006 | 0.006 | 0 | 0.52 | 0.44 | 0.44 | 0.18 | 0.66 | 0.57 |
| 10 | 4756587 | rs11252679 | 0.447 | 0.365 | -0.043 | -0.015 | 0.016 | 0.1 | 0.1 | 0 | 0.45 | 0.25 | 0.4 | 0.19 | 0.54 | 0.45 |
| 10 | 4758118 | rs17133449 | 0.856 | 0.262 | -0.276 | 0.054 | 0.041 | -0.073 | -0.073 | 0 | 0.33 | 0.05 | 0.32 | 0.41 | 0.27 | 0.22 |
| 10 | 4769379 | rs2397839 | -0.3 | 0.195 | -0.126 | -0.135 | 0.034 | 0.005 | 0.005 | 0.147 | 0.5 | 0.35 | 0.49 | 0.51 | 0.6 | 0.62 |
| 10 | 4772604 | rs10904328 | -0.981 | 0.036 | -0.161 | -0.137 | -0.017 | 0.003 | 0.003 | 0.009 | 0.63 | 0.54 | 0.64 | 0.57 | 0.73 | 0.78 |
| 10 | 4774565 | rs11252688 | 0.555 | 0.162 | -0.945 | 0.009 | 0.016 | -0.029 | -0.029 | 0 | 0.1 | 0.03 | 0.19 | 0.4 | 0.07 | 0.21 |
| 10 | 4775903 | rs11252690 | 1.101 | 0.833 | -1.155 | -0.294 | -0.077 | 0.021 | 0.021 | 0 | 0.25 | 0.02 | 0.19 | 0.11 | 0.3 | 0.23 |
| 10 | 4779211 | rs2123348 | -0.331 | 0.482 | -1.113 | -0.04 | -0.032 | 0.043 | 0.043 | 0 | 0.31 | 0.55 | 0.31 | 0.52 | 0.23 | 0.39 |
| 10 | 4780398 | rs12251644 | 1.19 | 0.697 | -1.356 | 0.077 | -0.004 | -0.002 | -0.002 | 0 | 0.08 | 0.06 | 0.18 | 0.1 | 0.11 | 0.13 |
| 10 | 4780818 | rs11596789 | 1.489 | 0.454 | -1.288 | 0.273 | 0.015 | -0.018 | -0.018 | 1.035 | 0.06 | 0.02 | 0.02 | 0.15 | 0.03 | 0.13 |
| 10 | 4781029 | rs10508288 | 0.583 | 0.833 | -1.248 | -0.48 | -0.072 | 0.025 | 0.025 | 0 | 0.33 | 0.1 | 0.38 | 0.22 | 0.41 | 0.34 |
| 10 | 4782841 | rs4146653 | 1.283 | 0.574 | -1.09 | -0.928 | 0.022 | 0.025 | 0.025 | 0 | 0.23 | 0.11 | 0.23 | 0.2 | 0.2 | 0.1 |
| 10 | 4786841 | rs7899916 | -1.094 | 0.935 | -0.944 | -0.633 | -0.043 | 0.04 | 0.04 | 0 | 0.48 | 0.84 | 0.42 | 0.56 | 0.44 | 0.64 |
| 10 | 4788438 | rs10904332 | -0.792 | 1.09 | -0.657 | -0.492 | 0 | -0.004 | -0.004 | 0 | 0.42 | 0.65 | 0.37 | 0.52 | 0.37 | 0.58 |
| 10 | 4791718 | rs4881340 | -0.629 | -0.12 | -0.361 | -0.189 | -0.003 | 0.001 | 0.001 | 0 | 0.34 | 0.66 | 0.31 | 0.51 | 0.23 | 0.46 |
| 10 | 4792577 | rs10904333 | -0.653 | -0.087 | -0.826 | -0.188 | 0.041 | -0.031 | -0.031 | 0 | 0.36 | 0.83 | 0.34 | 0.56 | 0.27 | 0.51 |
| 10 | 4797029 | rs7896112 | -0.804 | 0.079 | -1.106 | -0.57 | 0.006 | 0.068 | 0.068 | 0 | 0.68 | 0.87 | 0.55 | 0.74 | 0.72 | 0.79 |
| 10 | 4798756 | rs12357812 | -0.135 | 0.725 | -0.374 | -0.077 | -0.004 | 0.017 | 0.017 | 0 | 0.42 | 0.35 | 0.35 | 0.2 | 0.54 | 0.4 |
| 10 | 4799607 | rs2894969 | -0.736 | 0.72 | -0.427 | -0.094 | 0.036 | 0.043 | 0.043 | 0 | 0.5 | 0.35 | 0.45 | 0.32 | 0.64 | 0.52 |
| 10 | 4800054 | rs11252715 | 0.269 | 0.859 | -0.807 | -0.006 | -0.13 | 0.077 | 0.077 | 0 | 0.32 | 0.05 | 0.28 | 0.16 | 0.45 | 0.28 |
| 10 | 4800190 | rs11252717 | -0.152 | 0.899 | -0.531 | 0.329 | 0.001 | 0.029 | 0.029 | 0 | 0.47 | 0.54 | 0.37 | 0.2 | 0.54 | 0.4 |
| 10 | 4805703 | rs10508290 | -1.117 | 0.803 | -0.115 | 1.295 | 0.016 | 0.406 | 0.406 | 0 | 0.05 | 0.23 | 0.04 | 0.5 | 0.02 | 0.22 |
| 10 | 4808620 | rs1869210 | 0.147 | 1.13 | -0.815 | 0.079 | 0 | 0.001 | 0.001 | 0 | 0.61 | 0.42 | 0.68 | 0.86 | 0.58 | 0.78 |
| 10 | 4809075 | rs17133587 | -0.447 | 1.743 | -1.447 | 0.544 | -0.067 | -0.068 | -0.068 | 2.186 | 0.14 | 0.02 | 0.33 | 0.2 | 0.22 | 0.18 |
| 10 | 4811345 | rs10904337 | -0.112 | 1.098 | -1.066 | -0.078 | -0.003 | -0.009 | -0.009 | 0 | 0.33 | 0.32 | 0.25 | 0.19 | 0.31 | 0.21 |
| 10 | 4811703 | rs7893291 | 0.354 | 1.269 | -1.154 | -0.165 | -0.004 | -0.004 | -0.004 | 0 | 0.26 | 0.21 | 0.41 | 0.28 | 0.29 | 0.36 |
| 10 | 4813154 | rs11818567 | 0.503 | 1.343 | -1.23 | -0.587 | 0.001 | -0.013 | -0.013 | 0 | 0.1 | 0.07 | 0.05 | 0 | 0.09 | 0.06 |
| 10 | 4813211 | rs1901632 | 0.929 | 1.049 | -1.205 | -0.557 | -0.005 | -0.028 | -0.028 | 0 | 0.3 | 0.3 | 0.24 | 0.12 | 0.29 | 0.16 |
| 10 | 4814682 | rs7900146 | 0.612 | 1.555 | -1.121 | -0.383 | 0.015 | -0.037 | -0.037 | 0 | 0.19 | 0.12 | 0.21 | 0.53 | 0.16 | 0.24 |
| 10 | 4817965 | rs9423544 | 0.16 | 2.247 | -0.483 | -0.651 | 0.052 | -0.1 | -0.1 | 0 | 0.8 | 0.66 | 0.86 | 1 | 0.76 | 0.8 |
| 10 | 4822210 | rs7082636 | -0.363 | 2.082 | -0.5 | -0.31 | 0.076 | 0.007 | 0.007 | 0 | 0.06 | 0.18 | 0.06 | 0.04 | 0.12 | 0.06 |
| 10 | 4823252 | rs10904339 | -0.091 | 2.227 | -0.778 | -0.608 | -0.005 | -0.012 | -0.012 | 0 | 0.13 | 0.11 | 0.1 | 0 | 0.18 | 0.16 |
| 10 | 4823414 | rs4880687 | 0.482 | 1.51 | -0.684 | -0.953 | -0.007 | 0.007 | 0.007 | 0 | 0.61 | 0.42 | 0.58 | 0.87 | 0.55 | 0.62 |
| 10 | 4823794 | rs9423546 | 0.198 | 0.693 | -1.005 | 0.066 | -0.002 | 0.013 | 0.013 | 0.467 | 0.23 | 0.26 | 0.27 | 0.54 | 0.22 | 0.16 |
| 10 | 4824211 | rs9423547 | -0.456 | 0.623 | -0.937 | -0.143 | -0.01 | -0.011 | -0.011 | 0.262 | 0.33 | 0.44 | 0.42 | 0.77 | 0.35 | 0.4 |
| 10 | 4825336 | rs7084656 | -0.605 | 0.638 | -1.253 | -0.321 | 0.002 | 0.015 | 0.015 | 2.297 | 0.56 | 0.83 | 0.67 | 0.39 | 0.6 | 0.55 |
| 10 | 4827368 | rs9423548 | 0.193 | 0.692 | -0.943 | -0.472 | -0.011 | 0.043 | 0.043 | 1.035 | 0.46 | 0.37 | 0.44 | 0.31 | 0.5 | 0.52 |
| 10 | 4829103 | rs2397990 | -0.008 | 0.652 | -0.489 | -0.502 | 0 | -0.007 | -0.007 | 0 | 0.48 | 0.55 | 0.45 | 0.31 | 0.5 | 0.52 |
| 10 | 4830278 | rs1901649 | 0.761 | 0.647 | -0.59 | -0.555 | -0.008 | 0.003 | 0.003 | 0 | 0.24 | 0.32 | 0.26 | 0.28 | 0.23 | 0.29 |
| 10 | 4832920 | rs11595123 | 2.142 | 0.844 | -1.07 | -0.647 | -0.071 | 0.083 | 0.083 | 0 | 0.18 | 0.04 | 0.16 | 0.03 | 0.21 | 0.21 |
| 10 | 4833049 | rs4881344 | -0.216 | 0.789 | -0.927 | -0.698 | -0.001 | 0.003 | 0.003 | 1.552 | 0.49 | 0.62 | 0.46 | 0.31 | 0.5 | 0.53 |
| 10 | 4834556 | rs9423376 | -0.801 | 0.835 | -1.231 | -0.652 | 0.002 | 0.007 | 0.007 | 2.226 | 0.57 | 0.8 | 0.69 | 0.41 | 0.6 | 0.56 |
| 10 | 4840167 | rs12416000 | 1.072 | 1.03 | -1.602 | -0.565 | -0.029 | -0.049 | -0.049 | 0 | 0.08 | 0.02 | 0.12 | 0.19 | 0.11 | 0.23 |
| 10 | 4841905 | rs11252746 | 1.483 | 0.765 | -1.93 | -0.914 | 0.013 | 0.092 | 0.092 | 0.597 | 0.2 | 0.01 | 0.09 | 0.07 | 0.12 | 0.07 |
| 10 | 4842046 | rs10904346 | 0.152 | 0.906 | -1.895 | -0.822 | -0.039 | -0.06 | -0.06 | 0 | 0.25 | 0.05 | 0.45 | 0.66 | 0.31 | 0.34 |

| | | | | | | | | | | | | | | | | |
|---|---|---|---|---|---|---|---|---|---|---|---|---|---|---|---|---|
| 10 | 4842737 | rs4146754 | 0.556 | 0.774 | -1.92 | -0.847 | -0.037 | -0.138 | -0.138 | 0 | 0.22 | 0.03 | 0.25 | 0.55 | 0.23 | 0.31 |
| 10 | 4845062 | rs2167704 | 1.813 | 0.908 | -1.772 | -2.017 | -0.079 | 0.069 | 0.069 | 0 | 0.2 | 0.05 | 0.17 | 0.1 | 0.26 | 0.21 |
| 10 | 4845594 | rs11252752 | 1.416 | 0.443 | -1.42 | -2.143 | 0.01 | -0.005 | -0.005 | 0 | 0.17 | 0.11 | 0.09 | 0 | 0.12 | 0.14 |
| 10 | 4846866 | rs2397988 | 1.097 | 0.634 | -1.497 | -2.051 | -0.089 | 0.033 | 0.033 | 0 | 0.31 | 0.12 | 0.54 | 0.75 | 0.4 | 0.48 |
| 10 | 4847380 | rs17374031 | 1.164 | 0.92 | -1.387 | -1.959 | -0.03 | -0.023 | -0.023 | 0.722 | 0.07 | 0.02 | 0.11 | 0.12 | 0.14 | 0.15 |
| 10 | 4847924 | rs2961594 | 0.072 | 0.557 | -0.88 | -2.168 | -0.028 | -0.13 | -0.13 | 0 | 0.36 | 0.39 | 0.2 | 0.03 | 0.3 | 0.23 |
| 10 | 4850639 | rs2961618 | 0.746 | 1.29 | -0.465 | -1.873 | -0.001 | 0.056 | 0.056 | 0 | 0.56 | 0.3 | 0.69 | 0.86 | 0.58 | 0.57 |
| 10 | 4852869 | rs3110951 | 1.362 | 1.53 | 0.07 | -1.553 | -0.036 | -0.054 | -0.054 | 0.261 | 0.26 | 0.29 | 0.16 | 0.11 | 0.18 | 0.09 |
| 10 | 4854229 | rs3100196 | 1.474 | 1.494 | 0.005 | -1.55 | 0.007 | 0.004 | 0.004 | 0 | 0.25 | 0.15 | 0.14 | 0.11 | 0.17 | 0.08 |
| 10 | 4854369 | rs3100195 | -1.034 | 1.583 | 0.011 | -1.568 | 0.01 | -0.051 | -0.051 | 0.48 | 0.87 | 0.74 | 0.94 | 0.98 | 0.91 | 0.88 |
| 10 | 4855238 | rs2961609 | 0.398 | 1.494 | 0.037 | -1.508 | -0.02 | -0.033 | -0.033 | 0.02 | 0.34 | 0.4 | 0.26 | 0.23 | 0.31 | 0.24 |
| 10 | 4863351 | rs2961620 | 1.147 | 1.64 | -0.17 | -1.242 | 0.024 | -0.016 | -0.016 | 0.277 | 0.76 | 0.46 | 0.83 | 0.81 | 0.77 | 0.77 |
| 10 | 4864809 | rs2961559 | -0.348 | 1.524 | -0.271 | -1.376 | 0.009 | 0.028 | 0.028 | 0.064 | 0.51 | 0.57 | 0.36 | 0.29 | 0.46 | 0.27 |
| 10 | 4875894 | rs2961576 | 0.983 | 1.438 | -0.003 | -1.266 | 0.036 | 0.109 | 0.109 | 0.015 | 0.35 | 0.12 | 0.17 | 0.12 | 0.22 | 0.18 |
| 10 | 4884784 | rs2924269 | -0.513 | 1.447 | 0.199 | -1.288 | 0.044 | 0.025 | 0.025 | 0.047 | 0.94 | 0.66 | 0.96 | 0.98 | 0.97 | 0.9 |
| 10 | 4889403 | rs12240276 | 1.243 | 1.605 | 0.065 | -1.35 | 0.001 | -0.012 | -0.012 | 0 | 0.14 | 0.1 | 0.09 | 0 | 0.13 | 0.07 |
| 10 | 4889525 | rs2961597 | 0.365 | 1.646 | 0.163 | -1.341 | 0.055 | 0.067 | 0.067 | 0.367 | 0.91 | 0.5 | 0.93 | 0.91 | 0.94 | 0.88 |
| 10 | 4890145 | rs752534 | 1.249 | 1.588 | 0.118 | -1.34 | 0.001 | -0.012 | -0.012 | 0.002 | 0.14 | 0.1 | 0.09 | 0 | 0.13 | 0.07 |
| 10 | 4891221 | rs2924264 | 1.074 | 1.589 | 0.078 | -1.311 | 0.018 | 0.06 | 0.06 | 0 | 0.35 | 0.14 | 0.2 | 0.22 | 0.27 | 0.2 |
| 10 | 4893318 | rs10795202 | -0.378 | 1.626 | -0.041 | -1.288 | 0.024 | 0.051 | 0.051 | 0 | 0.55 | 0.33 | 0.73 | 0.69 | 0.67 | 0.68 |
| 10 | 4895192 | rs4363494 | 0.775 | 1.669 | -0.19 | -1.244 | -0.101 | 0.018 | 0.018 | 0.076 | 0.36 | 0.1 | 0.6 | 0.69 | 0.47 | 0.59 |
| 10 | 4899130 | rs10795205 | -0.543 | 1.586 | 0.071 | -1.549 | 0.049 | 0.02 | 0.02 | 0.014 | 0.94 | 0.67 | 0.96 | 0.98 | 0.97 | 0.92 |
| 10 | 4901798 | rs4359096 | 0.878 | 1.617 | -0.082 | -1.606 | 0.009 | 0.025 | 0.025 | 0 | 0.36 | 0.24 | 0.21 | 0.18 | 0.27 | 0.2 |
| 10 | 4909968 | rs10795206 | 0.733 | 1.559 | -0.367 | -1.579 | -0.061 | 0.074 | 0.074 | 0.079 | 0.37 | 0.18 | 0.59 | 0.73 | 0.47 | 0.6 |
| 10 | 4918182 | rs7092190 | -0.598 | 1.436 | -0.124 | -1.462 | 0.014 | 0.006 | 0.006 | 0 | 0.59 | 0.42 | 0.8 | 0.86 | 0.75 | 0.74 |
| 10 | 4919641 | rs4460716 | -0.363 | 1.452 | -0.129 | -1.451 | 0.019 | 0.029 | 0.029 | 0 | 0.57 | 0.42 | 0.78 | 0.79 | 0.73 | 0.71 |
| 10 | 4926008 | rs7071257 | -0.405 | 1.443 | -0.06 | -1.456 | 0 | -0.012 | -0.012 | 0 | 0.57 | 0.47 | 0.79 | 0.79 | 0.73 | 0.71 |
| 10 | 4929456 | rs3924662 | 0.042 | 1.169 | -0.01 | -1.383 | 0.054 | 0.02 | 0.02 | 0.305 | 0.92 | 0.56 | 0.96 | 0.98 | 0.97 | 0.92 |
| 10 | 4929527 | rs3924661 | -0.272 | 1.2 | -0.091 | -1.382 | 0.037 | 0.068 | 0.068 | 0 | 0.56 | 0.36 | 0.78 | 0.79 | 0.73 | 0.71 |
| 10 | 4932085 | rs10904368 | 0.702 | 1.257 | -0.499 | -1.341 | 0.043 | 0.098 | 0.098 | 0 | 0.34 | 0.14 | 0.16 | 0.12 | 0.21 | 0.18 |
| 10 | 4935482 | rs9888112 | -0.292 | 1.406 | 0.182 | -0.659 | 0.015 | -0.04 | -0.04 | 0 | 0.19 | 0.43 | 0.27 | 0.12 | 0.21 | 0.17 |
| 10 | 4943518 | rs4457653 | 1.416 | 1.393 | 0.269 | 0.126 | 0.034 | 0.12 | 0.12 | 0.053 | 0.82 | 0.59 | 0.66 | 0.37 | 0.79 | 0.74 |
| 10 | 4944816 | rs7090237 | 1.061 | 1.363 | 0.124 | 0.169 | 0.035 | 0.18 | 0.18 | 0.03 | 0.85 | 0.65 | 0.69 | 0.43 | 0.82 | 0.78 |
| 10 | 4960864 | rs7358080 | 0.053 | 1.058 | -0.783 | -0.702 | -0.017 | 0.142 | 0.142 | 0 | 0.41 | 0.2 | 0.25 | 0.12 | 0.41 | 0.25 |
| 10 | 4963327 | rs12774134 | 1.219 | 1.068 | -0.778 | -0.849 | -0.001 | 0.006 | 0.006 | 0 | 0.12 | 0.09 | 0.15 | 0.08 | 0.13 | 0.19 |
| 10 | 4965434 | rs4332428 | 1.195 | 1.059 | -0.698 | -0.852 | 0 | 0.001 | 0.001 | 0 | 0.12 | 0.12 | 0.15 | 0.08 | 0.13 | 0.19 |
| 10 | 4973690 | rs7901384 | -0.446 | 1.001 | -0.427 | -0.489 | 0.001 | 0.043 | 0.043 | 0.015 | 0.54 | 0.34 | 0.41 | 0.31 | 0.54 | 0.44 |
| 10 | 4976701 | rs7921514 | -0.421 | 0.95 | -0.442 | -0.533 | 0 | 0.035 | 0.035 | 0.1 | 0.55 | 0.37 | 0.4 | 0.2 | 0.55 | 0.44 |
| 10 | 4981696 | rs7076643 | -1.07 | 0.685 | 0.223 | -0.439 | -0.003 | 0.004 | 0.004 | 0 | 0.64 | 0.67 | 0.74 | 0.8 | 0.63 | 0.58 |
| 10 | 4982478 | rs7094041 | 0.403 | 0.675 | 0.055 | -0.464 | -0.005 | 0.1 | 0.1 | 0 | 0.43 | 0.25 | 0.25 | 0.12 | 0.41 | 0.25 |
| 10 | 4986044 | rs4339935 | -0.001 | 0.73 | -0.089 | -0.902 | 0 | 0.063 | 0.063 | 0 | 0.49 | 0.33 | 0.38 | 0.31 | 0.5 | 0.38 |
| 10 | 4990299 | rs7093917 | -0.675 | 0.601 | -0.03 | -0.761 | 0 | -0.007 | -0.007 | 0 | 0.59 | 0.42 | 0.41 | 0.36 | 0.56 | 0.49 |
| 10 | 4990339 | rs4323780 | -0.644 | 0.702 | -0.234 | -0.918 | 0.003 | -0.029 | -0.029 | 0 | 0.58 | 0.48 | 0.71 | 0.75 | 0.58 | 0.53 |
| 10 | 5008383 | rs7908994 | 1.314 | 0.612 | -0.627 | -0.917 | -0.005 | 0.02 | 0.02 | 0 | 0.12 | 0.07 | 0.16 | 0.06 | 0.13 | 0.19 |
| 10 | 5009364 | rs2904802 | -1.1 | 0.772 | -0.417 | -1.119 | -0.006 | 0.045 | 0.045 | 0 | 0.65 | 0.73 | 0.76 | 0.79 | 0.66 | 0.58 |
| 10 | 5009449 | rs7076886 | -0.13 | 1.103 | -0.293 | -1.277 | 0.019 | -0.067 | -0.067 | 0 | 0.22 | 0.61 | 0.28 | 0.06 | 0.31 | 0.24 |
| 10 | 5009759 | rs2904804 | -0.755 | 0.995 | -0.494 | -1.287 | -0.006 | 0.081 | 0.081 | 0.011 | 0.58 | 0.69 | 0.74 | 0.8 | 0.61 | 0.52 |
| 10 | 5011460 | rs2961611 | 0.441 | 1.38 | -0.34 | -0.629 | -0.019 | -0.049 | -0.049 | 0 | 0.1 | 0.02 | 0.32 | 0.49 | 0.11 | 0.16 |
| 10 | 5019320 | rs6650153 | 0.884 | 1.382 | 0.142 | -1.247 | -0.001 | -0.002 | -0.002 | 0.21 | 0.12 | 0.12 | 0.16 | 0.06 | 0.13 | 0.19 |
| 10 | 5021218 | rs6601882 | -0.781 | 1.001 | -0.018 | -1.352 | 0.004 | -0.073 | -0.073 | 0 | 0.28 | 0.55 | 0.3 | 0.09 | 0.35 | 0.3 |
| 10 | 5024394 | rs11252861 | -0.604 | 0.503 | -0.297 | -1.555 | 0.021 | -0.103 | -0.103 | 0 | 0.34 | 0.5 | 0.4 | 0.08 | 0.45 | 0.38 |
| 10 | 5026096 | rs7909151 | -0.303 | -0.268 | -0.199 | -1.561 | 0.048 | -0.111 | -0.111 | 0 | 0.68 | 0.56 | 0.57 | 0.93 | 0.47 | 0.68 |
| 10 | 5027660 | rs7915338 | -0.449 | -0.419 | -0.618 | -1.623 | 0.005 | -0.074 | -0.074 | 0 | 0.39 | 0.45 | 0.47 | 0.09 | 0.61 | 0.45 |
| 10 | 5035246 | rs10904387 | 0.312 | -0.457 | -1.265 | -0.922 | 0.022 | 0.002 | 0.002 | 0 | 0.35 | 0.19 | 0.43 | 0.62 | 0.27 | 0.54 |
| 10 | 5051532 | rs1937863 | -0.137 | -0.479 | -1.407 | -0.169 | 0.05 | -0.007 | -0.007 | 0 | 0.72 | 0.84 | 0.87 | 0.69 | 0.83 | 0.88 |
| 10 | 5053831 | rs2398174 | -0.145 | -0.474 | -1.451 | -0.166 | 0.05 | -0.009 | -0.009 | 0.433 | 0.72 | 0.86 | 0.87 | 0.69 | 0.83 | 0.88 |
| 10 | 5063398 | rs12414884 | -0.118 | 0.13 | -1.163 | -0.341 | -0.016 | 0.028 | 0.028 | 0 | 0.38 | 0.26 | 0.59 | 0.19 | 0.49 | 0.3 |
| 10 | 5063944 | rs4143630 | -0.036 | 0.219 | -0.875 | -0.43 | 0.005 | -0.011 | -0.011 | 0 | 0.41 | 0.57 | 0.53 | 0.72 | 0.53 | 0.49 |
| 10 | 5065248 | rs11252887 | 0.894 | -0.132 | -0.919 | -0.196 | 0.011 | 0.031 | 0.031 | 0 | 0.26 | 0.32 | 0.25 | 0.46 | 0.3 | 0.36 |
| 10 | 5065747 | rs2895059 | 0.393 | 0.02 | -1.125 | -0.04 | -0.026 | -0.022 | -0.022 | 0.855 | 0.08 | 0.03 | 0.29 | 0.13 | 0.2 | 0.05 |
| 10 | 5068113 | rs10904392 | 0.072 | -0.891 | -0.955 | -0.14 | -0.001 | 0 | 0 | 0.011 | 0.41 | 0.56 | 0.32 | 0.58 | 0.38 | 0.53 |
| 10 | 5075366 | rs17133974 | 1.007 | -1.154 | -1.192 | -0.237 | -0.004 | -0.048 | -0.048 | 0 | 0.22 | 0.02 | 0.09 | 0.23 | 0.14 | 0.17 |
| 10 | 5075493 | rs12249281 | -0.387 | -0.991 | -1.165 | -0.305 | 0.005 | -0.021 | -0.021 | 0 | 0.57 | 0.51 | 0.4 | 0.79 | 0.49 | 0.56 |
| 10 | 5076063 | rs7096045 | 0.948 | -1.042 | -1.04 | -0.183 | 0.01 | 0.034 | 0.034 | 0 | 0.29 | 0.38 | 0.28 | 0.43 | 0.31 | 0.36 |
| 10 | 5083884 | rs2801904 | -0.263 | -0.779 | -0.971 | -0.001 | -0.006 | 0.006 | 0.006 | 0 | 0.61 | 0.62 | 0.4 | 0.79 | 0.5 | 0.56 |
| 10 | 5084728 | rs11252900 | 0.9 | -0.622 | -1.032 | 0.074 | -0.007 | -0.014 | -0.014 | 0 | 0.06 | 0.02 | 0.24 | 0.13 | 0.1 | 0.05 |
| 10 | 5097405 | rs2096421 | -0.806 | -0.836 | -0.863 | 0.051 | -0.042 | 0.04 | 0.04 | 0.244 | 0.64 | 0.79 | 0.39 | 0.79 | 0.48 | 0.65 |
| 10 | 5104290 | rs10795241 | 0.686 | -0.641 | -1.161 | 0.158 | -0.006 | 0.021 | 0.021 | 0 | 0.33 | 0.41 | 0.27 | 0.42 | 0.29 | 0.36 |
| 10 | 5104383 | rs1937837 | -0.043 | -0.74 | -1.612 | 0.315 | -0.01 | 0.012 | 0.012 | 0 | 0.06 | 0.07 | 0.02 | 0.15 | 0.04 | 0.03 |

| | | | | | | | | | | | | | | | | |
|---|---|---|---|---|---|---|---|---|---|---|---|---|---|---|---|---|
| 10 | 5104908 | rs17395698 | -0.051 | -0.732 | -1.642 | 0.301 | 0.014 | -0.014 | -0.014 | 0.683 | 0.22 | 0.06 | 0.1 | 0.23 | 0.15 | 0.26 |
| 10 | 5107795 | rs12570598 | -0.501 | -0.417 | -1.517 | 0.458 | 0.019 | -0.016 | -0.016 | 0 | 0.28 | 0.2 | 0.12 | 0.38 | 0.18 | 0.29 |
| 10 | 5115854 | rs7903936 | -0.628 | -0.109 | -2.085 | 0.151 | -0.012 | 0.005 | 0.005 | 0 | 0.25 | 0.03 | 0.13 | 0.19 | 0.22 | 0.3 |
| 10 | 5115907 | rs2105450 | -0.358 | -0.577 | -3.11 | 0.014 | 0.106 | -0.008 | -0.008 | 0 | 0.57 | 0.06 | 0.57 | 0.34 | 0.7 | 0.71 |
| 10 | 5117424 | rs7923027 | 0.118 | -0.058 | -1.867 | 0.021 | 0.003 | 0.004 | 0.004 | 0.186 | 0.39 | 0.36 | 0.28 | 0.31 | 0.32 | 0.25 |
| 10 | 5118845 | rs10752001 | -0.249 | 0.943 | -1.915 | 0.174 | -0.025 | -0.015 | -0.015 | 0.821 | 0.89 | 0.92 | 0.85 | 0.88 | 0.85 | 0.83 |
| 10 | 5119544 | rs7901781 | -0.241 | 0.927 | -1.927 | 0.172 | -0.029 | -0.017 | -0.017 | 0 | 0.89 | 0.93 | 0.85 | 0.88 | 0.85 | 0.82 |
| 10 | 5121745 | rs12570303 | 0.145 | 0.461 | -1.55 | 0.082 | 0.001 | 0.001 | 0.001 | 0.284 | 0.4 | 0.4 | 0.29 | 0.3 | 0.31 | 0.26 |
| 10 | 5123136 | rs6601900 | -0.042 | 0.621 | -1.031 | 0.15 | -0.005 | 0.01 | 0.01 | 0 | 0.52 | 0.67 | 0.54 | 0.57 | 0.54 | 0.58 |
| 10 | 5130610 | rs1937905 | 0.623 | 0.393 | -1.051 | -0.279 | 0.018 | -0.039 | -0.039 | 0 | 0.22 | 0.13 | 0.32 | 0.65 | 0.14 | 0.46 |
| 10 | 5134416 | rs11252934 | 0.72 | 0.171 | -0.391 | -0.587 | 0.033 | -0.092 | -0.092 | 0.003 | 0.2 | 0.34 | 0.2 | 0 | 0.37 | 0.15 |
| 10 | 5134873 | rs11252935 | 0.813 | 0.153 | -0.345 | -0.591 | 0.032 | -0.08 | -0.08 | 0 | 0.19 | 0.3 | 0.19 | 0 | 0.37 | 0.14 |
| 10 | 5139003 | rs2801883 | 0.356 | 0.284 | -0.024 | -0.835 | -0.003 | -0.05 | -0.05 | 0 | 0.59 | 0.47 | 0.55 | 0.86 | 0.4 | 0.65 |
| 10 | 5139685 | rs12387 | -0.754 | 0.366 | 0.025 | -0.812 | 0.006 | 0.003 | 0.003 | 0.081 | 0.89 | 0.87 | 0.81 | 0.86 | 0.85 | 0.82 |
| 10 | 5143902 | rs2154306 | 1.362 | 0.586 | 0.034 | -0.876 | 0.002 | 0.002 | 0.002 | 1.433 | 0.36 | 0.31 | 0.21 | 0.22 | 0.26 | 0.2 |
| 10 | 5144037 | rs4881400 | -0.023 | 0.524 | -0.005 | -0.803 | 0.02 | -0.006 | -0.006 | 1.44 | 0.33 | 0.2 | 0.42 | 0.67 | 0.23 | 0.47 |
| 10 | 5147909 | rs2275928 | 0.514 | 0.542 | 0.158 | -0.854 | -0.001 | 0.015 | 0.015 | 0 | 0.56 | 0.62 | 0.41 | 0.22 | 0.63 | 0.36 |
| 10 | 5148387 | rs4512728 | 0.968 | 0.528 | 0.158 | -0.859 | 0.008 | -0.015 | -0.015 | 0.122 | 0.23 | 0.17 | 0.33 | 0.65 | 0.14 | 0.46 |
| 10 | 5149703 | rs3209896 | 0.497 | 0.517 | 0.156 | -0.893 | -0.001 | 0.015 | 0.015 | 0 | 0.56 | 0.62 | 0.41 | 0.22 | 0.63 | 0.34 |
| 10 | 5159453 | rs7921327 | -0.008 | 0.493 | 0.017 | -0.644 | -0.001 | 0.011 | 0.011 | 0.214 | 0.66 | 0.7 | 0.48 | 0.23 | 0.71 | 0.38 |
| 10 | 5161955 | rs1937920 | 1.144 | 0.578 | 0.353 | -0.715 | -0.022 | -0.028 | -0.028 | 0 | 0.37 | 0.41 | 0.22 | 0.23 | 0.26 | 0.22 |
| 10 | 5165820 | rs7086693 | 0.017 | 0.67 | 0.291 | -0.692 | 0.023 | -0.131 | -0.131 | 0.116 | 0.31 | 0.43 | 0.26 | 0 | 0.44 | 0.17 |
| 10 | 5172344 | rs2895064 | 0.981 | 0.651 | 0.473 | -0.697 | -0.019 | -0.003 | -0.003 | 0.099 | 0.19 | 0.17 | 0.18 | 0 | 0.36 | 0.15 |
| 10 | 5176727 | rs2186174 | 1.299 | 0.506 | 0.507 | -0.744 | 0.012 | 0.015 | 0.015 | 1.675 | 0.35 | 0.27 | 0.21 | 0.23 | 0.26 | 0.22 |
| 10 | 5176783 | rs2398203 | 1.451 | 0.484 | 0.529 | -0.754 | 0.034 | 0.049 | 0.049 | 2.526 | 0.34 | 0.14 | 0.2 | 0.23 | 0.26 | 0.21 |
| 10 | 5185582 | rs969819 | 0.912 | 0.513 | 0.608 | -0.714 | 0.008 | -0.015 | -0.015 | 0 | 0.24 | 0.17 | 0.33 | 0.64 | 0.15 | 0.45 |
| 10 | 5196273 | rs1781930 | -0.946 | 0.637 | 0.233 | -0.687 | -0.051 | -0.041 | -0.041 | 0.176 | 0.91 | 0.98 | 0.81 | 0.86 | 0.85 | 0.86 |
| 10 | 5197177 | rs17305952 | 1.1 | 0.605 | 0.26 | -0.677 | -0.016 | -0.007 | -0.007 | 0 | 0.19 | 0.19 | 0.18 | 0 | 0.36 | 0.15 |
| 10 | 5197881 | rs1781931 | 1.326 | 0.581 | 0.231 | -0.683 | 0.038 | 0.06 | 0.06 | 2.277 | 0.35 | 0.14 | 0.2 | 0.22 | 0.26 | 0.21 |
| 10 | 5201444 | rs1781938 | 0.271 | 1.598 | 0.385 | -0.676 | 0.018 | -0.01 | -0.01 | 0 | 0.8 | 0.73 | 0.73 | 0.86 | 0.77 | 0.82 |
| 10 | 5202925 | rs7067921 | 1.092 | 1.685 | -0.129 | -0.517 | -0.033 | -0.036 | -0.036 | 0 | 0.06 | 0 | 0.17 | 0.14 | 0.09 | 0.13 |
| 10 | 5203579 | rs11253020 | 0.602 | 1.501 | -0.004 | -0.637 | -0.015 | -0.01 | -0.01 | 0 | 0.19 | 0.21 | 0.19 | 0 | 0.36 | 0.15 |
| 10 | 5203864 | rs11253021 | 0.602 | 1.501 | -0.004 | -0.637 | -0.015 | -0.01 | -0.01 | 0 | 0.19 | 0.21 | 0.18 | 0 | 0.36 | 0.15 |
| 10 | 5227382 | rs2895055 | -0.699 | 1.132 | 0.186 | -0.588 | 0.005 | -0.064 | -0.064 | 0.018 | 0.39 | 0.51 | 0.46 | 0.1 | 0.58 | 0.33 |
| 10 | 5229002 | rs12269559 | -0.682 | 1.116 | 0.219 | -0.594 | 0.007 | -0.077 | -0.077 | 0.027 | 0.39 | 0.47 | 0.46 | 0.1 | 0.58 | 0.33 |
| 10 | 5229091 | rs11253038 | -0.71 | 1.068 | 0.207 | -0.615 | 0.006 | -0.078 | -0.078 | 0 | 0.39 | 0.47 | 0.46 | 0.1 | 0.58 | 0.33 |
| 10 | 5238196 | rs4880716 | 0.082 | 1.033 | 0.257 | -0.843 | 0.031 | -0.015 | -0.015 | 0 | 0.82 | 0.68 | 0.73 | 0.87 | 0.78 | 0.79 |
| 10 | 5238283 | rs3812617 | 0.73 | 1.107 | 0.098 | -0.824 | -0.043 | -0.031 | -0.031 | 0 | 0.08 | 0.02 | 0.2 | 0.12 | 0.13 | 0.11 |
| 10 | 5242637 | rs7101071 | -0.18 | 1.071 | 0.309 | -0.847 | -0.034 | 0.011 | 0.011 | 0.003 | 0.17 | 0.09 | 0.24 | 0.13 | 0.22 | 0.18 |
| 10 | 5243017 | rs7085249 | -0.18 | 1.071 | 0.307 | -0.847 | -0.036 | 0.011 | 0.011 | 0.003 | 0.17 | 0.08 | 0.24 | 0.13 | 0.22 | 0.18 |
| 10 | 5244295 | rs7083869 | 1.336 | 0.406 | 0.33 | -0.867 | 0.011 | 0.003 | 0.003 | 0 | 0.27 | 0.23 | 0.16 | 0.25 | 0.19 | 0.17 |
| 10 | 5246584 | rs11253042 | 0.622 | 0.936 | 0.004 | -1.325 | -0.021 | 0.114 | 0.114 | 0 | 0.46 | 0.14 | 0.32 | 0.26 | 0.52 | 0.28 |
| 10 | 5249213 | rs1889457 | -0.426 | 0.892 | 0.337 | -1.368 | -0.019 | -0.111 | -0.111 | 0 | 0.82 | 0.47 | 0.89 | 0.99 | 0.89 | 0.88 |
| 10 | 5250453 | rs1931679 | 2.025 | 1.093 | 0.22 | -1.294 | -0.02 | -0.023 | -0.023 | 0 | 0.05 | 0.01 | 0.17 | 0.11 | 0.07 | 0.1 |
| 10 | 5254804 | rs11253045 | 2.183 | 1.017 | 0.048 | -1.492 | -0.048 | 0.066 | 0.066 | 0 | 0.14 | 0.02 | 0.13 | 0 | 0.25 | 0.07 |
| 10 | 5257342 | rs11253046 | 0.633 | 0.114 | 0.211 | -1.396 | -0.037 | 0.037 | 0.037 | 0 | 0.32 | 0.21 | 0.26 | 0 | 0.49 | 0.21 |
| 10 | 5258571 | rs2050308 | 2.36 | -0.014 | 0.096 | -1.348 | 0 | -0.077 | -0.077 | 0.95 | 0.18 | 0.03 | 0.11 | 0.26 | 0.16 | 0.16 |
| 10 | 5261159 | rs11253048 | 0.122 | 0.13 | 0.534 | -1.056 | -0.011 | 0.001 | 0.001 | 0 | 0.14 | 0.12 | 0.25 | 0.11 | 0.14 | 0.13 |
| 10 | 5261703 | rs12775790 | 0.117 | -0.144 | 0.594 | -1.057 | 0.05 | 0.017 | 0.017 | 0 | 0.56 | 0.35 | 0.57 | 0.38 | 0.75 | 0.44 |
| 10 | 5262361 | rs10904442 | 0.988 | -0.097 | 0.292 | -0.997 | 0.035 | -0.056 | -0.056 | 0 | 0.23 | 0.1 | 0.31 | 0.61 | 0.12 | 0.41 |
| 10 | 5273767 | rs2104994 | 1.701 | -0.353 | 0.03 | -0.722 | 0.011 | -0.029 | -0.029 | 0 | 0.1 | 0.04 | 0.23 | 0.18 | 0.06 | 0.07 |
| 10 | 5275078 | rs11253059 | 0.132 | -0.235 | 0.273 | -0.731 | -0.026 | -0.097 | -0.097 | 0 | 0.18 | 0.34 | 0.09 | 0.01 | 0.11 | 0.12 |
| 10 | 5275762 | rs10904445 | 0.717 | -0.006 | -0.175 | -0.308 | 0.01 | -0.016 | -0.016 | 0 | 0.15 | 0.09 | 0.09 | 0.42 | 0.08 | 0.37 |
| 10 | 5286162 | rs9423393 | 0.592 | 0.345 | 0.145 | -0.616 | 0.035 | -0.063 | -0.063 | 0 | 0.71 | 0.46 | 0.64 | 0.88 | 0.64 | 0.69 |
| 10 | 5286600 | rs7895816 | 0.349 | 0.409 | 0.356 | -0.653 | 0 | -0.022 | -0.022 | 0.43 | 0.11 | 0.12 | 0.09 | 0 | 0.16 | 0.1 |
| 10 | 5294191 | rs2398152 | -0.716 | 0.418 | 0.681 | -0.682 | 0.017 | -0.128 | -0.128 | 0 | 0.22 | 0.39 | 0.15 | 0.01 | 0.24 | 0.2 |
| 10 | 5300727 | rs35814115 | 1.109 | 0.387 | 0.777 | -0.652 | 0.002 | -0.047 | -0.047 | 0 | 0.1 | 0.17 | 0.05 | 0.01 | 0.09 | 0.1 |
| 10 | 5300976 | rs11597152 | 1.253 | 0.385 | 0.776 | -0.653 | 0 | -0.002 | -0.002 | 0.472 | 0.09 | 0.08 | 0.04 | 0.01 | 0.09 | 0.04 |
| 10 | 5301811 | rs9423582 | 0.549 | 0.185 | 0.341 | -0.68 | 0.001 | -0.027 | -0.027 | 0 | 0.68 | 0.47 | 0.62 | 0.81 | 0.7 | 0.75 |
| 10 | 5302518 | rs12266035 | 0.189 | 0.183 | 0.376 | -0.692 | -0.017 | -0.094 | -0.094 | 14.219 | 0.16 | 0.29 | 0.08 | 0.01 | 0.11 | 0.12 |
| 10 | 5303742 | rs9423584 | -0.674 | 0.219 | 0.331 | -0.637 | 0.013 | 0.012 | 0.012 | 0 | 0.85 | 0.77 | 0.71 | 0.82 | 0.81 | 0.87 |
| 10 | 5307659 | rs2151898 | 0.946 | 0.017 | -0.173 | -0.637 | 0.005 | -0.006 | -0.006 | 0 | 0.61 | 0.45 | 0.43 | 0.7 | 0.58 | 0.63 |
| 10 | 5311075 | rs9423399 | 1.313 | -0.091 | -0.413 | -0.897 | -0.01 | 0.009 | 0.009 | 0 | 0.27 | 0.19 | 0.2 | 0 | 0.33 | 0.09 |
| 10 | 5313029 | rs869009 | 0.284 | 0.052 | -0.408 | -0.809 | 0.013 | -0.001 | -0.001 | 0 | 0.14 | 0.19 | 0.25 | 0.11 | 0.26 | 0.24 |
| 10 | 5313995 | rs1889460 | 0.284 | 0.052 | -0.408 | -0.809 | 0.013 | -0.001 | -0.001 | 0.695 | 0.14 | 0.19 | 0.25 | 0.11 | 0.26 | 0.24 |
| 10 | 5317870 | rs7082291 | 0.552 | 0.038 | -0.459 | -0.815 | -0.031 | 0.003 | 0.003 | 0.207 | 0.12 | 0.11 | 0.25 | 0.11 | 0.26 | 0.24 |
| 10 | 5320019 | rs11594521 | 1.614 | -0.053 | -0.433 | -0.847 | -0.001 | -0.001 | -0.001 | 0 | 0.09 | 0.07 | 0.04 | 0.01 | 0.09 | 0.04 |
| 10 | 5321569 | rs1931669 | -1.817 | -0.277 | -0.351 | -0.775 | -0.011 | 0.026 | 0.026 | 0 | 0.89 | 0.86 | 0.76 | 0.82 | 0.94 | 0.93 |
| 10 | 5324089 | rs9423588 | 1.377 | -0.341 | -0.369 | -0.842 | -0.016 | 0.049 | 0.049 | 0 | 0.26 | 0.11 | 0.19 | 0 | 0.32 | 0.09 |

| | | | | | | | | | | | | | | | | |
|---|---|---|---|---|---|---|---|---|---|---|---|---|---|---|---|---|
| 10 | 5325029 | rs10904449 | 1.031 | -0.555 | -0.405 | -0.589 | 0.008 | -0.01 | -0.01 | 0.291 | 0.16 | 0.1 | 0.05 | 0.41 | 0.08 | 0.38 |
| 10 | 5325154 | rs7097810 | 0.375 | -0.527 | -0.198 | -0.547 | 0.012 | -0.003 | -0.003 | 0 | 0.66 | 0.57 | 0.46 | 0.7 | 0.59 | 0.65 |
| 10 | 5326846 | rs7906894 | -0.094 | -0.504 | -0.474 | -0.579 | 0.006 | -0.01 | -0.01 | 0.06 | 0.25 | 0.26 | 0.31 | 0.12 | 0.37 | 0.31 |
| 10 | 5327477 | rs17155745 | 1.489 | -0.277 | -0.622 | -0.404 | -0.006 | 0.043 | 0.043 | 0 | 0.06 | 0.09 | 0.22 | 0.18 | 0.03 | 0.04 |
| 10 | 5327676 | rs9423591 | 0.703 | -0.293 | -0.878 | -0.239 | -0.031 | 0.072 | 0.072 | 0 | 0.32 | 0.13 | 0.22 | 0.19 | 0.38 | 0.2 |
| 10 | 5333008 | rs7076488 | -1.118 | -0.159 | -0.943 | 0.224 | 0 | 0.034 | 0.034 | 0 | 0.6 | 0.73 | 0.53 | 0.75 | 0.6 | 0.79 |
| 10 | 5333043 | rs7081029 | 0.471 | -0.354 | -0.643 | 0.629 | 0.012 | 0.005 | 0.005 | 0.413 | 0.31 | 0.31 | 0.17 | 0.33 | 0.26 | 0.25 |
| 10 | 5337743 | rs17134635 | 0.628 | -0.075 | -0.327 | 0.585 | 0.002 | -0.001 | -0.001 | 0 | 0.32 | 0.26 | 0.22 | 0.71 | 0.23 | 0.49 |
| 10 | 5339074 | rs1344354 | 0.493 | 0.052 | -1.24 | 0.676 | -0.001 | -0.001 | -0.001 | 0 | 0.34 | 0.32 | 0.19 | 0.63 | 0.22 | 0.47 |
| 10 | 5348337 | rs1987511 | -0.679 | 0.267 | -1.307 | 0.571 | 0.004 | 0.003 | 0.003 | 0.451 | 0.5 | 0.5 | 0.43 | 0.76 | 0.38 | 0.71 |
| 10 | 5349871 | rs7475343 | 0.643 | 0.274 | -1.406 | 0.613 | -0.01 | -0.008 | -0.008 | 0 | 0.11 | 0.09 | 0.23 | 0.11 | 0.14 | 0.2 |
| 10 | 5354677 | rs7896729 | -0.338 | -0.368 | -1.281 | 0.71 | -0.007 | 0 | 0 | 0 | 0.39 | 0.39 | 0.2 | 0.63 | 0.24 | 0.51 |
| 10 | 5355712 | rs9423407 | -0.61 | 0.66 | -1.492 | 0.398 | 0 | -0.002 | -0.002 | 0 | 0.43 | 0.37 | 0.28 | 0.5 | 0.44 | 0.51 |
| 10 | 5356862 | rs7894961 | -0.38 | -0.331 | -0.906 | 0.807 | -0.021 | 0.019 | 0.019 | 0.006 | 0.33 | 0.37 | 0.2 | 0.55 | 0.22 | 0.45 |
| 10 | 5365630 | rs7921890 | 0.117 | 0.627 | -1.312 | 0.441 | -0.012 | 0.015 | 0.015 | 0.774 | 0.42 | 0.23 | 0.64 | 0.53 | 0.46 | 0.35 |
| 10 | 5365776 | rs10159966 | 0.199 | 0.595 | -0.821 | -0.163 | -0.003 | 0.001 | 0.001 | 0.205 | 0.31 | 0.3 | 0.17 | 0.42 | 0.21 | 0.45 |
| 10 | 5368972 | rs4881433 | 0.93 | 0.312 | -0.886 | -0.2 | -0.005 | -0.017 | -0.017 | 0 | 0.16 | 0.1 | 0.13 | 0.27 | 0.16 | 0.19 |
| 10 | 5370986 | rs7477268 | 0.634 | 0.138 | -0.84 | -0.253 | -0.001 | 0.028 | 0.028 | 1.047 | 0.52 | 0.38 | 0.67 | 0.58 | 0.5 | 0.59 |
| 10 | 5375877 | rs6601945 | -0.625 | -0.26 | -1.039 | 0.116 | -0.05 | 0.057 | 0.057 | 0 | 0.28 | 0.49 | 0.2 | 0.4 | 0.16 | 0.38 |
| 10 | 5379362 | rs11253104 | -0.653 | -0.049 | -1.064 | 0.078 | -0.004 | 0.002 | 0.002 | 0 | 0.37 | 0.59 | 0.32 | 0.6 | 0.3 | 0.53 |
| 10 | 5380664 | rs11253111 | -1.298 | 0.225 | -1.355 | -0.186 | 0.002 | 0.041 | 0.041 | 0 | 0.52 | 0.66 | 0.49 | 0.7 | 0.46 | 0.64 |
| 10 | 5382676 | rs12570234 | 1.02 | 0.055 | -1.273 | -0.005 | -0.001 | 0.044 | 0.044 | 0 | 0.11 | 0.18 | 0.16 | 0.28 | 0.12 | 0.08 |
| 10 | 5383888 | rs12098412 | -0.112 | 0.01 | -1.213 | -0.114 | 0.019 | -0.016 | -0.016 | 0 | 0.33 | 0.27 | 0.28 | 0.64 | 0.26 | 0.29 |
| 10 | 5385029 | rs10904463 | -0.413 | 0.117 | -1.259 | -0.246 | 0 | -0.03 | -0.03 | 1.874 | 0.66 | 0.38 | 0.8 | 0.92 | 0.66 | 0.57 |
| 10 | 5385336 | rs11596042 | 1.015 | 0.462 | -1.471 | -0.087 | -0.054 | -0.02 | -0.02 | 0 | 0.33 | 0.09 | 0.4 | 0.27 | 0.38 | 0.28 |
| 10 | 5386067 | rs10904468 | 0.936 | 0.773 | -1.286 | -0.182 | -0.022 | -0.017 | -0.017 | 0 | 0.4 | 0.21 | 0.55 | 0.4 | 0.45 | 0.35 |
| 10 | 5388446 | rs10904469 | 0.215 | 0.707 | -1.261 | -0.239 | 0.006 | 0 | 0 | 0 | 0.49 | 0.38 | 0.58 | 0.5 | 0.46 | 0.38 |
| 10 | 5389209 | rs10450518 | 0.611 | 1.243 | -1.387 | -0.244 | -0.008 | -0.003 | -0.003 | 0 | 0.41 | 0.31 | 0.56 | 0.39 | 0.45 | 0.35 |
| 10 | 5390759 | rs10752010 | -0.148 | 1.497 | -1.475 | -0.456 | -0.019 | 0.021 | 0.021 | 0 | 0.85 | 0.87 | 0.88 | 0.97 | 0.76 | 0.91 |
| 10 | 5394166 | rs12770596 | -0.726 | 1.031 | -1.454 | -0.121 | -0.024 | 0.044 | 0.044 | 0.264 | 0.23 | 0.1 | 0.28 | 0.15 | 0.28 | 0.25 |
| 10 | 5395440 | rs11253122 | -1.161 | 0.328 | -1.372 | -0.123 | 0.002 | 0.016 | 0.016 | 0.258 | 0.27 | 0.77 | 0.2 | 0.19 | 0.28 | 0.25 |
| 10 | 5396328 | rs9423601 | -0.265 | -0.112 | -1.927 | -0.012 | 0.039 | 0.011 | 0.011 | 0.022 | 0.52 | 0.27 | 0.57 | 0.54 | 0.38 | 0.6 |
| 10 | 5401308 | rs7081234 | -0.373 | -0.293 | -1.366 | 0.101 | -0.021 | 0.034 | 0.034 | 0.419 | 0.55 | 0.69 | 0.66 | 0.44 | 0.51 | 0.62 |
| 10 | 5401566 | rs6601951 | -0.261 | -0.015 | -1.225 | 0.215 | 0.054 | -0.022 | -0.022 | 0 | 0.15 | 0.52 | 0.15 | 0.13 | 0.21 | 0.13 |
| 10 | 5404261 | rs12784234 | -0.021 | -0.023 | -0.893 | 0.204 | 0.075 | -0.053 | -0.053 | 0 | 0.14 | 0.39 | 0.15 | 0.03 | 0.2 | 0.1 |
| 10 | 5404347 | rs10904478 | -1.296 | -0.336 | -1.044 | 0.096 | -0.024 | -0.039 | -0.039 | 0 | 0.64 | 0.73 | 0.71 | 0.56 | 0.58 | 0.71 |
| 10 | 5405265 | rs12783734 | -0.005 | -0.103 | -0.834 | 0.21 | 0.071 | -0.051 | -0.051 | 0 | 0.14 | 0.39 | 0.15 | 0.03 | 0.2 | 0.1 |
| 10 | 5405315 | rs10904479 | 0.521 | -0.269 | -1.204 | 0.296 | -0.011 | 0.033 | 0.033 | 3.273 | 0.33 | 0.12 | 0.48 | 0.3 | 0.29 | 0.47 |
| 10 | 5408461 | rs10795268 | 0.464 | -0.073 | -1.003 | 0.441 | 0.017 | 0.04 | 0.04 | 2.13 | 0.35 | 0.17 | 0.5 | 0.31 | 0.28 | 0.49 |
| 10 | 5413634 | rs6601952 | -0.362 | -0.067 | -1.06 | 0.605 | 0.018 | -0.008 | -0.008 | 0 | 0.45 | 0.38 | 0.55 | 0.58 | 0.32 | 0.58 |
| 10 | 5415954 | rs10904481 | -0.303 | 0.001 | -1.243 | 0.549 | 0.033 | -0.007 | -0.007 | 0 | 0.44 | 0.33 | 0.55 | 0.55 | 0.31 | 0.56 |
| 10 | 5421135 | rs10795269 | -0.531 | -0.286 | -1.164 | 0.64 | 0.006 | 0.009 | 0.009 | 1.351 | 0.49 | 0.35 | 0.6 | 0.63 | 0.44 | 0.59 |
| 10 | 5422196 | rs11253141 | 0.032 | 0.285 | -1.135 | 0.599 | -0.021 | -0.003 | -0.003 | 0 | 0.38 | 0.26 | 0.55 | 0.48 | 0.4 | 0.46 |
| 10 | 5424341 | rs7069716 | -0.578 | 0.018 | -0.51 | 0.752 | -0.023 | 0.042 | 0.042 | 0 | 0.18 | 0.36 | 0.14 | 0.2 | 0.16 | 0.17 |
| 10 | 5424465 | rs10904487 | 1.018 | 0.856 | -0.986 | 0.121 | 0.03 | -0.112 | -0.112 | 0 | 0.72 | 0.61 | 0.78 | 0.97 | 0.63 | 0.78 |
| 10 | 5424727 | rs11253143 | 0.854 | 0.59 | -1.334 | 0.402 | 0.017 | -0.009 | -0.009 | 0.008 | 0.45 | 0.34 | 0.29 | 0.42 | 0.37 | 0.35 |
| 10 | 5432213 | rs7477986 | 0.471 | 0.681 | -0.459 | 0.637 | 0.007 | 0.007 | 0.007 | 0.007 | 0.11 | 0.16 | 0.12 | 0.1 | 0.16 | 0.12 |
| 10 | 5432565 | rs6422396 | 1.332 | 0.615 | -0.755 | 0.977 | -0.005 | 0.011 | 0.011 | 0 | 0.61 | 0.48 | 0.75 | 0.53 | 0.65 | 0.49 |
| 10 | 5433973 | rs12258526 | 0.696 | 0.42 | -0.763 | 0.68 | 0 | -0.011 | -0.011 | 0 | 0.06 | 0.06 | 0.03 | 0 | 0.03 | 0.08 |
| 10 | 5435918 | rs7097775 | 1.056 | 0.429 | -0.674 | 0.778 | 0.003 | 0.002 | 0.002 | 0.011 | 0.63 | 0.74 | 0.77 | 0.65 | 0.66 | 0.53 |
| 10 | 5436800 | rs7477292 | 0.507 | 0.376 | -0.731 | 0.719 | 0.018 | 0.03 | 0.03 | 0 | 0.7 | 0.82 | 0.82 | 0.83 | 0.74 | 0.77 |
| 10 | 5437365 | rs11253156 | 0.514 | 0.344 | -0.705 | 0.714 | 0.011 | 0.03 | 0.03 | 0 | 0.7 | 0.82 | 0.81 | 0.83 | 0.72 | 0.76 |
| 10 | 5441973 | rs10795273 | 1.213 | 0.369 | -0.14 | 0.774 | 0 | -0.002 | -0.002 | 0 | 0.64 | 0.66 | 0.77 | 0.75 | 0.65 | 0.55 |
| 10 | 5446794 | rs7477466 | 0.605 | 0.365 | -0.128 | 0.75 | 0 | -0.003 | -0.003 | 0 | 0.69 | 0.73 | 0.8 | 0.83 | 0.72 | 0.74 |
| 10 | 5456840 | rs3923348 | 0.844 | 0.364 | -0.086 | 0.755 | -0.01 | -0.005 | -0.005 | 0 | 0.7 | 0.65 | 0.8 | 0.72 | 0.72 | 0.75 |
| 10 | 5495157 | rs11253185 | -1.002 | 0.338 | -0.107 | 0.754 | 0.005 | 0.025 | 0.025 | 0 | 0.28 | 0.17 | 0.19 | 0.17 | 0.28 | 0.23 |
| 10 | 5496697 | rs10904506 | 1.125 | 0.169 | -0.204 | 0.691 | 0 | -0.002 | -0.002 | 0 | 0.65 | 0.64 | 0.77 | 0.65 | 0.65 | 0.56 |
| 10 | 5498248 | rs4242790 | 1.187 | 0.125 | -0.023 | 0.672 | -0.002 | -0.005 | -0.005 | 0 | 0.64 | 0.56 | 0.76 | 0.65 | 0.65 | 0.56 |
| 10 | 5499173 | rs11791 | 1.116 | 0.132 | -0.064 | 0.675 | 0 | -0.002 | -0.002 | 1.586 | 0.65 | 0.64 | 0.77 | 0.65 | 0.65 | 0.57 |
| 10 | 5503610 | rs10904508 | 1.021 | -0.04 | -0.469 | 0.611 | -0.094 | -0.034 | -0.034 | 1.122 | 0.23 | 0.04 | 0.43 | 0.29 | 0.33 | 0.24 |
| 10 | 5509387 | rs10737000 | -0.105 | -0.265 | -0.386 | 0.708 | 0.118 | 0.184 | 0.184 | 0 | 0.88 | 0.59 | 0.86 | 0.77 | 0.79 | 0.89 |
| 10 | 5511869 | rs10904511 | 1.062 | -0.448 | -0.644 | 0.428 | 0.01 | 0.005 | 0.005 | 0 | 0.68 | 0.59 | 0.79 | 0.65 | 0.66 | 0.61 |
| 10 | 5522332 | rs9423430 | -0.602 | -0.394 | -0.495 | 0.186 | 0.006 | -0.02 | -0.02 | 0 | 0.18 | 0.2 | 0.16 | 0.12 | 0.23 | 0.13 |
| 10 | 5524502 | rs7893671 | -1.037 | -0.924 | -0.694 | -0.047 | -0.003 | -0.02 | -0.02 | 0.915 | 0.24 | 0.26 | 0.17 | 0.13 | 0.25 | 0.15 |
| 10 | 5524609 | rs7913214 | -0.426 | -0.725 | -0.554 | -0.033 | -0.01 | -0.024 | -0.024 | 0 | 0.85 | 0.77 | 0.95 | 0.89 | 0.91 | 0.73 |
| 10 | 5526375 | rs11595157 | 0.67 | -0.803 | -0.938 | -0.24 | -0.007 | -0.008 | -0.008 | 0 | 0.18 | 0.34 | 0.15 | 0.13 | 0.16 | 0.18 |
| 10 | 5527716 | rs11253199 | -0.026 | -0.822 | -0.965 | -0.248 | -0.022 | -0.011 | -0.011 | 2.763 | 0.21 | 0.46 | 0.15 | 0.16 | 0.16 | 0.22 |
| 10 | 5529329 | rs11592540 | 0.337 | -0.628 | -1.021 | -0.21 | -0.026 | 0.108 | 0.108 | 2.373 | 0.26 | 0.07 | 0.14 | 0.06 | 0.24 | 0.35 |
| 10 | 5531392 | rs9423638 | 0.928 | -0.26 | -0.742 | 0.05 | 0.03 | 0.022 | 0.022 | 0 | 0.11 | 0.2 | 0.11 | 0.09 | 0.16 | 0.11 |

| | | | | | | | | | | | | | | | | |
|---|---|---|---|---|---|---|---|---|---|---|---|---|---|---|---|---|
| 10 | 5537569 | rs4471334 | -1.579 | -0.472 | -1.111 | 0.043 | -0.009 | 0.029 | 0.029 | 0 | 0.54 | 0.63 | 0.33 | 0.31 | 0.49 | 0.57 |
| 10 | 5541183 | rs10904517 | -0.438 | 0.504 | -1.548 | 0.027 | 0 | -0.002 | -0.002 | 0 | 0.35 | 0.52 | 0.23 | 0.31 | 0.32 | 0.31 |
| 10 | 5542109 | rs4881466 | -1.791 | 0.377 | -1.846 | 0.427 | 0.012 | -0.01 | -0.01 | 0.325 | 0.74 | 0.8 | 0.7 | 0.68 | 0.73 | 0.82 |
| 10 | 5543835 | rs4078294 | 1.086 | 0.442 | -1.831 | -0.078 | -0.055 | -0.028 | -0.028 | 0 | 0.37 | 0.14 | 0.63 | 0.63 | 0.47 | 0.34 |
| 10 | 5544093 | rs4075060 | -0.29 | 0.138 | -1.833 | -0.153 | 0.003 | -0.004 | -0.004 | 0 | 0.33 | 0.69 | 0.23 | 0.32 | 0.28 | 0.3 |
| 10 | 5544190 | rs4078293 | 1.004 | 0.618 | -1.8 | 0.095 | -0.051 | -0.045 | -0.045 | 0 | 0.35 | 0.12 | 0.61 | 0.63 | 0.44 | 0.34 |
| 10 | 5547309 | rs4881467 | 0.091 | 0.711 | -1.142 | 0.457 | -0.001 | -0.003 | -0.003 | 0 | 0.71 | 0.39 | 0.8 | 0.78 | 0.72 | 0.77 |
| 10 | 5550673 | rs12357686 | 0.684 | 1.405 | -1.441 | 0.748 | -0.046 | -0.038 | -0.038 | 0.035 | 0.15 | 0.06 | 0.34 | 0.29 | 0.22 | 0.19 |
| 10 | 5551181 | rs7919331 | -0.017 | 0.642 | -1.424 | 0.497 | -0.006 | 0.008 | 0.008 | 0.033 | 0.52 | 0.44 | 0.3 | 0.46 | 0.46 | 0.55 |
| 10 | 5551850 | rs11596364 | 1.028 | 0.241 | -1.26 | 0.248 | -0.001 | -0.003 | -0.003 | 0.192 | 0.12 | 0.13 | 0.07 | 0.08 | 0.1 | 0.13 |
| 10 | 5552475 | rs1133654 | -0.384 | 0.482 | -1.15 | 0.352 | -0.01 | -0.003 | -0.003 | 0 | 0.65 | 0.5 | 0.57 | 0.7 | 0.63 | 0.74 |
| 10 | 5553986 | rs4462243 | 0.619 | 0.325 | -1.29 | 0.214 | 0.012 | -0.018 | -0.018 | 0 | 0.44 | 0.25 | 0.39 | 0.48 | 0.42 | 0.5 |
| 10 | 5554141 | rs9423641 | 0.041 | 0.29 | -0.984 | 0.155 | 0.012 | 0.005 | 0.005 | 0 | 0.65 | 0.46 | 0.48 | 0.59 | 0.58 | 0.7 |
| 10 | 5554958 | rs7474671 | 0.209 | -0.024 | -0.885 | 0.21 | 0.001 | -0.001 | -0.001 | 1.096 | 0.21 | 0.2 | 0.13 | 0.23 | 0.19 | 0.26 |
| 10 | 5556594 | rs11595171 | 1.464 | 0.052 | -0.951 | 0.266 | -0.016 | -0.035 | -0.035 | 0 | 0.1 | 0.03 | 0.1 | 0.14 | 0.14 | 0.13 |
| 10 | 5558349 | rs4072067 | -0.313 | 0.116 | -0.608 | -0.057 | 0.001 | -0.029 | -0.029 | 0 | 0.85 | 0.83 | 0.88 | 0.97 | 0.8 | 0.9 |
| 10 | 5560844 | rs10904522 | 0.4 | 0.234 | -0.615 | -0.037 | 0 | 0.001 | 0.001 | 0 | 0.57 | 0.58 | 0.73 | 0.68 | 0.56 | 0.62 |
| 10 | 5565053 | rs4072069 | -0.504 | 0.6 | -0.092 | -0.043 | 0.005 | 0.003 | 0.003 | 1.126 | 0.23 | 0.48 | 0.25 | 0.21 | 0.2 | 0.19 |
| 10 | 5566269 | rs2231413 | -0.146 | 0.757 | -0.14 | -0.211 | 0.023 | 0.009 | 0.009 | 0 | 0.65 | 0.48 | 0.5 | 0.43 | 0.56 | 0.64 |
| 10 | 5567366 | rs1142825 | 0.032 | 1.074 | 0.334 | -0.145 | 0.017 | 0.06 | 0.06 | 0 | 0.23 | 0.51 | 0.32 | 0.61 | 0.25 | 0.35 |
| 10 | 5567695 | rs4242797 | -0.199 | 1.145 | 0.293 | -0.535 | 0.083 | 0.008 | 0.008 | 0 | 0.94 | 0.67 | 0.95 | 1 | 0.95 | 0.98 |
| 10 | 5567914 | rs4589188 | 0.512 | 0.601 | -0.648 | -0.819 | 0.005 | 0.014 | 0.014 | 0.96 | 0.37 | 0.3 | 0.22 | 0.12 | 0.34 | 0.4 |
| 10 | 5571354 | rs6601975 | -0.137 | 0.416 | -0.588 | 0.013 | -0.019 | -0.051 | -0.051 | 0 | 0.23 | 0.18 | 0.35 | 0.53 | 0.3 | 0.31 |
| 10 | 5571549 | rs10904523 | -0.037 | 0.22 | -0.428 | -0.585 | 0.012 | -0.005 | -0.005 | 0.574 | 0.3 | 0.37 | 0.42 | 0.71 | 0.41 | 0.41 |
| 10 | 5572893 | rs7095575 | -0.958 | 0.376 | -0.933 | -0.6 | 0.007 | 0.106 | 0.106 | 0 | 0.55 | 0.74 | 0.8 | 0.89 | 0.62 | 0.66 |
| 10 | 5578172 | rs4077511 | 0.453 | 0.894 | -1.018 | 0.021 | -0.042 | -0.035 | -0.035 | 0.653 | 0.11 | 0.01 | 0.31 | 0.15 | 0.16 | 0.18 |
| 10 | 5580628 | rs12572508 | -0.573 | 0.679 | -0.526 | 0.214 | -0.011 | 0.021 | 0.021 | 0 | 0.42 | 0.34 | 0.47 | 0.69 | 0.5 | 0.47 |
| 10 | 5591025 | rs4881475 | -0.349 | 0.422 | -0.545 | 0.276 | -0.027 | 0.034 | 0.034 | 0.039 | 0.26 | 0.48 | 0.17 | 0.34 | 0.22 | 0.32 |
| 10 | 5593361 | rs6601978 | -0.262 | 1.351 | -1.157 | -0.237 | -0.047 | 0.031 | 0.031 | 0 | 0.83 | 0.91 | 0.75 | 0.89 | 0.74 | 0.83 |
| 10 | 5594362 | rs11253227 | 0.143 | 0.573 | -1.236 | 0.02 | -0.003 | -0.042 | -0.042 | 2.82 | 0.29 | 0.1 | 0.42 | 0.69 | 0.3 | 0.43 |
| 10 | 5595293 | rs12415024 | -0.041 | 0.377 | -0.773 | -0.09 | -0.005 | 0.043 | 0.043 | 0 | 0.12 | 0.19 | 0.13 | 0.17 | 0.1 | 0.16 |
| 10 | 5595918 | rs7079395 | 0.155 | 0.492 | -0.71 | -0.277 | -0.005 | 0.017 | 0.017 | 0 | 0.37 | 0.25 | 0.52 | 0.73 | 0.4 | 0.48 |
| 10 | 5596392 | rs7922769 | -0.062 | 0.706 | -0.371 | -0.624 | -0.001 | -0.015 | -0.015 | 0 | 0.51 | 0.5 | 0.52 | 0.82 | 0.42 | 0.62 |
| 10 | 5600140 | rs9423652 | -0.105 | 0.369 | -0.816 | -0.842 | 0.028 | 0.066 | 0.066 | 3.323 | 0.75 | 0.89 | 0.9 | 0.98 | 0.83 | 0.85 |
| 10 | 5600202 | rs11253229 | -0.674 | 0.654 | -0.502 | -0.436 | -0.005 | -0.023 | -0.023 | 0.231 | 0.45 | 0.53 | 0.24 | 0.27 | 0.36 | 0.5 |
| 10 | 5602880 | rs4881479 | 0.143 | 0.651 | -0.476 | -0.033 | -0.002 | -0.004 | -0.004 | 0 | 0.69 | 0.64 | 0.72 | 0.7 | 0.72 | 0.72 |
| 10 | 5607678 | rs4529812 | 0.283 | 0.141 | -0.422 | -0.324 | -0.035 | -0.038 | -0.038 | 0 | 0.66 | 0.57 | 0.72 | 0.79 | 0.78 | 0.67 |
| 10 | 5607759 | rs4348801 | 0.896 | 0.641 | -0.905 | -0.81 | -0.004 | 0.004 | 0.004 | 0.464 | 0.39 | 0.35 | 0.42 | 0.74 | 0.41 | 0.5 |
| 10 | 5608202 | rs7899101 | -0.007 | 0.851 | -1.031 | -0.763 | -0.029 | 0.088 | 0.088 | 0 | 0.23 | 0.08 | 0.27 | 0.04 | 0.32 | 0.17 |
| 10 | 5610224 | rs4881482 | -0.008 | 0.738 | -0.08 | -0.703 | 0.029 | -0.006 | -0.006 | 0.111 | 0.95 | 0.74 | 0.96 | 0.99 | 0.96 | 0.98 |
| 10 | 5610349 | rs4881483 | 0.2 | 0.317 | -0.382 | -0.624 | -0.003 | 0.011 | 0.011 | 0 | 0.36 | 0.33 | 0.45 | 0.51 | 0.36 | 0.28 |
| 10 | 5612132 | rs9423451 | -0.166 | 0.203 | -0.275 | -0.68 | -0.014 | 0.017 | 0.017 | 0 | 0.63 | 0.49 | 0.56 | 0.48 | 0.7 | 0.72 |
| 10 | 5612864 | rs12769978 | 1.185 | 0.354 | -0.668 | -0.821 | 0.015 | 0.022 | 0.022 | 2.258 | 0.2 | 0.09 | 0.17 | 0.12 | 0.15 | 0.19 |
| 10 | 5617563 | rs4074889 | 0.416 | 0.36 | -0.719 | -0.715 | 0.016 | 0.014 | 0.014 | 0 | 0.39 | 0.24 | 0.49 | 0.64 | 0.3 | 0.43 |
| 10 | 5618776 | rs12262126 | -0.097 | 0.588 | -1.041 | -0.506 | 0.001 | 0.021 | 0.021 | 0 | 0.51 | 0.54 | 0.63 | 0.64 | 0.51 | 0.53 |
| 10 | 5621628 | rs11253241 | 0.425 | 1.007 | -1.285 | -0.043 | -0.025 | 0.019 | 0.019 | 0 | 0.24 | 0.13 | 0.16 | 0.24 | 0.3 | 0.26 |
| 10 | 5629111 | rs7898549 | 0.301 | 0.611 | -1.197 | 0.261 | 0.017 | -0.049 | -0.049 | 0 | 0.12 | 0 | 0.12 | 0.13 | 0.06 | 0.2 |
| 10 | 5631737 | rs11253250 | -0.84 | 1.378 | -0.162 | 1.229 | 0.032 | 0.189 | 0.189 | 0 | 0.18 | 0.38 | 0.15 | 0.46 | 0.14 | 0.28 |
| 10 | 5631773 | rs11597133 | 0.137 | 1.422 | -0.241 | 1.498 | -0.007 | -0.081 | -0.081 | 0.093 | 0.13 | 0.01 | 0.04 | 0.29 | 0.08 | 0.22 |
| 10 | 5632060 | rs6601981 | 0.437 | 1.668 | -1.245 | 1.109 | -0.05 | -0.044 | -0.044 | 0.001 | 0.27 | 0.03 | 0.41 | 0.24 | 0.28 | 0.29 |
| 10 | 5633371 | rs7074607 | 1.394 | 1.825 | -1.345 | 1.59 | -0.041 | -0.036 | -0.036 | 0.003 | 0.07 | 0.01 | 0.17 | 0.09 | 0.13 | 0.12 |
| 10 | 5633549 | rs7075917 | -0.884 | 1.884 | -0.912 | 1.385 | 0.004 | 0.019 | 0.019 | 0 | 0.52 | 0.33 | 0.53 | 0.54 | 0.5 | 0.63 |
| 10 | 5634775 | rs7392864 | -1.257 | 1.518 | -1.126 | 1.139 | -0.03 | -0.011 | -0.011 | 0 | 0.82 | 0.91 | 0.87 | 0.79 | 0.77 | 0.8 |
| 10 | 5636434 | rs28495843 | -1.254 | 1.512 | -1.137 | 1.136 | -0.032 | -0.012 | -0.012 | 3.686 | 0.82 | 0.91 | 0.87 | 0.79 | 0.77 | 0.8 |
| 10 | 5637650 | rs2380195 | -1.066 | 1.52 | -0.642 | 1.175 | 0.037 | 0.102 | 0.102 | 0 | 0.81 | 0.66 | 0.79 | 0.57 | 0.75 | 0.76 |
| 10 | 5637755 | rs2380196 | 0.321 | 1.476 | -0.655 | 1.013 | -0.003 | 0.017 | 0.017 | 0.856 | 0.33 | 0.25 | 0.45 | 0.2 | 0.33 | 0.31 |
| 10 | 5637772 | rs2380197 | 0.321 | 1.476 | -0.655 | 1.013 | -0.003 | 0.017 | 0.017 | 0.856 | 0.33 | 0.25 | 0.45 | 0.2 | 0.33 | 0.31 |
| 10 | 5642758 | rs4747897 | 0.951 | 1.052 | -0.453 | 0.972 | 0.011 | 0.076 | 0.076 | 0.016 | 0.41 | 0.18 | 0.3 | 0.26 | 0.36 | 0.23 |
| 10 | 5643044 | rs4747916 | 0.385 | 1.252 | -0.16 | 1.111 | 0.091 | 0.075 | 0.075 | 0 | 0.84 | 0.39 | 0.76 | 0.76 | 0.71 | 0.75 |
| 10 | 5643511 | rs7350438 | -0.024 | 1.211 | -0.241 | 1.109 | 0.032 | -0.04 | -0.04 | 0.271 | 0.43 | 0.18 | 0.46 | 0.5 | 0.35 | 0.52 |
| 10 | 5644867 | rs12242788 | 1.467 | 1.098 | -0.264 | 1.098 | -0.035 | -0.028 | -0.028 | 0.134 | 0.16 | 0.07 | 0.1 | 0.21 | 0.2 | 0.13 |
| 10 | 5644983 | rs7896306 | 0.22 | 1.292 | 0.328 | 1.4 | 0.172 | 0.258 | 0.258 | 0.089 | 0.93 | 0.46 | 0.84 | 0.79 | 0.87 | 0.87 |
| 10 | 5645477 | rs11595011 | -0.291 | 1.22 | 0.239 | 1.514 | -0.001 | -0.009 | -0.009 | 3.256 | 0.2 | 0.14 | 0.11 | 0.26 | 0.21 | 0.34 |
| 10 | 5645758 | rs4750500 | 0.076 | 1.517 | 0.398 | 1.621 | -0.001 | -0.007 | -0.007 | 0 | 0.62 | 0.3 | 0.42 | 0.59 | 0.6 | 0.58 |
| 10 | 5646999 | rs11259704 | 0.439 | 1.473 | 0.246 | 1.117 | 0.007 | 0.008 | 0.008 | 0.511 | 0.31 | 0.15 | 0.43 | 0.28 | 0.3 | 0.3 |
| 10 | 5647407 | rs17138235 | -0.436 | 1.453 | 0.124 | 1.117 | 0.013 | -0.009 | -0.009 | 1.441 | 0.44 | 0.15 | 0.52 | 0.43 | 0.43 | 0.52 |
| 10 | 5647942 | rs4748295 | -0.463 | 1.435 | 0.129 | 1.054 | 0.004 | -0.042 | -0.042 | 0 | 0.44 | 0.16 | 0.48 | 0.5 | 0.44 | 0.53 |
| 10 | 5651578 | rs1544216 | 0.091 | 1.528 | 0.399 | 0.999 | -0.001 | 0.028 | 0.028 | 0.792 | 0.33 | 0.24 | 0.44 | 0.2 | 0.34 | 0.3 |
| 10 | 5652277 | rs7896078 | -0.126 | 1.454 | 0.652 | 0.928 | -0.001 | -0.005 | -0.005 | 0 | 0.35 | 0.37 | 0.45 | 0.2 | 0.34 | 0.3 |

| | | | | | | | | | | | | | | | | |
|---|---|---|---|---|---|---|---|---|---|---|---|---|---|---|---|---|
| 10 | 5663188 | rs11255575 | 0.574 | 1.371 | 0.042 | 0.427 | 0 | 0.079 | 0.079 | 0.282 | 0.48 | 0.36 | 0.26 | 0.12 | 0.48 | 0.25 |
| 10 | 5672059 | rs2386660 | -0.431 | 1.291 | 0.384 | 1.115 | 0.146 | 0.386 | 0.386 | 0 | 0.08 | 0.3 | 0.16 | 0.4 | 0.19 | 0.12 |
| 10 | 5674342 | rs7088391 | -2.181 | 1.103 | 0.374 | 1.059 | -0.022 | -0.005 | -0.005 | 0.014 | 0.95 | 0.84 | 0.98 | 0.98 | 0.99 | 0.88 |
| 10 | 5675575 | rs4749861 | -2.276 | 1.103 | 0.363 | 1.059 | -0.022 | -0.007 | -0.007 | 2.522 | 0.95 | 0.88 | 0.98 | 0.98 | 0.99 | 0.88 |
| 10 | 5678058 | rs2386650 | 0.5 | 0.876 | 0.146 | 0.957 | 0.001 | 0.008 | 0.008 | 1.795 | 0.23 | 0.15 | 0.22 | 0.17 | 0.25 | 0.24 |
| 10 | 5679041 | rs7896605 | -1.713 | 1.096 | 0.594 | 1.526 | -0.01 | 0.026 | 0.026 | 0.588 | 0.94 | 0.84 | 0.96 | 0.92 | 0.98 | 0.88 |
| 10 | 5680842 | rs2386649 | -0.892 | 1.17 | 0.244 | 1.57 | 0.057 | 0.244 | 0.244 | 1.077 | 0.13 | 0.31 | 0.18 | 0.44 | 0.18 | 0.26 |
| 10 | 5680896 | rs1573 | -0.892 | 1.17 | 0.244 | 1.57 | 0.057 | 0.244 | 0.244 | 1.077 | 0.13 | 0.31 | 0.18 | 0.44 | 0.18 | 0.26 |
| 10 | 5681139 | rs12573500 | 0.408 | 1.663 | 0.455 | 1.764 | 0.021 | -0.019 | -0.019 | 2.523 | 0.07 | 0 | 0.27 | 0.06 | 0.03 | 0.06 |
| 10 | 5681783 | rs11256474 | -0.816 | 1.69 | 0.471 | 1.73 | 0.149 | 0.381 | 0.381 | 0 | 0.08 | 0.3 | 0.16 | 0.41 | 0.18 | 0.14 |
| 10 | 5682447 | rs3750644 | 0.246 | 1.38 | -0.657 | 1.51 | 0.003 | -0.039 | -0.039 | 0.406 | 0.07 | 0 | 0.28 | 0.11 | 0.05 | 0.06 |
| 10 | 5688085 | rs10905791 | -0.393 | 1.271 | -0.83 | 0.931 | -0.026 | 0.013 | 0.013 | 0.443 | 0.42 | 0.13 | 0.4 | 0.37 | 0.56 | 0.39 |
| 10 | 5689058 | rs11256831 | -1.163 | 1.368 | 0.283 | 1.808 | -0.011 | 0.261 | 0.261 | 0 | 0.14 | 0.34 | 0.44 | 0.5 | 0.11 | 0.25 |
| 10 | 5691135 | rs2386647 | -1.136 | 1.314 | 0.159 | 1.818 | -0.015 | 0.307 | 0.307 | 0 | 0.14 | 0.38 | 0.45 | 0.51 | 0.11 | 0.25 |
| 10 | 5693649 | rs7907734 | -0.809 | 2.016 | -1.052 | 1.172 | -0.017 | -0.009 | -0.009 | 0.02 | 0.17 | 0.07 | 0.13 | 0.16 | 0.21 | 0.11 |
| 10 | 5701761 | rs871747 | -1.277 | 2.267 | -0.648 | 1.781 | 0.016 | 0.124 | 0.124 | 0 | 0.1 | 0.17 | 0.25 | 0.51 | 0.15 | 0.29 |
| 10 | 5701795 | rs871748 | -1.967 | 1.995 | -0.211 | 1.49 | 0.01 | 0.139 | 0.139 | 0 | 0.18 | 0.35 | 0.36 | 0.51 | 0.19 | 0.34 |
| 10 | 5701833 | rs871749 | -0.632 | 3.004 | -0.727 | 1.339 | -0.067 | -0.029 | -0.029 | 0 | 0.11 | 0.03 | 0.14 | 0.1 | 0.24 | 0.08 |
| 10 | 5705084 | rs1539446 | -1.479 | 3.043 | -0.878 | 1.367 | -0.006 | -0.017 | -0.017 | 0 | 0.09 | 0.06 | 0.21 | 0.18 | 0.09 | 0.19 |
| 10 | 5705900 | rs4750217 | -0.855 | 2.631 | -0.741 | 0.519 | 0.016 | 0.001 | 0.001 | 2.105 | 0.5 | 0.39 | 0.5 | 0.46 | 0.57 | 0.55 |
| 10 | 5709909 | rs2386637 | 0.256 | 2.232 | -0.619 | 0.484 | -0.003 | -0.012 | -0.012 | 0.095 | 0.32 | 0.32 | 0.15 | 0.14 | 0.26 | 0.31 |
| 10 | 5721653 | rs2386626 | -1.336 | 2.342 | -0.616 | 0.707 | 0.006 | 0.009 | 0.009 | 2.531 | 0.59 | 0.54 | 0.69 | 0.47 | 0.53 | 0.62 |
| 10 | 5726253 | rs11596406 | -0.095 | 2.12 | -0.493 | 0.787 | 0.001 | 0.001 | 0.001 | 0 | 0.07 | 0.05 | 0.01 | 0.05 | 0.02 | 0.1 |
| 10 | 5734358 | rs9423936 | 0.19 | 2.373 | -0.577 | 0.738 | 0.001 | 0.003 | 0.003 | 0.013 | 0.3 | 0.28 | 0.37 | 0.21 | 0.29 | 0.31 |
| 10 | 5735648 | rs7096742 | 1.177 | 2.699 | -0.147 | 0.841 | -0.012 | -0.016 | -0.016 | 0 | 0.7 | 0.74 | 0.64 | 0.66 | 0.65 | 0.66 |
| 10 | 5739054 | rs7908235 | 1.398 | 2.716 | -0.084 | 0.904 | -0.002 | 0.004 | 0.004 | 3.089 | 0.4 | 0.4 | 0.37 | 0.48 | 0.37 | 0.34 |
| 10 | 5750078 | rs11254266 | 1.081 | 2.463 | -0.287 | 0.97 | -0.007 | -0.009 | -0.009 | 0 | 0.77 | 0.77 | 0.67 | 0.72 | 0.69 | 0.77 |
| 10 | 5751644 | rs10508295 | 0.992 | 2.196 | -0.452 | 0.896 | 0.007 | -0.011 | -0.011 | 0 | 0.21 | 0.01 | 0.08 | 0.16 | 0.14 | 0.21 |
| 10 | 5774915 | rs4748572 | -0.546 | 2.718 | 0.927 | 1.668 | -0.025 | -0.031 | -0.031 | 0.882 | 0.11 | 0.07 | 0.24 | 0.15 | 0.19 | 0.1 |
| 10 | 5794168 | rs2669139 | -1.13 | 2.662 | 1.192 | 1.677 | 0.013 | 0.445 | 0.445 | 0.721 | 0.06 | 0.34 | 0.08 | 0.43 | 0.03 | 0.23 |
| 10 | 5794475 | rs2669138 | 0.571 | 2.705 | 0.914 | 1.573 | 0 | 0 | 0 | 0 | 0.91 | 0.89 | 0.95 | 0.94 | 0.95 | 0.87 |
| 10 | 5799613 | rs2275774 | -0.18 | 2.725 | 0.898 | 1.582 | -0.04 | -0.003 | -0.003 | 0.469 | 0.1 | 0.02 | 0.1 | 0.06 | 0.18 | 0.08 |
| 10 | 5804531 | rs2797501 | 1.159 | 2.679 | 0.941 | 1.512 | 0.01 | 0.002 | 0.002 | 1.544 | 0.83 | 0.87 | 0.91 | 0.82 | 0.87 | 0.81 |
| 10 | 5805844 | rs1046393 | -0.105 | 2.688 | 0.935 | 1.545 | -0.023 | -0.027 | -0.027 | 0 | 0.1 | 0.07 | 0.24 | 0.15 | 0.19 | 0.1 |
| 10 | 5806248 | rs11255072 | 0.51 | 2.756 | 0.89 | 1.525 | -0.036 | -0.008 | -0.008 | 0.08 | 0.06 | 0.01 | 0.28 | 0.05 | 0.14 | 0.1 |
| 10 | 5808086 | rs1129614 | -0.286 | 1.404 | 1.186 | 1.73 | -0.023 | -0.026 | -0.026 | 0.149 | 0.11 | 0.07 | 0.24 | 0.14 | 0.19 | 0.1 |
| 10 | 5815108 | rs1567705 | 1.673 | 1.283 | 1.674 | 2.161 | -0.003 | 0.297 | 0.297 | 0 | 0.85 | 0.57 | 0.84 | 0.48 | 0.9 | 0.7 |
| 10 | 5820394 | rs4748916 | -2.589 | 1.435 | 1.11 | 1.612 | -0.009 | -0.025 | -0.025 | 0 | 0.29 | 0.22 | 0.35 | 0.36 | 0.32 | 0.34 |
| 10 | 5827074 | rs10795547 | 2.517 | 1.398 | 1.196 | 1.594 | -0.001 | -0.039 | -0.039 | 0 | 0.65 | 0.45 | 0.57 | 0.24 | 0.64 | 0.42 |
| 10 | 5852230 | rs1858447 | -2.436 | 1.758 | 0.484 | 0.911 | -0.017 | -0.026 | -0.026 | 0 | 0.3 | 0.23 | 0.36 | 0.36 | 0.35 | 0.34 |
| 10 | 5853767 | rs1567706 | -2.45 | 1.769 | 0.464 | 0.916 | -0.006 | -0.009 | -0.009 | 0 | 0.3 | 0.28 | 0.36 | 0.36 | 0.35 | 0.34 |
| 10 | 5856984 | rs7900361 | -2.002 | 1.408 | 0.14 | 0.744 | 0 | 0 | 0 | 0 | 0.19 | 0.19 | 0.13 | 0.2 | 0.15 | 0.23 |
| 10 | 5869839 | rs11596702 | -0.85 | 1.207 | -0.091 | 0.857 | 0.002 | -0.007 | -0.007 | 0 | 0.09 | 0.04 | 0.07 | 0.16 | 0.11 | 0.11 |
| 10 | 5874958 | rs11599388 | -1.576 | 1.467 | -0.17 | 0.886 | 0.004 | 0.014 | 0.014 | 0 | 0.29 | 0.38 | 0.36 | 0.35 | 0.34 | 0.3 |
| 10 | 5877213 | rs12220359 | -1.22 | 1.232 | -0.385 | 0.353 | -0.021 | -0.025 | -0.025 | 0 | 0.1 | 0.08 | 0.08 | 0.14 | 0.14 | 0.1 |
| 10 | 5880066 | rs4406717 | 1.381 | -0.049 | 0.21 | -0.084 | -0.059 | -0.019 | -0.019 | 0 | 0.71 | 0.55 | 0.82 | 0.77 | 0.82 | 0.67 |
| 10 | 5880488 | rs7904257 | -0.857 | 0.02 | 0.091 | -0.079 | 0.017 | -0.024 | -0.024 | 0 | 0.14 | 0.06 | 0.09 | 0.2 | 0.12 | 0.19 |
| 10 | 5881740 | rs11255499 | -0.082 | 0.068 | 0.134 | -0.884 | 0.012 | -0.003 | -0.003 | 0 | 0.18 | 0.25 | 0.18 | 0.18 | 0.21 | 0.18 |
| 10 | 5886149 | rs7098372 | 0.842 | -0.461 | -0.32 | -1.905 | -0.029 | 0.212 | 0.212 | 5.874 | 0.45 | 0.81 | 0.31 | 0.13 | 0.45 | 0.25 |
| 10 | 5900693 | rs907686 | 0.86 | -0.505 | -0.167 | -1.915 | -0.021 | 0.172 | 0.172 | 4.561 | 0.45 | 0.77 | 0.31 | 0.13 | 0.44 | 0.25 |
| 10 | 5916771 | rs1512694 | 0.619 | -0.473 | 0.357 | -2.02 | 0 | 0.057 | 0.057 | 0.933 | 0.51 | 0.39 | 0.32 | 0.19 | 0.47 | 0.27 |
| 10 | 5920121 | rs1052420 | 0.285 | -0.509 | 0.565 | -1.926 | -0.016 | -0.003 | -0.003 | 0.198 | 0.82 | 0.69 | 0.91 | 0.85 | 0.86 | 0.78 |
| 10 | 5926079 | rs3750657 | 0.453 | -0.535 | 0.57 | -1.988 | -0.004 | 0.014 | 0.014 | 0.71 | 0.54 | 0.61 | 0.35 | 0.19 | 0.48 | 0.28 |
| 10 | 5926216 | rs7919913 | 1.652 | -0.598 | 0.309 | -2.034 | -0.001 | -0.067 | -0.067 | 0 | 0.36 | 0.39 | 0.23 | 0.03 | 0.36 | 0.14 |
| 10 | 5930512 | rs2320 | 0.334 | -0.228 | -0.411 | -1.813 | -0.06 | -0.024 | -0.024 | 0.174 | 0.28 | 0.12 | 0.57 | 0.66 | 0.39 | 0.5 |
| 10 | 5951025 | rs595923 | 0.36 | -0.246 | -0.569 | -1.802 | -0.08 | -0.033 | -0.033 | 0 | 0.28 | 0.1 | 0.57 | 0.66 | 0.39 | 0.47 |
| 10 | 5951563 | rs593610 | 0.371 | -0.239 | -0.695 | -1.799 | -0.101 | -0.048 | -0.048 | 0.874 | 0.27 | 0.07 | 0.57 | 0.66 | 0.39 | 0.47 |
| 10 | 5957301 | rs11813102 | -1.186 | -0.753 | 0.203 | -1.008 | 0.052 | 0.158 | 0.158 | 0.524 | 0.06 | 0.26 | 0.05 | 0.22 | 0.04 | 0.14 |
| 10 | 5960499 | rs2274027 | -0.46 | -0.727 | -0.892 | -1.041 | 0.007 | 0.016 | 0.016 | 0.017 | 0.19 | 0.13 | 0.13 | 0.16 | 0.14 | 0.14 |
| 10 | 5961098 | rs593026 | 1.288 | -0.477 | -0.627 | -0.708 | 0.003 | 0.002 | 0.002 | 0 | 0.81 | 0.82 | 0.92 | 0.85 | 0.86 | 0.73 |
| 10 | 5967006 | rs2669127 | 0.141 | -0.436 | -0.716 | -0.692 | -0.008 | 0.022 | 0.022 | 0 | 0.88 | 0.98 | 0.94 | 0.95 | 0.88 | 0.84 |
| 10 | 5978411 | rs646406 | 0.693 | -0.34 | -0.667 | -0.663 | 0.002 | 0.001 | 0.001 | 0 | 0.81 | 0.82 | 0.92 | 0.85 | 0.86 | 0.73 |
| 10 | 5982577 | rs2669130 | -0.02 | -0.262 | -0.928 | -1.014 | 0.05 | 0.08 | 0.08 | 0 | 0.48 | 0.39 | 0.39 | 0.1 | 0.72 | 0.32 |
| 10 | 5988908 | rs9971043 | -0.771 | -0.251 | -0.435 | -0.675 | -0.018 | -0.043 | -0.043 | 0.313 | 0.08 | 0.15 | 0.03 | 0 | 0.03 | 0.08 |
| 10 | 5992119 | rs1998521 | -0.57 | 0.014 | -1.319 | -0.538 | -0.066 | -0.02 | -0.02 | 0 | 0.58 | 0.82 | 0.45 | 0.43 | 0.5 | 0.56 |
| 10 | 5992544 | rs17146980 | 0.863 | -0.636 | -2.009 | -0.969 | 0.018 | 0.048 | 0.048 | 0 | 0.18 | 0.02 | 0.12 | 0.08 | 0.13 | 0.16 |
| 10 | 5994694 | rs2296135 | -1.208 | -0.599 | -1.755 | -0.961 | -0.044 | -0.013 | -0.013 | 0 | 0.58 | 0.8 | 0.44 | 0.43 | 0.49 | 0.56 |
| 10 | 5995052 | rs2229135 | 1.006 | -0.129 | -1.372 | -1.149 | 0.001 | -0.01 | -0.01 | 0 | 0.09 | 0.11 | 0.14 | 0.05 | 0.08 | 0.05 |
| 10 | 5995492 | rs8177741 | 0.391 | -0.227 | -0.527 | -1.149 | -0.005 | 0.013 | 0.013 | 0 | 0.06 | 0.19 | 0.02 | 0.05 | 0.03 | 0.01 |

| | | | | | | | | | | | | | | | | |
|---|---|---|---|---|---|---|---|---|---|---|---|---|---|---|---|---|
| 10 | 5997820 | rs3136630 | -1.105 | -0.384 | 0.179 | -1.1 | 0.019 | 0.021 | 0.021 | 0 | 0.43 | 0.6 | 0.32 | 0.24 | 0.28 | 0.35 |
| 10 | 5998285 | rs3136627 | -1.303 | -0.231 | 0.294 | -1.038 | 0.016 | 0.037 | 0.037 | 0 | 0.51 | 0.57 | 0.41 | 0.31 | 0.43 | 0.46 |
| 10 | 5998659 | rs2296141 | 1.051 | -0.147 | 0.271 | -0.968 | 0.001 | -0.003 | -0.003 | 1.309 | 0.09 | 0.11 | 0.14 | 0.07 | 0.09 | 0.05 |
| 10 | 5998696 | rs2296140 | -1.078 | -0.776 | 0.103 | -1.284 | -0.019 | -0.037 | -0.037 | 0 | 0.26 | 0.27 | 0.17 | 0.07 | 0.12 | 0.13 |
| 10 | 6000259 | rs17320853 | 0.131 | -0.316 | 0.037 | -1.059 | 0.002 | -0.002 | -0.002 | 0.026 | 0.14 | 0.16 | 0.12 | 0.12 | 0.18 | 0.12 |
| 10 | 6001409 | rs3136621 | -1.196 | -0.291 | 0.147 | -1.076 | 0.001 | 0 | 0 | 0 | 0.27 | 0.65 | 0.28 | 0.3 | 0.28 | 0.36 |
| 10 | 6001442 | rs3136620 | -0.183 | -0.315 | 0.327 | -1.086 | 0.018 | -0.001 | -0.001 | 0.009 | 0.16 | 0.34 | 0.13 | 0.16 | 0.18 | 0.17 |
| 10 | 6001483 | rs3136619 | 0.31 | -0.348 | -0.354 | -1.16 | 0.037 | 0.044 | 0.044 | 0 | 0.24 | 0.1 | 0.21 | 0.13 | 0.18 | 0.2 |
| 10 | 6001696 | rs3136618 | -0.951 | -0.109 | -0.208 | -1.022 | 0.013 | 0.025 | 0.025 | 0 | 0.51 | 0.7 | 0.49 | 0.42 | 0.46 | 0.55 |
| 10 | 6002368 | rs2228059 | -1.018 | -0.067 | -0.465 | -1.008 | 0.009 | 0.023 | 0.023 | 0 | 0.51 | 0.75 | 0.49 | 0.43 | 0.46 | 0.53 |
| 10 | 6003451 | rs11574674 | -1.1 | 0.374 | -0.505 | -0.768 | -0.034 | -0.018 | -0.018 | 0 | 0.08 | 0.03 | 0.06 | 0.07 | 0.17 | 0.08 |
| 10 | 6004012 | rs8177692 | 0.016 | 0.726 | -0.851 | -0.238 | -0.017 | -0.038 | -0.038 | 0 | 0.24 | 0.12 | 0.31 | 0.35 | 0.31 | 0.18 |
| 10 | 6005674 | rs3136614 | -0.124 | 0.981 | -0.878 | -0.491 | -0.008 | 0.023 | 0.023 | 0 | 0.82 | 0.88 | 0.88 | 0.96 | 0.78 | 0.9 |
| 10 | 6008172 | rs2296139 | 0.31 | 1.208 | -0.779 | 0.011 | -0.001 | 0.003 | 0.003 | 0 | 0.14 | 0.15 | 0.24 | 0.27 | 0.14 | 0.13 |
| 10 | 6008613 | rs8177685 | -0.869 | 1.033 | -0.61 | 0.078 | -0.001 | 0.065 | 0.065 | 0.006 | 0.89 | 0.87 | 0.8 | 0.61 | 0.86 | 0.82 |
| 10 | 6009126 | rs7910212 | -0.27 | 0.937 | -0.076 | -0.098 | 0.012 | 0.227 | 0.227 | 0.143 | 0.86 | 0.67 | 0.73 | 0.52 | 0.85 | 0.74 |
| 10 | 6010227 | rs7097780 | 0.426 | 0.812 | -0.53 | -0.879 | 0.008 | -0.011 | -0.011 | 0 | 0.31 | 0.35 | 0.18 | 0.24 | 0.31 | 0.37 |
| 10 | 6011623 | rs8177667 | 0.46 | 0.913 | -1.058 | -1.074 | -0.024 | 0.06 | 0.06 | 0 | 0.19 | 0.08 | 0.09 | 0 | 0.2 | 0.12 |
| 10 | 6018594 | rs8177636 | 0.769 | 0.998 | -0.867 | -0.846 | -0.037 | 0.096 | 0.096 | 0 | 0.34 | 0.17 | 0.31 | 0.01 | 0.43 | 0.22 |
| 10 | 6024313 | rs4747826 | 0.673 | 0.641 | -0.945 | -0.284 | 0.022 | -0.094 | -0.094 | 0 | 0.2 | 0.06 | 0.28 | 0.47 | 0.18 | 0.2 |
| 10 | 6024331 | rs8177607 | 0.518 | 1.695 | -0.113 | 0.955 | 0 | 0 | 0 | 0.242 | 0.17 | 0.12 | 0.05 | 0.15 | 0.14 | 0.24 |
| 10 | 6027791 | rs8177585 | -0.225 | 1.493 | -0.407 | 1.064 | -0.002 | 0.001 | 0.001 | 0 | 0.3 | 0.24 | 0.17 | 0.24 | 0.32 | 0.36 |
| 10 | 6028491 | rs7904224 | 0.309 | 1.626 | -1.166 | 0.845 | -0.007 | -0.01 | -0.01 | 0 | 0.57 | 0.34 | 0.61 | 0.5 | 0.61 | 0.48 |
| 10 | 6029726 | rs6602360 | 0.638 | 2.133 | -1.357 | 1.344 | -0.03 | -0.047 | -0.047 | 0 | 0.78 | 0.89 | 0.72 | 0.59 | 0.66 | 0.64 |
| 10 | 6030194 | rs7088910 | -1.124 | 2.895 | -1.357 | 1.49 | -0.03 | -0.011 | -0.011 | 0 | 0.2 | 0.14 | 0.2 | 0.24 | 0.34 | 0.25 |
| 10 | 6030749 | rs10795726 | -1.181 | 2.48 | -1.187 | 1.663 | 0.011 | 0.009 | 0.009 | 0 | 0.24 | 0.29 | 0.33 | 0.59 | 0.34 | 0.49 |
| 10 | 6033415 | rs11256258 | -0.771 | 2.384 | -1.228 | 2.057 | 0 | 0.034 | 0.034 | 0 | 0.07 | 0.09 | 0.15 | 0.25 | 0.09 | 0.11 |
| 10 | 6035149 | rs11598704 | -0.924 | 2.361 | -1.345 | 1.763 | -0.003 | 0.086 | 0.086 | 0 | 0.25 | 0.1 | 0.3 | 0.12 | 0.22 | 0.29 |
| 10 | 6036144 | rs4749882 | 0.789 | 1.789 | -0.965 | 1.676 | 0.038 | 0.135 | 0.135 | 0 | 0.79 | 0.6 | 0.74 | 0.61 | 0.71 | 0.65 |
| 10 | 6037553 | rs7898880 | 1.931 | 1.8 | -2.059 | 0.661 | -0.085 | 0.124 | 0.124 | 0 | 0.19 | 0.02 | 0.12 | 0.01 | 0.26 | 0.13 |
| 10 | 6039058 | rs1323653 | 1.208 | 1.59 | -1.755 | 0.763 | -0.023 | 0.059 | 0.059 | 0.003 | 0.11 | 0 | 0.04 | 0.02 | 0.08 | 0.09 |
| 10 | 6041923 | rs10905599 | -0.331 | 1.643 | -1.499 | 1.072 | 0.022 | 0.068 | 0.068 | 0 | 0.13 | 0.18 | 0.21 | 0.36 | 0.22 | 0.31 |
| 10 | 6053866 | rs12722602 | 1.299 | 0.947 | -1.194 | 0.415 | 0.004 | 0 | 0 | 0.788 | 0.13 | 0.13 | 0.07 | 0.02 | 0.11 | 0.11 |
| 10 | 6054158 | rs12719919 | 1.95 | 0.844 | -1.246 | 0.489 | 0.012 | 0.014 | 0.014 | 0 | 0.07 | 0.04 | 0.03 | 0.02 | 0.05 | 0.05 |
| 10 | 6054765 | rs12722600 | 2.524 | 0.848 | -1.277 | 0.406 | 0.013 | 0.027 | 0.027 | 0 | 0.05 | 0.01 | 0.03 | 0 | 0.02 | 0.04 |
| 10 | 6055604 | rs9663421 | 0.954 | 0.769 | -1.5 | 0.082 | -0.047 | 0.12 | 0.12 | 0 | 0.25 | 0.09 | 0.33 | 0.05 | 0.31 | 0.15 |
| 10 | 6057732 | rs2386841 | -0.222 | 0.377 | -1.145 | 0.645 | 0.001 | 0.006 | 0.006 | 0 | 0.16 | 0.16 | 0.37 | 0.46 | 0.17 | 0.13 |
| 10 | 6059898 | rs7899538 | 0.55 | 0.419 | -1.148 | 0.689 | -0.006 | -0.018 | -0.018 | 1.289 | 0.1 | 0.05 | 0.07 | 0.24 | 0.11 | 0.11 |
| 10 | 6062108 | rs2274037 | 1.085 | 0.1 | -0.934 | 0.345 | 0.002 | 0.002 | 0.002 | 0 | 0.06 | 0.06 | 0.2 | 0.03 | 0.04 | 0.01 |
| 10 | 6063253 | rs2076846 | 0.937 | -0.248 | -0.758 | -0.342 | -0.002 | 0.019 | 0.019 | 0.462 | 0.36 | 0.1 | 0.19 | 0.26 | 0.33 | 0.3 |
| 10 | 6063508 | rs2228149 | 1.345 | -0.061 | -0.602 | -0.338 | 0.002 | 0.002 | 0.002 | 0 | 0.06 | 0.06 | 0.2 | 0.04 | 0.04 | 0.01 |
| 10 | 6063674 | rs942200 | -1.268 | -0.207 | -0.049 | -0.413 | 0.044 | 0.006 | 0.006 | 1.227 | 0.93 | 0.65 | 0.96 | 1 | 0.98 | 0.94 |
| 10 | 6065611 | rs7900744 | -0.098 | -0.279 | -0.548 | -0.396 | 0.008 | -0.035 | -0.035 | 0 | 0.21 | 0.46 | 0.13 | 0.15 | 0.19 | 0.22 |
| 10 | 6067688 | rs2025345 | 0.59 | 0.111 | -1.205 | -0.173 | -0.094 | -0.051 | -0.051 | 0 | 0.32 | 0.11 | 0.52 | 0.4 | 0.4 | 0.29 |
| 10 | 6069893 | rs12722561 | 1.293 | 0.281 | -0.635 | -0.359 | 0.087 | 0.079 | 0.079 | 0 | 0.05 | 0.13 | 0.06 | 0.14 | 0.13 | 0.14 |
| 10 | 6073374 | rs6602379 | -0.897 | 0.204 | -1.159 | -0.589 | -0.035 | 0.086 | 0.086 | 0 | 0.7 | 0.86 | 0.53 | 0.91 | 0.63 | 0.64 |
| 10 | 6077328 | rs12722527 | 1.772 | 0.3 | -0.958 | -0.421 | 0.041 | 0.034 | 0.034 | 0 | 0.05 | 0.06 | 0.06 | 0.15 | 0.14 | 0.15 |
| 10 | 6077796 | rs7910961 | -1.319 | 0.207 | -1.06 | -0.578 | -0.034 | 0.062 | 0.062 | 0 | 0.74 | 0.86 | 0.55 | 0.93 | 0.63 | 0.8 |
| 10 | 6077845 | rs11256433 | 1.173 | -0.197 | -1.289 | -0.581 | -0.002 | -0.028 | -0.028 | 0 | 0.22 | 0.12 | 0.15 | 0.26 | 0.22 | 0.21 |
| 10 | 6078079 | rs6602392 | 0.032 | -0.335 | -0.895 | -0.657 | -0.002 | 0.001 | 0.001 | 0 | 0.22 | 0.27 | 0.15 | 0.33 | 0.11 | 0.14 |
| 10 | 6079846 | rs7072398 | -0.417 | -0.199 | -0.917 | -0.649 | 0.004 | 0.038 | 0.038 | 0.13 | 0.5 | 0.68 | 0.61 | 0.4 | 0.5 | 0.4 |
| 10 | 6082953 | rs6602398 | 1.51 | 0.344 | -1.23 | -0.484 | -0.116 | 0.121 | 0.121 | 0.155 | 0.21 | 0.02 | 0.26 | 0.04 | 0.34 | 0.22 |
| 10 | 6085312 | rs4749926 | 0.376 | 0.087 | -1.006 | -0.414 | -0.015 | -0.006 | -0.006 | 0 | 0.32 | 0.29 | 0.54 | 0.39 | 0.41 | 0.31 |
| 10 | 6086093 | rs10905656 | 0.265 | 0.078 | -1.181 | -0.244 | -0.043 | -0.048 | -0.048 | 0.114 | 0.33 | 0.19 | 0.64 | 0.61 | 0.45 | 0.45 |
| 10 | 6087295 | rs1107345 | 1.125 | 0.068 | -1.624 | -0.122 | -0.033 | -0.079 | -0.079 | 0 | 0.21 | 0.05 | 0.17 | 0.27 | 0.22 | 0.21 |
| 10 | 6089571 | rs791589 | 0.561 | -0.668 | -1.503 | -0.264 | -0.001 | -0.001 | -0.001 | 0 | 0.76 | 0.3 | 0.67 | 0.67 | 0.86 | 0.84 |
| 10 | 6092093 | rs10905669 | 0.556 | -0.349 | -1.759 | -0.229 | -0.005 | -0.096 | -0.096 | 0 | 0.25 | 0.07 | 0.35 | 0.31 | 0.23 | 0.19 |
| 10 | 6096199 | rs10795763 | -0.312 | -0.568 | -1.61 | -0.172 | 0.002 | -0.02 | -0.02 | 0 | 0.56 | 0.17 | 0.34 | 0.4 | 0.54 | 0.53 |
| 10 | 6097165 | rs2256774 | 0.364 | -0.504 | -1.113 | -0.354 | -0.002 | -0.054 | -0.054 | 0.172 | 0.38 | 0.43 | 0.49 | 0.11 | 0.38 | 0.37 |
| 10 | 6099045 | rs2104286 | 1.215 | -0.446 | -1.808 | -0.289 | -0.067 | 0.025 | 0.025 | 0 | 0.18 | 0.03 | 0.1 | 0.13 | 0.22 | 0.19 |
| 10 | 6101713 | rs3118470 | 0.707 | -0.508 | -1.778 | -0.236 | -0.021 | -0.077 | -0.077 | 0 | 0.32 | 0.13 | 0.29 | 0.49 | 0.32 | 0.41 |
| 10 | 6102012 | rs12722489 | 1.725 | 0.034 | -1.543 | -0.067 | -0.032 | -0.029 | -0.029 | 0 | 0.05 | 0.01 | 0.06 | 0.13 | 0.14 | 0.13 |
| 10 | 6106266 | rs7072793 | 0.048 | -0.402 | -1.46 | -0.167 | 0.02 | 0.033 | 0.033 | 0 | 0.47 | 0.31 | 0.55 | 0.55 | 0.42 | 0.5 |
| 10 | 6110829 | rs7090512 | -0.832 | -0.549 | -1.385 | -0.224 | -0.005 | -0.013 | -0.013 | 0 | 0.67 | 0.59 | 0.66 | 0.69 | 0.65 | 0.69 |
| 10 | 6110875 | rs7090530 | -0.396 | -0.536 | -1.381 | -0.225 | 0.001 | 0.002 | 0.002 | 0 | 0.56 | 0.57 | 0.63 | 0.69 | 0.59 | 0.65 |
| 10 | 6113523 | rs12412095 | 0.926 | -0.452 | -1.491 | -0.071 | -0.018 | -0.019 | -0.019 | 0 | 0.32 | 0.11 | 0.19 | 0.28 | 0.31 | 0.29 |
| 10 | 6114010 | rs10905716 | 0.651 | -0.455 | -1.558 | -0.231 | -0.003 | 0.006 | 0.006 | 0 | 0.23 | 0.3 | 0.25 | 0.31 | 0.22 | 0.31 |
| 10 | 6114856 | rs10905718 | 0.622 | -0.247 | -1.613 | -0.187 | -0.019 | -0.019 | -0.019 | 0 | 0.32 | 0.11 | 0.19 | 0.28 | 0.31 | 0.29 |
| 10 | 6118966 | rs4749955 | 0.757 | -0.123 | -0.715 | 0.657 | 0.007 | 0.047 | 0.047 | 0 | 0.56 | 0.18 | 0.46 | 0.31 | 0.59 | 0.51 |

| | | | | | | | | | | | | | | | | |
|---|---|---|---|---|---|---|---|---|---|---|---|---|---|---|---|---|
| 10 | 6123495 | rs12251307 | -0.344 | -0.276 | -0.155 | 0.643 | -0.037 | 0.02 | 0.02 | 0 | 0.18 | 0.33 | 0.16 | 0.21 | 0.1 | 0.19 |
| 10 | 6130077 | rs6602437 | -0.404 | 0.102 | -0.561 | 0.747 | 0.006 | 0.005 | 0.005 | 0 | 0.48 | 0.61 | 0.48 | 0.45 | 0.54 | 0.37 |
| 10 | 6135025 | rs1571026 | -0.066 | -0.734 | 0.017 | 0.176 | -0.057 | -0.112 | -0.112 | 0.374 | 0.22 | 0.45 | 0.12 | 0.05 | 0.12 | 0.16 |
| 10 | 6143619 | rs3930531 | 0.8 | -1.244 | 0.421 | 0.465 | -0.004 | 0.003 | 0.003 | 0 | 0.4 | 0.35 | 0.45 | 0.33 | 0.41 | 0.45 |
| 10 | 6150501 | rs11256802 | 0.924 | -1.192 | 0.341 | 0.781 | -0.004 | 0.009 | 0.009 | 0 | 0.11 | 0.11 | 0.06 | 0.19 | 0.08 | 0.07 |
| 10 | 6153297 | rs6602450 | 1.386 | -1.256 | -0.12 | 0.945 | -0.011 | 0.021 | 0.021 | 0 | 0.34 | 0.15 | 0.33 | 0.22 | 0.33 | 0.3 |
| 10 | 6153868 | rs10905819 | 0.704 | -1.072 | -0.246 | 1.148 | -0.087 | 0.002 | 0.002 | 0 | 0.27 | 0.13 | 0.19 | 0.26 | 0.41 | 0.39 |
| 10 | 6158412 | rs8463 | 0.252 | -1.19 | 0.072 | 1.126 | -0.051 | -0.061 | -0.061 | 0 | 0.23 | 0.44 | 0.29 | 0.11 | 0.15 | 0.16 |
| 10 | 6161781 | rs4749997 | 0.993 | -1.022 | -0.161 | 1.145 | 0 | -0.027 | -0.027 | 0 | 0.4 | 0.26 | 0.6 | 0.39 | 0.39 | 0.45 |
| 10 | 6164310 | rs1073646 | 1.71 | -0.989 | -0.474 | 1.111 | -0.072 | 0.022 | 0.022 | 0.536 | 0.33 | 0.03 | 0.29 | 0.2 | 0.33 | 0.32 |
| 10 | 6165030 | rs1541020 | 0.797 | -0.72 | -0.204 | 1.208 | -0.076 | 0 | 0 | 0 | 0.24 | 0.13 | 0.2 | 0.24 | 0.39 | 0.26 |
| 10 | 6165525 | rs4750000 | 0.283 | -0.947 | 0.161 | 1.329 | -0.062 | 0.082 | 0.082 | 0 | 0.8 | 0.64 | 0.89 | 0.63 | 0.88 | 0.82 |
| 10 | 6167124 | rs10905856 | 0.127 | -0.925 | 0.015 | 1.174 | -0.004 | -0.002 | -0.002 | 0 | 0.12 | 0.1 | 0.06 | 0.17 | 0.06 | 0.14 |
| 10 | 6169272 | rs7905816 | 0.219 | -0.065 | 0.057 | 1.181 | 0.008 | 0.016 | 0.016 | 0 | 0.21 | 0.25 | 0.15 | 0.41 | 0.28 | 0.19 |
| 10 | 6169685 | rs4750005 | 0.451 | -0.181 | 0.241 | 0.668 | -0.003 | 0.004 | 0.004 | 0 | 0.59 | 0.37 | 0.45 | 0.28 | 0.47 | 0.49 |
| 10 | 6171924 | rs4310524 | -0.331 | -0.551 | -0.367 | 0.256 | 0 | 0.011 | 0.011 | 0 | 0.53 | 0.62 | 0.62 | 0.34 | 0.45 | 0.42 |
| 10 | 6173416 | rs7914915 | -1.355 | -0.576 | -0.376 | 0.364 | -0.04 | 0.014 | 0.014 | 0 | 0.91 | 0.83 | 0.93 | 0.92 | 0.97 | 0.89 |
| 10 | 6173465 | rs11256933 | 0.799 | -0.253 | -0.572 | 0.434 | -0.015 | -0.022 | -0.022 | 0 | 0.21 | 0.14 | 0.19 | 0.55 | 0.3 | 0.25 |
| 10 | 6176166 | rs4747886 | -0.223 | -0.458 | -0.529 | 0.336 | 0.007 | 0.007 | 0.007 | 0 | 0.48 | 0.53 | 0.39 | 0.37 | 0.39 | 0.38 |
| 10 | 6177083 | rs10905879 | 0.604 | -0.036 | -0.296 | 0.658 | -0.007 | -0.011 | -0.011 | 0.381 | 0.19 | 0.27 | 0.15 | 0.11 | 0.18 | 0.23 |
| 10 | 6177626 | rs12219630 | 0.45 | -0.033 | -0.566 | 0.673 | 0.002 | 0.019 | 0.019 | 0 | 0.53 | 0.39 | 0.55 | 0.35 | 0.46 | 0.38 |
| 10 | 6177682 | rs4747887 | 0.45 | -0.033 | -0.566 | 0.673 | 0.002 | 0.019 | 0.019 | 0 | 0.53 | 0.39 | 0.55 | 0.35 | 0.46 | 0.38 |
| 10 | 6178614 | rs1983890 | 0.763 | -0.119 | -0.608 | 0.565 | 0.013 | 0.083 | 0.083 | 0 | 0.44 | 0.31 | 0.41 | 0.09 | 0.4 | 0.4 |
| 10 | 6178941 | rs11598494 | 1.121 | -0.092 | -0.713 | 0.461 | 0.016 | 0.056 | 0.056 | 0 | 0.41 | 0.3 | 0.49 | 0.17 | 0.37 | 0.3 |
| 10 | 6178961 | rs12571538 | 1.202 | -0.029 | -0.565 | 0.648 | 0 | 0.004 | 0.004 | 0.09 | 0.11 | 0.1 | 0.15 | 0.26 | 0.09 | 0.09 |
| 10 | 6181709 | rs3763700 | 0.817 | 0.133 | -0.761 | 0.563 | 0.002 | 0.053 | 0.053 | 0 | 0.42 | 0.31 | 0.5 | 0.18 | 0.4 | 0.42 |
| 10 | 6182423 | rs3750671 | 0.099 | 0.711 | -0.797 | 1.01 | 0.001 | 0.004 | 0.004 | 0 | 0.75 | 0.78 | 0.8 | 0.55 | 0.68 | 0.73 |
| 10 | 6188543 | rs943119 | -0.08 | 0.491 | -0.642 | 0.923 | -0.006 | 0.017 | 0.017 | 0 | 0.13 | 0.15 | 0.08 | 0.25 | 0.12 | 0.13 |
| 10 | 6188591 | rs7074372 | -0.351 | 0.98 | -0.819 | 0.664 | 0.003 | 0.012 | 0.012 | 0.223 | 0.61 | 0.74 | 0.71 | 0.67 | 0.61 | 0.54 |
| 10 | 6190343 | rs725105 | -0.029 | 1.988 | -0.512 | 0.316 | 0.054 | -0.018 | -0.018 | 0 | 0.17 | 0.33 | 0.22 | 0.12 | 0.29 | 0.27 |
| 10 | 6194476 | rs7096344 | -0.157 | 0.869 | -0.741 | 0.538 | 0.008 | 0.01 | 0.01 | 0 | 0.34 | 0.25 | 0.18 | 0.25 | 0.25 | 0.31 |
| 10 | 6208465 | rs652318 | -0.271 | 1.46 | -1.225 | 0.691 | -0.041 | -0.023 | -0.023 | 0 | 0.21 | 0.09 | 0.23 | 0.25 | 0.23 | 0.18 |
| 10 | 6210655 | rs630204 | -0.284 | 0.485 | -1.723 | 0.219 | 0.057 | -0.009 | -0.009 | 0.709 | 0.63 | 0.29 | 0.61 | 0.39 | 0.76 | 0.69 |
| 10 | 6210740 | rs11599186 | 1.359 | 0.974 | -2.147 | -0.211 | -0.054 | 0.101 | 0.101 | 0 | 0.21 | 0.04 | 0.14 | 0.03 | 0.24 | 0.09 |
| 10 | 6213960 | rs3814195 | 0.814 | 0.185 | -1.479 | 0.435 | -0.069 | 0.072 | 0.072 | 0 | 0.23 | 0.39 | 0.15 | 0.33 | 0.14 | 0.19 |
| 10 | 6215091 | rs658386 | 0.904 | 0.359 | -1.614 | 0.441 | -0.05 | 0.026 | 0.026 | 0 | 0.21 | 0.08 | 0.15 | 0.15 | 0.23 | 0.17 |
| 10 | 6215257 | rs2387015 | 0.444 | 0.162 | -1.452 | 0.498 | -0.064 | 0.058 | 0.058 | 0 | 0.25 | 0.48 | 0.16 | 0.37 | 0.15 | 0.2 |
| 10 | 6218042 | rs616246 | -0.559 | 0.35 | -1.631 | 0.477 | 0.005 | 0.002 | 0.002 | 0 | 0.56 | 0.48 | 0.7 | 0.62 | 0.6 | 0.68 |
| 10 | 6218883 | rs602395 | 0.692 | 1.351 | -2.076 | 0.523 | -0.077 | 0.008 | 0.008 | 0 | 0.21 | 0.06 | 0.2 | 0.17 | 0.24 | 0.18 |
| 10 | 6219010 | rs12359393 | 0.682 | 1.252 | -1.978 | 0.341 | -0.053 | 0.041 | 0.041 | 0.015 | 0.2 | 0.03 | 0.23 | 0.1 | 0.24 | 0.42 |
| 10 | 6219037 | rs12777632 | 1.096 | 1.156 | -1.416 | 0.234 | -0.001 | -0.009 | -0.009 | 0.698 | 0.21 | 0.25 | 0.16 | 0.04 | 0.25 | 0.13 |
| 10 | 6220360 | rs674430 | -0.105 | 1.077 | -1.75 | 0.677 | 0.012 | 0.039 | 0.039 | 0 | 0.59 | 0.25 | 0.54 | 0.31 | 0.65 | 0.65 |
| 10 | 6220725 | rs11257090 | 0.424 | 1.062 | -1.924 | 0.575 | -0.015 | -0.025 | -0.025 | 0 | 0.52 | 0.22 | 0.56 | 0.5 | 0.57 | 0.51 |
| 10 | 6221318 | rs678395 | -0.616 | 1.423 | -1.261 | 0.579 | -0.013 | 0.065 | 0.065 | 0 | 0.16 | 0.07 | 0.08 | 0.03 | 0.11 | 0.14 |
| 10 | 6222819 | rs11257103 | -1.419 | 1.329 | -0.32 | 1.233 | 0.006 | 0.064 | 0.064 | 0.537 | 0.27 | 0.44 | 0.46 | 0.62 | 0.3 | 0.36 |
| 10 | 6224566 | rs17151529 | -1.425 | 1.923 | -0.547 | 1.338 | 0.008 | 0.003 | 0.003 | 0 | 0.14 | 0.17 | 0.2 | 0.21 | 0.29 | 0.2 |
| 10 | 6224765 | rs7083321 | -1.85 | 1.599 | -0.421 | 1.413 | 0.002 | 0.007 | 0.007 | 0 | 0.19 | 0.22 | 0.35 | 0.29 | 0.26 | 0.23 |
| 10 | 6228226 | rs666595 | 0.952 | 0.93 | -0.369 | 1.077 | 0 | -0.001 | -0.001 | 0 | 0.69 | 0.65 | 0.49 | 0.58 | 0.71 | 0.52 |
| 10 | 6228413 | rs647439 | 1.081 | 0.674 | -0.568 | 0.83 | 0.015 | 0.028 | 0.028 | 0 | 0.48 | 0.17 | 0.37 | 0.42 | 0.55 | 0.35 |
| 10 | 6228598 | rs646564 | 1.505 | 0.547 | -0.358 | 0.742 | -0.026 | -0.002 | -0.002 | 0 | 0.44 | 0.14 | 0.36 | 0.43 | 0.54 | 0.34 |
| 10 | 6228616 | rs6602502 | -0.364 | 0.55 | -0.938 | 0.237 | -0.02 | 0.094 | 0.094 | 0 | 0.22 | 0.09 | 0.18 | 0 | 0.19 | 0.13 |
| 10 | 6228878 | rs653659 | 1.39 | -0.035 | -0.323 | 0.829 | 0.019 | 0.029 | 0.029 | 0 | 0.49 | 0.23 | 0.38 | 0.43 | 0.56 | 0.35 |
| 10 | 6229574 | rs631902 | 0.538 | -0.023 | -0.308 | 0.687 | 0.003 | -0.016 | -0.016 | 0 | 0.59 | 0.31 | 0.5 | 0.5 | 0.61 | 0.38 |
| 10 | 6230178 | rs618982 | 0.999 | -0.203 | 0.104 | 0.126 | 0.006 | 0.005 | 0.005 | 0 | 0.65 | 0.29 | 0.7 | 0.79 | 0.79 | 0.71 |
| 10 | 6233300 | rs681966 | 0.703 | -0.007 | -0.064 | 0.313 | 0.003 | 0.003 | 0.003 | 0 | 0.65 | 0.34 | 0.75 | 0.69 | 0.73 | 0.58 |
| 10 | 6236332 | rs637452 | 1.573 | -0.284 | 0.621 | 0.045 | -0.022 | -0.069 | -0.069 | 0 | 0.83 | 0.64 | 0.77 | 0.93 | 0.87 | 0.94 |
| 10 | 6237156 | rs11257188 | -2.248 | -1.117 | 0.639 | 0.214 | -0.068 | -0.027 | -0.027 | 0.826 | 0.23 | 0.52 | 0.11 | 0.18 | 0.1 | 0.19 |
| 10 | 6237875 | rs11257192 | -1.27 | -0.884 | 0.682 | 0.437 | -0.025 | 0.031 | 0.031 | 0 | 0.05 | 0.13 | 0.04 | 0.09 | 0.01 | 0.1 |
| 10 | 6240195 | rs12219678 | -2.159 | -1.024 | 0.792 | 0.401 | -0.086 | 0.023 | 0.023 | 0.059 | 0.17 | 0.41 | 0.08 | 0.17 | 0.05 | 0.13 |
| 10 | 6241803 | rs7076428 | -1.72 | -0.89 | 0.598 | 0.395 | -0.03 | 0.001 | 0.001 | 0.375 | 0.14 | 0.28 | 0.18 | 0.16 | 0.07 | 0.18 |
| 10 | 6245317 | rs6602514 | -0.47 | -0.746 | 0.01 | -0.142 | -0.003 | -0.025 | -0.025 | 0 | 0.35 | 0.37 | 0.27 | 0.14 | 0.33 | 0.22 |
| 10 | 6251252 | rs2516614 | 0.347 | -0.493 | -0.033 | 0.476 | 0.049 | 0.05 | 0.05 | 0 | 0.26 | 0.34 | 0.52 | 0.56 | 0.37 | 0.5 |
| 10 | 6253651 | rs2516617 | -0.635 | -1.021 | 0.234 | 1.133 | -0.009 | 0.003 | 0.003 | 0.008 | 0.25 | 0.36 | 0.26 | 0.26 | 0.21 | 0.23 |
| 10 | 6253983 | rs2516618 | -0.093 | -0.911 | 0.607 | 1.496 | 0.019 | 0.035 | 0.035 | 0 | 0.16 | 0.34 | 0.23 | 0.17 | 0.17 | 0.22 |
| 10 | 6255931 | rs756567 | -1.339 | -1.752 | 0.011 | 1.135 | 0.024 | 0.009 | 0.009 | 0 | 0.38 | 0.69 | 0.29 | 0.33 | 0.25 | 0.31 |
| 10 | 6258052 | rs2148294 | 1.043 | -1.323 | -0.005 | 1.225 | -0.011 | 0.109 | 0.109 | 0 | 0.89 | 0.55 | 0.9 | 0.81 | 0.95 | 0.86 |
| 10 | 6259239 | rs2516624 | 0.651 | -1.013 | 0.199 | 0.906 | -0.04 | -0.072 | -0.072 | 0 | 0.88 | 0.57 | 0.96 | 0.98 | 0.97 | 0.93 |
| 10 | 6260717 | rs11257349 | -1.039 | -0.101 | -0.803 | 1.478 | -0.007 | 0.002 | 0.002 | 0.27 | 0.11 | 0.11 | 0.02 | 0.14 | 0.05 | 0.02 |
| 10 | 6269855 | rs7071157 | -0.247 | 0.425 | -1.28 | 1.001 | 0.031 | 0.035 | 0.035 | 0.016 | 0.43 | 0.23 | 0.52 | 0.68 | 0.43 | 0.59 |

| | | | | | | | | | | | | | | | | |
|---|---|---|---|---|---|---|---|---|---|---|---|---|---|---|---|---|
| 10 | 6270465 | rs3793721 | 0.057 | 0.426 | -1.192 | 0.587 | 0.021 | 0.028 | 0.028 | 0 | 0.51 | 0.38 | 0.47 | 0.62 | 0.6 | 0.56 |
| 10 | 6271291 | rs12255582 | -0.546 | 0.608 | -0.227 | 0.671 | 0.037 | 0.045 | 0.045 | 1.319 | 0.09 | 0.22 | 0.15 | 0.12 | 0.1 | 0.04 |
| 10 | 6272169 | rs7904792 | -1.132 | 0.282 | 0.094 | 1.193 | 0.012 | 0.118 | 0.118 | 0 | 0.19 | 0.42 | 0.13 | 0.49 | 0.19 | 0.25 |
| 10 | 6274511 | rs11257456 | -0.348 | 0.318 | -0.678 | 0.172 | -0.004 | -0.075 | -0.075 | 0 | 0.6 | 0.61 | 0.75 | 0.89 | 0.6 | 0.79 |
| 10 | 6275602 | rs3750667 | 1.327 | 0.552 | -1.339 | 0.306 | -0.08 | -0.149 | -0.149 | 4.142 | 0.25 | 0.03 | 0.39 | 0.35 | 0.31 | 0.34 |
| 10 | 6276574 | rs1064891 | 0.579 | 0.615 | -0.881 | 0.718 | -0.017 | 0.016 | 0.016 | 1.519 | 0.46 | 0.24 | 0.45 | 0.38 | 0.42 | 0.43 |
| 10 | 6277556 | rs11257470 | 0.686 | 0.722 | -0.544 | 1.013 | 0.009 | 0.1 | 0.1 | 3.063 | 0.47 | 0.23 | 0.44 | 0.27 | 0.4 | 0.42 |
| 10 | 6277661 | rs11257472 | 0.477 | 0.659 | -0.371 | 0.717 | -0.014 | -0.039 | -0.039 | 0 | 0.62 | 0.32 | 0.55 | 0.46 | 0.61 | 0.51 |
| 10 | 6278851 | rs11257482 | 0.493 | 0.664 | -0.383 | 0.724 | -0.015 | -0.05 | -0.05 | 0 | 0.62 | 0.31 | 0.55 | 0.47 | 0.61 | 0.51 |
| 10 | 6280493 | rs9733630 | 0.124 | 0.413 | 0.382 | 0.7 | 0.005 | 0.035 | 0.035 | 0.346 | 0.76 | 0.72 | 0.78 | 0.51 | 0.73 | 0.7 |
| 10 | 6284473 | rs17361911 | -0.527 | -0.281 | 0.807 | 0.783 | 0.007 | -0.042 | -0.042 | 0.382 | 0.4 | 0.45 | 0.43 | 0.13 | 0.48 | 0.4 |
| 10 | 6286606 | rs4750166 | -0.358 | 0.039 | 0.84 | 0.946 | -0.015 | 0.197 | 0.197 | 0 | 0.88 | 0.77 | 0.9 | 0.51 | 0.93 | 0.81 |
| 10 | 6286839 | rs7916923 | -0.33 | -0.023 | 0.83 | 0.846 | 0.007 | -0.059 | -0.059 | 3.681 | 0.39 | 0.37 | 0.3 | 0.09 | 0.49 | 0.29 |
| 10 | 6288042 | rs10795936 | 0.724 | -0.478 | 1.205 | 1.905 | -0.015 | 0.203 | 0.203 | 0 | 0.86 | 0.73 | 0.83 | 0.51 | 0.92 | 0.8 |
| 10 | 6292607 | rs2185691 | -0.078 | -0.581 | 1.546 | 1.546 | -0.011 | 0.02 | 0.02 | 0 | 0.11 | 0.16 | 0.19 | 0.13 | 0.06 | 0.24 |
| 10 | 6292805 | rs7897489 | -1.345 | -0.814 | 2.013 | 2.047 | 0.008 | 0.191 | 0.191 | 0.026 | 0.12 | 0.4 | 0.16 | 0.36 | 0.1 | 0.16 |
| 10 | 6292875 | rs2153754 | 0.319 | -1.23 | 1.52 | 1.835 | -0.007 | 0.001 | 0.001 | 0 | 0.68 | 0.66 | 0.77 | 0.68 | 0.73 | 0.76 |
| 10 | 6294533 | rs7094936 | -1.45 | -1.26 | 1.949 | 2.223 | -0.02 | 0.056 | 0.056 | 0 | 0.2 | 0.38 | 0.26 | 0.31 | 0.15 | 0.2 |
| 10 | 6295548 | rs10508301 | 1.698 | -1.307 | 1.006 | 2.005 | 0.029 | -0.017 | -0.017 | 0.027 | 0.71 | 0.23 | 0.65 | 0.4 | 0.83 | 0.58 |
| 10 | 6306289 | rs7092772 | -1.324 | -1.428 | 1.462 | 1.964 | -0.069 | 0.047 | 0.047 | 0.186 | 0.18 | 0.42 | 0.22 | 0.24 | 0.06 | 0.26 |
| 10 | 6307423 | rs11596955 | 1.789 | -1.565 | 1.223 | 1.801 | 0.062 | 0.178 | 0.178 | 0 | 0.5 | 0.26 | 0.36 | 0.15 | 0.58 | 0.35 |
| 10 | 6308484 | rs12780002 | -1.176 | -1.154 | 1.75 | 2.379 | -0.002 | 0.007 | 0.007 | 0.112 | 0.32 | 0.38 | 0.39 | 0.45 | 0.36 | 0.39 |
| 10 | 6309175 | rs10795949 | 1.006 | -1.074 | 1.355 | 2.323 | -0.003 | 0.001 | 0.001 | 0.073 | 0.66 | 0.4 | 0.55 | 0.39 | 0.63 | 0.61 |
| 10 | 6311749 | rs11257726 | -0.031 | -1.149 | 0.283 | 2.035 | 0.013 | 0.031 | 0.031 | 0 | 0.2 | 0.08 | 0.14 | 0.14 | 0.22 | 0.2 |
| 10 | 6313880 | rs12220579 | -0.044 | -1.116 | 0.174 | 2.073 | 0.008 | -0.038 | -0.038 | 0.502 | 0.07 | 0.01 | 0.11 | 0.17 | 0.05 | 0.21 |
| 10 | 6314221 | rs1544198 | 1.543 | -1.088 | 0.481 | 1.856 | 0.018 | 0.027 | 0.027 | 0.012 | 0.56 | 0.31 | 0.53 | 0.35 | 0.6 | 0.4 |
| 10 | 6314722 | rs11257741 | -0.401 | -0.502 | 0.289 | 1.409 | -0.009 | 0.024 | 0.024 | 0 | 0.2 | 0.03 | 0.14 | 0.13 | 0.21 | 0.19 |
| 10 | 6315794 | rs7077335 | 1.328 | 0.027 | 1.252 | 1.63 | 0.074 | 0.332 | 0.332 | 3.779 | 0.95 | 0.42 | 0.92 | 0.85 | 0.99 | 0.99 |
| 10 | 6318458 | rs17152987 | -1.357 | 0.37 | 0.142 | 1.032 | -0.027 | -0.023 | -0.023 | 0 | 0.07 | 0.03 | 0.09 | 0.12 | 0.08 | 0.04 |
| 10 | 6318937 | rs6602585 | 0.901 | -0.545 | 0.671 | 0.732 | -0.045 | -0.035 | -0.035 | 0 | 0.75 | 0.46 | 0.8 | 0.78 | 0.79 | 0.74 |
| 10 | 6335731 | rs17153204 | -1.39 | -0.841 | 1.663 | 0.881 | -0.008 | 0.003 | 0.003 | 0 | 0.17 | 0.16 | 0.14 | 0.24 | 0.13 | 0.26 |
| 10 | 6342636 | rs4747994 | 1.371 | -0.692 | 1.388 | 1.053 | 0.022 | 0.041 | 0.041 | 0.02 | 0.57 | 0.21 | 0.48 | 0.33 | 0.65 | 0.42 |
| 10 | 6342654 | rs4750246 | 0.877 | -0.657 | 1.634 | 1.039 | -0.033 | 0.049 | 0.049 | 0 | 0.83 | 0.57 | 0.85 | 0.75 | 0.91 | 0.76 |
| 10 | 6350590 | rs12262851 | -0.938 | -0.529 | 1.783 | 1.264 | -0.002 | -0.012 | -0.012 | 0.003 | 0.24 | 0.24 | 0.37 | 0.42 | 0.22 | 0.34 |
| 10 | 6352975 | rs7919234 | -1.56 | -0.676 | 1.71 | 1.193 | 0.017 | 0.009 | 0.009 | 0.054 | 0.29 | 0.2 | 0.18 | 0.26 | 0.16 | 0.23 |
| 10 | 6353355 | rs7901637 | 1.427 | -0.636 | 1.832 | 1.081 | -0.039 | 0.006 | 0.006 | 0.21 | 0.67 | 0.48 | 0.55 | 0.33 | 0.82 | 0.57 |
| 10 | 6356210 | rs764189 | 1.694 | -0.621 | 1.642 | 0.809 | 0.012 | 0.028 | 0.028 | 0 | 0.59 | 0.38 | 0.5 | 0.29 | 0.63 | 0.47 |
| 10 | 6357437 | rs7917326 | -1.139 | -0.537 | 1.499 | 0.941 | -0.003 | 0.022 | 0.022 | 0 | 0.14 | 0.12 | 0.34 | 0.68 | 0.12 | 0.38 |
| 10 | 6360293 | rs2387396 | 1.575 | -0.678 | 1.36 | 0.732 | 0.006 | 0.02 | 0.02 | 0 | 0.53 | 0.63 | 0.47 | 0.26 | 0.58 | 0.44 |
| 10 | 6363557 | rs2224781 | 0.582 | -0.66 | 1.261 | 0.745 | -0.007 | 0.02 | 0.02 | 0 | 0.87 | 0.79 | 0.86 | 0.78 | 0.89 | 0.8 |
| 10 | 6363948 | rs7919350 | -0.39 | -0.655 | 0.742 | 0.784 | -0.003 | 0.022 | 0.022 | 0 | 0.94 | 0.88 | 0.88 | 0.85 | 0.95 | 0.95 |
| 10 | 6364754 | rs12779447 | -0.841 | -0.605 | 0.682 | 0.749 | 0.052 | -0.01 | -0.01 | 0 | 0.07 | 0.11 | 0.08 | 0.02 | 0.18 | 0.08 |
| 10 | 6368760 | rs7087682 | 1.404 | -0.653 | 0.654 | 0.719 | 0.014 | 0.048 | 0.048 | 0 | 0.55 | 0.66 | 0.69 | 0.73 | 0.6 | 0.64 |
| 10 | 6370545 | rs7096921 | 1.558 | -0.489 | 0.504 | 0.281 | 0.008 | 0.02 | 0.02 | 0 | 0.51 | 0.45 | 0.47 | 0.26 | 0.58 | 0.44 |
| 10 | 6377100 | rs1557110 | 0.995 | 0.841 | 0.199 | 0.347 | -0.005 | 0.002 | 0.002 | 0.027 | 0.87 | 0.83 | 0.86 | 0.82 | 0.89 | 0.8 |
| 10 | 6387973 | rs7917852 | 1.841 | 0.913 | 0.01 | 0.356 | 0.023 | -0.01 | -0.01 | 0 | 0.79 | 0.71 | 0.77 | 0.8 | 0.69 | 0.71 |
| 10 | 6387992 | rs4750311 | 1.188 | -0.101 | -0.24 | 0.565 | -0.006 | 0.004 | 0.004 | 0 | 0.86 | 0.82 | 0.86 | 0.8 | 0.89 | 0.79 |
| 10 | 6388928 | rs17154230 | -1.296 | 0.459 | -0.65 | 0.452 | -0.038 | -0.018 | -0.018 | 0 | 0.09 | 0 | 0.06 | 0.1 | 0.09 | 0.14 |
| 10 | 6389605 | rs947470 | 0.227 | -0.332 | -0.81 | -0.431 | 0.009 | 0.036 | 0.036 | 0 | 0.59 | 0.4 | 0.73 | 0.84 | 0.65 | 0.64 |
| 10 | 6389914 | rs947473 | -0.27 | -0.747 | 0.004 | -0.694 | -0.004 | -0.059 | -0.059 | 0.084 | 0.74 | 0.71 | 0.85 | 0.97 | 0.76 | 0.81 |
| 10 | 6390107 | rs7917676 | 0.87 | -0.855 | 0.522 | -0.741 | -0.004 | -0.048 | -0.048 | 0 | 0.61 | 0.56 | 0.75 | 0.84 | 0.63 | 0.68 |
| 10 | 6390450 | rs947474 | 1.383 | -0.918 | 0.836 | -0.76 | 0.024 | 0.007 | 0.007 | 0.184 | 0.85 | 0.69 | 0.88 | 0.88 | 0.82 | 0.85 |
| 10 | 6391061 | rs2181622 | 1.722 | -1.641 | 0.586 | -0.724 | 0.042 | -0.005 | -0.005 | 0 | 0.66 | 0.22 | 0.74 | 0.7 | 0.73 | 0.64 |
| 10 | 6392848 | rs744254 | 1.632 | -1.693 | 1.185 | -0.72 | 0.013 | -0.003 | -0.003 | 0 | 0.67 | 0.28 | 0.75 | 0.7 | 0.73 | 0.65 |
| 10 | 6393009 | rs744253 | 1.526 | -1.684 | 1.197 | -0.686 | 0.006 | -0.004 | -0.004 | 1 | 0.7 | 0.28 | 0.76 | 0.7 | 0.74 | 0.67 |
| 10 | 6394995 | rs725206 | 1.91 | -0.947 | 0.493 | -1.307 | -0.063 | -0.053 | -0.053 | 0.46 | 0.32 | 0.06 | 0.53 | 0.66 | 0.34 | 0.41 |
| 10 | 6396795 | rs12570830 | 0.707 | -0.908 | 0.452 | -1.169 | 0.019 | -0.061 | -0.061 | 0 | 0.11 | 0 | 0.09 | 0.17 | 0.08 | 0.12 |
| 10 | 6401625 | rs4558075 | 0.725 | -0.501 | 0.798 | -1.356 | 0.009 | -0.035 | -0.035 | 0.931 | 0.86 | 0.78 | 0.9 | 0.98 | 0.83 | 0.88 |
| 10 | 6402741 | rs10796045 | 0.947 | -0.522 | 0.736 | -1.356 | 0.006 | -0.006 | -0.006 | 0.403 | 0.84 | 0.71 | 0.89 | 0.88 | 0.83 | 0.81 |
| 10 | 6405507 | rs10906340 | -0.583 | -0.307 | 0.293 | -1.37 | 0.005 | -0.015 | -0.015 | 1.064 | 0.1 | 0.11 | 0.05 | 0.02 | 0.09 | 0.1 |
| 10 | 6407737 | rs11258317 | -0.006 | -0.366 | 0.059 | -1.385 | -0.015 | 0.068 | 0.068 | 0 | 0.35 | 0.22 | 0.22 | 0.04 | 0.39 | 0.24 |
| 10 | 6409441 | rs7911141 | -1.39 | -0.55 | 0.916 | -0.561 | -0.023 | 0.169 | 0.169 | 0.118 | 0.1 | 0.48 | 0.07 | 0.17 | 0.03 | 0.12 |
| 10 | 6409502 | rs7915166 | -0.557 | -0.794 | 0.698 | -0.824 | -0.018 | -0.015 | -0.015 | 0 | 0.44 | 0.48 | 0.22 | 0.34 | 0.35 | 0.46 |
| 10 | 6410125 | rs4748026 | 1.523 | 0.053 | 0.53 | -0.496 | -0.045 | 0.064 | 0.064 | 0.077 | 0.54 | 0.17 | 0.7 | 0.69 | 0.53 | 0.48 |
| 10 | 6424490 | rs7904311 | -1.2 | 0.323 | 1.342 | -0.544 | 0.035 | 0.021 | 0.021 | 0 | 0.16 | 0.36 | 0.08 | 0.16 | 0.13 | 0.15 |
| 10 | 6426435 | rs1341929 | -0.109 | 0.365 | 0.78 | -0.667 | 0.007 | -0.051 | -0.051 | 0 | 0.84 | 0.79 | 0.87 | 0.99 | 0.77 | 0.86 |
| 10 | 6427952 | rs12763729 | 1.796 | 0.22 | 1.086 | -0.044 | -0.001 | 0 | 0 | 0 | 0.69 | 0.35 | 0.83 | 0.7 | 0.75 | 0.62 |
| 10 | 6430767 | rs10906402 | -0.619 | 0.12 | 1.096 | 0.153 | -0.018 | -0.059 | -0.059 | 0.181 | 0.13 | 0 | 0.05 | 0.15 | 0.12 | 0.21 |
| 10 | 6435119 | rs11258455 | -1.23 | 0.123 | 0.918 | -0.012 | -0.032 | 0.081 | 0.081 | 0.008 | 0.18 | 0.44 | 0.1 | 0.25 | 0.12 | 0.21 |

| | | | | | | | | | | | | | | | | |
|---|---|---|---|---|---|---|---|---|---|---|---|---|---|---|---|---|
| 10 | 6435374 | rs592456 | 1.098 | 0.689 | 0.169 | -0.384 | 0.006 | -0.026 | -0.026 | 0 | 0.9 | 0.89 | 0.94 | 0.97 | 0.91 | 0.86 |
| 10 | 6436034 | rs661985 | 1.429 | 0.753 | 0.113 | -0.328 | 0.01 | -0.022 | -0.022 | 0 | 0.89 | 0.82 | 0.93 | 0.93 | 0.9 | 0.8 |
| 10 | 6438081 | rs681604 | 1.397 | 0.234 | 0.243 | -0.376 | -0.028 | -0.024 | -0.024 | 0.458 | 0.83 | 0.33 | 0.89 | 0.83 | 0.92 | 0.88 |
| 10 | 6440676 | rs10906429 | -1.439 | 0.674 | -0.082 | -0.429 | -0.028 | -0.011 | -0.011 | 0.194 | 0.12 | 0.03 | 0.08 | 0.17 | 0.14 | 0.16 |
| 10 | 6443404 | rs12413588 | 1.513 | 1.537 | -0.974 | -1.251 | -0.019 | 0.225 | 0.225 | 0.723 | 0.55 | 0.14 | 0.54 | 0.79 | 0.46 | 0.61 |
| 10 | 6453415 | rs616490 | -0.411 | 1.661 | -0.048 | -1.18 | -0.02 | 0.058 | 0.058 | 0 | 0.21 | 0.07 | 0.11 | 0.08 | 0.26 | 0.16 |
| 10 | 6457597 | rs589142 | -2.212 | 1.659 | 0.053 | -1.25 | 0.019 | 0.179 | 0.179 | 0.942 | 0.48 | 0.78 | 0.45 | 0.25 | 0.55 | 0.4 |
| 10 | 6459414 | rs4750422 | -0.048 | 1.636 | 0.128 | -1.213 | -0.002 | -0.002 | -0.002 | 0 | 0.9 | 0.88 | 0.94 | 0.85 | 0.93 | 0.89 |
| 10 | 6459874 | rs10752330 | 0.79 | 1.589 | 0.921 | -1.233 | -0.015 | -0.019 | -0.019 | 0 | 0.87 | 0.53 | 0.9 | 0.85 | 0.93 | 0.88 |
| 10 | 6464156 | rs667899 | 0.459 | 1.794 | 1.01 | -1.262 | 0.144 | -0.051 | -0.051 | 0.011 | 0.87 | 0.47 | 0.85 | 0.97 | 0.78 | 0.87 |
| 10 | 6465273 | rs4750434 | -0.109 | 1.69 | 0.745 | -1.214 | -0.063 | 0.041 | 0.041 | 0.155 | 0.2 | 0.01 | 0.23 | 0.06 | 0.25 | 0.12 |
| 10 | 6469155 | rs582052 | -2.303 | 1.652 | 0.952 | -1.243 | 0.001 | 0.166 | 0.166 | 0.507 | 0.47 | 0.83 | 0.45 | 0.26 | 0.54 | 0.36 |
| 10 | 6469324 | rs4750439 | -0.33 | 1.662 | 1.01 | -1.247 | -0.007 | 0.044 | 0.044 | 0.319 | 0.22 | 0.08 | 0.23 | 0.07 | 0.26 | 0.13 |
| 10 | 6469673 | rs2453 | -0.093 | 1.663 | 0.99 | -1.247 | -0.063 | 0.042 | 0.042 | 0.147 | 0.2 | 0.01 | 0.23 | 0.06 | 0.25 | 0.12 |
| 10 | 6469817 | rs2236380 | 1.702 | 1.652 | 1.027 | -1.247 | -0.017 | 0.035 | 0.035 | 1.627 | 0.73 | 0.2 | 0.78 | 0.82 | 0.71 | 0.76 |
| 10 | 6470587 | rs7082071 | 0.25 | 1.675 | 1.099 | -1.242 | 0.054 | -0.022 | -0.022 | 0 | 0.1 | 0.13 | 0.11 | 0.03 | 0.22 | 0.12 |
| 10 | 6472247 | rs1889001 | 2.162 | 1.701 | 1.073 | -1.242 | -0.027 | 0.197 | 0.197 | 0.848 | 0.52 | 0.07 | 0.54 | 0.74 | 0.45 | 0.63 |
| 10 | 6472891 | rs11258747 | 0.062 | 1.702 | 1.12 | -1.21 | 0.069 | -0.021 | -0.021 | 0 | 0.09 | 0.13 | 0.11 | 0.03 | 0.22 | 0.12 |
| 10 | 6474027 | rs10796145 | 1.787 | 1.705 | 0.56 | -1.047 | -0.015 | 0.016 | 0.016 | 0.052 | 0.75 | 0.22 | 0.78 | 0.75 | 0.71 | 0.75 |
| 10 | 6474533 | rs17155850 | 0.019 | 1.349 | 0.502 | -0.884 | 0.008 | 0 | 0 | 0 | 0.09 | 0.02 | 0.05 | 0.14 | 0.06 | 0.09 |
| 10 | 6477267 | rs677986 | 0.551 | 1.497 | 0.577 | -0.923 | -0.001 | -0.003 | -0.003 | 0 | 0.77 | 0.75 | 0.75 | 0.87 | 0.74 | 0.87 |
| 10 | 6488790 | rs673964 | -0.785 | 1.401 | 0.266 | -0.975 | -0.017 | 0.054 | 0.054 | 0.233 | 0.23 | 0.13 | 0.28 | 0.08 | 0.3 | 0.15 |
| 10 | 6489652 | rs688391 | -1.824 | 1.315 | 0.57 | 0.162 | -0.003 | 0.005 | 0.005 | 0 | 0.37 | 0.45 | 0.36 | 0.54 | 0.36 | 0.33 |
| 10 | 6497307 | rs10508307 | -0.13 | 0.566 | -0.259 | -0.623 | 0.031 | -0.066 | -0.066 | 0.304 | 0.28 | 0.1 | 0.11 | 0.37 | 0.18 | 0.24 |
| 10 | 6497422 | rs586457 | -0.26 | 0.637 | -0.146 | -0.62 | -0.001 | -0.007 | -0.007 | 0 | 0.24 | 0.22 | 0.28 | 0.08 | 0.3 | 0.15 |
| 10 | 6500858 | rs11258960 | 0.269 | 0.697 | -0.509 | -0.586 | -0.078 | 0.058 | 0.058 | 0.194 | 0.21 | 0.01 | 0.27 | 0.07 | 0.29 | 0.13 |
| 10 | 6504126 | rs604663 | -1.365 | 0.754 | -0.376 | -0.679 | -0.004 | -0.011 | -0.011 | 4.469 | 0.54 | 0.39 | 0.4 | 0.52 | 0.48 | 0.36 |
| 10 | 6504676 | rs542817 | 0.334 | 0.651 | -0.273 | -0.66 | -0.012 | -0.038 | -0.038 | 0 | 0.17 | 0.25 | 0.05 | 0.08 | 0.09 | 0.06 |
| 10 | 6507603 | rs2026432 | 0.988 | 1.061 | -0.382 | -0.609 | 0.002 | 0.006 | 0.006 | 0 | 0.33 | 0.36 | 0.53 | 0.38 | 0.37 | 0.47 |
| 10 | 6508377 | rs3793727 | 0.365 | 1.467 | -0.734 | -0.758 | -0.052 | 0.009 | 0.009 | 0.285 | 0.23 | 0.11 | 0.25 | 0.21 | 0.33 | 0.13 |
| 10 | 6508563 | rs658230 | -0.804 | 1.24 | -0.79 | -0.886 | -0.023 | -0.021 | -0.021 | 0.314 | 0.55 | 0.26 | 0.38 | 0.54 | 0.53 | 0.41 |
| 10 | 6510454 | rs2026431 | 1.824 | 1.34 | -0.912 | -1.001 | -0.003 | 0.017 | 0.017 | 0 | 0.31 | 0.34 | 0.51 | 0.42 | 0.28 | 0.48 |
| 10 | 6511386 | rs591441 | 0.365 | 1.296 | -1.279 | -0.983 | 0.072 | -0.03 | -0.03 | 0 | 0.31 | 0.13 | 0.14 | 0.32 | 0.21 | 0.23 |
| 10 | 6517476 | rs12220818 | 0.526 | 2.028 | -0.748 | -0.534 | -0.024 | -0.013 | -0.013 | 0 | 0.07 | 0.02 | 0.07 | 0.08 | 0.14 | 0.1 |
| 10 | 6522506 | rs3793729 | 1.276 | 1.865 | -0.595 | -0.523 | -0.017 | 0.038 | 0.038 | 0 | 0.31 | 0.39 | 0.5 | 0.43 | 0.26 | 0.48 |
| 10 | 6522908 | rs494800 | 0.447 | 1.664 | -0.562 | -0.604 | 0.049 | -0.034 | -0.034 | 2.518 | 0.84 | 0.65 | 0.88 | 0.91 | 0.79 | 0.83 |
| 10 | 6526497 | rs501878 | -0.065 | 1.151 | -1.327 | -0.404 | 0.057 | 0.072 | 0.072 | 0.348 | 0.47 | 0.18 | 0.25 | 0.33 | 0.41 | 0.27 |
| 10 | 6530052 | rs790815 | -0.452 | 1.199 | -1.075 | -0.412 | -0.009 | -0.006 | -0.006 | 0 | 0.54 | 0.33 | 0.38 | 0.48 | 0.51 | 0.37 |
| 10 | 6531149 | rs587198 | -0.516 | 1.227 | -0.913 | -0.634 | -0.001 | -0.001 | -0.001 | 0 | 0.51 | 0.54 | 0.43 | 0.56 | 0.58 | 0.44 |
| 10 | 6532178 | rs472317 | 0.711 | 0.958 | -1.972 | -0.362 | -0.076 | -0.117 | -0.117 | 1.485 | 0.39 | 0.02 | 0.44 | 0.41 | 0.38 | 0.54 |
| 10 | 6533877 | rs3815975 | 0.029 | 1.025 | -1.461 | -0.498 | -0.002 | 0.002 | 0.002 | 3.651 | 0.41 | 0.55 | 0.44 | 0.44 | 0.42 | 0.3 |
| 10 | 6534384 | rs4750531 | -0.764 | 1.098 | -1.607 | -0.504 | 0.012 | -0.001 | -0.001 | 1.907 | 0.52 | 0.71 | 0.5 | 0.51 | 0.54 | 0.38 |
| 10 | 6534592 | rs648778 | 1.141 | 1.412 | -1.927 | -0.312 | -0.054 | -0.125 | -0.125 | 0 | 0.32 | 0.05 | 0.41 | 0.41 | 0.32 | 0.49 |
| 10 | 6536552 | rs6602745 | 0.149 | 1.503 | -1.242 | -0.018 | -0.038 | -0.004 | -0.004 | 0.035 | 0.09 | 0.02 | 0.06 | 0.07 | 0.13 | 0.1 |
| 10 | 6541923 | rs11259245 | 0.581 | 1.096 | -1.321 | 0.153 | -0.039 | -0.026 | -0.026 | 0.191 | 0.07 | 0.01 | 0.23 | 0.09 | 0.1 | 0.05 |
| 10 | 6543063 | rs1409876 | -0.631 | 0.264 | -1.418 | -0.174 | 0.012 | -0.012 | -0.012 | 0 | 0.45 | 0.72 | 0.27 | 0.48 | 0.39 | 0.43 |
| 10 | 6547909 | rs2387407 | -0.777 | -0.808 | -1.013 | -0.439 | -0.013 | -0.046 | -0.046 | 0 | 0.84 | 0.78 | 0.87 | 0.96 | 0.84 | 0.88 |
| 10 | 6550199 | rs596866 | 0.275 | -1.156 | -0.748 | -0.21 | -0.01 | 0.057 | 0.057 | 0 | 0.75 | 0.61 | 0.83 | 0.65 | 0.82 | 0.7 |
| 10 | 6550590 | rs500766 | -0.383 | -1.174 | -0.91 | -0.332 | -0.004 | -0.08 | -0.08 | 0 | 0.29 | 0.42 | 0.21 | 0.09 | 0.3 | 0.29 |
| 10 | 6551732 | rs7087802 | 1.546 | -1.22 | -1.177 | -0.216 | -0.141 | -0.116 | -0.116 | 0 | 0.43 | 0.05 | 0.6 | 0.47 | 0.5 | 0.35 |
| 10 | 6556492 | rs17137421 | 0.32 | -1.358 | -0.932 | -0.09 | 0.013 | -0.007 | -0.007 | 0 | 0.09 | 0.13 | 0.07 | 0.05 | 0.13 | 0.09 |
| 10 | 6556554 | rs571715 | -0.17 | -1.347 | -1.06 | 0.021 | -0.01 | 0.069 | 0.069 | 0 | 0.75 | 0.59 | 0.83 | 0.65 | 0.82 | 0.7 |
| 10 | 6565099 | rs521153 | -0.754 | -0.882 | -0.746 | -0.366 | -0.01 | -0.066 | -0.066 | 0 | 0.32 | 0.5 | 0.24 | 0.19 | 0.32 | 0.35 |
| 10 | 6566975 | rs4357599 | -0.918 | -1.075 | -1.179 | 0.021 | 0.006 | -0.008 | -0.008 | 0 | 0.52 | 0.31 | 0.29 | 0.44 | 0.43 | 0.49 |
| 10 | 6572225 | rs11259403 | 0.57 | -1.074 | -1.528 | 0.059 | -0.061 | -0.099 | -0.099 | 0 | 0.59 | 0.05 | 0.66 | 0.48 | 0.69 | 0.57 |
| 10 | 6575953 | rs7914917 | -0.042 | -0.858 | -0.813 | 0.227 | 0.002 | 0.009 | 0.009 | 1.538 | 0.14 | 0.31 | 0.08 | 0.14 | 0.11 | 0.13 |
| 10 | 6578628 | rs11259432 | 0.421 | -1.07 | -1.487 | -0.221 | 0.03 | -0.002 | -0.002 | 0 | 0.52 | 0.15 | 0.64 | 0.49 | 0.65 | 0.52 |
| 10 | 6582929 | rs4748138 | 0.085 | -0.669 | -0.816 | -0.082 | 0.003 | -0.018 | -0.018 | 0.941 | 0.94 | 0.78 | 0.95 | 1 | 0.98 | 0.95 |
| 10 | 6596242 | rs6602820 | 0.898 | -0.775 | -1.275 | 0.013 | -0.122 | -0.141 | -0.141 | 0 | 0.45 | 0.04 | 0.59 | 0.52 | 0.59 | 0.48 |
| 10 | 6597368 | rs2011261 | 0.435 | -0.323 | 0.02 | 0.092 | 0.004 | -0.016 | -0.016 | 0 | 0.94 | 0.79 | 0.95 | 1 | 0.97 | 0.95 |
| 10 | 6600833 | rs10752356 | 0.518 | -0.325 | -0.056 | 0.092 | 0.013 | -0.009 | -0.009 | 0.229 | 0.94 | 0.74 | 0.95 | 1 | 0.97 | 0.94 |
| 10 | 6606541 | rs4748153 | -0.179 | -0.48 | -1.2 | -0.068 | 0.02 | 0.047 | 0.047 | 3.603 | 0.17 | 0.11 | 0.08 | 0.01 | 0.11 | 0.08 |
| 10 | 6608313 | rs7072496 | 0.968 | -0.569 | -1.063 | -0.12 | 0.018 | 0.008 | 0.008 | 2.014 | 0.72 | 0.29 | 0.83 | 0.96 | 0.85 | 0.8 |
| 10 | 6615454 | rs943452 | -0.493 | -0.108 | -1.069 | 0.526 | -0.004 | -0.001 | -0.001 | 0.055 | 0.39 | 0.43 | 0.35 | 0.38 | 0.37 | 0.37 |
| 10 | 6616336 | rs4620621 | 0.445 | 0.148 | -2.173 | 0.554 | -0.092 | -0.113 | -0.113 | 0 | 0.52 | 0.04 | 0.6 | 0.54 | 0.61 | 0.54 |
| 10 | 6629225 | rs10906922 | 0.588 | 0.202 | -1.368 | 0.722 | 0.002 | -0.023 | -0.023 | 0 | 0.69 | 0.23 | 0.52 | 0.57 | 0.68 | 0.72 |
| 10 | 6630095 | rs755627 | 0.701 | 0.206 | -0.946 | 0.724 | 0 | 0.003 | 0.003 | 0 | 0.7 | 0.35 | 0.53 | 0.57 | 0.69 | 0.73 |
| 10 | 6632391 | rs2435588 | 0.795 | -0.143 | -0.383 | 0.375 | -0.016 | -0.035 | -0.035 | 1.219 | 0.86 | 0.74 | 0.82 | 0.96 | 0.88 | 0.85 |
| 10 | 6634620 | rs1556003 | 0.913 | -0.761 | -0.977 | 0.427 | 0.071 | 0.011 | 0.011 | 0 | 0.53 | 0.28 | 0.59 | 0.57 | 0.64 | 0.64 |

| | | | | | | | | | | | | | | | | |
|---|---|---|---|---|---|---|---|---|---|---|---|---|---|---|---|---|
| 10 | 6638856 | rs11253576 | -0.38 | -0.956 | -1.085 | 0.631 | 0.044 | -0.112 | -0.112 | 0 | 0.23 | 0.01 | 0.15 | 0.35 | 0.13 | 0.15 |
| 10 | 6639986 | rs1268331 | -0.809 | -0.423 | -1.301 | -0.314 | -0.001 | 0.079 | 0.079 | 0 | 0.64 | 0.76 | 0.8 | 0.89 | 0.62 | 0.78 |
| 10 | 6643803 | rs1041227 | -0.416 | 0.311 | -0.956 | 0.382 | 0.019 | 0.121 | 0.121 | 0 | 0.24 | 0.41 | 0.49 | 0.46 | 0.27 | 0.34 |
| 10 | 6643816 | rs1467681 | 0.426 | 0.453 | -1.869 | 0.158 | -0.053 | 0.038 | 0.038 | 0.01 | 0.26 | 0.14 | 0.17 | 0.13 | 0.28 | 0.3 |
| 10 | 6643851 | rs1041226 | 0.779 | 0.845 | -1.727 | 0.575 | -0.052 | -0.032 | -0.032 | 0.262 | 0.08 | 0.02 | 0.4 | 0.37 | 0.16 | 0.2 |
| 10 | 6644674 | rs12248647 | 1.061 | 0.203 | -1.793 | -0.267 | -0.022 | 0.099 | 0.099 | 0 | 0.27 | 0.05 | 0.14 | 0.13 | 0.27 | 0.11 |
| 10 | 6649132 | rs9423942 | 0.366 | -0.215 | -1.293 | -0.127 | 0 | -0.001 | -0.001 | 0 | 0.61 | 0.61 | 0.38 | 0.33 | 0.57 | 0.5 |
| 10 | 6649160 | rs11253614 | 0.318 | 0.529 | -1.613 | -0.381 | -0.038 | 0.03 | 0.03 | 0.226 | 0.22 | 0.14 | 0.18 | 0.15 | 0.22 | 0.36 |
| 10 | 6650426 | rs1409874 | 0.519 | 0.562 | -1.625 | 0.493 | -0.024 | -0.028 | -0.028 | 0 | 0.13 | 0.09 | 0.35 | 0.24 | 0.21 | 0.23 |
| 10 | 6650991 | rs975167 | -0.262 | -0.085 | -1.833 | 0.649 | -0.009 | -0.043 | -0.043 | 0 | 0.34 | 0.17 | 0.34 | 0.45 | 0.31 | 0.37 |
| 10 | 6651563 | rs1327249 | -0.429 | 0.226 | -1.795 | 0.588 | -0.068 | -0.068 | -0.068 | 0.477 | 0.33 | 0.18 | 0.52 | 0.43 | 0.43 | 0.48 |
| 10 | 6651941 | rs10904616 | 0.7 | 0.429 | -1.591 | 0.188 | -0.035 | 0.009 | 0.009 | 0 | 0.2 | 0.13 | 0.17 | 0.18 | 0.22 | 0.24 |
| 10 | 6655306 | rs1409875 | -1.688 | 0.053 | -0.947 | -0.206 | -0.001 | 0.002 | 0.002 | 0 | 0.73 | 0.78 | 0.85 | 0.87 | 0.71 | 0.79 |
| 10 | 6657775 | rs1576009 | -1.086 | -0.088 | -0.844 | -0.225 | 0 | -0.043 | -0.043 | 0.084 | 0.66 | 0.45 | 0.77 | 0.84 | 0.61 | 0.72 |
| 10 | 6662135 | rs1410135 | 0.89 | 0.216 | -1.214 | 0.049 | -0.018 | 0.01 | 0.01 | 0 | 0.18 | 0.11 | 0.19 | 0.18 | 0.17 | 0.23 |
| 10 | 6666308 | rs6602064 | 1.374 | 0.174 | -1.028 | -0.26 | -0.009 | 0.024 | 0.024 | 0 | 0.3 | 0.19 | 0.17 | 0.12 | 0.35 | 0.23 |
| 10 | 6666823 | rs7089639 | 0.28 | 0.106 | -0.951 | 0.019 | 0.002 | 0.012 | 0.012 | 0 | 0.61 | 0.62 | 0.72 | 0.69 | 0.65 | 0.61 |
| 10 | 6667616 | rs11253683 | 0.593 | 0.872 | -1.15 | 0.781 | -0.065 | -0.076 | -0.076 | 1.913 | 0.13 | 0.03 | 0.29 | 0.31 | 0.22 | 0.24 |
| 10 | 6668830 | rs11253689 | 0.326 | 0.896 | -1.038 | 0.373 | -0.029 | 0.006 | 0.006 | 0.047 | 0.14 | 0.12 | 0.17 | 0.15 | 0.2 | 0.25 |
| 10 | 6677767 | rs10795331 | 0.248 | 0.369 | -1.028 | 0.298 | -0.02 | -0.032 | -0.032 | 0.361 | 0.6 | 0.77 | 0.59 | 0.55 | 0.54 | 0.49 |
| 10 | 6685078 | rs6602077 | -0.092 | 0.147 | -0.353 | 0.28 | -0.034 | -0.014 | -0.014 | 0 | 0.77 | 0.58 | 0.87 | 0.8 | 0.85 | 0.87 |
| 10 | 6690147 | rs1475418 | -1.897 | 0.399 | -0.609 | -0.26 | -0.002 | -0.073 | -0.073 | 0.174 | 0.73 | 0.73 | 0.86 | 0.98 | 0.7 | 0.81 |
| 10 | 6691644 | rs10795350 | 0.791 | 0.733 | -0.791 | 0.018 | -0.04 | -0.048 | -0.048 | 1.06 | 0.15 | 0.08 | 0.27 | 0.21 | 0.25 | 0.3 |
| 10 | 6693011 | rs7916225 | 0.897 | 0.7 | -0.619 | 0.025 | -0.004 | -0.002 | -0.002 | 0.051 | 0.14 | 0.11 | 0.17 | 0.21 | 0.17 | 0.28 |
| 10 | 6695485 | rs4339947 | -2.209 | 0.696 | -0.567 | -0.034 | 0.003 | -0.055 | -0.055 | 0 | 0.73 | 0.73 | 0.85 | 0.98 | 0.68 | 0.8 |
| 10 | 6698096 | rs1475420 | 1.029 | 0.609 | -0.665 | 0.052 | -0.046 | -0.055 | -0.055 | 0.037 | 0.15 | 0.06 | 0.27 | 0.2 | 0.25 | 0.32 |
| 10 | 6699193 | rs17139562 | 1.365 | 0.252 | -0.697 | 0.089 | 0.034 | -0.034 | -0.034 | 0 | 0.15 | 0.04 | 0.08 | 0.3 | 0.08 | 0.1 |
| 10 | 6699236 | rs11253853 | 0.873 | 0.476 | -0.554 | -0.008 | -0.007 | -0.002 | -0.002 | 0.19 | 0.14 | 0.11 | 0.17 | 0.21 | 0.17 | 0.28 |
| 10 | 6700116 | rs11253857 | 0.943 | 0.471 | -0.612 | -0.01 | -0.044 | -0.017 | -0.017 | 0 | 0.14 | 0.05 | 0.17 | 0.21 | 0.17 | 0.28 |
| 10 | 6702598 | rs10904701 | -0.299 | 0.206 | -0.199 | 0.075 | -0.006 | -0.001 | -0.001 | 0 | 0.44 | 0.47 | 0.43 | 0.57 | 0.28 | 0.22 |
| 10 | 6709658 | rs2026628 | -1.067 | 0.26 | -0.322 | 0.075 | -0.001 | -0.006 | -0.006 | 0 | 0.57 | 0.44 | 0.67 | 0.77 | 0.53 | 0.52 |
| 10 | 6711880 | rs7098734 | -0.283 | 0.441 | -0.606 | 0.177 | 0.015 | -0.007 | -0.007 | 0 | 0.42 | 0.25 | 0.43 | 0.57 | 0.34 | 0.24 |
| 10 | 6714175 | rs7091048 | 0.823 | 0.447 | -0.844 | 0.341 | -0.019 | -0.061 | -0.061 | 0 | 0.15 | 0.05 | 0.27 | 0.25 | 0.16 | 0.07 |
| 10 | 6714257 | rs11253940 | 1.278 | 0.344 | -0.908 | 0.318 | 0.019 | -0.038 | -0.038 | 0 | 0.14 | 0.03 | 0.1 | 0.25 | 0.09 | 0.11 |
| 10 | 6714424 | rs10904722 | -0.169 | 0.578 | -0.727 | 0.163 | 0.008 | -0.006 | -0.006 | 0.282 | 0.4 | 0.31 | 0.3 | 0.53 | 0.28 | 0.43 |
| 10 | 6714979 | rs4237340 | -0.744 | 1.124 | -0.801 | 0.386 | 0.004 | 0.008 | 0.008 | 0.078 | 0.33 | 0.39 | 0.55 | 0.46 | 0.43 | 0.39 |
| 10 | 6718305 | rs17140077 | 0.321 | 1.092 | -0.891 | 0.396 | -0.024 | 0.003 | 0.003 | 0.136 | 0.24 | 0.11 | 0.18 | 0.28 | 0.19 | 0.32 |
| 10 | 6720799 | rs10508310 | 0.598 | 1.273 | -0.726 | 0.651 | 0 | -0.011 | -0.011 | 0.002 | 0.17 | 0.14 | 0.28 | 0.27 | 0.16 | 0.08 |
| 10 | 6724940 | rs12358653 | -0.225 | 1.26 | -0.553 | 0.437 | 0.057 | -0.016 | -0.016 | 0 | 0.05 | 0.2 | 0.04 | 0 | 0.06 | 0.03 |
| 10 | 6725587 | rs12242253 | 0.666 | 1.3 | -0.747 | 0.53 | -0.029 | -0.065 | -0.065 | 0.566 | 0.13 | 0.05 | 0.25 | 0.21 | 0.2 | 0.27 |
| 10 | 6727538 | rs2387595 | -0.302 | 1.092 | -0.501 | 0.432 | 0.007 | 0.007 | 0.007 | 0 | 0.42 | 0.49 | 0.49 | 0.53 | 0.39 | 0.41 |
| 10 | 6730443 | rs11254057 | -0.06 | 0.884 | -1.022 | 0.809 | -0.011 | -0.049 | -0.049 | 0 | 0.14 | 0.07 | 0.13 | 0.22 | 0.21 | 0.22 |
| 10 | 6730923 | rs2225145 | -0.394 | 0.464 | -0.657 | 0.342 | 0.001 | 0.004 | 0.004 | 0 | 0.51 | 0.54 | 0.64 | 0.65 | 0.58 | 0.69 |
| 10 | 6731071 | rs7893479 | -1.508 | 0.153 | -0.584 | -0.14 | -0.004 | -0.052 | -0.052 | 0 | 0.76 | 0.72 | 0.84 | 0.97 | 0.77 | 0.86 |
| 10 | 6731269 | rs7900106 | -1.575 | 0.146 | -0.495 | -0.066 | -0.007 | -0.032 | -0.032 | 0 | 0.77 | 0.72 | 0.85 | 0.94 | 0.8 | 0.85 |
| 10 | 6731376 | rs7896871 | -1.567 | 0.119 | -0.567 | -0.085 | -0.005 | -0.024 | -0.024 | 0 | 0.77 | 0.74 | 0.86 | 0.94 | 0.81 | 0.85 |
| 10 | 6731489 | rs7911217 | 0.476 | 0.117 | -0.98 | 0.337 | -0.007 | -0.033 | -0.033 | 0 | 0.16 | 0.07 | 0.15 | 0.25 | 0.15 | 0.08 |
| 10 | 6735768 | rs12768603 | 0.134 | -0.07 | -1.242 | -0.152 | 0.003 | 0.04 | 0.04 | 0.038 | 0.24 | 0.11 | 0.16 | 0.17 | 0.18 | 0.08 |
| 10 | 6736561 | rs10444205 | 0.469 | -0.586 | -1.295 | -0.322 | -0.031 | -0.009 | -0.009 | 0 | 0.31 | 0.25 | 0.51 | 0.64 | 0.45 | 0.57 |
| 10 | 6736921 | rs17394484 | 0.35 | -0.452 | -1.565 | -0.236 | 0.007 | 0.041 | 0.041 | 0.047 | 0.24 | 0.12 | 0.16 | 0.17 | 0.18 | 0.08 |
| 10 | 6737384 | rs4237341 | -2.055 | -0.348 | -1.451 | -0.239 | 0.002 | 0.006 | 0.006 | 0 | 0.81 | 0.83 | 0.87 | 0.95 | 0.82 | 0.87 |
| 10 | 6737505 | rs10795407 | 0.428 | -0.22 | -1.543 | -0.05 | -0.009 | -0.023 | -0.023 | 0 | 0.13 | 0.02 | 0.16 | 0.14 | 0.14 | 0.18 |
| 10 | 6737987 | rs10904780 | -0.191 | 0.329 | -1.369 | 0.133 | -0.005 | -0.006 | -0.006 | 0 | 0.24 | 0.23 | 0.53 | 0.57 | 0.39 | 0.46 |
| 10 | 6738313 | rs12573122 | 0.481 | 0.774 | -1.407 | 0.267 | -0.006 | -0.007 | -0.007 | 0.012 | 0.1 | 0.1 | 0.14 | 0.14 | 0.18 | 0.13 |
| 10 | 6738930 | rs4748306 | -0.051 | 0.257 | -1.405 | -0.025 | -0.038 | -0.03 | -0.03 | 0 | 0.29 | 0.22 | 0.55 | 0.61 | 0.42 | 0.54 |
| 10 | 6743597 | rs1327689 | -1.48 | -0.448 | -1.475 | -0.359 | 0 | -0.008 | -0.008 | 0 | 0.7 | 0.71 | 0.84 | 0.9 | 0.75 | 0.83 |
| 10 | 6743726 | rs11254134 | 0.251 | -0.333 | -1.321 | 0.215 | -0.003 | -0.023 | -0.023 | 0 | 0.13 | 0.02 | 0.15 | 0.15 | 0.14 | 0.16 |
| 10 | 6743781 | rs17141058 | 0.779 | 0.009 | -1.083 | 0.348 | 0.024 | -0.025 | -0.025 | 0 | 0.12 | 0.04 | 0.12 | 0.14 | 0.07 | 0.06 |
| 10 | 6750281 | rs7912136 | -0.999 | 0.274 | -0.686 | 0.235 | -0.012 | -0.051 | -0.051 | 0 | 0.36 | 0.47 | 0.27 | 0.18 | 0.29 | 0.26 |
| 10 | 6754079 | rs7909426 | 0.786 | 0.341 | -0.676 | 0.489 | 0.018 | -0.001 | -0.001 | 0 | 0.6 | 0.33 | 0.63 | 0.63 | 0.68 | 0.65 |
| 10 | 6756983 | rs7099454 | -1 | 0.291 | -0.609 | 0.419 | -0.015 | 0.015 | 0.015 | 5.345 | 0.26 | 0.35 | 0.31 | 0.35 | 0.2 | 0.29 |
| 10 | 6762403 | rs11597924 | 0.017 | 0.319 | -0.746 | 0.379 | -0.011 | -0.021 | -0.021 | 0 | 0.15 | 0.01 | 0.14 | 0.14 | 0.14 | 0.14 |
| 10 | 6768836 | rs11254272 | -0.3 | 0.474 | -0.876 | 0.285 | -0.028 | 0.004 | 0.004 | 0.167 | 0.19 | 0.01 | 0.18 | 0.14 | 0.19 | 0.15 |
| 10 | 6768893 | rs17141479 | 0.117 | 0.296 | -0.637 | 0.489 | 0.009 | -0.027 | -0.027 | 0.017 | 0.13 | 0.01 | 0.16 | 0.14 | 0.11 | 0.11 |
| 10 | 6769130 | rs1561875 | 0.268 | 1.026 | 0.191 | 0.708 | 0.027 | -0.117 | -0.117 | 0 | 0.83 | 0.68 | 0.91 | 1 | 0.84 | 0.91 |
| 10 | 6773835 | rs2119984 | -0.175 | 0.862 | 0.304 | 0.636 | -0.023 | 0.097 | 0.097 | 0 | 0.11 | 0.33 | 0.15 | 0.21 | 0.06 | 0.13 |
| 10 | 6774351 | rs4144205 | 0.129 | 1.499 | 0.27 | 0.259 | 0.017 | -0.084 | -0.084 | 0 | 0.86 | 0.76 | 0.94 | 1 | 0.87 | 0.94 |
| 10 | 6776447 | rs452121 | 0.469 | 1.813 | 0.226 | 0.914 | 0.035 | 0.026 | 0.026 | 0 | 0.8 | 0.61 | 0.63 | 0.76 | 0.77 | 0.75 |
| 10 | 6778087 | rs1992495 | -0.987 | 1.959 | -0.005 | 1.115 | 0.013 | 0.016 | 0.016 | 0.202 | 0.36 | 0.47 | 0.55 | 0.56 | 0.54 | 0.43 |

| | | | | | | | | | | | | | | | | |
|---|---|---|---|---|---|---|---|---|---|---|---|---|---|---|---|---|
| 10 | 6779443 | rs353212 | -0.645 | 1.244 | 0.123 | 1.282 | 0.047 | 0.131 | 0.131 | 0 | 0.13 | 0.32 | 0.33 | 0.32 | 0.18 | 0.18 |
| 10 | 6782072 | rs3763958 | 0.544 | 0.718 | -0.887 | 0.379 | -0.013 | 0.155 | 0.155 | 0 | 0.54 | 0.2 | 0.4 | 0.13 | 0.54 | 0.52 |
| 10 | 6782100 | rs353210 | -0.334 | 0.486 | -0.754 | 1.253 | -0.006 | -0.024 | -0.024 | 0 | 0.13 | 0.07 | 0.11 | 0.32 | 0.15 | 0.1 |
| 10 | 6782769 | rs353209 | 0.554 | 0.127 | -0.173 | 1.021 | -0.021 | 0.054 | 0.054 | 0.061 | 0.93 | 0.81 | 0.96 | 0.84 | 0.97 | 0.95 |
| 10 | 6783108 | rs353208 | -0.359 | -0.315 | -0.363 | 0.218 | -0.005 | -0.055 | -0.055 | 0.023 | 0.83 | 0.78 | 0.93 | 1 | 0.87 | 0.87 |
| 10 | 6784000 | rs353206 | 0.861 | -0.347 | -0.138 | 0.743 | -0.016 | 0.036 | 0.036 | 0 | 0.64 | 0.49 | 0.75 | 0.54 | 0.73 | 0.64 |
| 10 | 6786045 | rs353201 | -0.314 | -0.7 | 0.388 | 0.482 | -0.016 | -0.053 | -0.053 | 0.044 | 0.1 | 0.26 | 0.03 | 0.01 | 0.04 | 0.09 |
| 10 | 6788713 | rs1348307 | 0.868 | -0.358 | 0.11 | 0.611 | 0.01 | 0.013 | 0.013 | 0 | 0.11 | 0.1 | 0.31 | 0.23 | 0.24 | 0.21 |
| 10 | 6789167 | rs353197 | -0.884 | -1.985 | 0.243 | 0.538 | -0.017 | -0.104 | -0.104 | 0.692 | 0.2 | 0.35 | 0.06 | 0.01 | 0.12 | 0.11 |
| 10 | 6789463 | rs12570744 | 0.439 | -2.666 | -0.577 | 0.288 | 0.027 | -0.041 | -0.041 | 0.196 | 0.24 | 0.2 | 0.32 | 0.53 | 0.18 | 0.29 |
| 10 | 6791466 | rs17141695 | 0.962 | -2.832 | -0.837 | 0.245 | 0.023 | -0.035 | -0.035 | 0 | 0.18 | 0.01 | 0.15 | 0.18 | 0.14 | 0.17 |
| 10 | 6792431 | rs11254404 | -0.612 | -3.265 | -0.64 | 0.056 | 0.048 | 0.017 | 0.017 | 1.823 | 0.41 | 0.28 | 0.31 | 0.4 | 0.31 | 0.34 |
| 10 | 6793562 | rs11591887 | 1.791 | -3.353 | -0.577 | -0.037 | -0.023 | 0.014 | 0.014 | 0.319 | 0.34 | 0.21 | 0.5 | 0.28 | 0.53 | 0.39 |
| 10 | 6793861 | rs4748360 | -0.201 | -3.29 | -0.654 | 0.002 | 0.056 | 0 | 0 | 0 | 0.36 | 0.22 | 0.28 | 0.4 | 0.27 | 0.31 |
| 10 | 6795787 | rs2479108 | 1.222 | -3.367 | -0.424 | -0.089 | -0.032 | -0.033 | -0.033 | 0.561 | 0.07 | 0.17 | 0.01 | 0 | 0 | 0.01 |
| 10 | 6796079 | rs413721 | 0.809 | -3.322 | -0.434 | -0.106 | -0.007 | -0.012 | -0.012 | 0 | 0.45 | 0.48 | 0.54 | 0.3 | 0.57 | 0.48 |
| 10 | 6796743 | rs4747299 | -0.25 | -3.256 | -0.826 | -0.068 | 0.093 | 0.003 | 0.003 | 3.704 | 0.33 | 0.17 | 0.35 | 0.71 | 0.22 | 0.35 |
| 10 | 6797206 | rs1837976 | 0.368 | -2.365 | -0.166 | 0.106 | -0.009 | -0.173 | -0.173 | 0.032 | 0.72 | 0.53 | 0.89 | 0.99 | 0.79 | 0.76 |
| 10 | 6799269 | rs353189 | 0.037 | -2.249 | -0.542 | 0.297 | -0.017 | 0.002 | 0.002 | 4.691 | 0.85 | 0.8 | 0.88 | 0.77 | 0.94 | 0.88 |
| 10 | 6801162 | rs12218200 | -0.266 | -2.245 | -0.559 | 0.214 | 0.001 | 0 | 0 | 5.167 | 0.14 | 0.16 | 0.12 | 0.15 | 0.06 | 0.12 |
| 10 | 6801931 | rs7911544 | 1.416 | -2.297 | -0.657 | 0.149 | 0.068 | 0.019 | 0.019 | 0 | 0.51 | 0.26 | 0.65 | 0.44 | 0.67 | 0.55 |
| 10 | 6803655 | rs353194 | 0.438 | -2.138 | -0.737 | 0.591 | -0.011 | 0.027 | 0.027 | 4.181 | 0.84 | 0.79 | 0.79 | 0.45 | 0.91 | 0.82 |
| 10 | 6807941 | rs10904923 | -0.48 | -2.083 | -1.035 | 0.402 | 0.024 | -0.032 | -0.032 | 0.216 | 0.13 | 0.02 | 0.05 | 0.17 | 0.08 | 0.1 |
| 10 | 6811851 | rs11254538 | -0.829 | -2.219 | -0.826 | 0.484 | 0.005 | -0.007 | -0.007 | 0 | 0.15 | 0.14 | 0.12 | 0.29 | 0.09 | 0.13 |
| 10 | 6813002 | rs4747305 | 1.403 | -2.628 | -0.743 | 0.359 | 0.085 | 0.032 | 0.032 | 0 | 0.49 | 0.25 | 0.65 | 0.41 | 0.68 | 0.49 |
| 10 | 6813216 | rs2501685 | -1.8 | -2.643 | -0.662 | 0.35 | 0.072 | 0.025 | 0.025 | 0 | 0.5 | 0.7 | 0.35 | 0.58 | 0.32 | 0.51 |
| 10 | 6813564 | rs2387739 | 0.702 | -2.379 | -0.417 | 0.085 | -0.05 | -0.139 | -0.139 | 3.678 | 0.67 | 0.44 | 0.81 | 0.99 | 0.82 | 0.71 |
| 10 | 6816484 | rs894453 | -0.031 | -2.329 | -0.19 | 0.136 | 0.052 | 0.01 | 0.01 | 0 | 0.07 | 0.4 | 0.04 | 0.01 | 0.03 | 0.08 |
| 10 | 6819487 | rs2501686 | -0.643 | -2.345 | -0.223 | 0.291 | 0.023 | 0.19 | 0.19 | 0 | 0.15 | 0.43 | 0.15 | 0.31 | 0.12 | 0.22 |
| 10 | 6825715 | rs2439911 | -0.033 | -2.449 | -0.475 | 0.067 | 0.027 | 0.014 | 0.014 | 0.087 | 0.8 | 0.89 | 0.93 | 1 | 0.88 | 0.86 |
| 10 | 6829101 | rs2439909 | -0.343 | -2.386 | -0.428 | 0.114 | -0.008 | -0.033 | -0.033 | 2.898 | 0.9 | 0.89 | 0.95 | 1 | 0.92 | 0.96 |
| 10 | 6831860 | rs2439903 | 0.176 | -1.935 | -0.317 | 0.463 | -0.033 | 0.002 | 0.002 | 0.273 | 0.1 | 0.25 | 0.09 | 0.1 | 0.04 | 0.09 |
| 10 | 6832173 | rs10904971 | 0.977 | -1.853 | -0.536 | 0.665 | -0.03 | -0.028 | -0.028 | 0 | 0.07 | 0.01 | 0.07 | 0.29 | 0.09 | 0.14 |
| 10 | 6834239 | rs1372802 | 1.042 | -1.303 | -0.598 | 0.453 | -0.006 | 0.01 | 0.01 | 0 | 0.64 | 0.44 | 0.72 | 0.6 | 0.7 | 0.58 |
| 10 | 6845237 | rs10904989 | 1 | -2.257 | -0.894 | 0.781 | -0.023 | 0.007 | 0.007 | 0.689 | 0.6 | 0.52 | 0.76 | 0.49 | 0.78 | 0.49 |
| 10 | 6852426 | rs1899475 | 0.42 | -2.281 | -1.027 | 0.879 | -0.005 | 0.001 | 0.001 | 0 | 0.72 | 0.73 | 0.82 | 0.63 | 0.84 | 0.66 |
| 10 | 6852441 | rs11596811 | 0.967 | -2.056 | -1.026 | 0.983 | -0.028 | -0.043 | -0.043 | 2.543 | 0.08 | 0.01 | 0.07 | 0.27 | 0.08 | 0.16 |
| 10 | 6853878 | rs12783794 | 1.154 | -2.143 | -1.485 | 0.554 | -0.002 | 0.035 | 0.035 | 0 | 0.28 | 0.08 | 0.21 | 0.19 | 0.29 | 0.25 |
| 10 | 6856536 | rs7098362 | -0.206 | -2.301 | -0.923 | 0.85 | 0.059 | -0.019 | -0.019 | 0.077 | 0.21 | 0.1 | 0.07 | 0.23 | 0.1 | 0.22 |
| 10 | 6857219 | rs11254757 | -0.191 | -2.3 | -0.923 | 0.854 | 0.058 | -0.022 | -0.022 | 0 | 0.21 | 0.09 | 0.07 | 0.24 | 0.1 | 0.22 |
| 10 | 6859144 | rs10905018 | 0.712 | -2.334 | -0.843 | 0.764 | -0.018 | -0.001 | -0.001 | 0 | 0.69 | 0.65 | 0.84 | 0.67 | 0.87 | 0.69 |
| 10 | 6863484 | rs10458810 | -0.082 | -2.291 | -0.902 | 0.762 | 0.058 | -0.049 | -0.049 | 0 | 0.2 | 0.08 | 0.07 | 0.27 | 0.09 | 0.22 |
| 10 | 6864171 | rs7900770 | 0.609 | -2.282 | -0.839 | 0.759 | -0.016 | -0.002 | -0.002 | 0 | 0.69 | 0.65 | 0.84 | 0.72 | 0.87 | 0.69 |
| 10 | 6866042 | rs7083059 | 0.531 | -2.297 | -0.877 | 0.763 | 0.015 | 0.006 | 0.006 | 3.486 | 0.67 | 0.71 | 0.84 | 0.71 | 0.87 | 0.7 |
| 10 | 6880601 | rs920678 | 0.644 | -1.942 | -0.571 | 0.503 | -0.027 | -0.002 | -0.002 | 0 | 0.58 | 0.45 | 0.76 | 0.4 | 0.79 | 0.49 |
| 10 | 6883788 | rs10905035 | -0.638 | -1.375 | -0.866 | 0.638 | 0.058 | -0.071 | -0.071 | 0 | 0.32 | 0.2 | 0.16 | 0.55 | 0.18 | 0.45 |
| 10 | 6898219 | rs7091821 | -0.51 | -0.173 | -0.565 | 0.428 | -0.012 | 0.008 | 0.008 | 0 | 0.59 | 0.57 | 0.66 | 0.43 | 0.62 | 0.64 |
| 10 | 6904180 | rs7098268 | -0.284 | -0.366 | -0.728 | 0.498 | -0.007 | 0 | 0 | 0.998 | 0.54 | 0.56 | 0.63 | 0.49 | 0.59 | 0.57 |
| 10 | 6906476 | rs11254816 | -0.016 | 0.189 | -1.083 | 0.192 | -0.013 | 0.029 | 0.029 | 0.314 | 0.09 | 0.03 | 0.08 | 0 | 0.11 | 0.02 |
| 10 | 6908055 | rs7901626 | -0.669 | -0.071 | -0.845 | 0.064 | 0.017 | 0 | 0 | 0 | 0.53 | 0.4 | 0.63 | 0.49 | 0.6 | 0.64 |
| 10 | 6910884 | rs1470257 | 0.449 | 0.119 | -0.886 | 0.045 | -0.075 | -0.038 | -0.038 | 0 | 0.35 | 0.15 | 0.44 | 0.39 | 0.39 | 0.36 |
| 10 | 6911592 | rs17142082 | -0.966 | 0.086 | -0.583 | 0.068 | 0.002 | -0.015 | -0.015 | 0 | 0.15 | 0.24 | 0.17 | 0.07 | 0.18 | 0.08 |
| 10 | 6914957 | rs12258781 | 0.621 | 1.065 | -1.062 | -0.386 | -0.04 | -0.061 | -0.061 | 0 | 0.21 | 0.11 | 0.41 | 0.39 | 0.3 | 0.26 |
| 10 | 6915748 | rs11254835 | 0.624 | 1.249 | -1.229 | -0.562 | -0.039 | 0.045 | 0.045 | 0 | 0.13 | 0.04 | 0.08 | 0.01 | 0.17 | 0.01 |
| 10 | 6916628 | rs10508317 | -0.669 | 1.012 | -1.119 | -0.532 | -0.058 | -0.045 | -0.045 | 1.695 | 0.35 | 0.25 | 0.53 | 0.43 | 0.47 | 0.47 |
| 10 | 6916953 | rs17142104 | 0.674 | 0.887 | -0.896 | -0.568 | -0.002 | -0.019 | -0.019 | 4.327 | 0.06 | 0.05 | 0.01 | 0 | 0.04 | 0.05 |
| 10 | 6918209 | rs1340035 | -1.37 | 0.824 | -0.823 | -0.561 | -0.013 | -0.007 | -0.007 | 1.728 | 0.18 | 0.27 | 0.14 | 0.13 | 0.12 | 0.28 |
| 10 | 6918677 | rs10905055 | 2.144 | 0.519 | -1.427 | -0.837 | 0.017 | 0.132 | 0.132 | 1.15 | 0.3 | 0.06 | 0.21 | 0.09 | 0.28 | 0.2 |
| 10 | 6918775 | rs2804111 | -0.304 | 0.747 | -0.971 | -0.484 | -0.001 | -0.002 | -0.002 | 0 | 0.16 | 0.15 | 0.12 | 0.09 | 0.15 | 0.12 |
| 10 | 6920026 | rs11254850 | 0.811 | 1.054 | -1.082 | -0.36 | -0.022 | -0.035 | -0.035 | 0 | 0.16 | 0.1 | 0.32 | 0.37 | 0.21 | 0.25 |
| 10 | 6920619 | rs10795521 | 0.654 | 0.95 | -1.081 | -0.286 | 0.003 | -0.042 | -0.042 | 0.002 | 0.17 | 0.09 | 0.14 | 0.36 | 0.15 | 0.19 |
| 10 | 6921544 | rs11819078 | -0.124 | 0.92 | -0.579 | -0.442 | -0.001 | -0.045 | -0.045 | 0.04 | 0.09 | 0.17 | 0.08 | 0 | 0.08 | 0.06 |
| 10 | 6921783 | rs1417023 | 0.434 | 0.947 | -0.327 | -0.516 | 0.02 | -0.077 | -0.077 | 0 | 0.88 | 0.72 | 0.89 | 1 | 0.86 | 0.93 |
| 10 | 6925222 | rs7081563 | 1.175 | 1.069 | -0.65 | -0.727 | 0.001 | 0 | 0 | 0 | 0.5 | 0.49 | 0.58 | 0.54 | 0.48 | 0.58 |
| 10 | 6926362 | rs17142139 | -0.33 | 1.24 | -0.722 | -0.524 | 0 | -0.038 | -0.038 | 1.36 | 0.18 | 0.18 | 0.15 | 0 | 0.22 | 0.1 |
| 10 | 6928991 | rs9651532 | -0.021 | 0.805 | -0.916 | -0.186 | 0.007 | -0.02 | -0.02 | 0.402 | 0.21 | 0.19 | 0.14 | 0.44 | 0.16 | 0.22 |
| 10 | 6930655 | rs11254854 | -0.072 | 1.324 | -0.886 | 0.092 | -0.002 | 0 | 0 | 0 | 0.2 | 0.19 | 0.2 | 0.41 | 0.21 | 0.2 |
| 10 | 6930973 | rs10437542 | 0.409 | 1.243 | -0.665 | 0.137 | 0.006 | -0.031 | -0.031 | 0.065 | 0.06 | 0.1 | 0.03 | 0 | 0.07 | 0.05 |
| 10 | 6931440 | rs12220369 | 0.895 | 1.046 | -1.023 | 0.294 | -0.029 | -0.064 | -0.064 | 0 | 0.1 | 0 | 0.11 | 0.42 | 0.11 | 0.24 |

| | | | | | | | | | | | | | | | | |
|---|---|---|---|---|---|---|---|---|---|---|---|---|---|---|---|---|
| 10 | 6932438 | rs12774678 | 0.633 | 0.93 | -1.178 | -0.931 | -0.042 | 0.026 | 0.026 | 0.061 | 0.21 | 0.14 | 0.46 | 0.1 | 0.33 | 0.21 |
| 10 | 6934645 | rs10508315 | 0.494 | 0.171 | -0.809 | -0.906 | 0.007 | 0.034 | 0.034 | 0 | 0.43 | 0.31 | 0.34 | 0.75 | 0.41 | 0.56 |
| 10 | 6937239 | rs2501673 | -1.941 | 0.591 | 0.044 | -0.644 | 0.016 | -0.02 | -0.02 | 0 | 0.14 | 0.39 | 0.13 | 0.13 | 0.16 | 0.13 |
| 10 | 6940614 | rs7079649 | -1.26 | 0.627 | 0.076 | -0.628 | 0.015 | -0.048 | -0.048 | 0 | 0.1 | 0.39 | 0.06 | 0 | 0.08 | 0.03 |
| 10 | 6941553 | rs7907552 | 0.003 | 0.866 | -0.453 | -0.508 | 0.003 | -0.029 | -0.029 | 1.846 | 0.06 | 0.1 | 0.03 | 0 | 0.08 | 0.03 |
| 10 | 6941948 | rs11254864 | 0.663 | 0.962 | -0.457 | -0.333 | -0.027 | -0.028 | -0.028 | 0 | 0.13 | 0.1 | 0.42 | 0.22 | 0.27 | 0.18 |
| 10 | 6942275 | rs17317734 | 0.489 | 0.889 | -0.555 | -0.518 | -0.057 | -0.073 | -0.073 | 0.399 | 0.17 | 0.07 | 0.48 | 0.35 | 0.3 | 0.25 |
| 10 | 6944579 | rs10905060 | 0.503 | 0.581 | -0.562 | -0.648 | -0.065 | -0.106 | -0.106 | 0.004 | 0.2 | 0.04 | 0.24 | 0.29 | 0.26 | 0.23 |
| 10 | 6946407 | rs2490682 | -0.141 | 0.633 | -0.239 | -0.839 | -0.014 | -0.012 | -0.012 | 0 | 0.73 | 0.77 | 0.7 | 0.7 | 0.65 | 0.74 |
| 10 | 6946982 | rs9423750 | 1.078 | 0.082 | -0.173 | -1.226 | 0.001 | -0.014 | -0.014 | 0 | 0.36 | 0.27 | 0.49 | 0.53 | 0.4 | 0.53 |
| 10 | 6947860 | rs12243511 | 0.566 | 0.471 | -0.046 | -1.079 | 0.002 | -0.017 | -0.017 | 9.791 | 0.06 | 0.06 | 0.26 | 0 | 0.1 | 0.03 |
| 10 | 6951452 | rs10905061 | 0.738 | 0.338 | -0.03 | -1.099 | -0.017 | -0.039 | -0.039 | 0 | 0.21 | 0.13 | 0.25 | 0.31 | 0.27 | 0.25 |
| 10 | 6952227 | rs9423753 | -0.063 | -0.181 | 0.189 | -1.326 | -0.026 | -0.11 | -0.11 | 0 | 0.29 | 0.4 | 0.13 | 0 | 0.17 | 0.15 |
| 10 | 6952360 | rs10795522 | 1.985 | -0.038 | 0.031 | -1.184 | 0.001 | -0.047 | -0.047 | 0 | 0.29 | 0.15 | 0.22 | 0.54 | 0.3 | 0.48 |
| 10 | 6954586 | rs13377083 | 0.599 | -0.097 | 0.01 | -1.283 | -0.007 | -0.051 | -0.051 | 0 | 0.21 | 0.19 | 0.06 | 0 | 0.11 | 0.14 |
| 10 | 6954595 | rs4748519 | 0.186 | -0.057 | 0.059 | -1.287 | -0.002 | -0.006 | -0.006 | 0 | 0.6 | 0.61 | 0.35 | 0.54 | 0.46 | 0.63 |
| 10 | 6955401 | rs2387750 | -0.568 | 0.066 | 0.086 | -1.113 | 0.011 | 0.012 | 0.012 | 0.612 | 0.44 | 0.38 | 0.32 | 0.31 | 0.38 | 0.38 |
| 10 | 6956115 | rs9423993 | 0.459 | 0.113 | 0.199 | -1.016 | -0.001 | -0.017 | -0.017 | 0 | 0.54 | 0.59 | 0.5 | 0.84 | 0.57 | 0.73 |
| 10 | 6956478 | rs10795523 | -0.832 | 0.298 | 1.133 | -0.274 | 0.008 | 0.019 | 0.019 | 0 | 0.28 | 0.41 | 0.3 | 0.3 | 0.31 | 0.22 |
| 10 | 6957287 | rs2804116 | -0.737 | 0.285 | 0.93 | -0.316 | -0.022 | -0.033 | -0.033 | 0 | 0.93 | 0.8 | 0.98 | 1 | 0.99 | 1 |
| 10 | 6958808 | rs2804118 | 0.268 | 0.433 | 0.878 | -0.267 | 0.008 | 0.025 | 0.025 | 7.563 | 0.89 | 0.73 | 0.67 | 0.83 | 0.89 | 0.94 |
| 10 | 6959112 | rs2804121 | 1.266 | 0.391 | 0.889 | -0.329 | -0.009 | -0.038 | -0.038 | 0 | 0.6 | 0.22 | 0.36 | 0.54 | 0.58 | 0.7 |
| 10 | 6959853 | rs2501682 | 0.583 | 0.524 | 0.884 | -0.238 | 0.019 | 0.048 | 0.048 | 0 | 0.68 | 0.55 | 0.39 | 0.54 | 0.62 | 0.71 |
| 10 | 6962623 | rs10905064 | 0.551 | 0.51 | 0.442 | -0.204 | -0.009 | -0.009 | -0.009 | 0.001 | 0.17 | 0.16 | 0.26 | 0.31 | 0.24 | 0.17 |
| 10 | 6965134 | rs2501683 | 1.178 | 0.184 | 0.309 | -0.526 | -0.009 | 0.018 | 0.018 | 0.017 | 0.49 | 0.3 | 0.58 | 0.57 | 0.53 | 0.66 |
| 10 | 6968905 | rs3886013 | -0.123 | 0.549 | -0.372 | -0.338 | 0.001 | 0.004 | 0.004 | 0.004 | 0.25 | 0.28 | 0.26 | 0.29 | 0.28 | 0.2 |
| 10 | 6969816 | rs2490689 | 1.427 | 0.442 | -0.626 | -0.482 | -0.017 | -0.012 | -0.012 | 0 | 0.45 | 0.16 | 0.34 | 0.58 | 0.47 | 0.66 |
| 10 | 6972855 | rs7912259 | -0.473 | 1.011 | -0.651 | 0.204 | 0.003 | 0.009 | 0.009 | 0.019 | 0.24 | 0.29 | 0.27 | 0.28 | 0.29 | 0.2 |
| 10 | 6977250 | rs10905070 | -0.773 | 1.762 | -1.94 | 0.736 | -0.052 | 0.029 | 0.029 | 0 | 0.15 | 0.01 | 0.11 | 0.12 | 0.17 | 0.11 |
| 10 | 6978361 | rs17142281 | -0.637 | 1.369 | -1.563 | 1.069 | -0.025 | -0.056 | -0.056 | 0 | 0.19 | 0.1 | 0.25 | 0.29 | 0.25 | 0.23 |
| 10 | 6979595 | rs11254889 | -0.982 | 1.077 | -1.169 | 1.183 | -0.017 | -0.04 | -0.04 | 0 | 0.08 | 0.01 | 0.07 | 0.25 | 0.09 | 0.09 |
| 10 | 6983212 | rs11254895 | -1.012 | 1.076 | -1.177 | 1.328 | 0.001 | -0.025 | -0.025 | 1.563 | 0.14 | 0.07 | 0.12 | 0.27 | 0.12 | 0.06 |
| 10 | 6983811 | rs2490696 | -0.516 | 1.247 | -1.141 | 0.765 | 0 | 0 | 0 | 0.157 | 0.64 | 0.6 | 0.57 | 0.45 | 0.59 | 0.47 |
| 10 | 6984954 | rs7087785 | 0.871 | 0.622 | -1.383 | 0.145 | 0.054 | 0.042 | 0.042 | 0 | 0.3 | 0.11 | 0.17 | 0.2 | 0.19 | 0.26 |
| 10 | 6985007 | rs7086862 | -0.585 | 0.947 | -0.579 | 0.502 | 0.021 | -0.022 | -0.022 | 0 | 0.61 | 0.59 | 0.54 | 0.74 | 0.52 | 0.67 |
| 10 | 6993426 | rs11254900 | -0.028 | 0.861 | -0.187 | 0.626 | 0.002 | 0.017 | 0.017 | 2.419 | 0.16 | 0.26 | 0.15 | 0.24 | 0.15 | 0.1 |
| 10 | 6997213 | rs870871 | 0.287 | 1.147 | -0.356 | 0.498 | -0.021 | -0.007 | -0.007 | 0 | 0.37 | 0.29 | 0.54 | 0.56 | 0.47 | 0.54 |
| 10 | 6997298 | rs873814 | 0.555 | 1.354 | -0.411 | 0.645 | -0.001 | 0.013 | 0.013 | 0 | 0.13 | 0.17 | 0.32 | 0.33 | 0.2 | 0.36 |
| 10 | 6997861 | rs2387895 | -0.772 | 1.299 | -0.342 | 0.588 | 0.042 | 0.12 | 0.12 | 0 | 0.52 | 0.35 | 0.69 | 0.8 | 0.62 | 0.65 |
| 10 | 7000828 | rs4747368 | -1.203 | 1.155 | -0.45 | 0.42 | -0.001 | 0.043 | 0.043 | 0 | 0.63 | 0.76 | 0.73 | 0.81 | 0.65 | 0.73 |
| 10 | 7001118 | rs7070650 | -0.064 | 1.087 | -0.508 | 0.406 | -0.01 | 0.032 | 0.032 | 0 | 0.46 | 0.28 | 0.56 | 0.57 | 0.5 | 0.63 |
| 10 | 7001654 | rs946785 | -0.291 | 1.147 | -0.344 | 0.331 | 0.006 | 0.002 | 0.002 | 0 | 0.5 | 0.54 | 0.41 | 0.47 | 0.46 | 0.38 |
| 10 | 7015194 | rs7078447 | -0.255 | 1.722 | 0.232 | 0.382 | 0.036 | -0.036 | -0.036 | 0 | 0.1 | 0.24 | 0.16 | 0.02 | 0.12 | 0.05 |
| 10 | 7016064 | rs4146431 | 0.476 | 2.075 | 0.01 | 0.029 | 0.035 | -0.046 | -0.046 | 0 | 0.7 | 0.45 | 0.57 | 0.83 | 0.61 | 0.56 |
| 10 | 7021234 | rs7917304 | 1.067 | 1.688 | -0.748 | -0.211 | 0.033 | 0.057 | 0.057 | 0 | 0.31 | 0.16 | 0.2 | 0.17 | 0.23 | 0.24 |
| 10 | 7024059 | rs10905081 | 1.111 | 1.694 | -0.769 | -0.209 | 0.035 | 0.079 | 0.079 | 0 | 0.31 | 0.1 | 0.19 | 0.17 | 0.23 | 0.24 |
| 10 | 7025224 | rs2486872 | -0.888 | 1.862 | -0.642 | -0.373 | 0.003 | -0.029 | -0.029 | 1.222 | 0.9 | 0.9 | 0.83 | 1 | 0.87 | 0.97 |
| 10 | 7025379 | rs11254923 | 1.025 | 1.714 | -0.409 | -0.273 | -0.028 | -0.026 | -0.026 | 0 | 0.29 | 0.43 | 0.13 | 0.22 | 0.2 | 0.19 |
| 10 | 7028333 | rs2804143 | -0.239 | 1.926 | -0.402 | -0.249 | 0.008 | 0.011 | 0.011 | 0 | 0.51 | 0.59 | 0.5 | 0.41 | 0.56 | 0.6 |
| 10 | 7028723 | rs882778 | 0.886 | 1.539 | -0.625 | -0.269 | -0.013 | -0.011 | -0.011 | 0.074 | 0.28 | 0.31 | 0.13 | 0.22 | 0.21 | 0.22 |
| 10 | 7035823 | rs1544156 | 0.123 | 1.915 | -0.537 | -0.17 | -0.003 | 0.035 | 0.035 | 0 | 0.53 | 0.41 | 0.54 | 0.34 | 0.53 | 0.67 |
| 10 | 7039552 | rs2804133 | 0.217 | 2.183 | -0.7 | -0.224 | -0.007 | -0.003 | -0.003 | 0 | 0.16 | 0.14 | 0.2 | 0.17 | 0.21 | 0.35 |
| 10 | 7041980 | rs11254937 | 0.671 | 2.194 | -1.029 | -0.208 | -0.002 | 0.016 | 0.016 | 0 | 0.07 | 0.08 | 0.12 | 0.17 | 0.06 | 0.3 |
| 10 | 7042059 | rs7070721 | 1.144 | 2.156 | -1.328 | -0.984 | 0.017 | 0.06 | 0.06 | 0 | 0.37 | 0.27 | 0.35 | 0.18 | 0.31 | 0.34 |
| 10 | 7042270 | rs10905087 | -0.51 | 2.133 | -1.822 | -0.991 | -0.003 | -0.027 | -0.027 | 0 | 0.19 | 0.16 | 0.22 | 0 | 0.23 | 0.08 |
| 10 | 7042852 | rs2788628 | 0.502 | 1.76 | -1.733 | -0.24 | -0.033 | 0.038 | 0.038 | 0 | 0.09 | 0.04 | 0.09 | 0 | 0.17 | 0.06 |
| 10 | 7045019 | rs11254940 | 0.056 | 0.877 | -1.393 | 0.143 | -0.009 | -0.015 | -0.015 | 0 | 0.32 | 0.33 | 0.16 | 0.22 | 0.26 | 0.16 |
| 10 | 7048329 | rs8181411 | 0.328 | 0.67 | -1.915 | 0.29 | -0.02 | 0.022 | 0.022 | 0.942 | 0.27 | 0.02 | 0.19 | 0.18 | 0.2 | 0.12 |
| 10 | 7053911 | rs4747383 | -0.928 | 1.492 | -1.015 | 0.976 | 0.003 | 0.007 | 0.007 | 0 | 0.34 | 0.62 | 0.55 | 0.58 | 0.36 | 0.5 |
| 10 | 7057152 | rs11254946 | -0.938 | 1.691 | -1.124 | 1.021 | 0 | 0 | 0 | 0 | 0.37 | 0.63 | 0.53 | 0.54 | 0.39 | 0.54 |
| 10 | 7069403 | rs12264785 | -0.671 | 1.895 | -0.677 | 1.458 | -0.012 | 0.03 | 0.03 | 0 | 0.23 | 0.6 | 0.21 | 0.32 | 0.2 | 0.22 |
| 10 | 7071030 | rs11254958 | -0.579 | 1.963 | -0.915 | 1.372 | -0.065 | -0.056 | -0.056 | 3.429 | 0.13 | 0.03 | 0.33 | 0.22 | 0.19 | 0.33 |
| 10 | 7073365 | rs7079137 | 0.827 | 1.579 | -0.373 | 1.472 | -0.006 | 0.007 | 0.007 | 0 | 0.63 | 0.54 | 0.51 | 0.38 | 0.69 | 0.51 |
| 10 | 7073636 | rs10795528 | -1.08 | 2.556 | 0.381 | 1.805 | 0.092 | 0.044 | 0.044 | 0 | 0.18 | 0.46 | 0.3 | 0.21 | 0.29 | 0.28 |
| 10 | 7075633 | rs11254963 | 0.549 | 2.295 | 0.03 | 1.654 | -0.015 | 0.027 | 0.027 | 1.021 | 0.07 | 0.02 | 0.04 | 0 | 0.07 | 0.04 |
| 10 | 7083369 | rs12761228 | -1.469 | 2.238 | 0.851 | 2.525 | 0.034 | 0.263 | 0.263 | 0 | 0.16 | 0.38 | 0.24 | 0.46 | 0.15 | 0.21 |
| 10 | 7083438 | rs17419944 | -0.254 | 2.093 | 0.493 | 1.952 | 0.005 | 0.056 | 0.056 | 0 | 0.1 | 0.02 | 0.04 | 0 | 0.08 | 0.04 |
| 10 | 7084612 | rs867454 | -0.926 | 2.28 | 0.973 | 2.106 | 0.022 | -0.069 | -0.069 | 0.607 | 0.12 | 0.22 | 0.1 | 0 | 0.13 | 0.12 |
| 10 | 7085657 | rs2892380 | -1.347 | 2.397 | 0.686 | 2.451 | -0.033 | -0.004 | -0.004 | 0 | 0.06 | 0.01 | 0.2 | 0.26 | 0.12 | 0.19 |

| | | | | | | | | | | | | | | | | |
|---|---|---|---|---|---|---|---|---|---|---|---|---|---|---|---|---|
| 10 | 7086117 | rs12773886 | -0.438 | 2.579 | 0.927 | 1.998 | 0.042 | -0.053 | -0.053 | 2.377 | 0.09 | 0.2 | 0.08 | 0 | 0.12 | 0.11 |
| 10 | 7097117 | rs7080182 | 0.186 | 2.428 | 0.804 | 2.022 | -0.001 | -0.027 | -0.027 | 0 | 0.07 | 0.1 | 0.03 | 0 | 0.07 | 0.03 |
| 10 | 7111871 | rs4748667 | 1.353 | 2.428 | 0.645 | 2.213 | -0.013 | -0.018 | -0.018 | 0.084 | 0.6 | 0.27 | 0.41 | 0.49 | 0.47 | 0.4 |
| 10 | 7112076 | rs9423765 | -1.278 | 2.387 | 1.097 | 2.592 | 0.051 | 0.069 | 0.069 | 0 | 0.16 | 0.27 | 0.42 | 0.36 | 0.33 | 0.39 |
| 10 | 7116448 | rs9424004 | -0.382 | 2.185 | 0.643 | 2.66 | -0.03 | -0.022 | -0.022 | 0 | 0.05 | 0.01 | 0.17 | 0.19 | 0.18 | 0.13 |
| 10 | 7124442 | rs6602225 | -1.385 | 0.728 | 0.965 | 2.144 | -0.041 | 0.069 | 0.069 | 0 | 0.31 | 0.51 | 0.26 | 0.51 | 0.18 | 0.27 |
| 10 | 7131802 | rs10905104 | -1.08 | 0.888 | 0.271 | 1.812 | 0.018 | 0.047 | 0.047 | 0 | 0.25 | 0.13 | 0.09 | 0.02 | 0.16 | 0.14 |
| 10 | 7141150 | rs4748675 | -2.013 | 0.85 | 0.546 | 2.261 | 0.029 | 0.523 | 0.523 | 0 | 0.08 | 0.31 | 0.18 | 0.48 | 0.04 | 0.16 |
| 10 | 7143430 | rs10437543 | -0.29 | 0.673 | -0.322 | 2.124 | -0.015 | -0.014 | -0.014 | 0 | 0.1 | 0.18 | 0.09 | 0.06 | 0.06 | 0.11 |
| 10 | 7150098 | rs2388030 | 1.291 | 0.65 | -0.347 | 2.121 | 0.004 | -0.007 | -0.007 | 0 | 0.69 | 0.36 | 0.72 | 0.5 | 0.8 | 0.78 |
| 10 | 7150670 | rs2388031 | 0.285 | 0.661 | -0.325 | 2.018 | -0.033 | -0.099 | -0.099 | 0.115 | 0.83 | 0.43 | 0.92 | 0.99 | 0.93 | 0.95 |
| 10 | 7154147 | rs7906988 | 1.165 | 0.763 | -0.286 | 2.174 | -0.016 | 0.174 | 0.174 | 0 | 0.84 | 0.42 | 0.78 | 0.5 | 0.88 | 0.84 |
| 10 | 7154864 | rs10905106 | 1.709 | 0.892 | -1.123 | 1.73 | -0.033 | 0.137 | 0.137 | 0.061 | 0.31 | 0.05 | 0.23 | 0.1 | 0.32 | 0.14 |
| 10 | 7156076 | rs17142507 | -1.552 | 1.02 | -0.057 | 2.922 | -0.012 | 0.01 | 0.01 | 0 | 0.05 | 0 | 0.15 | 0.31 | 0.05 | 0.13 |
| 10 | 7157486 | rs2762576 | 1.175 | 1.093 | 0.052 | 2.087 | -0.03 | 0.057 | 0.057 | 0 | 0.8 | 0.53 | 0.87 | 0.78 | 0.88 | 0.76 |
| 10 | 7162204 | rs755647 | 1.262 | 1.419 | -1.001 | 1.954 | 0.004 | 0.007 | 0.007 | 0 | 0.87 | 0.81 | 0.82 | 0.79 | 0.82 | 0.81 |
| 10 | 7162312 | rs755648 | 1.717 | 1.415 | -0.965 | 1.903 | 0 | 0.019 | 0.019 | 0 | 0.83 | 0.8 | 0.8 | 0.63 | 0.8 | 0.68 |
| 10 | 7162564 | rs942432 | 2.033 | 1.687 | -1.115 | 1.8 | -0.016 | -0.03 | -0.03 | 0 | 0.67 | 0.73 | 0.58 | 0.53 | 0.63 | 0.56 |
| 10 | 7167104 | rs17331013 | -0.577 | 1.654 | -1.29 | 1.646 | -0.063 | -0.047 | -0.047 | 0 | 0.11 | 0.02 | 0.2 | 0.16 | 0.21 | 0.19 |
| 10 | 7167173 | rs2692781 | -1.64 | 0.665 | -0.826 | 1.382 | 0 | 0 | 0 | 0.566 | 0.19 | 0.19 | 0.22 | 0.18 | 0.17 | 0.2 |
| 10 | 7167907 | rs2762567 | -1.954 | 0.619 | -0.75 | 1.365 | -0.002 | 0.004 | 0.004 | 0.082 | 0.24 | 0.3 | 0.26 | 0.27 | 0.2 | 0.24 |
| 10 | 7168630 | rs17331068 | -0.99 | 1.053 | -1.444 | 0.72 | -0.026 | 0.039 | 0.039 | 0 | 0.16 | 0.06 | 0.22 | 0.1 | 0.17 | 0.11 |
| 10 | 7174215 | rs7094006 | 0.359 | 0.46 | -1.84 | 0.788 | 0.024 | -0.015 | -0.015 | 0.049 | 0.48 | 0.13 | 0.55 | 0.46 | 0.52 | 0.32 |
| 10 | 7187815 | rs2762559 | -0.371 | -0.365 | -1.782 | 1.155 | -0.004 | -0.042 | -0.042 | 0 | 0.28 | 0.1 | 0.22 | 0.37 | 0.31 | 0.51 |
| 10 | 7193364 | rs2247498 | -0.227 | -0.64 | -0.906 | 1.087 | -0.019 | 0.029 | 0.029 | 0.009 | 0.13 | 0.23 | 0.18 | 0.18 | 0.12 | 0.14 |
| 10 | 7194683 | rs1886283 | -0.497 | -0.93 | -1.215 | 0.966 | -0.011 | -0.041 | -0.041 | 0 | 0.34 | 0.09 | 0.22 | 0.39 | 0.34 | 0.51 |
| 10 | 7199054 | rs2497439 | -1.597 | -1.053 | 0.043 | 0.938 | -0.003 | -0.012 | -0.012 | 0 | 0.57 | 0.55 | 0.44 | 0.66 | 0.49 | 0.69 |
| 10 | 7202931 | rs2692759 | 0.089 | -1.045 | -0.638 | 0.806 | -0.026 | -0.043 | -0.043 | 0.012 | 0.09 | 0.01 | 0.04 | 0.11 | 0.09 | 0.07 |
| 10 | 7206424 | rs10905113 | 1.064 | -1.008 | -0.442 | 0.667 | 0.031 | 0.024 | 0.024 | 0 | 0.47 | 0.27 | 0.57 | 0.42 | 0.55 | 0.35 |
| 10 | 7221859 | rs10905115 | 1.016 | -0.735 | -0.395 | -0.435 | 0.028 | 0.089 | 0.089 | 0 | 0.49 | 0.15 | 0.65 | 0.63 | 0.61 | 0.4 |
| 10 | 7221938 | rs2762617 | -1.159 | -0.816 | -0.253 | -0.483 | -0.008 | -0.005 | -0.005 | 0 | 0.16 | 0.62 | 0.14 | 0.11 | 0.12 | 0.1 |
| 10 | 7223965 | rs10795531 | 1.226 | -0.831 | -0.597 | -0.536 | 0.042 | 0.109 | 0.109 | 0.237 | 0.51 | 0.13 | 0.65 | 0.65 | 0.62 | 0.38 |
| 10 | 7225045 | rs2762613 | -0.87 | -0.535 | -0.509 | -0.395 | 0.002 | 0.027 | 0.027 | 0 | 0.4 | 0.26 | 0.31 | 0.35 | 0.37 | 0.61 |
| 10 | 7230443 | rs2692795 | -0.508 | -0.064 | -1 | -1.496 | -0.021 | 0.169 | 0.169 | 0 | 0.43 | 0.16 | 0.3 | 0.16 | 0.42 | 0.45 |
| 10 | 7230524 | rs2462712 | -1.179 | 0.131 | -0.426 | -1.941 | 0.001 | 0.032 | 0.032 | 0 | 0.6 | 0.69 | 0.5 | 0.17 | 0.62 | 0.54 |
| 10 | 7231384 | rs2692796 | -0.986 | 0.109 | 0.103 | -1.965 | -0.003 | 0.007 | 0.007 | 0 | 0.56 | 0.39 | 0.39 | 0.17 | 0.53 | 0.51 |
| 10 | 7232269 | rs872542 | 0.029 | 0.133 | -0.322 | -1.987 | -0.022 | 0.153 | 0.153 | 0 | 0.4 | 0.05 | 0.24 | 0.16 | 0.38 | 0.43 |
| 10 | 7235805 | rs2692801 | -1.307 | 0.142 | -0.16 | -1.962 | -0.001 | 0.008 | 0.008 | 0 | 0.59 | 0.62 | 0.5 | 0.17 | 0.6 | 0.54 |
| 10 | 7239386 | rs2762604 | -0.18 | 0.147 | -0.892 | -1.917 | -0.032 | 0.184 | 0.184 | 0 | 0.42 | 0.06 | 0.25 | 0.16 | 0.38 | 0.46 |
| 10 | 7244207 | rs2147289 | 0.805 | 0.373 | -0.149 | -2.22 | -0.011 | 0.079 | 0.079 | 0 | 0.46 | 0.32 | 0.62 | 0.84 | 0.5 | 0.49 |
| 10 | 7249734 | rs10905117 | 1.177 | 0.608 | 0.52 | -2.265 | 0.15 | -0.005 | -0.005 | 0 | 0.92 | 0.61 | 0.84 | 1 | 0.86 | 0.97 |
| 10 | 7250148 | rs7920088 | 0.209 | 0.351 | -0.125 | -2.492 | -0.006 | 0.093 | 0.093 | 0 | 0.3 | 0.15 | 0.19 | 0.14 | 0.25 | 0.41 |
| 10 | 7251026 | rs2762598 | 1.079 | 0.372 | -0.084 | -2.221 | -0.006 | 0.085 | 0.085 | 0 | 0.42 | 0.25 | 0.53 | 0.84 | 0.42 | 0.44 |
| 10 | 7259446 | rs2692840 | 0.731 | 0.387 | 0.5 | -2.192 | 0.011 | 0.06 | 0.06 | 0.022 | 0.51 | 0.59 | 0.71 | 0.85 | 0.59 | 0.42 |
| 10 | 7264227 | rs2762583 | 0.91 | 0.156 | 0.918 | -2.426 | -0.005 | -0.038 | -0.038 | 0.376 | 0.81 | 0.68 | 0.83 | 0.91 | 0.82 | 0.82 |
| 10 | 7265468 | rs2692812 | 1.262 | 0.115 | 0.903 | -2.501 | -0.002 | -0.009 | -0.009 | 0.022 | 0.47 | 0.41 | 0.6 | 0.85 | 0.51 | 0.38 |
| 10 | 7266521 | rs10905119 | -0.103 | 0.14 | 0.985 | -2.473 | -0.005 | 0.097 | 0.097 | 0 | 0.34 | 0.18 | 0.22 | 0.06 | 0.3 | 0.43 |
| 10 | 7268383 | rs4747439 | -0.049 | 0.191 | 1.017 | -2.471 | -0.004 | 0.078 | 0.078 | 0 | 0.33 | 0.21 | 0.22 | 0.06 | 0.31 | 0.43 |
| 10 | 7273501 | rs2025465 | 0.289 | 0.156 | 1.209 | -2.497 | 0.002 | 0.018 | 0.018 | 0 | 0.31 | 0.23 | 0.22 | 0.06 | 0.28 | 0.42 |
| 10 | 7276215 | rs10905124 | 1.276 | 0.163 | 1.192 | -2.439 | -0.012 | 0.067 | 0.067 | 0.402 | 0.45 | 0.31 | 0.49 | 0.85 | 0.51 | 0.36 |
| 10 | 7280550 | rs10905129 | 1.218 | 0.254 | 1.205 | -2.35 | -0.018 | 0.124 | 0.124 | 0.957 | 0.46 | 0.26 | 0.5 | 0.85 | 0.51 | 0.37 |
| 10 | 7281467 | rs4997203 | 1.048 | 0.238 | 1.299 | -2.319 | 0.009 | 0.085 | 0.085 | 0.539 | 0.48 | 0.35 | 0.59 | 0.84 | 0.59 | 0.38 |
| 10 | 7285632 | rs2248474 | -0.053 | 0.23 | 1.339 | -2.358 | 0.005 | 0.009 | 0.009 | 0 | 0.87 | 0.89 | 0.96 | 0.91 | 0.92 | 0.86 |
| 10 | 7293894 | rs2762580 | 0.858 | 0.256 | 1.46 | -2.358 | 0.007 | -0.05 | -0.05 | 0 | 0.83 | 0.48 | 0.9 | 0.91 | 0.88 | 0.84 |
| 10 | 7295965 | rs1536494 | -0.235 | 0.11 | 0.613 | -2.338 | 0.02 | 0.092 | 0.092 | 0 | 0.35 | 0.18 | 0.22 | 0.08 | 0.29 | 0.47 |
| 10 | 7301591 | rs2767696 | -0.004 | 0.36 | 0.616 | -2.125 | -0.004 | -0.006 | -0.006 | 0.907 | 0.88 | 0.86 | 0.96 | 0.91 | 0.92 | 0.86 |
| 10 | 7304826 | rs2692776 | -0.277 | 0.424 | 0.521 | -2.128 | -0.013 | 0.069 | 0.069 | 0 | 0.32 | 0.18 | 0.29 | 0.11 | 0.35 | 0.46 |
| 10 | 7306383 | rs6602233 | 0.861 | 0.396 | 0.473 | -2.095 | -0.001 | 0.006 | 0.006 | 0.195 | 0.55 | 0.61 | 0.65 | 0.81 | 0.55 | 0.4 |
| 10 | 7309735 | rs1977427 | 0.59 | 0.712 | 0.476 | -1.855 | 0.03 | -0.051 | -0.051 | 0 | 0.82 | 0.63 | 0.83 | 0.91 | 0.79 | 0.83 |
| 10 | 7310320 | rs11255068 | 0.047 | 0.427 | -0.17 | -1.75 | 0.003 | 0.035 | 0.035 | 0 | 0.2 | 0.12 | 0.14 | 0.11 | 0.17 | 0.41 |
| 10 | 7315240 | rs10905143 | 0.057 | 1.278 | 0.09 | -1.916 | 0.009 | -0.045 | -0.045 | 0 | 0.38 | 0.53 | 0.68 | 0.78 | 0.5 | 0.32 |
| 10 | 7316963 | rs7076111 | -0.966 | 1.415 | -0.185 | -2.109 | -0.019 | 0.21 | 0.21 | 0.822 | 0.54 | 0.9 | 0.76 | 0.76 | 0.62 | 0.4 |
| 10 | 7317126 | rs6602234 | -0.186 | 1.341 | 0.438 | -2.153 | 0.037 | -0.062 | -0.062 | 0 | 0.1 | 0.29 | 0.05 | 0 | 0.09 | 0.05 |
| 10 | 7317647 | rs4748802 | -0.704 | 1.385 | 0.292 | -2.156 | 0.018 | 0.059 | 0.059 | 0 | 0.51 | 0.6 | 0.76 | 0.76 | 0.61 | 0.4 |
| 10 | 7317984 | rs7916739 | -0.456 | 1.278 | -0.154 | -1.971 | -0.024 | 0.157 | 0.157 | 1.31 | 0.46 | 0.77 | 0.76 | 0.76 | 0.53 | 0.36 |
| 10 | 7319088 | rs4611096 | -0.457 | 1.289 | -0.221 | -1.954 | -0.033 | 0.171 | 0.171 | 1.402 | 0.46 | 0.79 | 0.75 | 0.76 | 0.53 | 0.38 |
| 10 | 7321477 | rs4748803 | 0.02 | 1.968 | -0.233 | -1.548 | -0.039 | 0.038 | 0.038 | 0 | 0.29 | 0.14 | 0.59 | 0.72 | 0.41 | 0.28 |
| 10 | 7339814 | rs10905149 | -0.052 | 2.27 | 0.088 | -1.578 | -0.016 | 0.03 | 0.03 | 0.022 | 0.3 | 0.24 | 0.48 | 0.74 | 0.4 | 0.27 |
| 10 | 7340504 | rs12411681 | -0.587 | 2.024 | 0.415 | -1.6 | 0.001 | -0.012 | -0.012 | 0.625 | 0.4 | 0.42 | 0.63 | 0.74 | 0.52 | 0.34 |

| | | | | | | | | | | | | | | | | |
|---|---|---|---|---|---|---|---|---|---|---|---|---|---|---|---|---|
| 10 | 7353133 | rs7920106 | -0.828 | 1.926 | 0.582 | -1.434 | 0.01 | 0.037 | 0.037 | 0.012 | 0.46 | 0.64 | 0.7 | 0.7 | 0.62 | 0.36 |
| 10 | 7358878 | rs7900368 | -0.473 | 2.161 | 1.162 | -0.286 | 0.002 | 0.146 | 0.146 | 0.167 | 0.15 | 0.32 | 0.23 | 0.32 | 0.16 | 0.19 |
| 10 | 7377326 | rs4748830 | -1.385 | 2.596 | 0.705 | -0.291 | 0.081 | 0.116 | 0.116 | 0.164 | 0.2 | 0.6 | 0.39 | 0.33 | 0.3 | 0.22 |
| 10 | 7385798 | rs4748839 | -0.723 | 1.995 | 0.104 | -1.299 | 0.022 | 0.127 | 0.127 | 0 | 0.46 | 0.82 | 0.67 | 0.68 | 0.6 | 0.37 |
| 10 | 7392836 | rs10795542 | 0.208 | 1.952 | 0.326 | -1.286 | -0.007 | 0.006 | 0.006 | 0 | 0.32 | 0.28 | 0.46 | 0.68 | 0.44 | 0.31 |
| 10 | 7404033 | rs2764387 | -0.718 | 1.244 | 0.76 | -1.061 | 0.125 | -0.074 | -0.074 | 0 | 0.12 | 0.51 | 0.19 | 0 | 0.15 | 0.06 |
| 10 | 7411558 | rs10905155 | 0.508 | 0.644 | 0.29 | -1.342 | 0.092 | 0.111 | 0.111 | 0 | 0.38 | 0.08 | 0.27 | 0.25 | 0.25 | 0.47 |
| 10 | 7411642 | rs7099499 | 0.223 | 0.672 | 0.417 | -1.357 | 0.112 | -0.075 | -0.075 | 0 | 0.88 | 0.52 | 0.82 | 1 | 0.86 | 0.94 |
| 10 | 7412879 | rs11255094 | 0.467 | 0.579 | 0.018 | -1.321 | 0.097 | 0.138 | 0.138 | 0 | 0.38 | 0.08 | 0.27 | 0.2 | 0.25 | 0.47 |
| 10 | 7418147 | rs10453997 | 0.309 | 0.599 | 0.3 | -1.332 | 0.05 | -0.084 | -0.084 | 0 | 0.87 | 0.67 | 0.82 | 1 | 0.84 | 0.94 |
| 10 | 7425691 | rs4748899 | 0.329 | 0.578 | -0.139 | -1.334 | 0.022 | 0.076 | 0.076 | 0 | 0.47 | 0.37 | 0.53 | 0.75 | 0.59 | 0.47 |
| 10 | 7426644 | rs4083552 | 0.19 | 0.018 | -1.057 | -0.998 | 0.073 | 0.092 | 0.092 | 0 | 0.37 | 0.09 | 0.27 | 0.26 | 0.25 | 0.45 |
| 10 | 7434796 | rs11255106 | -1.036 | 0.146 | 0.177 | -0.685 | -0.045 | -0.079 | -0.079 | 0.22 | 0.14 | 0.26 | 0.04 | 0 | 0.08 | 0.11 |
| 10 | 7451241 | rs2249445 | 1.166 | -0.01 | 0.252 | -1.108 | -0.035 | -0.124 | -0.124 | 5.001 | 0.75 | 0.62 | 0.9 | 1 | 0.84 | 0.87 |
| 10 | 7460252 | rs719810 | -1.386 | -0.187 | 0.261 | -1.251 | 0.011 | -0.018 | -0.018 | 0 | 0.26 | 0.22 | 0.1 | 0 | 0.17 | 0.11 |
| 10 | 7463300 | rs2249455 | -1.885 | -0.352 | 0.747 | -1.394 | 0.013 | -0.023 | -0.023 | 0.663 | 0.31 | 0.69 | 0.14 | 0 | 0.2 | 0.14 |
| 10 | 7470868 | rs12773344 | -1.601 | -0.438 | 0.614 | -1.517 | 0.029 | 0.06 | 0.06 | 0.52 | 0.33 | 0.25 | 0.17 | 0.06 | 0.22 | 0.22 |
| 10 | 7473809 | rs2253543 | -1.31 | 0.205 | 0.543 | -1.049 | 0.033 | -0.013 | -0.013 | 0 | 0.07 | 0.59 | 0.07 | 0 | 0.04 | 0.03 |
| 10 | 7474760 | rs2253674 | -1.31 | 0.205 | 0.54 | -1.049 | 0.03 | -0.015 | -0.015 | 0 | 0.07 | 0.59 | 0.07 | 0 | 0.04 | 0.03 |
| 10 | 7475561 | rs2253805 | -1.177 | 0.474 | 0.684 | -0.793 | 0.104 | 0.047 | 0.047 | 0.309 | 0.05 | 0.53 | 0.07 | 0 | 0.04 | 0.03 |
| 10 | 7480692 | rs7077972 | -0.986 | -0.439 | -0.157 | -0.767 | 0.002 | 0.002 | 0.002 | 0 | 0.44 | 0.59 | 0.25 | 0.26 | 0.27 | 0.44 |
| 10 | 7491619 | rs2247230 | 0.158 | 0.541 | -0.239 | -1.193 | 0.004 | 0.009 | 0.009 | 0.194 | 0.5 | 0.53 | 0.52 | 0.35 | 0.57 | 0.48 |
| 10 | 7494290 | rs2247502 | 0.186 | 0.531 | -0.209 | -1.197 | 0.001 | 0.001 | 0.001 | 0.154 | 0.5 | 0.49 | 0.52 | 0.35 | 0.57 | 0.47 |
| 10 | 7497263 | rs11255141 | -0.092 | 0.242 | -0.482 | -1.32 | 0.018 | 0.021 | 0.021 | 0 | 0.13 | 0.07 | 0.06 | 0 | 0.07 | 0.08 |
| 10 | 7497817 | rs2247936 | -0.566 | 0.042 | -0.564 | -1.462 | -0.047 | -0.034 | -0.034 | 0 | 0.67 | 0.95 | 0.58 | 0.35 | 0.66 | 0.54 |
| 10 | 7498906 | rs11255142 | 1.361 | 0.397 | -0.847 | -1.798 | -0.01 | 0.028 | 0.028 | 0 | 0.27 | 0.17 | 0.16 | 0.18 | 0.29 | 0.18 |
| 10 | 7499528 | rs2248165 | -0.536 | 0.343 | -0.605 | -1.792 | 0.005 | 0.011 | 0.011 | 1.372 | 0.58 | 0.63 | 0.51 | 0.36 | 0.63 | 0.49 |
| 10 | 7506562 | rs4747484 | 0.58 | 0.413 | -1.058 | -1.813 | 0 | 0.053 | 0.053 | 0.086 | 0.36 | 0.25 | 0.18 | 0.18 | 0.37 | 0.22 |
| 10 | 7508117 | rs2251881 | -1.694 | 0.322 | -1.216 | -1.787 | -0.096 | -0.12 | -0.12 | 0 | 0.74 | 0.98 | 0.6 | 0.37 | 0.69 | 0.56 |
| 10 | 7511603 | rs11255149 | 0.967 | 0.372 | -1.102 | -1.865 | 0 | 0.068 | 0.068 | 0.078 | 0.12 | 0 | 0.06 | 0 | 0.08 | 0.08 |
| 10 | 7513248 | rs2265046 | -0.616 | 0.436 | -0.978 | -1.869 | 0.005 | 0.06 | 0.06 | 2.197 | 0.35 | 0.74 | 0.38 | 0.19 | 0.33 | 0.31 |
| 10 | 7514583 | rs2255576 | 0.012 | 0.519 | -0.839 | -1.783 | 0.017 | -0.037 | -0.037 | 0 | 0.24 | 0.37 | 0.34 | 0.19 | 0.25 | 0.27 |
| 10 | 7518411 | rs2255936 | -0.261 | 0.619 | -0.861 | -1.861 | -0.014 | -0.094 | -0.094 | 1.09 | 0.29 | 0.61 | 0.27 | 0.05 | 0.22 | 0.19 |
| 10 | 7523660 | rs2025573 | -0.948 | 0.797 | -1.37 | -1.822 | -0.015 | 0.013 | 0.013 | 0.579 | 0.64 | 0.69 | 0.51 | 0.33 | 0.59 | 0.5 |
| 10 | 7525161 | rs2025571 | -0.915 | 0.858 | -1.319 | -1.803 | -0.006 | 0.007 | 0.007 | 0.461 | 0.63 | 0.62 | 0.5 | 0.33 | 0.59 | 0.5 |
| 10 | 7526024 | rs1157042 | -0.055 | 0.815 | -1.336 | -1.842 | -0.002 | 0.062 | 0.062 | 1.2 | 0.43 | 0.26 | 0.27 | 0.25 | 0.43 | 0.34 |
| 10 | 7526078 | rs1157043 | 0.705 | 0.75 | -1.387 | -1.916 | -0.016 | 0.002 | 0.002 | 0.341 | 0.31 | 0.22 | 0.21 | 0.28 | 0.36 | 0.27 |
| 10 | 7526319 | rs1157044 | -1.657 | 0.691 | -1.368 | -1.816 | -0.055 | -0.036 | -0.036 | 0 | 0.72 | 0.9 | 0.54 | 0.34 | 0.65 | 0.54 |
| 10 | 7526447 | rs2245658 | 0.761 | 0.716 | -1.341 | -1.807 | -0.007 | -0.138 | -0.138 | 0.408 | 0.21 | 0.38 | 0.25 | 0.05 | 0.16 | 0.16 |
| 10 | 7527773 | rs2245841 | -1.393 | 0.942 | -1.756 | -1.517 | -0.044 | -0.028 | -0.028 | 0.683 | 0.72 | 0.88 | 0.53 | 0.34 | 0.65 | 0.51 |
| 10 | 7528721 | rs2245977 | -0.41 | 1.232 | -1.15 | -1.028 | 0.003 | 0.047 | 0.047 | 4.11 | 0.36 | 0.71 | 0.34 | 0.18 | 0.31 | 0.29 |
| 10 | 7529974 | rs873930 | -0.198 | 1.658 | -1.128 | -0.568 | -0.008 | 0.104 | 0.104 | 0.562 | 0.48 | 0.31 | 0.6 | 0.79 | 0.49 | 0.64 |
| 10 | 7530541 | rs2246218 | -0.298 | 1.474 | -1.445 | -0.349 | 0.005 | 0.119 | 0.119 | 0.12 | 0.57 | 0.75 | 0.45 | 0.21 | 0.59 | 0.35 |
| 10 | 7532999 | rs7099028 | -0.133 | 1.253 | -1.466 | 0.296 | 0.002 | 0.029 | 0.029 | 0 | 0.53 | 0.26 | 0.58 | 0.63 | 0.52 | 0.56 |
| 10 | 7547642 | rs4748964 | -0.303 | 1.076 | -1.123 | 0.633 | 0.003 | -0.007 | -0.007 | 1.033 | 0.41 | 0.32 | 0.37 | 0.44 | 0.57 | 0.43 |
| 10 | 7549849 | rs11591777 | -0.719 | 1.182 | -1.331 | 1.299 | -0.057 | -0.079 | -0.079 | 0.521 | 0.14 | 0.03 | 0.34 | 0.34 | 0.17 | 0.15 |
| 10 | 7560043 | rs17136510 | -1.421 | 0.578 | -0.339 | 1.168 | -0.015 | 0.066 | 0.066 | 0 | 0.16 | 0.44 | 0.24 | 0.27 | 0.11 | 0.12 |
| 10 | 7560704 | rs1537634 | -1.258 | 0.721 | -0.322 | 0.976 | -0.004 | -0.004 | -0.004 | 0.13 | 0.14 | 0.2 | 0.2 | 0.21 | 0.11 | 0.13 |
| 10 | 7565716 | rs11255178 | -0.474 | 1.142 | -0.294 | 0.998 | -0.002 | -0.001 | -0.001 | 0 | 0.25 | 0.3 | 0.32 | 0.35 | 0.32 | 0.26 |
| 10 | 7576231 | rs2050352 | -0.515 | 0.915 | -0.335 | 0.879 | 0.015 | -0.009 | -0.009 | 1.148 | 0.31 | 0.52 | 0.37 | 0.35 | 0.34 | 0.33 |
| 10 | 7578527 | rs2892445 | -0.803 | 0.852 | -0.368 | 0.837 | 0.003 | -0.01 | -0.01 | 0 | 0.33 | 0.61 | 0.37 | 0.34 | 0.34 | 0.33 |
| 10 | 7579089 | rs7083911 | -0.552 | 1.117 | -0.844 | 0.952 | 0.01 | -0.004 | -0.004 | 2.524 | 0.29 | 0.38 | 0.35 | 0.34 | 0.34 | 0.3 |
| 10 | 7581262 | rs17430079 | -0.425 | 0.697 | -1.584 | 0.985 | -0.019 | -0.051 | -0.051 | 0 | 0.14 | 0.01 | 0.1 | 0.17 | 0.13 | 0.25 |
| 10 | 7582855 | rs10795549 | -0.195 | 0.765 | -1.642 | 0.631 | -0.003 | 0.073 | 0.073 | 0 | 0.49 | 0.77 | 0.53 | 0.39 | 0.45 | 0.42 |
| 10 | 7583353 | rs12571901 | -0.487 | 0.336 | -1.214 | 0.858 | -0.003 | 0.048 | 0.048 | 0 | 0.16 | 0.19 | 0.24 | 0.48 | 0.13 | 0.22 |
| 10 | 7583364 | rs7897980 | -0.633 | 0.397 | -1.344 | -0.209 | -0.001 | 0.018 | 0.018 | 0 | 0.53 | 0.47 | 0.56 | 0.61 | 0.54 | 0.46 |
| 10 | 7597113 | rs7901010 | 0.271 | 0.074 | -0.927 | -0.633 | -0.059 | 0.083 | 0.083 | 0.083 | 0.28 | 0.08 | 0.18 | 0.2 | 0.31 | 0.31 |
| 10 | 7599350 | rs12413800 | 0.325 | 0.038 | -1.22 | -0.877 | -0.108 | 0.102 | 0.102 | 0 | 0.23 | 0.02 | 0.27 | 0.07 | 0.29 | 0.12 |
| 10 | 7601084 | rs3814696 | -0.223 | -0.805 | 0.027 | -0.236 | -0.025 | -0.007 | -0.007 | 0 | 0.17 | 0.24 | 0.13 | 0.12 | 0.1 | 0.2 |
| 10 | 7602519 | rs10508329 | 0.265 | -0.774 | -0.148 | -0.247 | 0.021 | 0.007 | 0.007 | 0 | 0.13 | 0.06 | 0.11 | 0.07 | 0.1 | 0.11 |
| 10 | 7602831 | rs10508330 | 0.17 | -0.812 | -0.006 | -0.253 | 0.016 | 0.009 | 0.009 | 0 | 0.13 | 0.1 | 0.11 | 0.07 | 0.09 | 0.11 |
| 10 | 7603104 | rs10752109 | 0.498 | -0.883 | 0.94 | -0.294 | -0.066 | -0.025 | -0.025 | 0 | 0.8 | 0.43 | 0.85 | 0.88 | 0.9 | 0.8 |
| 10 | 7603144 | rs7907148 | -0.069 | -0.8 | 0.569 | -0.258 | 0.007 | 0.002 | 0.002 | 0 | 0.16 | 0.18 | 0.13 | 0.12 | 0.1 | 0.2 |
| 10 | 7605067 | rs3824658 | 0.7 | -0.638 | 0.339 | 0.523 | -0.017 | 0.022 | 0.022 | 0 | 0.7 | 0.41 | 0.62 | 0.52 | 0.77 | 0.61 |
| 10 | 7611514 | rs12267936 | -0.668 | 0.172 | -0.184 | 0.949 | 0.014 | 0.258 | 0.258 | 0.621 | 0.14 | 0.35 | 0.3 | 0.4 | 0.13 | 0.23 |
| 10 | 7611998 | rs11599499 | -0.506 | 0.466 | -1.313 | -0.036 | -0.011 | 0.086 | 0.086 | 0 | 0.18 | 0.03 | 0.17 | 0.06 | 0.18 | 0.18 |
| 10 | 7617810 | rs1537632 | -0.298 | 0.874 | -0.539 | 0.469 | -0.004 | 0.012 | 0.012 | 0.774 | 0.18 | 0.28 | 0.11 | 0.19 | 0.16 | 0.26 |
| 10 | 7618686 | rs2275069 | -1.102 | 0.981 | -1.114 | 0.259 | -0.002 | -0.001 | -0.001 | 0 | 0.43 | 0.36 | 0.55 | 0.77 | 0.52 | 0.51 |
| 10 | 7618993 | rs10458802 | -1.102 | 0.986 | -1.112 | 0.262 | -0.002 | -0.001 | -0.001 | 0 | 0.43 | 0.36 | 0.54 | 0.77 | 0.52 | 0.51 |

| | | | | | | | | | | | | | | | | |
|---|---|---|---|---|---|---|---|---|---|---|---|---|---|---|---|---|
| 10 | 7619605 | rs10905190 | -0.246 | 1.292 | -1.581 | 0.529 | -0.083 | -0.107 | -0.107 | 0.109 | 0.25 | 0.06 | 0.44 | 0.57 | 0.35 | 0.24 |
| 10 | 7621578 | rs11255201 | 0.242 | 0.366 | -0.713 | 0.565 | -0.019 | 0.07 | 0.07 | 0 | 0.72 | 0.52 | 0.79 | 0.48 | 0.73 | 0.48 |
| 10 | 7622047 | rs12256183 | -0.294 | -0.035 | -0.993 | -0.551 | 0.015 | -0.012 | -0.012 | 0 | 0.77 | 0.77 | 0.94 | 1 | 0.9 | 0.87 |
| 10 | 7622904 | rs10905191 | -0.257 | 1.079 | -0.529 | -0.16 | 0.026 | -0.018 | -0.018 | 0 | 0.16 | 0.24 | 0.11 | 0.08 | 0.25 | 0.12 |
| 10 | 7623127 | rs12774458 | -0.186 | 0.875 | -0.318 | 0.065 | 0.007 | 0.001 | 0.001 | 0 | 0.7 | 0.6 | 0.51 | 0.74 | 0.66 | 0.73 |
| 10 | 7626689 | rs10905193 | 0.151 | 0.629 | -0.16 | 0.163 | -0.019 | 0.017 | 0.017 | 0 | 0.35 | 0.39 | 0.25 | 0.52 | 0.22 | 0.35 |
| 10 | 7628419 | rs885548 | 0.008 | 0.635 | -0.14 | 0.183 | -0.017 | 0.017 | 0.017 | 0 | 0.35 | 0.38 | 0.25 | 0.52 | 0.23 | 0.34 |
| 10 | 7628431 | rs11598825 | 0.835 | 0.779 | -0.541 | -0.113 | -0.019 | 0.05 | 0.05 | 0 | 0.23 | 0.14 | 0.19 | 0 | 0.28 | 0.18 |
| 10 | 7632504 | rs7904044 | -0.238 | 0.158 | 0.088 | 0.755 | -0.009 | 0.01 | 0.01 | 0 | 0.33 | 0.35 | 0.24 | 0.52 | 0.22 | 0.34 |
| 10 | 7634321 | rs871049 | 0.103 | 0.596 | -0.28 | 0.103 | 0.019 | 0.051 | 0.051 | 0 | 0.51 | 0.42 | 0.56 | 0.23 | 0.63 | 0.43 |
| 10 | 7642262 | rs11255225 | 0.779 | 0.666 | 0.154 | 0.75 | -0.001 | -0.001 | -0.001 | 0.049 | 0.7 | 0.3 | 0.77 | 0.67 | 0.72 | 0.71 |
| 10 | 7642284 | rs11255226 | -0.9 | 0.559 | 1.04 | 0.756 | 0.029 | 0.015 | 0.015 | 0 | 0.16 | 0.27 | 0.14 | 0.21 | 0.19 | 0.2 |
| 10 | 7646184 | rs11255232 | -0.498 | 0.597 | 1.082 | 0.801 | -0.049 | -0.04 | -0.04 | 0.018 | 0.08 | 0.01 | 0.11 | 0.2 | 0.15 | 0.11 |
| 10 | 7647289 | rs1572241 | 0.326 | 0.568 | 1.183 | 0.789 | -0.035 | 0.003 | 0.003 | 0 | 0.86 | 0.58 | 0.91 | 0.89 | 0.91 | 0.88 |
| 10 | 7648190 | rs10752113 | 0.738 | 0.574 | 1.057 | 0.84 | 0.003 | 0.004 | 0.004 | 0 | 0.79 | 0.71 | 0.82 | 0.69 | 0.76 | 0.77 |
| 10 | 7652410 | rs7083191 | -0.668 | 0.765 | 0.907 | 0.576 | 0 | 0.005 | 0.005 | 0 | 0.09 | 0.2 | 0.04 | 0.06 | 0.07 | 0.1 |
| 10 | 7653065 | rs1537628 | -1.075 | 0.91 | 0.305 | 0.748 | 0.004 | -0.004 | -0.004 | 0 | 0.2 | 0.29 | 0.15 | 0.23 | 0.24 | 0.15 |
| 10 | 7654501 | rs11255236 | -0.806 | 0.839 | -0.082 | 0.988 | 0 | 0.004 | 0.004 | 0 | 0.25 | 0.36 | 0.28 | 0.32 | 0.25 | 0.26 |
| 10 | 7654808 | rs7093342 | -0.525 | 0.22 | -0.474 | 1.001 | 0.006 | -0.006 | -0.006 | 0 | 0.33 | 0.31 | 0.31 | 0.39 | 0.25 | 0.33 |
| 10 | 7655352 | rs1931901 | -0.226 | -0.103 | -0.435 | 0.741 | -0.011 | 0.018 | 0.018 | 0.004 | 0.4 | 0.52 | 0.51 | 0.52 | 0.33 | 0.37 |
| 10 | 7655566 | rs1931902 | -0.454 | 0.33 | -0.916 | 0.142 | 0.012 | 0.033 | 0.033 | 0 | 0.61 | 0.67 | 0.81 | 0.91 | 0.67 | 0.69 |
| 10 | 7655665 | rs10508331 | 1.087 | -0.171 | -0.699 | 0.267 | -0.006 | -0.019 | -0.019 | 0 | 0.15 | 0.09 | 0.31 | 0.26 | 0.16 | 0.19 |
| 10 | 7655952 | rs7080791 | -0.213 | -0.591 | -0.562 | 0.175 | -0.009 | 0.013 | 0.013 | 0.093 | 0.37 | 0.55 | 0.47 | 0.5 | 0.27 | 0.34 |
| 10 | 7660490 | rs11255241 | 0.485 | -0.532 | -0.31 | 0.246 | -0.011 | 0.002 | 0.002 | 0 | 0.21 | 0.21 | 0.14 | 0.25 | 0.12 | 0.16 |
| 10 | 7661011 | rs7904081 | 0.866 | -0.05 | -0.75 | 0.232 | -0.065 | -0.078 | -0.078 | 1.187 | 0.23 | 0.08 | 0.33 | 0.37 | 0.37 | 0.29 |
| 10 | 7662452 | rs1022986 | 0.229 | -0.243 | -0.632 | -0.111 | 0.006 | 0.002 | 0.002 | 0.157 | 0.35 | 0.23 | 0.4 | 0.37 | 0.31 | 0.38 |
| 10 | 7668637 | rs10905203 | -1.078 | 0.063 | -0.308 | -0.248 | -0.001 | -0.059 | -0.059 | 0 | 0.84 | 0.75 | 0.92 | 0.98 | 0.86 | 0.88 |
| 10 | 7672986 | rs7921768 | 0.953 | 0.182 | -0.703 | -0.026 | -0.071 | 0.041 | 0.041 | 0.017 | 0.21 | 0.03 | 0.2 | 0.11 | 0.23 | 0.19 |
| 10 | 7673498 | rs17347867 | 0.37 | -0.27 | -0.957 | 0.122 | 0.02 | 0.015 | 0.015 | 0 | 0.23 | 0.07 | 0.1 | 0.2 | 0.19 | 0.3 |
| 10 | 7674300 | rs7086233 | 0.014 | -0.496 | -0.404 | 0.224 | 0 | 0 | 0 | 0 | 0.46 | 0.54 | 0.32 | 0.36 | 0.37 | 0.41 |
| 10 | 7677203 | rs10508332 | 0.554 | -0.33 | -0.128 | 0.439 | 0.002 | -0.019 | -0.019 | 0.031 | 0.29 | 0.44 | 0.46 | 0.22 | 0.28 | 0.25 |
| 10 | 7677416 | rs17142881 | 1.213 | -0.549 | -0.04 | 0.544 | 0 | 0 | 0 | 0.046 | 0.17 | 0.16 | 0.1 | 0.21 | 0.15 | 0.28 |
| 10 | 7678627 | rs7080582 | 0.005 | -0.579 | 0.168 | 0.297 | -0.021 | 0.014 | 0.014 | 0.197 | 0.38 | 0.49 | 0.41 | 0.59 | 0.31 | 0.49 |
| 10 | 7684323 | rs12769926 | -0.35 | -0.549 | 0.147 | 0.237 | 0.02 | 0.002 | 0.002 | 0 | 0.45 | 0.62 | 0.24 | 0.46 | 0.32 | 0.52 |
| 10 | 7685743 | rs17432951 | 1.374 | -0.308 | -0.145 | 0.113 | -0.053 | 0.074 | 0.074 | 0.002 | 0.24 | 0.05 | 0.22 | 0.09 | 0.26 | 0.21 |
| 10 | 7689194 | rs9329327 | 0.437 | 0.024 | 0.198 | 0.576 | 0.001 | 0.001 | 0.001 | 0 | 0.5 | 0.56 | 0.37 | 0.29 | 0.46 | 0.54 |
| 10 | 7692114 | rs7074758 | -1.243 | 0.535 | -0.082 | 0.563 | -0.008 | -0.019 | -0.019 | 1.105 | 0.81 | 0.9 | 0.77 | 0.74 | 0.76 | 0.78 |
| 10 | 7696641 | rs7091139 | 1.169 | 0.473 | -0.011 | 0.553 | 0.004 | -0.006 | -0.006 | 0.277 | 0.19 | 0.13 | 0.1 | 0.28 | 0.12 | 0.2 |
| 10 | 7697504 | rs11255267 | 1.333 | 0.505 | 0.078 | 0.511 | 0 | 0.021 | 0.021 | 0.752 | 0.35 | 0.31 | 0.27 | 0.11 | 0.31 | 0.26 |
| 10 | 7700353 | rs11255271 | 1.402 | 0.545 | -0.039 | 0.583 | -0.011 | -0.006 | -0.006 | 0.319 | 0.14 | 0.23 | 0.09 | 0.17 | 0.14 | 0.24 |
| 10 | 7700795 | rs11255272 | 1.441 | 0.565 | -0.288 | 0.551 | -0.005 | -0.004 | -0.004 | 0 | 0.14 | 0.16 | 0.08 | 0.17 | 0.13 | 0.24 |
| 10 | 7702343 | rs7094085 | 0.635 | 0.96 | -0.528 | 0.853 | 0 | -0.001 | -0.001 | 0 | 0.19 | 0.14 | 0.11 | 0.28 | 0.14 | 0.19 |
| 10 | 7704917 | rs927915 | 0.315 | 1.679 | -0.569 | 0.99 | -0.051 | -0.012 | -0.012 | 0 | 0.73 | 0.89 | 0.69 | 0.73 | 0.62 | 0.74 |
| 10 | 7707276 | rs10905207 | 0.115 | 1.416 | -0.656 | 0.672 | 0.005 | 0.046 | 0.046 | 0.041 | 0.75 | 0.64 | 0.53 | 0.55 | 0.7 | 0.71 |
| 10 | 7709909 | rs10905208 | 0.21 | 1.131 | -0.67 | 0.881 | 0.009 | 0.011 | 0.011 | 0.074 | 0.71 | 0.67 | 0.46 | 0.55 | 0.57 | 0.7 |
| 10 | 7716687 | rs7908119 | -0.229 | 0.246 | -1.012 | 0.561 | 0.029 | 0.012 | 0.012 | 0.026 | 0.61 | 0.73 | 0.67 | 0.63 | 0.71 | 0.56 |
| 10 | 7718053 | rs11255282 | 0.003 | 1.042 | -1.357 | 0.97 | -0.064 | -0.073 | -0.073 | 0.055 | 0.23 | 0.13 | 0.31 | 0.37 | 0.32 | 0.21 |
| 10 | 7722848 | rs7897216 | -0.609 | 0.482 | -0.839 | 0.385 | 0.002 | 0.011 | 0.011 | 0.529 | 0.45 | 0.42 | 0.5 | 0.61 | 0.56 | 0.45 |
| 10 | 7725787 | rs4749153 | -0.615 | 0.478 | -0.841 | 0.383 | 0.003 | 0.012 | 0.012 | 0.528 | 0.45 | 0.42 | 0.5 | 0.61 | 0.56 | 0.45 |
| 10 | 7730983 | rs11812109 | -0.995 | 0.435 | 0.006 | 0.574 | 0.059 | 0.064 | 0.064 | 0 | 0.09 | 0.25 | 0.12 | 0.12 | 0.14 | 0.07 |
| 10 | 7731885 | rs719748 | -0.701 | 0.261 | -0.408 | 0.634 | 0.001 | -0.002 | -0.002 | 0.071 | 0.44 | 0.64 | 0.66 | 0.45 | 0.58 | 0.41 |
| 10 | 7732323 | rs7917456 | 0.066 | 0.164 | -0.763 | 0.473 | -0.043 | -0.024 | -0.024 | 0 | 0.35 | 0.28 | 0.54 | 0.33 | 0.44 | 0.34 |
| 10 | 7739555 | rs10905220 | -0.587 | -0.135 | -0.67 | 0.456 | 0.004 | 0.001 | 0.001 | 0.391 | 0.49 | 0.39 | 0.65 | 0.4 | 0.54 | 0.4 |
| 10 | 7739760 | rs7918827 | -1.245 | -0.45 | -0.865 | -0.01 | 0.023 | 0.03 | 0.03 | 0.08 | 0.15 | 0.07 | 0.12 | 0.02 | 0.11 | 0.08 |
| 10 | 7739796 | rs7918844 | -0.027 | -0.026 | -0.656 | 0.176 | 0 | -0.008 | -0.008 | 0 | 0.08 | 0.08 | 0.1 | 0.02 | 0.06 | 0.05 |
| 10 | 7740248 | rs7921576 | -0.018 | -0.104 | -0.622 | 0.294 | 0 | 0.018 | 0.018 | 4.452 | 0.54 | 0.61 | 0.66 | 0.59 | 0.57 | 0.4 |
| 10 | 7744986 | rs1980857 | -1.26 | -0.723 | 0.111 | 0.806 | -0.005 | 0.097 | 0.097 | 0.241 | 0.17 | 0.45 | 0.35 | 0.27 | 0.15 | 0.2 |
| 10 | 7751626 | rs2026609 | -0.619 | -1.286 | 0.105 | 0.644 | 0.003 | 0.002 | 0.002 | 1.482 | 0.7 | 0.74 | 0.88 | 0.7 | 0.81 | 0.57 |
| 10 | 7755623 | rs4747574 | 0.982 | -1.449 | -0.09 | 0.368 | 0.032 | -0.072 | -0.072 | 0 | 0.45 | 0.1 | 0.5 | 0.43 | 0.6 | 0.34 |
| 10 | 7759595 | rs7072478 | -0.471 | -1.024 | 0.017 | 0.184 | 0.003 | 0.027 | 0.027 | 0 | 0.13 | 0.22 | 0.28 | 0.22 | 0.17 | 0.16 |
| 10 | 7759865 | rs7090433 | -2.12 | -1.079 | 0.096 | 0.143 | -0.002 | -0.002 | -0.002 | 0 | 0.29 | 0.61 | 0.42 | 0.27 | 0.3 | 0.24 |
| 10 | 7759926 | rs6602267 | -1.365 | -1.103 | 0.096 | 0.16 | -0.01 | 0.03 | 0.03 | 0 | 0.22 | 0.56 | 0.31 | 0.26 | 0.23 | 0.19 |
| 10 | 7760367 | rs1887332 | 0.352 | -1.069 | 0.013 | 0.257 | 0.004 | 0.02 | 0.02 | 0 | 0.06 | 0.07 | 0.23 | 0.16 | 0.12 | 0.11 |
| 10 | 7763421 | rs2277227 | 0.746 | -1.158 | -0.189 | 0.233 | 0.059 | -0.058 | -0.058 | 0 | 0.47 | 0.12 | 0.56 | 0.44 | 0.59 | 0.38 |
| 10 | 7764338 | rs927917 | -0.904 | -1.131 | -0.005 | 0.24 | 0.02 | 0.092 | 0.092 | 0 | 0.14 | 0.42 | 0.29 | 0.22 | 0.17 | 0.16 |
| 10 | 7767196 | rs4749197 | -1.054 | -1.377 | -0.097 | 0.237 | -0.008 | -0.002 | -0.002 | 0 | 0.7 | 0.73 | 0.87 | 0.7 | 0.81 | 0.58 |
| 10 | 7768221 | rs7912641 | 0.833 | -1.243 | 0.018 | 0.288 | -0.008 | -0.042 | -0.042 | 0.014 | 0.08 | 0.16 | 0.02 | 0 | 0.05 | 0.03 |
| 10 | 7770222 | rs3858256 | -1.736 | -1.201 | -0.411 | 0.313 | -0.009 | -0.001 | -0.001 | 0.135 | 0.28 | 0.61 | 0.41 | 0.27 | 0.27 | 0.24 |
| 10 | 7776831 | rs1926937 | 0.765 | -1.377 | -1.038 | 0.501 | 0.013 | -0.053 | -0.053 | 0 | 0.45 | 0.11 | 0.48 | 0.4 | 0.58 | 0.39 |

| | | | | | | | | | | | | | | | | |
|---|---|---|---|---|---|---|---|---|---|---|---|---|---|---|---|---|
| 10 | 7783546 | rs10508333 | 0.17 | -0.816 | -0.44 | 0.725 | 0.012 | 0.035 | 0.035 | 0 | 0.05 | 0.06 | 0.21 | 0.18 | 0.11 | 0.1 |
| 10 | 7785017 | rs3736965 | 0.912 | -1.67 | -0.065 | 0.545 | -0.016 | 0.04 | 0.04 | 0 | 0.74 | 0.44 | 0.7 | 0.66 | 0.75 | 0.61 |
| 10 | 7788495 | rs7912419 | 0.192 | -1.51 | -0.25 | 0.352 | 0.001 | 0.002 | 0.002 | 1.893 | 0.8 | 0.81 | 0.9 | 0.89 | 0.83 | 0.73 |
| 10 | 7788958 | rs2038993 | 0.865 | -1.516 | -0.073 | 0.42 | 0.008 | 0.013 | 0.013 | 0.838 | 0.73 | 0.65 | 0.69 | 0.67 | 0.71 | 0.62 |
| 10 | 7790894 | rs11255332 | 1.069 | -1.689 | -0.959 | 0.394 | -0.021 | -0.031 | -0.031 | 0 | 0.47 | 0.11 | 0.56 | 0.4 | 0.58 | 0.43 |
| 10 | 7793035 | rs2508 | -0.386 | -1.21 | -0.396 | 0.579 | 0.012 | 0.027 | 0.027 | 0 | 0.08 | 0.1 | 0.21 | 0.17 | 0.13 | 0.11 |
| 10 | 7794648 | rs7907897 | 0.749 | -1.792 | -0.837 | 0.444 | 0.028 | 0.024 | 0.024 | 0.031 | 0.5 | 0.29 | 0.49 | 0.44 | 0.58 | 0.33 |
| 10 | 7795911 | rs7342066 | -0.116 | -2.491 | -0.797 | 0.202 | 0.021 | 0.009 | 0.009 | 0 | 0.57 | 0.64 | 0.79 | 0.59 | 0.74 | 0.48 |
| 10 | 7800881 | rs11255344 | 0.769 | -2.593 | -0.553 | 0.416 | -0.006 | 0.001 | 0.001 | 0.426 | 0.21 | 0.14 | 0.28 | 0.18 | 0.24 | 0.23 |
| 10 | 7802785 | rs11255345 | 0.044 | -2.713 | -0.702 | 0.397 | 0.015 | -0.014 | -0.014 | 0 | 0.44 | 0.19 | 0.37 | 0.47 | 0.37 | 0.41 |
| 10 | 7805481 | rs2026612 | 2.115 | -2.783 | -0.65 | 0.388 | -0.059 | 0.023 | 0.023 | 0.346 | 0.25 | 0.15 | 0.39 | 0.15 | 0.44 | 0.11 |
| 10 | 7808086 | rs3736968 | 0.063 | -2.729 | -0.651 | 0.462 | 0.01 | -0.014 | -0.014 | 0 | 0.44 | 0.21 | 0.37 | 0.47 | 0.37 | 0.41 |
| 10 | 7813124 | rs12359107 | 0.779 | -2.492 | -0.454 | 0.345 | -0.016 | 0.002 | 0.002 | 0.526 | 0.2 | 0.11 | 0.28 | 0.18 | 0.25 | 0.23 |
| 10 | 7821794 | rs10905230 | -0.329 | -2.422 | -0.371 | 0.226 | 0.004 | 0.019 | 0.019 | 0 | 0.53 | 0.58 | 0.48 | 0.71 | 0.42 | 0.53 |
| 10 | 7825530 | rs4363498 | -0.128 | -2.333 | -0.188 | 0.26 | -0.009 | -0.013 | -0.013 | 0 | 0.81 | 0.76 | 0.88 | 0.86 | 0.86 | 0.66 |
| 10 | 7828752 | rs10795570 | 0.889 | -2.462 | -0.208 | 0.099 | 0.042 | -0.027 | -0.027 | 0 | 0.32 | 0.48 | 0.37 | 0.21 | 0.5 | 0.34 |
| 10 | 7829018 | rs11255367 | 2.237 | -2.409 | -0.675 | 0.138 | -0.016 | -0.059 | -0.059 | 1.172 | 0.17 | 0 | 0.07 | 0.26 | 0.11 | 0.12 |
| 10 | 7829037 | rs2802460 | 0.434 | -1.825 | -0.068 | -0.114 | -0.016 | 0.001 | 0.001 | 0.299 | 0.23 | 0.11 | 0.29 | 0.2 | 0.26 | 0.26 |
| 10 | 7836104 | rs1244414 | 0.396 | -1.615 | 0.119 | -0.116 | 0.003 | -0.033 | -0.033 | 0.249 | 0.11 | 0.21 | 0.09 | 0.07 | 0.1 | 0.25 |
| 10 | 7838932 | rs1244422 | -1.137 | -1.704 | 0.276 | -0.042 | -0.003 | -0.01 | -0.01 | 5.1 | 0.46 | 0.46 | 0.51 | 0.33 | 0.41 | 0.61 |
| 10 | 7842900 | rs12770829 | 1.795 | -1.712 | 0.289 | -0.035 | -0.03 | 0.007 | 0.007 | 0.028 | 0.22 | 0.15 | 0.29 | 0.14 | 0.41 | 0.09 |
| 10 | 7849688 | rs4655 | -0.346 | -1.701 | 0.302 | -0.047 | -0.001 | -0.001 | -0.001 | 2.013 | 0.35 | 0.33 | 0.37 | 0.27 | 0.36 | 0.5 |
| 10 | 7859156 | rs11255374 | 1.382 | -1.654 | 0.229 | -0.167 | -0.005 | 0.036 | 0.036 | 0 | 0.17 | 0.1 | 0.09 | 0.07 | 0.16 | 0.33 |
| 10 | 7860265 | rs2057604 | 1.735 | -1.454 | 0.108 | -0.044 | -0.035 | 0.008 | 0.008 | 0.035 | 0.22 | 0.14 | 0.29 | 0.14 | 0.41 | 0.09 |
| 10 | 7869372 | rs11255388 | 0.649 | -1.47 | -0.168 | -0.02 | -0.007 | -0.003 | -0.003 | 0.146 | 0.09 | 0.11 | 0.13 | 0.06 | 0.04 | 0.1 |
| 10 | 7872957 | rs1244507 | -0.911 | -1.542 | -0.199 | -0.039 | 0 | -0.007 | -0.007 | 0.443 | 0.43 | 0.51 | 0.5 | 0.34 | 0.4 | 0.6 |
| 10 | 7876830 | rs10508336 | -0.396 | -1.62 | -0.248 | -0.103 | 0.039 | 0.032 | 0.032 | 0 | 0.16 | 0.06 | 0.16 | 0.08 | 0.08 | 0.21 |
| 10 | 7886084 | rs2802457 | -1.197 | -1.368 | -0.238 | 0.048 | 0.009 | 0.025 | 0.025 | 0.01 | 0.5 | 0.53 | 0.54 | 0.35 | 0.44 | 0.7 |
| 10 | 7898502 | rs11255407 | 0.78 | -1.33 | -0.557 | 0.048 | 0 | 0.001 | 0.001 | 2.94 | 0.14 | 0.13 | 0.26 | 0.21 | 0.19 | 0.17 |
| 10 | 7898759 | rs1064134 | 0.055 | -1.132 | -0.588 | 0.335 | 0 | -0.007 | -0.007 | 1.113 | 0.39 | 0.39 | 0.4 | 0.28 | 0.39 | 0.59 |
| 10 | 7899231 | rs10905239 | 0.992 | -0.901 | -0.797 | 0.524 | 0.043 | -0.096 | -0.096 | 0 | 0.27 | 0.12 | 0.16 | 0.51 | 0.16 | 0.21 |
| 10 | 7903733 | rs6602275 | 0.523 | -0.609 | -0.731 | 0.303 | 0.011 | -0.003 | -0.003 | 1.627 | 0.08 | 0.01 | 0.12 | 0.06 | 0.04 | 0.1 |
| 10 | 7904358 | rs1312782 | -0.459 | -0.314 | -0.438 | 0.412 | 0.002 | 0.005 | 0.005 | 0 | 0.5 | 0.44 | 0.53 | 0.35 | 0.44 | 0.7 |
| 10 | 7906459 | rs1556479 | -0.435 | -0.655 | -0.378 | 0.721 | -0.013 | 0.001 | 0.001 | 0 | 0.69 | 0.6 | 0.73 | 0.65 | 0.74 | 0.75 |
| 10 | 7908552 | rs10905240 | 0.308 | 0.146 | -1.164 | 0.63 | -0.062 | -0.053 | -0.053 | 0 | 0.33 | 0.13 | 0.52 | 0.34 | 0.41 | 0.29 |
| 10 | 7922269 | rs1041541 | 0.596 | -0.308 | -0.855 | 0.446 | 0.01 | 0.015 | 0.015 | 0 | 0.29 | 0.19 | 0.23 | 0.28 | 0.23 | 0.26 |
| 10 | 7923508 | rs1244461 | 0.145 | -0.439 | -0.256 | 0.419 | 0.032 | 0.077 | 0.077 | 0 | 0.21 | 0.42 | 0.19 | 0.3 | 0.25 | 0.27 |
| 10 | 7928240 | rs1244459 | 0.77 | -0.392 | 0.048 | 0.329 | 0.013 | -0.022 | -0.022 | 0.437 | 0.12 | 0.14 | 0.11 | 0.05 | 0.16 | 0.21 |
| 10 | 7932563 | rs1244440 | -0.354 | -0.434 | -0.15 | 0.39 | 0.02 | 0.038 | 0.038 | 0 | 0.22 | 0.5 | 0.2 | 0.27 | 0.25 | 0.26 |
| 10 | 7935761 | rs10508338 | 0.472 | -0.572 | -0.23 | 0.266 | 0.004 | 0.009 | 0.009 | 0 | 0.63 | 0.35 | 0.77 | 0.72 | 0.65 | 0.58 |
| 10 | 7944469 | rs7069192 | -1.86 | -0.553 | -0.31 | 0.262 | 0 | -0.013 | -0.013 | 0 | 0.85 | 0.83 | 0.97 | 0.99 | 0.9 | 0.84 |
| 10 | 7950997 | rs12219547 | 1.471 | -0.504 | -0.616 | 0.294 | 0.002 | -0.008 | -0.008 | 1.374 | 0.25 | 0.14 | 0.21 | 0.31 | 0.22 | 0.24 |
| 10 | 7951445 | rs10508339 | 1.471 | -0.508 | -0.616 | 0.294 | 0.004 | -0.008 | -0.008 | 5.197 | 0.25 | 0.14 | 0.21 | 0.31 | 0.22 | 0.24 |
| 10 | 7954810 | rs4749347 | 0.678 | -0.561 | -0.6 | 0.28 | -0.047 | -0.043 | -0.043 | 0 | 0.36 | 0.17 | 0.56 | 0.4 | 0.43 | 0.32 |
| 10 | 7956230 | rs11255440 | 1.079 | -0.534 | -0.565 | 0.3 | 0.011 | 0.006 | 0.006 | 0 | 0.28 | 0.16 | 0.22 | 0.31 | 0.23 | 0.26 |
| 10 | 7961064 | rs4749353 | 0.514 | -0.497 | -0.428 | 0.302 | 0.014 | 0.023 | 0.023 | 0 | 0.61 | 0.28 | 0.75 | 0.72 | 0.64 | 0.56 |
| 10 | 7964341 | rs17143115 | 1.58 | -0.725 | -0.828 | 0.403 | 0.004 | -0.005 | -0.005 | 0.146 | 0.06 | 0.04 | 0.12 | 0.1 | 0.03 | 0.1 |
| 10 | 7972373 | rs10905257 | 0.266 | -0.626 | -0.75 | 0.329 | 0.007 | 0.023 | 0.023 | 0 | 0.62 | 0.27 | 0.76 | 0.72 | 0.63 | 0.57 |
| 10 | 7981522 | rs187821 | 0.128 | -0.441 | -0.203 | 0.518 | 0.016 | 0.01 | 0.01 | 0 | 0.19 | 0.3 | 0.17 | 0.2 | 0.24 | 0.26 |
| 10 | 7989049 | rs3858258 | 0.456 | -0.316 | -0.905 | 0.589 | -0.071 | -0.071 | -0.071 | 0 | 0.32 | 0.12 | 0.53 | 0.4 | 0.42 | 0.29 |
| 10 | 8002270 | rs9633771 | 0.802 | -0.601 | -0.859 | 0.542 | 0.02 | -0.019 | -0.019 | 0.25 | 0.31 | 0.19 | 0.32 | 0.41 | 0.24 | 0.33 |
| 10 | 8006798 | rs4747647 | 0.181 | -0.587 | -0.968 | 0.492 | 0.024 | 0.045 | 0.045 | 0 | 0.58 | 0.26 | 0.73 | 0.71 | 0.62 | 0.53 |
| 10 | 8007560 | rs1244229 | -0.289 | -0.713 | -0.376 | 0.414 | 0.038 | 0.06 | 0.06 | 0 | 0.79 | 0.55 | 0.8 | 0.73 | 0.74 | 0.72 |
| 10 | 8008411 | rs407644 | -1.264 | -0.58 | -0.399 | 0.338 | -0.001 | 0 | 0 | 0 | 0.91 | 0.94 | 0.94 | 0.96 | 0.89 | 0.83 |
| 10 | 8010498 | rs4747649 | 1.562 | -0.68 | -0.545 | 0.401 | -0.001 | -0.023 | -0.023 | 3.268 | 0.25 | 0.09 | 0.19 | 0.31 | 0.2 | 0.24 |
| 10 | 8011423 | rs10752122 | 0.432 | -0.686 | -0.286 | 0.356 | -0.049 | -0.065 | -0.065 | 0.044 | 0.32 | 0.17 | 0.55 | 0.48 | 0.44 | 0.32 |
| 10 | 8017823 | rs3898015 | 0.542 | -0.594 | -0.53 | 0.204 | -0.053 | -0.043 | -0.043 | 0 | 0.3 | 0.14 | 0.54 | 0.38 | 0.42 | 0.29 |
| 10 | 8018046 | rs11255468 | 1.062 | -0.633 | -0.627 | 0.203 | -0.142 | -0.077 | -0.077 | 0 | 0.24 | 0.04 | 0.43 | 0.27 | 0.41 | 0.19 |
| 10 | 8019124 | rs10795584 | -0.478 | -0.782 | -0.427 | 0.203 | 0.008 | 0.007 | 0.007 | 0 | 0.52 | 0.39 | 0.62 | 0.42 | 0.55 | 0.49 |
| 10 | 8025362 | rs12780500 | 0.192 | -0.562 | -0.584 | 0.137 | 0.01 | 0.001 | 0.001 | 0.177 | 0.28 | 0.21 | 0.33 | 0.27 | 0.24 | 0.32 |
| 10 | 8025394 | rs196528 | 0.944 | -1.191 | -0.715 | -0.13 | 0.02 | -0.006 | -0.006 | 0 | 0.42 | 0.32 | 0.27 | 0.49 | 0.36 | 0.34 |
| 10 | 8026609 | rs7894336 | -1.007 | -0.521 | -1.096 | 0.097 | -0.003 | -0.001 | -0.001 | 0 | 0.53 | 0.44 | 0.61 | 0.43 | 0.52 | 0.5 |
| 10 | 8028388 | rs4749442 | 0.638 | -0.59 | -1.254 | 0.143 | -0.148 | -0.096 | -0.096 | 0 | 0.24 | 0.02 | 0.43 | 0.29 | 0.4 | 0.2 |
| 10 | 8047550 | rs768166 | -0.054 | -0.925 | -0.768 | 0.178 | -0.044 | -0.026 | -0.026 | 0 | 0.28 | 0.38 | 0.19 | 0.15 | 0.13 | 0.3 |
| 10 | 8049837 | rs12245532 | -0.335 | -0.458 | -0.444 | 0.466 | -0.03 | 0.044 | 0.044 | 0.031 | 0.09 | 0.3 | 0.12 | 0.11 | 0.02 | 0.1 |
| 10 | 8064199 | rs10905269 | 0.12 | -0.119 | -1.267 | 0.287 | -0.01 | 0.024 | 0.024 | 0 | 0.43 | 0.25 | 0.45 | 0.3 | 0.45 | 0.38 |
| 10 | 8066178 | rs418704 | 0.268 | -0.202 | -1.89 | 0.202 | 0.005 | 0.014 | 0.014 | 0 | 0.66 | 0.22 | 0.56 | 0.73 | 0.67 | 0.66 |
| 10 | 8074021 | rs400431 | -0.628 | 0.383 | -1.521 | -0.037 | 0.009 | 0.049 | 0.049 | 0 | 0.48 | 0.63 | 0.34 | 0.26 | 0.48 | 0.33 |
| 10 | 8074437 | rs430753 | -1.005 | 0.615 | -1.525 | 0.005 | -0.007 | 0.006 | 0.006 | 0 | 0.69 | 0.72 | 0.64 | 0.81 | 0.66 | 0.64 |

| | | | | | | | | | | | | | | | | |
|---|---|---|---|---|---|---|---|---|---|---|---|---|---|---|---|---|
| 10 | 8077699 | rs406571 | -0.696 | 0.84 | -0.448 | 0.042 | -0.007 | 0.015 | 0.015 | 0.429 | 0.31 | 0.51 | 0.22 | 0.33 | 0.3 | 0.24 |
| 10 | 8083888 | rs578268 | -0.55 | 1.144 | -0.247 | -0.781 | 0 | -0.005 | -0.005 | 0 | 0.45 | 0.43 | 0.38 | 0.71 | 0.51 | 0.49 |
| 10 | 8086170 | rs10795588 | 1.159 | 1.265 | 0.389 | -1.289 | -0.017 | 0.17 | 0.17 | 0.924 | 0.43 | 0.21 | 0.66 | 0.94 | 0.46 | 0.67 |
| 10 | 8089136 | rs4143094 | 1.219 | -0.239 | 1.05 | -1.101 | 0.037 | -0.098 | -0.098 | 0 | 0.79 | 0.52 | 0.83 | 0.95 | 0.77 | 0.91 |
| 10 | 8092186 | rs1271899 | 1.176 | -0.175 | 0.652 | -1.107 | -0.005 | -0.003 | -0.003 | 0.301 | 0.81 | 0.78 | 0.85 | 0.96 | 0.77 | 0.91 |
| 10 | 8092683 | rs1244186 | 1.142 | -0.165 | 0.657 | -1.102 | -0.004 | -0.005 | -0.005 | 0.932 | 0.81 | 0.78 | 0.85 | 0.96 | 0.77 | 0.91 |
| 10 | 8093034 | rs501764 | 0.536 | -0.123 | 0.347 | -1.109 | -0.025 | 0.033 | 0.033 | 4.922 | 0.86 | 0.94 | 0.91 | 0.99 | 0.82 | 0.94 |
| 10 | 8093185 | rs485411 | 0.978 | -0.169 | 0.369 | -1.091 | -0.031 | 0.043 | 0.043 | 1.581 | 0.82 | 0.91 | 0.86 | 0.96 | 0.77 | 0.91 |
| 10 | 8097368 | rs10905277 | 1.228 | -0.285 | 0.217 | -0.884 | 0.002 | 0.048 | 0.048 | 0.009 | 0.43 | 0.21 | 0.43 | 0.67 | 0.54 | 0.67 |
| 10 | 8102272 | rs3802604 | 1.184 | -0.362 | 0.618 | -0.554 | 0.077 | 0.102 | 0.102 | 0 | 0.5 | 0.22 | 0.49 | 0.68 | 0.62 | 0.69 |
| 10 | 8104208 | rs3824662 | -0.168 | -0.545 | 0.409 | -0.56 | 0.087 | -0.009 | -0.009 | 0 | 0.29 | 0.08 | 0.36 | 0.27 | 0.19 | 0.17 |
| 10 | 8106701 | rs568727 | 0.576 | -0.409 | 0.954 | -0.574 | -0.009 | -0.022 | -0.022 | 0.105 | 0.59 | 0.51 | 0.66 | 0.73 | 0.62 | 0.76 |
| 10 | 8108592 | rs569421 | -0.649 | -0.639 | 0.957 | -0.639 | -0.039 | -0.026 | -0.026 | 0.005 | 0.37 | 0.46 | 0.18 | 0.26 | 0.25 | 0.24 |
| 10 | 8111409 | rs422628 | 1.14 | -0.429 | 0.563 | -0.618 | 0.011 | -0.039 | -0.039 | 0 | 0.81 | 0.74 | 0.87 | 0.95 | 0.78 | 0.84 |
| 10 | 8115362 | rs10905284 | 1.228 | -0.522 | 0.081 | -0.442 | 0.009 | 0.007 | 0.007 | 0 | 0.47 | 0.1 | 0.42 | 0.58 | 0.57 | 0.61 |
| 10 | 8116598 | rs1058240 | 0.535 | 0.296 | 0.518 | -0.335 | 0.064 | -0.034 | -0.034 | 0 | 0.9 | 0.74 | 0.89 | 0.98 | 0.82 | 0.93 |
| 10 | 8120143 | rs477461 | -0.525 | -1.062 | 0.022 | -0.163 | -0.023 | -0.007 | -0.007 | 0 | 0.27 | 0.29 | 0.12 | 0.24 | 0.18 | 0.16 |
| 10 | 8125819 | rs379568 | -1.059 | -0.346 | 0.556 | -0.03 | -0.032 | -0.002 | -0.002 | 0.101 | 0.15 | 0.41 | 0.17 | 0.11 | 0.09 | 0.11 |
| 10 | 8128498 | rs10508342 | -0.493 | 0.844 | -0.276 | -0.382 | -0.072 | 0.048 | 0.048 | 0.326 | 0.13 | 0.04 | 0.22 | 0.04 | 0.24 | 0.11 |
| 10 | 8129441 | rs7090113 | 0.089 | 0.11 | -0.408 | -0.531 | 0.012 | 0.006 | 0.006 | 0 | 0.09 | 0.06 | 0.03 | 0.06 | 0.02 | 0.09 |
| 10 | 8131840 | rs2664909 | 1.117 | -1.082 | -0.191 | -1.332 | -0.003 | 0.005 | 0.005 | 0.205 | 0.42 | 0.33 | 0.49 | 0.77 | 0.51 | 0.64 |
| 10 | 8132981 | rs2798842 | 0.989 | -1.162 | 0.101 | -1.303 | -0.005 | 0.022 | 0.022 | 0.449 | 0.43 | 0.34 | 0.49 | 0.77 | 0.52 | 0.64 |
| 10 | 8137383 | rs12776126 | -0.168 | -1.344 | 0.213 | -1.305 | -0.025 | 0.024 | 0.024 | 0.053 | 0.1 | 0.3 | 0.14 | 0.09 | 0.06 | 0.03 |
| 10 | 8138275 | rs1244159 | -0.266 | -1.516 | -0.261 | -0.864 | 0.006 | -0.007 | -0.007 | 0 | 0.85 | 0.89 | 0.88 | 0.75 | 0.87 | 0.95 |
| 10 | 8138906 | rs11255520 | 1.6 | -0.626 | -0.751 | -1.112 | -0.026 | -0.098 | -0.098 | 0 | 0.3 | 0.06 | 0.31 | 0.55 | 0.31 | 0.58 |
| 10 | 8141258 | rs11255522 | 0.758 | -0.251 | -0.667 | -1.098 | -0.051 | -0.016 | -0.016 | 0 | 0.11 | 0.03 | 0.23 | 0.08 | 0.21 | 0.08 |
| 10 | 8142012 | rs1514230 | 0.675 | -0.758 | -0.132 | -1.224 | 0.01 | 0.026 | 0.026 | 0.328 | 0.51 | 0.42 | 0.58 | 0.63 | 0.55 | 0.79 |
| 10 | 8142054 | rs1514231 | 0.439 | -0.525 | -0.41 | -1.254 | -0.03 | 0.025 | 0.025 | 0 | 0.1 | 0.08 | 0.23 | 0.03 | 0.23 | 0.14 |
| 10 | 8142617 | rs2671306 | -0.212 | -1.521 | -0.138 | -1.2 | -0.006 | -0.001 | -0.001 | 0 | 0.78 | 0.76 | 0.88 | 0.73 | 0.88 | 0.89 |
| 10 | 8142707 | rs1514232 | -0.419 | -0.981 | -0.359 | -1.059 | 0.006 | -0.02 | -0.02 | 0 | 0.26 | 0.3 | 0.3 | 0.1 | 0.34 | 0.1 |
| 10 | 8146142 | rs2798828 | 0.237 | -0.799 | -0.644 | -0.703 | -0.002 | -0.007 | -0.007 | 3.222 | 0.48 | 0.45 | 0.67 | 0.65 | 0.54 | 0.71 |
| 10 | 8153822 | rs1149908 | -0.522 | -1.548 | -0.977 | -0.163 | 0.068 | 0.024 | 0.024 | 0 | 0.37 | 0.72 | 0.35 | 0.27 | 0.15 | 0.4 |
| 10 | 8168485 | rs1704279 | 0.388 | -0.186 | -1.047 | -0.167 | 0.009 | 0.04 | 0.04 | 0.256 | 0.55 | 0.33 | 0.45 | 0.69 | 0.56 | 0.5 |
| 10 | 8171357 | rs4749641 | 0.863 | -0.444 | -1.363 | -0.388 | -0.012 | 0.078 | 0.078 | 0.474 | 0.13 | 0 | 0.1 | 0.01 | 0.1 | 0.1 |
| 10 | 8172575 | rs1618853 | 0.667 | -0.545 | -1.032 | -0.59 | 0.002 | 0.003 | 0.003 | 0 | 0.65 | 0.34 | 0.5 | 0.69 | 0.65 | 0.54 |
| 10 | 8177255 | rs17143250 | 0.182 | -0.548 | -0.192 | -0.46 | -0.005 | -0.017 | -0.017 | 0 | 0.11 | 0.16 | 0.29 | 0.09 | 0.09 | 0.13 |
| 10 | 8179864 | rs1243977 | -0.353 | -0.597 | 0.61 | -0.566 | 0.021 | 0.17 | 0.17 | 0.336 | 0.9 | 0.67 | 0.93 | 0.78 | 0.91 | 0.79 |
| 10 | 8184896 | rs490861 | 1.312 | -0.724 | 0.499 | -0.699 | 0.013 | 0.046 | 0.046 | 0.111 | 0.57 | 0.27 | 0.44 | 0.68 | 0.57 | 0.53 |
| 10 | 8186093 | rs2798832 | 0.031 | -0.707 | 0.762 | -0.697 | -0.001 | -0.024 | -0.024 | 0 | 0.77 | 0.7 | 0.56 | 0.89 | 0.76 | 0.76 |
| 10 | 8186946 | rs10905290 | 0.938 | -1.129 | -0.145 | -0.03 | 0 | -0.073 | -0.073 | 0 | 0.11 | 0.01 | 0.09 | 0.38 | 0.12 | 0.08 |
| 10 | 8187989 | rs2892530 | -0.855 | -0.976 | 0.004 | -0.19 | -0.003 | -0.039 | -0.039 | 0 | 0.4 | 0.48 | 0.31 | 0.16 | 0.42 | 0.33 |
| 10 | 8188595 | rs11255537 | 1.496 | -0.484 | -1.149 | -0.762 | -0.106 | -0.088 | -0.088 | 0 | 0.23 | 0.02 | 0.25 | 0.25 | 0.25 | 0.2 |
| 10 | 8192149 | rs541045 | 0.134 | -0.464 | -0.614 | -0.99 | 0 | -0.013 | -0.013 | 0 | 0.56 | 0.52 | 0.6 | 0.71 | 0.51 | 0.55 |
| 10 | 8194751 | rs10905291 | 0.943 | -0.583 | -0.161 | -0.87 | 0.003 | 0.178 | 0.178 | 0 | 0.09 | 0.29 | 0.07 | 0.25 | 0.07 | 0.18 |
| 10 | 8200390 | rs2664904 | -0.346 | -0.526 | -0.583 | -1.288 | 0.001 | -0.001 | -0.001 | 0 | 0.54 | 0.39 | 0.51 | 0.52 | 0.51 | 0.6 |
| 10 | 8200992 | rs526935 | -0.854 | -0.555 | -0.442 | -1.306 | -0.009 | 0.014 | 0.014 | 0 | 0.65 | 0.58 | 0.62 | 0.52 | 0.69 | 0.63 |
| 10 | 8203082 | rs11255542 | 1.961 | -0.34 | -0.711 | -1.426 | -0.079 | -0.083 | -0.083 | 0 | 0.19 | 0.02 | 0.21 | 0.21 | 0.18 | 0.17 |
| 10 | 8203475 | rs1999975 | -0.23 | -0.368 | -0.677 | -1.441 | 0.01 | -0.004 | -0.004 | 0 | 0.52 | 0.27 | 0.43 | 0.47 | 0.5 | 0.54 |
| 10 | 8206553 | rs479510 | -0.965 | -0.362 | -0.519 | -1.409 | 0.003 | -0.005 | -0.005 | 0 | 0.66 | 0.68 | 0.63 | 0.52 | 0.69 | 0.64 |
| 10 | 8209098 | rs10905304 | 1.919 | -0.06 | -0.624 | -1.354 | -0.068 | -0.085 | -0.085 | 0 | 0.19 | 0.02 | 0.2 | 0.2 | 0.17 | 0.17 |
| 10 | 8211676 | rs4749669 | 0.144 | -0.127 | -0.583 | -1.447 | -0.019 | -0.023 | -0.023 | 0.093 | 0.4 | 0.26 | 0.33 | 0.45 | 0.42 | 0.39 |
| 10 | 8213413 | rs548966 | -0.857 | -0.146 | -0.526 | -1.32 | -0.02 | 0.025 | 0.025 | 0 | 0.65 | 0.51 | 0.61 | 0.53 | 0.69 | 0.62 |
| 10 | 8216423 | rs17449616 | 1.354 | -0.191 | -0.578 | -1.484 | -0.003 | 0.001 | 0.001 | 0 | 0.13 | 0.12 | 0.17 | 0.06 | 0.19 | 0.1 |
| 10 | 8218069 | rs565371 | -0.595 | -0.492 | -0.846 | -1.848 | -0.001 | 0.036 | 0.036 | 0.154 | 0.47 | 0.38 | 0.4 | 0.07 | 0.51 | 0.33 |
| 10 | 8220230 | rs17449748 | 1.759 | -0.387 | -0.857 | -1.651 | -0.039 | 0.063 | 0.063 | 0 | 0.11 | 0.03 | 0.06 | 0 | 0.14 | 0.04 |
| 10 | 8223080 | rs1509968 | 2.57 | -0.418 | -0.876 | -1.655 | -0.009 | 0.066 | 0.066 | 0.052 | 0.12 | 0 | 0.1 | 0.01 | 0.09 | 0.09 |
| 10 | 8223674 | rs11814009 | 0.33 | -0.287 | -0.611 | -1.621 | -0.015 | 0.036 | 0.036 | 0 | 0.2 | 0.13 | 0.11 | 0.01 | 0.23 | 0.08 |
| 10 | 8224786 | rs10905311 | -0.049 | -0.441 | -0.494 | -1.396 | -0.022 | 0.047 | 0.047 | 0 | 0.19 | 0.35 | 0.11 | 0.27 | 0.17 | 0.26 |
| 10 | 8224850 | rs521331 | -1.374 | -0.366 | -0.554 | -1.313 | 0.002 | -0.022 | -0.022 | 0 | 0.74 | 0.8 | 0.67 | 0.53 | 0.76 | 0.66 |
| 10 | 8225401 | rs7078148 | 0.495 | 0.093 | -0.58 | -1.198 | -0.001 | -0.019 | -0.019 | 0.093 | 0.1 | 0.13 | 0.05 | 0 | 0.09 | 0.04 |
| 10 | 8229430 | rs2996520 | 0.049 | 0.161 | -0.526 | -1.006 | -0.002 | 0.017 | 0.017 | 0 | 0.24 | 0.27 | 0.15 | 0.31 | 0.25 | 0.22 |
| 10 | 8231126 | rs11255559 | 1.498 | 0.129 | -0.6 | -1.905 | 0 | -0.022 | -0.022 | 0 | 0.14 | 0.12 | 0.11 | 0.01 | 0.09 | 0.18 |
| 10 | 8231720 | rs1416560 | 0.518 | 0.203 | -0.706 | -1.924 | 0.01 | 0.061 | 0.061 | 0 | 0.39 | 0.2 | 0.32 | 0.19 | 0.3 | 0.34 |
| 10 | 8233615 | rs1779643 | 0.097 | 0.832 | -0.408 | -1.801 | -0.001 | -0.021 | -0.021 | 1.031 | 0.6 | 0.38 | 0.63 | 0.89 | 0.54 | 0.6 |
| 10 | 8235799 | rs532174 | 1.102 | 0.835 | -0.288 | -1.728 | 0.016 | -0.033 | -0.033 | 1.106 | 0.12 | 0.26 | 0.18 | 0.08 | 0.19 | 0.07 |
| 10 | 8237770 | rs2489253 | 0.287 | 0.802 | -0.61 | -1.858 | 0.032 | 0.051 | 0.051 | 0 | 0.49 | 0.3 | 0.45 | 0.71 | 0.41 | 0.42 |
| 10 | 8245150 | rs568057 | -0.753 | 0.964 | -0.329 | -2.011 | -0.001 | -0.037 | -0.037 | 0.029 | 0.74 | 0.74 | 0.81 | 0.97 | 0.74 | 0.7 |
| 10 | 8246475 | rs1629795 | -0.307 | 1.027 | -0.189 | -1.58 | 0.009 | -0.029 | -0.029 | 0 | 0.32 | 0.55 | 0.48 | 0.82 | 0.41 | 0.33 |
| 10 | 8247228 | rs1779640 | 0.349 | 0.728 | -0.14 | -1.238 | 0.005 | 0.023 | 0.023 | 0 | 0.21 | 0.29 | 0.29 | 0.74 | 0.22 | 0.26 |

| | | | | | | | | | | | | | | | | |
|---|---|---|---|---|---|---|---|---|---|---|---|---|---|---|---|---|
| 10 | 8251061 | rs11255579 | -0.1 | 0.555 | -0.403 | -1.682 | -0.002 | -0.076 | -0.076 | 0 | 0.44 | 0.51 | 0.35 | 0.04 | 0.41 | 0.45 |
| 10 | 8251307 | rs10905315 | 0.317 | 0.555 | -0.431 | -1.79 | -0.001 | -0.062 | -0.062 | 0.478 | 0.36 | 0.35 | 0.28 | 0.03 | 0.33 | 0.4 |
| 10 | 8255232 | rs1538512 | 0.853 | 0.607 | -0.48 | -1.789 | -0.003 | -0.076 | -0.076 | 0 | 0.17 | 0.31 | 0.18 | 0.02 | 0.16 | 0.16 |
| 10 | 8258675 | rs1416566 | -0.529 | 0.657 | -0.614 | -1.634 | -0.01 | -0.028 | -0.028 | 0 | 0.42 | 0.49 | 0.48 | 0.78 | 0.34 | 0.5 |
| 10 | 8263831 | rs10905324 | -0.007 | 0.869 | -1.12 | -1.364 | -0.008 | 0.01 | 0.01 | 0 | 0.2 | 0.27 | 0.37 | 0.73 | 0.17 | 0.26 |
| 10 | 8266151 | rs533919 | -0.44 | 1.968 | -1.911 | -1.474 | -0.083 | 0.011 | 0.011 | 0 | 0.8 | 0.92 | 0.76 | 0.89 | 0.62 | 0.85 |
| 10 | 8266475 | rs1243933 | 1.246 | 1.224 | -1.564 | -1.21 | -0.041 | 0.07 | 0.07 | 0.87 | 0.13 | 0 | 0.07 | 0.01 | 0.12 | 0.13 |
| 10 | 8270890 | rs494602 | -0.458 | 1.739 | -1.512 | -0.864 | -0.059 | -0.029 | -0.029 | 0.087 | 0.79 | 0.92 | 0.75 | 0.78 | 0.66 | 0.77 |
| 10 | 8273663 | rs4747760 | -0.48 | 0.876 | -1.252 | -0.797 | 0.008 | -0.01 | -0.01 | 0 | 0.18 | 0.15 | 0.34 | 0.74 | 0.14 | 0.26 |
| 10 | 8279913 | rs1936490 | 0.416 | 0.844 | -1.096 | -1.382 | 0.003 | -0.045 | -0.045 | 0 | 0.17 | 0.21 | 0.1 | 0.01 | 0.17 | 0.21 |
| 10 | 8280193 | rs1796858 | 0.789 | 0.83 | -1.155 | -1.388 | -0.053 | 0.083 | 0.083 | 0 | 0.15 | 0.01 | 0.09 | 0.01 | 0.16 | 0.21 |
| 10 | 8286284 | rs1779648 | -1.288 | 0.712 | -1.057 | -1.586 | 0 | -0.004 | -0.004 | 0 | 0.41 | 0.45 | 0.46 | 0.74 | 0.33 | 0.52 |
| 10 | 8286993 | rs11255598 | -0.916 | 0.688 | -1.01 | -1.494 | 0.003 | 0.001 | 0.001 | 1.348 | 0.24 | 0.22 | 0.37 | 0.73 | 0.19 | 0.31 |
| 10 | 8295136 | rs6602291 | -0.47 | 0.484 | -1.246 | -1.379 | 0.007 | -0.037 | -0.037 | 0 | 0.14 | 0.07 | 0.31 | 0.7 | 0.11 | 0.19 |
| 10 | 8300790 | rs7092384 | -1.008 | 0.341 | -1.035 | -0.738 | -0.006 | 0.003 | 0.003 | 0.763 | 0.23 | 0.35 | 0.14 | 0.22 | 0.16 | 0.21 |
| 10 | 8302481 | rs2298253 | -0.983 | 0.267 | -1.161 | -0.535 | 0.013 | 0.002 | 0.002 | 0 | 0.29 | 0.24 | 0.44 | 0.66 | 0.22 | 0.33 |
| 10 | 8303127 | rs7916633 | -0.614 | 0.244 | -1.156 | -0.485 | 0 | 0.003 | 0.003 | 0 | 0.15 | 0.18 | 0.06 | 0.3 | 0.09 | 0.12 |
| 10 | 8304149 | rs10905328 | -1.415 | 0.242 | -1.113 | -0.546 | 0.002 | -0.004 | -0.004 | 0 | 0.4 | 0.42 | 0.55 | 0.81 | 0.3 | 0.54 |
| 10 | 8307583 | rs11255607 | -0.662 | 0.295 | -1.07 | -0.463 | -0.001 | 0.004 | 0.004 | 0 | 0.15 | 0.19 | 0.07 | 0.3 | 0.09 | 0.12 |
| 10 | 8312604 | rs17143382 | 1.211 | 0.333 | -1.08 | -0.49 | 0.002 | -0.014 | -0.014 | 0 | 0.09 | 0.15 | 0.08 | 0.04 | 0.06 | 0.12 |
| 10 | 8314844 | rs12783630 | 1.18 | 0.333 | -1.08 | -0.49 | 0.001 | -0.015 | -0.015 | 0.294 | 0.09 | 0.15 | 0.08 | 0.04 | 0.06 | 0.12 |
| 10 | 8316529 | rs10905335 | 0.086 | 0.4 | -1.426 | -0.957 | -0.011 | -0.001 | -0.001 | 0 | 0.38 | 0.26 | 0.23 | 0.39 | 0.38 | 0.27 |
| 10 | 8318449 | rs10795607 | -1.546 | 0.644 | -1.206 | -1.198 | 0 | -0.113 | -0.113 | 0 | 0.66 | 0.6 | 0.72 | 0.97 | 0.61 | 0.74 |
| 10 | 8318517 | rs12780019 | 2.119 | 0.675 | -1.137 | -1.01 | 0.009 | -0.004 | -0.004 | 0 | 0.06 | 0.1 | 0.06 | 0.04 | 0.06 | 0.11 |
| 10 | 8318583 | rs17466327 | 1.113 | 0.66 | -1.208 | -1.013 | -0.022 | 0.035 | 0.035 | 0 | 0.25 | 0.12 | 0.17 | 0.16 | 0.3 | 0.18 |
| 10 | 8320405 | rs10795608 | -0.619 | 0.552 | -1.108 | -0.972 | -0.001 | -0.054 | -0.054 | 0.993 | 0.48 | 0.52 | 0.63 | 0.85 | 0.48 | 0.48 |
| 10 | 8323017 | rs11599385 | 0.454 | 0.056 | -1.184 | -0.62 | 0 | -0.005 | -0.005 | 0 | 0.2 | 0.2 | 0.12 | 0.04 | 0.16 | 0.26 |
| 10 | 8324047 | rs10737028 | -0.118 | -0.05 | -1.211 | -0.773 | 0.023 | 0.093 | 0.093 | 0 | 0.5 | 0.37 | 0.6 | 0.75 | 0.55 | 0.41 |
| 10 | 8324268 | rs6602294 | -1.122 | -0.137 | -1.188 | -0.763 | -0.022 | -0.059 | -0.059 | 0 | 0.7 | 0.58 | 0.72 | 0.79 | 0.72 | 0.67 |
| 10 | 8324716 | rs6602296 | -0.784 | -0.084 | -1.298 | -0.87 | -0.014 | -0.057 | -0.057 | 0.003 | 0.65 | 0.58 | 0.7 | 0.79 | 0.66 | 0.67 |
| 10 | 8325106 | rs11255616 | 0.512 | -0.061 | -1.328 | -0.686 | 0 | -0.004 | -0.004 | 0 | 0.12 | 0.1 | 0.05 | 0.11 | 0.08 | 0.08 |
| 10 | 8326554 | rs17466705 | 0.956 | -0.058 | -1.34 | -0.693 | -0.042 | 0.071 | 0.071 | 0.731 | 0.35 | 0.21 | 0.24 | 0.17 | 0.45 | 0.23 |
| 10 | 8331078 | rs4749728 | 1.442 | 0.025 | -1.289 | -0.717 | -0.025 | 0.039 | 0.039 | 0.178 | 0.31 | 0.21 | 0.22 | 0.17 | 0.39 | 0.22 |
| 10 | 8332014 | rs10752132 | 0.128 | -0.123 | -1.304 | -0.726 | 0.007 | 0.001 | 0.001 | 0 | 0.16 | 0.09 | 0.05 | 0.13 | 0.1 | 0.11 |
| 10 | 8334463 | rs10159927 | -1.221 | 0.263 | -1.061 | -0.622 | -0.015 | -0.09 | -0.09 | 1.315 | 0.72 | 0.58 | 0.73 | 0.9 | 0.72 | 0.71 |
| 10 | 8336932 | rs11255620 | -0.56 | 0.303 | -1.046 | -0.617 | -0.007 | -0.065 | -0.065 | 0.049 | 0.61 | 0.58 | 0.69 | 0.9 | 0.62 | 0.57 |
| 10 | 8337245 | rs1892063 | -0.199 | 0.341 | -0.983 | -0.536 | 0.005 | 0.027 | 0.027 | 0 | 0.53 | 0.42 | 0.63 | 0.87 | 0.56 | 0.44 |
| 10 | 8349075 | rs11595705 | 1.395 | 0.271 | -0.892 | -0.506 | 0.005 | 0.017 | 0.017 | 0 | 0.12 | 0.07 | 0.05 | 0 | 0.1 | 0.14 |
| 10 | 8350343 | rs11255628 | 0.028 | 0.24 | -0.991 | -0.45 | 0.016 | 0.047 | 0.047 | 0.451 | 0.48 | 0.33 | 0.3 | 0.32 | 0.52 | 0.27 |
| 10 | 8353101 | rs1999874 | -0.475 | -0.164 | -0.796 | -0.141 | -0.009 | -0.034 | -0.034 | 0 | 0.63 | 0.6 | 0.69 | 0.8 | 0.64 | 0.62 |
| 10 | 8354144 | rs10795618 | 0.116 | 0.109 | -0.909 | -0.533 | -0.001 | -0.001 | -0.001 | 0 | 0.12 | 0.12 | 0.05 | 0.13 | 0.08 | 0.08 |
| 10 | 8357299 | rs7896001 | -0.754 | 0.15 | -0.883 | -0.509 | -0.015 | 0.001 | 0.001 | 1.546 | 0.66 | 0.61 | 0.41 | 0.37 | 0.68 | 0.42 |
| 10 | 8362697 | rs7076802 | -0.655 | 0.68 | -1.094 | 0.07 | 0.019 | 0.031 | 0.031 | 0.175 | 0.49 | 0.38 | 0.33 | 0.37 | 0.53 | 0.3 |
| 10 | 8363061 | rs12777233 | 0.057 | 1.065 | -1.474 | -0.168 | -0.033 | 0.043 | 0.043 | 1.718 | 0.37 | 0.23 | 0.29 | 0.24 | 0.45 | 0.23 |
| 10 | 8367870 | rs1538246 | -0.879 | 0.937 | -1.291 | 0.72 | -0.004 | -0.025 | -0.025 | 0 | 0.18 | 0.14 | 0.2 | 0.6 | 0.19 | 0.14 |
| 10 | 8370629 | rs11255635 | 0.154 | 0.684 | -1.731 | -0.348 | 0.001 | 0.124 | 0.124 | 2.894 | 0.35 | 0.16 | 0.17 | 0.12 | 0.37 | 0.31 |
| 10 | 8378782 | rs10508350 | -0.063 | 0.453 | -1.132 | -0.565 | 0.015 | 0.061 | 0.061 | 5.743 | 0.46 | 0.37 | 0.22 | 0.14 | 0.41 | 0.35 |
| 10 | 8382415 | rs17136597 | 0.005 | 0.8 | -1.04 | -0.392 | -0.004 | -0.005 | -0.005 | 1.398 | 0.18 | 0.09 | 0.19 | 0.28 | 0.21 | 0.37 |
| 10 | 8383911 | rs2993391 | 0.368 | 0.58 | -0.517 | -0.055 | -0.01 | 0.025 | 0.025 | 0 | 0.92 | 0.89 | 0.75 | 0.85 | 0.97 | 0.94 |
| 10 | 8387006 | rs12771263 | 0.498 | 0.998 | -0.419 | -0.47 | 0.022 | 0.002 | 0.002 | 1.184 | 0.68 | 0.46 | 0.67 | 0.34 | 0.65 | 0.69 |
| 10 | 8388176 | rs7091759 | -0.298 | 0.66 | -0.701 | -1.542 | 0.003 | -0.017 | -0.017 | 0 | 0.32 | 0.29 | 0.21 | 0.01 | 0.32 | 0.28 |
| 10 | 8388512 | rs11814860 | 0.935 | 0.564 | -0.834 | -1.361 | 0.007 | 0.033 | 0.033 | 1.158 | 0.44 | 0.27 | 0.5 | 0.31 | 0.4 | 0.51 |
| 10 | 8390490 | rs4747790 | 0.421 | -0.404 | -0.782 | -1.692 | 0.049 | 0.003 | 0.003 | 0 | 0.56 | 0.24 | 0.67 | 0.45 | 0.68 | 0.67 |
| 10 | 8391429 | rs10508351 | 1.664 | 0.11 | -0.707 | -1.827 | -0.028 | -0.037 | -0.037 | 0.552 | 0.35 | 0.14 | 0.52 | 0.44 | 0.4 | 0.5 |
| 10 | 8394080 | rs10905349 | -1.464 | 0.278 | -0.493 | -1.565 | 0.095 | 0.303 | 0.303 | 0 | 0.88 | 0.64 | 0.8 | 0.48 | 0.84 | 0.87 |
| 10 | 8397438 | rs7074902 | -0.701 | -0.099 | -1.132 | -1.821 | 0.002 | 0.006 | 0.006 | 0.148 | 0.38 | 0.31 | 0.24 | 0.01 | 0.4 | 0.29 |
| 10 | 8398833 | rs2454059 | -1.037 | 0.059 | -0.772 | -1.826 | 0 | 0.001 | 0.001 | 0.112 | 0.46 | 0.48 | 0.42 | 0.52 | 0.51 | 0.39 |
| 10 | 8398914 | rs7081521 | 0.152 | 0.135 | -0.46 | -1.659 | -0.021 | 0.063 | 0.063 | 0 | 0.16 | 0.07 | 0.13 | 0 | 0.23 | 0.12 |
| 10 | 8399759 | rs2935686 | -1.465 | -0.02 | -0.703 | -1.799 | 0.008 | 0.026 | 0.026 | 0 | 0.76 | 0.66 | 0.65 | 0.61 | 0.76 | 0.62 |
| 10 | 8401112 | rs2787150 | 1.272 | -0.144 | -0.645 | -1.497 | -0.005 | -0.004 | -0.004 | 1.073 | 0.31 | 0.28 | 0.41 | 0.58 | 0.35 | 0.32 |
| 10 | 8405857 | rs11595087 | 0.434 | -0.255 | -0.359 | -1.302 | -0.025 | 0.074 | 0.074 | 0 | 0.12 | 0.01 | 0.06 | 0 | 0.14 | 0.1 |
| 10 | 8408629 | rs11255659 | -0.77 | -0.183 | -0.303 | -1.282 | -0.018 | 0.037 | 0.037 | 0 | 0.14 | 0.08 | 0.11 | 0.01 | 0.18 | 0.07 |
| 10 | 8411320 | rs9783189 | -1.742 | -0.214 | -0.169 | -1.266 | -0.007 | -0.06 | -0.06 | 0 | 0.4 | 0.42 | 0.22 | 0.07 | 0.35 | 0.26 |
| 10 | 8415364 | rs12264718 | -1.201 | 0.232 | 0.184 | -1.049 | -0.015 | -0.098 | -0.098 | 0 | 0.3 | 0.42 | 0.14 | 0.05 | 0.23 | 0.23 |
| 10 | 8416256 | rs12265137 | -0.143 | 0.433 | -0.221 | -0.609 | -0.001 | -0.044 | -0.044 | 0 | 0.12 | 0.17 | 0.07 | 0.01 | 0.13 | 0.1 |
| 10 | 8418090 | rs10905352 | -1.005 | 0.559 | 0.242 | -0.574 | -0.003 | -0.135 | -0.135 | 0 | 0.32 | 0.46 | 0.21 | 0.04 | 0.29 | 0.25 |
| 10 | 8418480 | rs11255669 | -0.813 | -0.066 | 0.639 | 0.763 | -0.053 | 0.176 | 0.176 | 0 | 0.15 | 0.4 | 0.3 | 0.38 | 0.1 | 0.18 |
| 10 | 8418546 | rs2787156 | 0.995 | 0.417 | 0.263 | 0.708 | -0.023 | -0.015 | -0.015 | 0.501 | 0.9 | 0.84 | 0.97 | 0.94 | 0.96 | 0.93 |
| 10 | 8420018 | rs12778305 | 0.627 | 0.873 | -0.25 | 0.712 | 0.066 | 0.074 | 0.074 | 0.768 | 0.52 | 0.21 | 0.44 | 0.31 | 0.4 | 0.44 |

| | | | | | | | | | | | | | | | | |
|---|---|---|---|---|---|---|---|---|---|---|---|---|---|---|---|---|
| 10 | 8425531 | rs998139 | -0.609 | 1.827 | -0.072 | 1.371 | -0.013 | -0.02 | -0.02 | 0 | 0.24 | 0.16 | 0.27 | 0.33 | 0.26 | 0.35 |
| 10 | 8434664 | rs2787131 | 1.06 | 1.521 | 0.794 | 1.573 | 0.023 | 0.014 | 0.014 | 0.342 | 0.54 | 0.4 | 0.42 | 0.41 | 0.36 | 0.42 |
| 10 | 8434884 | rs17384246 | -0.331 | 1.767 | 0.751 | 1.537 | -0.013 | -0.008 | -0.008 | 0 | 0.09 | 0.03 | 0.16 | 0.12 | 0.14 | 0.07 |
| 10 | 8436432 | rs17469061 | 0.116 | 2.388 | 0.984 | 1.52 | 0.043 | -0.015 | -0.015 | 0 | 0.11 | 0.13 | 0.13 | 0.05 | 0.25 | 0.16 |
| 10 | 8437546 | rs2787136 | 1.144 | 2.306 | 1.102 | 1.562 | 0 | 0 | 0 | 0 | 0.63 | 0.39 | 0.57 | 0.59 | 0.49 | 0.49 |
| 10 | 8439672 | rs11255680 | 0.132 | 2.166 | 0.235 | 1.405 | -0.048 | -0.071 | -0.071 | 0 | 0.13 | 0.01 | 0.2 | 0.2 | 0.16 | 0.17 |
| 10 | 8439744 | rs1415882 | -0.427 | 1.901 | 0.411 | 1.302 | -0.02 | -0.032 | -0.032 | 0.443 | 0.23 | 0.11 | 0.24 | 0.29 | 0.26 | 0.4 |
| 10 | 8442757 | rs2787138 | -1.152 | 1.834 | 0.378 | 0.843 | 0.032 | -0.008 | -0.008 | 1.828 | 0.25 | 0.28 | 0.22 | 0.2 | 0.35 | 0.27 |
| 10 | 8443306 | rs10795621 | 0.963 | 0.93 | 0.668 | 0.952 | 0.103 | 0.085 | 0.085 | 0.418 | 0.91 | 0.47 | 0.79 | 0.89 | 0.91 | 0.95 |
| 10 | 8444689 | rs1538245 | -0.945 | 0.49 | -0.535 | 1.056 | 0.008 | -0.02 | -0.02 | 0 | 0.18 | 0.09 | 0.17 | 0.26 | 0.12 | 0.21 |
| 10 | 8444810 | rs2787141 | 1.267 | 0.561 | -0.111 | 0.641 | -0.001 | -0.057 | -0.057 | 0 | 0.9 | 0.25 | 0.9 | 1 | 0.93 | 1 |
| 10 | 8445655 | rs12778770 | -0.636 | -0.21 | -0.766 | 0.79 | 0.02 | -0.005 | -0.005 | 0 | 0.12 | 0.06 | 0.04 | 0.11 | 0.07 | 0.1 |
| 10 | 8447916 | rs2152963 | 0.003 | 0.058 | -1.868 | 0.554 | -0.036 | -0.019 | -0.019 | 0 | 0.49 | 0.13 | 0.47 | 0.56 | 0.49 | 0.39 |
| 10 | 8449327 | rs10905357 | 0.257 | 0.013 | -1.507 | 0.27 | -0.008 | 0.004 | 0.004 | 1.904 | 0.23 | 0.18 | 0.22 | 0.22 | 0.3 | 0.21 |
| 10 | 8453142 | rs17143607 | 0.675 | -0.284 | -1.023 | 0.275 | 0.03 | -0.023 | -0.023 | 0 | 0.42 | 0.25 | 0.4 | 0.45 | 0.32 | 0.3 |
| 10 | 8455084 | rs17385118 | 0.173 | -0.168 | -1.102 | 0.166 | 0.001 | -0.017 | -0.017 | 0.48 | 0.08 | 0.06 | 0.07 | 0.16 | 0.08 | 0.16 |
| 10 | 8455919 | rs2454057 | 0.247 | -0.423 | -0.977 | 0.04 | -0.063 | 0.049 | 0.049 | 3.293 | 0.34 | 0.18 | 0.25 | 0.24 | 0.5 | 0.42 |
| 10 | 8456007 | rs11255686 | -0.132 | -0.575 | -0.109 | 0 | 0.001 | -0.008 | -0.008 | 0 | 0.62 | 0.6 | 0.71 | 0.76 | 0.45 | 0.53 |
| 10 | 8458317 | rs1361153 | 0.799 | -0.515 | -0.365 | 0.086 | 0.005 | -0.012 | -0.012 | 0 | 0.07 | 0.06 | 0.04 | 0.15 | 0.05 | 0.15 |
| 10 | 8462367 | rs7096950 | 0.879 | -0.732 | 0.023 | -0.096 | 0.039 | -0.023 | -0.023 | 0 | 0.4 | 0.28 | 0.4 | 0.46 | 0.28 | 0.28 |
| 10 | 8463078 | rs2787162 | 0.423 | -0.587 | 0.004 | -0.27 | -0.049 | 0.048 | 0.048 | 0.23 | 0.32 | 0.2 | 0.26 | 0.21 | 0.51 | 0.39 |
| 10 | 8463376 | rs2646422 | 0.426 | -0.595 | 0.021 | -0.274 | -0.048 | 0.046 | 0.046 | 0.213 | 0.32 | 0.2 | 0.26 | 0.21 | 0.51 | 0.39 |
| 10 | 8464847 | rs2454062 | -0.057 | -0.618 | 0 | -0.252 | -0.023 | 0.039 | 0.039 | 0 | 0.39 | 0.23 | 0.29 | 0.36 | 0.56 | 0.54 |
| 10 | 8465926 | rs2646419 | -1.747 | -1.368 | -0.335 | -0.634 | 0.004 | -0.038 | -0.038 | 0 | 0.74 | 0.81 | 0.66 | 0.55 | 0.78 | 0.75 |
| 10 | 8470387 | rs2646425 | -0.006 | -0.811 | -0.438 | -0.597 | 0.023 | 0.067 | 0.067 | 0.163 | 0.47 | 0.25 | 0.24 | 0.38 | 0.44 | 0.39 |
| 10 | 8479868 | rs7090925 | 1.001 | -0.324 | -0.268 | -0.241 | -0.013 | 0.009 | 0.009 | 0.355 | 0.23 | 0.14 | 0.14 | 0.22 | 0.32 | 0.19 |
| 10 | 8480488 | rs7921806 | 0.003 | -0.293 | -0.31 | -0.194 | -0.005 | -0.021 | -0.021 | 0 | 0.17 | 0.11 | 0.27 | 0.24 | 0.16 | 0.18 |
| 10 | 8483560 | rs17143655 | 0.003 | -0.185 | -0.443 | -0.133 | -0.013 | -0.043 | -0.043 | 0 | 0.16 | 0.06 | 0.26 | 0.24 | 0.14 | 0.21 |
| 10 | 8484113 | rs7096374 | 1.043 | -0.088 | -0.364 | -0.188 | -0.025 | 0.009 | 0.009 | 0.651 | 0.22 | 0.12 | 0.14 | 0.22 | 0.32 | 0.19 |
| 10 | 8489874 | rs7079593 | -0.931 | -0.601 | 0.102 | 0.016 | -0.002 | -0.002 | -0.002 | 0 | 0.27 | 0.26 | 0.34 | 0.34 | 0.28 | 0.3 |
| 10 | 8492791 | rs4747793 | -0.786 | -1.884 | 0.277 | 0 | -0.048 | 0.007 | 0.007 | 0 | 0.31 | 0.53 | 0.3 | 0.33 | 0.18 | 0.17 |
| 10 | 8496389 | rs11255703 | -1.028 | -1.911 | 0.274 | -0.474 | -0.009 | -0.004 | -0.004 | 0.254 | 0.23 | 0.27 | 0.26 | 0.2 | 0.14 | 0.35 |
| 10 | 8498892 | rs7079888 | 0.817 | -1.953 | 0.753 | -0.519 | -0.036 | -0.035 | -0.035 | 0 | 0.64 | 0.53 | 0.68 | 0.73 | 0.77 | 0.57 |
| 10 | 8500122 | rs4237397 | -1.072 | -1.957 | 0.686 | -0.521 | 0.036 | 0.034 | 0.034 | 0 | 0.34 | 0.25 | 0.31 | 0.27 | 0.22 | 0.43 |
| 10 | 8501583 | rs4619046 | -0.965 | -1.495 | 0.818 | -0.321 | -0.002 | -0.006 | -0.006 | 0 | 0.08 | 0.11 | 0.05 | 0.06 | 0.05 | 0.1 |
| 10 | 8507223 | rs7095711 | 1.766 | -1.42 | 1.126 | -0.317 | 0.041 | -0.03 | -0.03 | 0 | 0.88 | 0.67 | 0.91 | 0.94 | 0.88 | 0.83 |
| 10 | 8508246 | rs6602311 | 1.613 | -1.711 | 1.53 | -0.373 | -0.012 | -0.008 | -0.008 | 0 | 0.74 | 0.39 | 0.7 | 0.74 | 0.82 | 0.58 |
| 10 | 8508704 | rs17143770 | -1.528 | -1.875 | 1.546 | -0.341 | -0.035 | 0.002 | 0.002 | 0 | 0.22 | 0.37 | 0.26 | 0.26 | 0.13 | 0.36 |
| 10 | 8509339 | rs7072544 | -0.277 | -1.734 | 1.367 | -0.261 | -0.004 | 0 | 0 | 0.807 | 0.14 | 0.13 | 0.2 | 0.2 | 0.06 | 0.26 |
| 10 | 8511462 | rs11255711 | -0.676 | -1.905 | 0.592 | 0.005 | 0.009 | -0.015 | -0.015 | 0 | 0.3 | 0.24 | 0.33 | 0.36 | 0.21 | 0.23 |
| 10 | 8511627 | rs11255712 | -1.523 | -1.771 | 0.489 | -0.583 | 0 | 0 | 0 | 0 | 0.64 | 0.64 | 0.61 | 0.64 | 0.39 | 0.66 |
| 10 | 8512382 | rs10795631 | 0.472 | -1.568 | 0.202 | -0.439 | 0.006 | -0.001 | -0.001 | 2.218 | 0.58 | 0.32 | 0.45 | 0.37 | 0.68 | 0.43 |
| 10 | 8515348 | rs12785018 | -0.52 | -1.247 | -0.051 | -0.269 | 0.01 | -0.001 | -0.001 | 0 | 0.41 | 0.31 | 0.55 | 0.62 | 0.34 | 0.53 |
| 10 | 8517660 | rs7085991 | -0.175 | -1.293 | 0.037 | -0.18 | -0.002 | 0.001 | 0.001 | 0 | 0.31 | 0.29 | 0.5 | 0.62 | 0.25 | 0.45 |
| 10 | 8518880 | rs7090140 | 0.537 | -1.642 | -0.132 | -0.347 | 0.042 | 0.013 | 0.013 | 0 | 0.44 | 0.26 | 0.4 | 0.36 | 0.67 | 0.42 |
| 10 | 8519559 | rs4540753 | -0.225 | -2.088 | 0.313 | -0.468 | -0.041 | 0.017 | 0.017 | 0 | 0.35 | 0.47 | 0.51 | 0.61 | 0.23 | 0.45 |
| 10 | 8523708 | rs7904468 | 1.138 | -1.583 | -0.008 | -0.522 | -0.043 | -0.029 | -0.029 | 1.333 | 0.34 | 0.18 | 0.36 | 0.36 | 0.57 | 0.38 |
| 10 | 8523830 | rs17478850 | 1.423 | -0.42 | -0.316 | -0.261 | -0.045 | -0.039 | -0.039 | 0 | 0.07 | 0.01 | 0.13 | 0.25 | 0.14 | 0.16 |
| 10 | 8526255 | rs12266741 | 1.407 | -1.153 | -0.055 | -0.294 | -0.018 | 0.011 | 0.011 | 0 | 0.1 | 0.07 | 0.07 | 0 | 0.14 | 0.09 |
| 10 | 8529115 | rs4749765 | -0.557 | -1.546 | 0.189 | -0.143 | 0 | -0.001 | -0.001 | 0 | 0.43 | 0.38 | 0.57 | 0.6 | 0.39 | 0.53 |
| 10 | 8530042 | rs12569700 | 0.038 | -1.282 | 0.299 | 0.003 | -0.004 | 0.011 | 0.011 | 0 | 0.3 | 0.27 | 0.46 | 0.53 | 0.24 | 0.4 |
| 10 | 8531336 | rs11255716 | 0.183 | -1.282 | 0.281 | 0.003 | -0.011 | 0.029 | 0.029 | 0 | 0.3 | 0.32 | 0.46 | 0.53 | 0.24 | 0.4 |
| 10 | 8532635 | rs10795645 | 0.188 | -0.562 | 0.61 | 0.324 | -0.034 | 0.006 | 0.006 | 0 | 0.37 | 0.26 | 0.38 | 0.34 | 0.56 | 0.38 |
| 10 | 8533237 | rs10508354 | -0.115 | -0.647 | 0.674 | 0.476 | -0.002 | -0.01 | -0.01 | 0 | 0.58 | 0.54 | 0.56 | 0.68 | 0.41 | 0.57 |
| 10 | 8534603 | rs4747795 | 0.11 | -0.63 | 0.711 | 0.497 | 0.004 | 0.003 | 0.003 | 0 | 0.56 | 0.38 | 0.53 | 0.59 | 0.41 | 0.55 |
| 10 | 8534953 | rs7074625 | -0.028 | -0.578 | 0.601 | 0.576 | -0.003 | -0.01 | -0.01 | 0 | 0.57 | 0.53 | 0.55 | 0.67 | 0.41 | 0.57 |
| 10 | 8538492 | rs10752138 | -0.57 | -0.467 | -0.187 | 0.557 | -0.053 | 0.052 | 0.052 | 0 | 0.64 | 0.8 | 0.6 | 0.8 | 0.46 | 0.64 |
| 10 | 8545524 | rs10795650 | -0.2 | -0.625 | -0.45 | 0.478 | -0.002 | -0.002 | -0.002 | 0 | 0.57 | 0.47 | 0.55 | 0.59 | 0.42 | 0.54 |
| 10 | 8546906 | rs2388747 | -0.264 | -1.142 | -0.93 | 0.476 | -0.011 | -0.002 | -0.002 | 1.514 | 0.55 | 0.51 | 0.77 | 0.66 | 0.73 | 0.8 |
| 10 | 8547259 | rs2388746 | -0.763 | -1.186 | -1.496 | 0.126 | 0.053 | 0.078 | 0.078 | 0.007 | 0.33 | 0.18 | 0.28 | 0.2 | 0.23 | 0.22 |
| 10 | 8547572 | rs6602315 | -0.629 | -0.398 | -1.404 | 0.007 | -0.012 | 0.062 | 0.062 | 0 | 0.85 | 0.96 | 0.93 | 1 | 0.86 | 0.91 |
| 10 | 8548066 | rs4749774 | 0.014 | -0.525 | -1.595 | 0.277 | -0.023 | 0.028 | 0.028 | 0 | 0.8 | 0.95 | 0.89 | 0.87 | 0.81 | 0.83 |
| 10 | 8552196 | rs11255728 | 0.989 | -0.259 | -1.317 | 0.398 | -0.002 | 0.022 | 0.022 | 0 | 0.47 | 0.58 | 0.5 | 0.35 | 0.5 | 0.43 |
| 10 | 8553138 | rs11255731 | 0.228 | -0.175 | -1.209 | 0.772 | -0.006 | -0.007 | -0.007 | 2.028 | 0.15 | 0.11 | 0.15 | 0.22 | 0.16 | 0.25 |
| 10 | 8555318 | rs7922051 | 0.607 | -0.278 | -0.633 | 0.742 | -0.008 | 0.032 | 0.032 | 0 | 0.81 | 0.71 | 0.84 | 0.67 | 0.85 | 0.77 |
| 10 | 8558855 | rs7898388 | -0.997 | -0.47 | -0.365 | 0.671 | -0.007 | 0.007 | 0.007 | 0 | 0.3 | 0.35 | 0.3 | 0.31 | 0.24 | 0.33 |
| 10 | 8561142 | rs2646386 | -0.934 | -0.565 | 0.276 | 0.439 | -0.016 | 0.109 | 0.109 | 0 | 0.26 | 0.44 | 0.3 | 0.44 | 0.22 | 0.37 |
| 10 | 8561880 | rs2646387 | -0.907 | -0.719 | 0.898 | 0.505 | -0.012 | 0.05 | 0.05 | 0 | 0.26 | 0.58 | 0.32 | 0.5 | 0.22 | 0.35 |
| 10 | 8562348 | rs7095431 | 1.52 | -0.92 | 0.112 | -0.103 | -0.133 | 0.014 | 0.014 | 0 | 0.27 | 0.04 | 0.35 | 0.24 | 0.41 | 0.41 |

| | | | | | | | | | | | | | | | | |
|---|---|---|---|---|---|---|---|---|---|---|---|---|---|---|---|---|
| 10 | 8562425 | rs2765415 | -0.974 | -1.198 | 0.09 | -0.238 | 0.003 | 0 | 0 | 0 | 0.51 | 0.53 | 0.41 | 0.44 | 0.42 | 0.51 |
| 10 | 8563175 | rs7078855 | 1.616 | -0.99 | 0.111 | -0.159 | 0.003 | 0 | 0 | 0 | 0.26 | 0.28 | 0.34 | 0.31 | 0.36 | 0.38 |
| 10 | 8563593 | rs10905399 | 1.616 | -0.99 | 0.111 | -0.159 | 0.003 | 0 | 0 | 0.109 | 0.26 | 0.28 | 0.34 | 0.31 | 0.36 | 0.38 |
| 10 | 8565675 | rs2646438 | 0.025 | -1.173 | 0.22 | -0.368 | 0.03 | 0.081 | 0.081 | 0 | 0.51 | 0.64 | 0.64 | 0.74 | 0.57 | 0.71 |
| 10 | 8566311 | rs2646437 | 0.319 | -1.228 | 0.741 | -0.353 | -0.003 | 0.016 | 0.016 | 0 | 0.46 | 0.43 | 0.58 | 0.71 | 0.47 | 0.6 |
| 10 | 8566508 | rs2765399 | 0.109 | -1.315 | 0.722 | -0.464 | -0.004 | -0.007 | -0.007 | 0 | 0.61 | 0.42 | 0.66 | 0.71 | 0.63 | 0.7 |
| 10 | 8567129 | rs7093770 | 1.603 | -1.214 | 0.631 | -0.441 | -0.026 | 0.023 | 0.023 | 0 | 0.29 | 0.18 | 0.35 | 0.2 | 0.36 | 0.42 |
| 10 | 8567889 | rs12416202 | 1.706 | -1.219 | 0.564 | -0.536 | -0.106 | 0.076 | 0.076 | 0.035 | 0.29 | 0.05 | 0.33 | 0.14 | 0.37 | 0.41 |
| 10 | 8569242 | rs12146332 | 0.198 | -1.189 | 0.683 | -0.58 | 0.004 | -0.03 | -0.03 | 0 | 0.27 | 0.27 | 0.13 | 0 | 0.21 | 0.17 |
| 10 | 8572238 | rs2765410 | 0.326 | -1.248 | 0.733 | -0.483 | -0.03 | 0.095 | 0.095 | 0 | 0.16 | 0.25 | 0.23 | 0.53 | 0.09 | 0.17 |
| 10 | 8572572 | rs2646431 | -1.419 | -1.269 | 0.766 | -0.51 | -0.012 | -0.009 | -0.009 | 0 | 0.78 | 0.85 | 0.74 | 0.72 | 0.73 | 0.79 |
| 10 | 8572651 | rs7097313 | 0.52 | -1.273 | 0.829 | -0.5 | 0 | 0.015 | 0.015 | 0 | 0.15 | 0.17 | 0.22 | 0.53 | 0.09 | 0.17 |
| 10 | 8574074 | rs11255734 | 0.52 | -1.273 | 0.829 | -0.5 | 0 | 0.015 | 0.015 | 0 | 0.15 | 0.17 | 0.22 | 0.53 | 0.09 | 0.17 |
| 10 | 8575610 | rs2646397 | 0.347 | -1.265 | 0.729 | -0.478 | -0.025 | 0.079 | 0.079 | 0 | 0.16 | 0.24 | 0.23 | 0.53 | 0.09 | 0.17 |
| 10 | 8576161 | rs17143959 | 0.337 | -1.277 | 0.736 | -0.523 | -0.009 | -0.04 | -0.04 | 0.253 | 0.16 | 0.16 | 0.07 | 0 | 0.11 | 0.1 |
| 10 | 8576206 | rs4749779 | 0.889 | -0.563 | 1.078 | 0.02 | -0.042 | -0.036 | -0.036 | 0.366 | 0.21 | 0.06 | 0.25 | 0.28 | 0.27 | 0.2 |
| 10 | 8576629 | rs2646393 | -0.067 | -0.556 | 1.094 | 0.032 | -0.04 | 0.136 | 0.136 | 0 | 0.16 | 0.31 | 0.23 | 0.53 | 0.09 | 0.17 |
| 10 | 8576721 | rs17481499 | 1.424 | -0.387 | 1.023 | 0.014 | -0.03 | 0.055 | 0.055 | 0 | 0.16 | 0.07 | 0.09 | 0 | 0.18 | 0.1 |
| 10 | 8578342 | rs1830951 | 0.821 | -0.39 | 1.066 | 0.018 | -0.003 | 0.028 | 0.028 | 0 | 0.45 | 0.35 | 0.42 | 0.19 | 0.47 | 0.53 |
| 10 | 8578965 | rs7084731 | -0.08 | -0.363 | 0.981 | 0.14 | -0.044 | 0.215 | 0.215 | 0 | 0.84 | 0.58 | 0.76 | 0.47 | 0.91 | 0.83 |
| 10 | 8581226 | rs1409124 | 0.114 | -0.314 | 0.947 | 0.185 | 0.002 | -0.009 | -0.009 | 0 | 0.69 | 0.69 | 0.69 | 0.47 | 0.74 | 0.73 |
| 10 | 8583030 | rs1925932 | 0.197 | -0.314 | 0.893 | 0.186 | -0.012 | 0.038 | 0.038 | 0.182 | 0.68 | 0.53 | 0.67 | 0.47 | 0.74 | 0.73 |
| 10 | 8583805 | rs6602325 | 0.074 | -0.309 | 0.745 | 0.189 | -0.001 | 0.003 | 0.003 | 0 | 0.69 | 0.66 | 0.68 | 0.47 | 0.74 | 0.73 |
| 10 | 8591369 | rs7894791 | 1.01 | -0.377 | 0.736 | 0.152 | -0.001 | 0.066 | 0.066 | 0.04 | 0.42 | 0.25 | 0.37 | 0.2 | 0.41 | 0.5 |
| 10 | 8592643 | rs12265609 | 0.261 | -0.684 | 0.983 | 0.931 | 0 | 0 | 0 | 0 | 0.21 | 0.24 | 0.26 | 0.27 | 0.27 | 0.2 |
| 10 | 8592821 | rs4747803 | 0.75 | -0.631 | 0.458 | 1.046 | -0.045 | 0.138 | 0.138 | 0.087 | 0.83 | 0.64 | 0.76 | 0.47 | 0.91 | 0.83 |
| 10 | 8602170 | rs11255745 | -0.714 | -0.524 | -0.379 | 1.181 | -0.011 | 0.04 | 0.04 | 0 | 0.15 | 0.2 | 0.22 | 0.53 | 0.06 | 0.18 |
| 10 | 8606014 | rs17144046 | -0.336 | 0.262 | -0.513 | 1.068 | -0.005 | -0.016 | -0.016 | 0 | 0.25 | 0.22 | 0.31 | 0.45 | 0.29 | 0.24 |
| 10 | 8608790 | rs942930 | 0.003 | -0.249 | -0.562 | 0.478 | 0.007 | 0.048 | 0.048 | 0 | 0.5 | 0.34 | 0.55 | 0.35 | 0.47 | 0.45 |
| 10 | 8611014 | rs2031557 | 0.266 | -0.441 | -0.079 | -0.087 | -0.003 | 0.001 | 0.001 | 0 | 0.44 | 0.4 | 0.29 | 0.1 | 0.44 | 0.42 |
| 10 | 8611179 | rs7101305 | 0.426 | -0.784 | -0.116 | 0.009 | -0.037 | 0.019 | 0.019 | 0 | 0.37 | 0.26 | 0.6 | 0.66 | 0.43 | 0.37 |
| 10 | 8613894 | rs1325881 | 0.286 | -1.118 | 0.599 | 0.488 | -0.041 | 0.048 | 0.048 | 0.612 | 0.84 | 0.61 | 0.92 | 0.78 | 0.89 | 0.79 |
| 10 | 8616127 | rs10795659 | -0.587 | -1.072 | 0.287 | 0.511 | -0.023 | 0.016 | 0.016 | 0 | 0.16 | 0.19 | 0.1 | 0.23 | 0.13 | 0.22 |
| 10 | 8623589 | rs10905409 | -0.841 | -0.813 | -0.012 | 0.287 | -0.019 | 0.013 | 0.013 | 0 | 0.17 | 0.19 | 0.12 | 0.24 | 0.14 | 0.24 |
| 10 | 8628416 | rs7921734 | -0.932 | -1.449 | 0.152 | 0.174 | -0.018 | 0.03 | 0.03 | 0.411 | 0.17 | 0.28 | 0.11 | 0.25 | 0.13 | 0.32 |
| 10 | 8631309 | rs10508359 | -0.862 | -1.518 | 0.106 | 0.139 | -0.01 | -0.004 | -0.004 | 1.95 | 0.16 | 0.22 | 0.11 | 0.17 | 0.14 | 0.31 |
| 10 | 8633861 | rs4749791 | -1.395 | -2.159 | 0.09 | -0.025 | -0.029 | 0.002 | 0.002 | 0.366 | 0.38 | 0.47 | 0.41 | 0.4 | 0.25 | 0.53 |
| 10 | 8636739 | rs11255764 | 1.127 | -1.843 | -0.423 | -0.471 | -0.049 | 0.064 | 0.064 | 0 | 0.17 | 0.05 | 0.15 | 0.05 | 0.22 | 0.08 |
| 10 | 8638235 | rs4747806 | -0.291 | -2.056 | -0.55 | -0.759 | 0.013 | -0.099 | -0.099 | 0.14 | 0.32 | 0.51 | 0.26 | 0.14 | 0.35 | 0.21 |
| 10 | 8638679 | rs10905411 | -0.087 | -2.044 | -0.428 | -0.76 | 0.009 | -0.077 | -0.077 | 0.143 | 0.3 | 0.46 | 0.23 | 0.14 | 0.32 | 0.2 |
| 10 | 8638744 | rs12413367 | -0.148 | -2.249 | -0.44 | -0.705 | -0.006 | 0.001 | 0.001 | 1.784 | 0.18 | 0.17 | 0.3 | 0.21 | 0.17 | 0.44 |
| 10 | 8638851 | rs12411972 | -0.161 | -2.28 | -0.453 | -0.779 | -0.001 | 0.039 | 0.039 | 0.005 | 0.07 | 0.14 | 0.23 | 0.15 | 0.04 | 0.27 |
| 10 | 8644334 | rs17483588 | 0.259 | -2.378 | -0.361 | -0.846 | 0.011 | 0.036 | 0.036 | 0 | 0.16 | 0.07 | 0.1 | 0.05 | 0.14 | 0.15 |
| 10 | 8645163 | rs9787692 | 2.474 | -2.408 | -0.433 | -0.829 | -0.122 | -0.151 | -0.151 | 0 | 0.3 | 0.04 | 0.34 | 0.46 | 0.41 | 0.25 |
| 10 | 8646048 | rs10752143 | -0.252 | -2.44 | -0.341 | -0.811 | 0.003 | 0.002 | 0.002 | 0 | 0.73 | 0.74 | 0.83 | 0.83 | 0.83 | 0.82 |
| 10 | 8647256 | rs1409121 | 1.264 | -1.981 | -0.219 | -0.546 | -0.011 | 0.005 | 0.005 | 0 | 0.08 | 0.01 | 0.06 | 0.04 | 0.1 | 0.09 |
| 10 | 8651305 | rs12412087 | -1.079 | -1.947 | -0.082 | -0.527 | -0.017 | 0.046 | 0.046 | 0.252 | 0.11 | 0.2 | 0.24 | 0.23 | 0.07 | 0.36 |
| 10 | 8656947 | rs10905429 | 0.01 | -1.902 | -0.035 | -0.55 | 0.027 | 0.048 | 0.048 | 0 | 0.06 | 0.32 | 0.05 | 0.08 | 0.06 | 0.06 |
| 10 | 8660095 | rs6602329 | -0.511 | -1.755 | -0.62 | -0.496 | 0.01 | 0.038 | 0.038 | 0.484 | 0.5 | 0.62 | 0.36 | 0.27 | 0.43 | 0.31 |
| 10 | 8660732 | rs7907956 | 0.307 | -1.749 | -0.599 | -0.494 | -0.003 | -0.008 | -0.008 | 0.017 | 0.92 | 0.86 | 0.76 | 0.94 | 0.93 | 0.7 |
| 10 | 8663783 | rs11255779 | -0.29 | -1.76 | -0.604 | -0.483 | 0.009 | 0.033 | 0.033 | 0.578 | 0.48 | 0.63 | 0.33 | 0.27 | 0.4 | 0.3 |
| 10 | 8665147 | rs4454616 | -1.047 | -1.747 | -0.612 | -0.474 | -0.019 | 0.003 | 0.003 | 0.36 | 0.59 | 0.75 | 0.42 | 0.42 | 0.52 | 0.45 |
| 10 | 8666008 | rs7910940 | 1.76 | -1.69 | -0.791 | -0.435 | -0.116 | -0.139 | -0.139 | 1.052 | 0.29 | 0.04 | 0.33 | 0.45 | 0.4 | 0.24 |
| 10 | 8666609 | rs12218610 | 0.729 | -1.836 | -0.823 | -0.533 | -0.032 | 0.022 | 0.022 | 0 | 0.24 | 0.12 | 0.22 | 0.15 | 0.27 | 0.13 |
| 10 | 8668698 | rs4584486 | 0.639 | -1.861 | -0.734 | -0.482 | 0.004 | -0.003 | -0.003 | 0 | 0.11 | 0.1 | 0.07 | 0.14 | 0.1 | 0.15 |
| 10 | 8676366 | rs11255785 | 0.873 | -1.9 | -0.901 | -0.503 | 0.125 | 0.058 | 0.058 | 0.339 | 0.54 | 0.2 | 0.57 | 0.61 | 0.7 | 0.38 |
| 10 | 8678906 | rs7097369 | -0.315 | -1.778 | -0.65 | -0.376 | -0.009 | 0.005 | 0.005 | 0 | 0.05 | 0.09 | 0.02 | 0.08 | 0.04 | 0.09 |
| 10 | 8680784 | rs10905432 | -0.16 | -1.606 | -0.612 | -0.315 | -0.063 | -0.042 | -0.042 | 0 | 0.3 | 0.58 | 0.13 | 0.2 | 0.18 | 0.25 |
| 10 | 8682679 | rs7083359 | 0.462 | -1.494 | -0.738 | -0.283 | -0.049 | 0.027 | 0.027 | 0 | 0.24 | 0.1 | 0.24 | 0.15 | 0.29 | 0.14 |
| 10 | 8686095 | rs7098207 | 0.88 | -1.443 | -0.2 | -0.108 | 0.101 | 0.045 | 0.045 | 0.6 | 0.61 | 0.19 | 0.62 | 0.65 | 0.76 | 0.46 |
| 10 | 8686326 | rs827402 | 1.035 | -0.336 | 0.47 | 0.417 | 0.013 | 0.044 | 0.044 | 0.036 | 0.93 | 0.77 | 0.75 | 0.87 | 0.95 | 0.86 |
| 10 | 8689122 | rs827406 | -0.706 | -0.312 | 0.205 | 0.334 | 0.003 | 0.007 | 0.007 | 0 | 0.7 | 0.7 | 0.85 | 0.83 | 0.74 | 0.73 |
| 10 | 8693691 | rs827396 | -0.309 | -0.47 | 0.602 | 0.445 | -0.009 | -0.004 | -0.004 | 0 | 0.64 | 0.46 | 0.63 | 0.71 | 0.71 | 0.64 |
| 10 | 8693924 | rs827397 | 0.38 | -0.125 | 0.383 | 0.522 | -0.014 | -0.025 | -0.025 | 0 | 0.47 | 0.24 | 0.48 | 0.55 | 0.51 | 0.36 |
| 10 | 8694069 | rs7918220 | 0.4 | -0.076 | 0.375 | 0.539 | 0.007 | 0.003 | 0.003 | 0 | 0.17 | 0.23 | 0.15 | 0.16 | 0.2 | 0.28 |
| 10 | 8695795 | rs10905437 | -0.679 | -0.048 | 0.062 | 0.518 | 0.018 | -0.01 | -0.01 | 0.037 | 0.36 | 0.48 | 0.54 | 0.3 | 0.42 | 0.44 |
| 10 | 8696383 | rs706770 | -1.028 | -0.067 | 0.07 | 0.515 | -0.002 | -0.001 | -0.001 | 0.149 | 0.67 | 0.66 | 0.86 | 0.76 | 0.74 | 0.72 |
| 10 | 8696446 | rs706771 | 1.05 | -0.05 | 0.028 | 0.523 | -0.013 | -0.052 | -0.052 | 0.417 | 0.33 | 0.18 | 0.33 | 0.47 | 0.33 | 0.27 |
| 10 | 8697320 | rs2388822 | 0.534 | -0.03 | -0.111 | 0.529 | 0.003 | 0.001 | 0.001 | 0 | 0.17 | 0.23 | 0.16 | 0.16 | 0.21 | 0.28 |

| | | | | | | | | | | | | | | | | |
|---|---|---|---|---|---|---|---|---|---|---|---|---|---|---|---|---|
| 10 | 8701219 | rs10795668 | 1.298 | 0.123 | -0.799 | 0.543 | -0.072 | -0.103 | -0.103 | 0 | 0.29 | 0.02 | 0.31 | 0.37 | 0.32 | 0.21 |
| 10 | 8704983 | rs1334549 | 0.489 | 0.187 | -0.475 | 0.541 | -0.014 | 0.007 | 0.007 | 0 | 0.15 | 0.11 | 0.16 | 0.18 | 0.18 | 0.13 |
| 10 | 8707550 | rs10466295 | 0.489 | 0.187 | -0.475 | 0.541 | -0.014 | 0.007 | 0.007 | 0 | 0.15 | 0.11 | 0.16 | 0.18 | 0.18 | 0.13 |
| 10 | 8709338 | rs4508106 | -0.939 | 0.149 | -0.426 | 0.55 | 0.005 | 0.01 | 0.01 | 0 | 0.7 | 0.73 | 0.86 | 0.83 | 0.75 | 0.75 |
| 10 | 8711723 | rs4749806 | -0.837 | 0.272 | -0.397 | 0.506 | -0.003 | -0.003 | -0.003 | 0 | 0.68 | 0.66 | 0.85 | 0.76 | 0.74 | 0.73 |
| 10 | 8715144 | rs10905440 | -0.069 | 0.741 | -0.764 | 0.607 | -0.014 | 0.007 | 0.007 | 0 | 0.15 | 0.11 | 0.16 | 0.18 | 0.18 | 0.13 |
| 10 | 8734295 | rs1537603 | -0.209 | 0.481 | -1.103 | 0.668 | -0.027 | -0.001 | -0.001 | 0 | 0.46 | 0.21 | 0.46 | 0.44 | 0.5 | 0.3 |
| 10 | 8738908 | rs7898455 | 0.661 | 0.656 | -1.364 | 0.524 | -0.036 | -0.056 | -0.056 | 0 | 0.31 | 0.09 | 0.31 | 0.36 | 0.33 | 0.23 |
| 10 | 8739193 | rs11255840 | 0.259 | 0.714 | -1.233 | 0.52 | -0.008 | 0.018 | 0.018 | 0 | 0.15 | 0.11 | 0.16 | 0.08 | 0.18 | 0.07 |
| 10 | 8745420 | rs1970170 | 0.171 | 0.754 | -1.237 | 0.516 | -0.017 | 0.025 | 0.025 | 0 | 0.15 | 0.1 | 0.16 | 0.08 | 0.19 | 0.07 |
| 10 | 8745988 | rs11255847 | -0.287 | 0.616 | -0.968 | 0.463 | 0.002 | -0.015 | -0.015 | 0.185 | 0.18 | 0.23 | 0.18 | 0.08 | 0.2 | 0.09 |
| 10 | 8751545 | rs4112287 | 0.364 | 0.537 | -0.075 | 0.518 | -0.025 | 0.029 | 0.029 | 0 | 0.15 | 0.08 | 0.16 | 0.08 | 0.19 | 0.08 |
| 10 | 8754969 | rs17408580 | -0.32 | 0.502 | -0.021 | 0.489 | 0.002 | -0.013 | -0.013 | 0 | 0.18 | 0.23 | 0.19 | 0.08 | 0.2 | 0.1 |
| 10 | 8757444 | rs11255866 | -1.149 | 0.247 | -0.02 | 0.279 | 0.005 | 0.038 | 0.038 | 0 | 0.72 | 0.78 | 0.87 | 0.82 | 0.74 | 0.75 |
| 10 | 8762195 | rs12250379 | -0.851 | 0.321 | 0.295 | 0.419 | 0.01 | -0.015 | -0.015 | 0 | 0.36 | 0.48 | 0.33 | 0.28 | 0.39 | 0.37 |
| 10 | 8766942 | rs12259226 | 0.353 | 0.526 | 0.137 | 0.484 | 0.009 | -0.009 | -0.009 | 0.995 | 0.16 | 0.22 | 0.17 | 0.16 | 0.18 | 0.13 |
| 10 | 8767434 | rs958934 | 0.124 | 0.203 | 0.602 | 0.793 | 0.005 | 0.014 | 0.014 | 0 | 0.17 | 0.26 | 0.18 | 0.23 | 0.18 | 0.26 |
| 10 | 8767799 | rs11255875 | 0.04 | 0.213 | 0.607 | 0.834 | 0.004 | 0.014 | 0.014 | 0 | 0.18 | 0.26 | 0.19 | 0.23 | 0.19 | 0.27 |
| 10 | 8773143 | rs10905455 | -0.223 | -0.076 | -0.323 | 0.422 | 0.028 | 0.037 | 0.037 | 0 | 0.6 | 0.32 | 0.57 | 0.67 | 0.67 | 0.58 |
| 10 | 8773922 | rs17144396 | -0.917 | -0.344 | -0.026 | 0.591 | -0.005 | 0.044 | 0.044 | 0 | 0.08 | 0.11 | 0.09 | 0.26 | 0.07 | 0.1 |
| 10 | 8775730 | rs10752146 | 0.437 | -1.101 | 0.117 | -0.066 | -0.012 | -0.006 | -0.006 | 1.239 | 0.77 | 0.75 | 0.67 | 0.73 | 0.8 | 0.85 |
| 10 | 8776296 | rs17144411 | 0.528 | -1.321 | -0.855 | 0.1 | 0.021 | -0.135 | -0.135 | 0.054 | 0.22 | 0.02 | 0.18 | 0.49 | 0.17 | 0.32 |
| 10 | 8777692 | rs10508366 | 0.085 | -0.583 | -0.486 | -1.528 | 0.003 | -0.006 | -0.006 | 1.061 | 0.17 | 0.18 | 0.16 | 0.07 | 0.23 | 0.15 |
| 10 | 8780484 | rs7071899 | 0.411 | -1.183 | -0.136 | -1.658 | 0.064 | 0.047 | 0.047 | 0 | 0.38 | 0.21 | 0.39 | 0.73 | 0.25 | 0.46 |
| 10 | 8786636 | rs7908673 | -0.598 | -1.604 | 0.517 | -1.963 | -0.087 | -0.081 | -0.081 | 0 | 0.74 | 0.52 | 0.83 | 0.88 | 0.84 | 0.83 |
| 10 | 8786826 | rs7923068 | 0.164 | -1.622 | 0.568 | -2.006 | 0.01 | 0.027 | 0.027 | 0 | 0.36 | 0.29 | 0.44 | 0.14 | 0.59 | 0.37 |
| 10 | 8788259 | rs3897991 | 0.862 | -1.504 | 0.232 | -1.957 | -0.017 | -0.013 | -0.013 | 0.585 | 0.05 | 0.01 | 0.12 | 0.09 | 0.07 | 0.08 |
| 10 | 8788270 | rs3897990 | 0.85 | -1.671 | 0.339 | -1.945 | 0.062 | 0.046 | 0.046 | 0 | 0.38 | 0.22 | 0.39 | 0.73 | 0.25 | 0.46 |
| 10 | 8790819 | rs1334557 | 1.234 | -1.409 | 0.548 | -1.77 | 0.065 | 0.013 | 0.013 | 0 | 0.35 | 0.14 | 0.38 | 0.63 | 0.25 | 0.45 |
| 10 | 8795245 | rs8181367 | -0.603 | -1.4 | 0.833 | -1.871 | -0.045 | -0.055 | -0.055 | 0 | 0.78 | 0.68 | 0.88 | 0.89 | 0.85 | 0.93 |
| 10 | 8796596 | rs979534 | 0.703 | -1.266 | 0.958 | -1.905 | -0.038 | -0.027 | -0.027 | 0 | 0.42 | 0.56 | 0.45 | 0.73 | 0.26 | 0.55 |
| 10 | 8800398 | rs4749820 | 0.099 | -1.234 | 0.928 | -1.936 | 0.025 | 0.084 | 0.084 | 0 | 0.36 | 0.15 | 0.43 | 0.16 | 0.59 | 0.37 |
| 10 | 8808033 | rs983263 | 0.905 | -1.276 | 0.878 | -1.98 | 0.023 | 0.028 | 0.028 | 0 | 0.19 | 0.09 | 0.1 | 0.11 | 0.15 | 0.07 |
| 10 | 8811034 | rs9665552 | 0.226 | -1.268 | 0.942 | -2.179 | 0.019 | 0.069 | 0.069 | 0 | 0.35 | 0.16 | 0.43 | 0.16 | 0.59 | 0.38 |
| 10 | 8811751 | rs11255914 | 0.247 | -1.267 | 0.927 | -2.178 | 0.018 | 0.089 | 0.089 | 0 | 0.35 | 0.11 | 0.43 | 0.16 | 0.59 | 0.38 |
| 10 | 8819367 | rs982015 | -0.298 | -1.248 | 0.872 | -2.284 | -0.069 | -0.111 | -0.111 | 0 | 0.76 | 0.59 | 0.88 | 0.96 | 0.85 | 0.93 |
| 10 | 8826168 | rs2991810 | 0.637 | -1.25 | 0.909 | -2.279 | -0.009 | 0.003 | 0.003 | 0.03 | 0.92 | 0.58 | 0.93 | 0.96 | 0.99 | 0.92 |
| 10 | 8826990 | rs2243792 | 0.091 | -1.251 | 0.747 | -2.305 | -0.004 | -0.015 | -0.015 | 0 | 0.6 | 0.64 | 0.56 | 0.83 | 0.41 | 0.62 |
| 10 | 8831119 | rs420899 | 0.728 | -1.197 | 0.564 | -2.275 | 0.001 | -0.009 | -0.009 | 0.645 | 0.2 | 0.17 | 0.11 | 0.04 | 0.15 | 0.07 |
| 10 | 8834391 | rs17502419 | 1.171 | -1.095 | 0.424 | -2.156 | 0.074 | 0.053 | 0.053 | 0 | 0.36 | 0.14 | 0.38 | 0.76 | 0.25 | 0.46 |
| 10 | 8840705 | rs7069425 | 1.138 | -0.999 | 0.248 | -2.007 | 0.051 | 0.021 | 0.021 | 0 | 0.37 | 0.24 | 0.29 | 0.64 | 0.27 | 0.43 |
| 10 | 8841669 | rs2025758 | 0.027 | -0.433 | 0.083 | -2.074 | -0.026 | 0.015 | 0.015 | 0 | 0.32 | 0.28 | 0.51 | 0.29 | 0.44 | 0.49 |
| 10 | 8843949 | rs1334538 | 0.395 | 0.43 | -0.203 | -1.938 | -0.021 | -0.013 | -0.013 | 0.789 | 0.07 | 0.03 | 0.17 | 0.1 | 0.16 | 0.18 |
| 10 | 8845941 | rs406742 | 0.955 | 0.355 | -0.068 | -1.939 | 0.034 | 0.016 | 0.016 | 0 | 0.37 | 0.28 | 0.3 | 0.64 | 0.27 | 0.43 |
| 10 | 8847564 | rs17502745 | 0.006 | 0.686 | -0.135 | -1.739 | 0.027 | -0.027 | -0.027 | 0 | 0.11 | 0.26 | 0.15 | 0.07 | 0.16 | 0.11 |
| 10 | 8847765 | rs7913253 | 1.205 | 0.735 | -0.41 | -1.779 | -0.1 | 0.027 | 0.027 | 0 | 0.15 | 0.01 | 0.25 | 0.12 | 0.24 | 0.14 |
| 10 | 8851255 | rs2483935 | 0.584 | -0.019 | 0.12 | -1.483 | 0.053 | -0.082 | -0.082 | 0 | 0.84 | 0.48 | 0.83 | 0.95 | 0.84 | 0.89 |
| 10 | 8855244 | rs1577218 | -0.258 | -0.195 | -0.587 | -1.411 | -0.003 | -0.01 | -0.01 | 0 | 0.51 | 0.52 | 0.6 | 0.34 | 0.58 | 0.55 |
| 10 | 8857322 | rs10905471 | 0.46 | -0.457 | -0.548 | -1.307 | 0.011 | 0.02 | 0.02 | 0 | 0.52 | 0.46 | 0.57 | 0.65 | 0.58 | 0.55 |
| 10 | 8858346 | rs1018729 | -0.428 | -0.762 | -0.637 | -1.465 | -0.021 | 0.002 | 0.002 | 0.117 | 0.72 | 0.54 | 0.64 | 0.68 | 0.73 | 0.68 |
| 10 | 8866290 | rs12248916 | 0.705 | -0.669 | -0.439 | -1.418 | 0 | 0.004 | 0.004 | 0.135 | 0.22 | 0.16 | 0.32 | 0.32 | 0.25 | 0.31 |
| 10 | 8866733 | rs12249220 | 0.702 | -0.673 | -0.425 | -1.42 | 0 | 0.005 | 0.005 | 0.13 | 0.22 | 0.17 | 0.32 | 0.32 | 0.25 | 0.31 |
| 10 | 8872667 | rs12250687 | 0.63 | -0.564 | -0.759 | -1.381 | -0.006 | 0.009 | 0.009 | 0.286 | 0.25 | 0.27 | 0.38 | 0.34 | 0.27 | 0.39 |
| 10 | 8875926 | rs958866 | -0.201 | -0.729 | -0.703 | -1.405 | -0.037 | -0.04 | -0.04 | 0 | 0.92 | 0.71 | 0.97 | 1 | 0.99 | 0.93 |
| 10 | 8880984 | rs10737033 | -0.042 | -0.175 | -1.007 | -1.204 | -0.002 | -0.051 | -0.051 | 0 | 0.84 | 0.76 | 0.92 | 0.97 | 0.87 | 0.94 |
| 10 | 8887858 | rs729333 | 0.784 | -0.361 | -1.279 | -1.054 | -0.156 | -0.133 | -0.133 | 0 | 0.33 | 0.02 | 0.23 | 0.53 | 0.41 | 0.25 |
| 10 | 8903227 | rs17144704 | -0.033 | -0.481 | -0.504 | -0.611 | 0.013 | 0.007 | 0.007 | 0.354 | 0.1 | 0.05 | 0.03 | 0 | 0.04 | 0.04 |
| 10 | 8919297 | rs1373614 | 0.011 | 0.217 | -1.238 | -0.195 | 0.04 | 0.052 | 0.052 | 0.552 | 0.52 | 0.34 | 0.57 | 0.59 | 0.6 | 0.48 |
| 10 | 8926204 | rs4457655 | -0.165 | 0.227 | -1.074 | 0.088 | 0.021 | 0.039 | 0.039 | 0 | 0.5 | 0.39 | 0.62 | 0.68 | 0.46 | 0.58 |
| 10 | 8926700 | rs1444774 | 0.171 | -1.218 | -0.673 | 0.416 | -0.023 | 0.012 | 0.012 | 0 | 0.71 | 0.65 | 0.59 | 0.45 | 0.87 | 0.74 |
| 10 | 8933437 | rs290898 | 0.542 | -1.528 | -0.671 | 0.212 | -0.043 | 0.079 | 0.079 | 0 | 0.8 | 0.67 | 0.73 | 0.44 | 0.9 | 0.81 |
| 10 | 8934191 | rs7099087 | 0.381 | -1.55 | -0.63 | 0.229 | -0.031 | 0.07 | 0.07 | 0 | 0.12 | 0.15 | 0.23 | 0.53 | 0.02 | 0.16 |
| 10 | 8945541 | rs1444779 | -0.669 | -1.711 | -0.366 | -0.245 | -0.002 | -0.013 | -0.013 | 0 | 0.76 | 0.68 | 0.74 | 0.86 | 0.75 | 0.74 |
| 10 | 8951090 | rs1028827 | -0.642 | -1.715 | -0.287 | -0.247 | -0.002 | -0.014 | -0.014 | 0 | 0.76 | 0.68 | 0.74 | 0.86 | 0.75 | 0.74 |
| 10 | 8952873 | rs11255973 | 1.103 | -3.04 | -0.473 | -0.69 | -0.001 | -0.001 | -0.001 | 0 | 0.37 | 0.35 | 0.37 | 0.16 | 0.53 | 0.36 |
| 10 | 8961614 | rs1857231 | -0.368 | -3.009 | -0.56 | -0.726 | 0 | -0.01 | -0.01 | 0 | 0.66 | 0.61 | 0.75 | 0.83 | 0.63 | 0.68 |
| 10 | 8966406 | rs895068 | -0.18 | -3.176 | -0.614 | -0.776 | -0.001 | 0.028 | 0.028 | 0 | 0.38 | 0.31 | 0.28 | 0.19 | 0.4 | 0.34 |
| 10 | 8969083 | rs1243413 | -0.286 | -3.091 | -0.438 | -0.813 | 0 | 0.021 | 0.021 | 0 | 0.39 | 0.33 | 0.36 | 0.19 | 0.42 | 0.35 |
| 10 | 8969802 | rs1243411 | 0.431 | -3.132 | -0.465 | -0.818 | 0.005 | 0.015 | 0.015 | 0 | 0.29 | 0.25 | 0.21 | 0.16 | 0.3 | 0.29 |

| | | | | | | | | | | | | | | | | |
|---|---|---|---|---|---|---|---|---|---|---|---|---|---|---|---|---|
| 10 | 8972018 | rs10508372 | -0.159 | -3.407 | -0.483 | -0.88 | -0.001 | -0.015 | -0.015 | 0 | 0.26 | 0.21 | 0.28 | 0.65 | 0.06 | 0.3 |
| 10 | 8974161 | rs10508373 | 1.609 | -3.479 | -0.553 | -0.959 | -0.022 | -0.001 | -0.001 | 0.676 | 0.19 | 0.05 | 0.22 | 0.14 | 0.22 | 0.23 |
| 10 | 8974568 | rs1243403 | 1.803 | -3.434 | -0.504 | -0.954 | 0.007 | 0.008 | 0.008 | 0 | 0.06 | 0.04 | 0.04 | 0.02 | 0.04 | 0.05 |
| 10 | 8975723 | rs870376 | 1.609 | -3.607 | -0.18 | -1.554 | -0.021 | -0.074 | -0.074 | 1.961 | 0.13 | 0.28 | 0.08 | 0 | 0.08 | 0.06 |
| 10 | 8976707 | rs1243399 | -0.05 | -3.813 | -0.206 | -1.564 | 0.003 | 0.001 | 0.001 | 0 | 0.45 | 0.43 | 0.54 | 0.7 | 0.33 | 0.48 |
| 10 | 8978542 | rs7904957 | 1.777 | -3.802 | -0.447 | -1.611 | -0.02 | 0.015 | 0.015 | 0 | 0.22 | 0.06 | 0.24 | 0.14 | 0.24 | 0.18 |
| 10 | 8982241 | rs7902767 | 1.369 | -3.35 | -0.274 | -1.496 | -0.013 | 0 | 0 | 0 | 0.22 | 0.16 | 0.29 | 0.16 | 0.43 | 0.29 |
| 10 | 8988073 | rs10508374 | 1.591 | -3.159 | -0.194 | -1.386 | -0.014 | -0.051 | -0.051 | 0 | 0.13 | 0.2 | 0.08 | 0 | 0.08 | 0.06 |
| 10 | 8992281 | rs17505882 | 0.668 | -2.73 | -0.299 | -1.127 | -0.004 | 0.005 | 0.005 | 0 | 0.09 | 0.07 | 0.06 | 0.03 | 0.12 | 0.06 |
| 10 | 8996267 | rs17144879 | 0.279 | -2.857 | -0.197 | -0.895 | -0.025 | 0.036 | 0.036 | 0.062 | 0.17 | 0.19 | 0.23 | 0.51 | 0.07 | 0.22 |
| 10 | 9000841 | rs11255993 | 0.387 | -2.229 | -0.562 | -1.064 | -0.048 | 0.001 | 0.001 | 0 | 0.26 | 0.15 | 0.3 | 0.22 | 0.41 | 0.32 |
| 10 | 9003997 | rs827618 | 0.159 | -2.425 | -0.086 | -1.196 | 0.005 | -0.051 | -0.051 | 0.647 | 0.17 | 0.33 | 0.15 | 0.06 | 0.16 | 0.11 |
| 10 | 9010624 | rs827637 | -0.515 | -1.814 | -0.708 | -1.374 | -0.001 | -0.003 | -0.003 | 0 | 0.47 | 0.49 | 0.42 | 0.29 | 0.56 | 0.48 |
| 10 | 9011746 | rs6602348 | 1.743 | -1.382 | -1.081 | -0.629 | 0.008 | 0.012 | 0.012 | 2.098 | 0.22 | 0.1 | 0.26 | 0.16 | 0.22 | 0.07 |
| 10 | 9012053 | rs10905493 | 0.416 | -1.502 | -0.926 | -0.681 | -0.033 | 0.222 | 0.222 | 0.898 | 0.15 | 0.32 | 0.19 | 0.55 | 0.08 | 0.37 |
| 10 | 9013778 | rs827634 | -1.194 | -1.472 | -1.124 | -0.745 | -0.008 | 0.095 | 0.095 | 0.005 | 0.64 | 0.86 | 0.63 | 0.84 | 0.65 | 0.86 |
| 10 | 9013939 | rs11255999 | 1.789 | -1.266 | -1.012 | -0.665 | -0.049 | -0.019 | -0.019 | 1.363 | 0.22 | 0.12 | 0.23 | 0.22 | 0.36 | 0.31 |
| 10 | 9017944 | rs10795685 | -0.264 | -1.622 | -0.725 | -0.527 | -0.013 | 0.03 | 0.03 | 1.271 | 0.3 | 0.36 | 0.38 | 0.59 | 0.22 | 0.45 |
| 10 | 9020234 | rs827628 | -2.164 | -1.035 | -0.688 | -0.553 | -0.002 | 0.033 | 0.033 | 0.832 | 0.74 | 0.8 | 0.69 | 0.86 | 0.72 | 0.93 |
| 10 | 9020314 | rs827627 | 1.884 | -1.131 | -0.689 | -0.397 | -0.001 | 0.023 | 0.023 | 0 | 0.16 | 0.09 | 0.1 | 0.01 | 0.13 | 0.08 |
| 10 | 9022145 | rs4749828 | 1.95 | -0.88 | -0.761 | -0.321 | -0.036 | -0.027 | -0.027 | 0.009 | 0.21 | 0.12 | 0.24 | 0.22 | 0.33 | 0.3 |
| 10 | 9024136 | rs4747817 | 1.383 | -0.79 | -0.538 | -0.14 | -0.034 | -0.011 | -0.011 | 0 | 0.3 | 0.14 | 0.29 | 0.24 | 0.38 | 0.35 |
| 10 | 9028677 | rs1243438 | -0.433 | -0.763 | -0.49 | 0.061 | 0.004 | 0.008 | 0.008 | 0.331 | 0.54 | 0.54 | 0.64 | 0.67 | 0.58 | 0.69 |
| 10 | 9030765 | rs7920897 | 1.681 | 0.022 | -0.779 | 0.038 | -0.029 | 0.062 | 0.062 | 1.258 | 0.25 | 0.07 | 0.22 | 0.13 | 0.28 | 0.07 |
| 10 | 9034758 | rs7922444 | 0.469 | 0.075 | 0.087 | 0.718 | -0.004 | 0.02 | 0.02 | 0 | 0.74 | 0.71 | 0.73 | 0.55 | 0.8 | 0.6 |
| 10 | 9035618 | rs1243430 | 0.443 | 0.131 | -0.382 | 0.743 | 0.002 | -0.012 | -0.012 | 0.021 | 0.74 | 0.78 | 0.73 | 0.56 | 0.8 | 0.6 |
| 10 | 9035786 | rs12765699 | -0.555 | 0.122 | 0.222 | 1.435 | -0.008 | 0.114 | 0.114 | 0.003 | 0.1 | 0.17 | 0.17 | 0.4 | 0.07 | 0.28 |
| 10 | 9037492 | rs930579 | 0.002 | 0.184 | 0.182 | 1.082 | 0 | -0.015 | -0.015 | 0 | 0.13 | 0.22 | 0.12 | 0.05 | 0.13 | 0.12 |
| 10 | 9038134 | rs4749829 | -0.833 | 0.179 | 0.022 | 1.522 | -0.005 | 0.122 | 0.122 | 0 | 0.1 | 0.17 | 0.17 | 0.41 | 0.07 | 0.27 |
| 10 | 9038727 | rs11256013 | -0.904 | 0.11 | 0.46 | 1.551 | 0.015 | 0.286 | 0.286 | 0 | 0.09 | 0.24 | 0.25 | 0.53 | 0.07 | 0.23 |
| 10 | 9039506 | rs1705333 | -0.602 | -0.114 | 0.12 | 0.055 | -0.003 | -0.018 | -0.018 | 0 | 0.62 | 0.64 | 0.77 | 0.79 | 0.61 | 0.57 |
| 10 | 9041158 | rs1912584 | -0.03 | -0.491 | 0.089 | 0.183 | -0.014 | 0.1 | 0.1 | 0 | 0.1 | 0.21 | 0.34 | 0.65 | 0.06 | 0.21 |
| 10 | 9041345 | rs4749830 | 1.755 | -0.66 | -0.408 | -0.131 | -0.037 | 0.082 | 0.082 | 0.23 | 0.18 | 0.01 | 0.1 | 0.05 | 0.2 | 0.1 |
| 10 | 9041722 | rs4749831 | -0.112 | -0.38 | -0.066 | 0.138 | -0.004 | 0.049 | 0.049 | 0 | 0.1 | 0.16 | 0.34 | 0.65 | 0.06 | 0.21 |
| 10 | 9053043 | rs1775551 | -1 | -0.216 | -0.012 | 0.109 | -0.003 | -0.01 | -0.01 | 2.917 | 0.82 | 0.87 | 0.89 | 0.82 | 0.81 | 0.65 |
| 10 | 9061104 | rs7916897 | 0.691 | 0.028 | -0.92 | -0.68 | -0.083 | 0.064 | 0.064 | 0.001 | 0.23 | 0.07 | 0.17 | 0.07 | 0.31 | 0.12 |
| 10 | 9062856 | rs2197415 | -0.895 | -0.29 | -0.401 | -0.949 | 0.018 | 0.158 | 0.158 | 0 | 0.55 | 0.78 | 0.76 | 0.88 | 0.58 | 0.7 |
| 10 | 9064716 | rs1612986 | 0.644 | -0.32 | 0.01 | -0.759 | -0.007 | 0.014 | 0.014 | 0 | 0.21 | 0.36 | 0.17 | 0.18 | 0.2 | 0.31 |
| 10 | 9066242 | rs11256028 | 0.126 | -0.371 | -0.284 | -1.05 | -0.081 | -0.04 | -0.04 | 0 | 0.29 | 0.09 | 0.47 | 0.69 | 0.36 | 0.38 |
| 10 | 9074500 | rs1342766 | 0.755 | -0.39 | -0.136 | -1.175 | 0.013 | -0.005 | -0.005 | 0 | 0.22 | 0.31 | 0.15 | 0.19 | 0.24 | 0.3 |
| 10 | 9076230 | rs1623646 | -1.261 | -0.586 | -0.111 | -1.391 | 0.007 | 0.007 | 0.007 | 0 | 0.71 | 0.76 | 0.8 | 0.92 | 0.72 | 0.76 |
| 10 | 9076253 | rs11256031 | 0.23 | -0.564 | -0.083 | -1.379 | -0.016 | -0.004 | -0.004 | 0.061 | 0.29 | 0.22 | 0.46 | 0.68 | 0.34 | 0.36 |
| 10 | 9078999 | rs965307 | 1.509 | -0.655 | -0.082 | -1.455 | -0.001 | -0.007 | -0.007 | 0.421 | 0.12 | 0.12 | 0.14 | 0.05 | 0.11 | 0.03 |
| 10 | 9081709 | rs4747819 | 0.217 | -0.633 | -0.085 | -1.43 | -0.009 | -0.002 | -0.002 | 0 | 0.3 | 0.27 | 0.46 | 0.67 | 0.34 | 0.36 |
| 10 | 9091900 | rs860418 | -0.157 | -0.539 | 0.211 | -1.352 | 0.004 | -0.008 | -0.008 | 0 | 0.58 | 0.6 | 0.66 | 0.87 | 0.61 | 0.72 |
| 10 | 9092862 | rs860419 | 0.235 | -0.581 | -0.02 | -1.402 | 0.032 | 0.105 | 0.105 | 0 | 0.53 | 0.35 | 0.63 | 0.87 | 0.57 | 0.7 |
| 10 | 9095495 | rs1573110 | 1.106 | -0.247 | -0.587 | -1.108 | 0 | -0.002 | -0.002 | 0 | 0.12 | 0.11 | 0.14 | 0.05 | 0.11 | 0.03 |
| 10 | 9107135 | rs17145169 | 1.383 | -0.226 | -0.612 | -1.084 | 0.001 | 0.008 | 0.008 | 0 | 0.11 | 0.08 | 0.05 | 0.05 | 0.09 | 0.02 |
| 10 | 9108543 | rs10795690 | -0.754 | -0.094 | -0.613 | -1.039 | 0.002 | -0.013 | -0.013 | 0 | 0.71 | 0.77 | 0.81 | 0.92 | 0.71 | 0.75 |
| 10 | 9108558 | rs17434960 | 0.168 | 0.75 | -0.765 | -0.429 | -0.059 | 0.04 | 0.04 | 0 | 0.13 | 0.03 | 0.08 | 0.03 | 0.18 | 0.15 |
| 10 | 9121054 | rs2025289 | 0.621 | -0.327 | 0.118 | -0.706 | -0.056 | -0.077 | -0.077 | 0 | 0.22 | 0.42 | 0.07 | 0.09 | 0.11 | 0.06 |
| 10 | 9126643 | rs961945 | 0.301 | -0.321 | -0.048 | -0.96 | 0.044 | 0.16 | 0.16 | 0 | 0.5 | 0.27 | 0.54 | 0.75 | 0.53 | 0.66 |
| 10 | 9126840 | rs11256050 | 1.187 | -0.244 | -0.104 | -1.05 | 0.005 | -0.001 | -0.001 | 0 | 0.1 | 0.05 | 0.16 | 0.12 | 0.08 | 0.09 |
| 10 | 9127422 | rs10795693 | -1.409 | -0.247 | 0.006 | -1.075 | -0.023 | 0.04 | 0.04 | 0 | 0.85 | 0.93 | 0.88 | 0.96 | 0.75 | 0.82 |
| 10 | 9128153 | rs2013364 | 1.019 | -0.357 | 0.287 | -0.996 | -0.046 | -0.075 | -0.075 | 0.072 | 0.22 | 0.41 | 0.16 | 0.09 | 0.14 | 0.05 |
| 10 | 9128775 | rs2224696 | 1.08 | -0.386 | 0.254 | -0.95 | -0.04 | -0.052 | -0.052 | 2.537 | 0.21 | 0.36 | 0.15 | 0.11 | 0.14 | 0.05 |
| 10 | 9140415 | rs11256060 | 0.184 | -0.292 | 0.553 | -0.959 | -0.019 | -0.011 | -0.011 | 2.537 | 0.13 | 0.24 | 0.24 | 0.12 | 0.1 | 0.13 |
| 10 | 9142272 | rs11256063 | 1.148 | -0.249 | 0.301 | -0.929 | -0.003 | -0.023 | -0.023 | 0 | 0.09 | 0.01 | 0.14 | 0.12 | 0.08 | 0.09 |
| 10 | 9143333 | rs11256065 | 0.959 | -0.25 | 0.326 | -0.929 | -0.005 | -0.001 | -0.001 | 1.862 | 0.1 | 0.09 | 0.15 | 0.12 | 0.08 | 0.09 |
| 10 | 9151970 | rs391733 | -0.532 | -0.259 | -0.068 | -1.068 | 0.002 | 0.081 | 0.081 | 0 | 0.66 | 0.86 | 0.84 | 0.87 | 0.65 | 0.79 |
| 10 | 9155598 | rs441107 | 0.541 | -0.344 | -0.231 | -1.105 | 0.053 | 0.166 | 0.166 | 0 | 0.5 | 0.26 | 0.57 | 0.76 | 0.54 | 0.66 |
| 10 | 9164684 | rs7898051 | 1.242 | -0.247 | -0.385 | -1.018 | -0.055 | 0.047 | 0.047 | 0 | 0.14 | 0.01 | 0.11 | 0.05 | 0.24 | 0.16 |
| 10 | 9168739 | rs10905518 | 1.144 | -0.197 | -0.329 | -0.916 | 0 | -0.03 | -0.03 | 0 | 0.09 | 0.01 | 0.15 | 0.13 | 0.07 | 0.09 |
| 10 | 9172233 | rs1149915 | -1.838 | 0.032 | -0.842 | -1.01 | -0.051 | -0.05 | -0.05 | 0 | 0.82 | 0.97 | 0.9 | 0.81 | 0.83 | 0.65 |
| 10 | 9172291 | rs17145278 | 0.769 | -0.116 | -0.032 | -0.569 | -0.007 | 0 | 0 | 0 | 0.11 | 0.11 | 0.16 | 0.12 | 0.07 | 0.09 |
| 10 | 9174790 | rs9424040 | -0.286 | -0.125 | 0.164 | -0.653 | -0.012 | -0.073 | -0.073 | 0 | 0.38 | 0.44 | 0.28 | 0.12 | 0.4 | 0.23 |
| 10 | 9175519 | rs1149916 | -0.815 | -0.071 | -0.13 | -0.611 | 0.031 | 0.076 | 0.076 | 0 | 0.48 | 0.62 | 0.36 | 0.24 | 0.46 | 0.3 |
| 10 | 9191149 | rs1149927 | -0.948 | 0.418 | -0.439 | -1.075 | 0.047 | 0.121 | 0.121 | 0 | 0.49 | 0.7 | 0.45 | 0.24 | 0.46 | 0.31 |
| 10 | 9192570 | rs1149930 | -0.123 | 1.282 | -1.118 | -0.807 | -0.052 | 0.122 | 0.122 | 0 | 0.38 | 0.12 | 0.19 | 0.16 | 0.47 | 0.22 |

| | | | | | | | | | | | | | | | |
|---|---|---|---|---|---|---|---|---|---|---|---|---|---|---|---|
| 10 | 9196330 | rs1149936 | -0.146 | 1.298 | -1.1 | -0.802 | -0.031 | 0.104 | 0.104 | 0 | 0.38 | 0.16 | 0.19 | 0.16 | 0.46 | 0.21 |
| 10 | 9200102 | rs1149938 | -0.726 | 1.219 | -1.056 | -0.805 | -0.008 | 0.062 | 0.062 | 0 | 0.46 | 0.3 | 0.35 | 0.28 | 0.53 | 0.26 |
| 10 | 9201101 | rs10905522 | 0.791 | 0.612 | -0.999 | -0.545 | -0.008 | 0.008 | 0.008 | 0 | 0.09 | 0.15 | 0.15 | 0.12 | 0.07 | 0.05 |
| 10 | 9213854 | rs12778120 | 0.392 | 0.961 | -1.279 | -0.703 | 0.002 | 0.06 | 0.06 | 0 | 0.23 | 0.08 | 0.07 | 0.07 | 0.2 | 0.05 |
| 10 | 9216956 | rs1149952 | -0.405 | 1.529 | -1.104 | -0.463 | 0.003 | -0.004 | -0.004 | 0.868 | 0.24 | 0.26 | 0.3 | 0.21 | 0.33 | 0.32 |
| 10 | 9217212 | rs17485264 | -0.373 | 1.348 | -1.314 | -0.486 | 0.004 | -0.042 | -0.042 | 1.085 | 0.11 | 0.12 | 0.06 | 0 | 0.13 | 0.04 |
| 10 | 9221058 | rs12359419 | -0.148 | 1.254 | -2.013 | -0.5 | -0.034 | 0.107 | 0.107 | 0 | 0.29 | 0.04 | 0.11 | 0.11 | 0.26 | 0.06 |
| 10 | 9222418 | rs11256107 | 0.196 | 1.082 | -2.718 | 0.109 | -0.071 | -0.129 | -0.129 | 0 | 0.34 | 0.07 | 0.46 | 0.52 | 0.37 | 0.42 |
| 10 | 9223010 | rs1242988 | 0.28 | 1.305 | -2.279 | 0.415 | 0 | 0.005 | 0.005 | 0 | 0.87 | 0.83 | 0.58 | 0.71 | 0.8 | 0.75 |
| 10 | 9223424 | rs1242987 | 0.497 | 1.472 | -2.542 | 0.238 | 0.032 | 0.019 | 0.019 | 0 | 0.5 | 0.18 | 0.29 | 0.4 | 0.42 | 0.4 |
| 10 | 9233207 | rs7074400 | 0.549 | 1.162 | -1.284 | 0.097 | 0.026 | 0.014 | 0.014 | 0 | 0.05 | 0.26 | 0.03 | 0 | 0.01 | 0 |
| 10 | 9238010 | rs1242976 | 0.251 | 1.419 | -0.603 | 0.04 | -0.002 | -0.001 | -0.001 | 0 | 0.57 | 0.55 | 0.39 | 0.49 | 0.49 | 0.5 |
| 10 | 9242659 | rs1242967 | 1.389 | 1.245 | -0.715 | -0.144 | -0.025 | -0.082 | -0.082 | 0.144 | 0.23 | 0.08 | 0.21 | 0.3 | 0.23 | 0.36 |
| 10 | 9243785 | rs1001019 | 1.879 | 1.239 | -0.711 | -0.166 | 0.024 | 0.084 | 0.084 | 0.564 | 0.25 | 0.1 | 0.08 | 0.1 | 0.19 | 0.02 |
| 10 | 9247586 | rs1469993 | -0.081 | 1.277 | -0.654 | -0.163 | -0.041 | -0.035 | -0.035 | 0 | 0.62 | 0.76 | 0.43 | 0.49 | 0.51 | 0.52 |
| 10 | 9248331 | rs7911451 | 1.42 | 1.295 | -0.626 | -0.136 | -0.023 | -0.068 | -0.068 | 2.079 | 0.3 | 0.37 | 0.11 | 0.1 | 0.2 | 0.03 |
| 10 | 9248659 | rs12359319 | 0.475 | 1.285 | -0.612 | -0.131 | -0.003 | -0.003 | -0.003 | 0.024 | 0.54 | 0.5 | 0.32 | 0.4 | 0.44 | 0.41 |
| 10 | 9255870 | rs1242973 | 0.669 | 1.339 | -0.808 | -0.05 | -0.004 | -0.018 | -0.018 | 0.043 | 0.29 | 0.23 | 0.29 | 0.39 | 0.3 | 0.48 |
| 10 | 9258570 | rs7098546 | 1.115 | 1.425 | -0.98 | -0.067 | -0.036 | -0.072 | -0.072 | 0 | 0.27 | 0.08 | 0.23 | 0.3 | 0.27 | 0.36 |
| 10 | 9260741 | rs3098354 | -1.151 | 1.721 | -0.978 | -0.014 | 0.002 | 0.007 | 0.007 | 0 | 0.93 | 0.94 | 0.72 | 0.75 | 0.84 | 0.78 |
| 10 | 9260792 | rs1469994 | 1.021 | 1.359 | -0.869 | -0.141 | -0.001 | -0.004 | -0.004 | 0 | 0.28 | 0.24 | 0.25 | 0.3 | 0.27 | 0.36 |
| 10 | 9264824 | rs12774241 | 1.616 | 1.241 | -0.75 | -0.256 | -0.061 | -0.102 | -0.102 | 1.192 | 0.27 | 0.42 | 0.09 | 0.1 | 0.16 | 0.02 |
| 10 | 9266428 | rs2577969 | 0.129 | 1.429 | -0.765 | -0.028 | -0.001 | -0.002 | -0.002 | 0.718 | 0.37 | 0.37 | 0.33 | 0.46 | 0.35 | 0.53 |
| 10 | 9266677 | rs17407956 | 1.618 | 1.226 | -0.834 | -0.133 | -0.01 | 0.02 | 0.02 | 0 | 0.05 | 0.01 | 0.04 | 0 | 0.05 | 0.01 |
| 10 | 9268122 | rs10905530 | 0.996 | 1.292 | -0.947 | 0.044 | -0.008 | -0.036 | -0.036 | 0 | 0.27 | 0.15 | 0.24 | 0.3 | 0.26 | 0.37 |
| 10 | 9273578 | rs7089176 | -0.247 | 1.547 | -0.47 | -0.029 | 0 | -0.002 | -0.002 | 0 | 0.34 | 0.4 | 0.33 | 0.32 | 0.34 | 0.28 |
| 10 | 9277702 | rs4749842 | 0.024 | 1.364 | -0.509 | -0.042 | 0 | -0.017 | -0.017 | 0 | 0.1 | 0.23 | 0.19 | 0.03 | 0.1 | 0.04 |
| 10 | 9284374 | rs17145601 | 0.701 | 1.736 | -0.697 | -0.072 | -0.022 | -0.033 | -0.033 | 0.084 | 0.23 | 0.06 | 0.14 | 0.29 | 0.22 | 0.24 |
| 10 | 9286568 | rs4749844 | -0.269 | 1.85 | -0.317 | 0.011 | -0.01 | -0.021 | -0.021 | 0 | 0.36 | 0.54 | 0.36 | 0.31 | 0.36 | 0.29 |
| 10 | 9293290 | rs1143869 | -0.197 | 1.807 | -0.305 | 0.2 | 0.001 | -0.005 | -0.005 | 0.315 | 0.61 | 0.41 | 0.46 | 0.54 | 0.62 | 0.41 |
| 10 | 9296840 | rs17145627 | -0.061 | 2.123 | -0.143 | 0.523 | -0.005 | -0.007 | -0.007 | 0 | 0.1 | 0.06 | 0.07 | 0.2 | 0.12 | 0.16 |
| 10 | 9301624 | rs12245918 | -1.478 | 2.069 | 0.45 | 0.383 | 0.005 | -0.005 | -0.005 | 0 | 0.19 | 0.31 | 0.29 | 0.2 | 0.2 | 0.21 |
| 10 | 9301800 | rs2764399 | 1.704 | 2.418 | -0.214 | 0.523 | -0.065 | 0.004 | 0.004 | 0 | 0.45 | 0.05 | 0.3 | 0.29 | 0.4 | 0.35 |
| 10 | 9302625 | rs1242756 | -0.267 | 2.242 | 0.43 | 0.632 | -0.004 | 0.004 | 0.004 | 0 | 0.72 | 0.67 | 0.73 | 0.64 | 0.76 | 0.71 |
| 10 | 9317323 | rs10795702 | -0.293 | 2.171 | 0.535 | 0.653 | -0.011 | 0.011 | 0.011 | 0 | 0.72 | 0.5 | 0.7 | 0.65 | 0.76 | 0.59 |
| 10 | 9322527 | rs10795703 | 0.499 | 2.01 | -0.032 | 0.592 | -0.009 | -0.01 | -0.01 | 0 | 0.57 | 0.28 | 0.39 | 0.5 | 0.51 | 0.52 |
| 10 | 9325312 | rs10795704 | 2.023 | 2.252 | -0.841 | 0.619 | -0.045 | -0.043 | -0.043 | 0.888 | 0.39 | 0.04 | 0.27 | 0.29 | 0.34 | 0.35 |
| 10 | 9326101 | rs11256141 | -0.503 | 1.531 | -0.201 | 0.575 | -0.014 | -0.016 | -0.016 | 0 | 0.14 | 0.06 | 0.1 | 0.22 | 0.16 | 0.15 |
| 10 | 9328491 | rs17145704 | 0.465 | 1.162 | -0.068 | 0.426 | -0.001 | -0.009 | -0.009 | 1.884 | 0.08 | 0.13 | 0.22 | 0.04 | 0.1 | 0.06 |
| 10 | 9330087 | rs1149964 | 0.333 | 1.109 | 0.219 | 0.686 | -0.017 | 0.015 | 0.015 | 0 | 0.64 | 0.48 | 0.65 | 0.47 | 0.75 | 0.51 |
| 10 | 9330817 | rs1149966 | 1.903 | 1.759 | 0.061 | 0.447 | -0.004 | -0.05 | -0.05 | 0.621 | 0.39 | 0.11 | 0.31 | 0.41 | 0.43 | 0.36 |
| 10 | 9337376 | rs17558155 | 0.951 | 1.751 | 0.187 | 0.269 | -0.004 | 0.028 | 0.028 | 0 | 0.09 | 0.05 | 0.2 | 0.04 | 0.1 | 0.07 |
| 10 | 9337386 | rs17638790 | 0.047 | 1.989 | 0.385 | 0.351 | 0.02 | 0 | 0 | 0 | 0.07 | 0.09 | 0.13 | 0.1 | 0.19 | 0.13 |
| 10 | 9344681 | rs1149978 | 1.37 | 1.509 | 0.471 | 0.258 | 0.003 | 0.005 | 0.005 | 0.474 | 0.51 | 0.27 | 0.52 | 0.46 | 0.54 | 0.46 |
| 10 | 9354641 | rs17145800 | 0.037 | 1.455 | 0.831 | 0.266 | -0.003 | 0.002 | 0.002 | 0 | 0.09 | 0.1 | 0.09 | 0.09 | 0.1 | 0.1 |
| 10 | 9355284 | rs1149994 | 1.742 | 1.59 | 0.661 | 0.207 | 0.026 | -0.01 | -0.01 | 0 | 0.47 | 0.25 | 0.33 | 0.51 | 0.45 | 0.4 |
| 10 | 9355301 | rs1149995 | 1.231 | 1.595 | 1.023 | 0.458 | -0.016 | -0.009 | -0.009 | 0.007 | 0.57 | 0.33 | 0.54 | 0.55 | 0.56 | 0.49 |
| 10 | 9356003 | rs1149998 | -0.101 | 1.355 | 1.154 | 0.27 | -0.002 | 0 | 0 | 0 | 0.74 | 0.71 | 0.77 | 0.75 | 0.84 | 0.73 |
| 10 | 9357454 | rs1150007 | 0.964 | 1.262 | 1.262 | 0.281 | -0.015 | 0.003 | 0.003 | 0 | 0.64 | 0.46 | 0.68 | 0.65 | 0.75 | 0.64 |
| 10 | 9357543 | rs1886617 | -1.61 | 1.54 | 0.893 | 0.348 | -0.012 | -0.018 | -0.018 | 1.89 | 0.41 | 0.71 | 0.44 | 0.48 | 0.42 | 0.49 |
| 10 | 9357640 | rs1150009 | -0.934 | 0.273 | 0.883 | 0.198 | 0.004 | -0.008 | -0.008 | 0 | 0.86 | 0.89 | 0.94 | 0.76 | 0.92 | 0.8 |
| 10 | 9357949 | rs1150010 | 1.405 | 0.285 | 0.743 | 0.157 | 0.042 | -0.001 | -0.001 | 0 | 0.58 | 0.18 | 0.64 | 0.58 | 0.72 | 0.61 |
| 10 | 9358368 | rs1408405 | -0.17 | 0.335 | 0.877 | 0.255 | -0.004 | -0.005 | -0.005 | 0 | 0.09 | 0.16 | 0.15 | 0.04 | 0.06 | 0.04 |
| 10 | 9358638 | rs1150014 | 0.716 | 0.462 | 1.033 | 0.253 | -0.023 | 0.019 | 0.019 | 0 | 0.73 | 0.52 | 0.82 | 0.69 | 0.81 | 0.67 |
| 10 | 9360038 | rs1150018 | 0.166 | 0.407 | 0.644 | 0.292 | -0.006 | 0.015 | 0.015 | 0 | 0.78 | 0.76 | 0.86 | 0.63 | 0.82 | 0.65 |
| 10 | 9360823 | rs1150023 | 0.974 | 0.499 | 1.031 | 0.244 | -0.005 | 0.024 | 0.024 | 0 | 0.67 | 0.48 | 0.82 | 0.57 | 0.72 | 0.62 |
| 10 | 9368879 | rs1150034 | 1.051 | -0.371 | 0.335 | -0.488 | -0.023 | -0.048 | -0.048 | 0.331 | 0.42 | 0.16 | 0.57 | 0.49 | 0.57 | 0.58 |
| 10 | 9369313 | rs1150036 | 1.055 | -0.378 | 0.308 | -0.491 | -0.04 | -0.071 | -0.071 | 0 | 0.42 | 0.11 | 0.57 | 0.49 | 0.57 | 0.58 |
| 10 | 9371159 | rs1150038 | 2.444 | -0.185 | -0.408 | -0.867 | -0.058 | -0.117 | -0.117 | 0 | 0.15 | 0.01 | 0.17 | 0.42 | 0.2 | 0.34 |
| 10 | 9372367 | rs1150039 | -0.705 | -0.119 | 0.274 | -0.546 | 0.002 | 0.029 | 0.029 | 0 | 0.94 | 0.87 | 0.93 | 0.93 | 0.92 | 0.88 |
| 10 | 9372924 | rs12412664 | 0.497 | -0.172 | 0.154 | -0.577 | -0.015 | -0.047 | -0.047 | 0 | 0.22 | 0.12 | 0.09 | 0.37 | 0.19 | 0.26 |
| 10 | 9373946 | rs12773435 | 0.5 | -0.168 | 0.157 | -0.647 | -0.015 | -0.049 | -0.049 | 1.184 | 0.22 | 0.12 | 0.09 | 0.4 | 0.19 | 0.26 |
| 10 | 9377089 | rs1150045 | -0.623 | -0.136 | 0.695 | -0.765 | -0.035 | -0.046 | -0.046 | 0 | 0.64 | 0.44 | 0.68 | 0.87 | 0.76 | 0.84 |
| 10 | 9377513 | rs2147296 | 1.567 | -0.295 | 0.345 | -0.89 | 0.021 | 0.024 | 0.024 | 0.137 | 0.24 | 0.21 | 0.18 | 0.06 | 0.15 | 0.04 |
| 10 | 9381566 | rs7893180 | -0.628 | -0.15 | -0.028 | -0.876 | 0.037 | 0.034 | 0.034 | 0.815 | 0.51 | 0.85 | 0.4 | 0.44 | 0.36 | 0.37 |
| 10 | 9381652 | rs12784138 | -0.624 | -0.217 | -0.02 | -0.851 | 0.044 | 0.032 | 0.032 | 0.548 | 0.51 | 0.86 | 0.38 | 0.44 | 0.35 | 0.31 |
| 10 | 9383608 | rs11256160 | 2.973 | -0.44 | -0.181 | -0.994 | 0.032 | 0.084 | 0.084 | 0.171 | 0.14 | 0.02 | 0.05 | 0 | 0.1 | 0.01 |
| 10 | 9384033 | rs12775773 | 0.521 | -0.055 | 0.141 | -0.779 | -0.019 | -0.04 | -0.04 | 0 | 0.19 | 0.12 | 0.09 | 0.38 | 0.19 | 0.26 |
| 10 | 9384517 | rs10905540 | 0.412 | 0.083 | 0.005 | -0.726 | -0.058 | -0.085 | -0.085 | 0 | 0.44 | 0.07 | 0.57 | 0.5 | 0.59 | 0.63 |

| | | | | | | | | | | | | | | | | |
|---|---|---|---|---|---|---|---|---|---|---|---|---|---|---|---|---|
| 10 | 9394702 | rs1762782 | -0.734 | 0.318 | 0.422 | -0.935 | 0.036 | -0.061 | -0.061 | 0 | 0.22 | 0.51 | 0.23 | 0.13 | 0.26 | 0.24 |
| 10 | 9398565 | rs1747738 | -1.851 | -0.424 | 0.086 | -0.749 | -0.012 | -0.015 | -0.015 | 0.041 | 0.95 | 1 | 0.96 | 0.96 | 0.94 | 0.95 |
| 10 | 9399473 | rs1998596 | 0.033 | -0.378 | 0.141 | -0.689 | 0.082 | 0.092 | 0.092 | 0.619 | 0.57 | 0.09 | 0.62 | 0.77 | 0.71 | 0.75 |
| 10 | 9406724 | rs11256167 | -1.379 | -0.353 | 0.263 | -0.674 | 0.01 | 0.009 | 0.009 | 0.216 | 0.69 | 0.71 | 0.72 | 0.84 | 0.78 | 0.89 |
| 10 | 9409770 | rs1762757 | -1.37 | -0.346 | 0.247 | -0.672 | 0.01 | 0.01 | 0.01 | 0.267 | 0.69 | 0.71 | 0.72 | 0.84 | 0.78 | 0.88 |
| 10 | 9415189 | rs1932182 | 0.113 | -0.186 | 0.443 | -0.574 | 0 | 0 | 0 | 0 | 0.09 | 0.12 | 0.05 | 0.07 | 0.06 | 0.14 |
| 10 | 9419691 | rs2152221 | 0.651 | -0.179 | -0.166 | -0.592 | -0.019 | -0.032 | -0.032 | 0.452 | 0.87 | 0.39 | 0.9 | 0.93 | 0.94 | 0.85 |
| 10 | 9428807 | rs10905548 | -0.167 | -0.074 | -1.073 | -0.572 | 0.075 | 0.078 | 0.078 | 0.232 | 0.57 | 0.21 | 0.64 | 0.77 | 0.72 | 0.74 |
| 10 | 9429516 | rs11591404 | -1.125 | 0.682 | -0.917 | -0.354 | -0.002 | 0.003 | 0.003 | 0 | 0.14 | 0.15 | 0.27 | 0.07 | 0.13 | 0.26 |
| 10 | 9436362 | rs10905551 | 0.087 | 0.858 | -1.501 | -0.431 | -0.098 | -0.007 | -0.007 | 0 | 0.4 | 0.09 | 0.46 | 0.58 | 0.52 | 0.53 |
| 10 | 9440066 | rs10795712 | 0.02 | 0.804 | -1.085 | -0.454 | -0.05 | 0.015 | 0.015 | 1.601 | 0.41 | 0.19 | 0.48 | 0.58 | 0.53 | 0.53 |
| 10 | 9448569 | rs17146039 | -0.234 | 0.586 | -0.372 | -0.613 | -0.006 | -0.031 | -0.031 | 0 | 0.11 | 0.25 | 0.13 | 0.07 | 0.08 | 0.26 |
| 10 | 9459295 | rs12783041 | 0.618 | 0.455 | 0.131 | -0.596 | 0.013 | 0.035 | 0.035 | 0.726 | 0.44 | 0.28 | 0.36 | 0.36 | 0.39 | 0.21 |
| 10 | 9471974 | rs972701 | 0.169 | 0.677 | 0.12 | -0.594 | -0.036 | -0.004 | -0.004 | 0 | 0.39 | 0.19 | 0.38 | 0.57 | 0.5 | 0.48 |
| 10 | 9481178 | rs10795716 | 0.398 | 0.727 | -0.181 | -0.587 | -0.101 | -0.053 | -0.053 | 0 | 0.38 | 0.06 | 0.37 | 0.57 | 0.5 | 0.48 |
| 10 | 9481842 | rs10508394 | 0.32 | 0.762 | 0.046 | -0.59 | -0.036 | -0.009 | -0.009 | 0.333 | 0.38 | 0.15 | 0.37 | 0.56 | 0.5 | 0.48 |
| 10 | 9493837 | rs1572297 | -0.511 | 0.64 | 0.36 | -0.646 | 0.03 | -0.049 | -0.049 | 0 | 0.13 | 0.42 | 0.25 | 0.07 | 0.1 | 0.26 |
| 10 | 9511933 | rs915254 | -0.152 | 0.645 | -0.063 | -0.692 | 0.016 | 0.021 | 0.021 | 0 | 0.86 | 0.75 | 0.87 | 0.76 | 0.86 | 0.88 |
| 10 | 9516007 | rs10905565 | 2.286 | 0.417 | -0.245 | -0.943 | 0.009 | 0.085 | 0.085 | 7.677 | 0.2 | 0.08 | 0.14 | 0.03 | 0.16 | 0.02 |
| 10 | 9521776 | rs17515084 | 1.347 | 0.868 | -0.002 | -0.504 | 0.019 | 0.05 | 0.05 | 3.315 | 0.34 | 0.21 | 0.26 | 0.13 | 0.27 | 0.09 |
| 10 | 9533300 | rs4749872 | -0.235 | 1.007 | -0.558 | -0.382 | -0.098 | 0.011 | 0.011 | 2.03 | 0.44 | 0.08 | 0.45 | 0.61 | 0.55 | 0.65 |
| 10 | 9533566 | rs10430792 | -0.469 | 1.092 | -0.383 | -0.378 | 0.012 | 0.023 | 0.023 | 0 | 0.88 | 0.81 | 0.88 | 0.75 | 0.87 | 0.87 |
| 10 | 9540194 | rs17515236 | 1.358 | 1.518 | -0.71 | -0.361 | -0.042 | -0.006 | -0.006 | 0.249 | 0.07 | 0.01 | 0.08 | 0.06 | 0.15 | 0.05 |
| 10 | 9546541 | rs11528645 | 0.89 | 1.232 | -0.858 | -0.346 | -0.053 | -0.148 | -0.148 | 0.551 | 0.2 | 0.01 | 0.2 | 0.5 | 0.24 | 0.39 |
| 10 | 9550386 | rs7916168 | -0.912 | 0.881 | -0.389 | -0.328 | 0.037 | 0.085 | 0.085 | 2.653 | 0.56 | 0.27 | 0.56 | 0.87 | 0.67 | 0.75 |
| 10 | 9575095 | rs1324879 | -0.802 | 0.862 | -0.434 | -0.308 | 0.063 | 0.176 | 0.176 | 3.607 | 0.54 | 0.18 | 0.54 | 0.86 | 0.66 | 0.74 |
| 10 | 9577963 | rs2038813 | -1.248 | 0.625 | 0.249 | -0.261 | 0.008 | 0.006 | 0.006 | 0 | 0.2 | 0.53 | 0.24 | 0.24 | 0.15 | 0.28 |
| 10 | 9580829 | rs11817002 | -0.778 | 0.602 | 0.294 | -0.268 | 0.007 | -0.048 | -0.048 | 0.004 | 0.1 | 0.43 | 0.14 | 0 | 0.04 | 0.16 |
| 10 | 9590060 | rs1555732 | -0.35 | 0.803 | -1.089 | 0.002 | -0.049 | 0.011 | 0.011 | 0.774 | 0.46 | 0.1 | 0.5 | 0.57 | 0.58 | 0.59 |
| 10 | 9594872 | rs10905580 | -1.324 | 0.898 | -0.791 | -0.006 | -0.015 | -0.031 | -0.031 | 0.362 | 0.65 | 0.54 | 0.72 | 0.81 | 0.72 | 0.91 |
| 10 | 9601929 | rs7901214 | -1.2 | 1.238 | -0.754 | 0.302 | 0.009 | -0.006 | -0.006 | 0.107 | 0.62 | 0.53 | 0.64 | 0.71 | 0.55 | 0.88 |
| 10 | 9602307 | rs11256259 | 0.585 | 0.719 | -0.848 | 0.519 | -0.106 | -0.105 | -0.105 | 0 | 0.3 | 0.05 | 0.34 | 0.38 | 0.41 | 0.36 |
| 10 | 9604930 | rs1324882 | -0.817 | 0.748 | -0.568 | 0.101 | -0.011 | -0.009 | -0.009 | 1.077 | 0.31 | 0.37 | 0.37 | 0.34 | 0.26 | 0.51 |
| 10 | 9605649 | rs6602361 | -1.463 | 0.827 | -0.545 | 0.147 | -0.004 | -0.016 | -0.016 | 0 | 0.64 | 0.6 | 0.72 | 0.8 | 0.69 | 0.88 |
| 10 | 9610299 | rs4749880 | 0.444 | 0.92 | -1.239 | 0.255 | -0.058 | -0.093 | -0.093 | 0.121 | 0.33 | 0.14 | 0.35 | 0.45 | 0.43 | 0.37 |
| 10 | 9615467 | rs1324886 | 0.594 | 0.93 | -1.447 | 0.219 | -0.084 | -0.111 | -0.111 | 0 | 0.31 | 0.1 | 0.35 | 0.43 | 0.43 | 0.37 |
| 10 | 9617015 | rs10508393 | -0.242 | 0.872 | -1.425 | 0.129 | -0.016 | -0.082 | -0.082 | 0 | 0.19 | 0.29 | 0.24 | 0.02 | 0.16 | 0.29 |
| 10 | 9622626 | rs12769947 | -0.025 | 1.344 | -2.104 | 0.913 | 0.001 | -0.021 | -0.021 | 0 | 0.11 | 0.07 | 0.13 | 0.33 | 0.09 | 0.2 |
| 10 | 9622843 | rs17146207 | -0.53 | 1.563 | -1.9 | 0.776 | -0.079 | -0.097 | -0.097 | 0.454 | 0.34 | 0.12 | 0.38 | 0.43 | 0.46 | 0.38 |
| 10 | 9637947 | rs12249391 | -1.113 | 1.183 | -1.162 | 0.587 | 0.001 | -0.01 | -0.01 | 0.097 | 0.08 | 0.09 | 0.06 | 0.03 | 0.09 | 0.08 |
| 10 | 9644800 | rs4749884 | 0.267 | 1.986 | -2.18 | 0.889 | -0.131 | -0.176 | -0.176 | 0 | 0.7 | 0.97 | 0.58 | 0.54 | 0.61 | 0.54 |
| 10 | 9651244 | rs11592618 | 0.523 | 1.065 | -2.354 | -0.421 | 0.019 | 0.037 | 0.037 | 2.962 | 0.34 | 0.28 | 0.23 | 0.08 | 0.27 | 0.1 |
| 10 | 9652064 | rs7918989 | 0.601 | 0.747 | -2.524 | 0.12 | 0.022 | -0.074 | -0.074 | 0.264 | 0.19 | 0.08 | 0.17 | 0.46 | 0.17 | 0.34 |
| 10 | 9655895 | rs9651533 | 1.198 | 0.604 | -2.605 | -0.24 | 0.011 | 0 | 0 | 0 | 0.31 | 0.3 | 0.2 | 0.08 | 0.23 | 0.1 |
| 10 | 9670431 | rs7923998 | 0.458 | 0.877 | -2.112 | -0.451 | 0.003 | -0.007 | -0.007 | 0 | 0.44 | 0.46 | 0.34 | 0.18 | 0.35 | 0.18 |
| 10 | 9677810 | rs10905597 | 1.679 | 0.838 | -2.086 | -0.471 | 0.021 | 0.032 | 0.032 | 2.497 | 0.31 | 0.26 | 0.2 | 0.08 | 0.23 | 0.09 |
| 10 | 9685395 | rs10752170 | -0.797 | 1.817 | -2.145 | -0.718 | 0.074 | -0.103 | -0.103 | 0 | 0.81 | 0.6 | 0.82 | 0.94 | 0.78 | 0.91 |
| 10 | 9697754 | rs768365 | -1.158 | 1.786 | -2.138 | -0.77 | 0.003 | 0.088 | 0.088 | 1.32 | 0.47 | 0.32 | 0.6 | 0.87 | 0.52 | 0.78 |
| 10 | 9710675 | rs12570090 | 0.633 | 2.382 | -2.24 | -0.771 | 0.011 | 0.034 | 0.034 | 1.23 | 0.5 | 0.47 | 0.4 | 0.18 | 0.46 | 0.22 |
| 10 | 9717323 | rs960955 | -0.033 | 2.412 | -2.177 | -0.712 | -0.007 | -0.005 | -0.005 | 0 | 0.13 | 0.2 | 0.14 | 0.1 | 0.11 | 0.08 |
| 10 | 9720181 | rs12777207 | -0.804 | 2.389 | -2.18 | -0.71 | 0.003 | -0.007 | -0.007 | 0 | 0.18 | 0.2 | 0.19 | 0.1 | 0.22 | 0.11 |
| 10 | 9736439 | rs2027296 | -0.303 | 2.529 | -2.195 | -0.816 | 0.069 | -0.101 | -0.101 | 0 | 0.81 | 0.63 | 0.81 | 0.94 | 0.78 | 0.91 |
| 10 | 9749448 | rs7071857 | 0.729 | 2.336 | -2.252 | -0.651 | -0.032 | -0.07 | -0.07 | 0 | 0.11 | 0.01 | 0.23 | 0.28 | 0.13 | 0.33 |
| 10 | 9754068 | rs7088330 | 0.948 | 2.339 | -2.25 | -0.681 | -0.002 | -0.009 | -0.009 | 0 | 0.64 | 0.44 | 0.62 | 0.84 | 0.56 | 0.8 |
| 10 | 9776166 | rs10752172 | 1.566 | 2.478 | -2.286 | -0.668 | 0.033 | 0.039 | 0.039 | 0 | 0.51 | 0.36 | 0.39 | 0.56 | 0.44 | 0.45 |
| 10 | 9778195 | rs17146359 | 1.003 | 2.086 | -2.165 | -0.528 | -0.009 | 0.026 | 0.026 | 0 | 0.06 | 0.16 | 0.08 | 0.07 | 0.04 | 0.05 |
| 10 | 9790386 | rs12414761 | 1.46 | 2.119 | -2.257 | -0.501 | 0.014 | -0.049 | -0.049 | 1.395 | 0.18 | 0.09 | 0.17 | 0.48 | 0.16 | 0.33 |
| 10 | 9795653 | rs10905617 | 1.177 | 2.162 | -2.262 | -0.513 | 0 | -0.056 | -0.056 | 1.548 | 0.19 | 0.09 | 0.18 | 0.49 | 0.2 | 0.35 |
| 10 | 9801095 | rs3740219 | 0.342 | 2.181 | -2.306 | -0.556 | 0.015 | -0.062 | -0.062 | 0 | 0.25 | 0.09 | 0.23 | 0.5 | 0.23 | 0.37 |
| 10 | 9802634 | rs10795744 | 1.004 | 2.188 | -2.29 | -0.535 | 0.002 | -0.071 | -0.071 | 0 | 0.2 | 0.08 | 0.19 | 0.49 | 0.2 | 0.35 |
| 10 | 9804989 | rs913056 | 0.584 | 2.332 | -2.233 | -0.732 | 0.03 | -0.016 | -0.016 | 0.016 | 0.68 | 0.56 | 0.58 | 0.66 | 0.65 | 0.55 |
| 10 | 9807832 | rs10508388 | 0.61 | 2.267 | -2.251 | -0.633 | -0.007 | -0.059 | -0.059 | 0 | 0.1 | 0 | 0.21 | 0.26 | 0.1 | 0.32 |
| 10 | 9815411 | rs1926226 | -0.098 | 2.237 | -2.202 | -0.643 | 0.001 | -0.001 | -0.001 | 0 | 0.31 | 0.27 | 0.32 | 0.58 | 0.3 | 0.42 |
| 10 | 9823594 | rs11256338 | 1.941 | 2.233 | -2.259 | -0.696 | 0.015 | 0.063 | 0.063 | 1.283 | 0.34 | 0.27 | 0.25 | 0.08 | 0.31 | 0.11 |
| 10 | 9826653 | rs11256339 | -0.612 | 2.339 | -1.389 | -0.417 | -0.003 | -0.001 | -0.001 | 0 | 0.22 | 0.17 | 0.31 | 0.26 | 0.22 | 0.35 |
| 10 | 9830756 | rs2397779 | -0.255 | 2.065 | -1.437 | -0.26 | -0.008 | -0.034 | -0.034 | 0 | 0.11 | 0.05 | 0.23 | 0.25 | 0.12 | 0.32 |
| 10 | 9836771 | rs17441872 | -0.984 | 2.075 | -1.338 | -0.327 | 0 | -0.02 | -0.02 | 0 | 0.16 | 0.2 | 0.16 | 0.05 | 0.16 | 0.12 |
| 10 | 9844964 | rs7068776 | -0.451 | 2.332 | -0.912 | -0.412 | -0.013 | 0.013 | 0.013 | 0 | 0.83 | 0.88 | 0.79 | 0.93 | 0.81 | 0.9 |
| 10 | 9848073 | rs11256347 | 0.781 | 2.218 | -1.196 | -0.422 | -0.043 | -0.075 | -0.075 | 0 | 0.15 | 0.01 | 0.24 | 0.26 | 0.19 | 0.31 |

| | | | | | | | | | | | | | | | | |
|---|---|---|---|---|---|---|---|---|---|---|---|---|---|---|---|---|
| 10 | 9852429 | rs7081045 | -0.687 | 2.172 | -0.998 | -0.461 | -0.001 | -0.017 | -0.017 | 0.319 | 0.16 | 0.19 | 0.16 | 0.05 | 0.15 | 0.14 |
| 10 | 9863986 | rs10905625 | 1.923 | 1.062 | -1.543 | -1.244 | -0.055 | -0.016 | -0.016 | 0.816 | 0.37 | 0.09 | 0.31 | 0.62 | 0.38 | 0.4 |
| 10 | 9870680 | rs7910576 | 0.699 | 0.685 | -1.064 | -1.402 | -0.003 | -0.006 | -0.006 | 1.218 | 0.2 | 0.16 | 0.28 | 0.26 | 0.24 | 0.31 |
| 10 | 9870792 | rs11256351 | 2.738 | 0.465 | -1.106 | -0.937 | -0.047 | -0.067 | -0.067 | 0.162 | 0.35 | 0.09 | 0.3 | 0.57 | 0.37 | 0.4 |
| 10 | 9870869 | rs17647577 | 1.325 | 0.444 | -1.056 | -0.914 | -0.018 | 0.017 | 0.017 | 0 | 0.07 | 0.01 | 0.06 | 0.02 | 0.07 | 0.06 |
| 10 | 9891498 | rs7080250 | 0.633 | 0.38 | -0.756 | -0.95 | 0.009 | 0.005 | 0.005 | 0 | 0.62 | 0.65 | 0.57 | 0.64 | 0.62 | 0.56 |
| 10 | 9910342 | rs2153297 | -0.923 | 0.39 | -0.552 | -0.986 | -0.005 | 0.002 | 0.002 | 0 | 0.33 | 0.3 | 0.37 | 0.13 | 0.34 | 0.23 |
| 10 | 9917445 | rs10795749 | 0.993 | 0.377 | -0.549 | -0.986 | -0.008 | 0.117 | 0.117 | 0.28 | 0.57 | 0.25 | 0.57 | 0.82 | 0.61 | 0.74 |
| 10 | 9931024 | rs1538608 | -0.882 | 0.048 | -0.623 | -0.896 | -0.026 | -0.028 | -0.028 | 0.056 | 0.85 | 0.72 | 0.93 | 0.95 | 0.9 | 0.92 |
| 10 | 9931856 | rs10491043 | 1.544 | 0.149 | -0.906 | -0.842 | -0.035 | -0.06 | -0.06 | 0.031 | 0.15 | 0.01 | 0.22 | 0.24 | 0.17 | 0.29 |
| 10 | 9933991 | rs2050778 | -1.685 | 0.235 | -0.742 | -0.804 | -0.02 | 0 | 0 | 0 | 0.92 | 0.86 | 0.96 | 0.95 | 0.96 | 0.97 |
| 10 | 9937859 | rs10905633 | 2.43 | 0.436 | -0.827 | -0.174 | -0.045 | -0.099 | -0.099 | 0 | 0.35 | 0.1 | 0.31 | 0.49 | 0.39 | 0.39 |
| 10 | 9938980 | rs4142060 | 0.508 | 0.431 | -0.615 | -0.208 | 0.004 | -0.044 | -0.044 | 0 | 0.63 | 0.51 | 0.63 | 0.83 | 0.64 | 0.77 |
| 10 | 9943083 | rs7095367 | 0.241 | 0.526 | -0.574 | -0.137 | -0.005 | 0.049 | 0.049 | 0.376 | 0.25 | 0.38 | 0.31 | 0.34 | 0.23 | 0.35 |
| 10 | 9944159 | rs4749908 | 2.228 | 0.445 | -0.845 | -0.132 | -0.031 | -0.079 | -0.079 | 0 | 0.36 | 0.14 | 0.31 | 0.48 | 0.39 | 0.39 |
| 10 | 9944810 | rs10795751 | 1.846 | 0.47 | -0.816 | -0.08 | -0.03 | -0.065 | -0.065 | 0 | 0.39 | 0.15 | 0.33 | 0.48 | 0.41 | 0.42 |
| 10 | 9951762 | rs1339678 | 0.504 | 0.416 | -0.495 | -0.265 | -0.003 | 0.012 | 0.012 | 0.768 | 0.53 | 0.59 | 0.46 | 0.61 | 0.55 | 0.51 |
| 10 | 9963634 | rs10508381 | 0.417 | 0.395 | -0.434 | -0.196 | -0.006 | 0.011 | 0.011 | 0.197 | 0.23 | 0.29 | 0.28 | 0.27 | 0.19 | 0.36 |
| 10 | 9963835 | rs7090089 | 0.81 | -0.138 | -0.54 | -0.536 | 0.038 | 0.033 | 0.033 | 0 | 0.54 | 0.39 | 0.55 | 0.67 | 0.65 | 0.45 |
| 10 | 9963856 | rs4749918 | 0.372 | -0.135 | -0.622 | -0.708 | -0.005 | 0.03 | 0.03 | 0 | 0.69 | 0.74 | 0.72 | 0.87 | 0.71 | 0.81 |
| 10 | 9963973 | rs17146734 | 0.846 | -1.51 | -0.459 | -0.554 | -0.01 | -0.046 | -0.046 | 0 | 0.09 | 0.18 | 0.04 | 0 | 0.06 | 0.08 |
| 10 | 9964238 | rs7902229 | 1.279 | -1.893 | -1.499 | -0.266 | -0.104 | -0.059 | -0.059 | 2.609 | 0.39 | 0.11 | 0.39 | 0.39 | 0.4 | 0.37 |
| 10 | 9964251 | rs4747842 | 0.805 | -2.18 | -0.531 | -0.161 | -0.009 | 0.063 | 0.063 | 0.02 | 0.09 | 0.22 | 0.08 | 0.23 | 0.12 | 0.04 |
| 10 | 9964451 | rs4747843 | 0.304 | -2.007 | -0.686 | -0.526 | 0 | 0.011 | 0.011 | 0 | 0.24 | 0.21 | 0.18 | 0.24 | 0.3 | 0.21 |
| 10 | 9964571 | rs1934357 | -0.977 | -1.907 | -0.249 | -0.57 | -0.012 | 0.012 | 0.012 | 0.057 | 0.92 | 0.86 | 0.96 | 0.92 | 0.94 | 0.97 |
| 10 | 9964708 | rs1934358 | 0.625 | -1.769 | -0.374 | -0.716 | 0.001 | -0.031 | -0.031 | 0 | 0.23 | 0.09 | 0.33 | 0.3 | 0.21 | 0.42 |
| 10 | 9967519 | rs17146745 | 0.318 | -1.49 | 0.18 | -0.544 | -0.039 | -0.032 | -0.032 | 0 | 0.35 | 0.23 | 0.41 | 0.37 | 0.43 | 0.29 |
| 10 | 9971350 | rs1339693 | 1.103 | -1.159 | 0.524 | -0.399 | 0.003 | 0.013 | 0.013 | 0 | 0.07 | 0.02 | 0.04 | 0.02 | 0.06 | 0.08 |
| 10 | 9977037 | rs17146795 | 0.662 | -1.166 | 0.547 | -0.402 | -0.007 | -0.02 | -0.02 | 0 | 0.08 | 0.15 | 0.05 | 0.02 | 0.06 | 0.07 |
| 10 | 9987130 | rs10218944 | -0.45 | -1.218 | 0.849 | -0.457 | -0.013 | -0.061 | -0.061 | 0 | 0.13 | 0.36 | 0.1 | 0.02 | 0.1 | 0.08 |
| 10 | 9988716 | rs871158 | -0.787 | -1.478 | 0.684 | -0.567 | -0.009 | -0.04 | -0.04 | 0 | 0.56 | 0.84 | 0.52 | 0.38 | 0.55 | 0.42 |
| 10 | 10010389 | rs12262933 | 0.212 | -1.576 | 1.441 | -0.66 | -0.019 | -0.079 | -0.079 | 0 | 0.13 | 0.27 | 0.06 | 0 | 0.08 | 0.12 |
| 10 | 10020381 | rs7915729 | 0.039 | -1.618 | 1.336 | -0.646 | 0.002 | 0.001 | 0.001 | 0 | 0.42 | 0.46 | 0.51 | 0.41 | 0.48 | 0.32 |
| 10 | 10021054 | rs11499250 | 0.531 | -1.637 | 1.269 | -0.665 | -0.009 | -0.028 | -0.028 | 0 | 0.11 | 0.15 | 0.07 | 0.02 | 0.09 | 0.1 |
| 10 | 10021863 | rs1591890 | 0.17 | -1.641 | 1.247 | -0.662 | -0.017 | -0.013 | -0.013 | 0.251 | 0.41 | 0.33 | 0.5 | 0.41 | 0.48 | 0.32 |
| 10 | 10034211 | rs10508379 | -0.91 | -1.622 | 0.731 | -0.656 | -0.006 | -0.028 | -0.028 | 0.191 | 0.56 | 0.82 | 0.58 | 0.38 | 0.55 | 0.44 |
| 10 | 10034807 | rs6602387 | -0.532 | -1.617 | 0.729 | -0.641 | 0.012 | 0.017 | 0.017 | 0.672 | 0.49 | 0.69 | 0.54 | 0.37 | 0.52 | 0.37 |
| 10 | 10035760 | rs7071402 | 0.928 | -1.127 | 0.476 | -0.369 | -0.105 | -0.102 | -0.102 | 0 | 0.29 | 0.09 | 0.42 | 0.36 | 0.38 | 0.22 |
| 10 | 10042825 | rs7093588 | -0.912 | -1.337 | 0.304 | -0.461 | 0 | -0.001 | -0.001 | 0 | 0.52 | 0.83 | 0.56 | 0.39 | 0.53 | 0.36 |
| 10 | 10043822 | rs11256432 | -0.347 | -1.388 | 0.367 | -0.461 | -0.01 | -0.002 | -0.002 | 0.875 | 0.45 | 0.67 | 0.51 | 0.39 | 0.49 | 0.29 |
| 10 | 10045632 | rs1317706 | -0.398 | -1.371 | 0.465 | -0.458 | 0.043 | 0.01 | 0.01 | 0.084 | 0.07 | 0.4 | 0.06 | 0 | 0.04 | 0.01 |
| 10 | 10045642 | rs12570537 | 1.135 | -1.325 | -0.214 | -0.44 | -0.162 | -0.154 | -0.154 | 0 | 0.3 | 0.05 | 0.44 | 0.39 | 0.41 | 0.22 |
| 10 | 10049589 | rs17147020 | 1.092 | -1.354 | -0.217 | -0.277 | 0.028 | -0.078 | -0.078 | 0 | 0.13 | 0.01 | 0.1 | 0.29 | 0.09 | 0.16 |
| 10 | 10053013 | rs7095279 | -0.933 | -0.277 | 0.567 | -0.388 | 0 | -0.002 | -0.002 | 0.88 | 0.47 | 0.52 | 0.51 | 0.4 | 0.49 | 0.33 |
| 10 | 10061134 | rs11256450 | -1.227 | -0.198 | 0.007 | -0.353 | 0.006 | 0.009 | 0.009 | 1.122 | 0.52 | 0.65 | 0.54 | 0.4 | 0.53 | 0.36 |
| 10 | 10077160 | rs12240851 | -0.623 | -0.311 | 0.005 | -0.476 | 0.013 | -0.024 | -0.024 | 0 | 0.07 | 0.32 | 0.06 | 0 | 0.05 | 0.01 |
| 10 | 10079950 | rs7905144 | -0.487 | 0.086 | -0.18 | -0.291 | 0.002 | 0 | 0 | 0 | 0.41 | 0.57 | 0.49 | 0.38 | 0.46 | 0.26 |
| 10 | 10081005 | rs6602395 | -1.009 | 0.416 | -1.459 | -0.052 | 0.009 | 0.012 | 0.012 | 0.574 | 0.49 | 0.63 | 0.51 | 0.39 | 0.52 | 0.33 |
| 10 | 10081300 | rs7076367 | -0.307 | 1.101 | -2.215 | 0.736 | -0.164 | -0.147 | -0.147 | 2.871 | 0.31 | 0.04 | 0.4 | 0.39 | 0.4 | 0.18 |
| 10 | 10083583 | rs2762638 | 0.929 | 0.582 | -1.294 | 0.356 | -0.013 | -0.009 | -0.009 | 0 | 0.89 | 0.94 | 0.7 | 0.89 | 0.9 | 0.87 |
| 10 | 10085164 | rs2657506 | 0.937 | 0.529 | -1.644 | -0.091 | -0.064 | 0.107 | 0.107 | 0 | 0.83 | 0.99 | 0.91 | 1 | 0.83 | 0.85 |
| 10 | 10085984 | rs2657530 | 1.275 | 0.211 | -0.442 | 0.082 | 0.006 | -0.03 | -0.03 | 0 | 0.75 | 0.51 | 0.74 | 0.82 | 0.76 | 0.78 |
| 10 | 10087031 | rs1540956 | 0.913 | 0.476 | -0.029 | -0.086 | 0.003 | -0.022 | -0.022 | 0.02 | 0.92 | 0.87 | 0.95 | 1 | 0.9 | 0.94 |
| 10 | 10092440 | rs11256465 | 0.579 | 0.227 | -1.075 | -0.035 | -0.043 | -0.111 | -0.111 | 0 | 0.32 | 0.09 | 0.22 | 0.45 | 0.35 | 0.26 |
| 10 | 10094837 | rs2762637 | -0.571 | -0.324 | -1.075 | -0.186 | -0.024 | 0.089 | 0.089 | 0 | 0.55 | 0.79 | 0.45 | 0.7 | 0.53 | 0.51 |
| 10 | 10098119 | rs11256467 | 0.845 | -0.69 | -1.28 | -0.32 | 0.014 | 0.032 | 0.032 | 0 | 0.08 | 0.04 | 0.07 | 0.45 | 0.06 | 0.14 |
| 10 | 10098760 | rs17147166 | 0.37 | -0.712 | -0.872 | -0.303 | 0.021 | 0.021 | 0.021 | 0 | 0.06 | 0.15 | 0.04 | 0.08 | 0.08 | 0.05 |
| 10 | 10100700 | rs2025470 | -0.491 | -0.894 | -1.075 | -0.36 | -0.024 | -0.009 | -0.009 | 0.06 | 0.9 | 0.96 | 0.75 | 0.92 | 0.88 | 0.92 |
| 10 | 10101344 | rs11256470 | -1.175 | -0.612 | -0.762 | -0.212 | -0.003 | 0.002 | 0.002 | 0.045 | 0.24 | 0.66 | 0.23 | 0.24 | 0.18 | 0.28 |
| 10 | 10103503 | rs2657521 | -0.125 | -0.617 | -0.338 | -0.232 | -0.024 | -0.085 | -0.085 | 1.578 | 0.14 | 0.36 | 0.07 | 0 | 0.07 | 0.19 |
| 10 | 10104492 | rs2804070 | -0.343 | -0.672 | -0.358 | -0.265 | 0.006 | 0 | 0 | 0 | 0.47 | 0.49 | 0.3 | 0.46 | 0.43 | 0.48 |
| 10 | 10108651 | rs2209293 | 0.456 | -0.681 | -0.298 | -0.251 | 0.001 | -0.01 | -0.01 | 0 | 0.36 | 0.39 | 0.24 | 0.46 | 0.35 | 0.28 |
| 10 | 10108933 | rs6602399 | -1.369 | -0.614 | -0.262 | -0.258 | -0.003 | 0.003 | 0.003 | 0.052 | 0.23 | 0.65 | 0.23 | 0.24 | 0.18 | 0.27 |
| 10 | 10117897 | rs715814 | -0.451 | -0.541 | -0.301 | -0.214 | 0.125 | 0.303 | 0.303 | 0 | 0.08 | 0.33 | 0.16 | 0.23 | 0.11 | 0.09 |
| 10 | 10134319 | rs2657513 | 1.225 | -0.347 | -0.873 | -1.023 | -0.005 | 0.012 | 0.012 | 0 | 0.07 | 0.04 | 0.04 | 0 | 0.09 | 0.06 |
| 10 | 10136473 | rs12356435 | 0.939 | -0.275 | -0.662 | -0.743 | -0.107 | 0.165 | 0.165 | 0 | 0.26 | 0.05 | 0.19 | 0.01 | 0.34 | 0.26 |
| 10 | 10136486 | rs7089778 | -0.709 | -0.18 | -0.605 | -0.59 | 0.008 | 0.018 | 0.018 | 0 | 0.5 | 0.53 | 0.41 | 0.6 | 0.44 | 0.5 |
| 10 | 10136768 | rs7090158 | -0.864 | -0.547 | -0.98 | -0.349 | -0.031 | 0.04 | 0.04 | 0 | 0.24 | 0.22 | 0.24 | 0.6 | 0.15 | 0.37 |
| 10 | 10139363 | rs7073292 | 0.716 | -0.389 | -0.719 | -0.08 | -0.035 | 0.079 | 0.079 | 0 | 0.37 | 0.08 | 0.3 | 0.24 | 0.38 | 0.41 |

| | | | | | | | | | | | | | | | | |
|---|---|---|---|---|---|---|---|---|---|---|---|---|---|---|---|---|
| 10 | 10144990 | rs10905663 | -0.369 | -0.309 | -0.162 | 0.46 | -0.036 | 0.196 | 0.196 | 0 | 0.13 | 0.23 | 0.19 | 0.6 | 0.08 | 0.18 |
| 10 | 10155884 | rs1012131 | -0.517 | 0.16 | -1.098 | 0.205 | -0.028 | 0.087 | 0.087 | 0 | 0.56 | 0.8 | 0.45 | 0.69 | 0.54 | 0.59 |
| 10 | 10158656 | rs4749929 | -0.641 | -0.624 | -0.476 | 0.431 | -0.039 | 0.016 | 0.016 | 0 | 0.31 | 0.49 | 0.27 | 0.59 | 0.21 | 0.41 |
| 10 | 10160846 | rs4747847 | -1.523 | -0.117 | 0.417 | 1.001 | 0.069 | 0.192 | 0.192 | 0.025 | 0.1 | 0.36 | 0.09 | 0.24 | 0.07 | 0.11 |
| 10 | 10164309 | rs2762659 | -1.435 | -0.639 | 0.046 | 1.432 | 0.036 | -0.063 | -0.063 | 0 | 0.35 | 0.76 | 0.21 | 0.51 | 0.22 | 0.34 |
| 10 | 10164807 | rs4747849 | -0.72 | -0.583 | 0.338 | 1.483 | -0.005 | 0.065 | 0.065 | 0.189 | 0.07 | 0.17 | 0.06 | 0.25 | 0.04 | 0.11 |
| 10 | 10171642 | rs991956 | -0.759 | -0.597 | 0.465 | 1.356 | -0.036 | 0.014 | 0.014 | 0 | 0.24 | 0.29 | 0.1 | 0.27 | 0.15 | 0.23 |
| 10 | 10175411 | rs10490998 | -0.755 | -0.604 | 0.646 | 1.354 | -0.034 | 0.014 | 0.014 | 0 | 0.24 | 0.29 | 0.11 | 0.27 | 0.16 | 0.23 |
| 10 | 10179752 | rs7902516 | 0.135 | -0.133 | -0.661 | 1.647 | 0.014 | -0.058 | -0.058 | 2.744 | 0.73 | 0.88 | 0.73 | 0.61 | 0.8 | 0.77 |
| 10 | 10185315 | rs7090598 | 0.124 | -0.201 | -0.776 | 1.458 | 0.028 | -0.018 | -0.018 | 0 | 0.13 | 0.05 | 0.05 | 0.14 | 0.07 | 0.12 |
| 10 | 10188456 | rs11813494 | 0.043 | -0.731 | -1.104 | 0.985 | 0.023 | 0.022 | 0.022 | 0 | 0.33 | 0.24 | 0.16 | 0.19 | 0.24 | 0.26 |
| 10 | 10188827 | rs4611106 | 0.123 | 0.545 | -0.801 | 0.871 | 0.022 | 0.001 | 0.001 | 0 | 0.26 | 0.59 | 0.41 | 0.27 | 0.33 | 0.4 |
| 10 | 10189087 | rs4272705 | 0.047 | 0.682 | -0.753 | 0.818 | 0.007 | 0.006 | 0.006 | 0 | 0.55 | 0.66 | 0.72 | 0.66 | 0.62 | 0.58 |
| 10 | 10189194 | rs4622161 | 0.042 | 0.635 | -0.756 | 0.839 | 0.01 | 0.007 | 0.007 | 0 | 0.55 | 0.67 | 0.72 | 0.67 | 0.62 | 0.58 |
| 10 | 10190167 | rs6422406 | -1.405 | 1.048 | -0.701 | 0.777 | -0.011 | -0.03 | -0.03 | 0.42 | 0.73 | 0.79 | 0.7 | 0.61 | 0.73 | 0.82 |
| 10 | 10190595 | rs10905670 | 0.359 | 0.866 | -0.506 | 0.788 | 0.043 | -0.004 | -0.004 | 0 | 0.3 | 0.45 | 0.43 | 0.25 | 0.37 | 0.42 |
| 10 | 10190692 | rs11814585 | 0.164 | 0.434 | -0.541 | 0.631 | -0.027 | 0.036 | 0.036 | 0 | 0.24 | 0.26 | 0.16 | 0.36 | 0.16 | 0.28 |
| 10 | 10192591 | rs17147365 | 0.108 | 0.433 | -0.57 | 0.28 | -0.039 | -0.005 | -0.005 | 0 | 0.25 | 0.32 | 0.16 | 0.27 | 0.16 | 0.28 |
| 10 | 10194028 | rs927507 | 1.72 | 0.405 | -0.62 | 0.27 | 0.006 | -0.064 | -0.064 | 0.032 | 0.23 | 0.2 | 0.28 | 0.48 | 0.24 | 0.18 |
| 10 | 10194617 | rs7899795 | 1.707 | 0.406 | -0.619 | 0.27 | 0.006 | -0.062 | -0.062 | 2.498 | 0.24 | 0.2 | 0.28 | 0.48 | 0.24 | 0.18 |
| 10 | 10194737 | rs7903255 | 1.026 | 0.364 | -0.603 | 0.251 | -0.026 | -0.05 | -0.05 | 0 | 0.17 | 0.17 | 0.09 | 0.06 | 0.09 | 0.11 |
| 10 | 10195010 | rs9423818 | 0.707 | 0.549 | -0.538 | 0.309 | 0.011 | 0.046 | 0.046 | 1.117 | 0.1 | 0.16 | 0.12 | 0.21 | 0.1 | 0.18 |
| 10 | 10204589 | rs7097271 | -0.426 | 1.107 | -0.669 | 0.318 | -0.006 | 0.005 | 0.005 | 0.63 | 0.37 | 0.28 | 0.26 | 0.29 | 0.41 | 0.37 |
| 10 | 10216373 | rs10905673 | -0.001 | 1.004 | -0.755 | 0.021 | -0.067 | -0.005 | -0.005 | 0 | 0.11 | 0.03 | 0.07 | 0.09 | 0.2 | 0.09 |
| 10 | 10218843 | rs2759632 | 1.189 | 0.903 | -0.283 | 0.042 | 0.016 | -0.013 | -0.013 | 0 | 0.23 | 0.39 | 0.2 | 0.18 | 0.25 | 0.35 |
| 10 | 10228684 | rs7083994 | -0.015 | 0.517 | -0.28 | 0.081 | -0.044 | -0.024 | -0.024 | 0 | 0.25 | 0.38 | 0.24 | 0.23 | 0.16 | 0.26 |
| 10 | 10230151 | rs2819042 | -1.196 | 0.634 | -0.355 | 0.027 | -0.05 | -0.039 | -0.039 | 0 | 0.68 | 0.82 | 0.56 | 0.41 | 0.59 | 0.74 |
| 10 | 10230394 | rs2819041 | 1.279 | 0.605 | -0.612 | 0.017 | 0.023 | -0.014 | -0.014 | 0 | 0.23 | 0.4 | 0.21 | 0.17 | 0.26 | 0.26 |
| 10 | 10232363 | rs2819039 | -1.03 | 0.51 | -0.719 | 0.186 | -0.05 | -0.035 | -0.035 | 0.006 | 0.68 | 0.82 | 0.56 | 0.41 | 0.59 | 0.74 |
| 10 | 10234643 | rs2759635 | -2.048 | 0.594 | -0.848 | 0.185 | 0.01 | -0.059 | -0.059 | 0 | 0.78 | 0.86 | 0.63 | 0.49 | 0.78 | 0.83 |
| 10 | 10237192 | rs2819035 | -0.473 | 0.574 | -0.552 | 0.118 | 0.012 | -0.065 | -0.065 | 0 | 0.28 | 0.51 | 0.17 | 0.15 | 0.28 | 0.17 |
| 10 | 10237641 | rs2025276 | 0.637 | 0.508 | -1.054 | 0.094 | -0.006 | 0.084 | 0.084 | 0 | 0.41 | 0.28 | 0.31 | 0.15 | 0.43 | 0.37 |
| 10 | 10239143 | rs2025277 | -1.536 | -0.173 | -1.306 | 0.512 | 0.01 | -0.06 | -0.06 | 0 | 0.78 | 0.85 | 0.63 | 0.49 | 0.78 | 0.83 |
| 10 | 10239925 | rs7089164 | 0.27 | -0.071 | -0.795 | 0.272 | 0.026 | 0.029 | 0.029 | 0 | 0.15 | 0.08 | 0.07 | 0.06 | 0.08 | 0.08 |
| 10 | 10240837 | rs7902057 | -0.377 | 0.698 | -0.66 | 0.643 | 0 | -0.001 | -0.001 | 0.786 | 0.41 | 0.54 | 0.41 | 0.58 | 0.36 | 0.49 |
| 10 | 10240975 | rs1570411 | -0.82 | 0.866 | -0.821 | 0.499 | 0 | -0.023 | -0.023 | 0 | 0.58 | 0.55 | 0.6 | 0.8 | 0.51 | 0.72 |
| 10 | 10248165 | rs4749936 | -0.619 | 1.117 | -1.025 | 1.254 | -0.024 | -0.047 | -0.047 | 0 | 0.22 | 0.17 | 0.38 | 0.42 | 0.25 | 0.32 |
| 10 | 10249897 | rs11256530 | -0.956 | 0.318 | -0.453 | 0.966 | -0.018 | 0.026 | 0.026 | 0 | 0.21 | 0.31 | 0.14 | 0.31 | 0.16 | 0.09 |
| 10 | 10251910 | rs1885740 | -0.683 | -0.283 | -0.901 | 0.19 | 0.025 | 0.065 | 0.065 | 0 | 0.32 | 0.17 | 0.19 | 0.14 | 0.27 | 0.22 |
| 10 | 10254086 | rs7920248 | -0.519 | -0.405 | -0.576 | 0.145 | 0.022 | 0.032 | 0.032 | 0 | 0.28 | 0.22 | 0.17 | 0.13 | 0.21 | 0.2 |
| 10 | 10256332 | rs2210624 | -0.354 | -0.513 | -0.575 | -0.048 | 0.001 | 0.002 | 0.002 | 0 | 0.17 | 0.17 | 0.1 | 0.04 | 0.16 | 0.12 |
| 10 | 10260293 | rs2225655 | 0.219 | -0.716 | -0.17 | -0.174 | -0.025 | -0.056 | -0.056 | 0 | 0.85 | 0.67 | 0.91 | 0.98 | 0.91 | 0.91 |
| 10 | 10261530 | rs12257923 | -0.651 | -0.713 | -0.182 | -0.176 | 0.002 | 0.009 | 0.009 | 0 | 0.18 | 0.15 | 0.1 | 0.02 | 0.16 | 0.12 |
| 10 | 10263037 | rs7068043 | 0.194 | -0.239 | -0.278 | 0.009 | 0.005 | 0.024 | 0.024 | 0 | 0.32 | 0.36 | 0.51 | 0.51 | 0.36 | 0.38 |
| 10 | 10264651 | rs10905684 | 0.386 | -0.262 | -0.279 | -0.041 | 0.011 | 0.036 | 0.036 | 0.037 | 0.3 | 0.36 | 0.5 | 0.53 | 0.35 | 0.36 |
| 10 | 10270261 | rs7079940 | -0.047 | -0.371 | 0.002 | -0.195 | -0.028 | -0.042 | -0.042 | 0.425 | 0.85 | 0.67 | 0.91 | 0.96 | 0.91 | 0.91 |
| 10 | 10270642 | rs7915904 | -0.064 | -0.36 | -0.014 | -0.19 | -0.027 | -0.041 | -0.041 | 0 | 0.85 | 0.67 | 0.91 | 0.96 | 0.91 | 0.91 |
| 10 | 10270775 | rs7916069 | -0.064 | -0.36 | -0.014 | -0.166 | -0.027 | -0.035 | -0.035 | 0.409 | 0.85 | 0.67 | 0.91 | 0.96 | 0.91 | 0.9 |
| 10 | 10270836 | rs7917187 | -0.064 | -0.36 | -0.014 | -0.203 | -0.027 | -0.041 | -0.041 | 0 | 0.85 | 0.67 | 0.91 | 0.96 | 0.91 | 0.91 |
| 10 | 10277345 | rs17527069 | 0.288 | -0.356 | -0.531 | -0.393 | -0.026 | 0.041 | 0.041 | 0.022 | 0.23 | 0.02 | 0.19 | 0.1 | 0.19 | 0.25 |
| 10 | 10278847 | rs914987 | 0.521 | -0.143 | 0.313 | -0.013 | -0.043 | -0.064 | -0.064 | 0 | 0.82 | 0.52 | 0.88 | 0.96 | 0.89 | 0.89 |
| 10 | 10305616 | rs10905701 | 0.629 | -0.586 | 0.228 | -0.007 | -0.098 | -0.063 | -0.063 | 0 | 0.81 | 0.52 | 0.89 | 0.96 | 0.91 | 0.9 |
| 10 | 10310509 | rs4749944 | 0.028 | -0.574 | 0.434 | -0.019 | -0.054 | -0.03 | -0.03 | 0 | 0.87 | 0.67 | 0.93 | 0.96 | 0.94 | 0.93 |
| 10 | 10321874 | rs10905707 | 0.243 | 0.427 | 0.359 | 0.088 | 0 | 0.001 | 0.001 | 0 | 0.53 | 0.47 | 0.5 | 0.65 | 0.47 | 0.6 |
| 10 | 10324589 | rs17455682 | 0.086 | 1.167 | 0.269 | 0.076 | 0.003 | 0.079 | 0.079 | 0 | 0.34 | 0.21 | 0.25 | 0.09 | 0.35 | 0.26 |
| 10 | 10334623 | rs1889257 | -0.27 | 1.159 | 0.31 | 0.133 | -0.026 | 0.018 | 0.018 | 0 | 0.4 | 0.28 | 0.45 | 0.32 | 0.5 | 0.35 |
| 10 | 10342099 | rs7080874 | -0.17 | 1.041 | -0.337 | 0.186 | -0.044 | 0.106 | 0.106 | 0 | 0.63 | 0.84 | 0.67 | 0.85 | 0.55 | 0.66 |
| 10 | 10358556 | rs7917005 | -0.363 | 1.21 | -0.178 | 0.295 | -0.041 | -0.005 | -0.005 | 0.68 | 0.3 | 0.18 | 0.29 | 0.3 | 0.38 | 0.43 |
| 10 | 10364529 | rs4747852 | -1.013 | 0.883 | 0.097 | 0.176 | 0.003 | -0.003 | -0.003 | 0 | 0.42 | 0.46 | 0.37 | 0.34 | 0.47 | 0.52 |
| 10 | 10375562 | rs7905167 | -0.423 | 0.952 | 0.124 | 0.226 | -0.047 | -0.004 | -0.004 | 1.452 | 0.33 | 0.19 | 0.32 | 0.32 | 0.4 | 0.45 |
| 10 | 10385009 | rs7087371 | -0.147 | 0.949 | 0.122 | 0.044 | -0.071 | 0.122 | 0.122 | 0 | 0.3 | 0.07 | 0.22 | 0.08 | 0.37 | 0.37 |
| 10 | 10385660 | rs2398215 | -0.448 | 0.901 | 0.121 | 0.099 | -0.044 | 0.018 | 0.018 | 0 | 0.33 | 0.19 | 0.32 | 0.28 | 0.4 | 0.44 |
| 10 | 10395026 | rs7900897 | -1.549 | 0.671 | 0.687 | 0.706 | 0.002 | -0.017 | -0.017 | 0 | 0.06 | 0.24 | 0.03 | 0.01 | 0.04 | 0.03 |
| 10 | 10401012 | rs10905714 | -2.062 | 0.705 | 0.439 | 0.757 | 0.015 | 0.089 | 0.089 | 0.15 | 0.09 | 0.3 | 0.14 | 0.19 | 0.06 | 0.11 |
| 10 | 10401848 | rs1572119 | 1.378 | 0.659 | 0.59 | 0.802 | 0.066 | 0.222 | 0.222 | 0.798 | 0.92 | 0.55 | 0.92 | 0.8 | 0.97 | 0.92 |
| 10 | 10402541 | rs10905715 | -0.442 | 0.869 | -0.504 | 1.049 | -0.044 | -0.04 | -0.04 | 0.465 | 0.31 | 0.2 | 0.5 | 0.51 | 0.46 | 0.56 |
| 10 | 10403468 | rs7896031 | -0.329 | 0.174 | -0.589 | 1.126 | 0.003 | -0.004 | -0.004 | 0.165 | 0.1 | 0.09 | 0.15 | 0.19 | 0.06 | 0.07 |
| 10 | 10405808 | rs11256576 | 0.06 | 0.375 | -0.682 | 0.832 | -0.004 | 0.001 | 0.001 | 0.012 | 0.29 | 0.28 | 0.38 | 0.13 | 0.44 | 0.39 |
| 10 | 10408075 | rs10508403 | 0.419 | 0.254 | -0.801 | 1.302 | -0.001 | 0.025 | 0.025 | 0 | 0.07 | 0.11 | 0.16 | 0.22 | 0.09 | 0.2 |

| | | | | | | | | | | | | | | | | |
|---|---|---|---|---|---|---|---|---|---|---|---|---|---|---|---|---|
| 10 | 10409365 | rs3001719 | -1.425 | -0.441 | -1.109 | 0.455 | -0.001 | -0.008 | -0.008 | 0 | 0.67 | 0.66 | 0.78 | 0.74 | 0.68 | 0.83 |
| 10 | 10412208 | rs10795792 | 0.688 | -0.683 | -1.221 | 0.477 | 0.006 | -0.051 | -0.051 | 0 | 0.19 | 0.16 | 0.23 | 0.41 | 0.16 | 0.29 |
| 10 | 10412763 | rs10905717 | 0.802 | -0.379 | -1.027 | 0.605 | -0.006 | 0.007 | 0.007 | 0 | 0.1 | 0.12 | 0.15 | 0.2 | 0.06 | 0.08 |
| 10 | 10414194 | rs7098736 | -0.164 | -0.435 | -1.156 | 0.532 | 0.016 | -0.023 | -0.023 | 0 | 0.4 | 0.46 | 0.52 | 0.32 | 0.49 | 0.48 |
| 10 | 10416946 | rs2842139 | -0.774 | -0.099 | -1.606 | 0.13 | -0.009 | -0.021 | -0.021 | 0 | 0.63 | 0.57 | 0.78 | 0.73 | 0.69 | 0.82 |
| 10 | 10417509 | rs17147843 | 0.833 | 1.117 | -2.243 | 0.821 | -0.025 | -0.043 | -0.043 | 0 | 0.07 | 0.01 | 0.08 | 0.22 | 0.09 | 0.07 |
| 10 | 10423558 | rs11256583 | -0.248 | 1.204 | -1.873 | 0.271 | -0.074 | 0.055 | 0.055 | 0 | 0.27 | 0.11 | 0.41 | 0.13 | 0.44 | 0.35 |
| 10 | 10430827 | rs11256586 | -0.093 | 1.098 | -2.192 | 0.371 | -0.121 | -0.015 | -0.015 | 0.957 | 0.25 | 0.02 | 0.17 | 0.16 | 0.36 | 0.26 |
| 10 | 10436076 | rs2488166 | 0.552 | 0.233 | -2.979 | 0.628 | 0.024 | -0.087 | -0.087 | 0.176 | 0.26 | 0.09 | 0.22 | 0.38 | 0.19 | 0.25 |
| 10 | 10436135 | rs9424079 | 0.102 | -0.109 | -2.086 | 0.355 | -0.001 | -0.001 | -0.001 | 0 | 0.32 | 0.3 | 0.27 | 0.38 | 0.22 | 0.27 |
| 10 | 10444276 | rs1891399 | 0.259 | -0.271 | -1.647 | 0.393 | -0.001 | -0.001 | -0.001 | 0 | 0.43 | 0.59 | 0.5 | 0.42 | 0.42 | 0.5 |
| 10 | 10461038 | rs10905722 | 0.908 | -0.244 | -1.184 | 0.439 | 0.032 | 0.004 | 0.004 | 0.421 | 0.15 | 0.27 | 0.17 | 0.12 | 0.23 | 0.16 |
| 10 | 10463201 | rs7894041 | 0.195 | -0.636 | -1.278 | 0.479 | 0.023 | 0.003 | 0.003 | 0 | 0.33 | 0.46 | 0.47 | 0.35 | 0.39 | 0.36 |
| 10 | 10463425 | rs10905724 | 0.595 | -0.67 | -1.351 | 0.468 | 0 | -0.008 | -0.008 | 0.18 | 0.35 | 0.36 | 0.22 | 0.23 | 0.39 | 0.33 |
| 10 | 10467761 | rs2251525 | 0.783 | -0.776 | -1.607 | 0.521 | 0.01 | -0.113 | -0.113 | 0.695 | 0.28 | 0.04 | 0.21 | 0.39 | 0.19 | 0.25 |
| 10 | 10473907 | rs7895416 | -0.116 | -0.882 | -1.264 | 0.613 | 0 | 0 | 0 | 0 | 0.55 | 0.41 | 0.68 | 0.56 | 0.64 | 0.6 |
| 10 | 10480283 | rs2244673 | 0.363 | -0.323 | -0.985 | 0.726 | 0.013 | -0.008 | -0.008 | 0.005 | 0.31 | 0.42 | 0.25 | 0.33 | 0.31 | 0.34 |
| 10 | 10481352 | rs2244527 | -0.724 | -0.142 | -1.538 | 0.732 | 0.012 | 0.023 | 0.023 | 0.021 | 0.59 | 0.72 | 0.75 | 0.71 | 0.57 | 0.66 |
| 10 | 10486772 | rs2186063 | 0.429 | 0.42 | -1.69 | 1.085 | -0.032 | -0.039 | -0.039 | 0.407 | 0.19 | 0.1 | 0.27 | 0.23 | 0.25 | 0.25 |
| 10 | 10488292 | rs10905726 | 0.88 | 0.137 | -1.579 | 1.167 | -0.006 | -0.013 | -0.013 | 0 | 0.13 | 0.09 | 0.14 | 0.27 | 0.14 | 0.18 |
| 10 | 10495544 | rs2488172 | -0.284 | -0.089 | -1.375 | 0.784 | -0.003 | -0.006 | -0.006 | 0.017 | 0.57 | 0.51 | 0.69 | 0.56 | 0.58 | 0.64 |
| 10 | 10503251 | rs7895315 | 0.281 | -0.567 | -1.573 | 0.838 | -0.043 | 0.016 | 0.016 | 0 | 0.25 | 0.38 | 0.27 | 0.3 | 0.17 | 0.24 |
| 10 | 10506858 | rs2249834 | 0.372 | -0.577 | -1.506 | 0.792 | -0.045 | 0.011 | 0.011 | 0.876 | 0.35 | 0.16 | 0.44 | 0.26 | 0.42 | 0.4 |
| 10 | 10510613 | rs10905727 | 0.414 | -0.606 | -1.531 | 0.776 | -0.024 | 0.009 | 0.009 | 0 | 0.23 | 0.31 | 0.25 | 0.29 | 0.16 | 0.24 |
| 10 | 10521475 | rs1924507 | -0.535 | -0.524 | -1.687 | 0.759 | -0.001 | 0.004 | 0.004 | 0.099 | 0.56 | 0.56 | 0.69 | 0.57 | 0.57 | 0.62 |
| 10 | 10528970 | rs2255751 | 0.059 | -0.109 | -2.599 | 0.592 | -0.064 | 0.01 | 0.01 | 0.846 | 0.35 | 0.15 | 0.43 | 0.26 | 0.43 | 0.41 |
| 10 | 10530812 | rs11256624 | 1.202 | -0.057 | -3.004 | 0.313 | -0.048 | 0.048 | 0.048 | 0 | 0.1 | 0.02 | 0.1 | 0.01 | 0.21 | 0.11 |
| 10 | 10532725 | rs2477036 | -0.069 | -0.438 | -2.739 | 0.683 | -0.078 | -0.069 | -0.069 | 0 | 0.36 | 0.1 | 0.56 | 0.41 | 0.41 | 0.46 |
| 10 | 10539594 | rs4362053 | -0.076 | -0.065 | -2.54 | 0.712 | -0.019 | -0.08 | -0.08 | 0 | 0.8 | 0.95 | 0.68 | 0.73 | 0.81 | 0.64 |
| 10 | 10540164 | rs4747860 | 0.285 | -0.033 | -2.58 | 0.195 | 0.092 | 0.052 | 0.052 | 0.474 | 0.48 | 0.12 | 0.37 | 0.43 | 0.67 | 0.47 |
| 10 | 10542217 | rs2248503 | -0.691 | 1.16 | -1.661 | 0.468 | -0.008 | -0.015 | -0.015 | 0 | 0.24 | 0.21 | 0.45 | 0.38 | 0.27 | 0.4 |
| 10 | 10542851 | rs4747862 | -0.421 | 1.251 | -1.2 | -0.58 | 0.001 | -0.043 | -0.043 | 0 | 0.37 | 0.4 | 0.26 | 0.11 | 0.36 | 0.29 |
| 10 | 10546342 | rs2265025 | -0.44 | 1.336 | -1.787 | -0.273 | -0.13 | 0.059 | 0.059 | 0 | 0.33 | 0.08 | 0.29 | 0.27 | 0.46 | 0.32 |
| 10 | 10549627 | rs2247652 | 0.278 | 0.978 | -0.387 | -0.167 | 0.068 | 0.028 | 0.028 | 0 | 0.95 | 0.72 | 0.95 | 0.99 | 0.96 | 0.95 |
| 10 | 10552597 | rs2398243 | -0.572 | 0.308 | -0.158 | -0.926 | -0.006 | 0.042 | 0.042 | 0 | 0.47 | 0.66 | 0.34 | 0.3 | 0.47 | 0.41 |
| 10 | 10553889 | rs11256637 | -0.227 | 0.114 | 0.294 | -1.005 | 0.012 | -0.091 | -0.091 | 0 | 0.41 | 0.49 | 0.32 | 0.13 | 0.46 | 0.32 |
| 10 | 10558886 | rs2252120 | 0.416 | 0.201 | 0.306 | -1.017 | -0.007 | 0.077 | 0.077 | 0 | 0.52 | 0.3 | 0.66 | 0.7 | 0.52 | 0.59 |
| 10 | 10559149 | rs12570082 | 1.044 | -0.74 | 0.052 | -0.72 | 0.015 | -0.009 | -0.009 | 0 | 0.15 | 0.09 | 0.09 | 0.16 | 0.09 | 0.19 |
| 10 | 10559645 | rs12412511 | 1.28 | -0.67 | 0.2 | -1.035 | -0.006 | -0.028 | -0.028 | 0.151 | 0.21 | 0.21 | 0.21 | 0.03 | 0.31 | 0.12 |
| 10 | 10560189 | rs10795796 | -0.19 | -0.73 | 0.195 | -1.008 | 0 | -0.049 | -0.049 | 0 | 0.4 | 0.57 | 0.32 | 0.11 | 0.45 | 0.31 |
| 10 | 10562061 | rs4749960 | 0.037 | -0.721 | 0.15 | -0.405 | -0.004 | -0.006 | -0.006 | 0 | 0.55 | 0.48 | 0.64 | 0.63 | 0.57 | 0.67 |
| 10 | 10563782 | rs11817243 | -0.15 | -0.525 | 0.364 | -0.448 | 0 | -0.001 | -0.001 | 0.032 | 0.57 | 0.6 | 0.62 | 0.63 | 0.53 | 0.68 |
| 10 | 10564423 | rs12571685 | 0.974 | -0.804 | 0.444 | -0.417 | -0.044 | 0.048 | 0.048 | 0 | 0.15 | 0.24 | 0.11 | 0.23 | 0.11 | 0.19 |
| 10 | 10565427 | rs10508404 | 1.102 | -1.042 | 0.147 | -0.619 | -0.003 | 0.042 | 0.042 | 0.434 | 0.16 | 0.02 | 0.06 | 0.08 | 0.12 | 0.14 |
| 10 | 10565949 | rs1830809 | -0.094 | -0.936 | 0.875 | -0.405 | -0.046 | 0.045 | 0.045 | 0 | 0.79 | 0.66 | 0.85 | 0.66 | 0.86 | 0.79 |
| 10 | 10566117 | rs2224869 | 0.127 | -0.939 | 0.88 | -0.404 | -0.005 | 0.048 | 0.048 | 0 | 0.93 | 0.75 | 0.94 | 0.89 | 0.97 | 0.98 |
| 10 | 10566260 | rs12572320 | 1.001 | -0.923 | 0.86 | -0.402 | 0.019 | -0.021 | -0.021 | 0 | 0.14 | 0.08 | 0.1 | 0.23 | 0.11 | 0.19 |
| 10 | 10567159 | rs10905731 | 0.181 | -0.895 | 0.829 | -0.394 | -0.006 | 0.046 | 0.046 | 0.014 | 0.93 | 0.75 | 0.94 | 0.89 | 0.97 | 0.98 |
| 10 | 10570880 | rs2398245 | 0.394 | -0.813 | 0.976 | -0.398 | 0.002 | 0.036 | 0.036 | 0.262 | 0.53 | 0.32 | 0.6 | 0.63 | 0.52 | 0.66 |
| 10 | 10571906 | rs10905732 | 0.139 | -0.792 | 0.971 | -0.382 | -0.001 | 0.01 | 0.01 | 0.117 | 0.57 | 0.37 | 0.61 | 0.63 | 0.52 | 0.67 |
| 10 | 10572751 | rs1324322 | -0.729 | -0.627 | 0.789 | -0.484 | 0.016 | -0.035 | -0.035 | 0 | 0.73 | 0.67 | 0.73 | 0.86 | 0.63 | 0.86 |
| 10 | 10580437 | rs7094544 | 0.033 | -0.651 | 0.702 | -0.21 | 0.004 | -0.035 | -0.035 | 0 | 0.6 | 0.53 | 0.69 | 0.73 | 0.54 | 0.67 |
| 10 | 10581693 | rs12573727 | 1.202 | -0.698 | 0.325 | -0.262 | 0.009 | 0.002 | 0.002 | 0 | 0.12 | 0.08 | 0.06 | 0.14 | 0.1 | 0.19 |
| 10 | 10581764 | rs10752197 | 0.614 | -0.494 | 0.161 | -0.019 | 0.012 | 0.084 | 0.084 | 0 | 0.52 | 0.34 | 0.3 | 0.21 | 0.47 | 0.54 |
| 10 | 10582406 | rs17414384 | 1.087 | -0.307 | -0.221 | -0.025 | -0.064 | 0.077 | 0.077 | 1.88 | 0.2 | 0.07 | 0.2 | 0.03 | 0.32 | 0.11 |
| 10 | 10584734 | rs1324318 | 0.069 | -0.283 | -0.175 | 0.219 | -0.032 | 0.088 | 0.088 | 0 | 0.32 | 0.18 | 0.26 | 0.17 | 0.41 | 0.29 |
| 10 | 10586371 | rs2031340 | 0.034 | -0.407 | -0.212 | 0.208 | -0.034 | 0.095 | 0.095 | 0 | 0.33 | 0.17 | 0.26 | 0.17 | 0.41 | 0.29 |
| 10 | 10586835 | rs7071521 | 0.999 | -0.342 | -0.091 | 1.031 | 0.006 | -0.001 | -0.001 | 0.046 | 0.12 | 0.1 | 0.25 | 0.33 | 0.11 | 0.18 |
| 10 | 10587000 | rs7071686 | 1.267 | -0.232 | 0.007 | 0.99 | -0.009 | 0.034 | 0.034 | 0 | 0.08 | 0.1 | 0.1 | 0.18 | 0.08 | 0.16 |
| 10 | 10587027 | rs4749965 | 0.612 | -0.213 | -0.114 | 0.593 | -0.031 | 0.051 | 0.051 | 0 | 0.23 | 0.13 | 0.2 | 0.09 | 0.29 | 0.14 |
| 10 | 10587506 | rs10905739 | 0.011 | -0.419 | 0.197 | 0.794 | 0.007 | -0.001 | -0.001 | 0 | 0.28 | 0.37 | 0.18 | 0.24 | 0.25 | 0.3 |
| 10 | 10588705 | rs17148174 | 1.364 | 0.099 | -0.108 | 0.998 | 0.012 | -0.028 | -0.028 | 0.013 | 0.15 | 0.08 | 0.08 | 0.24 | 0.12 | 0.11 |
| 10 | 10593027 | rs2147290 | -0.889 | 0.206 | 0.248 | 1.261 | 0.009 | -0.003 | -0.003 | 3.414 | 0.5 | 0.72 | 0.64 | 0.56 | 0.48 | 0.55 |
| 10 | 10594364 | rs1324317 | 0.285 | 0.604 | 0.828 | 1.121 | 0.116 | 0.057 | 0.057 | 0.412 | 0.07 | 0.27 | 0.07 | 0.08 | 0.05 | 0.08 |
| 10 | 10597070 | rs7078952 | -0.41 | 0.287 | 0.493 | 1.286 | 0.005 | -0.009 | -0.009 | 0 | 0.32 | 0.68 | 0.29 | 0.53 | 0.2 | 0.35 |
| 10 | 10597399 | rs7073772 | 0.807 | 0.232 | 0.649 | 1.324 | 0.018 | -0.01 | -0.01 | 0 | 0.21 | 0.14 | 0.08 | 0.23 | 0.14 | 0.2 |
| 10 | 10601077 | rs1408386 | -0.874 | 0.418 | 0.459 | 1.179 | -0.01 | 0 | 0 | 0.15 | 0.41 | 0.71 | 0.41 | 0.58 | 0.37 | 0.4 |
| 10 | 10602446 | rs7085082 | -1.638 | 0.395 | 0.188 | 0.678 | -0.044 | 0.065 | 0.065 | 1.217 | 0.64 | 0.73 | 0.62 | 0.82 | 0.55 | 0.67 |
| 10 | 10604381 | rs1041044 | -1.722 | 0.381 | 0.224 | 0.666 | -0.049 | 0.082 | 0.082 | 1.11 | 0.64 | 0.74 | 0.69 | 0.89 | 0.54 | 0.72 |

| | | | | | | | | | | | | | | | | |
|---|---|---|---|---|---|---|---|---|---|---|---|---|---|---|---|---|
| 10 | 10606599 | rs11597484 | 0.708 | 0.493 | 0.835 | 1.366 | 0.014 | -0.018 | -0.018 | 0 | 0.22 | 0.17 | 0.16 | 0.31 | 0.14 | 0.21 |
| 10 | 10607323 | rs11594796 | 0.806 | 0.555 | 0.819 | 1.401 | -0.037 | -0.026 | -0.026 | 0 | 0.09 | 0.02 | 0.2 | 0.14 | 0.18 | 0.06 |
| 10 | 10607497 | rs7096494 | -1.671 | 0.418 | 0.799 | 1.377 | 0.019 | 0.053 | 0.053 | 0.41 | 0.53 | 0.74 | 0.42 | 0.62 | 0.43 | 0.53 |
| 10 | 10609578 | rs7896493 | 0.216 | 0.585 | 0.249 | 1.456 | 0.029 | 0.007 | 0.007 | 0 | 0.19 | 0.09 | 0.09 | 0.16 | 0.1 | 0.21 |
| 10 | 10610784 | rs12414807 | 0.908 | 0.761 | 0.34 | 1.595 | -0.04 | 0.006 | 0.006 | 0 | 0.08 | 0.01 | 0.1 | 0.05 | 0.13 | 0.05 |
| 10 | 10610969 | rs7918560 | -0.525 | 0.89 | 0.34 | 1.597 | 0.04 | -0.032 | -0.032 | 0 | 0.41 | 0.23 | 0.27 | 0.48 | 0.3 | 0.42 |
| 10 | 10611797 | rs11256679 | -1.206 | 0.922 | 0.323 | 1.613 | 0.013 | 0.012 | 0.012 | 0 | 0.51 | 0.37 | 0.3 | 0.54 | 0.44 | 0.54 |
| 10 | 10612816 | rs12569844 | 0.606 | 0.817 | 0.304 | 1.578 | 0.011 | -0.01 | -0.01 | 0.053 | 0.25 | 0.07 | 0.1 | 0.24 | 0.2 | 0.23 |
| 10 | 10613603 | rs12413166 | 0.964 | 0.854 | 0.302 | 1.568 | -0.038 | 0.006 | 0.006 | 0 | 0.08 | 0.02 | 0.1 | 0.05 | 0.13 | 0.05 |
| 10 | 10615416 | rs7084466 | -0.605 | 0.841 | 0.326 | 1.578 | -0.003 | 0.005 | 0.005 | 0 | 0.25 | 0.3 | 0.21 | 0.3 | 0.23 | 0.24 |
| 10 | 10616308 | rs2025467 | -0.462 | 0.824 | 0.513 | 1.592 | -0.001 | 0.007 | 0.007 | 0 | 0.36 | 0.41 | 0.32 | 0.48 | 0.34 | 0.36 |
| 10 | 10616485 | rs2025468 | 0.54 | 0.808 | 0.475 | 1.595 | 0.001 | -0.003 | -0.003 | 0.341 | 0.25 | 0.25 | 0.11 | 0.33 | 0.18 | 0.25 |
| 10 | 10618947 | rs11256681 | 0.69 | 0.512 | 0.243 | 1.353 | 0.012 | 0 | 0 | 0 | 0.17 | 0.11 | 0.08 | 0.15 | 0.09 | 0.18 |
| 10 | 10619227 | rs12251512 | -0.778 | 0.579 | 0.268 | 1.376 | -0.005 | -0.011 | -0.011 | 0 | 0.27 | 0.36 | 0.21 | 0.21 | 0.26 | 0.23 |
| 10 | 10624738 | rs4131374 | -0.007 | 0.854 | 0.388 | 1.276 | -0.005 | 0 | 0 | 0 | 0.42 | 0.36 | 0.27 | 0.57 | 0.27 | 0.44 |
| 10 | 10626134 | rs10905749 | 0.095 | 0.85 | 0.392 | 0.973 | 0.015 | 0 | 0 | 0.142 | 0.41 | 0.27 | 0.17 | 0.4 | 0.27 | 0.41 |
| 10 | 10628935 | rs11256691 | -0.488 | 0.502 | 0.445 | 0.333 | -0.002 | 0.019 | 0.019 | 0.692 | 0.57 | 0.62 | 0.4 | 0.72 | 0.51 | 0.62 |
| 10 | 10638836 | rs972186 | 0.413 | -0.071 | 0.583 | 1.431 | -0.026 | 0.155 | 0.155 | 0.027 | 0.82 | 0.6 | 0.71 | 0.59 | 0.9 | 0.89 |
| 10 | 10642515 | rs9329332 | 0.587 | -0.554 | -0.211 | 0.751 | 0.034 | 0.093 | 0.093 | 0 | 0.37 | 0.13 | 0.21 | 0.21 | 0.27 | 0.53 |
| 10 | 10644923 | rs7070851 | 0.92 | -0.776 | -0.377 | 0.62 | -0.016 | 0.048 | 0.048 | 0 | 0.28 | 0.04 | 0.17 | 0.19 | 0.22 | 0.46 |
| 10 | 10650389 | rs1327842 | 0.576 | -0.902 | -0.034 | 1.044 | 0.006 | -0.008 | -0.008 | 0 | 0.32 | 0.23 | 0.38 | 0.38 | 0.27 | 0.52 |
| 10 | 10650889 | rs7899028 | 0.15 | -0.52 | -0.235 | 0.774 | 0.007 | 0.003 | 0.003 | 0 | 0.58 | 0.47 | 0.61 | 0.44 | 0.51 | 0.75 |
| 10 | 10655038 | rs4749973 | -0.188 | -0.272 | -0.291 | 0.879 | -0.033 | 0.027 | 0.027 | 0 | 0.75 | 0.57 | 0.64 | 0.28 | 0.82 | 0.85 |
| 10 | 10655451 | rs10905767 | 0.422 | -0.317 | -0.314 | 0.816 | -0.001 | -0.008 | -0.008 | 0 | 0.56 | 0.44 | 0.39 | 0.2 | 0.46 | 0.72 |
| 10 | 10656351 | rs7086685 | -0.447 | -0.314 | -0.319 | 0.964 | -0.055 | 0.12 | 0.12 | 0.026 | 0.77 | 0.59 | 0.7 | 0.53 | 0.87 | 0.87 |
| 10 | 10657176 | rs4749974 | -0.168 | -0.3 | -0.318 | 0.901 | -0.036 | 0.028 | 0.028 | 0 | 0.75 | 0.56 | 0.64 | 0.27 | 0.82 | 0.84 |
| 10 | 10660838 | rs11597228 | 0.179 | -0.389 | -0.264 | 0.815 | 0.004 | -0.018 | -0.018 | 0.073 | 0.59 | 0.43 | 0.43 | 0.23 | 0.48 | 0.72 |
| 10 | 10666818 | rs11813043 | -0.631 | -0.364 | -0.241 | 0.818 | 0.021 | 0.005 | 0.005 | 0 | 0.08 | 0.28 | 0.07 | 0.05 | 0.05 | 0.1 |
| 10 | 10672067 | rs11256715 | 0.798 | -0.432 | -0.446 | 0.746 | 0.004 | 0.205 | 0.205 | 0 | 0.52 | 0.14 | 0.31 | 0.18 | 0.44 | 0.66 |
| 10 | 10679038 | rs2399547 | 0.581 | -0.241 | -0.387 | 0.574 | -0.06 | 0.037 | 0.037 | 0 | 0.2 | 0.12 | 0.3 | 0.07 | 0.36 | 0.15 |
| 10 | 10681795 | rs7920242 | 0.592 | -0.625 | -0.621 | 0.386 | 0.031 | 0.081 | 0.081 | 0 | 0.27 | 0.12 | 0.21 | 0.05 | 0.21 | 0.24 |
| 10 | 10686085 | rs4747873 | -0.007 | -0.437 | -0.477 | 0.476 | 0.002 | -0.013 | -0.013 | 0.492 | 0.59 | 0.42 | 0.38 | 0.23 | 0.49 | 0.73 |
| 10 | 10688361 | rs7910201 | 0.905 | -0.446 | -0.552 | 0.451 | -0.002 | 0.061 | 0.061 | 0.492 | 0.33 | 0.17 | 0.17 | 0.18 | 0.27 | 0.49 |
| 10 | 10689016 | rs11256725 | 1.527 | -0.538 | -0.551 | 0.478 | -0.039 | 0.073 | 0.073 | 0.48 | 0.27 | 0.05 | 0.12 | 0.14 | 0.24 | 0.45 |
| 10 | 10690346 | rs11256730 | 0.719 | -0.49 | -0.334 | 0.483 | 0.025 | 0.055 | 0.055 | 0 | 0.25 | 0.12 | 0.2 | 0.05 | 0.2 | 0.24 |
| 10 | 10690733 | rs7919635 | 0.719 | -0.49 | -0.334 | 0.483 | 0.025 | 0.055 | 0.055 | 0 | 0.25 | 0.12 | 0.2 | 0.05 | 0.2 | 0.24 |
| 10 | 10700036 | rs1359890 | 1.088 | 0.054 | -0.37 | 0.464 | -0.068 | 0.077 | 0.077 | 0 | 0.16 | 0.04 | 0.16 | 0 | 0.28 | 0.09 |
| 10 | 10702431 | rs2399551 | 0.867 | -0.009 | -0.233 | 0.509 | 0 | 0 | 0 | 0 | 0.34 | 0.31 | 0.19 | 0.18 | 0.28 | 0.48 |
| 10 | 10704323 | rs4749976 | 0.745 | -0.323 | -0.184 | 0.554 | -0.004 | -0.02 | -0.02 | 0 | 0.35 | 0.38 | 0.25 | 0.2 | 0.26 | 0.56 |
| 10 | 10707182 | rs6602448 | -0.537 | -0.323 | -0.171 | 0.695 | -0.028 | 0.065 | 0.065 | 0 | 0.76 | 0.54 | 0.66 | 0.34 | 0.82 | 0.84 |
| 10 | 10707200 | rs17148604 | 0.838 | -0.291 | -0.07 | 0.37 | -0.001 | -0.006 | -0.006 | 0 | 0.34 | 0.3 | 0.23 | 0.2 | 0.25 | 0.55 |
| 10 | 10707944 | rs10508415 | 1.212 | -0.288 | -0.071 | 0.371 | 0.001 | 0.008 | 0.008 | 0 | 0.31 | 0.22 | 0.13 | 0.17 | 0.24 | 0.46 |
| 10 | 10710599 | rs10905781 | -0.621 | -0.267 | -0.079 | 0.391 | -0.019 | -0.018 | -0.018 | 0 | 0.74 | 0.54 | 0.55 | 0.23 | 0.77 | 0.8 |
| 10 | 10718024 | rs10905782 | 0.826 | -0.27 | -0.074 | 0.388 | -0.005 | -0.035 | -0.035 | 0.004 | 0.33 | 0.4 | 0.15 | 0.18 | 0.26 | 0.55 |
| 10 | 10720800 | rs1327847 | 0.818 | -0.249 | -0.144 | 0.395 | -0.004 | -0.028 | -0.028 | 0.029 | 0.33 | 0.42 | 0.25 | 0.19 | 0.26 | 0.55 |
| 10 | 10723472 | rs763351 | 1.137 | -0.224 | -0.147 | 0.674 | -0.001 | -0.001 | -0.001 | 0 | 0.31 | 0.32 | 0.17 | 0.44 | 0.27 | 0.48 |
| 10 | 10725488 | rs7895677 | -0.722 | -0.235 | -0.188 | 0.688 | -0.032 | 0.123 | 0.123 | 0.096 | 0.75 | 0.53 | 0.48 | 0.5 | 0.8 | 0.83 |
| 10 | 10728076 | rs17517536 | 0.8 | -0.131 | -0.329 | 0.598 | -0.08 | 0.097 | 0.097 | 0 | 0.19 | 0.05 | 0.2 | 0 | 0.31 | 0.1 |
| 10 | 10729861 | rs10905788 | 0.958 | -0.208 | 0.016 | 0.875 | 0 | 0.02 | 0.02 | 0 | 0.29 | 0.22 | 0.13 | 0.16 | 0.24 | 0.45 |
| 10 | 10732112 | rs1327860 | 0.76 | -0.202 | 0.031 | 0.867 | -0.001 | -0.006 | -0.006 | 0.072 | 0.32 | 0.28 | 0.17 | 0.46 | 0.28 | 0.49 |
| 10 | 10743956 | rs1998432 | -0.044 | -0.247 | -0.052 | 0.883 | -0.01 | 0.005 | 0.005 | 0 | 0.46 | 0.32 | 0.44 | 0.44 | 0.55 | 0.38 |
| 10 | 10753513 | rs10905798 | 0.24 | -0.426 | 0.136 | 1.023 | 0.006 | -0.011 | -0.011 | 0 | 0.7 | 0.28 | 0.45 | 0.54 | 0.78 | 0.72 |
| 10 | 10756738 | rs7923994 | -0.101 | -0.16 | -0.429 | 1.1 | -0.018 | 0.064 | 0.064 | 0.001 | 0.76 | 0.62 | 0.48 | 0.58 | 0.81 | 0.84 |
| 10 | 10763509 | rs11256796 | 0.741 | 0.189 | -1.261 | 0.727 | -0.082 | -0.014 | -0.014 | 0 | 0.3 | 0.01 | 0.32 | 0.2 | 0.26 | 0.45 |
| 10 | 10769007 | rs10752199 | -0.359 | -0.075 | -0.796 | 0.896 | -0.019 | 0.015 | 0.015 | 0 | 0.76 | 0.65 | 0.78 | 0.68 | 0.82 | 0.83 |
| 10 | 10772031 | rs10752200 | 0.77 | -0.009 | -1.325 | 0.763 | -0.028 | 0.025 | 0.025 | 0 | 0.31 | 0.08 | 0.32 | 0.22 | 0.27 | 0.44 |
| 10 | 10773755 | rs11593549 | 0.124 | 0.625 | -0.977 | 1.157 | 0.017 | -0.045 | -0.045 | 0 | 0.22 | 0.09 | 0.13 | 0.29 | 0.21 | 0.17 |
| 10 | 10775958 | rs2399554 | 0.424 | 0.708 | -0.992 | 0.54 | 0 | 0.072 | 0.072 | 0 | 0.45 | 0.3 | 0.54 | 0.69 | 0.4 | 0.65 |
| 10 | 10776143 | rs11596743 | 0.501 | 1.278 | -0.868 | 0.031 | 0.009 | -0.039 | -0.039 | 0 | 0.13 | 0.15 | 0.12 | 0 | 0.21 | 0.09 |
| 10 | 10777085 | rs17148768 | 0.772 | 1.668 | -0.58 | -0.172 | -0.002 | -0.121 | -0.121 | 0.033 | 0.22 | 0.37 | 0.16 | 0.02 | 0.21 | 0.1 |
| 10 | 10777882 | rs17518702 | -0.439 | 1.683 | -1.297 | -0.068 | -0.028 | 0.031 | 0.031 | 0 | 0.21 | 0.15 | 0.16 | 0.13 | 0.29 | 0.22 |
| 10 | 10777966 | rs11256803 | -0.44 | 1.685 | -1.472 | -0.058 | -0.043 | 0.043 | 0.043 | 0 | 0.21 | 0.14 | 0.16 | 0.14 | 0.29 | 0.22 |
| 10 | 10780895 | rs12356893 | 0.957 | 1.236 | -1.354 | 0.374 | 0.046 | -0.101 | -0.101 | 0 | 0.38 | 0.12 | 0.24 | 0.44 | 0.23 | 0.52 |
| 10 | 10785439 | rs10752202 | 0.562 | 1.534 | -0.6 | 1.228 | 0.003 | 0.031 | 0.031 | 0 | 0.75 | 0.72 | 0.53 | 0.48 | 0.67 | 0.7 |
| 10 | 10788027 | rs10795819 | -0.795 | 1.725 | -1.102 | 1.327 | -0.055 | -0.066 | -0.066 | 0 | 0.1 | 0.01 | 0.07 | 0.27 | 0.18 | 0.15 |
| 10 | 10789590 | rs17519030 | 0.147 | 1.215 | -0.921 | -0.063 | -0.002 | -0.075 | -0.075 | 2.541 | 0.33 | 0.37 | 0.25 | 0.03 | 0.32 | 0.13 |
| 10 | 10791427 | rs10905807 | 0.256 | 0.998 | -0.836 | 0.556 | -0.001 | 0.021 | 0.021 | 0 | 0.11 | 0.1 | 0.29 | 0.26 | 0.11 | 0.16 |
| 10 | 10791450 | rs7922173 | -0.48 | 0.621 | -0.515 | 0.304 | 0.01 | 0.031 | 0.031 | 1.837 | 0.47 | 0.72 | 0.59 | 0.38 | 0.49 | 0.48 |
| 10 | 10796771 | rs10795820 | 0.376 | 0.546 | 0.011 | 0.375 | -0.006 | 0.087 | 0.087 | 0 | 0.12 | 0.23 | 0.3 | 0.3 | 0.11 | 0.16 |

| | | | | | | | | | | | | | | | | |
|---|---|---|---|---|---|---|---|---|---|---|---|---|---|---|---|---|
| 10 | 10797251 | rs7914208 | 0.225 | 0.37 | -0.009 | 0.048 | -0.009 | -0.091 | -0.091 | 0 | 0.38 | 0.48 | 0.29 | 0.08 | 0.38 | 0.31 |
| 10 | 10800157 | rs2224695 | 0.283 | 0.157 | -0.044 | 0.25 | -0.004 | 0.031 | 0.031 | 0 | 0.13 | 0.15 | 0.31 | 0.29 | 0.13 | 0.2 |
| 10 | 10800193 | rs2038662 | -1.23 | -0.308 | -0.544 | 0.51 | -0.004 | -0.003 | -0.003 | 0 | 0.56 | 0.74 | 0.62 | 0.54 | 0.51 | 0.54 |
| 10 | 10804089 | rs4363499 | -1.017 | -0.551 | -0.692 | 0.585 | -0.012 | 0.06 | 0.06 | 0 | 0.25 | 0.41 | 0.44 | 0.5 | 0.25 | 0.41 |
| 10 | 10806389 | rs9424093 | -0.125 | -0.532 | -0.651 | 0.609 | -0.001 | 0.085 | 0.085 | 0 | 0.13 | 0.35 | 0.11 | 0.19 | 0.12 | 0.2 |
| 10 | 10811867 | rs2399555 | -0.295 | -0.537 | -0.622 | 0.53 | -0.007 | 0.112 | 0.112 | 0 | 0.17 | 0.32 | 0.39 | 0.49 | 0.19 | 0.36 |
| 10 | 10813904 | rs9423832 | 1.012 | -0.573 | -1.06 | 0.425 | -0.001 | -0.029 | -0.029 | 0 | 0.08 | 0.09 | 0.04 | 0 | 0.07 | 0.06 |
| 10 | 10815196 | rs11599142 | 0.435 | -0.639 | -1.174 | 0.477 | 0.001 | -0.042 | -0.042 | 0 | 0.13 | 0.07 | 0.33 | 0.3 | 0.14 | 0.22 |
| 10 | 10818484 | rs2007504 | -1.715 | -0.587 | -1.778 | -0.177 | -0.028 | 0.114 | 0.114 | 0 | 0.8 | 0.98 | 0.87 | 0.97 | 0.79 | 0.88 |
| 10 | 10821168 | rs9423835 | 0.811 | -0.003 | -1.861 | -0.068 | -0.054 | 0.044 | 0.044 | 0.198 | 0.17 | 0.03 | 0.1 | 0.1 | 0.22 | 0.12 |
| 10 | 10834765 | rs11256834 | -0.863 | -0.284 | -1.566 | 0.139 | 0.004 | -0.009 | -0.009 | 0 | 0.22 | 0.26 | 0.4 | 0.51 | 0.22 | 0.41 |
| 10 | 10838596 | rs912083 | -0.002 | -0.256 | -1.442 | 0.267 | -0.015 | 0.051 | 0.051 | 0 | 0.09 | 0.2 | 0.08 | 0.18 | 0.08 | 0.16 |
| 10 | 10844011 | rs7921103 | -0.238 | -0.264 | -1.427 | 0.267 | 0 | 0.064 | 0.064 | 0 | 0.07 | 0.11 | 0.28 | 0.33 | 0.09 | 0.17 |
| 10 | 10846404 | rs2031142 | -1.037 | -0.517 | -1.807 | -0.112 | 0.009 | 0.027 | 0.027 | 0 | 0.69 | 0.8 | 0.73 | 0.79 | 0.67 | 0.64 |
| 10 | 10846843 | rs1475184 | -0.214 | -0.521 | -1.807 | -0.109 | 0 | 0.056 | 0.056 | 0 | 0.07 | 0.11 | 0.28 | 0.33 | 0.09 | 0.19 |
| 10 | 10848678 | rs7083333 | -0.257 | -0.544 | -1.819 | -0.121 | 0.012 | -0.021 | -0.021 | 0 | 0.62 | 0.69 | 0.46 | 0.46 | 0.58 | 0.48 |
| 10 | 10848955 | rs7083737 | -1.055 | -0.53 | -1.808 | -0.129 | 0.009 | 0.026 | 0.026 | 0 | 0.69 | 0.8 | 0.73 | 0.78 | 0.68 | 0.64 |
| 10 | 10855297 | rs867221 | 0.46 | -0.925 | -1.909 | -0.364 | 0.002 | 0.005 | 0.005 | 0 | 0.45 | 0.6 | 0.38 | 0.43 | 0.4 | 0.37 |
| 10 | 10859858 | rs10905835 | -0.336 | -0.83 | -1.928 | -1.309 | 0.014 | -0.026 | -0.026 | 0 | 0.61 | 0.7 | 0.62 | 0.54 | 0.59 | 0.45 |
| 10 | 10859974 | rs1885827 | -1.002 | -0.837 | -1.944 | -1.308 | 0.014 | 0.03 | 0.03 | 0 | 0.68 | 0.77 | 0.71 | 0.78 | 0.68 | 0.64 |
| 10 | 10861511 | rs11818010 | -0.015 | -0.571 | -1.728 | -1.074 | -0.001 | 0.001 | 0.001 | 0 | 0.06 | 0.05 | 0.1 | 0.24 | 0.09 | 0.19 |
| 10 | 10864227 | rs11813286 | 0.16 | -0.684 | -1.727 | -1.071 | 0.001 | 0.003 | 0.003 | 0 | 0.06 | 0.05 | 0.1 | 0.24 | 0.07 | 0.19 |
| 10 | 10874408 | rs4749995 | -0.053 | -0.728 | -1.763 | -1.168 | -0.003 | 0.112 | 0.112 | 0 | 0.5 | 0.68 | 0.61 | 0.76 | 0.47 | 0.53 |
| 10 | 10874535 | rs7085538 | 0.1 | 0.05 | -0.945 | -1.921 | 0.002 | 0.007 | 0.007 | 0 | 0.06 | 0.05 | 0.07 | 0.23 | 0.06 | 0.19 |
| 10 | 10878166 | rs12218816 | -0.757 | 0.08 | -0.955 | -1.983 | -0.015 | -0.004 | -0.004 | 0 | 0.2 | 0.18 | 0.12 | 0.24 | 0.25 | 0.3 |
| 10 | 10886244 | rs10752209 | -1.351 | -0.164 | -1.115 | -2.144 | 0.021 | 0.142 | 0.142 | 0 | 0.67 | 0.84 | 0.72 | 0.97 | 0.69 | 0.73 |
| 10 | 10888524 | rs11256861 | 0.185 | 0.095 | -0.825 | -1.835 | -0.065 | 0.076 | 0.076 | 1.625 | 0.14 | 0.02 | 0.06 | 0.02 | 0.18 | 0.08 |
| 10 | 10893317 | rs12221057 | -1.416 | 0.021 | -0.682 | -1.942 | -0.004 | 0.008 | 0.008 | 0 | 0.26 | 0.23 | 0.13 | 0.24 | 0.27 | 0.34 |
| 10 | 10895705 | rs11256872 | -0.89 | 0.087 | -0.609 | -1.867 | -0.017 | 0.019 | 0.019 | 0 | 0.12 | 0.2 | 0.06 | 0.22 | 0.08 | 0.24 |
| 10 | 10907971 | rs10905850 | -1.234 | 0.083 | -0.663 | -1.912 | -0.002 | 0.016 | 0.016 | 0 | 0.29 | 0.23 | 0.13 | 0.25 | 0.27 | 0.34 |
| 10 | 10912885 | rs1473737 | -1.302 | 0.329 | -0.232 | -2.155 | -0.015 | 0.079 | 0.079 | 0 | 0.56 | 0.77 | 0.58 | 0.98 | 0.5 | 0.65 |
| 10 | 10913845 | rs10905853 | 0.714 | 0.317 | -0.227 | -2.048 | 0.017 | -0.022 | -0.022 | 0 | 0.09 | 0.05 | 0.06 | 0.23 | 0.05 | 0.18 |
| 10 | 10914893 | rs11256885 | 0.389 | 0.333 | -0.206 | -2.11 | -0.003 | -0.012 | -0.012 | 0 | 0.29 | 0.59 | 0.45 | 0.73 | 0.25 | 0.34 |
| 10 | 10917971 | rs11817172 | 0.708 | 0.41 | -0.275 | -1.964 | 0.022 | -0.036 | -0.036 | 0 | 0.09 | 0.03 | 0.06 | 0.23 | 0.05 | 0.19 |
| 10 | 10918103 | rs7904753 | 0.831 | 0.474 | 0.135 | -1.958 | -0.004 | -0.001 | -0.001 | 1.329 | 0.25 | 0.46 | 0.41 | 0.73 | 0.21 | 0.34 |
| 10 | 10922068 | rs7098339 | -0.885 | 0.707 | -0.011 | -1.961 | 0.005 | 0.01 | 0.01 | 0 | 0.56 | 0.64 | 0.6 | 0.75 | 0.58 | 0.57 |
| 10 | 10926211 | rs7087274 | -1.114 | 0.419 | 0.437 | -1.274 | -0.011 | 0.135 | 0.135 | 0 | 0.48 | 0.78 | 0.55 | 0.76 | 0.46 | 0.46 |
| 10 | 10928268 | rs1018573 | -2.037 | 0.465 | 0.152 | -1.375 | -0.128 | 0.266 | 0.266 | 0 | 0.63 | 0.93 | 0.63 | 0.98 | 0.53 | 0.71 |
| 10 | 10930210 | rs7099477 | 0.037 | 0.722 | 0.544 | -0.827 | -0.075 | 0.001 | 0.001 | 0 | 0.85 | 0.98 | 0.92 | 0.91 | 0.79 | 0.91 |
| 10 | 10936611 | rs1570438 | 0.733 | 0.895 | 0.688 | -0.76 | -0.022 | 0.026 | 0.026 | 0 | 0.81 | 0.84 | 0.89 | 0.9 | 0.76 | 0.89 |
| 10 | 10943228 | rs11256894 | -0.093 | 0.357 | 1.148 | -0.415 | 0.003 | -0.002 | -0.002 | 0.141 | 0.11 | 0.08 | 0.08 | 0.1 | 0.16 | 0.08 |
| 10 | 10945351 | rs912069 | 0.08 | 0.345 | 1.465 | -0.381 | -0.001 | 0.011 | 0.011 | 0 | 0.24 | 0.26 | 0.43 | 0.62 | 0.23 | 0.33 |
| 10 | 10945862 | rs10905858 | 0.584 | 0.365 | 1.131 | -0.439 | -0.004 | 0.006 | 0.006 | 0 | 0.63 | 0.38 | 0.43 | 0.28 | 0.6 | 0.59 |
| 10 | 10948223 | rs1015087 | 1.423 | 0.549 | 0.527 | -0.79 | -0.021 | 0.105 | 0.105 | 0 | 0.45 | 0.23 | 0.34 | 0.02 | 0.5 | 0.32 |
| 10 | 10953351 | rs7067649 | 0.054 | 0.868 | 0.433 | -0.059 | -0.016 | 0.008 | 0.008 | 0 | 0.89 | 0.88 | 0.88 | 0.92 | 0.85 | 0.93 |
| 10 | 10956220 | rs7097329 | 1.393 | 0.804 | 0.445 | -0.058 | 0.011 | -0.057 | -0.057 | 0 | 0.8 | 0.63 | 0.88 | 0.92 | 0.8 | 0.93 |
| 10 | 10965003 | rs2399580 | -1.023 | 0.558 | 0.582 | 0.308 | -0.002 | -0.001 | -0.001 | 0.077 | 0.3 | 0.35 | 0.5 | 0.69 | 0.28 | 0.39 |
| 10 | 10968124 | rs734366 | -0.84 | 0.569 | 0.414 | 0.312 | -0.017 | -0.043 | -0.043 | 0 | 0.09 | 0.3 | 0.04 | 0 | 0.04 | 0.01 |
| 10 | 10968633 | rs731229 | -0.669 | 1.223 | -0.516 | 0.741 | 0 | 0.002 | 0.002 | 0 | 0.25 | 0.3 | 0.47 | 0.68 | 0.25 | 0.37 |
| 10 | 10973316 | rs992193 | 0.555 | 1.187 | -0.326 | 0.929 | 0.005 | 0.06 | 0.06 | 0 | 0.71 | 0.51 | 0.53 | 0.5 | 0.69 | 0.76 |
| 10 | 10974256 | rs10905868 | -0.679 | 2.089 | 0.104 | 0.569 | 0.057 | -0.003 | -0.003 | 0.014 | 0.17 | 0.32 | 0.17 | 0.16 | 0.24 | 0.26 |
| 10 | 10977601 | rs17149140 | -0.354 | 1.152 | -0.459 | -0.081 | 0.015 | 0.035 | 0.035 | 0 | 0.31 | 0.17 | 0.24 | 0.19 | 0.25 | 0.16 |
| 10 | 10978266 | rs2146553 | -0.149 | 1.614 | -1.083 | -0.829 | -0.102 | 0.133 | 0.133 | 3.967 | 0.68 | 0.91 | 0.78 | 0.92 | 0.55 | 0.66 |
| 10 | 10982410 | rs10752213 | 0.988 | 1.366 | -0.535 | -0.851 | 0.043 | 0.054 | 0.054 | 0 | 0.39 | 0.24 | 0.53 | 0.74 | 0.34 | 0.53 |
| 10 | 10982779 | rs10905872 | 0.978 | 1.297 | -0.567 | -0.863 | 0.043 | 0.052 | 0.052 | 0 | 0.39 | 0.23 | 0.53 | 0.74 | 0.34 | 0.53 |
| 10 | 10990421 | rs6602471 | 1.188 | 1.169 | -0.527 | -1.144 | 0.032 | 0.075 | 0.075 | 0 | 0.35 | 0.22 | 0.41 | 0.78 | 0.33 | 0.52 |
| 10 | 10990942 | rs7093099 | 0.381 | 1.025 | -0.508 | -1.348 | 0.001 | 0.032 | 0.032 | 0 | 0.2 | 0.08 | 0.14 | 0.09 | 0.18 | 0.11 |
| 10 | 10992865 | rs201070 | 0.013 | 0.976 | -0.471 | -1.374 | 0.042 | 0.166 | 0.166 | 0.246 | 0.54 | 0.85 | 0.7 | 0.81 | 0.45 | 0.59 |
| 10 | 10993826 | rs201071 | -0.909 | 0.937 | -0.052 | -1.388 | 0.05 | -0.006 | -0.006 | 0 | 0.09 | 0.48 | 0.25 | 0.02 | 0.06 | 0.04 |
| 10 | 10994808 | rs7922245 | 0.794 | 0.889 | -0.259 | -1.358 | 0.067 | 0.128 | 0.128 | 0.039 | 0.44 | 0.25 | 0.45 | 0.78 | 0.38 | 0.55 |
| 10 | 10996596 | rs7074477 | 0.999 | 0.141 | -0.648 | -1.536 | 0.044 | 0.026 | 0.026 | 0 | 0.31 | 0.21 | 0.45 | 0.76 | 0.27 | 0.54 |
| 10 | 11000339 | rs62209 | -1.006 | 0.397 | -0.145 | -1.375 | 0.023 | 0.049 | 0.049 | 2.474 | 0.27 | 0.78 | 0.19 | 0.08 | 0.17 | 0.14 |
| 10 | 11016076 | rs201086 | -1.286 | 0.301 | 0.061 | -1.366 | 0.048 | 0.094 | 0.094 | 1.442 | 0.3 | 0.84 | 0.21 | 0.09 | 0.18 | 0.15 |
| 10 | 11017020 | rs11256941 | 0.307 | 0.282 | 0.156 | -1.393 | 0.003 | 0.013 | 0.013 | 0 | 0.17 | 0.09 | 0.12 | 0.08 | 0.13 | 0.11 |
| 10 | 11024559 | rs201099 | -1.581 | 0.25 | 0.361 | -1.398 | 0.007 | -0.013 | -0.013 | 0 | 0.2 | 0.79 | 0.23 | 0.02 | 0.12 | 0.07 |
| 10 | 11038471 | rs201107 | -1.586 | 0.239 | 0.433 | -1.392 | -0.014 | -0.061 | -0.061 | 0 | 0.19 | 0.73 | 0.14 | 0 | 0.1 | 0.05 |
| 10 | 11041967 | rs201109 | -1.801 | 0.215 | 0.357 | -1.405 | 0.01 | -0.006 | -0.006 | 0 | 0.2 | 0.81 | 0.23 | 0.02 | 0.12 | 0.06 |
| 10 | 11048264 | rs201113 | -1.832 | 0.214 | 0.314 | -1.384 | 0.007 | -0.02 | -0.02 | 1.7 | 0.19 | 0.79 | 0.15 | 0 | 0.1 | 0.05 |
| 10 | 11048332 | rs201114 | -1.463 | 0.23 | 0.308 | -1.377 | -0.025 | -0.087 | -0.087 | 0.083 | 0.17 | 0.66 | 0.14 | 0 | 0.1 | 0.05 |

| | | | | | | | | | | | | | | | | |
|---|---|---|---|---|---|---|---|---|---|---|---|---|---|---|---|---|
| 10 | 11049977 | rs201119 | -1.568 | 0.218 | 0.134 | -1.358 | -0.012 | -0.057 | -0.057 | 0.104 | 0.18 | 0.75 | 0.15 | 0 | 0.11 | 0.05 |
| 10 | 11055589 | rs1547221 | -0.062 | 0.542 | 0.257 | -1.211 | 0.072 | -0.009 | -0.009 | 0.021 | 0.92 | 0.63 | 0.92 | 1 | 0.94 | 0.98 |
| 10 | 11057523 | rs7901515 | 0.461 | 0.987 | -1.059 | -1.097 | -0.137 | 0.053 | 0.053 | 1.835 | 0.27 | 0.03 | 0.19 | 0.16 | 0.37 | 0.3 |
| 10 | 11064241 | rs10795842 | 1.69 | 0.547 | -0.12 | -0.742 | -0.026 | -0.019 | -0.019 | 0 | 0.87 | 0.3 | 0.88 | 0.93 | 0.91 | 0.89 |
| 10 | 11067072 | rs10905883 | 2.214 | 0.711 | -0.325 | -0.797 | 0 | 0.006 | 0.006 | 0.341 | 0.8 | 0.18 | 0.73 | 0.9 | 0.8 | 0.84 |
| 10 | 11067117 | rs10905884 | 1.79 | 0.566 | -0.128 | -0.613 | -0.009 | -0.013 | -0.013 | 0.456 | 0.86 | 0.31 | 0.8 | 0.9 | 0.88 | 0.85 |
| 10 | 11075133 | rs7090372 | 1.186 | 0.704 | 0.15 | -0.578 | 0.035 | 0.026 | 0.026 | 0 | 0.89 | 0.44 | 0.82 | 0.9 | 0.91 | 0.83 |
| 10 | 11078390 | rs1999207 | -0.336 | 0.263 | -0.704 | -1.219 | 0.049 | 0.104 | 0.104 | 0 | 0.5 | 0.38 | 0.56 | 0.77 | 0.4 | 0.57 |
| 10 | 11082012 | rs10905889 | 0.266 | 0.219 | -0.549 | -0.79 | 0.006 | -0.012 | -0.012 | 0 | 0.19 | 0.12 | 0.38 | 0.67 | 0.15 | 0.42 |
| 10 | 11083239 | rs1291867 | 0.255 | 0.534 | 0.202 | -0.938 | 0.055 | -0.027 | -0.027 | 1.802 | 0.07 | 0.23 | 0.07 | 0 | 0.1 | 0.02 |
| 10 | 11088303 | rs1291875 | -1.113 | 0.044 | -0.105 | -1.305 | -0.011 | 0.034 | 0.034 | 0 | 0.31 | 0.76 | 0.25 | 0.02 | 0.32 | 0.16 |
| 10 | 11088921 | rs1291877 | -1.339 | 0.074 | 0.349 | -1.249 | 0.02 | -0.008 | -0.008 | 0 | 0.13 | 0.56 | 0.13 | 0.08 | 0.16 | 0.08 |
| 10 | 11089143 | rs1291878 | 1.138 | 0.098 | 0.331 | -1.241 | 0.031 | -0.029 | -0.029 | 0 | 0.06 | 0.14 | 0.06 | 0 | 0.09 | 0.02 |
| 10 | 11090637 | rs1291879 | -0.878 | 0.477 | 0.391 | -1.277 | 0.041 | 0.042 | 0.042 | 0 | 0.07 | 0.22 | 0.08 | 0.08 | 0.08 | 0.07 |
| 10 | 11090917 | rs976936 | 0.769 | 0.381 | 0.302 | -1.276 | 0.015 | -0.045 | -0.045 | 0.185 | 0.17 | 0.07 | 0.37 | 0.67 | 0.14 | 0.42 |
| 10 | 11093705 | rs1291886 | 0.035 | 0.634 | 0.604 | -1.554 | 0.123 | 0.008 | 0.008 | 0 | 0.07 | 0.43 | 0.09 | 0 | 0.1 | 0.02 |
| 10 | 11097512 | rs1291831 | -1.885 | 0.572 | 0.363 | -1.63 | -0.06 | 0.126 | 0.126 | 0.061 | 0.37 | 0.86 | 0.33 | 0.11 | 0.44 | 0.29 |
| 10 | 11101626 | rs6602475 | 0.036 | 0.276 | 0.423 | -1.571 | -0.016 | 0.026 | 0.026 | 0 | 0.21 | 0.19 | 0.15 | 0.02 | 0.23 | 0.14 |
| 10 | 11105845 | rs1291845 | -0.489 | 0.284 | 0.498 | -1.563 | 0.022 | -0.12 | -0.12 | 0 | 0.24 | 0.4 | 0.18 | 0.02 | 0.24 | 0.15 |
| 10 | 11107195 | rs1291846 | 0.156 | 0.291 | 0.539 | -1.56 | 0.05 | -0.025 | -0.025 | 0 | 0.07 | 0.24 | 0.07 | 0 | 0.09 | 0.02 |
| 10 | 11108377 | rs1291849 | -1.335 | 0.294 | 0.478 | -1.564 | 0.024 | -0.14 | -0.14 | 4.693 | 0.28 | 0.43 | 0.22 | 0.02 | 0.31 | 0.16 |
| 10 | 11115090 | rs1886281 | 0.299 | 0.389 | 0.438 | -1.497 | 0.067 | -0.023 | -0.023 | 0 | 0.07 | 0.21 | 0.08 | 0 | 0.1 | 0.02 |
| 10 | 11119519 | rs2773495 | -1.199 | 0.245 | 0.558 | -1.551 | 0.028 | -0.133 | -0.133 | 0 | 0.27 | 0.43 | 0.22 | 0.02 | 0.31 | 0.16 |
| 10 | 11119609 | rs11256967 | -0.59 | 0.241 | 0.543 | -1.554 | -0.003 | -0.033 | -0.033 | 0 | 0.25 | 0.26 | 0.19 | 0.02 | 0.28 | 0.15 |
| 10 | 11124499 | rs1291799 | -1.255 | 0.24 | 0.503 | -1.554 | 0.028 | -0.135 | -0.135 | 0.656 | 0.28 | 0.43 | 0.22 | 0.02 | 0.31 | 0.16 |
| 10 | 11128473 | rs10490956 | -0.393 | 0.262 | 0.04 | -1.539 | -0.009 | -0.015 | -0.015 | 0 | 0.24 | 0.22 | 0.19 | 0.01 | 0.28 | 0.15 |
| 10 | 11130833 | rs1291813 | -1.713 | 0.19 | 0.07 | -1.511 | -0.027 | 0.064 | 0.064 | 0 | 0.33 | 0.78 | 0.3 | 0.1 | 0.37 | 0.29 |
| 10 | 11132190 | rs1291818 | -1.622 | 0.117 | -0.115 | -1.539 | -0.001 | 0.004 | 0.004 | 0 | 0.48 | 0.59 | 0.61 | 0.8 | 0.49 | 0.64 |
| 10 | 11133119 | rs2224867 | -2.248 | -0.075 | 0.172 | -1.302 | 0.027 | -0.021 | -0.021 | 0 | 0.18 | 0.53 | 0.19 | 0.1 | 0.22 | 0.19 |
| 10 | 11138048 | rs10490954 | 0.593 | -0.245 | 0.078 | -1.161 | 0.003 | -0.005 | -0.005 | 0.229 | 0.17 | 0.17 | 0.38 | 0.73 | 0.11 | 0.37 |
| 10 | 11141426 | rs7083068 | -1.558 | -0.108 | 0.285 | -1.291 | 0.059 | -0.079 | -0.079 | 0.101 | 0.12 | 0.44 | 0.12 | 0.01 | 0.14 | 0.09 |
| 10 | 11141618 | rs7070530 | -1.852 | -0.116 | 0.206 | -1.295 | 0.021 | -0.087 | -0.087 | 0.091 | 0.14 | 0.56 | 0.16 | 0.01 | 0.16 | 0.1 |
| 10 | 11154164 | rs1750731 | -1.249 | 0.061 | 0.263 | -1.041 | -0.002 | -0.009 | -0.009 | 0.111 | 0.3 | 0.37 | 0.53 | 0.75 | 0.28 | 0.51 |
| 10 | 11154321 | rs7073060 | 0.097 | 0.206 | 0.099 | -1.19 | -0.02 | 0.025 | 0.025 | 0 | 0.16 | 0.1 | 0.14 | 0.05 | 0.21 | 0.15 |
| 10 | 11155332 | rs1750737 | 0.589 | 0.133 | 0.087 | -1.131 | 0.019 | -0.062 | -0.062 | 0.478 | 0.17 | 0.06 | 0.36 | 0.65 | 0.12 | 0.38 |
| 10 | 11156472 | rs1750738 | -0.121 | 0.13 | 0.158 | -1.136 | -0.004 | 0 | 0 | 0 | 0.22 | 0.3 | 0.4 | 0.72 | 0.17 | 0.47 |
| 10 | 11161978 | rs1750734 | -2.224 | 0.03 | 0.114 | -1.129 | 0.002 | 0.027 | 0.027 | 0 | 0.48 | 0.64 | 0.69 | 0.79 | 0.5 | 0.67 |
| 10 | 11162787 | rs17375715 | -1.378 | 0.028 | 0.061 | -1.159 | 0.008 | -0.012 | -0.012 | 1.461 | 0.25 | 0.31 | 0.27 | 0.15 | 0.31 | 0.27 |
| 10 | 11163048 | rs7083507 | -1.176 | -0.475 | -0.069 | -0.983 | 0.011 | -0.028 | -0.028 | 0 | 0.05 | 0.12 | 0.04 | 0 | 0.06 | 0.02 |
| 10 | 11163851 | rs7894817 | -0.098 | -0.483 | -0.23 | -0.95 | 0.003 | -0.019 | -0.019 | 0 | 0.12 | 0.2 | 0.12 | 0.07 | 0.11 | 0.15 |
| 10 | 11166455 | rs11256991 | -1.207 | -0.885 | -0.147 | -0.873 | 0.002 | -0.008 | -0.008 | 0.019 | 0.17 | 0.21 | 0.15 | 0.07 | 0.19 | 0.13 |
| 10 | 11166802 | rs1750730 | -0.462 | -1.306 | -0.187 | -0.928 | 0.006 | 0 | 0 | 0.217 | 0.3 | 0.28 | 0.45 | 0.72 | 0.21 | 0.48 |
| 10 | 11167624 | rs1747716 | 0.309 | -1.193 | -0.373 | -0.966 | 0.027 | -0.066 | -0.066 | 0.096 | 0.2 | 0.04 | 0.31 | 0.7 | 0.14 | 0.37 |
| 10 | 11167805 | rs1750726 | -2.085 | -1.511 | 0.215 | -1.37 | -0.001 | 0.104 | 0.104 | 0 | 0.53 | 0.72 | 0.68 | 0.85 | 0.48 | 0.72 |
| 10 | 11173303 | rs7915813 | -1.846 | -1.553 | 0.359 | -1.347 | 0.017 | 0.081 | 0.081 | 0.007 | 0.48 | 0.68 | 0.63 | 0.84 | 0.41 | 0.65 |
| 10 | 11173591 | rs1747718 | -0.946 | -1.578 | 0.561 | -1.367 | 0.021 | 0.018 | 0.018 | 0.013 | 0.33 | 0.24 | 0.4 | 0.77 | 0.22 | 0.47 |
| 10 | 11174777 | rs10905902 | 0.167 | -1.508 | 1.044 | -1.343 | -0.007 | -0.001 | -0.001 | 0 | 0.1 | 0.09 | 0.11 | 0.06 | 0.14 | 0.12 |
| 10 | 11177015 | rs12775751 | -0.071 | -2.384 | 0.56 | -1.676 | 0.029 | 0.023 | 0.023 | 0.903 | 0.29 | 0.21 | 0.15 | 0.08 | 0.15 | 0.14 |
| 10 | 11179447 | rs12245422 | -0.374 | -0.743 | 1.6 | -0.826 | 0.017 | -0.001 | -0.001 | 0 | 0.11 | 0.24 | 0.13 | 0.06 | 0.15 | 0.12 |
| 10 | 11185145 | rs7073803 | -1.231 | -0.652 | 1.524 | -0.748 | 0.003 | 0.003 | 0.003 | 0 | 0.33 | 0.28 | 0.41 | 0.78 | 0.22 | 0.47 |
| 10 | 11193951 | rs630840 | 0.079 | -0.643 | 1.199 | -0.708 | 0.037 | -0.052 | -0.052 | 0 | 0.2 | 0.04 | 0.31 | 0.71 | 0.14 | 0.36 |
| 10 | 11196369 | rs7896383 | -0.192 | -0.633 | 1.362 | -0.711 | 0.012 | 0.001 | 0.001 | 0 | 0.1 | 0.18 | 0.12 | 0.06 | 0.15 | 0.12 |
| 10 | 11196486 | rs17149655 | -1.48 | -0.549 | 1.474 | -0.796 | 0.001 | -0.067 | -0.067 | 0.252 | 0.16 | 0.29 | 0.2 | 0.07 | 0.13 | 0.16 |
| 10 | 11198255 | rs592033 | -1.789 | -0.561 | 1.25 | -0.741 | 0.023 | 0.099 | 0.099 | 0 | 0.47 | 0.7 | 0.62 | 0.82 | 0.4 | 0.69 |
| 10 | 11199917 | rs649882 | -2.047 | -0.693 | 1.39 | -0.687 | 0.004 | 0.162 | 0.162 | 0 | 0.52 | 0.76 | 0.64 | 0.85 | 0.47 | 0.71 |
| 10 | 11205224 | rs7909331 | -1.658 | -0.737 | 1.81 | -0.063 | -0.002 | -0.03 | -0.03 | 0 | 0.17 | 0.38 | 0.18 | 0.12 | 0.15 | 0.14 |
| 10 | 11210466 | rs688830 | -1.862 | -0.869 | 1.919 | 0.275 | 0.008 | 0.01 | 0.01 | 0 | 0.25 | 0.34 | 0.24 | 0.3 | 0.26 | 0.32 |
| 10 | 11212852 | rs693950 | -2.298 | -1.051 | 2.12 | 0.332 | 0.026 | 0.144 | 0.144 | 0 | 0.48 | 0.74 | 0.6 | 0.85 | 0.41 | 0.68 |
| 10 | 11215311 | rs660676 | -2.351 | -1.002 | 2.269 | 0.38 | 0 | 0 | 0 | 0.561 | 0.28 | 0.66 | 0.27 | 0.3 | 0.27 | 0.32 |
| 10 | 11217632 | rs583818 | -2.2 | -0.997 | 2.352 | 0.382 | 0.01 | 0.013 | 0.013 | 0 | 0.27 | 0.52 | 0.25 | 0.3 | 0.27 | 0.32 |
| 10 | 11221366 | rs658469 | -2.081 | -1.015 | 2.241 | 0.34 | 0.014 | 0.044 | 0.044 | 0 | 0.44 | 0.64 | 0.56 | 0.85 | 0.38 | 0.65 |
| 10 | 11222669 | rs10905909 | 0.319 | -0.956 | 2.117 | 0.511 | 0.039 | -0.11 | -0.11 | 0 | 0.18 | 0.02 | 0.3 | 0.62 | 0.12 | 0.33 |
| 10 | 11223837 | rs11257006 | -1.41 | -0.881 | 2.306 | 0.492 | -0.007 | -0.008 | -0.008 | 0.054 | 0.22 | 0.18 | 0.2 | 0.31 | 0.24 | 0.24 |
| 10 | 11224665 | rs596406 | -2.104 | -1.006 | 2.472 | 0.26 | 0.104 | 0.059 | 0.059 | 0 | 0.06 | 0.36 | 0.07 | 0.03 | 0.07 | 0.11 |
| 10 | 11225337 | rs625315 | -1.865 | -0.966 | 2.38 | 0.138 | 0.011 | 0.035 | 0.035 | 0 | 0.45 | 0.65 | 0.56 | 0.85 | 0.38 | 0.67 |
| 10 | 11227341 | rs638920 | 0.196 | -0.954 | 2.171 | 0.202 | 0.026 | -0.079 | -0.079 | 0 | 0.2 | 0.06 | 0.31 | 0.69 | 0.16 | 0.43 |
| 10 | 11228833 | rs4747894 | 0.278 | -0.956 | 2.155 | 0.209 | 0.018 | -0.123 | -0.123 | 0.105 | 0.19 | 0.03 | 0.31 | 0.58 | 0.15 | 0.43 |
| 10 | 11230987 | rs10905914 | -0.695 | -0.969 | 2.247 | 0.196 | -0.011 | -0.011 | -0.011 | 0 | 0.13 | 0.21 | 0.11 | 0.05 | 0.08 | 0.11 |
| 10 | 11231415 | rs7081571 | -0.401 | -0.959 | 2.23 | 0.199 | -0.005 | 0.011 | 0.011 | 0 | 0.12 | 0.15 | 0.1 | 0.16 | 0.08 | 0.11 |

| | | | | | | | | | | | | | | | | |
|---|---|---|---|---|---|---|---|---|---|---|---|---|---|---|---|---|
| 10 | 11231831 | rs624964 | -1.361 | -0.746 | 1.921 | 0.33 | -0.017 | -0.002 | -0.002 | 0.297 | 0.33 | 0.48 | 0.46 | 0.7 | 0.31 | 0.57 |
| 10 | 11237509 | rs4747895 | -0.205 | -0.728 | 1.577 | 0.041 | 0.02 | -0.036 | -0.036 | 0 | 0.79 | 0.87 | 0.66 | 0.3 | 0.81 | 0.55 |
| 10 | 11242780 | rs7085721 | 0.992 | -0.676 | 1.716 | -0.235 | -0.005 | -0.02 | -0.02 | 0 | 0.65 | 0.55 | 0.49 | 0.17 | 0.69 | 0.39 |
| 10 | 11246269 | rs1109861 | 1.569 | -0.692 | 1.23 | -0.288 | -0.084 | 0.229 | 0.229 | 0 | 0.42 | 0.07 | 0.36 | 0.12 | 0.54 | 0.27 |
| 10 | 11246337 | rs1109860 | 0.701 | -0.366 | 2.249 | 0.777 | -0.006 | -0.003 | -0.003 | 0 | 0.77 | 0.7 | 0.61 | 0.22 | 0.84 | 0.5 |
| 10 | 11249196 | rs10795849 | 0.972 | -0.33 | 2.193 | 0.838 | -0.043 | 0.019 | 0.019 | 0 | 0.72 | 0.47 | 0.56 | 0.31 | 0.79 | 0.5 |
| 10 | 11249240 | rs10795851 | 0.683 | -0.331 | 1.812 | 0.838 | 0.006 | -0.005 | -0.005 | 0 | 0.74 | 0.75 | 0.59 | 0.31 | 0.79 | 0.51 |
| 10 | 11251376 | rs2399667 | 0.81 | -0.301 | 1.425 | 0.862 | -0.033 | 0.016 | 0.016 | 0 | 0.72 | 0.58 | 0.57 | 0.32 | 0.79 | 0.51 |
| 10 | 11258812 | rs11596782 | -1.172 | 0.295 | 1.47 | 0.751 | 0.008 | -0.003 | -0.003 | 0 | 0.1 | 0.16 | 0.12 | 0.12 | 0.1 | 0.1 |
| 10 | 11258834 | rs4750029 | 2.061 | 0.142 | 1.616 | 1.037 | -0.005 | 0.005 | 0.005 | 0.377 | 0.74 | 0.28 | 0.69 | 0.62 | 0.8 | 0.65 |
| 10 | 11263204 | rs10160031 | 2.055 | 0.152 | 1.855 | 1.198 | -0.01 | 0.09 | 0.09 | 0.078 | 0.81 | 0.52 | 0.84 | 0.65 | 0.84 | 0.79 |
| 10 | 11263513 | rs2399669 | 1.675 | 0.169 | 1.75 | 1.205 | 0.003 | 0.11 | 0.11 | 0.223 | 0.84 | 0.55 | 0.85 | 0.65 | 0.85 | 0.79 |
| 10 | 11277613 | rs12098246 | -1.724 | 0.567 | 0.486 | 1.356 | -0.002 | 0.091 | 0.091 | 0 | 0.08 | 0.19 | 0.15 | 0.25 | 0.07 | 0.13 |
| 10 | 11287062 | rs4750033 | -1.247 | 1.224 | 0.21 | 1.2 | 0.005 | 0.012 | 0.012 | 0.79 | 0.13 | 0.14 | 0.25 | 0.23 | 0.19 | 0.35 |
| 10 | 11288418 | rs2765974 | -0.491 | 1.152 | 0.04 | 0.201 | 0.008 | 0.073 | 0.073 | 0 | 0.54 | 0.71 | 0.67 | 0.92 | 0.59 | 0.76 |
| 10 | 11290351 | rs2653521 | -0.486 | 1.214 | 0.143 | 0.608 | -0.002 | -0.016 | -0.016 | 0 | 0.53 | 0.49 | 0.58 | 0.82 | 0.6 | 0.75 |
| 10 | 11291657 | rs11257031 | -0.537 | 1.336 | -0.075 | 0.728 | -0.001 | 0 | 0 | 0 | 0.12 | 0.09 | 0.13 | 0.21 | 0.19 | 0.32 |
| 10 | 11292972 | rs2938021 | 0.144 | 1.025 | -0.149 | 0.701 | 0.004 | 0.008 | 0.008 | 0 | 0.43 | 0.39 | 0.42 | 0.65 | 0.42 | 0.43 |
| 10 | 11293778 | rs7086168 | -0.744 | 1.38 | -0.869 | -0.197 | -0.036 | 0.069 | 0.069 | 0 | 0.21 | 0.09 | 0.12 | 0.03 | 0.25 | 0.13 |
| 10 | 11293945 | rs3847420 | 0.081 | 0.768 | -0.965 | -0.53 | 0.014 | 0.018 | 0.018 | 0 | 0.43 | 0.66 | 0.49 | 0.34 | 0.39 | 0.28 |
| 10 | 11294546 | rs2765981 | -0.012 | -0.107 | -0.822 | -1.253 | -0.024 | -0.029 | -0.029 | 0.222 | 0.38 | 0.47 | 0.37 | 0.23 | 0.27 | 0.34 |
| 10 | 11294858 | rs2653517 | -0.53 | -0.202 | -0.819 | -1.193 | -0.016 | 0.032 | 0.032 | 0 | 0.76 | 0.68 | 0.7 | 0.46 | 0.79 | 0.68 |
| 10 | 11296726 | rs4750035 | 0.678 | -0.005 | -0.895 | -1.262 | -0.088 | 0.143 | 0.143 | 0 | 0.34 | 0.08 | 0.38 | 0.13 | 0.5 | 0.27 |
| 10 | 11296839 | rs11817658 | 0.312 | -0.554 | -0.665 | -1.149 | -0.027 | -0.028 | -0.028 | 1.984 | 0.4 | 0.48 | 0.36 | 0.26 | 0.27 | 0.36 |
| 10 | 11297309 | rs3740194 | -0.064 | -0.14 | -0.746 | -1.033 | -0.001 | 0.032 | 0.032 | 0 | 0.55 | 0.31 | 0.57 | 0.35 | 0.54 | 0.52 |
| 10 | 11298352 | rs2653514 | -0.217 | 0.474 | -0.1 | -0.712 | -0.006 | 0.003 | 0.003 | 0.014 | 0.11 | 0.3 | 0.12 | 0.09 | 0.1 | 0.07 |
| 10 | 11299011 | rs11257034 | 0.504 | 0.564 | -0.71 | -0.858 | -0.026 | 0.039 | 0.039 | 0 | 0.29 | 0.21 | 0.31 | 0.12 | 0.35 | 0.24 |
| 10 | 11302345 | rs1058348 | -0.19 | 0.944 | -0.822 | -0.052 | -0.003 | 0.002 | 0.002 | 0 | 0.12 | 0.13 | 0.13 | 0.07 | 0.15 | 0.09 |
| 10 | 11303759 | rs11257036 | 0.004 | 0.244 | -0.328 | 0.086 | 0.017 | -0.034 | -0.034 | 0 | 0.4 | 0.18 | 0.33 | 0.58 | 0.36 | 0.45 |
| 10 | 11305815 | rs2765989 | -1.049 | 0.009 | 0.376 | 0.285 | 0.006 | 0.023 | 0.023 | 0.077 | 0.14 | 0.33 | 0.16 | 0.14 | 0.14 | 0.19 |
| 10 | 11314769 | rs2765986 | -0.112 | -0.396 | -0.379 | 0.238 | 0.008 | -0.001 | -0.001 | 0 | 0.69 | 0.74 | 0.62 | 0.31 | 0.79 | 0.57 |
| 10 | 11315113 | rs3780993 | 0.288 | -0.71 | -0.927 | 0.035 | -0.007 | 0 | 0 | 0.479 | 0.13 | 0.02 | 0.13 | 0.07 | 0.09 | 0.09 |
| 10 | 11317357 | rs2243516 | -0.873 | -0.894 | -0.495 | -0.227 | 0.072 | 0.14 | 0.14 | 0.596 | 0.48 | 0.2 | 0.54 | 0.75 | 0.35 | 0.53 |
| 10 | 11327285 | rs11257046 | -0.369 | -1.201 | 0.135 | -0.247 | -0.002 | 0 | 0 | 3.185 | 0.39 | 0.38 | 0.45 | 0.62 | 0.28 | 0.43 |
| 10 | 11328131 | rs7099074 | 0.148 | -1.372 | 0.666 | -0.279 | 0.042 | 0.023 | 0.023 | 2.912 | 0.58 | 0.35 | 0.53 | 0.35 | 0.71 | 0.55 |
| 10 | 11329653 | rs10752217 | -0.229 | -1.256 | 0.906 | -0.334 | -0.008 | -0.03 | -0.03 | 0 | 0.84 | 0.78 | 0.86 | 0.92 | 0.88 | 0.91 |
| 10 | 11330576 | rs2277216 | -0.204 | -1.519 | 0.575 | -0.019 | -0.04 | 0.04 | 0.04 | 1.555 | 0.33 | 0.48 | 0.31 | 0.52 | 0.26 | 0.36 |
| 10 | 11331406 | rs17453876 | 0.078 | -1.053 | 0.068 | -0.345 | -0.001 | 0.036 | 0.036 | 1.67 | 0.14 | 0.07 | 0.12 | 0.06 | 0.12 | 0.07 |
| 10 | 11334678 | rs4747900 | 0.844 | -1.131 | -0.032 | -0.34 | 0.015 | 0.094 | 0.094 | 0 | 0.43 | 0.16 | 0.38 | 0.31 | 0.58 | 0.39 |
| 10 | 11335074 | rs1112520 | 0.878 | -0.407 | 0.332 | -0.624 | -0.006 | 0.004 | 0.004 | 0.071 | 0.37 | 0.35 | 0.32 | 0.3 | 0.45 | 0.26 |
| 10 | 11335395 | rs4129775 | 0.035 | -1.345 | 0.235 | -0.429 | 0.003 | 0.007 | 0.007 | 3.261 | 0.15 | 0.12 | 0.1 | 0.07 | 0.12 | 0.09 |
| 10 | 11337318 | rs17454012 | -0.291 | -1.32 | 0.292 | -0.298 | 0.006 | 0.008 | 0.008 | 0.66 | 0.2 | 0.2 | 0.15 | 0.11 | 0.15 | 0.13 |
| 10 | 11341596 | rs17150094 | -0.649 | -1.326 | 0.225 | -0.308 | -0.022 | -0.041 | -0.041 | 0 | 0.25 | 0.63 | 0.16 | 0.08 | 0.17 | 0.1 |
| 10 | 11342601 | rs1910231 | -0.418 | -1.392 | -0.185 | -0.276 | 0.041 | -0.012 | -0.012 | 0 | 0.66 | 0.82 | 0.54 | 0.39 | 0.75 | 0.47 |
| 10 | 11342887 | rs10905928 | 0.426 | -1.373 | -0.528 | -0.247 | 0.059 | -0.082 | -0.082 | 0 | 0.32 | 0.09 | 0.39 | 0.56 | 0.2 | 0.45 |
| 10 | 11345047 | rs10905929 | -0.645 | -0.688 | 0.658 | -0.229 | -0.051 | -0.08 | -0.08 | 0.054 | 0.21 | 0.4 | 0.12 | 0.06 | 0.11 | 0.1 |
| 10 | 11345345 | rs1875594 | 0.678 | -0.587 | -0.598 | -0.154 | 0.026 | 0.054 | 0.054 | 0.293 | 0.72 | 0.24 | 0.77 | 0.9 | 0.8 | 0.85 |
| 10 | 11355431 | rs11257055 | -0.562 | -0.816 | -0.228 | 0.643 | -0.032 | 0.035 | 0.035 | 0 | 0.23 | 0.36 | 0.15 | 0.39 | 0.16 | 0.36 |
| 10 | 11356160 | rs2246449 | -1.116 | -2.045 | -0.335 | 0.16 | 0.047 | -0.007 | -0.007 | 0.016 | 0.44 | 0.64 | 0.26 | 0.49 | 0.27 | 0.49 |
| 10 | 11357510 | rs2938015 | -1.153 | -2.181 | 0.013 | 0.101 | -0.017 | 0 | 0 | 0 | 0.44 | 0.56 | 0.26 | 0.48 | 0.27 | 0.48 |
| 10 | 11359469 | rs3781053 | 1.195 | -1.199 | -0.767 | 0.062 | 0 | -0.015 | -0.015 | 0 | 0.3 | 0.36 | 0.36 | 0.28 | 0.33 | 0.13 |
| 10 | 11360673 | rs2279809 | -0.231 | -0.683 | -0.819 | 0.075 | -0.006 | 0.005 | 0.005 | 0.081 | 0.57 | 0.59 | 0.53 | 0.41 | 0.52 | 0.3 |
| 10 | 11362317 | rs10795860 | -0.232 | -1.142 | -0.893 | 0.107 | -0.046 | 0.023 | 0.023 | 0.043 | 0.68 | 0.44 | 0.84 | 0.56 | 0.8 | 0.57 |
| 10 | 11367045 | rs12261260 | 1.221 | -0.916 | -0.919 | 0.046 | -0.022 | 0.008 | 0.008 | 0.852 | 0.28 | 0.13 | 0.37 | 0.3 | 0.33 | 0.12 |
| 10 | 11367447 | rs10905930 | -0.352 | -0.965 | -0.743 | 0.015 | 0.002 | 0.002 | 0.002 | 0 | 0.51 | 0.47 | 0.49 | 0.38 | 0.45 | 0.24 |
| 10 | 11368662 | rs12782642 | 1.075 | -0.968 | -0.798 | -0.003 | 0.004 | 0.074 | 0.074 | 0 | 0.12 | 0 | 0.05 | 0 | 0.1 | 0.08 |
| 10 | 11368747 | rs12268544 | 0.385 | -0.974 | -0.755 | 0.011 | -0.014 | -0.007 | -0.007 | 0 | 0.39 | 0.47 | 0.44 | 0.38 | 0.35 | 0.17 |
| 10 | 11369250 | rs2378991 | 0.005 | -0.437 | -0.947 | 0.124 | 0.015 | 0.059 | 0.059 | 0.183 | 0.43 | 0.26 | 0.44 | 0.32 | 0.42 | 0.2 |
| 10 | 11369870 | rs9329334 | 0.906 | -0.335 | -0.984 | 0.173 | -0.01 | -0.007 | -0.007 | 0 | 0.29 | 0.15 | 0.37 | 0.32 | 0.33 | 0.12 |
| 10 | 11375659 | rs17454839 | 0.233 | -0.39 | -0.888 | -0.009 | -0.007 | -0.04 | -0.04 | 0 | 0.15 | 0.17 | 0.09 | 0 | 0.12 | 0.08 |
| 10 | 11376999 | rs1051765 | 0.455 | -0.302 | -1.334 | 0.071 | 0.02 | 0.066 | 0.066 | 0 | 0.14 | 0.04 | 0.06 | 0 | 0.11 | 0.09 |
| 10 | 11384120 | rs12220081 | 0.727 | 0.125 | -1.256 | 0.623 | -0.051 | -0.05 | -0.05 | 0.011 | 0.27 | 0.09 | 0.23 | 0.33 | 0.32 | 0.17 |
| 10 | 11388290 | rs2440081 | 0.068 | -0.704 | -0.429 | 0.667 | -0.05 | 0.05 | 0.05 | 0.093 | 0.21 | 0.49 | 0.22 | 0.32 | 0.15 | 0.36 |
| 10 | 11388353 | rs2455584 | 0.077 | -0.71 | -0.275 | 0.664 | -0.048 | 0.051 | 0.051 | 0.088 | 0.21 | 0.45 | 0.22 | 0.32 | 0.15 | 0.36 |
| 10 | 11388843 | rs7098278 | 0.16 | -0.391 | -0.7 | 0.394 | -0.006 | 0.012 | 0.012 | 0 | 0.41 | 0.35 | 0.51 | 0.34 | 0.52 | 0.42 |
| 10 | 11389327 | rs3905635 | 0.02 | -0.276 | -0.635 | 0.167 | -0.015 | -0.01 | -0.01 | 0 | 0.66 | 0.46 | 0.74 | 0.69 | 0.75 | 0.62 |
| 10 | 11391939 | rs7090767 | 0.435 | 0.473 | -1.096 | 0.086 | -0.011 | 0.033 | 0.033 | 0 | 0.31 | 0.25 | 0.51 | 0.15 | 0.38 | 0.33 |
| 10 | 11392180 | rs6602495 | -0.49 | 0.412 | -0.633 | 0.138 | -0.009 | -0.001 | -0.001 | 0 | 0.72 | 0.66 | 0.8 | 0.69 | 0.77 | 0.59 |
| 10 | 11393873 | rs11257064 | 0.497 | 0.429 | -1.274 | 0.146 | -0.001 | 0.027 | 0.027 | 0 | 0.3 | 0.24 | 0.42 | 0.15 | 0.33 | 0.3 |

| | | | | | | | | | | | | | | | | |
|---|---|---|---|---|---|---|---|---|---|---|---|---|---|---|---|---|
| 10 | 11394809 | rs3847424 | 1.14 | 0.59 | -1.358 | 0.296 | -0.009 | 0.002 | 0.002 | 0 | 0.13 | 0.09 | 0.12 | 0.12 | 0.18 | 0.09 |
| 10 | 11395889 | rs4114417 | -0.153 | 0.509 | -1.346 | 0.25 | -0.005 | 0.016 | 0.016 | 0 | 0.44 | 0.41 | 0.55 | 0.28 | 0.5 | 0.39 |
| 10 | 11395977 | rs4114416 | 0.387 | 0.603 | -1.306 | 0.027 | -0.013 | 0.033 | 0.033 | 0 | 0.53 | 0.32 | 0.56 | 0.57 | 0.53 | 0.43 |
| 10 | 11396152 | rs12245515 | 0.634 | 1.101 | -1.168 | 0.113 | 0.003 | -0.002 | -0.002 | 0 | 0.13 | 0.13 | 0.12 | 0.12 | 0.17 | 0.09 |
| 10 | 11397752 | rs12775688 | 0.291 | 0.689 | -1.442 | 0.276 | -0.024 | -0.014 | -0.014 | 1.638 | 0.39 | 0.27 | 0.42 | 0.51 | 0.46 | 0.26 |
| 10 | 11402047 | rs3847427 | -0.123 | 0.352 | -1.618 | 0.166 | -0.02 | -0.005 | -0.005 | 0 | 0.61 | 0.47 | 0.66 | 0.64 | 0.72 | 0.51 |
| 10 | 11405290 | rs10752219 | -0.589 | 0.124 | -1.486 | 0.143 | -0.021 | 0.005 | 0.005 | 0 | 0.68 | 0.61 | 0.69 | 0.64 | 0.76 | 0.57 |
| 10 | 11412053 | rs11257069 | 1.067 | 0.288 | -2.324 | 0.572 | -0.038 | -0.117 | -0.117 | 0 | 0.23 | 0.03 | 0.15 | 0.35 | 0.22 | 0.16 |
| 10 | 11414182 | rs2455597 | 0.423 | 0.607 | -1.972 | -0.087 | -0.018 | -0.044 | -0.044 | 0 | 0.09 | 0.15 | 0.03 | 0 | 0.03 | 0.1 |
| 10 | 11417026 | rs7901534 | 0.705 | 0.692 | -2.212 | -0.17 | -0.068 | 0.146 | 0.146 | 0.027 | 0.32 | 0.11 | 0.36 | 0.09 | 0.43 | 0.36 |
| 10 | 11419738 | rs12254063 | -0.259 | 1.186 | -1.462 | 0.75 | -0.014 | -0.021 | -0.021 | 0 | 0.13 | 0.22 | 0.06 | 0.06 | 0.06 | 0.11 |
| 10 | 11421378 | rs7090793 | -0.185 | 1.854 | -1.821 | 1.224 | -0.076 | -0.065 | -0.065 | 0.195 | 0.13 | 0.01 | 0.25 | 0.19 | 0.22 | 0.08 |
| 10 | 11422431 | rs11257074 | -0.064 | 1.659 | -1.521 | 1.512 | -0.016 | -0.048 | -0.048 | 0 | 0.2 | 0.12 | 0.17 | 0.4 | 0.22 | 0.16 |
| 10 | 11422724 | rs10905940 | -0.648 | 1.732 | -1.247 | 1.166 | -0.005 | -0.004 | -0.004 | 0 | 0.28 | 0.26 | 0.57 | 0.6 | 0.37 | 0.34 |
| 10 | 11423434 | rs2454805 | -0.438 | 1.022 | -0.411 | 1.308 | -0.001 | 0.214 | 0.214 | 0 | 0.11 | 0.33 | 0.11 | 0.4 | 0.07 | 0.19 |
| 10 | 11423457 | rs4747905 | 0.452 | 1.06 | -1.527 | 0.344 | -0.098 | 0.052 | 0.052 | 0 | 0.41 | 0.11 | 0.47 | 0.32 | 0.43 | 0.49 |
| 10 | 11424447 | rs2454803 | 0.349 | 0.417 | -1.341 | 0.374 | 0.024 | 0.123 | 0.123 | 0 | 0.48 | 0.15 | 0.53 | 0.32 | 0.58 | 0.53 |
| 10 | 11430024 | rs2454800 | 0.19 | 0.063 | 0.019 | 0.496 | -0.033 | 0.15 | 0.15 | 0 | 0.81 | 0.65 | 0.84 | 0.55 | 0.87 | 0.76 |
| 10 | 11430558 | rs2938023 | 0.198 | -0.052 | 0.247 | 0.348 | 0.003 | 0.022 | 0.022 | 1.003 | 0.33 | 0.17 | 0.3 | 0.25 | 0.32 | 0.27 |
| 10 | 11431354 | rs4750051 | 0.206 | -0.054 | 0.363 | 0.347 | 0.002 | 0.022 | 0.022 | 0.854 | 0.33 | 0.18 | 0.3 | 0.25 | 0.32 | 0.27 |
| 10 | 11431693 | rs2454797 | 0.383 | -0.316 | 0.477 | 0.415 | -0.019 | 0.048 | 0.048 | 0.002 | 0.79 | 0.33 | 0.81 | 0.55 | 0.87 | 0.76 |
| 10 | 11433996 | rs7088943 | 0.064 | -0.648 | 0.492 | 0.172 | -0.043 | 0.058 | 0.058 | 1.222 | 0.11 | 0.45 | 0.13 | 0.09 | 0.04 | 0.04 |
| 10 | 11438205 | rs10905941 | 0.22 | -0.608 | 0.422 | 0.12 | 0 | 0.007 | 0.007 | 0 | 0.6 | 0.67 | 0.66 | 0.38 | 0.6 | 0.53 |
| 10 | 11438690 | rs10795865 | 0.257 | -0.594 | 0.224 | 0.14 | 0.002 | 0.03 | 0.03 | 0 | 0.57 | 0.17 | 0.62 | 0.38 | 0.6 | 0.53 |
| 10 | 11438833 | rs10905942 | 0.922 | -0.566 | 0.176 | 0.097 | -0.005 | 0.129 | 0.129 | 0.008 | 0.46 | 0.13 | 0.51 | 0.29 | 0.56 | 0.48 |
| 10 | 11439836 | rs2379113 | 0.947 | -0.378 | 0.13 | 0.071 | -0.036 | 0.124 | 0.124 | 0 | 0.45 | 0.13 | 0.51 | 0.3 | 0.55 | 0.48 |
| 10 | 11444870 | rs2440100 | -0.594 | -0.172 | 0.648 | 0.127 | -0.006 | 0.015 | 0.015 | 0 | 0.77 | 0.86 | 0.82 | 0.82 | 0.75 | 0.81 |
| 10 | 11448858 | rs2455600 | -0.794 | -0.212 | 0.824 | 0.146 | -0.048 | -0.008 | -0.008 | 0.023 | 0.14 | 0.39 | 0.13 | 0.09 | 0.05 | 0.06 |
| 10 | 11452260 | rs3847437 | -0.187 | -0.211 | 0.399 | 0.14 | -0.029 | -0.003 | -0.003 | 0.038 | 0.12 | 0.26 | 0.13 | 0.09 | 0.05 | 0.06 |
| 10 | 11455577 | rs4750054 | 0.262 | 0.069 | -0.523 | 0.149 | 0.012 | 0.017 | 0.017 | 0 | 0.63 | 0.37 | 0.68 | 0.74 | 0.7 | 0.75 |
| 10 | 11464063 | rs2440113 | -0.446 | 0.094 | -0.554 | 0.16 | 0.047 | 0.006 | 0.006 | 2.89 | 0.07 | 0.56 | 0.05 | 0 | 0.01 | 0.02 |
| 10 | 11465260 | rs7095714 | 0.037 | 0.281 | -0.898 | 0.282 | 0.014 | 0.021 | 0.021 | 0 | 0.63 | 0.34 | 0.68 | 0.74 | 0.71 | 0.75 |
| 10 | 11468441 | rs4750058 | 0.282 | 0.794 | -1.746 | -0.009 | -0.128 | 0.013 | 0.013 | 0 | 0.31 | 0.05 | 0.39 | 0.26 | 0.39 | 0.42 |
| 10 | 11473258 | rs2499902 | -0.462 | -0.144 | -1.147 | -0.553 | -0.007 | -0.051 | -0.051 | 2.922 | 0.34 | 0.63 | 0.24 | 0.08 | 0.25 | 0.17 |
| 10 | 11477584 | rs2440119 | -0.815 | -0.396 | -1.654 | 0.299 | 0.063 | -0.005 | -0.005 | 0 | 0.49 | 0.75 | 0.42 | 0.43 | 0.37 | 0.46 |
| 10 | 11479998 | rs7091016 | 0.868 | -0.348 | -2.169 | 0.156 | -0.143 | 0.14 | 0.14 | 0 | 0.35 | 0.06 | 0.38 | 0.17 | 0.47 | 0.34 |
| 10 | 11481180 | rs2454779 | -0.049 | -0.356 | -1.127 | 0.528 | -0.001 | 0.087 | 0.087 | 0 | 0.13 | 0.23 | 0.2 | 0.27 | 0.14 | 0.21 |
| 10 | 11482049 | rs2920412 | -0.079 | -0.214 | -1.105 | 1.043 | 0.039 | 0.142 | 0.142 | 0 | 0.13 | 0.22 | 0.29 | 0.42 | 0.21 | 0.27 |
| 10 | 11483103 | rs2499906 | -0.6 | -0.892 | -1.074 | 0.48 | -0.009 | 0.053 | 0.053 | 0 | 0.25 | 0.48 | 0.29 | 0.32 | 0.21 | 0.28 |
| 10 | 11483175 | rs4750060 | 0.754 | -0.745 | -1.234 | 0 | -0.007 | 0.016 | 0.016 | 0 | 0.1 | 0.06 | 0.06 | 0 | 0.12 | 0.06 |
| 10 | 11491077 | rs1530375 | -0.44 | -0.238 | -1.199 | -0.417 | 0 | -0.021 | -0.021 | 0.006 | 0.42 | 0.43 | 0.61 | 0.88 | 0.44 | 0.62 |
| 10 | 11492868 | rs11257101 | -0.567 | -0.168 | -1.207 | 0.294 | -0.002 | -0.016 | -0.016 | 0 | 0.29 | 0.27 | 0.36 | 0.38 | 0.3 | 0.4 |
| 10 | 11494666 | rs7070337 | 0.131 | -0.141 | -1.269 | 0.421 | -0.034 | -0.045 | -0.045 | 0 | 0.21 | 0.08 | 0.25 | 0.26 | 0.25 | 0.28 |
| 10 | 11498130 | rs11257104 | 0.756 | -0.25 | -1.196 | 0.309 | 0.007 | 0.009 | 0.009 | 0 | 0.16 | 0.14 | 0.08 | 0.09 | 0.11 | 0.21 |
| 10 | 11498486 | rs11257105 | 0.756 | -0.25 | -1.196 | 0.309 | 0.007 | 0.009 | 0.009 | 0 | 0.16 | 0.14 | 0.08 | 0.09 | 0.11 | 0.21 |
| 10 | 11505175 | rs3740204 | -0.943 | -1.281 | -0.7 | 0.013 | 0.006 | 0.009 | 0.009 | 0 | 0.42 | 0.58 | 0.3 | 0.74 | 0.3 | 0.49 |
| 10 | 11512648 | rs2440083 | 1.239 | -0.486 | -1.579 | -0.243 | -0.076 | 0.122 | 0.122 | 0.003 | 0.31 | 0.08 | 0.28 | 0.1 | 0.37 | 0.22 |
| 10 | 11530872 | rs11257127 | -0.431 | -0.114 | -0.792 | 0.342 | -0.005 | 0.035 | 0.035 | 0 | 0.16 | 0.18 | 0.31 | 0.61 | 0.17 | 0.22 |
| 10 | 11543661 | rs7085769 | -0.43 | -0.037 | -0.774 | 0.338 | 0.003 | 0.016 | 0.016 | 0 | 0.17 | 0.15 | 0.31 | 0.61 | 0.2 | 0.23 |
| 10 | 11546376 | rs12571504 | -0.75 | -0.223 | -0.832 | 0.16 | 0 | 0 | 0 | 0 | 0.31 | 0.27 | 0.38 | 0.7 | 0.29 | 0.43 |
| 10 | 11549584 | rs1550652 | 1.369 | -0.155 | -0.902 | 0.489 | 0.008 | -0.046 | -0.046 | 0 | 0.08 | 0 | 0.09 | 0.14 | 0.06 | 0.14 |
| 10 | 11564421 | rs4445554 | -0.375 | -0.08 | -0.822 | -0.131 | 0.003 | 0.015 | 0.015 | 0 | 0.17 | 0.16 | 0.32 | 0.61 | 0.19 | 0.22 |
| 10 | 11571368 | rs10905968 | 2.294 | -0.135 | -1.16 | -0.422 | -0.039 | -0.016 | -0.016 | 0 | 0.15 | 0.03 | 0.21 | 0.21 | 0.18 | 0.23 |
| 10 | 11574424 | rs12250042 | 0.789 | -0.38 | -0.774 | -0.727 | 0 | -0.012 | -0.012 | 0.843 | 0.46 | 0.43 | 0.35 | 0.1 | 0.49 | 0.29 |
| 10 | 11580354 | rs17150540 | 1.024 | -0.304 | -0.884 | -0.255 | -0.027 | 0.153 | 0.153 | 0 | 0.11 | 0.18 | 0.14 | 0.44 | 0.06 | 0.18 |
| 10 | 11582344 | rs10905969 | 1.581 | -0.435 | -0.991 | -0.467 | 0.006 | 0.014 | 0.014 | 0 | 0.09 | 0.07 | 0.05 | 0 | 0.09 | 0.08 |
| 10 | 11599041 | rs10905973 | 1.751 | -0.275 | -0.808 | -0.249 | 0.013 | 0.013 | 0.013 | 0 | 0.14 | 0.07 | 0.05 | 0.08 | 0.09 | 0.21 |
| 10 | 11623686 | rs11257179 | 0.917 | -0.123 | -0.832 | -0.209 | -0.036 | -0.033 | -0.033 | 0.154 | 0.06 | 0 | 0.18 | 0.17 | 0.14 | 0.04 |
| 10 | 11629403 | rs754426 | -0.697 | -0.255 | -0.66 | -0.17 | -0.025 | 0.219 | 0.219 | 0 | 0.87 | 0.71 | 0.85 | 0.56 | 0.93 | 0.8 |
| 10 | 11635029 | rs17136777 | 0.387 | -0.236 | -0.693 | -0.444 | -0.013 | 0.014 | 0.014 | 0 | 0.25 | 0.26 | 0.2 | 0.52 | 0.15 | 0.39 |
| 10 | 11663611 | rs11257199 | 1.728 | 0.329 | -0.962 | -0.755 | -0.047 | -0.008 | -0.008 | 0 | 0.15 | 0.02 | 0.21 | 0.19 | 0.18 | 0.22 |
| 10 | 11671268 | rs7094839 | 0.063 | 0.058 | -0.442 | -1.08 | 0 | -0.036 | -0.036 | 0 | 0.46 | 0.44 | 0.36 | 0.1 | 0.5 | 0.31 |
| 10 | 11680238 | rs7903301 | -1.055 | 0.199 | 0.069 | -0.828 | 0.044 | -0.007 | -0.007 | 0 | 0.15 | 0.46 | 0.17 | 0.11 | 0.14 | 0.1 |
| 10 | 11680543 | rs10795870 | 0.95 | 0.136 | -0.295 | -0.956 | -0.057 | 0.036 | 0.036 | 0.227 | 0.24 | 0.03 | 0.32 | 0.21 | 0.26 | 0.28 |
| 10 | 11681939 | rs4357603 | -0.412 | 0.424 | 0.252 | -0.716 | 0.109 | 0.115 | 0.115 | 0.124 | 0.05 | 0.31 | 0.11 | 0.11 | 0.06 | 0.03 |
| 10 | 11682396 | rs7101132 | -1.125 | 0.343 | 0.058 | -0.738 | 0.034 | -0.009 | -0.009 | 0 | 0.14 | 0.38 | 0.16 | 0.11 | 0.14 | 0.1 |
| 10 | 11682900 | rs6602510 | 0.185 | 0.057 | -0.188 | -0.849 | -0.016 | -0.045 | -0.045 | 0 | 0.08 | 0.16 | 0.04 | 0 | 0.05 | 0.06 |
| 10 | 11693166 | rs10905983 | 0.191 | -0.089 | -0.431 | -1.082 | -0.011 | -0.055 | -0.055 | 0 | 0.38 | 0.44 | 0.39 | 0.22 | 0.33 | 0.35 |
| 10 | 11694596 | rs6602512 | -1.419 | 0.896 | -0.81 | -1.094 | -0.024 | 0.024 | 0.024 | 0 | 0.9 | 0.93 | 0.9 | 0.96 | 0.88 | 0.89 |

| | | | | | | | | | | | | | | | | |
|---|---|---|---|---|---|---|---|---|---|---|---|---|---|---|---|---|
| 10 | 11697403 | rs11257215 | -0.414 | 0.726 | -0.562 | -1.355 | -0.002 | -0.083 | -0.083 | 0.019 | 0.31 | 0.35 | 0.26 | 0.04 | 0.29 | 0.2 |
| 10 | 11703136 | rs10905985 | 1.056 | 0.96 | -1.036 | -1.151 | -0.003 | 0.001 | 0.001 | 0 | 0.2 | 0.15 | 0.15 | 0.11 | 0.29 | 0.23 |
| 10 | 11703427 | rs7070729 | -0.147 | 0.279 | -1.193 | -0.74 | -0.038 | 0.022 | 0.022 | 0 | 0.65 | 0.47 | 0.69 | 0.48 | 0.78 | 0.67 |
| 10 | 11712575 | rs7084973 | -0.562 | 0.692 | -0.994 | -0.452 | -0.055 | 0.108 | 0.108 | 0 | 0.73 | 0.49 | 0.72 | 0.48 | 0.84 | 0.69 |
| 10 | 11712965 | rs12784561 | 0.046 | 1.118 | -1.105 | -0.424 | -0.004 | -0.006 | -0.006 | 0 | 0.05 | 0.05 | 0.04 | 0.11 | 0.1 | 0.03 |
| 10 | 11713277 | rs8181299 | -0.585 | 1.057 | -1.087 | -0.421 | -0.055 | 0.102 | 0.102 | 0 | 0.73 | 0.49 | 0.72 | 0.48 | 0.84 | 0.69 |
| 10 | 11713869 | rs11257231 | 0.799 | 1.387 | -1.64 | -0.466 | -0.002 | 0 | 0 | 0 | 0.21 | 0.17 | 0.15 | 0.1 | 0.29 | 0.23 |
| 10 | 11718331 | rs2011704 | 1.071 | 1.435 | -1.757 | -0.571 | -0.021 | 0.038 | 0.038 | 0.082 | 0.33 | 0.11 | 0.4 | 0.22 | 0.35 | 0.33 |
| 10 | 11719010 | rs12265836 | 1.004 | 1.489 | -1.761 | -0.545 | -0.003 | 0.002 | 0.002 | 0.674 | 0.21 | 0.15 | 0.14 | 0.09 | 0.28 | 0.22 |
| 10 | 11719074 | rs11257240 | 1.127 | 1.418 | -1.767 | -0.53 | -0.024 | 0.035 | 0.035 | 0 | 0.33 | 0.11 | 0.4 | 0.22 | 0.35 | 0.33 |
| 10 | 11722775 | rs3829155 | -0.315 | 2.049 | -0.932 | -0.35 | -0.013 | 0.006 | 0.006 | 3.086 | 0.25 | 0.16 | 0.19 | 0.21 | 0.4 | 0.26 |
| 10 | 11723616 | rs10508428 | -0.425 | 1.615 | -0.815 | -0.296 | -0.007 | 0.015 | 0.015 | 0 | 0.34 | 0.3 | 0.35 | 0.15 | 0.43 | 0.33 |
| 10 | 11727785 | rs1466696 | -1.327 | 1.659 | -0.571 | 0.163 | -0.019 | 0.003 | 0.003 | 0 | 0.13 | 0.1 | 0.2 | 0.1 | 0.21 | 0.2 |
| 10 | 11729374 | rs1466695 | -2.065 | 1.477 | 0.231 | -0.036 | 0.047 | -0.029 | -0.029 | 0 | 0.19 | 0.35 | 0.24 | 0.1 | 0.26 | 0.27 |
| 10 | 11730196 | rs17150715 | -1.007 | 1.612 | -0.011 | 0.048 | -0.001 | -0.01 | -0.01 | 0.656 | 0.08 | 0.09 | 0.15 | 0.02 | 0.09 | 0.16 |
| 10 | 11731124 | rs11257254 | -0.496 | 1.577 | -0.122 | 0.182 | -0.023 | -0.026 | -0.026 | 0 | 0.05 | 0.01 | 0.08 | 0.09 | 0.12 | 0.05 |
| 10 | 11736956 | rs2399726 | -0.297 | 1.071 | -0.255 | 0.26 | 0.005 | -0.012 | -0.012 | 1.095 | 0.39 | 0.25 | 0.24 | 0.46 | 0.38 | 0.41 |
| 10 | 11738432 | rs9730979 | -1.173 | 0.728 | 0.239 | -0.097 | 0.003 | 0.008 | 0.008 | 0 | 0.58 | 0.59 | 0.52 | 0.74 | 0.61 | 0.64 |
| 10 | 11740084 | rs10905996 | -1.034 | 0.745 | 0.335 | -0.082 | 0.006 | 0.007 | 0.007 | 0.249 | 0.54 | 0.42 | 0.5 | 0.75 | 0.62 | 0.68 |
| 10 | 11741083 | rs12356463 | -1.108 | 1.22 | 0.536 | 0.155 | 0.008 | -0.012 | -0.012 | 0 | 0.15 | 0.19 | 0.15 | 0.09 | 0.21 | 0.17 |
| 10 | 11743811 | rs12764425 | -0.915 | 1.409 | -0.09 | 0.213 | -0.069 | 0.026 | 0.026 | 0 | 0.15 | 0.04 | 0.15 | 0.08 | 0.23 | 0.2 |
| 10 | 11750055 | rs10795878 | -0.81 | 0.792 | -0.253 | 0.513 | 0.003 | 0.007 | 0.007 | 0 | 0.46 | 0.42 | 0.45 | 0.59 | 0.55 | 0.55 |
| 10 | 11751376 | rs11257269 | -0.39 | 0.99 | -0.672 | 0.658 | -0.022 | -0.032 | -0.032 | 0 | 0.22 | 0.13 | 0.35 | 0.3 | 0.23 | 0.22 |
| 10 | 11757180 | rs4750085 | 0.4 | 0.331 | -0.528 | -0.146 | 0.012 | 0.047 | 0.047 | 0 | 0.64 | 0.29 | 0.8 | 0.88 | 0.72 | 0.72 |
| 10 | 11767916 | rs10795879 | -0.599 | -0.276 | 0.06 | 0.162 | -0.003 | -0.001 | -0.001 | 0 | 0.55 | 0.58 | 0.51 | 0.59 | 0.61 | 0.62 |
| 10 | 11773389 | rs4747933 | -0.585 | -0.536 | 0.181 | 0.048 | 0.04 | 0.014 | 0.014 | 0 | 0.55 | 0.38 | 0.61 | 0.6 | 0.65 | 0.6 |
| 10 | 11779908 | rs718641 | -0.428 | -0.793 | -0.148 | 0.215 | -0.002 | 0.006 | 0.006 | 0.24 | 0.66 | 0.41 | 0.58 | 0.49 | 0.68 | 0.58 |
| 10 | 11780837 | rs2055040 | -0.492 | -1.209 | 0.075 | 0.056 | -0.03 | 0.039 | 0.039 | 0.242 | 0.67 | 0.6 | 0.65 | 0.5 | 0.71 | 0.6 |
| 10 | 11783873 | rs3814627 | 0.669 | -1.235 | 0.118 | 0.082 | 0.021 | -0.083 | -0.083 | 0.362 | 0.29 | 0.13 | 0.31 | 0.54 | 0.29 | 0.42 |
| 10 | 11789637 | rs4750093 | 0.227 | -0.891 | 0.028 | -0.225 | 0.005 | -0.031 | -0.031 | 0 | 0.37 | 0.31 | 0.35 | 0.55 | 0.36 | 0.46 |
| 10 | 11797675 | rs3750695 | 0.929 | -0.754 | -0.248 | -0.391 | -0.023 | -0.02 | -0.02 | 0 | 0.24 | 0.12 | 0.36 | 0.22 | 0.31 | 0.22 |
| 10 | 11803185 | rs10752230 | -0.959 | -0.506 | 0.27 | -0.424 | -0.017 | -0.002 | -0.002 | 0 | 0.67 | 0.55 | 0.69 | 0.69 | 0.7 | 0.67 |
| 10 | 11815455 | rs7101167 | -0.361 | -0.521 | 0.169 | -0.523 | 0.039 | -0.112 | -0.112 | 0.179 | 0.79 | 0.69 | 0.83 | 0.99 | 0.78 | 0.85 |
| 10 | 11817835 | rs2131265 | -0.449 | -0.055 | 0.409 | -0.254 | 0.015 | 0.04 | 0.04 | 0 | 0.49 | 0.66 | 0.55 | 0.68 | 0.47 | 0.51 |
| 10 | 11822949 | rs10795892 | 0.187 | -0.114 | 0.504 | -0.203 | 0.007 | -0.01 | -0.01 | 0 | 0.39 | 0.32 | 0.47 | 0.38 | 0.38 | 0.35 |
| 10 | 11824850 | rs7098491 | 0.572 | -0.152 | 0.917 | 0.145 | 0.086 | 0.264 | 0.264 | 0 | 0.92 | 0.57 | 0.91 | 0.74 | 0.92 | 0.84 |
| 10 | 11825696 | rs10508432 | 0.674 | -0.174 | 0.367 | -0.012 | 0.026 | 0.009 | 0.009 | 0 | 0.36 | 0.21 | 0.3 | 0.33 | 0.3 | 0.37 |
| 10 | 11828686 | rs4750112 | 1.31 | -0.165 | 0.089 | 0.005 | 0.052 | -0.024 | -0.024 | 0 | 0.28 | 0.12 | 0.29 | 0.31 | 0.21 | 0.3 |
| 10 | 11831111 | rs17150863 | 0.149 | 0.654 | 0.31 | 0.435 | 0.03 | 0.065 | 0.065 | 0 | 0.05 | 0.12 | 0.09 | 0.25 | 0.09 | 0.14 |
| 10 | 11837356 | rs11597793 | 0.429 | 0.58 | -0.064 | 0.284 | -0.029 | 0.037 | 0.037 | 0.373 | 0.08 | 0.01 | 0.04 | 0 | 0.08 | 0.06 |
| 10 | 11838400 | rs11257329 | 0.166 | 0.487 | 0.218 | 0.432 | 0.022 | 0.057 | 0.057 | 0 | 0.22 | 0.39 | 0.3 | 0.26 | 0.24 | 0.19 |
| 10 | 11843051 | rs942530 | -0.191 | 0.58 | -0.549 | 0.439 | -0.013 | -0.022 | -0.022 | 0.084 | 0.31 | 0.26 | 0.37 | 0.54 | 0.33 | 0.33 |
| 10 | 11846021 | rs1359363 | -0.266 | 0.554 | -0.74 | 0.394 | -0.025 | 0.006 | 0.006 | 0.348 | 0.34 | 0.28 | 0.34 | 0.25 | 0.34 | 0.32 |
| 10 | 11847227 | rs11257335 | 0.641 | 0.146 | -0.373 | 0.373 | -0.017 | 0.004 | 0.004 | 0 | 0.09 | 0.17 | 0.15 | 0.12 | 0.06 | 0.03 |
| 10 | 11848189 | rs942531 | -0.312 | 0.401 | -0.996 | 0.128 | -0.007 | 0.006 | 0.006 | 0 | 0.26 | 0.23 | 0.28 | 0.26 | 0.35 | 0.26 |
| 10 | 11850132 | rs1535976 | 0.359 | 0.542 | -0.831 | 0.399 | -0.003 | -0.001 | -0.001 | 0 | 0.43 | 0.35 | 0.38 | 0.5 | 0.42 | 0.44 |
| 10 | 11855599 | rs11599763 | 0.649 | 0.72 | -0.843 | 0.735 | 0 | -0.014 | -0.014 | 0 | 0.6 | 0.46 | 0.49 | 0.28 | 0.61 | 0.6 |
| 10 | 11856145 | rs10906021 | -0.1 | 1.178 | -1.518 | 0.955 | -0.037 | -0.066 | -0.066 | 0 | 0.21 | 0.09 | 0.26 | 0.3 | 0.27 | 0.21 |
| 10 | 11861758 | rs4750121 | -0.276 | 0.833 | -1.601 | 0.804 | 0.004 | -0.02 | -0.02 | 4.752 | 0.71 | 0.86 | 0.74 | 0.69 | 0.75 | 0.62 |
| 10 | 11869246 | rs10906025 | -0.138 | 1.357 | -1.284 | 0.585 | 0.011 | 0.006 | 0.006 | 0 | 0.29 | 0.34 | 0.53 | 0.54 | 0.46 | 0.43 |
| 10 | 11869521 | rs10508433 | 0.486 | 0.863 | -1.369 | -0.094 | 0 | -0.03 | -0.03 | 0 | 0.21 | 0.16 | 0.11 | 0 | 0.18 | 0.24 |
| 10 | 11870970 | rs6602542 | 0.376 | 0.579 | -1.707 | -0.33 | 0.063 | 0.185 | 0.185 | 0.222 | 0.5 | 0.21 | 0.65 | 0.73 | 0.6 | 0.66 |
| 10 | 11873528 | rs1998638 | 0.577 | 1.053 | -1.331 | 0.219 | -0.035 | -0.036 | -0.036 | 0 | 0.23 | 0.16 | 0.45 | 0.52 | 0.37 | 0.37 |
| 10 | 11874190 | rs7893993 | -0.805 | 0.552 | -0.595 | 0.421 | 0.004 | 0 | 0 | 0 | 0.54 | 0.55 | 0.66 | 0.51 | 0.69 | 0.68 |
| 10 | 11874754 | rs11257350 | -1.407 | 0.674 | 0.295 | 0.906 | 0 | 0.075 | 0.075 | 0 | 0.12 | 0.27 | 0.22 | 0.32 | 0.14 | 0.16 |
| 10 | 11877706 | rs1324646 | -0.119 | 0.786 | 0.586 | 0.013 | 0.033 | -0.062 | -0.062 | 0 | 0.88 | 0.8 | 0.9 | 1 | 0.86 | 0.91 |
| 10 | 11879615 | rs10795898 | -0.132 | 0.605 | 0.635 | 0.036 | -0.001 | 0.001 | 0.001 | 0 | 0.52 | 0.54 | 0.46 | 0.59 | 0.52 | 0.54 |
| 10 | 11879926 | rs942539 | -0.689 | 0.533 | 0.603 | 0.036 | 0.001 | -0.003 | -0.003 | 0 | 0.58 | 0.56 | 0.51 | 0.66 | 0.59 | 0.55 |
| 10 | 11881254 | rs7076760 | 0.444 | 0.549 | 0.592 | 0.041 | -0.019 | 0.012 | 0.012 | 0.167 | 0.11 | 0.06 | 0.1 | 0 | 0.14 | 0.11 |
| 10 | 11885961 | rs2031383 | -0.167 | 0.542 | 0.863 | 0.013 | -0.001 | 0.002 | 0.002 | 0 | 0.53 | 0.38 | 0.47 | 0.59 | 0.55 | 0.54 |
| 10 | 11892430 | rs10906027 | 1.405 | 0.568 | 0.892 | 0.057 | 0.018 | -0.031 | -0.031 | 0.231 | 0.29 | 0.23 | 0.35 | 0.39 | 0.21 | 0.34 |
| 10 | 11893971 | rs7079747 | -0.756 | 0.582 | 0.991 | -0.486 | -0.002 | -0.014 | -0.014 | 0 | 0.6 | 0.57 | 0.63 | 0.83 | 0.67 | 0.59 |
| 10 | 11904988 | rs7069533 | -1.005 | 0.48 | 1.045 | -0.447 | -0.019 | -0.027 | -0.027 | 0 | 0.64 | 0.47 | 0.61 | 0.76 | 0.66 | 0.67 |
| 10 | 11905880 | rs11257367 | -0.365 | 0.476 | 1.031 | -0.429 | 0.029 | 0.009 | 0.009 | 0.308 | 0.08 | 0.2 | 0.11 | 0.08 | 0.15 | 0.03 |
| 10 | 11909896 | rs12261792 | 1.651 | 0.454 | 1.01 | -0.436 | 0.02 | 0.017 | 0.017 | 4.094 | 0.27 | 0.22 | 0.24 | 0.16 | 0.19 | 0.3 |
| 10 | 11912431 | rs12253600 | -0.077 | 0.467 | 1.041 | -0.718 | 0.009 | 0.041 | 0.041 | 0.171 | 0.49 | 0.44 | 0.38 | 0.16 | 0.4 | 0.53 |
| 10 | 11915351 | rs7902251 | 0.408 | 0.432 | 0.614 | -0.711 | -0.007 | 0.02 | 0.02 | 0 | 0.4 | 0.29 | 0.5 | 0.75 | 0.44 | 0.44 |
| 10 | 11919872 | rs717800 | -0.126 | 0.477 | 0.62 | -0.717 | 0.008 | 0.081 | 0.081 | 0.093 | 0.48 | 0.3 | 0.65 | 0.83 | 0.59 | 0.47 |
| 10 | 11922731 | rs7074402 | -1.724 | 0.3 | 0.674 | -0.681 | -0.053 | -0.045 | -0.045 | 0.469 | 0.71 | 0.49 | 0.74 | 0.84 | 0.81 | 0.69 |

| | | | | | | | | | | | | | | | | |
|---|---|---|---|---|---|---|---|---|---|---|---|---|---|---|---|---|
| 10 | 11926406 | rs7093229 | -0.947 | 0.31 | 0.6 | -0.689 | -0.011 | -0.019 | -0.019 | 0 | 0.61 | 0.4 | 0.69 | 0.82 | 0.73 | 0.57 |
| 10 | 11931346 | rs11257386 | 0.403 | 0.419 | 0.414 | -0.698 | -0.005 | 0.031 | 0.031 | 0 | 0.41 | 0.3 | 0.48 | 0.73 | 0.44 | 0.43 |
| 10 | 11932234 | rs7900122 | -1.073 | 1.037 | 0.89 | -0.637 | 0.039 | -0.065 | -0.065 | 3.558 | 0.2 | 0.35 | 0.2 | 0.07 | 0.28 | 0.14 |
| 10 | 11932326 | rs12252853 | 0.294 | 0.628 | 0.755 | -0.34 | -0.004 | -0.04 | -0.04 | 0 | 0.1 | 0.16 | 0.05 | 0 | 0.08 | 0.11 |
| 10 | 11934634 | rs11257391 | 0.354 | 0.73 | 0.569 | -0.284 | -0.022 | -0.006 | -0.006 | 0 | 0.05 | 0.01 | 0.09 | 0.07 | 0.13 | 0.02 |
| 10 | 11936420 | rs12356904 | -1.115 | 0.646 | 0.688 | -0.344 | 0.01 | -0.094 | -0.094 | 0 | 0.21 | 0.51 | 0.17 | 0.07 | 0.24 | 0.16 |
| 10 | 11939481 | rs2031386 | 0.529 | 0.654 | 0.395 | -0.217 | 0.029 | -0.003 | -0.003 | 0.722 | 0.91 | 0.69 | 0.89 | 0.93 | 0.87 | 0.98 |
| 10 | 11943767 | rs1535975 | -0.264 | 0.726 | 0.235 | -0.042 | -0.023 | 0.007 | 0.007 | 0.638 | 0.09 | 0.04 | 0.14 | 0.07 | 0.16 | 0.03 |
| 10 | 11943885 | rs10795908 | -0.518 | 0.636 | 0.169 | -0.166 | 0.014 | 0.032 | 0.032 | 0 | 0.66 | 0.61 | 0.61 | 0.42 | 0.58 | 0.63 |
| 10 | 11944213 | rs11257408 | -1.193 | 1.047 | -0.136 | -0.037 | 0.018 | -0.078 | -0.078 | 0 | 0.21 | 0.42 | 0.22 | 0.08 | 0.25 | 0.12 |
| 10 | 11950934 | rs942537 | 0.556 | 0.436 | -0.59 | 0.404 | 0.003 | 0.003 | 0.003 | 0.195 | 0.51 | 0.49 | 0.42 | 0.23 | 0.38 | 0.52 |
| 10 | 11951896 | rs10795911 | 0.124 | 1.189 | -1.138 | 0.623 | -0.044 | -0.035 | -0.035 | 0 | 0.66 | 0.81 | 0.55 | 0.4 | 0.45 | 0.7 |
| 10 | 11966016 | rs11257429 | 0.551 | 0.175 | -1.748 | -0.453 | -0.019 | 0.108 | 0.108 | 3.09 | 0.4 | 0.17 | 0.32 | 0.17 | 0.57 | 0.34 |
| 10 | 11971155 | rs10430780 | 0.245 | -0.485 | -1.33 | -0.108 | -0.003 | 0.033 | 0.033 | 1.426 | 0.22 | 0.27 | 0.28 | 0.45 | 0.2 | 0.17 |
| 10 | 11975662 | rs7897567 | -1.672 | -0.127 | -1.385 | -0.303 | -0.046 | 0.07 | 0.07 | 6.082 | 0.65 | 0.81 | 0.71 | 0.84 | 0.52 | 0.69 |
| 10 | 11977300 | rs11257434 | 1.046 | -0.455 | -1.576 | -0.199 | -0.009 | -0.04 | -0.04 | 0 | 0.07 | 0.03 | 0.14 | 0.25 | 0.06 | 0.17 |
| 10 | 11978499 | rs7921794 | -1.492 | -0.449 | -1.572 | -0.205 | -0.037 | 0.115 | 0.115 | 3.708 | 0.61 | 0.83 | 0.7 | 0.84 | 0.47 | 0.66 |
| 10 | 11979002 | rs3740025 | -0.616 | -0.466 | -0.981 | -0.208 | 0.003 | -0.003 | -0.003 | 0 | 0.18 | 0.18 | 0.19 | 0.25 | 0.17 | 0.22 |
| 10 | 11981831 | rs12265381 | 0.421 | -0.427 | -0.701 | -0.191 | -0.005 | 0.04 | 0.04 | 1.864 | 0.21 | 0.27 | 0.25 | 0.45 | 0.17 | 0.17 |
| 10 | 12006358 | rs7080355 | 0.315 | -0.549 | -0.479 | -0.219 | -0.024 | -0.019 | -0.019 | 0 | 0.23 | 0.27 | 0.25 | 0.14 | 0.13 | 0.28 |
| 10 | 12020196 | rs10752250 | -0.108 | -0.501 | -0.398 | -0.3 | 0.121 | -0.028 | -0.028 | 0 | 0.92 | 0.68 | 0.94 | 1 | 0.88 | 0.9 |
| 10 | 12020268 | rs10795914 | 1.412 | -0.493 | -0.419 | -0.294 | -0.094 | -0.018 | -0.018 | 0 | 0.4 | 0.11 | 0.41 | 0.41 | 0.51 | 0.41 |
| 10 | 12028228 | rs6602555 | -1.021 | -0.538 | -0.447 | -0.307 | 0.043 | 0.031 | 0.031 | 5.312 | 0.52 | 0.64 | 0.54 | 0.59 | 0.36 | 0.48 |
| 10 | 12033741 | rs4750147 | 0.59 | -0.524 | -0.452 | -0.292 | -0.008 | 0.043 | 0.043 | 1.896 | 0.21 | 0.27 | 0.25 | 0.45 | 0.15 | 0.16 |
| 10 | 12043092 | rs4750149 | -0.458 | -0.423 | -0.717 | -0.29 | 0.036 | -0.041 | -0.041 | 0.025 | 0.92 | 0.87 | 0.95 | 1 | 0.87 | 0.89 |
| 10 | 12051574 | rs7899260 | 0.67 | -0.361 | -1.094 | -0.297 | -0.012 | 0.032 | 0.032 | 0 | 0.07 | 0.03 | 0.03 | 0 | 0.1 | 0.05 |
| 10 | 12062959 | rs7072007 | -0.586 | -0.543 | -1.047 | -0.238 | -0.025 | -0.017 | -0.017 | 0 | 0.77 | 0.75 | 0.75 | 0.86 | 0.89 | 0.72 |
| 10 | 12064869 | rs10795919 | 0.071 | -0.213 | -1.697 | 0.065 | 0.005 | 0.007 | 0.007 | 5.228 | 0.6 | 0.41 | 0.66 | 0.86 | 0.67 | 0.56 |
| 10 | 12075960 | rs10906052 | 0.322 | 0.368 | -2.348 | 0.291 | -0.18 | -0.072 | -0.072 | 2.814 | 0.39 | 0.05 | 0.4 | 0.41 | 0.51 | 0.41 |
| 10 | 12094866 | rs10508437 | -0.475 | -0.484 | -2.061 | 0.465 | 0.047 | 0.003 | 0.003 | 0.299 | 0.28 | 0.09 | 0.12 | 0.19 | 0.18 | 0.3 |
| 10 | 12094949 | rs10906055 | -0.019 | -0.871 | -1.725 | 0.835 | -0.011 | 0.056 | 0.056 | 3.756 | 0.13 | 0.19 | 0.17 | 0.28 | 0.1 | 0.07 |
| 10 | 12095729 | rs12782067 | 1.302 | 0.262 | -2.604 | 0.411 | -0.107 | 0.023 | 0.023 | 2.211 | 0.17 | 0.01 | 0.16 | 0.09 | 0.3 | 0.16 |
| 10 | 12099023 | rs11257506 | 0.117 | -0.8 | -1.474 | 0.067 | -0.02 | -0.028 | -0.028 | 0 | 0.26 | 0.35 | 0.21 | 0.15 | 0.22 | 0.2 |
| 10 | 12100735 | rs7908725 | 0.377 | -1.08 | -1.502 | 0.285 | 0.043 | -0.024 | -0.024 | 0 | 0.61 | 0.21 | 0.6 | 0.57 | 0.69 | 0.78 |
| 10 | 12108009 | rs7093150 | 0.404 | -1.19 | -1.808 | 0.097 | 0.017 | -0.006 | -0.006 | 0 | 0.53 | 0.19 | 0.53 | 0.53 | 0.54 | 0.69 |
| 10 | 12110518 | rs7894615 | 0.639 | -0.649 | -1.764 | 0.307 | -0.065 | -0.021 | -0.021 | 0 | 0.4 | 0.17 | 0.33 | 0.41 | 0.46 | 0.49 |
| 10 | 12122355 | rs10752254 | 0.47 | -0.597 | -1.674 | 0.35 | -0.022 | -0.056 | -0.056 | 2.137 | 0.62 | 0.2 | 0.47 | 0.48 | 0.59 | 0.76 |
| 10 | 12127376 | rs7081227 | -0.639 | -0.616 | -0.823 | 0.464 | 0.016 | 0.096 | 0.096 | 0 | 0.81 | 0.54 | 0.69 | 0.71 | 0.82 | 0.9 |
| 10 | 12130170 | rs11257529 | 0.941 | -1.112 | -1.204 | 0.522 | 0.11 | -0.004 | -0.004 | 0 | 0.29 | 0.03 | 0.11 | 0.2 | 0.12 | 0.38 |
| 10 | 12131081 | rs3740015 | 1.092 | -0.247 | -0.453 | 0.814 | -0.02 | -0.053 | -0.053 | 0 | 0.62 | 0.21 | 0.47 | 0.48 | 0.59 | 0.76 |
| 10 | 12140444 | rs2062986 | -0.505 | -0.244 | -0.181 | 0.825 | 0.021 | 0.111 | 0.111 | 0 | 0.81 | 0.56 | 0.69 | 0.7 | 0.82 | 0.9 |
| 10 | 12146858 | rs9424148 | -0.025 | -0.273 | 0.234 | 0.799 | 0.016 | 0.025 | 0.025 | 0 | 0.8 | 0.5 | 0.77 | 0.77 | 0.79 | 0.88 |
| 10 | 12154833 | rs1701470 | -0.329 | -0.247 | -0.141 | 0.543 | 0.003 | 0.054 | 0.054 | 0 | 0.29 | 0.6 | 0.48 | 0.5 | 0.3 | 0.16 |
| 10 | 12159472 | rs2658193 | -0.334 | -0.267 | 0.1 | 0.428 | 0.027 | -0.028 | -0.028 | 0 | 0.9 | 0.79 | 0.94 | 0.98 | 0.89 | 0.93 |
| 10 | 12170174 | rs6602563 | 0.042 | 0.63 | 0.189 | 0.815 | 0.018 | 0.004 | 0.004 | 0.094 | 0.63 | 0.55 | 0.47 | 0.57 | 0.58 | 0.81 |
| 10 | 12179270 | rs10906083 | 0.087 | 0.534 | 0.595 | 0.976 | 0.019 | 0.021 | 0.021 | 0.12 | 0.62 | 0.5 | 0.45 | 0.47 | 0.58 | 0.73 |
| 10 | 12186449 | rs7919751 | 0.173 | 0.531 | 0.434 | 0.969 | 0.002 | 0.006 | 0.006 | 0.039 | 0.58 | 0.48 | 0.43 | 0.44 | 0.51 | 0.7 |
| 10 | 12189712 | rs10906086 | 0.276 | 0.529 | 0.24 | 1.18 | -0.03 | 0.038 | 0.038 | 0.161 | 0.58 | 0.19 | 0.43 | 0.34 | 0.52 | 0.59 |
| 10 | 12191933 | rs10466280 | -0.114 | 0.55 | -0.038 | 1.143 | 0.015 | 0.015 | 0.015 | 0 | 0.35 | 0.41 | 0.5 | 0.53 | 0.41 | 0.31 |
| 10 | 12202408 | rs10906090 | -0.083 | 0.587 | -0.092 | 1.165 | 0.013 | 0.012 | 0.012 | 0 | 0.35 | 0.41 | 0.5 | 0.53 | 0.4 | 0.3 |
| 10 | 12208307 | rs1053403 | 0.009 | 0.597 | -0.063 | 1.166 | 0.01 | 0.01 | 0.01 | 0 | 0.35 | 0.4 | 0.5 | 0.53 | 0.41 | 0.31 |
| 10 | 12209752 | rs6686 | -0.501 | 0.641 | -0.059 | 1.164 | 0.004 | 0.001 | 0.001 | 0 | 0.39 | 0.43 | 0.52 | 0.56 | 0.48 | 0.35 |
| 10 | 12211590 | rs9633765 | 0.036 | 0.632 | -0.077 | 1.16 | 0.012 | 0.011 | 0.011 | 0 | 0.35 | 0.4 | 0.5 | 0.53 | 0.41 | 0.33 |
| 10 | 12215180 | rs2399786 | 0.262 | 0.683 | -0.103 | 1.146 | 0.008 | 0.003 | 0.003 | 0 | 0.61 | 0.54 | 0.48 | 0.44 | 0.52 | 0.64 |
| 10 | 12222316 | rs3780864 | -1.009 | 0.839 | -0.128 | 1.269 | 0.026 | 0.069 | 0.069 | 0 | 0.95 | 0.91 | 0.96 | 0.87 | 0.92 | 0.83 |
| 10 | 12227580 | rs3780866 | 2.328 | 0.601 | -1.13 | 0.787 | -0.076 | 0 | 0 | 0 | 0.15 | 0.01 | 0.11 | 0.09 | 0.24 | 0.13 |
| 10 | 12227895 | rs12260965 | -0.364 | 0.165 | -1.168 | 0.552 | -0.006 | -0.008 | -0.008 | 0 | 0.58 | 0.37 | 0.56 | 0.54 | 0.5 | 0.43 |
| 10 | 12230922 | rs12267330 | 0.705 | -1.422 | -0.674 | 0.369 | 0.012 | 0.01 | 0.01 | 1.242 | 0.27 | 0.22 | 0.1 | 0.11 | 0.12 | 0.25 |
| 10 | 12235711 | rs7072888 | 0.868 | -0.592 | -0.749 | 0.689 | -0.038 | 0.044 | 0.044 | 0 | 0.34 | 0.2 | 0.42 | 0.25 | 0.45 | 0.31 |
| 10 | 12236773 | rs3758379 | 0.29 | -0.028 | -0.509 | 0.99 | -0.023 | -0.032 | -0.032 | 0 | 0.1 | 0.02 | 0.06 | 0.1 | 0.13 | 0.17 |
| 10 | 12238610 | rs11257590 | 0.274 | -0.045 | -0.494 | 1.005 | -0.023 | -0.031 | -0.031 | 0 | 0.1 | 0.02 | 0.06 | 0.1 | 0.13 | 0.18 |
| 10 | 12239944 | rs17565905 | 0.356 | -0.204 | -0.301 | 1.149 | -0.018 | -0.002 | -0.002 | 0 | 0.18 | 0.14 | 0.25 | 0.21 | 0.23 | 0.13 |
| 10 | 12243821 | rs10906097 | 1.004 | -0.218 | 0.29 | 1.25 | -0.008 | 0.01 | 0.01 | 0 | 0.58 | 0.43 | 0.54 | 0.37 | 0.54 | 0.5 |
| 10 | 12244079 | rs10508438 | -0.736 | -0.251 | 0.78 | 1.391 | -0.003 | -0.003 | -0.003 | 0 | 0.28 | 0.28 | 0.37 | 0.5 | 0.33 | 0.29 |
| 10 | 12245520 | rs1317548 | 0.202 | -0.158 | 0.768 | 1.466 | 0.003 | 0.024 | 0.024 | 0.28 | 0.2 | 0.21 | 0.17 | 0.45 | 0.21 | 0.19 |
| 10 | 12245601 | rs938991 | -0.052 | -0.199 | 1.036 | 1.533 | 0.012 | 0.006 | 0.006 | 0.836 | 0.85 | 0.73 | 0.83 | 0.85 | 0.85 | 0.79 |
| 10 | 12249797 | rs4747968 | 1.134 | -0.257 | 0.864 | 1.217 | -0.049 | 0.071 | 0.071 | 0 | 0.57 | 0.18 | 0.57 | 0.33 | 0.57 | 0.52 |
| 10 | 12250861 | rs7100710 | -0.103 | -0.275 | 1.01 | 1.268 | 0.004 | 0.016 | 0.016 | 0 | 0.77 | 0.74 | 0.73 | 0.45 | 0.77 | 0.75 |
| 10 | 12253597 | rs12221133 | 0.031 | -0.285 | 1.026 | 1.301 | 0.006 | 0.022 | 0.022 | 0 | 0.22 | 0.23 | 0.27 | 0.53 | 0.26 | 0.19 |

| | | | | | | | | | | | | | | | | |
|---|---|---|---|---|---|---|---|---|---|---|---|---|---|---|---|---|
| 10 | 12258056 | rs10508440 | 0.291 | -0.258 | 0.986 | 1.179 | -0.038 | 0.056 | 0.056 | 0.456 | 0.14 | 0.02 | 0.15 | 0.05 | 0.19 | 0.19 |
| 10 | 12269343 | rs11257608 | 0.878 | -0.258 | 1.065 | 1.391 | -0.001 | 0.004 | 0.004 | 0.011 | 0.17 | 0.17 | 0.17 | 0.4 | 0.2 | 0.17 |
| 10 | 12280296 | rs10906104 | 0.416 | -0.315 | 1.129 | 1.352 | 0.002 | 0.007 | 0.007 | 0.001 | 0.72 | 0.38 | 0.68 | 0.41 | 0.74 | 0.7 |
| 10 | 12282461 | rs7075724 | 0.337 | -0.316 | 1.131 | 1.136 | 0.003 | 0.004 | 0.004 | 0 | 0.73 | 0.39 | 0.69 | 0.38 | 0.74 | 0.7 |
| 10 | 12283932 | rs11257612 | 0.089 | -0.345 | 1.011 | 1.077 | -0.034 | 0.065 | 0.065 | 0.352 | 0.15 | 0.02 | 0.16 | 0.05 | 0.18 | 0.2 |
| 10 | 12286251 | rs10906105 | -0.696 | -0.303 | 0.963 | 1.03 | -0.044 | 0.095 | 0.095 | 0.085 | 0.19 | 0.03 | 0.19 | 0.05 | 0.23 | 0.2 |
| 10 | 12288780 | rs4469792 | 1.241 | -0.182 | 1.043 | 1.24 | -0.065 | 0.078 | 0.078 | 0 | 0.57 | 0.19 | 0.51 | 0.33 | 0.55 | 0.51 |
| 10 | 12292344 | rs10951 | 0.686 | -0.174 | 0.931 | 1.23 | -0.029 | 0.001 | 0.001 | 0 | 0.64 | 0.73 | 0.61 | 0.61 | 0.6 | 0.69 |
| 10 | 12292454 | rs1051055 | 0.367 | -0.062 | 0.95 | 1.339 | -0.025 | -0.014 | -0.014 | 0 | 0.7 | 0.77 | 0.71 | 0.59 | 0.66 | 0.71 |
| 10 | 12295339 | rs11257622 | 0.965 | -0.024 | 0.783 | 1.407 | -0.003 | 0 | 0 | 0 | 0.17 | 0.14 | 0.21 | 0.35 | 0.19 | 0.1 |
| 10 | 12298194 | rs11819728 | 0.974 | -0.084 | 0.506 | 0.879 | -0.048 | 0.072 | 0.072 | 0 | 0.57 | 0.22 | 0.51 | 0.32 | 0.56 | 0.5 |
| 10 | 12300790 | rs7916879 | -0.723 | -0.016 | 0.26 | 0.811 | 0.034 | 0.023 | 0.023 | 0 | 0.32 | 0.46 | 0.42 | 0.59 | 0.37 | 0.36 |
| 10 | 12307894 | rs11257655 | 0.25 | 0.195 | -0.957 | 0.906 | 0.008 | 0.042 | 0.042 | 0 | 0.22 | 0.24 | 0.26 | 0.54 | 0.23 | 0.24 |
| 10 | 12307963 | rs11593567 | 0.749 | 0.213 | -1.484 | 1.042 | -0.069 | 0.071 | 0.071 | 0 | 0.56 | 0.17 | 0.5 | 0.33 | 0.54 | 0.49 |
| 10 | 12310737 | rs7895410 | 0.792 | 0.247 | -1.398 | 0.947 | -0.008 | -0.047 | -0.047 | 0 | 0.3 | 0.1 | 0.28 | 0.32 | 0.33 | 0.26 |
| 10 | 12314997 | rs10906115 | 0.181 | 0.624 | -1.776 | 0.721 | 0.008 | 0.008 | 0.008 | 0.329 | 0.39 | 0.3 | 0.4 | 0.4 | 0.37 | 0.49 |
| 10 | 12323464 | rs7069060 | -0.766 | 1.17 | -1.519 | 0.993 | -0.008 | 0.002 | 0.002 | 0 | 0.26 | 0.23 | 0.22 | 0.27 | 0.29 | 0.32 |
| 10 | 12335588 | rs2679246 | 0.459 | 0.843 | -1.739 | 1.256 | -0.005 | -0.071 | -0.071 | 0 | 0.72 | 0.86 | 0.68 | 0.64 | 0.7 | 0.67 |
| 10 | 12343629 | rs2096396 | -0.377 | 1.113 | -1.474 | 1.065 | -0.009 | -0.006 | -0.006 | 0 | 0.37 | 0.66 | 0.44 | 0.44 | 0.41 | 0.57 |
| 10 | 12344823 | rs4750199 | -1.002 | 1.563 | -1.253 | 1.203 | 0.002 | 0.001 | 0.001 | 0 | 0.22 | 0.24 | 0.22 | 0.32 | 0.31 | 0.23 |
| 10 | 12347361 | rs1573929 | 0.152 | 0.38 | -1.778 | 0.717 | 0.005 | 0.012 | 0.012 | 0.92 | 0.13 | 0.01 | 0.22 | 0.06 | 0.08 | 0.09 |
| 10 | 12347641 | rs1962249 | 0.496 | 0.324 | -1.009 | 1.362 | 0.018 | 0.152 | 0.152 | 0.112 | 0.85 | 0.72 | 0.76 | 0.59 | 0.81 | 0.8 |
| 10 | 12351036 | rs7081104 | 0.079 | -0.501 | -1.507 | 0.545 | 0.058 | 0.016 | 0.016 | 0.053 | 0.52 | 0.26 | 0.54 | 0.46 | 0.6 | 0.67 |
| 10 | 12351089 | rs7069309 | 0.147 | -0.468 | -1.506 | 0.312 | 0.013 | 0.045 | 0.045 | 0.019 | 0.46 | 0.2 | 0.52 | 0.4 | 0.54 | 0.64 |
| 10 | 12352028 | rs11593647 | 0.746 | -0.081 | -1.639 | 0.349 | -0.141 | -0.016 | -0.016 | 0 | 0.33 | 0.08 | 0.29 | 0.34 | 0.44 | 0.53 |
| 10 | 12353934 | rs4750201 | 0.999 | -0.78 | -1.109 | 0.064 | -0.019 | -0.039 | -0.039 | 0.044 | 0.21 | 0.22 | 0.11 | 0.09 | 0.16 | 0.1 |
| 10 | 12359441 | rs7098962 | 0.276 | -0.649 | -1.71 | 0.144 | 0.029 | 0.021 | 0.021 | 0.196 | 0.42 | 0.12 | 0.39 | 0.35 | 0.62 | 0.61 |
| 10 | 12364206 | rs11257707 | 0.431 | -1.038 | -1.093 | -0.189 | -0.045 | -0.057 | -0.057 | 0 | 0.15 | 0.24 | 0.12 | 0.04 | 0.07 | 0.09 |
| 10 | 12366615 | rs10906126 | 1.108 | -0.348 | -1.471 | -0.204 | -0.001 | 0.046 | 0.046 | 0 | 0.21 | 0.13 | 0.13 | 0.07 | 0.24 | 0.1 |
| 10 | 12369684 | rs10795950 | 0.953 | -0.868 | -1.332 | 0.284 | 0.002 | 0 | 0 | 0 | 0.41 | 0.43 | 0.4 | 0.47 | 0.57 | 0.47 |
| 10 | 12369861 | rs10906128 | -0.698 | -0.617 | -1.975 | 0.751 | 0.002 | -0.071 | -0.071 | 0.411 | 0.33 | 0.08 | 0.39 | 0.4 | 0.29 | 0.38 |
| 10 | 12376977 | rs7896794 | 0.535 | -0.958 | -1.57 | 0.514 | 0.042 | 0.002 | 0.002 | 0 | 0.74 | 0.88 | 0.83 | 0.75 | 0.84 | 0.71 |
| 10 | 12385344 | rs11257727 | -0.629 | -0.651 | -1.614 | 0.403 | 0.038 | 0.015 | 0.015 | 0 | 0.22 | 0.1 | 0.14 | 0.13 | 0.15 | 0.18 |
| 10 | 12389180 | rs2997063 | 0.828 | 0.021 | -1.29 | 0.468 | 0.022 | 0.087 | 0.087 | 0 | 0.51 | 0.25 | 0.54 | 0.27 | 0.62 | 0.47 |
| 10 | 12390116 | rs2997064 | 0.251 | -0.576 | -0.509 | 0.551 | -0.039 | 0.007 | 0.007 | 0 | 0.9 | 0.68 | 0.96 | 0.89 | 0.98 | 0.92 |
| 10 | 12396010 | rs2801499 | 0.576 | -0.48 | -0.655 | 0.603 | -0.017 | 0.034 | 0.034 | 0.835 | 0.68 | 0.44 | 0.65 | 0.46 | 0.75 | 0.58 |
| 10 | 12401096 | rs2801498 | 1.024 | -0.324 | -1.413 | 0.783 | -0.028 | 0.026 | 0.026 | 0 | 0.67 | 0.45 | 0.7 | 0.4 | 0.77 | 0.52 |
| 10 | 12402466 | rs7909593 | -0.008 | -0.6 | -1.423 | 1.161 | -0.016 | -0.034 | -0.034 | 0 | 0.09 | 0.03 | 0.09 | 0.23 | 0.09 | 0.29 |
| 10 | 12402701 | rs2997068 | 0.901 | -0.477 | -1.438 | 1.074 | -0.047 | 0.04 | 0.04 | 0 | 0.68 | 0.48 | 0.72 | 0.4 | 0.81 | 0.54 |
| 10 | 12403028 | rs17151584 | 0.19 | 0.207 | -2.254 | 0.975 | 0.017 | -0.061 | -0.061 | 1.24 | 0.12 | 0.03 | 0.14 | 0.26 | 0.08 | 0.09 |
| 10 | 12406899 | rs7074893 | -0.261 | 0.987 | -1.953 | 0.353 | -0.02 | -0.018 | -0.018 | 0 | 0.07 | 0.02 | 0.08 | 0.04 | 0.09 | 0.05 |
| 10 | 12412243 | rs10906138 | -0.525 | 1.18 | -2.012 | 0.111 | -0.025 | 0.016 | 0.016 | 0.185 | 0.21 | 0.08 | 0.15 | 0.17 | 0.26 | 0.35 |
| 10 | 12419475 | rs12261350 | -0.472 | 0.892 | -2.255 | -0.361 | -0.012 | 0.129 | 0.129 | 0 | 0.45 | 0.2 | 0.38 | 0.18 | 0.47 | 0.45 |
| 10 | 12423107 | rs11257754 | -0.233 | 1.141 | -2.249 | 0.064 | -0.004 | 0.011 | 0.011 | 1.016 | 0.16 | 0.1 | 0.15 | 0.13 | 0.16 | 0.21 |
| 10 | 12433942 | rs7073375 | -0.017 | 1.607 | -2.324 | 0.21 | -0.042 | -0.038 | -0.038 | 0 | 0.08 | 0.01 | 0.08 | 0.16 | 0.13 | 0.12 |
| 10 | 12434742 | rs11257765 | -0.748 | 2.309 | -2.445 | -0.113 | -0.051 | 0.012 | 0.012 | 0.481 | 0.2 | 0.07 | 0.26 | 0.2 | 0.3 | 0.3 |
| 10 | 12435379 | rs4750211 | 0.353 | 1.396 | -3.275 | 0.267 | 0.011 | -0.033 | -0.033 | 0 | 0.33 | 0.04 | 0.3 | 0.24 | 0.25 | 0.18 |
| 10 | 12436252 | rs4750212 | 0.373 | 1.363 | -3.313 | 0.662 | 0.002 | -0.108 | -0.108 | 2.367 | 0.33 | 0.05 | 0.34 | 0.34 | 0.25 | 0.19 |
| 10 | 12438783 | rs10906142 | 0.586 | 1.983 | -2.51 | 0.897 | 0.007 | 0.013 | 0.013 | 0 | 0.22 | 0.2 | 0.23 | 0.53 | 0.33 | 0.36 |
| 10 | 12440209 | rs11592475 | 0.119 | 1.274 | -2.978 | 0.254 | -0.031 | 0.019 | 0.019 | 2.317 | 0.21 | 0.03 | 0.27 | 0.16 | 0.25 | 0.18 |
| 10 | 12448088 | rs12254030 | 0.413 | 0.954 | -2.988 | 0.202 | 0.007 | -0.004 | -0.004 | 0 | 0.08 | 0.03 | 0.05 | 0.05 | 0.07 | 0.03 |
| 10 | 12448888 | rs1574114 | 0.105 | 1.115 | -2.096 | 0.281 | 0.03 | 0.031 | 0.031 | 0 | 0.49 | 0.4 | 0.36 | 0.43 | 0.43 | 0.45 |
| 10 | 12450495 | rs7100726 | 0.463 | 0.897 | -1.722 | -0.092 | 0.009 | 0.053 | 0.053 | 0 | 0.3 | 0.19 | 0.19 | 0.18 | 0.29 | 0.28 |
| 10 | 12451399 | rs2724786 | -2.013 | 0.81 | -0.994 | 0.095 | 0.038 | 0.043 | 0.043 | 0 | 0.81 | 0.74 | 0.8 | 0.71 | 0.72 | 0.7 |
| 10 | 12453032 | rs2724785 | -1.29 | 0.204 | -0.826 | 0.131 | -0.009 | -0.004 | -0.004 | 0 | 0.72 | 0.75 | 0.71 | 0.71 | 0.62 | 0.63 |
| 10 | 12458032 | rs2399860 | 0.592 | 0.115 | -0.261 | 0.095 | 0.001 | 0.004 | 0.004 | 0.384 | 0.22 | 0.14 | 0.38 | 0.22 | 0.2 | 0.21 |
| 10 | 12461773 | rs2801479 | -1.699 | 0.202 | 0.022 | 0.301 | -0.018 | -0.016 | -0.016 | 0.902 | 0.75 | 0.8 | 0.73 | 0.69 | 0.65 | 0.65 |
| 10 | 12464305 | rs10795953 | -0.728 | -0.035 | -0.004 | 0.337 | 0.005 | 0.025 | 0.025 | 0.271 | 0.46 | 0.35 | 0.55 | 0.39 | 0.41 | 0.39 |
| 10 | 12467228 | rs11592149 | -0.077 | -0.081 | -0.226 | 0.33 | -0.004 | 0.011 | 0.011 | 0 | 0.25 | 0.22 | 0.2 | 0.18 | 0.23 | 0.18 |
| 10 | 12468137 | rs1111056 | 1.053 | -0.224 | -0.027 | 0.352 | 0.006 | -0.004 | -0.004 | 0 | 0.48 | 0.31 | 0.38 | 0.51 | 0.55 | 0.58 |
| 10 | 12471105 | rs7918658 | 0.267 | -0.496 | 0.257 | 0.549 | -0.041 | 0.016 | 0.016 | 0 | 0.72 | 0.52 | 0.58 | 0.69 | 0.78 | 0.76 |
| 10 | 12475837 | rs1125791 | 0.387 | -0.567 | -0.593 | 0.508 | -0.003 | -0.012 | -0.012 | 0 | 0.2 | 0.03 | 0.35 | 0.21 | 0.17 | 0.21 |
| 10 | 12480633 | rs2801493 | 0.333 | -0.601 | -0.345 | 0.544 | 0.019 | 0 | 0 | 0 | 0.71 | 0.29 | 0.58 | 0.7 | 0.8 | 0.76 |
| 10 | 12483898 | rs2986651 | 0.394 | -0.002 | 0.066 | 0.563 | -0.006 | 0.025 | 0.025 | 0 | 0.27 | 0.48 | 0.29 | 0.39 | 0.28 | 0.26 |
| 10 | 12484556 | rs10906150 | -0.064 | -0.194 | -0.5 | 0.301 | 0.008 | -0.016 | -0.016 | 0 | 0.23 | 0.15 | 0.32 | 0.31 | 0.19 | 0.25 |
| 10 | 12487996 | rs6602588 | -1.53 | 0.547 | -0.153 | 0.266 | -0.025 | -0.027 | -0.027 | 2.195 | 0.71 | 0.77 | 0.63 | 0.67 | 0.65 | 0.58 |
| 10 | 12488714 | rs11257794 | -0.447 | 0.624 | 0.343 | 0.401 | 0.009 | 0.023 | 0.023 | 0 | 0.23 | 0.17 | 0.13 | 0.09 | 0.19 | 0.1 |
| 10 | 12491329 | rs11596121 | -0.298 | 0.745 | -0.114 | 0.862 | 0.006 | -0.001 | -0.001 | 0.762 | 0.1 | 0.04 | 0.1 | 0.12 | 0.06 | 0.08 |
| 10 | 12491347 | rs4750216 | 0.901 | 0.88 | 0.306 | 1.093 | 0.01 | 0.084 | 0.084 | 0.306 | 0.83 | 0.46 | 0.84 | 0.73 | 0.83 | 0.88 |

| | | | | | | | | | | | | | | | | |
|---|---|---|---|---|---|---|---|---|---|---|---|---|---|---|---|---|
| 10 | 12492379 | rs1077685 | -0.315 | 0.926 | 0.195 | 0.896 | 0.021 | 0.026 | 0.026 | 0 | 0.06 | 0.18 | 0.04 | 0.07 | 0.05 | 0.04 |
| 10 | 12492474 | rs7901627 | -0.765 | 1.468 | 0.122 | 1.62 | 0.014 | 0.088 | 0.088 | 0 | 0.06 | 0.08 | 0.07 | 0.34 | 0.11 | 0.1 |
| 10 | 12495536 | rs11257802 | 0.794 | 0.832 | -0.752 | 0.125 | 0.048 | 0.14 | 0.14 | 0.35 | 0.36 | 0.13 | 0.32 | 0.04 | 0.28 | 0.2 |
| 10 | 12496055 | rs11257804 | -0.072 | 1.553 | -1.193 | 0.373 | -0.021 | -0.042 | -0.042 | 0 | 0.22 | 0.12 | 0.24 | 0.51 | 0.3 | 0.22 |
| 10 | 12497747 | rs1108131 | 0.904 | 0.971 | -0.633 | -0.221 | 0.02 | -0.005 | -0.005 | 0 | 0.37 | 0.26 | 0.34 | 0.04 | 0.28 | 0.22 |
| 10 | 12497770 | rs2768367 | 0.016 | 1.156 | -0.159 | 0.457 | -0.003 | -0.012 | -0.012 | 1.027 | 0.84 | 0.8 | 0.91 | 0.87 | 0.88 | 0.74 |
| 10 | 12499860 | rs2399849 | -0.563 | 1.45 | 0.084 | 0.527 | 0.015 | -0.002 | -0.002 | 0 | 0.18 | 0.28 | 0.15 | 0.21 | 0.17 | 0.29 |
| 10 | 12501398 | rs2815611 | 0.277 | 1.449 | 0.359 | 0.644 | 0.007 | 0.012 | 0.012 | 0 | 0.16 | 0.3 | 0.2 | 0.15 | 0.19 | 0.14 |
| 10 | 12508956 | rs1571212 | 0.698 | 1.188 | -0.152 | 0.418 | -0.034 | 0.002 | 0.002 | 1.169 | 0.24 | 0.12 | 0.23 | 0.17 | 0.31 | 0.25 |
| 10 | 12509131 | rs7899150 | 0.144 | 1.166 | 0.193 | 1.022 | 0.018 | 0.185 | 0.185 | 0 | 0.06 | 0.13 | 0.16 | 0.44 | 0.09 | 0.07 |
| 10 | 12511729 | rs2768411 | 0.339 | 1.197 | 0.654 | 0.71 | 0.006 | 0.001 | 0.001 | 0.575 | 0.33 | 0.34 | 0.32 | 0.27 | 0.39 | 0.34 |
| 10 | 12511801 | rs7907938 | 0.46 | 1.202 | 0.628 | 0.712 | -0.027 | -0.011 | -0.011 | 0.115 | 0.32 | 0.21 | 0.31 | 0.27 | 0.39 | 0.34 |
| 10 | 12511933 | rs7894008 | 0.313 | 1.19 | 0.59 | 0.706 | -0.016 | -0.005 | -0.005 | 0 | 0.33 | 0.25 | 0.31 | 0.27 | 0.39 | 0.37 |
| 10 | 12512247 | rs7900817 | 0.033 | 0.951 | 0.407 | 0.496 | -0.021 | 0.012 | 0.012 | 1.242 | 0.41 | 0.33 | 0.37 | 0.28 | 0.47 | 0.4 |
| 10 | 12512388 | rs2815605 | -0.624 | 0.505 | -0.46 | 0.1 | -0.002 | -0.005 | -0.005 | 0 | 0.35 | 0.31 | 0.36 | 0.63 | 0.36 | 0.31 |
| 10 | 12515823 | rs1543595 | -1.962 | 0.237 | -0.702 | 0.107 | 0.018 | 0.05 | 0.05 | 0 | 0.75 | 0.86 | 0.72 | 0.96 | 0.78 | 0.71 |
| 10 | 12517891 | rs4620627 | -0.58 | 0.803 | -0.547 | 0.978 | 0 | 0.027 | 0.027 | 0 | 0.13 | 0.16 | 0.13 | 0.52 | 0.17 | 0.11 |
| 10 | 12520663 | rs12784955 | 0.013 | 0.629 | -0.677 | 0.635 | -0.01 | 0.001 | 0.001 | 0 | 0.17 | 0.1 | 0.14 | 0.12 | 0.17 | 0.24 |
| 10 | 12521907 | rs1536529 | -0.188 | 0.466 | -0.473 | 0.763 | 0.004 | 0.003 | 0.003 | 0 | 0.54 | 0.59 | 0.53 | 0.52 | 0.58 | 0.4 |
| 10 | 12522144 | rs1536531 | 0.926 | 0.546 | -0.427 | 0.958 | -0.006 | 0.027 | 0.027 | 0.141 | 0.13 | 0.17 | 0.09 | 0.15 | 0.11 | 0.1 |
| 10 | 12522713 | rs12781984 | 0.197 | 0.521 | -0.744 | 0.715 | -0.012 | 0.025 | 0.025 | 0 | 0.16 | 0.09 | 0.1 | 0.05 | 0.17 | 0.12 |
| 10 | 12524348 | rs2768429 | 0.677 | 0.252 | -0.613 | 1.175 | -0.003 | 0.013 | 0.013 | 0.035 | 0.15 | 0.14 | 0.15 | 0.21 | 0.12 | 0.1 |
| 10 | 12524637 | rs17491614 | 1.141 | 0.225 | -0.857 | 0.829 | -0.011 | 0.068 | 0.068 | 0 | 0.27 | 0.13 | 0.18 | 0.08 | 0.29 | 0.28 |
| 10 | 12524929 | rs12764478 | 0.005 | 0.579 | 0.115 | 0.934 | -0.004 | 0.009 | 0.009 | 0 | 0.17 | 0.13 | 0.08 | 0.05 | 0.18 | 0.12 |
| 10 | 12526550 | rs11257821 | -0.46 | 0.46 | -0.036 | 1.144 | 0.008 | 0.05 | 0.05 | 0 | 0.05 | 0.01 | 0.08 | 0.38 | 0.03 | 0.19 |
| 10 | 12527867 | rs2768431 | -0.125 | 0.484 | 0.316 | 0.966 | 0 | 0.03 | 0.03 | 0 | 0.26 | 0.33 | 0.25 | 0.39 | 0.25 | 0.15 |
| 10 | 12529195 | rs10906156 | 1.181 | 0.567 | -0.11 | 0.999 | -0.013 | 0.05 | 0.05 | 0 | 0.26 | 0.17 | 0.19 | 0.08 | 0.31 | 0.26 |
| 10 | 12531619 | rs10906157 | -0.833 | 0.584 | -0.063 | 1.097 | 0.001 | -0.043 | -0.043 | 0 | 0.23 | 0.16 | 0.16 | 0.44 | 0.19 | 0.36 |
| 10 | 12533288 | rs1998966 | -0.264 | 0.649 | -0.166 | 0.911 | -0.059 | 0.076 | 0.076 | 0 | 0.22 | 0.04 | 0.13 | 0.05 | 0.25 | 0.17 |
| 10 | 12534093 | rs2768436 | 0.038 | 0.573 | 0.226 | 0.436 | 0.02 | 0.015 | 0.015 | 0.045 | 0.44 | 0.7 | 0.45 | 0.56 | 0.42 | 0.41 |
| 10 | 12535056 | rs2768438 | -1.55 | 0.618 | 0.246 | 0.326 | -0.007 | 0.047 | 0.047 | 0 | 0.69 | 0.79 | 0.68 | 0.96 | 0.7 | 0.68 |
| 10 | 12537427 | rs17151786 | 1.434 | 0.158 | 0.65 | 0.58 | -0.003 | -0.011 | -0.011 | 1.443 | 0.22 | 0.27 | 0.17 | 0.1 | 0.23 | 0.17 |
| 10 | 12538491 | rs7078980 | 1.506 | 0.099 | 0.659 | 0.584 | 0.001 | 0.004 | 0.004 | 0 | 0.21 | 0.19 | 0.17 | 0.1 | 0.22 | 0.16 |
| 10 | 12538602 | rs7097560 | -1.281 | 0.052 | 0.995 | 0.539 | 0.019 | 0.085 | 0.085 | 0 | 0.49 | 0.37 | 0.46 | 0.81 | 0.54 | 0.41 |
| 10 | 12540800 | rs11257832 | 0.921 | 0.242 | 0.626 | 0.67 | -0.009 | -0.041 | -0.041 | 0 | 0.17 | 0.08 | 0.18 | 0.4 | 0.19 | 0.08 |
| 10 | 12542529 | rs2768444 | -0.51 | 0.263 | 0.835 | 0.646 | 0.012 | 0.035 | 0.035 | 0 | 0.33 | 0.45 | 0.37 | 0.48 | 0.36 | 0.35 |
| 10 | 12543729 | rs2815664 | -1.695 | 0.22 | 0.779 | 0.554 | -0.014 | -0.087 | -0.087 | 3.011 | 0.72 | 0.51 | 0.65 | 0.89 | 0.77 | 0.59 |
| 10 | 12544238 | rs1108959 | 0.064 | 0.41 | 0.348 | 0.81 | -0.013 | 0.003 | 0.003 | 0 | 0.26 | 0.19 | 0.24 | 0.2 | 0.28 | 0.2 |
| 10 | 12545786 | rs11257837 | -0.689 | 0.439 | 0.127 | 0.859 | 0.002 | -0.011 | -0.011 | 0 | 0.16 | 0.12 | 0.17 | 0.3 | 0.13 | 0.2 |
| 10 | 12547825 | rs12570530 | 0.784 | 0.284 | -0.113 | 0.435 | 0.003 | 0.173 | 0.173 | 0 | 0.34 | 0.11 | 0.32 | 0.03 | 0.29 | 0.35 |
| 10 | 12549612 | rs2068993 | 0.494 | 0.371 | 0.217 | 0.891 | 0.002 | 0.091 | 0.091 | 0 | 0.06 | 0.09 | 0.03 | 0.34 | 0.08 | 0.14 |
| 10 | 12549961 | rs2768447 | -0.572 | 0.461 | 0.466 | 0.858 | -0.006 | 0.007 | 0.007 | 0.293 | 0.59 | 0.5 | 0.63 | 0.56 | 0.58 | 0.76 |
| 10 | 12550970 | rs11257841 | 0.237 | 0.446 | 0.38 | 0.725 | -0.001 | 0.11 | 0.11 | 0.198 | 0.5 | 0.33 | 0.5 | 0.15 | 0.45 | 0.61 |
| 10 | 12551056 | rs11257842 | 0.233 | 0.459 | 0.364 | 0.741 | -0.001 | 0.094 | 0.094 | 0.188 | 0.5 | 0.36 | 0.5 | 0.15 | 0.44 | 0.61 |
| 10 | 12558701 | rs7078672 | 0.21 | -0.062 | 0.183 | 0.938 | 0.002 | 0.004 | 0.004 | 0 | 0.28 | 0.3 | 0.19 | 0.36 | 0.27 | 0.34 |
| 10 | 12566323 | rs11257857 | 0.031 | 0.015 | 0.139 | 1.02 | -0.002 | -0.006 | -0.006 | 0 | 0.55 | 0.49 | 0.47 | 0.38 | 0.52 | 0.63 |
| 10 | 12568166 | rs2815646 | -0.075 | 0.048 | 0.017 | 1 | 0 | 0.001 | 0.001 | 0.749 | 0.57 | 0.59 | 0.53 | 0.43 | 0.52 | 0.72 |
| 10 | 12570512 | rs12356332 | 1.135 | 0.151 | -0.808 | 0.98 | -0.012 | 0.03 | 0.03 | 0 | 0.21 | 0.02 | 0.14 | 0.1 | 0.21 | 0.16 |
| 10 | 12572496 | rs10906162 | 0.101 | 0.511 | -0.151 | 1.493 | 0.008 | 0.018 | 0.018 | 0 | 0.27 | 0.36 | 0.18 | 0.38 | 0.25 | 0.33 |
| 10 | 12573106 | rs2768463 | 0.096 | 0.602 | -0.089 | 1.601 | 0.003 | 0.007 | 0.007 | 0 | 0.35 | 0.29 | 0.4 | 0.32 | 0.34 | 0.42 |
| 10 | 12575817 | rs7079055 | 0.485 | 0.602 | -0.11 | 1.566 | 0.02 | 0.089 | 0.089 | 0 | 0.31 | 0.17 | 0.33 | 0.07 | 0.27 | 0.4 |
| 10 | 12576812 | rs1111060 | -1.076 | 0.699 | -0.195 | 1.664 | 0 | 0 | 0 | 0 | 0.63 | 0.61 | 0.57 | 0.65 | 0.6 | 0.75 |
| 10 | 12577793 | rs10458820 | -0.888 | 1.088 | 0.338 | 2.636 | 0.006 | 0.051 | 0.051 | 0 | 0.29 | 0.45 | 0.25 | 0.46 | 0.25 | 0.43 |
| 10 | 12577990 | rs6602593 | -1.311 | 1.126 | -0.036 | 2.615 | 0.005 | -0.004 | -0.004 | 0 | 0.37 | 0.6 | 0.41 | 0.72 | 0.39 | 0.47 |
| 10 | 12580515 | rs10906164 | -1.048 | 1.354 | -0.138 | 2.438 | -0.018 | 0.028 | 0.028 | 0 | 0.06 | 0.12 | 0.08 | 0.1 | 0.02 | 0.12 |
| 10 | 12582689 | rs7910874 | -1.119 | 1.823 | -0.455 | 2.347 | 0.006 | 0.067 | 0.067 | 0.262 | 0.07 | 0.09 | 0.19 | 0.33 | 0.14 | 0.03 |
| 10 | 12582784 | rs2815638 | -1.304 | 1.332 | -0.606 | 1.513 | 0.016 | -0.021 | -0.021 | 0 | 0.35 | 0.6 | 0.46 | 0.79 | 0.39 | 0.43 |
| 10 | 12583253 | rs1126368 | -1.307 | 0.572 | -1.01 | 1.974 | -0.005 | 0.088 | 0.088 | 0 | 0.07 | 0.12 | 0.18 | 0.29 | 0.1 | 0.13 |
| 10 | 12588117 | rs4750232 | -0.993 | 0.637 | -1.415 | 1.157 | -0.03 | -0.02 | -0.02 | 0 | 0.13 | 0.01 | 0.11 | 0.1 | 0.13 | 0.15 |
| 10 | 12589804 | rs2815633 | -0.706 | -0.171 | -1.977 | 1.396 | 0.037 | -0.066 | -0.066 | 0 | 0.21 | 0.06 | 0.3 | 0.28 | 0.15 | 0.11 |
| 10 | 12590175 | rs4747989 | -1.293 | -0.422 | -0.992 | 1.87 | -0.009 | 0.167 | 0.167 | 0 | 0.09 | 0.24 | 0.27 | 0.27 | 0.07 | 0.19 |
| 10 | 12591712 | rs7897059 | 0.976 | -0.12 | -0.76 | 1.17 | 0.017 | 0.006 | 0.006 | 0 | 0.51 | 0.57 | 0.42 | 0.47 | 0.6 | 0.49 |
| 10 | 12594436 | rs2768340 | 0.36 | -0.06 | -0.565 | 0.874 | -0.003 | -0.028 | -0.028 | 0 | 0.8 | 0.77 | 0.9 | 0.89 | 0.84 | 0.82 |
| 10 | 12595543 | rs1077746 | -0.197 | 0.162 | -0.952 | 0.875 | -0.018 | -0.02 | -0.02 | 0 | 0.46 | 0.25 | 0.39 | 0.59 | 0.42 | 0.48 |
| 10 | 12595815 | rs11592529 | 0.142 | -0.13 | -0.807 | 0.756 | -0.011 | -0.003 | -0.003 | 1.006 | 0.25 | 0.19 | 0.31 | 0.34 | 0.38 | 0.18 |
| 10 | 12597108 | rs2296754 | -0.012 | 0.623 | -0.564 | 0.767 | -0.034 | -0.007 | -0.007 | 0.305 | 0.25 | 0.13 | 0.28 | 0.34 | 0.42 | 0.17 |
| 10 | 12597458 | rs10906166 | 0.061 | 0.591 | -0.428 | 0.753 | 0.033 | 0.002 | 0.002 | 0 | 0.25 | 0.25 | 0.3 | 0.34 | 0.42 | 0.17 |
| 10 | 12598121 | rs10906169 | 1.027 | -0.433 | -0.59 | 0.454 | 0.002 | 0.041 | 0.041 | 0 | 0.33 | 0.29 | 0.32 | 0.11 | 0.34 | 0.37 |
| 10 | 12598255 | rs2768343 | -0.21 | -1.036 | -0.501 | 0.325 | -0.038 | -0.066 | -0.066 | 0 | 0.7 | 0.6 | 0.76 | 0.85 | 0.83 | 0.66 |

| | | | | | | | | | | | | | | | | |
|---|---|---|---|---|---|---|---|---|---|---|---|---|---|---|---|---|
| 10 | 12599694 | rs2815624 | -0.174 | -1.015 | -0.507 | 0.723 | -0.058 | 0.122 | 0.122 | 0 | 0.82 | 0.6 | 0.72 | 0.74 | 0.9 | 0.74 |
| 10 | 12600668 | rs2768348 | 0.913 | -1.115 | -0.767 | 0.635 | 0.027 | 0.008 | 0.008 | 0 | 0.38 | 0.36 | 0.37 | 0.35 | 0.32 | 0.33 |
| 10 | 12601504 | rs11257888 | -0.086 | -1.082 | -0.819 | 0.627 | 0.021 | 0.011 | 0.011 | 0 | 0.43 | 0.24 | 0.38 | 0.45 | 0.58 | 0.4 |
| 10 | 12603501 | rs12764138 | -0.084 | -1.072 | -0.811 | 0.587 | 0.02 | -0.012 | -0.012 | 0 | 0.43 | 0.24 | 0.42 | 0.51 | 0.58 | 0.4 |
| 10 | 12603912 | rs2648710 | 0.708 | -0.815 | -0.73 | 0.566 | 0.029 | 0.028 | 0.028 | 0 | 0.38 | 0.31 | 0.34 | 0.29 | 0.32 | 0.33 |
| 10 | 12608676 | rs1409961 | 0.242 | -0.312 | -0.871 | 0.443 | 0.003 | 0.088 | 0.088 | 0 | 0.47 | 0.39 | 0.42 | 0.19 | 0.45 | 0.51 |
| 10 | 12608850 | rs1112240 | 0.14 | 0.619 | -1.017 | 0.302 | 0.046 | 0.095 | 0.095 | 0 | 0.52 | 0.4 | 0.52 | 0.19 | 0.62 | 0.58 |
| 10 | 12612338 | rs12572894 | 1.086 | 0.272 | -0.957 | 0.648 | -0.05 | 0.007 | 0.007 | 0.338 | 0.16 | 0.02 | 0.09 | 0.09 | 0.16 | 0.15 |
| 10 | 12614891 | rs7081248 | 1.948 | 0.251 | -0.741 | 0.613 | -0.002 | 0.034 | 0.034 | 0 | 0.28 | 0.15 | 0.41 | 0.15 | 0.32 | 0.31 |
| 10 | 12622303 | rs952570 | -0.069 | 0.047 | -1.196 | 0.779 | -0.01 | -0.021 | -0.021 | 0 | 0.6 | 0.73 | 0.54 | 0.47 | 0.59 | 0.57 |
| 10 | 12622680 | rs7076395 | 0.451 | 0.034 | -1.106 | 0.626 | -0.04 | 0.194 | 0.194 | 0.021 | 0.14 | 0.54 | 0.14 | 0.35 | 0.07 | 0.11 |
| 10 | 12625436 | rs7074055 | 0.588 | 0.318 | -1.713 | 0.344 | -0.006 | 0.025 | 0.025 | 0 | 0.23 | 0.16 | 0.13 | 0.12 | 0.21 | 0.23 |
| 10 | 12626977 | rs2895527 | 0.259 | 0.149 | -1.83 | 0.447 | 0.05 | -0.009 | -0.009 | 0 | 0.48 | 0.78 | 0.58 | 0.49 | 0.43 | 0.42 |
| 10 | 12628501 | rs4364954 | 1.605 | 0.352 | -1.939 | -0.018 | -0.045 | 0.124 | 0.124 | 0 | 0.28 | 0.08 | 0.33 | 0.07 | 0.35 | 0.3 |
| 10 | 12629336 | rs11257908 | 0.879 | 0.298 | -1.745 | 0.316 | -0.051 | -0.03 | -0.03 | 0 | 0.1 | 0.01 | 0.1 | 0.12 | 0.13 | 0.13 |
| 10 | 12629368 | rs2895534 | -0.904 | 0.109 | -1.703 | 0.084 | 0.013 | 0.006 | 0.006 | 0.226 | 0.74 | 0.9 | 0.81 | 0.73 | 0.76 | 0.71 |
| 10 | 12631633 | rs7093225 | -1.006 | -0.285 | -1.045 | -0.198 | 0.012 | 0.046 | 0.046 | 0 | 0.56 | 0.66 | 0.64 | 0.78 | 0.56 | 0.56 |
| 10 | 12632156 | rs2399880 | 0.672 | -0.331 | -1.242 | -0.265 | 0 | 0.004 | 0.004 | 0 | 0.17 | 0.14 | 0.07 | 0.03 | 0.13 | 0.27 |
| 10 | 12633366 | rs10732936 | -1.704 | -0.367 | -1.545 | -0.256 | -0.04 | 0.085 | 0.085 | 0 | 0.73 | 0.92 | 0.72 | 0.88 | 0.69 | 0.76 |
| 10 | 12634257 | rs9787460 | 1.273 | -0.511 | -1.262 | 0.186 | -0.028 | 0.078 | 0.078 | 0.617 | 0.06 | 0.1 | 0.1 | 0.29 | 0.01 | 0.04 |
| 10 | 12634503 | rs9787594 | 0.838 | -0.601 | -1.233 | 0.136 | -0.04 | 0.031 | 0.031 | 0.801 | 0.1 | 0.15 | 0.03 | 0.28 | 0.01 | 0.04 |
| 10 | 12638147 | rs11257913 | 0.007 | -0.08 | -1.048 | 0.056 | 0.007 | -0.001 | -0.001 | 0 | 0.36 | 0.37 | 0.56 | 0.62 | 0.49 | 0.33 |
| 10 | 12638260 | rs10906179 | -1.087 | 0.364 | -1.36 | 0.056 | -0.007 | 0.083 | 0.083 | 0 | 0.72 | 0.86 | 0.72 | 0.91 | 0.68 | 0.76 |
| 10 | 12640797 | rs7911208 | -0.005 | 0.842 | -0.895 | 1.021 | -0.033 | -0.055 | -0.055 | 0.053 | 0.2 | 0.11 | 0.35 | 0.44 | 0.33 | 0.2 |
| 10 | 12641460 | rs10795966 | -0.589 | 0.558 | -0.642 | 0.957 | -0.002 | 0.003 | 0.003 | 1.578 | 0.23 | 0.22 | 0.4 | 0.44 | 0.34 | 0.21 |
| 10 | 12644580 | rs12414386 | 0.486 | -0.772 | -0.016 | 0.824 | -0.08 | 0.08 | 0.08 | 0 | 0.81 | 0.59 | 0.8 | 0.69 | 0.92 | 0.81 |
| 10 | 12646813 | rs17151970 | 0.214 | -0.834 | -0.432 | 0.348 | -0.004 | -0.029 | -0.029 | 0 | 0.16 | 0.22 | 0.09 | 0.07 | 0.12 | 0.1 |
| 10 | 12652657 | rs17151978 | 1.058 | -1.131 | -0.926 | 0.164 | 0.028 | 0.036 | 0.036 | 0 | 0.35 | 0.25 | 0.23 | 0.26 | 0.27 | 0.4 |
| 10 | 12654884 | rs4301695 | -0.25 | -1.179 | -1.562 | 0.263 | 0.014 | -0.006 | -0.006 | 0.64 | 0.41 | 0.24 | 0.47 | 0.58 | 0.58 | 0.4 |
| 10 | 12655800 | rs879753 | 1.445 | -0.906 | -1.84 | 0.344 | 0 | 0.032 | 0.032 | 0 | 0.3 | 0.09 | 0.2 | 0.19 | 0.26 | 0.38 |
| 10 | 12658889 | rs7912882 | 0.501 | -0.664 | -1.849 | 0.558 | -0.169 | -0.139 | -0.139 | 0.183 | 0.25 | 0.03 | 0.39 | 0.4 | 0.46 | 0.24 |
| 10 | 12661057 | rs12360308 | 1.459 | -1.049 | -1.875 | 0.405 | 0.019 | 0.022 | 0.022 | 0 | 0.29 | 0.1 | 0.18 | 0.2 | 0.25 | 0.37 |
| 10 | 12662552 | rs12355005 | 0.097 | -0.012 | -1.329 | 0.811 | -0.16 | -0.134 | -0.134 | 0.011 | 0.25 | 0.04 | 0.39 | 0.4 | 0.45 | 0.24 |
| 10 | 12669049 | rs11257922 | -0.487 | -0.394 | -1.111 | 0.655 | -0.066 | -0.052 | -0.052 | 0.01 | 0.35 | 0.13 | 0.57 | 0.55 | 0.54 | 0.33 |
| 10 | 12671886 | rs4480437 | 0.549 | -0.576 | -0.406 | 0.595 | -0.021 | -0.032 | -0.032 | 0 | 0.77 | 0.42 | 0.79 | 0.8 | 0.86 | 0.78 |
| 10 | 12672884 | rs12260523 | 1.64 | -0.967 | -0.529 | 0.387 | 0.052 | 0.045 | 0.045 | 0.267 | 0.37 | 0.25 | 0.2 | 0.22 | 0.26 | 0.43 |
| 10 | 12673160 | rs4750244 | 0.811 | -0.925 | -0.251 | 0.433 | 0.024 | 0.02 | 0.02 | 0.055 | 0.5 | 0.45 | 0.29 | 0.31 | 0.35 | 0.52 |
| 10 | 12673810 | rs17152029 | 1.945 | -0.984 | -0.355 | 0.395 | 0.059 | 0.088 | 0.088 | 0.477 | 0.33 | 0.12 | 0.19 | 0.18 | 0.26 | 0.36 |
| 10 | 12675061 | rs10906189 | 0.573 | 0.061 | -0.166 | 0.766 | 0.018 | 0.005 | 0.005 | 0 | 0.61 | 0.53 | 0.47 | 0.4 | 0.56 | 0.6 |
| 10 | 12676957 | rs11257926 | 1.666 | -0.171 | -0.383 | 0.707 | 0.085 | 0.052 | 0.052 | 0.432 | 0.38 | 0.18 | 0.2 | 0.26 | 0.27 | 0.42 |
| 10 | 12677614 | rs17152037 | 1.673 | -0.18 | -0.387 | 0.704 | 0.084 | 0.051 | 0.051 | 0.445 | 0.38 | 0.18 | 0.2 | 0.26 | 0.27 | 0.43 |
| 10 | 12679491 | rs2399866 | 0.512 | -0.069 | 0.371 | 0.796 | 0 | 0.001 | 0.001 | 0 | 0.58 | 0.58 | 0.4 | 0.57 | 0.44 | 0.68 |
| 10 | 12681113 | rs17580343 | 0.097 | 0.341 | 0.377 | 0.806 | 0.07 | 0.02 | 0.02 | 0 | 0.1 | 0.2 | 0.18 | 0.12 | 0.2 | 0.1 |
| 10 | 12686487 | rs1004247 | -0.665 | 0.068 | 0.264 | 0.752 | 0.017 | -0.026 | -0.026 | 0 | 0.11 | 0.2 | 0.08 | 0.07 | 0.1 | 0.08 |
| 10 | 12689914 | rs12780223 | -0.35 | 0.025 | -0.25 | 0.895 | -0.152 | -0.116 | -0.116 | 0 | 0.31 | 0.04 | 0.43 | 0.36 | 0.53 | 0.26 |
| 10 | 12692902 | rs10752269 | -1.301 | -0.036 | 0.441 | 1.092 | -0.001 | -0.001 | -0.001 | 0.309 | 0.42 | 0.51 | 0.57 | 0.58 | 0.54 | 0.41 |
| 10 | 12693760 | rs7922799 | -0.43 | 0.145 | 0.389 | 1.125 | -0.003 | 0.005 | 0.005 | 0.013 | 0.1 | 0.06 | 0.06 | 0.1 | 0.08 | 0.2 |
| 10 | 12695387 | rs7071404 | 1.956 | -0.218 | 0.68 | 1.12 | 0.071 | 0.027 | 0.027 | 0 | 0.42 | 0.19 | 0.29 | 0.32 | 0.32 | 0.45 |
| 10 | 12696147 | rs10906196 | 0.525 | 0.109 | 0.57 | 1.158 | -0.076 | -0.068 | -0.068 | 0 | 0.19 | 0.03 | 0.26 | 0.29 | 0.28 | 0.16 |
| 10 | 12702068 | rs11257933 | -0.191 | -0.174 | 0.637 | 1.143 | -0.007 | 0.006 | 0.006 | 0 | 0.1 | 0.06 | 0.06 | 0.09 | 0.08 | 0.21 |
| 10 | 12702818 | rs10508445 | -0.484 | -0.43 | 1.217 | 1.252 | 0.023 | 0.016 | 0.016 | 0.38 | 0.29 | 0.34 | 0.45 | 0.43 | 0.47 | 0.27 |
| 10 | 12703498 | rs10737061 | 0.265 | -0.441 | 1.454 | 1.146 | -0.034 | 0.033 | 0.033 | 0.528 | 0.82 | 0.63 | 0.8 | 0.76 | 0.88 | 0.92 |
| 10 | 12704191 | rs10752271 | -0.714 | -0.437 | 1.42 | 1.157 | -0.029 | 0.039 | 0.039 | 0 | 0.87 | 0.77 | 0.83 | 0.8 | 0.92 | 0.92 |
| 10 | 12706737 | rs4417163 | -0.478 | -0.525 | 1.25 | 1.159 | -0.027 | -0.02 | -0.02 | 0.447 | 0.32 | 0.25 | 0.5 | 0.44 | 0.48 | 0.28 |
| 10 | 12712070 | rs3995695 | 2.334 | -0.446 | 1.086 | 1.088 | 0.033 | 0.056 | 0.056 | 0 | 0.38 | 0.14 | 0.23 | 0.23 | 0.33 | 0.42 |
| 10 | 12714269 | rs12251362 | -1.203 | -0.455 | 1.536 | 1.119 | -0.003 | 0.001 | 0.001 | 0.076 | 0.41 | 0.45 | 0.51 | 0.44 | 0.55 | 0.46 |
| 10 | 12717333 | rs10795975 | -0.752 | -0.423 | 1.445 | 1.109 | -0.003 | -0.004 | -0.004 | 0 | 0.34 | 0.34 | 0.53 | 0.5 | 0.5 | 0.41 |
| 10 | 12718736 | rs10906202 | -0.707 | -0.503 | 1.405 | 1.06 | -0.022 | 0.023 | 0.023 | 0 | 0.87 | 0.78 | 0.83 | 0.8 | 0.92 | 0.93 |
| 10 | 12731447 | rs12220703 | 1.557 | -0.676 | 1.033 | 1.046 | 0.05 | 0.004 | 0.004 | 0 | 0.44 | 0.22 | 0.33 | 0.38 | 0.37 | 0.59 |
| 10 | 12740543 | rs10795977 | -1.131 | -0.436 | 1.267 | 1.114 | -0.004 | 0.001 | 0.001 | 0 | 0.41 | 0.44 | 0.53 | 0.45 | 0.55 | 0.45 |
| 10 | 12741744 | rs10906210 | 1.024 | -0.634 | 1.073 | 1.065 | 0.041 | -0.014 | -0.014 | 0 | 0.76 | 0.21 | 0.77 | 0.73 | 0.9 | 0.76 |
| 10 | 12743642 | rs2399864 | 1.509 | 0.043 | 1.391 | 1.206 | 0.012 | -0.042 | -0.042 | 1.307 | 0.6 | 0.24 | 0.55 | 0.53 | 0.65 | 0.58 |
| 10 | 12747672 | rs11257948 | -0.381 | 0.116 | 1.68 | 1.246 | -0.004 | -0.004 | -0.004 | 0 | 0.26 | 0.22 | 0.31 | 0.31 | 0.31 | 0.19 |
| 10 | 12748159 | rs7909195 | 0.712 | 0.117 | 1.703 | 0.986 | -0.013 | 0.018 | 0.018 | 0.057 | 0.94 | 0.76 | 0.95 | 0.94 | 0.97 | 0.8 |
| 10 | 12748816 | rs4750249 | 1.577 | 0.17 | 1.389 | 0.959 | 0.066 | -0.041 | -0.041 | 0.301 | 0.79 | 0.48 | 0.75 | 0.85 | 0.74 | 0.69 |
| 10 | 12752638 | rs11817741 | -1.268 | 0.391 | 1.425 | 1.081 | 0.046 | 0.059 | 0.059 | 0.015 | 0.06 | 0.34 | 0.03 | 0.01 | 0 | 0.02 |
| 10 | 12754862 | rs867764 | -0.74 | 0.547 | 1.171 | 1.16 | -0.163 | -0.147 | -0.147 | 3.269 | 0.28 | 0.02 | 0.43 | 0.41 | 0.5 | 0.28 |
| 10 | 12756149 | rs10906212 | 0.309 | 0.763 | 1.457 | 1.19 | 0.131 | 0.028 | 0.028 | 0 | 0.88 | 0.64 | 0.8 | 0.85 | 0.79 | 0.74 |
| 10 | 12758991 | rs11812913 | -1.378 | 0.067 | 1.51 | 1.151 | 0.001 | 0.128 | 0.128 | 0.182 | 0.06 | 0.28 | 0.08 | 0.11 | 0.01 | 0.05 |

| | | | | | | | | | | | | | | | | |
|---|---|---|---|---|---|---|---|---|---|---|---|---|---|---|---|---|
| 10 | 12762303 | rs723211 | 0.586 | 0.314 | 1.362 | 0.947 | 0.012 | 0.019 | 0.019 | 0.578 | 0.65 | 0.58 | 0.5 | 0.43 | 0.48 | 0.57 |
| 10 | 12762476 | rs723210 | 1.013 | 0.026 | 0.958 | 0.848 | -0.001 | 0.002 | 0.002 | 0 | 0.6 | 0.4 | 0.46 | 0.5 | 0.45 | 0.57 |
| 10 | 12771680 | rs7077423 | -1.296 | -0.352 | 1.093 | 0.796 | -0.022 | 0.017 | 0.017 | 0 | 0.36 | 0.56 | 0.36 | 0.38 | 0.3 | 0.29 |
| 10 | 12778614 | rs11257964 | 1.109 | -0.288 | 1.532 | 0.578 | -0.044 | -0.006 | -0.006 | 0 | 0.89 | 0.53 | 0.93 | 0.96 | 0.98 | 0.9 |
| 10 | 12783565 | rs17152177 | -0.896 | -0.01 | 0.889 | 0.544 | -0.025 | -0.027 | -0.027 | 0 | 0.08 | 0.18 | 0.04 | 0.01 | 0.03 | 0.08 |
| 10 | 12787065 | rs7894890 | -0.505 | 0.455 | 0.689 | 0.844 | -0.026 | 0.05 | 0.05 | 0 | 0.23 | 0.44 | 0.18 | 0.26 | 0.2 | 0.14 |
| 10 | 12794322 | rs4750255 | -0.038 | 0.855 | 0.019 | 0.61 | -0.052 | -0.002 | -0.002 | 0 | 0.27 | 0.14 | 0.29 | 0.26 | 0.31 | 0.17 |
| 10 | 12797007 | rs4072657 | -0.157 | 0.63 | 0.3 | 0.739 | -0.009 | 0.005 | 0.005 | 0.249 | 0.28 | 0.28 | 0.24 | 0.3 | 0.22 | 0.15 |
| 10 | 12797172 | rs4072656 | 0.249 | 0.782 | -0.278 | 0.705 | -0.044 | -0.012 | -0.012 | 0 | 0.23 | 0.16 | 0.29 | 0.26 | 0.31 | 0.16 |
| 10 | 12797285 | rs10906218 | 0.014 | 0.604 | 0.317 | 0.681 | 0.038 | -0.019 | -0.019 | 0.002 | 0.06 | 0.16 | 0.16 | 0.07 | 0.12 | 0.07 |
| 10 | 12797758 | rs4132050 | -0.883 | 0.522 | -0.225 | 0.76 | -0.006 | 0.004 | 0.004 | 0.336 | 0.34 | 0.56 | 0.4 | 0.37 | 0.34 | 0.23 |
| 10 | 12802455 | rs10508447 | -0.9 | 0.537 | -0.239 | 0.554 | 0 | -0.004 | -0.004 | 0 | 0.1 | 0.14 | 0.12 | 0.07 | 0.09 | 0.15 |
| 10 | 12805377 | rs3802572 | -0.021 | 0.251 | -0.48 | 0.396 | 0.005 | 0.004 | 0.004 | 4.774 | 0.44 | 0.31 | 0.52 | 0.59 | 0.51 | 0.45 |
| 10 | 12805430 | rs3802570 | 0.555 | -0.177 | -0.353 | 0.141 | 0.013 | 0.041 | 0.041 | 3.184 | 0.31 | 0.21 | 0.24 | 0.21 | 0.32 | 0.31 |
| 10 | 12810570 | rs2482052 | -0.066 | -0.096 | -0.268 | 0.648 | 0.007 | -0.017 | -0.017 | 0 | 0.75 | 0.83 | 0.69 | 0.62 | 0.77 | 0.62 |
| 10 | 12811171 | rs2482051 | -0.215 | 0.025 | 0.017 | 0.004 | -0.022 | -0.067 | -0.067 | 1.203 | 0.7 | 0.56 | 0.8 | 0.89 | 0.78 | 0.77 |
| 10 | 12826886 | rs3824654 | 0.314 | -0.092 | -0.561 | -0.035 | 0.013 | 0.123 | 0.123 | 1.191 | 0.32 | 0.1 | 0.21 | 0.16 | 0.28 | 0.29 |
| 10 | 12827660 | rs3802559 | 0.582 | 0.244 | -0.691 | 0.301 | -0.017 | 0.002 | 0.002 | 1.773 | 0.41 | 0.3 | 0.5 | 0.37 | 0.47 | 0.5 |
| 10 | 12827690 | rs3802558 | -0.529 | 0.639 | -0.923 | 0.526 | -0.019 | -0.044 | -0.044 | 0 | 0.64 | 0.57 | 0.77 | 0.8 | 0.71 | 0.71 |
| 10 | 12828290 | rs10906223 | -0.505 | 1.429 | -1.196 | 1.286 | -0.029 | -0.067 | -0.067 | 0.377 | 0.18 | 0.07 | 0.25 | 0.43 | 0.23 | 0.18 |
| 10 | 12830969 | rs12359004 | -0.361 | 1.227 | -0.894 | 0.779 | -0.028 | -0.015 | -0.015 | 0.683 | 0.05 | 0 | 0.02 | 0.05 | 0.05 | 0.03 |
| 10 | 12832593 | rs4747999 | -0.362 | 0.709 | -0.469 | 0.874 | -0.002 | -0.002 | -0.002 | 0 | 0.45 | 0.44 | 0.54 | 0.63 | 0.43 | 0.35 |
| 10 | 12833612 | rs11257991 | -0.228 | 0.023 | -0.353 | 1.082 | -0.027 | 0.214 | 0.214 | 0 | 0.16 | 0.36 | 0.19 | 0.49 | 0.09 | 0.18 |
| 10 | 12834926 | rs928336 | -1.048 | -0.007 | 0.489 | 0.962 | 0 | 0 | 0 | 1.229 | 0.46 | 0.54 | 0.57 | 0.63 | 0.44 | 0.37 |
| 10 | 12835045 | rs928337 | 1.209 | -0.329 | -0.205 | 0.255 | -0.094 | 0.174 | 0.174 | 0 | 0.3 | 0.06 | 0.28 | 0.01 | 0.39 | 0.19 |
| 10 | 12835383 | rs2493767 | -0.687 | -0.16 | 0.321 | 0.84 | -0.01 | 0.007 | 0.007 | 0 | 0.42 | 0.5 | 0.53 | 0.56 | 0.38 | 0.3 |
| 10 | 12837585 | rs11257997 | 0.241 | -0.097 | -0.163 | 0.977 | -0.031 | 0.04 | 0.04 | 1.34 | 0.3 | 0.11 | 0.35 | 0.17 | 0.34 | 0.18 |
| 10 | 12840842 | rs1556408 | -0.607 | -0.107 | 0.069 | 1.002 | 0.005 | -0.025 | -0.025 | 0.058 | 0.38 | 0.25 | 0.51 | 0.54 | 0.38 | 0.25 |
| 10 | 12847838 | rs4750267 | 0.916 | -0.215 | -0.148 | 1.257 | 0.022 | 0.029 | 0.029 | 0 | 0.58 | 0.25 | 0.43 | 0.31 | 0.63 | 0.71 |
| 10 | 12849721 | rs17583338 | 0.052 | -0.43 | -0.197 | 1.351 | -0.035 | -0.01 | -0.01 | 0.069 | 0.21 | 0.05 | 0.3 | 0.17 | 0.26 | 0.16 |
| 10 | 12849790 | rs10906228 | 0.069 | -0.463 | -0.184 | 1.627 | 0.017 | -0.011 | -0.011 | 0.078 | 0.07 | 0.03 | 0.12 | 0.33 | 0.02 | 0.03 |
| 10 | 12850236 | rs1999369 | -1.215 | -0.429 | -0.059 | 1.565 | -0.027 | 0.072 | 0.072 | 0.923 | 0.17 | 0.61 | 0.2 | 0.4 | 0.09 | 0.08 |
| 10 | 12851181 | rs10906230 | -1.254 | -0.318 | -0.236 | 1.499 | 0.016 | -0.026 | -0.026 | 0 | 0.4 | 0.69 | 0.52 | 0.58 | 0.36 | 0.27 |
| 10 | 12857635 | rs7080879 | 0.929 | -0.542 | -0.141 | 1.469 | 0.029 | 0.003 | 0.003 | 0.039 | 0.64 | 0.29 | 0.48 | 0.3 | 0.7 | 0.7 |
| 10 | 12858045 | rs3818675 | -1.302 | -0.665 | 0.425 | 1.623 | -0.076 | 0.154 | 0.154 | 0.751 | 0.14 | 0.53 | 0.17 | 0.41 | 0.04 | 0.11 |
| 10 | 12859068 | rs2431629 | -1.175 | -0.21 | 0.127 | 1.727 | -0.017 | 0.087 | 0.087 | 0 | 0.26 | 0.53 | 0.35 | 0.59 | 0.25 | 0.34 |
| 10 | 12860012 | rs17152297 | -0.895 | -0.341 | -0.2 | 1.944 | -0.007 | 0.027 | 0.027 | 0.048 | 0.14 | 0.16 | 0.1 | 0.31 | 0.1 | 0.2 |
| 10 | 12860373 | rs2493750 | 0.173 | -0.154 | -0.726 | 1.199 | -0.005 | 0.007 | 0.007 | 0 | 0.39 | 0.53 | 0.48 | 0.45 | 0.34 | 0.27 |
| 10 | 12860391 | rs2431628 | -0.559 | -0.061 | -1.191 | 0.757 | -0.004 | 0.011 | 0.011 | 0.88 | 0.33 | 0.36 | 0.25 | 0.54 | 0.3 | 0.53 |
| 10 | 12860747 | rs2161365 | -0.079 | -0.108 | -0.815 | 1.014 | -0.002 | -0.009 | -0.009 | 0 | 0.54 | 0.52 | 0.63 | 0.72 | 0.53 | 0.6 |
| 10 | 12860828 | rs766947 | -0.068 | -0.615 | -1.132 | 0.925 | 0.01 | 0.001 | 0.001 | 1.955 | 0.33 | 0.28 | 0.37 | 0.26 | 0.31 | 0.11 |
| 10 | 12861945 | rs1630635 | -0.608 | -0.81 | -0.752 | 1.35 | -0.004 | 0.039 | 0.039 | 0 | 0.08 | 0.1 | 0.17 | 0.33 | 0.06 | 0.2 |
| 10 | 12862628 | rs11812477 | -0.222 | -1.217 | -0.827 | 0.662 | 0.017 | 0.015 | 0.015 | 0.015 | 0.12 | 0.11 | 0.03 | 0 | 0.04 | 0.08 |
| 10 | 12863366 | rs1644426 | -0.319 | -1.51 | -0.305 | 0.593 | -0.026 | -0.026 | -0.026 | 0 | 0.66 | 0.43 | 0.69 | 0.82 | 0.73 | 0.81 |
| 10 | 12864310 | rs1644428 | -0.244 | -1.581 | 0.106 | 0.736 | -0.023 | -0.013 | -0.013 | 0 | 0.65 | 0.42 | 0.69 | 0.79 | 0.73 | 0.81 |
| 10 | 12866356 | rs2482023 | -0.766 | -1.65 | 0.282 | 0.652 | -0.007 | 0.013 | 0.013 | 0 | 0.87 | 0.81 | 0.79 | 0.86 | 0.85 | 0.86 |
| 10 | 12869000 | rs1644392 | 0.422 | -1.335 | -0.326 | 0.419 | -0.018 | -0.002 | -0.002 | 0 | 0.29 | 0.12 | 0.26 | 0.29 | 0.27 | 0.44 |
| 10 | 12869047 | rs1644393 | -1.048 | -1.551 | -0.249 | 0.401 | 0.021 | 0.039 | 0.039 | 1.404 | 0.74 | 0.84 | 0.74 | 0.92 | 0.76 | 0.85 |
| 10 | 12870334 | rs2431626 | 0.201 | -1.263 | 0.233 | 0.417 | 0.001 | 0.026 | 0.026 | 0 | 0.53 | 0.41 | 0.54 | 0.71 | 0.56 | 0.43 |
| 10 | 12873352 | rs1757063 | 0.053 | -1.574 | 0.506 | 0.504 | -0.001 | 0 | 0 | 0 | 0.32 | 0.31 | 0.24 | 0.28 | 0.29 | 0.44 |
| 10 | 12874421 | rs1757065 | 0.641 | -1.841 | 0.655 | 0.266 | -0.004 | -0.02 | -0.02 | 0 | 0.25 | 0.22 | 0.13 | 0.06 | 0.17 | 0.15 |
| 10 | 12874715 | rs2431621 | -1.523 | -1.977 | 0.174 | 0.402 | -0.155 | 0.085 | 0.085 | 0 | 0.65 | 0.95 | 0.64 | 0.84 | 0.49 | 0.76 |
| 10 | 12875285 | rs2431636 | 0.398 | -1.822 | 0.389 | 0.339 | -0.06 | -0.118 | -0.118 | 0.787 | 0.27 | 0.47 | 0.15 | 0.06 | 0.17 | 0.15 |
| 10 | 12877007 | rs15980 | -0.462 | -1.793 | -0.287 | 0.406 | 0.002 | -0.004 | -0.004 | 0.057 | 0.41 | 0.75 | 0.33 | 0.38 | 0.36 | 0.55 |
| 10 | 12877269 | rs13674 | 1.112 | -1.75 | -0.321 | 0.409 | 0.002 | 0 | 0 | 0 | 0.07 | 0.06 | 0.06 | 0.07 | 0.07 | 0.11 |
| 10 | 12878896 | rs1644406 | 1.725 | -1.676 | -0.416 | 0.279 | 0.032 | 0.095 | 0.095 | 0.984 | 0.17 | 0.02 | 0.06 | 0 | 0.1 | 0.04 |
| 10 | 12881855 | rs1627707 | -0.081 | -1.39 | -0.131 | 0.498 | -0.004 | 0.001 | 0.001 | 0.018 | 0.36 | 0.73 | 0.31 | 0.32 | 0.31 | 0.46 |
| 10 | 12889696 | rs2482069 | -0.293 | -1.625 | -0.096 | 0.645 | -0.025 | 0.001 | 0.001 | 0 | 0.37 | 0.43 | 0.33 | 0.43 | 0.22 | 0.32 |
| 10 | 12892282 | rs4750273 | -0.537 | -0.624 | 0.044 | 0.909 | 0.006 | 0.016 | 0.016 | 0 | 0.22 | 0.26 | 0.25 | 0.3 | 0.2 | 0.39 |
| 10 | 12897013 | rs2050292 | -0.798 | -0.464 | -0.461 | 0.446 | 0.007 | 0.107 | 0.107 | 0 | 0.59 | 0.77 | 0.62 | 0.89 | 0.6 | 0.75 |
| 10 | 12906431 | rs10906242 | 0.135 | -0.145 | -0.746 | 0.8 | -0.099 | -0.034 | -0.034 | 0.016 | 0.39 | 0.09 | 0.33 | 0.37 | 0.45 | 0.48 |
| 10 | 12913480 | rs17152403 | 0.944 | -0.356 | -0.114 | 0.635 | -0.007 | -0.008 | -0.008 | 0 | 0.06 | 0.01 | 0.06 | 0.04 | 0.07 | 0.08 |
| 10 | 12913754 | rs11258040 | 0.29 | -0.311 | -0.012 | 0.873 | -0.004 | -0.004 | -0.004 | 0 | 0.13 | 0.12 | 0.13 | 0.27 | 0.14 | 0.36 |
| 10 | 12914521 | rs4750277 | -0.701 | -0.807 | 0.258 | 0.573 | -0.017 | 0.03 | 0.03 | 0 | 0.78 | 0.64 | 0.71 | 0.65 | 0.84 | 0.77 |
| 10 | 12916042 | rs17152417 | -0.488 | -0.549 | 0.553 | 0.507 | -0.003 | -0.001 | -0.001 | 0 | 0.46 | 0.5 | 0.44 | 0.51 | 0.54 | 0.56 |
| 10 | 12918662 | rs11258041 | 1.222 | -0.707 | 0.393 | 0.557 | -0.038 | 0.009 | 0.009 | 0 | 0.24 | 0.12 | 0.22 | 0.2 | 0.32 | 0.12 |
| 10 | 12918764 | rs7909670 | -0.655 | -0.825 | 0.02 | 0.439 | -0.032 | -0.031 | -0.031 | 0 | 0.48 | 0.77 | 0.4 | 0.47 | 0.47 | 0.51 |
| 10 | 12923638 | rs1929394 | -0.419 | -0.768 | 0.351 | 0.522 | 0.001 | -0.005 | -0.005 | 0 | 0.18 | 0.15 | 0.12 | 0.25 | 0.14 | 0.38 |
| 10 | 12932822 | rs6602622 | -0.282 | -0.742 | 0.459 | 0.433 | 0 | 0 | 0 | 0 | 0.45 | 0.54 | 0.43 | 0.47 | 0.5 | 0.54 |

| | | | | | | | | | | | | | | | | |
|---|---|---|---|---|---|---|---|---|---|---|---|---|---|---|---|---|
| 10 | 12933926 | rs1888665 | 0.37 | -0.712 | -0.11 | 0.445 | -0.112 | -0.093 | -0.093 | 0.936 | 0.37 | 0.1 | 0.34 | 0.44 | 0.44 | 0.47 |
| 10 | 12934038 | rs1888667 | -0.621 | -0.483 | -0.052 | 0.514 | 0.009 | -0.006 | -0.006 | 0 | 0.53 | 0.77 | 0.46 | 0.51 | 0.55 | 0.58 |
| 10 | 12935808 | rs10906247 | -0.628 | -0.509 | -0.011 | 0.519 | 0.023 | -0.005 | -0.005 | 0 | 0.54 | 0.73 | 0.46 | 0.54 | 0.57 | 0.6 |
| 10 | 12936781 | rs2399915 | 0.309 | -0.613 | 0.066 | 0.578 | 0.004 | 0.052 | 0.052 | 0 | 0.46 | 0.21 | 0.55 | 0.36 | 0.45 | 0.39 |
| 10 | 12937365 | rs10906248 | 0.489 | -0.698 | 0.214 | 0.63 | -0.013 | 0.027 | 0.027 | 0 | 0.73 | 0.43 | 0.79 | 0.63 | 0.78 | 0.53 |
| 10 | 12938057 | rs4748009 | 0.348 | -0.607 | 0.736 | 0.652 | 0.01 | 0.074 | 0.074 | 0 | 0.13 | 0.24 | 0.12 | 0.24 | 0.13 | 0.37 |
| 10 | 12938754 | rs3232 | 1.23 | -0.591 | 0.582 | 0.608 | -0.107 | -0.02 | -0.02 | 0.075 | 0.23 | 0.03 | 0.22 | 0.23 | 0.32 | 0.11 |
| 10 | 12938961 | rs2280076 | 1.23 | -0.591 | 0.582 | 0.608 | -0.107 | -0.02 | -0.02 | 0.017 | 0.23 | 0.03 | 0.22 | 0.23 | 0.32 | 0.12 |
| 10 | 12939124 | rs10795991 | 0.173 | -0.583 | 0.988 | 0.768 | 0.009 | 0.074 | 0.074 | 0 | 0.13 | 0.24 | 0.12 | 0.23 | 0.13 | 0.37 |
| 10 | 12941157 | rs4748011 | -0.611 | -0.629 | 1.16 | 0.773 | 0.004 | 0.087 | 0.087 | 0.116 | 0.2 | 0.32 | 0.19 | 0.41 | 0.21 | 0.47 |
| 10 | 12942295 | rs7900033 | -0.008 | -0.354 | 1.177 | 0.661 | 0.016 | 0.057 | 0.057 | 0.163 | 0.83 | 0.72 | 0.83 | 0.73 | 0.8 | 0.6 |
| 10 | 12944980 | rs2026307 | -0.716 | -0.644 | 1.202 | 0.796 | -0.032 | 0.094 | 0.094 | 0 | 0.89 | 0.71 | 0.89 | 0.72 | 0.93 | 0.82 |
| 10 | 12945749 | rs11258055 | 0.736 | -0.874 | 0.572 | 0.28 | 0.003 | -0.018 | -0.018 | 0 | 0.19 | 0.16 | 0.08 | 0 | 0.15 | 0.11 |
| 10 | 12947995 | rs11258059 | 0.629 | -0.762 | 0.698 | 0.286 | -0.004 | -0.051 | -0.051 | 0 | 0.18 | 0.19 | 0.09 | 0 | 0.16 | 0.11 |
| 10 | 12948835 | rs7919134 | 0.285 | -0.766 | 0.495 | 0.282 | -0.013 | -0.014 | -0.014 | 0 | 0.44 | 0.24 | 0.55 | 0.51 | 0.46 | 0.56 |
| 10 | 12949079 | rs12266955 | 0.688 | -1.132 | 0.764 | 0.273 | -0.028 | 0.13 | 0.13 | 0 | 0.11 | 0.36 | 0.11 | 0.27 | 0.07 | 0.18 |
| 10 | 12950051 | rs7908635 | 0.789 | -1.131 | 0.694 | 0.289 | -0.029 | 0.107 | 0.107 | 0 | 0.11 | 0.25 | 0.12 | 0.3 | 0.07 | 0.19 |
| 10 | 12951727 | rs12243442 | 0.44 | -1.158 | 0.728 | 0.275 | -0.037 | 0.098 | 0.098 | 0 | 0.12 | 0.31 | 0.12 | 0.27 | 0.07 | 0.18 |
| 10 | 12952130 | rs12267985 | -0.212 | -0.791 | 0.363 | 0.133 | -0.022 | 0.108 | 0.108 | 0 | 0.18 | 0.49 | 0.19 | 0.31 | 0.13 | 0.21 |
| 10 | 12953137 | rs11813956 | 0.697 | -0.432 | -0.295 | 0.382 | 0.007 | 0.002 | 0.002 | 0 | 0.32 | 0.1 | 0.41 | 0.26 | 0.29 | 0.23 |
| 10 | 12953273 | rs10795995 | -0.166 | -0.424 | -0.364 | 0.355 | 0 | 0 | 0 | 0 | 0.36 | 0.65 | 0.25 | 0.31 | 0.29 | 0.32 |
| 10 | 12955138 | rs4748012 | -0.397 | 0.455 | -0.211 | 0.532 | -0.002 | -0.002 | -0.002 | 0 | 0.67 | 0.66 | 0.64 | 0.57 | 0.58 | 0.54 |
| 10 | 12959026 | rs281849 | -0.575 | -0.409 | -0.401 | 0.028 | -0.009 | -0.011 | -0.011 | 2.311 | 0.23 | 0.28 | 0.11 | 0.12 | 0.12 | 0.14 |
| 10 | 12966495 | rs281851 | 0.298 | 0.459 | -0.019 | 0.19 | -0.002 | -0.019 | -0.019 | 0 | 0.35 | 0.29 | 0.5 | 0.47 | 0.38 | 0.29 |
| 10 | 12974824 | rs1251015 | -0.815 | 0.523 | 0.176 | -0.314 | 0.015 | -0.089 | -0.089 | 0.161 | 0.67 | 0.58 | 0.77 | 0.9 | 0.65 | 0.77 |
| 10 | 12975556 | rs12767692 | 0.758 | 0.413 | 0.363 | -0.486 | 0.002 | -0.012 | -0.012 | 0.062 | 0.1 | 0.16 | 0.07 | 0.07 | 0.08 | 0.1 |
| 10 | 12976288 | rs1251007 | -1.137 | 0.378 | 0.43 | -0.539 | 0.004 | -0.034 | -0.034 | 0.357 | 0.75 | 0.75 | 0.8 | 0.93 | 0.72 | 0.85 |
| 10 | 12976696 | rs1251006 | 1.741 | 0.116 | 0.061 | -0.418 | -0.009 | 0.036 | 0.036 | 0 | 0.1 | 0.24 | 0.1 | 0.19 | 0.07 | 0.09 |
| 10 | 12976723 | rs10906258 | 0.112 | 0.236 | 0.107 | -0.475 | 0.017 | -0.03 | -0.03 | 0 | 0.32 | 0.21 | 0.28 | 0.42 | 0.25 | 0.49 |
| 10 | 12976933 | rs1620293 | -2.513 | 0.402 | -0.044 | -0.55 | -0.005 | -0.011 | -0.011 | 0.139 | 0.84 | 0.91 | 0.87 | 1 | 0.8 | 0.95 |
| 10 | 12977850 | rs7089944 | -2.145 | 0.404 | -0.184 | -0.549 | -0.036 | 0.049 | 0.049 | 0 | 0.83 | 0.94 | 0.86 | 0.97 | 0.74 | 0.93 |
| 10 | 12979679 | rs12254466 | 0.498 | 0.312 | -0.028 | -0.574 | -0.01 | -0.036 | -0.036 | 4.315 | 0.14 | 0.21 | 0.08 | 0.07 | 0.1 | 0.14 |
| 10 | 12979699 | rs1475610 | -2.572 | 0.353 | -0.043 | -0.59 | -0.01 | 0.009 | 0.009 | 0 | 0.84 | 0.89 | 0.87 | 1 | 0.8 | 0.96 |
| 10 | 12979866 | rs10906259 | 1.654 | 0.352 | -0.159 | -0.668 | -0.039 | 0.046 | 0.046 | 0.019 | 0.08 | 0 | 0.05 | 0 | 0.12 | 0.1 |
| 10 | 12981012 | rs17152523 | 0.06 | -0.036 | 0.053 | -0.597 | -0.01 | -0.052 | -0.052 | 0 | 0.18 | 0.25 | 0.1 | 0.07 | 0.12 | 0.15 |
| 10 | 12983576 | rs10906260 | -1.133 | -0.049 | -0.149 | -0.405 | -0.005 | -0.01 | -0.01 | 0 | 0.64 | 0.65 | 0.5 | 0.51 | 0.56 | 0.68 |
| 10 | 12985117 | rs281848 | 0.957 | -0.026 | -0.141 | -0.436 | 0 | 0.003 | 0.003 | 0 | 0.36 | 0.42 | 0.5 | 0.45 | 0.38 | 0.3 |
| 10 | 12987080 | rs11593289 | 1.232 | 0.205 | -0.49 | -0.909 | -0.099 | 0.167 | 0.167 | 0 | 0.26 | 0.04 | 0.18 | 0 | 0.34 | 0.16 |
| 10 | 12987096 | rs10906262 | -0.377 | 0.261 | -0.286 | -0.758 | -0.004 | 0 | 0 | 0 | 0.47 | 0.59 | 0.3 | 0.08 | 0.46 | 0.31 |
| 10 | 12987176 | rs10906263 | -0.377 | 0.261 | -0.416 | -0.758 | -0.005 | 0.028 | 0.028 | 0 | 0.47 | 0.6 | 0.3 | 0.08 | 0.46 | 0.31 |
| 10 | 12988942 | rs11258073 | -0.038 | 0.364 | -0.204 | 0.057 | -0.005 | -0.014 | -0.014 | 1.217 | 0.13 | 0.17 | 0.06 | 0.09 | 0.09 | 0.14 |
| 10 | 12989666 | rs2399892 | 0.15 | 0.595 | -0.563 | 0.418 | -0.006 | -0.005 | -0.005 | 1.721 | 0.26 | 0.26 | 0.41 | 0.28 | 0.37 | 0.19 |
| 10 | 12995888 | rs2580894 | -0.586 | 0.251 | -0.572 | 0.686 | -0.003 | -0.011 | -0.011 | 0 | 0.3 | 0.33 | 0.4 | 0.43 | 0.31 | 0.41 |
| 10 | 12997341 | rs10906264 | -0.069 | 0.113 | -0.864 | 0.826 | -0.014 | -0.04 | -0.04 | 1.849 | 0.1 | 0 | 0.27 | 0.16 | 0.1 | 0.12 |
| 10 | 12998574 | rs11258079 | 1.249 | 0.108 | -1.064 | 0.592 | -0.054 | -0.07 | -0.07 | 5.01 | 0.39 | 0.16 | 0.4 | 0.5 | 0.45 | 0.37 |
| 10 | 12999115 | rs11258082 | 0.799 | -0.187 | -1.022 | 0.451 | 0.055 | 0.04 | 0.04 | 0 | 0.53 | 0.17 | 0.51 | 0.57 | 0.64 | 0.52 |
| 10 | 13000495 | rs13377047 | 0.824 | -0.183 | -0.96 | 0.529 | 0.056 | 0.011 | 0.011 | 0 | 0.53 | 0.16 | 0.51 | 0.52 | 0.64 | 0.46 |
| 10 | 13001536 | rs2580887 | -0.428 | -0.237 | -0.205 | 0.222 | -0.068 | -0.132 | -0.132 | 0 | 0.82 | 0.61 | 0.93 | 1 | 0.93 | 0.89 |
| 10 | 13003442 | rs749062 | -0.528 | 0.078 | -0.361 | 0.553 | -0.002 | -0.002 | -0.002 | 0 | 0.39 | 0.47 | 0.3 | 0.48 | 0.45 | 0.51 |
| 10 | 13005288 | rs10508452 | -0.126 | 0.093 | -0.08 | 0.456 | -0.002 | 0.015 | 0.015 | 0 | 0.7 | 0.52 | 0.56 | 0.59 | 0.71 | 0.56 |
| 10 | 13005436 | rs7897133 | 0.705 | -0.032 | 0.329 | 0.658 | -0.032 | -0.019 | -0.019 | 0 | 0.39 | 0.2 | 0.39 | 0.55 | 0.45 | 0.39 |
| 10 | 13006444 | rs10906265 | -0.658 | -0.17 | 0.511 | 0.594 | -0.025 | -0.036 | -0.036 | 0 | 0.93 | 0.77 | 0.97 | 1 | 0.99 | 1 |
| 10 | 13007184 | rs2580912 | 0.213 | -0.169 | 0.201 | 0.583 | -0.067 | -0.125 | -0.125 | 0.032 | 0.81 | 0.61 | 0.93 | 1 | 0.93 | 0.91 |
| 10 | 13008083 | rs2580904 | 0.174 | -0.049 | 0.123 | 0.72 | -0.012 | 0.084 | 0.084 | 0 | 0.82 | 0.62 | 0.89 | 0.75 | 0.86 | 0.75 |
| 10 | 13008959 | rs10906273 | 0.998 | 0.396 | -0.525 | 0.803 | -0.038 | 0.007 | 0.007 | 0.006 | 0.33 | 0.12 | 0.34 | 0.31 | 0.38 | 0.23 |
| 10 | 13017220 | rs1866627 | -0.707 | 0.208 | -0.362 | 0.398 | 0.001 | 0 | 0 | 0 | 0.56 | 0.58 | 0.61 | 0.49 | 0.56 | 0.7 |
| 10 | 13022684 | rs557264 | -0.338 | 0.5 | -0.879 | -0.032 | -0.009 | 0.069 | 0.069 | 0 | 0.64 | 0.76 | 0.78 | 0.86 | 0.62 | 0.67 |
| 10 | 13028374 | rs10906276 | -1.009 | 0.626 | -1.125 | 0.395 | -0.003 | -0.002 | -0.002 | 0 | 0.15 | 0.15 | 0.28 | 0.16 | 0.14 | 0.21 |
| 10 | 13031971 | rs12763563 | -0.95 | 0.723 | -1.308 | 0.32 | -0.004 | -0.009 | -0.009 | 0 | 0.23 | 0.21 | 0.35 | 0.19 | 0.22 | 0.17 |
| 10 | 13033757 | rs920409 | 0.716 | 0.952 | -2.517 | 0.607 | -0.011 | -0.053 | -0.053 | 0.026 | 0.25 | 0.12 | 0.37 | 0.36 | 0.27 | 0.22 |
| 10 | 13033961 | rs11258111 | -0.158 | 1.452 | -2.069 | 0.461 | -0.066 | -0.045 | -0.045 | 0 | 0.12 | 0.01 | 0.05 | 0.13 | 0.19 | 0.16 |
| 10 | 13034057 | rs11597167 | -1.218 | 0.361 | -1.526 | 0.188 | -0.018 | -0.016 | -0.016 | 0 | 0.19 | 0.03 | 0.32 | 0.2 | 0.18 | 0.16 |
| 10 | 13039484 | rs7074830 | -0.774 | -0.053 | -1.248 | 0.292 | -0.003 | -0.033 | -0.033 | 0 | 0.11 | 0 | 0.18 | 0.16 | 0.08 | 0.06 |
| 10 | 13039569 | rs10906277 | 0.928 | -0.427 | -0.566 | 0.022 | -0.033 | -0.032 | -0.032 | 0 | 0.74 | 0.38 | 0.75 | 0.81 | 0.84 | 0.78 |
| 10 | 13049815 | rs11258120 | -1.177 | -0.246 | -0.79 | -0.156 | -0.002 | 0.031 | 0.031 | 0 | 0.17 | 0.05 | 0.26 | 0.1 | 0.17 | 0.17 |
| 10 | 13049885 | rs7921121 | -0.33 | -0.439 | -1.023 | -0.536 | -0.004 | 0.02 | 0.02 | 0 | 0.08 | 0.02 | 0.03 | 0.01 | 0.07 | 0.1 |
| 10 | 13050901 | rs7895277 | 0.115 | -0.001 | -1.266 | -1.139 | -0.056 | 0.066 | 0.066 | 0.496 | 0.28 | 0.14 | 0.19 | 0.1 | 0.37 | 0.28 |
| 10 | 13055619 | rs564022 | -0.865 | -1.424 | -1 | -0.945 | -0.062 | 0.003 | 0.003 | 0 | 0.75 | 0.6 | 0.87 | 0.83 | 0.86 | 0.76 |
| 10 | 13056587 | rs604772 | -0.918 | -1.45 | -0.833 | -0.96 | -0.061 | 0.003 | 0.003 | 0 | 0.76 | 0.64 | 0.87 | 0.83 | 0.88 | 0.76 |

| | | | | | | | | | | | | | | | | |
|---|---|---|---|---|---|---|---|---|---|---|---|---|---|---|---|---|
| 10 | 13057260 | rs17152660 | 1.67 | -1.691 | -1.117 | -0.685 | 0.009 | -0.011 | -0.011 | 0.091 | 0.12 | 0.06 | 0.06 | 0.11 | 0.09 | 0.05 |
| 10 | 13057323 | rs11258127 | 0.412 | -1.74 | -1.167 | -0.698 | -0.008 | 0.034 | 0.034 | 0 | 0.17 | 0.04 | 0.26 | 0.08 | 0.18 | 0.17 |
| 10 | 13057403 | rs618640 | -0.257 | -1.814 | -0.977 | -0.725 | -0.034 | -0.012 | -0.012 | 0 | 0.59 | 0.49 | 0.61 | 0.75 | 0.71 | 0.61 |
| 10 | 13057482 | rs513155 | 0.219 | -1.698 | -0.775 | -0.673 | 0.005 | 0.001 | 0.001 | 0 | 0.49 | 0.52 | 0.55 | 0.66 | 0.61 | 0.56 |
| 10 | 13058458 | rs11258128 | 1.341 | -1.731 | -0.786 | -0.534 | -0.013 | -0.029 | -0.029 | 0 | 0.16 | 0.11 | 0.34 | 0.5 | 0.24 | 0.18 |
| 10 | 13060410 | rs579882 | -0.088 | -1.816 | -0.651 | -0.633 | -0.035 | -0.008 | -0.008 | 0 | 0.62 | 0.51 | 0.62 | 0.77 | 0.72 | 0.62 |
| 10 | 13061445 | rs10906283 | 1.372 | -1.727 | -0.665 | -0.685 | -0.023 | 0.036 | 0.036 | 0 | 0.08 | 0.01 | 0.08 | 0 | 0.09 | 0.12 |
| 10 | 13063321 | rs2782278 | -1.095 | -1.804 | -0.659 | -0.719 | -0.058 | 0.002 | 0.002 | 0 | 0.75 | 0.62 | 0.87 | 0.83 | 0.87 | 0.77 |
| 10 | 13063627 | rs11594483 | 0.5 | -1.782 | -1.135 | -0.724 | -0.005 | 0.012 | 0.012 | 0 | 0.12 | 0.1 | 0.13 | 0.01 | 0.15 | 0.14 |
| 10 | 13064223 | rs565850 | -1.196 | -1.831 | -1.069 | -0.698 | -0.084 | 0.007 | 0.007 | 0 | 0.75 | 0.53 | 0.86 | 0.83 | 0.87 | 0.78 |
| 10 | 13070689 | rs4418694 | 0.461 | -1.637 | -1.287 | -0.697 | -0.003 | 0.001 | 0.001 | 0 | 0.12 | 0.09 | 0.12 | 0.01 | 0.15 | 0.14 |
| 10 | 13077232 | rs12765118 | 1.287 | -1.601 | -1.356 | -0.668 | -0.102 | 0.123 | 0.123 | 0.88 | 0.3 | 0.07 | 0.19 | 0.13 | 0.36 | 0.28 |
| 10 | 13077713 | rs7072705 | -0.24 | -1.675 | -1.276 | -0.572 | 0.004 | 0.004 | 0.004 | 0 | 0.3 | 0.31 | 0.33 | 0.22 | 0.27 | 0.31 |
| 10 | 13079319 | rs4409728 | 0.894 | -1.175 | -1.456 | -0.351 | -0.017 | -0.031 | -0.031 | 0 | 0.17 | 0.11 | 0.35 | 0.49 | 0.26 | 0.21 |
| 10 | 13080922 | rs10796006 | 0.328 | -0.932 | -1.422 | -0.597 | 0.009 | 0.04 | 0.04 | 0 | 0.21 | 0.05 | 0.15 | 0.12 | 0.18 | 0.17 |
| 10 | 13083201 | rs7082198 | 0.401 | -0.132 | -0.989 | -0.348 | -0.032 | 0.04 | 0.04 | 0.948 | 0.31 | 0.19 | 0.19 | 0.13 | 0.37 | 0.27 |
| 10 | 13090093 | rs1470093 | -0.807 | -0.421 | -0.715 | 0.536 | -0.027 | 0.035 | 0.035 | 0 | 0.29 | 0.43 | 0.27 | 0.39 | 0.23 | 0.41 |
| 10 | 13090534 | rs6602628 | -0.704 | 0.464 | -0.86 | 0.947 | 0.001 | 0.018 | 0.018 | 0 | 0.1 | 0.13 | 0.1 | 0.26 | 0.16 | 0.15 |
| 10 | 13091546 | rs663177 | 0.428 | 0.823 | -1.084 | 0.766 | -0.018 | -0.012 | -0.012 | 0 | 0.72 | 0.81 | 0.76 | 0.71 | 0.66 | 0.7 |
| 10 | 13093111 | rs553638 | 0.434 | 0.931 | -1.116 | 0.719 | -0.002 | 0.003 | 0.003 | 0 | 0.23 | 0.21 | 0.32 | 0.34 | 0.27 | 0.23 |
| 10 | 13093869 | rs17596659 | 0.585 | 0.86 | -2.001 | 0.205 | -0.017 | 0.072 | 0.072 | 1.362 | 0.39 | 0.06 | 0.24 | 0.22 | 0.31 | 0.32 |
| 10 | 13097242 | rs11258161 | 0.564 | 1.083 | -1.095 | 0.493 | 0.01 | -0.002 | -0.002 | 0 | 0.18 | 0.22 | 0.26 | 0.12 | 0.25 | 0.13 |
| 10 | 13097737 | rs1347979 | 0.552 | 1.082 | -1.096 | 0.492 | 0.009 | -0.002 | -0.002 | 0 | 0.18 | 0.22 | 0.26 | 0.12 | 0.25 | 0.13 |
| 10 | 13099421 | rs2065678 | -0.139 | 0.708 | -1.46 | 0.276 | 0.003 | -0.008 | -0.008 | 0.458 | 0.38 | 0.4 | 0.43 | 0.52 | 0.38 | 0.45 |
| 10 | 13100857 | rs17152780 | 1.821 | 1.065 | -1.505 | 0.155 | -0.033 | -0.036 | -0.036 | 0 | 0.06 | 0.01 | 0.17 | 0.12 | 0.14 | 0.03 |
| 10 | 13100999 | rs609445 | -0.173 | 0.882 | -1.496 | 0.119 | 0.028 | -0.033 | -0.033 | 0.094 | 0.29 | 0.41 | 0.34 | 0.16 | 0.35 | 0.27 |
| 10 | 13101370 | rs607742 | -1.103 | 0.522 | -1.449 | 0.202 | 0.019 | 0.011 | 0.011 | 0 | 0.55 | 0.78 | 0.55 | 0.33 | 0.55 | 0.4 |
| 10 | 13104362 | rs10906289 | -0.019 | 0.585 | -1.347 | 0.222 | -0.014 | -0.021 | -0.021 | 0 | 0.37 | 0.45 | 0.41 | 0.3 | 0.39 | 0.35 |
| 10 | 13104515 | rs2782292 | -1.218 | 0.293 | -0.977 | 0.464 | 0.009 | 0.001 | 0.001 | 0 | 0.69 | 0.8 | 0.78 | 0.68 | 0.73 | 0.72 |
| 10 | 13108440 | rs10796010 | -0.262 | 0.752 | -0.934 | 0.365 | 0.037 | -0.026 | -0.026 | 0.449 | 0.28 | 0.41 | 0.33 | 0.21 | 0.32 | 0.25 |
| 10 | 13114340 | rs10796011 | -0.36 | 0.37 | -1.288 | 0.877 | 0.001 | -0.005 | -0.005 | 0 | 0.84 | 0.84 | 0.85 | 0.8 | 0.83 | 0.89 |
| 10 | 13115887 | rs4750306 | 0.311 | 0.01 | -1.376 | 0.916 | -0.01 | -0.043 | -0.043 | 0 | 0.27 | 0.07 | 0.29 | 0.31 | 0.25 | 0.3 |
| 10 | 13121352 | rs7079869 | -0.384 | -0.023 | -1.306 | 0.941 | 0.005 | -0.007 | -0.007 | 0 | 0.72 | 0.78 | 0.7 | 0.62 | 0.75 | 0.61 |
| 10 | 13132786 | rs2083381 | -0.047 | 0.391 | -0.982 | 0.646 | -0.001 | 0.001 | 0.001 | 0 | 0.55 | 0.46 | 0.6 | 0.62 | 0.49 | 0.66 |
| 10 | 13135831 | rs1953314 | 0.144 | -0.607 | -0.769 | 0.642 | -0.054 | 0.029 | 0.029 | 0.127 | 0.77 | 0.54 | 0.7 | 0.76 | 0.85 | 0.83 |
| 10 | 13141144 | rs3829923 | -0.084 | 0.088 | -1.67 | 0.798 | -0.099 | -0.088 | -0.088 | 0 | 0.29 | 0.02 | 0.17 | 0.36 | 0.34 | 0.32 |
| 10 | 13148267 | rs660592 | 0.027 | -0.465 | -1.026 | 0.711 | -0.013 | 0.002 | 0.002 | 0.283 | 0.77 | 0.29 | 0.65 | 0.74 | 0.87 | 0.83 |
| 10 | 13151224 | rs2234968 | 0.645 | -0.142 | -1.608 | 0.278 | -0.058 | -0.011 | -0.011 | 0 | 0.25 | 0.01 | 0.23 | 0.19 | 0.27 | 0.26 |
| 10 | 13152666 | rs7921853 | 0.068 | 0.189 | -1.496 | 0.362 | -0.079 | 0.053 | 0.053 | 2.018 | 0.37 | 0.11 | 0.3 | 0.22 | 0.51 | 0.41 |
| 10 | 13155726 | rs1561570 | 0.078 | 0.999 | -0.209 | 0.509 | 0.008 | 0 | 0 | 0 | 0.49 | 0.25 | 0.54 | 0.53 | 0.5 | 0.43 |
| 10 | 13158262 | rs2244380 | -0.544 | 1.116 | 0.31 | 0.645 | 0.068 | 0.108 | 0.108 | 0 | 0.92 | 0.83 | 0.86 | 0.75 | 0.82 | 0.77 |
| 10 | 13161409 | rs7095146 | 0.066 | 1.142 | 0.268 | 0.625 | 0.005 | 0.005 | 0.005 | 1.458 | 0.49 | 0.24 | 0.55 | 0.53 | 0.52 | 0.44 |
| 10 | 13166875 | rs583277 | 0.144 | 1.156 | 0.326 | 0.644 | 0 | 0.007 | 0.007 | 2.772 | 0.19 | 0.23 | 0.16 | 0.3 | 0.23 | 0.29 |
| 10 | 13169066 | rs17512962 | 0.72 | 1.326 | 0.372 | 0.684 | 0.017 | 0.142 | 0.142 | 0.097 | 0.34 | 0.09 | 0.22 | 0.1 | 0.28 | 0.22 |
| 10 | 13169374 | rs825411 | -0.731 | 1.356 | 0.439 | 0.698 | 0.003 | -0.002 | -0.002 | 0 | 0.58 | 0.51 | 0.44 | 0.43 | 0.53 | 0.56 |
| 10 | 13169512 | rs10796028 | -0.232 | 1.681 | 0.576 | 0.945 | 0.02 | 0.067 | 0.067 | 3.08 | 0.2 | 0.35 | 0.21 | 0.34 | 0.24 | 0.33 |
| 10 | 13170467 | rs10906308 | 0.432 | 1.69 | 0.38 | 0.95 | -0.012 | -0.027 | -0.027 | 0 | 0.17 | 0.11 | 0.15 | 0.3 | 0.22 | 0.28 |
| 10 | 13175682 | rs7086894 | -0.621 | 1.766 | 0.206 | 0.962 | 0.029 | 0.049 | 0.049 | 2.893 | 0.23 | 0.4 | 0.23 | 0.34 | 0.3 | 0.35 |
| 10 | 13186875 | rs2184401 | 0.079 | 1.297 | -0.12 | 1.383 | -0.002 | -0.001 | -0.001 | 0.062 | 0.39 | 0.43 | 0.48 | 0.35 | 0.31 | 0.27 |
| 10 | 13193189 | rs10906313 | 0.001 | 1.546 | -0.741 | 1.264 | 0.085 | 0.105 | 0.105 | 0 | 0.87 | 0.7 | 0.83 | 0.71 | 0.77 | 0.72 |
| 10 | 13199611 | rs7899305 | -0.297 | 0.151 | -1.479 | 0.817 | 0.03 | 0.003 | 0.003 | 0.927 | 0.37 | 0.27 | 0.46 | 0.35 | 0.3 | 0.23 |
| 10 | 13208063 | rs11258226 | -0.597 | 0.715 | -1.685 | 1.112 | -0.013 | -0.07 | -0.07 | 0.662 | 0.28 | 0.13 | 0.3 | 0.41 | 0.28 | 0.32 |
| 10 | 13210175 | rs768255 | -0.917 | 0.565 | -1.21 | 0.818 | -0.001 | -0.001 | -0.001 | 0 | 0.54 | 0.56 | 0.5 | 0.49 | 0.56 | 0.54 |
| 10 | 13210325 | rs10508454 | 0.794 | 0.339 | -1.645 | 0.425 | -0.007 | 0.038 | 0.038 | 0 | 0.07 | 0 | 0.09 | 0 | 0.06 | 0.07 |
| 10 | 13212884 | rs2147279 | -1.083 | 0.832 | -1.476 | 0.213 | 0.006 | -0.01 | -0.01 | 0 | 0.72 | 0.77 | 0.59 | 0.58 | 0.77 | 0.75 |
| 10 | 13214160 | rs7073480 | -1.314 | 0.494 | -1.63 | 0.341 | -0.001 | 0.008 | 0.008 | 0 | 0.86 | 0.83 | 0.86 | 0.8 | 0.87 | 0.92 |
| 10 | 13217940 | rs1535856 | -1.169 | 0.451 | -1.615 | 0.326 | -0.002 | 0.021 | 0.021 | 0 | 0.86 | 0.8 | 0.77 | 0.79 | 0.87 | 0.92 |
| 10 | 13225119 | rs11258235 | 1.088 | 0.602 | -1.622 | 0.226 | -0.009 | -0.029 | -0.029 | 0 | 0.28 | 0.21 | 0.2 | 0.4 | 0.28 | 0.31 |
| 10 | 13226829 | rs942412 | -1.229 | 0.594 | -1.583 | 0.12 | -0.005 | 0.024 | 0.024 | 0.087 | 0.86 | 0.79 | 0.84 | 0.79 | 0.89 | 0.92 |
| 10 | 13227604 | rs6602640 | -0.868 | 0.525 | -1.497 | 0.147 | -0.01 | 0.028 | 0.028 | 0.266 | 0.83 | 0.72 | 0.75 | 0.76 | 0.88 | 0.9 |
| 10 | 13230950 | rs2296222 | 0.055 | 0.273 | -1.02 | -0.035 | 0.026 | 0.016 | 0.016 | 0 | 0.24 | 0.13 | 0.29 | 0.2 | 0.17 | 0.24 |
| 10 | 13231194 | rs2296221 | -0.701 | 0.194 | -0.61 | -0.042 | -0.001 | -0.002 | -0.002 | 0 | 0.55 | 0.44 | 0.54 | 0.36 | 0.6 | 0.59 |
| 10 | 13233867 | rs2274108 | 0.342 | 0.199 | -0.41 | -0.054 | 0.02 | 0.016 | 0.016 | 0 | 0.45 | 0.31 | 0.39 | 0.6 | 0.39 | 0.48 |
| 10 | 13239651 | rs2274110 | 1.648 | 0.252 | -0.796 | -0.341 | -0.033 | -0.017 | -0.017 | 1.199 | 0.15 | 0.01 | 0.13 | 0.1 | 0.19 | 0.14 |
| 10 | 13242379 | rs11258248 | 1.202 | 0.178 | -0.824 | -0.265 | 0.033 | -0.051 | -0.051 | 2.065 | 0.16 | 0.03 | 0.18 | 0.24 | 0.11 | 0.17 |
| 10 | 13242413 | rs4520495 | -0.016 | 0.375 | -0.592 | -0.335 | 0.232 | 0.022 | 0.022 | 0.319 | 0.07 | 0.32 | 0.09 | 0.02 | 0.18 | 0.03 |
| 10 | 13242785 | rs10906320 | -0.068 | -0.056 | -0.872 | -0.33 | 0.056 | -0.003 | -0.003 | 0.028 | 0.37 | 0.19 | 0.34 | 0.37 | 0.32 | 0.39 |
| 10 | 13243688 | rs11258249 | 1.087 | 0.026 | -0.754 | -0.274 | -0.018 | 0.041 | 0.041 | 0 | 0.08 | 0 | 0.11 | 0 | 0.07 | 0.07 |
| 10 | 13247476 | rs10796039 | -0.288 | -0.347 | -0.571 | -0.206 | -0.019 | 0.031 | 0.031 | 0 | 0.7 | 0.64 | 0.63 | 0.55 | 0.79 | 0.65 |

| | | | | | | | | | | | | | | | | |
|---|---|---|---|---|---|---|---|---|---|---|---|---|---|---|---|---|
| 10 | 13247611 | rs12414174 | 0.635 | -0.843 | -0.756 | -0.512 | 0.028 | -0.022 | -0.022 | 0.037 | 0.45 | 0.24 | 0.47 | 0.44 | 0.63 | 0.47 |
| 10 | 13248410 | rs3781081 | -0.718 | -0.673 | -0.46 | -0.212 | -0.008 | 0.017 | 0.017 | 0 | 0.72 | 0.7 | 0.65 | 0.55 | 0.79 | 0.65 |
| 10 | 13249379 | rs1886278 | -0.397 | -0.152 | -0.461 | -0.355 | 0 | 0.001 | 0.001 | 0 | 0.51 | 0.53 | 0.48 | 0.42 | 0.5 | 0.42 |
| 10 | 13254130 | rs9645605 | 1.082 | -0.115 | -0.664 | -0.298 | -0.019 | 0.04 | 0.04 | 0 | 0.08 | 0 | 0.1 | 0 | 0.07 | 0.07 |
| 10 | 13254480 | rs10508455 | -0.293 | 0.446 | -0.461 | -0.391 | 0 | 0.004 | 0.004 | 0 | 0.3 | 0.36 | 0.37 | 0.39 | 0.29 | 0.26 |
| 10 | 13255276 | rs11258253 | 0.103 | 0.271 | -0.555 | 0.047 | -0.002 | 0.005 | 0.005 | 0 | 0.26 | 0.34 | 0.4 | 0.56 | 0.23 | 0.34 |
| 10 | 13255513 | rs11814554 | 1.717 | 0.303 | -0.638 | 0.127 | -0.001 | 0.032 | 0.032 | 0 | 0.11 | 0.15 | 0.19 | 0.32 | 0.12 | 0.14 |
| 10 | 13258459 | rs17152941 | 1.639 | 0.307 | -0.694 | 0.102 | 0.006 | -0.037 | -0.037 | 0 | 0.14 | 0.06 | 0.2 | 0.24 | 0.12 | 0.2 |
| 10 | 13258742 | rs12778355 | 0.015 | 0.371 | -0.575 | -0.258 | 0.001 | -0.002 | -0.002 | 0.944 | 0.3 | 0.25 | 0.18 | 0.14 | 0.25 | 0.2 |
| 10 | 13262066 | rs727518 | -0.535 | 0.466 | -0.289 | -0.253 | -0.002 | -0.001 | -0.001 | 0 | 0.42 | 0.47 | 0.57 | 0.59 | 0.37 | 0.46 |
| 10 | 13262143 | rs1408363 | -1.519 | 0.152 | -0.466 | -0.258 | -0.008 | 0.001 | 0.001 | 0 | 0.65 | 0.7 | 0.68 | 0.71 | 0.56 | 0.62 |
| 10 | 13262912 | rs4589172 | -0.233 | -0.007 | -0.297 | -0.067 | -0.003 | 0.003 | 0.003 | 1.176 | 0.34 | 0.39 | 0.31 | 0.4 | 0.32 | 0.29 |
| 10 | 13263027 | rs4301691 | -0.831 | -0.032 | -0.651 | -0.114 | 0.003 | 0.021 | 0.021 | 0.129 | 0.52 | 0.4 | 0.54 | 0.66 | 0.45 | 0.51 |
| 10 | 13263558 | rs11815945 | 1.491 | 0.097 | -0.394 | 0.273 | 0.001 | 0.034 | 0.034 | 0 | 0.11 | 0.14 | 0.18 | 0.31 | 0.12 | 0.13 |
| 10 | 13264280 | rs2281797 | -0.397 | -0.38 | -0.573 | 0.21 | 0.004 | -0.014 | -0.014 | 3.202 | 0.32 | 0.3 | 0.44 | 0.61 | 0.27 | 0.39 |
| 10 | 13266987 | rs942413 | -0.91 | -0.226 | -0.996 | 0.088 | 0.007 | 0.019 | 0.019 | 0 | 0.54 | 0.56 | 0.48 | 0.4 | 0.45 | 0.46 |
| 10 | 13268140 | rs11258275 | -0.585 | -0.379 | -1.209 | 0.35 | 0 | 0 | 0 | 0 | 0.44 | 0.41 | 0.39 | 0.37 | 0.36 | 0.4 |
| 10 | 13268710 | rs4750319 | -0.185 | -0.332 | -1.052 | 0.544 | -0.019 | 0.003 | 0.003 | 0 | 0.28 | 0.36 | 0.3 | 0.31 | 0.23 | 0.34 |
| 10 | 13271834 | rs4750320 | -0.561 | 0.222 | -0.84 | 0.652 | 0.004 | 0.005 | 0.005 | 0 | 0.47 | 0.41 | 0.51 | 0.66 | 0.41 | 0.51 |
| 10 | 13275553 | rs3829925 | 0.017 | -0.15 | -0.578 | 1.101 | -0.018 | 0.039 | 0.039 | 0 | 0.87 | 0.72 | 0.92 | 0.84 | 0.89 | 0.71 |
| 10 | 13276534 | rs2093847 | 0.685 | -0.61 | -1.224 | 0.919 | 0.013 | -0.013 | -0.013 | 0 | 0.15 | 0.01 | 0.23 | 0.12 | 0.09 | 0.05 |
| 10 | 13277571 | rs2399955 | 0.216 | -0.569 | -1.675 | 0.769 | 0.057 | -0.054 | -0.054 | 0 | 0.53 | 0.15 | 0.45 | 0.49 | 0.62 | 0.73 |
| 10 | 13279827 | rs10906326 | 0.632 | -0.328 | -2.03 | 0.601 | -0.133 | -0.019 | -0.019 | 0 | 0.36 | 0.07 | 0.32 | 0.28 | 0.46 | 0.52 |
| 10 | 13281056 | rs2181841 | -0.008 | -0.493 | -0.744 | 0.761 | -0.008 | 0.019 | 0.019 | 0 | 0.3 | 0.39 | 0.46 | 0.45 | 0.26 | 0.16 |
| 10 | 13283807 | rs17152990 | 0.251 | -0.42 | -0.204 | 0.546 | -0.009 | 0.006 | 0.006 | 0 | 0.16 | 0.23 | 0.1 | 0.21 | 0.14 | 0.13 |
| 10 | 13283975 | rs2476981 | -0.614 | -0.896 | 0.042 | 0.183 | 0.004 | 0.001 | 0.001 | 0 | 0.41 | 0.6 | 0.32 | 0.39 | 0.32 | 0.34 |
| 10 | 13284032 | rs2476980 | -1.496 | -0.651 | -0.131 | 0.132 | -0.074 | -0.03 | -0.03 | 0 | 0.6 | 0.86 | 0.51 | 0.62 | 0.53 | 0.51 |
| 10 | 13284827 | rs10906331 | 0.463 | -0.629 | 0.431 | 0.391 | 0.019 | 0.167 | 0.167 | 0 | 0.06 | 0.17 | 0.07 | 0.23 | 0.06 | 0.08 |
| 10 | 13285299 | rs2025450 | -1.068 | -0.781 | 0.461 | 0.119 | 0.022 | 0.017 | 0.017 | 0 | 0.5 | 0.61 | 0.33 | 0.43 | 0.45 | 0.43 |
| 10 | 13285565 | rs2253447 | -0.147 | -0.663 | 0.507 | -0.142 | 0.023 | 0.052 | 0.052 | 0 | 0.1 | 0.26 | 0.18 | 0.17 | 0.09 | 0.08 |
| 10 | 13286331 | rs528320 | -0.478 | -0.816 | 0.215 | -0.272 | -0.003 | 0 | 0 | 0 | 0.17 | 0.22 | 0.2 | 0.2 | 0.16 | 0.18 |
| 10 | 13286601 | rs551389 | -0.365 | -0.806 | 0.032 | -0.296 | -0.002 | 0.012 | 0.012 | 0 | 0.22 | 0.14 | 0.24 | 0.17 | 0.19 | 0.17 |
| 10 | 13292404 | rs7920439 | 0.259 | -0.164 | 0.395 | -0.201 | 0.012 | -0.036 | -0.036 | 0.394 | 0.07 | 0.09 | 0.05 | 0 | 0.08 | 0.02 |
| 10 | 13294311 | rs472806 | 0.647 | -0.313 | 0.637 | -0.308 | -0.014 | -0.15 | -0.15 | 0 | 0.75 | 0.48 | 0.79 | 0.96 | 0.76 | 0.79 |
| 10 | 13303113 | rs7090352 | -0.952 | -0.131 | 0.285 | -0.01 | 0.015 | -0.001 | -0.001 | 0.572 | 0.24 | 0.31 | 0.22 | 0.2 | 0.29 | 0.24 |
| 10 | 13306566 | rs4465274 | -0.277 | -0.052 | -0.216 | -0.164 | 0 | -0.048 | -0.048 | 0 | 0.29 | 0.3 | 0.25 | 0.02 | 0.34 | 0.21 |
| 10 | 13310071 | rs10796051 | 1.099 | 0.049 | -1.341 | 0.068 | -0.083 | -0.098 | -0.098 | 0.104 | 0.4 | 0.05 | 0.47 | 0.42 | 0.42 | 0.44 |
| 10 | 13311227 | rs825632 | 0.3 | -0.023 | -0.378 | -0.064 | -0.013 | -0.014 | -0.014 | 0 | 0.77 | 0.66 | 0.88 | 0.81 | 0.85 | 0.8 |
| 10 | 13313949 | rs10508458 | -0.475 | 0.197 | -1.394 | -0.126 | -0.003 | 0.061 | 0.061 | 0 | 0.31 | 0.07 | 0.17 | 0.17 | 0.25 | 0.19 |
| 10 | 13314711 | rs4606378 | -0.51 | 0.432 | -1.365 | -0.152 | -0.034 | 0.07 | 0.07 | 0 | 0.3 | 0.07 | 0.19 | 0.18 | 0.28 | 0.24 |
| 10 | 13314732 | rs591670 | -0.509 | 0.32 | -0.873 | -0.046 | 0.006 | 0.029 | 0.029 | 0 | 0.32 | 0.19 | 0.21 | 0.22 | 0.29 | 0.3 |
| 10 | 13318060 | rs861136 | -0.644 | -0.065 | -0.733 | -0.244 | 0.02 | 0.039 | 0.039 | 0 | 0.39 | 0.27 | 0.23 | 0.26 | 0.35 | 0.45 |
| 10 | 13320848 | rs4750340 | 0.972 | 0.671 | -1.71 | 0.092 | -0.103 | -0.138 | -0.138 | 2.693 | 0.42 | 0.03 | 0.57 | 0.46 | 0.42 | 0.41 |
| 10 | 13326616 | rs615788 | -0.127 | 0.822 | -0.61 | -0.077 | -0.002 | -0.003 | -0.003 | 0 | 0.34 | 0.38 | 0.23 | 0.38 | 0.35 | 0.45 |
| 10 | 13330291 | rs7900830 | 0.175 | 0.764 | -0.567 | -0.056 | -0.008 | -0.006 | -0.006 | 0 | 0.31 | 0.36 | 0.18 | 0.31 | 0.29 | 0.37 |
| 10 | 13330402 | rs473407 | -0.816 | 0.833 | 0.066 | -0.162 | 0.032 | -0.038 | -0.038 | 0 | 0.21 | 0.43 | 0.17 | 0.12 | 0.23 | 0.16 |
| 10 | 13331621 | rs11258311 | 0.482 | 1.197 | 0.538 | -0.115 | 0.077 | -0.024 | -0.024 | 0 | 0.89 | 0.65 | 0.88 | 0.98 | 0.86 | 0.95 |
| 10 | 13332386 | rs9732147 | 0.213 | 1.233 | 0.833 | -0.149 | -0.006 | -0.006 | -0.006 | 0 | 0.32 | 0.34 | 0.17 | 0.31 | 0.29 | 0.37 |
| 10 | 13332690 | rs7087147 | -0.027 | 1.183 | 0.753 | -0.194 | 0.004 | 0.004 | 0.004 | 0.155 | 0.09 | 0.18 | 0.06 | 0.09 | 0.08 | 0.1 |
| 10 | 13334136 | rs1414395 | -0.41 | 1.222 | 0.791 | -0.045 | -0.001 | -0.009 | -0.009 | 0 | 0.39 | 0.55 | 0.27 | 0.35 | 0.42 | 0.42 |
| 10 | 13334321 | rs1414396 | -0.058 | 1.259 | 0.79 | -0.026 | 0.07 | -0.011 | -0.011 | 0 | 0.07 | 0.21 | 0.09 | 0.02 | 0.13 | 0.06 |
| 10 | 13336936 | rs471917 | -0.171 | 1.227 | 0.592 | -0.025 | 0.008 | -0.003 | -0.003 | 0 | 0.36 | 0.41 | 0.27 | 0.38 | 0.44 | 0.42 |
| 10 | 13338202 | rs505081 | 0.481 | 1.025 | 0.03 | -0.344 | -0.002 | 0.078 | 0.078 | 0 | 0.55 | 0.28 | 0.67 | 0.7 | 0.53 | 0.48 |
| 10 | 13341185 | rs12570229 | 1.071 | 1.186 | 0.41 | -0.248 | 0.041 | -0.028 | -0.028 | 0 | 0.74 | 0.43 | 0.73 | 0.78 | 0.7 | 0.79 |
| 10 | 13341731 | rs732701 | 0.554 | 0.327 | -0.095 | -0.679 | -0.01 | -0.036 | -0.036 | 0 | 0.35 | 0.36 | 0.16 | 0.1 | 0.28 | 0.31 |
| 10 | 13342654 | rs4615920 | 0.672 | 0.285 | -0.143 | -0.721 | -0.008 | -0.032 | -0.032 | 0 | 0.36 | 0.35 | 0.16 | 0.1 | 0.28 | 0.31 |
| 10 | 13347821 | rs10752292 | -0.049 | 0.459 | -0.714 | -0.732 | 0.063 | 0.133 | 0.133 | 0.018 | 0.52 | 0.15 | 0.7 | 0.72 | 0.65 | 0.58 |
| 10 | 13363717 | rs2254154 | -0.377 | 0.447 | -0.214 | -0.406 | 0.007 | 0.097 | 0.097 | 0 | 0.5 | 0.32 | 0.3 | 0.23 | 0.51 | 0.39 |
| 10 | 13366335 | rs10752294 | 0.372 | 0.435 | -0.186 | -0.411 | -0.028 | 0.118 | 0.118 | 0 | 0.45 | 0.16 | 0.66 | 0.76 | 0.49 | 0.62 |
| 10 | 13369506 | rs10752296 | 0.639 | 0.203 | -0.5 | -0.523 | -0.047 | 0.038 | 0.038 | 0.013 | 0.4 | 0.12 | 0.6 | 0.69 | 0.45 | 0.56 |
| 10 | 13387295 | rs1005089 | -0.311 | 0.463 | -0.609 | -0.483 | 0.008 | 0.083 | 0.083 | 0 | 0.54 | 0.71 | 0.74 | 0.76 | 0.55 | 0.65 |
| 10 | 13393621 | rs7923713 | 0.61 | 0.887 | -1.464 | -1.083 | -0.05 | 0.158 | 0.158 | 0.082 | 0.52 | 0.06 | 0.57 | 0.78 | 0.5 | 0.68 |
| 10 | 13398428 | rs17602403 | -0.076 | 0.851 | -1.037 | -1.078 | 0.015 | 0.037 | 0.037 | 0.389 | 0.35 | 0.23 | 0.23 | 0.16 | 0.3 | 0.19 |
| 10 | 13399789 | rs10906357 | -0.612 | 0.622 | -1.606 | -0.924 | -0.016 | 0.173 | 0.173 | 0 | 0.51 | 0.78 | 0.42 | 0.22 | 0.5 | 0.34 |
| 10 | 13410748 | rs10906359 | 0.044 | 0.725 | -1.278 | -1.626 | -0.002 | -0.075 | -0.075 | 0 | 0.64 | 0.41 | 0.75 | 0.97 | 0.68 | 0.82 |
| 10 | 13411415 | rs9787419 | 0.109 | 0.733 | -1.547 | -1.196 | -0.048 | 0.021 | 0.021 | 0 | 0.37 | 0.14 | 0.41 | 0.71 | 0.42 | 0.49 |
| 10 | 13413913 | rs10906361 | 0.165 | 0.893 | -1.531 | -1.198 | -0.017 | -0.003 | -0.003 | 3.303 | 0.35 | 0.15 | 0.34 | 0.68 | 0.36 | 0.41 |
| 10 | 13414068 | rs12217742 | 1.231 | 0.926 | -1.664 | -1.294 | -0.015 | -0.014 | -0.014 | 0 | 0.11 | 0.06 | 0.2 | 0.68 | 0.16 | 0.31 |
| 10 | 13415971 | rs1556992 | -0.188 | 0.892 | -1.631 | -1.381 | -0.013 | 0.038 | 0.038 | 0 | 0.37 | 0.31 | 0.52 | 0.84 | 0.41 | 0.54 |

| | | | | | | | | | | | | | | | | |
|---|---|---|---|---|---|---|---|---|---|---|---|---|---|---|---|---|
| 10 | 13417484 | rs11258351 | -0.232 | 1.053 | -1.679 | -1.154 | -0.002 | 0.002 | 0.002 | 0 | 0.29 | 0.29 | 0.49 | 0.84 | 0.36 | 0.5 |
| 10 | 13418659 | rs12775502 | -0.105 | 0.979 | -1.53 | -1.119 | 0 | -0.013 | -0.013 | 0.132 | 0.05 | 0.1 | 0.04 | 0.02 | 0.04 | 0.06 |
| 10 | 13418971 | rs10906364 | 0.135 | 0.983 | -1.511 | -1.022 | 0 | -0.004 | -0.004 | 0 | 0.26 | 0.28 | 0.31 | 0.82 | 0.3 | 0.41 |
| 10 | 13428593 | rs7087392 | -0.548 | 0.796 | -1.293 | -0.979 | -0.002 | 0.003 | 0.003 | 0 | 0.59 | 0.43 | 0.6 | 0.3 | 0.59 | 0.57 |
| 10 | 13432540 | rs12414133 | 0.002 | 0.878 | -0.989 | -0.578 | -0.012 | 0.122 | 0.122 | 0 | 0.35 | 0.12 | 0.39 | 0.14 | 0.38 | 0.32 |
| 10 | 13434975 | rs1414397 | 1.42 | 1.034 | -0.03 | 0.195 | 0.003 | 0.042 | 0.042 | 0.505 | 0.91 | 0.84 | 0.88 | 0.84 | 0.9 | 0.8 |
| 10 | 13438901 | rs7083156 | 2 | 0.89 | 0.179 | 0.303 | 0.006 | 0.093 | 0.093 | 0.083 | 0.89 | 0.66 | 0.83 | 0.78 | 0.9 | 0.72 |
| 10 | 13443995 | rs12768338 | -1.526 | 1.004 | 0.021 | 0.467 | 0.015 | 0.109 | 0.109 | 0 | 0.08 | 0.21 | 0.15 | 0.22 | 0.07 | 0.19 |
| 10 | 13448279 | rs17531080 | -0.079 | 1.097 | -0.566 | -0.094 | -0.033 | 0.034 | 0.034 | 0.13 | 0.12 | 0 | 0.08 | 0.06 | 0.12 | 0.07 |
| 10 | 13457100 | rs12570207 | -0.64 | 1.919 | -0.623 | -0.033 | -0.071 | 0.014 | 0.014 | 0 | 0.12 | 0.03 | 0.14 | 0.07 | 0.24 | 0.09 |
| 10 | 13472996 | rs4750355 | 1.541 | 1.288 | 1.348 | -0.03 | -0.024 | 0.039 | 0.039 | 1.677 | 0.87 | 0.38 | 0.82 | 0.78 | 0.91 | 0.8 |
| 10 | 13473126 | rs6602657 | 1.504 | 1.708 | 1.074 | 0.06 | 0.089 | 0.075 | 0.075 | 0 | 0.82 | 0.6 | 0.72 | 0.7 | 0.7 | 0.74 |
| 10 | 13473401 | rs6602658 | 0.849 | 0.96 | 0.73 | 0.182 | 0.02 | 0.107 | 0.107 | 0 | 0.92 | 0.78 | 0.85 | 0.78 | 0.91 | 0.86 |
| 10 | 13480789 | rs4750356 | 0.817 | 0.689 | -0.481 | -0.306 | -0.045 | -0.024 | -0.024 | 0 | 0.3 | 0.05 | 0.34 | 0.67 | 0.36 | 0.38 |
| 10 | 13482260 | rs12767849 | -0.126 | 0.334 | -0.305 | -0.932 | 0.038 | 0.069 | 0.069 | 0 | 0.42 | 0.69 | 0.42 | 0.21 | 0.32 | 0.44 |
| 10 | 13488453 | rs1105233 | -0.001 | 0.004 | 0.075 | -1.067 | -0.009 | -0.001 | -0.001 | 0.007 | 0.1 | 0.11 | 0.12 | 0.09 | 0.08 | 0.14 |
| 10 | 13494906 | rs11258392 | 0.352 | -0.092 | 0.778 | -1.15 | -0.014 | -0.061 | -0.061 | 0 | 0.3 | 0.39 | 0.29 | 0.11 | 0.24 | 0.31 |
| 10 | 13501202 | rs7922720 | 0.414 | -0.197 | 1.17 | -1.245 | 0.005 | 0.013 | 0.013 | 0.179 | 0.45 | 0.4 | 0.49 | 0.78 | 0.44 | 0.59 |
| 10 | 13501415 | rs1340261 | -0.312 | 0.025 | 1.584 | -0.962 | -0.048 | 0.054 | 0.054 | 0.082 | 0.76 | 0.86 | 0.78 | 0.88 | 0.67 | 0.85 |
| 10 | 13502070 | rs4750361 | -1.146 | 0.01 | 1.613 | -0.89 | -0.027 | 0.008 | 0.008 | 0.003 | 0.14 | 0.37 | 0.15 | 0.11 | 0.09 | 0.22 |
| 10 | 13503945 | rs7913260 | -0.355 | 0.017 | 1.441 | -0.866 | -0.002 | -0.001 | -0.001 | 0 | 0.11 | 0.1 | 0.12 | 0.09 | 0.08 | 0.14 |
| 10 | 13504706 | rs4237432 | -1.428 | 0.025 | 1.468 | -0.685 | -0.046 | 0.033 | 0.033 | 0.328 | 0.16 | 0.44 | 0.15 | 0.13 | 0.09 | 0.18 |
| 10 | 13511130 | rs2153442 | 0.858 | 0.055 | 1.627 | -0.724 | -0.004 | 0.011 | 0.011 | 0.637 | 0.6 | 0.39 | 0.62 | 0.75 | 0.58 | 0.67 |
| 10 | 13511986 | rs4748035 | 0.891 | 0.109 | 1.598 | -0.655 | 0.017 | -0.028 | -0.028 | 0 | 0.63 | 0.5 | 0.64 | 0.75 | 0.58 | 0.67 |
| 10 | 13516469 | rs12573227 | -0.354 | 0.13 | 1.219 | -0.617 | -0.009 | 0.007 | 0.007 | 0 | 0.11 | 0.13 | 0.12 | 0.11 | 0.09 | 0.14 |
| 10 | 13519743 | rs10508462 | 1.003 | 0.11 | 1.311 | -0.656 | 0.007 | -0.018 | -0.018 | 0.019 | 0.62 | 0.47 | 0.63 | 0.75 | 0.58 | 0.68 |
| 10 | 13523763 | rs4750369 | -0.318 | 0.162 | 1.248 | -0.634 | -0.05 | 0.048 | 0.048 | 0.148 | 0.77 | 0.87 | 0.78 | 0.87 | 0.67 | 0.86 |
| 10 | 13526000 | rs4748040 | 0.599 | 0.154 | 1.189 | -0.603 | 0.017 | -0.027 | -0.027 | 0 | 0.64 | 0.51 | 0.65 | 0.75 | 0.6 | 0.68 |
| 10 | 13531381 | rs7919627 | -0.983 | -0.049 | 0.978 | -0.485 | -0.005 | 0.004 | 0.004 | 0 | 0.24 | 0.29 | 0.2 | 0.25 | 0.2 | 0.24 |
| 10 | 13534851 | rs2251555 | -1.619 | -0.249 | 0.851 | -0.96 | -0.004 | -0.008 | -0.008 | 0.093 | 0.55 | 0.49 | 0.55 | 0.34 | 0.54 | 0.5 |
| 10 | 13537204 | rs12770479 | 0.011 | -0.13 | 0.896 | -0.969 | -0.033 | 0.059 | 0.059 | 0 | 0.3 | 0.18 | 0.35 | 0.1 | 0.34 | 0.14 |
| 10 | 13540037 | rs12764013 | 0.017 | -0.157 | 0.878 | -0.971 | -0.041 | 0.07 | 0.07 | 0 | 0.3 | 0.16 | 0.35 | 0.1 | 0.34 | 0.14 |
| 10 | 13540537 | rs7917804 | -0.109 | -0.159 | 0.882 | -0.971 | -0.009 | 0.014 | 0.014 | 0 | 0.32 | 0.23 | 0.36 | 0.12 | 0.34 | 0.27 |
| 10 | 13544854 | rs7904554 | 0.105 | -0.172 | 0.874 | -0.986 | -0.009 | 0.007 | 0.007 | 0.051 | 0.3 | 0.23 | 0.36 | 0.09 | 0.34 | 0.14 |
| 10 | 13553345 | rs11819227 | 0.522 | -0.415 | 0.878 | -1.148 | -0.024 | -0.071 | -0.071 | 0 | 0.13 | 0.32 | 0.09 | 0.01 | 0.1 | 0.12 |
| 10 | 13553391 | rs7074884 | 0.385 | -0.413 | 0.152 | -1.126 | 0.012 | 0.01 | 0.01 | 0 | 0.14 | 0.21 | 0.12 | 0.16 | 0.16 | 0.19 |
| 10 | 13557960 | rs10906399 | 0.876 | -0.092 | 0.106 | -1.227 | -0.006 | 0.003 | 0.003 | 0 | 0.08 | 0.07 | 0.07 | 0.07 | 0.08 | 0.06 |
| 10 | 13563816 | rs7910407 | 0.52 | -0.081 | 0.119 | -1.212 | -0.025 | 0.052 | 0.052 | 0 | 0.22 | 0.13 | 0.28 | 0.02 | 0.26 | 0.08 |
| 10 | 13573636 | rs4750377 | 0.28 | -0.225 | 0.248 | -0.996 | -0.002 | 0.137 | 0.137 | 0 | 0.55 | 0.25 | 0.57 | 0.86 | 0.58 | 0.69 |
| 10 | 13578863 | rs10796090 | 0.737 | -0.378 | 0.173 | -0.736 | 0.036 | 0.008 | 0.008 | 0.966 | 0.7 | 0.2 | 0.75 | 0.78 | 0.8 | 0.64 |
| 10 | 13580016 | rs10796091 | 0.408 | -0.371 | 0.468 | -0.698 | -0.033 | 0.011 | 0.011 | 0.12 | 0.75 | 0.61 | 0.8 | 0.79 | 0.82 | 0.77 |
| 10 | 13586627 | rs7913419 | 1.106 | -0.275 | -0.284 | -0.566 | -0.014 | -0.001 | -0.001 | 0 | 0.36 | 0.2 | 0.38 | 0.68 | 0.41 | 0.4 |
| 10 | 13588521 | rs4750379 | 0.605 | -0.284 | -0.073 | -1 | 0 | 0.03 | 0.03 | 0 | 0.43 | 0.35 | 0.4 | 0.77 | 0.45 | 0.52 |
| 10 | 13605368 | rs4279918 | -0.515 | -0.538 | -0.074 | -1.253 | -0.007 | -0.011 | -0.011 | 0.856 | 0.28 | 0.27 | 0.22 | 0.2 | 0.27 | 0.27 |
| 10 | 13606905 | rs7089579 | 0.985 | -1.072 | -0.163 | -1.676 | -0.06 | 0.083 | 0.083 | 0 | 0.44 | 0.12 | 0.43 | 0.72 | 0.5 | 0.43 |
| 10 | 13610149 | rs4750385 | -0.729 | -1.345 | 0.204 | -1.453 | -0.016 | -0.017 | -0.017 | 0 | 0.33 | 0.55 | 0.25 | 0.22 | 0.27 | 0.3 |
| 10 | 13615920 | rs2478116 | 0.232 | -1.002 | 0.38 | -1.25 | -0.031 | -0.023 | -0.023 | 0 | 0.73 | 0.52 | 0.77 | 0.78 | 0.74 | 0.71 |
| 10 | 13617554 | rs2502225 | 0.24 | -0.983 | 0.361 | -1.242 | -0.01 | -0.014 | -0.014 | 0 | 0.75 | 0.7 | 0.78 | 0.79 | 0.73 | 0.72 |
| 10 | 13618862 | rs12355018 | 0.69 | -0.975 | 0.188 | -1.31 | -0.025 | 0.038 | 0.038 | 0 | 0.14 | 0.04 | 0.13 | 0.04 | 0.23 | 0.11 |
| 10 | 13628528 | rs2478106 | -0.037 | -0.57 | 0.227 | -1.537 | 0.003 | -0.04 | -0.04 | 2.47 | 0.37 | 0.62 | 0.44 | 0.07 | 0.4 | 0.32 |
| 10 | 13636301 | rs11258459 | 1.873 | -0.512 | -0.064 | -1.002 | -0.018 | -0.122 | -0.122 | 0.115 | 0.21 | 0.04 | 0.26 | 0.48 | 0.22 | 0.09 |
| 10 | 13642086 | rs1409322 | -0.337 | -0.952 | 0.253 | -1.234 | -0.043 | -0.034 | -0.034 | 0 | 0.7 | 0.63 | 0.88 | 0.84 | 0.82 | 0.74 |
| 10 | 13643158 | rs9633780 | 0.487 | -0.953 | 0.243 | -1.268 | -0.026 | -0.005 | -0.005 | 0 | 0.06 | 0.09 | 0.01 | 0.04 | 0 | 0.07 |
| 10 | 13648003 | rs2502207 | -0.021 | -0.605 | 0.363 | -1.219 | -0.046 | -0.066 | -0.066 | 0 | 0.66 | 0.57 | 0.81 | 0.82 | 0.74 | 0.66 |
| 10 | 13650032 | rs2478118 | 0.746 | -0.47 | 0.365 | -1.285 | 0.015 | -0.082 | -0.082 | 0 | 0.33 | 0.43 | 0.42 | 0.04 | 0.39 | 0.3 |
| 10 | 13650236 | rs943088 | -1.21 | -0.434 | 0.379 | -1.259 | -0.013 | -0.028 | -0.028 | 0 | 0.77 | 0.75 | 0.88 | 0.9 | 0.81 | 0.81 |
| 10 | 13650853 | rs10796100 | 1.056 | -0.446 | 0.321 | -1.238 | -0.084 | 0.016 | 0.016 | 0 | 0.32 | 0.07 | 0.34 | 0.73 | 0.36 | 0.35 |
| 10 | 13653390 | rs7090752 | 0.687 | -0.461 | 0.395 | -1.252 | -0.017 | 0.071 | 0.071 | 0 | 0.35 | 0.2 | 0.39 | 0.83 | 0.35 | 0.43 |
| 10 | 13653653 | rs1135102 | 0.03 | -0.447 | 0.51 | -1.261 | 0.006 | -0.082 | -0.082 | 0 | 0.41 | 0.46 | 0.49 | 0.06 | 0.46 | 0.36 |
| 10 | 13659738 | rs2478126 | 0.127 | -0.469 | 0.666 | -1.223 | -0.012 | -0.047 | -0.047 | 0 | 0.11 | 0.32 | 0.1 | 0.02 | 0.08 | 0.09 |
| 10 | 13663011 | rs10906425 | -1.08 | -0.502 | 0.56 | -1.234 | -0.018 | -0.018 | -0.018 | 0.005 | 0.76 | 0.7 | 0.83 | 0.83 | 0.81 | 0.83 |
| 10 | 13664667 | rs11258480 | 1.764 | -0.385 | 0.454 | -1.111 | 0.004 | -0.141 | -0.141 | 0 | 0.25 | 0.05 | 0.26 | 0.47 | 0.21 | 0.09 |
| 10 | 13666931 | rs7098174 | -0.076 | -0.338 | 0.468 | -1.116 | -0.008 | -0.012 | -0.012 | 2.355 | 0.17 | 0.14 | 0.14 | 0.28 | 0.2 | 0.39 |
| 10 | 13667731 | rs7089227 | 0.521 | -0.41 | 0.505 | -1.14 | 0 | -0.001 | -0.001 | 0 | 0.56 | 0.45 | 0.66 | 0.51 | 0.61 | 0.37 |
| 10 | 13668716 | rs4750390 | 0.075 | -0.498 | 0.59 | -1.179 | 0.009 | -0.091 | -0.091 | 0 | 0.42 | 0.54 | 0.51 | 0.06 | 0.48 | 0.37 |
| 10 | 13670850 | rs7088328 | -1.193 | -0.628 | 0.43 | -1.124 | -0.037 | -0.036 | -0.036 | 0.649 | 0.77 | 0.66 | 0.83 | 0.84 | 0.82 | 0.83 |
| 10 | 13670971 | rs7088503 | -0.437 | -0.575 | 0.37 | -1.075 | 0.019 | 0.016 | 0.016 | 5.049 | 0.48 | 0.45 | 0.55 | 0.31 | 0.61 | 0.65 |
| 10 | 13680726 | rs4393223 | -1.094 | -0.538 | 0.35 | -1.26 | -0.011 | -0.011 | -0.011 | 0 | 0.77 | 0.74 | 0.83 | 0.84 | 0.82 | 0.83 |
| 10 | 13681108 | rs2399987 | 0.452 | -0.216 | 0.314 | -1.427 | 0.014 | 0.049 | 0.049 | 0 | 0.55 | 0.36 | 0.62 | 0.77 | 0.57 | 0.53 |

| | | | | | | | | | | | | | | | | |
|---|---|---|---|---|---|---|---|---|---|---|---|---|---|---|---|---|
| 10 | 13683905 | rs7897982 | 0.296 | -0.257 | 0.232 | -1.427 | -0.003 | -0.018 | -0.018 | 0 | 0.57 | 0.46 | 0.63 | 0.77 | 0.58 | 0.53 |
| 10 | 13684843 | rs10508463 | 0.302 | -0.241 | 0.233 | -1.42 | -0.005 | -0.025 | -0.025 | 1.335 | 0.57 | 0.44 | 0.63 | 0.77 | 0.58 | 0.53 |
| 10 | 13685766 | rs13005 | 0.284 | -0.242 | 0.232 | -1.42 | -0.004 | -0.023 | -0.023 | 0 | 0.57 | 0.44 | 0.63 | 0.77 | 0.58 | 0.53 |
| 10 | 13686642 | rs4748047 | -0.704 | -0.267 | 0.278 | -1.439 | 0 | -0.01 | -0.01 | 0 | 0.69 | 0.68 | 0.72 | 0.88 | 0.66 | 0.68 |
| 10 | 13687171 | rs9473 | 0.125 | -0.198 | 0.304 | -1.379 | -0.002 | -0.031 | -0.031 | 0.02 | 0.58 | 0.55 | 0.64 | 0.85 | 0.59 | 0.55 |
| 10 | 13687890 | rs17153667 | 0.817 | -0.711 | 0.584 | -0.824 | -0.02 | 0.047 | 0.047 | 0.187 | 0.29 | 0.35 | 0.19 | 0.57 | 0.22 | 0.16 |
| 10 | 13688051 | rs1060093 | 0.89 | -0.715 | 0.583 | -0.826 | -0.015 | 0.037 | 0.037 | 0.164 | 0.28 | 0.33 | 0.19 | 0.57 | 0.22 | 0.16 |
| 10 | 13688571 | rs729435 | 0.644 | -1.226 | 0.673 | -1.633 | 0.002 | -0.026 | -0.026 | 0 | 0.36 | 0.38 | 0.35 | 0.77 | 0.35 | 0.45 |
| 10 | 13696570 | rs10906442 | -0.203 | -1.06 | 0.512 | -1.614 | -0.036 | 0.161 | 0.161 | 0.818 | 0.47 | 0.83 | 0.54 | 0.18 | 0.49 | 0.48 |
| 10 | 13706408 | rs4748050 | -0.574 | -0.886 | 0.726 | -1.536 | -0.064 | -0.076 | -0.076 | 0 | 0.74 | 0.56 | 0.82 | 0.85 | 0.82 | 0.77 |
| 10 | 13708882 | rs7906964 | 0.383 | -0.694 | 0.025 | -1.526 | -0.005 | -0.029 | -0.029 | 0 | 0.38 | 0.36 | 0.44 | 0.04 | 0.43 | 0.32 |
| 10 | 13709426 | rs10796110 | 0.696 | -0.811 | -0.211 | -1.428 | -0.041 | 0.051 | 0.051 | 0 | 0.35 | 0.13 | 0.37 | 0.75 | 0.38 | 0.39 |
| 10 | 13710163 | rs9651540 | -0.311 | -0.877 | -0.162 | -1.44 | -0.03 | -0.014 | -0.014 | 0.328 | 0.69 | 0.37 | 0.8 | 0.79 | 0.82 | 0.68 |
| 10 | 13710897 | rs7093277 | 0.391 | -0.186 | -0.44 | -0.957 | -0.003 | 0.004 | 0.004 | 0 | 0.35 | 0.3 | 0.38 | 0.74 | 0.38 | 0.39 |
| 10 | 13716174 | rs11258513 | -0.181 | 0.644 | -0.441 | -0.944 | -0.023 | 0.051 | 0.051 | 0 | 0.4 | 0.34 | 0.46 | 0.08 | 0.49 | 0.36 |
| 10 | 13723773 | rs943097 | 0.067 | 0.828 | -1.174 | 0.32 | -0.065 | -0.016 | -0.016 | 0 | 0.18 | 0.01 | 0.19 | 0.14 | 0.21 | 0.15 |
| 10 | 13724841 | rs750976 | 0.202 | 0.138 | -1.581 | 0.484 | -0.02 | 0.026 | 0.026 | 0 | 0.68 | 0.95 | 0.57 | 0.28 | 0.72 | 0.61 |
| 10 | 13727937 | rs943099 | 0.712 | -0.309 | -1.309 | -0.431 | 0 | 0.04 | 0.04 | 0.225 | 0.42 | 0.32 | 0.4 | 0.11 | 0.59 | 0.34 |
| 10 | 13729426 | rs943101 | -0.565 | 0.315 | -0.389 | -0.354 | 0.003 | -0.029 | -0.029 | 0 | 0.24 | 0.53 | 0.22 | 0.19 | 0.28 | 0.18 |
| 10 | 13729505 | rs943102 | -0.721 | 0.337 | -0.265 | -0.395 | 0.003 | -0.023 | -0.023 | 0 | 0.26 | 0.6 | 0.22 | 0.2 | 0.28 | 0.18 |
| 10 | 13729819 | rs11258522 | 0.845 | 0.241 | 0.345 | -0.355 | -0.026 | 0.002 | 0.002 | 0.247 | 0.1 | 0.22 | 0.05 | 0.09 | 0.03 | 0.11 |
| 10 | 13730004 | rs10796113 | -0.997 | -0.003 | 0.158 | 0.452 | -0.001 | -0.009 | -0.009 | 1.729 | 0.8 | 0.83 | 0.72 | 0.63 | 0.79 | 0.76 |
| 10 | 13733923 | rs2182404 | -0.299 | 0.211 | 0.438 | 0.483 | -0.014 | 0.037 | 0.037 | 0.385 | 0.87 | 0.43 | 0.9 | 0.84 | 0.92 | 0.88 |
| 10 | 13734850 | rs2077150 | 1.468 | -0.097 | 0.091 | 0.377 | 0.004 | 0.011 | 0.011 | 0 | 0.43 | 0.21 | 0.44 | 0.41 | 0.41 | 0.48 |
| 10 | 13735405 | rs11258531 | 0.866 | -0.103 | 0.123 | 0.405 | -0.011 | -0.065 | -0.065 | 0.774 | 0.2 | 0.08 | 0.27 | 0.39 | 0.2 | 0.23 |
| 10 | 13735947 | rs2296596 | -0.169 | -0.088 | 0.505 | 0.372 | -0.009 | -0.01 | -0.01 | 0.018 | 0.15 | 0.2 | 0.07 | 0.11 | 0.11 | 0.12 |
| 10 | 13737029 | rs4314962 | 0.342 | -0.06 | 0.409 | 0.365 | -0.019 | -0.03 | -0.03 | 0 | 0.76 | 0.41 | 0.85 | 0.91 | 0.85 | 0.77 |
| 10 | 13737841 | rs10906449 | -1.257 | 0.255 | 0.513 | 0.559 | -0.006 | -0.002 | -0.002 | 0.972 | 0.46 | 0.65 | 0.49 | 0.5 | 0.49 | 0.4 |
| 10 | 13737972 | rs3750885 | 0.47 | 0.234 | 0.422 | 0.74 | -0.005 | -0.012 | -0.012 | 0.09 | 0.21 | 0.11 | 0.18 | 0.24 | 0.19 | 0.17 |
| 10 | 13738011 | rs3750884 | -0.206 | 0.237 | 0.582 | 0.73 | -0.031 | 0.161 | 0.161 | 0.019 | 0.14 | 0.35 | 0.17 | 0.32 | 0.06 | 0.18 |
| 10 | 13739528 | rs10906451 | 0.457 | 0.246 | 0.462 | 0.778 | -0.005 | -0.014 | -0.014 | 0 | 0.21 | 0.11 | 0.18 | 0.24 | 0.19 | 0.17 |
| 10 | 13740173 | rs7893899 | -1.166 | 0.537 | 0.576 | 0.626 | 0.029 | 0.051 | 0.051 | 0.387 | 0.26 | 0.53 | 0.32 | 0.34 | 0.31 | 0.24 |
| 10 | 13740314 | rs17627309 | 0.647 | 0.548 | 0.526 | 0.62 | -0.064 | 0.06 | 0.06 | 0 | 0.1 | 0.02 | 0.15 | 0.02 | 0.24 | 0.06 |
| 10 | 13740551 | rs2031609 | 0.457 | 0.557 | 0.56 | 0.624 | 0.045 | -0.047 | -0.047 | 0 | 0.11 | 0.14 | 0.15 | 0.02 | 0.24 | 0.06 |
| 10 | 13740917 | rs1887004 | -0.823 | 0.467 | 0.278 | 0.672 | 0.039 | 0.05 | 0.05 | 0.455 | 0.26 | 0.52 | 0.32 | 0.35 | 0.32 | 0.24 |
| 10 | 13742037 | rs12358118 | -0.644 | -0.335 | -0.231 | 0.257 | 0.001 | 0.003 | 0.003 | 0 | 0.31 | 0.28 | 0.26 | 0.21 | 0.27 | 0.22 |
| 10 | 13743517 | rs1541004 | -0.525 | 0.019 | -0.085 | 0.378 | 0 | -0.001 | -0.001 | 0 | 0.24 | 0.23 | 0.22 | 0.18 | 0.24 | 0.13 |
| 10 | 13743857 | rs1541006 | 0.353 | 0.22 | -0.591 | 0.511 | -0.062 | -0.039 | -0.039 | 0 | 0.17 | 0.01 | 0.19 | 0.15 | 0.22 | 0.11 |
| 10 | 13749228 | rs11258541 | 0.482 | 0.438 | -1.078 | 0.671 | -0.05 | -0.126 | -0.126 | 2.537 | 0.35 | 0.09 | 0.44 | 0.5 | 0.41 | 0.46 |
| 10 | 13752381 | rs2296599 | 0.22 | 0.236 | -0.303 | 0.592 | 0.01 | 0.011 | 0.011 | 0 | 0.2 | 0.4 | 0.2 | 0.24 | 0.21 | 0.17 |
| 10 | 13759976 | rs3814667 | -0.435 | 0.506 | -0.503 | 0.578 | 0.015 | 0.028 | 0.028 | 0.79 | 0.49 | 0.35 | 0.55 | 0.59 | 0.49 | 0.68 |
| 10 | 13765950 | rs12359348 | 0.328 | 0.268 | -1.165 | 0.863 | -0.048 | -0.13 | -0.13 | 0.051 | 0.34 | 0.09 | 0.41 | 0.48 | 0.4 | 0.45 |
| 10 | 13766692 | rs2015015 | 0.079 | 0.301 | -0.558 | 0.656 | 0.007 | -0.005 | -0.005 | 0 | 0.4 | 0.35 | 0.46 | 0.44 | 0.31 | 0.42 |
| 10 | 13771761 | rs756068 | 0.011 | 0.516 | -0.74 | 0.212 | 0.001 | 0.002 | 0.002 | 0.017 | 0.4 | 0.37 | 0.44 | 0.35 | 0.31 | 0.29 |
| 10 | 13776176 | rs10796117 | 0.297 | 0.905 | -0.903 | 0.101 | -0.012 | 0.024 | 0.024 | 1.039 | 0.59 | 0.68 | 0.43 | 0.27 | 0.56 | 0.34 |
| 10 | 13777194 | rs7900357 | 0.354 | 0.82 | -0.863 | 0.251 | -0.001 | 0 | 0 | 2.105 | 0.7 | 0.74 | 0.54 | 0.24 | 0.72 | 0.37 |
| 10 | 13780118 | rs9633782 | 0.507 | 0.754 | -1.095 | 0.168 | 0.006 | 0.019 | 0.019 | 0 | 0.66 | 0.75 | 0.57 | 0.26 | 0.68 | 0.4 |
| 10 | 13780362 | rs7074581 | 0.387 | 0.663 | -1.214 | 0.016 | -0.001 | -0.003 | -0.003 | 0.034 | 0.65 | 0.76 | 0.57 | 0.36 | 0.66 | 0.37 |
| 10 | 13780423 | rs11592088 | 0.329 | 0.954 | -1.941 | -0.57 | -0.07 | 0.021 | 0.021 | 0.101 | 0.22 | 0.06 | 0.25 | 0.15 | 0.31 | 0.13 |
| 10 | 13781195 | rs10906460 | 0.154 | 0.651 | -1.232 | -0.774 | 0.015 | -0.027 | -0.027 | 0 | 0.37 | 0.41 | 0.25 | 0.16 | 0.38 | 0.24 |
| 10 | 13782605 | rs6602679 | 0.112 | 0.204 | -1.675 | -1.333 | -0.02 | 0.112 | 0.112 | 0.061 | 0.62 | 0.79 | 0.71 | 0.96 | 0.61 | 0.77 |
| 10 | 13785400 | rs10737079 | 0.125 | 0.142 | -1.692 | -0.997 | -0.042 | 0.119 | 0.119 | 0.689 | 0.58 | 0.76 | 0.67 | 0.86 | 0.53 | 0.6 |
| 10 | 13787691 | rs7921755 | -0.205 | -0.254 | -1.74 | -0.416 | 0.051 | -0.014 | -0.014 | 0 | 0.52 | 0.3 | 0.66 | 0.55 | 0.61 | 0.58 |
| 10 | 13790265 | rs10906466 | -0.254 | -0.177 | -1.359 | -0.226 | 0.002 | -0.029 | -0.029 | 0.19 | 0.69 | 0.76 | 0.57 | 0.45 | 0.69 | 0.55 |
| 10 | 13791188 | rs12218115 | 1.043 | -0.503 | -0.624 | -0.284 | -0.018 | 0.071 | 0.071 | 0.556 | 0.25 | 0.4 | 0.21 | 0.34 | 0.23 | 0.18 |
| 10 | 13792522 | rs12257410 | 0.5 | -0.803 | -0.741 | -0.369 | -0.014 | 0.018 | 0.018 | 2.056 | 0.32 | 0.58 | 0.26 | 0.42 | 0.25 | 0.35 |
| 10 | 13795471 | rs4750407 | 0.206 | -0.672 | -1.084 | -0.134 | -0.021 | -0.036 | -0.036 | 0.255 | 0.18 | 0.09 | 0.32 | 0.32 | 0.27 | 0.36 |
| 10 | 13796225 | rs7095537 | 0.205 | -0.728 | -1.26 | -0.294 | -0.013 | -0.022 | -0.022 | 0.952 | 0.19 | 0.12 | 0.23 | 0.31 | 0.18 | 0.23 |
| 10 | 13798200 | rs4570478 | 1.302 | -0.73 | -1.031 | -0.341 | -0.02 | 0.024 | 0.024 | 1.109 | 0.27 | 0.51 | 0.23 | 0.3 | 0.24 | 0.17 |
| 10 | 13801269 | rs4424567 | 0.574 | -0.491 | -1.509 | -0.319 | -0.012 | -0.059 | -0.059 | 0 | 0.24 | 0.15 | 0.44 | 0.55 | 0.28 | 0.39 |
| 10 | 13802259 | rs4748054 | -0.527 | -0.429 | -1.511 | -0.912 | -0.082 | 0.229 | 0.229 | 0.228 | 0.63 | 0.91 | 0.77 | 0.96 | 0.61 | 0.77 |
| 10 | 13810059 | rs7910196 | -0.205 | -0.387 | -1.271 | -0.745 | 0.014 | 0.053 | 0.053 | 1.022 | 0.82 | 0.92 | 0.92 | 1 | 0.85 | 0.89 |
| 10 | 13817944 | rs2049745 | 0.311 | 0.088 | -0.862 | -0.874 | 0 | -0.023 | -0.023 | 0 | 0.76 | 0.3 | 0.81 | 0.95 | 0.75 | 0.88 |
| 10 | 13818934 | rs11258592 | 0.417 | 0.072 | -0.882 | -0.731 | -0.002 | 0.018 | 0.018 | 3.167 | 0.7 | 0.3 | 0.75 | 0.86 | 0.71 | 0.71 |
| 10 | 13821953 | rs2400015 | 0.134 | 0.198 | -0.521 | -0.592 | 0.023 | -0.043 | -0.043 | 0 | 0.75 | 0.61 | 0.77 | 0.86 | 0.72 | 0.71 |
| 10 | 13824009 | rs10906480 | -0.092 | 0.139 | -0.492 | -0.527 | 0.014 | -0.041 | -0.041 | 0 | 0.74 | 0.6 | 0.81 | 0.92 | 0.72 | 0.83 |
| 10 | 13825834 | rs4750414 | 0.525 | 0.108 | -0.847 | -0.224 | -0.014 | -0.055 | -0.055 | 0 | 0.34 | 0.16 | 0.46 | 0.53 | 0.34 | 0.52 |
| 10 | 13826018 | rs1919457 | -0.391 | 0.145 | -0.784 | -0.197 | 0.022 | -0.014 | -0.014 | 0 | 0.75 | 0.63 | 0.78 | 0.82 | 0.72 | 0.78 |
| 10 | 13826242 | rs10796127 | -0.024 | -0.106 | -0.883 | -0.134 | -0.003 | -0.004 | -0.004 | 0 | 0.39 | 0.41 | 0.3 | 0.29 | 0.37 | 0.25 |

| | | | | | | | | | | | | | | | | |
|---|---|---|---|---|---|---|---|---|---|---|---|---|---|---|---|---|
| 10 | 13831712 | rs1881584 | -0.373 | 0.255 | -1.246 | 0.181 | 0 | 0.006 | 0.006 | 0 | 0.29 | 0.32 | 0.43 | 0.54 | 0.33 | 0.51 |
| 10 | 13831873 | rs11258614 | -0.022 | 0.297 | -1.742 | 0.338 | -0.025 | -0.085 | -0.085 | 0 | 0.24 | 0.12 | 0.35 | 0.42 | 0.3 | 0.44 |
| 10 | 13832649 | rs11258616 | 0.499 | -0.028 | -1.814 | 0.127 | -0.014 | -0.041 | -0.041 | 1.755 | 0.13 | 0.01 | 0.15 | 0.19 | 0.16 | 0.09 |
| 10 | 13834516 | rs4750415 | 0.754 | -0.307 | -2.106 | -0.32 | 0.009 | 0.122 | 0.122 | 1.713 | 0.19 | 0 | 0.06 | 0 | 0.11 | 0.15 |
| 10 | 13834678 | rs4750416 | -0.457 | -0.452 | -1.613 | -0.493 | 0.031 | 0.05 | 0.05 | 0 | 0.45 | 0.63 | 0.36 | 0.26 | 0.36 | 0.3 |
| 10 | 13835616 | rs4145934 | 0.671 | 0.338 | -2.223 | -0.328 | -0.099 | -0.12 | -0.12 | 0 | 0.31 | 0.09 | 0.36 | 0.46 | 0.34 | 0.49 |
| 10 | 13836782 | rs3995590 | -0.261 | -0.544 | -1.508 | -0.146 | 0.012 | -0.007 | -0.007 | 0 | 0.64 | 0.38 | 0.63 | 0.57 | 0.66 | 0.75 |
| 10 | 13838604 | rs3740121 | 0.061 | 0.037 | -1.606 | 0.241 | -0.042 | -0.089 | -0.089 | 1.077 | 0.41 | 0.12 | 0.44 | 0.47 | 0.53 | 0.54 |
| 10 | 13841470 | rs7072101 | -1.04 | -0.135 | 0.041 | 0.614 | 0.069 | 0.168 | 0.168 | 0 | 0.08 | 0.44 | 0.2 | 0.23 | 0.07 | 0.19 |
| 10 | 13843530 | rs10906492 | 0.175 | -0.309 | -0.527 | 0.662 | -0.025 | 0.026 | 0.026 | 0 | 0.4 | 0.14 | 0.33 | 0.24 | 0.46 | 0.36 |
| 10 | 13850287 | rs11813129 | -0.228 | -0.18 | 0.163 | 0.615 | 0.024 | 0.013 | 0.013 | 0.061 | 0.08 | 0.16 | 0.13 | 0.12 | 0.11 | 0.1 |
| 10 | 13853852 | rs7088455 | 0.243 | -0.145 | 0.303 | 0.902 | -0.001 | 0.049 | 0.049 | 0 | 0.92 | 0.83 | 0.84 | 0.84 | 0.94 | 0.89 |
| 10 | 13854618 | rs10906496 | 0.96 | -0.499 | 0.3 | 0.936 | -0.005 | 0.004 | 0.004 | 1.245 | 0.37 | 0.34 | 0.29 | 0.35 | 0.37 | 0.33 |
| 10 | 13854997 | rs1343002 | -0.543 | -0.377 | 0.525 | 0.965 | 0.024 | 0.051 | 0.051 | 0 | 0.14 | 0.22 | 0.24 | 0.25 | 0.17 | 0.16 |
| 10 | 13856099 | rs12761607 | 0.843 | -0.306 | -0.321 | 0.904 | -0.011 | 0.008 | 0.008 | 0.949 | 0.38 | 0.26 | 0.34 | 0.35 | 0.37 | 0.33 |
| 10 | 13859410 | rs10906498 | -0.061 | -0.073 | -0.807 | 0.778 | -0.015 | 0.04 | 0.04 | 0 | 0.43 | 0.16 | 0.34 | 0.24 | 0.46 | 0.4 |
| 10 | 13863721 | rs7904993 | -0.611 | 0.398 | -0.277 | 1.471 | -0.002 | -0.008 | -0.008 | 0 | 0.14 | 0.13 | 0.13 | 0.33 | 0.14 | 0.35 |
| 10 | 13864263 | rs6602686 | -1.69 | 0.275 | 0.189 | 1.117 | 0 | 0 | 0 | 0 | 0.41 | 0.6 | 0.47 | 0.58 | 0.46 | 0.58 |
| 10 | 13865524 | rs11595591 | -0.495 | 0.756 | -0.338 | 1.062 | -0.028 | 0.045 | 0.045 | 1.933 | 0.2 | 0.1 | 0.16 | 0.1 | 0.27 | 0.14 |
| 10 | 13867888 | rs11258651 | 1.314 | 0.557 | -0.125 | 1.166 | -0.002 | -0.005 | -0.005 | 0.836 | 0.65 | 0.63 | 0.72 | 0.58 | 0.62 | 0.52 |
| 10 | 13868855 | rs11258652 | 0.007 | 0.557 | -0.152 | 0.773 | -0.031 | 0.04 | 0.04 | 1.611 | 0.17 | 0.09 | 0.13 | 0.09 | 0.25 | 0.12 |
| 10 | 13869240 | rs11258656 | 1.638 | 0.599 | 0.02 | 0.871 | 0.004 | -0.003 | -0.003 | 0 | 0.6 | 0.53 | 0.53 | 0.4 | 0.54 | 0.42 |
| 10 | 13869987 | rs10906503 | 0.571 | 0.695 | -0.243 | 0.439 | 0 | -0.012 | -0.012 | 0.019 | 0.8 | 0.77 | 0.86 | 0.91 | 0.76 | 0.89 |
| 10 | 13874376 | rs10906505 | -0.768 | 0.617 | -0.601 | 0.381 | -0.003 | 0.046 | 0.046 | 0.757 | 0.2 | 0.14 | 0.13 | 0.07 | 0.24 | 0.13 |
| 10 | 13875106 | rs10796134 | 1.215 | 0.788 | -0.474 | 0.794 | -0.008 | -0.011 | -0.011 | 0 | 0.65 | 0.68 | 0.63 | 0.62 | 0.58 | 0.52 |
| 10 | 13875590 | rs7899777 | -0.111 | 0.947 | -0.535 | 0.357 | -0.003 | -0.007 | -0.007 | 0 | 0.86 | 0.87 | 0.91 | 0.98 | 0.81 | 0.96 |
| 10 | 13875739 | rs7900034 | -1.061 | 1.15 | -0.353 | 0.408 | 0.006 | -0.004 | -0.004 | 5.696 | 0.39 | 0.48 | 0.47 | 0.61 | 0.5 | 0.51 |
| 10 | 13876469 | rs10796135 | 0.501 | 1.592 | -0.129 | 0.73 | 0.017 | 0.015 | 0.015 | 0 | 0.67 | 0.57 | 0.66 | 0.6 | 0.56 | 0.8 |
| 10 | 13879165 | rs7908696 | -0.11 | 1.498 | -0.199 | 0.686 | 0.001 | -0.019 | -0.019 | 0.036 | 0.83 | 0.84 | 0.89 | 0.96 | 0.78 | 0.89 |
| 10 | 13883460 | rs7894550 | -0.106 | 1.549 | -0.19 | 0.705 | 0.003 | -0.02 | -0.02 | 0 | 0.85 | 0.85 | 0.89 | 0.96 | 0.77 | 0.89 |
| 10 | 13884259 | rs7073646 | 1.202 | 1.327 | -0.359 | 0.337 | 0.049 | 0.019 | 0.019 | 0 | 0.44 | 0.32 | 0.42 | 0.37 | 0.27 | 0.37 |
| 10 | 13885487 | rs7097221 | -0.061 | 1.618 | -0.581 | 0.757 | 0.002 | 0.001 | 0.001 | 0 | 0.13 | 0.11 | 0.17 | 0.24 | 0.2 | 0.29 |
| 10 | 13887104 | rs10906515 | 0.703 | 0.573 | -0.565 | 0.267 | -0.004 | -0.003 | -0.003 | 0.068 | 0.51 | 0.51 | 0.49 | 0.39 | 0.41 | 0.4 |
| 10 | 13887165 | rs7920594 | 0.329 | 0.965 | -0.589 | 0.294 | 0.001 | -0.016 | -0.016 | 0 | 0.94 | 0.9 | 0.98 | 0.98 | 0.96 | 0.93 |
| 10 | 13898154 | rs10430781 | 0.405 | 0.738 | -0.758 | 0.235 | 0.002 | 0.001 | 0.001 | 0 | 0.54 | 0.47 | 0.48 | 0.64 | 0.45 | 0.56 |
| 10 | 13900071 | rs7096973 | 0.239 | 0.803 | -0.842 | 0.048 | 0.001 | 0.002 | 0.002 | 0 | 0.59 | 0.48 | 0.48 | 0.64 | 0.46 | 0.6 |
| 10 | 13902179 | rs4748063 | 0.884 | 0.261 | -1.196 | -0.277 | -0.045 | -0.001 | -0.001 | 0 | 0.43 | 0.47 | 0.37 | 0.64 | 0.2 | 0.53 |
| 10 | 13903948 | rs1881578 | -0.529 | 0.899 | -1.066 | -0.025 | -0.002 | 0.002 | 0.002 | 0 | 0.93 | 0.94 | 0.95 | 0.98 | 0.93 | 0.92 |
| 10 | 13911214 | rs10752332 | 0.812 | 0.738 | -1.016 | 0.345 | -0.002 | -0.001 | -0.001 | 0 | 0.52 | 0.49 | 0.47 | 0.53 | 0.42 | 0.53 |
| 10 | 13911267 | rs11258682 | -0.079 | 1.738 | -1.296 | 0.385 | -0.058 | -0.004 | -0.004 | 0 | 0.11 | 0.02 | 0.12 | 0.09 | 0.15 | 0.17 |
| 10 | 13913432 | rs12218323 | -0.643 | 1.162 | -1.17 | 0.622 | -0.036 | -0.019 | -0.019 | 0 | 0.38 | 0.23 | 0.34 | 0.31 | 0.48 | 0.29 |
| 10 | 13914363 | rs12777098 | 0.37 | 0.889 | -1.133 | 0.544 | 0 | -0.002 | -0.002 | 0 | 0.06 | 0.05 | 0.03 | 0.02 | 0.05 | 0.08 |
| 10 | 13914547 | rs7898937 | -0.889 | 0.862 | -1.005 | 0.542 | -0.016 | -0.011 | -0.011 | 0 | 0.42 | 0.35 | 0.42 | 0.42 | 0.47 | 0.32 |
| 10 | 13917675 | rs12778887 | 0.106 | 1.051 | -0.887 | 0.71 | -0.006 | -0.019 | -0.019 | 0 | 0.06 | 0.09 | 0.04 | 0.02 | 0.05 | 0.08 |
| 10 | 13917699 | rs11258686 | 0.039 | 1.225 | -0.993 | 0.724 | -0.036 | -0.035 | -0.035 | 0 | 0.21 | 0.16 | 0.27 | 0.3 | 0.34 | 0.19 |
| 10 | 13919625 | rs1881586 | 0.893 | 1.043 | -0.878 | 0.572 | 0.03 | 0.017 | 0.017 | 0 | 0.65 | 0.48 | 0.57 | 0.52 | 0.58 | 0.69 |
| 10 | 13926403 | rs4750436 | -0.902 | 0.891 | -1.145 | 0.415 | 0.004 | -0.006 | -0.006 | 0.259 | 0.35 | 0.46 | 0.44 | 0.68 | 0.39 | 0.48 |
| 10 | 13929130 | rs751498 | -0.8 | 0.992 | -1.031 | 0.542 | -0.009 | -0.01 | -0.01 | 0 | 0.38 | 0.3 | 0.38 | 0.31 | 0.52 | 0.28 |
| 10 | 13934962 | rs11258711 | 0.366 | 0.833 | -1.163 | 0.225 | -0.006 | -0.014 | -0.014 | 0.086 | 0.3 | 0.2 | 0.21 | 0.26 | 0.25 | 0.25 |
| 10 | 13936500 | rs1547833 | 0.631 | 1.074 | -1.139 | 0.012 | -0.005 | 0.005 | 0.005 | 0 | 0.42 | 0.45 | 0.53 | 0.54 | 0.41 | 0.52 |
| 10 | 13936554 | rs1547834 | -0.517 | 1.022 | -1.108 | -0.039 | -0.018 | 0.014 | 0.014 | 0 | 0.75 | 0.8 | 0.74 | 0.81 | 0.67 | 0.78 |
| 10 | 13937073 | rs2797877 | 0.831 | 1.057 | -1.222 | -0.046 | -0.007 | 0.004 | 0.004 | 0 | 0.38 | 0.44 | 0.5 | 0.49 | 0.38 | 0.47 |
| 10 | 13939530 | rs2610803 | 0.34 | 0.866 | -0.595 | 0.581 | -0.007 | 0.006 | 0.006 | 0 | 0.61 | 0.44 | 0.68 | 0.51 | 0.71 | 0.66 |
| 10 | 13942170 | rs10906535 | -0.055 | 0.489 | -0.727 | 0.393 | -0.011 | -0.005 | -0.005 | 0.107 | 0.35 | 0.42 | 0.23 | 0.25 | 0.23 | 0.25 |
| 10 | 13949609 | rs3235 | 0.481 | 0.541 | -1.031 | -0.127 | 0.013 | 0.026 | 0.026 | 0 | 0.6 | 0.67 | 0.72 | 0.73 | 0.67 | 0.71 |
| 10 | 13956618 | rs2698110 | 0.043 | 0.978 | -0.378 | 0.427 | 0.007 | 0.048 | 0.048 | 0 | 0.87 | 0.69 | 0.84 | 0.81 | 0.86 | 0.88 |
| 10 | 13962622 | rs2797866 | 0.892 | 1.178 | -0.036 | 0.339 | 0.042 | 0.058 | 0.058 | 0.656 | 0.84 | 0.6 | 0.8 | 0.76 | 0.8 | 0.91 |
| 10 | 13962655 | rs2610815 | 0.054 | 1.356 | -0.52 | 0.467 | 0 | 0.038 | 0.038 | 0 | 0.95 | 0.91 | 0.87 | 0.76 | 0.94 | 0.94 |
| 10 | 13965510 | rs2797862 | -0.196 | 1.352 | -0.433 | 0.498 | 0.008 | 0.023 | 0.023 | 0 | 0.92 | 0.86 | 0.86 | 0.8 | 0.86 | 0.94 |
| 10 | 13965860 | rs2797861 | -0.062 | 0.475 | -0.913 | -0.548 | 0.006 | 0.081 | 0.081 | 0 | 0.66 | 0.8 | 0.74 | 0.92 | 0.7 | 0.83 |
| 10 | 13967259 | rs2995648 | -0.915 | 1.416 | -0.638 | -0.438 | -0.005 | -0.003 | -0.003 | 0 | 0.22 | 0.17 | 0.21 | 0.27 | 0.3 | 0.2 |
| 10 | 13968125 | rs17632029 | 0.206 | 1.437 | -0.644 | -0.414 | -0.047 | 0.023 | 0.023 | 0.001 | 0.11 | 0.02 | 0.08 | 0.07 | 0.13 | 0.12 |
| 10 | 13968186 | rs1343005 | -0.476 | 0.868 | -0.671 | -1.15 | 0.022 | -0.139 | -0.139 | 0.584 | 0.66 | 0.53 | 0.65 | 0.98 | 0.55 | 0.73 |
| 10 | 13969539 | rs1534627 | 0.09 | 0.791 | -0.619 | -1.181 | 0.003 | 0.023 | 0.023 | 0 | 0.58 | 0.64 | 0.55 | 0.74 | 0.42 | 0.6 |
| 10 | 13969919 | rs4748064 | 0.37 | 0.793 | -0.543 | -1.468 | -0.004 | -0.034 | -0.034 | 0 | 0.3 | 0.29 | 0.33 | 0.02 | 0.45 | 0.26 |
| 10 | 13971726 | rs10906541 | 0.248 | 0.995 | -0.723 | -1.462 | -0.017 | -0.002 | -0.002 | 4.814 | 0.31 | 0.28 | 0.33 | 0.01 | 0.45 | 0.25 |
| 10 | 13973579 | rs7904694 | 1.149 | 0.813 | -0.724 | -0.587 | 0.019 | 0.219 | 0.219 | 0.261 | 0.07 | 0.23 | 0.1 | 0.27 | 0.06 | 0.08 |
| 10 | 13974185 | rs2797890 | -0.148 | 1.633 | -1.671 | -0.943 | -0.006 | -0.007 | -0.007 | 0.009 | 0.21 | 0.2 | 0.27 | 0.29 | 0.28 | 0.39 |
| 10 | 13975467 | rs10906543 | -0.011 | 1.694 | -1.957 | -1.168 | -0.01 | 0.063 | 0.063 | 0.001 | 0.19 | 0.04 | 0.15 | 0.03 | 0.23 | 0.1 |

| | | | | | | | | | | | | | | | | |
|---|---|---|---|---|---|---|---|---|---|---|---|---|---|---|---|---|
| 10 | 13976791 | rs7082056 | -0.846 | 1.698 | -1.95 | -1.127 | -0.045 | 0.081 | 0.081 | 0 | 0.37 | 0.17 | 0.39 | 0.19 | 0.5 | 0.26 |
| 10 | 13978235 | rs10906545 | 1.305 | 1.604 | -2.016 | -1.318 | -0.008 | 0.008 | 0.008 | 0.498 | 0.43 | 0.5 | 0.47 | 0.51 | 0.4 | 0.34 |
| 10 | 13980030 | rs1610163 | 0.676 | 1.678 | -2.03 | -1.379 | 0 | -0.008 | -0.008 | 0 | 0.12 | 0.13 | 0.08 | 0.03 | 0.12 | 0.11 |
| 10 | 13981162 | rs1026580 | 0.692 | 1.705 | -2.073 | -1.387 | -0.023 | 0.092 | 0.092 | 0 | 0.56 | 0.74 | 0.57 | 0.79 | 0.48 | 0.67 |
| 10 | 13981191 | rs11258736 | 0.454 | 1.781 | -2.093 | -1.411 | 0 | 0.021 | 0.021 | 0 | 0.14 | 0.06 | 0.13 | 0.03 | 0.15 | 0.1 |
| 10 | 13981491 | rs2614153 | 0.274 | 1.84 | -2.084 | -1.395 | -0.029 | 0.069 | 0.069 | 0.123 | 0.66 | 0.8 | 0.69 | 0.82 | 0.58 | 0.74 |
| 10 | 13983007 | rs10906548 | 0.598 | 1.617 | -1.839 | -1.153 | 0.004 | 0.015 | 0.015 | 0 | 0.12 | 0.06 | 0.12 | 0.05 | 0.13 | 0.08 |
| 10 | 13987999 | rs4553269 | 0.94 | 1.702 | -1.907 | -1.134 | -0.009 | -0.004 | -0.004 | 0 | 0.13 | 0.1 | 0.2 | 0.13 | 0.17 | 0.09 |
| 10 | 13988898 | rs12240404 | 0.79 | 1.827 | -1.87 | -1.146 | 0.004 | -0.015 | -0.015 | 0 | 0.11 | 0.14 | 0.07 | 0.02 | 0.11 | 0.09 |
| 10 | 13989027 | rs4354606 | 0.751 | 1.824 | -2.097 | -1.04 | -0.014 | 0.003 | 0.003 | 0.141 | 0.16 | 0.1 | 0.22 | 0.14 | 0.2 | 0.13 |
| 10 | 13989224 | rs1993742 | 1.022 | 1.797 | -2.219 | -1.257 | -0.004 | 0 | 0 | 0.621 | 0.53 | 0.63 | 0.52 | 0.5 | 0.47 | 0.43 |
| 10 | 13989299 | rs2926876 | 0.655 | 2.366 | -2.646 | -2.045 | 0.001 | 0.042 | 0.042 | 0 | 0.72 | 0.84 | 0.81 | 0.92 | 0.7 | 0.79 |
| 10 | 13990453 | rs10906550 | -0.065 | 2.063 | -2.446 | -1.886 | -0.036 | 0.057 | 0.057 | 0 | 0.19 | 0.07 | 0.19 | 0.04 | 0.31 | 0.13 |
| 10 | 13991865 | rs7081208 | -0.395 | 1.936 | -2.061 | -1.388 | -0.062 | 0.088 | 0.088 | 0 | 0.2 | 0.04 | 0.12 | 0.03 | 0.23 | 0.15 |
| 10 | 13994618 | rs12267038 | 0.346 | 1.704 | -1.611 | -1.106 | 0.001 | -0.007 | -0.007 | 0 | 0.16 | 0.13 | 0.17 | 0.03 | 0.16 | 0.15 |
| 10 | 13998736 | rs2457849 | 0.359 | 2.05 | -1.355 | -0.788 | 0 | -0.007 | -0.007 | 0.261 | 0.42 | 0.57 | 0.51 | 0.83 | 0.39 | 0.52 |
| 10 | 13998997 | rs2446574 | 0.26 | 2.047 | -1.369 | -0.784 | 0.01 | 0.028 | 0.028 | 0.151 | 0.43 | 0.64 | 0.51 | 0.83 | 0.39 | 0.52 |
| 10 | 14002143 | rs2447026 | -0.062 | 2.378 | -1.518 | -0.651 | -0.038 | 0.033 | 0.033 | 0.002 | 0.87 | 0.93 | 0.88 | 0.98 | 0.77 | 0.86 |
| 10 | 14003363 | rs2447027 | -0.076 | 1.7 | -1.204 | -0.338 | 0 | -0.001 | -0.001 | 1.329 | 0.42 | 0.58 | 0.52 | 0.77 | 0.4 | 0.51 |
| 10 | 14005536 | rs2446580 | -0.077 | 1.728 | -1.222 | -0.379 | 0 | -0.006 | -0.006 | 0 | 0.42 | 0.58 | 0.54 | 0.78 | 0.47 | 0.53 |
| 10 | 14006391 | rs2457861 | -0.366 | 0.983 | -1.261 | -0.014 | 0 | 0 | 0 | 0 | 0.41 | 0.63 | 0.58 | 0.54 | 0.4 | 0.55 |
| 10 | 14008445 | rs2446581 | -1.25 | -0.016 | -1.105 | 0.154 | -0.012 | 0.003 | 0.003 | 0.193 | 0.22 | 0.68 | 0.26 | 0.25 | 0.13 | 0.16 |
| 10 | 14010122 | rs2447014 | -0.23 | 0.096 | -1.756 | -0.019 | 0.041 | 0.028 | 0.028 | 0 | 0.53 | 0.91 | 0.57 | 0.34 | 0.62 | 0.49 |
| 10 | 14016290 | rs12358733 | 0.032 | 0.244 | -1.473 | 0.15 | 0.022 | -0.02 | -0.02 | 0 | 0.4 | 0.23 | 0.39 | 0.63 | 0.34 | 0.45 |
| 10 | 14016879 | rs10906555 | 0.027 | 0.209 | -1.203 | 0.16 | 0.017 | -0.002 | -0.002 | 0.308 | 0.41 | 0.34 | 0.4 | 0.62 | 0.34 | 0.45 |
| 10 | 14018232 | rs2926872 | -0.683 | 0.209 | -1.14 | 0.155 | -0.021 | 0.113 | 0.113 | 0.285 | 0.16 | 0.53 | 0.25 | 0.29 | 0.11 | 0.17 |
| 10 | 14018711 | rs4237437 | 0.02 | 0.221 | -1.12 | 0.153 | 0.012 | -0.001 | -0.001 | 0 | 0.41 | 0.37 | 0.4 | 0.62 | 0.34 | 0.47 |
| 10 | 14019456 | rs1963965 | 1.053 | 0.147 | -1.029 | 0.132 | -0.014 | 0.176 | 0.176 | 0 | 0.88 | 0.47 | 0.76 | 0.73 | 0.94 | 0.83 |
| 10 | 14021822 | rs10508470 | 0.195 | 0.152 | -1.179 | 0.245 | 0 | -0.082 | -0.082 | 0 | 0.38 | 0.07 | 0.37 | 0.62 | 0.32 | 0.44 |
| 10 | 14029701 | rs11258774 | -0.021 | 0.282 | -1.071 | 0.23 | 0.026 | 0.054 | 0.054 | 0 | 0.42 | 0.26 | 0.45 | 0.74 | 0.35 | 0.56 |
| 10 | 14029742 | rs11258775 | -0.021 | 0.282 | -1.071 | 0.23 | 0.026 | 0.054 | 0.054 | 0 | 0.42 | 0.26 | 0.45 | 0.74 | 0.35 | 0.56 |
| 10 | 14031722 | rs17314739 | -0.056 | 0.053 | -1.308 | 0.102 | -0.008 | 0.022 | 0.022 | 0.385 | 0.07 | 0.03 | 0.05 | 0.01 | 0.08 | 0.04 |
| 10 | 14032199 | rs2447011 | -0.451 | -0.046 | -1.164 | 0.005 | 0.05 | 0.195 | 0.195 | 0 | 0.48 | 0.74 | 0.61 | 0.92 | 0.39 | 0.61 |
| 10 | 14035266 | rs6602695 | -0.295 | -0.046 | -1.161 | 0.155 | -0.002 | -0.006 | -0.006 | 0 | 0.08 | 0.09 | 0.05 | 0.01 | 0.1 | 0.04 |
| 10 | 14038779 | rs1574584 | -1.117 | -0.038 | -1.44 | 0.146 | -0.069 | 0.223 | 0.223 | 0.262 | 0.58 | 0.87 | 0.66 | 0.94 | 0.48 | 0.66 |
| 10 | 14040425 | rs4329578 | -0.183 | -0.059 | -1.198 | 0.272 | 0.036 | 0.044 | 0.044 | 0 | 0.39 | 0.18 | 0.42 | 0.74 | 0.31 | 0.53 |
| 10 | 14047923 | rs1841557 | -0.997 | 0.111 | -0.217 | 0.736 | -0.004 | 0.049 | 0.049 | 0 | 0.21 | 0.5 | 0.28 | 0.28 | 0.16 | 0.28 |
| 10 | 14050695 | rs2292368 | 0.268 | -0.364 | -0.808 | 0.759 | 0.006 | -0.107 | -0.107 | 0.676 | 0.24 | 0.08 | 0.21 | 0.45 | 0.25 | 0.26 |
| 10 | 14051456 | rs2457837 | -1.188 | -0.135 | -0.81 | 0.597 | -0.004 | -0.025 | -0.025 | 0.458 | 0.24 | 0.77 | 0.4 | 0.38 | 0.18 | 0.28 |
| 10 | 14052850 | rs12777331 | 1.586 | -0.353 | -0.988 | 0.45 | -0.11 | 0.247 | 0.247 | 0 | 0.42 | 0.1 | 0.34 | 0.08 | 0.51 | 0.36 |
| 10 | 14053187 | rs2446588 | 1.436 | -0.365 | -0.217 | 0.458 | 0 | 0.211 | 0.211 | 1.735 | 0.83 | 0.5 | 0.63 | 0.62 | 0.87 | 0.77 |
| 10 | 14053253 | rs10906566 | 1.382 | -0.33 | -0.456 | 0.434 | -0.073 | 0.214 | 0.214 | 0 | 0.45 | 0.1 | 0.34 | 0.17 | 0.53 | 0.41 |
| 10 | 14054714 | rs1870745 | -0.864 | -0.271 | -0.111 | 0.487 | -0.002 | 0.019 | 0.019 | 0 | 0.22 | 0.62 | 0.3 | 0.29 | 0.17 | 0.25 |
| 10 | 14057498 | rs2447016 | 0.09 | -0.414 | -0.129 | 0.459 | 0.012 | -0.022 | -0.022 | 0 | 0.67 | 0.79 | 0.65 | 0.37 | 0.7 | 0.62 |
| 10 | 14060679 | rs2457857 | -0.994 | -0.381 | -0.027 | 0.502 | -0.004 | 0.015 | 0.015 | 0 | 0.23 | 0.64 | 0.3 | 0.29 | 0.18 | 0.26 |
| 10 | 14062413 | rs2447015 | 0.695 | -0.345 | -0.06 | 0.474 | 0 | 0 | 0 | 0.041 | 0.9 | 0.89 | 0.86 | 0.89 | 0.94 | 0.93 |
| 10 | 14064208 | rs2446571 | -0.085 | -0.18 | 0.06 | 0.487 | 0.023 | -0.097 | -0.097 | 0 | 0.14 | 0.35 | 0.06 | 0.01 | 0.09 | 0.09 |
| 10 | 14064519 | rs10906573 | 0.082 | -0.137 | -0.617 | 0.769 | 0.004 | -0.102 | -0.102 | 0.467 | 0.25 | 0.1 | 0.21 | 0.53 | 0.24 | 0.3 |
| 10 | 14070082 | rs10219103 | -0.062 | 0.019 | 0.296 | 0.322 | 0.01 | -0.07 | -0.07 | 0 | 0.13 | 0.3 | 0.07 | 0.01 | 0.09 | 0.08 |
| 10 | 14070693 | rs1599942 | 1.309 | -0.062 | 0.412 | 0.538 | -0.015 | 0.027 | 0.027 | 0 | 0.44 | 0.24 | 0.46 | 0.28 | 0.51 | 0.48 |
| 10 | 14070935 | rs1967380 | -0.143 | -0.092 | 0.835 | 0.618 | 0.013 | -0.041 | -0.041 | 0 | 0.15 | 0.46 | 0.13 | 0.09 | 0.09 | 0.11 |
| 10 | 14071061 | rs2447038 | 0.863 | -0.101 | 0.688 | 0.612 | -0.009 | 0.02 | 0.02 | 4.864 | 0.86 | 0.87 | 0.83 | 0.93 | 0.84 | 0.93 |
| 10 | 14076153 | rs12245506 | -1.049 | -0.114 | 0.717 | 0.713 | 0.014 | -0.032 | -0.032 | 0 | 0.23 | 0.34 | 0.23 | 0.15 | 0.22 | 0.16 |
| 10 | 14079996 | rs4508098 | 0.56 | -0.209 | 0.567 | 0.761 | 0.002 | -0.088 | -0.088 | 0 | 0.24 | 0.06 | 0.21 | 0.42 | 0.24 | 0.27 |
| 10 | 14085835 | rs922242 | -0.302 | -0.164 | 0.956 | 0.623 | -0.003 | -0.019 | -0.019 | 0.834 | 0.31 | 0.25 | 0.28 | 0.45 | 0.35 | 0.29 |
| 10 | 14086579 | rs7099081 | -0.654 | -0.203 | 1.113 | 0.625 | -0.002 | 0.029 | 0.029 | 0 | 0.34 | 0.42 | 0.31 | 0.57 | 0.35 | 0.33 |
| 10 | 14088837 | rs11258827 | -1.516 | -0.158 | 1.151 | 0.567 | 0.09 | 0.022 | 0.022 | 0.54 | 0.11 | 0.39 | 0.1 | 0.13 | 0.13 | 0.08 |
| 10 | 14089451 | rs12250709 | 0.227 | -0.168 | 1.071 | 0.549 | 0.004 | -0.066 | -0.066 | 2.309 | 0.14 | 0.29 | 0.15 | 0.03 | 0.1 | 0.08 |
| 10 | 14093180 | rs11258834 | 1.502 | -0.351 | 0.809 | 0.627 | -0.024 | 0.13 | 0.13 | 0 | 0.44 | 0.21 | 0.36 | 0.15 | 0.5 | 0.43 |
| 10 | 14093742 | rs7902073 | -0.561 | -0.354 | 0.837 | 0.732 | -0.019 | 0.017 | 0.017 | 0.255 | 0.06 | 0.12 | 0.04 | 0.14 | 0.02 | 0.06 |
| 10 | 14100104 | rs10906590 | 1.139 | 0.002 | 0.895 | 0.747 | -0.003 | 0.076 | 0.076 | 0 | 0.5 | 0.34 | 0.4 | 0.3 | 0.53 | 0.5 |
| 10 | 14101367 | rs4750458 | 0.177 | -0.024 | 0.983 | 0.72 | 0.012 | 0.056 | 0.056 | 0 | 0.82 | 0.6 | 0.69 | 0.77 | 0.86 | 0.78 |
| 10 | 14103607 | rs7099622 | -0.474 | -0.108 | 0.938 | 0.653 | -0.001 | -0.009 | -0.009 | 0 | 0.33 | 0.27 | 0.3 | 0.6 | 0.34 | 0.32 |
| 10 | 14103897 | rs11258844 | 0.255 | -0.197 | 0.811 | 0.649 | 0 | -0.066 | -0.066 | 0.604 | 0.14 | 0.29 | 0.16 | 0.03 | 0.1 | 0.09 |
| 10 | 14106314 | rs7079799 | 0.728 | -0.061 | 0.53 | 0.734 | -0.021 | -0.14 | -0.14 | 0 | 0.22 | 0.02 | 0.2 | 0.43 | 0.21 | 0.24 |
| 10 | 14106484 | rs7904460 | -0.705 | -0.065 | 0.657 | 0.727 | 0.015 | -0.034 | -0.034 | 0 | 0.75 | 0.83 | 0.78 | 0.55 | 0.79 | 0.74 |
| 10 | 14108711 | rs11258848 | 1.2 | 0.159 | -0.341 | -0.051 | -0.03 | 0.088 | 0.088 | 0.041 | 0.22 | 0.01 | 0.17 | 0.04 | 0.23 | 0.3 |
| 10 | 14117470 | rs11813322 | -0.254 | 0.165 | 0.193 | 0.461 | -0.018 | 0.019 | 0.019 | 0.082 | 0.05 | 0.1 | 0.03 | 0.14 | 0.02 | 0.06 |
| 10 | 14120334 | rs12257760 | 0.278 | 0.036 | -0.031 | 0.262 | -0.021 | -0.098 | -0.098 | 0 | 0.16 | 0.39 | 0.14 | 0.03 | 0.08 | 0.09 |

| | | | | | | | | | | | | | | | | |
|---|---|---|---|---|---|---|---|---|---|---|---|---|---|---|---|---|
| 10 | 14121089 | rs7920328 | -0.726 | -0.034 | -0.072 | 0.197 | -0.036 | -0.125 | -0.125 | 0.574 | 0.22 | 0.54 | 0.19 | 0.03 | 0.12 | 0.13 |
| 10 | 14123459 | rs2614143 | -1.048 | -0.26 | -0.419 | 0.103 | 0.044 | 0.021 | 0.021 | 0 | 0.47 | 0.7 | 0.49 | 0.63 | 0.36 | 0.41 |
| 10 | 14125252 | rs1359731 | -1.113 | -0.28 | -0.439 | 0.205 | 0.024 | 0.053 | 0.053 | 0 | 0.48 | 0.58 | 0.58 | 0.8 | 0.4 | 0.54 |
| 10 | 14125336 | rs898715 | -0.548 | -0.28 | -0.423 | 0.13 | 0.003 | -0.002 | -0.002 | 0 | 0.41 | 0.54 | 0.45 | 0.59 | 0.34 | 0.37 |
| 10 | 14125653 | rs898716 | 0.472 | -0.262 | -0.595 | 0.139 | 0.008 | -0.026 | -0.026 | 0 | 0.24 | 0.13 | 0.28 | 0.39 | 0.24 | 0.23 |
| 10 | 14129851 | rs2767288 | -1.006 | -0.058 | -0.561 | -0.581 | 0.031 | 0.103 | 0.103 | 0 | 0.49 | 0.64 | 0.58 | 0.85 | 0.41 | 0.55 |
| 10 | 14134826 | rs789767 | -0.41 | 0.16 | -0.942 | -0.414 | 0 | 0.011 | 0.011 | 0 | 0.41 | 0.39 | 0.52 | 0.72 | 0.43 | 0.53 |
| 10 | 14136905 | rs10796153 | 0.937 | -0.279 | -1.568 | -0.902 | 0.065 | 0.107 | 0.107 | 10.497 | 0.74 | 0.11 | 0.79 | 0.93 | 0.83 | 0.91 |
| 10 | 14137678 | rs789761 | 0.277 | 0.826 | -0.862 | -0.86 | 0.002 | -0.012 | -0.012 | 0 | 0.54 | 0.49 | 0.49 | 0.6 | 0.49 | 0.57 |
| 10 | 14137810 | rs789760 | -0.248 | 1.03 | -0.792 | -0.956 | 0.003 | -0.028 | -0.028 | 0.159 | 0.69 | 0.59 | 0.58 | 0.79 | 0.67 | 0.7 |
| 10 | 14145486 | rs9787650 | 1.24 | 1.182 | -0.919 | -0.922 | -0.071 | 0.002 | 0.002 | 0.506 | 0.41 | 0.16 | 0.46 | 0.57 | 0.47 | 0.51 |
| 10 | 14148412 | rs1327003 | -0.123 | 1.092 | -0.079 | -1.013 | 0 | 0 | 0 | 2.832 | 0.63 | 0.67 | 0.57 | 0.76 | 0.68 | 0.66 |
| 10 | 14148481 | rs1327002 | 0.185 | 1.079 | -0.224 | -0.949 | 0.045 | 0.068 | 0.068 | 0.086 | 0.58 | 0.21 | 0.53 | 0.73 | 0.68 | 0.65 |
| 10 | 14151277 | rs17154656 | -1.242 | 1.152 | -0.021 | -0.856 | 0.132 | 0.108 | 0.108 | 2.65 | 0.06 | 0.39 | 0.08 | 0.08 | 0.07 | 0.05 |
| 10 | 14155918 | rs1536450 | 1.221 | 1.164 | -0.285 | -0.909 | -0.005 | -0.009 | -0.009 | 0 | 0.41 | 0.18 | 0.35 | 0.5 | 0.43 | 0.49 |
| 10 | 14157261 | rs1327000 | 0.047 | 1.008 | -0.188 | -0.946 | -0.002 | -0.011 | -0.011 | 0 | 0.6 | 0.61 | 0.51 | 0.75 | 0.61 | 0.63 |
| 10 | 14161557 | rs1887241 | -0.14 | 0.041 | -0.018 | -0.995 | -0.023 | -0.051 | -0.051 | 0 | 0.2 | 0.33 | 0.18 | 0.05 | 0.13 | 0.11 |
| 10 | 14163007 | rs2104550 | -0.895 | 0.185 | -0.047 | -1.051 | -0.028 | 0.002 | 0.002 | 0.024 | 0.74 | 0.89 | 0.62 | 0.8 | 0.72 | 0.75 |
| 10 | 14163016 | rs7908576 | 0.136 | 0.203 | 0.238 | -1.1 | -0.021 | 0.025 | 0.025 | 1.048 | 0.57 | 0.71 | 0.45 | 0.61 | 0.51 | 0.61 |
| 10 | 14163942 | rs10796158 | 0.85 | 0.425 | 0.412 | -0.877 | -0.013 | 0.013 | 0.013 | 0.632 | 0.17 | 0.09 | 0.16 | 0.19 | 0.2 | 0.14 |
| 10 | 14166989 | rs11258878 | 1.359 | 0.369 | 0.764 | -0.934 | 0.01 | 0.027 | 0.027 | 2.65 | 0.33 | 0.52 | 0.27 | 0.54 | 0.35 | 0.47 |
| 10 | 14168048 | rs7906344 | -0.341 | 0.204 | 0.988 | -1.012 | -0.001 | 0.022 | 0.022 | 0.598 | 0.62 | 0.7 | 0.55 | 0.78 | 0.61 | 0.69 |
| 10 | 14168791 | rs12572893 | 1.743 | 0.264 | 1.201 | -0.979 | -0.002 | -0.02 | -0.02 | 1.035 | 0.31 | 0.22 | 0.22 | 0.49 | 0.31 | 0.43 |
| 10 | 14169524 | rs7903598 | -0.531 | 0.285 | 1.184 | -0.975 | -0.01 | 0.041 | 0.041 | 0.479 | 0.63 | 0.79 | 0.56 | 0.78 | 0.61 | 0.68 |
| 10 | 14172634 | rs11258880 | 1.829 | 0.297 | 1.184 | -0.959 | -0.001 | 0 | 0 | 2.362 | 0.31 | 0.27 | 0.23 | 0.49 | 0.31 | 0.42 |
| 10 | 14173658 | rs10906611 | -0.739 | 0.294 | 1.299 | -0.872 | -0.004 | -0.018 | -0.018 | 0 | 0.28 | 0.35 | 0.31 | 0.26 | 0.26 | 0.21 |
| 10 | 14182045 | rs9663251 | 0.113 | 0.036 | 1.526 | -0.773 | -0.002 | -0.014 | -0.014 | 0 | 0.58 | 0.45 | 0.54 | 0.75 | 0.58 | 0.67 |
| 10 | 14183105 | rs2400042 | 1.527 | 0.091 | 1.472 | -0.81 | -0.024 | -0.014 | -0.014 | 0 | 0.36 | 0.22 | 0.28 | 0.57 | 0.39 | 0.51 |
| 10 | 14183371 | rs11258894 | 0.46 | -0.054 | 1.374 | -0.884 | 0.001 | 0.009 | 0.009 | 0.57 | 0.48 | 0.58 | 0.38 | 0.6 | 0.42 | 0.56 |
| 10 | 14184348 | rs4748087 | -0.519 | -0.058 | 1.239 | -0.884 | -0.013 | 0.032 | 0.032 | 0.614 | 0.65 | 0.82 | 0.58 | 0.78 | 0.62 | 0.7 |
| 10 | 14184558 | rs7088074 | 1.209 | 0.059 | 0.938 | -0.824 | -0.007 | -0.034 | -0.034 | 0.599 | 0.38 | 0.25 | 0.25 | 0.49 | 0.36 | 0.48 |
| 10 | 14186840 | rs11258899 | -0.904 | 0.19 | 0.979 | -0.714 | -0.018 | -0.071 | -0.071 | 1.085 | 0.14 | 0.47 | 0.14 | 0.06 | 0.08 | 0.09 |
| 10 | 14193778 | rs12765358 | -0.349 | 0.446 | 0.184 | -0.594 | -0.001 | -0.005 | -0.005 | 1.454 | 0.23 | 0.21 | 0.27 | 0.21 | 0.22 | 0.19 |
| 10 | 14202285 | rs10752333 | 1.074 | 0.356 | -0.636 | -0.224 | -0.055 | -0.104 | -0.104 | 0 | 0.32 | 0.07 | 0.22 | 0.48 | 0.33 | 0.47 |
| 10 | 14208510 | rs10796164 | 0.437 | 0.567 | 0.031 | -0.229 | 0.021 | -0.028 | -0.028 | 0 | 0.64 | 0.5 | 0.58 | 0.72 | 0.54 | 0.71 |
| 10 | 14212066 | rs10906620 | -0.371 | 0.394 | 0.388 | 0.041 | -0.005 | 0.003 | 0.003 | 0 | 0.32 | 0.25 | 0.36 | 0.32 | 0.39 | 0.26 |
| 10 | 14218291 | rs11258918 | 0.75 | -0.096 | 0.87 | 0.129 | 0 | -0.016 | -0.016 | 0 | 0.58 | 0.2 | 0.55 | 0.53 | 0.57 | 0.6 |
| 10 | 14224687 | rs7901509 | -0.3 | -0.06 | 1.466 | 0.129 | 0.008 | -0.028 | -0.028 | 0.293 | 0.32 | 0.36 | 0.37 | 0.23 | 0.37 | 0.21 |
| 10 | 14225797 | rs10906624 | -0.659 | 0.088 | 1.309 | 0.128 | -0.01 | 0.008 | 0.008 | 0 | 0.35 | 0.35 | 0.38 | 0.46 | 0.31 | 0.38 |
| 10 | 14227908 | rs7905928 | -0.639 | 0.091 | 1.217 | 0.791 | -0.001 | 0.002 | 0.002 | 0 | 0.36 | 0.38 | 0.43 | 0.53 | 0.39 | 0.39 |
| 10 | 14230235 | rs6602713 | -1.033 | 0.158 | 0.944 | 0.779 | -0.003 | -0.043 | -0.043 | 0 | 0.41 | 0.78 | 0.48 | 0.53 | 0.42 | 0.41 |
| 10 | 14231954 | rs17154811 | 0.338 | -0.073 | 0.433 | 0.31 | 0.009 | 0.016 | 0.016 | 0.474 | 0.18 | 0.12 | 0.11 | 0.01 | 0.16 | 0.11 |
| 10 | 14234352 | rs7068441 | -0.05 | 0.081 | 0.72 | 0.701 | 0 | 0.008 | 0.008 | 0 | 0.29 | 0.21 | 0.34 | 0.23 | 0.3 | 0.21 |
| 10 | 14237503 | rs7080759 | 0.077 | -0.042 | 0.595 | 0.705 | 0.002 | 0.018 | 0.018 | 0.1 | 0.28 | 0.16 | 0.32 | 0.22 | 0.29 | 0.19 |
| 10 | 14238978 | rs10906625 | 0.499 | -0.061 | 0.204 | 0.833 | 0.009 | -0.033 | -0.033 | 0 | 0.6 | 0.18 | 0.57 | 0.42 | 0.59 | 0.63 |
| 10 | 14249417 | rs7078954 | -0.62 | 0.203 | 0.485 | 0.91 | -0.001 | 0.002 | 0.002 | 0 | 0.34 | 0.36 | 0.38 | 0.45 | 0.36 | 0.26 |
| 10 | 14251892 | rs4750489 | -0.236 | 0.22 | 0.459 | 0.948 | 0.001 | 0.002 | 0.002 | 0 | 0.29 | 0.28 | 0.36 | 0.24 | 0.29 | 0.2 |
| 10 | 14258278 | rs7095401 | -0.254 | 0.22 | 0.291 | 0.888 | -0.009 | 0.016 | 0.016 | 0 | 0.29 | 0.21 | 0.37 | 0.28 | 0.36 | 0.2 |
| 10 | 14260500 | rs11258947 | 1.442 | -0.1 | -0.187 | 0.711 | -0.121 | -0.073 | -0.073 | 0.249 | 0.4 | 0.09 | 0.41 | 0.42 | 0.41 | 0.52 |
| 10 | 14262019 | rs1983907 | -0.274 | 0.247 | 0.246 | 0.818 | -0.002 | 0.008 | 0.008 | 0 | 0.3 | 0.29 | 0.37 | 0.27 | 0.36 | 0.2 |
| 10 | 14266085 | rs2895596 | 1.865 | 0.481 | 0.99 | 1.138 | 0.091 | 0.271 | 0.271 | 1.162 | 0.9 | 0.52 | 0.91 | 0.77 | 0.84 | 0.88 |
| 10 | 14266643 | rs4750493 | -1.291 | 0.399 | 0.562 | 1.262 | -0.001 | 0.032 | 0.032 | 0 | 0.06 | 0.08 | 0.11 | 0.17 | 0.05 | 0.17 |
| 10 | 14266893 | rs4750494 | -0.984 | 0.485 | 0.636 | 1.096 | -0.001 | 0.001 | 0.001 | 0.215 | 0.53 | 0.48 | 0.51 | 0.51 | 0.51 | 0.52 |
| 10 | 14268106 | rs12219502 | -0.659 | 0.148 | 0.119 | 1 | 0.012 | -0.062 | -0.062 | 0 | 0.21 | 0.06 | 0.18 | 0.23 | 0.19 | 0.26 |
| 10 | 14272308 | rs11258957 | 1.299 | 0.325 | 0.676 | 1.086 | -0.043 | -0.004 | -0.004 | 0.018 | 0.38 | 0.24 | 0.44 | 0.4 | 0.44 | 0.46 |
| 10 | 14272502 | rs1879606 | 1.475 | 0.945 | 0.513 | 1.004 | -0.025 | -0.025 | -0.025 | 7.007 | 0.35 | 0.2 | 0.35 | 0.41 | 0.37 | 0.44 |
| 10 | 14275726 | rs7068964 | 1.4 | 0.033 | 1.007 | 0.904 | -0.056 | -0.028 | -0.028 | 0 | 0.35 | 0.2 | 0.42 | 0.39 | 0.43 | 0.45 |
| 10 | 14276055 | rs11258961 | -0.024 | -0.107 | 1.053 | 0.914 | 0.014 | -0.016 | -0.016 | 0 | 0.1 | 0.06 | 0.11 | 0.15 | 0.05 | 0.19 |
| 10 | 14276466 | rs10906633 | 1.55 | -0.072 | 1.198 | 0.888 | -0.089 | -0.05 | -0.05 | 0 | 0.34 | 0.15 | 0.41 | 0.39 | 0.43 | 0.45 |
| 10 | 14276664 | rs7902063 | 0.465 | -0.525 | 1.387 | 0.818 | 0.006 | 0.011 | 0.011 | 0.069 | 0.32 | 0.22 | 0.31 | 0.31 | 0.29 | 0.27 |
| 10 | 14277868 | rs7914612 | -0.53 | -0.512 | 1.291 | 0.823 | -0.014 | -0.024 | -0.024 | 0 | 0.66 | 0.53 | 0.68 | 0.7 | 0.7 | 0.72 |
| 10 | 14284219 | rs1218400 | 1.036 | -0.135 | 1.16 | 0.802 | -0.003 | -0.006 | -0.006 | 0 | 0.46 | 0.37 | 0.47 | 0.47 | 0.52 | 0.52 |
| 10 | 14284474 | rs1218397 | 1.218 | -0.149 | 1.205 | 0.796 | -0.017 | -0.022 | -0.022 | 0 | 0.44 | 0.26 | 0.46 | 0.47 | 0.51 | 0.52 |
| 10 | 14284566 | rs11258974 | 0.226 | -0.19 | 0.988 | 0.748 | 0.015 | -0.02 | -0.02 | 0 | 0.09 | 0.03 | 0.06 | 0.1 | 0.05 | 0.13 |
| 10 | 14285913 | rs1218386 | 0.682 | -0.188 | 1.233 | 0.742 | -0.017 | 0.007 | 0.007 | 0.003 | 0.8 | 0.68 | 0.86 | 0.8 | 0.86 | 0.8 |
| 10 | 14286606 | rs1218385 | 1.246 | -0.16 | 1.136 | 0.748 | -0.022 | -0.007 | -0.007 | 0.383 | 0.74 | 0.51 | 0.82 | 0.8 | 0.8 | 0.79 |
| 10 | 14288652 | rs1218379 | -0.136 | -0.081 | 0.876 | 0.816 | -0.002 | -0.001 | -0.001 | 0 | 0.37 | 0.32 | 0.4 | 0.39 | 0.38 | 0.33 |
| 10 | 14292871 | rs12260956 | 0.455 | -0.122 | 0.38 | 0.883 | 0.002 | -0.006 | -0.006 | 1.305 | 0.08 | 0.06 | 0.05 | 0.1 | 0.1 | 0.12 |
| 10 | 14293525 | rs10796175 | 0.489 | -0.24 | 0.839 | 0.838 | 0.001 | 0 | 0 | 0 | 0.3 | 0.26 | 0.35 | 0.31 | 0.27 | 0.27 |

| | | | | | | | | | | | | | | | | |
|---|---|---|---|---|---|---|---|---|---|---|---|---|---|---|---|---|
| 10 | 14299518 | rs1717516 | -1.567 | -0.315 | 0.983 | 0.791 | -0.02 | -0.022 | -0.022 | 0 | 0.25 | 0.54 | 0.19 | 0.18 | 0.21 | 0.15 |
| 10 | 14302366 | rs11258982 | -0.036 | -0.373 | 1.073 | 0.791 | 0.003 | -0.001 | -0.001 | 1.591 | 0.1 | 0.12 | 0.07 | 0.1 | 0.05 | 0.13 |
| 10 | 14306078 | rs1218335 | -2.042 | -0.354 | 1.202 | 0.778 | -0.021 | 0.003 | 0.003 | 2.418 | 0.3 | 0.6 | 0.2 | 0.28 | 0.24 | 0.27 |
| 10 | 14307461 | rs1218340 | -0.179 | -0.309 | 1.173 | 0.77 | 0.012 | 0.021 | 0.021 | 0 | 0.11 | 0.36 | 0.08 | 0.08 | 0.09 | 0.02 |
| 10 | 14307895 | rs4636552 | 1.357 | -0.303 | 1.066 | 0.782 | -0.015 | -0.023 | -0.023 | 0 | 0.07 | 0 | 0.05 | 0.08 | 0.1 | 0.07 |
| 10 | 14309999 | rs1218342 | -0.726 | -0.289 | 1.193 | 0.777 | 0.035 | -0.01 | -0.01 | 2.098 | 0.14 | 0.37 | 0.12 | 0.08 | 0.16 | 0.03 |
| 10 | 14311326 | rs7088164 | -0.325 | -0.328 | 1.099 | 0.784 | -0.023 | -0.002 | -0.002 | 0 | 0.11 | 0.2 | 0.08 | 0.1 | 0.05 | 0.12 |
| 10 | 14311615 | rs1218343 | -1.129 | -0.338 | 1.008 | 0.79 | -0.003 | 0.001 | 0.001 | 0.455 | 0.52 | 0.66 | 0.52 | 0.47 | 0.53 | 0.36 |
| 10 | 14312249 | rs1218345 | 0.053 | -0.314 | 1.03 | 0.81 | 0.004 | 0.002 | 0.002 | 0 | 0.34 | 0.62 | 0.39 | 0.32 | 0.3 | 0.27 |
| 10 | 14312775 | rs10906641 | -0.134 | -0.171 | 0.742 | 0.851 | 0.021 | -0.019 | -0.019 | 0 | 0.1 | 0.06 | 0.06 | 0.13 | 0.04 | 0.17 |
| 10 | 14318441 | rs1218353 | -1.197 | 0.047 | 0.885 | 0.751 | 0.02 | -0.012 | -0.012 | 0 | 0.21 | 0.43 | 0.16 | 0.15 | 0.25 | 0.09 |
| 10 | 14320952 | rs11258991 | 0.644 | -0.196 | 0.417 | 0.943 | -0.005 | -0.011 | -0.011 | 0 | 0.07 | 0.03 | 0.05 | 0.08 | 0.07 | 0.01 |
| 10 | 14321141 | rs12219057 | -0.356 | -0.359 | 0.252 | 0.791 | -0.005 | -0.055 | -0.055 | 0 | 0.42 | 0.17 | 0.44 | 0.47 | 0.44 | 0.34 |
| 10 | 14322668 | rs1218366 | -1.511 | -0.469 | 0.333 | 0.871 | -0.015 | -0.018 | -0.018 | 0.019 | 0.58 | 0.8 | 0.56 | 0.56 | 0.51 | 0.47 |
| 10 | 14322763 | rs10906645 | -0.208 | -0.707 | 0.344 | 0.627 | 0.016 | -0.019 | -0.019 | 0 | 0.09 | 0.02 | 0.06 | 0.1 | 0.05 | 0.12 |
| 10 | 14327174 | rs1218370 | -1.331 | -0.686 | 0.353 | 0.839 | 0.004 | 0.028 | 0.028 | 0.296 | 0.54 | 0.76 | 0.5 | 0.46 | 0.49 | 0.54 |
| 10 | 14327976 | rs1235022 | -1.099 | -0.721 | 0.496 | 0.777 | 0.014 | 0.074 | 0.074 | 0.162 | 0.52 | 0.72 | 0.5 | 0.39 | 0.49 | 0.47 |
| 10 | 14332431 | rs11258995 | -0.056 | -1.249 | 0.749 | 0.55 | -0.013 | 0.016 | 0.016 | 0 | 0.27 | 0.17 | 0.34 | 0.25 | 0.38 | 0.26 |
| 10 | 14333240 | rs12360009 | -0.243 | -1.269 | 0.848 | 0.546 | -0.005 | 0.006 | 0.006 | 0 | 0.28 | 0.22 | 0.36 | 0.28 | 0.38 | 0.28 |
| 10 | 14336842 | rs1218458 | -0.748 | -1.534 | 1.018 | 0.56 | -0.096 | -0.076 | -0.076 | 0 | 0.25 | 0.54 | 0.15 | 0.11 | 0.12 | 0.2 |
| 10 | 14340708 | rs867970 | -0.175 | -1.584 | 1.206 | 0.562 | 0.06 | 0.043 | 0.043 | 2.699 | 0.22 | 0.15 | 0.12 | 0.12 | 0.11 | 0.19 |
| 10 | 14342356 | rs4750506 | -0.231 | -1.508 | 1.376 | 0.608 | -0.011 | 0.001 | 0.001 | 0 | 0.28 | 0.19 | 0.35 | 0.35 | 0.37 | 0.35 |
| 10 | 14342616 | rs1755153 | -1.935 | -1.619 | 1.563 | 0.582 | 0.018 | 0.024 | 0.024 | 0.324 | 0.54 | 0.74 | 0.5 | 0.46 | 0.48 | 0.54 |
| 10 | 14343055 | rs10752336 | -0.202 | -1.587 | 1.367 | 0.594 | -0.009 | 0.001 | 0.001 | 0 | 0.28 | 0.19 | 0.35 | 0.35 | 0.37 | 0.35 |
| 10 | 14345658 | rs1218471 | -0.751 | -1.565 | 1.541 | 0.585 | -0.046 | -0.031 | -0.031 | 0.028 | 0.25 | 0.38 | 0.14 | 0.12 | 0.11 | 0.19 |
| 10 | 14349016 | rs7898492 | -0.942 | -1.563 | 1.412 | 0.591 | -0.003 | 0.001 | 0.001 | 0.545 | 0.27 | 0.29 | 0.15 | 0.21 | 0.15 | 0.32 |
| 10 | 14352808 | rs2480547 | 0.193 | -1.465 | 1.07 | 0.641 | 0.046 | 0.026 | 0.026 | 2.01 | 0.76 | 0.81 | 0.87 | 0.83 | 0.87 | 0.79 |
| 10 | 14353949 | rs10906649 | 1.494 | -1.531 | 1.128 | 0.683 | 0.013 | 0.012 | 0.012 | 0.249 | 0.45 | 0.27 | 0.49 | 0.52 | 0.51 | 0.46 |
| 10 | 14354960 | rs11259005 | 0.358 | -1.475 | 0.835 | 0.698 | -0.025 | 0.024 | 0.024 | 0 | 0.22 | 0.09 | 0.32 | 0.21 | 0.32 | 0.21 |
| 10 | 14361028 | rs1218427 | 0.925 | -1.497 | 0.882 | 0.705 | 0.048 | 0.036 | 0.036 | 0 | 0.52 | 0.72 | 0.54 | 0.55 | 0.55 | 0.53 |
| 10 | 14365113 | rs1218436 | 0.334 | -1.514 | 0.888 | 0.662 | 0.016 | 0.013 | 0.013 | 2.003 | 0.75 | 0.75 | 0.84 | 0.83 | 0.84 | 0.79 |
| 10 | 14365504 | rs1018664 | 1.484 | -1.43 | 1.177 | 0.649 | 0.01 | 0.012 | 0.012 | 0.093 | 0.46 | 0.28 | 0.5 | 0.52 | 0.51 | 0.46 |
| 10 | 14366146 | rs1218439 | 0.091 | -1.462 | 0.677 | 0.59 | 0.016 | 0.009 | 0.009 | 1.697 | 0.77 | 0.75 | 0.86 | 0.8 | 0.87 | 0.76 |
| 10 | 14367596 | rs17251862 | 0.531 | -1.157 | 0.359 | 0.673 | -0.085 | 0.029 | 0.029 | 0 | 0.21 | 0.02 | 0.28 | 0.16 | 0.3 | 0.16 |
| 10 | 14374370 | rs7077526 | 1.42 | -1.09 | 0.187 | 1.383 | -0.075 | 0.005 | 0.005 | 0.355 | 0.41 | 0.12 | 0.46 | 0.3 | 0.47 | 0.46 |
| 10 | 14375047 | rs7079603 | 0.714 | -0.927 | 0.008 | 1.505 | -0.03 | -0.046 | -0.046 | 0 | 0.08 | 0.01 | 0.06 | 0.21 | 0.11 | 0.03 |
| 10 | 14375259 | rs7082219 | -0.617 | -0.994 | 0.093 | 1.33 | 0.013 | -0.052 | -0.052 | 0 | 0.33 | 0.13 | 0.26 | 0.41 | 0.3 | 0.31 |
| 10 | 14377870 | rs1755117 | 0.735 | -0.987 | 0.472 | 1.067 | 0.064 | 0.109 | 0.109 | 0.797 | 0.95 | 0.58 | 0.95 | 0.95 | 1 | 0.96 |
| 10 | 14385164 | rs11259022 | 0.501 | -0.716 | -0.704 | 1.137 | -0.015 | -0.02 | -0.02 | 0.016 | 0.17 | 0.06 | 0.21 | 0.21 | 0.21 | 0.17 |
| 10 | 14391273 | rs4750511 | -0.888 | -0.531 | -0.592 | 1.391 | 0.021 | 0.046 | 0.046 | 0 | 0.51 | 0.32 | 0.48 | 0.67 | 0.52 | 0.47 |
| 10 | 14393233 | rs603261 | -0.748 | -1.255 | -1.321 | 0.908 | 0.069 | 0.014 | 0.014 | 0 | 0.55 | 0.76 | 0.7 | 0.57 | 0.69 | 0.54 |
| 10 | 14395411 | rs584933 | 0.638 | -0.898 | -1.683 | 1.028 | -0.002 | -0.012 | -0.012 | 0 | 0.09 | 0.04 | 0.07 | 0.15 | 0.11 | 0.02 |
| 10 | 14396207 | rs10906653 | 0.481 | -1.284 | -2.115 | 0.78 | -0.173 | 0.002 | 0.002 | 0 | 0.28 | 0.07 | 0.41 | 0.19 | 0.43 | 0.25 |
| 10 | 14396968 | rs11259027 | 1.778 | -1.325 | -1.907 | 0.614 | 0.011 | 0.031 | 0.031 | 0 | 0.15 | 0.07 | 0.08 | 0 | 0.14 | 0.13 |
| 10 | 14398225 | rs7083480 | 0.7 | -1.485 | -1.797 | 1.284 | -0.001 | -0.078 | -0.078 | 0 | 0.24 | 0.07 | 0.19 | 0.35 | 0.21 | 0.3 |
| 10 | 14401257 | rs684097 | -0.062 | -1.412 | -1.4 | 1.102 | 0.009 | -0.021 | -0.021 | 5.852 | 0.35 | 0.27 | 0.32 | 0.53 | 0.31 | 0.44 |
| 10 | 14404393 | rs11819533 | 0.483 | 0.125 | -1.668 | 0.834 | -0.038 | -0.025 | -0.025 | 0 | 0.15 | 0.1 | 0.25 | 0.17 | 0.27 | 0.2 |
| 10 | 14404442 | rs638707 | -0.483 | -0.359 | -1.538 | 0.633 | 0.006 | 0 | 0 | 1.408 | 0.57 | 0.44 | 0.66 | 0.56 | 0.75 | 0.43 |
| 10 | 14405718 | rs7908034 | 1.467 | -0.149 | -2.347 | 0.27 | 0.047 | -0.023 | -0.023 | 1.598 | 0.16 | 0.07 | 0.11 | 0.19 | 0.07 | 0.25 |
| 10 | 14405902 | rs17155131 | 1.1 | 0.041 | -2.395 | 0.093 | 0.012 | -0.05 | -0.05 | 0 | 0.13 | 0.2 | 0.13 | 0 | 0.19 | 0.03 |
| 10 | 14406135 | rs12411895 | 1.379 | -0.123 | -2.877 | 0.301 | 0.038 | -0.022 | -0.022 | 8.278 | 0.17 | 0.09 | 0.11 | 0.23 | 0.07 | 0.27 |
| 10 | 14409669 | rs11259035 | 0.687 | 0.717 | -2.828 | -0.101 | -0.027 | 0.078 | 0.078 | 0 | 0.22 | 0.03 | 0.2 | 0.1 | 0.19 | 0.16 |
| 10 | 14411279 | rs621649 | -0.169 | 1.14 | -2.146 | 0.504 | -0.028 | -0.079 | -0.079 | 0 | 0.27 | 0.1 | 0.28 | 0.38 | 0.32 | 0.21 |
| 10 | 14411956 | rs7899569 | -0.05 | 1.372 | -1.38 | 0.106 | -0.023 | 0.02 | 0.02 | 0 | 0.12 | 0.07 | 0.12 | 0 | 0.19 | 0.03 |
| 10 | 14412768 | rs582671 | -0.17 | 0.757 | -1.152 | 0.019 | -0.008 | -0.016 | -0.016 | 0 | 0.55 | 0.53 | 0.69 | 0.74 | 0.67 | 0.55 |
| 10 | 14413667 | rs1475539 | -0.462 | 0.737 | -1.131 | -0.207 | -0.001 | 0.002 | 0.002 | 0 | 0.32 | 0.3 | 0.22 | 0.22 | 0.34 | 0.28 |
| 10 | 14414508 | rs7101108 | -0.463 | 0.749 | -1.135 | -0.206 | -0.002 | 0.002 | 0.002 | 0 | 0.32 | 0.3 | 0.22 | 0.18 | 0.35 | 0.26 |
| 10 | 14414624 | rs7084966 | -0.546 | 0.647 | -0.946 | 0.353 | 0.001 | -0.006 | -0.006 | 0 | 0.33 | 0.38 | 0.27 | 0.26 | 0.35 | 0.28 |
| 10 | 14415617 | rs646646 | -0.752 | 0.507 | -1.639 | 0.75 | 0.002 | -0.011 | -0.011 | 0.02 | 0.6 | 0.36 | 0.61 | 0.58 | 0.61 | 0.64 |
| 10 | 14419282 | rs589345 | -0.382 | 0.483 | -1.653 | 0.992 | 0.008 | -0.009 | -0.009 | 0.182 | 0.55 | 0.31 | 0.55 | 0.5 | 0.59 | 0.58 |
| 10 | 14421577 | rs679295 | -0.436 | 0.461 | -1.289 | 0.99 | 0.005 | 0.006 | 0.006 | 0 | 0.56 | 0.4 | 0.6 | 0.59 | 0.58 | 0.59 |
| 10 | 14422809 | rs11259039 | 0.373 | 0.456 | -1.29 | 0.962 | -0.001 | 0 | 0 | 0 | 0.28 | 0.28 | 0.23 | 0.18 | 0.31 | 0.23 |
| 10 | 14423324 | rs621314 | -0.594 | -0.018 | -1.546 | 0.893 | 0 | 0.011 | 0.011 | 0 | 0.64 | 0.37 | 0.65 | 0.7 | 0.69 | 0.69 |
| 10 | 14426291 | rs685087 | 1.213 | 0.111 | -1.298 | 1.08 | -0.055 | -0.07 | -0.07 | 0.333 | 0.27 | 0.07 | 0.34 | 0.32 | 0.27 | 0.36 |
| 10 | 14426442 | rs10906656 | -0.582 | 0.061 | -1.162 | 0.676 | 0.006 | 0.004 | 0.004 | 0 | 0.43 | 0.39 | 0.34 | 0.38 | 0.44 | 0.37 |
| 10 | 14426467 | rs684202 | 0.565 | -0.044 | -0.908 | 0.713 | 0.001 | -0.018 | -0.018 | 0 | 0.33 | 0.27 | 0.42 | 0.48 | 0.29 | 0.42 |
| 10 | 14426616 | rs10906657 | -0.086 | -0.04 | -0.906 | 0.679 | 0.005 | 0.024 | 0.024 | 0.361 | 0.39 | 0.34 | 0.28 | 0.27 | 0.41 | 0.32 |
| 10 | 14426965 | rs673875 | -1.24 | -0.013 | -0.963 | 0.679 | 0.015 | -0.033 | -0.033 | 0 | 0.72 | 0.61 | 0.69 | 0.8 | 0.71 | 0.75 |
| 10 | 14427216 | rs670201 | 0.526 | -0.051 | -1.127 | 0.612 | -0.067 | -0.106 | -0.106 | 0 | 0.35 | 0.07 | 0.39 | 0.42 | 0.37 | 0.42 |

| | | | | | | | | | | | | | | | | |
|---|---|---|---|---|---|---|---|---|---|---|---|---|---|---|---|---|
| 10 | 14428263 | rs655463 | 1.175 | 0.046 | -1.062 | 0.659 | -0.054 | -0.121 | -0.121 | 0.021 | 0.27 | 0.07 | 0.35 | 0.41 | 0.27 | 0.38 |
| 10 | 14430522 | rs12259015 | 0.813 | -0.174 | -0.738 | 0.322 | -0.015 | -0.001 | -0.001 | 0 | 0.12 | 0.17 | 0.07 | 0.1 | 0.11 | 0.09 |
| 10 | 14430536 | rs10906662 | 0.495 | -0.129 | -0.733 | 0.286 | -0.001 | 0.004 | 0.004 | 0 | 0.28 | 0.27 | 0.21 | 0.17 | 0.31 | 0.22 |
| 10 | 14432890 | rs4268417 | -0.153 | -0.12 | -0.712 | 0.388 | -0.018 | -0.015 | -0.015 | 0 | 0.16 | 0.22 | 0.08 | 0.09 | 0.13 | 0.12 |
| 10 | 14433271 | rs672589 | -0.225 | 0.027 | -0.579 | 0.424 | -0.014 | -0.005 | -0.005 | 0 | 0.86 | 0.73 | 0.92 | 0.91 | 0.87 | 0.88 |
| 10 | 14433308 | rs7094560 | -0.23 | 0.043 | -0.998 | 0.337 | 0.014 | 0.016 | 0.016 | 0 | 0.6 | 0.71 | 0.5 | 0.34 | 0.6 | 0.46 |
| 10 | 14434447 | rs7095877 | 0.175 | 0.001 | -0.778 | 0.336 | -0.013 | -0.006 | -0.006 | 0.279 | 0.14 | 0.25 | 0.08 | 0.11 | 0.13 | 0.12 |
| 10 | 14436227 | rs11259056 | -1.614 | -0.025 | -1.011 | 0.502 | -0.041 | -0.12 | -0.12 | 0 | 0.8 | 0.93 | 0.63 | 0.55 | 0.75 | 0.65 |
| 10 | 14436836 | rs7911501 | 0.338 | -0.037 | -0.816 | 0.286 | 0.017 | -0.03 | -0.03 | 0.45 | 0.07 | 0.19 | 0.03 | 0.01 | 0.06 | 0.07 |
| 10 | 14436946 | rs7900809 | -0.434 | -0.125 | -0.847 | 0.254 | -0.021 | -0.03 | -0.03 | 0.147 | 0.2 | 0.3 | 0.1 | 0.11 | 0.16 | 0.16 |
| 10 | 14437100 | rs10906663 | 0.292 | -0.606 | -0.683 | -0.285 | -0.003 | -0.024 | -0.024 | 0.573 | 0.45 | 0.5 | 0.43 | 0.24 | 0.54 | 0.35 |
| 10 | 14438913 | rs789680 | -0.719 | -0.42 | -0.483 | -0.412 | 0.007 | -0.052 | -0.052 | 2.486 | 0.63 | 0.51 | 0.71 | 0.85 | 0.65 | 0.72 |
| 10 | 14439216 | rs1925935 | 1.041 | -0.423 | -0.442 | -0.364 | 0.012 | -0.004 | -0.004 | 0.984 | 0.14 | 0.15 | 0.16 | 0.11 | 0.23 | 0.13 |
| 10 | 14441716 | rs11259063 | -0.668 | -0.389 | -0.523 | -0.483 | 0.002 | 0.002 | 0.002 | 0.03 | 0.42 | 0.41 | 0.33 | 0.25 | 0.39 | 0.34 |
| 10 | 14444837 | rs590830 | -1.458 | -0.173 | -0.503 | -0.111 | -0.005 | 0.024 | 0.024 | 0 | 0.94 | 0.92 | 0.98 | 0.89 | 0.96 | 0.95 |
| 10 | 14445111 | rs10796185 | -0.896 | -0.199 | 0.001 | -0.214 | 0 | -0.002 | -0.002 | 0 | 0.71 | 0.71 | 0.58 | 0.41 | 0.7 | 0.58 |
| 10 | 14447060 | rs621480 | 0.157 | -0.229 | 0.073 | -0.292 | 0.012 | 0 | 0 | 0 | 0.39 | 0.47 | 0.56 | 0.59 | 0.49 | 0.52 |
| 10 | 14449211 | rs17155248 | 0.91 | -0.401 | 0.029 | -0.332 | -0.015 | -0.022 | -0.022 | 0.116 | 0.13 | 0.21 | 0.07 | 0.07 | 0.12 | 0.09 |
| 10 | 14452480 | rs2480536 | 0.446 | -0.433 | -0.135 | -0.318 | -0.006 | 0.01 | 0.01 | 0 | 0.34 | 0.43 | 0.45 | 0.52 | 0.35 | 0.45 |
| 10 | 14452543 | rs2480538 | 1.726 | -0.326 | -0.441 | -0.267 | -0.014 | -0.103 | -0.103 | 0 | 0.21 | 0.09 | 0.37 | 0.46 | 0.24 | 0.35 |
| 10 | 14458385 | rs613311 | -1.658 | -0.471 | -0.239 | -0.335 | -0.001 | -0.012 | -0.012 | 0 | 0.95 | 0.94 | 0.98 | 1 | 0.97 | 0.98 |
| 10 | 14458582 | rs880853 | -0.013 | -0.515 | -0.332 | -0.126 | 0.007 | 0.03 | 0.03 | 0.291 | 0.43 | 0.33 | 0.34 | 0.25 | 0.33 | 0.34 |
| 10 | 14464047 | rs1325896 | -0.023 | -0.497 | -0.333 | -0.145 | 0.002 | 0.036 | 0.036 | 0.267 | 0.43 | 0.33 | 0.36 | 0.25 | 0.36 | 0.33 |
| 10 | 14465110 | rs7080073 | 0.663 | -0.483 | -0.477 | 0.058 | -0.003 | 0 | 0 | 0.336 | 0.15 | 0.14 | 0.11 | 0.1 | 0.05 | 0.11 |
| 10 | 14467350 | rs1547219 | 0.739 | -0.269 | -0.482 | -0.035 | 0.002 | 0.003 | 0.003 | 0.039 | 0.11 | 0.1 | 0.08 | 0.07 | 0.12 | 0.09 |
| 10 | 14470284 | rs12570940 | 0.661 | -0.268 | -0.463 | -0.289 | 0.031 | -0.028 | -0.028 | 0.393 | 0.22 | 0.3 | 0.21 | 0.16 | 0.26 | 0.17 |
| 10 | 14470920 | rs7091633 | 0.749 | -0.534 | -0.431 | -0.099 | 0.002 | -0.002 | -0.002 | 0 | 0.18 | 0.15 | 0.12 | 0.19 | 0.05 | 0.15 |
| 10 | 14473162 | rs1325897 | -0.148 | -0.059 | -0.415 | -0.308 | 0.004 | -0.012 | -0.012 | 0 | 0.34 | 0.32 | 0.28 | 0.22 | 0.39 | 0.27 |
| 10 | 14475528 | rs10906673 | 0.274 | 0.258 | -0.968 | 0.02 | -0.016 | -0.013 | -0.013 | 0 | 0.09 | 0.03 | 0.12 | 0.1 | 0.14 | 0.13 |
| 10 | 14476631 | rs10906674 | 0.44 | 0.277 | -1.022 | 0.03 | -0.045 | -0.019 | -0.019 | 0 | 0.09 | 0.01 | 0.12 | 0.07 | 0.15 | 0.08 |
| 10 | 14477400 | rs7098908 | -0.56 | -0.15 | -0.946 | -0.192 | 0.017 | 0.05 | 0.05 | 0.224 | 0.51 | 0.56 | 0.47 | 0.34 | 0.5 | 0.47 |
| 10 | 14477442 | rs4750518 | -0.74 | -0.249 | -1.119 | -0.273 | 0.018 | 0.023 | 0.023 | 0 | 0.86 | 0.91 | 0.93 | 0.9 | 0.88 | 0.87 |
| 10 | 14478176 | rs10796190 | -1.075 | -0.322 | -1.12 | -0.649 | 0.012 | 0.042 | 0.042 | 0.648 | 0.51 | 0.41 | 0.35 | 0.27 | 0.43 | 0.39 |
| 10 | 14478286 | rs10906675 | -0.191 | -0.111 | -0.962 | -0.407 | -0.005 | -0.003 | -0.003 | 0 | 0.47 | 0.44 | 0.48 | 0.42 | 0.53 | 0.4 |
| 10 | 14478335 | rs10796191 | -0.255 | -0.136 | -0.843 | -0.456 | 0.004 | -0.013 | -0.013 | 0.496 | 0.84 | 0.78 | 0.83 | 0.85 | 0.82 | 0.82 |
| 10 | 14478462 | rs10906676 | -0.484 | -0.425 | -1.229 | -0.392 | -0.03 | -0.013 | -0.013 | 0 | 0.17 | 0.04 | 0.21 | 0.18 | 0.19 | 0.19 |
| 10 | 14478618 | rs11259096 | 0.598 | -0.558 | -0.618 | -0.339 | -0.019 | -0.073 | -0.073 | 0 | 0.12 | 0.29 | 0.03 | 0 | 0.05 | 0.07 |
| 10 | 14480073 | rs17155306 | 0.518 | -0.558 | -1.146 | -0.187 | 0.02 | -0.002 | -0.002 | 0 | 0.08 | 0.03 | 0.09 | 0.09 | 0.03 | 0.08 |
| 10 | 14484055 | rs10906678 | -1.04 | -0.49 | -1.212 | -0.208 | -0.01 | -0.015 | -0.015 | 0 | 0.4 | 0.42 | 0.3 | 0.24 | 0.33 | 0.4 |
| 10 | 14485205 | rs7911697 | -1.832 | -0.379 | -1.344 | -0.104 | -0.004 | -0.015 | -0.015 | 0 | 0.72 | 0.76 | 0.63 | 0.53 | 0.67 | 0.64 |
| 10 | 14487813 | rs12244056 | -0.587 | -0.458 | -1.37 | -0.142 | -0.01 | -0.034 | -0.034 | 0 | 0.29 | 0.38 | 0.26 | 0.2 | 0.24 | 0.28 |
| 10 | 14488967 | rs7905868 | 0.78 | -1.4 | -1.289 | -0.413 | 0 | -0.004 | -0.004 | 0.553 | 0.28 | 0.31 | 0.3 | 0.21 | 0.3 | 0.18 |
| 10 | 14489553 | rs17155309 | 0.321 | -1.048 | -1.303 | -0.193 | 0.032 | -0.018 | -0.018 | 0 | 0.13 | 0.04 | 0.04 | 0.08 | 0.08 | 0.14 |
| 10 | 14490169 | rs12249942 | -0.089 | -1.02 | -1.177 | -0.278 | 0.001 | 0.006 | 0.006 | 0.485 | 0.4 | 0.62 | 0.35 | 0.21 | 0.36 | 0.29 |
| 10 | 14491371 | rs12218877 | 0.261 | -0.883 | -0.899 | 0.319 | 0.015 | -0.002 | -0.002 | 0 | 0.1 | 0.04 | 0.04 | 0.04 | 0.08 | 0.12 |
| 10 | 14491517 | rs3889775 | -0.777 | -0.613 | -0.737 | 0.378 | -0.014 | -0.012 | -0.012 | 0 | 0.38 | 0.4 | 0.2 | 0.2 | 0.22 | 0.36 |
| 10 | 14493853 | rs10906681 | -0.9 | 0.375 | -0.678 | 1.047 | 0.002 | 0.014 | 0.014 | 1.012 | 0.11 | 0.1 | 0.15 | 0.26 | 0.13 | 0.22 |
| 10 | 14498050 | rs1325889 | 0.014 | 0.471 | -1.081 | 1.299 | -0.058 | -0.004 | -0.004 | 0.126 | 0.39 | 0.22 | 0.52 | 0.44 | 0.51 | 0.42 |
| 10 | 14498437 | rs2009555 | -0.597 | 0.941 | -1.171 | 1.556 | 0 | 0.003 | 0.003 | 0.471 | 0.08 | 0.1 | 0.17 | 0.2 | 0.09 | 0.17 |
| 10 | 14500618 | rs7914718 | -0.933 | -0.001 | -0.578 | 1.382 | -0.004 | -0.001 | -0.001 | 0 | 0.6 | 0.61 | 0.72 | 0.6 | 0.63 | 0.58 |
| 10 | 14500774 | rs10906682 | 0.331 | -0.04 | -0.593 | 1.364 | -0.022 | 0.003 | 0.003 | 0 | 0.21 | 0.39 | 0.21 | 0.15 | 0.12 | 0.17 |
| 10 | 14501236 | rs4750522 | 0.628 | -0.095 | -0.78 | 1.288 | -0.057 | -0.021 | -0.021 | 0 | 0.34 | 0.22 | 0.5 | 0.45 | 0.49 | 0.38 |
| 10 | 14502855 | rs4750524 | 1.756 | -0.054 | -0.791 | 1.286 | -0.076 | -0.001 | -0.001 | 0 | 0.26 | 0.11 | 0.32 | 0.25 | 0.4 | 0.21 |
| 10 | 14504402 | rs11259115 | 1.117 | -0.219 | -0.708 | 1.16 | 0.005 | -0.01 | -0.01 | 0 | 0.08 | 0.08 | 0.14 | 0.15 | 0.04 | 0.09 |
| 10 | 14505850 | rs7099780 | -1.048 | -0.401 | -0.334 | 1.055 | 0.002 | 0.002 | 0.002 | 0 | 0.62 | 0.68 | 0.73 | 0.68 | 0.66 | 0.66 |
| 10 | 14507883 | rs11259118 | 0.724 | 0.122 | 0.027 | 1.141 | -0.001 | -0.025 | -0.025 | 0.711 | 0.1 | 0.14 | 0.06 | 0 | 0.07 | 0.06 |
| 10 | 14509807 | rs873849 | 1.543 | 0.01 | -0.076 | 1.296 | -0.066 | -0.003 | -0.003 | 1.853 | 0.28 | 0.12 | 0.38 | 0.27 | 0.4 | 0.21 |
| 10 | 14510638 | rs1853878 | 0.92 | -0.089 | -0.096 | 1.205 | -0.072 | 0.029 | 0.029 | 0.043 | 0.36 | 0.16 | 0.42 | 0.28 | 0.46 | 0.29 |
| 10 | 14511347 | rs11259119 | -0.377 | -0.189 | 0.491 | 1.242 | 0.045 | 0.117 | 0.117 | 0.013 | 0.06 | 0.25 | 0.19 | 0.18 | 0.07 | 0.09 |
| 10 | 14511480 | rs11259120 | -0.09 | -0.187 | 0.491 | 1.259 | 0.05 | 0.127 | 0.127 | 0 | 0.05 | 0.21 | 0.19 | 0.18 | 0.07 | 0.09 |
| 10 | 14511505 | rs11259121 | 0.282 | -0.606 | 0.425 | 1.17 | 0.009 | -0.007 | -0.007 | 0 | 0.14 | 0.12 | 0.09 | 0.14 | 0.05 | 0.14 |
| 10 | 14512746 | rs7898971 | -0.534 | -0.629 | 0.443 | 1.123 | -0.004 | 0.011 | 0.011 | 0 | 0.16 | 0.31 | 0.23 | 0.18 | 0.13 | 0.16 |
| 10 | 14512831 | rs7077762 | 0.432 | -0.579 | 0.372 | 1.086 | -0.01 | -0.007 | -0.007 | 0.631 | 0.42 | 0.37 | 0.49 | 0.42 | 0.45 | 0.36 |
| 10 | 14514061 | rs11259131 | 0.563 | -0.599 | 0.322 | 1.128 | 0.006 | -0.006 | -0.006 | 0 | 0.13 | 0.14 | 0.09 | 0.14 | 0.05 | 0.14 |
| 10 | 14518009 | rs7089567 | -1.788 | -0.541 | 0.487 | 0.816 | -0.008 | 0.03 | 0.03 | 0.861 | 0.26 | 0.64 | 0.39 | 0.54 | 0.2 | 0.35 |
| 10 | 14518072 | rs7088537 | -0.716 | -0.54 | 0.684 | 0.968 | -0.026 | 0.099 | 0.099 | 0 | 0.18 | 0.34 | 0.27 | 0.36 | 0.11 | 0.24 |
| 10 | 14519492 | rs942932 | 0.973 | -0.347 | 0.442 | 0.841 | -0.076 | -0.011 | -0.011 | 0 | 0.39 | 0.15 | 0.43 | 0.38 | 0.48 | 0.33 |
| 10 | 14519623 | rs942933 | 0.173 | -0.327 | 0.518 | 0.796 | 0 | -0.001 | -0.001 | 2.307 | 0.46 | 0.44 | 0.54 | 0.56 | 0.56 | 0.43 |
| 10 | 14519892 | rs942935 | 0.236 | -0.354 | 0.34 | 0.794 | 0.001 | -0.009 | -0.009 | 3.103 | 0.44 | 0.34 | 0.53 | 0.56 | 0.55 | 0.43 |

| | | | | | | | | | | | | | | | | |
|---|---|---|---|---|---|---|---|---|---|---|---|---|---|---|---|---|
| 10 | 14520297 | rs942937 | 1.928 | -0.266 | 0.195 | 0.539 | -0.08 | -0.028 | -0.028 | 0 | 0.27 | 0.12 | 0.38 | 0.3 | 0.39 | 0.24 |
| 10 | 14523377 | rs7083394 | -0.491 | -0.248 | 0.483 | 0.576 | 0.066 | 0.056 | 0.056 | 0.103 | 0.08 | 0.32 | 0.19 | 0.11 | 0.08 | 0.08 |
| 10 | 14527223 | rs6602733 | -1.082 | -0.582 | 0.456 | 0.461 | -0.001 | -0.03 | -0.03 | 0 | 0.64 | 0.66 | 0.8 | 0.9 | 0.67 | 0.7 |
| 10 | 14531530 | rs10508474 | -0.294 | -0.241 | 0.484 | 0.515 | 0 | 0.019 | 0.019 | 0 | 0.86 | 0.87 | 0.83 | 0.57 | 0.88 | 0.75 |
| 10 | 14532894 | rs1998862 | -0.486 | 0.547 | 0.491 | 0.341 | 0.138 | 0.016 | 0.016 | 0 | 0.1 | 0.37 | 0.35 | 0.11 | 0.16 | 0.13 |
| 10 | 14533512 | rs7089824 | -0.401 | 0.55 | 0.501 | 0.342 | 0.14 | 0.018 | 0.018 | 0 | 0.1 | 0.35 | 0.35 | 0.12 | 0.16 | 0.13 |
| 10 | 14533738 | rs7897612 | 0.754 | 0.199 | 0.267 | 0.177 | 0.018 | 0.072 | 0.072 | 0 | 0.34 | 0.21 | 0.16 | 0.08 | 0.28 | 0.31 |
| 10 | 14535113 | rs1887091 | 0.705 | 0.356 | 0.301 | 0.362 | 0.008 | -0.002 | -0.002 | 0 | 0.41 | 0.45 | 0.62 | 0.42 | 0.52 | 0.33 |
| 10 | 14547448 | rs11259143 | -0.046 | 0.729 | 0.078 | 0.207 | 0.023 | 0.214 | 0.214 | 0 | 0.42 | 0.16 | 0.19 | 0.08 | 0.34 | 0.36 |
| 10 | 14547529 | rs7098808 | 1.095 | 0.611 | 0.237 | 0.242 | -0.013 | 0.023 | 0.023 | 0.112 | 0.06 | 0.13 | 0.04 | 0.08 | 0.04 | 0.08 |
| 10 | 14548864 | rs11259144 | -0.269 | 0.701 | 0.248 | 0.284 | 0.037 | 0.105 | 0.105 | 0 | 0.45 | 0.29 | 0.2 | 0.16 | 0.37 | 0.42 |
| 10 | 14563097 | rs3780915 | 0.363 | 0.873 | 0.68 | 0.416 | 0.124 | 0.022 | 0.022 | 0.025 | 0.9 | 0.68 | 0.66 | 0.88 | 0.85 | 0.87 |
| 10 | 14569422 | rs10752338 | 0.3 | 0.615 | 0.73 | 0.592 | -0.002 | 0.173 | 0.173 | 0 | 0.87 | 0.75 | 0.83 | 0.59 | 0.89 | 0.72 |
| 10 | 14574817 | rs11259159 | 0.265 | 0.535 | 0.406 | 0.218 | 0.002 | -0.002 | -0.002 | 0 | 0.55 | 0.44 | 0.66 | 0.5 | 0.64 | 0.48 |
| 10 | 14577404 | rs7082124 | -0.129 | 0.958 | 0.028 | 0.137 | -0.001 | -0.001 | -0.001 | 0 | 0.17 | 0.18 | 0.1 | 0.08 | 0.17 | 0.17 |
| 10 | 14578423 | rs7069426 | 0.466 | 0.972 | 0.01 | 0.132 | -0.018 | 0.028 | 0.028 | 1.064 | 0.13 | 0.07 | 0.07 | 0 | 0.15 | 0.11 |
| 10 | 14579015 | rs12359615 | 2.074 | 1.047 | 0.015 | 0.157 | -0.005 | -0.036 | -0.036 | 0 | 0.31 | 0.12 | 0.26 | 0.31 | 0.33 | 0.18 |
| 10 | 14589240 | rs11259176 | 0.375 | 1.018 | -0.001 | 0.127 | -0.022 | 0.032 | 0.032 | 1.142 | 0.13 | 0.07 | 0.07 | 0 | 0.15 | 0.1 |
| 10 | 14590542 | rs12767252 | -1.212 | 1.125 | 0.131 | 0.204 | -0.002 | -0.007 | -0.007 | 0 | 0.57 | 0.42 | 0.42 | 0.49 | 0.53 | 0.64 |
| 10 | 14596685 | rs7919989 | -1.112 | 0.923 | 0.217 | 0.309 | -0.007 | -0.004 | -0.004 | 0.239 | 0.55 | 0.42 | 0.39 | 0.52 | 0.5 | 0.61 |
| 10 | 14596739 | rs11259181 | 0.893 | 0.907 | 0.248 | 0.495 | -0.005 | 0.096 | 0.096 | 0 | 0.07 | 0.11 | 0.09 | 0.25 | 0.05 | 0.14 |
| 10 | 14603429 | rs11259186 | -1.372 | 0.911 | 0.278 | 0.459 | 0.009 | -0.004 | -0.004 | 0 | 0.58 | 0.5 | 0.41 | 0.61 | 0.52 | 0.67 |
| 10 | 14607610 | rs17155483 | -0.617 | 0.933 | 0.223 | 0.464 | 0.01 | 0.182 | 0.182 | 0 | 0.13 | 0.23 | 0.19 | 0.44 | 0.13 | 0.27 |
| 10 | 14609795 | rs7092057 | 0.523 | 0.928 | 0.216 | 0.43 | 0.095 | 0.051 | 0.051 | 0.143 | 0.87 | 0.62 | 0.66 | 0.8 | 0.83 | 0.81 |
| 10 | 14611258 | rs1571035 | -1.051 | 0.972 | 0.131 | 0.392 | 0.003 | 0.002 | 0.002 | 0.233 | 0.55 | 0.49 | 0.39 | 0.51 | 0.5 | 0.62 |
| 10 | 14618846 | rs7904177 | 1.287 | 1.28 | 0.6 | 0.926 | 0.048 | -0.035 | -0.035 | 0 | 0.75 | 0.58 | 0.59 | 0.8 | 0.69 | 0.71 |
| 10 | 14619045 | rs10906701 | -0.177 | 1.142 | 0.488 | 1.233 | -0.008 | 0.095 | 0.095 | 0 | 0.07 | 0.13 | 0.09 | 0.25 | 0.05 | 0.18 |
| 10 | 14625316 | rs7078538 | 0.718 | 1.413 | 0.373 | 0.342 | 0.09 | -0.015 | -0.015 | 0 | 0.88 | 0.6 | 0.69 | 0.92 | 0.86 | 0.84 |
| 10 | 14625801 | rs10508478 | -0.622 | 1.367 | -0.085 | -0.02 | -0.007 | -0.006 | -0.006 | 0.217 | 0.23 | 0.21 | 0.39 | 0.05 | 0.3 | 0.21 |
| 10 | 14625920 | rs12777920 | -0.174 | 1.439 | -0.447 | 0.37 | -0.025 | -0.025 | -0.025 | 0 | 0.05 | 0.01 | 0.07 | 0.18 | 0.1 | 0.12 |
| 10 | 14627383 | rs749232 | -0.83 | 1.2 | -0.547 | 0.395 | 0.013 | -0.03 | -0.03 | 0 | 0.94 | 0.84 | 0.94 | 1 | 0.92 | 0.99 |
| 10 | 14627825 | rs1409391 | 0.763 | 0.807 | -1.737 | 0.53 | 0.005 | 0.003 | 0.003 | 0 | 0.22 | 0.03 | 0.14 | 0.15 | 0.17 | 0.22 |
| 10 | 14627846 | rs1409390 | -0.055 | 0.691 | -1.8 | 0.651 | -0.01 | 0.055 | 0.055 | 0 | 0.14 | 0.26 | 0.14 | 0.26 | 0.1 | 0.19 |
| 10 | 14629401 | rs11259206 | 1.731 | 0.847 | -2.157 | 0.079 | -0.031 | -0.001 | -0.001 | 0.28 | 0.3 | 0.13 | 0.25 | 0.28 | 0.35 | 0.16 |
| 10 | 14632092 | rs7907299 | -0.709 | 0.561 | -2.045 | 0.142 | 0.017 | 0.025 | 0.025 | 0 | 0.42 | 0.33 | 0.32 | 0.65 | 0.35 | 0.57 |
| 10 | 14633723 | rs11259210 | -0.303 | 1.352 | -1.259 | 0.743 | 0.002 | 0.046 | 0.046 | 0 | 0.16 | 0.19 | 0.16 | 0.41 | 0.16 | 0.3 |
| 10 | 14641359 | rs11259213 | 0.542 | 1.147 | -1.835 | 0.491 | -0.02 | 0.032 | 0.032 | 0.032 | 0.51 | 0.15 | 0.38 | 0.43 | 0.5 | 0.38 |
| 10 | 14651050 | rs724177 | 0.753 | 0.825 | -1.696 | -0.006 | -0.005 | 0.203 | 0.203 | 0 | 0.39 | 0.11 | 0.26 | 0.13 | 0.41 | 0.18 |
| 10 | 14651346 | rs11259220 | -0.219 | 0.283 | -1.512 | 0.551 | 0.004 | -0.006 | -0.006 | 0.489 | 0.17 | 0.14 | 0.11 | 0.2 | 0.09 | 0.15 |
| 10 | 14652933 | rs717069 | 0.872 | 0.883 | -0.493 | 0.393 | -0.003 | 0.023 | 0.023 | 0 | 0.48 | 0.44 | 0.62 | 0.19 | 0.54 | 0.31 |
| 10 | 14655455 | rs943547 | -0.03 | 1.234 | -0.112 | 0.568 | 0.014 | 0.015 | 0.015 | 0.334 | 0.72 | 0.66 | 0.76 | 0.37 | 0.68 | 0.51 |
| 10 | 14658464 | rs7072976 | 0.128 | 1.144 | 0.07 | 0.5 | 0.002 | -0.049 | -0.049 | 0 | 0.1 | 0.16 | 0.05 | 0 | 0.09 | 0.07 |
| 10 | 14660199 | rs7921651 | -1.15 | 1.456 | 0.079 | 0.812 | 0.007 | -0.001 | -0.001 | 0 | 0.19 | 0.22 | 0.39 | 0.16 | 0.24 | 0.22 |
| 10 | 14661008 | rs11259230 | -0.076 | 1.46 | 0.186 | 0.862 | -0.018 | -0.047 | -0.047 | 0 | 0.24 | 0.13 | 0.21 | 0.52 | 0.27 | 0.43 |
| 10 | 14663384 | rs4750540 | 0.455 | 1.414 | 0.317 | 0.956 | -0.007 | 0.005 | 0.005 | 0 | 0.06 | 0.07 | 0.33 | 0.05 | 0.1 | 0.1 |
| 10 | 14664672 | rs17155589 | -0.552 | 0.919 | 0.293 | 0.865 | 0.045 | -0.03 | -0.03 | 1.059 | 0.2 | 0.06 | 0.11 | 0.17 | 0.1 | 0.2 |
| 10 | 14666221 | rs10906717 | -0.847 | 1.106 | 0.704 | 0.84 | 0.027 | 0.007 | 0.007 | 0.615 | 0.3 | 0.36 | 0.55 | 0.69 | 0.38 | 0.52 |
| 10 | 14667316 | rs11259235 | -0.58 | 0.801 | 0.57 | 0.998 | 0.009 | -0.012 | -0.012 | 0 | 0.17 | 0.13 | 0.1 | 0.29 | 0.1 | 0.19 |
| 10 | 14678962 | rs12218166 | 0.463 | 0.772 | 0.531 | 1.08 | 0.033 | -0.035 | -0.035 | 0 | 0.11 | 0.04 | 0.06 | 0.18 | 0.04 | 0.11 |
| 10 | 14679858 | rs8181403 | -0.38 | 0.979 | 0.598 | 1.116 | -0.01 | -0.025 | -0.025 | 0 | 0.26 | 0.16 | 0.21 | 0.5 | 0.28 | 0.43 |
| 10 | 14681979 | rs10906725 | 1.063 | 0.864 | 0.829 | 0.949 | -0.001 | 0.017 | 0.017 | 0 | 0.58 | 0.3 | 0.39 | 0.31 | 0.53 | 0.41 |
| 10 | 14682480 | rs1555985 | 1.94 | 0.924 | 0.658 | 0.654 | 0.012 | 0.129 | 0.129 | 0 | 0.43 | 0.2 | 0.3 | 0.13 | 0.48 | 0.23 |
| 10 | 14686790 | rs4463750 | -1.271 | 1.052 | 0.652 | 1.217 | -0.006 | -0.005 | -0.005 | 0.481 | 0.38 | 0.28 | 0.33 | 0.56 | 0.36 | 0.49 |
| 10 | 14689835 | rs10906729 | -1.061 | 1.009 | 0.62 | 1.354 | -0.004 | -0.002 | -0.002 | 1.043 | 0.3 | 0.27 | 0.19 | 0.34 | 0.33 | 0.39 |
| 10 | 14690743 | rs4489642 | -0.026 | 1.057 | 0.528 | 1.407 | -0.016 | 0.002 | 0.002 | 0.42 | 0.18 | 0.12 | 0.14 | 0.18 | 0.23 | 0.28 |
| 10 | 14691889 | rs10796216 | -1.959 | 1.109 | 0.919 | 1.321 | 0 | 0.002 | 0.002 | 0 | 0.46 | 0.59 | 0.57 | 0.58 | 0.45 | 0.57 |
| 10 | 14692912 | rs9630136 | 0.694 | 1.237 | 0.647 | 1.347 | -0.048 | -0.026 | -0.026 | 0 | 0.13 | 0.01 | 0.09 | 0.14 | 0.13 | 0.17 |
| 10 | 14693826 | rs10906731 | -1.461 | 1.169 | 1.052 | 1.308 | 0.009 | 0.041 | 0.041 | 0 | 0.23 | 0.49 | 0.44 | 0.25 | 0.18 | 0.24 |
| 10 | 14694058 | rs10796217 | -1.093 | 1.157 | 0.913 | 1.254 | 0.026 | 0.003 | 0.003 | 0 | 0.27 | 0.37 | 0.39 | 0.31 | 0.32 | 0.33 |
| 10 | 14695089 | rs3740114 | -0.693 | 1.208 | 0.569 | 1.311 | -0.006 | -0.004 | -0.004 | 0 | 0.23 | 0.25 | 0.36 | 0.31 | 0.27 | 0.32 |
| 10 | 14695682 | rs3740116 | -1.418 | 0.633 | 0.057 | 1.399 | -0.02 | 0.018 | 0.018 | 0 | 0.23 | 0.25 | 0.22 | 0.28 | 0.15 | 0.3 |
| 10 | 14696000 | rs7089802 | 0.955 | 0.548 | -0.389 | 1.517 | -0.009 | 0.02 | 0.02 | 0 | 0.81 | 0.84 | 0.76 | 0.48 | 0.9 | 0.75 |
| 10 | 14698054 | rs1870416 | 0.341 | 0.945 | -0.83 | 1.433 | -0.061 | -0.098 | -0.098 | 0 | 0.73 | 0.94 | 0.71 | 0.56 | 0.73 | 0.64 |
| 10 | 14698952 | rs7903263 | 0.882 | 0.499 | -0.123 | 1.21 | -0.014 | 0.036 | 0.036 | 0 | 0.71 | 0.48 | 0.5 | 0.38 | 0.76 | 0.57 |
| 10 | 14700158 | rs10906733 | 0.422 | 0.197 | -1.059 | 0.823 | 0.022 | 0.007 | 0.007 | 0 | 0.57 | 0.23 | 0.44 | 0.4 | 0.59 | 0.55 |
| 10 | 14700243 | rs10906734 | -0.6 | -0.441 | -1.122 | 0.729 | -0.008 | 0.006 | 0.006 | 0 | 0.36 | 0.42 | 0.4 | 0.52 | 0.3 | 0.52 |
| 10 | 14700470 | rs10508481 | -0.133 | -0.413 | -1.393 | 0.662 | 0.022 | -0.003 | -0.003 | 0 | 0.17 | 0.06 | 0.14 | 0.19 | 0.16 | 0.28 |
| 10 | 14700581 | rs12766190 | -0.962 | 0.199 | -0.592 | 0.664 | -0.003 | 0.004 | 0.004 | 0 | 0.21 | 0.26 | 0.14 | 0.27 | 0.2 | 0.33 |
| 10 | 14701678 | rs17155703 | -0.256 | 0.087 | -0.962 | 0.304 | 0.005 | 0.045 | 0.045 | 0 | 0.18 | 0.07 | 0.06 | 0.1 | 0.17 | 0.32 |

| | | | | | | | | | | | | | | | | |
|---|---|---|---|---|---|---|---|---|---|---|---|---|---|---|---|---|
| 10 | 14704727 | rs755277 | -0.503 | 0.288 | -1.293 | 0.183 | -0.044 | 0.034 | 0.034 | 0 | 0.23 | 0.07 | 0.16 | 0.15 | 0.26 | 0.28 |
| 10 | 14706454 | rs913295 | 0.495 | -0.036 | -0.601 | 0.099 | -0.021 | -0.019 | -0.019 | 0 | 0.37 | 0.54 | 0.36 | 0.24 | 0.26 | 0.31 |
| 10 | 14707693 | rs10906736 | 1.302 | -0.245 | -0.383 | -0.079 | 0.028 | 0.031 | 0.031 | 0.757 | 0.27 | 0.12 | 0.23 | 0.12 | 0.21 | 0.12 |
| 10 | 14711077 | rs4750545 | -0.409 | 0.587 | -0.192 | 0.349 | 0.004 | 0.011 | 0.011 | 0 | 0.14 | 0.15 | 0.15 | 0.3 | 0.16 | 0.22 |
| 10 | 14711514 | rs10906737 | 0.242 | 0.341 | 0.344 | 0.192 | 0.002 | -0.011 | -0.011 | 0 | 0.41 | 0.35 | 0.39 | 0.47 | 0.36 | 0.37 |
| 10 | 14712224 | rs11595636 | 0.864 | 0.37 | 0.198 | 0.242 | -0.002 | 0.011 | 0.011 | 0 | 0.07 | 0.04 | 0.11 | 0.06 | 0.06 | 0.16 |
| 10 | 14713077 | rs11259252 | 0.848 | 0.193 | 0.435 | 0.143 | -0.014 | -0.043 | -0.043 | 0.457 | 0.32 | 0.15 | 0.49 | 0.46 | 0.38 | 0.34 |
| 10 | 14714748 | rs10906742 | -0.152 | 0.17 | 0.935 | 0.119 | -0.001 | -0.006 | -0.006 | 0 | 0.56 | 0.39 | 0.36 | 0.48 | 0.53 | 0.5 |
| 10 | 14716516 | rs11259260 | 0.868 | 0.24 | 0.871 | 0.203 | 0.001 | -0.001 | -0.001 | 0 | 0.2 | 0.18 | 0.16 | 0.1 | 0.19 | 0.35 |
| 10 | 14717099 | rs9665534 | -0.537 | 0.276 | 0.96 | 0.247 | -0.006 | 0.005 | 0.005 | 0 | 0.61 | 0.39 | 0.7 | 0.59 | 0.7 | 0.76 |
| 10 | 14720899 | rs6602751 | 1.248 | 0.389 | 0.68 | 0.411 | -0.019 | -0.001 | -0.001 | 0 | 0.22 | 0.14 | 0.36 | 0.22 | 0.31 | 0.28 |
| 10 | 14720930 | rs7910586 | -1.745 | 0.252 | 0.909 | 0.497 | -0.014 | -0.009 | -0.009 | 0 | 0.72 | 0.65 | 0.77 | 0.86 | 0.8 | 0.89 |
| 10 | 14722720 | rs11259266 | 1.385 | 0.224 | 0.695 | 0.429 | 0.011 | 0.057 | 0.057 | 0 | 0.19 | 0.06 | 0.13 | 0.1 | 0.17 | 0.33 |
| 10 | 14722824 | rs11259267 | 1.385 | 0.224 | 0.695 | 0.429 | 0.011 | 0.057 | 0.057 | 0 | 0.19 | 0.06 | 0.13 | 0.1 | 0.18 | 0.33 |
| 10 | 14723053 | rs7919661 | -0.808 | 0.283 | 0.917 | 0.546 | -0.03 | 0.019 | 0.019 | 0 | 0.62 | 0.51 | 0.62 | 0.57 | 0.71 | 0.79 |
| 10 | 14723524 | rs7096784 | -0.431 | 0.244 | 0.586 | 0.379 | 0.016 | 0.035 | 0.035 | 0 | 0.55 | 0.34 | 0.56 | 0.35 | 0.62 | 0.73 |
| 10 | 14725088 | rs12767375 | -0.614 | 0.533 | 0.37 | 0.544 | -0.023 | -0.044 | -0.044 | 0 | 0.27 | 0.17 | 0.23 | 0.5 | 0.3 | 0.24 |
| 10 | 14725356 | rs12248918 | -0.145 | 0.592 | 0.739 | 0.668 | 0.021 | 0.142 | 0.142 | 0.044 | 0.17 | 0.31 | 0.2 | 0.51 | 0.18 | 0.16 |
| 10 | 14725454 | rs12264162 | -0.145 | 0.592 | 0.729 | 0.668 | 0.02 | 0.139 | 0.139 | 0.044 | 0.17 | 0.32 | 0.2 | 0.51 | 0.18 | 0.16 |
| 10 | 14725691 | rs4750551 | 0.83 | 0.63 | 0.278 | 0.559 | -0.051 | -0.008 | -0.008 | 1.76 | 0.22 | 0.07 | 0.36 | 0.22 | 0.32 | 0.28 |
| 10 | 14728558 | rs7922034 | -0.886 | 0.724 | 0.378 | 0.591 | 0.007 | -0.002 | -0.002 | 0 | 0.77 | 0.75 | 0.64 | 0.79 | 0.68 | 0.73 |
| 10 | 14730911 | rs2948719 | 0.54 | 0.733 | 0.41 | 0.594 | 0.014 | -0.003 | -0.003 | 0 | 0.24 | 0.31 | 0.36 | 0.21 | 0.33 | 0.27 |
| 10 | 14731449 | rs4750552 | 0.152 | 0.666 | -0.374 | 0.669 | -0.001 | -0.055 | -0.055 | 0.559 | 0.36 | 0.16 | 0.32 | 0.54 | 0.35 | 0.46 |
| 10 | 14733465 | rs3107803 | -0.24 | 0.643 | 0.028 | 0.887 | 0.008 | 0.032 | 0.032 | 0 | 0.32 | 0.53 | 0.5 | 0.43 | 0.37 | 0.34 |
| 10 | 14734417 | rs2609836 | -0.507 | 0.556 | -0.128 | 0.818 | -0.008 | 0.011 | 0.011 | 0 | 0.56 | 0.63 | 0.71 | 0.58 | 0.55 | 0.44 |
| 10 | 14734699 | rs1410077 | 0.722 | 0.341 | -0.038 | 0.945 | -0.007 | 0.042 | 0.042 | 0 | 0.07 | 0.09 | 0.11 | 0.22 | 0.04 | 0.06 |
| 10 | 14734972 | rs1410073 | -0.538 | 0.477 | -0.225 | 0.817 | -0.003 | 0.008 | 0.008 | 0 | 0.57 | 0.62 | 0.72 | 0.58 | 0.58 | 0.45 |
| 10 | 14736077 | rs2688841 | -0.629 | 1.03 | 0.258 | 0.973 | 0.004 | 0.014 | 0.014 | 0 | 0.32 | 0.57 | 0.5 | 0.43 | 0.37 | 0.35 |
| 10 | 14737912 | rs2688838 | -0.185 | 1.515 | 0.117 | 0.718 | 0.035 | 0.002 | 0.002 | 0 | 0.23 | 0.32 | 0.36 | 0.22 | 0.35 | 0.29 |
| 10 | 14740813 | rs11259283 | 0.567 | 0.569 | -0.99 | 0.067 | 0.055 | 0.196 | 0.196 | 0 | 0.42 | 0.15 | 0.2 | 0.13 | 0.32 | 0.36 |
| 10 | 14741040 | rs7911059 | -0.064 | 0.325 | -0.62 | 0.967 | -0.005 | 0.049 | 0.049 | 0.08 | 0.88 | 0.83 | 0.77 | 0.61 | 0.88 | 0.87 |
| 10 | 14741168 | rs10906748 | 0.187 | 1.083 | -0.249 | 1.103 | -0.001 | -0.002 | -0.002 | 1.35 | 0.21 | 0.19 | 0.35 | 0.26 | 0.35 | 0.25 |
| 10 | 14742856 | rs2601736 | -0.165 | 0.852 | -0.386 | 1.07 | 0.005 | -0.01 | -0.01 | 0 | 0.88 | 0.91 | 0.77 | 0.61 | 0.88 | 0.87 |
| 10 | 14743923 | rs11259286 | 1.351 | 1.015 | -0.551 | 0.9 | 0.07 | 0.181 | 0.181 | 0 | 0.4 | 0.14 | 0.19 | 0.12 | 0.31 | 0.36 |
| 10 | 14744006 | rs11259287 | 0.31 | 1.649 | -0.478 | 1.011 | -0.072 | -0.046 | -0.046 | 0 | 0.17 | 0.05 | 0.32 | 0.22 | 0.27 | 0.22 |
| 10 | 14753970 | rs7913807 | -0.055 | 1.478 | 0.092 | 0.936 | 0 | 0 | 0 | 0 | 0.31 | 0.62 | 0.26 | 0.25 | 0.28 | 0.25 |
| 10 | 14757013 | rs7341999 | 0.866 | 1.502 | -0.022 | 1.299 | -0.005 | -0.013 | -0.013 | 0 | 0.07 | 0.04 | 0.1 | 0.16 | 0.09 | 0.1 |
| 10 | 14757779 | rs4748109 | 0.351 | 1.599 | -0.028 | 1.415 | -0.023 | -0.003 | -0.003 | 0 | 0.17 | 0.12 | 0.31 | 0.19 | 0.28 | 0.22 |
| 10 | 14764993 | rs2688858 | -1.241 | 1.422 | -0.147 | 1.328 | 0.032 | 0.052 | 0.052 | 0.02 | 0.52 | 0.87 | 0.7 | 0.63 | 0.59 | 0.56 |
| 10 | 14765157 | rs7087109 | 0.729 | 1.199 | 0.269 | 1.186 | -0.007 | -0.028 | -0.028 | 0 | 0.05 | 0.12 | 0.02 | 0 | 0.02 | 0.08 |
| 10 | 14767687 | rs7897176 | 1.044 | 1.228 | 0.57 | 1.224 | 0.003 | 0.023 | 0.023 | 0 | 0.2 | 0.33 | 0.22 | 0.22 | 0.21 | 0.15 |
| 10 | 14769911 | rs883814 | -1.455 | 0.853 | 0.081 | 1.121 | 0.062 | 0.044 | 0.044 | 0 | 0.56 | 0.91 | 0.72 | 0.66 | 0.65 | 0.59 |
| 10 | 14777585 | rs2257172 | -1.383 | 1.001 | 0.157 | 1.216 | 0.065 | 0.147 | 0.147 | 0 | 0.52 | 0.9 | 0.68 | 0.71 | 0.63 | 0.57 |
| 10 | 14778101 | rs10906752 | 0.345 | 0.607 | 0.861 | 1.285 | 0.019 | 0.024 | 0.024 | 0 | 0.25 | 0.45 | 0.25 | 0.23 | 0.27 | 0.17 |
| 10 | 14778352 | rs7921961 | -0.481 | 0.637 | 0.913 | 1.349 | 0.006 | 0.022 | 0.022 | 0 | 0.33 | 0.6 | 0.35 | 0.38 | 0.35 | 0.28 |
| 10 | 14778920 | rs6602758 | 0.746 | 0.599 | 0.36 | 1.333 | -0.002 | -0.003 | -0.003 | 0 | 0.07 | 0.06 | 0.1 | 0.15 | 0.08 | 0.1 |
| 10 | 14779689 | rs2768702 | -0.979 | 0.527 | 0.189 | 1.186 | -0.036 | -0.055 | -0.055 | 0 | 0.41 | 0.9 | 0.4 | 0.54 | 0.39 | 0.42 |
| 10 | 14781173 | rs2601751 | -0.812 | 0.54 | 0.322 | 1.18 | -0.005 | -0.013 | -0.013 | 0 | 0.4 | 0.77 | 0.41 | 0.54 | 0.38 | 0.39 |
| 10 | 14793995 | rs12244530 | -1.098 | 0.168 | 0.741 | 1.244 | 0.017 | 0.139 | 0.139 | 0 | 0.08 | 0.35 | 0.05 | 0.15 | 0.02 | 0.19 |
| 10 | 14795628 | rs11259309 | 0.097 | 0.324 | 0.262 | 1.431 | 0 | 0.002 | 0.002 | 1.022 | 0.07 | 0.06 | 0.11 | 0.12 | 0.08 | 0.14 |
| 10 | 14806739 | rs2252188 | -0.051 | 0.144 | -1.254 | 1.09 | -0.084 | -0.097 | -0.097 | 0 | 0.77 | 0.99 | 0.61 | 0.7 | 0.72 | 0.8 |
| 10 | 14808696 | rs2609818 | -1.285 | -0.753 | -0.316 | 0.92 | -0.063 | 0.015 | 0.015 | 2.25 | 0.12 | 0.7 | 0.1 | 0.16 | 0.02 | 0.1 |
| 10 | 14813860 | rs2768713 | -1.224 | -0.731 | -0.411 | 0.929 | -0.06 | 0.015 | 0.015 | 1.718 | 0.12 | 0.71 | 0.11 | 0.16 | 0.02 | 0.11 |
| 10 | 14815729 | rs1974069 | -0.068 | -0.524 | -1.386 | 0.701 | 0.11 | 0.07 | 0.07 | 0 | 0.53 | 0.79 | 0.41 | 0.63 | 0.32 | 0.49 |
| 10 | 14815820 | rs1974068 | -0.036 | -0.89 | -1.492 | 0.652 | 0.174 | 0.101 | 0.101 | 0.135 | 0.52 | 0.81 | 0.4 | 0.65 | 0.3 | 0.48 |
| 10 | 14817067 | rs11259333 | -0.688 | 0.43 | -0.304 | 1.264 | 0.061 | 0.17 | 0.17 | 0.001 | 0.14 | 0.5 | 0.14 | 0.24 | 0.14 | 0.21 |
| 10 | 14818686 | rs2768715 | -0.386 | 1.173 | -0.974 | 0.455 | -0.009 | 0.005 | 0.005 | 0 | 0.25 | 0.21 | 0.35 | 0.06 | 0.34 | 0.26 |
| 10 | 14818870 | rs12256790 | 0.28 | 0.988 | -1.217 | 0.786 | 0.007 | 0.002 | 0.002 | 0 | 0.63 | 0.72 | 0.67 | 0.64 | 0.64 | 0.65 |
| 10 | 14820959 | rs2601744 | -0.731 | 0.888 | -1.699 | 0.396 | -0.097 | -0.07 | -0.07 | 0 | 0.66 | 0.92 | 0.42 | 0.38 | 0.54 | 0.52 |
| 10 | 14822491 | rs927935 | -0.632 | 0.862 | -1.622 | 0.402 | -0.071 | -0.039 | -0.039 | 1.026 | 0.65 | 0.87 | 0.41 | 0.37 | 0.53 | 0.52 |
| 10 | 14822827 | rs17773098 | 0.048 | 0.44 | -0.814 | 0.682 | -0.002 | -0.002 | -0.002 | 0 | 0.39 | 0.4 | 0.26 | 0.35 | 0.36 | 0.38 |
| 10 | 14839929 | rs10906761 | -0.634 | 0.163 | -0.42 | 0.859 | 0 | 0 | 0 | 0 | 0.51 | 0.5 | 0.55 | 0.4 | 0.44 | 0.44 |
| 10 | 14841026 | rs4750561 | -0.634 | 0.163 | -0.42 | 0.859 | 0 | 0 | 0 | 0 | 0.51 | 0.5 | 0.55 | 0.4 | 0.44 | 0.44 |
| 10 | 14843833 | rs2688848 | 0.24 | 0.194 | -0.568 | 1.262 | 0 | -0.025 | -0.025 | 0.281 | 0.08 | 0.03 | 0.09 | 0.23 | 0.07 | 0.11 |
| 10 | 14844884 | rs2688847 | -1.166 | -0.036 | -0.162 | 1.175 | -0.029 | 0.028 | 0.028 | 0 | 0.6 | 0.74 | 0.7 | 0.67 | 0.51 | 0.59 |
| 10 | 14845305 | rs11259350 | 0.455 | -0.639 | -0.142 | 0.74 | -0.005 | 0.056 | 0.056 | 0 | 0.26 | 0.11 | 0.26 | 0.02 | 0.31 | 0.1 |
| 10 | 14845687 | rs2400102 | -1.043 | -0.644 | 0.265 | 1.193 | -0.009 | 0.106 | 0.106 | 0 | 0.21 | 0.39 | 0.31 | 0.46 | 0.15 | 0.3 |
| 10 | 14846174 | rs2609839 | -0.379 | -0.616 | 0.054 | 1.174 | 0.006 | 0.015 | 0.015 | 0 | 0.38 | 0.31 | 0.39 | 0.2 | 0.37 | 0.24 |
| 10 | 14847149 | rs7091494 | -0.592 | -0.201 | 0.124 | 1.479 | -0.002 | 0.051 | 0.051 | 0 | 0.12 | 0.25 | 0.17 | 0.22 | 0.08 | 0.16 |

| | | | | | | | | | | | | | | | | |
|---|---|---|---|---|---|---|---|---|---|---|---|---|---|---|---|---|
| 10 | 14847220 | rs7079833 | -1.155 | -0.174 | 0.146 | 1.243 | -0.003 | 0.031 | 0.031 | 0 | 0.21 | 0.28 | 0.27 | 0.43 | 0.15 | 0.32 |
| 10 | 14861401 | rs2278871 | -0.737 | -0.888 | 0.698 | 1.049 | -0.04 | 0.075 | 0.075 | 0.126 | 0.16 | 0.34 | 0.13 | 0.3 | 0.08 | 0.37 |
| 10 | 14863864 | rs7900873 | -2.258 | -1.201 | 0.548 | 0.541 | 0.041 | 0.019 | 0.019 | 0 | 0.38 | 0.75 | 0.36 | 0.32 | 0.25 | 0.46 |
| 10 | 14864846 | rs7099185 | -2.444 | -1.406 | 0.407 | 0.688 | 0.075 | -0.04 | -0.04 | 3.451 | 0.41 | 0.82 | 0.4 | 0.43 | 0.26 | 0.5 |
| 10 | 14872427 | rs7094179 | -1.846 | -1.643 | -0.143 | 0.581 | 0.022 | 0.042 | 0.042 | 1.917 | 0.41 | 0.74 | 0.44 | 0.3 | 0.35 | 0.22 |
| 10 | 14874692 | rs6602768 | 0.077 | -0.874 | 0.471 | 0.472 | 0.026 | 0.047 | 0.047 | 0.112 | 0.13 | 0 | 0.08 | 0.05 | 0.06 | 0.22 |
| 10 | 14876885 | rs35258303 | -1.184 | -1.014 | 0.607 | 0.342 | 0 | 0.02 | 0.02 | 0 | 0.13 | 0.06 | 0.11 | 0.01 | 0.13 | 0.08 |
| 10 | 14884509 | rs12770830 | -0.63 | -1.001 | 0.425 | 0.349 | -0.018 | 0.069 | 0.069 | 0 | 0.12 | 0.01 | 0.08 | 0 | 0.09 | 0.06 |
| 10 | 14887979 | rs11593057 | -0.951 | -0.993 | 0.435 | 0.361 | -0.025 | 0.083 | 0.083 | 0 | 0.14 | 0.02 | 0.11 | 0 | 0.14 | 0.08 |
| 10 | 14891406 | rs17155992 | -0.19 | -0.998 | 0.475 | 0.349 | 0.003 | -0.001 | -0.001 | 0 | 0.14 | 0.11 | 0.09 | 0.05 | 0.06 | 0.22 |
| 10 | 14894901 | rs7894284 | -1.472 | -0.873 | 0.667 | 0.455 | -0.008 | 0.034 | 0.034 | 0 | 0.19 | 0.3 | 0.18 | 0.26 | 0.16 | 0.24 |
| 10 | 14920102 | rs12259906 | -2.083 | -0.889 | 0.152 | 0.255 | 0.005 | -0.007 | -0.007 | 0 | 0.3 | 0.68 | 0.29 | 0.42 | 0.16 | 0.45 |
| 10 | 14922023 | rs13377078 | -1.159 | -0.917 | 0.205 | 0.33 | -0.014 | 0.093 | 0.093 | 1.335 | 0.15 | 0.7 | 0.21 | 0.36 | 0.09 | 0.23 |
| 10 | 14941965 | rs10430783 | -2.014 | -0.333 | -0.418 | 0.282 | 0.011 | -0.004 | -0.004 | 0 | 0.39 | 0.65 | 0.39 | 0.4 | 0.27 | 0.49 |
| 10 | 14945406 | rs11594111 | -0.349 | -0.415 | -0.626 | 0.279 | -0.029 | 0.059 | 0.059 | 0 | 0.1 | 0.01 | 0.12 | 0 | 0.12 | 0.05 |
| 10 | 14954024 | rs12572872 | -1.515 | -0.354 | -0.464 | 0.417 | 0.024 | -0.003 | -0.003 | 0 | 0.37 | 0.7 | 0.3 | 0.38 | 0.24 | 0.48 |
| 10 | 14955178 | rs11517377 | 0.292 | 0.513 | -0.666 | 0.113 | -0.044 | 0.044 | 0.044 | 0 | 0.08 | 0.01 | 0.06 | 0 | 0.15 | 0.02 |
| 10 | 14978451 | rs12245497 | -0.735 | -0.298 | -0.446 | 0.29 | 0.004 | 0.009 | 0.009 | 0 | 0.11 | 0.1 | 0.14 | 0.01 | 0.1 | 0.05 |
| 10 | 14987440 | rs7920514 | -0.146 | -0.063 | -0.701 | 0.16 | -0.016 | 0.043 | 0.043 | 0 | 0.82 | 0.9 | 0.88 | 0.9 | 0.76 | 0.87 |
| 10 | 14988600 | rs11259405 | 0.955 | -0.585 | -0.703 | 0.665 | 0.035 | -0.02 | -0.02 | 0 | 0.54 | 0.27 | 0.65 | 0.51 | 0.59 | 0.51 |
| 10 | 14995149 | rs11593133 | 1.06 | -0.091 | -0.711 | 0.801 | 0.001 | 0.012 | 0.012 | 0.287 | 0.34 | 0.19 | 0.35 | 0.3 | 0.3 | 0.3 |
| 10 | 14996651 | rs1592159 | 0.91 | -0.049 | 0.528 | 1.375 | -0.007 | 0.046 | 0.046 | 0.101 | 0.69 | 0.43 | 0.72 | 0.46 | 0.66 | 0.6 |
| 10 | 15004452 | rs7085679 | 0.884 | 0.182 | 0.057 | 1.346 | 0.003 | 0.066 | 0.066 | 0.222 | 0.72 | 0.54 | 0.73 | 0.47 | 0.67 | 0.69 |
| 10 | 15008493 | rs4750568 | 0.853 | 0.215 | 0.072 | 1.364 | 0.003 | 0.07 | 0.07 | 0.382 | 0.72 | 0.54 | 0.73 | 0.48 | 0.67 | 0.69 |
| 10 | 15009109 | rs7087438 | -0.037 | 0.244 | -0.591 | 1.385 | -0.037 | -0.015 | -0.015 | 0 | 0.81 | 0.94 | 0.9 | 0.79 | 0.74 | 0.79 |
| 10 | 15014844 | rs5019235 | 0.413 | 0.3 | -0.596 | 1.018 | 0 | 0.02 | 0.02 | 0.319 | 0.72 | 0.64 | 0.74 | 0.48 | 0.67 | 0.69 |
| 10 | 15020847 | rs4276994 | -0.564 | 0.102 | -0.839 | 0.834 | 0.005 | -0.012 | -0.012 | 0 | 0.25 | 0.78 | 0.28 | 0.3 | 0.21 | 0.32 |
| 10 | 15026047 | rs12247446 | -0.664 | -0.375 | -0.196 | 0.452 | -0.037 | -0.049 | -0.049 | 2.02 | 0.14 | 0.21 | 0.05 | 0.04 | 0.06 | 0.1 |
| 10 | 15027214 | rs10752345 | -0.277 | -0.039 | -0.253 | 0.97 | -0.015 | 0.055 | 0.055 | 0.12 | 0.17 | 0.33 | 0.13 | 0.3 | 0.17 | 0.27 |
| 10 | 15037868 | rs2078000 | -0.029 | 0.79 | -1.189 | 0.48 | -0.045 | 0.03 | 0.03 | 0 | 0.8 | 0.9 | 0.76 | 0.86 | 0.73 | 0.81 |
| 10 | 15045021 | rs4750575 | 0.153 | 0.087 | -1.432 | 0.145 | -0.027 | 0.092 | 0.092 | 0 | 0.66 | 0.85 | 0.75 | 0.89 | 0.65 | 0.84 |
| 10 | 15045263 | rs4750578 | 0.522 | -0.139 | -1.109 | 0.264 | 0.006 | 0.007 | 0.007 | 0 | 0.36 | 0.48 | 0.47 | 0.68 | 0.39 | 0.5 |
| 10 | 15045634 | rs6602797 | -0.638 | -0.13 | -1.323 | 0.222 | -0.054 | 0.121 | 0.121 | 0.039 | 0.67 | 0.89 | 0.75 | 0.89 | 0.65 | 0.84 |
| 10 | 15057374 | rs2007712 | 0.768 | -0.711 | -1.349 | 0.323 | 0.007 | -0.011 | -0.011 | 0 | 0.17 | 0.05 | 0.13 | 0.15 | 0.16 | 0.23 |
| 10 | 15060276 | rs10906798 | 0.318 | -0.84 | -0.979 | 0.995 | -0.005 | 0.02 | 0.02 | 0.787 | 0.22 | 0.19 | 0.22 | 0.49 | 0.19 | 0.33 |
| 10 | 15060883 | rs7916087 | 1.195 | -0.713 | -0.898 | 0.948 | -0.002 | 0.043 | 0.043 | 0 | 0.16 | 0.17 | 0.2 | 0.42 | 0.18 | 0.31 |
| 10 | 15070615 | rs10906803 | -0.428 | -0.646 | -0.962 | 0.645 | -0.02 | 0.022 | 0.022 | 0.026 | 0.58 | 0.25 | 0.34 | 0.66 | 0.47 | 0.72 |
| 10 | 15071618 | rs11818063 | 0.398 | -0.89 | -1.134 | 0.516 | 0.05 | -0.047 | -0.047 | 0 | 0.39 | 0.13 | 0.2 | 0.51 | 0.3 | 0.39 |
| 10 | 15073244 | rs7918081 | -0.66 | -0.913 | -0.842 | 0.495 | -0.008 | 0.009 | 0.009 | 0 | 0.6 | 0.32 | 0.35 | 0.7 | 0.48 | 0.73 |
| 10 | 15079699 | rs17156239 | 0.907 | -0.852 | -0.946 | 0.493 | 0.006 | -0.01 | -0.01 | 0.136 | 0.18 | 0.09 | 0.14 | 0.18 | 0.15 | 0.27 |
| 10 | 15079983 | rs12269074 | 0.365 | -0.869 | -1.021 | 0.495 | 0.053 | -0.044 | -0.044 | 0 | 0.39 | 0.12 | 0.2 | 0.49 | 0.3 | 0.4 |
| 10 | 15081878 | rs4556439 | 0.309 | -0.944 | -0.743 | 0.467 | 0.049 | -0.013 | -0.013 | 0.501 | 0.4 | 0.22 | 0.22 | 0.5 | 0.31 | 0.41 |
| 10 | 15084568 | rs3763785 | 0.385 | -0.791 | -0.497 | 0.496 | -0.001 | 0 | 0 | 0 | 0.21 | 0.21 | 0.16 | 0.18 | 0.17 | 0.31 |
| 10 | 15086306 | rs3763787 | 0.317 | -0.82 | -0.315 | 0.158 | -0.008 | 0.002 | 0.002 | 0.065 | 0.36 | 0.37 | 0.25 | 0.44 | 0.3 | 0.34 |
| 10 | 15095122 | rs7099737 | -0.096 | -0.559 | -0.272 | -0.113 | 0.004 | 0.015 | 0.015 | 0 | 0.5 | 0.48 | 0.56 | 0.77 | 0.49 | 0.42 |
| 10 | 15095449 | rs12781344 | 2.141 | 0.192 | -0.129 | -0.584 | -0.001 | 0.065 | 0.065 | 3.504 | 0.16 | 0.04 | 0.04 | 0.04 | 0.09 | 0.05 |
| 10 | 15096448 | rs1935391 | -0.762 | 0.339 | -0.021 | -0.671 | -0.03 | 0.061 | 0.061 | 0 | 0.66 | 0.75 | 0.8 | 0.92 | 0.61 | 0.74 |
| 10 | 15097616 | rs10752349 | -0.915 | 0.381 | 0.054 | -0.637 | 0.026 | -0.06 | -0.06 | 0 | 0.78 | 0.74 | 0.71 | 0.98 | 0.64 | 0.77 |
| 10 | 15106441 | rs10906818 | 0.242 | 0.205 | 0.17 | -0.498 | 0.012 | 0.007 | 0.007 | 0 | 0.51 | 0.42 | 0.34 | 0.57 | 0.42 | 0.6 |
| 10 | 15108508 | rs17156272 | 0.751 | 0.435 | 0.434 | -0.427 | 0.062 | -0.017 | -0.017 | 0 | 0.06 | 0.24 | 0.16 | 0 | 0.07 | 0.07 |
| 10 | 15115700 | rs11259458 | 0.827 | 0.229 | 0.308 | -0.467 | -0.019 | -0.048 | -0.048 | 0 | 0.13 | 0.3 | 0.1 | 0.03 | 0.1 | 0.02 |
| 10 | 15117114 | rs7901579 | -1.121 | 0.467 | -0.196 | -0.609 | -0.079 | 0.151 | 0.151 | 1.791 | 0.7 | 0.91 | 0.82 | 0.95 | 0.65 | 0.82 |
| 10 | 15117672 | rs7902261 | -0.391 | 0.62 | 0.925 | 0.298 | 0.017 | 0.119 | 0.119 | 0 | 0.09 | 0.19 | 0.3 | 0.36 | 0.13 | 0.12 |
| 10 | 15125971 | rs1041158 | 0.404 | 1.157 | 1.386 | -0.008 | 0.053 | -0.028 | -0.028 | 0 | 0.93 | 0.74 | 0.92 | 1 | 0.9 | 0.95 |
| 10 | 15126323 | rs6602806 | -1.266 | 0.361 | 0.952 | -0.049 | 0.038 | 0.01 | 0.01 | 3.458 | 0.5 | 0.85 | 0.45 | 0.48 | 0.41 | 0.31 |
| 10 | 15127987 | rs10906823 | -0.609 | 0.245 | 1.813 | -0.56 | 0.082 | -0.013 | -0.013 | 0 | 0.08 | 0.37 | 0.19 | 0 | 0.07 | 0.07 |
| 10 | 15134222 | rs12254027 | -0.594 | -0.233 | 2.004 | -0.383 | -0.018 | -0.053 | -0.053 | 0 | 0.35 | 0.54 | 0.14 | 0.12 | 0.2 | 0.15 |
| 10 | 15137906 | rs6602811 | 0.39 | -0.221 | 2.183 | -0.414 | -0.02 | -0.08 | -0.08 | 0 | 0.25 | 0.31 | 0.11 | 0.05 | 0.14 | 0.1 |
| 10 | 15138615 | rs7896053 | -1.729 | -0.197 | 2.093 | -0.386 | -0.027 | -0.062 | -0.062 | 0.432 | 0.17 | 0.66 | 0.12 | 0.06 | 0.11 | 0.06 |
| 10 | 15148547 | rs913038 | -1.074 | -0.19 | 1.494 | -0.377 | 0.062 | 0.171 | 0.171 | 4.846 | 0.41 | 0.87 | 0.22 | 0.14 | 0.25 | 0.16 |
| 10 | 15156474 | rs6602815 | -1.836 | -0.015 | 2.073 | -0.244 | -0.011 | -0.059 | -0.059 | 0 | 0.1 | 0.71 | 0.11 | 0 | 0.07 | 0.01 |
| 10 | 15157771 | rs11259498 | -0.859 | -0.079 | 2.258 | -0.168 | 0.008 | 0.042 | 0.042 | 0 | 0.1 | 0.39 | 0.06 | 0.08 | 0.05 | 0.05 |
| 10 | 15158068 | rs7918717 | -0.66 | -0.172 | 1.775 | -0.227 | 0.039 | 0.096 | 0.096 | 6.843 | 0.34 | 0.79 | 0.21 | 0.08 | 0.21 | 0.13 |
| 10 | 15158087 | rs7921976 | 0.522 | -0.147 | 1.491 | -0.235 | 0.014 | 0.03 | 0.03 | 3.866 | 0.24 | 0.14 | 0.1 | 0.05 | 0.14 | 0.11 |
| 10 | 15158443 | rs11259499 | 0.263 | -0.101 | 1.224 | -0.198 | 0.004 | -0.005 | -0.005 | 0 | 0.07 | 0.05 | 0.02 | 0.09 | 0.04 | 0.05 |
| 10 | 15166555 | rs6602817 | -0.935 | -0.096 | 1.311 | -0.247 | 0.011 | 0.014 | 0.014 | 3.109 | 0.39 | 0.66 | 0.21 | 0.13 | 0.25 | 0.18 |
| 10 | 15166905 | rs1326207 | -1.094 | -0.11 | 0.394 | -0.244 | 0.054 | 0.152 | 0.152 | 6.476 | 0.42 | 0.9 | 0.23 | 0.16 | 0.26 | 0.18 |
| 10 | 15168559 | rs7091710 | -1.25 | 0.014 | 0.699 | -0.243 | -0.027 | -0.013 | -0.013 | 0 | 0.13 | 0.76 | 0.1 | 0.09 | 0.07 | 0.05 |
| 10 | 15176157 | rs12250830 | -0.937 | -0.026 | 0.334 | -0.268 | 0.075 | 0.143 | 0.143 | 6.466 | 0.39 | 0.85 | 0.2 | 0.14 | 0.21 | 0.17 |

| | | | | | | | | | | | | | | | | |
|---|---|---|---|---|---|---|---|---|---|---|---|---|---|---|---|---|
| 10 | 15176813 | rs12218990 | -1.049 | 0.418 | 0.885 | 0.355 | -0.015 | 0.027 | 0.027 | 0 | 0.11 | 0.43 | 0.08 | 0.1 | 0.06 | 0.05 |
| 10 | 15185503 | rs10737093 | 0.349 | 1.093 | -0.148 | 1.108 | 0.001 | 0.059 | 0.059 | 0.006 | 0.9 | 0.91 | 0.82 | 0.54 | 0.87 | 0.89 |
| 10 | 15199753 | rs7096608 | 0.272 | 0.295 | -0.249 | 0.011 | 0.02 | 0.04 | 0.04 | 0.019 | 0.2 | 0.1 | 0.08 | 0.06 | 0.12 | 0.06 |
| 10 | 15200560 | rs17356597 | -0.244 | 0.365 | -0.325 | 0.184 | 0.001 | -0.015 | -0.015 | 1.009 | 0.07 | 0.03 | 0.04 | 0.13 | 0.09 | 0.04 |
| 10 | 15203275 | rs10906841 | -0.882 | 0.638 | -0.041 | -0.111 | 0.003 | 0.014 | 0.014 | 0.049 | 0.35 | 0.32 | 0.2 | 0.22 | 0.34 | 0.13 |
| 10 | 15203310 | rs7902390 | -0.296 | 0.415 | 0.842 | -0.141 | 0.045 | 0.032 | 0.032 | 1.714 | 0.05 | 0.31 | 0.04 | 0.02 | 0.01 | 0.04 |
| 10 | 15205472 | rs10906842 | -1.24 | 0.499 | 0.492 | -0.134 | -0.003 | 0 | 0 | 0.462 | 0.13 | 0.2 | 0.09 | 0.16 | 0.15 | 0.06 |
| 10 | 15205655 | rs17156367 | 0.056 | 0.473 | 0.407 | -0.124 | 0.001 | -0.014 | -0.014 | 0.124 | 0.07 | 0.04 | 0.05 | 0.13 | 0.1 | 0.05 |
| 10 | 15206725 | rs11259527 | -0.799 | 0.425 | 0.677 | -0.152 | 0.011 | 0.004 | 0.004 | 0 | 0.06 | 0.22 | 0.05 | 0.03 | 0.05 | 0.02 |
| 10 | 15208011 | rs10906845 | 0.132 | 0.566 | 0.241 | -0.202 | -0.007 | 0.056 | 0.056 | 0.137 | 0.25 | 0.14 | 0.15 | 0.09 | 0.24 | 0.08 |
| 10 | 15212214 | rs3814170 | -0.964 | 0.524 | 0.723 | 0.104 | 0.002 | 0.039 | 0.039 | 0 | 0.11 | 0.31 | 0.06 | 0.15 | 0.11 | 0.09 |
| 10 | 15214583 | rs4750597 | -0.583 | 0.588 | -0.463 | 0.387 | 0.005 | 0.018 | 0.018 | 1.209 | 0.49 | 0.58 | 0.35 | 0.41 | 0.44 | 0.36 |
| 10 | 15219553 | rs4750598 | -0.098 | 0.514 | -0.997 | 0.265 | 0.012 | 0.016 | 0.016 | 0.363 | 0.7 | 0.29 | 0.59 | 0.85 | 0.77 | 0.56 |
| 10 | 15225054 | rs10906850 | -0.045 | 0.756 | -1.227 | 0.23 | -0.007 | 0.027 | 0.027 | 0.006 | 0.29 | 0.11 | 0.24 | 0.26 | 0.33 | 0.18 |
| 10 | 15225350 | rs10906851 | -0.2 | 0.517 | -0.851 | 0.532 | -0.014 | 0.019 | 0.019 | 0 | 0.32 | 0.46 | 0.22 | 0.33 | 0.26 | 0.38 |
| 10 | 15231633 | rs11259536 | 0.206 | 0.75 | -1.25 | 0.516 | -0.02 | 0.007 | 0.007 | 0.428 | 0.21 | 0.1 | 0.19 | 0.27 | 0.21 | 0.14 |
| 10 | 15231980 | rs7072433 | -0.402 | 0.766 | -0.938 | 0.691 | 0 | -0.001 | -0.001 | 0 | 0.58 | 0.63 | 0.67 | 0.51 | 0.56 | 0.62 |
| 10 | 15235954 | rs11259537 | 0.236 | 1.141 | -0.719 | 1.34 | 0.004 | 0.021 | 0.021 | 0 | 0.11 | 0.11 | 0.15 | 0.27 | 0.17 | 0.07 |
| 10 | 15237216 | rs10906855 | -0.121 | 0.896 | -0.324 | 1.381 | 0.016 | 0.043 | 0.043 | 0.085 | 0.33 | 0.49 | 0.32 | 0.47 | 0.38 | 0.3 |
| 10 | 15239498 | rs10906857 | -0.119 | 0.609 | -0.304 | 1.059 | 0.005 | -0.008 | -0.008 | 0 | 0.4 | 0.42 | 0.33 | 0.41 | 0.42 | 0.2 |
| 10 | 15240466 | rs4750603 | -1.745 | 0.566 | -0.422 | 1.173 | 0.002 | -0.005 | -0.005 | 0 | 0.78 | 0.8 | 0.6 | 0.81 | 0.79 | 0.57 |
| 10 | 15241923 | rs4237446 | 0.599 | 0.508 | 0.077 | 1.216 | 0.003 | 0.106 | 0.106 | 0.134 | 0.14 | 0.32 | 0.13 | 0.24 | 0.17 | 0.21 |
| 10 | 15242296 | rs3935268 | 0.863 | 0.506 | 0.084 | 1.221 | -0.003 | 0.071 | 0.071 | 0.139 | 0.13 | 0.23 | 0.11 | 0.23 | 0.15 | 0.21 |
| 10 | 15243205 | rs3935290 | -0.598 | 0.54 | 0.058 | 1.187 | -0.002 | 0.007 | 0.007 | 0 | 0.35 | 0.52 | 0.5 | 0.42 | 0.38 | 0.28 |
| 10 | 15243746 | rs4075162 | 0.427 | 0.306 | 0.089 | 1.123 | 0.001 | -0.013 | -0.013 | 0.926 | 0.2 | 0.16 | 0.14 | 0.14 | 0.21 | 0.06 |
| 10 | 15246346 | rs4237447 | -1.454 | 0.24 | -0.213 | 1.297 | 0.002 | -0.004 | -0.004 | 0 | 0.78 | 0.83 | 0.84 | 0.82 | 0.79 | 0.61 |
| 10 | 15256194 | rs3814165 | 0.755 | 0.235 | -0.253 | 1.457 | -0.029 | 0.013 | 0.013 | 0.302 | 0.22 | 0.01 | 0.13 | 0.14 | 0.19 | 0.19 |
| 10 | 15259495 | rs7085072 | -1.15 | 0.241 | -0.203 | 1.389 | 0.025 | 0.003 | 0.003 | 0.504 | 0.51 | 0.8 | 0.35 | 0.58 | 0.42 | 0.32 |
| 10 | 15260996 | rs11259554 | -0.315 | 0.228 | 0.059 | 1.361 | -0.002 | -0.026 | -0.026 | 0 | 0.22 | 0.35 | 0.15 | 0.17 | 0.21 | 0.08 |
| 10 | 15263100 | rs6602828 | 0.459 | 0.178 | -0.505 | 1.272 | 0.014 | -0.019 | -0.019 | 0.392 | 0.25 | 0.17 | 0.15 | 0.37 | 0.2 | 0.21 |
| 10 | 15265959 | rs7893353 | -0.836 | 0.645 | -0.61 | 1.241 | 0.016 | -0.008 | -0.008 | 1.194 | 0.51 | 0.8 | 0.55 | 0.57 | 0.43 | 0.32 |
| 10 | 15267411 | rs7921084 | -0.377 | 0.424 | -0.535 | 1.142 | 0.012 | -0.008 | -0.008 | 0.187 | 0.39 | 0.64 | 0.48 | 0.43 | 0.34 | 0.26 |
| 10 | 15268297 | rs11259556 | -0.546 | 0.442 | -0.385 | 1.103 | 0.025 | 0.03 | 0.03 | 0 | 0.13 | 0.3 | 0.13 | 0.17 | 0.14 | 0.07 |
| 10 | 15269846 | rs4750611 | 1.127 | 0.523 | -1.727 | 0.752 | -0.057 | -0.063 | -0.063 | 0 | 0.34 | 0.04 | 0.3 | 0.29 | 0.31 | 0.44 |
| 10 | 15270635 | rs4375345 | 0.633 | 0.732 | -1.514 | 0.302 | 0.016 | 0.011 | 0.011 | 0 | 0.2 | 0.27 | 0.38 | 0.29 | 0.3 | 0.15 |
| 10 | 15271162 | rs9423933 | 0.183 | -0.209 | -1.584 | -0.217 | 0.053 | 0.077 | 0.077 | 0 | 0.72 | 0.86 | 0.9 | 0.93 | 0.84 | 0.75 |
| 10 | 15274924 | rs10906867 | -0.117 | -0.015 | -1.43 | 0.204 | -0.001 | -0.001 | -0.001 | 1.431 | 0.1 | 0.09 | 0.09 | 0.16 | 0.14 | 0.09 |
| 10 | 15281660 | rs11816253 | -1.06 | -0.187 | -1.702 | 0.109 | 0.03 | 0.052 | 0.052 | 0 | 0.31 | 0.16 | 0.17 | 0.23 | 0.24 | 0.24 |
| 10 | 15285306 | rs11259573 | -0.171 | -0.485 | -1.738 | -0.101 | 0.066 | 0.071 | 0.071 | 0.033 | 0.2 | 0.05 | 0.07 | 0.07 | 0.09 | 0.15 |
| 10 | 15287493 | rs11259576 | -0.367 | 0.307 | -1.435 | 0.063 | -0.015 | 0.018 | 0.018 | 0 | 0.1 | 0.05 | 0.09 | 0.02 | 0.14 | 0.08 |
| 10 | 15292213 | rs11259577 | 0.671 | 0.01 | -2.153 | 0.49 | -0.058 | -0.176 | -0.176 | 0 | 0.28 | 0.01 | 0.4 | 0.48 | 0.23 | 0.38 |
| 10 | 15293903 | rs7915524 | 0.112 | 0.084 | -2.458 | -0.016 | -0.005 | 0.111 | 0.111 | 0 | 0.3 | 0.07 | 0.2 | 0.08 | 0.31 | 0.31 |
| 10 | 15294548 | rs2138929 | 0.216 | 0.317 | -1.337 | 0.221 | -0.01 | 0.009 | 0.009 | 0 | 0.33 | 0.31 | 0.3 | 0.22 | 0.43 | 0.23 |
| 10 | 15303527 | rs4750614 | 0.06 | 0.31 | -1.306 | 0.745 | 0.004 | -0.049 | -0.049 | 0 | 0.32 | 0.16 | 0.49 | 0.54 | 0.28 | 0.43 |
| 10 | 15304786 | rs7920144 | -0.741 | 0.033 | -0.865 | 0.665 | 0.002 | -0.003 | -0.003 | 0.003 | 0.14 | 0.14 | 0.05 | 0.14 | 0.09 | 0.1 |
| 10 | 15306347 | rs7073054 | -0.216 | 0.686 | -0.8 | 0.082 | -0.003 | 0.001 | 0.001 | 0 | 0.16 | 0.16 | 0.22 | 0.1 | 0.22 | 0.11 |
| 10 | 15308085 | rs12263544 | -0.386 | 0.536 | -0.349 | 0.131 | -0.012 | -0.004 | -0.004 | 0.104 | 0.14 | 0.19 | 0.05 | 0.11 | 0.09 | 0.11 |
| 10 | 15310265 | rs17156496 | -0.471 | 0.502 | -0.089 | 0.118 | -0.023 | -0.008 | -0.008 | 0 | 0.14 | 0.24 | 0.06 | 0.11 | 0.1 | 0.11 |
| 10 | 15310317 | rs11259586 | 1.525 | 0.588 | -0.877 | -0.291 | -0.069 | -0.077 | -0.077 | 0.06 | 0.39 | 0.06 | 0.31 | 0.62 | 0.39 | 0.62 |
| 10 | 15311580 | rs7897092 | -1 | 0.58 | -0.196 | -0.395 | 0.005 | 0.023 | 0.023 | 1.879 | 0.41 | 0.62 | 0.23 | 0.23 | 0.38 | 0.23 |
| 10 | 15318540 | rs6602830 | 0.259 | 0.634 | -0.257 | -0.41 | -0.018 | 0.009 | 0.009 | 0 | 0.21 | 0.17 | 0.45 | 0.13 | 0.26 | 0.15 |
| 10 | 15320660 | rs11259593 | 2.223 | 0.483 | 0.265 | -0.541 | -0.033 | -0.047 | -0.047 | 0 | 0.08 | 0.16 | 0.03 | 0.02 | 0.03 | 0.01 |
| 10 | 15322317 | rs7910208 | 0.188 | 0.538 | 0.05 | -0.277 | -0.008 | -0.005 | -0.005 | 0 | 0.22 | 0.21 | 0.26 | 0.25 | 0.26 | 0.16 |
| 10 | 15325014 | rs17156509 | 0.282 | 0.659 | -0.1 | -0.199 | -0.03 | -0.021 | -0.021 | 0 | 0.15 | 0.31 | 0.06 | 0.05 | 0.1 | 0.11 |
| 10 | 15327573 | rs12262545 | -0.69 | 0.721 | -0.24 | -0.24 | 0.001 | -0.005 | -0.005 | 0 | 0.28 | 0.32 | 0.17 | 0.21 | 0.28 | 0.19 |
| 10 | 15327869 | rs9787700 | 1.976 | 0.729 | -0.913 | -0.083 | -0.053 | -0.153 | -0.153 | 0 | 0.33 | 0.06 | 0.28 | 0.57 | 0.35 | 0.6 |
| 10 | 15333431 | rs2176489 | 1.823 | 0.903 | -0.788 | -0.496 | -0.046 | -0.02 | -0.02 | 0.022 | 0.34 | 0.05 | 0.5 | 0.68 | 0.35 | 0.59 |
| 10 | 15334823 | rs17358476 | 0.394 | 1.067 | -0.61 | -0.816 | -0.035 | 0.029 | 0.029 | 0 | 0.12 | 0.03 | 0.11 | 0.04 | 0.18 | 0.07 |
| 10 | 15338612 | rs6602832 | 0.197 | 0.82 | 0.014 | -0.953 | 0.004 | -0.032 | -0.032 | 0 | 0.71 | 0.71 | 0.69 | 0.91 | 0.64 | 0.82 |
| 10 | 15341575 | rs10796274 | 2.268 | 0.815 | 0.033 | -1.046 | -0.021 | 0.051 | 0.051 | 0.552 | 0.36 | 0.14 | 0.49 | 0.76 | 0.34 | 0.64 |
| 10 | 15345879 | rs1914547 | -0.554 | 0.746 | 0.243 | -1.104 | 0.002 | -0.012 | -0.012 | 0 | 0.79 | 0.78 | 0.73 | 0.92 | 0.73 | 0.87 |
| 10 | 15346448 | rs11812587 | -0.444 | 0.617 | 0.711 | -1.181 | 0.016 | -0.089 | -0.089 | 0.266 | 0.22 | 0.37 | 0.16 | 0.06 | 0.22 | 0.08 |
| 10 | 15346564 | rs4748157 | 2.424 | 0.655 | 0.639 | -1.157 | 0.003 | 0.039 | 0.039 | 0.471 | 0.35 | 0.12 | 0.48 | 0.76 | 0.32 | 0.64 |
| 10 | 15352058 | rs6602835 | -1.153 | 0.6 | 0.838 | -1.184 | -0.006 | -0.037 | -0.037 | 0 | 0.35 | 0.55 | 0.2 | 0.16 | 0.31 | 0.18 |
| 10 | 15355763 | rs17293194 | 0.283 | 0.638 | 0.835 | -1.174 | -0.005 | 0.002 | 0.002 | 0 | 0.21 | 0.2 | 0.27 | 0.08 | 0.27 | 0.13 |
| 10 | 15358362 | rs12777464 | -0.639 | 0.661 | 0.903 | -1.166 | 0.019 | -0.057 | -0.057 | 0 | 0.3 | 0.36 | 0.33 | 0.09 | 0.38 | 0.17 |
| 10 | 15358441 | rs10906884 | 1.356 | 0.517 | 0.801 | -1.216 | 0.003 | 0.08 | 0.08 | 0.72 | 0.56 | 0.35 | 0.75 | 0.84 | 0.59 | 0.78 |
| 10 | 15361751 | rs7080750 | -0.597 | 0.591 | 0.72 | -1.207 | 0.002 | -0.012 | -0.012 | 3.161 | 0.8 | 0.78 | 0.74 | 0.92 | 0.75 | 0.88 |
| 10 | 15362836 | rs7921541 | 0.583 | 0.533 | 0.723 | -1.233 | 0 | -0.023 | -0.023 | 0 | 0.66 | 0.58 | 0.7 | 0.81 | 0.66 | 0.78 |

| | | | | | | | | | | | | | | | | |
|---|---|---|---|---|---|---|---|---|---|---|---|---|---|---|---|---|
| 10 | 15364162 | rs1878624 | -0.605 | 0.472 | 0.972 | -1.177 | 0.002 | -0.007 | -0.007 | 0 | 0.26 | 0.29 | 0.16 | 0.15 | 0.26 | 0.17 |
| 10 | 15370036 | rs2352776 | 1.315 | 0.117 | 0.891 | -1.122 | 0.042 | 0.067 | 0.067 | 0 | 0.47 | 0.36 | 0.54 | 0.77 | 0.38 | 0.66 |
| 10 | 15376516 | rs7897087 | 2.154 | 0.174 | 0.41 | -0.939 | -0.025 | -0.09 | -0.09 | 0 | 0.35 | 0.04 | 0.48 | 0.64 | 0.33 | 0.66 |
| 10 | 15383768 | rs2138930 | -0.098 | 0.274 | 0.544 | -1.098 | 0.002 | -0.008 | -0.008 | 0 | 0.87 | 0.89 | 0.95 | 1 | 0.87 | 0.95 |
| 10 | 15383877 | rs4750625 | -0.96 | 0.15 | 0.649 | -1.148 | 0.011 | -0.033 | -0.033 | 0 | 0.39 | 0.48 | 0.41 | 0.22 | 0.5 | 0.28 |
| 10 | 15386588 | rs4750626 | 0.728 | 0.144 | 0.713 | -1.15 | 0 | -0.015 | -0.015 | 0 | 0.58 | 0.57 | 0.59 | 0.77 | 0.48 | 0.73 |
| 10 | 15391182 | rs4750628 | -0.831 | 0.152 | 0.747 | -1.148 | 0.009 | -0.02 | -0.02 | 0.018 | 0.38 | 0.42 | 0.4 | 0.24 | 0.5 | 0.26 |
| 10 | 15391610 | rs7904933 | -0.158 | 0.132 | 0.718 | -1.165 | -0.002 | -0.035 | -0.035 | 0 | 0.87 | 0.86 | 0.95 | 1 | 0.88 | 0.95 |
| 10 | 15393309 | rs10796285 | 1.365 | 0.289 | 0.759 | -1.167 | 0.033 | 0.057 | 0.057 | 0.025 | 0.48 | 0.36 | 0.54 | 0.76 | 0.39 | 0.68 |
| 10 | 15395181 | rs11599615 | -0.786 | 0.131 | 0.61 | -1.144 | -0.005 | 0.006 | 0.006 | 0 | 0.37 | 0.32 | 0.39 | 0.24 | 0.5 | 0.29 |
| 10 | 15396751 | rs7083529 | 0.431 | 0.38 | 0.438 | -1.104 | 0.012 | -0.023 | -0.023 | 0 | 0.15 | 0.23 | 0.23 | 0.08 | 0.17 | 0.08 |
| 10 | 15399458 | rs12264469 | -0.139 | 0.247 | 0.192 | -1.034 | 0.009 | 0.025 | 0.025 | 0.396 | 0.13 | 0.1 | 0.05 | 0 | 0.13 | 0.05 |
| 10 | 15406305 | rs7912836 | 1.958 | 0.228 | -0.07 | -0.967 | -0.077 | -0.085 | -0.085 | 0 | 0.38 | 0.05 | 0.51 | 0.63 | 0.41 | 0.66 |
| 10 | 15411829 | rs7912216 | -0.782 | 1.115 | 0.67 | -0.796 | -0.001 | -0.03 | -0.03 | 0 | 0.14 | 0.2 | 0.06 | 0 | 0.13 | 0.05 |
| 10 | 15412128 | rs7899526 | 0.933 | 1.176 | 0.758 | -0.839 | -0.01 | 0.015 | 0.015 | 0.003 | 0.6 | 0.36 | 0.6 | 0.76 | 0.55 | 0.73 |
| 10 | 15418175 | rs12774367 | -1.521 | 1.473 | 0.283 | -1.042 | -0.002 | 0.002 | 0.002 | 1.365 | 0.36 | 0.63 | 0.22 | 0.18 | 0.38 | 0.2 |
| 10 | 15422797 | rs12267785 | -1.55 | 1.475 | 0.177 | -0.963 | -0.006 | 0.009 | 0.009 | 1.753 | 0.36 | 0.66 | 0.22 | 0.26 | 0.38 | 0.21 |
| 10 | 15423620 | rs17360549 | 0.222 | 1.517 | 0.046 | -1.09 | -0.051 | 0.061 | 0.061 | 0 | 0.15 | 0.05 | 0.19 | 0.02 | 0.21 | 0.12 |
| 10 | 15427222 | rs1002598 | -0.703 | 1.328 | 0.206 | -1.018 | -0.003 | 0.002 | 0.002 | 0 | 0.23 | 0.23 | 0.15 | 0.16 | 0.27 | 0.16 |
| 10 | 15429075 | rs11259639 | -2.356 | 1.317 | 0.22 | -1.025 | 0.031 | 0.08 | 0.08 | 0.373 | 0.49 | 0.72 | 0.39 | 0.28 | 0.54 | 0.33 |
| 10 | 15443635 | rs4354607 | -0.886 | 1.211 | 0.156 | -0.916 | 0.007 | 0.011 | 0.011 | 2.063 | 0.13 | 0.47 | 0.08 | 0.1 | 0.11 | 0.05 |
| 10 | 15449578 | rs4748165 | -1.654 | 1.231 | -0.155 | -0.943 | -0.008 | 0.015 | 0.015 | 2.348 | 0.37 | 0.73 | 0.23 | 0.28 | 0.38 | 0.21 |
| 10 | 15470066 | rs7896716 | -0.754 | 1.387 | -0.292 | -0.921 | -0.041 | 0.003 | 0.003 | 0 | 0.9 | 0.97 | 0.9 | 0.93 | 0.8 | 0.93 |
| 10 | 15472860 | rs10906900 | -1.91 | 1.452 | -0.471 | -1.164 | 0.016 | 0.113 | 0.113 | 0.166 | 0.44 | 0.83 | 0.33 | 0.17 | 0.41 | 0.29 |
| 10 | 15476764 | rs10796290 | 2.02 | 1.41 | -0.41 | -1.032 | 0.052 | 0.043 | 0.043 | 0.625 | 0.45 | 0.18 | 0.56 | 0.74 | 0.38 | 0.64 |
| 10 | 15477626 | rs12246977 | 1.579 | 1.292 | -0.21 | -1.08 | -0.021 | -0.026 | -0.026 | 0 | 0.05 | 0.12 | 0.02 | 0 | 0.02 | 0.01 |
| 10 | 15479116 | rs7079961 | -1.612 | 1.293 | -0.056 | -1.126 | -0.001 | -0.005 | -0.005 | 0.611 | 0.41 | 0.43 | 0.3 | 0.2 | 0.43 | 0.35 |
| 10 | 15484342 | rs11259663 | -0.781 | 1.695 | -0.068 | -1.068 | -0.025 | 0.04 | 0.04 | 0.726 | 0.24 | 0.11 | 0.21 | 0.07 | 0.27 | 0.24 |
| 10 | 15494474 | rs4750646 | 0.862 | 1.449 | -0.149 | -1.138 | -0.002 | 0.005 | 0.005 | 0.393 | 0.68 | 0.3 | 0.8 | 0.83 | 0.69 | 0.81 |
| 10 | 15497132 | rs2057624 | 0.497 | 1.452 | -0.247 | -1.022 | 0.033 | -0.074 | -0.074 | 1.293 | 0.76 | 0.53 | 0.83 | 0.88 | 0.75 | 0.9 |
| 10 | 15497508 | rs7903131 | -0.346 | 1.556 | -0.45 | -1.201 | 0.021 | -0.093 | -0.093 | 0 | 0.21 | 0.38 | 0.15 | 0.03 | 0.21 | 0.07 |
| 10 | 15498463 | rs11259677 | -1.047 | 1.636 | -0.866 | -0.597 | -0.008 | 0.017 | 0.017 | 1.009 | 0.31 | 0.28 | 0.26 | 0.08 | 0.36 | 0.28 |
| 10 | 15505309 | rs10906902 | -1.18 | 1.44 | -1.027 | -0.224 | -0.01 | 0.005 | 0.005 | 0.209 | 0.16 | 0.09 | 0.18 | 0.09 | 0.22 | 0.18 |
| 10 | 15507240 | rs11599813 | -0.725 | 1.319 | -1.75 | -0.28 | -0.066 | 0.04 | 0.04 | 0.277 | 0.38 | 0.18 | 0.3 | 0.31 | 0.44 | 0.31 |
| 10 | 15507439 | rs10906903 | 1.487 | 1.074 | -1.721 | -0.514 | -0.047 | -0.025 | -0.025 | 0 | 0.43 | 0.11 | 0.41 | 0.57 | 0.48 | 0.56 |
| 10 | 15512600 | rs17156739 | 0.326 | 0.978 | -1.615 | -0.531 | 0.001 | -0.002 | -0.002 | 0 | 0.08 | 0.07 | 0.04 | 0.12 | 0.07 | 0.1 |
| 10 | 15513990 | rs7923056 | -0.685 | 1.182 | -0.867 | -0.667 | 0.016 | -0.059 | -0.059 | 0.011 | 0.37 | 0.57 | 0.3 | 0.2 | 0.38 | 0.21 |
| 10 | 15516402 | rs12260246 | 0.46 | 0.876 | -1.038 | -0.523 | -0.007 | 0.004 | 0.004 | 0 | 0.07 | 0.09 | 0.04 | 0.11 | 0.05 | 0.09 |
| 10 | 15524141 | rs10752384 | 1.171 | 0.896 | -0.877 | -0.556 | -0.025 | -0.013 | -0.013 | 0 | 0.86 | 0.38 | 0.92 | 0.92 | 0.94 | 0.93 |
| 10 | 15524162 | rs10508486 | 1.48 | 1.122 | -1.365 | -0.586 | -0.078 | 0.019 | 0.019 | 1.066 | 0.45 | 0.07 | 0.6 | 0.65 | 0.49 | 0.62 |
| 10 | 15525092 | rs4748169 | 1.505 | 1.116 | -1.367 | -0.564 | -0.08 | 0.012 | 0.012 | 1.072 | 0.44 | 0.07 | 0.6 | 0.63 | 0.49 | 0.61 |
| 10 | 15526255 | rs7893774 | 0.72 | 1.172 | -0.959 | -0.492 | 0 | 0.054 | 0.054 | 4.92 | 0.57 | 0.26 | 0.67 | 0.7 | 0.58 | 0.7 |
| 10 | 15529694 | rs12415975 | -1.075 | 1.512 | -0.911 | -0.352 | -0.016 | 0.004 | 0.004 | 0 | 0.11 | 0.1 | 0.21 | 0.13 | 0.21 | 0.13 |
| 10 | 15530101 | rs6602868 | 0.524 | 0.343 | -0.629 | -0.191 | 0.005 | 0.006 | 0.006 | 1.629 | 0.65 | 0.67 | 0.81 | 0.82 | 0.71 | 0.76 |
| 10 | 15533081 | rs4750656 | -0.606 | 0.151 | -0.696 | -0.294 | 0.027 | 0.025 | 0.025 | 2.906 | 0.82 | 0.9 | 0.93 | 1 | 0.9 | 0.94 |
| 10 | 15533137 | rs7072635 | 0.99 | 0.296 | -0.734 | -0.216 | 0.003 | 0.011 | 0.011 | 0.172 | 0.6 | 0.61 | 0.68 | 0.66 | 0.62 | 0.68 |
| 10 | 15533570 | rs4750657 | -0.935 | 0.596 | -0.98 | -0.267 | -0.017 | 0.017 | 0.017 | 0 | 0.2 | 0.15 | 0.23 | 0.19 | 0.27 | 0.16 |
| 10 | 15537270 | rs10752391 | 0.305 | 0.088 | -1.253 | -0.12 | -0.019 | 0.094 | 0.094 | 4.98 | 0.66 | 0.95 | 0.71 | 0.84 | 0.73 | 0.77 |
| 10 | 15537583 | rs7092715 | -0.409 | -0.211 | -1.188 | -0.286 | 0.039 | 0.097 | 0.097 | 2.57 | 0.76 | 0.95 | 0.91 | 1 | 0.88 | 0.9 |
| 10 | 15538218 | rs4750658 | 0.888 | -0.584 | -0.855 | 0.076 | 0.047 | 0.081 | 0.081 | 5.594 | 0.58 | 0.17 | 0.66 | 0.7 | 0.72 | 0.63 |
| 10 | 15539256 | rs17156833 | -0.337 | -0.622 | 0.414 | 0.012 | 0.065 | 0.103 | 0.103 | 5.052 | 0.05 | 0.54 | 0.05 | 0.05 | 0 | 0.01 |
| 10 | 15540058 | rs7070172 | -1.041 | -0.607 | 0.406 | 0.007 | 0.018 | 0.015 | 0.015 | 5.054 | 0.07 | 0.65 | 0.06 | 0.05 | 0.03 | 0.01 |
| 10 | 15540722 | rs1105180 | 0.91 | -0.618 | 0.336 | 0.079 | 0.043 | 0.077 | 0.077 | 5.132 | 0.58 | 0.19 | 0.67 | 0.66 | 0.7 | 0.63 |
| 10 | 15541140 | rs896433 | -1.503 | -0.618 | 0.341 | 0.06 | 0.001 | -0.015 | -0.015 | 2.273 | 0.19 | 0.71 | 0.25 | 0.15 | 0.2 | 0.13 |
| 10 | 15544077 | rs7901351 | 0.924 | -0.6 | 0.355 | 0.119 | 0.043 | 0.075 | 0.075 | 5.557 | 0.58 | 0.19 | 0.66 | 0.69 | 0.71 | 0.65 |
| 10 | 15545485 | rs17137176 | -1.6 | -0.34 | 0.485 | 0.167 | 0.001 | -0.025 | -0.025 | 0 | 0.19 | 0.71 | 0.25 | 0.1 | 0.2 | 0.12 |
| 10 | 15549080 | rs7910831 | -0.298 | -0.277 | 0.466 | 0.217 | 0.012 | -0.006 | -0.006 | 0 | 0.12 | 0.19 | 0.21 | 0.1 | 0.17 | 0.12 |
| 10 | 15552323 | rs4748177 | 0.705 | -0.695 | 0.3 | 0.997 | 0.031 | -0.082 | -0.082 | 0 | 0.59 | 0.12 | 0.65 | 0.44 | 0.71 | 0.63 |
| 10 | 15555232 | rs7085373 | 0.718 | -0.688 | 0.355 | 1.025 | 0.039 | -0.05 | -0.05 | 0 | 0.59 | 0.15 | 0.65 | 0.44 | 0.71 | 0.63 |
| 10 | 15557321 | rs878319 | -0.707 | -0.683 | 0.468 | 1.011 | 0.043 | 0.123 | 0.123 | 0.089 | 0.15 | 0.41 | 0.23 | 0.32 | 0.2 | 0.28 |
| 10 | 15561543 | rs7077361 | 0.816 | -0.654 | 0.595 | 0.962 | 0.027 | 0.064 | 0.064 | 0.057 | 0.2 | 0.07 | 0.07 | 0 | 0.1 | 0.08 |
| 10 | 15563450 | rs4750665 | 0.831 | -0.649 | 0.47 | 1.024 | 0.026 | -0.1 | -0.1 | 1.286 | 0.58 | 0.08 | 0.64 | 0.44 | 0.71 | 0.62 |
| 10 | 15573205 | rs9333240 | 0.057 | -0.389 | 0.488 | 0.792 | -0.005 | -0.004 | -0.004 | 0.003 | 0.12 | 0.09 | 0.2 | 0.13 | 0.2 | 0.14 |
| 10 | 15575030 | rs11818881 | 0.127 | -0.686 | 0.525 | 0.631 | -0.055 | -0.155 | -0.155 | 0 | 0.27 | 0.43 | 0.11 | 0 | 0.11 | 0.1 |
| 10 | 15595361 | rs11259736 | 1.598 | -0.561 | -0.462 | 0.079 | -0.043 | 0.108 | 0.108 | 9.436 | 0.17 | 0.01 | 0.1 | 0 | 0.18 | 0.05 |
| 10 | 15597263 | rs17137464 | 0.043 | -0.786 | 0.124 | 0.616 | 0 | -0.001 | -0.001 | 0 | 0.35 | 0.31 | 0.28 | 0.13 | 0.3 | 0.22 |
| 10 | 15597635 | rs10752399 | 0.62 | -0.585 | -0.729 | 0.977 | -0.033 | 0.005 | 0.005 | 6.253 | 0.45 | 0.07 | 0.57 | 0.37 | 0.57 | 0.53 |
| 10 | 15600737 | rs9333211 | -0.29 | -0.062 | -0.417 | 0.05 | 0 | -0.012 | -0.012 | 0 | 0.42 | 0.56 | 0.32 | 0.12 | 0.34 | 0.26 |
| 10 | 15604417 | rs7916526 | -0.266 | 0.003 | -0.587 | 0.092 | -0.006 | -0.046 | -0.046 | 0 | 0.42 | 0.53 | 0.31 | 0.12 | 0.34 | 0.26 |

| | | | | | | | | | | | | | | | | |
|---|---|---|---|---|---|---|---|---|---|---|---|---|---|---|---|---|
| 10 | 15609839 | rs12217558 | -1.01 | -0.029 | -0.647 | 0.73 | -0.004 | 0 | 0 | 0 | 0.07 | 0.09 | 0.08 | 0.15 | 0.05 | 0.07 |
| 10 | 15619054 | rs9333182 | -0.77 | -0.207 | -0.573 | 1.412 | 0.02 | 0.017 | 0.017 | 0 | 0.11 | 0.08 | 0.03 | 0.11 | 0.07 | 0.08 |
| 10 | 15620163 | rs1155200 | -1.213 | -0.275 | 0.378 | 1.372 | -0.037 | 0.16 | 0.16 | 0 | 0.15 | 0.39 | 0.27 | 0.39 | 0.08 | 0.27 |
| 10 | 15622887 | rs10906931 | 0.534 | 0.227 | -0.609 | 0.735 | -0.109 | 0.113 | 0.113 | 8.592 | 0.27 | 0.05 | 0.22 | 0.05 | 0.44 | 0.17 |
| 10 | 15626513 | rs1451668 | -0.941 | -0.072 | -0.621 | 1.083 | 0 | 0 | 0 | 0 | 0.5 | 0.47 | 0.57 | 0.55 | 0.45 | 0.53 |
| 10 | 15629360 | rs4750681 | -0.564 | -0.036 | -0.67 | 1.107 | -0.034 | 0.141 | 0.141 | 0.025 | 0.16 | 0.42 | 0.29 | 0.36 | 0.1 | 0.27 |
| 10 | 15633157 | rs1868996 | -0.035 | 0.672 | -0.974 | 0.803 | -0.161 | 0.051 | 0.051 | 0.514 | 0.33 | 0.02 | 0.29 | 0.18 | 0.36 | 0.25 |
| 10 | 15638751 | rs1057969 | 1.048 | 0.876 | -0.342 | 1.082 | 0.002 | 0.002 | 0.002 | 0 | 0.52 | 0.32 | 0.38 | 0.54 | 0.43 | 0.6 |
| 10 | 15640104 | rs980712 | 0.882 | 1.114 | -0.217 | 1.263 | -0.017 | 0.025 | 0.025 | 1.177 | 0.83 | 0.38 | 0.63 | 0.76 | 0.8 | 0.86 |
| 10 | 15645753 | rs1319475 | 0.102 | 1.424 | 0.71 | 1.287 | 0.046 | 0.005 | 0.005 | 0.391 | 0.87 | 0.76 | 0.67 | 0.85 | 0.78 | 0.9 |
| 10 | 15648104 | rs2282384 | -0.349 | 1.434 | 0.479 | 1.265 | -0.063 | -0.06 | -0.06 | 1 | 0.09 | 0.01 | 0.12 | 0.22 | 0.24 | 0.15 |
| 10 | 15650743 | rs9333156 | 1.445 | 1.333 | 0.608 | 1.147 | -0.001 | -0.001 | -0.001 | 0 | 0.57 | 0.58 | 0.41 | 0.54 | 0.44 | 0.64 |
| 10 | 15656674 | rs896431 | 0.966 | 0.438 | 1.122 | 0.766 | 0.031 | 0.055 | 0.055 | 0.262 | 0.83 | 0.43 | 0.61 | 0.75 | 0.74 | 0.88 |
| 10 | 15660502 | rs7092876 | -0.594 | 0.47 | 0.792 | 0.768 | 0.005 | 0.005 | 0.005 | 1.358 | 0.13 | 0.15 | 0.36 | 0.19 | 0.25 | 0.11 |
| 10 | 15662390 | rs10508489 | 0.741 | 0.272 | 0.852 | 0.701 | 0.024 | 0.041 | 0.041 | 0.658 | 0.16 | 0.05 | 0.09 | 0 | 0.07 | 0.07 |
| 10 | 15674416 | rs7082740 | -0.699 | 0.766 | 1.34 | 1.253 | 0.014 | 0.018 | 0.018 | 2.235 | 0.13 | 0.15 | 0.35 | 0.19 | 0.23 | 0.11 |
| 10 | 15682427 | rs1451665 | 1.327 | 0.745 | 1.378 | 1.254 | 0.013 | 0.043 | 0.043 | 0.43 | 0.83 | 0.41 | 0.6 | 0.76 | 0.77 | 0.87 |
| 10 | 15693659 | rs7089517 | -0.599 | 0.869 | 1.384 | 1.309 | 0.034 | 0.048 | 0.048 | 1.157 | 0.12 | 0.18 | 0.38 | 0.22 | 0.23 | 0.12 |
| 10 | 15702992 | rs9333114 | -0.666 | 1.205 | 1.201 | 1.619 | -0.009 | -0.012 | -0.012 | 0 | 0.08 | 0.03 | 0.12 | 0.31 | 0.07 | 0.09 |
| 10 | 15712828 | rs2039910 | 1.104 | 1.352 | 1.439 | 0.654 | 0.026 | 0.062 | 0.062 | 0.066 | 0.84 | 0.4 | 0.58 | 0.72 | 0.77 | 0.87 |
| 10 | 15713541 | rs6602051 | 1.228 | 1.344 | 1.009 | 0.574 | 0.062 | 0.047 | 0.047 | 0.819 | 0.8 | 0.54 | 0.56 | 0.71 | 0.67 | 0.81 |
| 10 | 15725723 | rs9333092 | -0.773 | 0.699 | 0.878 | 0.41 | -0.005 | -0.012 | -0.012 | 0 | 0.09 | 0.08 | 0.07 | 0.19 | 0.12 | 0.12 |
| 10 | 15729923 | rs1954181 | 1.766 | 0.519 | 1.557 | 0.325 | 0.012 | 0.061 | 0.061 | 7.3 | 0.86 | 0.52 | 0.89 | 0.79 | 0.86 | 0.85 |
| 10 | 15731185 | rs4748194 | 1.91 | 0.522 | 1.549 | 0.326 | 0.011 | 0.063 | 0.063 | 5.499 | 0.85 | 0.48 | 0.88 | 0.79 | 0.86 | 0.86 |
| 10 | 15731316 | rs9333077 | -1.071 | 0.58 | 0.998 | 0.349 | -0.003 | -0.014 | -0.014 | 0 | 0.11 | 0.1 | 0.08 | 0.18 | 0.12 | 0.12 |
| 10 | 15754920 | rs11575603 | -0.873 | 0.191 | 0.753 | -0.195 | 0.009 | 0.031 | 0.031 | 0.299 | 0.2 | 0.08 | 0.17 | 0.13 | 0.18 | 0.19 |
| 10 | 15760249 | rs9333075 | 1.583 | 0.195 | 2.09 | -0.39 | 0 | -0.005 | -0.005 | 0 | 0.7 | 0.33 | 0.72 | 0.83 | 0.71 | 0.77 |
| 10 | 15761209 | rs9333260 | -0.791 | 0.487 | 1.899 | -0.412 | -0.026 | 0.021 | 0.021 | 0 | 0.05 | 0.02 | 0.06 | 0.01 | 0.13 | 0.07 |
| 10 | 15769442 | rs1934937 | 1.19 | 0.143 | 2.007 | -0.468 | -0.004 | -0.044 | -0.044 | 0 | 0.69 | 0.4 | 0.72 | 0.86 | 0.68 | 0.75 |
| 10 | 15776239 | rs1592035 | 0.987 | -0.44 | 2.025 | -0.723 | -0.044 | -0.08 | -0.08 | 0 | 0.68 | 0.58 | 0.79 | 0.87 | 0.77 | 0.79 |
| 10 | 15779501 | rs17395099 | -1.16 | -0.485 | 1.921 | -0.727 | 0.043 | 0.073 | 0.073 | 3.913 | 0.29 | 0.16 | 0.19 | 0.12 | 0.22 | 0.23 |
| 10 | 15783765 | rs1341102 | 0.97 | -0.422 | 1.758 | -0.795 | -0.005 | -0.09 | -0.09 | 0.091 | 0.7 | 0.46 | 0.74 | 0.88 | 0.69 | 0.72 |
| 10 | 15786958 | rs1341103 | 1.045 | -0.837 | 1.834 | -0.657 | -0.033 | -0.085 | -0.085 | 0 | 0.67 | 0.45 | 0.78 | 0.91 | 0.77 | 0.76 |
| 10 | 15790632 | rs12242907 | -2.129 | -0.763 | 1.941 | -0.519 | 0.091 | 0.04 | 0.04 | 1.898 | 0.06 | 0.52 | 0.06 | 0 | 0.04 | 0.01 |
| 10 | 15798477 | rs7070312 | -1.178 | -1.067 | 2.021 | -0.647 | -0.006 | -0.016 | -0.016 | 0 | 0.28 | 0.28 | 0.17 | 0.1 | 0.17 | 0.26 |
| 10 | 15799150 | rs9943441 | -1.625 | -1.189 | 1.611 | -0.755 | 0.01 | 0 | 0 | 11.829 | 0.36 | 0.61 | 0.23 | 0.12 | 0.23 | 0.24 |
| 10 | 15801975 | rs11253625 | -0.819 | -0.948 | 1.465 | -0.643 | 0.028 | 0.077 | 0.077 | 15.294 | 0.24 | 0.07 | 0.17 | 0.11 | 0.18 | 0.22 |
| 10 | 15805171 | rs7909432 | -1.635 | -0.926 | 1.753 | -0.662 | -0.027 | -0.068 | -0.068 | 0 | 0.3 | 0.55 | 0.23 | 0.12 | 0.23 | 0.23 |
| 10 | 15810354 | rs11253637 | -1.281 | -1.118 | 2.001 | -0.58 | 0.023 | -0.006 | -0.006 | 0 | 0.08 | 0.48 | 0.07 | 0 | 0.05 | 0.01 |
| 10 | 15817004 | rs7904868 | -1.782 | -1.197 | 1.957 | -0.031 | -0.038 | 0.071 | 0.071 | 4.654 | 0.1 | 0.69 | 0.23 | 0.17 | 0.05 | 0.03 |
| 10 | 15835277 | rs12262809 | -1.549 | -1.318 | 1.708 | -0.443 | 0.043 | 0.057 | 0.057 | 10.353 | 0.32 | 0.84 | 0.33 | 0.16 | 0.26 | 0.24 |
| 10 | 15837431 | rs7904872 | -1.688 | -1.092 | 1.655 | -0.484 | -0.028 | -0.046 | -0.046 | 0 | 0.1 | 0.65 | 0.09 | 0.01 | 0.05 | 0.03 |
| 10 | 15843462 | rs10904627 | -0.221 | -0.683 | 1.336 | -0.478 | -0.032 | 0.047 | 0.047 | 6.505 | 0.23 | 0.03 | 0.25 | 0.13 | 0.23 | 0.26 |
| 10 | 15854752 | rs10904629 | -1.449 | -1.663 | 0.565 | -0.697 | -0.071 | -0.065 | -0.065 | 0.213 | 0.64 | 0.95 | 0.59 | 0.57 | 0.45 | 0.57 |
| 10 | 15861576 | rs10795322 | -0.224 | -1.196 | 1.511 | -1.193 | -0.024 | 0.048 | 0.048 | 6.528 | 0.22 | 0.01 | 0.21 | 0.09 | 0.2 | 0.25 |
| 10 | 15865076 | rs12258622 | -1.731 | -1.298 | 1.813 | -1.307 | -0.008 | -0.016 | -0.016 | 0 | 0.1 | 0.58 | 0.08 | 0.03 | 0.07 | 0.03 |
| 10 | 15872827 | rs7896076 | -1.34 | -1.468 | 1.834 | -1.433 | 0.004 | -0.022 | -0.022 | 0 | 0.08 | 0.43 | 0.06 | 0 | 0.05 | 0.01 |
| 10 | 15873288 | rs12265790 | -1.536 | -1.467 | 1.699 | -1.433 | -0.001 | -0.028 | -0.028 | 0 | 0.09 | 0.5 | 0.07 | 0 | 0.05 | 0.01 |
| 10 | 15895956 | rs12218978 | -0.591 | -1.073 | 0.933 | -1.279 | -0.006 | -0.029 | -0.029 | 0 | 0.25 | 0.26 | 0.24 | 0.05 | 0.19 | 0.23 |
| 10 | 15934921 | rs7079984 | -1.304 | -0.375 | 0.686 | -0.72 | 0.085 | 0.075 | 0.075 | 0.376 | 0.05 | 0.4 | 0.06 | 0.04 | 0.04 | 0.03 |
| 10 | 15943346 | rs11253684 | -0.68 | -0.537 | -0.476 | -0.844 | -0.018 | -0.031 | -0.031 | 0.276 | 0.12 | 0.77 | 0.09 | 0.03 | 0.06 | 0.07 |
| 10 | 15945316 | rs10752030 | -0.047 | -0.171 | -1.597 | -1.457 | -0.051 | 0.259 | 0.259 | 4.232 | 0.56 | 0.11 | 0.68 | 0.92 | 0.56 | 0.71 |
| 10 | 15950149 | rs1339474 | 0.175 | -0.157 | -1.699 | -1.506 | -0.039 | 0.293 | 0.293 | 1.064 | 0.47 | 0.08 | 0.55 | 0.9 | 0.45 | 0.63 |
| 10 | 15977878 | rs11253696 | 0.087 | 0.328 | -1.911 | -0.828 | 0.005 | 0.202 | 0.202 | 2.246 | 0.51 | 0.1 | 0.36 | 0.81 | 0.42 | 0.66 |
| 10 | 15982227 | rs2354501 | -0.735 | 0.735 | -1.52 | -0.793 | 0.016 | 0.156 | 0.156 | 0 | 0.57 | 0.85 | 0.69 | 0.88 | 0.58 | 0.68 |
| 10 | 15983045 | rs4748214 | -0.304 | 0.829 | -0.713 | -0.229 | 0.032 | 0.05 | 0.05 | 0.335 | 0.23 | 0.38 | 0.28 | 0.61 | 0.26 | 0.34 |
| 10 | 15984035 | rs10904645 | -0.359 | 0.148 | -0.434 | -0.21 | -0.04 | -0.028 | -0.028 | 0 | 0.33 | 0.47 | 0.37 | 0.26 | 0.21 | 0.34 |
| 10 | 15988902 | rs1538564 | -1.013 | 0.59 | 0.516 | 0.593 | 0.002 | -0.003 | -0.003 | 0.103 | 0.4 | 0.47 | 0.45 | 0.59 | 0.45 | 0.65 |
| 10 | 15996876 | rs10508496 | -0.924 | 0.535 | 1.796 | 0.77 | -0.011 | 0.071 | 0.071 | 0 | 0.24 | 0.39 | 0.17 | 0.4 | 0.22 | 0.41 |
| 10 | 15997352 | rs11253707 | -0.617 | 0.487 | 1.817 | 0.749 | -0.001 | 0.027 | 0.027 | 0.254 | 0.22 | 0.29 | 0.16 | 0.4 | 0.21 | 0.41 |
| 10 | 16000007 | rs867464 | -1.957 | 0.323 | 1.738 | 0.667 | -0.035 | 0.015 | 0.015 | 0 | 0.46 | 0.88 | 0.49 | 0.58 | 0.46 | 0.68 |
| 10 | 16005144 | rs11253709 | -1.221 | -0.129 | 2.299 | 0.766 | -0.035 | 0.12 | 0.12 | 0.109 | 0.25 | 0.47 | 0.16 | 0.41 | 0.17 | 0.43 |
| 10 | 16005841 | rs7907660 | -1.715 | 0.06 | 2.396 | 0.754 | -0.01 | 0.01 | 0.01 | 0 | 0.41 | 0.74 | 0.39 | 0.59 | 0.43 | 0.68 |
| 10 | 16013463 | rs1341558 | -1.172 | 0.107 | 2.444 | 0.753 | 0.024 | -0.024 | -0.024 | 0 | 0.22 | 0.5 | 0.33 | 0.18 | 0.26 | 0.26 |
| 10 | 16016468 | rs4748232 | -0.259 | 0.132 | 2.75 | 0.79 | 0.016 | -0.019 | -0.019 | 0 | 0.17 | 0.17 | 0.3 | 0.1 | 0.24 | 0.24 |
| 10 | 16018021 | rs10904656 | 1.947 | 0.05 | 2.661 | 0.769 | -0.022 | 0.039 | 0.039 | 0 | 0.53 | 0.12 | 0.53 | 0.37 | 0.57 | 0.3 |
| 10 | 16019793 | rs4748233 | -2.131 | 0.086 | 2.702 | 0.766 | -0.035 | 0.012 | 0.012 | 0 | 0.44 | 0.87 | 0.48 | 0.6 | 0.43 | 0.67 |
| 10 | 16021427 | rs11592119 | -0.656 | 0.268 | 2.854 | 0.77 | 0.038 | -0.006 | -0.006 | 0 | 0.18 | 0.3 | 0.31 | 0.18 | 0.26 | 0.24 |
| 10 | 16027818 | rs7079031 | -1.857 | 0.173 | 2.912 | 0.785 | 0.011 | 0.083 | 0.083 | 0.528 | 0.09 | 0.43 | 0.03 | 0.08 | 0.02 | 0.03 |

| | | | | | | | | | | | | | | | | |
|---|---|---|---|---|---|---|---|---|---|---|---|---|---|---|---|---|
| 10 | 16041011 | rs12773486 | -0.421 | 0.219 | 2.213 | 0.877 | 0.006 | -0.019 | -0.019 | 0 | 0.2 | 0.2 | 0.13 | 0.41 | 0.16 | 0.41 |
| 10 | 16043948 | rs11812670 | -0.554 | 0.334 | 1.962 | 0.967 | -0.016 | 0.071 | 0.071 | 0.015 | 0.23 | 0.37 | 0.14 | 0.42 | 0.17 | 0.41 |
| 10 | 16045536 | rs4747256 | -1.584 | 0.435 | 1.956 | 0.879 | -0.01 | -0.02 | -0.02 | 1.083 | 0.4 | 0.69 | 0.55 | 0.51 | 0.42 | 0.67 |
| 10 | 16050599 | rs12779523 | -0.547 | 0.526 | 1.548 | 0.977 | -0.025 | 0.099 | 0.099 | 0.154 | 0.22 | 0.41 | 0.14 | 0.41 | 0.16 | 0.41 |
| 10 | 16051758 | rs11253734 | -0.892 | 0.327 | 1.201 | 0.933 | 0.006 | 0.015 | 0.015 | 0 | 0.38 | 0.51 | 0.53 | 0.5 | 0.42 | 0.65 |
| 10 | 16061958 | rs870550 | 0.671 | 0.508 | 0.126 | 0.598 | 0 | -0.003 | -0.003 | 0 | 0.56 | 0.49 | 0.47 | 0.39 | 0.55 | 0.32 |
| 10 | 16065973 | rs17399470 | -1.039 | 0.173 | -0.122 | 0.369 | 0.001 | -0.006 | -0.006 | 0.024 | 0.1 | 0.06 | 0.07 | 0.02 | 0.09 | 0.09 |
| 10 | 16068563 | rs10904665 | 0.948 | -0.285 | -0.82 | 0.273 | 0.058 | 0.083 | 0.083 | 0.21 | 0.56 | 0.15 | 0.68 | 0.26 | 0.65 | 0.3 |
| 10 | 16070792 | rs947335 | 0.068 | -0.613 | 0.185 | 0.188 | 0.001 | -0.004 | -0.004 | 0 | 0.74 | 0.73 | 0.79 | 0.29 | 0.74 | 0.5 |
| 10 | 16073677 | rs10752039 | 0.476 | -0.658 | 1.295 | 0.161 | -0.018 | -0.004 | -0.004 | 0.25 | 0.69 | 0.42 | 0.77 | 0.28 | 0.74 | 0.46 |
| 10 | 16074308 | rs10795339 | 0.788 | -0.672 | 1.473 | 0.155 | -0.001 | -0.001 | -0.001 | 0 | 0.66 | 0.31 | 0.76 | 0.28 | 0.73 | 0.46 |
| 10 | 16078154 | rs7069992 | 0.14 | -0.574 | 1.405 | 0.247 | -0.001 | 0 | 0 | 0 | 0.73 | 0.72 | 0.8 | 0.29 | 0.74 | 0.49 |
| 10 | 16083443 | rs10904670 | 0.174 | -0.589 | 1.515 | 0.396 | -0.001 | -0.003 | -0.003 | 0 | 0.76 | 0.73 | 0.8 | 0.3 | 0.74 | 0.5 |
| 10 | 16099553 | rs2355247 | 0.129 | -0.642 | 1.604 | 0.372 | 0 | 0.002 | 0.002 | 0 | 0.77 | 0.72 | 0.8 | 0.29 | 0.74 | 0.5 |
| 10 | 16118555 | rs4748245 | 0.253 | -0.653 | 2.486 | 0.386 | 0.002 | 0.022 | 0.022 | 0 | 0.76 | 0.62 | 0.79 | 0.29 | 0.74 | 0.51 |
| 10 | 16134582 | rs12258951 | 1.438 | -0.943 | 2.477 | 0.235 | 0 | -0.001 | -0.001 | 0 | 0.61 | 0.41 | 0.72 | 0.26 | 0.66 | 0.32 |
| 10 | 16138489 | rs11253779 | 1.445 | -0.953 | 2.505 | 0.231 | -0.002 | -0.003 | -0.003 | 0.247 | 0.6 | 0.42 | 0.72 | 0.26 | 0.66 | 0.32 |
| 10 | 16139951 | rs4748251 | 1.275 | -0.921 | 2.579 | 0.246 | -0.012 | -0.019 | -0.019 | 0.016 | 0.62 | 0.48 | 0.73 | 0.26 | 0.67 | 0.32 |
| 10 | 16145787 | rs7098374 | 1.216 | -0.91 | 2.624 | 0.226 | 0.02 | 0.03 | 0.03 | 0 | 0.6 | 0.31 | 0.71 | 0.26 | 0.66 | 0.32 |
| 10 | 16148908 | rs1340151 | 0.05 | -0.885 | 2.665 | 0.394 | 0 | 0.005 | 0.005 | 0 | 0.77 | 0.7 | 0.81 | 0.29 | 0.74 | 0.51 |
| 10 | 16154458 | rs1340149 | 1.009 | -0.961 | 2.879 | 0.368 | -0.011 | -0.016 | -0.016 | 0 | 0.64 | 0.5 | 0.74 | 0.27 | 0.66 | 0.42 |
| 10 | 16154583 | rs12780183 | 1.113 | -0.984 | 2.773 | 0.368 | -0.008 | -0.012 | -0.012 | 0 | 0.64 | 0.44 | 0.73 | 0.27 | 0.66 | 0.42 |
| 10 | 16158880 | rs10795348 | 1.07 | -1.159 | 2.825 | 0.782 | -0.008 | -0.001 | -0.001 | 0.493 | 0.64 | 0.54 | 0.73 | 0.33 | 0.65 | 0.46 |
| 10 | 16160637 | rs1340144 | 1.756 | -0.385 | 1.888 | -0.019 | -0.03 | 0.248 | 0.248 | 0.837 | 0.44 | 0.06 | 0.55 | 0.13 | 0.38 | 0.24 |
| 10 | 16161797 | rs7914198 | 1.46 | -0.043 | 2.679 | 0.294 | -0.008 | -0.055 | -0.055 | 0.031 | 0.8 | 0.65 | 0.84 | 0.92 | 0.82 | 0.74 |
| 10 | 16174989 | rs1417088 | 0.707 | 0.058 | 2.31 | 0.322 | -0.007 | -0.038 | -0.038 | 0.71 | 0.87 | 0.79 | 0.91 | 0.97 | 0.9 | 0.88 |
| 10 | 16190790 | rs2991228 | -0.769 | 0.367 | 2.269 | 0.529 | 0.053 | 0.012 | 0.012 | 0.42 | 0.16 | 0.3 | 0.18 | 0.13 | 0.28 | 0.11 |
| 10 | 16196322 | rs12357007 | -0.182 | 0.402 | 1.873 | 0.513 | -0.043 | -0.027 | -0.027 | 1.012 | 0.12 | 0.07 | 0.15 | 0.12 | 0.26 | 0.11 |
| 10 | 16198980 | rs2226024 | -0.624 | 0.397 | 1.841 | 0.59 | -0.068 | -0.05 | -0.05 | 0 | 0.14 | 0.04 | 0.16 | 0.15 | 0.28 | 0.12 |
| 10 | 16200958 | rs946810 | 1.721 | -0.051 | 2.332 | 0.402 | -0.015 | -0.038 | -0.038 | 0.015 | 0.75 | 0.37 | 0.82 | 0.9 | 0.82 | 0.7 |
| 10 | 16204155 | rs4747262 | 1.438 | 0.032 | 2.049 | 0.413 | -0.013 | -0.052 | -0.052 | 0.021 | 0.78 | 0.71 | 0.85 | 0.91 | 0.82 | 0.7 |
| 10 | 16220828 | rs1934641 | 0.98 | 0.242 | 2.185 | 0.759 | 0.056 | 0.064 | 0.064 | 0.022 | 0.89 | 0.64 | 0.88 | 0.85 | 0.88 | 0.86 |
| 10 | 16221561 | rs9787585 | 2.042 | -0.056 | 2.188 | 0.476 | 0.011 | 0.03 | 0.03 | 0.341 | 0.73 | 0.25 | 0.78 | 0.84 | 0.79 | 0.68 |
| 10 | 16233175 | rs4748263 | -0.471 | 0.33 | 1.787 | 0.669 | -0.102 | -0.052 | -0.052 | 0.713 | 0.17 | 0.02 | 0.17 | 0.16 | 0.32 | 0.25 |
| 10 | 16241855 | rs10508502 | 2.337 | 0.252 | 1.974 | 0.727 | -0.033 | 0.014 | 0.014 | 1.371 | 0.57 | 0.22 | 0.61 | 0.64 | 0.47 | 0.46 |
| 10 | 16243408 | rs1340143 | 0.805 | 0.31 | 1.932 | 0.669 | -0.129 | -0.03 | -0.03 | 3.166 | 0.79 | 0.95 | 0.79 | 0.84 | 0.62 | 0.71 |
| 10 | 16244094 | rs7087787 | -1.291 | 0.219 | 2.41 | 0.593 | -0.022 | -0.078 | -0.078 | 0 | 0.14 | 0.35 | 0.09 | 0.03 | 0.1 | 0.17 |
| 10 | 16255006 | rs1340142 | 1.884 | 0.192 | 2.295 | 0.565 | 0.01 | 0.015 | 0.015 | 0.125 | 0.75 | 0.26 | 0.8 | 0.96 | 0.8 | 0.76 |
| 10 | 16256337 | rs4748264 | 1.738 | 0.19 | 2.534 | 0.655 | 0.004 | -0.04 | -0.04 | 1.943 | 0.77 | 0.56 | 0.82 | 0.84 | 0.79 | 0.73 |
| 10 | 16268673 | rs17136428 | -0.849 | 0.187 | 2.539 | 0.651 | -0.014 | -0.05 | -0.05 | 0 | 0.13 | 0.2 | 0.08 | 0.02 | 0.09 | 0.16 |
| 10 | 16269491 | rs7083560 | 2.656 | 0.157 | 2.465 | 0.704 | -0.01 | 0.013 | 0.013 | 1.101 | 0.58 | 0.37 | 0.62 | 0.66 | 0.48 | 0.48 |
| 10 | 16270884 | rs3858179 | 2.811 | 0.124 | 2.336 | 1.479 | -0.007 | -0.015 | -0.015 | 0 | 0.57 | 0.36 | 0.6 | 0.49 | 0.47 | 0.46 |
| 10 | 16271816 | rs17309402 | -0.971 | 0.084 | 2.295 | 1.106 | -0.084 | -0.028 | -0.028 | 0.195 | 0.19 | 0.07 | 0.19 | 0.16 | 0.32 | 0.27 |
| 10 | 16274905 | rs7092247 | 2.042 | 0.061 | 2.381 | 1.058 | -0.009 | -0.053 | -0.053 | 0 | 0.76 | 0.39 | 0.81 | 0.95 | 0.81 | 0.73 |
| 10 | 16279015 | rs1538761 | -0.197 | 0.091 | 2.356 | 1.07 | 0.017 | -0.027 | -0.027 | 0.803 | 0.93 | 0.86 | 0.93 | 0.99 | 0.91 | 0.93 |
| 10 | 16279342 | rs10904696 | -1.327 | 0.05 | 2.338 | 1.075 | -0.015 | -0.005 | -0.005 | 0 | 0.21 | 0.19 | 0.2 | 0.19 | 0.33 | 0.31 |
| 10 | 16281080 | rs7902512 | -0.76 | -0.078 | 2.354 | 0.762 | -0.011 | -0.04 | -0.04 | 0 | 0.12 | 0.18 | 0.08 | 0.02 | 0.09 | 0.19 |
| 10 | 16281483 | rs1413709 | -1.939 | -0.732 | 1.926 | 0.504 | -0.02 | 0.225 | 0.225 | 2.428 | 0.51 | 0.83 | 0.39 | 0.84 | 0.52 | 0.74 |
| 10 | 16284825 | rs11253863 | -1.064 | -1.045 | 2.343 | 0.732 | -0.013 | -0.021 | -0.021 | 3.319 | 0.13 | 0.19 | 0.08 | 0.08 | 0.1 | 0.19 |
| 10 | 16286345 | rs1417091 | 2.143 | -1.045 | 1.911 | 0.727 | 0.008 | 0.019 | 0.019 | 0.66 | 0.75 | 0.25 | 0.8 | 0.89 | 0.8 | 0.72 |
| 10 | 16297590 | rs1340133 | 0.191 | -0.929 | 1.919 | 0.757 | 0.024 | -0.033 | -0.033 | 0 | 0.92 | 0.84 | 0.93 | 0.99 | 0.9 | 0.93 |
| 10 | 16297598 | rs1340132 | 1.272 | -1.409 | 1.055 | 0.29 | 0.029 | 0.096 | 0.096 | 0 | 0.64 | 0.24 | 0.73 | 0.88 | 0.71 | 0.69 |
| 10 | 16302262 | rs10400069 | 0.704 | -1.423 | 1.289 | 0.483 | -0.01 | -0.037 | -0.037 | 2.04 | 0.68 | 0.64 | 0.77 | 0.81 | 0.73 | 0.7 |
| 10 | 16302893 | rs7921272 | -1.891 | -1.608 | 1.25 | 0.647 | -0.049 | 0.054 | 0.054 | 0 | 0.21 | 0.51 | 0.15 | 0.3 | 0.14 | 0.38 |
| 10 | 16303374 | rs10159475 | -0.631 | -1.664 | 1.433 | 0.588 | -0.024 | -0.031 | -0.031 | 0 | 0.24 | 0.32 | 0.17 | 0.17 | 0.18 | 0.22 |
| 10 | 16303877 | rs1340130 | 0.302 | -1.373 | 1.227 | 0.423 | 0.002 | -0.012 | -0.012 | 0 | 0.9 | 0.9 | 0.94 | 0.97 | 0.9 | 0.92 |
| 10 | 16305281 | rs872430 | -0.807 | -1.404 | 1.372 | 0.435 | -0.01 | 0.009 | 0.009 | 0 | 0.09 | 0.15 | 0.05 | 0.13 | 0.06 | 0.15 |
| 10 | 16308456 | rs963770 | 1.136 | -1.363 | 1.371 | 0.434 | -0.002 | 0.006 | 0.006 | 0.302 | 0.84 | 0.76 | 0.9 | 0.84 | 0.86 | 0.78 |
| 10 | 16310069 | rs10082530 | 0.597 | -1.423 | 1.666 | 0.629 | -0.014 | -0.063 | -0.063 | 0 | 0.15 | 0.34 | 0.15 | 0.03 | 0.12 | 0.07 |
| 10 | 16314717 | rs1340129 | 2.281 | -1.395 | 1.387 | 0.601 | -0.003 | 0.017 | 0.017 | 1.244 | 0.77 | 0.33 | 0.76 | 0.71 | 0.81 | 0.67 |
| 10 | 16317194 | rs10752046 | -0.112 | -1.391 | 1.39 | 0.584 | 0.003 | -0.007 | -0.007 | 0 | 0.91 | 0.91 | 0.94 | 0.97 | 0.9 | 0.92 |
| 10 | 16317571 | rs10508506 | 1.225 | -1.39 | 1.07 | 0.581 | 0.001 | 0 | 0 | 0.487 | 0.65 | 0.36 | 0.69 | 0.68 | 0.71 | 0.63 |
| 10 | 16322429 | rs10508507 | 1.713 | -1.293 | 1.056 | 0.704 | -0.009 | 0.007 | 0.007 | 2.435 | 0.78 | 0.34 | 0.8 | 0.75 | 0.83 | 0.71 |
| 10 | 16322710 | rs7075260 | 1.662 | -1.364 | 0.307 | 0.084 | -0.029 | 0.262 | 0.262 | 0.249 | 0.46 | 0.07 | 0.62 | 0.1 | 0.61 | 0.24 |
| 10 | 16324673 | rs10904706 | 0.939 | -0.922 | 1.055 | 0.562 | 0.02 | 0.015 | 0.015 | 1.811 | 0.82 | 0.59 | 0.83 | 0.8 | 0.83 | 0.79 |
| 10 | 16325878 | rs3802557 | 1.745 | -1.197 | 0.669 | 0.487 | 0.028 | -0.012 | -0.012 | 0 | 0.68 | 0.22 | 0.73 | 0.59 | 0.75 | 0.57 |
| 10 | 16342456 | rs1577221 | 0.575 | -0.993 | 0.442 | 0.536 | -0.01 | 0 | 0 | 0.469 | 0.83 | 0.89 | 0.87 | 0.82 | 0.82 | 0.79 |
| 10 | 16346741 | rs10904714 | 1.013 | -1.198 | 0.745 | 0.491 | 0.015 | 0.007 | 0.007 | 0 | 0.6 | 0.34 | 0.68 | 0.62 | 0.69 | 0.55 |
| 10 | 16355503 | rs10904716 | 0.179 | -1.145 | 0.368 | 0.704 | -0.009 | -0.107 | -0.107 | 0 | 0.24 | 0.09 | 0.13 | 0.48 | 0.2 | 0.32 |

| | | | | | | | | | | | | | | | | |
|---|---|---|---|---|---|---|---|---|---|---|---|---|---|---|---|---|
| 10 | 16358026 | rs6602094 | 1.265 | -1.147 | 0.839 | 0.706 | 0.051 | 0.042 | 0.042 | 0.298 | 0.57 | 0.21 | 0.65 | 0.63 | 0.67 | 0.54 |
| 10 | 16358599 | rs7088653 | 1.112 | -1.131 | 0.945 | 0.602 | 0.001 | 0.03 | 0.03 | 1.454 | 0.82 | 0.53 | 0.83 | 0.76 | 0.85 | 0.77 |
| 10 | 16378864 | rs2130227 | -0.274 | -0.923 | 0.436 | 0.593 | -0.053 | -0.13 | -0.13 | 0.12 | 0.26 | 0.46 | 0.16 | 0.05 | 0.18 | 0.19 |
| 10 | 16379231 | rs2130226 | 0.133 | -0.808 | -0.413 | 0.747 | -0.012 | -0.024 | -0.024 | 0 | 0.86 | 0.94 | 0.89 | 0.79 | 0.87 | 0.83 |
| 10 | 16386614 | rs7088589 | 1.108 | -0.809 | -0.464 | 0.685 | 0.051 | 0.044 | 0.044 | 0.224 | 0.57 | 0.21 | 0.65 | 0.65 | 0.66 | 0.55 |
| 10 | 16386748 | rs6602097 | 0.991 | -0.094 | -0.42 | 1.295 | -0.006 | -0.021 | -0.021 | 0 | 0.83 | 0.88 | 0.82 | 0.7 | 0.82 | 0.71 |
| 10 | 16389784 | rs2355638 | -0.083 | 0.037 | -1.08 | 1.424 | -0.074 | -0.146 | -0.146 | 0 | 0.8 | 0.99 | 0.91 | 0.58 | 0.8 | 0.79 |
| 10 | 16395866 | rs10904720 | 0.114 | -0.264 | -0.774 | 1.461 | 0.009 | -0.034 | -0.034 | 0.09 | 0.22 | 0.15 | 0.09 | 0.4 | 0.16 | 0.21 |
| 10 | 16396872 | rs17318083 | 1.079 | -0.272 | -1.522 | 1.036 | -0.174 | 0.107 | 0.107 | 0.532 | 0.39 | 0.03 | 0.54 | 0.18 | 0.52 | 0.32 |
| 10 | 16400072 | rs4748282 | 0.49 | -0.494 | -0.745 | 0.801 | 0.037 | 0.219 | 0.219 | 0.824 | 0.5 | 0.16 | 0.65 | 0.21 | 0.61 | 0.37 |
| 10 | 16408872 | rs11253941 | 0.884 | -0.322 | -0.345 | 1.332 | 0.002 | 0.004 | 0.004 | 0.028 | 0.13 | 0.14 | 0.1 | 0.34 | 0.12 | 0.13 |
| 10 | 16412510 | rs4748285 | 0.727 | -0.391 | -0.07 | 1.083 | -0.012 | 0.083 | 0.083 | 3.769 | 0.43 | 0.28 | 0.61 | 0.24 | 0.57 | 0.32 |
| 10 | 16421596 | rs7910788 | -0.633 | -0.442 | 0.18 | 1.094 | 0.024 | 0.016 | 0.016 | 0.468 | 0.63 | 0.83 | 0.82 | 0.68 | 0.74 | 0.6 |
| 10 | 16425017 | rs12261978 | 0.72 | -0.441 | 0.551 | 1.055 | -0.002 | 0.032 | 0.032 | 3.735 | 0.43 | 0.38 | 0.62 | 0.21 | 0.57 | 0.33 |
| 10 | 16427107 | rs7071889 | -0.094 | -0.45 | 0.457 | 1.05 | -0.006 | -0.003 | -0.003 | 0.065 | 0.56 | 0.55 | 0.72 | 0.59 | 0.68 | 0.46 |
| 10 | 16427375 | rs11253950 | 1.042 | -0.461 | 0.05 | 1.031 | 0.006 | -0.007 | -0.007 | 0 | 0.13 | 0.13 | 0.11 | 0.33 | 0.11 | 0.14 |
| 10 | 16427772 | rs4748292 | 0.985 | -0.461 | 0.041 | 1.054 | -0.056 | 0.076 | 0.076 | 4.789 | 0.41 | 0.13 | 0.59 | 0.27 | 0.57 | 0.33 |
| 10 | 16427927 | rs12218570 | 0.06 | -0.28 | 0.357 | 1.121 | -0.029 | 0.102 | 0.102 | 2.765 | 0.18 | 0.6 | 0.15 | 0.34 | 0.13 | 0.14 |
| 10 | 16430308 | rs11253965 | 0.571 | -0.196 | -0.14 | 1.114 | 0.005 | -0.023 | -0.023 | 0.903 | 0.16 | 0.15 | 0.11 | 0.33 | 0.14 | 0.14 |
| 10 | 16432797 | rs10795384 | -0.133 | -0.193 | -0.555 | 1.254 | 0.002 | -0.003 | -0.003 | 0 | 0.6 | 0.67 | 0.73 | 0.55 | 0.7 | 0.48 |
| 10 | 16435571 | rs7094646 | 0.651 | -0.029 | -0.663 | 0.947 | -0.015 | 0.156 | 0.156 | 0.348 | 0.17 | 0.52 | 0.41 | 0.34 | 0.15 | 0.15 |
| 10 | 16435727 | rs12572249 | 0.693 | 0.099 | -1.824 | 1.007 | -0.072 | 0.073 | 0.073 | 0.722 | 0.43 | 0.11 | 0.32 | 0.27 | 0.55 | 0.33 |
| 10 | 16438239 | rs1489494 | 0.016 | 0.513 | -1.468 | 0.637 | 0.018 | 0.022 | 0.022 | 0.914 | 0.64 | 0.84 | 0.81 | 0.71 | 0.73 | 0.63 |
| 10 | 16445703 | rs12774017 | -0.489 | 1.039 | -1.237 | -0.041 | -0.03 | 0.041 | 0.041 | 0.072 | 0.29 | 0.18 | 0.23 | 0.06 | 0.39 | 0.2 |
| 10 | 16447486 | rs10904728 | -0.344 | -0.023 | -1.13 | 0.76 | 0.02 | -0.035 | -0.035 | 0 | 0.32 | 0.18 | 0.26 | 0.49 | 0.25 | 0.37 |
| 10 | 16452812 | rs10904730 | -0.596 | 0.446 | -0.891 | 0.651 | -0.027 | 0.013 | 0.013 | 0 | 0.77 | 0.84 | 0.61 | 0.82 | 0.55 | 0.77 |
| 10 | 16458636 | rs12220516 | -0.568 | -0.256 | 0.104 | 1.405 | 0.072 | 0.289 | 0.289 | 3.231 | 0.09 | 0.48 | 0.13 | 0.19 | 0.09 | 0.23 |
| 10 | 16463310 | rs10795393 | -0.362 | -0.28 | 0.414 | 1.397 | 0.004 | 0.07 | 0.07 | 0.089 | 0.06 | 0.16 | 0.1 | 0.19 | 0.08 | 0.23 |
| 10 | 16469650 | rs7080624 | -0.388 | -0.552 | 1.197 | 1.619 | 0.104 | 0.043 | 0.043 | 0 | 0.72 | 0.53 | 0.49 | 0.63 | 0.49 | 0.72 |
| 10 | 16475766 | rs7075658 | 0.493 | -1.336 | 1.273 | 1.266 | -0.001 | 0 | 0 | 0.082 | 0.32 | 0.32 | 0.26 | 0.35 | 0.25 | 0.25 |
| 10 | 16477295 | rs7093030 | -1.01 | -1.257 | 1.388 | 1.242 | -0.038 | -0.021 | -0.021 | 0 | 0.74 | 0.8 | 0.51 | 0.62 | 0.49 | 0.72 |
| 10 | 16478278 | rs10904737 | -1.28 | -1.232 | 1.398 | 1.289 | -0.025 | 0.011 | 0.011 | 0 | 0.77 | 0.83 | 0.62 | 0.82 | 0.53 | 0.76 |
| 10 | 16488475 | rs11253996 | -0.161 | -1.419 | 1.879 | 1.655 | -0.003 | 0.001 | 0.001 | 0.635 | 0.39 | 0.6 | 0.35 | 0.47 | 0.31 | 0.47 |
| 10 | 16488505 | rs11253997 | -1.39 | -1.382 | 1.866 | 1.614 | -0.06 | 0.007 | 0.007 | 0 | 0.77 | 0.88 | 0.62 | 0.79 | 0.53 | 0.74 |
| 10 | 16506501 | rs7097885 | -0.084 | -1.431 | 2.023 | 1.668 | -0.004 | 0 | 0 | 0.056 | 0.43 | 0.52 | 0.36 | 0.5 | 0.28 | 0.49 |
| 10 | 16509209 | rs6602121 | 0.144 | -1.446 | 2.077 | 1.732 | 0.015 | -0.021 | -0.021 | 0 | 0.35 | 0.29 | 0.36 | 0.46 | 0.29 | 0.29 |
| 10 | 16509960 | rs2171730 | -0.589 | -1.218 | 2.065 | 1.846 | 0.044 | 0.249 | 0.249 | 1.933 | 0.07 | 0.33 | 0.06 | 0.18 | 0.05 | 0.23 |
| 10 | 16517383 | rs10904747 | -0.747 | -1.158 | 1.83 | 1.685 | -0.04 | 0.014 | 0.014 | 0 | 0.72 | 0.82 | 0.58 | 0.79 | 0.46 | 0.73 |
| 10 | 16519763 | rs7915486 | -0.586 | -1.369 | 1.712 | 1.753 | 0.055 | 0.022 | 0.022 | 0.21 | 0.46 | 0.81 | 0.44 | 0.61 | 0.36 | 0.54 |
| 10 | 16520784 | rs7919985 | -0.075 | -1.462 | 2.142 | 1.977 | 0.007 | 0.003 | 0.003 | 0 | 0.41 | 0.38 | 0.33 | 0.45 | 0.31 | 0.36 |
| 10 | 16521007 | rs7905643 | -1.398 | -1.445 | 2.083 | 1.991 | -0.012 | -0.001 | -0.001 | 0 | 0.76 | 0.82 | 0.61 | 0.78 | 0.52 | 0.77 |
| 10 | 16521820 | rs1571770 | 0.448 | -1.607 | 2.123 | 1.984 | 0.033 | -0.021 | -0.021 | 0 | 0.35 | 0.22 | 0.31 | 0.46 | 0.27 | 0.32 |
| 10 | 16522568 | rs2000200 | -0.507 | -1.411 | 2.436 | 2.151 | 0.042 | 0.038 | 0.038 | 0 | 0.69 | 0.48 | 0.51 | 0.62 | 0.45 | 0.55 |
| 10 | 16523309 | rs11254016 | -0.873 | -1.563 | 2.51 | 2.313 | 0.068 | 0.271 | 0.271 | 0.679 | 0.06 | 0.31 | 0.07 | 0.15 | 0.05 | 0.21 |
| 10 | 16523375 | rs902073 | 0.053 | -1.775 | 2.468 | 2.24 | -0.035 | 0.02 | 0.02 | 0 | 0.33 | 0.43 | 0.23 | 0.34 | 0.19 | 0.42 |
| 10 | 16523886 | rs7077602 | -1.948 | -1.65 | 2.459 | 2.32 | 0.021 | 0.309 | 0.309 | 1.118 | 0.09 | 0.41 | 0.18 | 0.29 | 0.08 | 0.25 |
| 10 | 16524175 | rs7081010 | -1.96 | -1.651 | 2.509 | 2.312 | 0.001 | 0.227 | 0.227 | 1.308 | 0.09 | 0.3 | 0.18 | 0.3 | 0.08 | 0.25 |
| 10 | 16526840 | rs2275731 | -0.048 | -1.848 | 2.074 | 1.959 | -0.002 | -0.001 | -0.001 | 0 | 0.6 | 0.38 | 0.39 | 0.45 | 0.39 | 0.5 |
| 10 | 16532574 | rs10508512 | -1.24 | -1.856 | 1.397 | 2.094 | -0.003 | 0.187 | 0.187 | 0 | 0.07 | 0.14 | 0.16 | 0.35 | 0.08 | 0.25 |
| 10 | 16535688 | rs7075143 | -2.615 | -1.971 | 1.403 | 1.877 | -0.035 | 0.239 | 0.239 | 2.593 | 0.13 | 0.56 | 0.18 | 0.36 | 0.09 | 0.29 |
| 10 | 16535888 | rs10795400 | -0.051 | -1.899 | 1.273 | 1.503 | 0.005 | -0.005 | -0.005 | 0 | 0.61 | 0.44 | 0.44 | 0.64 | 0.43 | 0.7 |
| 10 | 16536461 | rs7904568 | -0.863 | -1.61 | 0.871 | 1.568 | -0.07 | 0.059 | 0.059 | 0 | 0.69 | 0.87 | 0.59 | 0.83 | 0.48 | 0.74 |
| 10 | 16544654 | rs10466288 | -0.501 | -1.448 | 0.331 | 1.446 | 0.026 | -0.041 | -0.041 | 0.874 | 0.66 | 0.58 | 0.57 | 0.82 | 0.47 | 0.74 |
| 10 | 16545943 | rs12770890 | 0.278 | -1.584 | -0.215 | 1.505 | 0.027 | -0.002 | -0.002 | 1.351 | 0.35 | 0.19 | 0.29 | 0.34 | 0.31 | 0.38 |
| 10 | 16547058 | rs7904014 | 0.027 | -0.23 | -0.218 | 0.427 | 0.01 | -0.018 | -0.018 | 0 | 0.67 | 0.64 | 0.58 | 0.83 | 0.48 | 0.74 |
| 10 | 16548382 | rs10904757 | -0.216 | 0.034 | -1.448 | 0.541 | -0.095 | 0.06 | 0.06 | 0 | 0.7 | 0.92 | 0.6 | 0.83 | 0.48 | 0.74 |
| 10 | 16553355 | rs1052708 | -0.505 | -0.969 | -1.088 | 0.552 | 0.021 | -0.009 | -0.009 | 2.731 | 0.16 | 0.11 | 0.06 | 0.26 | 0.07 | 0.14 |
| 10 | 16555802 | rs4747277 | 0.139 | -0.656 | -2.052 | -0.146 | -0.034 | 0.074 | 0.074 | 0.117 | 0.3 | 0.11 | 0.37 | 0.15 | 0.3 | 0.34 |
| 10 | 16556710 | rs7909832 | -0.837 | -0.197 | -1.126 | -0.14 | -0.018 | -0.013 | -0.013 | 0.185 | 0.61 | 0.69 | 0.6 | 0.52 | 0.55 | 0.56 |
| 10 | 16558985 | rs2298126 | 0.004 | -0.419 | -0.72 | -0.341 | 0.006 | 0.003 | 0.003 | 0.226 | 0.4 | 0.3 | 0.4 | 0.41 | 0.32 | 0.46 |
| 10 | 16559381 | rs7913889 | -0.483 | -0.519 | 0.069 | -0.342 | -0.002 | -0.013 | -0.013 | 3.393 | 0.21 | 0.26 | 0.18 | 0.11 | 0.21 | 0.09 |
| 10 | 16566904 | rs10904764 | -0.698 | -0.144 | 0.181 | -0.396 | 0 | 0 | 0 | 0 | 0.34 | 0.57 | 0.21 | 0.29 | 0.26 | 0.2 |
| 10 | 16567300 | rs11254053 | -1.134 | -0.738 | 0.164 | -0.406 | 0.053 | 0.028 | 0.028 | 0 | 0.43 | 0.66 | 0.28 | 0.36 | 0.32 | 0.22 |
| 10 | 16570103 | rs11254055 | 0.28 | -0.74 | 0.55 | -0.413 | -0.001 | -0.001 | -0.001 | 0 | 0.12 | 0.12 | 0.1 | 0.07 | 0.13 | 0.04 |
| 10 | 16571980 | rs10904766 | 0.295 | -0.746 | 0.556 | -0.395 | -0.001 | -0.002 | -0.002 | 0 | 0.12 | 0.12 | 0.1 | 0.07 | 0.13 | 0.04 |
| 10 | 16572258 | rs11254059 | -0.263 | -0.799 | 0.693 | -0.406 | -0.039 | -0.013 | -0.013 | 0 | 0.3 | 0.51 | 0.18 | 0.28 | 0.19 | 0.19 |
| 10 | 16581629 | rs10904769 | 0.643 | -0.88 | 0.788 | -0.417 | 0.039 | -0.013 | -0.013 | 0 | 0.2 | 0.1 | 0.1 | 0.28 | 0.08 | 0.13 |
| 10 | 16581889 | rs7902458 | -1.206 | -0.908 | 0.874 | -0.429 | 0.057 | 0.032 | 0.032 | 0 | 0.43 | 0.71 | 0.28 | 0.35 | 0.31 | 0.22 |
| 10 | 16583249 | rs7086455 | -0.94 | -0.875 | 1.138 | -0.453 | 0.015 | -0.067 | -0.067 | 0 | 0.21 | 0.48 | 0.18 | 0.08 | 0.23 | 0.09 |

| | | | | | | | | | | | | | | | | |
|---|---|---|---|---|---|---|---|---|---|---|---|---|---|---|---|---|
| 10 | 16583945 | rs7087467 | -0.573 | -0.772 | 0.942 | -0.394 | -0.037 | -0.014 | -0.014 | 0 | 0.35 | 0.5 | 0.21 | 0.3 | 0.22 | 0.17 |
| 10 | 16584422 | rs10399997 | 0.049 | 0.127 | 1.119 | 0.001 | 0.002 | -0.031 | -0.031 | 0 | 0.07 | 0.11 | 0.06 | 0.01 | 0.09 | 0.05 |
| 10 | 16585522 | rs12412368 | -0.618 | 0.085 | 0.601 | -0.187 | -0.061 | -0.061 | -0.061 | 0 | 0.71 | 0.83 | 0.63 | 0.52 | 0.55 | 0.51 |
| 10 | 16590829 | rs7079565 | -1.604 | -0.395 | 0.601 | -0.495 | 0.016 | 0.039 | 0.039 | 0.149 | 0.29 | 0.72 | 0.22 | 0.09 | 0.25 | 0.1 |
| 10 | 16595396 | rs7084722 | -0.017 | 0.362 | 0.306 | -0.266 | -0.104 | -0.057 | -0.057 | 0 | 0.32 | 0.12 | 0.37 | 0.57 | 0.43 | 0.55 |
| 10 | 16605482 | rs11254079 | -0.323 | 0.63 | 1.221 | -0.045 | 0.002 | -0.017 | -0.017 | 0 | 0.07 | 0.07 | 0.09 | 0.01 | 0.12 | 0.05 |
| 10 | 16610962 | rs11254084 | 0.131 | 0.694 | 1.16 | 0.07 | -0.038 | -0.027 | -0.027 | 0 | 0.1 | 0.02 | 0.09 | 0.09 | 0.12 | 0.03 |
| 10 | 16611095 | rs7091082 | -1.595 | 0.469 | 1.8 | -0.058 | -0.013 | -0.076 | -0.076 | 1.421 | 0.12 | 0.65 | 0.13 | 0.01 | 0.12 | 0.05 |
| 10 | 16620884 | rs12221081 | -1.897 | 0.548 | 1.938 | 0.006 | 0 | -0.014 | -0.014 | 0 | 0.24 | 0.66 | 0.23 | 0.12 | 0.26 | 0.07 |
| 10 | 16632102 | rs7913156 | -2.203 | 0.366 | 3.131 | 0.897 | -0.018 | 0.091 | 0.091 | 0.127 | 0.15 | 0.56 | 0.15 | 0.19 | 0.13 | 0.07 |
| 10 | 16634299 | rs7068341 | -0.765 | 0.398 | 2.606 | 0.875 | 0 | 0.002 | 0.002 | 0 | 0.08 | 0.11 | 0.12 | 0.09 | 0.12 | 0.06 |
| 10 | 16635100 | rs6977 | -2.85 | 0.188 | 2.756 | 0.801 | -0.055 | 0.018 | 0.018 | 4.153 | 0.11 | 0.73 | 0.08 | 0.11 | 0.02 | 0.07 |
| 10 | 16645177 | rs7070681 | -2.667 | 0.342 | 2.698 | 0.55 | 0.013 | 0.01 | 0.01 | 3.704 | 0.19 | 0.83 | 0.19 | 0.1 | 0.17 | 0.13 |
| 10 | 16647000 | rs11254106 | -1.959 | 0.372 | 3.028 | 0.553 | 0 | -0.005 | -0.005 | 0 | 0.15 | 0.52 | 0.17 | 0.09 | 0.17 | 0.12 |
| 10 | 16649679 | rs6602135 | -2.247 | 0.372 | 2.368 | 0.557 | -0.017 | -0.024 | -0.024 | 0 | 0.17 | 0.72 | 0.18 | 0.08 | 0.17 | 0.12 |
| 10 | 16649790 | rs17138855 | 0.081 | 1.207 | 1.49 | 1.206 | -0.084 | -0.111 | -0.111 | 0 | 0.18 | 0.02 | 0.17 | 0.32 | 0.25 | 0.32 |
| 10 | 16653879 | rs12247824 | -0.878 | 0.511 | 1.834 | 0.912 | -0.004 | -0.003 | -0.003 | 0.681 | 0.1 | 0.06 | 0.13 | 0.08 | 0.16 | 0.09 |
| 10 | 16664475 | rs11254113 | -1.306 | 0.511 | 2.007 | 0.875 | 0.045 | 0.039 | 0.039 | 0 | 0.12 | 0.42 | 0.15 | 0.08 | 0.16 | 0.1 |
| 10 | 16670107 | rs17138883 | -0.819 | 0.536 | 0.441 | 0.887 | -0.003 | -0.002 | -0.002 | 0 | 0.1 | 0.08 | 0.12 | 0.09 | 0.16 | 0.09 |
| 10 | 16676551 | rs11254118 | -1.906 | 0.546 | 0.267 | 0.889 | -0.021 | -0.025 | -0.025 | 0 | 0.18 | 0.71 | 0.18 | 0.08 | 0.17 | 0.13 |
| 10 | 16677976 | rs12242999 | -1.913 | 0.552 | -0.658 | 0.9 | 0.029 | 0.025 | 0.025 | 1.951 | 0.2 | 0.85 | 0.2 | 0.09 | 0.18 | 0.15 |
| 10 | 16679279 | rs11254119 | -1.028 | 1.452 | -0.54 | 1.329 | -0.03 | -0.049 | -0.049 | 0 | 0.28 | 0.16 | 0.31 | 0.39 | 0.4 | 0.42 |
| 10 | 16682602 | rs17138910 | -0.977 | 0.75 | -0.849 | 1.591 | 0.014 | -0.038 | -0.038 | 0 | 0.26 | 0.12 | 0.3 | 0.31 | 0.23 | 0.43 |
| 10 | 16684232 | rs11254123 | -1.441 | 0.78 | 0.427 | 1.701 | -0.02 | -0.03 | -0.03 | 0 | 0.3 | 0.54 | 0.35 | 0.28 | 0.25 | 0.45 |
| 10 | 16685819 | rs7094368 | -1.271 | 0.805 | 0.731 | 1.685 | -0.008 | -0.012 | -0.012 | 0.021 | 0.3 | 0.65 | 0.36 | 0.28 | 0.25 | 0.46 |
| 10 | 16689400 | rs7087278 | -1.618 | 1.169 | 0.367 | 1.779 | -0.022 | -0.005 | -0.005 | 0 | 0.54 | 0.84 | 0.65 | 0.59 | 0.59 | 0.74 |
| 10 | 16689683 | rs7087965 | -0.224 | 1.184 | 0.748 | 1.652 | 0.026 | 0.051 | 0.051 | 0.056 | 0.26 | 0.46 | 0.31 | 0.31 | 0.32 | 0.28 |
| 10 | 16690571 | rs6602141 | -1.429 | 1.049 | 0.307 | 1.695 | -0.004 | -0.001 | -0.001 | 0.263 | 0.51 | 0.84 | 0.64 | 0.44 | 0.59 | 0.7 |
| 10 | 16695008 | rs7096268 | -0.05 | 1.089 | 0.239 | 1.687 | -0.021 | -0.032 | -0.032 | 0.4 | 0.25 | 0.11 | 0.26 | 0.3 | 0.32 | 0.26 |
| 10 | 16699689 | rs17421802 | -0.115 | 0.781 | 0.163 | 1.505 | -0.022 | 0.061 | 0.061 | 0 | 0.22 | 0.03 | 0.28 | 0.13 | 0.24 | 0.42 |
| 10 | 16702330 | rs7090316 | -0.6 | 0.887 | 0.4 | 1.81 | -0.013 | 0.012 | 0.012 | 0.78 | 0.26 | 0.62 | 0.35 | 0.3 | 0.23 | 0.43 |
| 10 | 16704314 | rs10795410 | -0.382 | 1.082 | 0.619 | 1.669 | 0.017 | 0.03 | 0.03 | 0.011 | 0.26 | 0.4 | 0.31 | 0.31 | 0.35 | 0.27 |
| 10 | 16705277 | rs7088676 | -0.798 | 1.198 | -0.095 | 1.464 | -0.042 | -0.031 | -0.031 | 0 | 0.38 | 0.2 | 0.38 | 0.45 | 0.46 | 0.39 |
| 10 | 16711372 | rs7092141 | -0.76 | 0.943 | 0.487 | 1.339 | -0.004 | -0.001 | -0.001 | 0 | 0.24 | 0.38 | 0.23 | 0.28 | 0.25 | 0.37 |
| 10 | 16711457 | rs7899419 | -0.379 | 0.806 | 0.596 | 1.384 | -0.01 | 0.038 | 0.038 | 0 | 0.06 | 0.18 | 0.05 | 0.07 | 0.03 | 0.05 |
| 10 | 16712610 | rs3847362 | -1.707 | 1.049 | -0.231 | 1.265 | -0.101 | -0.115 | -0.115 | 1.136 | 0.57 | 0.93 | 0.54 | 0.63 | 0.58 | 0.78 |
| 10 | 16714466 | rs10795412 | -0.922 | 0.719 | 1.076 | 1.13 | -0.001 | -0.013 | -0.013 | 0 | 0.21 | 0.22 | 0.22 | 0.15 | 0.22 | 0.3 |
| 10 | 16715194 | rs10752058 | -1.179 | 0.768 | 0.678 | 1.153 | -0.004 | -0.002 | -0.002 | 2.043 | 0.25 | 0.33 | 0.23 | 0.27 | 0.25 | 0.36 |
| 10 | 16718607 | rs10795415 | 0.641 | 1.099 | 0.69 | 1.387 | 0.003 | 0.004 | 0.004 | 0 | 0.64 | 0.38 | 0.5 | 0.5 | 0.58 | 0.42 |
| 10 | 16719035 | rs17139010 | 0.606 | 1.085 | 1.206 | 1.385 | 0.028 | 0.038 | 0.038 | 0 | 0.65 | 0.51 | 0.51 | 0.5 | 0.58 | 0.42 |
| 10 | 16721496 | rs3864821 | 0.215 | 1.132 | 1.702 | 1.533 | -0.001 | -0.002 | -0.002 | 0.313 | 0.73 | 0.69 | 0.61 | 0.86 | 0.67 | 0.66 |
| 10 | 16725247 | rs17331552 | -0.411 | 0.728 | 1.115 | 1.512 | -0.013 | 0.026 | 0.026 | 0 | 0.19 | 0.02 | 0.09 | 0.07 | 0.19 | 0.12 |
| 10 | 16728869 | rs7913448 | -1.271 | 0.502 | 1.885 | 1.52 | 0.042 | 0.304 | 0.304 | 0.635 | 0.09 | 0.38 | 0.13 | 0.38 | 0.11 | 0.21 |
| 10 | 16734349 | rs11254147 | -1.566 | 0.452 | 2.262 | 1.442 | -0.022 | 0.062 | 0.062 | 0 | 0.28 | 0.52 | 0.22 | 0.47 | 0.27 | 0.37 |
| 10 | 16736476 | rs4748311 | -1.355 | 0.458 | 2.467 | 1.439 | -0.016 | 0.051 | 0.051 | 0 | 0.26 | 0.34 | 0.2 | 0.45 | 0.26 | 0.32 |
| 10 | 16736638 | rs3864823 | 1.806 | 0.514 | 2.551 | 1.418 | 0.016 | 0.029 | 0.029 | 0 | 0.46 | 0.25 | 0.4 | 0.39 | 0.39 | 0.29 |
| 10 | 16737289 | rs3740172 | 1.522 | 0.532 | 2.722 | 1.425 | 0.003 | 0.002 | 0.002 | 0 | 0.49 | 0.51 | 0.44 | 0.43 | 0.42 | 0.37 |
| 10 | 16740784 | rs17139050 | -0.789 | 0.514 | 2.652 | 1.849 | 0.081 | 0.412 | 0.412 | 0.555 | 0.07 | 0.3 | 0.11 | 0.5 | 0.09 | 0.23 |
| 10 | 16742870 | rs10904794 | 1.808 | 0.427 | 2.382 | 1.774 | 0.019 | 0.061 | 0.061 | 0 | 0.47 | 0.23 | 0.42 | 0.3 | 0.41 | 0.32 |
| 10 | 16743365 | rs7093729 | -0.537 | 0.542 | 1.982 | 1.649 | -0.087 | 0.073 | 0.073 | 0.02 | 0.78 | 0.96 | 0.64 | 0.9 | 0.69 | 0.74 |
| 10 | 16743672 | rs10795417 | 1.515 | 0.54 | 2.162 | 1.778 | 0.003 | 0 | 0 | 0 | 0.49 | 0.54 | 0.43 | 0.47 | 0.43 | 0.38 |
| 10 | 16744523 | rs11254160 | -1.017 | 0.56 | 1.934 | 1.703 | 0.047 | 0.252 | 0.252 | 1.903 | 0.07 | 0.29 | 0.11 | 0.35 | 0.07 | 0.22 |
| 10 | 16748128 | rs7092024 | 0.792 | 0.525 | 1.319 | 0.916 | -0.099 | 0.083 | 0.083 | 0.001 | 0.59 | 0.91 | 0.56 | 0.8 | 0.5 | 0.58 |
| 10 | 16749526 | rs3951063 | 1.54 | 0.497 | 1.717 | 0.935 | 0.001 | 0.078 | 0.078 | 0 | 0.51 | 0.29 | 0.51 | 0.7 | 0.48 | 0.51 |
| 10 | 16760549 | rs12769960 | -0.511 | 0.697 | 1.68 | 0.961 | 0.014 | 0.055 | 0.055 | 0 | 0.18 | 0.02 | 0.07 | 0.04 | 0.17 | 0.11 |
| 10 | 16762523 | rs7084555 | 1.286 | 0.79 | 1.838 | 1.028 | -0.026 | 0.05 | 0.05 | 0 | 0.58 | 0.3 | 0.57 | 0.74 | 0.51 | 0.58 |
| 10 | 16763204 | rs11812463 | -1.486 | 0.874 | 2.358 | 1.053 | 0.086 | 0.159 | 0.159 | 0.91 | 0.06 | 0.45 | 0.04 | 0.12 | 0.02 | 0.04 |
| 10 | 16763882 | rs10732931 | 1.458 | 0.845 | 2.136 | 1.354 | -0.04 | -0.026 | -0.026 | 0 | 0.58 | 0.21 | 0.53 | 0.56 | 0.49 | 0.52 |
| 10 | 16764483 | rs6602151 | -0.36 | 0.802 | 1.969 | 0.913 | -0.007 | 0.004 | 0.004 | 0.173 | 0.8 | 0.84 | 0.69 | 0.9 | 0.71 | 0.75 |
| 10 | 16767461 | rs12261652 | -0.548 | 0.789 | 2.099 | 0.912 | 0.001 | 0.002 | 0.002 | 0 | 0.18 | 0.12 | 0.08 | 0.06 | 0.18 | 0.13 |
| 10 | 16771273 | rs7893556 | 0.68 | 0.732 | 1.786 | 1.007 | -0.068 | 0.079 | 0.079 | 0 | 0.64 | 0.88 | 0.61 | 0.85 | 0.52 | 0.61 |
| 10 | 16775663 | rs10795424 | 1.188 | 0.766 | 1.051 | 1.055 | -0.008 | 0.069 | 0.069 | 0 | 0.53 | 0.29 | 0.5 | 0.66 | 0.47 | 0.51 |
| 10 | 16779115 | rs12098816 | -1.199 | 0.324 | 1.576 | 1.395 | -0.017 | 0.057 | 0.057 | 0.049 | 0.2 | 0.44 | 0.21 | 0.26 | 0.19 | 0.15 |
| 10 | 16779918 | rs4748317 | -1.314 | 0.303 | 1.575 | 1.324 | -0.025 | 0.02 | 0.02 | 0 | 0.22 | 0.44 | 0.21 | 0.22 | 0.19 | 0.12 |
| 10 | 16782363 | rs4748319 | 0.233 | 0.915 | 1.575 | 1.23 | 0.099 | -0.004 | -0.004 | 0 | 0.75 | 0.51 | 0.66 | 0.81 | 0.54 | 0.68 |
| 10 | 16783674 | rs2124106 | -1.353 | 0.119 | 2.097 | 1.824 | -0.001 | -0.001 | -0.001 | 0 | 0.19 | 0.17 | 0.19 | 0.14 | 0.19 | 0.1 |
| 10 | 16784810 | rs7911748 | -1.47 | 0.113 | 1.997 | 1.755 | -0.016 | 0.011 | 0.011 | 0 | 0.2 | 0.32 | 0.2 | 0.21 | 0.19 | 0.11 |
| 10 | 16787560 | rs4748321 | -1.593 | 0.101 | 1.979 | 1.816 | -0.023 | 0.036 | 0.036 | 0 | 0.22 | 0.42 | 0.21 | 0.27 | 0.18 | 0.21 |
| 10 | 16790231 | rs11254169 | -1.715 | 0.015 | 2.29 | 1.784 | 0.012 | 0.068 | 0.068 | 0.017 | 0.12 | 0.41 | 0.11 | 0.17 | 0.1 | 0.19 |

| | | | | | | | | | | | | | | | | |
|---|---|---|---|---|---|---|---|---|---|---|---|---|---|---|---|---|
| 10 | 16806044 | rs3780981 | -2.127 | -0.322 | 2.198 | 2.426 | 0.039 | -0.1 | -0.1 | 0 | 0.32 | 0.89 | 0.35 | 0.44 | 0.24 | 0.29 |
| 10 | 16807745 | rs17156937 | -0.616 | -0.379 | 2.354 | 2.584 | -0.007 | 0.001 | 0.001 | 4.075 | 0.09 | 0.09 | 0.09 | 0.17 | 0.02 | 0.08 |
| 10 | 16820133 | rs3864817 | -1.268 | -0.355 | 3.179 | 2.768 | 0.013 | -0.02 | -0.02 | 0 | 0.17 | 0.31 | 0.17 | 0.08 | 0.19 | 0.12 |
| 10 | 16821021 | rs3780974 | -2.732 | -0.429 | 3.15 | 2.808 | 0.023 | -0.014 | -0.014 | 0 | 0.3 | 0.8 | 0.29 | 0.32 | 0.22 | 0.29 |
| 10 | 16821356 | rs2124107 | -2.813 | -0.447 | 3.15 | 2.799 | 0.03 | -0.017 | -0.017 | 0 | 0.31 | 0.83 | 0.3 | 0.32 | 0.22 | 0.29 |
| 10 | 16823657 | rs10904804 | -1.613 | -0.385 | 3.156 | 2.652 | 0.024 | -0.036 | -0.036 | 0.413 | 0.19 | 0.35 | 0.19 | 0.08 | 0.21 | 0.14 |
| 10 | 16825275 | rs17139162 | -2.169 | -0.376 | 3.156 | 2.705 | -0.051 | 0.176 | 0.176 | 3.489 | 0.13 | 0.35 | 0.1 | 0.32 | 0.06 | 0.19 |
| 10 | 16828829 | rs7088168 | -1.373 | -0.324 | 2.805 | 2.623 | -0.037 | 0.095 | 0.095 | 1.886 | 0.1 | 0.26 | 0.09 | 0.23 | 0.05 | 0.13 |
| 10 | 16831598 | rs10904815 | -2.243 | -0.316 | 2.87 | 2.781 | 0.004 | 0.001 | 0.001 | 0.147 | 0.22 | 0.63 | 0.21 | 0.21 | 0.23 | 0.2 |
| 10 | 16832561 | rs6602159 | -3.04 | -0.334 | 2.519 | 2.802 | -0.009 | -0.12 | -0.12 | 0 | 0.29 | 0.9 | 0.31 | 0.43 | 0.28 | 0.33 |
| 10 | 16834660 | rs3824633 | -1.135 | -0.255 | 2.604 | 2.709 | 0.004 | -0.021 | -0.021 | 0 | 0.16 | 0.23 | 0.16 | 0.07 | 0.19 | 0.12 |
| 10 | 16838417 | rs7068265 | -2.423 | -0.281 | 2.432 | 2.714 | -0.007 | 0.02 | 0.02 | 0.222 | 0.28 | 0.39 | 0.27 | 0.26 | 0.27 | 0.28 |
| 10 | 16838866 | rs7068881 | -2.092 | 0.132 | 1.678 | 2.493 | 0 | -0.03 | -0.03 | 0 | 0.26 | 0.79 | 0.23 | 0.29 | 0.25 | 0.24 |
| 10 | 16851934 | rs11254210 | -1.274 | 0.905 | 1.121 | 2.796 | -0.026 | 0.002 | 0.002 | 0.086 | 0.08 | 0.03 | 0.29 | 0.3 | 0.16 | 0.18 |
| 10 | 16852667 | rs7076779 | -1.332 | 0.864 | 1.957 | 2.396 | 0.019 | -0.063 | -0.063 | 0 | 0.16 | 0.35 | 0.11 | 0.05 | 0.17 | 0.16 |
| 10 | 16853415 | rs12253979 | -1.443 | 0.832 | 2.619 | 2.381 | 0.029 | -0.074 | -0.074 | 0 | 0.17 | 0.45 | 0.13 | 0.05 | 0.16 | 0.16 |
| 10 | 16853609 | rs7905776 | -2.041 | 0.8 | 2.666 | 2.357 | 0.005 | -0.058 | -0.058 | 2.558 | 0.2 | 0.63 | 0.16 | 0.05 | 0.2 | 0.18 |
| 10 | 16860239 | rs3808925 | -0.509 | -0.007 | 2.14 | 2.404 | 0.004 | -0.004 | -0.004 | 1.36 | 0.19 | 0.17 | 0.16 | 0.46 | 0.14 | 0.23 |
| 10 | 16864810 | rs7075036 | -2.012 | 0.03 | 2.172 | 2.473 | 0.023 | -0.013 | -0.013 | 0 | 0.37 | 0.75 | 0.31 | 0.47 | 0.33 | 0.38 |
| 10 | 16871456 | rs2603796 | -2.08 | 0.197 | 2.01 | 2.666 | -0.001 | -0.003 | -0.003 | 0.013 | 0.36 | 0.54 | 0.52 | 0.64 | 0.33 | 0.39 |
| 10 | 16872337 | rs7087360 | -1.265 | 1.08 | 2.329 | 3.042 | 0.019 | 0.002 | 0.002 | 0 | 0.17 | 0.43 | 0.13 | 0.1 | 0.16 | 0.16 |
| 10 | 16873744 | rs2603804 | -0.961 | 1.155 | 1.711 | 3.031 | 0.027 | 0.074 | 0.074 | 0.392 | 0.07 | 0.1 | 0.3 | 0.3 | 0.13 | 0.15 |
| 10 | 16876107 | rs7897625 | -1.271 | 0.249 | 1.589 | 2.379 | 0.01 | 0.043 | 0.043 | 0 | 0.07 | 0.16 | 0.05 | 0.14 | 0.1 | 0.11 |
| 10 | 16878204 | rs703062 | 1.469 | -0.201 | 1.692 | 2.433 | -0.016 | 0.036 | 0.036 | 1.285 | 0.74 | 0.66 | 0.76 | 0.49 | 0.81 | 0.63 |
| 10 | 16886806 | rs780827 | 1.836 | -0.114 | 2.054 | 2.336 | 0.006 | -0.013 | -0.013 | 0 | 0.69 | 0.3 | 0.71 | 0.47 | 0.75 | 0.62 |
| 10 | 16891317 | rs7094016 | -1.87 | 0.128 | 2.671 | 1.776 | 0.091 | 0.003 | 0.003 | 2.769 | 0.07 | 0.56 | 0.07 | 0 | 0.06 | 0.02 |
| 10 | 16891468 | rs12251746 | -1.631 | 0.057 | 2.076 | 2.105 | -0.008 | 0.064 | 0.064 | 0 | 0.06 | 0.11 | 0.09 | 0.21 | 0.05 | 0.07 |
| 10 | 16896387 | rs10904826 | -1.77 | 0.164 | 2.24 | 1.657 | 0.057 | -0.02 | -0.02 | 0.068 | 0.07 | 0.64 | 0.07 | 0 | 0.06 | 0.02 |
| 10 | 16905500 | rs7893395 | -1.578 | -0.152 | 1.078 | 2.202 | 0.008 | -0.02 | -0.02 | 0.713 | 0.21 | 0.17 | 0.39 | 0.6 | 0.19 | 0.37 |
| 10 | 16909589 | rs17139378 | -1.701 | -0.464 | 1.16 | 1.928 | -0.01 | 0.015 | 0.015 | 0 | 0.28 | 0.32 | 0.41 | 0.59 | 0.21 | 0.39 |
| 10 | 16911605 | rs703075 | -2.021 | 0.305 | 0.677 | 1.697 | -0.029 | -0.058 | -0.058 | 0 | 0.35 | 0.9 | 0.44 | 0.39 | 0.3 | 0.39 |
| 10 | 16912134 | rs780825 | -1.726 | 0.362 | 1.291 | 1.718 | -0.001 | 0.018 | 0.018 | 0.414 | 0.31 | 0.42 | 0.4 | 0.43 | 0.29 | 0.39 |
| 10 | 16918972 | rs1801241 | -1.846 | 0.055 | 1.906 | 1.771 | -0.038 | 0.137 | 0.137 | 0 | 0.13 | 0.35 | 0.29 | 0.31 | 0.09 | 0.18 |
| 10 | 16919052 | rs1801239 | -0.598 | -0.162 | 1.447 | 1.343 | 0.02 | 0.068 | 0.068 | 0 | 0.13 | 0.03 | 0.09 | 0 | 0.09 | 0.13 |
| 10 | 16921370 | rs780838 | 2.548 | 0.187 | 1.395 | 1.143 | 0.052 | -0.033 | -0.033 | 0 | 0.57 | 0.14 | 0.41 | 0.51 | 0.61 | 0.53 |
| 10 | 16923390 | rs780837 | 1.96 | 0.251 | 1.817 | 1.199 | 0 | -0.002 | -0.002 | 0.028 | 0.77 | 0.25 | 0.74 | 0.82 | 0.74 | 0.84 |
| 10 | 16925588 | rs2796833 | 1.425 | 0.279 | 1.6 | 1.211 | 0.026 | -0.018 | -0.018 | 0 | 0.8 | 0.58 | 0.77 | 0.82 | 0.74 | 0.83 |
| 10 | 16928435 | rs703065 | 1.987 | 0.279 | 1.268 | 1.197 | -0.013 | -0.081 | -0.081 | 0 | 0.75 | 0.89 | 0.55 | 0.51 | 0.71 | 0.67 |
| 10 | 16935506 | rs7068023 | 1.42 | 0.343 | 1.391 | 1.185 | -0.001 | -0.002 | -0.002 | 0 | 0.78 | 0.79 | 0.56 | 0.51 | 0.73 | 0.68 |
| 10 | 16938238 | rs780816 | 1.359 | 0.328 | 1.901 | 1.128 | 0.053 | -0.027 | -0.027 | 0 | 0.82 | 0.51 | 0.78 | 0.83 | 0.77 | 0.85 |
| 10 | 16939976 | rs10904833 | 2.656 | 1.22 | 1.321 | 0.763 | -0.039 | -0.008 | -0.008 | 0 | 0.52 | 0.13 | 0.34 | 0.53 | 0.48 | 0.48 |
| 10 | 16945193 | rs7099855 | 0.315 | 0.582 | 1.555 | 0.783 | 0.005 | 0.07 | 0.07 | 0.002 | 0.06 | 0.2 | 0.03 | 0.15 | 0.04 | 0.05 |
| 10 | 16948177 | rs780807 | 0.749 | 0.609 | 1.82 | 0.884 | 0.064 | 0.079 | 0.079 | 0.008 | 0.88 | 0.67 | 0.63 | 0.82 | 0.8 | 0.84 |
| 10 | 16948801 | rs780806 | 2.335 | 0.665 | 1.141 | 0.812 | 0.004 | 0.028 | 0.028 | 0 | 0.61 | 0.13 | 0.46 | 0.67 | 0.6 | 0.61 |
| 10 | 16949401 | rs780805 | 2.464 | 0.699 | 1.091 | 0.739 | 0.02 | -0.03 | -0.03 | 0 | 0.54 | 0.1 | 0.39 | 0.52 | 0.57 | 0.47 |
| 10 | 16962352 | rs11254275 | -1.501 | 0.861 | 1.259 | 0.573 | -0.001 | -0.011 | -0.011 | 0 | 0.27 | 0.27 | 0.33 | 0.08 | 0.31 | 0.31 |
| 10 | 16965227 | rs1810205 | -1.972 | 0.918 | 1.215 | 0.645 | -0.001 | 0 | 0 | 0 | 0.34 | 0.67 | 0.57 | 0.43 | 0.37 | 0.39 |
| 10 | 16968845 | rs1687716 | 1.409 | 1.735 | 1.253 | 0.492 | -0.001 | -0.023 | -0.023 | 0 | 0.77 | 0.79 | 0.78 | 0.95 | 0.73 | 0.72 |
| 10 | 16972011 | rs10904838 | -0.217 | 1.608 | 1.477 | 0.9 | 0.029 | 0.09 | 0.09 | 0.005 | 0.06 | 0.26 | 0.04 | 0.15 | 0.04 | 0.05 |
| 10 | 16973346 | rs7090658 | -1.897 | 2.35 | 0.623 | 0.798 | -0.022 | 0.023 | 0.023 | 4.286 | 0.44 | 0.84 | 0.65 | 0.37 | 0.52 | 0.41 |
| 10 | 16974133 | rs10904839 | -0.715 | 2.811 | 1.196 | 1.436 | 0.016 | 0.027 | 0.027 | 0 | 0.16 | 0.21 | 0.17 | 0.33 | 0.21 | 0.25 |
| 10 | 16975078 | rs2271461 | -0.54 | 2.829 | 1.042 | 1.432 | -0.019 | -0.022 | -0.022 | 0 | 0.14 | 0.09 | 0.16 | 0.32 | 0.2 | 0.25 |
| 10 | 16976226 | rs11591673 | -1.457 | 3.299 | 1.605 | 1.184 | 0.032 | -0.043 | -0.043 | 1.12 | 0.25 | 0.4 | 0.21 | 0.18 | 0.27 | 0.23 |
| 10 | 16976618 | rs2883972 | -0.129 | 3.323 | 1.537 | 1.181 | -0.026 | 0.023 | 0.023 | 1.067 | 0.11 | 0.07 | 0.13 | 0.04 | 0.16 | 0.12 |
| 10 | 16978429 | rs11254284 | -2.78 | 3.316 | 1.364 | 1.185 | 0.087 | 0.09 | 0.09 | 0.751 | 0.52 | 0.83 | 0.68 | 0.68 | 0.63 | 0.58 |
| 10 | 16980552 | rs3914238 | -2.585 | 3.34 | 2.074 | 1.337 | -0.017 | -0.039 | -0.039 | 0 | 0.35 | 0.75 | 0.53 | 0.49 | 0.37 | 0.41 |
| 10 | 16982244 | rs2271463 | -1.725 | 3.289 | 2.28 | 1.356 | 0.002 | 0.008 | 0.008 | 0 | 0.18 | 0.24 | 0.16 | 0.29 | 0.19 | 0.25 |
| 10 | 16985317 | rs7912716 | -1.073 | 3.325 | 2.257 | 1.341 | -0.001 | -0.003 | -0.003 | 0 | 0.12 | 0.12 | 0.11 | 0.21 | 0.13 | 0.23 |
| 10 | 16985766 | rs7913144 | -2.669 | 3.226 | 2.254 | 1.339 | -0.026 | -0.052 | -0.052 | 0 | 0.35 | 0.76 | 0.54 | 0.49 | 0.39 | 0.42 |
| 10 | 16986840 | rs11254288 | 0.055 | 3.06 | 2.135 | 1.326 | 0.031 | 0.032 | 0.032 | 0 | 0.07 | 0.23 | 0.05 | 0.09 | 0.07 | 0.03 |
| 10 | 16988186 | rs10904847 | -1.358 | 3.056 | 2.179 | 1.294 | 0.005 | 0.13 | 0.13 | 1.969 | 0.12 | 0.62 | 0.3 | 0.24 | 0.09 | 0.07 |
| 10 | 16990908 | rs11814420 | -1.881 | 3.173 | 1.967 | 1.262 | 0.031 | 0.012 | 0.012 | 2.123 | 0.34 | 0.49 | 0.53 | 0.36 | 0.5 | 0.35 |
| 10 | 16996953 | rs10752064 | -2.596 | 2.933 | 1.537 | 1.209 | 0.049 | 0.041 | 0.041 | 0 | 0.56 | 0.85 | 0.68 | 0.66 | 0.62 | 0.63 |
| 10 | 16997827 | rs10904851 | -0.547 | 3.006 | 1.501 | 1.214 | -0.062 | 0.039 | 0.039 | 0 | 0.23 | 0.09 | 0.17 | 0.16 | 0.32 | 0.26 |
| 10 | 16997891 | rs10508517 | -0.316 | 2.994 | 1.51 | 1.234 | 0.017 | -0.023 | -0.023 | 0 | 0.11 | 0.06 | 0.29 | 0.38 | 0.07 | 0.23 |
| 10 | 16999350 | rs7071576 | -0.693 | 3.04 | 1.591 | 0.968 | -0.013 | 0.014 | 0.014 | 0 | 0.24 | 0.19 | 0.18 | 0.16 | 0.32 | 0.26 |
| 10 | 17001812 | rs7906242 | -1.166 | 3.008 | 1.767 | 0.978 | 0.037 | -0.033 | -0.033 | 0 | 0.31 | 0.48 | 0.24 | 0.24 | 0.39 | 0.28 |
| 10 | 17004709 | rs1687713 | 2.16 | 2.867 | 1.546 | 1.019 | 0.047 | 0.047 | 0.047 | 0 | 0.44 | 0.19 | 0.32 | 0.34 | 0.38 | 0.37 |
| 10 | 17006385 | rs10904856 | 2.256 | 2.886 | 1.39 | 1.028 | 0.031 | 0.03 | 0.03 | 0 | 0.43 | 0.13 | 0.31 | 0.34 | 0.38 | 0.37 |

| | | | | | | | | | | | | | | | | |
|---|---|---|---|---|---|---|---|---|---|---|---|---|---|---|---|---|
| 10 | 17008183 | rs1687706 | 2.132 | 2.892 | 1.641 | 1.03 | 0.038 | 0.037 | 0.037 | 0 | 0.45 | 0.3 | 0.32 | 0.34 | 0.38 | 0.37 |
| 10 | 17008422 | rs2942359 | 0.761 | 3.106 | 2.007 | 1.022 | 0 | -0.003 | -0.003 | 0 | 0.68 | 0.7 | 0.55 | 0.45 | 0.7 | 0.55 |
| 10 | 17008873 | rs7913601 | 0.034 | 3.157 | 1.99 | 1.003 | -0.002 | -0.001 | -0.001 | 0.095 | 0.12 | 0.09 | 0.14 | 0.04 | 0.16 | 0.12 |
| 10 | 17009523 | rs12244831 | 0.024 | 3.128 | 1.968 | 1.026 | -0.053 | 0.013 | 0.013 | 0 | 0.2 | 0.05 | 0.14 | 0.16 | 0.24 | 0.23 |
| 10 | 17009736 | rs1707280 | 2.405 | 3.083 | 1.993 | 1.028 | 0.045 | 0.044 | 0.044 | 0 | 0.44 | 0.17 | 0.31 | 0.34 | 0.38 | 0.37 |
| 10 | 17010742 | rs1687703 | 2.354 | 2.774 | 1.996 | 1.087 | 0.028 | 0.048 | 0.048 | 0 | 0.45 | 0.26 | 0.35 | 0.34 | 0.41 | 0.38 |
| 10 | 17011175 | rs1398431 | 0.762 | 2.779 | 2.11 | 1.168 | -0.011 | 0.037 | 0.037 | 0.483 | 0.84 | 0.76 | 0.72 | 0.72 | 0.91 | 0.82 |
| 10 | 17012098 | rs9633766 | -1.536 | 2.784 | 2.186 | 1.169 | 0.005 | -0.001 | -0.001 | 0 | 0.38 | 0.38 | 0.35 | 0.38 | 0.48 | 0.44 |
| 10 | 17012113 | rs1687699 | 0.862 | 2.766 | 2.148 | 1.183 | -0.009 | 0.046 | 0.046 | 0.427 | 0.83 | 0.72 | 0.71 | 0.72 | 0.89 | 0.82 |
| 10 | 17015144 | rs2932908 | 1.76 | 3.021 | 1.719 | 1.152 | 0.02 | 0.008 | 0.008 | 0 | 0.68 | 0.49 | 0.56 | 0.69 | 0.66 | 0.67 |
| 10 | 17016258 | rs11254307 | -1.3 | 2.994 | 1.481 | 1.112 | -0.009 | -0.004 | -0.004 | 0 | 0.34 | 0.25 | 0.3 | 0.39 | 0.39 | 0.41 |
| 10 | 17016337 | rs11254308 | -1.324 | 2.966 | 1.468 | 1.1 | -0.009 | -0.003 | -0.003 | 0 | 0.34 | 0.25 | 0.3 | 0.39 | 0.39 | 0.43 |
| 10 | 17017492 | rs4366393 | -1.933 | 2.905 | 1.321 | 1.105 | 0.003 | 0.003 | 0.003 | 0 | 0.45 | 0.59 | 0.4 | 0.46 | 0.54 | 0.47 |
| 10 | 17019013 | rs3012499 | 0.175 | 2.418 | 1.651 | 1.246 | -0.011 | 0.11 | 0.11 | 0.145 | 0.9 | 0.69 | 0.75 | 0.81 | 0.95 | 0.85 |
| 10 | 17019845 | rs2932904 | 1.576 | 2.447 | 1.318 | 1.26 | -0.019 | 0.086 | 0.086 | 3.671 | 0.8 | 0.35 | 0.62 | 0.55 | 0.87 | 0.76 |
| 10 | 17023350 | rs12255119 | 0.103 | 2.556 | 1.123 | 1.142 | -0.011 | 0.063 | 0.063 | 0 | 0.18 | 0.05 | 0.13 | 0 | 0.2 | 0.19 |
| 10 | 17024503 | rs1801231 | 1.664 | 2.402 | 1.28 | 1.299 | -0.018 | 0.071 | 0.071 | 3.479 | 0.8 | 0.35 | 0.63 | 0.56 | 0.87 | 0.76 |
| 10 | 17025685 | rs3012492 | 1.666 | 2.479 | 1.247 | 1.266 | -0.014 | 0.06 | 0.06 | 3.032 | 0.79 | 0.35 | 0.62 | 0.56 | 0.85 | 0.73 |
| 10 | 17026384 | rs3012494 | 1.383 | 2.409 | 1.326 | 1.324 | -0.018 | 0.048 | 0.048 | 2.973 | 0.81 | 0.39 | 0.68 | 0.74 | 0.88 | 0.82 |
| 10 | 17029935 | rs7072049 | -2.033 | 2.509 | 0.301 | 1.145 | -0.025 | -0.019 | -0.019 | 0 | 0.45 | 0.82 | 0.42 | 0.6 | 0.45 | 0.49 |
| 10 | 17030854 | rs2942366 | 1.746 | 2.474 | 0.094 | 1.141 | -0.048 | -0.024 | -0.024 | 0.001 | 0.55 | 0.16 | 0.42 | 0.4 | 0.54 | 0.51 |
| 10 | 17031834 | rs1512706 | -0.256 | 2.275 | 0.135 | 1.187 | 0.008 | -0.044 | -0.044 | 0.033 | 0.91 | 0.97 | 0.76 | 0.81 | 0.95 | 0.85 |
| 10 | 17032281 | rs2291521 | -0.343 | 2.349 | 0.146 | 1.126 | -0.028 | 0.016 | 0.016 | 0 | 0.2 | 0.08 | 0.15 | 0.15 | 0.24 | 0.23 |
| 10 | 17035822 | rs3012479 | 1.241 | 1.91 | 0.43 | 1.127 | 0 | -0.002 | -0.002 | 4.277 | 0.64 | 0.44 | 0.51 | 0.46 | 0.66 | 0.55 |
| 10 | 17038701 | rs1276711 | -0.163 | 1.993 | 0.215 | 1.19 | 0.001 | -0.033 | -0.033 | 0.037 | 0.91 | 0.97 | 0.8 | 0.81 | 0.95 | 0.87 |
| 10 | 17039809 | rs11254325 | -0.717 | 2.147 | 0.94 | 1.075 | 0.036 | 0.105 | 0.105 | 0 | 0.11 | 0.33 | 0.13 | 0.17 | 0.1 | 0.1 |
| 10 | 17040116 | rs12258009 | 0.21 | 2.308 | 0.544 | 0.806 | 0.001 | 0.006 | 0.006 | 0 | 0.15 | 0.1 | 0.1 | 0.01 | 0.15 | 0.17 |
| 10 | 17040849 | rs7900486 | -1.214 | 2.728 | 0.816 | 1.21 | 0.051 | 0.14 | 0.14 | 0.073 | 0.13 | 0.27 | 0.34 | 0.37 | 0.2 | 0.13 |
| 10 | 17045711 | rs7922356 | -0.588 | 2.837 | 0.88 | 1.181 | 0.019 | 0.046 | 0.046 | 0 | 0.11 | 0.21 | 0.13 | 0.16 | 0.12 | 0.1 |
| 10 | 17047781 | rs4748341 | -1.542 | 2.793 | 0.866 | 1.345 | -0.003 | -0.003 | -0.003 | 0 | 0.31 | 0.29 | 0.51 | 0.57 | 0.41 | 0.36 |
| 10 | 17048022 | rs7084057 | -2.148 | 3.114 | 0.807 | 1.025 | 0.024 | 0.026 | 0.026 | 0 | 0.51 | 0.58 | 0.65 | 0.62 | 0.55 | 0.57 |
| 10 | 17052955 | rs11592014 | -1.226 | 3.092 | 0.133 | 1.069 | -0.014 | -0.019 | -0.019 | 0.011 | 0.31 | 0.25 | 0.51 | 0.56 | 0.39 | 0.35 |
| 10 | 17059462 | rs12246329 | -0.116 | 3.083 | 0.036 | 1.103 | 0.012 | -0.094 | -0.094 | 0 | 0.2 | 0.4 | 0.13 | 0.08 | 0.14 | 0.22 |
| 10 | 17063026 | rs4082518 | -0.741 | 3.05 | -0.331 | 1.214 | 0.083 | 0.206 | 0.206 | 0 | 0.11 | 0.27 | 0.33 | 0.37 | 0.18 | 0.09 |
| 10 | 17064594 | rs7896819 | -1.572 | 2.38 | -1.378 | 0.892 | 0.039 | 0.062 | 0.062 | 0 | 0.52 | 0.8 | 0.67 | 0.66 | 0.56 | 0.58 |
| 10 | 17064992 | rs7897550 | -0.145 | 2.366 | -1.305 | 0.934 | -0.056 | -0.101 | -0.101 | 0 | 0.21 | 0.07 | 0.37 | 0.41 | 0.28 | 0.26 |
| 10 | 17065067 | rs3902042 | -1.632 | 2.33 | -1.387 | 0.829 | 0.031 | 0.044 | 0.044 | 0 | 0.56 | 0.79 | 0.69 | 0.66 | 0.58 | 0.6 |
| 10 | 17065761 | rs17139621 | -0.827 | 2.437 | -0.983 | 1.085 | -0.051 | -0.084 | -0.084 | 0 | 0.21 | 0.06 | 0.38 | 0.38 | 0.28 | 0.32 |
| 10 | 17068112 | rs17431426 | -1.027 | 2.267 | -0.673 | 0.36 | 0.001 | -0.011 | -0.011 | 0.275 | 0.13 | 0.12 | 0.08 | 0.03 | 0.17 | 0.2 |
| 10 | 17068251 | rs12146414 | -0.93 | 1.188 | -0.362 | -0.205 | -0.018 | -0.045 | -0.045 | 0 | 0.19 | 0.27 | 0.08 | 0.01 | 0.09 | 0.11 |
| 10 | 17068275 | rs4525114 | 0.901 | 1.723 | -0.325 | 0.82 | -0.002 | 0.038 | 0.038 | 0 | 0.92 | 0.97 | 0.88 | 0.59 | 0.92 | 0.85 |
| 10 | 17071966 | rs12780809 | -0.829 | 1.679 | -0.488 | -0.369 | -0.011 | 0.046 | 0.046 | 0.113 | 0.26 | 0.14 | 0.21 | 0 | 0.3 | 0.09 |
| 10 | 17073135 | rs12766939 | -0.81 | 1.761 | -0.484 | -0.365 | -0.01 | 0.044 | 0.044 | 0.116 | 0.26 | 0.14 | 0.2 | 0 | 0.3 | 0.09 |
| 10 | 17073900 | rs2356215 | 1.013 | 1.427 | 0.442 | -0.554 | -0.019 | -0.041 | -0.041 | 0 | 0.81 | 0.33 | 0.84 | 0.94 | 0.87 | 0.74 |
| 10 | 17075573 | rs11254336 | 1.538 | 1.504 | 0.098 | -0.233 | -0.075 | 0.034 | 0.034 | 0.274 | 0.57 | 0.07 | 0.55 | 0.65 | 0.54 | 0.51 |
| 10 | 17077417 | rs17139663 | -0.436 | 1.479 | 0.279 | -0.109 | 0.006 | -0.011 | -0.011 | 0 | 0.13 | 0.14 | 0.15 | 0.3 | 0.11 | 0.37 |
| 10 | 17081153 | rs11254339 | -1.292 | 1.52 | 0.721 | -0.122 | -0.038 | 0.099 | 0.099 | 2.653 | 0.17 | 0.5 | 0.17 | 0.31 | 0.14 | 0.38 |
| 10 | 17086636 | rs7916057 | -0.435 | 1.596 | 0.703 | -0.262 | 0 | -0.139 | -0.139 | 0.575 | 0.26 | 0.4 | 0.23 | 0 | 0.27 | 0.07 |
| 10 | 17089942 | rs7082270 | 1.111 | 1.682 | 0.992 | 0.114 | 0.002 | -0.013 | -0.013 | 0 | 0.83 | 0.75 | 0.84 | 0.86 | 0.83 | 0.68 |
| 10 | 17098935 | rs7922682 | 0.494 | 2.928 | 0.822 | 0.08 | -0.003 | 0.011 | 0.011 | 0 | 0.93 | 0.97 | 0.86 | 0.96 | 0.81 | 0.95 |
| 10 | 17099016 | rs11254351 | 0.027 | 2.838 | 0.9 | 0.067 | -0.009 | 0 | 0 | 0 | 0.1 | 0.12 | 0.06 | 0.1 | 0.08 | 0.29 |
| 10 | 17102552 | rs4748345 | 0.517 | 2.877 | 0.894 | 0.139 | -0.006 | 0.005 | 0.005 | 0 | 0.93 | 0.97 | 0.86 | 0.95 | 0.82 | 0.95 |
| 10 | 17104173 | rs9665553 | 1.044 | 2.865 | 1.062 | 0.135 | 0.046 | -0.033 | -0.033 | 0 | 0.83 | 0.76 | 0.8 | 0.93 | 0.76 | 0.7 |
| 10 | 17105668 | rs1907360 | 1.635 | 2.826 | 0.758 | 0.008 | -0.086 | 0.036 | 0.036 | 0.015 | 0.55 | 0.07 | 0.53 | 0.64 | 0.47 | 0.51 |
| 10 | 17109730 | rs1907358 | 1.329 | 2.851 | 1.063 | 0.004 | 0.127 | -0.011 | -0.011 | 0.039 | 0.91 | 0.69 | 0.83 | 0.95 | 0.82 | 0.95 |
| 10 | 17110595 | rs10795440 | 0.408 | 2.871 | 0.912 | 0.014 | -0.002 | 0.001 | 0.001 | 0 | 0.93 | 0.97 | 0.85 | 0.95 | 0.81 | 0.95 |
| 10 | 17112417 | rs2172080 | -0.922 | 2.905 | 1.22 | 0.031 | -0.014 | -0.033 | -0.033 | 1.44 | 0.28 | 0.56 | 0.26 | 0.21 | 0.27 | 0.15 |
| 10 | 17119471 | rs7081149 | -0.095 | 3.232 | 0.948 | 0.282 | 0.002 | 0.011 | 0.011 | 0 | 0.22 | 0.14 | 0.14 | 0 | 0.22 | 0.03 |
| 10 | 17121910 | rs2130807 | 1.859 | 3.131 | 0.719 | 0.376 | 0.005 | -0.04 | -0.04 | 0 | 0.51 | 0.06 | 0.48 | 0.54 | 0.4 | 0.44 |
| 10 | 17122004 | rs7091854 | -1.452 | 3.121 | 0.947 | 0.462 | 0.062 | 0.252 | 0.252 | 0.8 | 0.05 | 0.38 | 0.11 | 0.21 | 0.05 | 0.12 |
| 10 | 17123463 | rs4748346 | 0.143 | 3.116 | 0.807 | 0.322 | 0.038 | -0.002 | -0.002 | 0.552 | 0.94 | 0.93 | 0.86 | 0.95 | 0.82 | 0.92 |
| 10 | 17133683 | rs1996316 | 1.546 | 2.547 | 0.066 | 0.286 | 0.013 | -0.073 | -0.073 | 0 | 0.49 | 0.04 | 0.48 | 0.51 | 0.38 | 0.42 |
| 10 | 17135164 | rs2356825 | 0.962 | 2.784 | 0.667 | 0.452 | -0.006 | 0.005 | 0.005 | 0 | 0.59 | 0.38 | 0.58 | 0.66 | 0.53 | 0.55 |
| 10 | 17136169 | rs2145939 | 1.027 | 3.031 | 0.939 | 0.432 | 0.162 | 0.2 | 0.2 | 0 | 0.9 | 0.64 | 0.75 | 0.68 | 0.8 | 0.81 |
| 10 | 17139931 | rs11816077 | 0.127 | 2.879 | 0.709 | 0.351 | -0.001 | 0.001 | 0.001 | 0 | 0.21 | 0.12 | 0.14 | 0 | 0.22 | 0.04 |
| 10 | 17141391 | rs10904876 | 1.038 | 3.012 | 0.298 | 0.355 | 0.002 | 0.004 | 0.004 | 0 | 0.6 | 0.39 | 0.62 | 0.68 | 0.63 | 0.56 |
| 10 | 17143000 | rs12261966 | -0.917 | 2.887 | 0.244 | 0.28 | 0.161 | 0.033 | 0.033 | 0 | 0.12 | 0.33 | 0.14 | 0.15 | 0.26 | 0.14 |
| 10 | 17143996 | rs10795445 | 0.344 | 2.736 | 0.16 | 0.11 | -0.014 | -0.027 | -0.027 | 0 | 0.81 | 0.67 | 0.84 | 0.91 | 0.85 | 0.69 |
| 10 | 17156151 | rs1801222 | 0.931 | 2.876 | -0.098 | 0.18 | 0.012 | -0.001 | -0.001 | 0.012 | 0.84 | 0.81 | 0.8 | 0.85 | 0.71 | 0.83 |

| | | | | | | | | | | | | | | | | |
|---|---|---|---|---|---|---|---|---|---|---|---|---|---|---|---|---|
| 10 | 17158132 | rs12243365 | -0.456 | 2.814 | -0.1 | 0.164 | 0.047 | 0.034 | 0.034 | 0.035 | 0.12 | 0.33 | 0.1 | 0.16 | 0.16 | 0.13 |
| 10 | 17158973 | rs12243895 | -0.712 | 2.262 | -0.124 | 0.126 | -0.019 | -0.029 | -0.029 | 0.15 | 0.37 | 0.48 | 0.31 | 0.27 | 0.36 | 0.4 |
| 10 | 17159192 | rs11254375 | -0.499 | 2.324 | -0.305 | 0.194 | 0.143 | 0.05 | 0.05 | 0 | 0.13 | 0.34 | 0.14 | 0.17 | 0.27 | 0.14 |
| 10 | 17161749 | rs10508521 | 0.767 | 2.217 | -0.686 | 0.115 | 0.001 | -0.001 | -0.001 | 1.058 | 0.16 | 0.09 | 0.11 | 0 | 0.18 | 0.03 |
| 10 | 17162503 | rs2277210 | -0.14 | 2.211 | -0.832 | 0.407 | 0.027 | 0.025 | 0.025 | 0 | 0.94 | 0.78 | 0.89 | 0.93 | 0.94 | 0.94 |
| 10 | 17163081 | rs10795449 | 0.11 | 2.024 | -1.484 | 0.438 | 0.002 | 0.009 | 0.009 | 0 | 0.57 | 0.38 | 0.59 | 0.65 | 0.57 | 0.53 |
| 10 | 17164221 | rs1914172 | 0.741 | 0.687 | -2.59 | 0.527 | 0.01 | -0.156 | -0.156 | 0 | 0.43 | 0.04 | 0.45 | 0.49 | 0.31 | 0.39 |
| 10 | 17167422 | rs7916688 | -0.29 | 0.262 | -2.64 | 0.671 | 0.01 | -0.02 | -0.02 | 0 | 0.39 | 0.93 | 0.29 | 0.38 | 0.27 | 0.41 |
| 10 | 17172952 | rs10904881 | 0.135 | 0.52 | -2.703 | 0.879 | -0.036 | -0.051 | -0.051 | 0 | 0.47 | 0.03 | 0.29 | 0.43 | 0.39 | 0.49 |
| 10 | 17185810 | rs10904886 | 0.306 | 0.549 | -0.717 | 0.457 | -0.002 | 0.013 | 0.013 | 0 | 0.14 | 0.1 | 0.23 | 0.03 | 0.15 | 0.11 |
| 10 | 17186625 | rs11254398 | 0.285 | 0.547 | -0.718 | 0.456 | -0.002 | 0.013 | 0.013 | 0 | 0.14 | 0.1 | 0.23 | 0.03 | 0.15 | 0.11 |
| 10 | 17188023 | rs7071287 | 0.325 | 0.545 | -0.707 | 0.456 | -0.002 | 0.012 | 0.012 | 0 | 0.13 | 0.08 | 0.23 | 0.03 | 0.15 | 0.11 |
| 10 | 17190561 | rs11254405 | 0.325 | 0.383 | -0.212 | 0.792 | -0.002 | 0.012 | 0.012 | 0 | 0.13 | 0.1 | 0.23 | 0.03 | 0.15 | 0.11 |
| 10 | 17192362 | rs6602180 | -0.294 | 0.877 | 0.246 | 0.804 | 0.002 | -0.006 | -0.006 | 0 | 0.4 | 0.38 | 0.43 | 0.26 | 0.49 | 0.35 |
| 10 | 17199778 | rs11254408 | 0.542 | 0.87 | 0.197 | 0.8 | -0.002 | 0.006 | 0.006 | 0 | 0.13 | 0.1 | 0.22 | 0.04 | 0.15 | 0.11 |
| 10 | 17204187 | rs11254413 | -0.394 | 0.864 | 0.303 | 0.8 | 0.007 | -0.015 | -0.015 | 0.413 | 0.41 | 0.4 | 0.43 | 0.26 | 0.49 | 0.35 |
| 10 | 17204944 | rs10904893 | 0.365 | 0.892 | 0.326 | 0.8 | -0.047 | -0.034 | -0.034 | 0 | 0.55 | 0.22 | 0.35 | 0.49 | 0.48 | 0.58 |
| 10 | 17210822 | rs17464824 | 0.421 | 0.851 | 0.357 | 0.998 | -0.018 | 0.007 | 0.007 | 0 | 0.55 | 0.32 | 0.37 | 0.45 | 0.5 | 0.58 |
| 10 | 17210881 | rs7096233 | 0.479 | 0.913 | 0.356 | 0.836 | -0.002 | 0.011 | 0.011 | 0 | 0.14 | 0.1 | 0.22 | 0.04 | 0.15 | 0.11 |
| 10 | 17211383 | rs7084476 | 0.595 | 0.924 | 0.447 | 0.875 | 0.037 | -0.013 | -0.013 | 0 | 0.28 | 0.45 | 0.22 | 0.22 | 0.34 | 0.24 |
| 10 | 17233450 | rs10490958 | 0.814 | 0.948 | 0.576 | 0.885 | -0.005 | 0 | 0 | 0.41 | 0.26 | 0.2 | 0.2 | 0.22 | 0.34 | 0.24 |
| 10 | 17234705 | rs7076638 | 0.476 | 0.817 | 0.551 | 0.852 | -0.002 | 0.011 | 0.011 | 0 | 0.14 | 0.1 | 0.22 | 0.04 | 0.15 | 0.1 |
| 10 | 17240367 | rs7071826 | -0.31 | 0.799 | 0.637 | 0.955 | -0.004 | 0.04 | 0.04 | 0 | 0.84 | 0.77 | 0.59 | 0.71 | 0.84 | 0.82 |
| 10 | 17243119 | rs7895248 | 0.488 | 0.811 | 0.564 | 0.942 | 0 | -0.011 | -0.011 | 3.726 | 0.14 | 0.12 | 0.22 | 0.04 | 0.15 | 0.1 |
| 10 | 17243351 | rs1885396 | 0.34 | 0.937 | 0.387 | 0.889 | -0.024 | -0.016 | -0.016 | 0 | 0.55 | 0.34 | 0.37 | 0.49 | 0.49 | 0.57 |
| 10 | 17244124 | rs10795459 | 0.285 | 0.937 | 0.396 | 0.889 | -0.011 | -0.007 | -0.007 | 1.229 | 0.56 | 0.41 | 0.37 | 0.49 | 0.49 | 0.58 |
| 10 | 17246355 | rs3758415 | -0.449 | 0.946 | 0.271 | 0.905 | -0.002 | 0.061 | 0.061 | 0 | 0.84 | 0.72 | 0.59 | 0.71 | 0.85 | 0.82 |
| 10 | 17247729 | rs4748363 | -0.23 | 0.987 | -0.007 | 0.938 | -0.008 | 0.085 | 0.085 | 0 | 0.83 | 0.61 | 0.58 | 0.71 | 0.85 | 0.82 |
| 10 | 17248287 | rs7084637 | 0.698 | 0.999 | -0.85 | 0.793 | -0.015 | -0.049 | -0.049 | 0.955 | 0.47 | 0.03 | 0.29 | 0.42 | 0.37 | 0.49 |
| 10 | 17249363 | rs10904907 | 0.8 | 0.751 | -0.756 | 0.609 | 0.008 | -0.051 | -0.051 | 0.253 | 0.29 | 0.01 | 0.16 | 0.3 | 0.19 | 0.29 |
| 10 | 17250900 | rs7095324 | -0.135 | 1.856 | 0.343 | 0.594 | 0.009 | -0.003 | -0.003 | 0 | 0.27 | 0.25 | 0.2 | 0.22 | 0.35 | 0.24 |
| 10 | 17259567 | rs7903137 | 0.932 | 1.998 | 0.822 | 0.757 | 0.015 | 0.008 | 0.008 | 1.27 | 0.67 | 0.42 | 0.36 | 0.65 | 0.45 | 0.63 |
| 10 | 17264486 | rs10752073 | 0.419 | 2.799 | 0.694 | 0.424 | -0.031 | 0.036 | 0.036 | 0 | 0.87 | 0.92 | 0.79 | 0.98 | 0.77 | 0.84 |
| 10 | 17270074 | rs6602186 | -1.831 | 2.325 | 1.689 | 0.517 | 0.069 | 0.055 | 0.055 | 3.05 | 0.05 | 0.54 | 0.04 | 0.01 | 0.02 | 0.04 |
| 10 | 17279517 | rs3249 | -0.301 | 2.756 | 1.755 | 0.576 | -0.039 | 0.003 | 0.003 | 0 | 0.12 | 0.06 | 0.13 | 0.08 | 0.24 | 0.08 |
| 10 | 17279562 | rs1049341 | -2.148 | 2.558 | 2.056 | 0.772 | 0.027 | 0.224 | 0.224 | 0.566 | 0.07 | 0.57 | 0.28 | 0.23 | 0.06 | 0.11 |
| 10 | 17283993 | rs12251583 | -1.84 | 2.6 | 2.025 | 0.79 | 0.061 | 0.284 | 0.284 | 0.056 | 0.06 | 0.32 | 0.26 | 0.22 | 0.06 | 0.1 |
| 10 | 17302891 | rs911607 | -0.327 | 2.83 | 1.344 | 0.71 | 0.04 | 0.005 | 0.005 | 0 | 0.12 | 0.16 | 0.14 | 0.1 | 0.25 | 0.08 |
| 10 | 17312690 | rs10904915 | -0.486 | 2.819 | 1.421 | 0.723 | 0.096 | 0.026 | 0.026 | 0 | 0.13 | 0.26 | 0.15 | 0.11 | 0.25 | 0.08 |
| 10 | 17319707 | rs359280 | -1.658 | 2.864 | 1.202 | 0.476 | 0.018 | -0.016 | -0.016 | 0 | 0.28 | 0.37 | 0.36 | 0.11 | 0.51 | 0.26 |
| 10 | 17327084 | rs359285 | -0.763 | 2.799 | 0.775 | 0.472 | -0.027 | 0.036 | 0.036 | 0.058 | 0.17 | 0.14 | 0.21 | 0.02 | 0.26 | 0.18 |
| 10 | 17333680 | rs359297 | -0.657 | 2.768 | 0.362 | 0.466 | -0.021 | 0.028 | 0.028 | 0 | 0.17 | 0.13 | 0.21 | 0.02 | 0.26 | 0.18 |
| 10 | 17334745 | rs10904919 | -1.189 | 1.926 | 0.194 | 0.505 | -0.008 | 0.015 | 0.015 | 0.115 | 0.33 | 0.72 | 0.42 | 0.13 | 0.54 | 0.3 |
| 10 | 17334935 | rs10795469 | -1.196 | 1.761 | 0.031 | 0.597 | -0.014 | -0.002 | -0.002 | 0 | 0.37 | 0.79 | 0.65 | 0.34 | 0.56 | 0.37 |
| 10 | 17337788 | rs359300 | 0.086 | 0.532 | -0.543 | 0.274 | 0.01 | 0.009 | 0.009 | 0.631 | 0.64 | 0.29 | 0.69 | 0.3 | 0.71 | 0.51 |
| 10 | 17340342 | rs11254509 | 0.161 | 0.852 | -1.015 | 0.333 | -0.009 | 0.055 | 0.055 | 3.501 | 0.48 | 0.23 | 0.36 | 0.34 | 0.49 | 0.35 |
| 10 | 17343390 | rs359308 | -0.7 | 0.659 | -0.323 | 0.984 | 0.002 | 0.325 | 0.325 | 0.086 | 0.06 | 0.19 | 0.36 | 0.51 | 0.08 | 0.12 |
| 10 | 17347404 | rs10904927 | 0.28 | 1.179 | -0.41 | 0.227 | 0 | 0 | 0 | 2.928 | 0.62 | 0.42 | 0.42 | 0.46 | 0.48 | 0.59 |
| 10 | 17349745 | rs384376 | 0.087 | 0.581 | -0.262 | -0.346 | 0.011 | -0.04 | -0.04 | 0 | 0.58 | 0.55 | 0.69 | 0.78 | 0.58 | 0.53 |
| 10 | 17351794 | rs400333 | -0.467 | -0.079 | -0.397 | -0.79 | -0.01 | -0.066 | -0.066 | 1.01 | 0.35 | 0.47 | 0.48 | 0.87 | 0.31 | 0.46 |
| 10 | 17351824 | rs415761 | -0.465 | -0.075 | -0.396 | -0.789 | -0.009 | -0.071 | -0.071 | 0 | 0.35 | 0.47 | 0.48 | 0.87 | 0.31 | 0.46 |
| 10 | 17352930 | rs452281 | 0.106 | -0.261 | 0.211 | -0.518 | 0.032 | 0.126 | 0.126 | 0.304 | 0.16 | 0.36 | 0.35 | 0.41 | 0.21 | 0.25 |
| 10 | 17353151 | rs11254514 | -0.047 | -0.054 | 0.061 | -0.536 | -0.023 | 0.089 | 0.089 | 0 | 0.18 | 0.25 | 0.14 | 0.47 | 0.11 | 0.22 |
| 10 | 17356447 | rs426357 | -0.739 | 0.388 | -0.242 | -0.504 | -0.008 | 0.012 | 0.012 | 0 | 0.26 | 0.25 | 0.43 | 0.77 | 0.27 | 0.42 |
| 10 | 17357756 | rs1055986 | -0.788 | 0.278 | -0.609 | -0.742 | -0.089 | 0.1 | 0.1 | 0.609 | 0.71 | 0.92 | 0.74 | 0.9 | 0.65 | 0.77 |
| 10 | 17362384 | rs7091114 | 0.461 | 0.26 | -0.215 | -0.849 | 0.071 | 0.238 | 0.238 | 0 | 0.48 | 0.72 | 0.31 | 0.12 | 0.37 | 0.4 |
| 10 | 17362784 | rs717768 | 0.468 | 0.257 | -0.229 | -0.85 | 0.053 | 0.133 | 0.133 | 0 | 0.48 | 0.61 | 0.3 | 0.12 | 0.37 | 0.4 |
| 10 | 17368394 | rs11254534 | 1.182 | 0.18 | 0.183 | -0.956 | -0.004 | -0.095 | -0.095 | 0.912 | 0.38 | 0.33 | 0.26 | 0.02 | 0.32 | 0.36 |
| 10 | 17376759 | rs12254144 | 1.127 | 0.218 | 0.085 | -0.949 | 0.004 | -0.062 | -0.062 | 0.807 | 0.39 | 0.55 | 0.27 | 0.02 | 0.31 | 0.36 |
| 10 | 17379520 | rs10795471 | -0.168 | -0.076 | -0.063 | -0.586 | 0.023 | -0.081 | -0.081 | 0 | 0.14 | 0.05 | 0.12 | 0.38 | 0.11 | 0.25 |
| 10 | 17380804 | rs359311 | 0.502 | 0.214 | 0.089 | -0.64 | -0.002 | 0.026 | 0.026 | 0.426 | 0.49 | 0.67 | 0.56 | 0.42 | 0.5 | 0.46 |
| 10 | 17384196 | rs10904931 | 0.667 | 0.197 | 0.063 | -0.674 | -0.004 | 0.011 | 0.011 | 0.136 | 0.48 | 0.58 | 0.55 | 0.41 | 0.49 | 0.44 |
| 10 | 17384422 | rs359315 | 0.079 | 0.208 | -0.101 | -0.682 | -0.043 | -0.085 | -0.085 | 0 | 0.57 | 0.84 | 0.59 | 0.5 | 0.56 | 0.47 |
| 10 | 17387716 | rs7067707 | -1.707 | 0.665 | 0.815 | -0.036 | 0.005 | 0.001 | 0.001 | 0 | 0.44 | 0.7 | 0.42 | 0.59 | 0.4 | 0.61 |
| 10 | 17387944 | rs11254539 | -0.464 | 0.897 | 0.74 | -0.103 | -0.004 | -0.018 | -0.018 | 0.012 | 0.21 | 0.24 | 0.18 | 0.09 | 0.18 | 0.2 |
| 10 | 17390943 | rs11254541 | -0.756 | 0.664 | 0.606 | 0.037 | -0.01 | -0.054 | -0.054 | 0 | 0.22 | 0.35 | 0.2 | 0.1 | 0.19 | 0.21 |
| 10 | 17394523 | rs11254543 | -0.44 | 0.599 | 0.55 | 0.117 | -0.001 | 0.023 | 0.023 | 0 | 0.18 | 0.22 | 0.15 | 0.51 | 0.15 | 0.3 |
| 10 | 17395500 | rs7896970 | -0.542 | 0.6 | 0.549 | 0.084 | -0.002 | -0.036 | -0.036 | 0 | 0.22 | 0.31 | 0.19 | 0.1 | 0.19 | 0.21 |
| 10 | 17396213 | rs4146638 | -0.285 | 0.598 | 0.474 | 0.097 | 0.011 | -0.062 | -0.062 | 0 | 0.17 | 0.13 | 0.15 | 0.51 | 0.15 | 0.3 |

| | | | | | | | | | | | | | | | | |
|---|---|---|---|---|---|---|---|---|---|---|---|---|---|---|---|---|
| 10 | 17396655 | rs4146637 | -0.399 | 0.592 | 0.479 | 0.094 | 0.012 | -0.04 | -0.04 | 0 | 0.18 | 0.18 | 0.15 | 0.51 | 0.15 | 0.29 |
| 10 | 17399616 | rs1020339 | -1.639 | 0.622 | 0.393 | 0.062 | -0.01 | 0.001 | 0.001 | 0 | 0.41 | 0.55 | 0.35 | 0.61 | 0.36 | 0.51 |
| 10 | 17400951 | rs932877 | -1.627 | 0.684 | -0.121 | 0.084 | 0.003 | 0 | 0 | 0.001 | 0.4 | 0.42 | 0.35 | 0.62 | 0.36 | 0.52 |
| 10 | 17401207 | rs10904935 | -0.631 | 0.704 | -0.146 | 0.123 | -0.005 | 0.039 | 0.039 | 0 | 0.2 | 0.27 | 0.16 | 0.52 | 0.17 | 0.31 |
| 10 | 17406855 | rs729169 | 0.256 | 0.782 | -0.275 | 0.032 | -0.017 | 0.056 | 0.056 | 0 | 0.14 | 0.06 | 0.16 | 0 | 0.15 | 0.19 |
| 10 | 17414971 | rs10904945 | -0.926 | 0.754 | -0.103 | 0.225 | 0.005 | -0.001 | -0.001 | 0.041 | 0.4 | 0.38 | 0.34 | 0.62 | 0.36 | 0.51 |
| 10 | 17426474 | rs2770197 | -1.018 | 1.029 | 0.164 | 0.478 | 0.003 | 0.016 | 0.016 | 0 | 0.18 | 0.25 | 0.38 | 0.64 | 0.22 | 0.21 |
| 10 | 17431776 | rs359322 | -1.079 | 0.951 | 0.822 | 0.456 | 0.044 | 0.147 | 0.147 | 0 | 0.18 | 0.42 | 0.37 | 0.59 | 0.22 | 0.24 |
| 10 | 17439103 | rs10159766 | -1.087 | 0.974 | 1.039 | 0.585 | 0.046 | 0.151 | 0.151 | 0 | 0.18 | 0.45 | 0.37 | 0.59 | 0.22 | 0.22 |
| 10 | 17439939 | rs10904949 | -1.814 | 0.862 | 0.794 | 0.174 | 0.003 | 0.006 | 0.006 | 0 | 0.31 | 0.58 | 0.48 | 0.59 | 0.32 | 0.35 |
| 10 | 17451308 | rs746570 | -1.579 | 1.221 | 0.702 | -0.087 | 0.018 | 0.017 | 0.017 | 0 | 0.29 | 0.55 | 0.48 | 0.59 | 0.35 | 0.37 |
| 10 | 17451324 | rs746571 | 0.367 | 1.114 | 0.641 | -0.114 | 0.008 | -0.035 | -0.035 | 0 | 0.1 | 0.13 | 0.11 | 0.01 | 0.13 | 0.14 |
| 10 | 17456574 | rs4561106 | -1.942 | 0.992 | 0.35 | -0.12 | -0.04 | 0.069 | 0.069 | 0 | 0.43 | 0.75 | 0.6 | 0.68 | 0.46 | 0.53 |
| 10 | 17474443 | rs7911857 | -1.196 | 0.561 | 1.121 | 0.104 | -0.009 | -0.013 | -0.013 | 0 | 0.14 | 0.51 | 0.08 | 0.1 | 0.08 | 0.12 |
| 10 | 17474469 | rs7911736 | -2.198 | 0.561 | 1.091 | 0.035 | -0.001 | -0.011 | -0.011 | 0 | 0.25 | 0.63 | 0.19 | 0.1 | 0.21 | 0.3 |
| 10 | 17474594 | rs7897269 | -1.094 | 0.565 | 1.24 | 0.042 | 0.108 | 0.183 | 0.183 | 0 | 0.05 | 0.32 | 0.03 | 0.09 | 0.02 | 0.06 |
| 10 | 17476686 | rs12246530 | -1.18 | 0.57 | 1.015 | 0.066 | -0.009 | -0.009 | -0.009 | 0 | 0.14 | 0.51 | 0.08 | 0.1 | 0.08 | 0.12 |
| 10 | 17477507 | rs4748376 | -2.099 | 0.545 | 0.61 | 0.064 | 0.002 | 0.026 | 0.026 | 1.284 | 0.26 | 0.76 | 0.27 | 0.16 | 0.22 | 0.35 |
| 10 | 17486038 | rs11254602 | -2.232 | 0.571 | 0.526 | -0.009 | -0.014 | 0.073 | 0.073 | 1.124 | 0.28 | 0.8 | 0.25 | 0.1 | 0.25 | 0.35 |
| 10 | 17491652 | rs3928773 | -1.688 | 0.654 | 0.931 | 0.039 | 0.033 | -0.039 | -0.039 | 0 | 0.19 | 0.54 | 0.21 | 0.1 | 0.19 | 0.3 |
| 10 | 17493857 | rs12259924 | -0.465 | 0.54 | 0.884 | 0.365 | 0.005 | 0.028 | 0.028 | 0 | 0.13 | 0.15 | 0.14 | 0.3 | 0.15 | 0.24 |
| 10 | 17498867 | rs7077568 | -1.017 | 0.412 | 0.818 | 0.326 | -0.004 | -0.014 | -0.014 | 0 | 0.16 | 0.13 | 0.19 | 0.3 | 0.16 | 0.28 |
| 10 | 17504935 | rs7088825 | -0.793 | 0.484 | 0.946 | 0.312 | 0.017 | -0.057 | -0.057 | 0.058 | 0.14 | 0.25 | 0.17 | 0.01 | 0.16 | 0.23 |
| 10 | 17507136 | rs12266595 | -2.476 | 0.31 | 0.666 | 0.245 | -0.001 | -0.041 | -0.041 | 0 | 0.32 | 0.75 | 0.41 | 0.44 | 0.28 | 0.52 |
| 10 | 17513477 | rs7089753 | -0.365 | 0.319 | 0.73 | 0.21 | -0.007 | -0.039 | -0.039 | 0.154 | 0.11 | 0.25 | 0.18 | 0.04 | 0.09 | 0.14 |
| 10 | 17518204 | rs7069823 | -1.804 | 0.301 | 0.629 | 0.236 | 0.002 | 0.019 | 0.019 | 0.175 | 0.23 | 0.35 | 0.31 | 0.34 | 0.24 | 0.38 |
| 10 | 17519718 | rs10508540 | -0.398 | 0.273 | 0.611 | 0.266 | 0.018 | 0.103 | 0.103 | 0.017 | 0.12 | 0.3 | 0.14 | 0.3 | 0.13 | 0.23 |
| 10 | 17520288 | rs7920635 | -1.153 | 0.276 | 0.606 | 0.275 | 0 | 0.021 | 0.021 | 0.02 | 0.19 | 0.3 | 0.19 | 0.3 | 0.19 | 0.34 |
| 10 | 17521395 | rs1277443 | -0.701 | -0.296 | -0.168 | -0.645 | 0.024 | 0.034 | 0.034 | 4.093 | 0.43 | 0.3 | 0.24 | 0.3 | 0.33 | 0.55 |
| 10 | 17521991 | rs1290099 | 0.568 | -0.769 | -0.519 | -1.006 | 0.034 | 0.102 | 0.102 | 0.358 | 0.27 | 0.09 | 0.11 | 0 | 0.17 | 0.31 |
| 10 | 17522699 | rs1556056 | -0.846 | -0.729 | -0.436 | -0.938 | 0.027 | 0.049 | 0.049 | 2.86 | 0.46 | 0.32 | 0.26 | 0.29 | 0.33 | 0.59 |
| 10 | 17523041 | rs12572065 | 0.865 | -0.807 | -1.221 | -0.299 | 0.03 | -0.057 | -0.057 | 1.96 | 0.15 | 0.05 | 0.13 | 0.25 | 0.09 | 0.12 |
| 10 | 17531322 | rs9423961 | -0.212 | -0.052 | -1.374 | -0.52 | -0.057 | -0.07 | -0.07 | 0 | 0.73 | 0.82 | 0.6 | 0.45 | 0.65 | 0.71 |
| 10 | 17531569 | rs2226248 | 0.797 | -0.259 | -0.943 | -0.691 | -0.015 | 0.003 | 0.003 | 0 | 0.58 | 0.35 | 0.38 | 0.41 | 0.52 | 0.56 |
| 10 | 17532277 | rs2479893 | 0.249 | -0.118 | -0.915 | -0.464 | 0.023 | 0.02 | 0.02 | 2.543 | 0.38 | 0.19 | 0.19 | 0.32 | 0.27 | 0.53 |
| 10 | 17534158 | rs1753375 | -1.418 | 0.61 | -0.747 | 0.051 | 0.019 | 0.048 | 0.048 | 0 | 0.94 | 0.93 | 0.81 | 0.74 | 0.85 | 0.94 |
| 10 | 17534259 | rs1781408 | -1.617 | -0.288 | -1.016 | -0.125 | -0.072 | -0.053 | -0.053 | 0 | 0.6 | 0.86 | 0.52 | 0.54 | 0.49 | 0.77 |
| 10 | 17535270 | rs4633352 | 0.499 | -0.227 | -1.13 | -0.169 | -0.018 | -0.047 | -0.047 | 3.3 | 0.14 | 0.08 | 0.15 | 0.33 | 0.16 | 0.25 |
| 10 | 17537366 | rs1753371 | 0.442 | -0.333 | -1.263 | -0.343 | 0.025 | 0.219 | 0.219 | 0.823 | 0.3 | 0.01 | 0.12 | 0 | 0.19 | 0.32 |
| 10 | 17538889 | rs4748378 | -0.973 | -0.289 | -0.666 | -0.3 | 0.019 | 0.05 | 0.05 | 0 | 0.28 | 0.54 | 0.37 | 0.41 | 0.3 | 0.42 |
| 10 | 17539140 | rs997946 | 0.993 | -0.276 | -0.979 | -0.326 | -0.006 | 0.166 | 0.166 | 0.255 | 0.24 | 0 | 0.1 | 0 | 0.16 | 0.29 |
| 10 | 17539320 | rs12357059 | 0.693 | -0.003 | -0.567 | -0.128 | 0 | 0.031 | 0.031 | 0.036 | 0.11 | 0.15 | 0.11 | 0.36 | 0.13 | 0.16 |
| 10 | 17540410 | rs1753370 | -1.433 | -0.004 | -0.584 | -0.053 | -0.02 | -0.016 | -0.016 | 7.352 | 0.6 | 0.72 | 0.55 | 0.54 | 0.5 | 0.76 |
| 10 | 17544529 | rs11593604 | 0.277 | 0.368 | -0.73 | 0.188 | -0.015 | 0.034 | 0.034 | 0 | 0.14 | 0.08 | 0.16 | 0.01 | 0.16 | 0.23 |
| 10 | 17547113 | rs7075034 | -0.999 | 0.367 | -0.517 | 0.42 | 0.001 | 0.001 | 0.001 | 0 | 0.5 | 0.5 | 0.49 | 0.5 | 0.46 | 0.71 |
| 10 | 17548038 | rs12773877 | -0.354 | 0.481 | -0.581 | 0.381 | 0 | -0.006 | -0.006 | 0 | 0.37 | 0.37 | 0.3 | 0.3 | 0.32 | 0.5 |
| 10 | 17554259 | rs2884126 | -0.256 | 0.98 | -0.507 | 1.276 | 0 | 0.025 | 0.025 | 0 | 0.13 | 0.12 | 0.11 | 0.35 | 0.15 | 0.18 |
| 10 | 17559867 | rs7074210 | -0.144 | 1.005 | -0.558 | 1.271 | -0.005 | 0.007 | 0.007 | 0 | 0.12 | 0.09 | 0.11 | 0.34 | 0.15 | 0.22 |
| 10 | 17565864 | rs10904962 | -0.402 | 1.08 | -0.454 | 1.425 | -0.003 | 0.023 | 0.023 | 0 | 0.11 | 0.08 | 0.1 | 0.35 | 0.14 | 0.24 |
| 10 | 17566940 | rs7899306 | 0.367 | 0.791 | -1.074 | 1.105 | -0.009 | 0.027 | 0.027 | 0 | 0.35 | 0.06 | 0.32 | 0.28 | 0.36 | 0.28 |
| 10 | 17567062 | rs6602200 | -0.36 | 0.778 | -0.077 | 1.145 | 0.107 | 0.145 | 0.145 | 0.829 | 0.91 | 0.45 | 0.77 | 0.83 | 0.86 | 0.97 |
| 10 | 17575053 | rs7911087 | -0.331 | -0.009 | -0.558 | 1.018 | 0.007 | 0.014 | 0.014 | 0.275 | 0.56 | 0.4 | 0.43 | 0.25 | 0.57 | 0.57 |
| 10 | 17576155 | rs11254652 | 1.31 | 0.039 | -0.99 | 1.014 | -0.037 | -0.043 | -0.043 | 0.007 | 0.14 | 0.02 | 0.13 | 0.17 | 0.14 | 0.17 |
| 10 | 17576227 | rs9787557 | 1.538 | -0.025 | -0.951 | 1.068 | 0.013 | -0.034 | -0.034 | 0 | 0.21 | 0.04 | 0.17 | 0.28 | 0.21 | 0.09 |
| 10 | 17578161 | rs2884127 | -0.29 | -0.034 | -0.828 | 0.874 | -0.033 | -0.021 | -0.021 | 0.022 | 0.64 | 0.8 | 0.56 | 0.62 | 0.63 | 0.73 |
| 10 | 17579441 | rs10752086 | -1.041 | 0.133 | -0.15 | 1.05 | -0.006 | 0.03 | 0.03 | 0.048 | 0.59 | 0.65 | 0.56 | 0.72 | 0.56 | 0.86 |
| 10 | 17580108 | rs17141297 | -0.056 | 0.129 | -0.446 | 0.923 | -0.012 | -0.023 | -0.023 | 0 | 0.36 | 0.43 | 0.33 | 0.26 | 0.32 | 0.52 |
| 10 | 17580890 | rs7090660 | -1.05 | 0.385 | -0.473 | 1.638 | 0 | 0.003 | 0.003 | 0 | 0.33 | 0.38 | 0.38 | 0.69 | 0.32 | 0.58 |
| 10 | 17581157 | rs2884128 | -0.274 | 0.521 | -1.532 | 1.131 | -0.065 | -0.137 | -0.137 | 0 | 0.22 | 0.01 | 0.18 | 0.38 | 0.22 | 0.33 |
| 10 | 17590042 | rs10795496 | -0.383 | 0.304 | -0.677 | 0.966 | 0 | 0.013 | 0.013 | 0.296 | 0.3 | 0.66 | 0.51 | 0.58 | 0.28 | 0.27 |
| 10 | 17590681 | rs4748380 | 0.619 | -0.519 | -0.57 | 0.878 | -0.012 | -0.002 | -0.002 | 0 | 0.82 | 0.79 | 0.87 | 0.81 | 0.9 | 0.84 |
| 10 | 17591060 | rs11254660 | -0.851 | 0.045 | -0.327 | 0.959 | -0.026 | 0.05 | 0.05 | 0 | 0.26 | 0.47 | 0.25 | 0.37 | 0.23 | 0.33 |
| 10 | 18314067 | rs2357782 | -0.386 | -0.525 | -1.107 | 0.798 | 0.008 | -0.076 | -0.076 | 0.04 | 0.65 | 0.93 | 0.62 | 0.65 | 0.74 | 0.5 |
| 10 | 18316940 | rs1935506 | 0.308 | -0.475 | -1.067 | 0.741 | 0.015 | -0.001 | -0.001 | 0.021 | 0.55 | 0.41 | 0.52 | 0.45 | 0.68 | 0.47 |
| 10 | 18334521 | rs2488136 | -0.149 | -0.382 | -0.837 | 0.662 | 0.005 | 0.004 | 0.004 | 1.021 | 0.61 | 0.68 | 0.63 | 0.7 | 0.73 | 0.48 |
| 10 | 18336330 | rs2488137 | 0.481 | -0.48 | -0.93 | 0.573 | 0.022 | 0.306 | 0.306 | 0.746 | 0.12 | 0.42 | 0.23 | 0.3 | 0.13 | 0.2 |
| 10 | 18338262 | rs10828143 | -0.688 | -0.659 | -1.057 | 0.398 | -0.005 | -0.013 | -0.013 | 6.721 | 0.21 | 0.44 | 0.11 | 0.23 | 0.16 | 0.08 |
| 10 | 18338686 | rs2488140 | -0.53 | -0.579 | -1.054 | 0.426 | 0.002 | 0.128 | 0.128 | 0 | 0.22 | 0.5 | 0.28 | 0.33 | 0.22 | 0.23 |
| 10 | 18340587 | rs2497791 | -0.455 | -0.576 | -1.036 | 0.429 | 0 | 0.112 | 0.112 | 0 | 0.22 | 0.43 | 0.27 | 0.31 | 0.22 | 0.23 |

| | | | | | | | | | | | | | | | | |
|---|---|---|---|---|---|---|---|---|---|---|---|---|---|---|---|---|
| 10 | 18342172 | rs2497793 | -1.334 | -0.582 | -1.149 | 0.353 | -0.008 | -0.061 | -0.061 | 0 | 0.72 | 0.94 | 0.67 | 0.69 | 0.77 | 0.6 |
| 10 | 18343319 | rs4748438 | 0.36 | -0.122 | -0.788 | -0.196 | -0.001 | 0.001 | 0.001 | 0.684 | 0.42 | 0.4 | 0.35 | 0.36 | 0.5 | 0.25 |
| 10 | 18345137 | rs7920075 | 0.049 | 0.146 | -0.864 | -0.265 | -0.001 | -0.027 | -0.027 | 0 | 0.45 | 0.46 | 0.39 | 0.18 | 0.55 | 0.28 |
| 10 | 18351509 | rs10764296 | -0.048 | 0.31 | -1.132 | 0.288 | -0.021 | 0.075 | 0.075 | 0.257 | 0.47 | 0.19 | 0.28 | 0.3 | 0.52 | 0.38 |
| 10 | 18352728 | rs2488151 | -0.254 | 0.483 | -1.153 | 0.716 | 0.061 | -0.013 | -0.013 | 0.119 | 0.53 | 0.28 | 0.44 | 0.37 | 0.64 | 0.48 |
| 10 | 18358107 | rs7072642 | -0.137 | 0.259 | -0.269 | 0.502 | -0.001 | 0.026 | 0.026 | 3.791 | 0.55 | 0.37 | 0.31 | 0.28 | 0.54 | 0.39 |
| 10 | 18361329 | rs2488155 | -0.931 | -0.159 | -0.583 | 0.564 | -0.059 | -0.133 | -0.133 | 0 | 0.7 | 0.92 | 0.57 | 0.43 | 0.67 | 0.55 |
| 10 | 18365424 | rs2497816 | -0.965 | -0.338 | 0.153 | 0.878 | 0.004 | 0.094 | 0.094 | 0 | 0.19 | 0.43 | 0.23 | 0.25 | 0.18 | 0.17 |
| 10 | 18370324 | rs2488158 | -1.498 | -0.467 | 0.067 | 0.819 | -0.025 | 0.099 | 0.099 | 0.23 | 0.25 | 0.41 | 0.28 | 0.4 | 0.19 | 0.24 |
| 10 | 18373035 | rs7903081 | 0.686 | -0.121 | -1.415 | -0.092 | -0.102 | 0.161 | 0.161 | 2.451 | 0.29 | 0.04 | 0.19 | 0.03 | 0.33 | 0.2 |
| 10 | 18375174 | rs1935499 | -0.315 | -0.522 | -1.399 | -0.298 | -0.001 | -0.004 | -0.004 | 0 | 0.57 | 0.51 | 0.38 | 0.41 | 0.55 | 0.4 |
| 10 | 18377343 | rs7079408 | 1.337 | -0.07 | -1.642 | -0.396 | -0.109 | 0.112 | 0.112 | 0.061 | 0.21 | 0.03 | 0.16 | 0.03 | 0.28 | 0.21 |
| 10 | 18381086 | rs6415964 | -0.687 | -0.821 | -1.138 | -0.204 | 0.001 | -0.031 | -0.031 | 0 | 0.81 | 0.85 | 0.86 | 1 | 0.81 | 0.83 |
| 10 | 18383222 | rs7087949 | -0.712 | -0.904 | -0.796 | 1.055 | 0.001 | 0.136 | 0.136 | 0 | 0.16 | 0.25 | 0.2 | 0.43 | 0.16 | 0.16 |
| 10 | 18383929 | rs11012659 | 0.899 | -0.888 | -1.205 | 1.169 | 0.002 | -0.035 | -0.035 | 0.271 | 0.19 | 0.05 | 0.31 | 0.28 | 0.17 | 0.27 |
| 10 | 18387689 | rs11012674 | -0.367 | -0.462 | -0.354 | 1.724 | -0.002 | 0.053 | 0.053 | 0 | 0.12 | 0.18 | 0.1 | 0.44 | 0.1 | 0.08 |
| 10 | 18387746 | rs4573589 | -1.819 | -0.516 | -0.165 | 1.801 | 0 | 0.001 | 0.001 | 0 | 0.22 | 0.28 | 0.27 | 0.58 | 0.22 | 0.21 |
| 10 | 18388277 | rs10508557 | -0.374 | -0.507 | -0.076 | 1.726 | 0 | -0.005 | -0.005 | 0 | 0.09 | 0.07 | 0.17 | 0.07 | 0.13 | 0.09 |
| 10 | 18388434 | rs10508558 | -0.854 | -0.504 | -0.03 | 1.991 | 0.001 | 0.099 | 0.099 | 0 | 0.07 | 0.15 | 0.07 | 0.34 | 0.05 | 0.1 |
| 10 | 18391562 | rs11012691 | -0.191 | -0.565 | -0.136 | 1.963 | 0.012 | -0.059 | -0.059 | 0.383 | 0.12 | 0.03 | 0.09 | 0.25 | 0.08 | 0.08 |
| 10 | 18392980 | rs10828242 | -1.933 | -0.411 | -0.047 | 1.895 | 0.001 | -0.005 | -0.005 | 0 | 0.26 | 0.3 | 0.29 | 0.58 | 0.25 | 0.25 |
| 10 | 18397565 | rs12769993 | -0.344 | -0.692 | 0.091 | 1.696 | -0.001 | 0.008 | 0.008 | 0 | 0.09 | 0.16 | 0.18 | 0.07 | 0.12 | 0.09 |
| 10 | 18398954 | rs11599314 | -0.248 | -0.69 | 0.055 | 1.697 | -0.001 | -0.005 | -0.005 | 1.466 | 0.09 | 0.07 | 0.17 | 0.07 | 0.12 | 0.09 |
| 10 | 18399519 | rs4748442 | 0.871 | -0.716 | 0.444 | 1.621 | -0.013 | -0.031 | -0.031 | 1.215 | 0.69 | 0.6 | 0.65 | 0.82 | 0.66 | 0.69 |
| 10 | 18402179 | rs7904976 | -2.026 | -0.856 | 0.563 | 1.766 | -0.015 | -0.006 | -0.006 | 0.03 | 0.41 | 0.54 | 0.45 | 0.58 | 0.42 | 0.42 |
| 10 | 18409137 | rs10828254 | -2.338 | -0.637 | 0.556 | 1.738 | -0.012 | 0.05 | 0.05 | 0 | 0.29 | 0.58 | 0.4 | 0.42 | 0.31 | 0.28 |
| 10 | 18410168 | rs7478172 | -0.876 | -0.584 | 0.506 | 1.617 | 0 | 0.018 | 0.018 | 0.667 | 0.12 | 0.14 | 0.22 | 0.25 | 0.17 | 0.21 |
| 10 | 18414220 | rs7897163 | -1.58 | -0.573 | 0.608 | 1.216 | -0.018 | -0.03 | -0.03 | 0.018 | 0.19 | 0.47 | 0.33 | 0.07 | 0.22 | 0.16 |
| 10 | 18414466 | rs7897452 | -1.415 | -0.41 | 0.678 | 1.311 | 0.035 | 0.042 | 0.042 | 0.393 | 0.13 | 0.41 | 0.29 | 0.07 | 0.19 | 0.15 |
| 10 | 18415963 | rs10764318 | 0.194 | -0.265 | 0.577 | 1.415 | 0.062 | 0.094 | 0.094 | 0.026 | 0.07 | 0.23 | 0.17 | 0.06 | 0.12 | 0.1 |
| 10 | 18418180 | rs7897594 | -1.37 | -0.72 | 0.342 | 1.427 | -0.029 | 0.018 | 0.018 | 0 | 0.29 | 0.37 | 0.28 | 0.31 | 0.29 | 0.3 |
| 10 | 18420326 | rs11596960 | -2.13 | -0.9 | 0.218 | 1.404 | 0.008 | 0.002 | 0.002 | 0 | 0.46 | 0.61 | 0.51 | 0.5 | 0.47 | 0.44 |
| 10 | 18420384 | rs11596974 | -0.753 | -0.961 | 0.362 | 1.54 | 0.011 | -0.013 | -0.013 | 0 | 0.26 | 0.22 | 0.31 | 0.34 | 0.28 | 0.2 |
| 10 | 18425519 | rs12570727 | -2.011 | -0.924 | 0.39 | 1.46 | 0.012 | -0.041 | -0.041 | 0.695 | 0.38 | 0.21 | 0.4 | 0.5 | 0.42 | 0.41 |
| 10 | 18427430 | rs1891393 | -1.68 | -0.995 | 0.441 | 1.46 | -0.019 | 0.051 | 0.051 | 0 | 0.32 | 0.52 | 0.26 | 0.45 | 0.31 | 0.32 |
| 10 | 18428415 | rs10764319 | -1.568 | -0.996 | 0.629 | 1.452 | -0.019 | 0.053 | 0.053 | 0 | 0.32 | 0.44 | 0.25 | 0.44 | 0.31 | 0.31 |
| 10 | 18433986 | rs1571787 | -1.656 | -0.675 | 0.605 | 1.675 | -0.001 | -0.001 | -0.001 | 0.189 | 0.24 | 0.26 | 0.35 | 0.43 | 0.29 | 0.29 |
| 10 | 18434591 | rs12247369 | -0.403 | -0.741 | 0.592 | 1.574 | -0.018 | -0.018 | -0.018 | 0 | 0.19 | 0.26 | 0.18 | 0.11 | 0.18 | 0.12 |
| 10 | 18442482 | rs10740993 | -2.573 | -0.779 | 0.531 | 1.635 | 0.01 | 0.012 | 0.012 | 0 | 0.45 | 0.64 | 0.44 | 0.58 | 0.48 | 0.45 |
| 10 | 18442940 | rs17604757 | 0.129 | -0.731 | 0.449 | 1.539 | -0.001 | -0.019 | -0.019 | 0.011 | 0.08 | 0.08 | 0.04 | 0.01 | 0.07 | 0.1 |
| 10 | 18444535 | rs11012829 | -0.596 | -0.766 | 0.439 | 1.564 | 0.014 | 0.021 | 0.021 | 0 | 0.19 | 0.12 | 0.08 | 0.11 | 0.18 | 0.18 |
| 10 | 18445483 | rs11012832 | -0.621 | -0.768 | 0.554 | 1.563 | -0.006 | -0.01 | -0.01 | 0 | 0.19 | 0.19 | 0.09 | 0.11 | 0.19 | 0.18 |
| 10 | 18447295 | rs10828278 | -0.775 | -0.648 | 0.457 | 1.769 | 0 | 0.018 | 0.018 | 0 | 0.11 | 0.16 | 0.12 | 0.26 | 0.12 | 0.1 |
| 10 | 18449161 | rs2068356 | -0.512 | -0.738 | 0.452 | 1.765 | -0.001 | 0.031 | 0.031 | 0 | 0.09 | 0.12 | 0.12 | 0.25 | 0.09 | 0.09 |
| 10 | 18450043 | rs12358164 | -2.106 | -0.828 | 0.427 | 1.576 | 0.014 | -0.012 | -0.012 | 0 | 0.34 | 0.24 | 0.32 | 0.36 | 0.34 | 0.31 |
| 10 | 18451380 | rs11012843 | -0.651 | -0.881 | 0.598 | 1.588 | 0.004 | 0.072 | 0.072 | 0 | 0.16 | 0.4 | 0.24 | 0.21 | 0.17 | 0.17 |
| 10 | 18458707 | rs16916916 | 0.351 | -0.765 | 0.501 | 1.703 | 0.004 | -0.019 | -0.019 | 0 | 0.06 | 0.09 | 0.04 | 0 | 0.07 | 0.08 |
| 10 | 18459282 | rs7090118 | -1.949 | -0.782 | 0.617 | 1.738 | -0.015 | 0.033 | 0.033 | 0 | 0.34 | 0.5 | 0.35 | 0.4 | 0.37 | 0.33 |
| 10 | 18465479 | rs17662793 | 1.045 | -0.413 | 0.443 | 0.973 | -0.02 | 0.002 | 0.002 | 0 | 0.22 | 0.15 | 0.17 | 0.2 | 0.28 | 0.26 |
| 10 | 18466094 | rs10828295 | -1.926 | -0.868 | 0.879 | 1.392 | -0.017 | 0.049 | 0.049 | 0 | 0.27 | 0.49 | 0.32 | 0.3 | 0.26 | 0.33 |
| 10 | 18468124 | rs11012883 | -0.268 | -0.842 | 0.859 | 1.505 | -0.001 | 0.07 | 0.07 | 0 | 0.07 | 0.09 | 0.07 | 0.25 | 0.06 | 0.12 |
| 10 | 18468589 | rs7911644 | -2.2 | -0.912 | 0.847 | 1.483 | -0.006 | 0.018 | 0.018 | 0.2 | 0.34 | 0.6 | 0.31 | 0.42 | 0.33 | 0.4 |
| 10 | 18469522 | rs12569914 | 0.865 | -0.847 | 0.419 | 1.386 | -0.001 | -0.002 | -0.002 | 0.065 | 0.06 | 0.04 | 0.09 | 0.05 | 0.1 | 0.12 |
| 10 | 18470153 | rs16916928 | -1.141 | -0.855 | 0.507 | 1.381 | -0.005 | -0.002 | -0.002 | 0 | 0.25 | 0.29 | 0.25 | 0.19 | 0.27 | 0.26 |
| 10 | 18471961 | rs1624525 | -1.256 | -0.867 | 0.372 | 1.346 | -0.003 | 0.001 | 0.001 | 0 | 0.57 | 0.45 | 0.47 | 0.43 | 0.67 | 0.55 |
| 10 | 18474727 | rs1779228 | 1.116 | -1.305 | 0.836 | 1.918 | -0.004 | -0.012 | -0.012 | 0 | 0.81 | 0.88 | 0.81 | 0.82 | 0.77 | 0.75 |
| 10 | 18475947 | rs1779237 | 1.681 | -1.32 | 0.842 | 1.938 | -0.008 | 0.034 | 0.034 | 0.107 | 0.68 | 0.61 | 0.7 | 0.58 | 0.67 | 0.62 |
| 10 | 18479198 | rs11593318 | -0.781 | -0.866 | 0.404 | 1.907 | -0.004 | -0.018 | -0.018 | 0 | 0.12 | 0.05 | 0.1 | 0.13 | 0.13 | 0.13 |
| 10 | 18481044 | rs16916944 | -0.761 | -0.849 | 0.417 | 1.859 | -0.005 | -0.012 | -0.012 | 0 | 0.12 | 0.05 | 0.1 | 0.12 | 0.13 | 0.12 |
| 10 | 18481080 | rs2482100 | 1.126 | -0.992 | 1.415 | 2.183 | -0.002 | 0.091 | 0.091 | 0 | 0.85 | 0.73 | 0.84 | 0.73 | 0.84 | 0.76 |
| 10 | 18482691 | rs1277729 | 1.383 | -1.166 | 1.706 | 1.925 | -0.001 | 0 | 0 | 0.015 | 0.72 | 0.34 | 0.76 | 0.51 | 0.8 | 0.72 |
| 10 | 18486612 | rs1277726 | -0.062 | -1.18 | 2.066 | 2.062 | 0.028 | -0.039 | -0.039 | 0 | 0.84 | 0.92 | 0.9 | 0.69 | 0.88 | 0.78 |
| 10 | 18488465 | rs1573221 | 0.638 | -1.266 | 2.364 | 2.037 | -0.082 | 0.116 | 0.116 | 0 | 0.79 | 0.61 | 0.83 | 0.49 | 0.87 | 0.78 |
| 10 | 18497295 | rs1277765 | 1.063 | -1.284 | 2.701 | 2.018 | -0.052 | 0.076 | 0.076 | 0 | 0.74 | 0.6 | 0.81 | 0.55 | 0.83 | 0.78 |
| 10 | 18499676 | rs1277768 | 1.341 | -1.286 | 2.618 | 2.009 | -0.008 | 0.002 | 0.002 | 0 | 0.72 | 0.34 | 0.79 | 0.49 | 0.83 | 0.78 |
| 10 | 18499691 | rs16916984 | -1.569 | -1.281 | 2.698 | 1.991 | 0.065 | 0.066 | 0.066 | 2.197 | 0.08 | 0.41 | 0.07 | 0.06 | 0.04 | 0 |
| 10 | 18499794 | rs1277769 | 0.393 | -1.227 | 2.436 | 2.102 | -0.015 | 0.013 | 0.013 | 0 | 0.81 | 0.8 | 0.88 | 0.67 | 0.88 | 0.78 |
| 10 | 18504830 | rs11013044 | -1.865 | -0.996 | 1.696 | 1.842 | 0.021 | 0.147 | 0.147 | 0.593 | 0.11 | 0.24 | 0.13 | 0.3 | 0.1 | 0.08 |
| 10 | 18505937 | rs7901394 | -0.596 | -0.91 | 1.387 | 1.465 | 0.014 | -0.027 | -0.027 | 0 | 0.05 | 0.14 | 0.03 | 0 | 0.04 | 0 |

| | | | | | | | | | | | | | | | | |
|---|---|---|---|---|---|---|---|---|---|---|---|---|---|---|---|---|
| 10 | 18506017 | rs2482117 | 0.847 | -0.949 | 0.972 | 1.545 | -0.014 | -0.001 | -0.001 | 0.095 | 0.69 | 0.4 | 0.75 | 0.38 | 0.78 | 0.71 |
| 10 | 18506911 | rs1779246 | 0.343 | -0.439 | 0.328 | 1.503 | 0.022 | -0.027 | -0.027 | 0 | 0.75 | 0.9 | 0.86 | 0.74 | 0.77 | 0.76 |
| 10 | 18510996 | rs1779217 | 1.14 | -0.159 | 0.713 | 1.054 | 0.026 | 0.035 | 0.035 | 0 | 0.6 | 0.27 | 0.68 | 0.35 | 0.63 | 0.62 |
| 10 | 18511918 | rs7896607 | 0.693 | -0.161 | 0.719 | 1.128 | 0.003 | -0.001 | -0.001 | 0 | 0.65 | 0.39 | 0.73 | 0.44 | 0.69 | 0.67 |
| 10 | 18515834 | rs1613804 | -0.018 | -0.363 | 0.807 | 1.343 | -0.023 | 0.027 | 0.027 | 0 | 0.75 | 0.73 | 0.85 | 0.58 | 0.81 | 0.76 |
| 10 | 18518934 | rs11013093 | -1.827 | -0.711 | 0.179 | 1.178 | -0.001 | -0.001 | -0.001 | 0.706 | 0.18 | 0.81 | 0.21 | 0.27 | 0.15 | 0.15 |
| 10 | 18520535 | rs1277748 | 0.987 | -0.697 | 0.184 | 1.146 | 0.06 | 0.058 | 0.058 | 0 | 0.58 | 0.18 | 0.68 | 0.36 | 0.66 | 0.62 |
| 10 | 18522975 | rs1277753 | -0.331 | -0.708 | 0.012 | 1.23 | 0.066 | -0.135 | -0.135 | 0.021 | 0.75 | 0.98 | 0.88 | 0.64 | 0.81 | 0.77 |
| 10 | 18526110 | rs7098242 | 0.966 | -0.104 | -0.322 | 0.521 | -0.074 | 0.077 | 0.077 | 0 | 0.4 | 0.14 | 0.44 | 0.31 | 0.49 | 0.45 |
| 10 | 18526451 | rs1757234 | 0.832 | -0.159 | -0.041 | 0.309 | -0.006 | -0.018 | -0.018 | 0 | 0.15 | 0.19 | 0.21 | 0.06 | 0.1 | 0.09 |
| 10 | 18527039 | rs1757235 | 0.474 | 0.336 | -0.043 | 0.296 | -0.001 | 0.029 | 0.029 | 0 | 0.53 | 0.39 | 0.63 | 0.41 | 0.51 | 0.51 |
| 10 | 18527454 | rs1757236 | 0.165 | 0.211 | -0.195 | 0.187 | -0.004 | -0.022 | -0.022 | 0 | 0.21 | 0.24 | 0.23 | 0.07 | 0.14 | 0.14 |
| 10 | 18528245 | rs11013126 | 1.563 | 0.123 | -0.818 | 0.149 | -0.129 | -0.006 | -0.006 | 1.116 | 0.33 | 0.03 | 0.41 | 0.31 | 0.44 | 0.37 |
| 10 | 18529419 | rs1779229 | -0.545 | 0.124 | -0.476 | 0.215 | -0.006 | 0.026 | 0.026 | 0 | 0.67 | 0.61 | 0.78 | 0.6 | 0.68 | 0.59 |
| 10 | 18529560 | rs914363 | 0.443 | 0.14 | -0.553 | 0.144 | -0.038 | 0.07 | 0.07 | 0.479 | 0.46 | 0.18 | 0.48 | 0.4 | 0.52 | 0.45 |
| 10 | 18531619 | rs11013137 | 1.275 | 0.062 | -0.445 | 0.234 | -0.088 | 0.031 | 0.031 | 0 | 0.37 | 0.09 | 0.48 | 0.36 | 0.52 | 0.48 |
| 10 | 18532229 | rs11812617 | -0.26 | 0.754 | 0.458 | 0.313 | -0.008 | -0.001 | -0.001 | 0 | 0.07 | 0.07 | 0.06 | 0.08 | 0.08 | 0.03 |
| 10 | 18532672 | rs1331328 | -0.451 | 0.733 | 0.559 | 0.31 | -0.024 | -0.076 | -0.076 | 0.064 | 0.21 | 0.37 | 0.25 | 0.06 | 0.14 | 0.13 |
| 10 | 18533377 | rs16917091 | -0.965 | 0.745 | 0.625 | 0.496 | 0.006 | -0.019 | -0.019 | 0 | 0.15 | 0.26 | 0.11 | 0.11 | 0.12 | 0.12 |
| 10 | 18534850 | rs1757214 | -0.477 | 0.982 | 0.663 | 0.673 | 0.019 | 0.024 | 0.024 | 0 | 0.18 | 0.46 | 0.3 | 0.15 | 0.15 | 0.11 |
| 10 | 18536423 | rs1034139 | -0.957 | 1.131 | 0.476 | 0.785 | 0.021 | 0.019 | 0.019 | 0 | 0.24 | 0.6 | 0.39 | 0.26 | 0.25 | 0.25 |
| 10 | 18538659 | rs12245847 | -0.283 | 0.849 | 0.14 | 0.459 | -0.001 | 0.009 | 0.009 | 0.988 | 0.39 | 0.52 | 0.46 | 0.63 | 0.41 | 0.44 |
| 10 | 18539795 | rs993864 | 0.778 | 0.855 | -0.659 | -0.353 | -0.042 | 0.099 | 0.099 | 0.117 | 0.21 | 0.08 | 0.23 | 0.01 | 0.23 | 0.32 |
| 10 | 18542339 | rs17681709 | 0.795 | 0.797 | -0.94 | -0.661 | -0.031 | 0.036 | 0.036 | 0 | 0.1 | 0.04 | 0.09 | 0 | 0.17 | 0.05 |
| 10 | 18543846 | rs10764365 | -0.794 | 0.557 | -0.534 | -0.647 | -0.005 | 0.006 | 0.006 | 0.703 | 0.75 | 0.77 | 0.66 | 0.95 | 0.74 | 0.57 |
| 10 | 18549016 | rs7099380 | -0.472 | 0.56 | -0.741 | -0.421 | -0.02 | 0.006 | 0.006 | 1.742 | 0.64 | 0.68 | 0.47 | 0.68 | 0.56 | 0.42 |
| 10 | 18549115 | rs1106380 | 1.111 | 0.132 | -0.401 | -0.56 | 0.012 | -0.007 | -0.007 | 0 | 0.25 | 0.15 | 0.15 | 0.37 | 0.23 | 0.25 |
| 10 | 18549641 | rs2357928 | -0.405 | 0.373 | -0.237 | -1.17 | -0.03 | 0.012 | 0.012 | 0.014 | 0.62 | 0.71 | 0.48 | 0.68 | 0.54 | 0.42 |
| 10 | 18552276 | rs2489219 | -0.909 | 0.355 | -0.386 | -1.18 | -0.053 | -0.064 | -0.064 | 0 | 0.84 | 0.95 | 0.71 | 0.73 | 0.77 | 0.79 |
| 10 | 18553968 | rs10828399 | -0.525 | 0.252 | -0.312 | -1.219 | -0.024 | 0.015 | 0.015 | 0 | 0.63 | 0.69 | 0.48 | 0.72 | 0.54 | 0.49 |
| 10 | 18555510 | rs12254400 | 1.485 | -0.142 | -0.327 | -1.343 | -0.006 | -0.036 | -0.036 | 0 | 0.14 | 0.2 | 0.16 | 0.05 | 0.09 | 0.08 |
| 10 | 18558798 | rs1411930 | -0.39 | -0.577 | -0.464 | -1.53 | -0.011 | -0.066 | -0.066 | 0 | 0.34 | 0.54 | 0.32 | 0.12 | 0.28 | 0.14 |
| 10 | 18558867 | rs1411929 | -0.319 | -0.575 | -0.463 | -1.528 | -0.012 | -0.069 | -0.069 | 0 | 0.34 | 0.53 | 0.32 | 0.13 | 0.28 | 0.14 |
| 10 | 18562538 | rs1277733 | -1.18 | -0.371 | -0.448 | -1.329 | 0.004 | 0.006 | 0.006 | 0 | 0.84 | 0.83 | 0.77 | 0.75 | 0.78 | 0.8 |
| 10 | 18563985 | rs1277731 | 2.298 | -0.436 | -0.64 | -1.484 | -0.034 | 0.07 | 0.07 | 0 | 0.2 | 0.12 | 0.22 | 0.01 | 0.23 | 0.29 |
| 10 | 18564095 | rs16917138 | 1.118 | -0.453 | -0.576 | -1.497 | -0.009 | 0.019 | 0.019 | 0.558 | 0.13 | 0.08 | 0.13 | 0.08 | 0.18 | 0.05 |
| 10 | 18564922 | rs9633646 | -0.814 | -0.456 | -0.598 | -1.502 | 0.002 | -0.053 | -0.053 | 0 | 0.63 | 0.56 | 0.63 | 0.91 | 0.59 | 0.65 |
| 10 | 18565470 | rs9633647 | 0.804 | -0.492 | -1.019 | -1.328 | 0.016 | -0.054 | -0.054 | 2.76 | 0.3 | 0.14 | 0.25 | 0.57 | 0.3 | 0.29 |
| 10 | 18567397 | rs10741019 | -0.956 | -0.43 | -1.048 | -1.373 | -0.001 | 0.012 | 0.012 | 0 | 0.62 | 0.67 | 0.61 | 0.95 | 0.66 | 0.61 |
| 10 | 18571215 | rs7072759 | -0.594 | -0.342 | -1.401 | -1.199 | -0.013 | 0.035 | 0.035 | 0 | 0.5 | 0.75 | 0.62 | 0.43 | 0.47 | 0.42 |
| 10 | 18571411 | rs11013317 | -0.431 | -0.336 | -1.104 | -1.074 | -0.062 | 0.195 | 0.195 | 0 | 0.44 | 0.14 | 0.45 | 0.83 | 0.52 | 0.49 |
| 10 | 18577425 | rs7914038 | -1.13 | -0.665 | -0.456 | -0.863 | -0.034 | -0.017 | -0.017 | 0 | 0.28 | 0.53 | 0.25 | 0.26 | 0.16 | 0.25 |
| 10 | 18582514 | rs16917151 | -0.173 | -0.329 | -0.725 | -0.852 | -0.023 | 0.046 | 0.046 | 0 | 0.18 | 0.05 | 0.16 | 0.12 | 0.18 | 0.2 |
| 10 | 18585219 | rs7089228 | -0.983 | -0.234 | -0.509 | -1.034 | -0.001 | -0.007 | -0.007 | 0.229 | 0.38 | 0.36 | 0.3 | 0.23 | 0.34 | 0.32 |
| 10 | 18588285 | rs11013421 | -0.407 | -0.034 | -0.823 | -1.166 | -0.002 | 0.046 | 0.046 | 1.184 | 0.26 | 0.18 | 0.2 | 0.09 | 0.24 | 0.23 |
| 10 | 18591861 | rs4748447 | -0.023 | 0.302 | -0.623 | -0.972 | 0.007 | -0.01 | -0.01 | 0 | 0.77 | 0.73 | 0.77 | 0.2 | 0.73 | 0.61 |
| 10 | 18594604 | rs4748448 | 0.412 | 0.017 | -1.136 | -1.304 | 0.007 | 0.043 | 0.043 | 0.397 | 0.65 | 0.77 | 0.63 | 0.19 | 0.65 | 0.58 |
| 10 | 18600263 | rs1547168 | 1.465 | 0.109 | -1.001 | -1.755 | -0.014 | 0.004 | 0.004 | 0 | 0.31 | 0.25 | 0.36 | 0.04 | 0.38 | 0.32 |
| 10 | 18600905 | rs1750477 | 0.408 | -0.674 | -1.091 | -1.487 | 0.015 | 0.014 | 0.014 | 0 | 0.62 | 0.65 | 0.7 | 0.24 | 0.76 | 0.5 |
| 10 | 18602417 | rs2489214 | 0.446 | -0.223 | -1.299 | -1.144 | -0.011 | 0.023 | 0.023 | 0.122 | 0.66 | 0.87 | 0.66 | 0.29 | 0.67 | 0.68 |
| 10 | 18605942 | rs10828479 | 0.589 | 0.223 | -1.02 | -1.486 | -0.007 | -0.058 | -0.058 | 0 | 0.6 | 0.52 | 0.36 | 0.18 | 0.59 | 0.34 |
| 10 | 18611677 | rs41383652 | 1.375 | 0.209 | -1.058 | -1.677 | 0.002 | -0.044 | -0.044 | 1.692 | 0.39 | 0.43 | 0.23 | 0.03 | 0.34 | 0.23 |
| 10 | 18614995 | rs11013624 | 0.796 | 0.608 | -1.123 | -1.409 | -0.04 | 0.017 | 0.017 | 0 | 0.17 | 0.08 | 0.12 | 0.13 | 0.25 | 0.11 |
| 10 | 18620950 | rs9645525 | 0.413 | 0.403 | -1.016 | -1.033 | -0.039 | 0.05 | 0.05 | 0 | 0.59 | 0.78 | 0.51 | 0.29 | 0.51 | 0.54 |
| 10 | 18623463 | rs7093791 | -0.469 | 0.375 | -0.877 | -0.603 | -0.006 | -0.05 | -0.05 | 0 | 0.83 | 0.89 | 0.65 | 0.5 | 0.79 | 0.71 |
| 10 | 18626566 | rs11013700 | 0.571 | 0.426 | -0.287 | -0.718 | 0.034 | 0.109 | 0.109 | 0 | 0.15 | 0.34 | 0.15 | 0.27 | 0.17 | 0.26 |
| 10 | 18629603 | rs10828545 | 1.052 | 0.649 | -0.479 | -0.82 | 0.005 | 0.016 | 0.016 | 0 | 0.44 | 0.55 | 0.37 | 0.31 | 0.38 | 0.48 |
| 10 | 18630884 | rs10458703 | 0.952 | 1.192 | 0.083 | -0.415 | 0 | 0.009 | 0.009 | 0 | 0.11 | 0.1 | 0.12 | 0.2 | 0.15 | 0.19 |
| 10 | 18632601 | rs11013723 | -0.3 | 1.025 | 0.035 | -0.734 | 0.001 | -0.001 | -0.001 | 0 | 0.13 | 0.17 | 0.19 | 0.04 | 0.1 | 0.08 |
| 10 | 18632799 | rs4453118 | 1.745 | 1.188 | 0.262 | -0.943 | -0.004 | -0.091 | -0.091 | 0 | 0.36 | 0.34 | 0.19 | 0.05 | 0.29 | 0.33 |
| 10 | 18633777 | rs7919793 | 0.238 | 1.161 | -0.162 | -0.794 | -0.019 | 0.038 | 0.038 | 0.841 | 0.21 | 0.09 | 0.1 | 0.11 | 0.22 | 0.1 |
| 10 | 18634862 | rs11013738 | 1.31 | 1.065 | -0.02 | -0.801 | -0.022 | -0.033 | -0.033 | 0 | 0.06 | 0.18 | 0.04 | 0 | 0.01 | 0 |
| 10 | 18636887 | rs10764427 | 0.346 | 1.167 | -0.288 | -0.817 | -0.021 | 0.041 | 0.041 | 0.48 | 0.19 | 0.09 | 0.1 | 0.1 | 0.21 | 0.09 |
| 10 | 18639242 | rs11013768 | 2.115 | 0.996 | -0.212 | -0.996 | 0.028 | 0.031 | 0.031 | 0 | 0.34 | 0.19 | 0.15 | 0.02 | 0.21 | 0.25 |
| 10 | 18642730 | rs11594922 | -0.112 | 1.242 | -0.198 | -0.461 | -0.005 | 0.011 | 0.011 | 0.569 | 0.21 | 0.15 | 0.1 | 0.11 | 0.2 | 0.09 |
| 10 | 18643513 | rs1298103 | -0.839 | 1.272 | -0.145 | 0.027 | 0.001 | 0.057 | 0.057 | 0 | 0.9 | 0.89 | 0.93 | 0.5 | 0.93 | 0.82 |
| 10 | 18644913 | rs2799571 | -0.03 | 1.28 | -0.118 | -0.093 | -0.013 | 0.053 | 0.053 | 0 | 0.84 | 0.8 | 0.9 | 0.51 | 0.91 | 0.82 |
| 10 | 18646986 | rs4748453 | -0.477 | 1.54 | -0.239 | -0.28 | -0.036 | -0.035 | -0.035 | 0 | 0.19 | 0.15 | 0.23 | 0.24 | 0.31 | 0.24 |
| 10 | 18649635 | rs10764440 | -1.939 | 1.151 | -0.125 | -0.444 | 0.004 | 0.003 | 0.003 | 0 | 0.54 | 0.46 | 0.54 | 0.93 | 0.59 | 0.57 |

| | | | | | | | | | | | | | | | | |
|---|---|---|---|---|---|---|---|---|---|---|---|---|---|---|---|---|
| 10 | 18651860 | rs6482355 | -1.672 | 1.147 | -0.07 | -0.41 | 0.012 | 0.049 | 0.049 | 0 | 0.49 | 0.45 | 0.41 | 0.89 | 0.56 | 0.52 |
| 10 | 18652077 | rs11013849 | -0.39 | 1.102 | -0.13 | -0.271 | 0.001 | -0.014 | -0.014 | 0 | 0.07 | 0.11 | 0.09 | 0.01 | 0.14 | 0.03 |
| 10 | 18653986 | rs11013859 | -0.855 | 1.045 | -0.14 | -0.219 | 0.017 | -0.006 | -0.006 | 0 | 0.35 | 0.26 | 0.2 | 0.65 | 0.29 | 0.29 |
| 10 | 18654027 | rs11013860 | 0.823 | 1.006 | -0.166 | -0.199 | 0 | 0.048 | 0.048 | 0 | 0.1 | 0.12 | 0.08 | 0.56 | 0.08 | 0.21 |
| 10 | 18656953 | rs7901640 | -1.281 | 1.151 | -0.059 | -0.24 | 0 | 0.005 | 0.005 | 0.12 | 0.4 | 0.4 | 0.38 | 0.67 | 0.41 | 0.35 |
| 10 | 18656963 | rs7920123 | -0.902 | 1.08 | -0.1 | -0.22 | 0.011 | -0.002 | -0.002 | 0 | 0.36 | 0.29 | 0.28 | 0.65 | 0.28 | 0.29 |
| 10 | 18662873 | rs1325992 | 0.117 | 1.124 | -0.125 | -0.22 | -0.015 | -0.026 | -0.026 | 1.463 | 0.14 | 0.11 | 0.18 | 0.23 | 0.21 | 0.21 |
| 10 | 18670017 | rs10828616 | -0.705 | 1.111 | -0.131 | -0.29 | 0.007 | -0.02 | -0.02 | 0 | 0.3 | 0.22 | 0.24 | 0.64 | 0.25 | 0.28 |
| 10 | 18671971 | rs4747344 | -1.188 | 1.151 | -0.004 | -0.34 | -0.001 | -0.001 | -0.001 | 0.534 | 0.25 | 0.3 | 0.31 | 0.27 | 0.36 | 0.28 |
| 10 | 18672145 | rs4748457 | -0.318 | 1.397 | 0.071 | -0.827 | 0.03 | -0.037 | -0.037 | 0 | 0.07 | 0.17 | 0.09 | 0 | 0.14 | 0.03 |
| 10 | 18673100 | rs17611696 | 0.647 | 1.325 | -0.068 | -0.773 | -0.014 | 0.026 | 0.026 | 0.088 | 0.14 | 0.06 | 0.08 | 0.09 | 0.16 | 0.07 |
| 10 | 18674224 | rs4748458 | -1.066 | 1.282 | 0.107 | -0.716 | 0.002 | 0.003 | 0.003 | 0 | 0.48 | 0.48 | 0.5 | 0.92 | 0.56 | 0.55 |
| 10 | 18675148 | rs7078741 | -1.166 | 1.22 | 0.21 | -0.729 | 0.003 | -0.039 | -0.039 | 0 | 0.49 | 0.52 | 0.52 | 0.93 | 0.58 | 0.56 |
| 10 | 18680935 | rs10828623 | 0.761 | 1.282 | 1.058 | -0.215 | 0.104 | 0.007 | 0.007 | 1.069 | 0.91 | 0.62 | 0.72 | 0.98 | 0.85 | 0.91 |
| 10 | 18682539 | rs10828625 | 0.503 | 0.945 | 0.819 | 0.084 | -0.005 | -0.002 | -0.002 | 0 | 0.72 | 0.73 | 0.56 | 0.72 | 0.65 | 0.71 |
| 10 | 18683267 | rs7069531 | 1.168 | 1.17 | 1.032 | -0.026 | 0.007 | 0.059 | 0.059 | 0 | 0.48 | 0.37 | 0.29 | 0.13 | 0.44 | 0.34 |
| 10 | 18685765 | rs1570940 | 0.472 | 1.136 | 1.106 | 0.311 | -0.001 | 0.013 | 0.013 | 0.078 | 0.21 | 0.17 | 0.22 | 0.51 | 0.2 | 0.2 |
| 10 | 18685891 | rs1570939 | 0.531 | 1.096 | 1.098 | 0.28 | 0.015 | 0.019 | 0.019 | 0 | 0.71 | 0.56 | 0.51 | 0.69 | 0.64 | 0.58 |
| 10 | 18686452 | rs11014049 | 1.622 | 1.178 | 0.776 | 0.101 | 0.026 | 0.098 | 0.098 | 0 | 0.42 | 0.23 | 0.22 | 0.12 | 0.32 | 0.31 |
| 10 | 18691153 | rs3750591 | -0.681 | 0.876 | 0.651 | 0.519 | 0.002 | 0.021 | 0.021 | 1.283 | 0.3 | 0.62 | 0.33 | 0.48 | 0.32 | 0.25 |
| 10 | 18695892 | rs7909027 | -0.889 | 0.902 | 0.522 | 0.305 | -0.007 | -0.034 | -0.034 | 0.029 | 0.33 | 0.75 | 0.35 | 0.55 | 0.34 | 0.28 |
| 10 | 18697657 | rs11819196 | -0.06 | 0.925 | 0.814 | 0.302 | -0.002 | 0.012 | 0.012 | 0 | 0.24 | 0.24 | 0.26 | 0.6 | 0.24 | 0.37 |
| 10 | 18700219 | rs12267833 | -0.681 | 1.019 | 0.978 | 0.288 | 0.123 | -0.012 | -0.012 | 0.246 | 0.08 | 0.35 | 0.09 | 0 | 0.11 | 0.03 |
| 10 | 18703097 | rs10828662 | 0.982 | 1.068 | 0.525 | 0.249 | 0.018 | 0.015 | 0.015 | 0 | 0.64 | 0.5 | 0.47 | 0.62 | 0.54 | 0.57 |
| 10 | 18704776 | rs11014144 | 0.397 | 1.345 | 0.102 | 0.204 | -0.041 | -0.101 | -0.101 | 0.06 | 0.18 | 0.01 | 0.15 | 0.28 | 0.19 | 0.2 |
| 10 | 18705120 | rs11593522 | -0.783 | 1.295 | 0.269 | -0.199 | 0.004 | 0.05 | 0.05 | 0 | 0.43 | 0.29 | 0.41 | 0.77 | 0.43 | 0.46 |
| 10 | 18707296 | rs982003 | 0.7 | 1.434 | 0.316 | -0.14 | 0.162 | -0.022 | -0.022 | 0.025 | 0.88 | 0.54 | 0.66 | 0.94 | 0.77 | 0.91 |
| 10 | 18707995 | rs6482406 | -1.214 | 1.268 | -0.767 | -0.011 | -0.009 | 0 | 0 | 0 | 0.95 | 0.96 | 0.76 | 0.94 | 0.88 | 0.92 |
| 10 | 18711195 | rs11014171 | 1.317 | 1.189 | -0.809 | -0.067 | 0.018 | 0.17 | 0.17 | 0 | 0.43 | 0.15 | 0.24 | 0.17 | 0.34 | 0.42 |
| 10 | 18711288 | rs10828679 | -0.779 | 1.201 | -0.818 | -0.065 | -0.001 | 0 | 0 | 0 | 0.94 | 0.89 | 0.76 | 0.94 | 0.88 | 0.92 |
| 10 | 18712250 | rs4748463 | -1.236 | 1.205 | -0.733 | -0.047 | 0.006 | 0.107 | 0.107 | 0 | 0.52 | 0.66 | 0.5 | 0.75 | 0.54 | 0.52 |
| 10 | 18712285 | rs11014175 | 0.166 | 1.306 | -0.969 | -0.028 | -0.044 | -0.054 | -0.054 | 0.086 | 0.2 | 0.03 | 0.17 | 0.22 | 0.21 | 0.19 |
| 10 | 18712352 | rs4747345 | -0.759 | 1.017 | -0.647 | -0.166 | 0.009 | 0.152 | 0.152 | 0 | 0.51 | 0.7 | 0.51 | 0.77 | 0.54 | 0.5 |
| 10 | 18716145 | rs10741058 | -0.448 | 1.254 | -0.379 | 0.374 | -0.007 | 0.002 | 0.002 | 0 | 0.95 | 0.95 | 0.76 | 0.96 | 0.88 | 0.93 |
| 10 | 18733293 | rs4748466 | -0.184 | 1.129 | -0.382 | 0.43 | -0.004 | -0.008 | -0.008 | 0 | 0.07 | 0.03 | 0.22 | 0.13 | 0.12 | 0.24 |
| 10 | 18734235 | rs10508561 | -0.224 | 1.434 | -0.486 | 0.525 | -0.023 | -0.023 | -0.023 | 0.998 | 0.18 | 0.1 | 0.19 | 0.23 | 0.25 | 0.19 |
| 10 | 18735355 | rs10828705 | 0.161 | 1.002 | 0.033 | 0.35 | 0 | 0.042 | 0.042 | 0.977 | 0.71 | 0.4 | 0.68 | 0.49 | 0.72 | 0.74 |
| 10 | 18736474 | rs11014262 | 0.274 | 0.941 | 0.066 | 0.324 | -0.002 | 0.018 | 0.018 | 1.355 | 0.7 | 0.35 | 0.68 | 0.49 | 0.72 | 0.74 |
| 10 | 18741794 | rs729245 | 0.05 | 0.965 | 0.468 | 0.339 | 0.007 | 0.087 | 0.087 | 0.619 | 0.72 | 0.49 | 0.68 | 0.48 | 0.72 | 0.74 |
| 10 | 18755609 | rs12219393 | -0.341 | 0.659 | 0.232 | 0.485 | -0.003 | 0.052 | 0.052 | 0 | 0.13 | 0.22 | 0.13 | 0.5 | 0.1 | 0.17 |
| 10 | 18756881 | rs10828749 | 0.618 | 0.728 | 0.198 | 0.073 | -0.008 | -0.004 | -0.004 | 0 | 0.58 | 0.35 | 0.38 | 0.51 | 0.43 | 0.43 |
| 10 | 18760635 | rs11598027 | 0.749 | 1.446 | 0.3 | 0.123 | 0.014 | 0 | 0 | 0 | 0.06 | 0.07 | 0.2 | 0.09 | 0.13 | 0.12 |
| 10 | 18762948 | rs7895467 | -0.471 | 1.105 | 0.547 | -0.168 | 0.018 | -0.023 | -0.023 | 0.972 | 0.28 | 0.29 | 0.23 | 0.24 | 0.32 | 0.18 |
| 10 | 18763854 | rs4418698 | 0.845 | 0.718 | 0.159 | -0.309 | 0.003 | 0.052 | 0.052 | 0.054 | 0.54 | 0.29 | 0.41 | 0.58 | 0.45 | 0.66 |
| 10 | 18776197 | rs4748472 | -1.236 | 0.506 | -0.765 | -0.871 | -0.062 | 0.144 | 0.144 | 0 | 0.68 | 0.92 | 0.66 | 0.89 | 0.68 | 0.77 |
| 10 | 18782880 | rs8181477 | -0.246 | 0.552 | -0.082 | -0.45 | -0.001 | 0.027 | 0.027 | 0 | 0.46 | 0.39 | 0.38 | 0.68 | 0.47 | 0.48 |
| 10 | 18789075 | rs12416030 | 0.242 | 1.096 | 0.187 | 0.109 | 0.033 | 0.042 | 0.042 | 0.226 | 0.12 | 0.23 | 0.37 | 0.24 | 0.19 | 0.34 |
| 10 | 18789498 | rs10764540 | 0.198 | 1.418 | 0.123 | -0.317 | 0.029 | -0.069 | -0.069 | 0.492 | 0.22 | 0.3 | 0.21 | 0.05 | 0.28 | 0.11 |
| 10 | 18790858 | rs10741083 | -0.288 | 2.015 | 0.002 | 0.321 | 0.08 | 0.091 | 0.091 | 0 | 0.25 | 0.4 | 0.48 | 0.53 | 0.37 | 0.3 |
| 10 | 18794073 | rs10828803 | 0.786 | 0.975 | -0.459 | -0.047 | 0.006 | -0.008 | -0.008 | 0.84 | 0.24 | 0.2 | 0.09 | 0.35 | 0.22 | 0.22 |
| 10 | 18804094 | rs7088091 | 0.351 | 1.282 | -0.339 | -0.065 | -0.004 | 0.003 | 0.003 | 0 | 0.36 | 0.29 | 0.3 | 0.71 | 0.37 | 0.38 |
| 10 | 18804689 | rs12784387 | 1.138 | 1.326 | -0.325 | -0.203 | 0.031 | 0.071 | 0.071 | 0.021 | 0.12 | 0.25 | 0.3 | 0.24 | 0.21 | 0.28 |
| 10 | 18807057 | rs7910506 | -1.473 | 0.957 | -0.598 | -0.717 | 0.021 | 0.047 | 0.047 | 0.087 | 0.69 | 0.84 | 0.77 | 0.99 | 0.75 | 0.83 |
| 10 | 18819782 | rs17621077 | 1.407 | 0.957 | -0.731 | -0.697 | -0.038 | 0.099 | 0.099 | 0.129 | 0.19 | 0.03 | 0.11 | 0 | 0.2 | 0.09 |
| 10 | 18821074 | rs12779641 | 1.967 | 1.081 | -0.529 | -0.419 | 0.017 | 0.033 | 0.033 | 0 | 0.1 | 0.16 | 0.25 | 0.18 | 0.15 | 0.13 |
| 10 | 18825694 | rs7082235 | -1.277 | 0.892 | -0.529 | -0.746 | 0.026 | 0.051 | 0.051 | 0 | 0.67 | 0.82 | 0.76 | 0.98 | 0.75 | 0.82 |
| 10 | 18827828 | rs6482515 | 0.63 | 0.956 | -0.294 | -0.669 | 0.006 | -0.001 | -0.001 | 0.132 | 0.29 | 0.33 | 0.42 | 0.24 | 0.32 | 0.38 |
| 10 | 18828371 | rs2228645 | 1.875 | 1.007 | -0.276 | -0.636 | 0.019 | 0.065 | 0.065 | 0 | 0.11 | 0.19 | 0.33 | 0.24 | 0.16 | 0.31 |
| 10 | 18835354 | rs10828879 | -1.104 | 0.874 | -0.314 | -0.619 | -0.013 | -0.077 | -0.077 | 0 | 0.66 | 0.62 | 0.76 | 0.96 | 0.76 | 0.78 |
| 10 | 18837285 | rs3817494 | 0.658 | 0.853 | -0.322 | -0.509 | -0.012 | 0.005 | 0.005 | 0 | 0.35 | 0.27 | 0.32 | 0.72 | 0.43 | 0.38 |
| 10 | 18841795 | rs10828906 | -1.156 | 0.768 | -0.304 | -0.507 | 0.018 | 0.029 | 0.029 | 0 | 0.67 | 0.79 | 0.76 | 0.96 | 0.75 | 0.77 |
| 10 | 18872699 | rs7081271 | -2.451 | 0.6 | -0.913 | -0.675 | -0.026 | 0.094 | 0.094 | 0 | 0.85 | 0.99 | 0.89 | 0.99 | 0.87 | 0.87 |
| 10 | 18886261 | rs10828975 | 0.409 | 1.097 | -0.588 | -0.429 | -0.004 | 0.004 | 0.004 | 0 | 0.34 | 0.26 | 0.3 | 0.72 | 0.37 | 0.38 |
| 10 | 18892382 | rs7077786 | -1.3 | 1.2 | -0.688 | -0.425 | 0.019 | 0.015 | 0.015 | 0 | 0.69 | 0.8 | 0.8 | 0.98 | 0.78 | 0.83 |
| 10 | 18896035 | rs7081972 | 0.532 | 1.488 | -0.939 | 0.058 | -0.05 | -0.058 | -0.058 | 0 | 0.15 | 0.04 | 0.16 | 0.35 | 0.24 | 0.17 |
| 10 | 18922659 | rs4468242 | 1.164 | 1.318 | -0.575 | 0.02 | 0.02 | 0.066 | 0.066 | 0.098 | 0.13 | 0.24 | 0.35 | 0.24 | 0.16 | 0.32 |
| 10 | 18948072 | rs11015435 | 1.142 | 1.325 | -0.579 | 0.049 | 0.02 | 0.068 | 0.068 | 0.108 | 0.13 | 0.24 | 0.34 | 0.24 | 0.17 | 0.32 |
| 10 | 18950555 | rs10829156 | -1.348 | 1.1 | -0.736 | -0.567 | 0.005 | -0.038 | -0.038 | 1.927 | 0.69 | 0.71 | 0.79 | 0.98 | 0.8 | 0.83 |
| 10 | 18951085 | rs17622920 | 1.172 | 1.14 | -1.139 | -0.435 | 0.001 | 0.038 | 0.038 | 0.063 | 0.22 | 0.13 | 0.13 | 0.02 | 0.21 | 0.13 |

| | | | | | | | | | | | | | | | | |
|---|---|---|---|---|---|---|---|---|---|---|---|---|---|---|---|---|
| 10 | 18962541 | rs10829186 | 0.797 | 1.396 | -1.27 | -0.023 | -0.065 | -0.07 | -0.07 | 0 | 0.11 | 0.02 | 0.14 | 0.31 | 0.19 | 0.15 |
| 10 | 18964840 | rs6482597 | 0.016 | 1.102 | -1.034 | -0.26 | -0.006 | 0.003 | 0.003 | 0 | 0.34 | 0.25 | 0.3 | 0.73 | 0.37 | 0.38 |
| 10 | 18972378 | rs7896918 | -0.503 | 0.945 | -0.807 | -0.105 | -0.002 | -0.004 | -0.004 | 0 | 0.56 | 0.45 | 0.44 | 0.75 | 0.64 | 0.51 |
| 10 | 18974986 | rs12359581 | 0.402 | 0.845 | -1.021 | 0.042 | 0.019 | -0.001 | -0.001 | 6.888 | 0.39 | 0.23 | 0.32 | 0.45 | 0.38 | 0.29 |
| 10 | 18975192 | rs10159584 | -0.46 | 0.88 | -0.384 | 0.117 | -0.002 | 0.018 | 0.018 | 1.674 | 0.43 | 0.64 | 0.62 | 0.7 | 0.52 | 0.57 |
| 10 | 18977879 | rs7067650 | -1.236 | 0.52 | 0.39 | 0.48 | 0.063 | 0.253 | 0.253 | 0.131 | 0.08 | 0.46 | 0.1 | 0.18 | 0.07 | 0.19 |
| 10 | 18979469 | rs11015633 | 0.749 | 0.522 | -0.246 | 0.083 | -0.005 | 0.01 | 0.01 | 0 | 0.55 | 0.22 | 0.46 | 0.38 | 0.56 | 0.56 |
| 10 | 18982244 | rs4293025 | -0.573 | 0.242 | -0.296 | -0.314 | 0.011 | 0.068 | 0.068 | 0 | 0.74 | 0.84 | 0.74 | 0.96 | 0.79 | 0.82 |
| 10 | 18986052 | rs10829247 | -0.066 | 0.448 | 0.166 | -0.028 | 0.046 | -0.028 | -0.028 | 0 | 0.08 | 0.31 | 0.14 | 0.03 | 0.09 | 0.03 |
| 10 | 18987158 | rs7092524 | 0.197 | 0.57 | -0.441 | 0.236 | 0.02 | -0.01 | -0.01 | 0.044 | 0.57 | 0.24 | 0.48 | 0.4 | 0.62 | 0.6 |
| 10 | 18988807 | rs984292 | 0.346 | 0.553 | -0.368 | 0.241 | -0.044 | 0.015 | 0.015 | 0 | 0.41 | 0.17 | 0.45 | 0.63 | 0.5 | 0.44 |
| 10 | 18990315 | rs1542727 | -0.954 | -0.11 | -0.073 | 0.406 | -0.002 | 0 | 0 | 0 | 0.76 | 0.75 | 0.83 | 0.74 | 0.83 | 0.8 |
| 10 | 18990816 | rs12266596 | 0.744 | 0.467 | 0.155 | 0.704 | -0.001 | -0.005 | -0.005 | 0 | 0.18 | 0.24 | 0.25 | 0.1 | 0.17 | 0.2 |
| 10 | 18991696 | rs7086190 | -0.383 | -0.127 | 0.074 | 0.749 | -0.024 | -0.021 | -0.021 | 0 | 0.63 | 0.58 | 0.79 | 0.73 | 0.77 | 0.65 |
| 10 | 18993303 | rs10764694 | -0.488 | -0.031 | -0.707 | 0.714 | 0.015 | 0.014 | 0.014 | 0 | 0.64 | 0.72 | 0.8 | 0.74 | 0.77 | 0.64 |
| 10 | 18996761 | rs12573176 | -0.055 | 0.352 | -0.969 | 1.016 | 0.025 | -0.031 | -0.031 | 0 | 0.23 | 0.1 | 0.25 | 0.37 | 0.18 | 0.16 |
| 10 | 19002550 | rs7098949 | -0.662 | 1.696 | -0.161 | 1.165 | 0.101 | 0.091 | 0.091 | 3.499 | 0.23 | 0.4 | 0.38 | 0.4 | 0.36 | 0.22 |
| 10 | 19002710 | rs16917693 | -0.011 | 2.342 | 0.138 | 1.08 | 0.034 | -0.036 | -0.036 | 0.106 | 0.07 | 0.14 | 0.08 | 0 | 0.1 | 0.03 |
| 10 | 19002887 | rs11015839 | -0.238 | 2.279 | 0.294 | 1.07 | 0.034 | 0.066 | 0.066 | 0 | 0.16 | 0.19 | 0.3 | 0.4 | 0.26 | 0.19 |
| 10 | 19007722 | rs7087861 | -0.802 | 1.586 | -0.411 | 0.507 | 0.028 | 0.09 | 0.09 | 0 | 0.72 | 0.96 | 0.76 | 0.93 | 0.79 | 0.67 |
| 10 | 19007922 | rs10764712 | 0.357 | 1.584 | 0.151 | 0.474 | -0.001 | -0.018 | -0.018 | 0 | 0.55 | 0.41 | 0.52 | 0.76 | 0.56 | 0.47 |
| 10 | 19009698 | rs7097370 | -1.197 | 1.945 | 0.366 | 0.863 | 0.077 | 0.013 | 0.013 | 1.552 | 0.28 | 0.58 | 0.39 | 0.37 | 0.44 | 0.34 |
| 10 | 19012514 | rs7098497 | -0.72 | 1.942 | 0.382 | 0.858 | 0.1 | -0.028 | -0.028 | 0 | 0.17 | 0.54 | 0.24 | 0.16 | 0.23 | 0.21 |
| 10 | 19013062 | rs12260987 | -0.664 | 1.941 | 0.378 | 0.858 | 0.101 | -0.026 | -0.026 | 0 | 0.17 | 0.53 | 0.24 | 0.16 | 0.23 | 0.21 |
| 10 | 19013396 | rs11015915 | 0.491 | 2.078 | 0.709 | 0.849 | 0.002 | 0.016 | 0.016 | 0 | 0.08 | 0.17 | 0.12 | 0.16 | 0.07 | 0.17 |
| 10 | 19013738 | rs7894279 | 0.924 | 1.974 | 0.581 | 0.841 | 0.024 | 0.003 | 0.003 | 1.933 | 0.44 | 0.31 | 0.36 | 0.56 | 0.35 | 0.34 |
| 10 | 19025197 | rs11593491 | -0.66 | 2 | 0.572 | 0.855 | 0.082 | -0.036 | -0.036 | 0.03 | 0.17 | 0.44 | 0.24 | 0.1 | 0.23 | 0.2 |
| 10 | 19026307 | rs16917736 | -0.017 | 1.935 | 0.599 | 0.91 | 0.031 | 0.179 | 0.179 | 0.09 | 0.06 | 0.22 | 0.18 | 0.22 | 0.05 | 0.1 |
| 10 | 19026841 | rs7096907 | -1.227 | 1.921 | 0.561 | 0.885 | 0.009 | 0.124 | 0.124 | 0.017 | 0.73 | 0.98 | 0.78 | 0.95 | 0.79 | 0.73 |
| 10 | 19029312 | rs11006713 | 1.027 | 1.893 | 0.599 | 0.878 | -0.005 | -0.006 | -0.006 | 0 | 0.11 | 0.06 | 0.15 | 0.21 | 0.21 | 0.14 |
| 10 | 19032354 | rs11006727 | 0.043 | 1.902 | 0.61 | 0.972 | 0.062 | 0.089 | 0.089 | 0 | 0.17 | 0.28 | 0.28 | 0.35 | 0.26 | 0.2 |
| 10 | 19033920 | rs7899514 | -1.22 | 1.878 | 0.558 | 0.938 | 0.01 | 0.075 | 0.075 | 0.024 | 0.73 | 0.98 | 0.75 | 0.9 | 0.79 | 0.72 |
| 10 | 19034276 | rs7901044 | -0.691 | 1.995 | 0.685 | 0.947 | 0.08 | -0.023 | -0.023 | 0 | 0.17 | 0.44 | 0.23 | 0.16 | 0.23 | 0.21 |
| 10 | 19034951 | rs10763625 | 0.938 | 1.95 | 0.611 | 0.992 | 0.006 | 0.004 | 0.004 | 0 | 0.11 | 0.09 | 0.15 | 0.21 | 0.21 | 0.14 |
| 10 | 19052600 | rs10826382 | -0.324 | 1.937 | 0.644 | 0.996 | 0.064 | 0.142 | 0.142 | 0 | 0.18 | 0.31 | 0.33 | 0.42 | 0.26 | 0.23 |
| 10 | 19054576 | rs2495839 | -0.306 | 1.946 | 0.648 | 0.977 | 0.065 | 0.141 | 0.141 | 0 | 0.18 | 0.31 | 0.33 | 0.42 | 0.26 | 0.23 |
| 10 | 19061889 | rs16917797 | -0.197 | 1.876 | 0.659 | 0.984 | 0.03 | 0.181 | 0.181 | 0.157 | 0.06 | 0.22 | 0.18 | 0.22 | 0.05 | 0.09 |
| 10 | 19061898 | rs17703879 | 1.132 | 1.962 | 0.679 | 0.91 | 0.01 | -0.014 | -0.014 | 2.429 | 0.06 | 0.04 | 0.07 | 0 | 0.1 | 0.03 |
| 10 | 19073973 | rs2495833 | -0.234 | 1.966 | 0.823 | 0.97 | 0.074 | 0.145 | 0.145 | 0 | 0.17 | 0.33 | 0.34 | 0.42 | 0.26 | 0.24 |
| 10 | 19074461 | rs2495832 | 0.849 | 1.979 | 0.818 | 0.99 | 0.009 | 0.004 | 0.004 | 0 | 0.12 | 0.08 | 0.16 | 0.2 | 0.21 | 0.14 |
| 10 | 19074958 | rs16917823 | 0.293 | 1.908 | 0.791 | 0.963 | 0.018 | 0.141 | 0.141 | 0 | 0.05 | 0.16 | 0.18 | 0.22 | 0.05 | 0.1 |
| 10 | 19078873 | rs10826440 | 1.407 | 1.776 | 0.69 | 0.789 | 0.032 | -0.001 | -0.001 | 0 | 0.38 | 0.25 | 0.2 | 0.37 | 0.28 | 0.26 |
| 10 | 19081595 | rs2451518 | -0.217 | 1.783 | 0.509 | 0.725 | 0.057 | 0.037 | 0.037 | 0 | 0.2 | 0.35 | 0.28 | 0.34 | 0.29 | 0.29 |
| 10 | 19086041 | rs2495826 | -0.363 | 1.771 | 0.386 | 0.721 | 0.027 | 0.05 | 0.05 | 0.983 | 0.64 | 0.73 | 0.65 | 0.8 | 0.71 | 0.53 |
| 10 | 19090812 | rs913655 | -0.477 | 1.788 | 0.415 | 0.73 | 0.062 | 0.038 | 0.038 | 0 | 0.22 | 0.46 | 0.28 | 0.35 | 0.29 | 0.29 |
| 10 | 19092008 | rs10763663 | -0.88 | 1.817 | 0.437 | 0.761 | 0.062 | 0.094 | 0.094 | 4.504 | 0.14 | 0.34 | 0.26 | 0.23 | 0.15 | 0.13 |
| 10 | 19095467 | rs2480317 | -1.138 | 1.806 | 0.359 | 0.745 | 0 | 0.112 | 0.112 | 0.05 | 0.73 | 0.97 | 0.77 | 0.95 | 0.78 | 0.7 |
| 10 | 19096597 | rs7094458 | -0.582 | 1.783 | 0.419 | 0.785 | 0.056 | 0.084 | 0.084 | 4.295 | 0.12 | 0.27 | 0.25 | 0.23 | 0.15 | 0.13 |
| 10 | 19103954 | rs1116767 | 1.36 | 1.47 | 0.043 | 0.543 | -0.017 | 0.05 | 0.05 | 0 | 0.18 | 0.02 | 0.21 | 0.05 | 0.18 | 0.15 |
| 10 | 19118287 | rs11007193 | 0.161 | 1.452 | 0.16 | 0.597 | -0.005 | -0.001 | -0.001 | 0 | 0.62 | 0.65 | 0.51 | 0.57 | 0.53 | 0.52 |
| 10 | 19120169 | rs7073789 | 0.207 | 1.498 | 0.183 | 0.548 | 0.005 | -0.003 | -0.003 | 0 | 0.62 | 0.59 | 0.49 | 0.6 | 0.47 | 0.54 |
| 10 | 19122016 | rs7083302 | 0.078 | 1.492 | 0.177 | 0.548 | -0.006 | 0.002 | 0.002 | 0.276 | 0.63 | 0.67 | 0.52 | 0.61 | 0.53 | 0.55 |
| 10 | 19123809 | rs12775126 | 0.974 | 1.213 | -0.078 | 0.627 | 0.114 | 0.041 | 0.041 | 4.256 | 0.47 | 0.16 | 0.22 | 0.37 | 0.32 | 0.41 |
| 10 | 19133713 | rs1329245 | 0.134 | 1.409 | 0.073 | 0.813 | 0.025 | 0.186 | 0.186 | 0.31 | 0.06 | 0.21 | 0.18 | 0.24 | 0.05 | 0.1 |
| 10 | 19160029 | rs16917919 | 0.923 | 1.362 | -0.001 | 0.659 | 0.036 | 0.009 | 0.009 | 2.14 | 0.43 | 0.31 | 0.24 | 0.38 | 0.34 | 0.29 |
| 10 | 19160986 | rs4748537 | 0.029 | 1.423 | -0.281 | 0.67 | -0.027 | 0.055 | 0.055 | 0 | 0.61 | 0.85 | 0.62 | 0.75 | 0.57 | 0.6 |
| 10 | 19190079 | rs9417699 | 0.836 | 1.165 | -1.129 | 0.41 | 0.011 | 0.083 | 0.083 | 0.162 | 0.22 | 0.02 | 0.2 | 0.05 | 0.18 | 0.25 |
| 10 | 19192853 | rs3847372 | 0.002 | 1.194 | -1.103 | 0.431 | -0.009 | 0.038 | 0.038 | 0 | 0.62 | 0.79 | 0.62 | 0.76 | 0.6 | 0.57 |
| 10 | 19197265 | rs1339061 | 0.479 | 0.831 | -1.289 | 0.276 | 0.035 | 0.106 | 0.106 | 0.655 | 0.25 | 0.02 | 0.21 | 0.05 | 0.18 | 0.3 |
| 10 | 19198427 | rs3737265 | -0.787 | 1.357 | -0.417 | 0.913 | 0.059 | 0.075 | 0.075 | 2.239 | 0.22 | 0.62 | 0.39 | 0.38 | 0.28 | 0.28 |
| 10 | 19204242 | rs11007642 | 0.203 | 0.254 | -1.207 | 0.406 | 0.021 | 0.064 | 0.064 | 0 | 0.28 | 0.1 | 0.15 | 0.12 | 0.26 | 0.17 |
| 10 | 19212287 | rs9888029 | -0.148 | -0.56 | -1.192 | 0.117 | 0.01 | 0.016 | 0.016 | 0 | 0.31 | 0.31 | 0.32 | 0.16 | 0.29 | 0.33 |
| 10 | 19228730 | rs10763749 | -0.563 | -1.247 | -0.673 | 0.172 | 0.024 | 0.012 | 0.012 | 0 | 0.47 | 0.6 | 0.41 | 0.35 | 0.4 | 0.52 |
| 10 | 19248555 | rs1890507 | -1.386 | -0.694 | -1.101 | 0.503 | -0.079 | 0.005 | 0.005 | 0.622 | 0.72 | 0.92 | 0.7 | 0.76 | 0.66 | 0.76 |
| 10 | 19249300 | rs11007843 | 0.354 | -0.933 | -0.839 | 0.492 | -0.001 | -0.002 | -0.002 | 0 | 0.43 | 0.42 | 0.34 | 0.24 | 0.38 | 0.46 |
| 10 | 19257664 | rs10740814 | -1.084 | -0.861 | -0.715 | 0.567 | -0.031 | -0.018 | -0.018 | 0.146 | 0.68 | 0.81 | 0.65 | 0.65 | 0.62 | 0.7 |
| 10 | 19272870 | rs2813777 | -0.325 | -0.916 | -0.554 | 0.538 | -0.006 | -0.002 | -0.002 | 0.223 | 0.54 | 0.46 | 0.52 | 0.65 | 0.59 | 0.47 |
| 10 | 19274291 | rs1934053 | -1.754 | -0.631 | -0.955 | 0.524 | -0.083 | 0.043 | 0.043 | 1.373 | 0.78 | 0.97 | 0.77 | 0.9 | 0.73 | 0.8 |
| 10 | 19291847 | rs1339057 | 0.511 | -0.99 | -0.547 | 0.79 | 0.03 | 0.038 | 0.038 | 0.014 | 0.43 | 0.34 | 0.33 | 0.24 | 0.38 | 0.46 |

| | | | | | | | | | | | | | | | | |
|---|---|---|---|---|---|---|---|---|---|---|---|---|---|---|---|---|
| 10 | 19297989 | rs4284312 | 0.251 | -1.002 | -0.459 | 0.736 | -0.014 | 0.083 | 0.083 | 0.669 | 0.08 | 0.25 | 0.12 | 0.15 | 0.02 | 0.1 |
| 10 | 19302567 | rs11008081 | 0.944 | -1.033 | -0.803 | 0.47 | 0.016 | 0.083 | 0.083 | 0 | 0.28 | 0.17 | 0.28 | 0.06 | 0.29 | 0.29 |
| 10 | 19307536 | rs12356231 | 1.856 | -1.009 | -0.916 | 0.418 | -0.057 | 0.069 | 0.069 | 2.621 | 0.12 | 0.02 | 0.1 | 0 | 0.17 | 0.03 |
| 10 | 19315274 | rs4145924 | 0.987 | -1.19 | -0.758 | 0.585 | 0.014 | 0.087 | 0.087 | 0 | 0.27 | 0.15 | 0.27 | 0.05 | 0.28 | 0.27 |
| 10 | 19320865 | rs10740828 | -0.005 | -1.263 | -0.347 | 0.925 | 0.013 | -0.003 | -0.003 | 0 | 0.47 | 0.57 | 0.38 | 0.48 | 0.4 | 0.51 |
| 10 | 19321154 | rs7079204 | 0.403 | -1.173 | -0.721 | 0.886 | 0.014 | 0.005 | 0.005 | 0.09 | 0.43 | 0.36 | 0.33 | 0.37 | 0.37 | 0.45 |
| 10 | 19327404 | rs2813783 | 0.647 | -1.034 | -0.964 | 0.835 | 0.015 | 0.066 | 0.066 | 0 | 0.32 | 0.19 | 0.3 | 0.18 | 0.31 | 0.31 |
| 10 | 19328435 | rs2789310 | 1.509 | -1.372 | -1.145 | 0.714 | 0.001 | 0.002 | 0.002 | 0 | 0.09 | 0.07 | 0.07 | 0 | 0.1 | 0.02 |
| 10 | 19328816 | rs12781326 | 2.051 | -1.367 | -1.158 | 0.961 | -0.019 | 0.029 | 0.029 | 0 | 0.15 | 0.18 | 0.08 | 0.27 | 0.1 | 0.23 |
| 10 | 19330039 | rs7073111 | 1.035 | -1.005 | -1.619 | 0.799 | 0.023 | 0.01 | 0.01 | 0 | 0.23 | 0.04 | 0.2 | 0.18 | 0.2 | 0.29 |
| 10 | 19333762 | rs2050739 | -1.371 | -0.284 | -1.451 | 0.993 | -0.084 | -0.02 | -0.02 | 0 | 0.75 | 0.94 | 0.7 | 0.72 | 0.68 | 0.8 |
| 10 | 19334637 | rs1999956 | -0.877 | -0.523 | -1.302 | 0.991 | -0.041 | -0.041 | -0.041 | 0.249 | 0.59 | 0.8 | 0.65 | 0.56 | 0.57 | 0.62 |
| 10 | 19336118 | rs952630 | 0.458 | -0.453 | -0.955 | 0.897 | 0.01 | 0.016 | 0.016 | 0 | 0.35 | 0.28 | 0.42 | 0.29 | 0.35 | 0.37 |
| 10 | 19340096 | rs11008298 | -0.096 | -0.161 | -0.911 | 0.753 | 0.023 | 0.005 | 0.005 | 0 | 0.42 | 0.33 | 0.45 | 0.44 | 0.37 | 0.49 |
| 10 | 19341911 | rs7084034 | -0.836 | 0.483 | -1.037 | 0.942 | -0.08 | -0.003 | -0.003 | 0 | 0.74 | 0.91 | 0.67 | 0.75 | 0.61 | 0.78 |
| 10 | 19343525 | rs10826891 | -0.565 | 0.391 | -1.246 | 0.93 | -0.084 | -0.01 | -0.01 | 0 | 0.72 | 0.9 | 0.58 | 0.72 | 0.6 | 0.75 |
| 10 | 19343732 | rs7475679 | 0 | 0.06 | -0.837 | 0.96 | 0.011 | 0.008 | 0.008 | 0.203 | 0.43 | 0.66 | 0.32 | 0.32 | 0.41 | 0.31 |
| 10 | 19344979 | rs10826893 | -0.588 | -0.102 | -0.128 | 0.94 | 0.053 | 0.05 | 0.05 | 0 | 0.2 | 0.28 | 0.41 | 0.36 | 0.3 | 0.15 |
| 10 | 19349377 | rs10763854 | -1.091 | -0.125 | 0.202 | 0.246 | 0.011 | -0.032 | -0.032 | 0.122 | 0.25 | 0.37 | 0.19 | 0.11 | 0.31 | 0.2 |
| 10 | 19357411 | rs2358276 | -1.475 | -0.652 | 0.022 | 0.152 | -0.023 | -0.009 | -0.009 | 0.891 | 0.58 | 0.71 | 0.6 | 0.53 | 0.52 | 0.52 |
| 10 | 19357737 | rs2358277 | 0.001 | -0.766 | 0.079 | 0.174 | -0.041 | 0.038 | 0.038 | 0.75 | 0.34 | 0.46 | 0.41 | 0.44 | 0.21 | 0.33 |
| 10 | 19365648 | rs2884423 | -0.485 | -0.699 | -0.07 | 0.276 | -0.011 | -0.01 | -0.01 | 2.928 | 0.22 | 0.16 | 0.21 | 0.37 | 0.28 | 0.22 |
| 10 | 19369100 | rs12242254 | -0.5 | -0.711 | -0.098 | 0.259 | -0.083 | 0.077 | 0.077 | 0 | 0.22 | 0.05 | 0.17 | 0.09 | 0.31 | 0.18 |
| 10 | 19373472 | rs10740840 | -1.523 | -0.815 | 0.184 | 0.233 | -0.015 | 0.002 | 0.002 | 0 | 0.57 | 0.63 | 0.6 | 0.56 | 0.52 | 0.51 |
| 10 | 19374137 | rs11008420 | 0.007 | -0.811 | 0.007 | 0.201 | -0.006 | -0.003 | -0.003 | 0 | 0.34 | 0.31 | 0.38 | 0.17 | 0.24 | 0.29 |
| 10 | 19375392 | rs2358300 | -1.677 | -0.809 | -0.032 | 0.202 | -0.057 | -0.021 | -0.021 | 0.201 | 0.58 | 0.84 | 0.61 | 0.56 | 0.53 | 0.52 |
| 10 | 19375491 | rs11008423 | 0.232 | -0.752 | -0.247 | -0.249 | -0.026 | -0.01 | -0.01 | 0 | 0.29 | 0.3 | 0.14 | 0.16 | 0.18 | 0.24 |
| 10 | 19384232 | rs10826935 | -1.005 | -0.576 | -0.288 | -0.243 | 0.004 | 0.046 | 0.046 | 0 | 0.51 | 0.35 | 0.53 | 0.26 | 0.49 | 0.42 |
| 10 | 19385420 | rs4748553 | -1.005 | -0.576 | -0.302 | -0.276 | 0.004 | 0.045 | 0.045 | 0 | 0.51 | 0.35 | 0.53 | 0.26 | 0.49 | 0.42 |
| 10 | 19386040 | rs10826941 | -0.515 | 0.195 | -0.665 | -0.207 | -0.069 | 0.056 | 0.056 | 0 | 0.18 | 0.05 | 0.14 | 0.08 | 0.24 | 0.13 |
| 10 | 19397980 | rs11008507 | 0.823 | 0.309 | -0.524 | -0.157 | -0.061 | 0.06 | 0.06 | 0 | 0.1 | 0.01 | 0.08 | 0 | 0.19 | 0.02 |
| 10 | 19403598 | rs12218913 | -1.181 | -1.296 | -0.719 | 0.178 | 0.024 | -0.011 | -0.011 | 0.339 | 0.54 | 0.78 | 0.55 | 0.58 | 0.41 | 0.49 |
| 10 | 19405531 | rs16918253 | -0.057 | -1.207 | -0.393 | 0.15 | -0.008 | -0.027 | -0.027 | 0 | 0.08 | 0.14 | 0.03 | 0.01 | 0.06 | 0.14 |
| 10 | 19406306 | rs12252665 | -0.191 | -1.306 | -0.187 | 0.086 | -0.026 | -0.104 | -0.104 | 0 | 0.21 | 0.45 | 0.34 | 0.03 | 0.17 | 0.16 |
| 10 | 19413368 | rs10450317 | -0.586 | -1.204 | -0.323 | 0.328 | -0.011 | -0.006 | -0.006 | 0.944 | 0.41 | 0.46 | 0.45 | 0.24 | 0.29 | 0.31 |
| 10 | 19417181 | rs11008573 | 0.05 | -0.862 | -0.279 | 0.401 | -0.054 | 0.043 | 0.043 | 0 | 0.17 | 0.02 | 0.14 | 0.11 | 0.2 | 0.17 |
| 10 | 19418567 | rs12252510 | -0.351 | -0.767 | -0.004 | 0.406 | -0.012 | -0.012 | -0.012 | 0.426 | 0.3 | 0.38 | 0.14 | 0.11 | 0.21 | 0.24 |
| 10 | 19420181 | rs11008597 | -0.732 | 0.044 | -1.176 | 0.346 | -0.001 | -0.007 | -0.007 | 0 | 0.55 | 0.77 | 0.58 | 0.56 | 0.46 | 0.5 |
| 10 | 19422749 | rs7922413 | -0.887 | 0.054 | -0.619 | 0 | -0.012 | -0.03 | -0.03 | 2.414 | 0.17 | 0.25 | 0.13 | 0.06 | 0.1 | 0.08 |
| 10 | 19424694 | rs2884429 | -0.836 | 1.217 | -0.744 | 0.206 | -0.023 | -0.003 | -0.003 | 0 | 0.29 | 0.22 | 0.47 | 0.31 | 0.38 | 0.16 |
| 10 | 19425704 | rs10826986 | 0.156 | 1.801 | -0.383 | -0.001 | 0.008 | -0.042 | -0.042 | 0 | 0.12 | 0.12 | 0.06 | 0 | 0.11 | 0.03 |
| 10 | 19428643 | rs12775323 | -1.531 | 1.749 | -0.406 | 0.067 | -0.018 | 0.019 | 0.019 | 0 | 0.3 | 0.24 | 0.41 | 0.14 | 0.39 | 0.13 |
| 10 | 19431373 | rs12769501 | -0.599 | 1.733 | -0.36 | 0.068 | -0.05 | 0.036 | 0.036 | 0 | 0.17 | 0.06 | 0.35 | 0.14 | 0.27 | 0.1 |
| 10 | 19432352 | rs10826998 | -1.582 | 1.312 | -0.351 | -0.308 | -0.011 | 0.014 | 0.014 | 0 | 0.38 | 0.32 | 0.45 | 0.14 | 0.42 | 0.15 |
| 10 | 19435382 | rs17646341 | 1.122 | 1.085 | -0.829 | -0.779 | -0.007 | 0.04 | 0.04 | 0 | 0.09 | 0.02 | 0.04 | 0 | 0.09 | 0.05 |
| 10 | 19436404 | rs12775097 | 1.105 | 0.796 | -1.233 | -0.293 | 0.002 | -0.041 | -0.041 | 0 | 0.07 | 0 | 0.02 | 0.13 | 0.05 | 0.14 |
| 10 | 19436862 | rs11008674 | 0.609 | 1.146 | -1.161 | 0.213 | -0.001 | 0.034 | 0.034 | 2.915 | 0.05 | 0.03 | 0.07 | 0.22 | 0.07 | 0.06 |
| 10 | 19437915 | rs7080782 | -1.755 | 1.164 | -1.595 | -0.291 | -0.041 | 0.05 | 0.05 | 0 | 0.91 | 0.99 | 0.93 | 1 | 0.86 | 0.92 |
| 10 | 19444182 | rs9417900 | 1.843 | 0.594 | -0.84 | -0.282 | -0.016 | -0.002 | -0.002 | 0 | 0.29 | 0.37 | 0.26 | 0.33 | 0.23 | 0.47 |
| 10 | 19446660 | rs12783745 | 0.343 | 1.386 | -0.839 | -0.155 | -0.03 | -0.055 | -0.055 | 0 | 0.15 | 0.07 | 0.36 | 0.31 | 0.22 | 0.12 |
| 10 | 19463367 | rs6481773 | 0.605 | 1.563 | -0.498 | -0.076 | -0.005 | 0.003 | 0.003 | 0 | 0.13 | 0.1 | 0.3 | 0.14 | 0.21 | 0.11 |
| 10 | 19466829 | rs10827018 | 1.417 | 1.645 | -0.44 | -0.123 | 0 | 0.002 | 0.002 | 0 | 0.11 | 0.13 | 0.12 | 0.17 | 0.11 | 0.04 |
| 10 | 19467440 | rs4748559 | -0.196 | 1.602 | -0.419 | -0.131 | 0 | 0 | 0 | 0 | 0.24 | 0.24 | 0.42 | 0.31 | 0.32 | 0.16 |
| 10 | 19471662 | rs10827030 | -0.174 | 1.602 | -0.419 | -0.131 | 0 | 0 | 0 | 0.076 | 0.24 | 0.24 | 0.42 | 0.31 | 0.32 | 0.15 |
| 10 | 19471726 | rs16918323 | 0.755 | 1.553 | -0.367 | -0.264 | -0.001 | -0.039 | -0.039 | 0.006 | 0.14 | 0.14 | 0.1 | 0 | 0.14 | 0.04 |
| 10 | 19471894 | rs4748560 | -0.061 | 1.553 | -0.371 | -0.26 | 0.001 | 0 | 0 | 0 | 0.24 | 0.24 | 0.42 | 0.31 | 0.32 | 0.15 |
| 10 | 19472350 | rs12357450 | 1.825 | 1.321 | -0.271 | -0.17 | 0.002 | 0.001 | 0.001 | 0 | 0.34 | 0.34 | 0.21 | 0.32 | 0.24 | 0.52 |
| 10 | 19478110 | rs10827035 | -0.704 | 1.449 | -0.299 | -0.322 | -0.002 | 0.001 | 0.001 | 0 | 0.38 | 0.33 | 0.52 | 0.31 | 0.46 | 0.2 |
| 10 | 19480296 | rs7094873 | -0.855 | 1.431 | -0.163 | -0.33 | 0.017 | -0.017 | -0.017 | 0 | 0.39 | 0.48 | 0.53 | 0.31 | 0.46 | 0.2 |
| 10 | 19482108 | rs11008874 | 0.77 | 1.481 | -0.229 | -0.339 | -0.022 | -0.011 | -0.011 | 0 | 0.19 | 0.11 | 0.16 | 0.17 | 0.2 | 0.09 |
| 10 | 19482577 | rs2884432 | -0.8 | 1.495 | -0.165 | -0.351 | -0.001 | 0.001 | 0.001 | 0 | 0.38 | 0.34 | 0.51 | 0.31 | 0.46 | 0.21 |
| 10 | 19483105 | rs10827041 | 0.746 | 1.49 | -0.229 | -0.317 | -0.021 | -0.01 | -0.01 | 0.109 | 0.19 | 0.11 | 0.16 | 0.17 | 0.2 | 0.09 |
| 10 | 19488366 | rs7916072 | -0.732 | 1.25 | -0.138 | -0.209 | 0.017 | 0.028 | 0.028 | 0.153 | 0.34 | 0.48 | 0.59 | 0.53 | 0.4 | 0.2 |
| 10 | 19490727 | rs10508577 | -0.901 | 1.24 | -0.26 | -0.265 | -0.04 | -0.016 | -0.016 | 0 | 0.8 | 0.91 | 0.81 | 0.83 | 0.74 | 0.86 |
| 10 | 19492702 | rs16918344 | 0.83 | 1.225 | -0.232 | -0.288 | -0.001 | -0.024 | -0.024 | 0 | 0.14 | 0.11 | 0.1 | 0 | 0.14 | 0.04 |
| 10 | 19493309 | rs10508578 | 2.102 | 1.225 | -0.254 | -0.286 | -0.01 | 0.034 | 0.034 | 0 | 0.08 | 0.01 | 0.04 | 0 | 0.09 | 0.03 |
| 10 | 19494514 | rs10763883 | 0.371 | 1.149 | -0.43 | -0.408 | -0.027 | -0.016 | -0.016 | 0 | 0.64 | 0.81 | 0.67 | 0.7 | 0.57 | 0.67 |
| 10 | 19495188 | rs7081985 | 0.128 | 1.124 | -0.443 | -0.421 | -0.014 | -0.021 | -0.021 | 0.808 | 0.67 | 0.81 | 0.74 | 0.7 | 0.64 | 0.69 |
| 10 | 19496816 | rs11816914 | 1.295 | 1.083 | -0.303 | -0.414 | -0.016 | 0.007 | 0.007 | 0 | 0.4 | 0.61 | 0.32 | 0.56 | 0.26 | 0.53 |

| | | | | | | | | | | | | | | | | |
|---|---|---|---|---|---|---|---|---|---|---|---|---|---|---|---|---|
| 10 | 19497686 | rs2358352 | -1.809 | 1.32 | -0.335 | -0.398 | -0.048 | -0.053 | -0.053 | 0.065 | 0.88 | 0.96 | 0.86 | 0.83 | 0.83 | 0.89 |
| 10 | 19497776 | rs2884433 | 1.468 | 1.275 | -0.31 | -0.421 | 0.002 | -0.003 | -0.003 | 0 | 0.4 | 0.37 | 0.23 | 0.47 | 0.29 | 0.67 |
| 10 | 19498722 | rs2245136 | -1.039 | 1.565 | -0.465 | -0.234 | -0.03 | -0.043 | -0.043 | 0.272 | 0.86 | 0.9 | 0.73 | 0.61 | 0.78 | 0.86 |
| 10 | 19501994 | rs2245456 | -0.333 | 1.556 | -0.469 | -0.314 | -0.011 | 0.005 | 0.005 | 0 | 0.75 | 0.81 | 0.76 | 0.83 | 0.69 | 0.85 |
| 10 | 19502571 | rs2583646 | -0.021 | 1.393 | -0.532 | -0.286 | -0.061 | 0.11 | 0.11 | 0 | 0.66 | 0.88 | 0.56 | 0.86 | 0.51 | 0.77 |
| 10 | 19509564 | rs1889515 | 1.465 | 1.048 | -0.298 | 0.095 | 0 | 0 | 0 | 0 | 0.08 | 0.07 | 0.04 | 0 | 0.09 | 0.03 |
| 10 | 19511109 | rs4278424 | 0.72 | 1.166 | -0.33 | 0.151 | 0.001 | 0.007 | 0.007 | 0 | 0.08 | 0.05 | 0.32 | 0.31 | 0.08 | 0.11 |
| 10 | 19517528 | rs10827096 | 0.432 | 1.326 | -0.201 | 0.151 | 0.008 | 0.098 | 0.098 | 0 | 0.13 | 0.24 | 0.24 | 0.36 | 0.16 | 0.1 |
| 10 | 19520364 | rs756308 | 0.247 | 1.32 | -0.041 | 0.191 | 0.028 | 0.177 | 0.177 | 0 | 0.14 | 0.39 | 0.25 | 0.35 | 0.16 | 0.1 |
| 10 | 19520750 | rs7088424 | -1.089 | 1.563 | -0.096 | -0.056 | 0.007 | 0.023 | 0.023 | 0 | 0.37 | 0.54 | 0.61 | 0.5 | 0.37 | 0.23 |
| 10 | 19521707 | rs10763889 | -0.29 | 1.504 | -0.107 | -0.087 | -0.002 | 0.001 | 0.001 | 0 | 0.22 | 0.19 | 0.37 | 0.14 | 0.21 | 0.13 |
| 10 | 19534443 | rs10437417 | -0.433 | 1.483 | -0.131 | -0.096 | -0.001 | -0.005 | -0.005 | 0 | 0.23 | 0.23 | 0.37 | 0.14 | 0.21 | 0.13 |
| 10 | 19535852 | rs10437418 | -1.203 | 1.547 | -0.104 | -0.076 | 0.004 | 0.014 | 0.014 | 0 | 0.38 | 0.57 | 0.62 | 0.49 | 0.38 | 0.23 |
| 10 | 19536320 | rs10128464 | 0.029 | 1.608 | -0.084 | -0.07 | 0.029 | 0.081 | 0.081 | 0 | 0.1 | 0.31 | 0.19 | 0.19 | 0.12 | 0.05 |
| 10 | 19536557 | rs10763899 | -1.145 | 1.602 | -0.145 | -0.069 | 0.011 | -0.025 | -0.025 | 0 | 0.37 | 0.51 | 0.56 | 0.33 | 0.38 | 0.21 |
| 10 | 19540391 | rs10827154 | -0.175 | 1.625 | -0.178 | -0.057 | -0.006 | -0.012 | -0.012 | 0 | 0.2 | 0.17 | 0.44 | 0.33 | 0.21 | 0.16 |
| 10 | 19541168 | rs10827159 | 0.448 | 1.567 | -0.197 | -0.095 | 0.004 | -0.074 | -0.074 | 4.243 | 0.16 | 0.24 | 0.11 | 0 | 0.17 | 0.05 |
| 10 | 19543516 | rs11009148 | 0.547 | 1.569 | -0.219 | -0.082 | -0.019 | 0.028 | 0.028 | 0 | 0.16 | 0.04 | 0.31 | 0.14 | 0.15 | 0.12 |
| 10 | 19543985 | rs11009149 | 1.824 | 1.583 | -0.222 | -0.086 | -0.025 | 0.041 | 0.041 | 0 | 0.08 | 0.02 | 0.05 | 0 | 0.08 | 0.02 |
| 10 | 19547472 | rs1889513 | -0.251 | 1.516 | -0.109 | 0.017 | 0.014 | -0.003 | -0.003 | 0.53 | 0.2 | 0.38 | 0.24 | 0.19 | 0.23 | 0.08 |
| 10 | 19549734 | rs2484086 | 0.997 | 1.522 | -0.306 | 0.012 | -0.018 | -0.047 | -0.047 | 0 | 0.1 | 0 | 0.09 | 0.12 | 0.11 | 0.17 |
| 10 | 19550078 | rs10827168 | -0.85 | 1.719 | 0.04 | -0.291 | 0 | -0.028 | -0.028 | 0.154 | 0.34 | 0.34 | 0.43 | 0.14 | 0.32 | 0.19 |
| 10 | 19550156 | rs12259609 | 0.567 | 1.743 | -0.175 | -0.256 | 0.005 | -0.038 | -0.038 | 0 | 0.07 | 0.19 | 0.06 | 0 | 0.06 | 0.01 |
| 10 | 19554984 | rs2484089 | 1.334 | 1.623 | -0.354 | -0.29 | -0.006 | -0.075 | -0.075 | 3.142 | 0.41 | 0.15 | 0.24 | 0.51 | 0.42 | 0.69 |
| 10 | 19558475 | rs17667025 | 0.933 | 2.208 | -0.325 | 0.17 | -0.022 | 0.046 | 0.046 | 0 | 0.1 | 0.02 | 0.05 | 0 | 0.11 | 0.04 |
| 10 | 19560251 | rs2499073 | 1.076 | 2.25 | -0.205 | 0.184 | 0.008 | 0.002 | 0.002 | 0 | 0.5 | 0.43 | 0.32 | 0.49 | 0.46 | 0.69 |
| 10 | 19560960 | rs8181385 | -1.049 | 1.993 | -0.075 | 0.941 | 0.005 | -0.005 | -0.005 | 0 | 0.31 | 0.4 | 0.5 | 0.33 | 0.33 | 0.21 |
| 10 | 19567647 | rs16918457 | -0.239 | 1.517 | -0.917 | 0.735 | -0.006 | 0.006 | 0.006 | 0 | 0.11 | 0.05 | 0.05 | 0 | 0.12 | 0.07 |
| 10 | 19572729 | rs2499065 | -0.615 | 1.168 | -0.454 | 1.298 | -0.005 | -0.012 | -0.012 | 0 | 0.33 | 0.33 | 0.47 | 0.45 | 0.4 | 0.51 |
| 10 | 19578706 | rs7912589 | -0.327 | 0.754 | 0.009 | 1.193 | -0.011 | -0.026 | -0.026 | 0 | 0.06 | 0.08 | 0.03 | 0 | 0.02 | 0.01 |
| 10 | 19579258 | rs16918470 | -0.118 | 0.754 | 0.102 | 1.194 | -0.012 | -0.029 | -0.029 | 0.173 | 0.06 | 0.09 | 0.03 | 0 | 0.02 | 0.01 |
| 10 | 19579806 | rs1609614 | -1.296 | 0.891 | 0.091 | 1.353 | 0.005 | 0.003 | 0.003 | 0 | 0.16 | 0.21 | 0.12 | 0.13 | 0.19 | 0.18 |
| 10 | 19587093 | rs2496053 | -1.692 | 0.139 | 0.047 | 1.249 | 0 | 0 | 0 | 0 | 0.39 | 0.39 | 0.42 | 0.31 | 0.3 | 0.57 |
| 10 | 19587993 | rs2496050 | 0.43 | 0.009 | -0.058 | 1.152 | 0.022 | 0.001 | 0.001 | 1.068 | 0.14 | 0.08 | 0.28 | 0.16 | 0.09 | 0.29 |
| 10 | 19591645 | rs1889517 | -0.264 | 0.042 | -0.046 | 1.157 | 0.048 | 0.015 | 0.015 | 0.884 | 0.18 | 0.09 | 0.28 | 0.17 | 0.1 | 0.31 |
| 10 | 19592431 | rs2499082 | -0.207 | 0.038 | -0.047 | 1.156 | 0.044 | 0.013 | 0.013 | 1.303 | 0.18 | 0.09 | 0.28 | 0.17 | 0.1 | 0.29 |
| 10 | 19596177 | rs2496096 | -2.096 | 0.13 | 0.053 | 1.246 | -0.003 | 0.004 | 0.004 | 0 | 0.42 | 0.47 | 0.5 | 0.47 | 0.41 | 0.55 |
| 10 | 19603579 | rs1591331 | 1.469 | 0.124 | 0.018 | 1.246 | -0.002 | -0.011 | -0.011 | 0 | 0.64 | 0.65 | 0.54 | 0.52 | 0.61 | 0.44 |
| 10 | 19605298 | rs7894610 | -1.126 | 0.012 | 0.035 | 1.178 | -0.033 | -0.057 | -0.057 | 0 | 0.11 | 0.22 | 0.04 | 0 | 0.02 | 0.02 |
| 10 | 19613431 | rs16918492 | -1.098 | 0.065 | 0.052 | 1.184 | -0.015 | -0.06 | -0.06 | 0 | 0.11 | 0.21 | 0.04 | 0 | 0.04 | 0.03 |
| 10 | 19615429 | rs16918494 | -1.118 | 0.065 | 0.05 | 1.184 | -0.015 | -0.062 | -0.062 | 0 | 0.11 | 0.23 | 0.04 | 0 | 0.04 | 0.03 |
| 10 | 19618011 | rs7071861 | 0.183 | 0.09 | 0.03 | 1.232 | 0.011 | -0.036 | -0.036 | 0 | 0.8 | 0.88 | 0.64 | 0.68 | 0.8 | 0.63 |
| 10 | 19619962 | rs16918512 | 0.196 | 0.065 | 0.034 | 1.156 | -0.013 | -0.03 | -0.03 | 0.007 | 0.06 | 0.08 | 0.03 | 0 | 0.02 | 0.01 |
| 10 | 19620407 | rs7100382 | 0.921 | 0.084 | 0.022 | 1.196 | -0.007 | -0.018 | -0.018 | 0.158 | 0.69 | 0.72 | 0.54 | 0.52 | 0.62 | 0.45 |
| 10 | 19620439 | rs7100403 | 1.485 | 0.152 | -0.045 | 1.239 | -0.004 | 0.007 | 0.007 | 0 | 0.58 | 0.49 | 0.5 | 0.52 | 0.59 | 0.43 |
| 10 | 19636869 | rs1609746 | 1.498 | 0.087 | 0.025 | 1.2 | -0.005 | 0.008 | 0.008 | 0 | 0.58 | 0.48 | 0.51 | 0.52 | 0.59 | 0.42 |
| 10 | 19640125 | rs6481882 | 0.29 | 0.054 | -0.073 | 1.242 | -0.004 | -0.052 | -0.052 | 0 | 0.79 | 0.92 | 0.65 | 0.68 | 0.78 | 0.65 |
| 10 | 19663737 | rs1889520 | 1.135 | 0.032 | 0.016 | 1.252 | -0.012 | -0.04 | -0.04 | 0 | 0.68 | 0.75 | 0.55 | 0.52 | 0.62 | 0.46 |
| 10 | 19663851 | rs1889521 | 1.747 | 0.019 | 0.067 | 1.245 | -0.001 | -0.005 | -0.005 | 0 | 0.62 | 0.62 | 0.51 | 0.52 | 0.59 | 0.44 |
| 10 | 19664446 | rs12761627 | -0.49 | 0.038 | 0.069 | 1.246 | 0.006 | 0.019 | 0.019 | 0.022 | 0.12 | 0.18 | 0.1 | 0.16 | 0.17 | 0.19 |
| 10 | 19668100 | rs10763972 | 1.816 | 0.03 | 0.068 | 1.241 | -0.001 | -0.001 | -0.001 | 0.101 | 0.58 | 0.57 | 0.51 | 0.52 | 0.59 | 0.43 |
| 10 | 19669367 | rs11009669 | -1.011 | 0.102 | -0.133 | 1.333 | -0.004 | 0.006 | 0.006 | 0.005 | 0.15 | 0.18 | 0.18 | 0.31 | 0.27 | 0.25 |
| 10 | 19676200 | rs7897577 | -0.38 | -0.014 | -0.128 | 1.269 | 0.025 | 0.002 | 0.002 | 0.215 | 0.84 | 0.93 | 0.73 | 0.83 | 0.89 | 0.71 |
| 10 | 19676561 | rs12771333 | -0.101 | 0.003 | -0.113 | 1.29 | -0.006 | -0.007 | -0.007 | 0 | 0.1 | 0.06 | 0.08 | 0.16 | 0.15 | 0.2 |
| 10 | 19679886 | rs4276997 | -0.696 | -0.019 | -0.042 | 1.319 | 0.017 | -0.011 | -0.011 | 0.1 | 0.87 | 0.96 | 0.73 | 0.84 | 0.9 | 0.74 |
| 10 | 19682953 | rs7917396 | 1.147 | -0.556 | -0.78 | 0.442 | 0.012 | 0.061 | 0.061 | 7.415 | 0.15 | 0.07 | 0.07 | 0 | 0.08 | 0.02 |
| 10 | 19684858 | rs10763980 | -0.323 | -0.24 | -0.497 | 0.721 | -0.006 | -0.005 | -0.005 | 0 | 0.83 | 0.8 | 0.64 | 0.85 | 0.87 | 0.74 |
| 10 | 19689148 | rs4433484 | -0.956 | -0.237 | -0.499 | 0.724 | -0.007 | -0.002 | -0.002 | 2.152 | 0.87 | 0.82 | 0.66 | 0.85 | 0.9 | 0.76 |
| 10 | 19690815 | rs12415964 | -0.518 | -0.257 | -0.747 | 1.154 | -0.014 | -0.022 | -0.022 | 0 | 0.06 | 0.03 | 0.06 | 0.13 | 0.09 | 0.1 |
| 10 | 19691164 | rs12415710 | -0.944 | -0.419 | -0.819 | 1.145 | -0.013 | -0.025 | -0.025 | 0 | 0.12 | 0.11 | 0.11 | 0.26 | 0.16 | 0.17 |
| 10 | 19692463 | rs10827364 | 1.511 | -0.655 | -0.119 | 1.032 | -0.022 | 0.016 | 0.016 | 0 | 0.63 | 0.53 | 0.51 | 0.6 | 0.72 | 0.56 |
| 10 | 19693587 | rs7907773 | -0.392 | -0.678 | -0.153 | 1.017 | 0.008 | 0.007 | 0.007 | 0 | 0.83 | 0.87 | 0.65 | 0.85 | 0.88 | 0.74 |
| 10 | 19695156 | rs10763994 | 1.458 | -0.623 | -0.235 | 1.049 | -0.015 | 0.009 | 0.009 | 0 | 0.63 | 0.43 | 0.5 | 0.6 | 0.72 | 0.55 |
| 10 | 19695260 | rs11009788 | 1.417 | -0.554 | -1.166 | 0.58 | -0.096 | -0.105 | -0.105 | 0.91 | 0.23 | 0.04 | 0.25 | 0.45 | 0.36 | 0.32 |
| 10 | 19696007 | rs4274131 | -1.002 | -0.969 | -0.304 | 0.811 | 0.007 | 0 | 0 | 0.25 | 0.87 | 0.93 | 0.68 | 0.85 | 0.89 | 0.75 |
| 10 | 19699004 | rs7910052 | 0.686 | -0.968 | -0.31 | 0.814 | 0.003 | -0.066 | -0.066 | 1.299 | 0.77 | 0.88 | 0.59 | 0.61 | 0.77 | 0.6 |
| 10 | 19703136 | rs7908470 | 0.444 | -0.767 | 0.074 | 0.452 | -0.054 | -0.056 | -0.056 | 0.593 | 0.73 | 0.49 | 0.59 | 0.82 | 0.84 | 0.71 |
| 10 | 19710332 | rs11009827 | -0.561 | 0.173 | 0.048 | 0.806 | 0.001 | 0.028 | 0.028 | 0 | 0.05 | 0.05 | 0.07 | 0.15 | 0.07 | 0.03 |
| 10 | 19714510 | rs12267414 | -0.922 | 0.062 | 0.326 | 0.612 | 0.001 | -0.036 | -0.036 | 0 | 0.1 | 0.17 | 0.06 | 0.01 | 0.07 | 0.05 |

| | | | | | | | | | | | | | | | | |
|---|---|---|---|---|---|---|---|---|---|---|---|---|---|---|---|---|
| 10 | 19714554 | rs12267418 | -0.758 | -0.33 | 0.283 | 0.366 | -0.041 | -0.141 | -0.141 | 0 | 0.21 | 0.44 | 0.11 | 0.01 | 0.11 | 0.07 |
| 10 | 19715913 | rs7094477 | -1.114 | -0.305 | -0.252 | 0.459 | -0.005 | -0.006 | -0.006 | 0 | 0.27 | 0.62 | 0.19 | 0.15 | 0.18 | 0.11 |
| 10 | 19719841 | rs2486053 | -0.154 | 0.569 | -1.226 | 0.922 | -0.045 | -0.094 | -0.094 | 2.034 | 0.8 | 0.95 | 0.59 | 0.66 | 0.77 | 0.64 |
| 10 | 19721013 | rs2358395 | -0.101 | 0.896 | -2.18 | -0.198 | -0.039 | 0.129 | 0.129 | 1.135 | 0.25 | 0.06 | 0.15 | 0 | 0.26 | 0.15 |
| 10 | 19722515 | rs2993646 | -0.053 | 0.054 | -2.171 | -0.381 | 0.011 | -0.031 | -0.031 | 0 | 0.59 | 0.26 | 0.41 | 0.51 | 0.64 | 0.48 |
| 10 | 19722747 | rs10827401 | 0.738 | -0.205 | -0.862 | -0.495 | 0 | 0.011 | 0.011 | 0 | 0.11 | 0.34 | 0.12 | 0.1 | 0.08 | 0.12 |
| 10 | 19724287 | rs10827409 | 0.845 | -0.217 | -0.248 | -0.478 | -0.017 | -0.025 | -0.025 | 0.01 | 0.08 | 0.12 | 0.04 | 0.05 | 0.05 | 0.03 |
| 10 | 19725008 | rs2486056 | -1.229 | -0.28 | -0.219 | -0.476 | 0.006 | 0.035 | 0.035 | 0.409 | 0.77 | 0.69 | 0.56 | 0.66 | 0.77 | 0.62 |
| 10 | 19728251 | rs2496101 | -1.627 | -0.191 | -0.414 | -0.372 | -0.051 | -0.083 | -0.083 | 0.581 | 0.8 | 0.95 | 0.62 | 0.68 | 0.77 | 0.63 |
| 10 | 19739459 | rs6415957 | -1.525 | -0.61 | -0.447 | -0.683 | -0.006 | -0.024 | -0.024 | 0.451 | 0.89 | 0.92 | 0.97 | 1 | 0.96 | 0.99 |
| 10 | 19751780 | rs2358398 | -0.602 | -0.206 | -0.567 | -0.867 | -0.001 | 0.035 | 0.035 | 1.626 | 0.82 | 0.92 | 0.89 | 1 | 0.83 | 0.93 |
| 10 | 19753117 | rs10764023 | 0.855 | -0.597 | -0.269 | -0.691 | -0.001 | 0.007 | 0.007 | 0.458 | 0.31 | 0.3 | 0.44 | 0.42 | 0.32 | 0.42 |
| 10 | 19754883 | rs7089570 | 0.582 | -0.766 | -0.046 | -0.766 | -0.002 | -0.069 | -0.069 | 0 | 0.7 | 0.7 | 0.82 | 0.99 | 0.74 | 0.82 |
| 10 | 19756569 | rs12245764 | 0.193 | -0.657 | 0.176 | -0.776 | -0.001 | 0.003 | 0.003 | 0 | 0.48 | 0.6 | 0.44 | 0.58 | 0.49 | 0.49 |
| 10 | 19758593 | rs12358346 | 1.168 | -0.586 | -0.013 | -0.744 | 0.004 | 0.01 | 0.01 | 0 | 0.07 | 0.03 | 0.04 | 0 | 0.07 | 0.03 |
| 10 | 19765768 | rs17200443 | 2.537 | -0.767 | -0.551 | -0.745 | -0.005 | -0.021 | -0.021 | 0 | 0.1 | 0 | 0.04 | 0.14 | 0.07 | 0.13 |
| 10 | 19767583 | rs12265190 | 0.325 | -1.026 | -0.675 | -0.814 | -0.029 | -0.029 | -0.029 | 0 | 0.33 | 0.25 | 0.3 | 0.47 | 0.43 | 0.38 |
| 10 | 19769058 | rs11592961 | 0.523 | -1.03 | -0.632 | -0.801 | -0.054 | -0.036 | -0.036 | 0.006 | 0.32 | 0.16 | 0.28 | 0.44 | 0.42 | 0.38 |
| 10 | 19769510 | rs10764037 | 0.383 | -1.062 | -0.623 | -0.882 | 0.012 | 0.003 | 0.003 | 0.27 | 0.47 | 0.57 | 0.56 | 0.53 | 0.4 | 0.55 |
| 10 | 19769651 | rs10764038 | -1.175 | -0.934 | -0.702 | -0.854 | 0 | 0.026 | 0.026 | 1.435 | 0.82 | 0.91 | 0.89 | 1 | 0.83 | 0.93 |
| 10 | 19775692 | rs1537346 | -0.277 | -1.051 | -0.715 | -0.872 | -0.01 | 0.088 | 0.088 | 1.018 | 0.73 | 0.9 | 0.85 | 1 | 0.74 | 0.84 |
| 10 | 19776636 | rs1332926 | 0.535 | -1.145 | -0.546 | -0.852 | -0.001 | -0.022 | -0.022 | 0 | 0.56 | 0.54 | 0.73 | 0.89 | 0.66 | 0.71 |
| 10 | 19777189 | rs9663503 | 1.905 | -1.317 | -0.466 | -0.656 | -0.001 | -0.006 | -0.006 | 2.939 | 0.26 | 0.23 | 0.41 | 0.37 | 0.28 | 0.4 |
| 10 | 19778056 | rs2151234 | -0.211 | -1.504 | -0.666 | -0.553 | -0.005 | -0.012 | -0.012 | 0.473 | 0.45 | 0.46 | 0.42 | 0.34 | 0.59 | 0.42 |
| 10 | 19778072 | rs2184035 | -2.157 | -1.545 | -0.625 | -0.628 | 0.015 | 0.005 | 0.005 | 4.739 | 0.89 | 0.91 | 0.97 | 1 | 0.96 | 0.99 |
| 10 | 19778596 | rs16906129 | 0.529 | -1.334 | -0.483 | -0.482 | -0.028 | -0.016 | -0.016 | 1.945 | 0.16 | 0.27 | 0.11 | 0.11 | 0.09 | 0.12 |
| 10 | 19780463 | rs16918863 | -0.565 | -1.492 | -0.399 | -0.416 | -0.041 | 0.056 | 0.056 | 0.723 | 0.13 | 0.59 | 0.12 | 0.13 | 0.06 | 0.05 |
| 10 | 19780543 | rs10827499 | 2.122 | -1.432 | -0.834 | -0.363 | -0.082 | -0.092 | -0.092 | 0 | 0.24 | 0.02 | 0.4 | 0.3 | 0.29 | 0.38 |
| 10 | 19783593 | rs10827500 | -0.554 | -1.353 | -0.707 | -0.335 | -0.032 | 0.052 | 0.052 | 0.335 | 0.14 | 0.58 | 0.13 | 0.14 | 0.08 | 0.06 |
| 10 | 19785754 | rs7092838 | -1.499 | -1.537 | -0.812 | -0.463 | -0.016 | -0.008 | -0.008 | 0.078 | 0.66 | 0.88 | 0.57 | 0.7 | 0.67 | 0.61 |
| 10 | 19787418 | rs12256835 | -0.94 | -1.691 | -0.979 | -0.542 | -0.028 | -0.003 | -0.003 | 0 | 0.3 | 0.63 | 0.21 | 0.25 | 0.17 | 0.19 |
| 10 | 19791144 | rs2026465 | -0.162 | -1.698 | -0.996 | -0.547 | -0.006 | 0.011 | 0.011 | 0 | 0.43 | 0.26 | 0.38 | 0.45 | 0.56 | 0.45 |
| 10 | 19795045 | rs6481955 | -0.153 | -1.69 | -0.823 | -0.516 | -0.064 | -0.025 | -0.025 | 0 | 0.22 | 0.51 | 0.18 | 0.13 | 0.1 | 0.15 |
| 10 | 19796290 | rs3852464 | -0.166 | -1.638 | -0.834 | -0.525 | -0.061 | -0.025 | -0.025 | 0.117 | 0.22 | 0.51 | 0.18 | 0.13 | 0.1 | 0.15 |
| 10 | 19796594 | rs7474936 | -0.205 | -1.678 | -0.841 | -0.525 | -0.066 | -0.01 | -0.01 | 0.026 | 0.22 | 0.51 | 0.18 | 0.2 | 0.1 | 0.15 |
| 10 | 19796958 | rs1332925 | 0.314 | -1.463 | -0.963 | -0.587 | -0.023 | 0.046 | 0.046 | 0.015 | 0.14 | 0.07 | 0.11 | 0.02 | 0.18 | 0.15 |
| 10 | 19800492 | rs3852468 | 0.866 | -1.514 | -0.792 | -0.474 | -0.004 | 0.004 | 0.004 | 0 | 0.15 | 0.17 | 0.11 | 0.22 | 0.09 | 0.12 |
| 10 | 19802238 | rs6481957 | -1.838 | -1.536 | -0.955 | -0.507 | -0.026 | -0.043 | -0.043 | 0 | 0.67 | 0.88 | 0.53 | 0.67 | 0.67 | 0.61 |
| 10 | 19807623 | rs11813491 | -0.047 | -1.566 | -0.835 | -0.464 | -0.06 | -0.015 | -0.015 | 2.364 | 0.2 | 0.52 | 0.15 | 0.19 | 0.09 | 0.18 |
| 10 | 19807875 | rs10494868 | 1.269 | -1.602 | -0.855 | -0.479 | 0.015 | 0 | 0 | 0 | 0.13 | 0.04 | 0.09 | 0.11 | 0.08 | 0.11 |
| 10 | 19809674 | rs10494867 | -0.137 | -1.545 | -0.833 | -0.418 | -0.048 | 0.017 | 0.017 | 0.52 | 0.21 | 0.63 | 0.15 | 0.24 | 0.1 | 0.19 |
| 10 | 19811046 | rs3852470 | 2.143 | -1.519 | -0.988 | -0.509 | -0.106 | -0.132 | -0.132 | 0 | 0.25 | 0.02 | 0.44 | 0.32 | 0.3 | 0.38 |
| 10 | 19811364 | rs13377040 | 0.11 | -1.482 | -0.837 | -0.429 | -0.059 | -0.034 | -0.034 | 0 | 0.2 | 0.52 | 0.14 | 0.13 | 0.09 | 0.18 |
| 10 | 19812767 | rs12766985 | -0.047 | -1.188 | -1.194 | -0.48 | -0.035 | 0.003 | 0.003 | 0 | 0.39 | 0.24 | 0.35 | 0.43 | 0.5 | 0.38 |
| 10 | 19813583 | rs12776128 | 0.945 | -0.971 | -1.651 | -0.42 | -0.146 | -0.162 | -0.162 | 6.808 | 0.3 | 0.04 | 0.48 | 0.4 | 0.46 | 0.52 |
| 10 | 19814454 | rs2358415 | 0.439 | -1.508 | -0.323 | -0.512 | -0.023 | -0.018 | -0.018 | 0 | 0.65 | 0.39 | 0.77 | 0.74 | 0.76 | 0.72 |
| 10 | 19820560 | rs7097568 | -0.506 | -1.458 | 0.059 | -0.307 | 0.028 | 0.126 | 0.126 | 0.946 | 0.06 | 0.36 | 0.03 | 0.11 | 0.01 | 0.1 |
| 10 | 19823467 | rs10764052 | 0.699 | -1.502 | -0.414 | -0.492 | -0.091 | 0.004 | 0.004 | 0.039 | 0.35 | 0.1 | 0.31 | 0.38 | 0.51 | 0.29 |
| 10 | 19824906 | rs11010274 | 0.047 | -1.793 | 0.106 | -0.554 | 0.016 | 0.008 | 0.008 | 2.842 | 0.69 | 0.28 | 0.81 | 0.74 | 0.85 | 0.72 |
| 10 | 19825052 | rs11010275 | 0.12 | -1.813 | 0.154 | -0.558 | 0.008 | 0.008 | 0.008 | 2.68 | 0.69 | 0.3 | 0.81 | 0.82 | 0.85 | 0.72 |
| 10 | 19826238 | rs7076844 | 0.065 | -1.89 | 0.193 | -0.615 | -0.025 | -0.02 | -0.02 | 0.966 | 0.68 | 0.58 | 0.81 | 0.77 | 0.81 | 0.79 |
| 10 | 19827911 | rs6481968 | 1.478 | -1.776 | -0.082 | -0.588 | 0.004 | 0.008 | 0.008 | 0 | 0.08 | 0.08 | 0.04 | 0 | 0.07 | 0.03 |
| 10 | 19830436 | rs7078866 | 0.681 | -1.801 | -0.041 | -0.607 | 0.077 | 0.064 | 0.064 | 1.136 | 0.6 | 0.21 | 0.77 | 0.74 | 0.78 | 0.66 |
| 10 | 19834795 | rs3844357 | -0.712 | -1.646 | 0.136 | -0.372 | 0.028 | 0.024 | 0.024 | 0 | 0.77 | 0.85 | 0.88 | 0.85 | 0.86 | 0.82 |
| 10 | 19839161 | rs6481973 | -0.695 | -1.638 | 0.109 | -0.373 | 0.023 | 0.006 | 0.006 | 0 | 0.76 | 0.79 | 0.87 | 0.77 | 0.86 | 0.82 |
| 10 | 19846068 | rs10740916 | -0.039 | -1.665 | 0.11 | -0.375 | 0.01 | 0.005 | 0.005 | 1.809 | 0.7 | 0.26 | 0.82 | 0.74 | 0.85 | 0.72 |
| 10 | 19846368 | rs3904894 | -0.959 | -1.644 | 0.077 | -0.418 | 0.029 | 0.035 | 0.035 | 0 | 0.78 | 0.9 | 0.88 | 0.88 | 0.87 | 0.83 |
| 10 | 19858231 | rs10740921 | 0.212 | -1.606 | 0.107 | -0.425 | 0.043 | 0.026 | 0.026 | 2.804 | 0.68 | 0.21 | 0.81 | 0.74 | 0.85 | 0.72 |
| 10 | 19863970 | rs17729873 | 0.442 | -1.028 | -0.16 | -0.394 | -0.072 | -0.019 | -0.019 | 0 | 0.33 | 0.16 | 0.3 | 0.41 | 0.47 | 0.25 |
| 10 | 19864706 | rs10827607 | -0.284 | -1.111 | -0.027 | -0.527 | 0 | -0.023 | -0.023 | 0 | 0.16 | 0.2 | 0.08 | 0 | 0.11 | 0.13 |
| 10 | 19865757 | rs11010520 | -0.325 | -1.084 | -0.006 | -0.51 | -0.003 | -0.034 | -0.034 | 0 | 0.08 | 0.2 | 0.04 | 0 | 0.05 | 0.08 |
| 10 | 19867761 | rs10827613 | -0.234 | -1.047 | 0.217 | -0.517 | 0.006 | -0.01 | -0.01 | 0 | 0.43 | 0.52 | 0.38 | 0.38 | 0.52 | 0.36 |
| 10 | 19884264 | rs10827628 | -0.552 | -1.067 | 0.42 | -0.445 | -0.002 | -0.007 | -0.007 | 0.718 | 0.47 | 0.53 | 0.39 | 0.38 | 0.56 | 0.38 |
| 10 | 19886330 | rs1409737 | -0.545 | -1.113 | 0.422 | -0.442 | -0.002 | -0.008 | -0.008 | 1.351 | 0.47 | 0.53 | 0.39 | 0.38 | 0.56 | 0.38 |
| 10 | 19888896 | rs3844358 | -0.864 | -1.213 | 0.486 | -0.408 | 0.013 | 0.027 | 0.027 | 0 | 0.51 | 0.65 | 0.41 | 0.41 | 0.58 | 0.38 |
| 10 | 19893332 | rs3852479 | 1.583 | -0.714 | 0.188 | -0.265 | -0.087 | -0.053 | -0.053 | 0 | 0.13 | 0.01 | 0.15 | 0.19 | 0.27 | 0.12 |
| 10 | 19899961 | rs3852483 | 0.081 | -1.737 | 0.77 | -0.289 | -0.057 | -0.036 | -0.036 | 0 | 0.76 | 0.49 | 0.9 | 0.82 | 0.89 | 0.83 |
| 10 | 19907943 | rs11010707 | -0.513 | -1.668 | 0.609 | -0.122 | 0.056 | 0.093 | 0.093 | 0.782 | 0.05 | 0.39 | 0.04 | 0.03 | 0.01 | 0.08 |
| 10 | 19909770 | rs10827649 | -0.168 | -1.79 | 0.286 | -0.141 | 0.005 | -0.004 | -0.004 | 0.109 | 0.44 | 0.42 | 0.58 | 0.42 | 0.39 | 0.57 |

| | | | | | | | | | | | | | | | | |
|---|---|---|---|---|---|---|---|---|---|---|---|---|---|---|---|---|
| 10 | 19921009 | rs7906048 | -0.711 | -1.61 | 0.477 | -0.008 | -0.048 | -0.009 | -0.009 | 1.34 | 0.15 | 0.37 | 0.07 | 0.14 | 0.06 | 0.11 |
| 10 | 19921103 | rs7906180 | -0.89 | -1.607 | 0.495 | -0.009 | -0.022 | 0.019 | 0.019 | 0.021 | 0.08 | 0.31 | 0.03 | 0.06 | 0.01 | 0.08 |
| 10 | 19930367 | rs10827728 | 1.093 | -1.555 | 0.433 | -0.001 | -0.008 | -0.006 | -0.006 | 0.202 | 0.07 | 0.11 | 0.05 | 0.08 | 0.05 | 0.03 |
| 10 | 19931174 | rs11010886 | 1.093 | -1.555 | 0.406 | 0 | -0.005 | -0.003 | -0.003 | 0.209 | 0.07 | 0.08 | 0.04 | 0.08 | 0.05 | 0.03 |
| 10 | 19935793 | rs12413789 | 0.983 | -1.792 | -0.017 | -0.034 | -0.047 | -0.116 | -0.116 | 0.035 | 0.29 | 0.04 | 0.48 | 0.35 | 0.31 | 0.44 |
| 10 | 19936411 | rs1326983 | -0.45 | -1.893 | 0.363 | -0.154 | 0.01 | 0.003 | 0.003 | 0 | 0.53 | 0.6 | 0.44 | 0.6 | 0.61 | 0.45 |
| 10 | 19941027 | rs7087709 | -1.061 | -1.699 | 0.455 | -0.076 | -0.019 | 0.079 | 0.079 | 0.11 | 0.08 | 0.37 | 0.04 | 0.14 | 0.01 | 0.08 |
| 10 | 19948827 | rs11812122 | -0.86 | -1.678 | 0.36 | -0.069 | -0.012 | 0.079 | 0.079 | 0.039 | 0.08 | 0.36 | 0.03 | 0.14 | 0.01 | 0.08 |
| 10 | 19954929 | rs1831047 | -0.918 | -1.937 | -0.07 | -0.078 | 0.018 | 0.009 | 0.009 | 1.797 | 0.85 | 0.9 | 0.95 | 0.87 | 0.92 | 0.9 |
| 10 | 19955587 | rs7086414 | -0.498 | -1.966 | -0.244 | -0.065 | 0.027 | -0.033 | -0.033 | 0 | 0.54 | 0.86 | 0.47 | 0.51 | 0.6 | 0.45 |
| 10 | 19957829 | rs1926820 | 0.704 | -1.79 | -0.2 | -0.265 | -0.026 | -0.045 | -0.045 | 0 | 0.38 | 0.16 | 0.36 | 0.48 | 0.55 | 0.34 |
| 10 | 19970957 | rs2148617 | 0.998 | -1.664 | -0.315 | -0.141 | -0.017 | -0.077 | -0.077 | 2.648 | 0.3 | 0.11 | 0.49 | 0.37 | 0.32 | 0.45 |
| 10 | 19973096 | rs1326971 | -0.699 | -1.796 | 0.023 | -0.307 | 0.006 | -0.001 | -0.001 | 0 | 0.86 | 0.84 | 0.96 | 0.95 | 0.93 | 0.91 |
| 10 | 19975850 | rs7924093 | 0.168 | -1.848 | 0.183 | -0.296 | -0.036 | -0.031 | -0.031 | 0 | 0.75 | 0.65 | 0.9 | 0.84 | 0.88 | 0.81 |
| 10 | 19981007 | rs10827765 | 0.439 | -1.641 | -0.479 | -0.217 | -0.027 | -0.066 | -0.066 | 4.613 | 0.38 | 0.13 | 0.52 | 0.37 | 0.4 | 0.48 |
| 10 | 20002736 | rs2025789 | -0.799 | -1.496 | -0.211 | -0.322 | -0.029 | 0.032 | 0.032 | 1.414 | 0.58 | 0.79 | 0.6 | 0.64 | 0.49 | 0.69 |
| 10 | 20002757 | rs2025788 | -0.799 | -1.496 | -0.211 | -0.322 | -0.029 | 0.032 | 0.032 | 1.414 | 0.58 | 0.79 | 0.6 | 0.64 | 0.49 | 0.69 |
| 10 | 20004996 | rs595118 | -0.534 | -1.548 | -0.236 | -0.317 | 0.001 | 0.032 | 0.032 | 2.294 | 0.53 | 0.77 | 0.6 | 0.59 | 0.48 | 0.65 |
| 10 | 20015932 | rs9417254 | 0.589 | -1.606 | -0.353 | -0.353 | -0.054 | -0.083 | -0.083 | 0.351 | 0.36 | 0.1 | 0.52 | 0.4 | 0.41 | 0.44 |
| 10 | 20029770 | rs7073601 | -0.978 | -1.715 | -0.416 | -0.367 | -0.081 | -0.144 | -0.144 | 0 | 0.67 | 0.97 | 0.52 | 0.6 | 0.62 | 0.55 |
| 10 | 20042199 | rs7071564 | 0.377 | -1.629 | -0.046 | -0.304 | -0.004 | -0.006 | -0.006 | 0 | 0.08 | 0.1 | 0.04 | 0.04 | 0.05 | 0.06 |
| 10 | 20051083 | rs11011420 | 0.051 | -1.544 | 0.204 | -0.354 | 0.004 | 0.007 | 0.007 | 0 | 0.53 | 0.4 | 0.42 | 0.47 | 0.55 | 0.39 |
| 10 | 20066869 | rs4748610 | -0.869 | -1.298 | 0.12 | -0.087 | -0.02 | -0.053 | -0.053 | 0 | 0.67 | 0.84 | 0.5 | 0.53 | 0.62 | 0.51 |
| 10 | 20071857 | rs12775805 | -0.828 | -1.049 | 0.033 | -0.216 | -0.005 | -0.001 | -0.001 | 0 | 0.2 | 0.24 | 0.17 | 0.23 | 0.14 | 0.19 |
| 10 | 20076614 | rs7908480 | -1.215 | -1.118 | -0.335 | -0.145 | -0.024 | -0.087 | -0.087 | 0.308 | 0.7 | 0.87 | 0.48 | 0.56 | 0.64 | 0.56 |
| 10 | 20077441 | rs1411823 | 0.175 | -1.207 | 0.103 | -0.239 | 0.002 | 0.008 | 0.008 | 2.266 | 0.49 | 0.45 | 0.35 | 0.21 | 0.51 | 0.36 |
| 10 | 20088820 | rs7910169 | -0.45 | -1.142 | -0.079 | -0.145 | -0.002 | 0.003 | 0.003 | 1.184 | 0.63 | 0.68 | 0.4 | 0.37 | 0.59 | 0.48 |
| 10 | 20088859 | rs2150599 | -1.461 | -1.002 | -0.279 | -0.151 | -0.005 | -0.087 | -0.087 | 0 | 0.74 | 0.88 | 0.5 | 0.52 | 0.7 | 0.58 |
| 10 | 20098492 | rs7086438 | -1.435 | -1.006 | -0.291 | -0.148 | -0.007 | -0.091 | -0.091 | 0 | 0.73 | 0.88 | 0.5 | 0.52 | 0.7 | 0.58 |
| 10 | 20104426 | rs7922577 | -0.46 | -1.004 | -0.28 | -0.129 | -0.016 | -0.047 | -0.047 | 0.019 | 0.64 | 0.79 | 0.47 | 0.45 | 0.6 | 0.52 |
| 10 | 20106794 | rs12257233 | -1.058 | -1.08 | -0.344 | -0.198 | 0.008 | -0.091 | -0.091 | 0.052 | 0.71 | 0.88 | 0.55 | 0.54 | 0.71 | 0.55 |
| 10 | 20110335 | rs11011636 | 0.773 | -1.05 | -0.167 | -0.331 | -0.006 | -0.031 | -0.031 | 0 | 0.08 | 0.11 | 0.04 | 0 | 0.05 | 0.05 |
| 10 | 20118838 | rs12263557 | 1.434 | -1.313 | -0.586 | -0.543 | -0.012 | 0.003 | 0.003 | 0 | 0.27 | 0.22 | 0.25 | 0.22 | 0.39 | 0.24 |
| 10 | 20121156 | rs379242 | -1.385 | -0.622 | 0.055 | 0.026 | -0.019 | 0.051 | 0.051 | 5.768 | 0.57 | 0.72 | 0.7 | 0.77 | 0.52 | 0.73 |
| 10 | 20121877 | rs2478143 | -0.501 | 0.06 | -0.15 | 0.52 | -0.004 | 0.001 | 0.001 | 2.109 | 0.37 | 0.33 | 0.54 | 0.62 | 0.41 | 0.53 |
| 10 | 20128601 | rs333688 | -0.984 | 0.183 | -0.186 | 0.524 | 0 | 0.003 | 0.003 | 4.526 | 0.46 | 0.57 | 0.59 | 0.61 | 0.46 | 0.55 |
| 10 | 20129463 | rs333692 | -0.022 | 0.349 | -1.409 | 0.949 | 0.002 | -0.079 | -0.079 | 0.232 | 0.26 | 0.14 | 0.42 | 0.46 | 0.28 | 0.39 |
| 10 | 20132665 | rs4748613 | -0.532 | 0.735 | -1.125 | 0.963 | 0.002 | -0.008 | -0.008 | 0 | 0.35 | 0.29 | 0.56 | 0.63 | 0.35 | 0.41 |
| 10 | 20135727 | rs6482067 | -0.173 | 1.254 | -1.446 | 1.163 | -0.044 | -0.046 | -0.046 | 0 | 0.87 | 0.99 | 0.73 | 0.81 | 0.89 | 0.81 |
| 10 | 20137094 | rs1326243 | 0.121 | 0.863 | -1.561 | 0.947 | 0.006 | 0.007 | 0.007 | 0.578 | 0.08 | 0.06 | 0.08 | 0.05 | 0.07 | 0.03 |
| 10 | 20137692 | rs11594168 | 0.104 | 0.72 | -1.17 | 2.084 | 0.007 | 0 | 0 | 0 | 0.08 | 0.06 | 0.08 | 0.08 | 0.07 | 0.03 |
| 10 | 20137907 | rs1555872 | 0.018 | 0.946 | -0.479 | 2.067 | 0.002 | 0 | 0 | 1.744 | 0.86 | 0.85 | 0.85 | 0.84 | 0.88 | 0.81 |
| 10 | 20138208 | rs17677605 | 0.157 | 1.503 | -0.989 | 2.174 | -0.015 | -0.027 | -0.027 | 0 | 0.18 | 0.08 | 0.14 | 0.2 | 0.22 | 0.2 |
| 10 | 20139722 | rs2481940 | -1.432 | 1.441 | -1.042 | 1.866 | -0.039 | -0.031 | -0.031 | 0.118 | 0.92 | 1 | 0.94 | 0.87 | 0.91 | 0.86 |
| 10 | 20140808 | rs12766994 | -0.421 | 1.325 | -0.744 | 1.37 | -0.001 | 0.004 | 0.004 | 0 | 0.31 | 0.29 | 0.27 | 0.47 | 0.3 | 0.31 |
| 10 | 20141709 | rs10508593 | 1.311 | 1.205 | -0.408 | 1.336 | -0.003 | 0.002 | 0.002 | 3.495 | 0.06 | 0.2 | 0.04 | 0.03 | 0.02 | 0.01 |
| 10 | 20141731 | rs16919426 | -0.341 | 1.234 | -0.524 | 1.347 | -0.001 | 0.01 | 0.01 | 0 | 0.3 | 0.29 | 0.26 | 0.47 | 0.29 | 0.31 |
| 10 | 20142655 | rs1331148 | -0.329 | 1.261 | -0.567 | 1.342 | -0.002 | 0.027 | 0.027 | 0 | 0.27 | 0.29 | 0.24 | 0.46 | 0.27 | 0.29 |
| 10 | 20144397 | rs7080901 | -0.123 | 0.67 | -1.651 | 0.461 | 0.03 | 0.156 | 0.156 | 2.674 | 0.43 | 0.89 | 0.28 | 0.21 | 0.36 | 0.15 |
| 10 | 20147680 | rs10764170 | 0.071 | 1.009 | -1.624 | 0.304 | -0.01 | -0.02 | -0.02 | 0.077 | 0.31 | 0.26 | 0.23 | 0.56 | 0.37 | 0.29 |
| 10 | 20150639 | rs1326242 | -1.331 | 0.529 | -2.007 | 0.058 | -0.052 | 0.032 | 0.032 | 0 | 0.57 | 0.93 | 0.64 | 0.7 | 0.58 | 0.43 |
| 10 | 20150985 | rs12266731 | 1.239 | 0.607 | -2.035 | 0.154 | 0.003 | 0 | 0 | 0 | 0.12 | 0.19 | 0.29 | 0.2 | 0.16 | 0.1 |
| 10 | 20151213 | rs11011649 | -1.409 | 0.517 | -2.249 | 0.045 | -0.068 | -0.01 | -0.01 | 0 | 0.6 | 0.97 | 0.65 | 0.7 | 0.61 | 0.46 |
| 10 | 20152558 | rs12572398 | 1.081 | 0.308 | -1.786 | 0.191 | 0.006 | -0.022 | -0.022 | 0 | 0.12 | 0.01 | 0.07 | 0.12 | 0.08 | 0.12 |
| 10 | 20154249 | rs10508599 | 0.076 | 0.259 | -1.623 | -0.168 | 0.009 | 0.011 | 0.011 | 0 | 0.44 | 0.36 | 0.28 | 0.13 | 0.31 | 0.29 |
| 10 | 20159948 | rs4747386 | -0.729 | 0.585 | -1.573 | 0.301 | -0.032 | -0.023 | -0.023 | 0.197 | 0.76 | 0.8 | 0.73 | 0.53 | 0.63 | 0.7 |
| 10 | 20161663 | rs4748619 | 1.87 | 0.255 | -1.661 | 0.241 | 0.021 | -0.05 | -0.05 | 2.061 | 0.09 | 0.01 | 0.2 | 0.18 | 0.04 | 0.11 |
| 10 | 20163833 | rs752318 | 2.438 | 0.012 | -1.759 | 0.079 | -0.019 | 0.035 | 0.035 | 0.446 | 0.07 | 0 | 0.05 | 0.01 | 0.07 | 0.01 |
| 10 | 20167823 | rs10508600 | -0.227 | -0.681 | -1.646 | -0.272 | -0.018 | -0.022 | -0.022 | 0.6 | 0.35 | 0.44 | 0.36 | 0.21 | 0.22 | 0.16 |
| 10 | 20169529 | rs1475361 | -0.183 | -0.585 | -1.642 | -0.215 | 0.003 | -0.002 | -0.002 | 0 | 0.45 | 0.57 | 0.51 | 0.43 | 0.41 | 0.43 |
| 10 | 20170037 | rs12779437 | 0.926 | -0.58 | -1.609 | 0.066 | -0.021 | 0.041 | 0.041 | 1.986 | 0.23 | 0.31 | 0.23 | 0.39 | 0.19 | 0.3 |
| 10 | 20170754 | rs1830994 | 2.215 | -0.606 | -1.629 | 0.095 | 0.008 | -0.048 | -0.048 | 0 | 0.09 | 0.01 | 0.25 | 0.19 | 0.05 | 0.12 |
| 10 | 20171770 | rs7087684 | 0.517 | -0.254 | -1.656 | -0.038 | -0.008 | 0.023 | 0.023 | 1.465 | 0.28 | 0.32 | 0.23 | 0.4 | 0.25 | 0.31 |
| 10 | 20175790 | rs7097965 | 0.93 | -0.272 | -1.551 | 0.054 | -0.027 | 0.046 | 0.046 | 1.959 | 0.23 | 0.33 | 0.24 | 0.39 | 0.19 | 0.29 |
| 10 | 20176132 | rs10827894 | 0.108 | -0.246 | -1.624 | 0 | -0.029 | 0 | 0 | 0 | 0.4 | 0.46 | 0.49 | 0.45 | 0.3 | 0.37 |
| 10 | 20177944 | rs17679459 | 2.37 | 0.161 | -1.749 | 0.18 | -0.004 | -0.029 | -0.029 | 0 | 0.18 | 0.06 | 0.15 | 0.22 | 0.22 | 0.24 |
| 10 | 20178703 | rs10764175 | -1.431 | 0.775 | -0.968 | 0.698 | -0.062 | -0.009 | -0.009 | 0 | 0.86 | 0.94 | 0.79 | 0.84 | 0.75 | 0.77 |
| 10 | 20179223 | rs1467658 | -0.703 | 0.925 | -0.577 | 0.697 | 0.047 | -0.04 | -0.04 | 1.083 | 0.88 | 0.7 | 0.92 | 0.98 | 0.83 | 0.96 |
| 10 | 20179266 | rs12415168 | 1.447 | 0.745 | -0.876 | 0.935 | 0.011 | -0.042 | -0.042 | 5.33 | 0.1 | 0.01 | 0.22 | 0.19 | 0.06 | 0.13 |

| | | | | | | | | | | | | | | | | |
|---|---|---|---|---|---|---|---|---|---|---|---|---|---|---|---|---|
| 10 | 20179337 | rs1555871 | 0.644 | 0.774 | -0.957 | 0.698 | -0.047 | 0.043 | 0.043 | 0.078 | 0.25 | 0.09 | 0.24 | 0.17 | 0.26 | 0.3 |
| 10 | 20180184 | rs12572892 | 1.469 | 0.58 | -0.648 | 0.985 | 0.013 | -0.027 | -0.027 | 0 | 0.1 | 0.01 | 0.03 | 0.13 | 0.05 | 0.15 |
| 10 | 20180225 | rs11011661 | 1.415 | 0.911 | -0.414 | 1.15 | -0.019 | -0.02 | -0.02 | 0 | 0.05 | 0.01 | 0.1 | 0.1 | 0.07 | 0.15 |
| 10 | 20180660 | rs1467659 | -0.539 | 0.818 | -0.149 | 1.211 | 0.037 | 0.217 | 0.217 | 0.715 | 0.13 | 0.41 | 0.22 | 0.39 | 0.14 | 0.21 |
| 10 | 20187467 | rs7904103 | -0.669 | 0.985 | -0.193 | 1.005 | 0.014 | 0.022 | 0.022 | 0 | 0.89 | 0.84 | 0.65 | 0.84 | 0.85 | 0.87 |
| 10 | 20193797 | rs11592239 | 0.825 | 0.966 | -0.153 | 0.866 | 0.038 | -0.022 | -0.022 | 0 | 0.09 | 0.2 | 0.06 | 0.03 | 0.12 | 0.05 |
| 10 | 20198442 | rs17685390 | 2.034 | 0.78 | -0.664 | 0.931 | -0.007 | 0.011 | 0.011 | 1.919 | 0.28 | 0.05 | 0.18 | 0.18 | 0.29 | 0.33 |
| 10 | 20204247 | rs11011671 | 0.307 | 0.795 | -0.036 | 1.12 | 0.004 | 0.106 | 0.106 | 0.239 | 0.07 | 0.23 | 0.08 | 0.29 | 0.07 | 0.03 |
| 10 | 20204325 | rs1326235 | -0.671 | 0.794 | 0.048 | 1.077 | 0.045 | 0.15 | 0.15 | 1.522 | 0.1 | 0.55 | 0.12 | 0.28 | 0.1 | 0.06 |
| 10 | 20204628 | rs1326236 | -0.307 | 0.795 | 0.059 | 1.078 | 0.067 | 0.182 | 0.182 | 0.535 | 0.09 | 0.44 | 0.1 | 0.28 | 0.1 | 0.06 |
| 10 | 20204915 | rs16919551 | 0.064 | 1.361 | -0.079 | 1.176 | 0.07 | 0.071 | 0.071 | 0 | 0.09 | 0.26 | 0.2 | 0.13 | 0.16 | 0.15 |
| 10 | 20205463 | rs12778053 | 1.407 | 1.368 | -0.394 | 1.149 | 0.023 | 0.032 | 0.032 | 0 | 0.06 | 0.08 | 0.15 | 0.1 | 0.1 | 0.13 |
| 10 | 20207349 | rs2038912 | 0.683 | 1.112 | -0.757 | 0.899 | -0.008 | -0.01 | -0.01 | 0 | 0.58 | 0.18 | 0.4 | 0.53 | 0.56 | 0.74 |
| 10 | 20207400 | rs2038913 | -1.544 | 1.049 | -0.712 | 0.672 | 0.019 | 0.026 | 0.026 | 3.859 | 0.84 | 0.89 | 0.75 | 0.94 | 0.89 | 0.95 |
| 10 | 20208336 | rs11011674 | 1.452 | 1.231 | -0.666 | 0.8 | 0.033 | 0.048 | 0.048 | 0 | 0.06 | 0.1 | 0.15 | 0.1 | 0.1 | 0.14 |
| 10 | 20208483 | rs11596680 | 1.67 | 1.002 | -0.99 | 0.742 | 0.002 | 0.025 | 0.025 | 0.2 | 0.35 | 0.09 | 0.19 | 0.21 | 0.34 | 0.35 |
| 10 | 20208995 | rs1887035 | -0.64 | 0.874 | -0.973 | 0.641 | -0.019 | 0.028 | 0.028 | 0.789 | 0.72 | 0.84 | 0.57 | 0.77 | 0.71 | 0.71 |
| 10 | 20211425 | rs7075937 | 0.691 | 1.431 | -0.957 | 0.814 | -0.001 | -0.003 | -0.003 | 0 | 0.14 | 0.12 | 0.25 | 0.21 | 0.22 | 0.24 |
| 10 | 20216174 | rs12765810 | 1.692 | 0.938 | -1.109 | 0.754 | -0.013 | 0.018 | 0.018 | 1.45 | 0.33 | 0.05 | 0.2 | 0.18 | 0.31 | 0.36 |
| 10 | 20217660 | rs11011683 | 0.259 | 0.808 | -1.081 | 0.985 | -0.012 | -0.06 | -0.06 | 0 | 0.3 | 0.05 | 0.22 | 0.35 | 0.24 | 0.39 |
| 10 | 20222092 | rs10827906 | -0.39 | 0.997 | -0.037 | 1.095 | 0.043 | 0.175 | 0.175 | 0.223 | 0.17 | 0.42 | 0.22 | 0.41 | 0.21 | 0.17 |
| 10 | 20223198 | rs9651366 | 0.784 | 1.195 | 0.077 | 1.168 | 0.015 | 0.086 | 0.086 | 0.976 | 0.1 | 0.26 | 0.09 | 0.27 | 0.1 | 0.06 |
| 10 | 20223450 | rs9651367 | -0.638 | 1.083 | -0.094 | 1.123 | -0.036 | 0.028 | 0.028 | 2.18 | 0.72 | 0.88 | 0.58 | 0.78 | 0.71 | 0.74 |
| 10 | 20225200 | rs7081201 | -0.493 | 0.974 | 0.184 | 1.139 | 0.017 | 0.014 | 0.014 | 0 | 0.3 | 0.38 | 0.28 | 0.3 | 0.34 | 0.32 |
| 10 | 20225688 | rs2148300 | -1.187 | 0.959 | 0.198 | 1.15 | -0.016 | -0.022 | -0.022 | 0 | 0.39 | 0.66 | 0.37 | 0.58 | 0.44 | 0.39 |
| 10 | 20225914 | rs7897805 | 0.739 | 1.009 | 0.17 | 1.033 | 0.021 | 0.099 | 0.099 | 0.994 | 0.09 | 0.28 | 0.1 | 0.27 | 0.1 | 0.06 |
| 10 | 20229749 | rs6482071 | -1.623 | 1.042 | -0.311 | 0.827 | 0.001 | -0.003 | -0.003 | 0 | 0.94 | 0.9 | 0.77 | 0.95 | 0.93 | 0.97 |
| 10 | 20237290 | rs11011694 | -0.984 | 1.051 | -0.28 | 1.018 | -0.008 | -0.017 | -0.017 | 1.016 | 0.37 | 0.65 | 0.37 | 0.59 | 0.4 | 0.38 |
| 10 | 20243009 | rs11011698 | 1.528 | 1.013 | -0.079 | 1.155 | 0.003 | 0.022 | 0.022 | 0 | 0.1 | 0.15 | 0.12 | 0.14 | 0.11 | 0.12 |
| 10 | 20246814 | rs7909155 | -0.134 | 0.943 | 0.064 | 1.22 | -0.022 | 0.132 | 0.132 | 0.298 | 0.19 | 0.4 | 0.15 | 0.47 | 0.15 | 0.22 |
| 10 | 20250530 | rs6482072 | -0.042 | 1.05 | -0.323 | 0.9 | -0.005 | -0.006 | -0.006 | 0.687 | 0.65 | 0.73 | 0.5 | 0.51 | 0.64 | 0.65 |
| 10 | 20250984 | rs1409341 | 0.141 | 1.095 | -0.36 | 1.013 | 0 | 0.019 | 0.019 | 0 | 0.2 | 0.22 | 0.2 | 0.35 | 0.21 | 0.26 |
| 10 | 20253995 | rs4143249 | 0.868 | 1.056 | -0.381 | 1.017 | 0 | -0.041 | -0.041 | 0 | 0.17 | 0.08 | 0.16 | 0.33 | 0.15 | 0.25 |
| 10 | 20254725 | rs11011705 | -0.132 | 1.134 | -0.401 | 0.787 | -0.001 | -0.004 | -0.004 | 0 | 0.24 | 0.22 | 0.22 | 0.31 | 0.24 | 0.27 |
| 10 | 20256243 | rs10827912 | 0.574 | 1.152 | -0.38 | 0.714 | 0.011 | -0.004 | -0.004 | 0 | 0.75 | 0.37 | 0.58 | 0.68 | 0.7 | 0.88 |
| 10 | 20258993 | rs11011707 | 0.85 | 0.997 | -0.623 | 0.21 | -0.042 | 0.015 | 0.015 | 1.119 | 0.22 | 0.04 | 0.18 | 0.17 | 0.21 | 0.25 |
| 10 | 20259437 | rs17687258 | 2.216 | 1.051 | -0.59 | 0.232 | -0.027 | 0.034 | 0.034 | 0.681 | 0.07 | 0.01 | 0.05 | 0.01 | 0.09 | 0.02 |
| 10 | 20267705 | rs716829 | 2.176 | 0.968 | -0.598 | 0.224 | 0.026 | 0.013 | 0.013 | 2.006 | 0.32 | 0.16 | 0.19 | 0.19 | 0.28 | 0.34 |
| 10 | 20268754 | rs7075311 | 0.955 | 1.002 | -0.516 | 0.238 | -0.018 | 0.019 | 0.019 | 0.482 | 0.21 | 0.08 | 0.18 | 0.17 | 0.2 | 0.25 |
| 10 | 20269831 | rs10047272 | 1.153 | 1.123 | -0.432 | 0.36 | -0.002 | 0.01 | 0.01 | 0 | 0.1 | 0.17 | 0.09 | 0.17 | 0.11 | 0.15 |
| 10 | 20272455 | rs12263607 | 0.224 | 1.171 | -1.062 | 0.427 | 0.004 | 0.012 | 0.012 | 3.996 | 0.23 | 0.25 | 0.25 | 0.41 | 0.28 | 0.17 |
| 10 | 20272662 | rs7070003 | -0.328 | 1.015 | -1.126 | 0.487 | -0.002 | 0.021 | 0.021 | 2.845 | 0.72 | 0.83 | 0.58 | 0.77 | 0.73 | 0.72 |
| 10 | 20273583 | rs1111367 | 0.016 | 0.909 | -1.36 | 0.663 | 0.001 | 0.002 | 0.002 | 2.23 | 0.62 | 0.41 | 0.55 | 0.76 | 0.65 | 0.76 |
| 10 | 20275034 | rs7900457 | -0.197 | 1.365 | -1.086 | 1.34 | 0.004 | 0.03 | 0.03 | 0.69 | 0.16 | 0.17 | 0.19 | 0.4 | 0.2 | 0.17 |
| 10 | 20277083 | rs4311960 | 0.077 | 1.138 | -1.439 | 0.789 | -0.004 | -0.008 | -0.008 | 0 | 0.86 | 0.9 | 0.71 | 0.81 | 0.82 | 0.92 |
| 10 | 20278286 | rs10827919 | -0.469 | 1.252 | -1.593 | 0.766 | -0.006 | -0.026 | -0.026 | 0 | 0.3 | 0.25 | 0.33 | 0.51 | 0.35 | 0.22 |
| 10 | 20279125 | rs10764181 | 0.003 | 1.149 | -2.583 | -0.052 | -0.007 | 0.13 | 0.13 | 1.326 | 0.55 | 0.91 | 0.51 | 0.75 | 0.61 | 0.58 |
| 10 | 20282977 | rs12779340 | 1.819 | 1.244 | -2.537 | -0.02 | -0.017 | 0.042 | 0.042 | 1.31 | 0.08 | 0.01 | 0.06 | 0.01 | 0.1 | 0.04 |
| 10 | 20284949 | rs7079519 | 0.418 | 0.991 | -2.419 | -0.091 | -0.003 | 0.05 | 0.05 | 0 | 0.46 | 0.62 | 0.34 | 0.22 | 0.47 | 0.45 |
| 10 | 20288812 | rs1980820 | -0.193 | 1.7 | -2.005 | 0.208 | 0.004 | 0.043 | 0.043 | 0 | 0.23 | 0.3 | 0.28 | 0.52 | 0.25 | 0.2 |
| 10 | 20299308 | rs7912767 | -0.948 | 1.523 | -2.113 | 0.112 | -0.017 | -0.017 | -0.017 | 0.008 | 0.35 | 0.3 | 0.43 | 0.55 | 0.43 | 0.33 |
| 10 | 20301670 | rs7921016 | -0.369 | 1.646 | -2.571 | -0.251 | -0.018 | -0.008 | -0.008 | 0.097 | 0.41 | 0.31 | 0.42 | 0.59 | 0.5 | 0.31 |
| 10 | 20305643 | rs11011739 | 1.08 | 1.646 | -2.435 | 0.136 | -0.052 | -0.026 | -0.026 | 0 | 0.11 | 0.03 | 0.16 | 0.14 | 0.15 | 0.16 |
| 10 | 20305784 | rs10827926 | -0.512 | 1.566 | -2.094 | 0.09 | 0.009 | 0.022 | 0.022 | 0 | 0.29 | 0.34 | 0.37 | 0.55 | 0.33 | 0.3 |
| 10 | 20305855 | rs11011740 | 1.179 | 1.566 | -2.115 | 0.114 | 0.002 | 0.061 | 0.061 | 0 | 0.15 | 0.21 | 0.19 | 0.41 | 0.18 | 0.14 |
| 10 | 20305941 | rs10764184 | -0.357 | 1.132 | -2.166 | -0.308 | 0.005 | 0.005 | 0.005 | 0.299 | 0.38 | 0.43 | 0.44 | 0.56 | 0.44 | 0.34 |
| 10 | 20309308 | rs7077535 | 0.155 | 0.951 | -2.297 | -0.596 | -0.087 | 0.103 | 0.103 | 0 | 0.25 | 0.02 | 0.19 | 0.08 | 0.31 | 0.19 |
| 10 | 20311913 | rs4311961 | -0.092 | 0.031 | -1.67 | -0.598 | 0.001 | -0.001 | -0.001 | 0.123 | 0.76 | 0.82 | 0.69 | 0.87 | 0.83 | 0.73 |
| 10 | 20312209 | rs4636557 | 0.342 | -0.727 | -1.319 | -1.273 | 0.004 | -0.011 | -0.011 | 0.634 | 0.57 | 0.45 | 0.6 | 0.88 | 0.76 | 0.57 |
| 10 | 20313068 | rs4748630 | 0.501 | 0.514 | -1.51 | -0.704 | -0.048 | -0.04 | -0.04 | 0 | 0.09 | 0.03 | 0.37 | 0.3 | 0.21 | 0.34 |
| 10 | 20315373 | rs2358660 | 0.261 | 0.016 | -1.269 | -0.971 | -0.026 | -0.074 | -0.074 | 0 | 0.57 | 0.54 | 0.79 | 0.91 | 0.75 | 0.57 |
| 10 | 20316168 | rs10764186 | -0.33 | -0.336 | -1.466 | -1.165 | 0.047 | 0.115 | 0.115 | 6.03 | 0.72 | 0.93 | 0.87 | 0.91 | 0.82 | 0.67 |
| 10 | 20319192 | rs7907793 | 0.22 | -0.409 | -1.006 | -1.203 | -0.007 | -0.029 | -0.029 | 1.341 | 0.65 | 0.68 | 0.83 | 0.91 | 0.8 | 0.66 |
| 10 | 20319892 | rs10740960 | 0.334 | -0.425 | -0.943 | -1.21 | -0.042 | -0.09 | -0.09 | 0 | 0.64 | 0.53 | 0.81 | 0.91 | 0.79 | 0.65 |
| 10 | 20326159 | rs17688735 | 0.935 | -0.363 | -1.566 | -1.697 | -0.073 | 0.254 | 0.254 | 0 | 0.38 | 0.06 | 0.26 | 0 | 0.44 | 0.2 |
| 10 | 20328816 | rs4748633 | 0.534 | -0.832 | -1.416 | -1.539 | 0.063 | 0.049 | 0.049 | 5.329 | 0.23 | 0.09 | 0.27 | 0.14 | 0.1 | 0.31 |
| 10 | 20342316 | rs7923873 | -0.63 | -0.805 | -1.318 | -1.604 | -0.021 | -0.043 | -0.043 | 0 | 0.38 | 0.53 | 0.18 | 0.1 | 0.16 | 0.39 |
| 10 | 20349877 | rs10827940 | 0.241 | -0.709 | -1.421 | -1.651 | 0.037 | 0.048 | 0.048 | 4.888 | 0.63 | 0.72 | 0.82 | 0.91 | 0.78 | 0.59 |
| 10 | 20361929 | rs6482086 | -0.269 | -0.623 | -1.342 | -1.532 | 0.02 | -0.025 | -0.025 | 1.01 | 0.3 | 0.75 | 0.26 | 0.61 | 0.17 | 0.21 |

| | | | | | | | | | | | | | | | | |
|---|---|---|---|---|---|---|---|---|---|---|---|---|---|---|---|---|
| 10 | 20368998 | rs4336923 | -0.721 | -0.625 | -1.159 | -1.588 | -0.002 | -0.003 | -0.003 | 0.177 | 0.49 | 0.56 | 0.32 | 0.7 | 0.29 | 0.51 |
| 10 | 20374678 | rs7090101 | 1.346 | -0.595 | -1.21 | -1.514 | -0.017 | 0.178 | 0.178 | 1.64 | 0.17 | 0.43 | 0.19 | 0.61 | 0.11 | 0.12 |
| 10 | 20375875 | rs11011788 | -0.208 | -0.761 | -1.277 | -1.641 | -0.05 | -0.072 | -0.072 | 0 | 0.31 | 0.36 | 0.13 | 0.09 | 0.13 | 0.32 |
| 10 | 20377401 | rs11011789 | 1.301 | -0.687 | -1.255 | -1.6 | 0.003 | 0.167 | 0.167 | 1.841 | 0.17 | 0.43 | 0.2 | 0.62 | 0.14 | 0.13 |
| 10 | 20378035 | rs6482088 | -1.258 | -0.203 | -1.405 | -1.748 | -0.064 | 0.208 | 0.208 | 0 | 0.57 | 0.85 | 0.69 | 1 | 0.47 | 0.76 |
| 10 | 20378446 | rs10764200 | -1.708 | -0.207 | -1.438 | -1.762 | -0.134 | 0.274 | 0.274 | 0 | 0.64 | 0.98 | 0.73 | 1 | 0.52 | 0.84 |
| 10 | 20378627 | rs12264569 | 1.987 | -0.276 | -1.331 | -1.725 | 0.076 | 0.039 | 0.039 | 0.91 | 0.05 | 0.29 | 0.04 | 0 | 0.02 | 0.01 |
| 10 | 20379812 | rs7101197 | -1.164 | -0.26 | -1.193 | -1.742 | -0.016 | 0.037 | 0.037 | 0 | 0.56 | 0.7 | 0.68 | 1 | 0.47 | 0.76 |
| 10 | 20380428 | rs10827963 | -0.684 | -0.407 | -1.219 | -1.601 | 0.003 | -0.007 | -0.007 | 0.055 | 0.39 | 0.27 | 0.49 | 0.39 | 0.34 | 0.64 |
| 10 | 20389236 | rs7920683 | -1.078 | -0.519 | -1.331 | -1.645 | 0.004 | 0.003 | 0.003 | 0 | 0.48 | 0.4 | 0.53 | 0.4 | 0.4 | 0.74 |
| 10 | 20390641 | rs2358839 | -0.422 | -0.589 | -1.209 | -1.399 | 0.001 | -0.03 | -0.03 | 0 | 0.34 | 0.21 | 0.49 | 0.6 | 0.32 | 0.65 |
| 10 | 20396171 | rs16919862 | 0.302 | -0.731 | -1.184 | -1.498 | -0.017 | -0.046 | -0.046 | 0.346 | 0.11 | 0.14 | 0.05 | 0 | 0.05 | 0.09 |
| 10 | 20397548 | rs2255452 | 1.561 | -0.906 | -1.432 | -1.708 | -0.161 | 0.238 | 0.238 | 1.027 | 0.33 | 0.03 | 0.27 | 0 | 0.49 | 0.14 |
| 10 | 20399179 | rs2461943 | -2.07 | -1.041 | -1.443 | -1.701 | 0.023 | 0.013 | 0.013 | 1.052 | 0.87 | 0.92 | 0.94 | 0.91 | 0.94 | 0.78 |
| 10 | 20405138 | rs2461941 | 1.36 | -0.984 | -1.326 | -1.728 | -0.061 | 0.094 | 0.094 | 0.028 | 0.34 | 0.17 | 0.29 | 0 | 0.5 | 0.14 |
| 10 | 20405650 | rs11011808 | 1.298 | -1.085 | -1.312 | -1.756 | -0.003 | -0.017 | -0.017 | 0 | 0.08 | 0.09 | 0.03 | 0 | 0.04 | 0.08 |
| 10 | 20407180 | rs10458694 | -0.272 | -1.034 | -1.367 | -1.669 | 0.044 | -0.009 | -0.009 | 0 | 0.38 | 0.69 | 0.24 | 0.4 | 0.18 | 0.29 |
| 10 | 20407490 | rs2461935 | 0.533 | -0.829 | -1.232 | -1.512 | 0.007 | -0.003 | -0.003 | 0 | 0.6 | 0.73 | 0.49 | 0.4 | 0.65 | 0.33 |
| 10 | 20416814 | rs11812627 | 1.407 | -0.836 | -1.474 | -1.358 | -0.041 | 0.17 | 0.17 | 1.691 | 0.2 | 0.49 | 0.17 | 0.4 | 0.12 | 0.12 |
| 10 | 20419343 | rs4412661 | 1.069 | -0.819 | -1.668 | -1.359 | -0.021 | 0.07 | 0.07 | 0.549 | 0.22 | 0.62 | 0.19 | 0.39 | 0.14 | 0.12 |
| 10 | 20419531 | rs2461926 | 0.683 | -0.511 | -1.483 | -1.163 | 0.008 | -0.004 | -0.004 | 0 | 0.61 | 0.69 | 0.48 | 0.4 | 0.66 | 0.34 |
| 10 | 20428197 | rs4237353 | 0.063 | -0.297 | -1.52 | -1.029 | -0.021 | -0.036 | -0.036 | 0 | 0.18 | 0.13 | 0.42 | 0.5 | 0.27 | 0.36 |
| 10 | 20431361 | rs2460594 | -1.372 | -0.463 | -1.466 | -1.101 | -0.032 | -0.02 | -0.02 | 1.168 | 0.85 | 0.79 | 0.93 | 0.91 | 0.94 | 0.77 |
| 10 | 20431639 | rs2461920 | 0.771 | -0.528 | -1.303 | -1.178 | -0.002 | 0.001 | 0.001 | 0 | 0.61 | 0.58 | 0.47 | 0.4 | 0.64 | 0.33 |
| 10 | 20437311 | rs7097590 | -1.598 | -0.582 | -1.469 | -1.145 | -0.017 | -0.01 | -0.01 | 0.088 | 0.87 | 0.86 | 0.94 | 0.91 | 0.94 | 0.77 |
| 10 | 20443553 | rs7896864 | 0.55 | -0.718 | -1.635 | -1.247 | 0.007 | 0 | 0 | 0 | 0.6 | 0.69 | 0.5 | 0.38 | 0.65 | 0.33 |
| 10 | 20445601 | rs4518993 | -0.624 | -0.764 | -1.687 | -1.253 | -0.042 | -0.108 | -0.108 | 0 | 0.27 | 0.14 | 0.45 | 0.53 | 0.31 | 0.45 |
| 10 | 20447821 | rs7086733 | -1.657 | -0.767 | -1.643 | -1.295 | 0.002 | 0.002 | 0.002 | 1.009 | 0.87 | 0.91 | 0.94 | 0.91 | 0.94 | 0.78 |
| 10 | 20449266 | rs7074575 | 1.078 | -0.836 | -1.681 | -1.326 | 0.015 | 0.022 | 0.022 | 0.47 | 0.53 | 0.71 | 0.47 | 0.38 | 0.6 | 0.25 |
| 10 | 20450560 | rs2884566 | -2.06 | -0.798 | -1.632 | -1.372 | -0.009 | -0.004 | -0.004 | 0 | 0.89 | 0.89 | 0.94 | 0.91 | 0.94 | 0.78 |
| 10 | 20451608 | rs10827977 | -0.6 | -0.845 | -1.559 | -1.395 | -0.009 | -0.011 | -0.011 | 0.071 | 0.8 | 0.8 | 0.9 | 0.91 | 0.9 | 0.7 |
| 10 | 20457389 | rs2358859 | 1.341 | -0.912 | -1.562 | -1.654 | -0.035 | 0.04 | 0.04 | 0 | 0.34 | 0.25 | 0.3 | 0 | 0.47 | 0.14 |
| 10 | 20457686 | rs4748640 | -1.072 | -0.76 | -1.418 | -1.52 | -0.01 | -0.012 | -0.012 | 0 | 0.82 | 0.83 | 0.91 | 0.91 | 0.91 | 0.7 |
| 10 | 20464904 | rs11593311 | 0.404 | -0.715 | -1.433 | -1.452 | -0.063 | -0.095 | -0.095 | 0.021 | 0.18 | 0.07 | 0.42 | 0.51 | 0.29 | 0.36 |
| 10 | 20467173 | rs11011841 | 1.109 | -0.613 | -1.287 | -1.371 | -0.039 | 0.153 | 0.153 | 2.8 | 0.21 | 0.47 | 0.18 | 0.39 | 0.15 | 0.13 |
| 10 | 20476083 | rs2358873 | 1.387 | -0.602 | -1.396 | -1.372 | -0.026 | 0.17 | 0.17 | 1.268 | 0.19 | 0.4 | 0.18 | 0.4 | 0.15 | 0.13 |
| 10 | 20481262 | rs10827993 | 0.593 | -0.051 | -1.641 | -1.566 | 0.001 | 0.002 | 0.002 | 0 | 0.49 | 0.52 | 0.36 | 0.4 | 0.49 | 0.25 |
| 10 | 20493901 | rs7895512 | -0.373 | -0.12 | -1.837 | -1.619 | -0.102 | -0.101 | -0.101 | 0 | 0.37 | 0.12 | 0.56 | 0.51 | 0.45 | 0.53 |
| 10 | 20495893 | rs11011869 | 1.358 | 0 | -1.889 | -1.718 | -0.093 | 0.198 | 0.198 | 0 | 0.28 | 0.02 | 0.17 | 0 | 0.36 | 0.13 |
| 10 | 20498082 | rs11011872 | 0.457 | 0.398 | -1.618 | -1.537 | 0.002 | -0.006 | -0.006 | 0 | 0.4 | 0.65 | 0.33 | 0.42 | 0.41 | 0.23 |
| 10 | 20500592 | rs16920092 | 1.505 | 0.363 | -1.542 | -1.526 | -0.034 | 0.155 | 0.155 | 2.209 | 0.2 | 0.43 | 0.18 | 0.42 | 0.14 | 0.13 |
| 10 | 20500710 | rs12259741 | 0.481 | 0.359 | -1.775 | -1.523 | 0.003 | -0.005 | -0.005 | 0.041 | 0.39 | 0.65 | 0.32 | 0.42 | 0.38 | 0.22 |
| 10 | 20500743 | rs10827997 | 0.94 | 0.602 | -1.589 | -1.255 | -0.081 | -0.09 | -0.09 | 0.137 | 0.12 | 0.03 | 0.39 | 0.53 | 0.27 | 0.35 |
| 10 | 20501152 | rs10508619 | 0.902 | 0.407 | -1.458 | -1.159 | -0.042 | 0.162 | 0.162 | 0 | 0.24 | 0.03 | 0.17 | 0.01 | 0.23 | 0.13 |
| 10 | 20501730 | rs12572697 | 0.791 | 0.675 | -0.646 | -0.189 | -0.033 | -0.018 | -0.018 | 0 | 0.06 | 0.02 | 0.1 | 0.32 | 0.17 | 0.14 |
| 10 | 20501874 | rs2358866 | 0.285 | -0.706 | -0.31 | -0.41 | 0.058 | -0.062 | -0.062 | 0.012 | 0.67 | 0.23 | 0.75 | 0.54 | 0.82 | 0.67 |
| 10 | 20503946 | rs11011876 | -0.086 | 0.028 | -0.076 | 0.072 | -0.038 | 0.116 | 0.116 | 2.925 | 0.21 | 0.61 | 0.19 | 0.42 | 0.12 | 0.13 |
| 10 | 20504157 | rs16920113 | 0.192 | 0.24 | -0.483 | 0.378 | -0.026 | 0.115 | 0.115 | 0.731 | 0.18 | 0.3 | 0.16 | 0.42 | 0.12 | 0.13 |
| 10 | 20505081 | rs10764214 | -0.126 | 0.749 | -0.719 | -0.334 | 0 | -0.023 | -0.023 | 0.798 | 0.61 | 0.61 | 0.74 | 0.91 | 0.67 | 0.6 |
| 10 | 20516326 | rs11011890 | -0.417 | 2.274 | 0.146 | 1.305 | 0.054 | 0.099 | 0.099 | 0 | 0.15 | 0.26 | 0.24 | 0.25 | 0.19 | 0.19 |
| 10 | 20516738 | rs7085600 | 0.683 | 2.3 | -0.119 | 0.817 | 0.036 | -0.007 | -0.007 | 0 | 0.66 | 0.6 | 0.36 | 0.31 | 0.56 | 0.54 |
| 10 | 20516978 | rs7074175 | -0.857 | 2.294 | -0.115 | 0.788 | 0.018 | 0.007 | 0.007 | 0 | 0.29 | 0.3 | 0.31 | 0.25 | 0.34 | 0.23 |
| 10 | 20518371 | rs16920135 | 0.207 | 2.091 | 0.351 | 0.724 | 0.007 | -0.027 | -0.027 | 0 | 0.09 | 0.24 | 0.09 | 0.04 | 0.09 | 0.1 |
| 10 | 20521825 | rs2249520 | 0.57 | 1.996 | 0.319 | 0.728 | 0.024 | 0.123 | 0.123 | 0 | 0.85 | 0.67 | 0.58 | 0.52 | 0.8 | 0.67 |
| 10 | 20523233 | rs2681936 | -0.483 | 2.059 | 0.208 | 0.813 | 0.024 | 0.143 | 0.143 | 0 | 0.95 | 0.9 | 0.67 | 0.56 | 0.89 | 0.79 |
| 10 | 20538901 | rs11011905 | -0.031 | 2.121 | 0.698 | 0.784 | 0.04 | 0.104 | 0.104 | 0 | 0.16 | 0.32 | 0.24 | 0.25 | 0.19 | 0.19 |
| 10 | 20565765 | rs12355139 | 0.54 | 1.558 | 0.11 | 0.203 | 0.029 | 0.131 | 0.131 | 0 | 0.29 | 0.08 | 0.12 | 0.01 | 0.22 | 0.22 |
| 10 | 20569460 | rs2778958 | 0.21 | 1.982 | 0.926 | 1.064 | 0.034 | 0.151 | 0.151 | 0.014 | 0.94 | 0.85 | 0.67 | 0.56 | 0.88 | 0.76 |
| 10 | 20575991 | rs1106674 | -1.238 | 2.056 | 1.133 | 0.998 | 0.022 | 0.07 | 0.07 | 0 | 0.17 | 0.28 | 0.42 | 0.48 | 0.22 | 0.33 |
| 10 | 20577412 | rs2297490 | -0.072 | 2.083 | 1.301 | 1.086 | -0.003 | 0.022 | 0.022 | 0 | 0.91 | 0.85 | 0.96 | 0.84 | 0.94 | 0.93 |
| 10 | 20578530 | rs955429 | -0.222 | 2.081 | 1.24 | 1.098 | 0.051 | 0.186 | 0.186 | 0 | 0.06 | 0.17 | 0.33 | 0.44 | 0.12 | 0.24 |
| 10 | 20586743 | rs10740971 | -0.243 | 1.845 | 1.109 | 0.99 | 0.002 | -0.005 | -0.005 | 0.901 | 0.92 | 0.9 | 0.96 | 0.85 | 0.94 | 0.94 |
| 10 | 20593087 | rs12219125 | 0.192 | 1.988 | 1.269 | 1.128 | 0.071 | 0.045 | 0.045 | 0 | 0.07 | 0.15 | 0.16 | 0.09 | 0.14 | 0.05 |
| 10 | 20595489 | rs985477 | 1.121 | 1.733 | 1.281 | 1.107 | 0.017 | 0.075 | 0.075 | 0 | 0.73 | 0.48 | 0.52 | 0.36 | 0.72 | 0.59 |
| 10 | 20599021 | rs987548 | 0.456 | 1.028 | 1.137 | 1.256 | -0.01 | 0.111 | 0.111 | 0 | 0.9 | 0.66 | 0.92 | 0.82 | 0.95 | 0.92 |
| 10 | 20604377 | rs11011957 | 0.698 | 1.061 | 0.407 | 1.1 | -0.007 | -0.027 | -0.027 | 0 | 0.07 | 0 | 0.03 | 0.06 | 0.05 | 0.09 |
| 10 | 20604722 | rs12218734 | -0.407 | 1.323 | 0.866 | 1.021 | 0.079 | 0.054 | 0.054 | 0 | 0.07 | 0.19 | 0.16 | 0.09 | 0.15 | 0.06 |
| 10 | 20611613 | rs1888979 | 0.563 | 0.502 | 0.367 | 0.865 | -0.016 | 0.058 | 0.058 | 0 | 0.9 | 0.73 | 0.94 | 0.84 | 0.95 | 0.92 |
| 10 | 20612660 | rs6482112 | 0.639 | 0.482 | 0.327 | 0.822 | -0.021 | 0.057 | 0.057 | 0 | 0.89 | 0.7 | 0.95 | 0.85 | 0.95 | 0.92 |

| | | | | | | | | | | | | | | | | |
|---|---|---|---|---|---|---|---|---|---|---|---|---|---|---|---|---|
| 10 | 20622826 | rs12781696 | -0.086 | 1.049 | -0.568 | 0.857 | -0.033 | -0.054 | -0.054 | 0.18 | 0.33 | 0.2 | 0.45 | 0.48 | 0.38 | 0.37 |
| 10 | 20623564 | rs7098524 | -0.502 | 1.014 | -0.551 | 0.667 | 0.009 | 0.024 | 0.024 | 0 | 0.46 | 0.38 | 0.65 | 0.67 | 0.57 | 0.56 |
| 10 | 20629321 | rs12777951 | 0.337 | 1.009 | -0.528 | 0.855 | -0.034 | -0.054 | -0.054 | 1.147 | 0.12 | 0.02 | 0.16 | 0.24 | 0.16 | 0.14 |
| 10 | 20631413 | rs7070298 | -0.481 | 0.59 | 0.374 | 0.667 | 0.004 | 0.038 | 0.038 | 0 | 0.16 | 0.38 | 0.2 | 0.24 | 0.17 | 0.15 |
| 10 | 20637499 | rs10828029 | 0.306 | 0.356 | 0.258 | 0.671 | -0.009 | -0.053 | -0.053 | 0 | 0.12 | 0.02 | 0.14 | 0.24 | 0.11 | 0.17 |
| 10 | 20638885 | rs7081455 | -0.815 | 0.324 | 0.255 | 0.606 | 0.028 | 0.11 | 0.11 | 0.003 | 0.51 | 0.75 | 0.68 | 0.8 | 0.57 | 0.6 |
| 10 | 20639359 | rs4518994 | 0.348 | 0.316 | 0.294 | 0.626 | -0.005 | -0.004 | -0.004 | 0.475 | 0.35 | 0.33 | 0.5 | 0.56 | 0.46 | 0.43 |
| 10 | 20642974 | rs11011992 | 0.525 | 0.137 | 0.466 | 0.423 | -0.012 | -0.01 | -0.01 | 0 | 0.34 | 0.27 | 0.53 | 0.53 | 0.42 | 0.37 |
| 10 | 20644506 | rs7090902 | 0.634 | 0.138 | 0.343 | 0.419 | -0.001 | 0.002 | 0.002 | 0 | 0.11 | 0.02 | 0.14 | 0.1 | 0.08 | 0.12 |
| 10 | 20644979 | rs4501895 | 0.623 | 0.155 | 0.425 | 0.439 | -0.024 | -0.02 | -0.02 | 0 | 0.33 | 0.23 | 0.53 | 0.53 | 0.42 | 0.38 |
| 10 | 20646220 | rs4748669 | -0.236 | -0.603 | 0.09 | -0.226 | -0.002 | 0.135 | 0.135 | 0 | 0.48 | 0.25 | 0.7 | 0.75 | 0.52 | 0.49 |
| 10 | 20647896 | rs4237357 | 1.175 | -0.799 | -0.223 | -0.235 | -0.096 | -0.05 | -0.05 | 0 | 0.3 | 0.08 | 0.58 | 0.61 | 0.41 | 0.35 |
| 10 | 20648238 | rs944697 | -0.549 | -0.894 | 0.248 | -0.235 | 0.003 | 0.004 | 0.004 | 0 | 0.5 | 0.52 | 0.73 | 0.83 | 0.57 | 0.52 |
| 10 | 20648302 | rs1154689 | -0.333 | -0.87 | 0.265 | -0.27 | 0.001 | 0.013 | 0.013 | 0.023 | 0.48 | 0.5 | 0.69 | 0.65 | 0.52 | 0.45 |
| 10 | 20648411 | rs883580 | 1.028 | -0.849 | 0.226 | -0.262 | 0.009 | 0.04 | 0.04 | 0 | 0.11 | 0.04 | 0.07 | 0.04 | 0.09 | 0.1 |
| 10 | 20650216 | rs12359855 | 0.972 | -1.317 | -0.344 | -0.817 | -0.054 | 0.13 | 0.13 | 0 | 0.2 | 0.03 | 0.12 | 0.01 | 0.23 | 0.21 |
| 10 | 20650878 | rs1332553 | 0.102 | -1.025 | 0.087 | -0.568 | 0.001 | 0.015 | 0.015 | 0.033 | 0.49 | 0.5 | 0.69 | 0.65 | 0.52 | 0.46 |
| 10 | 20658827 | rs10764228 | -0.227 | -1.056 | 0.093 | -0.584 | 0.004 | 0.03 | 0.03 | 0 | 0.54 | 0.7 | 0.76 | 0.86 | 0.58 | 0.57 |
| 10 | 20661390 | rs10740972 | -0.082 | -0.995 | -0.057 | -0.561 | 0 | 0.015 | 0.015 | 0 | 0.54 | 0.43 | 0.7 | 0.67 | 0.53 | 0.53 |
| 10 | 20665695 | rs10828037 | 0.139 | -1.032 | -0.345 | -0.404 | 0.028 | 0.103 | 0.103 | 0 | 0.5 | 0.28 | 0.57 | 0.69 | 0.56 | 0.56 |
| 10 | 20666974 | rs11012011 | 1.27 | -1.042 | -0.441 | -0.234 | -0.094 | -0.083 | -0.083 | 0 | 0.34 | 0.08 | 0.44 | 0.5 | 0.42 | 0.4 |
| 10 | 20668209 | rs12218939 | 0.171 | -0.991 | -0.415 | -0.181 | 0.015 | 0.045 | 0.045 | 0.221 | 0.49 | 0.26 | 0.53 | 0.6 | 0.52 | 0.52 |
| 10 | 20669511 | rs7923550 | 0.842 | -1.012 | -0.415 | -0.539 | 0.012 | 0.027 | 0.027 | 0.003 | 0.13 | 0.09 | 0.07 | 0.03 | 0.1 | 0.11 |
| 10 | 20670316 | rs17709943 | 1.065 | -1.003 | -0.466 | -0.536 | 0.01 | 0.042 | 0.042 | 0 | 0.12 | 0.05 | 0.06 | 0.03 | 0.1 | 0.11 |
| 10 | 20672656 | rs10828042 | -0.863 | -1.291 | -0.295 | -0.772 | 0.01 | 0.034 | 0.034 | 0 | 0.72 | 0.83 | 0.8 | 0.88 | 0.72 | 0.7 |
| 10 | 20673914 | rs7917108 | -0.269 | -0.918 | -0.407 | -0.546 | 0.005 | 0.015 | 0.015 | 0 | 0.54 | 0.44 | 0.68 | 0.87 | 0.59 | 0.58 |
| 10 | 20674653 | rs12772794 | 1.154 | -0.93 | -0.449 | -0.451 | -0.09 | -0.049 | -0.049 | 0 | 0.34 | 0.08 | 0.42 | 0.56 | 0.45 | 0.44 |
| 10 | 20676808 | rs1332551 | -0.234 | -0.859 | -0.368 | -0.414 | 0.001 | -0.008 | -0.008 | 0 | 0.54 | 0.45 | 0.68 | 0.86 | 0.58 | 0.58 |
| 10 | 20676986 | rs1332550 | -0.043 | -0.851 | -0.474 | -0.416 | 0.022 | 0.059 | 0.059 | 0 | 0.53 | 0.31 | 0.52 | 0.67 | 0.59 | 0.58 |
| 10 | 20677661 | rs1412563 | 0.384 | -0.44 | -0.311 | -0.599 | -0.003 | -0.004 | -0.004 | 0 | 0.4 | 0.4 | 0.6 | 0.83 | 0.47 | 0.47 |
| 10 | 20679473 | rs1984058 | 0.266 | -0.078 | -0.421 | -0.378 | -0.008 | 0.046 | 0.046 | 0 | 0.18 | 0.08 | 0.08 | 0.1 | 0.15 | 0.2 |
| 10 | 20679495 | rs11012022 | -0.289 | -0.043 | -0.51 | -0.219 | -0.022 | -0.004 | -0.004 | 0 | 0.22 | 0.1 | 0.15 | 0.25 | 0.23 | 0.21 |
| 10 | 20684324 | rs7918915 | -0.179 | -0.249 | 0.43 | -0.279 | -0.015 | 0.001 | 0.001 | 0 | 0.21 | 0.41 | 0.13 | 0.24 | 0.15 | 0.2 |
| 10 | 20697852 | rs1339658 | -0.905 | -0.148 | 0.202 | -0.33 | 0.01 | 0.003 | 0.003 | 1.003 | 0.83 | 0.88 | 0.7 | 0.86 | 0.86 | 0.91 |
| 10 | 20698281 | rs12570227 | 0.718 | -0.149 | 0.258 | -0.291 | -0.001 | -0.013 | -0.013 | 0.242 | 0.18 | 0.11 | 0.1 | 0.25 | 0.15 | 0.21 |
| 10 | 20698518 | rs10508623 | 0.77 | -0.147 | 0.255 | -0.306 | -0.004 | 0.037 | 0.037 | 0 | 0.18 | 0.09 | 0.08 | 0.1 | 0.15 | 0.2 |
| 10 | 20701000 | rs3925284 | 1.949 | -0.189 | 0.448 | -0.446 | -0.034 | -0.045 | -0.045 | 0 | 0.38 | 0.15 | 0.35 | 0.51 | 0.44 | 0.55 |
| 10 | 20707062 | rs16920492 | 1.114 | -0.09 | 0.391 | -0.501 | -0.001 | 0.032 | 0.032 | 0 | 0.18 | 0.11 | 0.08 | 0.07 | 0.15 | 0.2 |
| 10 | 20717789 | rs10508626 | 0.747 | -0.133 | 0.606 | -0.606 | 0.002 | -0.004 | -0.004 | 0.558 | 0.11 | 0.16 | 0.09 | 0.11 | 0.09 | 0.09 |
| 10 | 20720261 | rs7085871 | 0.135 | -0.102 | 0.763 | -0.63 | -0.004 | -0.01 | -0.01 | 0 | 0.23 | 0.5 | 0.14 | 0.22 | 0.18 | 0.21 |
| 10 | 20722097 | rs11012039 | 0.858 | -0.022 | 0.406 | -0.639 | 0.003 | -0.039 | -0.039 | 0 | 0.1 | 0.16 | 0.11 | 0.02 | 0.14 | 0.05 |
| 10 | 20740273 | rs6482121 | -1.322 | 0.017 | 0.294 | -0.626 | 0.001 | -0.002 | -0.002 | 0.291 | 0.84 | 0.89 | 0.67 | 0.84 | 0.83 | 0.9 |
| 10 | 20743148 | rs10828054 | -0.923 | -0.126 | 0.391 | -0.553 | -0.005 | 0.008 | 0.008 | 0.115 | 0.39 | 0.3 | 0.28 | 0.37 | 0.38 | 0.35 |
| 10 | 20746200 | rs12769394 | 0.978 | -0.124 | 0.259 | -0.567 | -0.032 | 0.002 | 0.002 | 0 | 0.1 | 0.01 | 0.06 | 0.07 | 0.09 | 0.09 |
| 10 | 20747422 | rs10828055 | -0.216 | -0.046 | 0.362 | -0.557 | 0 | 0 | 0 | 0 | 0.27 | 0.25 | 0.2 | 0.25 | 0.28 | 0.25 |
| 10 | 20761187 | rs12243166 | 0.572 | 0.003 | 0.569 | -0.483 | 0.005 | 0.005 | 0.005 | 0.055 | 0.12 | 0.19 | 0.09 | 0.13 | 0.08 | 0.07 |
| 10 | 20761804 | rs6482125 | -0.699 | 0.092 | 0.709 | -0.48 | -0.001 | -0.035 | -0.035 | 0.014 | 0.32 | 0.48 | 0.23 | 0.24 | 0.33 | 0.26 |
| 10 | 20762314 | rs11012067 | 0.076 | 0.09 | 0.685 | -0.506 | -0.002 | -0.002 | -0.002 | 0 | 0.23 | 0.27 | 0.15 | 0.24 | 0.21 | 0.24 |
| 10 | 20764149 | rs11012071 | 0.269 | 0.133 | 0.445 | -0.489 | 0.006 | -0.043 | -0.043 | 0 | 0.08 | 0.16 | 0.08 | 0 | 0.12 | 0.02 |
| 10 | 20764339 | rs7100193 | -0.66 | 0.169 | 0.004 | -0.457 | 0 | 0.001 | 0.001 | 0.703 | 0.34 | 0.65 | 0.25 | 0.24 | 0.33 | 0.26 |
| 10 | 20766268 | rs5025968 | 0.145 | 0.399 | 0.582 | -0.024 | 0 | 0 | 0 | 0 | 0.73 | 0.75 | 0.61 | 0.75 | 0.79 | 0.82 |
| 10 | 20768098 | rs11012077 | 0.608 | 0.784 | 0.548 | 0.117 | 0.027 | -0.012 | -0.012 | 0 | 0.06 | 0.18 | 0.05 | 0.02 | 0.07 | 0.04 |
| 10 | 20774281 | rs1416802 | 1.021 | 0.823 | 0.418 | 0.306 | 0.011 | 0.083 | 0.083 | 0.592 | 0.8 | 0.38 | 0.57 | 0.62 | 0.77 | 0.78 |
| 10 | 20775973 | rs7911952 | 0.088 | 0.465 | -0.47 | -0.624 | 0.003 | 0.092 | 0.092 | 0 | 0.29 | 0.11 | 0.16 | 0.06 | 0.23 | 0.25 |
| 10 | 20778051 | rs12355916 | -0.225 | 0.966 | 0.548 | -0.614 | 0.009 | -0.054 | -0.054 | 0 | 0.13 | 0.25 | 0.1 | 0.02 | 0.16 | 0.08 |
| 10 | 20779606 | rs1339661 | -0.084 | 1.05 | 0.553 | -0.539 | 0.01 | -0.049 | -0.049 | 0 | 0.12 | 0.22 | 0.09 | 0.02 | 0.16 | 0.08 |
| 10 | 20783841 | rs10828065 | -1.524 | 1.067 | 0.758 | -0.527 | 0.029 | 0 | 0 | 0 | 0.21 | 0.48 | 0.21 | 0.21 | 0.25 | 0.15 |
| 10 | 20792026 | rs11012097 | -0.217 | 0.956 | 0.787 | -0.55 | 0.019 | -0.062 | -0.062 | 0 | 0.12 | 0.32 | 0.12 | 0.03 | 0.14 | 0.07 |
| 10 | 20795467 | rs12245757 | -0.828 | 1.033 | 0.587 | -0.406 | 0.035 | 0.055 | 0.055 | 0 | 0.08 | 0.25 | 0.13 | 0.11 | 0.1 | 0.07 |
| 10 | 20795797 | rs1361360 | 0.955 | 0.547 | 0.031 | -0.739 | -0.054 | -0.064 | -0.064 | 0 | 0.27 | 0.14 | 0.31 | 0.52 | 0.42 | 0.4 |
| 10 | 20796639 | rs16920576 | 0.446 | 0.121 | 0.147 | -1.076 | 0.029 | 0.065 | 0.065 | 0 | 0.05 | 0.15 | 0.1 | 0.11 | 0.08 | 0.07 |
| 10 | 20807211 | rs10764242 | -0.433 | 0.015 | -0.168 | -1.6 | 0.004 | -0.023 | -0.023 | 0 | 0.56 | 0.61 | 0.29 | 0.1 | 0.39 | 0.36 |
| 10 | 20810525 | rs16920624 | 0.789 | -0.765 | -0.295 | -1.675 | -0.004 | -0.032 | -0.032 | 0 | 0.24 | 0.21 | 0.06 | 0 | 0.07 | 0.13 |
| 10 | 20820829 | rs11012119 | 0.299 | -0.842 | -0.076 | -1.643 | -0.103 | -0.075 | -0.075 | 0.422 | 0.28 | 0.45 | 0.1 | 0.11 | 0.1 | 0.14 |
| 10 | 20822253 | rs2496621 | -1.201 | -0.507 | -0.167 | -1.727 | -0.018 | 0.041 | 0.041 | 0 | 0.6 | 0.77 | 0.42 | 0.28 | 0.44 | 0.54 |
| 10 | 20822606 | rs1339666 | 0.346 | -0.437 | 0.048 | -1.585 | 0.002 | -0.001 | -0.001 | 0 | 0.07 | 0.09 | 0.15 | 0.06 | 0.12 | 0.16 |
| 10 | 20824223 | rs2496622 | -0.724 | -0.584 | 0.058 | -1.725 | 0.029 | 0.033 | 0.033 | 0.338 | 0.41 | 0.73 | 0.32 | 0.27 | 0.29 | 0.4 |
| 10 | 20827105 | rs953093 | 0.489 | -0.6 | 0.102 | -1.732 | -0.096 | -0.159 | -0.159 | 0 | 0.27 | 0.4 | 0.1 | 0.01 | 0.1 | 0.14 |
| 10 | 20836426 | rs2066124 | -0.872 | -0.606 | 0.003 | -1.73 | 0.035 | 0.038 | 0.038 | 0.955 | 0.42 | 0.74 | 0.33 | 0.27 | 0.29 | 0.4 |

| | | | | | | | | | | | | | | | | |
|---|---|---|---|---|---|---|---|---|---|---|---|---|---|---|---|---|
| 10 | 20836995 | rs11012133 | -0.649 | -0.746 | 0.031 | -1.703 | 0.01 | 0.073 | 0.073 | 0.028 | 0.11 | 0.3 | 0.11 | 0.21 | 0.1 | 0.11 |
| 10 | 20842031 | rs7081156 | 0.713 | -0.739 | -0.097 | -1.715 | -0.072 | -0.113 | -0.113 | 0 | 0.25 | 0.34 | 0.07 | 0 | 0.07 | 0.13 |
| 10 | 20863245 | rs12355783 | 0.941 | 0.155 | -0.554 | -1.819 | -0.023 | 0.059 | 0.059 | 0 | 0.28 | 0.18 | 0.2 | 0.09 | 0.29 | 0.16 |
| 10 | 20865157 | rs11819199 | 0.999 | -0.478 | 0.057 | -1.414 | -0.048 | -0.088 | -0.088 | 0 | 0.22 | 0.33 | 0.07 | 0 | 0.07 | 0.11 |
| 10 | 20866328 | rs7099358 | -0.743 | -0.341 | 0.157 | -1.378 | -0.013 | -0.014 | -0.014 | 0 | 0.4 | 0.59 | 0.32 | 0.26 | 0.29 | 0.4 |
| 10 | 20872496 | rs7094536 | 0.042 | -0.398 | 0.047 | -1.359 | -0.006 | 0 | 0 | 0 | 0.31 | 0.34 | 0.14 | 0.21 | 0.13 | 0.24 |
| 10 | 20878450 | rs11595313 | -0.014 | -0.226 | 0.118 | -1.414 | -0.065 | -0.08 | -0.08 | 0 | 0.31 | 0.5 | 0.2 | 0.12 | 0.19 | 0.21 |
| 10 | 20882399 | rs11012178 | -0.326 | 0.181 | -0.011 | -1.273 | -0.013 | -0.001 | -0.001 | 0 | 0.1 | 0.07 | 0.15 | 0.1 | 0.15 | 0.09 |
| 10 | 20892178 | rs10828088 | -0.341 | 0.137 | 0.002 | -1.275 | -0.008 | 0.001 | 0.001 | 0 | 0.1 | 0.09 | 0.16 | 0.1 | 0.15 | 0.1 |
| 10 | 20894107 | rs7079857 | -0.619 | -0.089 | 0.027 | -1.551 | 0.044 | 0.057 | 0.057 | 0.063 | 0.49 | 0.61 | 0.24 | 0.14 | 0.29 | 0.39 |
| 10 | 20896023 | rs7921871 | 1.202 | -0.272 | -0.305 | -1.646 | 0.002 | 0.034 | 0.034 | 0 | 0.17 | 0.09 | 0.08 | 0 | 0.15 | 0.13 |
| 10 | 20897903 | rs11598854 | 0.606 | -0.299 | 0.086 | -1.087 | -0.011 | -0.006 | -0.006 | 0 | 0.25 | 0.22 | 0.07 | 0.08 | 0.07 | 0.14 |
| 10 | 20903394 | rs11012190 | -0.136 | -0.286 | 0.092 | -1.092 | -0.041 | -0.04 | -0.04 | 0 | 0.3 | 0.4 | 0.15 | 0.14 | 0.15 | 0.29 |
| 10 | 20905395 | rs10828089 | 0.438 | 0.036 | -0.116 | -0.991 | -0.007 | -0.006 | -0.006 | 0.043 | 0.06 | 0.05 | 0.06 | 0.09 | 0.07 | 0.08 |
| 10 | 20907665 | rs2359537 | 0.778 | -0.289 | -0.066 | -1.075 | -0.038 | -0.072 | -0.072 | 0 | 0.23 | 0.24 | 0.06 | 0 | 0.07 | 0.12 |
| 10 | 20910166 | rs10828090 | -0.769 | -0.032 | -0.355 | -1.036 | -0.007 | -0.014 | -0.014 | 0 | 0.39 | 0.55 | 0.32 | 0.22 | 0.3 | 0.37 |
| 10 | 20911600 | rs1925530 | -0.918 | 1.231 | -0.685 | -0.535 | 0.031 | 0.004 | 0.004 | 0 | 0.13 | 0.2 | 0.22 | 0.14 | 0.23 | 0.23 |
| 10 | 20918345 | rs12573031 | 0.493 | 0.745 | -1.608 | -0.076 | 0.002 | -0.058 | -0.058 | 0.692 | 0.4 | 0.05 | 0.21 | 0.42 | 0.34 | 0.42 |
| 10 | 20921881 | rs1925533 | 0.068 | 0.992 | -1.616 | -0.414 | -0.012 | 0.034 | 0.034 | 0 | 0.54 | 0.19 | 0.39 | 0.41 | 0.51 | 0.52 |
| 10 | 20923412 | rs11012211 | -0.052 | 1.531 | -0.557 | -0.215 | 0.057 | 0.039 | 0.039 | 0.068 | 0.11 | 0.33 | 0.14 | 0.15 | 0.15 | 0.13 |
| 10 | 20924940 | rs12415669 | 1.194 | 1.42 | -0.822 | -0.276 | -0.029 | 0.074 | 0.074 | 0 | 0.12 | 0.01 | 0.17 | 0 | 0.17 | 0.1 |
| 10 | 20926396 | rs4748677 | 1.434 | 0.929 | -0.735 | -0.439 | 0 | -0.113 | -0.113 | 0.661 | 0.3 | 0.02 | 0.15 | 0.4 | 0.22 | 0.35 |
| 10 | 20928181 | rs11012215 | 0.436 | 1.633 | 0.695 | 0.164 | -0.007 | -0.026 | -0.026 | 0.005 | 0.23 | 0.26 | 0.14 | 0.09 | 0.16 | 0.2 |
| 10 | 20928365 | rs12219402 | 0.172 | 1.609 | 1.069 | 0.158 | -0.027 | -0.029 | -0.029 | 0 | 0.26 | 0.52 | 0.18 | 0.2 | 0.17 | 0.21 |
| 10 | 20929319 | rs1925537 | 0.028 | 1.698 | 0.869 | 0.181 | 0.087 | 0.232 | 0.232 | 2.605 | 0.85 | 0.58 | 0.67 | 0.64 | 0.82 | 0.86 |
| 10 | 20929522 | rs1925538 | 0.394 | 1.497 | 0.194 | 0.111 | -0.025 | -0.02 | -0.02 | 0 | 0.62 | 0.23 | 0.5 | 0.44 | 0.65 | 0.65 |
| 10 | 20933096 | rs7898772 | 0.628 | 1.352 | 0.142 | -0.078 | -0.012 | -0.055 | -0.055 | 0.076 | 0.6 | 0.21 | 0.49 | 0.51 | 0.64 | 0.64 |
| 10 | 20934045 | rs2359579 | 1.074 | 1.465 | -0.098 | -0.345 | -0.018 | 0.044 | 0.044 | 0 | 0.13 | 0.08 | 0.17 | 0 | 0.17 | 0.1 |
| 10 | 20934310 | rs2359580 | 0.086 | 1.673 | 0.706 | -0.191 | 0.049 | 0.128 | 0.128 | 0.744 | 0.84 | 0.64 | 0.68 | 0.67 | 0.8 | 0.85 |
| 10 | 20935127 | rs7916534 | 1.007 | 1.573 | 0.357 | -0.048 | -0.008 | 0.013 | 0.013 | 0 | 0.48 | 0.13 | 0.33 | 0.48 | 0.48 | 0.53 |
| 10 | 20936755 | rs4748682 | 1.553 | 1.671 | 0.321 | -0.151 | 0.04 | -0.025 | -0.025 | 0 | 0.41 | 0.16 | 0.3 | 0.51 | 0.36 | 0.42 |
| 10 | 20938264 | rs11012223 | 0.592 | 1.733 | 0.245 | -0.095 | -0.048 | 0.01 | 0.01 | 0 | 0.09 | 0 | 0.11 | 0.03 | 0.15 | 0.14 |
| 10 | 20945813 | rs10828096 | -0.675 | 1.83 | 0.855 | -0.065 | 0.045 | 0.102 | 0.102 | 1.721 | 0.89 | 0.84 | 0.71 | 0.73 | 0.83 | 0.9 |
| 10 | 20949214 | rs7084463 | 1.159 | 1.764 | 1.142 | -0.127 | -0.011 | -0.002 | -0.002 | 0 | 0.45 | 0.57 | 0.33 | 0.51 | 0.36 | 0.45 |
| 10 | 20949825 | rs2147871 | 1.315 | 1.803 | 1.041 | -0.148 | 0 | -0.03 | -0.03 | 0 | 0.13 | 0.11 | 0.11 | 0 | 0.17 | 0.1 |
| 10 | 20955012 | rs4747415 | -1.132 | 1.799 | 1.023 | -0.103 | 0.053 | 0.118 | 0.118 | 0 | 0.93 | 0.91 | 0.72 | 0.73 | 0.83 | 0.91 |
| 10 | 20960724 | rs6482136 | 1.536 | 1.669 | 1.197 | -0.17 | 0.031 | -0.003 | -0.003 | 0 | 0.42 | 0.29 | 0.31 | 0.49 | 0.36 | 0.45 |
| 10 | 20962633 | rs12249320 | -0.635 | 1.689 | 1.395 | -0.16 | 0.026 | -0.019 | -0.019 | 0 | 0.14 | 0.34 | 0.15 | 0.12 | 0.15 | 0.15 |
| 10 | 20963377 | rs7074040 | 0.98 | 1.281 | 1.248 | -0.336 | 0 | -0.002 | -0.002 | 0 | 0.52 | 0.53 | 0.43 | 0.5 | 0.53 | 0.56 |
| 10 | 20964156 | rs1570844 | 0.459 | 1.605 | 1.346 | -0.198 | -0.02 | -0.012 | -0.012 | 0 | 0.25 | 0.36 | 0.16 | 0.2 | 0.15 | 0.21 |
| 10 | 20964434 | rs17726796 | 1.564 | 1.658 | 1.31 | -0.228 | -0.001 | -0.023 | -0.023 | 0 | 0.12 | 0.1 | 0.11 | 0 | 0.17 | 0.07 |
| 10 | 20965953 | rs6415962 | -0.203 | 1.667 | 1.342 | -0.189 | 0.073 | 0.159 | 0.159 | 0 | 0.89 | 0.8 | 0.7 | 0.72 | 0.82 | 0.9 |
| 10 | 20965972 | rs10828101 | 0.498 | 1.397 | 1.221 | -0.194 | -0.017 | -0.011 | -0.011 | 0 | 0.25 | 0.35 | 0.16 | 0.19 | 0.16 | 0.21 |
| 10 | 20966815 | rs1925546 | 0.789 | 1.428 | 1.174 | -0.205 | 0.036 | 0.012 | 0.012 | 0.157 | 0.66 | 0.5 | 0.45 | 0.68 | 0.51 | 0.69 |
| 10 | 20970307 | rs10828105 | 0.714 | 1.265 | 0.841 | -0.334 | 0.026 | 0.032 | 0.032 | 0.015 | 0.23 | 0.09 | 0.14 | 0.11 | 0.15 | 0.2 |
| 10 | 20975241 | rs4748690 | 1.845 | 1.161 | 0.494 | -0.466 | 0.073 | -0.057 | -0.057 | 0 | 0.33 | 0.1 | 0.24 | 0.43 | 0.25 | 0.41 |
| 10 | 20975801 | rs10764262 | -1.417 | 1.327 | 0.678 | -0.463 | 0.019 | 0.058 | 0.058 | 0 | 0.58 | 0.8 | 0.5 | 0.29 | 0.6 | 0.51 |
| 10 | 20977925 | rs10764265 | -0.359 | 1.308 | 0.689 | -0.513 | 0.02 | 0.039 | 0.039 | 0.028 | 0.39 | 0.74 | 0.32 | 0.26 | 0.34 | 0.34 |
| 10 | 20978879 | rs11012272 | 0.641 | 1.336 | 0.655 | -0.504 | -0.057 | 0.061 | 0.061 | 0 | 0.17 | 0.04 | 0.16 | 0.02 | 0.24 | 0.16 |
| 10 | 20980016 | rs9645521 | -0.323 | 1.043 | 1.217 | -0.28 | -0.011 | -0.021 | -0.021 | 0 | 0.38 | 0.56 | 0.3 | 0.26 | 0.34 | 0.35 |
| 10 | 20983854 | rs16920893 | 1.045 | 1.179 | 1.218 | -0.234 | 0.017 | -0.078 | -0.078 | 0.234 | 0.14 | 0.27 | 0.13 | 0 | 0.18 | 0.07 |
| 10 | 20984907 | rs4748699 | 0.549 | 1.164 | 1.156 | -0.214 | 0.031 | -0.004 | -0.004 | 0 | 0.23 | 0.06 | 0.17 | 0.18 | 0.15 | 0.28 |
| 10 | 20984912 | rs7895134 | -0.766 | 1.411 | 1.179 | -0.159 | 0.049 | 0.101 | 0.101 | 0 | 0.91 | 0.9 | 0.69 | 0.73 | 0.8 | 0.91 |
| 10 | 20988778 | rs7075945 | -1.115 | 1.096 | 1.165 | -0.15 | 0.021 | 0.073 | 0.073 | 0 | 0.92 | 0.92 | 0.74 | 0.74 | 0.85 | 0.93 |
| 10 | 20989087 | rs10828110 | 0.365 | 0.68 | 1.195 | -0.554 | 0.009 | 0.003 | 0.003 | 0 | 0.23 | 0.19 | 0.18 | 0.18 | 0.16 | 0.28 |
| 10 | 20993102 | rs7095613 | -0.635 | 1.052 | 1.409 | -0.28 | 0.006 | 0.099 | 0.099 | 0 | 0.14 | 0.4 | 0.17 | 0.26 | 0.1 | 0.28 |
| 10 | 20994847 | rs6482146 | -1.071 | 1.063 | 1.146 | -0.316 | 0 | -0.003 | -0.003 | 0 | 0.28 | 0.67 | 0.29 | 0.27 | 0.28 | 0.35 |
| 10 | 20996285 | rs7097699 | 0.255 | 1.154 | 1.256 | -0.453 | 0.02 | -0.072 | -0.072 | 0.005 | 0.16 | 0.62 | 0.16 | 0.08 | 0.19 | 0.08 |
| 10 | 20998112 | rs16920937 | 0.63 | 1.278 | 0.764 | -0.478 | 0.025 | -0.08 | -0.08 | 0 | 0.13 | 0.29 | 0.13 | 0 | 0.18 | 0.07 |
| 10 | 21002760 | rs7095517 | -1.257 | 0.922 | 0.968 | -0.264 | 0.023 | -0.029 | -0.029 | 0 | 0.25 | 0.5 | 0.32 | 0.21 | 0.26 | 0.3 |
| 10 | 21003993 | rs10828113 | -0.131 | 0.958 | 0.711 | -0.247 | -0.002 | 0.014 | 0.014 | 0 | 0.11 | 0.13 | 0.15 | 0.15 | 0.09 | 0.16 |
| 10 | 21011704 | rs7080393 | -0.733 | 0.805 | 0.638 | -0.182 | 0.001 | 0.001 | 0.001 | 0 | 0.63 | 0.59 | 0.59 | 0.68 | 0.59 | 0.81 |
| 10 | 21013845 | rs4748702 | 0.984 | 0.841 | 0.635 | -0.238 | 0.022 | -0.055 | -0.055 | 0 | 0.4 | 0.14 | 0.29 | 0.5 | 0.35 | 0.54 |
| 10 | 21015343 | rs7070319 | 0.661 | 0.828 | 0.41 | -0.208 | -0.059 | 0.062 | 0.062 | 0 | 0.11 | 0.02 | 0.13 | 0.01 | 0.16 | 0.13 |
| 10 | 21024318 | rs970758 | -0.307 | -0.077 | 0.407 | 0.168 | -0.001 | 0.005 | 0.005 | 0 | 0.11 | 0.09 | 0.15 | 0.14 | 0.09 | 0.16 |
| 10 | 21024607 | rs10828118 | -0.137 | -0.072 | 0.351 | 0.17 | 0 | -0.004 | -0.004 | 0.105 | 0.11 | 0.05 | 0.14 | 0.14 | 0.09 | 0.16 |
| 10 | 21026889 | rs7075837 | -0.087 | -0.462 | 0.143 | 0.145 | 0.082 | 0.012 | 0.012 | 0 | 0.46 | 0.77 | 0.35 | 0.47 | 0.32 | 0.6 |
| 10 | 21030580 | rs7079544 | -0.192 | -0.754 | 0.38 | -0.194 | 0.004 | -0.056 | -0.056 | 0 | 0.1 | 0.3 | 0.07 | 0 | 0.06 | 0.07 |
| 10 | 21030854 | rs1326248 | -0.539 | -0.818 | -0.009 | -0.075 | 0.116 | 0.022 | 0.022 | 0 | 0.49 | 0.79 | 0.43 | 0.68 | 0.33 | 0.66 |

| | | | | | | | | | | | | | | | | |
|---|---|---|---|---|---|---|---|---|---|---|---|---|---|---|---|---|
| 10 | 21033698 | rs7478154 | 0.7 | -0.539 | -0.217 | 0.352 | 0.045 | -0.014 | -0.014 | 0 | 0.4 | 0.2 | 0.27 | 0.48 | 0.32 | 0.47 |
| 10 | 21033932 | rs11012309 | -0.486 | -0.503 | -0.21 | 0.373 | -0.018 | 0.061 | 0.061 | 1.551 | 0.07 | 0.2 | 0.1 | 0.17 | 0.03 | 0.14 |
| 10 | 21034954 | rs4748711 | 0.667 | -0.493 | -0.232 | 0.377 | 0.042 | -0.004 | -0.004 | 0 | 0.41 | 0.23 | 0.28 | 0.48 | 0.32 | 0.47 |
| 10 | 21036074 | rs11012310 | -0.799 | 0.023 | -0.346 | 0.682 | -0.017 | 0.061 | 0.061 | 0.831 | 0.06 | 0.19 | 0.1 | 0.17 | 0.03 | 0.14 |
| 10 | 21046758 | rs10458682 | -0.306 | 0.275 | -1.383 | 0.383 | -0.042 | -0.066 | -0.066 | 0 | 0.59 | 0.92 | 0.48 | 0.69 | 0.41 | 0.68 |
| 10 | 21048682 | rs4748713 | 0.01 | -0.257 | -0.526 | 0.062 | 0.003 | 0 | 0 | 0 | 0.49 | 0.51 | 0.42 | 0.63 | 0.36 | 0.61 |
| 10 | 21050514 | rs11012326 | -0.298 | -0.133 | -0.909 | -0.234 | -0.023 | 0.057 | 0.057 | 0 | 0.1 | 0.01 | 0.06 | 0 | 0.08 | 0.09 |
| 10 | 21050688 | rs2765839 | 0.938 | 0.248 | -1.231 | -0.559 | -0.026 | 0.036 | 0.036 | 0 | 0.1 | 0.02 | 0.08 | 0 | 0.16 | 0.08 |
| 10 | 21052023 | rs4748718 | 0.185 | -0.296 | -0.571 | -0.428 | 0.041 | 0.028 | 0.028 | 0 | 0.49 | 0.68 | 0.36 | 0.52 | 0.38 | 0.52 |
| 10 | 21054582 | rs10828129 | 0.111 | -0.322 | -0.512 | -0.421 | 0.036 | 0.027 | 0.027 | 0 | 0.51 | 0.71 | 0.37 | 0.52 | 0.38 | 0.54 |
| 10 | 21056587 | rs4747421 | -0.406 | -0.591 | 0.056 | -0.39 | -0.019 | -0.069 | -0.069 | 2.921 | 0.17 | 0.45 | 0.16 | 0.09 | 0.1 | 0.14 |
| 10 | 21058067 | rs10764284 | 0.36 | -0.585 | 0.139 | -0.366 | 0 | 0 | 0 | 0 | 0.47 | 0.52 | 0.35 | 0.47 | 0.38 | 0.51 |
| 10 | 21058174 | rs10740984 | 0.64 | -0.598 | 0.418 | -0.369 | 0.019 | 0.023 | 0.023 | 0 | 0.44 | 0.32 | 0.33 | 0.44 | 0.37 | 0.51 |
| 10 | 21063061 | rs7070846 | -1.827 | -0.528 | 0.499 | -0.326 | 0.005 | 0.054 | 0.054 | 2.104 | 0.07 | 0.37 | 0.04 | 0.08 | 0.01 | 0.02 |
| 10 | 21066981 | rs2765841 | 0.02 | -0.535 | 0.394 | -0.43 | 0.02 | -0.039 | -0.039 | 0 | 0.13 | 0.31 | 0.11 | 0.08 | 0.17 | 0.06 |
| 10 | 21067683 | rs2765843 | -0.8 | -0.615 | 0.386 | -0.373 | -0.016 | -0.016 | -0.016 | 0 | 0.66 | 0.76 | 0.55 | 0.69 | 0.59 | 0.75 |
| 10 | 21072601 | rs7913231 | -0.579 | -0.815 | 0.637 | -0.2 | -0.003 | 0.115 | 0.115 | 1.514 | 0.06 | 0.27 | 0.11 | 0.18 | 0.03 | 0.11 |
| 10 | 21073521 | rs4748722 | 0.346 | -0.848 | 0.608 | -0.294 | 0.029 | 0.024 | 0.024 | 0.023 | 0.47 | 0.32 | 0.34 | 0.5 | 0.37 | 0.55 |
| 10 | 21074344 | rs2296612 | -0.31 | -0.891 | 0.672 | -0.202 | 0.014 | 0.005 | 0.005 | 0 | 0.55 | 0.76 | 0.46 | 0.7 | 0.41 | 0.68 |
| 10 | 21074455 | rs2296613 | 1.635 | -0.872 | 0.569 | -0.191 | 0.024 | -0.024 | -0.024 | 0 | 0.33 | 0.17 | 0.2 | 0.44 | 0.27 | 0.4 |
| 10 | 21076100 | rs2296606 | -1.404 | -0.804 | 0.648 | -0.194 | 0.011 | 0.204 | 0.204 | 6.907 | 0.08 | 0.49 | 0.13 | 0.22 | 0.03 | 0.12 |
| 10 | 21078422 | rs11012340 | -0.55 | -0.71 | 0.599 | -0.133 | 0.005 | 0.108 | 0.108 | 2.129 | 0.06 | 0.28 | 0.08 | 0.16 | 0.03 | 0.13 |
| 10 | 21081843 | rs3781490 | 1.152 | -0.742 | 0.281 | -0.053 | 0.024 | 0.009 | 0.009 | 0 | 0.38 | 0.2 | 0.22 | 0.41 | 0.32 | 0.47 |
| 10 | 21083773 | rs11012341 | 1.105 | -0.493 | 0.104 | -0.127 | -0.002 | -0.033 | -0.033 | 0 | 0.07 | 0.11 | 0.11 | 0 | 0.05 | 0.05 |
| 10 | 21086468 | rs10491054 | 1.105 | -0.493 | 0.089 | -0.127 | -0.002 | -0.032 | -0.032 | 0.038 | 0.07 | 0.11 | 0.11 | 0 | 0.05 | 0.05 |
| 10 | 21097769 | rs3758601 | 0.182 | -0.362 | -0.081 | -0.045 | 0.05 | 0.012 | 0.012 | 0 | 0.51 | 0.74 | 0.41 | 0.64 | 0.37 | 0.61 |
| 10 | 21108377 | rs1006363 | 0.69 | 0.399 | -0.657 | 0.429 | -0.088 | -0.078 | -0.078 | 0 | 0.13 | 0.02 | 0.22 | 0.25 | 0.3 | 0.19 |
| 10 | 21120116 | rs10491056 | 0.671 | -0.914 | -0.562 | 0.223 | 0.052 | 0.004 | 0.004 | 0 | 0.5 | 0.64 | 0.29 | 0.53 | 0.35 | 0.43 |
| 10 | 21131710 | rs788999 | 0.98 | -0.758 | -0.653 | 0.047 | -0.027 | 0.031 | 0.031 | 0 | 0.1 | 0.03 | 0.11 | 0 | 0.18 | 0.08 |
| 10 | 21137746 | rs10828141 | 0.757 | -1.035 | -0.625 | 0.109 | 0.051 | 0.018 | 0.018 | 0 | 0.53 | 0.72 | 0.35 | 0.64 | 0.38 | 0.54 |
| 10 | 21139389 | rs4025981 | 1.738 | -1.565 | -0.975 | -0.095 | 0.046 | 0.073 | 0.073 | 1.42 | 0.18 | 0.01 | 0.05 | 0.03 | 0.08 | 0.05 |
| 10 | 21145121 | rs3791198 | -0.197 | -1.247 | -0.428 | 0.016 | -0.127 | -0.006 | -0.006 | 0.04 | 0.34 | 0.09 | 0.41 | 0.26 | 0.44 | 0.36 |
| 10 | 21147418 | rs703093 | 2.155 | -1.63 | 0.028 | 0.256 | -0.007 | -0.05 | -0.05 | 0 | 0.26 | 0.1 | 0.14 | 0.4 | 0.26 | 0.37 |
| 10 | 21147520 | rs1409272 | -1.956 | -1.64 | 0.455 | 0.251 | -0.011 | 0.079 | 0.079 | 0 | 0.71 | 0.49 | 0.65 | 0.48 | 0.74 | 0.62 |
| 10 | 21157028 | rs1475354 | 1.195 | -0.793 | -0.114 | 0.041 | -0.075 | 0.07 | 0.07 | 1.603 | 0.13 | 0 | 0.12 | 0.02 | 0.21 | 0.21 |
| 10 | 21181703 | rs788965 | 1.942 | -1.807 | 0.227 | 0.553 | -0.004 | -0.045 | -0.045 | 0 | 0.26 | 0.1 | 0.15 | 0.39 | 0.26 | 0.35 |
| 10 | 21185661 | rs788970 | 1.735 | -1.799 | 0.334 | 0.578 | 0.002 | -0.011 | -0.011 | 0.122 | 0.28 | 0.26 | 0.16 | 0.45 | 0.26 | 0.38 |
| 10 | 21191984 | rs7907121 | -1.12 | -1.726 | 0.706 | 0.553 | 0.059 | 0.021 | 0.021 | 0 | 0.5 | 0.33 | 0.59 | 0.4 | 0.65 | 0.57 |
| 10 | 21195712 | rs10828148 | -1.206 | -2.061 | 0.489 | 0.461 | 0.035 | -0.003 | -0.003 | 0 | 0.54 | 0.65 | 0.76 | 0.43 | 0.66 | 0.47 |
| 10 | 21196566 | rs16921240 | 0.676 | -1.87 | 0.417 | 0.432 | -0.022 | 0.016 | 0.016 | 0.136 | 0.12 | 0.15 | 0.21 | 0.16 | 0.05 | 0.04 |
| 10 | 21198380 | rs11012384 | -1.097 | -1.981 | 0.153 | 0.306 | 0.004 | 0.002 | 0.002 | 0 | 0.47 | 0.53 | 0.71 | 0.53 | 0.64 | 0.54 |
| 10 | 21201658 | rs7071220 | 0.115 | -1.695 | 0.075 | 0.378 | -0.015 | -0.009 | -0.009 | 0 | 0.3 | 0.29 | 0.46 | 0.34 | 0.55 | 0.4 |
| 10 | 21203132 | rs12259450 | 1.221 | -1.678 | -0.07 | 0.412 | 0.011 | -0.009 | -0.009 | 0 | 0.1 | 0.06 | 0.2 | 0.11 | 0.05 | 0.04 |
| 10 | 21206340 | rs11012389 | 0.057 | -1.632 | 0.067 | 0.408 | -0.001 | -0.001 | -0.001 | 1.257 | 0.3 | 0.35 | 0.46 | 0.34 | 0.55 | 0.4 |
| 10 | 21207377 | rs11012391 | -1.224 | -2.124 | -0.012 | 0.349 | -0.015 | 0 | 0 | 0 | 0.56 | 0.55 | 0.8 | 0.53 | 0.67 | 0.49 |
| 10 | 21208182 | rs11012392 | 0.601 | -2.228 | -0.458 | 0.148 | 0.052 | -0.002 | -0.002 | 0 | 0.18 | 0.06 | 0.21 | 0.14 | 0.06 | 0.04 |
| 10 | 21209194 | rs10764292 | -1.515 | -2.479 | -0.529 | 0.264 | 0.029 | -0.028 | -0.028 | 0 | 0.66 | 0.88 | 0.87 | 0.59 | 0.72 | 0.61 |
| 10 | 21214677 | rs7086221 | -2.293 | -2.471 | -0.268 | 0.279 | -0.001 | 0.004 | 0.004 | 0 | 0.75 | 0.79 | 0.87 | 0.6 | 0.76 | 0.64 |
| 10 | 21216368 | rs10508635 | -1.486 | -2.46 | -0.115 | 0.316 | 0.003 | -0.002 | -0.002 | 0.321 | 0.65 | 0.74 | 0.85 | 0.54 | 0.72 | 0.62 |
| 10 | 21218979 | rs4748737 | -0.715 | -2.65 | -0.126 | 0.267 | 0.006 | 0.001 | 0.001 | 0 | 0.53 | 0.59 | 0.67 | 0.4 | 0.68 | 0.59 |
| 10 | 21219406 | rs4748738 | -0.858 | -2.637 | -0.122 | 0.271 | 0.004 | 0 | 0 | 0 | 0.54 | 0.59 | 0.67 | 0.41 | 0.69 | 0.6 |
| 10 | 21220134 | rs11596224 | 1.43 | -2.543 | -0.175 | 0.309 | 0.037 | -0.037 | -0.037 | 1.493 | 0.1 | 0.02 | 0.18 | 0.14 | 0.03 | 0.03 |
| 10 | 21224119 | rs952738 | 0.731 | -2.73 | -0.046 | 0.126 | -0.013 | -0.005 | -0.005 | 0 | 0.32 | 0.25 | 0.49 | 0.32 | 0.58 | 0.36 |
| 10 | 21224986 | rs7921413 | 0.134 | -2.663 | -0.329 | 0.186 | -0.032 | -0.029 | -0.029 | 0 | 0.21 | 0.29 | 0.18 | 0.09 | 0.1 | 0.21 |
| 10 | 21225243 | rs7098616 | 0.02 | -2.659 | -0.379 | 0.187 | -0.039 | -0.022 | -0.022 | 0 | 0.21 | 0.36 | 0.17 | 0.14 | 0.11 | 0.21 |
| 10 | 21236328 | rs1539211 | 0.499 | -1.91 | -0.817 | 0.17 | -0.016 | -0.003 | -0.003 | 0 | 0.34 | 0.28 | 0.49 | 0.32 | 0.53 | 0.38 |
| 10 | 21239877 | rs1856762 | -1.988 | -1.648 | -1.398 | -0.012 | -0.002 | 0.027 | 0.027 | 0 | 0.74 | 0.7 | 0.86 | 0.55 | 0.75 | 0.65 |
| 10 | 21240532 | rs4543885 | 0.182 | -1.441 | -1.717 | 0.259 | 0.078 | -0.113 | -0.113 | 0 | 0.18 | 0.03 | 0.22 | 0.36 | 0.07 | 0.05 |
| 10 | 21240818 | rs16905928 | -0.042 | -0.632 | -1.225 | 0.584 | 0.01 | 0.006 | 0.006 | 0.166 | 0.13 | 0.08 | 0.12 | 0 | 0.07 | 0.08 |
| 10 | 21242841 | rs11012413 | -0.282 | 0.263 | -1.172 | 0.937 | -0.03 | -0.072 | -0.072 | 0 | 0.15 | 0.04 | 0.31 | 0.35 | 0.23 | 0.08 |
| 10 | 21244087 | rs33932343 | -0.431 | 0.568 | -0.862 | 0.605 | 0.001 | -0.018 | -0.018 | 0 | 0.44 | 0.32 | 0.62 | 0.51 | 0.48 | 0.55 |
| 10 | 21252195 | rs1417184 | -1.455 | 0.134 | 0.413 | 1.094 | -0.021 | 0.271 | 0.271 | 2.63 | 0.14 | 0.5 | 0.1 | 0.39 | 0.1 | 0.09 |
| 10 | 21262579 | rs11813945 | -0.24 | 0.276 | -0.229 | 0.979 | -0.061 | -0.042 | -0.042 | 0 | 0.2 | 0.03 | 0.15 | 0.26 | 0.21 | 0.31 |
| 10 | 21270480 | rs1832677 | -1.416 | -0.04 | 0.963 | 0.877 | -0.029 | 0.273 | 0.273 | 0.407 | 0.12 | 0.42 | 0.07 | 0.41 | 0.06 | 0.09 |
| 10 | 21271153 | rs960941 | -2.137 | -0.108 | 1.407 | 0.847 | -0.053 | 0.157 | 0.157 | 3.942 | 0.15 | 0.62 | 0.08 | 0.4 | 0.06 | 0.09 |
| 10 | 21272444 | rs11012457 | -0.775 | 0.001 | 1.313 | 0.87 | -0.016 | 0.195 | 0.195 | 0.104 | 0.08 | 0.26 | 0.05 | 0.41 | 0.05 | 0.09 |
| 10 | 21273438 | rs10508637 | -0.04 | -0.368 | 1.073 | 0.748 | 0.02 | -0.057 | -0.057 | 1.222 | 0.25 | 0.1 | 0.19 | 0.45 | 0.2 | 0.4 |
| 10 | 21274956 | rs11012460 | -1.34 | 0.428 | 1.598 | 1.081 | 0.002 | 0.299 | 0.299 | 0.967 | 0.09 | 0.34 | 0.06 | 0.41 | 0.06 | 0.08 |
| 10 | 21276403 | rs7893616 | -1.444 | 0.441 | 1.577 | 1.087 | -0.012 | 0.288 | 0.288 | 0.832 | 0.12 | 0.43 | 0.07 | 0.4 | 0.07 | 0.1 |

| | | | | | | | | | | | | | | | | |
|---|---|---|---|---|---|---|---|---|---|---|---|---|---|---|---|---|
| 10 | 21287267 | rs7898090 | -0.963 | 0.361 | 1.237 | 1.159 | -0.045 | 0.199 | 0.199 | 0.839 | 0.13 | 0.58 | 0.08 | 0.4 | 0.07 | 0.09 |
| 10 | 21288921 | rs608788 | -0.283 | 1.812 | -0.588 | 0.911 | -0.106 | 0.044 | 0.044 | 1.088 | 0.17 | 0.01 | 0.29 | 0.06 | 0.29 | 0.28 |
| 10 | 21291331 | rs640056 | -0.03 | 0.581 | 0.168 | 1.134 | -0.032 | 0.172 | 0.172 | 0 | 0.15 | 0.34 | 0.21 | 0.46 | 0.09 | 0.08 |
| 10 | 21291924 | rs11595730 | 0.245 | 0.707 | -0.337 | -0.329 | -0.028 | 0.161 | 0.161 | 0 | 0.45 | 0.23 | 0.37 | 0.06 | 0.49 | 0.42 |
| 10 | 21294051 | rs10828173 | -0.218 | 0.662 | 1.002 | 0.927 | -0.023 | 0.109 | 0.109 | 0.978 | 0.17 | 0.66 | 0.09 | 0.55 | 0.1 | 0.15 |
| 10 | 21296519 | rs599087 | 0.376 | 0.765 | 0.569 | 1.055 | -0.029 | -0.061 | -0.061 | 0.099 | 0.11 | 0.01 | 0.29 | 0.2 | 0.11 | 0.25 |
| 10 | 21301630 | rs11012485 | 0.147 | 0.587 | 1.233 | 1.091 | -0.008 | 0.245 | 0.245 | 0 | 0.15 | 0.37 | 0.07 | 0.54 | 0.1 | 0.15 |
| 10 | 21301918 | rs6482158 | -0.183 | 0.538 | 0.71 | 1.095 | -0.006 | 0.033 | 0.033 | 1.162 | 0.18 | 0.74 | 0.1 | 0.56 | 0.1 | 0.15 |
| 10 | 21302821 | rs597837 | -0.56 | 1.304 | 0.348 | 0.736 | -0.074 | 0.005 | 0.005 | 0.051 | 0.28 | 0.09 | 0.53 | 0.25 | 0.38 | 0.47 |
| 10 | 21304646 | rs7920067 | -0.453 | 0.814 | 0.966 | 1.124 | -0.018 | 0.085 | 0.085 | 0.771 | 0.17 | 0.68 | 0.1 | 0.54 | 0.1 | 0.14 |
| 10 | 21311248 | rs10828176 | -0.519 | 0.861 | 0.273 | 1.085 | -0.008 | 0.044 | 0.044 | 1.06 | 0.18 | 0.74 | 0.1 | 0.56 | 0.11 | 0.16 |
| 10 | 21313556 | rs661924 | 0.272 | 1.319 | 0.191 | 0.915 | 0.023 | 0.032 | 0.032 | 0 | 0.69 | 0.39 | 0.66 | 0.47 | 0.62 | 0.73 |
| 10 | 21314935 | rs10764302 | -0.827 | 0.094 | 0.375 | 1.238 | -0.017 | 0.021 | 0.021 | 0 | 0.23 | 0.33 | 0.23 | 0.36 | 0.18 | 0.25 |
| 10 | 21315152 | rs3858201 | -0.045 | -0.201 | 0.363 | 1.343 | -0.008 | -0.009 | -0.009 | 0 | 0.59 | 0.54 | 0.63 | 0.67 | 0.62 | 0.67 |
| 10 | 21315553 | rs3858202 | 0.59 | -0.054 | 0.085 | 1.659 | 0.005 | -0.017 | -0.017 | 0.01 | 0.6 | 0.17 | 0.49 | 0.59 | 0.59 | 0.67 |
| 10 | 21317504 | rs7898342 | 0.435 | -0.049 | 0.68 | 1.54 | -0.001 | -0.01 | -0.01 | 0 | 0.54 | 0.49 | 0.56 | 0.73 | 0.56 | 0.67 |
| 10 | 21318625 | rs638929 | 0.279 | 0.034 | 0.794 | 1.476 | 0.001 | -0.002 | -0.002 | 0 | 0.57 | 0.56 | 0.68 | 0.73 | 0.6 | 0.67 |
| 10 | 21319705 | rs561415 | 0.298 | 0.058 | 0.791 | 1.474 | 0.001 | -0.002 | -0.002 | 0 | 0.57 | 0.56 | 0.68 | 0.74 | 0.59 | 0.67 |
| 10 | 21332250 | rs2986673 | 0.43 | 0.012 | 0.884 | 1.756 | -0.005 | -0.005 | -0.005 | 0 | 0.56 | 0.51 | 0.67 | 0.63 | 0.59 | 0.59 |
| 10 | 21344747 | rs2119243 | -1.078 | -0.064 | 0.899 | 1.874 | 0.052 | 0.013 | 0.013 | 0.018 | 0.48 | 0.86 | 0.39 | 0.46 | 0.42 | 0.42 |
| 10 | 21349368 | rs893881 | -0.749 | -0.095 | 0.922 | 1.986 | -0.008 | -0.02 | -0.02 | 0 | 0.44 | 0.48 | 0.32 | 0.31 | 0.42 | 0.41 |
| 10 | 21352013 | rs893880 | 0.58 | 0.034 | -0.028 | 2.209 | -0.015 | -0.03 | -0.03 | 0 | 0.91 | 0.96 | 0.82 | 0.73 | 0.92 | 0.89 |
| 10 | 21354398 | rs1440295 | 1.278 | 0.047 | -0.009 | 1.817 | -0.149 | 0.054 | 0.054 | 0.054 | 0.41 | 0.02 | 0.43 | 0.25 | 0.51 | 0.39 |
| 10 | 21358142 | rs7918237 | 1.748 | 0.641 | 1.188 | 2.175 | -0.001 | 0.016 | 0.016 | 0 | 0.61 | 0.3 | 0.58 | 0.28 | 0.63 | 0.43 |
| 10 | 21363291 | rs11012506 | -1.033 | 0.917 | 1.815 | 2.578 | 0.02 | 0.253 | 0.253 | 0 | 0.06 | 0.21 | 0.08 | 0.25 | 0.03 | 0.13 |
| 10 | 21366778 | rs9633625 | -2.55 | 0.85 | 2.054 | 2.645 | -0.021 | 0.078 | 0.078 | 0.1 | 0.19 | 0.66 | 0.18 | 0.36 | 0.15 | 0.34 |
| 10 | 21370420 | rs7097035 | -1.261 | 0.888 | 2.377 | 2.512 | 0.012 | 0.024 | 0.024 | 0.006 | 0.1 | 0.28 | 0.09 | 0.11 | 0.12 | 0.21 |
| 10 | 21370995 | rs4631770 | 1.489 | 0.954 | 2.394 | 2.707 | 0.028 | 0.109 | 0.109 | 0 | 0.89 | 0.83 | 0.75 | 0.75 | 0.91 | 0.8 |
| 10 | 21371016 | rs1123796 | 2.965 | 0.95 | 2.286 | 2.642 | 0 | 0.016 | 0.016 | 0 | 0.75 | 0.27 | 0.65 | 0.6 | 0.78 | 0.6 |
| 10 | 21373696 | rs10828185 | -2.868 | 0.33 | 2.466 | 2.836 | -0.017 | 0.093 | 0.093 | 0.117 | 0.19 | 0.66 | 0.23 | 0.4 | 0.15 | 0.35 |
| 10 | 21376689 | rs621429 | -1.505 | 0.354 | 2.523 | 2.796 | -0.005 | 0.02 | 0.02 | 0.037 | 0.21 | 0.4 | 0.18 | 0.34 | 0.19 | 0.22 |
| 10 | 21378085 | rs515249 | -1.339 | 0.386 | 2.472 | 2.782 | 0.002 | 0.028 | 0.028 | 0.046 | 0.19 | 0.32 | 0.17 | 0.36 | 0.2 | 0.22 |
| 10 | 21378412 | rs11012519 | -0.431 | 0.389 | 2.389 | 2.674 | 0.014 | 0.035 | 0.035 | 0 | 0.05 | 0.1 | 0.06 | 0.11 | 0.1 | 0.15 |
| 10 | 21404530 | rs11012530 | -0.228 | 0.396 | 2.345 | 2.646 | 0.007 | 0.018 | 0.018 | 0 | 0.05 | 0.09 | 0.06 | 0.12 | 0.11 | 0.14 |
| 10 | 21406933 | rs591652 | -1.142 | 0.403 | 2.469 | 2.715 | -0.001 | 0.011 | 0.011 | 0 | 0.18 | 0.23 | 0.17 | 0.33 | 0.18 | 0.23 |
| 10 | 21407139 | rs522736 | -1.245 | 0.397 | 2.5 | 2.71 | -0.002 | 0.014 | 0.014 | 0 | 0.19 | 0.27 | 0.18 | 0.33 | 0.18 | 0.23 |
| 10 | 21409757 | rs10764307 | -1.269 | 0.406 | 2.498 | 2.738 | 0.05 | 0.346 | 0.346 | 0 | 0.07 | 0.29 | 0.14 | 0.28 | 0.03 | 0.14 |
| 10 | 21419271 | rs10828196 | -0.433 | 0.453 | 1.864 | 2.709 | 0.001 | 0.035 | 0.035 | 0 | 0.05 | 0.06 | 0.11 | 0.22 | 0.03 | 0.13 |
| 10 | 21421272 | rs188764 | -1.932 | 0.534 | 2.151 | 2.532 | 0.002 | 0.05 | 0.05 | 0.037 | 0.27 | 0.54 | 0.24 | 0.48 | 0.31 | 0.4 |
| 10 | 21444603 | rs1751915 | -0.867 | 0.5 | 2.091 | 2.41 | 0 | 0.009 | 0.009 | 0 | 0.18 | 0.23 | 0.16 | 0.32 | 0.18 | 0.29 |
| 10 | 21444900 | rs7090013 | -2.371 | 0.438 | 1.844 | 2.257 | 0.018 | 0.05 | 0.05 | 0.122 | 0.16 | 0.4 | 0.21 | 0.22 | 0.16 | 0.25 |
| 10 | 21449777 | rs11599589 | -0.561 | 0.39 | 1.469 | 2.176 | 0 | -0.008 | -0.008 | 0 | 0.16 | 0.12 | 0.12 | 0.35 | 0.15 | 0.21 |
| 10 | 21453797 | rs1527042 | -2.551 | 0.513 | 0.934 | 2.268 | 0.02 | 0.181 | 0.181 | 0.783 | 0.17 | 0.5 | 0.22 | 0.35 | 0.16 | 0.25 |
| 10 | 21461860 | rs10828201 | -0.797 | -0.029 | 0.584 | 2.292 | 0.012 | -0.015 | -0.015 | 0.715 | 0.08 | 0.06 | 0.11 | 0.21 | 0.03 | 0.14 |
| 10 | 21463833 | rs2279311 | -1.409 | -0.142 | 0.58 | 2.165 | -0.018 | 0.029 | 0.029 | 0.01 | 0.31 | 0.58 | 0.28 | 0.58 | 0.2 | 0.41 |
| 10 | 21470357 | rs7095405 | -0.937 | -0.084 | 0.09 | 1.527 | -0.015 | 0.048 | 0.048 | 0 | 0.09 | 0.15 | 0.08 | 0.26 | 0.03 | 0.17 |
| 10 | 21470801 | rs7099403 | -0.902 | 0.362 | -0.316 | 0.695 | 0.022 | 0.017 | 0.017 | 0.46 | 0.36 | 0.78 | 0.3 | 0.76 | 0.28 | 0.48 |
| 10 | 21477482 | rs11012566 | -1.223 | 1.108 | 0.312 | 0.587 | 0.098 | 0.097 | 0.097 | 0 | 0.06 | 0.18 | 0.12 | 0.12 | 0.13 | 0.08 |
| 10 | 21487254 | rs3911716 | -2.084 | 0.926 | 1.187 | 0.065 | 0.139 | 0.002 | 0.002 | 0 | 0.08 | 0.42 | 0.1 | 0.02 | 0.12 | 0.08 |
| 10 | 21495362 | rs12412833 | -0.449 | 0.992 | 0.117 | -0.329 | -0.039 | 0.009 | 0.009 | 0 | 0.13 | 0.01 | 0.12 | 0.09 | 0.11 | 0.12 |
| 10 | 21502882 | rs7900067 | -1.803 | 0.388 | 0.564 | -0.707 | -0.005 | -0.068 | -0.068 | 0.388 | 0.16 | 0.65 | 0.14 | 0.01 | 0.13 | 0.1 |
| 10 | 21506018 | rs10828212 | -1.947 | 0.034 | 0.161 | -0.896 | 0.014 | 0.021 | 0.021 | 0 | 0.19 | 0.75 | 0.15 | 0.01 | 0.12 | 0.12 |
| 10 | 21510840 | rs10828213 | -1.822 | 0.044 | 0.336 | -0.889 | 0.003 | -0.012 | -0.012 | 1.045 | 0.18 | 0.71 | 0.14 | 0.01 | 0.12 | 0.12 |
| 10 | 21523325 | rs10828224 | -1.376 | -0.714 | -0.035 | -1.668 | 0.02 | 0.05 | 0.05 | 0.483 | 0.38 | 0.67 | 0.3 | 0.05 | 0.23 | 0.38 |
| 10 | 21530561 | rs12762419 | -1.359 | -0.976 | 0.49 | -1.636 | -0.013 | -0.075 | -0.075 | 0 | 0.33 | 0.51 | 0.25 | 0.11 | 0.19 | 0.31 |
| 10 | 21532085 | rs7092555 | -0.753 | -1.035 | 0.653 | -1.716 | 0.059 | -0.092 | -0.092 | 0.018 | 0.15 | 0.31 | 0.2 | 0 | 0.15 | 0.12 |
| 10 | 21532263 | rs7080918 | -1.762 | -1.177 | -0.57 | -2.61 | -0.125 | 0.166 | 0.166 | 0 | 0.62 | 0.89 | 0.54 | 0.15 | 0.5 | 0.56 |
| 10 | 21542327 | rs12269257 | 0.084 | -0.677 | 0.175 | -2.134 | -0.014 | -0.015 | -0.015 | 0.963 | 0.17 | 0.2 | 0.04 | 0.07 | 0.04 | 0.18 |
| 10 | 21544231 | rs7090954 | -0.513 | -0.661 | 0.231 | -2.298 | 0.067 | -0.09 | -0.09 | 0.005 | 0.15 | 0.37 | 0.21 | 0.01 | 0.16 | 0.12 |
| 10 | 21544357 | rs6482170 | -1.582 | -0.729 | -0.436 | -2.334 | -0.002 | -0.088 | -0.088 | 1.047 | 0.29 | 0.6 | 0.25 | 0.02 | 0.2 | 0.3 |
| 10 | 21546536 | rs7076532 | -0.953 | -0.148 | -0.706 | -1.9 | 0 | -0.033 | -0.033 | 0 | 0.21 | 0.71 | 0.26 | 0.01 | 0.17 | 0.1 |
| 10 | 21552094 | rs7896069 | -0.902 | -0.324 | -1.001 | -2.147 | 0.038 | 0.153 | 0.153 | 0 | 0.47 | 0.74 | 0.3 | 0.06 | 0.36 | 0.4 |
| 10 | 21567436 | rs12358291 | -0.873 | -0.08 | -0.047 | -1.881 | 0.118 | 0.036 | 0.036 | 0 | 0.05 | 0.29 | 0.11 | 0.01 | 0.06 | 0.06 |
| 10 | 21567724 | rs3895938 | -0.45 | 0.205 | -1.253 | -1.581 | -0.061 | -0.039 | -0.039 | 0.038 | 0.64 | 0.86 | 0.67 | 0.56 | 0.61 | 0.66 |
| 10 | 21569443 | rs11012639 | -0.008 | 1.042 | -0.924 | -1.666 | -0.049 | 0.063 | 0.063 | 0 | 0.88 | 0.98 | 0.88 | 0.99 | 0.82 | 0.88 |
| 10 | 21575812 | rs10828230 | 0.302 | 0.657 | -0.25 | -1.815 | 0.008 | -0.109 | -0.109 | 0 | 0.74 | 0.66 | 0.82 | 0.97 | 0.74 | 0.7 |
| 10 | 21581432 | rs2293438 | -1.035 | -0.181 | 0.156 | -2.004 | -0.044 | -0.108 | -0.108 | 0 | 0.29 | 0.48 | 0.2 | 0.03 | 0.17 | 0.25 |
| 10 | 21583368 | rs11012645 | 0.683 | -0.133 | 0.148 | -1.975 | -0.001 | -0.025 | -0.025 | 0 | 0.16 | 0.14 | 0.07 | 0.02 | 0.12 | 0.2 |
| 10 | 21586367 | rs7910000 | 0.633 | -0.137 | 0.161 | -1.977 | 0.008 | 0.014 | 0.014 | 0 | 0.15 | 0.09 | 0.07 | 0.02 | 0.12 | 0.2 |

| | | | | | | | | | | | | | | | | |
|---|---|---|---|---|---|---|---|---|---|---|---|---|---|---|---|---|
| 10 | 21587780 | rs10828232 | -0.737 | -0.273 | 0.249 | -2.042 | -0.004 | -0.033 | -0.033 | 5.62 | 0.27 | 0.27 | 0.12 | 0.03 | 0.18 | 0.24 |
| 10 | 21595291 | rs7922927 | 0.773 | -0.258 | 0.238 | -2.001 | 0.003 | -0.011 | -0.011 | 0 | 0.14 | 0.12 | 0.05 | 0.02 | 0.09 | 0.18 |
| 10 | 21599726 | rs12264555 | -0.276 | -0.111 | 0.598 | -2.167 | 0.042 | -0.072 | -0.072 | 0 | 0.12 | 0.49 | 0.17 | 0.01 | 0.08 | 0.06 |
| 10 | 21602923 | rs12241514 | -1.55 | -0.249 | 0.491 | -1.994 | 0.061 | 0.125 | 0.125 | 10.449 | 0.38 | 0.78 | 0.27 | 0.08 | 0.23 | 0.19 |
| 10 | 21603187 | rs16921721 | -0.398 | -0.287 | 0.459 | -1.911 | -0.001 | -0.009 | -0.009 | 0 | 0.26 | 0.25 | 0.09 | 0.07 | 0.16 | 0.13 |
| 10 | 21616815 | rs2049587 | 0.75 | -0.059 | 0.115 | -1.308 | 0.015 | 0 | 0 | 0.004 | 0.51 | 0.58 | 0.49 | 0.56 | 0.44 | 0.35 |
| 10 | 21628352 | rs7898226 | -0.612 | -0.2 | 0.488 | -1.322 | 0.025 | -0.081 | -0.081 | 0 | 0.13 | 0.49 | 0.11 | 0.01 | 0.08 | 0.06 |
| 10 | 21638971 | rs2039028 | -1.104 | -0.635 | 0.245 | -1.559 | 0.055 | 0.073 | 0.073 | 11.222 | 0.35 | 0.76 | 0.18 | 0.07 | 0.19 | 0.16 |
| 10 | 21639081 | rs2039029 | -0.139 | -0.288 | 0.581 | -1.367 | 0.014 | -0.048 | -0.048 | 0 | 0.11 | 0.44 | 0.09 | 0.01 | 0.05 | 0.06 |
| 10 | 21644741 | rs7893639 | -0.086 | -0.281 | 0.614 | -1.365 | 0.011 | -0.05 | -0.05 | 0 | 0.11 | 0.39 | 0.07 | 0.01 | 0.04 | 0.06 |
| 10 | 21646236 | rs12413399 | 0.498 | -0.27 | 0.593 | -1.359 | -0.001 | -0.034 | -0.034 | 0 | 0.08 | 0.16 | 0.05 | 0.01 | 0.04 | 0.06 |
| 10 | 21646353 | rs7915882 | -0.808 | 0.236 | 0.113 | -1.444 | -0.041 | 0.039 | 0.039 | 0 | 0.91 | 0.98 | 0.9 | 1 | 0.82 | 0.86 |
| 10 | 21653816 | rs3858205 | -0.688 | -0.472 | -0.276 | -1.069 | -0.141 | -0.141 | -0.141 | 0.04 | 0.66 | 0.95 | 0.48 | 0.56 | 0.47 | 0.39 |
| 10 | 21657474 | rs1556002 | -0.39 | -0.667 | 0.258 | -1.21 | -0.052 | -0.124 | -0.124 | 2.066 | 0.24 | 0.56 | 0.15 | 0.02 | 0.13 | 0.18 |
| 10 | 21668199 | rs12245880 | -0.479 | -0.889 | 0.093 | -1.161 | -0.024 | -0.053 | -0.053 | 0.022 | 0.26 | 0.64 | 0.17 | 0.06 | 0.16 | 0.18 |
| 10 | 21679527 | rs10828239 | 1.213 | -0.56 | -0.084 | -0.754 | 0.001 | -0.038 | -0.038 | 1.468 | 0.29 | 0.17 | 0.25 | 0.42 | 0.26 | 0.2 |
| 10 | 21683824 | rs16921796 | 0.304 | 0.056 | -0.675 | -0.621 | -0.001 | -0.009 | -0.009 | 0 | 0.07 | 0.13 | 0.04 | 0.04 | 0.04 | 0.06 |
| 10 | 21687123 | rs7923552 | -0.533 | -0.123 | -1.506 | -0.389 | -0.139 | -0.093 | -0.093 | 0.165 | 0.6 | 0.97 | 0.46 | 0.55 | 0.48 | 0.36 |
| 10 | 21701220 | rs12775448 | 0.483 | 0.064 | -1.169 | -0.752 | -0.021 | 0.069 | 0.069 | 0 | 0.17 | 0.03 | 0.12 | 0.01 | 0.18 | 0.12 |
| 10 | 21701447 | rs12776687 | -0.961 | -0.652 | -0.548 | -1.011 | 0.021 | 0.027 | 0.027 | 2.624 | 0.35 | 0.71 | 0.19 | 0.12 | 0.24 | 0.17 |
| 10 | 21709301 | rs12770588 | 0.734 | -0.549 | -0.407 | -0.995 | -0.005 | 0.013 | 0.013 | 0 | 0.17 | 0.08 | 0.12 | 0.01 | 0.18 | 0.11 |
| 10 | 21721758 | rs959652 | -1.005 | -0.688 | -0.365 | -1.039 | 0.047 | 0.079 | 0.079 | 2.266 | 0.36 | 0.79 | 0.2 | 0.11 | 0.23 | 0.17 |
| 10 | 21725452 | rs7899812 | -1.078 | -0.777 | -0.139 | -1.066 | 0.04 | 0.059 | 0.059 | 1.971 | 0.37 | 0.72 | 0.19 | 0.11 | 0.24 | 0.18 |
| 10 | 21734014 | rs12767504 | 0.334 | -0.739 | 0.461 | -1.081 | 0.004 | 0.024 | 0.024 | 0 | 0.2 | 0.11 | 0.12 | 0.01 | 0.18 | 0.13 |
| 10 | 21735144 | rs11593040 | 1.572 | -0.549 | -0.046 | -0.733 | -0.069 | -0.119 | -0.119 | 0.001 | 0.26 | 0.01 | 0.22 | 0.31 | 0.26 | 0.19 |
| 10 | 21762267 | rs11012716 | -0.823 | -0.752 | 0.764 | -0.838 | -0.017 | -0.034 | -0.034 | 0 | 0.33 | 0.42 | 0.17 | 0.17 | 0.23 | 0.17 |
| 10 | 21768926 | rs11595807 | -0.012 | -0.258 | 1.161 | -0.69 | -0.042 | -0.051 | -0.051 | 0 | 0.16 | 0.35 | 0.07 | 0.08 | 0.08 | 0.06 |
| 10 | 21838380 | rs4553272 | -1.509 | 1.016 | 0.394 | -0.622 | -0.168 | -0.148 | -0.148 | 0 | 0.69 | 0.97 | 0.47 | 0.37 | 0.53 | 0.38 |
| 10 | 21869609 | rs1243191 | -2.362 | 1.705 | 0.277 | -0.733 | -0.034 | 0.033 | 0.033 | 0 | 0.94 | 0.99 | 0.91 | 1 | 0.85 | 0.97 |
| 10 | 21874370 | rs11012746 | -0.73 | 1.35 | 0.43 | -0.867 | -0.005 | 0.095 | 0.095 | 0 | 0.23 | 0.51 | 0.24 | 0.31 | 0.19 | 0.15 |
| 10 | 21908803 | rs1243188 | 1.016 | 1.064 | 0.487 | -1.018 | -0.015 | -0.107 | -0.107 | 0.527 | 0.44 | 0.4 | 0.34 | 0.03 | 0.32 | 0.21 |
| 10 | 21916728 | rs1243182 | 0.22 | 1.15 | 0.553 | -1.001 | 0.026 | 0.015 | 0.015 | 0.319 | 0.5 | 0.55 | 0.34 | 0.07 | 0.33 | 0.24 |
| 10 | 21928049 | rs9418419 | -0.371 | 1.498 | 0.503 | -0.801 | 0.012 | 0.039 | 0.039 | 0 | 0.12 | 0.12 | 0.2 | 0.36 | 0.23 | 0.15 |
| 10 | 21936665 | rs1243190 | -1.687 | 1.258 | 0.543 | -0.838 | -0.037 | -0.064 | -0.064 | 0 | 0.69 | 0.83 | 0.59 | 0.42 | 0.59 | 0.41 |
| 10 | 21957229 | rs2183271 | -0.018 | 1.297 | 0.589 | -0.908 | 0.023 | 0.05 | 0.05 | 0.127 | 0.53 | 0.62 | 0.26 | 0.05 | 0.38 | 0.29 |
| 10 | 21970077 | rs1243197 | -0.724 | 1.167 | 0.656 | -0.768 | 0.001 | -0.004 | -0.004 | 0 | 0.56 | 0.54 | 0.55 | 0.39 | 0.55 | 0.37 |
| 10 | 21998445 | rs7477335 | 0.433 | 1.24 | 0.789 | -0.783 | 0 | -0.001 | -0.001 | 0 | 0.43 | 0.48 | 0.45 | 0.64 | 0.45 | 0.63 |
| 10 | 22030023 | rs4747437 | 1.163 | 1.206 | 0.744 | -0.926 | 0.001 | 0.017 | 0.017 | 1.504 | 0.33 | 0.41 | 0.41 | 0.57 | 0.36 | 0.61 |
| 10 | 22032942 | rs7072776 | -1.308 | 0.087 | 0.456 | -1.346 | 0.005 | -0.036 | -0.036 | 0 | 0.59 | 0.44 | 0.65 | 0.95 | 0.71 | 0.79 |
| 10 | 22067207 | rs7895753 | 1.993 | 0.201 | 0.218 | -1.12 | -0.007 | -0.017 | -0.017 | 0 | 0.09 | 0.09 | 0.02 | 0 | 0.02 | 0 |
| 10 | 22095737 | rs1970467 | 0.532 | 0.535 | -0.024 | -1.014 | -0.059 | -0.06 | -0.06 | 0 | 0.1 | 0.01 | 0.15 | 0.33 | 0.17 | 0.17 |
| 10 | 22219556 | rs2666750 | 0.188 | -0.491 | 0.251 | -2.098 | 0.075 | 0.178 | 0.178 | 2.453 | 0.46 | 0.22 | 0.62 | 0.95 | 0.67 | 0.76 |
| 10 | 22278857 | rs16921976 | -0.052 | -0.035 | 0.402 | -1.816 | 0.021 | 0.018 | 0.018 | 0 | 0.06 | 0.31 | 0.05 | 0.03 | 0.02 | 0.03 |
| 10 | 22280429 | rs6650129 | -0.515 | -0.048 | 0.324 | -1.856 | 0.014 | -0.011 | -0.011 | 0 | 0.59 | 0.63 | 0.66 | 0.95 | 0.7 | 0.79 |
| 10 | 22295747 | rs4304660 | 0.34 | 0.721 | -0.732 | -1.095 | -0.017 | 0.129 | 0.129 | 0.05 | 0.71 | 0.94 | 0.75 | 0.96 | 0.72 | 0.83 |
| 10 | 22299503 | rs7898685 | 1.014 | 0.593 | -0.37 | -1.213 | 0.024 | 0.048 | 0.048 | 0.321 | 0.6 | 0.72 | 0.7 | 0.93 | 0.69 | 0.83 |
| 10 | 22311305 | rs10740998 | 0.609 | 1.387 | 1.038 | -0.759 | -0.029 | 0.025 | 0.025 | 0 | 0.87 | 0.93 | 0.74 | 0.95 | 0.76 | 0.9 |
| 10 | 22318101 | rs4529807 | 0.39 | 1.384 | 0.962 | -0.757 | -0.076 | 0.036 | 0.036 | 0.007 | 0.88 | 0.99 | 0.74 | 0.95 | 0.76 | 0.88 |
| 10 | 22319508 | rs12248406 | -2.625 | 1.068 | 1.585 | -0.845 | -0.047 | -0.057 | -0.057 | 1.231 | 0.11 | 0.73 | 0.07 | 0.01 | 0.02 | 0.03 |
| 10 | 22326472 | rs11012847 | 2.643 | 1.387 | 1.328 | -0.702 | -0.133 | 0.115 | 0.115 | 0.393 | 0.55 | 0.04 | 0.5 | 0.8 | 0.54 | 0.64 |
| 10 | 22336151 | rs6482198 | 0.477 | 1.545 | 2.106 | -0.735 | -0.02 | 0.017 | 0.017 | 0 | 0.88 | 0.92 | 0.75 | 0.95 | 0.78 | 0.89 |
| 10 | 22360446 | rs2986341 | 2.315 | 1.549 | 2.54 | -0.816 | -0.023 | 0.129 | 0.129 | 0.834 | 0.68 | 0.14 | 0.65 | 0.95 | 0.7 | 0.86 |
| 10 | 22362410 | rs3011631 | -1.21 | 1.712 | 2.527 | -0.754 | -0.043 | 0.043 | 0.043 | 0 | 0.92 | 0.99 | 0.9 | 1 | 0.82 | 0.96 |
| 10 | 22362612 | rs11818094 | -3.022 | 0.743 | 3.594 | -0.154 | 0.053 | 0.05 | 0.05 | 2.001 | 0.05 | 0.34 | 0.04 | 0.01 | 0.02 | 0.01 |
| 10 | 22394843 | rs3011642 | -2.488 | 0.816 | 3.534 | 0.026 | 0.002 | -0.01 | -0.01 | 0 | 0.22 | 0.69 | 0.32 | 0.22 | 0.22 | 0.13 |
| 10 | 22397995 | rs3011644 | -0.978 | 0.826 | 3.5 | 0.031 | -0.029 | -0.01 | -0.01 | 0 | 0.13 | 0.07 | 0.26 | 0.2 | 0.21 | 0.11 |
| 10 | 22427400 | rs13376918 | -2.946 | 0.694 | 3.683 | 0.078 | -0.021 | 0.003 | 0.003 | 3.702 | 0.17 | 0.69 | 0.25 | 0.22 | 0.1 | 0.1 |
| 10 | 22441887 | rs11012878 | -0.401 | 0.743 | 3.445 | 0.102 | 0 | 0.009 | 0.009 | 0 | 0.07 | 0.07 | 0.19 | 0.21 | 0.09 | 0.07 |
| 10 | 22460830 | rs10828296 | -1.976 | 0.637 | 3.807 | 0.114 | 0.009 | -0.011 | -0.011 | 0 | 0.22 | 0.62 | 0.32 | 0.22 | 0.23 | 0.14 |
| 10 | 22471741 | rs1926690 | -1.384 | 0.633 | 3.93 | 0.102 | 0.034 | 0.002 | 0.002 | 0.001 | 0.18 | 0.33 | 0.29 | 0.22 | 0.24 | 0.14 |
| 10 | 22474979 | rs4748792 | -2.126 | 0.614 | 3.831 | 0.102 | 0.003 | -0.011 | -0.011 | 0 | 0.23 | 0.66 | 0.32 | 0.22 | 0.24 | 0.15 |
| 10 | 22477498 | rs11593316 | -0.829 | 0.614 | 3.828 | 0.101 | 0.01 | 0.002 | 0.002 | 0 | 0.16 | 0.21 | 0.27 | 0.21 | 0.23 | 0.13 |
| 10 | 22479936 | rs6482203 | -1.895 | 0.614 | 3.857 | 0.108 | 0.019 | -0.018 | -0.018 | 2.813 | 0.22 | 0.53 | 0.31 | 0.22 | 0.23 | 0.14 |
| 10 | 22496104 | rs913110 | -2.35 | 0.596 | 3.57 | 0.091 | -0.001 | -0.006 | -0.006 | 0 | 0.27 | 0.64 | 0.32 | 0.22 | 0.25 | 0.14 |
| 10 | 22496669 | rs1097960 | -0.542 | 0.598 | 3.587 | 0.105 | 0 | -0.001 | -0.001 | 0.483 | 0.14 | 0.12 | 0.27 | 0.21 | 0.21 | 0.11 |
| 10 | 22498209 | rs2243897 | -2.169 | 0.595 | 3.807 | 0.079 | 0.002 | -0.03 | -0.03 | 0.303 | 0.26 | 0.56 | 0.31 | 0.22 | 0.25 | 0.14 |
| 10 | 22498468 | rs838759 | -1.355 | 0.604 | 3.478 | 0.083 | 0.024 | 0 | 0 | 0 | 0.18 | 0.27 | 0.28 | 0.21 | 0.23 | 0.13 |
| 10 | 22501329 | rs1555922 | -0.64 | 0.59 | 3.202 | 0.065 | 0 | -0.001 | -0.001 | 0.255 | 0.14 | 0.13 | 0.27 | 0.21 | 0.21 | 0.11 |
| 10 | 22504189 | rs2148959 | -2.002 | 0.577 | 3.088 | 0.105 | 0.001 | 0.011 | 0.011 | 0.373 | 0.27 | 0.71 | 0.33 | 0.21 | 0.25 | 0.14 |

| | | | | | | | | | | | | | | | | |
|---|---|---|---|---|---|---|---|---|---|---|---|---|---|---|---|---|
| 10 | 22505064 | rs11012894 | -1.342 | 0.559 | 2.664 | 0.091 | -0.009 | -0.025 | -0.025 | 0.38 | 0.09 | 0.44 | 0.04 | 0.01 | 0.03 | 0.01 |
| 10 | 22509765 | rs4539216 | 0.794 | 0.625 | 1.216 | 0.442 | -0.026 | -0.128 | -0.128 | 0 | 0.19 | 0.01 | 0.28 | 0.41 | 0.17 | 0.28 |
| 10 | 22517518 | rs1556010 | -2.149 | 0.913 | 2.194 | 0.166 | -0.086 | 0.036 | 0.036 | 0.189 | 0.45 | 0.97 | 0.49 | 0.34 | 0.42 | 0.37 |
| 10 | 22531317 | rs12250660 | -2.688 | 1.649 | 3.637 | 0.141 | -0.05 | -0.071 | -0.071 | 0 | 0.11 | 0.52 | 0.06 | 0.01 | 0.03 | 0.02 |
| 10 | 22532959 | rs2478473 | -1.69 | 2.005 | 3.642 | 0.387 | -0.054 | -0.001 | -0.001 | 0 | 0.17 | 0.07 | 0.37 | 0.18 | 0.3 | 0.16 |
| 10 | 22538174 | rs7067636 | 2.537 | 1.995 | 3.821 | 0.37 | 0.005 | -0.012 | -0.012 | 0 | 0.75 | 0.3 | 0.58 | 0.81 | 0.7 | 0.83 |
| 10 | 22539138 | rs6482206 | 1.624 | 1.426 | 3.464 | 0.39 | -0.04 | 0 | 0 | 0.01 | 0.79 | 0.89 | 0.6 | 0.8 | 0.71 | 0.82 |
| 10 | 22568695 | rs7083590 | -3.974 | 1.068 | 4.29 | 0.366 | -0.021 | -0.029 | -0.029 | 0 | 0.07 | 0.69 | 0.06 | 0 | 0.01 | 0.01 |
| 10 | 22594765 | rs11012937 | -2.315 | 1.026 | 4.368 | 0.335 | -0.048 | -0.07 | -0.07 | 2.761 | 0.11 | 0.61 | 0.08 | 0 | 0.02 | 0.01 |
| 10 | 22658281 | rs1170549 | 0.805 | 0.605 | 4.028 | 0.468 | -0.054 | -0.057 | -0.057 | 0 | 0.88 | 0.99 | 0.77 | 0.78 | 0.82 | 0.85 |
| 10 | 22683351 | rs16922235 | -1.819 | 0.12 | 4.668 | 0.031 | -0.018 | -0.034 | -0.034 | 0 | 0.08 | 0.25 | 0.05 | 0 | 0.02 | 0.01 |
| 10 | 22712097 | rs7909597 | -2.539 | 0.118 | 4.511 | 0.005 | -0.049 | -0.07 | -0.07 | 0 | 0.11 | 0.63 | 0.08 | 0 | 0.02 | 0.01 |
| 10 | 22723311 | rs7080269 | -2.871 | -0.355 | 3.663 | 0.108 | -0.058 | -0.075 | -0.075 | 0 | 0.13 | 0.62 | 0.09 | 0.01 | 0.04 | 0.01 |
| 10 | 22724200 | rs6482214 | -1.91 | -0.354 | 3.711 | 0.109 | -0.004 | -0.02 | -0.02 | 0 | 0.09 | 0.45 | 0.07 | 0.01 | 0.02 | 0.01 |
| 10 | 22727983 | rs11012993 | -2.056 | -0.315 | 3.122 | 0.127 | 0.038 | 0.04 | 0.04 | 5.075 | 0.06 | 0.6 | 0.06 | 0.01 | 0 | 0 |
| 10 | 22734088 | rs11012997 | -2.056 | -0.315 | 3.122 | 0.127 | 0.038 | 0.04 | 0.04 | 5.075 | 0.06 | 0.6 | 0.06 | 0.01 | 0 | 0 |
| 10 | 22734795 | rs7918856 | -2.056 | -0.315 | 3.122 | 0.127 | 0.038 | 0.04 | 0.04 | 5.075 | 0.06 | 0.6 | 0.06 | 0.01 | 0 | 0 |
| 10 | 22743447 | rs12243038 | -2.02 | -0.273 | 2.145 | 0.139 | -0.054 | -0.077 | -0.077 | 0.058 | 0.14 | 0.65 | 0.1 | 0.01 | 0.06 | 0.02 |
| 10 | 22770341 | rs7087192 | -1.89 | 0.17 | 1.065 | 0.398 | -0.028 | -0.031 | -0.031 | 0.012 | 0.12 | 0.4 | 0.15 | 0.08 | 0.06 | 0.06 |
| 10 | 22777875 | rs16922359 | -1.888 | -0.157 | 1.412 | 0.149 | -0.037 | -0.087 | -0.087 | 3.709 | 0.15 | 0.32 | 0.12 | 0 | 0.09 | 0.08 |
| 10 | 22779693 | rs12570045 | -0.763 | 0.485 | 0.831 | 1.191 | -0.047 | -0.072 | -0.072 | 0 | 0.12 | 0.02 | 0.11 | 0.4 | 0.19 | 0.16 |
| 10 | 22781927 | rs16922373 | -2.114 | 0.219 | 1.596 | 1.017 | 0.044 | 0.079 | 0.079 | 1.054 | 0.06 | 0.38 | 0.1 | 0.08 | 0.03 | 0.02 |
| 10 | 22790884 | rs11593715 | -1.904 | 0.2 | 2.184 | 0.89 | -0.036 | -0.084 | -0.084 | 0 | 0.13 | 0.44 | 0.11 | 0 | 0.08 | 0.08 |
| 10 | 22796761 | rs7092910 | -1.756 | 0.969 | 1.703 | 1.426 | -0.024 | -0.052 | -0.052 | 0 | 0.16 | 0.12 | 0.17 | 0.4 | 0.21 | 0.21 |
| 10 | 22822524 | rs7899156 | 1.98 | 1.161 | 2.025 | 1.317 | 0.015 | 0.021 | 0.021 | 0 | 0.74 | 0.64 | 0.81 | 0.65 | 0.68 | 0.78 |
| 10 | 22826093 | rs1053454 | 1.997 | 1.608 | 1.706 | 1.078 | -0.044 | -0.037 | -0.037 | 0 | 0.77 | 0.85 | 0.81 | 0.65 | 0.64 | 0.78 |
| 10 | 22837014 | rs4532931 | 2.451 | 1.807 | 2.259 | 1.469 | 0.002 | 0.004 | 0.004 | 0 | 0.67 | 0.39 | 0.64 | 0.57 | 0.63 | 0.78 |
| 10 | 22847439 | rs7094131 | 2.662 | 1.718 | 2.451 | 1.335 | -0.036 | -0.042 | -0.042 | 0 | 0.65 | 0.21 | 0.63 | 0.55 | 0.61 | 0.77 |
| 10 | 22847893 | rs7081744 | 2.651 | 1.719 | 2.452 | 1.263 | -0.036 | -0.018 | -0.018 | 0 | 0.65 | 0.21 | 0.75 | 0.63 | 0.61 | 0.77 |
| 10 | 22851609 | rs7078635 | 1.084 | 1.801 | 2.567 | 1.226 | 0.001 | -0.001 | -0.001 | 0.067 | 0.93 | 0.95 | 0.95 | 1 | 0.94 | 0.96 |
| 10 | 22852948 | rs7088318 | 2.659 | 1.777 | 2.662 | 1.252 | -0.013 | -0.007 | -0.007 | 0.655 | 0.65 | 0.34 | 0.75 | 0.61 | 0.6 | 0.77 |
| 10 | 22859530 | rs3793753 | 2.226 | 3.034 | 3.4 | 1.213 | 0.036 | -0.013 | -0.013 | 0 | 0.81 | 0.55 | 0.77 | 0.86 | 0.81 | 0.91 |
| 10 | 22859656 | rs12253847 | -2.012 | 1.939 | 3.22 | 0.398 | -0.002 | -0.035 | -0.035 | 0 | 0.38 | 0.38 | 0.2 | 0.04 | 0.42 | 0.36 |
| 10 | 22860068 | rs7919839 | 0.53 | 1.002 | 2.295 | 0.127 | 0.05 | 0.053 | 0.053 | 5.74 | 0.8 | 0.98 | 0.9 | 0.94 | 0.88 | 0.83 |
| 10 | 22863004 | rs16922479 | -0.587 | 1.374 | 2.561 | 0.047 | -0.019 | 0.005 | 0.005 | 0.429 | 0.15 | 0.34 | 0.11 | 0.1 | 0.11 | 0.13 |
| 10 | 22864209 | rs11013053 | -1.908 | 2.028 | 2.592 | -0.11 | 0.035 | -0.047 | -0.047 | 0 | 0.3 | 0.56 | 0.2 | 0.11 | 0.35 | 0.22 |
| 10 | 22866892 | rs10764338 | 0.908 | 1.95 | 2.557 | -0.13 | -0.016 | -0.017 | -0.017 | 1.364 | 0.77 | 0.72 | 0.87 | 0.9 | 0.86 | 0.83 |
| 10 | 22867451 | rs7071450 | 2.08 | 2.172 | 2.609 | -0.158 | 0 | 0.021 | 0.021 | 0.094 | 0.55 | 0.39 | 0.66 | 0.86 | 0.51 | 0.59 |
| 10 | 22869502 | rs12357384 | -1.047 | 2.208 | 2.435 | -0.162 | -0.107 | 0.076 | 0.076 | 0.031 | 0.19 | 0.04 | 0.17 | 0.03 | 0.35 | 0.22 |
| 10 | 22873207 | rs7087066 | 1.605 | 1.913 | 2.555 | -0.147 | -0.007 | 0.016 | 0.016 | 0.346 | 0.59 | 0.65 | 0.76 | 0.95 | 0.59 | 0.63 |
| 10 | 22874216 | rs7912144 | 0.119 | 1.786 | 2.606 | -0.148 | 0.001 | 0.016 | 0.016 | 0.02 | 0.11 | 0.2 | 0.09 | 0.1 | 0.1 | 0.12 |
| 10 | 22878239 | rs7921549 | 1.679 | 1.78 | 2.394 | -0.288 | -0.002 | -0.026 | -0.026 | 1.524 | 0.66 | 0.66 | 0.81 | 0.97 | 0.64 | 0.71 |
| 10 | 22882081 | rs10764342 | 2.061 | 1.768 | 2.549 | -0.284 | 0 | -0.066 | -0.066 | 0 | 0.55 | 0.45 | 0.76 | 0.95 | 0.58 | 0.62 |
| 10 | 22885232 | rs10764344 | 1.45 | 1.806 | 2.437 | -0.259 | -0.007 | 0 | 0 | 0 | 0.68 | 0.71 | 0.83 | 0.97 | 0.66 | 0.71 |
| 10 | 22889530 | rs2765993 | 0.664 | 1.491 | 2.401 | -0.2 | 0.004 | 0.003 | 0.003 | 0.65 | 0.84 | 0.84 | 0.9 | 0.98 | 0.89 | 0.85 |
| 10 | 22890965 | rs2913109 | 1.029 | 1.486 | 2.571 | -0.181 | -0.011 | -0.009 | -0.009 | 0.676 | 0.83 | 0.56 | 0.86 | 0.9 | 0.88 | 0.84 |
| 10 | 22892152 | rs2559524 | 1.52 | 1.469 | 2.378 | -0.189 | -0.023 | -0.022 | -0.022 | 0.095 | 0.8 | 0.32 | 0.84 | 0.9 | 0.89 | 0.85 |
| 10 | 22892263 | rs7093543 | 0.515 | 1.487 | 2.183 | -0.185 | -0.023 | 0.023 | 0.023 | 1.796 | 0.09 | 0.01 | 0.07 | 0.02 | 0.09 | 0.11 |
| 10 | 22894491 | rs2765996 | 0.236 | 1.489 | 2.085 | -0.182 | 0.004 | 0.039 | 0.039 | 0.698 | 0.87 | 0.94 | 0.9 | 0.97 | 0.9 | 0.85 |
| 10 | 22898017 | rs2559527 | 2.66 | 1.582 | 2.045 | -0.213 | -0.009 | 0.107 | 0.107 | 0.408 | 0.56 | 0.23 | 0.73 | 0.87 | 0.6 | 0.62 |
| 10 | 22905234 | rs1778353 | -2.196 | 1.662 | 2.111 | -0.149 | -0.015 | 0.053 | 0.053 | 0 | 0.37 | 0.77 | 0.25 | 0.13 | 0.36 | 0.36 |
| 10 | 22908266 | rs1778356 | -1.308 | 1.526 | 2.221 | -0.091 | -0.016 | -0.016 | -0.016 | 0 | 0.18 | 0.65 | 0.16 | 0.1 | 0.11 | 0.15 |
| 10 | 22912326 | rs11013069 | -1.415 | 1.576 | 2.059 | -0.208 | 0.022 | -0.003 | -0.003 | 0.255 | 0.17 | 0.37 | 0.12 | 0.1 | 0.2 | 0.18 |
| 10 | 22912425 | rs1750761 | -0.19 | 0.437 | 0.481 | -0.581 | 0.015 | 0.022 | 0.022 | 5.193 | 0.17 | 0.05 | 0.11 | 0.13 | 0.15 | 0.12 |
| 10 | 22913051 | rs1627983 | -1.661 | 0.814 | 1.098 | -0.677 | -0.007 | -0.017 | -0.017 | 0.024 | 0.23 | 0.7 | 0.15 | 0.1 | 0.16 | 0.16 |
| 10 | 22932491 | rs1409395 | -1.873 | 0.983 | 0.98 | -0.755 | -0.021 | 0.035 | 0.035 | 0 | 0.24 | 0.18 | 0.12 | 0.03 | 0.3 | 0.29 |
| 10 | 22941536 | rs16922560 | -0.4 | 0.954 | 1.221 | -0.768 | -0.002 | -0.02 | -0.02 | 0 | 0.14 | 0.17 | 0.07 | 0.02 | 0.11 | 0.11 |
| 10 | 22945961 | rs1171496 | -2.606 | 0.969 | 1.497 | -0.758 | -0.006 | 0.008 | 0.008 | 0 | 0.38 | 0.64 | 0.23 | 0.12 | 0.41 | 0.35 |
| 10 | 22949035 | rs12098721 | -0.401 | 0.994 | 1.473 | -0.714 | -0.005 | -0.04 | -0.04 | 0 | 0.14 | 0.22 | 0.08 | 0.02 | 0.12 | 0.1 |
| 10 | 22961707 | rs16922578 | -0.366 | 1.204 | 1.443 | -0.72 | -0.001 | -0.017 | -0.017 | 0.107 | 0.14 | 0.16 | 0.08 | 0.02 | 0.12 | 0.1 |
| 10 | 22966606 | rs10159847 | -1.226 | 1.185 | 1.614 | -0.741 | -0.015 | -0.072 | -0.072 | 0 | 0.21 | 0.33 | 0.11 | 0.02 | 0.15 | 0.14 |
| 10 | 22971463 | rs11013095 | -2.433 | 1.272 | 1.554 | -0.792 | -0.018 | 0.031 | 0.031 | 0 | 0.37 | 0.31 | 0.19 | 0.04 | 0.42 | 0.36 |
| 10 | 22974261 | rs6482228 | -2.618 | 0.347 | 1.417 | -0.825 | 0.005 | 0.093 | 0.093 | 0 | 0.38 | 0.24 | 0.16 | 0.05 | 0.32 | 0.38 |
| 10 | 22978029 | rs1891877 | -3.139 | 0.469 | 0.971 | -1.038 | -0.101 | 0.23 | 0.23 | 1.591 | 0.52 | 0.97 | 0.37 | 0.16 | 0.45 | 0.49 |
| 10 | 22984875 | rs11013103 | -1.055 | 0.45 | 1.171 | -1.019 | -0.001 | -0.023 | -0.023 | 1.925 | 0.21 | 0.25 | 0.09 | 0.02 | 0.16 | 0.16 |
| 10 | 22985068 | rs11013104 | -1.925 | 0.448 | 1.434 | -1.021 | -0.002 | -0.006 | -0.006 | 0 | 0.23 | 0.72 | 0.15 | 0.11 | 0.16 | 0.16 |
| 10 | 22990428 | rs7079078 | -2.484 | 0.512 | 1.48 | -0.937 | -0.003 | 0.072 | 0.072 | 0 | 0.4 | 0.73 | 0.22 | 0.13 | 0.34 | 0.4 |
| 10 | 22990528 | rs12255136 | -2.192 | 0.501 | 1.407 | -0.97 | -0.003 | 0.032 | 0.032 | 0 | 0.38 | 0.3 | 0.16 | 0.03 | 0.33 | 0.4 |
| 10 | 22991590 | rs7084115 | -2.542 | 0.611 | 1.314 | -0.93 | 0.016 | 0.136 | 0.136 | 1.132 | 0.46 | 0.72 | 0.22 | 0.06 | 0.36 | 0.36 |

| | | | | | | | | | | | | | | | | |
|---|---|---|---|---|---|---|---|---|---|---|---|---|---|---|---|---|
| 10 | 22995449 | rs12259546 | -2.747 | -0.084 | 1.514 | -0.843 | 0.016 | 0.219 | 0.219 | 1.04 | 0.46 | 0.85 | 0.31 | 0.07 | 0.38 | 0.37 |
| 10 | 23020917 | rs6482232 | -2.292 | -0.18 | 1.527 | -0.836 | -0.004 | -0.034 | -0.034 | 0.012 | 0.34 | 0.55 | 0.19 | 0.1 | 0.28 | 0.4 |
| 10 | 23022966 | rs6482235 | -2.466 | -0.202 | 1.499 | -0.849 | 0.023 | 0.076 | 0.076 | 0 | 0.41 | 0.72 | 0.22 | 0.11 | 0.3 | 0.4 |
| 10 | 23027581 | rs17527491 | -1.382 | -0.242 | 1.248 | -0.637 | -0.059 | 0.059 | 0.059 | 0 | 0.23 | 0.02 | 0.14 | 0.08 | 0.23 | 0.32 |
| 10 | 23030282 | rs7091875 | -2.442 | -0.185 | 1.344 | -0.582 | 0.004 | 0.007 | 0.007 | 0 | 0.4 | 0.57 | 0.24 | 0.15 | 0.3 | 0.4 |
| 10 | 23038083 | rs7092729 | 1.125 | -0.146 | 1.273 | -0.501 | 0.021 | 0.081 | 0.081 | 0.601 | 0.08 | 0.4 | 0.06 | 0.07 | 0.03 | 0.01 |
| 10 | 23049139 | rs7922014 | -2.495 | -0.093 | 0.414 | -0.005 | -0.055 | -0.045 | -0.045 | 0.008 | 0.47 | 0.96 | 0.41 | 0.42 | 0.38 | 0.44 |
| 10 | 23054884 | rs12248702 | 0.939 | -0.622 | 1.398 | -0.178 | 0.02 | 0.006 | 0.006 | 0.504 | 0.08 | 0.39 | 0.05 | 0.02 | 0.03 | 0.01 |
| 10 | 23062951 | rs7093771 | -2.08 | -0.616 | 0.661 | -0.108 | 0.029 | 0.064 | 0.064 | 0 | 0.32 | 0.8 | 0.22 | 0.12 | 0.2 | 0.32 |
| 10 | 23071128 | rs10828334 | -1.994 | -0.68 | 0.961 | -0.227 | -0.012 | -0.062 | -0.062 | 0.183 | 0.29 | 0.6 | 0.21 | 0.07 | 0.22 | 0.29 |
| 10 | 23079385 | rs7922263 | -0.647 | -0.744 | 1.503 | -0.155 | -0.036 | -0.101 | -0.101 | 0.05 | 0.17 | 0.36 | 0.08 | 0.01 | 0.1 | 0.05 |
| 10 | 23082827 | rs11013131 | -0.53 | -0.706 | 1.295 | -0.177 | -0.037 | 0.002 | 0.002 | 0 | 0.1 | 0.01 | 0.1 | 0.05 | 0.14 | 0.2 |
| 10 | 23090535 | rs2222871 | -2.731 | -1.491 | 1.186 | 0.3 | 0.096 | 0.005 | 0.005 | 0.01 | 0.5 | 0.84 | 0.43 | 0.56 | 0.3 | 0.4 |
| 10 | 23097537 | rs11013139 | -1.328 | -1.541 | 1.51 | 0.443 | -0.007 | -0.011 | -0.011 | 0 | 0.22 | 0.23 | 0.26 | 0.16 | 0.17 | 0.28 |
| 10 | 23102588 | rs7074731 | -0.205 | -1.567 | 1.367 | 0.431 | -0.001 | -0.001 | -0.001 | 0 | 0.17 | 0.18 | 0.23 | 0.08 | 0.17 | 0.23 |
| 10 | 23106191 | rs7083521 | -1.563 | -1.6 | 1.442 | 0.494 | -0.072 | 0.173 | 0.173 | 0.325 | 0.19 | 0.51 | 0.16 | 0.48 | 0.05 | 0.14 |
| 10 | 23107822 | rs7093538 | -0.946 | -1.661 | 1.418 | 0.403 | -0.081 | -0.038 | -0.038 | 0 | 0.15 | 0.39 | 0.07 | 0.09 | 0.03 | 0.08 |
| 10 | 23110328 | rs10764345 | -1.961 | -1.535 | 1.374 | 0.429 | 0.024 | 0.038 | 0.038 | 0 | 0.42 | 0.61 | 0.33 | 0.17 | 0.28 | 0.34 |
| 10 | 23110797 | rs7091463 | -1.421 | -1.532 | 1.338 | 0.529 | -0.045 | 0.065 | 0.065 | 1.362 | 0.25 | 0.42 | 0.15 | 0.4 | 0.13 | 0.11 |
| 10 | 23113841 | rs1890950 | -2.131 | -1.097 | 0.593 | 0.242 | 0.047 | 0.225 | 0.225 | 0 | 0.5 | 0.8 | 0.38 | 0.17 | 0.38 | 0.37 |
| 10 | 23121820 | rs7084892 | -1.687 | -0.935 | 1.215 | 0.603 | -0.008 | 0.012 | 0.012 | 0 | 0.36 | 0.44 | 0.37 | 0.58 | 0.3 | 0.41 |
| 10 | 23128475 | rs1417374 | -1.112 | -0.817 | 1.304 | 0.446 | -0.019 | 0.029 | 0.029 | 0.129 | 0.27 | 0.1 | 0.26 | 0.18 | 0.29 | 0.35 |
| 10 | 23128862 | rs10734042 | -2.673 | -1.067 | 1.161 | 0.566 | 0.014 | 0.058 | 0.058 | 0 | 0.5 | 0.87 | 0.44 | 0.6 | 0.4 | 0.45 |
| 10 | 23130720 | rs10741010 | -2.648 | -1.072 | 1.486 | 0.6 | 0.029 | 0.055 | 0.055 | 0 | 0.49 | 0.71 | 0.43 | 0.6 | 0.39 | 0.44 |
| 10 | 23137611 | rs11013152 | -1.337 | -0.902 | 1.693 | 0.585 | 0 | -0.005 | -0.005 | 0 | 0.29 | 0.25 | 0.27 | 0.18 | 0.29 | 0.34 |
| 10 | 23137898 | rs11013153 | 0.006 | -1.169 | 1.795 | 0.726 | -0.007 | 0.195 | 0.195 | 0 | 0.06 | 0.12 | 0.09 | 0.42 | 0.02 | 0.07 |
| 10 | 23140318 | rs10828350 | -1.791 | -1.19 | 1.821 | 0.488 | 0.002 | 0.003 | 0.003 | 0 | 0.39 | 0.35 | 0.32 | 0.19 | 0.34 | 0.37 |
| 10 | 23140731 | rs10828351 | -1.101 | -1.21 | 1.75 | 0.479 | -0.003 | -0.008 | -0.008 | 0 | 0.26 | 0.24 | 0.26 | 0.18 | 0.25 | 0.32 |
| 10 | 23151064 | rs2222870 | -1.2 | -1.259 | 1.74 | 0.687 | -0.029 | 0.085 | 0.085 | 0 | 0.2 | 0.29 | 0.14 | 0.38 | 0.1 | 0.1 |
| 10 | 23153350 | rs7070284 | -2.46 | -1.242 | 1.446 | 0.64 | 0.059 | 0.063 | 0.063 | 0 | 0.49 | 0.81 | 0.44 | 0.57 | 0.37 | 0.44 |
| 10 | 23157013 | rs7097449 | -1.307 | -1.188 | 1.777 | 0.71 | -0.037 | 0.122 | 0.122 | 0 | 0.21 | 0.35 | 0.12 | 0.38 | 0.1 | 0.1 |
| 10 | 23165066 | rs3898027 | -1.085 | -1.148 | 1.913 | 0.753 | -0.018 | 0.235 | 0.235 | 0 | 0.09 | 0.21 | 0.07 | 0.38 | 0.02 | 0.08 |
| 10 | 23199531 | rs4747449 | -2.333 | -0.977 | 1.358 | 0.508 | 0.023 | -0.004 | -0.004 | 0 | 0.49 | 0.88 | 0.51 | 0.52 | 0.39 | 0.45 |
| 10 | 23206001 | rs6482244 | -2.09 | -1.041 | 1.224 | 0.46 | 0.107 | -0.088 | -0.088 | 0 | 0.43 | 0.87 | 0.42 | 0.48 | 0.25 | 0.39 |
| 10 | 23206446 | rs1922124 | -2.557 | -1.018 | 1.335 | 0.453 | 0.041 | 0.018 | 0.018 | 0 | 0.49 | 0.81 | 0.5 | 0.52 | 0.39 | 0.45 |
| 10 | 23214761 | rs7098947 | -0.756 | -1.314 | 1.836 | 0.289 | -0.03 | -0.023 | -0.023 | 0.036 | 0.18 | 0.23 | 0.18 | 0.14 | 0.12 | 0.26 |
| 10 | 23217699 | rs7067598 | -2.273 | -1.339 | 1.562 | 0.283 | 0.104 | -0.06 | -0.06 | 0 | 0.43 | 0.81 | 0.42 | 0.47 | 0.25 | 0.38 |
| 10 | 23222070 | rs16922812 | -1.479 | -1.087 | 1.905 | 0.412 | -0.053 | 0.138 | 0.138 | 0 | 0.23 | 0.37 | 0.33 | 0.47 | 0.13 | 0.33 |
| 10 | 23225414 | rs2448132 | -1.265 | -1.1 | 1.894 | 0.408 | -0.023 | 0.065 | 0.065 | 0 | 0.21 | 0.26 | 0.31 | 0.45 | 0.12 | 0.32 |
| 10 | 23230827 | rs10764366 | -1.686 | -0.696 | 1.925 | 0.325 | 0.012 | -0.059 | -0.059 | 0.015 | 0.28 | 0.59 | 0.17 | 0.13 | 0.32 | 0.27 |
| 10 | 23234144 | rs1148301 | -2.311 | -0.751 | 1.774 | 0.405 | -0.026 | -0.119 | -0.119 | 3.574 | 0.37 | 0.9 | 0.44 | 0.46 | 0.33 | 0.36 |
| 10 | 23247377 | rs10828382 | -1.753 | -0.653 | 1.965 | 0.211 | -0.008 | 0.022 | 0.022 | 0 | 0.3 | 0.71 | 0.2 | 0.13 | 0.36 | 0.29 |
| 10 | 23249109 | rs11013210 | -0.655 | -0.615 | 2.211 | 0.207 | 0.014 | -0.01 | -0.01 | 0 | 0.15 | 0.16 | 0.1 | 0.12 | 0.25 | 0.24 |
| 10 | 23249878 | rs7100326 | 0.095 | -0.946 | 2.309 | 0.234 | -0.007 | -0.047 | -0.047 | 0 | 0.11 | 0.18 | 0.06 | 0 | 0.09 | 0.03 |
| 10 | 23258966 | rs7907625 | -0.665 | -0.802 | 2.182 | 0.264 | 0.005 | -0.012 | -0.012 | 0 | 0.15 | 0.16 | 0.08 | 0.13 | 0.18 | 0.21 |
| 10 | 23263945 | rs3101795 | -2.051 | -1.017 | 1.945 | 0.246 | 0 | -0.002 | -0.002 | 0.021 | 0.4 | 0.6 | 0.43 | 0.5 | 0.36 | 0.38 |
| 10 | 23273110 | rs1060761 | -2.294 | -0.994 | 1.572 | 0.146 | 0.008 | -0.03 | -0.03 | 3.676 | 0.44 | 0.73 | 0.45 | 0.47 | 0.38 | 0.41 |
| 10 | 23274996 | rs1170665 | -0.439 | -1.486 | 0.709 | -0.253 | -0.019 | -0.138 | -0.138 | 1.279 | 0.23 | 0.02 | 0.34 | 0.35 | 0.21 | 0.16 |
| 10 | 23283308 | rs10828392 | -0.419 | -0.994 | 1.498 | -0.028 | 0.012 | -0.026 | -0.026 | 0.03 | 0.16 | 0.26 | 0.09 | 0.11 | 0.18 | 0.21 |
| 10 | 23286466 | rs11013228 | 1.943 | -1.089 | 1.439 | 0.021 | 0.013 | -0.025 | -0.025 | 0 | 0.55 | 0.25 | 0.55 | 0.55 | 0.62 | 0.58 |
| 10 | 23289062 | rs10828393 | -0.03 | -0.886 | 1.437 | 0.189 | -0.018 | -0.082 | -0.082 | 0.63 | 0.87 | 0.99 | 0.72 | 0.67 | 0.89 | 0.87 |
| 10 | 23290952 | rs12259839 | 1.292 | -0.882 | 1.587 | 0.079 | -0.001 | -0.039 | -0.039 | 0 | 0.09 | 0.16 | 0.03 | 0 | 0.06 | 0.04 |
| 10 | 23297252 | rs10828395 | -0.722 | -0.982 | 1.833 | 0.022 | 0.011 | -0.031 | -0.031 | 0.439 | 0.17 | 0.28 | 0.1 | 0.12 | 0.19 | 0.2 |
| 10 | 23297301 | rs11817610 | 2.144 | -1.061 | 1.836 | 0.045 | 0.017 | -0.02 | -0.02 | 0 | 0.55 | 0.3 | 0.56 | 0.55 | 0.62 | 0.56 |
| 10 | 23298069 | rs4747451 | -0.6 | -1.038 | 1.847 | 0.058 | -0.001 | -0.016 | -0.016 | 0 | 0.87 | 0.91 | 0.73 | 0.67 | 0.91 | 0.85 |
| 10 | 23299017 | rs7914172 | -0.43 | -1.04 | 1.942 | 0.067 | -0.002 | 0.073 | 0.073 | 0 | 0.87 | 0.82 | 0.72 | 0.67 | 0.91 | 0.85 |
| 10 | 23300214 | rs10764371 | 2.051 | -1.078 | 1.889 | 0.053 | 0.008 | -0.028 | -0.028 | 0.459 | 0.64 | 0.3 | 0.6 | 0.55 | 0.7 | 0.61 |
| 10 | 23301436 | rs7100499 | -1.389 | -1.037 | 1.949 | 0.059 | -0.01 | -0.07 | -0.07 | 0 | 0.22 | 0.43 | 0.11 | 0.12 | 0.21 | 0.23 |
| 10 | 23306089 | rs11814322 | 2.081 | -1.013 | 1.832 | 0.071 | 0.017 | -0.022 | -0.022 | 0 | 0.55 | 0.3 | 0.56 | 0.55 | 0.61 | 0.57 |
| 10 | 23307046 | rs10764372 | 1.257 | -0.869 | 1.964 | 0.259 | -0.012 | 0.051 | 0.051 | 0 | 0.75 | 0.64 | 0.66 | 0.67 | 0.82 | 0.8 |
| 10 | 23307206 | rs993352 | 0.652 | -0.843 | 1.792 | 0.339 | -0.008 | 0.019 | 0.019 | 0.104 | 0.79 | 0.81 | 0.69 | 0.7 | 0.86 | 0.82 |
| 10 | 23310769 | rs12257453 | -0.021 | -0.526 | 1.602 | 0.358 | -0.015 | -0.076 | -0.076 | 0 | 0.14 | 0.28 | 0.04 | 0 | 0.08 | 0.09 |
| 10 | 23317917 | rs12261088 | 0.012 | -0.466 | 1.385 | 0.403 | -0.015 | -0.07 | -0.07 | 0.528 | 0.14 | 0.29 | 0.05 | 0 | 0.08 | 0.09 |
| 10 | 23319824 | rs11013262 | 0.184 | -0.408 | 1.108 | 0.412 | -0.011 | -0.059 | -0.059 | 0 | 0.13 | 0.22 | 0.04 | 0 | 0.08 | 0.09 |
| 10 | 23324361 | rs7898656 | -0.543 | -0.075 | 1.235 | 0.862 | 0.002 | -0.021 | -0.021 | 0 | 0.11 | 0.15 | 0.07 | 0.04 | 0.15 | 0.18 |
| 10 | 23332946 | rs1382579 | 0.832 | 0.122 | 0.238 | 1.372 | 0.004 | 0.174 | 0.174 | 0 | 0.95 | 0.91 | 0.75 | 0.55 | 0.97 | 0.88 |
| 10 | 23357212 | rs6482252 | -0.028 | 0.25 | -0.103 | 0.371 | 0.001 | 0.003 | 0.003 | 0 | 0.57 | 0.69 | 0.46 | 0.62 | 0.56 | 0.62 |
| 10 | 23364720 | rs7898113 | 0.066 | 0.357 | -0.594 | 0.249 | 0.035 | 0.031 | 0.031 | 0 | 0.49 | 0.3 | 0.44 | 0.45 | 0.6 | 0.55 |
| 10 | 23369421 | rs722258 | -0.45 | 0.506 | -0.661 | 0.049 | 0.014 | -0.032 | -0.032 | 0 | 0.32 | 0.57 | 0.47 | 0.34 | 0.42 | 0.4 |

| | | | | | | | | | | | | | | | | |
|---|---|---|---|---|---|---|---|---|---|---|---|---|---|---|---|---|
| 10 | 23372022 | rs7073142 | 0.307 | 0.273 | -1.473 | -0.063 | 0.008 | 0.056 | 0.056 | 0 | 0.27 | 0.06 | 0.17 | 0.15 | 0.18 | 0.25 |
| 10 | 23372056 | rs7073159 | 0.27 | 0.311 | -1.262 | -0.032 | 0.011 | 0.023 | 0.023 | 0 | 0.27 | 0.14 | 0.19 | 0.21 | 0.18 | 0.25 |
| 10 | 23374433 | rs11592906 | 0.19 | 0.357 | -0.898 | -0.073 | -0.001 | -0.01 | -0.01 | 0.034 | 0.12 | 0.14 | 0.18 | 0.05 | 0.14 | 0.09 |
| 10 | 23379444 | rs11013284 | 1.205 | 0.752 | -1.122 | 0.376 | -0.038 | -0.034 | -0.034 | 0 | 0.11 | 0.04 | 0.17 | 0.18 | 0.18 | 0.15 |
| 10 | 23383522 | rs2886428 | -0.405 | -0.226 | -0.638 | 0.106 | -0.009 | -0.036 | -0.036 | 0.038 | 0.36 | 0.55 | 0.39 | 0.35 | 0.39 | 0.41 |
| 10 | 23387293 | rs1975958 | 0.79 | -0.335 | -0.623 | -0.008 | 0.007 | 0.019 | 0.019 | 0.057 | 0.19 | 0.33 | 0.31 | 0.33 | 0.25 | 0.3 |
| 10 | 23390612 | rs11591816 | 0.504 | -0.334 | -0.601 | -0.025 | 0.001 | -0.025 | -0.025 | 0 | 0.17 | 0.17 | 0.07 | 0.03 | 0.13 | 0.11 |
| 10 | 23394012 | rs12769723 | 0.475 | -0.337 | -0.74 | -0.056 | -0.003 | -0.043 | -0.043 | 2.054 | 0.17 | 0.19 | 0.07 | 0.03 | 0.13 | 0.11 |
| 10 | 23394654 | rs7094823 | 1.337 | -0.275 | -1.055 | -0.001 | -0.06 | -0.058 | -0.058 | 0 | 0.16 | 0.02 | 0.18 | 0.26 | 0.25 | 0.28 |
| 10 | 23394876 | rs2147941 | -0.012 | -0.149 | -0.759 | 0.057 | -0.032 | -0.093 | -0.093 | 0 | 0.2 | 0.49 | 0.2 | 0.1 | 0.13 | 0.12 |
| 10 | 23396191 | rs6482253 | -0.019 | -0.343 | -0.792 | 0.168 | -0.007 | -0.019 | -0.019 | 1.219 | 0.36 | 0.63 | 0.39 | 0.36 | 0.38 | 0.4 |
| 10 | 23396688 | rs10828409 | 0.406 | -0.628 | -1.228 | -0.256 | 0.013 | 0.058 | 0.058 | 0.774 | 0.31 | 0.15 | 0.22 | 0.13 | 0.24 | 0.24 |
| 10 | 23400191 | rs11013291 | 0.19 | -0.491 | -1.105 | -0.051 | 0.008 | -0.001 | -0.001 | 0.007 | 0.32 | 0.27 | 0.22 | 0.3 | 0.45 | 0.37 |
| 10 | 23400393 | rs11013292 | -0.072 | -0.819 | -1.449 | 0.15 | 0.017 | 0.009 | 0.009 | 0 | 0.62 | 0.73 | 0.66 | 0.35 | 0.66 | 0.53 |
| 10 | 23403403 | rs1983880 | -0.088 | -0.881 | -1.406 | 0.317 | 0.017 | -0.023 | -0.023 | 0 | 0.82 | 0.82 | 0.83 | 0.51 | 0.95 | 0.8 |
| 10 | 23404861 | rs17540863 | 0.893 | 0.111 | -1.226 | -0.012 | -0.026 | 0.025 | 0.025 | 0 | 0.23 | 0.08 | 0.18 | 0.19 | 0.29 | 0.28 |
| 10 | 23408278 | rs7427 | 0.026 | 0.242 | -0.769 | -0.042 | 0.018 | -0.017 | -0.017 | 0 | 0.23 | 0.23 | 0.16 | 0.12 | 0.33 | 0.27 |
| 10 | 23422210 | rs6482254 | -1.169 | -0.187 | -0.64 | 0.509 | -0.034 | -0.098 | -0.098 | 0 | 0.78 | 0.91 | 0.54 | 0.53 | 0.69 | 0.68 |
| 10 | 23424277 | rs7090625 | -0.076 | -0.129 | 0.225 | 0.717 | -0.026 | 0.231 | 0.231 | 0 | 0.85 | 0.65 | 0.64 | 0.54 | 0.88 | 0.87 |
| 10 | 23431618 | rs2057599 | 0.591 | -0.364 | 0.415 | 0.598 | -0.036 | 0.073 | 0.073 | 2.204 | 0.78 | 0.32 | 0.58 | 0.53 | 0.85 | 0.82 |
| 10 | 23435524 | rs1467646 | 0.53 | 0.023 | 0.232 | -0.34 | -0.007 | 0.002 | 0.002 | 0 | 0.12 | 0.21 | 0.06 | 0.06 | 0.12 | 0.12 |
| 10 | 23444095 | rs744107 | 0.53 | 0.023 | 0.232 | -0.381 | -0.007 | -0.001 | -0.001 | 0.052 | 0.12 | 0.21 | 0.06 | 0.06 | 0.12 | 0.12 |
| 10 | 23466734 | rs7916519 | -0.433 | -0.083 | -0.198 | -0.808 | 0.014 | 0.088 | 0.088 | 0 | 0.46 | 0.66 | 0.44 | 0.18 | 0.42 | 0.48 |
| 10 | 23477062 | rs1408986 | -0.882 | -0.319 | -0.037 | -0.971 | -0.022 | 0.097 | 0.097 | 0 | 0.58 | 0.74 | 0.65 | 0.87 | 0.48 | 0.55 |
| 10 | 23480318 | rs12415184 | -0.718 | -0.361 | 0.073 | -0.978 | -0.003 | 0.003 | 0.003 | 0 | 0.56 | 0.61 | 0.63 | 0.84 | 0.47 | 0.5 |
| 10 | 23482635 | rs7918487 | -0.808 | -0.304 | 0.092 | -0.955 | -0.001 | -0.023 | -0.023 | 0 | 0.58 | 0.6 | 0.63 | 0.87 | 0.47 | 0.56 |
| 10 | 23486297 | rs1886764 | -0.769 | -0.284 | 0.132 | -0.947 | -0.003 | -0.003 | -0.003 | 0 | 0.57 | 0.6 | 0.64 | 0.84 | 0.48 | 0.51 |
| 10 | 23506529 | rs927751 | -0.869 | -0.062 | 0.049 | -0.973 | -0.034 | 0.123 | 0.123 | 0 | 0.58 | 0.75 | 0.66 | 0.87 | 0.49 | 0.6 |
| 10 | 23530278 | rs1325591 | 1.633 | -0.132 | -0.157 | -1.089 | -0.076 | 0.089 | 0.089 | 0 | 0.24 | 0.02 | 0.17 | 0.08 | 0.3 | 0.29 |
| 10 | 23538351 | rs17543414 | 0.859 | -0.308 | -0.209 | -1.094 | -0.074 | -0.01 | -0.01 | 0 | 0.16 | 0.02 | 0.13 | 0.14 | 0.26 | 0.22 |
| 10 | 23539203 | rs17543456 | 1.238 | -0.986 | -0.01 | -1.202 | 0.009 | 0.063 | 0.063 | 0 | 0.22 | 0.1 | 0.2 | 0 | 0.17 | 0.18 |
| 10 | 23547519 | rs4237360 | -0.48 | -0.954 | -0.061 | -1.179 | 0.087 | 0.091 | 0.091 | 0 | 0.44 | 0.74 | 0.56 | 0.74 | 0.25 | 0.41 |
| 10 | 23552058 | rs7070758 | 0.885 | -0.973 | 0.224 | -1.175 | -0.036 | -0.16 | -0.16 | 0 | 0.26 | 0.47 | 0.4 | 0.02 | 0.17 | 0.21 |
| 10 | 23559052 | rs7078858 | 1.172 | -0.966 | 0.377 | -1.172 | 0.007 | -0.009 | -0.009 | 0 | 0.23 | 0.2 | 0.38 | 0.01 | 0.17 | 0.21 |
| 10 | 23566105 | rs10764398 | 0.211 | -0.888 | 0.284 | -1.107 | -0.036 | 0.019 | 0.019 | 0 | 0.18 | 0.24 | 0.15 | 0.72 | 0.08 | 0.2 |
| 10 | 23570792 | rs11013364 | 0.244 | -0.89 | 0.282 | -1.059 | -0.033 | 0.031 | 0.031 | 0 | 0.18 | 0.23 | 0.15 | 0.67 | 0.08 | 0.15 |
| 10 | 23572382 | rs4454613 | 0.084 | -0.88 | 0.247 | -1.105 | -0.017 | 0.004 | 0.004 | 0.232 | 0.76 | 0.76 | 0.83 | 0.3 | 0.88 | 0.84 |
| 10 | 23572471 | rs4396189 | -1.972 | -0.8 | 0.248 | -1.087 | -0.01 | -0.026 | -0.026 | 0 | 0.94 | 0.89 | 0.97 | 0.97 | 0.96 | 0.99 |
| 10 | 23591365 | rs7089898 | -0.454 | -0.883 | 0.371 | -1.012 | 0.068 | 0.07 | 0.07 | 0 | 0.44 | 0.7 | 0.55 | 0.73 | 0.25 | 0.41 |
| 10 | 23596420 | rs12266732 | 0.484 | -0.877 | 0.447 | -0.828 | 0.015 | 0.09 | 0.09 | 0 | 0.06 | 0.22 | 0.04 | 0.08 | 0.03 | 0.08 |
| 10 | 23598327 | rs12217944 | 1.122 | -0.868 | 0.017 | -0.823 | 0.039 | -0.075 | -0.075 | 0 | 0.13 | 0.02 | 0.1 | 0.63 | 0.04 | 0.12 |
| 10 | 23600800 | rs4747461 | 0.998 | -0.87 | 0.086 | -0.824 | 0.023 | -0.027 | -0.027 | 0 | 0.13 | 0.08 | 0.11 | 0.64 | 0.04 | 0.12 |
| 10 | 23605930 | rs4326695 | -1.108 | -0.691 | -0.228 | -1.056 | -0.031 | 0.097 | 0.097 | 0 | 0.6 | 0.75 | 0.67 | 0.87 | 0.46 | 0.61 |
| 10 | 23607192 | rs4603206 | -0.335 | -0.755 | -0.245 | -1.042 | 0.065 | 0.066 | 0.066 | 0 | 0.44 | 0.7 | 0.55 | 0.73 | 0.25 | 0.41 |
| 10 | 23622202 | rs4259746 | 0.177 | -0.407 | -0.194 | -0.799 | -0.001 | -0.007 | -0.007 | 0 | 0.42 | 0.45 | 0.51 | 0.19 | 0.38 | 0.47 |
| 10 | 23623523 | rs10828431 | 0.167 | -0.443 | -0.348 | -0.666 | -0.037 | 0.057 | 0.057 | 0 | 0.18 | 0.23 | 0.15 | 0.63 | 0.09 | 0.14 |
| 10 | 23640882 | rs4748867 | -0.865 | -0.349 | -0.334 | -0.715 | -0.029 | 0.085 | 0.085 | 0 | 0.6 | 0.74 | 0.66 | 0.86 | 0.47 | 0.61 |
| 10 | 23647619 | rs2366163 | -1.857 | -0.061 | -0.753 | -0.759 | -0.102 | 0.101 | 0.101 | 0 | 0.74 | 0.97 | 0.8 | 0.91 | 0.64 | 0.7 |
| 10 | 23649989 | rs7906070 | -0.205 | -0.09 | -0.339 | -0.525 | -0.002 | -0.029 | -0.029 | 0.193 | 0.42 | 0.5 | 0.51 | 0.2 | 0.38 | 0.47 |
| 10 | 23661291 | rs12414475 | 0.037 | -0.57 | -0.507 | 0.108 | 0.034 | 0.04 | 0.04 | 0.403 | 0.32 | 0.82 | 0.52 | 0.18 | 0.22 | 0.25 |
| 10 | 23664096 | rs10734046 | -1.529 | -0.523 | -0.861 | -0.061 | -0.121 | 0.153 | 0.153 | 0 | 0.65 | 0.96 | 0.78 | 0.86 | 0.52 | 0.65 |
| 10 | 23665438 | rs1398024 | -0.04 | -1.644 | 0.523 | 0.514 | -0.023 | -0.092 | -0.092 | 0 | 0.3 | 0.55 | 0.48 | 0.05 | 0.21 | 0.23 |
| 10 | 23670597 | rs4287251 | -0.249 | -1.637 | 0.381 | 0.473 | 0.005 | -0.044 | -0.044 | 0 | 0.32 | 0.68 | 0.41 | 0.01 | 0.2 | 0.17 |
| 10 | 23673298 | rs4019731 | -1.101 | -1.63 | -0.182 | 0.709 | 0.101 | 0.111 | 0.111 | 0.895 | 0.47 | 0.88 | 0.54 | 0.69 | 0.27 | 0.4 |
| 10 | 23681762 | rs2012456 | -1.11 | -1.622 | -0.15 | 0.786 | 0.113 | 0.102 | 0.102 | 2.043 | 0.47 | 0.88 | 0.52 | 0.66 | 0.26 | 0.36 |
| 10 | 23685456 | rs10828441 | -1.081 | -1.47 | -0.21 | 0.795 | 0.083 | 0.103 | 0.103 | 0.668 | 0.47 | 0.78 | 0.52 | 0.7 | 0.26 | 0.37 |
| 10 | 23688651 | rs10458699 | -0.165 | -1.31 | -0.603 | 0.817 | 0.017 | -0.012 | -0.012 | 0 | 0.14 | 0.09 | 0.09 | 0.52 | 0.06 | 0.17 |
| 10 | 23691638 | rs10828450 | -0.54 | -0.849 | -0.311 | 0.651 | -0.017 | -0.01 | -0.01 | 0.077 | 0.45 | 0.55 | 0.48 | 0.65 | 0.26 | 0.37 |
| 10 | 23708307 | rs6482274 | -0.965 | -0.882 | 0.159 | 1.45 | -0.024 | 0.178 | 0.178 | 0.118 | 0.14 | 0.29 | 0.09 | 0.35 | 0.09 | 0.19 |
| 10 | 23708636 | rs7909628 | -0.955 | -1.784 | -0.859 | 0.968 | 0.126 | 0.043 | 0.043 | 0.47 | 0.5 | 0.88 | 0.29 | 0.56 | 0.29 | 0.37 |
| 10 | 23721412 | rs956007 | -0.492 | -2.499 | -0.666 | 0.589 | 0.023 | 0.084 | 0.084 | 0 | 0.33 | 0.17 | 0.3 | 0.14 | 0.26 | 0.29 |
| 10 | 23723000 | rs907451 | -1.384 | -2.982 | 0.253 | 1.173 | 0.123 | -0.04 | -0.04 | 2.966 | 0.52 | 0.87 | 0.44 | 0.46 | 0.3 | 0.43 |
| 10 | 23725892 | rs2137157 | 1.378 | -2.263 | -0.297 | 0.822 | -0.169 | 0.003 | 0.003 | 0.261 | 0.33 | 0.02 | 0.36 | 0.29 | 0.46 | 0.41 |
| 10 | 23726000 | rs1511825 | 0.046 | -2.353 | 0.526 | 0.641 | -0.063 | -0.038 | -0.038 | 0.424 | 0.12 | 0.53 | 0.09 | 0.02 | 0.02 | 0.01 |
| 10 | 23726421 | rs7080648 | 0.552 | -2.272 | 0.294 | 0.63 | -0.028 | -0.046 | -0.046 | 1.718 | 0.09 | 0.34 | 0.06 | 0 | 0.02 | 0.01 |
| 10 | 23745857 | rs11013503 | 1.048 | -2.166 | -0.115 | 0.605 | -0.03 | 0.041 | 0.041 | 0 | 0.08 | 0.02 | 0.06 | 0 | 0.12 | 0.01 |
| 10 | 23755800 | rs7918459 | -0.328 | -2.232 | 0.361 | 0.8 | -0.01 | -0.011 | -0.011 | 0 | 0.41 | 0.43 | 0.35 | 0.22 | 0.28 | 0.4 |
| 10 | 23771164 | rs12266508 | -0.791 | -1.603 | 0.055 | 1.044 | 0.089 | -0.038 | -0.038 | 0 | 0.51 | 0.87 | 0.45 | 0.46 | 0.33 | 0.44 |
| 10 | 23776811 | rs10828462 | -0.33 | -1.713 | 0.548 | 0.943 | 0.009 | 0.011 | 0.011 | 0 | 0.45 | 0.45 | 0.38 | 0.19 | 0.33 | 0.41 |

| | | | | | | | | | | | | | | | | |
|---|---|---|---|---|---|---|---|---|---|---|---|---|---|---|---|---|
| 10 | 23781152 | rs10828463 | -0.704 | -1.785 | 0.749 | 0.915 | 0.059 | 0.094 | 0.094 | 0.244 | 0.5 | 0.66 | 0.4 | 0.19 | 0.33 | 0.4 |
| 10 | 23781797 | rs6482290 | -0.89 | -1.807 | 0.809 | 0.94 | 0.085 | 0.147 | 0.147 | 1.657 | 0.52 | 0.84 | 0.43 | 0.21 | 0.33 | 0.4 |
| 10 | 23783728 | rs4612701 | -0.965 | -1.83 | 0.805 | 1.075 | 0.067 | -0.066 | -0.066 | 0 | 0.53 | 0.9 | 0.46 | 0.46 | 0.33 | 0.43 |
| 10 | 23785362 | rs4748902 | -0.903 | -1.82 | 0.789 | 1.097 | 0.084 | -0.028 | -0.028 | 0 | 0.53 | 0.84 | 0.48 | 0.47 | 0.33 | 0.43 |
| 10 | 23788819 | rs7901238 | -0.37 | -1.808 | 1.04 | 0.752 | 0.031 | 0.05 | 0.05 | 0 | 0.47 | 0.39 | 0.37 | 0.19 | 0.33 | 0.4 |
| 10 | 23794804 | rs1877325 | -0.636 | -1.725 | 0.296 | 0.806 | 0.06 | 0.093 | 0.093 | 0.272 | 0.49 | 0.66 | 0.4 | 0.19 | 0.32 | 0.4 |
| 10 | 23806382 | rs983990 | -0.632 | -1.797 | 0.237 | 0.757 | 0.075 | 0.131 | 0.131 | 0.975 | 0.51 | 0.72 | 0.4 | 0.18 | 0.32 | 0.4 |
| 10 | 23809840 | rs4570482 | -0.698 | -1.341 | -0.514 | 0.883 | 0.044 | 0.198 | 0.198 | 1.234 | 0.53 | 0.91 | 0.31 | 0.17 | 0.37 | 0.4 |
| 10 | 23810151 | rs11013549 | -0.222 | -1.518 | -0.385 | 0.797 | 0.073 | 0.118 | 0.118 | 0.004 | 0.46 | 0.69 | 0.27 | 0.12 | 0.31 | 0.3 |
| 10 | 23820137 | rs9417385 | -1.292 | -0.992 | -1.068 | 1.227 | -0.17 | 0.025 | 0.025 | 0.273 | 0.69 | 0.99 | 0.74 | 0.78 | 0.59 | 0.68 |
| 10 | 23837290 | rs10828478 | -0.854 | -1.211 | -0.888 | 0.882 | 0.002 | -0.079 | -0.079 | 0 | 0.55 | 0.95 | 0.33 | 0.41 | 0.37 | 0.45 |
| 10 | 23838779 | rs1337312 | -0.486 | -1.44 | 0.533 | 1.064 | -0.016 | 0.202 | 0.202 | 1.01 | 0.09 | 0.33 | 0.09 | 0.23 | 0.04 | 0.03 |
| 10 | 23846829 | rs10764403 | -0.445 | -1.796 | 0.681 | -0.17 | -0.017 | -0.098 | -0.098 | 0 | 0.26 | 0.41 | 0.22 | 0.04 | 0.23 | 0.18 |
| 10 | 23854039 | rs7901530 | 0.316 | -2.045 | 1.124 | -0.249 | -0.05 | -0.051 | -0.051 | 0 | 0.09 | 0.17 | 0.04 | 0 | 0.01 | 0.01 |
| 10 | 23860231 | rs10828495 | -1.377 | -2.13 | 1.11 | -0.369 | 0.089 | 0.196 | 0.196 | 0.931 | 0.4 | 0.8 | 0.36 | 0.07 | 0.26 | 0.23 |
| 10 | 23861418 | rs10764406 | -1.194 | -2.129 | 1.414 | -0.369 | 0.026 | 0.019 | 0.019 | 0.713 | 0.38 | 0.63 | 0.35 | 0.04 | 0.26 | 0.22 |
| 10 | 23863296 | rs12260319 | -0.696 | -2.269 | 1.538 | -0.547 | -0.031 | -0.051 | -0.051 | 0 | 0.1 | 0.28 | 0.11 | 0 | 0.02 | 0.1 |
| 10 | 23891240 | rs10828509 | 0.274 | -2.222 | 1.303 | -0.525 | 0 | -0.017 | -0.017 | 0 | 0.19 | 0.17 | 0.21 | 0.03 | 0.22 | 0.11 |
| 10 | 23895866 | rs2050610 | -1.631 | -2.255 | 1.095 | -0.534 | 0.084 | 0.189 | 0.189 | 1.102 | 0.39 | 0.79 | 0.36 | 0.08 | 0.26 | 0.23 |
| 10 | 23896359 | rs16923535 | -0.996 | -2.148 | 1.453 | -0.494 | -0.028 | -0.106 | -0.106 | 0 | 0.29 | 0.5 | 0.25 | 0.07 | 0.23 | 0.12 |
| 10 | 23897514 | rs7075982 | -0.902 | -2.142 | 1.429 | -0.491 | -0.029 | -0.117 | -0.117 | 0 | 0.29 | 0.39 | 0.24 | 0.07 | 0.23 | 0.12 |
| 10 | 23907200 | rs16923552 | 0.364 | -2.015 | 1.385 | -0.382 | -0.032 | -0.05 | -0.05 | 0.175 | 0.09 | 0.19 | 0.04 | 0 | 0.02 | 0 |
| 10 | 23916761 | rs12261593 | -0.083 | -2.102 | 1.638 | -0.386 | -0.042 | -0.052 | -0.052 | 0.903 | 0.11 | 0.43 | 0.07 | 0.02 | 0.01 | 0.02 |
| 10 | 23919919 | rs1577815 | 0.237 | -2.351 | 1.517 | -0.236 | 0.004 | 0.009 | 0.009 | 0.591 | 0.19 | 0.14 | 0.16 | 0.15 | 0.18 | 0.1 |
| 10 | 23927266 | rs7097486 | -1.603 | -2.612 | 0.883 | -0.371 | 0.13 | 0.163 | 0.163 | 4.569 | 0.42 | 0.81 | 0.33 | 0.17 | 0.23 | 0.24 |
| 10 | 23936020 | rs1337322 | 0.24 | -1.959 | 0.189 | -0.333 | -0.002 | -0.001 | -0.001 | 0 | 0.39 | 0.62 | 0.34 | 0.45 | 0.43 | 0.3 |
| 10 | 23938791 | rs12260863 | 0.176 | -2.596 | 0.872 | -0.408 | -0.041 | -0.051 | -0.051 | 0 | 0.11 | 0.4 | 0.07 | 0.02 | 0.01 | 0.02 |
| 10 | 23939405 | rs7096031 | -1.515 | -2.718 | 0.775 | -0.422 | 0.136 | 0.165 | 0.165 | 5.105 | 0.42 | 0.89 | 0.34 | 0.19 | 0.22 | 0.24 |
| 10 | 23940363 | rs11013633 | -0.827 | -2.518 | 0.807 | -0.33 | -0.041 | -0.052 | -0.052 | 0 | 0.31 | 0.49 | 0.27 | 0.16 | 0.21 | 0.22 |
| 10 | 23943468 | rs7081315 | -0.831 | -2.328 | 0.82 | -0.261 | -0.053 | -0.073 | -0.073 | 0 | 0.12 | 0.29 | 0.11 | 0.01 | 0.03 | 0.12 |
| 10 | 23952687 | rs7091045 | -1.522 | -2.666 | 0.419 | -0.285 | 0.157 | 0.142 | 0.142 | 3.191 | 0.45 | 0.89 | 0.38 | 0.22 | 0.26 | 0.33 |
| 10 | 23954214 | rs10764422 | -1.193 | -2.705 | 1.005 | -0.303 | 0.029 | 0.03 | 0.03 | 3.031 | 0.4 | 0.68 | 0.31 | 0.17 | 0.23 | 0.22 |
| 10 | 23985407 | rs7900989 | -1.312 | -2.411 | 1.347 | -0.267 | -0.08 | -0.056 | -0.056 | 0 | 0.16 | 0.42 | 0.15 | 0.07 | 0.06 | 0.22 |
| 10 | 23991074 | rs6482316 | -1.343 | -2.405 | 1.364 | -0.264 | -0.079 | -0.057 | -0.057 | 0 | 0.17 | 0.48 | 0.16 | 0.07 | 0.06 | 0.22 |
| 10 | 24001598 | rs7477699 | -0.091 | -2.217 | 0.636 | -0.061 | 0.032 | 0.002 | 0.002 | 2.339 | 0.24 | 0.09 | 0.17 | 0.16 | 0.2 | 0.09 |
| 10 | 24003567 | rs10508661 | -0.186 | -2.242 | 0.906 | -0.076 | 0.021 | 0.011 | 0.011 | 2.5 | 0.25 | 0.18 | 0.19 | 0.16 | 0.2 | 0.09 |
| 10 | 24025126 | rs7909219 | -0.27 | -2.316 | 1.609 | -0.108 | 0.002 | 0.002 | 0.002 | 2.354 | 0.26 | 0.25 | 0.19 | 0.16 | 0.2 | 0.09 |
| 10 | 24029291 | rs6482335 | -0.452 | -2.32 | 1.679 | -0.11 | -0.024 | -0.014 | -0.014 | 0.855 | 0.27 | 0.34 | 0.2 | 0.16 | 0.2 | 0.09 |
| 10 | 24030759 | rs10508662 | -0.346 | -2.406 | 1.824 | -0.187 | -0.003 | -0.027 | -0.027 | 16.484 | 0.06 | 0.14 | 0.11 | 0.02 | 0.02 | 0.05 |
| 10 | 24031397 | rs12782051 | -0.181 | -2.415 | 1.834 | -0.188 | -0.002 | -0.001 | -0.001 | 0.015 | 0.26 | 0.25 | 0.19 | 0.16 | 0.2 | 0.09 |
| 10 | 24034646 | rs10508663 | -1.034 | -2.292 | 1.753 | -0.19 | -0.043 | -0.043 | -0.043 | 0 | 0.33 | 0.46 | 0.31 | 0.18 | 0.25 | 0.14 |
| 10 | 24046158 | rs12146296 | -0.009 | -2.323 | 1.791 | -0.185 | 0.017 | 0.006 | 0.006 | 2.02 | 0.23 | 0.17 | 0.18 | 0.16 | 0.2 | 0.08 |
| 10 | 24055066 | rs6482343 | -0.464 | -2.257 | 1.916 | -0.197 | -0.038 | -0.027 | -0.027 | 0.11 | 0.27 | 0.47 | 0.22 | 0.18 | 0.21 | 0.12 |
| 10 | 24056662 | rs6482344 | -1.3 | -2.324 | 2.027 | -0.234 | 0.026 | 0.026 | 0.026 | 2.725 | 0.35 | 0.7 | 0.34 | 0.18 | 0.23 | 0.17 |
| 10 | 24058804 | rs10828544 | -0.858 | -2.569 | 1.95 | -0.317 | -0.008 | -0.005 | -0.005 | 1.251 | 0.32 | 0.63 | 0.24 | 0.19 | 0.21 | 0.14 |
| 10 | 24059607 | rs11812868 | 1.639 | -2.158 | 1.246 | -0.289 | -0.156 | -0.118 | -0.118 | 0 | 0.32 | 0.03 | 0.36 | 0.42 | 0.44 | 0.32 |
| 10 | 24064899 | rs7898294 | -2.456 | -2.961 | 2.107 | -0.243 | 0.071 | 0.067 | 0.067 | 3.308 | 0.05 | 0.4 | 0.05 | 0.02 | 0 | 0.03 |
| 10 | 24068099 | rs7087962 | -1.146 | -2.985 | 2.048 | -0.25 | 0.013 | 0.012 | 0.012 | 2.944 | 0.35 | 0.66 | 0.34 | 0.19 | 0.23 | 0.16 |
| 10 | 24077166 | rs999749 | -0.78 | -3.009 | 1.956 | -0.256 | -0.042 | -0.034 | -0.034 | 0.083 | 0.32 | 0.42 | 0.31 | 0.19 | 0.23 | 0.18 |
| 10 | 24083384 | rs12412791 | -1.159 | -2.922 | 1.741 | -0.232 | -0.018 | -0.032 | -0.032 | 0 | 0.08 | 0.14 | 0.1 | 0.02 | 0.02 | 0.09 |
| 10 | 24091546 | rs1034287 | -0.978 | -2.94 | 1.901 | -0.254 | -0.02 | -0.022 | -0.022 | 0 | 0.34 | 0.58 | 0.33 | 0.19 | 0.25 | 0.2 |
| 10 | 24104679 | rs7907848 | -1.064 | -3.024 | 1.951 | -0.248 | 0.038 | 0.031 | 0.031 | 4.531 | 0.35 | 0.72 | 0.25 | 0.19 | 0.22 | 0.17 |
| 10 | 24107651 | rs11013728 | -0.816 | -2.999 | 1.946 | -0.237 | -0.021 | -0.016 | -0.016 | 0 | 0.33 | 0.61 | 0.24 | 0.19 | 0.22 | 0.17 |
| 10 | 24108358 | rs2366591 | -0.5 | -2.998 | 2.015 | -0.236 | -0.031 | -0.02 | -0.02 | 0 | 0.29 | 0.39 | 0.2 | 0.18 | 0.22 | 0.16 |
| 10 | 24115017 | rs1414443 | -0.99 | -3.047 | 1.711 | -0.25 | -0.043 | -0.03 | -0.03 | 0 | 0.35 | 0.54 | 0.24 | 0.19 | 0.25 | 0.28 |
| 10 | 24120093 | rs1335813 | -0.245 | -3.059 | 1.374 | -0.255 | 0.022 | 0.008 | 0.008 | 0.372 | 0.28 | 0.16 | 0.2 | 0.19 | 0.24 | 0.21 |
| 10 | 24121297 | rs11013736 | -0.927 | -2.976 | 1.316 | -0.218 | 0.019 | 0.015 | 0.015 | 1.689 | 0.37 | 0.68 | 0.26 | 0.21 | 0.25 | 0.29 |
| 10 | 24128433 | rs11013742 | 0.511 | -2.884 | 1.013 | -0.352 | 0.007 | 0.01 | 0.01 | 0 | 0.18 | 0.12 | 0.15 | 0.11 | 0.2 | 0.11 |
| 10 | 24132033 | rs9418444 | 1.687 | -3.551 | 0.432 | -0.333 | -0.025 | -0.109 | -0.109 | 2.53 | 0.4 | 0.04 | 0.36 | 0.43 | 0.6 | 0.35 |
| 10 | 24133028 | rs11013746 | -2.167 | -3.172 | 1.461 | -0.627 | -0.046 | -0.039 | -0.039 | 0 | 0.09 | 0.68 | 0.13 | 0.03 | 0 | 0.02 |
| 10 | 24154216 | rs3906205 | -0.213 | -3.326 | 1.701 | -0.619 | 0.018 | 0.006 | 0.006 | 0 | 0.23 | 0.15 | 0.16 | 0.17 | 0.2 | 0.13 |
| 10 | 24160236 | rs7919049 | -1.072 | -3.021 | 1.735 | -0.414 | 0.043 | 0.012 | 0.012 | 0.351 | 0.28 | 0.79 | 0.29 | 0.24 | 0.2 | 0.13 |
| 10 | 24165579 | rs16924005 | -0.007 | -2.89 | 1.637 | -0.365 | -0.015 | -0.009 | -0.009 | 0 | 0.21 | 0.28 | 0.16 | 0.15 | 0.2 | 0.13 |
| 10 | 24165775 | rs12247225 | -0.068 | -2.884 | 1.597 | -0.362 | -0.023 | -0.014 | -0.014 | 0 | 0.21 | 0.32 | 0.17 | 0.15 | 0.2 | 0.13 |
| 10 | 24172108 | rs9299784 | 0.361 | -2.864 | 1.513 | -0.345 | -0.003 | 0.001 | 0.001 | 0 | 0.19 | 0.18 | 0.16 | 0.21 | 0.2 | 0.11 |
| 10 | 24176704 | rs9971218 | -0.177 | -2.716 | 1.444 | -0.72 | -0.016 | -0.056 | -0.056 | 0.556 | 0.28 | 0.39 | 0.31 | 0.1 | 0.29 | 0.23 |
| 10 | 24181848 | rs10828568 | 0.214 | -2.662 | 1.056 | -0.701 | 0.012 | 0.02 | 0.02 | 0 | 0.25 | 0.17 | 0.2 | 0.13 | 0.27 | 0.2 |
| 10 | 24187822 | rs7081769 | 0.157 | -2.689 | 1.048 | -0.708 | 0.01 | 0.015 | 0.015 | 0 | 0.26 | 0.19 | 0.2 | 0.13 | 0.27 | 0.21 |
| 10 | 24187867 | rs11013778 | -0.369 | -2.647 | 0.812 | -0.684 | -0.004 | -0.008 | -0.008 | 0 | 0.32 | 0.67 | 0.33 | 0.16 | 0.28 | 0.24 |

| | | | | | | | | | | | | | | | | |
|---|---|---|---|---|---|---|---|---|---|---|---|---|---|---|---|---|
| 10 | 24199672 | rs3935858 | -0.835 | -2.682 | 0.717 | -0.727 | 0.03 | 0.038 | 0.038 | 0.23 | 0.42 | 0.69 | 0.37 | 0.16 | 0.33 | 0.28 |
| 10 | 24206076 | rs1938048 | -0.706 | -2.48 | -0.195 | -0.877 | 0.067 | 0.121 | 0.121 | 0.884 | 0.41 | 0.77 | 0.38 | 0.1 | 0.32 | 0.22 |
| 10 | 24207529 | rs11013784 | -0.511 | -2.492 | -0.682 | -0.805 | 0.078 | 0.142 | 0.142 | 1.053 | 0.35 | 0.86 | 0.4 | 0.12 | 0.27 | 0.29 |
| 10 | 24215163 | rs7893219 | -0.556 | -2.3 | 0.233 | -0.53 | -0.044 | -0.052 | -0.052 | 0 | 0.15 | 0.51 | 0.11 | 0.05 | 0.06 | 0.05 |
| 10 | 24216156 | rs4529808 | -1.089 | -2.394 | -0.386 | -0.833 | 0.08 | 0.211 | 0.211 | 4.774 | 0.46 | 0.89 | 0.41 | 0.18 | 0.33 | 0.28 |
| 10 | 24217196 | rs4474347 | -1.065 | -2.385 | -0.374 | -0.837 | 0.088 | 0.208 | 0.208 | 4.586 | 0.46 | 0.9 | 0.41 | 0.18 | 0.33 | 0.28 |
| 10 | 24219417 | rs12248960 | -0.915 | -2.467 | -0.155 | -0.873 | -0.011 | -0.009 | -0.009 | 0 | 0.18 | 0.69 | 0.23 | 0.11 | 0.07 | 0.08 |
| 10 | 24221752 | rs7083271 | 0.051 | -2.484 | 0.315 | -0.888 | -0.046 | -0.096 | -0.096 | 0 | 0.14 | 0.47 | 0.1 | 0 | 0.05 | 0.01 |
| 10 | 24225035 | rs11013798 | -0.905 | -2.494 | 0.069 | -0.892 | 0.041 | 0.077 | 0.077 | 1.638 | 0.43 | 0.67 | 0.28 | 0.13 | 0.32 | 0.25 |
| 10 | 24230240 | rs11013804 | 0.345 | -2.489 | 0.012 | -0.884 | 0.014 | 0.095 | 0.095 | 0.047 | 0.27 | 0.13 | 0.17 | 0.08 | 0.26 | 0.2 |
| 10 | 24233614 | rs4420167 | -1.138 | -2.512 | 0.017 | -0.89 | 0.077 | 0.181 | 0.181 | 2.183 | 0.45 | 0.81 | 0.4 | 0.19 | 0.33 | 0.28 |
| 10 | 24234777 | rs4748918 | -0.023 | -2.519 | 0.225 | -0.889 | -0.009 | -0.034 | -0.034 | 0 | 0.31 | 0.41 | 0.31 | 0.13 | 0.26 | 0.23 |
| 10 | 24239466 | rs1938047 | -1.089 | -2.51 | 0.067 | -0.884 | 0.077 | 0.184 | 0.184 | 3.607 | 0.44 | 0.87 | 0.4 | 0.19 | 0.33 | 0.27 |
| 10 | 24244556 | rs7905904 | 0.143 | -2.432 | 0.124 | -0.876 | -0.026 | -0.035 | -0.035 | 2.443 | 0.14 | 0.47 | 0.1 | 0.05 | 0.06 | 0.04 |
| 10 | 24247356 | rs11013811 | 0.312 | -2.263 | -0.81 | -0.876 | 0.012 | 0.098 | 0.098 | 0.223 | 0.26 | 0.1 | 0.18 | 0.08 | 0.25 | 0.21 |
| 10 | 24250080 | rs4466724 | -1.136 | -2.115 | -1.273 | -0.19 | -0.112 | -0.028 | -0.028 | 0 | 0.62 | 0.96 | 0.71 | 0.66 | 0.46 | 0.6 |
| 10 | 24262896 | rs17578320 | 0.283 | -1.647 | -0.857 | -0.591 | 0.01 | 0.063 | 0.063 | 0 | 0.29 | 0.16 | 0.21 | 0.07 | 0.27 | 0.26 |
| 10 | 24264334 | rs7915067 | 0.407 | -1.654 | -1.076 | -0.577 | 0.008 | 0.138 | 0.138 | 0 | 0.3 | 0.09 | 0.18 | 0.07 | 0.27 | 0.29 |
| 10 | 24264843 | rs7912455 | -1.368 | -1.606 | -1.761 | -0.372 | -0.171 | -0.047 | -0.047 | 3.652 | 0.7 | 0.96 | 0.7 | 0.64 | 0.52 | 0.61 |
| 10 | 24267337 | rs11013821 | 0.859 | -1.756 | -1.262 | -0.469 | 0.006 | 0.143 | 0.143 | 0 | 0.3 | 0.09 | 0.19 | 0.07 | 0.27 | 0.29 |
| 10 | 24269640 | rs1832311 | -1.057 | -1.053 | -0.813 | 0.17 | -0.119 | -0.081 | -0.081 | 0 | 0.7 | 0.94 | 0.69 | 0.52 | 0.52 | 0.6 |
| 10 | 24274773 | rs12782619 | 0.242 | -1.23 | -1.353 | 0.121 | 0.072 | 0.103 | 0.103 | 0.82 | 0.46 | 0.75 | 0.44 | 0.19 | 0.32 | 0.32 |
| 10 | 24278748 | rs10828584 | -0.555 | -0.906 | -0.911 | 0.558 | 0.011 | -0.039 | -0.039 | 0.663 | 0.25 | 0.19 | 0.26 | 0.46 | 0.21 | 0.31 |
| 10 | 24280011 | rs7101307 | -0.797 | 0.045 | -1.597 | -0.496 | -0.035 | 0.122 | 0.122 | 3.963 | 0.46 | 0.92 | 0.62 | 0.68 | 0.44 | 0.49 |
| 10 | 24281017 | rs10508667 | 0.231 | 0.104 | -1.088 | -0.903 | -0.01 | 0.003 | 0.003 | 0.241 | 0.28 | 0.22 | 0.27 | 0.3 | 0.37 | 0.33 |
| 10 | 24283192 | rs2252484 | -0.358 | 0.02 | 0.178 | -0.837 | 0.005 | 0.004 | 0.004 | 0.476 | 0.34 | 0.62 | 0.35 | 0.42 | 0.39 | 0.35 |
| 10 | 24285290 | rs10764442 | 0.43 | 0.769 | 0.525 | -0.611 | -0.005 | -0.005 | -0.005 | 0 | 0.64 | 0.32 | 0.48 | 0.43 | 0.65 | 0.59 |
| 10 | 24287260 | rs10741035 | -0.066 | 0.693 | 0.759 | -0.692 | 0.049 | 0.019 | 0.019 | 0 | 0.74 | 0.57 | 0.52 | 0.34 | 0.69 | 0.64 |
| 10 | 24294067 | rs2027238 | -0.743 | 0.887 | 0.478 | -0.756 | 0 | -0.001 | -0.001 | 0 | 0.31 | 0.3 | 0.3 | 0.32 | 0.35 | 0.38 |
| 10 | 24294791 | rs4336925 | -0.304 | 0.945 | 0.238 | -0.759 | 0.003 | -0.014 | -0.014 | 0 | 0.12 | 0.09 | 0.06 | 0 | 0.08 | 0.11 |
| 10 | 24295328 | rs1832312 | 0.99 | 0.978 | 0.204 | -0.651 | -0.071 | -0.088 | -0.088 | 0 | 0.16 | 0.03 | 0.23 | 0.38 | 0.29 | 0.25 |
| 10 | 24295992 | rs4748923 | 1 | 0.966 | 0.207 | -0.688 | -0.067 | -0.049 | -0.049 | 0 | 0.16 | 0.03 | 0.18 | 0.26 | 0.28 | 0.24 |
| 10 | 24302480 | rs10764445 | -0.577 | 1.007 | 0.723 | -0.604 | 0 | -0.059 | -0.059 | 0 | 0.12 | 0.17 | 0.08 | 0 | 0.11 | 0.11 |
| 10 | 24308545 | rs10741036 | -0.081 | 0.952 | 0.786 | -0.325 | 0.03 | 0.07 | 0.07 | 0.018 | 0.75 | 0.6 | 0.55 | 0.45 | 0.71 | 0.66 |
| 10 | 24316134 | rs4354609 | -0.087 | 1.032 | 0.505 | -0.361 | 0.008 | 0.016 | 0.016 | 0 | 0.73 | 0.7 | 0.52 | 0.43 | 0.68 | 0.69 |
| 10 | 24318765 | rs6482357 | -0.631 | 0.699 | 0.631 | -0.411 | 0.037 | 0.015 | 0.015 | 0 | 0.23 | 0.41 | 0.22 | 0.29 | 0.3 | 0.28 |
| 10 | 24326623 | rs2067701 | 0.133 | 0.417 | 0.479 | -1.109 | 0 | -0.03 | -0.03 | 0.006 | 0.61 | 0.6 | 0.37 | 0.07 | 0.59 | 0.42 |
| 10 | 24333063 | rs10828593 | -0.072 | 0.466 | 1.048 | -1.21 | -0.014 | 0.052 | 0.052 | 0 | 0.63 | 0.76 | 0.38 | 0.08 | 0.58 | 0.42 |
| 10 | 24334928 | rs10828594 | 0.632 | -0.373 | 1.073 | -1.68 | -0.038 | 0.195 | 0.195 | 0 | 0.46 | 0.2 | 0.29 | 0.06 | 0.49 | 0.35 |
| 10 | 24343063 | rs11013866 | -0.759 | -0.67 | 2.167 | -1.249 | -0.06 | -0.095 | -0.095 | 5.476 | 0.17 | 0.45 | 0.1 | 0.02 | 0.09 | 0.07 |
| 10 | 24350577 | rs12262912 | -2.183 | -0.807 | 2.089 | -1.366 | 0.056 | 0.09 | 0.09 | 6.508 | 0.29 | 0.82 | 0.19 | 0.04 | 0.18 | 0.14 |
| 10 | 24350775 | rs12256944 | -1.585 | -1.327 | 2.316 | -1.372 | -0.027 | -0.038 | -0.038 | 2.698 | 0.25 | 0.66 | 0.13 | 0.03 | 0.12 | 0.12 |
| 10 | 24365207 | rs12242424 | -0.843 | -0.979 | 2.614 | -1.118 | -0.089 | -0.075 | -0.075 | 0 | 0.17 | 0.47 | 0.09 | 0.03 | 0.07 | 0.05 |
| 10 | 24370937 | rs7097422 | -1.397 | -0.937 | 2.573 | -1.131 | -0.055 | -0.076 | -0.076 | 0.213 | 0.19 | 0.57 | 0.12 | 0.04 | 0.1 | 0.09 |
| 10 | 24372953 | rs6482365 | -1.452 | -0.76 | 2.113 | -1.032 | -0.078 | -0.071 | -0.071 | 0 | 0.22 | 0.59 | 0.11 | 0.04 | 0.09 | 0.09 |
| 10 | 24381792 | rs11013893 | -1.488 | -0.755 | 2.129 | -1.02 | -0.076 | -0.069 | -0.069 | 0.774 | 0.22 | 0.58 | 0.11 | 0.04 | 0.1 | 0.09 |
| 10 | 24395147 | rs1326644 | -1.632 | -0.653 | 2.05 | -1.041 | -0.045 | -0.077 | -0.077 | 0 | 0.24 | 0.59 | 0.13 | 0.05 | 0.14 | 0.12 |
| 10 | 24399674 | rs11013898 | -0.822 | -0.396 | 1.115 | -1.047 | 0.013 | 0.023 | 0.023 | 2.052 | 0.16 | 0.11 | 0.07 | 0 | 0.13 | 0.1 |
| 10 | 24407947 | rs11013906 | -1.131 | 0.638 | 0.754 | 0.062 | 0 | 0.01 | 0.01 | 0 | 0.24 | 0.24 | 0.24 | 0.45 | 0.21 | 0.29 |
| 10 | 24417442 | rs7902917 | -0.873 | 0.646 | 1.017 | 0.043 | 0.004 | 0.032 | 0.032 | 0 | 0.21 | 0.22 | 0.25 | 0.46 | 0.2 | 0.29 |
| 10 | 24419413 | rs7911785 | -0.928 | 0.56 | 0.467 | -0.561 | 0.008 | -0.028 | -0.028 | 0 | 0.64 | 0.71 | 0.59 | 0.5 | 0.64 | 0.78 |
| 10 | 24425167 | rs10741045 | -0.933 | 0.569 | 0.388 | -0.54 | 0.001 | -0.009 | -0.009 | 0.428 | 0.64 | 0.67 | 0.54 | 0.47 | 0.63 | 0.78 |
| 10 | 24426693 | rs10764449 | -0.634 | 0.229 | 0.49 | -0.702 | 0.004 | 0.082 | 0.082 | 0.069 | 0.25 | 0.43 | 0.23 | 0.47 | 0.25 | 0.29 |
| 10 | 24430388 | rs11814429 | 0.521 | 0.271 | 0.112 | -1.163 | -0.008 | 0.106 | 0.106 | 0.485 | 0.37 | 0.17 | 0.28 | 0.02 | 0.37 | 0.41 |
| 10 | 24447600 | rs1529529 | 0.667 | 0.193 | -0.014 | -0.869 | -0.015 | 0.018 | 0.018 | 0.545 | 0.66 | 0.52 | 0.61 | 0.4 | 0.74 | 0.69 |
| 10 | 24447877 | rs10828610 | 0.233 | 0.196 | -0.004 | -0.848 | 0.015 | 0.079 | 0.079 | 0.086 | 0.72 | 0.54 | 0.63 | 0.43 | 0.75 | 0.74 |
| 10 | 24451240 | rs2031648 | -0.015 | 0.184 | 0.096 | -0.851 | 0.002 | 0.02 | 0.02 | 0 | 0.74 | 0.71 | 0.64 | 0.43 | 0.76 | 0.74 |
| 10 | 24456486 | rs10764451 | 0.005 | 0.189 | 0.076 | -0.85 | 0.013 | 0.051 | 0.051 | 0 | 0.74 | 0.66 | 0.63 | 0.42 | 0.75 | 0.74 |
| 10 | 24457776 | rs4086430 | 0.765 | 0.316 | -0.035 | -0.956 | -0.021 | 0.021 | 0.021 | 2.673 | 0.3 | 0.21 | 0.26 | 0.02 | 0.37 | 0.44 |
| 10 | 24467344 | rs1326641 | 1.13 | 0.312 | -0.088 | -0.904 | 0.004 | -0.003 | -0.003 | 0.016 | 0.36 | 0.3 | 0.35 | 0.38 | 0.37 | 0.25 |
| 10 | 24475352 | rs11013944 | 0.248 | 0.611 | -0.243 | -0.882 | -0.003 | -0.02 | -0.02 | 1.121 | 0.32 | 0.3 | 0.28 | 0.06 | 0.38 | 0.45 |
| 10 | 24484239 | rs7083993 | 0.669 | 0.559 | -0.198 | 0.025 | 0.007 | 0.041 | 0.041 | 0 | 0.74 | 0.64 | 0.52 | 0.38 | 0.74 | 0.73 |
| 10 | 24485269 | rs10741048 | 0.559 | 0.625 | -0.638 | 0.254 | 0 | -0.024 | -0.024 | 0 | 0.83 | 0.87 | 0.67 | 0.38 | 0.86 | 0.73 |
| 10 | 24491058 | rs954729 | 0.39 | 0.632 | -0.928 | 0.391 | 0 | 0.001 | 0.001 | 0 | 0.65 | 0.65 | 0.48 | 0.38 | 0.66 | 0.64 |
| 10 | 24493326 | rs6482373 | -0.308 | 0.223 | -1.244 | 0.195 | 0.011 | 0.041 | 0.041 | 0.8 | 0.9 | 1 | 0.76 | 0.97 | 0.94 | 0.97 |
| 10 | 24502070 | rs4748930 | -0.162 | 0.402 | -1.284 | 0.334 | -0.008 | 0.008 | 0.008 | 0 | 0.64 | 0.71 | 0.5 | 0.69 | 0.59 | 0.62 |
| 10 | 24503255 | rs3936155 | -0.121 | 0.345 | -1.31 | 0.591 | -0.008 | 0.002 | 0.002 | 0 | 0.64 | 0.71 | 0.5 | 0.62 | 0.59 | 0.58 |
| 10 | 24506475 | rs11013979 | 1.147 | 0.193 | -2.173 | 0.958 | -0.093 | -0.085 | -0.085 | 0 | 0.28 | 0.02 | 0.29 | 0.35 | 0.34 | 0.13 |
| 10 | 24513629 | rs7907546 | -0.508 | -0.375 | -1.422 | 0.864 | 0 | -0.001 | -0.001 | 0 | 0.62 | 0.63 | 0.56 | 0.68 | 0.58 | 0.6 |

| | | | | | | | | | | | | | | | | |
|---|---|---|---|---|---|---|---|---|---|---|---|---|---|---|---|---|
| 10 | 24518094 | rs7081091 | 0.281 | -0.351 | -1.391 | 0.748 | 0.002 | -0.027 | -0.027 | 0 | 0.32 | 0.33 | 0.29 | 0.15 | 0.39 | 0.28 |
| 10 | 24518180 | rs7077598 | -0.3 | -0.497 | -1.075 | 0.854 | -0.016 | 0.075 | 0.075 | 0 | 0.92 | 0.81 | 0.77 | 0.83 | 0.97 | 0.85 |
| 10 | 24523789 | rs12413189 | 1.338 | -0.385 | -1.099 | 1.142 | -0.041 | -0.077 | -0.077 | 0 | 0.29 | 0.05 | 0.34 | 0.36 | 0.32 | 0.18 |
| 10 | 24524433 | rs11013986 | 0.789 | -0.413 | -0.936 | 1.024 | -0.006 | 0.002 | 0.002 | 0 | 0.28 | 0.21 | 0.22 | 0.15 | 0.38 | 0.23 |
| 10 | 24529308 | rs1543740 | -0.255 | -0.43 | -0.917 | 1.057 | 0.002 | -0.006 | -0.006 | 0 | 0.38 | 0.38 | 0.51 | 0.33 | 0.41 | 0.4 |
| 10 | 24534056 | rs10508671 | -0.818 | -0.495 | -0.924 | 1.117 | -0.031 | 0.093 | 0.093 | 0.21 | 0.1 | 0.29 | 0.24 | 0.19 | 0.03 | 0.17 |
| 10 | 24534194 | rs1543739 | 0.628 | -0.412 | -0.397 | 0.9 | 0.02 | -0.048 | -0.048 | 3.053 | 0.3 | 0.43 | 0.23 | 0.22 | 0.38 | 0.25 |
| 10 | 24537108 | rs6482377 | 0.457 | -0.399 | -0.493 | 1.002 | 0.008 | -0.004 | -0.004 | 0.301 | 0.4 | 0.35 | 0.59 | 0.55 | 0.37 | 0.36 |
| 10 | 24539039 | rs3814686 | -0.088 | -0.755 | -0.32 | 1.251 | 0.002 | -0.007 | -0.007 | 0 | 0.66 | 0.65 | 0.81 | 0.61 | 0.72 | 0.49 |
| 10 | 24542427 | rs1138000 | -0.116 | -0.773 | -0.34 | 1.161 | 0.002 | -0.004 | -0.004 | 0 | 0.66 | 0.67 | 0.81 | 0.63 | 0.72 | 0.49 |
| 10 | 24542941 | rs17583120 | 1.69 | -0.362 | -0.629 | 1.294 | -0.065 | 0.044 | 0.044 | 2.135 | 0.29 | 0.06 | 0.29 | 0.18 | 0.35 | 0.15 |
| 10 | 24545194 | rs2280171 | -0.406 | -0.339 | -0.548 | 1.29 | 0.002 | -0.012 | -0.012 | 0.279 | 0.67 | 0.74 | 0.81 | 0.63 | 0.73 | 0.48 |
| 10 | 24546608 | rs12777541 | 1.616 | -0.248 | -0.64 | 1.13 | -0.05 | 0.043 | 0.043 | 1.357 | 0.31 | 0.08 | 0.3 | 0.21 | 0.36 | 0.18 |
| 10 | 24546741 | rs12782581 | 1.616 | -0.248 | -0.64 | 1.13 | -0.05 | 0.043 | 0.043 | 1.357 | 0.31 | 0.08 | 0.3 | 0.21 | 0.36 | 0.18 |
| 10 | 24553625 | rs11013998 | 0.228 | -0.096 | -0.425 | 0.924 | 0.012 | -0.053 | -0.053 | 0.084 | 0.26 | 0.37 | 0.26 | 0.17 | 0.33 | 0.21 |
| 10 | 24554196 | rs11597573 | 0.178 | 0.034 | -0.442 | 0.901 | -0.004 | -0.055 | -0.055 | 0 | 0.24 | 0.36 | 0.24 | 0.16 | 0.29 | 0.2 |
| 10 | 24558972 | rs6482382 | -0.118 | 0.027 | -0.542 | 1.179 | -0.001 | -0.001 | -0.001 | 0 | 0.68 | 0.68 | 0.81 | 0.75 | 0.73 | 0.56 |
| 10 | 24564653 | rs11014002 | 0.128 | -0.057 | -0.495 | 1.794 | -0.01 | -0.029 | -0.029 | 0 | 0.07 | 0 | 0.04 | 0.22 | 0.05 | 0.13 |
| 10 | 24565181 | rs17505709 | 1.427 | -0.167 | -0.762 | 1.631 | 0.011 | 0.12 | 0.12 | 3.635 | 0.42 | 0.16 | 0.5 | 0.21 | 0.39 | 0.22 |
| 10 | 24565347 | rs4748935 | -0.168 | -0.385 | -0.906 | 1.794 | 0.047 | -0.045 | -0.045 | 0 | 0.86 | 0.98 | 0.91 | 0.78 | 0.93 | 0.64 |
| 10 | 24567595 | rs220341 | 0.48 | -0.348 | -0.479 | 1.713 | 0.007 | -0.005 | -0.005 | 0 | 0.82 | 0.87 | 0.84 | 0.76 | 0.88 | 0.61 |
| 10 | 24571677 | rs373648 | -1.138 | -0.018 | 0.744 | 1.837 | 0.035 | -0.03 | -0.03 | 0 | 0.34 | 0.48 | 0.34 | 0.23 | 0.51 | 0.3 |
| 10 | 24574409 | rs220358 | -0.657 | 0.044 | 1.034 | 1.709 | 0.066 | -0.071 | -0.071 | 0 | 0.29 | 0.4 | 0.28 | 0.13 | 0.47 | 0.19 |
| 10 | 24575653 | rs16924466 | -0.746 | -0.038 | 0.896 | 1.756 | -0.005 | 0.014 | 0.014 | 0 | 0.08 | 0.18 | 0.07 | 0.16 | 0.05 | 0.06 |
| 10 | 24576622 | rs220355 | -0.38 | 0.146 | 0.599 | 1.708 | -0.011 | 0.006 | 0.006 | 1.07 | 0.27 | 0.19 | 0.26 | 0.13 | 0.45 | 0.19 |
| 10 | 24584362 | rs387690 | 1.091 | -0.195 | 0.676 | 2.131 | -0.059 | 0.074 | 0.074 | 0.36 | 0.75 | 0.57 | 0.78 | 0.4 | 0.86 | 0.51 |
| 10 | 24586929 | rs220371 | -0.402 | -0.04 | 0.362 | 1.955 | 0.055 | -0.06 | -0.06 | 0 | 0.88 | 0.98 | 0.92 | 0.77 | 0.96 | 0.66 |
| 10 | 24588732 | rs7350447 | 1.623 | 0.142 | 0.275 | 1.842 | -0.044 | 0.127 | 0.127 | 4.171 | 0.51 | 0.14 | 0.6 | 0.36 | 0.48 | 0.28 |
| 10 | 24590566 | rs7913679 | -1.087 | -0.281 | 0.585 | 2.119 | 0.015 | 0.246 | 0.246 | 0.464 | 0.07 | 0.24 | 0.07 | 0.28 | 0.06 | 0.17 |
| 10 | 24592764 | rs220366 | 1.018 | -0.392 | 0.782 | 1.57 | -0.073 | 0.107 | 0.107 | 0.14 | 0.78 | 0.5 | 0.85 | 0.48 | 0.89 | 0.48 |
| 10 | 24595755 | rs17383772 | -0.039 | -0.057 | 0.233 | 1.325 | 0 | 0.035 | 0.035 | 0 | 0.19 | 0.13 | 0.15 | 0.11 | 0.18 | 0.14 |
| 10 | 24595889 | rs12220707 | -0.038 | -0.003 | 0.286 | 1.646 | -0.001 | -0.001 | -0.001 | 0.188 | 0.14 | 0.13 | 0.06 | 0.29 | 0.06 | 0.35 |
| 10 | 24595983 | rs10828624 | -1.033 | 0.004 | 0.481 | 1.65 | -0.072 | 0.123 | 0.123 | 0.405 | 0.2 | 0.49 | 0.09 | 0.36 | 0.08 | 0.43 |
| 10 | 24598415 | rs11014020 | -1.179 | 0.34 | 0.168 | 1.293 | 0.012 | -0.026 | -0.026 | 0 | 0.31 | 0.8 | 0.25 | 0.33 | 0.26 | 0.32 |
| 10 | 24598722 | rs7894356 | -0.592 | 0.618 | 0.19 | 1.202 | 0.022 | 0.146 | 0.146 | 0.357 | 0.05 | 0.21 | 0.06 | 0.16 | 0.05 | 0.1 |
| 10 | 24600008 | rs12258036 | 0.91 | 0.759 | -0.137 | 0.935 | -0.002 | 0.016 | 0.016 | 0.367 | 0.46 | 0.39 | 0.59 | 0.25 | 0.48 | 0.32 |
| 10 | 24600547 | rs10741050 | 1.232 | 1.044 | 0.218 | 1.33 | 0.033 | 0.021 | 0.021 | 0.527 | 0.71 | 0.45 | 0.72 | 0.67 | 0.61 | 0.68 |
| 10 | 24600918 | rs7095243 | -0.194 | 0.91 | -0.273 | 1.587 | 0.004 | -0.105 | -0.105 | 0.017 | 0.2 | 0.05 | 0.12 | 0.4 | 0.16 | 0.35 |
| 10 | 24605003 | rs10828627 | -0.825 | 1.232 | 0.63 | 1.677 | 0.033 | 0.219 | 0.219 | 2.454 | 0.07 | 0.33 | 0.07 | 0.22 | 0.05 | 0.1 |
| 10 | 24605328 | rs11014023 | -0.563 | 1.384 | 0.206 | 1.387 | -0.004 | -0.011 | -0.011 | 0 | 0.16 | 0.17 | 0.15 | 0.14 | 0.14 | 0.13 |
| 10 | 24607648 | rs7098629 | -0.611 | 1.383 | 0.234 | 1.384 | -0.006 | -0.017 | -0.017 | 0 | 0.16 | 0.19 | 0.15 | 0.13 | 0.14 | 0.13 |
| 10 | 24610724 | rs1080524 | -0.232 | 1.804 | -0.019 | 1.446 | 0.028 | 0.001 | 0.001 | 0 | 0.9 | 0.88 | 0.86 | 0.9 | 0.75 | 0.91 |
| 10 | 24612478 | rs12354978 | 0.636 | 1.737 | -0.134 | 1.274 | -0.011 | 0.01 | 0.01 | 0.015 | 0.06 | 0 | 0.07 | 0.04 | 0.06 | 0.08 |
| 10 | 24614315 | rs6482383 | -0.964 | 1.584 | 0.055 | 1.338 | -0.04 | 0.002 | 0.002 | 0 | 0.21 | 0.46 | 0.21 | 0.22 | 0.15 | 0.18 |
| 10 | 24614504 | rs10443985 | 0.678 | 1.799 | -0.002 | 1.258 | -0.017 | -0.033 | -0.033 | 0.269 | 0.13 | 0.01 | 0.08 | 0.13 | 0.11 | 0.1 |
| 10 | 24614557 | rs442463 | -0.206 | 1.892 | 0.151 | 0.837 | 0.045 | -0.009 | -0.009 | 0 | 0.11 | 0.2 | 0.11 | 0.05 | 0.25 | 0.09 |
| 10 | 24616847 | rs7087182 | -1.02 | 0.462 | 0.166 | 1.028 | -0.03 | -0.066 | -0.066 | 0 | 0.19 | 0.43 | 0.17 | 0.14 | 0.14 | 0.12 |
| 10 | 24620265 | rs11014033 | -0.685 | 0.52 | -0.551 | 1.303 | 0.008 | -0.082 | -0.082 | 0 | 0.19 | 0.04 | 0.12 | 0.41 | 0.15 | 0.37 |
| 10 | 24622926 | rs714326 | -0.93 | 0.916 | 0.079 | 0.701 | 0.02 | 0.044 | 0.044 | 0 | 0.08 | 0.14 | 0.08 | 0.17 | 0.1 | 0.12 |
| 10 | 24624059 | rs11014034 | -0.905 | 0.55 | -0.204 | 0.262 | -0.016 | -0.055 | -0.055 | 0.216 | 0.12 | 0.16 | 0.08 | 0 | 0.08 | 0.03 |
| 10 | 24629653 | rs11592861 | -0.181 | 1.073 | -0.347 | 0.361 | 0.003 | -0.022 | -0.022 | 0 | 0.37 | 0.39 | 0.25 | 0.22 | 0.38 | 0.28 |
| 10 | 24632135 | rs10828629 | -0.281 | 0.89 | -0.231 | 0.762 | -0.003 | -0.007 | -0.007 | 0.014 | 0.55 | 0.5 | 0.36 | 0.62 | 0.6 | 0.72 |
| 10 | 24632537 | rs220345 | -0.042 | 1.387 | -0.078 | 1.234 | -0.008 | -0.008 | -0.008 | 0 | 0.41 | 0.32 | 0.4 | 0.41 | 0.41 | 0.24 |
| 10 | 24636556 | rs1414684 | 0.052 | 0.983 | 0.189 | 0.965 | 0.001 | -0.002 | -0.002 | 0 | 0.73 | 0.69 | 0.46 | 0.79 | 0.71 | 0.8 |
| 10 | 24642865 | rs220350 | -0.762 | 1.062 | 0.212 | 1.03 | 0.017 | 0.044 | 0.044 | 0 | 0.43 | 0.3 | 0.34 | 0.33 | 0.38 | 0.53 |
| 10 | 24644589 | rs11014051 | -0.187 | 1.01 | 0.548 | 1.014 | 0.032 | 0.169 | 0.169 | 0.011 | 0.13 | 0.37 | 0.18 | 0.28 | 0.12 | 0.1 |
| 10 | 24647899 | rs11014053 | 0.879 | 1.242 | 0.341 | 0.999 | -0.024 | -0.027 | -0.027 | 1.466 | 0.48 | 0.19 | 0.3 | 0.44 | 0.51 | 0.42 |
| 10 | 24651563 | rs2214172 | -0.005 | 0.731 | 0.183 | 0.77 | 0.006 | 0.024 | 0.024 | 0 | 0.21 | 0.07 | 0.11 | 0.15 | 0.18 | 0.39 |
| 10 | 24652905 | rs220338 | -0.734 | 0.649 | 0.493 | 1.002 | -0.006 | -0.025 | -0.025 | 0 | 0.27 | 0.36 | 0.26 | 0.15 | 0.23 | 0.41 |
| 10 | 24653883 | rs220335 | -0.768 | 0.384 | 0.368 | 1.125 | -0.005 | 0.002 | 0.002 | 0 | 0.37 | 0.45 | 0.4 | 0.37 | 0.34 | 0.54 |
| 10 | 24654757 | rs10828638 | -1.718 | 0.218 | 0.471 | 1.103 | 0.03 | 0.048 | 0.048 | 0 | 0.08 | 0.23 | 0.16 | 0.11 | 0.07 | 0.05 |
| 10 | 24658034 | rs16924656 | -0.471 | -0.413 | 0.533 | 1.047 | -0.021 | -0.026 | -0.026 | 0 | 0.19 | 0.28 | 0.12 | 0.11 | 0.1 | 0.34 |
| 10 | 24658539 | rs1123509 | -0.098 | -0.91 | 0.009 | 0.855 | -0.012 | -0.037 | -0.037 | 5.598 | 0.33 | 0.5 | 0.2 | 0.22 | 0.27 | 0.18 |
| 10 | 24662302 | rs1889874 | -0.07 | -0.947 | -0.031 | 0.937 | -0.018 | 0.006 | 0.006 | 0 | 0.19 | 0.26 | 0.15 | 0.21 | 0.1 | 0.34 |
| 10 | 24663585 | rs10828639 | -0.13 | -0.958 | -0.036 | 1.402 | -0.018 | 0.049 | 0.049 | 0 | 0.19 | 0.27 | 0.21 | 0.4 | 0.1 | 0.4 |
| 10 | 24663615 | rs10828640 | -0.13 | -0.958 | -0.036 | 1.402 | -0.018 | 0.049 | 0.049 | 0 | 0.19 | 0.27 | 0.2 | 0.39 | 0.1 | 0.4 |
| 10 | 24665266 | rs7083113 | -0.208 | -0.98 | -0.062 | 1.427 | -0.009 | -0.02 | -0.02 | 0 | 0.34 | 0.53 | 0.2 | 0.23 | 0.28 | 0.19 |
| 10 | 24666826 | rs16924711 | 0.569 | -0.954 | -0.115 | 1.441 | 0.014 | -0.019 | -0.019 | 0.166 | 0.16 | 0.11 | 0.14 | 0.21 | 0.1 | 0.34 |
| 10 | 24667091 | rs11014072 | -1.095 | -0.988 | -0.073 | 1.443 | 0.039 | 0.02 | 0.02 | 0 | 0.5 | 0.67 | 0.35 | 0.44 | 0.37 | 0.53 |

| | | | | | | | | | | | | | | | | |
|---|---|---|---|---|---|---|---|---|---|---|---|---|---|---|---|---|
| 10 | 24670973 | rs7081094 | -0.89 | -1.197 | 0.146 | 1.292 | 0.011 | 0.001 | 0.001 | 0 | 0.47 | 0.4 | 0.35 | 0.43 | 0.36 | 0.58 |
| 10 | 24671365 | rs7902724 | -0.421 | -0.999 | 0.157 | 1.319 | 0.013 | 0.024 | 0.024 | 0.02 | 0.35 | 0.63 | 0.21 | 0.22 | 0.28 | 0.18 |
| 10 | 24674934 | rs7896427 | -1.25 | -0.993 | 0.026 | 1.315 | 0.036 | 0.057 | 0.057 | 0.062 | 0.52 | 0.84 | 0.42 | 0.63 | 0.37 | 0.59 |
| 10 | 24675337 | rs6482390 | -0.954 | -0.645 | 0.55 | 1.518 | 0.034 | -0.041 | -0.041 | 0 | 0.37 | 0.74 | 0.31 | 0.52 | 0.28 | 0.25 |
| 10 | 24682940 | rs6482393 | -0.929 | -0.405 | 0.26 | 0.642 | -0.004 | -0.01 | -0.01 | 0 | 0.34 | 0.56 | 0.2 | 0.23 | 0.28 | 0.18 |
| 10 | 24686022 | rs7068383 | -0.781 | -0.414 | 0.274 | 0.608 | -0.016 | -0.044 | -0.044 | 0.371 | 0.33 | 0.43 | 0.21 | 0.22 | 0.28 | 0.19 |
| 10 | 24691079 | rs7912789 | -0.554 | 0.227 | -1.183 | 0.837 | -0.004 | -0.029 | -0.029 | 0.127 | 0.29 | 0.08 | 0.2 | 0.32 | 0.27 | 0.19 |
| 10 | 24693878 | rs4601665 | -0.459 | 0.414 | -0.834 | 0.548 | -0.004 | -0.007 | -0.007 | 0 | 0.43 | 0.44 | 0.4 | 0.35 | 0.34 | 0.34 |
| 10 | 24694715 | rs1556401 | -0.832 | 0.15 | -0.336 | 0.271 | -0.036 | -0.061 | -0.061 | 1.79 | 0.25 | 0.39 | 0.22 | 0.1 | 0.13 | 0.13 |
| 10 | 24694909 | rs17513725 | 0.19 | 1.214 | -1.106 | 0.229 | -0.034 | 0.019 | 0.019 | 0.51 | 0.15 | 0.11 | 0.22 | 0.12 | 0.27 | 0.25 |
| 10 | 24696277 | rs7915144 | -0.917 | 1.284 | -1.655 | 0.625 | -0.068 | -0.086 | -0.086 | 2.616 | 0.77 | 0.96 | 0.75 | 0.69 | 0.77 | 0.71 |
| 10 | 24697860 | rs7096689 | -0.209 | 0.992 | -0.435 | 0.727 | -0.062 | -0.025 | -0.025 | 0 | 0.12 | 0.29 | 0.05 | 0.05 | 0.02 | 0.01 |
| 10 | 24702109 | rs4748946 | 0.261 | 1.158 | -0.757 | 0.867 | -0.008 | -0.015 | -0.015 | 0 | 0.06 | 0.01 | 0.06 | 0.05 | 0.06 | 0.1 |
| 10 | 24706072 | rs3858219 | 0.239 | 1.101 | -0.762 | 0.778 | -0.067 | -0.008 | -0.008 | 0 | 0.2 | 0.02 | 0.22 | 0.17 | 0.29 | 0.29 |
| 10 | 24706471 | rs3858220 | 0.186 | 1.229 | -0.505 | 0.955 | 0.039 | 0.046 | 0.046 | 0 | 0.06 | 0.1 | 0.17 | 0.2 | 0.18 | 0.08 |
| 10 | 24707669 | rs2366915 | -1.266 | 1.035 | -0.317 | 0.715 | -0.009 | 0.012 | 0.012 | 0.02 | 0.25 | 0.49 | 0.28 | 0.26 | 0.26 | 0.19 |
| 10 | 24711413 | rs17513940 | -0.73 | 1.653 | -0.069 | 1.017 | 0.01 | 0.01 | 0.01 | 0 | 0.4 | 0.52 | 0.49 | 0.49 | 0.51 | 0.46 |
| 10 | 24712864 | rs16924739 | -0.447 | 1.718 | 0.172 | 1.109 | 0.021 | 0.008 | 0.008 | 0 | 0.05 | 0.13 | 0.07 | 0.07 | 0.08 | 0.03 |
| 10 | 24720098 | rs7087652 | -1.033 | 2.06 | -0.165 | 1.745 | 0.025 | 0.044 | 0.044 | 1.102 | 0.21 | 0.34 | 0.2 | 0.34 | 0.31 | 0.17 |
| 10 | 24720444 | rs3762085 | -1.744 | 1.67 | 0.278 | 1.212 | 0.019 | 0.048 | 0.048 | 0.007 | 0.3 | 0.55 | 0.28 | 0.36 | 0.37 | 0.26 |
| 10 | 24720488 | rs3762086 | -0.334 | 1.824 | 0.332 | 1.442 | -0.002 | 0.032 | 0.032 | 0.978 | 0.13 | 0.23 | 0.11 | 0.19 | 0.15 | 0.1 |
| 10 | 24720825 | rs10508674 | -1.841 | 0.699 | -0.255 | 1.366 | -0.074 | 0.033 | 0.033 | 0 | 0.58 | 0.87 | 0.52 | 0.69 | 0.52 | 0.65 |
| 10 | 24721946 | rs2274831 | 0.106 | 0.68 | 0.085 | 1.377 | -0.002 | 0.051 | 0.051 | 1.203 | 0.14 | 0.31 | 0.12 | 0.19 | 0.15 | 0.1 |
| 10 | 24723566 | rs1331291 | 0.096 | 0.309 | 0.01 | 1.322 | -0.013 | 0.003 | 0.003 | 0.856 | 0.28 | 0.24 | 0.23 | 0.37 | 0.15 | 0.44 |
| 10 | 24724335 | rs16924760 | -0.898 | 0.416 | 0.169 | 1.147 | 0.019 | 0.014 | 0.014 | 0 | 0.2 | 0.43 | 0.2 | 0.19 | 0.22 | 0.17 |
| 10 | 24729218 | rs1537170 | 0.174 | 0.536 | 0.175 | 1.227 | -0.007 | -0.008 | -0.008 | 0.032 | 0.12 | 0.16 | 0.05 | 0.09 | 0.1 | 0.13 |
| 10 | 24742828 | rs11014094 | -0.023 | 0.485 | 0.003 | 1.215 | -0.011 | 0.022 | 0.022 | 1.864 | 0.11 | 0.04 | 0.05 | 0.04 | 0.14 | 0.11 |
| 10 | 24748968 | rs11014097 | 0.175 | -0.075 | -0.074 | 1.392 | 0.048 | -0.032 | -0.032 | 0.317 | 0.22 | 0.05 | 0.18 | 0.3 | 0.14 | 0.34 |
| 10 | 24753146 | rs10764472 | -0.197 | 0.212 | -0.11 | 1.364 | 0.048 | -0.025 | -0.025 | 0 | 0.27 | 0.05 | 0.2 | 0.32 | 0.18 | 0.38 |
| 10 | 24777635 | rs2067887 | -0.683 | 1.068 | 0.489 | 1.093 | 0.004 | 0.002 | 0.002 | 0 | 0.44 | 0.57 | 0.29 | 0.37 | 0.35 | 0.54 |
| 10 | 24779757 | rs10764473 | -0.882 | 0.927 | 0.535 | 1.033 | 0.003 | 0.003 | 0.003 | 0 | 0.47 | 0.58 | 0.4 | 0.5 | 0.49 | 0.56 |
| 10 | 24780659 | rs12252729 | 0.336 | 0.733 | 0.229 | 0.856 | 0.02 | -0.014 | -0.014 | 0 | 0.28 | 0.21 | 0.2 | 0.35 | 0.19 | 0.41 |
| 10 | 24781090 | rs11014114 | -0.352 | 0.729 | 0.179 | 0.792 | -0.039 | 0.069 | 0.069 | 1.667 | 0.13 | 0.02 | 0.06 | 0.01 | 0.17 | 0.11 |
| 10 | 24782246 | rs12265701 | 0.099 | 0.428 | 0.464 | 1.035 | 0.004 | -0.004 | -0.004 | 0 | 0.29 | 0.26 | 0.21 | 0.36 | 0.2 | 0.41 |
| 10 | 24786291 | rs12782580 | 0.348 | 0.308 | 0.442 | 1.101 | 0.024 | -0.02 | -0.02 | 0 | 0.27 | 0.16 | 0.2 | 0.33 | 0.19 | 0.36 |
| 10 | 24786510 | rs11014115 | -0.143 | 0.524 | 0.627 | 1.056 | -0.006 | 0.015 | 0.015 | 0 | 0.32 | 0.37 | 0.33 | 0.51 | 0.33 | 0.45 |
| 10 | 24790393 | rs1112284 | -0.423 | 0.71 | 0.415 | 0.329 | 0.039 | 0.079 | 0.079 | 3.082 | 0.46 | 0.68 | 0.3 | 0.21 | 0.34 | 0.34 |
| 10 | 24793880 | rs10508679 | -1.869 | 0.844 | 0.584 | 0.507 | 0.039 | 0.135 | 0.135 | 0 | 0.16 | 0.46 | 0.22 | 0.29 | 0.23 | 0.16 |
| 10 | 24798368 | rs12252802 | -0.787 | 0.887 | 0.841 | 0.734 | 0.006 | -0.013 | -0.013 | 0 | 0.08 | 0.29 | 0.04 | 0.06 | 0.05 | 0.07 |
| 10 | 24799175 | rs10828661 | 0.223 | 0.89 | 0.672 | 0.698 | -0.001 | -0.021 | -0.021 | 0 | 0.36 | 0.3 | 0.24 | 0.11 | 0.29 | 0.26 |
| 10 | 24805665 | rs7910985 | 0.075 | 0.86 | 0.347 | 0.576 | -0.005 | -0.025 | -0.025 | 0 | 0.37 | 0.36 | 0.25 | 0.15 | 0.29 | 0.27 |
| 10 | 24806631 | rs7090837 | -0.854 | 0.884 | -0.326 | 0.624 | 0.019 | 0.006 | 0.006 | 3.217 | 0.5 | 0.82 | 0.45 | 0.43 | 0.48 | 0.42 |
| 10 | 24810663 | rs2297330 | 0.264 | 0.928 | -0.257 | 0.227 | 0.004 | 0.012 | 0.012 | 8.891 | 0.26 | 0.19 | 0.17 | 0.09 | 0.22 | 0.15 |
| 10 | 24813327 | rs2297328 | -1.22 | 0.934 | 0.394 | 0.724 | 0.048 | 0.153 | 0.153 | 0.508 | 0.07 | 0.31 | 0.09 | 0.17 | 0.04 | 0.11 |
| 10 | 24813454 | rs10828663 | 0.237 | 1.009 | -0.463 | 0.351 | 0.001 | 0.022 | 0.022 | 2.232 | 0.29 | 0.21 | 0.21 | 0.11 | 0.29 | 0.11 |
| 10 | 24814503 | rs10828664 | -0.591 | 1.436 | -0.718 | 0.644 | 0.002 | 0.002 | 0.002 | 1.337 | 0.55 | 0.67 | 0.49 | 0.37 | 0.6 | 0.41 |
| 10 | 24818152 | rs10828665 | -0.525 | 1.108 | -0.558 | 0.857 | -0.001 | -0.003 | -0.003 | 0 | 0.21 | 0.27 | 0.21 | 0.24 | 0.25 | 0.2 |
| 10 | 24823076 | rs2297325 | -0.158 | 0.872 | -0.363 | 0.606 | 0.009 | 0.027 | 0.027 | 0.079 | 0.65 | 0.59 | 0.56 | 0.41 | 0.64 | 0.49 |
| 10 | 24827151 | rs7100799 | 0.512 | 0.379 | -0.248 | 0.304 | 0.034 | -0.01 | -0.01 | 0.244 | 0.4 | 0.33 | 0.33 | 0.38 | 0.36 | 0.21 |
| 10 | 24828744 | rs7904181 | 0.787 | 0.071 | -0.303 | 0.074 | 0.042 | 0.055 | 0.055 | 0 | 0.35 | 0.26 | 0.21 | 0.14 | 0.25 | 0.25 |
| 10 | 24830350 | rs1537169 | -1.057 | 0.058 | -0.282 | 0.328 | -0.002 | 0.088 | 0.088 | 0 | 0.09 | 0.23 | 0.17 | 0.17 | 0.1 | 0.05 |
| 10 | 24832885 | rs3748218 | -1.66 | -0.161 | -0.343 | 0.533 | -0.02 | 0.128 | 0.128 | 0 | 0.17 | 0.33 | 0.22 | 0.4 | 0.15 | 0.25 |
| 10 | 24833137 | rs2250411 | -0.44 | 0.351 | -0.695 | 0.174 | -0.012 | 0.036 | 0.036 | 0 | 0.33 | 0.25 | 0.31 | 0.14 | 0.4 | 0.22 |
| 10 | 24833614 | rs1888657 | 0.189 | 0.382 | -0.391 | 0.3 | 0.034 | 0.026 | 0.026 | 1.986 | 0.28 | 0.35 | 0.45 | 0.5 | 0.4 | 0.4 |
| 10 | 24833705 | rs1888656 | -0.776 | -0.084 | -0.594 | 0.526 | 0.01 | -0.016 | -0.016 | 0 | 0.09 | 0.03 | 0.17 | 0.27 | 0.07 | 0.21 |
| 10 | 24837691 | rs11014147 | 0.43 | -0.109 | -0.263 | 0.135 | 0.001 | -0.002 | -0.002 | 0 | 0.57 | 0.45 | 0.56 | 0.7 | 0.51 | 0.52 |
| 10 | 24837715 | rs2249911 | -0.773 | 0.198 | -1.133 | 0.052 | -0.032 | 0.026 | 0.026 | 0 | 0.76 | 0.88 | 0.79 | 0.98 | 0.67 | 0.82 |
| 10 | 24842278 | rs11014155 | 0.454 | 0.486 | -1.274 | 0.463 | -0.02 | 0.028 | 0.028 | 0 | 0.49 | 0.19 | 0.49 | 0.58 | 0.5 | 0.49 |
| 10 | 24842340 | rs2246603 | -0.964 | 0.605 | -0.927 | 1.07 | 0.005 | 0.068 | 0.068 | 0 | 0.07 | 0.08 | 0.19 | 0.29 | 0.14 | 0.18 |
| 10 | 24846116 | rs7901814 | 0.168 | -0.027 | -1.462 | 0.293 | 0.032 | 0.105 | 0.105 | 0 | 0.34 | 0.05 | 0.22 | 0.11 | 0.29 | 0.25 |
| 10 | 24847361 | rs10828672 | -0.363 | -0.326 | -0.847 | 0.263 | 0.035 | 0.052 | 0.052 | 0 | 0.44 | 0.61 | 0.31 | 0.19 | 0.37 | 0.27 |
| 10 | 24848149 | rs914320 | -0.363 | -0.326 | -0.847 | 0.263 | 0.035 | 0.052 | 0.052 | 0 | 0.44 | 0.61 | 0.31 | 0.19 | 0.37 | 0.27 |
| 10 | 24851663 | rs1556398 | 0.073 | -0.389 | -0.613 | 0.462 | 0.014 | -0.005 | -0.005 | 0 | 0.37 | 0.23 | 0.36 | 0.36 | 0.39 | 0.35 |
| 10 | 24852365 | rs10764478 | 0.363 | 0.127 | -1.111 | 0.531 | 0.003 | 0.048 | 0.048 | 0 | 0.22 | 0.31 | 0.38 | 0.56 | 0.22 | 0.27 |
| 10 | 24852400 | rs10764479 | 0.37 | 0.102 | -1.121 | 0.643 | 0.003 | 0.058 | 0.058 | 0 | 0.22 | 0.32 | 0.38 | 0.52 | 0.22 | 0.26 |
| 10 | 24854876 | rs2265265 | -0.404 | 0.306 | -1.295 | 0.308 | 0.015 | 0.004 | 0.004 | 0 | 0.51 | 0.6 | 0.44 | 0.46 | 0.48 | 0.38 |
| 10 | 24856314 | rs2256220 | 0.452 | 0.594 | -1.436 | 0.447 | -0.043 | -0.062 | -0.062 | 0 | 0.32 | 0.09 | 0.35 | 0.41 | 0.43 | 0.29 |
| 10 | 24858390 | rs4332427 | 0.592 | 0.87 | -1.383 | -0.068 | -0.005 | 0.087 | 0.087 | 0.892 | 0.28 | 0.12 | 0.22 | 0.09 | 0.29 | 0.28 |
| 10 | 24863020 | rs2001237 | 0.239 | 0.9 | -0.971 | 0.367 | 0.009 | 0.041 | 0.041 | 0.061 | 0.2 | 0.25 | 0.4 | 0.55 | 0.25 | 0.32 |

| | | | | | | | | | | | | | | | | |
|---|---|---|---|---|---|---|---|---|---|---|---|---|---|---|---|---|
| 10 | 24865081 | rs1962279 | -0.353 | 0.631 | -0.729 | 0.125 | 0.002 | 0.002 | 0.002 | 0.031 | 0.52 | 0.55 | 0.59 | 0.65 | 0.52 | 0.49 |
| 10 | 24870596 | rs10764480 | -0.396 | 0.603 | -0.741 | 0.12 | 0.001 | 0.001 | 0.001 | 0 | 0.53 | 0.55 | 0.59 | 0.65 | 0.51 | 0.49 |
| 10 | 24878834 | rs7894175 | -0.078 | 0.417 | -0.631 | 0.483 | 0.01 | 0.026 | 0.026 | 0 | 0.48 | 0.42 | 0.46 | 0.37 | 0.51 | 0.39 |
| 10 | 24887932 | rs2367063 | 0.992 | 0.479 | -0.698 | 0.416 | -0.005 | 0.102 | 0.102 | 0 | 0.31 | 0.15 | 0.18 | 0.06 | 0.27 | 0.17 |
| 10 | 24927447 | rs7082393 | 1.185 | 0.501 | -0.543 | 0.497 | -0.004 | -0.003 | -0.003 | 0.234 | 0.16 | 0.14 | 0.28 | 0.31 | 0.24 | 0.23 |
| 10 | 24933700 | rs10764482 | 0.615 | 0.603 | -0.765 | 0.002 | -0.027 | 0.056 | 0.056 | 0 | 0.73 | 0.58 | 0.62 | 0.44 | 0.74 | 0.63 |
| 10 | 24933956 | rs10828681 | 0.434 | 0.599 | -0.742 | -0.002 | 0.006 | 0.013 | 0.013 | 0 | 0.48 | 0.44 | 0.47 | 0.41 | 0.51 | 0.4 |
| 10 | 24948072 | rs10764484 | 0.445 | 0.665 | -0.912 | 0.008 | 0.016 | 0.03 | 0.03 | 0 | 0.49 | 0.41 | 0.47 | 0.39 | 0.51 | 0.4 |
| 10 | 24953538 | rs7911476 | -0.997 | 0.602 | -0.605 | 0.226 | -0.003 | 0.011 | 0.011 | 0 | 0.89 | 0.92 | 0.77 | 0.77 | 0.77 | 0.78 |
| 10 | 24975152 | rs2886854 | 0.41 | 0.491 | -0.125 | 0.14 | 0.014 | 0.037 | 0.037 | 0 | 0.52 | 0.63 | 0.48 | 0.38 | 0.51 | 0.39 |
| 10 | 24980584 | rs12161720 | 0.091 | 0.47 | -0.16 | 0.17 | 0.003 | 0.002 | 0.002 | 0 | 0.37 | 0.34 | 0.29 | 0.36 | 0.26 | 0.39 |
| 10 | 24986146 | rs6482414 | -0.922 | 0.562 | -0.154 | 0.185 | 0.002 | 0.035 | 0.035 | 0 | 0.89 | 0.89 | 0.77 | 0.73 | 0.77 | 0.77 |
| 10 | 24996710 | rs10734055 | 0.652 | 0.219 | -0.184 | 0.209 | 0.012 | 0.028 | 0.028 | 0 | 0.48 | 0.4 | 0.47 | 0.37 | 0.51 | 0.39 |
| 10 | 24999668 | rs7067743 | -0.383 | 0.3 | -0.137 | 0.256 | 0.021 | 0.112 | 0.112 | 0 | 0.87 | 0.77 | 0.76 | 0.73 | 0.77 | 0.77 |
| 10 | 25004441 | rs10764490 | 0.708 | -0.099 | -0.79 | -0.207 | 0.003 | -0.002 | -0.002 | 0 | 0.36 | 0.4 | 0.41 | 0.37 | 0.41 | 0.34 |
| 10 | 25007306 | rs12219181 | 0.319 | -0.323 | -1.011 | 0.142 | 0.033 | -0.032 | -0.032 | 0 | 0.21 | 0.12 | 0.17 | 0.31 | 0.12 | 0.19 |
| 10 | 25015683 | rs16924943 | 0.21 | 0.179 | -0.565 | 0.575 | -0.001 | -0.004 | -0.004 | 0.749 | 0.11 | 0.07 | 0.21 | 0.24 | 0.16 | 0.22 |
| 10 | 25025919 | rs7073062 | -0.17 | -0.299 | -0.453 | -0.004 | 0.013 | 0.005 | 0.005 | 0 | 0.41 | 0.4 | 0.46 | 0.6 | 0.37 | 0.56 |
| 10 | 25028611 | rs1414456 | -0.588 | -0.411 | -0.219 | -0.521 | -0.013 | 0.09 | 0.09 | 0 | 0.61 | 0.77 | 0.76 | 0.93 | 0.63 | 0.83 |
| 10 | 25031272 | rs10828692 | -0.633 | -0.448 | 0.146 | -0.537 | 0 | -0.096 | -0.096 | 0 | 0.59 | 0.6 | 0.75 | 0.92 | 0.63 | 0.82 |
| 10 | 25035311 | rs7916054 | -0.569 | -0.472 | 0.2 | -0.561 | -0.001 | -0.103 | -0.103 | 0.094 | 0.58 | 0.59 | 0.74 | 0.93 | 0.62 | 0.82 |
| 10 | 25040622 | rs10466055 | -0.525 | -0.414 | 0.042 | -0.55 | -0.01 | 0.075 | 0.075 | 0 | 0.6 | 0.77 | 0.75 | 0.93 | 0.62 | 0.82 |
| 10 | 25048987 | rs2152432 | -0.664 | -0.443 | 0.077 | -0.494 | -0.003 | 0.031 | 0.031 | 0 | 0.59 | 0.68 | 0.75 | 0.89 | 0.62 | 0.82 |
| 10 | 25055592 | rs951857 | -0.845 | -0.676 | 0.257 | -0.558 | 0 | -0.058 | -0.058 | 4.937 | 0.64 | 0.63 | 0.79 | 0.95 | 0.66 | 0.87 |
| 10 | 25057052 | rs7903504 | 1.046 | -0.865 | 0.306 | -0.289 | -0.022 | 0.006 | 0.006 | 0.059 | 0.07 | 0.13 | 0.02 | 0.08 | 0.01 | 0.01 |
| 10 | 25059038 | rs16925018 | 0.139 | -1.128 | 0.341 | 0.44 | -0.029 | 0.246 | 0.246 | 0 | 0.07 | 0.2 | 0.11 | 0.37 | 0.01 | 0.19 |
| 10 | 25063291 | rs11014225 | 1.33 | -0.776 | -0.116 | 0.203 | -0.021 | 0.01 | 0.01 | 0 | 0.06 | 0.13 | 0.02 | 0.08 | 0.01 | 0.01 |
| 10 | 25066353 | rs4747500 | -0.503 | -0.732 | -0.084 | 0.12 | 0.001 | 0.002 | 0.002 | 1.959 | 0.61 | 0.61 | 0.77 | 0.82 | 0.66 | 0.85 |
| 10 | 25069865 | rs10741064 | -0.477 | -0.697 | -0.009 | 0.137 | -0.002 | -0.027 | -0.027 | 0 | 0.6 | 0.58 | 0.77 | 0.82 | 0.66 | 0.85 |
| 10 | 25078878 | rs1537900 | -0.268 | -0.535 | -0.508 | 0.344 | -0.006 | 0.003 | 0.003 | 1.096 | 0.53 | 0.74 | 0.61 | 0.6 | 0.52 | 0.63 |
| 10 | 25081855 | rs11593521 | -0.185 | 0.189 | -0.234 | 0.853 | -0.015 | -0.051 | -0.051 | 0 | 0.13 | 0.07 | 0.29 | 0.34 | 0.18 | 0.21 |
| 10 | 25095648 | rs12414747 | -0.127 | 0.376 | -0.086 | 0.476 | -0.005 | 0.003 | 0.003 | 0.303 | 0.26 | 0.31 | 0.3 | 0.18 | 0.39 | 0.33 |
| 10 | 25099184 | rs4237366 | -1.462 | 0.166 | 0.459 | 0.569 | 0.035 | 0.03 | 0.03 | 0.035 | 0.5 | 0.66 | 0.62 | 0.59 | 0.6 | 0.56 |
| 10 | 25101407 | rs7910171 | -0.298 | 0.285 | 0.376 | 0.98 | 0.017 | 0.134 | 0.134 | 0 | 0.14 | 0.23 | 0.31 | 0.36 | 0.2 | 0.22 |
| 10 | 25116923 | rs1314611 | 0.311 | 0.03 | 0.352 | 0.806 | -0.011 | 0.104 | 0.104 | 0 | 0.82 | 0.71 | 0.72 | 0.63 | 0.85 | 0.79 |
| 10 | 25117861 | rs4553267 | 0.277 | 0.361 | -0.12 | 0.559 | -0.138 | 0.032 | 0.032 | 1.778 | 0.23 | 0.03 | 0.27 | 0.21 | 0.4 | 0.26 |
| 10 | 25120376 | rs10741067 | 1.247 | 0.146 | 0.363 | 0.479 | 0.004 | 0.005 | 0.005 | 0 | 0.53 | 0.43 | 0.42 | 0.37 | 0.44 | 0.52 |
| 10 | 25122898 | rs4453120 | 0.012 | 0.213 | 0.257 | 0.575 | -0.014 | 0.098 | 0.098 | 0 | 0.82 | 0.7 | 0.72 | 0.64 | 0.85 | 0.8 |
| 10 | 25130623 | rs2478085 | -0.019 | 0.446 | -0.036 | 0.477 | -0.125 | 0.011 | 0.011 | 0.719 | 0.26 | 0.07 | 0.28 | 0.27 | 0.41 | 0.28 |
| 10 | 25137051 | rs2484648 | 0.258 | 0.546 | -0.04 | 0.387 | -0.125 | 0.04 | 0.04 | 2.579 | 0.23 | 0.03 | 0.34 | 0.19 | 0.37 | 0.29 |
| 10 | 25137772 | rs1045873 | 0.269 | 0.514 | -0.038 | 0.389 | -0.127 | 0.04 | 0.04 | 2.506 | 0.23 | 0.03 | 0.34 | 0.19 | 0.37 | 0.29 |
| 10 | 25144730 | rs10741068 | 0.567 | 0.291 | 0.605 | 0.545 | -0.007 | -0.008 | -0.008 | 0 | 0.59 | 0.67 | 0.39 | 0.51 | 0.48 | 0.54 |
| 10 | 25152336 | rs2152433 | 0.555 | 0.236 | 0.711 | 0.901 | -0.027 | -0.035 | -0.035 | 0 | 0.13 | 0.03 | 0.26 | 0.28 | 0.18 | 0.21 |
| 10 | 25152621 | rs7903234 | 0.319 | -0.015 | 0.864 | 0.964 | -0.002 | 0.027 | 0.027 | 0 | 0.81 | 0.81 | 0.74 | 0.6 | 0.83 | 0.83 |
| 10 | 25155407 | rs9418462 | 0.835 | 0.112 | 0.986 | 0.945 | -0.009 | 0.019 | 0.019 | 0.7 | 0.66 | 0.45 | 0.46 | 0.52 | 0.71 | 0.6 |
| 10 | 25158024 | rs4748990 | 0.049 | 0.102 | 0.815 | 0.884 | -0.004 | -0.019 | -0.019 | 0 | 0.18 | 0.41 | 0.27 | 0.18 | 0.14 | 0.22 |
| 10 | 25158221 | rs1971470 | 0.802 | 0.059 | 0.062 | 0.738 | -0.019 | 0.001 | 0.001 | 0 | 0.43 | 0.26 | 0.32 | 0.39 | 0.5 | 0.44 |
| 10 | 25161334 | rs2478094 | 0.207 | 0.513 | 0.139 | 0.975 | -0.008 | -0.052 | -0.052 | 0 | 0.85 | 0.93 | 0.75 | 0.71 | 0.84 | 0.88 |
| 10 | 25164979 | rs1932422 | 0.256 | 0.614 | -0.336 | 0.995 | -0.001 | -0.016 | -0.016 | 0 | 0.85 | 0.89 | 0.75 | 0.72 | 0.84 | 0.85 |
| 10 | 25177744 | rs7916535 | -0.152 | 0.326 | -0.644 | 0.65 | -0.002 | 0.005 | 0.005 | 0 | 0.55 | 0.62 | 0.53 | 0.62 | 0.46 | 0.68 |
| 10 | 25177970 | rs881487 | -0.372 | 0.346 | -0.858 | 0.676 | -0.022 | 0.03 | 0.03 | 0 | 0.6 | 0.72 | 0.58 | 0.76 | 0.49 | 0.71 |
| 10 | 25178864 | rs11014285 | 0.513 | 0.486 | -1.181 | 0.503 | -0.055 | 0.026 | 0.026 | 0 | 0.1 | 0.01 | 0.08 | 0.03 | 0.16 | 0.14 |
| 10 | 25180919 | rs10828718 | -0.239 | 0.355 | -0.901 | 0.61 | -0.006 | 0.011 | 0.011 | 0 | 0.55 | 0.68 | 0.54 | 0.62 | 0.46 | 0.69 |
| 10 | 25182708 | rs2148270 | 0.07 | -0.404 | -0.629 | 0.546 | 0.017 | 0.018 | 0.018 | 0.391 | 0.41 | 0.35 | 0.32 | 0.31 | 0.26 | 0.26 |
| 10 | 25188285 | rs41408444 | 0.184 | -0.756 | -0.809 | 0.475 | 0 | 0.005 | 0.005 | 0 | 0.11 | 0.12 | 0.21 | 0.3 | 0.1 | 0.15 |
| 10 | 25197073 | rs6482454 | 0.845 | -1.267 | -0.116 | 0.12 | -0.06 | -0.116 | -0.116 | 0 | 0.23 | 0.31 | 0.09 | 0.01 | 0.1 | 0.09 |
| 10 | 25197134 | rs2148271 | -1.08 | -0.775 | -0.813 | 0.288 | -0.032 | -0.046 | -0.046 | 4.699 | 0.69 | 0.82 | 0.59 | 0.55 | 0.62 | 0.47 |
| 10 | 25198782 | rs16925145 | 1.146 | -0.505 | -0.635 | -0.082 | -0.075 | -0.156 | -0.156 | 0.211 | 0.25 | 0.4 | 0.11 | 0.01 | 0.12 | 0.09 |
| 10 | 25206554 | rs7908585 | 1.091 | -0.598 | -0.481 | -0.163 | -0.05 | 0.053 | 0.053 | 0 | 0.2 | 0.1 | 0.17 | 0.04 | 0.35 | 0.19 |
| 10 | 25216806 | rs1033962 | 0.161 | -0.656 | -0.536 | -0.1 | -0.001 | 0 | 0 | 0.003 | 0.4 | 0.58 | 0.32 | 0.29 | 0.24 | 0.24 |
| 10 | 25219544 | rs4748991 | 0.126 | -0.644 | -0.958 | 0.066 | 0.038 | 0.001 | 0.001 | 0.052 | 0.41 | 0.66 | 0.33 | 0.39 | 0.24 | 0.24 |
| 10 | 25219669 | rs4748992 | 0.419 | -0.659 | -0.955 | 0.059 | -0.001 | 0.004 | 0.004 | 3.433 | 0.15 | 0.18 | 0.22 | 0.28 | 0.12 | 0.15 |
| 10 | 25219838 | rs1409307 | -0.604 | -0.784 | -1.047 | 0.18 | -0.067 | -0.079 | -0.079 | 0 | 0.63 | 0.89 | 0.51 | 0.43 | 0.59 | 0.43 |
| 10 | 25224462 | rs11014306 | 0.451 | -0.933 | -0.279 | 0.003 | -0.078 | -0.058 | -0.058 | 0 | 0.25 | 0.48 | 0.11 | 0.11 | 0.12 | 0.09 |
| 10 | 25227988 | rs10828727 | 0.323 | 0.145 | -0.725 | 0.14 | -0.004 | -0.034 | -0.034 | 0.021 | 0.25 | 0.15 | 0.3 | 0.4 | 0.27 | 0.38 |
| 10 | 25231171 | rs3748223 | 0.957 | 0.996 | -0.558 | 0.244 | 0.001 | -0.004 | -0.004 | 0 | 0.51 | 0.16 | 0.34 | 0.51 | 0.46 | 0.53 |
| 10 | 25231515 | rs10828731 | 0.919 | 1.01 | -0.534 | 0.221 | 0.003 | -0.022 | -0.022 | 0 | 0.51 | 0.16 | 0.45 | 0.53 | 0.46 | 0.53 |
| 10 | 25235781 | rs10764514 | -0.153 | 1.288 | 0.297 | -0.033 | -0.018 | -0.081 | -0.081 | 0.3 | 0.21 | 0.32 | 0.13 | 0.03 | 0.17 | 0.2 |
| 10 | 25236223 | rs274296 | -0.499 | 1.825 | 0.654 | -0.009 | -0.003 | 0.011 | 0.011 | 0 | 0.17 | 0.18 | 0.29 | 0.44 | 0.2 | 0.23 |

| | | | | | | | | | | | | | | | | |
|---|---|---|---|---|---|---|---|---|---|---|---|---|---|---|---|---|
| 10 | 25237193 | rs10741071 | 1.498 | 1.799 | 0.573 | 0.027 | 0.05 | 0.089 | 0.089 | 0 | 0.72 | 0.49 | 0.49 | 0.55 | 0.63 | 0.74 |
| 10 | 25242922 | rs3808907 | -1.932 | 1.093 | 0.433 | -0.052 | -0.011 | -0.004 | -0.004 | 0 | 0.16 | 0.81 | 0.21 | 0.03 | 0.02 | 0.03 |
| 10 | 25244919 | rs274311 | -2.177 | 1.339 | 0.317 | -0.015 | 0.004 | -0.037 | -0.037 | 1.061 | 0.45 | 0.85 | 0.59 | 0.48 | 0.38 | 0.46 |
| 10 | 25247006 | rs274304 | -1.799 | 1.345 | 0.92 | -0.016 | 0.001 | 0.012 | 0.012 | 0.115 | 0.39 | 0.55 | 0.43 | 0.45 | 0.37 | 0.44 |
| 10 | 25250053 | rs6482465 | -1.932 | 1.517 | 1.079 | -0.158 | -0.003 | -0.094 | -0.094 | 0 | 0.13 | 0.52 | 0.1 | 0.01 | 0.07 | 0.04 |
| 10 | 25254151 | rs274288 | -2.26 | 1.757 | 1.011 | -0.113 | -0.028 | -0.029 | -0.029 | 0.964 | 0.43 | 0.83 | 0.58 | 0.48 | 0.4 | 0.44 |
| 10 | 25271270 | rs2307047 | -1.264 | 2.191 | 1.349 | 0.064 | 0.005 | 0.012 | 0.012 | 0.532 | 0.3 | 0.29 | 0.48 | 0.47 | 0.31 | 0.4 |
| 10 | 25273988 | rs2279547 | -2.169 | 2.171 | 1.3 | 0.053 | -0.014 | -0.014 | -0.014 | 7.983 | 0.39 | 0.7 | 0.57 | 0.49 | 0.4 | 0.44 |
| 10 | 25291250 | rs11014341 | 0.913 | 1.607 | 0.82 | 0.026 | -0.028 | 0.038 | 0.038 | 0 | 0.1 | 0.01 | 0.05 | 0.03 | 0.12 | 0.09 |
| 10 | 25305371 | rs2291583 | -1.32 | 1.666 | 0.929 | 0.07 | -0.014 | -0.063 | -0.063 | 0 | 0.28 | 0.16 | 0.46 | 0.48 | 0.29 | 0.4 |
| 10 | 25314538 | rs12249889 | -0.8 | 1.673 | 0.886 | 0.074 | -0.044 | -0.107 | -0.107 | 0 | 0.24 | 0.08 | 0.43 | 0.49 | 0.27 | 0.39 |
| 10 | 25321244 | rs10764519 | -2.014 | 1.681 | 1.015 | 0.05 | 0 | 0.003 | 0.003 | 0.218 | 0.36 | 0.38 | 0.48 | 0.49 | 0.33 | 0.42 |
| 10 | 25325937 | rs10508687 | -1.632 | 1.715 | 0.998 | 0.039 | 0.049 | -0.058 | -0.058 | 0 | 0.11 | 0.45 | 0.1 | 0 | 0.08 | 0.04 |
| 10 | 25336332 | rs4749000 | -1.399 | 1.781 | 0.652 | 0.186 | 0.004 | 0.033 | 0.033 | 0.447 | 0.3 | 0.38 | 0.49 | 0.46 | 0.29 | 0.39 |
| 10 | 25342819 | rs12243641 | -0.099 | 1.842 | -0.305 | 0.232 | -0.029 | 0.005 | 0.005 | 0.052 | 0.08 | 0.01 | 0.31 | 0.42 | 0.1 | 0.17 |
| 10 | 25357584 | rs923757 | -0.351 | 1.728 | -0.209 | 0.278 | -0.054 | -0.169 | -0.169 | 0.034 | 0.24 | 0.03 | 0.43 | 0.46 | 0.24 | 0.38 |
| 10 | 25359690 | rs10828753 | 0.045 | 1.873 | 0.329 | 0.668 | -0.012 | -0.015 | -0.015 | 0 | 0.88 | 0.91 | 0.79 | 0.57 | 0.8 | 0.7 |
| 10 | 25370348 | rs7901590 | -0.977 | 1.351 | 0.909 | 0.42 | 0.088 | -0.018 | -0.018 | 1.377 | 0.09 | 0.44 | 0.08 | 0 | 0.07 | 0.05 |
| 10 | 25375783 | rs10508683 | -1.503 | 1.31 | 0.426 | 0.4 | -0.008 | -0.068 | -0.068 | 0 | 0.12 | 0.66 | 0.1 | 0 | 0.07 | 0.05 |
| 10 | 25385858 | rs1388925 | 0.312 | 1.89 | 0.033 | 0.722 | -0.03 | -0.038 | -0.038 | 0 | 0.86 | 0.91 | 0.75 | 0.57 | 0.73 | 0.7 |
| 10 | 25394337 | rs943043 | -0.405 | 0.823 | 0.5 | 0.646 | -0.054 | -0.136 | -0.136 | 0 | 0.26 | 0.07 | 0.49 | 0.41 | 0.31 | 0.34 |
| 10 | 25399644 | rs12258541 | -1.175 | 0.738 | 0.384 | 0.507 | 0.011 | -0.063 | -0.063 | 2.242 | 0.1 | 0.62 | 0.11 | 0.02 | 0.08 | 0.04 |
| 10 | 25406857 | rs7072278 | -0.75 | 0.585 | 0.372 | 0.545 | -0.002 | -0.022 | -0.022 | 0.008 | 0.08 | 0.53 | 0.06 | 0.01 | 0.02 | 0 |
| 10 | 25408128 | rs11014404 | -0.568 | 0.892 | -0.338 | 0.721 | -0.042 | -0.128 | -0.128 | 3.191 | 0.26 | 0.09 | 0.48 | 0.43 | 0.31 | 0.35 |
| 10 | 25417521 | rs10764522 | -1.073 | 0.956 | -0.054 | 0.762 | -0.01 | -0.029 | -0.029 | 0 | 0.34 | 0.7 | 0.58 | 0.45 | 0.4 | 0.37 |
| 10 | 25417900 | rs4747512 | -0.857 | 0.935 | 0.604 | 0.759 | 0.012 | 0.031 | 0.031 | 0 | 0.32 | 0.4 | 0.56 | 0.44 | 0.4 | 0.37 |
| 10 | 25418407 | rs1926050 | -1.073 | 0.716 | 0.129 | 0.898 | 0.002 | 0.028 | 0.028 | 0 | 0.33 | 0.55 | 0.44 | 0.45 | 0.38 | 0.31 |
| 10 | 25419620 | rs16925405 | -0.206 | -0.027 | -0.951 | 0.986 | -0.042 | -0.172 | -0.172 | 0 | 0.25 | 0.01 | 0.26 | 0.45 | 0.25 | 0.18 |
| 10 | 25428658 | rs10828759 | -1.005 | -0.105 | -0.16 | 0.55 | -0.004 | 0 | 0 | 0 | 0.4 | 0.46 | 0.46 | 0.52 | 0.36 | 0.32 |
| 10 | 25433725 | rs11014425 | -0.88 | -0.142 | -0.029 | 0.534 | -0.004 | 0 | 0 | 0.07 | 0.4 | 0.46 | 0.46 | 0.53 | 0.36 | 0.32 |
| 10 | 25437862 | rs2887041 | -0.043 | -0.194 | -0.07 | 0.58 | -0.053 | -0.148 | -0.148 | 0 | 0.27 | 0.04 | 0.28 | 0.43 | 0.29 | 0.19 |
| 10 | 25438640 | rs7912264 | -1.37 | -0.574 | -0.218 | 0.274 | -0.047 | -0.055 | -0.055 | 0 | 0.17 | 0.45 | 0.19 | 0.1 | 0.09 | 0.17 |
| 10 | 25457920 | rs12783891 | -0.03 | -0.404 | -0.508 | 0.671 | -0.066 | -0.119 | -0.119 | 0.162 | 0.27 | 0.04 | 0.19 | 0.37 | 0.3 | 0.2 |
| 10 | 25459750 | rs11014431 | -0.302 | -0.751 | -0.374 | 0.792 | -0.014 | 0.053 | 0.053 | 0 | 0.09 | 0.14 | 0.15 | 0.28 | 0.07 | 0.14 |
| 10 | 25465327 | rs11014433 | 0.73 | -1.132 | -1.261 | -0.009 | -0.041 | 0.085 | 0.085 | 0.098 | 0.22 | 0.03 | 0.15 | 0.07 | 0.22 | 0.22 |
| 10 | 25466693 | rs1001948 | -0.496 | -1.655 | -0.555 | -0.005 | 0.035 | 0.045 | 0.045 | 0 | 0.53 | 0.71 | 0.44 | 0.69 | 0.4 | 0.42 |
| 10 | 25470573 | rs11014435 | -1.851 | -2.201 | 0.084 | -0.183 | -0.077 | 0.047 | 0.047 | 2.614 | 0.18 | 0.46 | 0.22 | 0.27 | 0.1 | 0.22 |
| 10 | 25474028 | rs7903171 | -0.883 | -1.968 | 0.33 | -0.162 | 0.035 | 0.032 | 0.032 | 3.932 | 0.05 | 0.29 | 0.05 | 0 | 0.01 | 0.04 |
| 10 | 25515067 | rs2807231 | -1.011 | -2.118 | -0.028 | 0.01 | -0.036 | 0.035 | 0.035 | 1.31 | 0.15 | 0.24 | 0.19 | 0.31 | 0.09 | 0.22 |
| 10 | 25522280 | rs10508689 | -0.347 | -2.072 | -0.081 | 0 | -0.02 | 0.026 | 0.026 | 0 | 0.12 | 0.17 | 0.17 | 0.31 | 0.09 | 0.18 |
| 10 | 25523366 | rs16925478 | -0.784 | -2.081 | -0.074 | -0.066 | -0.036 | 0.054 | 0.054 | 1.476 | 0.14 | 0.23 | 0.19 | 0.34 | 0.09 | 0.22 |
| 10 | 25531636 | rs915028 | -0.571 | -2.121 | -0.14 | -0.114 | -0.007 | 0.007 | 0.007 | 0 | 0.14 | 0.13 | 0.16 | 0.32 | 0.08 | 0.18 |
| 10 | 25532099 | rs1339999 | -0.018 | -2.11 | -0.317 | -0.104 | 0.036 | -0.051 | -0.051 | 0.196 | 0.12 | 0.03 | 0.14 | 0.27 | 0.08 | 0.17 |
| 10 | 25533536 | rs1416995 | -0.319 | -2.029 | 0.149 | -0.27 | -0.085 | 0.102 | 0.102 | 1.424 | 0.17 | 0.47 | 0.2 | 0.32 | 0.08 | 0.22 |
| 10 | 25546418 | rs10828769 | 0.617 | -1.598 | -0.429 | -0.307 | -0.076 | 0.001 | 0.001 | 0.039 | 0.2 | 0.03 | 0.14 | 0.14 | 0.25 | 0.19 |
| 10 | 25555314 | rs2807256 | 0.477 | -1.386 | -0.473 | -0.021 | -0.045 | 0.13 | 0.13 | 0 | 0.37 | 0.14 | 0.24 | 0.15 | 0.41 | 0.3 |
| 10 | 25559939 | rs2491198 | -0.041 | 0.424 | -0.49 | 0.498 | -0.097 | 0.051 | 0.051 | 0 | 0.74 | 0.87 | 0.53 | 0.84 | 0.57 | 0.59 |
| 10 | 25571944 | rs12263037 | -1.169 | -0.517 | 0.334 | 0.992 | -0.05 | 0.19 | 0.19 | 0.276 | 0.13 | 0.39 | 0.18 | 0.32 | 0.07 | 0.22 |
| 10 | 25582637 | rs2807253 | 0.076 | -0.175 | -0.412 | -0.028 | 0.028 | 0.053 | 0.053 | 0 | 0.49 | 0.39 | 0.41 | 0.23 | 0.62 | 0.47 |
| 10 | 25590161 | rs532538 | 0.24 | -0.906 | -0.013 | -0.513 | 0.013 | 0.014 | 0.014 | 0.047 | 0.62 | 0.37 | 0.46 | 0.23 | 0.71 | 0.53 |
| 10 | 25591909 | rs564371 | -0.283 | -1.462 | -0.055 | -0.362 | 0.018 | -0.05 | -0.05 | 0 | 0.79 | 0.9 | 0.58 | 0.37 | 0.81 | 0.74 |
| 10 | 25594793 | rs558947 | 1.351 | -0.196 | -0.636 | -0.958 | -0.096 | 0.235 | 0.235 | 0.05 | 0.48 | 0.04 | 0.33 | 0.22 | 0.5 | 0.45 |
| 10 | 25599875 | rs824147 | 1.309 | 0.12 | -0.646 | -0.962 | -0.005 | 0.239 | 0.239 | 0.146 | 0.49 | 0.04 | 0.34 | 0.22 | 0.55 | 0.46 |
| 10 | 25616383 | rs16925561 | -0.201 | -0.172 | -0.084 | -0.939 | -0.056 | 0.033 | 0.033 | 3.077 | 0.14 | 0.42 | 0.08 | 0.13 | 0.05 | 0.18 |
| 10 | 25616473 | rs2781256 | 0.894 | -0.217 | -0.278 | -0.234 | -0.021 | -0.034 | -0.034 | 0 | 0.06 | 0 | 0.09 | 0.15 | 0.1 | 0.17 |
| 10 | 25624290 | rs1340002 | 1.241 | -0.656 | -0.794 | -0.959 | 0.003 | -0.035 | -0.035 | 0 | 0.1 | 0.02 | 0.02 | 0.17 | 0.08 | 0.1 |
| 10 | 25626272 | rs539357 | -0.519 | -0.738 | -0.766 | -0.937 | 0.016 | -0.05 | -0.05 | 0 | 0.83 | 0.9 | 0.6 | 0.37 | 0.84 | 0.76 |
| 10 | 25630924 | rs543850 | 0.712 | -0.898 | -0.798 | -1.125 | -0.027 | -0.024 | -0.024 | 0 | 0.67 | 0.4 | 0.51 | 0.24 | 0.78 | 0.59 |
| 10 | 25638181 | rs2795044 | 0.766 | -0.678 | -0.346 | -1.021 | -0.007 | -0.007 | -0.007 | 1.226 | 0.73 | 0.65 | 0.52 | 0.25 | 0.75 | 0.58 |
| 10 | 25641990 | rs1413396 | -0.047 | -0.831 | 0.777 | -0.812 | -0.026 | 0.081 | 0.081 | 0 | 0.82 | 0.63 | 0.69 | 0.4 | 0.85 | 0.81 |
| 10 | 25653453 | rs10828778 | 1.67 | -1.006 | 1.02 | -1.041 | -0.025 | -0.044 | -0.044 | 0 | 0.61 | 0.45 | 0.46 | 0.22 | 0.69 | 0.52 |
| 10 | 25656838 | rs959751 | -1.206 | -0.833 | 1.08 | -0.872 | -0.011 | 0.01 | 0.01 | 0.161 | 0.13 | 0.23 | 0.09 | 0.13 | 0.1 | 0.24 |
| 10 | 25664256 | rs1773634 | 2.417 | -0.746 | 0.862 | -1.124 | -0.103 | 0.32 | 0.32 | 0.374 | 0.42 | 0.02 | 0.32 | 0.05 | 0.43 | 0.38 |
| 10 | 25665449 | rs1773631 | -0.231 | -0.545 | 0.973 | -1.008 | 0.012 | -0.02 | -0.02 | 0 | 0.82 | 0.83 | 0.6 | 0.35 | 0.83 | 0.79 |
| 10 | 25668687 | rs1773625 | 1.051 | -0.536 | 0.969 | -1.028 | -0.02 | -0.03 | -0.03 | 0 | 0.69 | 0.59 | 0.51 | 0.22 | 0.73 | 0.55 |
| 10 | 25670435 | rs12265226 | 0.132 | -0.529 | 1.011 | -1.056 | 0.053 | -0.059 | -0.059 | 0 | 0.12 | 0.21 | 0.14 | 0 | 0.21 | 0.07 |
| 10 | 25672136 | rs1760763 | 1.297 | -0.679 | 0.83 | -0.94 | -0.014 | -0.026 | -0.026 | 0 | 0.67 | 0.59 | 0.5 | 0.22 | 0.71 | 0.54 |
| 10 | 25674081 | rs1591250 | 0.953 | -0.581 | 0.451 | -0.89 | 0.012 | 0.014 | 0.014 | 0 | 0.72 | 0.77 | 0.53 | 0.24 | 0.73 | 0.55 |
| 10 | 25681590 | rs2009148 | -0.198 | -0.385 | 0.404 | -0.78 | -0.001 | 0.004 | 0.004 | 0 | 0.1 | 0.06 | 0.06 | 0.1 | 0.11 | 0.24 |

| | | | | | | | | | | | | | | | | |
|---|---|---|---|---|---|---|---|---|---|---|---|---|---|---|---|---|
| 10 | 25686900 | rs11818261 | 0.794 | -0.438 | 0.448 | -0.783 | -0.002 | 0.001 | 0.001 | 0 | 0.1 | 0.13 | 0.04 | 0.16 | 0.08 | 0.08 |
| 10 | 25693426 | rs10508693 | 0.111 | -0.265 | 0.832 | -0.93 | 0.067 | -0.066 | -0.066 | 0 | 0.12 | 0.23 | 0.13 | 0 | 0.2 | 0.08 |
| 10 | 25702556 | rs1773691 | 1.7 | -0.537 | 0.648 | -1.109 | 0.066 | 0.123 | 0.123 | 0.06 | 0.54 | 0.25 | 0.45 | 0.06 | 0.63 | 0.46 |
| 10 | 25705149 | rs1754277 | 0.782 | -0.631 | 0.73 | -1.024 | -0.025 | -0.004 | -0.004 | 0 | 0.73 | 0.59 | 0.54 | 0.32 | 0.77 | 0.74 |
| 10 | 25706232 | rs956088 | -0.595 | -0.678 | 0.79 | -1.022 | 0.011 | -0.035 | -0.035 | 0 | 0.84 | 0.88 | 0.61 | 0.34 | 0.85 | 0.81 |
| 10 | 25706976 | rs1773681 | -0.503 | -0.679 | 0.81 | -1.022 | 0.009 | -0.02 | -0.02 | 0 | 0.83 | 0.84 | 0.6 | 0.34 | 0.85 | 0.81 |
| 10 | 25708501 | rs4749023 | -0.145 | -0.506 | 0.785 | -1.023 | 0.017 | -0.084 | -0.084 | 0 | 0.2 | 0.43 | 0.17 | 0.1 | 0.2 | 0.26 |
| 10 | 25711579 | rs1773678 | 1.176 | -0.148 | 1.023 | -0.47 | -0.023 | -0.004 | -0.004 | 0 | 0.73 | 0.61 | 0.54 | 0.32 | 0.77 | 0.74 |
| 10 | 25712004 | rs7392687 | 1.188 | -0.149 | 1.022 | -0.47 | -0.023 | -0.004 | -0.004 | 0 | 0.73 | 0.61 | 0.54 | 0.32 | 0.77 | 0.74 |
| 10 | 25714745 | rs10764548 | 1.177 | -0.142 | 1.025 | -0.467 | -0.023 | -0.004 | -0.004 | 0 | 0.73 | 0.61 | 0.54 | 0.32 | 0.77 | 0.74 |
| 10 | 25725491 | rs16925850 | 0.594 | -0.136 | 1.037 | -0.423 | -0.033 | -0.011 | -0.011 | 0 | 0.1 | 0.04 | 0.03 | 0.13 | 0.13 | 0.08 |
| 10 | 25726938 | rs1773658 | 1.197 | -0.196 | 1.019 | -0.41 | -0.025 | -0.005 | -0.005 | 0 | 0.73 | 0.61 | 0.54 | 0.32 | 0.77 | 0.74 |
| 10 | 25727031 | rs1754248 | 1.223 | -0.168 | 1.017 | -0.412 | -0.022 | -0.005 | -0.005 | 0 | 0.73 | 0.61 | 0.54 | 0.32 | 0.76 | 0.74 |
| 10 | 25728493 | rs11014558 | -0.779 | 0.239 | 1.116 | -0.102 | 0.05 | 0.036 | 0.036 | 0 | 0.14 | 0.29 | 0.07 | 0.18 | 0.16 | 0.09 |
| 10 | 25729604 | rs1773654 | 0.41 | 0.065 | 0.685 | -0.153 | 0.005 | -0.027 | -0.027 | 0.639 | 0.86 | 0.88 | 0.61 | 0.3 | 0.86 | 0.8 |
| 10 | 25733319 | rs11014565 | 1.391 | 0.215 | 1.014 | 0.279 | 0.007 | 0.043 | 0.043 | 0 | 0.78 | 0.62 | 0.62 | 0.36 | 0.8 | 0.84 |
| 10 | 25734412 | rs12770990 | -1.155 | 0.396 | 1.027 | 0.046 | 0.038 | -0.047 | -0.047 | 0 | 0.15 | 0.33 | 0.08 | 0.08 | 0.16 | 0.09 |
| 10 | 25735631 | rs12356315 | 0.04 | 0.678 | 0.509 | 0.264 | -0.024 | -0.014 | -0.014 | 0 | 0.07 | 0.04 | 0.05 | 0.1 | 0.14 | 0.08 |
| 10 | 25737340 | rs4747520 | 0.506 | 0.405 | 0.067 | 0.069 | 0.007 | 0.015 | 0.015 | 0 | 0.51 | 0.21 | 0.34 | 0.41 | 0.47 | 0.55 |
| 10 | 25740053 | rs1329249 | -0.909 | 0.318 | 0.114 | -0.121 | -0.051 | 0.168 | 0.168 | 0 | 0.7 | 0.94 | 0.72 | 1 | 0.62 | 0.74 |
| 10 | 25747511 | rs12415532 | -2.342 | 1.183 | 2.112 | 0.692 | 0.077 | 0.047 | 0.047 | 0.795 | 0.12 | 0.52 | 0.25 | 0.17 | 0.14 | 0.18 |
| 10 | 25747588 | rs1418127 | -0.082 | 1.065 | 2.017 | 1.011 | -0.006 | 0.098 | 0.098 | 1.185 | 0.06 | 0.07 | 0.07 | 0.57 | 0.01 | 0.05 |
| 10 | 25756895 | rs1410931 | 0.984 | 2.042 | 1.976 | 0.239 | -0.035 | 0.134 | 0.134 | 0.443 | 0.58 | 0.24 | 0.46 | 0.82 | 0.5 | 0.53 |
| 10 | 25757095 | rs12763436 | -1.16 | 2.049 | 2.091 | 0.239 | 0.059 | 0.042 | 0.042 | 0 | 0.09 | 0.22 | 0.2 | 0.17 | 0.13 | 0.17 |
| 10 | 25764141 | rs1928364 | -0.512 | 2.017 | 1.854 | 0.219 | -0.05 | 0.105 | 0.105 | 0.013 | 0.74 | 0.92 | 0.73 | 0.99 | 0.66 | 0.75 |
| 10 | 25764413 | rs10508695 | -2.556 | 2.029 | 1.982 | 0.229 | 0.042 | 0.02 | 0.02 | 0.176 | 0.14 | 0.53 | 0.25 | 0.17 | 0.16 | 0.19 |
| 10 | 25765688 | rs1773636 | 1.035 | 2.022 | 1.816 | 0.4 | -0.035 | 0.104 | 0.104 | 0.382 | 0.58 | 0.24 | 0.42 | 0.75 | 0.5 | 0.52 |
| 10 | 25773608 | rs2987833 | 0.942 | 2.027 | 1.858 | 0.041 | -0.027 | 0.107 | 0.107 | 0.416 | 0.58 | 0.26 | 0.46 | 0.82 | 0.5 | 0.54 |
| 10 | 25779135 | rs3005188 | -0.195 | 2.208 | 1.905 | 0.108 | -0.014 | 0.025 | 0.025 | 0 | 0.73 | 0.85 | 0.72 | 0.99 | 0.66 | 0.75 |
| 10 | 25781407 | rs1329255 | 0.713 | 1.957 | 1.623 | 0.258 | -0.027 | 0.114 | 0.114 | 0.342 | 0.59 | 0.24 | 0.49 | 0.79 | 0.53 | 0.57 |
| 10 | 25784552 | rs3005182 | 0.854 | 1.969 | 1.668 | 0.263 | -0.011 | 0.03 | 0.03 | 0.043 | 0.6 | 0.37 | 0.5 | 0.79 | 0.53 | 0.61 |
| 10 | 25786532 | rs4097490 | -1.648 | 2.101 | 1.651 | 0.526 | -0.018 | 0.066 | 0.066 | 2.265 | 0.49 | 0.86 | 0.57 | 0.71 | 0.51 | 0.49 |
| 10 | 25787306 | rs4749039 | -1.514 | 1.68 | 2.127 | 0.576 | 0.013 | 0.051 | 0.051 | 3.052 | 0.48 | 0.84 | 0.44 | 0.71 | 0.45 | 0.48 |
| 10 | 25789096 | rs10508697 | -2.184 | 1.662 | 2.275 | 0.575 | 0.038 | 0.145 | 0.145 | 2.707 | 0.06 | 0.38 | 0.04 | 0.14 | 0 | 0.12 |
| 10 | 25791017 | rs4749042 | -0.714 | 1.707 | 2.125 | 0.492 | -0.037 | -0.097 | -0.097 | 0 | 0.35 | 0.14 | 0.29 | 0.5 | 0.39 | 0.31 |
| 10 | 25791843 | rs1329252 | -1.929 | 1.987 | 1.993 | 0.419 | -0.055 | -0.063 | -0.063 | 0.443 | 0.51 | 0.95 | 0.49 | 0.57 | 0.53 | 0.45 |
| 10 | 25811435 | rs7086249 | -1.391 | 1.808 | 1.972 | 0.508 | 0.009 | -0.008 | -0.008 | 0.044 | 0.44 | 0.69 | 0.44 | 0.57 | 0.44 | 0.41 |
| 10 | 25812375 | rs11591807 | -0.825 | 1.923 | 1.193 | 0.539 | -0.007 | 0.003 | 0.003 | 0 | 0.06 | 0.05 | 0.11 | 0.07 | 0.14 | 0.06 |
| 10 | 25818777 | rs4459183 | -0.695 | 1.548 | 0.623 | 0.591 | -0.036 | -0.116 | -0.116 | 0 | 0.34 | 0.1 | 0.29 | 0.5 | 0.36 | 0.31 |
| 10 | 25823753 | rs7100942 | -1.102 | 1.369 | 0.767 | 0.586 | 0.006 | -0.005 | -0.005 | 0 | 0.42 | 0.37 | 0.4 | 0.57 | 0.42 | 0.37 |
| 10 | 25828840 | rs2152214 | -0.657 | 2.105 | 0.399 | 0.45 | -0.134 | 0.117 | 0.117 | 0 | 0.27 | 0.04 | 0.32 | 0.09 | 0.39 | 0.3 |
| 10 | 25831178 | rs11014598 | -0.947 | 1.104 | 0.875 | 0.846 | -0.026 | 0.207 | 0.207 | 0.07 | 0.07 | 0.14 | 0.03 | 0.47 | 0.03 | 0.05 |
| 10 | 25832727 | rs9919420 | -1.859 | 0.994 | 0.87 | 0.74 | 0.036 | 0.01 | 0.01 | 1.148 | 0.43 | 0.83 | 0.43 | 0.7 | 0.4 | 0.62 |
| 10 | 25839112 | rs11014603 | -0.634 | 0.75 | 0.913 | 0.858 | -0.023 | 0.201 | 0.201 | 0 | 0.06 | 0.1 | 0.03 | 0.48 | 0.03 | 0.05 |
| 10 | 25843033 | rs10828822 | -1.254 | 0.753 | 1.086 | 0.785 | -0.029 | 0.288 | 0.288 | 0.482 | 0.07 | 0.2 | 0.04 | 0.47 | 0.03 | 0.05 |
| 10 | 25847478 | rs7903336 | -1.781 | 0.947 | 1.493 | 0.739 | 0.023 | 0.014 | 0.014 | 0.878 | 0.42 | 0.81 | 0.42 | 0.7 | 0.4 | 0.61 |
| 10 | 25849053 | rs7087655 | -0.827 | 1.384 | 1.762 | 0.498 | 0.02 | 0.096 | 0.096 | 0 | 0.07 | 0.33 | 0.06 | 0.13 | 0.03 | 0.21 |
| 10 | 25856412 | rs11014611 | -0.597 | 1.377 | 1.384 | 0.481 | -0.044 | 0.091 | 0.091 | 0 | 0.26 | 0.13 | 0.31 | 0.08 | 0.34 | 0.28 |
| 10 | 25862212 | rs11815379 | -1.204 | 1.91 | 1.714 | 0.644 | 0.082 | 0.093 | 0.093 | 0.255 | 0.08 | 0.42 | 0.07 | 0.13 | 0.06 | 0.22 |
| 10 | 25864482 | rs4749049 | -1.753 | 1.894 | 1.292 | 0.575 | -0.054 | 0.065 | 0.065 | 3.019 | 0.36 | 0.83 | 0.45 | 0.24 | 0.45 | 0.59 |
| 10 | 25865513 | rs4747531 | -0.761 | 1.831 | 1.171 | 0.465 | 0.008 | -0.012 | -0.012 | 0 | 0.22 | 0.2 | 0.31 | 0.09 | 0.34 | 0.28 |
| 10 | 25866387 | rs6482497 | -0.985 | 1.731 | 1.079 | 0.329 | -0.016 | 0.002 | 0.002 | 0 | 0.23 | 0.16 | 0.27 | 0.01 | 0.32 | 0.28 |
| 10 | 25868585 | rs12357174 | -0.992 | 1.733 | 1.08 | 0.33 | -0.016 | 0.002 | 0.002 | 0 | 0.23 | 0.16 | 0.27 | 0.01 | 0.32 | 0.28 |
| 10 | 25894777 | rs10764567 | -0.84 | 1.638 | 0.902 | 0.994 | -0.014 | -0.03 | -0.03 | 0 | 0.16 | 0.13 | 0.25 | 0.5 | 0.24 | 0.21 |
| 10 | 25917909 | rs11014640 | -0.418 | 1.284 | 0.345 | 0.136 | -0.014 | 0.033 | 0.033 | 0 | 0.15 | 0.05 | 0.16 | 0 | 0.18 | 0.14 |
| 10 | 25926559 | rs10741097 | -1.83 | 1.171 | -0.001 | 0.204 | -0.061 | 0.02 | 0.02 | 3.388 | 0.46 | 0.92 | 0.5 | 0.64 | 0.46 | 0.45 |
| 10 | 25932400 | rs1004845 | -1.194 | 1.23 | 0.463 | 0.132 | -0.008 | -0.019 | -0.019 | 0 | 0.3 | 0.21 | 0.32 | 0.64 | 0.29 | 0.35 |
| 10 | 25936459 | rs10508704 | -0.471 | 1.233 | 0.922 | 0.128 | 0.12 | 0.019 | 0.019 | 0 | 0.06 | 0.27 | 0.08 | 0 | 0.1 | 0.04 |
| 10 | 25958020 | rs11014665 | -0.786 | 1.185 | 0.31 | 0.362 | 0.075 | 0.059 | 0.059 | 0.319 | 0.12 | 0.42 | 0.2 | 0.11 | 0.19 | 0.06 |
| 10 | 25958966 | rs11014666 | -0.573 | 1.165 | 0.339 | 0.32 | 0.012 | 0.009 | 0.009 | 1.803 | 0.23 | 0.16 | 0.17 | 0.01 | 0.18 | 0.14 |
| 10 | 25958982 | rs6482510 | -1.609 | 1.19 | -0.033 | 0.316 | -0.003 | -0.027 | -0.027 | 0.522 | 0.35 | 0.58 | 0.37 | 0.12 | 0.37 | 0.2 |
| 10 | 25961933 | rs7897614 | -2.022 | 0.435 | -0.338 | -0.621 | 0.011 | 0.181 | 0.181 | 3.158 | 0.43 | 0.91 | 0.42 | 0.14 | 0.39 | 0.25 |
| 10 | 25968240 | rs7080019 | -1.412 | 0.436 | -0.338 | -0.602 | 0.002 | 0.009 | 0.009 | 4.145 | 0.35 | 0.62 | 0.37 | 0.12 | 0.38 | 0.2 |
| 10 | 25970198 | rs16926251 | -0.271 | 0.541 | -0.56 | -0.574 | -0.018 | -0.031 | -0.031 | 0 | 0.06 | 0.18 | 0.04 | 0 | 0.03 | 0.02 |
| 10 | 25972010 | rs2290167 | -1.641 | 0.647 | -1.302 | -0.51 | 0.026 | 0.144 | 0.144 | 1.875 | 0.42 | 0.82 | 0.41 | 0.15 | 0.41 | 0.25 |
| 10 | 25979743 | rs11014680 | -0.934 | 0.659 | -1.226 | -0.446 | 0.015 | 0.065 | 0.065 | 1.312 | 0.22 | 0.07 | 0.16 | 0 | 0.17 | 0.14 |
| 10 | 25988086 | rs2505500 | 0.222 | 0.388 | -1.564 | -0.53 | 0.009 | 0.093 | 0.093 | 3.653 | 0.16 | 0.01 | 0.08 | 0 | 0.12 | 0.18 |
| 10 | 25988527 | rs2505502 | -0.002 | 0.921 | -2.138 | -0.551 | -0.032 | -0.008 | -0.008 | 0 | 0.81 | 0.95 | 0.86 | 0.82 | 0.77 | 0.85 |
| 10 | 25988830 | rs7091819 | -0.09 | 1.335 | -1.673 | -0.166 | 0.008 | 0.027 | 0.027 | 0.004 | 0.19 | 0.25 | 0.33 | 0.31 | 0.24 | 0.32 |

| | | | | | | | | | | | | | | | | |
|---|---|---|---|---|---|---|---|---|---|---|---|---|---|---|---|---|
| 10 | 25991772 | rs2442826 | -0.013 | 1.334 | -1.819 | -1.045 | -0.013 | 0.093 | 0.093 | 0.129 | 0.38 | 0.21 | 0.23 | 0.04 | 0.39 | 0.32 |
| 10 | 25996101 | rs10764581 | -0.179 | 1.544 | -0.999 | -0.459 | 0.017 | 0.002 | 0.002 | 0 | 0.17 | 0.28 | 0.17 | 0.19 | 0.22 | 0.14 |
| 10 | 25997959 | rs10828853 | -0.999 | 1.666 | -0.973 | -0.517 | 0.033 | -0.019 | -0.019 | 1.056 | 0.24 | 0.37 | 0.24 | 0.19 | 0.34 | 0.23 |
| 10 | 25998924 | rs10828856 | -0.785 | 1.393 | -1.469 | -0.148 | -0.03 | -0.046 | -0.046 | 0 | 0.26 | 0.13 | 0.2 | 0.38 | 0.31 | 0.51 |
| 10 | 26002103 | rs11014704 | -0.252 | 1.431 | -1.436 | -0.525 | -0.021 | 0.021 | 0.021 | 0 | 0.12 | 0.07 | 0.06 | 0.03 | 0.15 | 0.03 |
| 10 | 26003851 | rs2367972 | 1.389 | 0.725 | -0.952 | -0.545 | -0.001 | -0.001 | -0.001 | 4.549 | 0.67 | 0.36 | 0.69 | 0.67 | 0.7 | 0.53 |
| 10 | 26014312 | rs4749074 | 0.718 | 0.65 | -0.64 | -0.451 | -0.07 | 0.095 | 0.095 | 0.28 | 0.82 | 0.49 | 0.76 | 0.71 | 0.89 | 0.63 |
| 10 | 26015203 | rs2442871 | -0.009 | 0.657 | -0.552 | -0.43 | -0.018 | 0.223 | 0.223 | 0 | 0.89 | 0.58 | 0.84 | 0.73 | 0.95 | 0.73 |
| 10 | 26019204 | rs872974 | -0.162 | 0.732 | -1.61 | -1.1 | -0.041 | 0.065 | 0.065 | 0.674 | 0.43 | 0.16 | 0.54 | 0.7 | 0.47 | 0.3 |
| 10 | 26019753 | rs2250803 | 0.516 | 0.584 | -1.7 | -0.982 | -0.005 | 0.004 | 0.004 | 0 | 0.71 | 0.39 | 0.7 | 0.7 | 0.74 | 0.61 |
| 10 | 26020424 | rs10828869 | 0.164 | 0.934 | -2.143 | -0.042 | -0.014 | -0.034 | -0.034 | 0 | 0.31 | 0.07 | 0.4 | 0.67 | 0.33 | 0.27 |
| 10 | 26023828 | rs2505453 | 0.041 | 0.994 | -1.78 | 0.306 | 0 | -0.038 | -0.038 | 0 | 0.8 | 0.91 | 0.75 | 0.68 | 0.78 | 0.65 |
| 10 | 26026515 | rs2442867 | 0.596 | 0.869 | -2.139 | -0.002 | -0.022 | 0.049 | 0.049 | 0 | 0.45 | 0.14 | 0.42 | 0.62 | 0.43 | 0.47 |
| 10 | 26026785 | rs2263000 | 0.344 | 1.159 | -0.992 | 0.488 | 0.006 | 0.011 | 0.011 | 0.121 | 0.77 | 0.72 | 0.7 | 0.68 | 0.72 | 0.64 |
| 10 | 26028530 | rs2442865 | 0.557 | 1.111 | -1.194 | 1.366 | -0.003 | 0.067 | 0.067 | 0 | 0.92 | 0.93 | 0.72 | 0.49 | 0.88 | 0.72 |
| 10 | 26028813 | rs4256890 | -0.237 | 1.015 | -1.653 | 0.165 | -0.031 | 0.009 | 0.009 | 0 | 0.2 | 0.05 | 0.09 | 0.14 | 0.19 | 0.11 |
| 10 | 26039061 | rs2442855 | -0.05 | 0.807 | -0.929 | -0.002 | 0 | -0.001 | -0.001 | 0.013 | 0.61 | 0.65 | 0.48 | 0.49 | 0.59 | 0.57 |
| 10 | 26047663 | rs2505470 | 0.451 | 0.805 | -0.598 | -0.579 | -0.008 | 0.008 | 0.008 | 0 | 0.09 | 0.04 | 0.06 | 0 | 0.1 | 0.1 |
| 10 | 26053548 | rs2442846 | 0.053 | 0.674 | 0.182 | -0.649 | 0.018 | -0.064 | -0.064 | 0.019 | 0.11 | 0.24 | 0.07 | 0 | 0.11 | 0.11 |
| 10 | 26054210 | rs2442845 | 0.078 | 0.677 | 0.184 | -0.648 | 0.018 | -0.064 | -0.064 | 0 | 0.11 | 0.24 | 0.07 | 0 | 0.11 | 0.11 |
| 10 | 26055070 | rs2505478 | -0.742 | 0.285 | 0.482 | -1.375 | 0.022 | 0.092 | 0.092 | 0 | 0.43 | 0.73 | 0.3 | 0.04 | 0.39 | 0.23 |
| 10 | 26055631 | rs2437382 | 0.591 | 0.24 | 0.477 | -1.368 | 0.005 | -0.043 | -0.043 | 7.071 | 0.1 | 0.17 | 0.06 | 0 | 0.1 | 0.1 |
| 10 | 26056683 | rs2442843 | -0.733 | 0.276 | 0.469 | -1.372 | 0.024 | 0.105 | 0.105 | 0 | 0.43 | 0.75 | 0.3 | 0.04 | 0.39 | 0.23 |
| 10 | 26066590 | rs12261856 | 0.219 | -0.007 | 1.009 | -1.347 | -0.032 | -0.07 | -0.07 | 0 | 0.15 | 0.34 | 0.11 | 0.03 | 0.12 | 0.09 |
| 10 | 26072818 | rs17664916 | 1.036 | 0.068 | 0.552 | -1.327 | -0.002 | -0.043 | -0.043 | 0.091 | 0.17 | 0.18 | 0.12 | 0.01 | 0.17 | 0.03 |
| 10 | 26073673 | rs16926353 | 0.349 | 0.126 | 0.388 | -1.081 | 0.016 | 0.031 | 0.031 | 0.084 | 0.13 | 0.08 | 0.08 | 0.03 | 0.12 | 0.09 |
| 10 | 26093881 | rs17812509 | -0.039 | -0.004 | 0.488 | -0.84 | 0.008 | -0.005 | -0.005 | 0.127 | 0.16 | 0.12 | 0.08 | 0.14 | 0.11 | 0.1 |
| 10 | 26096852 | rs7071770 | 0.098 | 0.364 | 0.487 | -0.885 | -0.003 | 0 | 0 | 0 | 0.53 | 0.45 | 0.38 | 0.5 | 0.41 | 0.5 |
| 10 | 26101453 | rs2917191 | -1.385 | 0.989 | -0.196 | -0.687 | -0.079 | -0.104 | -0.104 | 0.248 | 0.87 | 0.99 | 0.62 | 0.51 | 0.76 | 0.75 |
| 10 | 26116844 | rs2437401 | -1.075 | 0.448 | 0.519 | -0.471 | -0.028 | -0.046 | -0.046 | 1.184 | 0.65 | 0.81 | 0.48 | 0.51 | 0.56 | 0.71 |
| 10 | 26119044 | rs2437399 | 0.347 | 0.541 | 0.139 | -0.457 | -0.013 | -0.052 | -0.052 | 0 | 0.14 | 0.08 | 0.12 | 0.31 | 0.13 | 0.24 |
| 10 | 26122732 | rs4498875 | 0.257 | 0.58 | 0.109 | -0.872 | 0 | -0.054 | -0.054 | 0.153 | 0.33 | 0.3 | 0.25 | 0.05 | 0.29 | 0.25 |
| 10 | 26148710 | rs11014790 | 0.173 | 0.708 | -0.957 | -0.787 | -0.002 | -0.025 | -0.025 | 0 | 0.35 | 0.32 | 0.29 | 0.06 | 0.33 | 0.32 |
| 10 | 26156382 | rs2367987 | 0.018 | 0.126 | -1.106 | -0.68 | 0.027 | -0.019 | -0.019 | 0 | 0.15 | 0.07 | 0.05 | 0.13 | 0.1 | 0.15 |
| 10 | 26156728 | rs4749079 | -0.373 | 0.861 | -0.811 | -0.903 | 0.003 | -0.01 | -0.01 | 0 | 0.34 | 0.29 | 0.27 | 0.05 | 0.29 | 0.25 |
| 10 | 26158125 | rs11014799 | -1.061 | 1.315 | -0.615 | -0.191 | 0 | -0.02 | -0.02 | 0 | 0.09 | 0.1 | 0.09 | 0 | 0.08 | 0.04 |
| 10 | 26158386 | rs10741102 | 0.267 | 1.421 | -0.556 | 0.213 | 0.004 | -0.008 | -0.008 | 0 | 0.59 | 0.45 | 0.53 | 0.69 | 0.53 | 0.63 |
| 10 | 26162766 | rs4749080 | -0.59 | 1.365 | -0.623 | 0.929 | -0.003 | -0.022 | -0.022 | 0.029 | 0.11 | 0.04 | 0.21 | 0.27 | 0.12 | 0.16 |
| 10 | 26164407 | rs7078573 | 0.405 | 1.09 | -0.987 | 0.665 | 0.026 | 0.011 | 0.011 | 0 | 0.41 | 0.24 | 0.3 | 0.35 | 0.38 | 0.4 |
| 10 | 26167135 | rs7100959 | -0.401 | 1.218 | 0.104 | 0.188 | 0.012 | -0.095 | -0.095 | 0 | 0.15 | 0.39 | 0.14 | 0.01 | 0.17 | 0.05 |
| 10 | 26171121 | rs11014824 | 0.195 | 1.227 | 0.089 | 0.168 | -0.015 | 0.015 | 0.015 | 0 | 0.06 | 0.01 | 0.07 | 0 | 0.11 | 0.03 |
| 10 | 26173364 | rs12247783 | -0.13 | 1.143 | 0.263 | 0.191 | 0.003 | -0.026 | -0.026 | 0 | 0.09 | 0.37 | 0.07 | 0.01 | 0.06 | 0.02 |
| 10 | 26173695 | rs12413953 | 0.186 | 1.108 | 0.597 | 0.343 | 0.005 | 0.107 | 0.107 | 0 | 0.11 | 0.18 | 0.23 | 0.32 | 0.11 | 0.15 |
| 10 | 26174841 | rs7907646 | 0.863 | 1.284 | 0.527 | 0.222 | 0.046 | 0.042 | 0.042 | 0 | 0.42 | 0.17 | 0.28 | 0.31 | 0.39 | 0.43 |
| 10 | 26176051 | rs7100118 | -0.848 | 1.327 | 0.562 | 0.196 | -0.062 | -0.006 | -0.006 | 0 | 0.66 | 0.83 | 0.62 | 0.63 | 0.64 | 0.61 |
| 10 | 26177672 | rs16926449 | 1.017 | 1.286 | 0.948 | 0.18 | -0.002 | -0.011 | -0.011 | 1.559 | 0.06 | 0.06 | 0.03 | 0 | 0.06 | 0.02 |
| 10 | 26178787 | rs12248876 | -0.223 | 1.287 | 0.946 | 0.18 | 0.005 | -0.035 | -0.035 | 0.743 | 0.09 | 0.37 | 0.07 | 0.01 | 0.06 | 0.02 |
| 10 | 26179166 | rs7922301 | 0.215 | 1.298 | 0.894 | 0.185 | 0.012 | 0.015 | 0.015 | 0 | 0.51 | 0.55 | 0.34 | 0.31 | 0.45 | 0.45 |
| 10 | 26181956 | rs7083201 | -0.335 | 1.309 | 1.309 | 0.195 | -0.004 | 0.004 | 0.004 | 0 | 0.58 | 0.6 | 0.58 | 0.63 | 0.6 | 0.6 |
| 10 | 26186577 | rs10828899 | 0.8 | 1.276 | 1.073 | 0.196 | 0.028 | 0.025 | 0.025 | 0 | 0.42 | 0.32 | 0.27 | 0.3 | 0.38 | 0.43 |
| 10 | 26191114 | rs7920650 | -0.747 | 1.333 | 1.248 | 0.127 | -0.026 | -0.001 | -0.001 | 0 | 0.66 | 0.77 | 0.62 | 0.64 | 0.67 | 0.62 |
| 10 | 26196544 | rs4749083 | 0.166 | 1.213 | 1.375 | 0.084 | 0.008 | 0.043 | 0.043 | 0 | 0.54 | 0.26 | 0.48 | 0.63 | 0.5 | 0.57 |
| 10 | 26197355 | rs4328138 | 0.093 | 1.993 | 1.393 | 0.07 | -0.003 | 0.036 | 0.036 | 0 | 0.54 | 0.32 | 0.52 | 0.63 | 0.56 | 0.58 |
| 10 | 26203228 | rs11819262 | 0.328 | 1.983 | 1.523 | 0.056 | 0.036 | -0.036 | -0.036 | 0.258 | 0.08 | 0.17 | 0.08 | 0.01 | 0.13 | 0.02 |
| 10 | 26204979 | rs7904360 | -0.513 | 1.927 | 1.542 | 0.07 | -0.004 | -0.004 | -0.004 | 0 | 0.64 | 0.43 | 0.6 | 0.63 | 0.72 | 0.62 |
| 10 | 26210316 | rs7073941 | 0.092 | 1.923 | 1.544 | 0.083 | -0.002 | 0.03 | 0.03 | 0 | 0.54 | 0.32 | 0.52 | 0.63 | 0.57 | 0.58 |
| 10 | 26212094 | rs10828912 | 0.15 | 1.889 | 1.803 | 0.077 | 0 | 0.016 | 0.016 | 0 | 0.54 | 0.41 | 0.52 | 0.62 | 0.56 | 0.58 |
| 10 | 26214318 | rs6482520 | -0.265 | 1.854 | 1.821 | 0.058 | 0.008 | -0.012 | -0.012 | 0.36 | 0.62 | 0.49 | 0.56 | 0.63 | 0.63 | 0.6 |
| 10 | 26221227 | rs11014863 | 0.3 | 1.922 | 1.773 | 0.066 | 0.013 | 0.062 | 0.062 | 0 | 0.53 | 0.22 | 0.47 | 0.62 | 0.49 | 0.58 |
| 10 | 26224097 | rs11014866 | 0.197 | 1.953 | 1.719 | 0.093 | 0.04 | -0.04 | -0.04 | 0 | 0.08 | 0.19 | 0.08 | 0 | 0.13 | 0.03 |
| 10 | 26241456 | rs7923691 | -0.75 | 1.892 | 1.777 | 0.08 | -0.001 | -0.005 | -0.005 | 0 | 0.67 | 0.82 | 0.63 | 0.63 | 0.72 | 0.63 |
| 10 | 26252005 | rs9787562 | 0.405 | 2.004 | 1.864 | 0.246 | -0.002 | 0.023 | 0.023 | 0 | 0.1 | 0.08 | 0.22 | 0.33 | 0.13 | 0.15 |
| 10 | 26286159 | rs12257119 | 0.343 | 1.981 | 1.948 | 0.211 | -0.005 | -0.036 | -0.036 | 0 | 0.08 | 0.25 | 0.05 | 0 | 0.06 | 0.02 |
| 10 | 26286941 | rs993398 | -0.839 | 2.003 | 1.945 | 0.196 | -0.044 | -0.009 | -0.009 | 0.205 | 0.67 | 0.84 | 0.61 | 0.62 | 0.66 | 0.62 |
| 10 | 26293048 | rs11014897 | 0.335 | 1.991 | 1.812 | 0.203 | -0.004 | -0.035 | -0.035 | 0 | 0.08 | 0.25 | 0.05 | 0 | 0.06 | 0.02 |
| 10 | 26307291 | rs11014914 | -0.629 | 2.021 | 1.817 | 0.203 | 0.018 | -0.065 | -0.065 | 0 | 0.12 | 0.3 | 0.11 | 0.01 | 0.16 | 0.04 |
| 10 | 26310410 | rs3737275 | 0.231 | 1.997 | 1.671 | 0.181 | 0.012 | 0.054 | 0.054 | 0 | 0.53 | 0.25 | 0.48 | 0.62 | 0.5 | 0.58 |
| 10 | 26310679 | rs17666079 | 0.572 | 1.99 | 1.594 | 0.208 | 0.03 | 0.075 | 0.075 | 0 | 0.46 | 0.18 | 0.3 | 0.29 | 0.47 | 0.46 |
| 10 | 26311217 | rs7077072 | -0.937 | 2.036 | 1.677 | 0.127 | 0 | -0.001 | -0.001 | 0 | 0.69 | 0.71 | 0.66 | 0.62 | 0.75 | 0.63 |

| | | | | | | | | | | | | | | | | |
|---|---|---|---|---|---|---|---|---|---|---|---|---|---|---|---|---|
| 10 | 26311675 | rs17738738 | -0.352 | 1.945 | 1.687 | 0.15 | 0.004 | -0.017 | -0.017 | 0 | 0.11 | 0.1 | 0.12 | 0.01 | 0.22 | 0.05 |
| 10 | 26312425 | rs1339816 | 0.969 | 2.05 | 1.685 | 0.164 | 0.059 | 0.048 | 0.048 | 0 | 0.42 | 0.21 | 0.23 | 0.29 | 0.34 | 0.43 |
| 10 | 26314305 | rs12573019 | 0.463 | 2.082 | 1.672 | 0.181 | -0.051 | -0.061 | -0.061 | 2.327 | 0.1 | 0.01 | 0.24 | 0.33 | 0.16 | 0.15 |
| 10 | 26318337 | rs1339811 | 0.324 | 2.086 | 1.691 | 0.179 | -0.011 | -0.006 | -0.006 | 0 | 0.11 | 0.08 | 0.25 | 0.32 | 0.16 | 0.15 |
| 10 | 26337526 | rs1521037 | -0.337 | 2.112 | 1.699 | 0.172 | -0.002 | -0.009 | -0.009 | 0.406 | 0.11 | 0.09 | 0.12 | 0.01 | 0.22 | 0.05 |
| 10 | 26344939 | rs11014929 | 0.082 | 1.879 | 1.71 | 0.192 | -0.007 | 0.041 | 0.041 | 0 | 0.54 | 0.3 | 0.35 | 0.3 | 0.56 | 0.48 |
| 10 | 26344948 | rs11014930 | 0.321 | 1.895 | 1.712 | 0.224 | -0.018 | -0.025 | -0.025 | 0.261 | 0.11 | 0.07 | 0.25 | 0.32 | 0.16 | 0.15 |
| 10 | 26350660 | rs4747545 | -0.627 | 1.895 | 1.741 | 0.222 | -0.003 | -0.001 | -0.001 | 0 | 0.66 | 0.43 | 0.6 | 0.62 | 0.72 | 0.63 |
| 10 | 26352676 | rs4631780 | 0.105 | 1.902 | 1.664 | 0.224 | -0.007 | 0.041 | 0.041 | 0 | 0.54 | 0.3 | 0.35 | 0.3 | 0.56 | 0.48 |
| 10 | 26352952 | rs11593128 | 0.105 | 1.902 | 1.658 | 0.224 | -0.006 | 0.036 | 0.036 | 0 | 0.54 | 0.31 | 0.35 | 0.3 | 0.56 | 0.48 |
| 10 | 26353592 | rs1521032 | 0.099 | 1.909 | 1.696 | 0.228 | -0.007 | 0.039 | 0.039 | 0 | 0.54 | 0.31 | 0.35 | 0.3 | 0.56 | 0.48 |
| 10 | 26355906 | rs3824700 | 0.099 | 1.909 | 1.691 | 0.228 | -0.006 | 0.037 | 0.037 | 0 | 0.54 | 0.31 | 0.35 | 0.29 | 0.55 | 0.48 |
| 10 | 26355992 | rs3824699 | -0.799 | 1.89 | 1.7 | 0.221 | -0.001 | -0.001 | -0.001 | 0.618 | 0.67 | 0.69 | 0.63 | 0.62 | 0.72 | 0.63 |
| 10 | 26379661 | rs16926578 | 0.845 | 1.81 | 1.289 | 0.097 | -0.006 | -0.022 | -0.022 | 0 | 0.07 | 0.08 | 0.03 | 0 | 0.06 | 0.02 |
| 10 | 26381812 | rs7916320 | 0.042 | 1.738 | 1.33 | 0.109 | -0.004 | 0.03 | 0.03 | 0 | 0.54 | 0.35 | 0.35 | 0.3 | 0.56 | 0.48 |
| 10 | 26392824 | rs11014959 | 0.474 | 1.74 | 1.261 | 0.107 | 0.038 | 0.064 | 0.064 | 0 | 0.47 | 0.25 | 0.28 | 0.29 | 0.44 | 0.45 |
| 10 | 26393354 | rs12573080 | -0.632 | 1.985 | 1.469 | 0.28 | -0.01 | 0.007 | 0.007 | 0 | 0.66 | 0.48 | 0.6 | 0.58 | 0.72 | 0.62 |
| 10 | 26411093 | rs4749092 | 0.409 | 2.076 | 1.161 | 0.309 | -0.056 | -0.073 | -0.073 | 0 | 0.11 | 0.01 | 0.23 | 0.28 | 0.16 | 0.15 |
| 10 | 26413547 | rs1891382 | -0.806 | 1.957 | 1.454 | 0.275 | 0.004 | -0.025 | -0.025 | 0 | 0.67 | 0.86 | 0.64 | 0.58 | 0.73 | 0.62 |
| 10 | 26418298 | rs17739436 | 0.684 | 1.998 | 1.473 | 0.158 | 0.005 | -0.012 | -0.012 | 0 | 0.07 | 0.07 | 0.07 | 0 | 0.12 | 0.03 |
| 10 | 26422301 | rs17739453 | 0.684 | 1.998 | 1.473 | 0.158 | 0.005 | -0.012 | -0.012 | 0 | 0.07 | 0.07 | 0.07 | 0 | 0.12 | 0.03 |
| 10 | 26433366 | rs2096176 | 0.613 | 1.954 | 1.643 | 0.377 | 0.02 | 0.019 | 0.019 | 0 | 0.47 | 0.35 | 0.38 | 0.35 | 0.44 | 0.46 |
| 10 | 26440886 | rs1935483 | 0.184 | 1.957 | 1.645 | 0.406 | -0.019 | -0.028 | -0.028 | 0 | 0.12 | 0.07 | 0.24 | 0.29 | 0.16 | 0.15 |
| 10 | 26443646 | rs3818956 | -0.684 | 1.956 | 1.752 | 0.409 | 0.001 | 0.001 | 0.001 | 0 | 0.66 | 0.71 | 0.71 | 0.64 | 0.73 | 0.63 |
| 10 | 26446312 | rs3758449 | 0.161 | 1.959 | 1.762 | 0.408 | -0.001 | 0.004 | 0.004 | 0 | 0.54 | 0.43 | 0.45 | 0.36 | 0.56 | 0.48 |
| 10 | 26452144 | rs4749093 | -0.595 | 1.931 | 1.872 | 0.394 | -0.005 | -0.005 | -0.005 | 0 | 0.66 | 0.62 | 0.7 | 0.64 | 0.72 | 0.63 |
| 10 | 26456397 | rs1935485 | -0.342 | 2.111 | 1.831 | 0.349 | 0.046 | -0.05 | -0.05 | 0 | 0.11 | 0.18 | 0.13 | 0.01 | 0.22 | 0.05 |
| 10 | 26457637 | rs16926628 | 0.505 | 2.118 | 1.827 | 0.35 | 0.043 | -0.037 | -0.037 | 0 | 0.07 | 0.18 | 0.08 | 0 | 0.12 | 0.03 |
| 10 | 26460246 | rs1888252 | -0.265 | 2.058 | 1.787 | 0.366 | 0.004 | -0.013 | -0.013 | 0.246 | 0.59 | 0.55 | 0.63 | 0.63 | 0.6 | 0.61 |
| 10 | 26461232 | rs12266309 | 0.629 | 2.066 | 1.877 | 0.377 | 0.022 | 0.02 | 0.02 | 0 | 0.46 | 0.3 | 0.37 | 0.35 | 0.44 | 0.46 |
| 10 | 26462790 | rs3740232 | 0.99 | 2.028 | 1.859 | 0.374 | 0.053 | 0.012 | 0.012 | 0 | 0.42 | 0.23 | 0.32 | 0.35 | 0.34 | 0.44 |
| 10 | 26463130 | rs1999240 | -0.277 | 2.094 | 1.811 | 0.371 | 0.005 | -0.014 | -0.014 | 2.428 | 0.59 | 0.55 | 0.63 | 0.63 | 0.6 | 0.61 |
| 10 | 26480204 | rs10828962 | -0.847 | 2.12 | 1.949 | 0.465 | 0.079 | 0.116 | 0.116 | 0 | 0.17 | 0.43 | 0.3 | 0.28 | 0.25 | 0.18 |
| 10 | 26483183 | rs11014993 | 1.037 | 1.532 | 1.426 | 0.103 | 0.009 | -0.003 | -0.003 | 0 | 0.29 | 0.25 | 0.27 | 0.29 | 0.21 | 0.29 |
| 10 | 26483265 | rs16926660 | 1.051 | 1.555 | 1.326 | 0.084 | 0.005 | -0.027 | -0.027 | 0 | 0.07 | 0.11 | 0.06 | 0 | 0.09 | 0.02 |
| 10 | 26483504 | rs4749097 | -0.617 | 1.46 | 1.323 | 0.217 | 0.013 | 0.087 | 0.087 | 0 | 0.54 | 0.84 | 0.66 | 0.63 | 0.58 | 0.52 |
| 10 | 26483710 | rs4749098 | 0.235 | 0.917 | 1.686 | 0.554 | -0.009 | -0.004 | -0.004 | 0 | 0.11 | 0.09 | 0.22 | 0.28 | 0.15 | 0.17 |
| 10 | 26500166 | rs994502 | 0.284 | -0.26 | 1.438 | 0.256 | -0.009 | -0.001 | -0.001 | 0 | 0.33 | 0.67 | 0.3 | 0.35 | 0.17 | 0.24 |
| 10 | 26500658 | rs10508715 | 0.771 | -0.29 | 1.56 | 0.24 | 0.025 | -0.01 | -0.01 | 0 | 0.28 | 0.2 | 0.24 | 0.35 | 0.17 | 0.24 |
| 10 | 26505496 | rs2236418 | 0.081 | -0.336 | 1.455 | 0.226 | 0.1 | -0.006 | -0.006 | 0.691 | 0.34 | 0.87 | 0.31 | 0.35 | 0.17 | 0.24 |
| 10 | 26512375 | rs8190612 | 0.652 | -0.147 | 1.21 | 0.057 | -0.015 | 0.052 | 0.052 | 0 | 0.13 | 0.01 | 0.06 | 0.02 | 0.15 | 0.11 |
| 10 | 26514831 | rs7079102 | -1.824 | -0.036 | 1.606 | 0.156 | 0.073 | 0.068 | 0.068 | 6.993 | 0.05 | 0.51 | 0.05 | 0 | 0 | 0 |
| 10 | 26527590 | rs11015008 | -0.022 | -0.082 | 0.536 | 0.21 | -0.011 | -0.051 | -0.051 | 1.406 | 0.23 | 0.02 | 0.16 | 0.24 | 0.23 | 0.22 |
| 10 | 26528835 | rs1330581 | 0.075 | -0.133 | 0.952 | 0.313 | 0.064 | -0.023 | -0.023 | 0 | 0.37 | 0.87 | 0.32 | 0.4 | 0.27 | 0.29 |
| 10 | 26534809 | rs1805398 | -0.52 | 0.171 | 1.559 | 0.244 | 0.149 | 0.056 | 0.056 | 0 | 0.06 | 0.39 | 0.09 | 0.04 | 0.1 | 0.02 |
| 10 | 26537170 | rs11015012 | -1.087 | 0.155 | 1.466 | 0.237 | 0.142 | 0.044 | 0.044 | 0.025 | 0.07 | 0.51 | 0.1 | 0.05 | 0.11 | 0.03 |
| 10 | 26552723 | rs4749102 | 0.273 | 0.011 | 0.792 | 0.31 | 0.016 | -0.003 | -0.003 | 0 | 0.35 | 0.66 | 0.31 | 0.38 | 0.28 | 0.29 |
| 10 | 26564433 | rs7900976 | 0.644 | -0.067 | 1.198 | 0.289 | -0.056 | 0.013 | 0.013 | 0.512 | 0.3 | 0.38 | 0.23 | 0.34 | 0.17 | 0.26 |
| 10 | 26565634 | rs7067571 | 0.028 | -0.014 | 1.001 | 0.29 | 0.047 | -0.013 | -0.013 | 0 | 0.37 | 0.86 | 0.33 | 0.38 | 0.28 | 0.29 |
| 10 | 26566456 | rs11015025 | 1.124 | -0.046 | 0.772 | 0.433 | -0.01 | -0.062 | -0.062 | 0 | 0.1 | 0 | 0.09 | 0.23 | 0.09 | 0.13 |
| 10 | 26570721 | rs8190754 | 0.674 | -0.01 | 1.16 | 0.554 | -0.03 | 0.009 | 0.009 | 0 | 0.28 | 0.33 | 0.22 | 0.34 | 0.17 | 0.26 |
| 10 | 26574893 | rs7902107 | 0.058 | 0.011 | 1.263 | 0.549 | 0.019 | -0.002 | -0.002 | 0 | 0.36 | 0.74 | 0.33 | 0.38 | 0.28 | 0.28 |
| 10 | 26602427 | rs12248143 | 0.671 | -0.03 | 0.911 | 0.617 | -0.014 | 0.003 | 0.003 | 0 | 0.27 | 0.28 | 0.21 | 0.34 | 0.17 | 0.25 |
| 10 | 26612058 | rs1148158 | 0.487 | 0.495 | 0.286 | 0.795 | -0.031 | -0.028 | -0.028 | 0.03 | 0.09 | 0.02 | 0.27 | 0.34 | 0.14 | 0.16 |
| 10 | 26624887 | rs11015040 | 0.4 | 0.156 | 0.155 | 0.005 | 0.002 | 0.002 | 0.002 | 0 | 0.36 | 0.34 | 0.29 | 0.3 | 0.31 | 0.32 |
| 10 | 26625316 | rs7077123 | 0.081 | 0.204 | -0.889 | 0.075 | 0.014 | 0.009 | 0.009 | 0 | 0.4 | 0.67 | 0.34 | 0.34 | 0.34 | 0.34 |
| 10 | 26630518 | rs11015043 | -0.136 | 0.714 | -1.524 | 0.221 | 0.022 | 0.08 | 0.08 | 0.275 | 0.53 | 0.86 | 0.4 | 0.33 | 0.43 | 0.4 |
| 10 | 26631987 | rs6482556 | -0.203 | 0.651 | -1.519 | 0.217 | 0.003 | 0.05 | 0.05 | 1.105 | 0.54 | 0.86 | 0.41 | 0.37 | 0.45 | 0.42 |
| 10 | 26633775 | rs7073826 | -0.065 | 0.707 | -1.143 | -0.389 | 0.014 | 0.052 | 0.052 | 0.31 | 0.51 | 0.59 | 0.45 | 0.2 | 0.58 | 0.51 |
| 10 | 26636901 | rs3006842 | -0.079 | 1.1 | -1.247 | -0.295 | -0.008 | 0.11 | 0.11 | 1.25 | 0.51 | 0.3 | 0.47 | 0.75 | 0.55 | 0.51 |
| 10 | 26636925 | rs3006843 | 0.088 | 1.405 | -1.346 | -0.19 | 0.028 | 0.073 | 0.073 | 1.918 | 0.46 | 0.29 | 0.43 | 0.69 | 0.46 | 0.43 |
| 10 | 26643343 | rs1335545 | -0.213 | 1.221 | -0.699 | -0.237 | -0.011 | 0.124 | 0.124 | 0.7 | 0.47 | 0.24 | 0.47 | 0.75 | 0.5 | 0.5 |
| 10 | 26654660 | rs3006801 | 0.61 | 0.812 | -0.663 | 0.37 | 0.035 | -0.047 | -0.047 | 0 | 0.37 | 0.23 | 0.17 | 0.45 | 0.3 | 0.29 |
| 10 | 26660774 | rs2992274 | -0.467 | 0.798 | -0.453 | 0.341 | -0.001 | -0.033 | -0.033 | 0.666 | 0.51 | 0.52 | 0.59 | 0.84 | 0.48 | 0.52 |
| 10 | 26660883 | rs4749118 | -1.522 | 0.609 | -0.087 | 0.788 | 0.02 | 0.115 | 0.115 | 0 | 0.15 | 0.29 | 0.42 | 0.38 | 0.18 | 0.23 |
| 10 | 26664506 | rs11015068 | -0.821 | 0.085 | 0.018 | 0.976 | 0.002 | 0.207 | 0.207 | 0.455 | 0.06 | 0.18 | 0.34 | 0.32 | 0.06 | 0.14 |
| 10 | 26673486 | rs3006783 | 0.235 | -0.083 | 0.063 | 0.764 | -0.021 | 0.056 | 0.056 | 0 | 0.68 | 0.54 | 0.59 | 0.46 | 0.72 | 0.49 |
| 10 | 26681484 | rs2152305 | 0.792 | -0.011 | 0.805 | 1.014 | -0.018 | 0.162 | 0.162 | 0.146 | 0.88 | 0.64 | 0.75 | 0.68 | 0.91 | 0.74 |
| 10 | 26685333 | rs12412360 | -0.053 | 0.021 | 0.744 | 0.882 | -0.003 | 0.023 | 0.023 | 0 | 0.08 | 0.11 | 0.21 | 0.22 | 0.09 | 0.18 |

| | | | | | | | | | | | | | | | | |
|---|---|---|---|---|---|---|---|---|---|---|---|---|---|---|---|---|
| 10 | 26688228 | rs17746517 | 0.489 | 1.1 | 0.687 | 0.59 | -0.019 | -0.008 | -0.008 | 0 | 0.05 | 0.01 | 0.1 | 0.05 | 0.13 | 0.06 |
| 10 | 26695750 | rs867354 | 0.315 | 0.373 | 0.309 | 0.072 | -0.013 | 0.01 | 0.01 | 1.148 | 0.66 | 0.7 | 0.43 | 0.76 | 0.53 | 0.67 |
| 10 | 26697294 | rs945294 | -0.487 | 0.356 | 0.35 | 0.037 | -0.008 | -0.003 | -0.003 | 0 | 0.75 | 0.78 | 0.64 | 0.95 | 0.61 | 0.84 |
| 10 | 26700784 | rs2992299 | 0.352 | -0.313 | -0.179 | -0.271 | -0.049 | -0.106 | -0.106 | 0 | 0.27 | 0.1 | 0.2 | 0.49 | 0.38 | 0.38 |
| 10 | 26702050 | rs4749123 | 0.752 | -1.052 | 0.252 | -0.09 | 0.034 | 0.035 | 0.035 | 2.855 | 0.41 | 0.31 | 0.25 | 0.23 | 0.27 | 0.28 |
| 10 | 26704514 | rs945296 | -0.749 | -0.895 | -0.068 | -0.016 | -0.01 | -0.007 | -0.007 | 0 | 0.55 | 0.46 | 0.71 | 0.73 | 0.73 | 0.69 |
| 10 | 26705832 | rs868093 | -0.02 | -0.769 | -0.3 | -0.339 | -0.002 | -0.001 | -0.001 | 0 | 0.35 | 0.32 | 0.58 | 0.24 | 0.48 | 0.33 |
| 10 | 26706338 | rs11015094 | 0.428 | 0.203 | -0.639 | 0.107 | -0.015 | -0.016 | -0.016 | 0.054 | 0.05 | 0.01 | 0.06 | 0.08 | 0.08 | 0.09 |
| 10 | 26713667 | rs3006824 | 0.66 | 0.366 | -0.351 | 0.159 | -0.005 | -0.003 | -0.003 | 0 | 0.2 | 0.17 | 0.15 | 0.48 | 0.26 | 0.34 |
| 10 | 26716863 | rs7895480 | 0.214 | 0.032 | -0.329 | 0.02 | 0.041 | 0.015 | 0.015 | 0.473 | 0.5 | 0.42 | 0.47 | 0.45 | 0.36 | 0.48 |
| 10 | 26718561 | rs3006819 | -0.528 | 0.128 | -0.323 | -0.028 | 0.041 | -0.098 | -0.098 | 1.36 | 0.7 | 0.59 | 0.62 | 0.93 | 0.61 | 0.82 |
| 10 | 26726606 | rs1537746 | 0.225 | 0.084 | -0.493 | 0.097 | 0.062 | 0.004 | 0.004 | 0.409 | 0.5 | 0.35 | 0.47 | 0.5 | 0.36 | 0.49 |
| 10 | 26727454 | rs2992333 | -0.676 | 0.016 | -0.475 | -0.239 | 0.056 | -0.123 | -0.123 | 0 | 0.7 | 0.5 | 0.6 | 0.94 | 0.6 | 0.82 |
| 10 | 26730316 | rs2050447 | 0.723 | 0.022 | -0.57 | -0.203 | -0.007 | -0.016 | -0.016 | 0.021 | 0.2 | 0.15 | 0.15 | 0.48 | 0.26 | 0.34 |
| 10 | 26734485 | rs10764613 | 0.107 | 0.323 | -0.792 | -0.026 | 0.014 | 0.006 | 0.006 | 0.271 | 0.51 | 0.47 | 0.47 | 0.44 | 0.35 | 0.49 |
| 10 | 26740751 | rs7904878 | 0.237 | 0.61 | -1.469 | -0.53 | 0.006 | -0.021 | -0.021 | 0 | 0.17 | 0.01 | 0.13 | 0.15 | 0.15 | 0.14 |
| 10 | 26746468 | rs3006835 | 0.265 | 1.034 | -0.773 | -0.618 | -0.018 | 0.013 | 0.013 | 0 | 0.71 | 0.79 | 0.63 | 0.94 | 0.62 | 0.84 |
| 10 | 26764577 | rs7085145 | -0.068 | 0.475 | -1.621 | 0.214 | -0.088 | 0.045 | 0.045 | 0 | 0.42 | 0.08 | 0.36 | 0.64 | 0.51 | 0.46 |
| 10 | 26774694 | rs786870 | -0.34 | 0.009 | -0.899 | -0.228 | -0.031 | 0.042 | 0.042 | 0.252 | 0.81 | 0.94 | 0.8 | 0.92 | 0.77 | 0.8 |
| 10 | 26778606 | rs12412826 | -0.103 | -0.729 | -0.845 | -0.152 | 0 | 0.027 | 0.027 | 0 | 0.1 | 0.01 | 0.07 | 0.07 | 0.08 | 0.11 |
| 10 | 26788462 | rs10764627 | -0.417 | -0.937 | 0.026 | 0.439 | 0.021 | -0.022 | -0.022 | 0 | 0.36 | 0.27 | 0.36 | 0.52 | 0.26 | 0.48 |
| 10 | 26789281 | rs4749159 | 0.059 | -0.999 | -0.388 | 0.863 | -0.002 | 0.017 | 0.017 | 0 | 0.07 | 0.09 | 0.2 | 0.22 | 0.06 | 0.19 |
| 10 | 26795922 | rs787034 | 0.62 | -0.913 | -0.387 | 0.419 | -0.011 | -0.001 | -0.001 | 0 | 0.14 | 0.22 | 0.09 | 0.16 | 0.13 | 0.09 |
| 10 | 26796810 | rs787033 | -0.524 | -0.387 | -0.715 | 0.259 | 0.061 | 0.087 | 0.087 | 0.9 | 0.54 | 0.85 | 0.48 | 0.69 | 0.37 | 0.57 |
| 10 | 26801721 | rs7096662 | -1.122 | 0.157 | 0.367 | 0.506 | 0.027 | 0.032 | 0.032 | 0 | 0.1 | 0.3 | 0.12 | 0.1 | 0.07 | 0.09 |
| 10 | 26802361 | rs12783959 | -1.118 | 0.648 | 0.245 | 0.96 | 0.014 | 0.05 | 0.05 | 0 | 0.2 | 0.33 | 0.38 | 0.4 | 0.21 | 0.29 |
| 10 | 26804638 | rs11015146 | -0.226 | 0.964 | -0.549 | 0.95 | -0.025 | -0.019 | -0.019 | 0 | 0.18 | 0.11 | 0.14 | 0.24 | 0.18 | 0.19 |
| 10 | 26806629 | rs1775230 | -0.576 | 0.343 | -0.513 | 0.357 | -0.015 | -0.025 | -0.025 | 0 | 0.81 | 0.78 | 0.89 | 0.93 | 0.85 | 0.82 |
| 10 | 26808561 | rs787068 | 1.453 | 0.615 | -0.718 | 0.29 | 0.009 | 0.019 | 0.019 | 0 | 0.38 | 0.25 | 0.24 | 0.28 | 0.35 | 0.21 |
| 10 | 26812175 | rs11015149 | 2.069 | 1.275 | -0.77 | 0.298 | -0.016 | -0.015 | -0.015 | 0.603 | 0.05 | 0.02 | 0.08 | 0.07 | 0.11 | 0.05 |
| 10 | 26814452 | rs702988 | -1.629 | 0.852 | -0.585 | 0.17 | 0.007 | 0.002 | 0.002 | 0 | 0.82 | 0.86 | 0.91 | 0.96 | 0.86 | 0.87 |
| 10 | 26816541 | rs1013471 | 0.553 | 0.711 | 0.044 | 0.132 | 0.002 | 0.013 | 0.013 | 0.263 | 0.22 | 0.34 | 0.18 | 0.27 | 0.19 | 0.2 |
| 10 | 26818574 | rs787057 | -0.089 | 0.429 | 0.267 | 0.102 | -0.015 | -0.013 | -0.013 | 0 | 0.73 | 0.7 | 0.86 | 0.89 | 0.84 | 0.8 |
| 10 | 26820960 | rs1318662 | 1.658 | 0.29 | -0.193 | 0.126 | -0.01 | -0.006 | -0.006 | 0 | 0.21 | 0.13 | 0.15 | 0.26 | 0.2 | 0.2 |
| 10 | 26822825 | rs787052 | -1.171 | 0.213 | -0.097 | 0.105 | -0.027 | -0.008 | -0.008 | 0 | 0.88 | 0.8 | 0.93 | 0.93 | 0.92 | 0.83 |
| 10 | 26823045 | rs10741112 | 1.062 | 0.447 | -0.044 | -0.019 | 0.006 | 0.058 | 0.058 | 0 | 0.38 | 0.19 | 0.52 | 0.24 | 0.4 | 0.34 |
| 10 | 26823941 | rs787050 | -0.92 | 0.498 | -0.085 | -0.035 | 0.008 | -0.004 | -0.004 | 1.553 | 0.36 | 0.57 | 0.51 | 0.64 | 0.4 | 0.48 |
| 10 | 26826118 | rs7903201 | 1.565 | 0.454 | -0.232 | -0.097 | -0.01 | 0.151 | 0.151 | 0.441 | 0.3 | 0.09 | 0.27 | 0.03 | 0.32 | 0.14 |
| 10 | 26827600 | rs10764631 | -0.239 | 0.597 | -0.059 | -0.057 | -0.001 | -0.003 | -0.003 | 1.023 | 0.55 | 0.54 | 0.69 | 0.33 | 0.6 | 0.5 |
| 10 | 26828889 | rs7902011 | -0.858 | 0.455 | -0.628 | -0.234 | 0.003 | 0.01 | 0.01 | 0.028 | 0.3 | 0.54 | 0.44 | 0.31 | 0.3 | 0.36 |
| 10 | 26829338 | rs11015160 | 1.327 | 0.802 | -0.696 | 0.029 | -0.01 | 0.106 | 0.106 | 0 | 0.27 | 0.11 | 0.28 | 0.06 | 0.31 | 0.19 |
| 10 | 26830113 | rs7910019 | 1.601 | 0.837 | -0.689 | 0.016 | -0.022 | 0.11 | 0.11 | 0 | 0.25 | 0.09 | 0.27 | 0.03 | 0.31 | 0.14 |
| 10 | 26833501 | rs787043 | 0.943 | 0.952 | -0.665 | 0.106 | -0.002 | 0.016 | 0.016 | 0 | 0.11 | 0.1 | 0.09 | 0.34 | 0.11 | 0.13 |
| 10 | 26836009 | rs11015165 | 0.358 | 1.094 | -0.241 | -0.546 | -0.029 | -0.004 | -0.004 | 0 | 0.11 | 0.08 | 0.11 | 0.07 | 0.18 | 0.14 |
| 10 | 26836125 | rs10764632 | -0.706 | 1.193 | -0.277 | -0.593 | 0 | 0.002 | 0.002 | 0 | 0.51 | 0.57 | 0.69 | 0.37 | 0.57 | 0.54 |
| 10 | 26839856 | rs7895809 | 0.848 | 1.185 | -1.374 | -0.69 | 0 | 0.007 | 0.007 | 0 | 0.07 | 0.05 | 0.25 | 0.2 | 0.08 | 0.19 |
| 10 | 26845653 | rs10764633 | 0.289 | 1.154 | -1.453 | -1.037 | -0.011 | 0.087 | 0.087 | 0 | 0.26 | 0.13 | 0.28 | 0.03 | 0.31 | 0.14 |
| 10 | 26857061 | rs1753336 | 0.131 | 1.422 | -1.249 | -0.311 | -0.081 | -0.011 | -0.011 | 0 | 0.42 | 0.08 | 0.41 | 0.4 | 0.49 | 0.47 |
| 10 | 26858860 | rs12219836 | 1.041 | 1.586 | -1.11 | -0.16 | -0.029 | -0.027 | -0.027 | 0 | 0.05 | 0.01 | 0.1 | 0.1 | 0.15 | 0.02 |
| 10 | 26861229 | rs787016 | -0.612 | 0.444 | -0.491 | -0.096 | 0.011 | 0.011 | 0.011 | 0 | 0.69 | 0.78 | 0.64 | 0.92 | 0.76 | 0.78 |
| 10 | 26867795 | rs11815280 | 1.023 | 0.445 | -1.077 | -0.103 | 0.014 | 0.056 | 0.056 | 1.173 | 0.21 | 0.05 | 0.34 | 0.07 | 0.2 | 0.08 |
| 10 | 26873049 | rs796820 | -0.076 | 1.192 | -0.895 | 0.324 | 0.058 | 0.043 | 0.043 | 0.353 | 0.5 | 0.22 | 0.48 | 0.43 | 0.62 | 0.53 |
| 10 | 26875066 | rs4749165 | -1.091 | 1.273 | -0.89 | 0.431 | -0.033 | -0.071 | -0.071 | 0.016 | 0.67 | 0.55 | 0.62 | 0.9 | 0.76 | 0.78 |
| 10 | 26876272 | rs6482566 | -1.429 | 1.804 | -0.668 | 1.496 | 0.001 | -0.004 | -0.004 | 0 | 0.38 | 0.36 | 0.33 | 0.72 | 0.38 | 0.56 |
| 10 | 26876769 | rs12784646 | -0.884 | 1.506 | -1.289 | 1.271 | -0.031 | -0.043 | -0.043 | 0 | 0.21 | 0.05 | 0.2 | 0.25 | 0.24 | 0.31 |
| 10 | 26883359 | rs12772391 | -0.82 | 1.398 | -1.266 | 1.287 | -0.032 | -0.048 | -0.048 | 0 | 0.2 | 0.05 | 0.2 | 0.26 | 0.23 | 0.28 |
| 10 | 26900228 | rs7074426 | -1.108 | 1.498 | -0.262 | 1.895 | 0.072 | 0.256 | 0.256 | 0.036 | 0.06 | 0.28 | 0.1 | 0.32 | 0.07 | 0.06 |
| 10 | 26901421 | rs7075871 | -0.44 | 1.268 | -0.79 | 1.014 | -0.012 | 0.123 | 0.123 | 0 | 0.55 | 0.75 | 0.7 | 0.82 | 0.53 | 0.58 |
| 10 | 26908475 | rs7911815 | -1.417 | 0.733 | -0.297 | 1.759 | -0.018 | 0.206 | 0.206 | 1.048 | 0.11 | 0.37 | 0.14 | 0.67 | 0.06 | 0.19 |
| 10 | 26929728 | rs2015190 | -0.82 | 0.442 | -0.357 | 1.427 | -0.001 | -0.004 | -0.004 | 1.785 | 0.27 | 0.25 | 0.39 | 0.4 | 0.26 | 0.26 |
| 10 | 26931729 | rs1408529 | 1.012 | 0.637 | -0.49 | 1.397 | -0.019 | -0.058 | -0.058 | 1.51 | 0.27 | 0.13 | 0.23 | 0.38 | 0.34 | 0.41 |
| 10 | 26933522 | rs1748333 | 1.247 | 0.565 | -0.47 | 1.463 | -0.052 | -0.047 | -0.047 | 0.522 | 0.24 | 0.05 | 0.21 | 0.31 | 0.33 | 0.24 |
| 10 | 26934152 | rs1677722 | 1.233 | 0.503 | -0.455 | 1.436 | -0.033 | -0.03 | -0.03 | 0 | 0.25 | 0.09 | 0.22 | 0.31 | 0.34 | 0.25 |
| 10 | 26946998 | rs1748329 | -0.989 | 0.456 | -0.152 | 1.35 | 0.003 | -0.032 | -0.032 | 0 | 0.93 | 0.86 | 0.98 | 1 | 0.96 | 0.9 |
| 10 | 26948937 | rs1748326 | 0.456 | -0.305 | -0.326 | 0.829 | -0.006 | -0.01 | -0.01 | 2.55 | 0.46 | 0.51 | 0.45 | 0.6 | 0.45 | 0.5 |
| 10 | 26949370 | rs10829043 | 0.974 | -0.4 | -0.263 | 0.761 | 0.016 | -0.016 | -0.016 | 0 | 0.16 | 0.11 | 0.19 | 0.22 | 0.11 | 0.09 |
| 10 | 26950217 | rs1853682 | 0.626 | -0.282 | -0.281 | 0.754 | -0.005 | 0.005 | 0.005 | 0 | 0.43 | 0.49 | 0.43 | 0.52 | 0.44 | 0.33 |
| 10 | 26956056 | rs11015211 | 1.061 | -0.354 | -0.609 | 0.502 | -0.066 | 0.102 | 0.102 | 0.039 | 0.19 | 0.01 | 0.13 | 0 | 0.24 | 0.14 |
| 10 | 26956129 | rs2252096 | 1.948 | -0.398 | -0.61 | 0.598 | -0.054 | -0.039 | -0.039 | 0 | 0.25 | 0.05 | 0.22 | 0.31 | 0.34 | 0.23 |

| | | | | | | | | | | | | | | | | |
|---|---|---|---|---|---|---|---|---|---|---|---|---|---|---|---|---|
| 10 | 26959202 | rs2489556 | 0.516 | -0.628 | -0.419 | 0.682 | 0.028 | 0.128 | 0.128 | 0 | 0.46 | 0.2 | 0.43 | 0.82 | 0.45 | 0.48 |
| 10 | 26971879 | rs12415035 | -0.338 | -0.737 | 0.103 | 0.853 | -0.009 | -0.074 | -0.074 | 2.099 | 0.36 | 0.14 | 0.31 | 0.52 | 0.32 | 0.38 |
| 10 | 26974263 | rs1748343 | -0.181 | -0.689 | 0.496 | 0.81 | 0.005 | -0.022 | -0.022 | 0 | 0.68 | 0.49 | 0.58 | 0.82 | 0.69 | 0.62 |
| 10 | 26985971 | rs2489585 | -0.423 | -0.554 | 0.456 | 0.668 | 0.017 | -0.092 | -0.092 | 0.078 | 0.7 | 0.54 | 0.59 | 0.9 | 0.7 | 0.78 |
| 10 | 26993080 | rs11015236 | -0.265 | -0.763 | 0.517 | 1.049 | -0.031 | 0.032 | 0.032 | 0 | 0.22 | 0.3 | 0.18 | 0.57 | 0.12 | 0.42 |
| 10 | 26997889 | rs1780179 | 0.733 | -0.672 | 0.835 | 0.795 | -0.001 | -0.002 | -0.002 | 0 | 0.56 | 0.47 | 0.64 | 0.4 | 0.62 | 0.34 |
| 10 | 26997925 | rs1780180 | 0.628 | -0.68 | 0.831 | 0.924 | -0.002 | 0.002 | 0.002 | 0 | 0.57 | 0.47 | 0.65 | 0.48 | 0.62 | 0.53 |
| 10 | 26997972 | rs1748358 | -0.422 | -0.547 | 0.647 | 0.708 | 0.004 | -0.049 | -0.049 | 0 | 0.78 | 0.77 | 0.87 | 1 | 0.74 | 0.76 |
| 10 | 27004841 | rs11015244 | 0.871 | -0.651 | 0.445 | 1.068 | 0.023 | -0.052 | -0.052 | 0 | 0.15 | 0.03 | 0.13 | 0.25 | 0.12 | 0.11 |
| 10 | 27009334 | rs1780196 | -0.453 | -0.498 | 0.562 | 0.682 | -0.013 | 0.022 | 0.022 | 1.785 | 0.86 | 0.93 | 0.92 | 0.93 | 0.87 | 0.87 |
| 10 | 27011112 | rs1564271 | 1.908 | -0.048 | -0.691 | 0.447 | 0.002 | 0.012 | 0.012 | 0 | 0.29 | 0.14 | 0.29 | 0.27 | 0.29 | 0.21 |
| 10 | 27011863 | rs11015249 | -0.315 | 0.027 | -0.422 | 0.397 | -0.008 | -0.011 | -0.011 | 0 | 0.42 | 0.46 | 0.37 | 0.62 | 0.4 | 0.61 |
| 10 | 27014467 | rs10508720 | 0.91 | 0.173 | -0.412 | 0.327 | 0.003 | -0.015 | -0.015 | 0 | 0.14 | 0.16 | 0.25 | 0.04 | 0.18 | 0.05 |
| 10 | 27016958 | rs3118153 | -0.393 | 0.22 | -0.415 | 0.337 | 0.003 | -0.014 | -0.014 | 0 | 0.86 | 0.84 | 0.92 | 0.94 | 0.87 | 0.87 |
| 10 | 27019578 | rs4369307 | 0.936 | 0.245 | -0.542 | 0.339 | 0.003 | -0.012 | -0.012 | 0 | 0.14 | 0.16 | 0.26 | 0.04 | 0.18 | 0.04 |
| 10 | 27021006 | rs2448111 | 0.25 | 0.006 | -0.477 | 0.734 | 0.016 | -0.001 | -0.001 | 0.26 | 0.82 | 0.59 | 0.84 | 0.84 | 0.87 | 0.85 |
| 10 | 27023402 | rs16927305 | 0.456 | -0.558 | -1.021 | 0.656 | -0.022 | 0.022 | 0.022 | 0 | 0.15 | 0.2 | 0.09 | 0.24 | 0.09 | 0.29 |
| 10 | 27025659 | rs3802609 | -1.435 | -0.271 | -1.97 | -0.214 | -0.092 | 0.123 | 0.123 | 3.411 | 0.64 | 0.88 | 0.61 | 0.8 | 0.54 | 0.67 |
| 10 | 27033193 | rs6482573 | -0.909 | -0.937 | -1.893 | 0.338 | 0.039 | 0.025 | 0.025 | 1.606 | 0.44 | 0.81 | 0.46 | 0.46 | 0.4 | 0.54 |
| 10 | 27034362 | rs11015262 | 0.594 | -1.168 | -0.89 | 0.039 | 0.008 | -0.073 | -0.073 | 0.447 | 0.12 | 0.29 | 0.07 | 0 | 0.13 | 0.12 |
| 10 | 27044460 | rs12220461 | 1.396 | -1.575 | -1.146 | -0.026 | 0.029 | 0.004 | 0.004 | 0 | 0.13 | 0.04 | 0.09 | 0.15 | 0.09 | 0.05 |
| 10 | 27045387 | rs1031858 | -1.857 | -1.84 | -1.265 | 0.087 | -0.018 | 0.095 | 0.095 | 3.563 | 0.58 | 0.88 | 0.55 | 0.62 | 0.49 | 0.62 |
| 10 | 27055896 | rs6482574 | 0.785 | -2.018 | -0.763 | 0.879 | -0.005 | 0.008 | 0.008 | 3.487 | 0.3 | 0.36 | 0.37 | 0.31 | 0.26 | 0.21 |
| 10 | 27058273 | rs3818409 | -0.247 | -2.298 | -0.98 | 0.982 | 0.003 | -0.005 | -0.005 | 0 | 0.33 | 0.71 | 0.36 | 0.28 | 0.32 | 0.43 |
| 10 | 27059608 | rs1324637 | -2.16 | -2.269 | -0.988 | 0.963 | -0.028 | 0.103 | 0.103 | 3.02 | 0.57 | 0.91 | 0.55 | 0.62 | 0.49 | 0.61 |
| 10 | 27064648 | rs11015279 | 1.456 | -2.215 | -0.956 | 0.994 | 0.021 | -0.052 | -0.052 | 0 | 0.11 | 0.02 | 0.08 | 0.2 | 0.07 | 0.12 |
| 10 | 27068310 | rs7094825 | 1.874 | -2.169 | -0.854 | 0.925 | 0.008 | -0.04 | -0.04 | 0 | 0.19 | 0.36 | 0.28 | 0.12 | 0.18 | 0.09 |
| 10 | 27073535 | rs17816509 | 0.876 | -2.61 | -1.224 | 0.791 | 0.041 | -0.075 | -0.075 | 0.133 | 0.21 | 0.04 | 0.16 | 0.29 | 0.17 | 0.28 |
| 10 | 27103056 | rs10159978 | 0.534 | -2.42 | -1.054 | 0.881 | -0.008 | 0.092 | 0.092 | 0.025 | 0.13 | 0.29 | 0.08 | 0.16 | 0.14 | 0.35 |
| 10 | 27115039 | rs4749190 | 2.134 | -2.409 | -1.112 | 0.892 | 0.021 | -0.005 | -0.005 | 0 | 0.12 | 0.04 | 0.09 | 0.15 | 0.09 | 0.05 |
| 10 | 27119069 | rs2182294 | 2.706 | -2.485 | -0.775 | 0.803 | -0.003 | 0.003 | 0.003 | 0.422 | 0.16 | 0.16 | 0.27 | 0.12 | 0.18 | 0.07 |
| 10 | 27121133 | rs7082503 | -0.296 | -2.6 | -0.482 | 0.932 | -0.014 | 0.12 | 0.12 | 0.091 | 0.14 | 0.39 | 0.09 | 0.17 | 0.14 | 0.37 |
| 10 | 27125541 | rs2505971 | -0.4 | -2.492 | -0.386 | 0.626 | -0.044 | -0.048 | -0.048 | 0 | 0.78 | 0.41 | 0.85 | 0.85 | 0.86 | 0.77 |
| 10 | 27125592 | rs17816696 | -0.835 | -3.159 | -0.587 | 1.044 | 0.026 | -0.065 | -0.065 | 0 | 0.27 | 0.13 | 0.2 | 0.48 | 0.23 | 0.43 |
| 10 | 27126196 | rs10829089 | -0.914 | -2.58 | 0.012 | 1.005 | -0.025 | 0.044 | 0.044 | 0 | 0.27 | 0.39 | 0.17 | 0.36 | 0.21 | 0.49 |
| 10 | 27150093 | rs1409128 | -0.898 | -2.334 | -0.156 | 0.931 | -0.014 | -0.003 | -0.003 | 0 | 0.41 | 0.53 | 0.27 | 0.51 | 0.31 | 0.54 |
| 10 | 27155368 | rs7914045 | -1.075 | -1.513 | 0.495 | 1.378 | 0.011 | 0.004 | 0.004 | 0 | 0.44 | 0.7 | 0.48 | 0.36 | 0.42 | 0.37 |
| 10 | 27157722 | rs7082748 | 1.203 | -1.505 | 0.759 | 1.584 | 0.006 | 0.009 | 0.009 | 2.373 | 0.21 | 0.27 | 0.32 | 0.22 | 0.25 | 0.11 |
| 10 | 27160932 | rs2505963 | 1.548 | -1.682 | 0.729 | 1.682 | 0.034 | 0.111 | 0.111 | 0.22 | 0.5 | 0.08 | 0.47 | 0.45 | 0.57 | 0.39 |
| 10 | 27165095 | rs11015358 | -0.44 | -1.801 | 0.999 | 1.583 | -0.013 | -0.008 | -0.008 | 0 | 0.2 | 0.23 | 0.11 | 0.16 | 0.16 | 0.25 |
| 10 | 27168322 | rs4287252 | -1.564 | -1.797 | 0.82 | 1.613 | 0.046 | -0.028 | -0.028 | 0.103 | 0.48 | 0.84 | 0.5 | 0.44 | 0.42 | 0.38 |
| 10 | 27168850 | rs7076236 | -1.025 | -1.769 | 0.83 | 1.499 | -0.067 | -0.024 | -0.024 | 0 | 0.25 | 0.45 | 0.17 | 0.19 | 0.16 | 0.27 |
| 10 | 27170960 | rs7086043 | 1.014 | -1.405 | 0.58 | 1.419 | 0.003 | -0.007 | -0.007 | 0 | 0.21 | 0.27 | 0.32 | 0.15 | 0.25 | 0.13 |
| 10 | 27177212 | rs2477926 | 0.404 | -1.289 | 0.767 | 1.319 | -0.02 | 0.313 | 0.313 | 3.971 | 0.94 | 0.75 | 0.95 | 0.73 | 0.99 | 0.82 |
| 10 | 27177245 | rs10829131 | -0.755 | -1.452 | 0.36 | 1.239 | -0.02 | -0.011 | -0.011 | 0 | 0.2 | 0.25 | 0.14 | 0.15 | 0.15 | 0.22 |
| 10 | 27178756 | rs10159883 | -1.617 | -1.34 | -0.539 | 0.913 | -0.036 | 0.028 | 0.028 | 0.687 | 0.29 | 0.62 | 0.2 | 0.43 | 0.18 | 0.42 |
| 10 | 27185286 | rs2505956 | 0.215 | -1.002 | -0.776 | 0.248 | 0.029 | -0.002 | -0.002 | 0 | 0.64 | 0.24 | 0.6 | 0.72 | 0.74 | 0.67 |
| 10 | 27192279 | rs2182296 | -0.725 | -0.766 | -0.75 | 0.079 | -0.032 | 0.027 | 0.027 | 0 | 0.73 | 0.89 | 0.62 | 0.84 | 0.68 | 0.89 |
| 10 | 27193821 | rs7902288 | -0.714 | -0.677 | -0.385 | 0.299 | -0.047 | 0.089 | 0.089 | 3.643 | 0.1 | 0.56 | 0.05 | 0.11 | 0.02 | 0.22 |
| 10 | 27200291 | rs787728 | -1.178 | -0.662 | -0.905 | 0.125 | 0.029 | 0.08 | 0.08 | 0.495 | 0.48 | 0.82 | 0.49 | 0.27 | 0.45 | 0.43 |
| 10 | 27200881 | rs11015373 | -1.296 | -0.61 | 0.004 | 0.597 | -0.053 | 0.081 | 0.081 | 4.343 | 0.1 | 0.61 | 0.06 | 0.12 | 0.02 | 0.2 |
| 10 | 27201386 | rs12251191 | -1.296 | -0.61 | 0.004 | 0.597 | -0.053 | 0.081 | 0.081 | 4.343 | 0.1 | 0.61 | 0.06 | 0.12 | 0.02 | 0.2 |
| 10 | 27204531 | rs1815323 | -1.372 | -0.234 | -0.264 | 0.185 | -0.005 | -0.035 | -0.035 | 0 | 0.13 | 0.18 | 0.11 | 0.02 | 0.11 | 0.07 |
| 10 | 27210569 | rs2797930 | -1.297 | -0.335 | -0.021 | 0.198 | -0.02 | -0.01 | -0.01 | 0 | 0.14 | 0.36 | 0.1 | 0.08 | 0.1 | 0.04 |
| 10 | 27210616 | rs11015383 | -0.312 | -0.524 | -0.362 | 0.862 | -0.009 | -0.037 | -0.037 | 0 | 0.1 | 0.01 | 0.11 | 0.44 | 0.13 | 0.36 |
| 10 | 27213946 | rs2148117 | -0.971 | -0.517 | -0.304 | 0.826 | 0.008 | 0.06 | 0.06 | 3.047 | 0.08 | 0.36 | 0.07 | 0.07 | 0.06 | 0.03 |
| 10 | 27221632 | rs7909447 | 0.063 | -0.162 | -1.714 | -0.229 | -0.016 | -0.006 | -0.006 | 0 | 0.43 | 0.21 | 0.57 | 0.48 | 0.42 | 0.35 |
| 10 | 27227168 | rs3928123 | 0.465 | -0.174 | -1.727 | 0.037 | -0.003 | -0.008 | -0.008 | 0 | 0.08 | 0.01 | 0.15 | 0.46 | 0.11 | 0.35 |
| 10 | 27241780 | rs11015418 | 0.192 | -0.224 | -2.232 | -0.599 | -0.087 | -0.089 | -0.089 | 0 | 0.47 | 0.08 | 0.39 | 0.47 | 0.58 | 0.55 |
| 10 | 27253308 | rs9703520 | -0.592 | 0.031 | -2.028 | -0.59 | -0.026 | -0.008 | -0.008 | 0.001 | 0.59 | 0.83 | 0.46 | 0.54 | 0.61 | 0.6 |
| 10 | 27255751 | rs7896781 | -0.907 | 0.085 | -2.164 | -0.544 | -0.054 | -0.045 | -0.045 | 0 | 0.66 | 0.93 | 0.67 | 0.57 | 0.66 | 0.63 |
| 10 | 27265641 | rs7914382 | -0.51 | 0.108 | -2.486 | -0.646 | -0.028 | -0.009 | -0.009 | 1.287 | 0.8 | 0.96 | 0.9 | 0.85 | 0.8 | 0.72 |
| 10 | 27266501 | rs10829152 | 0.594 | 0.249 | -2.73 | -0.682 | -0.015 | 0.016 | 0.016 | 1.454 | 0.43 | 0.15 | 0.41 | 0.45 | 0.41 | 0.43 |
| 10 | 27277310 | rs10829153 | 0.976 | -0.182 | -2.708 | -0.719 | 0.047 | 0.004 | 0.004 | 0.66 | 0.56 | 0.16 | 0.47 | 0.45 | 0.58 | 0.46 |
| 10 | 27288925 | rs10741125 | -0.282 | -0.278 | -2.367 | -0.838 | 0.001 | 0.045 | 0.045 | 0.638 | 0.76 | 0.87 | 0.71 | 0.99 | 0.78 | 0.88 |
| 10 | 27290161 | rs11015445 | -0.753 | -0.406 | -1.99 | -0.579 | -0.013 | 0.066 | 0.066 | 0 | 0.09 | 0.18 | 0.06 | 0.27 | 0.07 | 0.24 |
| 10 | 27293344 | rs10829158 | -0.753 | -0.408 | -1.989 | -0.571 | -0.013 | 0.066 | 0.066 | 0 | 0.09 | 0.18 | 0.06 | 0.27 | 0.07 | 0.24 |
| 10 | 27302772 | rs1411283 | 1.481 | -0.412 | -2.016 | -0.584 | 0.068 | 0.034 | 0.034 | 0.177 | 0.52 | 0.17 | 0.49 | 0.44 | 0.58 | 0.44 |
| 10 | 27314444 | rs7088171 | -0.777 | -0.5 | -1.578 | -0.614 | 0.002 | 0.003 | 0.003 | 0.585 | 0.18 | 0.24 | 0.18 | 0.55 | 0.19 | 0.42 |

| | | | | | | | | | | | | | | | | |
|---|---|---|---|---|---|---|---|---|---|---|---|---|---|---|---|---|
| 10 | 27317840 | rs10829163 | -0.777 | -0.5 | -1.578 | -0.614 | 0.002 | 0.003 | 0.003 | 0.585 | 0.18 | 0.24 | 0.18 | 0.55 | 0.19 | 0.42 |
| 10 | 27319258 | rs11015463 | -0.22 | -0.47 | -1.584 | -0.625 | 0.005 | 0.008 | 0.008 | 0.449 | 0.74 | 0.82 | 0.7 | 0.99 | 0.77 | 0.86 |
| 10 | 27333284 | rs7091830 | 1.631 | -0.103 | -1.823 | -0.663 | -0.113 | -0.113 | -0.113 | 0.615 | 0.22 | 0.02 | 0.32 | 0.34 | 0.3 | 0.29 |
| 10 | 27334674 | rs7097009 | 1.494 | -0.511 | -1.045 | -0.268 | 0.096 | 0.039 | 0.039 | 0.644 | 0.52 | 0.18 | 0.5 | 0.44 | 0.6 | 0.46 |
| 10 | 27343232 | rs4749219 | 0.362 | -0.456 | -1.177 | -0.267 | 0.03 | 0.019 | 0.019 | 0.051 | 0.64 | 0.82 | 0.59 | 0.71 | 0.68 | 0.7 |
| 10 | 27343834 | rs4749220 | -0.203 | -0.223 | -1.085 | -0.07 | -0.007 | 0.017 | 0.017 | 0 | 0.07 | 0.15 | 0.04 | 0.13 | 0.06 | 0.07 |
| 10 | 27348423 | rs11015492 | -0.42 | -0.224 | -1.079 | -0.071 | -0.011 | 0.031 | 0.031 | 0 | 0.07 | 0.19 | 0.04 | 0.13 | 0.06 | 0.08 |
| 10 | 27353250 | rs11015501 | -0.205 | -0.215 | -1.096 | -0.067 | -0.008 | 0.025 | 0.025 | 0 | 0.07 | 0.17 | 0.04 | 0.13 | 0.06 | 0.07 |
| 10 | 27368126 | rs3737055 | 1.458 | -0.266 | -1.265 | -0.022 | 0.078 | 0.036 | 0.036 | 2.242 | 0.51 | 0.13 | 0.5 | 0.44 | 0.6 | 0.44 |
| 10 | 27368325 | rs3737054 | -0.788 | -0.316 | -1.143 | 0.007 | -0.004 | 0.034 | 0.034 | 0.928 | 0.18 | 0.31 | 0.17 | 0.4 | 0.2 | 0.27 |
| 10 | 27369987 | rs11015508 | -1.203 | -0.287 | -1.241 | -0.015 | -0.007 | 0.071 | 0.071 | 0.14 | 0.11 | 0.28 | 0.06 | 0.27 | 0.11 | 0.26 |
| 10 | 27371126 | rs10829172 | 1.505 | -0.295 | -1.315 | -0.048 | 0.078 | 0.04 | 0.04 | 2.464 | 0.52 | 0.13 | 0.5 | 0.44 | 0.6 | 0.45 |
| 10 | 27379044 | rs10764670 | -0.944 | -0.271 | -1.118 | 0.036 | -0.001 | 0.109 | 0.109 | 0.108 | 0.08 | 0.24 | 0.04 | 0.26 | 0.06 | 0.23 |
| 10 | 27381349 | rs2297145 | -1.042 | -0.316 | -1.152 | 0.018 | -0.008 | 0.036 | 0.036 | 1.252 | 0.2 | 0.32 | 0.19 | 0.54 | 0.21 | 0.43 |
| 10 | 27389314 | rs3737056 | 1.326 | -0.236 | -1.132 | -0.004 | -0.015 | -0.049 | -0.049 | 0 | 0.08 | 0.01 | 0.12 | 0.28 | 0.11 | 0.17 |
| 10 | 27392541 | rs10764672 | 0.107 | -0.31 | -1.147 | -0.161 | -0.017 | -0.082 | -0.082 | 0 | 0.71 | 0.42 | 0.68 | 0.99 | 0.82 | 0.87 |
| 10 | 27395102 | rs1753415 | -0.367 | -0.326 | -1.189 | -0.161 | 0.023 | 0.069 | 0.069 | 1.171 | 0.76 | 0.89 | 0.72 | 0.99 | 0.83 | 0.88 |
| 10 | 27403635 | rs12782501 | 1.335 | -0.3 | -1.188 | -0.048 | -0.015 | -0.049 | -0.049 | 0 | 0.08 | 0.01 | 0.12 | 0.28 | 0.11 | 0.17 |
| 10 | 27409555 | rs10829190 | -0.297 | -0.355 | -0.994 | -0.161 | 0.014 | 0.01 | 0.01 | 0.641 | 0.75 | 0.83 | 0.71 | 0.99 | 0.83 | 0.87 |
| 10 | 27415906 | rs10829192 | -1.103 | -0.401 | -0.907 | -0.092 | -0.007 | 0.028 | 0.028 | 0.805 | 0.21 | 0.31 | 0.19 | 0.55 | 0.22 | 0.43 |
| 10 | 27418065 | rs11015558 | -0.987 | -0.386 | -0.9 | -0.086 | -0.007 | 0.034 | 0.034 | 0.793 | 0.2 | 0.31 | 0.19 | 0.55 | 0.22 | 0.43 |
| 10 | 27422413 | rs7910688 | -0.987 | -0.386 | -0.898 | -0.086 | -0.007 | 0.036 | 0.036 | 0.793 | 0.2 | 0.31 | 0.19 | 0.55 | 0.22 | 0.43 |
| 10 | 27434483 | rs2274634 | -0.999 | -0.386 | -0.912 | -0.086 | -0.007 | 0.035 | 0.035 | 0.735 | 0.2 | 0.31 | 0.19 | 0.55 | 0.22 | 0.43 |
| 10 | 27441973 | rs1775350 | 0.158 | -0.402 | -0.97 | -0.149 | 0.02 | 0.069 | 0.069 | 1.216 | 0.24 | 0.11 | 0.28 | 0.01 | 0.17 | 0.12 |
| 10 | 27443012 | rs2274636 | 1.439 | -0.401 | -0.994 | -0.1 | -0.015 | -0.048 | -0.048 | 0 | 0.08 | 0.01 | 0.12 | 0.28 | 0.11 | 0.17 |
| 10 | 27444344 | rs7907988 | -0.274 | -0.42 | -0.958 | -0.175 | 0.013 | 0.013 | 0.013 | 0.597 | 0.75 | 0.84 | 0.71 | 0.99 | 0.83 | 0.87 |
| 10 | 27459670 | rs1981296 | 1.098 | -0.276 | -0.815 | -0.052 | 0.001 | 0.001 | 0.001 | 0.238 | 0.54 | 0.52 | 0.52 | 0.44 | 0.6 | 0.43 |
| 10 | 27464600 | rs6482601 | 1.321 | 0.326 | -0.931 | -0.135 | 0 | 0.008 | 0.008 | 1.201 | 0.52 | 0.52 | 0.48 | 0.44 | 0.5 | 0.42 |
| 10 | 27474078 | rs1557169 | 1.065 | -0.212 | -0.852 | -0.081 | 0.003 | 0.001 | 0.001 | 0.238 | 0.54 | 0.53 | 0.52 | 0.46 | 0.6 | 0.43 |
| 10 | 27475494 | rs10764686 | 1.42 | -0.245 | -0.948 | -0.095 | 0.085 | 0.042 | 0.042 | 2 | 0.51 | 0.13 | 0.49 | 0.46 | 0.6 | 0.43 |
| 10 | 27475579 | rs10764687 | -0.096 | -0.034 | -1.07 | -0.082 | 0.01 | 0.04 | 0.04 | 0 | 0.06 | 0.2 | 0.02 | 0.11 | 0.05 | 0.06 |
| 10 | 27477821 | rs882475 | 0.585 | -0.051 | -1.073 | -0.07 | 0.012 | -0.007 | -0.007 | 2.637 | 0.6 | 0.64 | 0.57 | 0.59 | 0.67 | 0.63 |
| 10 | 27483972 | rs788230 | 0.093 | -0.074 | -1.279 | -0.077 | -0.024 | -0.078 | -0.078 | 0.428 | 0.27 | 0.02 | 0.2 | 0.29 | 0.25 | 0.26 |
| 10 | 27484837 | rs11015603 | -0.33 | -0.141 | -1.213 | -0.137 | -0.009 | -0.025 | -0.025 | 0.708 | 0.06 | 0.1 | 0.03 | 0.01 | 0.06 | 0.06 |
| 10 | 27490233 | rs788201 | 0.033 | 0.016 | -1.167 | 0.019 | -0.021 | -0.085 | -0.085 | 0 | 0.26 | 0.03 | 0.21 | 0.29 | 0.25 | 0.26 |
| 10 | 27491870 | rs788203 | 0.046 | 0.017 | -1.166 | 0.019 | -0.021 | -0.085 | -0.085 | 0.121 | 0.26 | 0.02 | 0.2 | 0.29 | 0.25 | 0.26 |
| 10 | 27497191 | rs7918793 | 1.206 | 0.215 | -0.979 | -0.037 | -0.001 | -0.017 | -0.017 | 0.077 | 0.06 | 0.1 | 0.21 | 0.01 | 0.05 | 0.02 |
| 10 | 27508369 | rs7902034 | -0.035 | 0.16 | -1.169 | 0.071 | 0.026 | -0.032 | -0.032 | 0 | 0.71 | 0.88 | 0.76 | 0.67 | 0.73 | 0.7 |
| 10 | 27508758 | rs10764690 | 0.673 | 0.103 | -1.502 | -0.315 | -0.128 | -0.053 | -0.053 | 0 | 0.33 | 0.07 | 0.39 | 0.41 | 0.47 | 0.35 |
| 10 | 27516588 | rs2800403 | 0.021 | 0.385 | -0.919 | 0.148 | 0.021 | -0.039 | -0.039 | 0 | 0.69 | 0.75 | 0.76 | 0.56 | 0.72 | 0.66 |
| 10 | 27519221 | rs2780670 | -0.016 | 0.396 | -1.012 | 0.139 | 0.026 | -0.03 | -0.03 | 0 | 0.71 | 0.87 | 0.76 | 0.67 | 0.72 | 0.7 |
| 10 | 27524480 | rs2780667 | 0.648 | 0.464 | -0.652 | 0.082 | -0.007 | -0.004 | -0.004 | 0 | 0.58 | 0.51 | 0.52 | 0.42 | 0.63 | 0.43 |
| 10 | 27540461 | rs11015625 | 0.728 | 0.713 | -0.519 | 0.107 | -0.038 | 0.034 | 0.034 | 0 | 0.06 | 0 | 0.07 | 0 | 0.12 | 0.04 |
| 10 | 27549961 | rs651134 | 0.519 | 0.31 | 0.034 | 0.309 | -0.033 | 0.213 | 0.213 | 2.183 | 0.9 | 0.37 | 0.94 | 0.76 | 0.96 | 0.73 |
| 10 | 27560120 | rs10829219 | 1.107 | 0.579 | -0.865 | 0.093 | -0.027 | 0.071 | 0.071 | 0.968 | 0.51 | 0.13 | 0.43 | 0.41 | 0.51 | 0.41 |
| 10 | 27563961 | rs12217470 | 1.19 | 0.814 | -1.094 | 0.198 | -0.036 | -0.086 | -0.086 | 1.014 | 0.2 | 0.02 | 0.21 | 0.26 | 0.19 | 0.25 |
| 10 | 27566971 | rs532680 | 0.251 | 1.438 | -0.285 | 0.522 | -0.002 | 0.083 | 0.083 | 1.923 | 0.16 | 0.03 | 0.09 | 0 | 0.14 | 0.1 |
| 10 | 27569521 | rs2483499 | 0.228 | 1.943 | -0.017 | 1.058 | -0.017 | -0.059 | -0.059 | 0.019 | 0.1 | 0.01 | 0.13 | 0.33 | 0.13 | 0.16 |
| 10 | 27572440 | rs17748862 | 0.218 | 2.213 | 0.28 | 0.383 | 0.05 | -0.033 | -0.033 | 0 | 0.1 | 0.16 | 0.26 | 0.02 | 0.17 | 0.04 |
| 10 | 27577568 | rs2780686 | 0.264 | 2.117 | -0.174 | 0.253 | -0.04 | 0.055 | 0.055 | 0 | 0.88 | 0.97 | 0.89 | 1 | 0.82 | 0.92 |
| 10 | 27586972 | rs11015657 | 1.371 | 1.306 | 0.155 | 0.626 | 0.003 | 0.069 | 0.069 | 0 | 0.54 | 0.28 | 0.35 | 0.4 | 0.44 | 0.45 |
| 10 | 27616543 | rs11015696 | -1.156 | 0.678 | 0.499 | 1.361 | -0.013 | 0.056 | 0.056 | 0 | 0.11 | 0.13 | 0.14 | 0.3 | 0.09 | 0.24 |
| 10 | 27617789 | rs237591 | 0.844 | 1.07 | 0.116 | 0.789 | -0.051 | 0.059 | 0.059 | 0 | 0.87 | 0.97 | 0.91 | 1 | 0.81 | 0.89 |
| 10 | 27624562 | rs11015701 | -1.49 | 0.95 | 0.719 | 0.986 | -0.031 | 0.063 | 0.063 | 0 | 0.15 | 0.37 | 0.18 | 0.21 | 0.13 | 0.23 |
| 10 | 27629880 | rs237597 | -0.077 | 1.171 | 0.571 | 0.938 | -0.041 | 0.034 | 0.034 | 0 | 0.93 | 0.99 | 0.93 | 1 | 0.84 | 0.93 |
| 10 | 27632128 | rs7078512 | -1.64 | 0.767 | 1.159 | 1.038 | -0.03 | 0.095 | 0.095 | 0 | 0.16 | 0.5 | 0.18 | 0.21 | 0.13 | 0.23 |
| 10 | 27633772 | rs10490914 | -1.338 | 0.809 | 0.863 | 0.685 | -0.015 | -0.077 | -0.077 | 0 | 0.13 | 0.3 | 0.14 | 0.01 | 0.12 | 0.06 |
| 10 | 27637918 | rs500981 | 1.119 | 0.887 | 0.677 | 0.822 | 0.018 | -0.055 | -0.055 | 0.364 | 0.64 | 0.53 | 0.45 | 0.22 | 0.61 | 0.38 |
| 10 | 27648069 | rs10829248 | -0.183 | 0.871 | -0.038 | 1.548 | -0.002 | 0.012 | 0.012 | 0 | 0.06 | 0 | 0.1 | 0.39 | 0.04 | 0.12 |
| 10 | 27648950 | rs11015719 | -0.727 | 0.835 | 0.131 | 1.532 | -0.004 | 0.069 | 0.069 | 0 | 0.08 | 0.09 | 0.1 | 0.39 | 0.04 | 0.12 |
| 10 | 27654517 | rs237568 | 1.308 | 0.796 | 0.249 | 1.518 | -0.026 | -0.008 | -0.008 | 0 | 0.63 | 0.74 | 0.58 | 0.41 | 0.57 | 0.52 |
| 10 | 27659316 | rs118917 | -0.406 | 0.7 | 0.612 | 1.248 | 0.005 | -0.033 | -0.033 | 0.015 | 0.08 | 0.25 | 0.07 | 0.01 | 0.08 | 0.05 |
| 10 | 27661533 | rs237577 | 0.414 | 1.117 | 0.046 | 1.512 | -0.021 | -0.002 | -0.002 | 0 | 0.77 | 0.83 | 0.77 | 0.77 | 0.7 | 0.74 |
| 10 | 27664646 | rs237579 | 0.596 | 1.072 | 0.025 | 1.339 | -0.023 | -0.043 | -0.043 | 0 | 0.75 | 0.83 | 0.7 | 0.58 | 0.7 | 0.57 |
| 10 | 27667087 | rs10490915 | -1.01 | 0.629 | 0.215 | 1.639 | -0.005 | 0.08 | 0.08 | 0 | 0.07 | 0.09 | 0.1 | 0.38 | 0.03 | 0.12 |
| 10 | 27673128 | rs16927796 | -0.854 | 0.33 | 0.701 | 0.926 | -0.004 | -0.04 | -0.04 | 0.245 | 0.08 | 0.29 | 0.18 | 0.01 | 0.03 | 0.03 |
| 10 | 27674288 | rs493965 | 1.151 | 1.346 | 0.695 | 0.99 | 0.047 | 0.02 | 0.02 | 0 | 0.74 | 0.51 | 0.51 | 0.38 | 0.68 | 0.64 |
| 10 | 27678534 | rs506659 | 1.295 | 1.28 | 1.266 | 1.428 | 0.052 | 0.074 | 0.074 | 0 | 0.75 | 0.49 | 0.55 | 0.6 | 0.69 | 0.84 |
| 10 | 27679816 | rs663263 | 1.281 | 1.216 | 1.456 | 1.436 | 0.039 | 0.017 | 0.017 | 0 | 0.73 | 0.49 | 0.51 | 0.37 | 0.68 | 0.65 |

| | | | | | | | | | | | | | | | | |
|---|---|---|---|---|---|---|---|---|---|---|---|---|---|---|---|---|
| 10 | 27681526 | rs7100677 | -1.435 | 1.368 | 1.595 | 1.441 | 0.013 | 0.04 | 0.04 | 0 | 0.24 | 0.33 | 0.42 | 0.4 | 0.27 | 0.15 |
| 10 | 27682273 | rs603872 | 0.761 | 0.976 | 0.46 | 0.147 | 0 | -0.005 | -0.005 | 0 | 0.33 | 0.32 | 0.32 | 0.17 | 0.37 | 0.33 |
| 10 | 27683028 | rs617297 | 0.693 | 0.991 | 0.542 | 0.226 | 0.027 | 0.01 | 0.01 | 0 | 0.73 | 0.46 | 0.52 | 0.37 | 0.68 | 0.65 |
| 10 | 27687225 | rs2505323 | 0.594 | 0.995 | 0.535 | 0.228 | 0.043 | 0.019 | 0.019 | 0 | 0.73 | 0.53 | 0.52 | 0.37 | 0.68 | 0.65 |
| 10 | 27687965 | rs2505327 | 0.594 | 0.978 | 0.567 | 0.217 | 0.043 | 0.019 | 0.019 | 0 | 0.73 | 0.53 | 0.52 | 0.37 | 0.68 | 0.65 |
| 10 | 27687994 | rs2484173 | 0.594 | 0.978 | 0.567 | 0.217 | 0.043 | 0.019 | 0.019 | 0 | 0.73 | 0.53 | 0.52 | 0.37 | 0.68 | 0.65 |
| 10 | 27688109 | rs2429485 | 0.594 | 0.978 | 0.567 | 0.217 | 0.043 | 0.019 | 0.019 | 0 | 0.73 | 0.53 | 0.52 | 0.37 | 0.68 | 0.65 |
| 10 | 27688460 | rs2505328 | 0.15 | 0.675 | 0.066 | -0.164 | -0.004 | 0.038 | 0.038 | 0 | 0.52 | 0.39 | 0.4 | 0.37 | 0.54 | 0.59 |
| 10 | 27694888 | rs2429498 | 0.494 | 0.241 | 0.203 | -0.195 | -0.008 | 0.019 | 0.019 | 0 | 0.38 | 0.42 | 0.41 | 0.54 | 0.39 | 0.39 |
| 10 | 27696561 | rs12359622 | 1.805 | 0.46 | -0.046 | -0.32 | -0.031 | 0.114 | 0.114 | 0.385 | 0.23 | 0.05 | 0.12 | 0 | 0.2 | 0.15 |
| 10 | 27700887 | rs2429499 | 0.035 | 0.455 | 0.235 | -0.271 | 0.048 | -0.122 | -0.122 | 0 | 0.79 | 0.58 | 0.67 | 0.98 | 0.72 | 0.89 |
| 10 | 27701920 | rs11015752 | 0.655 | 0.439 | 0.1 | -0.058 | 0 | 0.036 | 0.036 | 0 | 0.06 | 0.02 | 0.11 | 0.36 | 0.04 | 0.09 |
| 10 | 27704360 | rs7915825 | 1.005 | 0.294 | 0.377 | -0.23 | 0.013 | -0.029 | -0.029 | 0.611 | 0.11 | 0.12 | 0.25 | 0.01 | 0.2 | 0.05 |
| 10 | 27713525 | rs2477341 | 0.672 | 0.343 | 0.251 | -0.187 | 0.001 | 0.001 | 0.001 | 0 | 0.38 | 0.38 | 0.42 | 0.75 | 0.39 | 0.41 |
| 10 | 27723033 | rs7342136 | 0.454 | 0.315 | -0.018 | -0.141 | 0.003 | -0.054 | -0.054 | 0 | 0.41 | 0.17 | 0.27 | 0.44 | 0.34 | 0.51 |
| 10 | 27734691 | rs17681969 | -0.116 | 0.383 | 0.588 | 0.095 | 0.011 | -0.059 | -0.059 | 1.435 | 0.17 | 0.26 | 0.27 | 0.01 | 0.22 | 0.08 |
| 10 | 27734720 | rs12572763 | 0.006 | 0.241 | 0.366 | 0.241 | 0.002 | 0.048 | 0.048 | 0 | 0.06 | 0.03 | 0.1 | 0.37 | 0.05 | 0.1 |
| 10 | 27738002 | rs2484204 | 1.279 | 0.128 | 0.637 | -0.175 | -0.007 | 0.045 | 0.045 | 1.532 | 0.55 | 0.26 | 0.43 | 0.39 | 0.55 | 0.48 |
| 10 | 27740959 | rs660498 | -1.046 | 0.376 | 0.503 | -0.461 | 0.016 | -0.131 | -0.131 | 1.335 | 0.35 | 0.48 | 0.39 | 0.02 | 0.39 | 0.25 |
| 10 | 27751896 | rs652645 | -0.638 | 0.224 | 0.217 | -0.22 | 0.005 | 0.01 | 0.01 | 0 | 0.85 | 0.87 | 0.93 | 0.98 | 0.87 | 0.84 |
| 10 | 27768083 | rs661882 | -0.147 | 0.344 | 0.471 | -0.341 | -0.001 | -0.026 | -0.026 | 0 | 0.18 | 0.22 | 0.28 | 0.01 | 0.24 | 0.08 |
| 10 | 27775770 | rs7475332 | -0.544 | 0.262 | 0.949 | 0.735 | 0.028 | 0.053 | 0.053 | 0.134 | 0.19 | 0.39 | 0.31 | 0.23 | 0.24 | 0.19 |
| 10 | 27776411 | rs2505331 | -0.12 | 0.028 | 1.193 | 0.337 | -0.014 | 0.016 | 0.016 | 0 | 0.77 | 0.7 | 0.79 | 0.65 | 0.81 | 0.74 |
| 10 | 27783076 | rs2477333 | 0.96 | 0.103 | 0.99 | 0.444 | 0.003 | 0.031 | 0.031 | 2.816 | 0.56 | 0.3 | 0.45 | 0.37 | 0.56 | 0.53 |
| 10 | 27792089 | rs10764698 | 0.61 | -0.127 | 0.709 | 0.158 | -0.002 | 0 | 0 | 1.346 | 0.48 | 0.5 | 0.39 | 0.44 | 0.49 | 0.49 |
| 10 | 27812418 | rs10829268 | 0.596 | -0.115 | 0.754 | 0.179 | -0.001 | 0 | 0 | 0.632 | 0.48 | 0.48 | 0.39 | 0.46 | 0.49 | 0.49 |
| 10 | 27815665 | rs2642273 | 0.134 | -0.082 | 0.56 | 0.179 | -0.029 | 0.053 | 0.053 | 0 | 0.28 | 0.16 | 0.4 | 0.21 | 0.32 | 0.24 |
| 10 | 27829355 | rs2477317 | 1.987 | -0.161 | 0.428 | 0.28 | 0.012 | -0.018 | -0.018 | 0 | 0.12 | 0.02 | 0.13 | 0.19 | 0.09 | 0.16 |
| 10 | 27831551 | rs7074813 | 0.023 | -0.062 | 0.846 | 0.413 | -0.006 | 0.006 | 0.006 | 0 | 0.23 | 0.23 | 0.21 | 0.34 | 0.19 | 0.27 |
| 10 | 27840836 | rs2763301 | 0.487 | -0.018 | 0.99 | 0.281 | -0.004 | 0.008 | 0.008 | 0 | 0.29 | 0.28 | 0.41 | 0.2 | 0.32 | 0.24 |
| 10 | 27847941 | rs1874085 | -1.088 | 0.024 | 0.846 | 0.224 | -0.013 | 0.066 | 0.066 | 1.486 | 0.55 | 0.68 | 0.66 | 0.81 | 0.52 | 0.67 |
| 10 | 27851166 | rs2763299 | 0.706 | -0.2 | 0.876 | 0.364 | -0.011 | 0.01 | 0.01 | 0 | 0.36 | 0.33 | 0.51 | 0.21 | 0.49 | 0.28 |
| 10 | 27855762 | rs2763298 | -0.883 | -0.209 | 0.391 | 0.217 | 0.006 | 0.025 | 0.025 | 0.227 | 0.63 | 0.71 | 0.76 | 0.81 | 0.68 | 0.7 |
| 10 | 27857574 | rs903281 | -0.021 | 0.005 | 0.001 | 0.348 | 0 | -0.001 | -0.001 | 0 | 0.23 | 0.21 | 0.21 | 0.35 | 0.19 | 0.24 |
| 10 | 27860061 | rs12358998 | 1.56 | -0.28 | -0.416 | 0.235 | -0.074 | 0.128 | 0.128 | 0.292 | 0.21 | 0.01 | 0.14 | 0.02 | 0.21 | 0.15 |
| 10 | 27862968 | rs2254027 | -0.506 | -0.639 | 0.146 | 0.341 | 0.001 | -0.001 | -0.001 | 0 | 0.64 | 0.68 | 0.78 | 0.84 | 0.66 | 0.71 |
| 10 | 27865573 | rs4598563 | 0.486 | -0.63 | 0.293 | 0.618 | 0.005 | 0.021 | 0.021 | 0.002 | 0.3 | 0.34 | 0.3 | 0.49 | 0.34 | 0.28 |
| 10 | 27867209 | rs2763309 | 0.074 | -0.717 | 0.15 | 0.533 | -0.009 | 0.008 | 0.008 | 0 | 0.68 | 0.42 | 0.7 | 0.54 | 0.76 | 0.53 |
| 10 | 27873733 | rs12771357 | 0.08 | -0.506 | 0.093 | 0.596 | -0.027 | -0.035 | -0.035 | 0 | 0.26 | 0.41 | 0.19 | 0.17 | 0.21 | 0.25 |
| 10 | 27875006 | rs1494209 | -1.065 | -0.487 | -0.203 | 0.594 | -0.086 | 0.176 | 0.176 | 0 | 0.58 | 0.85 | 0.57 | 0.78 | 0.55 | 0.68 |
| 10 | 27883558 | rs10734066 | -1.064 | -0.436 | -0.182 | 0.642 | -0.083 | 0.176 | 0.176 | 0 | 0.58 | 0.84 | 0.57 | 0.78 | 0.55 | 0.67 |
| 10 | 27888928 | rs7075864 | 0.831 | -0.443 | -0.139 | 0.672 | -0.016 | -0.016 | -0.016 | 0.835 | 0.28 | 0.17 | 0.27 | 0.34 | 0.33 | 0.26 |
| 10 | 27889818 | rs1494201 | -1.01 | -0.297 | -0.498 | 0.351 | -0.082 | 0.178 | 0.178 | 0 | 0.58 | 0.84 | 0.57 | 0.78 | 0.55 | 0.67 |
| 10 | 27890831 | rs7069985 | -0.216 | -0.191 | -0.015 | 0.354 | -0.029 | -0.038 | -0.038 | 0 | 0.26 | 0.43 | 0.2 | 0.17 | 0.21 | 0.25 |
| 10 | 27898930 | rs749333 | 0.091 | 0.101 | 0.236 | 0.686 | 0.003 | 0.073 | 0.073 | 0.165 | 0.3 | 0.48 | 0.28 | 0.38 | 0.3 | 0.38 |
| 10 | 27901470 | rs10829290 | 0.407 | 0.106 | 0.185 | 0.78 | -0.007 | -0.022 | -0.022 | 0 | 0.27 | 0.18 | 0.23 | 0.26 | 0.29 | 0.35 |
| 10 | 27903074 | rs10458709 | 0.412 | 0.077 | 0.269 | 0.743 | -0.013 | -0.033 | -0.033 | 0 | 0.27 | 0.17 | 0.23 | 0.26 | 0.28 | 0.34 |
| 10 | 27903259 | rs10764725 | 0.614 | 0.062 | 0.225 | 0.676 | -0.005 | -0.021 | -0.021 | 0 | 0.23 | 0.17 | 0.21 | 0.26 | 0.24 | 0.31 |
| 10 | 27904052 | rs1690690 | 0.156 | -0.167 | 0.528 | 0.164 | -0.001 | -0.003 | -0.003 | 0 | 0.59 | 0.44 | 0.68 | 0.75 | 0.59 | 0.57 |
| 10 | 27904646 | rs2815572 | 0.808 | -0.311 | 0.153 | -0.028 | -0.001 | -0.008 | -0.008 | 0 | 0.36 | 0.32 | 0.41 | 0.5 | 0.38 | 0.33 |
| 10 | 27906212 | rs1908274 | 0.402 | -0.431 | 0.264 | 0.057 | -0.002 | 0.163 | 0.163 | 0 | 0.06 | 0.15 | 0.22 | 0.34 | 0.06 | 0.12 |
| 10 | 27910782 | rs2642306 | 1.245 | -0.481 | 0.237 | 0.033 | -0.012 | 0.048 | 0.048 | 0 | 0.3 | 0.19 | 0.19 | 0.16 | 0.32 | 0.21 |
| 10 | 27913243 | rs1168579 | 0.077 | -0.565 | 0.311 | -0.045 | -0.01 | 0.025 | 0.025 | 0 | 0.49 | 0.25 | 0.49 | 0.5 | 0.5 | 0.57 |
| 10 | 27914696 | rs1168581 | -0.085 | -0.542 | 0.695 | 0.023 | -0.004 | -0.026 | -0.026 | 0 | 0.81 | 0.55 | 0.87 | 0.88 | 0.84 | 0.86 |
| 10 | 27917124 | rs10829295 | -1.03 | -0.519 | 0.672 | -0.165 | 0.013 | 0.317 | 0.317 | 0 | 0.11 | 0.37 | 0.26 | 0.47 | 0.1 | 0.17 |
| 10 | 27919931 | rs997750 | 0.074 | -0.408 | 0.533 | -0.227 | 0 | -0.001 | -0.001 | 0 | 0.43 | 0.57 | 0.46 | 0.62 | 0.42 | 0.38 |
| 10 | 27923457 | rs2815523 | 0.065 | -0.408 | 0.533 | -0.227 | 0 | -0.001 | -0.001 | 0.278 | 0.43 | 0.57 | 0.46 | 0.62 | 0.42 | 0.38 |
| 10 | 27928326 | rs1168587 | 0.728 | -0.482 | 0.283 | -0.216 | -0.044 | 0.173 | 0.173 | 0 | 0.42 | 0.19 | 0.38 | 0.14 | 0.45 | 0.37 |
| 10 | 27930818 | rs1690678 | -0.558 | -0.562 | 0.42 | -0.216 | -0.019 | 0.082 | 0.082 | 0 | 0.83 | 0.71 | 0.83 | 0.77 | 0.87 | 0.74 |
| 10 | 27934698 | rs2815538 | -0.073 | -0.526 | 0.423 | -0.221 | -0.001 | -0.031 | -0.031 | 0 | 0.44 | 0.53 | 0.53 | 0.86 | 0.43 | 0.55 |
| 10 | 27934905 | rs11015933 | 0.628 | -0.527 | 0.422 | -0.182 | -0.004 | 0.059 | 0.059 | 0 | 0.06 | 0.02 | 0.21 | 0.47 | 0.07 | 0.13 |
| 10 | 27936242 | rs2642279 | -0.066 | -0.508 | 0.44 | -0.647 | -0.001 | -0.031 | -0.031 | 0 | 0.44 | 0.53 | 0.53 | 0.86 | 0.43 | 0.55 |
| 10 | 27942642 | rs1687724 | -1.145 | -0.469 | 0.415 | -0.655 | 0.003 | -0.085 | -0.085 | 0 | 0.86 | 0.73 | 0.9 | 1 | 0.87 | 0.92 |
| 10 | 27944386 | rs1687721 | 0.721 | -0.441 | 0.056 | -0.635 | -0.041 | 0.171 | 0.171 | 0 | 0.42 | 0.17 | 0.36 | 0.14 | 0.44 | 0.37 |
| 10 | 27950367 | rs2007516 | 1.063 | -0.512 | 0.162 | -0.603 | -0.004 | 0.025 | 0.025 | 0 | 0.32 | 0.25 | 0.2 | 0.15 | 0.33 | 0.21 |
| 10 | 27957785 | rs2172086 | 0.42 | -0.757 | -0.238 | -0.772 | 0.027 | 0.079 | 0.079 | 0 | 0.4 | 0.3 | 0.36 | 0.13 | 0.34 | 0.34 |
| 10 | 27959429 | rs2815545 | 1.221 | -0.705 | -0.157 | -0.802 | -0.01 | 0.016 | 0.016 | 0.881 | 0.32 | 0.25 | 0.24 | 0.2 | 0.37 | 0.25 |
| 10 | 27960532 | rs2815548 | 0.218 | -0.684 | -0.138 | -0.772 | 0.007 | -0.028 | -0.028 | 0 | 0.72 | 0.55 | 0.6 | 0.31 | 0.71 | 0.59 |
| 10 | 27960822 | rs2637301 | -1.377 | -0.685 | 0.147 | -0.803 | 0.016 | -0.082 | -0.082 | 0.34 | 0.83 | 0.76 | 0.89 | 1 | 0.81 | 0.91 |

| | | | | | | | | | | | | | | | | |
|---|---|---|---|---|---|---|---|---|---|---|---|---|---|---|---|---|
| 10 | 27962714 | rs2253055 | 0.6 | -1.352 | -0.222 | -0.482 | 0.006 | 0.099 | 0.099 | 0 | 0.27 | 0.12 | 0.26 | 0.1 | 0.21 | 0.27 |
| 10 | 27963246 | rs2492909 | -0.574 | -0.951 | -0.02 | -0.225 | 0.008 | -0.004 | -0.004 | 0.59 | 0.49 | 0.49 | 0.64 | 0.81 | 0.42 | 0.64 |
| 10 | 27963892 | rs2130809 | 0.172 | -0.859 | -0.708 | -0.26 | 0.03 | 0.094 | 0.094 | 0 | 0.4 | 0.25 | 0.34 | 0.12 | 0.34 | 0.34 |
| 10 | 27964470 | rs2253230 | 0.237 | -0.813 | -0.721 | -0.241 | 0.031 | 0.086 | 0.086 | 0 | 0.4 | 0.3 | 0.35 | 0.12 | 0.34 | 0.34 |
| 10 | 27965984 | rs2815552 | -0.045 | -0.563 | -1.417 | 0.307 | 0.086 | -0.005 | -0.005 | 0 | 0.52 | 0.15 | 0.52 | 0.63 | 0.65 | 0.45 |
| 10 | 27968383 | rs2815554 | 0.488 | 0.462 | -1.424 | 0.054 | -0.182 | 0.136 | 0.136 | 1.009 | 0.35 | 0.03 | 0.24 | 0.21 | 0.43 | 0.28 |
| 10 | 27974779 | rs2368251 | 0.663 | 0.546 | 0.267 | 0.778 | 0.036 | 0.015 | 0.015 | 0 | 0.94 | 0.77 | 0.96 | 0.98 | 0.95 | 0.91 |
| 10 | 27978030 | rs2815560 | 1.095 | 0.897 | 0.178 | 1.444 | 0.019 | 0.061 | 0.061 | 0 | 0.86 | 0.81 | 0.85 | 0.67 | 0.82 | 0.76 |
| 10 | 27987627 | rs2815569 | 1.379 | 0.62 | 0.859 | 1.746 | 0.018 | 0.065 | 0.065 | 0 | 0.83 | 0.74 | 0.84 | 0.67 | 0.81 | 0.76 |
| 10 | 27988298 | rs5024546 | -0.649 | 0.688 | 0.874 | 1.781 | 0.003 | 0.054 | 0.054 | 0 | 0.08 | 0.07 | 0.11 | 0.33 | 0.11 | 0.18 |
| 10 | 28001001 | rs11015961 | -1.854 | 0.155 | 1.008 | 1.69 | -0.029 | 0.229 | 0.229 | 0 | 0.19 | 0.54 | 0.18 | 0.5 | 0.12 | 0.32 |
| 10 | 28003559 | rs2815477 | 2.181 | 0.216 | 1.224 | 1.668 | -0.007 | 0.07 | 0.07 | 0.21 | 0.77 | 0.36 | 0.78 | 0.5 | 0.8 | 0.63 |
| 10 | 28004725 | rs11015964 | -0.583 | 0.218 | 1.118 | 1.748 | -0.002 | 0.014 | 0.014 | 0 | 0.1 | 0.08 | 0.11 | 0.33 | 0.11 | 0.2 |
| 10 | 28005782 | rs2815478 | 2.363 | 0.215 | 1.287 | 1.739 | -0.005 | 0.068 | 0.068 | 0.117 | 0.77 | 0.39 | 0.75 | 0.42 | 0.8 | 0.62 |
| 10 | 28016082 | rs12358501 | 0.91 | 1.145 | 0.836 | 1.249 | -0.054 | 0.003 | 0.003 | 0 | 0.1 | 0.01 | 0.12 | 0.08 | 0.18 | 0.02 |
| 10 | 28017572 | rs7358151 | -1.395 | 1.161 | 1.053 | 1.307 | 0.026 | 0.294 | 0.294 | 4.372 | 0.06 | 0.41 | 0.11 | 0.17 | 0.01 | 0.13 |
| 10 | 28020768 | rs2637306 | -0.123 | 1.288 | 0.67 | 1.193 | 0.008 | -0.032 | -0.032 | 0 | 0.94 | 0.9 | 0.97 | 1 | 0.92 | 0.95 |
| 10 | 28027163 | rs10508728 | 0.82 | 1.512 | 0.049 | 0.651 | -0.079 | 0.133 | 0.133 | 1.457 | 0.2 | 0.02 | 0.14 | 0 | 0.23 | 0.07 |
| 10 | 28040823 | rs2637317 | -0.024 | 1.224 | 0.915 | 1.402 | 0.011 | -0.033 | -0.033 | 0 | 0.94 | 0.89 | 0.97 | 1 | 0.92 | 0.95 |
| 10 | 28050840 | rs11006706 | -0.489 | 1.258 | 0.732 | 1.703 | -0.006 | 0.012 | 0.012 | 0 | 0.1 | 0.05 | 0.18 | 0.37 | 0.11 | 0.21 |
| 10 | 28054040 | rs2246842 | 1.505 | 1.184 | 0.92 | 1.641 | 0.012 | 0.11 | 0.11 | 0 | 0.81 | 0.68 | 0.78 | 0.56 | 0.81 | 0.75 |
| 10 | 28071715 | rs2815492 | 1.883 | 1.123 | 0.833 | 1.55 | -0.005 | 0.067 | 0.067 | 0 | 0.77 | 0.48 | 0.64 | 0.38 | 0.8 | 0.61 |
| 10 | 28085825 | rs10826348 | -0.529 | 1.185 | 0.71 | 1.646 | -0.004 | 0.02 | 0.02 | 0 | 0.09 | 0.05 | 0.18 | 0.37 | 0.11 | 0.22 |
| 10 | 28087929 | rs2782336 | 1.647 | 1.139 | 0.863 | 1.586 | 0.01 | 0.123 | 0.123 | 0 | 0.81 | 0.65 | 0.77 | 0.55 | 0.81 | 0.75 |
| 10 | 28093718 | rs2782334 | 1.579 | 1.021 | 0.762 | 1.683 | -0.003 | 0.099 | 0.099 | 0 | 0.82 | 0.7 | 0.83 | 0.56 | 0.83 | 0.77 |
| 10 | 28100491 | rs9651357 | -0.832 | 1.004 | 0.771 | 1.709 | -0.005 | 0.158 | 0.158 | 0 | 0.1 | 0.21 | 0.15 | 0.38 | 0.1 | 0.22 |
| 10 | 28101311 | rs1061577 | 0.39 | 1.225 | 0.298 | 1.217 | -0.04 | 0.042 | 0.042 | 0 | 0.09 | 0 | 0.08 | 0 | 0.15 | 0.07 |
| 10 | 28107057 | rs1931888 | 1.032 | 1.079 | 0.611 | 1.473 | -0.002 | 0.106 | 0.106 | 0 | 0.83 | 0.72 | 0.83 | 0.55 | 0.83 | 0.77 |
| 10 | 28121137 | rs11006736 | 1.162 | 1.563 | 0.59 | 1.327 | 0.03 | 0.1 | 0.1 | 0 | 0.81 | 0.71 | 0.76 | 0.51 | 0.74 | 0.71 |
| 10 | 28133529 | rs915174 | 1.064 | 1.411 | 0.719 | 1.539 | 0.035 | 0.089 | 0.089 | 0 | 0.8 | 0.7 | 0.69 | 0.56 | 0.73 | 0.69 |
| 10 | 28133687 | rs915175 | 1.146 | 1.379 | 0.723 | 1.527 | 0.032 | 0.088 | 0.088 | 0 | 0.79 | 0.69 | 0.68 | 0.55 | 0.73 | 0.68 |
| 10 | 28141107 | rs11006744 | 1.353 | 0.99 | 0.772 | 1.162 | 0.001 | 0.007 | 0.007 | 0 | 0.69 | 0.4 | 0.57 | 0.41 | 0.66 | 0.57 |
| 10 | 28141445 | rs10826354 | 0.496 | 0.729 | 0.239 | 1.106 | -0.011 | -0.047 | -0.047 | 4.163 | 0.08 | 0.01 | 0.1 | 0.28 | 0.1 | 0.08 |
| 10 | 28150935 | rs11006747 | 1.393 | 0.82 | 0.88 | 1.024 | 0.003 | -0.015 | -0.015 | 0 | 0.67 | 0.33 | 0.6 | 0.42 | 0.64 | 0.57 |
| 10 | 28156285 | rs2152019 | 1.114 | 0.893 | 1.127 | 1.129 | 0.013 | 0.085 | 0.085 | 0 | 0.77 | 0.66 | 0.73 | 0.53 | 0.72 | 0.69 |
| 10 | 28186106 | rs1413772 | 1.119 | 0.925 | 1.179 | 1.038 | 0.024 | 0.114 | 0.114 | 0 | 0.77 | 0.59 | 0.73 | 0.51 | 0.72 | 0.7 |
| 10 | 28193276 | rs2368274 | 0.124 | 0.954 | 1.248 | 0.99 | -0.002 | 0.024 | 0.024 | 0 | 0.09 | 0.05 | 0.16 | 0.49 | 0.1 | 0.21 |
| 10 | 28193887 | rs880706 | 1.235 | 0.937 | 1.161 | 1.013 | 0.025 | 0.115 | 0.115 | 0 | 0.77 | 0.59 | 0.72 | 0.51 | 0.72 | 0.7 |
| 10 | 28201966 | rs7897553 | 0.53 | 0.705 | 0.831 | 0.595 | 0.009 | 0.012 | 0.012 | 0 | 0.67 | 0.65 | 0.59 | 0.51 | 0.55 | 0.62 |
| 10 | 28208262 | rs6481493 | -0.64 | 0.404 | 1.067 | 0.595 | -0.023 | 0.084 | 0.084 | 0 | 0.12 | 0.39 | 0.15 | 0.19 | 0.09 | 0.15 |
| 10 | 28224563 | rs11006776 | 1.57 | -0.186 | 0.197 | -0.336 | -0.009 | -0.029 | -0.029 | 0.321 | 0.23 | 0.06 | 0.15 | 0.27 | 0.21 | 0.23 |
| 10 | 28225897 | rs2297646 | -0.544 | 0.009 | 0.748 | -0.106 | 0.001 | 0.013 | 0.013 | 0 | 0.74 | 0.75 | 0.65 | 0.51 | 0.68 | 0.7 |
| 10 | 28228037 | rs11006778 | 0.879 | -0.449 | 0.719 | -0.238 | -0.017 | 0.045 | 0.045 | 1.054 | 0.56 | 0.28 | 0.45 | 0.33 | 0.46 | 0.49 |
| 10 | 28228865 | rs7893462 | 0.262 | -0.441 | 0.785 | -0.238 | 0.018 | 0.027 | 0.027 | 0.113 | 0.64 | 0.47 | 0.58 | 0.5 | 0.55 | 0.61 |
| 10 | 28230586 | rs11006779 | 0.556 | -0.436 | 0.67 | -0.236 | -0.014 | -0.023 | -0.023 | 0 | 0.61 | 0.29 | 0.56 | 0.5 | 0.55 | 0.61 |
| 10 | 28238836 | rs7074562 | 0.082 | -0.41 | 0.51 | -0.224 | 0.016 | 0.023 | 0.023 | 0.011 | 0.65 | 0.53 | 0.58 | 0.5 | 0.55 | 0.62 |
| 10 | 28244079 | rs4367844 | -0.721 | -0.48 | 0.526 | -0.243 | -0.035 | 0.069 | 0.069 | 0.035 | 0.13 | 0.38 | 0.16 | 0.19 | 0.09 | 0.15 |
| 10 | 28245890 | rs11596934 | 1.112 | -0.42 | 0.216 | -0.761 | -0.029 | 0.09 | 0.09 | 1.696 | 0.53 | 0.15 | 0.42 | 0.31 | 0.47 | 0.46 |
| 10 | 28254487 | rs4749286 | -0.011 | -0.381 | 0.333 | -0.644 | 0.008 | 0.006 | 0.006 | 0.009 | 0.67 | 0.63 | 0.54 | 0.41 | 0.55 | 0.62 |
| 10 | 28259142 | rs4749290 | 0.733 | -1.472 | 0.284 | -0.867 | -0.025 | 0.069 | 0.069 | 0 | 0.27 | 0.13 | 0.25 | 0.05 | 0.25 | 0.19 |
| 10 | 28270909 | rs12779067 | 0.889 | -1.48 | 0.271 | -0.871 | -0.045 | 0.099 | 0.099 | 0 | 0.26 | 0.08 | 0.25 | 0.05 | 0.25 | 0.18 |
| 10 | 28271283 | rs12415087 | 0.416 | -1.511 | 0.436 | -0.851 | -0.007 | 0.023 | 0.023 | 0 | 0.29 | 0.22 | 0.26 | 0.08 | 0.25 | 0.2 |
| 10 | 28273575 | rs10826379 | -0.948 | -1.939 | 0.525 | -1.011 | 0.036 | 0.04 | 0.04 | 0 | 0.51 | 0.65 | 0.46 | 0.43 | 0.36 | 0.48 |
| 10 | 28286740 | rs7915784 | -0.421 | -2.051 | 0.461 | -0.953 | -0.04 | -0.031 | -0.031 | 0 | 0.13 | 0.36 | 0.11 | 0.1 | 0.06 | 0.13 |
| 10 | 28288057 | rs7097367 | 0.259 | -1.634 | 0.424 | -0.78 | 0.002 | 0.022 | 0.022 | 0 | 0.36 | 0.27 | 0.34 | 0.14 | 0.3 | 0.29 |
| 10 | 28296076 | rs7076276 | -0.33 | -1.569 | 0.425 | -0.673 | -0.014 | 0.041 | 0.041 | 0.003 | 0.12 | 0.2 | 0.1 | 0.29 | 0.06 | 0.2 |
| 10 | 28296673 | rs7095042 | -0.538 | -1.55 | 0.418 | -0.761 | 0.008 | 0.007 | 0.007 | 0 | 0.45 | 0.37 | 0.43 | 0.42 | 0.34 | 0.47 |
| 10 | 28297248 | rs1419365 | 0.227 | -1.492 | 0.059 | -0.97 | 0.001 | -0.063 | -0.063 | 0 | 0.34 | 0.38 | 0.36 | 0.12 | 0.3 | 0.29 |
| 10 | 28298688 | rs4747610 | -0.448 | -1.164 | -0.153 | -0.849 | 0.011 | 0.02 | 0.02 | 0 | 0.48 | 0.57 | 0.49 | 0.4 | 0.35 | 0.48 |
| 10 | 28311369 | rs2275881 | 0.479 | -0.789 | 0.508 | -0.473 | -0.002 | 0.026 | 0.026 | 0 | 0.22 | 0.39 | 0.26 | 0.26 | 0.18 | 0.21 |
| 10 | 28313132 | rs7098777 | 0.584 | -0.766 | 0.575 | -0.463 | 0.001 | 0.027 | 0.027 | 0 | 0.21 | 0.37 | 0.26 | 0.26 | 0.18 | 0.21 |
| 10 | 28314473 | rs10763636 | -0.521 | -0.942 | 0.468 | -0.856 | -0.003 | 0.008 | 0.008 | 0 | 0.25 | 0.38 | 0.29 | 0.72 | 0.26 | 0.48 |
| 10 | 28314957 | rs7906846 | 0.91 | -0.832 | 0.435 | -0.821 | 0.008 | 0.026 | 0.026 | 0 | 0.21 | 0.37 | 0.28 | 0.26 | 0.19 | 0.21 |
| 10 | 28317101 | rs12257232 | -0.394 | -0.799 | 0.243 | -0.922 | -0.002 | -0.001 | -0.001 | 0 | 0.34 | 0.63 | 0.38 | 0.38 | 0.34 | 0.36 |
| 10 | 28319692 | rs2490059 | -0.751 | -0.344 | 0.171 | -0.5 | -0.003 | -0.014 | -0.014 | 0 | 0.26 | 0.27 | 0.17 | 0.04 | 0.33 | 0.15 |
| 10 | 28322859 | rs11006802 | 0.217 | -0.429 | -0.191 | -0.233 | 0.001 | 0.01 | 0.01 | 0 | 0.07 | 0.02 | 0.16 | 0.58 | 0.07 | 0.26 |
| 10 | 28326647 | rs10826385 | 0.369 | -0.39 | -0.283 | -0.215 | -0.001 | 0.019 | 0.019 | 0 | 0.06 | 0.02 | 0.15 | 0.58 | 0.07 | 0.26 |
| 10 | 28326775 | rs2249120 | 0.296 | -0.18 | -0.069 | -0.866 | -0.021 | 0.048 | 0.048 | 0 | 0.15 | 0.07 | 0.09 | 0.02 | 0.18 | 0.11 |
| 10 | 28337017 | rs2253165 | 0.264 | -0.46 | 0.528 | -0.975 | 0 | -0.033 | -0.033 | 0 | 0.45 | 0.42 | 0.37 | 0.12 | 0.43 | 0.4 |

| | | | | | | | | | | | | | | | | |
|---|---|---|---|---|---|---|---|---|---|---|---|---|---|---|---|---|
| 10 | 28337570 | rs12245062 | -2.344 | -0.209 | 1.219 | -0.262 | 0.03 | 0.118 | 0.118 | 0.124 | 0.06 | 0.44 | 0.06 | 0.11 | 0.01 | 0.11 |
| 10 | 28342740 | rs12249305 | -2.23 | -0.192 | 0.988 | -0.258 | 0.017 | 0.091 | 0.091 | 0.01 | 0.05 | 0.38 | 0.06 | 0.11 | 0.01 | 0.11 |
| 10 | 28349519 | rs7919932 | -1.125 | -0.025 | 0.934 | -0.381 | 0.022 | -0.014 | -0.014 | 0 | 0.13 | 0.36 | 0.13 | 0.12 | 0.17 | 0.11 |
| 10 | 28349623 | rs2255018 | -0.168 | -0.551 | 0.743 | 0.112 | -0.009 | -0.008 | -0.008 | 0 | 0.58 | 0.34 | 0.49 | 0.41 | 0.52 | 0.52 |
| 10 | 28356087 | rs11006830 | -0.787 | -0.787 | 1.192 | 0.766 | -0.016 | 0.093 | 0.093 | 0.155 | 0.11 | 0.19 | 0.29 | 0.63 | 0.09 | 0.34 |
| 10 | 28359713 | rs1781837 | -0.349 | -0.768 | 1.061 | 0.676 | -0.004 | 0.009 | 0.009 | 0 | 0.89 | 0.93 | 0.95 | 0.96 | 0.87 | 0.92 |
| 10 | 28375931 | rs7920402 | -0.41 | -0.925 | 1.239 | 0.669 | 0.006 | 0.031 | 0.031 | 0 | 0.06 | 0.26 | 0.1 | 0.07 | 0.02 | 0.05 |
| 10 | 28378613 | rs11006853 | -0.689 | -0.708 | 0.694 | 0.877 | -0.007 | 0.164 | 0.164 | 0 | 0.05 | 0.17 | 0.13 | 0.25 | 0.05 | 0.15 |
| 10 | 28378758 | rs2997211 | -0.454 | -0.657 | 0.526 | 0.857 | -0.012 | 0.015 | 0.015 | 0 | 0.89 | 0.95 | 0.95 | 0.96 | 0.88 | 0.92 |
| 10 | 28379604 | rs1317790 | -0.371 | -0.692 | 0.493 | 0.883 | 0.013 | 0.007 | 0.007 | 0 | 0.09 | 0.07 | 0.26 | 0.51 | 0.08 | 0.28 |
| 10 | 28382756 | rs2801848 | -0.376 | -0.566 | 0.606 | 0.894 | -0.014 | 0.015 | 0.015 | 0 | 0.89 | 0.95 | 0.95 | 0.96 | 0.88 | 0.88 |
| 10 | 28397812 | rs11006874 | -1.644 | -0.666 | 0.896 | 1.105 | 0.004 | 0.31 | 0.31 | 0.388 | 0.07 | 0.34 | 0.17 | 0.37 | 0.06 | 0.21 |
| 10 | 28410989 | rs11006885 | -0.974 | -0.568 | 0.47 | 1.137 | -0.005 | 0.204 | 0.204 | 0.016 | 0.06 | 0.2 | 0.13 | 0.3 | 0.05 | 0.15 |
| 10 | 28414791 | rs7080796 | -0.907 | -1.137 | 0.395 | 0.986 | -0.039 | 0.096 | 0.096 | 0 | 0.18 | 0.29 | 0.31 | 0.45 | 0.09 | 0.29 |
| 10 | 28417828 | rs12411923 | -0.27 | -1.037 | 0.492 | 1.042 | -0.026 | -0.034 | -0.034 | 0 | 0.1 | 0.22 | 0.07 | 0.04 | 0.04 | 0.05 |
| 10 | 28419760 | rs2997200 | 0.983 | -1.174 | 0.407 | 0.73 | -0.028 | 0.097 | 0.097 | 0 | 0.75 | 0.46 | 0.63 | 0.46 | 0.78 | 0.66 |
| 10 | 28425895 | rs12268470 | -0.587 | -0.397 | 0.192 | 0.787 | 0.017 | 0.138 | 0.138 | 0.634 | 0.06 | 0.15 | 0.25 | 0.35 | 0.12 | 0.16 |
| 10 | 28430660 | rs2985525 | -0.692 | -0.536 | 0.158 | 0.695 | -0.013 | 0.005 | 0.005 | 0 | 0.9 | 0.96 | 0.93 | 0.93 | 0.88 | 0.92 |
| 10 | 28433708 | rs10826406 | -0.893 | -0.965 | 0.488 | 0.524 | -0.044 | 0.091 | 0.091 | 0 | 0.19 | 0.25 | 0.21 | 0.51 | 0.1 | 0.38 |
| 10 | 28440867 | rs10826408 | -0.858 | -0.924 | 0.496 | 0.542 | -0.001 | 0.021 | 0.021 | 0 | 0.14 | 0.19 | 0.29 | 0.36 | 0.16 | 0.19 |
| 10 | 28445503 | rs2997203 | -0.776 | -0.755 | 0.644 | 0.614 | -0.01 | 0.003 | 0.003 | 0 | 0.9 | 0.96 | 0.93 | 0.93 | 0.88 | 0.94 |
| 10 | 28447052 | rs7100675 | -0.057 | -0.828 | 0.804 | 0.665 | -0.031 | 0.247 | 0.247 | 0 | 0.09 | 0.16 | 0.17 | 0.45 | 0.05 | 0.32 |
| 10 | 28453046 | rs2763316 | 0.62 | -0.816 | 0.791 | 0.648 | 0.008 | 0.046 | 0.046 | 0 | 0.83 | 0.75 | 0.67 | 0.57 | 0.75 | 0.8 |
| 10 | 28458902 | rs2763321 | 1.403 | -0.695 | 0.897 | 0.58 | -0.007 | 0.009 | 0.009 | 0 | 0.73 | 0.47 | 0.64 | 0.37 | 0.78 | 0.53 |
| 10 | 28460483 | rs6481505 | -1.084 | -0.446 | 0.707 | 0.598 | 0.04 | 0.115 | 0.115 | 0 | 0.08 | 0.34 | 0.12 | 0.17 | 0.05 | 0.1 |
| 10 | 28462019 | rs7903957 | -0.537 | -0.245 | 0.01 | 0.605 | -0.008 | -0.003 | -0.003 | 0.365 | 0.89 | 0.93 | 0.9 | 0.89 | 0.88 | 0.94 |
| 10 | 28498435 | rs11006920 | -0.158 | 0.357 | -0.292 | 0.359 | 0.01 | 0.088 | 0.088 | 0.361 | 0.05 | 0.1 | 0.25 | 0.31 | 0.13 | 0.14 |
| 10 | 28501345 | rs7081651 | -0.641 | -0.748 | -0.238 | -0.212 | -0.011 | -0.004 | -0.004 | 0.813 | 0.44 | 0.42 | 0.49 | 0.6 | 0.34 | 0.5 |
| 10 | 28501356 | rs7081654 | -0.084 | -0.985 | -0.275 | -0.549 | 0.024 | 0.026 | 0.026 | 0 | 0.28 | 0.23 | 0.15 | 0.07 | 0.17 | 0.13 |
| 10 | 28520798 | rs6481514 | -0.878 | -0.468 | 0.219 | -0.315 | 0.005 | -0.015 | -0.015 | 1.074 | 0.18 | 0.29 | 0.11 | 0.13 | 0.16 | 0.19 |
| 10 | 28525971 | rs11006940 | -0.878 | -0.468 | 0.219 | -0.315 | 0.005 | -0.015 | -0.015 | 0 | 0.18 | 0.29 | 0.11 | 0.13 | 0.16 | 0.19 |
| 10 | 28527917 | rs7916525 | 0.012 | -0.429 | -0.063 | -0.341 | -0.018 | 0.026 | 0.026 | 0 | 0.13 | 0.04 | 0.06 | 0.06 | 0.13 | 0.08 |
| 10 | 28540638 | rs10763654 | -1.409 | -0.814 | 0.202 | -0.25 | -0.037 | 0.039 | 0.039 | 0 | 0.31 | 0.4 | 0.3 | 0.56 | 0.22 | 0.48 |
| 10 | 28552168 | rs11006970 | -0.446 | -0.836 | 0.07 | -0.228 | 0.042 | -0.057 | -0.057 | 0.28 | 0.2 | 0.12 | 0.21 | 0.53 | 0.11 | 0.41 |
| 10 | 28559892 | rs3847384 | -0.975 | -0.783 | 0.344 | -0.256 | -0.004 | -0.031 | -0.031 | 0.908 | 0.21 | 0.37 | 0.12 | 0.13 | 0.17 | 0.21 |
| 10 | 28563732 | rs11006976 | -0.754 | -0.745 | 0.364 | -0.24 | -0.001 | -0.006 | -0.006 | 0 | 0.19 | 0.17 | 0.1 | 0.13 | 0.17 | 0.21 |
| 10 | 28564732 | rs7100335 | -0.006 | -0.68 | -0.048 | -0.182 | -0.033 | 0.033 | 0.033 | 0 | 0.14 | 0.01 | 0.07 | 0.06 | 0.14 | 0.1 |
| 10 | 28568378 | rs11006981 | -0.191 | -0.685 | -0.051 | -0.183 | -0.027 | -0.019 | -0.019 | 0 | 0.16 | 0.01 | 0.07 | 0.13 | 0.15 | 0.17 |
| 10 | 28569959 | rs12256403 | 0.682 | 0.023 | -0.166 | -0.435 | -0.006 | -0.007 | -0.007 | 0 | 0.26 | 0.25 | 0.38 | 0.38 | 0.37 | 0.31 |
| 10 | 28579969 | rs3936497 | -0.165 | -0.142 | -0.442 | -0.374 | -0.017 | 0.075 | 0.075 | 0 | 0.14 | 0.18 | 0.27 | 0.53 | 0.08 | 0.3 |
| 10 | 28584597 | rs1148174 | -1.213 | 0.028 | -0.769 | -0.515 | -0.059 | -0.06 | -0.06 | 0 | 0.77 | 0.9 | 0.68 | 0.67 | 0.7 | 0.71 |
| 10 | 28585496 | rs11006995 | -0.223 | -0.044 | -0.586 | -0.369 | -0.009 | 0.03 | 0.03 | 0 | 0.14 | 0.14 | 0.18 | 0.5 | 0.06 | 0.34 |
| 10 | 28592380 | rs7903242 | 0.063 | -0.02 | -0.575 | -0.369 | -0.012 | 0.04 | 0.04 | 0 | 0.13 | 0.14 | 0.18 | 0.43 | 0.05 | 0.28 |
| 10 | 28596995 | rs3847388 | -0.404 | 0.37 | -0.493 | -0.806 | 0.004 | 0.001 | 0.001 | 0 | 0.46 | 0.39 | 0.37 | 0.55 | 0.35 | 0.55 |
| 10 | 28597192 | rs1148176 | 1.635 | 0.87 | -0.91 | -1.097 | 0.003 | 0.056 | 0.056 | 0 | 0.2 | 0.09 | 0.12 | 0 | 0.19 | 0.11 |
| 10 | 28605526 | rs786399 | -0.075 | 0.95 | -0.62 | -1.145 | 0 | -0.039 | -0.039 | 0 | 0.41 | 0.57 | 0.27 | 0.07 | 0.36 | 0.22 |
| 10 | 28606902 | rs11007018 | 0.786 | 0.888 | -0.804 | -1.291 | -0.053 | 0.057 | 0.057 | 0 | 0.12 | 0.02 | 0.11 | 0.02 | 0.18 | 0.07 |
| 10 | 28609141 | rs786394 | 0.977 | 0.75 | -0.833 | -1.064 | 0.023 | -0.077 | -0.077 | 0 | 0.29 | 0.13 | 0.28 | 0.51 | 0.26 | 0.48 |
| 10 | 28609690 | rs786391 | 0.977 | 0.75 | -0.833 | -1.064 | 0.023 | -0.077 | -0.077 | 0 | 0.29 | 0.13 | 0.28 | 0.51 | 0.26 | 0.48 |
| 10 | 28611563 | rs12254001 | 0.176 | 1.045 | -0.959 | -1.046 | 0.035 | 0.073 | 0.073 | 0 | 0.18 | 0.29 | 0.45 | 0.42 | 0.29 | 0.2 |
| 10 | 28613779 | rs12098807 | 1.151 | 0.404 | -0.291 | -0.887 | -0.03 | -0.047 | -0.047 | 1.708 | 0.23 | 0.25 | 0.09 | 0.05 | 0.09 | 0.11 |
| 10 | 28615573 | rs1148178 | 0.413 | 0.514 | -0.685 | -0.815 | -0.043 | -0.123 | -0.123 | 0 | 0.29 | 0.51 | 0.16 | 0.05 | 0.19 | 0.15 |
| 10 | 28617268 | rs1148184 | -0.529 | 0.765 | -0.935 | -0.655 | 0 | -0.004 | -0.004 | 0 | 0.59 | 0.7 | 0.44 | 0.55 | 0.45 | 0.63 |
| 10 | 28619487 | rs811496 | 0.171 | 0.584 | -0.713 | -0.913 | -0.017 | -0.075 | -0.075 | 0 | 0.3 | 0.55 | 0.17 | 0.05 | 0.2 | 0.18 |
| 10 | 28624009 | rs4747618 | 0.021 | 0.93 | -1.094 | -0.648 | -0.037 | -0.115 | -0.115 | 0 | 0.18 | 0.04 | 0.27 | 0.51 | 0.22 | 0.34 |
| 10 | 28624136 | rs1144518 | -1.198 | 1.579 | -1.07 | -0.774 | 0.001 | -0.002 | -0.002 | 0 | 0.71 | 0.69 | 0.62 | 0.62 | 0.74 | 0.74 |
| 10 | 28632464 | rs1776118 | 1.459 | 1.659 | -0.888 | -0.921 | 0.001 | 0.015 | 0.015 | 0 | 0.22 | 0.16 | 0.12 | 0 | 0.2 | 0.12 |
| 10 | 28633039 | rs1249330 | -0.413 | 1.733 | -0.51 | -0.983 | 0.03 | 0.106 | 0.106 | 0 | 0.23 | 0.35 | 0.34 | 0.52 | 0.31 | 0.43 |
| 10 | 28633450 | rs1776127 | 1.356 | 1.701 | -0.65 | -0.93 | 0.003 | 0.021 | 0.021 | 0 | 0.23 | 0.17 | 0.12 | 0.04 | 0.21 | 0.16 |
| 10 | 28634926 | rs1249328 | -0.737 | 1.842 | -0.389 | -0.395 | -0.011 | -0.01 | -0.01 | 0 | 0.69 | 0.7 | 0.55 | 0.61 | 0.63 | 0.72 |
| 10 | 28637660 | rs7900823 | 0.29 | 1.661 | -0.717 | -0.383 | -0.042 | 0.028 | 0.028 | 0 | 0.08 | 0.01 | 0.09 | 0.03 | 0.16 | 0.06 |
| 10 | 28640915 | rs703027 | -0.346 | 1.671 | -0.434 | -0.25 | -0.001 | 0.015 | 0.015 | 0 | 0.19 | 0.18 | 0.34 | 0.5 | 0.22 | 0.37 |
| 10 | 28641805 | rs703014 | -0.369 | 1.377 | -0.462 | -0.215 | -0.003 | -0.002 | -0.002 | 0 | 0.65 | 0.57 | 0.61 | 0.63 | 0.66 | 0.66 |
| 10 | 28646618 | rs4082298 | -0.046 | 1.132 | -0.646 | 0.049 | 0.013 | 0.029 | 0.029 | 0 | 0.65 | 0.52 | 0.52 | 0.49 | 0.64 | 0.46 |
| 10 | 28649830 | rs10763660 | -0.273 | 1.057 | -0.918 | 0.286 | 0 | -0.002 | -0.002 | 0 | 0.28 | 0.28 | 0.25 | 0.37 | 0.29 | 0.19 |
| 10 | 28651538 | rs1776099 | 0.125 | 1.133 | -0.791 | 0.841 | -0.003 | -0.057 | -0.057 | 0 | 0.18 | 0.09 | 0.22 | 0.39 | 0.2 | 0.18 |
| 10 | 28652865 | rs1313019 | -0.317 | 1.091 | -0.154 | 0.572 | -0.013 | 0.121 | 0.121 | 0.023 | 0.05 | 0.11 | 0.13 | 0.2 | 0.04 | 0.11 |
| 10 | 28652892 | rs11007056 | -0.571 | 1.437 | -0.62 | -0.326 | 0.004 | -0.049 | -0.049 | 0.19 | 0.22 | 0.26 | 0.19 | 0.01 | 0.33 | 0.16 |
| 10 | 28654384 | rs10826459 | -0.582 | 1.612 | -0.492 | -0.101 | 0.012 | -0.051 | -0.051 | 0 | 0.18 | 0.22 | 0.16 | 0.01 | 0.29 | 0.16 |

| | | | | | | | | | | | | | | | | |
|---|---|---|---|---|---|---|---|---|---|---|---|---|---|---|---|---|
| 10 | 28668098 | rs4595437 | -1.013 | 0.51 | -0.143 | 0.527 | -0.026 | 0.017 | 0.017 | 0 | 0.21 | 0.34 | 0.21 | 0.31 | 0.13 | 0.29 |
| 10 | 28671644 | rs10508738 | -1.001 | 0.438 | 0.701 | 0.402 | 0.01 | -0.055 | -0.055 | 0 | 0.15 | 0.48 | 0.11 | 0.1 | 0.1 | 0.13 |
| 10 | 28672502 | rs11007074 | -2.197 | 0.401 | 0.93 | 0.589 | 0.002 | -0.007 | -0.007 | 0.546 | 0.26 | 0.73 | 0.29 | 0.4 | 0.21 | 0.41 |
| 10 | 28672541 | rs11007075 | -0.582 | 0.401 | 1.285 | 0.588 | 0.017 | -0.034 | -0.034 | 0.254 | 0.13 | 0.38 | 0.1 | 0.1 | 0.09 | 0.1 |
| 10 | 28672643 | rs10826461 | -1.83 | 0.468 | 1.362 | 0.668 | 0.012 | 0.11 | 0.11 | 0.15 | 0.13 | 0.32 | 0.18 | 0.3 | 0.13 | 0.31 |
| 10 | 28674644 | rs12249943 | -0.786 | 0.474 | 1.039 | 0.712 | 0.001 | 0.014 | 0.014 | 0 | 0.09 | 0.14 | 0.16 | 0.19 | 0.11 | 0.14 |
| 10 | 28680096 | rs10826464 | -1.894 | 0.579 | 0.639 | 0.551 | -0.003 | 0.015 | 0.015 | 2.961 | 0.18 | 0.7 | 0.24 | 0.3 | 0.17 | 0.31 |
| 10 | 28682973 | rs1984273 | -0.959 | 0.359 | -0.075 | 0.119 | -0.002 | 0.014 | 0.014 | 0.09 | 0.55 | 0.7 | 0.45 | 0.42 | 0.51 | 0.5 |
| 10 | 28688678 | rs734210 | 0.534 | 0.112 | -0.495 | -0.021 | 0.001 | 0.041 | 0.041 | 0 | 0.12 | 0.03 | 0.06 | 0.05 | 0.11 | 0.11 |
| 10 | 28698034 | rs9971071 | -1.168 | 0.466 | -0.558 | 0.163 | -0.098 | -0.073 | -0.073 | 0 | 0.57 | 0.94 | 0.49 | 0.45 | 0.54 | 0.58 |
| 10 | 28702779 | rs7087410 | -0.705 | 0.174 | -0.118 | 0.004 | -0.002 | 0.049 | 0.049 | 0.949 | 0.42 | 0.68 | 0.31 | 0.2 | 0.47 | 0.46 |
| 10 | 28704213 | rs10733990 | -0.573 | 0.211 | -0.597 | 0.463 | 0.026 | -0.027 | -0.027 | 0.046 | 0.61 | 0.7 | 0.62 | 0.52 | 0.71 | 0.72 |
| 10 | 28707263 | rs1249285 | 0.625 | 0.88 | -0.586 | 0.562 | 0.01 | -0.044 | -0.044 | 0.495 | 0.36 | 0.06 | 0.34 | 0.31 | 0.33 | 0.24 |
| 10 | 28709327 | rs9416794 | 0.066 | 1.11 | 0.056 | 0.581 | -0.005 | 0.002 | 0.002 | 0 | 0.43 | 0.53 | 0.53 | 0.46 | 0.42 | 0.41 |
| 10 | 28709550 | rs9416795 | -1.154 | 1.358 | -0.551 | 0.596 | 0.018 | -0.001 | -0.001 | 0.264 | 0.7 | 0.78 | 0.87 | 0.71 | 0.83 | 0.77 |
| 10 | 28716336 | rs1249280 | 0.392 | 1.132 | -0.924 | 0.717 | -0.001 | -0.004 | -0.004 | 0.217 | 0.38 | 0.12 | 0.39 | 0.34 | 0.41 | 0.27 |
| 10 | 28722502 | rs11007107 | 0.074 | 2.197 | -0.515 | 0.655 | -0.026 | -0.017 | -0.017 | 0.057 | 0.05 | 0.01 | 0.17 | 0.25 | 0.15 | 0.2 |
| 10 | 28734355 | rs7085106 | -0.142 | 1.865 | -0.475 | 0.402 | -0.012 | 0.018 | 0.018 | 0 | 0.2 | 0.15 | 0.17 | 0.07 | 0.27 | 0.16 |
| 10 | 28734655 | rs16928895 | -0.878 | 1.495 | 0.34 | 0.731 | 0.085 | 0.048 | 0.048 | 0.064 | 0.11 | 0.35 | 0.21 | 0.16 | 0.14 | 0.24 |
| 10 | 28738098 | rs6481523 | 0.302 | 0.925 | -0.087 | 1.081 | -0.015 | 0.034 | 0.034 | 0 | 0.78 | 0.69 | 0.72 | 0.68 | 0.82 | 0.88 |
| 10 | 28748263 | rs4554782 | 0.392 | 1.633 | -0.804 | 0.847 | -0.101 | -0.111 | -0.111 | 0.366 | 0.7 | 0.92 | 0.57 | 0.58 | 0.57 | 0.7 |
| 10 | 28748824 | rs12762979 | 1.538 | 1.423 | -1.067 | 0.502 | 0.017 | -0.014 | -0.014 | 0 | 0.32 | 0.22 | 0.35 | 0.37 | 0.23 | 0.44 |
| 10 | 28752106 | rs2153299 | 0.86 | 1.499 | -0.949 | 0.361 | -0.016 | -0.012 | -0.012 | 0 | 0.57 | 0.31 | 0.47 | 0.51 | 0.46 | 0.61 |
| 10 | 28757370 | rs2807749 | 0.614 | 1.631 | -0.856 | 0.645 | -0.014 | -0.009 | -0.009 | 0.276 | 0.14 | 0.11 | 0.19 | 0.17 | 0.27 | 0.2 |
| 10 | 28761159 | rs2790441 | 0.724 | 1.551 | -0.93 | 0.593 | -0.019 | -0.015 | -0.015 | 0 | 0.13 | 0.09 | 0.17 | 0.15 | 0.24 | 0.2 |
| 10 | 28761901 | rs2251068 | -0.996 | 1.638 | -0.794 | 0.384 | 0.034 | 0.046 | 0.046 | 0 | 0.52 | 0.72 | 0.46 | 0.36 | 0.63 | 0.43 |
| 10 | 28770018 | rs2790453 | 0.419 | 1.239 | -0.734 | 0.355 | -0.003 | 0.077 | 0.077 | 0.544 | 0.51 | 0.33 | 0.75 | 0.77 | 0.54 | 0.66 |
| 10 | 28775881 | rs11007129 | 0.762 | 1.263 | -0.986 | 1.088 | -0.025 | -0.048 | -0.048 | 0.589 | 0.09 | 0.01 | 0.34 | 0.32 | 0.11 | 0.08 |
| 10 | 28783350 | rs2807741 | 0.467 | 1.205 | -0.732 | 0.438 | 0.003 | -0.005 | -0.005 | 0 | 0.61 | 0.4 | 0.59 | 0.66 | 0.49 | 0.63 |
| 10 | 28786795 | rs11007131 | 0.671 | 1.534 | -1.151 | 0.425 | -0.023 | -0.027 | -0.027 | 0 | 0.17 | 0.14 | 0.26 | 0.26 | 0.28 | 0.26 |
| 10 | 28787794 | rs2790430 | -1.23 | 1.435 | -1.169 | 0.239 | -0.047 | 0.047 | 0.047 | 0 | 0.89 | 0.96 | 0.93 | 0.95 | 0.84 | 0.94 |
| 10 | 28793239 | rs2998285 | -0.217 | 0.506 | -0.992 | 0.206 | -0.009 | 0.01 | 0.01 | 0 | 0.83 | 0.85 | 0.89 | 0.94 | 0.78 | 0.89 |
| 10 | 28800202 | rs11007134 | 0.38 | -0.011 | -1.382 | -0.122 | -0.021 | 0.04 | 0.04 | 0 | 0.23 | 0.08 | 0.12 | 0.13 | 0.21 | 0.23 |
| 10 | 28804787 | rs2790450 | 0.062 | -0.081 | -1.242 | 0.012 | -0.008 | 0.003 | 0.003 | 0 | 0.65 | 0.64 | 0.63 | 0.69 | 0.48 | 0.66 |
| 10 | 28819406 | rs1072523 | 0.054 | -0.074 | -1.232 | 0.088 | -0.012 | 0.003 | 0.003 | 0 | 0.65 | 0.67 | 0.63 | 0.68 | 0.48 | 0.66 |
| 10 | 28821216 | rs2807761 | 0.154 | -0.093 | -1.129 | 0.021 | -0.003 | 0.001 | 0.001 | 0 | 0.64 | 0.6 | 0.63 | 0.69 | 0.49 | 0.66 |
| 10 | 28830108 | rs11007143 | 0.342 | -0.162 | -1.374 | 0.022 | -0.042 | 0.048 | 0.048 | 0 | 0.24 | 0.06 | 0.13 | 0.13 | 0.25 | 0.19 |
| 10 | 28840124 | rs332120 | 0.592 | 0.016 | -1.457 | -0.047 | -0.056 | 0.146 | 0.146 | 0 | 0.22 | 0.01 | 0.09 | 0 | 0.19 | 0.1 |
| 10 | 28848789 | rs2807755 | 0.828 | -0.324 | -1.905 | 0.13 | -0.013 | -0.094 | -0.094 | 0 | 0.13 | 0.01 | 0.3 | 0.32 | 0.12 | 0.16 |
| 10 | 28858889 | rs10490898 | 1.515 | -0.226 | -1.819 | -0.045 | -0.061 | -0.066 | -0.066 | 0 | 0.15 | 0.03 | 0.24 | 0.24 | 0.29 | 0.22 |
| 10 | 28867397 | rs6481534 | 1.533 | -0.227 | -1.82 | -0.046 | -0.061 | -0.066 | -0.066 | 0.842 | 0.15 | 0.03 | 0.24 | 0.24 | 0.29 | 0.22 |
| 10 | 28880160 | rs332153 | 0.331 | -0.233 | -1.745 | -0.046 | -0.035 | 0.047 | 0.047 | 0 | 0.24 | 0.07 | 0.13 | 0.13 | 0.25 | 0.19 |
| 10 | 28883068 | rs332157 | -0.153 | -0.272 | -1.704 | -0.061 | -0.016 | 0.007 | 0.007 | 0 | 0.66 | 0.7 | 0.65 | 0.69 | 0.51 | 0.68 |
| 10 | 28887192 | rs2807754 | 1.092 | -0.293 | -1.54 | -0.075 | -0.024 | 0.005 | 0.005 | 0 | 0.4 | 0.49 | 0.51 | 0.56 | 0.26 | 0.49 |
| 10 | 28910715 | rs332182 | -0.119 | -0.208 | -1.671 | -0.045 | -0.017 | 0.007 | 0.007 | 0 | 0.66 | 0.7 | 0.65 | 0.69 | 0.51 | 0.68 |
| 10 | 28913309 | rs332185 | 0.249 | -0.084 | -1.745 | -0.587 | -0.027 | 0.053 | 0.053 | 0 | 0.26 | 0.08 | 0.13 | 0.13 | 0.26 | 0.2 |
| 10 | 28925701 | rs332114 | -0.556 | 0.257 | -1.009 | -0.506 | 0.001 | -0.022 | -0.022 | 0 | 0.83 | 0.82 | 0.9 | 0.93 | 0.81 | 0.91 |
| 10 | 28928158 | rs332118 | 1.445 | -0.271 | -1.053 | -0.266 | -0.049 | -0.061 | -0.061 | 0.383 | 0.2 | 0.04 | 0.25 | 0.24 | 0.3 | 0.32 |
| 10 | 28929097 | rs1265846 | 0.701 | -0.444 | -0.826 | -0.219 | -0.017 | 0.014 | 0.014 | 0 | 0.4 | 0.48 | 0.51 | 0.5 | 0.26 | 0.41 |
| 10 | 28936753 | rs1780221 | -0.359 | -0.452 | -1.128 | -0.494 | -0.058 | 0.101 | 0.101 | 1.565 | 0.7 | 0.86 | 0.82 | 0.97 | 0.63 | 0.86 |
| 10 | 28939852 | rs11007183 | 0.484 | -0.508 | -1.177 | -0.347 | 0.007 | -0.021 | -0.021 | 0.05 | 0.13 | 0.11 | 0.31 | 0.43 | 0.11 | 0.19 |
| 10 | 28941227 | rs12411515 | 1.371 | -0.553 | -1.256 | -0.378 | -0.062 | -0.065 | -0.065 | 0 | 0.22 | 0.04 | 0.27 | 0.27 | 0.34 | 0.36 |
| 10 | 28943648 | rs1265837 | 1.746 | -0.595 | -1.158 | -0.403 | -0.036 | -0.038 | -0.038 | 0 | 0.27 | 0.48 | 0.22 | 0.21 | 0.15 | 0.26 |
| 10 | 28945223 | rs628590 | 1.284 | -0.332 | -0.637 | -0.207 | 0.002 | 0.002 | 0.002 | 2.06 | 0.49 | 0.51 | 0.48 | 0.49 | 0.49 | 0.62 |
| 10 | 28948379 | rs12571273 | 0.24 | 0.148 | -1.03 | -0.058 | -0.02 | -0.085 | -0.085 | 0 | 0.13 | 0.01 | 0.29 | 0.26 | 0.11 | 0.12 |
| 10 | 28948902 | rs678642 | 0.123 | 0.179 | -1.045 | -0.376 | -0.077 | 0.154 | 0.154 | 2.311 | 0.71 | 0.9 | 0.84 | 0.97 | 0.66 | 0.86 |
| 10 | 28950841 | rs10826486 | -0.873 | 0.342 | -0.294 | 0.529 | 0 | 0.036 | 0.036 | 0 | 0.15 | 0.18 | 0.33 | 0.43 | 0.13 | 0.2 |
| 10 | 28957854 | rs11007191 | -0.664 | 0.337 | -0.326 | 0.526 | 0.003 | 0.131 | 0.131 | 0 | 0.13 | 0.25 | 0.32 | 0.44 | 0.12 | 0.21 |
| 10 | 28961432 | rs10826489 | -0.773 | 0.38 | -0.398 | 0.546 | -0.003 | 0.061 | 0.061 | 0 | 0.16 | 0.24 | 0.33 | 0.44 | 0.13 | 0.22 |
| 10 | 28975214 | rs663378 | -0.788 | 1.029 | -0.695 | -0.096 | -0.032 | 0.032 | 0.032 | 0 | 0.91 | 0.95 | 0.93 | 0.97 | 0.84 | 0.95 |
| 10 | 28975522 | rs17832466 | 0.77 | 1.008 | -0.872 | -0.173 | -0.01 | -0.04 | -0.04 | 0 | 0.27 | 0.13 | 0.32 | 0.4 | 0.34 | 0.38 |
| 10 | 28975865 | rs660371 | 1.431 | 0.662 | -1.106 | -0.384 | -0.007 | 0.031 | 0.031 | 0.305 | 0.27 | 0.1 | 0.2 | 0.21 | 0.26 | 0.31 |
| 10 | 28977683 | rs9663912 | 0.379 | 0.501 | -0.383 | -0.394 | -0.001 | -0.054 | -0.054 | 0 | 0.19 | 0.23 | 0.09 | 0 | 0.17 | 0.07 |
| 10 | 28978553 | rs12570151 | 0.299 | 0.255 | -0.644 | 0.135 | 0.005 | 0.056 | 0.056 | 0 | 0.08 | 0.1 | 0.26 | 0.25 | 0.08 | 0.06 |
| 10 | 28983584 | rs4749335 | -0.402 | 0.239 | -0.5 | 0.115 | 0.026 | 0.144 | 0.144 | 0.192 | 0.1 | 0.24 | 0.27 | 0.25 | 0.08 | 0.07 |
| 10 | 28990162 | rs786880 | 0.367 | 0.365 | -1.517 | 0.113 | -0.001 | 0.012 | 0.012 | 0.48 | 0.54 | 0.32 | 0.52 | 0.62 | 0.59 | 0.69 |
| 10 | 28990644 | rs10826496 | -0.676 | 0.325 | -1.302 | 0.77 | 0.002 | 0.035 | 0.035 | 0 | 0.09 | 0.1 | 0.27 | 0.25 | 0.09 | 0.07 |
| 10 | 28994376 | rs11592762 | 0.118 | 0.363 | -1.331 | 0.82 | -0.036 | -0.072 | -0.072 | 0 | 0.26 | 0.1 | 0.32 | 0.41 | 0.35 | 0.38 |
| 10 | 28995563 | rs12569910 | -0.256 | 0.568 | -1.075 | 0.148 | 0.009 | 0.022 | 0.022 | 0 | 0.41 | 0.25 | 0.58 | 0.66 | 0.43 | 0.5 |

| | | | | | | | | | | | | | | | | |
|---|---|---|---|---|---|---|---|---|---|---|---|---|---|---|---|---|
| 10 | 29001607 | rs7914314 | 0.738 | 0.541 | -0.168 | -0.1 | -0.008 | -0.093 | -0.093 | 0.011 | 0.84 | 0.42 | 0.88 | 0.97 | 0.86 | 0.85 |
| 10 | 29003943 | rs4749338 | 0.852 | 0.953 | -0.323 | 0.089 | -0.017 | 0.001 | 0.001 | 0 | 0.61 | 0.35 | 0.53 | 0.36 | 0.48 | 0.48 |
| 10 | 29005087 | rs1249012 | -1.159 | 0.921 | -0.01 | 0.067 | -0.011 | 0.001 | 0.001 | 0 | 0.39 | 0.64 | 0.46 | 0.64 | 0.52 | 0.54 |
| 10 | 29005257 | rs10826501 | -0.05 | 0.953 | -0.041 | 0.053 | -0.056 | -0.079 | -0.079 | 0 | 0.25 | 0.11 | 0.35 | 0.61 | 0.38 | 0.4 |
| 10 | 29008583 | rs2782389 | -0.937 | 0.95 | -0.029 | 0.053 | 0.024 | 0.008 | 0.008 | 0 | 0.37 | 0.5 | 0.45 | 0.62 | 0.52 | 0.47 |
| 10 | 29010588 | rs10826503 | 0.757 | 0.796 | -0.429 | -0.16 | 0.014 | 0.021 | 0.021 | 0 | 0.48 | 0.3 | 0.51 | 0.36 | 0.38 | 0.42 |
| 10 | 29010863 | rs2782398 | -0.61 | 0.941 | 0.118 | -0.091 | 0.041 | -0.081 | -0.081 | 0 | 0.12 | 0.38 | 0.1 | 0 | 0.14 | 0.08 |
| 10 | 29015323 | rs1762004 | 0.636 | 1.745 | 0.034 | 0.507 | 0.054 | 0.045 | 0.045 | 0 | 0.75 | 0.67 | 0.67 | 0.65 | 0.6 | 0.75 |
| 10 | 29016918 | rs1623555 | 0.535 | 2.358 | 0.134 | 0.513 | 0.007 | 0.074 | 0.074 | 0 | 0.83 | 0.75 | 0.78 | 0.68 | 0.81 | 0.85 |
| 10 | 29017304 | rs11594758 | -0.112 | 2.403 | 0.104 | 0.463 | -0.025 | 0.002 | 0.002 | 0 | 0.06 | 0.01 | 0.1 | 0.02 | 0.2 | 0.09 |
| 10 | 29018935 | rs2489633 | 1.294 | 2.378 | 0.129 | 0.605 | 0.045 | 0.034 | 0.034 | 0 | 0.77 | 0.73 | 0.68 | 0.64 | 0.59 | 0.75 |
| 10 | 29024130 | rs1014607 | -0.091 | 1.413 | -0.945 | -0.735 | 0.001 | -0.094 | -0.094 | 0 | 0.34 | 0.61 | 0.21 | 0.01 | 0.31 | 0.13 |
| 10 | 29025678 | rs1248990 | -0.136 | 0.488 | -0.688 | -0.7 | -0.001 | -0.001 | -0.001 | 0.416 | 0.3 | 0.32 | 0.26 | 0.35 | 0.18 | 0.18 |
| 10 | 29025823 | rs2050361 | -0.206 | 0.225 | 0.247 | -0.898 | 0.015 | -0.147 | -0.147 | 0 | 0.21 | 0.46 | 0.17 | 0.01 | 0.19 | 0.09 |
| 10 | 29026214 | rs1334754 | -0.349 | 0.181 | 0.229 | -0.917 | 0.006 | -0.137 | -0.137 | 0 | 0.21 | 0.58 | 0.17 | 0.01 | 0.19 | 0.09 |
| 10 | 29026933 | rs951753 | -0.126 | 0.179 | 0.353 | -0.922 | 0.024 | -0.144 | -0.144 | 0 | 0.2 | 0.5 | 0.16 | 0.01 | 0.18 | 0.08 |
| 10 | 29027223 | rs1248993 | 0.499 | 0.099 | 0.342 | -0.997 | -0.017 | -0.083 | -0.083 | 0.009 | 0.18 | 0.31 | 0.06 | 0 | 0.11 | 0.07 |
| 10 | 29027618 | rs1248994 | -2.18 | 0.228 | 0.023 | -0.76 | -0.005 | 0.005 | 0.005 | 0.322 | 0.6 | 0.85 | 0.58 | 0.43 | 0.58 | 0.39 |
| 10 | 29029711 | rs11591945 | 0.741 | 0.419 | -0.101 | -0.697 | -0.047 | -0.016 | -0.016 | 0 | 0.07 | 0.01 | 0.1 | 0.05 | 0.2 | 0.11 |
| 10 | 29029736 | rs1248999 | -0.278 | 0.341 | 0.058 | -0.751 | -0.006 | 0.001 | 0.001 | 1.127 | 0.27 | 0.27 | 0.15 | 0.05 | 0.31 | 0.18 |
| 10 | 29029776 | rs11596179 | 0.76 | 0.365 | -0.045 | -0.731 | -0.047 | -0.016 | -0.016 | 0.502 | 0.07 | 0.01 | 0.1 | 0.05 | 0.2 | 0.11 |
| 10 | 29030691 | rs1249003 | 0.497 | 0.227 | 0.132 | -0.739 | -0.023 | -0.091 | -0.091 | 0 | 0.2 | 0.32 | 0.06 | 0 | 0.11 | 0.08 |
| 10 | 29031926 | rs1572248 | -0.257 | 0.436 | 0.152 | -0.492 | 0.113 | 0.075 | 0.075 | 0.421 | 0.2 | 0.38 | 0.33 | 0.37 | 0.37 | 0.21 |
| 10 | 29031951 | rs1249006 | -2.764 | 0.284 | -0.373 | -0.518 | 0.003 | -0.017 | -0.017 | 0.182 | 0.73 | 0.97 | 0.88 | 0.83 | 0.78 | 0.54 |
| 10 | 29034873 | rs787116 | 0.49 | 0.291 | -0.04 | -0.642 | 0.004 | -0.042 | -0.042 | 0.747 | 0.16 | 0.19 | 0.1 | 0.03 | 0.18 | 0.1 |
| 10 | 29034954 | rs7915909 | 1.286 | 0.315 | 0.196 | -0.624 | -0.001 | 0.001 | 0.001 | 0 | 0.41 | 0.49 | 0.28 | 0.25 | 0.45 | 0.6 |
| 10 | 29035016 | rs1265787 | -0.507 | 1.101 | 0.627 | -0.886 | 0 | -0.037 | -0.037 | 0 | 0.2 | 0.2 | 0.12 | 0.03 | 0.19 | 0.11 |
| 10 | 29043820 | rs11007230 | 1.399 | 1.268 | 0.572 | -0.716 | -0.015 | -0.01 | -0.01 | 0 | 0.44 | 0.28 | 0.49 | 0.57 | 0.45 | 0.6 |
| 10 | 29045858 | rs1775921 | -0.853 | 1.299 | 0.932 | -0.734 | -0.012 | -0.066 | -0.066 | 0.586 | 0.82 | 0.8 | 0.92 | 1 | 0.89 | 0.96 |
| 10 | 29050284 | rs1762024 | 0.562 | 1.263 | 1.075 | -0.775 | 0 | -0.002 | -0.002 | 0 | 0.64 | 0.4 | 0.61 | 0.6 | 0.69 | 0.72 |
| 10 | 29052912 | rs17692054 | -0.585 | 1.475 | 0.823 | -0.78 | 0.079 | -0.014 | -0.014 | 0.392 | 0.07 | 0.18 | 0.12 | 0.03 | 0.18 | 0.07 |
| 10 | 29053205 | rs7085567 | 1.071 | 1.371 | 0.848 | -0.753 | -0.021 | 0.015 | 0.015 | 0 | 0.06 | 0.13 | 0.02 | 0.06 | 0.03 | 0.05 |
| 10 | 29054609 | rs1265786 | 0.915 | 1.366 | 0.814 | -0.762 | -0.011 | -0.007 | -0.007 | 0 | 0.58 | 0.33 | 0.51 | 0.62 | 0.51 | 0.66 |
| 10 | 29054941 | rs17759689 | 2.015 | 1.324 | 0.625 | -0.786 | 0.088 | -0.108 | -0.108 | 2.641 | 0.39 | 0.11 | 0.37 | 0.55 | 0.27 | 0.53 |
| 10 | 29056628 | rs787111 | -0.239 | 1.269 | 0.729 | -0.725 | 0.01 | -0.077 | -0.077 | 0.326 | 0.21 | 0.38 | 0.16 | 0.08 | 0.23 | 0.13 |
| 10 | 29056775 | rs11007233 | 0.328 | 1.459 | -0.027 | -0.499 | 0.026 | 0.073 | 0.073 | 0.113 | 0.12 | 0.14 | 0.27 | 0.3 | 0.18 | 0.18 |
| 10 | 29058083 | rs11007234 | 0.574 | 0.925 | -0.56 | -0.312 | 0.015 | 0.049 | 0.049 | 0 | 0.5 | 0.41 | 0.5 | 0.62 | 0.49 | 0.65 |
| 10 | 29067529 | rs1002931 | 1.181 | 0.208 | -1.165 | -0.062 | -0.002 | -0.183 | -0.183 | 2.383 | 0.38 | 0.02 | 0.38 | 0.52 | 0.32 | 0.51 |
| 10 | 29071586 | rs7087796 | 1.196 | -0.177 | -0.589 | 0.007 | 0.011 | 0.005 | 0.005 | 0.87 | 0.5 | 0.1 | 0.42 | 0.54 | 0.56 | 0.57 |
| 10 | 29073571 | rs1105572 | 1.174 | -0.263 | -0.29 | -0.026 | 0.032 | 0.026 | 0.026 | 0.177 | 0.51 | 0.17 | 0.43 | 0.54 | 0.57 | 0.57 |
| 10 | 29075122 | rs11007241 | -0.969 | -0.289 | 0.414 | 0.031 | 0.002 | 0.013 | 0.013 | 0 | 0.2 | 0.24 | 0.3 | 0.39 | 0.23 | 0.24 |
| 10 | 29084646 | rs1761972 | -1.059 | -0.591 | 1.099 | -0.725 | -0.005 | -0.044 | -0.044 | 0 | 0.3 | 0.31 | 0.14 | 0.05 | 0.25 | 0.24 |
| 10 | 29091723 | rs7893940 | 1.563 | -0.668 | 1.098 | -0.602 | 0 | -0.002 | -0.002 | 0 | 0.56 | 0.42 | 0.51 | 0.68 | 0.65 | 0.63 |
| 10 | 29093379 | rs12766414 | 1.912 | -0.739 | 0.576 | -0.56 | 0.006 | 0.074 | 0.074 | 1.233 | 0.49 | 0.1 | 0.43 | 0.59 | 0.56 | 0.49 |
| 10 | 29093914 | rs1775914 | 0.756 | -0.813 | 0.503 | -0.595 | -0.009 | 0.072 | 0.072 | 0 | 0.16 | 0.05 | 0.06 | 0.06 | 0.12 | 0.08 |
| 10 | 29093958 | rs1761984 | 1.877 | -0.613 | 1.572 | 0.095 | -0.008 | -0.003 | -0.003 | 0 | 0.81 | 0.7 | 0.75 | 0.77 | 0.84 | 0.79 |
| 10 | 29094608 | rs1761985 | 2.684 | -0.594 | 1.783 | 0.103 | 0.011 | 0.002 | 0.002 | 2.81 | 0.66 | 0.31 | 0.5 | 0.65 | 0.68 | 0.57 |
| 10 | 29098011 | rs1627701 | 2.586 | -0.577 | 1.741 | 0.09 | -0.049 | -0.007 | -0.007 | 0 | 0.82 | 0.48 | 0.76 | 0.88 | 0.85 | 0.78 |
| 10 | 29098958 | rs1775908 | 2.757 | -0.554 | 1.734 | 0.091 | -0.001 | 0.031 | 0.031 | 1.302 | 0.75 | 0.48 | 0.54 | 0.68 | 0.73 | 0.62 |
| 10 | 29099016 | rs12415919 | -0.367 | -0.592 | 1.398 | 0.11 | -0.01 | 0.003 | 0.003 | 0 | 0.06 | 0.11 | 0.15 | 0.05 | 0.05 | 0.03 |
| 10 | 29099655 | rs10826527 | -0.686 | -0.527 | 1.4 | 0.103 | -0.044 | -0.045 | -0.045 | 0 | 0.08 | 0.01 | 0.21 | 0.2 | 0.12 | 0.16 |
| 10 | 29101875 | rs2065695 | -1.294 | -0.631 | 1.427 | 0.088 | -0.001 | 0 | 0 | 0 | 0.19 | 0.19 | 0.42 | 0.32 | 0.25 | 0.33 |
| 10 | 29106668 | rs1775983 | 1.274 | -0.611 | 1.108 | 0.124 | -0.004 | 0.036 | 0.036 | 0.093 | 0.76 | 0.57 | 0.56 | 0.68 | 0.75 | 0.67 |
| 10 | 29110316 | rs787349 | 0.895 | -0.543 | 0.598 | 0.305 | -0.025 | 0.011 | 0.011 | 0.12 | 0.71 | 0.52 | 0.54 | 0.66 | 0.71 | 0.63 |
| 10 | 29112699 | rs787346 | 0.921 | -0.446 | -0.679 | 0.322 | 0.032 | 0.001 | 0.001 | 4.006 | 0.66 | 0.24 | 0.51 | 0.66 | 0.7 | 0.57 |
| 10 | 29115095 | rs1532393 | 0.942 | -0.814 | -0.021 | 0.395 | -0.037 | 0.003 | 0.003 | 1.453 | 0.72 | 0.36 | 0.67 | 0.67 | 0.87 | 0.56 |
| 10 | 29115431 | rs2643949 | 1.604 | -0.411 | -0.123 | 0.251 | 0.033 | 0.005 | 0.005 | 0.597 | 0.61 | 0.21 | 0.58 | 0.64 | 0.75 | 0.51 |
| 10 | 29123532 | rs1691908 | 1.221 | -0.704 | 0.936 | 0 | -0.005 | -0.005 | -0.005 | 2.647 | 0.68 | 0.38 | 0.61 | 0.72 | 0.78 | 0.58 |
| 10 | 29124168 | rs7078596 | -0.825 | -0.796 | 1.263 | 0.035 | -0.036 | 0.013 | 0.013 | 0.289 | 0.2 | 0.41 | 0.24 | 0.22 | 0.11 | 0.33 |
| 10 | 29125050 | rs12783403 | -0.716 | -0.802 | 1.638 | 0.032 | -0.026 | 0.012 | 0.012 | 0 | 0.19 | 0.33 | 0.23 | 0.22 | 0.11 | 0.33 |
| 10 | 29135359 | rs787338 | 1.932 | -0.77 | 1.591 | 0.002 | 0.032 | 0.04 | 0.04 | 0.515 | 0.63 | 0.21 | 0.47 | 0.73 | 0.77 | 0.57 |
| 10 | 29154573 | rs787343 | 1.998 | -0.897 | 2.069 | 0.144 | 0.047 | 0.011 | 0.011 | 0.624 | 0.65 | 0.16 | 0.56 | 0.67 | 0.79 | 0.57 |
| 10 | 29155832 | rs11007273 | -0.544 | -0.942 | 2.05 | 0.165 | -0.003 | 0.011 | 0.011 | 0 | 0.06 | 0.1 | 0.02 | 0.19 | 0.01 | 0.1 |
| 10 | 29156072 | rs4747633 | -0.941 | -0.967 | 2.358 | 0.143 | 0.005 | 0.003 | 0.003 | 0 | 0.13 | 0.13 | 0.28 | 0.1 | 0.08 | 0.26 |
| 10 | 29156687 | rs7901873 | 0.858 | -0.975 | 2.536 | 0.103 | 0.017 | -0.044 | -0.044 | 0.174 | 0.9 | 0.85 | 0.91 | 1 | 0.88 | 0.95 |
| 10 | 29161199 | rs1631827 | 2.005 | -0.959 | 2.378 | 0.121 | -0.002 | 0.018 | 0.018 | 0 | 0.81 | 0.53 | 0.86 | 0.8 | 0.87 | 0.84 |
| 10 | 29163413 | rs1691940 | 1.931 | -0.968 | 1.902 | 0.2 | 0.004 | 0 | 0 | 0 | 0.67 | 0.31 | 0.57 | 0.67 | 0.8 | 0.55 |
| 10 | 29169492 | rs810894 | 2.022 | -0.954 | 1.639 | 0.183 | 0.027 | 0.011 | 0.011 | 0.401 | 0.66 | 0.25 | 0.57 | 0.68 | 0.81 | 0.53 |
| 10 | 29172222 | rs12256310 | -1.538 | -0.641 | 1.357 | 0.345 | -0.022 | 0.092 | 0.092 | 0.222 | 0.08 | 0.27 | 0.05 | 0.19 | 0.01 | 0.1 |

| | | | | | | | | | | | | | | | | |
|---|---|---|---|---|---|---|---|---|---|---|---|---|---|---|---|---|
| 10 | 29183272 | rs1773881 | -0.172 | 0.123 | -0.006 | -0.848 | -0.037 | 0.078 | 0.078 | 0 | 0.74 | 0.83 | 0.82 | 1 | 0.69 | 0.91 |
| 10 | 29183362 | rs1691935 | -0.055 | 0.113 | 0.633 | -1.457 | 0.053 | -0.094 | -0.094 | 5.765 | 0.17 | 0.35 | 0.22 | 0.02 | 0.22 | 0.24 |
| 10 | 29183994 | rs1773883 | -0.815 | -0.537 | 0.076 | -1.253 | -0.004 | 0.019 | 0.019 | 0 | 0.31 | 0.7 | 0.37 | 0.3 | 0.33 | 0.32 |
| 10 | 29193826 | rs703039 | 0.336 | -0.686 | 0.149 | -1.381 | 0.031 | -0.076 | -0.076 | 2.665 | 0.16 | 0.23 | 0.18 | 0.02 | 0.2 | 0.24 |
| 10 | 29196847 | rs11007308 | 0.92 | -0.403 | -0.16 | -1.528 | -0.056 | 0.112 | 0.112 | 0.117 | 0.25 | 0.07 | 0.15 | 0 | 0.29 | 0.1 |
| 10 | 29197207 | rs1343992 | 1.03 | -0.656 | -0.419 | -1.458 | -0.063 | 0.074 | 0.074 | 0.832 | 0.42 | 0.03 | 0.44 | 0.69 | 0.42 | 0.59 |
| 10 | 29199966 | rs11596869 | 0.296 | -0.416 | 0.389 | -0.974 | -0.005 | 0.006 | 0.006 | 0 | 0.12 | 0.2 | 0.19 | 0.24 | 0.1 | 0.03 |
| 10 | 29203987 | rs703031 | -0.025 | -0.246 | 0.457 | -1.084 | 0.049 | -0.074 | -0.074 | 0.064 | 0.14 | 0.25 | 0.18 | 0.02 | 0.19 | 0.21 |
| 10 | 29204907 | rs703030 | 0.811 | -0.735 | 0.205 | -1.131 | -0.046 | -0.073 | -0.073 | 2.418 | 0.81 | 0.54 | 0.7 | 0.87 | 0.9 | 0.86 |
| 10 | 29205665 | rs11598943 | 0.061 | -0.573 | -0.585 | -0.699 | 0.014 | -0.01 | -0.01 | 0 | 0.27 | 0.2 | 0.12 | 0.27 | 0.26 | 0.23 |
| 10 | 29206762 | rs4749364 | -1.059 | -0.44 | -1.037 | -1.344 | -0.039 | 0.104 | 0.104 | 0.257 | 0.61 | 0.76 | 0.75 | 0.97 | 0.56 | 0.75 |
| 10 | 29207121 | rs12762778 | 1.104 | -0.482 | -1.114 | -1.02 | 0.043 | -0.024 | -0.024 | 0.107 | 0.24 | 0.03 | 0.09 | 0.19 | 0.17 | 0.21 |
| 10 | 29211753 | rs10826541 | -0.092 | -0.648 | -1.456 | -1.418 | 0.02 | 0.165 | 0.165 | 1.394 | 0.46 | 0.23 | 0.49 | 0.82 | 0.45 | 0.57 |
| 10 | 29211930 | rs10763675 | 2.234 | -0.596 | -1.411 | -1.265 | -0.04 | -0.118 | -0.118 | 0.671 | 0.16 | 0.04 | 0.36 | 0.52 | 0.18 | 0.34 |
| 10 | 29214224 | rs7912457 | -0.158 | -0.717 | -0.643 | -1.458 | 0.021 | 0.037 | 0.037 | 1.201 | 0.07 | 0.24 | 0.18 | 0.1 | 0.05 | 0.04 |
| 10 | 29216834 | rs11592131 | 0.302 | -0.7 | -0.589 | -1.451 | 0.039 | 0.055 | 0.055 | 0 | 0.06 | 0.19 | 0.17 | 0.1 | 0.05 | 0.04 |
| 10 | 29217961 | rs11599254 | 1.434 | -1.095 | -0.969 | -1.511 | 0.025 | 0.038 | 0.038 | 0 | 0.06 | 0.14 | 0.04 | 0.1 | 0.1 | 0.04 |
| 10 | 29220291 | rs12767457 | 1.29 | -1.174 | -1.168 | -1.5 | 0.036 | 0.004 | 0.004 | 0 | 0.24 | 0.01 | 0.09 | 0.18 | 0.17 | 0.19 |
| 10 | 29221648 | rs12259250 | -1.24 | -1.089 | -0.901 | -1.487 | -0.007 | -0.004 | -0.004 | 0 | 0.25 | 0.58 | 0.29 | 0.25 | 0.22 | 0.24 |
| 10 | 29230913 | rs1908573 | -1.248 | -1.174 | -1.08 | -1.604 | -0.005 | -0.029 | -0.029 | 0 | 0.6 | 0.6 | 0.73 | 0.97 | 0.55 | 0.76 |
| 10 | 29233160 | rs1832845 | -0.046 | -1.187 | -1.449 | -1.442 | -0.002 | 0.013 | 0.013 | 0 | 0.31 | 0.34 | 0.51 | 0.64 | 0.29 | 0.53 |
| 10 | 29237182 | rs970257 | -1.254 | -1.146 | -1.375 | -1.661 | -0.064 | 0.148 | 0.148 | 0.794 | 0.62 | 0.83 | 0.76 | 0.97 | 0.55 | 0.75 |
| 10 | 29237685 | rs16929394 | -0.629 | -0.944 | -1.178 | -1.255 | -0.012 | -0.023 | -0.023 | 0 | 0.17 | 0.32 | 0.24 | 0.15 | 0.12 | 0.19 |
| 10 | 29240402 | rs12774206 | 1.428 | -0.715 | -1.275 | -1.208 | 0.043 | -0.028 | -0.028 | 1.131 | 0.21 | 0.02 | 0.18 | 0.22 | 0.16 | 0.18 |
| 10 | 29242724 | rs1624345 | -2.304 | -0.172 | -1.112 | -1.159 | -0.013 | 0.011 | 0.011 | 3.293 | 0.88 | 0.91 | 0.91 | 0.96 | 0.78 | 0.87 |
| 10 | 29244111 | rs1557118 | -0.411 | -0.927 | -0.888 | -0.96 | 0.015 | 0.018 | 0.018 | 0 | 0.45 | 0.31 | 0.46 | 0.37 | 0.37 | 0.39 |
| 10 | 29252565 | rs788065 | 0.647 | -0.61 | -0.896 | -0.584 | -0.025 | -0.006 | -0.006 | 0.666 | 0.4 | 0.32 | 0.5 | 0.55 | 0.47 | 0.46 |
| 10 | 29258204 | rs7081328 | -0.101 | -0.78 | -0.43 | -0.73 | -0.045 | -0.025 | -0.025 | 0.002 | 0.13 | 0.48 | 0.16 | 0.1 | 0.06 | 0.09 |
| 10 | 29258268 | rs813716 | 0.9 | -0.983 | -0.75 | -0.928 | 0.02 | 0.11 | 0.11 | 0.046 | 0.57 | 0.24 | 0.64 | 0.8 | 0.6 | 0.64 |
| 10 | 29259450 | rs17755119 | 0.976 | -0.913 | -0.874 | -0.951 | -0.032 | 0.043 | 0.043 | 0 | 0.14 | 0.03 | 0.08 | 0.05 | 0.19 | 0.11 |
| 10 | 29261497 | rs788063 | -1.537 | -0.931 | -0.79 | -0.981 | 0.006 | -0.048 | -0.048 | 0.187 | 0.89 | 0.85 | 0.92 | 1 | 0.9 | 0.97 |
| 10 | 29261978 | rs1248416 | 0.508 | -0.936 | -1.295 | -0.934 | 0.028 | 0.058 | 0.058 | 2.018 | 0.7 | 0.21 | 0.71 | 0.85 | 0.79 | 0.76 |
| 10 | 29262074 | rs1248417 | 0.977 | -0.929 | -1.326 | -0.95 | 0.017 | 0.153 | 0.153 | 0.122 | 0.56 | 0.17 | 0.64 | 0.8 | 0.6 | 0.64 |
| 10 | 29262803 | rs11814043 | -0.114 | -0.728 | -1.232 | -0.76 | 0.005 | 0.004 | 0.004 | 0.452 | 0.37 | 0.31 | 0.31 | 0.34 | 0.39 | 0.43 |
| 10 | 29265375 | rs11593724 | 0.541 | -0.103 | -1.466 | -0.652 | -0.032 | 0.044 | 0.044 | 0 | 0.14 | 0.03 | 0.08 | 0.04 | 0.19 | 0.11 |
| 10 | 29267139 | rs788046 | 0.428 | -0.37 | -1.355 | -0.464 | 0.045 | 0.094 | 0.094 | 0.299 | 0.52 | 0.1 | 0.53 | 0.65 | 0.63 | 0.66 |
| 10 | 29269836 | rs788049 | 0.355 | -0.328 | -0.127 | 0.441 | -0.002 | 0.113 | 0.113 | 0.082 | 0.94 | 0.67 | 0.89 | 0.85 | 0.99 | 0.95 |
| 10 | 29272115 | rs7089095 | -0.733 | 0.269 | -0.949 | 0.378 | -0.001 | 0.012 | 0.012 | 0 | 0.48 | 0.36 | 0.52 | 0.56 | 0.5 | 0.47 |
| 10 | 29274739 | rs16929457 | -0.566 | -0.102 | -0.18 | 0.03 | -0.009 | -0.044 | -0.044 | 0.494 | 0.08 | 0.18 | 0.06 | 0 | 0.06 | 0.04 |
| 10 | 29276863 | rs11007338 | -0.347 | 0.328 | -1.386 | 0.616 | 0.004 | -0.03 | -0.03 | 0 | 0.41 | 0.25 | 0.38 | 0.57 | 0.4 | 0.41 |
| 10 | 29277541 | rs10826549 | -0.796 | 0.217 | -1.199 | 0.113 | 0.008 | 0.041 | 0.041 | 0 | 0.3 | 0.19 | 0.2 | 0.15 | 0.27 | 0.22 |
| 10 | 29277762 | rs10826550 | -0.5 | 0.095 | -1.152 | 0.164 | 0.009 | -0.005 | -0.005 | 1.621 | 0.13 | 0.08 | 0.09 | 0.11 | 0.11 | 0.07 |
| 10 | 29278237 | rs2484510 | 0.968 | 0.312 | -0.752 | 0.26 | -0.001 | 0.004 | 0.004 | 0.347 | 0.38 | 0.37 | 0.49 | 0.4 | 0.38 | 0.33 |
| 10 | 29278939 | rs2487854 | 0.192 | -0.058 | -0.045 | 0.114 | -0.023 | 0.024 | 0.024 | 0.013 | 0.68 | 0.52 | 0.7 | 0.57 | 0.77 | 0.55 |
| 10 | 29282337 | rs1248424 | 1.656 | -0.041 | -0.096 | -0.022 | -0.01 | -0.045 | -0.045 | 0 | 0.34 | 0.13 | 0.45 | 0.4 | 0.35 | 0.34 |
| 10 | 29289031 | rs11007350 | -0.714 | -0.161 | 0.044 | -0.287 | 0.006 | 0.083 | 0.083 | 1.899 | 0.47 | 0.33 | 0.32 | 0.21 | 0.54 | 0.32 |
| 10 | 29290203 | rs6481557 | -0.075 | -0.156 | 0.31 | -0.323 | -0.003 | 0.011 | 0.011 | 4.742 | 0.33 | 0.34 | 0.23 | 0.16 | 0.38 | 0.2 |
| 10 | 29290958 | rs11007352 | 1.165 | -0.186 | 0.204 | -0.362 | 0.005 | -0.016 | -0.016 | 0 | 0.08 | 0.04 | 0.04 | 0.11 | 0.08 | 0.06 |
| 10 | 29291556 | rs1773860 | 0.923 | -0.22 | 0.483 | -0.146 | 0.001 | 0.018 | 0.018 | 1.442 | 0.52 | 0.34 | 0.45 | 0.34 | 0.51 | 0.47 |
| 10 | 29295203 | rs2783592 | 1.909 | -0.171 | 0.49 | -0.12 | -0.007 | -0.008 | -0.008 | 0.968 | 0.34 | 0.13 | 0.33 | 0.3 | 0.34 | 0.34 |
| 10 | 29295577 | rs2783590 | 0.238 | -0.209 | 0.535 | -0.088 | -0.004 | 0.126 | 0.126 | 0 | 0.87 | 0.79 | 0.69 | 0.53 | 0.9 | 0.75 |
| 10 | 29296678 | rs7920971 | -0.099 | -0.321 | 0.229 | -0.217 | -0.001 | 0.007 | 0.007 | 1.511 | 0.33 | 0.32 | 0.21 | 0.16 | 0.37 | 0.22 |
| 10 | 29297998 | rs10740793 | -1.385 | -0.295 | 0.428 | -0.201 | 0.002 | 0.012 | 0.012 | 0.189 | 0.5 | 0.47 | 0.54 | 0.65 | 0.51 | 0.57 |
| 10 | 29300299 | rs1773879 | 1.909 | -0.367 | 0.404 | -0.208 | -0.003 | -0.001 | -0.001 | 0 | 0.35 | 0.2 | 0.33 | 0.31 | 0.35 | 0.34 |
| 10 | 29300987 | rs10826553 | -0.398 | -0.188 | 0.489 | -0.121 | -0.001 | 0.045 | 0.045 | 0 | 0.12 | 0.14 | 0.3 | 0.47 | 0.09 | 0.26 |
| 10 | 29302946 | rs1773877 | 1.03 | -0.315 | 0.462 | -0.202 | -0.003 | 0.037 | 0.037 | 0 | 0.67 | 0.46 | 0.54 | 0.47 | 0.7 | 0.56 |
| 10 | 29303059 | rs7093231 | 0.282 | -0.32 | 0.469 | -0.193 | -0.009 | -0.022 | -0.022 | 0.279 | 0.06 | 0.12 | 0.04 | 0.02 | 0.04 | 0.09 |
| 10 | 29306940 | rs11599857 | -0.393 | -0.306 | 0.456 | -0.169 | 0 | 0.035 | 0.035 | 0 | 0.12 | 0.13 | 0.3 | 0.47 | 0.09 | 0.26 |
| 10 | 29308480 | rs1763382 | 0.46 | -0.303 | 0.479 | -0.191 | 0.008 | 0.078 | 0.078 | 0 | 0.73 | 0.58 | 0.58 | 0.49 | 0.74 | 0.65 |
| 10 | 29312679 | rs10466058 | -1.403 | -0.192 | 0.102 | -0.089 | -0.009 | 0.047 | 0.047 | 0 | 0.18 | 0.27 | 0.34 | 0.48 | 0.13 | 0.35 |
| 10 | 29312885 | rs7081804 | -1.141 | -0.124 | 0.233 | 0.009 | -0.014 | 0.1 | 0.1 | 0 | 0.14 | 0.26 | 0.31 | 0.46 | 0.09 | 0.26 |
| 10 | 29313072 | rs2487848 | -0.994 | 0.041 | 0.379 | -0.839 | 0.01 | -0.057 | -0.057 | 0 | 0.79 | 0.74 | 0.84 | 0.97 | 0.73 | 0.88 |
| 10 | 29314226 | rs11007391 | -0.469 | 0.11 | 0.174 | -0.35 | 0.005 | -0.001 | -0.001 | 0 | 0.08 | 0.15 | 0.07 | 0.08 | 0.07 | 0.1 |
| 10 | 29314775 | rs16929500 | 0.12 | 0.077 | 0.134 | -0.423 | -0.024 | -0.029 | -0.029 | 0.053 | 0.06 | 0.12 | 0.02 | 0 | 0.02 | 0.03 |
| 10 | 29315161 | rs12779321 | 0.041 | 0.505 | -0.208 | -0.281 | -0.044 | 0.005 | 0.005 | 0 | 0.21 | 0.05 | 0.14 | 0.16 | 0.24 | 0.17 |
| 10 | 29315331 | rs11007392 | -1.549 | 0.4 | 0.623 | 0.003 | 0.002 | -0.004 | -0.004 | 0 | 0.09 | 0.21 | 0.08 | 0.08 | 0.06 | 0.09 |
| 10 | 29315965 | rs11007395 | -1.188 | 0.126 | 0.535 | -0.181 | 0.002 | -0.028 | -0.028 | 0 | 0.14 | 0.25 | 0.18 | 0.07 | 0.16 | 0.19 |
| 10 | 29316537 | rs2887372 | -1.163 | -0.137 | 0.599 | -0.484 | 0 | -0.071 | -0.071 | 0 | 0.36 | 0.44 | 0.39 | 0.14 | 0.35 | 0.4 |
| 10 | 29319000 | rs2783425 | 1.288 | -0.133 | 0.557 | -0.166 | -0.002 | -0.002 | -0.002 | 0 | 0.55 | 0.37 | 0.5 | 0.51 | 0.52 | 0.5 |

| | | | | | | | | | | | | | | | | |
|---|---|---|---|---|---|---|---|---|---|---|---|---|---|---|---|---|
| 10 | 29320403 | rs4411195 | -0.501 | 0.093 | 0.593 | -0.007 | 0 | -0.003 | -0.003 | 0 | 0.26 | 0.26 | 0.19 | 0.21 | 0.3 | 0.3 |
| 10 | 29321793 | rs703044 | 0.813 | -0.018 | 0.476 | 0.16 | 0.005 | 0.007 | 0.007 | 0 | 0.59 | 0.45 | 0.52 | 0.5 | 0.53 | 0.5 |
| 10 | 29321843 | rs1023310 | -1.006 | 0.022 | 0.247 | 0.166 | 0.001 | -0.006 | -0.006 | 0.114 | 0.37 | 0.45 | 0.41 | 0.27 | 0.44 | 0.4 |
| 10 | 29323568 | rs788053 | -1.271 | -0.561 | 0.35 | -0.156 | -0.014 | -0.075 | -0.075 | 0 | 0.31 | 0.55 | 0.22 | 0.21 | 0.3 | 0.26 |
| 10 | 29323774 | rs7899901 | 1.531 | -1.841 | 0.922 | -0.243 | -0.044 | -0.043 | -0.043 | 0 | 0.72 | 0.56 | 0.72 | 0.76 | 0.85 | 0.68 |
| 10 | 29323844 | rs2039978 | 1.604 | -1.677 | 1.443 | -0.289 | -0.046 | -0.083 | -0.083 | 0 | 0.8 | 0.58 | 0.74 | 0.93 | 0.9 | 0.81 |
| 10 | 29324846 | rs1041701 | 1.928 | -1.814 | 1.005 | -0.344 | 0.027 | 0.023 | 0.023 | 0 | 0.45 | 0.2 | 0.65 | 0.6 | 0.63 | 0.49 |
| 10 | 29324879 | rs11007399 | 0.411 | -1.649 | 1.388 | -0.155 | -0.001 | 0.026 | 0.026 | 0 | 0.08 | 0.09 | 0.09 | 0.22 | 0.08 | 0.13 |
| 10 | 29327163 | rs7074969 | -0.373 | -1.715 | 1.624 | -0.189 | -0.002 | 0.098 | 0.098 | 0 | 0.11 | 0.18 | 0.1 | 0.35 | 0.09 | 0.16 |
| 10 | 29327405 | rs1248280 | 1.604 | -1.87 | 1.968 | -0.295 | 0.008 | 0.001 | 0.001 | 0.435 | 0.6 | 0.34 | 0.72 | 0.62 | 0.74 | 0.57 |
| 10 | 29330089 | rs2887370 | 2.463 | -1.881 | 2.014 | -0.309 | 0.084 | 0.048 | 0.048 | 0.471 | 0.52 | 0.16 | 0.67 | 0.6 | 0.66 | 0.55 |
| 10 | 29330277 | rs1342011 | 1.708 | -1.852 | 2.105 | -0.296 | -0.027 | 0 | 0 | 0 | 0.62 | 0.48 | 0.73 | 0.62 | 0.74 | 0.57 |
| 10 | 29332168 | rs2804558 | 0.779 | -1.941 | 2.203 | -0.358 | -0.01 | -0.04 | -0.04 | 0.289 | 0.88 | 0.82 | 0.93 | 1 | 0.93 | 0.89 |
| 10 | 29332324 | rs1763371 | 1.84 | -1.993 | 2.235 | -0.368 | -0.033 | -0.044 | -0.044 | 0 | 0.81 | 0.66 | 0.9 | 0.86 | 0.9 | 0.82 |
| 10 | 29334226 | rs7074390 | -1.295 | -1.989 | 2.103 | -0.34 | 0.018 | -0.019 | -0.019 | 0 | 0.26 | 0.19 | 0.19 | 0.26 | 0.21 | 0.36 |
| 10 | 29336849 | rs788076 | 0.602 | -1.924 | 2.116 | -0.307 | -0.006 | -0.022 | -0.022 | 0.449 | 0.94 | 0.77 | 0.95 | 1 | 0.98 | 1 |
| 10 | 29337888 | rs7893339 | -0.914 | -1.845 | 1.739 | -0.268 | -0.006 | 0.01 | 0.01 | 0 | 0.07 | 0.15 | 0.03 | 0.13 | 0.03 | 0.07 |
| 10 | 29338209 | rs2368945 | 2.155 | -1.794 | 1.36 | -0.258 | 0.044 | 0.028 | 0.028 | 0.095 | 0.53 | 0.31 | 0.68 | 0.6 | 0.66 | 0.55 |
| 10 | 29342810 | rs4749381 | 1.75 | -1.556 | 1.258 | -0.39 | -0.011 | -0.042 | -0.042 | 0 | 0.77 | 0.72 | 0.88 | 0.85 | 0.84 | 0.83 |
| 10 | 29342976 | rs4749382 | 1.75 | -1.556 | 1.258 | -0.39 | -0.011 | -0.042 | -0.042 | 0 | 0.77 | 0.72 | 0.88 | 0.85 | 0.84 | 0.83 |
| 10 | 29343457 | rs1763403 | -1.08 | -1.568 | 1.17 | -0.403 | 0.003 | -0.002 | -0.002 | 0 | 0.1 | 0.05 | 0.05 | 0.01 | 0.07 | 0.02 |
| 10 | 29343825 | rs1763404 | 1.194 | -1.574 | 1.489 | -0.363 | -0.011 | -0.03 | -0.03 | 0 | 0.87 | 0.7 | 0.92 | 0.87 | 0.91 | 0.86 |
| 10 | 29344360 | rs4749384 | 0.201 | -1.427 | 1.072 | -0.094 | -0.001 | 0.005 | 0.005 | 0 | 0.08 | 0.09 | 0.08 | 0.18 | 0.08 | 0.13 |
| 10 | 29346895 | rs2804553 | -1.523 | -1.639 | 1.017 | -0.43 | 0.013 | -0.001 | -0.001 | 0 | 0.21 | 0.14 | 0.13 | 0.03 | 0.17 | 0.12 |
| 10 | 29347384 | rs2804552 | -1.678 | -1.572 | 1.239 | -0.374 | -0.017 | -0.074 | -0.074 | 4.533 | 0.14 | 0.23 | 0.09 | 0.01 | 0.1 | 0.02 |
| 10 | 29348259 | rs2783436 | -1.927 | -1.602 | 1.418 | -0.388 | -0.023 | -0.087 | -0.087 | 0.105 | 0.15 | 0.28 | 0.1 | 0.01 | 0.1 | 0.02 |
| 10 | 29349286 | rs1773842 | -0.029 | -1.553 | 1.635 | -0.173 | 0.006 | 0.038 | 0.038 | 0 | 0.9 | 0.87 | 0.95 | 0.76 | 0.9 | 0.85 |
| 10 | 29349540 | rs1773841 | -1.642 | -1.6 | 1.536 | -0.353 | -0.017 | -0.076 | -0.076 | 0 | 0.14 | 0.24 | 0.09 | 0.01 | 0.1 | 0.02 |
| 10 | 29356437 | rs4749385 | 1.884 | -1.486 | 1.737 | -0.067 | -0.003 | -0.003 | -0.003 | 0 | 0.65 | 0.4 | 0.78 | 0.62 | 0.77 | 0.66 |
| 10 | 29356716 | rs1763430 | -1.529 | -1.554 | 1.865 | -0.111 | 0 | 0.002 | 0.002 | 0.017 | 0.22 | 0.22 | 0.14 | 0.37 | 0.19 | 0.25 |
| 10 | 29357507 | rs12767581 | 2.525 | -1.708 | 1.467 | 0.315 | -0.002 | -0.029 | -0.029 | 0.28 | 0.5 | 0.04 | 0.61 | 0.53 | 0.64 | 0.52 |
| 10 | 29361017 | rs10763683 | 2.552 | -1.585 | 1.935 | 0.172 | 0.049 | 0.111 | 0.111 | 0.219 | 0.52 | 0.2 | 0.68 | 0.61 | 0.64 | 0.55 |
| 10 | 29362371 | rs10763685 | 0.046 | -1.597 | 1.899 | 0.165 | -0.002 | -0.004 | -0.004 | 0 | 0.93 | 0.97 | 0.96 | 0.99 | 0.93 | 0.91 |
| 10 | 29364393 | rs573192 | 0.051 | -1.599 | 1.937 | 0.163 | -0.004 | -0.042 | -0.042 | 0 | 0.12 | 0.17 | 0.09 | 0.01 | 0.13 | 0.12 |
| 10 | 29364697 | rs10826566 | -1.711 | -1.567 | 1.68 | 0.185 | -0.019 | 0.005 | 0.005 | 0 | 0.25 | 0.35 | 0.16 | 0.36 | 0.16 | 0.25 |
| 10 | 29365730 | rs1889735 | 2.04 | -1.403 | 0.886 | 0.282 | 0.049 | 0.101 | 0.101 | 0.199 | 0.53 | 0.25 | 0.68 | 0.61 | 0.64 | 0.55 |
| 10 | 29367763 | rs501561 | 1.155 | -0.621 | 0.89 | 1.022 | 0 | 0.004 | 0.004 | 0 | 0.82 | 0.88 | 0.79 | 0.63 | 0.82 | 0.74 |
| 10 | 29370208 | rs985079 | -1.456 | -0.785 | 0.912 | 1.018 | -0.015 | 0.004 | 0.004 | 0.591 | 0.21 | 0.29 | 0.13 | 0.27 | 0.15 | 0.25 |
| 10 | 29370430 | rs534145 | 1.036 | -0.298 | 0.81 | 1.545 | 0.026 | 0.26 | 0.26 | 0 | 0.95 | 0.83 | 0.9 | 0.73 | 0.96 | 0.94 |
| 10 | 29371517 | rs10826567 | 0.293 | 0.47 | -0.229 | 1.273 | -0.044 | -0.062 | -0.062 | 0.306 | 0.12 | 0.05 | 0.2 | 0.29 | 0.23 | 0.17 |
| 10 | 29378718 | rs1338541 | 0.748 | -0.024 | 0.034 | 0.46 | -0.005 | -0.041 | -0.041 | 0 | 0.55 | 0.49 | 0.64 | 0.81 | 0.64 | 0.54 |
| 10 | 29378926 | rs6481566 | 0.661 | -0.195 | 0.125 | 0.321 | -0.004 | -0.04 | -0.04 | 0 | 0.62 | 0.56 | 0.67 | 0.87 | 0.68 | 0.6 |
| 10 | 29379116 | rs7095662 | 0.556 | 0.074 | 0.398 | 0.421 | 0.007 | 0.029 | 0.029 | 0.638 | 0.14 | 0.2 | 0.22 | 0.3 | 0.23 | 0.18 |
| 10 | 29381502 | rs1144522 | 0.343 | -0.022 | 0.088 | 0.252 | -0.066 | -0.001 | -0.001 | 0 | 0.16 | 0.01 | 0.18 | 0.1 | 0.16 | 0.23 |
| 10 | 29382968 | rs544736 | -1.653 | -0.116 | 0.676 | 0.247 | -0.004 | 0.001 | 0.001 | 0 | 0.54 | 0.5 | 0.55 | 0.49 | 0.6 | 0.64 |
| 10 | 29384015 | rs7069400 | 1.748 | -0.034 | 0.396 | 0.284 | -0.002 | -0.014 | -0.014 | 0.935 | 0.4 | 0.31 | 0.42 | 0.51 | 0.4 | 0.36 |
| 10 | 29386668 | rs10763691 | 1.754 | -0.03 | 0.398 | 0.294 | -0.002 | -0.018 | -0.018 | 0 | 0.4 | 0.3 | 0.42 | 0.5 | 0.4 | 0.36 |
| 10 | 29386824 | rs568206 | 0.171 | 0.065 | 0.534 | 0.156 | 0 | -0.02 | -0.02 | 0.025 | 0.11 | 0.08 | 0.1 | 0 | 0.12 | 0.09 |
| 10 | 29388289 | rs2018838 | 1.171 | 0.059 | 0.546 | 0.143 | -0.027 | -0.112 | -0.112 | 0 | 0.79 | 0.61 | 0.85 | 0.97 | 0.84 | 0.83 |
| 10 | 29393640 | rs494650 | -0.078 | 0.518 | -0.195 | 0.367 | 0 | -0.038 | -0.038 | 0 | 0.11 | 0.12 | 0.1 | 0 | 0.13 | 0.09 |
| 10 | 29399839 | rs7901884 | -0.264 | 0.658 | -0.457 | 0.967 | -0.018 | -0.032 | -0.032 | 0 | 0.14 | 0.08 | 0.23 | 0.25 | 0.24 | 0.17 |
| 10 | 29402761 | rs10763693 | 1.163 | 0.857 | -0.361 | 0.684 | 0.005 | 0.01 | 0.01 | 0 | 0.51 | 0.43 | 0.48 | 0.4 | 0.45 | 0.58 |
| 10 | 29402944 | rs4749397 | 1.163 | 0.857 | -0.275 | 0.678 | 0.001 | 0.003 | 0.003 | 0 | 0.51 | 0.44 | 0.48 | 0.4 | 0.45 | 0.58 |
| 10 | 29405387 | rs573036 | 0.227 | 0.883 | -0.279 | 0.683 | -0.006 | 0.024 | 0.024 | 0 | 0.16 | 0.03 | 0.12 | 0.12 | 0.15 | 0.11 |
| 10 | 29409192 | rs2793345 | -0.636 | 1.121 | 0.053 | 0.599 | -0.004 | -0.004 | -0.004 | 0.06 | 0.22 | 0.23 | 0.3 | 0.47 | 0.26 | 0.19 |
| 10 | 29415626 | rs10443969 | 1.947 | 1.112 | -0.024 | 0.604 | 0.003 | 0.009 | 0.009 | 0 | 0.51 | 0.44 | 0.48 | 0.36 | 0.46 | 0.58 |
| 10 | 29416667 | rs2793346 | -0.501 | 1.18 | -0.375 | 0.709 | -0.022 | -0.044 | -0.044 | 0 | 0.21 | 0.14 | 0.3 | 0.5 | 0.25 | 0.19 |
| 10 | 29423049 | rs2783596 | 0.12 | 1.379 | -0.412 | 0.807 | -0.031 | -0.006 | -0.006 | 0.241 | 0.13 | 0.04 | 0.09 | 0.14 | 0.15 | 0.18 |
| 10 | 29424176 | rs10508742 | -0.186 | 1.403 | -0.322 | 0.885 | -0.01 | -0.007 | -0.007 | 0.947 | 0.08 | 0.06 | 0.09 | 0.13 | 0.14 | 0.09 |
| 10 | 29424298 | rs10508743 | -1.218 | 1.395 | -0.302 | 0.944 | -0.01 | -0.005 | -0.005 | 0.618 | 0.16 | 0.17 | 0.31 | 0.49 | 0.25 | 0.2 |
| 10 | 29425176 | rs2788451 | 0.004 | 1.247 | -0.1 | 0.785 | -0.004 | -0.051 | -0.051 | 0 | 0.81 | 0.83 | 0.89 | 1 | 0.85 | 0.93 |
| 10 | 29426092 | rs2783419 | 0.043 | 1.249 | -0.12 | 0.786 | -0.009 | -0.069 | -0.069 | 0 | 0.81 | 0.8 | 0.88 | 1 | 0.85 | 0.93 |
| 10 | 29428123 | rs2804569 | -1.678 | 1.329 | -0.281 | 0.765 | -0.034 | -0.014 | -0.014 | 0 | 0.29 | 0.24 | 0.4 | 0.63 | 0.4 | 0.36 |
| 10 | 29434036 | rs478229 | 1.35 | 0.962 | -0.647 | 0.523 | -0.004 | 0.038 | 0.038 | 0 | 0.52 | 0.68 | 0.49 | 0.39 | 0.49 | 0.55 |
| 10 | 29437892 | rs533844 | 0.758 | 0.9 | -0.296 | 0.599 | 0.001 | 0.061 | 0.061 | 0 | 0.75 | 0.66 | 0.65 | 0.53 | 0.72 | 0.77 |
| 10 | 29437974 | rs548193 | 1.528 | 0.861 | -0.386 | 0.548 | -0.001 | 0.006 | 0.006 | 0 | 0.52 | 0.58 | 0.49 | 0.41 | 0.5 | 0.64 |
| 10 | 29438786 | rs549758 | -0.818 | 0.686 | -0.294 | 0.984 | 0.003 | 0.033 | 0.033 | 0 | 0.17 | 0.16 | 0.29 | 0.36 | 0.22 | 0.08 |
| 10 | 29441347 | rs564281 | 1.433 | -0.032 | -0.844 | 0.559 | -0.045 | 0.078 | 0.078 | 0 | 0.42 | 0.16 | 0.35 | 0.25 | 0.47 | 0.43 |
| 10 | 29441360 | rs534371 | 1.001 | 0.048 | -0.656 | 0.681 | 0.007 | 0.096 | 0.096 | 0 | 0.49 | 0.25 | 0.38 | 0.26 | 0.52 | 0.44 |

| | | | | | | | | | | | | | | | | |
|---|---|---|---|---|---|---|---|---|---|---|---|---|---|---|---|---|
| 10 | 29441518 | rs12413557 | -0.373 | -0.1 | -0.26 | 1.184 | 0.016 | 0.011 | 0.011 | 0 | 0.07 | 0.16 | 0.18 | 0.07 | 0.06 | 0.04 |
| 10 | 29446722 | rs1014083 | -0.472 | 0.148 | -0.664 | 1.344 | 0.001 | -0.024 | -0.024 | 0 | 0.12 | 0.05 | 0.11 | 0.29 | 0.13 | 0.08 |
| 10 | 29449051 | rs7902675 | -1.13 | -0.022 | 0.151 | 1.068 | 0.035 | -0.008 | -0.008 | 0 | 0.17 | 0.52 | 0.17 | 0.12 | 0.2 | 0.09 |
| 10 | 29449129 | rs511991 | 0.72 | -0.536 | -0.187 | 1.114 | -0.003 | 0.081 | 0.081 | 1.164 | 0.72 | 0.55 | 0.55 | 0.48 | 0.71 | 0.77 |
| 10 | 29450498 | rs1591773 | 1.366 | -0.547 | -0.16 | 1.066 | 0.036 | -0.029 | -0.029 | 0 | 0.57 | 0.31 | 0.48 | 0.54 | 0.59 | 0.72 |
| 10 | 29453842 | rs526182 | -1.125 | -0.771 | 0.597 | 1.297 | -0.057 | 0.093 | 0.093 | 0.015 | 0.18 | 0.34 | 0.16 | 0.33 | 0.14 | 0.24 |
| 10 | 29457642 | rs571214 | -0.007 | -0.349 | 0.782 | 1.381 | 0.052 | 0.002 | 0.002 | 0 | 0.93 | 0.88 | 0.73 | 0.95 | 0.87 | 0.93 |
| 10 | 29460122 | rs1118649 | 1.85 | -0.56 | 1.116 | 1.303 | 0.015 | 0.019 | 0.019 | 0 | 0.74 | 0.54 | 0.62 | 0.69 | 0.68 | 0.66 |
| 10 | 29461164 | rs2788453 | -0.16 | -0.39 | 0.588 | 1.452 | -0.033 | -0.041 | -0.041 | 0 | 0.07 | 0 | 0.11 | 0.2 | 0.07 | 0.19 |
| 10 | 29463797 | rs12256237 | -1.567 | -0.48 | 1.114 | 1.251 | -0.015 | -0.037 | -0.037 | 0 | 0.18 | 0.26 | 0.1 | 0.1 | 0.15 | 0.12 |
| 10 | 29463830 | rs11007455 | 2.537 | -0.782 | 0.59 | 1.268 | 0.045 | -0.022 | -0.022 | 0.035 | 0.5 | 0.08 | 0.52 | 0.4 | 0.56 | 0.48 |
| 10 | 29466058 | rs495219 | -2.035 | -0.866 | 0.84 | 1.318 | 0.003 | -0.004 | -0.004 | 0 | 0.37 | 0.64 | 0.37 | 0.45 | 0.24 | 0.42 |
| 10 | 29466492 | rs1932273 | 2.303 | -0.844 | 0.949 | 1.339 | 0.068 | 0.001 | 0.001 | 3.504 | 0.54 | 0.2 | 0.55 | 0.4 | 0.61 | 0.49 |
| 10 | 29467377 | rs478463 | 1.704 | -0.849 | 0.961 | 1.319 | 0.013 | -0.006 | -0.006 | 0 | 0.6 | 0.36 | 0.6 | 0.54 | 0.72 | 0.56 |
| 10 | 29468229 | rs2532750 | -0.529 | -0.821 | 0.768 | 1.377 | -0.03 | -0.044 | -0.044 | 0 | 0.07 | 0.01 | 0.1 | 0.19 | 0.07 | 0.27 |
| 10 | 29468276 | rs557493 | 1.337 | -0.862 | 1.159 | 1.339 | 0.001 | 0.01 | 0.01 | 0 | 0.78 | 0.65 | 0.77 | 0.72 | 0.78 | 0.64 |
| 10 | 29470176 | rs1271994 | 0.035 | -0.924 | 1.044 | 1.276 | -0.067 | 0.065 | 0.065 | 0.475 | 0.13 | 0.33 | 0.14 | 0.17 | 0.04 | 0.06 |
| 10 | 29472548 | rs570942 | 0.618 | -0.833 | 0.995 | 1.287 | 0.029 | 0.043 | 0.043 | 0.051 | 0.05 | 0.13 | 0.05 | 0.14 | 0.11 | 0.07 |
| 10 | 29473292 | rs4265509 | 2.489 | -0.771 | 0.479 | 1.231 | -0.115 | 0.009 | 0.009 | 1.494 | 0.47 | 0.03 | 0.34 | 0.31 | 0.5 | 0.42 |
| 10 | 29473575 | rs16929852 | 0.003 | -0.89 | 1.135 | 0.97 | -0.048 | 0.034 | 0.034 | 0.063 | 0.12 | 0.22 | 0.14 | 0.17 | 0.04 | 0.06 |
| 10 | 29474877 | rs16929855 | -1.993 | -0.877 | 1.163 | 0.942 | -0.073 | -0.026 | -0.026 | 0 | 0.28 | 0.5 | 0.25 | 0.17 | 0.18 | 0.1 |
| 10 | 29475869 | rs2532747 | -2.03 | -0.781 | 0.905 | 1.097 | -0.02 | 0.011 | 0.011 | 1.304 | 0.29 | 0.35 | 0.32 | 0.32 | 0.2 | 0.31 |
| 10 | 29481588 | rs10490901 | -1.003 | -0.625 | 0.505 | 0.919 | -0.004 | -0.034 | -0.034 | 0 | 0.12 | 0.11 | 0.07 | 0 | 0.11 | 0.03 |
| 10 | 29483846 | rs9971350 | -0.328 | -0.757 | 0.953 | 1.245 | -0.035 | 0.035 | 0.035 | 1.021 | 0.15 | 0.33 | 0.17 | 0.16 | 0.09 | 0.07 |
| 10 | 29484595 | rs576692 | -0.661 | -0.632 | 0.517 | 1.588 | -0.001 | 0.009 | 0.009 | 0.57 | 0.11 | 0.1 | 0.12 | 0.33 | 0.14 | 0.28 |
| 10 | 29486078 | rs7893901 | -0.504 | -0.591 | 0.441 | 1.471 | -0.041 | 0.03 | 0.03 | 2.273 | 0.17 | 0.44 | 0.18 | 0.16 | 0.1 | 0.07 |
| 10 | 29489709 | rs501138 | -1.209 | -0.411 | -0.474 | 1.14 | 0.007 | -0.012 | -0.012 | 1.496 | 0.32 | 0.7 | 0.33 | 0.49 | 0.24 | 0.37 |
| 10 | 29490619 | rs521753 | -1.221 | 0.193 | -1.324 | 1.091 | 0.019 | -0.062 | -0.062 | 0 | 0.43 | 0.88 | 0.38 | 0.56 | 0.35 | 0.49 |
| 10 | 29493230 | rs532405 | -0.676 | 1.091 | -1.134 | 1.144 | -0.076 | -0.095 | -0.095 | 0 | 0.13 | 0.01 | 0.34 | 0.52 | 0.22 | 0.27 |
| 10 | 29493922 | rs11007465 | -0.595 | 1.07 | -0.417 | 0.151 | 0.015 | -0.132 | -0.132 | 0 | 0.17 | 0.4 | 0.11 | 0.01 | 0.18 | 0.17 |
| 10 | 29496472 | rs511920 | -1.168 | 0.4 | -0.513 | -0.204 | 0.001 | 0.003 | 0.003 | 0.226 | 0.34 | 0.42 | 0.51 | 0.62 | 0.37 | 0.42 |
| 10 | 29503772 | rs16929930 | 1.477 | 0.332 | -1.569 | -1.004 | -0.014 | -0.066 | -0.066 | 0.153 | 0.33 | 0.07 | 0.26 | 0.37 | 0.29 | 0.36 |
| 10 | 29508589 | rs555438 | -1.125 | 0.448 | -1.418 | -1.208 | -0.033 | 0.082 | 0.082 | 0 | 0.79 | 0.91 | 0.86 | 0.99 | 0.76 | 0.88 |
| 10 | 29509590 | rs7094704 | 0.668 | 0.208 | -1.071 | -1.06 | 0.016 | 0.016 | 0.016 | 0 | 0.2 | 0.19 | 0.08 | 0.07 | 0.1 | 0.08 |
| 10 | 29512320 | rs2532774 | -0.614 | 0.51 | -0.826 | -0.938 | -0.001 | 0 | 0 | 0 | 0.41 | 0.42 | 0.47 | 0.63 | 0.44 | 0.47 |
| 10 | 29515295 | rs493933 | -1.27 | 0.075 | -1.051 | -1.257 | -0.004 | 0.006 | 0.006 | 0 | 0.7 | 0.76 | 0.64 | 0.8 | 0.66 | 0.73 |
| 10 | 29515598 | rs496776 | -0.907 | -0.076 | -1.204 | -1.267 | -0.014 | 0.108 | 0.108 | 0 | 0.53 | 0.76 | 0.57 | 0.78 | 0.52 | 0.61 |
| 10 | 29516516 | rs501298 | -0.081 | -0.17 | -1.7 | -1.022 | 0 | -0.016 | -0.016 | 0 | 0.33 | 0.28 | 0.43 | 0.63 | 0.34 | 0.4 |
| 10 | 29519230 | rs12267702 | 0.857 | -0.27 | -2.41 | -1.041 | -0.007 | -0.009 | -0.009 | 0 | 0.22 | 0.27 | 0.1 | 0.07 | 0.12 | 0.09 |
| 10 | 29521744 | rs4747646 | 0.471 | 0.474 | -2.126 | -0.758 | 0.054 | -0.013 | -0.013 | 0 | 0.43 | 0.29 | 0.37 | 0.42 | 0.35 | 0.41 |
| 10 | 29523746 | rs17392645 | 0.843 | 0.766 | -2.167 | 0.026 | 0.018 | 0.066 | 0.066 | 0 | 0.17 | 0.05 | 0.07 | 0.03 | 0.13 | 0.12 |
| 10 | 29524450 | rs581090 | -0.244 | 1.419 | -1.824 | 0.608 | -0.066 | -0.076 | -0.076 | 0 | 0.11 | 0.02 | 0.32 | 0.54 | 0.22 | 0.31 |
| 10 | 29532110 | rs561406 | -0.45 | 0.67 | -1.834 | -0.274 | 0.016 | 0.072 | 0.072 | 0 | 0.34 | 0.2 | 0.2 | 0.15 | 0.32 | 0.32 |
| 10 | 29536990 | rs6481572 | 0.397 | 0.662 | -1.414 | -0.021 | -0.006 | -0.008 | -0.008 | 1.276 | 0.41 | 0.53 | 0.31 | 0.33 | 0.32 | 0.32 |
| 10 | 29541789 | rs558951 | 0.041 | 0.339 | -2.059 | -0.758 | 0.024 | 0.095 | 0.095 | 0.414 | 0.46 | 0.73 | 0.29 | 0.16 | 0.39 | 0.36 |
| 10 | 29542629 | rs7909464 | 0.205 | -0.077 | -2.18 | -1.025 | 0.023 | 0.028 | 0.028 | 0.224 | 0.4 | 0.66 | 0.17 | 0.16 | 0.23 | 0.33 |
| 10 | 29545201 | rs2532772 | -0.907 | 0.15 | -2.356 | -1.293 | -0.047 | 0.109 | 0.109 | 0.589 | 0.6 | 0.84 | 0.37 | 0.11 | 0.48 | 0.4 |
| 10 | 29546473 | rs1860403 | -1.383 | 0.426 | -2.853 | -0.914 | -0.103 | -0.165 | -0.165 | 1.133 | 0.76 | 0.99 | 0.52 | 0.42 | 0.65 | 0.71 |
| 10 | 29546557 | rs2532770 | 2.233 | 0.529 | -2.659 | -1.03 | -0.022 | 0.031 | 0.031 | 1.983 | 0.11 | 0.05 | 0.11 | 0 | 0.16 | 0.04 |
| 10 | 29549209 | rs9943444 | 0.698 | -0.265 | -2.107 | -1.142 | 0.012 | 0.019 | 0.019 | 0 | 0.36 | 0.6 | 0.18 | 0.1 | 0.24 | 0.21 |
| 10 | 29550249 | rs4622160 | 0.279 | -0.35 | -2.372 | -1.162 | 0.066 | 0.173 | 0.173 | 0.257 | 0.4 | 0.88 | 0.21 | 0.1 | 0.24 | 0.21 |
| 10 | 29551255 | rs11007489 | 0.438 | -0.126 | -2.371 | -0.92 | -0.028 | -0.069 | -0.069 | 0.14 | 0.21 | 0.04 | 0.14 | 0.28 | 0.18 | 0.27 |
| 10 | 29554384 | rs11007491 | 1.032 | -0.394 | -1.431 | -1.164 | -0.005 | -0.017 | -0.017 | 0 | 0.29 | 0.3 | 0.13 | 0.1 | 0.22 | 0.21 |
| 10 | 29556150 | rs2796323 | -0.378 | -0.057 | -1.458 | -1.158 | 0.005 | -0.011 | -0.011 | 0 | 0.61 | 0.58 | 0.38 | 0.28 | 0.51 | 0.48 |
| 10 | 29562039 | rs2532759 | -0.561 | -0.113 | -1.511 | -1.142 | 0.007 | -0.005 | -0.005 | 0.098 | 0.63 | 0.61 | 0.39 | 0.3 | 0.51 | 0.53 |
| 10 | 29562460 | rs10826586 | 0.638 | -0.237 | -2.179 | -1.297 | -0.011 | 0.011 | 0.011 | 0.51 | 0.37 | 0.08 | 0.21 | 0.28 | 0.27 | 0.37 |
| 10 | 29567747 | rs2817834 | -0.753 | -0.289 | -1.856 | -1.273 | 0.021 | 0.071 | 0.071 | 0 | 0.46 | 0.72 | 0.33 | 0.24 | 0.39 | 0.47 |
| 10 | 29568777 | rs2493340 | 1.928 | 0.041 | -1.602 | -1.276 | 0.021 | -0.057 | -0.057 | 0 | 0.13 | 0.2 | 0.14 | 0 | 0.19 | 0.06 |
| 10 | 29569272 | rs7069887 | 1.407 | 0.147 | -1.608 | -1.034 | -0.018 | -0.027 | -0.027 | 0 | 0.12 | 0.07 | 0.14 | 0.17 | 0.14 | 0.19 |
| 10 | 29569355 | rs2488264 | -1.979 | 0.695 | -1.835 | -0.959 | -0.042 | 0.01 | 0.01 | 0 | 0.86 | 0.99 | 0.82 | 0.9 | 0.77 | 0.82 |
| 10 | 29569541 | rs2488265 | 0.106 | 0.004 | -1.202 | -1.031 | 0.017 | 0.042 | 0.042 | 0 | 0.46 | 0.59 | 0.26 | 0.21 | 0.36 | 0.37 |
| 10 | 29575046 | rs2817811 | 1.003 | 0.12 | -1.62 | -0.989 | -0.002 | 0.008 | 0.008 | 0.129 | 0.34 | 0.31 | 0.23 | 0.18 | 0.33 | 0.23 |
| 10 | 29576177 | rs2796298 | 0.828 | 0.074 | -1.564 | -0.959 | -0.007 | 0.017 | 0.017 | 0 | 0.38 | 0.31 | 0.25 | 0.21 | 0.37 | 0.32 |
| 10 | 29576414 | rs2532766 | 0.313 | 0.727 | -1.408 | -0.6 | -0.029 | -0.01 | -0.01 | 0.151 | 0.2 | 0.14 | 0.22 | 0.21 | 0.31 | 0.26 |
| 10 | 29576828 | rs16930100 | 0.838 | 0.629 | -0.733 | -0.253 | -0.004 | -0.012 | -0.012 | 0.204 | 0.07 | 0.08 | 0.01 | 0 | 0.02 | 0 |
| 10 | 29579482 | rs2479061 | 0.061 | 0.945 | -0.791 | -0.188 | -0.007 | -0.027 | -0.027 | 0 | 0.67 | 0.75 | 0.49 | 0.42 | 0.66 | 0.55 |
| 10 | 29579603 | rs10826591 | 0.502 | 0.836 | -0.619 | -0.445 | 0.003 | 0.011 | 0.011 | 0 | 0.51 | 0.43 | 0.36 | 0.4 | 0.49 | 0.44 |
| 10 | 29581998 | rs1751956 | 1.487 | 0.81 | -0.057 | -0.699 | 0.005 | -0.049 | -0.049 | 0.657 | 0.1 | 0.15 | 0.13 | 0.01 | 0.14 | 0.12 |
| 10 | 29582248 | rs1774950 | 0.261 | 0.692 | -0.328 | -0.712 | -0.006 | 0.004 | 0.004 | 0.946 | 0.53 | 0.33 | 0.58 | 0.53 | 0.55 | 0.54 |

| | | | | | | | | | | | | | | | | |
|---|---|---|---|---|---|---|---|---|---|---|---|---|---|---|---|---|
| 10 | 29583431 | rs10733993 | -1.254 | 0.731 | -0.051 | -0.582 | -0.001 | -0.016 | -0.016 | 0.824 | 0.72 | 0.73 | 0.81 | 0.9 | 0.72 | 0.85 |
| 10 | 29585480 | rs10763703 | -0.401 | 0.355 | -0.28 | -0.71 | -0.005 | -0.021 | -0.021 | 0 | 0.58 | 0.59 | 0.73 | 0.83 | 0.64 | 0.69 |
| 10 | 29585987 | rs10826596 | 0.603 | 0.476 | -0.011 | -0.737 | 0.006 | 0.008 | 0.008 | 0.021 | 0.2 | 0.25 | 0.28 | 0.27 | 0.22 | 0.26 |
| 10 | 29586483 | rs10763704 | 0.598 | 0.45 | 0.04 | -0.71 | -0.009 | 0.013 | 0.013 | 0.486 | 0.42 | 0.34 | 0.46 | 0.61 | 0.45 | 0.49 |
| 10 | 29589009 | rs1327566 | -0.733 | 0.564 | -0.031 | -0.639 | 0.024 | 0.037 | 0.037 | 0.304 | 0.28 | 0.48 | 0.34 | 0.37 | 0.29 | 0.43 |
| 10 | 29591701 | rs2148899 | 0.681 | 0.671 | -0.079 | -0.569 | 0.008 | 0.014 | 0.014 | 0 | 0.18 | 0.25 | 0.26 | 0.27 | 0.2 | 0.26 |
| 10 | 29594791 | rs10763707 | -0.452 | 0.626 | -0.101 | -0.594 | 0.001 | -0.002 | -0.002 | 0.047 | 0.3 | 0.29 | 0.32 | 0.34 | 0.26 | 0.41 |
| 10 | 29595135 | rs10826606 | 0.643 | 0.467 | -0.473 | -0.158 | -0.074 | -0.126 | -0.126 | 0.342 | 0.34 | 0.06 | 0.41 | 0.51 | 0.36 | 0.37 |
| 10 | 29597554 | rs7094590 | -0.78 | 0.427 | -0.064 | -0.375 | -0.025 | -0.035 | -0.035 | 2.079 | 0.18 | 0.37 | 0.12 | 0.11 | 0.13 | 0.19 |
| 10 | 29600240 | rs16930144 | 0.262 | 0.511 | -0.46 | -0.282 | 0.008 | 0.006 | 0.006 | 0 | 0.12 | 0.08 | 0.06 | 0.07 | 0.06 | 0.15 |
| 10 | 29606974 | rs11007524 | -0.065 | 0.776 | -1.06 | 0.005 | 0.028 | 0.067 | 0.067 | 0.073 | 0.45 | 0.2 | 0.47 | 0.63 | 0.43 | 0.52 |
| 10 | 29609234 | rs7076385 | -0.837 | 1.073 | -0.285 | -0.084 | 0.016 | -0.014 | -0.014 | 0 | 0.17 | 0.24 | 0.14 | 0.11 | 0.22 | 0.22 |
| 10 | 29612818 | rs789930 | 0.703 | 0.506 | 0.616 | -0.187 | 0.062 | 0.058 | 0.058 | 1.744 | 0.95 | 0.6 | 0.95 | 1 | 1 | 1 |
| 10 | 29615837 | rs2256713 | -0.008 | 0.109 | 0.012 | -0.444 | -0.036 | -0.042 | -0.042 | 0.387 | 0.71 | 0.5 | 0.77 | 0.86 | 0.79 | 0.68 |
| 10 | 29617150 | rs10491049 | 0.688 | 0.064 | -0.064 | -0.466 | -0.004 | 0.037 | 0.037 | 0 | 0.22 | 0.05 | 0.18 | 0.1 | 0.2 | 0.23 |
| 10 | 29621419 | rs932838 | 0.463 | 0.233 | -0.032 | -0.707 | -0.045 | 0.052 | 0.052 | 0.039 | 0.14 | 0.05 | 0.15 | 0.05 | 0.18 | 0.11 |
| 10 | 29622943 | rs12779048 | -0.151 | 0.153 | 0.264 | -0.758 | -0.032 | -0.042 | -0.042 | 0 | 0.3 | 0.47 | 0.24 | 0.13 | 0.21 | 0.31 |
| 10 | 29628094 | rs789920 | 0.908 | -0.266 | -0.075 | -1.113 | -0.002 | 0.03 | 0.03 | 0 | 0.44 | 0.32 | 0.56 | 0.81 | 0.54 | 0.44 |
| 10 | 29628552 | rs11598403 | 1.439 | -0.824 | -1.071 | -0.886 | -0.008 | -0.054 | -0.054 | 0 | 0.19 | 0.01 | 0.11 | 0.21 | 0.13 | 0.08 |
| 10 | 29632242 | rs7094201 | 1.042 | -0.483 | -0.317 | -0.719 | -0.01 | -0.043 | -0.043 | 0 | 0.11 | 0.16 | 0.05 | 0.01 | 0.08 | 0.12 |
| 10 | 29635856 | rs1751960 | 0.885 | -0.005 | -0.118 | -0.303 | -0.002 | 0.035 | 0.035 | 1.185 | 0.36 | 0.48 | 0.29 | 0.5 | 0.37 | 0.26 |
| 10 | 29637337 | rs10826611 | -0.236 | 0.586 | -0.161 | 0.165 | 0.01 | 0.018 | 0.018 | 0 | 0.27 | 0.34 | 0.48 | 0.42 | 0.33 | 0.38 |
| 10 | 29642160 | rs4747651 | 0.079 | 0.917 | -0.978 | 0.309 | -0.034 | -0.047 | -0.047 | 0.362 | 0.24 | 0.13 | 0.45 | 0.42 | 0.33 | 0.32 |
| 10 | 29648177 | rs2182758 | 0.085 | 0.869 | -1.265 | 0.101 | -0.065 | -0.112 | -0.112 | 0 | 0.24 | 0.05 | 0.41 | 0.46 | 0.29 | 0.32 |
| 10 | 29656633 | rs11007540 | 0.217 | 0.729 | -0.802 | 0.149 | -0.077 | -0.091 | -0.091 | 0 | 0.26 | 0.06 | 0.47 | 0.39 | 0.31 | 0.43 |
| 10 | 29666463 | rs913290 | 0.053 | 0.293 | -0.464 | -0.053 | -0.001 | -0.002 | -0.002 | 0 | 0.58 | 0.39 | 0.78 | 0.8 | 0.72 | 0.7 |
| 10 | 29666617 | rs1751963 | -0.015 | 0.862 | -0.436 | -0.246 | -0.019 | -0.097 | -0.097 | 0 | 0.26 | 0.29 | 0.13 | 0.01 | 0.19 | 0.19 |
| 10 | 29670331 | rs4749414 | -0.572 | 1.593 | -0.696 | 0.668 | -0.027 | -0.033 | -0.033 | 0 | 0.26 | 0.15 | 0.48 | 0.39 | 0.31 | 0.43 |
| 10 | 29671014 | rs7902673 | -0.565 | 1.601 | -0.722 | 0.66 | -0.034 | -0.04 | -0.04 | 0 | 0.26 | 0.12 | 0.47 | 0.39 | 0.31 | 0.43 |
| 10 | 29682041 | rs7896382 | -0.806 | 1.762 | -0.205 | 0.48 | 0.009 | -0.011 | -0.011 | 0 | 0.06 | 0.11 | 0.15 | 0.03 | 0.08 | 0.06 |
| 10 | 29697227 | rs4237392 | -0.811 | 1.721 | -0.044 | 0.835 | -0.002 | -0.014 | -0.014 | 0 | 0.23 | 0.19 | 0.36 | 0.56 | 0.25 | 0.35 |
| 10 | 29698286 | rs11007559 | -1.171 | 2.424 | 0.328 | 0.673 | 0.068 | -0.036 | -0.036 | 0 | 0.11 | 0.25 | 0.22 | 0.04 | 0.22 | 0.09 |
| 10 | 29701247 | rs7902800 | -1.92 | 2.352 | 1.111 | 0.63 | 0.024 | 0.013 | 0.013 | 0.637 | 0.34 | 0.59 | 0.6 | 0.59 | 0.47 | 0.45 |
| 10 | 29713509 | rs10763717 | -0.239 | 2.451 | 1.04 | 0.938 | -0.003 | 0.046 | 0.046 | 0.337 | 0.1 | 0.11 | 0.21 | 0.24 | 0.09 | 0.17 |
| 10 | 29757491 | rs4747659 | -1.898 | 2.29 | 1.137 | 0.882 | -0.001 | 0.021 | 0.021 | 2.537 | 0.31 | 0.4 | 0.54 | 0.59 | 0.33 | 0.42 |
| 10 | 29758346 | rs16930246 | -0.124 | 2.284 | 1.15 | 0.877 | 0.006 | -0.011 | -0.011 | 0 | 0.06 | 0.08 | 0.16 | 0.03 | 0.08 | 0.06 |
| 10 | 29759225 | rs11007607 | -1.36 | 2.294 | 1.144 | 0.886 | 0.038 | 0.041 | 0.041 | 0.004 | 0.16 | 0.28 | 0.28 | 0.24 | 0.23 | 0.2 |
| 10 | 29759782 | rs7906814 | -1.977 | 2.293 | 1.115 | 0.878 | 0.004 | 0.005 | 0.005 | 0.018 | 0.29 | 0.35 | 0.43 | 0.56 | 0.38 | 0.39 |
| 10 | 29761314 | rs10826629 | -1.155 | 2.299 | 1.146 | 0.882 | 0 | 0.012 | 0.012 | 0.431 | 0.23 | 0.28 | 0.37 | 0.56 | 0.25 | 0.36 |
| 10 | 29765730 | rs6481596 | -1.891 | 1.82 | 1.272 | 0.446 | -0.006 | 0.012 | 0.012 | 0.089 | 0.46 | 0.67 | 0.52 | 0.6 | 0.46 | 0.6 |
| 10 | 29768499 | rs7096453 | -2.321 | 1.756 | 1.284 | 0.405 | 0.007 | 0.016 | 0.016 | 1.486 | 0.5 | 0.47 | 0.64 | 0.63 | 0.55 | 0.65 |
| 10 | 29773036 | rs11007614 | 0.794 | 1.716 | 0.949 | 0.431 | -0.043 | -0.077 | -0.077 | 0.812 | 0.12 | 0.01 | 0.14 | 0.32 | 0.15 | 0.19 |
| 10 | 29777306 | rs10826644 | -1.398 | 1.674 | 0.981 | 0.346 | -0.015 | 0.042 | 0.042 | 1.253 | 0.29 | 0.62 | 0.42 | 0.56 | 0.25 | 0.46 |
| 10 | 29779199 | rs3780853 | -1.787 | 1.639 | 0.824 | 0.351 | -0.002 | -0.003 | -0.003 | 0 | 0.34 | 0.71 | 0.47 | 0.56 | 0.39 | 0.49 |
| 10 | 29780468 | rs11007622 | -0.575 | 1.762 | 0.662 | -0.332 | 0.036 | -0.026 | -0.026 | 1.249 | 0.11 | 0.17 | 0.22 | 0.04 | 0.22 | 0.09 |
| 10 | 29781479 | rs10826648 | 1.848 | 1.979 | 0.46 | -0.445 | 0.039 | 0.057 | 0.057 | 2.839 | 0.44 | 0.11 | 0.28 | 0.62 | 0.39 | 0.38 |
| 10 | 29783692 | rs754865 | 0.395 | 1.879 | 0.316 | -0.366 | -0.013 | -0.054 | -0.054 | 0 | 0.15 | 0.07 | 0.16 | 0.34 | 0.16 | 0.2 |
| 10 | 29784489 | rs12775988 | -1.266 | 1.994 | 0.515 | -0.508 | -0.01 | -0.003 | -0.003 | 0 | 0.22 | 0.2 | 0.44 | 0.03 | 0.32 | 0.31 |
| 10 | 29785191 | rs11007625 | -1.768 | 1.884 | 0.826 | -0.48 | -0.009 | 0.008 | 0.008 | 0.535 | 0.4 | 0.64 | 0.63 | 0.37 | 0.48 | 0.51 |
| 10 | 29786463 | rs6481610 | -1.394 | 1.43 | 0.786 | -0.42 | -0.018 | -0.02 | -0.02 | 0 | 0.06 | 0.3 | 0.04 | 0.01 | 0.01 | 0.11 |
| 10 | 29789094 | rs4747661 | 0.474 | 1.741 | 0.368 | -0.371 | -0.024 | -0.052 | -0.052 | 0.009 | 0.15 | 0.07 | 0.15 | 0.32 | 0.17 | 0.21 |
| 10 | 29789648 | rs2025853 | -2.084 | 1.737 | 0.431 | -0.415 | 0.002 | 0.001 | 0.001 | 1.647 | 0.55 | 0.64 | 0.7 | 0.4 | 0.6 | 0.62 |
| 10 | 29792539 | rs10826649 | -1.616 | 1.847 | 0.804 | -0.711 | 0.006 | -0.07 | -0.07 | 0 | 0.37 | 0.56 | 0.4 | 0.08 | 0.35 | 0.39 |
| 10 | 29803661 | rs7910521 | -1.56 | 1.908 | 0.951 | -0.568 | 0.019 | -0.073 | -0.073 | 0 | 0.37 | 0.57 | 0.44 | 0.08 | 0.43 | 0.37 |
| 10 | 29803731 | rs7913801 | -1.547 | 1.908 | 0.957 | -0.569 | 0.021 | -0.078 | -0.078 | 0 | 0.37 | 0.56 | 0.44 | 0.08 | 0.43 | 0.37 |
| 10 | 29808562 | rs2368395 | -1.567 | 1.926 | 0.842 | -0.561 | 0.036 | -0.089 | -0.089 | 5.476 | 0.35 | 0.55 | 0.43 | 0.08 | 0.43 | 0.32 |
| 10 | 29809291 | rs7089591 | -0.713 | 1.991 | 0.876 | -0.563 | -0.003 | -0.066 | -0.066 | 0 | 0.11 | 0.34 | 0.14 | 0 | 0.09 | 0.18 |
| 10 | 29812422 | rs7070135 | -0.641 | 1.993 | 0.84 | -0.562 | -0.002 | -0.062 | -0.062 | 0 | 0.11 | 0.3 | 0.14 | 0 | 0.09 | 0.18 |
| 10 | 29812602 | rs7070678 | -1.225 | 1.968 | 0.496 | -0.476 | 0.019 | -0.069 | -0.069 | 0.019 | 0.37 | 0.56 | 0.44 | 0.08 | 0.43 | 0.36 |
| 10 | 29821089 | rs17756919 | -1.024 | 2.039 | 0.156 | -0.363 | 0.016 | -0.038 | -0.038 | 22.448 | 0.29 | 0.33 | 0.36 | 0.08 | 0.39 | 0.28 |
| 10 | 29826069 | rs7904705 | -1.121 | 1.95 | -0.422 | -0.109 | -0.032 | 0.031 | 0.031 | 0 | 0.41 | 0.76 | 0.57 | 0.36 | 0.46 | 0.43 |
| 10 | 29839513 | rs10159964 | -0.829 | 2.467 | 0.775 | -0.251 | 0.082 | 0.114 | 0.114 | 0 | 0.17 | 0.4 | 0.26 | 0.35 | 0.24 | 0.21 |
| 10 | 29839787 | rs10160013 | -0.848 | 2.462 | 0.87 | -0.253 | 0.09 | 0.124 | 0.124 | 0 | 0.17 | 0.42 | 0.27 | 0.35 | 0.24 | 0.21 |
| 10 | 29839864 | rs1270874 | -1.467 | 1.38 | 0.054 | -0.33 | 0.017 | 0.018 | 0.018 | 0 | 0.63 | 0.63 | 0.71 | 0.82 | 0.73 | 0.77 |
| 10 | 29840164 | rs3740002 | -0.046 | 1.603 | 0.391 | -0.255 | 0.037 | 0.056 | 0.056 | 0 | 0.16 | 0.23 | 0.25 | 0.35 | 0.24 | 0.21 |
| 10 | 29840566 | rs2887238 | -1.107 | 1.564 | 0.427 | 0.214 | -0.011 | -0.012 | -0.012 | 0 | 0.4 | 0.59 | 0.55 | 0.51 | 0.46 | 0.53 |
| 10 | 29841556 | rs4749456 | -0.101 | 1.602 | 0.287 | 0.249 | 0.065 | 0.096 | 0.096 | 0 | 0.16 | 0.32 | 0.25 | 0.36 | 0.24 | 0.21 |
| 10 | 29842681 | rs12413724 | 1.186 | 1.571 | 0.181 | 0.212 | -0.04 | 0.005 | 0.005 | 0 | 0.14 | 0.02 | 0.13 | 0.11 | 0.15 | 0.11 |
| 10 | 29843833 | rs1547169 | 0.005 | 1.565 | 0.215 | 0.224 | 0.041 | 0.063 | 0.063 | 0 | 0.16 | 0.24 | 0.25 | 0.36 | 0.24 | 0.21 |

| | | | | | | | | | | | | | | | | |
|---|---|---|---|---|---|---|---|---|---|---|---|---|---|---|---|---|
| 10 | 29851213 | rs12267267 | -0.704 | 1.518 | 0.092 | 0.219 | 0.055 | 0.092 | 0.092 | 0 | 0.2 | 0.4 | 0.27 | 0.36 | 0.24 | 0.36 |
| 10 | 29853972 | rs7077610 | -1.017 | 1.5 | 0.129 | 0.205 | 0.041 | 0.031 | 0.031 | 0 | 0.34 | 0.46 | 0.41 | 0.47 | 0.4 | 0.47 |
| 10 | 29855186 | rs7091361 | -1.475 | 1.242 | 0.111 | 0.15 | -0.024 | -0.029 | -0.029 | 0 | 0.43 | 0.72 | 0.59 | 0.51 | 0.48 | 0.54 |
| 10 | 29859879 | rs1328326 | -1.466 | 1.247 | 0.096 | 0.151 | -0.019 | -0.023 | -0.023 | 0 | 0.43 | 0.67 | 0.59 | 0.51 | 0.48 | 0.53 |
| 10 | 29860704 | rs11599713 | 0.847 | 1.559 | 0.031 | 0.179 | -0.026 | 0.015 | 0.015 | 0 | 0.09 | 0.04 | 0.09 | 0.07 | 0.15 | 0.13 |
| 10 | 29861341 | rs1247697 | -1.989 | 1.373 | -0.126 | 0.181 | -0.081 | -0.105 | -0.105 | 3.034 | 0.79 | 0.99 | 0.85 | 0.69 | 0.73 | 0.81 |
| 10 | 29864370 | rs1328327 | -1.911 | 2.046 | 0.542 | 0.238 | -0.024 | -0.028 | -0.028 | 0 | 0.43 | 0.7 | 0.59 | 0.52 | 0.47 | 0.53 |
| 10 | 29864741 | rs7903861 | -0.938 | 2.09 | 0.527 | 0.275 | 0.071 | 0.11 | 0.11 | 0 | 0.17 | 0.37 | 0.26 | 0.37 | 0.23 | 0.22 |
| 10 | 29865078 | rs973286 | -1.459 | 1.907 | 0.565 | 0.239 | 0.012 | 0.014 | 0.014 | 0 | 0.39 | 0.44 | 0.56 | 0.52 | 0.47 | 0.4 |
| 10 | 29865286 | rs973285 | -1.493 | 1.305 | 0.605 | 0.661 | 0.005 | 0.156 | 0.156 | 1.011 | 0.12 | 0.54 | 0.19 | 0.29 | 0.08 | 0.22 |
| 10 | 29874502 | rs2368369 | -0.87 | 1.866 | 0.138 | 0.47 | 0.072 | 0.073 | 0.073 | 0 | 0.16 | 0.26 | 0.24 | 0.33 | 0.25 | 0.22 |
| 10 | 29881488 | rs4083633 | 0.618 | 1.904 | -0.533 | 0.124 | -0.084 | 0.085 | 0.085 | 0 | 0.17 | 0.02 | 0.22 | 0.03 | 0.27 | 0.14 |
| 10 | 29882153 | rs2505924 | 0.069 | 0.873 | -0.723 | -0.244 | -0.008 | -0.006 | -0.006 | 0.038 | 0.38 | 0.21 | 0.25 | 0.4 | 0.41 | 0.23 |
| 10 | 29882991 | rs2488684 | -1.044 | -0.075 | -0.346 | -0.622 | -0.003 | 0.065 | 0.065 | 0 | 0.52 | 0.58 | 0.63 | 0.27 | 0.53 | 0.55 |
| 10 | 29889441 | rs2488682 | -0.967 | 0.575 | -0.274 | -0.551 | -0.001 | -0.004 | -0.004 | 0 | 0.48 | 0.37 | 0.36 | 0.21 | 0.5 | 0.42 |
| 10 | 29891260 | rs12416605 | 0.675 | 0.72 | -1.067 | -0.546 | -0.085 | 0.099 | 0.099 | 0 | 0.18 | 0.02 | 0.22 | 0.03 | 0.27 | 0.14 |
| 10 | 29892003 | rs2479691 | 0.232 | 0.526 | -0.928 | -0.456 | 0.019 | 0.019 | 0.019 | 0 | 0.27 | 0.13 | 0.13 | 0.18 | 0.22 | 0.16 |
| 10 | 29892296 | rs2505903 | 0.182 | 0.526 | -0.897 | -0.456 | 0.011 | 0.011 | 0.011 | 0 | 0.27 | 0.19 | 0.13 | 0.19 | 0.22 | 0.16 |
| 10 | 29894001 | rs2479693 | -0.473 | 0.451 | -0.69 | -0.145 | -0.004 | 0.068 | 0.068 | 0 | 0.48 | 0.29 | 0.48 | 0.23 | 0.52 | 0.39 |
| 10 | 29896243 | rs1247421 | -0.601 | -0.635 | -0.615 | 0.13 | 0.028 | -0.006 | -0.006 | 0 | 0.56 | 0.67 | 0.59 | 0.47 | 0.64 | 0.58 |
| 10 | 29896899 | rs1247422 | -0.198 | -0.79 | -0.239 | -0.185 | 0.065 | 0.022 | 0.022 | 0 | 0.3 | 0.4 | 0.45 | 0.36 | 0.41 | 0.22 |
| 10 | 29897181 | rs1247423 | 0.805 | -0.55 | 0.132 | -0.334 | 0.013 | -0.045 | -0.045 | 0.905 | 0.1 | 0.13 | 0.2 | 0 | 0.13 | 0.09 |
| 10 | 29898439 | rs1247434 | -0.562 | -0.591 | 0.111 | -0.336 | 0.058 | 0.018 | 0.018 | 0 | 0.3 | 0.47 | 0.46 | 0.36 | 0.41 | 0.22 |
| 10 | 29899946 | rs1247442 | -0.738 | -0.59 | 0.116 | -0.335 | 0.065 | 0.023 | 0.023 | 0 | 0.29 | 0.47 | 0.46 | 0.36 | 0.41 | 0.22 |
| 10 | 29900664 | rs2505907 | 1 | -0.701 | 0.018 | -0.402 | 0.016 | 0.034 | 0.034 | 0.017 | 0.18 | 0.12 | 0.1 | 0.06 | 0.14 | 0.1 |
| 10 | 29902184 | rs7904156 | 1.027 | -1.53 | 0.006 | -0.501 | 0.033 | 0.041 | 0.041 | 4.547 | 0.48 | 0.34 | 0.47 | 0.65 | 0.65 | 0.43 |
| 10 | 29902804 | rs2479686 | 0.751 | -1.073 | -0.073 | -0.419 | 0.013 | 0.006 | 0.006 | 0 | 0.17 | 0.11 | 0.1 | 0.02 | 0.13 | 0.05 |
| 10 | 29905358 | rs1777319 | 0.476 | -1.036 | -0.119 | -0.251 | -0.011 | 0.021 | 0.021 | 0.009 | 0.1 | 0.07 | 0.19 | 0 | 0.13 | 0.09 |
| 10 | 29909555 | rs1247420 | 0.478 | -1.04 | -0.089 | -0.252 | -0.004 | -0.001 | -0.001 | 0 | 0.1 | 0.09 | 0.19 | 0 | 0.13 | 0.09 |
| 10 | 29917425 | rs2505910 | 1.012 | -1.193 | 0.022 | -0.144 | -0.01 | -0.02 | -0.02 | 0 | 0.15 | 0.12 | 0.1 | 0.05 | 0.1 | 0.05 |
| 10 | 29933714 | rs2488688 | 0.141 | -1.139 | 0.087 | 0.027 | 0.044 | 0.031 | 0.031 | 0.337 | 0.49 | 0.67 | 0.3 | 0.37 | 0.43 | 0.39 |
| 10 | 29934922 | rs1247459 | -0.3 | -1.2 | 0.054 | -0.147 | 0.003 | 0.002 | 0.002 | 0 | 0.34 | 0.35 | 0.29 | 0.39 | 0.31 | 0.48 |
| 10 | 29936332 | rs12573620 | 0.675 | -1.229 | -0.023 | -0.104 | 0.015 | 0.007 | 0.007 | 0 | 0.23 | 0.17 | 0.14 | 0.21 | 0.15 | 0.12 |
| 10 | 29940175 | rs2505917 | -1 | -1.034 | -0.295 | -0.03 | -0.028 | -0.015 | -0.015 | 0.057 | 0.7 | 0.84 | 0.65 | 0.69 | 0.65 | 0.84 |
| 10 | 29947754 | rs1247456 | 0.474 | -0.874 | -0.173 | 0.011 | 0.001 | -0.001 | -0.001 | 0 | 0.13 | 0.2 | 0.2 | 0.19 | 0.17 | 0.37 |
| 10 | 29952872 | rs10826675 | 0.189 | -1.162 | 0.309 | -0.068 | -0.104 | -0.022 | -0.022 | 0 | 0.26 | 0.43 | 0.24 | 0.23 | 0.14 | 0.2 |
| 10 | 29955649 | rs9888055 | 0.609 | -0.555 | -0.192 | 0.032 | -0.082 | -0.05 | -0.05 | 0 | 0.3 | 0.42 | 0.15 | 0.19 | 0.18 | 0.21 |
| 10 | 29958333 | rs3886283 | 0.913 | 0.111 | -0.456 | 0.399 | -0.015 | -0.024 | -0.024 | 2.658 | 0.24 | 0.15 | 0.17 | 0.3 | 0.29 | 0.21 |
| 10 | 29959765 | rs1148214 | -0.265 | 0.268 | -1.004 | 0.661 | -0.064 | -0.071 | -0.071 | 0 | 0.77 | 0.89 | 0.71 | 0.66 | 0.7 | 0.86 |
| 10 | 29963887 | rs7098565 | -0.216 | -0.995 | -0.464 | 0.135 | -0.024 | -0.014 | -0.014 | 0.021 | 0.31 | 0.33 | 0.14 | 0.19 | 0.18 | 0.21 |
| 10 | 29965262 | rs1148212 | -0.056 | -0.067 | -0.049 | 0.646 | 0.001 | 0.019 | 0.019 | 0.06 | 0.2 | 0.22 | 0.24 | 0.27 | 0.21 | 0.3 |
| 10 | 29970252 | rs1148226 | -0.463 | 0.077 | 0.134 | 0.461 | -0.006 | 0.013 | 0.013 | 0.275 | 0.6 | 0.67 | 0.52 | 0.82 | 0.55 | 0.71 |
| 10 | 29970592 | rs1148204 | 0.549 | 0.108 | 0.361 | 0.488 | -0.003 | -0.007 | -0.007 | 0 | 0.2 | 0.21 | 0.25 | 0.27 | 0.23 | 0.3 |
| 10 | 29972550 | rs11007696 | 0.252 | 0.07 | 0.501 | 0.522 | -0.016 | -0.001 | -0.001 | 0.328 | 0.4 | 0.42 | 0.25 | 0.45 | 0.31 | 0.38 |
| 10 | 29972943 | rs17294592 | 0.662 | 0.507 | 0.361 | 0.264 | -0.012 | 0.01 | 0.01 | 0 | 0.1 | 0.05 | 0.29 | 0 | 0.14 | 0.06 |
| 10 | 29973953 | rs3858235 | -0.866 | 0.175 | 0.481 | 0.183 | 0.006 | -0.019 | -0.019 | 0.101 | 0.19 | 0.35 | 0.37 | 0.18 | 0.19 | 0.23 |
| 10 | 29974861 | rs4385793 | 0.982 | 0.287 | 0.449 | 0.264 | -0.001 | -0.058 | -0.058 | 0.294 | 0.3 | 0.08 | 0.17 | 0.37 | 0.26 | 0.24 |
| 10 | 29977344 | rs7901774 | -0.235 | 0.353 | 0.444 | 0.265 | 0.021 | -0.066 | -0.066 | 0.235 | 0.15 | 0.28 | 0.36 | 0.08 | 0.17 | 0.18 |
| 10 | 29977431 | rs4749479 | 0.784 | 0.384 | 0.081 | 0.281 | -0.01 | 0.006 | 0.006 | 0 | 0.09 | 0.05 | 0.29 | 0 | 0.13 | 0.06 |
| 10 | 29979708 | rs4747673 | 0.625 | 0.387 | 0.091 | 0.297 | 0.011 | 0.002 | 0.002 | 0 | 0.1 | 0.12 | 0.3 | 0.11 | 0.15 | 0.08 |
| 10 | 29982954 | rs6481619 | 0.845 | 0.42 | -0.083 | 0.373 | 0.011 | -0.009 | -0.009 | 0.43 | 0.31 | 0.19 | 0.18 | 0.37 | 0.27 | 0.24 |
| 10 | 29989182 | rs1624281 | 0.383 | 1.104 | 0.102 | 0.098 | -0.001 | 0.001 | 0.001 | 1.092 | 0.56 | 0.44 | 0.44 | 0.74 | 0.49 | 0.56 |
| 10 | 29991935 | rs1752582 | -0.761 | 0.718 | 0.008 | 0.088 | -0.001 | -0.001 | -0.001 | 0 | 0.72 | 0.74 | 0.8 | 0.82 | 0.68 | 0.75 |
| 10 | 29993344 | rs1777334 | -0.003 | 0.705 | -0.44 | 0.154 | 0 | -0.001 | -0.001 | 1.097 | 0.56 | 0.43 | 0.46 | 0.64 | 0.5 | 0.55 |
| 10 | 30000399 | rs1247086 | -0.283 | 0.395 | -0.478 | 0.129 | -0.01 | -0.029 | -0.029 | 0 | 0.4 | 0.49 | 0.45 | 0.27 | 0.35 | 0.22 |
| 10 | 30000415 | rs1247087 | -0.522 | 0.458 | -0.577 | -0.009 | -0.001 | -0.007 | -0.007 | 0 | 0.46 | 0.55 | 0.52 | 0.34 | 0.43 | 0.38 |
| 10 | 30000738 | rs1247089 | -0.33 | 0.143 | -0.856 | 0.113 | -0.02 | -0.009 | -0.009 | 0 | 0.59 | 0.76 | 0.72 | 0.54 | 0.58 | 0.38 |
| 10 | 30001433 | rs7898486 | -0.567 | 0.068 | -0.742 | 0.108 | -0.039 | 0.043 | 0.043 | 0 | 0.57 | 0.79 | 0.58 | 0.71 | 0.52 | 0.51 |
| 10 | 30005534 | rs7101204 | -1.048 | 0.04 | -0.15 | 0.162 | -0.023 | -0.027 | -0.027 | 1.168 | 0.18 | 0.39 | 0.15 | 0.21 | 0.13 | 0.22 |
| 10 | 30008609 | rs1927456 | -0.21 | 0.244 | -0.598 | 0.272 | -0.006 | -0.001 | -0.001 | 0 | 0.45 | 0.49 | 0.35 | 0.49 | 0.39 | 0.46 |
| 10 | 30008663 | rs1927457 | -0.298 | 0.16 | -0.656 | 0.215 | 0.009 | 0 | 0 | 0 | 0.45 | 0.4 | 0.31 | 0.48 | 0.32 | 0.44 |
| 10 | 30009270 | rs10740808 | 0.101 | 1.373 | -0.95 | -0.347 | -0.033 | 0.031 | 0.031 | 0 | 0.74 | 0.84 | 0.63 | 0.94 | 0.63 | 0.75 |
| 10 | 30009340 | rs11007712 | 0.022 | 1.732 | -1.279 | -0.227 | -0.057 | 0.001 | 0.001 | 0 | 0.12 | 0.04 | 0.28 | 0.11 | 0.25 | 0.16 |
| 10 | 30011304 | rs12773173 | 0.688 | 0.204 | -1.108 | -0.354 | 0.017 | 0.015 | 0.015 | 0.794 | 0.3 | 0.24 | 0.13 | 0.21 | 0.19 | 0.23 |
| 10 | 30012334 | rs7100694 | -0.934 | 0.971 | -1.252 | -0.546 | -0.028 | 0.043 | 0.043 | 0 | 0.8 | 0.93 | 0.74 | 0.94 | 0.77 | 0.81 |
| 10 | 30012843 | rs11007714 | 1.78 | 0.19 | -0.894 | -0.088 | 0 | -0.002 | -0.002 | 0 | 0.17 | 0.15 | 0.07 | 0.23 | 0.1 | 0.24 |
| 10 | 30014073 | rs10826682 | 2.002 | 0.344 | -1.135 | 0.231 | -0.074 | -0.089 | -0.089 | 0.72 | 0.16 | 0.02 | 0.19 | 0.27 | 0.24 | 0.09 |
| 10 | 30014423 | rs11007716 | 0.761 | 0.298 | -1.08 | 0.188 | 0.008 | -0.017 | -0.017 | 0 | 0.46 | 0.26 | 0.34 | 0.48 | 0.42 | 0.32 |
| 10 | 30014723 | rs1539536 | -1.268 | 0.264 | -1.057 | 0.179 | -0.021 | 0.037 | 0.037 | 0 | 0.8 | 0.91 | 0.73 | 0.94 | 0.77 | 0.81 |

| | | | | | | | | | | | | | | | | |
|---|---|---|---|---|---|---|---|---|---|---|---|---|---|---|---|---|
| 10 | 30015800 | rs7477689 | 1.113 | 0.147 | -0.31 | 0.251 | -0.075 | 0.063 | 0.063 | 0 | 0.2 | 0.45 | 0.11 | 0.3 | 0.1 | 0.29 |
| 10 | 30017048 | rs7098831 | -0.173 | 0.46 | -0.379 | 0.171 | -0.061 | 0.055 | 0.055 | 0 | 0.68 | 0.84 | 0.47 | 0.85 | 0.54 | 0.62 |
| 10 | 30018443 | rs7070379 | -0.321 | 0.132 | -0.176 | 0.202 | -0.031 | -0.017 | -0.017 | 0 | 0.61 | 0.75 | 0.48 | 0.6 | 0.52 | 0.67 |
| 10 | 30019507 | rs11007719 | 0.704 | 0.888 | 0.07 | 0.391 | -0.047 | 0.01 | 0.01 | 0 | 0.11 | 0.05 | 0.26 | 0.09 | 0.23 | 0.19 |
| 10 | 30019868 | rs12359769 | 0.73 | 0.975 | 0.073 | 0.395 | -0.045 | 0.012 | 0.012 | 0 | 0.11 | 0.05 | 0.26 | 0.09 | 0.22 | 0.19 |
| 10 | 30028267 | rs2369194 | 0.296 | 0.908 | 0.329 | 0.398 | 0.023 | -0.009 | -0.009 | 0 | 0.12 | 0.2 | 0.29 | 0.09 | 0.23 | 0.2 |
| 10 | 30032940 | rs733212 | -0.744 | 0.708 | 0.432 | 0.346 | -0.011 | 0.002 | 0.002 | 0 | 0.1 | 0.37 | 0.18 | 0.09 | 0.05 | 0.06 |
| 10 | 30034377 | rs4749487 | 0.727 | 0.845 | 0.067 | 0.402 | -0.016 | -0.012 | -0.012 | 0.516 | 0.08 | 0.05 | 0.2 | 0.14 | 0.21 | 0.17 |
| 10 | 30041106 | rs1247098 | -1.361 | -0.053 | 0.04 | 0.218 | 0.066 | 0.069 | 0.069 | 3.942 | 0.64 | 0.84 | 0.86 | 0.78 | 0.8 | 0.76 |
| 10 | 30044314 | rs1927461 | -0.124 | -0.094 | 0.421 | 0.419 | -0.062 | 0.012 | 0.012 | 0 | 0.27 | 0.48 | 0.23 | 0.32 | 0.16 | 0.34 |
| 10 | 30046498 | rs11007728 | 0.924 | -0.08 | 0.158 | 0.457 | -0.01 | 0.001 | 0.001 | 0 | 0.19 | 0.22 | 0.08 | 0.23 | 0.12 | 0.26 |
| 10 | 30054255 | rs2096249 | -0.521 | 0.174 | 0.592 | 0.338 | 0.018 | 0.019 | 0.019 | 0 | 0.1 | 0.38 | 0.19 | 0.09 | 0.06 | 0.08 |
| 10 | 30054560 | rs2105374 | -0.493 | 0.572 | 0.103 | 0.383 | 0.001 | 0.011 | 0.011 | 0 | 0.32 | 0.56 | 0.43 | 0.36 | 0.32 | 0.21 |
| 10 | 30054852 | rs1752584 | 1.363 | 1.347 | -0.619 | 0.471 | -0.032 | 0.027 | 0.027 | 0.012 | 0.05 | 0.01 | 0.17 | 0 | 0.12 | 0.05 |
| 10 | 30056318 | rs12357935 | 1.423 | 0.979 | -0.529 | 0.566 | -0.077 | -0.092 | -0.092 | 1.81 | 0.18 | 0.02 | 0.23 | 0.26 | 0.26 | 0.12 |
| 10 | 30074800 | rs1927450 | 0.402 | 1.198 | -0.245 | 0.543 | -0.008 | -0.002 | -0.002 | 0 | 0.09 | 0.06 | 0.18 | 0.15 | 0.18 | 0.16 |
| 10 | 30075215 | rs1927446 | -0.196 | 1.448 | 0.347 | 0.597 | 0.1 | 0.033 | 0.033 | 0.028 | 0.09 | 0.24 | 0.2 | 0.07 | 0.19 | 0.12 |
| 10 | 30076326 | rs11007742 | 1.579 | 0.869 | 0.229 | 0.672 | 0.033 | -0.019 | -0.019 | 0 | 0.17 | 0.1 | 0.05 | 0.22 | 0.11 | 0.26 |
| 10 | 30076923 | rs1927443 | -0.777 | 0.812 | 0.308 | 0.47 | 0.004 | 0.001 | 0.001 | 1.066 | 0.48 | 0.47 | 0.68 | 0.54 | 0.71 | 0.45 |
| 10 | 30078483 | rs11007747 | 1.254 | 0.898 | -0.34 | 0.502 | -0.098 | -0.087 | -0.087 | 0.596 | 0.26 | 0.02 | 0.24 | 0.25 | 0.3 | 0.15 |
| 10 | 30080681 | rs10826698 | -0.519 | 1.059 | -0.106 | 0.545 | 0.019 | 0.002 | 0.002 | 0.172 | 0.17 | 0.17 | 0.38 | 0.22 | 0.37 | 0.29 |
| 10 | 30080696 | rs10826699 | 0.712 | 1.047 | -0.124 | 0.545 | -0.008 | -0.003 | -0.003 | 0 | 0.09 | 0.06 | 0.18 | 0.15 | 0.18 | 0.16 |
| 10 | 30084049 | rs10826700 | 0.4 | 0.993 | -0.049 | 0.54 | 0.034 | 0.007 | 0.007 | 0.262 | 0.08 | 0.11 | 0.19 | 0.07 | 0.18 | 0.13 |
| 10 | 30084908 | rs3006538 | -0.982 | 0.807 | 0.232 | 0.488 | 0.009 | 0.003 | 0.003 | 0.555 | 0.5 | 0.52 | 0.69 | 0.54 | 0.71 | 0.45 |
| 10 | 30085845 | rs2986957 | 0.092 | 0.825 | 0.224 | 0.491 | -0.021 | -0.007 | -0.007 | 0 | 0.37 | 0.28 | 0.44 | 0.4 | 0.49 | 0.32 |
| 10 | 30087346 | rs3006546 | -1.124 | -0.086 | 0.164 | 0.388 | 0.028 | 0.003 | 0.003 | 1.504 | 0.55 | 0.63 | 0.81 | 0.56 | 0.73 | 0.54 |
| 10 | 30087359 | rs2986961 | -0.836 | 0.078 | -0.014 | 0.454 | 0.008 | 0.003 | 0.003 | 0.579 | 0.49 | 0.52 | 0.69 | 0.54 | 0.71 | 0.45 |
| 10 | 30090748 | rs4397737 | -0.055 | 0.277 | -0.165 | 0.481 | -0.025 | -0.004 | -0.004 | 0.528 | 0.17 | 0.11 | 0.34 | 0.19 | 0.36 | 0.29 |
| 10 | 30093367 | rs2986968 | 0.64 | 0.015 | -0.695 | 0.502 | -0.005 | 0 | 0 | 0 | 0.29 | 0.22 | 0.28 | 0.25 | 0.33 | 0.16 |
| 10 | 30094779 | rs12219543 | 0.784 | -0.405 | -1.022 | 0.507 | 0.027 | -0.006 | -0.006 | 0.007 | 0.22 | 0.18 | 0.15 | 0.25 | 0.11 | 0.34 |
| 10 | 30096628 | rs2986971 | -0.142 | 0 | -1.16 | 0.443 | -0.001 | -0.001 | -0.001 | 0 | 0.55 | 0.54 | 0.4 | 0.55 | 0.48 | 0.48 |
| 10 | 30097481 | rs2986972 | -1.178 | 0.164 | -1.757 | 0.297 | 0.024 | 0.044 | 0.044 | 4.85 | 0.75 | 0.91 | 0.87 | 0.8 | 0.84 | 0.8 |
| 10 | 30098281 | rs1010625 | 0.722 | 0.47 | -1.877 | 0.424 | -0.078 | -0.043 | -0.043 | 0.6 | 0.25 | 0.03 | 0.23 | 0.21 | 0.3 | 0.12 |
| 10 | 30099114 | rs3006536 | -0.182 | 0.406 | -1.294 | 0.225 | 0.008 | 0.009 | 0.009 | 0.339 | 0.55 | 0.38 | 0.5 | 0.66 | 0.68 | 0.59 |
| 10 | 30099518 | rs1927455 | -0.675 | -0.413 | -1.738 | -0.142 | 0.024 | 0.024 | 0.024 | 9.063 | 0.77 | 0.84 | 0.9 | 0.92 | 0.89 | 0.84 |
| 10 | 30099672 | rs1343735 | -0.618 | 1.195 | -1.871 | 0.368 | 0.04 | 0.01 | 0.01 | 0 | 0.25 | 0.36 | 0.28 | 0.33 | 0.37 | 0.27 |
| 10 | 30099778 | rs2986974 | 0.365 | 1.095 | -2.595 | 0.63 | -0.111 | -0.061 | -0.061 | 0.985 | 0.44 | 0.06 | 0.52 | 0.38 | 0.54 | 0.33 |
| 10 | 30101219 | rs7070647 | -0.232 | 1.643 | -1.539 | 1.166 | 0.018 | 0.039 | 0.039 | 0 | 0.11 | 0.16 | 0.15 | 0.27 | 0.19 | 0.23 |
| 10 | 30102047 | rs874697 | -0.702 | 0.891 | -2.057 | 0.782 | 0.001 | 0.039 | 0.039 | 0 | 0.54 | 0.85 | 0.48 | 0.64 | 0.55 | 0.64 |
| 10 | 30108560 | rs6481636 | -0.414 | 0.592 | -0.522 | 1.239 | 0.002 | 0 | 0 | 0 | 0.48 | 0.53 | 0.49 | 0.52 | 0.5 | 0.42 |
| 10 | 30110095 | rs10826705 | 0.158 | 0.522 | -0.154 | 1.472 | 0.01 | 0.015 | 0.015 | 0 | 0.39 | 0.23 | 0.44 | 0.33 | 0.43 | 0.37 |
| 10 | 30116428 | rs11007765 | -0.419 | 0.727 | 0.174 | 1.621 | 0.055 | 0.25 | 0.25 | 0 | 0.06 | 0.18 | 0.14 | 0.39 | 0.12 | 0.22 |
| 10 | 30118622 | rs11593046 | 1.699 | 0.763 | 0.018 | 1.467 | -0.018 | 0.004 | 0.004 | 0.039 | 0.06 | 0.01 | 0.05 | 0.03 | 0.07 | 0.06 |
| 10 | 30123871 | rs4749496 | 0.426 | 0.742 | 0.154 | 1.439 | -0.001 | 0.011 | 0.011 | 0 | 0.39 | 0.15 | 0.43 | 0.33 | 0.43 | 0.36 |
| 10 | 30124617 | rs1929239 | -0.672 | 0.863 | 0.289 | 1.41 | 0.002 | 0.02 | 0.02 | 0 | 0.54 | 0.58 | 0.63 | 0.91 | 0.62 | 0.66 |
| 10 | 30125633 | rs11007773 | -0.449 | 1.029 | -0.042 | 1.693 | 0.058 | 0.257 | 0.257 | 0.013 | 0.06 | 0.17 | 0.14 | 0.39 | 0.12 | 0.22 |
| 10 | 30128609 | rs11595141 | 0.541 | 0.663 | -0.308 | 1.303 | -0.027 | 0.021 | 0.021 | 0 | 0.09 | 0.02 | 0.09 | 0.03 | 0.13 | 0.07 |
| 10 | 30138411 | rs6481643 | -1.192 | 0.594 | -0.35 | 1.363 | -0.01 | 0.084 | 0.084 | 0 | 0.64 | 0.91 | 0.73 | 0.8 | 0.69 | 0.67 |
| 10 | 30143168 | rs7096754 | -0.789 | 1.035 | 0.408 | 1.646 | 0.024 | 0.24 | 0.24 | 0.747 | 0.11 | 0.3 | 0.13 | 0.36 | 0.1 | 0.22 |
| 10 | 30149457 | rs11007783 | 0.124 | 1.159 | -0.18 | 1.173 | -0.03 | 0.031 | 0.031 | 0 | 0.1 | 0.04 | 0.1 | 0.02 | 0.14 | 0.07 |
| 10 | 30161387 | rs4749499 | -0.669 | 1.017 | 0.08 | 1.558 | 0.005 | 0.133 | 0.133 | 0.004 | 0.1 | 0.19 | 0.12 | 0.41 | 0.09 | 0.27 |
| 10 | 30164762 | rs2989404 | -0.112 | 0.636 | -0.083 | 1.249 | 0.008 | -0.002 | -0.002 | 0.423 | 0.53 | 0.55 | 0.36 | 0.46 | 0.4 | 0.53 |
| 10 | 30165983 | rs7096778 | -0.968 | 1.277 | 0.568 | 1.538 | 0.009 | 0.046 | 0.046 | 0 | 0.51 | 0.7 | 0.66 | 0.67 | 0.59 | 0.62 |
| 10 | 30167754 | rs10740811 | -0.992 | 1.285 | 0.533 | 1.491 | 0.009 | 0.02 | 0.02 | 0 | 0.51 | 0.72 | 0.66 | 0.59 | 0.59 | 0.59 |
| 10 | 30168844 | rs10508757 | -0.273 | 1.452 | 0.025 | 1.389 | -0.012 | 0.017 | 0.017 | 0 | 0.39 | 0.29 | 0.5 | 0.31 | 0.5 | 0.34 |
| 10 | 30170992 | rs1109609 | -1.104 | 1.703 | -0.41 | 1.468 | 0.001 | 0.052 | 0.052 | 0.35 | 0.22 | 0.67 | 0.23 | 0.48 | 0.21 | 0.36 |
| 10 | 30171504 | rs914278 | -0.381 | 2.545 | -0.884 | 1.053 | 0.002 | -0.001 | -0.001 | 0 | 0.32 | 0.26 | 0.45 | 0.32 | 0.43 | 0.27 |
| 10 | 30172754 | rs867515 | -0.313 | 1.893 | -0.956 | 0.992 | -0.002 | -0.002 | -0.002 | 0.03 | 0.32 | 0.28 | 0.41 | 0.39 | 0.36 | 0.27 |
| 10 | 30175340 | rs7096905 | -0.508 | 0.862 | -1.327 | 0.5 | 0.025 | 0.037 | 0.037 | 1.923 | 0.6 | 0.72 | 0.67 | 0.73 | 0.69 | 0.72 |
| 10 | 30177888 | rs3006564 | -0.4 | 0.748 | -1.279 | 0.179 | 0.022 | 0.044 | 0.044 | 0 | 0.44 | 0.74 | 0.54 | 0.42 | 0.57 | 0.44 |
| 10 | 30180967 | rs7075330 | -0.096 | 0.172 | -0.358 | -0.1 | -0.04 | -0.054 | -0.054 | 0 | 0.1 | 0.21 | 0.04 | 0 | 0.02 | 0.01 |
| 10 | 30181616 | rs2989412 | -0.118 | 0.396 | -0.409 | -0.131 | 0.019 | -0.002 | -0.002 | 0.037 | 0.34 | 0.52 | 0.49 | 0.42 | 0.52 | 0.43 |
| 10 | 30182564 | rs17294753 | 0.871 | -0.385 | -1.129 | -0.294 | -0.048 | -0.004 | -0.004 | 0 | 0.17 | 0.03 | 0.09 | 0.14 | 0.2 | 0.28 |
| 10 | 30182960 | rs2989415 | 0.839 | 0.663 | -0.698 | -0.493 | 0.027 | 0.043 | 0.043 | 0 | 0.14 | 0.25 | 0.18 | 0.36 | 0.27 | 0.16 |
| 10 | 30185933 | rs2150558 | -0.731 | 0.634 | -0.712 | -0.507 | -0.006 | 0.04 | 0.04 | 0.038 | 0.43 | 0.73 | 0.53 | 0.42 | 0.54 | 0.43 |
| 10 | 30186233 | rs1929254 | 0.495 | 0.571 | -0.53 | -0.539 | 0.056 | -0.105 | -0.105 | 0 | 0.22 | 0.34 | 0.33 | 0.06 | 0.25 | 0.26 |
| 10 | 30188995 | rs2989426 | -0.048 | 0.549 | -0.512 | -0.427 | 0.02 | 0.035 | 0.035 | 0 | 0.21 | 0.38 | 0.19 | 0.36 | 0.29 | 0.16 |
| 10 | 30192326 | rs3119568 | -0.572 | 0.687 | 0 | -0.243 | 0.033 | 0.036 | 0.036 | 0.011 | 0.34 | 0.47 | 0.29 | 0.56 | 0.49 | 0.54 |
| 10 | 30192591 | rs10826723 | 0.639 | 0.711 | -0.076 | -0.233 | 0 | 0 | 0 | 0 | 0.19 | 0.16 | 0.11 | 0.19 | 0.21 | 0.35 |

| | | | | | | | | | | | | | | | | |
|---|---|---|---|---|---|---|---|---|---|---|---|---|---|---|---|---|
| 10 | 30194164 | rs2989430 | -1.083 | 0.522 | 0.009 | -0.046 | 0.093 | -0.066 | -0.066 | 0 | 0.58 | 0.84 | 0.63 | 0.56 | 0.73 | 0.74 |
| 10 | 30194393 | rs1610357 | -0.929 | 0.551 | 0.121 | -0.033 | 0.058 | -0.017 | -0.017 | 0.573 | 0.56 | 0.68 | 0.62 | 0.56 | 0.73 | 0.74 |
| 10 | 30195110 | rs12356319 | -0.152 | 0.645 | -0.226 | -0.039 | 0.01 | -0.04 | -0.04 | 0 | 0.42 | 0.45 | 0.44 | 0.2 | 0.45 | 0.55 |
| 10 | 30201227 | rs2481916 | -0.8 | 0.558 | -0.49 | 0.095 | 0.021 | -0.003 | -0.003 | 0 | 0.54 | 0.39 | 0.59 | 0.58 | 0.71 | 0.73 |
| 10 | 30201353 | rs1573027 | 0.364 | 1.19 | -0.703 | 0.314 | -0.06 | -0.069 | -0.069 | 4.493 | 0.13 | 0.06 | 0.16 | 0.36 | 0.27 | 0.2 |
| 10 | 30204072 | rs1756740 | -0.816 | 0.2 | -0.746 | 0.235 | -0.007 | -0.006 | -0.006 | 0 | 0.7 | 0.92 | 0.77 | 0.75 | 0.74 | 0.79 |
| 10 | 30207693 | rs3119572 | -0.9 | 0.086 | -0.745 | 0.199 | -0.002 | -0.051 | -0.051 | 0 | 0.7 | 0.93 | 0.79 | 0.66 | 0.76 | 0.79 |
| 10 | 30210147 | rs1756737 | -0.553 | -0.164 | -0.003 | 0.134 | -0.011 | 0.008 | 0.008 | 1.62 | 0.59 | 0.6 | 0.67 | 0.53 | 0.68 | 0.68 |
| 10 | 30212034 | rs17294774 | 1.248 | -0.092 | -0.644 | -0.016 | 0 | -0.002 | -0.002 | 0 | 0.17 | 0.06 | 0.1 | 0.14 | 0.18 | 0.21 |
| 10 | 30212086 | rs2776633 | -1.512 | -0.056 | -0.609 | -0.108 | -0.007 | -0.032 | -0.032 | 0.975 | 0.68 | 0.9 | 0.78 | 0.69 | 0.72 | 0.76 |
| 10 | 30212755 | rs2776632 | -0.985 | -0.076 | -0.608 | -0.111 | 0.019 | -0.028 | -0.028 | 0.44 | 0.54 | 0.84 | 0.7 | 0.55 | 0.58 | 0.58 |
| 10 | 30212930 | rs928325 | -1.7 | 0.006 | -0.518 | 0.055 | 0.008 | -0.041 | -0.041 | 0 | 0.72 | 0.9 | 0.8 | 0.69 | 0.76 | 0.79 |
| 10 | 30214703 | rs1246758 | 0.51 | 0.176 | -0.115 | -0.046 | 0.044 | 0.061 | 0.061 | 0 | 0.19 | 0.41 | 0.27 | 0.29 | 0.25 | 0.21 |
| 10 | 30215238 | rs2795591 | -0.89 | 0.026 | -0.242 | -0.063 | 0.008 | -0.003 | -0.003 | 0 | 0.59 | 0.64 | 0.68 | 0.45 | 0.68 | 0.64 |
| 10 | 30218346 | rs7922737 | 0.504 | 0.033 | -0.175 | 0.002 | -0.016 | 0.009 | 0.009 | 0.276 | 0.19 | 0.28 | 0.14 | 0.23 | 0.13 | 0.17 |
| 10 | 30220668 | rs2487930 | -0.817 | 0.339 | -0.071 | 0.094 | 0.016 | 0.005 | 0.005 | 0.218 | 0.54 | 0.67 | 0.66 | 0.45 | 0.64 | 0.62 |
| 10 | 30222053 | rs11007820 | 0.147 | 0.407 | -0.721 | -0.186 | -0.013 | 0.06 | 0.06 | 0.472 | 0.35 | 0.21 | 0.4 | 0.16 | 0.38 | 0.42 |
| 10 | 30223380 | rs11007823 | -0.82 | 0.454 | -0.876 | -0.084 | -0.016 | 0.007 | 0.007 | 0.874 | 0.58 | 0.64 | 0.54 | 0.39 | 0.53 | 0.6 |
| 10 | 30223537 | rs11007825 | -0.533 | 0.128 | -0.641 | -0.477 | 0.001 | 0.004 | 0.004 | 0.399 | 0.38 | 0.57 | 0.43 | 0.19 | 0.33 | 0.34 |
| 10 | 30224987 | rs1774230 | -1.399 | 0.344 | -1.255 | -0.544 | -0.003 | -0.061 | -0.061 | 0.372 | 0.72 | 0.88 | 0.76 | 0.59 | 0.74 | 0.81 |
| 10 | 30225724 | rs2776650 | 0.356 | 1.158 | -1.278 | -0.142 | -0.021 | -0.052 | -0.052 | 0.957 | 0.32 | 0.25 | 0.34 | 0.48 | 0.43 | 0.39 |
| 10 | 30230749 | rs10763769 | -0.346 | 1.193 | -0.646 | -0.62 | -0.002 | -0.011 | -0.011 | 0.18 | 0.13 | 0.18 | 0.11 | 0.09 | 0.12 | 0.13 |
| 10 | 30230765 | rs7078452 | -0.563 | 1.034 | -1.029 | -0.504 | -0.004 | -0.02 | -0.02 | 0.043 | 0.15 | 0.12 | 0.25 | 0.01 | 0.19 | 0.17 |
| 10 | 30233296 | rs1929247 | -0.974 | 0.58 | -1.36 | -0.59 | -0.016 | 0.087 | 0.087 | 0.28 | 0.24 | 0.15 | 0.26 | 0.05 | 0.24 | 0.11 |
| 10 | 30233374 | rs6481648 | -1.162 | 0.167 | -0.738 | 0.469 | -0.015 | 0.038 | 0.038 | 2.709 | 0.19 | 0.31 | 0.26 | 0.37 | 0.1 | 0.18 |
| 10 | 30233437 | rs2795593 | 0.355 | 0.34 | -1.601 | -0.02 | -0.076 | -0.16 | -0.16 | 0 | 0.31 | 0.05 | 0.35 | 0.52 | 0.36 | 0.39 |
| 10 | 30234246 | rs17294794 | -0.54 | 0.161 | -0.978 | -0.06 | 0.007 | 0.005 | 0.005 | 0.062 | 0.09 | 0.15 | 0.16 | 0.11 | 0.12 | 0.07 |
| 10 | 30242543 | rs2776641 | 0.275 | -0.083 | -0.628 | -0.13 | 0.006 | -0.043 | -0.043 | 0 | 0.37 | 0.1 | 0.45 | 0.6 | 0.36 | 0.42 |
| 10 | 30243844 | rs2795610 | -1.053 | -0.725 | 0.015 | -0.409 | -0.019 | 0.058 | 0.058 | 0 | 0.57 | 0.76 | 0.67 | 0.72 | 0.5 | 0.57 |
| 10 | 30245561 | rs12243869 | -0.535 | -0.754 | 0.585 | -0.413 | -0.036 | -0.035 | -0.035 | 0 | 0.1 | 0.39 | 0.05 | 0.01 | 0.01 | 0.08 |
| 10 | 30246232 | rs1246762 | -1.143 | -0.926 | 0.171 | -0.513 | -0.07 | 0.058 | 0.058 | 0 | 0.61 | 0.86 | 0.69 | 0.72 | 0.51 | 0.66 |
| 10 | 30249705 | rs2795617 | 0.628 | -0.862 | 0.073 | -0.542 | -0.025 | -0.079 | -0.079 | 0.226 | 0.37 | 0.07 | 0.44 | 0.6 | 0.36 | 0.43 |
| 10 | 30250420 | rs2795622 | 0.092 | -0.757 | 0.701 | -0.896 | -0.006 | 0.079 | 0.079 | 0 | 0.47 | 0.36 | 0.62 | 0.76 | 0.49 | 0.49 |
| 10 | 30253179 | rs1617192 | -0.695 | -0.681 | 0.574 | -0.831 | -0.071 | 0.105 | 0.105 | 0 | 0.61 | 0.86 | 0.69 | 0.77 | 0.5 | 0.66 |
| 10 | 30254563 | rs1774221 | -0.646 | -0.67 | 0.795 | -0.827 | -0.039 | 0.08 | 0.08 | 0 | 0.6 | 0.81 | 0.68 | 0.77 | 0.5 | 0.66 |
| 10 | 30264822 | rs3847396 | -0.658 | -0.823 | 0.789 | -0.899 | -0.028 | 0.095 | 0.095 | 0 | 0.6 | 0.82 | 0.68 | 0.83 | 0.47 | 0.69 |
| 10 | 30265093 | rs11007837 | -0.67 | -0.946 | 1.181 | -0.946 | -0.023 | -0.016 | -0.016 | 0 | 0.09 | 0.42 | 0.06 | 0.01 | 0.01 | 0.08 |
| 10 | 30266409 | rs10763770 | -0.703 | -0.932 | 1.171 | -0.942 | -0.028 | 0.096 | 0.096 | 0 | 0.6 | 0.82 | 0.68 | 0.83 | 0.47 | 0.69 |
| 10 | 30267084 | rs4747692 | -0.025 | -0.953 | 1.14 | -0.95 | 0.017 | 0.097 | 0.097 | 0.046 | 0.49 | 0.36 | 0.61 | 0.82 | 0.46 | 0.61 |
| 10 | 30270596 | rs11007842 | -0.405 | -0.896 | 0.994 | -1.128 | -0.018 | -0.011 | -0.011 | 0 | 0.08 | 0.26 | 0.04 | 0.01 | 0.01 | 0.07 |
| 10 | 30273031 | rs1571764 | -0.688 | -0.88 | 0.872 | -1.118 | -0.027 | 0.12 | 0.12 | 0 | 0.6 | 0.82 | 0.68 | 0.88 | 0.47 | 0.69 |
| 10 | 30274368 | rs2369341 | 0.086 | -0.894 | 0.796 | -1.124 | 0.014 | 0.141 | 0.141 | 0.492 | 0.48 | 0.29 | 0.6 | 0.88 | 0.46 | 0.62 |
| 10 | 30278236 | rs4747695 | -0.744 | -0.551 | 0.491 | -0.974 | -0.012 | 0.059 | 0.059 | 0 | 0.6 | 0.72 | 0.67 | 0.88 | 0.47 | 0.69 |
| 10 | 30282715 | rs11007847 | -0.943 | -0.473 | 0.266 | -0.477 | -0.016 | -0.009 | -0.009 | 0 | 0.08 | 0.27 | 0.04 | 0.01 | 0.01 | 0.07 |
| 10 | 30285513 | rs11007849 | -0.318 | -0.357 | -0.901 | -0.748 | 0.036 | 0.185 | 0.185 | 0 | 0.52 | 0.8 | 0.64 | 0.84 | 0.41 | 0.58 |
| 10 | 30286263 | rs12764479 | -1.124 | -1.109 | -0.711 | -0.529 | -0.027 | -0.008 | -0.008 | 0 | 0.14 | 0.21 | 0.11 | 0.12 | 0.11 | 0.1 |
| 10 | 30287398 | rs7917431 | 0.16 | -0.079 | -1.833 | -1.042 | -0.141 | 0.136 | 0.136 | 0 | 0.3 | 0.03 | 0.18 | 0.12 | 0.4 | 0.14 |
| 10 | 30288622 | rs2795624 | -0.346 | -1.634 | -2.078 | -0.485 | 0.087 | -0.022 | -0.022 | 0.191 | 0.35 | 0.83 | 0.4 | 0.59 | 0.18 | 0.51 |
| 10 | 30291963 | rs1774240 | -0.952 | -1.312 | -2.371 | -0.639 | -0.089 | 0.012 | 0.012 | 3.413 | 0.66 | 0.94 | 0.59 | 0.73 | 0.58 | 0.66 |
| 10 | 30292597 | rs11007851 | 0.181 | -1.323 | -1.089 | -0.685 | 0.013 | -0.032 | -0.032 | 0 | 0.37 | 0.48 | 0.27 | 0.2 | 0.46 | 0.22 |
| 10 | 30295538 | rs11597994 | -0.214 | -1.412 | -0.748 | -0.687 | 0.012 | 0.025 | 0.025 | 0.05 | 0.06 | 0.22 | 0.05 | 0.05 | 0.05 | 0.03 |
| 10 | 30296648 | rs4749521 | 0.815 | -1.473 | -0.869 | -0.415 | -0.108 | 0.076 | 0.076 | 0 | 0.3 | 0.12 | 0.26 | 0.18 | 0.41 | 0.13 |
| 10 | 30297987 | rs1418276 | -0.023 | -1.421 | -0.757 | -0.402 | 0.004 | -0.009 | -0.009 | 0 | 0.38 | 0.49 | 0.32 | 0.24 | 0.47 | 0.23 |
| 10 | 30300420 | rs17294968 | 0.776 | -1.534 | -1.336 | -0.347 | -0.155 | 0.045 | 0.045 | 0 | 0.31 | 0.04 | 0.27 | 0.22 | 0.42 | 0.15 |
| 10 | 30300996 | rs2505132 | 0.759 | -1.869 | -1.609 | -0.235 | -0.095 | -0.005 | -0.005 | 0 | 0.35 | 0.06 | 0.41 | 0.28 | 0.43 | 0.35 |
| 10 | 30304918 | rs1063205 | 1.106 | -1.834 | -1.069 | -0.239 | -0.044 | 0.008 | 0.008 | 0 | 0.27 | 0.14 | 0.26 | 0.3 | 0.38 | 0.21 |
| 10 | 30306804 | rs2478839 | 0.945 | -2.051 | -0.451 | 0.141 | -0.08 | 0.065 | 0.065 | 1.317 | 0.35 | 0.09 | 0.45 | 0.2 | 0.43 | 0.27 |
| 10 | 30308675 | rs11598974 | -0.241 | -2.128 | 0.106 | 0.1 | 0.058 | 0.06 | 0.06 | 0.071 | 0.06 | 0.36 | 0.06 | 0.03 | 0.05 | 0.06 |
| 10 | 30310120 | rs2014144 | -0.501 | -2.591 | -0.945 | -0.024 | -0.05 | 0.103 | 0.103 | 6.567 | 0.49 | 0.92 | 0.39 | 0.34 | 0.48 | 0.37 |
| 10 | 30316208 | rs2185724 | 0.302 | -2.772 | -0.844 | -0.101 | 0 | -0.01 | -0.01 | 0 | 0.4 | 0.37 | 0.3 | 0.3 | 0.43 | 0.26 |
| 10 | 30316872 | rs3739996 | 1.262 | -2.629 | -1.823 | -0.075 | -0.056 | 0.034 | 0.034 | 0 | 0.29 | 0.11 | 0.24 | 0.22 | 0.39 | 0.13 |
| 10 | 30317826 | rs7920682 | -0.351 | -2.222 | -1.984 | 0.06 | -0.032 | 0.117 | 0.117 | 6.202 | 0.5 | 0.91 | 0.38 | 0.34 | 0.48 | 0.37 |
| 10 | 30317949 | rs2478835 | 0.807 | -1.699 | -1.649 | -0.027 | -0.095 | 0.037 | 0.037 | 0.963 | 0.33 | 0.06 | 0.45 | 0.19 | 0.43 | 0.27 |
| 10 | 30331829 | rs1342150 | 0.664 | -1.449 | -1.734 | 0.104 | -0.094 | 0.03 | 0.03 | 3.392 | 0.34 | 0.07 | 0.39 | 0.2 | 0.42 | 0.28 |
| 10 | 30332445 | rs4749523 | 0.617 | -1.679 | -1.643 | -0.083 | -0.063 | 0.038 | 0.038 | 0 | 0.27 | 0.12 | 0.27 | 0.19 | 0.37 | 0.21 |
| 10 | 30340018 | rs2478833 | 0.601 | -1.559 | -1.565 | -0.465 | -0.036 | 0.114 | 0.114 | 0.393 | 0.4 | 0.19 | 0.35 | 0.08 | 0.51 | 0.41 |
| 10 | 30346833 | rs2478832 | 1.377 | -1.499 | -1.938 | -0.558 | -0.158 | 0.167 | 0.167 | 0.309 | 0.32 | 0.02 | 0.27 | 0.04 | 0.42 | 0.27 |
| 10 | 30355052 | rs1122730 | -0.255 | -1.852 | -2.181 | 0.108 | 0.007 | -0.082 | -0.082 | 0.922 | 0.42 | 0.9 | 0.48 | 0.59 | 0.34 | 0.27 |
| 10 | 30355193 | rs2255515 | -0.542 | -2.703 | -2.162 | 0.188 | -0.003 | -0.109 | -0.109 | 0 | 0.81 | 0.98 | 0.82 | 0.66 | 0.85 | 0.7 |

| | | | | | | | | | | | | | | | | |
|---|---|---|---|---|---|---|---|---|---|---|---|---|---|---|---|---|
| 10 | 30359559 | rs10826757 | 0.095 | -2.992 | -0.885 | 0.701 | -0.011 | -0.009 | -0.009 | 0 | 0.31 | 0.32 | 0.26 | 0.27 | 0.25 | 0.15 |
| 10 | 30366521 | rs2478830 | 1.541 | -3.109 | -1.168 | 0.472 | -0.001 | 0.237 | 0.237 | 0.381 | 0.44 | 0.11 | 0.45 | 0.08 | 0.58 | 0.47 |
| 10 | 30367323 | rs10826758 | 0.369 | -3.021 | -0.788 | 0.697 | 0.003 | 0.029 | 0.029 | 0 | 0.06 | 0.06 | 0.14 | 0.31 | 0.03 | 0.1 |
| 10 | 30369860 | rs2505127 | -0.691 | -3.018 | -0.623 | 0.758 | 0.003 | -0.093 | -0.093 | 0.876 | 0.85 | 0.99 | 0.92 | 0.72 | 0.88 | 0.73 |
| 10 | 30376771 | rs4749528 | -0.742 | -2.857 | -0.257 | 0.854 | 0.037 | -0.043 | -0.043 | 0 | 0.4 | 0.73 | 0.39 | 0.55 | 0.31 | 0.23 |
| 10 | 30378092 | rs1935436 | -0.848 | -2.859 | 0.132 | 0.853 | 0.039 | -0.035 | -0.035 | 0 | 0.41 | 0.74 | 0.39 | 0.55 | 0.31 | 0.23 |
| 10 | 30387015 | rs16931056 | 0.519 | -2.83 | -0.028 | 0.826 | 0.002 | 0.006 | 0.006 | 0 | 0.06 | 0.06 | 0.06 | 0.22 | 0.03 | 0.06 |
| 10 | 30389213 | rs3847404 | -0.876 | -2.735 | 0.322 | 0.881 | 0.036 | -0.033 | -0.033 | 0.19 | 0.41 | 0.74 | 0.39 | 0.53 | 0.33 | 0.22 |
| 10 | 30389836 | rs7072215 | -1.101 | -2.751 | 0.518 | 0.769 | 0.001 | 0.008 | 0.008 | 0.801 | 0.06 | 0.24 | 0.03 | 0.02 | 0.01 | 0.06 |
| 10 | 30397456 | rs17295031 | 0.187 | -2.75 | 0.485 | 0.796 | -0.017 | -0.02 | -0.02 | 0 | 0.28 | 0.12 | 0.26 | 0.32 | 0.29 | 0.11 |
| 10 | 30397839 | rs12220219 | 0.465 | -2.762 | 0.13 | 0.722 | -0.087 | -0.067 | -0.067 | 0 | 0.25 | 0.01 | 0.23 | 0.3 | 0.28 | 0.11 |
| 10 | 30398791 | rs2505115 | -0.358 | -2.396 | 0.146 | 0.965 | 0.005 | -0.035 | -0.035 | 0 | 0.86 | 0.93 | 0.91 | 0.68 | 0.86 | 0.73 |
| 10 | 30398869 | rs9988732 | -1.291 | -2.854 | 0.379 | 0.828 | 0.001 | -0.01 | -0.01 | 0 | 0.39 | 0.72 | 0.48 | 0.63 | 0.39 | 0.31 |
| 10 | 30400396 | rs12266240 | -0.459 | -2.337 | 0.718 | 1.235 | -0.017 | 0.012 | 0.012 | 0 | 0.08 | 0.16 | 0.12 | 0.1 | 0.05 | 0.1 |
| 10 | 30407527 | rs2505082 | 0.143 | -2.376 | 0.72 | 1.301 | -0.007 | 0.025 | 0.025 | 0 | 0.8 | 0.71 | 0.85 | 0.7 | 0.8 | 0.72 |
| 10 | 30407984 | rs10826765 | -0.563 | -2.661 | 0.574 | 1.122 | -0.023 | 0.032 | 0.032 | 0.575 | 0.36 | 0.4 | 0.34 | 0.55 | 0.29 | 0.33 |
| 10 | 30408814 | rs2887510 | -1.033 | -2.75 | 0.151 | 1.068 | 0.033 | 0.003 | 0.003 | 0.204 | 0.41 | 0.62 | 0.44 | 0.65 | 0.3 | 0.41 |
| 10 | 30411253 | rs2066341 | 0.309 | -2.614 | -0.808 | 1.022 | 0.003 | -0.053 | -0.053 | 0 | 0.26 | 0.06 | 0.21 | 0.32 | 0.25 | 0.22 |
| 10 | 30413194 | rs2478837 | 1.616 | -2.332 | -0.971 | 0.638 | -0.111 | 0.237 | 0.237 | 0.003 | 0.39 | 0.03 | 0.4 | 0.05 | 0.5 | 0.29 |
| 10 | 30420194 | rs3121293 | 1.634 | -2.505 | -1.144 | 0.679 | -0.1 | 0.229 | 0.229 | 0.049 | 0.39 | 0.03 | 0.44 | 0.07 | 0.51 | 0.29 |
| 10 | 30423885 | rs11007913 | 0.088 | -2.238 | -0.934 | 1.285 | -0.009 | -0.131 | -0.131 | 0 | 0.26 | 0.01 | 0.18 | 0.36 | 0.23 | 0.22 |
| 10 | 30430255 | rs2505093 | 0.685 | -1.517 | 0.06 | 1.797 | -0.013 | 0.03 | 0.03 | 1.802 | 0.9 | 0.8 | 0.82 | 0.75 | 0.93 | 0.75 |
| 10 | 30433280 | rs2482839 | -0.424 | -1.515 | 0.026 | 1.827 | 0.002 | -0.018 | -0.018 | 0 | 0.94 | 0.97 | 0.91 | 0.94 | 0.94 | 0.86 |
| 10 | 30433291 | rs2482838 | 0.706 | -1.521 | 0.057 | 1.839 | -0.011 | 0.029 | 0.029 | 1.873 | 0.91 | 0.82 | 0.82 | 0.76 | 0.93 | 0.81 |
| 10 | 30433467 | rs3006509 | 0.081 | -1.618 | 0.218 | 1.873 | -0.002 | 0.002 | 0.002 | 0 | 0.14 | 0.12 | 0.12 | 0.31 | 0.17 | 0.21 |
| 10 | 30433590 | rs2505098 | 0.11 | -1.62 | 0.588 | 1.91 | 0.007 | -0.008 | -0.008 | 0 | 0.94 | 0.95 | 0.91 | 0.94 | 0.94 | 0.86 |
| 10 | 30442069 | rs532904 | 0.867 | -1.561 | 0.889 | 1.998 | -0.018 | 0.042 | 0.042 | 1.631 | 0.9 | 0.78 | 0.82 | 0.75 | 0.92 | 0.75 |
| 10 | 30444255 | rs581385 | 0.798 | -1.556 | 0.875 | 1.986 | -0.016 | 0.038 | 0.038 | 1.677 | 0.91 | 0.8 | 0.82 | 0.75 | 0.93 | 0.75 |
| 10 | 30454233 | rs2488034 | 1.237 | -1.632 | 1.224 | 1.672 | -0.001 | 0 | 0 | 0.253 | 0.62 | 0.62 | 0.62 | 0.31 | 0.67 | 0.57 |
| 10 | 30468830 | rs586286 | 0.626 | -1.591 | 1.325 | 2.031 | -0.02 | 0.101 | 0.101 | 1.296 | 0.9 | 0.82 | 0.88 | 0.69 | 0.92 | 0.79 |
| 10 | 30469934 | rs1571957 | 1.104 | -1.594 | 1.642 | 1.91 | -0.007 | 0 | 0 | 4.958 | 0.64 | 0.59 | 0.68 | 0.37 | 0.67 | 0.62 |
| 10 | 30470826 | rs473195 | -0.448 | -1.536 | 1.119 | 1.998 | 0.001 | -0.002 | -0.002 | 0 | 0.25 | 0.14 | 0.2 | 0.27 | 0.25 | 0.17 |
| 10 | 30472392 | rs1977583 | 1.287 | -1.631 | 0.88 | 1.794 | 0.048 | 0.055 | 0.055 | 0 | 0.59 | 0.13 | 0.54 | 0.28 | 0.67 | 0.55 |
| 10 | 30477204 | rs2505112 | 1.441 | -1.672 | 0.925 | 1.682 | 0.066 | 0.092 | 0.092 | 0.1 | 0.58 | 0.14 | 0.53 | 0.26 | 0.67 | 0.46 |
| 10 | 30481446 | rs2488005 | 1.612 | -1.796 | 1.414 | 1.858 | -0.047 | 0.142 | 0.142 | 1.456 | 0.85 | 0.33 | 0.81 | 0.63 | 0.92 | 0.77 |
| 10 | 30488590 | rs2689220 | 1.074 | -1.782 | 1.763 | 1.865 | -0.023 | 0.333 | 0.333 | 0.571 | 0.88 | 0.56 | 0.84 | 0.63 | 0.92 | 0.79 |
| 10 | 30490704 | rs2462021 | 1.469 | -1.763 | 1.705 | 1.862 | 0.059 | 0.082 | 0.082 | 0 | 0.59 | 0.14 | 0.54 | 0.26 | 0.67 | 0.46 |
| 10 | 30492605 | rs2151114 | -0.94 | -1.759 | 1.751 | 1.865 | -0.013 | 0.01 | 0.01 | 0 | 0.29 | 0.39 | 0.28 | 0.35 | 0.26 | 0.23 |
| 10 | 30496543 | rs2488011 | 1.887 | -1.742 | 1.764 | 1.852 | -0.052 | 0.112 | 0.112 | 0.781 | 0.84 | 0.32 | 0.8 | 0.61 | 0.92 | 0.71 |
| 10 | 30501007 | rs2488015 | -0.253 | -1.745 | 1.863 | 1.633 | 0.001 | 0.09 | 0.09 | 1.206 | 0.93 | 0.96 | 0.9 | 0.61 | 0.93 | 0.79 |
| 10 | 30502532 | rs7895257 | -1.393 | -1.736 | 1.923 | 1.389 | 0.057 | 0.053 | 0.053 | 4.476 | 0.06 | 0.43 | 0.06 | 0 | 0.01 | 0 |
| 10 | 30503579 | rs2779068 | 1.452 | -1.725 | 1.627 | 1.407 | 0.059 | 0.086 | 0.086 | 0.004 | 0.58 | 0.13 | 0.54 | 0.27 | 0.66 | 0.48 |
| 10 | 30509608 | rs671123 | 0.283 | -1.628 | 1.716 | 1.472 | -0.008 | 0.207 | 0.207 | 0.576 | 0.92 | 0.84 | 0.82 | 0.49 | 0.92 | 0.71 |
| 10 | 30511016 | rs553395 | -0.1 | -1.612 | 1.45 | 1.441 | 0.002 | 0.081 | 0.081 | 1.265 | 0.93 | 0.96 | 0.9 | 0.55 | 0.93 | 0.75 |
| 10 | 30512247 | rs11591768 | -0.563 | -1.586 | 1.802 | 1.269 | 0.002 | 0.004 | 0.004 | 0 | 0.25 | 0.22 | 0.2 | 0.23 | 0.25 | 0.19 |
| 10 | 30512258 | rs638572 | -2.018 | -1.391 | 2.35 | 1.596 | 0.002 | 0.021 | 0.021 | 5.241 | 0.08 | 0.52 | 0.13 | 0.02 | 0.01 | 0.03 |
| 10 | 30530796 | rs577377 | 1.163 | -1.523 | 1.923 | 1.508 | -0.048 | 0.173 | 0.173 | 0 | 0.83 | 0.65 | 0.81 | 0.46 | 0.86 | 0.66 |
| 10 | 30533660 | rs2689208 | 1.64 | -1.355 | 0.822 | 1.557 | -0.016 | 0.015 | 0.015 | 0.069 | 0.78 | 0.29 | 0.77 | 0.46 | 0.88 | 0.65 |
| 10 | 30537797 | rs7906520 | -0.572 | -0.82 | 0.216 | 1.612 | -0.001 | -0.012 | -0.012 | 0.095 | 0.13 | 0.07 | 0.11 | 0.17 | 0.15 | 0.11 |
| 10 | 30560230 | rs531707 | 0.957 | -0.887 | 0.607 | 1.779 | -0.017 | 0.092 | 0.092 | 0 | 0.85 | 0.8 | 0.84 | 0.46 | 0.88 | 0.7 |
| 10 | 30570045 | rs3006499 | -0.975 | -0.568 | 0.298 | 1.357 | -0.002 | 0.002 | 0.002 | 0.152 | 0.19 | 0.16 | 0.08 | 0.23 | 0.13 | 0.23 |
| 10 | 30572687 | rs612507 | -1.544 | -0.961 | 0.969 | 1.352 | -0.005 | 0.001 | 0.001 | 4.413 | 0.25 | 0.68 | 0.11 | 0.27 | 0.14 | 0.32 |
| 10 | 30574681 | rs2990498 | -1.823 | -0.22 | 1.605 | 1.275 | 0.013 | -0.019 | -0.019 | 0 | 0.18 | 0.46 | 0.17 | 0.13 | 0.17 | 0.19 |
| 10 | 30584513 | rs12355619 | -0.009 | -0.242 | 1.349 | 1.542 | -0.017 | 0 | 0 | 0 | 0.06 | 0 | 0.06 | 0.05 | 0.08 | 0.1 |
| 10 | 30588673 | rs586594 | 0.094 | -0.342 | 0.974 | 1.174 | 0.001 | 0.006 | 0.006 | 0 | 0.83 | 0.93 | 0.95 | 0.9 | 0.87 | 0.75 |
| 10 | 30609810 | rs2480296 | -1.381 | -0.018 | 1.339 | 1.02 | 0.003 | 0.042 | 0.042 | 3.524 | 0.4 | 0.71 | 0.32 | 0.79 | 0.31 | 0.48 |
| 10 | 30610365 | rs4749552 | -0.979 | 0.125 | 1.502 | 1.247 | 0.032 | 0.337 | 0.337 | 0 | 0.07 | 0.2 | 0.15 | 0.52 | 0.05 | 0.17 |
| 10 | 30620795 | rs2247081 | 0.778 | 0.313 | 1.191 | 1.046 | 0.009 | 0.019 | 0.019 | 0 | 0.88 | 0.75 | 0.89 | 0.88 | 0.85 | 0.87 |
| 10 | 30628478 | rs2254706 | -0.279 | 0.451 | 1.199 | 0.91 | -0.008 | 0 | 0 | 0 | 0.26 | 0.2 | 0.12 | 0.21 | 0.25 | 0.3 |
| 10 | 30629226 | rs1047991 | -0.279 | 0.451 | 1.149 | 0.917 | -0.011 | -0.002 | -0.002 | 0.498 | 0.26 | 0.19 | 0.12 | 0.21 | 0.25 | 0.3 |
| 10 | 30639220 | rs11008005 | -0.925 | 0.387 | 1.202 | 0.826 | 0.03 | 0.345 | 0.345 | 0 | 0.08 | 0.21 | 0.16 | 0.54 | 0.05 | 0.17 |
| 10 | 30641430 | rs1762598 | -0.308 | 0.392 | 1.041 | 0.751 | -0.007 | 0.012 | 0.012 | 4.022 | 0.26 | 0.23 | 0.13 | 0.17 | 0.24 | 0.3 |
| 10 | 30643087 | rs10826787 | -0.18 | 0.288 | 0.673 | -0.439 | -0.037 | -0.029 | -0.029 | 0.296 | 0.08 | 0 | 0.08 | 0.09 | 0.14 | 0.1 |
| 10 | 30643872 | rs10826790 | 0.977 | 0.278 | 0.749 | -0.446 | -0.017 | 0.075 | 0.075 | 0.544 | 0.56 | 0.32 | 0.61 | 0.19 | 0.56 | 0.41 |
| 10 | 30650152 | rs2480289 | 1.283 | 0.291 | 0.707 | -0.393 | 0.003 | -0.003 | -0.003 | 0.278 | 0.85 | 0.53 | 0.86 | 0.89 | 0.85 | 0.87 |
| 10 | 30651481 | rs2489857 | -0.831 | 0.668 | 0.209 | -0.28 | -0.001 | -0.001 | -0.001 | 0 | 0.37 | 0.67 | 0.32 | 0.7 | 0.29 | 0.52 |
| 10 | 30653146 | rs7093966 | -1.477 | 0.662 | 0.302 | -0.022 | 0.006 | -0.01 | -0.01 | 0 | 0.15 | 0.48 | 0.14 | 0.09 | 0.15 | 0.13 |
| 10 | 30654528 | rs4749554 | -0.935 | 1.379 | -0.713 | -0.083 | -0.031 | 0.022 | 0.022 | 0 | 0.05 | 0.01 | 0.29 | 0 | 0.16 | 0.1 |
| 10 | 30658147 | rs2252396 | -0.313 | 0.464 | -0.674 | -0.402 | -0.002 | -0.008 | -0.008 | 0.564 | 0.18 | 0.16 | 0.11 | 0.06 | 0.17 | 0.16 |

| | | | | | | | | | | | | | | | | |
|---|---|---|---|---|---|---|---|---|---|---|---|---|---|---|---|---|
| 10 | 30661341 | rs10826795 | 0.621 | 0.516 | -1.802 | -1.249 | -0.1 | 0.174 | 0.174 | 0.739 | 0.29 | 0.02 | 0.25 | 0.04 | 0.35 | 0.26 |
| 10 | 30663275 | rs2297040 | 0.317 | 0.35 | -1.254 | -0.844 | -0.001 | 0.004 | 0.004 | 0 | 0.36 | 0.52 | 0.43 | 0.63 | 0.34 | 0.28 |
| 10 | 30663368 | rs2297039 | 0.317 | 0.35 | -1.254 | -0.844 | -0.001 | 0.004 | 0.004 | 0 | 0.36 | 0.52 | 0.43 | 0.63 | 0.34 | 0.28 |
| 10 | 30669430 | rs1041677 | -0.954 | 0.406 | -1.782 | -1.372 | 0.017 | 0.031 | 0.031 | 0.12 | 0.85 | 0.92 | 0.95 | 0.91 | 0.93 | 0.88 |
| 10 | 30671658 | rs1831249 | 0.576 | 0.274 | -1.537 | -1.403 | -0.003 | -0.001 | -0.001 | 0 | 0.23 | 0.23 | 0.44 | 0.72 | 0.29 | 0.33 |
| 10 | 30673598 | rs2265749 | -0.385 | 0.558 | -1.132 | -1.261 | -0.004 | -0.019 | -0.019 | 0 | 0.33 | 0.61 | 0.27 | 0.15 | 0.28 | 0.28 |
| 10 | 30678440 | rs2244677 | 0.062 | 0.232 | -0.981 | -0.474 | 0.002 | 0.042 | 0.042 | 0.008 | 0.5 | 0.56 | 0.37 | 0.18 | 0.53 | 0.46 |
| 10 | 30687815 | rs1754239 | 0.899 | -0.322 | -1.004 | 0.187 | 0.008 | 0 | 0 | 2.55 | 0.76 | 0.25 | 0.71 | 0.75 | 0.81 | 0.67 |
| 10 | 30699265 | rs1536833 | -0.265 | -1.011 | -1.746 | 0.394 | 0.043 | -0.036 | -0.036 | 0.219 | 0.4 | 0.13 | 0.24 | 0.46 | 0.29 | 0.57 |
| 10 | 30703513 | rs2480277 | 0.751 | -2.011 | -1.83 | 0.456 | -0.005 | -0.028 | -0.028 | 0.559 | 0.38 | 0.13 | 0.53 | 0.35 | 0.6 | 0.32 |
| 10 | 30706590 | rs12770958 | 1.061 | -2.425 | -2.172 | 0.481 | 0.019 | -0.089 | -0.089 | 1.241 | 0.21 | 0.02 | 0.16 | 0.36 | 0.16 | 0.37 |
| 10 | 30711929 | rs694346 | 0 | -2.39 | -2.104 | 0.324 | 0.038 | -0.034 | -0.034 | 0 | 0.6 | 0.84 | 0.72 | 0.53 | 0.7 | 0.43 |
| 10 | 30716397 | rs11008047 | 0.668 | -2.537 | -1.424 | 0.31 | 0.002 | -0.003 | -0.003 | 0 | 0.13 | 0.14 | 0.15 | 0.17 | 0.07 | 0.1 |
| 10 | 30723706 | rs1360119 | -0.25 | -2.566 | -1.053 | 0.256 | 0.056 | 0.042 | 0.042 | 4.665 | 0.05 | 0.56 | 0.05 | 0 | 0.02 | 0 |
| 10 | 30727084 | rs303448 | 1.38 | -2.701 | -1.179 | 0.274 | -0.008 | -0.019 | -0.019 | 0 | 0.38 | 0.14 | 0.52 | 0.35 | 0.6 | 0.32 |
| 10 | 30727454 | rs928620 | -0.828 | -2.761 | -1.124 | 0.276 | -0.054 | -0.011 | -0.011 | 0.439 | 0.22 | 0.68 | 0.19 | 0.17 | 0.1 | 0.1 |
| 10 | 30728101 | rs1042058 | 1.357 | -2.66 | -1.132 | 0.307 | -0.022 | -0.034 | -0.034 | 0 | 0.38 | 0.1 | 0.52 | 0.35 | 0.6 | 0.33 |
| 10 | 30731491 | rs303435 | 0.064 | -2.69 | -0.423 | 0.302 | 0.06 | -0.01 | -0.01 | 0 | 0.56 | 0.78 | 0.71 | 0.54 | 0.7 | 0.43 |
| 10 | 30731660 | rs8176984 | 0.186 | -2.57 | 0.262 | 0.283 | -0.044 | 0.021 | 0.021 | 2.307 | 0.15 | 0.37 | 0.17 | 0.19 | 0.09 | 0.1 |
| 10 | 30734883 | rs598009 | -0.095 | -2.779 | -0.148 | 0.263 | 0.049 | 0.032 | 0.032 | 0 | 0.69 | 0.94 | 0.79 | 0.8 | 0.82 | 0.63 |
| 10 | 30735609 | rs1751648 | 0.404 | -2.587 | -0.098 | 0.312 | 0.067 | 0.026 | 0.026 | 0.021 | 0.48 | 0.86 | 0.61 | 0.4 | 0.63 | 0.35 |
| 10 | 30739058 | rs303428 | 1.143 | -2.792 | -0.05 | 0.308 | 0.03 | -0.017 | -0.017 | 0 | 0.4 | 0.16 | 0.56 | 0.39 | 0.65 | 0.35 |
| 10 | 30742388 | rs598672 | 0.723 | -2.47 | 0.332 | 0.344 | 0.064 | 0.021 | 0.021 | 0 | 0.51 | 0.35 | 0.68 | 0.54 | 0.69 | 0.43 |
| 10 | 30749895 | rs3034 | -0.501 | -2.524 | -0.017 | 0.171 | 0.026 | 0.102 | 0.102 | 0 | 0.78 | 0.95 | 0.9 | 0.95 | 0.87 | 0.72 |
| 10 | 30756188 | rs590855 | -0.48 | -2.531 | 0.038 | 0.155 | 0.033 | 0.075 | 0.075 | 0 | 0.78 | 0.91 | 0.9 | 0.95 | 0.87 | 0.71 |
| 10 | 30762088 | rs593400 | 0.766 | -2.363 | 0.231 | 0.375 | 0.067 | 0.033 | 0.033 | 0 | 0.51 | 0.32 | 0.67 | 0.56 | 0.68 | 0.42 |
| 10 | 30762973 | rs2771242 | 0.334 | -2.065 | -0.515 | 0.484 | 0.037 | 0.013 | 0.013 | 0.328 | 0.56 | 0.73 | 0.71 | 0.57 | 0.69 | 0.41 |
| 10 | 30778738 | rs753173 | 0.75 | -1.682 | -1.319 | 0.613 | -0.002 | -0.087 | -0.087 | 0 | 0.21 | 0.03 | 0.18 | 0.38 | 0.19 | 0.29 |
| 10 | 30781197 | rs2767634 | 0.004 | -1.269 | -1.283 | 0.563 | -0.009 | 0.027 | 0.027 | 0 | 0.21 | 0.03 | 0.11 | 0.14 | 0.15 | 0.23 |
| 10 | 30782289 | rs603805 | 0.086 | -0.194 | -1.105 | 0.85 | -0.035 | -0.028 | -0.028 | 0 | 0.07 | 0 | 0.07 | 0.22 | 0.09 | 0.08 |
| 10 | 30789588 | rs3124199 | -0.137 | -0.111 | -1.41 | -0.191 | -0.052 | -0.018 | -0.018 | 0 | 0.62 | 0.78 | 0.65 | 0.5 | 0.55 | 0.53 |
| 10 | 30790264 | rs745290 | 0.653 | -0.171 | -0.919 | -0.548 | -0.028 | -0.013 | -0.013 | 0.378 | 0.35 | 0.36 | 0.24 | 0.37 | 0.26 | 0.34 |
| 10 | 30790361 | rs12779604 | 1.442 | 0.439 | -1.233 | -0.854 | -0.02 | -0.065 | -0.065 | 0.499 | 0.16 | 0.06 | 0.29 | 0.41 | 0.17 | 0.23 |
| 10 | 30791045 | rs3124197 | 0.378 | 0.406 | -1.046 | -0.967 | -0.007 | 0.101 | 0.101 | 0 | 0.29 | 0.11 | 0.26 | 0.08 | 0.28 | 0.24 |
| 10 | 30791941 | rs745453 | -0.852 | 0.441 | -1.036 | -0.962 | -0.015 | 0.01 | 0.01 | 0 | 0.82 | 0.8 | 0.82 | 0.87 | 0.71 | 0.81 |
| 10 | 30792776 | rs728975 | 0.23 | 0.44 | -0.906 | -0.934 | -0.001 | -0.042 | -0.042 | 0.08 | 0.3 | 0.33 | 0.29 | 0.08 | 0.28 | 0.24 |
| 10 | 30798293 | rs3118956 | 1.144 | 0.591 | -0.957 | -0.964 | -0.019 | 0.04 | 0.04 | 0 | 0.15 | 0.05 | 0.18 | 0 | 0.19 | 0.11 |
| 10 | 30799639 | rs3118957 | 0.153 | 0.162 | -0.42 | -0.498 | 0 | -0.002 | -0.002 | 0.943 | 0.49 | 0.5 | 0.58 | 0.53 | 0.45 | 0.48 |
| 10 | 30802799 | rs11008080 | 0.018 | -0.492 | -0.143 | -0.372 | -0.008 | -0.004 | -0.004 | 0 | 0.37 | 0.33 | 0.25 | 0.36 | 0.26 | 0.34 |
| 10 | 30813618 | rs4301703 | -0.432 | 1.173 | -0.023 | -0.08 | 0.046 | -0.035 | -0.035 | 1.44 | 0.2 | 0.28 | 0.26 | 0.13 | 0.28 | 0.18 |
| 10 | 30815163 | rs10826802 | 0.577 | 0.81 | -0.784 | 0.279 | 0.008 | -0.044 | -0.044 | 0.161 | 0.13 | 0.04 | 0.25 | 0.57 | 0.1 | 0.23 |
| 10 | 30816383 | rs7905629 | -1.208 | 0.405 | -1.001 | -0.365 | -0.027 | 0.14 | 0.14 | 1.005 | 0.46 | 0.8 | 0.61 | 0.7 | 0.46 | 0.44 |
| 10 | 30817129 | rs7085583 | -0.15 | 0.549 | -0.476 | -0.506 | 0.002 | -0.007 | -0.007 | 0 | 0.18 | 0.21 | 0.24 | 0.13 | 0.24 | 0.17 |
| 10 | 30822317 | rs10826804 | 1.061 | 0.21 | -1.258 | -1.054 | -0.071 | 0.126 | 0.126 | 0 | 0.37 | 0.1 | 0.27 | 0.2 | 0.49 | 0.41 |
| 10 | 30825119 | rs16931605 | -0.735 | 0.255 | 0.064 | -0.658 | 0.028 | 0.099 | 0.099 | 6.954 | 0.05 | 0.29 | 0.02 | 0.06 | 0.01 | 0.07 |
| 10 | 30828754 | rs17295289 | -0.126 | 0.136 | -0.536 | -0.699 | -0.075 | 0.054 | 0.054 | 0.689 | 0.26 | 0.05 | 0.27 | 0.18 | 0.32 | 0.26 |
| 10 | 30829020 | rs6481694 | 0.929 | -0.049 | 0.456 | -0.5 | -0.002 | -0.055 | -0.055 | 0 | 0.11 | 0.21 | 0.08 | 0 | 0.07 | 0.07 |
| 10 | 30829330 | rs6481696 | -1.138 | 0.16 | 0.246 | -0.456 | -0.006 | 0.005 | 0.005 | 1.35 | 0.29 | 0.47 | 0.18 | 0.31 | 0.24 | 0.33 |
| 10 | 30829554 | rs3124181 | 0.032 | 0.108 | -0.314 | -0.638 | -0.039 | 0.085 | 0.085 | 0 | 0.4 | 0.18 | 0.31 | 0.24 | 0.43 | 0.36 |
| 10 | 30830592 | rs3105630 | 0.961 | 0.204 | -0.062 | -0.404 | 0.018 | 0.019 | 0.019 | 0.003 | 0.38 | 0.23 | 0.26 | 0.31 | 0.31 | 0.29 |
| 10 | 30831940 | rs906232 | 0.925 | 0.233 | 0.1 | -0.391 | 0.014 | 0.015 | 0.015 | 0 | 0.39 | 0.27 | 0.26 | 0.31 | 0.31 | 0.29 |
| 10 | 30831958 | rs906233 | -1.213 | 0.505 | 0.05 | -0.474 | -0.027 | 0.081 | 0.081 | 0 | 0.71 | 0.86 | 0.75 | 0.98 | 0.67 | 0.82 |
| 10 | 30834632 | rs11008099 | -0.344 | 0.784 | 0.821 | 0.908 | -0.003 | -0.018 | -0.018 | 0 | 0.08 | 0.04 | 0.06 | 0.23 | 0.08 | 0.19 |
| 10 | 30834897 | rs12244483 | 0.972 | 0.222 | 0.895 | 0.782 | 0.003 | -0.001 | -0.001 | 0 | 0.07 | 0.09 | 0.07 | 0.01 | 0.05 | 0.02 |
| 10 | 30838729 | rs11008100 | -1.353 | 0.403 | 1.181 | 0.905 | -0.004 | -0.033 | -0.033 | 0 | 0.28 | 0.22 | 0.4 | 0.42 | 0.31 | 0.34 |
| 10 | 30838846 | rs7100253 | -1.432 | 0.389 | 1.549 | 0.87 | -0.002 | 0.069 | 0.069 | 0 | 0.3 | 0.53 | 0.44 | 0.42 | 0.31 | 0.33 |
| 10 | 30838906 | rs7082831 | -1.294 | -0.09 | 1.604 | 0.781 | 0.017 | -0.017 | -0.017 | 0.627 | 0.1 | 0.33 | 0.07 | 0.05 | 0.1 | 0.04 |
| 10 | 30840142 | rs8181390 | -0.354 | -0.048 | 0.686 | 0.817 | -0.029 | 0.034 | 0.034 | 0 | 0.09 | 0.13 | 0.05 | 0.24 | 0.04 | 0.12 |
| 10 | 30842791 | rs7067585 | -1.736 | -0.056 | 0.623 | 0.807 | -0.03 | 0.012 | 0.012 | 0 | 0.18 | 0.28 | 0.11 | 0.27 | 0.12 | 0.17 |
| 10 | 30843141 | rs7067941 | -0.926 | 0.497 | 0.189 | 0.674 | 0.002 | 0.009 | 0.009 | 0 | 0.08 | 0.07 | 0.04 | 0.03 | 0.08 | 0.04 |
| 10 | 30851219 | rs906236 | -0.864 | 0.514 | 0.525 | 0.972 | -0.011 | 0.071 | 0.071 | 0 | 0.22 | 0.33 | 0.36 | 0.37 | 0.2 | 0.3 |
| 10 | 30853414 | rs7090979 | -1.274 | 1.022 | 0.861 | 0.698 | 0.018 | 0.076 | 0.076 | 0 | 0.29 | 0.48 | 0.34 | 0.41 | 0.26 | 0.3 |
| 10 | 30860535 | rs2369932 | -0.023 | 0.67 | 0.646 | 0.662 | 0.017 | -0.011 | -0.011 | 0 | 0.85 | 0.91 | 0.93 | 0.76 | 0.9 | 0.86 |
| 10 | 30862180 | rs7092615 | 0.227 | 0.616 | 1.197 | 0.639 | -0.059 | 0.053 | 0.053 | 0 | 0.84 | 0.53 | 0.89 | 0.76 | 0.9 | 0.86 |
| 10 | 30867140 | rs4747715 | 0.872 | 0.898 | 0.581 | 0.17 | -0.003 | -0.005 | -0.005 | 0 | 0.58 | 0.55 | 0.6 | 0.35 | 0.64 | 0.56 |
| 10 | 30877649 | rs3123276 | 1.011 | 1.489 | 0.908 | 0.084 | 0.071 | -0.07 | -0.07 | 0 | 0.85 | 0.61 | 0.88 | 0.95 | 0.83 | 0.93 |
| 10 | 30883179 | rs11008138 | -0.708 | 0.869 | 0.412 | 0.324 | -0.006 | -0.006 | -0.006 | 0 | 0.35 | 0.39 | 0.3 | 0.26 | 0.27 | 0.41 |
| 10 | 30887952 | rs16931765 | -0.474 | 0.812 | 0.685 | 0.182 | -0.008 | -0.054 | -0.054 | 0.035 | 0.21 | 0.24 | 0.13 | 0.02 | 0.14 | 0.22 |
| 10 | 30888135 | rs12248774 | -1.26 | 0.743 | 0.809 | 0.165 | -0.014 | -0.014 | -0.014 | 0.014 | 0.36 | 0.42 | 0.3 | 0.26 | 0.28 | 0.42 |

| | | | | | | | | | | | | | | | | |
|---|---|---|---|---|---|---|---|---|---|---|---|---|---|---|---|---|
| 10 | 30890572 | rs623635 | 0.796 | 1.027 | 0.562 | 0.448 | -0.007 | 0.003 | 0.003 | 0.672 | 0.86 | 0.91 | 0.71 | 0.52 | 0.82 | 0.67 |
| 10 | 30895478 | rs661893 | 1.764 | 0.782 | 0.961 | 0.168 | 0.004 | 0.048 | 0.048 | 0 | 0.49 | 0.42 | 0.42 | 0.26 | 0.55 | 0.31 |
| 10 | 30896831 | rs448247 | 1.558 | 0.785 | 1.207 | 0.161 | -0.005 | 0.014 | 0.014 | 0 | 0.62 | 0.49 | 0.57 | 0.5 | 0.68 | 0.48 |
| 10 | 30898322 | rs7902622 | -0.745 | 0.709 | 1.211 | 0.155 | -0.038 | -0.131 | -0.131 | 0 | 0.23 | 0.38 | 0.13 | 0.01 | 0.14 | 0.2 |
| 10 | 30898870 | rs366620 | -0.544 | 0.748 | 0.99 | 0.186 | -0.022 | 0.019 | 0.019 | 0 | 0.92 | 0.97 | 0.75 | 0.52 | 0.87 | 0.73 |
| 10 | 30899851 | rs16931784 | 0.473 | 0.74 | 1.053 | 0.168 | 0.002 | -0.03 | -0.03 | 0.271 | 0.14 | 0.07 | 0.16 | 0.24 | 0.13 | 0.19 |
| 10 | 30902048 | rs2814637 | 0.713 | 0.797 | 0.923 | 0.228 | -0.009 | -0.006 | -0.006 | 0 | 0.86 | 0.89 | 0.71 | 0.52 | 0.82 | 0.68 |
| 10 | 30902141 | rs10159940 | 0.482 | 0.815 | 0.669 | 0.222 | -0.009 | -0.044 | -0.044 | 0.086 | 0.15 | 0.19 | 0.1 | 0.01 | 0.1 | 0.18 |
| 10 | 30904892 | rs389258 | 0.583 | 0.824 | 0.668 | 0.252 | -0.018 | -0.032 | -0.032 | 0 | 0.86 | 0.93 | 0.72 | 0.52 | 0.82 | 0.69 |
| 10 | 30905676 | rs16931785 | 0.377 | 0.643 | 0.727 | -0.446 | -0.009 | -0.044 | -0.044 | 0 | 0.15 | 0.18 | 0.1 | 0.01 | 0.1 | 0.17 |
| 10 | 30905934 | rs1798571 | 0.692 | 0.709 | 0.499 | -0.409 | -0.001 | -0.013 | -0.013 | 0.454 | 0.61 | 0.59 | 0.48 | 0.27 | 0.64 | 0.34 |
| 10 | 30909900 | rs1506784 | -1.329 | 1.039 | 0.036 | 0.066 | -0.014 | -0.05 | -0.05 | 0 | 0.13 | 0.18 | 0.05 | 0.01 | 0.09 | 0.05 |
| 10 | 30912550 | rs406896 | 0.407 | 1.409 | -0.215 | 0.515 | -0.005 | -0.003 | -0.003 | 0 | 0.73 | 0.78 | 0.56 | 0.34 | 0.7 | 0.52 |
| 10 | 30912963 | rs10826830 | -0.27 | 1.706 | -0.323 | 0.463 | -0.041 | 0.037 | 0.037 | 0 | 0.07 | 0.01 | 0.06 | 0 | 0.15 | 0.01 |
| 10 | 30921287 | rs1603005 | 1.04 | 1.429 | -0.005 | 0.849 | -0.002 | -0.001 | -0.001 | 0 | 0.74 | 0.28 | 0.6 | 0.69 | 0.73 | 0.61 |
| 10 | 30936214 | rs10826834 | -1.478 | 1.338 | 1.642 | 0.951 | 0.073 | 0.143 | 0.143 | 0.038 | 0.06 | 0.36 | 0.26 | 0.18 | 0.07 | 0.11 |
| 10 | 30937904 | rs3104001 | 1.012 | 1.302 | 1.643 | 0.978 | -0.018 | 0.003 | 0.003 | 0 | 0.83 | 0.89 | 0.88 | 0.86 | 0.8 | 0.73 |
| 10 | 30937956 | rs2015920 | 2.059 | 1.282 | 1.801 | 0.949 | 0.019 | 0.021 | 0.021 | 1.647 | 0.75 | 0.42 | 0.61 | 0.67 | 0.73 | 0.62 |
| 10 | 30952268 | rs1494131 | -1.122 | 1.41 | 1.486 | 0.846 | -0.006 | 0.002 | 0.002 | 0 | 0.09 | 0.09 | 0.06 | 0.01 | 0.11 | 0.19 |
| 10 | 30960206 | rs7097635 | 1.442 | 1.308 | 1.182 | 1.192 | -0.009 | 0.018 | 0.018 | 2.45 | 0.54 | 0.29 | 0.43 | 0.39 | 0.5 | 0.48 |
| 10 | 30960307 | rs7097789 | 0.304 | 1.291 | 1.354 | 1.456 | -0.016 | -0.063 | -0.063 | 0 | 0.72 | 0.81 | 0.77 | 0.58 | 0.68 | 0.8 |
| 10 | 30965085 | rs11008171 | 0.038 | 1.308 | 1.266 | 1.637 | -0.036 | 0.035 | 0.035 | 0 | 0.21 | 0.06 | 0.13 | 0.12 | 0.24 | 0.16 |
| 10 | 30975675 | rs11008178 | -1.251 | 1.106 | 1.463 | 1.608 | -0.039 | 0.048 | 0.048 | 0 | 0.37 | 0.23 | 0.41 | 0.27 | 0.4 | 0.49 |
| 10 | 30976768 | rs11008181 | -0.543 | 1.15 | 1.333 | 1.605 | -0.047 | -0.003 | -0.003 | 0 | 0.27 | 0.08 | 0.34 | 0.26 | 0.3 | 0.31 |
| 10 | 30976818 | rs10826843 | 1.752 | 1.075 | 1.58 | 1.739 | -0.007 | 0.02 | 0.02 | 0.734 | 0.54 | 0.32 | 0.44 | 0.4 | 0.51 | 0.48 |
| 10 | 30984306 | rs7078126 | 1.018 | 0.791 | 1.975 | 2.014 | 0.006 | 0.156 | 0.156 | 0 | 0.93 | 0.81 | 0.89 | 0.66 | 0.92 | 0.97 |
| 10 | 30985081 | rs10826846 | 2.168 | 0.774 | 1.915 | 2.007 | 0.005 | 0.251 | 0.251 | 4.651 | 0.89 | 0.38 | 0.85 | 0.66 | 0.91 | 0.97 |
| 10 | 30986700 | rs4749589 | -0.586 | 0.705 | 1.645 | 1.968 | -0.032 | -0.006 | -0.006 | 0 | 0.25 | 0.1 | 0.33 | 0.26 | 0.29 | 0.31 |
| 10 | 30990074 | rs10763819 | 1.184 | 0.76 | 1.739 | 1.928 | -0.002 | 0.005 | 0.005 | 0.02 | 0.65 | 0.37 | 0.52 | 0.41 | 0.64 | 0.66 |
| 10 | 30993448 | rs10826848 | 1.081 | 0.908 | 1.296 | 2.239 | -0.005 | 0.074 | 0.074 | 0 | 0.93 | 0.88 | 0.9 | 0.73 | 0.92 | 0.95 |
| 10 | 31000901 | rs10763821 | 2.147 | 0.784 | 1.712 | 2.063 | -0.001 | 0.303 | 0.303 | 0 | 0.85 | 0.54 | 0.64 | 0.5 | 0.85 | 0.83 |
| 10 | 31010783 | rs4749600 | -1.847 | 0.776 | 2.035 | 2.318 | 0.024 | 0.264 | 0.264 | 0 | 0.07 | 0.22 | 0.3 | 0.35 | 0.08 | 0.12 |
| 10 | 31010850 | rs2029747 | -1.754 | 0.776 | 2.064 | 2.318 | 0.016 | 0.226 | 0.226 | 0 | 0.07 | 0.19 | 0.29 | 0.35 | 0.08 | 0.12 |
| 10 | 31015069 | rs10740830 | 2.601 | 0.864 | 2.331 | 2.337 | 0.017 | 0.227 | 0.227 | 0 | 0.86 | 0.67 | 0.64 | 0.52 | 0.84 | 0.83 |
| 10 | 31015182 | rs10740831 | 0.712 | 0.87 | 2.297 | 2.341 | -0.002 | -0.003 | -0.003 | 0.206 | 0.94 | 0.95 | 0.95 | 0.87 | 0.91 | 0.95 |
| 10 | 31015533 | rs4080824 | 2.519 | 0.865 | 2.28 | 2.337 | 0.012 | 0.189 | 0.189 | 0 | 0.87 | 0.73 | 0.65 | 0.52 | 0.84 | 0.86 |
| 10 | 31019935 | rs7920155 | 1.174 | 1.022 | 1.984 | 2.871 | 0.001 | 0.018 | 0.018 | 0 | 0.92 | 0.88 | 0.95 | 0.86 | 0.91 | 0.95 |
| 10 | 31022598 | rs11008211 | -0.122 | 0.962 | 1.484 | 2.58 | -0.054 | 0.016 | 0.016 | 0 | 0.18 | 0.04 | 0.1 | 0.09 | 0.2 | 0.13 |
| 10 | 31024081 | rs2047142 | 0.838 | 1.032 | 1.575 | 2.443 | 0.011 | 0.035 | 0.035 | 0 | 0.93 | 0.84 | 0.94 | 0.86 | 0.91 | 0.95 |
| 10 | 31024135 | rs2047141 | 1.474 | 0.929 | 1.705 | 2.353 | -0.002 | -0.014 | -0.014 | 0.751 | 0.65 | 0.27 | 0.5 | 0.42 | 0.64 | 0.74 |
| 10 | 31030449 | rs7918466 | -2.287 | 0.801 | 1.956 | 2.581 | 0.05 | 0.423 | 0.423 | 0.098 | 0.07 | 0.35 | 0.34 | 0.36 | 0.05 | 0.09 |
| 10 | 31038752 | rs7077990 | 1.098 | 0.795 | 0.832 | 1.275 | 0.017 | 0.153 | 0.153 | 0.536 | 0.46 | 0.21 | 0.31 | 0.17 | 0.4 | 0.36 |
| 10 | 31040092 | rs7924137 | 0.702 | 0.967 | 1.176 | 1.809 | 0 | -0.001 | -0.001 | 0 | 0.6 | 0.53 | 0.68 | 0.58 | 0.62 | 0.49 |
| 10 | 31040363 | rs903270 | 0.747 | 1.015 | 1.171 | 1.873 | -0.027 | 0.031 | 0.031 | 0 | 0.63 | 0.71 | 0.73 | 0.69 | 0.62 | 0.49 |
| 10 | 31041134 | rs10826862 | 1.403 | 1.354 | 1.369 | 1.631 | 0 | -0.006 | -0.006 | 0 | 0.58 | 0.38 | 0.39 | 0.26 | 0.47 | 0.46 |
| 10 | 31042981 | rs7097542 | 0.955 | 1.4 | 1.518 | 1.946 | 0.002 | 0.004 | 0.004 | 0.52 | 0.77 | 0.71 | 0.77 | 0.61 | 0.71 | 0.57 |
| 10 | 31043732 | rs10826863 | 1.472 | 1.509 | 1.428 | 1.929 | 0.031 | 0.025 | 0.025 | 0 | 0.66 | 0.6 | 0.48 | 0.65 | 0.55 | 0.74 |
| 10 | 31044294 | rs6481711 | 0.185 | 1.517 | 1.399 | 1.898 | -0.039 | 0.042 | 0.042 | 0 | 0.85 | 0.92 | 0.85 | 0.99 | 0.77 | 0.86 |
| 10 | 31045442 | rs11008227 | -0.707 | 1.488 | 1.427 | 1.944 | -0.005 | 0.086 | 0.086 | 0.004 | 0.06 | 0.15 | 0.08 | 0.38 | 0.06 | 0.28 |
| 10 | 31046041 | rs12247732 | -0.629 | 1.502 | 1.371 | 1.993 | 0.017 | 0.143 | 0.143 | 0 | 0.19 | 0.29 | 0.35 | 0.34 | 0.22 | 0.1 |
| 10 | 31046403 | rs16931887 | -0.635 | 1.483 | 1.328 | 2.008 | 0.004 | 0.039 | 0.039 | 0.018 | 0.06 | 0.08 | 0.07 | 0.38 | 0.06 | 0.28 |
| 10 | 31050678 | rs11008232 | -0.635 | 1.483 | 1.328 | 2.008 | 0.004 | 0.039 | 0.039 | 0.018 | 0.06 | 0.08 | 0.07 | 0.38 | 0.06 | 0.28 |
| 10 | 31054186 | rs947091 | 1.489 | 1.482 | 1.27 | 1.091 | 0.011 | -0.036 | -0.036 | 0.709 | 0.59 | 0.55 | 0.41 | 0.27 | 0.46 | 0.5 |
| 10 | 31057094 | rs7080202 | 0.504 | 1.527 | 1.083 | 1.065 | -0.033 | 0.028 | 0.028 | 0 | 0.87 | 0.93 | 0.86 | 0.99 | 0.77 | 0.88 |
| 10 | 31059336 | rs947095 | 0.633 | 1.526 | 1.369 | 1.067 | 0.037 | -0.057 | -0.057 | 0 | 0.86 | 0.81 | 0.85 | 0.99 | 0.78 | 0.88 |
| 10 | 31061216 | rs7099648 | 0.885 | 1.506 | 1.305 | 1.058 | 0.085 | -0.099 | -0.099 | 0.014 | 0.84 | 0.63 | 0.83 | 0.99 | 0.77 | 0.88 |
| 10 | 31077396 | rs10763832 | -1.477 | 1.825 | 1.058 | 1.3 | 0.017 | 0.025 | 0.025 | 0 | 0.25 | 0.38 | 0.49 | 0.62 | 0.31 | 0.43 |
| 10 | 31081548 | rs2050877 | -0.828 | 1.814 | 0.909 | 0.987 | 0 | -0.016 | -0.016 | 0 | 0.07 | 0.1 | 0.05 | 0.01 | 0.06 | 0.12 |
| 10 | 31082661 | rs7100064 | -0.681 | 1.906 | 1.043 | 1.503 | -0.007 | 0 | 0 | 0 | 0.17 | 0.14 | 0.41 | 0.16 | 0.25 | 0.16 |
| 10 | 31086001 | rs7916670 | -1.849 | 1.421 | 0.923 | 1.807 | -0.027 | 0.113 | 0.113 | 0.092 | 0.22 | 0.44 | 0.15 | 0.47 | 0.18 | 0.45 |
| 10 | 31099139 | rs1340949 | -2.271 | 1.669 | 0.058 | 1.043 | -0.039 | 0.07 | 0.07 | 0 | 0.49 | 0.88 | 0.64 | 0.66 | 0.53 | 0.64 |
| 10 | 31101541 | rs7088786 | -1.934 | 1.619 | 0.589 | 1.03 | 0.005 | -0.008 | -0.008 | 0 | 0.44 | 0.51 | 0.61 | 0.65 | 0.53 | 0.64 |
| 10 | 31101743 | rs1417569 | -3.015 | 1.1 | -0.231 | 0.372 | -0.039 | 0.186 | 0.186 | 0.636 | 0.64 | 0.96 | 0.73 | 0.89 | 0.65 | 0.76 |
| 10 | 31102222 | rs7092692 | -0.294 | -0.096 | -0.92 | -0.501 | -0.006 | -0.002 | -0.002 | 0 | 0.38 | 0.4 | 0.25 | 0.34 | 0.26 | 0.31 |
| 10 | 31102564 | rs7093320 | 0.162 | -0.09 | -1.004 | -0.498 | 0.014 | -0.008 | -0.008 | 0 | 0.34 | 0.2 | 0.24 | 0.34 | 0.25 | 0.31 |
| 10 | 31108819 | rs10763839 | 0.569 | 0.546 | -1.33 | -0.26 | -0.049 | -0.118 | -0.118 | 1.911 | 0.15 | 0.01 | 0.23 | 0.44 | 0.16 | 0.33 |
| 10 | 31109503 | rs1538955 | -0.42 | 0.438 | -0.8 | -0.152 | -0.014 | 0.075 | 0.075 | 1.806 | 0.23 | 0.57 | 0.2 | 0.36 | 0.13 | 0.22 |
| 10 | 31113388 | rs4747731 | -0.957 | 0.636 | -1.222 | -0.214 | -0.024 | 0.082 | 0.082 | 6.062 | 0.84 | 0.98 | 0.87 | 0.98 | 0.8 | 0.86 |
| 10 | 31117269 | rs7921109 | -1.223 | 0.49 | -1.019 | -0.219 | 0.001 | -0.008 | -0.008 | 0 | 0.86 | 0.89 | 0.9 | 0.99 | 0.85 | 0.92 |

| | | | | | | | | | | | | | | | | |
|---|---|---|---|---|---|---|---|---|---|---|---|---|---|---|---|---|
| 10 | 31118934 | rs4749618 | -1.726 | 0.542 | -1.33 | -0.21 | -0.013 | 0.058 | 0.058 | 6.985 | 0.87 | 0.98 | 0.91 | 0.99 | 0.85 | 0.92 |
| 10 | 31119770 | rs12572052 | 0.39 | 0.715 | -1.411 | 0.064 | -0.02 | -0.041 | -0.041 | 0.016 | 0.14 | 0.08 | 0.25 | 0.44 | 0.19 | 0.36 |
| 10 | 31120198 | rs12770855 | 0.885 | 0.772 | -1.468 | -0.025 | -0.002 | -0.001 | -0.001 | 0 | 0.07 | 0.07 | 0.04 | 0.07 | 0.08 | 0.01 |
| 10 | 31124111 | rs2994652 | 0.4 | 0.626 | -1.499 | -0.136 | -0.001 | -0.003 | -0.003 | 2.252 | 0.64 | 0.37 | 0.58 | 0.44 | 0.61 | 0.45 |
| 10 | 31125508 | rs2994655 | 0.681 | 0.604 | -1.4 | -0.089 | -0.003 | 0.002 | 0.002 | 3.299 | 0.58 | 0.38 | 0.54 | 0.37 | 0.53 | 0.44 |
| 10 | 31129883 | rs947099 | 1.841 | 0.781 | -1.705 | -0.166 | -0.008 | 0.009 | 0.009 | 5.301 | 0.38 | 0.13 | 0.48 | 0.33 | 0.34 | 0.34 |
| 10 | 31130983 | rs11008285 | -0.141 | 1.137 | -1.137 | 0.365 | 0.003 | -0.021 | -0.021 | 0 | 0.18 | 0.2 | 0.13 | 0.04 | 0.2 | 0.11 |
| 10 | 31132475 | rs7894787 | -0.259 | 1.026 | -1.335 | 1.004 | -0.018 | -0.04 | -0.04 | 0 | 0.14 | 0.08 | 0.18 | 0.44 | 0.19 | 0.35 |
| 10 | 31134214 | rs3740236 | -0.649 | 0.558 | -1.5 | 1.388 | 0.001 | -0.068 | -0.068 | 0 | 0.12 | 0.02 | 0.3 | 0.33 | 0.1 | 0.08 |
| 10 | 31134557 | rs17229657 | -0.457 | 0.671 | -0.808 | 0.924 | 0.003 | 0.033 | 0.033 | 1.008 | 0.31 | 0.59 | 0.39 | 0.49 | 0.36 | 0.42 |
| 10 | 31135019 | rs4749619 | -1.56 | 0.661 | -1.372 | 0.69 | -0.065 | -0.067 | -0.067 | 0.507 | 0.5 | 0.93 | 0.59 | 0.47 | 0.53 | 0.49 |
| 10 | 31138817 | rs10160116 | 0.03 | 0.569 | -0.411 | 0.645 | 0.002 | 0.008 | 0.008 | 0.931 | 0.29 | 0.33 | 0.43 | 0.4 | 0.35 | 0.41 |
| 10 | 31144430 | rs4747735 | -1.658 | 0.557 | -0.396 | 0.627 | -0.053 | -0.06 | -0.06 | 0 | 0.5 | 0.92 | 0.59 | 0.46 | 0.53 | 0.49 |
| 10 | 31146615 | rs16932010 | -0.255 | 0.546 | -0.131 | 0.621 | 0.005 | -0.056 | -0.056 | 0 | 0.2 | 0.31 | 0.14 | 0.07 | 0.18 | 0.08 |
| 10 | 31147607 | rs7090992 | -1.758 | 0.545 | 0.132 | 0.621 | -0.001 | -0.008 | -0.008 | 0 | 0.5 | 0.83 | 0.58 | 0.46 | 0.53 | 0.49 |
| 10 | 31149437 | rs994464 | 0.049 | 0.546 | 0.127 | 0.622 | 0.002 | 0.008 | 0.008 | 0.931 | 0.29 | 0.33 | 0.43 | 0.39 | 0.35 | 0.41 |
| 10 | 31149545 | rs17295437 | 0.249 | 0.497 | 0.104 | 0.591 | -0.028 | 0.051 | 0.051 | 0 | 0.17 | 0.04 | 0.11 | 0.07 | 0.18 | 0.08 |
| 10 | 31152916 | rs4749628 | -1.668 | 0.493 | 0.13 | 0.595 | -0.017 | -0.025 | -0.025 | 0.103 | 0.5 | 0.87 | 0.59 | 0.46 | 0.53 | 0.49 |
| 10 | 31158415 | rs10826883 | 0.108 | 0.024 | -0.599 | -0.13 | 0.004 | 0.018 | 0.018 | 1.23 | 0.32 | 0.59 | 0.46 | 0.39 | 0.36 | 0.41 |
| 10 | 31166206 | rs10763849 | -1.179 | -0.021 | -0.607 | -0.135 | 0.003 | 0 | 0 | 1.254 | 0.49 | 0.64 | 0.57 | 0.46 | 0.53 | 0.49 |
| 10 | 31170360 | rs1587006 | -0.599 | 0.073 | -0.546 | -0.167 | -0.04 | -0.153 | -0.153 | 0 | 0.71 | 0.95 | 0.71 | 0.46 | 0.73 | 0.58 |
| 10 | 31171998 | rs2994613 | 1.098 | 0.147 | -0.48 | -0.267 | 0 | -0.004 | -0.004 | 0.625 | 0.4 | 0.33 | 0.25 | 0.07 | 0.38 | 0.17 |
| 10 | 31184125 | rs3006598 | 2.682 | 0.155 | -0.229 | -0.268 | -0.006 | -0.082 | -0.082 | 0 | 0.23 | 0.28 | 0.14 | 0 | 0.2 | 0.1 |
| 10 | 31193339 | rs2934637 | 1.327 | 0.094 | -0.312 | -0.328 | 0.004 | -0.028 | -0.028 | 0 | 0.37 | 0.62 | 0.23 | 0.01 | 0.31 | 0.17 |
| 10 | 31203092 | rs1038972 | -1.529 | 0.417 | -0.822 | -0.158 | -0.041 | -0.017 | -0.017 | 0 | 0.93 | 0.99 | 0.94 | 0.96 | 0.85 | 0.95 |
| 10 | 31210609 | rs3006603 | -0.558 | 0.131 | -0.963 | -0.065 | -0.055 | -0.058 | -0.058 | 0 | 0.78 | 0.97 | 0.78 | 0.74 | 0.75 | 0.7 |
| 10 | 31216112 | rs17229725 | 1.855 | 0.151 | -0.91 | -0.16 | -0.012 | 0.021 | 0.021 | 0 | 0.1 | 0.03 | 0.06 | 0.01 | 0.11 | 0.07 |
| 10 | 31233525 | rs3006574 | -0.053 | 0.088 | -0.85 | 0.071 | -0.011 | -0.069 | -0.069 | 0.174 | 0.69 | 0.91 | 0.7 | 0.41 | 0.69 | 0.58 |
| 10 | 31245835 | rs3011603 | -0.448 | 0.111 | -0.858 | 0.109 | -0.049 | -0.063 | -0.063 | 0 | 0.76 | 0.96 | 0.77 | 0.69 | 0.72 | 0.7 |
| 10 | 31259248 | rs1494119 | -1.131 | 0.19 | -0.877 | -0.146 | -0.076 | -0.009 | -0.009 | 0 | 0.49 | 0.91 | 0.58 | 0.4 | 0.49 | 0.48 |
| 10 | 31275977 | rs2994647 | 0.531 | 0.267 | -0.467 | -0.023 | -0.006 | -0.005 | -0.005 | 0.614 | 0.59 | 0.77 | 0.62 | 0.4 | 0.58 | 0.51 |
| 10 | 31282422 | rs2887738 | 1.167 | 0.296 | -0.024 | 0.149 | 0.007 | 0.019 | 0.019 | 0 | 0.08 | 0.2 | 0.07 | 0.08 | 0.07 | 0.01 |
| 10 | 31289921 | rs12219452 | 2.107 | 0.206 | -0.397 | 0.158 | -0.012 | -0.037 | -0.037 | 0 | 0.07 | 0 | 0.02 | 0.16 | 0.04 | 0.14 |
| 10 | 31294981 | rs1690627 | 0.732 | 0.077 | -0.33 | 0.048 | -0.053 | -0.045 | -0.045 | 0 | 0.24 | 0.09 | 0.4 | 0.32 | 0.34 | 0.43 |
| 10 | 31297648 | rs2642192 | 1.011 | 0.023 | 0.32 | 0.03 | 0.002 | 0.021 | 0.021 | 0.022 | 0.51 | 0.64 | 0.58 | 0.32 | 0.54 | 0.52 |
| 10 | 31298325 | rs1690624 | 0.437 | 0.019 | 0.249 | 0.014 | 0.003 | 0.006 | 0.006 | 0 | 0.62 | 0.72 | 0.64 | 0.33 | 0.66 | 0.58 |
| 10 | 31307374 | rs2793104 | -0.209 | 0.018 | 0.131 | 0.092 | 0.009 | -0.035 | -0.035 | 0 | 0.69 | 0.86 | 0.7 | 0.41 | 0.73 | 0.58 |
| 10 | 31343788 | rs2487457 | 0.966 | 0.007 | -0.132 | 0.046 | -0.008 | 0.085 | 0.085 | 0.072 | 0.52 | 0.83 | 0.6 | 0.32 | 0.54 | 0.53 |
| 10 | 31350494 | rs1776616 | 0.017 | -0.226 | -0.311 | 0.044 | 0.02 | -0.052 | -0.052 | 0 | 0.67 | 0.9 | 0.7 | 0.4 | 0.73 | 0.58 |
| 10 | 31361464 | rs7095095 | 2.434 | -0.189 | -0.129 | -0.095 | -0.002 | 0.024 | 0.024 | 0.324 | 0.21 | 0.17 | 0.23 | 0.07 | 0.2 | 0.13 |
| 10 | 31361749 | rs7098952 | 3.005 | -0.226 | -0.159 | -0.111 | -0.049 | 0.027 | 0.027 | 0.388 | 0.19 | 0.02 | 0.21 | 0.07 | 0.2 | 0.13 |
| 10 | 31363143 | rs793086 | -0.605 | -0.244 | -0.112 | -0.165 | 0.021 | 0.032 | 0.032 | 0.253 | 0.73 | 0.87 | 0.88 | 0.77 | 0.79 | 0.73 |
| 10 | 31364621 | rs793088 | 1.435 | 0.103 | -0.308 | -0.23 | -0.014 | -0.025 | -0.025 | 0.833 | 0.21 | 0.12 | 0.37 | 0.3 | 0.29 | 0.28 |
| 10 | 31367031 | rs11008337 | 1.286 | 0.15 | -0.281 | -0.393 | -0.012 | 0.053 | 0.053 | 0.001 | 0.36 | 0.29 | 0.33 | 0.15 | 0.37 | 0.18 |
| 10 | 31374166 | rs12244754 | 2.652 | 0.187 | -0.172 | -0.208 | -0.01 | 0.053 | 0.053 | 0.332 | 0.2 | 0.12 | 0.13 | 0 | 0.21 | 0.09 |
| 10 | 31374935 | rs16932182 | 0.464 | 0.225 | -0.129 | -0.041 | 0.005 | 0.002 | 0.002 | 0 | 0.05 | 0.07 | 0.15 | 0.08 | 0.09 | 0.16 |
| 10 | 31379105 | rs2793108 | -0.038 | 0.151 | 0.07 | -0.007 | -0.003 | -0.014 | -0.014 | 0.145 | 0.41 | 0.34 | 0.62 | 0.47 | 0.56 | 0.51 |
| 10 | 31381616 | rs12248722 | 0.919 | 0.03 | 0.215 | 0.087 | -0.036 | 0.141 | 0.141 | 0.195 | 0.12 | 0.31 | 0.11 | 0.3 | 0.04 | 0.14 |
| 10 | 31386822 | rs793094 | 1.011 | 0.145 | -0.788 | 0.05 | -0.066 | -0.049 | -0.049 | 1.003 | 0.3 | 0.09 | 0.41 | 0.32 | 0.39 | 0.33 |
| 10 | 31387374 | rs12573099 | 1.512 | -0.146 | -0.594 | 0.072 | 0.008 | 0.001 | 0.001 | 0 | 0.1 | 0.08 | 0.08 | 0.3 | 0.04 | 0.14 |
| 10 | 31388250 | rs7080444 | 3.162 | 0.143 | -0.644 | -0.135 | -0.042 | 0.117 | 0.117 | 0.4 | 0.18 | 0.02 | 0.12 | 0 | 0.2 | 0.09 |
| 10 | 31389265 | rs867991 | 0.547 | 0.207 | -0.628 | -0.065 | -0.095 | -0.071 | -0.071 | 0.176 | 0.35 | 0.1 | 0.45 | 0.39 | 0.47 | 0.34 |
| 10 | 31389575 | rs867993 | 0.118 | 0.199 | -0.169 | 0.121 | 0.015 | -0.013 | -0.013 | 0.048 | 0.64 | 0.75 | 0.79 | 0.47 | 0.77 | 0.59 |
| 10 | 31389909 | rs12256012 | -0.107 | 0.624 | -0.444 | 0.172 | 0.054 | 0.041 | 0.041 | 0.186 | 0.06 | 0.17 | 0.16 | 0.08 | 0.09 | 0.17 |
| 10 | 31402767 | rs793098 | 0.255 | 0.433 | -0.898 | 0.101 | 0.019 | -0.006 | -0.006 | 0.01 | 0.59 | 0.7 | 0.75 | 0.4 | 0.66 | 0.55 |
| 10 | 31411355 | rs1538981 | -0.119 | 0.448 | -1.401 | 0.071 | -0.115 | -0.069 | -0.069 | 0 | 0.39 | 0.09 | 0.57 | 0.38 | 0.49 | 0.36 |
| 10 | 31412936 | rs2066250 | -1.553 | 0.179 | -1.164 | 0.228 | 0.005 | -0.034 | -0.034 | 0 | 0.69 | 0.83 | 0.78 | 0.64 | 0.72 | 0.71 |
| 10 | 31414304 | rs867768 | 0.783 | 0.675 | -1.44 | -0.007 | -0.072 | -0.113 | -0.113 | 0 | 0.21 | 0.02 | 0.45 | 0.31 | 0.25 | 0.27 |
| 10 | 31418463 | rs1340991 | -2.648 | 0.522 | -1.203 | -0.337 | -0.026 | 0.073 | 0.073 | 0.1 | 0.81 | 0.95 | 0.86 | 1 | 0.77 | 0.91 |
| 10 | 31424353 | rs1250302 | 0.744 | 0.631 | -0.746 | -0.189 | -0.001 | -0.008 | -0.008 | 0 | 0.18 | 0.18 | 0.31 | 0.07 | 0.19 | 0.11 |
| 10 | 31425683 | rs11008356 | -0.732 | 0.737 | 0.145 | -0.24 | 0.143 | 0.064 | 0.064 | 2.352 | 0.07 | 0.45 | 0.09 | 0.03 | 0.07 | 0.14 |
| 10 | 31428177 | rs2050888 | 1.393 | 0.813 | -0.12 | 0.055 | -0.02 | -0.016 | -0.016 | 4.47 | 0.06 | 0.01 | 0.17 | 0.22 | 0.1 | 0.17 |
| 10 | 31429305 | rs16932218 | -0.587 | 0.685 | 0.603 | 0.103 | 0.098 | 0.034 | 0.034 | 1.322 | 0.06 | 0.33 | 0.08 | 0.03 | 0.07 | 0.14 |
| 10 | 31429850 | rs1272045 | -0.663 | 0.476 | 0.556 | 0.009 | 0.005 | -0.021 | -0.021 | 0 | 0.37 | 0.5 | 0.44 | 0.33 | 0.36 | 0.42 |
| 10 | 31431419 | rs2797914 | 0.568 | 0.039 | 0.807 | 0.354 | -0.007 | 0.002 | 0.002 | 3.225 | 0.15 | 0.13 | 0.06 | 0.23 | 0.1 | 0.18 |
| 10 | 31435153 | rs1624116 | 0.441 | 0.034 | 0.584 | 0.306 | -0.082 | -0.026 | -0.026 | 0 | 0.43 | 0.1 | 0.58 | 0.38 | 0.54 | 0.41 |
| 10 | 31435410 | rs11008362 | 0.865 | -0.037 | 0.619 | 0.283 | 0.009 | 0.028 | 0.028 | 0 | 0.07 | 0.06 | 0.05 | 0.23 | 0.09 | 0.17 |
| 10 | 31438487 | rs1250317 | 0.481 | -0.051 | 0.563 | 0.288 | -0.084 | -0.03 | -0.03 | 0 | 0.43 | 0.1 | 0.58 | 0.38 | 0.54 | 0.4 |
| 10 | 31442435 | rs12571605 | 0.444 | -0.611 | 1.445 | 0.73 | -0.022 | 0.043 | 0.043 | 0.713 | 0.13 | 0.24 | 0.1 | 0.25 | 0.05 | 0.19 |

| | | | | | | | | | | | | | | | | |
|---|---|---|---|---|---|---|---|---|---|---|---|---|---|---|---|---|
| 10 | 31446322 | rs1764443 | -0.971 | -0.596 | 1.435 | 0.736 | -0.028 | 0.129 | 0.129 | 1.117 | 0.22 | 0.49 | 0.17 | 0.48 | 0.15 | 0.36 |
| 10 | 31448076 | rs11008376 | 0.128 | -0.574 | 1.388 | 0.729 | 0.007 | 0.029 | 0.029 | 0 | 0.08 | 0.12 | 0.05 | 0.21 | 0.09 | 0.16 |
| 10 | 31453736 | rs7908936 | 0.551 | -0.566 | 1.367 | 0.733 | 0.009 | 0.033 | 0.033 | 0 | 0.07 | 0.09 | 0.05 | 0.22 | 0.09 | 0.16 |
| 10 | 31457279 | rs1250294 | 0.033 | -0.87 | 1.271 | 0.468 | -0.047 | 0.106 | 0.106 | 2.187 | 0.15 | 0.43 | 0.12 | 0.25 | 0.06 | 0.2 |
| 10 | 31465307 | rs793081 | -1.687 | -0.229 | 0.818 | -0.337 | -0.039 | 0.092 | 0.092 | 0.871 | 0.74 | 0.91 | 0.79 | 0.92 | 0.66 | 0.9 |
| 10 | 31465875 | rs793082 | 0.203 | -0.391 | 0.958 | -0.17 | -0.009 | 0.073 | 0.073 | 0.523 | 0.49 | 0.27 | 0.63 | 0.61 | 0.49 | 0.57 |
| 10 | 31466155 | rs7091901 | -0.504 | 0.004 | 0.914 | 0.168 | 0.015 | 0.058 | 0.058 | 0 | 0.09 | 0.27 | 0.07 | 0.17 | 0.09 | 0.19 |
| 10 | 31466397 | rs10826912 | -0.287 | 0.003 | 0.945 | 0.141 | 0.003 | 0.03 | 0.03 | 0 | 0.08 | 0.16 | 0.06 | 0.16 | 0.09 | 0.19 |
| 10 | 31476778 | rs11008410 | -0.311 | 0.071 | 0.633 | -0.106 | 0.002 | 0 | 0 | 0 | 0.07 | 0.2 | 0.05 | 0.08 | 0.06 | 0.17 |
| 10 | 31478422 | rs7899488 | -0.045 | 0.001 | 0.5 | -0.112 | -0.014 | 0.066 | 0.066 | 0.267 | 0.43 | 0.31 | 0.33 | 0.17 | 0.47 | 0.39 |
| 10 | 31479103 | rs2995653 | -0.099 | 0.119 | 0.274 | -0.027 | -0.001 | 0.001 | 0.001 | 1.682 | 0.54 | 0.47 | 0.66 | 0.72 | 0.53 | 0.62 |
| 10 | 31516419 | rs2505731 | 0.679 | -0.961 | 1.437 | 0.506 | -0.058 | 0.087 | 0.087 | 3.437 | 0.89 | 0.68 | 0.95 | 0.83 | 0.99 | 0.9 |
| 10 | 31521891 | rs2505730 | 1.776 | -1.135 | 1.401 | 0.419 | -0.05 | 0.01 | 0.01 | 12.24 | 0.86 | 0.26 | 0.91 | 0.83 | 0.99 | 0.9 |
| 10 | 31534411 | rs11008425 | -0.183 | -1.049 | 0.973 | 0.447 | 0.002 | -0.006 | -0.006 | 0 | 0.08 | 0.07 | 0.03 | 0.16 | 0.01 | 0.1 |
| 10 | 31544026 | rs1314004 | -0.149 | -1.344 | -0.07 | -0.126 | -0.053 | 0.097 | 0.097 | 7.725 | 0.72 | 0.97 | 0.69 | 0.9 | 0.71 | 0.89 |
| 10 | 31557545 | rs1314007 | -1.241 | -1.626 | 0.29 | -0.242 | -0.015 | -0.024 | -0.024 | 0 | 0.19 | 0.74 | 0.12 | 0.02 | 0.07 | 0.17 |
| 10 | 31565307 | rs1314011 | -0.363 | -1.655 | 0.994 | 0.071 | 0.03 | 0.187 | 0.187 | 0.186 | 0.09 | 0.41 | 0.07 | 0.16 | 0.06 | 0.1 |
| 10 | 31578918 | rs12242798 | -0.44 | -1.682 | 1.068 | 0.044 | -0.061 | -0.03 | -0.03 | 0.025 | 0.11 | 0.43 | 0.06 | 0.02 | 0.02 | 0.09 |
| 10 | 31586372 | rs9417869 | 0.775 | -1.424 | 0.122 | 0.408 | -0.086 | -0.093 | -0.093 | 0 | 0.19 | 0.01 | 0.14 | 0.28 | 0.26 | 0.25 |
| 10 | 31607215 | rs3737178 | -1.504 | -1.295 | 0.508 | 0.425 | 0.015 | -0.005 | -0.005 | 0 | 0.2 | 0.8 | 0.12 | 0.18 | 0.07 | 0.19 |
| 10 | 31662486 | rs10826945 | 0.268 | -1.306 | 0.664 | 0.424 | -0.042 | -0.018 | -0.018 | 0 | 0.09 | 0.18 | 0.04 | 0.02 | 0.02 | 0.09 |
| 10 | 31670562 | rs161229 | -1.469 | -1.29 | 0.079 | 0.376 | 0.005 | -0.004 | -0.004 | 0 | 0.2 | 0.79 | 0.12 | 0.18 | 0.07 | 0.18 |
| 10 | 31677206 | rs431073 | -1.005 | -1.149 | -0.099 | 0.307 | -0.057 | 0.005 | 0.005 | 0 | 0.88 | 0.98 | 0.63 | 0.93 | 0.8 | 0.95 |
| 10 | 31740091 | rs223531 | 0.399 | -1.193 | 0.241 | 0.389 | -0.003 | -0.039 | -0.039 | 0 | 0.07 | 0.17 | 0.04 | 0 | 0.06 | 0.08 |
| 10 | 31741273 | rs12217563 | 0.806 | -1.357 | -0.043 | 0.478 | -0.056 | -0.145 | -0.145 | 2.048 | 0.31 | 0.01 | 0.17 | 0.43 | 0.29 | 0.45 |
| 10 | 31781004 | rs220065 | -1.232 | -1.104 | 0.588 | 0.774 | -0.052 | 0.063 | 0.063 | 3.288 | 0.15 | 0.59 | 0.09 | 0.18 | 0.07 | 0.18 |
| 10 | 31795212 | rs1618092 | 0.418 | -1.124 | 0.453 | 0.765 | -0.131 | -0.126 | -0.126 | 0.766 | 0.47 | 0.04 | 0.55 | 0.5 | 0.54 | 0.52 |
| 10 | 31813958 | rs220072 | -0.199 | -1.333 | 1.616 | 0.836 | -0.001 | -0.014 | -0.014 | 0 | 0.06 | 0.07 | 0.03 | 0 | 0.06 | 0.08 |
| 10 | 31817905 | rs7349 | -2.691 | -1.355 | 1.421 | 0.829 | 0.073 | -0.064 | -0.064 | 0 | 0.19 | 0.94 | 0.12 | 0.18 | 0.07 | 0.18 |
| 10 | 31818563 | rs220071 | -0.503 | -1.182 | 1.522 | 0.893 | -0.003 | -0.039 | -0.039 | 0 | 0.07 | 0.17 | 0.04 | 0 | 0.06 | 0.08 |
| 10 | 31830736 | rs1592078 | -2.101 | -1.179 | 1.165 | 0.904 | -0.024 | 0.006 | 0.006 | 4.597 | 0.17 | 0.75 | 0.13 | 0.18 | 0.07 | 0.19 |
| 10 | 31831830 | rs7075638 | -0.997 | -1.134 | 1.033 | 0.747 | 0.006 | -0.042 | -0.042 | 0 | 0.09 | 0.33 | 0.06 | 0 | 0.06 | 0.08 |
| 10 | 31860345 | rs10826966 | -1.066 | -0.963 | -0.227 | 0.847 | -0.035 | 0.007 | 0.007 | 4.227 | 0.16 | 0.75 | 0.14 | 0.15 | 0.06 | 0.18 |
| 10 | 31872826 | rs12781751 | -0.002 | -0.397 | -0.471 | 0.837 | 0 | -0.001 | -0.001 | 0 | 0.45 | 0.44 | 0.57 | 0.44 | 0.52 | 0.34 |
| 10 | 31874969 | rs10763867 | 0.196 | -0.86 | -0.883 | 1.124 | 0.037 | -0.029 | -0.029 | 0 | 0.45 | 0.08 | 0.57 | 0.39 | 0.61 | 0.37 |
| 10 | 31883612 | rs7899550 | -0.722 | -1.667 | -0.812 | 0.859 | 0.028 | -0.04 | -0.04 | 0 | 0.66 | 0.79 | 0.68 | 0.5 | 0.71 | 0.55 |
| 10 | 31886363 | rs7920301 | -0.869 | -1.747 | -0.973 | 0.822 | -0.016 | -0.127 | -0.127 | 0.929 | 0.67 | 0.94 | 0.69 | 0.5 | 0.69 | 0.55 |
| 10 | 31894818 | rs11008540 | -0.066 | -1.669 | -0.246 | 0.614 | -0.088 | -0.092 | -0.092 | 0 | 0.17 | 0.49 | 0.09 | 0.02 | 0.07 | 0.16 |
| 10 | 31899073 | rs727345 | 0.586 | -1.777 | -1.134 | 0.792 | 0.043 | -0.026 | -0.026 | 0 | 0.43 | 0.08 | 0.56 | 0.36 | 0.62 | 0.39 |
| 10 | 31901584 | rs7917824 | -0.683 | -1.796 | -1.193 | 0.785 | 0.042 | -0.011 | -0.011 | 0 | 0.57 | 0.73 | 0.64 | 0.54 | 0.66 | 0.59 |
| 10 | 31905494 | rs11008546 | 0.376 | -0.994 | -1.521 | 0.812 | -0.044 | -0.083 | -0.083 | 0.016 | 0.36 | 0.08 | 0.53 | 0.4 | 0.57 | 0.41 |
| 10 | 31908498 | rs796076 | -0.421 | -0.557 | -1.17 | 0.536 | 0.008 | 0.017 | 0.017 | 2.844 | 0.55 | 0.6 | 0.58 | 0.62 | 0.57 | 0.53 |
| 10 | 31909119 | rs703067 | 0.203 | -0.277 | -0.907 | 0.217 | -0.031 | 0.04 | 0.04 | 0.848 | 0.34 | 0.22 | 0.47 | 0.13 | 0.45 | 0.31 |
| 10 | 31912104 | rs16932483 | -0.63 | -0.712 | -1.399 | 0.181 | 0.033 | 0.075 | 0.075 | 0.452 | 0.44 | 0.77 | 0.45 | 0.33 | 0.4 | 0.25 |
| 10 | 31916689 | rs4313476 | 0.914 | -0.015 | -2.121 | -0.199 | -0.137 | 0.09 | 0.09 | 0.483 | 0.29 | 0.03 | 0.28 | 0.11 | 0.35 | 0.3 |
| 10 | 31935360 | rs11815055 | 0.757 | -0.262 | -1.352 | -0.548 | -0.006 | 0.038 | 0.038 | 0 | 0.15 | 0.07 | 0.25 | 0.08 | 0.19 | 0.08 |
| 10 | 31937671 | rs796149 | -1.046 | -0.414 | -1.473 | -0.683 | -0.053 | 0.051 | 0.051 | 0 | 0.91 | 0.99 | 0.92 | 1 | 0.85 | 0.92 |
| 10 | 31939485 | rs2797129 | 0.273 | -0.504 | -1.124 | -0.675 | 0.001 | 0.001 | 0.001 | 1.592 | 0.71 | 0.34 | 0.8 | 0.74 | 0.79 | 0.82 |
| 10 | 31950753 | rs796143 | -0.114 | -0.767 | -0.659 | -0.824 | 0.016 | -0.145 | -0.145 | 0 | 0.81 | 0.57 | 0.86 | 1 | 0.83 | 0.9 |
| 10 | 31957790 | rs796140 | 0.352 | -0.789 | -0.611 | -0.808 | -0.016 | -0.006 | -0.006 | 0 | 0.74 | 0.49 | 0.82 | 0.74 | 0.79 | 0.81 |
| 10 | 31966507 | rs796130 | 1.066 | -0.809 | -0.712 | -0.811 | 0.031 | 0.069 | 0.069 | 0.186 | 0.55 | 0.26 | 0.55 | 0.65 | 0.59 | 0.73 |
| 10 | 31968759 | rs796126 | 1.066 | -0.809 | -0.712 | -0.811 | 0.031 | 0.069 | 0.069 | 0.186 | 0.55 | 0.26 | 0.55 | 0.65 | 0.59 | 0.73 |
| 10 | 31968783 | rs10508765 | 0.607 | -0.824 | -0.671 | -0.807 | -0.001 | -0.01 | -0.01 | 0 | 0.17 | 0.16 | 0.26 | 0.08 | 0.2 | 0.08 |
| 10 | 31974332 | rs4747759 | -0.092 | -0.743 | -0.547 | -0.875 | 0.027 | -0.143 | -0.143 | 0 | 0.79 | 0.61 | 0.84 | 1 | 0.79 | 0.87 |
| 10 | 31975163 | rs1418635 | 1.184 | -0.889 | -0.628 | -0.7 | 0.003 | 0.118 | 0.118 | 0.144 | 0.52 | 0.17 | 0.53 | 0.66 | 0.54 | 0.7 |
| 10 | 31975192 | rs1418634 | 0.327 | -0.819 | -0.587 | -0.823 | 0.008 | -0.109 | -0.109 | 0.93 | 0.63 | 0.54 | 0.59 | 0.91 | 0.59 | 0.78 |
| 10 | 31979399 | rs796156 | 1.083 | -0.592 | -1.276 | -0.242 | -0.078 | -0.063 | -0.063 | 0 | 0.47 | 0.07 | 0.49 | 0.43 | 0.51 | 0.63 |
| 10 | 31982953 | rs796159 | -1.385 | -0.071 | -1.181 | -0.239 | -0.023 | -0.026 | -0.026 | 0.842 | 0.55 | 0.8 | 0.52 | 0.53 | 0.54 | 0.42 |
| 10 | 31985404 | rs10508766 | 1.585 | 0.097 | -1.412 | -0.381 | -0.045 | -0.154 | -0.154 | 0 | 0.19 | 0.01 | 0.32 | 0.41 | 0.19 | 0.26 |
| 10 | 31985518 | rs796153 | 0.484 | 0.312 | -0.335 | -0.578 | -0.001 | 0.006 | 0.006 | 0 | 0.66 | 0.34 | 0.7 | 0.72 | 0.63 | 0.76 |
| 10 | 31987246 | rs10826979 | 0.711 | 0.186 | 0.368 | -0.624 | -0.02 | -0.056 | -0.056 | 0.312 | 0.14 | 0.31 | 0.18 | 0.05 | 0.1 | 0.13 |
| 10 | 31987319 | rs12245995 | 0.19 | 0.591 | 0.133 | -0.358 | -0.001 | 0.029 | 0.029 | 0 | 0.07 | 0.11 | 0.1 | 0.21 | 0.09 | 0.03 |
| 10 | 31989021 | rs11008599 | 0.251 | 0.789 | 0.058 | -0.794 | 0.021 | -0.053 | -0.053 | 0.968 | 0.12 | 0.19 | 0.1 | 0.01 | 0.15 | 0.07 |
| 10 | 31996242 | rs2277251 | -0.101 | 0.28 | -0.065 | -1.163 | 0.001 | -0.04 | -0.04 | 0 | 0.46 | 0.49 | 0.47 | 0.9 | 0.46 | 0.64 |
| 10 | 31996352 | rs2277253 | -1.017 | 0.093 | -0.142 | -1.086 | 0.011 | 0.035 | 0.035 | 0 | 0.62 | 0.68 | 0.64 | 0.92 | 0.67 | 0.72 |
| 10 | 31997937 | rs12219561 | 0.354 | 0.135 | 0.351 | -0.526 | 0.002 | 0.022 | 0.022 | 0 | 0.1 | 0.13 | 0.08 | 0.19 | 0.09 | 0.19 |
| 10 | 31998017 | rs3858244 | -1.823 | 0.125 | -0.132 | -0.648 | -0.004 | -0.016 | -0.016 | 0 | 0.45 | 0.92 | 0.35 | 0.42 | 0.39 | 0.43 |
| 10 | 32001010 | rs3898094 | -1.353 | 0.192 | 0.757 | -0.568 | -0.005 | -0.02 | -0.02 | 0 | 0.34 | 0.41 | 0.21 | 0.29 | 0.32 | 0.38 |
| 10 | 32012831 | rs17295738 | -0.414 | 0.195 | 0.79 | -0.58 | -0.002 | -0.033 | -0.033 | 2.747 | 0.24 | 0.25 | 0.13 | 0.1 | 0.23 | 0.19 |

| | | | | | | | | | | | | | | | | |
|---|---|---|---|---|---|---|---|---|---|---|---|---|---|---|---|---|
| 10 | 32016090 | rs11008612 | 0.555 | 0.213 | 1.09 | -0.569 | 0.007 | 0.051 | 0.051 | 0 | 0.1 | 0.23 | 0.09 | 0.19 | 0.09 | 0.2 |
| 10 | 32018561 | rs11008614 | 0.581 | 0.168 | 0.951 | -0.581 | -0.014 | 0.005 | 0.005 | 0.059 | 0.15 | 0.18 | 0.07 | 0.19 | 0.11 | 0.17 |
| 10 | 32022199 | rs3908708 | -1.652 | 0.17 | 1.129 | -0.642 | 0.009 | 0 | 0 | 0.407 | 0.38 | 0.69 | 0.26 | 0.39 | 0.32 | 0.43 |
| 10 | 32029550 | rs12218350 | -1.729 | 0.14 | 1.097 | -0.654 | 0.014 | 0.002 | 0.002 | 0 | 0.39 | 0.71 | 0.26 | 0.39 | 0.32 | 0.43 |
| 10 | 32030344 | rs6481756 | -1.76 | 0.117 | 0.912 | -0.669 | 0.014 | -0.025 | -0.025 | 0.035 | 0.41 | 0.9 | 0.27 | 0.39 | 0.32 | 0.43 |
| 10 | 32031788 | rs7896634 | 0.224 | 0.202 | 1.067 | -0.64 | 0.021 | -0.063 | -0.063 | 0 | 0.11 | 0.27 | 0.1 | 0.01 | 0.12 | 0.06 |
| 10 | 32035898 | rs11008623 | 0.672 | 0.211 | 0.701 | -0.575 | 0.017 | 0 | 0 | 0 | 0.14 | 0.06 | 0.06 | 0.17 | 0.12 | 0.17 |
| 10 | 32038452 | rs6481758 | -0.906 | 0.423 | 0.941 | -0.432 | 0.03 | 0.032 | 0.032 | 0.033 | 0.2 | 0.4 | 0.17 | 0.21 | 0.21 | 0.23 |
| 10 | 32040764 | rs2808068 | 0.923 | 0.844 | 0.06 | -0.788 | 0.023 | 0.042 | 0.042 | 0 | 0.48 | 0.26 | 0.48 | 0.61 | 0.39 | 0.47 |
| 10 | 32044641 | rs2778646 | -0.462 | 1.179 | -0.241 | -0.909 | -0.019 | 0.038 | 0.038 | 0 | 0.68 | 0.77 | 0.68 | 0.82 | 0.57 | 0.65 |
| 10 | 32045932 | rs2808067 | -0.037 | 1.227 | -0.339 | -0.983 | -0.041 | 0.045 | 0.045 | 0 | 0.78 | 0.88 | 0.65 | 0.95 | 0.63 | 0.75 |
| 10 | 32047523 | rs10826989 | 0.349 | 0.707 | -1.411 | -0.754 | 0.03 | -0.056 | -0.056 | 1.103 | 0.26 | 0.09 | 0.11 | 0.35 | 0.22 | 0.26 |
| 10 | 32050944 | rs2797128 | 1.127 | 0.857 | -1.275 | -0.503 | 0.017 | -0.034 | -0.034 | 0 | 0.37 | 0.1 | 0.4 | 0.58 | 0.29 | 0.44 |
| 10 | 32053818 | rs2778668 | -0.144 | 1.508 | -0.515 | -0.537 | 0.027 | -0.028 | -0.028 | 0 | 0.84 | 0.8 | 0.75 | 0.96 | 0.72 | 0.79 |
| 10 | 32054404 | rs2808062 | 0.359 | 0.481 | -1.116 | -0.323 | -0.033 | -0.059 | -0.059 | 1.079 | 0.58 | 0.87 | 0.56 | 0.5 | 0.54 | 0.52 |
| 10 | 32057106 | rs2778665 | 1.153 | 0.098 | -1.176 | -0.478 | -0.005 | -0.034 | -0.034 | 1.075 | 0.46 | 0.12 | 0.33 | 0.5 | 0.42 | 0.47 |
| 10 | 32067305 | rs2808055 | 0.652 | 0.35 | 0.304 | -0.938 | -0.002 | -0.01 | -0.01 | 0 | 0.66 | 0.36 | 0.66 | 0.93 | 0.64 | 0.74 |
| 10 | 32070331 | rs2808052 | 0.616 | 0.666 | 0.646 | -0.672 | 0.021 | -0.032 | -0.032 | 0 | 0.73 | 0.58 | 0.65 | 0.83 | 0.65 | 0.72 |
| 10 | 32071859 | rs2797142 | 0.711 | 0.658 | 0.359 | -0.725 | 0.002 | -0.009 | -0.009 | 0 | 0.69 | 0.37 | 0.59 | 0.83 | 0.6 | 0.69 |
| 10 | 32074960 | rs7921886 | -0.247 | 0.793 | 0.297 | -0.773 | 0.008 | -0.035 | -0.035 | 0 | 0.08 | 0.18 | 0.09 | 0 | 0.08 | 0.01 |
| 10 | 32080912 | rs2799016 | 0.521 | 1.019 | 0.123 | -1.068 | 0.017 | -0.041 | -0.041 | 0 | 0.71 | 0.43 | 0.63 | 0.86 | 0.64 | 0.7 |
| 10 | 32081924 | rs2256582 | 0.33 | 0.838 | -0.234 | -1.127 | -0.088 | 0.131 | 0.131 | 1.436 | 0.67 | 0.88 | 0.64 | 0.86 | 0.58 | 0.67 |
| 10 | 32085248 | rs11008641 | 0.614 | 0.436 | -0.668 | -0.858 | 0.023 | -0.095 | -0.095 | 0 | 0.25 | 0.1 | 0.18 | 0.45 | 0.2 | 0.26 |
| 10 | 32089191 | rs2799003 | 1.084 | 0.582 | -0.797 | -1.33 | -0.029 | 0.264 | 0.264 | 1.077 | 0.45 | 0.11 | 0.47 | 0.86 | 0.4 | 0.52 |
| 10 | 32091885 | rs2808029 | -0.008 | 0.62 | 0.108 | -1.346 | -0.001 | -0.012 | -0.012 | 0.098 | 0.65 | 0.68 | 0.63 | 0.86 | 0.57 | 0.65 |
| 10 | 32092377 | rs10508768 | 0.28 | 0.702 | 0.371 | -1.321 | -0.002 | -0.117 | -0.117 | 0.121 | 0.18 | 0.56 | 0.15 | 0 | 0.17 | 0.12 |
| 10 | 32094751 | rs1044798 | 2.405 | 0.699 | -0.272 | -1.168 | -0.016 | -0.083 | -0.083 | 0.023 | 0.21 | 0.09 | 0.3 | 0.41 | 0.2 | 0.27 |
| 10 | 32095249 | rs1044795 | 0.58 | 0.615 | -0.145 | -1.219 | 0.001 | -0.003 | -0.003 | 0 | 0.27 | 0.32 | 0.22 | 0.45 | 0.19 | 0.24 |
| 10 | 32095666 | rs2799002 | 2.504 | 0.662 | -0.133 | -1.16 | -0.028 | -0.104 | -0.104 | 1.117 | 0.21 | 0.05 | 0.3 | 0.41 | 0.21 | 0.27 |
| 10 | 32110907 | rs2808105 | 0.268 | 0.762 | -0.234 | -1.275 | 0.011 | 0.015 | 0.015 | 0 | 0.51 | 0.6 | 0.54 | 0.42 | 0.5 | 0.46 |
| 10 | 32114950 | rs2255555 | 2.627 | 0.779 | -0.348 | -1.324 | -0.016 | -0.082 | -0.082 | 1.623 | 0.21 | 0.09 | 0.3 | 0.41 | 0.2 | 0.27 |
| 10 | 32128023 | rs2808097 | -0.694 | 0.773 | -0.412 | -1.348 | -0.038 | 0.048 | 0.048 | 1.622 | 0.78 | 0.92 | 0.76 | 0.87 | 0.68 | 0.71 |
| 10 | 32128611 | rs2808096 | 2.752 | 0.789 | -0.45 | -1.325 | -0.028 | -0.104 | -0.104 | 0 | 0.21 | 0.05 | 0.3 | 0.41 | 0.2 | 0.27 |
| 10 | 32133667 | rs7091876 | 1.099 | 0.808 | -0.457 | -1.341 | -0.026 | 0.054 | 0.054 | 0 | 0.12 | 0.03 | 0.11 | 0.01 | 0.12 | 0.06 |
| 10 | 32133756 | rs2808094 | 0.241 | 0.787 | -0.335 | -1.283 | 0.005 | 0.008 | 0.008 | 0 | 0.5 | 0.4 | 0.51 | 0.42 | 0.49 | 0.46 |
| 10 | 32141460 | rs2799021 | 2.675 | 0.793 | -0.575 | -1.237 | -0.028 | -0.104 | -0.104 | 0 | 0.21 | 0.05 | 0.3 | 0.41 | 0.2 | 0.27 |
| 10 | 32161486 | rs10508769 | 0.772 | 0.765 | -0.483 | -1.243 | 0.002 | -0.005 | -0.005 | 0 | 0.26 | 0.29 | 0.22 | 0.45 | 0.19 | 0.24 |
| 10 | 32167385 | rs2808079 | 0.327 | 0.875 | -0.804 | -1.245 | 0.011 | 0.019 | 0.019 | 0 | 0.49 | 0.29 | 0.5 | 0.42 | 0.49 | 0.46 |
| 10 | 32169634 | rs2799027 | -0.697 | 1.026 | -0.889 | -1.291 | -0.068 | -0.029 | -0.029 | 0 | 0.79 | 0.97 | 0.75 | 0.81 | 0.69 | 0.7 |
| 10 | 32196106 | rs2797403 | -0.736 | 0.821 | -0.919 | -1.644 | -0.08 | 0.035 | 0.035 | 1.703 | 0.79 | 0.98 | 0.76 | 0.87 | 0.68 | 0.71 |
| 10 | 32214122 | rs16932785 | -0.393 | 0.761 | -0.555 | -1.53 | -0.008 | -0.016 | -0.016 | 0 | 0.27 | 0.2 | 0.2 | 0.01 | 0.29 | 0.19 |
| 10 | 32264033 | rs1775725 | 0.677 | 0.791 | -0.695 | -1.481 | -0.071 | 0.011 | 0.011 | 0.912 | 0.18 | 0.02 | 0.25 | 0.13 | 0.29 | 0.26 |
| 10 | 32273277 | rs11818697 | -0.401 | 0.901 | -0.429 | -1.402 | -0.019 | 0.04 | 0.04 | 0 | 0.27 | 0.15 | 0.19 | 0.01 | 0.29 | 0.19 |
| 10 | 32278603 | rs806811 | 0.215 | 0.731 | -0.187 | -1.389 | 0.001 | -0.082 | -0.082 | 0 | 0.73 | 0.69 | 0.82 | 1 | 0.77 | 0.82 |
| 10 | 32285574 | rs211268 | 0.905 | 0.753 | -0.205 | -1.383 | -0.001 | 0.02 | 0.02 | 0 | 0.13 | 0.05 | 0.05 | 0 | 0.11 | 0.12 |
| 10 | 32298840 | rs13470 | -0.563 | 0.761 | -0.302 | -1.404 | -0.004 | -0.002 | -0.002 | 0 | 0.26 | 0.19 | 0.16 | 0.01 | 0.23 | 0.17 |
| 10 | 32303758 | rs172432 | -0.575 | 0.757 | -0.303 | -1.405 | -0.004 | -0.001 | -0.001 | 0.027 | 0.26 | 0.19 | 0.16 | 0.01 | 0.23 | 0.17 |
| 10 | 32308394 | rs2286746 | 1.935 | 0.731 | -0.567 | -1.232 | -0.024 | 0.034 | 0.034 | 0 | 0.06 | 0.01 | 0.11 | 0.45 | 0.08 | 0.19 |
| 10 | 32309005 | rs1775715 | -0.364 | 0.719 | -0.299 | -1.291 | 0.005 | 0.03 | 0.03 | 0 | 0.47 | 0.33 | 0.51 | 0.58 | 0.53 | 0.52 |
| 10 | 32333847 | rs17487587 | 0.889 | 0.767 | -0.41 | -1.198 | -0.008 | 0.006 | 0.006 | 0 | 0.12 | 0.08 | 0.11 | 0 | 0.12 | 0.05 |
| 10 | 32338355 | rs11008750 | 1.971 | 0.644 | -0.299 | -1.083 | 0.003 | 0.026 | 0.026 | 0 | 0.26 | 0.29 | 0.31 | 0.41 | 0.23 | 0.31 |
| 10 | 32339792 | rs11594506 | 2.059 | 0.652 | -0.853 | -1.08 | -0.074 | -0.134 | -0.134 | 0.706 | 0.24 | 0.02 | 0.28 | 0.42 | 0.23 | 0.31 |
| 10 | 32352929 | rs7921970 | 0.258 | 0.476 | -0.725 | -1.09 | 0.034 | -0.088 | -0.088 | 0 | 0.28 | 0.19 | 0.22 | 0.46 | 0.21 | 0.25 |
| 10 | 32353837 | rs806813 | 0.829 | 0.709 | -0.587 | -1.232 | -0.041 | 0.027 | 0.027 | 0.243 | 0.18 | 0.07 | 0.28 | 0.13 | 0.33 | 0.27 |
| 10 | 32358438 | rs16932874 | 0.4 | 0.569 | -0.013 | -1.293 | 0.028 | -0.108 | -0.108 | 0 | 0.16 | 0.34 | 0.14 | 0.01 | 0.13 | 0.07 |
| 10 | 32360188 | rs211315 | 0.16 | 0.463 | -0.036 | -1.247 | 0.001 | -0.004 | -0.004 | 0 | 0.57 | 0.43 | 0.62 | 0.52 | 0.65 | 0.7 |
| 10 | 32361215 | rs806818 | -0.527 | 0.397 | -0.081 | -1.359 | -0.024 | -0.081 | -0.081 | 0 | 0.84 | 0.73 | 0.93 | 0.99 | 0.88 | 0.88 |
| 10 | 32361839 | rs740859 | 0.886 | 0.465 | -0.284 | -1.335 | -0.033 | 0.059 | 0.059 | 0 | 0.14 | 0.03 | 0.11 | 0.01 | 0.13 | 0.07 |
| 10 | 32364670 | rs806822 | 1.152 | 0.487 | -0.348 | -1.285 | -0.054 | -0.044 | -0.044 | 0 | 0.41 | 0.21 | 0.55 | 0.51 | 0.54 | 0.58 |
| 10 | 32366761 | rs1251370 | 0.615 | 0.425 | -0.117 | -1.336 | 0.012 | 0.054 | 0.054 | 0 | 0.67 | 0.27 | 0.77 | 0.99 | 0.76 | 0.81 |
| 10 | 32389424 | rs2887830 | -0.137 | 0.391 | -0.219 | -1.108 | -0.02 | 0.031 | 0.031 | 0 | 0.45 | 0.33 | 0.62 | 0.64 | 0.55 | 0.5 |
| 10 | 32390950 | rs211402 | 0.81 | 0.314 | -0.17 | -1.156 | 0.011 | 0.025 | 0.025 | 0 | 0.15 | 0.06 | 0.05 | 0 | 0.12 | 0.12 |
| 10 | 32393142 | rs211408 | -1.099 | 0.336 | -0.007 | -1.162 | -0.001 | -0.01 | -0.01 | 0 | 0.6 | 0.51 | 0.68 | 0.64 | 0.67 | 0.62 |
| 10 | 32393213 | rs211409 | -0.866 | 0.3 | 0.066 | -1.239 | 0.014 | -0.098 | -0.098 | 0 | 0.85 | 0.67 | 0.86 | 0.99 | 0.9 | 0.93 |
| 10 | 32396400 | rs211412 | 0.645 | 0.285 | 0.027 | -1.172 | -0.014 | -0.071 | -0.071 | 0.123 | 0.16 | 0.23 | 0.07 | 0 | 0.12 | 0.12 |
| 10 | 32400396 | rs73477 | -0.397 | 0.33 | -0.266 | -1.154 | 0.036 | -0.02 | -0.02 | 0 | 0.43 | 0.29 | 0.3 | 0.47 | 0.34 | 0.35 |
| 10 | 32404028 | rs2988089 | 0.94 | 0.341 | -0.274 | -1.172 | 0.01 | 0.008 | 0.008 | 0 | 0.15 | 0.1 | 0.06 | 0 | 0.12 | 0.12 |
| 10 | 32406441 | rs425411 | -0.068 | 0.361 | -0.269 | -1.226 | 0.014 | -0.042 | -0.042 | 0 | 0.68 | 0.43 | 0.48 | 0.84 | 0.57 | 0.66 |
| 10 | 32409150 | rs2907801 | 0.816 | 0.323 | -0.327 | -1.185 | 0.012 | 0.047 | 0.047 | 0 | 0.15 | 0.05 | 0.05 | 0 | 0.12 | 0.11 |

| | | | | | | | | | | | | | | | | |
|---|---|---|---|---|---|---|---|---|---|---|---|---|---|---|---|---|
| 10 | 32409654 | rs2907803 | 0.45 | 0.525 | -0.164 | -1.067 | 0.002 | 0.006 | 0.006 | 0 | 0.42 | 0.62 | 0.33 | 0.38 | 0.33 | 0.39 |
| 10 | 32413670 | rs3095259 | 1.749 | 0.525 | -0.173 | -1.069 | -0.028 | -0.076 | -0.076 | 2.257 | 0.25 | 0.08 | 0.17 | 0.37 | 0.23 | 0.31 |
| 10 | 32414174 | rs3006628 | 1.753 | 0.531 | -0.172 | -1.066 | -0.008 | -0.047 | -0.047 | 0 | 0.26 | 0.14 | 0.18 | 0.37 | 0.23 | 0.31 |
| 10 | 32415805 | rs3006633 | 0.618 | 0.718 | -0.119 | -0.923 | 0.002 | 0.006 | 0.006 | 0.007 | 0.42 | 0.62 | 0.33 | 0.38 | 0.33 | 0.39 |
| 10 | 32420091 | rs4749705 | 0.62 | 0.617 | -0.41 | -1.12 | 0.017 | -0.073 | -0.073 | 0 | 0.28 | 0.15 | 0.23 | 0.42 | 0.21 | 0.28 |
| 10 | 32420282 | rs3006650 | 0.602 | 0.701 | -0.433 | -1.084 | -0.002 | 0.001 | 0.001 | 0.054 | 0.42 | 0.52 | 0.35 | 0.45 | 0.35 | 0.4 |
| 10 | 32422064 | rs3006653 | 0.618 | 1.042 | -0.751 | -0.887 | 0 | -0.001 | -0.001 | 0.088 | 0.46 | 0.54 | 0.41 | 0.48 | 0.4 | 0.41 |
| 10 | 32428236 | rs2998074 | -1.115 | 1.145 | -0.86 | -0.872 | -0.014 | 0.012 | 0.012 | 0 | 0.83 | 0.88 | 0.64 | 0.87 | 0.67 | 0.7 |
| 10 | 32431923 | rs11594690 | 1.138 | 1.216 | -0.981 | -0.969 | -0.019 | 0.056 | 0.056 | 0 | 0.12 | 0.02 | 0.05 | 0 | 0.13 | 0.08 |
| 10 | 32432000 | rs10827014 | -0.619 | 1.249 | -0.924 | -0.954 | -0.006 | 0.004 | 0.004 | 0.087 | 0.8 | 0.82 | 0.62 | 0.87 | 0.64 | 0.69 |
| 10 | 32434312 | rs10763879 | 1.709 | 1.082 | -0.84 | -0.948 | 0 | -0.003 | -0.003 | 0 | 0.29 | 0.3 | 0.26 | 0.48 | 0.26 | 0.32 |
| 10 | 32435853 | rs9299674 | -0.608 | 1.088 | -0.831 | -0.438 | 0.004 | -0.003 | -0.003 | 0 | 0.41 | 0.41 | 0.24 | 0.39 | 0.29 | 0.3 |
| 10 | 32450499 | rs2998029 | -0.359 | 0.912 | -0.681 | 0.08 | -0.006 | 0.005 | 0.005 | 0 | 0.82 | 0.86 | 0.64 | 0.87 | 0.67 | 0.72 |
| 10 | 32450551 | rs7083956 | -0.512 | -0.073 | -0.578 | 0.474 | -0.024 | 0.037 | 0.037 | 0 | 0.25 | 0.35 | 0.18 | 0.38 | 0.13 | 0.22 |
| 10 | 32456826 | rs2998033 | 0.029 | 0.634 | -0.968 | 0.428 | -0.015 | -0.012 | -0.012 | 0 | 0.68 | 0.75 | 0.58 | 0.65 | 0.65 | 0.57 |
| 10 | 32460069 | rs3006679 | 0.861 | 0.202 | -1.929 | 0.123 | -0.033 | -0.135 | -0.135 | 0 | 0.32 | 0.06 | 0.23 | 0.54 | 0.34 | 0.29 |
| 10 | 32460326 | rs2998036 | -0.006 | -0.097 | -1.454 | -0.085 | 0.004 | 0.042 | 0.042 | 0 | 0.5 | 0.26 | 0.46 | 0.61 | 0.51 | 0.41 |
| 10 | 32467111 | rs2998050 | -0.284 | -0.285 | -1.405 | -0.145 | -0.009 | 0.005 | 0.005 | 0 | 0.57 | 0.3 | 0.5 | 0.61 | 0.59 | 0.48 |
| 10 | 32469529 | rs2998056 | 0.446 | -0.36 | -1.357 | -0.114 | -0.017 | -0.062 | -0.062 | 0.673 | 0.31 | 0.16 | 0.23 | 0.48 | 0.36 | 0.34 |
| 10 | 32476236 | rs3006702 | -0.683 | -0.742 | -1.585 | -0.42 | -0.119 | 0.025 | 0.025 | 0.01 | 0.54 | 0.96 | 0.53 | 0.65 | 0.56 | 0.51 |
| 10 | 32487433 | rs12256995 | 0.48 | -0.792 | 0.024 | -0.568 | -0.005 | -0.07 | -0.07 | 0 | 0.16 | 0.24 | 0.14 | 0 | 0.14 | 0.09 |
| 10 | 32492118 | rs3006608 | 0.758 | -0.709 | -0.142 | -0.555 | -0.013 | -0.021 | -0.021 | 0.082 | 0.31 | 0.26 | 0.32 | 0.59 | 0.38 | 0.33 |
| 10 | 32493733 | rs11595812 | -0.346 | -0.828 | -0.033 | -0.563 | -0.024 | -0.07 | -0.07 | 0.514 | 0.22 | 0.54 | 0.19 | 0.07 | 0.18 | 0.19 |
| 10 | 32499938 | rs2806150 | -0.68 | -0.737 | -0.427 | -0.407 | -0.062 | 0.055 | 0.055 | 0 | 0.55 | 0.89 | 0.53 | 0.65 | 0.56 | 0.53 |
| 10 | 32503359 | rs2806154 | -0.168 | -1.164 | 0.082 | -0.376 | -0.031 | 0.019 | 0.019 | 0 | 0.26 | 0.39 | 0.15 | 0.27 | 0.2 | 0.21 |
| 10 | 32509660 | rs3891660 | 0.842 | -0.69 | -0.252 | -0.336 | -0.037 | -0.034 | -0.034 | 0 | 0.06 | 0.01 | 0.12 | 0.07 | 0.13 | 0.04 |
| 10 | 32510010 | rs1080543 | -1.763 | -1.125 | -0.533 | -0.688 | -0.086 | -0.063 | -0.063 | 0.151 | 0.59 | 0.89 | 0.46 | 0.35 | 0.5 | 0.38 |
| 10 | 32513318 | rs12775713 | -0.688 | -1.256 | -0.072 | -0.749 | 0.008 | -0.005 | -0.005 | 0 | 0.39 | 0.37 | 0.28 | 0.35 | 0.34 | 0.27 |
| 10 | 32515093 | rs2769214 | -1.453 | -1.203 | 0.101 | -0.702 | -0.003 | 0 | 0 | 0.058 | 0.56 | 0.64 | 0.43 | 0.35 | 0.5 | 0.37 |
| 10 | 32524898 | rs2769202 | -0.971 | -0.841 | 0.185 | -0.571 | 0 | -0.007 | -0.007 | 0 | 0.34 | 0.39 | 0.33 | 0.14 | 0.35 | 0.23 |
| 10 | 32531607 | rs2031791 | -1.007 | -0.857 | -0.038 | -0.333 | 0.003 | 0.022 | 0.022 | 0.021 | 0.36 | 0.69 | 0.35 | 0.2 | 0.35 | 0.24 |
| 10 | 32534087 | rs16915609 | 0.548 | -0.968 | 0.064 | -0.3 | 0.001 | -0.002 | -0.002 | 0 | 0.2 | 0.19 | 0.15 | 0.21 | 0.15 | 0.12 |
| 10 | 32538176 | rs12268212 | -0.731 | -0.623 | 0.32 | -0.203 | -0.007 | 0.003 | 0.003 | 1.829 | 0.21 | 0.62 | 0.22 | 0.17 | 0.2 | 0.14 |
| 10 | 32547292 | rs12261062 | 0.772 | -0.351 | -0.156 | -0.158 | -0.003 | 0.003 | 0.003 | 2.315 | 0.13 | 0.12 | 0.14 | 0.05 | 0.15 | 0.07 |
| 10 | 32552782 | rs2769240 | -1.436 | -0.722 | -0.745 | -0.145 | -0.139 | -0.157 | -0.157 | 0 | 0.58 | 0.96 | 0.5 | 0.39 | 0.5 | 0.39 |
| 10 | 32554586 | rs2806144 | -0.9 | -0.994 | 0.525 | -0.366 | 0.009 | 0.03 | 0.03 | 0 | 0.47 | 0.58 | 0.44 | 0.24 | 0.44 | 0.28 |
| 10 | 32563922 | rs2986936 | 0.171 | -0.339 | 0.64 | -0.229 | 0.04 | 0.016 | 0.016 | 0 | 0.74 | 0.57 | 0.62 | 0.76 | 0.69 | 0.58 |
| 10 | 32574842 | rs11008865 | 0.43 | -0.344 | 0.568 | -0.226 | -0.008 | -0.015 | -0.015 | 0 | 0.13 | 0.08 | 0.21 | 0.21 | 0.15 | 0.36 |
| 10 | 32575444 | rs10508773 | 0.477 | -0.338 | 0.571 | -0.223 | -0.008 | -0.014 | -0.014 | 0.667 | 0.13 | 0.08 | 0.21 | 0.21 | 0.15 | 0.36 |
| 10 | 32584506 | rs2479358 | 0.184 | -0.349 | 0.833 | -0.237 | 0.012 | 0.004 | 0.004 | 0 | 0.74 | 0.69 | 0.62 | 0.76 | 0.69 | 0.58 |
| 10 | 32587500 | rs713267 | 0.582 | -0.379 | 0.988 | -0.251 | 0.007 | 0 | 0 | 0.226 | 0.71 | 0.41 | 0.6 | 0.76 | 0.69 | 0.58 |
| 10 | 32593393 | rs2479362 | -1.494 | -0.359 | 0.794 | -0.233 | 0.017 | -0.026 | -0.026 | 0 | 0.87 | 0.78 | 0.83 | 0.96 | 0.84 | 0.94 |
| 10 | 32610543 | rs1475435 | 1.057 | -0.615 | 0.019 | -0.415 | -0.094 | 0.091 | 0.091 | 0.787 | 0.53 | 0.06 | 0.43 | 0.75 | 0.54 | 0.48 |
| 10 | 32610703 | rs1475436 | 1.921 | -0.516 | 0.015 | -0.348 | -0.111 | -0.158 | -0.158 | 0.517 | 0.36 | 0.04 | 0.31 | 0.55 | 0.4 | 0.36 |
| 10 | 32619572 | rs11592754 | 0.833 | -0.187 | 0.612 | -0.573 | -0.045 | -0.039 | -0.039 | 0.577 | 0.1 | 0 | 0.18 | 0.11 | 0.12 | 0.27 |
| 10 | 32623818 | rs2490508 | -0.907 | -0.189 | 0.626 | -0.627 | 0.045 | -0.048 | -0.048 | 0 | 0.85 | 0.58 | 0.82 | 0.96 | 0.84 | 0.94 |
| 10 | 32634972 | rs2370759 | 0.383 | -0.17 | 0.586 | -0.6 | -0.015 | -0.024 | -0.024 | 0 | 0.12 | 0.06 | 0.21 | 0.21 | 0.15 | 0.36 |
| 10 | 32639215 | rs2505374 | 1.018 | -0.284 | 0.231 | -0.617 | -0.094 | 0.082 | 0.082 | 0.865 | 0.54 | 0.08 | 0.44 | 0.75 | 0.54 | 0.49 |
| 10 | 32640139 | rs4291577 | 1.21 | -0.285 | 0.248 | -0.622 | -0.01 | 0.025 | 0.025 | 0 | 0.11 | 0.03 | 0.11 | 0 | 0.14 | 0.08 |
| 10 | 32641653 | rs1126369 | 0.528 | -0.292 | 0.803 | -0.188 | 0.023 | 0.002 | 0.002 | 0 | 0.71 | 0.5 | 0.6 | 0.75 | 0.68 | 0.57 |
| 10 | 32644899 | rs2479351 | 0.825 | -0.207 | -0.108 | -0.154 | -0.023 | 0.01 | 0.01 | 3.183 | 0.66 | 0.12 | 0.56 | 0.75 | 0.68 | 0.57 |
| 10 | 32659387 | rs2505366 | 0.543 | 0.097 | 0.154 | 0.046 | -0.004 | 0.02 | 0.02 | 0 | 0.59 | 0.19 | 0.5 | 0.72 | 0.63 | 0.55 |
| 10 | 32668717 | rs7086385 | 0.206 | -0.068 | 0.82 | -0.001 | -0.01 | -0.017 | -0.017 | 0 | 0.23 | 0.3 | 0.22 | 0.14 | 0.21 | 0.2 |
| 10 | 32671117 | rs2490527 | 1.04 | 0.002 | 0.692 | -0.029 | 0.007 | 0.03 | 0.03 | 0 | 0.56 | 0.2 | 0.48 | 0.69 | 0.62 | 0.53 |
| 10 | 32672577 | rs2251013 | 0.252 | 0.129 | 0.844 | 0.169 | 0.054 | 0.049 | 0.049 | 0.15 | 0.85 | 0.64 | 0.72 | 0.83 | 0.84 | 0.73 |
| 10 | 32678925 | rs7075248 | -0.448 | 0.005 | 0.406 | 0.069 | -0.006 | -0.009 | -0.009 | 0 | 0.34 | 0.57 | 0.35 | 0.3 | 0.33 | 0.29 |
| 10 | 32684126 | rs7922515 | 0.838 | -0.054 | 0.479 | 0.071 | 0.005 | 0.007 | 0.007 | 0.34 | 0.21 | 0.2 | 0.2 | 0.15 | 0.18 | 0.18 |
| 10 | 32684427 | rs7099897 | 0.933 | -0.056 | 0.434 | 0.07 | 0.007 | 0.011 | 0.011 | 0.524 | 0.2 | 0.16 | 0.19 | 0.15 | 0.18 | 0.18 |
| 10 | 32687497 | rs7072959 | 1.066 | 0.063 | 0.473 | 0.09 | 0.004 | 0.014 | 0.014 | 0 | 0.08 | 0.09 | 0.09 | 0.15 | 0.09 | 0.09 |
| 10 | 32690521 | rs2490472 | 1.62 | -0.128 | -0.082 | -0.036 | -0.159 | -0.141 | -0.141 | 0 | 0.43 | 0.03 | 0.37 | 0.56 | 0.5 | 0.43 |
| 10 | 32693485 | rs17296114 | -0.565 | 0.108 | 0.374 | 0.021 | -0.029 | -0.009 | -0.009 | 0 | 0.18 | 0.08 | 0.27 | 0.14 | 0.21 | 0.3 |
| 10 | 32696496 | rs4394741 | 0.039 | 0.116 | 0.224 | 0.03 | -0.057 | -0.047 | -0.047 | 0.655 | 0.13 | 0.02 | 0.25 | 0.14 | 0.18 | 0.28 |
| 10 | 32700343 | rs12242183 | -1.313 | 0.064 | 0.223 | -0.027 | 0.006 | -0.01 | -0.01 | 0.293 | 0.32 | 0.53 | 0.41 | 0.28 | 0.32 | 0.4 |
| 10 | 32702456 | rs2370771 | -1.499 | -0.339 | -0.239 | 0.112 | 0.014 | 0.009 | 0.009 | 0.643 | 0.3 | 0.76 | 0.4 | 0.27 | 0.23 | 0.39 |
| 10 | 32704465 | rs2370772 | 0.315 | -0.437 | -0.252 | 0.075 | -0.004 | -0.002 | -0.002 | 0 | 0.23 | 0.26 | 0.22 | 0.16 | 0.18 | 0.19 |
| 10 | 32714338 | rs2505389 | 0.729 | -0.617 | -0.878 | -0.22 | -0.036 | 0.106 | 0.106 | 1.317 | 0.43 | 0.14 | 0.55 | 0.27 | 0.55 | 0.58 |
| 10 | 32714392 | rs2505390 | -0.962 | -0.927 | -0.312 | 0.159 | -0.014 | 0.165 | 0.165 | 2.082 | 0.83 | 0.6 | 0.89 | 0.68 | 0.88 | 0.89 |
| 10 | 32724827 | rs11594190 | 0.887 | -1.018 | -0.018 | 0.04 | -0.003 | 0.032 | 0.032 | 0 | 0.12 | 0.28 | 0.11 | 0.16 | 0.1 | 0.09 |
| 10 | 32743146 | rs2243926 | 1.724 | -0.876 | -0.297 | -0.155 | -0.092 | 0.132 | 0.132 | 0.501 | 0.33 | 0.13 | 0.33 | 0.1 | 0.48 | 0.3 |

| | | | | | | | | | | | | | | | | |
|---|---|---|---|---|---|---|---|---|---|---|---|---|---|---|---|---|
| 10 | 32798947 | rs10740855 | -0.743 | -1.091 | -0.283 | -0.081 | 0.01 | 0.002 | 0.002 | 0 | 0.47 | 0.59 | 0.56 | 0.59 | 0.37 | 0.57 |
| 10 | 32804398 | rs7075342 | 1.058 | -1.072 | -0.376 | -0.04 | -0.004 | 0.03 | 0.03 | 0 | 0.12 | 0.28 | 0.11 | 0.16 | 0.1 | 0.09 |
| 10 | 32815921 | rs10827071 | 2.006 | -1.136 | -0.673 | -0.037 | -0.023 | -0.058 | -0.058 | 0.053 | 0.19 | 0.03 | 0.19 | 0.25 | 0.17 | 0.18 |
| 10 | 32834243 | rs11008969 | 2.269 | -1.157 | -0.704 | -0.038 | -0.032 | -0.074 | -0.074 | 0.089 | 0.11 | 0.01 | 0.05 | 0.32 | 0.12 | 0.1 |
| 10 | 32836249 | rs2990987 | 1.945 | -1.135 | -0.674 | -0.043 | -0.022 | -0.058 | -0.058 | 0 | 0.2 | 0.03 | 0.19 | 0.25 | 0.17 | 0.19 |
| 10 | 32846254 | rs2370782 | 0.134 | -1.166 | -0.646 | -0.068 | 0.004 | -0.048 | -0.048 | 0 | 0.33 | 0.12 | 0.44 | 0.43 | 0.27 | 0.48 |
| 10 | 32856445 | rs12266925 | 0.755 | -1.182 | -0.634 | -0.094 | 0 | -0.002 | -0.002 | 0 | 0.11 | 0.09 | 0.24 | 0.17 | 0.09 | 0.29 |
| 10 | 32856746 | rs12268559 | 0.929 | -1.163 | -0.618 | -0.087 | 0 | 0.001 | 0.001 | 0 | 0.1 | 0.09 | 0.24 | 0.17 | 0.09 | 0.29 |
| 10 | 32864730 | rs10508775 | 1.947 | -1.186 | -0.702 | -0.178 | -0.173 | 0.154 | 0.154 | 0.645 | 0.32 | 0.02 | 0.32 | 0.1 | 0.46 | 0.3 |
| 10 | 32888475 | rs9663143 | -0.577 | -1.219 | -0.504 | -0.118 | 0 | 0 | 0 | 0 | 0.45 | 0.44 | 0.55 | 0.59 | 0.37 | 0.57 |
| 10 | 32919407 | rs1759666 | -1.479 | -1.196 | -0.37 | -0.072 | -0.007 | 0.134 | 0.134 | 0.652 | 0.85 | 0.69 | 0.9 | 0.68 | 0.88 | 0.9 |
| 10 | 32919749 | rs12416352 | 1.049 | -1.129 | -0.391 | -0.058 | 0 | 0.003 | 0.003 | 0 | 0.09 | 0.09 | 0.24 | 0.17 | 0.09 | 0.29 |
| 10 | 32952029 | rs10218979 | 0.303 | -1.184 | -0.398 | -0.064 | 0.001 | -0.006 | -0.006 | 0 | 0.13 | 0.09 | 0.25 | 0.17 | 0.1 | 0.3 |
| 10 | 32952768 | rs7089887 | -0.677 | -1.303 | -0.452 | -0.14 | 0.003 | 0 | 0 | 0 | 0.46 | 0.44 | 0.55 | 0.59 | 0.37 | 0.58 |
| 10 | 32959844 | rs10827098 | 0.168 | -1.319 | -0.527 | -0.137 | 0.004 | -0.051 | -0.051 | 0 | 0.33 | 0.12 | 0.44 | 0.43 | 0.27 | 0.48 |
| 10 | 32960793 | rs11009027 | 2.066 | -1.31 | -0.533 | -0.139 | -0.022 | -0.058 | -0.058 | 0.104 | 0.2 | 0.03 | 0.19 | 0.25 | 0.17 | 0.19 |
| 10 | 32963988 | rs1627683 | 1.966 | -1.312 | -0.544 | -0.17 | -0.173 | 0.155 | 0.155 | 0.599 | 0.32 | 0.03 | 0.32 | 0.09 | 0.47 | 0.3 |
| 10 | 33010304 | rs7075648 | 0.4 | -1.058 | -0.615 | -0.278 | -0.014 | 0.037 | 0.037 | 0.033 | 0.15 | 0.51 | 0.13 | 0.16 | 0.1 | 0.1 |
| 10 | 33013011 | rs1541135 | -0.254 | -1.121 | -0.702 | -0.307 | 0.001 | -0.001 | -0.001 | 0 | 0.61 | 0.63 | 0.7 | 0.43 | 0.67 | 0.69 |
| 10 | 33016633 | rs4145862 | 1.977 | -0.964 | -0.676 | -0.221 | -0.022 | -0.058 | -0.058 | 0 | 0.2 | 0.03 | 0.19 | 0.25 | 0.17 | 0.19 |
| 10 | 33050681 | rs11598580 | 0.82 | -1.075 | -0.356 | -0.118 | -0.004 | 0.029 | 0.029 | 0 | 0.13 | 0.32 | 0.12 | 0.15 | 0.1 | 0.1 |
| 10 | 33080104 | rs7914502 | -0.753 | -1.081 | -0.706 | -0.184 | 0.02 | 0.007 | 0.007 | 0 | 0.48 | 0.63 | 0.57 | 0.59 | 0.37 | 0.58 |
| 10 | 33092798 | rs11009088 | 1.8 | -1.043 | -0.744 | -0.162 | -0.005 | -0.039 | -0.039 | 0 | 0.12 | 0.03 | 0.06 | 0.32 | 0.12 | 0.1 |
| 10 | 33123774 | rs11009107 | 1.763 | -0.999 | -0.93 | -0.317 | -0.174 | 0.15 | 0.15 | 0.677 | 0.32 | 0.02 | 0.31 | 0.1 | 0.46 | 0.29 |
| 10 | 33137551 | rs2504011 | 1.744 | -1.003 | -0.932 | -0.319 | -0.174 | 0.151 | 0.151 | 0.585 | 0.32 | 0.02 | 0.31 | 0.1 | 0.46 | 0.3 |
| 10 | 33166816 | rs10827150 | 0.903 | -1.094 | -0.567 | -0.143 | -0.001 | -0.002 | -0.002 | 0.377 | 0.1 | 0.09 | 0.1 | 0.19 | 0.09 | 0.29 |
| 10 | 33170951 | rs2504004 | -0.218 | -1.181 | -0.566 | -0.092 | -0.025 | 0.079 | 0.079 | 0.026 | 0.78 | 0.59 | 0.87 | 0.69 | 0.83 | 0.87 |
| 10 | 33181994 | rs2000217 | 1.368 | -1.245 | -0.589 | -0.108 | 0 | -0.017 | -0.017 | 0 | 0.1 | 0.05 | 0.24 | 0.16 | 0.1 | 0.1 |
| 10 | 33187619 | rs11009132 | 0.623 | -1.355 | -0.608 | -0.22 | -0.02 | -0.108 | -0.108 | 0 | 0.3 | 0.04 | 0.42 | 0.41 | 0.27 | 0.27 |
| 10 | 33188102 | rs1342548 | -1.321 | -1.394 | -0.47 | -0.169 | -0.01 | 0.128 | 0.128 | 0.703 | 0.84 | 0.66 | 0.9 | 0.69 | 0.88 | 0.88 |
| 10 | 33190567 | rs2153875 | 0.815 | -1.462 | -0.595 | -0.249 | 0.026 | -0.061 | -0.061 | 2.432 | 0.66 | 0.19 | 0.76 | 0.5 | 0.74 | 0.55 |
| 10 | 33195877 | rs3824602 | 1.423 | -1.429 | -0.479 | -0.221 | -0.002 | 0.106 | 0.106 | 0.021 | 0.14 | 0.25 | 0.09 | 0.31 | 0.12 | 0.13 |
| 10 | 33197197 | rs1316757 | 2.165 | -1.434 | -0.617 | -0.223 | -0.011 | -0.042 | -0.042 | 0 | 0.11 | 0.02 | 0.06 | 0.31 | 0.12 | 0.13 |
| 10 | 33198911 | rs2488320 | 1.111 | -1.46 | -0.795 | -0.318 | -0.029 | 0.096 | 0.096 | 0.353 | 0.42 | 0.15 | 0.56 | 0.26 | 0.56 | 0.36 |
| 10 | 33209266 | rs2230396 | -1.553 | -1.437 | -0.688 | -0.246 | -0.008 | 0.131 | 0.131 | 0.572 | 0.85 | 0.66 | 0.9 | 0.69 | 0.88 | 0.87 |
| 10 | 33213680 | rs3780873 | 1.22 | -1.358 | -1.132 | -0.231 | -0.022 | -0.048 | -0.048 | 0 | 0.09 | 0.01 | 0.09 | 0.19 | 0.07 | 0.29 |
| 10 | 33214802 | rs2298141 | 1.723 | -1.443 | -1.085 | -0.206 | 0 | -0.046 | -0.046 | 0.004 | 0.21 | 0.04 | 0.19 | 0.25 | 0.17 | 0.19 |
| 10 | 33215725 | rs2503997 | 0.111 | -1.442 | -1.078 | -0.23 | 0.066 | 0.006 | 0.006 | 1.45 | 0.55 | 0.17 | 0.66 | 0.44 | 0.66 | 0.66 |
| 10 | 33218076 | rs10763902 | 0.514 | -1.32 | -1.065 | -0.217 | 0.007 | -0.07 | -0.07 | 0 | 0.31 | 0.09 | 0.42 | 0.41 | 0.27 | 0.27 |
| 10 | 33226552 | rs10827163 | 1.618 | -1.273 | -1.042 | -0.209 | 0 | -0.046 | -0.046 | 2.683 | 0.21 | 0.04 | 0.19 | 0.25 | 0.17 | 0.19 |
| 10 | 33227477 | rs16933819 | 1.618 | -1.273 | -1.042 | -0.224 | 0 | -0.044 | -0.044 | 0 | 0.21 | 0.04 | 0.19 | 0.24 | 0.17 | 0.19 |
| 10 | 33232628 | rs10827164 | 0.404 | -1.677 | -1.654 | -0.511 | 0.007 | -0.053 | -0.053 | 0.375 | 0.37 | 0.12 | 0.44 | 0.44 | 0.35 | 0.33 |
| 10 | 33236901 | rs1009002 | 1.328 | -1.662 | -1.785 | -0.592 | -0.165 | 0.163 | 0.163 | 0 | 0.34 | 0.03 | 0.33 | 0.1 | 0.45 | 0.26 |
| 10 | 33242401 | rs11009157 | 0.496 | -1.744 | -1.517 | -0.462 | 0.016 | -0.016 | -0.016 | 0.71 | 0.34 | 0.25 | 0.26 | 0.44 | 0.25 | 0.5 |
| 10 | 33245272 | rs1187073 | -1.923 | -1.611 | -1.279 | -0.363 | -0.002 | 0.12 | 0.12 | 0.038 | 0.86 | 0.71 | 0.9 | 0.7 | 0.88 | 0.88 |
| 10 | 33249133 | rs1187078 | 0.986 | -1.581 | -1.511 | -0.423 | -0.073 | 0.077 | 0.077 | 0.001 | 0.39 | 0.13 | 0.54 | 0.26 | 0.54 | 0.38 |
| 10 | 33255172 | rs1775114 | 1.112 | -1.335 | -1.483 | -0.417 | -0.089 | 0.158 | 0.158 | 0 | 0.35 | 0.11 | 0.35 | 0.1 | 0.46 | 0.3 |
| 10 | 33260699 | rs17296289 | 1.221 | -1.259 | -1.824 | -0.347 | -0.017 | -0.052 | -0.052 | 6.228 | 0.09 | 0 | 0.07 | 0.2 | 0.07 | 0.28 |
| 10 | 33261418 | rs2457705 | 1.472 | -1.113 | -1.706 | -0.423 | -0.151 | 0.165 | 0.165 | 0 | 0.31 | 0.03 | 0.3 | 0.09 | 0.43 | 0.28 |
| 10 | 33264180 | rs11009162 | 1.659 | -1.08 | -1.398 | -0.222 | -0.004 | 0.002 | 0.002 | 1.065 | 0.2 | 0.24 | 0.18 | 0.24 | 0.16 | 0.18 |
| 10 | 33265555 | rs1187098 | -1.568 | -0.742 | -1.361 | -0.143 | 0 | 0.131 | 0.131 | 0 | 0.86 | 0.72 | 0.89 | 0.69 | 0.87 | 0.87 |
| 10 | 33266888 | rs1187095 | 0.679 | -0.365 | -1.823 | -0.49 | -0.088 | 0.137 | 0.137 | 0 | 0.3 | 0.06 | 0.32 | 0.1 | 0.35 | 0.25 |
| 10 | 33272133 | rs2475193 | 0.639 | -0.71 | -1.255 | 0.084 | -0.001 | -0.001 | -0.001 | 0.438 | 0.19 | 0.19 | 0.14 | 0.23 | 0.25 | 0.22 |
| 10 | 33275354 | rs11009164 | 0.629 | -1.376 | -0.569 | 0.167 | -0.001 | -0.01 | -0.01 | 0.133 | 0.07 | 0.05 | 0.04 | 0 | 0.04 | 0.02 |
| 10 | 33276335 | rs16933853 | 0.486 | -1.697 | -0.57 | 0.373 | 0.003 | -0.002 | -0.002 | 0 | 0.08 | 0.05 | 0.22 | 0.18 | 0.03 | 0.04 |
| 10 | 33277782 | rs10827167 | -0.324 | -1.807 | -0.135 | 0.33 | -0.003 | 0.006 | 0.006 | 1.422 | 0.37 | 0.45 | 0.26 | 0.44 | 0.3 | 0.45 |
| 10 | 33281942 | rs1187086 | 1.306 | -1.819 | -0.35 | 0.308 | 0.014 | 0.023 | 0.023 | 0.154 | 0.44 | 0.21 | 0.47 | 0.38 | 0.63 | 0.49 |
| 10 | 33284653 | rs11591508 | 0.55 | -2.239 | -0.13 | 0.55 | -0.002 | 0.033 | 0.033 | 0 | 0.11 | 0.14 | 0.09 | 0.26 | 0.08 | 0.25 |
| 10 | 33296362 | rs16933862 | -0.975 | -2.23 | 0.191 | 0.764 | 0.031 | -0.011 | -0.011 | 0 | 0.16 | 0.08 | 0.24 | 0.18 | 0.08 | 0.07 |
| 10 | 33297667 | rs1418540 | -0.705 | -1.986 | 0.355 | 0.747 | -0.01 | 0.01 | 0.01 | 6.298 | 0.32 | 0.64 | 0.24 | 0.44 | 0.22 | 0.38 |
| 10 | 33299459 | rs6481819 | -0.658 | -1.973 | 0.473 | 0.75 | -0.028 | 0.046 | 0.046 | 5.643 | 0.3 | 0.55 | 0.23 | 0.44 | 0.21 | 0.37 |
| 10 | 33311032 | rs12146170 | 0.978 | -1.864 | -0.109 | 0.752 | -0.012 | -0.014 | -0.014 | 1.467 | 0.17 | 0.03 | 0.1 | 0.16 | 0.13 | 0.15 |
| 10 | 33317051 | rs1984867 | -0.969 | -1.91 | 0.028 | 0.843 | 0.021 | -0.021 | -0.021 | 0 | 0.33 | 0.22 | 0.23 | 0.43 | 0.24 | 0.37 |
| 10 | 33324516 | rs941831 | 0.301 | -1.945 | -0.327 | 0.943 | 0.015 | -0.014 | -0.014 | 0 | 0.08 | 0.02 | 0.21 | 0.21 | 0.03 | 0.04 |
| 10 | 33325916 | rs1557148 | -1.194 | -1.495 | 0.061 | 0.836 | -0.02 | 0.012 | 0.012 | 0.148 | 0.35 | 0.42 | 0.25 | 0.44 | 0.26 | 0.38 |
| 10 | 33334273 | rs11009195 | -0.312 | -1.328 | -0.771 | 0.496 | 0.002 | 0.02 | 0.02 | 2.046 | 0.23 | 0.04 | 0.13 | 0.17 | 0.17 | 0.18 |
| 10 | 33342328 | rs11009199 | 0.236 | -0.849 | -0.483 | 0.829 | -0.014 | -0.041 | -0.041 | 1.357 | 0.18 | 0.03 | 0.11 | 0.18 | 0.14 | 0.15 |
| 10 | 33353843 | rs2666229 | -1.031 | -1.014 | 0.251 | 0.887 | 0.011 | -0.006 | -0.006 | 0 | 0.38 | 0.3 | 0.4 | 0.38 | 0.28 | 0.29 |
| 10 | 33356139 | rs2666261 | -2.05 | -0.951 | -0.012 | 0.862 | -0.079 | 0.182 | 0.182 | 0.736 | 0.56 | 0.92 | 0.53 | 0.77 | 0.43 | 0.61 |

| | | | | | | | | | | | | | | | | |
|---|---|---|---|---|---|---|---|---|---|---|---|---|---|---|---|---|
| 10 | 33357552 | rs2666260 | -0.181 | -1.059 | 0.058 | 0.846 | -0.001 | 0.006 | 0.006 | 0 | 0.18 | 0.2 | 0.29 | 0.2 | 0.16 | 0.12 |
| 10 | 33358070 | rs12359953 | 0.964 | -1.179 | -0.067 | 0.771 | -0.003 | -0.046 | -0.046 | 0 | 0.17 | 0.01 | 0.06 | 0.17 | 0.11 | 0.17 |
| 10 | 33361337 | rs2666282 | -2.133 | -0.931 | 0.403 | 1.06 | -0.015 | 0.014 | 0.014 | 0.39 | 0.54 | 0.76 | 0.52 | 0.52 | 0.44 | 0.45 |
| 10 | 33361371 | rs2756003 | -0.607 | -1.027 | 0.804 | 0.999 | 0.002 | -0.018 | -0.018 | 0 | 0.27 | 0.34 | 0.2 | 0.16 | 0.24 | 0.2 |
| 10 | 33363021 | rs752251 | -1.316 | -1.021 | 0.966 | 1.014 | 0.019 | 0.01 | 0.01 | 1.018 | 0.41 | 0.74 | 0.29 | 0.34 | 0.33 | 0.35 |
| 10 | 33368504 | rs1923272 | -2.22 | -1.1 | 0.979 | 0.827 | -0.001 | 0.015 | 0.015 | 0.364 | 0.54 | 0.75 | 0.52 | 0.54 | 0.43 | 0.45 |
| 10 | 33368644 | rs7088801 | 0.877 | -1.263 | 1.065 | 0.849 | -0.013 | 0.016 | 0.016 | 0 | 0.11 | 0.22 | 0.05 | 0.18 | 0.09 | 0.15 |
| 10 | 33372725 | rs10827185 | 0.346 | -1.233 | 0.937 | 0.873 | -0.016 | -0.073 | -0.073 | 0 | 0.13 | 0.01 | 0.24 | 0.26 | 0.11 | 0.11 |
| 10 | 33373750 | rs10827186 | 0.153 | -1.068 | 1.158 | 0.88 | 0.007 | -0.026 | -0.026 | 0 | 0.21 | 0.28 | 0.18 | 0.11 | 0.19 | 0.18 |
| 10 | 33374808 | rs1923268 | -1.435 | -1.289 | 1.178 | 1.017 | -0.014 | -0.007 | -0.007 | 0.842 | 0.35 | 0.62 | 0.37 | 0.67 | 0.24 | 0.44 |
| 10 | 33379800 | rs6481830 | -1.194 | -1.341 | 1.125 | 1.022 | -0.04 | 0.021 | 0.021 | 1.048 | 0.33 | 0.6 | 0.2 | 0.54 | 0.18 | 0.44 |
| 10 | 33383439 | rs2750066 | 0.227 | -1.184 | 0.736 | 0.995 | 0.004 | -0.011 | -0.011 | 3.128 | 0.2 | 0.1 | 0.28 | 0.26 | 0.18 | 0.17 |
| 10 | 33391568 | rs2666242 | -1.137 | -1.228 | 0.743 | 1.034 | -0.033 | 0.049 | 0.049 | 0 | 0.31 | 0.52 | 0.36 | 0.42 | 0.24 | 0.27 |
| 10 | 33393443 | rs2057253 | -1.724 | -1.237 | 0.705 | 1.042 | 0.006 | 0.003 | 0.003 | 0.724 | 0.49 | 0.49 | 0.46 | 0.52 | 0.35 | 0.43 |
| 10 | 33395124 | rs16934017 | -1.144 | -1.1 | 0.801 | 1.124 | -0.037 | 0.007 | 0.007 | 1.481 | 0.3 | 0.46 | 0.18 | 0.26 | 0.17 | 0.26 |
| 10 | 33398251 | rs7070343 | -0.65 | -0.808 | 0.392 | 0.538 | -0.025 | -0.044 | -0.044 | 0.375 | 0.24 | 0.33 | 0.16 | 0.1 | 0.16 | 0.18 |
| 10 | 33402524 | rs11009231 | 0.763 | -0.527 | -0.52 | -0.153 | -0.015 | -0.011 | -0.011 | 0.096 | 0.16 | 0.01 | 0.1 | 0.1 | 0.11 | 0.17 |
| 10 | 33405575 | rs2750055 | -1.11 | -0.14 | 0.013 | -0.051 | 0.016 | 0.017 | 0.017 | 0 | 0.37 | 0.75 | 0.45 | 0.67 | 0.3 | 0.44 |
| 10 | 33411435 | rs2070302 | -0.474 | 0.022 | 0.246 | 0.07 | 0.007 | -0.011 | -0.011 | 0 | 0.26 | 0.27 | 0.32 | 0.62 | 0.21 | 0.4 |
| 10 | 33420095 | rs2750052 | -1.831 | -0.373 | 0.118 | -0.367 | 0.022 | 0.212 | 0.212 | 0 | 0.5 | 0.81 | 0.53 | 0.75 | 0.43 | 0.59 |
| 10 | 33421548 | rs11009235 | -1.012 | -0.469 | 0.535 | -0.482 | -0.002 | -0.001 | -0.001 | 0 | 0.36 | 0.64 | 0.36 | 0.63 | 0.3 | 0.42 |
| 10 | 33424448 | rs2755987 | -1.672 | -0.489 | 1.111 | -0.5 | 0.027 | -0.044 | -0.044 | 0.477 | 0.27 | 0.39 | 0.28 | 0.12 | 0.31 | 0.21 |
| 10 | 33426123 | rs722986 | 0.267 | -0.441 | 1.187 | -0.515 | 0.004 | -0.017 | -0.017 | 0 | 0.07 | 0.09 | 0.06 | 0.01 | 0.11 | 0.03 |
| 10 | 33426147 | rs722988 | -1.649 | -0.507 | 1.228 | -0.543 | -0.003 | -0.027 | -0.027 | 0.272 | 0.44 | 0.56 | 0.47 | 0.75 | 0.39 | 0.56 |
| 10 | 33427023 | rs881423 | -0.089 | -0.521 | 1.223 | -0.508 | -0.059 | 0.05 | 0.05 | 0 | 0.23 | 0.42 | 0.26 | 0.62 | 0.12 | 0.37 |
| 10 | 33427220 | rs11009243 | 0.195 | -0.499 | 1.108 | -0.492 | -0.017 | -0.021 | -0.021 | 0 | 0.13 | 0.04 | 0.15 | 0.11 | 0.14 | 0.16 |
| 10 | 33429181 | rs2750051 | -0.393 | -0.492 | 1.263 | -0.506 | 0.044 | -0.017 | -0.017 | 0 | 0.07 | 0.26 | 0.07 | 0.01 | 0.06 | 0.02 |
| 10 | 33429922 | rs2750050 | -1.742 | -0.497 | 1.16 | -0.533 | 0.019 | -0.04 | -0.04 | 1.515 | 0.26 | 0.37 | 0.26 | 0.12 | 0.28 | 0.2 |
| 10 | 33430409 | rs11009247 | 0.46 | -0.483 | 0.655 | -0.498 | -0.031 | -0.048 | -0.048 | 3.837 | 0.11 | 0.02 | 0.12 | 0.11 | 0.11 | 0.15 |
| 10 | 33434232 | rs11009251 | 0.018 | -0.523 | 0.836 | -0.476 | 0.014 | -0.014 | -0.014 | 0 | 0.22 | 0.24 | 0.26 | 0.63 | 0.13 | 0.37 |
| 10 | 33435249 | rs17230662 | -0.145 | -0.255 | 0.378 | -0.783 | -0.019 | 0.009 | 0.009 | 0 | 0.11 | 0.07 | 0.09 | 0 | 0.18 | 0.04 |
| 10 | 33437213 | rs7097848 | -0.145 | -0.255 | 0.382 | -0.783 | -0.014 | 0.006 | 0.006 | 0 | 0.11 | 0.08 | 0.09 | 0.01 | 0.18 | 0.04 |
| 10 | 33442404 | rs4551704 | -1.608 | -0.293 | -0.16 | -0.83 | 0.022 | 0.245 | 0.245 | 0 | 0.51 | 0.85 | 0.54 | 0.75 | 0.41 | 0.56 |
| 10 | 33442904 | rs6481836 | -0.968 | -0.898 | -0.132 | -0.812 | 0.092 | 0.113 | 0.113 | 0 | 0.4 | 0.77 | 0.44 | 0.75 | 0.23 | 0.53 |
| 10 | 33444312 | rs927096 | -0.069 | -0.968 | -0.209 | -0.781 | -0.007 | 0.007 | 0.007 | 0 | 0.1 | 0.07 | 0.08 | 0 | 0.14 | 0.03 |
| 10 | 33445364 | rs6481837 | -1.255 | -0.526 | -0.814 | -0.585 | 0.065 | 0.236 | 0.236 | 0 | 0.5 | 0.89 | 0.52 | 0.73 | 0.37 | 0.54 |
| 10 | 33446557 | rs7923059 | -1.236 | -0.671 | -0.789 | -0.618 | 0.084 | 0.215 | 0.215 | 0 | 0.51 | 0.83 | 0.51 | 0.74 | 0.38 | 0.55 |
| 10 | 33450242 | rs2506128 | -1.356 | -0.206 | -0.974 | -0.724 | -0.034 | 0.163 | 0.163 | 0 | 0.56 | 0.84 | 0.55 | 0.74 | 0.46 | 0.61 |
| 10 | 33450700 | rs11009265 | -1.164 | -0.177 | 0.009 | -0.485 | -0.001 | -0.035 | -0.035 | 0 | 0.3 | 0.6 | 0.28 | 0.12 | 0.32 | 0.24 |
| 10 | 33451379 | rs2474710 | -1.416 | -0.449 | 0.328 | -0.012 | -0.005 | 0.006 | 0.006 | 0.376 | 0.33 | 0.34 | 0.17 | 0.47 | 0.28 | 0.35 |
| 10 | 33454715 | rs2474741 | -1.302 | -0.49 | 0.614 | -0.032 | -0.014 | 0.019 | 0.019 | 0.193 | 0.32 | 0.39 | 0.18 | 0.47 | 0.26 | 0.36 |
| 10 | 33460744 | rs6481838 | -1.878 | -0.637 | 0.675 | -0.146 | 0.039 | 0.049 | 0.049 | 2.609 | 0.43 | 0.82 | 0.38 | 0.58 | 0.34 | 0.41 |
| 10 | 33462578 | rs7099537 | -1.473 | -0.497 | 1.105 | -0.059 | 0.035 | 0.102 | 0.102 | 3.428 | 0.1 | 0.41 | 0.19 | 0.11 | 0.07 | 0.06 |
| 10 | 33463601 | rs7912484 | -0.971 | -0.512 | 1.111 | -0.053 | 0.034 | 0.147 | 0.147 | 3.697 | 0.08 | 0.4 | 0.19 | 0.11 | 0.05 | 0.05 |
| 10 | 33464952 | rs2506134 | -1.995 | -0.022 | 0.823 | -0.453 | 0.014 | 0.088 | 0.088 | 0.497 | 0.48 | 0.7 | 0.5 | 0.73 | 0.43 | 0.5 |
| 10 | 33465037 | rs2506135 | -1.591 | -0.168 | 1.173 | -0.487 | -0.022 | -0.014 | -0.014 | 0 | 0.38 | 0.54 | 0.23 | 0.6 | 0.31 | 0.37 |
| 10 | 33466566 | rs1044268 | -0.361 | -0.153 | 1 | -0.47 | -0.005 | 0.016 | 0.016 | 0.017 | 0.18 | 0.19 | 0.11 | 0.49 | 0.12 | 0.27 |
| 10 | 33467108 | rs10080 | -2.551 | -0.133 | 0.959 | -0.493 | 0.013 | 0.085 | 0.085 | 0.444 | 0.47 | 0.7 | 0.5 | 0.73 | 0.43 | 0.5 |
| 10 | 33467312 | rs10827206 | -0.35 | -0.142 | 0.978 | -0.487 | -0.02 | 0.043 | 0.043 | 0.051 | 0.18 | 0.23 | 0.12 | 0.49 | 0.12 | 0.27 |
| 10 | 33467321 | rs1044210 | -1.831 | -0.141 | 0.997 | -0.489 | -0.02 | -0.012 | -0.012 | 0.228 | 0.36 | 0.57 | 0.23 | 0.6 | 0.31 | 0.36 |
| 10 | 33467539 | rs2506140 | -1.002 | -0.094 | 0.678 | -0.493 | 0.008 | -0.022 | -0.022 | 2.94 | 0.18 | 0.33 | 0.11 | 0.11 | 0.18 | 0.09 |
| 10 | 33468014 | rs2506141 | -2.642 | -0.053 | 0.775 | -0.527 | -0.021 | 0.213 | 0.213 | 0.997 | 0.51 | 0.88 | 0.53 | 0.73 | 0.49 | 0.54 |
| 10 | 33468169 | rs2506143 | -1.122 | 0.185 | 0.825 | -0.463 | 0.008 | -0.022 | -0.022 | 1.8 | 0.18 | 0.33 | 0.11 | 0.11 | 0.18 | 0.09 |
| 10 | 33468456 | rs2506144 | -1.122 | 0.185 | 0.825 | -0.463 | 0.008 | -0.022 | -0.022 | 0 | 0.18 | 0.33 | 0.11 | 0.11 | 0.18 | 0.09 |
| 10 | 33471130 | rs2506145 | -0.947 | -0.228 | 0.916 | -0.36 | -0.01 | 0.004 | 0.004 | 0.163 | 0.17 | 0.35 | 0.1 | 0.13 | 0.13 | 0.09 |
| 10 | 33473330 | rs2474730 | -0.797 | -0.244 | 0.898 | -0.355 | -0.02 | 0.008 | 0.008 | 0.235 | 0.16 | 0.33 | 0.09 | 0.13 | 0.12 | 0.09 |
| 10 | 33475282 | rs2228638 | 0.707 | -0.105 | 0.183 | -0.386 | -0.041 | -0.034 | -0.034 | 0 | 0.07 | 0.01 | 0.25 | 0.13 | 0.11 | 0.13 |
| 10 | 33484696 | rs734187 | -1.509 | -0.423 | 0.322 | -0.536 | 0.044 | 0.025 | 0.025 | 0 | 0.33 | 0.24 | 0.2 | 0.64 | 0.24 | 0.36 |
| 10 | 33484829 | rs734186 | -2.048 | -0.409 | 0.544 | -0.552 | 0.037 | 0.095 | 0.095 | 0 | 0.45 | 0.74 | 0.49 | 0.74 | 0.37 | 0.49 |
| 10 | 33485962 | rs2474723 | -1.564 | -0.383 | 0.317 | -0.43 | -0.022 | -0.014 | -0.014 | 0.045 | 0.32 | 0.41 | 0.21 | 0.64 | 0.26 | 0.39 |
| 10 | 33488814 | rs11009281 | -0.241 | -0.375 | -0.007 | -0.271 | 0.017 | -0.016 | -0.016 | 4.479 | 0.13 | 0.09 | 0.08 | 0.51 | 0.08 | 0.24 |
| 10 | 33489889 | rs1555321 | -0.865 | -0.282 | -0.049 | -0.361 | 0.008 | -0.047 | -0.047 | 0 | 0.25 | 0.1 | 0.35 | 0.6 | 0.23 | 0.39 |
| 10 | 33492093 | rs2247715 | -1.984 | -0.129 | 0.163 | -0.478 | 0.009 | 0.047 | 0.047 | 0 | 0.45 | 0.38 | 0.56 | 0.74 | 0.44 | 0.53 |
| 10 | 33492543 | rs16934171 | -0.84 | -0.134 | 0.125 | -0.395 | 0.002 | -0.049 | -0.049 | 2.409 | 0.25 | 0.09 | 0.43 | 0.6 | 0.25 | 0.38 |
| 10 | 33493070 | rs10827211 | -0.853 | -0.135 | 0.128 | -0.375 | 0.004 | -0.044 | -0.044 | 0 | 0.25 | 0.11 | 0.43 | 0.6 | 0.25 | 0.38 |
| 10 | 33495983 | rs2474714 | -2.001 | -0.132 | 0.149 | -0.443 | 0.015 | 0.075 | 0.075 | 0 | 0.44 | 0.32 | 0.56 | 0.74 | 0.43 | 0.52 |
| 10 | 33503179 | rs2506155 | -0.458 | -0.303 | 0.052 | -0.433 | 0.006 | 0.007 | 0.007 | 0 | 0.17 | 0.11 | 0.1 | 0.12 | 0.16 | 0.14 |
| 10 | 33504465 | rs2254826 | -2.02 | -0.034 | 0.108 | -0.442 | 0.005 | 0.049 | 0.049 | 0 | 0.44 | 0.36 | 0.56 | 0.74 | 0.45 | 0.53 |
| 10 | 33504784 | rs10827214 | -0.941 | 0.021 | 0.048 | -0.272 | 0.006 | -0.013 | -0.013 | 0 | 0.26 | 0.18 | 0.44 | 0.62 | 0.25 | 0.39 |

| | | | | | | | | | | | | | | | | |
|---|---|---|---|---|---|---|---|---|---|---|---|---|---|---|---|---|
| 10 | 33504883 | rs743168 | -1.104 | 0.021 | 0.048 | -0.272 | 0.002 | -0.008 | -0.008 | 0 | 0.27 | 0.23 | 0.44 | 0.62 | 0.26 | 0.4 |
| 10 | 33506928 | rs1360456 | -0.941 | 0.021 | 0.02 | -0.271 | 0.004 | -0.01 | -0.01 | 0 | 0.26 | 0.18 | 0.43 | 0.62 | 0.25 | 0.39 |
| 10 | 33508142 | rs2269096 | -1.083 | 0.011 | 0.012 | -0.268 | 0.001 | -0.004 | -0.004 | 0.097 | 0.27 | 0.23 | 0.44 | 0.62 | 0.26 | 0.4 |
| 10 | 33517647 | rs1331317 | -0.93 | -0.419 | -0.388 | -0.548 | -0.005 | -0.005 | -0.005 | 0 | 0.38 | 0.36 | 0.49 | 0.63 | 0.31 | 0.41 |
| 10 | 33520354 | rs11009311 | -0.419 | -0.581 | -0.435 | -0.448 | -0.031 | 0.033 | 0.033 | 0 | 0.25 | 0.27 | 0.19 | 0.52 | 0.15 | 0.25 |
| 10 | 33525836 | rs927099 | -2.242 | -0.596 | -0.583 | -0.724 | -0.049 | 0.158 | 0.158 | 0 | 0.59 | 0.88 | 0.65 | 0.76 | 0.5 | 0.55 |
| 10 | 33526391 | rs11009313 | 0.298 | -0.598 | -0.096 | -0.694 | 0.012 | -0.017 | -0.017 | 0 | 0.14 | 0.33 | 0.31 | 0.11 | 0.15 | 0.16 |
| 10 | 33526827 | rs12765284 | 0.083 | -0.534 | -0.37 | -0.575 | 0.016 | -0.023 | -0.023 | 0.705 | 0.14 | 0.08 | 0.15 | 0.49 | 0.09 | 0.22 |
| 10 | 33529028 | rs2269091 | -0.259 | -0.571 | -0.274 | -0.506 | -0.013 | -0.022 | -0.022 | 0.014 | 0.24 | 0.29 | 0.33 | 0.15 | 0.2 | 0.16 |
| 10 | 33530050 | rs2269088 | 0.879 | -0.278 | -0.708 | -0.42 | -0.042 | -0.025 | -0.025 | 0.023 | 0.11 | 0.01 | 0.27 | 0.13 | 0.15 | 0.15 |
| 10 | 33530297 | rs12762312 | -1.282 | -0.46 | -0.526 | -0.384 | -0.006 | -0.011 | -0.011 | 0.304 | 0.44 | 0.47 | 0.31 | 0.65 | 0.35 | 0.38 |
| 10 | 33532755 | rs17413155 | -0.264 | -0.038 | -0.819 | -0.098 | -0.012 | -0.001 | -0.001 | 0 | 0.16 | 0.07 | 0.09 | 0.14 | 0.18 | 0.14 |
| 10 | 33533148 | rs17413169 | -0.412 | 0.55 | -0.96 | -0.222 | -0.049 | 0.042 | 0.042 | 0 | 0.2 | 0.08 | 0.13 | 0.12 | 0.27 | 0.14 |
| 10 | 33534880 | rs10490939 | 0.722 | 0.783 | -1.023 | -0.083 | -0.032 | -0.015 | -0.015 | 0.322 | 0.11 | 0.02 | 0.28 | 0.11 | 0.15 | 0.16 |
| 10 | 33535907 | rs1888688 | -0.248 | 0.907 | -0.874 | -0.127 | 0.001 | 0.11 | 0.11 | 0 | 0.14 | 0.21 | 0.17 | 0.5 | 0.13 | 0.22 |
| 10 | 33538621 | rs11009323 | -0.574 | 0.601 | -0.93 | -0.362 | -0.012 | 0.012 | 0.012 | 0.186 | 0.29 | 0.34 | 0.25 | 0.53 | 0.21 | 0.25 |
| 10 | 33540448 | rs11598565 | -0.484 | 0.803 | -1.217 | -0.472 | -0.092 | 0.047 | 0.047 | 0 | 0.31 | 0.11 | 0.4 | 0.22 | 0.42 | 0.3 |
| 10 | 33543702 | rs4934583 | -2.19 | 0.559 | -0.928 | -0.573 | -0.017 | 0.084 | 0.084 | 0.611 | 0.63 | 0.86 | 0.69 | 0.74 | 0.64 | 0.54 |
| 10 | 33543929 | rs1319013 | -1.327 | 0.706 | -0.712 | -0.394 | -0.007 | 0.031 | 0.031 | 0 | 0.44 | 0.37 | 0.56 | 0.7 | 0.53 | 0.52 |
| 10 | 33545081 | rs11593943 | -0.499 | 1.004 | -0.702 | -0.316 | -0.044 | 0.091 | 0.091 | 0.017 | 0.3 | 0.18 | 0.23 | 0.13 | 0.39 | 0.27 |
| 10 | 33547465 | rs3780869 | 0.405 | 1.483 | -0.475 | -0.242 | 0.007 | 0.022 | 0.022 | 0.82 | 0.19 | 0.16 | 0.1 | 0.04 | 0.15 | 0.04 |
| 10 | 33549468 | rs16934292 | 1.179 | 1.419 | -0.456 | -0.209 | 0.006 | 0.007 | 0.007 | 2.177 | 0.15 | 0.16 | 0.08 | 0.04 | 0.11 | 0.03 |
| 10 | 33552695 | rs2070296 | -0.352 | 1.885 | 0.077 | 0.366 | 0.097 | 0.33 | 0.33 | 0 | 0.12 | 0.32 | 0.2 | 0.46 | 0.16 | 0.23 |
| 10 | 33553257 | rs2073320 | -1.753 | 2.126 | -0.587 | 0.267 | 0.039 | 0.029 | 0.029 | 0.021 | 0.56 | 0.79 | 0.51 | 0.64 | 0.62 | 0.52 |
| 10 | 33557046 | rs10740868 | -1.144 | 2.253 | -0.6 | 0.368 | -0.007 | 0.003 | 0.003 | 0 | 0.42 | 0.67 | 0.39 | 0.59 | 0.49 | 0.36 |
| 10 | 33557279 | rs10827224 | -0.39 | 2.362 | -0.622 | 0.912 | 0.012 | 0.091 | 0.091 | 0.015 | 0.1 | 0.09 | 0.17 | 0.46 | 0.16 | 0.23 |
| 10 | 33560552 | rs4934584 | -0.252 | 2.001 | -0.784 | 0.133 | -0.004 | 0.01 | 0.01 | 0 | 0.15 | 0.14 | 0.15 | 0.06 | 0.17 | 0.17 |
| 10 | 33561560 | rs17296436 | -0.448 | 2.234 | -1.082 | -0.026 | -0.035 | 0.054 | 0.054 | 0 | 0.18 | 0.06 | 0.18 | 0.06 | 0.24 | 0.18 |
| 10 | 33561574 | rs17296443 | -0.131 | 2.842 | -1.11 | -0.243 | -0.034 | 0.034 | 0.034 | 0 | 0.12 | 0.07 | 0.08 | 0.04 | 0.17 | 0.11 |
| 10 | 33561788 | rs10827227 | -0.726 | 2.154 | -1.502 | -0.242 | -0.024 | -0.047 | -0.047 | 0 | 0.44 | 0.85 | 0.36 | 0.51 | 0.43 | 0.41 |
| 10 | 33567133 | rs10827228 | 0.757 | 1.577 | -0.296 | -0.192 | -0.001 | 0 | 0 | 0 | 0.49 | 0.44 | 0.44 | 0.53 | 0.5 | 0.55 |
| 10 | 33568935 | rs1952983 | 1.035 | 1.327 | -0.1 | -0.216 | -0.056 | -0.055 | -0.055 | 0.474 | 0.46 | 0.12 | 0.4 | 0.53 | 0.5 | 0.55 |
| 10 | 33572039 | rs869636 | 0.317 | 1.258 | 0.019 | -0.073 | -0.002 | -0.013 | -0.013 | 0 | 0.58 | 0.47 | 0.57 | 0.82 | 0.62 | 0.79 |
| 10 | 33572836 | rs870087 | 0.857 | 1.276 | 0.03 | -0.259 | -0.013 | -0.011 | -0.011 | 0 | 0.61 | 0.5 | 0.55 | 0.68 | 0.65 | 0.65 |
| 10 | 33573630 | rs2804470 | -0.314 | 1.297 | -0.074 | -0.232 | -0.009 | -0.021 | -0.021 | 0.039 | 0.09 | 0.06 | 0.13 | 0.29 | 0.11 | 0.25 |
| 10 | 33573806 | rs7079372 | 0.087 | 1.162 | 0.137 | -0.342 | -0.018 | 0.01 | 0.01 | 0 | 0.72 | 0.65 | 0.64 | 0.68 | 0.78 | 0.71 |
| 10 | 33574405 | rs2776928 | -1.224 | 1.091 | 0.146 | -0.359 | -0.043 | -0.053 | -0.053 | 0 | 0.81 | 0.71 | 0.78 | 0.97 | 0.9 | 0.95 |
| 10 | 33579890 | rs1331326 | 0.82 | 1.207 | 0.019 | -0.271 | -0.007 | -0.008 | -0.008 | 0.173 | 0.62 | 0.53 | 0.55 | 0.68 | 0.65 | 0.65 |
| 10 | 33584131 | rs6481844 | 1.371 | 0.854 | -0.212 | -0.323 | -0.051 | -0.029 | -0.029 | 1.49 | 0.45 | 0.11 | 0.54 | 0.56 | 0.47 | 0.47 |
| 10 | 33586035 | rs7910405 | 0.67 | 0.702 | 0.288 | -0.299 | 0.009 | 0 | 0 | 0 | 0.16 | 0.36 | 0.14 | 0.11 | 0.15 | 0.1 |
| 10 | 33587881 | rs2776930 | -0.978 | 0.669 | 0.563 | -0.351 | -0.017 | -0.085 | -0.085 | 0 | 0.8 | 0.67 | 0.9 | 0.97 | 0.85 | 0.9 |
| 10 | 33592406 | rs2776932 | -0.133 | 0.697 | 0.543 | -0.308 | -0.038 | -0.061 | -0.061 | 1.187 | 0.1 | 0.01 | 0.13 | 0.3 | 0.14 | 0.28 |
| 10 | 33594002 | rs2065364 | 0.251 | 0.658 | 0.681 | -0.324 | -0.005 | -0.002 | -0.002 | 0.272 | 0.69 | 0.64 | 0.78 | 0.67 | 0.72 | 0.62 |
| 10 | 33602046 | rs2804492 | 1.031 | 0.55 | 0.186 | -0.328 | 0.019 | 0.012 | 0.012 | 1.748 | 0.52 | 0.29 | 0.62 | 0.55 | 0.58 | 0.52 |
| 10 | 33605336 | rs11009337 | -0.354 | 0.184 | 0.782 | 0.172 | -0.006 | -0.003 | -0.003 | 0.017 | 0.07 | 0.05 | 0.1 | 0.3 | 0.09 | 0.26 |
| 10 | 33605632 | rs2768420 | 1.651 | 0.123 | 0.666 | 0.057 | 0.018 | -0.039 | -0.039 | 0 | 0.61 | 0.1 | 0.7 | 0.6 | 0.71 | 0.56 |
| 10 | 33605676 | rs2804493 | -0.037 | 0.114 | 0.94 | 0.032 | -0.022 | -0.111 | -0.111 | 0 | 0.76 | 0.39 | 0.84 | 0.99 | 0.83 | 0.87 |
| 10 | 33606189 | rs2776937 | 1.392 | 0.1 | 0.765 | 0.068 | 0.043 | -0.039 | -0.039 | 0 | 0.63 | 0.13 | 0.74 | 0.6 | 0.75 | 0.59 |
| 10 | 33607828 | rs11492493 | -0.919 | 0.379 | 0.812 | -0.239 | 0.013 | 0.116 | 0.116 | 1.35 | 0.09 | 0.22 | 0.12 | 0.3 | 0.09 | 0.26 |
| 10 | 33614721 | rs4934597 | -0.704 | 0.475 | 0.657 | -0.215 | 0.007 | 0.064 | 0.064 | 0.065 | 0.08 | 0.15 | 0.1 | 0.3 | 0.09 | 0.25 |
| 10 | 33616085 | rs11009344 | -1.418 | 0.457 | 0.769 | -0.243 | 0.007 | 0.047 | 0.047 | 0 | 0.13 | 0.22 | 0.13 | 0.29 | 0.12 | 0.27 |
| 10 | 33616929 | rs1360457 | -0.936 | 0.423 | 0.945 | -0.481 | -0.006 | -0.05 | -0.05 | 0.014 | 0.83 | 0.8 | 0.93 | 0.99 | 0.88 | 0.91 |
| 10 | 33620707 | rs2804498 | 1.301 | 0.423 | 0.973 | -0.461 | 0.01 | 0.044 | 0.044 | 0.206 | 0.65 | 0.24 | 0.72 | 0.77 | 0.71 | 0.64 |
| 10 | 33622977 | rs1888684 | -0.838 | 0.464 | 0.795 | -0.489 | 0.003 | -0.023 | -0.023 | 0 | 0.82 | 0.82 | 0.92 | 0.99 | 0.88 | 0.92 |
| 10 | 33637243 | rs11812929 | 1.635 | 0.452 | 0.786 | -0.501 | -0.001 | -0.025 | -0.025 | 1.512 | 0.12 | 0.15 | 0.05 | 0 | 0.08 | 0.01 |
| 10 | 33638672 | rs2768399 | -0.865 | 0.544 | 0.873 | -0.451 | -0.006 | -0.057 | -0.057 | 0 | 0.81 | 0.75 | 0.91 | 0.99 | 0.86 | 0.91 |
| 10 | 33639488 | rs10827245 | -0.027 | 0.613 | 0.815 | -0.362 | -0.013 | -0.018 | -0.018 | 0 | 0.06 | 0.04 | 0.08 | 0.22 | 0.09 | 0.22 |
| 10 | 33642408 | rs2804891 | -0.121 | 0.447 | 0.617 | -0.492 | -0.03 | -0.132 | -0.132 | 0.815 | 0.72 | 0.41 | 0.86 | 0.99 | 0.83 | 0.89 |
| 10 | 33645685 | rs11009356 | 0.418 | 0.944 | -0.018 | -0.379 | -0.064 | 0.019 | 0.019 | 0 | 0.13 | 0.03 | 0.16 | 0.1 | 0.22 | 0.14 |
| 10 | 33647091 | rs2804894 | 0.914 | -0.048 | 0.608 | -0.183 | 0 | 0 | 0 | 0 | 0.64 | 0.32 | 0.59 | 0.64 | 0.74 | 0.62 |
| 10 | 33648832 | rs2254822 | 1.317 | -0.032 | 0.495 | -0.208 | 0.018 | 0.016 | 0.016 | 1.842 | 0.6 | 0.25 | 0.55 | 0.64 | 0.7 | 0.58 |
| 10 | 33649023 | rs2150686 | -0.166 | -0.059 | 0.509 | -0.834 | 0.03 | 0.077 | 0.077 | 0 | 0.07 | 0.16 | 0.09 | 0.22 | 0.09 | 0.22 |
| 10 | 33649860 | rs2804502 | 0.547 | -0.044 | 0.214 | -1.03 | -0.036 | -0.178 | -0.178 | 0.794 | 0.68 | 0.45 | 0.81 | 0.99 | 0.79 | 0.85 |
| 10 | 33655450 | rs11009360 | -1.059 | 0.35 | 0.167 | -0.382 | 0.034 | 0.077 | 0.077 | 0.482 | 0.13 | 0.34 | 0.18 | 0.23 | 0.13 | 0.28 |
| 10 | 33656035 | rs1320485 | 0.292 | -0.27 | -0.672 | -1.122 | -0.028 | -0.092 | -0.092 | 2.168 | 0.66 | 0.52 | 0.78 | 0.93 | 0.75 | 0.81 |
| 10 | 33657364 | rs10740870 | -0.409 | -0.192 | -1.935 | -1.564 | -0.063 | 0.251 | 0.251 | 1.684 | 0.58 | 0.91 | 0.62 | 0.9 | 0.49 | 0.54 |
| 10 | 33657994 | rs9633635 | 0.218 | -0.842 | -1.131 | -1.282 | 0.015 | 0.017 | 0.017 | 0.743 | 0.42 | 0.58 | 0.31 | 0.73 | 0.36 | 0.41 |
| 10 | 33667278 | rs10763930 | -0.089 | -0.772 | -0.181 | -1.009 | 0.02 | 0.193 | 0.193 | 2.317 | 0.07 | 0.39 | 0.21 | 0.19 | 0.03 | 0.06 |
| 10 | 33667617 | rs2804887 | 0.975 | -0.127 | -0.678 | -1.35 | -0.115 | 0.078 | 0.078 | 0.195 | 0.26 | 0.02 | 0.23 | 0.11 | 0.31 | 0.34 |

| | | | | | | | | | | | | | | | | |
|---|---|---|---|---|---|---|---|---|---|---|---|---|---|---|---|---|
| 10 | 33670320 | rs2776947 | 0.444 | -0.118 | -0.473 | -1.55 | 0 | -0.03 | -0.03 | 0.784 | 0.85 | 0.86 | 0.86 | 1 | 0.84 | 0.87 |
| 10 | 33672884 | rs10827252 | 0.112 | -0.105 | -0.396 | -1.471 | 0.004 | 0.032 | 0.032 | 0.297 | 0.54 | 0.54 | 0.43 | 0.71 | 0.5 | 0.48 |
| 10 | 33675053 | rs2804456 | 0.475 | -0.281 | -0.354 | -1.577 | -0.135 | 0.145 | 0.145 | 0.624 | 0.32 | 0.03 | 0.27 | 0.11 | 0.37 | 0.38 |
| 10 | 33677323 | rs10763931 | 0.613 | -0.674 | -0.266 | -1.713 | 0.008 | 0.01 | 0.01 | 0 | 0.3 | 0.25 | 0.41 | 0.89 | 0.26 | 0.47 |
| 10 | 33679777 | rs7097345 | 0.505 | -0.681 | -0.213 | -1.716 | -0.038 | -0.102 | -0.102 | 9.113 | 0.32 | 0.48 | 0.43 | 0.89 | 0.26 | 0.47 |
| 10 | 33682553 | rs7898847 | 0.658 | -0.553 | -0.288 | -1.577 | 0.029 | 0.002 | 0.002 | 1.154 | 0.06 | 0.3 | 0.04 | 0 | 0.03 | 0.01 |
| 10 | 33683090 | rs2804884 | -0.499 | -0.251 | -0.776 | -1.493 | -0.119 | 0.198 | 0.198 | 1.875 | 0.62 | 0.98 | 0.71 | 0.89 | 0.6 | 0.63 |
| 10 | 33687297 | rs3851080 | -0.465 | -0.238 | -0.579 | -1.493 | -0.015 | 0.096 | 0.096 | 0.338 | 0.6 | 0.79 | 0.69 | 0.89 | 0.6 | 0.61 |
| 10 | 33689743 | rs2785264 | -0.238 | 0.178 | -0.421 | -0.786 | -0.01 | 0.048 | 0.048 | 1.18 | 0.34 | 0.8 | 0.55 | 0.17 | 0.34 | 0.21 |
| 10 | 33690581 | rs7910887 | -0.715 | -0.555 | 0.332 | -0.131 | 0.032 | 0.034 | 0.034 | 0 | 0.51 | 0.61 | 0.34 | 0.73 | 0.38 | 0.5 |
| 10 | 33691106 | rs2785267 | -1.328 | -0.644 | 0.393 | -0.444 | 0.053 | 0.049 | 0.049 | 7.49 | 0.05 | 0.43 | 0.05 | 0 | 0 | 0 |
| 10 | 33699673 | rs2785273 | -0.655 | -0.371 | -0.534 | -0.244 | 0.066 | 0.184 | 0.184 | 0.21 | 0.51 | 0.74 | 0.64 | 0.88 | 0.41 | 0.56 |
| 10 | 33701628 | rs2785277 | -0.639 | -0.346 | -0.476 | -0.264 | 0.013 | 0.003 | 0.003 | 0 | 0.49 | 0.54 | 0.61 | 0.87 | 0.4 | 0.55 |
| 10 | 33705496 | rs11593792 | -0.62 | 0.388 | -0.933 | 1.424 | -0.009 | 0.054 | 0.054 | 0 | 0.15 | 0.17 | 0.16 | 0.36 | 0.08 | 0.12 |
| 10 | 33708150 | rs17230787 | -1.067 | 1.3 | -1.285 | 0.932 | -0.061 | -0.055 | -0.055 | 0 | 0.12 | 0.01 | 0.16 | 0.15 | 0.18 | 0.23 |
| 10 | 33710224 | rs2804473 | -0.449 | 0.528 | -1.639 | 0.244 | 0.016 | 0.071 | 0.071 | 0.449 | 0.41 | 0.73 | 0.31 | 0.21 | 0.3 | 0.33 |
| 10 | 33711828 | rs2804898 | -0.578 | 0.298 | -1.756 | 0.132 | 0.021 | 0.132 | 0.132 | 1.587 | 0.43 | 0.86 | 0.29 | 0.2 | 0.3 | 0.32 |
| 10 | 33717004 | rs11009376 | -0.335 | -0.204 | -0.569 | -0.152 | -0.057 | -0.046 | -0.046 | 0 | 0.31 | 0.52 | 0.18 | 0.15 | 0.15 | 0.12 |
| 10 | 33722748 | rs1125432 | -0.384 | -0.362 | 0.769 | -0.205 | -0.055 | -0.05 | -0.05 | 0 | 0.31 | 0.53 | 0.19 | 0.15 | 0.16 | 0.12 |
| 10 | 33725829 | rs2804479 | -1.114 | 0.107 | 0.963 | -0.656 | 0.028 | -0.041 | -0.041 | 1.252 | 0.09 | 0.27 | 0.16 | 0.01 | 0.11 | 0.1 |
| 10 | 33728704 | rs2785285 | -0.961 | -0.164 | 1.032 | -0.736 | -0.019 | -0.064 | -0.064 | 0.631 | 0.34 | 0.44 | 0.3 | 0.1 | 0.25 | 0.21 |
| 10 | 33730855 | rs2804904 | -1.783 | -0.609 | 0.37 | -1.058 | -0.069 | 0.195 | 0.195 | 0 | 0.55 | 0.88 | 0.43 | 0.17 | 0.48 | 0.41 |
| 10 | 33733364 | rs2804486 | 0.005 | -0.485 | 0.485 | -0.903 | -0.005 | -0.018 | -0.018 | 0 | 0.27 | 0.3 | 0.17 | 0.09 | 0.17 | 0.12 |
| 10 | 33733976 | rs10508782 | 0.202 | -0.411 | 0.275 | -0.842 | 0.028 | -0.023 | -0.023 | 1.974 | 0.12 | 0.19 | 0.12 | 0.08 | 0.15 | 0.16 |
| 10 | 33735664 | rs17230822 | 0.36 | -0.628 | -0.034 | -0.739 | 0.037 | 0.069 | 0.069 | 0 | 0.23 | 0.07 | 0.13 | 0.09 | 0.13 | 0.11 |
| 10 | 33746806 | rs2245294 | -0.969 | -0.269 | 0.693 | -0.841 | 0.028 | -0.051 | -0.051 | 0 | 0.3 | 0.58 | 0.33 | 0.13 | 0.36 | 0.34 |
| 10 | 33746909 | rs11009395 | -0.267 | -0.223 | 1.32 | -0.674 | 0.041 | 0.03 | 0.03 | 0 | 0.13 | 0.22 | 0.16 | 0.19 | 0.16 | 0.19 |
| 10 | 33748515 | rs945587 | 0.282 | -0.782 | 0.799 | -1.237 | 0.027 | 0.063 | 0.063 | 0 | 0.55 | 0.32 | 0.63 | 0.92 | 0.64 | 0.7 |
| 10 | 33748685 | rs945589 | 1.307 | -0.961 | 0.571 | -0.431 | -0.106 | -0.064 | -0.064 | 0 | 0.4 | 0.05 | 0.28 | 0.62 | 0.46 | 0.44 |
| 10 | 33758430 | rs7900617 | -0.888 | -1.248 | 0.681 | -0.536 | 0.012 | 0.012 | 0.012 | 0 | 0.69 | 0.74 | 0.64 | 0.81 | 0.78 | 0.76 |
| 10 | 33763923 | rs7070107 | -1.053 | -1.215 | 0.713 | -0.658 | -0.002 | -0.051 | -0.051 | 0.924 | 0.72 | 0.72 | 0.88 | 0.97 | 0.8 | 0.84 |
| 10 | 33763999 | rs10763943 | 0.537 | -1.227 | 0.773 | -0.669 | 0.004 | 0.03 | 0.03 | 0 | 0.49 | 0.43 | 0.67 | 0.78 | 0.57 | 0.61 |
| 10 | 33764406 | rs12247780 | -0.095 | -1.231 | 0.756 | -0.67 | -0.017 | -0.011 | -0.011 | 0 | 0.13 | 0.17 | 0.1 | 0.07 | 0.07 | 0.06 |
| 10 | 33764769 | rs1336546 | -0.253 | -1.204 | 0.742 | -0.664 | -0.001 | -0.006 | -0.006 | 4.421 | 0.62 | 0.63 | 0.78 | 0.85 | 0.64 | 0.67 |
| 10 | 33767547 | rs12569831 | 0.796 | -1.028 | 0.637 | -0.556 | 0.01 | 0 | 0 | 1.084 | 0.11 | 0.11 | 0.11 | 0.12 | 0.16 | 0.17 |
| 10 | 33770092 | rs10827266 | -0.081 | -1.213 | 0.618 | -0.601 | -0.018 | -0.062 | -0.062 | 0 | 0.58 | 0.47 | 0.78 | 0.91 | 0.73 | 0.78 |
| 10 | 33774663 | rs11009405 | 1.644 | -0.196 | -0.556 | 0.187 | -0.011 | -0.087 | -0.087 | 0.119 | 0.3 | 0.08 | 0.21 | 0.38 | 0.28 | 0.29 |
| 10 | 33774731 | rs2804860 | -1.382 | 0.016 | -0.499 | 0.439 | 0.012 | -0.029 | -0.029 | 0.365 | 0.64 | 0.75 | 0.72 | 0.57 | 0.65 | 0.61 |
| 10 | 33775261 | rs10508785 | -0.198 | -0.114 | 0.21 | 0.354 | 0.043 | 0.054 | 0.054 | 1.125 | 0.07 | 0.34 | 0.19 | 0.07 | 0.06 | 0.06 |
| 10 | 33780972 | rs7908846 | -1.021 | -0.218 | -0.461 | 0.139 | 0.008 | 0.021 | 0.021 | 0.136 | 0.7 | 0.75 | 0.84 | 0.92 | 0.74 | 0.74 |
| 10 | 33781045 | rs7923589 | 0.043 | -0.273 | -0.426 | 0.097 | 0.013 | 0.003 | 0.003 | 0.054 | 0.57 | 0.63 | 0.79 | 0.92 | 0.67 | 0.69 |
| 10 | 33781428 | rs4934972 | 0.043 | -0.273 | -0.426 | 0.097 | 0.013 | 0.003 | 0.003 | 0.054 | 0.57 | 0.63 | 0.79 | 0.92 | 0.67 | 0.69 |
| 10 | 33782231 | rs12263033 | -1.181 | -0.255 | -0.435 | 0.106 | 0.009 | 0.023 | 0.023 | 0.136 | 0.7 | 0.74 | 0.83 | 0.92 | 0.74 | 0.74 |
| 10 | 33782618 | rs16934579 | 0.215 | -0.227 | -0.422 | 0.013 | -0.017 | -0.034 | -0.034 | 0 | 0.12 | 0.11 | 0.04 | 0 | 0.05 | 0.04 |
| 10 | 33786642 | rs11009407 | 0.156 | -0.22 | -0.44 | 0.27 | 0.004 | 0.004 | 0.004 | 0 | 0.55 | 0.57 | 0.56 | 0.75 | 0.67 | 0.64 |
| 10 | 33788096 | rs1414992 | 0.856 | 0.38 | -1.72 | 0.368 | -0.053 | -0.02 | -0.02 | 0 | 0.23 | 0.08 | 0.2 | 0.29 | 0.36 | 0.16 |
| 10 | 33788915 | rs4623845 | -0.522 | 0.407 | -1.388 | 0.128 | 0.019 | -0.001 | -0.001 | 0 | 0.69 | 0.77 | 0.61 | 0.75 | 0.74 | 0.69 |
| 10 | 33794668 | rs1993186 | -0.317 | 1.215 | -0.936 | 0.016 | -0.014 | -0.009 | -0.009 | 0.602 | 0.4 | 0.33 | 0.65 | 0.52 | 0.51 | 0.31 |
| 10 | 33795480 | rs2785245 | 0.521 | 0.852 | -1.293 | -0.874 | 0.04 | 0.006 | 0.006 | 0 | 0.29 | 0.12 | 0.41 | 0.74 | 0.2 | 0.37 |
| 10 | 33795913 | rs2804864 | 0.686 | 0.724 | -1.31 | -0.852 | 0.061 | -0.027 | -0.027 | 1.9 | 0.27 | 0.09 | 0.39 | 0.7 | 0.17 | 0.36 |
| 10 | 33798381 | rs11592507 | 0.197 | 1.499 | -0.596 | -1.379 | 0.059 | -0.153 | -0.153 | 0 | 0.31 | 0.42 | 0.32 | 0.03 | 0.42 | 0.17 |
| 10 | 33798746 | rs2605506 | 0.702 | 1.418 | -0.552 | -1.573 | 0.035 | 0.014 | 0.014 | 0 | 0.33 | 0.22 | 0.43 | 0.69 | 0.25 | 0.39 |
| 10 | 33806159 | rs11009411 | 1.806 | 1.297 | -0.604 | -1.766 | 0.014 | 0.046 | 0.046 | 1.506 | 0.2 | 0.1 | 0.07 | 0 | 0.14 | 0.11 |
| 10 | 33806255 | rs1574514 | -1.1 | 1.345 | -0.54 | -1.777 | -0.004 | -0.006 | -0.006 | 0.002 | 0.89 | 0.9 | 0.85 | 0.72 | 0.84 | 0.68 |
| 10 | 33808216 | rs2601768 | 0.133 | 1.269 | -0.381 | -1.864 | 0.024 | 0.068 | 0.068 | 0 | 0.43 | 0.3 | 0.59 | 0.97 | 0.4 | 0.7 |
| 10 | 33814932 | rs11009415 | 0.422 | 1.86 | -0.678 | -2.04 | 0.01 | 0.104 | 0.104 | 0 | 0.39 | 0.22 | 0.21 | 0 | 0.34 | 0.22 |
| 10 | 33820356 | rs1855990 | -1.158 | 1.477 | -0.542 | -1.742 | 0.003 | -0.039 | -0.039 | 6.886 | 0.74 | 0.72 | 0.9 | 1 | 0.82 | 0.87 |
| 10 | 33821216 | rs2785252 | 1.534 | 1.223 | -0.665 | -1.535 | 0.068 | -0.132 | -0.132 | 2.343 | 0.23 | 0.02 | 0.36 | 0.61 | 0.13 | 0.21 |
| 10 | 33836041 | rs1869237 | 0.359 | 1.559 | -0.246 | -2.168 | 0.022 | 0.196 | 0.196 | 0 | 0.43 | 0.2 | 0.57 | 0.97 | 0.41 | 0.7 |
| 10 | 33837510 | rs1414984 | 0.254 | 1.569 | -0.177 | -2.189 | 0.04 | -0.116 | -0.116 | 0 | 0.31 | 0.45 | 0.32 | 0.03 | 0.42 | 0.17 |
| 10 | 33841791 | rs2785255 | 0.35 | 1.499 | -0.483 | -2.152 | 0.024 | 0.161 | 0.161 | 0 | 0.43 | 0.24 | 0.58 | 0.97 | 0.41 | 0.7 |
| 10 | 33843547 | rs2804875 | -0.542 | 1.682 | -0.081 | -1.735 | 0.039 | -0.144 | -0.144 | 0.392 | 0.74 | 0.6 | 0.76 | 0.98 | 0.68 | 0.85 |
| 10 | 33844273 | rs2249415 | -0.523 | 1.7 | -0.454 | -1.728 | 0.024 | -0.111 | -0.111 | 0.517 | 0.73 | 0.51 | 0.75 | 0.98 | 0.68 | 0.85 |
| 10 | 33847448 | rs2785260 | 0.767 | 1.365 | -0.828 | -1.527 | -0.028 | 0.089 | 0.089 | 0 | 0.41 | 0.17 | 0.56 | 0.72 | 0.44 | 0.44 |
| 10 | 33848021 | rs2601758 | 0.041 | 1.36 | -0.479 | -1.572 | -0.003 | -0.009 | -0.009 | 0 | 0.64 | 0.4 | 0.74 | 0.74 | 0.7 | 0.57 |
| 10 | 33849179 | rs11009437 | 1.011 | 1.585 | -1.131 | -1.505 | 0 | 0.109 | 0.109 | 1.576 | 0.2 | 0.05 | 0.08 | 0 | 0.14 | 0.12 |
| 10 | 33850159 | rs11009438 | -0.409 | 1.882 | -0.131 | -0.636 | 0.004 | 0.028 | 0.028 | 0 | 0.14 | 0.24 | 0.16 | 0.26 | 0.16 | 0.31 |
| 10 | 33850943 | rs11597016 | -0.445 | 1.874 | -0.603 | -0.771 | -0.001 | -0.025 | -0.025 | 1.027 | 0.23 | 0.23 | 0.18 | 0.02 | 0.27 | 0.14 |
| 10 | 33855128 | rs12257625 | -0.552 | 2.035 | -0.563 | -0.804 | 0 | -0.037 | -0.037 | 0 | 0.26 | 0.24 | 0.24 | 0.02 | 0.36 | 0.15 |

| | | | | | | | | | | | | | | | | |
|---|---|---|---|---|---|---|---|---|---|---|---|---|---|---|---|---|
| 10 | 33859760 | rs1901670 | 0.192 | 1.762 | -0.404 | 0.116 | -0.001 | 0.027 | 0.027 | 0 | 0.92 | 0.91 | 0.87 | 0.66 | 0.9 | 0.69 |
| 10 | 33863906 | rs6481857 | 0.795 | 1.198 | 0.227 | -0.442 | -0.018 | -0.009 | -0.009 | 0 | 0.66 | 0.74 | 0.65 | 0.57 | 0.61 | 0.52 |
| 10 | 33865825 | rs2084882 | 0.801 | 1.174 | 0.222 | -0.694 | -0.016 | 0.01 | 0.01 | 0 | 0.65 | 0.74 | 0.7 | 0.66 | 0.61 | 0.53 |
| 10 | 33867286 | rs11009449 | -0.455 | 1.177 | 0.218 | -0.702 | -0.015 | 0.021 | 0.021 | 0 | 0.26 | 0.17 | 0.17 | 0 | 0.29 | 0.15 |
| 10 | 33868055 | rs1336540 | 1.14 | 0.666 | -0.004 | -1.261 | -0.007 | 0.012 | 0.012 | 0.092 | 0.41 | 0.17 | 0.52 | 0.58 | 0.42 | 0.35 |
| 10 | 33869945 | rs7071708 | -0.718 | 0.714 | -0.075 | -1.224 | -0.001 | 0.024 | 0.024 | 0 | 0.92 | 0.91 | 0.87 | 0.66 | 0.9 | 0.68 |
| 10 | 33872211 | rs12776899 | 1.154 | 0.57 | -0.129 | -1.338 | -0.055 | -0.007 | -0.007 | 0.192 | 0.39 | 0.1 | 0.52 | 0.58 | 0.45 | 0.33 |
| 10 | 33874231 | rs12357869 | 0.293 | 0.607 | -0.092 | -1.516 | -0.015 | 0.01 | 0.01 | 0 | 0.23 | 0.18 | 0.15 | 0 | 0.27 | 0.13 |
| 10 | 33879069 | rs2383992 | 1.269 | 0.46 | 0.361 | -1.46 | -0.006 | 0.001 | 0.001 | 0 | 0.38 | 0.46 | 0.5 | 0.56 | 0.37 | 0.31 |
| 10 | 33884138 | rs16915736 | 0.912 | 0.295 | -0.049 | -1.507 | 0 | -0.002 | -0.002 | 0 | 0.54 | 0.45 | 0.55 | 0.56 | 0.49 | 0.44 |
| 10 | 33885828 | rs11009463 | 0.214 | 0.429 | 0.263 | -1.221 | -0.004 | 0.144 | 0.144 | 0 | 0.87 | 0.66 | 0.8 | 0.67 | 0.88 | 0.73 |
| 10 | 33885939 | rs7097479 | 1.369 | 0.444 | -0.038 | -1.233 | -0.002 | 0.084 | 0.084 | 2.935 | 0.77 | 0.55 | 0.7 | 0.56 | 0.77 | 0.58 |
| 10 | 33887875 | rs2167712 | -0.525 | 0.306 | -0.463 | -0.986 | -0.006 | 0.003 | 0.003 | 0.733 | 0.08 | 0.05 | 0.09 | 0.11 | 0.09 | 0.13 |
| 10 | 33891964 | rs1902403 | -0.63 | 0.302 | -0.232 | -0.676 | -0.019 | 0.106 | 0.106 | 0 | 0.94 | 0.87 | 0.97 | 0.68 | 0.96 | 0.83 |
| 10 | 33901538 | rs10827274 | 0.569 | 0.477 | -0.676 | -0.668 | 0.006 | 0.035 | 0.035 | 0.189 | 0.49 | 0.23 | 0.65 | 0.56 | 0.51 | 0.42 |
| 10 | 33907172 | rs11594539 | -0.207 | 0.224 | -0.361 | -0.684 | -0.01 | -0.056 | -0.056 | 0.041 | 0.17 | 0.22 | 0.06 | 0 | 0.11 | 0.12 |
| 10 | 33911780 | rs6481863 | -1.352 | 0.57 | -0.351 | -0.405 | -0.001 | -0.003 | -0.003 | 0 | 0.28 | 0.69 | 0.25 | 0.26 | 0.3 | 0.32 |
| 10 | 33913412 | rs7900449 | -0.777 | 0.564 | -0.168 | -0.469 | -0.015 | -0.027 | -0.027 | 2.315 | 0.16 | 0.1 | 0.14 | 0.25 | 0.25 | 0.26 |
| 10 | 33913854 | rs11009475 | 1.049 | 0.656 | -0.714 | -0.882 | -0.042 | -0.06 | -0.06 | 0 | 0.28 | 0.08 | 0.36 | 0.59 | 0.32 | 0.32 |
| 10 | 33916527 | rs17296817 | 0.6 | 0.138 | -0.674 | -1.024 | 0.015 | 0.038 | 0.038 | 1.166 | 0.15 | 0.05 | 0.09 | 0.09 | 0.12 | 0.14 |
| 10 | 33917458 | rs11009478 | 0.286 | 0.575 | -0.494 | -0.817 | -0.007 | -0.025 | -0.025 | 0.044 | 0.07 | 0.01 | 0.05 | 0.25 | 0.09 | 0.18 |
| 10 | 33920586 | rs7908016 | -0.199 | 0.574 | -0.271 | -1.031 | 0.001 | -0.02 | -0.02 | 0 | 0.09 | 0.13 | 0.09 | 0 | 0.15 | 0.08 |
| 10 | 33925036 | rs3862575 | -0.539 | 0.247 | 0.718 | -0.995 | 0.027 | -0.03 | -0.03 | 0 | 0.09 | 0.31 | 0.07 | 0.02 | 0.05 | 0.05 |
| 10 | 33940796 | rs1902409 | -1.66 | 0.282 | 0.734 | -0.779 | 0.029 | 0.043 | 0.043 | 0 | 0.15 | 0.39 | 0.13 | 0.17 | 0.14 | 0.14 |
| 10 | 33951730 | rs17231072 | -1.381 | 0.537 | 0.561 | -0.651 | 0.006 | 0.036 | 0.036 | 0 | 0.13 | 0.16 | 0.18 | 0.22 | 0.17 | 0.23 |
| 10 | 33953263 | rs12415658 | -0.932 | 0.538 | 0.557 | -0.651 | 0.015 | 0.002 | 0.002 | 0 | 0.05 | 0.1 | 0.12 | 0.06 | 0.09 | 0.14 |
| 10 | 33955680 | rs7919494 | -2.687 | 0.437 | 1.095 | -0.731 | 0.018 | 0.047 | 0.047 | 0 | 0.18 | 0.35 | 0.23 | 0.22 | 0.24 | 0.25 |
| 10 | 33956346 | rs2891376 | -1.529 | 0.458 | 0.994 | -0.726 | 0.013 | -0.006 | -0.006 | 0 | 0.08 | 0.14 | 0.14 | 0.06 | 0.13 | 0.15 |
| 10 | 33969931 | rs12762034 | -0.282 | 0.443 | 0.96 | -0.702 | 0.004 | -0.006 | -0.006 | 0 | 0.07 | 0.04 | 0.06 | 0.14 | 0.08 | 0.09 |
| 10 | 33970134 | rs7070395 | -2.226 | 0.44 | 1.025 | -0.71 | 0.011 | 0.034 | 0.034 | 0 | 0.16 | 0.28 | 0.22 | 0.22 | 0.21 | 0.24 |
| 10 | 33973093 | rs12357304 | 0.626 | 0.415 | 0.344 | -1.259 | -0.036 | 0.083 | 0.083 | 1.079 | 0.14 | 0 | 0.03 | 0 | 0.1 | 0.03 |
| 10 | 33990654 | rs12777908 | -2.227 | 0.423 | 0.696 | -1 | 0.011 | 0.052 | 0.052 | 0 | 0.18 | 0.35 | 0.23 | 0.32 | 0.24 | 0.24 |
| 10 | 33993436 | rs3923643 | -2.173 | 0.428 | 0.693 | -0.997 | 0.013 | 0.056 | 0.056 | 0 | 0.18 | 0.36 | 0.23 | 0.33 | 0.24 | 0.24 |
| 10 | 33998632 | rs4129050 | -1.305 | 0.423 | 0.672 | -1.006 | -0.002 | 0.035 | 0.035 | 0 | 0.13 | 0.22 | 0.11 | 0.25 | 0.16 | 0.11 |
| 10 | 34003178 | rs11009501 | 0.01 | -0.2 | 0.571 | -1.587 | 0.011 | 0.064 | 0.064 | 1.311 | 0.31 | 0.16 | 0.26 | 0.07 | 0.22 | 0.16 |
| 10 | 34010790 | rs4306251 | -0.813 | 0.002 | 0.628 | -1.11 | -0.006 | 0.061 | 0.061 | 0 | 0.1 | 0.22 | 0.1 | 0.24 | 0.11 | 0.1 |
| 10 | 34014264 | rs7096272 | -1.611 | 0.019 | 0.368 | -1.124 | 0.006 | 0.023 | 0.023 | 0 | 0.18 | 0.29 | 0.23 | 0.32 | 0.24 | 0.23 |
| 10 | 34023497 | rs11009506 | -1.023 | -0.84 | -0.007 | -1.765 | -0.02 | -0.087 | -0.087 | 0 | 0.36 | 0.49 | 0.37 | 0.18 | 0.31 | 0.22 |
| 10 | 34028228 | rs11009508 | 0.563 | -0.796 | -0.58 | -1.557 | 0.013 | 0.004 | 0.004 | 0 | 0.09 | 0.12 | 0.15 | 0.07 | 0.11 | 0.21 |
| 10 | 34028762 | rs7073701 | -0.91 | -1.073 | -0.644 | -1.634 | 0.036 | 0.05 | 0.05 | 0 | 0.29 | 0.21 | 0.29 | 0.18 | 0.21 | 0.21 |
| 10 | 34033366 | rs12414563 | 0.076 | -1.098 | -0.671 | -1.673 | 0.028 | 0.037 | 0.037 | 0 | 0.15 | 0.1 | 0.21 | 0.03 | 0.1 | 0.1 |
| 10 | 34035796 | rs2812668 | -1.414 | -1.161 | -0.547 | -1.581 | -0.003 | -0.018 | -0.018 | 0.107 | 0.21 | 0.3 | 0.15 | 0.16 | 0.2 | 0.14 |
| 10 | 34036098 | rs2812669 | -0.762 | -1.17 | -0.62 | -1.585 | -0.003 | -0.002 | -0.002 | 0 | 0.14 | 0.21 | 0.08 | 0.16 | 0.11 | 0.11 |
| 10 | 34037557 | rs2812672 | -1.256 | -1.158 | -0.883 | -1.585 | -0.01 | 0.026 | 0.026 | 0 | 0.19 | 0.09 | 0.12 | 0.16 | 0.2 | 0.14 |
| 10 | 34039348 | rs1537207 | -0.208 | -1.586 | -1.125 | -1.782 | -0.003 | -0.023 | -0.023 | 0 | 0.6 | 0.64 | 0.56 | 0.9 | 0.67 | 0.5 |
| 10 | 34047975 | rs11009512 | 1.317 | -1.553 | -1.751 | -1.732 | -0.108 | 0.055 | 0.055 | 0 | 0.35 | 0.08 | 0.47 | 0.75 | 0.48 | 0.33 |
| 10 | 34063173 | rs2812612 | 0.166 | -1.555 | -1.556 | -1.709 | 0.008 | -0.012 | -0.012 | 0 | 0.11 | 0.02 | 0.05 | 0.14 | 0.1 | 0.1 |
| 10 | 34064066 | rs10827283 | 0.414 | -1.922 | -1.393 | -1.957 | 0.004 | 0.002 | 0.002 | 3.265 | 0.72 | 0.26 | 0.82 | 0.86 | 0.81 | 0.74 |
| 10 | 34066789 | rs2812613 | -0.989 | -1.783 | -1.227 | -1.872 | -0.053 | -0.106 | -0.106 | 0.015 | 0.84 | 0.37 | 0.89 | 1 | 0.91 | 0.83 |
| 10 | 34074562 | rs7911429 | 0.952 | -1.647 | -1.481 | -1.844 | 0.011 | 0.029 | 0.029 | 0 | 0.15 | 0.07 | 0.22 | 0.03 | 0.12 | 0.12 |
| 10 | 34078973 | rs17231134 | 1.073 | -1.582 | -1.509 | -1.824 | 0.011 | 0.038 | 0.038 | 0 | 0.14 | 0.04 | 0.21 | 0.03 | 0.12 | 0.11 |
| 10 | 34079819 | rs2777515 | 0.499 | -1.932 | -1.487 | -1.811 | 0 | -0.001 | -0.001 | 0.76 | 0.08 | 0.06 | 0.05 | 0.14 | 0.07 | 0.09 |
| 10 | 34083447 | rs7896638 | 0.228 | -2.073 | -1.6 | -1.265 | 0.019 | -0.036 | -0.036 | 0 | 0.65 | 0.22 | 0.68 | 0.53 | 0.72 | 0.47 |
| 10 | 34083871 | rs7900289 | 1.22 | -2.112 | -1.536 | -1.149 | 0.02 | 0.034 | 0.034 | 0 | 0.14 | 0.04 | 0.2 | 0.03 | 0.11 | 0.11 |
| 10 | 34091207 | rs2777525 | 0.374 | -1.791 | -1.366 | -0.948 | -0.002 | 0.001 | 0.001 | 0 | 0.08 | 0.08 | 0.05 | 0.15 | 0.07 | 0.09 |
| 10 | 34097009 | rs2777531 | -0.444 | -1.045 | -1.362 | -0.95 | 0.005 | 0.009 | 0.009 | 1.401 | 0.7 | 0.34 | 0.7 | 0.67 | 0.69 | 0.53 |
| 10 | 34098326 | rs10827290 | 1.524 | -0.878 | -1.789 | -0.781 | -0.121 | -0.172 | -0.172 | 0.99 | 0.27 | 0.03 | 0.33 | 0.49 | 0.38 | 0.27 |
| 10 | 34105728 | rs1331690 | -0.027 | -0.894 | -0.864 | -0.732 | 0.002 | -0.006 | -0.006 | 0 | 0.57 | 0.54 | 0.49 | 0.69 | 0.55 | 0.44 |
| 10 | 34109601 | rs2777534 | 0.318 | -0.972 | -0.884 | -0.674 | 0.011 | 0.01 | 0.01 | 5.776 | 0.16 | 0.11 | 0.12 | 0.15 | 0.13 | 0.11 |
| 10 | 34112672 | rs2812640 | 0.264 | -1.009 | -0.628 | -0.673 | -0.005 | 0.018 | 0.018 | 0 | 0.05 | 0.09 | 0.06 | 0.15 | 0.06 | 0.08 |
| 10 | 34113989 | rs2812645 | 0.234 | -1.335 | -0.747 | -0.855 | 0 | 0.009 | 0.009 | 0.119 | 0.51 | 0.48 | 0.46 | 0.68 | 0.51 | 0.38 |
| 10 | 34116863 | rs2812656 | -0.314 | -1.308 | -0.658 | -0.62 | -0.034 | 0.017 | 0.017 | 4.521 | 0.19 | 0.44 | 0.14 | 0.24 | 0.13 | 0.14 |
| 10 | 34119814 | rs2812661 | -0.459 | -1.277 | -0.323 | -0.627 | -0.007 | 0.016 | 0.016 | 1.19 | 0.55 | 0.64 | 0.47 | 0.68 | 0.52 | 0.38 |
| 10 | 34120563 | rs2777493 | -0.345 | -1.555 | -0.228 | -0.808 | -0.077 | -0.052 | -0.052 | 0.026 | 0.24 | 0.44 | 0.26 | 0.19 | 0.16 | 0.21 |
| 10 | 34127825 | rs1331683 | -0.939 | -1.247 | -0.154 | -0.747 | -0.01 | -0.006 | -0.006 | 0.011 | 0.69 | 0.74 | 0.68 | 0.7 | 0.64 | 0.51 |
| 10 | 34128582 | rs7068747 | 1.219 | -1.3 | -0.368 | -0.814 | -0.001 | -0.001 | -0.001 | 0 | 0.41 | 0.43 | 0.39 | 0.58 | 0.43 | 0.33 |
| 10 | 34135567 | rs7901673 | -0.749 | -0.961 | -0.213 | -0.674 | 0.008 | 0.003 | 0.003 | 0.456 | 0.68 | 0.66 | 0.67 | 0.7 | 0.64 | 0.51 |
| 10 | 34147295 | rs10508791 | 1.237 | -0.785 | -0.838 | -0.727 | 0.023 | 0.038 | 0.038 | 0 | 0.18 | 0.1 | 0.21 | 0.03 | 0.14 | 0.13 |
| 10 | 34148613 | rs2001177 | 0.824 | 0.015 | -0.096 | -0.011 | -0.005 | 0.009 | 0.009 | 0 | 0.55 | 0.43 | 0.58 | 0.71 | 0.55 | 0.48 |

| | | | | | | | | | | | | | | | | |
|---|---|---|---|---|---|---|---|---|---|---|---|---|---|---|---|---|
| 10 | 34148669 | rs867214 | 0.98 | 0.026 | -0.177 | -0.127 | -0.017 | 0.016 | 0.016 | 0 | 0.53 | 0.28 | 0.56 | 0.62 | 0.54 | 0.45 |
| 10 | 34151392 | rs1757469 | 0.201 | 0.294 | -0.626 | 0.088 | -0.046 | -0.058 | -0.058 | 0.162 | 0.7 | 0.84 | 0.68 | 0.62 | 0.63 | 0.5 |
| 10 | 34151735 | rs1757470 | -0.306 | -0.097 | -0.14 | 0.049 | 0 | -0.013 | -0.013 | 0.751 | 0.81 | 0.85 | 0.85 | 0.77 | 0.79 | 0.78 |
| 10 | 34151815 | rs1740727 | -0.958 | 0.098 | 0.735 | 0.01 | -0.012 | 0.037 | 0.037 | 7.518 | 0.14 | 0.45 | 0.09 | 0.16 | 0.07 | 0.11 |
| 10 | 34160241 | rs16934885 | -0.795 | 0.751 | -0.267 | 0.12 | 0.005 | -0.001 | -0.001 | 0 | 0.06 | 0.1 | 0.17 | 0.06 | 0.1 | 0.17 |
| 10 | 34161499 | rs1780434 | -0.822 | 0.929 | -0.708 | 0.073 | 0.004 | 0.029 | 0.029 | 1.152 | 0.54 | 0.72 | 0.72 | 0.58 | 0.55 | 0.55 |
| 10 | 34162083 | rs2800812 | -0.379 | 1.018 | -0.735 | 0.134 | 0.007 | -0.014 | -0.014 | 0 | 0.44 | 0.69 | 0.45 | 0.48 | 0.4 | 0.35 |
| 10 | 34165004 | rs10827299 | 0.548 | 1.42 | -1.154 | 0.363 | -0.029 | -0.11 | -0.11 | 0.289 | 0.14 | 0.01 | 0.12 | 0.41 | 0.15 | 0.19 |
| 10 | 34168090 | rs1757471 | -0.794 | 1.358 | -0.803 | -0.247 | 0 | 0.005 | 0.005 | 2.162 | 0.51 | 0.52 | 0.49 | 0.84 | 0.52 | 0.48 |
| 10 | 34173438 | rs7905447 | -0.047 | 1.385 | -1.206 | 0.046 | -0.018 | -0.041 | -0.041 | 0.004 | 0.26 | 0.06 | 0.18 | 0.74 | 0.26 | 0.35 |
| 10 | 34178427 | rs876644 | -0.795 | 1.512 | -0.769 | 0.222 | -0.008 | 0.029 | 0.029 | 0.085 | 0.2 | 0.1 | 0.3 | 0.13 | 0.23 | 0.23 |
| 10 | 34184063 | rs1757476 | -1.125 | 1.438 | -0.526 | 0.076 | -0.012 | -0.044 | -0.044 | 0 | 0.66 | 0.59 | 0.75 | 0.91 | 0.7 | 0.69 |
| 10 | 34186405 | rs1740720 | -1.046 | 1.515 | -0.359 | 0.07 | -0.011 | -0.051 | -0.051 | 0 | 0.65 | 0.47 | 0.72 | 0.91 | 0.68 | 0.69 |
| 10 | 34189978 | rs1740721 | -0.84 | 0.988 | -0.308 | 0.442 | -0.002 | 0.001 | 0.001 | 0.567 | 0.53 | 0.6 | 0.68 | 0.55 | 0.5 | 0.56 |
| 10 | 34192113 | rs12254866 | 0.023 | 1.174 | -0.892 | 0.488 | -0.011 | -0.036 | -0.036 | 0 | 0.34 | 0.2 | 0.24 | 0.4 | 0.29 | 0.37 |
| 10 | 34194532 | rs4934622 | -1.236 | 1.256 | -0.163 | 0.428 | 0.004 | -0.024 | -0.024 | 0 | 0.1 | 0.2 | 0.24 | 0.07 | 0.11 | 0.14 |
| 10 | 34201851 | rs2057227 | 0.206 | 1.781 | -0.603 | 0.612 | -0.011 | 0 | 0 | 0 | 0.7 | 0.74 | 0.75 | 0.65 | 0.61 | 0.84 |
| 10 | 34201924 | rs1615127 | -0.697 | 1.4 | -0.1 | 1.362 | -0.005 | 0.108 | 0.108 | 0.002 | 0.18 | 0.3 | 0.15 | 0.49 | 0.13 | 0.23 |
| 10 | 34202500 | rs10763961 | -0.667 | 2.18 | -0.37 | 1.1 | -0.022 | -0.022 | -0.022 | 0 | 0.16 | 0.09 | 0.33 | 0.45 | 0.26 | 0.25 |
| 10 | 34203029 | rs1740723 | -0.749 | 1.532 | -0.176 | -0.172 | 0.002 | -0.057 | -0.057 | 0.756 | 0.1 | 0.2 | 0.05 | 0 | 0.09 | 0.03 |
| 10 | 34203669 | rs12252306 | 0.696 | 0.781 | -0.387 | -1.012 | 0.018 | 0.05 | 0.05 | 0 | 0.39 | 0.3 | 0.21 | 0.06 | 0.28 | 0.3 |
| 10 | 34206497 | rs17505533 | 0.628 | 0.627 | -0.284 | -1.259 | 0.013 | 0.13 | 0.13 | 0 | 0.41 | 0.21 | 0.22 | 0.05 | 0.33 | 0.33 |
| 10 | 34207049 | rs927043 | 0.432 | -0.021 | -0.093 | -1.2 | -0.01 | -0.002 | -0.002 | 0 | 0.21 | 0.16 | 0.11 | 0.09 | 0.11 | 0.1 |
| 10 | 34207459 | rs743106 | -1.089 | 0.062 | -0.144 | -1.309 | -0.008 | 0.011 | 0.011 | 0 | 0.95 | 0.96 | 0.85 | 1 | 0.88 | 0.96 |
| 10 | 34209459 | rs743107 | -0.088 | 0.001 | -0.156 | -1.17 | 0.011 | 0.007 | 0.007 | 1.601 | 0.25 | 0.18 | 0.14 | 0.14 | 0.18 | 0.12 |
| 10 | 34210538 | rs1160769 | 0.622 | -0.037 | -0.561 | -1.163 | 0.066 | -0.037 | -0.037 | 0 | 0.2 | 0.01 | 0.09 | 0.14 | 0.11 | 0.1 |
| 10 | 34216438 | rs10827315 | 0.237 | 0.349 | -0.546 | -1.094 | -0.083 | -0.049 | -0.049 | 2.694 | 0.18 | 0.07 | 0.38 | 0.75 | 0.35 | 0.41 |
| 10 | 34216806 | rs10734012 | -0.802 | 0.01 | 0.126 | -1.272 | 0.002 | -0.079 | -0.079 | 0.438 | 0.44 | 0.43 | 0.53 | 0.9 | 0.49 | 0.51 |
| 10 | 34217265 | rs1002386 | -0.68 | 0.018 | 0.204 | -1.271 | 0.004 | -0.075 | -0.075 | 0 | 0.42 | 0.38 | 0.52 | 0.9 | 0.49 | 0.47 |
| 10 | 34217624 | rs10763963 | -0.048 | -0.092 | -0.162 | -1.201 | -0.05 | -0.03 | -0.03 | 0 | 0.3 | 0.39 | 0.23 | 0.17 | 0.25 | 0.17 |
| 10 | 34218093 | rs11815649 | 1.297 | -0.177 | -0.736 | -1.581 | -0.003 | 0.121 | 0.121 | 0 | 0.43 | 0.25 | 0.22 | 0.05 | 0.36 | 0.42 |
| 10 | 34219485 | rs7091026 | 0.861 | -0.523 | -0.058 | -1.108 | -0.025 | -0.039 | -0.039 | 0.963 | 0.84 | 0.33 | 0.72 | 0.93 | 0.85 | 0.89 |
| 10 | 34226309 | rs224731 | -0.324 | -0.248 | -0.803 | -0.999 | -0.002 | 0.032 | 0.032 | 0.012 | 0.68 | 0.96 | 0.61 | 0.77 | 0.67 | 0.79 |
| 10 | 34226551 | rs7091513 | 1.072 | -0.682 | -0.929 | -1.167 | 0.01 | 0.074 | 0.074 | 0 | 0.19 | 0.09 | 0.1 | 0.03 | 0.13 | 0.19 |
| 10 | 34230325 | rs224747 | -1.018 | -0.316 | -1.417 | -0.525 | 0.052 | 0.065 | 0.065 | 0 | 0.45 | 0.75 | 0.64 | 0.3 | 0.4 | 0.26 |
| 10 | 34233002 | rs11009597 | -0.343 | 0.132 | -1.63 | -0.187 | 0.002 | 0.002 | 0.002 | 0.613 | 0.29 | 0.04 | 0.48 | 0.26 | 0.29 | 0.2 |
| 10 | 34247385 | rs546254 | -0.228 | 0.652 | -0.718 | -0.257 | -0.002 | 0 | 0 | 0.674 | 0.37 | 0.36 | 0.33 | 0.26 | 0.45 | 0.3 |
| 10 | 34253008 | rs1617306 | -0.27 | 0.125 | -0.581 | -0.296 | 0.006 | 0.047 | 0.047 | 1.684 | 0.56 | 0.31 | 0.45 | 0.36 | 0.61 | 0.44 |
| 10 | 34256352 | rs11009606 | 0.4 | 0.395 | -0.343 | 0.423 | -0.001 | 0.025 | 0.025 | 0 | 0.05 | 0.07 | 0.17 | 0.17 | 0.03 | 0.18 |
| 10 | 34256803 | rs1780437 | 0.43 | 0.51 | -0.415 | 0.181 | -0.011 | -0.001 | -0.001 | 0 | 0.28 | 0.2 | 0.3 | 0.26 | 0.38 | 0.31 |
| 10 | 34258488 | rs11009607 | -0.214 | 0.301 | -0.686 | 0.583 | -0.011 | -0.111 | -0.111 | 0.759 | 0.27 | 0.08 | 0.28 | 0.5 | 0.21 | 0.4 |
| 10 | 34259148 | rs11009608 | 0.312 | 0.247 | -0.742 | 0.462 | -0.032 | -0.094 | -0.094 | 0 | 0.22 | 0.02 | 0.11 | 0.33 | 0.19 | 0.22 |
| 10 | 34259248 | rs11009609 | 0.316 | 0.278 | -0.678 | 0.476 | -0.032 | -0.094 | -0.094 | 0.242 | 0.23 | 0.02 | 0.11 | 0.33 | 0.19 | 0.22 |
| 10 | 34263354 | rs2136785 | -0.802 | 0.237 | 0.561 | 0.51 | 0.032 | 0.145 | 0.145 | 0.002 | 0.07 | 0.33 | 0.19 | 0.14 | 0.03 | 0.16 |
| 10 | 34266223 | rs224714 | -0.217 | 0.195 | 0.916 | 0.28 | -0.009 | -0.037 | -0.037 | 0 | 0.27 | 0.57 | 0.24 | 0.19 | 0.28 | 0.27 |
| 10 | 34275113 | rs10827318 | 0.042 | 0.232 | 0.628 | 0.297 | 0 | -0.001 | -0.001 | 1.201 | 0.25 | 0.25 | 0.21 | 0.19 | 0.27 | 0.27 |
| 10 | 34279023 | rs224762 | -0.29 | 0.331 | 0.632 | 0.264 | -0.011 | -0.004 | -0.004 | 0 | 0.32 | 0.56 | 0.4 | 0.33 | 0.3 | 0.43 |
| 10 | 34279238 | rs224763 | -0.218 | 0.508 | -0.169 | 0.341 | 0.001 | -0.001 | -0.001 | 0.776 | 0.3 | 0.27 | 0.38 | 0.33 | 0.3 | 0.43 |
| 10 | 34289996 | rs1740713 | -0.089 | 0.736 | 0.145 | 0.256 | 0.013 | 0.006 | 0.006 | 0 | 0.32 | 0.34 | 0.33 | 0.36 | 0.4 | 0.31 |
| 10 | 34302891 | rs127822 | 1.102 | -0.055 | 1.216 | -0.415 | -0.059 | -0.103 | -0.103 | 0 | 0.85 | 0.43 | 0.88 | 0.99 | 0.93 | 0.96 |
| 10 | 34315821 | rs323003 | 1.123 | 0.012 | 2.343 | -0.398 | 0.027 | -0.013 | -0.013 | 0.095 | 0.92 | 0.51 | 0.9 | 1 | 0.95 | 1 |
| 10 | 34315931 | rs2665896 | -0.459 | 0.377 | 1.734 | -0.833 | -0.009 | 0.032 | 0.032 | 0.085 | 0.18 | 0.09 | 0.12 | 0.12 | 0.19 | 0.12 |
| 10 | 34316633 | rs322999 | 0.51 | 0.11 | 2.14 | -0.818 | 0.033 | 0.007 | 0.007 | 1.25 | 0.95 | 0.68 | 0.92 | 1 | 0.95 | 1 |
| 10 | 34316972 | rs1705007 | -1.059 | 0.436 | 1.725 | -0.799 | -0.007 | 0.037 | 0.037 | 0 | 0.25 | 0.16 | 0.18 | 0.17 | 0.29 | 0.18 |
| 10 | 34317035 | rs322998 | 1.61 | 0.431 | 1.41 | -0.795 | 0.013 | 0.053 | 0.053 | 0.228 | 0.64 | 0.25 | 0.7 | 0.83 | 0.66 | 0.81 |
| 10 | 34319506 | rs1705015 | -1.262 | 0.883 | 1.178 | -0.641 | -0.01 | 0.012 | 0.012 | 0 | 0.27 | 0.18 | 0.21 | 0.17 | 0.39 | 0.2 |
| 10 | 34323378 | rs1705013 | -1.323 | 0.848 | 1.653 | -0.184 | 0.041 | -0.006 | -0.006 | 0 | 0.26 | 0.41 | 0.29 | 0.27 | 0.35 | 0.33 |
| 10 | 34334898 | rs10508795 | -1.649 | 0.29 | 2.165 | -0.208 | -0.029 | 0.035 | 0.035 | 2.28 | 0.09 | 0.53 | 0.14 | 0.05 | 0.01 | 0.13 |
| 10 | 34342143 | rs7090872 | -0.779 | -0.269 | 1.61 | -0.477 | -0.068 | -0.017 | -0.017 | 0 | 0.21 | 0.52 | 0.25 | 0.18 | 0.07 | 0.14 |
| 10 | 34369037 | rs10827324 | -0.515 | 0.741 | 1.405 | 0.124 | 0.014 | 0.002 | 0.002 | 1.374 | 0.2 | 0.16 | 0.13 | 0.24 | 0.32 | 0.16 |
| 10 | 34371333 | rs7893965 | -1.875 | 0.306 | 1.823 | 0.115 | -0.006 | 0.174 | 0.174 | 0.578 | 0.06 | 0.3 | 0.22 | 0.18 | 0.01 | 0.16 |
| 10 | 34380866 | rs7074105 | -0.677 | 0.227 | 1.309 | -0.217 | -0.05 | 0.005 | 0.005 | 0.033 | 0.23 | 0.07 | 0.23 | 0.21 | 0.33 | 0.19 |
| 10 | 34383172 | rs1078386 | 1.196 | 0.309 | 0.349 | -0.883 | -0.048 | -0.013 | -0.013 | 0 | 0.49 | 0.05 | 0.46 | 0.59 | 0.46 | 0.51 |
| 10 | 34393732 | rs7076988 | -1.136 | 0.395 | 1.292 | -0.748 | 0.01 | 0.019 | 0.019 | 0.206 | 0.3 | 0.58 | 0.37 | 0.39 | 0.34 | 0.35 |
| 10 | 34397063 | rs7082440 | -2.276 | 0.344 | 1.68 | -0.828 | 0.078 | 0.166 | 0.166 | 3.783 | 0.05 | 0.48 | 0.05 | 0.11 | 0.01 | 0.02 |
| 10 | 34398520 | rs12415499 | -2.101 | 0.347 | 1.582 | -0.827 | 0.044 | 0.251 | 0.251 | 4.67 | 0.06 | 0.45 | 0.14 | 0.18 | 0.01 | 0.16 |
| 10 | 34401632 | rs11009644 | -0.196 | 0.677 | 0.971 | -1.51 | 0 | -0.062 | -0.062 | 0 | 0.22 | 0.3 | 0.21 | 0.01 | 0.21 | 0.14 |
| 10 | 34408636 | rs11009651 | -0.384 | 0.343 | 1.051 | -1.655 | -0.009 | -0.085 | -0.085 | 0.551 | 0.3 | 0.36 | 0.19 | 0 | 0.25 | 0.12 |
| 10 | 34408745 | rs3004928 | 0.981 | 0.678 | 1.221 | -1.869 | 0.004 | 0.002 | 0.002 | 0.044 | 0.5 | 0.44 | 0.62 | 0.86 | 0.46 | 0.71 |

| | | | | | | | | | | | | | | | | |
|---|---|---|---|---|---|---|---|---|---|---|---|---|---|---|---|---|
| 10 | 34415490 | rs4126706 | -0.18 | 0.683 | 1.163 | -1.866 | -0.013 | -0.105 | -0.105 | 0 | 0.3 | 0.4 | 0.2 | 0 | 0.25 | 0.11 |
| 10 | 34417835 | rs1415546 | -0.747 | 0.589 | 1.126 | -1.878 | 0.031 | -0.047 | -0.047 | 0 | 0.24 | 0.36 | 0.2 | 0.13 | 0.31 | 0.2 |
| 10 | 34417876 | rs1415547 | -0.328 | 0.606 | 0.847 | -1.871 | -0.028 | 0.027 | 0.027 | 0.042 | 0.21 | 0.14 | 0.18 | 0.13 | 0.29 | 0.2 |
| 10 | 34423253 | rs1692712 | -1.311 | 0.451 | 0.452 | -1.882 | 0.025 | 0.116 | 0.116 | 0 | 0.52 | 0.34 | 0.36 | 0.13 | 0.56 | 0.33 |
| 10 | 34428128 | rs1753583 | -0.236 | 0.537 | 0.294 | -1.827 | -0.052 | 0.044 | 0.044 | 0.015 | 0.21 | 0.1 | 0.17 | 0.13 | 0.29 | 0.2 |
| 10 | 34428832 | rs1536771 | -0.271 | 0.615 | 0.331 | -1.817 | -0.003 | 0.001 | 0.001 | 0.085 | 0.21 | 0.21 | 0.19 | 0.13 | 0.3 | 0.18 |
| 10 | 34433123 | rs10827332 | 0.044 | 0.15 | -0.044 | -1.798 | -0.022 | 0.154 | 0.154 | 0 | 0.28 | 0.07 | 0.16 | 0 | 0.26 | 0.14 |
| 10 | 34439852 | rs920515 | -1.172 | 0.39 | 0.385 | -1.674 | 0.01 | 0.054 | 0.054 | 0.021 | 0.53 | 0.42 | 0.35 | 0.14 | 0.58 | 0.31 |
| 10 | 34441217 | rs1148242 | 0.002 | 0.751 | -0.092 | -1.467 | -0.094 | 0.019 | 0.019 | 0 | 0.17 | 0.03 | 0.15 | 0.14 | 0.31 | 0.18 |
| 10 | 34443115 | rs686993 | -0.515 | 0.352 | 0.702 | -1.245 | -0.01 | -0.016 | -0.016 | 0 | 0.37 | 0.5 | 0.29 | 0.29 | 0.3 | 0.2 |
| 10 | 34444434 | rs1822126 | -1.061 | 0.445 | 1.041 | -1.34 | 0.026 | -0.035 | -0.035 | 0 | 0.07 | 0.23 | 0.05 | 0 | 0.07 | 0.03 |
| 10 | 34444709 | rs1977894 | -0.577 | 0.398 | 1.144 | -1.392 | -0.002 | -0.075 | -0.075 | 0 | 0.37 | 0.37 | 0.2 | 0 | 0.29 | 0.17 |
| 10 | 34451109 | rs1660619 | -1.314 | 0.618 | 1.41 | -1.433 | 0.04 | 0.026 | 0.026 | 0.273 | 0.25 | 0.55 | 0.3 | 0.41 | 0.33 | 0.23 |
| 10 | 34452072 | rs2248375 | 0.221 | 0.534 | 1.38 | -1.474 | 0.027 | 0.124 | 0.124 | 0 | 0.28 | 0.11 | 0.15 | 0 | 0.23 | 0.14 |
| 10 | 34452698 | rs2786555 | 0.11 | 0.513 | 1.378 | -1.474 | 0.032 | 0.098 | 0.098 | 0 | 0.29 | 0.13 | 0.15 | 0 | 0.23 | 0.14 |
| 10 | 34453509 | rs16935155 | -0.858 | 0.516 | 1.546 | -1.439 | 0.013 | -0.014 | -0.014 | 0 | 0.07 | 0.3 | 0.04 | 0 | 0.04 | 0 |
| 10 | 34459326 | rs748104 | -0.89 | 0.479 | 1.194 | -1.451 | -0.013 | 0.058 | 0.058 | 0.236 | 0.46 | 0.31 | 0.37 | 0.36 | 0.49 | 0.36 |
| 10 | 34459407 | rs748103 | 0.109 | 0.485 | 1.186 | -1.453 | 0 | -0.002 | -0.002 | 0.005 | 0.3 | 0.29 | 0.24 | 0.23 | 0.23 | 0.18 |
| 10 | 34461345 | rs1822127 | -0.829 | 0.481 | 1.188 | -1.452 | -0.016 | 0.048 | 0.048 | 0.304 | 0.45 | 0.31 | 0.37 | 0.35 | 0.49 | 0.33 |
| 10 | 34466656 | rs1290002 | 0.247 | 0.54 | 0.982 | -1.526 | -0.091 | 0.014 | 0.014 | 0.608 | 0.16 | 0.02 | 0.14 | 0.11 | 0.28 | 0.17 |
| 10 | 34467741 | rs1036731 | 0.25 | 0.53 | 0.975 | -1.524 | -0.093 | -0.005 | -0.005 | 0.013 | 0.15 | 0.01 | 0.12 | 0.11 | 0.26 | 0.15 |
| 10 | 34470204 | rs1274466 | -1.46 | 0.422 | 1.098 | -1.551 | 0.007 | 0.075 | 0.075 | 0.318 | 0.53 | 0.67 | 0.43 | 0.33 | 0.54 | 0.35 |
| 10 | 34470304 | rs1660627 | -0.398 | 0.433 | 1.079 | -1.56 | 0.012 | 0.014 | 0.014 | 0.009 | 0.37 | 0.62 | 0.29 | 0.22 | 0.26 | 0.19 |
| 10 | 34471937 | rs1296521 | -1.434 | 0.468 | 1.084 | -1.522 | 0.004 | 0.046 | 0.046 | 0.339 | 0.53 | 0.65 | 0.43 | 0.39 | 0.54 | 0.36 |
| 10 | 34473335 | rs1660630 | -1.649 | 0.353 | 0.916 | -1.53 | -0.054 | 0.046 | 0.046 | 0.235 | 0.56 | 0.82 | 0.44 | 0.39 | 0.52 | 0.37 |
| 10 | 34475080 | rs1660632 | -0.122 | 0.208 | 1.141 | -1.726 | -0.03 | -0.023 | -0.023 | 0 | 0.32 | 0.52 | 0.27 | 0.28 | 0.23 | 0.19 |
| 10 | 34477824 | rs1274470 | -0.823 | 0.393 | 0.336 | -1.921 | 0.025 | -0.086 | -0.086 | 0.353 | 0.23 | 0.57 | 0.2 | 0.03 | 0.28 | 0.13 |
| 10 | 34478675 | rs2799459 | -0.052 | 0.304 | 0.154 | -1.677 | 0.019 | 0.026 | 0.026 | 0 | 0.31 | 0.22 | 0.23 | 0.2 | 0.23 | 0.18 |
| 10 | 34490854 | rs647758 | -1.518 | 0.817 | 0.043 | -1.945 | 0.016 | 0.081 | 0.081 | 0 | 0.58 | 0.74 | 0.52 | 0.25 | 0.62 | 0.38 |
| 10 | 34491850 | rs10827337 | 0.104 | 0.753 | 0.18 | -2.079 | 0.027 | 0.035 | 0.035 | 0 | 0.29 | 0.19 | 0.2 | 0.15 | 0.2 | 0.16 |
| 10 | 34492188 | rs1121682 | 0.372 | 0.869 | -0.276 | -2.173 | -0.053 | 0.045 | 0.045 | 0.148 | 0.17 | 0.06 | 0.17 | 0.09 | 0.26 | 0.16 |
| 10 | 34495289 | rs678188 | -0.582 | 0.651 | -0.233 | -2.173 | -0.043 | 0.043 | 0.043 | 0 | 0.23 | 0.11 | 0.25 | 0.11 | 0.39 | 0.19 |
| 10 | 34498716 | rs1765169 | -1.808 | 0.627 | -0.577 | -2.176 | 0.014 | 0.115 | 0.115 | 0 | 0.58 | 0.78 | 0.52 | 0.23 | 0.62 | 0.38 |
| 10 | 34499647 | rs10827338 | -0.334 | 0.645 | -0.561 | -2.163 | 0.041 | 0.066 | 0.066 | 0 | 0.35 | 0.71 | 0.28 | 0.13 | 0.24 | 0.2 |
| 10 | 34505150 | rs622408 | -1.853 | 0.544 | -0.632 | -2.169 | 0.003 | 0.118 | 0.118 | 0 | 0.59 | 0.83 | 0.53 | 0.23 | 0.63 | 0.38 |
| 10 | 34505864 | rs12778087 | 0.011 | 0.467 | -0.429 | -2.002 | 0.023 | 0.168 | 0.168 | 0 | 0.28 | 0.07 | 0.15 | 0 | 0.21 | 0.12 |
| 10 | 34509032 | rs671228 | -1.235 | 0.598 | -0.425 | -1.926 | 0.013 | 0.156 | 0.156 | 0 | 0.51 | 0.72 | 0.45 | 0.22 | 0.52 | 0.36 |
| 10 | 34513830 | rs2570326 | -1.148 | 0.406 | -2.367 | -2.038 | -0.052 | 0.195 | 0.195 | 0.326 | 0.47 | 0.82 | 0.5 | 0.2 | 0.47 | 0.36 |
| 10 | 34520359 | rs1545214 | -0.04 | 1.588 | -2.214 | -1.784 | -0.077 | 0.038 | 0.038 | 0.848 | 0.15 | 0.03 | 0.23 | 0.09 | 0.25 | 0.16 |
| 10 | 34524792 | rs11592647 | -0.314 | 1.155 | -2.068 | -1.663 | -0.005 | 0.207 | 0.207 | 0 | 0.29 | 0.03 | 0.14 | 0 | 0.22 | 0.11 |
| 10 | 34527387 | rs1111267 | -0.775 | 1.964 | -1.045 | -0.642 | -0.058 | 0.025 | 0.025 | 0 | 0.12 | 0.05 | 0.2 | 0.09 | 0.23 | 0.13 |
| 10 | 34534237 | rs11009690 | -1.393 | 0.961 | 0.289 | -0.496 | 0.079 | 0.153 | 0.153 | 1.908 | 0.05 | 0.39 | 0.04 | 0.07 | 0.01 | 0.04 |
| 10 | 34536049 | rs593992 | -1.244 | 0.276 | 1.068 | -0.909 | -0.04 | -0.016 | -0.016 | 0 | 0.11 | 0.66 | 0.09 | 0.07 | 0.02 | 0.03 |
| 10 | 34538563 | rs11009696 | -1.648 | 0.981 | 1.244 | -1.103 | 0.108 | 0.033 | 0.033 | 0 | 0.09 | 0.34 | 0.1 | 0.07 | 0.11 | 0.1 |
| 10 | 34539038 | rs2248686 | -0.548 | 0.93 | 1.269 | -1.106 | -0.009 | -0.037 | -0.037 | 0 | 0.09 | 0.43 | 0.07 | 0 | 0.02 | 0 |
| 10 | 34544880 | rs1348106 | 0.182 | 0.944 | 0 | -0.834 | 0.008 | -0.007 | -0.007 | 0 | 0.61 | 0.62 | 0.47 | 0.37 | 0.51 | 0.55 |
| 10 | 34547562 | rs1095687 | -0.107 | 0.964 | 0.004 | -0.837 | -0.022 | 0.005 | 0.005 | 0.336 | 0.64 | 0.76 | 0.49 | 0.37 | 0.52 | 0.56 |
| 10 | 34552061 | rs7901148 | 0.484 | 0.97 | 0.199 | -0.859 | 0.004 | 0.035 | 0.035 | 0.117 | 0.55 | 0.29 | 0.41 | 0.37 | 0.5 | 0.56 |
| 10 | 34564546 | rs2570339 | -0.046 | 0.974 | 0.507 | -0.833 | 0.003 | -0.034 | -0.034 | 0 | 0.09 | 0.49 | 0.08 | 0 | 0.02 | 0 |
| 10 | 34566190 | rs1660614 | -0.02 | 0.927 | 0.384 | -0.897 | -0.018 | 0.009 | 0.009 | 0.646 | 0.62 | 0.75 | 0.48 | 0.37 | 0.51 | 0.56 |
| 10 | 34569584 | rs10827343 | -0.558 | 1.054 | 0.517 | -0.887 | 0.062 | 0.006 | 0.006 | 0 | 0.08 | 0.23 | 0.09 | 0.07 | 0.12 | 0.09 |
| 10 | 34570702 | rs16935275 | 0.112 | 1.053 | 0.495 | -0.889 | 0 | -0.011 | -0.011 | 0.143 | 0.07 | 0.08 | 0.08 | 0 | 0.12 | 0.06 |
| 10 | 34576089 | rs584581 | -0.54 | 0.634 | 0.221 | -0.952 | -0.039 | -0.029 | -0.029 | 0 | 0.65 | 0.82 | 0.55 | 0.47 | 0.6 | 0.66 |
| 10 | 34580880 | rs1985313 | 1.229 | 0.062 | -0.595 | -1.727 | -0.051 | -0.099 | -0.099 | 0.084 | 0.29 | 0.02 | 0.3 | 0.4 | 0.32 | 0.47 |
| 10 | 34582534 | rs2582860 | 0.303 | 0.15 | -0.245 | -1.646 | 0.086 | 0.021 | 0.021 | 0 | 0.08 | 0.28 | 0.09 | 0.08 | 0.12 | 0.09 |
| 10 | 34583697 | rs1441027 | -0.507 | 0.105 | -0.192 | -1.65 | 0 | -0.005 | -0.005 | 0 | 0.57 | 0.47 | 0.46 | 0.4 | 0.48 | 0.56 |
| 10 | 34588112 | rs1936426 | -0.667 | 0.344 | -0.251 | -1.652 | -0.003 | 0.005 | 0.005 | 0.763 | 0.58 | 0.64 | 0.51 | 0.39 | 0.5 | 0.57 |
| 10 | 34588467 | rs16935300 | 0.36 | 0.368 | -0.239 | -1.646 | 0.084 | 0.02 | 0.02 | 0 | 0.08 | 0.26 | 0.09 | 0.08 | 0.12 | 0.09 |
| 10 | 34591402 | rs2570334 | -1.201 | 0.32 | -0.352 | -1.616 | -0.063 | -0.056 | -0.056 | 0 | 0.66 | 0.9 | 0.59 | 0.47 | 0.62 | 0.66 |
| 10 | 34599342 | rs12241064 | 0.305 | 0.272 | 0.478 | -1.441 | -0.02 | -0.055 | -0.055 | 0.71 | 0.09 | 0.32 | 0.06 | 0 | 0.03 | 0.03 |
| 10 | 34602653 | rs10763978 | -0.811 | 0.131 | 0.285 | -1.548 | -0.041 | 0.029 | 0.029 | 1.99 | 0.51 | 0.91 | 0.53 | 0.47 | 0.5 | 0.61 |
| 10 | 34620220 | rs3781128 | -0.819 | 0.17 | 0.493 | -1.571 | 0.013 | 0.06 | 0.06 | 1.054 | 0.5 | 0.73 | 0.52 | 0.39 | 0.48 | 0.57 |
| 10 | 34627039 | rs10827349 | 0.377 | 0.117 | 0.655 | -1.577 | -0.03 | -0.056 | -0.056 | 0.571 | 0.1 | 0.28 | 0.06 | 0 | 0.03 | 0.02 |
| 10 | 34628649 | rs2002773 | -0.784 | 0.095 | 0.573 | -1.614 | 0.011 | 0.068 | 0.068 | 1.042 | 0.51 | 0.75 | 0.52 | 0.39 | 0.48 | 0.59 |
| 10 | 34629060 | rs7922046 | -0.588 | 0.103 | 0.509 | -1.589 | 0 | 0 | 0 | 0.898 | 0.48 | 0.54 | 0.5 | 0.39 | 0.48 | 0.57 |
| 10 | 34629851 | rs2384210 | -0.288 | 0.157 | 0.481 | -1.598 | 0.009 | 0.009 | 0.009 | 1.728 | 0.44 | 0.42 | 0.42 | 0.39 | 0.38 | 0.53 |
| 10 | 34631239 | rs2384211 | -0.896 | 0.165 | 0.439 | -1.595 | -0.002 | 0.04 | 0.04 | 0.033 | 0.52 | 0.74 | 0.52 | 0.47 | 0.51 | 0.62 |
| 10 | 34635529 | rs12247156 | -0.736 | 0.157 | 0.401 | -1.618 | 0.015 | -0.097 | -0.097 | 1.573 | 0.17 | 0.35 | 0.18 | 0 | 0.17 | 0.09 |
| 10 | 34640064 | rs10763981 | -1.035 | 0.285 | -0.078 | -1.349 | -0.036 | -0.019 | -0.019 | 0 | 0.53 | 0.91 | 0.53 | 0.51 | 0.49 | 0.63 |

| | | | | | | | | | | | | | | | | |
|---|---|---|---|---|---|---|---|---|---|---|---|---|---|---|---|---|
| 10 | 34659351 | rs10763985 | 1.066 | 0.301 | -0.355 | -1.015 | -0.037 | -0.115 | -0.115 | 0.69 | 0.28 | 0.03 | 0.27 | 0.43 | 0.28 | 0.5 |
| 10 | 34672463 | rs16935349 | -1.065 | 0.308 | 0.196 | -1 | -0.034 | -0.056 | -0.056 | 0 | 0.17 | 0.64 | 0.13 | 0.08 | 0.1 | 0.05 |
| 10 | 34692235 | rs4113505 | 0.277 | 0.29 | 0.373 | -0.92 | -0.016 | -0.005 | -0.005 | 0 | 0.1 | 0.35 | 0.06 | 0.07 | 0.03 | 0.03 |
| 10 | 34693172 | rs4934633 | -0.824 | 0.321 | -0.081 | -0.919 | -0.031 | -0.064 | -0.064 | 0.31 | 0.17 | 0.64 | 0.13 | 0.07 | 0.1 | 0.05 |
| 10 | 34694931 | rs3905150 | -0.786 | 0.403 | -0.776 | -0.737 | 0.017 | 0.081 | 0.081 | 1.024 | 0.51 | 0.87 | 0.52 | 0.44 | 0.48 | 0.56 |
| 10 | 34697974 | rs3851066 | -1.022 | 0.46 | -0.908 | -0.718 | -0.025 | 0.027 | 0.027 | 0.648 | 0.53 | 0.92 | 0.53 | 0.47 | 0.48 | 0.58 |
| 10 | 34705559 | rs2252655 | -0.848 | 0.441 | -1.066 | -0.489 | 0.01 | -0.044 | -0.044 | 3.484 | 0.49 | 0.95 | 0.47 | 0.53 | 0.41 | 0.44 |
| 10 | 34714686 | rs7079002 | -0.147 | 0.193 | -0.177 | -0.446 | 0.01 | -0.005 | -0.005 | 0.043 | 0.59 | 0.37 | 0.6 | 0.51 | 0.63 | 0.58 |
| 10 | 34731264 | rs773964 | -0.984 | 0.104 | 0.144 | -0.527 | -0.013 | -0.049 | -0.049 | 0 | 0.71 | 0.92 | 0.7 | 0.61 | 0.68 | 0.61 |
| 10 | 34735753 | rs17475284 | 0.636 | 0.552 | -0.51 | -0.802 | -0.102 | 0.064 | 0.064 | 0.123 | 0.26 | 0.04 | 0.32 | 0.13 | 0.32 | 0.15 |
| 10 | 34739166 | rs982687 | 1.118 | 0.324 | 0.015 | -0.708 | 0.008 | -0.031 | -0.031 | 0.563 | 0.07 | 0.11 | 0.14 | 0 | 0.07 | 0.02 |
| 10 | 34743703 | rs1469874 | 0.025 | -0.246 | 0.326 | -0.55 | 0.054 | -0.032 | -0.032 | 0.622 | 0.37 | 0.84 | 0.38 | 0.45 | 0.3 | 0.42 |
| 10 | 34750875 | rs773944 | 0.15 | -0.124 | 0.629 | -0.508 | 0.007 | -0.004 | -0.004 | 0 | 0.36 | 0.3 | 0.36 | 0.44 | 0.34 | 0.42 |
| 10 | 34786887 | rs773990 | 0.061 | -0.061 | 0.706 | -0.503 | -0.003 | -0.029 | -0.029 | 1.907 | 0.06 | 0.13 | 0.05 | 0 | 0.05 | 0.01 |
| 10 | 34797585 | rs7477871 | -0.212 | -0.195 | 0.696 | -0.6 | 0.005 | -0.028 | -0.028 | 0.1 | 0.07 | 0.28 | 0.06 | 0 | 0.05 | 0.04 |
| 10 | 34802967 | rs4934638 | 0.959 | -0.171 | 0.286 | -0.526 | 0.006 | -0.012 | -0.012 | 0 | 0.26 | 0.2 | 0.28 | 0.42 | 0.24 | 0.37 |
| 10 | 34803585 | rs1025637 | 1.045 | -0.17 | 0.232 | -0.543 | 0.009 | -0.023 | -0.023 | 0 | 0.26 | 0.16 | 0.28 | 0.42 | 0.25 | 0.37 |
| 10 | 34806744 | rs1780578 | -1.204 | -0.029 | 0.681 | -0.529 | -0.013 | -0.063 | -0.063 | 0.122 | 0.11 | 0.63 | 0.1 | 0 | 0.05 | 0.01 |
| 10 | 34813742 | rs773970 | -0.389 | -0.131 | 0.235 | -0.466 | 0.033 | -0.016 | -0.016 | 0.64 | 0.39 | 0.84 | 0.43 | 0.43 | 0.36 | 0.43 |
| 10 | 34821063 | rs773946 | -0.616 | -0.263 | 0.05 | -0.421 | -0.002 | -0.04 | -0.04 | 0 | 0.44 | 0.94 | 0.44 | 0.43 | 0.39 | 0.46 |
| 10 | 34842837 | rs10508807 | 0.281 | -0.323 | 0.414 | -0.458 | -0.005 | 0.009 | 0.009 | 0 | 0.07 | 0.04 | 0.05 | 0 | 0.07 | 0.04 |
| 10 | 34851940 | rs7071602 | -0.572 | -0.329 | 0.531 | -0.448 | 0.041 | 0.007 | 0.007 | 0.526 | 0.43 | 0.82 | 0.44 | 0.43 | 0.38 | 0.46 |
| 10 | 34855095 | rs1540953 | -0.128 | -0.227 | 0.869 | -0.399 | -0.004 | 0.001 | 0.001 | 0 | 0.37 | 0.39 | 0.38 | 0.43 | 0.38 | 0.46 |
| 10 | 34895329 | rs2496720 | 1.808 | -0.419 | -0.091 | -0.46 | 0.015 | -0.128 | -0.128 | 0 | 0.23 | 0.01 | 0.14 | 0.3 | 0.12 | 0.33 |
| 10 | 34898549 | rs7098206 | 0.396 | 0.373 | 0.694 | -0.257 | -0.002 | -0.013 | -0.013 | 0.109 | 0.23 | 0.2 | 0.31 | 0.43 | 0.29 | 0.4 |
| 10 | 34941134 | rs2490817 | -0.223 | 0.162 | 0.318 | -0.174 | 0 | -0.013 | -0.013 | 0 | 0.28 | 0.25 | 0.3 | 0.43 | 0.31 | 0.44 |
| 10 | 34971389 | rs11598578 | 0.403 | 0.325 | -0.025 | -0.65 | -0.132 | 0.083 | 0.083 | 0.022 | 0.27 | 0.05 | 0.34 | 0.14 | 0.39 | 0.13 |
| 10 | 34972844 | rs2496737 | -1.393 | 0.155 | -0.21 | -0.63 | -0.045 | -0.142 | -0.142 | 1.058 | 0.6 | 0.99 | 0.77 | 0.51 | 0.68 | 0.53 |
| 10 | 34984884 | rs2496742 | -0.948 | 0.26 | 0.156 | -0.582 | 0.033 | 0.002 | 0.002 | 0.474 | 0.53 | 0.37 | 0.69 | 0.51 | 0.68 | 0.53 |
| 10 | 34984959 | rs2477009 | -0.966 | 0.262 | 0.248 | -0.582 | 0.007 | 0 | 0 | 0 | 0.53 | 0.45 | 0.7 | 0.51 | 0.68 | 0.53 |
| 10 | 34989041 | rs2496745 | 0.06 | 0.258 | 0.258 | -0.338 | 0.001 | -0.006 | -0.006 | 0 | 0.26 | 0.25 | 0.34 | 0.37 | 0.28 | 0.4 |
| 10 | 34999031 | rs10508810 | -1.056 | 1.023 | 0.789 | -0.371 | 0.183 | -0.006 | -0.006 | 0.306 | 0.08 | 0.41 | 0.1 | 0 | 0.1 | 0.01 |
| 10 | 35016575 | rs11009892 | 0.284 | -0.359 | 0.706 | -0.228 | 0 | -0.012 | -0.012 | 0 | 0.23 | 0.18 | 0.31 | 0.35 | 0.29 | 0.39 |
| 10 | 35026141 | rs10450314 | 0.941 | -0.485 | 0.102 | -0.479 | 0.003 | -0.127 | -0.127 | 1.924 | 0.39 | 0.02 | 0.23 | 0.52 | 0.28 | 0.47 |
| 10 | 35028713 | rs7075263 | 0.612 | -0.244 | 0.618 | -0.431 | 0.029 | -0.023 | -0.023 | 0 | 0.25 | 0.28 | 0.36 | 0.16 | 0.4 | 0.13 |
| 10 | 35031758 | rs17493513 | 0.511 | -0.69 | 0.589 | -0.358 | -0.136 | 0.077 | 0.077 | 0 | 0.28 | 0.09 | 0.38 | 0.16 | 0.46 | 0.15 |
| 10 | 35053262 | rs10764002 | -0.281 | -0.842 | 0.685 | -0.362 | -0.011 | -0.005 | -0.005 | 0.8 | 0.69 | 0.61 | 0.65 | 0.7 | 0.73 | 0.58 |
| 10 | 35058930 | rs9417142 | 0.779 | -0.843 | 0.524 | -0.329 | 0.013 | -0.005 | -0.005 | 0 | 0.39 | 0.28 | 0.25 | 0.53 | 0.26 | 0.44 |
| 10 | 35075817 | rs7897283 | -0.155 | -0.79 | 0.614 | -0.267 | -0.009 | -0.002 | -0.002 | 0 | 0.3 | 0.61 | 0.35 | 0.3 | 0.26 | 0.38 |
| 10 | 35099799 | rs7342064 | -0.249 | -0.781 | 0.674 | -0.263 | -0.023 | -0.001 | -0.001 | 0.021 | 0.29 | 0.5 | 0.33 | 0.3 | 0.26 | 0.39 |
| 10 | 35102818 | rs4934671 | 0.756 | -0.872 | 0.452 | -0.183 | 0.011 | -0.03 | -0.03 | 0 | 0.21 | 0.1 | 0.26 | 0.31 | 0.21 | 0.37 |
| 10 | 35107733 | rs4078277 | -0.577 | -0.87 | 0.434 | -0.02 | 0.013 | 0.017 | 0.017 | 0 | 0.9 | 0.81 | 0.92 | 0.87 | 0.88 | 0.93 |
| 10 | 35111495 | rs10827437 | -0.46 | -0.884 | 0.661 | -0.026 | 0.043 | 0.05 | 0.05 | 0 | 0.9 | 0.72 | 0.92 | 0.87 | 0.88 | 0.93 |
| 10 | 35114524 | rs11009966 | 1.616 | -1.166 | 0.26 | -0.142 | 0.03 | -0.15 | -0.15 | 1.125 | 0.27 | 0.01 | 0.17 | 0.38 | 0.15 | 0.36 |
| 10 | 35116998 | rs4638261 | 0.183 | -1.134 | 0.547 | -0.165 | -0.007 | -0.001 | -0.001 | 0.191 | 0.31 | 0.36 | 0.33 | 0.32 | 0.23 | 0.38 |
| 10 | 35118276 | rs7911088 | 0.034 | -1.011 | 0.044 | -0.152 | -0.012 | 0.003 | 0.003 | 0 | 0.59 | 0.46 | 0.6 | 0.55 | 0.65 | 0.55 |
| 10 | 35123628 | rs4934517 | 0.398 | -0.775 | -0.636 | -0.183 | 0.026 | 0.02 | 0.02 | 0.002 | 0.53 | 0.28 | 0.53 | 0.54 | 0.59 | 0.5 |
| 10 | 35130901 | rs10827443 | 0.268 | -0.608 | -0.577 | -0.236 | -0.113 | 0.035 | 0.035 | 0 | 0.29 | 0.13 | 0.38 | 0.22 | 0.47 | 0.16 |
| 10 | 35147414 | rs2492459 | 0.406 | -1.057 | -0.317 | -0.377 | 0.004 | 0.004 | 0.004 | 0 | 0.5 | 0.5 | 0.48 | 0.65 | 0.35 | 0.7 |
| 10 | 35148739 | rs2492460 | 0.745 | -1.041 | -0.057 | -0.232 | 0.002 | 0.012 | 0.012 | 0.004 | 0.8 | 0.85 | 0.87 | 0.87 | 0.83 | 0.88 |
| 10 | 35150975 | rs11598558 | -0.934 | -1.203 | -0.014 | -0.293 | -0.021 | -0.056 | -0.056 | 0.71 | 0.15 | 0.27 | 0.08 | 0.01 | 0.07 | 0.06 |
| 10 | 35154346 | rs7915550 | 0.514 | -0.864 | -0.06 | 0.183 | 0.02 | -0.014 | -0.014 | 0 | 0.3 | 0.22 | 0.2 | 0.31 | 0.18 | 0.42 |
| 10 | 35158707 | rs2492461 | 0.526 | -0.877 | -0.294 | 0.19 | 0.052 | -0.009 | -0.009 | 0 | 0.52 | 0.31 | 0.65 | 0.57 | 0.68 | 0.5 |
| 10 | 35169778 | rs10827457 | 0.591 | -1.012 | -0.464 | 0.228 | 0.027 | -0.021 | -0.021 | 0.14 | 0.29 | 0.22 | 0.21 | 0.3 | 0.18 | 0.42 |
| 10 | 35173730 | rs16935780 | -1.281 | -0.064 | 0.37 | 0.194 | 0.032 | 0.009 | 0.009 | 5.487 | 0.07 | 0.44 | 0.07 | 0 | 0.02 | 0.01 |
| 10 | 35174347 | rs2504249 | 0.336 | 0.518 | -0.839 | 0.502 | -0.005 | -0.01 | -0.01 | 0 | 0.9 | 0.96 | 0.93 | 0.88 | 0.88 | 0.92 |
| 10 | 35194388 | rs4469833 | -0.676 | 0.852 | -0.783 | 0.465 | -0.007 | -0.005 | -0.005 | 0 | 0.37 | 0.33 | 0.34 | 0.43 | 0.39 | 0.26 |
| 10 | 35195406 | rs2492448 | -0.046 | 0.374 | -0.574 | 0.369 | -0.006 | -0.007 | -0.007 | 0.196 | 0.28 | 0.39 | 0.34 | 0.26 | 0.29 | 0.37 |
| 10 | 35196538 | rs2492450 | -0.735 | 0.574 | -0.376 | -0.189 | 0.001 | 0.003 | 0.003 | 0 | 0.33 | 0.32 | 0.42 | 0.26 | 0.37 | 0.46 |
| 10 | 35207153 | rs11591754 | 0.366 | 0.772 | -0.329 | -0.047 | 0.006 | -0.022 | -0.022 | 0 | 0.19 | 0.09 | 0.19 | 0.26 | 0.18 | 0.24 |
| 10 | 35207822 | rs2504262 | -0.304 | 0.696 | -0.04 | -0.048 | 0 | 0.002 | 0.002 | 0 | 0.26 | 0.23 | 0.27 | 0.27 | 0.3 | 0.39 |
| 10 | 35208068 | rs17496856 | 0.451 | 0.675 | 0.235 | 0.136 | 0 | 0 | 0 | 0 | 0.09 | 0.06 | 0.05 | 0.11 | 0.11 | 0.09 |
| 10 | 35212149 | rs11597065 | -0.551 | 0.573 | 0.607 | 0.132 | 0 | 0.01 | 0.01 | 0 | 0.28 | 0.4 | 0.3 | 0.29 | 0.26 | 0.16 |
| 10 | 35212572 | rs10764013 | -0.483 | 0.578 | 0.515 | 0.135 | -0.001 | -0.002 | -0.002 | 1.427 | 0.28 | 0.27 | 0.29 | 0.29 | 0.26 | 0.17 |
| 10 | 35220151 | rs7078774 | 0.301 | 0.718 | 0.419 | 0.198 | 0.003 | -0.009 | -0.009 | 0.256 | 0.1 | 0.13 | 0.07 | 0.07 | 0.13 | 0.09 |
| 10 | 35220175 | rs10764014 | -1.215 | 0.602 | 0.44 | 0.095 | 0.004 | 0.015 | 0.015 | 0 | 0.49 | 0.63 | 0.53 | 0.59 | 0.49 | 0.55 |
| 10 | 35222113 | rs7895855 | -2.157 | 0.563 | 0.169 | 0.097 | -0.118 | -0.093 | -0.093 | 1.196 | 0.62 | 0.98 | 0.62 | 0.66 | 0.62 | 0.64 |
| 10 | 35229637 | rs12255903 | -0.049 | 0.229 | 0.948 | 0.083 | -0.004 | 0.003 | 0.003 | 0 | 0.23 | 0.33 | 0.26 | 0.32 | 0.24 | 0.38 |
| 10 | 35244427 | rs12254380 | -0.383 | 0.222 | 0.645 | -0.023 | -0.006 | -0.005 | -0.005 | 0 | 0.26 | 0.66 | 0.29 | 0.29 | 0.25 | 0.38 |

| | | | | | | | | | | | | | | | | |
|---|---|---|---|---|---|---|---|---|---|---|---|---|---|---|---|---|
| 10 | 35246612 | rs7086375 | -0.796 | 0.23 | 0.319 | -0.543 | 0.001 | 0.002 | 0.002 | 0 | 0.47 | 0.54 | 0.52 | 0.18 | 0.52 | 0.54 |
| 10 | 35246623 | rs12573778 | -0.485 | 0.098 | 0.059 | -0.321 | 0 | -0.001 | -0.001 | 3.185 | 0.39 | 0.42 | 0.26 | 0.26 | 0.41 | 0.21 |
| 10 | 35247587 | rs7091165 | -1.741 | -0.25 | 0.679 | 0.557 | 0.031 | -0.011 | -0.011 | 0 | 0.52 | 0.79 | 0.54 | 0.53 | 0.46 | 0.56 |
| 10 | 35250069 | rs11010055 | -0.448 | -0.092 | 0.672 | 0.549 | -0.016 | -0.016 | -0.016 | 3.057 | 0.24 | 0.37 | 0.25 | 0.24 | 0.2 | 0.4 |
| 10 | 35251225 | rs10827475 | -0.744 | -0.095 | 0.621 | 0.692 | -0.013 | 0.035 | 0.035 | 0 | 0.26 | 0.4 | 0.28 | 0.33 | 0.23 | 0.16 |
| 10 | 35256179 | rs11010057 | -0.986 | 0.689 | 1.034 | 1.095 | -0.02 | 0.016 | 0.016 | 0 | 0.24 | 0.49 | 0.27 | 0.28 | 0.2 | 0.38 |
| 10 | 35257823 | rs4007287 | -1.225 | 0.791 | 0.289 | 1.041 | -0.047 | -0.049 | -0.049 | 0 | 0.23 | 0.07 | 0.28 | 0.31 | 0.32 | 0.17 |
| 10 | 35300623 | rs2001893 | -2.022 | 0.674 | 0.841 | 0.958 | 0.001 | -0.002 | -0.002 | 0.054 | 0.34 | 0.42 | 0.37 | 0.33 | 0.36 | 0.33 |
| 10 | 35323130 | rs12184386 | 2.182 | 0.853 | 0.7 | 0.926 | -0.107 | -0.035 | -0.035 | 0 | 0.48 | 0.02 | 0.42 | 0.39 | 0.47 | 0.43 |
| 10 | 35341301 | rs2148483 | -0.581 | 0.781 | 0.85 | 0.854 | -0.017 | 0.031 | 0.031 | 0 | 0.16 | 0.05 | 0.13 | 0.06 | 0.17 | 0.2 |
| 10 | 35344969 | rs1926554 | -1.814 | 0.775 | 0.99 | 0.859 | -0.01 | -0.003 | -0.003 | 0 | 0.32 | 0.2 | 0.33 | 0.31 | 0.36 | 0.33 |
| 10 | 35359475 | rs12240347 | -2.234 | 0.774 | 1.013 | 0.859 | 0.001 | -0.001 | -0.001 | 0.416 | 0.34 | 0.42 | 0.36 | 0.33 | 0.36 | 0.33 |
| 10 | 35362977 | rs12248333 | -2.234 | 0.774 | 1.013 | 0.859 | 0.001 | -0.001 | -0.001 | 3.718 | 0.34 | 0.42 | 0.36 | 0.33 | 0.36 | 0.33 |
| 10 | 35399668 | rs12769189 | -1.675 | 0.761 | 1.006 | 0.858 | -0.024 | -0.012 | -0.012 | 0 | 0.31 | 0.15 | 0.33 | 0.31 | 0.36 | 0.33 |
| 10 | 35414524 | rs11595640 | -0.576 | 0.762 | 0.987 | 0.88 | -0.019 | 0.024 | 0.024 | 0 | 0.16 | 0.05 | 0.13 | 0.07 | 0.18 | 0.2 |
| 10 | 35415555 | rs4934730 | -1.688 | 0.764 | 0.989 | 0.819 | -0.025 | -0.013 | -0.013 | 0 | 0.31 | 0.15 | 0.33 | 0.31 | 0.36 | 0.33 |
| 10 | 35426755 | rs17499247 | -2.087 | 0.764 | 1.084 | 0.819 | 0.001 | -0.001 | -0.001 | 2.159 | 0.33 | 0.33 | 0.35 | 0.31 | 0.36 | 0.33 |
| 10 | 35432405 | rs16935888 | -0.295 | 0.766 | 1.062 | 0.825 | -0.009 | -0.025 | -0.025 | 0 | 0.15 | 0.07 | 0.2 | 0.26 | 0.18 | 0.13 |
| 10 | 35459976 | rs10508816 | -2.251 | 0.764 | 1.042 | 0.821 | 0.003 | -0.002 | -0.002 | 0 | 0.34 | 0.45 | 0.36 | 0.33 | 0.36 | 0.33 |
| 10 | 35464778 | rs1531550 | -0.352 | 0.764 | 0.851 | 0.821 | -0.007 | -0.024 | -0.024 | 0 | 0.15 | 0.09 | 0.2 | 0.26 | 0.18 | 0.13 |
| 10 | 35474421 | rs2505639 | 1.832 | 0.775 | 0.986 | 0.825 | 0 | -0.001 | -0.001 | 0.368 | 0.65 | 0.41 | 0.63 | 0.67 | 0.64 | 0.67 |
| 10 | 35474699 | rs4934540 | -2.031 | 0.792 | 0.503 | 0.818 | -0.001 | 0 | 0 | 0 | 0.33 | 0.3 | 0.35 | 0.31 | 0.36 | 0.33 |
| 10 | 35484949 | rs1057108 | -2.022 | 0.806 | 0.492 | 0.824 | -0.001 | 0 | 0 | 0 | 0.33 | 0.31 | 0.35 | 0.31 | 0.36 | 0.33 |
| 10 | 35496946 | rs1148247 | 2.301 | 0.626 | 0.253 | 0.931 | -0.119 | 0.003 | 0.003 | 0 | 0.45 | 0.02 | 0.38 | 0.32 | 0.43 | 0.42 |
| 10 | 35502533 | rs3740082 | -1.105 | 0.509 | 1.127 | 0.77 | 0.001 | 0.019 | 0.019 | 0 | 0.17 | 0.24 | 0.22 | 0.26 | 0.18 | 0.13 |
| 10 | 35526574 | rs7070384 | -1.474 | 0.703 | 0.986 | 0.715 | 0.001 | -0.052 | -0.052 | 0.023 | 0.18 | 0.4 | 0.13 | 0.07 | 0.16 | 0.2 |
| 10 | 35535597 | rs17500653 | -0.702 | 0.718 | 0.904 | 0.782 | -0.01 | 0.015 | 0.015 | 0 | 0.12 | 0.05 | 0.1 | 0.05 | 0.14 | 0.04 |
| 10 | 35535695 | rs12242110 | -1.711 | 0.794 | 1.183 | 0.913 | 0.008 | 0.008 | 0.008 | 0 | 0.28 | 0.37 | 0.33 | 0.32 | 0.32 | 0.3 |
| 10 | 35541026 | rs12777960 | -1.634 | 0.846 | 0.966 | 0.908 | 0.014 | 0.011 | 0.011 | 0 | 0.3 | 0.52 | 0.35 | 0.33 | 0.33 | 0.3 |
| 10 | 35549257 | rs3936503 | -1.865 | 0.856 | 0.975 | 0.917 | 0.013 | 0.012 | 0.012 | 0 | 0.3 | 0.54 | 0.34 | 0.33 | 0.32 | 0.3 |
| 10 | 35554054 | rs12261843 | -1.744 | 0.741 | 1.031 | 0.937 | 0.002 | 0.018 | 0.018 | 0 | 0.28 | 0.56 | 0.34 | 0.33 | 0.29 | 0.29 |
| 10 | 35554947 | rs7921820 | -1.023 | 0.354 | 0.842 | 0.864 | 0.001 | -0.002 | -0.002 | 0 | 0.21 | 0.24 | 0.26 | 0.26 | 0.2 | 0.27 |
| 10 | 35567127 | rs11010152 | -1.524 | 0.193 | 0.877 | 1.049 | -0.007 | 0 | 0 | 0 | 0.41 | 0.6 | 0.47 | 0.42 | 0.34 | 0.44 |
| 10 | 35577194 | rs4934744 | -1.527 | 0.192 | 0.876 | 1.061 | -0.007 | -0.001 | -0.001 | 0.437 | 0.41 | 0.6 | 0.47 | 0.42 | 0.34 | 0.44 |
| 10 | 35597715 | rs6481952 | -1.294 | 0.173 | 0.949 | 1.056 | 0.003 | 0.003 | 0.003 | 0 | 0.37 | 0.37 | 0.44 | 0.36 | 0.34 | 0.31 |
| 10 | 35641571 | rs2474519 | -1.504 | 0.146 | 1.173 | 1.086 | 0.003 | -0.002 | -0.002 | 0.605 | 0.39 | 0.42 | 0.45 | 0.42 | 0.35 | 0.32 |
| 10 | 35662640 | rs11813258 | -1.136 | 0.136 | 1.124 | 1.076 | 0.003 | -0.004 | -0.004 | 1.015 | 0.33 | 0.21 | 0.39 | 0.36 | 0.32 | 0.31 |
| 10 | 35671151 | rs2504352 | -1.746 | 0.119 | 1.501 | 1.084 | 0.005 | 0 | 0 | 0 | 0.41 | 0.65 | 0.47 | 0.42 | 0.35 | 0.37 |
| 10 | 35699170 | rs4934750 | -0.728 | 0.104 | 1.356 | 1.045 | -0.048 | -0.079 | -0.079 | 0 | 0.28 | 0.03 | 0.33 | 0.34 | 0.25 | 0.2 |
| 10 | 35718212 | rs3003981 | -1.712 | 0.056 | 1.451 | 1.04 | -0.004 | -0.001 | -0.001 | 0.294 | 0.41 | 0.62 | 0.47 | 0.42 | 0.35 | 0.32 |
| 10 | 35719339 | rs3013361 | -0.887 | 0.09 | 1.295 | 1.019 | -0.003 | 0.041 | 0.041 | 0 | 0.05 | 0.19 | 0.05 | 0.06 | 0.03 | 0.01 |
| 10 | 35724454 | rs10827503 | 1.963 | 0.01 | 0.674 | 1.207 | -0.134 | 0.099 | 0.099 | 0 | 0.43 | 0.02 | 0.35 | 0.23 | 0.51 | 0.36 |
| 10 | 35755345 | rs4934753 | -0.868 | 0.115 | 0.628 | 1.433 | 0.005 | 0 | 0 | 0 | 0.33 | 0.21 | 0.39 | 0.36 | 0.31 | 0.31 |
| 10 | 35815727 | rs10827512 | -0.108 | 0.052 | 0.608 | 1.522 | -0.026 | -0.083 | -0.083 | 0.008 | 0.12 | 0.01 | 0.14 | 0.34 | 0.13 | 0.25 |
| 10 | 35858665 | rs1043583 | -1.345 | 0.2 | 0.898 | 1.5 | -0.012 | 0.005 | 0.005 | 0 | 0.37 | 0.53 | 0.45 | 0.42 | 0.33 | 0.31 |
| 10 | 35870714 | rs12770412 | -1.079 | 0.173 | 0.79 | 1.68 | -0.001 | 0 | 0 | 0 | 0.36 | 0.41 | 0.47 | 0.45 | 0.32 | 0.31 |
| 10 | 35874895 | rs2474755 | 1.372 | 0.334 | 1.461 | 1.466 | 0.058 | 0.087 | 0.087 | 0 | 0.93 | 0.64 | 0.91 | 0.91 | 0.94 | 0.9 |
| 10 | 35878792 | rs7899248 | -1.657 | 0.436 | 0.976 | 1.588 | 0 | 0.057 | 0.057 | 0.393 | 0.27 | 0.44 | 0.33 | 0.5 | 0.28 | 0.45 |
| 10 | 35883547 | rs7094532 | -1.609 | 0.709 | 1.075 | 1.68 | 0.098 | 0.289 | 0.289 | 1.491 | 0.07 | 0.38 | 0.09 | 0.17 | 0.07 | 0.13 |
| 10 | 35887109 | rs1895119 | -2.056 | 0.668 | 1.184 | 1.646 | 0.061 | 0.234 | 0.234 | 1.688 | 0.09 | 0.48 | 0.1 | 0.16 | 0.07 | 0.13 |
| 10 | 35901715 | rs640827 | -1.452 | 0.862 | -0.095 | 1.279 | -0.045 | -0.097 | -0.097 | 0 | 0.23 | 0.07 | 0.29 | 0.33 | 0.28 | 0.33 |
| 10 | 35902836 | rs615271 | -1.148 | 0.817 | -0.031 | 1.391 | -0.04 | -0.109 | -0.109 | 1.549 | 0.2 | 0.03 | 0.27 | 0.32 | 0.24 | 0.3 |
| 10 | 35912391 | rs10827519 | -0.813 | 0.573 | 0.149 | 1.225 | 0.004 | -0.09 | -0.09 | 0 | 0.19 | 0.03 | 0.14 | 0.31 | 0.17 | 0.18 |
| 10 | 35913205 | rs618644 | 0.374 | 0.469 | 1.071 | 1.2 | 0.026 | 0.044 | 0.044 | 2.428 | 0.95 | 0.68 | 0.94 | 0.99 | 0.99 | 0.94 |
| 10 | 35915111 | rs619319 | 1.394 | 0.259 | 1.284 | 1.284 | 0.042 | -0.031 | -0.031 | 0 | 0.68 | 0.19 | 0.67 | 0.49 | 0.79 | 0.67 |
| 10 | 35919451 | rs10827522 | -1.274 | 0.313 | 1.536 | 1.172 | 0.001 | -0.002 | -0.002 | 0 | 0.26 | 0.3 | 0.2 | 0.35 | 0.21 | 0.22 |
| 10 | 35923186 | rs674991 | 1.297 | 0.32 | 1.266 | 1.122 | 0.04 | -0.034 | -0.034 | 0 | 0.68 | 0.18 | 0.66 | 0.5 | 0.79 | 0.66 |
| 10 | 35936753 | rs596642 | 1.454 | 0.314 | 1.263 | 1.152 | 0.047 | -0.042 | -0.042 | 0 | 0.66 | 0.18 | 0.64 | 0.5 | 0.77 | 0.66 |
| 10 | 35948710 | rs618443 | 1.849 | 0.489 | 1.49 | 1.019 | 0.038 | 0.137 | 0.137 | 2.417 | 0.88 | 0.54 | 0.83 | 0.85 | 0.87 | 0.85 |
| 10 | 35962887 | rs596815 | 1.421 | 0.441 | 1.125 | 0.868 | 0.029 | -0.017 | -0.017 | 0 | 0.6 | 0.25 | 0.55 | 0.51 | 0.66 | 0.59 |
| 10 | 35964183 | rs10764049 | -1.453 | 0.434 | 1.232 | 0.871 | -0.006 | 0.018 | 0.018 | 0 | 0.35 | 0.46 | 0.4 | 0.47 | 0.32 | 0.35 |
| 10 | 35973308 | rs661752 | 0.179 | 0.375 | 1.333 | 0.879 | 0.055 | 0.08 | 0.08 | 1.791 | 0.94 | 0.64 | 0.96 | 0.98 | 0.98 | 0.94 |
| 10 | 35983852 | rs612766 | 0.962 | 0.321 | 1.323 | 0.867 | 0.002 | 0.019 | 0.019 | 3.444 | 0.92 | 0.48 | 0.9 | 0.96 | 0.98 | 0.9 |
| 10 | 35991767 | rs641297 | 0.036 | 0.329 | 1.21 | 0.848 | 0.029 | 0.031 | 0.031 | 0.986 | 0.94 | 0.72 | 0.94 | 0.99 | 0.98 | 0.9 |
| 10 | 35993394 | rs2486509 | 1.496 | 0.362 | 0.485 | 0.909 | 0.027 | -0.025 | -0.025 | 8.664 | 0.63 | 0.27 | 0.62 | 0.55 | 0.74 | 0.61 |
| 10 | 35993853 | rs2486507 | 0.636 | 0.478 | 0.634 | 0.782 | 0.026 | 0.027 | 0.027 | 2.806 | 0.92 | 0.56 | 0.92 | 0.99 | 0.98 | 0.9 |
| 10 | 36000671 | rs654221 | -0.778 | 1.101 | -0.989 | 0.615 | -0.043 | -0.001 | -0.001 | 0 | 0.29 | 0.13 | 0.22 | 0.28 | 0.34 | 0.34 |
| 10 | 36003170 | rs686040 | 0.892 | 0.796 | -0.285 | 0.915 | -0.007 | 0.023 | 0.023 | 2.473 | 0.82 | 0.37 | 0.7 | 0.81 | 0.83 | 0.75 |
| 10 | 36007848 | rs10827533 | 0.638 | 1.047 | -1.099 | 0.647 | -0.01 | -0.011 | -0.011 | 0 | 0.53 | 0.19 | 0.49 | 0.46 | 0.55 | 0.47 |

| | | | | | | | | | | | | | | | | |
|---|---|---|---|---|---|---|---|---|---|---|---|---|---|---|---|---|
| 10 | 36011038 | rs603401 | 0.754 | 1.165 | -0.11 | 0.583 | -0.001 | 0.015 | 0.015 | 0 | 0.84 | 0.8 | 0.83 | 0.79 | 0.84 | 0.84 |
| 10 | 36017327 | rs586934 | 1.239 | 1.482 | 0.249 | 0.212 | -0.008 | 0.02 | 0.02 | 0 | 0.64 | 0.32 | 0.61 | 0.72 | 0.59 | 0.57 |
| 10 | 36017815 | rs599516 | 1.019 | 1.143 | 0.932 | 0.575 | 0.082 | 0.202 | 0.202 | 0.791 | 0.94 | 0.66 | 0.86 | 0.85 | 0.95 | 0.93 |
| 10 | 36026685 | rs2790254 | -0.555 | 1.221 | 0.329 | 0.36 | -0.001 | 0.023 | 0.023 | 0 | 0.12 | 0.08 | 0.08 | 0.05 | 0.11 | 0.09 |
| 10 | 36029626 | rs875655 | 1.221 | 1.118 | 0.213 | 0.079 | 0.007 | 0.029 | 0.029 | 0.702 | 0.53 | 0.38 | 0.56 | 0.6 | 0.57 | 0.47 |
| 10 | 36035048 | rs946558 | 0.264 | 1.103 | 0.777 | 0.229 | -0.011 | 0.049 | 0.049 | 0 | 0.84 | 0.76 | 0.78 | 0.71 | 0.88 | 0.82 |
| 10 | 36036100 | rs945495 | 0.599 | 0.876 | 0.74 | 0.144 | -0.031 | 0.074 | 0.074 | 0.619 | 0.82 | 0.52 | 0.75 | 0.73 | 0.88 | 0.84 |
| 10 | 36036780 | rs17511194 | -0.513 | 1.041 | 0.609 | 0.074 | -0.001 | -0.001 | -0.001 | 0 | 0.16 | 0.14 | 0.13 | 0.1 | 0.2 | 0.21 |
| 10 | 36037466 | rs12411410 | -0.373 | 1.156 | 0.37 | 0.163 | -0.04 | 0 | 0 | 0 | 0.11 | 0.03 | 0.1 | 0.11 | 0.15 | 0.18 |
| 10 | 36043507 | rs654218 | -0.336 | 1.665 | 0.825 | 0.671 | 0.019 | -0.019 | -0.019 | 0 | 0.22 | 0.27 | 0.26 | 0.19 | 0.29 | 0.16 |
| 10 | 36045994 | rs2152541 | 1.144 | 1.326 | 1.034 | 0.857 | 0.023 | 0.001 | 0.001 | 0 | 0.61 | 0.46 | 0.45 | 0.41 | 0.52 | 0.63 |
| 10 | 36046482 | rs587003 | 0.357 | 0.933 | 1.238 | 0.803 | -0.031 | 0.1 | 0.1 | 0.255 | 0.87 | 0.65 | 0.92 | 0.82 | 0.92 | 0.82 |
| 10 | 36057038 | rs7922611 | 1.946 | 0.653 | 0.529 | 0.293 | 0.017 | -0.001 | -0.001 | 0.882 | 0.37 | 0.32 | 0.35 | 0.39 | 0.28 | 0.35 |
| 10 | 36058844 | rs7899614 | 1.305 | 0.994 | 0.863 | 0.31 | 0.004 | 0.004 | 0.004 | 0 | 0.56 | 0.54 | 0.59 | 0.58 | 0.53 | 0.48 |
| 10 | 36059053 | rs2804818 | 1.374 | 0.992 | 0.913 | 0.293 | -0.002 | -0.002 | -0.002 | 0 | 0.55 | 0.43 | 0.57 | 0.58 | 0.53 | 0.48 |
| 10 | 36059240 | rs661100 | 1.322 | 0.952 | 0.841 | 0.277 | 0.003 | 0.003 | 0.003 | 0 | 0.56 | 0.56 | 0.6 | 0.58 | 0.53 | 0.48 |
| 10 | 36062442 | rs606815 | 1.071 | 0.956 | 0.815 | 0.385 | 0.007 | -0.004 | -0.004 | 0 | 0.67 | 0.63 | 0.69 | 0.7 | 0.65 | 0.58 |
| 10 | 36067367 | rs7897664 | 1.045 | 1.452 | 0.48 | 0.375 | -0.003 | -0.003 | -0.003 | 0.81 | 0.74 | 0.76 | 0.71 | 0.73 | 0.69 | 0.79 |
| 10 | 36067407 | rs2784750 | -0.016 | 1.468 | 0.443 | 0.399 | 0.012 | 0.052 | 0.052 | 0 | 0.88 | 0.82 | 0.81 | 0.75 | 0.86 | 0.87 |
| 10 | 36070546 | rs16936333 | 0.134 | 1.409 | 0.027 | 0.01 | -0.031 | -0.036 | -0.036 | 0 | 0.11 | 0.05 | 0.1 | 0.15 | 0.15 | 0.26 |
| 10 | 36071555 | rs2804814 | 0.008 | 1.039 | 0.424 | -0.077 | -0.035 | 0.04 | 0.04 | 0 | 0.12 | 0.02 | 0.09 | 0.02 | 0.17 | 0.08 |
| 10 | 36072829 | rs7906986 | -0.37 | 1.104 | 0.538 | 0.006 | -0.025 | 0.02 | 0.02 | 0.058 | 0.09 | 0.02 | 0.07 | 0.02 | 0.13 | 0.02 |
| 10 | 36074474 | rs4934765 | -2.191 | 0.972 | 0.828 | -0.101 | -0.011 | -0.075 | -0.075 | 0 | 0.25 | 0.43 | 0.17 | 0.08 | 0.23 | 0.14 |
| 10 | 36076774 | rs10827540 | 0.004 | 1.016 | 1.014 | -0.016 | -0.002 | -0.024 | -0.024 | 8.247 | 0.09 | 0.07 | 0.04 | 0 | 0.07 | 0.08 |
| 10 | 36079145 | rs10218989 | -0.221 | 1.008 | 1.014 | -0.019 | -0.006 | -0.063 | -0.063 | 0 | 0.1 | 0.18 | 0.05 | 0 | 0.07 | 0.08 |
| 10 | 36086651 | rs10827543 | -1.818 | 0.966 | 0.927 | -0.015 | -0.038 | -0.069 | -0.069 | 0.018 | 0.17 | 0.53 | 0.13 | 0.06 | 0.09 | 0.12 |
| 10 | 36101809 | rs945494 | -0.397 | 1.057 | 1.097 | -0.139 | -0.002 | -0.069 | -0.069 | 0.009 | 0.11 | 0.27 | 0.05 | 0 | 0.07 | 0.08 |
| 10 | 36104056 | rs11010301 | -1.111 | 1.039 | 0.995 | -0.147 | -0.013 | -0.092 | -0.092 | 0.052 | 0.13 | 0.44 | 0.08 | 0 | 0.07 | 0.08 |
| 10 | 36111597 | rs11596679 | -0.51 | 1.434 | 0.417 | 0.059 | -0.025 | 0.023 | 0.023 | 1.528 | 0.11 | 0.01 | 0.07 | 0.02 | 0.14 | 0.03 |
| 10 | 36116230 | rs16936389 | 0.685 | 1.38 | 0.583 | 0.313 | -0.049 | -0.043 | -0.043 | 2.463 | 0.11 | 0.01 | 0.19 | 0.21 | 0.12 | 0.16 |
| 10 | 36121546 | rs2767063 | -1.402 | 1.063 | 1.013 | 0.163 | -0.003 | 0.021 | 0.021 | 0 | 0.17 | 0.12 | 0.16 | 0.08 | 0.18 | 0.17 |
| 10 | 36121611 | rs2767062 | -1.856 | 1 | 1.447 | 0.136 | -0.011 | -0.069 | -0.069 | 0 | 0.21 | 0.59 | 0.21 | 0.08 | 0.18 | 0.17 |
| 10 | 36161439 | rs2247156 | -1.423 | 1.221 | 0.924 | 0.293 | -0.011 | 0.028 | 0.028 | 0 | 0.15 | 0.08 | 0.17 | 0.07 | 0.18 | 0.12 |
| 10 | 36171734 | rs601807 | -1.929 | 1.532 | 1.189 | 0.499 | 0.049 | 0.002 | 0.002 | 0 | 0.22 | 0.48 | 0.33 | 0.24 | 0.27 | 0.17 |
| 10 | 36176901 | rs10827561 | -1.631 | 1.444 | 0.123 | 0.537 | 0.011 | -0.002 | -0.002 | 0 | 0.21 | 0.39 | 0.27 | 0.18 | 0.22 | 0.17 |
| 10 | 36190616 | rs2804808 | 1.613 | 1.098 | 0.31 | 0.69 | 0.007 | 0.026 | 0.026 | 0.027 | 0.77 | 0.39 | 0.65 | 0.72 | 0.76 | 0.82 |
| 10 | 36191613 | rs708424 | -1.063 | 0.864 | -0.63 | 0.531 | -0.001 | 0.005 | 0.005 | 0 | 0.16 | 0.25 | 0.17 | 0.16 | 0.16 | 0.14 |
| 10 | 36198869 | rs2804805 | -0.686 | 0.495 | -1.642 | 0.335 | 0.014 | 0.005 | 0.005 | 0 | 0.16 | 0 | 0.1 | 0.06 | 0.13 | 0.22 |
| 10 | 36198981 | rs2767070 | -0.904 | 0.773 | -1.729 | 0.354 | -0.03 | -0.005 | -0.005 | 0 | 0.21 | 0.02 | 0.26 | 0.12 | 0.21 | 0.29 |
| 10 | 36201437 | rs11010324 | 0.577 | 0.994 | -1.556 | 0.403 | -0.021 | 0.07 | 0.07 | 0.558 | 0.45 | 0.18 | 0.35 | 0.32 | 0.46 | 0.31 |
| 10 | 36204758 | rs708431 | 0.633 | 0.634 | -0.586 | 0.286 | -0.014 | -0.034 | -0.034 | 0.749 | 0.7 | 0.48 | 0.79 | 0.91 | 0.76 | 0.67 |
| 10 | 36205466 | rs2797921 | -0.877 | 1.213 | -0.801 | 0.739 | -0.025 | -0.025 | -0.025 | 2.555 | 0.06 | 0 | 0.07 | 0.06 | 0.09 | 0.22 |
| 10 | 36206905 | rs708433 | 0.696 | 1.119 | -0.244 | 0.66 | -0.008 | -0.023 | -0.023 | 0.503 | 0.79 | 0.66 | 0.83 | 0.85 | 0.79 | 0.76 |
| 10 | 36209003 | rs10764062 | 1.176 | 2.296 | -0.246 | 1.718 | 0.001 | -0.01 | -0.01 | 0.421 | 0.57 | 0.45 | 0.47 | 0.31 | 0.54 | 0.43 |
| 10 | 36211761 | rs1343030 | 1.059 | 2.342 | -0.238 | 1.498 | -0.002 | 0.008 | 0.008 | 0.088 | 0.65 | 0.57 | 0.7 | 0.61 | 0.63 | 0.49 |
| 10 | 36215550 | rs10827563 | 1.476 | 2.426 | -0.132 | 1.71 | 0.011 | -0.005 | -0.005 | 2.087 | 0.58 | 0.5 | 0.56 | 0.37 | 0.5 | 0.37 |
| 10 | 36217248 | rs772072 | 0.196 | 2.188 | -0.039 | 1.833 | -0.011 | -0.016 | -0.016 | 0.032 | 0.84 | 0.91 | 0.87 | 0.86 | 0.8 | 0.75 |
| 10 | 36218364 | rs7069154 | 0.006 | 2.028 | -0.062 | 1.718 | -0.006 | -0.004 | -0.004 | 0 | 0.16 | 0.16 | 0.16 | 0.18 | 0.14 | 0.23 |
| 10 | 36218722 | rs772067 | 0.106 | 2.055 | -0.178 | 1.725 | -0.012 | -0.015 | -0.015 | 0.027 | 0.85 | 0.91 | 0.87 | 0.87 | 0.8 | 0.78 |
| 10 | 36219717 | rs1976962 | 1.275 | 2.03 | -0.112 | 1.756 | -0.004 | 0.006 | 0.006 | 0 | 0.62 | 0.6 | 0.66 | 0.56 | 0.61 | 0.48 |
| 10 | 36222010 | rs2696310 | 1.417 | 2.093 | -0.082 | 1.605 | 0 | -0.003 | -0.003 | 0.373 | 0.57 | 0.48 | 0.58 | 0.37 | 0.54 | 0.43 |
| 10 | 36227101 | rs2696309 | 1.157 | 2.693 | -0.311 | 1.771 | 0.009 | 0.036 | 0.036 | 0 | 0.75 | 0.66 | 0.68 | 0.53 | 0.69 | 0.67 |
| 10 | 36230317 | rs708438 | 0.901 | 1.792 | -0.233 | 2.018 | -0.001 | -0.002 | -0.002 | 0.461 | 0.8 | 0.8 | 0.85 | 0.8 | 0.77 | 0.74 |
| 10 | 36244805 | rs11594750 | -1.089 | 1.305 | -0.45 | 1.414 | -0.002 | 0.009 | 0.009 | 0 | 0.35 | 0.37 | 0.41 | 0.44 | 0.38 | 0.31 |
| 10 | 36244899 | rs7921289 | 1.154 | 0.888 | -0.873 | 1.195 | -0.053 | 0.043 | 0.043 | 1.22 | 0.46 | 0.15 | 0.39 | 0.33 | 0.48 | 0.44 |
| 10 | 36246259 | rs708412 | -0.087 | 0.71 | -0.132 | 1.202 | -0.011 | 0.096 | 0.096 | 0 | 0.91 | 0.76 | 0.84 | 0.78 | 0.94 | 0.83 |
| 10 | 36247208 | rs10764063 | -0.368 | 0.642 | 0.519 | 1.564 | 0.026 | 0.249 | 0.249 | 0.303 | 0.07 | 0.25 | 0.2 | 0.34 | 0.08 | 0.05 |
| 10 | 36253686 | rs771872 | 0.227 | 0.666 | 0.463 | 1.501 | -0.019 | 0.062 | 0.062 | 0 | 0.89 | 0.78 | 0.84 | 0.78 | 0.93 | 0.79 |
| 10 | 36262488 | rs708419 | 1.504 | 0.71 | -0.017 | 1.278 | -0.009 | 0.063 | 0.063 | 0.014 | 0.49 | 0.17 | 0.38 | 0.34 | 0.49 | 0.48 |
| 10 | 36277984 | rs11010349 | -0.182 | 1.184 | 0.02 | 1.284 | -0.022 | 0.009 | 0.009 | 0.005 | 0.07 | 0.03 | 0.2 | 0.34 | 0.12 | 0.06 |
| 10 | 36284854 | rs16936512 | -0.218 | 0.888 | 0.196 | 1.186 | 0.002 | 0.006 | 0.006 | 0 | 0.31 | 0.26 | 0.24 | 0.09 | 0.33 | 0.26 |
| 10 | 36296842 | rs16936535 | -0.127 | 0.623 | 0.783 | 1.355 | -0.009 | 0.023 | 0.023 | 0 | 0.09 | 0.12 | 0.15 | 0.22 | 0.07 | 0.18 |
| 10 | 36306640 | rs10740920 | 1.662 | 0.661 | 0.651 | 1.397 | -0.002 | 0.149 | 0.149 | 0.334 | 0.8 | 0.43 | 0.63 | 0.43 | 0.81 | 0.77 |
| 10 | 36306742 | rs12218305 | -0.763 | 0.671 | 0.287 | 1.37 | -0.001 | 0.04 | 0.04 | 0 | 0.08 | 0.08 | 0.18 | 0.33 | 0.09 | 0.05 |
| 10 | 36322229 | rs3904594 | 0.966 | 0.37 | -0.272 | 0.864 | 0.037 | 0.038 | 0.038 | 0.784 | 0.48 | 0.16 | 0.56 | 0.43 | 0.61 | 0.62 |
| 10 | 36328422 | rs11010375 | 0.605 | -0.105 | 0.431 | 0.884 | -0.011 | -0.016 | -0.016 | 0 | 0.58 | 0.54 | 0.66 | 0.7 | 0.68 | 0.82 |
| 10 | 36331363 | rs2891528 | -0.487 | -0.472 | 0.523 | 0.621 | -0.018 | -0.05 | -0.05 | 0 | 0.8 | 0.79 | 0.91 | 0.98 | 0.92 | 0.91 |
| 10 | 36333217 | rs12249188 | 0.117 | -0.476 | 0.555 | 0.664 | -0.019 | 0.155 | 0.155 | 0.018 | 0.1 | 0.32 | 0.1 | 0.29 | 0.06 | 0.19 |
| 10 | 36333349 | rs11527676 | 1.658 | -0.429 | 0.301 | 0.654 | 0.035 | 0.038 | 0.038 | 0.345 | 0.48 | 0.22 | 0.57 | 0.41 | 0.62 | 0.63 |

| | | | | | | | | | | | | | | | | |
|---|---|---|---|---|---|---|---|---|---|---|---|---|---|---|---|---|
| 10 | 36337678 | rs3851074 | 1.731 | -0.529 | 0.183 | 0.62 | 0.022 | 0.02 | 0.02 | 0.538 | 0.47 | 0.15 | 0.56 | 0.43 | 0.61 | 0.63 |
| 10 | 36344376 | rs10508828 | 0.25 | -0.474 | 0.144 | 0.598 | -0.006 | -0.061 | -0.061 | 0 | 0.14 | 0.04 | 0.18 | 0.26 | 0.14 | 0.08 |
| 10 | 36345362 | rs7920455 | 1.191 | -0.466 | 0.405 | 0.584 | -0.004 | -0.002 | -0.002 | 0.581 | 0.61 | 0.62 | 0.7 | 0.72 | 0.75 | 0.77 |
| 10 | 36346582 | rs12782605 | 1.878 | -0.445 | 0.298 | 0.531 | 0.006 | 0.008 | 0.008 | 1.036 | 0.46 | 0.12 | 0.54 | 0.43 | 0.6 | 0.62 |
| 10 | 36351747 | rs12146388 | -0.781 | -0.178 | 0.576 | 0.52 | 0.03 | 0.162 | 0.162 | 0.899 | 0.15 | 0.47 | 0.14 | 0.3 | 0.14 | 0.2 |
| 10 | 36359231 | rs7088865 | -1.076 | -0.259 | 0.565 | 0.491 | 0.009 | 0.114 | 0.114 | 1.221 | 0.17 | 0.53 | 0.15 | 0.3 | 0.16 | 0.21 |
| 10 | 36361666 | rs16936633 | -0.068 | -0.323 | 0.584 | 0.311 | -0.012 | 0.143 | 0.143 | 1.294 | 0.11 | 0.47 | 0.1 | 0.18 | 0.07 | 0.18 |
| 10 | 36373516 | rs7920214 | -0.201 | -0.269 | 0.095 | 0.296 | -0.006 | -0.081 | -0.081 | 0 | 0.86 | 0.97 | 0.82 | 0.65 | 0.86 | 0.91 |
| 10 | 36379503 | rs7906685 | 1.343 | -0.261 | 0.4 | 0.188 | 0.025 | 0.011 | 0.011 | 0 | 0.53 | 0.37 | 0.55 | 0.44 | 0.66 | 0.63 |
| 10 | 36383244 | rs16936653 | 0.038 | -0.281 | 0.555 | 0.059 | -0.02 | -0.062 | -0.062 | 0 | 0.22 | 0.23 | 0.12 | 0.01 | 0.12 | 0.08 |
| 10 | 36385519 | rs10827596 | 1.616 | -0.4 | -0.169 | -0.475 | -0.011 | 0.013 | 0.013 | 0.235 | 0.42 | 0.19 | 0.48 | 0.42 | 0.58 | 0.53 |
| 10 | 36395066 | rs16936674 | 0.499 | -0.54 | -0.473 | -0.559 | -0.03 | -0.114 | -0.114 | 0.496 | 0.21 | 0.32 | 0.12 | 0 | 0.13 | 0.06 |
| 10 | 36399558 | rs10827602 | 2.354 | 0.095 | -0.824 | -0.565 | -0.108 | -0.037 | -0.037 | 0.021 | 0.31 | 0.03 | 0.36 | 0.31 | 0.36 | 0.44 |
| 10 | 36401997 | rs12248963 | -0.074 | -0.8 | -0.18 | -0.419 | 0.001 | 0.014 | 0.014 | 0 | 0.13 | 0.18 | 0.14 | 0.2 | 0.14 | 0.21 |
| 10 | 36402390 | rs12249110 | 0.421 | -0.808 | -0.182 | -0.423 | -0.014 | 0.038 | 0.038 | 0.011 | 0.1 | 0.18 | 0.1 | 0.2 | 0.06 | 0.19 |
| 10 | 36403798 | rs991629 | -1.698 | -0.82 | -0.203 | -0.452 | 0.017 | -0.086 | -0.086 | 0 | 0.84 | 0.97 | 0.84 | 0.64 | 0.88 | 0.85 |
| 10 | 36405541 | rs915134 | 0.463 | -0.823 | -0.146 | -0.468 | -0.03 | -0.153 | -0.153 | 2.294 | 0.21 | 0.39 | 0.17 | 0 | 0.14 | 0.06 |
| 10 | 36406944 | rs10764081 | -1.493 | -0.527 | -0.331 | -0.312 | 0.017 | -0.034 | -0.034 | 0 | 0.84 | 0.89 | 0.83 | 0.64 | 0.88 | 0.85 |
| 10 | 36408181 | rs11010462 | -0.392 | -0.061 | -0.123 | -0.121 | 0 | 0.014 | 0.014 | 0 | 0.13 | 0.18 | 0.14 | 0.21 | 0.14 | 0.2 |
| 10 | 36411104 | rs994980 | 1.389 | -0.077 | -0.117 | -0.123 | -0.009 | 0.017 | 0.017 | 0.162 | 0.41 | 0.2 | 0.48 | 0.41 | 0.58 | 0.54 |
| 10 | 36414157 | rs7072062 | -0.321 | 0.825 | 0.485 | 0.123 | -0.014 | 0.045 | 0.045 | 0.022 | 0.09 | 0.18 | 0.1 | 0.21 | 0.07 | 0.19 |
| 10 | 36421016 | rs11010488 | -0.934 | 0.855 | 0.166 | 0.13 | 0.003 | 0.039 | 0.039 | 0 | 0.14 | 0.25 | 0.15 | 0.21 | 0.14 | 0.21 |
| 10 | 36455895 | rs315644 | 0.347 | 0.114 | -0.111 | -0.138 | -0.032 | -0.022 | -0.022 | 6.612 | 0.67 | 0.45 | 0.76 | 0.79 | 0.77 | 0.77 |
| 10 | 36461567 | rs1451193 | 1.328 | 0.535 | -0.782 | -0.084 | -0.096 | 0.012 | 0.012 | 0 | 0.39 | 0.13 | 0.4 | 0.36 | 0.49 | 0.53 |
| 10 | 36466891 | rs1413611 | 1.576 | 0.652 | -0.933 | -0.109 | -0.079 | -0.046 | -0.046 | 0 | 0.35 | 0.05 | 0.37 | 0.38 | 0.39 | 0.48 |
| 10 | 36467648 | rs315634 | -0.497 | 0.299 | -0.759 | -0.104 | -0.038 | 0.027 | 0.027 | 0 | 0.87 | 0.81 | 0.91 | 0.8 | 0.94 | 0.81 |
| 10 | 36469499 | rs7092191 | 0.42 | 1.064 | -1.245 | 0.118 | 0.003 | 0.024 | 0.024 | 0 | 0.46 | 0.22 | 0.51 | 0.42 | 0.52 | 0.55 |
| 10 | 36472611 | rs315632 | 0.364 | 0.756 | -1.522 | 0.451 | 0.004 | -0.014 | -0.014 | 0 | 0.17 | 0.1 | 0.21 | 0.4 | 0.14 | 0.17 |
| 10 | 36473044 | rs899865 | 0.296 | 1.086 | -1.594 | 0.146 | -0.087 | -0.035 | -0.035 | 0 | 0.36 | 0.05 | 0.37 | 0.35 | 0.4 | 0.5 |
| 10 | 36474705 | rs315631 | 1.187 | 0.915 | -0.704 | -0.277 | 0.013 | 0.056 | 0.056 | 1.126 | 0.7 | 0.23 | 0.69 | 0.95 | 0.73 | 0.85 |
| 10 | 36474822 | rs315630 | 1.011 | 1.252 | -0.145 | -0.16 | -0.002 | -0.004 | -0.004 | 0 | 0.89 | 0.87 | 0.85 | 0.95 | 0.89 | 0.94 |
| 10 | 36474879 | rs7078826 | 1.772 | 1.69 | 0.715 | 0.142 | 0.06 | -0.005 | -0.005 | 0.009 | 0.77 | 0.48 | 0.68 | 0.76 | 0.67 | 0.83 |
| 10 | 36484336 | rs315580 | 1.129 | 1.56 | 1.035 | -0.121 | 0.048 | -0.042 | -0.042 | 0 | 0.92 | 0.78 | 0.93 | 1 | 0.87 | 0.95 |
| 10 | 36490707 | rs11010560 | 1.849 | 1.529 | 0.689 | -0.261 | -0.023 | 0.061 | 0.061 | 4.023 | 0.69 | 0.18 | 0.72 | 0.85 | 0.64 | 0.72 |
| 10 | 36502489 | rs315620 | -1.287 | 1.351 | 0.939 | -0.263 | -0.004 | -0.031 | -0.031 | 0.248 | 0.16 | 0.16 | 0.09 | 0.04 | 0.17 | 0.12 |
| 10 | 36511451 | rs11010572 | -1.928 | 1.631 | 1.382 | -0.082 | 0.106 | -0.003 | -0.003 | 0.586 | 0.15 | 0.49 | 0.18 | 0.11 | 0.19 | 0.16 |
| 10 | 36514772 | rs11815387 | -1.616 | 1.797 | 1.027 | -0.205 | -0.02 | 0.01 | 0.01 | 0 | 0.1 | 0.05 | 0.09 | 0 | 0.19 | 0.07 |
| 10 | 36525137 | rs1219564 | 0.269 | 0.959 | 0.784 | 0.15 | 0.007 | -0.011 | -0.011 | 0 | 0.71 | 0.72 | 0.64 | 0.63 | 0.77 | 0.71 |
| 10 | 36526084 | rs1219563 | 0.945 | 0.806 | 1.456 | -0.156 | -0.019 | 0.003 | 0.003 | 0 | 0.61 | 0.41 | 0.57 | 0.63 | 0.67 | 0.63 |
| 10 | 36528096 | rs1219562 | 1.419 | 0.853 | 1.744 | -0.151 | 0.019 | -0.004 | -0.004 | 0 | 0.52 | 0.25 | 0.46 | 0.54 | 0.51 | 0.45 |
| 10 | 36529124 | rs7085738 | -0.929 | 0.832 | 1.885 | -0.148 | -0.011 | -0.055 | -0.055 | 0 | 0.1 | 0.26 | 0.06 | 0 | 0.09 | 0.08 |
| 10 | 36536174 | rs2598582 | 1.454 | 1.001 | 1.707 | -0.17 | 0.005 | -0.012 | -0.012 | 0 | 0.49 | 0.18 | 0.44 | 0.52 | 0.5 | 0.44 |
| 10 | 36537486 | rs4934799 | -0.389 | 0.936 | 1.838 | -0.098 | 0.004 | -0.01 | -0.01 | 0.048 | 0.28 | 0.25 | 0.39 | 0.45 | 0.23 | 0.38 |
| 10 | 36542535 | rs1219604 | 0.328 | 0.945 | 2.037 | -0.104 | 0.029 | 0.006 | 0.006 | 0.276 | 0.94 | 0.66 | 0.95 | 1 | 0.97 | 0.94 |
| 10 | 36547975 | rs1219591 | 1.01 | 0.959 | 1.949 | -0.101 | 0.019 | -0.028 | -0.028 | 0 | 0.91 | 0.59 | 0.92 | 0.98 | 0.95 | 0.93 |
| 10 | 36551131 | rs2122723 | -0.422 | 0.938 | 1.944 | -0.104 | 0.003 | -0.009 | -0.009 | 0 | 0.28 | 0.25 | 0.39 | 0.45 | 0.23 | 0.37 |
| 10 | 36553587 | rs1219582 | 0.853 | 0.965 | 1.584 | -0.104 | -0.001 | -0.001 | -0.001 | 0 | 0.63 | 0.34 | 0.53 | 0.52 | 0.72 | 0.56 |
| 10 | 36584286 | rs1597615 | -0.25 | 1.045 | 1.074 | 0.306 | 0.01 | -0.023 | -0.023 | 0.341 | 0.25 | 0.21 | 0.39 | 0.37 | 0.2 | 0.24 |
| 10 | 36587545 | rs2796780 | -0.663 | 1.09 | 1.084 | 0.294 | -0.008 | -0.027 | -0.027 | 0 | 0.31 | 0.22 | 0.45 | 0.36 | 0.34 | 0.29 |
| 10 | 36590152 | rs1219615 | -0.005 | 0.859 | 1.153 | 0.371 | -0.013 | 0.062 | 0.062 | 0 | 0.83 | 0.75 | 0.66 | 0.62 | 0.88 | 0.82 |
| 10 | 36592064 | rs1219610 | 1.372 | 0.936 | 1.169 | 0.249 | -0.004 | -0.01 | -0.01 | 0.906 | 0.65 | 0.66 | 0.47 | 0.48 | 0.63 | 0.57 |
| 10 | 36599534 | rs1668775 | -0.676 | 0.681 | 1.478 | 0.392 | -0.001 | -0.001 | -0.001 | 0 | 0.16 | 0.22 | 0.28 | 0.16 | 0.12 | 0.15 |
| 10 | 36600257 | rs1779874 | -0.315 | 0.657 | 1.698 | 0.39 | -0.017 | 0.007 | 0.007 | 0 | 0.14 | 0.25 | 0.25 | 0.14 | 0.1 | 0.14 |
| 10 | 36606260 | rs1857579 | -1.598 | 0.569 | 2.413 | 0.606 | -0.02 | 0.139 | 0.139 | 0 | 0.23 | 0.46 | 0.27 | 0.41 | 0.19 | 0.31 |
| 10 | 36612604 | rs12573241 | -0.884 | 0.562 | 2.485 | 0.822 | -0.003 | 0.055 | 0.055 | 0 | 0.16 | 0.31 | 0.2 | 0.21 | 0.15 | 0.21 |
| 10 | 36613554 | rs2800092 | -2.835 | 0.635 | 2.429 | 0.831 | -0.014 | 0.031 | 0.031 | 0.582 | 0.47 | 0.88 | 0.63 | 0.57 | 0.47 | 0.51 |
| 10 | 36614838 | rs713052 | -2.663 | 0.606 | 2.351 | 0.861 | 0.002 | 0.01 | 0.01 | 0.419 | 0.43 | 0.79 | 0.6 | 0.55 | 0.44 | 0.5 |
| 10 | 36615382 | rs10508843 | -0.771 | 0.621 | 2.157 | 0.871 | -0.004 | 0.042 | 0.042 | 0 | 0.15 | 0.25 | 0.19 | 0.21 | 0.16 | 0.21 |
| 10 | 36618434 | rs2796787 | -1.906 | 0.639 | 2.107 | 0.873 | 0.002 | 0.029 | 0.029 | 0 | 0.28 | 0.48 | 0.4 | 0.34 | 0.29 | 0.29 |
| 10 | 36620416 | rs1125556 | 0.017 | 0.56 | 2.007 | 0.896 | -0.001 | 0 | 0 | 0.388 | 0.13 | 0.17 | 0.25 | 0.14 | 0.1 | 0.14 |
| 10 | 36621986 | rs16936947 | -0.137 | 0.609 | 1.958 | 0.918 | -0.008 | 0.047 | 0.047 | 0 | 0.08 | 0.15 | 0.07 | 0.23 | 0.04 | 0.1 |
| 10 | 36626060 | rs12571499 | -0.915 | 0.687 | 1.957 | 0.718 | -0.005 | 0.026 | 0.026 | 0 | 0.14 | 0.18 | 0.18 | 0.21 | 0.15 | 0.21 |
| 10 | 36631915 | rs2796792 | -1.486 | 0.824 | 0.285 | 0.218 | -0.031 | -0.061 | -0.061 | 0.048 | 0.64 | 0.83 | 0.65 | 0.51 | 0.61 | 0.71 |
| 10 | 36632539 | rs7905625 | -0.144 | 0.85 | 0.373 | 0.211 | -0.001 | -0.003 | -0.003 | 1.164 | 0.15 | 0.12 | 0.17 | 0.2 | 0.17 | 0.22 |
| 10 | 36632635 | rs7905986 | -0.039 | 0.851 | 0.363 | 0.211 | -0.001 | -0.007 | -0.007 | 0 | 0.14 | 0.1 | 0.16 | 0.2 | 0.17 | 0.22 |
| 10 | 36633732 | rs1936561 | -1.591 | 1.179 | 0.266 | 0.139 | 0.029 | -0.054 | -0.054 | 0 | 0.66 | 0.8 | 0.69 | 0.51 | 0.74 | 0.74 |
| 10 | 36634124 | rs2796793 | -1.335 | 1.407 | -0.229 | 0.172 | -0.018 | -0.012 | -0.012 | 0 | 0.34 | 0.28 | 0.5 | 0.34 | 0.46 | 0.42 |
| 10 | 36638274 | rs1632588 | -0.883 | 0.999 | -0.029 | 0.346 | 0 | -0.003 | -0.003 | 0 | 0.26 | 0.27 | 0.42 | 0.34 | 0.26 | 0.33 |
| 10 | 36638425 | rs1631619 | -0.875 | 0.997 | -0.029 | 0.345 | 0 | -0.003 | -0.003 | 0 | 0.26 | 0.27 | 0.42 | 0.34 | 0.26 | 0.32 |

| | | | | | | | | | | | | | | | | |
|---|---|---|---|---|---|---|---|---|---|---|---|---|---|---|---|---|
| 10 | 36643454 | rs7916089 | 0.684 | 0.729 | 0.218 | -0.005 | -0.002 | 0.002 | 0.002 | 0 | 0.3 | 0.26 | 0.15 | 0.16 | 0.29 | 0.35 |
| 10 | 36644263 | rs2796806 | -0.268 | 0.873 | 0.42 | 0.203 | -0.025 | 0.008 | 0.008 | 0.011 | 0.16 | 0.27 | 0.25 | 0.14 | 0.09 | 0.11 |
| 10 | 36647717 | rs11010639 | 0.382 | 1.178 | -0.136 | 0.15 | 0 | -0.009 | -0.009 | 0 | 0.26 | 0.25 | 0.16 | 0.16 | 0.29 | 0.3 |
| 10 | 36648040 | rs1779876 | 0.038 | 0.89 | -0.439 | 0.525 | -0.003 | 0.005 | 0.005 | 0 | 0.11 | 0.16 | 0.24 | 0.14 | 0.08 | 0.1 |
| 10 | 36649186 | rs2796809 | -0.357 | 0.713 | -0.688 | 0.445 | 0.003 | -0.004 | -0.004 | 0 | 0.14 | 0.17 | 0.28 | 0.14 | 0.13 | 0.11 |
| 10 | 36651214 | rs7911097 | -0.078 | 0.816 | -0.821 | -0.027 | 0.009 | 0.028 | 0.028 | 0.269 | 0.48 | 0.4 | 0.36 | 0.2 | 0.43 | 0.51 |
| 10 | 36651245 | rs2800086 | -0.651 | 1.452 | -0.54 | 0.759 | 0.002 | -0.011 | -0.011 | 0 | 0.63 | 0.74 | 0.6 | 0.55 | 0.65 | 0.67 |
| 10 | 36651355 | rs7911503 | -0.637 | 1.171 | -0.57 | 0.829 | 0.007 | -0.002 | -0.002 | 0 | 0.59 | 0.72 | 0.59 | 0.46 | 0.65 | 0.7 |
| 10 | 36652055 | rs12415815 | 0.355 | 1.094 | -0.241 | 0.493 | -0.009 | -0.005 | -0.005 | 0 | 0.39 | 0.5 | 0.28 | 0.34 | 0.29 | 0.44 |
| 10 | 36654368 | rs2800089 | -2.005 | 0.973 | -0.319 | 0.796 | 0.012 | -0.013 | -0.013 | 0 | 0.76 | 0.86 | 0.77 | 0.66 | 0.81 | 0.81 |
| 10 | 36655226 | rs2796765 | -0.227 | 0.969 | -0.237 | 0.763 | -0.009 | 0.003 | 0.003 | 0 | 0.29 | 0.22 | 0.39 | 0.22 | 0.36 | 0.28 |
| 10 | 36659020 | rs2995470 | -2.079 | 0.946 | -0.229 | 0.807 | 0.018 | -0.026 | -0.026 | 0 | 0.76 | 0.89 | 0.77 | 0.64 | 0.81 | 0.81 |
| 10 | 36663024 | rs11010651 | 1.406 | 0.635 | -0.172 | 0.636 | -0.008 | -0.036 | -0.036 | 0.483 | 0.28 | 0.36 | 0.15 | 0.14 | 0.23 | 0.31 |
| 10 | 36666512 | rs2384598 | 1.091 | 0.312 | -0.347 | 0.657 | -0.015 | -0.018 | -0.018 | 0 | 0.14 | 0.09 | 0.18 | 0.16 | 0.18 | 0.12 |
| 10 | 36669344 | rs7898044 | -1.068 | 0.245 | -0.135 | 0.405 | 0.003 | -0.007 | -0.007 | 0 | 0.61 | 0.69 | 0.58 | 0.49 | 0.64 | 0.69 |
| 10 | 36686162 | rs7085749 | 1.383 | -0.751 | -0.255 | 0.34 | -0.004 | -0.005 | -0.005 | 0.702 | 0.26 | 0.25 | 0.11 | 0.14 | 0.16 | 0.29 |
| 10 | 36687210 | rs17627563 | 1.288 | -0.583 | -0.578 | 0.297 | -0.031 | 0.025 | 0.025 | 0 | 0.14 | 0.01 | 0.18 | 0.07 | 0.18 | 0.17 |
| 10 | 36688414 | rs17627587 | 0.491 | -0.506 | -0.247 | 0.292 | 0.026 | -0.007 | -0.007 | 0 | 0.09 | 0.12 | 0.11 | 0.09 | 0.16 | 0.09 |
| 10 | 36688528 | rs10827637 | -0.393 | -0.441 | -0.408 | 0.343 | 0.038 | 0.016 | 0.016 | 0.192 | 0.46 | 0.7 | 0.35 | 0.42 | 0.34 | 0.5 |
| 10 | 36700944 | rs10508845 | 0.88 | -0.65 | 0.112 | 0.194 | -0.014 | -0.025 | -0.025 | 0.047 | 0.24 | 0.26 | 0.11 | 0.12 | 0.17 | 0.29 |
| 10 | 36702570 | rs2796771 | -1.919 | -0.644 | -0.079 | 0.303 | -0.026 | -0.055 | -0.055 | 0.012 | 0.73 | 0.89 | 0.73 | 0.59 | 0.69 | 0.72 |
| 10 | 36714690 | rs4934813 | -0.86 | -0.493 | 0.007 | 0.346 | -0.001 | 0.002 | 0.002 | 0 | 0.46 | 0.6 | 0.58 | 0.38 | 0.48 | 0.39 |
| 10 | 36717061 | rs1857576 | 1.108 | -0.438 | -0.091 | 0.12 | -0.005 | -0.01 | -0.01 | 2.109 | 0.24 | 0.22 | 0.1 | 0.12 | 0.17 | 0.3 |
| 10 | 36718126 | rs12771570 | -0.433 | -0.425 | -0.328 | 0.357 | -0.054 | 0.048 | 0.048 | 0 | 0.33 | 0.18 | 0.42 | 0.18 | 0.45 | 0.32 |
| 10 | 36718300 | rs7095178 | 0.898 | -0.396 | -0.097 | 0.334 | 0 | -0.017 | -0.017 | 0 | 0.15 | 0.13 | 0.2 | 0.03 | 0.2 | 0.14 |
| 10 | 36725466 | rs2891552 | 1.111 | -0.392 | -0.16 | 0.329 | 0.001 | 0.004 | 0.004 | 0 | 0.22 | 0.22 | 0.13 | 0.12 | 0.2 | 0.29 |
| 10 | 36726471 | rs2245720 | 1.045 | -0.42 | -0.358 | 0.507 | -0.055 | -0.078 | -0.078 | 0.456 | 0.36 | 0.13 | 0.33 | 0.5 | 0.39 | 0.34 |
| 10 | 36730835 | rs2503048 | 1.108 | -0.446 | -0.37 | 0.451 | -0.043 | -0.09 | -0.09 | 0 | 0.34 | 0.11 | 0.39 | 0.45 | 0.35 | 0.3 |
| 10 | 36734783 | rs2503044 | 1.246 | -0.28 | -0.379 | 0.542 | -0.073 | -0.154 | -0.154 | 0 | 0.34 | 0.03 | 0.36 | 0.46 | 0.32 | 0.28 |
| 10 | 36737862 | rs10827641 | -0.352 | -0.039 | 0.612 | 0.532 | 0.022 | 0.019 | 0.019 | 2.166 | 0.05 | 0.29 | 0.03 | 0 | 0 | 0 |
| 10 | 36738839 | rs7087074 | -0.755 | 0.067 | 0.635 | 0.564 | 0.002 | 0.004 | 0.004 | 0 | 0.4 | 0.37 | 0.39 | 0.27 | 0.39 | 0.37 |
| 10 | 36746633 | rs2503040 | 1.746 | 0.071 | 0.402 | 0.793 | -0.073 | -0.093 | -0.093 | 0 | 0.26 | 0.05 | 0.26 | 0.34 | 0.3 | 0.26 |
| 10 | 36753404 | rs12772289 | 0.373 | 0.051 | 1.531 | 0.79 | -0.02 | 0.002 | 0.002 | 0 | 0.12 | 0.06 | 0.13 | 0.06 | 0.15 | 0.15 |
| 10 | 36755806 | rs12779724 | 0.373 | 0.065 | 1.531 | 0.79 | -0.028 | 0.002 | 0.002 | 0 | 0.12 | 0.05 | 0.13 | 0.06 | 0.18 | 0.15 |
| 10 | 36765473 | rs2503035 | 1.449 | -0.035 | 1.575 | 0.807 | 0.017 | 0.03 | 0.03 | 0 | 0.52 | 0.28 | 0.49 | 0.6 | 0.54 | 0.57 |
| 10 | 36788039 | rs2505629 | 1.401 | -0.351 | 1.71 | 0.725 | 0.029 | 0.027 | 0.027 | 0.007 | 0.55 | 0.23 | 0.63 | 0.59 | 0.59 | 0.62 |
| 10 | 36793742 | rs7911626 | -1.253 | -0.311 | 1.786 | 0.701 | -0.005 | -0.017 | -0.017 | 1.578 | 0.38 | 0.39 | 0.27 | 0.24 | 0.37 | 0.33 |
| 10 | 36800702 | rs2505562 | 1.378 | -0.263 | 1.693 | 0.728 | 0.017 | 0.016 | 0.016 | 0.009 | 0.56 | 0.3 | 0.64 | 0.59 | 0.59 | 0.65 |
| 10 | 36814121 | rs16937058 | -1.613 | 0.502 | 1.719 | 0.883 | 0.022 | 0.108 | 0.108 | 0.32 | 0.07 | 0.22 | 0.09 | 0.17 | 0.04 | 0.07 |
| 10 | 36832921 | rs11010701 | -0.054 | 0.532 | 1.496 | 0.723 | -0.001 | -0.008 | -0.008 | 0 | 0.12 | 0.1 | 0.1 | 0.05 | 0.18 | 0.08 |
| 10 | 36837670 | rs2483025 | 1.62 | 0.542 | 1.822 | 0.82 | 0.06 | 0.18 | 0.18 | 1.133 | 0.93 | 0.6 | 0.88 | 0.83 | 0.96 | 0.92 |
| 10 | 36842708 | rs17629831 | 0.177 | 0.562 | 1.393 | 0.768 | -0.035 | 0.042 | 0.042 | 0 | 0.11 | 0.02 | 0.08 | 0.05 | 0.18 | 0.08 |
| 10 | 36847928 | rs2095107 | 1.742 | 0.528 | 1.516 | 0.989 | -0.018 | -0.032 | -0.032 | 1.741 | 0.61 | 0.17 | 0.52 | 0.58 | 0.59 | 0.65 |
| 10 | 36848789 | rs2505583 | 1.388 | 0.589 | 1.651 | 1.016 | 0 | 0 | 0 | 0 | 0.64 | 0.34 | 0.54 | 0.58 | 0.59 | 0.65 |
| 10 | 36855630 | rs4934823 | 0.078 | 0.633 | 1.616 | 1.022 | -0.012 | -0.025 | -0.025 | 0.619 | 0.12 | 0.07 | 0.2 | 0.2 | 0.13 | 0.15 |
| 10 | 36856115 | rs2503066 | 0.394 | 0.65 | 1.645 | 1.029 | 0.007 | 0.042 | 0.042 | 0 | 0.85 | 0.79 | 0.72 | 0.62 | 0.83 | 0.78 |
| 10 | 36856595 | rs2505590 | -0.031 | 0.653 | 1.647 | 1.03 | 0.003 | 0.021 | 0.021 | 0 | 0.87 | 0.8 | 0.79 | 0.8 | 0.87 | 0.85 |
| 10 | 36858817 | rs3006958 | 1.616 | 0.686 | 1.353 | 0.908 | -0.014 | -0.026 | -0.026 | 0 | 0.62 | 0.21 | 0.53 | 0.58 | 0.59 | 0.65 |
| 10 | 36862892 | rs1332770 | -1.066 | 0.669 | 1.371 | 0.39 | 0.002 | -0.105 | -0.105 | 0 | 0.2 | 0.41 | 0.17 | 0.05 | 0.24 | 0.13 |
| 10 | 36877054 | rs7082532 | -1.08 | 0.701 | 0.543 | 0.432 | 0.003 | -0.087 | -0.087 | 0 | 0.21 | 0.43 | 0.18 | 0.1 | 0.25 | 0.14 |
| 10 | 36885921 | rs1360573 | -1.242 | 0.508 | 0.37 | 0.36 | -0.003 | -0.041 | -0.041 | 0 | 0.23 | 0.28 | 0.15 | 0.11 | 0.24 | 0.16 |
| 10 | 36886920 | rs2503032 | -0.909 | 0.704 | -0.322 | 0.524 | -0.011 | -0.007 | -0.007 | 0 | 0.28 | 0.12 | 0.26 | 0.38 | 0.28 | 0.22 |
| 10 | 36887434 | rs11010724 | -0.059 | 0.834 | -0.457 | 0.975 | 0.001 | -0.008 | -0.008 | 0 | 0.09 | 0.08 | 0.12 | 0.13 | 0.12 | 0.28 |
| 10 | 36890536 | rs4934826 | 0.524 | 1.184 | 0.594 | 0.85 | -0.004 | -0.008 | -0.008 | 0.25 | 0.67 | 0.57 | 0.68 | 0.61 | 0.67 | 0.74 |
| 10 | 36890663 | rs2483003 | 0.393 | 1.195 | 0.43 | 0.867 | -0.057 | 0.01 | 0.01 | 0.756 | 0.13 | 0.01 | 0.18 | 0.07 | 0.19 | 0.11 |
| 10 | 36891028 | rs1332772 | 0.937 | 1.649 | 0.405 | 0.826 | -0.005 | 0.002 | 0.002 | 0 | 0.66 | 0.71 | 0.67 | 0.59 | 0.62 | 0.72 |
| 10 | 36892217 | rs1332773 | 1.664 | 1.775 | 0.226 | 0.651 | 0.006 | 0.005 | 0.005 | 0 | 0.36 | 0.33 | 0.32 | 0.32 | 0.3 | 0.21 |
| 10 | 36893210 | rs7082117 | 0.591 | 1.771 | 0.204 | 0.657 | 0.007 | -0.017 | -0.017 | 0 | 0.18 | 0.11 | 0.1 | 0.32 | 0.15 | 0.13 |
| 10 | 36895284 | rs10827651 | 1.583 | 1.617 | 0.469 | 0.643 | 0.01 | 0.008 | 0.008 | 0 | 0.35 | 0.27 | 0.32 | 0.32 | 0.3 | 0.21 |
| 10 | 36896437 | rs11010727 | 1.503 | 1.493 | 0.44 | 0.683 | -0.011 | -0.009 | -0.009 | 0 | 0.37 | 0.4 | 0.32 | 0.31 | 0.29 | 0.21 |
| 10 | 36897650 | rs12265071 | 1.688 | 1.227 | 0.237 | 0.762 | 0.012 | 0.022 | 0.022 | 0 | 0.33 | 0.29 | 0.23 | 0.13 | 0.24 | 0.16 |
| 10 | 36898886 | rs11593196 | 0.975 | 1.092 | -0.108 | 0.811 | 0.011 | -0.021 | -0.021 | 0.201 | 0.16 | 0.1 | 0.08 | 0.32 | 0.12 | 0.12 |
| 10 | 36901232 | rs944733 | 1.21 | 1.629 | 0.187 | 0.819 | -0.002 | -0.033 | -0.033 | 0.332 | 0.34 | 0.43 | 0.25 | 0.13 | 0.28 | 0.17 |
| 10 | 36905424 | rs4934574 | 0.313 | 1.566 | 0.095 | 0.866 | -0.001 | -0.013 | -0.013 | 0 | 0.13 | 0.08 | 0.22 | 0.14 | 0.16 | 0.33 |
| 10 | 36909064 | rs2990536 | 0.326 | 1.647 | 0.042 | 0.91 | 0.002 | 0.067 | 0.067 | 0 | 0.68 | 0.55 | 0.54 | 0.54 | 0.65 | 0.45 |
| 10 | 36911213 | rs7067849 | -0.523 | 1.693 | -0.311 | 0.934 | 0 | -0.002 | -0.002 | 0 | 0.29 | 0.28 | 0.32 | 0.28 | 0.29 | 0.51 |
| 10 | 36920321 | rs10827678 | 1.847 | 0.977 | -0.826 | 0.257 | 0 | 0 | 0 | 0 | 0.2 | 0.2 | 0.19 | 0.14 | 0.18 | 0.12 |
| 10 | 36921049 | rs885161 | -0.389 | 1.039 | -1.255 | 0.569 | -0.001 | 0.003 | 0.003 | 0 | 0.29 | 0.28 | 0.32 | 0.28 | 0.29 | 0.51 |
| 10 | 36922991 | rs16937197 | 0.965 | 0.959 | -1.623 | 0.765 | 0 | -0.006 | -0.006 | 0 | 0.14 | 0.1 | 0.1 | 0.14 | 0.13 | 0.18 |

| | | | | | | | | | | | | | | | | |
|---|---|---|---|---|---|---|---|---|---|---|---|---|---|---|---|---|
| 10 | 36927691 | rs3006968 | -0.491 | 0.79 | -1.603 | 0.713 | -0.045 | -0.067 | -0.067 | 0 | 0.37 | 0.14 | 0.39 | 0.58 | 0.43 | 0.33 |
| 10 | 36949132 | rs2782619 | 0.096 | -0.007 | -1.048 | 0.276 | 0.007 | 0.025 | 0.025 | 0 | 0.64 | 0.77 | 0.67 | 0.77 | 0.67 | 0.75 |
| 10 | 36951018 | rs2782618 | 0.373 | -0.341 | -0.921 | 0.06 | -0.032 | 0.003 | 0.003 | 0 | 0.37 | 0.5 | 0.38 | 0.65 | 0.24 | 0.42 |
| 10 | 36952850 | rs2779332 | 0.057 | -1.125 | -0.489 | -0.006 | 0.008 | -0.016 | -0.016 | 4.999 | 0.72 | 0.5 | 0.71 | 0.72 | 0.74 | 0.76 |
| 10 | 36955908 | rs2483032 | -0.695 | -1.17 | -0.588 | 0.066 | 0.006 | -0.02 | -0.02 | 0 | 0.8 | 0.74 | 0.76 | 0.86 | 0.79 | 0.86 |
| 10 | 36956358 | rs2779331 | 0.87 | -1.253 | -0.706 | 0.112 | -0.005 | 0.004 | 0.004 | 0 | 0.28 | 0.26 | 0.33 | 0.51 | 0.19 | 0.31 |
| 10 | 36964639 | rs884263 | 0.341 | -0.747 | -0.645 | -0.262 | 0.04 | -0.063 | -0.063 | 0 | 0.34 | 0.55 | 0.37 | 0.13 | 0.49 | 0.38 |
| 10 | 36969777 | rs1815507 | -0.846 | -0.77 | -0.93 | 0.027 | -0.033 | -0.039 | -0.039 | 0 | 0.7 | 0.92 | 0.6 | 0.69 | 0.69 | 0.71 |
| 10 | 36978306 | rs1412702 | 0.383 | -1.022 | -0.562 | 0.056 | -0.027 | 0.016 | 0.016 | 0 | 0.36 | 0.38 | 0.22 | 0.56 | 0.21 | 0.33 |
| 10 | 36978385 | rs1412701 | 0.94 | -0.645 | -0.508 | -0.113 | -0.022 | 0.019 | 0.019 | 0 | 0.25 | 0.26 | 0.3 | 0.09 | 0.44 | 0.28 |
| 10 | 36978597 | rs7068831 | 0.905 | -0.65 | -0.505 | -0.116 | -0.007 | 0.003 | 0.003 | 0 | 0.25 | 0.29 | 0.3 | 0.09 | 0.44 | 0.28 |
| 10 | 37004038 | rs4442492 | 0.81 | -0.718 | -0.247 | -0.305 | 0.03 | -0.048 | -0.048 | 0 | 0.25 | 0.34 | 0.26 | 0.05 | 0.43 | 0.25 |
| 10 | 37010297 | rs7918988 | 0.837 | -1.348 | -0.205 | -0.39 | 0.029 | -0.036 | -0.036 | 0 | 0.27 | 0.18 | 0.21 | 0.6 | 0.17 | 0.34 |
| 10 | 37011209 | rs2782614 | -1.108 | -1.441 | -0.194 | -0.515 | -0.026 | -0.032 | -0.032 | 0 | 0.69 | 0.91 | 0.6 | 0.69 | 0.7 | 0.71 |
| 10 | 37011284 | rs1537325 | 1.071 | -1.275 | -0.017 | -0.574 | -0.017 | 0.011 | 0.011 | 0 | 0.24 | 0.25 | 0.25 | 0.06 | 0.43 | 0.25 |
| 10 | 37014308 | rs16937227 | -0.12 | -1.402 | 0.674 | -0.498 | 0.031 | 0.001 | 0.001 | 0 | 0.07 | 0.32 | 0.07 | 0.03 | 0.05 | 0.11 |
| 10 | 37030676 | rs10437422 | -0.881 | -1.293 | -0.07 | -0.31 | -0.012 | -0.034 | -0.034 | 0.085 | 0.69 | 0.89 | 0.6 | 0.68 | 0.7 | 0.68 |
| 10 | 37035679 | rs10827697 | -0.765 | -1.286 | 0.045 | -0.305 | 0 | -0.006 | -0.006 | 0 | 0.68 | 0.79 | 0.57 | 0.68 | 0.7 | 0.68 |
| 10 | 37046656 | rs2027129 | 0.233 | -1.444 | -0.253 | -0.485 | 0.009 | -0.052 | -0.052 | 0 | 0.39 | 0.47 | 0.33 | 0.08 | 0.52 | 0.34 |
| 10 | 37047826 | rs7098998 | 1.433 | -1.962 | -0.517 | -0.022 | 0.031 | -0.089 | -0.089 | 0.625 | 0.18 | 0.03 | 0.13 | 0.35 | 0.1 | 0.22 |
| 10 | 37048430 | rs7907995 | -1.133 | -1.517 | -0.558 | -0.085 | -0.031 | -0.024 | -0.024 | 0.038 | 0.66 | 0.88 | 0.51 | 0.68 | 0.65 | 0.66 |
| 10 | 37049251 | rs2891569 | -0.079 | -1.239 | -0.316 | -0.364 | -0.026 | 0.062 | 0.062 | 0 | 0.12 | 0.33 | 0.12 | 0.24 | 0.07 | 0.12 |
| 10 | 37053548 | rs10466051 | 0.207 | -0.902 | -0.421 | -0.779 | 0.017 | -0.063 | -0.063 | 1.485 | 0.37 | 0.46 | 0.25 | 0.08 | 0.48 | 0.32 |
| 10 | 37056102 | rs10827707 | 0.173 | -0.997 | -0.175 | -0.53 | 0.027 | -0.036 | -0.036 | 0 | 0.14 | 0.41 | 0.11 | 0.04 | 0.14 | 0.11 |
| 10 | 37060221 | rs10827721 | 0.173 | -0.997 | -0.175 | -0.53 | 0.027 | -0.036 | -0.036 | 0.057 | 0.14 | 0.41 | 0.11 | 0.04 | 0.13 | 0.11 |
| 10 | 37062991 | rs2804974 | 0.758 | -1.27 | -0.729 | -0.703 | -0.097 | 0.166 | 0.166 | 0 | 0.4 | 0.06 | 0.34 | 0.13 | 0.54 | 0.3 |
| 10 | 37063692 | rs2185863 | 0.152 | -1.291 | -0.655 | -0.679 | -0.004 | -0.009 | -0.009 | 0 | 0.63 | 0.43 | 0.57 | 0.75 | 0.66 | 0.6 |
| 10 | 37066341 | rs11010872 | -0.416 | -1.238 | -0.35 | -0.449 | -0.039 | 0.02 | 0.02 | 0 | 0.31 | 0.48 | 0.25 | 0.66 | 0.2 | 0.39 |
| 10 | 37069380 | rs10508853 | -0.278 | -1.584 | 0.413 | -0.712 | -0.031 | -0.097 | -0.097 | 0 | 0.14 | 0.55 | 0.12 | 0 | 0.08 | 0.03 |
| 10 | 37070799 | rs10508855 | 0.878 | -1.538 | -0.241 | -0.587 | 0.06 | -0.096 | -0.096 | 0 | 0.17 | 0.03 | 0.13 | 0.63 | 0.06 | 0.28 |
| 10 | 37083125 | rs1573156 | -0.667 | -1.14 | 0.056 | -1.015 | 0.009 | -0.064 | -0.064 | 0 | 0.29 | 0.45 | 0.3 | 0.09 | 0.28 | 0.18 |
| 10 | 37083715 | rs16937276 | 0.654 | -1.122 | -0.108 | -1.011 | -0.009 | -0.04 | -0.04 | 0 | 0.09 | 0.14 | 0.06 | 0 | 0.07 | 0.03 |
| 10 | 37083941 | rs2782637 | 0.752 | -0.897 | -0.23 | -0.884 | -0.001 | -0.002 | -0.002 | 0.485 | 0.15 | 0.19 | 0.2 | 0.07 | 0.15 | 0.07 |
| 10 | 37084210 | rs2782636 | -1.299 | -0.524 | -0.87 | -1.119 | -0.102 | 0.042 | 0.042 | 0 | 0.78 | 0.95 | 0.82 | 0.87 | 0.59 | 0.73 |
| 10 | 37093411 | rs4934833 | -0.369 | -0.679 | -0.382 | -1.147 | 0.007 | -0.022 | -0.022 | 0 | 0.61 | 0.48 | 0.59 | 0.77 | 0.46 | 0.64 |
| 10 | 37096338 | rs12415110 | 1.205 | -0.991 | -0.605 | -1.129 | 0.057 | -0.093 | -0.093 | 0 | 0.18 | 0.04 | 0.12 | 0.63 | 0.07 | 0.29 |
| 10 | 37097830 | rs1855640 | -0.938 | -0.855 | -0.351 | -1.136 | -0.06 | 0.059 | 0.059 | 0 | 0.66 | 0.83 | 0.65 | 0.8 | 0.52 | 0.72 |
| 10 | 37099723 | rs12571505 | 0.572 | -0.987 | -0.39 | -1.237 | -0.001 | -0.003 | -0.003 | 0.184 | 0.33 | 0.28 | 0.4 | 0.14 | 0.32 | 0.33 |
| 10 | 37100768 | rs2490846 | -0.481 | -0.345 | -0.12 | -1.003 | 0.021 | -0.035 | -0.035 | 0 | 0.64 | 0.55 | 0.62 | 0.8 | 0.52 | 0.71 |
| 10 | 37105314 | rs10827730 | -0.373 | -0.463 | -0.201 | -0.982 | 0.005 | -0.024 | -0.024 | 0 | 0.6 | 0.52 | 0.57 | 0.77 | 0.46 | 0.63 |
| 10 | 37107000 | rs12257945 | -0.382 | -0.449 | -0.25 | -0.976 | 0.007 | -0.032 | -0.032 | 0 | 0.6 | 0.52 | 0.57 | 0.77 | 0.46 | 0.65 |
| 10 | 37122315 | rs1777589 | 0.47 | -0.152 | -0.426 | -1 | -0.034 | 0.087 | 0.087 | 0.86 | 0.4 | 0.16 | 0.36 | 0.2 | 0.45 | 0.29 |
| 10 | 37130138 | rs1192701 | -0.273 | -0.63 | 0.302 | -1.106 | -0.035 | 0.039 | 0.039 | 0 | 0.74 | 0.58 | 0.81 | 0.34 | 0.83 | 0.6 |
| 10 | 37142348 | rs12258125 | 0.185 | -0.61 | 0.499 | -1.127 | 0.016 | -0.056 | -0.056 | 0 | 0.33 | 0.5 | 0.48 | 0.13 | 0.38 | 0.32 |
| 10 | 37146971 | rs11010918 | 1.315 | -0.674 | 0.331 | -1.031 | -0.024 | -0.008 | -0.008 | 0 | 0.05 | 0.01 | 0.04 | 0.04 | 0.05 | 0.1 |
| 10 | 37155513 | rs11010921 | 1.694 | -0.663 | 0.044 | -1.281 | -0.096 | 0.05 | 0.05 | 0 | 0.19 | 0.03 | 0.15 | 0.07 | 0.29 | 0.19 |
| 10 | 37158006 | rs1192658 | -0.733 | -0.927 | 0.782 | -0.631 | 0.045 | -0.066 | -0.066 | 0.938 | 0.82 | 0.93 | 0.89 | 0.39 | 0.92 | 0.73 |
| 10 | 37162907 | rs1192655 | -0.738 | -0.901 | 0.656 | -0.619 | 0.044 | -0.065 | -0.065 | 0.059 | 0.82 | 0.93 | 0.89 | 0.39 | 0.92 | 0.73 |
| 10 | 37169295 | rs1192691 | -0.747 | -0.922 | 0.636 | -0.649 | 0.002 | 0.002 | 0.002 | 0 | 0.83 | 0.86 | 0.89 | 0.39 | 0.92 | 0.73 |
| 10 | 37170407 | rs1192690 | 1.186 | -0.853 | 0.507 | -0.813 | -0.028 | 0.08 | 0.08 | 0.161 | 0.43 | 0.22 | 0.36 | 0.2 | 0.47 | 0.29 |
| 10 | 37173697 | rs12269644 | 0.093 | -0.805 | 0.63 | -0.827 | -0.001 | 0 | 0 | 0 | 0.32 | 0.3 | 0.44 | 0.14 | 0.37 | 0.34 |
| 10 | 37175719 | rs1192673 | 1.074 | -0.785 | 0.262 | -0.863 | -0.03 | 0.087 | 0.087 | 0.15 | 0.43 | 0.21 | 0.36 | 0.2 | 0.46 | 0.28 |
| 10 | 37188470 | rs2488750 | -0.664 | -1.015 | 0.88 | -0.267 | 0.024 | -0.034 | -0.034 | 0 | 0.22 | 0.13 | 0.23 | 0.7 | 0.13 | 0.41 |
| 10 | 37194703 | rs2488663 | 0.137 | -1.03 | 0.87 | -0.262 | 0.053 | -0.085 | -0.085 | 0.014 | 0.17 | 0.04 | 0.18 | 0.67 | 0.08 | 0.3 |
| 10 | 37203527 | rs2504401 | -0.702 | -1.025 | 0.875 | -0.267 | 0.024 | -0.034 | -0.034 | 0.129 | 0.22 | 0.13 | 0.23 | 0.7 | 0.13 | 0.41 |
| 10 | 37207701 | rs1192644 | 1.447 | -1.087 | 1.063 | -0.417 | -0.027 | -0.009 | -0.009 | 1.39 | 0.63 | 0.43 | 0.61 | 0.3 | 0.74 | 0.56 |
| 10 | 37213409 | rs1192650 | 1.45 | -1.05 | 1.076 | -0.403 | -0.028 | -0.009 | -0.009 | 0.063 | 0.63 | 0.43 | 0.62 | 0.3 | 0.75 | 0.56 |
| 10 | 37214419 | rs2504418 | 0.297 | -1.059 | 0.859 | -0.396 | 0.049 | -0.106 | -0.106 | 1.752 | 0.16 | 0.03 | 0.17 | 0.66 | 0.08 | 0.28 |
| 10 | 37219159 | rs1192635 | 1.503 | -1.055 | 0.775 | -0.404 | -0.007 | -0.002 | -0.002 | 0 | 0.63 | 0.33 | 0.6 | 0.3 | 0.74 | 0.56 |
| 10 | 37222063 | rs2488736 | -0.069 | -0.98 | -0.139 | -0.139 | 0.06 | -0.085 | -0.085 | 0.039 | 0.18 | 0.05 | 0.18 | 0.67 | 0.08 | 0.29 |
| 10 | 37231470 | rs1405639 | 0.536 | -0.58 | 0.471 | -0.401 | -0.006 | -0.093 | -0.093 | 0 | 0.86 | 0.69 | 0.88 | 1 | 0.87 | 0.97 |
| 10 | 37236853 | rs2488741 | 0.151 | -0.949 | -0.316 | 0.171 | 0.047 | -0.086 | -0.086 | 6.054 | 0.16 | 0.04 | 0.09 | 0.56 | 0.08 | 0.3 |
| 10 | 37239565 | rs10764116 | 1.661 | -1.221 | -0.343 | 0.179 | 0.091 | -0.023 | -0.023 | 0 | 0.62 | 0.15 | 0.73 | 0.35 | 0.77 | 0.54 |
| 10 | 37245609 | rs2995757 | 0.334 | -1.2 | -0.315 | 0.212 | 0.045 | -0.08 | -0.08 | 0 | 0.16 | 0.04 | 0.09 | 0.56 | 0.08 | 0.3 |
| 10 | 37245706 | rs1830206 | 1.558 | -1.173 | -0.393 | 0.184 | 0.091 | -0.021 | -0.021 | 0 | 0.62 | 0.15 | 0.73 | 0.35 | 0.77 | 0.54 |
| 10 | 37250388 | rs10827747 | -0.377 | -0.84 | 0.019 | 0.152 | 0.017 | -0.018 | -0.018 | 0 | 0.08 | 0.2 | 0.07 | 0.03 | 0.07 | 0.12 |
| 10 | 37254098 | rs1852491 | -0.124 | -0.541 | -0.251 | 0.198 | 0.02 | -0.065 | -0.065 | 0.048 | 0.19 | 0.07 | 0.15 | 0.57 | 0.15 | 0.31 |
| 10 | 37256635 | rs2488747 | 1.276 | -0.512 | -0.372 | 0.019 | 0.039 | 0.04 | 0.04 | 0.256 | 0.59 | 0.24 | 0.69 | 0.32 | 0.67 | 0.51 |
| 10 | 37267174 | rs16905836 | -0.31 | -0.222 | -0.651 | 0.413 | 0.03 | -0.113 | -0.113 | 0.669 | 0.2 | 0.06 | 0.14 | 0.58 | 0.13 | 0.31 |

| | | | | | | | | | | | | | | | | |
|---|---|---|---|---|---|---|---|---|---|---|---|---|---|---|---|---|
| 10 | 37274522 | rs12358845 | 0.148 | 0.152 | -0.5 | 0.281 | -0.034 | -0.001 | -0.001 | 0 | 0.06 | 0 | 0.04 | 0.04 | 0.09 | 0.13 |
| 10 | 37278930 | rs2505150 | -1.239 | 0.432 | -0.97 | -0.398 | 0.001 | 0.035 | 0.035 | 0.335 | 0.69 | 0.83 | 0.69 | 0.79 | 0.68 | 0.78 |
| 10 | 37280354 | rs7358066 | 0.982 | 0.426 | -0.736 | -0.796 | -0.008 | 0.007 | 0.007 | 0 | 0.24 | 0.2 | 0.36 | 0.11 | 0.32 | 0.27 |
| 10 | 37280842 | rs2504408 | -0.472 | 0.227 | -0.699 | -0.992 | -0.005 | 0.112 | 0.112 | 0.793 | 0.49 | 0.3 | 0.55 | 0.75 | 0.51 | 0.62 |
| 10 | 37283349 | rs12416632 | 0.285 | 0.143 | -0.77 | -0.849 | 0.035 | -0.153 | -0.153 | 0 | 0.24 | 0.01 | 0.17 | 0.56 | 0.17 | 0.34 |
| 10 | 37286763 | rs12773606 | 1.158 | 0.389 | -0.48 | -0.941 | -0.034 | -0.002 | -0.002 | 0 | 0.06 | 0 | 0.04 | 0.04 | 0.1 | 0.14 |
| 10 | 37287260 | rs11010985 | -0.547 | 0.326 | 0.144 | -0.966 | 0.013 | -0.058 | -0.058 | 0 | 0.17 | 0.31 | 0.11 | 0.05 | 0.16 | 0.16 |
| 10 | 37296499 | rs11010990 | 1.126 | 0.434 | 0.123 | -0.896 | -0.034 | -0.002 | -0.002 | 0 | 0.06 | 0 | 0.04 | 0.05 | 0.1 | 0.14 |
| 10 | 37300026 | rs2459418 | -0.133 | 0.302 | 0.172 | -1.005 | -0.004 | 0.109 | 0.109 | 0.938 | 0.51 | 0.71 | 0.45 | 0.25 | 0.49 | 0.37 |
| 10 | 37302621 | rs2459420 | -0.126 | 0.375 | 0.229 | -1.004 | 0 | 0.033 | 0.033 | 0.111 | 0.5 | 0.58 | 0.44 | 0.25 | 0.47 | 0.37 |
| 10 | 37309400 | rs2505160 | 0.158 | 0.433 | 0.231 | -0.78 | -0.006 | 0.031 | 0.031 | 0 | 0.45 | 0.38 | 0.42 | 0.27 | 0.45 | 0.31 |
| 10 | 37312705 | rs1914159 | -0.909 | 0.604 | -0.046 | -0.578 | -0.006 | -0.015 | -0.015 | 0 | 0.65 | 0.8 | 0.54 | 0.4 | 0.61 | 0.51 |
| 10 | 37323786 | rs2505171 | 0.063 | 0.691 | -0.053 | -0.578 | 0.005 | 0.042 | 0.042 | 1.542 | 0.49 | 0.64 | 0.47 | 0.33 | 0.49 | 0.33 |
| 10 | 37341389 | rs2505176 | -0.782 | 0.773 | -0.1 | -0.34 | 0.002 | -0.018 | -0.018 | 0 | 0.65 | 0.76 | 0.55 | 0.47 | 0.64 | 0.53 |
| 10 | 37342776 | rs1914170 | 0.615 | 0.812 | -0.074 | -0.273 | -0.001 | -0.01 | -0.01 | 0 | 0.07 | 0.07 | 0.04 | 0 | 0.06 | 0.03 |
| 10 | 37347942 | rs7917864 | -0.225 | 0.905 | 0.137 | -0.226 | 0.078 | -0.002 | -0.002 | 0.683 | 0.1 | 0.33 | 0.12 | 0.06 | 0.17 | 0.05 |
| 10 | 37358339 | rs2051563 | -0.977 | 0.634 | -0.147 | -0.389 | 0 | -0.016 | -0.016 | 2.581 | 0.3 | 0.35 | 0.2 | 0.23 | 0.27 | 0.31 |
| 10 | 37364068 | rs1200909 | -1.269 | -0.221 | -0.423 | -0.749 | -0.007 | -0.029 | -0.029 | 0 | 0.34 | 0.34 | 0.16 | 0.13 | 0.19 | 0.24 |
| 10 | 37364259 | rs1200910 | -0.643 | -0.16 | -0.286 | -0.669 | -0.022 | -0.034 | -0.034 | 0.178 | 0.27 | 0.31 | 0.14 | 0.13 | 0.11 | 0.22 |
| 10 | 37380969 | rs1767347 | 0.882 | 0.245 | -0.185 | -0.69 | -0.011 | 0.012 | 0.012 | 0.143 | 0.31 | 0.2 | 0.33 | 0.2 | 0.41 | 0.22 |
| 10 | 37383596 | rs2459444 | 0.796 | 0.276 | -0.112 | -0.706 | 0 | -0.002 | -0.002 | 0.399 | 0.32 | 0.25 | 0.34 | 0.2 | 0.42 | 0.22 |
| 10 | 37384046 | rs2486118 | -0.853 | 0.134 | 0.066 | -0.814 | 0.002 | 0.005 | 0.005 | 0 | 0.58 | 0.56 | 0.5 | 0.32 | 0.58 | 0.49 |
| 10 | 37386777 | rs2505140 | 0.634 | 0.521 | 0.036 | -0.589 | 0 | -0.003 | -0.003 | 0 | 0.32 | 0.26 | 0.34 | 0.2 | 0.42 | 0.23 |
| 10 | 37411550 | rs1200931 | -0.78 | 0.56 | 0.284 | -0.637 | 0.032 | -0.047 | -0.047 | 0 | 0.67 | 0.56 | 0.61 | 0.8 | 0.56 | 0.85 |
| 10 | 37416376 | rs1878249 | -0.293 | 0.637 | 0.024 | -0.891 | -0.003 | -0.033 | -0.033 | 0 | 0.57 | 0.52 | 0.53 | 0.76 | 0.42 | 0.75 |
| 10 | 37425238 | rs1200907 | -0.254 | 0.601 | -0.072 | -0.761 | -0.002 | -0.006 | -0.006 | 0 | 0.44 | 0.53 | 0.48 | 0.33 | 0.59 | 0.27 |
| 10 | 37479884 | rs1782076 | 0.934 | 0.614 | 0.043 | -0.432 | -0.023 | 0.022 | 0.022 | 0.031 | 0.31 | 0.18 | 0.37 | 0.2 | 0.43 | 0.15 |
| 10 | 37487831 | rs1200894 | 0.214 | 0.653 | -0.048 | -0.428 | 0.019 | -0.017 | -0.017 | 0 | 0.37 | 0.48 | 0.43 | 0.32 | 0.5 | 0.22 |
| 10 | 37497698 | rs1623321 | 0.766 | 0.626 | -0.041 | -0.482 | 0.007 | -0.011 | -0.011 | 0.058 | 0.32 | 0.29 | 0.37 | 0.2 | 0.43 | 0.15 |
| 10 | 37505192 | rs1200875 | 1.584 | 0.756 | -0.514 | -0.599 | -0.086 | 0.024 | 0.024 | 0.417 | 0.19 | 0.02 | 0.27 | 0.08 | 0.26 | 0.09 |
| 10 | 37520625 | rs1926127 | 0.218 | 0.609 | -0.24 | -0.435 | 0.092 | -0.009 | -0.009 | 0.123 | 0.83 | 0.56 | 0.84 | 0.85 | 0.78 | 0.89 |
| 10 | 37558907 | rs1200817 | 0.294 | 0.824 | -0.66 | -0.238 | 0.006 | -0.008 | -0.008 | 4.785 | 0.61 | 0.61 | 0.58 | 0.66 | 0.46 | 0.74 |
| 10 | 37559338 | rs1200818 | 1.535 | 0.55 | -1.562 | -0.175 | -0.079 | -0.163 | -0.163 | 0.002 | 0.28 | 0.02 | 0.28 | 0.41 | 0.34 | 0.52 |
| 10 | 37582939 | rs7905906 | -0.458 | 0.902 | -0.968 | -0.273 | -0.012 | 0.013 | 0.013 | 0 | 0.7 | 0.67 | 0.66 | 0.79 | 0.64 | 0.81 |
| 10 | 37583900 | rs1926131 | -0.45 | 0.895 | -0.971 | -0.276 | -0.013 | 0.013 | 0.013 | 0 | 0.7 | 0.67 | 0.66 | 0.79 | 0.64 | 0.8 |
| 10 | 37602764 | rs11011112 | -0.579 | 0.824 | -0.814 | -0.258 | 0.016 | 0.008 | 0.008 | 0 | 0.06 | 0.17 | 0.06 | 0.09 | 0.07 | 0.07 |
| 10 | 37627810 | rs35829275 | -0.576 | 0.824 | -0.814 | -0.258 | 0.016 | 0.008 | 0.008 | 0 | 0.06 | 0.17 | 0.06 | 0.09 | 0.07 | 0.07 |
| 10 | 37628199 | rs7078879 | -0.576 | 0.824 | -0.814 | -0.258 | 0.016 | 0.008 | 0.008 | 0 | 0.06 | 0.17 | 0.06 | 0.09 | 0.07 | 0.07 |
| 10 | 37628244 | rs7096958 | -0.576 | 0.824 | -0.814 | -0.258 | 0.016 | 0.008 | 0.008 | 0 | 0.06 | 0.17 | 0.06 | 0.09 | 0.07 | 0.07 |
| 10 | 37662915 | rs1938421 | -0.647 | 0.87 | -0.995 | -0.463 | -0.082 | 0.128 | 0.128 | 0 | 0.71 | 0.97 | 0.72 | 0.91 | 0.67 | 0.88 |
| 10 | 37665513 | rs11011159 | -0.602 | 0.944 | -0.964 | -0.432 | 0.012 | 0.005 | 0.005 | 0.79 | 0.07 | 0.18 | 0.06 | 0.09 | 0.07 | 0.07 |
| 10 | 37671202 | rs1938416 | -0.219 | 0.495 | -1.074 | -0.547 | -0.002 | 0.119 | 0.119 | 0 | 0.65 | 0.84 | 0.71 | 0.91 | 0.67 | 0.88 |
| 10 | 37681073 | rs10764131 | -0.43 | 0.843 | -0.399 | 0.009 | 0.036 | 0.093 | 0.093 | 0.047 | 0.08 | 0.16 | 0.2 | 0.19 | 0.13 | 0.06 |
| 10 | 37682903 | rs10764132 | -0.491 | 0.841 | -0.334 | 0.011 | 0.06 | 0.131 | 0.131 | 0.096 | 0.09 | 0.22 | 0.2 | 0.18 | 0.13 | 0.06 |
| 10 | 37703778 | rs12774006 | -0.919 | 0.858 | -0.531 | -0.115 | 0.011 | 0.007 | 0.007 | 1.191 | 0.07 | 0.19 | 0.07 | 0.09 | 0.07 | 0.07 |
| 10 | 37704970 | rs2505693 | -0.942 | 0.825 | -0.783 | -0.473 | -0.01 | 0.04 | 0.04 | 0.19 | 0.77 | 0.85 | 0.84 | 0.94 | 0.74 | 0.92 |
| 10 | 37706195 | rs2253294 | -0.203 | 1.021 | -0.736 | -0.506 | 0.017 | 0.022 | 0.022 | 0.004 | 0.16 | 0.21 | 0.22 | 0.19 | 0.22 | 0.11 |
| 10 | 37712642 | rs1938419 | -0.33 | 1.016 | -0.794 | -0.244 | -0.003 | 0.005 | 0.005 | 0.205 | 0.7 | 0.64 | 0.78 | 0.81 | 0.67 | 0.85 |
| 10 | 37721225 | rs12250475 | 0.663 | 0.649 | -0.934 | -0.437 | -0.008 | -0.003 | -0.003 | 5.123 | 0.16 | 0.16 | 0.06 | 0.18 | 0.09 | 0.17 |
| 10 | 37733381 | rs16936517 | -0.617 | 0.941 | -1.169 | -0.271 | 0.009 | 0.027 | 0.027 | 1.191 | 0.07 | 0.19 | 0.07 | 0.13 | 0.07 | 0.07 |
| 10 | 37736944 | rs11011186 | -0.577 | 0.947 | -1.168 | -0.285 | 0.011 | 0.03 | 0.03 | 1.265 | 0.07 | 0.19 | 0.07 | 0.13 | 0.07 | 0.07 |
| 10 | 37769019 | rs2057360 | -0.633 | 0.735 | -1.123 | -0.254 | 0.012 | 0.032 | 0.032 | 1.324 | 0.07 | 0.2 | 0.07 | 0.13 | 0.07 | 0.09 |
| 10 | 37777277 | rs1208802 | 0.991 | 0.684 | -1.267 | -0.298 | -0.067 | 0.075 | 0.075 | 0 | 0.21 | 0.03 | 0.24 | 0.08 | 0.26 | 0.07 |
| 10 | 37790656 | rs2475204 | -0.074 | 0.72 | -1.13 | -0.2 | 0.026 | 0.069 | 0.069 | 0 | 0.08 | 0.13 | 0.16 | 0.18 | 0.15 | 0.06 |
| 10 | 37802532 | rs2475206 | -1.178 | 0.519 | -1.248 | -0.291 | 0.001 | -0.001 | -0.001 | 0.087 | 0.48 | 0.58 | 0.55 | 0.46 | 0.56 | 0.3 |
| 10 | 37807847 | rs4934856 | 0.845 | 0.536 | -1.229 | -0.284 | 0 | -0.001 | -0.001 | 0 | 0.53 | 0.48 | 0.45 | 0.53 | 0.44 | 0.69 |
| 10 | 37808117 | rs2474876 | 0.212 | 0.668 | -1.618 | -0.32 | 0.035 | 0.003 | 0.003 | 0 | 0.08 | 0.13 | 0.06 | 0.08 | 0.16 | 0.05 |
| 10 | 37812532 | rs11011225 | -0.396 | 0.552 | -1.678 | -0.435 | -0.001 | 0.002 | 0.002 | 0 | 0.16 | 0.18 | 0.11 | 0.13 | 0.15 | 0.14 |
| 10 | 37815014 | rs11011230 | -0.566 | 0.94 | -1.812 | -0.451 | -0.006 | 0.01 | 0.01 | 0 | 0.24 | 0.23 | 0.17 | 0.14 | 0.26 | 0.15 |
| 10 | 37818247 | rs11011236 | -0.658 | 0.921 | -1.908 | -0.459 | -0.006 | 0.014 | 0.014 | 0 | 0.25 | 0.23 | 0.18 | 0.13 | 0.26 | 0.15 |
| 10 | 37819709 | rs10827791 | -0.036 | 1.103 | -1.942 | -0.217 | 0 | -0.003 | -0.003 | 0.801 | 0.08 | 0.07 | 0.04 | 0.05 | 0.07 | 0.1 |
| 10 | 37826322 | rs9417245 | 2.257 | 1.123 | -2.375 | -0.285 | -0.061 | -0.168 | -0.168 | 0 | 0.27 | 0.02 | 0.32 | 0.44 | 0.31 | 0.51 |
| 10 | 37826492 | rs2505702 | -0.363 | 1.621 | -1.3 | 0.595 | 0.078 | 0.188 | 0.188 | 0.014 | 0.06 | 0.16 | 0.17 | 0.25 | 0.14 | 0.1 |
| 10 | 37835654 | rs17653445 | 0.551 | 0.852 | -1.955 | 0.036 | -0.028 | 0.018 | 0.018 | 0 | 0.14 | 0.01 | 0.18 | 0.09 | 0.14 | 0.06 |
| 10 | 37837513 | rs10827797 | -0.86 | 0.882 | -1.716 | 0.151 | -0.001 | 0.003 | 0.003 | 0 | 0.1 | 0.13 | 0.06 | 0.13 | 0.08 | 0.1 |
| 10 | 37843457 | rs16937511 | 0.611 | 0.941 | -1.936 | -0.048 | -0.007 | -0.011 | -0.011 | 0.52 | 0.08 | 0.01 | 0.02 | 0.1 | 0.05 | 0.12 |
| 10 | 37846835 | rs11011254 | -0.286 | 1.267 | -1.911 | 0.014 | -0.065 | 0.019 | 0.019 | 0 | 0.19 | 0.01 | 0.2 | 0.12 | 0.21 | 0.07 |
| 10 | 37879010 | rs2224371 | 0.053 | 1.762 | -1.778 | 0.394 | 0.008 | 0.018 | 0.018 | 0 | 0.86 | 0.8 | 0.87 | 0.72 | 0.76 | 0.85 |
| 10 | 37883937 | rs7081464 | 1.506 | 1.211 | -1.593 | 0.349 | 0.012 | 0.011 | 0.011 | 0 | 0.51 | 0.56 | 0.46 | 0.44 | 0.42 | 0.7 |

| | | | | | | | | | | | | | | | | |
|---|---|---|---|---|---|---|---|---|---|---|---|---|---|---|---|---|
| 10 | 37898267 | rs7082477 | 0.911 | 1.125 | -1.551 | 0.353 | 0.012 | 0.027 | 0.027 | 0 | 0.73 | 0.63 | 0.67 | 0.6 | 0.64 | 0.83 |
| 10 | 37899173 | rs4934879 | 0.911 | 1.125 | -1.551 | 0.353 | 0.012 | 0.027 | 0.027 | 0 | 0.73 | 0.63 | 0.67 | 0.6 | 0.64 | 0.83 |
| 10 | 37905848 | rs11011284 | 0.31 | 1.114 | -1.553 | 0.351 | -0.045 | 0.034 | 0.034 | 0 | 0.2 | 0.02 | 0.2 | 0.12 | 0.21 | 0.07 |
| 10 | 37908064 | rs7086522 | 0.822 | 0.829 | -0.831 | 0.135 | 0.038 | -0.023 | -0.023 | 0 | 0.07 | 0.21 | 0.06 | 0 | 0.03 | 0.04 |
| 10 | 37909407 | rs10734027 | -0.436 | 1.048 | -1.629 | 0.203 | 0.066 | 0.147 | 0.147 | 0 | 0.92 | 0.74 | 0.9 | 0.72 | 0.83 | 0.89 |
| 10 | 37910494 | rs7080970 | 1.374 | 1.033 | -1.652 | 0.199 | 0.012 | 0.037 | 0.037 | 0 | 0.71 | 0.58 | 0.66 | 0.55 | 0.63 | 0.77 |
| 10 | 37912214 | rs3920554 | 0.762 | 1.027 | -1.653 | 0.193 | 0.022 | 0.051 | 0.051 | 0.423 | 0.82 | 0.63 | 0.86 | 0.72 | 0.75 | 0.85 |
| 10 | 37923183 | rs289637 | -1.485 | 1.014 | -1.641 | 0.184 | 0.012 | 0.027 | 0.027 | 0 | 0.27 | 0.37 | 0.33 | 0.4 | 0.36 | 0.17 |
| 10 | 37923731 | rs289638 | -1.834 | 1.077 | -1.778 | 0.013 | -0.005 | 0.002 | 0.002 | 0 | 0.45 | 0.44 | 0.51 | 0.56 | 0.5 | 0.28 |
| 10 | 37932791 | rs11592503 | 0.34 | 1.029 | -1.878 | -0.085 | -0.019 | 0.039 | 0.039 | 0 | 0.14 | 0.07 | 0.1 | 0 | 0.17 | 0.06 |
| 10 | 37938147 | rs12775925 | -1.268 | 0.95 | -1.884 | -0.039 | 0.006 | 0.037 | 0.037 | 0 | 0.36 | 0.2 | 0.42 | 0.23 | 0.32 | 0.11 |
| 10 | 37946651 | rs11011311 | 0.356 | 0.859 | -1.732 | -0.003 | -0.022 | -0.027 | -0.027 | 0.157 | 0.05 | 0.09 | 0.01 | 0 | 0.01 | 0 |
| 10 | 37958940 | rs11011319 | 0.49 | 1.184 | -2.054 | 0.025 | -0.057 | -0.001 | -0.001 | 0.523 | 0.13 | 0.05 | 0.07 | 0.1 | 0.17 | 0.17 |
| 10 | 37961832 | rs12411757 | 0.258 | 1.074 | -1.897 | 0.149 | 0.004 | -0.013 | -0.013 | 0.404 | 0.09 | 0.07 | 0.16 | 0.11 | 0.08 | 0.06 |
| 10 | 37975505 | rs289650 | -1.521 | 1.046 | -1.972 | 0.069 | -0.005 | 0 | 0 | 0 | 0.42 | 0.48 | 0.51 | 0.58 | 0.5 | 0.27 |
| 10 | 38000856 | rs10827818 | -0.599 | 1.149 | -1.958 | 0.127 | -0.007 | -0.004 | -0.004 | 0 | 0.25 | 0.18 | 0.38 | 0.23 | 0.25 | 0.07 |
| 10 | 38030639 | rs16937569 | -0.332 | 1.168 | -1.753 | 0.181 | 0.039 | 0.185 | 0.185 | 1.007 | 0.1 | 0.26 | 0.22 | 0.41 | 0.19 | 0.12 |
| 10 | 38032830 | rs2295647 | 0.963 | 1.209 | -1.873 | 0.248 | -0.011 | -0.011 | -0.011 | 2.898 | 0.06 | 0.05 | 0.07 | 0.14 | 0.17 | 0.06 |
| 10 | 43281089 | rs7074877 | -0.19 | -0.626 | -1.044 | -0.243 | 0.022 | 0.028 | 0.028 | 0 | 0.49 | 0.63 | 0.49 | 0.64 | 0.42 | 0.59 |
| 10 | 43341976 | rs2744085 | -1.302 | -0.004 | -0.882 | -0.331 | -0.008 | 0 | 0 | 0 | 0.86 | 0.93 | 0.85 | 0.86 | 0.83 | 0.82 |
| 10 | 43343989 | rs11239794 | 1.266 | 0.237 | -0.908 | -0.448 | -0.013 | 0.009 | 0.009 | 0.351 | 0.16 | 0.15 | 0.1 | 0.03 | 0.23 | 0.06 |
| 10 | 43369124 | rs2744074 | -0.721 | -0.018 | -0.824 | -0.225 | -0.002 | 0.003 | 0.003 | 0.475 | 0.6 | 0.64 | 0.65 | 0.76 | 0.62 | 0.75 |
| 10 | 43369916 | rs2744072 | 0.033 | -0.217 | -1.535 | -0.079 | -0.003 | -0.076 | -0.076 | 0 | 0.39 | 0.16 | 0.31 | 0.55 | 0.4 | 0.47 |
| 10 | 43373200 | rs2744070 | 0.109 | -0.198 | -1.512 | -0.176 | -0.006 | -0.078 | -0.078 | 0.588 | 0.38 | 0.16 | 0.31 | 0.55 | 0.41 | 0.48 |
| 10 | 43373847 | rs3128720 | 0.023 | -0.2 | -1.513 | -0.177 | -0.002 | -0.075 | -0.075 | 1.157 | 0.39 | 0.16 | 0.31 | 0.55 | 0.4 | 0.47 |
| 10 | 43374226 | rs2744069 | -0.486 | -0.17 | -1.506 | -0.211 | -0.008 | -0.017 | -0.017 | 0 | 0.47 | 0.24 | 0.36 | 0.55 | 0.51 | 0.5 |
| 10 | 43374908 | rs2744065 | -0.079 | -0.26 | -1.5 | -0.037 | -0.001 | -0.071 | -0.071 | 0 | 0.39 | 0.18 | 0.31 | 0.55 | 0.4 | 0.47 |
| 10 | 43377166 | rs3121323 | 0.462 | -0.125 | -1.131 | -0.11 | -0.009 | -0.001 | -0.001 | 0 | 0.13 | 0.08 | 0.15 | 0.15 | 0.17 | 0.18 |
| 10 | 43379515 | rs3121324 | -0.112 | -0.164 | -1.133 | -0.118 | 0.001 | 0.011 | 0.011 | 0 | 0.17 | 0.14 | 0.19 | 0.15 | 0.19 | 0.19 |
| 10 | 43385069 | rs7081599 | -0.039 | -0.123 | -0.542 | -0.07 | -0.014 | 0.049 | 0.049 | 12.946 | 0.16 | 0.3 | 0.29 | 0.23 | 0.13 | 0.23 |
| 10 | 43388285 | rs3121327 | 0.424 | -0.073 | -0.807 | -0.185 | -0.006 | 0.026 | 0.026 | 0 | 0.14 | 0.09 | 0.15 | 0.11 | 0.18 | 0.16 |
| 10 | 43389133 | rs788254 | 1.386 | -0.49 | -1.123 | -0.44 | 0.053 | -0.019 | -0.019 | 0.019 | 0.24 | 0.1 | 0.23 | 0.24 | 0.11 | 0.26 |
| 10 | 43398194 | rs11239821 | 0.609 | 0.045 | -0.945 | -0.652 | -0.02 | 0.015 | 0.015 | 0 | 0.2 | 0.18 | 0.16 | 0.04 | 0.3 | 0.08 |
| 10 | 43407238 | rs2796545 | 1.439 | -0.175 | -0.852 | -0.531 | 0.032 | -0.006 | -0.006 | 0 | 0.24 | 0.14 | 0.23 | 0.23 | 0.11 | 0.26 |
| 10 | 43414098 | rs2796546 | 1.616 | -0.214 | -0.973 | -0.491 | 0.03 | -0.076 | -0.076 | 0 | 0.23 | 0.01 | 0.22 | 0.24 | 0.11 | 0.26 |
| 10 | 43416861 | rs7067682 | 0.398 | -0.303 | -0.561 | -0.495 | -0.018 | 0.058 | 0.058 | 12.747 | 0.17 | 0.36 | 0.31 | 0.23 | 0.13 | 0.24 |
| 10 | 43417874 | rs1774203 | 0.991 | 0.1 | -0.837 | -0.616 | 0.025 | -0.001 | -0.001 | 0.502 | 0.3 | 0.11 | 0.27 | 0.24 | 0.19 | 0.28 |
| 10 | 43418668 | rs2796569 | 0.999 | 0.103 | -0.839 | -0.599 | 0.027 | 0.004 | 0.004 | 0.761 | 0.3 | 0.12 | 0.27 | 0.24 | 0.19 | 0.28 |
| 10 | 43419094 | rs11239831 | 1.226 | 0.103 | -0.504 | -0.721 | -0.014 | -0.048 | -0.048 | 0.78 | 0.1 | 0.19 | 0.11 | 0 | 0.06 | 0.11 |
| 10 | 43420922 | rs1775091 | -0.197 | 0.284 | -0.589 | -0.591 | 0.033 | -0.001 | -0.001 | 0.814 | 0.48 | 0.62 | 0.59 | 0.47 | 0.32 | 0.52 |
| 10 | 43434757 | rs788269 | -0.491 | 1.239 | -0.368 | -0.79 | 0.036 | -0.027 | -0.027 | 0 | 0.32 | 0.52 | 0.27 | 0.28 | 0.41 | 0.29 |
| 10 | 43439353 | rs11239840 | 1.491 | 1.231 | -0.354 | -0.857 | -0.002 | 0.023 | 0.023 | 0 | 0.11 | 0.06 | 0.08 | 0.09 | 0.1 | 0.11 |
| 10 | 43440869 | rs3121313 | 0.032 | 1.066 | -0.436 | -1.017 | -0.004 | -0.121 | -0.121 | 0 | 0.2 | 0.39 | 0.27 | 0.02 | 0.2 | 0.15 |
| 10 | 43441729 | rs1582227 | 0.304 | 1.11 | -0.404 | -1.043 | 0.017 | -0.048 | -0.048 | 3.183 | 0.25 | 0.35 | 0.2 | 0.13 | 0.32 | 0.18 |
| 10 | 43444167 | rs12355806 | 0.864 | 1.324 | -0.399 | -1.031 | 0.014 | -0.068 | -0.068 | 0.14 | 0.14 | 0.23 | 0.18 | 0 | 0.15 | 0.12 |
| 10 | 43444534 | rs2506028 | 0.819 | 1.171 | -0.694 | -0.915 | 0.047 | 0.169 | 0.169 | 1.583 | 0.49 | 0.13 | 0.47 | 0.7 | 0.39 | 0.56 |
| 10 | 43448012 | rs3004255 | -0.522 | 1.193 | -0.42 | -0.681 | 0 | -0.001 | -0.001 | 0 | 0.55 | 0.58 | 0.49 | 0.58 | 0.62 | 0.48 |
| 10 | 43449749 | rs2506031 | 2.23 | 1.385 | -1.036 | -0.789 | -0.031 | 0.002 | 0.002 | 0.207 | 0.13 | 0.01 | 0.15 | 0.08 | 0.16 | 0.14 |
| 10 | 43451934 | rs947692 | 1.066 | 1.14 | -0.83 | -0.891 | 0.017 | 0.02 | 0.02 | 0.288 | 0.37 | 0.24 | 0.39 | 0.26 | 0.25 | 0.37 |
| 10 | 43452436 | rs2506034 | 1.037 | 1.245 | -1.164 | -0.834 | -0.057 | -0.14 | -0.14 | 1.176 | 0.22 | 0.03 | 0.24 | 0.45 | 0.29 | 0.3 |
| 10 | 43455339 | rs947696 | 0.254 | 1.133 | -0.592 | -0.961 | -0.004 | -0.055 | -0.055 | 0 | 0.23 | 0.34 | 0.18 | 0.11 | 0.23 | 0.14 |
| 10 | 43460062 | rs11239845 | 1.646 | 1.282 | -0.822 | -0.773 | -0.002 | 0.001 | 0.001 | 0 | 0.11 | 0.09 | 0.08 | 0.03 | 0.15 | 0.04 |
| 10 | 43467392 | rs17431892 | 0.728 | 1.204 | -0.802 | -0.793 | 0.013 | -0.058 | -0.058 | 0 | 0.13 | 0.23 | 0.18 | 0 | 0.14 | 0.12 |
| 10 | 43471575 | rs10900290 | 1.046 | 0.607 | -1.492 | -0.791 | 0.041 | -0.022 | -0.022 | 0.034 | 0.33 | 0.07 | 0.29 | 0.32 | 0.19 | 0.33 |
| 10 | 43474623 | rs2256432 | 1.457 | 0.407 | -1.868 | -1.024 | 0.013 | -0.085 | -0.085 | 0 | 0.3 | 0.02 | 0.34 | 0.32 | 0.2 | 0.33 |
| 10 | 43474694 | rs10900291 | 0.549 | 0.954 | -1.257 | -1.255 | 0.014 | -0.091 | -0.091 | 0 | 0.23 | 0.35 | 0.23 | 0 | 0.26 | 0.16 |
| 10 | 43480529 | rs3004257 | 1.335 | 0.851 | -1.38 | -1.206 | 0.004 | 0.026 | 0.026 | 0 | 0.12 | 0.06 | 0.09 | 0.08 | 0.09 | 0.12 |
| 10 | 43481956 | rs2066394 | 1.302 | 0.944 | -1.315 | -1.287 | 0.001 | -0.051 | -0.051 | 0.002 | 0.19 | 0.22 | 0.19 | 0 | 0.19 | 0.14 |
| 10 | 43483879 | rs11239855 | 1.802 | 0.904 | -1.594 | -1.071 | -0.013 | 0.035 | 0.035 | 0 | 0.09 | 0.01 | 0.04 | 0.06 | 0.07 | 0.11 |
| 10 | 43483971 | rs3123728 | 0.963 | 1.191 | -1.177 | -1.17 | 0.049 | -0.013 | -0.013 | 0 | 0.07 | 0.19 | 0.13 | 0.02 | 0.14 | 0.05 |
| 10 | 43484163 | rs3004258 | -0.728 | 1.234 | -1.895 | -1.286 | -0.001 | -0.027 | -0.027 | 2.736 | 0.76 | 0.71 | 0.8 | 0.89 | 0.76 | 0.84 |
| 10 | 43485964 | rs4949062 | 1.235 | 0.903 | -1.915 | -1.128 | -0.006 | -0.006 | -0.006 | 0 | 0.26 | 0.26 | 0.25 | 0.03 | 0.31 | 0.17 |
| 10 | 43486041 | rs3004259 | -1.693 | 0.81 | -1.947 | -0.955 | 0.002 | 0.02 | 0.02 | 0 | 0.88 | 0.94 | 0.91 | 0.89 | 0.9 | 0.88 |
| 10 | 43494298 | rs746995 | 2.906 | 0.901 | -1.992 | -0.943 | -0.014 | 0.042 | 0.042 | 0 | 0.08 | 0 | 0.02 | 0 | 0.07 | 0.08 |
| 10 | 43499398 | rs2506039 | 2.203 | 0.898 | -1.997 | -0.828 | -0.02 | -0.006 | -0.006 | 0 | 0.13 | 0 | 0.14 | 0.1 | 0.12 | 0.14 |
| 10 | 43499481 | rs1254986 | 0.383 | 1.053 | -1.703 | -0.822 | -0.007 | 0.019 | 0.019 | 0 | 0.38 | 0.34 | 0.34 | 0.13 | 0.4 | 0.29 |
| 10 | 43505098 | rs1271546 | -2.652 | 0.972 | -1.839 | -0.704 | 0.006 | -0.089 | -0.089 | 0 | 0.84 | 0.99 | 0.8 | 0.76 | 0.92 | 0.81 |
| 10 | 43518525 | rs2419089 | -0.5 | 1.499 | -1.246 | -0.812 | 0.016 | 0.014 | 0.014 | 0 | 0.5 | 0.4 | 0.43 | 0.47 | 0.56 | 0.46 |
| 10 | 43525148 | rs7068646 | 1.207 | 1.526 | -1.271 | -0.792 | -0.009 | -0.009 | -0.009 | 2.353 | 0.12 | 0.06 | 0.09 | 0.33 | 0.16 | 0.16 |

| | | | | | | | | | | | | | | | | |
|---|---|---|---|---|---|---|---|---|---|---|---|---|---|---|---|---|
| 10 | 43526023 | rs17432714 | 1.317 | 1.104 | -0.987 | -0.531 | -0.005 | 0.009 | 0.009 | 0 | 0.07 | 0.06 | 0.04 | 0 | 0.07 | 0.02 |
| 10 | 43530071 | rs2565190 | -0.372 | 1.188 | -0.953 | -0.324 | 0.016 | 0.007 | 0.007 | 0 | 0.5 | 0.41 | 0.43 | 0.48 | 0.56 | 0.46 |
| 10 | 43537155 | rs873300 | -0.065 | 1.224 | -0.85 | -0.143 | -0.002 | 0.002 | 0.002 | 0 | 0.09 | 0.06 | 0.09 | 0.12 | 0.09 | 0.1 |
| 10 | 43549708 | rs7088902 | -0.001 | 1.262 | -0.433 | -0.232 | 0.01 | -0.041 | -0.041 | 0 | 0.15 | 0.35 | 0.19 | 0.11 | 0.17 | 0.08 |
| 10 | 43549935 | rs7078220 | -0.598 | 1.323 | -0.208 | -0.149 | 0.029 | -0.028 | -0.028 | 0 | 0.18 | 0.5 | 0.23 | 0.14 | 0.2 | 0.09 |
| 10 | 43550567 | rs4948701 | 0.088 | 1.352 | -0.187 | -0.144 | 0.013 | -0.034 | -0.034 | 0 | 0.15 | 0.35 | 0.19 | 0.09 | 0.17 | 0.08 |
| 10 | 43552895 | rs7069590 | -0.82 | 1.461 | -0.522 | 0.082 | 0.033 | 0.027 | 0.027 | 0 | 0.22 | 0.51 | 0.25 | 0.28 | 0.25 | 0.18 |
| 10 | 43562340 | rs2505988 | 0.319 | 1.556 | -1.952 | -0.57 | -0.048 | 0.093 | 0.093 | 0 | 0.28 | 0.11 | 0.34 | 0.09 | 0.37 | 0.33 |
| 10 | 43563854 | rs3026720 | 0.088 | 1.39 | -1.479 | -0.486 | 0.013 | -0.036 | -0.036 | 0 | 0.18 | 0.62 | 0.24 | 0.11 | 0.2 | 0.08 |
| 10 | 43571447 | rs2505999 | -0.13 | 1.133 | -2.236 | -0.104 | -0.014 | 0.017 | 0.017 | 0 | 0.38 | 0.2 | 0.4 | 0.3 | 0.39 | 0.41 |
| 10 | 43573558 | rs2506010 | -0.14 | 1.132 | -2.237 | -0.105 | -0.014 | 0.017 | 0.017 | 0 | 0.38 | 0.2 | 0.4 | 0.3 | 0.39 | 0.41 |
| 10 | 43584148 | rs2506021 | -0.507 | 0.623 | -2.345 | -0.333 | 0.034 | 0.056 | 0.056 | 2.243 | 0.48 | 0.3 | 0.43 | 0.31 | 0.42 | 0.49 |
| 10 | 43585874 | rs2435344 | -1.611 | 0.907 | -2.763 | -0.093 | -0.035 | -0.167 | -0.167 | 0 | 0.7 | 0.98 | 0.75 | 0.53 | 0.74 | 0.69 |
| 10 | 43589056 | rs4606438 | 0.413 | 0.993 | -2.631 | -0.228 | 0.002 | -0.023 | -0.023 | 0 | 0.27 | 0.64 | 0.31 | 0.1 | 0.33 | 0.18 |
| 10 | 43590280 | rs17158508 | 1.015 | 1.234 | -2.879 | -0.137 | -0.003 | -0.034 | -0.034 | 0 | 0.12 | 0.08 | 0.12 | 0.19 | 0.12 | 0.12 |
| 10 | 43594545 | rs2505532 | -0.525 | 1.241 | -2.902 | -0.163 | -0.005 | -0.009 | -0.009 | 0 | 0.42 | 0.24 | 0.42 | 0.43 | 0.41 | 0.51 |
| 10 | 43596728 | rs3026737 | -0.262 | 0.872 | -3.426 | -0.147 | -0.004 | -0.043 | -0.043 | 0.184 | 0.31 | 0.15 | 0.26 | 0.38 | 0.29 | 0.35 |
| 10 | 43598195 | rs2472739 | 0.199 | 1.254 | -2.9 | -0.392 | -0.023 | 0.085 | 0.085 | 0 | 0.25 | 0.06 | 0.28 | 0.12 | 0.26 | 0.29 |
| 10 | 43605860 | rs1864402 | -0.926 | 1.07 | -2.381 | 0.138 | -0.015 | -0.109 | -0.109 | 2.668 | 0.73 | 0.93 | 0.75 | 0.53 | 0.75 | 0.7 |
| 10 | 43606687 | rs1800860 | 0.365 | 1.747 | -2.086 | -0.218 | -0.038 | 0.005 | 0.005 | 0 | 0.82 | 0.89 | 0.62 | 0.8 | 0.72 | 0.75 |
| 10 | 43610119 | rs1799939 | -0.243 | 0.845 | -2.242 | -0.719 | 0.031 | 0.073 | 0.073 | 0 | 0.26 | 0.09 | 0.26 | 0.12 | 0.19 | 0.24 |
| 10 | 43612609 | rs2742234 | -1.004 | 0.158 | -2.221 | -0.88 | 0.03 | -0.04 | -0.04 | 0.096 | 0.67 | 0.84 | 0.74 | 0.47 | 0.76 | 0.6 |
| 10 | 43622217 | rs2075912 | -1.693 | -0.259 | -2.411 | -1.088 | 0.022 | -0.101 | -0.101 | 0 | 0.77 | 0.94 | 0.78 | 0.48 | 0.83 | 0.7 |
| 10 | 43623812 | rs17028 | 2.277 | 0.385 | -2.704 | -1.229 | -0.06 | -0.077 | -0.077 | 0 | 0.14 | 0.01 | 0.16 | 0.2 | 0.21 | 0.15 |
| 10 | 43624833 | rs2435355 | 1.749 | 0.404 | -2.118 | -1.305 | -0.009 | -0.011 | -0.011 | 0.426 | 0.18 | 0.15 | 0.21 | 0.2 | 0.24 | 0.17 |
| 10 | 43649563 | rs7476512 | 1.037 | 0.572 | -1.497 | -1.391 | 0.002 | -0.003 | -0.003 | 0 | 0.15 | 0.18 | 0.21 | 0.06 | 0.29 | 0.12 |
| 10 | 43659536 | rs2503865 | -1.048 | -0.144 | -1.685 | -1.341 | 0.048 | -0.035 | -0.035 | 3.95 | 0.66 | 0.88 | 0.75 | 0.38 | 0.76 | 0.53 |
| 10 | 43663260 | rs7910805 | 0.47 | -0.156 | -1.563 | -1.393 | 0.026 | 0.048 | 0.048 | 0 | 0.27 | 0.14 | 0.26 | 0.12 | 0.19 | 0.23 |
| 10 | 43666322 | rs12773930 | 2.002 | -0.322 | -1.773 | -1.518 | 0.016 | 0.008 | 0.008 | 0.591 | 0.12 | 0.01 | 0.13 | 0.09 | 0.09 | 0.11 |
| 10 | 43666623 | rs2505556 | 0.326 | -0.261 | -0.97 | -1.51 | 0.001 | -0.012 | -0.012 | 0.684 | 0.39 | 0.64 | 0.48 | 0.26 | 0.57 | 0.29 |
| 10 | 43691691 | rs1254968 | 1.817 | -0.473 | -1.308 | -1.422 | 0.007 | -0.149 | -0.149 | 0 | 0.23 | 0.02 | 0.21 | 0.52 | 0.17 | 0.32 |
| 10 | 43695484 | rs1272142 | 0.902 | -0.461 | -1.278 | -1.419 | 0.049 | -0.082 | -0.082 | 0.008 | 0.34 | 0.05 | 0.24 | 0.62 | 0.24 | 0.48 |
| 10 | 43707762 | rs17158616 | 1.171 | -0.276 | -1.163 | -1.434 | -0.004 | -0.009 | -0.009 | 0 | 0.07 | 0.08 | 0.1 | 0.03 | 0.05 | 0.1 |
| 10 | 43713047 | rs12241923 | 0.483 | -0.138 | -0.447 | -1.338 | -0.007 | 0 | 0 | 0 | 0.36 | 0.39 | 0.29 | 0.62 | 0.25 | 0.47 |
| 10 | 43724215 | rs12775743 | 0.438 | -0.116 | -0.716 | -1.374 | 0.039 | 0.078 | 0.078 | 0 | 0.26 | 0.1 | 0.25 | 0.12 | 0.19 | 0.22 |
| 10 | 43730794 | rs2503846 | -1.19 | 0.564 | -0.496 | -1.309 | 0.023 | -0.021 | -0.021 | 0.636 | 0.85 | 0.78 | 0.81 | 0.96 | 0.73 | 0.88 |
| 10 | 43731521 | rs2503850 | -0.189 | 0.454 | -0.445 | -1.322 | 0.011 | -0.021 | -0.021 | 0 | 0.74 | 0.67 | 0.78 | 0.86 | 0.66 | 0.71 |
| 10 | 43746702 | rs2460554 | 1.473 | 0.43 | -0.701 | -1.277 | -0.02 | -0.006 | -0.006 | 0.789 | 0.22 | 0.16 | 0.26 | 0.21 | 0.3 | 0.18 |
| 10 | 43747330 | rs12244413 | 1.236 | -0.052 | -0.549 | -1.213 | -0.009 | -0.003 | -0.003 | 0.506 | 0.11 | 0.12 | 0.04 | 0.1 | 0.07 | 0.16 |
| 10 | 43750069 | rs1879298 | 0.009 | -0.247 | -0.24 | -1.223 | -0.011 | -0.031 | -0.031 | 0 | 0.28 | 0.27 | 0.27 | 0.09 | 0.18 | 0.22 |
| 10 | 43750260 | rs2503853 | 0.999 | -0.111 | -0.19 | -0.846 | 0.016 | -0.051 | -0.051 | 0 | 0.24 | 0.14 | 0.26 | 0.55 | 0.18 | 0.32 |
| 10 | 43752285 | rs2460557 | -1.028 | -0.331 | -0.158 | -0.695 | -0.003 | 0.006 | 0.006 | 0 | 0.75 | 0.77 | 0.74 | 0.49 | 0.85 | 0.7 |
| 10 | 43753071 | rs12766919 | 0.217 | -0.133 | -0.411 | -1.033 | -0.01 | 0.012 | 0.012 | 0 | 0.26 | 0.23 | 0.24 | 0.04 | 0.34 | 0.19 |
| 10 | 43754308 | rs1879302 | -0.231 | -0.27 | -0.453 | -1.157 | 0.056 | 0.045 | 0.045 | 0 | 0.54 | 0.35 | 0.51 | 0.26 | 0.7 | 0.47 |
| 10 | 43754540 | rs17158702 | 0.774 | 0.062 | -0.202 | -0.914 | 0.006 | -0.002 | -0.002 | 0 | 0.18 | 0.15 | 0.19 | 0.22 | 0.15 | 0.23 |
| 10 | 43755620 | rs1879304 | 0.172 | 0.17 | -0.226 | -0.919 | -0.002 | -0.019 | -0.019 | 0.947 | 0.26 | 0.29 | 0.25 | 0.04 | 0.34 | 0.19 |
| 10 | 43757943 | rs1879307 | 1.746 | -0.137 | -0.608 | -0.808 | 0.032 | -0.123 | -0.123 | 0 | 0.18 | 0.02 | 0.23 | 0.51 | 0.12 | 0.27 |
| 10 | 43767816 | rs11816932 | 0.474 | 0.509 | -0.045 | -0.682 | -0.006 | -0.015 | -0.015 | 0 | 0.18 | 0.23 | 0.2 | 0.12 | 0.16 | 0.14 |
| 10 | 43769560 | rs2503857 | 0.045 | 1.051 | 0.252 | -0.433 | 0 | -0.002 | -0.002 | 0 | 0.34 | 0.31 | 0.31 | 0.22 | 0.37 | 0.3 |
| 10 | 43772876 | rs1879314 | 0.674 | 0.731 | 0.912 | 0.023 | -0.007 | -0.045 | -0.045 | 0 | 0.91 | 0.37 | 0.91 | 1 | 0.97 | 0.99 |
| 10 | 43793625 | rs2503872 | 0.131 | 1.541 | -1.716 | 0.455 | -0.005 | 0.016 | 0.016 | 1.01 | 0.36 | 0.19 | 0.22 | 0.31 | 0.34 | 0.3 |
| 10 | 43794257 | rs11238505 | -0.092 | 2.097 | -1.439 | 1.21 | -0.047 | -0.033 | -0.033 | 0 | 0.1 | 0.02 | 0.22 | 0.4 | 0.26 | 0.16 |
| 10 | 43796180 | rs2503870 | -0.176 | 0.953 | -1.218 | 0.338 | 0.001 | 0.019 | 0.019 | 4.359 | 0.34 | 0.26 | 0.24 | 0.22 | 0.33 | 0.25 |
| 10 | 43799588 | rs2505523 | 0.147 | 0.501 | -1.527 | 0.633 | -0.023 | 0.074 | 0.074 | 0.78 | 0.46 | 0.21 | 0.4 | 0.34 | 0.55 | 0.39 |
| 10 | 43804617 | rs2505521 | 0.276 | -0.09 | -0.924 | 1.185 | -0.023 | 0.042 | 0.042 | 0.108 | 0.77 | 0.43 | 0.77 | 0.65 | 0.83 | 0.77 |
| 10 | 43809570 | rs2460540 | 0.119 | -0.19 | -1.397 | 1.424 | -0.085 | -0.171 | -0.171 | 0.016 | 0.27 | 0.03 | 0.36 | 0.5 | 0.29 | 0.44 |
| 10 | 43809825 | rs2505516 | -0.222 | -0.407 | -0.6 | 1.443 | 0.044 | 0.29 | 0.29 | 2.549 | 0.93 | 0.59 | 0.92 | 0.84 | 0.99 | 0.94 |
| 10 | 43814049 | rs2503875 | 0.188 | -0.354 | -0.622 | 1.503 | -0.085 | -0.142 | -0.142 | 0.253 | 0.27 | 0.03 | 0.36 | 0.43 | 0.29 | 0.43 |
| 10 | 43814896 | rs2503876 | 0.578 | -0.306 | -0.42 | 1.105 | 0.028 | 0.061 | 0.061 | 0.348 | 0.53 | 0.31 | 0.44 | 0.35 | 0.6 | 0.4 |
| 10 | 43820626 | rs2493655 | -0.769 | -0.883 | -0.118 | 0.966 | 0 | 0 | 0 | 0 | 0.45 | 0.5 | 0.4 | 0.52 | 0.38 | 0.55 |
| 10 | 43821380 | rs2460535 | 1.058 | -1.415 | 0.588 | 1.068 | -0.035 | 0.146 | 0.146 | 0.034 | 0.81 | 0.49 | 0.87 | 0.67 | 0.89 | 0.73 |
| 10 | 43828231 | rs7087184 | -0.527 | -1.602 | 0.667 | 0.993 | 0.006 | -0.011 | -0.011 | 0.022 | 0.21 | 0.22 | 0.2 | 0.26 | 0.19 | 0.19 |
| 10 | 43831280 | rs2264603 | -0.661 | -1.613 | 0.875 | 0.956 | -0.018 | -0.063 | -0.063 | 1.124 | 0.84 | 0.81 | 0.88 | 1 | 0.88 | 0.9 |
| 10 | 43834545 | rs2252761 | 0.875 | -1.534 | -0.018 | 0.274 | -0.053 | -0.096 | -0.096 | 0 | 0.43 | 0.1 | 0.58 | 0.54 | 0.55 | 0.48 |
| 10 | 43849580 | rs11238518 | 1.489 | -1.633 | 0.206 | -0.016 | 0 | -0.053 | -0.053 | 0 | 0.21 | 0.21 | 0.13 | 0 | 0.19 | 0.15 |
| 10 | 43850568 | rs2245413 | 0.723 | -1.57 | 0.348 | 0.303 | -0.03 | 0.184 | 0.184 | 0 | 0.81 | 0.53 | 0.84 | 0.58 | 0.89 | 0.74 |
| 10 | 43855778 | rs2493665 | 0.747 | -1.469 | 0.201 | 0.28 | -0.009 | -0.012 | -0.012 | 1.118 | 0.09 | 0.23 | 0.03 | 0.03 | 0.03 | 0.15 |
| 10 | 43869097 | rs10899795 | -0.823 | -1.624 | 0.053 | 0.233 | -0.013 | 0.009 | 0.009 | 0.092 | 0.33 | 0.55 | 0.26 | 0.43 | 0.22 | 0.35 |
| 10 | 43871999 | rs7094556 | 1.42 | -1.537 | -0.845 | -0.305 | -0.113 | -0.033 | -0.033 | 0 | 0.23 | 0.07 | 0.42 | 0.29 | 0.39 | 0.3 |

| | | | | | | | | | | | | | | | | |
|---|---|---|---|---|---|---|---|---|---|---|---|---|---|---|---|---|
| 10 | 43874817 | rs4519046 | -2.097 | -1.489 | -0.929 | -0.408 | -0.045 | -0.118 | -0.118 | 2.735 | 0.84 | 0.99 | 0.79 | 0.6 | 0.81 | 0.81 |
| 10 | 43879573 | rs10899797 | -0.249 | -1.548 | -0.411 | -1.125 | 0.034 | 0.037 | 0.037 | 0.163 | 0.38 | 0.76 | 0.25 | 0.26 | 0.26 | 0.36 |
| 10 | 43890151 | rs17153622 | 0.679 | -1.603 | -0.095 | -1.125 | -0.013 | -0.017 | -0.017 | 0 | 0.11 | 0.43 | 0.04 | 0.03 | 0.04 | 0.14 |
| 10 | 43895101 | rs7070769 | -1.247 | -1.682 | -0.254 | -1.211 | -0.028 | 0.077 | 0.077 | 0.649 | 0.55 | 0.89 | 0.37 | 0.3 | 0.41 | 0.5 |
| 10 | 43907495 | rs2252783 | 0.457 | -1.633 | -0.1 | -1.161 | 0.01 | 0.003 | 0.003 | 0.272 | 0.67 | 0.33 | 0.77 | 0.7 | 0.77 | 0.65 |
| 10 | 43923728 | rs2247979 | 0.157 | -1.542 | -0.611 | -1.148 | -0.014 | -0.005 | -0.005 | 0 | 0.69 | 0.36 | 0.78 | 0.74 | 0.81 | 0.7 |
| 10 | 43940898 | rs10899803 | 0.824 | -1.5 | -0.673 | -1.085 | -0.011 | 0.064 | 0.064 | 0 | 0.06 | 0.12 | 0.08 | 0.22 | 0.03 | 0.07 |
| 10 | 43989935 | rs7100846 | 1.281 | -0.969 | -0.392 | -1.197 | 0.001 | -0.024 | -0.024 | 0 | 0.2 | 0.19 | 0.13 | 0 | 0.18 | 0.15 |
| 10 | 44002994 | rs3006399 | 0.488 | -0.858 | -0.083 | -0.767 | -0.022 | -0.074 | -0.074 | 2.374 | 0.73 | 0.47 | 0.84 | 0.92 | 0.81 | 0.72 |
| 10 | 44003499 | rs1106215 | -1.183 | -1 | -0.674 | -0.73 | 0.001 | -0.017 | -0.017 | 0 | 0.86 | 0.92 | 0.9 | 1 | 0.89 | 0.91 |
| 10 | 44013866 | rs12240317 | -0.794 | -1.036 | -0.44 | -0.404 | -0.066 | 0.067 | 0.067 | 0.193 | 0.19 | 0.57 | 0.13 | 0.28 | 0.07 | 0.25 |
| 10 | 44014299 | rs2393949 | -0.554 | -1.087 | -0.829 | -0.945 | -0.001 | -0.063 | -0.063 | 0.079 | 0.67 | 0.72 | 0.76 | 0.99 | 0.69 | 0.76 |
| 10 | 44014339 | rs4948737 | -0.726 | -0.988 | -0.574 | -0.536 | -0.066 | 0.076 | 0.076 | 0.193 | 0.19 | 0.57 | 0.13 | 0.28 | 0.07 | 0.25 |
| 10 | 44015353 | rs7068859 | -0.172 | -0.704 | -0.896 | -1.535 | 0.008 | 0.071 | 0.071 | 0.019 | 0.37 | 0.23 | 0.23 | 0.01 | 0.38 | 0.29 |
| 10 | 44015746 | rs7084621 | -0.025 | -1.169 | -0.885 | -1.571 | 0.012 | -0.013 | -0.013 | 0 | 0.24 | 0.2 | 0.09 | 0.01 | 0.16 | 0.17 |
| 10 | 44015891 | rs7085915 | -1.446 | -1.069 | -1.223 | -1.407 | -0.038 | 0.084 | 0.084 | 0 | 0.55 | 0.88 | 0.37 | 0.36 | 0.49 | 0.38 |
| 10 | 44015907 | rs11238575 | -0.186 | -0.844 | 0.162 | -1.716 | -0.004 | -0.106 | -0.106 | 0 | 0.33 | 0.4 | 0.22 | 0.01 | 0.28 | 0.22 |
| 10 | 44021769 | rs2999241 | 0.264 | -0.914 | 0.018 | -1.631 | -0.003 | -0.092 | -0.092 | 0 | 0.64 | 0.64 | 0.72 | 1 | 0.68 | 0.77 |
| 10 | 44021832 | rs2999242 | 0.416 | -0.903 | 0.021 | -1.625 | -0.003 | -0.094 | -0.094 | 0 | 0.62 | 0.62 | 0.72 | 1 | 0.68 | 0.77 |
| 10 | 44030617 | rs1338559 | 0.375 | -0.269 | 0.378 | -1.293 | 0.005 | -0.118 | -0.118 | 0.454 | 0.67 | 0.61 | 0.72 | 1 | 0.69 | 0.83 |
| 10 | 44037408 | rs7910230 | -0.431 | -0.283 | 0.431 | -1.302 | -0.017 | 0.022 | 0.022 | 0 | 0.29 | 0.2 | 0.26 | 0.01 | 0.3 | 0.17 |
| 10 | 44046676 | rs2860340 | 1.019 | -0.196 | 0.615 | -0.804 | 0.003 | -0.001 | -0.001 | 0 | 0.56 | 0.49 | 0.51 | 0.55 | 0.53 | 0.64 |
| 10 | 44050715 | rs7893169 | -0.48 | -0.196 | 0.716 | -0.813 | 0.017 | 0.124 | 0.124 | 0 | 0.11 | 0.19 | 0.21 | 0.44 | 0.16 | 0.19 |
| 10 | 44051819 | rs10793442 | -0.064 | -0.19 | 0.694 | -0.811 | 0 | 0.045 | 0.045 | 0 | 0.1 | 0.11 | 0.21 | 0.44 | 0.16 | 0.19 |
| 10 | 44052903 | rs2230661 | 0.977 | -0.198 | 0.644 | -0.788 | 0.003 | -0.001 | -0.001 | 0 | 0.56 | 0.49 | 0.51 | 0.55 | 0.53 | 0.65 |
| 10 | 44063812 | rs2393938 | -0.416 | -0.096 | 0.36 | -0.56 | 0.004 | 0.063 | 0.063 | 0 | 0.11 | 0.15 | 0.21 | 0.45 | 0.16 | 0.18 |
| 10 | 44064778 | rs9422541 | 1.179 | -0.219 | 0.103 | -0.812 | -0.034 | 0.002 | 0.002 | 0.162 | 0.54 | 0.2 | 0.51 | 0.54 | 0.56 | 0.58 |
| 10 | 44070587 | rs7075288 | -0.022 | -0.315 | 0.337 | -0.859 | 0 | -0.059 | -0.059 | 0 | 0.69 | 0.69 | 0.73 | 1 | 0.69 | 0.83 |
| 10 | 44071499 | rs9422546 | 0.896 | -0.15 | -0.055 | -0.715 | -0.016 | 0.011 | 0.011 | 0 | 0.54 | 0.21 | 0.49 | 0.55 | 0.53 | 0.65 |
| 10 | 44087671 | rs10793446 | -0.75 | 0.159 | 0.185 | -0.82 | -0.016 | -0.054 | -0.054 | 0 | 0.36 | 0.44 | 0.46 | 0.78 | 0.31 | 0.37 |
| 10 | 44094186 | rs10793447 | -0.438 | 0.256 | 0.258 | -0.78 | -0.004 | -0.004 | -0.004 | 0.906 | 0.27 | 0.24 | 0.41 | 0.75 | 0.24 | 0.31 |
| 10 | 44101391 | rs6593362 | -1.134 | 0.417 | -0.148 | -0.816 | -0.005 | -0.017 | -0.017 | 0.031 | 0.43 | 0.44 | 0.5 | 0.75 | 0.38 | 0.39 |
| 10 | 44104513 | rs7342073 | -2.041 | 0.424 | -0.33 | -0.922 | 0.003 | 0.031 | 0.031 | 0 | 0.65 | 0.8 | 0.72 | 0.76 | 0.63 | 0.65 |
| 10 | 44105044 | rs10899836 | 0.744 | 0.657 | -0.305 | -0.151 | 0.005 | -0.029 | -0.029 | 1.533 | 0.17 | 0.08 | 0.2 | 0.36 | 0.15 | 0.17 |
| 10 | 44109024 | rs2039723 | 0.242 | 0.697 | -0.344 | -1.171 | -0.003 | 0.002 | 0.002 | 0 | 0.23 | 0.2 | 0.37 | 0.74 | 0.2 | 0.31 |
| 10 | 44109759 | rs10899838 | -1.026 | 0.703 | -0.337 | -1.189 | 0.004 | -0.021 | -0.021 | 0.512 | 0.43 | 0.51 | 0.58 | 0.74 | 0.43 | 0.44 |
| 10 | 44110651 | rs17154024 | -0.893 | 1.025 | -0.359 | -1.072 | 0.001 | -0.04 | -0.04 | 0.006 | 0.21 | 0.31 | 0.21 | 0 | 0.24 | 0.17 |
| 10 | 44111806 | rs10899839 | -1.384 | 1.125 | -0.192 | -0.854 | -0.002 | -0.097 | -0.097 | 0.011 | 0.3 | 0.53 | 0.3 | 0.04 | 0.27 | 0.31 |
| 10 | 44112245 | rs12354886 | -0.901 | 1.709 | -0.641 | -0.394 | 0 | -0.039 | -0.039 | 0 | 0.2 | 0.29 | 0.21 | 0 | 0.22 | 0.17 |
| 10 | 44115660 | rs3123678 | 0.314 | 1.431 | -0.695 | 0.176 | -0.002 | 0.003 | 0.003 | 0.477 | 0.57 | 0.69 | 0.53 | 0.45 | 0.46 | 0.56 |
| 10 | 44116906 | rs10899841 | -0.558 | 2.037 | -0.586 | 0.058 | 0.008 | 0.026 | 0.026 | 0.215 | 0.24 | 0.26 | 0.34 | 0.52 | 0.29 | 0.31 |
| 10 | 44133991 | rs11238609 | -0.812 | 1.999 | -0.379 | 0.256 | 0.017 | 0 | 0 | 0 | 0.27 | 0.36 | 0.39 | 0.66 | 0.31 | 0.41 |
| 10 | 44137293 | rs10751327 | -0.793 | 1.891 | 0.242 | 0.21 | 0.018 | 0.001 | 0.001 | 0.62 | 0.27 | 0.36 | 0.39 | 0.67 | 0.31 | 0.41 |
| 10 | 44138691 | rs10736822 | -0.582 | 1.894 | 0.24 | 0.236 | 0.019 | 0.065 | 0.065 | 0.352 | 0.26 | 0.35 | 0.36 | 0.52 | 0.29 | 0.31 |
| 10 | 44142466 | rs10751328 | -0.802 | 1.891 | 0.241 | 0.235 | 0.016 | 0.001 | 0.001 | 0.595 | 0.27 | 0.36 | 0.39 | 0.67 | 0.3 | 0.41 |
| 10 | 44143154 | rs10793464 | -0.535 | 1.885 | 0.262 | 0.234 | 0.004 | 0.016 | 0.016 | 0.334 | 0.25 | 0.25 | 0.34 | 0.52 | 0.28 | 0.31 |
| 10 | 44145822 | rs3128247 | 1.247 | 2.042 | 0.181 | 0.131 | 0 | 0 | 0 | 0 | 0.54 | 0.59 | 0.49 | 0.3 | 0.46 | 0.46 |
| 10 | 44147007 | rs3128248 | -0.425 | 2.167 | -0.253 | 0.084 | -0.015 | 0.042 | 0.042 | 0 | 0.87 | 0.98 | 0.95 | 0.97 | 0.87 | 0.92 |
| 10 | 44160627 | rs3123665 | 0.343 | 2.117 | -0.232 | 0.102 | -0.039 | 0.071 | 0.071 | 0 | 0.81 | 0.95 | 0.88 | 0.96 | 0.75 | 0.86 |
| 10 | 44175098 | rs2505644 | 1.108 | 1.899 | 0.232 | 0.012 | 0.002 | 0.025 | 0.025 | 0.801 | 0.53 | 0.47 | 0.48 | 0.29 | 0.45 | 0.46 |
| 10 | 44185247 | rs12358200 | -0.392 | 1.983 | -0.279 | 0.102 | -0.033 | 0.031 | 0.031 | 0 | 0.06 | 0.01 | 0.05 | 0 | 0.1 | 0.05 |
| 10 | 44191192 | rs10899857 | -1.163 | 1.679 | -0.529 | 0.235 | -0.009 | -0.071 | -0.071 | 0 | 0.25 | 0.11 | 0.36 | 0.55 | 0.31 | 0.22 |
| 10 | 44193355 | rs10899858 | 0.044 | 1.306 | -0.475 | 0.039 | -0.002 | -0.005 | -0.005 | 0 | 0.08 | 0.06 | 0.05 | 0 | 0.11 | 0.06 |
| 10 | 44195467 | rs17154173 | -0.802 | 0.961 | -0.572 | 0.108 | 0.006 | -0.01 | -0.01 | 2.106 | 0.06 | 0.02 | 0.03 | 0.15 | 0.04 | 0.1 |
| 10 | 44196014 | rs17154178 | -1.061 | 0.923 | -0.589 | 0.091 | 0.009 | -0.016 | -0.016 | 0 | 0.07 | 0.02 | 0.03 | 0.15 | 0.04 | 0.1 |
| 10 | 44196101 | rs10751332 | -1.491 | 0.879 | -0.093 | 0.076 | 0 | 0.137 | 0.137 | 0 | 0.19 | 0.28 | 0.29 | 0.5 | 0.21 | 0.19 |
| 10 | 44199230 | rs7094994 | -2.206 | 0.821 | 0.092 | 0.027 | -0.01 | 0.019 | 0.019 | 0 | 0.25 | 0.35 | 0.33 | 0.66 | 0.25 | 0.29 |
| 10 | 44203201 | rs2050502 | -1.513 | 0.801 | 0.117 | 0.043 | -0.01 | 0.041 | 0.041 | 0 | 0.15 | 0.2 | 0.27 | 0.66 | 0.14 | 0.24 |
| 10 | 44207259 | rs11238645 | -0.506 | 0.879 | 0.1 | -0.186 | -0.02 | 0.027 | 0.027 | 0 | 0.11 | 0.06 | 0.1 | 0 | 0.17 | 0.06 |
| 10 | 44207375 | rs11238646 | -1.299 | 0.874 | 0.122 | -0.158 | -0.008 | -0.009 | -0.009 | 0 | 0.15 | 0.11 | 0.13 | 0.15 | 0.2 | 0.16 |
| 10 | 44207709 | rs10899867 | -2.593 | 0.781 | 0.296 | -0.197 | -0.006 | -0.001 | -0.001 | 0 | 0.3 | 0.46 | 0.39 | 0.66 | 0.32 | 0.3 |
| 10 | 44208251 | rs2505647 | 2.233 | 0.499 | -0.492 | -1.076 | -0.055 | 0.028 | 0.028 | 0 | 0.29 | 0.04 | 0.28 | 0.27 | 0.3 | 0.38 |
| 10 | 44210600 | rs868471 | -2.107 | 0.551 | -0.212 | -1.021 | -0.006 | -0.002 | -0.002 | 0.198 | 0.31 | 0.5 | 0.39 | 0.66 | 0.32 | 0.3 |
| 10 | 44223180 | rs7903170 | -0.187 | 0.55 | -0.178 | -1.012 | -0.01 | 0.273 | 0.273 | 0 | 0.12 | 0.28 | 0.25 | 0.51 | 0.1 | 0.14 |
| 10 | 44223771 | rs10751337 | -1.821 | 0.553 | -0.197 | -1.015 | -0.002 | 0.119 | 0.119 | 0 | 0.27 | 0.45 | 0.37 | 0.51 | 0.28 | 0.21 |
| 10 | 44223790 | rs2505649 | 2.026 | 0.489 | 0.058 | -1.098 | 0 | -0.005 | -0.005 | 0 | 0.32 | 0.37 | 0.32 | 0.28 | 0.32 | 0.39 |
| 10 | 44233503 | rs12258494 | -0.493 | 0.547 | 0.235 | -0.881 | -0.012 | 0.295 | 0.295 | 0 | 0.12 | 0.32 | 0.25 | 0.51 | 0.1 | 0.14 |
| 10 | 44234466 | rs11238680 | -1.119 | 0.545 | 0.219 | -0.898 | -0.023 | 0.112 | 0.112 | 0 | 0.16 | 0.36 | 0.28 | 0.66 | 0.13 | 0.24 |
| 10 | 44248758 | rs2505655 | -0.649 | 0.472 | -0.01 | -0.193 | -0.034 | 0.117 | 0.117 | 1.644 | 0.68 | 0.88 | 0.72 | 0.93 | 0.65 | 0.7 |

| | | | | | | | | | | | | | | | |
|---|---|---|---|---|---|---|---|---|---|---|---|---|---|---|---|
| 10 | 44249237 | rs11238689 | 0.695 | 0.699 | 0.5 | 0.517 | -0.036 | 0.039 | 0.039 | 0 | 0.07 | 0 | 0.05 | 0 | 0.11 | 0.06 |
| 10 | 44256350 | rs4536162 | -2.458 | 0.635 | 0.556 | 0.552 | -0.005 | -0.002 | -0.002 | 0 | 0.3 | 0.5 | 0.39 | 0.66 | 0.31 | 0.3 |
| 10 | 44268654 | rs4080021 | -0.529 | 0.717 | 0.632 | 0.648 | -0.009 | 0.243 | 0.243 | 0 | 0.11 | 0.23 | 0.24 | 0.5 | 0.1 | 0.14 |
| 10 | 44277067 | rs4245609 | -2.109 | 0.72 | 0.616 | 0.86 | -0.008 | 0.069 | 0.069 | 0 | 0.26 | 0.42 | 0.34 | 0.6 | 0.26 | 0.29 |
| 10 | 44278838 | rs10751344 | -1.814 | 0.723 | 0.621 | 0.862 | 0.003 | 0.085 | 0.085 | 0 | 0.23 | 0.38 | 0.33 | 0.6 | 0.24 | 0.29 |
| 10 | 44285921 | rs3750725 | -1.483 | 0.821 | 0.445 | 0.847 | -0.013 | 0.151 | 0.151 | 0 | 0.15 | 0.28 | 0.27 | 0.47 | 0.14 | 0.12 |
| 10 | 44291108 | rs11238711 | 0.387 | 0.749 | 0.407 | 0.846 | -0.009 | 0.015 | 0.015 | 0 | 0.08 | 0.02 | 0.05 | 0 | 0.11 | 0.06 |
| 10 | 44292364 | rs751464 | -1.811 | 0.885 | 0.408 | 0.823 | -0.001 | 0.16 | 0.16 | 0 | 0.17 | 0.36 | 0.32 | 0.47 | 0.19 | 0.12 |
| 10 | 44293839 | rs10899911 | -2.104 | 0.835 | 0.47 | 0.86 | 0.003 | 0.074 | 0.074 | 0 | 0.24 | 0.38 | 0.35 | 0.47 | 0.27 | 0.17 |
| 10 | 44295539 | rs11238713 | -0.714 | 0.784 | 0.529 | 0.896 | -0.011 | 0.128 | 0.128 | 0 | 0.11 | 0.18 | 0.23 | 0.48 | 0.1 | 0.11 |
| 10 | 44296893 | rs10899912 | -0.577 | 0.795 | 0.52 | 0.927 | -0.003 | 0.077 | 0.077 | 0 | 0.1 | 0.12 | 0.22 | 0.48 | 0.1 | 0.11 |
| 10 | 44298644 | rs12357048 | -1.14 | 1.023 | 0.677 | 0.851 | 0.017 | 0.008 | 0.008 | 0.322 | 0.16 | 0.19 | 0.12 | 0.16 | 0.22 | 0.18 |
| 10 | 44298868 | rs12360468 | -0.463 | 1.008 | 0.685 | 0.789 | 0.001 | -0.021 | -0.021 | 0 | 0.13 | 0.1 | 0.09 | 0 | 0.18 | 0.08 |
| 10 | 44303219 | rs7922745 | -2.771 | 1.08 | 0.676 | 0.86 | 0.019 | -0.001 | -0.001 | 0 | 0.33 | 0.52 | 0.43 | 0.65 | 0.38 | 0.29 |
| 10 | 44303930 | rs1540967 | -0.916 | 0.994 | 0.718 | 0.636 | 0.042 | -0.024 | -0.024 | 0 | 0.05 | 0.16 | 0.07 | 0 | 0.07 | 0.01 |
| 10 | 44307607 | rs7906426 | -1.236 | 0.978 | 0.647 | 0.783 | 0.003 | 0.074 | 0.074 | 0 | 0.12 | 0.29 | 0.14 | 0.18 | 0.1 | 0.06 |
| 10 | 44308000 | rs7900896 | -2.145 | 1.008 | 0.419 | 0.722 | 0.023 | 0.056 | 0.056 | 0 | 0.28 | 0.51 | 0.26 | 0.34 | 0.31 | 0.23 |
| 10 | 44314431 | rs7913725 | -1.999 | 0.802 | 1.119 | 0.815 | 0.028 | 0.057 | 0.057 | 2.148 | 0.14 | 0.63 | 0.16 | 0.17 | 0.14 | 0.1 |
| 10 | 44315371 | rs11238722 | 1.558 | 0.554 | 0.258 | 0.555 | -0.045 | 0.092 | 0.092 | 0 | 0.34 | 0.15 | 0.32 | 0.23 | 0.42 | 0.47 |
| 10 | 44319891 | rs7075076 | 0.106 | 0.68 | 0.997 | 0.641 | 0.03 | -0.034 | -0.034 | 0 | 0.07 | 0.17 | 0.05 | 0 | 0.07 | 0.02 |
| 10 | 44321351 | rs10899918 | 0.108 | 0.279 | 0.81 | 0.652 | 0 | -0.002 | -0.002 | 0 | 0.6 | 0.62 | 0.68 | 0.89 | 0.62 | 0.7 |
| 10 | 44321629 | rs12570694 | 0.218 | 0.302 | 0.57 | 1.203 | -0.004 | 0.095 | 0.095 | 0 | 0.05 | 0.13 | 0.19 | 0.33 | 0.06 | 0.09 |
| 10 | 44334916 | rs11238734 | -0.869 | 0.241 | 0.467 | 1.185 | -0.019 | -0.06 | -0.06 | 0 | 0.13 | 0.05 | 0.36 | 0.54 | 0.14 | 0.14 |
| 10 | 44336374 | rs7901526 | 0.165 | 0.072 | 0.768 | 1.059 | -0.022 | -0.03 | -0.03 | 0 | 0.56 | 0.78 | 0.43 | 0.51 | 0.53 | 0.58 |
| 10 | 44341964 | rs10899922 | 0.622 | 0.06 | 0.896 | 1.055 | 0.006 | -0.003 | -0.003 | 0 | 0.48 | 0.37 | 0.34 | 0.51 | 0.48 | 0.57 |
| 10 | 44344738 | rs10793503 | 0.623 | 0.205 | 0.944 | 1.09 | 0.015 | -0.002 | -0.002 | 0.272 | 0.51 | 0.4 | 0.34 | 0.52 | 0.47 | 0.57 |
| 10 | 44358219 | rs2169200 | 1.923 | -0.2 | 0.904 | 0.982 | -0.053 | 0.189 | 0.189 | 0 | 0.42 | 0.13 | 0.27 | 0.19 | 0.47 | 0.51 |
| 10 | 44358300 | rs1008803 | 1.591 | -0.199 | 0.903 | 0.981 | -0.026 | 0.054 | 0.054 | 0 | 0.45 | 0.27 | 0.31 | 0.37 | 0.49 | 0.57 |
| 10 | 44362266 | rs11238750 | 1.914 | -0.187 | 0.805 | 0.809 | -0.053 | 0.189 | 0.189 | 0 | 0.42 | 0.13 | 0.27 | 0.19 | 0.47 | 0.51 |
| 10 | 44373858 | rs10899930 | -0.51 | -0.03 | 1.113 | 0.968 | 0.049 | 0.09 | 0.09 | 1.798 | 0.06 | 0.28 | 0.09 | 0.07 | 0.04 | 0.07 |
| 10 | 44374899 | rs7920511 | 0.71 | -0.082 | 0.975 | 1.029 | -0.006 | -0.008 | -0.008 | 0 | 0.6 | 0.7 | 0.47 | 0.57 | 0.56 | 0.68 |
| 10 | 44375239 | rs2613102 | -0.668 | -0.319 | 0.944 | 0.991 | -0.004 | -0.011 | -0.011 | 0 | 0.9 | 0.89 | 0.92 | 0.98 | 0.89 | 0.94 |
| 10 | 44375635 | rs10899933 | -0.462 | -0.235 | 1.315 | 1.129 | 0.001 | 0.137 | 0.137 | 0 | 0.1 | 0.25 | 0.11 | 0.22 | 0.05 | 0.14 |
| 10 | 44376756 | rs10899937 | 1.791 | -0.252 | 1.201 | 1.028 | -0.025 | 0.104 | 0.104 | 0 | 0.46 | 0.25 | 0.34 | 0.34 | 0.5 | 0.54 |
| 10 | 44378740 | rs7097453 | 0.138 | 0.206 | 1.573 | 1.292 | 0.003 | 0.031 | 0.031 | 0 | 0.07 | 0.17 | 0.06 | 0.06 | 0.03 | 0.07 |
| 10 | 44383053 | rs10899938 | -0.58 | 0.198 | 1.535 | 1.287 | 0.009 | 0.038 | 0.038 | 0.279 | 0.1 | 0.32 | 0.09 | 0.06 | 0.04 | 0.07 |
| 10 | 44384191 | rs11238766 | 1.889 | 0.134 | 1.057 | 1.231 | -0.033 | 0.124 | 0.124 | 0 | 0.52 | 0.17 | 0.35 | 0.35 | 0.52 | 0.57 |
| 10 | 44390863 | rs10899942 | 0.278 | 0.505 | 0.899 | 1.122 | -0.002 | 0.021 | 0.021 | 0 | 0.07 | 0.16 | 0.06 | 0.06 | 0.03 | 0.07 |
| 10 | 44391001 | rs1578854 | 0.873 | 0.647 | 0.778 | 1.157 | -0.008 | -0.014 | -0.014 | 3.013 | 0.67 | 0.74 | 0.5 | 0.57 | 0.6 | 0.71 |
| 10 | 44392068 | rs7895918 | 1.103 | 0.589 | 0.73 | 1.161 | -0.013 | -0.013 | -0.013 | 0.1 | 0.64 | 0.73 | 0.46 | 0.57 | 0.55 | 0.7 |
| 10 | 44396631 | rs898548 | 0.442 | 0.743 | 0.685 | 1.239 | -0.008 | -0.012 | -0.012 | 0.905 | 0.73 | 0.79 | 0.6 | 0.62 | 0.69 | 0.7 |
| 10 | 44405258 | rs882902 | 0.989 | 0.643 | 0.918 | 1.178 | 0.02 | 0.011 | 0.011 | 0.405 | 0.67 | 0.5 | 0.54 | 0.62 | 0.62 | 0.69 |
| 10 | 44418891 | rs1147902 | 0.437 | 0.717 | 0.78 | 1.226 | -0.011 | -0.013 | -0.013 | 0.007 | 0.73 | 0.79 | 0.6 | 0.63 | 0.67 | 0.71 |
| 10 | 44427264 | rs1408939 | 1.376 | 0.718 | 0.961 | 1.288 | -0.017 | -0.001 | -0.001 | 7.948 | 0.6 | 0.3 | 0.42 | 0.44 | 0.57 | 0.55 |
| 10 | 44428806 | rs1749798 | 0.591 | 0.719 | 1.041 | 1.279 | 0.007 | 0.018 | 0.018 | 0 | 0.7 | 0.68 | 0.57 | 0.5 | 0.66 | 0.63 |
| 10 | 44429319 | rs984001 | -0.868 | 0.713 | 0.416 | 1.058 | 0.014 | -0.031 | -0.031 | 0 | 0.06 | 0.15 | 0.05 | 0 | 0.05 | 0.01 |
| 10 | 44430746 | rs898550 | 0.997 | 0.631 | -0.108 | 1.156 | -0.024 | -0.001 | -0.001 | 0 | 0.6 | 0.25 | 0.41 | 0.43 | 0.57 | 0.55 |
| 10 | 44432455 | rs12572433 | -0.441 | -0.021 | -0.522 | 1.213 | 0.01 | -0.063 | -0.063 | 0 | 0.13 | 0.01 | 0.09 | 0.2 | 0.09 | 0.07 |
| 10 | 44435246 | rs4948582 | 1.055 | 0.244 | -0.444 | 1.289 | 0.017 | -0.019 | -0.019 | 0 | 0.58 | 0.19 | 0.43 | 0.59 | 0.69 | 0.53 |
| 10 | 44437232 | rs2863230 | -0.79 | 0.212 | 0.418 | 1.29 | -0.004 | -0.01 | -0.01 | 0 | 0.16 | 0.37 | 0.2 | 0.11 | 0.13 | 0.15 |
| 10 | 44437946 | rs10793509 | 0.399 | 0.163 | 0.857 | 1.437 | 0.015 | -0.022 | -0.022 | 0 | 0.81 | 0.63 | 0.73 | 0.88 | 0.83 | 0.84 |
| 10 | 44438863 | rs11238782 | 2.314 | 0.069 | 0.593 | 1.416 | 0.001 | 0.047 | 0.047 | 2.957 | 0.53 | 0.15 | 0.33 | 0.4 | 0.59 | 0.47 |
| 10 | 44443729 | rs2798997 | -0.726 | -0.014 | 0.952 | 1.348 | -0.014 | -0.037 | -0.037 | 0.882 | 0.92 | 0.88 | 0.98 | 1 | 0.97 | 0.96 |
| 10 | 44446878 | rs6593374 | -1.76 | 0.045 | 0.912 | 1.455 | 0.003 | -0.01 | -0.01 | 0.05 | 0.24 | 0.27 | 0.35 | 0.44 | 0.21 | 0.27 |
| 10 | 44448664 | rs4948794 | 0.082 | 0.187 | 0.838 | 1.572 | 0.003 | -0.032 | -0.032 | 0 | 0.1 | 0.04 | 0.1 | 0.19 | 0.1 | 0.07 |
| 10 | 44448756 | rs1325488 | 1.738 | 1.036 | 0.31 | 1.388 | -0.058 | -0.031 | -0.031 | 0 | 0.09 | 0.01 | 0.11 | 0.16 | 0.21 | 0.17 |
| 10 | 44454186 | rs10508876 | 0.401 | 0.054 | 0.828 | 1.453 | -0.026 | -0.03 | -0.03 | 0 | 0.05 | 0 | 0.12 | 0.07 | 0.07 | 0.07 |
| 10 | 44456029 | rs7924201 | 2.756 | -0.072 | 0.834 | 1.408 | 0.018 | 0.07 | 0.07 | 0.467 | 0.5 | 0.13 | 0.34 | 0.39 | 0.59 | 0.47 |
| 10 | 44456716 | rs1532232 | 0.159 | -0.032 | 0.761 | 1.35 | 0.002 | -0.036 | -0.036 | 0 | 0.1 | 0.03 | 0.1 | 0.19 | 0.1 | 0.07 |
| 10 | 44458714 | rs898549 | 2.581 | -0.09 | 0.815 | 1.263 | -0.018 | 0.032 | 0.032 | 0.424 | 0.5 | 0.07 | 0.33 | 0.39 | 0.59 | 0.45 |
| 10 | 44463444 | rs7901081 | -1.862 | 0.119 | 1.432 | 1.365 | -0.003 | 0.023 | 0.023 | 0.668 | 0.22 | 0.31 | 0.34 | 0.44 | 0.2 | 0.29 |
| 10 | 44469148 | rs6593376 | -1.867 | 0.121 | 1.432 | 1.365 | -0.003 | 0.02 | 0.02 | 0.701 | 0.23 | 0.31 | 0.34 | 0.45 | 0.2 | 0.29 |
| 10 | 44470909 | rs1147909 | 0.401 | 0.232 | 1.563 | 1 | -0.015 | 0.008 | 0.008 | 0 | 0.85 | 0.71 | 0.76 | 0.84 | 0.89 | 0.82 |
| 10 | 44471109 | rs1460542 | 1.067 | 0.164 | 1.367 | 1.02 | 0.013 | -0.011 | -0.011 | 0 | 0.8 | 0.61 | 0.71 | 0.84 | 0.83 | 0.79 |
| 10 | 44473036 | rs10899958 | -1.201 | 0.194 | 0.868 | 0.997 | -0.001 | 0.003 | 0.003 | 0.117 | 0.1 | 0.2 | 0.16 | 0.11 | 0.09 | 0.12 |
| 10 | 44479236 | rs4948803 | 1.094 | 0.16 | 1.195 | 0.72 | 0.003 | -0.068 | -0.068 | 0 | 0.74 | 0.55 | 0.69 | 0.86 | 0.79 | 0.77 |
| 10 | 44482885 | rs10508877 | -0.461 | -0.065 | 0.993 | 0.531 | -0.003 | 0.004 | 0.004 | 0 | 0.14 | 0.15 | 0.16 | 0.1 | 0.13 | 0.17 |
| 10 | 44483397 | rs2862856 | -0.411 | -0.108 | 1.293 | 0.475 | 0.007 | -0.044 | -0.044 | 0 | 0.88 | 0.86 | 0.95 | 1 | 0.9 | 0.91 |
| 10 | 44489563 | rs4948806 | 0.473 | -0.155 | 1.231 | 0.495 | 0.044 | -0.021 | -0.021 | 0.032 | 0.82 | 0.63 | 0.72 | 0.86 | 0.83 | 0.78 |

| | | | | | | | | | | | | | | | | |
|---|---|---|---|---|---|---|---|---|---|---|---|---|---|---|---|---|
| 10 | 44490414 | rs4948591 | 2.897 | -0.35 | 0.703 | 0.582 | -0.096 | -0.087 | -0.087 | 3.228 | 0.41 | 0.08 | 0.3 | 0.46 | 0.52 | 0.4 |
| 10 | 44490532 | rs17154827 | 0.898 | -0.331 | 0.623 | 0.546 | -0.006 | -0.044 | -0.044 | 0 | 0.07 | 0 | 0.09 | 0.18 | 0.09 | 0.07 |
| 10 | 44494546 | rs10793513 | 1.757 | -0.297 | 0.746 | 0.093 | 0.004 | 0.025 | 0.025 | 0 | 0.57 | 0.38 | 0.52 | 0.76 | 0.66 | 0.6 |
| 10 | 44501460 | rs11238807 | -0.563 | -0.198 | 0.464 | 0.143 | -0.001 | -0.002 | -0.002 | 0 | 0.15 | 0.21 | 0.17 | 0.11 | 0.13 | 0.17 |
| 10 | 44501813 | rs10899964 | -0.547 | -0.198 | 0.443 | 0.144 | 0 | -0.002 | -0.002 | 0 | 0.14 | 0.2 | 0.17 | 0.11 | 0.13 | 0.17 |
| 10 | 44511905 | rs4948819 | 0.803 | -0.21 | 0.559 | 0.11 | -0.001 | -0.026 | -0.026 | 0 | 0.76 | 0.71 | 0.72 | 0.85 | 0.8 | 0.78 |
| 10 | 44513645 | rs4948822 | -1.317 | -0.343 | 0.792 | 0.185 | -0.008 | -0.009 | -0.009 | 0.102 | 0.25 | 0.39 | 0.3 | 0.24 | 0.24 | 0.25 |
| 10 | 44515716 | rs10899970 | 2.882 | -0.327 | 0.819 | 0.094 | -0.041 | 0.026 | 0.026 | 0.505 | 0.42 | 0.19 | 0.39 | 0.62 | 0.51 | 0.47 |
| 10 | 44517536 | rs7908204 | -2.052 | -0.32 | 0.706 | 0.107 | -0.023 | -0.042 | -0.042 | 0.594 | 0.34 | 0.53 | 0.33 | 0.24 | 0.28 | 0.3 |
| 10 | 44520403 | rs6593391 | -0.439 | -0.246 | 0.722 | 0.177 | -0.005 | -0.025 | -0.025 | 0 | 0.06 | 0.15 | 0.03 | 0 | 0.04 | 0.05 |
| 10 | 44532138 | rs1492712 | -1.479 | -0.22 | 0.687 | 0.218 | -0.015 | -0.034 | -0.034 | 0.168 | 0.3 | 0.52 | 0.33 | 0.24 | 0.28 | 0.3 |
| 10 | 44539913 | rs2047009 | 2.697 | -0.174 | 0.546 | 0.25 | -0.058 | -0.06 | -0.06 | 0 | 0.4 | 0.09 | 0.3 | 0.5 | 0.49 | 0.38 |
| 10 | 44541565 | rs10508881 | 2.085 | -0.186 | 0.639 | 0.159 | -0.001 | 0.002 | 0.002 | 0.658 | 0.47 | 0.36 | 0.38 | 0.51 | 0.56 | 0.45 |
| 10 | 44544935 | rs2639469 | 1.024 | -0.134 | 0.751 | 0.249 | -0.002 | -0.006 | -0.006 | 0.235 | 0.72 | 0.61 | 0.63 | 0.74 | 0.78 | 0.7 |
| 10 | 44553342 | rs268325 | -1.656 | -0.273 | 0.68 | 0.01 | 0.001 | 0.01 | 0.01 | 0 | 0.34 | 0.38 | 0.41 | 0.2 | 0.3 | 0.36 |
| 10 | 44553583 | rs2802493 | -1.97 | -0.287 | 0.688 | 0.305 | 0.002 | 0.001 | 0.001 | 0 | 0.4 | 0.69 | 0.55 | 0.39 | 0.34 | 0.45 |
| 10 | 44557502 | rs187901 | -1.395 | -0.078 | 0.941 | 0.487 | 0.1 | 0.244 | 0.244 | 0.056 | 0.09 | 0.39 | 0.18 | 0.25 | 0.11 | 0.15 |
| 10 | 44558945 | rs268319 | -0.9 | -0.194 | 0.958 | 0.425 | 0.026 | 0.114 | 0.114 | 0 | 0.12 | 0.3 | 0.19 | 0.25 | 0.12 | 0.15 |
| 10 | 44559434 | rs17155030 | 0.598 | -0.162 | 0.713 | 0.448 | -0.018 | -0.018 | -0.018 | 0.116 | 0.08 | 0.01 | 0.11 | 0.13 | 0.12 | 0.09 |
| 10 | 44561651 | rs268314 | -1.785 | -0.225 | 0.77 | 0.382 | 0.007 | 0.08 | 0.08 | 1.82 | 0.23 | 0.52 | 0.32 | 0.38 | 0.24 | 0.23 |
| 10 | 44562687 | rs7898629 | 0.261 | -0.158 | 0.649 | 0.378 | 0 | -0.003 | -0.003 | 4.568 | 0.08 | 0.1 | 0.05 | 0.06 | 0.08 | 0.06 |
| 10 | 44564178 | rs12572132 | 0.221 | -0.155 | 0.491 | 0.326 | -0.039 | -0.015 | -0.015 | 2.094 | 0.09 | 0.01 | 0.13 | 0.07 | 0.1 | 0.11 |
| 10 | 44567765 | rs187900 | -1.544 | -0.451 | 0.718 | 0.287 | -0.021 | 0.055 | 0.055 | 0 | 0.19 | 0.45 | 0.22 | 0.25 | 0.14 | 0.18 |
| 10 | 44579062 | rs1254531 | -1.359 | -0.44 | 0.684 | 0.292 | -0.015 | 0.041 | 0.041 | 0 | 0.18 | 0.35 | 0.2 | 0.25 | 0.14 | 0.18 |
| 10 | 44580839 | rs268300 | -0.203 | -0.424 | 0.708 | 0.313 | 0.028 | 0.191 | 0.191 | 0.11 | 0.05 | 0.21 | 0.13 | 0.19 | 0.05 | 0.11 |
| 10 | 44581683 | rs268304 | -2.172 | -0.662 | 0.797 | 0.11 | -0.003 | -0.001 | -0.001 | 0.724 | 0.35 | 0.62 | 0.36 | 0.37 | 0.3 | 0.29 |
| 10 | 44581989 | rs268305 | -0.808 | -0.646 | 0.782 | 0.127 | -0.012 | 0.058 | 0.058 | 0.202 | 0.21 | 0.37 | 0.27 | 0.31 | 0.21 | 0.22 |
| 10 | 44583027 | rs2818912 | -2.141 | -0.689 | 0.341 | 0.012 | 0.011 | 0.035 | 0.035 | 0 | 0.45 | 0.66 | 0.48 | 0.23 | 0.34 | 0.41 |
| 10 | 44597898 | rs11238850 | 0.252 | -0.92 | 0.275 | 0.108 | 0 | 0 | 0 | 0 | 0.12 | 0.12 | 0.05 | 0.07 | 0.08 | 0.07 |
| 10 | 44598058 | rs10508882 | -0.23 | -0.94 | 0.265 | 0.137 | -0.003 | 0.016 | 0.016 | 0.002 | 0.18 | 0.19 | 0.23 | 0.31 | 0.18 | 0.21 |
| 10 | 44598146 | rs268289 | -0.865 | -0.991 | 0.247 | 0.103 | -0.01 | 0.029 | 0.029 | 0.151 | 0.25 | 0.15 | 0.34 | 0.14 | 0.22 | 0.29 |
| 10 | 44599013 | rs2802490 | -3.177 | -0.864 | 0.184 | 0.16 | -0.09 | -0.071 | -0.071 | 0 | 0.63 | 0.9 | 0.7 | 0.55 | 0.51 | 0.62 |
| 10 | 44615859 | rs174849 | 0.151 | -0.648 | 0.186 | 0.268 | 0 | 0.005 | 0.005 | 0 | 0.12 | 0.09 | 0.18 | 0.07 | 0.1 | 0.14 |
| 10 | 44618925 | rs4948844 | -1.657 | -0.556 | 0.111 | 0.361 | -0.007 | 0.004 | 0.004 | 0 | 0.31 | 0.32 | 0.29 | 0.38 | 0.26 | 0.27 |
| 10 | 44619943 | rs7895805 | -0.404 | -0.061 | 0.111 | 0.3 | -0.004 | -0.01 | -0.01 | 0 | 0.16 | 0.17 | 0.14 | 0.12 | 0.16 | 0.11 |
| 10 | 44631156 | rs2047008 | -0.474 | 0.296 | 0.062 | 0.204 | 0.001 | -0.017 | -0.017 | 0 | 0.18 | 0.22 | 0.23 | 0.11 | 0.14 | 0.17 |
| 10 | 44632262 | rs268274 | -0.655 | 0.276 | 0.013 | 0.183 | -0.001 | -0.01 | -0.01 | 0 | 0.22 | 0.21 | 0.24 | 0.12 | 0.2 | 0.22 |
| 10 | 44640797 | rs10899990 | -0.379 | 0.519 | -0.287 | 0.102 | 0.003 | -0.044 | -0.044 | 0 | 0.14 | 0.18 | 0.15 | 0 | 0.13 | 0.12 |
| 10 | 44664524 | rs2802487 | -2.042 | 0.28 | -0.608 | 0.508 | -0.039 | -0.039 | -0.039 | 1.452 | 0.59 | 0.89 | 0.7 | 0.53 | 0.53 | 0.6 |
| 10 | 44673148 | rs1680635 | -0.911 | -0.221 | -0.257 | 0.575 | 0 | 0.032 | 0.032 | 0 | 0.32 | 0.14 | 0.17 | 0.19 | 0.29 | 0.3 |
| 10 | 44675056 | rs17155437 | 0.136 | -0.272 | -0.344 | 0.517 | -0.024 | -0.016 | -0.016 | 2.86 | 0.15 | 0.04 | 0.14 | 0.13 | 0.19 | 0.12 |
| 10 | 44678454 | rs17155478 | -0.37 | -0.634 | 0.745 | 0.366 | -0.005 | -0.018 | -0.018 | 0 | 0.07 | 0.33 | 0.05 | 0 | 0.01 | 0.01 |
| 10 | 44678775 | rs1704227 | -1.555 | -0.644 | 0.672 | 0.444 | -0.063 | 0.187 | 0.187 | 0.384 | 0.12 | 0.53 | 0.2 | 0.28 | 0.03 | 0.15 |
| 10 | 44680405 | rs2804036 | -1.699 | -0.651 | 0.561 | 0.394 | -0.008 | 0 | 0 | 0 | 0.3 | 0.66 | 0.24 | 0.35 | 0.13 | 0.33 |
| 10 | 44691241 | rs1623851 | -1.643 | -0.406 | 0.361 | 0.341 | -0.032 | 0.008 | 0.008 | 0 | 0.32 | 0.54 | 0.25 | 0.35 | 0.18 | 0.35 |
| 10 | 44694201 | rs11597731 | -0.714 | -0.314 | -0.017 | 0.387 | -0.002 | 0.001 | 0.001 | 0 | 0.18 | 0.26 | 0.16 | 0.16 | 0.19 | 0.13 |
| 10 | 44694606 | rs12570491 | -1.469 | -0.118 | -0.036 | 0.42 | -0.002 | 0 | 0 | 0.086 | 0.24 | 0.23 | 0.21 | 0.24 | 0.27 | 0.19 |
| 10 | 44695308 | rs7902040 | -0.248 | -0.443 | -0.088 | 0.217 | 0.012 | 0.017 | 0.017 | 0 | 0.2 | 0.15 | 0.04 | 0.08 | 0.1 | 0.27 |
| 10 | 44696034 | rs2802477 | -2.531 | -0.01 | -0.625 | -0.038 | -0.11 | -0.106 | -0.106 | 0 | 0.65 | 0.97 | 0.76 | 0.56 | 0.62 | 0.79 |
| 10 | 44697616 | rs7894238 | 0.935 | -0.327 | -0.301 | -0.021 | -0.048 | -0.03 | -0.03 | 0.066 | 0.11 | 0.02 | 0.13 | 0.12 | 0.18 | 0.1 |
| 10 | 44698075 | rs2054620 | -1.715 | -0.33 | -0.215 | -0.014 | 0.002 | -0.003 | -0.003 | 0 | 0.42 | 0.43 | 0.34 | 0.48 | 0.35 | 0.51 |
| 10 | 44702276 | rs1680640 | -1.605 | -0.276 | -0.195 | 0.01 | 0 | -0.001 | -0.001 | 0.365 | 0.39 | 0.42 | 0.34 | 0.48 | 0.35 | 0.51 |
| 10 | 44713311 | rs1905150 | -0.833 | -0.286 | -0.122 | 0.058 | -0.001 | 0.002 | 0.002 | 0 | 0.25 | 0.32 | 0.2 | 0.35 | 0.16 | 0.36 |
| 10 | 44730995 | rs7907961 | -0.652 | -0.145 | -0.132 | 0.077 | 0.009 | 0.055 | 0.055 | 0 | 0.18 | 0.25 | 0.3 | 0.41 | 0.24 | 0.22 |
| 10 | 44732917 | rs677465 | -0.264 | -0.466 | -0.011 | 0.107 | -0.004 | -0.016 | -0.016 | 0 | 0.2 | 0.26 | 0.05 | 0.08 | 0.11 | 0.25 |
| 10 | 44736389 | rs736136 | 0.047 | -0.396 | 0.128 | 0.121 | 0.005 | -0.001 | -0.001 | 0 | 0.14 | 0.22 | 0.15 | 0.14 | 0.17 | 0.12 |
| 10 | 44740698 | rs661697 | -0.693 | -0.671 | 0.257 | 0.103 | 0.017 | 0.017 | 0.017 | 0 | 0.23 | 0.19 | 0.04 | 0.06 | 0.1 | 0.24 |
| 10 | 44746395 | rs541483 | -1.215 | -0.671 | 0.162 | 0.164 | -0.007 | 0.001 | 0.001 | 0 | 0.31 | 0.32 | 0.21 | 0.35 | 0.15 | 0.36 |
| 10 | 44747059 | rs535176 | -1.262 | -0.667 | 0.078 | 0.166 | -0.059 | 0.023 | 0.023 | 0 | 0.32 | 0.49 | 0.22 | 0.35 | 0.15 | 0.36 |
| 10 | 44749854 | rs11238956 | 1.938 | -1.038 | -0.527 | 0.106 | -0.081 | -0.011 | -0.011 | 0.254 | 0.26 | 0.02 | 0.2 | 0.24 | 0.3 | 0.16 |
| 10 | 44753867 | rs501120 | -1.487 | -1.407 | 0.501 | 0.308 | -0.07 | 0.019 | 0.019 | 0 | 0.32 | 0.5 | 0.22 | 0.34 | 0.15 | 0.36 |
| 10 | 44761623 | rs607363 | -1.475 | -1.371 | 0.463 | 0.316 | -0.068 | 0.021 | 0.021 | 0 | 0.32 | 0.54 | 0.23 | 0.34 | 0.15 | 0.35 |
| 10 | 44773984 | rs1632484 | -1.335 | -1.303 | 0.233 | -0.249 | -0.059 | -0.099 | -0.099 | 0 | 0.31 | 0.44 | 0.09 | 0.07 | 0.14 | 0.27 |
| 10 | 44786364 | rs800314 | -0.374 | -1.011 | 0.191 | -0.055 | -0.022 | -0.029 | -0.029 | 0 | 0.17 | 0.23 | 0.04 | 0.06 | 0.07 | 0.22 |
| 10 | 44789602 | rs809601 | 0.572 | -1.251 | 0.652 | -0.222 | -0.022 | -0.038 | -0.038 | 0 | 0.92 | 0.71 | 0.93 | 1 | 0.98 | 0.95 |
| 10 | 44793299 | rs2505734 | 1.761 | -1.231 | 0.339 | 0.029 | 0.088 | -0.019 | -0.019 | 0 | 0.51 | 0.17 | 0.58 | 0.54 | 0.66 | 0.47 |
| 10 | 44794277 | rs1436927 | 0.054 | -1.342 | 0.493 | 0.188 | 0.004 | 0 | 0 | 0.034 | 0.15 | 0.16 | 0.15 | 0.14 | 0.22 | 0.15 |
| 10 | 44797087 | rs754713 | 1.376 | -1.394 | 0.58 | 0.184 | -0.011 | 0.046 | 0.046 | 3.032 | 0.71 | 0.4 | 0.62 | 0.57 | 0.73 | 0.68 |
| 10 | 44801673 | rs800323 | 1.357 | -1.408 | 0.584 | 0.146 | -0.021 | -0.062 | -0.062 | 0.013 | 0.69 | 0.47 | 0.76 | 0.85 | 0.73 | 0.78 |

| | | | | | | | | | | | | | | | | |
|---|---|---|---|---|---|---|---|---|---|---|---|---|---|---|---|---|
| 10 | 44806054 | rs2209065 | 1.159 | -1.683 | 0.693 | 0.196 | -0.06 | 0.017 | 0.017 | 0 | 0.72 | 0.44 | 0.75 | 0.68 | 0.89 | 0.63 |
| 10 | 44810205 | rs11238983 | -1.083 | -1.296 | 0.405 | 0.081 | -0.022 | -0.071 | -0.071 | 0 | 0.23 | 0.36 | 0.22 | 0.06 | 0.15 | 0.23 |
| 10 | 44814336 | rs7082209 | -1.342 | -1.314 | 0.397 | 0.184 | -0.025 | 0.059 | 0.059 | 0 | 0.24 | 0.37 | 0.35 | 0.39 | 0.15 | 0.34 |
| 10 | 44815709 | rs2505735 | 0.934 | -1.457 | 0.569 | -0.505 | -0.034 | -0.115 | -0.115 | 0 | 0.69 | 0.5 | 0.75 | 0.94 | 0.82 | 0.73 |
| 10 | 44820513 | rs11238986 | 1.137 | -1.248 | 0.477 | -0.377 | -0.021 | -0.035 | -0.035 | 0.3 | 0.07 | 0.11 | 0.04 | 0 | 0.03 | 0.01 |
| 10 | 44821387 | rs10508884 | -1.16 | -1.217 | 0.335 | -0.518 | 0.003 | 0.008 | 0.008 | 0 | 0.29 | 0.31 | 0.22 | 0.13 | 0.28 | 0.18 |
| 10 | 44828212 | rs982098 | 1.993 | -1.226 | 0.411 | -0.229 | -0.036 | -0.03 | -0.03 | 0 | 0.44 | 0.19 | 0.46 | 0.57 | 0.55 | 0.45 |
| 10 | 44828234 | rs982097 | 1.48 | -1.226 | 0.542 | -0.341 | 0.009 | 0.026 | 0.026 | 0 | 0.52 | 0.27 | 0.47 | 0.64 | 0.56 | 0.6 |
| 10 | 44829205 | rs11238994 | 1.239 | -1.292 | 0.415 | -0.519 | 0 | -0.022 | -0.022 | 0.106 | 0.12 | 0.11 | 0.06 | 0 | 0.1 | 0.02 |
| 10 | 44829517 | rs1144484 | 0.299 | -1.336 | 0.397 | -0.519 | -0.024 | 0.002 | 0.002 | 0 | 0.74 | 0.53 | 0.67 | 0.77 | 0.82 | 0.74 |
| 10 | 44830727 | rs7918046 | -0.114 | -1.226 | 0.378 | -0.511 | -0.007 | 0.003 | 0.003 | 0 | 0.24 | 0.22 | 0.33 | 0.16 | 0.3 | 0.16 |
| 10 | 44833031 | rs11599561 | 1.546 | -1.225 | 0.293 | -0.508 | 0.001 | -0.006 | -0.006 | 0 | 0.11 | 0.07 | 0.04 | 0 | 0.1 | 0.02 |
| 10 | 44835963 | rs1144482 | 2.199 | -1.442 | 0.497 | -0.569 | -0.07 | -0.033 | -0.033 | 0 | 0.4 | 0.16 | 0.45 | 0.62 | 0.51 | 0.45 |
| 10 | 44838019 | rs10900025 | -0.826 | -1.316 | 0.617 | -0.649 | 0 | -0.007 | -0.007 | 0 | 0.37 | 0.43 | 0.4 | 0.17 | 0.4 | 0.19 |
| 10 | 44838530 | rs77839 | 0.748 | -1.308 | 0.538 | -0.656 | 0 | 0.018 | 0.018 | 0.272 | 0.59 | 0.35 | 0.56 | 0.83 | 0.59 | 0.77 |
| 10 | 44838832 | rs12360225 | 1.413 | -1.143 | 0.085 | -0.563 | 0.001 | -0.01 | -0.01 | 0 | 0.11 | 0.09 | 0.05 | 0 | 0.1 | 0.02 |
| 10 | 44845097 | rs17156139 | 1.014 | -1.057 | -0.085 | -0.509 | -0.034 | -0.023 | -0.023 | 0 | 0.13 | 0.06 | 0.14 | 0.13 | 0.19 | 0.12 |
| 10 | 44845164 | rs751425 | -0.524 | -0.934 | 0.262 | -0.32 | 0.002 | -0.036 | -0.036 | 0 | 0.41 | 0.52 | 0.42 | 0.19 | 0.41 | 0.34 |
| 10 | 44849350 | rs1144479 | 0.973 | -0.128 | -0.551 | 0.145 | -0.053 | -0.032 | -0.032 | 0 | 0.38 | 0.2 | 0.44 | 0.62 | 0.49 | 0.43 |
| 10 | 44850424 | rs266109 | -0.863 | -0.022 | -0.224 | 0.206 | 0.026 | -0.045 | -0.045 | 0 | 0.89 | 0.72 | 0.79 | 0.98 | 0.89 | 0.95 |
| 10 | 44853297 | rs1147877 | 0.662 | 0.039 | -1.024 | 0.239 | -0.062 | -0.025 | -0.025 | 0 | 0.41 | 0.22 | 0.46 | 0.63 | 0.51 | 0.44 |
| 10 | 44853606 | rs874304 | 0.675 | 0.073 | -1.028 | 0.269 | -0.06 | -0.032 | -0.032 | 0.346 | 0.41 | 0.21 | 0.45 | 0.62 | 0.51 | 0.44 |
| 10 | 44855663 | rs266105 | -0.397 | -0.25 | -0.353 | 0.463 | -0.01 | 0 | 0 | 0 | 0.86 | 0.77 | 0.77 | 0.87 | 0.89 | 0.75 |
| 10 | 44855740 | rs11595588 | 0.598 | -0.071 | -0.786 | 0.385 | 0.012 | 0.003 | 0.003 | 0 | 0.57 | 0.25 | 0.47 | 0.57 | 0.6 | 0.4 |
| 10 | 44855927 | rs17391002 | 0.114 | 0.271 | -0.483 | 0.446 | 0.052 | 0.01 | 0.01 | 0 | 0.14 | 0.34 | 0.26 | 0.15 | 0.23 | 0.17 |
| 10 | 44856863 | rs4948877 | 0.73 | -0.251 | -1.33 | 0.612 | -0.032 | -0.053 | -0.053 | 0.167 | 0.15 | 0.04 | 0.16 | 0.16 | 0.17 | 0.18 |
| 10 | 44859347 | rs17156191 | 0.776 | -0.448 | -1.505 | 0.581 | 0.041 | -0.033 | -0.033 | 0 | 0.27 | 0.16 | 0.16 | 0.36 | 0.17 | 0.25 |
| 10 | 44859618 | rs17482472 | 0.719 | -0.15 | -1.437 | 0.497 | 0 | 0 | 0 | 0.013 | 0.07 | 0.07 | 0.09 | 0.09 | 0.1 | 0.06 |
| 10 | 44862220 | rs10900029 | 1.191 | -0.308 | -1.56 | 0.412 | 0.019 | -0.067 | -0.067 | 0 | 0.3 | 0.1 | 0.21 | 0.43 | 0.24 | 0.31 |
| 10 | 44865841 | rs1804429 | 1.833 | -0.228 | -1.371 | 0.371 | 0 | 0 | 0 | 0 | 0.06 | 0.09 | 0.03 | 0.06 | 0.03 | 0.06 |
| 10 | 44865976 | rs1065297 | 0.632 | -0.368 | -1.399 | 0.316 | -0.004 | -0.005 | -0.005 | 0.255 | 0.08 | 0.12 | 0.04 | 0.06 | 0.03 | 0.06 |
| 10 | 44866208 | rs266093 | 0.024 | -0.248 | -1.15 | 0.089 | 0.002 | -0.031 | -0.031 | 0 | 0.65 | 0.4 | 0.78 | 0.82 | 0.66 | 0.73 |
| 10 | 44866247 | rs17883887 | 0.85 | -0.111 | -0.756 | -0.111 | 0.053 | -0.092 | -0.092 | 0 | 0.14 | 0.36 | 0.11 | 0 | 0.2 | 0.04 |
| 10 | 44866275 | rs266092 | 1.892 | -0.217 | -0.988 | -0.127 | 0.006 | 0.034 | 0.034 | 0 | 0.08 | 0.02 | 0.02 | 0 | 0.05 | 0.08 |
| 10 | 44866628 | rs2839696 | 2.279 | -0.196 | -0.971 | -0.108 | 0.001 | 0 | 0 | 0 | 0.05 | 0.09 | 0.03 | 0.06 | 0.03 | 0.06 |
| 10 | 44867146 | rs1029153 | 0.755 | -0.207 | -1.001 | -0.034 | -0.027 | 0 | 0 | 0 | 0.26 | 0.14 | 0.47 | 0.19 | 0.3 | 0.26 |
| 10 | 44867304 | rs2522 | 2.227 | -0.203 | -0.965 | -0.023 | 0 | 0 | 0 | 0 | 0.06 | 0.09 | 0.03 | 0.06 | 0.03 | 0.06 |
| 10 | 44868257 | rs1801157 | 2.419 | -0.497 | -1.081 | 0.138 | -0.026 | -0.043 | -0.043 | 0 | 0.25 | 0.05 | 0.19 | 0.27 | 0.23 | 0.24 |
| 10 | 44868465 | rs12258838 | 2.183 | 0.281 | -0.749 | -0.16 | 0.001 | 0 | 0 | 2.715 | 0.05 | 0.09 | 0.03 | 0.06 | 0.03 | 0.06 |
| 10 | 44869427 | rs266089 | -1.481 | 0.264 | -0.749 | -0.147 | -0.016 | -0.006 | -0.006 | 0.76 | 0.83 | 0.7 | 0.91 | 0.85 | 0.89 | 0.83 |
| 10 | 44870015 | rs266088 | 0.446 | 0.448 | -1.428 | -0.303 | 0.003 | 0.011 | 0.011 | 0 | 0.15 | 0.1 | 0.12 | 0.11 | 0.15 | 0.14 |
| 10 | 44870705 | rs17881412 | 2.106 | 0.518 | -1.47 | -0.255 | 0.001 | 0 | 0 | 0 | 0.05 | 0.09 | 0.03 | 0.06 | 0.03 | 0.06 |
| 10 | 44871003 | rs12413068 | 1.528 | 0.661 | -1.569 | -0.23 | -0.064 | -0.03 | -0.03 | 0.714 | 0.18 | 0.05 | 0.44 | 0.13 | 0.26 | 0.2 |
| 10 | 44871062 | rs266087 | 0.9 | 0.529 | -1.57 | -0.228 | 0.032 | -0.03 | -0.03 | 0 | 0.45 | 0.21 | 0.32 | 0.53 | 0.39 | 0.43 |
| 10 | 44871346 | rs17881575 | 1.302 | 0.751 | -1.583 | -0.189 | -0.025 | -0.028 | -0.028 | 0 | 0.07 | 0 | 0.07 | 0.09 | 0.1 | 0.06 |
| 10 | 44871548 | rs2297630 | 1.501 | 0.762 | -1.556 | -0.194 | -0.073 | -0.04 | -0.04 | 0.759 | 0.17 | 0.02 | 0.43 | 0.13 | 0.26 | 0.2 |
| 10 | 44873849 | rs2839695 | 0.432 | 0.792 | -1.843 | 0.007 | 0.003 | -0.039 | -0.039 | 2.609 | 0.15 | 0.2 | 0.1 | 0 | 0.2 | 0.05 |
| 10 | 44881455 | rs2839686 | -0.537 | 0.999 | -1.792 | 0.085 | -0.008 | 0.012 | 0.012 | 0 | 0.22 | 0.22 | 0.17 | 0.09 | 0.3 | 0.11 |
| 10 | 44886206 | rs754618 | 0.784 | 1.131 | -1.854 | 0.137 | -0.043 | 0.019 | 0.019 | 0 | 0.26 | 0.08 | 0.46 | 0.13 | 0.31 | 0.29 |
| 10 | 44887235 | rs7900257 | 1.252 | 0.711 | -1.59 | 0.27 | 0.022 | -0.025 | -0.025 | 0 | 0.37 | 0.22 | 0.26 | 0.5 | 0.29 | 0.37 |
| 10 | 44888130 | rs6593412 | -0.896 | 1.25 | -2.101 | 0.085 | 0.021 | 0.083 | 0.083 | 0.061 | 0.47 | 0.31 | 0.64 | 0.22 | 0.6 | 0.39 |
| 10 | 44889652 | rs1855531 | -0.372 | 0.956 | -2.267 | 0.666 | -0.06 | 0 | 0 | 0.328 | 0.33 | 0.1 | 0.54 | 0.22 | 0.4 | 0.35 |
| 10 | 44894310 | rs2781546 | -0.823 | 0.534 | -2.03 | 1.024 | -0.03 | -0.124 | -0.124 | 0.422 | 0.27 | 0.08 | 0.4 | 0.5 | 0.24 | 0.44 |
| 10 | 44894443 | rs1779384 | -1.041 | 0.164 | -1.243 | 0.606 | -0.011 | 0.004 | 0.004 | 0 | 0.29 | 0.31 | 0.17 | 0.33 | 0.21 | 0.26 |
| 10 | 44895371 | rs1761316 | -0.314 | 0.199 | -1.288 | 0.788 | -0.011 | -0.065 | -0.065 | 0.065 | 0.19 | 0.07 | 0.13 | 0.35 | 0.18 | 0.34 |
| 10 | 44896035 | rs4948881 | 0.943 | -0.044 | -1.781 | 0.205 | -0.055 | 0.062 | 0.062 | 0 | 0.41 | 0.13 | 0.54 | 0.23 | 0.46 | 0.26 |
| 10 | 44896989 | rs1761325 | 0.226 | -0.072 | -1.711 | 0.343 | -0.005 | -0.039 | -0.039 | 0 | 0.21 | 0.04 | 0.13 | 0.25 | 0.2 | 0.24 |
| 10 | 44922040 | rs17156360 | 1.155 | -0.127 | -0.739 | 0.16 | -0.001 | -0.01 | -0.01 | 0.23 | 0.14 | 0.13 | 0.15 | 0.22 | 0.13 | 0.21 |
| 10 | 44923802 | rs4948627 | 0.982 | -0.159 | -0.216 | -0.146 | -0.017 | 0.037 | 0.037 | 0 | 0.4 | 0.27 | 0.55 | 0.22 | 0.43 | 0.26 |
| 10 | 44931721 | rs11239043 | 0.352 | -0.194 | -0.104 | -0.253 | 0.007 | 0.024 | 0.024 | 0 | 0.49 | 0.43 | 0.62 | 0.22 | 0.57 | 0.28 |
| 10 | 44932080 | rs11591534 | 1.173 | -0.14 | -0.437 | -0.232 | -0.002 | -0.013 | -0.013 | 0 | 0.09 | 0.09 | 0.06 | 0 | 0.14 | 0.02 |
| 10 | 44934817 | rs10900037 | 1.078 | -0.055 | -0.505 | -0.083 | -0.056 | 0.053 | 0.053 | 0 | 0.39 | 0.13 | 0.54 | 0.22 | 0.43 | 0.26 |
| 10 | 44935055 | rs1147937 | -0.481 | 0.101 | -0.212 | 0.263 | 0 | 0.023 | 0.023 | 0 | 0.75 | 0.67 | 0.77 | 0.56 | 0.76 | 0.62 |
| 10 | 44953571 | rs10900040 | 1.171 | 0.801 | -0.861 | -0.039 | -0.057 | 0.066 | 0.066 | 0 | 0.11 | 0.01 | 0.14 | 0.02 | 0.19 | 0.08 |
| 10 | 44961811 | rs1147949 | -0.754 | -0.311 | -0.15 | 0.198 | -0.002 | -0.014 | -0.014 | 0 | 0.44 | 0.54 | 0.38 | 0.77 | 0.42 | 0.7 |
| 10 | 44972848 | rs1147963 | 0.051 | -0.597 | -0.021 | 0.319 | 0.007 | -0.023 | -0.023 | 1.594 | 0.7 | 0.28 | 0.7 | 0.48 | 0.76 | 0.52 |
| 10 | 44974143 | rs1573237 | 0.46 | -0.513 | -0.123 | 0.451 | 0.011 | 0.087 | 0.087 | 0 | 0.23 | 0.39 | 0.24 | 0.45 | 0.23 | 0.37 |
| 10 | 44987921 | rs10900048 | 0.855 | -0.45 | -0.421 | 0.289 | -0.004 | 0.127 | 0.127 | 0 | 0.49 | 0.09 | 0.57 | 0.23 | 0.58 | 0.3 |
| 10 | 45002370 | rs11598086 | 0.707 | -0.441 | -0.339 | 0.296 | 0.013 | 0.093 | 0.093 | 0 | 0.54 | 0.23 | 0.6 | 0.23 | 0.61 | 0.3 |

| | | | | | | | | | | | | | | | | |
|---|---|---|---|---|---|---|---|---|---|---|---|---|---|---|---|---|
| 10 | 45002620 | rs10900054 | 0.441 | -0.564 | -0.192 | 0.473 | 0.014 | 0.085 | 0.085 | 0 | 0.23 | 0.38 | 0.24 | 0.46 | 0.22 | 0.37 |
| 10 | 45003179 | rs915085 | 0.127 | -0.19 | -0.343 | 0.642 | -0.028 | 0.039 | 0.039 | 0 | 0.84 | 0.76 | 0.9 | 0.76 | 0.92 | 0.8 |
| 10 | 45005002 | rs4948636 | 0.545 | -0.363 | -1.045 | 0.389 | -0.025 | 0.06 | 0.06 | 0.383 | 0.42 | 0.15 | 0.57 | 0.23 | 0.56 | 0.28 |
| 10 | 45005077 | rs11239100 | 0.226 | -0.35 | -1.372 | 0.421 | 0.006 | 0.002 | 0.002 | 1.833 | 0.41 | 0.36 | 0.27 | 0.4 | 0.33 | 0.48 |
| 10 | 45005988 | rs10900055 | 0.249 | 0.255 | -2.165 | 0.281 | -0.119 | -0.004 | -0.004 | 0 | 0.39 | 0.06 | 0.55 | 0.25 | 0.53 | 0.27 |
| 10 | 45006398 | rs1889923 | 0.123 | 0.673 | -2.218 | 0.218 | -0.124 | 0.005 | 0.005 | 1.655 | 0.42 | 0.05 | 0.55 | 0.27 | 0.54 | 0.31 |
| 10 | 45007373 | rs17406863 | 0.38 | 0.947 | -2.005 | 0.009 | -0.021 | 0.03 | 0.03 | 0 | 0.08 | 0.04 | 0.08 | 0 | 0.15 | 0.02 |
| 10 | 45010296 | rs11239104 | -0.553 | 0.969 | -1.613 | 0.246 | 0.002 | 0.025 | 0.025 | 0.369 | 0.2 | 0.22 | 0.15 | 0.31 | 0.22 | 0.27 |
| 10 | 45012008 | rs6593427 | 0.135 | 0.214 | -2.077 | 0.11 | 0.003 | 0.028 | 0.028 | 0 | 0.26 | 0.1 | 0.21 | 0.23 | 0.26 | 0.31 |
| 10 | 45012043 | rs12265667 | 0.739 | 0.19 | -2.095 | 0.281 | -0.014 | 0.025 | 0.025 | 0 | 0.1 | 0.2 | 0.03 | 0.12 | 0.06 | 0.11 |
| 10 | 45012946 | rs10900056 | 1.979 | 0.42 | -2.631 | -0.244 | -0.036 | 0.078 | 0.078 | 1.347 | 0.15 | 0.02 | 0.07 | 0 | 0.15 | 0.08 |
| 10 | 45026907 | rs10900058 | -1.293 | 0.027 | -1.514 | 0.365 | 0.004 | 0.003 | 0.003 | 0.336 | 0.48 | 0.56 | 0.39 | 0.62 | 0.47 | 0.6 |
| 10 | 45032962 | rs12359952 | 0.774 | 0.057 | -1.467 | 0.389 | -0.016 | 0.023 | 0.023 | 0 | 0.11 | 0.2 | 0.03 | 0.12 | 0.05 | 0.11 |
| 10 | 45034672 | rs7072122 | 0.851 | 0.047 | -1.31 | 0.364 | 0.002 | -0.002 | -0.002 | 0 | 0.1 | 0.05 | 0.02 | 0.12 | 0.05 | 0.11 |
| 10 | 45047674 | rs10437521 | -0.95 | -0.052 | -1.356 | 0.332 | -0.021 | 0.098 | 0.098 | 0 | 0.24 | 0.46 | 0.18 | 0.39 | 0.22 | 0.29 |
| 10 | 45049498 | rs7076575 | -1.354 | -0.09 | -1.457 | 0.318 | 0.001 | 0.001 | 0.001 | 0 | 0.47 | 0.4 | 0.36 | 0.62 | 0.47 | 0.62 |
| 10 | 45052047 | rs11239130 | -0.827 | -0.179 | -1.492 | 0.317 | 0.007 | -0.002 | -0.002 | 0 | 0.38 | 0.3 | 0.25 | 0.43 | 0.32 | 0.45 |
| 10 | 45052108 | rs11239131 | 0.737 | -0.047 | -1.485 | 0.291 | -0.013 | 0.016 | 0.016 | 0 | 0.11 | 0.19 | 0.03 | 0.12 | 0.06 | 0.11 |
| 10 | 45053522 | rs11597734 | 0.347 | 0.162 | -1.678 | 0.265 | -0.033 | -0.008 | -0.008 | 0 | 0.22 | 0.02 | 0.19 | 0.23 | 0.24 | 0.31 |
| 10 | 45059905 | rs10159976 | -0.923 | 0.246 | -1.715 | 0.267 | -0.005 | -0.012 | -0.012 | 0 | 0.45 | 0.27 | 0.51 | 0.51 | 0.52 | 0.57 |
| 10 | 45062495 | rs11239139 | -0.912 | 0.228 | -1.688 | 0.252 | -0.041 | 0.022 | 0.022 | 0 | 0.45 | 0.14 | 0.5 | 0.43 | 0.52 | 0.55 |
| 10 | 45063198 | rs12415991 | 0.996 | 0.339 | -1.769 | 0.302 | -0.037 | -0.048 | -0.048 | 0.359 | 0.08 | 0.01 | 0.11 | 0.18 | 0.15 | 0.15 |
| 10 | 45066922 | rs10900072 | -0.821 | 0.169 | -1.595 | 0.247 | -0.008 | 0.015 | 0.015 | 0 | 0.43 | 0.33 | 0.51 | 0.43 | 0.51 | 0.55 |
| 10 | 45072819 | rs899889 | -0.582 | 0.267 | -1.403 | 0.331 | -0.035 | -0.006 | -0.006 | 1.963 | 0.38 | 0.19 | 0.5 | 0.43 | 0.48 | 0.53 |
| 10 | 45078170 | rs11239157 | -0.577 | 0.272 | -1.404 | 0.333 | -0.036 | -0.007 | -0.007 | 0.05 | 0.37 | 0.19 | 0.5 | 0.43 | 0.48 | 0.53 |
| 10 | 45102420 | rs7918194 | 1.267 | 0.056 | -0.885 | 0.339 | -0.032 | 0.058 | 0.058 | 0 | 0.43 | 0.24 | 0.4 | 0.23 | 0.44 | 0.26 |
| 10 | 45107555 | rs10900079 | -0.8 | 0.45 | -0.811 | 0.123 | 0.014 | 0.007 | 0.007 | 0 | 0.27 | 0.4 | 0.3 | 0.28 | 0.39 | 0.27 |
| 10 | 45117466 | rs1904837 | -1.157 | 0.387 | -0.701 | -0.122 | -0.001 | 0.006 | 0.006 | 0 | 0.42 | 0.42 | 0.52 | 0.34 | 0.48 | 0.49 |
| 10 | 45118949 | rs17156689 | 0.868 | 0.366 | -0.734 | -0.132 | 0.024 | 0.013 | 0.013 | 0 | 0.17 | 0.06 | 0.25 | 0.18 | 0.11 | 0.29 |
| 10 | 45123379 | rs11239177 | -1.218 | 0.418 | -0.732 | -0.117 | -0.001 | -0.005 | -0.005 | 0.679 | 0.43 | 0.48 | 0.54 | 0.34 | 0.48 | 0.49 |
| 10 | 45124638 | rs999639 | 0.151 | 0.52 | -0.982 | -0.154 | -0.037 | 0.032 | 0.032 | 0 | 0.14 | 0.08 | 0.17 | 0.08 | 0.21 | 0.14 |
| 10 | 45125211 | rs17156715 | 0.971 | 0.631 | -0.993 | -0.163 | -0.009 | 0.012 | 0.012 | 0 | 0.08 | 0.07 | 0.07 | 0.03 | 0.09 | 0.1 |
| 10 | 45126887 | rs1002589 | -1.255 | 0.52 | -0.921 | -0.154 | -0.001 | 0.019 | 0.019 | 0 | 0.43 | 0.42 | 0.53 | 0.26 | 0.47 | 0.48 |
| 10 | 45126974 | rs1002588 | -0.484 | 0.504 | -0.906 | -0.132 | -0.001 | -0.001 | -0.001 | 0.028 | 0.28 | 0.29 | 0.35 | 0.26 | 0.26 | 0.35 |
| 10 | 45133277 | rs11239187 | -0.729 | 1.105 | -0.985 | 0.003 | -0.045 | 0.018 | 0.018 | 0 | 0.23 | 0.14 | 0.26 | 0.16 | 0.37 | 0.2 |
| 10 | 45135344 | rs17488134 | 0.677 | 1.278 | -0.982 | -0.053 | -0.037 | 0.067 | 0.067 | 0 | 0.21 | 0.08 | 0.13 | 0 | 0.23 | 0.11 |
| 10 | 45141957 | rs7922897 | -0.554 | 1.07 | -0.637 | 0.289 | -0.009 | 0.026 | 0.026 | 0.027 | 0.38 | 0.7 | 0.32 | 0.26 | 0.37 | 0.29 |
| 10 | 45161109 | rs1480597 | -0.074 | 0.937 | -0.289 | 0.801 | -0.023 | 0.028 | 0.028 | 1.585 | 0.11 | 0.09 | 0.1 | 0 | 0.19 | 0.05 |
| 10 | 45163756 | rs4035631 | -0.074 | 0.937 | -0.289 | 0.801 | -0.023 | 0.028 | 0.028 | 1.585 | 0.11 | 0.09 | 0.1 | 0 | 0.19 | 0.05 |
| 10 | 45169414 | rs12266496 | -1.865 | 0.794 | -0.03 | 1.184 | 0 | 0.018 | 0.018 | 0.019 | 0.27 | 0.66 | 0.23 | 0.45 | 0.28 | 0.23 |
| 10 | 45176155 | rs11813028 | 0.237 | 0.847 | -0.035 | 1.175 | -0.019 | 0.014 | 0.014 | 0.403 | 0.09 | 0.09 | 0.1 | 0 | 0.18 | 0.05 |
| 10 | 45181550 | rs899888 | -0.613 | 0.81 | 0.157 | 1.213 | -0.019 | 0.241 | 0.241 | 0 | 0.14 | 0.49 | 0.11 | 0.45 | 0.08 | 0.17 |
| 10 | 45182316 | rs999473 | -1.68 | 0.8 | 0.118 | 1.334 | 0 | 0.018 | 0.018 | 0.062 | 0.27 | 0.71 | 0.23 | 0.5 | 0.28 | 0.23 |
| 10 | 45193587 | rs11239211 | -1.408 | 0.785 | 0.06 | 1.339 | 0.004 | -0.003 | -0.003 | 0 | 0.24 | 0.35 | 0.2 | 0.24 | 0.28 | 0.21 |
| 10 | 45209441 | rs12772796 | 0.123 | 0.765 | 0.016 | 1.059 | 0.002 | -0.002 | -0.002 | 0 | 0.11 | 0.08 | 0.06 | 0.01 | 0.08 | 0.12 |
| 10 | 45216112 | rs11239219 | 0.24 | 0.979 | 0.286 | 1.521 | 0.001 | -0.001 | -0.001 | 0 | 0.06 | 0.03 | 0.02 | 0.04 | 0.05 | 0.07 |
| 10 | 45230151 | rs960442 | -0.06 | 1.168 | 0.289 | 1.631 | -0.028 | -0.016 | -0.016 | 0.017 | 0.1 | 0 | 0.06 | 0.1 | 0.11 | 0.14 |
| 10 | 45237208 | rs7895327 | -1.289 | 1.002 | 0.828 | 1.671 | 0.008 | 0.093 | 0.093 | 1.39 | 0.19 | 0.66 | 0.19 | 0.45 | 0.2 | 0.12 |
| 10 | 45248254 | rs10900108 | 0.357 | 1.464 | -0.007 | 0.667 | 0.024 | -0.032 | -0.032 | 4.749 | 0.92 | 0.89 | 0.91 | 1 | 0.82 | 0.95 |
| 10 | 45249394 | rs7912740 | -0.148 | 1.887 | -0.722 | 0.655 | -0.054 | -0.016 | -0.016 | 0 | 0.25 | 0.1 | 0.37 | 0.32 | 0.3 | 0.39 |
| 10 | 45252001 | rs2985704 | -0.451 | 1.325 | -0.486 | 0.557 | -0.001 | 0.002 | 0.002 | 0.063 | 0.32 | 0.73 | 0.3 | 0.31 | 0.35 | 0.19 |
| 10 | 45254830 | rs2487504 | -0.266 | 1.241 | -0.179 | 0.508 | 0.019 | -0.027 | -0.027 | 0 | 0.11 | 0.18 | 0.09 | 0.03 | 0.17 | 0.06 |
| 10 | 45262491 | rs1408815 | -1.047 | 0.666 | -0.565 | 0.615 | 0.016 | 0.019 | 0.019 | 0.131 | 0.61 | 0.75 | 0.67 | 0.72 | 0.67 | 0.63 |
| 10 | 45264421 | rs2256688 | -0.374 | 0.646 | -0.001 | 0.71 | 0.008 | -0.007 | -0.007 | 0.062 | 0.29 | 0.48 | 0.24 | 0.28 | 0.3 | 0.18 |
| 10 | 45266317 | rs1325159 | 1.508 | 0.604 | 0.318 | 0.664 | -0.003 | -0.068 | -0.068 | 0.507 | 0.16 | 0.24 | 0.11 | 0 | 0.16 | 0.09 |
| 10 | 45270373 | rs2611371 | -0.414 | 0.596 | 0.266 | 0.678 | 0.003 | -0.007 | -0.007 | 0 | 0.3 | 0.57 | 0.25 | 0.27 | 0.3 | 0.18 |
| 10 | 45273349 | rs2793304 | -1.205 | 0.403 | 0.156 | 0.746 | 0.028 | 0.039 | 0.039 | 0 | 0.57 | 0.75 | 0.65 | 0.73 | 0.63 | 0.63 |
| 10 | 45274144 | rs12268505 | 1.178 | 0.528 | 0.523 | 0.812 | -0.005 | 0.046 | 0.046 | 0 | 0.17 | 0.33 | 0.17 | 0.27 | 0.16 | 0.12 |
| 10 | 45280577 | rs2793296 | -0.143 | 0.535 | 0.534 | 0.825 | 0.007 | -0.018 | -0.018 | 0 | 0.12 | 0.15 | 0.07 | 0.03 | 0.14 | 0.07 |
| 10 | 45282703 | rs7071453 | 0.949 | 0.601 | 0.494 | 0.82 | -0.003 | 0.101 | 0.101 | 0 | 0.18 | 0.46 | 0.18 | 0.3 | 0.16 | 0.12 |
| 10 | 45300460 | rs1158153 | -0.669 | 0.598 | 0.648 | 0.775 | -0.001 | -0.001 | -0.001 | 0 | 0.3 | 0.71 | 0.26 | 0.3 | 0.29 | 0.17 |
| 10 | 45303318 | rs7918692 | 1.397 | 0.625 | 0.604 | 0.568 | -0.008 | -0.066 | -0.066 | 0 | 0.16 | 0.2 | 0.11 | 0 | 0.15 | 0.09 |
| 10 | 45307205 | rs10900114 | -0.339 | 0.733 | 0.447 | 0.646 | 0.044 | -0.059 | -0.059 | 0.953 | 0.11 | 0.29 | 0.08 | 0 | 0.14 | 0.06 |
| 10 | 45317052 | rs1963172 | 1.541 | 1.373 | -0.104 | 0.812 | -0.064 | -0.061 | -0.061 | 0 | 0.12 | 0.01 | 0.32 | 0.24 | 0.21 | 0.2 |
| 10 | 45317971 | rs7070244 | 0.715 | 1.036 | 0.46 | 0.866 | -0.03 | 0.075 | 0.075 | 0 | 0.2 | 0.54 | 0.19 | 0.29 | 0.15 | 0.11 |
| 10 | 45319654 | rs7911712 | 0.717 | 1.097 | 0.262 | 0.84 | 0.008 | -0.011 | -0.011 | 0.677 | 0.07 | 0.08 | 0.05 | 0 | 0.14 | 0.06 |
| 10 | 45320830 | rs11239252 | -0.323 | 0.803 | -0.169 | 0.755 | -0.039 | -0.093 | -0.093 | 0 | 0.33 | 0.07 | 0.38 | 0.44 | 0.32 | 0.45 |
| 10 | 45329849 | rs11239257 | 0.827 | 1.059 | -0.46 | 0.077 | -0.03 | -0.02 | -0.02 | 0 | 0.13 | 0 | 0.05 | 0.11 | 0.11 | 0.2 |
| 10 | 45339904 | rs1830888 | 0.977 | 1.225 | -0.44 | -0.452 | -0.052 | 0.033 | 0.033 | 0.974 | 0.25 | 0.03 | 0.35 | 0.2 | 0.28 | 0.25 |

| | | | | | | | | | | | | | | | | |
|---|---|---|---|---|---|---|---|---|---|---|---|---|---|---|---|---|
| 10 | 45340044 | rs7068975 | 0.516 | 1.479 | 0.69 | -0.137 | -0.002 | -0.029 | -0.029 | 0 | 0.17 | 0.2 | 0.1 | 0 | 0.1 | 0.09 |
| 10 | 45351530 | rs11239266 | -0.676 | 1.465 | 0.652 | -0.152 | -0.004 | -0.022 | -0.022 | 0 | 0.24 | 0.22 | 0.17 | 0 | 0.29 | 0.15 |
| 10 | 45352419 | rs7067780 | -0.178 | 0.298 | 0.859 | 0.242 | -0.032 | 0.047 | 0.047 | 0 | 0.2 | 0.5 | 0.19 | 0.27 | 0.15 | 0.1 |
| 10 | 45355020 | rs12098754 | -0.303 | 0.378 | 0.729 | 0.234 | -0.02 | 0.019 | 0.019 | 0 | 0.22 | 0.46 | 0.2 | 0.27 | 0.17 | 0.12 |
| 10 | 45357291 | rs912739 | -0.407 | 0.503 | 0.332 | 0.196 | -0.024 | 0.029 | 0.029 | 0 | 0.94 | 1 | 0.95 | 1 | 0.9 | 0.93 |
| 10 | 45359483 | rs6593445 | -0.389 | 0.106 | 0.597 | 0.412 | -0.023 | 0.032 | 0.032 | 0 | 0.22 | 0.56 | 0.2 | 0.27 | 0.17 | 0.13 |
| 10 | 45368956 | rs7910103 | -0.82 | -0.205 | 0.365 | 0.118 | -0.029 | -0.021 | -0.021 | 0 | 0.26 | 0.6 | 0.18 | 0.17 | 0.17 | 0.11 |
| 10 | 45371418 | rs1018702 | -0.81 | -0.208 | 0.373 | 0.127 | -0.031 | -0.022 | -0.022 | 0.431 | 0.26 | 0.6 | 0.18 | 0.17 | 0.17 | 0.11 |
| 10 | 45375296 | rs3740086 | 0.449 | 0.116 | 0.031 | 0.134 | -0.003 | 0.005 | 0.005 | 0 | 0.71 | 0.38 | 0.66 | 0.37 | 0.72 | 0.59 |
| 10 | 45380472 | rs10793562 | 0.083 | 0.077 | 0.993 | 0.265 | 0.001 | 0.016 | 0.016 | 0 | 0.75 | 0.67 | 0.71 | 0.45 | 0.72 | 0.61 |
| 10 | 45384047 | rs2065466 | 0.788 | -0.07 | 0.668 | -0.071 | -0.001 | -0.001 | -0.001 | 0.434 | 0.57 | 0.51 | 0.58 | 0.45 | 0.57 | 0.56 |
| 10 | 45384409 | rs1570985 | 0.167 | 0.317 | 0.279 | -0.526 | 0.038 | -0.057 | -0.057 | 0 | 0.16 | 0.38 | 0.17 | 0.06 | 0.2 | 0.18 |
| 10 | 45384811 | rs4506594 | -0.288 | -0.07 | 0.57 | -0.425 | -0.007 | -0.027 | -0.027 | 0 | 0.76 | 0.82 | 0.72 | 0.45 | 0.71 | 0.64 |
| 10 | 45390103 | rs10900127 | 1.379 | -0.108 | 0.415 | -0.478 | -0.041 | 0.091 | 0.091 | 0.278 | 0.46 | 0.12 | 0.47 | 0.32 | 0.48 | 0.47 |
| 10 | 45395839 | rs7909074 | 0.745 | 0.167 | 0.57 | -0.16 | -0.002 | -0.008 | -0.008 | 0 | 0.58 | 0.42 | 0.56 | 0.37 | 0.57 | 0.55 |
| 10 | 45399088 | rs1408814 | 0.604 | 0.251 | 0.127 | -0.124 | -0.001 | 0.006 | 0.006 | 0 | 0.59 | 0.55 | 0.57 | 0.37 | 0.58 | 0.55 |
| 10 | 45402198 | rs1332626 | 0.638 | 0.222 | -0.334 | -0.141 | 0.002 | 0.018 | 0.018 | 0.036 | 0.16 | 0.05 | 0.13 | 0.07 | 0.13 | 0.09 |
| 10 | 45402325 | rs1332624 | 0.517 | 0.178 | -0.383 | -0.159 | -0.001 | 0.018 | 0.018 | 0.008 | 0.17 | 0.02 | 0.13 | 0.07 | 0.13 | 0.09 |
| 10 | 45407333 | rs4147126 | 0.235 | 0.57 | 0.173 | 0.261 | 0 | 0.008 | 0.008 | 0 | 0.74 | 0.65 | 0.75 | 0.5 | 0.7 | 0.61 |
| 10 | 45408011 | rs7904676 | -0.16 | 0.662 | 0.03 | 0.279 | -0.004 | -0.001 | -0.001 | 0 | 0.88 | 0.87 | 0.81 | 0.48 | 0.88 | 0.66 |
| 10 | 45409244 | rs6593449 | 0.782 | 0.411 | -0.25 | -0.069 | -0.047 | -0.082 | -0.082 | 0.044 | 0.76 | 0.86 | 0.74 | 0.45 | 0.7 | 0.53 |
| 10 | 45410971 | rs4543933 | 1.362 | 0.324 | 0.266 | -0.387 | 0.003 | -0.007 | -0.007 | 0 | 0.64 | 0.56 | 0.63 | 0.35 | 0.59 | 0.36 |
| 10 | 45412777 | rs17417060 | 0.32 | 0.317 | 0.034 | -0.404 | -0.005 | 0.035 | 0.035 | 0 | 0.1 | 0.03 | 0.05 | 0.05 | 0.1 | 0.16 |
| 10 | 45412787 | rs7923073 | 0.038 | 0.357 | 0.014 | -0.402 | -0.052 | -0.102 | -0.102 | 0.18 | 0.81 | 0.9 | 0.79 | 0.47 | 0.76 | 0.56 |
| 10 | 45415345 | rs932986 | 1.225 | 0.306 | 0.296 | -0.403 | 0.013 | -0.015 | -0.015 | 0 | 0.65 | 0.55 | 0.62 | 0.35 | 0.59 | 0.38 |
| 10 | 45420076 | rs1332622 | 1.495 | 0.132 | 0.58 | -0.446 | -0.005 | 0.018 | 0.018 | 3.331 | 0.62 | 0.33 | 0.6 | 0.32 | 0.59 | 0.36 |
| 10 | 45420416 | rs10793569 | 0.195 | 0.168 | 0.752 | -0.436 | 0.009 | 0.052 | 0.052 | 0 | 0.8 | 0.72 | 0.78 | 0.46 | 0.77 | 0.56 |
| 10 | 45424516 | rs11239282 | 1.347 | 0.109 | 0.149 | -0.596 | -0.006 | 0.048 | 0.048 | 2.669 | 0.55 | 0.34 | 0.56 | 0.31 | 0.53 | 0.33 |
| 10 | 45424779 | rs7098274 | 1.103 | 0.091 | 0.163 | -0.224 | -0.004 | 0 | 0 | 2.627 | 0.63 | 0.34 | 0.62 | 0.42 | 0.6 | 0.41 |
| 10 | 45426616 | rs1109423 | 0.108 | -0.237 | -0.49 | -0.891 | 0.007 | 0.109 | 0.109 | 0 | 0.21 | 0.04 | 0.14 | 0 | 0.17 | 0.12 |
| 10 | 45430462 | rs17417442 | 1.731 | -0.288 | -0.277 | -0.925 | -0.004 | 0.024 | 0.024 | 1.015 | 0.4 | 0.31 | 0.44 | 0.33 | 0.4 | 0.23 |
| 10 | 45430699 | rs7067692 | -0.698 | -0.102 | -1.171 | -0.807 | -0.03 | -0.055 | -0.055 | 0.049 | 0.83 | 0.91 | 0.77 | 0.47 | 0.75 | 0.55 |
| 10 | 45430956 | rs7084511 | 0.806 | -0.147 | -1.176 | -0.826 | 0.01 | 0.06 | 0.06 | 0 | 0.15 | 0.04 | 0.11 | 0 | 0.11 | 0.1 |
| 10 | 45431321 | rs10751362 | -0.101 | -0.117 | -1.107 | -0.764 | -0.013 | -0.028 | -0.028 | 0 | 0.79 | 0.84 | 0.76 | 0.4 | 0.75 | 0.54 |
| 10 | 45431416 | rs10900132 | -0.655 | -0.016 | -1.265 | -0.746 | -0.028 | -0.052 | -0.052 | 3.543 | 0.84 | 0.91 | 0.77 | 0.47 | 0.76 | 0.56 |
| 10 | 45432386 | rs2297494 | -0.709 | -0.683 | -0.932 | -0.872 | -0.004 | 0.046 | 0.046 | 0 | 0.22 | 0.14 | 0.16 | 0.06 | 0.23 | 0.18 |
| 10 | 45442303 | rs11597898 | -1.083 | -0.657 | -0.898 | -0.841 | 0.01 | 0.117 | 0.117 | 0 | 0.36 | 0.2 | 0.27 | 0.06 | 0.34 | 0.28 |
| 10 | 45444490 | rs1332628 | -1.103 | -0.626 | -0.973 | -0.695 | 0.002 | 0.128 | 0.128 | 0 | 0.3 | 0.12 | 0.21 | 0.05 | 0.27 | 0.25 |
| 10 | 45446048 | rs7908664 | -0.092 | -0.561 | -0.816 | -0.546 | 0.018 | 0.039 | 0.039 | 0 | 0.16 | 0.06 | 0.15 | 0 | 0.12 | 0.12 |
| 10 | 45446791 | rs10900135 | -0.632 | -0.565 | -0.156 | -0.281 | 0.018 | 0.055 | 0.055 | 0.304 | 0.12 | 0.33 | 0.08 | 0.15 | 0.11 | 0.16 |
| 10 | 45446917 | rs10900136 | -0.827 | -1.17 | -0.601 | -0.281 | 0.028 | 0.005 | 0.005 | 0 | 0.42 | 0.64 | 0.28 | 0.4 | 0.31 | 0.3 |
| 10 | 45451178 | rs6593451 | -0.725 | -0.888 | -1.078 | -0.813 | -0.018 | 0.064 | 0.064 | 0.326 | 0.61 | 0.79 | 0.46 | 0.97 | 0.41 | 0.65 |
| 10 | 45453950 | rs10900143 | 0.092 | -1.143 | -0.606 | 0.003 | -0.026 | 0.073 | 0.073 | 0.199 | 0.26 | 0.4 | 0.25 | 0.51 | 0.17 | 0.37 |
| 10 | 45455227 | rs12773699 | 0.03 | -1.131 | -0.606 | 0.038 | -0.028 | 0.086 | 0.086 | 0.298 | 0.26 | 0.45 | 0.25 | 0.5 | 0.17 | 0.37 |
| 10 | 45464035 | rs3814566 | -0.05 | -1.307 | -0.669 | 0.361 | -0.017 | 0.007 | 0.007 | 0.015 | 0.59 | 0.52 | 0.65 | 0.47 | 0.72 | 0.48 |
| 10 | 45464100 | rs3829908 | 0.065 | -1.368 | -0.696 | 0.206 | -0.015 | -0.015 | -0.015 | 0 | 0.57 | 0.5 | 0.65 | 0.39 | 0.72 | 0.47 |
| 10 | 45467703 | rs914700 | 0.213 | -1.31 | -0.996 | 0.321 | 0.036 | 0.101 | 0.101 | 0 | 0.55 | 0.18 | 0.59 | 0.35 | 0.65 | 0.45 |
| 10 | 45471222 | rs3740098 | 0.237 | -1.317 | -1.039 | 0.315 | 0.04 | 0.107 | 0.107 | 0 | 0.55 | 0.16 | 0.59 | 0.35 | 0.65 | 0.45 |
| 10 | 45475917 | rs3758402 | 0.059 | -1.493 | -0.747 | 1.325 | 0.02 | -0.015 | -0.015 | 0.003 | 0.62 | 0.78 | 0.79 | 0.62 | 0.68 | 0.56 |
| 10 | 45478092 | rs870957 | -0.475 | -1.732 | -0.744 | 1.596 | -0.024 | 0.066 | 0.066 | 0 | 0.21 | 0.38 | 0.14 | 0.32 | 0.17 | 0.32 |
| 10 | 45481452 | rs7896801 | 0.281 | -1.612 | -0.941 | 1.613 | 0.014 | -0.026 | -0.026 | 1.374 | 0.66 | 0.78 | 0.7 | 0.58 | 0.74 | 0.71 |
| 10 | 45491371 | rs2297492 | 0.189 | -1.571 | -1.262 | 1.71 | 0.016 | -0.058 | -0.058 | 0.095 | 0.16 | 0.02 | 0.09 | 0.26 | 0.09 | 0.1 |
| 10 | 45495555 | rs12219891 | 0.253 | -1.579 | -1.26 | 1.822 | 0.015 | -0.058 | -0.058 | 0 | 0.15 | 0.02 | 0.09 | 0.26 | 0.09 | 0.1 |
| 10 | 45497915 | rs7068660 | 1.647 | -1.118 | -1.816 | 1.157 | -0.128 | 0.101 | 0.101 | 0.396 | 0.23 | 0.03 | 0.43 | 0.05 | 0.44 | 0.23 |
| 10 | 45499799 | rs11471 | -0.241 | -1.408 | -1.44 | 1.52 | 0.013 | -0.016 | -0.016 | 0 | 0.67 | 0.78 | 0.81 | 0.62 | 0.76 | 0.76 |
| 10 | 45500221 | rs11494 | -0.497 | -1.631 | -1.029 | 1.338 | 0.028 | 0.257 | 0.257 | 4.685 | 0.1 | 0.52 | 0.23 | 0.33 | 0.06 | 0.21 |
| 10 | 45504012 | rs6593454 | -0.236 | -1.631 | -0.868 | 1.303 | 0.056 | 0.134 | 0.134 | 1.786 | 0.08 | 0.45 | 0.09 | 0.15 | 0.06 | 0.16 |
| 10 | 45511408 | rs6593456 | -0.217 | -1.8 | -1.089 | 1.239 | 0.009 | -0.017 | -0.017 | 0 | 0.65 | 0.73 | 0.8 | 0.53 | 0.73 | 0.73 |
| 10 | 45511577 | rs4948652 | 0.737 | -1.804 | -1.1 | 1.242 | 0.046 | 0.152 | 0.152 | 0 | 0.54 | 0.21 | 0.56 | 0.2 | 0.66 | 0.5 |
| 10 | 45515053 | rs7902918 | -0.676 | -1.661 | -0.927 | 1.407 | 0.056 | 0.299 | 0.299 | 4.545 | 0.09 | 0.49 | 0.22 | 0.33 | 0.06 | 0.21 |
| 10 | 45548029 | rs1936395 | 0.088 | -2.325 | -0.775 | 1.449 | -0.063 | 0.046 | 0.046 | 0.862 | 0.2 | 0.55 | 0.21 | 0.31 | 0.09 | 0.22 |
| 10 | 45548068 | rs1936396 | -0.299 | -2.415 | -0.89 | 1.528 | 0.021 | -0.033 | -0.033 | 0 | 0.67 | 0.82 | 0.77 | 0.6 | 0.77 | 0.76 |
| 10 | 45548366 | rs7478211 | -1.144 | -2.462 | -0.936 | 1.688 | -0.001 | 0.006 | 0.006 | 0 | 0.79 | 0.77 | 0.84 | 0.77 | 0.82 | 0.81 |
| 10 | 45550750 | rs7905526 | -0.176 | -2.287 | -0.844 | 1.439 | 0.008 | -0.014 | -0.014 | 0 | 0.66 | 0.73 | 0.76 | 0.51 | 0.73 | 0.73 |
| 10 | 45552865 | rs10793581 | -0.289 | -2.288 | -0.845 | 1.437 | 0.015 | -0.022 | -0.022 | 0 | 0.67 | 0.76 | 0.77 | 0.6 | 0.77 | 0.76 |
| 10 | 45553282 | rs10751369 | 1.057 | -2.183 | -0.832 | 1.035 | 0.027 | 0.153 | 0.153 | 1.407 | 0.45 | 0.16 | 0.54 | 0.2 | 0.63 | 0.49 |
| 10 | 45559980 | rs7077032 | 0.032 | -2.105 | -0.968 | 1.332 | 0.01 | -0.013 | -0.013 | 0 | 0.66 | 0.75 | 0.74 | 0.44 | 0.73 | 0.64 |
| 10 | 45565290 | rs7913550 | 0.72 | -1.774 | -0.738 | 0.847 | 0.031 | 0.042 | 0.042 | 0.984 | 0.51 | 0.41 | 0.59 | 0.2 | 0.7 | 0.55 |
| 10 | 45581813 | rs10793583 | -0.333 | -1.583 | -0.56 | 1.453 | 0.008 | -0.015 | -0.015 | 0 | 0.76 | 0.81 | 0.83 | 0.7 | 0.81 | 0.69 |

| | | | | | | | | | | | | | | | | |
|---|---|---|---|---|---|---|---|---|---|---|---|---|---|---|---|---|
| 10 | 45589783 | rs2002497 | 0.48 | -1.928 | -0.513 | 1.208 | 0.022 | 0.004 | 0.004 | 2.079 | 0.55 | 0.4 | 0.66 | 0.46 | 0.75 | 0.61 |
| 10 | 45598618 | rs1856591 | 0.65 | -1.088 | -0.625 | 1.025 | 0.004 | 0.005 | 0.005 | 0 | 0.41 | 0.43 | 0.61 | 0.52 | 0.51 | 0.38 |
| 10 | 45610044 | rs12413391 | 0.723 | -1.101 | -0.745 | 0.957 | 0.01 | -0.004 | -0.004 | 0.156 | 0.39 | 0.44 | 0.59 | 0.42 | 0.51 | 0.38 |
| 10 | 45613231 | rs2311745 | -0.429 | -0.982 | -0.713 | 0.803 | -0.01 | 0.023 | 0.023 | 0 | 0.74 | 0.67 | 0.8 | 0.72 | 0.81 | 0.67 |
| 10 | 45648517 | rs10751374 | -0.051 | -1.226 | -0.701 | 0.852 | -0.005 | -0.004 | -0.004 | 0 | 0.76 | 0.3 | 0.82 | 0.89 | 0.85 | 0.84 |
| 10 | 45658078 | rs3851262 | 0.365 | -1.358 | -0.616 | 0.997 | 0.019 | -0.006 | -0.006 | 0 | 0.73 | 0.2 | 0.74 | 0.79 | 0.85 | 0.81 |
| 10 | 48428267 | rs1870174 | 0.949 | -2.133 | 0.067 | 0.454 | -0.009 | 0.061 | 0.061 | 1.886 | 0.91 | 0.55 | 0.89 | 0.93 | 0.98 | 0.9 |
| 10 | 48429650 | rs2236723 | 1.045 | -2.133 | -0.077 | 0.48 | -0.057 | 0.081 | 0.081 | 0 | 0.83 | 0.46 | 0.8 | 0.74 | 0.94 | 0.8 |
| 10 | 48431110 | rs2853838 | 1.409 | -2.118 | -0.195 | 0.435 | -0.019 | -0.011 | -0.011 | 0 | 0.7 | 0.33 | 0.78 | 0.78 | 0.79 | 0.66 |
| 10 | 48438147 | rs1902724 | 1.578 | -1.891 | -0.631 | 0.563 | 0.065 | 0.018 | 0.018 | 0 | 0.58 | 0.19 | 0.65 | 0.6 | 0.69 | 0.54 |
| 10 | 48442146 | rs2853840 | 1.898 | -1.91 | -0.491 | 0.636 | 0.007 | 0.002 | 0.002 | 0 | 0.6 | 0.37 | 0.66 | 0.59 | 0.68 | 0.56 |
| 10 | 48442894 | rs17537178 | -0.568 | -1.911 | -0.313 | 0.383 | -0.006 | 0.036 | 0.036 | 0 | 0.22 | 0.17 | 0.16 | 0.11 | 0.26 | 0.24 |
| 10 | 48443093 | rs11593867 | -1.301 | -1.904 | -0.303 | 0.754 | 0.025 | -0.038 | -0.038 | 0.187 | 0.29 | 0.17 | 0.28 | 0.33 | 0.27 | 0.34 |
| 10 | 48445554 | rs1993488 | 1.783 | -1.366 | 0.44 | 1.218 | 0.003 | 0.292 | 0.292 | 1.758 | 0.91 | 0.55 | 0.84 | 0.78 | 0.98 | 0.92 |
| 10 | 48456176 | rs878263 | 0.067 | -0.144 | -0.28 | 0.865 | 0.011 | 0.006 | 0.006 | 0 | 0.47 | 0.44 | 0.56 | 0.51 | 0.59 | 0.53 |
| 10 | 48471020 | rs12414155 | 0.497 | -0.595 | -0.191 | 1.234 | 0.048 | 0.032 | 0.032 | 0 | 0.5 | 0.36 | 0.56 | 0.58 | 0.63 | 0.57 |
| 10 | 48473929 | rs7076160 | 0.167 | -0.304 | -0.162 | 1.167 | -0.005 | -0.003 | -0.003 | 0 | 0.25 | 0.21 | 0.3 | 0.18 | 0.22 | 0.29 |
| 10 | 48480228 | rs10788705 | -0.677 | 0.549 | -0.295 | 1.037 | 0.008 | 0.018 | 0.018 | 0 | 0.37 | 0.29 | 0.55 | 0.76 | 0.34 | 0.58 |
| 10 | 48480248 | rs2376611 | -0.875 | 0.864 | -0.208 | 1.44 | 0.009 | -0.001 | -0.001 | 0 | 0.48 | 0.34 | 0.56 | 0.49 | 0.42 | 0.56 |
| 10 | 48480371 | rs10788706 | 0.244 | 0.937 | -0.338 | 1.141 | -0.002 | -0.001 | -0.001 | 2.141 | 0.25 | 0.22 | 0.3 | 0.18 | 0.22 | 0.24 |
| 10 | 48480480 | rs9421818 | 1.118 | 1.326 | -0.156 | 1.167 | -0.004 | 0.011 | 0.011 | 0 | 0.43 | 0.36 | 0.35 | 0.42 | 0.48 | 0.31 |
| 10 | 48482465 | rs11204232 | 0.714 | 0.561 | 0.056 | 1.425 | -0.001 | 0 | 0 | 1.762 | 0.2 | 0.13 | 0.25 | 0.18 | 0.14 | 0.17 |
| 10 | 48483044 | rs9421742 | 0.389 | 0.687 | 0.131 | 1.373 | -0.001 | -0.004 | -0.004 | 0 | 0.6 | 0.64 | 0.62 | 0.78 | 0.67 | 0.72 |
| 10 | 48483653 | rs9421743 | 0.757 | 0.76 | -0.167 | 1.139 | 0.005 | 0.027 | 0.027 | 0 | 0.48 | 0.39 | 0.38 | 0.42 | 0.56 | 0.4 |
| 10 | 48485956 | rs4075854 | -0.088 | 0.797 | -0.245 | 1.187 | -0.001 | -0.027 | -0.027 | 0.233 | 0.8 | 0.77 | 0.87 | 0.96 | 0.81 | 0.89 |
| 10 | 48497360 | rs4397779 | -0.714 | 1.34 | -0.374 | 1.011 | -0.01 | 0.054 | 0.054 | 0.013 | 0.46 | 0.24 | 0.4 | 0.29 | 0.5 | 0.36 |
| 10 | 48499234 | rs9421747 | 0.175 | 1.619 | -0.536 | 0.847 | 0.006 | 0.028 | 0.028 | 0 | 0.23 | 0.11 | 0.18 | 0.13 | 0.21 | 0.19 |
| 10 | 48501475 | rs9421750 | -0.995 | 1.579 | -0.491 | 0.922 | -0.01 | 0.053 | 0.053 | 0.018 | 0.46 | 0.24 | 0.4 | 0.29 | 0.5 | 0.36 |
| 10 | 48508137 | rs7895237 | -1.165 | 1.535 | -0.689 | 0.893 | 0.006 | 0.052 | 0.052 | 0 | 0.51 | 0.6 | 0.45 | 0.29 | 0.51 | 0.36 |
| 10 | 48511877 | rs4598610 | -0.909 | 1.628 | -0.742 | 0.827 | 0.003 | 0.031 | 0.031 | 0 | 0.45 | 0.59 | 0.4 | 0.3 | 0.44 | 0.29 |
| 10 | 48512114 | rs4601698 | -1.264 | 1.091 | -0.733 | 0.832 | 0.009 | 0.008 | 0.008 | 0.132 | 0.49 | 0.6 | 0.41 | 0.49 | 0.45 | 0.35 |
| 10 | 48516636 | rs7901713 | 0.996 | 1.703 | -0.61 | 0.986 | -0.014 | -0.014 | -0.014 | 0 | 0.69 | 0.27 | 0.69 | 0.61 | 0.64 | 0.81 |
| 10 | 48521126 | rs4341471 | -0.585 | 1.847 | -0.852 | 0.352 | -0.073 | -0.039 | -0.039 | 0.183 | 0.16 | 0.04 | 0.22 | 0.2 | 0.31 | 0.14 |
| 10 | 48524900 | rs4255475 | 0.814 | 0.798 | -0.153 | 0.073 | -0.031 | -0.087 | -0.087 | 0 | 0.86 | 0.43 | 0.93 | 1 | 0.95 | 0.96 |
| 10 | 48529987 | rs12262083 | -0.613 | 1.211 | -1.11 | 0.002 | -0.046 | 0.058 | 0.058 | 0 | 0.38 | 0.27 | 0.3 | 0.28 | 0.46 | 0.28 |
| 10 | 49659559 | rs10776601 | 1.431 | -0.843 | 1.332 | -0.758 | -0.055 | -0.092 | -0.092 | 0 | 0.8 | 0.64 | 0.7 | 0.97 | 0.91 | 0.75 |
| 10 | 49661665 | rs4838592 | 1.322 | -0.86 | 1.338 | -0.611 | -0.084 | -0.118 | -0.118 | 0 | 0.82 | 0.55 | 0.93 | 0.97 | 0.96 | 0.82 |
| 10 | 49683897 | rs3844600 | 1.178 | -0.401 | 0.654 | -0.31 | 0.01 | -0.01 | -0.01 | 0 | 0.69 | 0.28 | 0.74 | 0.38 | 0.76 | 0.75 |
| 10 | 49686384 | rs10857580 | -1.092 | -0.752 | 0.868 | -0.227 | -0.046 | 0.081 | 0.081 | 0 | 0.21 | 0.33 | 0.21 | 0.61 | 0.1 | 0.33 |
| 10 | 49696844 | rs2377354 | 1.697 | -0.784 | 0.139 | -0.255 | 0.067 | -0.008 | -0.008 | 0 | 0.63 | 0.12 | 0.66 | 0.32 | 0.77 | 0.56 |
| 10 | 49698113 | rs1077960 | -1.065 | -0.368 | 0.862 | 0.088 | -0.029 | 0.088 | 0.088 | 1.658 | 0.17 | 0.29 | 0.21 | 0.62 | 0.09 | 0.31 |
| 10 | 49701384 | rs11812546 | -0.744 | -0.51 | 0.779 | -0.405 | -0.005 | -0.035 | -0.035 | 2.798 | 0.22 | 0.28 | 0.16 | 0.05 | 0.21 | 0.1 |
| 10 | 49702383 | rs7913593 | -0.458 | 0.528 | -0.028 | -0.342 | 0.001 | 0.008 | 0.008 | 0 | 0.18 | 0.21 | 0.19 | 0.34 | 0.24 | 0.4 |
| 10 | 49703242 | rs10776610 | -0.399 | 0.527 | -0.012 | -0.398 | 0.004 | 0.012 | 0.012 | 0 | 0.18 | 0.21 | 0.19 | 0.33 | 0.24 | 0.39 |
| 10 | 49705636 | rs9663790 | -1.203 | 0.309 | 0.085 | -0.376 | 0.008 | -0.007 | -0.007 | 0.457 | 0.32 | 0.67 | 0.34 | 0.63 | 0.32 | 0.26 |
| 10 | 49730920 | rs11101368 | -0.205 | 0.594 | 0.544 | -0.421 | 0.004 | -0.069 | -0.069 | 0 | 0.21 | 0.32 | 0.21 | 0.06 | 0.24 | 0.13 |
| 10 | 49735098 | rs11101371 | 0.093 | 1.213 | 0.455 | -0.242 | -0.004 | 0.009 | 0.009 | 0 | 0.12 | 0.14 | 0.14 | 0.36 | 0.17 | 0.35 |
| 10 | 49735563 | rs10776612 | -1.284 | 1.076 | 0.45 | -0.276 | -0.004 | -0.006 | -0.006 | 0.212 | 0.36 | 0.62 | 0.37 | 0.42 | 0.4 | 0.48 |
| 10 | 49741135 | rs1947075 | 0.477 | 0.757 | 0.338 | -0.306 | -0.006 | 0.005 | 0.005 | 0 | 0.65 | 0.48 | 0.67 | 0.37 | 0.63 | 0.64 |
| 10 | 49743779 | rs11101376 | -0.329 | 0.964 | 0.583 | 0.195 | 0.019 | 0.144 | 0.144 | 0 | 0.13 | 0.28 | 0.16 | 0.46 | 0.13 | 0.34 |
| 10 | 49749918 | rs6537567 | -1.348 | 1.13 | 0.472 | -0.721 | -0.007 | 0.162 | 0.162 | 0.692 | 0.51 | 0.73 | 0.44 | 0.98 | 0.49 | 0.75 |
| 10 | 49753654 | rs7096053 | -0.162 | 0.985 | 0.699 | -0.626 | -0.013 | -0.077 | -0.077 | 0 | 0.27 | 0.37 | 0.19 | 0.08 | 0.25 | 0.15 |
| 10 | 49756258 | rs747533 | -0.101 | 1.17 | 0.701 | -0.629 | -0.002 | -0.048 | -0.048 | 2.337 | 0.27 | 0.27 | 0.2 | 0.07 | 0.27 | 0.15 |
| 10 | 49756362 | rs747532 | -1.578 | 0.961 | 0.568 | -0.853 | -0.012 | 0.1 | 0.1 | 1.364 | 0.54 | 0.7 | 0.46 | 0.98 | 0.52 | 0.77 |
| 10 | 49757251 | rs935270 | -1.217 | 1.124 | 0.354 | -1.24 | 0.002 | -0.06 | -0.06 | 0 | 0.49 | 0.49 | 0.46 | 0.96 | 0.56 | 0.83 |
| 10 | 49759708 | rs1838571 | -0.82 | 0.842 | 0.154 | -1.311 | -0.004 | -0.029 | -0.029 | 0 | 0.43 | 0.42 | 0.39 | 0.96 | 0.44 | 0.76 |
| 10 | 49760869 | rs4838419 | -1.645 | 0.727 | 0.062 | -1.371 | -0.042 | 0.163 | 0.163 | 0.059 | 0.57 | 0.78 | 0.67 | 0.97 | 0.52 | 0.79 |
| 10 | 49760981 | rs7895981 | -1.021 | 0.305 | -0.716 | -1.923 | -0.033 | 0.062 | 0.062 | 0.193 | 0.79 | 0.86 | 0.8 | 0.97 | 0.73 | 0.86 |
| 10 | 49761146 | rs11101380 | 1.294 | 0.224 | -0.594 | -1.23 | 0.008 | -0.021 | -0.021 | 0 | 0.1 | 0.03 | 0.05 | 0.43 | 0.08 | 0.29 |
| 10 | 49761403 | rs749377 | -0.29 | 0.362 | -0.364 | -1.977 | 0.002 | 0.021 | 0.021 | 0.366 | 0.28 | 0.29 | 0.26 | 0.89 | 0.24 | 0.64 |
| 10 | 49762077 | rs2004665 | -1.036 | 0.345 | -0.341 | -1.908 | -0.01 | 0.09 | 0.09 | 0.205 | 0.55 | 0.66 | 0.65 | 0.97 | 0.53 | 0.79 |
| 10 | 49763717 | rs7098068 | 1.716 | 0.315 | -0.169 | -1.701 | -0.001 | 0.006 | 0.006 | 0 | 0.16 | 0.12 | 0.32 | 0.07 | 0.16 | 0.12 |
| 10 | 49764050 | rs7896644 | -0.396 | 0.128 | 0.204 | -1.84 | -0.028 | -0.102 | -0.102 | 0 | 0.74 | 0.57 | 0.62 | 0.92 | 0.79 | 0.86 |
| 10 | 49764131 | rs7085335 | 1.829 | 0.135 | -0.001 | -1.822 | -0.002 | 0.009 | 0.009 | 0 | 0.16 | 0.12 | 0.32 | 0.07 | 0.16 | 0.12 |
| 10 | 49766885 | rs935264 | -0.494 | 0.047 | 0.124 | -1.849 | -0.026 | -0.071 | -0.071 | 2.499 | 0.89 | 0.59 | 0.94 | 0.99 | 0.94 | 0.97 |
| 10 | 49767546 | rs1838579 | -0.075 | 0.066 | 0.107 | -1.841 | -0.026 | -0.088 | -0.088 | 0 | 0.27 | 0.45 | 0.38 | 0.08 | 0.21 | 0.15 |
| 10 | 49773295 | rs2120905 | -0.915 | 0.176 | -0.358 | -1.912 | 0.01 | 0.023 | 0.023 | 0.426 | 0.41 | 0.64 | 0.33 | 0.9 | 0.35 | 0.66 |
| 10 | 49778422 | rs1838577 | 0.71 | 0.203 | -0.153 | -1.75 | -0.016 | -0.038 | -0.038 | 2.412 | 0.09 | 0.12 | 0.04 | 0.01 | 0.06 | 0.03 |
| 10 | 49782417 | rs10491040 | 1.816 | 0.372 | -0.411 | -1.753 | -0.011 | 0.034 | 0.034 | 0.29 | 0.16 | 0.06 | 0.33 | 0.07 | 0.16 | 0.1 |

| | | | | | | | | | | | | | | | |
|---|---|---|---|---|---|---|---|---|---|---|---|---|---|---|---|
| 10 | 49783708 | rs7906475 | 1.017 | 0.388 | -0.38 | -1.853 | -0.035 | 0.029 | 0.029 | 0 | 0.06 | 0 | 0.07 | 0 | 0.12 | 0.08 |
| 10 | 49783859 | rs7919520 | 0.366 | 0.241 | -0.36 | -2.01 | -0.026 | 0.029 | 0.029 | 0.019 | 0.22 | 0.15 | 0.23 | 0.02 | 0.3 | 0.15 |
| 10 | 49786443 | rs935291 | -1.52 | -0.122 | -0.255 | -2.107 | 0.016 | -0.008 | -0.008 | 0 | 0.69 | 0.75 | 0.83 | 0.99 | 0.76 | 0.9 |
| 10 | 49787078 | rs7098855 | -0.014 | -0.141 | -0.275 | -1.604 | -0.016 | -0.016 | -0.016 | 0 | 0.36 | 0.53 | 0.48 | 0.69 | 0.29 | 0.54 |
| 10 | 49790972 | rs2289808 | 0.403 | -0.015 | -0.545 | -1.699 | 0.012 | 0.023 | 0.023 | 0 | 0.29 | 0.22 | 0.45 | 0.73 | 0.28 | 0.5 |
| 10 | 49792382 | rs10857606 | 0.905 | -0.058 | -0.516 | -1.533 | 0.004 | -0.032 | -0.032 | 0.868 | 0.25 | 0.14 | 0.4 | 0.61 | 0.26 | 0.38 |
| 10 | 49793554 | rs4320911 | 0.946 | -0.21 | -0.319 | -1.889 | -0.015 | -0.044 | -0.044 | 0 | 0.12 | 0.12 | 0.05 | 0.01 | 0.07 | 0.05 |
| 10 | 49794481 | rs1345107 | -2.148 | -0.193 | -0.147 | -2 | -0.02 | -0.125 | -0.125 | 0 | 0.76 | 0.68 | 0.85 | 1 | 0.81 | 0.92 |
| 10 | 49795636 | rs10491035 | -0.042 | 0.019 | -0.241 | -2.025 | -0.017 | 0.06 | 0.06 | 0 | 0.39 | 0.3 | 0.57 | 0.96 | 0.44 | 0.71 |
| 10 | 49797216 | rs1814474 | -0.935 | -0.812 | 0.09 | -1.746 | -0.035 | -0.168 | -0.168 | 0.153 | 0.64 | 0.58 | 0.81 | 0.99 | 0.74 | 0.87 |
| 10 | 49797472 | rs870353 | 0.813 | -0.504 | -0.334 | -1.448 | 0.005 | -0.011 | -0.011 | 0 | 0.12 | 0.09 | 0.04 | 0.01 | 0.07 | 0.05 |
| 10 | 49798463 | rs1806437 | -0.552 | 0.038 | -0.144 | -0.966 | 0.003 | -0.009 | -0.009 | 0.437 | 0.31 | 0.37 | 0.3 | 0.23 | 0.34 | 0.32 |
| 10 | 49799453 | rs3851552 | -0.909 | 0.729 | -0.096 | -0.079 | -0.004 | -0.003 | -0.003 | 0 | 0.25 | 0.24 | 0.29 | 0.39 | 0.35 | 0.4 |
| 10 | 49800427 | rs1445164 | -0.052 | 0.526 | 0.15 | 0.41 | -0.013 | 0.06 | 0.06 | 0.758 | 0.13 | 0.16 | 0.07 | 0.53 | 0.09 | 0.28 |
| 10 | 49801239 | rs12218054 | -0.252 | 1.51 | 0.401 | 0.332 | 0.084 | 0.084 | 0.084 | 0.006 | 0.09 | 0.19 | 0.11 | 0.25 | 0.19 | 0.21 |
| 10 | 49807631 | rs7914253 | 0.41 | 1.132 | 0.069 | 0.145 | 0.008 | 0.005 | 0.005 | 0.324 | 0.76 | 0.31 | 0.77 | 0.68 | 0.72 | 0.67 |
| 10 | 49810977 | rs7915707 | -0.121 | 1.094 | -0.799 | 0.326 | 0.008 | 0.043 | 0.043 | 0 | 0.49 | 0.33 | 0.59 | 0.65 | 0.46 | 0.5 |
| 10 | 49818668 | rs4838632 | -0.881 | 1.824 | -0.628 | 0.802 | -0.001 | -0.002 | -0.002 | 0 | 0.18 | 0.17 | 0.19 | 0.27 | 0.25 | 0.24 |
| 10 | 49822801 | rs3844492 | 0.009 | 0.856 | -0.743 | 0.273 | 0.019 | 0.023 | 0.023 | 3.611 | 0.4 | 0.59 | 0.45 | 0.27 | 0.3 | 0.27 |
| 10 | 49834326 | rs1822861 | -0.535 | 0.809 | 0.338 | 0.225 | -0.002 | 0.007 | 0.007 | 0.416 | 0.5 | 0.3 | 0.57 | 0.55 | 0.5 | 0.51 |
| 10 | 49853274 | rs7081000 | -1.441 | 0.567 | 0.893 | 0.087 | 0.068 | 0.064 | 0.064 | 1.058 | 0.06 | 0.43 | 0.04 | 0 | 0 | 0 |
| 10 | 49856349 | rs2377595 | -0.709 | 0.973 | 0.263 | 0.428 | 0.009 | 0.062 | 0.062 | 0.474 | 0.51 | 0.71 | 0.61 | 0.63 | 0.46 | 0.5 |
| 10 | 49858748 | rs11101413 | 0.863 | 0.754 | -0.137 | 0.405 | -0.018 | -0.078 | -0.078 | 1.546 | 0.23 | 0.03 | 0.36 | 0.3 | 0.21 | 0.18 |
| 10 | 49860040 | rs1002196 | 1.021 | 0.741 | -0.366 | -0.363 | -0.019 | -0.025 | -0.025 | 0 | 0.23 | 0.1 | 0.21 | 0.28 | 0.29 | 0.18 |
| 10 | 49863781 | rs12257645 | -0.507 | 1.024 | 0.009 | -0.434 | -0.01 | -0.012 | -0.012 | 0.179 | 0.58 | 0.44 | 0.69 | 0.64 | 0.63 | 0.58 |
| 10 | 49865727 | rs11101416 | -0.483 | 1.08 | -0.178 | -0.406 | 0.008 | 0.009 | 0.009 | 0.13 | 0.57 | 0.54 | 0.7 | 0.63 | 0.63 | 0.58 |
| 10 | 49868035 | rs6537570 | -0.579 | 1.218 | -0.239 | -0.778 | -0.018 | 0.017 | 0.017 | 0 | 0.13 | 0.05 | 0.1 | 0.01 | 0.2 | 0.15 |
| 10 | 49871496 | rs10219012 | -0.843 | 1.21 | -0.158 | -0.781 | 0.017 | -0.066 | -0.066 | 0 | 0.16 | 0.19 | 0.12 | 0.01 | 0.21 | 0.16 |
| 10 | 49886813 | rs6537571 | -0.325 | 0.299 | -0.685 | -0.825 | 0.007 | 0.008 | 0.008 | 0 | 0.44 | 0.55 | 0.32 | 0.29 | 0.33 | 0.29 |
| 10 | 49897219 | rs950275 | -0.004 | 0.035 | -0.353 | -0.932 | -0.001 | -0.002 | -0.002 | 0 | 0.34 | 0.25 | 0.24 | 0.26 | 0.28 | 0.2 |
| 10 | 49900195 | rs10857618 | -0.263 | 0.408 | -0.853 | -0.794 | -0.016 | -0.024 | -0.024 | 0 | 0.77 | 0.81 | 0.54 | 0.38 | 0.75 | 0.61 |
| 10 | 49902845 | rs10458720 | 0.064 | 0.252 | -1.149 | -0.885 | -0.107 | -0.113 | -0.113 | 0 | 0.16 | 0.01 | 0.45 | 0.63 | 0.27 | 0.32 |
| 10 | 49907313 | rs10857619 | -0.472 | -0.616 | -1.078 | -1.464 | 0.029 | 0.065 | 0.065 | 0 | 0.5 | 0.42 | 0.63 | 0.67 | 0.54 | 0.62 |
| 10 | 49910758 | rs4255465 | -0.601 | -0.707 | -0.895 | -1.482 | -0.024 | -0.027 | -0.027 | 0 | 0.87 | 0.62 | 0.9 | 0.93 | 0.93 | 0.88 |
| 10 | 49911701 | rs17011106 | 0.822 | -0.702 | -0.845 | -1.48 | -0.014 | -0.015 | -0.015 | 0 | 0.11 | 0.24 | 0.09 | 0.07 | 0.07 | 0.12 |
| 10 | 49911737 | rs2377623 | -0.747 | -0.672 | -1.032 | -1.467 | -0.024 | -0.027 | -0.027 | 0 | 0.87 | 0.62 | 0.9 | 0.93 | 0.93 | 0.88 |
| 10 | 49918587 | rs7912944 | 0.602 | -1.161 | -0.86 | -1.792 | 0.006 | -0.02 | -0.02 | 0 | 0.27 | 0.17 | 0.14 | 0.01 | 0.23 | 0.17 |
| 10 | 49919801 | rs10776637 | 0.712 | -1.061 | -1.383 | -1.626 | 0.01 | 0.018 | 0.018 | 0 | 0.21 | 0.24 | 0.46 | 0.62 | 0.26 | 0.33 |
| 10 | 49920608 | rs1904601 | -0.283 | -1.078 | -1.439 | -1.638 | -0.013 | -0.06 | -0.06 | 0 | 0.6 | 0.45 | 0.76 | 0.91 | 0.69 | 0.7 |
| 10 | 49923404 | rs11101437 | 0.415 | 0.045 | -1.55 | -1.039 | -0.044 | -0.056 | -0.056 | 0 | 0.2 | 0.12 | 0.45 | 0.62 | 0.26 | 0.34 |
| 10 | 49925611 | rs17771443 | 1.107 | 0.085 | -1.658 | -1.054 | -0.033 | 0.115 | 0.115 | 0 | 0.19 | 0.02 | 0.09 | 0.01 | 0.21 | 0.17 |
| 10 | 49927627 | rs7100103 | -0.059 | 0.378 | -1.151 | -0.595 | 0.013 | 0.021 | 0.021 | 0 | 0.22 | 0.3 | 0.46 | 0.62 | 0.27 | 0.3 |
| 10 | 49930509 | rs10508904 | -0.391 | -0.359 | -1.326 | -0.438 | -0.03 | -0.007 | -0.007 | 0 | 0.36 | 0.51 | 0.25 | 0.33 | 0.28 | 0.45 |
| 10 | 49930541 | rs17781278 | 0.335 | 0.192 | -1.854 | -0.226 | 0.013 | -0.024 | -0.024 | 0 | 0.17 | 0.1 | 0.08 | 0.2 | 0.13 | 0.26 |
| 10 | 49931149 | rs10508906 | 0.006 | 0.73 | -1.587 | -0.367 | -0.001 | 0.024 | 0.024 | 3.36 | 0.7 | 0.8 | 0.79 | 0.84 | 0.71 | 0.86 |
| 10 | 49933488 | rs10776641 | -0.33 | 1.1 | -2.253 | 0.292 | -0.188 | -0.029 | -0.029 | 0 | 0.36 | 0.03 | 0.57 | 0.71 | 0.52 | 0.45 |
| 10 | 49933974 | rs7072606 | -0.387 | 0.15 | -1.506 | 0.581 | -0.019 | 0.036 | 0.036 | 0 | 0.11 | 0.15 | 0.11 | 0.18 | 0.08 | 0.17 |
| 10 | 49936336 | rs11101442 | 0.668 | -0.233 | -1.927 | 1.023 | 0.006 | 0 | 0 | 0 | 0.39 | 0.13 | 0.24 | 0.41 | 0.34 | 0.44 |
| 10 | 49944249 | rs4838645 | 0.138 | -0.163 | -1.036 | 0.863 | 0.002 | 0.006 | 0.006 | 0.155 | 0.49 | 0.59 | 0.36 | 0.4 | 0.47 | 0.47 |
| 10 | 49944770 | rs7075318 | -0.028 | -0.154 | -1.029 | 0.876 | -0.003 | 0.005 | 0.005 | 1.191 | 0.54 | 0.61 | 0.5 | 0.64 | 0.54 | 0.57 |
| 10 | 49945594 | rs4838646 | 0.345 | -0.203 | -0.344 | 0.823 | 0.009 | 0.016 | 0.016 | 0.109 | 0.48 | 0.39 | 0.34 | 0.43 | 0.47 | 0.44 |
| 10 | 49947197 | rs11101446 | 0.245 | -0.32 | -0.181 | 0.748 | -0.049 | -0.026 | -0.026 | 0 | 0.37 | 0.23 | 0.52 | 0.41 | 0.44 | 0.38 |
| 10 | 49948011 | rs1871599 | 0.565 | -0.437 | 0.092 | 0.733 | 0.011 | 0.027 | 0.027 | 0.075 | 0.47 | 0.32 | 0.33 | 0.41 | 0.47 | 0.44 |
| 10 | 49950133 | rs11101448 | 0.222 | -0.371 | 0.116 | 0.782 | -0.046 | -0.025 | -0.025 | 0 | 0.37 | 0.2 | 0.52 | 0.4 | 0.45 | 0.38 |
| 10 | 49953604 | rs12242299 | 1.288 | -0.44 | 0.137 | 0.677 | 0.012 | -0.007 | -0.007 | 0 | 0.37 | 0.19 | 0.24 | 0.41 | 0.33 | 0.41 |
| 10 | 49955077 | rs10857630 | -0.788 | -0.222 | 0.128 | 0.747 | 0.007 | 0.039 | 0.039 | 0.005 | 0.14 | 0.31 | 0.24 | 0.21 | 0.19 | 0.16 |
| 10 | 49959702 | rs17011160 | 0.168 | -0.24 | -0.009 | 0.303 | 0 | -0.044 | -0.044 | 0.02 | 0.09 | 0.21 | 0.09 | 0 | 0.13 | 0.03 |
| 10 | 49963131 | rs2170127 | 0.499 | -0.227 | -0.347 | 0.431 | -0.093 | 0.061 | 0.061 | 1.932 | 0.31 | 0.05 | 0.38 | 0.19 | 0.39 | 0.25 |
| 10 | 49968596 | rs11101460 | -0.184 | -1.06 | 0.23 | 0.565 | -0.015 | 0.01 | 0.01 | 0 | 0.12 | 0.13 | 0.11 | 0.16 | 0.08 | 0.2 |
| 10 | 49973323 | rs11101462 | -1.124 | -1.506 | 0.433 | 0.38 | -0.035 | 0.004 | 0.004 | 0 | 0.13 | 0.26 | 0.12 | 0.15 | 0.07 | 0.13 |
| 10 | 49973858 | rs10508908 | -0.193 | -1.999 | 0.283 | 0.422 | -0.006 | 0.008 | 0.008 | 0 | 0.58 | 0.51 | 0.4 | 0.55 | 0.46 | 0.45 |
| 10 | 49974148 | rs4838647 | 1.477 | -2.304 | -0.18 | 0.082 | 0.028 | 0.051 | 0.051 | 0.057 | 0.31 | 0.16 | 0.19 | 0.01 | 0.26 | 0.15 |
| 10 | 49976229 | rs17836440 | 0.265 | -2.346 | -0.171 | 0.151 | 0.055 | 0.085 | 0.085 | 0.183 | 0.49 | 0.22 | 0.34 | 0.37 | 0.44 | 0.35 |
| 10 | 49977882 | rs11595065 | -0.292 | -2.408 | 0.008 | 0.145 | -0.003 | 0.004 | 0.004 | 0 | 0.58 | 0.44 | 0.39 | 0.55 | 0.47 | 0.45 |
| 10 | 49978113 | rs2620888 | 0.597 | -2.509 | -0.465 | 0.361 | -0.114 | 0.066 | 0.066 | 0 | 0.31 | 0.07 | 0.41 | 0.17 | 0.49 | 0.41 |
| 10 | 49978696 | rs11598288 | 0.319 | -2.358 | -0.096 | 0.669 | -0.005 | -0.071 | -0.071 | 3.414 | 0.17 | 0.04 | 0.13 | 0.37 | 0.17 | 0.19 |
| 10 | 49979699 | rs6537575 | -0.986 | -2.597 | 0.155 | 0.804 | -0.028 | 0.033 | 0.033 | 0.358 | 0.28 | 0.35 | 0.21 | 0.6 | 0.2 | 0.31 |
| 10 | 49983172 | rs2683611 | -0.14 | -2.526 | 0.458 | 0.372 | -0.017 | -0.068 | -0.068 | 0.044 | 0.88 | 0.69 | 0.93 | 1 | 0.94 | 0.98 |
| 10 | 49983506 | rs12217548 | 0.25 | -2.565 | 0.537 | 0.52 | -0.017 | 0.01 | 0.01 | 0 | 0.08 | 0.09 | 0.04 | 0.16 | 0.03 | 0.1 |

| | | | | | | | | | | | | | | | | |
|---|---|---|---|---|---|---|---|---|---|---|---|---|---|---|---|---|
| 10 | 49984302 | rs12412647 | 0.815 | -2.582 | 0.428 | 0.721 | -0.002 | 0.033 | 0.033 | 0 | 0.05 | 0.01 | 0.14 | 0.31 | 0.04 | 0.14 |
| 10 | 49984690 | rs2947344 | 1.31 | -2.676 | 0.739 | 0.609 | 0.058 | -0.026 | -0.026 | 0.069 | 0.6 | 0.22 | 0.73 | 0.48 | 0.74 | 0.69 |
| 10 | 49990081 | rs10776644 | -0.336 | -2.625 | 0.975 | 0.066 | -0.037 | -0.01 | -0.01 | 0 | 0.17 | 0.31 | 0.09 | 0.16 | 0.09 | 0.12 |
| 10 | 49994270 | rs4838650 | -0.787 | -3.648 | 0.356 | -0.629 | -0.042 | -0.035 | -0.035 | 0 | 0.28 | 0.33 | 0.13 | 0.16 | 0.16 | 0.16 |
| 10 | 49997272 | rs2928391 | -0.518 | -3.251 | 0.539 | -0.315 | -0.03 | 0.015 | 0.015 | 0.542 | 0.27 | 0.47 | 0.31 | 0.67 | 0.21 | 0.32 |
| 10 | 50008287 | rs2448544 | -0.354 | -3.505 | 0.382 | -0.369 | -0.019 | 0.011 | 0.011 | 0.268 | 0.27 | 0.39 | 0.29 | 0.67 | 0.21 | 0.32 |
| 10 | 50009291 | rs10857647 | -0.409 | -3.603 | 0.428 | -0.428 | -0.055 | -0.026 | -0.026 | 0 | 0.2 | 0.4 | 0.11 | 0.16 | 0.09 | 0.12 |
| 10 | 50013402 | rs2170132 | -1.293 | -3.32 | -0.221 | -0.624 | 0.089 | 0.138 | 0.138 | 2.594 | 0.47 | 0.79 | 0.4 | 0.83 | 0.3 | 0.44 |
| 10 | 50014273 | rs1871611 | 0.574 | -3.16 | -0.51 | -0.546 | 0.032 | -0.01 | -0.01 | 0 | 0.42 | 0.21 | 0.37 | 0.65 | 0.38 | 0.42 |
| 10 | 50018165 | rs2928402 | -0.984 | -2.969 | 0.004 | -1.026 | -0.024 | -0.035 | -0.035 | 0 | 0.24 | 0.68 | 0.22 | 0.1 | 0.14 | 0.14 |
| 10 | 50024030 | rs2620896 | -1.347 | -3.162 | -0.396 | -1.271 | 0.056 | 0.219 | 0.219 | 2.555 | 0.52 | 0.92 | 0.5 | 0.81 | 0.39 | 0.47 |
| 10 | 50026020 | rs7919656 | -0.773 | -2.886 | -0.084 | -1.461 | -0.006 | 0.002 | 0.002 | 0 | 0.63 | 0.59 | 0.52 | 0.82 | 0.51 | 0.54 |
| 10 | 50029461 | rs17699451 | 0.59 | -2.984 | 0.186 | -1.431 | -0.009 | -0.005 | -0.005 | 0 | 0.17 | 0.3 | 0.16 | 0.1 | 0.12 | 0.13 |
| 10 | 50030023 | rs10857650 | 0.638 | -3.172 | -0.017 | -1.511 | 0.011 | -0.036 | -0.036 | 0 | 0.23 | 0.1 | 0.26 | 0.71 | 0.24 | 0.32 |
| 10 | 50034833 | rs6537579 | 0.029 | -3.16 | 0.243 | -1.527 | -0.021 | -0.014 | -0.014 | 0 | 0.19 | 0.4 | 0.17 | 0.1 | 0.12 | 0.13 |
| 10 | 50035284 | rs12248218 | 0.173 | -3.145 | 0.167 | -1.537 | 0.013 | 0.018 | 0.018 | 0 | 0.42 | 0.6 | 0.28 | 0.1 | 0.27 | 0.29 |
| 10 | 50038800 | rs3747874 | -1.022 | -2.937 | 0.121 | -1.526 | -0.036 | 0.046 | 0.046 | 0 | 0.66 | 0.72 | 0.53 | 0.81 | 0.51 | 0.61 |
| 10 | 50040610 | rs3747872 | -0.706 | -3.004 | 0.131 | -1.563 | -0.013 | -0.045 | -0.045 | 0 | 0.43 | 0.53 | 0.43 | 0.81 | 0.37 | 0.45 |
| 10 | 50041276 | rs2620898 | -1.454 | -2.686 | -0.158 | -1.388 | -0.111 | 0.07 | 0.07 | 0 | 0.7 | 0.93 | 0.58 | 0.81 | 0.54 | 0.63 |
| 10 | 50057325 | rs7094742 | -0.037 | -2.875 | 0.115 | -1.426 | 0.007 | -0.003 | -0.003 | 0 | 0.31 | 0.15 | 0.31 | 0.71 | 0.33 | 0.37 |
| 10 | 50064431 | rs11101535 | 1.311 | -3.005 | -0.174 | -1.276 | -0.031 | -0.097 | -0.097 | 0.115 | 0.14 | 0.02 | 0.18 | 0.59 | 0.17 | 0.17 |
| 10 | 50069395 | rs2663052 | -1.02 | -3.196 | -0.01 | -1.443 | -0.04 | 0.039 | 0.039 | 0 | 0.6 | 0.82 | 0.48 | 0.71 | 0.49 | 0.55 |
| 10 | 50075461 | rs2663019 | -1.144 | -3.075 | 0.414 | -1.305 | -0.004 | -0.004 | -0.004 | 0 | 0.38 | 0.66 | 0.39 | 0.71 | 0.35 | 0.39 |
| 10 | 50076325 | rs11101537 | 1.464 | -2.931 | 0.295 | -1.211 | -0.035 | -0.094 | -0.094 | 0.309 | 0.13 | 0.01 | 0.18 | 0.59 | 0.17 | 0.17 |
| 10 | 50087458 | rs2663030 | -0.234 | -2.64 | 0.384 | -1.433 | 0.002 | 0.003 | 0.003 | 0.377 | 0.36 | 0.39 | 0.35 | 0.72 | 0.35 | 0.4 |
| 10 | 50091045 | rs2671712 | 0.324 | -2.951 | 0.201 | -1.577 | -0.001 | -0.001 | -0.001 | 0 | 0.59 | 0.53 | 0.63 | 0.28 | 0.66 | 0.6 |
| 10 | 50097819 | rs2671692 | -0.066 | -2.706 | 0.206 | -1.556 | 0.027 | 0.038 | 0.038 | 0 | 0.46 | 0.39 | 0.59 | 0.27 | 0.6 | 0.44 |
| 10 | 50103690 | rs17011395 | 0.18 | -2.259 | 0.741 | -1.324 | -0.015 | -0.043 | -0.043 | 0.107 | 0.08 | 0.18 | 0.05 | 0 | 0.02 | 0.02 |
| 10 | 50107537 | rs2663041 | 0.877 | -2.483 | 0.083 | -1.574 | -0.148 | -0.07 | -0.07 | 0 | 0.23 | 0.05 | 0.46 | 0.29 | 0.44 | 0.35 |
| 10 | 50109330 | rs7098403 | -0.58 | -2.529 | 0.49 | -1.617 | -0.011 | -0.053 | -0.053 | 0.055 | 0.1 | 0.32 | 0.07 | 0 | 0.02 | 0.02 |
| 10 | 50111346 | rs11101557 | -0.147 | -2.502 | 0.452 | -1.6 | -0.005 | -0.04 | -0.04 | 0 | 0.09 | 0.28 | 0.06 | 0 | 0.02 | 0.02 |
| 10 | 50114814 | rs12257105 | -0.419 | -2.516 | -0.003 | -1.605 | -0.028 | -0.073 | -0.073 | 0.387 | 0.12 | 0.43 | 0.08 | 0 | 0.02 | 0.02 |
| 10 | 50119054 | rs1913517 | -0.754 | -2.24 | -0.985 | -1.51 | -0.019 | 0.002 | 0.002 | 0 | 0.69 | 0.68 | 0.48 | 0.72 | 0.49 | 0.64 |
| 10 | 50120095 | rs4838657 | -0.121 | -1.946 | -1.434 | -1.047 | 0.04 | -0.011 | -0.011 | 0 | 0.35 | 0.07 | 0.29 | 0.72 | 0.32 | 0.45 |
| 10 | 50122181 | rs7094610 | 0.038 | -1.728 | -1.428 | -1.059 | 0.032 | 0.002 | 0.002 | 0 | 0.35 | 0.08 | 0.29 | 0.72 | 0.34 | 0.46 |
| 10 | 50125972 | rs2663056 | 0.752 | 0.774 | -0.749 | -1.108 | 0.072 | -0.037 | -0.037 | 0.018 | 0.08 | 0.13 | 0.15 | 0 | 0.26 | 0.08 |
| 10 | 50126875 | rs11101565 | 0.308 | 0.43 | -1.316 | -1.019 | 0.008 | -0.082 | -0.082 | 0 | 0.32 | 0.08 | 0.28 | 0.61 | 0.31 | 0.35 |
| 10 | 50127136 | rs2943248 | 0.466 | 0.101 | -1.142 | -1.387 | 0.011 | -0.086 | -0.086 | 0 | 0.29 | 0.32 | 0.23 | 0 | 0.38 | 0.25 |
| 10 | 50133442 | rs17782114 | 0.838 | 0.085 | -1.599 | -1.368 | -0.077 | -0.16 | -0.16 | 0 | 0.28 | 0.03 | 0.36 | 0.38 | 0.29 | 0.36 |
| 10 | 50135158 | rs2671702 | -0.388 | -0.997 | -0.701 | -1.121 | -0.014 | 0.021 | 0.021 | 0.176 | 0.63 | 0.49 | 0.61 | 0.39 | 0.67 | 0.66 |
| 10 | 50137109 | rs2377694 | -0.196 | -1.17 | 0.072 | -1.344 | -0.053 | -0.149 | -0.149 | 0.412 | 0.74 | 0.58 | 0.84 | 1 | 0.87 | 0.79 |
| 10 | 50138459 | rs7911009 | 0.458 | -0.993 | -1.478 | -1.457 | 0.03 | 0.08 | 0.08 | 1.512 | 0.2 | 0.09 | 0.07 | 0 | 0.13 | 0.21 |
| 10 | 50140588 | rs2940708 | 0.472 | -1.176 | -0.997 | -1.047 | -0.032 | -0.092 | -0.092 | 0.118 | 0.84 | 0.34 | 0.86 | 1 | 0.9 | 0.99 |
| 10 | 50140873 | rs2377631 | -0.373 | -1.038 | -1.947 | -0.4 | 0.064 | -0.065 | -0.065 | 0 | 0.36 | 0.13 | 0.22 | 0.52 | 0.27 | 0.33 |
| 10 | 50143591 | rs10776658 | 0.007 | -0.398 | -1.894 | -1.121 | -0.076 | 0.051 | 0.051 | 0.14 | 0.4 | 0.1 | 0.49 | 0.36 | 0.5 | 0.5 |
| 10 | 50145820 | rs2457466 | 0.044 | -0.753 | -0.936 | -1.459 | -0.008 | 0.035 | 0.035 | 0 | 0.54 | 0.65 | 0.46 | 0.64 | 0.47 | 0.5 |
| 10 | 50147561 | rs2671716 | -0.503 | -0.603 | -0.725 | -1.387 | -0.009 | -0.03 | -0.03 | 1.939 | 0.94 | 0.82 | 0.95 | 1 | 0.96 | 0.99 |
| 10 | 50155130 | rs2663067 | 0.29 | -1.072 | 0.118 | -1.518 | 0 | 0.086 | 0.086 | 0 | 0.5 | 0.29 | 0.43 | 0.62 | 0.46 | 0.49 |
| 10 | 50157552 | rs2663028 | 0.215 | -1.069 | 0.207 | -1.517 | 0 | 0.07 | 0.07 | 0 | 0.51 | 0.31 | 0.44 | 0.62 | 0.46 | 0.49 |
| 10 | 50162809 | rs7098224 | 0.187 | -1.029 | 0.339 | -1.495 | -0.028 | 0.046 | 0.046 | 0 | 0.4 | 0.24 | 0.49 | 0.35 | 0.5 | 0.5 |
| 10 | 50165128 | rs3829206 | 0.224 | -1.025 | 0.335 | -1.494 | -0.02 | 0.04 | 0.04 | 0 | 0.41 | 0.28 | 0.5 | 0.35 | 0.5 | 0.49 |
| 10 | 50173551 | rs7091429 | -0.482 | -1.009 | 0.246 | -1.487 | -0.001 | -0.005 | -0.005 | 0 | 0.49 | 0.51 | 0.55 | 0.38 | 0.54 | 0.51 |
| 10 | 50176407 | rs17011563 | 0.15 | -1.33 | 0.034 | -1.846 | -0.037 | -0.064 | -0.064 | 0 | 0.17 | 0.22 | 0.06 | 0.01 | 0.06 | 0.11 |
| 10 | 50177420 | rs7895907 | 1.894 | -0.853 | -0.316 | -1.958 | -0.031 | 0.03 | 0.03 | 0 | 0.06 | 0.01 | 0.07 | 0 | 0.11 | 0.01 |
| 10 | 50181364 | rs12573522 | 0.937 | -1.121 | -0.551 | -2.073 | -0.063 | -0.061 | -0.061 | 0 | 0.15 | 0.01 | 0.2 | 0.23 | 0.2 | 0.29 |
| 10 | 50186317 | rs2292585 | -0.616 | -1.278 | 0.15 | -1.802 | 0.002 | -0.013 | -0.013 | 0 | 0.58 | 0.4 | 0.54 | 0.62 | 0.51 | 0.52 |
| 10 | 50189949 | rs7093656 | 0.381 | -1.222 | -0.184 | -1.77 | -0.008 | 0.015 | 0.015 | 0 | 0.4 | 0.38 | 0.46 | 0.35 | 0.48 | 0.48 |
| 10 | 50192120 | rs2252811 | 0.606 | -1.111 | -0.713 | -1.728 | -0.06 | 0.047 | 0.047 | 0.833 | 0.4 | 0.15 | 0.45 | 0.6 | 0.45 | 0.34 |
| 10 | 50197435 | rs7086917 | -0.069 | -0.076 | -0.741 | -1.195 | -0.02 | 0.047 | 0.047 | 0 | 0.41 | 0.27 | 0.46 | 0.35 | 0.49 | 0.55 |
| 10 | 50210819 | rs2245162 | 0.175 | -0.498 | -0.319 | -1.504 | -0.001 | -0.004 | -0.004 | 0.17 | 0.55 | 0.44 | 0.61 | 0.64 | 0.62 | 0.34 |
| 10 | 50211170 | rs2245147 | 0.176 | -0.592 | -0.166 | -1.545 | 0.011 | 0.021 | 0.021 | 2.111 | 0.55 | 0.38 | 0.6 | 0.64 | 0.62 | 0.34 |
| 10 | 50217438 | rs883017 | 0.77 | -0.954 | -0.41 | -2.149 | 0.025 | 0.065 | 0.065 | 0.235 | 0.5 | 0.33 | 0.53 | 0.64 | 0.57 | 0.42 |
| 10 | 50218092 | rs17700388 | 1.445 | -0.957 | -0.494 | -2.263 | -0.046 | 0.041 | 0.041 | 0.321 | 0.1 | 0.03 | 0.17 | 0 | 0.22 | 0.03 |
| 10 | 50218184 | rs2255486 | -0.397 | -1.211 | -0.145 | -2.24 | 0.053 | 0.138 | 0.138 | 1.712 | 0.94 | 0.58 | 0.94 | 0.91 | 1 | 0.92 |
| 10 | 50218813 | rs12775969 | 0.123 | -1.24 | -0.183 | -2.255 | 0.022 | -0.009 | -0.009 | 0 | 0.19 | 0.16 | 0.2 | 0.27 | 0.15 | 0.4 |
| 10 | 50222585 | rs2254863 | 0.305 | -1.245 | -0.313 | -2.261 | -0.043 | 0.006 | 0.006 | 0.045 | 0.74 | 0.38 | 0.74 | 0.64 | 0.84 | 0.52 |
| 10 | 50226804 | rs11814277 | 1.562 | -1.058 | -0.368 | -2.254 | -0.041 | 0.036 | 0.036 | 0 | 0.1 | 0.03 | 0.17 | 0 | 0.21 | 0.03 |
| 10 | 50227129 | rs3810954 | 0.07 | -1.23 | -0.348 | -2.248 | 0.029 | -0.01 | -0.01 | 0.557 | 0.2 | 0.15 | 0.2 | 0.24 | 0.15 | 0.4 |
| 10 | 50227323 | rs3763776 | 0.6 | -1.375 | -0.439 | -2.321 | 0.001 | -0.053 | -0.053 | 0 | 0.12 | 0.02 | 0.18 | 0.23 | 0.13 | 0.26 |

| | | | | | | | | | | | | | | | | |
|---|---|---|---|---|---|---|---|---|---|---|---|---|---|---|---|---|
| 10 | 50231840 | rs2175918 | 1.064 | -1.54 | -0.438 | -2.379 | -0.015 | -0.004 | -0.004 | 0 | 0.62 | 0.38 | 0.71 | 0.67 | 0.79 | 0.45 |
| 10 | 50232180 | rs2244085 | -0.888 | -1.588 | -0.291 | -2.396 | -0.073 | -0.107 | -0.107 | 0.111 | 0.83 | 0.67 | 0.93 | 0.97 | 0.94 | 0.92 |
| 10 | 50235619 | rs12765128 | 0.698 | -1.555 | -0.401 | -2.384 | 0.003 | -0.053 | -0.053 | 0 | 0.12 | 0.02 | 0.18 | 0.23 | 0.13 | 0.25 |
| 10 | 50238874 | rs2254393 | 1.075 | -1.574 | -0.274 | -2.392 | 0.005 | 0.006 | 0.006 | 0 | 0.62 | 0.35 | 0.71 | 0.64 | 0.79 | 0.44 |
| 10 | 50243539 | rs1125517 | -0.41 | -1.453 | -0.42 | -2.349 | 0.017 | -0.003 | -0.003 | 0 | 0.2 | 0.18 | 0.21 | 0.24 | 0.15 | 0.4 |
| 10 | 50244148 | rs2943252 | 0.074 | -1.313 | -0.215 | -2.254 | 0.006 | -0.107 | -0.107 | 0.6 | 0.72 | 0.51 | 0.75 | 0.88 | 0.73 | 0.81 |
| 10 | 50249026 | rs12776211 | 0.393 | -1.16 | -0.466 | -2.062 | 0.006 | -0.049 | -0.049 | 0 | 0.1 | 0.03 | 0.21 | 0.21 | 0.11 | 0.22 |
| 10 | 50255568 | rs17011692 | 1.359 | -1 | -0.317 | -2.118 | 0.025 | -0.033 | -0.033 | 0 | 0.1 | 0.13 | 0.16 | 0 | 0.19 | 0.03 |
| 10 | 50262144 | rs9804403 | -1.581 | -1.255 | -0.061 | -2.187 | 0.003 | -0.002 | -0.002 | 0 | 0.26 | 0.63 | 0.41 | 0.28 | 0.28 | 0.31 |
| 10 | 50263201 | rs4272720 | -0.246 | -1.257 | -0.012 | -2.185 | 0.043 | -0.076 | -0.076 | 0.737 | 0.18 | 0.56 | 0.21 | 0.07 | 0.2 | 0.11 |
| 10 | 50271800 | rs4550151 | -1.356 | -1.419 | 0.005 | -2.115 | -0.042 | 0.099 | 0.099 | 0 | 0.15 | 0.48 | 0.24 | 0.28 | 0.09 | 0.29 |
| 10 | 50285084 | rs17782649 | 0.388 | -1.372 | -0.15 | -2.114 | 0.002 | -0.011 | -0.011 | 0 | 0.08 | 0.06 | 0.2 | 0.21 | 0.08 | 0.2 |
| 10 | 50288261 | rs12098770 | 0.016 | -1.604 | -0.05 | -2.214 | -0.036 | -0.037 | -0.037 | 2.593 | 0.1 | 0.18 | 0.02 | 0.01 | 0.01 | 0.15 |
| 10 | 50288931 | rs12776460 | -0.384 | -1.477 | -0.037 | -2.274 | 0.006 | 0.002 | 0.002 | 0 | 0.2 | 0.23 | 0.36 | 0.28 | 0.27 | 0.31 |
| 10 | 50293456 | rs12269023 | -1.985 | -1.801 | -0.182 | -2.185 | 0.056 | 0.098 | 0.098 | 0.165 | 0.46 | 0.72 | 0.45 | 0.32 | 0.34 | 0.54 |
| 10 | 50296622 | rs4445564 | -1.581 | -1.764 | -0.154 | -2.167 | 0 | 0.001 | 0.001 | 0 | 0.29 | 0.66 | 0.41 | 0.22 | 0.28 | 0.38 |
| 10 | 50300179 | rs4240498 | -0.696 | -1.643 | -0.192 | -2.06 | -0.072 | -0.001 | -0.001 | 0 | 0.26 | 0.39 | 0.27 | 0.31 | 0.14 | 0.37 |
| 10 | 50300505 | rs6537495 | 0.542 | -1.577 | -0.146 | -2.335 | -0.079 | -0.074 | -0.074 | 0 | 0.18 | 0.32 | 0.07 | 0.1 | 0.06 | 0.16 |
| 10 | 50306386 | rs10508911 | 0.579 | -1.304 | -0.289 | -2.309 | -0.016 | 0.005 | 0.005 | 0 | 0.14 | 0.11 | 0.15 | 0.01 | 0.19 | 0.16 |
| 10 | 50308491 | rs7080003 | 0.007 | -1.507 | -0.206 | -2.467 | -0.059 | -0.118 | -0.118 | 0 | 0.2 | 0.47 | 0.11 | 0.03 | 0.12 | 0.11 |
| 10 | 50309905 | rs4488117 | 0.625 | -1.396 | -0.247 | -2.282 | 0.003 | -0.01 | -0.01 | 0 | 0.08 | 0.07 | 0.2 | 0.2 | 0.08 | 0.22 |
| 10 | 50313621 | rs4509682 | -1.835 | -1.569 | -0.364 | -2.394 | 0.04 | 0.178 | 0.178 | 0 | 0.47 | 0.77 | 0.48 | 0.23 | 0.39 | 0.5 |
| 10 | 50315893 | rs13088 | -1.833 | -1.612 | -0.384 | -2.405 | 0.04 | 0.178 | 0.178 | 0 | 0.47 | 0.78 | 0.48 | 0.24 | 0.39 | 0.5 |
| 10 | 50318484 | rs12255055 | -1.254 | -1.646 | -0.316 | -2.08 | 0.006 | 0.026 | 0.026 | 0 | 0.4 | 0.65 | 0.47 | 0.3 | 0.39 | 0.37 |
| 10 | 50326124 | rs4417190 | -1.688 | -1.651 | -0.319 | -2.065 | 0.027 | 0.051 | 0.051 | 0 | 0.45 | 0.65 | 0.47 | 0.35 | 0.39 | 0.51 |
| 10 | 50326725 | rs4620641 | -1.769 | -1.655 | -0.317 | -2.066 | 0.037 | 0.072 | 0.072 | 0 | 0.46 | 0.74 | 0.48 | 0.35 | 0.39 | 0.51 |
| 10 | 50329746 | rs4838368 | 0.729 | -1.772 | -0.262 | -2.134 | 0.031 | 0.017 | 0.017 | 0.709 | 0.18 | 0.08 | 0.04 | 0 | 0.06 | 0.08 |
| 10 | 50332539 | rs4603218 | -1.144 | -2.03 | -0.926 | -2.756 | 0.091 | 0.232 | 0.232 | 0 | 0.53 | 0.75 | 0.25 | 0.09 | 0.32 | 0.47 |
| 10 | 50332930 | rs4550152 | 0.13 | -2.049 | -0.88 | -2.764 | -0.051 | -0.103 | -0.103 | 4.438 | 0.23 | 0.52 | 0.12 | 0.02 | 0.11 | 0.11 |
| 10 | 50336672 | rs4284329 | 0.496 | -1.946 | -0.913 | -2.71 | 0.024 | 0.046 | 0.046 | 0 | 0.31 | 0.23 | 0.14 | 0.07 | 0.21 | 0.36 |
| 10 | 50343022 | rs10218995 | -0.611 | -1.884 | -0.816 | -2.695 | 0 | -0.003 | -0.003 | 0.291 | 0.45 | 0.58 | 0.5 | 0.39 | 0.47 | 0.48 |
| 10 | 50348719 | rs4073201 | 0.751 | -1.759 | -0.769 | -2.63 | 0.003 | -0.017 | -0.017 | 0 | 0.26 | 0.3 | 0.14 | 0.06 | 0.2 | 0.22 |
| 10 | 50355418 | rs2466744 | -2.026 | -1.705 | -0.807 | -2.553 | 0.021 | 0.008 | 0.008 | 0 | 0.7 | 0.7 | 0.57 | 0.39 | 0.59 | 0.72 |
| 10 | 50357938 | rs7902240 | 0.924 | -1.719 | -1.216 | -2.56 | 0.033 | 0.019 | 0.019 | 0.254 | 0.19 | 0.1 | 0.05 | 0 | 0.06 | 0.08 |
| 10 | 50362930 | rs12198 | -1.58 | -1.206 | -0.983 | -2.699 | 0.034 | 0.005 | 0.005 | 0 | 0.68 | 0.56 | 0.56 | 0.37 | 0.59 | 0.73 |
| 10 | 50368339 | rs11101001 | -0.281 | -0.915 | -1.085 | -2.409 | 0.005 | 0.005 | 0.005 | 0 | 0.39 | 0.62 | 0.27 | 0.09 | 0.38 | 0.3 |
| 10 | 50374171 | rs2725200 | -1.298 | -0.662 | -0.372 | -1.644 | -0.016 | 0.023 | 0.023 | 0 | 0.2 | 0.27 | 0.25 | 0.34 | 0.16 | 0.3 |
| 10 | 50381921 | rs10857437 | -0.578 | -0.56 | -0.638 | -1.893 | -0.002 | 0.02 | 0.02 | 0.405 | 0.14 | 0.07 | 0.11 | 0.04 | 0.11 | 0.2 |
| 10 | 50382038 | rs6537501 | -1.198 | -0.459 | -0.663 | -1.819 | -0.006 | 0.03 | 0.03 | 0.002 | 0.43 | 0.17 | 0.49 | 0.32 | 0.54 | 0.47 |
| 10 | 50382478 | rs7910476 | -0.708 | -0.471 | -0.37 | -1.413 | 0.002 | -0.016 | -0.016 | 1.781 | 0.08 | 0.03 | 0.21 | 0.33 | 0.08 | 0.27 |
| 10 | 50386498 | rs10776544 | -0.812 | -0.637 | 0.006 | -1.661 | 0.005 | -0.003 | -0.003 | 0 | 0.36 | 0.23 | 0.33 | 0.4 | 0.33 | 0.47 |
| 10 | 50393013 | rs1962336 | -0.861 | -0.686 | 0.089 | -1.729 | 0.007 | 0.001 | 0.001 | 0 | 0.37 | 0.24 | 0.35 | 0.4 | 0.33 | 0.47 |
| 10 | 50415005 | rs11101015 | -1.488 | -1.02 | -0.025 | -1.877 | 0.048 | 0.037 | 0.037 | 0 | 0.45 | 0.69 | 0.4 | 0.4 | 0.33 | 0.52 |
| 10 | 50416034 | rs11101017 | -0.625 | -0.897 | -0.138 | -1.748 | 0.002 | -0.024 | -0.024 | 0.004 | 0.09 | 0.03 | 0.21 | 0.33 | 0.09 | 0.27 |
| 10 | 50418087 | rs12571488 | 0.144 | -0.964 | -0.117 | -1.808 | -0.001 | -0.013 | -0.013 | 0 | 0.25 | 0.21 | 0.12 | 0.09 | 0.22 | 0.17 |
| 10 | 50418949 | rs12249370 | -0.194 | -0.412 | -0.183 | -1.939 | 0.037 | -0.057 | -0.057 | 0 | 0.85 | 0.75 | 0.84 | 0.99 | 0.75 | 0.96 |
| 10 | 50419622 | rs10857441 | 1.671 | -0.393 | -0.259 | -1.97 | -0.011 | -0.006 | -0.006 | 0 | 0.43 | 0.48 | 0.47 | 0.57 | 0.38 | 0.48 |
| 10 | 50423250 | rs12770471 | 0.379 | -0.704 | -0.137 | -1.802 | -0.002 | -0.017 | -0.017 | 0 | 0.23 | 0.21 | 0.11 | 0.09 | 0.21 | 0.17 |
| 10 | 50424674 | rs10857443 | 1.26 | 0.135 | 0.168 | -1.418 | 0.007 | -0.071 | -0.071 | 0.007 | 0.78 | 0.27 | 0.75 | 0.99 | 0.75 | 0.93 |
| 10 | 50425252 | rs2377991 | 1.034 | 0.11 | 0.196 | -1.387 | 0.01 | -0.071 | -0.071 | 0 | 0.78 | 0.27 | 0.76 | 0.99 | 0.74 | 0.93 |
| 10 | 50430027 | rs12253763 | -1.151 | -1.219 | 0.768 | -0.747 | 0.022 | 0.019 | 0.019 | 0 | 0.08 | 0.21 | 0.23 | 0.13 | 0.1 | 0.22 |
| 10 | 50433556 | rs9645532 | 0.196 | -1.25 | 0.75 | -0.748 | -0.016 | -0.015 | -0.015 | 0 | 0.92 | 0.54 | 0.94 | 1 | 0.99 | 0.99 |
| 10 | 50443840 | rs7072265 | -0.097 | -1.348 | 0.685 | -0.784 | 0.017 | 0.017 | 0.017 | 0 | 0.36 | 0.25 | 0.16 | 0.3 | 0.31 | 0.32 |
| 10 | 50445299 | rs2377987 | 0.553 | -1.257 | 0.217 | -0.648 | -0.048 | -0.046 | -0.046 | 0 | 0.91 | 0.69 | 0.96 | 1 | 0.99 | 0.99 |
| 10 | 50449175 | rs2002392 | -0.047 | -1.121 | -1.343 | -0.582 | -0.04 | -0.037 | -0.037 | 0 | 0.93 | 0.78 | 0.97 | 1 | 0.99 | 0.98 |
| 10 | 50451667 | rs4838470 | 0.966 | -0.755 | -1.881 | -0.456 | -0.059 | -0.051 | -0.051 | 0.016 | 0.44 | 0.03 | 0.53 | 0.57 | 0.56 | 0.46 |
| 10 | 50451843 | rs11101060 | -1.524 | -0.469 | -0.398 | -0.117 | -0.008 | 0.01 | 0.01 | 0 | 0.08 | 0.15 | 0.11 | 0.1 | 0.06 | 0.2 |
| 10 | 50453668 | rs12776010 | -0.732 | -0.383 | -0.82 | -0.088 | 0.02 | 0.031 | 0.031 | 0 | 0.12 | 0.04 | 0.05 | 0.05 | 0.09 | 0.04 |
| 10 | 50453933 | rs17009617 | -1.277 | -0.113 | -0.456 | 0.155 | -0.008 | -0.004 | -0.004 | 0 | 0.1 | 0.14 | 0.22 | 0.11 | 0.06 | 0.12 |
| 10 | 50463693 | rs7906545 | -0.841 | 0.168 | -0.069 | 0.078 | 0.026 | -0.023 | -0.023 | 0 | 0.05 | 0.14 | 0.04 | 0 | 0.08 | 0.02 |
| 10 | 50467224 | rs7914160 | 1.89 | -0.067 | 0.986 | -0.088 | -0.001 | -0.051 | -0.051 | 0 | 0.83 | 0.51 | 0.86 | 0.89 | 0.85 | 0.81 |
| 10 | 50470330 | rs12411581 | -0.543 | -1.115 | 1.195 | 0.232 | -0.036 | 0.017 | 0.017 | 0 | 0.17 | 0.23 | 0.09 | 0.24 | 0.09 | 0.06 |
| 10 | 50471176 | rs10857453 | 1.567 | -0.913 | 1.354 | -0.033 | 0.001 | -0.025 | -0.025 | 2.414 | 0.74 | 0.35 | 0.67 | 0.85 | 0.75 | 0.71 |
| 10 | 50479493 | rs2377976 | 0.431 | -0.92 | 1.513 | -0.089 | 0.061 | -0.025 | -0.025 | 2.276 | 0.89 | 0.65 | 0.79 | 0.96 | 0.89 | 0.87 |
| 10 | 50479824 | rs6537519 | -0.091 | -0.986 | 1.477 | -0.067 | 0.071 | -0.006 | -0.006 | 0 | 0.91 | 0.66 | 0.8 | 0.96 | 0.9 | 0.88 |
| 10 | 50481447 | rs17009755 | 0.33 | -0.98 | 1.385 | -0.065 | -0.003 | 0.002 | 0.002 | 0 | 0.07 | 0.05 | 0.17 | 0.04 | 0.09 | 0.12 |
| 10 | 50482252 | rs4838484 | -0.562 | -1.036 | 1.329 | -0.077 | 0.001 | 0 | 0 | 0 | 0.11 | 0.12 | 0.09 | 0.11 | 0.07 | 0.14 |
| 10 | 50486727 | rs7082468 | 1.749 | -1.328 | 0.731 | 0.095 | 0.042 | -0.029 | -0.029 | 0.014 | 0.56 | 0.11 | 0.56 | 0.58 | 0.66 | 0.64 |
| 10 | 50488542 | rs10857460 | 0.12 | -1.002 | 1.186 | 0.008 | 0.071 | -0.005 | -0.005 | 0 | 0.91 | 0.66 | 0.79 | 0.96 | 0.9 | 0.88 |

| | | | | | | | | | | | | | | | | |
|---|---|---|---|---|---|---|---|---|---|---|---|---|---|---|---|---|
| 10 | 50496662 | rs6537521 | 1.225 | -1.149 | 1.376 | -0.03 | -0.027 | -0.018 | -0.018 | 0.505 | 0.86 | 0.56 | 0.87 | 0.9 | 0.93 | 0.85 |
| 10 | 50501467 | rs7893248 | 1.538 | -1.098 | 1.489 | -0.014 | 0 | -0.045 | -0.045 | 0 | 0.79 | 0.48 | 0.71 | 0.85 | 0.84 | 0.73 |
| 10 | 50503524 | rs7919151 | 0.645 | -1.062 | 1.352 | 0.019 | 0.038 | 0.053 | 0.053 | 0.037 | 0.93 | 0.73 | 0.9 | 0.9 | 0.94 | 0.84 |
| 10 | 50506398 | rs6537523 | 0.912 | -0.869 | 0.807 | -0.163 | -0.001 | -0.01 | -0.01 | 0 | 0.49 | 0.44 | 0.34 | 0.21 | 0.55 | 0.52 |
| 10 | 50506558 | rs2292200 | -1.643 | -1.07 | 0.963 | 0.531 | 0 | -0.002 | -0.002 | 0 | 0.19 | 0.23 | 0.26 | 0.26 | 0.13 | 0.12 |
| 10 | 50506613 | rs2292201 | -0.415 | -0.844 | 1.251 | 0.819 | 0.052 | 0.012 | 0.012 | 0.723 | 0.12 | 0.31 | 0.21 | 0.1 | 0.13 | 0.1 |
| 10 | 50507362 | rs11101081 | 1.181 | -1.014 | 0.524 | 0.922 | -0.019 | -0.027 | -0.027 | 0 | 0.11 | 0.01 | 0.06 | 0.12 | 0.12 | 0.12 |
| 10 | 50507454 | rs7897237 | 1.362 | -1.042 | 0.701 | 0.888 | 0.012 | 0.062 | 0.062 | 0.069 | 0.5 | 0.32 | 0.35 | 0.3 | 0.57 | 0.64 |
| 10 | 50508228 | rs7900404 | -0.672 | -1.194 | 0.449 | 0.856 | -0.002 | 0.043 | 0.043 | 0 | 0.19 | 0.05 | 0.14 | 0.11 | 0.2 | 0.23 |
| 10 | 50510522 | rs2377965 | -2.105 | -1.249 | 0.783 | 0.861 | 0.007 | -0.039 | -0.039 | 0 | 0.6 | 0.51 | 0.59 | 0.81 | 0.55 | 0.52 |
| 10 | 50514253 | rs11101083 | 0.234 | -1.233 | 0.685 | 1.015 | -0.007 | -0.038 | -0.038 | 0 | 0.18 | 0.1 | 0.16 | 0.34 | 0.16 | 0.16 |
| 10 | 50515246 | rs12572137 | 1.075 | -1.264 | 0.707 | 0.987 | 0.001 | 0.001 | 0.001 | 0.186 | 0.15 | 0.14 | 0.1 | 0.27 | 0.09 | 0.06 |
| 10 | 50516184 | rs2889802 | 1.972 | -1.121 | 0.465 | 0.904 | -0.028 | 0.072 | 0.072 | 0.085 | 0.38 | 0.23 | 0.38 | 0.19 | 0.45 | 0.48 |
| 10 | 50518351 | rs10857465 | 1.958 | -1.093 | 0.422 | 0.916 | -0.017 | 0.045 | 0.045 | 0 | 0.38 | 0.28 | 0.39 | 0.19 | 0.46 | 0.48 |
| 10 | 50533237 | rs12217617 | 0.053 | -0.802 | 0.236 | 1.117 | -0.002 | 0.008 | 0.008 | 0 | 0.11 | 0.09 | 0.26 | 0.31 | 0.15 | 0.11 |
| 10 | 50534862 | rs12411843 | -1.398 | -0.92 | 0.179 | 1.041 | -0.03 | -0.039 | -0.039 | 0.644 | 0.31 | 0.13 | 0.38 | 0.41 | 0.33 | 0.34 |
| 10 | 50536578 | rs2377941 | -1.411 | -0.963 | 0.191 | 1.036 | -0.007 | -0.04 | -0.04 | 0 | 0.29 | 0.14 | 0.35 | 0.42 | 0.26 | 0.3 |
| 10 | 50540023 | rs10745259 | -0.24 | -1.246 | 0.221 | 0.812 | 0 | -0.002 | -0.002 | 0 | 0.82 | 0.85 | 0.87 | 0.78 | 0.83 | 0.73 |
| 10 | 50544378 | rs953866 | 0.052 | -1.279 | 0.311 | 0.909 | 0 | 0.03 | 0.03 | 0.57 | 0.23 | 0.3 | 0.28 | 0.37 | 0.22 | 0.14 |
| 10 | 50546999 | rs10857475 | 1.161 | -1.006 | -0.268 | 0.595 | 0.003 | 0.033 | 0.033 | 0 | 0.44 | 0.33 | 0.46 | 0.33 | 0.55 | 0.53 |
| 10 | 50547459 | rs11101099 | 1.059 | -0.959 | -0.587 | 0.682 | -0.018 | -0.027 | -0.027 | 0 | 0.12 | 0.01 | 0.06 | 0.12 | 0.12 | 0.12 |
| 10 | 50551420 | rs10776562 | 0.787 | -0.711 | -0.219 | 0.593 | -0.016 | 0.053 | 0.053 | 0 | 0.44 | 0.27 | 0.43 | 0.33 | 0.49 | 0.5 |
| 10 | 50558099 | rs4562729 | -0.717 | -0.405 | 0.195 | 0.595 | -0.026 | -0.031 | -0.031 | 0 | 0.06 | 0.14 | 0.03 | 0 | 0.01 | 0 |
| 10 | 50558280 | rs4494243 | 0.269 | -0.769 | -0.13 | 0.716 | 0.003 | -0.015 | -0.015 | 0.428 | 0.16 | 0.03 | 0.09 | 0.19 | 0.17 | 0.16 |
| 10 | 50559202 | rs7915976 | 0.48 | -0.62 | 0.341 | 0.737 | 0.04 | 0.02 | 0.02 | 0 | 0.08 | 0.15 | 0.19 | 0.1 | 0.15 | 0.09 |
| 10 | 50561190 | rs2889780 | -1.046 | -0.354 | 0.763 | 1.083 | -0.006 | 0.028 | 0.028 | 0 | 0.28 | 0.33 | 0.24 | 0.48 | 0.25 | 0.22 |
| 10 | 50568273 | rs11599426 | 1.326 | -0.855 | 0.906 | 0.851 | -0.024 | -0.046 | -0.046 | 0 | 0.56 | 0.11 | 0.52 | 0.42 | 0.59 | 0.7 |
| 10 | 50569286 | rs11101114 | -1.146 | -0.866 | 1.047 | 0.846 | 0.005 | 0.009 | 0.009 | 0 | 0.37 | 0.58 | 0.43 | 0.58 | 0.4 | 0.3 |
| 10 | 50570421 | rs7909260 | 2.01 | -1.113 | 0.498 | 0.825 | -0.067 | -0.042 | -0.042 | 0.016 | 0.43 | 0.09 | 0.47 | 0.39 | 0.49 | 0.64 |
| 10 | 50572862 | rs17009950 | 0.523 | -1.075 | 0.479 | 0.517 | 0.008 | 0.019 | 0.019 | 0 | 0.13 | 0.11 | 0.06 | 0.05 | 0.1 | 0.06 |
| 10 | 50588903 | rs7900263 | -0.328 | -0.998 | 0.503 | 0.556 | -0.027 | -0.032 | -0.032 | 0 | 0.06 | 0.2 | 0.03 | 0 | 0.01 | 0 |
| 10 | 50588990 | rs7903692 | -1.242 | -0.49 | 0.697 | 0.644 | -0.005 | 0.001 | 0.001 | 0 | 0.43 | 0.52 | 0.3 | 0.53 | 0.35 | 0.26 |
| 10 | 50591739 | rs10857481 | -0.11 | -0.284 | -0.074 | 0.741 | 0 | 0.014 | 0.014 | 0.015 | 0.21 | 0.24 | 0.13 | 0.47 | 0.23 | 0.2 |
| 10 | 50595930 | rs10857482 | 1.007 | 0 | -0.594 | 0.411 | -0.034 | 0.057 | 0.057 | 0.307 | 0.43 | 0.13 | 0.44 | 0.33 | 0.49 | 0.65 |
| 10 | 50615806 | rs7086360 | 0.657 | 0.565 | -0.421 | 0.396 | 0.019 | 0.085 | 0.085 | 0 | 0.48 | 0.22 | 0.39 | 0.26 | 0.47 | 0.55 |
| 10 | 50617166 | rs10776571 | 0.605 | 0.566 | -0.026 | 0.341 | 0.006 | -0.01 | -0.01 | 0 | 0.61 | 0.4 | 0.46 | 0.27 | 0.55 | 0.62 |
| 10 | 50621859 | rs4838513 | 0.536 | 0.748 | 0.202 | 0.864 | 0.046 | 0.142 | 0.142 | 0.604 | 0.88 | 0.62 | 0.78 | 0.73 | 0.91 | 0.91 |
| 10 | 50623895 | rs10857488 | 0.887 | 0.667 | 0.268 | 0.785 | 0.003 | -0.009 | -0.009 | 0.158 | 0.59 | 0.43 | 0.46 | 0.27 | 0.55 | 0.62 |
| 10 | 50634771 | rs17783727 | 0.044 | 0.876 | 0.067 | 1.167 | -0.011 | -0.062 | -0.062 | 0 | 0.26 | 0.11 | 0.17 | 0.35 | 0.3 | 0.24 |
| 10 | 50634956 | rs3945802 | 1.241 | 0.612 | 0.142 | 0.905 | 0 | -0.001 | -0.001 | 0.065 | 0.45 | 0.4 | 0.42 | 0.23 | 0.46 | 0.54 |
| 10 | 50635226 | rs2025448 | -1.058 | 0.796 | 0.532 | 1.113 | 0.013 | 0.081 | 0.081 | 0 | 0.1 | 0.24 | 0.16 | 0.26 | 0.09 | 0.09 |
| 10 | 50640272 | rs10857493 | -0.657 | 0.665 | 0.499 | 0.778 | -0.013 | 0.049 | 0.049 | 0.006 | 0.32 | 0.53 | 0.44 | 0.45 | 0.28 | 0.44 |
| 10 | 50643498 | rs1924490 | -0.168 | 1.504 | 0.487 | 0.46 | 0.007 | -0.026 | -0.026 | 0 | 0.91 | 0.87 | 0.94 | 1 | 0.87 | 0.95 |
| 10 | 50652373 | rs7906303 | 1.199 | 1.375 | 0.015 | 0.222 | -0.001 | 0 | 0 | 0 | 0.48 | 0.32 | 0.46 | 0.46 | 0.5 | 0.48 |
| 10 | 50652603 | rs12572848 | 0.465 | 1.609 | -0.246 | 0.16 | -0.015 | -0.023 | -0.023 | 0 | 0.08 | 0.03 | 0.13 | 0.15 | 0.1 | 0.22 |
| 10 | 50660509 | rs7907557 | 1.121 | 1.544 | -0.088 | 0.115 | -0.016 | -0.017 | -0.017 | 0 | 0.46 | 0.19 | 0.44 | 0.46 | 0.5 | 0.48 |
| 10 | 50666808 | rs4253231 | 0.29 | 1.534 | 0.029 | 0.126 | 0.002 | 0.009 | 0.009 | 0 | 0.08 | 0.11 | 0.14 | 0.15 | 0.1 | 0.22 |
| 10 | 50667105 | rs2228529 | 0.725 | 1.446 | 0.274 | 0.046 | -0.003 | 0.045 | 0.045 | 0.747 | 0.21 | 0.12 | 0.21 | 0.04 | 0.21 | 0.34 |
| 10 | 50678717 | rs2228526 | 0.74 | 1.447 | 0.283 | 0.046 | -0.003 | 0.041 | 0.041 | 0.815 | 0.21 | 0.14 | 0.21 | 0.04 | 0.21 | 0.34 |
| 10 | 50681033 | rs2229760 | 2.186 | 1.317 | 0.327 | 0.039 | -0.018 | 0.013 | 0.013 | 0.195 | 0.37 | 0.09 | 0.3 | 0.32 | 0.39 | 0.24 |
| 10 | 50684228 | rs3750749 | 0.19 | 1.275 | 1.309 | 0.152 | -0.005 | 0.16 | 0.16 | 0 | 0.08 | 0.13 | 0.21 | 0.41 | 0.07 | 0.1 |
| 10 | 50701156 | rs4253132 | -0.106 | 1.311 | 1.371 | 0.061 | 0.036 | -0.063 | -0.063 | 0 | 0.89 | 0.71 | 0.93 | 1 | 0.86 | 0.95 |
| 10 | 50702190 | rs4253126 | 0.934 | 1.308 | 1.256 | 0.052 | -0.027 | 0.02 | 0.02 | 0.013 | 0.12 | 0.01 | 0.03 | 0.08 | 0.09 | 0.04 |
| 10 | 50710123 | rs4838524 | -0.206 | 1.317 | 1.399 | 0.054 | 0.016 | -0.053 | -0.053 | 0 | 0.89 | 0.8 | 0.93 | 1 | 0.86 | 0.95 |
| 10 | 50717586 | rs4253082 | -0.793 | 1.275 | 1.489 | 0.112 | -0.013 | 0.095 | 0.095 | 0.036 | 0.22 | 0.38 | 0.27 | 0.49 | 0.16 | 0.14 |
| 10 | 50727787 | rs4253060 | 0.248 | 1.248 | 1.537 | 0.122 | -0.005 | 0.141 | 0.141 | 0 | 0.08 | 0.12 | 0.21 | 0.41 | 0.07 | 0.1 |
| 10 | 50728371 | rs1018603 | -0.125 | 1.311 | 1.564 | 0.094 | 0.039 | -0.063 | -0.063 | 0.016 | 0.89 | 0.71 | 0.93 | 1 | 0.86 | 0.95 |
| 10 | 50732280 | rs2228528 | -0.385 | 1.263 | 1.197 | 0.101 | 0.001 | -0.003 | -0.003 | 0.15 | 0.2 | 0.18 | 0.24 | 0.49 | 0.16 | 0.14 |
| 10 | 50740143 | rs1012553 | 1.337 | 1.413 | 1.007 | 0.104 | -0.001 | 0 | 0 | 1.394 | 0.67 | 0.33 | 0.67 | 0.51 | 0.73 | 0.82 |
| 10 | 50757504 | rs2377902 | 0.609 | 1.278 | 0.951 | 0.03 | 0 | -0.026 | -0.026 | 0 | 0.59 | 0.47 | 0.57 | 0.8 | 0.59 | 0.38 |
| 10 | 50762747 | rs1917822 | -1.131 | 1.255 | 0.979 | 0.03 | -0.001 | 0.006 | 0.006 | 0 | 0.43 | 0.54 | 0.41 | 0.5 | 0.4 | 0.49 |
| 10 | 50763795 | rs2222638 | -0.433 | 1.25 | 1.069 | 0.053 | -0.002 | -0.009 | -0.009 | 0 | 0.21 | 0.17 | 0.26 | 0.48 | 0.19 | 0.15 |
| 10 | 50764246 | rs12161717 | -0.618 | 1.249 | 0.976 | 0.053 | 0.004 | 0.1 | 0.1 | 0 | 0.22 | 0.4 | 0.28 | 0.48 | 0.19 | 0.15 |
| 10 | 50765354 | rs17784218 | 0.901 | 1.273 | 0.721 | 0.05 | -0.047 | 0.022 | 0.022 | 0 | 0.12 | 0.01 | 0.05 | 0.08 | 0.12 | 0.05 |
| 10 | 50768203 | rs1917821 | 2.124 | 1.277 | 0.787 | 0.088 | -0.025 | 0.01 | 0.01 | 0 | 0.37 | 0.09 | 0.3 | 0.32 | 0.4 | 0.23 |
| 10 | 50769606 | rs7081904 | 0.732 | 1.261 | 0.887 | 0.029 | 0.005 | 0.004 | 0.004 | 0.123 | 0.81 | 0.7 | 0.72 | 0.82 | 0.8 | 0.72 |
| 10 | 50770315 | rs2103201 | -0.444 | 1.25 | 0.811 | 0.006 | 0 | -0.037 | -0.037 | 0.177 | 0.91 | 0.84 | 0.95 | 1 | 0.9 | 0.96 |
| 10 | 50771419 | rs1949884 | -0.646 | 1.243 | 0.759 | -0.023 | -0.003 | -0.027 | -0.027 | 3.664 | 0.33 | 0.45 | 0.38 | 0.2 | 0.31 | 0.58 |
| 10 | 50771462 | rs1949883 | -0.3 | 1.264 | 0.658 | 0.176 | 0.016 | 0.073 | 0.073 | 0 | 0.12 | 0.31 | 0.25 | 0.18 | 0.1 | 0.24 |

| | | | | | | | | | | | | | | | | |
|---|---|---|---|---|---|---|---|---|---|---|---|---|---|---|---|---|
| 10 | 50772362 | rs1917820 | 1.433 | 1.264 | 0.526 | 0.247 | -0.013 | 0.043 | 0.043 | 0 | 0.4 | 0.18 | 0.48 | 0.69 | 0.44 | 0.32 |
| 10 | 50786698 | rs12259435 | -1.229 | 1.158 | 0.616 | 0.867 | 0.042 | 0.352 | 0.352 | 0.09 | 0.05 | 0.21 | 0.22 | 0.37 | 0.05 | 0.09 |
| 10 | 50789851 | rs1917817 | -0.129 | 1.27 | 0.062 | 0.805 | -0.006 | -0.016 | -0.016 | 0.411 | 0.09 | 0.05 | 0.22 | 0.19 | 0.1 | 0.24 |
| 10 | 50797262 | rs1917803 | -1.447 | 1.195 | 0.045 | 0.407 | 0.004 | 0.004 | 0.004 | 0 | 0.32 | 0.5 | 0.51 | 0.67 | 0.29 | 0.4 |
| 10 | 50801368 | rs11595012 | 0.013 | 1.299 | -0.625 | 0.279 | -0.025 | 0.021 | 0.021 | 0 | 0.13 | 0.02 | 0.09 | 0.08 | 0.14 | 0.08 |
| 10 | 50804578 | rs1880674 | 0.361 | 1.523 | -0.509 | 0.039 | 0 | -0.043 | -0.043 | 0.603 | 0.87 | 0.81 | 0.92 | 0.97 | 0.88 | 0.95 |
| 10 | 50810891 | rs11101179 | 2.029 | 1.478 | -1.968 | -0.105 | -0.043 | 0.003 | 0.003 | 1.134 | 0.28 | 0.01 | 0.18 | 0.21 | 0.31 | 0.14 |
| 10 | 50815512 | rs885834 | 0.891 | 1.509 | -1.618 | -0.064 | -0.023 | 0.131 | 0.131 | 0.278 | 0.56 | 0.18 | 0.34 | 0.24 | 0.61 | 0.53 |
| 10 | 50815696 | rs7067651 | 0.507 | 2.087 | -1.097 | 0.279 | -0.005 | -0.007 | -0.007 | 0 | 0.16 | 0.1 | 0.1 | 0.01 | 0.18 | 0.25 |
| 10 | 50816943 | rs733722 | -1.445 | 1.904 | -0.801 | 1.398 | -0.007 | -0.004 | -0.004 | 0 | 0.17 | 0.19 | 0.46 | 0.43 | 0.22 | 0.26 |
| 10 | 50818595 | rs3750752 | -0.486 | 0.988 | -0.678 | 1.194 | -0.001 | 0.025 | 0.025 | 0 | 0.07 | 0.07 | 0.3 | 0.2 | 0.05 | 0.09 |
| 10 | 50821834 | rs10857517 | -0.215 | 0.968 | -0.646 | 1.182 | -0.004 | 0.044 | 0.044 | 0 | 0.06 | 0.08 | 0.3 | 0.19 | 0.04 | 0.09 |
| 10 | 50826163 | rs7091005 | 0.554 | 1.178 | -0.673 | 0.464 | 0.005 | -0.035 | -0.035 | 0 | 0.91 | 0.89 | 0.94 | 1 | 0.89 | 0.93 |
| 10 | 50833839 | rs868750 | 0.179 | 1.357 | -1.459 | 0.2 | -0.047 | 0.087 | 0.087 | 0 | 0.17 | 0.01 | 0.12 | 0.06 | 0.18 | 0.16 |
| 10 | 50834978 | rs4838392 | -0.744 | 0.978 | -1.508 | 0.86 | -0.049 | -0.024 | -0.024 | 0 | 0.37 | 0.14 | 0.33 | 0.56 | 0.45 | 0.28 |
| 10 | 50835264 | rs12246528 | -0.29 | 0.91 | -1.222 | 0.905 | -0.022 | 0.058 | 0.058 | 0 | 0.29 | 0.17 | 0.26 | 0.13 | 0.36 | 0.2 |
| 10 | 50836466 | rs2377877 | 0.935 | 0.125 | 0.11 | 1.391 | -0.029 | 0.112 | 0.112 | 0.37 | 0.86 | 0.48 | 0.76 | 0.81 | 0.94 | 0.89 |
| 10 | 50838874 | rs2177370 | -0.303 | 0.459 | -0.53 | 1.156 | -0.002 | -0.032 | -0.032 | 0 | 0.4 | 0.51 | 0.6 | 0.75 | 0.35 | 0.43 |
| 10 | 50841704 | rs3793791 | 1.191 | 0.504 | -0.023 | 2.107 | 0.006 | 0.115 | 0.115 | 0 | 0.88 | 0.76 | 0.68 | 0.78 | 0.93 | 0.82 |
| 10 | 50846882 | rs11101190 | -1.086 | 0.547 | -0.116 | 2.815 | -0.004 | 0.038 | 0.038 | 0 | 0.06 | 0.07 | 0.18 | 0.18 | 0.06 | 0.17 |
| 10 | 50853192 | rs11101191 | -1.337 | -0.045 | 0.872 | 2.028 | -0.001 | 0.083 | 0.083 | 0 | 0.1 | 0.11 | 0.2 | 0.45 | 0.09 | 0.21 |
| 10 | 50853526 | rs1917814 | 1.38 | -0.05 | 1.037 | 2.026 | 0 | 0.253 | 0.253 | 0 | 0.89 | 0.76 | 0.79 | 0.55 | 0.91 | 0.79 |
| 10 | 51795395 | rs10824914 | 0.334 | -0.641 | -1.342 | 0.452 | -0.003 | -0.023 | -0.023 | 0 | 0.21 | 0.07 | 0.36 | 0.29 | 0.19 | 0.2 |
| 10 | 51797239 | rs4042937 | 0.397 | -0.99 | -1.698 | -0.615 | 0.059 | 0.34 | 0.34 | 0.278 | 0.48 | 0.13 | 0.49 | 0.94 | 0.62 | 0.71 |
| 10 | 51822906 | rs3964873 | 0.268 | -1.356 | -0.653 | -0.1 | -0.034 | -0.097 | -0.097 | 1.392 | 0.76 | 0.38 | 0.84 | 0.97 | 0.84 | 0.91 |
| 10 | 51832220 | rs12411414 | -0.142 | -0.992 | -0.997 | 0.531 | -0.02 | -0.077 | -0.077 | 0.365 | 0.22 | 0.09 | 0.2 | 0.62 | 0.2 | 0.36 |
| 10 | 52000536 | rs7902744 | 0.895 | -1.127 | -0.872 | 0.357 | -0.148 | -0.1 | -0.1 | 3.943 | 0.31 | 0.03 | 0.42 | 0.3 | 0.48 | 0.25 |
| 10 | 52002204 | rs7076242 | -0.732 | -1.497 | 0.11 | 0.5 | -0.057 | 0.064 | 0.064 | 1.427 | 0.25 | 0.56 | 0.18 | 0.43 | 0.09 | 0.26 |
| 10 | 52002800 | rs1436216 | -0.634 | -1.498 | 0.396 | 0.499 | -0.08 | 0.117 | 0.117 | 0.712 | 0.24 | 0.48 | 0.17 | 0.43 | 0.09 | 0.26 |
| 10 | 52002980 | rs3739979 | -0.634 | -1.498 | 0.403 | 0.499 | -0.079 | 0.115 | 0.115 | 0.712 | 0.24 | 0.48 | 0.17 | 0.43 | 0.09 | 0.26 |
| 10 | 52005293 | rs7077415 | 0.269 | -1.412 | 0.524 | 0.634 | -0.028 | 0.112 | 0.112 | 0 | 0.14 | 0.26 | 0.1 | 0.35 | 0.07 | 0.2 |
| 10 | 52008730 | rs10825492 | 0.535 | -1.418 | 0.373 | 0.631 | -0.009 | 0.041 | 0.041 | 0 | 0.13 | 0.15 | 0.09 | 0.35 | 0.07 | 0.2 |
| 10 | 52008748 | rs2196463 | 0.398 | -1.502 | 0.37 | 0.444 | -0.002 | -0.001 | -0.001 | 0 | 0.31 | 0.29 | 0.22 | 0.11 | 0.29 | 0.24 |
| 10 | 52012044 | rs2813302 | 0.222 | -1.47 | 0.459 | 0.532 | 0.024 | -0.015 | -0.015 | 0.864 | 0.61 | 0.33 | 0.64 | 0.41 | 0.77 | 0.48 |
| 10 | 52013434 | rs1865749 | -1.429 | -1.215 | 0.368 | 0.468 | 0.009 | 0.006 | 0.006 | 0.014 | 0.8 | 0.84 | 0.78 | 0.79 | 0.87 | 0.84 |
| 10 | 52014345 | rs1583522 | -0.872 | -1.202 | 0.407 | 0.467 | -0.002 | -0.002 | -0.002 | 0.85 | 0.77 | 0.7 | 0.76 | 0.76 | 0.84 | 0.68 |
| 10 | 52019288 | rs2574983 | -1.371 | -1.19 | 0.14 | 0.421 | -0.002 | -0.001 | -0.001 | 0.835 | 0.79 | 0.77 | 0.77 | 0.73 | 0.86 | 0.84 |
| 10 | 52019352 | rs7908522 | 0.219 | -1.219 | -0.146 | 0.373 | -0.024 | 0.043 | 0.043 | 0 | 0.45 | 0.24 | 0.51 | 0.59 | 0.56 | 0.44 |
| 10 | 52020424 | rs2008644 | 1.239 | -1.051 | -0.493 | 0.419 | -0.05 | 0.069 | 0.069 | 0 | 0.2 | 0.08 | 0.16 | 0.03 | 0.27 | 0.19 |
| 10 | 52020651 | rs16917064 | -0.504 | -1.925 | 0.216 | 1.257 | -0.026 | 0.009 | 0.009 | 0.578 | 0.11 | 0.22 | 0.08 | 0.11 | 0.04 | 0.22 |
| 10 | 52020743 | rs11813798 | -0.259 | -0.98 | -0.178 | 1.398 | -0.004 | 0.022 | 0.022 | 0 | 0.16 | 0.23 | 0.11 | 0.35 | 0.11 | 0.36 |
| 10 | 52025167 | rs2574943 | 0.074 | -0.561 | -0.342 | 1.088 | -0.018 | -0.004 | -0.004 | 0 | 0.78 | 0.7 | 0.76 | 0.73 | 0.86 | 0.84 |
| 10 | 52032275 | rs11597071 | -0.587 | 0.51 | -1.602 | 1.361 | -0.133 | 0.019 | 0.019 | 0 | 0.24 | 0.02 | 0.3 | 0.12 | 0.37 | 0.18 |
| 10 | 52032872 | rs930324 | -0.635 | 0.489 | -0.969 | 1.289 | -0.013 | 0.008 | 0.008 | 0 | 0.22 | 0.23 | 0.28 | 0.13 | 0.32 | 0.16 |
| 10 | 52033901 | rs1436222 | -0.896 | -0.877 | -1.362 | 0.75 | -0.065 | 0.104 | 0.104 | 0.014 | 0.61 | 0.84 | 0.61 | 0.84 | 0.51 | 0.65 |
| 10 | 52039157 | rs2813323 | -0.075 | -1.225 | -0.298 | 0.8 | 0.036 | 0.062 | 0.062 | 0 | 0.93 | 0.69 | 0.92 | 0.92 | 0.97 | 0.83 |
| 10 | 52043336 | rs2842131 | 0.293 | -1.171 | -0.233 | 0.791 | 0.016 | -0.015 | -0.015 | 0 | 0.7 | 0.57 | 0.66 | 0.79 | 0.66 | 0.66 |
| 10 | 52043966 | rs12357549 | 1.286 | -1.879 | -0.622 | 0.852 | 0.029 | -0.057 | -0.057 | 0.128 | 0.16 | 0.03 | 0.18 | 0.31 | 0.11 | 0.11 |
| 10 | 52043987 | rs2574951 | 0.482 | -1.774 | -0.018 | 0.901 | -0.061 | 0.067 | 0.067 | 1.601 | 0.76 | 0.34 | 0.86 | 0.6 | 0.92 | 0.63 |
| 10 | 52045632 | rs1008716 | 0.468 | -1.235 | -0.145 | 0.908 | -0.003 | 0.013 | 0.013 | 0 | 0.64 | 0.52 | 0.59 | 0.49 | 0.69 | 0.46 |
| 10 | 52046187 | rs2574952 | 0.805 | -1.524 | 0.671 | 0.617 | -0.026 | -0.062 | -0.062 | 0 | 0.89 | 0.57 | 0.91 | 1 | 0.96 | 0.97 |
| 10 | 52046803 | rs6481165 | 0.035 | -1.298 | 0.369 | 0.711 | 0.018 | -0.012 | -0.012 | 0 | 0.18 | 0.21 | 0.23 | 0.09 | 0.25 | 0.12 |
| 10 | 52047351 | rs10825563 | 0.007 | -2.059 | -0.225 | 0.55 | 0.03 | 0.016 | 0.016 | 0.32 | 0.37 | 0.28 | 0.26 | 0.35 | 0.27 | 0.28 |
| 10 | 52051645 | rs4935597 | 1.048 | -1.987 | -0.285 | 0.367 | -0.08 | -0.011 | -0.011 | 0 | 0.16 | 0.01 | 0.21 | 0.09 | 0.25 | 0.12 |
| 10 | 52052292 | rs1369861 | -0.199 | -2.08 | -0.256 | 0.632 | 0.013 | 0.002 | 0.002 | 0.15 | 0.41 | 0.37 | 0.32 | 0.69 | 0.28 | 0.5 |
| 10 | 52053115 | rs892987 | -1.539 | -1.962 | -0.202 | 0.628 | -0.098 | 0.055 | 0.055 | 0 | 0.63 | 0.89 | 0.58 | 0.78 | 0.53 | 0.67 |
| 10 | 52054343 | rs2082929 | -1.022 | -1.645 | -0.253 | 0.615 | -0.008 | 0.02 | 0.02 | 0 | 0.57 | 0.39 | 0.54 | 0.79 | 0.53 | 0.62 |
| 10 | 52061783 | rs10159676 | -0.111 | -1.925 | -0.201 | 0.619 | -0.043 | 0.008 | 0.008 | 0.7 | 0.37 | 0.5 | 0.34 | 0.65 | 0.25 | 0.43 |
| 10 | 52065890 | rs14327 | -1.473 | -1.604 | -0.693 | 0.591 | -0.108 | 0.067 | 0.067 | 0 | 0.63 | 0.9 | 0.6 | 0.8 | 0.53 | 0.67 |
| 10 | 52068298 | rs2842084 | -1.44 | -1.593 | -0.689 | 0.612 | -0.107 | 0.054 | 0.054 | 0 | 0.63 | 0.9 | 0.6 | 0.79 | 0.53 | 0.67 |
| 10 | 52069663 | rs17721080 | 0.099 | -1.19 | -0.633 | 0.782 | -0.031 | 0.035 | 0.035 | 0 | 0.22 | 0.09 | 0.25 | 0.09 | 0.29 | 0.14 |
| 10 | 52070947 | rs2251601 | -0.201 | -1.265 | -0.477 | 1.058 | -0.011 | -0.029 | -0.029 | 0 | 0.25 | 0.19 | 0.31 | 0.48 | 0.31 | 0.37 |
| 10 | 52073479 | rs4935605 | -0.074 | -1.336 | -0.502 | 0.832 | -0.042 | 0.059 | 0.059 | 0 | 0.25 | 0.08 | 0.27 | 0.09 | 0.31 | 0.13 |
| 10 | 52081612 | rs7070330 | -0.093 | -1.409 | -0.272 | 0.519 | 0.006 | 0 | 0 | 2.788 | 0.3 | 0.23 | 0.24 | 0.32 | 0.23 | 0.24 |
| 10 | 52085411 | rs7078238 | -0.19 | -1.342 | -0.411 | 0.566 | -0.017 | 0 | 0 | 0.113 | 0.3 | 0.32 | 0.25 | 0.32 | 0.22 | 0.24 |
| 10 | 52088733 | rs2177371 | -0.687 | 0.056 | -0.774 | -0.148 | -0.008 | 0.004 | 0.004 | 0.437 | 0.6 | 0.64 | 0.68 | 0.86 | 0.52 | 0.66 |
| 10 | 52093995 | rs1344848 | 1.571 | -0.061 | -1.376 | 0.06 | -0.039 | -0.068 | -0.068 | 1.996 | 0.17 | 0.01 | 0.24 | 0.24 | 0.16 | 0.19 |
| 10 | 52097415 | rs17785945 | 1.088 | -0.088 | -1.344 | 0.153 | -0.023 | -0.076 | -0.076 | 0 | 0.19 | 0.05 | 0.26 | 0.27 | 0.18 | 0.24 |
| 10 | 52099505 | rs7905772 | 0.844 | -0.105 | -0.921 | 0.441 | 0.002 | 0.008 | 0.008 | 0 | 0.21 | 0.33 | 0.25 | 0.21 | 0.2 | 0.07 |

| | | | | | | | | | | | | | | | | |
|---|---|---|---|---|---|---|---|---|---|---|---|---|---|---|---|---|
| 10 | 52103707 | rs10763354 | -0.917 | -0.037 | -1.125 | 0.433 | 0.005 | 0.001 | 0.001 | 1.075 | 0.56 | 0.52 | 0.63 | 0.49 | 0.53 | 0.37 |
| 10 | 52104339 | rs11005278 | 1.041 | 0.29 | -1.552 | 0.782 | -0.051 | -0.061 | -0.061 | 0 | 0.17 | 0.04 | 0.22 | 0.21 | 0.19 | 0.07 |
| 10 | 52108830 | rs7070574 | 0.341 | -0.025 | -1.516 | 0.419 | -0.018 | -0.072 | -0.072 | 0.267 | 0.37 | 0.15 | 0.47 | 0.44 | 0.35 | 0.27 |
| 10 | 52109375 | rs10825780 | -0.055 | 0.329 | -1.044 | -0.334 | 0.002 | -0.018 | -0.018 | 0.114 | 0.23 | 0.48 | 0.19 | 0.16 | 0.21 | 0.14 |
| 10 | 52112030 | rs7069860 | -0.706 | 0.428 | -1.121 | -0.721 | 0.001 | -0.024 | -0.024 | 0 | 0.76 | 0.77 | 0.85 | 1 | 0.8 | 0.75 |
| 10 | 52117268 | rs2574976 | 0.31 | 0.116 | -1.055 | -0.702 | -0.004 | -0.041 | -0.041 | 0 | 0.21 | 0.24 | 0.1 | 0.01 | 0.13 | 0.2 |
| 10 | 52119549 | rs10825789 | 1.353 | 0.471 | -1.208 | -0.296 | -0.012 | -0.06 | -0.06 | 0 | 0.19 | 0.09 | 0.26 | 0.31 | 0.21 | 0.19 |
| 10 | 52120612 | rs1436206 | -0.218 | 0.702 | -0.999 | -0.503 | 0.008 | -0.013 | -0.013 | 0 | 0.61 | 0.57 | 0.5 | 0.26 | 0.54 | 0.5 |
| 10 | 52127483 | rs2060560 | 0.542 | 0.57 | -1.115 | -0.806 | 0.026 | 0.077 | 0.077 | 0 | 0.43 | 0.29 | 0.37 | 0.2 | 0.37 | 0.39 |
| 10 | 52131032 | rs2688875 | 0.658 | 0.488 | -1.124 | -0.801 | 0.027 | 0.122 | 0.122 | 0 | 0.42 | 0.19 | 0.36 | 0.21 | 0.36 | 0.39 |
| 10 | 52135215 | rs2842110 | 1.326 | 0.382 | -1.222 | -0.936 | 0.025 | 0.065 | 0.065 | 0 | 0.2 | 0.08 | 0.08 | 0.01 | 0.15 | 0.2 |
| 10 | 52139069 | rs768185 | 1.692 | 0.407 | -0.984 | -0.753 | 0 | -0.005 | -0.005 | 2.763 | 0.2 | 0.19 | 0.29 | 0.32 | 0.22 | 0.2 |
| 10 | 52141744 | rs2339400 | 0.268 | 0.433 | -0.951 | -0.753 | -0.001 | 0 | 0 | 0 | 0.37 | 0.38 | 0.41 | 0.32 | 0.38 | 0.27 |
| 10 | 52160862 | rs10825903 | 0.297 | 0.417 | -0.776 | -0.764 | 0.004 | -0.036 | -0.036 | 2.08 | 0.17 | 0.26 | 0.13 | 0.05 | 0.17 | 0.12 |
| 10 | 52162253 | rs17721658 | 1.393 | 0.411 | -0.639 | -0.576 | 0 | 0.001 | 0.001 | 0.07 | 0.22 | 0.22 | 0.28 | 0.2 | 0.22 | 0.19 |
| 10 | 52163991 | rs884880 | 1.624 | 0.414 | -0.638 | -0.574 | -0.012 | 0.003 | 0.003 | 0.739 | 0.21 | 0.09 | 0.27 | 0.2 | 0.21 | 0.19 |
| 10 | 52188013 | rs11005626 | 1.443 | 0.385 | -0.298 | -0.597 | 0 | 0.004 | 0.004 | 1.079 | 0.22 | 0.2 | 0.29 | 0.2 | 0.21 | 0.19 |
| 10 | 52193777 | rs1203412 | -1.284 | 0.336 | -0.27 | -0.879 | 0.004 | -0.019 | -0.019 | 0.116 | 0.79 | 0.82 | 0.9 | 0.99 | 0.82 | 0.79 |
| 10 | 52197398 | rs1203408 | -0.176 | -0.012 | -0.05 | -0.677 | 0.007 | 0.007 | 0.007 | 0 | 0.49 | 0.46 | 0.53 | 0.36 | 0.58 | 0.3 |
| 10 | 52199196 | rs10825996 | 0.821 | 0.141 | -0.114 | -0.735 | -0.001 | 0.006 | 0.006 | 0 | 0.35 | 0.25 | 0.38 | 0.26 | 0.36 | 0.22 |
| 10 | 52201167 | rs10082515 | -0.241 | 0.115 | -0.18 | -0.67 | -0.025 | 0.333 | 0.333 | 0.871 | 0.09 | 0.32 | 0.11 | 0.44 | 0.03 | 0.3 |
| 10 | 52209389 | rs10826013 | 0.707 | 0.138 | -0.172 | -0.713 | -0.002 | -0.005 | -0.005 | 0.163 | 0.36 | 0.36 | 0.39 | 0.26 | 0.36 | 0.22 |
| 10 | 52222605 | rs910819 | -1.553 | 0.122 | -0.262 | -0.746 | 0.007 | 0.074 | 0.074 | 0.028 | 0.81 | 0.93 | 0.91 | 0.99 | 0.82 | 0.79 |
| 10 | 52231729 | rs1203380 | -0.327 | 0.084 | -0.1 | -0.698 | -0.013 | 0.016 | 0.016 | 0.05 | 0.72 | 0.66 | 0.81 | 0.56 | 0.79 | 0.49 |
| 10 | 52232694 | rs10763500 | 0.689 | 0.045 | 0.104 | -0.686 | -0.004 | -0.013 | -0.013 | 0.943 | 0.37 | 0.42 | 0.4 | 0.26 | 0.36 | 0.22 |
| 10 | 52237841 | rs16937491 | 0.449 | 0.089 | 0.14 | -0.533 | -0.031 | 0.165 | 0.165 | 0 | 0.06 | 0.14 | 0.07 | 0.37 | 0.01 | 0.26 |
| 10 | 52247531 | rs1011881 | 0.306 | 0.237 | -0.035 | -0.639 | -0.003 | -0.011 | -0.011 | 0.123 | 0.58 | 0.61 | 0.68 | 0.47 | 0.58 | 0.41 |
| 10 | 52258698 | rs2339506 | 0.489 | 0.214 | 0.005 | -0.651 | -0.001 | 0.005 | 0.005 | 0.266 | 0.57 | 0.45 | 0.66 | 0.47 | 0.57 | 0.4 |
| 10 | 52262468 | rs10826118 | 0.339 | 0.202 | 0.12 | -0.671 | -0.003 | -0.003 | -0.003 | 0 | 0.6 | 0.64 | 0.69 | 0.52 | 0.6 | 0.45 |
| 10 | 52263910 | rs2339507 | 0.792 | 0.173 | 0.032 | -0.721 | 0 | 0.005 | 0.005 | 0 | 0.36 | 0.31 | 0.39 | 0.27 | 0.37 | 0.22 |
| 10 | 52265265 | rs7920818 | 1.907 | 0.12 | -0.065 | -0.868 | 0.001 | -0.006 | -0.006 | 0 | 0.2 | 0.22 | 0.28 | 0.26 | 0.2 | 0.15 |
| 10 | 52268049 | rs16937786 | 0.41 | 0.131 | 0.061 | -0.822 | -0.029 | 0.204 | 0.204 | 0.411 | 0.06 | 0.18 | 0.07 | 0.37 | 0.01 | 0.26 |
| 10 | 52269914 | rs2064760 | -0.545 | 0.166 | -0.125 | -0.887 | -0.052 | 0.037 | 0.037 | 0.079 | 0.73 | 0.48 | 0.81 | 0.62 | 0.81 | 0.54 |
| 10 | 52271385 | rs3011770 | 1.002 | 0.169 | -0.151 | -0.924 | -0.032 | -0.005 | -0.005 | 0 | 0.16 | 0.08 | 0.14 | 0.15 | 0.24 | 0.12 |
| 10 | 52272007 | rs7071603 | 0.422 | 0.203 | -0.129 | -0.918 | -0.001 | -0.004 | -0.004 | 0 | 0.57 | 0.4 | 0.66 | 0.46 | 0.57 | 0.41 |
| 10 | 52277400 | rs4935727 | 0.328 | 0.235 | -0.071 | -0.904 | -0.001 | -0.003 | -0.003 | 0.49 | 0.58 | 0.41 | 0.66 | 0.45 | 0.57 | 0.41 |
| 10 | 52279428 | rs7904833 | 1.004 | 0.215 | -0.119 | -0.939 | 0.006 | -0.001 | -0.001 | 0.489 | 0.44 | 0.34 | 0.56 | 0.51 | 0.44 | 0.39 |
| 10 | 52279502 | rs2983348 | -1.358 | 0.316 | -0.051 | -0.803 | 0.005 | -0.009 | -0.009 | 0 | 0.8 | 0.88 | 0.9 | 0.99 | 0.82 | 0.8 |
| 10 | 52281609 | rs2092996 | 0.316 | 0.386 | -0.21 | -0.713 | -0.001 | -0.004 | -0.004 | 0 | 0.58 | 0.38 | 0.66 | 0.46 | 0.57 | 0.41 |
| 10 | 52296835 | rs10826161 | 1.283 | 0.163 | -0.461 | -0.734 | 0.019 | -0.048 | -0.048 | 0.106 | 0.23 | 0.08 | 0.31 | 0.64 | 0.21 | 0.38 |
| 10 | 52306053 | rs2983360 | 0.312 | 0.283 | -0.453 | -0.869 | -0.033 | 0.036 | 0.036 | 0.03 | 0.38 | 0.19 | 0.45 | 0.74 | 0.44 | 0.47 |
| 10 | 52315912 | rs11006119 | 0.589 | 0.108 | -0.676 | -0.958 | -0.058 | 0.035 | 0.035 | 3.051 | 0.35 | 0.07 | 0.37 | 0.24 | 0.37 | 0.31 |
| 10 | 52321378 | rs9919413 | 1.327 | -0.042 | -0.626 | -0.833 | 0.01 | -0.025 | -0.025 | 0.241 | 0.23 | 0.11 | 0.31 | 0.64 | 0.21 | 0.38 |
| 10 | 52322047 | rs11006141 | 0.364 | -0.034 | -0.873 | -0.922 | 0.005 | -0.011 | -0.011 | 0 | 0.36 | 0.11 | 0.41 | 0.64 | 0.36 | 0.44 |
| 10 | 52323909 | rs12219692 | 2.227 | 0.038 | -0.884 | -0.629 | -0.03 | -0.098 | -0.098 | 0 | 0.19 | 0.02 | 0.25 | 0.27 | 0.2 | 0.09 |
| 10 | 52325713 | rs2983362 | -1.395 | 0.192 | -0.487 | -0.64 | 0.001 | -0.021 | -0.021 | 0 | 0.81 | 0.85 | 0.89 | 1 | 0.82 | 0.84 |
| 10 | 52326285 | rs11006155 | -0.114 | 0.603 | -0.603 | -0.666 | -0.005 | 0.09 | 0.09 | 0 | 0.47 | 0.2 | 0.5 | 0.74 | 0.52 | 0.51 |
| 10 | 52337406 | rs7072826 | 1.092 | 0.863 | -0.562 | -0.575 | -0.021 | 0.003 | 0.003 | 0 | 0.11 | 0.05 | 0.11 | 0.1 | 0.18 | 0.07 |
| 10 | 52338020 | rs11006190 | 1.197 | 0.865 | -0.594 | -0.574 | -0.061 | -0.014 | -0.014 | 0.23 | 0.1 | 0.01 | 0.1 | 0.1 | 0.18 | 0.07 |
| 10 | 52338100 | rs10508930 | -0.055 | 0.71 | -0.644 | -0.516 | -0.002 | -0.001 | -0.001 | 0 | 0.13 | 0.08 | 0.11 | 0.35 | 0.13 | 0.31 |
| 10 | 52348894 | rs3011710 | 0.009 | 1.004 | -0.254 | -0.782 | -0.014 | 0.054 | 0.054 | 0 | 0.43 | 0.22 | 0.49 | 0.74 | 0.5 | 0.53 |
| 10 | 52350006 | rs11006229 | 1.021 | 0.987 | -0.261 | -0.683 | -0.027 | 0.005 | 0.005 | 0 | 0.11 | 0.04 | 0.1 | 0.1 | 0.18 | 0.07 |
| 10 | 52350814 | rs17496403 | 0.93 | 0.986 | -0.266 | -0.683 | -0.002 | 0.001 | 0.001 | 0 | 0.11 | 0.09 | 0.11 | 0.1 | 0.18 | 0.07 |
| 10 | 52354605 | rs10763585 | -0.699 | 0.88 | -0.543 | -0.82 | -0.024 | 0.039 | 0.039 | 0.618 | 0.64 | 0.78 | 0.6 | 0.75 | 0.61 | 0.66 |
| 10 | 52355506 | rs10826234 | -0.038 | 0.895 | -0.617 | -0.733 | -0.003 | -0.019 | -0.019 | 0.664 | 0.43 | 0.51 | 0.49 | 0.75 | 0.44 | 0.49 |
| 10 | 52356264 | rs10763587 | -1.571 | 0.522 | -0.518 | -0.465 | 0.006 | 0.014 | 0.014 | 0.855 | 0.3 | 0.6 | 0.26 | 0.44 | 0.31 | 0.38 |
| 10 | 52357891 | rs11006259 | -0.065 | 0.487 | -1.172 | -0.34 | -0.048 | 0.087 | 0.087 | 0.306 | 0.44 | 0.17 | 0.42 | 0.32 | 0.49 | 0.36 |
| 10 | 52359689 | rs17581567 | 1.062 | 0.642 | -1.269 | -0.379 | -0.045 | 0.009 | 0.009 | 0 | 0.11 | 0.03 | 0.1 | 0.1 | 0.18 | 0.07 |
| 10 | 52361844 | rs10740762 | -1.113 | 0.638 | -1.005 | -0.472 | -0.002 | -0.002 | -0.002 | 0 | 0.67 | 0.64 | 0.59 | 0.33 | 0.69 | 0.53 |
| 10 | 52370610 | rs7897345 | 0.899 | 0.614 | -0.976 | -0.504 | -0.023 | -0.14 | -0.14 | 0 | 0.22 | 0.43 | 0.17 | 0.01 | 0.2 | 0.17 |
| 10 | 52370630 | rs4935738 | 0.517 | 0.614 | -0.939 | -0.308 | 0.009 | -0.022 | -0.022 | 0 | 0.36 | 0.23 | 0.36 | 0.58 | 0.32 | 0.59 |
| 10 | 52371188 | rs6481442 | -1.619 | 0.631 | -0.969 | -0.312 | -0.003 | -0.01 | -0.01 | 0 | 0.71 | 0.75 | 0.64 | 0.69 | 0.7 | 0.83 |
| 10 | 52381154 | rs13376734 | 0.945 | 0.675 | -0.983 | -0.305 | -0.023 | -0.14 | -0.14 | 0 | 0.22 | 0.43 | 0.17 | 0.01 | 0.2 | 0.17 |
| 10 | 52388236 | rs4475874 | 1.056 | 0.81 | -1.669 | -0.257 | -0.031 | 0.045 | 0.045 | 0.505 | 0.32 | 0.1 | 0.31 | 0.22 | 0.3 | 0.29 |
| 10 | 52393270 | rs7093195 | -0.711 | 0.785 | -1.697 | -0.264 | 0.001 | -0.014 | -0.014 | 0 | 0.4 | 0.65 | 0.33 | 0.47 | 0.39 | 0.54 |
| 10 | 52404195 | rs4636605 | 0.573 | 1.135 | -1.64 | -0.379 | -0.017 | 0.049 | 0.049 | 0.639 | 0.32 | 0.09 | 0.31 | 0.22 | 0.31 | 0.29 |
| 10 | 52414456 | rs7904306 | -0.31 | 1.132 | -0.611 | -0.086 | -0.032 | 0.023 | 0.023 | 0 | 0.14 | 0.06 | 0.11 | 0.1 | 0.18 | 0.07 |
| 10 | 52484993 | rs4934999 | 1.383 | 0.599 | -0.811 | 0.144 | -0.074 | -0.044 | -0.044 | 0.315 | 0.47 | 0.07 | 0.6 | 0.43 | 0.48 | 0.32 |
| 10 | 52487724 | rs10821669 | 0.569 | 0.494 | -0.203 | -0.052 | 0 | -0.008 | -0.008 | 0 | 0.66 | 0.37 | 0.8 | 0.85 | 0.71 | 0.6 |

| | | | | | | | | | | | | | | | | |
|---|---|---|---|---|---|---|---|---|---|---|---|---|---|---|---|---|
| 10 | 52489148 | rs10994191 | -0.061 | 0.438 | -0.483 | -0.216 | -0.067 | 0.052 | 0.052 | 0 | 0.15 | 0.01 | 0.11 | 0.09 | 0.22 | 0.15 |
| 10 | 52499880 | rs4497350 | 0.651 | 0.277 | -0.311 | -0.226 | 0.052 | 0.107 | 0.107 | 0.724 | 0.67 | 0.16 | 0.81 | 0.88 | 0.75 | 0.64 |
| 10 | 52550014 | rs10821825 | 1.714 | 0.374 | -0.44 | -0.097 | -0.061 | -0.035 | -0.035 | 0.057 | 0.48 | 0.08 | 0.61 | 0.42 | 0.49 | 0.33 |
| 10 | 52554416 | rs12358745 | -1.216 | 0.313 | 0.296 | -0.173 | -0.004 | -0.032 | -0.032 | 0 | 0.33 | 0.22 | 0.23 | 0.55 | 0.35 | 0.45 |
| 10 | 52558492 | rs7895606 | 1.975 | 0.269 | 0.894 | -0.204 | -0.019 | 0.002 | 0.002 | 0.057 | 0.49 | 0.14 | 0.62 | 0.42 | 0.49 | 0.33 |
| 10 | 52559634 | rs10994507 | 0.594 | 0.229 | 0.922 | -0.221 | -0.031 | -0.05 | -0.05 | 1.076 | 0.15 | 0.33 | 0.07 | 0.09 | 0.09 | 0.14 |
| 10 | 52565940 | rs4078160 | 0.486 | 0.491 | 0.92 | -0.062 | -0.01 | -0.083 | -0.083 | 0 | 0.83 | 0.54 | 0.88 | 0.97 | 0.84 | 0.78 |
| 10 | 52566975 | rs4382847 | 0.258 | 0.499 | 1.095 | -0.061 | -0.002 | -0.071 | -0.071 | 0.014 | 0.86 | 0.74 | 0.9 | 0.99 | 0.84 | 0.81 |
| 10 | 52568616 | rs4256922 | 0.455 | 0.513 | 1.024 | -0.053 | 0.001 | -0.084 | -0.084 | 0.023 | 0.84 | 0.56 | 0.88 | 0.97 | 0.84 | 0.78 |
| 10 | 52571924 | rs4460764 | -0.65 | 0.527 | 0.745 | -0.032 | -0.075 | -0.142 | -0.142 | 0 | 0.2 | 0.03 | 0.19 | 0.46 | 0.26 | 0.32 |
| 10 | 52576068 | rs4245008 | 0.612 | 0.048 | 1.131 | -0.248 | -0.003 | -0.075 | -0.075 | 0 | 0.84 | 0.59 | 0.89 | 0.97 | 0.84 | 0.78 |
| 10 | 52584110 | rs10821871 | -1.104 | 0.045 | 1.105 | -0.273 | -0.004 | -0.03 | -0.03 | 0 | 0.22 | 0.29 | 0.14 | 0.07 | 0.23 | 0.24 |
| 10 | 52608657 | rs4935194 | 1.939 | 0.673 | 1.157 | -0.156 | -0.016 | -0.012 | -0.012 | 0 | 0.62 | 0.3 | 0.53 | 0.49 | 0.52 | 0.41 |
| 10 | 52612101 | rs10994720 | -0.883 | 0.7 | 1.031 | -0.145 | -0.003 | -0.028 | -0.028 | 0 | 0.21 | 0.28 | 0.13 | 0.06 | 0.22 | 0.22 |
| 10 | 52612662 | rs12769066 | -0.081 | 0.715 | 1.045 | -0.138 | -0.074 | -0.091 | -0.091 | 0 | 0.14 | 0.04 | 0.29 | 0.45 | 0.26 | 0.34 |
| 10 | 52630185 | rs12768916 | -1.422 | 0.782 | 1.091 | -0.124 | 0 | 0.003 | 0.003 | 0 | 0.26 | 0.3 | 0.32 | 0.46 | 0.32 | 0.4 |
| 10 | 52633459 | rs4317935 | -0.606 | 1.029 | 1.123 | 0.07 | 0.054 | 0.125 | 0.125 | 0 | 0.17 | 0.25 | 0.32 | 0.49 | 0.26 | 0.37 |
| 10 | 52635240 | rs17591964 | 0.545 | -0.155 | 1.114 | -0.915 | -0.036 | 0.022 | 0.022 | 0 | 0.11 | 0.02 | 0.11 | 0.09 | 0.16 | 0.18 |
| 10 | 52638933 | rs6479768 | 0.069 | -0.166 | 1.263 | -0.92 | 0.024 | -0.039 | -0.039 | 2.831 | 0.86 | 0.73 | 0.86 | 0.91 | 0.83 | 0.81 |
| 10 | 52642384 | rs17500846 | 0.112 | -0.325 | 1.062 | -0.963 | -0.058 | 0.064 | 0.064 | 0 | 0.24 | 0.03 | 0.16 | 0.17 | 0.28 | 0.33 |
| 10 | 52643179 | rs12244405 | 0.072 | -0.325 | 1.079 | -0.963 | -0.03 | 0.065 | 0.065 | 0 | 0.24 | 0.07 | 0.17 | 0.17 | 0.28 | 0.33 |
| 10 | 52645248 | rs10994856 | -0.819 | -0.248 | 0.499 | -0.957 | 0 | -0.01 | -0.01 | 0 | 0.18 | 0.22 | 0.11 | 0.05 | 0.2 | 0.2 |
| 10 | 52645424 | rs10994860 | -0.81 | -0.245 | 0.5 | -0.957 | 0 | -0.01 | -0.01 | 0 | 0.18 | 0.22 | 0.11 | 0.05 | 0.2 | 0.2 |
| 10 | 52646093 | rs10821905 | -0.996 | -0.227 | 0.345 | -0.949 | -0.002 | -0.039 | -0.039 | 0 | 0.19 | 0.3 | 0.12 | 0.05 | 0.2 | 0.2 |
| 10 | 52646344 | rs3808919 | 0.95 | -0.178 | -0.29 | -0.726 | 0.009 | 0.188 | 0.188 | 0 | 0.52 | 0.2 | 0.65 | 0.78 | 0.51 | 0.46 |
| 10 | 52650502 | rs7918123 | 0.776 | -0.501 | -0.354 | -0.449 | -0.004 | 0.087 | 0.087 | 0 | 0.45 | 0.16 | 0.53 | 0.69 | 0.56 | 0.49 |
| 10 | 52681585 | rs10995011 | -0.978 | -0.499 | 0.064 | -0.784 | -0.023 | -0.037 | -0.037 | 0 | 0.15 | 0.42 | 0.09 | 0.11 | 0.1 | 0.16 |
| 10 | 52685197 | rs2339624 | -0.581 | -0.272 | -0.671 | -0.578 | -0.006 | -0.002 | -0.002 | 0 | 0.09 | 0.18 | 0.04 | 0.11 | 0.07 | 0.14 |
| 10 | 52702677 | rs10490967 | -0.313 | -0.282 | -1.309 | -0.756 | -0.037 | 0.233 | 0.233 | 0.754 | 0.36 | 0.02 | 0.2 | 0.08 | 0.34 | 0.32 |
| 10 | 52714759 | rs2816838 | -0.211 | -0.586 | -1.074 | -0.26 | 0.057 | 0.097 | 0.097 | 0 | 0.74 | 0.94 | 0.87 | 0.91 | 0.85 | 0.76 |
| 10 | 52717885 | rs7097894 | 0.788 | -0.549 | -1.745 | -0.609 | -0.002 | 0.16 | 0.16 | 0.643 | 0.47 | 0.14 | 0.51 | 0.75 | 0.6 | 0.65 |
| 10 | 52724547 | rs4336980 | 0.524 | -0.092 | -1.95 | -0.604 | -0.056 | -0.002 | -0.002 | 0 | 0.12 | 0.02 | 0.16 | 0.12 | 0.15 | 0.23 |
| 10 | 52724799 | rs6479812 | 0.602 | -0.353 | -0.439 | -0.829 | 0.065 | 0.193 | 0.193 | 0 | 0.5 | 0.24 | 0.62 | 0.8 | 0.67 | 0.5 |
| 10 | 52727674 | rs2666538 | -0.801 | -0.69 | 0.06 | -0.751 | 0.058 | 0.116 | 0.116 | 0 | 0.75 | 0.97 | 0.88 | 0.93 | 0.87 | 0.71 |
| 10 | 52728613 | rs7914943 | 0.056 | -0.598 | 0.551 | -0.719 | 0.022 | -0.013 | -0.013 | 0.254 | 0.08 | 0.36 | 0.05 | 0 | 0.02 | 0.01 |
| 10 | 52730716 | rs10995179 | 0.341 | -0.823 | 0.104 | 0.184 | 0.11 | 0.04 | 0.04 | 0 | 0.58 | 0.21 | 0.78 | 0.74 | 0.81 | 0.6 |
| 10 | 52733719 | rs6479825 | 0.082 | -0.498 | 0.307 | 0.29 | 0.046 | 0.037 | 0.037 | 1.041 | 0.05 | 0.31 | 0.04 | 0 | 0.01 | 0.02 |
| 10 | 52736525 | rs7100341 | 0.895 | 0.002 | -0.155 | 0.304 | 0.004 | 0.02 | 0.02 | 0 | 0.28 | 0.31 | 0.32 | 0.44 | 0.33 | 0.24 |
| 10 | 52738475 | rs7076050 | 0.895 | 0.002 | -0.155 | 0.304 | 0.004 | 0.02 | 0.02 | 0 | 0.28 | 0.31 | 0.32 | 0.44 | 0.33 | 0.24 |
| 10 | 52752634 | rs2816835 | -1.214 | -0.308 | -0.496 | -0.167 | 0.018 | 0.057 | 0.057 | 0.022 | 0.8 | 0.91 | 0.9 | 0.95 | 0.87 | 0.82 |
| 10 | 52761458 | rs7893612 | -1.467 | -0.336 | -0.64 | -0.173 | 0.011 | 0.064 | 0.064 | 0 | 0.82 | 0.97 | 0.9 | 0.95 | 0.88 | 0.83 |
| 10 | 52762887 | rs10995319 | 0.988 | -0.177 | -0.759 | -0.221 | -0.061 | 0.094 | 0.094 | 0.048 | 0.26 | 0.02 | 0.23 | 0.11 | 0.22 | 0.31 |
| 10 | 52771503 | rs10128514 | 1.177 | 0.071 | -0.248 | 0.142 | 0.008 | 0.028 | 0.028 | 0 | 0.26 | 0.31 | 0.3 | 0.51 | 0.31 | 0.28 |
| 10 | 52785053 | rs10822114 | 0.677 | -0.022 | -0.123 | 0.093 | 0.004 | 0.005 | 0.005 | 0 | 0.19 | 0.22 | 0.16 | 0.24 | 0.28 | 0.13 |
| 10 | 52789016 | rs10822131 | 0.9 | -0.019 | -0.11 | 0.094 | -0.007 | -0.008 | -0.008 | 0.065 | 0.17 | 0.14 | 0.15 | 0.24 | 0.28 | 0.13 |
| 10 | 52792194 | rs10822139 | 0.48 | 0.107 | -0.119 | -0.002 | 0 | -0.016 | -0.016 | 0.692 | 0.32 | 0.35 | 0.28 | 0.18 | 0.28 | 0.34 |
| 10 | 52792499 | rs2339681 | -1.227 | 0.134 | -0.108 | 0.16 | 0.026 | 0.05 | 0.05 | 0 | 0.8 | 0.93 | 0.9 | 0.93 | 0.87 | 0.8 |
| 10 | 52792744 | rs10822142 | 0.61 | 0.368 | 0.126 | 0.378 | 0.006 | 0.006 | 0.006 | 0 | 0.19 | 0.22 | 0.16 | 0.24 | 0.28 | 0.13 |
| 10 | 52793177 | rs2339680 | 1.061 | 0.197 | -0.051 | 0.563 | 0 | 0.019 | 0.019 | 0 | 0.26 | 0.29 | 0.45 | 0.45 | 0.3 | 0.29 |
| 10 | 52809354 | rs10761738 | 0.101 | 0.086 | -0.051 | 0.507 | 0.004 | -0.024 | -0.024 | 0 | 0.2 | 0.27 | 0.26 | 0.09 | 0.28 | 0.1 |
| 10 | 52809783 | rs2666536 | -0.982 | 0.022 | -0.548 | 0.57 | -0.007 | -0.045 | -0.045 | 0 | 0.83 | 0.95 | 0.91 | 0.76 | 0.86 | 0.79 |
| 10 | 52813733 | rs16913257 | 0.339 | 0.036 | -0.537 | 0.557 | 0.001 | -0.009 | -0.009 | 0 | 0.19 | 0.22 | 0.25 | 0.09 | 0.28 | 0.1 |
| 10 | 52814442 | rs2660204 | -1.568 | 0.036 | -0.655 | 0.625 | -0.036 | -0.001 | -0.001 | 0 | 0.86 | 1 | 0.91 | 0.9 | 0.88 | 0.87 |
| 10 | 52822348 | rs4935216 | 0.91 | -0.127 | -0.618 | 0.472 | 0.009 | 0.029 | 0.029 | 0.685 | 0.54 | 0.15 | 0.59 | 0.67 | 0.56 | 0.67 |
| 10 | 52828115 | rs16913371 | 1.568 | -0.12 | -0.764 | 0.4 | -0.093 | -0.143 | -0.143 | 0.244 | 0.27 | 0.03 | 0.42 | 0.41 | 0.35 | 0.26 |
| 10 | 52829340 | rs10995547 | -0.733 | 0.01 | -0.164 | -0.054 | -0.011 | -0.035 | -0.035 | 0 | 0.1 | 0.44 | 0.06 | 0.01 | 0.04 | 0.04 |
| 10 | 52831004 | rs10740130 | 0.457 | 0.066 | -0.246 | -0.046 | -0.003 | 0.003 | 0.003 | 0.024 | 0.18 | 0.16 | 0.24 | 0.06 | 0.27 | 0.1 |
| 10 | 52835691 | rs9414804 | -1.137 | -0.003 | -0.148 | 0.011 | -0.031 | 0.09 | 0.09 | 0.607 | 0.12 | 0.57 | 0.07 | 0.15 | 0.04 | 0.1 |
| 10 | 52835732 | rs12355844 | 0.676 | 0.091 | -0.253 | -0.018 | 0.007 | 0.059 | 0.059 | 0.47 | 0.55 | 0.16 | 0.59 | 0.69 | 0.56 | 0.67 |
| 10 | 52841372 | rs7097311 | 0.051 | 0.235 | -0.087 | 0.003 | 0.01 | -0.069 | -0.069 | 0 | 0.2 | 0.32 | 0.25 | 0.07 | 0.27 | 0.11 |
| 10 | 52842575 | rs16913603 | -0.281 | 0.535 | -0.372 | 0.202 | -0.001 | 0.022 | 0.022 | 0 | 0.06 | 0.07 | 0.01 | 0.15 | 0.03 | 0.1 |
| 10 | 52852610 | rs2339675 | 0.276 | 0.599 | -0.517 | -0.343 | -0.002 | -0.001 | -0.001 | 0 | 0.18 | 0.17 | 0.24 | 0.07 | 0.27 | 0.1 |
| 10 | 52856168 | rs7910047 | -0.878 | 0.476 | -0.535 | -0.306 | -0.013 | -0.036 | -0.036 | 0.032 | 0.8 | 0.64 | 0.84 | 0.87 | 0.85 | 0.87 |
| 10 | 52860815 | rs16913900 | -0.537 | 0.423 | -0.535 | -0.346 | -0.004 | -0.028 | -0.028 | 0.022 | 0.26 | 0.37 | 0.28 | 0.15 | 0.3 | 0.19 |
| 10 | 52863364 | rs4303185 | -0.426 | 0.499 | -0.687 | -0.427 | -0.012 | -0.045 | -0.045 | 1.259 | 0.77 | 0.6 | 0.84 | 0.86 | 0.82 | 0.83 |
| 10 | 52877778 | rs2879523 | 0.688 | 0.514 | -0.932 | -0.442 | -0.009 | 0.123 | 0.123 | 0.487 | 0.54 | 0.12 | 0.54 | 0.72 | 0.54 | 0.67 |
| 10 | 52878398 | rs4935225 | 1.374 | 0.577 | -1.354 | -0.498 | -0.113 | -0.136 | -0.136 | 0.002 | 0.28 | 0.03 | 0.39 | 0.53 | 0.34 | 0.27 |
| 10 | 52878970 | rs2339628 | 0.773 | 0.755 | -0.249 | -0.111 | -0.004 | -0.032 | -0.032 | 0 | 0.62 | 0.43 | 0.59 | 0.8 | 0.59 | 0.77 |
| 10 | 52881906 | rs10761830 | -0.402 | 0.749 | -0.688 | -0.051 | -0.012 | -0.038 | -0.038 | 0 | 0.8 | 0.63 | 0.84 | 0.87 | 0.85 | 0.87 |

| | | | | | | | | | | | | | | | | |
|---|---|---|---|---|---|---|---|---|---|---|---|---|---|---|---|---|
| 10 | 52883081 | rs4935227 | 0.37 | 1.154 | -1.697 | 0.247 | 0.001 | 0.063 | 0.063 | 0.142 | 0.55 | 0.26 | 0.55 | 0.69 | 0.52 | 0.64 |
| 10 | 52897900 | rs10995831 | -0.404 | 1.351 | -1.936 | 0.427 | -0.058 | -0.04 | -0.04 | 0.375 | 0.1 | 0.01 | 0.15 | 0.11 | 0.23 | 0.08 |
| 10 | 52899265 | rs10822288 | 0.441 | 1.019 | -2.491 | 0.119 | -0.073 | 0.091 | 0.091 | 0.426 | 0.5 | 0.07 | 0.6 | 0.69 | 0.52 | 0.66 |
| 10 | 52905494 | rs1904694 | -0.091 | 0.682 | -1.649 | 0.168 | -0.002 | -0.002 | -0.002 | 0 | 0.46 | 0.54 | 0.35 | 0.36 | 0.43 | 0.43 |
| 10 | 52908166 | rs2127610 | -0.249 | 0.431 | -1.47 | -0.039 | -0.01 | -0.043 | -0.043 | 0 | 0.19 | 0.35 | 0.16 | 0.11 | 0.16 | 0.12 |
| 10 | 52909710 | rs1904699 | -0.087 | 0.682 | -1.425 | -0.16 | 0.032 | 0.079 | 0.079 | 0 | 0.51 | 0.79 | 0.46 | 0.31 | 0.47 | 0.44 |
| 10 | 52911474 | rs1904696 | 0.189 | 0.401 | -0.884 | -0.061 | -0.001 | -0.002 | -0.002 | 0 | 0.46 | 0.53 | 0.41 | 0.33 | 0.47 | 0.45 |
| 10 | 52913970 | rs10508937 | 0.62 | 0.386 | -1.126 | 0.043 | -0.017 | -0.027 | -0.027 | 0 | 0.08 | 0 | 0.08 | 0.11 | 0.07 | 0.11 |
| 10 | 52916173 | rs7893773 | -0.11 | 0.314 | -0.956 | -0.002 | -0.015 | 0.053 | 0.053 | 0 | 0.51 | 0.8 | 0.46 | 0.34 | 0.5 | 0.44 |
| 10 | 52927242 | rs2339732 | 0.188 | -0.045 | -0.759 | -0.016 | 0.006 | 0.007 | 0.007 | 0 | 0.46 | 0.57 | 0.4 | 0.34 | 0.44 | 0.43 |
| 10 | 52929610 | rs9414844 | -0.41 | -0.175 | -0.85 | -0.092 | -0.029 | 0.011 | 0.011 | 0 | 0.54 | 0.9 | 0.45 | 0.34 | 0.47 | 0.43 |
| 10 | 52931142 | rs1904677 | -0.012 | -0.191 | -0.715 | -0.1 | 0.016 | 0.051 | 0.051 | 0 | 0.51 | 0.84 | 0.44 | 0.35 | 0.47 | 0.43 |
| 10 | 52936070 | rs7100315 | -0.95 | -0.191 | -0.686 | -0.099 | 0.004 | -0.023 | -0.023 | 0 | 0.07 | 0.24 | 0.04 | 0 | 0.03 | 0 |
| 10 | 52936188 | rs6479991 | 0.048 | -0.164 | -0.716 | -0.086 | 0.041 | 0.051 | 0.051 | 0 | 0.5 | 0.7 | 0.42 | 0.34 | 0.44 | 0.43 |
| 10 | 52939040 | rs7085697 | -0.352 | -0.157 | -0.722 | -0.083 | -0.006 | 0.029 | 0.029 | 0 | 0.54 | 0.85 | 0.45 | 0.35 | 0.47 | 0.44 |
| 10 | 52949162 | rs1904683 | -0.001 | -0.14 | -0.676 | -0.07 | 0.027 | 0.035 | 0.035 | 0 | 0.49 | 0.66 | 0.41 | 0.34 | 0.44 | 0.43 |
| 10 | 52957721 | rs7897633 | -0.675 | 0.355 | -1.068 | -0.09 | -0.093 | -0.085 | -0.085 | 0 | 0.6 | 0.88 | 0.54 | 0.46 | 0.57 | 0.55 |
| 10 | 52964590 | rs7905063 | -0.649 | -0.054 | -1.012 | -0.035 | -0.105 | -0.092 | -0.092 | 0.016 | 0.62 | 0.88 | 0.53 | 0.46 | 0.55 | 0.55 |
| 10 | 52991510 | rs1865645 | -0.603 | -0.719 | -0.939 | -0.065 | -0.063 | -0.04 | -0.04 | 0 | 0.63 | 0.82 | 0.52 | 0.42 | 0.49 | 0.52 |
| 10 | 52994866 | rs10490978 | 0.333 | -1.288 | 0.014 | -0.311 | 0.012 | 0.01 | 0.01 | 0.115 | 0.46 | 0.29 | 0.38 | 0.43 | 0.41 | 0.42 |
| 10 | 52995974 | rs1915672 | -0.276 | -1.324 | 0.569 | -0.327 | -0.001 | -0.001 | -0.001 | 0 | 0.53 | 0.51 | 0.43 | 0.43 | 0.44 | 0.44 |
| 10 | 53001872 | rs10822481 | 0.371 | -1.293 | 0.491 | -0.319 | 0.012 | 0.012 | 0.012 | 0.252 | 0.45 | 0.24 | 0.38 | 0.42 | 0.4 | 0.41 |
| 10 | 53009688 | rs10996325 | -0.424 | -1.28 | 0.7 | -0.325 | -0.017 | -0.033 | -0.033 | 0 | 0.11 | 0.15 | 0.08 | 0.01 | 0.08 | 0.09 |
| 10 | 53010887 | rs10996331 | -0.912 | -1.298 | 0.721 | -0.332 | -0.023 | -0.041 | -0.041 | 0 | 0.12 | 0.19 | 0.09 | 0.01 | 0.08 | 0.09 |
| 10 | 53012950 | rs16915897 | -1.523 | -1.332 | 1.039 | -0.347 | -0.052 | -0.087 | -0.087 | 0 | 0.15 | 0.49 | 0.14 | 0.01 | 0.09 | 0.09 |
| 10 | 53013479 | rs10822496 | 0.268 | -1.345 | 0.998 | -0.318 | 0.014 | 0.019 | 0.019 | 1.006 | 0.47 | 0.24 | 0.37 | 0.43 | 0.41 | 0.46 |
| 10 | 53015492 | rs9943368 | -0.631 | -1.286 | 0.989 | -0.302 | -0.024 | -0.014 | -0.014 | 0.191 | 0.6 | 0.68 | 0.5 | 0.44 | 0.5 | 0.55 |
| 10 | 53020936 | rs10490975 | -2.01 | -1.281 | 1.343 | -0.238 | -0.053 | -0.09 | -0.09 | 0 | 0.15 | 0.55 | 0.15 | 0.01 | 0.09 | 0.09 |
| 10 | 53025206 | rs12240684 | -1.465 | -1.211 | 0.846 | -0.208 | -0.036 | -0.064 | -0.064 | 0 | 0.13 | 0.3 | 0.1 | 0.01 | 0.08 | 0.09 |
| 10 | 53026775 | rs1561039 | -0.336 | -1.33 | 0.682 | -0.423 | -0.012 | 0.029 | 0.029 | 0 | 0.17 | 0.11 | 0.1 | 0.02 | 0.2 | 0.13 |
| 10 | 53029510 | rs7077723 | -1.456 | -1.305 | 0.768 | -0.408 | -0.005 | -0.076 | -0.076 | 0 | 0.28 | 0.37 | 0.19 | 0.03 | 0.28 | 0.22 |
| 10 | 53035096 | rs3844131 | -1.123 | -1.545 | 0.534 | 0.343 | 0.005 | -0.005 | -0.005 | 0.854 | 0.17 | 0.18 | 0.12 | 0.18 | 0.12 | 0.16 |
| 10 | 53035622 | rs11594220 | -0.999 | -1.538 | 0.537 | 0.346 | 0.004 | -0.005 | -0.005 | 0.823 | 0.17 | 0.18 | 0.12 | 0.18 | 0.12 | 0.15 |
| 10 | 53042298 | rs10996555 | -1.946 | -2.205 | 0.593 | 0.55 | 0.061 | -0.002 | -0.002 | 0.189 | 0.46 | 0.74 | 0.53 | 0.44 | 0.33 | 0.43 |
| 10 | 53047303 | rs12264442 | -0.621 | -2.13 | 1.096 | 0.542 | -0.02 | 0.026 | 0.026 | 0.054 | 0.16 | 0.3 | 0.13 | 0.18 | 0.12 | 0.16 |
| 10 | 53048551 | rs2339710 | -1.389 | -2.165 | 1.337 | 0.503 | -0.044 | 0.021 | 0.021 | 0.703 | 0.19 | 0.6 | 0.18 | 0.18 | 0.12 | 0.17 |
| 10 | 53058057 | rs1915681 | -0.706 | -2.072 | 0.421 | 0.575 | 0.013 | -0.012 | -0.012 | 1.644 | 0.27 | 0.2 | 0.37 | 0.27 | 0.21 | 0.26 |
| 10 | 53058385 | rs12251902 | -0.452 | -2.052 | 0.443 | 0.589 | -0.004 | 0.004 | 0.004 | 0.007 | 0.15 | 0.19 | 0.11 | 0.18 | 0.12 | 0.17 |
| 10 | 53059406 | rs2043556 | -0.93 | -1.744 | 0.243 | 0.56 | 0.005 | -0.005 | -0.005 | 0 | 0.28 | 0.23 | 0.37 | 0.27 | 0.22 | 0.25 |
| 10 | 53060249 | rs10822646 | -0.417 | -1.637 | -0.002 | 0.673 | 0.003 | 0.008 | 0.008 | 0 | 0.21 | 0.17 | 0.18 | 0.09 | 0.17 | 0.18 |
| 10 | 53060905 | rs16916483 | 0.385 | -1.9 | 0.142 | 0.885 | -0.026 | 0.02 | 0.02 | 0.181 | 0.12 | 0.12 | 0.2 | 0.17 | 0.06 | 0.06 |
| 10 | 53062052 | rs10490969 | -0.961 | -1.672 | -0.023 | 0.801 | -0.019 | 0.009 | 0.009 | 1.27 | 0.14 | 0.21 | 0.09 | 0.17 | 0.06 | 0.21 |
| 10 | 53064997 | rs7100331 | -0.203 | -1.446 | -0.157 | 0.821 | -0.008 | 0.007 | 0.007 | 0.398 | 0.08 | 0.08 | 0.05 | 0.14 | 0.03 | 0.11 |
| 10 | 53068604 | rs12261736 | -0.245 | -0.994 | -0.532 | 0.716 | 0 | 0 | 0 | 0 | 0.34 | 0.29 | 0.31 | 0.38 | 0.36 | 0.25 |
| 10 | 53070556 | rs3862580 | -0.96 | -0.268 | -0.7 | 0.213 | -0.013 | 0.005 | 0.005 | 0 | 0.73 | 0.72 | 0.67 | 0.73 | 0.71 | 0.66 |
| 10 | 53070953 | rs7898516 | -0.425 | -0.283 | -0.733 | 0.028 | 0.001 | 0.002 | 0.002 | 0 | 0.49 | 0.56 | 0.52 | 0.53 | 0.52 | 0.58 |
| 10 | 53072168 | rs7091469 | 0.611 | 0.232 | -0.89 | 0.045 | -0.006 | -0.002 | -0.002 | 0 | 0.13 | 0.11 | 0.16 | 0.17 | 0.19 | 0.16 |
| 10 | 53084089 | rs4935240 | 1.207 | -0.259 | -1.158 | -0.065 | -0.076 | -0.133 | -0.133 | 3.375 | 0.19 | 0.01 | 0.25 | 0.37 | 0.25 | 0.32 |
| 10 | 53092302 | rs211068 | -0.462 | -0.324 | -0.618 | -0.22 | 0 | -0.001 | -0.001 | 0.832 | 0.49 | 0.53 | 0.55 | 0.51 | 0.56 | 0.58 |
| 10 | 53093126 | rs211067 | -0.704 | -0.314 | -0.897 | -0.287 | -0.001 | -0.009 | -0.009 | 0 | 0.56 | 0.73 | 0.6 | 0.53 | 0.6 | 0.59 |
| 10 | 53093557 | rs10996926 | 1.695 | -0.04 | -1.248 | -0.622 | -0.041 | 0.075 | 0.075 | 0 | 0.14 | 0.02 | 0.06 | 0.02 | 0.1 | 0.12 |
| 10 | 53095119 | rs1822690 | -0.547 | 0.075 | -1.039 | -0.391 | 0.001 | -0.009 | -0.009 | 0 | 0.56 | 0.73 | 0.6 | 0.53 | 0.6 | 0.59 |
| 10 | 53104260 | rs211083 | 0.332 | 0.286 | -0.557 | -0.311 | -0.008 | -0.029 | -0.029 | 0 | 0.12 | 0.16 | 0.08 | 0 | 0.08 | 0.09 |
| 10 | 53109723 | rs7099142 | 0.508 | 1.099 | -0.542 | -0.016 | -0.017 | -0.007 | -0.007 | 0.324 | 0.11 | 0.08 | 0.16 | 0.17 | 0.21 | 0.14 |
| 10 | 53114079 | rs1189311 | 0.135 | 1.207 | -0.114 | 0.236 | 0.017 | 0.013 | 0.013 | 0 | 0.71 | 0.64 | 0.56 | 0.39 | 0.68 | 0.71 |
| 10 | 53114847 | rs1189310 | -0.531 | 1.38 | 0.028 | 0.59 | 0.018 | 0.079 | 0.079 | 0 | 0.85 | 0.82 | 0.69 | 0.51 | 0.76 | 0.81 |
| 10 | 53131006 | rs1550718 | 0.327 | 0.691 | -1.001 | -0.335 | -0.046 | 0.121 | 0.121 | 0 | 0.44 | 0.09 | 0.31 | 0.23 | 0.44 | 0.45 |
| 10 | 53131697 | rs1189326 | 0.295 | 0.698 | -0.705 | -0.189 | 0.003 | 0.003 | 0.003 | 0 | 0.47 | 0.52 | 0.33 | 0.24 | 0.39 | 0.48 |
| 10 | 53134556 | rs1189322 | -0.364 | 0.924 | -1.124 | 0.128 | -0.03 | -0.026 | -0.026 | 0 | 0.73 | 0.83 | 0.55 | 0.36 | 0.71 | 0.67 |
| 10 | 53137136 | rs1189320 | -0.271 | 0.829 | -0.958 | 0.374 | -0.006 | 0.008 | 0.008 | 0 | 0.36 | 0.44 | 0.35 | 0.46 | 0.34 | 0.61 |
| 10 | 53139938 | rs1373497 | 0.466 | 1.077 | -1.05 | 0.671 | -0.016 | -0.04 | -0.04 | 2.293 | 0.15 | 0.08 | 0.17 | 0.35 | 0.19 | 0.37 |
| 10 | 53144956 | rs10740244 | -0.147 | 0.754 | -1.166 | -0.032 | -0.021 | 0.021 | 0.021 | 2.657 | 0.34 | 0.18 | 0.28 | 0.27 | 0.37 | 0.32 |
| 10 | 53145705 | rs10822867 | 1.029 | 0.482 | -1.045 | 0.074 | -0.017 | 0.012 | 0.012 | 5.65 | 0.16 | 0.04 | 0.13 | 0.09 | 0.15 | 0.12 |
| 10 | 53148778 | rs4935244 | 0.139 | 0.169 | -0.231 | 0.176 | -0.001 | 0.066 | 0.066 | 0.948 | 0.71 | 0.46 | 0.45 | 0.43 | 0.68 | 0.62 |
| 10 | 53154782 | rs10822909 | 1.226 | -0.096 | 0.184 | -0.472 | -0.008 | -0.072 | -0.072 | 0 | 0.32 | 0.34 | 0.13 | 0.03 | 0.24 | 0.17 |
| 10 | 53156299 | rs11815733 | 0.931 | 0.068 | -0.232 | -0.568 | -0.051 | -0.021 | -0.021 | 0 | 0.19 | 0.01 | 0.11 | 0.17 | 0.2 | 0.16 |
| 10 | 53159076 | rs293233 | -0.291 | -0.269 | 0.037 | -0.557 | -0.002 | 0.002 | 0.002 | 0.465 | 0.7 | 0.63 | 0.43 | 0.35 | 0.67 | 0.5 |
| 10 | 53159566 | rs10997388 | 0.014 | -0.168 | -0.207 | -0.528 | -0.027 | 0.022 | 0.022 | 0 | 0.38 | 0.2 | 0.29 | 0.33 | 0.44 | 0.27 |
| 10 | 53163763 | rs10822922 | 0.78 | -0.064 | 0.796 | 0.049 | -0.009 | 0.007 | 0.007 | 0 | 0.64 | 0.54 | 0.41 | 0.39 | 0.65 | 0.59 |

| | | | | | | | | | | | | | | | | |
|---|---|---|---|---|---|---|---|---|---|---|---|---|---|---|---|---|
| 10 | 53164356 | rs10762113 | -0.585 | 0.15 | 0.304 | 0.314 | 0 | 0.028 | 0.028 | 0.039 | 0.83 | 0.83 | 0.75 | 0.54 | 0.81 | 0.7 |
| 10 | 53165265 | rs1550720 | -0.145 | 0.161 | 0.335 | 0.298 | -0.001 | 0.023 | 0.023 | 0 | 0.8 | 0.76 | 0.7 | 0.49 | 0.79 | 0.68 |
| 10 | 53172877 | rs7902518 | 0.183 | 0.168 | 0.437 | 0.25 | -0.001 | -0.001 | -0.001 | 0 | 0.11 | 0.19 | 0.29 | 0.14 | 0.13 | 0.24 |
| 10 | 53175181 | rs293242 | 0.526 | 0.222 | 0.135 | 0.316 | 0 | 0.002 | 0.002 | 0 | 0.74 | 0.75 | 0.68 | 0.49 | 0.76 | 0.67 |
| 10 | 53175263 | rs293243 | -1.083 | 0.602 | -0.216 | 0.551 | -0.02 | -0.04 | -0.04 | 0 | 0.88 | 0.95 | 0.79 | 0.54 | 0.85 | 0.74 |
| 10 | 53184548 | rs293326 | -0.945 | 0.615 | -0.193 | 0.544 | -0.014 | -0.032 | -0.032 | 0 | 0.88 | 0.92 | 0.79 | 0.53 | 0.85 | 0.74 |
| 10 | 53186479 | rs293330 | -1.2 | 0.576 | -0.028 | 0.114 | -0.002 | -0.004 | -0.004 | 0 | 0.48 | 0.61 | 0.62 | 0.51 | 0.56 | 0.5 |
| 10 | 53192833 | rs12248673 | -0.386 | 0.87 | 0.023 | 0.123 | -0.023 | -0.017 | -0.017 | 0.057 | 0.33 | 0.22 | 0.49 | 0.44 | 0.48 | 0.3 |
| 10 | 53197117 | rs10997547 | 1.366 | 0.752 | 0.174 | -0.25 | -0.007 | -0.101 | -0.101 | 0.849 | 0.4 | 0.45 | 0.18 | 0.02 | 0.29 | 0.24 |
| 10 | 53197242 | rs10997549 | 0.013 | 0.801 | -0.244 | 0.108 | -0.051 | -0.052 | -0.052 | 0.5 | 0.22 | 0.1 | 0.21 | 0.31 | 0.34 | 0.21 |
| 10 | 53200276 | rs17531066 | 1.179 | 1.021 | -0.323 | 0.101 | -0.029 | -0.016 | -0.016 | 0.014 | 0.1 | 0.06 | 0.14 | 0.13 | 0.2 | 0.1 |
| 10 | 53201249 | rs10997575 | 1.272 | 0.605 | 0.044 | -0.331 | 0.006 | -0.017 | -0.017 | 0 | 0.43 | 0.38 | 0.19 | 0.02 | 0.32 | 0.26 |
| 10 | 53201604 | rs293245 | 0.029 | 0.533 | 0.106 | -0.27 | 0.003 | -0.016 | -0.016 | 0.036 | 0.7 | 0.8 | 0.49 | 0.37 | 0.69 | 0.58 |
| 10 | 53207634 | rs293257 | 0.877 | 0.573 | 0.307 | -0.65 | -0.009 | -0.114 | -0.114 | 0 | 0.44 | 0.53 | 0.21 | 0.02 | 0.32 | 0.26 |
| 10 | 53208813 | rs293259 | 0.584 | 0.574 | 0.296 | -0.643 | 0.027 | 0.116 | 0.116 | 2.923 | 0.48 | 0.73 | 0.28 | 0.07 | 0.34 | 0.32 |
| 10 | 53211449 | rs293266 | -0.08 | 0.563 | 0.122 | -0.656 | -0.037 | 0.085 | 0.085 | 0.166 | 0.59 | 0.78 | 0.57 | 0.18 | 0.48 | 0.4 |
| 10 | 53219368 | rs10823039 | 0.29 | 0.642 | -0.19 | -0.536 | 0.003 | 0.005 | 0.005 | 0 | 0.15 | 0.24 | 0.36 | 0.16 | 0.17 | 0.14 |
| 10 | 53220753 | rs293290 | 0.295 | 0.759 | -0.911 | -0.174 | -0.019 | 0.087 | 0.087 | 0 | 0.58 | 0.77 | 0.57 | 0.18 | 0.5 | 0.42 |
| 10 | 53223351 | rs10997638 | -0.39 | 0.542 | -0.747 | 0.444 | 0.003 | -0.001 | -0.001 | 1.515 | 0.2 | 0.16 | 0.26 | 0.46 | 0.16 | 0.28 |
| 10 | 53223446 | rs10823041 | 0.731 | 0.619 | -0.388 | -0.462 | -0.003 | -0.074 | -0.074 | 0 | 0.44 | 0.48 | 0.26 | 0.06 | 0.36 | 0.35 |
| 10 | 53223663 | rs16918343 | 0.508 | 0.751 | -0.193 | -0.326 | 0 | 0 | 0 | 0 | 0.11 | 0.18 | 0.23 | 0.14 | 0.12 | 0.07 |
| 10 | 53223673 | rs293296 | 0.063 | 0.926 | 0.217 | -0.462 | 0.026 | 0.017 | 0.017 | 0 | 0.7 | 0.62 | 0.65 | 0.6 | 0.61 | 0.82 |
| 10 | 53224750 | rs11819049 | 0.211 | 0.608 | 0.004 | -2.212 | 0 | -0.04 | -0.04 | 0 | 0.45 | 0.54 | 0.31 | 0.09 | 0.46 | 0.39 |
| 10 | 53234155 | rs1444400 | -1.081 | 0.857 | -0.174 | -2.343 | -0.011 | 0.116 | 0.116 | 0 | 0.68 | 0.87 | 0.63 | 0.13 | 0.67 | 0.6 |
| 10 | 53235068 | rs12359399 | 1.519 | 0.645 | -0.807 | -2.836 | -0.056 | 0.143 | 0.143 | 2.001 | 0.25 | 0.07 | 0.16 | 0.01 | 0.24 | 0.19 |
| 10 | 53236023 | rs10997716 | 0.166 | 0.637 | -0.634 | -2.757 | 0.005 | -0.065 | -0.065 | 0 | 0.22 | 0.31 | 0.32 | 0.05 | 0.21 | 0.2 |
| 10 | 53237452 | rs6480276 | -1.42 | 0.52 | -0.893 | -2.709 | -0.01 | 0.098 | 0.098 | 0 | 0.68 | 0.86 | 0.68 | 0.18 | 0.67 | 0.61 |
| 10 | 53237647 | rs6480277 | 0.718 | 0.508 | -0.432 | -2.894 | -0.002 | -0.005 | -0.005 | 0 | 0.37 | 0.32 | 0.27 | 0.09 | 0.38 | 0.39 |
| 10 | 53238950 | rs10823071 | 1.432 | 0.781 | -0.658 | -3.431 | -0.021 | 0.163 | 0.163 | 0.18 | 0.38 | 0.12 | 0.34 | 0.83 | 0.37 | 0.48 |
| 10 | 53240022 | rs10508942 | 0.761 | 0.762 | -0.587 | -3.456 | 0.009 | 0.278 | 0.278 | 0.407 | 0.48 | 0.18 | 0.41 | 0.91 | 0.48 | 0.65 |
| 10 | 53240189 | rs4935017 | 1.379 | 0.661 | -0.439 | -3.404 | -0.005 | -0.009 | -0.009 | 0 | 0.12 | 0.16 | 0.24 | 0.08 | 0.09 | 0.13 |
| 10 | 53240557 | rs293305 | 1.973 | 0.699 | -0.626 | -3.373 | -0.037 | 0.089 | 0.089 | 4.797 | 0.18 | 0.03 | 0.11 | 0.01 | 0.15 | 0.09 |
| 10 | 53241687 | rs293309 | 0.015 | 0.708 | -0.373 | -3.37 | -0.002 | 0.007 | 0.007 | 0.369 | 0.38 | 0.34 | 0.43 | 0.09 | 0.34 | 0.22 |
| 10 | 53242283 | rs293312 | 0.933 | 0.708 | -0.12 | -3.129 | 0.007 | -0.068 | -0.068 | 0.03 | 0.77 | 0.46 | 0.72 | 0.87 | 0.72 | 0.71 |
| 10 | 53243107 | rs293319 | -0.483 | 0.506 | -0.085 | -2.799 | 0.016 | -0.008 | -0.008 | 0 | 0.88 | 0.72 | 0.82 | 0.87 | 0.81 | 0.79 |
| 10 | 53244090 | rs10762176 | 0.375 | 0.647 | -0.146 | -2.817 | -0.001 | 0.2 | 0.2 | 0.868 | 0.54 | 0.21 | 0.42 | 0.83 | 0.54 | 0.54 |
| 10 | 53244911 | rs1326250 | -0.56 | 0.823 | -0.268 | -2.771 | 0.001 | -0.022 | -0.022 | 0 | 0.7 | 0.69 | 0.7 | 0.86 | 0.66 | 0.7 |
| 10 | 53245022 | rs7091396 | -0.238 | 0.821 | 0.098 | -2.17 | -0.01 | -0.078 | -0.078 | 0 | 0.15 | 0.36 | 0.26 | 0.03 | 0.12 | 0.16 |
| 10 | 53245152 | rs766208 | 0.01 | 0.786 | 0.099 | -2.169 | 0.028 | -0.082 | -0.082 | 0 | 0.85 | 0.64 | 0.7 | 0.97 | 0.81 | 0.8 |
| 10 | 53248191 | rs1926208 | 1.208 | 0.39 | -0.295 | -2.32 | -0.001 | -0.007 | -0.007 | 0 | 0.78 | 0.3 | 0.69 | 0.87 | 0.75 | 0.76 |
| 10 | 53248566 | rs1326251 | -0.799 | 0.32 | 0.06 | -2.339 | -0.008 | -0.095 | -0.095 | 0 | 0.15 | 0.59 | 0.12 | 0 | 0.09 | 0.02 |
| 10 | 53250698 | rs10823078 | 0.158 | 0.354 | -0.4 | -2.874 | -0.002 | -0.029 | -0.029 | 0 | 0.29 | 0.31 | 0.35 | 0.02 | 0.27 | 0.3 |
| 10 | 53250879 | rs10508945 | 1.146 | 0.167 | -0.773 | -3.219 | -0.006 | 0.039 | 0.039 | 0 | 0.2 | 0.17 | 0.11 | 0.06 | 0.22 | 0.06 |
| 10 | 53251706 | rs986401 | 0.66 | 0.192 | -0.715 | -3.171 | -0.003 | -0.022 | -0.022 | 0.927 | 0.29 | 0.31 | 0.32 | 0.02 | 0.27 | 0.27 |
| 10 | 53252315 | rs7921499 | -0.086 | 0.123 | -0.299 | -3.166 | -0.013 | 0.076 | 0.076 | 0.013 | 0.61 | 0.23 | 0.46 | 0.85 | 0.57 | 0.51 |
| 10 | 53260053 | rs12772194 | 1.594 | 0.178 | -0.301 | -3.086 | -0.026 | 0.06 | 0.06 | 0 | 0.18 | 0.07 | 0.1 | 0.06 | 0.22 | 0.06 |
| 10 | 53261988 | rs1926212 | 0.402 | 0.003 | -0.361 | -3.128 | 0.015 | 0.109 | 0.109 | 0 | 0.43 | 0.16 | 0.35 | 0.79 | 0.35 | 0.45 |
| 10 | 53263809 | rs12257749 | 0.188 | 0.028 | -0.384 | -3.138 | -0.019 | 0.03 | 0.03 | 0 | 0.28 | 0.23 | 0.3 | 0.19 | 0.37 | 0.24 |
| 10 | 53276530 | rs10997890 | 0.592 | -0.019 | -0.638 | -3.169 | 0.025 | 0.053 | 0.053 | 0 | 0.39 | 0.22 | 0.36 | 0.79 | 0.28 | 0.46 |
| 10 | 53276747 | rs10128493 | 1.061 | 0.029 | -0.688 | -3.165 | -0.029 | 0.009 | 0.009 | 0 | 0.22 | 0.1 | 0.19 | 0.19 | 0.31 | 0.2 |
| 10 | 53281191 | rs10823125 | 1.347 | -0.214 | -0.86 | -3.209 | 0.006 | 0.037 | 0.037 | 0 | 0.3 | 0.08 | 0.28 | 0.78 | 0.24 | 0.38 |
| 10 | 53282106 | rs10997909 | 0.674 | -0.064 | -0.659 | -3.042 | -0.015 | 0.053 | 0.053 | 0 | 0.29 | 0.2 | 0.32 | 0.08 | 0.27 | 0.27 |
| 10 | 53282308 | rs10740281 | -0.34 | 0.01 | -0.667 | -2.957 | 0.003 | -0.044 | -0.044 | 0.275 | 0.72 | 0.37 | 0.72 | 0.87 | 0.62 | 0.75 |
| 10 | 53286396 | rs10823129 | 0.227 | -0.18 | -1.333 | -2.958 | 0.038 | 0.064 | 0.064 | 0 | 0.44 | 0.64 | 0.38 | 0.8 | 0.29 | 0.43 |
| 10 | 53289428 | rs9988728 | 0.101 | 0.418 | -1.023 | -2.746 | -0.004 | -0.006 | -0.006 | 0 | 0.31 | 0.27 | 0.34 | 0.02 | 0.28 | 0.32 |
| 10 | 53291813 | rs2038918 | 0.368 | -0.189 | -0.763 | -2.413 | 0.002 | 0.059 | 0.059 | 0 | 0.28 | 0.12 | 0.34 | 0.85 | 0.29 | 0.42 |
| 10 | 53292516 | rs2148309 | 0.656 | -0.309 | -0.785 | -2.553 | -0.029 | -0.124 | -0.124 | 1.232 | 0.75 | 0.57 | 0.79 | 0.98 | 0.78 | 0.78 |
| 10 | 53293729 | rs7342004 | 0.233 | -0.345 | -0.739 | -2.547 | -0.05 | -0.097 | -0.097 | 0 | 0.81 | 0.64 | 0.85 | 0.98 | 0.86 | 0.78 |
| 10 | 53295014 | rs7100765 | 0.351 | -0.345 | -0.747 | -2.548 | -0.056 | -0.105 | -0.105 | 1.124 | 0.8 | 0.61 | 0.84 | 0.98 | 0.86 | 0.78 |
| 10 | 53303397 | rs10998031 | 0.404 | -0.297 | -0.847 | -2.593 | 0.004 | -0.017 | -0.017 | 0 | 0.48 | 0.56 | 0.5 | 0.13 | 0.54 | 0.33 |
| 10 | 53304989 | rs1326257 | 0.859 | -0.293 | -0.719 | -2.495 | -0.023 | -0.119 | -0.119 | 0.616 | 0.76 | 0.61 | 0.8 | 0.98 | 0.77 | 0.78 |
| 10 | 53309898 | rs10823172 | 0.212 | -0.33 | -0.688 | -2.462 | -0.051 | -0.099 | -0.099 | 0.078 | 0.81 | 0.65 | 0.84 | 0.98 | 0.86 | 0.78 |
| 10 | 53317506 | rs10823184 | -0.543 | -0.076 | -0.772 | -2.483 | 0.026 | 0.114 | 0.114 | 0 | 0.51 | 0.32 | 0.47 | 0.88 | 0.43 | 0.66 |
| 10 | 53317666 | rs10998097 | -0.126 | -0.124 | -0.822 | -2.525 | -0.023 | -0.061 | -0.061 | 0 | 0.2 | 0.28 | 0.13 | 0.02 | 0.12 | 0.2 |
| 10 | 53318238 | rs4492772 | -0.859 | -0.126 | -0.536 | -2.562 | -0.014 | -0.093 | -0.093 | 0.094 | 0.26 | 0.62 | 0.21 | 0.02 | 0.21 | 0.21 |
| 10 | 53326226 | rs7897063 | 0.75 | -0.115 | -0.659 | -2.519 | -0.05 | 0.091 | 0.091 | 0.069 | 0.31 | 0.04 | 0.34 | 0.85 | 0.31 | 0.46 |
| 10 | 53330663 | rs11817932 | 0.218 | -0.146 | -0.489 | -2.441 | -0.013 | -0.044 | -0.044 | 0.091 | 0.17 | 0.23 | 0.12 | 0.02 | 0.12 | 0.2 |
| 10 | 53336732 | rs4935018 | 1.401 | -0.306 | -0.817 | -2.645 | -0.106 | 0.15 | 0.15 | 1.973 | 0.34 | 0.09 | 0.36 | 0.13 | 0.39 | 0.3 |
| 10 | 53337255 | rs10823230 | 0.191 | -0.312 | -0.838 | -2.668 | -0.068 | 0.155 | 0.155 | 1.159 | 0.38 | 0.08 | 0.39 | 0.85 | 0.4 | 0.46 |

| | | | | | | | | | | | | | | | | |
|---|---|---|---|---|---|---|---|---|---|---|---|---|---|---|---|---|
| 10 | 53342024 | rs10823235 | 0.852 | -0.289 | -0.906 | -2.679 | -0.05 | 0.091 | 0.091 | 0.069 | 0.31 | 0.04 | 0.34 | 0.85 | 0.31 | 0.46 |
| 10 | 53344284 | rs10823243 | 1.086 | -0.255 | -0.815 | -2.646 | -0.031 | 0.103 | 0.103 | 0.123 | 0.39 | 0.22 | 0.39 | 0.13 | 0.41 | 0.3 |
| 10 | 53346871 | rs12262193 | 0.071 | -0.376 | -0.778 | -2.645 | -0.001 | -0.018 | -0.018 | 0 | 0.56 | 0.45 | 0.51 | 0.15 | 0.51 | 0.47 |
| 10 | 53348048 | rs1922127 | 0.267 | 0.025 | -1.296 | -2.464 | -0.004 | -0.044 | -0.044 | 0 | 0.17 | 0.23 | 0.13 | 0.02 | 0.14 | 0.18 |
| 10 | 53350411 | rs7923153 | 0.423 | 0.697 | -1.814 | -2.13 | 0.001 | -0.009 | -0.009 | 0.405 | 0.9 | 0.89 | 0.9 | 0.94 | 0.84 | 0.95 |
| 10 | 53356764 | rs10823257 | 0.194 | 0.295 | -2.125 | -1.93 | -0.039 | 0.079 | 0.079 | 0.296 | 0.29 | 0.08 | 0.35 | 0.85 | 0.32 | 0.44 |
| 10 | 53356952 | rs1917841 | 0.194 | 0.583 | -1.936 | -1.548 | -0.106 | 0.266 | 0.266 | 0.789 | 0.43 | 0.09 | 0.4 | 0.07 | 0.46 | 0.41 |
| 10 | 53363713 | rs1922139 | 0.305 | 0.575 | -0.98 | -0.099 | -0.005 | 0.024 | 0.024 | 0.398 | 0.71 | 0.51 | 0.73 | 0.61 | 0.69 | 0.62 |
| 10 | 53365163 | rs7078670 | 0.316 | 0.675 | -1.733 | -0.227 | -0.034 | 0.055 | 0.055 | 0.854 | 0.28 | 0.11 | 0.18 | 0.15 | 0.26 | 0.23 |
| 10 | 53365208 | rs7077665 | 0.194 | 0.651 | -1.806 | -0.277 | -0.047 | 0.061 | 0.061 | 0.872 | 0.29 | 0.08 | 0.2 | 0.16 | 0.27 | 0.24 |
| 10 | 53366180 | rs10998465 | 0.544 | 0.815 | -1.789 | 0.179 | -0.04 | -0.056 | -0.056 | 1.812 | 0.09 | 0.01 | 0.12 | 0.16 | 0.13 | 0.19 |
| 10 | 53375420 | rs2339794 | -0.152 | -0.244 | -0.48 | -0.042 | -0.029 | 0.015 | 0.015 | 0.185 | 0.87 | 0.55 | 0.86 | 0.86 | 0.87 | 0.9 |
| 10 | 53378000 | rs4935274 | 0.645 | -0.099 | -1.005 | -0.268 | -0.032 | 0.054 | 0.054 | 0 | 0.15 | 0.04 | 0.15 | 0.05 | 0.18 | 0.07 |
| 10 | 53382116 | rs6480375 | 0.469 | 0.134 | -1.039 | -0.294 | 0.075 | 0.064 | 0.064 | 0 | 0.88 | 0.59 | 0.75 | 0.8 | 0.81 | 0.86 |
| 10 | 53385460 | rs12414562 | -0.729 | -0.266 | -0.859 | -0.451 | -0.005 | 0.016 | 0.016 | 0 | 0.27 | 0.24 | 0.41 | 0.14 | 0.29 | 0.28 |
| 10 | 53388223 | rs7915186 | 1.115 | -0.46 | -0.665 | -0.521 | 0.015 | 0.082 | 0.082 | 2.553 | 0.58 | 0.19 | 0.46 | 0.71 | 0.58 | 0.64 |
| 10 | 53393469 | rs10998594 | -0.229 | -0.41 | -0.49 | -0.514 | -0.01 | 0.02 | 0.02 | 0 | 0.24 | 0.22 | 0.36 | 0.11 | 0.29 | 0.14 |
| 10 | 53393629 | rs10733849 | 0.053 | -0.565 | -0.413 | -0.491 | -0.046 | -0.004 | -0.004 | 0 | 0.85 | 0.54 | 0.87 | 0.85 | 0.87 | 0.93 |
| 10 | 53404013 | rs7079618 | 0.554 | -0.54 | -0.373 | -0.447 | -0.055 | -0.003 | -0.003 | 0.239 | 0.83 | 0.43 | 0.83 | 0.81 | 0.87 | 0.8 |
| 10 | 53407953 | rs1922137 | -0.376 | -0.346 | -0.317 | -0.34 | 0.081 | 0.18 | 0.18 | 0.421 | 0.94 | 0.62 | 0.93 | 0.84 | 0.92 | 0.95 |
| 10 | 53411242 | rs7895839 | 0.885 | -0.243 | -0.654 | -0.412 | -0.006 | 0.018 | 0.018 | 2.473 | 0.66 | 0.22 | 0.51 | 0.7 | 0.61 | 0.66 |
| 10 | 53419753 | rs12412862 | 0.896 | 0.021 | -1.213 | -0.547 | -0.01 | 0.03 | 0.03 | 0 | 0.15 | 0.08 | 0.19 | 0.05 | 0.18 | 0.06 |
| 10 | 53425328 | rs4575213 | 0.893 | 0.207 | -1.476 | -0.279 | -0.105 | 0.05 | 0.05 | 2.491 | 0.4 | 0.08 | 0.33 | 0.27 | 0.42 | 0.44 |
| 10 | 53435774 | rs10128113 | -0.366 | 0.536 | -0.677 | -0.289 | 0.009 | -0.01 | -0.01 | 2.932 | 0.12 | 0.15 | 0.19 | 0.08 | 0.14 | 0.21 |
| 10 | 53442559 | rs6480412 | -0.573 | 0.041 | -0.614 | -0.48 | -0.006 | -0.046 | -0.046 | 0.182 | 0.43 | 0.49 | 0.47 | 0.14 | 0.38 | 0.33 |
| 10 | 53445667 | rs10740321 | -0.752 | 0.229 | -0.392 | 0.009 | 0.007 | 0.014 | 0.014 | 0.862 | 0.55 | 0.69 | 0.66 | 0.4 | 0.6 | 0.57 |
| 10 | 53455408 | rs1045767 | -1.23 | 0.229 | 1.683 | 0.239 | -0.006 | 0.041 | 0.041 | 0 | 0.1 | 0.3 | 0.06 | 0.14 | 0.08 | 0.12 |
| 10 | 53455862 | rs16921605 | 0.133 | 0.495 | 1.924 | 0.435 | 0.024 | 0.027 | 0.027 | 0 | 0.07 | 0.14 | 0.06 | 0.08 | 0.08 | 0.08 |
| 10 | 53456599 | rs16921608 | -1.855 | 0.494 | 1.859 | 0.435 | 0.011 | 0.067 | 0.067 | 0 | 0.18 | 0.45 | 0.12 | 0.21 | 0.16 | 0.21 |
| 10 | 53458047 | rs3740228 | -1.845 | 0.493 | 1.866 | 0.434 | 0.011 | 0.066 | 0.066 | 0 | 0.18 | 0.45 | 0.12 | 0.22 | 0.16 | 0.2 |
| 10 | 53458167 | rs3824686 | -0.666 | 0.496 | 1.777 | 0.432 | -0.002 | 0.004 | 0.004 | 0 | 0.08 | 0.08 | 0.05 | 0.14 | 0.07 | 0.11 |
| 10 | 53467198 | rs12255839 | 0.342 | 0.541 | 1.757 | 0.448 | 0.001 | 0.001 | 0.001 | 0 | 0.07 | 0.05 | 0.05 | 0.07 | 0.07 | 0.08 |
| 10 | 53467979 | rs10823447 | -2.038 | 0.369 | 1.871 | 0.41 | -0.011 | 0.019 | 0.019 | 0.199 | 0.2 | 0.57 | 0.14 | 0.21 | 0.16 | 0.2 |
| 10 | 53468025 | rs7089114 | -2.325 | 0.413 | 1.851 | 0.432 | -0.014 | 0.006 | 0.006 | 1.104 | 0.21 | 0.68 | 0.15 | 0.21 | 0.16 | 0.21 |
| 10 | 53478713 | rs4400745 | -0.169 | 0.644 | 1.347 | 0.026 | 0.003 | 0.022 | 0.022 | 1.549 | 0.58 | 0.7 | 0.43 | 0.26 | 0.59 | 0.51 |
| 10 | 53482734 | rs11594325 | 0.784 | 0.852 | 1.242 | -0.02 | 0.003 | 0.045 | 0.045 | 0.053 | 0.46 | 0.4 | 0.35 | 0.19 | 0.48 | 0.41 |
| 10 | 53494326 | rs10999157 | 0.898 | 0.889 | 1.056 | -0.024 | -0.008 | -0.003 | -0.003 | 1.111 | 0.09 | 0.05 | 0.06 | 0.08 | 0.1 | 0.08 |
| 10 | 53497295 | rs10762342 | -0.745 | 0.817 | 1.281 | -0.083 | 0.01 | 0.01 | 0.01 | 0.072 | 0.18 | 0.21 | 0.12 | 0.21 | 0.17 | 0.2 |
| 10 | 53498108 | rs10999171 | 1.454 | 0.752 | 0.859 | -0.159 | 0.001 | 0.15 | 0.15 | 0 | 0.39 | 0.2 | 0.29 | 0.05 | 0.42 | 0.29 |
| 10 | 53507053 | rs4466778 | -0.434 | 0.73 | 1.065 | -0.098 | -0.006 | 0.025 | 0.025 | 3.212 | 0.63 | 0.78 | 0.44 | 0.26 | 0.59 | 0.51 |
| 10 | 53510461 | rs4568954 | 0.414 | 0.826 | 0.771 | -0.2 | 0.011 | 0.072 | 0.072 | 0.125 | 0.51 | 0.39 | 0.36 | 0.16 | 0.52 | 0.38 |
| 10 | 53512766 | rs10999270 | 0.379 | 0.831 | 0.775 | -0.19 | 0.005 | 0.028 | 0.028 | 0.128 | 0.51 | 0.46 | 0.37 | 0.16 | 0.51 | 0.37 |
| 10 | 53525356 | rs1937723 | 0.658 | 0.811 | 0.776 | -0.299 | -0.007 | -0.085 | -0.085 | 0 | 0.44 | 0.59 | 0.34 | 0.08 | 0.42 | 0.29 |
| 10 | 53525737 | rs10999342 | 0.721 | 0.82 | 0.787 | -0.296 | 0 | -0.042 | -0.042 | 0 | 0.44 | 0.53 | 0.33 | 0.08 | 0.42 | 0.28 |
| 10 | 53528622 | rs16922836 | -0.042 | 0.334 | 0.36 | 0.177 | -0.018 | -0.156 | -0.156 | 0.9 | 0.26 | 0.01 | 0.19 | 0.36 | 0.21 | 0.19 |
| 10 | 53541632 | rs2211614 | -0.249 | 0.581 | 0.949 | 0.204 | 0.044 | -0.021 | -0.021 | 0 | 0.13 | 0.38 | 0.1 | 0.07 | 0.11 | 0.12 |
| 10 | 53545982 | rs17629265 | 1.066 | 0.443 | 0.872 | 0.07 | -0.071 | 0.227 | 0.227 | 0 | 0.39 | 0.07 | 0.28 | 0.09 | 0.45 | 0.3 |
| 10 | 53546023 | rs12266397 | -0.182 | 0.43 | 1.072 | 0.065 | 0.005 | 0.035 | 0.035 | 0 | 0.55 | 0.67 | 0.4 | 0.15 | 0.56 | 0.41 |
| 10 | 53557567 | rs11592752 | 0.915 | 0.105 | 0.954 | -0.373 | -0.002 | 0.067 | 0.067 | 0 | 0.41 | 0.29 | 0.31 | 0.09 | 0.45 | 0.34 |
| 10 | 53561613 | rs4415704 | 0.384 | 0.293 | 1.365 | 0.048 | 0.004 | 0.004 | 0.004 | 0 | 0.56 | 0.56 | 0.47 | 0.49 | 0.56 | 0.49 |
| 10 | 53562468 | rs11813335 | -0.276 | 0.313 | 1.343 | 0.056 | 0 | 0.008 | 0.008 | 0 | 0.66 | 0.65 | 0.53 | 0.55 | 0.66 | 0.57 |
| 10 | 53563270 | rs10999561 | -0.071 | 0.797 | 1.29 | 0.216 | -0.004 | 0.002 | 0.002 | 0 | 0.1 | 0.07 | 0.06 | 0.06 | 0.1 | 0.08 |
| 10 | 53572544 | rs1937688 | -0.539 | 0.8 | 1.223 | 0.381 | 0 | -0.006 | -0.006 | 0 | 0.66 | 0.71 | 0.5 | 0.59 | 0.68 | 0.64 |
| 10 | 53581387 | rs16923587 | -0.236 | 0.933 | 1.209 | -0.268 | 0.028 | -0.024 | -0.024 | 0 | 0.13 | 0.28 | 0.08 | 0.07 | 0.1 | 0.08 |
| 10 | 53584234 | rs1937690 | -0.561 | 0.954 | 0.692 | -0.261 | 0.001 | 0.001 | 0.001 | 0 | 0.65 | 0.7 | 0.53 | 0.66 | 0.69 | 0.59 |
| 10 | 53592537 | rs1937697 | -0.513 | 0.961 | -0.133 | -0.191 | 0.033 | 0.055 | 0.055 | 1.133 | 0.17 | 0.53 | 0.12 | 0.18 | 0.12 | 0.17 |
| 10 | 53593979 | rs6480499 | -0.023 | 0.68 | -1.227 | -0.257 | -0.063 | -0.077 | -0.077 | 0 | 0.44 | 0.1 | 0.37 | 0.46 | 0.52 | 0.41 |
| 10 | 53602190 | rs12247820 | -1.394 | 0.703 | 0.142 | -0.554 | 0.024 | -0.055 | -0.055 | 0 | 0.11 | 0.3 | 0.09 | 0.01 | 0.1 | 0.04 |
| 10 | 53608977 | rs1937701 | -1.544 | 0.405 | -0.049 | -0.837 | 0.001 | -0.018 | -0.018 | 0.011 | 0.23 | 0.71 | 0.21 | 0.02 | 0.18 | 0.12 |
| 10 | 53609086 | rs17549153 | -0.526 | 0.085 | -0.522 | -0.899 | -0.015 | -0.005 | -0.005 | 0.832 | 0.14 | 0.02 | 0.13 | 0.11 | 0.13 | 0.19 |
| 10 | 53613913 | rs7080306 | -0.895 | 0.299 | 0.678 | -0.926 | 0.052 | 0.025 | 0.025 | 0.007 | 0.08 | 0.35 | 0.09 | 0.01 | 0.06 | 0.07 |
| 10 | 53614863 | rs1937707 | -1.317 | 0.203 | 0.366 | -1.409 | -0.001 | -0.045 | -0.045 | 1.624 | 0.39 | 0.41 | 0.36 | 0.08 | 0.31 | 0.36 |
| 10 | 53615000 | rs1937710 | -1.095 | 0.301 | 0.442 | -1.032 | -0.002 | -0.065 | -0.065 | 0.052 | 0.18 | 0.34 | 0.17 | 0.06 | 0.15 | 0.14 |
| 10 | 53626998 | rs6480524 | -0.852 | 0.898 | 0.552 | -1.206 | 0.024 | -0.084 | -0.084 | 0 | 0.25 | 0.42 | 0.28 | 0.1 | 0.23 | 0.21 |
| 10 | 53627572 | rs7907601 | -0.354 | 1.758 | 0.059 | -1.69 | -0.047 | 0.05 | 0.05 | 0 | 0.12 | 0.04 | 0.13 | 0 | 0.24 | 0.04 |
| 10 | 53627833 | rs7071831 | -0.712 | 1.193 | 0.889 | -1.905 | 0.005 | 0.017 | 0.017 | 1.079 | 0.09 | 0.45 | 0.22 | 0.08 | 0.03 | 0.05 |
| 10 | 53629958 | rs7917131 | -0.922 | 1.208 | 0.871 | -1.907 | -0.016 | 0.025 | 0.025 | 0 | 0.18 | 0.13 | 0.32 | 0.08 | 0.27 | 0.09 |
| 10 | 53633984 | rs16924439 | 0.615 | 1.515 | 0.864 | -1.99 | -0.004 | 0.002 | 0.002 | 0 | 0.06 | 0.04 | 0.06 | 0.01 | 0.06 | 0.02 |
| 10 | 53637163 | rs10999888 | 0.151 | 1.508 | 0.843 | -2.059 | -0.042 | 0.021 | 0.021 | 0 | 0.15 | 0.03 | 0.18 | 0.08 | 0.17 | 0.18 |

| | | | | | | | | | | | | | | | | |
|---|---|---|---|---|---|---|---|---|---|---|---|---|---|---|---|---|
| 10 | 53637549 | rs17631875 | -0.087 | 1.536 | 1.019 | -2.076 | -0.008 | -0.011 | -0.011 | 0 | 0.07 | 0.14 | 0.19 | 0.07 | 0.04 | 0.05 |
| 10 | 53638306 | rs7921498 | -2.065 | 0.708 | -0.087 | -1.727 | -0.002 | -0.089 | -0.089 | 0 | 0.72 | 0.89 | 0.73 | 0.61 | 0.73 | 0.68 |
| 10 | 53639784 | rs7914093 | -0.569 | 0.732 | 0.481 | -1.77 | 0.059 | -0.146 | -0.146 | 1.07 | 0.22 | 0.5 | 0.22 | 0 | 0.3 | 0.05 |
| 10 | 53640416 | rs10823798 | 0.012 | 0.74 | 0.45 | -1.683 | 0.009 | -0.017 | -0.017 | 0 | 0.43 | 0.25 | 0.31 | 0.52 | 0.37 | 0.58 |
| 10 | 53644408 | rs17632249 | 1.455 | 0.808 | 0.434 | -1.675 | -0.002 | -0.007 | -0.007 | 0 | 0.06 | 0.04 | 0.06 | 0 | 0.06 | 0.01 |
| 10 | 53645308 | rs7908692 | -0.511 | 0.804 | 0.777 | -1.683 | 0.009 | 0.013 | 0.013 | 0.015 | 0.49 | 0.37 | 0.5 | 0.6 | 0.42 | 0.62 |
| 10 | 53645972 | rs6480542 | 0.189 | 0.776 | 1.193 | -1.703 | 0.042 | -0.085 | -0.085 | 0 | 0.14 | 0.3 | 0.15 | 0 | 0.24 | 0.04 |
| 10 | 53646127 | rs6480543 | -0.688 | 0.77 | 1.099 | -1.696 | 0.06 | -0.16 | -0.16 | 0.172 | 0.22 | 0.52 | 0.22 | 0 | 0.3 | 0.05 |
| 10 | 53648350 | rs6480545 | -2.031 | 0.762 | 1.071 | -1.691 | 0.009 | -0.039 | -0.039 | 0.03 | 0.71 | 0.8 | 0.72 | 0.6 | 0.72 | 0.67 |
| 10 | 53648544 | rs6480546 | -0.583 | 0.771 | 1.11 | -1.696 | 0.042 | -0.125 | -0.125 | 0.556 | 0.22 | 0.4 | 0.22 | 0 | 0.3 | 0.05 |
| 10 | 53650135 | rs6480547 | -1.97 | 0.762 | 1.058 | -1.666 | 0.007 | -0.026 | -0.026 | 0.021 | 0.71 | 0.77 | 0.72 | 0.6 | 0.71 | 0.67 |
| 10 | 53651912 | rs1937671 | 0.078 | 0.795 | 1.227 | -1.617 | 0.003 | -0.011 | -0.011 | 1.069 | 0.22 | 0.24 | 0.39 | 0.16 | 0.21 | 0.22 |
| 10 | 53652218 | rs1937670 | -0.194 | 0.766 | 1.145 | -1.64 | 0.001 | -0.014 | -0.014 | 0 | 0.23 | 0.27 | 0.39 | 0.17 | 0.21 | 0.22 |
| 10 | 53657648 | rs10762479 | -1.467 | 0.775 | 0.909 | -1.585 | -0.014 | -0.043 | -0.043 | 0.11 | 0.65 | 0.83 | 0.66 | 0.6 | 0.66 | 0.66 |
| 10 | 53660836 | rs1937662 | 1.095 | 0.747 | 0.917 | -1.595 | 0.014 | -0.063 | -0.063 | 0 | 0.26 | 0.1 | 0.11 | 0.44 | 0.21 | 0.4 |
| 10 | 53664992 | rs1937672 | -0.786 | 0.765 | 1.02 | -1.498 | 0.001 | -0.002 | -0.002 | 0 | 0.56 | 0.48 | 0.57 | 0.58 | 0.48 | 0.63 |
| 10 | 53667885 | rs6480565 | -1.941 | 0.686 | 0.384 | -1.468 | -0.004 | -0.086 | -0.086 | 0.29 | 0.71 | 0.89 | 0.73 | 0.6 | 0.72 | 0.67 |
| 10 | 53673286 | rs4595495 | -0.632 | 0.67 | 0.726 | -1.466 | 0.026 | -0.049 | -0.049 | 0 | 0.28 | 0.38 | 0.31 | 0.09 | 0.4 | 0.22 |
| 10 | 53677635 | rs2154281 | 0.341 | 0.396 | -0.118 | -1.499 | 0.01 | -0.054 | -0.054 | 0.803 | 0.32 | 0.16 | 0.22 | 0.46 | 0.27 | 0.42 |
| 10 | 53679060 | rs1937683 | -1.673 | 0.448 | -0.596 | -1.366 | 0.008 | -0.074 | -0.074 | 0.13 | 0.65 | 0.88 | 0.56 | 0.54 | 0.66 | 0.63 |
| 10 | 53682131 | rs2879629 | -0.985 | 0.35 | -0.649 | -1.378 | -0.06 | -0.066 | -0.066 | 0 | 0.55 | 0.87 | 0.48 | 0.46 | 0.53 | 0.52 |
| 10 | 53688297 | rs2339893 | -0.37 | 0.018 | -0.292 | -1.097 | 0.041 | -0.031 | -0.031 | 1.112 | 0.43 | 0.68 | 0.33 | 0.48 | 0.3 | 0.48 |
| 10 | 53689774 | rs7903366 | -0.378 | 0.018 | -0.253 | -1.099 | 0.039 | -0.025 | -0.025 | 0.271 | 0.43 | 0.67 | 0.33 | 0.48 | 0.3 | 0.48 |
| 10 | 53692081 | rs11000178 | -0.325 | 0.076 | -0.03 | -1.081 | 0.016 | -0.01 | -0.01 | 0.341 | 0.43 | 0.61 | 0.33 | 0.47 | 0.3 | 0.48 |
| 10 | 53692726 | rs17638883 | -0.408 | -0.137 | -0.691 | -1.319 | -0.031 | 0.081 | 0.081 | 0.109 | 0.25 | 0.05 | 0.16 | 0.11 | 0.29 | 0.21 |
| 10 | 53694448 | rs11000189 | 1.276 | -0.397 | -1.014 | -0.745 | 0.005 | -0.017 | -0.017 | 0.062 | 0.12 | 0.05 | 0.16 | 0.3 | 0.12 | 0.21 |
| 10 | 53694967 | rs11000191 | 0.148 | -0.378 | -1.213 | -0.885 | -0.068 | 0.068 | 0.068 | 0 | 0.23 | 0.01 | 0.15 | 0.1 | 0.28 | 0.2 |
| 10 | 53700409 | rs17638971 | 1.408 | -0.187 | -1.3 | -0.952 | -0.031 | 0.03 | 0.03 | 0 | 0.08 | 0.03 | 0.1 | 0 | 0.1 | 0.02 |
| 10 | 53708732 | rs2339899 | 0.391 | -0.53 | -1.421 | -1.234 | 0 | 0.002 | 0.002 | 0 | 0.22 | 0.19 | 0.28 | 0.3 | 0.21 | 0.29 |
| 10 | 53709352 | rs2339898 | -0.795 | -1.011 | -1.708 | -0.832 | 0.07 | 0.069 | 0.069 | 0.364 | 0.46 | 0.85 | 0.42 | 0.64 | 0.35 | 0.56 |
| 10 | 53710325 | rs7905683 | 0.507 | -1.023 | -1.284 | -0.781 | -0.064 | 0.081 | 0.081 | 0.459 | 0.23 | 0.57 | 0.13 | 0.34 | 0.14 | 0.27 |
| 10 | 53710540 | rs7905966 | 1.459 | -1.243 | -0.549 | -0.241 | 0.009 | 0.179 | 0.179 | 0.546 | 0.06 | 0.27 | 0.03 | 0.13 | 0.03 | 0.01 |
| 10 | 53711220 | rs10508952 | 1.092 | -1.28 | -0.393 | -0.342 | 0.019 | -0.034 | -0.034 | 3.136 | 0.15 | 0.09 | 0.07 | 0.34 | 0.1 | 0.26 |
| 10 | 53711836 | rs12255275 | -0.147 | -1.287 | -0.371 | -0.347 | -0.065 | 0.082 | 0.082 | 0.459 | 0.23 | 0.57 | 0.13 | 0.34 | 0.14 | 0.27 |
| 10 | 53712759 | rs11000264 | 0.57 | -1.129 | -0.667 | -0.598 | 0.034 | -0.035 | -0.035 | 0.93 | 0.26 | 0.1 | 0.22 | 0.64 | 0.22 | 0.47 |
| 10 | 53713462 | rs10466028 | 0.162 | -1.186 | -0.638 | -0.618 | 0.01 | -0.004 | -0.004 | 0.584 | 0.29 | 0.29 | 0.24 | 0.64 | 0.22 | 0.47 |
| 10 | 53714654 | rs10740403 | -0.381 | -1.018 | -0.48 | -0.577 | 0.001 | 0.001 | 0.001 | 0 | 0.38 | 0.38 | 0.37 | 0.64 | 0.32 | 0.56 |
| 10 | 53721176 | rs1903960 | 1.833 | -0.187 | -0.263 | -0.279 | -0.002 | 0.01 | 0.01 | 0 | 0.28 | 0.21 | 0.34 | 0.25 | 0.27 | 0.21 |
| 10 | 53721665 | rs1903961 | -1.672 | 0.137 | -0.141 | -0.159 | 0.014 | -0.03 | -0.03 | 0 | 0.92 | 0.88 | 0.89 | 1 | 0.9 | 0.98 |
| 10 | 53727464 | rs10508951 | 0.822 | 0.092 | -0.021 | -0.114 | 0.017 | -0.044 | -0.044 | 0.474 | 0.09 | 0.15 | 0.13 | 0 | 0.1 | 0.08 |
| 10 | 53730326 | rs1904034 | -0.296 | -0.429 | -0.131 | 0.174 | 0 | -0.026 | -0.026 | 0 | 0.76 | 0.8 | 0.81 | 0.67 | 0.79 | 0.7 |
| 10 | 53731491 | rs16925357 | 1.362 | -0.551 | 0.28 | 0.203 | 0.033 | 0.231 | 0.231 | 0.915 | 0.06 | 0.32 | 0.03 | 0.13 | 0.03 | 0.01 |
| 10 | 53733545 | rs2454566 | 1.664 | -0.592 | 0.066 | -0.24 | -0.041 | 0.031 | 0.031 | 0.614 | 0.32 | 0.2 | 0.43 | 0.25 | 0.38 | 0.29 |
| 10 | 53734733 | rs1904036 | 0.681 | -0.778 | 0.037 | -0.284 | -0.003 | -0.001 | -0.001 | 0.001 | 0.46 | 0.4 | 0.62 | 0.57 | 0.49 | 0.49 |
| 10 | 53736306 | rs2454564 | -0.568 | -0.798 | 0.02 | -0.301 | 0 | -0.024 | -0.024 | 0 | 0.76 | 0.79 | 0.81 | 0.67 | 0.79 | 0.7 |
| 10 | 53740094 | rs1904042 | 0.676 | -0.736 | -0.09 | -0.284 | -0.003 | -0.001 | -0.001 | 0 | 0.46 | 0.4 | 0.61 | 0.57 | 0.49 | 0.49 |
| 10 | 53742120 | rs1904046 | 0.007 | -0.612 | 0.018 | -0.168 | -0.018 | 0.032 | 0.032 | 0 | 0.7 | 0.47 | 0.77 | 0.67 | 0.77 | 0.69 |
| 10 | 53747846 | rs10740406 | -0.991 | -0.581 | -0.073 | -0.334 | -0.001 | 0.038 | 0.038 | 0.395 | 0.59 | 0.66 | 0.46 | 0.76 | 0.52 | 0.68 |
| 10 | 53748419 | rs1993860 | -1.64 | -0.462 | 0.076 | -0.061 | 0.017 | -0.034 | -0.034 | 0 | 0.91 | 0.87 | 0.89 | 1 | 0.9 | 0.98 |
| 10 | 53752475 | rs11000400 | -0.141 | -0.801 | 0.122 | 0.14 | -0.08 | 0.1 | 0.1 | 0 | 0.23 | 0.5 | 0.12 | 0.34 | 0.13 | 0.28 |
| 10 | 53753066 | rs11000404 | 1.536 | -0.789 | -0.006 | 0.164 | -0.013 | 0.002 | 0.002 | 0 | 0.05 | 0.01 | 0.14 | 0.32 | 0.07 | 0.11 |
| 10 | 53754335 | rs1904050 | -0.508 | -0.816 | 0.051 | 0.102 | -0.063 | 0.02 | 0.02 | 0 | 0.27 | 0.47 | 0.26 | 0.66 | 0.2 | 0.39 |
| 10 | 53758289 | rs4275567 | 1.179 | -0.712 | -0.199 | 0.168 | 0.021 | -0.031 | -0.031 | 2.92 | 0.15 | 0.09 | 0.08 | 0.34 | 0.1 | 0.27 |
| 10 | 53762669 | rs2447643 | -0.206 | -0.615 | -0.359 | -0.023 | -0.041 | -0.013 | -0.013 | 0.463 | 0.63 | 0.83 | 0.61 | 0.58 | 0.55 | 0.63 |
| 10 | 53762921 | rs2454539 | 0.724 | -0.24 | 0.038 | -0.195 | -0.008 | 0.022 | 0.022 | 0.197 | 0.41 | 0.4 | 0.49 | 0.23 | 0.43 | 0.36 |
| 10 | 53764919 | rs2454535 | 1.516 | -0.002 | -0.148 | -0.148 | -0.05 | -0.025 | -0.025 | 0 | 0.24 | 0.08 | 0.36 | 0.27 | 0.29 | 0.21 |
| 10 | 53765301 | rs2246714 | -0.511 | -0.171 | -0.014 | -0.466 | 0 | -0.015 | -0.015 | 0 | 0.68 | 0.68 | 0.79 | 0.91 | 0.68 | 0.7 |
| 10 | 53767465 | rs11000432 | 0.578 | -0.183 | -0.072 | -0.472 | -0.013 | 0.099 | 0.099 | 0 | 0.3 | 0.13 | 0.2 | 0.09 | 0.32 | 0.3 |
| 10 | 53770239 | rs1903989 | -0.066 | -0.301 | 0.445 | -0.489 | 0.001 | 0.002 | 0.002 | 0 | 0.38 | 0.38 | 0.4 | 0.64 | 0.37 | 0.48 |
| 10 | 53771085 | rs726195 | -0.09 | -0.28 | 0.351 | -0.27 | 0.008 | 0.035 | 0.035 | 0.163 | 0.21 | 0.28 | 0.28 | 0.34 | 0.21 | 0.2 |
| 10 | 53774826 | rs1903979 | 1.188 | -0.197 | 0.14 | -0.39 | -0.052 | 0.027 | 0.027 | 0 | 0.34 | 0.21 | 0.47 | 0.27 | 0.4 | 0.24 |
| 10 | 53776070 | rs10823959 | 2.021 | -0.2 | 0.132 | -0.389 | -0.027 | -0.011 | -0.011 | 0 | 0.24 | 0.09 | 0.36 | 0.27 | 0.29 | 0.21 |
| 10 | 53776666 | rs10823962 | 0.575 | -0.589 | 0.314 | -0.285 | -0.001 | 0.022 | 0.022 | 0 | 0.31 | 0.24 | 0.21 | 0.09 | 0.32 | 0.3 |
| 10 | 53778213 | rs10740409 | 0.812 | -0.575 | 0.276 | -0.189 | -0.041 | 0 | 0 | 0 | 0.4 | 0.23 | 0.61 | 0.61 | 0.47 | 0.41 |
| 10 | 53781310 | rs9664844 | 0.213 | -0.357 | -0.026 | -0.052 | 0 | -0.004 | -0.004 | 0 | 0.62 | 0.68 | 0.63 | 0.57 | 0.61 | 0.53 |
| 10 | 53781397 | rs1903953 | -0.255 | -0.285 | -0.343 | -0.019 | -0.009 | -0.031 | -0.031 | 0 | 0.37 | 0.18 | 0.34 | 0.43 | 0.39 | 0.47 |
| 10 | 53786295 | rs4935306 | -0.721 | -0.365 | -0.276 | -0.297 | -0.003 | 0.036 | 0.036 | 0 | 0.7 | 0.81 | 0.8 | 0.91 | 0.68 | 0.7 |
| 10 | 53787149 | rs12243919 | 0.537 | -0.374 | -0.497 | 0.077 | -0.014 | 0.014 | 0.014 | 0 | 0.22 | 0.3 | 0.14 | 0.3 | 0.19 | 0.29 |
| 10 | 53788132 | rs12763079 | 0.431 | 0.084 | -1.026 | 0.201 | -0.006 | 0.053 | 0.053 | 0 | 0.18 | 0.03 | 0.11 | 0.09 | 0.2 | 0.16 |

| | | | | | | | | | | | | | | | | |
|---|---|---|---|---|---|---|---|---|---|---|---|---|---|---|---|---|
| 10 | 53789388 | rs7901487 | -0.322 | 0.636 | -0.511 | 0.541 | -0.016 | 0 | 0 | 0 | 0.44 | 0.63 | 0.6 | 0.56 | 0.46 | 0.38 |
| 10 | 53790034 | rs10733891 | -0.183 | 0.802 | -0.397 | 0.612 | 0.018 | 0.007 | 0.007 | 0 | 0.4 | 0.49 | 0.58 | 0.56 | 0.44 | 0.37 |
| 10 | 53790625 | rs10740411 | -0.777 | 0.911 | -0.221 | 0.893 | 0.019 | 0.091 | 0.091 | 0 | 0.74 | 0.48 | 0.61 | 0.62 | 0.72 | 0.81 |
| 10 | 53791217 | rs7898220 | -0.57 | 0.963 | -0.311 | 0.934 | 0.017 | 0.066 | 0.066 | 0 | 0.72 | 0.42 | 0.56 | 0.58 | 0.66 | 0.81 |
| 10 | 53792463 | rs10762542 | 0.059 | 0.643 | -0.76 | 0.429 | -0.011 | -0.001 | -0.001 | 3.031 | 0.59 | 0.67 | 0.59 | 0.6 | 0.55 | 0.54 |
| 10 | 53792576 | rs10762543 | -0.18 | 0.983 | -0.731 | 0.586 | 0.036 | 0.058 | 0.058 | 0 | 0.82 | 0.68 | 0.73 | 0.73 | 0.8 | 0.8 |
| 10 | 53792671 | rs10762544 | -0.222 | 0.985 | -0.704 | 0.587 | 0.029 | 0.047 | 0.047 | 0 | 0.82 | 0.72 | 0.73 | 0.73 | 0.8 | 0.79 |
| 10 | 53799707 | rs1903999 | -1.102 | 1.408 | -0.747 | 0.618 | 0.006 | -0.034 | -0.034 | 0 | 0.89 | 0.89 | 0.89 | 1 | 0.89 | 0.98 |
| 10 | 53803755 | rs17560636 | 0.845 | 1.047 | -0.636 | -0.118 | 0 | -0.02 | -0.02 | 0 | 0.25 | 0.25 | 0.15 | 0.11 | 0.26 | 0.33 |
| 10 | 53804712 | rs10823999 | 0.351 | 0.006 | -0.39 | -0.283 | -0.018 | 0.022 | 0.022 | 0.365 | 0.2 | 0.2 | 0.15 | 0.26 | 0.14 | 0.19 |
| 10 | 53805360 | rs6480687 | -1.11 | 0.136 | -0.631 | -0.837 | -0.003 | -0.019 | -0.019 | 0 | 0.64 | 0.68 | 0.74 | 0.9 | 0.63 | 0.65 |
| 10 | 53806602 | rs17641137 | 1.63 | 0.337 | -1.244 | -0.732 | -0.022 | 0.083 | 0.083 | 0 | 0.19 | 0.05 | 0.34 | 0.04 | 0.22 | 0.16 |
| 10 | 53809027 | rs1904006 | 1.134 | 0.434 | -1.233 | -0.607 | -0.062 | 0.017 | 0.017 | 0 | 0.2 | 0.01 | 0.17 | 0.15 | 0.25 | 0.26 |
| 10 | 53809894 | rs17641202 | 0.939 | -0.065 | -0.975 | -0.759 | 0.004 | 0.067 | 0.067 | 0 | 0.23 | 0.11 | 0.37 | 0.05 | 0.22 | 0.19 |
| 10 | 53811431 | rs10824002 | -0.511 | -0.288 | -0.182 | -0.48 | -0.04 | 0.002 | 0.002 | 0.952 | 0.23 | 0.44 | 0.23 | 0.72 | 0.18 | 0.26 |
| 10 | 53814383 | rs12778630 | 0.477 | -0.318 | -0.272 | -0.559 | -0.009 | 0.019 | 0.019 | 0 | 0.39 | 0.3 | 0.33 | 0.26 | 0.42 | 0.38 |
| 10 | 53816171 | rs1903997 | 0.429 | -0.45 | -0.29 | -0.539 | -0.05 | 0.116 | 0.116 | 0.008 | 0.39 | 0.16 | 0.5 | 0.16 | 0.47 | 0.37 |
| 10 | 53818156 | rs10824006 | 0.68 | -0.015 | -0.287 | -0.365 | -0.006 | 0.027 | 0.027 | 0 | 0.37 | 0.15 | 0.26 | 0.25 | 0.38 | 0.38 |
| 10 | 53821238 | rs4411238 | -0.048 | -0.456 | -0.137 | -0.486 | -0.009 | 0.002 | 0.002 | 2.485 | 0.4 | 0.63 | 0.48 | 0.62 | 0.32 | 0.35 |
| 10 | 53821988 | rs7077500 | 0.838 | -0.232 | -0.314 | -0.354 | -0.025 | 0.021 | 0.021 | 0.326 | 0.2 | 0.09 | 0.21 | 0.13 | 0.26 | 0.25 |
| 10 | 53825576 | rs1903998 | -0.642 | -0.47 | -0.137 | -0.093 | -0.02 | 0.306 | 0.306 | 0.431 | 0.9 | 0.67 | 0.88 | 0.56 | 0.96 | 0.89 |
| 10 | 53826682 | rs10824010 | -0.266 | -0.434 | -0.391 | -0.318 | 0 | 0 | 0 | 0 | 0.33 | 0.36 | 0.34 | 0.67 | 0.36 | 0.42 |
| 10 | 53833671 | rs11000703 | 0.7 | -0.403 | -0.899 | -0.33 | -0.007 | 0.022 | 0.022 | 0 | 0.37 | 0.12 | 0.24 | 0.24 | 0.37 | 0.36 |
| 10 | 53838761 | rs4935027 | -0.273 | -0.367 | -0.325 | -0.103 | -0.005 | 0 | 0 | 0 | 0.41 | 0.58 | 0.49 | 0.63 | 0.35 | 0.35 |
| 10 | 53840971 | rs10762553 | 0.567 | -0.779 | -0.034 | -0.307 | 0.012 | 0.073 | 0.073 | 0.397 | 0.55 | 0.31 | 0.69 | 0.82 | 0.62 | 0.78 |
| 10 | 53841105 | rs12255651 | 0.848 | -0.725 | 0.273 | 0.148 | 0.001 | -0.037 | -0.037 | 1.673 | 0.16 | 0.12 | 0.13 | 0 | 0.19 | 0.08 |
| 10 | 53841337 | rs1904024 | 0.512 | -0.824 | 0.407 | 0.297 | -0.037 | -0.03 | -0.03 | 0 | 0.79 | 0.37 | 0.85 | 0.83 | 0.87 | 0.88 |
| 10 | 53843374 | rs7072999 | -0.418 | -0.71 | -0.145 | 0.51 | -0.055 | 0 | 0 | 0 | 0.37 | 0.13 | 0.29 | 0.29 | 0.41 | 0.37 |
| 10 | 53851164 | rs7477839 | -0.776 | -1.102 | 0.969 | 0.608 | 0.003 | 0 | 0 | 0 | 0.3 | 0.45 | 0.29 | 0.31 | 0.28 | 0.2 |
| 10 | 53855492 | rs10762562 | 1.187 | -1.459 | 0.982 | 0.648 | -0.04 | 0.022 | 0.022 | 0 | 0.42 | 0.14 | 0.54 | 0.36 | 0.54 | 0.43 |
| 10 | 53857848 | rs10824034 | 0.04 | -1.588 | 1.302 | 0.635 | 0.015 | -0.009 | -0.009 | 0.228 | 0.28 | 0.25 | 0.16 | 0.33 | 0.18 | 0.37 |
| 10 | 53859440 | rs1904013 | -0.529 | -1.563 | 1.363 | 0.735 | -0.004 | 0.166 | 0.166 | 0 | 0.1 | 0.3 | 0.15 | 0.31 | 0.08 | 0.1 |
| 10 | 53864402 | rs11000792 | -0.649 | -1.27 | 1.478 | 0.64 | 0.014 | 0.016 | 0.016 | 0.346 | 0.45 | 0.36 | 0.31 | 0.34 | 0.37 | 0.48 |
| 10 | 53871455 | rs11594206 | 0.468 | -1.196 | 0.954 | 0.587 | 0.015 | 0.154 | 0.154 | 2.997 | 0.26 | 0.06 | 0.14 | 0.01 | 0.18 | 0.28 |
| 10 | 53877006 | rs10762569 | 1.14 | -1.235 | 0.958 | 0.619 | -0.04 | 0.067 | 0.067 | 0.201 | 0.44 | 0.13 | 0.53 | 0.33 | 0.53 | 0.42 |
| 10 | 53877294 | rs9651311 | 0.356 | -1.216 | 1.096 | 0.667 | 0.01 | 0.005 | 0.005 | 2.483 | 0.63 | 0.4 | 0.66 | 0.34 | 0.7 | 0.49 |
| 10 | 53887029 | rs16927026 | -0.091 | -1.154 | -0.094 | 0.696 | 0.032 | 0.24 | 0.24 | 0 | 0.09 | 0.37 | 0.16 | 0.28 | 0.08 | 0.09 |
| 10 | 53890697 | rs9971270 | -0.36 | -1.1 | -0.675 | 0.744 | -0.015 | -0.003 | -0.003 | 0 | 0.35 | 0.45 | 0.3 | 0.34 | 0.26 | 0.4 |
| 10 | 53895744 | rs2001283 | -0.298 | -1.039 | -0.904 | 0.6 | -0.006 | -0.117 | -0.117 | 0 | 0.8 | 0.53 | 0.87 | 0.95 | 0.83 | 0.9 |
| 10 | 53896150 | rs12413303 | -0.499 | -0.409 | -0.566 | 0.851 | -0.019 | -0.007 | -0.007 | 0 | 0.36 | 0.5 | 0.35 | 0.63 | 0.29 | 0.48 |
| 10 | 53904166 | rs7910428 | -0.705 | 0.091 | -0.147 | 1.878 | -0.001 | 0.007 | 0.007 | 0 | 0.24 | 0.36 | 0.25 | 0.23 | 0.22 | 0.14 |
| 10 | 53910521 | rs1910534 | 0.362 | 0.093 | -0.378 | 1.006 | -0.001 | 0.028 | 0.028 | 0 | 0.74 | 0.56 | 0.69 | 0.44 | 0.74 | 0.76 |
| 10 | 53923411 | rs10437352 | -0.238 | -0.423 | -1.163 | 0.869 | 0.061 | -0.029 | -0.029 | 0 | 0.55 | 0.75 | 0.36 | 0.49 | 0.38 | 0.58 |
| 10 | 53929658 | rs10762627 | 0.085 | -1.771 | -0.632 | 1.223 | -0.089 | 0.129 | 0.129 | 0 | 0.8 | 0.53 | 0.84 | 0.41 | 0.89 | 0.56 |
| 10 | 53929791 | rs11001056 | 0.542 | -1.945 | -1.253 | 0.823 | -0.004 | 0.126 | 0.126 | 0.45 | 0.4 | 0.07 | 0.59 | 0.23 | 0.58 | 0.35 |
| 10 | 53930786 | rs7910470 | -0.391 | -2.406 | -0.722 | 0.831 | 0.001 | -0.004 | -0.004 | 0 | 0.27 | 0.25 | 0.15 | 0.39 | 0.16 | 0.46 |
| 10 | 53932566 | rs7902949 | -0.91 | -2.725 | -1.253 | 0.065 | 0.034 | 0.1 | 0.1 | 0.377 | 0.7 | 0.93 | 0.81 | 0.88 | 0.75 | 0.86 |
| 10 | 53932625 | rs7095803 | 0.841 | -2.319 | -1.039 | 0.122 | -0.037 | 0.074 | 0.074 | 0.135 | 0.34 | 0.1 | 0.57 | 0.23 | 0.56 | 0.3 |
| 10 | 53934284 | rs10824201 | -0.241 | -0.862 | -0.979 | -0.132 | -0.02 | 0.02 | 0.02 | 0 | 0.3 | 0.22 | 0.46 | 0.22 | 0.32 | 0.29 |
| 10 | 53935082 | rs7903745 | 0.934 | -0.786 | -0.543 | -0.142 | 0.004 | -0.01 | -0.01 | 0 | 0.22 | 0.23 | 0.2 | 0.12 | 0.31 | 0.19 |
| 10 | 53943122 | rs10824214 | -0.136 | -0.796 | -0.455 | 0.269 | -0.004 | -0.003 | -0.003 | 0 | 0.42 | 0.52 | 0.27 | 0.36 | 0.36 | 0.46 |
| 10 | 53943887 | rs955779 | -0.284 | -0.848 | -0.382 | 0.246 | 0.005 | 0.003 | 0.003 | 1.094 | 0.43 | 0.61 | 0.27 | 0.36 | 0.36 | 0.46 |
| 10 | 53944877 | rs10128399 | -0.332 | -1.213 | -0.536 | 0.226 | -0.073 | -0.005 | -0.005 | 1.294 | 0.25 | 0.59 | 0.15 | 0.2 | 0.11 | 0.29 |
| 10 | 53945382 | rs7921595 | -0.332 | -1.213 | -0.536 | 0.232 | -0.073 | -0.006 | -0.006 | 0 | 0.25 | 0.59 | 0.15 | 0.2 | 0.11 | 0.29 |
| 10 | 53946150 | rs11001100 | -0.02 | -1.201 | -0.364 | 0.238 | -0.096 | 0.005 | 0.005 | 0 | 0.23 | 0.46 | 0.14 | 0.2 | 0.11 | 0.29 |
| 10 | 53958511 | rs11001157 | 1.207 | -0.823 | -0.735 | 0.317 | -0.044 | 0.005 | 0.005 | 0 | 0.19 | 0.1 | 0.14 | 0.16 | 0.24 | 0.16 |
| 10 | 53959455 | rs2174257 | 0.297 | -1.16 | -0.331 | 0.782 | 0.007 | 0 | 0 | 0.335 | 0.44 | 0.36 | 0.41 | 0.56 | 0.56 | 0.4 |
| 10 | 53966300 | rs1194492 | 0.809 | -1.155 | -0.892 | 0.743 | -0.055 | 0.054 | 0.054 | 0 | 0.33 | 0.05 | 0.42 | 0.23 | 0.32 | 0.29 |
| 10 | 53966802 | rs7898960 | -0.226 | -1.314 | 0.498 | 1.155 | -0.037 | 0.006 | 0.006 | 0 | 0.19 | 0.24 | 0.12 | 0.2 | 0.11 | 0.29 |
| 10 | 53972465 | rs1875789 | -0.209 | -1.346 | 0.565 | 1.351 | -0.028 | 0.228 | 0.228 | 0 | 0.12 | 0.27 | 0.17 | 0.47 | 0.07 | 0.11 |
| 10 | 53974062 | rs11001213 | -0.168 | -1.34 | 0.541 | 1.302 | -0.027 | 0.226 | 0.226 | 0 | 0.12 | 0.26 | 0.18 | 0.5 | 0.08 | 0.11 |
| 10 | 53974269 | rs12256137 | -0.686 | -0.649 | 0.733 | 0.856 | -0.023 | -0.039 | -0.039 | 0 | 0.14 | 0.29 | 0.07 | 0.04 | 0.11 | 0.3 |
| 10 | 53975769 | rs1406476 | -1.066 | -0.423 | 0.521 | 0.865 | 0.01 | -0.056 | -0.056 | 0.1 | 0.78 | 0.68 | 0.73 | 0.94 | 0.73 | 0.82 |
| 10 | 53983258 | rs1729578 | 0.69 | -0.333 | 0.151 | 0.946 | -0.001 | 0.004 | 0.004 | 0 | 0.26 | 0.23 | 0.27 | 0.16 | 0.3 | 0.29 |
| 10 | 54002021 | rs1194501 | 0.649 | -0.625 | 0.803 | 0.851 | 0.001 | 0.031 | 0.031 | 0 | 0.27 | 0.18 | 0.37 | 0.16 | 0.3 | 0.26 |
| 10 | 54003008 | rs1875794 | 0.671 | -1.106 | 0.783 | 0.836 | 0.055 | 0.023 | 0.023 | 0 | 0.49 | 0.14 | 0.36 | 0.42 | 0.59 | 0.41 |
| 10 | 54004847 | rs11001343 | 0.525 | -1.132 | 0.806 | 0.844 | 0.052 | 0.127 | 0.127 | 0 | 0.52 | 0.21 | 0.48 | 0.77 | 0.6 | 0.53 |
| 10 | 54005644 | rs1194505 | 0.269 | -1.216 | 0.802 | 0.799 | 0.012 | 0.029 | 0.029 | 0 | 0.34 | 0.22 | 0.41 | 0.23 | 0.34 | 0.43 |
| 10 | 54007174 | rs1201585 | -0.327 | -1.116 | 0.861 | 0.777 | -0.007 | -0.08 | -0.08 | 0 | 0.87 | 0.62 | 0.91 | 1 | 0.94 | 0.96 |

| | | | | | | | | | | | | | | | | |
|---|---|---|---|---|---|---|---|---|---|---|---|---|---|---|---|---|
| 10 | 54009116 | rs7907681 | 0.671 | -1.189 | 0.6 | 0.868 | 0.058 | 0.047 | 0.047 | 0 | 0.49 | 0.17 | 0.36 | 0.4 | 0.59 | 0.4 |
| 10 | 54011665 | rs10824300 | 0.687 | -1.177 | 0.571 | 0.889 | 0.054 | -0.026 | -0.026 | 1.126 | 0.49 | 0.16 | 0.36 | 0.51 | 0.59 | 0.41 |
| 10 | 54028464 | rs10824310 | 0.676 | -1.146 | 0.664 | 0.663 | 0.02 | -0.014 | -0.014 | 0 | 0.08 | 0.03 | 0.04 | 0.09 | 0.04 | 0.24 |
| 10 | 54031443 | rs1194516 | -0.779 | -1.193 | 0.853 | 0.605 | -0.004 | -0.061 | -0.061 | 0.013 | 0.89 | 0.76 | 0.93 | 1 | 0.94 | 0.96 |
| 10 | 54032702 | rs1406473 | -0.1 | -1.181 | 0.779 | 0.808 | -0.023 | 0.061 | 0.061 | 0 | 0.14 | 0.3 | 0.18 | 0.23 | 0.07 | 0.13 |
| 10 | 54037188 | rs1919460 | 0.805 | -1.336 | 0.767 | 0.747 | 0.051 | -0.031 | -0.031 | 0 | 0.48 | 0.21 | 0.35 | 0.51 | 0.59 | 0.43 |
| 10 | 54039559 | rs7083967 | 0.822 | -1.335 | 0.768 | 0.723 | 0.05 | -0.029 | -0.029 | 0 | 0.48 | 0.21 | 0.35 | 0.51 | 0.59 | 0.43 |
| 10 | 54042556 | rs11001444 | -1.419 | -1.221 | 0.675 | 0.769 | -0.061 | 0.056 | 0.056 | 0 | 0.23 | 0.44 | 0.24 | 0.32 | 0.11 | 0.37 |
| 10 | 54042782 | rs16928871 | 0.116 | -1.07 | 0.726 | 0.624 | -0.02 | 0.002 | 0.002 | 0 | 0.09 | 0.14 | 0.06 | 0.09 | 0.04 | 0.24 |
| 10 | 54044262 | rs3829169 | 0.116 | -1.07 | 0.726 | 0.624 | -0.02 | 0.002 | 0.002 | 0 | 0.09 | 0.14 | 0.06 | 0.09 | 0.04 | 0.24 |
| 10 | 54048234 | rs7916242 | 0.022 | -0.981 | 0.203 | 0.563 | -0.003 | 0 | 0 | 0.029 | 0.61 | 0.59 | 0.44 | 0.61 | 0.63 | 0.73 |
| 10 | 54053817 | rs1881597 | 0.243 | -0.675 | -0.379 | -0.002 | 0.019 | 0.003 | 0.003 | 0 | 0.34 | 0.21 | 0.23 | 0.3 | 0.3 | 0.45 |
| 10 | 54059350 | rs7916906 | -0.71 | -0.53 | -0.518 | -0.285 | 0.009 | 0.035 | 0.035 | 0 | 0.54 | 0.66 | 0.38 | 0.28 | 0.41 | 0.5 |
| 10 | 54060101 | rs11815205 | 1.217 | 0.272 | -1.286 | -0.361 | -0.002 | -0.019 | -0.019 | 0.081 | 0.13 | 0.1 | 0.08 | 0 | 0.16 | 0.05 |
| 10 | 54063162 | rs12773082 | 0.968 | 0.084 | -1.523 | -0.096 | -0.07 | -0.009 | -0.009 | 0 | 0.3 | 0.02 | 0.2 | 0.2 | 0.3 | 0.17 |
| 10 | 54065821 | rs7100461 | -0.143 | 0.626 | -0.563 | 0.268 | -0.001 | -0.012 | -0.012 | 0 | 0.06 | 0.06 | 0.05 | 0 | 0.06 | 0.04 |
| 10 | 54068904 | rs1896368 | 0.026 | 0.627 | -0.571 | 0.587 | -0.029 | -0.006 | -0.006 | 0 | 0.59 | 0.28 | 0.37 | 0.39 | 0.52 | 0.5 |
| 10 | 54072901 | rs11001553 | -1.491 | 1.062 | -0.083 | 0.875 | 0.01 | -0.005 | -0.005 | 0 | 0.11 | 0.2 | 0.08 | 0.1 | 0.09 | 0.26 |
| 10 | 54079054 | rs2288335 | -0.818 | 0.914 | -0.079 | 0.904 | 0.002 | 0.003 | 0.003 | 0.467 | 0.08 | 0.07 | 0.04 | 0.09 | 0.06 | 0.18 |
| 10 | 54080807 | rs10824351 | -1.852 | 0.828 | 0.528 | 0.791 | -0.01 | -0.03 | -0.03 | 0 | 0.16 | 0.56 | 0.13 | 0.09 | 0.08 | 0.18 |
| 10 | 54086453 | rs11001604 | -1.794 | 0.821 | 0.532 | 0.799 | 0.009 | -0.015 | -0.015 | 0 | 0.15 | 0.44 | 0.12 | 0.09 | 0.08 | 0.18 |
| 10 | 54087203 | rs1991392 | -1.538 | 0.967 | 0.328 | 0.885 | -0.002 | 0.001 | 0.001 | 0.098 | 0.09 | 0.12 | 0.06 | 0.1 | 0.06 | 0.25 |
| 10 | 54101139 | rs1441123 | 0.279 | 1.225 | 0 | 1.217 | -0.099 | -0.047 | -0.047 | 0 | 0.69 | 0.92 | 0.6 | 0.56 | 0.57 | 0.6 |
| 10 | 54103425 | rs11001702 | -0.923 | 1.063 | 0.712 | 1.614 | 0.059 | 0.41 | 0.41 | 0.148 | 0.08 | 0.35 | 0.19 | 0.38 | 0.08 | 0.26 |
| 10 | 54116123 | rs16929708 | 0.12 | 0.986 | 0.684 | 0.616 | 0.024 | -0.007 | -0.007 | 0 | 0.06 | 0.23 | 0.06 | 0 | 0.03 | 0 |
| 10 | 54116418 | rs10762715 | 0.313 | 1.501 | -0.151 | 0.852 | -0.038 | -0.01 | -0.01 | 1.093 | 0.79 | 0.9 | 0.67 | 0.72 | 0.65 | 0.74 |
| 10 | 54128413 | rs1194683 | -0.214 | 0.387 | -1.082 | 0.939 | -0.115 | -0.082 | -0.082 | 0 | 0.74 | 0.92 | 0.51 | 0.37 | 0.52 | 0.56 |
| 10 | 54153591 | rs12769610 | -0.773 | -0.155 | -0.485 | 0.693 | 0.028 | 0.024 | 0.024 | 0.212 | 0.15 | 0.04 | 0.05 | 0.09 | 0.08 | 0.22 |
| 10 | 54171706 | rs7893873 | -1.081 | -0.455 | 0.405 | 0.835 | -0.018 | 0.01 | 0.01 | 0.215 | 0.3 | 0.6 | 0.23 | 0.63 | 0.23 | 0.35 |
| 10 | 54175474 | rs12250292 | -1.062 | -0.505 | 1.086 | 0.992 | 0.033 | 0.276 | 0.276 | 0 | 0.06 | 0.32 | 0.14 | 0.26 | 0.02 | 0.19 |
| 10 | 54175591 | rs11001943 | -0.928 | -0.519 | 1.116 | 0.7 | 0.013 | 0.17 | 0.17 | 0 | 0.06 | 0.25 | 0.13 | 0.23 | 0.02 | 0.19 |
| 10 | 54196773 | rs1194716 | -1.435 | -0.433 | 1.178 | 0.463 | 0.017 | 0.027 | 0.027 | 0.407 | 0.39 | 0.67 | 0.4 | 0.83 | 0.29 | 0.54 |
| 10 | 54207465 | rs11002107 | -0.674 | -0.518 | 1.364 | 0.471 | -0.01 | 0.012 | 0.012 | 0 | 0.28 | 0.36 | 0.2 | 0.63 | 0.24 | 0.37 |
| 10 | 54212597 | rs1194743 | -1.704 | -0.597 | 0.693 | -0.087 | -0.032 | 0.145 | 0.145 | 0.307 | 0.69 | 0.99 | 0.79 | 0.95 | 0.75 | 0.79 |
| 10 | 54213550 | rs1194742 | -1.3 | -0.653 | 1.862 | 0.935 | -0.021 | 0.056 | 0.056 | 0 | 0.11 | 0.25 | 0.19 | 0.2 | 0.05 | 0.19 |
| 10 | 54225116 | rs12411572 | -0.354 | -0.63 | 1.874 | 0.878 | -0.001 | 0.017 | 0.017 | 0 | 0.05 | 0.05 | 0.11 | 0.19 | 0.02 | 0.19 |
| 10 | 54234187 | rs7906127 | -0.951 | -0.638 | 1.758 | 0.896 | -0.015 | 0.013 | 0.013 | 0 | 0.28 | 0.4 | 0.2 | 0.64 | 0.23 | 0.37 |
| 10 | 54236752 | rs1618625 | -1.226 | -0.459 | 1.796 | 0.898 | 0.003 | 0.046 | 0.046 | 0 | 0.08 | 0.22 | 0.18 | 0.15 | 0.05 | 0.19 |
| 10 | 54240004 | rs4935325 | -0.421 | -0.456 | 1.752 | 0.861 | 0 | 0.012 | 0.012 | 0 | 0.05 | 0.04 | 0.12 | 0.2 | 0.02 | 0.19 |
| 10 | 54243993 | rs10824558 | -1.301 | -0.138 | 1.861 | 0.969 | -0.03 | 0.004 | 0.004 | 0.205 | 0.3 | 0.5 | 0.22 | 0.68 | 0.23 | 0.37 |
| 10 | 54244585 | rs11002230 | -1.151 | 0.532 | 2.014 | 0.828 | -0.009 | -0.01 | -0.01 | 7.519 | 0.23 | 0.37 | 0.13 | 0.19 | 0.21 | 0.15 |
| 10 | 54315112 | rs11002517 | -0.917 | 0.589 | 0.967 | 0.577 | 0.003 | 0.014 | 0.014 | 0 | 0.18 | 0.32 | 0.11 | 0.22 | 0.17 | 0.15 |
| 10 | 54335093 | rs6480894 | -0.53 | 0.959 | -0.064 | -0.396 | 0.003 | -0.039 | -0.039 | 0.019 | 0.38 | 0.4 | 0.27 | 0.17 | 0.41 | 0.29 |
| 10 | 54335375 | rs2204758 | 0.554 | 1.505 | 0.08 | -0.478 | 0.013 | -0.019 | -0.019 | 0 | 0.72 | 0.61 | 0.58 | 0.24 | 0.64 | 0.54 |
| 10 | 54336509 | rs11002604 | -0.514 | 0.95 | -0.562 | -1.189 | 0.003 | 0.058 | 0.058 | 0 | 0.18 | 0.06 | 0.17 | 0.04 | 0.16 | 0.1 |
| 10 | 54337021 | rs10824621 | -0.124 | 0.741 | -0.862 | -0.559 | -0.042 | -0.057 | -0.057 | 0.593 | 0.23 | 0.11 | 0.35 | 0.39 | 0.32 | 0.19 |
| 10 | 54344288 | rs1919763 | 1.166 | 0.448 | 0.002 | -0.792 | 0 | -0.001 | -0.001 | 0.263 | 0.52 | 0.46 | 0.45 | 0.52 | 0.5 | 0.68 |
| 10 | 54345990 | rs1919764 | 0.213 | 0.375 | 0.104 | -0.673 | 0.03 | 0.057 | 0.057 | 0.151 | 0.7 | 0.51 | 0.61 | 0.55 | 0.65 | 0.77 |
| 10 | 54346781 | rs1919765 | 0.252 | 0.372 | 0.244 | -0.671 | 0.016 | 0.027 | 0.027 | 0.003 | 0.71 | 0.61 | 0.63 | 0.58 | 0.65 | 0.78 |
| 10 | 54353199 | rs4617536 | 1.38 | 0.1 | -0.208 | -1.525 | -0.005 | 0.152 | 0.152 | 0 | 0.34 | 0.08 | 0.19 | 0.03 | 0.33 | 0.16 |
| 10 | 54354784 | rs4620668 | 0.631 | -0.474 | -0.097 | -1.338 | -0.001 | -0.005 | -0.005 | 3.939 | 0.44 | 0.49 | 0.44 | 0.35 | 0.45 | 0.58 |
| 10 | 54355982 | rs10762799 | 0.514 | -0.502 | 0.373 | -1.35 | 0.003 | 0.006 | 0.006 | 0.227 | 0.46 | 0.4 | 0.43 | 0.35 | 0.45 | 0.54 |
| 10 | 54358549 | rs10762804 | 0.554 | -0.523 | 0.523 | -1.357 | -0.002 | -0.004 | -0.004 | 0 | 0.46 | 0.48 | 0.45 | 0.36 | 0.45 | 0.59 |
| 10 | 54361898 | rs10824650 | -0.062 | -0.673 | 0.264 | -1.454 | -0.05 | -0.06 | -0.06 | 0 | 0.57 | 0.11 | 0.44 | 0.52 | 0.55 | 0.72 |
| 10 | 54363515 | rs1919768 | 0.398 | -0.83 | 1.01 | -1.412 | -0.001 | -0.005 | -0.005 | 4.711 | 0.44 | 0.47 | 0.45 | 0.35 | 0.45 | 0.55 |
| 10 | 54372284 | rs10824668 | 0.175 | -0.789 | 0.991 | -1.394 | 0.002 | 0.003 | 0.003 | 0 | 0.47 | 0.44 | 0.44 | 0.36 | 0.45 | 0.55 |
| 10 | 54372305 | rs11002735 | 1.525 | -0.793 | 0.93 | -1.415 | -0.072 | 0.228 | 0.228 | 0.273 | 0.31 | 0.01 | 0.17 | 0.01 | 0.32 | 0.16 |
| 10 | 54372399 | rs10824669 | -0.324 | -0.759 | 1.027 | -1.392 | -0.001 | -0.001 | -0.001 | 0.178 | 0.6 | 0.6 | 0.58 | 0.39 | 0.55 | 0.63 |
| 10 | 54373291 | rs10740487 | 0.334 | -0.759 | 0.992 | -1.385 | 0.01 | 0.017 | 0.017 | 0.077 | 0.47 | 0.31 | 0.42 | 0.36 | 0.45 | 0.55 |
| 10 | 54373493 | rs10733910 | -0.281 | -0.69 | 0.879 | -1.348 | 0.004 | -0.001 | -0.001 | 0.181 | 0.58 | 0.54 | 0.57 | 0.39 | 0.55 | 0.63 |
| 10 | 54383916 | rs16931762 | 0.107 | -0.58 | 0.581 | -1.253 | -0.038 | -0.081 | -0.081 | 0 | 0.18 | 0.45 | 0.2 | 0.05 | 0.11 | 0.08 |
| 10 | 54384757 | rs10733912 | 0.542 | -0.654 | 0.486 | -1.209 | 0 | 0.004 | 0.004 | 0 | 0.44 | 0.22 | 0.4 | 0.37 | 0.44 | 0.56 |
| 10 | 54385382 | rs11002799 | 1.48 | -0.732 | 0.398 | -1.269 | -0.016 | 0.129 | 0.129 | 0.376 | 0.31 | 0.08 | 0.18 | 0.04 | 0.32 | 0.16 |
| 10 | 54386443 | rs12220324 | 0.742 | -0.703 | 0.289 | -1.243 | -0.008 | -0.045 | -0.045 | 0 | 0.11 | 0.03 | 0.21 | 0.33 | 0.12 | 0.4 |
| 10 | 54386713 | rs12416338 | 0.145 | -0.689 | 0.602 | -1.064 | -0.035 | -0.073 | -0.073 | 0 | 0.18 | 0.42 | 0.2 | 0.05 | 0.11 | 0.08 |
| 10 | 54391014 | rs11002822 | -0.296 | -0.58 | 0.453 | -0.991 | 0.009 | 0.003 | 0.003 | 0 | 0.6 | 0.5 | 0.58 | 0.39 | 0.55 | 0.64 |
| 10 | 54391792 | rs1919780 | -0.175 | -0.558 | 0.356 | -0.981 | -0.001 | -0.001 | -0.001 | 0 | 0.58 | 0.37 | 0.57 | 0.39 | 0.55 | 0.64 |
| 10 | 54405487 | rs7909247 | 0.006 | -0.46 | -0.412 | -0.966 | -0.021 | -0.068 | -0.068 | 0 | 0.8 | 0.49 | 0.77 | 0.91 | 0.85 | 0.92 |
| 10 | 54408006 | rs10740495 | -0.119 | -0.249 | -1.382 | -0.819 | -0.011 | -0.013 | -0.013 | 0 | 0.59 | 0.29 | 0.59 | 0.43 | 0.58 | 0.65 |

| | | | | | | | | | | | | | | | | |
|---|---|---|---|---|---|---|---|---|---|---|---|---|---|---|---|---|
| 10 | 54410645 | rs17662737 | 0.62 | -0.831 | -2.182 | -0.413 | 0.004 | 0.107 | 0.107 | 0 | 0.31 | 0.02 | 0.12 | 0.16 | 0.24 | 0.23 |
| 10 | 54410927 | rs11002920 | -0.204 | -0.237 | -2.183 | 0.005 | -0.058 | -0.025 | -0.025 | 0 | 0.3 | 0.13 | 0.35 | 0.3 | 0.35 | 0.47 |
| 10 | 54411117 | rs7071467 | 0.898 | 0.224 | -1.879 | -0.044 | -0.01 | 0.023 | 0.023 | 0.124 | 0.15 | 0.07 | 0.08 | 0.04 | 0.16 | 0.07 |
| 10 | 54411308 | rs10762857 | -0.603 | -0.013 | -1.739 | 0.591 | -0.041 | -0.074 | -0.074 | 0 | 0.73 | 0.84 | 0.63 | 0.59 | 0.64 | 0.78 |
| 10 | 54412493 | rs1159798 | -0.493 | -0.351 | -1.092 | 0.258 | -0.042 | -0.035 | -0.035 | 3.428 | 0.65 | 0.52 | 0.81 | 0.75 | 0.73 | 0.66 |
| 10 | 54413554 | rs991681 | -0.26 | 0.659 | -1.187 | 0.831 | -0.035 | -0.025 | -0.025 | 0 | 0.34 | 0.28 | 0.37 | 0.43 | 0.43 | 0.45 |
| 10 | 54417882 | rs1160630 | 0.405 | 0.582 | -0.869 | 0.789 | -0.011 | -0.003 | -0.003 | 0.296 | 0.62 | 0.28 | 0.53 | 0.59 | 0.53 | 0.68 |
| 10 | 54422338 | rs10762864 | 0.169 | 1.215 | -1.057 | 0.892 | -0.09 | -0.007 | -0.007 | 0 | 0.21 | 0.05 | 0.28 | 0.21 | 0.33 | 0.19 |
| 10 | 54423398 | rs7898709 | -0.614 | 1.09 | 0.029 | 0.844 | 0.016 | 0.094 | 0.094 | 0.03 | 0.12 | 0.41 | 0.16 | 0.2 | 0.12 | 0.13 |
| 10 | 54425442 | rs7095115 | -0.56 | 1.08 | 0.156 | 0.839 | 0.016 | 0.087 | 0.087 | 0 | 0.12 | 0.38 | 0.16 | 0.19 | 0.12 | 0.12 |
| 10 | 54427594 | rs1949095 | 0.684 | 0.499 | -0.682 | 0.769 | -0.027 | 0.006 | 0.006 | 0.722 | 0.25 | 0.01 | 0.11 | 0.15 | 0.23 | 0.29 |
| 10 | 54429118 | rs10762869 | 0.864 | 0.66 | 0.132 | 0.875 | 0.01 | 0.096 | 0.096 | 0.633 | 0.86 | 0.4 | 0.81 | 0.8 | 0.88 | 0.87 |
| 10 | 54430631 | rs12416665 | -0.404 | 0.724 | 0.173 | 0.903 | 0.006 | 0.039 | 0.039 | 0 | 0.1 | 0.18 | 0.14 | 0.19 | 0.12 | 0.13 |
| 10 | 54434511 | rs11003047 | -0.845 | 0.75 | 0.261 | 0.903 | 0.018 | 0.088 | 0.088 | 0.012 | 0.12 | 0.43 | 0.17 | 0.2 | 0.12 | 0.13 |
| 10 | 54435422 | rs6480949 | -0.625 | 0.809 | 0.078 | 0.945 | 0.014 | 0.075 | 0.075 | 0.007 | 0.11 | 0.36 | 0.16 | 0.2 | 0.12 | 0.13 |
| 10 | 54440485 | rs2218330 | -0.15 | 0.645 | -0.36 | 0.701 | 0.005 | 0.012 | 0.012 | 0 | 0.55 | 0.39 | 0.49 | 0.66 | 0.58 | 0.63 |
| 10 | 54442734 | rs1000219 | -0.201 | 0.888 | -0.476 | 0.667 | 0.001 | 0.001 | 0.001 | 0.213 | 0.35 | 0.34 | 0.4 | 0.54 | 0.4 | 0.36 |
| 10 | 54446999 | rs1443136 | 0.4 | 0.191 | -0.908 | 0.182 | 0.05 | 0.052 | 0.052 | 1.677 | 0.32 | 0.19 | 0.11 | 0.22 | 0.2 | 0.46 |
| 10 | 54447346 | rs4935336 | -0.175 | 0.856 | -0.372 | 0.161 | -0.005 | -0.005 | -0.005 | 0 | 0.46 | 0.47 | 0.58 | 0.57 | 0.6 | 0.38 |
| 10 | 54453387 | rs7917895 | -0.768 | 1.202 | 0.465 | 0.556 | 0.073 | 0.155 | 0.155 | 0.251 | 0.11 | 0.32 | 0.15 | 0.23 | 0.15 | 0.15 |
| 10 | 54469155 | rs4597018 | 0.576 | 0.696 | -0.382 | 0.007 | -0.079 | -0.011 | -0.011 | 0 | 0.49 | 0.05 | 0.55 | 0.54 | 0.51 | 0.49 |
| 10 | 54475551 | rs10824774 | 1.038 | 0.662 | -0.011 | -0.075 | 0.049 | 0.062 | 0.062 | 4.438 | 0.26 | 0.11 | 0.12 | 0.17 | 0.18 | 0.3 |
| 10 | 54477048 | rs11003065 | -0.763 | 0.877 | 0.333 | 0.052 | 0.068 | 0.077 | 0.077 | 0.273 | 0.14 | 0.24 | 0.17 | 0.24 | 0.22 | 0.17 |
| 10 | 54498390 | rs11816856 | -0.515 | 0.643 | -0.222 | -0.133 | 0.054 | -0.016 | -0.016 | 0 | 0.06 | 0.23 | 0.06 | 0.01 | 0.06 | 0.01 |
| 10 | 54498496 | rs10824783 | 0.691 | 0.363 | -0.226 | -0.208 | 0.029 | 0.032 | 0.032 | 3.739 | 0.28 | 0.23 | 0.13 | 0.17 | 0.18 | 0.3 |
| 10 | 54498900 | rs12385766 | -0.931 | 1.091 | -0.16 | 0.168 | 0.024 | -0.003 | -0.003 | 0 | 0.13 | 0.23 | 0.16 | 0.14 | 0.17 | 0.16 |
| 10 | 54499040 | rs12385751 | -0.862 | 1.15 | -0.174 | 0.183 | 0.027 | 0.002 | 0.002 | 0.424 | 0.12 | 0.23 | 0.16 | 0.15 | 0.17 | 0.16 |
| 10 | 54499639 | rs11003100 | -0.688 | 1.138 | -0.25 | 0.191 | 0.018 | 0 | 0 | 0 | 0.12 | 0.22 | 0.17 | 0.14 | 0.17 | 0.16 |
| 10 | 54501332 | rs12571712 | 0.343 | 0.774 | -0.74 | 0.257 | 0.004 | 0.03 | 0.03 | 5.245 | 0.49 | 0.22 | 0.56 | 0.54 | 0.49 | 0.47 |
| 10 | 54501892 | rs1975459 | 0.581 | 0.68 | -0.857 | 0.242 | -0.023 | 0.06 | 0.06 | 4.639 | 0.47 | 0.17 | 0.56 | 0.58 | 0.49 | 0.47 |
| 10 | 54504049 | rs1992529 | 0.481 | 0.731 | -0.745 | 0.336 | -0.003 | 0.051 | 0.051 | 4.247 | 0.51 | 0.11 | 0.59 | 0.59 | 0.57 | 0.51 |
| 10 | 54504396 | rs4935044 | 0.441 | 0.735 | -0.737 | 0.337 | 0.013 | 0.056 | 0.056 | 3.858 | 0.51 | 0.17 | 0.6 | 0.59 | 0.57 | 0.52 |
| 10 | 54505972 | rs2891402 | -0.819 | 0.673 | -0.205 | 0.254 | 0.005 | -0.012 | -0.012 | 0 | 0.2 | 0.25 | 0.22 | 0.16 | 0.22 | 0.17 |
| 10 | 54507871 | rs10824787 | -0.767 | 0.602 | 0.367 | 0.158 | 0.015 | -0.01 | -0.01 | 0 | 0.19 | 0.29 | 0.23 | 0.16 | 0.23 | 0.17 |
| 10 | 54515590 | rs3829168 | -0.292 | 0.642 | 0.578 | 0.166 | -0.02 | -0.004 | -0.004 | 0.137 | 0.14 | 0.08 | 0.19 | 0.16 | 0.23 | 0.17 |
| 10 | 54517687 | rs10740519 | -0.988 | 0.273 | 0.582 | 0.009 | -0.039 | -0.01 | -0.01 | 0 | 0.73 | 0.62 | 0.85 | 0.75 | 0.82 | 0.74 |
| 10 | 54518968 | rs920727 | 0.882 | 0.461 | 0.631 | 0.095 | 0.018 | 0.021 | 0.021 | 0.555 | 0.8 | 0.31 | 0.7 | 0.74 | 0.76 | 0.79 |
| 10 | 54525509 | rs2506 | -1.03 | 0.477 | 0.583 | 0.102 | 0.043 | 0.048 | 0.048 | 0.182 | 0.19 | 0.6 | 0.28 | 0.26 | 0.24 | 0.21 |
| 10 | 54525678 | rs2083771 | -1.041 | 0.474 | 0.582 | 0.101 | 0.042 | 0.047 | 0.047 | 0.259 | 0.2 | 0.6 | 0.28 | 0.26 | 0.24 | 0.21 |
| 10 | 54525857 | rs2099903 | -0.615 | 0.455 | 0.645 | 0.096 | 0.07 | 0 | 0 | 0 | 0.17 | 0.45 | 0.23 | 0.16 | 0.24 | 0.21 |
| 10 | 54526018 | rs2120131 | -0.603 | 0.465 | 0.665 | 0.101 | 0.071 | 0 | 0 | 0 | 0.17 | 0.45 | 0.24 | 0.16 | 0.24 | 0.21 |
| 10 | 54526206 | rs10824792 | 0.684 | 0.417 | 0.273 | 0.053 | 0.003 | 0.03 | 0.03 | 2.958 | 0.53 | 0.13 | 0.6 | 0.57 | 0.58 | 0.53 |
| 10 | 54526622 | rs10082466 | -0.734 | 0.749 | 0.449 | 0.07 | 0.073 | 0 | 0 | 0 | 0.17 | 0.46 | 0.24 | 0.16 | 0.24 | 0.21 |
| 10 | 54526762 | rs11595876 | 1.161 | 0.699 | -0.58 | -0.063 | -0.022 | 0.038 | 0.038 | 0 | 0.07 | 0.01 | 0.03 | 0 | 0.08 | 0.01 |
| 10 | 54529257 | rs1838065 | 0.196 | 0.607 | -1.019 | -0.024 | -0.015 | -0.083 | -0.083 | 0.034 | 0.39 | 0.08 | 0.4 | 0.45 | 0.39 | 0.3 |
| 10 | 54530636 | rs1982266 | -0.462 | 0.606 | -0.909 | -0.121 | 0.024 | 0.069 | 0.069 | 0 | 0.51 | 0.21 | 0.66 | 0.61 | 0.58 | 0.46 |
| 10 | 54531235 | rs1800450 | 1.11 | 0.824 | -0.921 | -0.085 | -0.036 | -0.03 | -0.03 | 0 | 0.11 | 0.01 | 0.22 | 0.15 | 0.14 | 0.15 |
| 10 | 54531534 | rs11003123 | 0.781 | 0.639 | -0.5 | -0.18 | 0.018 | -0.023 | -0.023 | 0.892 | 0.2 | 0.54 | 0.17 | 0.14 | 0.2 | 0.26 |
| 10 | 54536839 | rs7899547 | -0.806 | 0.502 | -0.457 | -0.153 | -0.009 | 0.027 | 0.027 | 1.602 | 0.65 | 0.85 | 0.66 | 0.66 | 0.65 | 0.6 |
| 10 | 54537089 | rs10824799 | 0.369 | 0.315 | -0.329 | -0.252 | -0.009 | -0.041 | -0.041 | 0.07 | 0.23 | 0.61 | 0.19 | 0.14 | 0.22 | 0.3 |
| 10 | 54538159 | rs11003134 | 0.245 | 0.659 | -0.644 | -0.263 | -0.004 | 0.004 | 0.004 | 0 | 0.13 | 0.12 | 0.16 | 0.1 | 0.15 | 0.16 |
| 10 | 54540783 | rs4935347 | -1.47 | 0.523 | -0.494 | -0.378 | 0.018 | 0.036 | 0.036 | 3.54 | 0.74 | 0.85 | 0.85 | 0.81 | 0.8 | 0.74 |
| 10 | 54542138 | rs7477069 | -0.086 | 0.148 | -0.113 | 0.238 | -0.019 | -0.039 | -0.039 | 2.922 | 0.3 | 0.19 | 0.36 | 0.37 | 0.35 | 0.43 |
| 10 | 54544665 | rs7083840 | 0.854 | 0.242 | 0.136 | 0.03 | 0.024 | 0.001 | 0.001 | 0 | 0.1 | 0.17 | 0.09 | 0.08 | 0.14 | 0.04 |
| 10 | 54544685 | rs12247509 | 1.14 | 0.239 | 0.141 | 0.028 | 0.026 | 0.003 | 0.003 | 0 | 0.09 | 0.13 | 0.08 | 0.08 | 0.14 | 0.04 |
| 10 | 54545202 | rs7474747 | -1.304 | 0.13 | 0.283 | -0.113 | -0.019 | -0.062 | -0.062 | 2.761 | 0.65 | 0.54 | 0.78 | 0.82 | 0.71 | 0.69 |
| 10 | 54545590 | rs10762885 | -0.564 | 0.142 | 0.214 | -0.106 | 0.002 | 0.031 | 0.031 | 2.516 | 0.55 | 0.39 | 0.69 | 0.73 | 0.57 | 0.65 |
| 10 | 54547156 | rs12248290 | 1.093 | 0.135 | 0.126 | -0.031 | 0.024 | 0.002 | 0.002 | 0 | 0.09 | 0.13 | 0.08 | 0.08 | 0.14 | 0.04 |
| 10 | 54547173 | rs11003138 | -0.027 | 0.121 | -0.244 | 0.024 | 0.002 | 0.045 | 0.045 | 2.575 | 0.43 | 0.26 | 0.56 | 0.66 | 0.43 | 0.49 |
| 10 | 54552805 | rs10824804 | 0.528 | 0.091 | -0.227 | 0.29 | 0.018 | -0.005 | -0.005 | 0 | 0.48 | 0.24 | 0.36 | 0.53 | 0.48 | 0.52 |
| 10 | 54554024 | rs4935349 | 0.154 | 0.257 | 0.613 | 0.594 | 0.043 | 0.001 | 0.001 | 0 | 0.86 | 0.61 | 0.86 | 0.9 | 0.86 | 0.96 |
| 10 | 54557568 | rs7923443 | 0.05 | 0.276 | 0.621 | 0.57 | 0.045 | -0.012 | -0.012 | 0 | 0.86 | 0.64 | 0.88 | 0.92 | 0.86 | 0.96 |
| 10 | 54561593 | rs12771797 | 0.053 | 0.271 | 0.461 | 0.566 | 0 | -0.001 | -0.001 | 0 | 0.3 | 0.25 | 0.32 | 0.32 | 0.26 | 0.26 |
| 10 | 54561643 | rs11003158 | 0.081 | 0.141 | 0.448 | 0.583 | -0.004 | -0.017 | -0.017 | 0.936 | 0.26 | 0.26 | 0.27 | 0.4 | 0.3 | 0.3 |
| 10 | 54563015 | rs12254353 | 0.984 | 0.136 | 0.559 | 0.558 | -0.005 | 0.077 | 0.077 | 3.444 | 0.42 | 0.23 | 0.36 | 0.27 | 0.43 | 0.45 |
| 10 | 54563148 | rs11003159 | -0.249 | 0.369 | 0.719 | 0.492 | 0 | -0.001 | -0.001 | 0.061 | 0.3 | 0.25 | 0.32 | 0.32 | 0.26 | 0.26 |
| 10 | 54564407 | rs11003173 | -0.206 | -0.117 | 0.633 | 0.542 | -0.002 | -0.001 | -0.001 | 0 | 0.29 | 0.25 | 0.17 | 0.33 | 0.22 | 0.25 |
| 10 | 54564689 | rs7474987 | -0.068 | -0.003 | 0.124 | 0.604 | -0.033 | -0.006 | -0.006 | 0.061 | 0.68 | 0.52 | 0.8 | 0.67 | 0.77 | 0.76 |
| 10 | 54566151 | rs7095605 | -0.343 | 0.756 | 0.185 | 0.668 | 0.015 | -0.004 | -0.004 | 0 | 0.14 | 0.27 | 0.11 | 0.11 | 0.13 | 0.04 |

| | | | | | | | | | | | | | | | | |
|---|---|---|---|---|---|---|---|---|---|---|---|---|---|---|---|---|
| 10 | 54570851 | rs7917334 | 0.449 | 0.562 | -0.146 | 0.986 | 0.003 | 0.005 | 0.005 | 0 | 0.63 | 0.32 | 0.36 | 0.67 | 0.63 | 0.54 |
| 10 | 54570951 | rs7918389 | 0.486 | 0.598 | -0.2 | 1.174 | -0.03 | -0.074 | -0.074 | 0 | 0.59 | 0.11 | 0.32 | 0.55 | 0.59 | 0.52 |
| 10 | 54571607 | rs11003177 | 0.717 | 0.578 | -0.191 | 1.166 | -0.024 | -0.067 | -0.067 | 0 | 0.6 | 0.15 | 0.32 | 0.58 | 0.59 | 0.53 |
| 10 | 54572138 | rs11003182 | 0.717 | 0.578 | -0.191 | 1.166 | -0.024 | -0.067 | -0.067 | 0.188 | 0.6 | 0.14 | 0.32 | 0.58 | 0.59 | 0.53 |
| 10 | 54574756 | rs6480974 | 0.511 | 0.452 | 0.102 | 1.082 | 0.002 | 0.003 | 0.003 | 0 | 0.62 | 0.39 | 0.39 | 0.67 | 0.64 | 0.55 |
| 10 | 54575721 | rs7068367 | -0.103 | 0.548 | 0.434 | 1.107 | -0.009 | -0.015 | -0.015 | 0.014 | 0.81 | 0.64 | 0.77 | 0.88 | 0.86 | 0.75 |
| 10 | 54583109 | rs11817079 | 0.522 | 0.536 | 0.269 | 1.1 | -0.015 | -0.025 | -0.025 | 0 | 0.16 | 0.08 | 0.36 | 0.24 | 0.22 | 0.21 |
| 10 | 54583602 | rs11003189 | 1.024 | 0.323 | -0.335 | 0.98 | -0.012 | 0.063 | 0.063 | 0 | 0.28 | 0.02 | 0.13 | 0.14 | 0.26 | 0.23 |
| 10 | 54585401 | rs3897993 | -0.39 | 0.521 | 0.227 | 1.184 | -0.012 | -0.066 | -0.066 | 0 | 0.82 | 0.64 | 0.76 | 0.95 | 0.86 | 0.76 |
| 10 | 54586048 | rs4098805 | 0.425 | 0.586 | -0.553 | 1.052 | 0.004 | 0.011 | 0.011 | 0.618 | 0.18 | 0.24 | 0.39 | 0.29 | 0.23 | 0.22 |
| 10 | 54588090 | rs1935924 | 0.916 | 0.586 | -0.609 | 1.106 | -0.026 | -0.081 | -0.081 | 0.144 | 0.61 | 0.14 | 0.32 | 0.54 | 0.59 | 0.54 |
| 10 | 54597592 | rs12764383 | 0.341 | 0.822 | -0.455 | 1.122 | -0.008 | -0.017 | -0.017 | 0 | 0.17 | 0.11 | 0.36 | 0.27 | 0.22 | 0.21 |
| 10 | 54617058 | rs12777905 | 0.424 | 0.838 | -0.849 | 1.089 | -0.039 | -0.051 | -0.051 | 0.18 | 0.16 | 0.04 | 0.35 | 0.24 | 0.22 | 0.21 |
| 10 | 54629374 | rs11003231 | 0.499 | 0.537 | -0.804 | 1.003 | -0.016 | 0.008 | 0.008 | 0.176 | 0.38 | 0.12 | 0.33 | 0.35 | 0.35 | 0.31 |
| 10 | 54635124 | rs7920685 | 0.029 | 0.47 | 0.121 | 1.158 | -0.006 | 0.057 | 0.057 | 0.828 | 0.78 | 0.37 | 0.75 | 0.7 | 0.83 | 0.71 |
| 10 | 54635791 | rs10824826 | 0.644 | 0.433 | -0.107 | 0.925 | 0.007 | 0.021 | 0.021 | 0.321 | 0.39 | 0.2 | 0.36 | 0.36 | 0.35 | 0.32 |
| 10 | 54640441 | rs4147265 | 0.726 | 0.419 | -0.17 | 0.915 | -0.013 | 0.018 | 0.018 | 0.374 | 0.39 | 0.12 | 0.33 | 0.34 | 0.35 | 0.31 |
| 10 | 54642295 | rs10824829 | 0.301 | 0.594 | -0.149 | 1.174 | -0.102 | -0.048 | -0.048 | 0 | 0.38 | 0.05 | 0.37 | 0.39 | 0.48 | 0.4 |
| 10 | 54647063 | rs4935371 | -0.524 | 0.564 | 0.25 | 1.064 | 0.012 | 0.045 | 0.045 | 0 | 0.49 | 0.62 | 0.6 | 0.64 | 0.57 | 0.44 |
| 10 | 54653454 | rs10824831 | -0.337 | 0.466 | 1.031 | 0.812 | 0.044 | -0.014 | -0.014 | 0 | 0.08 | 0.38 | 0.08 | 0.01 | 0.06 | 0.02 |
| 10 | 54653845 | rs7083678 | -0.651 | 0.487 | 0.756 | 0.85 | 0.074 | 0.286 | 0.286 | 0.807 | 0.11 | 0.5 | 0.19 | 0.23 | 0.12 | 0.06 |
| 10 | 54668597 | rs2222547 | -0.317 | 0.207 | -0.1 | 0.135 | -0.005 | -0.034 | -0.034 | 1.348 | 0.12 | 0.16 | 0.06 | 0.01 | 0.07 | 0.26 |
| 10 | 54669514 | rs17590583 | 0.403 | 0.343 | -0.624 | -0.154 | -0.019 | 0.037 | 0.037 | 0.003 | 0.21 | 0.1 | 0.15 | 0.15 | 0.25 | 0.19 |
| 10 | 54669848 | rs7087469 | -0.2 | 0.206 | -0.131 | -0.159 | -0.015 | -0.054 | -0.054 | 1.404 | 0.13 | 0.24 | 0.09 | 0.03 | 0.09 | 0.27 |
| 10 | 54671011 | rs1578778 | 0.688 | 0.442 | -0.55 | 0.028 | 0.001 | 0.028 | 0.028 | 0.122 | 0.52 | 0.24 | 0.62 | 0.56 | 0.49 | 0.42 |
| 10 | 54671654 | rs7093457 | -0.65 | -0.144 | -0.974 | -0.31 | -0.038 | -0.026 | -0.026 | 0.006 | 0.59 | 0.78 | 0.42 | 0.49 | 0.55 | 0.4 |
| 10 | 54672175 | rs2204345 | -0.03 | -0.602 | -1.307 | -0.592 | 0.016 | 0.05 | 0.05 | 0.462 | 0.51 | 0.2 | 0.61 | 0.58 | 0.53 | 0.59 |
| 10 | 54672184 | rs2204344 | -0.378 | -0.407 | -0.514 | -0.404 | 0.068 | 0.151 | 0.151 | 0 | 0.87 | 0.6 | 0.87 | 0.76 | 0.82 | 0.84 |
| 10 | 54675600 | rs17590772 | 0.684 | -0.516 | -1.05 | -0.737 | 0.014 | 0.02 | 0.02 | 0 | 0.11 | 0.02 | 0.1 | 0.07 | 0.08 | 0.06 |
| 10 | 54676411 | rs1343065 | -0.107 | -0.473 | -0.48 | -0.838 | -0.007 | -0.021 | -0.021 | 0.3 | 0.53 | 0.42 | 0.68 | 0.75 | 0.64 | 0.74 |
| 10 | 54680621 | rs1917182 | -1.004 | -0.873 | -0.383 | -0.708 | -0.011 | -0.014 | -0.014 | 0 | 0.93 | 0.67 | 0.97 | 1 | 1 | 1 |
| 10 | 54682395 | rs11003258 | 0.333 | -0.352 | -1.071 | -1.034 | -0.052 | 0.071 | 0.071 | 0 | 0.13 | 0.02 | 0.2 | 0.01 | 0.16 | 0.18 |
| 10 | 54684217 | rs10508977 | 0.44 | -0.554 | -1.417 | -1.259 | 0.01 | 0.037 | 0.037 | 0 | 0.19 | 0.03 | 0.07 | 0.07 | 0.13 | 0.15 |
| 10 | 54689409 | rs10824839 | -0.234 | -0.213 | -1.018 | -1.049 | -0.014 | -0.097 | -0.097 | 0 | 0.59 | 0.5 | 0.68 | 0.94 | 0.71 | 0.67 |
| 10 | 54692621 | rs16935459 | -0.136 | 0.267 | -1.115 | 0.525 | -0.004 | -0.043 | -0.043 | 0 | 0.15 | 0.07 | 0.2 | 0.37 | 0.18 | 0.21 |
| 10 | 54695338 | rs1578777 | -0.227 | 0.022 | -0.709 | -0.157 | -0.002 | -0.038 | -0.038 | 0 | 0.41 | 0.48 | 0.43 | 0.15 | 0.39 | 0.31 |
| 10 | 54695965 | rs16935489 | -0.654 | -0.145 | -0.28 | 0.53 | 0.006 | 0.01 | 0.01 | 0.012 | 0.51 | 0.64 | 0.5 | 0.52 | 0.44 | 0.52 |
| 10 | 54696181 | rs11003269 | -0.093 | 0.046 | -0.485 | 0.444 | -0.011 | -0.007 | -0.007 | 1.346 | 0.27 | 0.16 | 0.3 | 0.21 | 0.27 | 0.22 |
| 10 | 54697623 | rs12571703 | -0.383 | -0.157 | -0.238 | 0.623 | 0.004 | -0.003 | -0.003 | 0 | 0.33 | 0.33 | 0.21 | 0.39 | 0.26 | 0.39 |
| 10 | 54697730 | rs10740529 | 0.137 | -0.064 | -0.514 | 0.479 | 0.02 | -0.04 | -0.04 | 0.099 | 0.85 | 0.61 | 0.69 | 0.92 | 0.85 | 0.93 |
| 10 | 54699867 | rs10824841 | 1.311 | -0.105 | -1.166 | 0.58 | -0.014 | -0.008 | -0.008 | 0 | 0.31 | 0.21 | 0.22 | 0.37 | 0.37 | 0.35 |
| 10 | 54703555 | rs17603971 | -0.3 | -0.403 | -1.149 | 0.859 | 0.021 | -0.009 | -0.009 | 3.904 | 0.27 | 0.16 | 0.18 | 0.32 | 0.23 | 0.39 |
| 10 | 54706817 | rs10824842 | 0.358 | 0.466 | -1.177 | 0.76 | -0.026 | 0.014 | 0.014 | 0 | 0.17 | 0.06 | 0.22 | 0.1 | 0.19 | 0.14 |
| 10 | 54715100 | rs10824843 | 0.054 | -0.006 | -1.057 | 1.07 | 0.015 | -0.041 | -0.041 | 0 | 0.65 | 0.24 | 0.68 | 0.58 | 0.69 | 0.59 |
| 10 | 54716416 | rs11003281 | 0.08 | 0.357 | -0.32 | 1.117 | -0.001 | -0.004 | -0.004 | 0 | 0.78 | 0.78 | 0.75 | 0.87 | 0.78 | 0.8 |
| 10 | 54717446 | rs7077492 | -0.951 | 0.471 | -0.304 | 1.68 | 0.005 | 0.134 | 0.134 | 0 | 0.11 | 0.23 | 0.32 | 0.35 | 0.12 | 0.1 |
| 10 | 54725776 | rs7902775 | -0.266 | 0.018 | -0.796 | 1.125 | 0.046 | 0 | 0 | 0 | 0.53 | 0.69 | 0.41 | 0.48 | 0.35 | 0.31 |
| 10 | 54725850 | rs7916982 | -0.118 | -0.25 | -0.12 | 1.289 | -0.005 | -0.003 | -0.003 | 0 | 0.47 | 0.5 | 0.37 | 0.37 | 0.34 | 0.31 |
| 10 | 54726056 | rs6480983 | 0.018 | -0.443 | 0.157 | 1.387 | 0.017 | 0.01 | 0.01 | 0.042 | 0.44 | 0.39 | 0.33 | 0.35 | 0.32 | 0.3 |
| 10 | 54731281 | rs1343045 | -0.301 | -1.184 | 0.936 | 1.037 | -0.016 | 0.013 | 0.013 | 0 | 0.68 | 0.42 | 0.71 | 0.6 | 0.76 | 0.78 |
| 10 | 54732463 | rs2384101 | -0.126 | -1.319 | 1.32 | 0.919 | -0.016 | -0.001 | -0.001 | 3.355 | 0.17 | 0.24 | 0.09 | 0.16 | 0.13 | 0.14 |
| 10 | 54739291 | rs1539512 | 0.534 | -1.22 | 1.279 | 0.906 | 0.058 | 0.001 | 0.001 | 0.3 | 0.53 | 0.23 | 0.63 | 0.53 | 0.63 | 0.64 |
| 10 | 54744574 | rs10824854 | 0.73 | -1.157 | 1.343 | 0.92 | 0.007 | -0.005 | -0.005 | 0 | 0.26 | 0.17 | 0.24 | 0.28 | 0.22 | 0.17 |
| 10 | 54747748 | rs11003306 | 0.493 | -1.226 | 1.252 | 0.909 | -0.006 | -0.011 | -0.011 | 0.235 | 0.13 | 0.18 | 0.07 | 0.07 | 0.12 | 0.13 |
| 10 | 54753677 | rs1935936 | 0.273 | -0.693 | 1.326 | 0.891 | 0.003 | -0.001 | -0.001 | 0.147 | 0.56 | 0.39 | 0.64 | 0.56 | 0.59 | 0.67 |
| 10 | 54758648 | rs7894017 | 0.238 | -0.729 | 1.304 | 0.898 | -0.012 | 0.005 | 0.005 | 0 | 0.15 | 0.23 | 0.09 | 0.16 | 0.13 | 0.15 |
| 10 | 54761717 | rs7915183 | 0.199 | -0.731 | 1.486 | 0.897 | -0.015 | 0.008 | 0.008 | 0 | 0.15 | 0.27 | 0.09 | 0.15 | 0.13 | 0.16 |
| 10 | 54762533 | rs1917205 | -0.91 | -0.734 | 1.539 | 0.897 | 0 | 0 | 0 | 0.002 | 0.73 | 0.71 | 0.74 | 0.71 | 0.75 | 0.82 |
| 10 | 54765036 | rs2248899 | 0.309 | -0.733 | 1.09 | 0.832 | 0.017 | -0.003 | -0.003 | 0 | 0.56 | 0.29 | 0.62 | 0.56 | 0.6 | 0.67 |
| 10 | 54765608 | rs2249003 | 1.131 | 0.36 | -0.082 | 1.142 | -0.066 | -0.041 | -0.041 | 0 | 0.34 | 0.04 | 0.46 | 0.33 | 0.38 | 0.36 |
| 10 | 54770299 | rs7074652 | 0.585 | 0.455 | 0.02 | 1.095 | -0.007 | -0.013 | -0.013 | 0.345 | 0.14 | 0.18 | 0.07 | 0.07 | 0.12 | 0.13 |
| 10 | 54777403 | rs2253169 | -0.182 | 1.103 | -0.707 | 0.832 | -0.076 | 0.033 | 0.033 | 4.953 | 0.32 | 0.09 | 0.22 | 0.23 | 0.42 | 0.35 |
| 10 | 54789631 | rs12360114 | 0.719 | 1.024 | -0.279 | 1.43 | -0.022 | -0.051 | -0.051 | 0 | 0.25 | 0.07 | 0.26 | 0.3 | 0.25 | 0.17 |
| 10 | 54790431 | rs2265884 | 0.761 | 1.333 | 0.018 | 1.409 | -0.003 | -0.023 | -0.023 | 0.103 | 0.46 | 0.47 | 0.42 | 0.25 | 0.34 | 0.57 |
| 10 | 54794208 | rs1917190 | 0.966 | 1.246 | -0.097 | 1.462 | -0.009 | -0.034 | -0.034 | 0.439 | 0.25 | 0.1 | 0.26 | 0.3 | 0.25 | 0.17 |
| 10 | 54794763 | rs11003342 | -0.771 | 1.19 | 0.246 | 1.926 | -0.002 | -0.002 | -0.002 | 0 | 0.18 | 0.16 | 0.22 | 0.29 | 0.25 | 0.2 |
| 10 | 54798006 | rs2812335 | 0.683 | 1.244 | -0.2 | 1.646 | 0.003 | -0.01 | -0.01 | 1.765 | 0.23 | 0.18 | 0.25 | 0.35 | 0.2 | 0.15 |
| 10 | 54798085 | rs2812336 | -0.65 | 1.19 | -0.275 | 0.911 | -0.003 | -0.004 | -0.004 | 0 | 0.38 | 0.48 | 0.43 | 0.56 | 0.39 | 0.32 |
| 10 | 54799931 | rs1917186 | -1.387 | 1.413 | -0.371 | 0.699 | 0.001 | 0 | 0 | 0 | 0.19 | 0.24 | 0.16 | 0.19 | 0.24 | 0.14 |

| | | | | | | | | | | | | | | | | |
|---|---|---|---|---|---|---|---|---|---|---|---|---|---|---|---|---|
| 10 | 54801847 | rs2812338 | -1.777 | 1.321 | -0.226 | 0.456 | 0.005 | 0.035 | 0.035 | 0 | 0.55 | 0.7 | 0.59 | 0.81 | 0.63 | 0.47 |
| 10 | 54802372 | rs16936758 | -0.636 | 1.343 | -0.215 | 0.53 | -0.027 | -0.033 | -0.033 | 0 | 0.13 | 0.09 | 0.15 | 0.23 | 0.22 | 0.13 |
| 10 | 54802561 | rs7096396 | 0.289 | 1.443 | -0.208 | 0.615 | -0.056 | -0.028 | -0.028 | 0 | 0.11 | 0.01 | 0.13 | 0.1 | 0.19 | 0.18 |
| 10 | 54803809 | rs1319032 | -1.175 | 1.453 | -0.188 | 0.626 | -0.001 | 0.029 | 0.029 | 0 | 0.46 | 0.66 | 0.53 | 0.78 | 0.55 | 0.42 |
| 10 | 54804747 | rs2177610 | -0.172 | 1.46 | -0.067 | 0.55 | -0.013 | -0.006 | -0.006 | 0.086 | 0.32 | 0.23 | 0.4 | 0.61 | 0.38 | 0.34 |
| 10 | 54809993 | rs10824865 | -0.102 | 1.504 | -0.133 | 0.515 | -0.018 | -0.028 | -0.028 | 0 | 0.32 | 0.19 | 0.39 | 0.57 | 0.38 | 0.33 |
| 10 | 54810396 | rs1917173 | 0.165 | 1.477 | -0.172 | 0.525 | -0.028 | -0.035 | -0.035 | 0 | 0.16 | 0.09 | 0.16 | 0.23 | 0.24 | 0.14 |
| 10 | 54810578 | rs1917172 | 1.153 | 1.496 | -0.161 | 0.504 | 0.005 | -0.038 | -0.038 | 0 | 0.21 | 0.16 | 0.25 | 0.48 | 0.19 | 0.15 |
| 10 | 54824741 | rs4142734 | -1.346 | 1.277 | -0.26 | 0.351 | 0.007 | 0.04 | 0.04 | 0 | 0.53 | 0.46 | 0.57 | 0.79 | 0.62 | 0.46 |
| 10 | 54828992 | rs10824875 | -1.349 | 1.297 | -0.371 | 0.361 | 0.001 | -0.009 | -0.009 | 0 | 0.53 | 0.52 | 0.57 | 0.79 | 0.62 | 0.46 |
| 10 | 54831160 | rs7090905 | -0.624 | 1.395 | -0.707 | 0.438 | 0.012 | 0.005 | 0.005 | 2.948 | 0.16 | 0.31 | 0.18 | 0.13 | 0.18 | 0.18 |
| 10 | 54832274 | rs11003362 | 0.342 | 0.701 | -1.208 | 0.495 | -0.037 | 0.1 | 0.1 | 0 | 0.27 | 0.57 | 0.29 | 0.47 | 0.19 | 0.15 |
| 10 | 54837838 | rs4437945 | -0.411 | 1.105 | -0.942 | -0.259 | -0.004 | 0.028 | 0.028 | 0 | 0.55 | 0.69 | 0.54 | 0.79 | 0.54 | 0.42 |
| 10 | 54843499 | rs10762919 | -0.104 | 1.111 | -1.247 | -0.035 | -0.062 | 0.075 | 0.075 | 0 | 0.65 | 0.84 | 0.75 | 0.85 | 0.59 | 0.67 |
| 10 | 54845270 | rs11813961 | 0.772 | -0.152 | -1.402 | 0.462 | 0.03 | -0.031 | -0.031 | 0 | 0.19 | 0.1 | 0.24 | 0.35 | 0.12 | 0.16 |
| 10 | 54850707 | rs2384148 | -0.248 | -0.448 | -0.423 | 0.656 | 0.005 | -0.012 | -0.012 | 0.188 | 0.6 | 0.65 | 0.71 | 0.76 | 0.59 | 0.67 |
| 10 | 54854441 | rs7093161 | 0.196 | -0.235 | -0.495 | 0.654 | -0.029 | 0.032 | 0.032 | 0 | 0.47 | 0.8 | 0.51 | 0.41 | 0.51 | 0.55 |
| 10 | 54856226 | rs4268426 | 1.06 | -0.333 | -0.544 | 0.68 | 0.063 | -0.072 | -0.072 | 0 | 0.18 | 0.04 | 0.24 | 0.34 | 0.11 | 0.13 |
| 10 | 54859973 | rs10824884 | -0.552 | -0.137 | 0.086 | 0.607 | -0.002 | 0.001 | 0.001 | 0 | 0.65 | 0.65 | 0.75 | 0.78 | 0.65 | 0.72 |
| 10 | 54860616 | rs11003383 | 0.896 | 0.408 | -0.058 | 0.763 | 0 | 0.001 | 0.001 | 0.336 | 0.08 | 0.05 | 0.11 | 0.22 | 0.21 | 0.09 |
| 10 | 54861336 | rs12241649 | 0.846 | 0.455 | -0.014 | 0.737 | -0.027 | 0.029 | 0.029 | 0 | 0.32 | 0.19 | 0.23 | 0.22 | 0.34 | 0.28 |
| 10 | 54862560 | rs7914633 | -1.19 | 0.331 | -0.021 | 0.727 | 0.044 | -0.064 | -0.064 | 0 | 0.79 | 0.95 | 0.73 | 0.66 | 0.85 | 0.83 |
| 10 | 54863538 | rs11003386 | 1.229 | 0.097 | 0.216 | 0.545 | 0.012 | 0.007 | 0.007 | 0 | 0.29 | 0.22 | 0.31 | 0.24 | 0.26 | 0.42 |
| 10 | 54864327 | rs10824886 | -0.481 | 0.102 | 0.462 | 0.546 | 0.067 | 0.047 | 0.047 | 0.379 | 0.16 | 0.34 | 0.17 | 0.22 | 0.26 | 0.12 |
| 10 | 54864975 | rs10218945 | -0.142 | 0.085 | 0.396 | 0.505 | -0.005 | -0.005 | -0.005 | 0 | 0.46 | 0.63 | 0.49 | 0.46 | 0.51 | 0.55 |
| 10 | 54865321 | rs10218930 | 1.011 | 0.09 | 0.348 | 0.497 | -0.011 | 0.013 | 0.013 | 0 | 0.32 | 0.24 | 0.23 | 0.21 | 0.33 | 0.28 |
| 10 | 54867617 | rs1904056 | -1.019 | -0.361 | 0.192 | 0.543 | 0.028 | -0.02 | -0.02 | 0 | 0.78 | 0.83 | 0.73 | 0.66 | 0.86 | 0.83 |
| 10 | 54869231 | rs2126756 | 0.989 | -0.148 | 0.193 | 0.6 | 0.013 | 0.007 | 0.007 | 0 | 0.29 | 0.22 | 0.31 | 0.24 | 0.25 | 0.45 |
| 10 | 54879218 | rs16937219 | -0.064 | 0.083 | 0.318 | 0.566 | 0.002 | -0.026 | -0.026 | 0 | 0.05 | 0.12 | 0.05 | 0 | 0.05 | 0.03 |
| 10 | 54879944 | rs2384130 | 0.501 | -0.057 | 0.313 | 0.548 | -0.003 | -0.017 | -0.017 | 0.436 | 0.39 | 0.24 | 0.32 | 0.43 | 0.53 | 0.33 |
| 10 | 54882943 | rs1006678 | -0.064 | -0.085 | 0.372 | 0.567 | 0.014 | 0.001 | 0.001 | 0 | 0.43 | 0.32 | 0.37 | 0.42 | 0.59 | 0.35 |
| 10 | 54883866 | rs11003396 | -1.274 | -0.244 | 0.484 | 0.588 | 0.046 | -0.038 | -0.038 | 0 | 0.79 | 0.91 | 0.73 | 0.66 | 0.86 | 0.82 |
| 10 | 54890625 | rs10824895 | 0.737 | 0.351 | 0.133 | 0.279 | -0.001 | 0 | 0 | 0 | 0.27 | 0.24 | 0.19 | 0.24 | 0.34 | 0.2 |
| 10 | 54901002 | rs1871072 | 1.162 | 0.109 | 0.418 | 0.301 | -0.018 | -0.012 | -0.012 | 0 | 0.32 | 0.41 | 0.32 | 0.25 | 0.26 | 0.43 |
| 10 | 54903766 | rs2384123 | -0.593 | 0.084 | 0.375 | 0.308 | 0.007 | 0.001 | 0.001 | 0 | 0.45 | 0.42 | 0.4 | 0.41 | 0.6 | 0.39 |
| 10 | 54909599 | rs1903927 | 1.157 | 0.235 | 0.313 | 0.312 | -0.045 | -0.03 | -0.03 | 3.028 | 0.14 | 0.04 | 0.16 | 0.18 | 0.21 | 0.13 |
| 10 | 54909607 | rs1903928 | -1.554 | 0.141 | 0.344 | 0.31 | 0.047 | -0.038 | -0.038 | 0 | 0.79 | 0.91 | 0.73 | 0.66 | 0.86 | 0.82 |
| 10 | 54911319 | rs6481001 | 0.128 | 0.304 | 0.608 | 0.281 | 0.024 | 0.005 | 0.005 | 0 | 0.2 | 0.38 | 0.19 | 0.18 | 0.21 | 0.13 |
| 10 | 54914951 | rs1962474 | -0.671 | 0.333 | 0.426 | 0.319 | 0.013 | 0 | 0 | 0 | 0.45 | 0.4 | 0.39 | 0.41 | 0.6 | 0.4 |
| 10 | 54917339 | rs11003430 | 0.368 | 0.43 | 0.463 | 0.318 | 0.031 | -0.018 | -0.018 | 0 | 0.31 | 0.4 | 0.24 | 0.24 | 0.39 | 0.27 |
| 10 | 54924612 | rs1903942 | -1.612 | 0.107 | 0.278 | 0.224 | 0.023 | -0.027 | -0.027 | 0.015 | 0.81 | 0.89 | 0.71 | 0.65 | 0.85 | 0.84 |
| 10 | 54927678 | rs11003436 | 1.668 | -0.09 | 0.271 | 0.019 | 0.001 | 0 | 0 | 0 | 0.26 | 0.24 | 0.16 | 0.25 | 0.2 | 0.41 |
| 10 | 54929003 | rs12782349 | 0.625 | 0.129 | 0.111 | 0.049 | 0.02 | -0.015 | -0.015 | 0 | 0.31 | 0.38 | 0.23 | 0.23 | 0.38 | 0.29 |
| 10 | 54930114 | rs1903935 | -1.399 | 0.159 | -0.414 | 0.074 | 0.07 | -0.081 | -0.081 | 0 | 0.77 | 0.95 | 0.72 | 0.65 | 0.84 | 0.84 |
| 10 | 54930122 | rs12765822 | 0.194 | 0.495 | -0.134 | -0.165 | 0.011 | -0.008 | -0.008 | 0 | 0.31 | 0.3 | 0.23 | 0.23 | 0.38 | 0.29 |
| 10 | 54933879 | rs10762941 | -0.418 | 0.483 | -0.598 | 0.024 | 0.011 | -0.028 | -0.028 | 0 | 0.62 | 0.73 | 0.43 | 0.48 | 0.63 | 0.72 |
| 10 | 54939153 | rs1903916 | 0.574 | 0.426 | -0.866 | -0.906 | -0.011 | 0.191 | 0.191 | 0 | 0.44 | 0.12 | 0.63 | 0.82 | 0.56 | 0.31 |
| 10 | 54940738 | rs951769 | 0.913 | 0.929 | -0.689 | -0.995 | -0.028 | 0.098 | 0.098 | 0 | 0.4 | 0.22 | 0.58 | 0.81 | 0.46 | 0.29 |
| 10 | 54942130 | rs1903922 | -0.117 | 0.637 | -0.346 | -1.236 | 0.028 | 0.076 | 0.076 | 0 | 0.54 | 0.34 | 0.68 | 0.82 | 0.62 | 0.34 |
| 10 | 54942566 | rs1903923 | 0.231 | 0.528 | -0.231 | -1.206 | 0.017 | 0.177 | 0.177 | 0.223 | 0.5 | 0.22 | 0.62 | 0.82 | 0.54 | 0.32 |
| 10 | 54944056 | rs1903926 | 0.744 | 0.583 | -0.38 | -1.16 | -0.035 | 0.145 | 0.145 | 0 | 0.44 | 0.21 | 0.6 | 0.81 | 0.5 | 0.3 |
| 10 | 54944868 | rs4504976 | -0.009 | 0.745 | -0.281 | -1.188 | 0 | -0.004 | -0.004 | 0.703 | 0.34 | 0.35 | 0.28 | 0.18 | 0.34 | 0.55 |
| 10 | 54945433 | rs1903968 | 1.06 | 0.888 | 0.54 | -0.959 | 0.014 | -0.095 | -0.095 | 0 | 0.81 | 0.75 | 0.88 | 1 | 0.81 | 0.83 |
| 10 | 54950997 | rs7477874 | -0.141 | 1.28 | 0.014 | -0.395 | -0.043 | -0.049 | -0.049 | 0.887 | 0.25 | 0.09 | 0.18 | 0.33 | 0.32 | 0.24 |
| 10 | 54955143 | rs4935400 | -0.617 | 1.149 | 0.908 | -0.487 | 0.005 | 0.003 | 0.003 | 0 | 0.29 | 0.25 | 0.2 | 0.32 | 0.32 | 0.24 |
| 10 | 54955296 | rs2384167 | -0.259 | 1.069 | 0.956 | -0.814 | 0.018 | -0.096 | -0.096 | 0 | 0.65 | 0.49 | 0.7 | 0.87 | 0.61 | 0.48 |
| 10 | 54961731 | rs1903920 | 1.483 | 1.112 | 0.992 | -0.75 | 0.039 | -0.042 | -0.042 | 0 | 0.33 | 0.24 | 0.47 | 0.54 | 0.26 | 0.21 |
| 10 | 54962573 | rs10740542 | -0.179 | 1.142 | 1.353 | -0.841 | 0.009 | -0.053 | -0.053 | 0.151 | 0.9 | 0.77 | 0.92 | 1 | 0.91 | 0.89 |
| 10 | 54964412 | rs10824945 | 0.424 | 1.142 | 1.392 | -0.883 | -0.021 | 0.055 | 0.055 | 0 | 0.29 | 0.18 | 0.23 | 0.14 | 0.34 | 0.45 |
| 10 | 54964568 | rs11003448 | 0.321 | 1.156 | 1.449 | -0.833 | 0.01 | 0.013 | 0.013 | 0.744 | 0.24 | 0.23 | 0.17 | 0.32 | 0.29 | 0.22 |
| 10 | 54968630 | rs11003453 | 1.776 | 0.495 | 1.144 | -0.872 | 0.021 | -0.036 | -0.036 | 3.205 | 0.3 | 0.26 | 0.48 | 0.55 | 0.25 | 0.2 |
| 10 | 54981512 | rs1903933 | 0.265 | 0.681 | 1.384 | -0.524 | 0.003 | -0.008 | -0.008 | 0 | 0.58 | 0.46 | 0.48 | 0.73 | 0.56 | 0.46 |
| 10 | 54982181 | rs11003466 | 0.646 | 0.646 | 1.329 | -0.535 | 0 | -0.012 | -0.012 | 0.039 | 0.54 | 0.47 | 0.44 | 0.72 | 0.5 | 0.43 |
| 10 | 54990473 | rs2891447 | 0.471 | 0.512 | 1.418 | -0.838 | 0.005 | -0.054 | -0.054 | 0 | 0.87 | 0.77 | 0.9 | 1 | 0.87 | 0.88 |
| 10 | 54991244 | rs11814008 | 1.527 | 0.445 | 1.16 | -0.857 | 0.042 | -0.072 | -0.072 | 0 | 0.34 | 0.11 | 0.46 | 0.54 | 0.26 | 0.25 |
| 10 | 54991869 | rs10824952 | 0.804 | 0.666 | 1.214 | -0.668 | -0.034 | -0.047 | -0.047 | 0 | 0.2 | 0.09 | 0.17 | 0.32 | 0.29 | 0.22 |
| 10 | 54993618 | rs11003476 | 0.556 | 0.714 | 1.436 | -1.47 | -0.005 | 0.005 | 0.005 | 0 | 0.55 | 0.61 | 0.64 | 0.86 | 0.52 | 0.46 |
| 10 | 54993785 | rs12412557 | 0.243 | 0.585 | 1.047 | -1.387 | -0.035 | 0.087 | 0.087 | 0 | 0.31 | 0.15 | 0.26 | 0.14 | 0.35 | 0.43 |
| 10 | 54995173 | rs1903950 | 0.969 | 0.458 | 1.148 | -1.346 | -0.002 | 0.009 | 0.009 | 0.217 | 0.48 | 0.44 | 0.62 | 0.83 | 0.5 | 0.33 |

| | | | | | | | | | | | | | | | | |
|---|---|---|---|---|---|---|---|---|---|---|---|---|---|---|---|---|
| 10 | 54999514 | rs1871071 | 1.385 | 0.517 | 0.938 | -1.169 | -0.054 | -0.089 | -0.089 | 0.459 | 0.18 | 0.09 | 0.34 | 0.45 | 0.29 | 0.19 |
| 10 | 55009632 | rs7893174 | 2.106 | 0.408 | 1.325 | -1.213 | -0.008 | 0.012 | 0.012 | 0.351 | 0.28 | 0.28 | 0.29 | 0.39 | 0.21 | 0.1 |
| 10 | 55013346 | rs10740544 | 0.143 | 0.336 | 1.355 | -0.981 | -0.012 | -0.028 | -0.028 | 0 | 0.89 | 0.86 | 0.95 | 0.98 | 0.93 | 0.84 |
| 10 | 55021657 | rs7100812 | 1.427 | 0.505 | 1.479 | -0.937 | -0.005 | 0 | 0 | 0.215 | 0.38 | 0.41 | 0.36 | 0.39 | 0.31 | 0.15 |
| 10 | 55022209 | rs7071141 | 0.78 | 0.564 | 0.752 | -0.847 | 0.012 | 0.043 | 0.043 | 2.62 | 0.19 | 0.26 | 0.36 | 0.45 | 0.29 | 0.19 |
| 10 | 55027700 | rs7918231 | 0.13 | 0.719 | 0.619 | -0.943 | 0.031 | 0.074 | 0.074 | 0 | 0.3 | 0.12 | 0.18 | 0.13 | 0.21 | 0.39 |
| 10 | 55030113 | rs10824967 | 0.707 | 0.498 | 0.66 | -0.902 | -0.031 | -0.06 | -0.06 | 0.167 | 0.2 | 0.14 | 0.35 | 0.45 | 0.29 | 0.18 |
| 10 | 55031469 | rs10824971 | 0.577 | 0.528 | 0.676 | -0.889 | 0.011 | 0.044 | 0.044 | 2.881 | 0.2 | 0.27 | 0.36 | 0.45 | 0.29 | 0.19 |
| 10 | 55034244 | rs4935411 | 0.215 | 0.611 | -0.188 | -0.862 | 0.006 | 0.079 | 0.079 | 0 | 0.3 | 0.13 | 0.21 | 0.13 | 0.28 | 0.4 |
| 10 | 55036091 | rs4935412 | -0.455 | 0.524 | -0.007 | -0.989 | 0.039 | 0.044 | 0.044 | 0 | 0.5 | 0.23 | 0.57 | 0.59 | 0.59 | 0.6 |
| 10 | 55037733 | rs12764429 | -0.355 | 0.857 | 0.267 | -0.904 | -0.018 | 0.029 | 0.029 | 0 | 0.07 | 0.01 | 0.05 | 0 | 0.09 | 0.1 |
| 10 | 55044886 | rs7095212 | 0.02 | 1.061 | 0.504 | -0.813 | 0.011 | 0.019 | 0.019 | 2.053 | 0.2 | 0.23 | 0.33 | 0.56 | 0.29 | 0.2 |
| 10 | 55046704 | rs4144618 | 0.314 | 1.054 | 0.465 | -0.816 | -0.017 | -0.02 | -0.02 | 0 | 0.18 | 0.11 | 0.32 | 0.56 | 0.29 | 0.2 |
| 10 | 55047264 | rs10762960 | 1.568 | 0.828 | 0.434 | -1.16 | -0.008 | 0.062 | 0.062 | 1.477 | 0.43 | 0.19 | 0.35 | 0.36 | 0.42 | 0.21 |
| 10 | 55048345 | rs7901256 | 1.3 | 0.69 | 0.616 | -0.745 | -0.002 | 0.034 | 0.034 | 1.41 | 0.52 | 0.39 | 0.41 | 0.36 | 0.48 | 0.29 |
| 10 | 55054817 | rs10824989 | 1.356 | 0.695 | 0.618 | -0.749 | -0.001 | 0.034 | 0.034 | 1.031 | 0.52 | 0.4 | 0.41 | 0.35 | 0.48 | 0.29 |
| 10 | 55055218 | rs4935417 | 0.617 | 0.745 | 0.211 | -0.786 | -0.016 | -0.083 | -0.083 | 0 | 0.71 | 0.6 | 0.75 | 0.9 | 0.77 | 0.48 |
| 10 | 55057721 | rs4261195 | -0.135 | 1.009 | -0.303 | -0.536 | -0.037 | -0.045 | -0.045 | 0 | 0.18 | 0.08 | 0.32 | 0.56 | 0.29 | 0.19 |
| 10 | 55080874 | rs7899729 | -0.31 | 0.365 | -0.229 | -0.56 | 0.005 | 0.065 | 0.065 | 0 | 0.19 | 0.07 | 0.2 | 0.08 | 0.15 | 0.32 |
| 10 | 55093531 | rs2891450 | -1.272 | 0.568 | -0.806 | -0.176 | -0.025 | 0.012 | 0.012 | 1.513 | 0.34 | 0.14 | 0.46 | 0.63 | 0.38 | 0.5 |
| 10 | 55094843 | rs2384205 | -0.843 | 0.001 | -0.822 | -0.45 | 0.019 | 0.059 | 0.059 | 0 | 0.18 | 0.07 | 0.17 | 0.1 | 0.14 | 0.32 |
| 10 | 55104497 | rs9664298 | -0.778 | -0.29 | -0.491 | -0.389 | -0.002 | -0.003 | -0.003 | 0.454 | 0.17 | 0.18 | 0.17 | 0.1 | 0.09 | 0.29 |
| 10 | 55107805 | rs7100563 | -1.405 | -0.443 | 0.069 | -0.924 | -0.012 | -0.051 | -0.051 | 0 | 0.27 | 0.63 | 0.4 | 0.12 | 0.23 | 0.38 |
| 10 | 55109173 | rs12571715 | -1.167 | -0.631 | 0.45 | -0.99 | 0.006 | 0.006 | 0.006 | 0 | 0.13 | 0.1 | 0.27 | 0.11 | 0.1 | 0.31 |
| 10 | 55109587 | rs10825004 | -1.94 | -0.695 | 0.638 | -1.025 | -0.05 | -0.057 | -0.057 | 0 | 0.18 | 0.56 | 0.32 | 0.11 | 0.11 | 0.3 |
| 10 | 55115414 | rs12415081 | -1.92 | -0.719 | 1.168 | -1.037 | -0.045 | -0.05 | -0.05 | 0 | 0.17 | 0.48 | 0.3 | 0.11 | 0.1 | 0.32 |
| 10 | 55116060 | rs12570447 | -1.208 | -0.685 | 1.128 | -1.024 | -0.018 | -0.015 | -0.015 | 0 | 0.13 | 0.23 | 0.26 | 0.11 | 0.08 | 0.3 |
| 10 | 55117633 | rs10508994 | -1.485 | -0.671 | 1.168 | -1.018 | -0.014 | -0.014 | -0.014 | 0.723 | 0.15 | 0.23 | 0.28 | 0.12 | 0.1 | 0.31 |
| 10 | 55121585 | rs11003595 | -2.115 | -0.814 | 1.19 | -1 | -0.031 | -0.071 | -0.071 | 0 | 0.27 | 0.57 | 0.41 | 0.17 | 0.21 | 0.41 |
| 10 | 55124591 | rs1909409 | -0.11 | 0.147 | 0.645 | -1.294 | 0.001 | -0.003 | -0.003 | 0.248 | 0.87 | 0.84 | 0.89 | 0.95 | 0.85 | 0.9 |
| 10 | 55127060 | rs11003596 | 0.059 | 0.171 | 0.641 | -1.289 | 0.007 | -0.005 | -0.005 | 0 | 0.12 | 0.15 | 0.1 | 0.04 | 0.15 | 0.1 |
| 10 | 55146357 | rs1909419 | 0.146 | 0.21 | -0.025 | -2.042 | -0.059 | 0.064 | 0.064 | 0 | 0.79 | 0.91 | 0.84 | 0.94 | 0.73 | 0.81 |
| 10 | 55147591 | rs885507 | 1.181 | 0.007 | 0.271 | -2.167 | -0.026 | 0.059 | 0.059 | 0 | 0.63 | 0.72 | 0.54 | 0.77 | 0.63 | 0.46 |
| 10 | 55156862 | rs2173520 | -0.543 | -0.047 | 0.279 | -2.151 | 0.002 | 0.002 | 0.002 | 0.146 | 0.16 | 0.21 | 0.3 | 0.15 | 0.1 | 0.35 |
| 10 | 55161298 | rs2133471 | 1.763 | 0.198 | 0.247 | -2.528 | 0.039 | 0.188 | 0.188 | 0 | 0.48 | 0.14 | 0.38 | 0.79 | 0.44 | 0.4 |
| 10 | 55162850 | rs12411918 | -0.077 | 0.176 | 0.158 | -2.516 | 0.014 | 0.001 | 0.001 | 0.34 | 0.13 | 0.05 | 0.27 | 0.15 | 0.1 | 0.31 |
| 10 | 55167790 | rs7916768 | -0.136 | 0.504 | 0.173 | -2.475 | -0.061 | 0.052 | 0.052 | 0 | 0.8 | 0.94 | 0.87 | 0.93 | 0.75 | 0.79 |
| 10 | 55173321 | rs17692775 | 2.038 | -0.023 | -0.761 | -2.844 | -0.031 | 0.172 | 0.172 | 0 | 0.31 | 0.03 | 0.16 | 0.09 | 0.3 | 0.13 |
| 10 | 55184722 | rs10762979 | 0.698 | -0.027 | -0.616 | -2.27 | -0.003 | -0.077 | -0.077 | 3.125 | 0.17 | 0.02 | 0.24 | 0.61 | 0.17 | 0.25 |
| 10 | 55186807 | rs12411335 | 0.096 | -0.041 | -0.438 | -2.322 | 0.005 | -0.003 | -0.003 | 1.676 | 0.14 | 0.11 | 0.29 | 0.2 | 0.13 | 0.32 |
| 10 | 55188322 | rs12266210 | 1.578 | -0.196 | 0.299 | -2.456 | -0.006 | -0.057 | -0.057 | 3.805 | 0.38 | 0.34 | 0.2 | 0.09 | 0.31 | 0.13 |
| 10 | 55189979 | rs1198121 | 0.786 | 0.136 | 0.538 | -2.302 | -0.001 | -0.011 | -0.011 | 0 | 0.07 | 0.08 | 0.1 | 0 | 0.14 | 0.07 |
| 10 | 55190288 | rs1198120 | -0.723 | -0.072 | 0.716 | -2.356 | -0.001 | -0.031 | -0.031 | 0 | 0.87 | 0.74 | 0.92 | 0.94 | 0.88 | 0.9 |
| 10 | 55190325 | rs1198119 | -0.732 | 0.114 | 0.509 | -2.19 | -0.034 | 0.018 | 0.018 | 0 | 0.3 | 0.27 | 0.46 | 0.25 | 0.4 | 0.51 |
| 10 | 55195348 | rs11003625 | -0.086 | -0.266 | 0.916 | -1.874 | 0.072 | 0.049 | 0.049 | 0.499 | 0.06 | 0.38 | 0.04 | 0 | 0.01 | 0 |
| 10 | 55196871 | rs7078885 | 0.422 | -0.77 | -0.525 | -2.19 | 0.009 | 0.011 | 0.011 | 0 | 0.39 | 0.27 | 0.34 | 0.25 | 0.28 | 0.36 |
| 10 | 55198707 | rs405588 | -0.77 | 0.949 | -0.903 | -2.445 | -0.055 | 0.052 | 0.052 | 0.044 | 0.9 | 0.98 | 0.85 | 0.99 | 0.72 | 0.87 |
| 10 | 55199473 | rs10825023 | 0.755 | 0.708 | -0.557 | -1.715 | -0.023 | -0.031 | -0.031 | 0 | 0.4 | 0.43 | 0.36 | 0.24 | 0.28 | 0.35 |
| 10 | 55200488 | rs1487469 | 1.812 | 0.165 | -0.956 | -1.99 | -0.062 | -0.047 | -0.047 | 0 | 0.23 | 0.25 | 0.08 | 0.08 | 0.09 | 0.09 |
| 10 | 55201785 | rs12251298 | 1.636 | 0.177 | -0.964 | -2 | -0.056 | -0.045 | -0.045 | 0.071 | 0.24 | 0.25 | 0.08 | 0.08 | 0.1 | 0.09 |
| 10 | 55202802 | rs7903405 | 1.155 | 0.415 | -0.794 | -2.044 | 0.062 | -0.056 | -0.056 | 0 | 0.68 | 0.48 | 0.53 | 0.82 | 0.53 | 0.61 |
| 10 | 55205997 | rs17620281 | 0.767 | 0.377 | -1.109 | -1.852 | -0.026 | -0.021 | -0.021 | 0.003 | 0.13 | 0.03 | 0.25 | 0.16 | 0.16 | 0.25 |
| 10 | 55216633 | rs1198114 | -0.702 | 0.092 | -1.468 | -0.461 | -0.094 | -0.114 | -0.114 | 0 | 0.7 | 0.91 | 0.56 | 0.46 | 0.55 | 0.67 |
| 10 | 55216772 | rs1198113 | 0.182 | 0.122 | -1.073 | -0.679 | 0 | -0.003 | -0.003 | 0 | 0.19 | 0.27 | 0.29 | 0.15 | 0.19 | 0.27 |
| 10 | 55219934 | rs351842 | -1.687 | 0.372 | -1.133 | -0.652 | -0.025 | 0.025 | 0.025 | 0.218 | 0.9 | 0.97 | 0.84 | 0.99 | 0.71 | 0.88 |
| 10 | 55221282 | rs10825025 | 1.969 | -0.35 | -0.94 | -0.764 | -0.037 | -0.043 | -0.043 | 1.333 | 0.25 | 0.24 | 0.09 | 0.08 | 0.12 | 0.11 |
| 10 | 55227059 | rs351833 | 0.189 | 0.254 | -1.109 | -0.519 | 0.008 | -0.01 | -0.01 | 0.695 | 0.17 | 0.37 | 0.28 | 0.15 | 0.16 | 0.26 |
| 10 | 55227588 | rs16937445 | 0.51 | 0.268 | -1.148 | -0.516 | 0 | -0.001 | -0.001 | 0 | 0.15 | 0.2 | 0.25 | 0.15 | 0.16 | 0.25 |
| 10 | 55229956 | rs10825026 | 1.231 | 0.231 | -1.301 | -0.441 | 0.017 | -0.086 | -0.086 | 0 | 0.21 | 0.08 | 0.33 | 0.54 | 0.18 | 0.21 |
| 10 | 55233826 | rs17620681 | 1.639 | 0.588 | -1.501 | -0.404 | -0.049 | 0.039 | 0.039 | 0 | 0.19 | 0.06 | 0.12 | 0.06 | 0.24 | 0.23 |
| 10 | 55235962 | rs370244 | -1.482 | 1.195 | -1.062 | -0.063 | -0.043 | 0.043 | 0.043 | 0.171 | 0.91 | 0.98 | 0.85 | 0.99 | 0.73 | 0.89 |
| 10 | 55236073 | rs16937446 | 0.131 | 0.782 | -0.927 | 0.137 | -0.001 | 0.001 | 0.001 | 0 | 0.14 | 0.18 | 0.25 | 0.24 | 0.15 | 0.26 |
| 10 | 55241540 | rs7916009 | 0.628 | 0.538 | -0.8 | 0.294 | -0.004 | -0.007 | -0.007 | 0 | 0.54 | 0.45 | 0.27 | 0.33 | 0.4 | 0.44 |
| 10 | 55247985 | rs1685295 | -0.242 | 0.215 | -0.55 | 0.245 | 0.016 | 0.035 | 0.035 | 0 | 0.16 | 0.33 | 0.26 | 0.24 | 0.18 | 0.25 |
| 10 | 55248100 | rs626723 | 0.064 | 0.253 | -0.618 | 0.261 | 0.015 | 0.028 | 0.028 | 0 | 0.14 | 0.23 | 0.26 | 0.25 | 0.18 | 0.25 |
| 10 | 55249035 | rs445637 | -0.605 | 0.116 | -0.782 | 0.095 | -0.008 | -0.011 | -0.011 | 1.207 | 0.36 | 0.49 | 0.6 | 0.74 | 0.33 | 0.45 |
| 10 | 55249446 | rs11003639 | 0.775 | -0.67 | -1.711 | -0.439 | -0.054 | 0.109 | 0.109 | 0 | 0.31 | 0.01 | 0.16 | 0.07 | 0.29 | 0.26 |
| 10 | 55250151 | rs7913433 | 0.397 | -1.131 | -0.861 | -0.119 | -0.027 | 0.072 | 0.072 | 0.007 | 0.2 | 0.35 | 0.32 | 0.5 | 0.12 | 0.21 |
| 10 | 55252387 | rs12779337 | -0.714 | -0.739 | -0.664 | -0.316 | 0.048 | -0.015 | -0.015 | 0.011 | 0.07 | 0.34 | 0.09 | 0.01 | 0.08 | 0.01 |

| | | | | | | | | | | | | | | | | |
|---|---|---|---|---|---|---|---|---|---|---|---|---|---|---|---|---|
| 10 | 55268006 | rs10825038 | 0.984 | -1.085 | -0.209 | -0.166 | -0.019 | 0.043 | 0.043 | 0 | 0.16 | 0.16 | 0.26 | 0.39 | 0.09 | 0.2 |
| 10 | 55269440 | rs12775493 | 0.406 | -1.101 | -0.223 | -0.158 | -0.05 | 0.083 | 0.083 | 0 | 0.19 | 0.31 | 0.27 | 0.39 | 0.1 | 0.19 |
| 10 | 55277005 | rs10825046 | 0.862 | -0.991 | -0.75 | -0.405 | -0.01 | -0.004 | -0.004 | 0.181 | 0.26 | 0.23 | 0.21 | 0.26 | 0.34 | 0.29 |
| 10 | 55278234 | rs10825047 | 0.529 | -1.085 | -0.983 | -0.671 | -0.002 | 0 | 0 | 0 | 0.3 | 0.27 | 0.27 | 0.4 | 0.22 | 0.36 |
| 10 | 55293890 | rs4935444 | -0.078 | -1.109 | -0.99 | -0.685 | 0 | 0 | 0 | 0 | 0.54 | 0.55 | 0.5 | 0.65 | 0.57 | 0.66 |
| 10 | 55294187 | rs4935078 | 0.146 | -1.184 | -1.04 | -0.835 | -0.023 | -0.015 | -0.015 | 0 | 0.34 | 0.48 | 0.36 | 0.33 | 0.29 | 0.37 |
| 10 | 55294874 | rs583012 | -1.151 | -1.083 | -1.308 | -0.898 | -0.12 | 0.171 | 0.171 | 0 | 0.75 | 0.97 | 0.78 | 0.98 | 0.66 | 0.86 |
| 10 | 55294933 | rs10825060 | 1.218 | -1.099 | -1.296 | -0.886 | -0.017 | -0.012 | -0.012 | 0.022 | 0.27 | 0.23 | 0.22 | 0.27 | 0.36 | 0.31 |
| 10 | 55295059 | rs422296 | -0.503 | -1.192 | -1.239 | -0.815 | 0.108 | 0.071 | 0.071 | 0 | 0.48 | 0.74 | 0.56 | 0.71 | 0.3 | 0.55 |
| 10 | 55296717 | rs445739 | -0.833 | -0.928 | -1.599 | -0.695 | 0.028 | 0.045 | 0.045 | 0 | 0.55 | 0.93 | 0.63 | 0.72 | 0.39 | 0.57 |
| 10 | 55301336 | rs7896942 | 1.196 | -1.356 | -1.048 | -0.388 | -0.027 | 0.104 | 0.104 | 0 | 0.17 | 0.26 | 0.26 | 0.4 | 0.1 | 0.2 |
| 10 | 55301663 | rs421811 | 1.244 | -1.104 | -1.563 | -0.261 | -0.065 | -0.07 | -0.07 | 1.047 | 0.25 | 0.08 | 0.17 | 0.26 | 0.3 | 0.3 |
| 10 | 55301827 | rs12255740 | 0.305 | -1.22 | -1.061 | -0.268 | 0.005 | -0.001 | -0.001 | 0 | 0.27 | 0.19 | 0.26 | 0.32 | 0.19 | 0.33 |
| 10 | 55308157 | rs406255 | -0.357 | -1.286 | -0.875 | 0.031 | -0.026 | 0.04 | 0.04 | 2.804 | 0.21 | 0.72 | 0.32 | 0.37 | 0.08 | 0.23 |
| 10 | 55318856 | rs689581 | -0.579 | -1.001 | -1.133 | -0.305 | -0.048 | 0.063 | 0.063 | 0 | 0.67 | 0.78 | 0.67 | 0.81 | 0.5 | 0.71 |
| 10 | 55331574 | rs1118489 | 0.802 | -1.351 | -1.023 | -0.059 | -0.007 | 0.031 | 0.031 | 0 | 0.16 | 0.14 | 0.35 | 0.35 | 0.09 | 0.36 |
| 10 | 55333463 | rs16937581 | 1.139 | -1.503 | -1.561 | -0.137 | -0.004 | 0.001 | 0.001 | 1.165 | 0.18 | 0.12 | 0.06 | 0.12 | 0.08 | 0.07 |
| 10 | 55333926 | rs391865 | 0.019 | -0.85 | -1.84 | -0.312 | 0.1 | -0.019 | -0.019 | 0 | 0.44 | 0.72 | 0.52 | 0.5 | 0.21 | 0.44 |
| 10 | 55334005 | rs10825066 | 0.109 | 0.289 | -1.979 | 0.025 | -0.104 | -0.115 | -0.115 | 0.591 | 0.25 | 0.02 | 0.19 | 0.31 | 0.33 | 0.28 |
| 10 | 55338875 | rs448075 | -0.315 | 0.253 | -1.512 | -0.995 | -0.03 | 0.043 | 0.043 | 0.237 | 0.69 | 0.76 | 0.64 | 0.93 | 0.58 | 0.8 |
| 10 | 55346502 | rs4935450 | 0.208 | -0.694 | -1.865 | -1.602 | -0.025 | 0.206 | 0.206 | 0 | 0.44 | 0.15 | 0.48 | 0.88 | 0.45 | 0.74 |
| 10 | 55355543 | rs11003694 | -0.212 | -0.961 | -2.216 | -1.482 | 0.092 | 0.127 | 0.127 | 2.048 | 0.52 | 0.76 | 0.33 | 0.29 | 0.36 | 0.46 |
| 10 | 55365816 | rs10825072 | 0.322 | -0.993 | -1.488 | -1.57 | -0.003 | -0.014 | -0.014 | 0 | 0.4 | 0.49 | 0.44 | 0.7 | 0.35 | 0.42 |
| 10 | 55367569 | rs1489843 | 0.734 | -1.027 | -1.521 | -1.611 | -0.051 | 0.083 | 0.083 | 0 | 0.32 | 0.19 | 0.37 | 0.07 | 0.43 | 0.2 |
| 10 | 55369981 | rs2130373 | -0.038 | -1.082 | -1.13 | -1.571 | 0.053 | 0.083 | 0.083 | 2.797 | 0.48 | 0.3 | 0.67 | 0.7 | 0.63 | 0.54 |
| 10 | 55371601 | rs10825073 | 0.765 | -1.071 | -1.032 | -1.52 | 0.011 | 0.01 | 0.01 | 0 | 0.27 | 0.26 | 0.18 | 0.23 | 0.21 | 0.38 |
| 10 | 55384743 | rs12256830 | 0.539 | -1.411 | -0.988 | -1.509 | -0.024 | -0.01 | -0.01 | 0 | 0.09 | 0.13 | 0.03 | 0.05 | 0.03 | 0.05 |
| 10 | 55389030 | rs9416286 | -1.369 | -1.032 | -0.783 | -1.282 | -0.026 | -0.044 | -0.044 | 0 | 0.84 | 0.91 | 0.7 | 0.36 | 0.79 | 0.66 |
| 10 | 55390656 | rs16937609 | 0.85 | -1.158 | -0.556 | -1.394 | -0.046 | -0.087 | -0.087 | 0.097 | 0.25 | 0.35 | 0.14 | 0.06 | 0.15 | 0.08 |
| 10 | 55391418 | rs12778875 | 0.71 | -1.114 | -0.514 | -1.359 | 0.008 | 0.024 | 0.024 | 0.188 | 0.19 | 0.29 | 0.32 | 0.65 | 0.21 | 0.34 |
| 10 | 55391609 | rs7906396 | -0.818 | -1.115 | -0.525 | -1.36 | 0.022 | 0.061 | 0.061 | 0.268 | 0.81 | 0.6 | 0.66 | 0.35 | 0.79 | 0.66 |
| 10 | 55394423 | rs16937617 | 0.951 | -1.098 | -0.557 | -1.353 | -0.015 | -0.027 | -0.027 | 0 | 0.18 | 0.13 | 0.31 | 0.65 | 0.21 | 0.34 |
| 10 | 55394615 | rs11003722 | -0.214 | -1.132 | -0.834 | -1.429 | -0.008 | -0.027 | -0.027 | 0 | 0.43 | 0.47 | 0.44 | 0.72 | 0.36 | 0.42 |
| 10 | 55402784 | rs10762999 | -0.888 | -1.019 | -0.858 | -1.37 | 0.011 | 0.029 | 0.029 | 0.322 | 0.81 | 0.68 | 0.66 | 0.34 | 0.78 | 0.66 |
| 10 | 55405246 | rs17696599 | 1.412 | -1.236 | -0.506 | -1.47 | 0.021 | 0.044 | 0.044 | 5.422 | 0.2 | 0.08 | 0.11 | 0.05 | 0.14 | 0.08 |
| 10 | 55405696 | rs7067829 | -0.863 | -1.185 | -0.481 | -1.34 | 0.011 | 0.023 | 0.023 | 0.293 | 0.81 | 0.67 | 0.66 | 0.33 | 0.78 | 0.66 |
| 10 | 55407368 | rs1080097 | -0.014 | -1.447 | -0.656 | -1.544 | 0.001 | 0.013 | 0.013 | 0 | 0.59 | 0.57 | 0.52 | 0.18 | 0.61 | 0.31 |
| 10 | 55407502 | rs1080098 | 0.126 | -1.434 | -0.675 | -1.54 | 0 | 0.013 | 0.013 | 0 | 0.6 | 0.58 | 0.52 | 0.18 | 0.61 | 0.31 |
| 10 | 55410001 | rs7100273 | -0.299 | -1.377 | -0.699 | -1.529 | -0.023 | 0.128 | 0.128 | 0.092 | 0.4 | 0.16 | 0.46 | 0.82 | 0.4 | 0.69 |
| 10 | 55417490 | rs7923617 | 0.706 | -1.392 | -0.878 | -1.493 | 0.017 | 0.037 | 0.037 | 0.064 | 0.22 | 0.07 | 0.14 | 0.16 | 0.18 | 0.34 |
| 10 | 55418087 | rs11003740 | 1.054 | -1.222 | -0.779 | -1.37 | -0.035 | -0.055 | -0.055 | 0.218 | 0.17 | 0.09 | 0.32 | 0.66 | 0.22 | 0.34 |
| 10 | 55439071 | rs12355616 | 0.729 | -1.476 | -0.59 | -1.551 | -0.07 | -0.123 | -0.123 | 0 | 0.24 | 0.4 | 0.13 | 0.06 | 0.13 | 0.09 |
| 10 | 55446058 | rs1686681 | -0.812 | -1.428 | -0.846 | -1.659 | -0.007 | 0.01 | 0.01 | 3.506 | 0.61 | 0.44 | 0.58 | 0.82 | 0.51 | 0.73 |
| 10 | 55447975 | rs1489825 | 1.187 | -1.353 | -1.059 | -1.603 | -0.063 | -0.086 | -0.086 | 0 | 0.17 | 0.04 | 0.3 | 0.66 | 0.22 | 0.35 |
| 10 | 55461257 | rs17697532 | 0.592 | -1.341 | -1.034 | -1.596 | 0.019 | 0.047 | 0.047 | 0.109 | 0.23 | 0.06 | 0.15 | 0.15 | 0.17 | 0.34 |
| 10 | 55470411 | rs12775516 | 1.482 | -1.412 | -1.022 | -1.657 | 0.033 | 0.083 | 0.083 | 5.464 | 0.2 | 0.06 | 0.1 | 0 | 0.13 | 0.04 |
| 10 | 55473675 | rs1489838 | 0.417 | -1.387 | -1.064 | -1.661 | 0.027 | 0.047 | 0.047 | 0.024 | 0.24 | 0.11 | 0.16 | 0.15 | 0.18 | 0.34 |
| 10 | 55477195 | rs1954343 | 0.129 | -1.231 | -0.924 | -1.675 | -0.007 | -0.009 | -0.009 | 0 | 0.4 | 0.39 | 0.4 | 0.68 | 0.33 | 0.39 |
| 10 | 55479314 | rs1906928 | -0.917 | -0.975 | -0.88 | -1.754 | 0.012 | -0.023 | -0.023 | 0 | 0.65 | 0.58 | 0.59 | 0.83 | 0.53 | 0.73 |
| 10 | 55480316 | rs12784193 | 1.271 | -0.815 | -1.197 | -1.774 | 0.017 | 0.114 | 0.114 | 6.354 | 0.19 | 0.01 | 0.09 | 0 | 0.13 | 0.03 |
| 10 | 55483713 | rs825786 | -0.88 | -0.548 | -0.48 | -1.496 | 0.005 | 0.025 | 0.025 | 0 | 0.47 | 0.58 | 0.54 | 0.86 | 0.43 | 0.69 |
| 10 | 55499548 | rs1418889 | 0.143 | -0.207 | 0.012 | -1.25 | -0.027 | 0.023 | 0.023 | 1.104 | 0.79 | 0.94 | 0.82 | 0.81 | 0.77 | 0.7 |
| 10 | 55520279 | rs12251255 | 0.082 | -0.023 | -0.501 | -1.234 | -0.001 | -0.023 | -0.023 | 0 | 0.15 | 0.15 | 0.12 | 0.04 | 0.16 | 0.14 |
| 10 | 55526557 | rs7922420 | -1.172 | 0.771 | -1.195 | -1.365 | -0.086 | 0.137 | 0.137 | 0 | 0.78 | 0.98 | 0.82 | 1 | 0.7 | 0.93 |
| 10 | 55535920 | rs7914477 | 0.02 | 0.738 | -0.362 | -0.179 | -0.066 | -0.086 | -0.086 | 0.666 | 0.13 | 0.02 | 0.3 | 0.65 | 0.17 | 0.35 |
| 10 | 55539871 | rs12241129 | -1.151 | 0.469 | 0.237 | -0.118 | 0.012 | -0.006 | -0.006 | 0 | 0.37 | 0.7 | 0.3 | 0.45 | 0.3 | 0.37 |
| 10 | 55539954 | rs12783120 | -1.151 | 0.403 | 1.106 | 0.049 | -0.023 | 0.114 | 0.114 | 0 | 0.17 | 0.38 | 0.14 | 0.4 | 0.1 | 0.22 |
| 10 | 55540207 | rs16937734 | -1.134 | 0.406 | 1.107 | 0.05 | -0.022 | 0.116 | 0.116 | 0 | 0.17 | 0.38 | 0.14 | 0.4 | 0.1 | 0.22 |
| 10 | 55542453 | rs11003834 | -1.183 | 0.365 | 1.271 | 0.104 | -0.021 | 0.099 | 0.099 | 0 | 0.16 | 0.41 | 0.15 | 0.39 | 0.09 | 0.22 |
| 10 | 55545455 | rs12250304 | -0.748 | 0.369 | 1.213 | -0.29 | -0.021 | 0.01 | 0.01 | 0 | 0.13 | 0.27 | 0.09 | 0.11 | 0.07 | 0.11 |
| 10 | 55546323 | rs12253911 | -0.748 | 0.369 | 1.213 | -0.308 | -0.021 | 0.008 | 0.008 | 0 | 0.13 | 0.28 | 0.09 | 0.11 | 0.07 | 0.11 |
| 10 | 55548958 | rs11003842 | -0.307 | 0.619 | 0.881 | -0.31 | -0.05 | -0.038 | -0.038 | 0 | 0.16 | 0.04 | 0.33 | 0.84 | 0.18 | 0.48 |
| 10 | 55559110 | rs7478212 | -0.278 | 0.285 | 1.198 | -0.323 | -0.05 | -0.024 | -0.024 | 0 | 0.16 | 0.05 | 0.34 | 0.83 | 0.18 | 0.45 |
| 10 | 55563013 | rs11003860 | -0.296 | 0.355 | 0.736 | -0.236 | -0.05 | -0.042 | -0.042 | 0 | 0.16 | 0.07 | 0.34 | 0.81 | 0.18 | 0.45 |
| 10 | 55564846 | rs11003862 | -1.668 | 0.248 | 0.792 | -0.347 | 0.007 | 0.063 | 0.063 | 0 | 0.28 | 0.27 | 0.41 | 0.95 | 0.24 | 0.56 |
| 10 | 55575155 | rs12783799 | -0.347 | 0.286 | 0.623 | -0.244 | 0.011 | -0.015 | -0.015 | 0 | 0.1 | 0.07 | 0.06 | 0.13 | 0.06 | 0.12 |
| 10 | 55577022 | rs16937774 | -0.123 | 0.29 | 0.739 | -0.092 | -0.008 | -0.047 | -0.047 | 0 | 0.15 | 0.06 | 0.29 | 0.65 | 0.16 | 0.35 |
| 10 | 55579294 | rs10763009 | -1.086 | 0.232 | 0.947 | -0.247 | 0.008 | -0.018 | -0.018 | 1.443 | 0.27 | 0.51 | 0.22 | 0.22 | 0.26 | 0.27 |
| 10 | 55590238 | rs12763311 | -0.397 | 0.232 | 0.431 | -0.179 | 0.008 | -0.012 | -0.012 | 7.338 | 0.1 | 0.06 | 0.07 | 0.11 | 0.07 | 0.11 |

| | | | | | | | | | | | | | | | | |
|---|---|---|---|---|---|---|---|---|---|---|---|---|---|---|---|---|
| 10 | 55590722 | rs11003872 | -1.222 | 0.393 | 0.88 | -0.085 | -0.006 | 0.007 | 0.007 | 0 | 0.34 | 0.25 | 0.43 | 0.72 | 0.37 | 0.5 |
| 10 | 55592455 | rs1900416 | -0.308 | 0.366 | 0.916 | -0.095 | -0.013 | 0.032 | 0.032 | 0 | 0.2 | 0.13 | 0.15 | 0.06 | 0.24 | 0.15 |
| 10 | 55594780 | rs4310525 | -1.19 | 0.39 | 1.125 | -0.135 | 0.002 | -0.004 | -0.004 | 0 | 0.36 | 0.35 | 0.45 | 0.72 | 0.39 | 0.52 |
| 10 | 55623229 | rs11003889 | -0.029 | 0.829 | 1.07 | 0.101 | -0.018 | 0.011 | 0.011 | 0 | 0.16 | 0.09 | 0.13 | 0.01 | 0.23 | 0.14 |
| 10 | 55623309 | rs10825130 | -0.922 | 0.465 | 1.278 | 0.143 | -0.032 | 0.037 | 0.037 | 0.717 | 0.23 | 0.46 | 0.34 | 0.74 | 0.16 | 0.38 |
| 10 | 55632374 | rs12762435 | -0.556 | 0.7 | 1.106 | 0.261 | -0.007 | -0.005 | -0.005 | 1.12 | 0.09 | 0.05 | 0.04 | 0.11 | 0.08 | 0.09 |
| 10 | 55634290 | rs11003901 | -1.177 | 0.548 | 1.799 | 0.641 | 0.015 | 0.137 | 0.137 | 4.881 | 0.05 | 0.25 | 0.06 | 0.11 | 0 | 0.01 |
| 10 | 55634794 | rs4478892 | 1.02 | 1.22 | 1.951 | 0.476 | 0.058 | 0.008 | 0.008 | 0 | 0.73 | 0.48 | 0.73 | 0.75 | 0.67 | 0.71 |
| 10 | 55635024 | rs4478893 | 0.342 | 1.224 | 1.999 | 0.457 | -0.045 | 0.013 | 0.013 | 0 | 0.79 | 0.83 | 0.8 | 0.85 | 0.67 | 0.72 |
| 10 | 55635719 | rs7097365 | 0.402 | 1.224 | 1.993 | 0.457 | 0.012 | -0.004 | -0.004 | 0 | 0.78 | 0.73 | 0.79 | 0.85 | 0.67 | 0.72 |
| 10 | 55645842 | rs11003908 | -1.815 | 1.028 | 1.863 | 0.562 | 0.02 | 0.319 | 0.319 | 3.455 | 0.07 | 0.34 | 0.09 | 0.25 | 0.02 | 0.11 |
| 10 | 55657419 | rs10825157 | 0.842 | 1.027 | 1.793 | 0.483 | 0.058 | -0.057 | -0.057 | 0.042 | 0.74 | 0.44 | 0.76 | 0.89 | 0.68 | 0.82 |
| 10 | 55659824 | rs16913796 | -0.883 | 0.894 | 1.806 | 0.434 | -0.001 | -0.001 | -0.001 | 0 | 0.11 | 0.13 | 0.06 | 0.1 | 0.09 | 0.11 |
| 10 | 55661245 | rs2593126 | -2.515 | 0.866 | 1.984 | 0.47 | -0.028 | 0.09 | 0.09 | 0.013 | 0.2 | 0.65 | 0.18 | 0.36 | 0.11 | 0.23 |
| 10 | 55662156 | rs11003924 | -2.509 | 0.873 | 2.039 | 0.523 | -0.024 | 0.15 | 0.15 | 0 | 0.19 | 0.54 | 0.16 | 0.36 | 0.11 | 0.23 |
| 10 | 55667101 | rs2593103 | 0.125 | 0.821 | 1.793 | 0.396 | -0.044 | 0.03 | 0.03 | 0 | 0.81 | 0.87 | 0.83 | 1 | 0.68 | 0.83 |
| 10 | 55667263 | rs2254115 | -1.172 | 0.748 | 1.815 | 0.447 | -0.003 | -0.003 | -0.003 | 0 | 0.12 | 0.2 | 0.07 | 0.1 | 0.09 | 0.11 |
| 10 | 55668753 | rs7088140 | 1.133 | 0.728 | 1.852 | 0.6 | -0.02 | 0.018 | 0.018 | 0 | 0.69 | 0.68 | 0.75 | 0.87 | 0.6 | 0.74 |
| 10 | 55685073 | rs12412902 | -0.804 | 0.743 | 1.837 | 0.61 | -0.001 | -0.002 | -0.002 | 0 | 0.1 | 0.09 | 0.08 | 0.13 | 0.09 | 0.07 |
| 10 | 55695655 | rs2384337 | 2.128 | 0.708 | 2.009 | 0.668 | -0.024 | -0.004 | -0.004 | 0 | 0.61 | 0.23 | 0.67 | 0.63 | 0.59 | 0.63 |
| 10 | 55698362 | rs724434 | -0.64 | 0.714 | 1.964 | 0.659 | -0.003 | -0.005 | -0.005 | 0 | 0.09 | 0.05 | 0.07 | 0.12 | 0.08 | 0.08 |
| 10 | 55700204 | rs2610837 | 0.581 | 0.743 | 2.227 | 0.641 | -0.012 | -0.019 | -0.019 | 1.948 | 0.76 | 0.79 | 0.82 | 1 | 0.69 | 0.83 |
| 10 | 55701068 | rs10825169 | 1.584 | 0.721 | 2.287 | 0.672 | 0.03 | -0.02 | -0.02 | 0.059 | 0.64 | 0.48 | 0.71 | 0.74 | 0.59 | 0.65 |
| 10 | 55703451 | rs1900444 | 1.116 | 0.659 | 2.012 | 0.574 | 0.021 | -0.032 | -0.032 | 0.014 | 0.71 | 0.57 | 0.78 | 0.86 | 0.67 | 0.73 |
| 10 | 55704488 | rs10740560 | 1.075 | 0.682 | 1.993 | 0.584 | 0.022 | -0.031 | -0.031 | 0 | 0.71 | 0.58 | 0.79 | 0.86 | 0.67 | 0.73 |
| 10 | 55717918 | rs2610919 | 0.533 | 0.764 | 1.842 | 0.57 | -0.005 | -0.032 | -0.032 | 0 | 0.77 | 0.78 | 0.84 | 1 | 0.71 | 0.85 |
| 10 | 55720024 | rs2446609 | 0.871 | 0.875 | 1.809 | 0.592 | 0.007 | -0.013 | -0.013 | 0 | 0.73 | 0.67 | 0.82 | 0.86 | 0.7 | 0.74 |
| 10 | 55735463 | rs2660175 | 1.605 | 0.859 | 1.614 | 0.513 | 0.019 | 0.047 | 0.047 | 0 | 0.53 | 0.44 | 0.4 | 0.14 | 0.46 | 0.28 |
| 10 | 55736655 | rs2610895 | 1.364 | 0.588 | 1.652 | 0.521 | 0.003 | 0.01 | 0.01 | 0 | 0.55 | 0.49 | 0.44 | 0.24 | 0.5 | 0.39 |
| 10 | 55744510 | rs10825183 | -0.099 | 0.68 | 1.829 | 1.402 | 0.007 | 0.078 | 0.078 | 0 | 0.17 | 0.22 | 0.15 | 0.38 | 0.19 | 0.4 |
| 10 | 55745211 | rs2660181 | 1.266 | 0.642 | 1.855 | 1.305 | 0 | 0.002 | 0.002 | 0 | 0.58 | 0.67 | 0.46 | 0.24 | 0.51 | 0.4 |
| 10 | 55746740 | rs7077514 | -1.084 | 0.627 | 1.913 | 1.312 | 0.018 | 0.172 | 0.172 | 2.363 | 0.05 | 0.32 | 0.03 | 0.11 | 0.01 | 0.01 |
| 10 | 55747792 | rs7094237 | -1.211 | 0.634 | 1.868 | 1.302 | -0.006 | 0.037 | 0.037 | 0.009 | 0.24 | 0.58 | 0.21 | 0.6 | 0.21 | 0.46 |
| 10 | 55752601 | rs1911420 | -0.093 | 0.687 | 1.699 | 1.317 | -0.083 | 0.085 | 0.085 | 0.962 | 0.81 | 0.96 | 0.77 | 0.94 | 0.74 | 0.9 |
| 10 | 55752717 | rs11003974 | -1.83 | 0.59 | 1.841 | 1.364 | 0.007 | 0.166 | 0.166 | 4.475 | 0.07 | 0.42 | 0.04 | 0.11 | 0.01 | 0.01 |
| 10 | 55754940 | rs6481053 | -0.236 | 0.627 | 1.717 | 1.37 | 0.008 | 0.093 | 0.093 | 0 | 0.18 | 0.23 | 0.19 | 0.5 | 0.2 | 0.45 |
| 10 | 55755491 | rs2135720 | -0.213 | 0.627 | 1.713 | 1.37 | 0.01 | 0.11 | 0.11 | 0 | 0.18 | 0.24 | 0.19 | 0.5 | 0.2 | 0.45 |
| 10 | 55757502 | rs1911413 | -0.204 | 0.657 | 1.718 | 1.417 | 0.011 | 0.089 | 0.089 | 0 | 0.18 | 0.23 | 0.19 | 0.44 | 0.2 | 0.44 |
| 10 | 55766507 | rs2660178 | 0.743 | 0.627 | 1.636 | 1.52 | -0.065 | 0.003 | 0.003 | 0.061 | 0.75 | 0.93 | 0.65 | 0.7 | 0.7 | 0.84 |
| 10 | 55766609 | rs7918590 | -1.313 | 0.437 | 1.849 | 1.566 | 0.021 | 0.207 | 0.207 | 2.468 | 0.05 | 0.34 | 0.03 | 0.14 | 0.01 | 0.02 |
| 10 | 55768822 | rs2660149 | -2.229 | 0.448 | 1.584 | 1.539 | -0.04 | 0.247 | 0.247 | 4.385 | 0.12 | 0.57 | 0.08 | 0.25 | 0.05 | 0.12 |
| 10 | 55770929 | rs11598849 | -0.007 | 0.835 | 0.94 | 1.238 | -0.057 | 0.06 | 0.06 | 0 | 0.16 | 0.04 | 0.22 | 0.06 | 0.24 | 0.12 |
| 10 | 55774649 | rs2660141 | 0.253 | 0.806 | 1.369 | 1.574 | 0.005 | 0.032 | 0.032 | 0.175 | 0.82 | 0.8 | 0.79 | 0.57 | 0.77 | 0.58 |
| 10 | 55798725 | rs16937872 | -1.729 | 0.625 | 1.545 | 1.764 | -0.023 | 0.255 | 0.255 | 0.007 | 0.09 | 0.42 | 0.06 | 0.25 | 0.03 | 0.17 |
| 10 | 55801554 | rs1911402 | -0.156 | 0.864 | 1.289 | 1.5 | -0.015 | 0.013 | 0.013 | 0 | 0.15 | 0.12 | 0.22 | 0.06 | 0.22 | 0.1 |
| 10 | 55806316 | rs1911409 | 0.922 | 0.888 | 1.447 | 1.697 | 0.025 | 0.08 | 0.08 | 0.356 | 0.78 | 0.62 | 0.75 | 0.45 | 0.74 | 0.43 |
| 10 | 55819700 | rs1911382 | 1.026 | 0.86 | 1.35 | 1.706 | -0.036 | -0.021 | -0.021 | 0 | 0.7 | 0.76 | 0.63 | 0.62 | 0.56 | 0.54 |
| 10 | 55824973 | rs17643564 | 0.126 | 0.892 | 1.327 | 1.446 | 0.017 | -0.014 | -0.014 | 0 | 0.13 | 0.13 | 0.23 | 0.06 | 0.23 | 0.1 |
| 10 | 55825382 | rs7917052 | 2.26 | 0.64 | 1.413 | 1.381 | 0.026 | 0.075 | 0.075 | 0 | 0.5 | 0.33 | 0.41 | 0.21 | 0.48 | 0.3 |
| 10 | 55825695 | rs2221567 | 2.15 | 0.623 | 1.394 | 1.36 | 0.004 | 0.009 | 0.009 | 0 | 0.51 | 0.44 | 0.45 | 0.21 | 0.49 | 0.31 |
| 10 | 55828241 | rs11004034 | -0.022 | 0.945 | 0.788 | 1.358 | 0.019 | -0.017 | -0.017 | 0.022 | 0.14 | 0.17 | 0.23 | 0.06 | 0.23 | 0.1 |
| 10 | 55832500 | rs4465289 | -0.437 | 0.933 | 0.461 | 1.668 | -0.046 | -0.058 | -0.058 | 1.949 | 0.88 | 0.96 | 0.87 | 0.62 | 0.82 | 0.67 |
| 10 | 55838466 | rs12246412 | -0.172 | 0.643 | 1.079 | 1.577 | 0.016 | 0.066 | 0.066 | 0 | 0.07 | 0.28 | 0.06 | 0.07 | 0.03 | 0.05 |
| 10 | 55840376 | rs10825219 | -0.689 | 0.354 | 0.592 | 1.493 | 0.001 | -0.003 | -0.003 | 0 | 0.11 | 0.08 | 0.13 | 0.26 | 0.04 | 0.08 |
| 10 | 55841321 | rs7921240 | -0.44 | 0.444 | 0.374 | 1.428 | -0.061 | -0.091 | -0.091 | 0 | 0.85 | 0.96 | 0.84 | 0.62 | 0.77 | 0.67 |
| 10 | 55841776 | rs7921902 | 0.5 | 0.478 | 0.348 | 1.349 | -0.115 | -0.115 | -0.115 | 0.57 | 0.72 | 0.9 | 0.62 | 0.57 | 0.53 | 0.58 |
| 10 | 55842152 | rs16937893 | -1.118 | 0.362 | 0.582 | 0.681 | -0.032 | 0.04 | 0.04 | 0 | 0.13 | 0.55 | 0.15 | 0.16 | 0.08 | 0.23 |
| 10 | 55843303 | rs10825223 | 1.204 | 0.383 | 0.55 | 0.677 | -0.028 | 0.021 | 0.021 | 0 | 0.61 | 0.82 | 0.49 | 0.31 | 0.49 | 0.48 |
| 10 | 55847948 | rs1911424 | 1.99 | 0.489 | 0.27 | 0.547 | 0.029 | 0.164 | 0.164 | 0 | 0.45 | 0.19 | 0.33 | 0.17 | 0.41 | 0.24 |
| 10 | 55848977 | rs17644321 | -0.201 | 0.986 | 0.255 | 0.689 | 0.001 | -0.003 | -0.003 | 0 | 0.15 | 0.12 | 0.24 | 0.09 | 0.24 | 0.11 |
| 10 | 55852282 | rs7893476 | 0.509 | 0.454 | 0.227 | 1.118 | -0.057 | -0.107 | -0.107 | 2.296 | 0.8 | 0.92 | 0.73 | 0.41 | 0.75 | 0.54 |
| 10 | 55856732 | rs1911432 | 0.487 | 0.397 | 0.167 | 0.864 | -0.065 | -0.125 | -0.125 | 0 | 0.8 | 0.94 | 0.75 | 0.43 | 0.76 | 0.53 |
| 10 | 55859752 | rs7094268 | 1.024 | 0.596 | 0.313 | -0.03 | -0.004 | -0.026 | -0.026 | 0 | 0.66 | 0.47 | 0.62 | 0.25 | 0.67 | 0.32 |
| 10 | 55866536 | rs1876325 | 0.93 | 0.417 | 0.1 | -0.107 | -0.008 | -0.027 | -0.027 | 0 | 0.64 | 0.47 | 0.64 | 0.25 | 0.67 | 0.31 |
| 10 | 55867735 | rs7080145 | 0.917 | 0.509 | 0.082 | -0.57 | -0.005 | -0.053 | -0.053 | 0.016 | 0.64 | 0.5 | 0.63 | 0.21 | 0.66 | 0.3 |
| 10 | 55870356 | rs12413011 | -0.443 | 0.336 | -0.092 | -1.385 | -0.001 | -0.007 | -0.007 | 0 | 0.26 | 0.23 | 0.27 | 0.86 | 0.21 | 0.45 |
| 10 | 55871271 | rs7074276 | -1.003 | 0.836 | -1.376 | -2.525 | -0.115 | 0.214 | 0.214 | 0.001 | 0.7 | 0.96 | 0.73 | 1 | 0.66 | 0.78 |
| 10 | 55871323 | rs7095441 | -0.358 | 0.943 | -0.763 | -2.847 | -0.046 | 0.096 | 0.096 | 0.013 | 0.61 | 0.75 | 0.66 | 1 | 0.57 | 0.69 |
| 10 | 55876860 | rs4007264 | 1.575 | 0.539 | -0.697 | -3.109 | -0.025 | 0.038 | 0.038 | 0 | 0.22 | 0.14 | 0.31 | 0.12 | 0.31 | 0.13 |

| | | | | | | | | | | | | | | | | |
|---|---|---|---|---|---|---|---|---|---|---|---|---|---|---|---|---|
| 10 | 55878046 | rs11004081 | 0.752 | -0.129 | -0.015 | -3.123 | -0.027 | -0.042 | -0.042 | 0 | 0.09 | 0.14 | 0.05 | 0.01 | 0.03 | 0.11 |
| 10 | 55880531 | rs4935499 | -0.698 | 0.203 | 0.214 | -3.3 | -0.008 | 0.02 | 0.02 | 0 | 0.63 | 0.74 | 0.7 | 0.99 | 0.66 | 0.71 |
| 10 | 55884408 | rs11004085 | -0.035 | 0.397 | 0.177 | -3.204 | 0.004 | -0.055 | -0.055 | 0.051 | 0.52 | 0.45 | 0.63 | 0.98 | 0.63 | 0.59 |
| 10 | 55884789 | rs10825242 | 0.654 | 0.554 | 0.345 | -3.256 | 0.01 | 0.026 | 0.026 | 0 | 0.25 | 0.12 | 0.27 | 0.86 | 0.21 | 0.45 |
| 10 | 55885524 | rs12360522 | 1.663 | 0.647 | 0.227 | -3.238 | -0.056 | 0.058 | 0.058 | 0 | 0.25 | 0.14 | 0.35 | 0.12 | 0.41 | 0.15 |
| 10 | 55886676 | rs7074636 | -0.295 | 0.367 | 0.274 | -3.194 | 0.006 | -0.058 | -0.058 | 0 | 0.47 | 0.55 | 0.36 | 0.02 | 0.34 | 0.39 |
| 10 | 55889237 | rs4935500 | -1.635 | 0.565 | -0.142 | -3.266 | -0.08 | 0.066 | 0.066 | 0 | 0.73 | 0.86 | 0.64 | 0.88 | 0.55 | 0.84 |
| 10 | 55890367 | rs7904409 | -1.398 | 0.212 | -0.23 | -3.276 | -0.097 | 0.089 | 0.089 | 0 | 0.7 | 0.85 | 0.6 | 0.88 | 0.51 | 0.83 |
| 10 | 55892389 | rs4272709 | 0.535 | 0.076 | 0.244 | -3.535 | 0.001 | -0.02 | -0.02 | 0 | 0.24 | 0.23 | 0.29 | 0.88 | 0.25 | 0.39 |
| 10 | 55892402 | rs7914232 | 0.243 | -0.258 | 0.429 | -3.712 | -0.019 | -0.11 | -0.11 | 0.017 | 0.38 | 0.55 | 0.3 | 0.02 | 0.23 | 0.38 |
| 10 | 55898144 | rs4447073 | 0.604 | -0.248 | 0.402 | -3.705 | -0.001 | 0.007 | 0.007 | 0 | 0.25 | 0.19 | 0.28 | 0.88 | 0.25 | 0.39 |
| 10 | 55903108 | rs7907790 | -1.808 | -0.213 | 0.275 | -3.702 | -0.1 | 0.096 | 0.096 | 0 | 0.71 | 0.86 | 0.6 | 0.89 | 0.51 | 0.83 |
| 10 | 55913771 | rs11004102 | 0.763 | -0.283 | 0.548 | -3.705 | -0.026 | -0.041 | -0.041 | 0 | 0.08 | 0.23 | 0.04 | 0.01 | 0.02 | 0.15 |
| 10 | 55914673 | rs6481063 | 0.302 | -0.322 | 0.463 | -3.724 | -0.036 | -0.158 | -0.158 | 0 | 0.36 | 0.44 | 0.29 | 0.02 | 0.23 | 0.39 |
| 10 | 55919780 | rs4539220 | 1.365 | -0.553 | 0.274 | -3.713 | -0.048 | 0.139 | 0.139 | 0 | 0.26 | 0.05 | 0.24 | 0 | 0.26 | 0.16 |
| 10 | 55921305 | rs11004106 | 0.848 | -0.386 | 0.372 | -3.633 | 0.016 | 0.033 | 0.033 | 0 | 0.23 | 0.13 | 0.26 | 0.87 | 0.19 | 0.38 |
| 10 | 55921316 | rs11004107 | 1.339 | -0.548 | 0.29 | -3.605 | 0.004 | 0.003 | 0.003 | 0 | 0.18 | 0.11 | 0.23 | 0.81 | 0.17 | 0.37 |
| 10 | 55922706 | rs10509006 | 0.507 | -1.016 | 0.34 | -3.551 | -0.005 | -0.099 | -0.099 | 2.685 | 0.35 | 0.38 | 0.29 | 0.01 | 0.28 | 0.31 |
| 10 | 55925040 | rs4415663 | 0.437 | -1.012 | 0.046 | -3.549 | -0.013 | -0.156 | -0.156 | 0 | 0.36 | 0.45 | 0.29 | 0.01 | 0.28 | 0.31 |
| 10 | 55935394 | rs10763079 | 0.659 | -0.739 | -0.598 | -3.483 | -0.014 | 0.038 | 0.038 | 0 | 0.35 | 0.69 | 0.3 | 0.05 | 0.41 | 0.26 |
| 10 | 55935933 | rs6481064 | 0.161 | -0.748 | -0.711 | -3.491 | -0.011 | 0.157 | 0.157 | 0 | 0.43 | 0.79 | 0.32 | 0.05 | 0.44 | 0.31 |
| 10 | 55943184 | rs7093302 | -0.536 | -0.672 | -0.69 | -3.309 | 0.002 | 0.049 | 0.049 | 0 | 0.75 | 0.87 | 0.69 | 0.13 | 0.77 | 0.52 |
| 10 | 55948714 | rs1561674 | -0.889 | -1.084 | -0.89 | -3.643 | 0.124 | 0.226 | 0.226 | 0 | 0.45 | 0.74 | 0.49 | 0.93 | 0.3 | 0.66 |
| 10 | 55949151 | rs16905686 | 1.365 | -1.155 | -0.694 | -3.653 | 0.013 | 0.005 | 0.005 | 0 | 0.18 | 0.13 | 0.25 | 0.84 | 0.13 | 0.4 |
| 10 | 55949899 | rs16905691 | 0.336 | -1.204 | -0.452 | -3.698 | 0.004 | 0.004 | 0.004 | 0 | 0.25 | 0.24 | 0.27 | 0.84 | 0.16 | 0.45 |
| 10 | 55949937 | rs10825265 | -0.984 | -1.098 | -0.75 | -3.804 | -0.093 | 0.177 | 0.177 | 0 | 0.69 | 0.9 | 0.7 | 0.97 | 0.66 | 0.76 |
| 10 | 55951332 | rs7082778 | -1.075 | -1.367 | -0.317 | -3.483 | 0.113 | 0.192 | 0.192 | 0 | 0.45 | 0.73 | 0.48 | 0.93 | 0.29 | 0.66 |
| 10 | 55955444 | rs4935502 | 1.224 | -1.312 | -0.238 | -3.408 | 0.012 | 0.004 | 0.004 | 0 | 0.18 | 0.13 | 0.25 | 0.84 | 0.13 | 0.4 |
| 10 | 55955485 | rs7921598 | -0.454 | -1.258 | -0.086 | -3.346 | -0.04 | -0.043 | -0.043 | 0 | 0.2 | 0.53 | 0.21 | 0.09 | 0.14 | 0.21 |
| 10 | 55955610 | rs10825269 | -0.415 | -1.258 | 0.064 | -3.346 | -0.043 | -0.046 | -0.046 | 0 | 0.2 | 0.49 | 0.21 | 0.09 | 0.14 | 0.21 |
| 10 | 55960818 | rs11004134 | -0.369 | -1.256 | 0.201 | -3.344 | -0.042 | -0.033 | -0.033 | 0 | 0.19 | 0.48 | 0.2 | 0.11 | 0.13 | 0.2 |
| 10 | 55964008 | rs2384414 | 0.989 | -1.204 | 0.098 | -3.44 | -0.003 | -0.014 | -0.014 | 0 | 0.71 | 0.41 | 0.71 | 0.86 | 0.76 | 0.71 |
| 10 | 55964451 | rs4082042 | -0.498 | -1.166 | 0.07 | -3.349 | -0.045 | 0.037 | 0.037 | 0.323 | 0.91 | 0.99 | 0.92 | 0.97 | 0.88 | 0.91 |
| 10 | 55970177 | rs10825274 | -0.906 | -1.417 | -0.036 | -3.5 | 0.126 | 0.146 | 0.146 | 0 | 0.42 | 0.78 | 0.36 | 0.84 | 0.22 | 0.62 |
| 10 | 55970290 | rs10825275 | -0.897 | -1.433 | 0.488 | -3.497 | -0.056 | -0.106 | -0.106 | 0.268 | 0.23 | 0.6 | 0.11 | 0.02 | 0.09 | 0.21 |
| 10 | 55971020 | rs11004139 | 0.931 | -1.392 | 0.987 | -3.476 | -0.026 | -0.06 | -0.06 | 0.864 | 0.12 | 0.19 | 0.05 | 0.02 | 0.06 | 0.16 |
| 10 | 55971427 | rs11004141 | 1.237 | -1.429 | 0.743 | -3.504 | 0.019 | 0.002 | 0.002 | 0 | 0.18 | 0.1 | 0.25 | 0.82 | 0.13 | 0.4 |
| 10 | 55971921 | rs10825278 | 0.332 | -1.466 | 1.031 | -3.44 | -0.043 | -0.094 | -0.094 | 0.014 | 0.14 | 0.35 | 0.07 | 0.02 | 0.06 | 0.16 |
| 10 | 55973889 | rs10763086 | 0.087 | -1.29 | 0.613 | -3.38 | 0.009 | 0.052 | 0.052 | 0 | 0.67 | 0.79 | 0.64 | 0.15 | 0.72 | 0.46 |
| 10 | 55985621 | rs7068210 | -0.671 | -0.822 | 0.984 | -3.085 | -0.011 | 0.015 | 0.015 | 0 | 0.77 | 0.89 | 0.7 | 0.16 | 0.77 | 0.6 |
| 10 | 55988979 | rs11004153 | 0.845 | -0.979 | 1.004 | -3.157 | 0.057 | 0.18 | 0.18 | 0 | 0.55 | 0.19 | 0.56 | 0.14 | 0.68 | 0.34 |
| 10 | 55996761 | rs721825 | -0.528 | -0.808 | 0.991 | -2.986 | -0.005 | 0.011 | 0.011 | 0 | 0.77 | 0.86 | 0.7 | 0.16 | 0.77 | 0.57 |
| 10 | 56018508 | rs1020207 | -0.293 | -0.888 | 0.782 | -2.891 | 0.002 | -0.019 | -0.019 | 0.165 | 0.74 | 0.61 | 0.66 | 0.19 | 0.75 | 0.59 |
| 10 | 56019051 | rs2043994 | 1.132 | -0.967 | 0.235 | -3.228 | 0.004 | 0.163 | 0.163 | 0 | 0.26 | 0.01 | 0.19 | 0.01 | 0.17 | 0.19 |
| 10 | 56022809 | rs1975407 | 1.701 | -0.507 | 0.381 | -3.127 | -0.079 | 0.109 | 0.109 | 0 | 0.24 | 0.08 | 0.21 | 0.06 | 0.38 | 0.11 |
| 10 | 56025604 | rs11004189 | -0.448 | -0.433 | 0.682 | -2.82 | 0.008 | -0.009 | -0.009 | 0 | 0.76 | 0.7 | 0.67 | 0.19 | 0.76 | 0.61 |
| 10 | 56035204 | rs966910 | 0.131 | -0.578 | 0.121 | -2.83 | 0.038 | 0.202 | 0.202 | 0 | 0.51 | 0.21 | 0.44 | 0.16 | 0.61 | 0.37 |
| 10 | 56037808 | rs11591584 | -0.094 | -0.606 | 0.593 | -2.842 | 0.007 | 0.022 | 0.022 | 0 | 0.52 | 0.41 | 0.45 | 0.15 | 0.61 | 0.37 |
| 10 | 56038831 | rs2044002 | 0 | -0.601 | 0.609 | -2.84 | 0.014 | 0.047 | 0.047 | 0 | 0.52 | 0.37 | 0.45 | 0.15 | 0.61 | 0.37 |
| 10 | 56042135 | rs12262364 | -0.349 | -0.585 | 0.358 | -2.905 | 0.002 | 0.014 | 0.014 | 0 | 0.24 | 0.2 | 0.24 | 0.12 | 0.24 | 0.22 |
| 10 | 56054752 | rs920526 | 0.695 | -0.691 | 0.642 | -3.018 | -0.005 | 0.053 | 0.053 | 0 | 0.52 | 0.59 | 0.42 | 0.11 | 0.5 | 0.28 |
| 10 | 56062045 | rs17725634 | 0.724 | -0.715 | 0.599 | -3.067 | 0 | -0.035 | -0.035 | 0.178 | 0.13 | 0.14 | 0.07 | 0.01 | 0.12 | 0.14 |
| 10 | 56062317 | rs10825299 | 0.94 | -0.778 | 0.87 | -3.09 | -0.008 | -0.028 | -0.028 | 0 | 0.24 | 0.27 | 0.2 | 0.07 | 0.15 | 0.22 |
| 10 | 56063948 | rs7919725 | -1.031 | -0.717 | 0.765 | -3.055 | -0.012 | -0.007 | -0.007 | 0 | 0.78 | 0.86 | 0.67 | 0.23 | 0.76 | 0.58 |
| 10 | 56071606 | rs9416306 | 0.595 | -0.707 | 0.847 | -3.045 | -0.007 | 0.047 | 0.047 | 0 | 0.53 | 0.63 | 0.44 | 0.11 | 0.51 | 0.38 |
| 10 | 56072270 | rs857369 | 0.588 | -0.7 | 0.851 | -3.059 | -0.007 | 0.046 | 0.046 | 0 | 0.53 | 0.63 | 0.44 | 0.11 | 0.51 | 0.38 |
| 10 | 56072949 | rs857373 | -0.924 | -0.679 | 0.85 | -3.048 | -0.005 | -0.002 | -0.002 | 7.258 | 0.77 | 0.83 | 0.67 | 0.23 | 0.75 | 0.59 |
| 10 | 56075990 | rs16905888 | -0.248 | -0.633 | 0.774 | -3.018 | 0.001 | 0.015 | 0.015 | 0 | 0.24 | 0.2 | 0.23 | 0.12 | 0.23 | 0.21 |
| 10 | 56076004 | rs11004222 | 1.868 | -0.676 | 0.767 | -3.054 | -0.074 | 0.137 | 0.137 | 0 | 0.27 | 0.06 | 0.21 | 0.05 | 0.36 | 0.14 |
| 10 | 56079721 | rs12255969 | -0.462 | -0.679 | 0.899 | -3.051 | -0.008 | -0.041 | -0.041 | 0 | 0.26 | 0.39 | 0.25 | 0.12 | 0.24 | 0.21 |
| 10 | 56079770 | rs11004230 | 0.078 | -0.683 | 0.899 | -3.058 | 0.032 | 0.16 | 0.16 | 0 | 0.51 | 0.26 | 0.44 | 0.16 | 0.6 | 0.35 |
| 10 | 56083605 | rs1342303 | -0.609 | -0.445 | 1.082 | -2.84 | -0.007 | -0.041 | -0.041 | 0.036 | 0.26 | 0.39 | 0.25 | 0.12 | 0.24 | 0.21 |
| 10 | 56087474 | rs965266 | -0.564 | -0.369 | 1.091 | -2.772 | 0.011 | 0 | 0 | 0 | 0.75 | 0.57 | 0.64 | 0.23 | 0.74 | 0.59 |
| 10 | 56087484 | rs965267 | -0.671 | -0.396 | 1.17 | -2.774 | 0.005 | 0.001 | 0.001 | 0 | 0.77 | 0.75 | 0.66 | 0.23 | 0.75 | 0.59 |
| 10 | 56087659 | rs12571998 | -0.413 | -0.458 | 1.001 | -2.889 | -0.001 | -0.011 | -0.011 | 0.273 | 0.24 | 0.25 | 0.23 | 0.12 | 0.23 | 0.21 |
| 10 | 56088367 | rs857392 | -0.824 | -0.462 | 0.981 | -2.856 | -0.015 | -0.008 | -0.008 | 0.248 | 0.77 | 0.87 | 0.67 | 0.23 | 0.75 | 0.59 |
| 10 | 56088627 | rs1319836 | -0.545 | -0.489 | 0.981 | -2.868 | 0.005 | -0.002 | -0.002 | 0 | 0.75 | 0.63 | 0.64 | 0.23 | 0.74 | 0.59 |
| 10 | 56102910 | rs2795927 | -0.619 | -0.368 | 1.243 | -2.872 | 0.006 | -0.049 | -0.049 | 0.684 | 0.24 | 0.42 | 0.24 | 0.12 | 0.23 | 0.21 |
| 10 | 56104514 | rs2583024 | -1.069 | -0.373 | 1.098 | -2.874 | 0.002 | -0.022 | -0.022 | 0 | 0.28 | 0.56 | 0.36 | 0.17 | 0.27 | 0.3 |

| | | | | | | | | | | | | | | | | |
|---|---|---|---|---|---|---|---|---|---|---|---|---|---|---|---|---|
| 10 | 56108631 | rs2583026 | 0.565 | -0.236 | 1.123 | -2.937 | -0.009 | 0.047 | 0.047 | 0 | 0.55 | 0.68 | 0.45 | 0.08 | 0.55 | 0.38 |
| 10 | 56114857 | rs11594789 | -0.511 | -0.104 | 1.108 | -2.563 | 0 | 0.001 | 0.001 | 0 | 0.25 | 0.22 | 0.22 | 0.09 | 0.21 | 0.22 |
| 10 | 56123140 | rs17654584 | 0.74 | -0.755 | 1.301 | -2.112 | 0.017 | 0.01 | 0.01 | 0 | 0.22 | 0.15 | 0.08 | 0.01 | 0.12 | 0.14 |
| 10 | 56126202 | rs2028440 | -0.877 | -0.737 | 1.169 | -2.115 | -0.021 | -0.009 | -0.009 | 0 | 0.82 | 0.9 | 0.68 | 0.18 | 0.77 | 0.59 |
| 10 | 56131587 | rs10763103 | 0.197 | -0.651 | 1.397 | -2.177 | -0.022 | -0.067 | -0.067 | 0 | 0.24 | 0.31 | 0.1 | 0.01 | 0.12 | 0.14 |
| 10 | 56132046 | rs4639844 | 0.233 | -0.616 | 0.938 | -2.168 | 0.005 | -0.019 | -0.019 | 0 | 0.24 | 0.24 | 0.09 | 0.01 | 0.12 | 0.14 |
| 10 | 56151359 | rs10763105 | -0.6 | -0.623 | 0.834 | -2.155 | -0.02 | -0.065 | -0.065 | 0.186 | 0.77 | 0.76 | 0.92 | 0.99 | 0.88 | 0.86 |
| 10 | 56173800 | rs4935107 | 0.749 | -0.128 | -0.035 | -2.048 | 0.001 | -0.063 | -0.063 | 0 | 0.43 | 0.43 | 0.42 | 0.07 | 0.55 | 0.31 |
| 10 | 56174108 | rs4935109 | -0.54 | -0.206 | -0.169 | -2.07 | -0.042 | -0.104 | -0.104 | 0 | 0.7 | 0.6 | 0.87 | 0.98 | 0.86 | 0.82 |
| 10 | 56190305 | rs1219862 | -0.286 | -0.334 | -0.536 | -1.816 | -0.031 | 0.071 | 0.071 | 0.265 | 0.73 | 0.86 | 0.44 | 0.1 | 0.68 | 0.49 |
| 10 | 56200276 | rs1930171 | 0.255 | 0.105 | -0.371 | -1.503 | 0.003 | -0.021 | -0.021 | 0 | 0.46 | 0.41 | 0.34 | 0.07 | 0.6 | 0.31 |
| 10 | 56207497 | rs1231248 | 0.192 | 0.324 | -0.167 | -0.028 | -0.008 | 0.013 | 0.013 | 0 | 0.65 | 0.5 | 0.79 | 0.58 | 0.73 | 0.58 |
| 10 | 56208736 | rs2026414 | 0.616 | 0.385 | -0.462 | 0.313 | 0.005 | 0.068 | 0.068 | 0.453 | 0.55 | 0.21 | 0.63 | 0.32 | 0.61 | 0.37 |
| 10 | 56208967 | rs1930149 | 0.281 | 0.714 | -0.138 | 0.175 | -0.038 | -0.003 | -0.003 | 0.059 | 0.66 | 0.75 | 0.65 | 0.32 | 0.55 | 0.38 |
| 10 | 56212367 | rs2891522 | 0.792 | 0.627 | 0.51 | 0.2 | -0.002 | 0.003 | 0.003 | 1.168 | 0.56 | 0.44 | 0.6 | 0.28 | 0.52 | 0.29 |
| 10 | 56218932 | rs6481099 | -1.556 | 0.513 | 0.686 | 0.719 | 0.011 | 0.075 | 0.075 | 0 | 0.11 | 0.22 | 0.17 | 0.3 | 0.13 | 0.29 |
| 10 | 56219586 | rs1234676 | -0.783 | 0.563 | 0.692 | 0.697 | -0.015 | -0.015 | -0.015 | 0 | 0.84 | 0.91 | 0.85 | 0.63 | 0.76 | 0.73 |
| 10 | 56224597 | rs4145278 | -0.274 | -0.769 | 0.306 | 0.185 | 0 | -0.003 | -0.003 | 0 | 0.37 | 0.34 | 0.58 | 0.56 | 0.36 | 0.35 |
| 10 | 56225300 | rs7911134 | -0.775 | -0.742 | 0.456 | 0.331 | 0.008 | -0.006 | -0.006 | 0 | 0.44 | 0.68 | 0.63 | 0.56 | 0.4 | 0.5 |
| 10 | 56234349 | rs10825335 | -0.695 | -0.658 | 0.376 | 0.547 | 0.033 | 0.132 | 0.132 | 0 | 0.06 | 0.12 | 0.15 | 0.28 | 0.11 | 0.15 |
| 10 | 56237639 | rs11004331 | 0.505 | -0.829 | 0.054 | 0.379 | 0.028 | 0.029 | 0.029 | 0.016 | 0.14 | 0.05 | 0.03 | 0.04 | 0.06 | 0.13 |
| 10 | 56245351 | rs16906192 | -0.274 | -1.048 | -0.219 | 0.424 | 0.039 | 0.035 | 0.035 | 1.249 | 0.27 | 0.09 | 0.11 | 0.15 | 0.16 | 0.19 |
| 10 | 56245667 | rs1342309 | -1.138 | -1.343 | 0.345 | 0.357 | 0.062 | 0.215 | 0.215 | 0.974 | 0.13 | 0.49 | 0.2 | 0.31 | 0.15 | 0.3 |
| 10 | 56245720 | rs11004335 | 1.016 | -1.172 | -0.244 | 0.043 | 0.019 | 0.01 | 0.01 | 4.574 | 0.17 | 0.06 | 0.34 | 0.14 | 0.13 | 0.13 |
| 10 | 56247631 | rs1930177 | -0.998 | -0.712 | 0.214 | 0.192 | 0.017 | -0.009 | -0.009 | 0.139 | 0.54 | 0.68 | 0.61 | 0.51 | 0.38 | 0.56 |
| 10 | 56258202 | rs6481100 | -0.925 | -0.588 | -0.582 | 0.23 | 0.022 | 0.018 | 0.018 | 0 | 0.56 | 0.71 | 0.63 | 0.59 | 0.37 | 0.58 |
| 10 | 56261099 | rs11004340 | -0.314 | -0.177 | -0.61 | 0.609 | 0.002 | -0.055 | -0.055 | 0 | 0.18 | 0.08 | 0.19 | 0.33 | 0.17 | 0.34 |
| 10 | 56262826 | rs1418366 | 0.323 | -0.128 | -0.584 | 0.71 | -0.012 | -0.004 | -0.004 | 0 | 0.06 | 0.01 | 0.12 | 0.28 | 0.07 | 0.22 |
| 10 | 56270331 | rs10825340 | -0.857 | 0.049 | -0.476 | -0.409 | 0.006 | -0.022 | -0.022 | 1.29 | 0.54 | 0.6 | 0.69 | 0.97 | 0.53 | 0.72 |
| 10 | 56271032 | rs16906230 | 0.73 | 0.455 | -0.108 | 0.014 | 0.085 | 0.137 | 0.137 | 0 | 0.08 | 0.19 | 0.13 | 0.33 | 0.17 | 0.3 |
| 10 | 56271258 | rs10825341 | 0.592 | 0.092 | -0.307 | 0.04 | 0.052 | -0.098 | -0.098 | 0.044 | 0.17 | 0.04 | 0.16 | 0.35 | 0.08 | 0.25 |
| 10 | 56284213 | rs16906240 | 0.912 | 0.258 | -0.184 | 0.044 | 0.07 | 0.113 | 0.113 | 0 | 0.08 | 0.17 | 0.12 | 0.32 | 0.17 | 0.29 |
| 10 | 56286068 | rs1930145 | -0.223 | 0.086 | -0.11 | -0.012 | 0 | 0.001 | 0.001 | 0 | 0.24 | 0.19 | 0.22 | 0.43 | 0.11 | 0.27 |
| 10 | 56286556 | rs1930146 | -1.705 | 0.055 | -0.181 | -0.217 | 0.018 | 0.111 | 0.111 | 0 | 0.58 | 0.77 | 0.72 | 0.97 | 0.57 | 0.77 |
| 10 | 56287101 | rs10509013 | 1.128 | 0.073 | -0.358 | -0.185 | -0.054 | -0.022 | -0.022 | 0 | 0.2 | 0.05 | 0.33 | 0.22 | 0.24 | 0.15 |
| 10 | 56287266 | rs17730358 | 1.859 | 0.271 | -0.353 | -0.183 | -0.028 | 0.115 | 0.115 | 0.055 | 0.24 | 0.06 | 0.19 | 0.03 | 0.3 | 0.11 |
| 10 | 56287689 | rs10825347 | -1.9 | 0.446 | 0.578 | 0.771 | -0.005 | 0.061 | 0.061 | 3.296 | 0.41 | 0.72 | 0.42 | 0.76 | 0.4 | 0.64 |
| 10 | 56288866 | rs12772008 | 0.155 | 0.799 | 0.716 | 1.072 | 0.117 | 0.187 | 0.187 | 0 | 0.08 | 0.23 | 0.13 | 0.33 | 0.17 | 0.3 |
| 10 | 56295820 | rs1758827 | -1.599 | -0.323 | 0.083 | 0.786 | -0.044 | 0.088 | 0.088 | 0.822 | 0.27 | 0.45 | 0.28 | 0.43 | 0.19 | 0.3 |
| 10 | 56295951 | rs1930167 | -0.405 | 0.669 | -0.062 | 0.147 | -0.009 | -0.008 | -0.008 | 0 | 0.07 | 0.02 | 0.06 | 0.09 | 0.09 | 0.04 |
| 10 | 56296300 | rs1781451 | -1.151 | 0.487 | 0.176 | 0.065 | 0.044 | -0.01 | -0.01 | 0.88 | 0.07 | 0.38 | 0.06 | 0 | 0.06 | 0.05 |
| 10 | 56301930 | rs1930165 | -0.572 | 0.752 | -0.976 | -0.116 | 0 | 0.059 | 0.059 | 0.333 | 0.44 | 0.33 | 0.39 | 0.75 | 0.44 | 0.62 |
| 10 | 56302037 | rs1930164 | -0.15 | 1.367 | -1.494 | 0.704 | -0.091 | -0.078 | -0.078 | 0.41 | 0.24 | 0.12 | 0.21 | 0.41 | 0.36 | 0.4 |
| 10 | 56302374 | rs11004362 | 0.145 | 1.267 | -1.254 | 0.725 | 0.006 | -0.048 | -0.048 | 0.059 | 0.18 | 0.07 | 0.18 | 0.34 | 0.11 | 0.23 |
| 10 | 56302543 | rs11004365 | 0.147 | 1.257 | -1.259 | 0.721 | 0.005 | -0.047 | -0.047 | 4.259 | 0.17 | 0.08 | 0.18 | 0.34 | 0.11 | 0.23 |
| 10 | 56303319 | rs1219786 | -0.162 | 1.299 | -1.342 | 0.706 | 0.003 | -0.045 | -0.045 | 0.002 | 0.83 | 0.91 | 0.82 | 0.64 | 0.89 | 0.76 |
| 10 | 56304066 | rs10825350 | -0.357 | 1.45 | -1.275 | 0.565 | -0.007 | -0.006 | -0.006 | 0 | 0.19 | 0.22 | 0.21 | 0.39 | 0.31 | 0.36 |
| 10 | 56304174 | rs2482814 | -0.022 | 1.421 | -1.346 | 0.561 | -0.003 | -0.052 | -0.052 | 0 | 0.81 | 0.92 | 0.81 | 0.64 | 0.85 | 0.74 |
| 10 | 56304265 | rs7099825 | 1.041 | 0.228 | -1.826 | -0.344 | 0.044 | 0.107 | 0.107 | 5.504 | 0.41 | 0.16 | 0.42 | 0.23 | 0.36 | 0.28 |
| 10 | 56304815 | rs9787725 | -0.209 | 1.314 | -1.802 | -0.107 | -0.047 | -0.009 | -0.009 | 0 | 0.23 | 0.12 | 0.31 | 0.74 | 0.27 | 0.5 |
| 10 | 56305644 | rs12257225 | 0.461 | 1.385 | -1.725 | 0.204 | -0.02 | 0 | 0 | 0 | 0.07 | 0.05 | 0.14 | 0.36 | 0.17 | 0.25 |
| 10 | 56313403 | rs10825356 | 0.013 | -0.158 | -2.045 | 0.145 | -0.027 | -0.034 | -0.034 | 1.22 | 0.31 | 0.26 | 0.34 | 0.52 | 0.48 | 0.48 |
| 10 | 56317742 | rs4587642 | 0.739 | -0.197 | -2.135 | -0.214 | 0.031 | 0.055 | 0.055 | 7.137 | 0.34 | 0.14 | 0.35 | 0.18 | 0.28 | 0.19 |
| 10 | 56319852 | rs1937389 | -1.061 | -0.142 | -1.486 | -0.295 | -0.028 | -0.139 | -0.139 | 0.42 | 0.8 | 0.54 | 0.85 | 0.99 | 0.83 | 0.88 |
| 10 | 56323564 | rs10825357 | -0.438 | -0.03 | -1.609 | 0.316 | 0.005 | -0.01 | -0.01 | 0 | 0.24 | 0.21 | 0.23 | 0.32 | 0.18 | 0.26 |
| 10 | 56324127 | rs12777241 | -0.366 | -0.149 | -2.034 | 0.434 | -0.05 | -0.146 | -0.146 | 0.403 | 0.3 | 0.04 | 0.32 | 0.4 | 0.25 | 0.21 |
| 10 | 56331169 | rs12769874 | 0.059 | 0.071 | 0.239 | 0.417 | -0.001 | -0.002 | -0.002 | 0 | 0.13 | 0.16 | 0.14 | 0.1 | 0.17 | 0.1 |
| 10 | 56345511 | rs2026410 | 0.019 | 0.039 | 0.231 | 0.391 | 0.005 | 0.011 | 0.011 | 0 | 0.85 | 0.73 | 0.78 | 0.82 | 0.81 | 0.83 |
| 10 | 56349666 | rs10763119 | 0.426 | 0.317 | 0.514 | 0.333 | 0.016 | 0.007 | 0.007 | 0 | 0.83 | 0.65 | 0.75 | 0.82 | 0.78 | 0.87 |
| 10 | 56361499 | rs11004407 | 0.242 | -0.424 | 0 | 0.302 | 0.027 | -0.051 | -0.051 | 0 | 0.26 | 0.14 | 0.26 | 0.36 | 0.19 | 0.23 |
| 10 | 56362285 | rs10825366 | 0.37 | 0.198 | 0.377 | 0.033 | -0.006 | 0.015 | 0.015 | 0 | 0.12 | 0.07 | 0.1 | 0 | 0.17 | 0.05 |
| 10 | 56366138 | rs1937411 | 1.006 | 1.28 | 0.791 | -0.064 | 0.027 | -0.001 | -0.001 | 0.12 | 0.06 | 0.09 | 0.11 | 0.07 | 0.18 | 0.16 |
| 10 | 56366524 | rs7913327 | 0.71 | 0.956 | 0.767 | -0.362 | 0.018 | -0.038 | -0.038 | 0.003 | 0.63 | 0.48 | 0.59 | 0.79 | 0.52 | 0.7 |
| 10 | 56370710 | rs11004411 | 0.452 | 1.506 | 0.702 | 0.315 | -0.009 | 0.021 | 0.021 | 0 | 0.12 | 0.05 | 0.1 | 0 | 0.17 | 0.05 |
| 10 | 56371176 | rs1937409 | 0.766 | 1.47 | 0.762 | 0.342 | 0.087 | 0 | 0 | 0 | 0.77 | 0.59 | 0.67 | 0.76 | 0.62 | 0.71 |
| 10 | 56375357 | rs10763120 | -0.038 | 0.992 | 0.284 | 0.434 | 0.004 | 0.002 | 0.002 | 0 | 0.85 | 0.83 | 0.79 | 0.83 | 0.8 | 0.85 |
| 10 | 56386858 | rs11004420 | 0.201 | 0.454 | 0.409 | 0.638 | 0.006 | 0.005 | 0.005 | 0 | 0.58 | 0.5 | 0.54 | 0.52 | 0.58 | 0.49 |
| 10 | 56388664 | rs10740588 | -0.423 | 0.524 | 0.081 | 0.649 | -0.007 | 0.045 | 0.045 | 0 | 0.78 | 0.66 | 0.7 | 0.61 | 0.82 | 0.7 |
| 10 | 56389870 | rs12770729 | 0.326 | 0.373 | -0.32 | 0.638 | -0.02 | 0.038 | 0.038 | 0.274 | 0.54 | 0.23 | 0.47 | 0.45 | 0.57 | 0.43 |

| | | | | | | | | | | | | | | | | |
|---|---|---|---|---|---|---|---|---|---|---|---|---|---|---|---|---|
| 10 | 56399124 | rs10825369 | 0.338 | 0.373 | -0.447 | 0.505 | -0.003 | 0.111 | 0.111 | 0.533 | 0.49 | 0.22 | 0.37 | 0.35 | 0.54 | 0.41 |
| 10 | 56401270 | rs4935116 | 0.408 | 0.168 | 0.286 | 1.095 | -0.011 | 0.074 | 0.074 | 1.251 | 0.78 | 0.44 | 0.71 | 0.61 | 0.83 | 0.68 |
| 10 | 56401411 | rs4935117 | -0.03 | 0.232 | 0.087 | 0.526 | -0.001 | -0.013 | -0.013 | 0.153 | 0.18 | 0.13 | 0.15 | 0.08 | 0.15 | 0.09 |
| 10 | 56411394 | rs10509016 | -0.165 | 0.285 | 0.313 | 0.978 | -0.006 | 0.041 | 0.041 | 0 | 0.07 | 0.14 | 0.04 | 0.13 | 0.06 | 0.16 |
| 10 | 56411969 | rs1414688 | 0.077 | 0.283 | 0.313 | 0.977 | 0.003 | -0.008 | -0.008 | 0.276 | 0.07 | 0.05 | 0.03 | 0.11 | 0.06 | 0.16 |
| 10 | 56413404 | rs11004434 | -1.294 | 0.315 | 0.568 | 1.019 | -0.006 | 0.048 | 0.048 | 0 | 0.12 | 0.27 | 0.1 | 0.23 | 0.12 | 0.25 |
| 10 | 56417062 | rs11004437 | 0.331 | 0.408 | 0.136 | 0.804 | -0.007 | 0.01 | 0.01 | 0 | 0.06 | 0.03 | 0.04 | 0.01 | 0.06 | 0.09 |
| 10 | 56423968 | rs11004439 | -0.122 | 0.221 | -0.019 | 0.464 | 0.007 | 0.047 | 0.047 | 0 | 0.29 | 0.12 | 0.27 | 0.14 | 0.25 | 0.13 |
| 10 | 56428277 | rs7902184 | -0.871 | 0.078 | 0.721 | 0.563 | -0.007 | -0.015 | -0.015 | 0.566 | 0.4 | 0.53 | 0.35 | 0.24 | 0.32 | 0.28 |
| 10 | 56435989 | rs7898511 | 0.57 | 0.293 | 0.939 | 0.904 | 0.002 | 0.015 | 0.015 | 0 | 0.56 | 0.55 | 0.39 | 0.27 | 0.6 | 0.45 |
| 10 | 56441751 | rs2384520 | 0.119 | 0.222 | 0.762 | 0.893 | 0.003 | 0.101 | 0.101 | 0 | 0.31 | 0.11 | 0.28 | 0.08 | 0.26 | 0.12 |
| 10 | 56445616 | rs7095795 | 0.803 | 0.243 | 0.641 | 0.952 | 0.014 | 0.037 | 0.037 | 0 | 0.54 | 0.34 | 0.36 | 0.32 | 0.6 | 0.45 |
| 10 | 56448886 | rs16906408 | 0.689 | 0.245 | 0.687 | 0.992 | -0.003 | 0.012 | 0.012 | 0 | 0.07 | 0.1 | 0.04 | 0.14 | 0.06 | 0.16 |
| 10 | 56456576 | rs7098216 | 0.74 | 0.249 | 0.611 | 0.946 | 0.012 | 0.034 | 0.034 | 0 | 0.54 | 0.34 | 0.36 | 0.32 | 0.6 | 0.45 |
| 10 | 56462471 | rs10763126 | 1.127 | 0.25 | 0.44 | 0.817 | 0.019 | 0.155 | 0.155 | 0 | 0.5 | 0.24 | 0.31 | 0.18 | 0.54 | 0.35 |
| 10 | 56464007 | rs10763127 | -0.514 | 0.44 | 0.208 | 1.083 | 0.035 | 0.3 | 0.3 | 0 | 0.94 | 0.76 | 0.7 | 0.54 | 0.92 | 0.74 |
| 10 | 56466912 | rs17500902 | 0.635 | 0.273 | -0.45 | 0.552 | -0.026 | 0.112 | 0.112 | 0 | 0.18 | 0.01 | 0.1 | 0 | 0.16 | 0.04 |
| 10 | 56467563 | rs12413003 | -0.434 | 0.381 | -0.056 | 1.001 | 0 | 0.002 | 0.002 | 0 | 0.11 | 0.14 | 0.09 | 0.26 | 0.11 | 0.22 |
| 10 | 56478284 | rs12764105 | -0.2 | 0.399 | -0.145 | 1.008 | -0.002 | -0.028 | -0.028 | 0 | 0.1 | 0.07 | 0.08 | 0.26 | 0.11 | 0.22 |
| 10 | 56497659 | rs1336197 | -1.234 | 1.095 | 0.248 | 0.687 | 0.012 | 0.048 | 0.048 | 1.913 | 0.3 | 0.5 | 0.36 | 0.43 | 0.33 | 0.37 |
| 10 | 56500064 | rs10825391 | 1.357 | 1.118 | 0.197 | 0.433 | 0.013 | 0.105 | 0.105 | 0 | 0.45 | 0.25 | 0.25 | 0.12 | 0.42 | 0.32 |
| 10 | 56502293 | rs1336199 | -0.501 | 1.233 | 0.719 | 0.543 | -0.016 | -0.026 | -0.026 | 0 | 0.16 | 0.13 | 0.16 | 0.36 | 0.22 | 0.29 |
| 10 | 56502386 | rs953128 | 0.534 | 1.19 | 0.986 | 0.512 | -0.004 | -0.01 | -0.01 | 0 | 0.12 | 0.2 | 0.18 | 0.07 | 0.11 | 0.08 |
| 10 | 56504373 | rs1832185 | -1.198 | 1.257 | 0.778 | 0.464 | -0.001 | -0.024 | -0.024 | 0 | 0.89 | 0.91 | 0.85 | 1 | 0.89 | 0.94 |
| 10 | 56507333 | rs2384529 | -0.604 | 1.217 | 0.688 | 0.542 | -0.015 | -0.024 | -0.024 | 0 | 0.16 | 0.13 | 0.16 | 0.36 | 0.23 | 0.28 |
| 10 | 56517410 | rs10763141 | 0.414 | 1.136 | 0.831 | 0.687 | 0 | 0.05 | 0.05 | 0 | 0.77 | 0.67 | 0.52 | 0.47 | 0.76 | 0.5 |
| 10 | 56529521 | rs11004564 | -1.014 | 1.163 | 0.462 | 0.575 | 0.004 | 0.006 | 0.006 | 0 | 0.44 | 0.63 | 0.62 | 0.61 | 0.41 | 0.55 |
| 10 | 56542240 | rs16906534 | 0.332 | 1.402 | 0.167 | 0.663 | -0.03 | -0.029 | -0.029 | 0 | 0.06 | 0.01 | 0.07 | 0.15 | 0.11 | 0.08 |
| 10 | 56545395 | rs7071691 | -0.2 | 1.131 | 0.379 | 0.611 | 0.017 | 0.014 | 0.014 | 0 | 0.22 | 0.42 | 0.18 | 0.18 | 0.24 | 0.21 |
| 10 | 56547552 | rs2384533 | -0.649 | 0.403 | -0.543 | -0.152 | -0.001 | 0.027 | 0.027 | 0 | 0.44 | 0.33 | 0.61 | 0.71 | 0.43 | 0.53 |
| 10 | 56550427 | rs11004579 | 1.575 | 0.154 | -0.826 | -0.605 | -0.006 | 0.038 | 0.038 | 0 | 0.24 | 0.12 | 0.14 | 0.07 | 0.23 | 0.14 |
| 10 | 56554529 | rs11004581 | 1.518 | 0.582 | -0.503 | -0.037 | 0.025 | 0.185 | 0.185 | 0 | 0.05 | 0.12 | 0.28 | 0.45 | 0.08 | 0.28 |
| 10 | 56554972 | rs16906543 | 0.978 | 0.365 | -0.591 | -0.163 | 0.004 | -0.014 | -0.014 | 2.749 | 0.07 | 0.05 | 0.04 | 0.12 | 0.08 | 0.16 |
| 10 | 56573906 | rs10825402 | 0.219 | 1.095 | -0.791 | -0.236 | -0.009 | -0.014 | -0.014 | 0.142 | 0.2 | 0.17 | 0.5 | 0.47 | 0.25 | 0.39 |
| 10 | 56575182 | rs10825403 | -0.294 | 1.045 | -0.377 | -0.333 | 0.018 | 0.035 | 0.035 | 0 | 0.75 | 0.66 | 0.82 | 0.7 | 0.7 | 0.74 |
| 10 | 56577441 | rs11004585 | -0.057 | 0.79 | -0.602 | -0.446 | -0.005 | -0.018 | -0.018 | 1.782 | 0.4 | 0.37 | 0.58 | 0.54 | 0.39 | 0.49 |
| 10 | 56589230 | rs7081418 | 0.989 | 0.766 | -0.743 | -0.514 | 0.016 | 0.045 | 0.045 | 0 | 0.34 | 0.22 | 0.21 | 0.14 | 0.32 | 0.25 |
| 10 | 56595219 | rs10430482 | 1.1 | 0.947 | -0.807 | -0.213 | -0.065 | -0.075 | -0.075 | 0 | 0.16 | 0.01 | 0.16 | 0.24 | 0.2 | 0.12 |
| 10 | 56595890 | rs10763150 | 0.106 | 0.883 | -0.525 | -0.235 | -0.013 | 0.006 | 0.006 | 0.046 | 0.67 | 0.55 | 0.76 | 0.65 | 0.72 | 0.74 |
| 10 | 56606168 | rs16906592 | 0.873 | 1.593 | -0.657 | 0.337 | 0.055 | 0.201 | 0.201 | 0 | 0.06 | 0.16 | 0.32 | 0.42 | 0.11 | 0.31 |
| 10 | 56614745 | rs1840335 | -0.362 | 1.413 | -1.235 | -0.098 | 0 | 0.105 | 0.105 | 0 | 0.55 | 0.78 | 0.67 | 0.72 | 0.55 | 0.63 |
| 10 | 56616917 | rs2061048 | 0.995 | 1.375 | -0.72 | 0.013 | 0.022 | 0.011 | 0.011 | 0 | 0.16 | 0.27 | 0.16 | 0.16 | 0.21 | 0.12 |
| 10 | 56618052 | rs3902590 | 0.971 | 1.362 | -0.673 | 0.046 | 0.013 | 0.008 | 0.008 | 0 | 0.17 | 0.23 | 0.16 | 0.2 | 0.22 | 0.11 |
| 10 | 56619339 | rs12415187 | 0.581 | 1.508 | -0.854 | -0.234 | -0.002 | -0.001 | -0.001 | 0 | 0.16 | 0.16 | 0.1 | 0.18 | 0.18 | 0.15 |
| 10 | 56619592 | rs7085479 | -0.239 | 1.419 | -0.829 | -0.236 | -0.008 | -0.002 | -0.002 | 0.076 | 0.33 | 0.29 | 0.24 | 0.34 | 0.39 | 0.26 |
| 10 | 56622131 | rs7078039 | -1.031 | 1.056 | -1.165 | -0.611 | 0.033 | 0.09 | 0.09 | 0.45 | 0.68 | 0.83 | 0.77 | 0.89 | 0.73 | 0.76 |
| 10 | 56624092 | rs7087645 | -0.732 | 1.127 | -0.865 | -0.566 | -0.001 | -0.027 | -0.027 | 0.05 | 0.44 | 0.48 | 0.57 | 0.84 | 0.51 | 0.6 |
| 10 | 56626553 | rs10825411 | 1.195 | 1.037 | -1.048 | -0.499 | -0.009 | -0.055 | -0.055 | 0 | 0.25 | 0.27 | 0.19 | 0.05 | 0.22 | 0.15 |
| 10 | 56627225 | rs7913159 | 0.205 | 1.075 | -1.256 | -0.21 | -0.001 | -0.014 | -0.014 | 0.098 | 0.47 | 0.46 | 0.34 | 0.21 | 0.46 | 0.27 |
| 10 | 56627577 | rs10825412 | -0.219 | 0.83 | -1.051 | 0.091 | -0.005 | -0.002 | -0.002 | 0 | 0.59 | 0.52 | 0.5 | 0.38 | 0.64 | 0.42 |
| 10 | 56628289 | rs11004610 | 0.074 | 0.647 | -1.696 | -0.35 | 0.04 | 0.147 | 0.147 | 0 | 0.52 | 0.22 | 0.61 | 0.73 | 0.4 | 0.58 |
| 10 | 56641728 | rs11004618 | 0.487 | 0.325 | -1.783 | -0.011 | -0.011 | -0.034 | -0.034 | 0 | 0.18 | 0.14 | 0.34 | 0.52 | 0.19 | 0.44 |
| 10 | 56644401 | rs7350463 | -0.324 | 0.589 | -1.887 | -0.565 | -0.005 | 0.005 | 0.005 | 0 | 0.8 | 0.8 | 0.86 | 0.87 | 0.71 | 0.79 |
| 10 | 56644738 | rs4935569 | -1.209 | 0.281 | -2.01 | -0.718 | -0.016 | -0.026 | -0.026 | 0 | 0.94 | 0.88 | 0.98 | 0.99 | 0.98 | 0.98 |
| 10 | 56648924 | rs4935126 | -0.587 | -0.116 | -1.289 | -0.836 | 0.015 | 0.014 | 0.014 | 0 | 0.53 | 0.54 | 0.63 | 0.64 | 0.58 | 0.68 |
| 10 | 56649668 | rs11813604 | 0.527 | 0.018 | -1.334 | -0.818 | -0.035 | 0.04 | 0.04 | 0 | 0.18 | 0.08 | 0.12 | 0.13 | 0.29 | 0.21 |
| 10 | 56662603 | rs1454471 | -0.585 | 0.013 | -1.005 | -0.814 | 0.059 | 0.083 | 0.083 | 0 | 0.51 | 0.76 | 0.63 | 0.64 | 0.55 | 0.7 |
| 10 | 56670968 | rs17596680 | 1.322 | 0.353 | -1.105 | -0.823 | -0.03 | 0.032 | 0.032 | 0 | 0.13 | 0.06 | 0.09 | 0.02 | 0.23 | 0.09 |
| 10 | 56673719 | rs12767364 | 1.087 | 0.388 | -1.082 | -0.709 | -0.082 | 0.088 | 0.088 | 0.744 | 0.23 | 0.07 | 0.19 | 0.07 | 0.28 | 0.13 |
| 10 | 56691305 | rs1733795 | 1.38 | -0.566 | -0.874 | -0.205 | 0.089 | -0.007 | -0.007 | 1.035 | 0.26 | 0.1 | 0.22 | 0.23 | 0.13 | 0.11 |
| 10 | 56701951 | rs1349893 | -1.096 | -0.301 | -0.791 | -0.133 | -0.024 | -0.041 | -0.041 | 0.509 | 0.83 | 0.9 | 0.67 | 0.48 | 0.79 | 0.55 |
| 10 | 56707648 | rs1733774 | 0.608 | -0.801 | -0.57 | -0.196 | 0.039 | 0.012 | 0.012 | 0 | 0.29 | 0.18 | 0.25 | 0.23 | 0.17 | 0.12 |
| 10 | 56709224 | rs1670837 | 0.703 | -1.006 | -0.4 | -0.133 | -0.081 | -0.012 | -0.012 | 0.871 | 0.28 | 0.43 | 0.25 | 0.23 | 0.13 | 0.11 |
| 10 | 56711013 | rs10763154 | -0.032 | -0.846 | -0.463 | -0.156 | 0.019 | 0.013 | 0.013 | 0 | 0.43 | 0.37 | 0.44 | 0.67 | 0.31 | 0.58 |
| 10 | 56716972 | rs2045145 | 1.328 | -0.978 | -1.057 | -0.202 | 0.085 | 0.084 | 0.084 | 0 | 0.24 | 0 | 0.18 | 0.07 | 0.12 | 0.1 |
| 10 | 56723297 | rs897116 | 1.629 | -0.483 | -0.807 | 0.336 | -0.007 | 0.008 | 0.008 | 0 | 0.09 | 0.04 | 0.19 | 0.55 | 0.13 | 0.37 |
| 10 | 56726940 | rs3911311 | 0.33 | -0.57 | -0.318 | 0.106 | -0.007 | -0.003 | -0.003 | 2.303 | 0.43 | 0.39 | 0.44 | 0.63 | 0.3 | 0.58 |
| 10 | 56728229 | rs1237360 | -0.952 | 0.231 | -1.273 | -0.435 | -0.003 | 0.006 | 0.006 | 0 | 0.72 | 0.74 | 0.7 | 0.89 | 0.72 | 0.83 |
| 10 | 56731004 | rs1219535 | -0.1 | 0.352 | -1.205 | -0.364 | -0.07 | 0.048 | 0.048 | 0 | 0.27 | 0.14 | 0.24 | 0.22 | 0.41 | 0.23 |

| | | | | | | | | | | | | | | | | |
|---|---|---|---|---|---|---|---|---|---|---|---|---|---|---|---|---|
| 10 | 56734168 | rs1219531 | -0.228 | 0.333 | -0.711 | -0.31 | -0.009 | 0.011 | 0.011 | 0 | 0.71 | 0.74 | 0.69 | 0.89 | 0.68 | 0.83 |
| 10 | 56740556 | rs11004654 | 0.132 | -0.819 | -0.699 | 0.068 | -0.024 | -0.031 | -0.031 | 0.03 | 0.37 | 0.24 | 0.4 | 0.53 | 0.46 | 0.43 |
| 10 | 56742013 | rs1219525 | 0.237 | -1.341 | -0.346 | -0.152 | -0.02 | -0.028 | -0.028 | 0 | 0.37 | 0.44 | 0.28 | 0.26 | 0.25 | 0.4 |
| 10 | 56745512 | rs11004656 | 0.721 | -1.62 | -0.313 | -0.248 | 0.001 | 0 | 0 | 0 | 0.28 | 0.23 | 0.24 | 0.13 | 0.21 | 0.28 |
| 10 | 56751980 | rs10763161 | -1.248 | -1.555 | -0.802 | -0.248 | -0.1 | -0.053 | -0.053 | 0 | 0.78 | 0.99 | 0.73 | 0.78 | 0.76 | 0.84 |
| 10 | 56754578 | rs7096888 | -0.1 | -1.484 | -0.174 | -0.22 | -0.011 | -0.016 | -0.016 | 2.09 | 0.36 | 0.57 | 0.29 | 0.26 | 0.26 | 0.41 |
| 10 | 56759380 | rs4317895 | -0.671 | -1.443 | -0.084 | -0.195 | -0.002 | 0.02 | 0.02 | 0 | 0.67 | 0.73 | 0.72 | 0.87 | 0.72 | 0.82 |
| 10 | 56763586 | rs6481134 | 0.557 | -1.568 | 0.023 | -0.221 | -0.021 | -0.044 | -0.044 | 0 | 0.29 | 0.35 | 0.24 | 0.12 | 0.18 | 0.26 |
| 10 | 56764321 | rs10763165 | -1.142 | -1.567 | -0.085 | -0.233 | -0.004 | 0.024 | 0.024 | 0 | 0.7 | 0.81 | 0.74 | 0.87 | 0.75 | 0.83 |
| 10 | 56764751 | rs10825427 | -0.782 | -1.564 | -0.187 | -0.232 | -0.002 | 0.021 | 0.021 | 0 | 0.67 | 0.74 | 0.72 | 0.87 | 0.72 | 0.82 |
| 10 | 56768101 | rs2141841 | -1.143 | -1.527 | -0.212 | -0.23 | -0.023 | 0.042 | 0.042 | 0 | 0.71 | 0.88 | 0.74 | 0.84 | 0.75 | 0.8 |
| 10 | 56770436 | rs1882469 | 0.272 | -1.524 | -0.22 | -0.23 | -0.053 | 0.039 | 0.039 | 0 | 0.38 | 0.19 | 0.46 | 0.75 | 0.53 | 0.55 |
| 10 | 56775106 | rs16906812 | 0.99 | -1.486 | -0.079 | -0.298 | 0.013 | 0.044 | 0.044 | 0 | 0.22 | 0.09 | 0.2 | 0.08 | 0.15 | 0.2 |
| 10 | 56775306 | rs2204866 | 1.081 | -1.782 | 0.099 | -0.22 | 0.011 | 0.014 | 0.014 | 0 | 0.21 | 0.12 | 0.19 | 0.08 | 0.13 | 0.18 |
| 10 | 56777024 | rs10763170 | -0.618 | -0.979 | 0.203 | 0.188 | -0.013 | 0.047 | 0.047 | 0 | 0.44 | 0.32 | 0.48 | 0.76 | 0.58 | 0.6 |
| 10 | 56777654 | rs4363501 | 0.548 | -1.073 | 0.015 | 0.344 | -0.087 | -0.115 | -0.115 | 2.209 | 0.27 | 0.1 | 0.39 | 0.49 | 0.48 | 0.43 |
| 10 | 56784302 | rs2384610 | -0.325 | -1.071 | -0.016 | 0.347 | -0.053 | 0.036 | 0.036 | 0 | 0.38 | 0.19 | 0.45 | 0.74 | 0.53 | 0.55 |
| 10 | 56793962 | rs1922163 | -1.255 | -1.075 | -0.408 | 0.344 | 0.012 | 0.105 | 0.105 | 0 | 0.49 | 0.76 | 0.54 | 0.76 | 0.62 | 0.61 |
| 10 | 56795078 | rs1922166 | -1.01 | -0.756 | -1.409 | 0.529 | 0 | -0.008 | -0.008 | 0 | 0.46 | 0.5 | 0.51 | 0.75 | 0.58 | 0.59 |
| 10 | 56795648 | rs11004690 | 0.196 | -0.765 | -1.452 | 0.557 | 0.015 | -0.016 | -0.016 | 0 | 0.11 | 0.09 | 0.07 | 0.26 | 0.05 | 0.13 |
| 10 | 56795696 | rs10825446 | -0.991 | -0.732 | -1.457 | 0.206 | 0 | -0.007 | -0.007 | 0 | 0.46 | 0.49 | 0.51 | 0.76 | 0.58 | 0.59 |
| 10 | 56817533 | rs17528196 | 0.237 | -0.21 | -2.044 | 0.787 | -0.087 | -0.008 | -0.008 | 0.856 | 0.27 | 0.1 | 0.29 | 0.28 | 0.46 | 0.42 |
| 10 | 56824247 | rs10430541 | -0.122 | -0.328 | -1.6 | 0.38 | -0.019 | -0.01 | -0.01 | 0 | 0.27 | 0.22 | 0.39 | 0.74 | 0.38 | 0.48 |
| 10 | 56826443 | rs10825455 | 0.212 | -0.842 | -0.903 | 0.516 | 0.015 | -0.027 | -0.027 | 0 | 0.06 | 0.11 | 0.03 | 0.01 | 0.06 | 0.07 |
| 10 | 56840255 | rs2250775 | -0.561 | -1.367 | -1.832 | 0.57 | 0.088 | 0.11 | 0.11 | 0.022 | 0.45 | 0.82 | 0.26 | 0.3 | 0.3 | 0.4 |
| 10 | 56846298 | rs16906891 | 0.374 | -1.688 | -1.98 | 0.814 | -0.002 | -0.004 | -0.004 | 1.387 | 0.07 | 0.06 | 0.14 | 0.13 | 0.01 | 0.04 |
| 10 | 56847319 | rs7920748 | 0.292 | -2.244 | -2.106 | 0.67 | 0.059 | -0.022 | -0.022 | 0.02 | 0.19 | 0.07 | 0.05 | 0.23 | 0.08 | 0.22 |
| 10 | 56847522 | rs1733743 | -1.337 | -2.263 | -2.135 | 0.596 | 0.108 | 0.009 | 0.009 | 0.011 | 0.52 | 0.88 | 0.39 | 0.41 | 0.31 | 0.5 |
| 10 | 56849669 | rs11004735 | 0.287 | -2.255 | -2.094 | 0.638 | 0.059 | -0.031 | -0.031 | 0 | 0.19 | 0.07 | 0.05 | 0.26 | 0.08 | 0.22 |
| 10 | 56875956 | rs16906966 | 0.523 | -2.174 | -2.205 | 0.652 | -0.002 | -0.004 | -0.004 | 1.075 | 0.07 | 0.06 | 0.14 | 0.13 | 0.01 | 0.04 |
| 10 | 56877796 | rs7899461 | -0.083 | -1.914 | -2.307 | 0.7 | 0.069 | -0.028 | -0.028 | 0.444 | 0.2 | 0.07 | 0.07 | 0.26 | 0.09 | 0.23 |
| 10 | 56880137 | rs1857449 | -1.103 | -1.851 | -2.243 | 0.797 | 0.116 | 0.05 | 0.05 | 0 | 0.52 | 0.86 | 0.48 | 0.62 | 0.33 | 0.51 |
| 10 | 56880393 | rs11004740 | 0.218 | -0.605 | -1.534 | 0.272 | 0.026 | 0.058 | 0.058 | 0 | 0.67 | 0.25 | 0.67 | 0.85 | 0.77 | 0.72 |
| 10 | 56886783 | rs7088769 | -0.009 | 0.316 | -1.516 | 0.363 | 0.005 | 0.028 | 0.028 | 0 | 0.48 | 0.31 | 0.46 | 0.43 | 0.48 | 0.54 |
| 10 | 56886920 | rs7092223 | 0.033 | 0.603 | -0.634 | 0.242 | 0.002 | 0.014 | 0.014 | 0.185 | 0.66 | 0.52 | 0.67 | 0.61 | 0.65 | 0.6 |
| 10 | 56887138 | rs16907003 | -0.08 | 0.831 | -0.057 | -0.106 | 0.044 | -0.086 | -0.086 | 0 | 0.15 | 0.42 | 0.14 | 0.02 | 0.18 | 0.08 |
| 10 | 56894035 | rs10825469 | 0.386 | 0.331 | -0.279 | -0.083 | 0.012 | 0.004 | 0.004 | 0 | 0.33 | 0.14 | 0.18 | 0.34 | 0.26 | 0.34 |
| 10 | 56896643 | rs7916117 | -0.134 | 0.654 | -0.212 | -0.013 | 0.015 | 0.025 | 0.025 | 0.005 | 0.45 | 0.58 | 0.27 | 0.36 | 0.33 | 0.42 |
| 10 | 56897671 | rs4935576 | -0.211 | 1.028 | 0.088 | 0.231 | 0.028 | 0.018 | 0.018 | 0.213 | 0.19 | 0.34 | 0.21 | 0.22 | 0.27 | 0.31 |
| 10 | 56898863 | rs7081462 | 0.057 | 1.261 | -0.185 | 0.328 | -0.029 | -0.026 | -0.026 | 0.016 | 0.15 | 0.07 | 0.2 | 0.27 | 0.29 | 0.26 |
| 10 | 56900638 | rs10825471 | 0.122 | 0.952 | 0.306 | 0.376 | 0.008 | 0.061 | 0.061 | 0 | 0.41 | 0.3 | 0.47 | 0.16 | 0.4 | 0.31 |
| 10 | 56902502 | rs2799610 | -0.174 | 0.953 | 0.166 | 0.373 | 0 | 0 | 0 | 0 | 0.55 | 0.41 | 0.4 | 0.63 | 0.57 | 0.67 |
| 10 | 56904461 | rs10509025 | 1.157 | 0.599 | -0.764 | -0.013 | -0.057 | 0.089 | 0.089 | 0 | 0.23 | 0.05 | 0.27 | 0.05 | 0.3 | 0.12 |
| 10 | 56905947 | rs7905584 | 0.316 | 0.87 | -0.465 | 0.277 | 0.007 | 0.07 | 0.07 | 0 | 0.41 | 0.28 | 0.47 | 0.16 | 0.4 | 0.31 |
| 10 | 56914138 | rs2799580 | -0.512 | 1.328 | -0.111 | 0.879 | -0.095 | -0.086 | -0.086 | 0.042 | 0.15 | 0.02 | 0.2 | 0.26 | 0.29 | 0.27 |
| 10 | 56916347 | rs11004759 | 1.348 | 0.848 | -0.031 | 0.785 | -0.015 | -0.034 | -0.034 | 0 | 0.67 | 0.24 | 0.59 | 0.49 | 0.63 | 0.67 |
| 10 | 56918363 | rs12357313 | 0.66 | -0.05 | 0.932 | 0.914 | -0.022 | -0.09 | -0.09 | 0 | 0.85 | 0.49 | 0.91 | 0.97 | 0.93 | 0.93 |
| 10 | 56918888 | rs2589444 | 1.148 | -0.046 | 0.432 | 1.038 | -0.035 | 0.026 | 0.026 | 1.041 | 0.81 | 0.33 | 0.8 | 0.75 | 0.92 | 0.87 |
| 10 | 56920229 | rs2680316 | 0.893 | -0.246 | 0.298 | 1.015 | -0.036 | 0.005 | 0.005 | 0.185 | 0.74 | 0.42 | 0.65 | 0.78 | 0.83 | 0.8 |
| 10 | 56920839 | rs2589452 | -0.096 | 0.262 | -0.493 | 0.97 | -0.004 | 0.011 | 0.011 | 0 | 0.59 | 0.71 | 0.62 | 0.34 | 0.59 | 0.58 |
| 10 | 56920872 | rs2589453 | 0.091 | 0.493 | -0.473 | 1.269 | 0.001 | -0.008 | -0.008 | 0 | 0.6 | 0.69 | 0.63 | 0.41 | 0.62 | 0.62 |
| 10 | 56927420 | rs2680312 | -0.057 | 1.289 | -0.724 | 1.258 | 0.005 | 0.037 | 0.037 | 0 | 0.59 | 0.72 | 0.65 | 0.7 | 0.6 | 0.73 |
| 10 | 56929835 | rs16907085 | -0.422 | 1.224 | -0.535 | 1.372 | 0.017 | 0.036 | 0.036 | 0 | 0.09 | 0.1 | 0.19 | 0.25 | 0.15 | 0.13 |
| 10 | 56933271 | rs12572873 | 0.233 | 1.227 | -0.53 | 1.42 | 0.005 | 0.027 | 0.027 | 0 | 0.07 | 0.07 | 0.1 | 0.24 | 0.12 | 0.11 |
| 10 | 56937626 | rs7086823 | -1.057 | 0.814 | -0.778 | 0.868 | -0.009 | -0.013 | -0.013 | 0 | 0.58 | 0.5 | 0.53 | 0.69 | 0.66 | 0.54 |
| 10 | 56937693 | rs7899074 | 1.223 | 0.844 | -0.884 | 0.792 | 0 | -0.002 | -0.002 | 0 | 0.1 | 0.11 | 0.06 | 0.14 | 0.12 | 0.09 |
| 10 | 56947195 | rs2589423 | 1.525 | 0.379 | -1.341 | 0.464 | -0.074 | -0.05 | -0.05 | 1.709 | 0.26 | 0.02 | 0.35 | 0.25 | 0.25 | 0.28 |
| 10 | 56947230 | rs12265891 | -0.603 | 1.055 | -0.901 | 0.56 | -0.003 | 0.048 | 0.048 | 0.339 | 0.49 | 0.34 | 0.43 | 0.69 | 0.51 | 0.49 |
| 10 | 56957207 | rs7091255 | -0.603 | 1.23 | -0.816 | 0.636 | -0.002 | 0.02 | 0.02 | 1.603 | 0.49 | 0.41 | 0.43 | 0.69 | 0.51 | 0.5 |
| 10 | 56960923 | rs7100925 | -1.272 | 1.217 | -0.948 | 0.621 | -0.018 | -0.008 | -0.008 | 0 | 0.6 | 0.5 | 0.56 | 0.66 | 0.68 | 0.53 |
| 10 | 56961505 | rs2166153 | 0.006 | 1.268 | -1.034 | 0.675 | -0.003 | -0.007 | -0.007 | 0.082 | 0.43 | 0.19 | 0.34 | 0.43 | 0.41 | 0.36 |
| 10 | 56970011 | rs2589450 | -0.518 | 1.259 | -1.072 | 0.569 | 0.011 | -0.01 | -0.01 | 0 | 0.59 | 0.8 | 0.67 | 0.61 | 0.65 | 0.66 |
| 10 | 56970216 | rs10509027 | 0.47 | 1.325 | -1.09 | 0.533 | -0.051 | 0.057 | 0.057 | 0 | 0.11 | 0.02 | 0.12 | 0 | 0.22 | 0.06 |
| 10 | 56971092 | rs16907120 | 0.521 | 1.344 | -1.038 | 0.6 | 0.013 | 0.057 | 0.057 | 0 | 0.07 | 0.1 | 0.12 | 0.27 | 0.12 | 0.11 |
| 10 | 56972812 | rs2247977 | -1.363 | 1.358 | -1.176 | 0.575 | 0.001 | -0.004 | -0.004 | 0.274 | 0.93 | 0.93 | 0.96 | 0.89 | 0.94 | 0.92 |
| 10 | 56974190 | rs11004778 | 0.566 | 1.362 | -1.205 | 0.556 | 0.004 | 0.008 | 0.008 | 0 | 0.34 | 0.14 | 0.29 | 0.28 | 0.29 | 0.26 |
| 10 | 56974951 | rs11004779 | -0.21 | 1.349 | -1.197 | 0.56 | -0.042 | 0.077 | 0.077 | 0 | 0.46 | 0.18 | 0.41 | 0.28 | 0.51 | 0.35 |
| 10 | 56989733 | rs16907126 | -0.488 | 1.024 | -0.576 | 0.959 | 0.051 | 0.137 | 0.137 | 0 | 0.09 | 0.23 | 0.15 | 0.27 | 0.13 | 0.14 |
| 10 | 56993816 | rs2680327 | 1.141 | 1.029 | -0.592 | 0.913 | 0 | 0.003 | 0.003 | 2.131 | 0.43 | 0.41 | 0.47 | 0.33 | 0.43 | 0.45 |

| | | | | | | | | | | | | | | | | |
|---|---|---|---|---|---|---|---|---|---|---|---|---|---|---|---|---|
| 10 | 56994603 | rs2680328 | 0.698 | 0.994 | -0.702 | 0.945 | 0.029 | 0.134 | 0.134 | 0.036 | 0.83 | 0.67 | 0.81 | 0.62 | 0.82 | 0.79 |
| 10 | 56996554 | rs17611966 | 0.168 | 1.04 | -1.25 | 0.857 | 0.007 | 0.01 | 0.01 | 0 | 0.35 | 0.17 | 0.29 | 0.29 | 0.29 | 0.27 |
| 10 | 57000058 | rs17611988 | -0.196 | 1.102 | -1.42 | 0.874 | -0.009 | 0.043 | 0.043 | 0 | 0.4 | 0.17 | 0.34 | 0.27 | 0.38 | 0.32 |
| 10 | 57001859 | rs16907157 | -0.623 | 0.842 | -1.087 | 1.115 | 0.045 | 0.135 | 0.135 | 0 | 0.08 | 0.18 | 0.14 | 0.27 | 0.12 | 0.16 |
| 10 | 57004253 | rs10509029 | 1.375 | 0.891 | -1.427 | 0.635 | -0.022 | 0.068 | 0.068 | 0 | 0.18 | 0.02 | 0.11 | 0.08 | 0.15 | 0.15 |
| 10 | 57007337 | rs7083999 | -0.007 | 0.987 | -1.24 | 0.741 | 0.022 | 0.042 | 0.042 | 0.024 | 0.37 | 0.17 | 0.32 | 0.26 | 0.31 | 0.28 |
| 10 | 57008142 | rs12252515 | 0.421 | 1.473 | -0.562 | 1.083 | 0 | 0 | 0 | 0 | 0.06 | 0.06 | 0.04 | 0.12 | 0.06 | 0.06 |
| 10 | 57012515 | rs10509031 | -0.34 | 1.499 | -0.501 | 1.242 | 0.043 | 0.173 | 0.173 | 0 | 0.05 | 0.12 | 0.11 | 0.35 | 0.11 | 0.12 |
| 10 | 57014225 | rs10763189 | 0.919 | 1.47 | -0.53 | 0.411 | 0.018 | -0.002 | -0.002 | 0 | 0.61 | 0.46 | 0.54 | 0.41 | 0.53 | 0.58 |
| 10 | 57018033 | rs10763190 | 0.52 | 0.739 | -1.024 | 0.09 | 0.001 | 0.004 | 0.004 | 0 | 0.66 | 0.64 | 0.73 | 0.84 | 0.65 | 0.73 |
| 10 | 57019821 | rs2249500 | 0.336 | 0.147 | -0.615 | 0.393 | -0.018 | -0.047 | -0.047 | 0 | 0.82 | 0.78 | 0.89 | 0.97 | 0.89 | 0.89 |
| 10 | 57025991 | rs9971326 | -0.092 | 0.373 | -0.914 | 0.554 | 0.001 | 0.069 | 0.069 | 0 | 0.14 | 0.01 | 0.07 | 0.02 | 0.1 | 0.09 |
| 10 | 57026067 | rs12416366 | -0.521 | 0.71 | -0.212 | 1.045 | 0.034 | 0.383 | 0.383 | 0.017 | 0.1 | 0.37 | 0.26 | 0.37 | 0.09 | 0.15 |
| 10 | 57029123 | rs11004794 | -0.353 | 0.836 | -1.488 | -0.317 | -0.032 | 0.138 | 0.138 | 0.409 | 0.54 | 0.75 | 0.65 | 0.76 | 0.5 | 0.69 |
| 10 | 57034040 | rs1237728 | -0.045 | 0.664 | -1.472 | -0.626 | 0.002 | 0.035 | 0.035 | 0 | 0.81 | 0.89 | 0.89 | 1 | 0.79 | 0.82 |
| 10 | 57035385 | rs12412156 | -0.356 | 0.015 | -0.468 | -0.553 | -0.023 | 0.049 | 0.049 | 0.28 | 0.11 | 0.36 | 0.09 | 0.09 | 0.04 | 0.11 |
| 10 | 57039062 | rs11004796 | 0.191 | 0.065 | -0.993 | -0.921 | -0.005 | 0.02 | 0.02 | 0 | 0.24 | 0.16 | 0.3 | 0.16 | 0.22 | 0.15 |
| 10 | 57045932 | rs2680310 | -1.071 | -0.366 | -1.189 | -1.133 | -0.053 | 0.025 | 0.025 | 0.171 | 0.63 | 0.8 | 0.52 | 0.3 | 0.49 | 0.48 |
| 10 | 57057172 | rs2441771 | -1.559 | 0.314 | -1.408 | -0.649 | -0.03 | -0.104 | -0.104 | 0 | 0.73 | 0.93 | 0.71 | 0.42 | 0.69 | 0.62 |
| 10 | 57059690 | rs16907220 | -0.412 | 0.471 | -0.843 | -0.737 | -0.057 | -0.009 | -0.009 | 0 | 0.18 | 0.51 | 0.14 | 0.13 | 0.09 | 0.18 |
| 10 | 57060014 | rs17541119 | 0.517 | 0.617 | -0.983 | -0.87 | -0.087 | 0.048 | 0.048 | 0 | 0.21 | 0.03 | 0.39 | 0.09 | 0.28 | 0.18 |
| 10 | 57061057 | rs1073665 | -0.647 | -0.009 | -0.496 | -0.461 | -0.06 | -0.005 | -0.005 | 0 | 0.18 | 0.42 | 0.13 | 0.13 | 0.09 | 0.18 |
| 10 | 57077430 | rs16907255 | -0.839 | 0.502 | -0.549 | -0.193 | 0.003 | 0.007 | 0.007 | 0 | 0.12 | 0.17 | 0.12 | 0.21 | 0.15 | 0.27 |
| 10 | 57077635 | rs2441760 | -0.917 | 0.313 | -1.064 | -0.757 | 0.063 | 0.017 | 0.017 | 1.242 | 0.63 | 0.85 | 0.7 | 0.31 | 0.73 | 0.61 |
| 10 | 57089420 | rs2488825 | 1.029 | 0.594 | -1.148 | -1.139 | 0.003 | 0.043 | 0.043 | 0 | 0.21 | 0.16 | 0.14 | 0 | 0.22 | 0.15 |
| 10 | 57095400 | rs1900474 | -0.792 | 0.984 | -0.101 | -0.752 | -0.001 | 0.005 | 0.005 | 0 | 0.41 | 0.59 | 0.56 | 0.31 | 0.51 | 0.46 |
| 10 | 57099471 | rs6481157 | -0.97 | 0.95 | -0.197 | -0.228 | -0.001 | 0.003 | 0.003 | 0.268 | 0.43 | 0.7 | 0.57 | 0.41 | 0.51 | 0.46 |
| 10 | 57108195 | rs1900482 | -0.85 | 1.022 | -0.129 | -0.512 | -0.004 | 0.021 | 0.021 | 0.035 | 0.42 | 0.63 | 0.57 | 0.31 | 0.51 | 0.46 |
| 10 | 57117502 | rs7897491 | -0.954 | 1.018 | -0.205 | -0.515 | -0.003 | 0.04 | 0.04 | 0.23 | 0.42 | 0.69 | 0.57 | 0.29 | 0.51 | 0.46 |
| 10 | 57120283 | rs7918437 | 0.173 | 1.373 | 0.262 | -0.644 | -0.011 | 0.01 | 0.01 | 0 | 0.27 | 0.23 | 0.4 | 0.11 | 0.33 | 0.18 |
| 10 | 57128289 | rs1992821 | -0.808 | 0.979 | -0.059 | -0.534 | -0.004 | 0.051 | 0.051 | 0.207 | 0.43 | 0.68 | 0.55 | 0.3 | 0.51 | 0.44 |
| 10 | 57135857 | rs12784587 | 0.284 | 0.878 | 0.182 | -0.263 | -0.008 | 0.023 | 0.023 | 0 | 0.35 | 0.31 | 0.3 | 0.25 | 0.39 | 0.34 |
| 10 | 57139743 | rs1777675 | -0.699 | 0.963 | 0.433 | -0.574 | 0.002 | 0.04 | 0.04 | 0.072 | 0.61 | 0.74 | 0.47 | 0.29 | 0.62 | 0.49 |
| 10 | 57141578 | rs2441764 | 0.059 | 0.801 | 0.568 | -0.635 | -0.002 | -0.083 | -0.083 | 0 | 0.24 | 0.3 | 0.15 | 0.04 | 0.23 | 0.15 |
| 10 | 57155507 | rs2488834 | -0.108 | 0.788 | 0.653 | -0.451 | -0.01 | -0.132 | -0.132 | 0.369 | 0.26 | 0.43 | 0.17 | 0.04 | 0.23 | 0.15 |
| 10 | 57164148 | rs1777674 | -0.73 | 0.855 | 0.626 | -0.279 | -0.003 | 0.042 | 0.042 | 0.009 | 0.61 | 0.8 | 0.45 | 0.29 | 0.61 | 0.49 |
| 10 | 57166573 | rs10160121 | -0.681 | 0.841 | 0.58 | -0.519 | -0.015 | 0.074 | 0.074 | 0 | 0.57 | 0.69 | 0.35 | 0.04 | 0.53 | 0.38 |
| 10 | 57170638 | rs10763198 | -0.554 | 0.903 | 0.619 | -0.492 | 0 | -0.075 | -0.075 | 0 | 0.55 | 0.53 | 0.33 | 0.04 | 0.53 | 0.37 |
| 10 | 57180675 | rs7916325 | -0.175 | 0.603 | 0.392 | -0.729 | -0.009 | -0.11 | -0.11 | 0 | 0.26 | 0.51 | 0.17 | 0.03 | 0.23 | 0.14 |
| 10 | 57183077 | rs11004845 | 0.973 | 0.572 | -0.18 | -0.768 | -0.063 | 0.233 | 0.233 | 0.028 | 0.31 | 0.02 | 0.17 | 0 | 0.3 | 0.22 |
| 10 | 57183203 | rs11004846 | 0.973 | 0.572 | -0.18 | -0.768 | -0.063 | 0.233 | 0.233 | 0.028 | 0.31 | 0.02 | 0.17 | 0 | 0.3 | 0.22 |
| 10 | 57184517 | rs11004848 | -0.072 | 0.688 | 0.123 | -0.531 | -0.001 | -0.056 | -0.056 | 0.593 | 0.24 | 0.27 | 0.15 | 0.04 | 0.23 | 0.14 |
| 10 | 57188995 | rs11004852 | -0.346 | 0.935 | -0.297 | -0.231 | -0.014 | 0.055 | 0.055 | 0 | 0.59 | 0.68 | 0.38 | 0.19 | 0.53 | 0.38 |
| 10 | 57191812 | rs16907428 | -0.682 | 0.715 | 0.037 | -0.464 | 0 | -0.008 | -0.008 | 1.257 | 0.37 | 0.57 | 0.32 | 0.16 | 0.34 | 0.25 |
| 10 | 57208210 | rs7073734 | -0.381 | 0.846 | 0.505 | -0.795 | 0 | -0.005 | -0.005 | 0 | 0.36 | 0.64 | 0.32 | 0.18 | 0.33 | 0.21 |
| 10 | 57213026 | rs7908118 | 0.766 | 0.824 | 0.541 | -0.179 | -0.013 | -0.048 | -0.048 | 0 | 0.26 | 0.16 | 0.35 | 0.5 | 0.31 | 0.46 |
| 10 | 57213690 | rs9633612 | 0.213 | 0.819 | 0.788 | -0.385 | -0.001 | 0.024 | 0.024 | 0.114 | 0.27 | 0.15 | 0.23 | 0.17 | 0.26 | 0.16 |
| 10 | 57237203 | rs10763206 | 0.413 | 0.804 | 0.599 | -0.274 | 0 | -0.008 | -0.008 | 0 | 0.29 | 0.34 | 0.5 | 0.79 | 0.33 | 0.51 |
| 10 | 57244379 | rs4935594 | -0.449 | 0.834 | 0.543 | -0.29 | -0.001 | -0.006 | -0.006 | 0 | 0.34 | 0.34 | 0.28 | 0.2 | 0.33 | 0.25 |
| 10 | 57263910 | rs2050319 | -1.524 | 0.835 | 0.233 | -0.264 | -0.038 | 0.196 | 0.196 | 0.07 | 0.65 | 0.9 | 0.81 | 0.99 | 0.66 | 0.76 |
| 10 | 57268819 | rs10763215 | 0.791 | 0.874 | 0.268 | -0.178 | -0.012 | -0.048 | -0.048 | 0 | 0.26 | 0.16 | 0.35 | 0.5 | 0.32 | 0.46 |
| 10 | 57272557 | rs10825534 | 0.765 | 0.877 | 0.27 | -0.176 | -0.006 | -0.027 | -0.027 | 0.003 | 0.26 | 0.22 | 0.36 | 0.5 | 0.32 | 0.46 |
| 10 | 57272932 | rs10825535 | 0.765 | 0.877 | 0.26 | -0.176 | -0.006 | -0.029 | -0.029 | 3.189 | 0.26 | 0.21 | 0.36 | 0.5 | 0.32 | 0.46 |
| 10 | 57292460 | rs10509034 | 1.112 | 0.82 | -0.038 | -0.406 | -0.024 | -0.025 | -0.025 | 0 | 0.1 | 0 | 0.14 | 0.11 | 0.12 | 0.11 |
| 10 | 57295962 | rs1413676 | -0.41 | 0.727 | 0.294 | -0.341 | -0.002 | -0.021 | -0.021 | 0.585 | 0.36 | 0.38 | 0.29 | 0.21 | 0.33 | 0.26 |
| 10 | 57306282 | rs1413666 | -1.319 | 0.74 | -0.073 | -0.482 | -0.03 | 0.186 | 0.186 | 0.093 | 0.65 | 0.89 | 0.81 | 0.99 | 0.67 | 0.76 |
| 10 | 57313274 | rs10825541 | 0.677 | 0.699 | 0.382 | -0.18 | 0.002 | 0.016 | 0.016 | 0 | 0.27 | 0.26 | 0.37 | 0.49 | 0.32 | 0.44 |
| 10 | 57335764 | rs16907523 | -0.402 | 0.65 | 0.55 | -0.155 | -0.001 | -0.038 | -0.038 | 0 | 0.35 | 0.42 | 0.29 | 0.21 | 0.33 | 0.27 |
| 10 | 57369117 | rs4336940 | 0.271 | 0.642 | 0.156 | 0.114 | -0.001 | -0.02 | -0.02 | 2.569 | 0.28 | 0.21 | 0.37 | 0.49 | 0.33 | 0.44 |
| 10 | 57390762 | rs2461906 | -0.654 | 0.641 | 0.029 | 0.14 | -0.006 | -0.044 | -0.044 | 0 | 0.73 | 0.81 | 0.63 | 0.52 | 0.67 | 0.56 |
| 10 | 57396751 | rs1916530 | 0.167 | 0.547 | 0.075 | -0.052 | 0 | 0.013 | 0.013 | 0 | 0.16 | 0.09 | 0.26 | 0.12 | 0.15 | 0.16 |
| 10 | 57401749 | rs1916527 | 0.739 | 0.284 | 0.202 | -0.483 | 0 | -0.001 | -0.001 | 0 | 0.36 | 0.34 | 0.27 | 0.41 | 0.36 | 0.34 |
| 10 | 57405689 | rs11004942 | 0.789 | 0.295 | 0.228 | -0.493 | 0.001 | 0.009 | 0.009 | 0 | 0.16 | 0.1 | 0.26 | 0.12 | 0.15 | 0.17 |
| 10 | 57409695 | rs1922135 | -0.653 | 0.12 | 0.432 | -0.556 | -0.016 | -0.02 | -0.02 | 0 | 0.56 | 0.68 | 0.56 | 0.53 | 0.52 | 0.5 |
| 10 | 57416961 | rs1916521 | 0.784 | 0.122 | 0.511 | -0.552 | -0.001 | -0.001 | -0.001 | 0 | 0.37 | 0.37 | 0.27 | 0.41 | 0.36 | 0.34 |
| 10 | 57419530 | rs2461899 | 1.531 | 0.113 | 0.052 | -0.738 | -0.057 | -0.105 | -0.105 | 1.955 | 0.26 | 0.02 | 0.19 | 0.34 | 0.26 | 0.21 |
| 10 | 57420417 | rs2463951 | -1.732 | 0.401 | 0.131 | -0.347 | -0.036 | -0.093 | -0.093 | 0.404 | 0.82 | 0.95 | 0.73 | 0.67 | 0.77 | 0.68 |
| 10 | 57424219 | rs12261063 | 0.255 | 0.327 | 0.34 | -0.514 | -0.007 | -0.03 | -0.03 | 1.133 | 0.19 | 0.3 | 0.29 | 0.12 | 0.16 | 0.18 |
| 10 | 57427515 | rs10763222 | 0.467 | 0.383 | 0.237 | -0.628 | 0.001 | 0.018 | 0.018 | 0 | 0.49 | 0.38 | 0.36 | 0.41 | 0.51 | 0.37 |

| | | | | | | | | | | | | | | | | |
|---|---|---|---|---|---|---|---|---|---|---|---|---|---|---|---|---|
| 10 | 57429570 | rs7898669 | 0.432 | 0.441 | 0.369 | -0.603 | -0.001 | 0.012 | 0.012 | 0 | 0.51 | 0.5 | 0.37 | 0.42 | 0.51 | 0.37 |
| 10 | 57430146 | rs7922805 | 0.573 | 0.442 | 0.388 | -0.603 | 0 | 0.001 | 0.001 | 0 | 0.5 | 0.41 | 0.36 | 0.42 | 0.51 | 0.37 |
| 10 | 57430517 | rs11004954 | 0.466 | 0.438 | 0.76 | -0.609 | 0 | 0.052 | 0.052 | 0 | 0.5 | 0.24 | 0.35 | 0.42 | 0.51 | 0.37 |
| 10 | 57432506 | rs10763223 | -0.616 | 0.399 | 0.663 | -0.609 | 0 | 0.008 | 0.008 | 0.022 | 0.68 | 0.59 | 0.64 | 0.54 | 0.65 | 0.55 |
| 10 | 57439221 | rs2392951 | -0.85 | 0.392 | 0.716 | -0.612 | -0.006 | -0.034 | -0.034 | 0.069 | 0.71 | 0.73 | 0.66 | 0.54 | 0.66 | 0.55 |
| 10 | 57442231 | rs2392955 | -0.618 | 0.395 | 0.724 | -0.608 | -0.001 | -0.003 | -0.003 | 0 | 0.68 | 0.62 | 0.64 | 0.54 | 0.65 | 0.55 |
| 10 | 57449086 | rs12245354 | 0.721 | 0.38 | 0.765 | -0.634 | -0.004 | -0.02 | -0.02 | 0 | 0.17 | 0.26 | 0.28 | 0.12 | 0.14 | 0.18 |
| 10 | 57456261 | rs7902447 | -0.492 | 0.377 | 0.784 | -0.632 | 0 | 0.029 | 0.029 | 0.047 | 0.67 | 0.5 | 0.63 | 0.54 | 0.65 | 0.55 |
| 10 | 57458863 | rs3936639 | 0.585 | 0.386 | 0.725 | -0.631 | 0 | 0.028 | 0.028 | 0 | 0.51 | 0.34 | 0.36 | 0.42 | 0.51 | 0.37 |
| 10 | 57460416 | rs6481176 | -0.668 | 0.396 | 0.543 | -0.603 | -0.002 | -0.017 | -0.017 | 0 | 0.68 | 0.65 | 0.65 | 0.54 | 0.65 | 0.55 |
| 10 | 57476292 | rs6481177 | 0.388 | 0.419 | 0.578 | -0.592 | -0.001 | 0.023 | 0.023 | 0 | 0.52 | 0.54 | 0.38 | 0.42 | 0.51 | 0.37 |
| 10 | 57494790 | rs2384675 | -0.816 | 0.386 | 0.429 | -0.588 | 0.009 | 0.054 | 0.054 | 0.166 | 0.7 | 0.49 | 0.64 | 0.54 | 0.66 | 0.54 |
| 10 | 57497070 | rs1917824 | -0.719 | 0.405 | 0.464 | -0.581 | 0.002 | 0.014 | 0.014 | 0.328 | 0.7 | 0.59 | 0.65 | 0.54 | 0.66 | 0.54 |
| 10 | 57497638 | rs10733938 | -0.356 | 0.366 | 0.458 | -0.603 | 0.002 | 0.049 | 0.049 | 0.017 | 0.66 | 0.38 | 0.64 | 0.54 | 0.65 | 0.54 |
| 10 | 57499265 | rs10740630 | -0.417 | 0.331 | 0.343 | -0.599 | 0.003 | 0.037 | 0.037 | 0 | 0.67 | 0.44 | 0.64 | 0.54 | 0.65 | 0.54 |
| 10 | 57499807 | rs2177367 | -0.725 | 0.437 | 0.11 | -0.516 | -0.001 | -0.001 | -0.001 | 0 | 0.71 | 0.63 | 0.66 | 0.54 | 0.66 | 0.54 |
| 10 | 57504582 | rs10509041 | 0.107 | 0.246 | 0.598 | -0.045 | 0.004 | 0.018 | 0.018 | 0 | 0.17 | 0.09 | 0.27 | 0.12 | 0.14 | 0.19 |
| 10 | 57521574 | rs7915549 | -1.144 | 0.216 | 0.58 | -0.017 | -0.01 | 0.045 | 0.045 | 0 | 0.3 | 0.49 | 0.41 | 0.52 | 0.25 | 0.36 |
| 10 | 57524666 | rs11005008 | 0.01 | 0.218 | 0.538 | 0.003 | 0.006 | 0.127 | 0.127 | 0 | 0.12 | 0.19 | 0.11 | 0.41 | 0.11 | 0.17 |
| 10 | 57537162 | rs11005017 | -0.961 | 0.205 | -0.056 | -0.043 | 0.001 | -0.004 | -0.004 | 0 | 0.31 | 0.59 | 0.41 | 0.52 | 0.25 | 0.36 |
| 10 | 57545246 | rs7897503 | 0.17 | 0.29 | -0.036 | -0.082 | -0.016 | -0.006 | -0.006 | 0.116 | 0.15 | 0.29 | 0.16 | 0.11 | 0.09 | 0.13 |
| 10 | 57547092 | rs11005023 | 1.007 | 0.557 | -0.907 | 0.199 | -0.059 | -0.075 | -0.075 | 0 | 0.25 | 0.06 | 0.22 | 0.33 | 0.3 | 0.31 |
| 10 | 57549296 | rs16907840 | 0.88 | 0.336 | -0.825 | -0.509 | 0.001 | -0.017 | -0.017 | 0 | 0.23 | 0.15 | 0.15 | 0 | 0.23 | 0.05 |
| 10 | 57557708 | rs1916487 | 1.266 | 0.122 | -0.566 | -0.585 | -0.019 | -0.007 | -0.007 | 0.047 | 0.15 | 0.29 | 0.17 | 0.11 | 0.09 | 0.13 |
| 10 | 57560172 | rs7898484 | -0.272 | 0.072 | -0.406 | -0.521 | -0.003 | 0.036 | 0.036 | 0 | 0.53 | 0.76 | 0.57 | 0.62 | 0.52 | 0.55 |
| 10 | 57563408 | rs2893661 | -2.267 | -0.073 | -0.543 | -0.491 | 0.01 | -0.016 | -0.016 | 0 | 0.89 | 0.91 | 0.83 | 0.85 | 0.92 | 0.83 |
| 10 | 57570418 | rs1933909 | 0.992 | -0.078 | -0.703 | -0.561 | -0.021 | -0.025 | -0.025 | 0 | 0.2 | 0.06 | 0.17 | 0.23 | 0.25 | 0.25 |
| 10 | 57571135 | rs10763263 | -1.268 | 0.074 | 0.004 | -0.51 | 0 | -0.001 | -0.001 | 0 | 0.58 | 0.47 | 0.59 | 0.56 | 0.61 | 0.58 |
| 10 | 57574054 | rs1856438 | -1.746 | 0.063 | 0.131 | -0.697 | -0.011 | -0.006 | -0.006 | 0 | 0.84 | 0.76 | 0.84 | 0.89 | 0.9 | 0.87 |
| 10 | 57575119 | rs2392962 | -0.268 | 0.056 | 0.221 | -0.705 | 0.006 | 0.036 | 0.036 | 0 | 0.47 | 0.32 | 0.43 | 0.38 | 0.48 | 0.45 |
| 10 | 57584038 | rs2153136 | 0.316 | 0.069 | 0.209 | -0.709 | 0.005 | -0.002 | -0.002 | 3.18 | 0.53 | 0.47 | 0.5 | 0.51 | 0.58 | 0.45 |
| 10 | 57586185 | rs4304686 | 0.363 | 0.112 | 0.157 | -0.66 | 0.002 | -0.039 | -0.039 | 0 | 0.31 | 0.17 | 0.33 | 0.38 | 0.33 | 0.42 |
| 10 | 57587064 | rs11005046 | 1.947 | 0.241 | 0.218 | -0.532 | 0.006 | 0.015 | 0.015 | 0 | 0.26 | 0.23 | 0.25 | 0.39 | 0.3 | 0.29 |
| 10 | 57594500 | rs4587661 | 1.079 | 0.238 | 0.138 | -0.53 | 0.008 | 0 | 0 | 0 | 0.11 | 0.16 | 0.15 | 0.13 | 0.13 | 0.14 |
| 10 | 57595769 | rs1361258 | 1.801 | -0.052 | 0.194 | -0.514 | -0.003 | -0.006 | -0.006 | 0.873 | 0.26 | 0.17 | 0.24 | 0.38 | 0.3 | 0.3 |
| 10 | 57597140 | rs10740636 | 0.477 | -0.342 | 0.37 | -0.471 | -0.002 | -0.001 | -0.001 | 0 | 0.54 | 0.46 | 0.49 | 0.58 | 0.62 | 0.61 |
| 10 | 57601709 | rs10763269 | 0.151 | -0.082 | 0.439 | -0.637 | 0.02 | 0.033 | 0.033 | 0 | 0.29 | 0.19 | 0.15 | 0.01 | 0.22 | 0.16 |
| 10 | 57605472 | rs7086413 | 0.686 | -0.003 | 0.519 | -0.492 | -0.008 | -0.002 | -0.002 | 0 | 0.6 | 0.32 | 0.42 | 0.41 | 0.56 | 0.49 |
| 10 | 57611683 | rs1538424 | -0.203 | -0.055 | 0.711 | -0.5 | -0.022 | 0.087 | 0.087 | 0 | 0.72 | 0.53 | 0.59 | 0.54 | 0.77 | 0.64 |
| 10 | 57616711 | rs10733940 | 0.625 | -0.029 | 0.72 | -0.499 | -0.001 | -0.001 | -0.001 | 0 | 0.61 | 0.37 | 0.42 | 0.41 | 0.56 | 0.5 |
| 10 | 57621292 | rs12414696 | 1.577 | -0.087 | 0.375 | -0.547 | 0 | 0.027 | 0.027 | 0.114 | 0.16 | 0.08 | 0.1 | 0 | 0.15 | 0.03 |
| 10 | 57624771 | rs12770927 | 1.144 | -0.005 | 0.413 | -0.504 | 0.03 | 0.001 | 0.001 | 0 | 0.11 | 0.17 | 0.17 | 0.13 | 0.21 | 0.15 |
| 10 | 57625173 | rs12774407 | 1.197 | -0.004 | 0.412 | -0.504 | 0.018 | 0 | 0 | 0 | 0.11 | 0.13 | 0.17 | 0.13 | 0.21 | 0.15 |
| 10 | 57628056 | rs17551858 | 1.727 | -0.024 | 0.211 | -0.484 | -0.002 | 0.022 | 0.022 | 0.048 | 0.15 | 0.08 | 0.1 | 0 | 0.15 | 0.03 |
| 10 | 57630800 | rs1113253 | 1.538 | -0.067 | 0.153 | -0.445 | 0.012 | 0 | 0 | 0.592 | 0.14 | 0.11 | 0.05 | 0.01 | 0.06 | 0.14 |
| 10 | 57634812 | rs7902926 | -0.207 | 0.215 | 0.493 | -0.287 | -0.004 | 0.014 | 0.014 | 0 | 0.73 | 0.68 | 0.61 | 0.54 | 0.78 | 0.64 |
| 10 | 57634920 | rs7909547 | -0.025 | 0.285 | -0.383 | -0.24 | -0.021 | 0.052 | 0.052 | 0 | 0.72 | 0.46 | 0.59 | 0.58 | 0.77 | 0.69 |
| 10 | 57638469 | rs2211082 | 1.286 | 0.597 | -1.155 | 0.082 | -0.017 | -0.053 | -0.053 | 0 | 0.11 | 0.01 | 0.12 | 0.21 | 0.12 | 0.14 |
| 10 | 57647581 | rs7908845 | 1.21 | 0.709 | -0.903 | 0.08 | -0.047 | -0.064 | -0.064 | 0 | 0.31 | 0.12 | 0.27 | 0.4 | 0.35 | 0.33 |
| 10 | 57667668 | rs12764655 | 0.277 | 0.068 | -0.944 | -0.11 | 0.007 | -0.001 | -0.001 | 1.305 | 0.13 | 0.15 | 0.15 | 0.13 | 0.18 | 0.2 |
| 10 | 57669027 | rs1338803 | 1.382 | 0.337 | -1.149 | -0.039 | -0.079 | -0.146 | -0.146 | 2.806 | 0.29 | 0.03 | 0.25 | 0.41 | 0.32 | 0.32 |
| 10 | 57671424 | rs11005106 | 0.53 | 0.193 | -0.487 | -0.587 | 0.009 | -0.008 | -0.008 | 2.01 | 0.16 | 0.13 | 0.06 | 0.01 | 0.07 | 0.14 |
| 10 | 57678522 | rs10825663 | -0.99 | 0.225 | -0.557 | -0.584 | 0.001 | -0.003 | -0.003 | 0 | 0.29 | 0.27 | 0.21 | 0.15 | 0.25 | 0.34 |
| 10 | 57683893 | rs12782822 | -0.126 | 1.014 | -0.791 | 0.007 | -0.009 | 0 | 0 | 1.036 | 0.12 | 0.07 | 0.15 | 0.13 | 0.18 | 0.2 |
| 10 | 57690158 | rs1572688 | -0.198 | 0.875 | -0.814 | -0.178 | 0.009 | 0.097 | 0.097 | 0 | 0.5 | 0.35 | 0.6 | 0.81 | 0.57 | 0.56 |
| 10 | 57696970 | rs16908017 | -0.176 | 0.892 | -0.828 | 0.021 | 0.024 | 0.048 | 0.048 | 0 | 0.13 | 0.22 | 0.26 | 0.27 | 0.19 | 0.21 |
| 10 | 57696982 | rs4515893 | 0.837 | 0.867 | -0.941 | 0.043 | -0.023 | -0.074 | -0.074 | 0 | 0.36 | 0.12 | 0.34 | 0.55 | 0.38 | 0.35 |
| 10 | 57706974 | rs17639747 | -0.445 | 1.179 | -0.679 | 0.139 | 0.008 | 0.014 | 0.014 | 0 | 0.14 | 0.14 | 0.26 | 0.27 | 0.21 | 0.22 |
| 10 | 57707708 | rs16908031 | -0.238 | 1.235 | -0.458 | 0.264 | 0.086 | 0.08 | 0.08 | 0 | 0.1 | 0.3 | 0.19 | 0.16 | 0.14 | 0.16 |
| 10 | 57710042 | rs1908345 | 0.056 | 0.995 | -0.515 | 0.152 | -0.004 | 0.026 | 0.026 | 0 | 0.46 | 0.38 | 0.51 | 0.7 | 0.53 | 0.5 |
| 10 | 57710416 | rs7904250 | 0.862 | 0.998 | -0.671 | 0.168 | -0.028 | -0.106 | -0.106 | 0.037 | 0.37 | 0.08 | 0.32 | 0.54 | 0.38 | 0.33 |
| 10 | 57714991 | rs11005123 | 0.877 | 0.904 | -0.657 | 0.172 | -0.023 | -0.099 | -0.099 | 0 | 0.37 | 0.08 | 0.32 | 0.54 | 0.38 | 0.34 |
| 10 | 57718804 | rs10430551 | 0.208 | 0.9 | -0.579 | 0.073 | -0.023 | 0.072 | 0.072 | 0 | 0.45 | 0.22 | 0.5 | 0.7 | 0.53 | 0.5 |
| 10 | 57722576 | rs1908348 | 0.38 | 0.212 | -0.637 | 0.002 | 0.012 | 0.066 | 0.066 | 0.138 | 0.43 | 0.27 | 0.43 | 0.71 | 0.39 | 0.46 |
| 10 | 57724824 | rs4297390 | 0.868 | 0.204 | -0.643 | 0.005 | -0.003 | 0.001 | 0.001 | 0 | 0.06 | 0.06 | 0.03 | 0.06 | 0.02 | 0.08 |
| 10 | 57724936 | rs4485029 | 0.419 | 0.272 | -0.621 | -0.007 | 0.012 | 0.064 | 0.064 | 0.138 | 0.43 | 0.27 | 0.44 | 0.71 | 0.39 | 0.46 |
| 10 | 57726702 | rs7101054 | 0.437 | 0.266 | -0.623 | -0.028 | 0.01 | 0.064 | 0.064 | 0.072 | 0.43 | 0.27 | 0.44 | 0.71 | 0.39 | 0.45 |
| 10 | 57727193 | rs1908363 | -0.126 | 0.027 | -0.557 | -0.259 | -0.001 | -0.02 | -0.02 | 0.256 | 0.56 | 0.47 | 0.57 | 0.84 | 0.48 | 0.61 |
| 10 | 57736009 | rs12248518 | -0.754 | 0.003 | -0.176 | 0.019 | -0.033 | 0.046 | 0.046 | 0 | 0.26 | 0.43 | 0.32 | 0.37 | 0.22 | 0.39 |

| | | | | | | | | | | | | | | | | |
|---|---|---|---|---|---|---|---|---|---|---|---|---|---|---|---|---|
| 10 | 57747976 | rs11005161 | 0.448 | -0.068 | -0.644 | -0.273 | 0.012 | 0.158 | 0.158 | 0 | 0.49 | 0.15 | 0.42 | 0.73 | 0.45 | 0.5 |
| 10 | 57750714 | rs1874155 | 0.17 | 0.093 | 0.106 | -0.221 | 0.013 | -0.013 | -0.013 | 0 | 0.12 | 0.32 | 0.11 | 0.08 | 0.12 | 0.14 |
| 10 | 57757228 | rs7093287 | 1.16 | -0.114 | -0.034 | -0.156 | -0.018 | -0.017 | -0.017 | 1.816 | 0.38 | 0.16 | 0.33 | 0.59 | 0.39 | 0.34 |
| 10 | 57762772 | rs10763292 | 1.16 | -0.114 | -0.034 | -0.156 | -0.018 | -0.017 | -0.017 | 0 | 0.38 | 0.16 | 0.33 | 0.59 | 0.39 | 0.34 |
| 10 | 57766407 | rs10825720 | 1.135 | -0.147 | 0.284 | -0.152 | -0.007 | -0.006 | -0.006 | 0 | 0.38 | 0.21 | 0.33 | 0.6 | 0.39 | 0.34 |
| 10 | 57769143 | rs10825723 | 1.165 | -0.153 | 0.525 | -0.155 | -0.017 | -0.016 | -0.016 | 0 | 0.38 | 0.16 | 0.33 | 0.59 | 0.39 | 0.34 |
| 10 | 57770652 | rs10825726 | 1.22 | -0.167 | 0.536 | -0.117 | -0.029 | -0.051 | -0.051 | 2.1 | 0.37 | 0.12 | 0.33 | 0.55 | 0.39 | 0.31 |
| 10 | 57776775 | rs7067574 | 1.144 | -0.178 | 0.529 | -0.122 | -0.017 | -0.016 | -0.016 | 0.056 | 0.38 | 0.16 | 0.33 | 0.59 | 0.39 | 0.34 |
| 10 | 57787290 | rs10740647 | 0.063 | -0.097 | 0.547 | -0.25 | -0.002 | 0.024 | 0.024 | 0 | 0.55 | 0.36 | 0.45 | 0.69 | 0.56 | 0.53 |
| 10 | 57792128 | rs9415368 | -0.008 | -0.094 | 0.518 | -0.255 | 0 | -0.026 | -0.026 | 2.295 | 0.56 | 0.48 | 0.47 | 0.73 | 0.56 | 0.57 |
| 10 | 57794104 | rs9415372 | -0.007 | -0.153 | 0.88 | -0.28 | -0.001 | 0.008 | 0.008 | 0 | 0.55 | 0.39 | 0.44 | 0.73 | 0.56 | 0.56 |
| 10 | 57795477 | rs1587516 | 1.379 | -0.137 | 0.899 | -0.259 | -0.03 | -0.044 | -0.044 | 0.153 | 0.37 | 0.11 | 0.31 | 0.55 | 0.39 | 0.29 |
| 10 | 57800284 | rs11005194 | 1.267 | -0.117 | 0.905 | -0.262 | 0.005 | -0.005 | -0.005 | 0.924 | 0.05 | 0.01 | 0.03 | 0.06 | 0.04 | 0.08 |
| 10 | 57808398 | rs4586075 | -0.03 | -0.131 | 0.926 | -0.276 | -0.004 | 0.031 | 0.031 | 0 | 0.56 | 0.6 | 0.56 | 0.82 | 0.57 | 0.62 |
| 10 | 57809570 | rs9415380 | -0.172 | -0.194 | 1.137 | -0.303 | -0.017 | 0.099 | 0.099 | 0 | 0.57 | 0.76 | 0.57 | 0.82 | 0.57 | 0.62 |
| 10 | 57819376 | rs7078466 | 0.022 | -0.161 | 0.985 | -0.289 | -0.001 | -0.005 | -0.005 | 0 | 0.55 | 0.54 | 0.56 | 0.83 | 0.57 | 0.62 |
| 10 | 57843879 | rs1875290 | 0.424 | -0.131 | 1.054 | -0.259 | 0 | 0.013 | 0.013 | 0 | 0.5 | 0.52 | 0.52 | 0.77 | 0.53 | 0.54 |
| 10 | 57844640 | rs1908354 | 0.415 | -0.139 | 1.063 | -0.253 | 0 | 0.038 | 0.038 | 0 | 0.49 | 0.34 | 0.41 | 0.63 | 0.53 | 0.44 |
| 10 | 57850451 | rs9415410 | 0.377 | -0.132 | 1.059 | -0.257 | 0.001 | 0.05 | 0.05 | 0 | 0.51 | 0.58 | 0.53 | 0.77 | 0.53 | 0.54 |
| 10 | 57852543 | rs7910631 | 0.423 | -0.129 | 1.014 | -0.256 | 0 | 0.013 | 0.013 | 0 | 0.5 | 0.52 | 0.52 | 0.77 | 0.53 | 0.54 |
| 10 | 57868529 | rs11005218 | 0.465 | 0.018 | 1.041 | -0.155 | 0.012 | -0.008 | -0.008 | 0 | 0.11 | 0.16 | 0.1 | 0.06 | 0.13 | 0.14 |
| 10 | 57892777 | rs2454994 | -0.689 | -0.23 | 0.303 | -0.259 | -0.041 | 0.019 | 0.019 | 0 | 0.84 | 0.74 | 0.77 | 0.71 | 0.94 | 0.72 |
| 10 | 57914433 | rs12785159 | 0.336 | -0.132 | -0.106 | -0.557 | -0.016 | 0.006 | 0.006 | 0 | 0.2 | 0.15 | 0.19 | 0.06 | 0.33 | 0.14 |
| 10 | 57916382 | rs10733945 | -1.052 | -0.144 | -0.116 | -0.526 | 0.053 | -0.024 | -0.024 | 0 | 0.85 | 0.95 | 0.89 | 0.81 | 0.94 | 0.78 |
| 10 | 57932693 | rs7350417 | -0.298 | -0.181 | -0.047 | -0.134 | 0.01 | -0.005 | -0.005 | 0 | 0.26 | 0.28 | 0.23 | 0.16 | 0.36 | 0.23 |
| 10 | 57941502 | rs2392998 | -1.022 | -0.316 | -0.268 | -0.07 | 0.048 | -0.02 | -0.02 | 0 | 0.85 | 0.95 | 0.89 | 0.81 | 0.94 | 0.78 |
| 10 | 57982870 | rs7070470 | 0.295 | 0.357 | -0.262 | -0.035 | -0.005 | -0.001 | -0.001 | 0.614 | 0.17 | 0.11 | 0.14 | 0.11 | 0.23 | 0.14 |
| 10 | 57996884 | rs2435852 | -0.395 | 0.326 | -0.266 | -0.082 | -0.025 | 0.013 | 0.013 | 0 | 0.27 | 0.17 | 0.23 | 0.16 | 0.37 | 0.25 |
| 10 | 57997348 | rs10825774 | 1.394 | -0.011 | -1.014 | -0.201 | -0.09 | -0.174 | -0.174 | 0.121 | 0.34 | 0.04 | 0.28 | 0.47 | 0.35 | 0.27 |
| 10 | 57999830 | rs2435858 | 1.052 | -0.132 | -0.843 | -0.201 | 0.049 | -0.007 | -0.007 | 0.012 | 0.18 | 0.07 | 0.13 | 0.19 | 0.11 | 0.24 |
| 10 | 58005331 | rs2893680 | 1.547 | 0.254 | -1.148 | -0.156 | -0.028 | -0.153 | -0.153 | 0 | 0.22 | 0.02 | 0.18 | 0.4 | 0.2 | 0.14 |
| 10 | 58015268 | rs4631808 | 1.385 | 0.188 | -0.792 | 0.431 | -0.09 | -0.083 | -0.083 | 0.083 | 0.34 | 0.04 | 0.28 | 0.34 | 0.35 | 0.27 |
| 10 | 58023584 | rs2245618 | -0.085 | 0.199 | -0.303 | -0.355 | 0.005 | -0.003 | -0.003 | 0.871 | 0.73 | 0.73 | 0.77 | 0.83 | 0.63 | 0.76 |
| 10 | 58032727 | rs2462476 | -0.195 | 0.081 | 0.067 | -0.42 | -0.014 | -0.018 | -0.018 | 0.056 | 0.37 | 0.42 | 0.36 | 0.25 | 0.27 | 0.45 |
| 10 | 58037493 | rs2462456 | -1.139 | 0.387 | -0.227 | -0.359 | -0.011 | 0.002 | 0.002 | 0 | 0.88 | 0.93 | 0.89 | 1 | 0.81 | 0.96 |
| 10 | 58040286 | rs6481203 | -0.354 | -0.563 | -0.022 | -0.249 | -0.03 | -0.018 | -0.018 | 1.299 | 0.38 | 0.53 | 0.46 | 0.34 | 0.28 | 0.49 |
| 10 | 58060764 | rs7923504 | 0.763 | -0.633 | 0.593 | -0.323 | -0.009 | -0.003 | -0.003 | 1.283 | 0.18 | 0.19 | 0.15 | 0.18 | 0.12 | 0.25 |
| 10 | 58063346 | rs10825782 | 1.505 | -0.583 | 0.317 | -0.369 | -0.064 | -0.148 | -0.148 | 0 | 0.34 | 0.06 | 0.29 | 0.48 | 0.35 | 0.27 |
| 10 | 58067127 | rs7099218 | 0.424 | -0.597 | 0.44 | -0.382 | -0.008 | -0.053 | -0.053 | 0 | 0.19 | 0.33 | 0.22 | 0.07 | 0.17 | 0.22 |
| 10 | 58074938 | rs16908708 | 1.433 | -0.617 | 0.39 | -0.379 | -0.011 | -0.035 | -0.035 | 0 | 0.34 | 0.22 | 0.4 | 0.58 | 0.35 | 0.28 |
| 10 | 58077694 | rs7909329 | -0.452 | -0.633 | 0.429 | -0.386 | 0.002 | 0.001 | 0.001 | 0 | 0.37 | 0.62 | 0.47 | 0.35 | 0.29 | 0.51 |
| 10 | 58079353 | rs12249018 | -0.354 | -0.656 | 0.528 | -0.395 | -0.021 | -0.036 | -0.036 | 0 | 0.37 | 0.5 | 0.37 | 0.25 | 0.28 | 0.47 |
| 10 | 58082333 | rs11005311 | 1.57 | -0.644 | 0.417 | -0.395 | -0.052 | -0.128 | -0.128 | 1.543 | 0.33 | 0.08 | 0.29 | 0.49 | 0.35 | 0.24 |
| 10 | 58083638 | rs2393059 | -0.372 | -0.591 | 0.49 | -0.365 | -0.006 | 0.006 | 0.006 | 0 | 0.71 | 0.75 | 0.77 | 0.84 | 0.64 | 0.75 |
| 10 | 58086141 | rs4144609 | 0.698 | -0.651 | 0.485 | -0.382 | -0.011 | 0.083 | 0.083 | 0.71 | 0.51 | 0.19 | 0.43 | 0.67 | 0.46 | 0.49 |
| 10 | 58086503 | rs12263637 | 0.191 | -0.641 | 0.513 | -0.419 | -0.015 | -0.027 | -0.027 | 0 | 0.2 | 0.55 | 0.34 | 0.16 | 0.18 | 0.27 |
| 10 | 58090264 | rs9971280 | 0.014 | -0.644 | 0.599 | -0.666 | -0.018 | -0.051 | -0.051 | 1.188 | 0.21 | 0.62 | 0.35 | 0.1 | 0.17 | 0.22 |
| 10 | 58096806 | rs7918738 | 1.507 | -0.666 | 0.527 | -0.673 | -0.052 | -0.053 | -0.053 | 0 | 0.34 | 0.13 | 0.39 | 0.65 | 0.35 | 0.32 |
| 10 | 58099085 | rs2393068 | -0.386 | -0.686 | 0.692 | -0.692 | 0.023 | -0.004 | -0.004 | 0 | 0.7 | 0.49 | 0.65 | 0.73 | 0.63 | 0.71 |
| 10 | 58102205 | rs4144608 | -0.518 | -0.687 | 0.706 | -0.694 | -0.012 | 0.009 | 0.009 | 0 | 0.72 | 0.79 | 0.78 | 0.83 | 0.64 | 0.75 |
| 10 | 58116156 | rs10825784 | 1.623 | -0.643 | 0.694 | -0.699 | -0.099 | -0.152 | -0.152 | 0.184 | 0.33 | 0.04 | 0.28 | 0.55 | 0.35 | 0.28 |
| 10 | 58117124 | rs11005326 | 0.305 | -0.666 | 0.717 | -0.715 | -0.012 | -0.112 | -0.112 | 0 | 0.19 | 0.38 | 0.23 | 0 | 0.17 | 0.18 |
| 10 | 58118630 | rs2241666 | -0.514 | -0.642 | 0.717 | -0.7 | -0.013 | 0.01 | 0.01 | 0 | 0.72 | 0.8 | 0.78 | 0.83 | 0.64 | 0.75 |
| 10 | 58138121 | rs3849972 | 1.512 | -0.644 | 0.813 | -0.63 | -0.065 | -0.118 | -0.118 | 0.185 | 0.34 | 0.06 | 0.29 | 0.54 | 0.35 | 0.28 |
| 10 | 58201703 | rs10825795 | -0.923 | -0.731 | 0.521 | -0.767 | -0.016 | 0.115 | 0.115 | 0.5 | 0.49 | 0.8 | 0.56 | 0.27 | 0.54 | 0.47 |
| 10 | 58239664 | rs4935649 | -0.795 | -0.82 | 0.1 | -0.502 | 0.013 | 0.005 | 0.005 | 0 | 0.81 | 0.87 | 0.85 | 0.81 | 0.89 | 0.74 |
| 10 | 58249479 | rs12761806 | -0.174 | -0.424 | 0.416 | -0.466 | -0.022 | 0.014 | 0.014 | 0 | 0.27 | 0.18 | 0.22 | 0.17 | 0.36 | 0.25 |
| 10 | 58250470 | rs7086393 | 0.522 | -0.615 | 0.511 | -0.611 | 0.001 | 0.032 | 0.032 | 0.105 | 0.59 | 0.3 | 0.5 | 0.73 | 0.58 | 0.63 |
| 10 | 58266235 | rs1907912 | 0.55 | -0.35 | 0.551 | -0.519 | -0.003 | 0.004 | 0.004 | 0.975 | 0.65 | 0.34 | 0.52 | 0.73 | 0.61 | 0.64 |
| 10 | 58276545 | rs11005382 | 0.661 | -0.373 | 0.602 | -0.524 | -0.04 | -0.065 | -0.065 | 0 | 0.12 | 0.27 | 0.15 | 0 | 0.05 | 0.08 |
| 10 | 58281746 | rs1907893 | 0.073 | 0.189 | 0.238 | -0.208 | 0.041 | 0.016 | 0.016 | 0.073 | 0.77 | 0.63 | 0.66 | 0.73 | 0.66 | 0.72 |
| 10 | 58285762 | rs1907895 | 0.402 | -0.572 | -0.556 | -0.239 | 0.043 | 0.012 | 0.012 | 0.402 | 0.47 | 0.28 | 0.48 | 0.54 | 0.37 | 0.43 |
| 10 | 58291800 | rs12220494 | 0.538 | -1.505 | 0.26 | -0.164 | -0.034 | 0.039 | 0.039 | 1.397 | 0.82 | 0.44 | 0.78 | 0.8 | 0.88 | 0.79 |
| 10 | 58296254 | rs10825819 | 0.987 | -1.775 | -0.276 | -0.428 | 0.047 | 0.032 | 0.032 | 2.751 | 0.69 | 0.14 | 0.6 | 0.8 | 0.78 | 0.7 |
| 10 | 58317415 | rs11005401 | 1.109 | -1.846 | 0 | -0.57 | 0.003 | -0.003 | -0.003 | 0 | 0.08 | 0.07 | 0.02 | 0 | 0.04 | 0.02 |
| 10 | 58326495 | rs7070694 | -0.583 | -1.908 | 0.595 | -0.403 | -0.022 | -0.1 | -0.1 | 0.31 | 0.17 | 0.38 | 0.21 | 0 | 0.13 | 0.1 |
| 10 | 58329209 | rs7080327 | 0.448 | -2.121 | -0.014 | -0.552 | 0.008 | 0.025 | 0.025 | 0 | 0.2 | 0.07 | 0.17 | 0.15 | 0.18 | 0.21 |
| 10 | 58338839 | rs1907928 | -0.65 | -2.075 | 0.121 | -0.568 | -0.022 | -0.125 | -0.125 | 0.132 | 0.18 | 0.52 | 0.23 | 0 | 0.14 | 0.1 |
| 10 | 58340060 | rs1907925 | -1.073 | -1.969 | -0.634 | -0.414 | 0.035 | 0.059 | 0.059 | 0 | 0.39 | 0.7 | 0.39 | 0.22 | 0.31 | 0.36 |

| | | | | | | | | | | | | | | | | |
|---|---|---|---|---|---|---|---|---|---|---|---|---|---|---|---|---|
| 10 | 58340191 | rs7474843 | -1.079 | -1.964 | -0.632 | -0.424 | 0.035 | 0.056 | 0.056 | 0 | 0.39 | 0.7 | 0.39 | 0.24 | 0.31 | 0.36 |
| 10 | 58340947 | rs11005411 | -0.795 | -2.403 | -0.648 | -0.322 | 0.016 | 0.006 | 0.006 | 0 | 0.26 | 0.22 | 0.3 | 0.21 | 0.12 | 0.22 |
| 10 | 58341064 | rs1948426 | -0.865 | -2.382 | -0.873 | -0.317 | 0.072 | 0.045 | 0.045 | 0 | 0.26 | 0.14 | 0.2 | 0.13 | 0.13 | 0.2 |
| 10 | 58341455 | rs7090782 | 0.591 | -1.799 | -0.575 | -0.061 | 0.035 | 0.009 | 0.009 | 0.004 | 0.56 | 0.69 | 0.61 | 0.63 | 0.67 | 0.54 |
| 10 | 58342829 | rs4586076 | 0.517 | -1.302 | 0.068 | 0.456 | -0.024 | 0.047 | 0.047 | 0 | 0.87 | 0.81 | 0.81 | 0.75 | 0.93 | 0.85 |
| 10 | 58343819 | rs7909510 | 1.3 | -1.297 | -0.067 | 0.368 | -0.006 | 0.011 | 0.011 | 0 | 0.82 | 0.8 | 0.78 | 0.75 | 0.86 | 0.81 |
| 10 | 58365415 | rs1873800 | -0.803 | -1.091 | 0.143 | 0.109 | 0.002 | -0.008 | -0.008 | 0.312 | 0.39 | 0.59 | 0.3 | 0.3 | 0.39 | 0.38 |
| 10 | 58391125 | rs16909125 | 0.196 | -1.02 | 0.107 | -0.177 | 0.016 | -0.023 | -0.023 | 0 | 0.12 | 0.05 | 0.17 | 0.26 | 0.05 | 0.14 |
| 10 | 58394334 | rs10825827 | -0.107 | -1.005 | 0.662 | -0.339 | 0.018 | -0.097 | -0.097 | 0 | 0.72 | 0.55 | 0.82 | 0.92 | 0.71 | 0.8 |
| 10 | 58424161 | rs10825843 | 0.822 | -1.083 | 0.668 | -0.289 | 0.003 | -0.026 | -0.026 | 0 | 0.62 | 0.41 | 0.7 | 0.78 | 0.61 | 0.63 |
| 10 | 58457011 | rs4622179 | 0.043 | -0.93 | 0.765 | -0.354 | 0.048 | -0.029 | -0.029 | 0 | 0.88 | 0.69 | 0.89 | 0.92 | 0.82 | 0.91 |
| 10 | 58468167 | rs7916845 | 1.407 | -0.957 | 0.807 | -0.36 | 0.044 | 0.002 | 0.002 | 0 | 0.79 | 0.53 | 0.78 | 0.79 | 0.72 | 0.81 |
| 10 | 58479428 | rs7099123 | -0.003 | -0.893 | 0.838 | -0.361 | 0.054 | -0.023 | -0.023 | 0.687 | 0.88 | 0.72 | 0.89 | 0.92 | 0.82 | 0.91 |
| 10 | 58481762 | rs7911138 | 0.839 | -0.985 | 0.785 | -0.393 | 0.013 | -0.03 | -0.03 | 0 | 0.65 | 0.52 | 0.72 | 0.78 | 0.61 | 0.69 |
| 10 | 58486549 | rs10763400 | 1.474 | -1.236 | 0.782 | -0.665 | -0.011 | 0.113 | 0.113 | 0 | 0.54 | 0.21 | 0.65 | 0.77 | 0.56 | 0.56 |
| 10 | 58491092 | rs16909295 | 1.081 | -1.207 | 0.884 | -0.81 | -0.002 | -0.017 | -0.017 | 0 | 0.07 | 0.06 | 0.02 | 0 | 0.05 | 0.06 |
| 10 | 58494646 | rs2203751 | -0.727 | -1.178 | 0.952 | -0.828 | -0.017 | 0 | 0 | 0 | 0.93 | 0.94 | 0.93 | 0.92 | 0.85 | 0.93 |
| 10 | 58497890 | rs12219855 | 1.21 | -1.216 | 1.014 | -0.817 | -0.004 | 0.013 | 0.013 | 0 | 0.57 | 0.36 | 0.68 | 0.8 | 0.55 | 0.63 |
| 10 | 58499813 | rs10825863 | -0.996 | -1.225 | 1.007 | -0.805 | 0 | 0.014 | 0.014 | 0 | 0.1 | 0.2 | 0.13 | 0.12 | 0.12 | 0.11 |
| 10 | 58500557 | rs12776363 | 1.505 | -1.219 | 1.018 | -0.844 | -0.009 | 0.065 | 0.065 | 0 | 0.16 | 0.03 | 0.07 | 0 | 0.11 | 0.11 |
| 10 | 58501439 | rs1914568 | 0.314 | -1.456 | 0.895 | -0.807 | 0.012 | -0.015 | -0.015 | 0 | 0.12 | 0.07 | 0.18 | 0.26 | 0.04 | 0.15 |
| 10 | 58503111 | rs12265565 | -0.946 | -1.406 | 0.835 | -0.785 | -0.001 | 0.012 | 0.012 | 0 | 0.1 | 0.2 | 0.14 | 0.12 | 0.12 | 0.12 |
| 10 | 58504123 | rs1914564 | -0.02 | -1.405 | 0.899 | -0.795 | 0.059 | -0.02 | -0.02 | 1.341 | 0.89 | 0.68 | 0.9 | 0.92 | 0.83 | 0.91 |
| 10 | 58505135 | rs7924028 | 2.354 | -1.411 | 0.783 | -0.8 | -0.048 | -0.04 | -0.04 | 0 | 0.42 | 0.13 | 0.47 | 0.52 | 0.51 | 0.4 |
| 10 | 58508377 | rs10763402 | 1.483 | -1.338 | 0.787 | -0.799 | 0.01 | 0.037 | 0.037 | 0 | 0.52 | 0.33 | 0.61 | 0.64 | 0.63 | 0.52 |
| 10 | 58537877 | rs12355708 | 1.207 | -1.251 | 0.677 | -0.746 | 0 | 0.034 | 0.034 | 0.077 | 0.06 | 0.01 | 0.02 | 0 | 0.05 | 0.06 |
| 10 | 58539202 | rs2393120 | -1.559 | -1.291 | 0.793 | -0.721 | -0.005 | -0.003 | -0.003 | 0 | 0.47 | 0.52 | 0.43 | 0.39 | 0.32 | 0.51 |
| 10 | 58541887 | rs10509061 | 2.115 | -1.266 | 0.669 | -0.716 | -0.048 | -0.031 | -0.031 | 0 | 0.42 | 0.13 | 0.47 | 0.53 | 0.51 | 0.4 |
| 10 | 58552182 | rs1914596 | -1.035 | -1.153 | 0.838 | -0.966 | -0.027 | -0.028 | -0.028 | 0 | 0.39 | 0.49 | 0.32 | 0.27 | 0.21 | 0.39 |
| 10 | 58560560 | rs10825892 | 1.011 | -1.153 | 0.628 | -1.03 | -0.018 | 0.119 | 0.119 | 0 | 0.61 | 0.2 | 0.73 | 0.91 | 0.66 | 0.67 |
| 10 | 58560654 | rs10740686 | 1.041 | -1.162 | 0.693 | -1.045 | 0.024 | -0.042 | -0.042 | 0 | 0.85 | 0.36 | 0.86 | 0.91 | 0.84 | 0.85 |
| 10 | 58562211 | rs10825898 | -0.358 | -1.165 | 0.866 | -1.048 | 0.024 | -0.053 | -0.053 | 0 | 0.1 | 0.34 | 0.08 | 0 | 0.08 | 0.06 |
| 10 | 58563729 | rs11005540 | 0.944 | -1.16 | 0.711 | -1.045 | -0.002 | 0.031 | 0.031 | 0.077 | 0.07 | 0.02 | 0.03 | 0 | 0.06 | 0.06 |
| 10 | 58564731 | rs10763419 | 1.65 | -1.257 | 0.648 | -1.029 | -0.054 | 0.068 | 0.068 | 0 | 0.49 | 0.07 | 0.55 | 0.66 | 0.61 | 0.52 |
| 10 | 58573238 | rs981155 | 0.023 | -1.226 | 0.713 | -0.999 | -0.005 | 0.007 | 0.007 | 0 | 0.13 | 0.12 | 0.19 | 0.25 | 0.06 | 0.16 |
| 10 | 58577802 | rs10763422 | 1.673 | -1.232 | 0.721 | -1.021 | 0.01 | 0.093 | 0.093 | 0 | 0.5 | 0.2 | 0.57 | 0.66 | 0.61 | 0.52 |
| 10 | 58580335 | rs7098642 | 1.368 | -1.241 | 0.892 | -1.024 | -0.008 | 0.008 | 0.008 | 0 | 0.69 | 0.55 | 0.68 | 0.67 | 0.72 | 0.69 |
| 10 | 58584083 | rs10740687 | 1.686 | -1.233 | 0.69 | -1.043 | 0.011 | 0.082 | 0.082 | 0 | 0.5 | 0.21 | 0.57 | 0.64 | 0.61 | 0.52 |
| 10 | 58593295 | rs7907512 | 1.583 | -1.221 | 0.709 | -1.038 | 0.002 | -0.001 | -0.001 | 0 | 0.67 | 0.35 | 0.65 | 0.65 | 0.72 | 0.69 |
| 10 | 58594360 | rs16909567 | 1.414 | -1.156 | 0.71 | -0.996 | -0.002 | 0.004 | 0.004 | 0 | 0.15 | 0.11 | 0.07 | 0 | 0.11 | 0.11 |
| 10 | 58607217 | rs1980776 | 1.163 | -1.178 | 0.758 | -0.976 | -0.005 | 0.011 | 0.011 | 0 | 0.74 | 0.38 | 0.69 | 0.65 | 0.8 | 0.76 |
| 10 | 58612274 | rs11005554 | -0.8 | -1.232 | 0.804 | -0.998 | -0.022 | -0.009 | -0.009 | 0 | 0.32 | 0.56 | 0.33 | 0.28 | 0.17 | 0.33 |
| 10 | 58617298 | rs7893434 | 1.505 | -1.2 | 0.636 | -0.993 | -0.033 | 0.09 | 0.09 | 0 | 0.15 | 0.01 | 0.06 | 0 | 0.11 | 0.11 |
| 10 | 58622012 | rs12355683 | 0.871 | -1.205 | 0.649 | -0.979 | -0.001 | 0.032 | 0.032 | 0.108 | 0.07 | 0.02 | 0.03 | 0 | 0.06 | 0.06 |
| 10 | 58622841 | rs4935672 | -0.364 | -1.199 | 0.755 | -0.95 | -0.047 | 0.1 | 0.1 | 0 | 0.14 | 0.3 | 0.2 | 0.27 | 0.04 | 0.16 |
| 10 | 58632384 | rs10763432 | -0.813 | -1.129 | 0.616 | -0.967 | -0.02 | -0.003 | -0.003 | 0.225 | 0.94 | 0.96 | 0.93 | 0.93 | 0.86 | 0.92 |
| 10 | 58633205 | rs7917259 | 0.789 | -1.357 | 0.62 | -1.066 | 0 | -0.034 | -0.034 | 0.114 | 0.63 | 0.39 | 0.78 | 0.93 | 0.67 | 0.72 |
| 10 | 58634596 | rs16909599 | -0.468 | -1.37 | 0.59 | -1.063 | -0.01 | 0.058 | 0.058 | 0 | 0.16 | 0.29 | 0.25 | 0.27 | 0.12 | 0.24 |
| 10 | 58636572 | rs10825918 | 1.085 | -1.386 | 0.626 | -1.079 | 0.012 | -0.083 | -0.083 | 0.174 | 0.78 | 0.45 | 0.84 | 0.93 | 0.78 | 0.83 |
| 10 | 58638739 | rs16909609 | 0.133 | -1.391 | 0.615 | -1.077 | 0.007 | -0.032 | -0.032 | 0 | 0.13 | 0.06 | 0.2 | 0.26 | 0.09 | 0.19 |
| 10 | 58640650 | rs6481237 | 2.166 | -1.406 | 0.578 | -1.085 | -0.012 | 0.005 | 0.005 | 0 | 0.62 | 0.13 | 0.59 | 0.65 | 0.67 | 0.6 |
| 10 | 58646990 | rs4935676 | -0.762 | -1.48 | 0.576 | -1.112 | -0.016 | 0.044 | 0.044 | 0 | 0.17 | 0.32 | 0.26 | 0.27 | 0.12 | 0.24 |
| 10 | 58658969 | rs10825929 | 0.678 | -1.419 | 0.703 | -1.1 | 0.002 | -0.055 | -0.055 | 0.942 | 0.64 | 0.43 | 0.77 | 0.91 | 0.66 | 0.69 |
| 10 | 58665441 | rs10763437 | 1.693 | -1.367 | 0.548 | -1.063 | -0.107 | 0.036 | 0.036 | 0 | 0.46 | 0.07 | 0.53 | 0.65 | 0.56 | 0.49 |
| 10 | 58670516 | rs2818836 | -0.827 | -1.433 | 0.661 | -1.048 | -0.018 | -0.016 | -0.016 | 1.231 | 0.35 | 0.5 | 0.35 | 0.28 | 0.23 | 0.35 |
| 10 | 58670856 | rs3000612 | -0.816 | -1.433 | 0.669 | -1.048 | -0.02 | -0.017 | -0.017 | 0 | 0.35 | 0.47 | 0.35 | 0.28 | 0.23 | 0.35 |
| 10 | 58672533 | rs2255696 | 0.748 | -1.306 | 0.682 | -1.021 | 0.029 | -0.06 | -0.06 | 0 | 0.82 | 0.57 | 0.88 | 0.92 | 0.8 | 0.85 |
| 10 | 58677382 | rs2818849 | 0.426 | -1.338 | 0.666 | -1.048 | 0.006 | -0.106 | -0.106 | 0 | 0.67 | 0.5 | 0.82 | 0.92 | 0.7 | 0.73 |
| 10 | 58681058 | rs2393151 | 1.388 | -1.318 | 0.567 | -1.035 | -0.009 | 0.052 | 0.052 | 0 | 0.15 | 0.07 | 0.06 | 0 | 0.11 | 0.12 |
| 10 | 58684269 | rs2578084 | 1.351 | -1.331 | 0.67 | -0.957 | 0.004 | 0.029 | 0.029 | 0 | 0.51 | 0.35 | 0.59 | 0.66 | 0.57 | 0.51 |
| 10 | 58687200 | rs2251096 | 0.647 | -1.332 | 0.651 | -0.992 | -0.001 | -0.003 | -0.003 | 0 | 0.65 | 0.3 | 0.78 | 0.91 | 0.69 | 0.72 |
| 10 | 58687410 | rs2393158 | 0.324 | -1.386 | 1.06 | -1.032 | -0.008 | -0.08 | -0.08 | 0 | 0.21 | 0.28 | 0.09 | 0 | 0.12 | 0.15 |
| 10 | 58693527 | rs7895488 | -1.136 | -1.31 | 1.036 | -1.029 | -0.001 | -0.009 | -0.009 | 0 | 0.35 | 0.69 | 0.22 | 0.09 | 0.3 | 0.28 |
| 10 | 58696144 | rs11005583 | 0.018 | -1.274 | 0.974 | -0.996 | 0.024 | -0.009 | -0.009 | 0 | 0.11 | 0.2 | 0.09 | 0.07 | 0.18 | 0.12 |
| 10 | 58697216 | rs12414237 | 0.186 | -1.328 | 0.796 | -0.98 | 0.01 | -0.053 | -0.053 | 0 | 0.15 | 0.05 | 0.2 | 0.27 | 0.11 | 0.22 |
| 10 | 58702033 | rs12762470 | -0.873 | -1.371 | 0.865 | -0.989 | 0.012 | 0.01 | 0.01 | 0 | 0.38 | 0.31 | 0.29 | 0.27 | 0.23 | 0.37 |
| 10 | 58702440 | rs2393179 | -0.878 | -1.38 | 0.86 | -0.993 | 0.012 | 0.011 | 0.011 | 0 | 0.38 | 0.31 | 0.29 | 0.27 | 0.23 | 0.37 |
| 10 | 58721707 | rs12772977 | 0.474 | -2.321 | 0.794 | -1.339 | 0.015 | 0.168 | 0.168 | 0.02 | 0.27 | 0.04 | 0.09 | 0 | 0.13 | 0.13 |
| 10 | 58723878 | rs2393190 | 1.855 | -1.31 | 1.429 | -0.548 | -0.049 | -0.088 | -0.088 | 0 | 0.55 | 0.04 | 0.63 | 0.55 | 0.62 | 0.55 |

| | | | | | | | | | | | | | | | | |
|---|---|---|---|---|---|---|---|---|---|---|---|---|---|---|---|---|
| 10 | 58725991 | rs10763440 | 1.647 | -0.897 | 1.602 | -1.42 | 0.002 | 0.009 | 0.009 | 0.064 | 0.81 | 0.35 | 0.76 | 0.79 | 0.77 | 0.8 |
| 10 | 58730014 | rs11816606 | -0.838 | -0.964 | 1.689 | -1.44 | -0.008 | 0.032 | 0.032 | 0 | 0.05 | 0.1 | 0.09 | 0.13 | 0.06 | 0.07 |
| 10 | 58736121 | rs7068823 | -1.531 | -0.943 | 1.711 | -1.43 | 0.05 | 0.023 | 0.023 | 0 | 0.19 | 0.53 | 0.24 | 0.21 | 0.26 | 0.21 |
| 10 | 58739433 | rs10763441 | 1.486 | -0.98 | 1.697 | -1.46 | -0.041 | 0.137 | 0.137 | 0 | 0.58 | 0.08 | 0.65 | 0.79 | 0.62 | 0.65 |
| 10 | 58742985 | rs11005592 | -1.473 | -0.862 | 1.838 | -1.351 | 0.043 | 0.027 | 0.027 | 0 | 0.18 | 0.46 | 0.22 | 0.21 | 0.24 | 0.21 |
| 10 | 58746084 | rs7095023 | -0.59 | -0.857 | 1.698 | -1.342 | -0.008 | -0.028 | -0.028 | 0.039 | 0.06 | 0.21 | 0.03 | 0 | 0.02 | 0.03 |
| 10 | 58749365 | rs11005600 | -1.144 | -0.773 | 1.603 | -1.302 | 0.009 | 0.006 | 0.006 | 0 | 0.16 | 0.25 | 0.18 | 0.2 | 0.24 | 0.2 |
| 10 | 58750203 | rs4545461 | 1.524 | -0.806 | 1.889 | -1.34 | 0.005 | -0.002 | -0.002 | 0 | 0.8 | 0.32 | 0.74 | 0.79 | 0.76 | 0.79 |
| 10 | 58750903 | rs7922854 | -1.655 | -0.713 | 1.865 | -1.328 | 0.033 | 0.014 | 0.014 | 0.014 | 0.19 | 0.42 | 0.22 | 0.19 | 0.25 | 0.21 |
| 10 | 58760115 | rs4935683 | -1.043 | -0.311 | 1.953 | -1.262 | 0.005 | -0.008 | -0.008 | 0 | 0.31 | 0.36 | 0.27 | 0.19 | 0.37 | 0.29 |
| 10 | 58760917 | rs2393163 | 1.452 | -0.374 | 2.127 | -1.294 | -0.009 | 0.035 | 0.035 | 0 | 0.62 | 0.28 | 0.69 | 0.81 | 0.62 | 0.69 |
| 10 | 58763382 | rs4630213 | 0.756 | -0.376 | 2.114 | -1.293 | 0.002 | -0.01 | -0.01 | 0 | 0.94 | 0.75 | 0.97 | 1 | 0.99 | 0.98 |
| 10 | 58764472 | rs10740696 | 1.638 | -0.383 | 2.051 | -1.324 | -0.067 | 0.119 | 0.119 | 0 | 0.6 | 0.09 | 0.57 | 0.79 | 0.62 | 0.68 |
| 10 | 58768319 | rs7912714 | -1.086 | -0.366 | 2.098 | -1.329 | 0.021 | 0.015 | 0.015 | 0 | 0.15 | 0.27 | 0.18 | 0.19 | 0.22 | 0.16 |
| 10 | 58772800 | rs11817737 | 0.836 | -0.363 | 2.079 | -1.329 | -0.006 | 0.014 | 0.014 | 0 | 0.16 | 0.07 | 0.08 | 0 | 0.14 | 0.13 |
| 10 | 58773025 | rs10733970 | 1.371 | -0.364 | 2.105 | -1.31 | -0.001 | -0.01 | -0.01 | 0.964 | 0.93 | 0.43 | 0.92 | 0.98 | 1 | 0.98 |
| 10 | 58781082 | rs12359387 | 2.013 | -0.362 | 2.087 | -1.311 | -0.004 | 0.007 | 0.007 | 1.066 | 0.76 | 0.17 | 0.73 | 0.82 | 0.76 | 0.81 |
| 10 | 58794959 | rs10825969 | -1.003 | -0.317 | 1.965 | -1.269 | 0.016 | -0.015 | -0.015 | 0 | 0.14 | 0.25 | 0.11 | 0.07 | 0.18 | 0.1 |
| 10 | 58801996 | rs16910039 | -0.969 | -0.336 | 1.578 | -1.792 | 0.018 | -0.062 | -0.062 | 0 | 0.14 | 0.25 | 0.11 | 0 | 0.18 | 0.09 |
| 10 | 58804299 | rs16910063 | 0.178 | -0.86 | 0.869 | -1.3 | -0.001 | -0.026 | -0.026 | 0 | 0.17 | 0.14 | 0.08 | 0 | 0.13 | 0.12 |
| 10 | 58809636 | rs10825976 | -1.165 | -0.854 | -0.043 | -1.153 | 0.056 | 0.143 | 0.143 | 0 | 0.4 | 0.86 | 0.28 | 0.17 | 0.25 | 0.36 |
| 10 | 58822244 | rs3104660 | -1.067 | -0.756 | 1.49 | -1.472 | 0.035 | 0.073 | 0.073 | 0 | 0.39 | 0.78 | 0.25 | 0 | 0.27 | 0.21 |
| 10 | 58823631 | rs7081489 | -0.059 | -0.854 | 1.459 | -1.534 | -0.015 | 0.165 | 0.165 | 0 | 0.27 | 0.06 | 0.13 | 0 | 0.23 | 0.17 |
| 10 | 58825350 | rs3098360 | -1.408 | -0.894 | 1.496 | -1.522 | 0.045 | 0.227 | 0.227 | 0.186 | 0.43 | 0.88 | 0.26 | 0.06 | 0.29 | 0.28 |
| 10 | 58828890 | rs11812237 | -0.677 | -0.87 | 1.775 | -1.548 | 0.002 | -0.035 | -0.035 | 0.548 | 0.07 | 0.19 | 0.05 | 0 | 0.03 | 0.04 |
| 10 | 58836950 | rs3104656 | -1.061 | -0.862 | 1.836 | -1.552 | 0.009 | -0.039 | -0.039 | 0.074 | 0.38 | 0.66 | 0.25 | 0 | 0.27 | 0.21 |
| 10 | 58842551 | rs3105324 | -0.489 | -0.829 | 1.802 | -1.525 | -0.019 | -0.139 | -0.139 | 0 | 0.32 | 0.56 | 0.19 | 0 | 0.24 | 0.16 |
| 10 | 58847690 | rs3105330 | -1.283 | -0.869 | 1.829 | -1.271 | -0.009 | 0.03 | 0.03 | 0.536 | 0.11 | 0.24 | 0.13 | 0.15 | 0.06 | 0.19 |
| 10 | 58847806 | rs3104645 | -1.136 | -0.925 | 1.868 | -1.381 | 0.009 | 0.016 | 0.016 | 0 | 0.41 | 0.39 | 0.26 | 0.15 | 0.3 | 0.36 |
| 10 | 58849853 | rs3105341 | -1.293 | -0.894 | 1.875 | -1.355 | -0.007 | 0.029 | 0.029 | 0.583 | 0.11 | 0.24 | 0.13 | 0.15 | 0.06 | 0.19 |
| 10 | 58853258 | rs3104641 | -1.049 | -0.922 | 1.912 | -1.381 | -0.01 | 0.015 | 0.015 | 0.291 | 0.1 | 0.19 | 0.13 | 0.14 | 0.06 | 0.18 |
| 10 | 58853784 | rs4589208 | -0.247 | -0.935 | 1.894 | -1.405 | 0.015 | 0.095 | 0.095 | 0.325 | 0.3 | 0.14 | 0.13 | 0 | 0.24 | 0.16 |
| 10 | 58871052 | rs12253767 | -0.188 | -0.946 | 2.411 | -1.41 | -0.001 | 0.161 | 0.161 | 0.49 | 0.29 | 0.08 | 0.13 | 0 | 0.24 | 0.16 |
| 10 | 58874198 | rs995236 | -1.057 | -0.969 | 2.499 | -1.416 | -0.01 | 0.013 | 0.013 | 0.216 | 0.1 | 0.19 | 0.12 | 0.14 | 0.06 | 0.18 |
| 10 | 58877088 | rs12767717 | -0.168 | -0.969 | 2.489 | -1.419 | -0.004 | 0.182 | 0.182 | 0.404 | 0.29 | 0.06 | 0.12 | 0 | 0.23 | 0.17 |
| 10 | 58879797 | rs3105307 | -1.076 | -0.962 | 2.521 | -1.412 | -0.014 | -0.057 | -0.057 | 2.843 | 0.41 | 0.49 | 0.27 | 0.14 | 0.3 | 0.34 |
| 10 | 58885646 | rs924792 | -1.104 | -0.96 | 2.409 | -1.411 | 0.017 | 0.043 | 0.043 | 0 | 0.4 | 0.33 | 0.26 | 0.14 | 0.3 | 0.34 |
| 10 | 58892275 | rs3104678 | -1.085 | -0.978 | 2.409 | -1.411 | 0.018 | 0.041 | 0.041 | 0 | 0.4 | 0.32 | 0.26 | 0.14 | 0.3 | 0.34 |
| 10 | 58905069 | rs1503914 | 0.881 | -0.928 | 2.483 | -1.413 | 0.018 | 0.031 | 0.031 | 0 | 0.58 | 0.33 | 0.72 | 0.86 | 0.7 | 0.67 |
| 10 | 58914777 | rs11005667 | -0.715 | -0.964 | 2.481 | -1.443 | -0.026 | -0.158 | -0.158 | 0.025 | 0.34 | 0.5 | 0.16 | 0.01 | 0.25 | 0.19 |
| 10 | 58916759 | rs6481254 | -0.568 | -0.94 | 2.483 | -1.439 | -0.022 | -0.157 | -0.157 | 0 | 0.33 | 0.44 | 0.16 | 0.01 | 0.25 | 0.19 |
| 10 | 58922421 | rs1394112 | -0.64 | -0.941 | 2.46 | -1.439 | -0.025 | -0.165 | -0.165 | 1.202 | 0.34 | 0.5 | 0.16 | 0.01 | 0.25 | 0.19 |
| 10 | 58922857 | rs12247842 | -0.64 | -0.941 | 2.46 | -1.439 | -0.025 | -0.165 | -0.165 | 0.385 | 0.34 | 0.5 | 0.16 | 0.01 | 0.25 | 0.19 |
| 10 | 58923590 | rs12248271 | -0.245 | -0.946 | 2.396 | -1.441 | 0.02 | 0.07 | 0.07 | 0.43 | 0.29 | 0.14 | 0.12 | 0 | 0.23 | 0.16 |
| 10 | 58936492 | rs7082897 | -0.56 | -0.94 | 2.555 | -1.436 | -0.013 | -0.118 | -0.118 | 0 | 0.33 | 0.42 | 0.16 | 0.01 | 0.25 | 0.19 |
| 10 | 58936553 | rs7081958 | -1.52 | -0.955 | 2.638 | -1.441 | 0.043 | 0.123 | 0.123 | 0 | 0.45 | 0.75 | 0.29 | 0.15 | 0.32 | 0.36 |
| 10 | 58936939 | rs10160131 | -0.557 | -0.947 | 2.565 | -1.438 | -0.013 | -0.118 | -0.118 | 0.027 | 0.33 | 0.42 | 0.16 | 0.01 | 0.25 | 0.19 |
| 10 | 58937708 | rs4935688 | -1.148 | -0.92 | 2.463 | -1.396 | -0.01 | 0.017 | 0.017 | 0.301 | 0.1 | 0.23 | 0.12 | 0.14 | 0.06 | 0.17 |
| 10 | 58938552 | rs6481257 | -1.676 | -1.016 | 2.327 | -1.446 | 0.062 | 0.227 | 0.227 | 0.167 | 0.48 | 0.9 | 0.31 | 0.18 | 0.32 | 0.4 |
| 10 | 58943359 | rs10825991 | -1.374 | -0.944 | 2.319 | -1.419 | -0.003 | -0.025 | -0.025 | 0 | 0.44 | 0.59 | 0.28 | 0.14 | 0.31 | 0.36 |
| 10 | 58945586 | rs12247721 | -0.213 | -0.898 | 1.413 | -1.397 | -0.004 | 0.182 | 0.182 | 0.39 | 0.29 | 0.06 | 0.12 | 0 | 0.23 | 0.17 |
| 10 | 58948297 | rs10825993 | -0.891 | -0.426 | 1.679 | -1.321 | 0.029 | 0.114 | 0.114 | 0 | 0.43 | 0.28 | 0.27 | 0.15 | 0.34 | 0.34 |
| 10 | 58953960 | rs2893721 | -1.489 | -0.445 | 1.715 | -1.325 | 0.045 | 0.228 | 0.228 | 0 | 0.49 | 0.89 | 0.33 | 0.19 | 0.36 | 0.38 |
| 10 | 58954023 | rs2393225 | -0.803 | -0.38 | 1.595 | -1.313 | 0.014 | 0.045 | 0.045 | 0 | 0.42 | 0.35 | 0.27 | 0.14 | 0.34 | 0.33 |
| 10 | 58986586 | rs12766354 | -0.177 | -0.406 | 1.266 | -1.355 | 0.025 | 0.066 | 0.066 | 0 | 0.33 | 0.19 | 0.14 | 0 | 0.24 | 0.2 |
| 10 | 58995327 | rs3907689 | -1.342 | -0.188 | 1.452 | -1.343 | 0.007 | 0.03 | 0.03 | 0 | 0.48 | 0.61 | 0.32 | 0.19 | 0.38 | 0.39 |
| 10 | 59006901 | rs2938357 | -1.342 | -0.178 | 1.309 | -1.338 | 0.01 | 0.048 | 0.048 | 0 | 0.48 | 0.62 | 0.31 | 0.19 | 0.38 | 0.39 |
| 10 | 59007742 | rs1614490 | -1.422 | -0.212 | 1.113 | -1.256 | 0.027 | 0.04 | 0.04 | 0.027 | 0.13 | 0.38 | 0.17 | 0.19 | 0.13 | 0.17 |
| 10 | 59010858 | rs1717543 | -1.205 | -0.172 | 0.363 | -1.235 | 0.026 | 0.107 | 0.107 | 0 | 0.49 | 0.69 | 0.32 | 0.19 | 0.38 | 0.4 |
| 10 | 59013036 | rs7072438 | -0.204 | 0.039 | -0.199 | -1.534 | -0.014 | 0.132 | 0.132 | 0 | 0.29 | 0.1 | 0.13 | 0 | 0.25 | 0.19 |
| 10 | 59018855 | rs1679489 | -1.13 | -0.143 | 0.292 | -1.073 | -0.004 | 0.062 | 0.062 | 0 | 0.13 | 0.3 | 0.26 | 0.21 | 0.08 | 0.25 |
| 10 | 59020450 | rs1679495 | -1.283 | -0.214 | -0.518 | -1.016 | -0.026 | 0.105 | 0.105 | 0.23 | 0.49 | 0.96 | 0.45 | 0.29 | 0.36 | 0.54 |
| 10 | 59028417 | rs2393230 | -1.142 | -0.182 | -0.4 | -1.294 | -0.052 | 0.063 | 0.063 | 1.679 | 0.58 | 0.78 | 0.62 | 0.27 | 0.48 | 0.6 |
| 10 | 59054965 | rs1717524 | -0.451 | -0.234 | 0.708 | -1.305 | -0.009 | 0.004 | 0.004 | 0 | 0.07 | 0.22 | 0.19 | 0.09 | 0.04 | 0.1 |
| 10 | 59079675 | rs4568922 | -0.451 | -0.51 | 1.235 | -1.582 | -0.01 | 0.004 | 0.004 | 1.88 | 0.21 | 0.26 | 0.26 | 0.18 | 0.18 | 0.3 |
| 10 | 59084209 | rs7915278 | -1.3 | -0.549 | 1.309 | -1.57 | 0.013 | 0.019 | 0.019 | 1.234 | 0.1 | 0.51 | 0.22 | 0.1 | 0.04 | 0.1 |
| 10 | 59089930 | rs10826017 | -0.296 | -0.557 | 1.533 | -1.543 | 0.002 | 0.027 | 0.027 | 0 | 0.07 | 0.1 | 0.19 | 0.14 | 0.06 | 0.17 |
| 10 | 59091105 | rs11005753 | -0.604 | -0.55 | 1.579 | -1.626 | 0.008 | 0.027 | 0.027 | 0.169 | 0.08 | 0.4 | 0.2 | 0.09 | 0.03 | 0.09 |
| 10 | 59092325 | rs4935699 | -0.837 | -0.554 | 1.586 | -1.628 | 0 | 0.019 | 0.019 | 0.509 | 0.09 | 0.4 | 0.2 | 0.09 | 0.03 | 0.09 |

| | | | | | | | | | | | | | | | | |
|---|---|---|---|---|---|---|---|---|---|---|---|---|---|---|---|---|
| 10 | 59092351 | rs1904761 | -1.832 | -0.156 | 1.378 | -1.786 | -0.12 | 0.029 | 0.029 | 1.569 | 0.6 | 0.97 | 0.66 | 0.27 | 0.53 | 0.61 |
| 10 | 59094568 | rs16913959 | -0.379 | -0.103 | 1.585 | -1.871 | -0.011 | 0.003 | 0.003 | 0.004 | 0.08 | 0.24 | 0.19 | 0.09 | 0.03 | 0.09 |
| 10 | 59111940 | rs1904757 | 0.238 | -0.083 | 1.47 | -1.889 | 0.001 | -0.015 | -0.015 | 0.579 | 0.16 | 0.17 | 0.21 | 0.04 | 0.17 | 0.15 |
| 10 | 59113403 | rs7911894 | 0.509 | -0.217 | 2.064 | -1.657 | -0.013 | 0.056 | 0.056 | 0 | 0.26 | 0.14 | 0.15 | 0 | 0.27 | 0.2 |
| 10 | 59113708 | rs1011077 | -1.146 | -0.294 | 2.712 | -0.976 | 0.016 | 0.036 | 0.036 | 0 | 0.08 | 0.39 | 0.21 | 0.09 | 0.04 | 0.09 |
| 10 | 59130616 | rs3008777 | -1.702 | -0.358 | 2.725 | -0.963 | 0.003 | 0.023 | 0.023 | 0.122 | 0.1 | 0.41 | 0.21 | 0.1 | 0.04 | 0.1 |
| 10 | 59143964 | rs7900471 | -0.875 | -0.269 | 2.359 | -0.911 | -0.001 | 0.01 | 0.01 | 0 | 0.07 | 0.1 | 0.07 | 0.11 | 0.05 | 0.18 |
| 10 | 59146776 | rs2928445 | -2.3 | -0.282 | 2.27 | -0.908 | -0.004 | 0.044 | 0.044 | 0.07 | 0.15 | 0.54 | 0.3 | 0.21 | 0.09 | 0.28 |
| 10 | 59146948 | rs2928446 | -0.975 | -0.24 | 1.718 | -0.876 | -0.008 | 0.014 | 0.014 | 0 | 0.07 | 0.18 | 0.2 | 0.09 | 0.03 | 0.09 |
| 10 | 59154057 | rs2127615 | -1.089 | -0.24 | 1.69 | -0.846 | 0.01 | 0.052 | 0.052 | 0 | 0.08 | 0.3 | 0.21 | 0.1 | 0.03 | 0.09 |
| 10 | 59158985 | rs12359659 | -0.874 | 0.049 | 0.519 | -1.254 | -0.004 | 0.014 | 0.014 | 0 | 0.28 | 0.25 | 0.37 | 0.15 | 0.34 | 0.4 |
| 10 | 59165924 | rs4412689 | -0.345 | -0.251 | 0.212 | -1.648 | 0.051 | 0.104 | 0.104 | 0 | 0.41 | 0.77 | 0.33 | 0.18 | 0.24 | 0.43 |
| 10 | 59172666 | rs4445565 | 0.108 | -0.43 | 0.551 | -1.772 | 0.024 | 0.04 | 0.04 | 0 | 0.31 | 0.19 | 0.11 | 0.05 | 0.18 | 0.31 |
| 10 | 59172837 | rs4436472 | 0.223 | -0.438 | 0.66 | -1.953 | 0.003 | 0.031 | 0.031 | 0 | 0.57 | 0.64 | 0.66 | 0.95 | 0.62 | 0.58 |
| 10 | 59176243 | rs7077068 | 0.406 | -0.45 | 1.287 | -1.958 | 0.004 | -0.006 | -0.006 | 0.014 | 0.55 | 0.34 | 0.63 | 0.94 | 0.62 | 0.58 |
| 10 | 59181959 | rs4275536 | -0.494 | -0.41 | 1.203 | -1.842 | 0.051 | 0.129 | 0.129 | 0.184 | 0.05 | 0.31 | 0.16 | 0.14 | 0.03 | 0.08 |
| 10 | 59189178 | rs4462262 | 0.465 | -0.469 | 1.104 | -1.887 | 0.004 | 0.001 | 0.001 | 0 | 0.57 | 0.33 | 0.63 | 0.94 | 0.62 | 0.58 |
| 10 | 59213174 | rs12573259 | 1.676 | -0.686 | 0.666 | -1.877 | -0.064 | 0.041 | 0.041 | 0 | 0.37 | 0.1 | 0.37 | 0.69 | 0.46 | 0.18 |
| 10 | 59215019 | rs4497328 | -0.424 | -0.379 | 0.825 | -1.824 | -0.004 | -0.012 | -0.012 | 0 | 0.87 | 0.84 | 0.78 | 1 | 0.81 | 0.89 |
| 10 | 59217371 | rs11005804 | 0.496 | -0.429 | 0.453 | -1.866 | 0.009 | 0.066 | 0.066 | 0 | 0.22 | 0.08 | 0.08 | 0 | 0.16 | 0.23 |
| 10 | 59219261 | rs11005805 | 0.403 | -0.242 | 0.351 | -1.698 | 0.001 | -0.036 | -0.036 | 0.011 | 0.58 | 0.42 | 0.66 | 0.95 | 0.64 | 0.59 |
| 10 | 59242718 | rs7081956 | -0.071 | -0.323 | 0.288 | -1.571 | -0.014 | -0.061 | -0.061 | 0 | 0.27 | 0.3 | 0.11 | 0.04 | 0.15 | 0.29 |
| 10 | 59243976 | rs2393289 | -0.394 | -0.086 | 0.765 | -1.695 | 0.027 | -0.065 | -0.065 | 1.814 | 0.87 | 0.72 | 0.77 | 1 | 0.81 | 0.9 |
| 10 | 59245816 | rs7095419 | -0.394 | -0.086 | 0.765 | -1.695 | 0.027 | -0.065 | -0.065 | 0 | 0.87 | 0.73 | 0.77 | 1 | 0.81 | 0.9 |
| 10 | 59258746 | rs2393287 | -0.468 | -0.086 | 0.746 | -1.695 | 0.019 | -0.055 | -0.055 | 0 | 0.88 | 0.77 | 0.78 | 1 | 0.81 | 0.9 |
| 10 | 59268458 | rs11005826 | 0.085 | -0.429 | 0.871 | -1.849 | -0.001 | -0.023 | -0.023 | 2.458 | 0.24 | 0.22 | 0.1 | 0.04 | 0.15 | 0.28 |
| 10 | 59271322 | rs6481286 | 1.238 | -0.737 | 0.647 | -1.986 | 0.001 | -0.025 | -0.025 | 0 | 0.46 | 0.46 | 0.55 | 0.82 | 0.51 | 0.33 |
| 10 | 59272405 | rs10763490 | 0.424 | -0.742 | 0.628 | -1.979 | -0.001 | 0.009 | 0.009 | 0 | 0.7 | 0.67 | 0.64 | 0.86 | 0.66 | 0.6 |
| 10 | 59276436 | rs867860 | 0.291 | -0.741 | 0.267 | -1.939 | 0.016 | 0.006 | 0.006 | 0 | 0.18 | 0.11 | 0.14 | 0.13 | 0.15 | 0.29 |
| 10 | 59286797 | rs12354504 | 0.455 | -0.661 | 0.248 | -1.931 | 0.006 | 0.008 | 0.008 | 0 | 0.23 | 0.19 | 0.09 | 0.04 | 0.15 | 0.28 |
| 10 | 59298937 | rs17693954 | 0.259 | -0.624 | 0.176 | -1.883 | 0.018 | 0.002 | 0.002 | 0 | 0.18 | 0.09 | 0.14 | 0.13 | 0.15 | 0.29 |
| 10 | 59299296 | rs11005844 | 1.718 | -0.817 | 0.343 | -1.874 | -0.061 | 0.028 | 0.028 | 0 | 0.39 | 0.13 | 0.36 | 0.68 | 0.47 | 0.23 |
| 10 | 59300009 | rs2393283 | 0.09 | -1.006 | 0.396 | -1.715 | -0.003 | -0.008 | -0.008 | 0 | 0.26 | 0.28 | 0.24 | 0.18 | 0.18 | 0.37 |
| 10 | 59300303 | rs11005847 | 0.304 | -1.011 | 0.364 | -1.708 | 0.018 | 0.003 | 0.003 | 0 | 0.18 | 0.09 | 0.14 | 0.13 | 0.15 | 0.29 |
| 10 | 59302685 | rs10826079 | -0.939 | -0.849 | 0.42 | -1.626 | 0.017 | 0.021 | 0.021 | 0 | 0.44 | 0.39 | 0.39 | 0.32 | 0.33 | 0.66 |
| 10 | 59305355 | rs2393369 | 0.264 | -1.01 | 0.478 | -1.924 | -0.009 | -0.067 | -0.067 | 0 | 0.63 | 0.49 | 0.68 | 0.84 | 0.68 | 0.57 |
| 10 | 59312333 | rs7923928 | -0.442 | -1.499 | 0.613 | -1.848 | -0.012 | 0.003 | 0.003 | 0 | 0.17 | 0.2 | 0.3 | 0.19 | 0.08 | 0.29 |
| 10 | 59313571 | rs12414228 | 0.135 | -1.552 | 0.651 | -1.775 | 0.013 | -0.01 | -0.01 | 0.367 | 0.14 | 0.07 | 0.23 | 0.14 | 0.08 | 0.24 |
| 10 | 59317730 | rs10826086 | 1.946 | -1.59 | 0.488 | -2.16 | -0.045 | 0.026 | 0.026 | 0 | 0.36 | 0.19 | 0.34 | 0.68 | 0.48 | 0.24 |
| 10 | 59318263 | rs7899724 | 0.179 | -1.662 | 0.63 | -2.248 | 0.004 | 0.048 | 0.048 | 0.086 | 0.32 | 0.18 | 0.12 | 0.01 | 0.26 | 0.27 |
| 10 | 59324591 | rs12416395 | -0.103 | -1.664 | 0.512 | -2.097 | -0.011 | 0.003 | 0.003 | 0 | 0.17 | 0.2 | 0.3 | 0.19 | 0.08 | 0.29 |
| 10 | 59340263 | rs4333949 | -0.415 | -1.645 | 0.011 | -2.29 | -0.037 | 0.197 | 0.197 | 1.594 | 0.72 | 0.99 | 0.91 | 0.99 | 0.78 | 0.76 |
| 10 | 59342621 | rs11005865 | -0.627 | -1.463 | 0.536 | -1.891 | -0.014 | -0.035 | -0.035 | 0 | 0.08 | 0.14 | 0.04 | 0 | 0.05 | 0.08 |
| 10 | 59353945 | rs10826091 | -0.64 | -1.525 | 0.677 | -1.821 | -0.002 | 0.002 | 0.002 | 0 | 0.2 | 0.28 | 0.41 | 0.23 | 0.19 | 0.3 |
| 10 | 59358541 | rs4586071 | -2.022 | -1.6 | 0.546 | -1.893 | 0.028 | 0.033 | 0.033 | 0 | 0.41 | 0.76 | 0.52 | 0.33 | 0.35 | 0.51 |
| 10 | 59359706 | rs2393261 | -0.965 | -1.171 | 0.524 | -1.473 | -0.001 | -0.023 | -0.023 | 0 | 0.12 | 0.33 | 0.07 | 0.1 | 0.11 | 0.13 |
| 10 | 59362526 | rs7088834 | -0.465 | -1.076 | -0.105 | -1.432 | 0.001 | 0.008 | 0.008 | 2.822 | 0.1 | 0.05 | 0.06 | 0.1 | 0.11 | 0.13 |
| 10 | 59369677 | rs2393281 | 0.535 | -1.148 | 0.198 | -1.184 | -0.024 | 0.021 | 0.021 | 0 | 0.81 | 0.7 | 0.7 | 0.76 | 0.86 | 0.75 |
| 10 | 59371321 | rs7894146 | -1.154 | -1.07 | 0.691 | -1.381 | 0.002 | -0.025 | -0.025 | 0.247 | 0.11 | 0.58 | 0.07 | 0.07 | 0.06 | 0.17 |
| 10 | 59372482 | rs2393368 | -0.065 | -0.777 | -0.033 | -1.761 | 0.02 | 0.076 | 0.076 | 0 | 0.5 | 0.69 | 0.45 | 0.67 | 0.54 | 0.45 |
| 10 | 59379460 | rs7909667 | 1.157 | -1.059 | -0.342 | -2.08 | 0.014 | 0.031 | 0.031 | 0 | 0.29 | 0.13 | 0.11 | 0 | 0.2 | 0.17 |
| 10 | 59384883 | rs9415477 | 1.158 | -1.023 | -0.397 | -2.065 | -0.055 | 0.018 | 0.018 | 0 | 0.39 | 0.19 | 0.37 | 0.6 | 0.48 | 0.28 |
| 10 | 59386322 | rs9416575 | 0.289 | -1.17 | -0.393 | -2.133 | 0.003 | -0.015 | -0.015 | 0.087 | 0.77 | 0.81 | 0.56 | 0.67 | 0.74 | 0.64 |
| 10 | 59388098 | rs4501916 | 0.099 | -1.176 | -0.414 | -2.142 | -0.005 | -0.043 | -0.043 | 0.007 | 0.79 | 0.92 | 0.57 | 0.67 | 0.75 | 0.65 |
| 10 | 59389388 | rs4548550 | 1.081 | -1.26 | -0.391 | -2.136 | -0.055 | 0.017 | 0.017 | 0 | 0.39 | 0.19 | 0.38 | 0.6 | 0.48 | 0.28 |
| 10 | 59392180 | rs12411458 | 1.343 | -1.052 | -0.587 | -2.023 | -0.011 | 0.057 | 0.057 | 1.769 | 0.11 | 0.01 | 0.06 | 0 | 0.12 | 0.04 |
| 10 | 59392443 | rs11591507 | 1.002 | -1.059 | 0.088 | -1.725 | 0.009 | 0.032 | 0.032 | 0 | 0.31 | 0.19 | 0.14 | 0.1 | 0.23 | 0.21 |
| 10 | 59392718 | rs7100584 | -0.963 | -1.073 | 0.05 | -1.709 | 0.003 | -0.036 | -0.036 | 0 | 0.88 | 0.96 | 0.8 | 0.76 | 0.87 | 0.69 |
| 10 | 59394044 | rs10826094 | 0.1 | -1.027 | 0.462 | -1.644 | 0.015 | -0.009 | -0.009 | 0.394 | 0.09 | 0.3 | 0.05 | 0.07 | 0.05 | 0.17 |
| 10 | 59395309 | rs9416577 | 0.032 | -0.982 | 0.082 | -1.617 | 0.007 | -0.005 | -0.005 | 0 | 0.78 | 0.85 | 0.57 | 0.76 | 0.76 | 0.64 |
| 10 | 59404400 | rs4617511 | 1.053 | -0.513 | -0.065 | -1.495 | -0.013 | 0.006 | 0.006 | 0 | 0.39 | 0.33 | 0.39 | 0.61 | 0.48 | 0.27 |
| 10 | 59410897 | rs12777563 | 0.697 | -0.475 | -0.126 | -1.671 | 0.009 | 0.033 | 0.033 | 0 | 0.3 | 0.17 | 0.13 | 0.09 | 0.23 | 0.18 |
| 10 | 59417476 | rs7474675 | 0.589 | -0.339 | -0.461 | -1.624 | -0.009 | -0.065 | -0.065 | 0 | 0.32 | 0.28 | 0.14 | 0.1 | 0.24 | 0.19 |
| 10 | 59420427 | rs4443999 | 0.619 | -0.219 | -0.267 | -1.192 | -0.009 | -0.013 | -0.013 | 2.4 | 0.69 | 0.55 | 0.52 | 0.74 | 0.71 | 0.46 |
| 10 | 59420647 | rs4428995 | 0.626 | -0.217 | -0.302 | -1.199 | -0.011 | -0.017 | -0.017 | 0 | 0.69 | 0.54 | 0.52 | 0.74 | 0.71 | 0.46 |
| 10 | 59423191 | rs7089259 | -0.588 | -0.222 | -0.507 | -1.114 | 0.008 | 0.012 | 0.012 | 0 | 0.5 | 0.48 | 0.46 | 0.19 | 0.38 | 0.34 |
| 10 | 59425399 | rs4935168 | 0.338 | 0.284 | -0.135 | -0.714 | 0 | -0.001 | -0.001 | 0.063 | 0.59 | 0.4 | 0.6 | 0.57 | 0.59 | 0.47 |
| 10 | 59426642 | rs4394757 | 1.115 | 0.22 | -0.193 | -0.819 | -0.001 | -0.002 | -0.002 | 0 | 0.38 | 0.26 | 0.34 | 0.59 | 0.39 | 0.35 |
| 10 | 59430088 | rs1505907 | -1.465 | 0.506 | 0.718 | -0.805 | 0.007 | -0.01 | -0.01 | 0 | 0.28 | 0.31 | 0.25 | 0.22 | 0.31 | 0.42 |

| | | | | | | | | | | | | | | | | |
|---|---|---|---|---|---|---|---|---|---|---|---|---|---|---|---|---|
| 10 | 59433399 | rs1505908 | -1.444 | 0.505 | 0.718 | -0.828 | 0.007 | -0.011 | -0.011 | 0 | 0.28 | 0.31 | 0.25 | 0.22 | 0.31 | 0.42 |
| 10 | 59441244 | rs6481309 | -1.964 | 0.326 | 1.229 | -1.068 | -0.001 | 0.03 | 0.03 | 0.206 | 0.43 | 0.57 | 0.33 | 0.22 | 0.42 | 0.5 |
| 10 | 59443530 | rs1553571 | -1.435 | 0.357 | 1.428 | -1.15 | -0.002 | -0.118 | -0.118 | 1.313 | 0.32 | 0.33 | 0.19 | 0.01 | 0.31 | 0.22 |
| 10 | 59443875 | rs7087313 | -2.533 | 0.35 | 1.547 | -1.032 | -0.008 | 0.074 | 0.074 | 1.073 | 0.48 | 0.74 | 0.39 | 0.35 | 0.46 | 0.61 |
| 10 | 59450117 | rs7907840 | -0.052 | 0.382 | 1.438 | -0.997 | -0.001 | 0.004 | 0.004 | 0 | 0.1 | 0.1 | 0.13 | 0.21 | 0.11 | 0.28 |
| 10 | 59452554 | rs7900293 | -2.024 | 0.369 | 1.5 | -1.02 | 0.002 | -0.038 | -0.038 | 0 | 0.41 | 0.44 | 0.31 | 0.22 | 0.42 | 0.5 |
| 10 | 59456787 | rs12253675 | -1.376 | 0.392 | 1.576 | -0.948 | -0.009 | -0.013 | -0.013 | 0 | 0.25 | 0.37 | 0.21 | 0.22 | 0.22 | 0.41 |
| 10 | 59472250 | rs9416608 | -1.864 | 0.345 | 1.734 | -0.883 | -0.009 | 0.017 | 0.017 | 3.852 | 0.28 | 0.44 | 0.26 | 0.35 | 0.25 | 0.52 |
| 10 | 59472580 | rs9416609 | -1.54 | 0.338 | 1.808 | -0.892 | 0.007 | 0.051 | 0.051 | 0.572 | 0.14 | 0.22 | 0.18 | 0.35 | 0.15 | 0.44 |
| 10 | 59472805 | rs9416610 | -1.54 | 0.338 | 1.808 | -0.892 | 0.007 | 0.051 | 0.051 | 0.572 | 0.14 | 0.22 | 0.18 | 0.35 | 0.15 | 0.44 |
| 10 | 59484180 | rs10509077 | 0.156 | 0.334 | 1.765 | -0.895 | -0.018 | -0.071 | -0.071 | 0 | 0.14 | 0.21 | 0.07 | 0 | 0.09 | 0.08 |
| 10 | 59485326 | rs17623893 | -1.35 | 0.334 | 1.775 | -0.884 | -0.009 | -0.006 | -0.006 | 0 | 0.24 | 0.31 | 0.19 | 0.22 | 0.2 | 0.38 |
| 10 | 59486084 | rs7902910 | -1.384 | 0.332 | 1.755 | -0.881 | -0.006 | -0.002 | -0.002 | 0.008 | 0.24 | 0.31 | 0.19 | 0.22 | 0.2 | 0.39 |
| 10 | 59487008 | rs7476817 | 2.09 | 0.327 | 1.771 | -0.954 | 0.001 | -0.009 | -0.009 | 0 | 0.56 | 0.49 | 0.64 | 0.65 | 0.56 | 0.39 |
| 10 | 59487363 | rs1505929 | -0.238 | 0.534 | 1.649 | -1.18 | -0.025 | 0.035 | 0.035 | 0 | 0.16 | 0.06 | 0.12 | 0 | 0.22 | 0.09 |
| 10 | 59496520 | rs11005930 | -0.146 | 0.21 | 1.689 | -1.101 | -0.003 | -0.005 | -0.005 | 2.293 | 0.1 | 0.1 | 0.13 | 0.18 | 0.11 | 0.29 |
| 10 | 59500344 | rs11005931 | 1.166 | 0.206 | 1.688 | -1.103 | 0.002 | 0.013 | 0.013 | 0.552 | 0.86 | 0.8 | 0.83 | 0.69 | 0.85 | 0.56 |
| 10 | 59529513 | rs10826110 | 2.306 | -0.394 | 1.82 | -1.27 | 0.029 | 0.121 | 0.121 | 2.011 | 0.5 | 0.23 | 0.62 | 0.69 | 0.57 | 0.38 |
| 10 | 59538180 | rs10826113 | 1.374 | -0.396 | 1.796 | -1.265 | 0 | 0.049 | 0.049 | 0.905 | 0.84 | 0.69 | 0.82 | 0.69 | 0.86 | 0.56 |
| 10 | 59538196 | rs12247460 | 0.131 | -0.387 | 1.741 | -1.261 | -0.006 | -0.011 | -0.011 | 0 | 0.09 | 0.08 | 0.12 | 0.18 | 0.11 | 0.23 |
| 10 | 59549120 | rs7475733 | 1.32 | -0.376 | 1.733 | -1.282 | -0.021 | -0.01 | -0.01 | 0 | 0.69 | 0.46 | 0.74 | 0.69 | 0.77 | 0.48 |
| 10 | 59557630 | rs7097813 | 2.308 | -0.382 | 1.762 | -1.292 | 0.013 | 0.052 | 0.052 | 1.933 | 0.52 | 0.39 | 0.64 | 0.68 | 0.57 | 0.38 |
| 10 | 59558639 | rs1910975 | 1.182 | -0.12 | 1.682 | -1.299 | -0.004 | 0.045 | 0.045 | 1.396 | 0.84 | 0.7 | 0.82 | 0.69 | 0.86 | 0.53 |
| 10 | 59565280 | rs1505919 | 1.218 | -0.076 | 1.583 | -1.294 | -0.001 | 0.067 | 0.067 | 1.331 | 0.84 | 0.64 | 0.81 | 0.69 | 0.87 | 0.53 |
| 10 | 59565698 | rs1505920 | -0.186 | -0.259 | 1.917 | -1.79 | -0.018 | -0.083 | -0.083 | 0 | 0.14 | 0.24 | 0.07 | 0 | 0.09 | 0.06 |
| 10 | 59572370 | rs7909889 | -1.039 | -0.052 | 1.442 | -1.545 | 0.007 | -0.095 | -0.095 | 0 | 0.18 | 0.33 | 0.13 | 0 | 0.19 | 0.1 |
| 10 | 59572975 | rs7393803 | 1.758 | 0.014 | 1.555 | -1.098 | -0.011 | 0.006 | 0.006 | 0.287 | 0.87 | 0.59 | 0.87 | 0.83 | 0.9 | 0.68 |
| 10 | 59574735 | rs7094817 | -1.815 | 0.243 | 1.67 | -1.504 | -0.005 | -0.091 | -0.091 | 0 | 0.25 | 0.54 | 0.18 | 0.09 | 0.25 | 0.2 |
| 10 | 59576183 | rs10733977 | 2.089 | 0.213 | 1.509 | -1.747 | -0.022 | 0.216 | 0.216 | 1.594 | 0.59 | 0.16 | 0.7 | 0.91 | 0.59 | 0.7 |
| 10 | 59577377 | rs2393390 | -0.489 | 0.128 | 2.015 | -1.757 | 0.003 | -0.034 | -0.034 | 0.189 | 0.08 | 0.13 | 0.05 | 0.01 | 0.07 | 0.15 |
| 10 | 59578949 | rs12572355 | -0.886 | 0.093 | 2.003 | -1.744 | 0.014 | -0.09 | -0.09 | 1.561 | 0.17 | 0.39 | 0.13 | 0.07 | 0.18 | 0.05 |
| 10 | 59582520 | rs9416634 | 2.162 | 0.109 | 1.745 | -1.793 | -0.015 | 0.206 | 0.206 | 1.749 | 0.58 | 0.18 | 0.71 | 0.94 | 0.59 | 0.68 |
| 10 | 59582901 | rs7077489 | -0.333 | 0.119 | 1.949 | -1.78 | -0.002 | -0.023 | -0.023 | 0 | 0.08 | 0.07 | 0.05 | 0.01 | 0.07 | 0.16 |
| 10 | 59588583 | rs9416637 | 1.594 | 0.11 | 2.164 | -1.791 | 0.008 | -0.148 | -0.148 | 0 | 0.68 | 0.48 | 0.77 | 0.95 | 0.67 | 0.85 |
| 10 | 59590078 | rs7474890 | 1.28 | 0.139 | 2.5 | -1.786 | -0.002 | -0.068 | -0.068 | 0 | 0.73 | 0.71 | 0.84 | 1 | 0.75 | 0.87 |
| 10 | 59604302 | rs17625635 | -0.051 | 0.141 | 2.507 | -1.796 | -0.005 | -0.04 | -0.04 | 0.017 | 0.12 | 0.11 | 0.05 | 0 | 0.09 | 0.03 |
| 10 | 59606236 | rs16911135 | -0.634 | 0.182 | 2.603 | -1.785 | 0 | -0.043 | -0.043 | 0 | 0.08 | 0.15 | 0.05 | 0.01 | 0.07 | 0.18 |
| 10 | 59607602 | rs6481369 | 0.545 | 0.182 | 2.333 | -1.771 | -0.001 | -0.025 | -0.025 | 0 | 0.84 | 0.82 | 0.89 | 1 | 0.84 | 0.9 |
| 10 | 59631611 | rs2537632 | 2.222 | -0.282 | 2.202 | -1.849 | 0.012 | 0.221 | 0.221 | 2.008 | 0.58 | 0.16 | 0.73 | 0.94 | 0.64 | 0.71 |
| 10 | 59636838 | rs2252080 | -0.138 | -0.275 | 2.395 | -1.854 | 0 | -0.02 | -0.02 | 0 | 0.87 | 0.89 | 0.94 | 1 | 0.9 | 0.96 |
| 10 | 59642500 | rs17704490 | -0.71 | -0.205 | 2.368 | -1.61 | 0.001 | -0.011 | -0.011 | 1.089 | 0.13 | 0.11 | 0.07 | 0.01 | 0.11 | 0.08 |
| 10 | 59648851 | rs11005971 | -1.342 | -0.303 | 2.398 | -1.568 | 0.041 | -0.061 | -0.061 | 0 | 0.13 | 0.38 | 0.11 | 0.05 | 0.15 | 0.17 |
| 10 | 59668649 | rs12258613 | -2.814 | -0.358 | 2.125 | -1.345 | 0.009 | 0.037 | 0.037 | 1.497 | 0.29 | 0.77 | 0.21 | 0.13 | 0.25 | 0.26 |
| 10 | 59668737 | rs16911174 | -1.084 | -0.355 | 2.191 | -1.354 | -0.021 | -0.092 | -0.092 | 0.405 | 0.15 | 0.34 | 0.09 | 0.01 | 0.09 | 0.08 |
| 10 | 59671798 | rs7101032 | -1.354 | -0.262 | 1.818 | -1.307 | 0.028 | 0.008 | 0.008 | 0 | 0.14 | 0.29 | 0.11 | 0.13 | 0.17 | 0.17 |
| 10 | 59682064 | rs16911207 | -1.124 | -0.557 | 0.956 | -1.105 | -0.005 | -0.037 | -0.037 | 0 | 0.14 | 0.16 | 0.07 | 0.01 | 0.09 | 0.08 |
| 10 | 59686104 | rs2243952 | 1.026 | -0.582 | 0.933 | -1.147 | -0.031 | -0.064 | -0.064 | 0 | 0.71 | 0.42 | 0.87 | 0.91 | 0.83 | 0.78 |
| 10 | 59697552 | rs12764204 | -0.391 | -0.492 | 0.498 | -1.198 | 0.018 | 0.009 | 0.009 | 0.019 | 0.2 | 0.11 | 0.07 | 0 | 0.13 | 0.1 |
| 10 | 59700476 | rs16911231 | -1.028 | -0.491 | 0.675 | -1.073 | 0.004 | -0.002 | -0.002 | 0 | 0.06 | 0.19 | 0.03 | 0.05 | 0.05 | 0.06 |
| 10 | 59701343 | rs2430076 | 0.107 | -0.229 | 0.433 | -1.036 | 0.012 | 0.02 | 0.02 | 0 | 0.81 | 0.89 | 0.92 | 1 | 0.86 | 0.91 |
| 10 | 59707476 | rs2159862 | -0.34 | -0.567 | 0.501 | -0.994 | 0.002 | -0.026 | -0.026 | 0.086 | 0.83 | 0.86 | 0.94 | 1 | 0.9 | 0.91 |
| 10 | 59712446 | rs2537668 | 0.93 | -0.422 | -0.009 | -1.5 | -0.012 | 0.166 | 0.166 | 0 | 0.46 | 0.16 | 0.67 | 0.8 | 0.55 | 0.69 |
| 10 | 59715050 | rs10826129 | 0.225 | -0.491 | 0.626 | -1.593 | 0.022 | -0.094 | -0.094 | 0 | 0.24 | 0.36 | 0.18 | 0.08 | 0.27 | 0.06 |
| 10 | 59715472 | rs2658618 | 1.052 | -0.466 | 0.584 | -1.575 | -0.02 | 0.182 | 0.182 | 0 | 0.47 | 0.13 | 0.67 | 0.8 | 0.55 | 0.69 |
| 10 | 59716360 | rs10763523 | 0.404 | -0.472 | 0.688 | -1.645 | 0.021 | 0.077 | 0.077 | 0 | 0.56 | 0.28 | 0.74 | 0.92 | 0.62 | 0.84 |
| 10 | 59716853 | rs2537665 | 0.25 | -0.479 | 0.754 | -1.648 | 0.003 | -0.023 | -0.023 | 2.227 | 0.57 | 0.43 | 0.75 | 0.92 | 0.62 | 0.84 |
| 10 | 59718844 | rs2658636 | 0.818 | -0.464 | 0.686 | -1.633 | 0.016 | 0.193 | 0.193 | 0 | 0.49 | 0.21 | 0.68 | 0.88 | 0.56 | 0.76 |
| 10 | 59719339 | rs2537662 | 1.928 | -0.535 | 0.076 | -1.311 | -0.121 | -0.159 | -0.159 | 1.01 | 0.24 | 0.02 | 0.29 | 0.55 | 0.37 | 0.2 |
| 10 | 59724431 | rs2658641 | 0.775 | -0.872 | 1.301 | -1.48 | 0.013 | -0.004 | -0.004 | 0 | 0.69 | 0.28 | 0.82 | 0.96 | 0.82 | 0.82 |
| 10 | 59727834 | rs2658645 | 0.47 | -0.96 | 1.29 | -1.522 | -0.06 | -0.159 | -0.159 | 0.931 | 0.73 | 0.51 | 0.88 | 1 | 0.86 | 0.81 |
| 10 | 59729698 | rs2537650 | 1.716 | -0.874 | 1.271 | -1.497 | -0.046 | 0.15 | 0.15 | 0.012 | 0.43 | 0.15 | 0.66 | 0.81 | 0.53 | 0.63 |
| 10 | 59731757 | rs10509081 | -0.469 | -0.959 | 1.289 | -1.452 | 0.002 | -0.018 | -0.018 | 3.949 | 0.08 | 0.21 | 0.07 | 0.03 | 0.07 | 0.06 |
| 10 | 59739454 | rs1930447 | -0.103 | -1 | 1.332 | -1.474 | 0.008 | -0.025 | -0.025 | 0 | 0.77 | 0.78 | 0.92 | 1 | 0.87 | 0.82 |
| 10 | 59740012 | rs2110020 | 1.208 | -1.01 | 1.383 | -1.485 | 0.022 | 0.007 | 0.007 | 0.37 | 0.64 | 0.3 | 0.8 | 0.96 | 0.79 | 0.75 |
| 10 | 59740431 | rs2779630 | 0.69 | -1.011 | 1.415 | -1.485 | -0.012 | -0.06 | -0.06 | 0 | 0.69 | 0.57 | 0.85 | 0.96 | 0.8 | 0.76 |
| 10 | 59740636 | rs1930449 | 1.61 | -0.857 | 1.385 | -1.421 | -0.04 | 0.148 | 0.148 | 0.02 | 0.43 | 0.15 | 0.66 | 0.81 | 0.54 | 0.64 |
| 10 | 59743252 | rs1999505 | -1.557 | -0.829 | 1.34 | -1.386 | 0.013 | -0.051 | -0.051 | 0 | 0.13 | 0.48 | 0.11 | 0.03 | 0.09 | 0.06 |
| 10 | 59749408 | rs1930458 | -2.18 | -0.828 | 1.314 | -1.387 | -0.02 | -0.097 | -0.097 | 0 | 0.18 | 0.56 | 0.13 | 0.04 | 0.11 | 0.13 |
| 10 | 59758221 | rs1930462 | 0.086 | -0.765 | 1.168 | -1.402 | 0.007 | 0.047 | 0.047 | 0 | 0.63 | 0.76 | 0.79 | 0.92 | 0.65 | 0.83 |

| | | | | | | | | | | | | | | | |
|---|---|---|---|---|---|---|---|---|---|---|---|---|---|---|---|
| 10 | 59759251 | rs11006003 | -1.276 | -1.031 | 1.307 | -1.163 | -0.006 | 0 | 0 | 0 | 0.08 | 0.12 | 0.03 | 0.08 | 0.02 | 0.15 |
| 10 | 59760871 | rs10509084 | -1.788 | -1.033 | 1.309 | -1.198 | 0 | 0.001 | 0.001 | 0 | 0.14 | 0.19 | 0.1 | 0.11 | 0.1 | 0.2 |
| 10 | 59761078 | rs11006004 | -2.468 | -1.052 | 1.424 | -1.21 | -0.033 | -0.046 | -0.046 | 0.416 | 0.2 | 0.58 | 0.14 | 0.11 | 0.11 | 0.21 |
| 10 | 59763232 | rs13376839 | -1.229 | -1.019 | 1.473 | -1.213 | 0.058 | 0.037 | 0.037 | 1.858 | 0.06 | 0.39 | 0.05 | 0 | 0.01 | 0.01 |
| 10 | 59763468 | rs7906925 | 0.108 | -0.992 | 1.29 | -1.267 | -0.013 | -0.085 | -0.085 | 0 | 0.65 | 0.62 | 0.82 | 0.92 | 0.72 | 0.84 |
| 10 | 59766998 | rs1930465 | -0.043 | -1.001 | 1.034 | -1.258 | 0.011 | 0.043 | 0.043 | 0 | 0.67 | 0.78 | 0.82 | 0.92 | 0.72 | 0.84 |
| 10 | 59767845 | rs1930470 | 0.553 | -1.04 | 1.097 | -1.266 | -0.005 | -0.05 | -0.05 | 1.276 | 0.59 | 0.57 | 0.8 | 0.92 | 0.71 | 0.76 |
| 10 | 59768050 | rs1930472 | 0.553 | -1.04 | 1.097 | -1.266 | -0.005 | -0.05 | -0.05 | 0 | 0.59 | 0.57 | 0.8 | 0.92 | 0.71 | 0.76 |
| 10 | 59770449 | rs11006011 | 1.623 | -1.025 | 0.905 | -1.259 | -0.057 | 0.184 | 0.184 | 0.012 | 0.45 | 0.04 | 0.68 | 0.81 | 0.62 | 0.63 |
| 10 | 59772188 | rs11006013 | -0.416 | -1.019 | 1.025 | -1.248 | 0 | -0.009 | -0.009 | 0.817 | 0.06 | 0.09 | 0.02 | 0 | 0.02 | 0.07 |
| 10 | 59776781 | rs1889066 | -0.008 | -0.859 | 0.977 | -1.236 | -0.005 | 0.005 | 0.005 | 0 | 0.07 | 0.06 | 0.07 | 0.03 | 0.1 | 0.06 |
| 10 | 59790055 | rs10740725 | 1.337 | -1.074 | 1.028 | -1.282 | 0.014 | 0.285 | 0.285 | 0.18 | 0.48 | 0.1 | 0.7 | 0.88 | 0.62 | 0.75 |
| 10 | 59799112 | rs1953092 | -0.631 | -1.084 | 1.098 | -1.299 | -0.003 | -0.036 | -0.036 | 0 | 0.82 | 0.85 | 0.93 | 1 | 0.89 | 0.91 |
| 10 | 59799718 | rs11006023 | -0.935 | -0.805 | 0.858 | -1.126 | 0.031 | -0.022 | -0.022 | 0 | 0.08 | 0.42 | 0.05 | 0 | 0.03 | 0.06 |
| 10 | 59805449 | rs12411438 | 1.141 | -0.577 | -0.458 | -0.895 | -0.066 | 0.146 | 0.146 | 0.013 | 0.45 | 0.04 | 0.68 | 0.77 | 0.62 | 0.61 |
| 10 | 59808144 | rs12777369 | 0.964 | -0.49 | -0.394 | -0.857 | 0.022 | 0.14 | 0.14 | 0 | 0.47 | 0.23 | 0.69 | 0.77 | 0.62 | 0.61 |
| 10 | 59819581 | rs10826146 | 0.54 | -0.333 | -0.57 | -0.789 | 0 | -0.005 | -0.005 | 0.085 | 0.18 | 0.25 | 0.11 | 0.12 | 0.17 | 0.09 |
| 10 | 59819898 | rs1910220 | -1.383 | -0.382 | -0.586 | -0.826 | 0.021 | 0.154 | 0.154 | 0 | 0.53 | 0.8 | 0.29 | 0.23 | 0.38 | 0.38 |
| 10 | 59820496 | rs1759365 | -0.988 | -0.345 | -0.089 | -0.411 | -0.007 | -0.026 | -0.026 | 1.456 | 0.35 | 0.53 | 0.18 | 0.11 | 0.21 | 0.29 |
| 10 | 59823894 | rs2184033 | 1.469 | 0.2 | -0.467 | 0.715 | -0.034 | -0.133 | -0.133 | 0 | 0.67 | 0.06 | 0.68 | 0.53 | 0.7 | 0.72 |
| 10 | 59825583 | rs1698417 | 1.521 | 0.164 | 1.041 | 0.674 | 0.004 | 0.028 | 0.028 | 0.447 | 0.83 | 0.56 | 0.87 | 0.82 | 0.83 | 0.85 |
| 10 | 59829407 | rs2221296 | -0.436 | 0.159 | 1.358 | 0.675 | -0.002 | 0.026 | 0.026 | 0.072 | 0.08 | 0.11 | 0.03 | 0.14 | 0.05 | 0.1 |
| 10 | 59836753 | rs1698425 | 1.807 | 0.152 | 1.714 | 0.67 | -0.002 | 0.008 | 0.008 | 0.39 | 0.84 | 0.71 | 0.88 | 0.82 | 0.83 | 0.87 |
| 10 | 59837172 | rs1759320 | -1.174 | 0.314 | 2.182 | 0.711 | 0.047 | 0.019 | 0.019 | 0 | 0.13 | 0.37 | 0.11 | 0.14 | 0.15 | 0.13 |
| 10 | 59838496 | rs1759325 | 1.879 | 0.31 | 2.121 | 0.636 | 0 | 0.016 | 0.016 | 0.404 | 0.84 | 0.65 | 0.88 | 0.82 | 0.83 | 0.87 |
| 10 | 59842423 | rs1625223 | 1.667 | 0.305 | 1.994 | 0.634 | -0.009 | -0.019 | -0.019 | 3.942 | 0.86 | 0.9 | 0.9 | 0.82 | 0.83 | 0.86 |
| 10 | 59846118 | rs1759339 | 1.397 | 0.321 | 2.087 | 0.609 | 0.015 | -0.035 | -0.035 | 0 | 0.92 | 0.78 | 0.91 | 0.96 | 0.89 | 0.97 |
| 10 | 59849685 | rs1759345 | 0.68 | 0.341 | 1.836 | 0.617 | -0.006 | 0.009 | 0.009 | 0 | 0.94 | 0.96 | 0.93 | 0.96 | 0.89 | 0.96 |
| 10 | 59850445 | rs1698455 | 0.917 | 0.381 | 1.947 | 0.616 | 0.012 | -0.024 | -0.024 | 0 | 0.93 | 0.9 | 0.93 | 0.96 | 0.89 | 0.96 |
| 10 | 59854651 | rs1698469 | 2.247 | 0.252 | 2.024 | 0.623 | -0.007 | 0.005 | 0.005 | 0.356 | 0.79 | 0.38 | 0.82 | 0.79 | 0.81 | 0.82 |
| 10 | 59855010 | rs1698471 | 1.206 | 0.472 | 1.371 | 0.467 | 0.009 | -0.026 | -0.026 | 0 | 0.91 | 0.86 | 0.89 | 0.96 | 0.86 | 0.94 |
| 10 | 59882663 | rs1698372 | -1.099 | 0.356 | 1.135 | 0.428 | -0.009 | 0.01 | 0.01 | 0 | 0.14 | 0.15 | 0.13 | 0.09 | 0.21 | 0.14 |
| 10 | 59887998 | rs10826158 | -2.3 | 0.125 | 1.377 | 1.018 | -0.001 | 0.165 | 0.165 | 0.093 | 0.12 | 0.52 | 0.2 | 0.25 | 0.08 | 0.14 |
| 10 | 59892940 | rs11817349 | -1.243 | 0.315 | 0.729 | 0.732 | 0.008 | -0.034 | -0.034 | 1.338 | 0.06 | 0.14 | 0.09 | 0 | 0.06 | 0.04 |
| 10 | 59893930 | rs2247911 | -1.984 | 0.049 | 0.298 | 0.939 | 0.001 | -0.086 | -0.086 | 0.037 | 0.34 | 0.86 | 0.36 | 0.48 | 0.32 | 0.34 |
| 10 | 59897533 | rs17635621 | -0.962 | -0.563 | 0.266 | 0.481 | 0.006 | 0.007 | 0.007 | 0 | 0.15 | 0.13 | 0.1 | 0.08 | 0.12 | 0.12 |
| 10 | 59905757 | rs7079625 | -1.069 | -0.435 | 0.215 | 0.564 | 0.011 | 0.013 | 0.013 | 0 | 0.15 | 0.05 | 0.12 | 0.1 | 0.13 | 0.11 |
| 10 | 59907497 | rs17635794 | 1.35 | -1.168 | -0.744 | 0.297 | 0.01 | -0.023 | -0.023 | 0 | 0.17 | 0.01 | 0.08 | 0.14 | 0.09 | 0.2 |
| 10 | 59908845 | rs11006067 | -0.643 | -1.035 | -0.343 | 0.277 | 0.013 | 0.005 | 0.005 | 0 | 0.16 | 0.04 | 0.13 | 0.11 | 0.14 | 0.11 |
| 10 | 59909204 | rs1199088 | -2.201 | -1.144 | 0.102 | 0.472 | 0.013 | -0.034 | -0.034 | 0.107 | 0.33 | 0.8 | 0.37 | 0.47 | 0.32 | 0.3 |
| 10 | 59912870 | rs3862864 | -0.351 | -0.987 | 1.067 | 0.29 | 0.077 | -0.014 | -0.014 | 0 | 0.05 | 0.22 | 0.06 | 0 | 0.09 | 0.01 |
| 10 | 59915056 | rs7083712 | -1.257 | -0.931 | 1.547 | 0.328 | -0.007 | 0.097 | 0.097 | 0 | 0.14 | 0.25 | 0.19 | 0.44 | 0.11 | 0.23 |
| 10 | 59918833 | rs11815860 | -0.384 | -0.946 | 1.53 | 0.285 | -0.003 | -0.023 | -0.023 | 0.33 | 0.05 | 0.1 | 0.07 | 0 | 0.05 | 0.03 |
| 10 | 59925048 | rs7072206 | 0.601 | -0.916 | 1.496 | 0.388 | -0.006 | -0.021 | -0.021 | 0 | 0.07 | 0.01 | 0.04 | 0.14 | 0.05 | 0.11 |
| 10 | 59941452 | rs11006074 | -0.454 | -1.07 | 1.863 | 0.233 | 0.009 | 0.015 | 0.015 | 0 | 0.15 | 0.08 | 0.13 | 0.1 | 0.13 | 0.12 |
| 10 | 59943234 | rs7073090 | -0.792 | -1.021 | 1.805 | 0.263 | 0.053 | 0.168 | 0.168 | 0 | 0.06 | 0.18 | 0.12 | 0.28 | 0.1 | 0.1 |
| 10 | 59944924 | rs1199106 | -2.408 | -1.048 | 1.919 | 0.223 | -0.003 | 0.023 | 0.023 | 0 | 0.24 | 0.69 | 0.25 | 0.43 | 0.21 | 0.28 |
| 10 | 59946672 | rs1199105 | -1.4 | -1.093 | 2.125 | 0.218 | 0.03 | 0.024 | 0.024 | 0 | 0.17 | 0.39 | 0.11 | 0.15 | 0.15 | 0.15 |
| 10 | 59972453 | rs2153281 | -2.417 | -0.785 | 2.017 | 0.141 | -0.003 | 0.023 | 0.023 | 0 | 0.24 | 0.69 | 0.25 | 0.43 | 0.21 | 0.27 |
| 10 | 59985671 | rs2790232 | -0.338 | -0.799 | 2.047 | 0.132 | 0.028 | 0.015 | 0.015 | 0 | 0.06 | 0.29 | 0.03 | 0 | 0.02 | 0.05 |
| 10 | 60002345 | rs6481383 | -2.899 | -0.838 | 1.984 | 0.109 | 0.023 | -0.074 | -0.074 | 1.518 | 0.4 | 0.84 | 0.39 | 0.53 | 0.34 | 0.38 |
| 10 | 60026425 | rs7899961 | -0.377 | -0.592 | 2.528 | 0.31 | 0.003 | 0.225 | 0.225 | 0 | 0.09 | 0.2 | 0.11 | 0.43 | 0.06 | 0.18 |
| 10 | 60028894 | rs2251039 | -2.5 | -0.599 | 2.213 | 0.306 | -0.003 | 0.026 | 0.026 | 0 | 0.24 | 0.69 | 0.25 | 0.43 | 0.21 | 0.26 |
| 10 | 60029163 | rs1416763 | -2.552 | -0.602 | 2.224 | 0.305 | -0.002 | 0.011 | 0.011 | 0 | 0.25 | 0.7 | 0.25 | 0.43 | 0.21 | 0.26 |
| 10 | 60035152 | rs2275442 | 0.393 | -0.592 | 2.112 | 0.304 | 0.011 | 0.075 | 0.075 | 0 | 0.07 | 0.11 | 0.04 | 0.15 | 0.05 | 0.1 |
| 10 | 60042625 | rs11812882 | -0.573 | -0.56 | 1.876 | 0.315 | -0.006 | -0.021 | -0.021 | 0 | 0.05 | 0.09 | 0.06 | 0 | 0.04 | 0.02 |
| 10 | 60049524 | rs16912085 | -1.298 | -0.558 | 1.889 | 0.316 | -0.002 | -0.031 | -0.031 | 0 | 0.07 | 0.3 | 0.08 | 0 | 0.04 | 0.03 |
| 10 | 60050453 | rs17620218 | 1.08 | -0.493 | 1.081 | -0.043 | -0.006 | 0.021 | 0.021 | 0 | 0.15 | 0.06 | 0.11 | 0 | 0.18 | 0.07 |
| 10 | 60052999 | rs12264744 | 0.841 | -0.219 | 0.935 | 0.294 | -0.006 | -0.017 | -0.017 | 0 | 0.06 | 0.01 | 0.03 | 0.15 | 0.05 | 0.1 |
| 10 | 60059641 | rs2590289 | -0.926 | -0.236 | 1 | 0.253 | 0.016 | 0.087 | 0.087 | 0 | 0.1 | 0.26 | 0.13 | 0.28 | 0.12 | 0.14 |
| 10 | 60061246 | rs965867 | -1.828 | -0.261 | 0.914 | 0.241 | 0.003 | 0.094 | 0.094 | 0 | 0.16 | 0.5 | 0.19 | 0.28 | 0.15 | 0.16 |
| 10 | 60074817 | rs2590303 | -0.897 | -0.243 | 0.619 | 0.35 | -0.013 | 0.092 | 0.092 | 0 | 0.09 | 0.26 | 0.11 | 0.28 | 0.06 | 0.17 |
| 10 | 60100089 | rs4948291 | -0.664 | -0.464 | -0.301 | -0.037 | 0.012 | 0.104 | 0.104 | 1.382 | 0.38 | 0.11 | 0.23 | 0.15 | 0.35 | 0.25 |
| 10 | 60110335 | rs11599360 | 0.458 | -0.485 | -0.424 | -0.094 | -0.019 | 0.145 | 0.145 | 0.008 | 0.22 | 0.02 | 0.13 | 0 | 0.21 | 0.07 |
| 10 | 60115223 | rs1905455 | -1.13 | -0.267 | 0.006 | 0.352 | 0.026 | -0.005 | -0.005 | 0 | 0.52 | 0.75 | 0.45 | 0.52 | 0.45 | 0.45 |
| 10 | 60118681 | rs10826172 | -1.027 | -0.245 | -0.308 | 0.163 | 0.012 | 0.004 | 0.004 | 0.656 | 0.49 | 0.62 | 0.39 | 0.44 | 0.4 | 0.42 |
| 10 | 60121414 | rs11593650 | -0.136 | 0.383 | 0.367 | 0.524 | -0.043 | 0.047 | 0.047 | 0 | 0.08 | 0.01 | 0.06 | 0 | 0.12 | 0.06 |
| 10 | 60134096 | rs1994174 | -1.131 | -0.259 | 0.844 | 0.951 | -0.014 | 0.205 | 0.205 | 0 | 0.11 | 0.52 | 0.15 | 0.28 | 0.06 | 0.17 |
| 10 | 60140709 | rs12355138 | -0.574 | -0.058 | 0.118 | -0.096 | 0.001 | 0.005 | 0.005 | 0 | 0.17 | 0.13 | 0.14 | 0 | 0.2 | 0.05 |

| | | | | | | | | | | | | | | | | |
|---|---|---|---|---|---|---|---|---|---|---|---|---|---|---|---|---|
| 10 | 60141537 | rs10826176 | -1.684 | -0.748 | 0.97 | 0.195 | -0.027 | -0.014 | -0.014 | 0 | 0.32 | 0.51 | 0.31 | 0.29 | 0.29 | 0.23 |
| 10 | 60142800 | rs9971104 | 0.267 | -0.599 | 0.534 | 0.015 | 0.006 | -0.011 | -0.011 | 0 | 0.21 | 0.35 | 0.18 | 0.18 | 0.18 | 0.2 |
| 10 | 60143021 | rs11006127 | -0.058 | -0.683 | 0.667 | -0.012 | 0.007 | -0.007 | -0.007 | 0.021 | 0.08 | 0.04 | 0.12 | 0.3 | 0.06 | 0.15 |
| 10 | 60144039 | rs10826177 | -0.461 | -0.678 | 0.984 | -0.079 | -0.019 | -0.129 | -0.129 | 0.287 | 0.23 | 0.37 | 0.19 | 0 | 0.22 | 0.08 |
| 10 | 60144884 | rs2279340 | -0.49 | -0.562 | 1.142 | 0.018 | 0 | -0.002 | -0.002 | 0.023 | 0.26 | 0.29 | 0.21 | 0.18 | 0.23 | 0.21 |
| 10 | 60144998 | rs2279339 | 0.272 | -0.595 | 1.392 | 0.007 | -0.005 | 0.003 | 0.003 | 0 | 0.2 | 0.21 | 0.15 | 0.18 | 0.18 | 0.18 |
| 10 | 60150445 | rs11006130 | 0.43 | -0.662 | 1.224 | -0.079 | 0 | -0.03 | -0.03 | 0 | 0.14 | 0.17 | 0.12 | 0 | 0.17 | 0.06 |
| 10 | 60151026 | rs10826179 | -2.034 | -0.814 | 1.205 | -0.077 | 0.008 | 0.003 | 0.003 | 0 | 0.49 | 0.86 | 0.45 | 0.47 | 0.44 | 0.41 |
| 10 | 60155120 | rs1049432 | 0.269 | -0.729 | 1.41 | 0.062 | 0.007 | -0.01 | -0.01 | 0 | 0.2 | 0.31 | 0.17 | 0.18 | 0.18 | 0.18 |
| 10 | 60156166 | rs11006132 | -0.435 | -0.726 | 1.418 | 0.078 | 0.003 | -0.016 | -0.016 | 0 | 0.26 | 0.32 | 0.21 | 0.18 | 0.23 | 0.2 |
| 10 | 60159362 | rs12245545 | -1.34 | -0.631 | 0.773 | -0.009 | 0.006 | -0.011 | -0.011 | 0 | 0.35 | 0.64 | 0.33 | 0.47 | 0.27 | 0.35 |
| 10 | 60167882 | rs6481387 | -1.194 | -0.489 | 0.452 | 0.144 | -0.012 | -0.062 | -0.062 | 0 | 0.41 | 0.88 | 0.38 | 0.49 | 0.36 | 0.31 |
| 10 | 60201928 | rs7916849 | -0.234 | -1.15 | 0.494 | 0.12 | -0.041 | 0.007 | 0.007 | 0 | 0.69 | 0.46 | 0.77 | 0.67 | 0.74 | 0.59 |
| 10 | 60205817 | rs2114563 | -0.853 | -1.323 | -0.033 | 0.156 | 0.025 | -0.03 | -0.03 | 0.026 | 0.74 | 0.87 | 0.82 | 0.65 | 0.76 | 0.57 |
| 10 | 60219560 | rs1125138 | 1.438 | -1.1 | -0.395 | 0.249 | -0.151 | 0.033 | 0.033 | 0.268 | 0.32 | 0.02 | 0.45 | 0.21 | 0.41 | 0.31 |
| 10 | 60225935 | rs2114561 | -0.306 | -1.403 | 0.369 | 0.419 | -0.044 | -0.02 | -0.02 | 0 | 0.69 | 0.52 | 0.79 | 0.74 | 0.74 | 0.57 |
| 10 | 60230128 | rs10082474 | -1.184 | -0.523 | -0.273 | 0.357 | -0.091 | -0.055 | -0.055 | 0.203 | 0.68 | 0.85 | 0.59 | 0.61 | 0.51 | 0.68 |
| 10 | 60236950 | rs12262404 | 0.159 | -0.891 | -0.052 | 0.174 | -0.016 | -0.002 | -0.002 | 0 | 0.43 | 0.54 | 0.4 | 0.41 | 0.31 | 0.47 |
| 10 | 60240354 | rs17644676 | -0.68 | -0.942 | -0.301 | -0.014 | 0.006 | 0.02 | 0.02 | 1.142 | 0.28 | 0.18 | 0.16 | 0.16 | 0.22 | 0.21 |
| 10 | 60242641 | rs10826197 | -1.22 | -0.702 | -0.345 | 0.068 | -0.035 | -0.021 | -0.021 | 0 | 0.67 | 0.75 | 0.58 | 0.61 | 0.53 | 0.68 |
| 10 | 60245684 | rs10763548 | -1.684 | -0.733 | -0.629 | -0.092 | -0.172 | -0.004 | -0.004 | 0.111 | 0.72 | 0.98 | 0.68 | 0.79 | 0.57 | 0.72 |
| 10 | 60255969 | rs1427213 | -0.191 | -0.754 | -0.065 | -0.468 | -0.001 | -0.001 | -0.001 | 0 | 0.49 | 0.5 | 0.49 | 0.59 | 0.43 | 0.6 |
| 10 | 60258297 | rs1346303 | -0.501 | -0.78 | -0.207 | -0.498 | -0.012 | -0.06 | -0.06 | 0 | 0.19 | 0.27 | 0.08 | 0 | 0.11 | 0.08 |
| 10 | 60261697 | rs6481397 | -1.129 | -0.219 | -0.073 | -0.127 | -0.016 | -0.009 | -0.009 | 0 | 0.68 | 0.77 | 0.59 | 0.59 | 0.54 | 0.68 |
| 10 | 60271824 | rs7069916 | -0.188 | 0.37 | 0.047 | -0.34 | 0.015 | -0.009 | -0.009 | 0 | 0.45 | 0.61 | 0.27 | 0.48 | 0.37 | 0.49 |
| 10 | 60274606 | rs7897298 | -0.813 | 0.463 | -0.025 | -0.501 | -0.006 | -0.005 | -0.005 | 0.712 | 0.54 | 0.64 | 0.33 | 0.49 | 0.5 | 0.56 |
| 10 | 60279742 | rs1649031 | -0.855 | 0.456 | 0.139 | -0.503 | -0.023 | -0.02 | -0.02 | 0 | 0.54 | 0.75 | 0.34 | 0.49 | 0.5 | 0.56 |
| 10 | 60280153 | rs1658495 | -0.915 | 0.44 | 0.19 | -0.51 | -0.057 | -0.053 | -0.053 | 0 | 0.55 | 0.86 | 0.35 | 0.49 | 0.5 | 0.56 |
| 10 | 60312599 | rs1649059 | 0.483 | 0.429 | 0.149 | -0.515 | -0.022 | -0.02 | -0.02 | 0 | 0.46 | 0.25 | 0.66 | 0.51 | 0.5 | 0.44 |
| 10 | 60314017 | rs16912382 | -0.154 | 0.396 | 0.409 | -0.538 | 0.013 | -0.055 | -0.055 | 0 | 0.11 | 0.16 | 0.06 | 0 | 0.14 | 0.08 |
| 10 | 60316211 | rs7099881 | -0.706 | 0.468 | 0.048 | -0.467 | -0.002 | -0.002 | -0.002 | 0 | 0.53 | 0.57 | 0.31 | 0.49 | 0.49 | 0.56 |
| 10 | 60317087 | rs1649058 | 0.444 | 0.419 | -0.098 | -0.462 | -0.022 | -0.019 | -0.019 | 1.213 | 0.46 | 0.25 | 0.66 | 0.51 | 0.51 | 0.44 |
| 10 | 60324282 | rs1649048 | 0.503 | 0.431 | -0.327 | -0.457 | -0.057 | -0.053 | -0.053 | 0 | 0.45 | 0.14 | 0.65 | 0.51 | 0.5 | 0.44 |
| 10 | 60334002 | rs1982173 | 1.078 | 0.697 | -0.597 | -0.363 | -0.063 | -0.058 | -0.058 | 0 | 0.35 | 0.14 | 0.6 | 0.42 | 0.4 | 0.31 |
| 10 | 60335090 | rs6481407 | -1.187 | 0.693 | -0.473 | -0.364 | -0.009 | -0.008 | -0.008 | 0.089 | 0.64 | 0.67 | 0.37 | 0.58 | 0.59 | 0.69 |
| 10 | 60351663 | rs7091956 | 0.41 | 0.731 | -0.464 | -0.186 | -0.002 | 0.032 | 0.032 | 0.02 | 0.27 | 0.32 | 0.18 | 0.46 | 0.25 | 0.34 |
| 10 | 60365975 | rs12258523 | -0.474 | 0.863 | -1.052 | -0.16 | -0.013 | -0.063 | -0.063 | 0 | 0.38 | 0.18 | 0.21 | 0.46 | 0.37 | 0.49 |
| 10 | 60369477 | rs12244874 | -0.071 | 0.423 | -0.782 | 0.186 | 0.02 | -0.058 | -0.058 | 0 | 0.28 | 0.16 | 0.16 | 0.46 | 0.24 | 0.39 |
| 10 | 60370141 | rs7097457 | -0.315 | 0.979 | -0.355 | -0.091 | 0.012 | -0.02 | -0.02 | 0 | 0.06 | 0.1 | 0.07 | 0.02 | 0.1 | 0.06 |
| 10 | 60383526 | rs11006209 | 0.413 | 0.89 | -0.763 | 0.056 | -0.044 | -0.058 | -0.058 | 0 | 0.33 | 0.16 | 0.3 | 0.47 | 0.44 | 0.36 |
| 10 | 60389001 | rs2028205 | -0.564 | 0.748 | -0.122 | -0.004 | -0.007 | -0.01 | -0.01 | 0 | 0.65 | 0.7 | 0.69 | 0.54 | 0.56 | 0.64 |
| 10 | 60396065 | rs920259 | 0.075 | 0.43 | 0.474 | 0.107 | 0.001 | -0.002 | -0.002 | 0 | 0.67 | 0.32 | 0.62 | 0.64 | 0.76 | 0.65 |
| 10 | 60408485 | rs7085920 | -0.267 | 0.634 | 0.834 | -0.219 | 0.009 | -0.004 | -0.004 | 0 | 0.13 | 0.18 | 0.1 | 0.12 | 0.14 | 0.15 |
| 10 | 60421828 | rs4948308 | -1.051 | 0.393 | 0.481 | -0.298 | -0.008 | -0.033 | -0.033 | 0 | 0.22 | 0.52 | 0.2 | 0.12 | 0.17 | 0.22 |
| 10 | 60422453 | rs4948535 | 1.048 | 0.201 | 0.763 | -0.335 | 0.001 | 0 | 0 | 3.07 | 0.76 | 0.23 | 0.78 | 0.88 | 0.83 | 0.79 |
| 10 | 60431371 | rs7910029 | 0.989 | 0.208 | 0.933 | -0.376 | -0.003 | -0.006 | -0.006 | 0.04 | 0.77 | 0.28 | 0.78 | 0.9 | 0.83 | 0.81 |
| 10 | 60459795 | rs11006227 | -1.121 | 0.234 | 0.926 | -0.379 | -0.016 | -0.036 | -0.036 | 0.195 | 0.23 | 0.6 | 0.2 | 0.1 | 0.17 | 0.23 |
| 10 | 60472605 | rs7896348 | -1.147 | 0.441 | 0.938 | -0.435 | 0.004 | -0.053 | -0.053 | 0 | 0.24 | 0.53 | 0.22 | 0.1 | 0.24 | 0.25 |
| 10 | 60510598 | rs2393481 | 0.745 | 0.249 | 0.11 | -0.729 | -0.006 | -0.037 | -0.037 | 1.109 | 0.62 | 0.42 | 0.75 | 0.9 | 0.69 | 0.66 |
| 10 | 60512403 | rs10826233 | 0.856 | 0.345 | -0.191 | -0.712 | 0.012 | 0.03 | 0.03 | 0.692 | 0.6 | 0.31 | 0.73 | 0.9 | 0.68 | 0.66 |
| 10 | 60516335 | rs11815410 | 0.507 | 0.819 | -0.574 | -0.284 | 0.009 | 0.005 | 0.005 | 1.617 | 0.49 | 0.17 | 0.59 | 0.52 | 0.43 | 0.45 |
| 10 | 60523769 | rs10509093 | -0.026 | 1.088 | -0.445 | -0.442 | -0.018 | -0.033 | -0.033 | 0 | 0.25 | 0.19 | 0.28 | 0.48 | 0.37 | 0.39 |
| 10 | 60531282 | rs2393479 | -0.07 | 0.944 | 0.221 | -0.582 | 0.004 | -0.133 | -0.133 | 0.158 | 0.79 | 0.58 | 0.89 | 1 | 0.81 | 0.86 |
| 10 | 60535920 | rs999845 | -0.177 | 0.915 | 0.142 | -0.577 | -0.003 | -0.104 | -0.104 | 0 | 0.8 | 0.65 | 0.89 | 1 | 0.82 | 0.86 |
| 10 | 60548276 | rs11812567 | 0.409 | 0.694 | 0.222 | -0.263 | -0.002 | -0.001 | -0.001 | 0.776 | 0.56 | 0.59 | 0.63 | 0.53 | 0.45 | 0.47 |
| 10 | 60555578 | rs11006263 | -0.947 | 0.931 | 0.261 | -0.142 | -0.001 | -0.003 | -0.003 | 0 | 0.08 | 0.1 | 0.02 | 0.04 | 0.05 | 0.07 |
| 10 | 60556751 | rs2893790 | -0.344 | 1.166 | 0.499 | 0.048 | 0.001 | 0.007 | 0.007 | 0 | 0.36 | 0.4 | 0.31 | 0.56 | 0.36 | 0.26 |
| 10 | 60565959 | rs3740452 | 1.121 | 1.319 | 1.584 | -0.009 | 0.011 | -0.028 | -0.028 | 0 | 0.72 | 0.48 | 0.75 | 0.77 | 0.7 | 0.68 |
| 10 | 60569646 | rs9416745 | 1.304 | 1.349 | 1.79 | 0.043 | 0.002 | -0.01 | -0.01 | 1.243 | 0.7 | 0.42 | 0.74 | 0.73 | 0.7 | 0.68 |
| 10 | 60570927 | rs4948548 | 1.17 | 1.294 | 1.858 | 0.065 | 0.005 | -0.009 | -0.009 | 0.177 | 0.71 | 0.63 | 0.75 | 0.73 | 0.7 | 0.67 |
| 10 | 60577393 | rs4344442 | 0.67 | 1.307 | 1.888 | 0.064 | 0.002 | -0.002 | -0.002 | 0 | 0.75 | 0.77 | 0.78 | 0.76 | 0.7 | 0.69 |
| 10 | 60580635 | rs1899979 | 0.872 | 1.243 | 1.822 | 0.064 | 0.015 | -0.017 | -0.017 | 0 | 0.74 | 0.66 | 0.76 | 0.74 | 0.7 | 0.68 |
| 10 | 60581614 | rs1992657 | 0.897 | 1.245 | 1.697 | 0.064 | 0.007 | -0.006 | -0.006 | 0 | 0.75 | 0.71 | 0.77 | 0.75 | 0.7 | 0.69 |
| 10 | 60585616 | rs9416747 | 1.247 | 1.256 | 1.529 | 0.069 | 0.006 | -0.011 | -0.011 | 0.658 | 0.7 | 0.6 | 0.75 | 0.72 | 0.69 | 0.62 |
| 10 | 60588553 | rs4948550 | 1.075 | 1.246 | 0.995 | 0.073 | 0.001 | -0.007 | -0.007 | 0 | 0.69 | 0.38 | 0.73 | 0.73 | 0.69 | 0.62 |
| 10 | 60590203 | rs9416748 | 1.603 | 0.748 | 1.889 | 0.505 | -0.015 | 0.032 | 0.032 | 8.905 | 0.87 | 0.39 | 0.88 | 0.86 | 0.92 | 0.8 |
| 10 | 60604883 | rs1867551 | -1.097 | 1.334 | 0.013 | 0.481 | -0.001 | -0.001 | -0.001 | 0 | 0.1 | 0.11 | 0.1 | 0.16 | 0.09 | 0.2 |
| 10 | 60605408 | rs3099152 | -1.556 | 1.838 | 0.25 | 0.604 | 0.02 | 0.018 | 0.018 | 0 | 0.1 | 0.18 | 0.09 | 0.15 | 0.16 | 0.31 |
| 10 | 60606271 | rs12218563 | -0.693 | 1.827 | 0.189 | 0.6 | -0.012 | -0.01 | -0.01 | 0 | 0.09 | 0.06 | 0.09 | 0.15 | 0.09 | 0.11 |

| | | | | | | | | | | | | | | | | |
|---|---|---|---|---|---|---|---|---|---|---|---|---|---|---|---|---|
| 10 | 60606990 | rs3125452 | 0.26 | 1.921 | 0.019 | 0.383 | -0.005 | 0.03 | 0.03 | 0 | 0.77 | 0.85 | 0.75 | 0.86 | 0.81 | 0.83 |
| 10 | 60610919 | rs7910260 | 1.73 | 1.528 | 0.737 | 0.136 | -0.007 | 0.011 | 0.011 | 0.142 | 0.55 | 0.34 | 0.53 | 0.57 | 0.56 | 0.42 |
| 10 | 60614717 | rs7350384 | 1.776 | 1.418 | 0.457 | -0.25 | -0.037 | -0.042 | -0.042 | 0.595 | 0.42 | 0.07 | 0.46 | 0.56 | 0.43 | 0.38 |
| 10 | 60615095 | rs11006280 | 1.731 | 1.507 | 1.144 | 0.272 | -0.001 | 0 | 0 | 0 | 0.63 | 0.37 | 0.61 | 0.74 | 0.68 | 0.57 |
| 10 | 60619002 | rs12261770 | -0.615 | 0.71 | 1.517 | 0.34 | 0.012 | 0.032 | 0.032 | 0 | 0.05 | 0.13 | 0.08 | 0.15 | 0.07 | 0.1 |
| 10 | 60620540 | rs3099157 | -2.298 | 0.713 | 1.867 | 0.326 | 0.061 | 0.032 | 0.032 | 0 | 0.12 | 0.39 | 0.13 | 0.15 | 0.15 | 0.26 |
| 10 | 60630041 | rs16912758 | -0.771 | 0.766 | 1.649 | 0.221 | -0.011 | -0.075 | -0.075 | 0 | 0.21 | 0.28 | 0.06 | 0 | 0.11 | 0.12 |
| 10 | 60653783 | rs7094854 | -0.783 | 0.84 | 1.061 | 0.276 | -0.023 | -0.107 | -0.107 | 0 | 0.22 | 0.38 | 0.08 | 0.01 | 0.12 | 0.22 |
| 10 | 60658833 | rs282687 | 0.573 | 0.505 | 0.005 | -0.089 | -0.013 | -0.028 | -0.028 | 0.782 | 0.71 | 0.34 | 0.83 | 0.88 | 0.84 | 0.64 |
| 10 | 60664514 | rs282682 | 0.421 | 0.526 | -0.047 | -0.069 | -0.015 | -0.046 | -0.046 | 0.511 | 0.7 | 0.38 | 0.82 | 0.88 | 0.81 | 0.64 |
| 10 | 60668561 | rs10826248 | -0.752 | 0.818 | -0.27 | 0.732 | -0.004 | -0.023 | -0.023 | 0.918 | 0.23 | 0.16 | 0.41 | 0.38 | 0.24 | 0.28 |
| 10 | 60671286 | rs3910870 | -0.752 | 0.836 | -0.883 | 0.689 | -0.043 | -0.064 | -0.064 | 0 | 0.16 | 0.04 | 0.23 | 0.29 | 0.23 | 0.06 |
| 10 | 60683190 | rs7906427 | -1.548 | 0.902 | 0.07 | 0.938 | 0 | 0.002 | 0.002 | 1.77 | 0.37 | 0.37 | 0.52 | 0.53 | 0.41 | 0.31 |
| 10 | 60693417 | rs3851256 | -0.353 | 0.885 | -0.006 | 0.95 | -0.013 | -0.075 | -0.075 | 0 | 0.21 | 0.09 | 0.4 | 0.44 | 0.22 | 0.28 |
| 10 | 60694657 | rs10826255 | -0.794 | 1.026 | 0.432 | 1.137 | 0.001 | -0.006 | -0.006 | 0 | 0.15 | 0.14 | 0.11 | 0.08 | 0.19 | 0.03 |
| 10 | 60711370 | rs12219375 | -0.903 | 1.404 | 0.278 | 1.277 | 0.006 | 0.132 | 0.132 | 0 | 0.06 | 0.1 | 0.29 | 0.44 | 0.1 | 0.26 |
| 10 | 60724482 | rs10826259 | -1.423 | 1.386 | 0.299 | 1.276 | 0.008 | 0.035 | 0.035 | 0 | 0.12 | 0.15 | 0.32 | 0.4 | 0.25 | 0.26 |
| 10 | 60725392 | rs3844157 | -1.345 | 1.415 | 0.309 | 1.265 | 0.008 | 0.036 | 0.036 | 0 | 0.12 | 0.15 | 0.32 | 0.4 | 0.24 | 0.27 |
| 10 | 60727887 | rs3851241 | -2.456 | 1.313 | 0.694 | 1.405 | 0.036 | 0.118 | 0.118 | 0 | 0.19 | 0.37 | 0.45 | 0.55 | 0.28 | 0.3 |
| 10 | 60732753 | rs10763593 | -1.731 | 1.757 | 0.714 | 1.451 | 0.059 | 0.112 | 0.112 | 0 | 0.14 | 0.24 | 0.33 | 0.41 | 0.3 | 0.28 |
| 10 | 60733376 | rs3844159 | -1.877 | 1.56 | 0.684 | 1.42 | 0.026 | 0.098 | 0.098 | 0 | 0.14 | 0.32 | 0.32 | 0.3 | 0.17 | 0.09 |
| 10 | 60740316 | rs204940 | -0.696 | 1.38 | 0.76 | 1.277 | 0.013 | 0.165 | 0.165 | 0 | 0.07 | 0.21 | 0.11 | 0.24 | 0.06 | 0.22 |
| 10 | 60741387 | rs282661 | -1.733 | 1.189 | 0.499 | 1.237 | -0.033 | 0.289 | 0.289 | 0.074 | 0.15 | 0.48 | 0.26 | 0.44 | 0.1 | 0.33 |
| 10 | 60754780 | rs2447112 | -0.814 | 0.797 | -0.092 | 0.602 | 0.009 | -0.021 | -0.021 | 0.011 | 0.22 | 0.56 | 0.29 | 0.15 | 0.25 | 0.15 |
| 10 | 60759141 | rs11597180 | 0.786 | 0.935 | -0.404 | 0.381 | -0.037 | 0.097 | 0.097 | 0.171 | 0.16 | 0 | 0.11 | 0 | 0.14 | 0.02 |
| 10 | 60761865 | rs1198056 | -1.26 | 0.937 | 0.176 | 0.901 | 0.005 | -0.023 | -0.023 | 0 | 0.24 | 0.57 | 0.3 | 0.15 | 0.25 | 0.15 |
| 10 | 60763470 | rs10826264 | -1.396 | 0.987 | 0.012 | 0.641 | -0.04 | -0.027 | -0.027 | 1.177 | 0.42 | 0.22 | 0.51 | 0.53 | 0.47 | 0.53 |
| 10 | 60765799 | rs10159574 | -1.299 | 0.876 | -0.049 | 0.605 | -0.027 | -0.028 | -0.028 | 0 | 0.4 | 0.22 | 0.5 | 0.53 | 0.47 | 0.53 |
| 10 | 60787802 | rs11006349 | -1.401 | 1.049 | -0.526 | 0.561 | -0.024 | -0.031 | -0.031 | 1.387 | 0.4 | 0.27 | 0.5 | 0.52 | 0.47 | 0.52 |
| 10 | 60790128 | rs17829274 | -0.685 | 1.319 | -1.948 | 0.323 | -0.053 | -0.03 | -0.03 | 0 | 0.21 | 0.04 | 0.23 | 0.2 | 0.22 | 0.39 |
| 10 | 60804875 | rs12779873 | -0.726 | 1.58 | -2.015 | -0.185 | -0.092 | 0.12 | 0.12 | 0 | 0.19 | 0.01 | 0.15 | 0 | 0.24 | 0.05 |
| 10 | 60818997 | rs10491007 | -0.79 | 0.457 | -0.241 | 0.155 | -0.032 | 0.13 | 0.13 | 0.96 | 0.11 | 0.46 | 0.08 | 0.15 | 0.02 | 0.09 |
| 10 | 60820345 | rs1992668 | -0.353 | 0.33 | -0.633 | -0.045 | -0.007 | 0.008 | 0.008 | 0 | 0.93 | 1 | 0.96 | 0.97 | 0.93 | 0.91 |
| 10 | 60821456 | rs16913019 | -0.27 | 0.304 | -0.348 | -0.005 | -0.008 | -0.003 | -0.003 | 4.239 | 0.06 | 0.06 | 0.03 | 0.03 | 0.01 | 0.06 |
| 10 | 60832582 | rs7911331 | -1.741 | 0.095 | -0.111 | -0.061 | -0.051 | 0.11 | 0.11 | 2.417 | 0.12 | 0.47 | 0.08 | 0.15 | 0.02 | 0.09 |
| 10 | 60839167 | rs16913073 | -0.681 | -0.028 | -0.4 | -0.449 | -0.017 | 0.021 | 0.021 | 0 | 0.08 | 0.11 | 0.04 | 0.15 | 0.02 | 0.08 |
| 10 | 60845241 | rs7068712 | -1.985 | -0.086 | -0.634 | -0.292 | -0.043 | 0.03 | 0.03 | 0.351 | 0.23 | 0.5 | 0.21 | 0.27 | 0.14 | 0.25 |
| 10 | 60851998 | rs12411643 | -0.063 | -0.059 | -0.232 | -0.322 | -0.037 | -0.015 | -0.015 | 0.102 | 0.1 | 0 | 0.11 | 0.12 | 0.13 | 0.16 |
| 10 | 60853419 | rs11006367 | -1.319 | -0.102 | 0.488 | -0.338 | 0.086 | 0.082 | 0.082 | 3.724 | 0.05 | 0.47 | 0.04 | 0 | 0 | 0 |
| 10 | 60861901 | rs7900300 | -2.068 | -0.161 | 0.858 | -0.324 | -0.038 | 0.017 | 0.017 | 0.417 | 0.23 | 0.53 | 0.2 | 0.27 | 0.14 | 0.24 |
| 10 | 60865856 | rs6481458 | -2.169 | -0.114 | 0.954 | -0.329 | -0.025 | 0.006 | 0.006 | 0.705 | 0.24 | 0.63 | 0.21 | 0.28 | 0.14 | 0.25 |
| 10 | 60869017 | rs11006373 | -2.157 | -0.197 | 0.925 | -0.366 | -0.037 | 0.012 | 0.012 | 0.494 | 0.24 | 0.54 | 0.2 | 0.28 | 0.14 | 0.25 |
| 10 | 60875922 | rs16913153 | -0.715 | -0.191 | 0.935 | -0.366 | 0.004 | -0.012 | -0.012 | 0.016 | 0.13 | 0.25 | 0.14 | 0.13 | 0.13 | 0.17 |
| 10 | 60879016 | rs892250 | -2.192 | -0.211 | 1.377 | -0.392 | -0.032 | 0.009 | 0.009 | 1.059 | 0.23 | 0.57 | 0.2 | 0.27 | 0.14 | 0.25 |
| 10 | 60880409 | rs12412348 | -0.617 | -0.2 | 1.328 | -0.39 | 0 | -0.007 | -0.007 | 0 | 0.13 | 0.15 | 0.13 | 0.13 | 0.13 | 0.17 |
| 10 | 60887354 | rs10826271 | 1.555 | -0.652 | 0.108 | -0.147 | -0.073 | -0.121 | -0.121 | 0 | 0.25 | 0.03 | 0.33 | 0.34 | 0.32 | 0.35 |
| 10 | 60888567 | rs1836034 | 0.818 | -0.679 | 0.173 | -0.19 | -0.037 | -0.016 | -0.016 | 0 | 0.1 | 0 | 0.11 | 0.13 | 0.13 | 0.17 |
| 10 | 60892390 | rs996355 | -1.63 | -0.699 | 0.293 | -0.192 | -0.013 | 0.001 | 0.001 | 0.671 | 0.24 | 0.66 | 0.21 | 0.28 | 0.14 | 0.25 |
| 10 | 60894724 | rs7894911 | -1.537 | -0.698 | 0.289 | -0.192 | -0.017 | 0.003 | 0.003 | 0.969 | 0.24 | 0.65 | 0.21 | 0.27 | 0.14 | 0.25 |
| 10 | 60896265 | rs1897620 | -0.83 | -0.876 | 0.299 | -0.542 | -0.024 | 0.071 | 0.071 | 2.274 | 0.63 | 0.79 | 0.75 | 0.82 | 0.65 | 0.75 |
| 10 | 60901336 | rs12268935 | -0.226 | -0.923 | 0.642 | -0.416 | -0.036 | -0.027 | -0.027 | 0 | 0.09 | 0.21 | 0.05 | 0.03 | 0.01 | 0.08 |
| 10 | 60901603 | rs7902312 | -1.65 | -0.981 | 0.555 | -0.46 | -0.009 | 0.01 | 0.01 | 0 | 0.93 | 0.99 | 0.96 | 0.97 | 0.94 | 0.92 |
| 10 | 60908468 | rs987059 | -1.485 | -0.966 | 0.8 | -0.313 | -0.031 | -0.026 | -0.026 | 0 | 0.24 | 0.58 | 0.2 | 0.18 | 0.14 | 0.25 |
| 10 | 60909461 | rs12411934 | 0.376 | -0.895 | 0.671 | -0.283 | -0.016 | -0.013 | -0.013 | 1.446 | 0.12 | 0.04 | 0.12 | 0.15 | 0.13 | 0.17 |
| 10 | 60912886 | rs11815411 | 0.815 | -0.858 | 0.663 | -0.257 | -0.037 | -0.034 | -0.034 | 0.235 | 0.1 | 0 | 0.12 | 0.15 | 0.13 | 0.17 |
| 10 | 60925428 | rs9299588 | -0.327 | -0.964 | 0.676 | -0.513 | -0.039 | -0.025 | -0.025 | 0 | 0.1 | 0.15 | 0.04 | 0.03 | 0.01 | 0.05 |
| 10 | 60928800 | rs7915977 | -1.488 | -1.071 | 0.889 | -0.542 | -0.044 | -0.077 | -0.077 | 0 | 0.24 | 0.45 | 0.12 | 0.11 | 0.13 | 0.28 |
| 10 | 60929485 | rs11006390 | 0.241 | -1.042 | 0.918 | -0.531 | 0 | -0.013 | -0.013 | 0.019 | 0.12 | 0.17 | 0.05 | 0.07 | 0.11 | 0.2 |
| 10 | 60942534 | rs4336984 | -2.073 | -0.908 | 0.789 | -0.589 | 0.027 | 0.04 | 0.04 | 0.193 | 0.3 | 0.76 | 0.23 | 0.17 | 0.19 | 0.39 |
| 10 | 60954419 | rs2452512 | -1.723 | -0.958 | 0.686 | -0.518 | -0.04 | -0.062 | -0.062 | 0 | 0.25 | 0.57 | 0.13 | 0.11 | 0.13 | 0.28 |
| 10 | 60966706 | rs10509097 | -1.225 | -0.959 | 0.593 | -0.518 | -0.027 | -0.046 | -0.046 | 0.79 | 0.23 | 0.35 | 0.1 | 0.11 | 0.13 | 0.26 |
| 10 | 60981423 | rs10509098 | -1.293 | -0.977 | 0.637 | -0.518 | -0.035 | -0.055 | -0.055 | 1.805 | 0.23 | 0.37 | 0.11 | 0.12 | 0.13 | 0.26 |
| 10 | 60981954 | rs12412923 | 0.687 | -0.981 | 0.606 | -0.521 | -0.032 | 0.013 | 0.013 | 0 | 0.11 | 0.02 | 0.04 | 0.08 | 0.11 | 0.2 |
| 10 | 60995585 | rs11006408 | -0.317 | -0.978 | 0.643 | -0.516 | -0.031 | -0.019 | -0.019 | 0.006 | 0.1 | 0.13 | 0.04 | 0.03 | 0.01 | 0.05 |
| 10 | 61003189 | rs11006413 | -1.427 | -0.974 | 0.534 | -0.517 | -0.053 | -0.08 | -0.08 | 0.026 | 0.24 | 0.45 | 0.11 | 0.12 | 0.13 | 0.26 |
| 10 | 61006410 | rs11815693 | 0.707 | -0.954 | 0.461 | -0.505 | -0.032 | 0.008 | 0.008 | 0 | 0.11 | 0.01 | 0.04 | 0.08 | 0.12 | 0.21 |
| 10 | 61007534 | rs1459996 | 1.135 | -0.915 | 0.512 | -0.485 | -0.049 | -0.073 | -0.073 | 0 | 0.75 | 0.46 | 0.87 | 0.88 | 0.87 | 0.72 |
| 10 | 61035570 | rs11819163 | -1.18 | -0.811 | 0.303 | -0.443 | -0.05 | -0.079 | -0.079 | 0 | 0.24 | 0.48 | 0.11 | 0.12 | 0.13 | 0.28 |
| 10 | 61050653 | rs11006428 | 1.263 | -0.364 | -0.331 | -0.148 | -0.065 | -0.153 | -0.153 | 0 | 0.25 | 0.02 | 0.37 | 0.56 | 0.31 | 0.32 |

| | | | | | | | | | | | | | | | | |
|---|---|---|---|---|---|---|---|---|---|---|---|---|---|---|---|---|
| 10 | 61057736 | rs284592 | 1.306 | -0.507 | 0.586 | -0.165 | 0.004 | -0.003 | -0.003 | 0.222 | 0.65 | 0.56 | 0.72 | 0.65 | 0.64 | 0.59 |
| 10 | 61060828 | rs284598 | 1.633 | -0.555 | 0.815 | -0.187 | 0 | 0.001 | 0.001 | 0.139 | 0.62 | 0.35 | 0.68 | 0.64 | 0.63 | 0.57 |
| 10 | 61063157 | rs415267 | 0.537 | -0.471 | 0.842 | -0.214 | -0.003 | 0.011 | 0.011 | 0 | 0.79 | 0.71 | 0.77 | 0.74 | 0.77 | 0.8 |
| 10 | 61064655 | rs12264332 | -0.294 | -0.563 | 1.073 | -0.281 | -0.008 | -0.006 | -0.006 | 0 | 0.06 | 0.18 | 0.03 | 0.03 | 0.01 | 0.05 |
| 10 | 61067088 | rs284583 | 1.495 | -0.533 | 0.811 | -0.286 | -0.005 | -0.067 | -0.067 | 1.863 | 0.8 | 0.46 | 0.86 | 0.9 | 0.84 | 0.76 |
| 10 | 61068522 | rs10509100 | 0.208 | -0.203 | 0.18 | -0.137 | -0.015 | 0.007 | 0.007 | 0 | 0.08 | 0.06 | 0.04 | 0.07 | 0.11 | 0.17 |
| 10 | 61071899 | rs2169127 | -1.788 | -0.368 | 0.284 | 0.012 | 0.001 | -0.001 | -0.001 | 0 | 0.36 | 0.62 | 0.28 | 0.41 | 0.31 | 0.47 |
| 10 | 61073257 | rs10826276 | 1.008 | -0.188 | 0.352 | -0.236 | -0.035 | -0.041 | -0.041 | 0 | 0.89 | 0.78 | 0.92 | 0.96 | 0.96 | 0.86 |
| 10 | 61073614 | rs11812286 | -0.375 | 0.279 | -0.078 | -0.031 | 0.007 | -0.007 | -0.007 | 0.312 | 0.1 | 0.17 | 0.05 | 0.07 | 0.12 | 0.17 |
| 10 | 61075275 | rs11815025 | -0.252 | 0.142 | -0.063 | -0.174 | -0.014 | 0.031 | 0.031 | 0 | 0.11 | 0.07 | 0.08 | 0.01 | 0.13 | 0.14 |
| 10 | 61082343 | rs284640 | -1.196 | 0.709 | 0.538 | 0.341 | 0 | 0.01 | 0.01 | 0 | 0.36 | 0.5 | 0.39 | 0.49 | 0.37 | 0.38 |
| 10 | 61085082 | rs1999668 | 1.405 | 0.393 | 0.583 | 0.253 | 0.021 | -0.044 | -0.044 | 0 | 0.57 | 0.21 | 0.57 | 0.5 | 0.63 | 0.52 |
| 10 | 61086444 | rs388516 | -1.569 | 0.407 | 0.634 | 0.259 | 0.011 | -0.034 | -0.034 | 2.241 | 0.41 | 0.69 | 0.41 | 0.49 | 0.37 | 0.39 |
| 10 | 61087914 | rs436207 | -1.613 | 0.42 | 0.641 | 0.265 | 0.012 | -0.034 | -0.034 | 0 | 0.41 | 0.69 | 0.41 | 0.5 | 0.37 | 0.48 |
| 10 | 61091607 | rs284630 | -1.185 | 0.63 | 0.979 | 0.314 | 0.003 | 0.026 | 0.026 | 0 | 0.37 | 0.48 | 0.39 | 0.48 | 0.38 | 0.39 |
| 10 | 61097761 | rs16913417 | -0.587 | 0.555 | 1.207 | 0.427 | -0.004 | 0.058 | 0.058 | 0 | 0.14 | 0.21 | 0.13 | 0.33 | 0.13 | 0.2 |
| 10 | 61102036 | rs1380558 | 0.626 | 0.542 | 1.492 | 0.386 | 0.008 | -0.033 | -0.033 | 1.622 | 0.76 | 0.57 | 0.72 | 0.83 | 0.75 | 0.71 |
| 10 | 61102489 | rs6481463 | 0.398 | 0.54 | 1.937 | 0.369 | 0.012 | -0.017 | -0.017 | 0 | 0.78 | 0.67 | 0.79 | 0.86 | 0.76 | 0.83 |
| 10 | 61105475 | rs10763603 | 1.897 | 0.469 | 1.761 | 0.372 | 0.002 | -0.021 | -0.021 | 0 | 0.62 | 0.35 | 0.59 | 0.51 | 0.63 | 0.51 |
| 10 | 61113123 | rs7912407 | -0.212 | 0.518 | 1.886 | 0.369 | 0 | 0 | 0 | 0.022 | 0.16 | 0.14 | 0.13 | 0.13 | 0.14 | 0.13 |
| 10 | 61113279 | rs16913479 | -0.446 | 0.352 | 1.639 | 0.288 | -0.01 | -0.087 | -0.087 | 0 | 0.15 | 0.03 | 0.11 | 0.33 | 0.13 | 0.2 |
| 10 | 61113993 | rs284609 | -1.629 | 0.438 | 2.059 | -0.078 | -0.012 | -0.017 | -0.017 | 0 | 0.26 | 0.53 | 0.45 | 0.18 | 0.23 | 0.32 |
| 10 | 61115114 | rs7098034 | -0.636 | 0.683 | 2.104 | -0.137 | -0.005 | -0.058 | -0.058 | 0 | 0.1 | 0.23 | 0.08 | 0 | 0.07 | 0.07 |
| 10 | 61123193 | rs453670 | -0.159 | 0.612 | 2.086 | -0.161 | 0.008 | -0.013 | -0.013 | 1.03 | 0.06 | 0.07 | 0.06 | 0.01 | 0.1 | 0.03 |
| 10 | 61124829 | rs1931973 | -1.042 | 0.794 | 2.252 | -0.166 | 0.043 | -0.072 | -0.072 | 0.089 | 0.12 | 0.32 | 0.1 | 0 | 0.12 | 0.07 |
| 10 | 61136215 | rs384626 | -2.627 | 0.4 | 1.826 | -0.166 | -0.05 | -0.046 | -0.046 | 1.029 | 0.52 | 0.83 | 0.68 | 0.46 | 0.49 | 0.55 |
| 10 | 61140376 | rs513817 | -2.485 | 0.42 | 1.907 | -0.168 | 0.006 | -0.014 | -0.014 | 0.065 | 0.51 | 0.76 | 0.6 | 0.43 | 0.47 | 0.41 |
| 10 | 61141321 | rs7902722 | -0.484 | 0.462 | 1.872 | -0.05 | 0.029 | -0.049 | -0.049 | 0 | 0.11 | 0.23 | 0.09 | 0.02 | 0.12 | 0.07 |
| 10 | 61150322 | rs284616 | -1.461 | 0.689 | 1.777 | -0.049 | -0.016 | 0.048 | 0.048 | 0 | 0.31 | 0.42 | 0.53 | 0.43 | 0.29 | 0.44 |
| 10 | 61157502 | rs10740769 | 2.196 | 0.93 | 1.911 | -0.105 | -0.005 | -0.024 | -0.024 | 5.744 | 0.49 | 0.21 | 0.32 | 0.51 | 0.46 | 0.44 |
| 10 | 61162106 | rs12356480 | -0.015 | 0.987 | 2.117 | -0.055 | 0.02 | -0.029 | -0.029 | 0 | 0.1 | 0.15 | 0.09 | 0.04 | 0.12 | 0.07 |
| 10 | 61181517 | rs1414627 | -1.001 | 0.943 | 2.024 | -0.069 | -0.005 | 0.021 | 0.021 | 0.299 | 0.25 | 0.27 | 0.41 | 0.4 | 0.25 | 0.3 |
| 10 | 61190125 | rs1450774 | -1.983 | 0.943 | 1.904 | -0.059 | 0.003 | -0.023 | -0.023 | 0 | 0.4 | 0.64 | 0.59 | 0.47 | 0.4 | 0.51 |
| 10 | 61195106 | rs4462284 | 0.117 | 1.07 | 1.712 | -0.013 | 0.064 | -0.04 | -0.04 | 0 | 0.91 | 0.83 | 0.86 | 0.96 | 0.86 | 0.8 |
| 10 | 61208539 | rs2060690 | 0.923 | 0.961 | 1.66 | 0.036 | 0.037 | 0.003 | 0.003 | 1.571 | 0.78 | 0.59 | 0.55 | 0.78 | 0.76 | 0.75 |
| 10 | 61208949 | rs11006494 | -0.322 | 0.957 | 1.23 | 0.121 | -0.023 | -0.075 | -0.075 | 0 | 0.12 | 0.02 | 0.11 | 0.22 | 0.15 | 0.24 |
| 10 | 61227930 | rs10826293 | -1.577 | 0.826 | 1.532 | 0.064 | 0.018 | 0.048 | 0.048 | 0 | 0.23 | 0.42 | 0.2 | 0.26 | 0.26 | 0.3 |
| 10 | 61237072 | rs2393556 | -1.684 | 0.825 | 1.568 | 0.066 | 0.008 | 0.034 | 0.034 | 0 | 0.24 | 0.39 | 0.19 | 0.26 | 0.26 | 0.31 |
| 10 | 61239474 | rs1977684 | 0.361 | 0.861 | 1.608 | 0.053 | 0.047 | 0.03 | 0.03 | 0 | 0.85 | 0.73 | 0.59 | 0.8 | 0.77 | 0.83 |
| 10 | 61242961 | rs10826297 | 2.418 | 0.724 | 1.552 | -0.019 | -0.025 | -0.032 | -0.032 | 0 | 0.49 | 0.16 | 0.33 | 0.52 | 0.47 | 0.44 |
| 10 | 61243709 | rs10826299 | 2.418 | 0.725 | 1.517 | -0.013 | -0.041 | -0.045 | -0.045 | 0 | 0.49 | 0.15 | 0.33 | 0.52 | 0.47 | 0.44 |
| 10 | 61250658 | rs7916869 | 0.599 | 0.96 | 1.744 | 0.17 | 0.04 | 0.043 | 0.043 | 0 | 0.86 | 0.73 | 0.6 | 0.8 | 0.81 | 0.82 |
| 10 | 61254813 | rs7915236 | 1.291 | 0.962 | 1.636 | 0.183 | 0.002 | -0.009 | -0.009 | 0 | 0.73 | 0.58 | 0.53 | 0.79 | 0.76 | 0.75 |
| 10 | 61259602 | rs12356684 | -0.787 | 1.373 | 1.625 | 0.189 | 0.015 | -0.061 | -0.061 | 0 | 0.19 | 0.23 | 0.13 | 0.05 | 0.21 | 0.09 |
| 10 | 61260801 | rs6481467 | -0.804 | 1.223 | 2.149 | 0.73 | 0.007 | 0.07 | 0.07 | 0 | 0.12 | 0.23 | 0.13 | 0.22 | 0.16 | 0.23 |
| 10 | 61266568 | rs7898048 | 0.081 | 1.448 | 2.029 | 0.812 | -0.002 | 0.071 | 0.071 | 0.247 | 0.07 | 0.14 | 0.27 | 0.18 | 0.06 | 0.05 |
| 10 | 61273488 | rs7919742 | -1.512 | 1.343 | 1.894 | 0.308 | 0.06 | 0.084 | 0.084 | 0 | 0.19 | 0.39 | 0.18 | 0.26 | 0.26 | 0.3 |
| 10 | 61284378 | rs1763891 | 1.61 | 0.845 | 1.244 | 0.266 | -0.003 | 0.004 | 0.004 | 2.112 | 0.58 | 0.29 | 0.48 | 0.63 | 0.59 | 0.62 |
| 10 | 61286319 | rs1763894 | 1.62 | 0.844 | 1.244 | 0.279 | -0.003 | 0.003 | 0.003 | 2.118 | 0.58 | 0.29 | 0.48 | 0.63 | 0.59 | 0.62 |
| 10 | 61294973 | rs1763885 | 0.663 | 0.902 | 0.949 | 0.287 | 0 | -0.021 | -0.021 | 0 | 0.77 | 0.66 | 0.63 | 0.82 | 0.78 | 0.88 |
| 10 | 61297547 | rs1763887 | 1.183 | 0.687 | -0.013 | 0.397 | -0.002 | 0.005 | 0.005 | 2.209 | 0.58 | 0.31 | 0.48 | 0.63 | 0.59 | 0.62 |
| 10 | 61302380 | rs16913702 | -0.269 | 0.616 | -0.467 | 0.416 | -0.024 | -0.052 | -0.052 | 0.028 | 0.08 | 0.01 | 0.08 | 0.15 | 0.11 | 0.21 |
| 10 | 61303642 | rs1981138 | 1.166 | 0.769 | -0.399 | 0.424 | -0.062 | -0.051 | -0.051 | 0 | 0.56 | 0.16 | 0.4 | 0.59 | 0.56 | 0.55 |
| 10 | 61303675 | rs1763895 | 0.597 | 1.292 | 0.166 | 0.077 | 0.04 | -0.027 | -0.027 | 0 | 0.75 | 0.49 | 0.59 | 0.78 | 0.71 | 0.84 |
| 10 | 61322015 | rs11006576 | 0.454 | 0.595 | -1.234 | 0.234 | -0.063 | -0.038 | -0.038 | 0 | 0.44 | 0.11 | 0.58 | 0.52 | 0.51 | 0.43 |
| 10 | 61324243 | rs1909646 | -0.271 | 0.359 | -0.249 | 0.009 | 0.042 | 0.022 | 0.022 | 0 | 0.12 | 0.36 | 0.16 | 0.11 | 0.13 | 0.15 |
| 10 | 61324576 | rs10509107 | 0.817 | 0.54 | 0.229 | 0.022 | 0.024 | 0.029 | 0.029 | 0 | 0.47 | 0.35 | 0.65 | 0.61 | 0.58 | 0.52 |
| 10 | 61331409 | rs7087516 | -0.459 | 0.62 | 0.213 | -0.007 | 0.056 | 0.046 | 0.046 | 0.191 | 0.11 | 0.37 | 0.15 | 0.11 | 0.13 | 0.15 |
| 10 | 61336292 | rs11813512 | -1.002 | 0.612 | 0.348 | -0.012 | 0.035 | 0.011 | 0.011 | 0 | 0.14 | 0.36 | 0.16 | 0.11 | 0.14 | 0.16 |
| 10 | 61337787 | rs7082987 | -1.222 | 0.149 | 0.329 | -0.281 | -0.011 | -0.005 | -0.005 | 0 | 0.35 | 0.39 | 0.29 | 0.33 | 0.31 | 0.38 |
| 10 | 61339147 | rs4948334 | 1.378 | -0.025 | 0.112 | -0.358 | 0.082 | 0.083 | 0.083 | 0 | 0.5 | 0.28 | 0.73 | 0.63 | 0.66 | 0.62 |
| 10 | 61339897 | rs10826320 | -0.149 | -0.119 | 0.082 | -0.401 | 0.025 | -0.009 | -0.009 | 0 | 0.23 | 0.06 | 0.14 | 0.23 | 0.17 | 0.23 |
| 10 | 61340114 | rs4948337 | -0.394 | -0.129 | 0.17 | -0.403 | -0.002 | -0.008 | -0.008 | 0 | 0.83 | 0.8 | 0.95 | 0.94 | 0.89 | 0.9 |
| 10 | 61340313 | rs12264752 | -0.117 | -0.127 | 0.17 | -0.418 | 0.047 | -0.005 | -0.005 | 0 | 0.07 | 0.34 | 0.05 | 0 | 0.06 | 0.04 |
| 10 | 61341519 | rs10826322 | 1.874 | -0.057 | 0.153 | -0.399 | -0.043 | -0.01 | -0.01 | 0 | 0.45 | 0.16 | 0.59 | 0.53 | 0.52 | 0.46 |
| 10 | 61346166 | rs12570492 | 0.135 | -0.043 | 0.163 | -0.36 | 0.028 | -0.022 | -0.022 | 1.66 | 0.2 | 0.05 | 0.12 | 0.23 | 0.14 | 0.2 |
| 10 | 61346942 | rs1909637 | -0.717 | 0.402 | 0.344 | -0.122 | -0.004 | 0.002 | 0.002 | 0 | 0.93 | 0.94 | 0.99 | 0.94 | 0.99 | 0.98 |
| 10 | 61354446 | rs6481469 | 0.746 | 0.419 | 0.365 | -0.119 | -0.019 | -0.021 | -0.021 | 0 | 0.81 | 0.71 | 0.94 | 0.86 | 0.89 | 0.88 |
| 10 | 61360110 | rs11006597 | 2.049 | 0.217 | 0.195 | -0.217 | -0.027 | -0.007 | -0.007 | 0 | 0.55 | 0.17 | 0.61 | 0.52 | 0.56 | 0.57 |

| | | | | | | | | | | | | | | | | |
|---|---|---|---|---|---|---|---|---|---|---|---|---|---|---|---|---|
| 10 | 61367131 | rs12268265 | -1.096 | -0.091 | 0.307 | -0.135 | -0.008 | 0.015 | 0.015 | 0 | 0.2 | 0.38 | 0.17 | 0.23 | 0.18 | 0.16 |
| 10 | 61376156 | rs11006606 | -1.171 | -0.232 | 0.206 | -0.352 | 0.009 | 0.008 | 0.008 | 0.049 | 0.31 | 0.55 | 0.34 | 0.41 | 0.35 | 0.34 |
| 10 | 61378404 | rs4948234 | -0.329 | 0.085 | -0.035 | -0.434 | -0.001 | -0.001 | -0.001 | 0 | 0.9 | 0.88 | 0.93 | 0.93 | 0.92 | 0.96 |
| 10 | 61379398 | rs10509111 | 0.435 | 0.075 | -0.04 | -0.451 | -0.005 | -0.032 | -0.032 | 0 | 0.09 | 0.14 | 0.04 | 0 | 0.08 | 0.07 |
| 10 | 61380392 | rs2169685 | -0.064 | 0.166 | -0.146 | -0.235 | -0.056 | -0.016 | -0.016 | 0 | 0.19 | 0.11 | 0.37 | 0.17 | 0.28 | 0.23 |
| 10 | 61380753 | rs2126771 | -0.08 | 0.111 | 0.128 | -0.287 | 0.001 | 0.06 | 0.06 | 0 | 0.11 | 0.35 | 0.1 | 0.18 | 0.07 | 0.1 |
| 10 | 61383345 | rs10826335 | 0.523 | 0.085 | 0.05 | -0.256 | -0.019 | 0.008 | 0.008 | 0 | 0.71 | 0.78 | 0.58 | 0.78 | 0.66 | 0.73 |
| 10 | 61384326 | rs11006609 | 0.931 | 0.301 | -0.213 | -0.199 | -0.002 | 0 | 0 | 0 | 0.61 | 0.41 | 0.62 | 0.6 | 0.62 | 0.64 |
| 10 | 61385731 | rs4948235 | 1.085 | 0.499 | -1.085 | -0.114 | -0.083 | -0.041 | -0.041 | 0 | 0.48 | 0.07 | 0.59 | 0.52 | 0.51 | 0.55 |
| 10 | 61385774 | rs4948236 | 0.59 | -0.168 | -0.286 | -0.06 | -0.01 | -0.005 | -0.005 | 0.61 | 0.69 | 0.36 | 0.82 | 0.68 | 0.79 | 0.78 |
| 10 | 61388368 | rs16913885 | -1.122 | 0.42 | -0.159 | 0.265 | 0.001 | 0.012 | 0.012 | 0 | 0.14 | 0.21 | 0.13 | 0.19 | 0.17 | 0.08 |
| 10 | 61396958 | rs1948703 | -0.817 | 0.313 | 0.585 | 0.302 | -0.003 | 0.002 | 0.002 | 0 | 0.08 | 0.06 | 0.09 | 0.19 | 0.11 | 0.08 |
| 10 | 61403940 | rs11006635 | 0.568 | 0.38 | -0.522 | -0.947 | 0.003 | 0.151 | 0.151 | 0 | 0.42 | 0.22 | 0.28 | 0.08 | 0.34 | 0.23 |
| 10 | 61413353 | rs12356193 | 0.578 | 0.7 | -0.056 | -1.117 | -0.023 | 0.042 | 0.042 | 0.01 | 0.14 | 0.06 | 0.09 | 0 | 0.17 | 0.11 |
| 10 | 61414011 | rs2242206 | 0.921 | 0.509 | 0.054 | -0.445 | -0.02 | -0.1 | -0.1 | 0.01 | 0.22 | 0.07 | 0.38 | 0.58 | 0.27 | 0.53 |
| 10 | 61424666 | rs11006670 | 1.619 | 0.414 | -0.578 | -1.042 | -0.041 | 0.125 | 0.125 | 0 | 0.24 | 0.03 | 0.18 | 0.06 | 0.2 | 0.19 |
| 10 | 61425189 | rs4948351 | 1.668 | 0.543 | -0.077 | -0.082 | -0.043 | -0.143 | -0.143 | 0.017 | 0.18 | 0.01 | 0.31 | 0.47 | 0.21 | 0.4 |
| 10 | 61433292 | rs10826342 | -0.901 | 0.697 | 1.132 | -0.461 | -0.006 | 0.084 | 0.084 | 1.967 | 0.75 | 0.57 | 0.58 | 0.41 | 0.73 | 0.47 |
| 10 | 61434519 | rs1171606 | 0.78 | 0.698 | 1.139 | -0.461 | -0.001 | 0.001 | 0.001 | 2.111 | 0.24 | 0.22 | 0.4 | 0.59 | 0.27 | 0.53 |
| 10 | 61438710 | rs1177697 | -1.071 | 0.824 | 0.956 | -0.589 | 0.002 | 0.007 | 0.007 | 0 | 0.51 | 0.51 | 0.62 | 0.8 | 0.6 | 0.71 |
| 10 | 61444364 | rs1171659 | 0.546 | 0.683 | 0.949 | -0.351 | -0.001 | -0.001 | -0.001 | 0 | 0.25 | 0.26 | 0.41 | 0.69 | 0.27 | 0.54 |
| 10 | 61449512 | rs1171652 | 0.834 | 0.713 | 0.904 | -0.391 | -0.001 | 0.005 | 0.005 | 1.953 | 0.23 | 0.24 | 0.38 | 0.62 | 0.22 | 0.53 |
| 10 | 61449564 | rs7094971 | 0.857 | 0.822 | 0.712 | -0.463 | -0.002 | -0.014 | -0.014 | 0.153 | 0.13 | 0.15 | 0.09 | 0 | 0.16 | 0.1 |
| 10 | 61450153 | rs1171650 | 0.751 | 0.791 | 0.798 | -0.465 | -0.011 | 0.038 | 0.038 | 1.583 | 0.24 | 0.35 | 0.38 | 0.62 | 0.23 | 0.53 |
| 10 | 61452116 | rs1171646 | -0.231 | 0.728 | 0.655 | -0.415 | 0 | 0.002 | 0.002 | 0 | 0.4 | 0.66 | 0.5 | 0.62 | 0.43 | 0.64 |
| 10 | 61460975 | rs1171627 | 0.652 | 0.705 | 0.633 | -0.806 | 0.082 | 0.057 | 0.057 | 0.056 | 0.93 | 0.73 | 0.94 | 0.94 | 0.9 | 0.94 |
| 10 | 61465713 | rs12772745 | 0.819 | 0.758 | -0.617 | -1.141 | -0.049 | 0.186 | 0.186 | 0 | 0.41 | 0.14 | 0.29 | 0.08 | 0.37 | 0.24 |
| 10 | 61465838 | rs1171619 | 0.305 | 0.9 | -0.111 | -0.32 | -0.003 | -0.066 | -0.066 | 0 | 0.81 | 0.78 | 0.85 | 1 | 0.77 | 0.87 |
| 10 | 61467182 | rs1171617 | 0.428 | 0.886 | -0.105 | -0.307 | -0.001 | -0.065 | -0.065 | 0.012 | 0.79 | 0.77 | 0.85 | 1 | 0.76 | 0.86 |
| 10 | 61468589 | rs1171616 | 0.453 | 0.893 | -0.102 | -0.303 | -0.002 | -0.065 | -0.065 | 1.748 | 0.79 | 0.76 | 0.85 | 1 | 0.76 | 0.86 |
| 10 | 61478755 | rs2137906 | -1.182 | 0.916 | -0.071 | 0.949 | 0 | 0.126 | 0.126 | 0 | 0.05 | 0.11 | 0.07 | 0.34 | 0.07 | 0.22 |
| 10 | 61479385 | rs1171609 | 0.461 | 0.777 | -0.403 | 0.336 | 0.003 | -0.026 | -0.026 | 0 | 0.57 | 0.27 | 0.68 | 0.52 | 0.59 | 0.48 |
| 10 | 61485660 | rs2616664 | 0.733 | 0.769 | -0.165 | 0.415 | 0.003 | 0.192 | 0.192 | 0 | 0.93 | 0.82 | 0.92 | 0.66 | 0.93 | 0.78 |
| 10 | 61494037 | rs1171582 | 1.295 | 0.776 | -0.888 | 0.008 | -0.102 | -0.171 | -0.171 | 0 | 0.39 | 0.03 | 0.54 | 0.52 | 0.4 | 0.38 |
| 10 | 61498947 | rs7896470 | -1.039 | 0.66 | -0.236 | 0.194 | 0.003 | 0.086 | 0.086 | 0 | 0.09 | 0.14 | 0.13 | 0.33 | 0.13 | 0.25 |
| 10 | 61507014 | rs10993988 | 0.393 | 0.679 | -0.541 | -0.031 | -0.056 | 0.062 | 0.062 | 0 | 0.28 | 0.07 | 0.17 | 0.14 | 0.27 | 0.28 |
| 10 | 61508784 | rs10128501 | -1.231 | 0.58 | -0.005 | -0.069 | 0.017 | -0.005 | -0.005 | 0 | 0.49 | 0.62 | 0.36 | 0.48 | 0.46 | 0.55 |
| 10 | 61512057 | rs1171594 | 1.001 | 0.585 | -0.55 | 0.227 | -0.024 | -0.058 | -0.058 | 0.048 | 0.48 | 0.13 | 0.6 | 0.52 | 0.52 | 0.46 |
| 10 | 61521278 | rs4948359 | -0.153 | 0.844 | -0.189 | -0.147 | -0.009 | 0.044 | 0.044 | 0 | 0.29 | 0.18 | 0.19 | 0.15 | 0.27 | 0.28 |
| 10 | 61531043 | rs2137907 | 1.626 | 0.777 | 0.457 | 0.358 | -0.013 | 0.036 | 0.036 | 0 | 0.77 | 0.4 | 0.76 | 0.64 | 0.75 | 0.67 |
| 10 | 61533567 | rs10994002 | -0.059 | 1.257 | 0.195 | -0.143 | -0.02 | 0.034 | 0.034 | 0 | 0.28 | 0.23 | 0.17 | 0.05 | 0.31 | 0.23 |
| 10 | 61533605 | rs10821592 | -1.243 | 1.183 | 0.635 | -0.018 | 0.006 | 0.103 | 0.103 | 0.452 | 0.14 | 0.47 | 0.18 | 0.25 | 0.17 | 0.29 |
| 10 | 61546009 | rs4948364 | -1.298 | 1.276 | 0.768 | -0.045 | -0.017 | 0.059 | 0.059 | 0.951 | 0.43 | 0.77 | 0.36 | 0.3 | 0.47 | 0.49 |
| 10 | 61546775 | rs16914097 | 0.045 | 1.244 | 0.761 | 0.006 | 0.004 | -0.012 | -0.012 | 0.277 | 0.06 | 0.03 | 0.1 | 0.15 | 0.11 | 0.16 |
| 10 | 61550057 | rs3802695 | -0.33 | 1.206 | 0.871 | 0.003 | -0.02 | -0.037 | -0.037 | 0 | 0.93 | 1 | 0.96 | 0.88 | 0.92 | 0.95 |
| 10 | 61552692 | rs1053266 | 1.326 | 1.241 | 0.984 | -0.065 | 0.009 | 0.058 | 0.058 | 0 | 0.5 | 0.2 | 0.6 | 0.61 | 0.45 | 0.44 |
| 10 | 61556986 | rs10761414 | -1.401 | 0.899 | 1.311 | 0.084 | -0.004 | 0.01 | 0.01 | 1.254 | 0.44 | 0.73 | 0.36 | 0.39 | 0.47 | 0.53 |
| 10 | 61562977 | rs10994023 | -1.985 | 0.767 | 1.494 | 0.08 | 0.045 | 0.099 | 0.099 | 0 | 0.05 | 0.24 | 0.06 | 0.08 | 0.07 | 0.05 |
| 10 | 61566000 | rs10509112 | -1.14 | 0.916 | 1.471 | -0.172 | 0.003 | -0.009 | -0.009 | 0 | 0.41 | 0.46 | 0.33 | 0.3 | 0.43 | 0.49 |
| 10 | 61575119 | rs7098890 | -2.649 | 0.842 | 1.964 | -0.188 | 0.097 | 0.014 | 0.014 | 0.015 | 0.07 | 0.4 | 0.06 | 0 | 0.08 | 0.02 |
| 10 | 61575320 | rs7099156 | -1.162 | 1.065 | 2.189 | -0.177 | 0.014 | -0.007 | -0.007 | 0 | 0.39 | 0.55 | 0.47 | 0.34 | 0.43 | 0.5 |
| 10 | 61576850 | rs7088886 | -1.671 | 1.024 | 2.221 | -0.262 | 0.005 | 0.058 | 0.058 | 1.906 | 0.48 | 0.71 | 0.36 | 0.3 | 0.51 | 0.55 |
| 10 | 61580949 | rs10994030 | -1.446 | 1.044 | 2.215 | -0.231 | 0.004 | 0.117 | 0.117 | 0.009 | 0.12 | 0.4 | 0.15 | 0.25 | 0.13 | 0.25 |
| 10 | 61588347 | rs10458645 | -1.662 | 1.034 | 2.137 | -0.254 | 0.001 | 0.047 | 0.047 | 1.685 | 0.47 | 0.72 | 0.37 | 0.3 | 0.51 | 0.54 |
| 10 | 61595110 | rs1822913 | -0.595 | 1.06 | 2.062 | -0.268 | -0.025 | 0.04 | 0.04 | 0.049 | 0.34 | 0.31 | 0.22 | 0.05 | 0.38 | 0.3 |
| 10 | 61598118 | rs7090532 | -2.386 | 0.97 | 2.214 | -0.305 | 0.061 | -0.011 | -0.011 | 0.375 | 0.07 | 0.39 | 0.04 | 0.01 | 0.04 | 0.01 |
| 10 | 61600910 | rs7079588 | -1.47 | 0.948 | 2.237 | -0.272 | -0.001 | 0.163 | 0.163 | 0.188 | 0.12 | 0.47 | 0.16 | 0.25 | 0.15 | 0.26 |
| 10 | 61600952 | rs4948372 | -1.607 | 0.927 | 2.118 | -0.31 | -0.034 | 0.097 | 0.097 | 1.777 | 0.46 | 0.82 | 0.35 | 0.28 | 0.48 | 0.49 |
| 10 | 61612119 | rs2271562 | -0.795 | 0.949 | 2.077 | -0.245 | 0.004 | 0.172 | 0.172 | 0.083 | 0.09 | 0.35 | 0.13 | 0.25 | 0.11 | 0.25 |
| 10 | 61612159 | rs17791484 | 0.124 | 1.058 | 1.801 | -0.286 | -0.137 | 0.141 | 0.141 | 0.081 | 0.27 | 0.02 | 0.15 | 0.03 | 0.3 | 0.22 |
| 10 | 61614418 | rs1709342 | -1.856 | 0.914 | 1.972 | -0.194 | -0.012 | 0.117 | 0.117 | 0.006 | 0.14 | 0.47 | 0.15 | 0.26 | 0.13 | 0.25 |
| 10 | 61616606 | rs1664284 | -1.043 | 1.004 | 1.771 | -0.192 | -0.005 | 0.011 | 0.011 | 0.574 | 0.4 | 0.63 | 0.31 | 0.28 | 0.41 | 0.46 |
| 10 | 61624697 | rs1177701 | -1.523 | 0.803 | 1.565 | 0.059 | -0.008 | 0.029 | 0.029 | 1.028 | 0.47 | 0.64 | 0.37 | 0.36 | 0.48 | 0.48 |
| 10 | 61630250 | rs1684898 | -0.17 | 0.784 | 1.753 | -0.333 | -0.003 | 0.012 | 0.012 | 0 | 0.71 | 0.74 | 0.83 | 0.88 | 0.74 | 0.72 |
| 10 | 61631994 | rs16914297 | -1.263 | 0.744 | 1.964 | -0.321 | -0.026 | 0.012 | 0.012 | 0 | 0.15 | 0.32 | 0.14 | 0.15 | 0.13 | 0.18 |
| 10 | 61636147 | rs2440906 | -0.086 | 0.779 | 1.846 | -0.388 | -0.02 | 0.05 | 0.05 | 0.029 | 0.71 | 0.8 | 0.82 | 0.87 | 0.71 | 0.72 |
| 10 | 61640320 | rs1626088 | 1.32 | 0.771 | 1.943 | -0.133 | 0.002 | 0.031 | 0.031 | 0.905 | 0.54 | 0.33 | 0.58 | 0.67 | 0.54 | 0.53 |
| 10 | 61645590 | rs1184089 | 0.042 | 0.867 | 1.899 | -0.125 | -0.003 | 0.002 | 0.002 | 0.098 | 0.68 | 0.67 | 0.77 | 0.84 | 0.66 | 0.66 |
| 10 | 61651701 | rs1180657 | 0.562 | 0.843 | 1.931 | -0.129 | 0.02 | -0.054 | -0.054 | 0 | 0.63 | 0.52 | 0.73 | 0.84 | 0.6 | 0.65 |

| | | | | | | | | | | | | | | | | |
|---|---|---|---|---|---|---|---|---|---|---|---|---|---|---|---|---|
| 10 | 61661564 | rs7914022 | -0.986 | 0.768 | 2.165 | -0.069 | -0.019 | 0.054 | 0.054 | 0 | 0.14 | 0.32 | 0.21 | 0.22 | 0.13 | 0.18 |
| 10 | 61663377 | rs3793865 | -1.308 | 0.796 | 2.063 | -0.293 | 0.048 | 0.029 | 0.029 | 0 | 0.07 | 0.21 | 0.08 | 0.04 | 0.11 | 0.07 |
| 10 | 61663637 | rs1171826 | 0.293 | 0.74 | 1.914 | -0.353 | 0.014 | -0.039 | -0.039 | 0 | 0.66 | 0.56 | 0.74 | 0.84 | 0.62 | 0.65 |
| 10 | 61665886 | rs1171830 | 1.734 | 0.647 | 1.311 | -0.256 | -0.126 | -0.06 | -0.06 | 0 | 0.45 | 0.04 | 0.47 | 0.63 | 0.45 | 0.42 |
| 10 | 61693239 | rs1664276 | -0.106 | 0.622 | 1.505 | -0.349 | -0.019 | 0.074 | 0.074 | 0 | 0.7 | 0.83 | 0.81 | 0.98 | 0.71 | 0.77 |
| 10 | 61694304 | rs1664278 | 0.084 | 0.625 | 1.662 | -0.317 | 0 | -0.022 | -0.022 | 0 | 0.68 | 0.65 | 0.8 | 0.88 | 0.71 | 0.69 |
| 10 | 61696682 | rs3099371 | 1.098 | 0.569 | 1.807 | -0.3 | 0.007 | 0.028 | 0.028 | 0.452 | 0.54 | 0.34 | 0.6 | 0.69 | 0.57 | 0.5 |
| 10 | 61698103 | rs7073136 | -1.129 | 0.542 | 2.29 | -0.166 | -0.008 | 0.036 | 0.036 | 0 | 0.14 | 0.22 | 0.21 | 0.2 | 0.15 | 0.17 |
| 10 | 61702152 | rs6479673 | -1.796 | 0.54 | 2.24 | -0.159 | -0.014 | 0.042 | 0.042 | 0.241 | 0.18 | 0.41 | 0.22 | 0.2 | 0.17 | 0.21 |
| 10 | 61707047 | rs1171849 | 2.28 | 0.162 | 1.896 | -0.356 | -0.006 | 0.002 | 0.002 | 1.568 | 0.75 | 0.28 | 0.72 | 0.7 | 0.78 | 0.68 |
| 10 | 61707879 | rs16914411 | -0.404 | 0.323 | 1.673 | -0.27 | -0.004 | 0.01 | 0.01 | 0 | 0.11 | 0.11 | 0.17 | 0.2 | 0.12 | 0.19 |
| 10 | 61709195 | rs2166271 | -0.205 | 0.402 | 1.399 | -0.488 | -0.01 | 0.069 | 0.069 | 0 | 0.33 | 0.18 | 0.21 | 0.12 | 0.34 | 0.34 |
| 10 | 61711410 | rs1684927 | 1.041 | 0.122 | 1.584 | -0.346 | -0.008 | 0.036 | 0.036 | 0 | 0.85 | 0.71 | 0.79 | 0.79 | 0.84 | 0.8 |
| 10 | 61715836 | rs12784462 | -0.366 | 0.409 | 1.271 | -0.415 | -0.02 | 0.05 | 0.05 | 0 | 0.33 | 0.21 | 0.22 | 0.1 | 0.36 | 0.33 |
| 10 | 61716887 | rs10821617 | -0.796 | 0.366 | 1.254 | -0.439 | -0.001 | -0.002 | -0.002 | 0 | 0.09 | 0.14 | 0.05 | 0.06 | 0.09 | 0.08 |
| 10 | 61722273 | rs7078432 | -1.576 | 0.402 | 1.275 | -0.395 | -0.002 | -0.003 | -0.003 | 0.287 | 0.54 | 0.48 | 0.43 | 0.36 | 0.57 | 0.6 |
| 10 | 61726870 | rs1684895 | 0.389 | 0.461 | 1.112 | -0.68 | 0.012 | -0.054 | -0.054 | 0.149 | 0.59 | 0.44 | 0.7 | 0.84 | 0.57 | 0.61 |
| 10 | 61728196 | rs10994094 | -0.558 | 0.343 | 0.979 | -0.514 | -0.016 | 0.017 | 0.017 | 0 | 0.13 | 0.18 | 0.15 | 0.14 | 0.14 | 0.19 |
| 10 | 61730651 | rs10994097 | -0.822 | 0.237 | 0.772 | -0.709 | -0.016 | 0.048 | 0.048 | 0 | 0.42 | 0.28 | 0.36 | 0.24 | 0.46 | 0.5 |
| 10 | 61744247 | rs10821629 | -1.909 | -0.212 | 1.144 | -0.655 | -0.014 | -0.01 | -0.01 | 0 | 0.32 | 0.58 | 0.23 | 0.22 | 0.28 | 0.31 |
| 10 | 61745573 | rs12219264 | -0.54 | -0.278 | 1.384 | -0.67 | -0.045 | 0.025 | 0.025 | 0 | 0.16 | 0.36 | 0.16 | 0.15 | 0.13 | 0.19 |
| 10 | 61749766 | rs10821633 | -2.071 | -0.287 | 1.361 | -0.69 | -0.023 | -0.015 | -0.015 | 2.536 | 0.32 | 0.57 | 0.23 | 0.22 | 0.27 | 0.31 |
| 10 | 61750470 | rs10994118 | -0.538 | -0.268 | 1.421 | -0.673 | -0.044 | 0.023 | 0.023 | 0 | 0.16 | 0.34 | 0.16 | 0.15 | 0.13 | 0.2 |
| 10 | 61752055 | rs10821636 | -1.375 | -0.27 | 1.354 | -0.687 | -0.039 | 0 | 0 | 0 | 0.2 | 0.48 | 0.19 | 0.15 | 0.18 | 0.19 |
| 10 | 61754080 | rs10509116 | -1.43 | -0.218 | 1.295 | -0.687 | -0.002 | -0.014 | -0.014 | 0.401 | 0.16 | 0.21 | 0.07 | 0.07 | 0.14 | 0.1 |
| 10 | 61754849 | rs7080898 | -0.442 | -0.235 | 1.154 | -0.678 | 0 | 0.014 | 0.014 | 0 | 0.11 | 0.02 | 0.05 | 0.06 | 0.1 | 0.1 |
| 10 | 61759870 | rs10821644 | -1.387 | -0.256 | 1.351 | -0.687 | -0.017 | -0.039 | -0.039 | 0 | 0.14 | 0.39 | 0.07 | 0.06 | 0.11 | 0.1 |
| 10 | 61769337 | rs10821653 | -0.568 | 0.167 | 1.266 | -0.663 | -0.039 | 0.016 | 0.016 | 0 | 0.16 | 0.28 | 0.16 | 0.15 | 0.13 | 0.18 |
| 10 | 61780209 | rs10761438 | -2.466 | 0.038 | 1.158 | -0.813 | 0.009 | 0.007 | 0.007 | 1.409 | 0.38 | 0.7 | 0.31 | 0.31 | 0.38 | 0.35 |
| 10 | 61781721 | rs2279421 | -0.257 | 0.015 | 1.186 | -0.805 | -0.038 | 0.023 | 0.023 | 0 | 0.15 | 0.28 | 0.15 | 0.15 | 0.12 | 0.18 |
| 10 | 61781947 | rs2279420 | -0.515 | 0.032 | 0.978 | -0.798 | 0.001 | 0.022 | 0.022 | 0 | 0.12 | 0.04 | 0.06 | 0.06 | 0.11 | 0.1 |
| 10 | 61784928 | rs7068231 | -2.366 | 0.118 | 0.737 | -1.059 | 0.006 | 0.025 | 0.025 | 1.075 | 0.58 | 0.77 | 0.43 | 0.31 | 0.6 | 0.54 |
| 10 | 61786965 | rs11816586 | 0.54 | -0.379 | 0.642 | -1.091 | -0.055 | 0.095 | 0.095 | 0 | 0.2 | 0.07 | 0.12 | 0 | 0.21 | 0.19 |
| 10 | 61787249 | rs1050745 | -1.367 | -0.684 | 1.1 | -0.787 | -0.04 | -0.012 | -0.012 | 0 | 0.28 | 0.5 | 0.23 | 0.21 | 0.24 | 0.29 |
| 10 | 61790331 | rs4528269 | 0.245 | -1.006 | -0.211 | -1.186 | -0.022 | -0.008 | -0.008 | 2.08 | 0.69 | 0.41 | 0.54 | 0.28 | 0.82 | 0.66 |
| 10 | 61790383 | rs12357206 | 1.967 | -0.894 | -1.234 | -1.255 | -0.166 | 0.059 | 0.059 | 0.007 | 0.37 | 0.03 | 0.32 | 0.22 | 0.5 | 0.39 |
| 10 | 61791039 | rs7911953 | -1.246 | -1.065 | -0.597 | -0.789 | 0.016 | -0.018 | -0.018 | 0 | 0.85 | 0.88 | 0.63 | 0.3 | 0.92 | 0.74 |
| 10 | 61793424 | rs10821659 | 0.941 | -0.554 | -0.282 | -0.806 | 0.001 | -0.001 | -0.001 | 0 | 0.48 | 0.5 | 0.41 | 0.24 | 0.61 | 0.41 |
| 10 | 61793446 | rs16914571 | -0.306 | 0.112 | -0.337 | -0.459 | 0.001 | 0.007 | 0.007 | 0 | 0.13 | 0.13 | 0.06 | 0.04 | 0.08 | 0.12 |
| 10 | 61795908 | rs10761445 | -0.9 | 0.667 | -0.381 | 0.136 | 0 | 0 | 0 | 0 | 0.13 | 0.14 | 0.32 | 0.2 | 0.19 | 0.18 |
| 10 | 61800984 | rs2893823 | -0.916 | 0.624 | -0.335 | 0.175 | -0.005 | -0.005 | -0.005 | 0.6 | 0.13 | 0.1 | 0.28 | 0.19 | 0.19 | 0.2 |
| 10 | 61801020 | rs2393576 | 1.445 | 0.581 | -1.566 | 0.044 | -0.122 | -0.023 | -0.023 | 0 | 0.35 | 0.05 | 0.29 | 0.23 | 0.45 | 0.41 |
| 10 | 61801121 | rs2393578 | 0.2 | 0.06 | -1.292 | -0.472 | -0.01 | -0.036 | -0.036 | 0.88 | 0.78 | 0.75 | 0.87 | 0.94 | 0.86 | 0.81 |
| 10 | 61801142 | rs2393579 | 0.323 | 0.061 | -1.291 | -0.442 | -0.01 | -0.031 | -0.031 | 0 | 0.78 | 0.75 | 0.87 | 0.94 | 0.86 | 0.8 |
| 10 | 61801282 | rs2393581 | 0.63 | 0.017 | -1.314 | -0.484 | -0.005 | 0.004 | 0.004 | 0 | 0.75 | 0.8 | 0.68 | 0.79 | 0.72 | 0.68 |
| 10 | 61801402 | rs2393582 | 0.641 | 0.015 | -1.315 | -0.485 | -0.005 | 0.004 | 0.004 | 0 | 0.75 | 0.8 | 0.68 | 0.79 | 0.72 | 0.68 |
| 10 | 61801523 | rs4431977 | 0.967 | 0.055 | -1.257 | -0.46 | 0.001 | 0.009 | 0.009 | 0.016 | 0.67 | 0.7 | 0.6 | 0.78 | 0.67 | 0.61 |
| 10 | 61801614 | rs10740004 | 0.982 | 0.066 | -1.265 | -0.456 | 0.001 | 0.014 | 0.014 | 0.021 | 0.67 | 0.71 | 0.6 | 0.78 | 0.67 | 0.62 |
| 10 | 61801739 | rs10821662 | 0.369 | 0.035 | -1.289 | -0.471 | -0.001 | -0.012 | -0.012 | 0 | 0.78 | 0.71 | 0.64 | 0.83 | 0.77 | 0.74 |
| 10 | 61802242 | rs2393602 | 0.964 | -0.13 | -1.235 | -0.641 | 0.001 | -0.021 | -0.021 | 0 | 0.68 | 0.63 | 0.57 | 0.81 | 0.64 | 0.69 |
| 10 | 61802342 | rs2393603 | 0.636 | -0.158 | -1.262 | -0.666 | -0.008 | -0.028 | -0.028 | 0.405 | 0.73 | 0.64 | 0.62 | 0.81 | 0.76 | 0.71 |
| 10 | 61802726 | rs9888034 | 1.589 | -0.159 | -1.264 | -0.666 | -0.002 | 0.02 | 0.02 | 0.144 | 0.57 | 0.61 | 0.53 | 0.76 | 0.55 | 0.56 |
| 10 | 61803062 | rs1002442 | -0.033 | -0.141 | -1.336 | -0.657 | -0.056 | 0.048 | 0.048 | 0 | 0.17 | 0.02 | 0.09 | 0.06 | 0.21 | 0.15 |
| 10 | 61804282 | rs7900212 | 1.591 | -0.379 | -0.63 | -0.628 | 0 | 0.133 | 0.133 | 0.126 | 0.54 | 0.24 | 0.5 | 0.76 | 0.55 | 0.56 |
| 10 | 61808738 | rs16914606 | -0.398 | -0.254 | -0.071 | -0.066 | 0.004 | 0.011 | 0.011 | 0 | 0.15 | 0.11 | 0.05 | 0.08 | 0.1 | 0.16 |
| 10 | 61813046 | rs9665327 | -0.063 | -0.333 | -0.08 | 0.029 | -0.041 | 0.097 | 0.097 | 3.289 | 0.23 | 0.57 | 0.24 | 0.57 | 0.11 | 0.16 |
| 10 | 61813685 | rs4948383 | -0.788 | -0.187 | -0.154 | -0.116 | 0.029 | -0.008 | -0.008 | 0 | 0.37 | 0.7 | 0.29 | 0.64 | 0.21 | 0.33 |
| 10 | 61815048 | rs10994173 | -0.934 | 0.153 | 0.366 | -0.291 | 0.04 | 0.018 | 0.018 | 0 | 0.07 | 0.41 | 0.04 | 0.01 | 0.01 | 0.03 |
| 10 | 61815367 | rs4948384 | -1.395 | 0.854 | -0.553 | -0.412 | -0.018 | 0.043 | 0.043 | 0.02 | 0.18 | 0.11 | 0.11 | 0.05 | 0.19 | 0.13 |
| 10 | 61816736 | rs4372392 | -0.174 | 0.948 | -0.865 | -0.594 | -0.052 | 0.093 | 0.093 | 0 | 0.17 | 0.02 | 0.19 | 0.04 | 0.23 | 0.14 |
| 10 | 61820631 | rs7893313 | 0.177 | 1.534 | -0.569 | -0.42 | 0.012 | 0.007 | 0.007 | 0 | 0.6 | 0.56 | 0.71 | 0.41 | 0.63 | 0.67 |
| 10 | 61823109 | rs4568956 | 0.721 | 1.734 | -0.217 | -0.519 | -0.005 | 0.011 | 0.011 | 0 | 0.71 | 0.73 | 0.74 | 0.84 | 0.65 | 0.78 |
| 10 | 61826411 | rs7902905 | 0.72 | 1.785 | -0.316 | -0.781 | -0.002 | -0.035 | -0.035 | 0 | 0.71 | 0.71 | 0.74 | 0.95 | 0.64 | 0.83 |
| 10 | 61831271 | rs10821668 | -0.135 | 1.943 | -0.88 | -0.797 | -0.09 | 0.132 | 0.132 | 0 | 0.22 | 0.02 | 0.21 | 0.03 | 0.27 | 0.15 |
| 10 | 61839831 | rs9804190 | -1.268 | 1.292 | 0.137 | -0.758 | 0.013 | -0.126 | -0.126 | 2.728 | 0.23 | 0.56 | 0.16 | 0.06 | 0.23 | 0.15 |
| 10 | 61839971 | rs2241538 | -0.124 | 0.981 | -0.202 | -0.64 | -0.006 | 0 | 0 | 0 | 0.3 | 0.32 | 0.35 | 0.68 | 0.25 | 0.39 |
| 10 | 61840167 | rs7917800 | -0.043 | 0.676 | 0.055 | -0.656 | 0.042 | -0.045 | -0.045 | 0 | 0.08 | 0.24 | 0.07 | 0 | 0.11 | 0.01 |
| 10 | 61843567 | rs2393596 | -0.385 | 0.366 | 0.285 | -0.366 | -0.031 | 0.007 | 0.007 | 0 | 0.29 | 0.37 | 0.34 | 0.72 | 0.2 | 0.35 |
| 10 | 61847040 | rs11596260 | 0.285 | 0.343 | -0.043 | -0.76 | 0.001 | 0.087 | 0.087 | 2.074 | 0.37 | 0.24 | 0.29 | 0.08 | 0.37 | 0.3 |

| | | | | | | | | | | | | | | | | |
|---|---|---|---|---|---|---|---|---|---|---|---|---|---|---|---|---|
| 10 | 61847439 | rs10761450 | 0.361 | 0.193 | -0.014 | -0.758 | 0.033 | 0.046 | 0.046 | 0.496 | 0.64 | 0.23 | 0.59 | 0.23 | 0.74 | 0.57 |
| 10 | 61853056 | rs3793854 | 1.113 | 0.385 | 0.28 | -0.416 | 0.014 | -0.006 | -0.006 | 0 | 0.31 | 0.44 | 0.34 | 0.22 | 0.37 | 0.33 |
| 10 | 61853486 | rs10821673 | 1.453 | 0.381 | 0.287 | -0.445 | -0.018 | 0.018 | 0.018 | 0 | 0.29 | 0.23 | 0.32 | 0.16 | 0.37 | 0.27 |
| 10 | 61854292 | rs3793856 | 0.106 | -0.016 | 0.399 | -0.352 | -0.04 | 0.057 | 0.057 | 1.166 | 0.23 | 0.35 | 0.3 | 0.64 | 0.14 | 0.24 |
| 10 | 61855027 | rs10761453 | -0.611 | 0.076 | 0.166 | -0.446 | -0.043 | 0.197 | 0.197 | 1.585 | 0.64 | 0.92 | 0.73 | 0.93 | 0.63 | 0.64 |
| 10 | 61856035 | rs10994198 | 0.064 | 0.08 | 0.212 | -0.442 | 0.008 | 0.086 | 0.086 | 2.302 | 0.55 | 0.71 | 0.67 | 0.79 | 0.56 | 0.51 |
| 10 | 61856861 | rs2393599 | 0.707 | 0.154 | 0.241 | -0.37 | -0.005 | 0.033 | 0.033 | 0 | 0.08 | 0.19 | 0.05 | 0.13 | 0.07 | 0.14 |
| 10 | 61861872 | rs3793859 | 0.463 | 0.172 | -0.154 | -0.454 | -0.061 | 0.211 | 0.211 | 2.602 | 0.35 | 0.05 | 0.23 | 0.07 | 0.37 | 0.36 |
| 10 | 61863033 | rs10994199 | -0.258 | -0.084 | 0.392 | -0.188 | 0.02 | 0.137 | 0.137 | 0 | 0.11 | 0.43 | 0.12 | 0.2 | 0.08 | 0.21 |
| 10 | 61868716 | rs3750800 | 0.457 | 0.085 | -0.151 | -0.37 | -0.076 | 0.175 | 0.175 | 3.109 | 0.33 | 0.03 | 0.23 | 0.07 | 0.32 | 0.32 |
| 10 | 61870135 | rs10761454 | -0.267 | 0.078 | 0.243 | -0.386 | -0.011 | 0.102 | 0.102 | 1.744 | 0.62 | 0.79 | 0.68 | 0.92 | 0.62 | 0.67 |
| 10 | 61870669 | rs12049756 | 0.247 | -0.175 | 0.402 | -0.123 | -0.017 | 0.028 | 0.028 | 0.55 | 0.24 | 0.23 | 0.28 | 0.5 | 0.13 | 0.2 |
| 10 | 61871598 | rs4369346 | 1.31 | -0.132 | 0.41 | -0.159 | -0.005 | -0.001 | -0.001 | 0 | 0.32 | 0.31 | 0.34 | 0.29 | 0.4 | 0.31 |
| 10 | 61876720 | rs12779035 | 1.022 | -0.142 | 0.256 | -0.297 | -0.025 | 0.142 | 0.142 | 3.038 | 0.28 | 0.04 | 0.19 | 0.07 | 0.27 | 0.28 |
| 10 | 61877706 | rs10994209 | 1.421 | -0.214 | 0.589 | -0.006 | -0.017 | 0.004 | 0.004 | 0 | 0.29 | 0.24 | 0.34 | 0.22 | 0.4 | 0.26 |
| 10 | 61878214 | rs10994210 | -0.731 | -0.211 | 0.622 | -0.156 | -0.017 | 0.129 | 0.129 | 1.935 | 0.63 | 0.83 | 0.73 | 0.93 | 0.62 | 0.68 |
| 10 | 61881937 | rs2393612 | 1.253 | -0.187 | 0.553 | -0.12 | -0.026 | -0.011 | -0.011 | 0.411 | 0.3 | 0.24 | 0.33 | 0.28 | 0.4 | 0.32 |
| 10 | 61882020 | rs12220169 | 0.183 | -0.132 | 0.623 | 0.095 | 0.008 | -0.012 | -0.012 | 0 | 0.22 | 0.14 | 0.27 | 0.52 | 0.13 | 0.2 |
| 10 | 61882414 | rs10994212 | -0.237 | -0.054 | 0.691 | -0.076 | -0.023 | 0.022 | 0.022 | 0 | 0.31 | 0.33 | 0.32 | 0.65 | 0.22 | 0.35 |
| 10 | 61882484 | rs10761459 | -0.36 | -0.048 | 0.521 | -0.074 | -0.023 | 0.018 | 0.018 | 0.001 | 0.31 | 0.54 | 0.35 | 0.65 | 0.22 | 0.35 |
| 10 | 61883364 | rs10761461 | -0.198 | 0.008 | -0.44 | -0.022 | 0.007 | -0.007 | -0.007 | 0 | 0.31 | 0.27 | 0.3 | 0.65 | 0.2 | 0.35 |
| 10 | 61886610 | rs10994214 | -0.049 | 0.227 | -0.725 | -0.475 | -0.026 | 0.112 | 0.112 | 1.689 | 0.37 | 0.18 | 0.29 | 0.07 | 0.4 | 0.33 |
| 10 | 61890937 | rs10740011 | 0.407 | -0.203 | -0.593 | -0.061 | 0.006 | 0.019 | 0.019 | 0.056 | 0.46 | 0.34 | 0.57 | 0.69 | 0.55 | 0.44 |
| 10 | 61891032 | rs7070364 | -0.037 | -0.251 | -0.27 | -0.281 | -0.008 | -0.057 | -0.057 | 0 | 0.36 | 0.55 | 0.32 | 0.14 | 0.34 | 0.35 |
| 10 | 61893145 | rs10509123 | 0.192 | 0.375 | -0.68 | -0.241 | -0.046 | 0.04 | 0.04 | 0 | 0.26 | 0.16 | 0.26 | 0.14 | 0.35 | 0.27 |
| 10 | 61893898 | rs3213829 | -0.349 | -0.08 | -0.348 | -0.274 | 0.012 | 0.046 | 0.046 | 0.106 | 0.47 | 0.69 | 0.42 | 0.25 | 0.52 | 0.37 |
| 10 | 61895547 | rs16914791 | -0.233 | -0.526 | -0.427 | 0.666 | 0.06 | -0.056 | -0.056 | 6.009 | 0.13 | 0.02 | 0.14 | 0.21 | 0.05 | 0.14 |
| 10 | 61902023 | rs2288359 | 0.774 | -0.616 | 0.269 | 0.537 | -0.076 | 0.077 | 0.077 | 0.022 | 0.83 | 0.66 | 0.83 | 0.71 | 0.94 | 0.83 |
| 10 | 61907907 | rs12357972 | -0.11 | -0.627 | -0.038 | 0.583 | 0.037 | -0.035 | -0.035 | 0 | 0.13 | 0.05 | 0.15 | 0.21 | 0.05 | 0.14 |
| 10 | 61915955 | rs2393614 | -1.031 | -0.141 | -0.541 | 0.254 | -0.055 | 0.046 | 0.046 | 0.148 | 0.79 | 0.91 | 0.76 | 0.87 | 0.68 | 0.8 |
| 10 | 61926866 | rs10509125 | 0.508 | -0.178 | -0.28 | 0.307 | 0.006 | 0.003 | 0.003 | 0 | 0.61 | 0.47 | 0.57 | 0.57 | 0.61 | 0.61 |
| 10 | 61938252 | rs4948255 | 0.953 | -0.197 | -1.244 | 0.248 | -0.019 | -0.149 | -0.149 | 0 | 0.36 | 0.04 | 0.24 | 0.46 | 0.37 | 0.32 |
| 10 | 61939519 | rs10821695 | -0.617 | 0.233 | -0.87 | 0.189 | -0.005 | 0 | 0 | 0.292 | 0.27 | 0.35 | 0.31 | 0.24 | 0.24 | 0.25 |
| 10 | 61940899 | rs2288358 | 1.03 | 0.066 | -1.438 | 0.356 | -0.039 | -0.154 | -0.154 | 0 | 0.35 | 0.03 | 0.24 | 0.46 | 0.37 | 0.31 |
| 10 | 61941250 | rs10994246 | -0.196 | -0.116 | -0.667 | 0.67 | 0.004 | -0.007 | -0.007 | 0 | 0.12 | 0.12 | 0.15 | 0.2 | 0.04 | 0.14 |
| 10 | 61941275 | rs16914582 | 0.627 | 0.44 | -0.191 | 0.69 | -0.005 | 0.009 | 0.009 | 0 | 0.07 | 0.09 | 0.14 | 0.11 | 0.05 | 0.12 |
| 10 | 61944112 | rs16914585 | 0.151 | 0.42 | -0.039 | 0.688 | 0.001 | 0.033 | 0.033 | 0 | 0.08 | 0.22 | 0.15 | 0.11 | 0.06 | 0.13 |
| 10 | 61945643 | rs16914587 | 0.151 | 0.42 | 0.007 | 0.688 | 0.003 | 0.038 | 0.038 | 0 | 0.08 | 0.23 | 0.16 | 0.11 | 0.06 | 0.13 |
| 10 | 61945931 | rs7087489 | -0.776 | 1.035 | -0.22 | 0.269 | -0.019 | 0.019 | 0.019 | 1.2 | 0.78 | 0.82 | 0.75 | 0.87 | 0.71 | 0.8 |
| 10 | 61954476 | rs4310561 | -0.029 | 0.781 | -0.373 | 0.419 | -0.026 | -0.006 | -0.006 | 0 | 0.7 | 0.81 | 0.63 | 0.66 | 0.68 | 0.68 |
| 10 | 61957168 | rs10821699 | 1.165 | 0.754 | -0.746 | 0.51 | -0.066 | -0.082 | -0.082 | 0.135 | 0.44 | 0.09 | 0.28 | 0.43 | 0.45 | 0.38 |
| 10 | 61960135 | rs16914619 | -0.019 | 0.556 | -0.534 | 0.313 | 0.024 | -0.043 | -0.043 | 1.256 | 0.14 | 0.08 | 0.25 | 0.3 | 0.09 | 0.26 |
| 10 | 61961795 | rs10994250 | 0.382 | 0.679 | 0.082 | 0.246 | 0.017 | -0.02 | -0.02 | 0 | 0.13 | 0.18 | 0.14 | 0.07 | 0.19 | 0.18 |
| 10 | 61964752 | rs10994253 | 0.132 | 0.72 | -0.315 | 0.285 | 0.02 | -0.071 | -0.071 | 0 | 0.14 | 0.03 | 0.25 | 0.3 | 0.09 | 0.26 |
| 10 | 61965188 | rs10732408 | -0.808 | 0.74 | 0.124 | 0.069 | 0.01 | -0.076 | -0.076 | 0 | 0.76 | 0.63 | 0.71 | 0.87 | 0.75 | 0.84 |
| 10 | 61966755 | rs4948398 | 0.283 | 0.662 | 0.092 | 0.083 | -0.03 | 0.129 | 0.129 | 1.352 | 0.6 | 0.25 | 0.55 | 0.8 | 0.56 | 0.65 |
| 10 | 61981247 | rs1340655 | 0.141 | 0.676 | 0.085 | 0.088 | 0.008 | -0.038 | -0.038 | 0 | 0.63 | 0.51 | 0.58 | 0.8 | 0.57 | 0.66 |
| 10 | 61984604 | rs10994259 | 0.266 | 0.772 | -0.799 | 0.116 | -0.041 | 0.15 | 0.15 | 1.372 | 0.59 | 0.22 | 0.54 | 0.8 | 0.57 | 0.65 |
| 10 | 61988660 | rs10821704 | 1.057 | 0.839 | -0.905 | 0.243 | -0.08 | -0.032 | -0.032 | 0.008 | 0.45 | 0.13 | 0.3 | 0.5 | 0.48 | 0.39 |
| 10 | 61998060 | rs1340654 | -0.457 | 1.025 | -0.325 | 0.284 | -0.001 | -0.004 | -0.004 | 0.886 | 0.78 | 0.79 | 0.73 | 0.79 | 0.75 | 0.84 |
| 10 | 62008708 | rs6479706 | 0.155 | 0.929 | -0.23 | 0.126 | 0 | -0.049 | -0.049 | 0 | 0.71 | 0.63 | 0.7 | 0.87 | 0.73 | 0.76 |
| 10 | 62009518 | rs16914651 | 0.748 | 0.48 | -1.116 | 0.578 | 0.027 | -0.063 | -0.063 | 0 | 0.25 | 0.15 | 0.16 | 0.34 | 0.19 | 0.19 |
| 10 | 62010523 | rs12767186 | -1.047 | 1.524 | -0.288 | 0.965 | 0.033 | 0.012 | 0.012 | 1.044 | 0.82 | 0.73 | 0.69 | 0.76 | 0.74 | 0.89 |
| 10 | 62014629 | rs16914660 | 0.89 | 1.653 | -0.665 | 0.977 | 0.021 | -0.032 | -0.032 | 0 | 0.35 | 0.14 | 0.18 | 0.41 | 0.28 | 0.29 |
| 10 | 62015782 | rs2018783 | 0.209 | 1.867 | -0.195 | 1.045 | -0.008 | -0.001 | -0.001 | 0 | 0.53 | 0.28 | 0.33 | 0.52 | 0.54 | 0.61 |
| 10 | 62023034 | rs16914677 | 0.781 | 1.805 | 0.097 | 0.948 | -0.003 | -0.024 | -0.024 | 0 | 0.11 | 0.01 | 0.06 | 0.08 | 0.11 | 0.15 |
| 10 | 62028797 | rs1459731 | -1.445 | 1.689 | 0.132 | 0.977 | 0.001 | 0.008 | 0.008 | 2.008 | 0.78 | 0.74 | 0.71 | 0.72 | 0.79 | 0.92 |
| 10 | 62034355 | rs1380453 | 0.708 | 1.619 | 0.188 | 0.892 | 0.013 | 0.011 | 0.011 | 0 | 0.5 | 0.41 | 0.52 | 0.62 | 0.46 | 0.62 |
| 10 | 62041469 | rs922116 | -0.649 | 1.522 | 0.094 | 0.828 | -0.01 | 0.009 | 0.009 | 0.015 | 0.26 | 0.24 | 0.17 | 0.13 | 0.28 | 0.28 |
| 10 | 62042160 | rs12780624 | -0.591 | 1.526 | 0.096 | 0.539 | -0.009 | 0.009 | 0.009 | 0.069 | 0.25 | 0.24 | 0.17 | 0.08 | 0.28 | 0.27 |
| 10 | 62045698 | rs10994293 | 0.369 | 1.244 | 0.196 | 0.671 | -0.031 | 0.037 | 0.037 | 0.122 | 0.09 | 0.19 | 0.25 | 0.18 | 0.05 | 0.16 |
| 10 | 62048219 | rs7095717 | 0.379 | 1.558 | 0.086 | 0.59 | -0.008 | 0.006 | 0.006 | 0 | 0.13 | 0.14 | 0.09 | 0.03 | 0.16 | 0.13 |
| 10 | 62055781 | rs10821713 | 1.231 | 1.652 | 0.073 | 0.709 | 0.012 | -0.044 | -0.044 | 6.358 | 0.42 | 0.19 | 0.26 | 0.44 | 0.4 | 0.48 |
| 10 | 62056468 | rs9633531 | -0.307 | 1.967 | 0.415 | 0.625 | 0.019 | -0.009 | -0.009 | 0 | 0.66 | 0.58 | 0.64 | 0.7 | 0.62 | 0.81 |
| 10 | 62063473 | rs12771158 | 0.548 | 2.02 | 0.582 | 0.66 | 0.015 | -0.029 | -0.029 | 0.05 | 0.11 | 0.15 | 0.1 | 0.01 | 0.14 | 0.11 |
| 10 | 62067448 | rs997238 | -0.392 | 1.957 | 0.578 | 0.665 | 0.035 | -0.011 | -0.011 | 2.639 | 0.67 | 0.52 | 0.64 | 0.7 | 0.62 | 0.82 |
| 10 | 62072030 | rs10821717 | -0.106 | 1.945 | 0.624 | 0.654 | 0.014 | -0.008 | -0.008 | 0 | 0.66 | 0.42 | 0.63 | 0.7 | 0.61 | 0.8 |
| 10 | 62078203 | rs1340652 | -1.43 | 1.365 | 0.655 | 0.705 | 0.01 | 0.013 | 0.013 | 1.579 | 0.78 | 0.7 | 0.74 | 0.71 | 0.76 | 0.92 |
| 10 | 62091912 | rs12217173 | 1.224 | 1.349 | 0.281 | 0.716 | -0.021 | -0.032 | -0.032 | 0.842 | 0.43 | 0.19 | 0.28 | 0.43 | 0.44 | 0.48 |

| | | | | | | | | | | | | | | | | |
|---|---|---|---|---|---|---|---|---|---|---|---|---|---|---|---|---|
| 10 | 62093251 | rs11816516 | -0.484 | 1.413 | 0.293 | 0.632 | -0.035 | 0.089 | 0.089 | 0.833 | 0.23 | 0.1 | 0.14 | 0.05 | 0.25 | 0.27 |
| 10 | 62094202 | rs991405 | -1.584 | 1.419 | 0.565 | 0.69 | 0.015 | 0.026 | 0.026 | 1.703 | 0.78 | 0.66 | 0.74 | 0.72 | 0.77 | 0.93 |
| 10 | 62094752 | rs2199210 | -1.131 | 1.419 | 0.404 | 0.691 | 0.004 | 0.012 | 0.012 | 1.936 | 0.75 | 0.47 | 0.71 | 0.72 | 0.76 | 0.91 |
| 10 | 62102842 | rs2061487 | -1.274 | 1.401 | 0.364 | 0.663 | 0.002 | -0.002 | -0.002 | 0 | 0.76 | 0.43 | 0.71 | 0.74 | 0.77 | 0.93 |
| 10 | 62104776 | rs4582919 | 0.667 | 1.414 | 0.351 | 0.669 | -0.001 | 0.043 | 0.043 | 0 | 0.51 | 0.36 | 0.56 | 0.66 | 0.51 | 0.65 |
| 10 | 62105053 | rs10821736 | 0.4 | 1.314 | 0.35 | 0.744 | -0.009 | 0.078 | 0.078 | 0 | 0.08 | 0.16 | 0.28 | 0.23 | 0.07 | 0.16 |
| 10 | 62136206 | rs10821745 | 0.783 | 1.332 | 0.269 | 0.815 | -0.005 | 0.048 | 0.048 | 0 | 0.07 | 0.08 | 0.27 | 0.25 | 0.07 | 0.15 |
| 10 | 62137665 | rs4147263 | 1.33 | 1.372 | 0.253 | 0.938 | -0.009 | 0.008 | 0.008 | 0 | 0.42 | 0.24 | 0.27 | 0.37 | 0.43 | 0.52 |
| 10 | 62143348 | rs7895653 | -0.458 | 1.38 | 0.38 | 0.91 | -0.003 | 0.032 | 0.032 | 0.149 | 0.2 | 0.27 | 0.36 | 0.33 | 0.22 | 0.3 |
| 10 | 62143794 | rs7896287 | 0.712 | 1.405 | 0.362 | 0.862 | -0.007 | 0.008 | 0.008 | 0 | 0.11 | 0.08 | 0.08 | 0.01 | 0.13 | 0.14 |
| 10 | 62149543 | rs16914968 | 0.722 | 1.403 | 0.355 | 0.886 | -0.002 | 0.001 | 0.001 | 0.62 | 0.13 | 0.11 | 0.09 | 0.08 | 0.14 | 0.14 |
| 10 | 62150848 | rs3808942 | -1.118 | 1.353 | 0.441 | 0.917 | -0.007 | 0.004 | 0.004 | 2.654 | 0.74 | 0.61 | 0.72 | 0.72 | 0.76 | 0.91 |
| 10 | 62161618 | rs4948417 | 0.78 | 1.38 | 0.354 | 0.948 | -0.006 | 0.052 | 0.052 | 0 | 0.07 | 0.09 | 0.27 | 0.25 | 0.07 | 0.15 |
| 10 | 62170256 | rs7099869 | 0.342 | 1.413 | 0.276 | 0.831 | -0.002 | -0.006 | -0.006 | 0 | 0.12 | 0.13 | 0.09 | 0.01 | 0.12 | 0.13 |
| 10 | 62171796 | rs10761495 | -0.004 | 1.409 | 0.348 | 0.929 | -0.001 | 0.003 | 0.003 | 0.202 | 0.58 | 0.56 | 0.38 | 0.46 | 0.58 | 0.64 |
| 10 | 62176933 | rs10821759 | 0.449 | 1.471 | 0.002 | 0.934 | 0.003 | 0.006 | 0.006 | 0.63 | 0.13 | 0.18 | 0.13 | 0.17 | 0.17 | 0.14 |
| 10 | 62177839 | rs10761497 | 0.076 | 1.631 | 0.3 | 0.921 | -0.005 | 0.007 | 0.007 | 1.484 | 0.57 | 0.69 | 0.4 | 0.58 | 0.56 | 0.61 |
| 10 | 62179812 | rs10994336 | 1.012 | 1.262 | 0.207 | 1.06 | -0.015 | -0.033 | -0.033 | 0 | 0.06 | 0.01 | 0.27 | 0.19 | 0.07 | 0.15 |
| 10 | 62184600 | rs12257109 | -0.097 | 1.228 | 0.356 | 1.039 | 0.011 | 0.026 | 0.026 | 0.737 | 0.16 | 0.37 | 0.14 | 0.17 | 0.17 | 0.15 |
| 10 | 62185494 | rs4948418 | 0.91 | 1.291 | 0.119 | 0.992 | -0.022 | -0.033 | -0.033 | 0 | 0.06 | 0.01 | 0.27 | 0.19 | 0.08 | 0.15 |
| 10 | 62200882 | rs4948420 | 2.059 | 1.06 | -0.025 | 0.562 | 0.017 | 0.026 | 0.026 | 6.463 | 0.15 | 0.08 | 0.16 | 0 | 0.1 | 0.02 |
| 10 | 62215071 | rs10994357 | -1.115 | 1.13 | 0.378 | 0.904 | -0.003 | 0.022 | 0.022 | 0.735 | 0.31 | 0.41 | 0.43 | 0.41 | 0.32 | 0.45 |
| 10 | 62220244 | rs1934750 | 0.97 | 1.296 | 0.49 | 0.949 | -0.01 | 0.009 | 0.009 | 0 | 0.44 | 0.28 | 0.29 | 0.39 | 0.44 | 0.47 |
| 10 | 62223362 | rs4948422 | -1.062 | 1.12 | 0.557 | 0.991 | -0.001 | 0.001 | 0.001 | 0.998 | 0.32 | 0.34 | 0.44 | 0.5 | 0.31 | 0.47 |
| 10 | 62235173 | rs17810349 | 0.284 | 1.147 | 0.524 | 0.954 | 0 | 0.011 | 0.011 | 0 | 0.14 | 0.08 | 0.08 | 0.05 | 0.1 | 0.17 |
| 10 | 62235443 | rs7090243 | -0.104 | 1.179 | 0.586 | 0.943 | -0.001 | -0.003 | -0.003 | 0 | 0.58 | 0.49 | 0.36 | 0.35 | 0.54 | 0.58 |
| 10 | 62243232 | rs10761507 | -1.679 | 1.175 | 0.574 | 0.932 | 0.002 | -0.015 | -0.015 | 0 | 0.76 | 0.76 | 0.72 | 0.8 | 0.76 | 0.88 |
| 10 | 62243554 | rs17816232 | 0.233 | 1.044 | 0.504 | 0.941 | 0.001 | 0.016 | 0.016 | 0 | 0.14 | 0.08 | 0.08 | 0.05 | 0.1 | 0.17 |
| 10 | 62244379 | rs7898622 | -0.054 | 1.146 | 0.532 | 0.927 | 0 | 0.006 | 0.006 | 0 | 0.56 | 0.37 | 0.34 | 0.35 | 0.52 | 0.58 |
| 10 | 62246634 | rs10761508 | 1.159 | 1.173 | 0.459 | 0.999 | -0.01 | 0.004 | 0.004 | 0 | 0.41 | 0.26 | 0.25 | 0.33 | 0.42 | 0.42 |
| 10 | 62255455 | rs7077572 | 0.186 | 1.191 | 0.411 | 0.836 | -0.004 | -0.022 | -0.022 | 0.118 | 0.15 | 0.22 | 0.09 | 0.05 | 0.1 | 0.17 |
| 10 | 62259224 | rs2393638 | -0.338 | 1.185 | 0.328 | 0.875 | 0 | -0.035 | -0.035 | 0 | 0.59 | 0.51 | 0.61 | 0.76 | 0.63 | 0.71 |
| 10 | 62260544 | rs10994386 | 0.685 | 1.144 | 0.16 | 1.122 | -0.001 | 0.007 | 0.007 | 1.497 | 0.11 | 0.13 | 0.07 | 0.17 | 0.12 | 0.12 |
| 10 | 62276317 | rs10761514 | -0.782 | 1.077 | 0.263 | 1.151 | -0.005 | 0.038 | 0.038 | 0.075 | 0.19 | 0.23 | 0.35 | 0.45 | 0.2 | 0.29 |
| 10 | 62277679 | rs7098650 | -1.574 | 1.161 | 0.355 | 1.067 | 0.007 | -0.018 | -0.018 | 0 | 0.75 | 0.73 | 0.69 | 0.8 | 0.73 | 0.88 |
| 10 | 62278812 | rs2153522 | -0.654 | 1.202 | -0.01 | 1.091 | 0.001 | -0.007 | -0.007 | 0 | 0.21 | 0.19 | 0.35 | 0.32 | 0.19 | 0.35 |
| 10 | 62278842 | rs12146207 | 1.157 | 1.17 | -0.095 | 1.068 | -0.006 | 0.015 | 0.015 | 0 | 0.4 | 0.34 | 0.26 | 0.31 | 0.42 | 0.41 |
| 10 | 62279124 | rs10994397 | 0.647 | 1.318 | -0.114 | 0.763 | -0.001 | 0.042 | 0.042 | 0.137 | 0.07 | 0.08 | 0.27 | 0.27 | 0.07 | 0.17 |
| 10 | 62281234 | rs10994398 | 0.829 | 1.685 | -0.516 | 0.44 | -0.021 | 0.026 | 0.026 | 1.495 | 0.1 | 0.06 | 0.07 | 0.01 | 0.16 | 0.19 |
| 10 | 62286187 | rs7906431 | 0.396 | 1.275 | -0.234 | 0.65 | 0.001 | 0.004 | 0.004 | 0.758 | 0.51 | 0.44 | 0.36 | 0.48 | 0.54 | 0.53 |
| 10 | 62294097 | rs17817819 | 0.404 | 1.048 | -0.31 | 0.616 | -0.005 | 0.037 | 0.037 | 0 | 0.13 | 0 | 0.06 | 0.05 | 0.09 | 0.17 |
| 10 | 62298616 | rs10821792 | 1.026 | 1.1 | -0.27 | 0.697 | -0.021 | -0.03 | -0.03 | 0.078 | 0.07 | 0.01 | 0.27 | 0.27 | 0.08 | 0.18 |
| 10 | 62305640 | rs7072878 | -0.24 | 1.059 | -0.167 | 0.527 | 0 | -0.003 | -0.003 | 0 | 0.57 | 0.44 | 0.63 | 0.75 | 0.63 | 0.71 |
| 10 | 62313241 | rs7085315 | -0.16 | 0.943 | -0.549 | 0.582 | 0.012 | 0.034 | 0.034 | 0 | 0.56 | 0.39 | 0.65 | 0.73 | 0.62 | 0.68 |
| 10 | 62319434 | rs2393646 | -1.449 | 0.94 | -0.545 | 0.53 | -0.003 | -0.033 | -0.033 | 0 | 0.71 | 0.48 | 0.72 | 0.8 | 0.73 | 0.88 |
| 10 | 62320330 | rs2068043 | -0.809 | 0.965 | -0.606 | 0.729 | 0.007 | -0.071 | -0.071 | 0.763 | 0.22 | 0.1 | 0.35 | 0.36 | 0.17 | 0.38 |
| 10 | 62327427 | rs1596194 | 0.377 | 0.966 | -0.618 | 0.725 | 0.01 | -0.002 | -0.002 | 0 | 0.49 | 0.37 | 0.37 | 0.44 | 0.55 | 0.49 |
| 10 | 62330759 | rs17818833 | 0.226 | 1.03 | -0.873 | 0.763 | -0.006 | 0.026 | 0.026 | 0 | 0.14 | 0 | 0.07 | 0.07 | 0.1 | 0.19 |
| 10 | 62334468 | rs1442550 | -0.695 | 1.061 | -0.844 | 0.773 | 0.003 | -0.083 | -0.083 | 0 | 0.22 | 0.09 | 0.35 | 0.36 | 0.17 | 0.37 |
| 10 | 62354797 | rs1009410 | -1.208 | 0.976 | -0.364 | 0.602 | -0.002 | -0.049 | -0.049 | 0 | 0.72 | 0.53 | 0.72 | 0.8 | 0.73 | 0.88 |
| 10 | 62366430 | rs7894641 | -0.169 | 1.058 | -1.141 | 0.556 | -0.029 | 0.035 | 0.035 | 0.928 | 0.21 | 0.01 | 0.12 | 0.12 | 0.2 | 0.25 |
| 10 | 62368339 | rs1006955 | 0.011 | 0.76 | -0.858 | 0.406 | -0.001 | -0.004 | -0.004 | 0.018 | 0.48 | 0.52 | 0.6 | 0.66 | 0.52 | 0.62 |
| 10 | 62370693 | rs6479720 | -1.59 | 0.767 | -0.632 | 0.496 | -0.007 | -0.032 | -0.032 | 0 | 0.71 | 0.6 | 0.73 | 0.79 | 0.72 | 0.87 |
| 10 | 62374041 | rs4143095 | -0.823 | 1.089 | -0.685 | 0.575 | 0.016 | 0.011 | 0.011 | 3.287 | 0.62 | 0.57 | 0.46 | 0.56 | 0.6 | 0.64 |
| 10 | 62384304 | rs10821803 | 0.455 | 1.041 | -0.732 | 0.545 | -0.006 | 0.013 | 0.013 | 0 | 0.4 | 0.46 | 0.32 | 0.46 | 0.4 | 0.38 |
| 10 | 62386654 | rs10821805 | -0.782 | 1.135 | -0.89 | -0.082 | -0.094 | -0.088 | -0.088 | 1.345 | 0.3 | 0.06 | 0.4 | 0.35 | 0.3 | 0.5 |
| 10 | 62390594 | rs10821807 | 0.636 | 0.925 | -0.835 | 0.075 | -0.003 | -0.022 | -0.022 | 0.181 | 0.35 | 0.42 | 0.25 | 0.19 | 0.37 | 0.26 |
| 10 | 62390646 | rs10821808 | -0.354 | 1.051 | -0.861 | 0.078 | 0 | -0.001 | -0.001 | 0 | 0.48 | 0.53 | 0.56 | 0.42 | 0.51 | 0.57 |
| 10 | 62396248 | rs7068934 | -1.839 | 0.897 | -0.907 | 0.072 | -0.006 | 0.058 | 0.058 | 2.286 | 0.7 | 0.55 | 0.68 | 0.55 | 0.7 | 0.82 |
| 10 | 62409221 | rs10761538 | -1.922 | 1.02 | -0.913 | 0.026 | -0.004 | 0.055 | 0.055 | 2.538 | 0.71 | 0.6 | 0.69 | 0.55 | 0.7 | 0.83 |
| 10 | 62412705 | rs10761540 | 0.801 | 0.984 | -0.92 | 0.011 | -0.002 | -0.018 | -0.018 | 0.016 | 0.34 | 0.4 | 0.25 | 0.19 | 0.37 | 0.26 |
| 10 | 62412920 | rs12779419 | 0.225 | 0.985 | -0.956 | 0.021 | -0.029 | 0.031 | 0.031 | 0.826 | 0.22 | 0.01 | 0.12 | 0.13 | 0.2 | 0.25 |
| 10 | 62418569 | rs10761545 | -1.257 | 0.991 | -0.842 | 0.044 | -0.012 | 0.03 | 0.03 | 0.786 | 0.65 | 0.53 | 0.65 | 0.55 | 0.68 | 0.77 |
| 10 | 62418615 | rs10821819 | -1.806 | 0.997 | -0.825 | 0.055 | -0.006 | 0.051 | 0.051 | 2.071 | 0.7 | 0.6 | 0.69 | 0.55 | 0.71 | 0.83 |
| 10 | 62419251 | rs12785023 | -0.986 | 0.884 | -0.987 | 0.038 | -0.019 | -0.004 | -0.004 | 0 | 0.36 | 0.21 | 0.44 | 0.34 | 0.34 | 0.56 |
| 10 | 62420405 | rs10994465 | 0.854 | 0.949 | -0.975 | 0.027 | -0.044 | -0.042 | -0.042 | 0.193 | 0.08 | 0.02 | 0.27 | 0.23 | 0.11 | 0.25 |
| 10 | 62421191 | rs16915297 | 1.424 | 0.9 | -0.889 | -0.047 | -0.008 | -0.024 | -0.024 | 0 | 0.06 | 0.11 | 0.04 | 0 | 0.03 | 0.07 |
| 10 | 62428196 | rs10509137 | 1.787 | 0.907 | -0.889 | -0.044 | 0.003 | 0.002 | 0.002 | 0 | 0.05 | 0.06 | 0.03 | 0 | 0.03 | 0.06 |
| 10 | 62437648 | rs7080822 | 0.651 | 0.955 | -0.866 | -0.044 | -0.001 | -0.008 | -0.008 | 0 | 0.34 | 0.38 | 0.25 | 0.2 | 0.37 | 0.27 |

| | | | | | | | | | | | | | | | | |
|---|---|---|---|---|---|---|---|---|---|---|---|---|---|---|---|---|
| 10 | 62478197 | rs975262 | -0.666 | 1.06 | -0.68 | 0.118 | -0.04 | -0.039 | -0.039 | 0.276 | 0.3 | 0.17 | 0.39 | 0.35 | 0.3 | 0.49 |
| 10 | 62481380 | rs1442539 | -0.713 | 1.051 | -0.578 | 0.114 | -0.014 | -0.014 | -0.014 | 0 | 0.31 | 0.24 | 0.41 | 0.35 | 0.31 | 0.5 |
| 10 | 62493950 | rs1904398 | -0.779 | 1.059 | -0.446 | 0.006 | 0.002 | 0.002 | 0.002 | 0 | 0.53 | 0.65 | 0.61 | 0.43 | 0.51 | 0.58 |
| 10 | 62495498 | rs11813034 | -0.129 | 1.121 | -0.948 | -0.05 | -0.033 | 0.034 | 0.034 | 0.44 | 0.19 | 0.03 | 0.08 | 0.11 | 0.19 | 0.22 |
| 10 | 62498474 | rs16915451 | 0.432 | 1.363 | -0.298 | -0.081 | 0.053 | -0.047 | -0.047 | 0 | 0.1 | 0.27 | 0.11 | 0.01 | 0.17 | 0.11 |
| 10 | 62501766 | rs11817236 | -0.859 | 1.033 | -0.54 | -0.21 | -0.007 | -0.023 | -0.023 | 1.549 | 0.53 | 0.19 | 0.53 | 0.53 | 0.51 | 0.62 |
| 10 | 62501994 | rs1904402 | 0.183 | 0.965 | -0.349 | -0.301 | -0.007 | -0.037 | -0.037 | 0 | 0.6 | 0.51 | 0.64 | 0.81 | 0.63 | 0.67 |
| 10 | 62509300 | rs10994493 | 0.806 | 0.684 | -0.35 | 0.024 | 0.01 | 0.002 | 0.002 | 0 | 0.23 | 0.07 | 0.21 | 0.19 | 0.18 | 0.15 |
| 10 | 62512484 | rs7092606 | 0.485 | 0.801 | 0.122 | 0.316 | 0.013 | 0.029 | 0.029 | 0 | 0.85 | 0.74 | 0.83 | 0.76 | 0.88 | 0.74 |
| 10 | 62514053 | rs7893206 | -0.737 | 0.745 | 0.059 | 0.259 | -0.015 | -0.037 | -0.037 | 0 | 0.92 | 0.79 | 0.95 | 1 | 0.98 | 0.94 |
| 10 | 62518655 | rs7087628 | -0.696 | 0.781 | 0.08 | 0.304 | -0.005 | 0.016 | 0.016 | 0.641 | 0.54 | 0.4 | 0.49 | 0.3 | 0.55 | 0.51 |
| 10 | 62519731 | rs720600 | 1.378 | 0.71 | -0.053 | 0.442 | -0.001 | 0.008 | 0.008 | 2.037 | 0.31 | 0.33 | 0.34 | 0.46 | 0.34 | 0.23 |
| 10 | 62523470 | rs10761558 | -0.119 | 1.002 | -0.136 | -0.402 | -0.019 | 0.026 | 0.026 | 3.084 | 0.28 | 0.17 | 0.35 | 0.21 | 0.36 | 0.3 |
| 10 | 62524139 | rs10761559 | -0.141 | 1.001 | -0.134 | -0.404 | -0.015 | 0.022 | 0.022 | 2.657 | 0.28 | 0.19 | 0.36 | 0.21 | 0.36 | 0.3 |
| 10 | 62525906 | rs2448339 | 1.282 | 0.748 | -0.031 | -0.398 | 0.014 | 0.012 | 0.012 | 2.138 | 0.36 | 0.26 | 0.45 | 0.62 | 0.35 | 0.3 |
| 10 | 62526660 | rs2448340 | 1.028 | 0.748 | 0.03 | -0.398 | -0.003 | -0.003 | -0.003 | 0 | 0.39 | 0.39 | 0.46 | 0.62 | 0.35 | 0.3 |
| 10 | 62535957 | rs2448341 | 1.306 | 0.786 | -0.139 | -0.416 | 0.024 | -0.004 | -0.004 | 0.061 | 0.36 | 0.23 | 0.43 | 0.62 | 0.33 | 0.36 |
| 10 | 62536778 | rs744383 | -0.798 | 0.949 | 0.108 | -0.538 | 0.015 | -0.01 | -0.01 | 0.496 | 0.79 | 0.69 | 0.8 | 0.85 | 0.81 | 0.82 |
| 10 | 62540453 | rs1904415 | -0.52 | 0.981 | -0.032 | -0.529 | 0.013 | -0.023 | -0.023 | 0 | 0.76 | 0.58 | 0.78 | 0.85 | 0.8 | 0.82 |
| 10 | 62547124 | rs2448348 | -0.64 | 0.968 | -0.018 | -0.521 | 0.01 | -0.012 | -0.012 | 0 | 0.77 | 0.67 | 0.79 | 0.84 | 0.79 | 0.79 |
| 10 | 62548674 | rs1871445 | 1.795 | 0.977 | -0.093 | -0.621 | 0.016 | -0.015 | -0.015 | 0 | 0.32 | 0.23 | 0.42 | 0.61 | 0.29 | 0.34 |
| 10 | 62551276 | rs3213075 | 0.016 | 1.098 | -0.299 | -0.653 | -0.017 | 0.026 | 0.026 | 3.62 | 0.27 | 0.17 | 0.21 | 0.14 | 0.36 | 0.28 |
| 10 | 62552732 | rs3213077 | 0.024 | 0.969 | 0.149 | -0.494 | 0.002 | -0.027 | -0.027 | 0 | 0.14 | 0.14 | 0.11 | 0.01 | 0.2 | 0.15 |
| 10 | 62554500 | rs1060373 | -0.56 | 0.811 | 0.171 | 0.097 | 0.004 | 0.007 | 0.007 | 0 | 0.13 | 0.09 | 0.14 | 0.09 | 0.13 | 0.13 |
| 10 | 62557260 | rs2448358 | 0.41 | 1.114 | 0.269 | 0.354 | 0.01 | -0.022 | -0.022 | 0 | 0.75 | 0.62 | 0.78 | 0.84 | 0.79 | 0.79 |
| 10 | 62565043 | rs7093708 | -1.292 | 1.596 | 0.549 | 1.13 | 0.062 | 0.079 | 0.079 | 0 | 0.24 | 0.42 | 0.3 | 0.39 | 0.35 | 0.3 |
| 10 | 62570821 | rs7077462 | -0.978 | 1.563 | 0.662 | 1.157 | 0.023 | 0.032 | 0.032 | 0 | 0.21 | 0.26 | 0.25 | 0.34 | 0.33 | 0.27 |
| 10 | 62588168 | rs7909618 | -1.426 | 1.595 | 0.783 | 1.169 | 0.068 | 0.085 | 0.085 | 0 | 0.24 | 0.44 | 0.29 | 0.34 | 0.34 | 0.28 |
| 10 | 62593918 | rs6479734 | -1.504 | 1.556 | 0.859 | 1.152 | 0.067 | 0.084 | 0.084 | 0 | 0.24 | 0.49 | 0.29 | 0.34 | 0.34 | 0.28 |
| 10 | 62596062 | rs4948428 | -0.912 | 0.608 | 0.198 | 0.983 | -0.001 | -0.002 | -0.002 | 0.269 | 0.45 | 0.51 | 0.58 | 0.67 | 0.46 | 0.38 |
| 10 | 62600239 | rs12242535 | 0.042 | 0.908 | 0.02 | 1.39 | 0.009 | 0.075 | 0.075 | 0 | 0.1 | 0.15 | 0.17 | 0.32 | 0.12 | 0.11 |
| 10 | 62600287 | rs4132554 | -1.043 | 0.606 | -0.254 | 0.852 | -0.003 | 0.083 | 0.083 | 0 | 0.48 | 0.66 | 0.61 | 0.67 | 0.49 | 0.39 |
| 10 | 62604856 | rs2393702 | -0.448 | 0.356 | -0.002 | 1.331 | 0.003 | -0.024 | -0.024 | 4.71 | 0.17 | 0.13 | 0.1 | 0.28 | 0.13 | 0.24 |
| 10 | 62606784 | rs2893863 | -0.058 | -0.096 | -0.118 | 0.897 | 0.006 | 0.012 | 0.012 | 2.624 | 0.22 | 0.17 | 0.28 | 0.02 | 0.16 | 0.24 |
| 10 | 62607532 | rs10821845 | 0.143 | -0.031 | 0.113 | 1.305 | 0.011 | 0.07 | 0.07 | 0.117 | 0.07 | 0.12 | 0.06 | 0.19 | 0.08 | 0.11 |
| 10 | 62612714 | rs7091284 | 0.079 | -0.379 | 0 | 0.929 | 0.006 | 0.017 | 0.017 | 2.683 | 0.21 | 0.13 | 0.26 | 0.08 | 0.15 | 0.25 |
| 10 | 62614516 | rs1002074 | -1.552 | -0.546 | 0.453 | 0.87 | 0.031 | 0.043 | 0.043 | 0.085 | 0.05 | 0.34 | 0.05 | 0.03 | 0.02 | 0.02 |
| 10 | 62614958 | rs10994538 | 0.068 | -0.507 | 0.204 | 1.048 | 0.001 | -0.015 | -0.015 | 0 | 0.15 | 0.12 | 0.1 | 0.25 | 0.12 | 0.23 |
| 10 | 62616883 | rs954488 | -0.682 | -0.513 | 0.368 | 1.049 | -0.013 | 0.021 | 0.021 | 0 | 0.21 | 0.35 | 0.13 | 0.28 | 0.15 | 0.25 |
| 10 | 62618398 | rs7085788 | -1.452 | -0.582 | 0.254 | 1.02 | 0.006 | -0.001 | -0.001 | 0 | 0.45 | 0.62 | 0.43 | 0.39 | 0.31 | 0.52 |
| 10 | 62619842 | rs10994544 | -0.918 | -0.236 | 0.126 | 1.033 | -0.011 | 0.011 | 0.011 | 0 | 0.19 | 0.31 | 0.12 | 0.25 | 0.13 | 0.23 |
| 10 | 62623814 | rs4948267 | -0.918 | -0.236 | 0.126 | 1.033 | -0.011 | 0.011 | 0.011 | 0 | 0.19 | 0.31 | 0.12 | 0.25 | 0.13 | 0.23 |
| 10 | 62626454 | rs10821847 | -1.188 | -0.486 | -0.267 | 1.206 | 0.025 | -0.032 | -0.032 | 0 | 0.22 | 0.16 | 0.26 | 0.28 | 0.12 | 0.23 |
| 10 | 62626579 | rs12245333 | -1.409 | -0.456 | 0.46 | 0.908 | -0.04 | -0.008 | -0.008 | 2.282 | 0.23 | 0.37 | 0.35 | 0.13 | 0.1 | 0.12 |
| 10 | 62627865 | rs10994553 | -1.434 | 0.019 | 0.524 | 0.819 | 0.027 | -0.02 | -0.02 | 0.189 | 0.07 | 0.16 | 0.09 | 0.06 | 0.09 | 0.08 |
| 10 | 62630615 | rs3763742 | -1.827 | 0.236 | 0.658 | 0.782 | 0.057 | -0.025 | -0.025 | 0 | 0.08 | 0.22 | 0.11 | 0.06 | 0.12 | 0.08 |
| 10 | 62634839 | rs12251212 | -1.151 | 0.092 | 0.783 | 0.982 | -0.012 | 0.002 | 0.002 | 10.738 | 0.14 | 0.2 | 0.15 | 0.14 | 0.09 | 0.04 |
| 10 | 62638036 | rs12266720 | -1.978 | 0.015 | 0.795 | 0.655 | -0.022 | -0.032 | -0.032 | 0.02 | 0.28 | 0.39 | 0.24 | 0.16 | 0.2 | 0.24 |
| 10 | 62640488 | rs7081336 | -2.179 | -0.018 | 1.085 | 0.687 | -0.011 | -0.031 | -0.031 | 0 | 0.31 | 0.48 | 0.32 | 0.23 | 0.28 | 0.3 |
| 10 | 62647306 | rs16915675 | -1.921 | -0.134 | 1.125 | 1.623 | 0 | 0.005 | 0.005 | 0.855 | 0.27 | 0.23 | 0.29 | 0.51 | 0.21 | 0.32 |
| 10 | 62650130 | rs16915679 | -1.184 | -0.158 | 1.004 | 1.604 | -0.029 | -0.089 | -0.089 | 1.349 | 0.14 | 0.01 | 0.16 | 0.34 | 0.13 | 0.13 |
| 10 | 62652165 | rs7898435 | -2.289 | -0.198 | 1.221 | 1.325 | -0.018 | 0.039 | 0.039 | 0.02 | 0.33 | 0.54 | 0.39 | 0.6 | 0.25 | 0.37 |
| 10 | 62653374 | rs12265809 | -1.929 | -0.144 | 0.803 | 1.357 | 0.003 | -0.01 | -0.01 | 0 | 0.29 | 0.24 | 0.29 | 0.49 | 0.22 | 0.31 |
| 10 | 62658304 | rs7908589 | -1.965 | -0.127 | 0.756 | 1.317 | -0.002 | 0.006 | 0.006 | 0.157 | 0.32 | 0.36 | 0.39 | 0.59 | 0.25 | 0.31 |
| 10 | 62659093 | rs7097615 | -1.481 | -0.132 | 0.686 | 1.307 | -0.01 | 0.075 | 0.075 | 0.589 | 0.25 | 0.41 | 0.36 | 0.47 | 0.19 | 0.25 |
| 10 | 62660292 | rs10509141 | -1.551 | -0.603 | 0.879 | 0.926 | -0.039 | 0.024 | 0.024 | 0 | 0.2 | 0.36 | 0.2 | 0.24 | 0.11 | 0.16 |
| 10 | 62661961 | rs2893869 | -2.02 | -0.504 | 1.326 | 0.752 | -0.003 | -0.002 | -0.002 | 0.401 | 0.27 | 0.69 | 0.25 | 0.24 | 0.16 | 0.18 |
| 10 | 62662781 | rs2893871 | -1.771 | -0.457 | 1.495 | 0.746 | -0.01 | -0.009 | -0.009 | 0 | 0.23 | 0.3 | 0.21 | 0.22 | 0.15 | 0.17 |
| 10 | 62663124 | rs12252953 | -2.015 | -0.242 | 1.984 | 0.843 | 0.011 | 0.056 | 0.056 | 1.35 | 0.11 | 0.37 | 0.13 | 0.14 | 0.1 | 0.08 |
| 10 | 62664013 | rs7087599 | -0.91 | 0.224 | 0.442 | -0.234 | -0.089 | 0.117 | 0.117 | 0 | 0.77 | 0.99 | 0.84 | 0.96 | 0.7 | 0.87 |
| 10 | 62664755 | rs10821855 | -1.347 | -0.3 | 1.029 | -0.128 | -0.011 | -0.001 | -0.001 | 11.912 | 0.17 | 0.35 | 0.18 | 0.18 | 0.12 | 0.15 |
| 10 | 62665460 | rs10994572 | -1.183 | -0.122 | 1.025 | -0.133 | -0.002 | -0.002 | -0.002 | 0 | 0.16 | 0.2 | 0.17 | 0.18 | 0.12 | 0.15 |
| 10 | 62671305 | rs2271126 | -1.789 | -0.275 | 0.85 | -0.028 | -0.027 | -0.016 | -0.016 | 0 | 0.24 | 0.6 | 0.22 | 0.22 | 0.14 | 0.21 |
| 10 | 62676054 | rs11595278 | 0.979 | -0.112 | 0.129 | 0.668 | -0.044 | -0.077 | -0.077 | 0.097 | 0.13 | 0.02 | 0.06 | 0.33 | 0.15 | 0.23 |
| 10 | 62682169 | rs7913287 | 0.141 | -0.379 | -0.046 | 0.072 | 0.004 | 0.076 | 0.076 | 0.568 | 0.49 | 0.32 | 0.59 | 0.76 | 0.5 | 0.58 |
| 10 | 62682308 | rs7913431 | -0.265 | -0.479 | 0.305 | -0.018 | 0.006 | -0.029 | -0.029 | 0 | 0.58 | 0.48 | 0.63 | 0.8 | 0.53 | 0.61 |
| 10 | 62697642 | rs957854 | -0.305 | -0.491 | 0.462 | -0.023 | -0.019 | 0.05 | 0.05 | 0 | 0.58 | 0.66 | 0.64 | 0.8 | 0.53 | 0.61 |
| 10 | 62730841 | rs6479743 | -0.099 | -0.555 | 0.352 | -0.046 | 0.005 | -0.038 | -0.038 | 0.063 | 0.56 | 0.44 | 0.54 | 0.78 | 0.52 | 0.61 |
| 10 | 62733317 | rs4948437 | -0.618 | -0.635 | 0.366 | -0.081 | -0.093 | 0.185 | 0.185 | 0 | 0.62 | 0.89 | 0.63 | 0.93 | 0.54 | 0.65 |

| | | | | | | | | | | | | | | | | |
|---|---|---|---|---|---|---|---|---|---|---|---|---|---|---|---|---|
| 10 | 62749393 | rs6479744 | 0.267 | -0.603 | 0.431 | -0.027 | 0.001 | 0.03 | 0.03 | 1.053 | 0.49 | 0.37 | 0.51 | 0.76 | 0.5 | 0.59 |
| 10 | 62751135 | rs10821866 | -0.193 | -0.531 | 0.108 | 0.082 | 0.004 | -0.02 | -0.02 | 0 | 0.56 | 0.48 | 0.54 | 0.75 | 0.53 | 0.61 |
| 10 | 62755890 | rs10994604 | 0.571 | -0.57 | 0.167 | 0.269 | -0.004 | 0.006 | 0.006 | 0 | 0.42 | 0.39 | 0.54 | 0.43 | 0.39 | 0.42 |
| 10 | 62773645 | rs10509142 | -0.052 | -0.475 | -0.46 | -0.016 | 0.03 | 0.049 | 0.049 | 0.377 | 0.51 | 0.67 | 0.61 | 0.59 | 0.43 | 0.52 |
| 10 | 62780127 | rs10994613 | 2.069 | -0.503 | -0.505 | 0.023 | 0.014 | 0.015 | 0.015 | 27.422 | 0.08 | 0.05 | 0.02 | 0.04 | 0.04 | 0.09 |
| 10 | 62788795 | rs10437343 | -0.081 | -0.182 | -0.364 | 0.045 | 0.03 | 0.048 | 0.048 | 0.34 | 0.52 | 0.7 | 0.61 | 0.59 | 0.43 | 0.52 |
| 10 | 62803417 | rs1864758 | -0.057 | -0.129 | -0.45 | 0.068 | 0.025 | 0.038 | 0.038 | 0.307 | 0.52 | 0.66 | 0.62 | 0.59 | 0.43 | 0.52 |
| 10 | 62804925 | rs12772906 | 0.796 | 0.16 | -1.033 | -0.229 | -0.042 | 0.038 | 0.038 | 0 | 0.16 | 0.02 | 0.11 | 0.08 | 0.14 | 0.15 |
| 10 | 62826952 | rs7922793 | -0.742 | 0.652 | -1.044 | -0.165 | -0.036 | 0.009 | 0.009 | 0.292 | 0.73 | 0.87 | 0.65 | 0.78 | 0.63 | 0.73 |
| 10 | 62842602 | rs7087951 | 0.033 | 0.348 | -0.013 | -0.376 | 0.001 | -0.012 | -0.012 | 0 | 0.63 | 0.36 | 0.57 | 0.75 | 0.57 | 0.6 |
| 10 | 62856612 | rs16915839 | 0.739 | -0.448 | -0.344 | -0.709 | 0.048 | 0.111 | 0.111 | 0.97 | 0.37 | 0.16 | 0.25 | 0.12 | 0.24 | 0.2 |
| 10 | 62866831 | rs4131334 | 0.925 | -0.038 | -0.859 | -0.552 | -0.078 | -0.112 | -0.112 | 0 | 0.15 | 0.02 | 0.25 | 0.61 | 0.22 | 0.2 |
| 10 | 62869980 | rs7911472 | -0.275 | -0.018 | -0.223 | -0.617 | 0.005 | -0.022 | -0.022 | 0 | 0.58 | 0.47 | 0.54 | 0.75 | 0.5 | 0.56 |
| 10 | 62896874 | rs1906461 | -0.369 | 0.06 | -0.477 | -0.563 | -0.006 | 0.016 | 0.016 | 0.138 | 0.59 | 0.63 | 0.63 | 0.75 | 0.51 | 0.56 |
| 10 | 62917396 | rs2807691 | -0.916 | 0.457 | -1.239 | -0.538 | -0.071 | 0.044 | 0.044 | 0.206 | 0.68 | 0.9 | 0.68 | 0.79 | 0.57 | 0.69 |
| 10 | 62936427 | rs12253075 | 0.682 | 0.176 | -0.91 | -0.569 | -0.004 | -0.029 | -0.029 | 0 | 0.07 | 0.07 | 0.04 | 0 | 0.04 | 0.01 |
| 10 | 62969458 | rs2633295 | -0.939 | 0.035 | -1.588 | -0.734 | -0.086 | 0.085 | 0.085 | 0.239 | 0.64 | 0.91 | 0.69 | 0.8 | 0.57 | 0.72 |
| 10 | 63004505 | rs4948446 | -0.572 | 0.022 | -1.303 | -0.581 | -0.057 | 0.071 | 0.071 | 0.569 | 0.6 | 0.84 | 0.66 | 0.76 | 0.54 | 0.72 |
| 10 | 63018988 | rs11596067 | 0.314 | 0.017 | -1.427 | -0.965 | -0.003 | 0.016 | 0.016 | 0 | 0.11 | 0.09 | 0.07 | 0.04 | 0.12 | 0.14 |
| 10 | 63023314 | rs4145320 | -1.016 | -0.107 | -1.569 | -0.994 | -0.085 | 0.113 | 0.113 | 0.21 | 0.62 | 0.89 | 0.69 | 0.84 | 0.56 | 0.73 |
| 10 | 63024071 | rs1906468 | -0.176 | -0.315 | -0.454 | -0.985 | -0.004 | -0.027 | -0.027 | 0 | 0.41 | 0.4 | 0.55 | 0.78 | 0.37 | 0.54 |
| 10 | 63032694 | rs4643025 | 0.125 | -0.167 | -1.121 | -0.947 | -0.061 | -0.008 | -0.008 | 0 | 0.14 | 0.01 | 0.15 | 0.1 | 0.16 | 0.14 |
| 10 | 63035242 | rs10994701 | 1.086 | -0.129 | -0.618 | -1.285 | 0.006 | -0.041 | -0.041 | 0.533 | 0.38 | 0.44 | 0.34 | 0.09 | 0.43 | 0.26 |
| 10 | 63038236 | rs3107461 | 0.18 | -0.284 | -0.973 | -0.867 | 0.034 | -0.002 | -0.002 | 0 | 0.64 | 0.24 | 0.54 | 0.32 | 0.73 | 0.54 |
| 10 | 63043987 | rs3107457 | -0.169 | -0.306 | -0.926 | -0.654 | 0.026 | 0.058 | 0.058 | 0 | 0.39 | 0.06 | 0.25 | 0.26 | 0.32 | 0.31 |
| 10 | 63053788 | rs3125326 | 0.165 | -0.312 | 0.1 | -0.547 | 0.002 | 0.007 | 0.007 | 0.953 | 0.58 | 0.67 | 0.5 | 0.34 | 0.57 | 0.52 |
| 10 | 63057855 | rs3107450 | 0.273 | -0.3 | 0.276 | -0.358 | 0.044 | 0.087 | 0.087 | 0 | 0.4 | 0.13 | 0.27 | 0.26 | 0.33 | 0.33 |
| 10 | 63061633 | rs2139783 | 2.091 | -0.17 | 0.259 | -0.382 | -0.009 | 0.025 | 0.025 | 0 | 0.06 | 0 | 0.02 | 0 | 0.04 | 0.04 |
| 10 | 63063557 | rs1915448 | 0.198 | -0.166 | 0.582 | -0.256 | 0.032 | 0.046 | 0.046 | 0 | 0.41 | 0.26 | 0.36 | 0.33 | 0.36 | 0.34 |
| 10 | 63070228 | rs3125338 | 0.192 | -0.154 | 0.953 | -0.094 | 0.007 | 0.11 | 0.11 | 0 | 0.82 | 0.73 | 0.68 | 0.47 | 0.79 | 0.65 |
| 10 | 63081373 | rs4948449 | 0.362 | -0.186 | 1.241 | -0.058 | 0.015 | 0.201 | 0.201 | 0 | 0.81 | 0.59 | 0.67 | 0.49 | 0.79 | 0.65 |
| 10 | 63096664 | rs12411660 | 1.287 | 0.018 | 0.775 | 0.01 | -0.023 | 0.074 | 0.074 | 0 | 0.05 | 0.01 | 0.23 | 0.49 | 0.09 | 0.16 |
| 10 | 63099550 | rs10509145 | 0.909 | -0.085 | 0.789 | -0.294 | -0.063 | 0.124 | 0.124 | 0 | 0.38 | 0.2 | 0.3 | 0.1 | 0.45 | 0.31 |
| 10 | 63112808 | rs10821885 | 0.45 | 0.115 | 0.911 | 0.441 | 0.026 | 0.178 | 0.178 | 0.335 | 0.81 | 0.62 | 0.66 | 0.48 | 0.76 | 0.66 |
| 10 | 63127295 | rs12243939 | -0.779 | -0.245 | 0.978 | 0.247 | 0.027 | -0.003 | -0.003 | 0 | 0.06 | 0.32 | 0.05 | 0.01 | 0.05 | 0.04 |
| 10 | 63127639 | rs10994740 | -0.719 | -0.667 | 0.408 | -0.026 | 0.067 | 0.048 | 0.048 | 0 | 0.47 | 0.81 | 0.46 | 0.39 | 0.4 | 0.34 |
| 10 | 63146547 | rs16916041 | 1.446 | -0.713 | 0.232 | -0.088 | -0.007 | -0.045 | -0.045 | 0 | 0.12 | 0.01 | 0.19 | 0.21 | 0.11 | 0.15 |
| 10 | 63149922 | rs10994747 | -0.929 | -1.184 | 0.278 | -0.234 | 0.072 | 0.071 | 0.071 | 0 | 0.47 | 0.79 | 0.44 | 0.37 | 0.39 | 0.32 |
| 10 | 63157465 | rs10994750 | -0.504 | -1.197 | 0.762 | -0.199 | -0.026 | 0.034 | 0.034 | 3.104 | 0.19 | 0.43 | 0.26 | 0.27 | 0.14 | 0.21 |
| 10 | 63163526 | rs12573098 | 1.764 | -1.178 | 0.732 | -0.19 | 0.004 | -0.007 | -0.007 | 0 | 0.12 | 0.08 | 0.2 | 0.21 | 0.09 | 0.15 |
| 10 | 63166773 | rs10994754 | -0.9 | -1.182 | 0.709 | -0.228 | 0.083 | 0.066 | 0.066 | 0 | 0.47 | 0.79 | 0.42 | 0.37 | 0.35 | 0.33 |
| 10 | 63167132 | rs2176929 | 0.436 | -1.149 | 0.815 | -0.243 | -0.03 | -0.058 | -0.058 | 0.159 | 0.28 | 0.38 | 0.16 | 0.09 | 0.21 | 0.12 |
| 10 | 63167262 | rs2139779 | 0.769 | -1.239 | 1.142 | -0.261 | 0.029 | 0.052 | 0.052 | 0.128 | 0.26 | 0.14 | 0.14 | 0.09 | 0.21 | 0.11 |
| 10 | 63167404 | rs2139780 | 0.512 | -1.218 | 1.055 | -0.251 | -0.022 | -0.05 | -0.05 | 0.008 | 0.28 | 0.34 | 0.16 | 0.09 | 0.21 | 0.11 |
| 10 | 63168488 | rs16916104 | 0.512 | -1.218 | 1.054 | -0.251 | -0.02 | -0.046 | -0.046 | 0 | 0.28 | 0.33 | 0.16 | 0.09 | 0.21 | 0.11 |
| 10 | 63169896 | rs10509148 | -0.14 | -1.164 | 1.099 | -0.186 | 0.032 | -0.003 | -0.003 | 0 | 0.06 | 0.31 | 0.05 | 0.01 | 0.05 | 0.04 |
| 10 | 63172966 | rs1907217 | -0.389 | -1.184 | 1.087 | -0.182 | -0.021 | 0.01 | 0.01 | 2.347 | 0.18 | 0.42 | 0.28 | 0.22 | 0.13 | 0.2 |
| 10 | 63176708 | rs10994757 | -0.24 | -1.174 | 1.097 | -0.175 | 0.042 | 0.058 | 0.058 | 0.135 | 0.07 | 0.33 | 0.08 | 0.07 | 0.05 | 0.05 |
| 10 | 63183583 | rs1915433 | -0.914 | -0.849 | 0.808 | -0.186 | 0.058 | 0.062 | 0.062 | 0.162 | 0.47 | 0.76 | 0.45 | 0.38 | 0.38 | 0.32 |
| 10 | 63186686 | rs12572590 | 1.259 | -0.642 | 0.812 | -0.087 | 0 | 0.001 | 0.001 | 4.753 | 0.13 | 0.12 | 0.25 | 0.22 | 0.14 | 0.19 |
| 10 | 63221167 | rs10994774 | -0.908 | -0.846 | 0.819 | -0.131 | 0.057 | 0.045 | 0.045 | 0 | 0.47 | 0.79 | 0.46 | 0.39 | 0.41 | 0.3 |
| 10 | 63223455 | rs11814548 | 0.717 | -0.973 | 1.043 | -0.217 | -0.014 | -0.036 | -0.036 | 0 | 0.27 | 0.3 | 0.15 | 0.1 | 0.21 | 0.11 |
| 10 | 63236711 | rs2650703 | 1.544 | -1.123 | 0.555 | -0.324 | -0.135 | 0.072 | 0.072 | 0 | 0.33 | 0.07 | 0.34 | 0.15 | 0.43 | 0.32 |
| 10 | 63237480 | rs2650704 | 1.544 | -1.129 | 0.555 | -0.316 | -0.132 | 0.068 | 0.068 | 0 | 0.33 | 0.07 | 0.34 | 0.15 | 0.43 | 0.32 |
| 10 | 63237913 | rs2993425 | 0.824 | -1.116 | 0.61 | -0.286 | 0.004 | 0.008 | 0.008 | 1.23 | 0.6 | 0.37 | 0.49 | 0.24 | 0.63 | 0.42 |
| 10 | 63240300 | rs2650706 | 0.816 | -1.117 | 0.61 | -0.287 | 0.004 | 0.008 | 0.008 | 1.275 | 0.6 | 0.37 | 0.48 | 0.24 | 0.63 | 0.42 |
| 10 | 63243436 | rs2650711 | 1.511 | -1.158 | 0.45 | -0.306 | -0.132 | 0.068 | 0.068 | 0 | 0.33 | 0.07 | 0.34 | 0.15 | 0.43 | 0.32 |
| 10 | 63248358 | rs4948281 | 1.113 | -1.296 | 0.539 | -0.31 | -0.002 | -0.005 | -0.005 | 0.102 | 0.14 | 0.12 | 0.23 | 0.2 | 0.16 | 0.16 |
| 10 | 63248540 | rs2249689 | 1.534 | -1.239 | 0.501 | -0.286 | -0.132 | 0.068 | 0.068 | 0 | 0.33 | 0.07 | 0.34 | 0.15 | 0.43 | 0.32 |
| 10 | 63259595 | rs2650753 | -0.67 | -1.182 | 0.834 | -0.148 | 0.005 | -0.017 | -0.017 | 0 | 0.83 | 0.89 | 0.84 | 0.49 | 0.86 | 0.64 |
| 10 | 63264872 | rs10994785 | 1.164 | -1.201 | 0.733 | -0.054 | -0.022 | 0.005 | 0.005 | 0 | 0.07 | 0 | 0.12 | 0.49 | 0.09 | 0.17 |
| 10 | 63279936 | rs7070380 | 0.167 | -1.161 | 0.93 | -0.388 | 0.02 | 0.004 | 0.004 | 0.269 | 0.05 | 0.3 | 0.05 | 0 | 0.03 | 0.01 |
| 10 | 63300506 | rs16916259 | 1.762 | -1.115 | 0.689 | -0.307 | 0.001 | -0.006 | -0.006 | 0 | 0.12 | 0.08 | 0.21 | 0.26 | 0.12 | 0.16 |
| 10 | 63301072 | rs2606083 | 0.34 | -1.199 | 0.745 | -0.354 | 0.007 | 0.012 | 0.012 | 3.576 | 0.69 | 0.78 | 0.62 | 0.17 | 0.72 | 0.47 |
| 10 | 63303731 | rs2606086 | 0.428 | -1.207 | 0.973 | -0.357 | -0.007 | -0.013 | -0.013 | 2.08 | 0.69 | 0.67 | 0.61 | 0.17 | 0.72 | 0.47 |
| 10 | 63306445 | rs10740046 | -0.389 | -1.141 | 0.885 | -0.25 | 0.002 | 0.002 | 0.002 | 0 | 0.82 | 0.85 | 0.83 | 0.44 | 0.84 | 0.65 |
| 10 | 63307297 | rs12267116 | -1.287 | -1.138 | 1.037 | -0.475 | -0.004 | -0.027 | -0.027 | 0.04 | 0.12 | 0.64 | 0.15 | 0.03 | 0.11 | 0.09 |
| 10 | 63307475 | rs2787702 | -0.27 | -1.187 | 0.895 | -0.584 | 0.016 | 0.004 | 0.004 | 0 | 0.59 | 0.74 | 0.68 | 0.35 | 0.64 | 0.56 |
| 10 | 63308159 | rs2254096 | 1.536 | -1.2 | 0.711 | -0.608 | -0.106 | 0.129 | 0.129 | 0 | 0.34 | 0.06 | 0.32 | 0.09 | 0.41 | 0.3 |

| | | | | | | | | | | | | | | | | |
|---|---|---|---|---|---|---|---|---|---|---|---|---|---|---|---|---|
| 10 | 63324847 | rs2606094 | -0.36 | -1.189 | 0.787 | -0.585 | 0.011 | -0.001 | -0.001 | 0 | 0.59 | 0.7 | 0.68 | 0.36 | 0.64 | 0.56 |
| 10 | 63332067 | rs7914692 | -1.021 | -1.109 | 0.791 | -0.609 | 0.018 | 0.06 | 0.06 | 1.465 | 0.12 | 0.46 | 0.13 | 0.09 | 0.12 | 0.11 |
| 10 | 63342112 | rs2814027 | 1.39 | -1.051 | 0.501 | -0.667 | -0.05 | 0.084 | 0.084 | 0 | 0.34 | 0.16 | 0.33 | 0.09 | 0.41 | 0.3 |
| 10 | 63342764 | rs11594617 | -0.35 | -0.91 | 0.523 | -0.463 | 0.031 | 0.025 | 0.025 | 0.21 | 0.07 | 0.25 | 0.08 | 0.04 | 0.08 | 0.08 |
| 10 | 63350295 | rs7894852 | 1.106 | -0.909 | -0.144 | -0.534 | 0.003 | 0.057 | 0.057 | 0 | 0.23 | 0.12 | 0.13 | 0.04 | 0.2 | 0.1 |
| 10 | 63352842 | rs16916326 | 1.663 | -0.49 | 0.079 | -0.229 | -0.019 | -0.037 | -0.037 | 4.244 | 0.11 | 0.01 | 0.21 | 0.17 | 0.12 | 0.15 |
| 10 | 63354442 | rs7088920 | 0.883 | -0.708 | 0.15 | -0.484 | 0.004 | -0.016 | -0.016 | 0 | 0.11 | 0.13 | 0.1 | 0.05 | 0.15 | 0.08 |
| 10 | 63355099 | rs4454676 | -1.322 | -0.596 | 0.981 | -0.123 | 0.009 | 0.067 | 0.067 | 1.834 | 0.12 | 0.56 | 0.12 | 0.09 | 0.11 | 0.11 |
| 10 | 63356349 | rs2650714 | -0.072 | -0.591 | 0.489 | -0.194 | -0.001 | -0.008 | -0.008 | 0 | 0.8 | 0.86 | 0.78 | 0.39 | 0.79 | 0.55 |
| 10 | 63361987 | rs2814021 | -0.022 | -0.621 | 0.478 | 0.142 | -0.001 | -0.007 | -0.007 | 0 | 0.79 | 0.85 | 0.8 | 0.52 | 0.79 | 0.55 |
| 10 | 63369632 | rs1456285 | 0.569 | -0.698 | 0.846 | 0.101 | 0.009 | 0.094 | 0.094 | 0 | 0.73 | 0.56 | 0.72 | 0.48 | 0.72 | 0.49 |
| 10 | 63371375 | rs10509154 | 1.124 | -0.8 | 0.488 | -0.209 | -0.117 | 0.07 | 0.07 | 0.01 | 0.31 | 0.07 | 0.33 | 0.11 | 0.42 | 0.24 |
| 10 | 63377651 | rs16916421 | 1.592 | -0.763 | 1.07 | -0.046 | -0.023 | 0.017 | 0.017 | 7.34 | 0.12 | 0.23 | 0.21 | 0.15 | 0.08 | 0.11 |
| 10 | 63377998 | rs17279244 | 1.49 | -0.716 | 0.856 | -0.058 | 0.01 | 0.093 | 0.093 | 0.223 | 0.26 | 0.12 | 0.13 | 0.05 | 0.2 | 0.09 |
| 10 | 63381954 | rs1456292 | 0.916 | -0.315 | 1.761 | 0.815 | 0.046 | 0.271 | 0.271 | 0 | 0.84 | 0.57 | 0.77 | 0.52 | 0.77 | 0.65 |
| 10 | 63382567 | rs1869928 | 0.047 | -0.325 | 1.771 | 0.845 | 0.029 | 0.265 | 0.265 | 0 | 0.88 | 0.7 | 0.83 | 0.55 | 0.85 | 0.73 |
| 10 | 63394596 | rs1456282 | 1.166 | -0.312 | 1.532 | 0.782 | -0.02 | 0.017 | 0.017 | 8.972 | 0.11 | 0.21 | 0.2 | 0.15 | 0.08 | 0.11 |
| 10 | 63415264 | rs4144119 | -0.414 | 0.1 | 1.153 | 0.937 | 0.008 | 0.111 | 0.111 | 0 | 0.9 | 0.89 | 0.86 | 0.56 | 0.85 | 0.73 |
| 10 | 63421242 | rs12414729 | -0.709 | 0.081 | 1.034 | 0.415 | 0.013 | 0.051 | 0.051 | 0.003 | 0.05 | 0.12 | 0.06 | 0.11 | 0.07 | 0.07 |
| 10 | 63426226 | rs11815699 | 1.01 | -0.076 | 1.231 | 0.315 | -0.013 | 0.021 | 0.021 | 8.596 | 0.12 | 0.19 | 0.21 | 0.18 | 0.09 | 0.13 |
| 10 | 63429315 | rs10994867 | -0.22 | 0.126 | 1.116 | 0.429 | 0.028 | 0.203 | 0.203 | 0 | 0.89 | 0.77 | 0.81 | 0.41 | 0.84 | 0.72 |
| 10 | 63438578 | rs9629894 | -0.349 | 0.139 | 0.731 | 0.497 | -0.012 | 0.011 | 0.011 | 0 | 0.9 | 0.95 | 0.82 | 0.44 | 0.84 | 0.72 |
| 10 | 63452956 | rs16916504 | -0.523 | 0.51 | 0.411 | 0.114 | 0.003 | -0.026 | -0.026 | 0 | 0.22 | 0.34 | 0.16 | 0.09 | 0.22 | 0.15 |
| 10 | 63462365 | rs7922049 | 0.976 | 0.52 | -0.174 | 0.008 | -0.022 | 0.036 | 0.036 | 0.022 | 0.12 | 0.08 | 0.07 | 0 | 0.14 | 0.05 |
| 10 | 63464225 | rs10994884 | 0.784 | 0.412 | -0.099 | 0.093 | -0.002 | -0.002 | -0.002 | 0.064 | 0.66 | 0.29 | 0.65 | 0.35 | 0.62 | 0.58 |
| 10 | 63476983 | rs17281372 | 1.357 | 0.441 | -0.767 | -0.256 | -0.084 | 0.208 | 0.208 | 0.267 | 0.43 | 0.07 | 0.38 | 0.11 | 0.43 | 0.32 |
| 10 | 63493724 | rs10821916 | 0.58 | 0.408 | 0.308 | 0.148 | 0 | 0.068 | 0.068 | 0 | 0.1 | 0.09 | 0.17 | 0.55 | 0.15 | 0.27 |
| 10 | 63495205 | rs16916549 | 1.298 | 0.196 | 0.443 | 0.083 | -0.03 | 0.029 | 0.029 | 8.37 | 0.12 | 0.21 | 0.22 | 0.2 | 0.07 | 0.11 |
| 10 | 63502431 | rs10994906 | 0.066 | 0.241 | 0.504 | 0.064 | 0.036 | 0.278 | 0.278 | 0 | 0.12 | 0.31 | 0.18 | 0.55 | 0.15 | 0.27 |
| 10 | 63507669 | rs10509158 | -0.554 | 0.228 | 0.416 | -0.096 | -0.018 | -0.033 | -0.033 | 0 | 0.25 | 0.42 | 0.28 | 0.18 | 0.2 | 0.14 |
| 10 | 63507847 | rs7902688 | -1.609 | 0.149 | 0.417 | -0.168 | 0.011 | 0.066 | 0.066 | 0 | 0.42 | 0.78 | 0.5 | 0.77 | 0.39 | 0.48 |
| 10 | 63507915 | rs10761592 | 0.909 | 0.348 | 0.549 | -0.086 | -0.001 | 0.003 | 0.003 | 0.299 | 0.71 | 0.44 | 0.65 | 0.69 | 0.71 | 0.77 |
| 10 | 63509111 | rs16916568 | -0.358 | 0.344 | 0.448 | -0.092 | -0.001 | -0.001 | -0.001 | 0.011 | 0.23 | 0.22 | 0.28 | 0.21 | 0.22 | 0.16 |
| 10 | 63518232 | rs12761488 | 1.049 | 0.364 | 0.388 | -0.057 | -0.018 | 0.039 | 0.039 | 7.841 | 0.13 | 0.21 | 0.23 | 0.23 | 0.08 | 0.13 |
| 10 | 63520698 | rs1992625 | -0.499 | 0.556 | 0.335 | 0.038 | 0.007 | -0.003 | -0.003 | 0 | 0.87 | 0.86 | 0.89 | 0.91 | 0.8 | 0.9 |
| 10 | 63523266 | rs2675612 | 0.765 | 0.524 | 0.465 | 0.089 | -0.001 | 0 | 0 | 0.678 | 0.7 | 0.39 | 0.66 | 0.69 | 0.72 | 0.77 |
| 10 | 63524979 | rs2588917 | 1.629 | 0.296 | -0.116 | 0.071 | -0.127 | 0.233 | 0.233 | 0.823 | 0.45 | 0.07 | 0.42 | 0.11 | 0.45 | 0.31 |
| 10 | 63525301 | rs973237 | 0.819 | 0.825 | 0.777 | 0.926 | 0.002 | 0.016 | 0.016 | 0.253 | 0.76 | 0.69 | 0.7 | 0.68 | 0.74 | 0.75 |
| 10 | 63530322 | rs2675616 | 1.168 | 0.936 | 0.253 | 0.558 | 0.009 | -0.028 | -0.028 | 0.207 | 0.68 | 0.6 | 0.53 | 0.14 | 0.59 | 0.51 |
| 10 | 63541723 | rs751891 | -0.043 | 1.005 | 0.168 | 0.734 | -0.005 | -0.017 | -0.017 | 0 | 0.18 | 0.12 | 0.24 | 0.22 | 0.19 | 0.15 |
| 10 | 63547718 | rs2675626 | 0.57 | 0.895 | 0.072 | 0.402 | 0.013 | -0.021 | -0.021 | 0 | 0.69 | 0.6 | 0.53 | 0.18 | 0.6 | 0.52 |
| 10 | 63553849 | rs7904544 | -1.075 | 0.258 | -0.083 | 0.521 | -0.015 | 0.005 | 0.005 | 0.305 | 0.31 | 0.41 | 0.34 | 0.31 | 0.27 | 0.25 |
| 10 | 63557049 | rs920869 | -0.637 | 0.66 | 0.134 | 0.752 | -0.006 | -0.003 | -0.003 | 1.05 | 0.27 | 0.2 | 0.27 | 0.37 | 0.34 | 0.37 |
| 10 | 63565555 | rs2263272 | -0.803 | 0.22 | 0.415 | 0.707 | -0.028 | 0.031 | 0.031 | 2.119 | 0.22 | 0.34 | 0.17 | 0.28 | 0.15 | 0.21 |
| 10 | 63568544 | rs12257215 | -0.265 | 0.164 | 0.368 | 0.438 | -0.03 | -0.088 | -0.088 | 0 | 0.17 | 0.33 | 0.09 | 0.01 | 0.1 | 0.1 |
| 10 | 63576286 | rs2675638 | -0.967 | 0.018 | 0.49 | 0.419 | -0.012 | 0.004 | 0.004 | 0 | 0.36 | 0.68 | 0.38 | 0.69 | 0.4 | 0.48 |
| 10 | 63576349 | rs12247913 | 0.032 | 0.002 | 0.49 | 0.279 | 0.019 | 0.033 | 0.033 | 3.975 | 0.16 | 0.1 | 0.07 | 0.01 | 0.1 | 0.1 |
| 10 | 63583730 | rs12266520 | -0.569 | 0.303 | 0.427 | 0.232 | 0.004 | -0.06 | -0.06 | 0 | 0.26 | 0.23 | 0.19 | 0.01 | 0.29 | 0.21 |
| 10 | 63590396 | rs2675645 | 0.622 | 0.439 | 0.603 | 0.203 | -0.001 | -0.014 | -0.014 | 0 | 0.71 | 0.34 | 0.76 | 0.92 | 0.7 | 0.74 |
| 10 | 63590712 | rs2588944 | -0.123 | 0.21 | 0.411 | 0.447 | 0.002 | -0.003 | -0.003 | 0.001 | 0.16 | 0.16 | 0.15 | 0.29 | 0.13 | 0.21 |
| 10 | 63594669 | rs12248902 | 2.295 | 0.31 | 0.294 | 0.581 | -0.018 | 0.083 | 0.083 | 0.199 | 0.44 | 0.18 | 0.47 | 0.26 | 0.43 | 0.34 |
| 10 | 63595025 | rs16916781 | 1.466 | 0.367 | 0.303 | 0.591 | 0 | -0.016 | -0.016 | 0 | 0.05 | 0.07 | 0.02 | 0 | 0.05 | 0.02 |
| 10 | 63595191 | rs10994944 | -0.6 | 0.337 | 0.377 | 0.645 | 0.002 | 0.027 | 0.027 | 0 | 0.13 | 0.14 | 0.15 | 0.35 | 0.14 | 0.24 |
| 10 | 63596968 | rs17284864 | 0.292 | 0.353 | 0.468 | 0.526 | 0 | -0.028 | -0.028 | 0.619 | 0.2 | 0.17 | 0.16 | 0.01 | 0.24 | 0.19 |
| 10 | 63600048 | rs10994948 | 2.073 | -0.098 | 0.567 | 0.707 | -0.013 | 0.066 | 0.066 | 0.154 | 0.44 | 0.17 | 0.47 | 0.27 | 0.43 | 0.31 |
| 10 | 63604654 | rs1899888 | -1.276 | -0.132 | 0.741 | 0.692 | -0.002 | 0.001 | 0.001 | 0.337 | 0.44 | 0.4 | 0.39 | 0.43 | 0.46 | 0.45 |
| 10 | 63604701 | rs1444466 | -1.793 | -0.112 | 0.524 | 0.703 | -0.012 | 0.04 | 0.04 | 0.372 | 0.47 | 0.74 | 0.41 | 0.44 | 0.47 | 0.45 |
| 10 | 63606148 | rs2675651 | 0.139 | -0.082 | 0.699 | 0.894 | -0.005 | 0.055 | 0.055 | 0 | 0.92 | 0.9 | 0.88 | 0.71 | 0.9 | 0.77 |
| 10 | 63607143 | rs2588963 | -1.332 | 0.084 | 0.927 | 0.491 | 0.033 | 0.009 | 0.009 | 0.062 | 0.33 | 0.49 | 0.35 | 0.42 | 0.39 | 0.41 |
| 10 | 63609810 | rs2588966 | -1.279 | 0.104 | 0.276 | 0.578 | 0.016 | 0.014 | 0.014 | 0 | 0.36 | 0.41 | 0.62 | 0.5 | 0.41 | 0.45 |
| 10 | 63617300 | rs3851252 | 0.72 | -0.082 | -0.158 | 0.569 | -0.079 | -0.032 | -0.032 | 0 | 0.36 | 0.08 | 0.31 | 0.39 | 0.41 | 0.42 |
| 10 | 63619709 | rs2588935 | -1.211 | -0.037 | 0.272 | 0.419 | 0.011 | -0.032 | -0.032 | 1.791 | 0.14 | 0.56 | 0.11 | 0.08 | 0.12 | 0.07 |
| 10 | 63625277 | rs2675598 | -0.599 | -0.025 | 0.117 | 0.582 | 0.008 | -0.002 | -0.002 | 0 | 0.55 | 0.65 | 0.71 | 0.59 | 0.58 | 0.55 |
| 10 | 63629411 | rs1992626 | -0.6 | -0.021 | 0.118 | 0.583 | 0.008 | -0.002 | -0.002 | 0 | 0.55 | 0.65 | 0.71 | 0.59 | 0.58 | 0.55 |
| 10 | 63633467 | rs2675607 | -0.368 | 0.035 | 0.356 | 0.516 | -0.001 | -0.001 | -0.001 | 0.008 | 0.52 | 0.53 | 0.43 | 0.49 | 0.56 | 0.54 |
| 10 | 63636254 | rs10994959 | 0.438 | 0.17 | -0.435 | 0.569 | -0.098 | -0.049 | -0.049 | 0 | 0.37 | 0.08 | 0.3 | 0.41 | 0.43 | 0.41 |
| 10 | 63638766 | rs10509163 | -1.005 | 0.147 | 0.57 | 0.304 | -0.012 | -0.008 | -0.008 | 0 | 0.15 | 0.38 | 0.34 | 0.13 | 0.09 | 0.11 |
| 10 | 63645959 | rs1530439 | 0.741 | 0.18 | 0.146 | 0.394 | -0.012 | -0.042 | -0.042 | 0 | 0.3 | 0.16 | 0.22 | 0.44 | 0.28 | 0.4 |
| 10 | 63646610 | rs4948486 | 0.909 | 0.933 | 0.21 | -0.175 | -0.064 | 0.094 | 0.094 | 0 | 0.28 | 0.06 | 0.19 | 0.11 | 0.31 | 0.18 |

| | | | | | | | | | | | | | | | | |
|---|---|---|---|---|---|---|---|---|---|---|---|---|---|---|---|---|
| 10 | 63648027 | rs12784097 | -0.411 | 0.797 | 0.644 | -0.056 | 0.017 | 0.019 | 0.019 | 0 | 0.08 | 0.18 | 0.17 | 0.09 | 0.09 | 0.09 |
| 10 | 63649483 | rs7093142 | 0.43 | 0.774 | 1.112 | -0.117 | -0.006 | 0.016 | 0.016 | 0 | 0.35 | 0.32 | 0.26 | 0.11 | 0.38 | 0.24 |
| 10 | 63655873 | rs7091928 | -0.655 | 0.838 | 0.901 | -0.097 | 0.004 | 0.014 | 0.014 | 0.319 | 0.49 | 0.57 | 0.47 | 0.2 | 0.54 | 0.38 |
| 10 | 63660689 | rs6415872 | -0.303 | 0.796 | 0.658 | 0.019 | -0.005 | 0.015 | 0.015 | 0 | 0.45 | 0.6 | 0.5 | 0.27 | 0.5 | 0.36 |
| 10 | 63661340 | rs4509706 | -0.135 | 0.93 | 0.357 | 0.158 | -0.009 | -0.009 | -0.009 | 0 | 0.07 | 0.04 | 0.16 | 0.13 | 0.09 | 0.1 |
| 10 | 63669865 | rs4948487 | -0.224 | 0.836 | 0.326 | 0.13 | -0.009 | 0.017 | 0.017 | 0 | 0.43 | 0.6 | 0.49 | 0.27 | 0.49 | 0.34 |
| 10 | 63671348 | rs2393781 | 0.086 | 0.922 | -0.014 | 0.187 | -0.021 | 0.021 | 0.021 | 0 | 0.39 | 0.32 | 0.44 | 0.27 | 0.46 | 0.33 |
| 10 | 63674312 | rs10509164 | 0.134 | 0.622 | -0.501 | 0.464 | -0.032 | -0.034 | -0.034 | 0 | 0.06 | 0 | 0.16 | 0.12 | 0.08 | 0.1 |
| 10 | 63679281 | rs7901348 | -0.155 | 0.142 | -0.274 | 0.485 | 0.002 | 0.008 | 0.008 | 0 | 0.48 | 0.53 | 0.52 | 0.26 | 0.56 | 0.39 |
| 10 | 63685154 | rs4948488 | 0.226 | -0.715 | -0.016 | 1.068 | -0.041 | 0.048 | 0.048 | 0 | 0.75 | 0.7 | 0.6 | 0.59 | 0.82 | 0.75 |
| 10 | 63688672 | rs2893881 | 0.086 | -0.603 | 0.077 | 1.251 | -0.034 | 0.022 | 0.022 | 0 | 0.78 | 0.74 | 0.72 | 0.68 | 0.85 | 0.81 |
| 10 | 63688728 | rs17215180 | 1.047 | -0.651 | -0.257 | 0.596 | -0.03 | 0.05 | 0.05 | 0.069 | 0.37 | 0.32 | 0.32 | 0.12 | 0.48 | 0.25 |
| 10 | 63689077 | rs6479778 | -0.511 | -0.555 | -0.464 | 0.497 | 0.048 | -0.048 | -0.048 | 0 | 0.79 | 0.9 | 0.73 | 0.68 | 0.85 | 0.81 |
| 10 | 63691908 | rs10821932 | 0.113 | -0.338 | -0.144 | 0.184 | 0.002 | 0.014 | 0.014 | 0 | 0.26 | 0.21 | 0.15 | 0.08 | 0.2 | 0.34 |
| 10 | 63696889 | rs4948491 | 0.329 | -0.519 | 0.083 | 0.505 | -0.001 | 0 | 0 | 0.013 | 0.64 | 0.65 | 0.52 | 0.59 | 0.69 | 0.67 |
| 10 | 63699895 | rs7073837 | 0.064 | -0.288 | 0.17 | 0.393 | 0.03 | 0.037 | 0.037 | 0 | 0.55 | 0.74 | 0.49 | 0.64 | 0.61 | 0.37 |
| 10 | 63708007 | rs10994981 | 0.692 | -0.477 | 0.513 | 0.481 | -0.001 | -0.004 | -0.004 | 0 | 0.44 | 0.39 | 0.43 | 0.5 | 0.52 | 0.3 |
| 10 | 63721176 | rs7090445 | -0.328 | -0.492 | 0.412 | 0.447 | 0.006 | 0.011 | 0.011 | 0 | 0.65 | 0.81 | 0.53 | 0.64 | 0.67 | 0.48 |
| 10 | 63724773 | rs10821938 | 0.399 | -0.508 | 0.502 | 0.445 | -0.003 | 0.002 | 0.002 | 0 | 0.55 | 0.49 | 0.46 | 0.49 | 0.59 | 0.43 |
| 10 | 63727187 | rs4506592 | -0.136 | -0.409 | 0.435 | 0.489 | 0.003 | 0.01 | 0.01 | 0 | 0.63 | 0.7 | 0.51 | 0.65 | 0.65 | 0.48 |
| 10 | 63730012 | rs12249208 | -0.022 | -0.345 | 0.375 | 0.646 | 0.007 | 0.067 | 0.067 | 0 | 0.06 | 0.12 | 0.04 | 0.16 | 0.06 | 0.05 |
| 10 | 63752159 | rs7089424 | -0.089 | -0.253 | 0.016 | 0.469 | 0.007 | 0.009 | 0.009 | 0 | 0.35 | 0.24 | 0.48 | 0.36 | 0.33 | 0.51 |
| 10 | 63763369 | rs7922024 | 0.282 | 0.362 | 0.306 | 0.559 | 0.002 | -0.015 | -0.015 | 0 | 0.05 | 0.07 | 0.04 | 0 | 0.04 | 0.04 |
| 10 | 63769839 | rs7894504 | -0.567 | 0.137 | -0.216 | 0.573 | -0.03 | -0.001 | -0.001 | 0 | 0.55 | 0.77 | 0.71 | 0.62 | 0.49 | 0.69 |
| 10 | 63773039 | rs7087125 | -0.168 | 0.277 | -1.459 | 0.456 | -0.047 | -0.029 | -0.029 | 0 | 0.43 | 0.13 | 0.48 | 0.54 | 0.45 | 0.56 |
| 10 | 63786554 | rs11593907 | -0.301 | -0.08 | -1.589 | -0.073 | 0.005 | 0.118 | 0.118 | 1.192 | 0.3 | 0.09 | 0.17 | 0.15 | 0.24 | 0.21 |
| 10 | 63801030 | rs7902146 | 0.249 | -0.037 | -0.42 | 0.276 | -0.001 | -0.001 | -0.001 | 0 | 0.65 | 0.59 | 0.57 | 0.72 | 0.72 | 0.67 |
| 10 | 63803472 | rs12357548 | 0.566 | -0.429 | -0.588 | 0.459 | -0.032 | -0.014 | -0.014 | 0.431 | 0.32 | 0.2 | 0.4 | 0.37 | 0.5 | 0.39 |
| 10 | 63805617 | rs4948496 | -0.783 | -0.441 | -0.469 | 0.452 | -0.003 | -0.001 | -0.001 | 0 | 0.67 | 0.71 | 0.59 | 0.63 | 0.5 | 0.61 |
| 10 | 63806809 | rs6479782 | -0.621 | -0.437 | -0.49 | 0.498 | -0.004 | -0.006 | -0.006 | 0.479 | 0.66 | 0.7 | 0.39 | 0.5 | 0.46 | 0.48 |
| 10 | 63808666 | rs7902719 | -0.663 | -0.433 | -0.105 | 0.472 | 0.01 | 0.004 | 0.004 | 0 | 0.66 | 0.62 | 0.56 | 0.63 | 0.47 | 0.59 |
| 10 | 63809624 | rs10821948 | 0.147 | -1.09 | 0.063 | 0.13 | 0.038 | 0.005 | 0.005 | 0 | 0.52 | 0.31 | 0.31 | 0.5 | 0.38 | 0.43 |
| 10 | 63811678 | rs10821949 | 0.81 | -0.912 | 0.098 | 0.184 | -0.04 | -0.017 | -0.017 | 0 | 0.32 | 0.21 | 0.4 | 0.36 | 0.5 | 0.39 |
| 10 | 63814914 | rs10821951 | 1.207 | -0.907 | 0.474 | 0.427 | 0.045 | 0.197 | 0.197 | 1.171 | 0.91 | 0.46 | 0.75 | 0.83 | 0.95 | 0.85 |
| 10 | 63830189 | rs16916945 | 1.138 | -1.124 | -0.498 | 0.312 | -0.025 | -0.026 | -0.026 | 0 | 0.18 | 0.01 | 0.11 | 0.22 | 0.12 | 0.1 |
| 10 | 63831638 | rs7078374 | -0.344 | -1.258 | -0.222 | -0.128 | -0.022 | -0.082 | -0.082 | 0 | 0.83 | 0.74 | 0.92 | 1 | 0.86 | 0.9 |
| 10 | 63835692 | rs3740352 | 1.224 | -0.561 | -1.146 | 0.012 | -0.031 | -0.022 | -0.022 | 0 | 0.28 | 0.15 | 0.35 | 0.37 | 0.38 | 0.38 |
| 10 | 63835891 | rs7101171 | 1.017 | -0.572 | -0.995 | -0.153 | 0.026 | -0.055 | -0.055 | 0 | 0.1 | 0.23 | 0.08 | 0 | 0.13 | 0.06 |
| 10 | 63837006 | rs2393730 | 1.319 | -0.519 | -1.31 | -0.037 | -0.044 | -0.026 | -0.026 | 0 | 0.28 | 0.14 | 0.35 | 0.36 | 0.39 | 0.36 |
| 10 | 63840062 | rs4948503 | -0.337 | -0.59 | -1.364 | 0.293 | 0.039 | -0.052 | -0.052 | 0.955 | 0.6 | 0.84 | 0.73 | 0.52 | 0.69 | 0.69 |
| 10 | 63843112 | rs2278305 | -0.605 | -0.288 | -0.952 | 0.558 | 0.01 | -0.067 | -0.067 | 0 | 0.8 | 0.93 | 0.84 | 0.73 | 0.81 | 0.81 |
| 10 | 63848656 | rs9414761 | 1.072 | 0.243 | -0.213 | 0.816 | 0.006 | 0.086 | 0.086 | 0.603 | 0.8 | 0.59 | 0.47 | 0.53 | 0.74 | 0.63 |
| 10 | 63849790 | rs7913907 | 0.797 | 0.091 | -0.584 | 1.042 | 0 | 0.041 | 0.041 | 0.634 | 0.88 | 0.79 | 0.55 | 0.54 | 0.86 | 0.7 |
| 10 | 63861600 | rs1122157 | 0.052 | 0.61 | -0.512 | 0.083 | 0.009 | -0.09 | -0.09 | 0 | 0.26 | 0.27 | 0.16 | 0.05 | 0.28 | 0.18 |
| 10 | 63870456 | rs10821958 | 0.027 | 0.735 | -0.487 | -0.198 | 0.003 | -0.034 | -0.034 | 0 | 0.15 | 0.17 | 0.12 | 0 | 0.19 | 0.1 |
| 10 | 63875960 | rs7923071 | -0.176 | 1.044 | -0.313 | 0.575 | 0.007 | 0.02 | 0.02 | 0 | 0.71 | 0.7 | 0.79 | 0.65 | 0.66 | 0.72 |
| 10 | 63883113 | rs7915967 | 0.806 | 0.779 | -0.817 | -0.38 | -0.004 | -0.043 | -0.043 | 0 | 0.23 | 0.14 | 0.17 | 0.52 | 0.28 | 0.28 |
| 10 | 63894285 | rs10761610 | 0.172 | 0.623 | -1.042 | -1.1 | -0.007 | 0.048 | 0.048 | 0.102 | 0.67 | 0.76 | 0.48 | 0.85 | 0.65 | 0.58 |
| 10 | 63897452 | rs4948508 | -0.548 | 0.438 | -0.916 | -1.557 | 0.011 | 0.109 | 0.109 | 0 | 0.54 | 0.76 | 0.68 | 0.73 | 0.59 | 0.7 |
| 10 | 63905893 | rs10995054 | 0.484 | 0.43 | -0.615 | -1.683 | 0.005 | -0.096 | -0.096 | 0 | 0.28 | 0.3 | 0.18 | 0.03 | 0.3 | 0.19 |
| 10 | 63907602 | rs4256883 | 1.566 | 0.416 | -0.528 | -1.633 | -0.005 | -0.006 | -0.006 | 0.002 | 0.21 | 0.17 | 0.4 | 0.64 | 0.28 | 0.29 |
| 10 | 63912835 | rs10821962 | -0.646 | -0.191 | 0.274 | -2.037 | 0.007 | 0.053 | 0.053 | 0 | 0.34 | 0.74 | 0.29 | 0.08 | 0.31 | 0.4 |
| 10 | 63926234 | rs10995070 | 0.724 | 0.009 | 0.327 | -2.006 | -0.005 | 0 | 0 | 0 | 0.08 | 0.05 | 0.07 | 0.01 | 0.14 | 0.07 |
| 10 | 63935516 | rs6415873 | -0.326 | -0.137 | 0.509 | -2.173 | -0.004 | -0.15 | -0.15 | 0 | 0.21 | 0.41 | 0.15 | 0.01 | 0.23 | 0.13 |
| 10 | 63935987 | rs2116583 | -0.473 | -0.13 | 0.343 | -2.17 | -0.009 | -0.135 | -0.135 | 0 | 0.22 | 0.52 | 0.17 | 0.01 | 0.23 | 0.13 |
| 10 | 63941939 | rs10821967 | 1.465 | -0.555 | -0.258 | -2.34 | 0.063 | 0.271 | 0.271 | 0 | 0.5 | 0.2 | 0.64 | 0.89 | 0.62 | 0.51 |
| 10 | 63943292 | rs12767995 | 0.054 | -0.337 | 0.105 | -2.129 | 0.005 | -0.007 | -0.007 | 0 | 0.18 | 0.12 | 0.11 | 0 | 0.18 | 0.08 |
| 10 | 63944188 | rs7903510 | -0.871 | -0.341 | -0.047 | -2.146 | -0.012 | -0.083 | -0.083 | 0.345 | 0.3 | 0.56 | 0.2 | 0.1 | 0.25 | 0.36 |
| 10 | 63944984 | rs7908029 | -0.955 | -0.342 | -0.005 | -2.147 | -0.015 | -0.117 | -0.117 | 0.025 | 0.29 | 0.48 | 0.2 | 0.1 | 0.25 | 0.36 |
| 10 | 63952639 | rs1432411 | -0.96 | -0.347 | -0.079 | -2.149 | -0.012 | -0.074 | -0.074 | 0 | 0.31 | 0.56 | 0.2 | 0.1 | 0.26 | 0.38 |
| 10 | 63954188 | rs10995080 | -0.824 | -0.341 | -0.098 | -2.146 | -0.017 | -0.111 | -0.111 | 0.175 | 0.3 | 0.48 | 0.2 | 0.1 | 0.26 | 0.37 |
| 10 | 63955526 | rs12253181 | -0.734 | -0.333 | -0.447 | -2.154 | -0.03 | -0.129 | -0.129 | 0 | 0.24 | 0.56 | 0.15 | 0 | 0.18 | 0.09 |
| 10 | 63956698 | rs7088302 | -0.602 | -0.094 | -0.52 | -1.878 | -0.006 | -0.038 | -0.038 | 0.835 | 0.28 | 0.65 | 0.2 | 0.1 | 0.22 | 0.36 |
| 10 | 63958112 | rs3125734 | 0.658 | 0.055 | -0.491 | -1.788 | 0.02 | 0.059 | 0.059 | 0 | 0.57 | 0.31 | 0.7 | 0.9 | 0.64 | 0.51 |
| 10 | 63962026 | rs7090884 | -0.216 | 0.285 | -0.434 | -1.366 | -0.001 | 0.013 | 0.013 | 0 | 0.46 | 0.38 | 0.47 | 0.58 | 0.52 | 0.52 |
| 10 | 63966387 | rs10995086 | -0.605 | -0.478 | 0.242 | -2.319 | -0.013 | -0.063 | -0.063 | 3.189 | 0.16 | 0.21 | 0.1 | 0.01 | 0.16 | 0.08 |
| 10 | 63974229 | rs10995090 | 0.058 | -0.492 | 0.186 | -2.324 | -0.002 | -0.025 | -0.025 | 0 | 0.12 | 0.09 | 0.09 | 0 | 0.12 | 0.06 |
| 10 | 63974255 | rs10733769 | 1.321 | -0.588 | 0.446 | -2.713 | -0.001 | -0.062 | -0.062 | 0.767 | 0.58 | 0.46 | 0.7 | 0.89 | 0.59 | 0.49 |
| 10 | 63982984 | rs10740069 | 0.255 | -0.529 | 0.479 | -2.651 | -0.002 | -0.01 | -0.01 | 0 | 0.75 | 0.75 | 0.81 | 0.9 | 0.75 | 0.57 |

| | | | | | | | | | | | | | | | | |
|---|---|---|---|---|---|---|---|---|---|---|---|---|---|---|---|---|
| 10 | 63984957 | rs2195851 | -0.327 | -0.549 | 0.425 | -2.673 | -0.005 | 0.04 | 0.04 | 0.959 | 0.85 | 0.93 | 0.9 | 1 | 0.86 | 0.84 |
| 10 | 63995983 | rs3852448 | -0.218 | -0.538 | 0.429 | -2.663 | -0.005 | 0.042 | 0.042 | 1.003 | 0.85 | 0.93 | 0.9 | 0.99 | 0.86 | 0.84 |
| 10 | 64005711 | rs3815999 | 0.14 | -0.679 | 0.801 | -2.595 | 0.006 | -0.072 | -0.072 | 0 | 0.74 | 0.63 | 0.8 | 0.89 | 0.75 | 0.57 |
| 10 | 64007604 | rs10995093 | -0.238 | -0.764 | 0.838 | -2.662 | -0.024 | -0.097 | -0.097 | 0.256 | 0.17 | 0.29 | 0.11 | 0.01 | 0.16 | 0.07 |
| 10 | 64014095 | rs2116581 | 0.004 | -0.665 | 0.851 | -2.599 | 0.002 | -0.031 | -0.031 | 0 | 0.75 | 0.72 | 0.81 | 0.89 | 0.75 | 0.57 |
| 10 | 64024648 | rs7923556 | 1.311 | -0.689 | 0.655 | -2.634 | 0.014 | 0.145 | 0.145 | 0.617 | 0.55 | 0.24 | 0.68 | 0.88 | 0.59 | 0.49 |
| 10 | 64026725 | rs7076551 | 0.317 | -0.278 | 0.729 | -2.459 | 0.004 | -0.074 | -0.074 | 0 | 0.73 | 0.61 | 0.79 | 0.89 | 0.75 | 0.57 |
| 10 | 64029895 | rs4979766 | 0.076 | -0.229 | 0.005 | -2.438 | 0.004 | -0.049 | -0.049 | 0 | 0.74 | 0.69 | 0.8 | 0.89 | 0.75 | 0.57 |
| 10 | 64056068 | rs1835862 | 0.788 | -0.098 | -1.009 | -2.358 | -0.006 | 0.109 | 0.109 | 0.293 | 0.59 | 0.25 | 0.68 | 0.89 | 0.59 | 0.49 |
| 10 | 64059261 | rs6479805 | 0.605 | 0.193 | -1.12 | -2.196 | 0.002 | 0.028 | 0.028 | 0.673 | 0.62 | 0.27 | 0.7 | 0.92 | 0.66 | 0.47 |
| 10 | 64063345 | rs17291653 | -0.3 | 0.81 | -1.325 | -1.37 | -0.002 | -0.017 | -0.017 | 0.19 | 0.12 | 0.08 | 0.07 | 0 | 0.13 | 0.15 |
| 10 | 64068343 | rs3998448 | 0.159 | 0.903 | -1.752 | -0.913 | -0.108 | -0.052 | -0.052 | 0.16 | 0.25 | 0.02 | 0.52 | 0.69 | 0.39 | 0.3 |
| 10 | 64069979 | rs3852421 | 0.465 | -0.194 | -0.862 | -1.445 | -0.016 | -0.083 | -0.083 | 0 | 0.76 | 0.69 | 0.87 | 0.92 | 0.85 | 0.88 |
| 10 | 64070219 | rs3864807 | -0.146 | -0.692 | -0.852 | -1.368 | -0.012 | -0.034 | -0.034 | 0 | 0.39 | 0.47 | 0.33 | 0.13 | 0.27 | 0.3 |
| 10 | 64074412 | rs10995114 | -0.816 | -0.894 | -0.156 | -1.117 | -0.014 | -0.032 | -0.032 | 1.561 | 0.06 | 0.25 | 0.05 | 0 | 0.03 | 0.01 |
| 10 | 64075136 | rs10821978 | -0.113 | -0.822 | 0.124 | -1.204 | 0.007 | -0.013 | -0.013 | 0 | 0.93 | 0.88 | 0.96 | 0.94 | 0.95 | 0.9 |
| 10 | 64088654 | rs7919462 | 0.835 | -1.193 | -0.381 | -1.583 | -0.042 | 0.095 | 0.095 | 0 | 0.4 | 0.11 | 0.58 | 0.78 | 0.62 | 0.67 |
| 10 | 64092543 | rs12249008 | -0.134 | -1.7 | 0.422 | -1.811 | 0.01 | 0.01 | 0.01 | 0 | 0.44 | 0.37 | 0.36 | 0.13 | 0.31 | 0.2 |
| 10 | 64095849 | rs4093501 | 0.664 | -1.67 | 0.358 | -1.793 | -0.025 | -0.023 | -0.023 | 0 | 0.07 | 0 | 0.03 | 0.09 | 0.05 | 0.11 |
| 10 | 64117558 | rs3999187 | 1.125 | -1.69 | 0.49 | -1.797 | -0.063 | 0.089 | 0.089 | 0 | 0.4 | 0.08 | 0.58 | 0.78 | 0.62 | 0.67 |
| 10 | 64128133 | rs10995132 | -0.254 | -1.703 | 0.769 | -1.799 | -0.008 | -0.031 | -0.031 | 0 | 0.44 | 0.45 | 0.35 | 0.13 | 0.3 | 0.2 |
| 10 | 64128913 | rs7080900 | -0.217 | -1.708 | 0.78 | -1.801 | 0 | -0.013 | -0.013 | 0.388 | 0.44 | 0.42 | 0.36 | 0.13 | 0.3 | 0.2 |
| 10 | 64128954 | rs7096931 | 0.902 | -2.077 | 0.729 | -2.028 | -0.005 | -0.008 | -0.008 | 0 | 0.23 | 0.24 | 0.26 | 0.13 | 0.17 | 0.16 |
| 10 | 64136875 | rs1873687 | 0.668 | -1.678 | 0.645 | -1.776 | 0.02 | 0.092 | 0.092 | 0 | 0.46 | 0.24 | 0.59 | 0.78 | 0.63 | 0.69 |
| 10 | 64145543 | rs10761628 | -0.743 | -1.652 | 0.76 | -1.779 | 0.002 | 0 | 0 | 0 | 0.51 | 0.49 | 0.4 | 0.23 | 0.37 | 0.31 |
| 10 | 64149481 | rs3852425 | 1.291 | -1.732 | 0.417 | -1.398 | -0.067 | 0.006 | 0.006 | 0 | 0.38 | 0.1 | 0.56 | 0.71 | 0.59 | 0.48 |
| 10 | 64156110 | rs10740076 | 0.332 | -1.799 | 0.49 | -1.396 | 0.066 | 0.096 | 0.096 | 0 | 0.51 | 0.27 | 0.61 | 0.8 | 0.67 | 0.61 |
| 10 | 64159333 | rs3758490 | 1.408 | -1.622 | 0.272 | -1.244 | -0.103 | -0.04 | -0.04 | 0 | 0.37 | 0.06 | 0.56 | 0.69 | 0.6 | 0.46 |
| 10 | 64165285 | rs10995147 | 0.759 | -1.474 | 0.551 | -1.203 | 0.002 | 0.047 | 0.047 | 0 | 0.44 | 0.31 | 0.6 | 0.77 | 0.63 | 0.67 |
| 10 | 64166207 | rs4746718 | -1.069 | -1.475 | 0.558 | -1.246 | -0.001 | -0.015 | -0.015 | 0 | 0.13 | 0.24 | 0.05 | 0.09 | 0.07 | 0.13 |
| 10 | 64169941 | rs12779966 | 0.445 | -1.376 | 0.362 | -1.058 | -0.001 | -0.003 | -0.003 | 0 | 0.48 | 0.45 | 0.61 | 0.78 | 0.63 | 0.55 |
| 10 | 64176173 | rs4745969 | 0.316 | -1.262 | 0.476 | -1.015 | -0.002 | -0.019 | -0.019 | 0 | 0.49 | 0.49 | 0.61 | 0.85 | 0.65 | 0.72 |
| 10 | 64182727 | rs4636539 | 0.511 | -1.141 | 0.612 | -1.126 | 0.01 | 0.042 | 0.042 | 0 | 0.47 | 0.62 | 0.61 | 0.78 | 0.63 | 0.54 |
| 10 | 64185908 | rs10995154 | 0.832 | -1.126 | 0.557 | -1.12 | 0.001 | 0.029 | 0.029 | 0 | 0.44 | 0.29 | 0.59 | 0.78 | 0.63 | 0.54 |
| 10 | 64191520 | rs7907542 | 0.359 | -0.972 | 0.311 | -1.118 | 0.003 | -0.004 | -0.004 | 0 | 0.49 | 0.51 | 0.62 | 0.85 | 0.65 | 0.72 |
| 10 | 64195778 | rs17312431 | -0.001 | -1.001 | -0.099 | -1.167 | 0.006 | 0.003 | 0.003 | 0.766 | 0.19 | 0.13 | 0.09 | 0 | 0.14 | 0.06 |
| 10 | 64195873 | rs10128203 | 0.014 | -0.984 | 0.067 | -0.984 | -0.012 | -0.011 | -0.011 | 0.012 | 0.64 | 0.53 | 0.68 | 0.77 | 0.76 | 0.63 |
| 10 | 64199222 | rs7904487 | 0.792 | -1.105 | 0.026 | -0.918 | 0.011 | -0.01 | -0.01 | 0 | 0.75 | 0.62 | 0.7 | 0.77 | 0.77 | 0.79 |
| 10 | 64200060 | rs6479810 | -0.039 | -1.454 | -0.296 | -0.602 | -0.012 | -0.001 | -0.001 | 0 | 0.66 | 0.45 | 0.7 | 0.69 | 0.8 | 0.7 |
| 10 | 64200492 | rs17312550 | -0.362 | -1.207 | -0.898 | -0.838 | -0.016 | 0.11 | 0.11 | 2.226 | 0.22 | 0.05 | 0.12 | 0 | 0.19 | 0.09 |
| 10 | 64205327 | rs7074870 | -0.061 | -1.193 | -0.391 | -0.632 | -0.022 | -0.003 | -0.003 | 1.196 | 0.65 | 0.54 | 0.66 | 0.66 | 0.75 | 0.7 |
| 10 | 64206712 | rs17221319 | 1.051 | -0.667 | -1.232 | -0.412 | -0.109 | -0.116 | -0.116 | 0 | 0.34 | 0.07 | 0.48 | 0.57 | 0.49 | 0.46 |
| 10 | 64208221 | rs10822007 | 0.975 | -1.105 | -1.203 | 0.028 | -0.062 | -0.061 | -0.061 | 0 | 0.44 | 0.05 | 0.57 | 0.51 | 0.58 | 0.54 |
| 10 | 64208433 | rs3847337 | 0.123 | -1.593 | -0.635 | -0.143 | 0.004 | 0.005 | 0.005 | 0.001 | 0.28 | 0.26 | 0.3 | 0.26 | 0.25 | 0.3 |
| 10 | 64211640 | rs11598789 | -0.107 | -1.548 | -0.623 | -0.174 | 0.003 | 0.002 | 0.002 | 0.742 | 0.09 | 0.07 | 0.03 | 0 | 0.1 | 0.03 |
| 10 | 64219000 | rs12220488 | 1.227 | -1.178 | -0.88 | 0.066 | -0.088 | -0.103 | -0.103 | 0.088 | 0.43 | 0.04 | 0.55 | 0.54 | 0.54 | 0.53 |
| 10 | 64220494 | rs2138555 | -1.23 | -1.242 | -0.107 | -0.028 | 0.047 | 0.038 | 0.038 | 0 | 0.49 | 0.76 | 0.43 | 0.5 | 0.42 | 0.5 |
| 10 | 64220831 | rs11818767 | 2.534 | -1.183 | -0.319 | 0.101 | 0.011 | -0.004 | -0.004 | 0 | 0.06 | 0.01 | 0.04 | 0.2 | 0.04 | 0.04 |
| 10 | 64224100 | rs6479815 | 0.394 | -1.195 | 0.061 | -0.139 | -0.001 | -0.008 | -0.008 | 0.497 | 0.26 | 0.27 | 0.3 | 0.31 | 0.3 | 0.35 |
| 10 | 64230159 | rs7923418 | -0.988 | -1.238 | 0.265 | -0.091 | -0.043 | 0.018 | 0.018 | 0.618 | 0.21 | 0.38 | 0.12 | 0.18 | 0.12 | 0.13 |
| 10 | 64231895 | rs1955332 | -0.595 | -1.085 | 0.39 | 0.056 | -0.008 | -0.011 | -0.011 | 0 | 0.42 | 0.57 | 0.4 | 0.31 | 0.42 | 0.42 |
| 10 | 64235506 | rs753518 | 0.188 | -1.128 | 0.474 | 0.026 | -0.001 | -0.001 | -0.001 | 2.047 | 0.32 | 0.43 | 0.36 | 0.31 | 0.37 | 0.39 |
| 10 | 64238165 | rs9804184 | 0.553 | -1.347 | 0.147 | -0.129 | 0.01 | -0.011 | -0.011 | 0 | 0.83 | 0.88 | 0.94 | 1 | 0.92 | 0.94 |
| 10 | 64243828 | rs4282885 | 0.752 | -0.935 | 0.035 | 0.177 | 0.006 | 0.001 | 0.001 | 0.358 | 0.46 | 0.28 | 0.52 | 0.51 | 0.49 | 0.51 |
| 10 | 64252883 | rs6479821 | -0.124 | -0.937 | 0.146 | 0.328 | 0.016 | 0.171 | 0.171 | 0 | 0.06 | 0.16 | 0.06 | 0.18 | 0.05 | 0.05 |
| 10 | 64257828 | rs10509168 | 0.98 | -0.941 | -0.561 | 0.166 | -0.035 | -0.052 | -0.052 | 0 | 0.42 | 0.15 | 0.53 | 0.51 | 0.51 | 0.5 |
| 10 | 64260847 | rs12245332 | -0.089 | -1.016 | -0.264 | 0.17 | -0.004 | -0.008 | -0.008 | 0 | 0.4 | 0.45 | 0.35 | 0.3 | 0.37 | 0.42 |
| 10 | 64261198 | rs16917302 | -1.245 | -0.895 | 0.383 | 0.476 | -0.069 | 0.063 | 0.063 | 0.429 | 0.18 | 0.39 | 0.13 | 0.19 | 0.12 | 0.08 |
| 10 | 64270701 | rs16917312 | 0.26 | 0.206 | -0.575 | 0.283 | 0.007 | -0.008 | -0.008 | 0.585 | 0.53 | 0.29 | 0.58 | 0.5 | 0.55 | 0.49 |
| 10 | 64271051 | rs10995184 | 1.179 | 0.136 | -0.895 | -0.1 | -0.045 | -0.066 | -0.066 | 0 | 0.37 | 0.13 | 0.47 | 0.48 | 0.41 | 0.42 |
| 10 | 64275082 | rs12255773 | 0.313 | -0.281 | -0.149 | -0.268 | -0.025 | -0.072 | -0.072 | 0.026 | 0.12 | 0.32 | 0.05 | 0 | 0.04 | 0.04 |
| 10 | 64278181 | rs10995189 | 0.087 | -0.241 | -0.306 | -0.245 | -0.001 | -0.023 | -0.023 | 0 | 0.16 | 0.17 | 0.11 | 0.02 | 0.15 | 0.08 |
| 10 | 64278634 | rs10509170 | 1.444 | -0.225 | -0.394 | -0.165 | -0.054 | -0.078 | -0.078 | 0 | 0.37 | 0.11 | 0.47 | 0.48 | 0.4 | 0.42 |
| 10 | 64286975 | rs10995192 | -0.627 | -0.234 | 0.091 | -0.443 | 0.002 | 0.008 | 0.008 | 0 | 0.43 | 0.6 | 0.37 | 0.27 | 0.38 | 0.46 |
| 10 | 64288592 | rs2393892 | 0.212 | -0.073 | -0.092 | -0.363 | -0.003 | 0.016 | 0.016 | 0 | 0.26 | 0.2 | 0.26 | 0.15 | 0.26 | 0.29 |
| 10 | 64301817 | rs16917398 | -0.52 | 0.375 | 0.247 | -0.139 | -0.007 | -0.03 | -0.03 | 0.208 | 0.13 | 0.21 | 0.06 | 0.05 | 0.11 | 0.04 |
| 10 | 64301941 | rs12249410 | -0.265 | 0.333 | 0.329 | -0.056 | -0.008 | -0.005 | -0.005 | 0 | 0.15 | 0.2 | 0.08 | 0.11 | 0.1 | 0.17 |
| 10 | 64303351 | rs10761643 | -0.557 | -0.004 | 0.172 | -0.22 | -0.004 | -0.015 | -0.015 | 0 | 0.39 | 0.55 | 0.27 | 0.27 | 0.3 | 0.49 |
| 10 | 64303915 | rs10822019 | -0.877 | 0.04 | 0.117 | 0.177 | 0.009 | -0.009 | -0.009 | 0.262 | 0.39 | 0.64 | 0.34 | 0.48 | 0.31 | 0.31 |

| | | | | | | | | | | | | | | | | |
|---|---|---|---|---|---|---|---|---|---|---|---|---|---|---|---|---|
| 10 | 64304083 | rs10995205 | -0.003 | 0.304 | 0.04 | -0.033 | 0.012 | -0.058 | -0.058 | 0 | 0.27 | 0.36 | 0.13 | 0.1 | 0.25 | 0.29 |
| 10 | 64307317 | rs2138564 | -1.565 | 0.416 | -0.048 | -0.092 | -0.082 | -0.045 | -0.045 | 0.023 | 0.61 | 0.9 | 0.44 | 0.6 | 0.5 | 0.62 |
| 10 | 64307474 | rs2138563 | -0.458 | 0.284 | 0.321 | -0.115 | 0.002 | -0.067 | -0.067 | 1.309 | 0.35 | 0.5 | 0.18 | 0.14 | 0.3 | 0.31 |
| 10 | 64311230 | rs10995208 | 0.891 | 0.265 | 0.53 | 0.142 | -0.005 | 0 | 0 | 0.481 | 0.21 | 0.23 | 0.17 | 0.25 | 0.12 | 0.26 |
| 10 | 64312533 | rs7921139 | -0.595 | 0.328 | 0.509 | 0.112 | 0.004 | -0.049 | -0.049 | 2.52 | 0.35 | 0.44 | 0.18 | 0.14 | 0.31 | 0.31 |
| 10 | 64313875 | rs4745849 | 1.132 | 0.365 | 0.617 | 0.158 | 0.011 | -0.004 | -0.004 | 0 | 0.19 | 0.16 | 0.16 | 0.23 | 0.12 | 0.26 |
| 10 | 64315820 | rs7903566 | -0.133 | 0.384 | 0.507 | 0.168 | -0.042 | 0.077 | 0.077 | 0.139 | 0.27 | 0.46 | 0.27 | 0.46 | 0.2 | 0.31 |
| 10 | 64317700 | rs6479827 | -0.048 | 0.44 | 0.458 | 0.151 | 0.006 | -0.026 | -0.026 | 0 | 0.27 | 0.31 | 0.14 | 0.12 | 0.26 | 0.29 |
| 10 | 64321478 | rs4746447 | -0.02 | 0.355 | 0.889 | 0.137 | -0.03 | 0.006 | 0.006 | 1.542 | 0.21 | 0.38 | 0.19 | 0.26 | 0.13 | 0.26 |
| 10 | 64329247 | rs4508090 | 0.174 | 0.38 | 0.411 | 0.128 | -0.04 | 0.006 | 0.006 | 0.182 | 0.2 | 0.38 | 0.19 | 0.23 | 0.12 | 0.26 |
| 10 | 64344371 | rs10822027 | -0.818 | 0.995 | -0.099 | -0.133 | 0.003 | -0.002 | -0.002 | 0 | 0.41 | 0.54 | 0.37 | 0.42 | 0.43 | 0.37 |
| 10 | 64350533 | rs2176289 | 0.541 | 0.91 | -0.445 | -0.711 | 0.009 | 0.013 | 0.013 | 0 | 0.49 | 0.43 | 0.38 | 0.55 | 0.52 | 0.41 |
| 10 | 64355339 | rs2138566 | 0.352 | 0.823 | -0.469 | -0.763 | 0.01 | 0.005 | 0.005 | 0 | 0.54 | 0.43 | 0.4 | 0.55 | 0.58 | 0.47 |
| 10 | 64356993 | rs1949359 | 1.132 | 0.885 | -0.477 | -0.707 | -0.003 | 0 | 0 | 0 | 0.07 | 0.05 | 0.1 | 0.19 | 0.12 | 0.06 |
| 10 | 64358445 | rs2944537 | -0.8 | 0.812 | -0.433 | -0.785 | -0.028 | -0.055 | -0.055 | 0 | 0.83 | 0.73 | 0.69 | 0.96 | 0.89 | 0.78 |
| 10 | 64360090 | rs17316874 | 0.465 | 1.327 | -0.433 | -0.669 | 0.001 | -0.009 | -0.009 | 0 | 0.11 | 0.11 | 0.1 | 0.01 | 0.15 | 0.06 |
| 10 | 64361700 | rs10822031 | -0.067 | 1.143 | -0.44 | -0.59 | -0.002 | 0.008 | 0.008 | 0.286 | 0.28 | 0.3 | 0.29 | 0.41 | 0.3 | 0.31 |
| 10 | 64362223 | rs1914191 | 0.997 | 1.176 | -0.753 | -0.467 | -0.031 | -0.043 | -0.043 | 0.274 | 0.07 | 0 | 0.09 | 0.19 | 0.12 | 0.06 |
| 10 | 64369001 | rs2393902 | -0.336 | 1.095 | -0.551 | -0.626 | 0.009 | -0.023 | -0.023 | 0 | 0.32 | 0.39 | 0.29 | 0.23 | 0.34 | 0.31 |
| 10 | 64369924 | rs2944540 | -0.903 | 0.896 | -0.711 | -0.696 | 0.006 | 0.077 | 0.077 | 0 | 0.49 | 0.65 | 0.6 | 0.26 | 0.45 | 0.53 |
| 10 | 64374360 | rs10822036 | 0.746 | 0.788 | -1.1 | -0.555 | -0.005 | -0.003 | -0.003 | 0 | 0.21 | 0.24 | 0.19 | 0.22 | 0.18 | 0.25 |
| 10 | 64374948 | rs4745856 | -1.351 | 1.179 | -1.618 | -0.316 | -0.081 | -0.128 | -0.128 | 0 | 0.66 | 0.96 | 0.74 | 0.45 | 0.64 | 0.65 |
| 10 | 64379326 | rs2393904 | -0.894 | 1.596 | -0.815 | -0.523 | 0.061 | -0.042 | -0.042 | 0 | 0.21 | 0.35 | 0.23 | 0.15 | 0.3 | 0.13 |
| 10 | 64380336 | rs2393903 | -1.263 | 0.967 | -1.927 | -0.444 | -0.053 | -0.047 | -0.047 | 0 | 0.52 | 0.91 | 0.72 | 0.47 | 0.58 | 0.58 |
| 10 | 64387934 | rs2893906 | -1.224 | -0.109 | -1.971 | -0.609 | 0.019 | 0.018 | 0.018 | 11.201 | 0.73 | 0.97 | 0.85 | 0.81 | 0.78 | 0.88 |
| 10 | 64388365 | rs10995239 | 1.127 | 0.246 | -1.98 | -0.58 | -0.104 | -0.108 | -0.108 | 0 | 0.27 | 0.09 | 0.2 | 0.53 | 0.35 | 0.36 |
| 10 | 64396790 | rs12098699 | -0.106 | -0.257 | -1.232 | -1.321 | -0.02 | -0.079 | -0.079 | 0.006 | 0.24 | 0.55 | 0.23 | 0.07 | 0.21 | 0.25 |
| 10 | 64398466 | rs10995251 | 1.318 | -0.07 | -1.335 | -1.148 | -0.108 | -0.066 | -0.066 | 0 | 0.28 | 0.09 | 0.2 | 0.6 | 0.36 | 0.36 |
| 10 | 64401650 | rs10995253 | 0.872 | -0.209 | -1.19 | -1.119 | -0.001 | -0.002 | -0.002 | 0 | 0.13 | 0.13 | 0.38 | 0.23 | 0.14 | 0.26 |
| 10 | 64401836 | rs10995254 | 0.506 | -0.242 | -0.924 | -1.13 | -0.002 | 0.036 | 0.036 | 0 | 0.14 | 0.25 | 0.44 | 0.26 | 0.14 | 0.27 |
| 10 | 64406802 | rs4537656 | 0.423 | -0.321 | -0.438 | -1.154 | -0.003 | 0.041 | 0.041 | 0 | 0.14 | 0.27 | 0.38 | 0.23 | 0.14 | 0.26 |
| 10 | 64408367 | rs7068361 | -0.011 | -0.431 | -0.582 | -1.282 | 0.018 | 0.035 | 0.035 | 3.454 | 0.5 | 0.56 | 0.43 | 0.67 | 0.57 | 0.6 |
| 10 | 64415184 | rs7076156 | -1.352 | -0.322 | -1.314 | -1.255 | -0.006 | 0.163 | 0.163 | 9.674 | 0.67 | 0.98 | 0.83 | 0.9 | 0.73 | 0.87 |
| 10 | 64424284 | rs10822047 | -1.352 | -0.318 | -1.37 | -1.253 | -0.006 | 0.163 | 0.163 | 9.674 | 0.67 | 0.98 | 0.83 | 0.89 | 0.73 | 0.87 |
| 10 | 64427649 | rs7075349 | 0.641 | -0.427 | -1.18 | -1.278 | 0.003 | 0.005 | 0.005 | 0 | 0.65 | 0.37 | 0.43 | 0.71 | 0.67 | 0.49 |
| 10 | 64431206 | rs12413946 | 0.517 | -0.445 | -1.193 | -1.327 | 0.007 | -0.009 | -0.009 | 0.5 | 0.05 | 0.12 | 0.09 | 0.04 | 0.03 | 0.03 |
| 10 | 64441204 | rs12766391 | 1.457 | -0.401 | -1.567 | -1.163 | -0.114 | -0.076 | -0.076 | 0.203 | 0.28 | 0.08 | 0.24 | 0.54 | 0.38 | 0.35 |
| 10 | 64441802 | rs224112 | 0.037 | -0.46 | -1.206 | -1.002 | -0.036 | -0.13 | -0.13 | 0 | 0.2 | 0.48 | 0.11 | 0.02 | 0.17 | 0.22 |
| 10 | 64443119 | rs224116 | -0.803 | -0.033 | -2.039 | -0.531 | -0.026 | -0.054 | -0.054 | 0 | 0.9 | 0.98 | 0.63 | 0.78 | 0.88 | 0.74 |
| 10 | 64445760 | rs224120 | -0.67 | -0.347 | -2.01 | -0.734 | -0.003 | 0.083 | 0.083 | 9.033 | 0.66 | 0.98 | 0.83 | 0.8 | 0.69 | 0.87 |
| 10 | 64448420 | rs17703956 | -0.89 | -0.942 | -1.327 | -0.598 | 0 | 0.013 | 0.013 | 0 | 0.14 | 0.09 | 0.12 | 0.04 | 0.1 | 0.11 |
| 10 | 64454061 | rs10995276 | 0.49 | -1.384 | -2.018 | -0.597 | -0.1 | 0.155 | 0.155 | 4.322 | 0.34 | 0.02 | 0.26 | 0.14 | 0.36 | 0.34 |
| 10 | 64456179 | rs10490965 | -0.479 | -1.498 | -1.033 | -0.618 | 0.024 | 0.012 | 0.012 | 0 | 0.06 | 0.24 | 0.05 | 0.02 | 0.06 | 0.09 |
| 10 | 64460929 | rs224126 | -1.605 | -1.624 | -0.702 | -0.549 | -0.004 | 0.017 | 0.017 | 0.115 | 0.13 | 0.43 | 0.15 | 0.13 | 0.09 | 0.21 |
| 10 | 64466651 | rs11599754 | 0.271 | -1.725 | -0.668 | -0.631 | -0.061 | 0.224 | 0.224 | 5.419 | 0.41 | 0.1 | 0.28 | 0.15 | 0.41 | 0.41 |
| 10 | 64481684 | rs17470703 | 0.937 | -1.632 | -0.574 | -0.555 | -0.005 | 0.036 | 0.036 | 0 | 0.07 | 0.01 | 0.02 | 0.01 | 0.05 | 0.07 |
| 10 | 64487073 | rs224149 | -1.084 | -1.927 | -0.446 | -0.647 | 0.003 | -0.017 | -0.017 | 0 | 0.59 | 0.51 | 0.44 | 0.27 | 0.52 | 0.65 |
| 10 | 64493222 | rs224050 | -1.51 | -2.09 | -0.052 | -0.505 | -0.031 | -0.005 | -0.005 | 0 | 0.66 | 0.82 | 0.49 | 0.45 | 0.59 | 0.72 |
| 10 | 64496870 | rs224055 | -1.154 | -2.097 | -0.125 | -0.509 | 0.012 | -0.007 | -0.007 | 0 | 0.62 | 0.56 | 0.47 | 0.45 | 0.58 | 0.72 |
| 10 | 64508475 | rs224071 | -0.973 | -2.096 | -0.172 | -0.441 | 0.003 | -0.017 | -0.017 | 0 | 0.59 | 0.52 | 0.44 | 0.3 | 0.52 | 0.65 |
| 10 | 64514111 | rs10822056 | -0.671 | -2.305 | -0.544 | -0.503 | -0.009 | 0.033 | 0.033 | 1.355 | 0.56 | 0.61 | 0.35 | 0.39 | 0.5 | 0.51 |
| 10 | 64516306 | rs10761660 | 0.947 | -2.128 | -0.504 | -0.454 | -0.052 | 0.132 | 0.132 | 4.31 | 0.34 | 0.1 | 0.25 | 0.18 | 0.36 | 0.33 |
| 10 | 64523481 | rs224039 | -1.681 | -1.656 | 0.33 | -0.044 | -0.015 | -0.002 | -0.002 | 0.025 | 0.13 | 0.34 | 0.15 | 0.1 | 0.1 | 0.21 |
| 10 | 64525547 | rs224045 | -0.899 | -1.872 | 0.336 | -0.101 | 0 | -0.004 | -0.004 | 0 | 0.55 | 0.51 | 0.46 | 0.48 | 0.58 | 0.7 |
| 10 | 64532193 | rs224087 | -1.972 | -1.457 | 0.72 | 0.022 | -0.022 | -0.001 | -0.001 | 0.257 | 0.14 | 0.46 | 0.16 | 0.11 | 0.09 | 0.2 |
| 10 | 64552242 | rs10995307 | -0.24 | -1.41 | 0.221 | 0.082 | -0.003 | 0.116 | 0.116 | 0.7 | 0.49 | 0.31 | 0.37 | 0.28 | 0.47 | 0.55 |
| 10 | 64560470 | rs1444418 | -1.731 | -1.585 | 0.475 | 0.929 | -0.021 | 0.143 | 0.143 | 0.704 | 0.16 | 0.4 | 0.4 | 0.33 | 0.14 | 0.16 |
| 10 | 64563702 | rs9971363 | -2.412 | -1.635 | 0.436 | 0.937 | -0.033 | 0.161 | 0.161 | 0.426 | 0.18 | 0.43 | 0.43 | 0.42 | 0.15 | 0.26 |
| 10 | 64564227 | rs7074164 | -1.541 | -1.575 | 0.507 | 1.052 | -0.018 | 0.072 | 0.072 | 0.002 | 0.1 | 0.2 | 0.35 | 0.23 | 0.07 | 0.2 |
| 10 | 64566572 | rs10995312 | -2.074 | -1.58 | 0.245 | 1.034 | -0.025 | 0.149 | 0.149 | 0.513 | 0.17 | 0.38 | 0.42 | 0.43 | 0.15 | 0.26 |
| 10 | 64567364 | rs1509964 | -2.056 | -1.575 | 0.248 | 1.036 | -0.025 | 0.151 | 0.151 | 0.447 | 0.17 | 0.38 | 0.42 | 0.43 | 0.15 | 0.26 |
| 10 | 64567938 | rs9990 | -2.056 | -1.575 | 0.248 | 1.036 | -0.025 | 0.151 | 0.151 | 0.447 | 0.17 | 0.38 | 0.42 | 0.43 | 0.15 | 0.26 |
| 10 | 64570038 | rs10128333 | -2.129 | -1.693 | 0.163 | 0.92 | -0.024 | 0.076 | 0.076 | 0.021 | 0.23 | 0.41 | 0.44 | 0.44 | 0.19 | 0.34 |
| 10 | 64580061 | rs224277 | 0.424 | -1.457 | -0.393 | 0.985 | -0.004 | 0.007 | 0.007 | 0 | 0.42 | 0.18 | 0.55 | 0.56 | 0.43 | 0.45 |
| 10 | 64588424 | rs224292 | 1.114 | -1.458 | -0.369 | 1.108 | -0.042 | -0.019 | -0.019 | 0 | 0.33 | 0.08 | 0.23 | 0.31 | 0.36 | 0.24 |
| 10 | 64590819 | rs224299 | -0.067 | -1.258 | -0.219 | 0.96 | 0.008 | 0.065 | 0.065 | 0 | 0.49 | 0.23 | 0.6 | 0.58 | 0.49 | 0.53 |
| 10 | 64594164 | rs2136613 | -0.328 | -1.129 | -0.162 | 1.034 | 0.004 | 0.014 | 0.014 | 0 | 0.52 | 0.45 | 0.67 | 0.42 | 0.51 | 0.46 |
| 10 | 64597189 | rs10822061 | 0.248 | -1.234 | -0.36 | 1.06 | -0.005 | -0.01 | -0.01 | 0 | 0.06 | 0.02 | 0.33 | 0.24 | 0.08 | 0.07 |
| 10 | 64602767 | rs224308 | 0.324 | -1.167 | -0.217 | 1.276 | -0.003 | -0.002 | -0.002 | 0.41 | 0.45 | 0.5 | 0.31 | 0.4 | 0.43 | 0.43 |

| | | | | | | | | | | | | | | | | |
|---|---|---|---|---|---|---|---|---|---|---|---|---|---|---|---|---|
| 10 | 64610718 | rs1509957 | -0.274 | -1.142 | -0.387 | 1.205 | 0.008 | 0.03 | 0.03 | 0 | 0.5 | 0.41 | 0.35 | 0.37 | 0.49 | 0.5 |
| 10 | 64610932 | rs10822065 | 0.123 | -1.166 | -0.393 | 1.207 | -0.01 | 0.061 | 0.061 | 0 | 0.46 | 0.34 | 0.3 | 0.19 | 0.44 | 0.41 |
| 10 | 64615451 | rs10761673 | -0.268 | -1.156 | -0.389 | 1.214 | 0.008 | 0.03 | 0.03 | 0 | 0.5 | 0.4 | 0.35 | 0.37 | 0.49 | 0.5 |
| 10 | 64620042 | rs911610 | 1.066 | -0.985 | -1.019 | 1.231 | -0.161 | 0.02 | 0.02 | 0 | 0.26 | 0.04 | 0.23 | 0.18 | 0.38 | 0.36 |
| 10 | 64626026 | rs10995337 | -0.161 | -0.391 | -0.199 | 1.431 | -0.014 | -0.024 | -0.024 | 0.229 | 0.05 | 0.01 | 0.34 | 0.22 | 0.09 | 0.07 |
| 10 | 64644428 | rs4746697 | -0.254 | -0.351 | -0.27 | 1.162 | -0.017 | -0.029 | -0.029 | 1.732 | 0.08 | 0.01 | 0.11 | 0.42 | 0.11 | 0.13 |
| 10 | 64644619 | rs12246498 | 0.06 | -0.456 | 0.201 | 1.025 | 0.028 | -0.037 | -0.037 | 0 | 0.11 | 0.37 | 0.07 | 0.03 | 0.08 | 0.05 |
| 10 | 64646868 | rs12252317 | 0.06 | -0.456 | 0.201 | 1.025 | 0.028 | -0.037 | -0.037 | 0.321 | 0.11 | 0.37 | 0.07 | 0.03 | 0.08 | 0.05 |
| 10 | 64647504 | rs9633562 | 0.145 | -0.79 | -0.054 | 0.913 | -0.006 | -0.001 | -0.001 | 0 | 0.57 | 0.55 | 0.72 | 0.42 | 0.63 | 0.67 |
| 10 | 64654109 | rs953027 | -1.964 | -0.704 | -0.078 | 0.87 | 0.025 | -0.061 | -0.061 | 0.289 | 0.36 | 0.8 | 0.29 | 0.53 | 0.3 | 0.29 |
| 10 | 64655913 | rs10995356 | 1.449 | -0.683 | -0.083 | 0.869 | -0.081 | 0.018 | 0.018 | 1.611 | 0.4 | 0.07 | 0.57 | 0.32 | 0.45 | 0.51 |
| 10 | 64667404 | rs950214 | -1.05 | -0.692 | 0.068 | 0.916 | -0.003 | 0.128 | 0.128 | 0 | 0.21 | 0.44 | 0.19 | 0.46 | 0.19 | 0.18 |
| 10 | 64671764 | rs9651304 | -1.574 | -0.716 | 0.068 | 0.731 | -0.013 | 0.112 | 0.112 | 1.73 | 0.24 | 0.37 | 0.21 | 0.5 | 0.22 | 0.24 |
| 10 | 64673225 | rs10761678 | 0.645 | -0.533 | 0.484 | 0.841 | 0.051 | 0.137 | 0.137 | 1.403 | 0.54 | 0.21 | 0.76 | 0.81 | 0.66 | 0.75 |
| 10 | 64673830 | rs4746725 | 0.044 | -0.388 | 0.397 | 0.906 | -0.002 | -0.034 | -0.034 | 0 | 0.64 | 0.5 | 0.78 | 0.81 | 0.67 | 0.75 |
| 10 | 64676376 | rs944684 | -0.864 | -0.163 | -0.767 | 1.049 | -0.018 | 0.031 | 0.031 | 0 | 0.75 | 0.89 | 0.86 | 0.85 | 0.75 | 0.8 |
| 10 | 64701797 | rs1546479 | -0.473 | -0.093 | -0.754 | 1.198 | -0.014 | -0.047 | -0.047 | 0 | 0.71 | 0.83 | 0.77 | 0.54 | 0.67 | 0.73 |
| 10 | 64703368 | rs1888968 | -0.977 | -0.101 | -0.78 | 1.206 | -0.012 | 0.022 | 0.022 | 0 | 0.76 | 0.86 | 0.86 | 0.85 | 0.75 | 0.8 |
| 10 | 64704741 | rs10822092 | -0.021 | -0.052 | -0.754 | 1.262 | -0.016 | -0.005 | -0.005 | 0.019 | 0.66 | 0.71 | 0.69 | 0.36 | 0.57 | 0.62 |
| 10 | 64709041 | rs2065449 | 1.044 | -0.075 | -0.752 | 1.225 | -0.052 | 0.055 | 0.055 | 0.745 | 0.45 | 0.11 | 0.59 | 0.32 | 0.49 | 0.57 |
| 10 | 64719454 | rs7082792 | 0.063 | -0.051 | -0.646 | 1.226 | 0.014 | -0.054 | -0.054 | 0.014 | 0.13 | 0.44 | 0.08 | 0.03 | 0.08 | 0.05 |
| 10 | 64745589 | rs4746781 | 0.163 | 0.056 | -1.295 | 0.694 | 0.001 | 0.099 | 0.099 | 0 | 0.16 | 0.21 | 0.19 | 0.49 | 0.18 | 0.17 |
| 10 | 64745865 | rs6479860 | -0.455 | 0.177 | -0.909 | 0.073 | -0.019 | 0.025 | 0.025 | 0 | 0.76 | 0.88 | 0.87 | 0.85 | 0.75 | 0.79 |
| 10 | 64747544 | rs7080482 | 0.108 | 0.433 | -1.146 | 0.237 | 0.003 | 0.041 | 0.041 | 0 | 0.62 | 0.26 | 0.78 | 0.82 | 0.67 | 0.74 |
| 10 | 64760686 | rs10995388 | -0.574 | 0.48 | -1.38 | 0.77 | -0.023 | -0.051 | -0.051 | 0 | 0.13 | 0.01 | 0.1 | 0.17 | 0.13 | 0.06 |
| 10 | 64761091 | rs11597905 | -1.036 | 1.167 | -2.134 | 1.103 | -0.072 | -0.045 | -0.045 | 0.228 | 0.37 | 0.12 | 0.56 | 0.59 | 0.48 | 0.35 |
| 10 | 64762369 | rs2664317 | -0.415 | 0.41 | -1.357 | 0.451 | -0.041 | -0.076 | -0.076 | 0 | 0.81 | 0.59 | 0.83 | 0.95 | 0.9 | 0.77 |
| 10 | 64763125 | rs12413575 | 0.433 | 0.628 | -1.401 | 0.85 | -0.044 | -0.077 | -0.077 | 0 | 0.22 | 0.09 | 0.45 | 0.39 | 0.34 | 0.28 |
| 10 | 64765220 | rs2842282 | -0.212 | 0.243 | -0.675 | 0.671 | -0.005 | -0.018 | -0.018 | 0 | 0.74 | 0.74 | 0.54 | 0.58 | 0.66 | 0.7 |
| 10 | 64765654 | rs10822109 | -0.706 | 0.252 | -0.705 | 0.672 | -0.074 | -0.007 | -0.007 | 0.022 | 0.38 | 0.12 | 0.56 | 0.62 | 0.48 | 0.37 |
| 10 | 64770248 | rs2664328 | -0.369 | 0.169 | -0.666 | 0.645 | -0.004 | -0.037 | -0.037 | 1.8 | 0.84 | 0.94 | 0.89 | 0.8 | 0.86 | 0.93 |
| 10 | 64802127 | rs10995404 | 0.546 | 0.129 | -0.423 | 0.557 | -0.017 | -0.071 | -0.071 | 0.036 | 0.16 | 0.25 | 0.17 | 0.02 | 0.1 | 0.23 |
| 10 | 64805235 | rs1155233 | 0.523 | 0.392 | -0.589 | 1.162 | -0.008 | -0.019 | -0.019 | 0 | 0.09 | 0.03 | 0.1 | 0.22 | 0.14 | 0.1 |
| 10 | 64811552 | rs2842269 | 1.497 | 0.462 | -0.236 | 1.169 | -0.018 | 0.051 | 0.051 | 0.466 | 0.45 | 0.25 | 0.26 | 0.35 | 0.43 | 0.43 |
| 10 | 64812131 | rs2664339 | 1.658 | 0.474 | -0.319 | 1.174 | -0.089 | 0.037 | 0.037 | 1.238 | 0.43 | 0.1 | 0.25 | 0.35 | 0.43 | 0.43 |
| 10 | 64816444 | rs2495012 | 0.287 | 0.383 | -0.331 | 1.069 | 0.005 | 0.006 | 0.006 | 0.238 | 0.67 | 0.68 | 0.72 | 0.78 | 0.76 | 0.7 |
| 10 | 64819367 | rs11813180 | -0.04 | 0.781 | -0.202 | 1.39 | -0.009 | -0.017 | -0.017 | 0 | 0.09 | 0.05 | 0.1 | 0.24 | 0.14 | 0.12 |
| 10 | 64819996 | rs6479868 | 0.261 | 0.761 | 0.044 | 1.144 | 0.038 | 0.037 | 0.037 | 0 | 0.14 | 0.32 | 0.36 | 0.17 | 0.19 | 0.17 |
| 10 | 64823212 | rs2842276 | 0.576 | 0.639 | 0.424 | 0.877 | -0.017 | -0.023 | -0.023 | 0 | 0.77 | 0.62 | 0.83 | 0.83 | 0.82 | 0.87 |
| 10 | 64829830 | rs16918155 | -0.226 | 0.753 | 0.035 | 0.888 | -0.011 | -0.048 | -0.048 | 0 | 0.15 | 0.06 | 0.14 | 0.27 | 0.19 | 0.22 |
| 10 | 64833459 | rs10733783 | 0.3 | 0.836 | 0.475 | 0.748 | 0.031 | -0.013 | -0.013 | 6.142 | 0.95 | 0.84 | 0.94 | 1 | 0.94 | 1 |
| 10 | 64839442 | rs1010889 | 1.114 | 1.16 | 0.155 | 0.698 | 0.022 | 0.003 | 0.003 | 0.045 | 0.5 | 0.38 | 0.59 | 0.54 | 0.41 | 0.56 |
| 10 | 64846332 | rs10733785 | 0.328 | 1.509 | 0.184 | 0.408 | 0.029 | -0.054 | -0.054 | 0.173 | 0.81 | 0.78 | 0.86 | 0.97 | 0.71 | 0.88 |
| 10 | 64848773 | rs12253611 | -1.746 | 1.117 | 0.881 | 1.132 | 0.071 | 0.181 | 0.181 | 0.427 | 0.07 | 0.25 | 0.15 | 0.23 | 0.09 | 0.04 |
| 10 | 64855558 | rs10761702 | -0.172 | 0.224 | 0.439 | 0.513 | -0.014 | -0.052 | -0.052 | 0.01 | 0.71 | 0.37 | 0.53 | 0.88 | 0.75 | 0.64 |
| 10 | 64859984 | rs10509185 | -0.183 | 0.217 | 0.418 | 0.51 | -0.021 | -0.078 | -0.078 | 0.148 | 0.71 | 0.43 | 0.54 | 0.88 | 0.75 | 0.64 |
| 10 | 64862962 | rs10761703 | 0.394 | -0.55 | 0.08 | 0.672 | 0.002 | 0 | 0 | 0 | 0.64 | 0.39 | 0.69 | 0.36 | 0.76 | 0.67 |
| 10 | 64868040 | rs10761708 | 0.301 | -0.573 | 0.162 | 0.691 | -0.049 | 0.026 | 0.026 | 0.933 | 0.66 | 0.52 | 0.78 | 0.63 | 0.79 | 0.71 |
| 10 | 64869239 | rs10995441 | 0.43 | -0.606 | 0.112 | 0.655 | -0.035 | 0.017 | 0.017 | 1.859 | 0.64 | 0.52 | 0.76 | 0.63 | 0.75 | 0.7 |
| 10 | 64875546 | rs4414112 | 0.319 | -0.843 | -0.431 | 0.276 | 0.012 | -0.011 | -0.011 | 0 | 0.68 | 0.31 | 0.53 | 0.64 | 0.79 | 0.67 |
| 10 | 64875745 | rs4545434 | 0.093 | -0.932 | -0.49 | 0.243 | 0.029 | -0.083 | -0.083 | 0.145 | 0.23 | 0.11 | 0.41 | 0.35 | 0.2 | 0.29 |
| 10 | 64879229 | rs1553789 | 0.706 | -0.931 | -0.469 | 0.25 | 0.034 | -0.005 | -0.005 | 0 | 0.64 | 0.29 | 0.46 | 0.38 | 0.77 | 0.66 |
| 10 | 64893433 | rs2175197 | 0.649 | -0.42 | -1.083 | -0.348 | -0.019 | 0.078 | 0.078 | 1.82 | 0.43 | 0.3 | 0.28 | 0.09 | 0.47 | 0.43 |
| 10 | 64894047 | rs871689 | 1.697 | -0.469 | -1.241 | -0.529 | -0.099 | 0.125 | 0.125 | 0.27 | 0.22 | 0.02 | 0.15 | 0.01 | 0.29 | 0.15 |
| 10 | 64894059 | rs871690 | 0.017 | -0.652 | -1.357 | -0.541 | 0.014 | -0.005 | -0.005 | 0.143 | 0.67 | 0.31 | 0.49 | 0.37 | 0.81 | 0.67 |
| 10 | 64897433 | rs6479884 | -0.771 | -0.731 | -1.105 | -0.606 | -0.003 | -0.048 | -0.048 | 0 | 0.11 | 0.49 | 0.07 | 0.01 | 0.05 | 0.04 |
| 10 | 64914467 | rs1046521 | 1.825 | -0.742 | -1.189 | -0.594 | -0.099 | 0.125 | 0.125 | 0.288 | 0.22 | 0.03 | 0.15 | 0.01 | 0.29 | 0.15 |
| 10 | 64916469 | rs1009984 | -0.17 | -0.812 | -0.748 | -0.496 | -0.066 | 0.072 | 0.072 | 0.191 | 0.71 | 0.48 | 0.58 | 0.64 | 0.83 | 0.69 |
| 10 | 64919511 | rs12413415 | 1.478 | -0.737 | -0.858 | -0.47 | 0.009 | -0.147 | -0.147 | 0 | 0.2 | 0.01 | 0.37 | 0.35 | 0.15 | 0.27 |
| 10 | 64920588 | rs7100693 | 2.608 | -0.587 | -1.044 | -0.626 | -0.063 | 0.063 | 0.063 | 0 | 0.11 | 0.01 | 0.12 | 0 | 0.18 | 0.1 |
| 10 | 64923353 | rs7080760 | 0.779 | -0.938 | -0.499 | -0.311 | 0.025 | 0.001 | 0.001 | 1.762 | 0.06 | 0.31 | 0.03 | 0 | 0.01 | 0 |
| 10 | 64926482 | rs4747045 | 1.336 | -0.928 | -0.651 | -0.299 | 0.009 | -0.147 | -0.147 | 0.939 | 0.2 | 0.01 | 0.37 | 0.35 | 0.15 | 0.27 |
| 10 | 64927823 | rs1935 | -1.027 | -0.902 | -0.627 | -0.28 | -0.015 | 0.034 | 0.034 | 0.249 | 0.55 | 0.29 | 0.31 | 0.36 | 0.49 | 0.53 |
| 10 | 64945364 | rs3211105 | 0.165 | -1.15 | -0.336 | -0.044 | 0.01 | -0.011 | -0.011 | 0 | 0.68 | 0.31 | 0.54 | 0.64 | 0.8 | 0.69 |
| 10 | 64993101 | rs12259115 | -0.864 | -0.868 | -0.115 | -0.222 | 0.018 | -0.032 | -0.032 | 0 | 0.1 | 0.51 | 0.08 | 0.01 | 0.05 | 0.03 |
| 10 | 65047395 | rs10822152 | 2.147 | -1.088 | -0.715 | -0.368 | -0.005 | 0.076 | 0.076 | 8.286 | 0.17 | 0.02 | 0.08 | 0.05 | 0.15 | 0.28 |
| 10 | 65058224 | rs12773283 | -0.95 | -1.11 | -0.614 | -0.176 | 0.007 | -0.011 | -0.011 | 0 | 0.78 | 0.78 | 0.61 | 0.65 | 0.84 | 0.73 |
| 10 | 65101800 | rs7091664 | -1.286 | -1.063 | -0.721 | -0.018 | 0.014 | -0.142 | -0.142 | 0.248 | 0.8 | 0.98 | 0.63 | 0.65 | 0.85 | 0.73 |
| 10 | 65104894 | rs10761746 | 0.195 | -1.136 | -0.599 | -0.082 | 0.01 | -0.011 | -0.011 | 0.079 | 0.68 | 0.31 | 0.54 | 0.64 | 0.8 | 0.69 |

| | | | | | | | | | | | | | | | | |
|---|---|---|---|---|---|---|---|---|---|---|---|---|---|---|---|---|
| 10 | 65129917 | rs4746119 | 1.131 | -1.077 | -0.633 | -0.03 | 0.01 | -0.148 | -0.148 | 0.779 | 0.2 | 0.01 | 0.37 | 0.35 | 0.15 | 0.27 |
| 10 | 65133156 | rs2393967 | 1.365 | -0.858 | -0.787 | -0.231 | -0.124 | -0.076 | -0.076 | 0 | 0.24 | 0.02 | 0.2 | 0.28 | 0.34 | 0.24 |
| 10 | 65133822 | rs7923609 | 1.057 | -0.653 | -0.004 | 0.216 | -0.017 | 0.05 | 0.05 | 0.255 | 0.45 | 0.29 | 0.31 | 0.34 | 0.49 | 0.52 |
| 10 | 65162153 | rs7896783 | 1.02 | -0.629 | -0.013 | 0.14 | -0.016 | 0.025 | 0.025 | 0.347 | 0.45 | 0.26 | 0.3 | 0.4 | 0.49 | 0.52 |
| 10 | 65212240 | rs7906819 | -0.798 | -0.691 | 0.175 | 0.094 | 0.031 | -0.018 | -0.018 | 0 | 0.09 | 0.47 | 0.07 | 0.01 | 0.05 | 0.03 |
| 10 | 65214700 | rs12257258 | -0.769 | -0.691 | 0.171 | 0.094 | 0.031 | -0.017 | -0.017 | 0 | 0.09 | 0.47 | 0.07 | 0.01 | 0.05 | 0.03 |
| 10 | 65216244 | rs12246496 | -1.028 | -0.715 | 0.16 | 0.084 | 0.02 | -0.028 | -0.028 | 0 | 0.1 | 0.47 | 0.07 | 0.01 | 0.05 | 0.03 |
| 10 | 65234512 | rs17814528 | 1.001 | -0.649 | -0.031 | 0.148 | -0.105 | -0.052 | -0.052 | 0 | 0.23 | 0.03 | 0.23 | 0.3 | 0.31 | 0.17 |
| 10 | 65253700 | rs7910662 | 0.324 | -0.674 | -0.007 | 0.156 | 0.015 | -0.008 | -0.008 | 0 | 0.68 | 0.29 | 0.54 | 0.7 | 0.8 | 0.69 |
| 10 | 65274927 | rs10761779 | 0.937 | -0.511 | 0.101 | 0.203 | -0.015 | 0.026 | 0.026 | 0.326 | 0.45 | 0.26 | 0.3 | 0.4 | 0.49 | 0.52 |
| 10 | 65318766 | rs10761785 | 0.462 | -0.364 | 0.184 | 0.131 | -0.003 | 0.022 | 0.022 | 0 | 0.48 | 0.42 | 0.63 | 0.71 | 0.51 | 0.55 |
| 10 | 65325126 | rs10822181 | -0.687 | -0.574 | 0.258 | 0.148 | 0.002 | 0.015 | 0.015 | 0 | 0.51 | 0.53 | 0.37 | 0.28 | 0.48 | 0.44 |
| 10 | 65325478 | rs10822182 | 0.927 | -0.581 | 0.241 | 0.192 | -0.016 | 0.014 | 0.014 | 0.119 | 0.46 | 0.31 | 0.54 | 0.44 | 0.49 | 0.53 |
| 10 | 65333946 | rs6479909 | 0.395 | -0.477 | 0.252 | 0.144 | 0.035 | -0.137 | -0.137 | 0.053 | 0.27 | 0.43 | 0.22 | 0.01 | 0.32 | 0.14 |
| 10 | 65350383 | rs10822186 | 0.505 | -0.54 | 0.322 | 0.22 | 0.002 | 0.019 | 0.019 | 0.162 | 0.5 | 0.38 | 0.55 | 0.45 | 0.51 | 0.54 |
| 10 | 65351581 | rs6479911 | -0.823 | -0.557 | 0.034 | -0.004 | 0.016 | -0.103 | -0.103 | 2.724 | 0.74 | 0.65 | 0.82 | 0.99 | 0.72 | 0.86 |
| 10 | 65359701 | rs4746244 | 0.413 | -0.799 | -0.146 | 0.153 | 0.027 | -0.083 | -0.083 | 1.002 | 0.2 | 0.05 | 0.15 | 0.25 | 0.15 | 0.27 |
| 10 | 65376395 | rs7076601 | 0.039 | -0.626 | 0.169 | -0.07 | 0.03 | -0.025 | -0.025 | 0 | 0.27 | 0.51 | 0.25 | 0.22 | 0.3 | 0.17 |
| 10 | 65381740 | rs7075302 | 0.809 | -0.683 | 0.116 | 0.038 | 0.005 | -0.004 | -0.004 | 0 | 0.53 | 0.39 | 0.6 | 0.57 | 0.56 | 0.61 |
| 10 | 65384567 | rs7089698 | 0.824 | -0.694 | 0.111 | 0.034 | 0.005 | -0.003 | -0.003 | 0 | 0.52 | 0.39 | 0.6 | 0.57 | 0.56 | 0.61 |
| 10 | 65392260 | rs12246748 | 0.291 | -0.582 | 0.255 | -0.038 | 0.023 | -0.128 | -0.128 | 0 | 0.25 | 0.4 | 0.2 | 0.01 | 0.27 | 0.16 |
| 10 | 65393613 | rs1255496 | -0.231 | -0.578 | 0.047 | -0.023 | 0.021 | -0.031 | -0.031 | 0 | 0.29 | 0.56 | 0.28 | 0.21 | 0.32 | 0.19 |
| 10 | 65399851 | rs3858120 | 0.244 | -0.697 | -0.154 | 0.393 | -0.018 | 0.042 | 0.042 | 0 | 0.76 | 0.68 | 0.78 | 0.6 | 0.81 | 0.76 |
| 10 | 65414090 | rs2393992 | 0.392 | -0.299 | -0.274 | -0.046 | 0.005 | -0.003 | -0.003 | 0 | 0.42 | 0.51 | 0.57 | 0.65 | 0.49 | 0.54 |
| 10 | 65417520 | rs10509192 | 0.406 | -0.548 | -0.168 | -0.18 | -0.013 | -0.023 | -0.023 | 0 | 0.07 | 0.22 | 0.03 | 0.03 | 0.03 | 0.03 |
| 10 | 65425682 | rs4369293 | 0.581 | -0.483 | -0.049 | -0.134 | 0.007 | -0.002 | -0.002 | 0 | 0.41 | 0.43 | 0.56 | 0.65 | 0.49 | 0.54 |
| 10 | 65452546 | rs1255476 | 0.138 | -0.625 | -0.49 | -0.148 | 0.027 | 0.065 | 0.065 | 0 | 0.57 | 0.72 | 0.69 | 0.82 | 0.61 | 0.64 |
| 10 | 65461640 | rs2619601 | -0.158 | -0.571 | -0.755 | -0.2 | -0.009 | 0.081 | 0.081 | 0 | 0.66 | 0.9 | 0.71 | 0.87 | 0.64 | 0.7 |
| 10 | 65461793 | rs10995587 | 0.549 | -0.471 | -0.414 | 0.056 | -0.006 | -0.006 | -0.006 | 0.058 | 0.55 | 0.44 | 0.61 | 0.66 | 0.56 | 0.66 |
| 10 | 65483118 | rs1255485 | -0.848 | -0.452 | -0.559 | -0.715 | -0.008 | -0.065 | -0.065 | 0 | 0.33 | 0.4 | 0.18 | 0.07 | 0.24 | 0.15 |
| 10 | 65483262 | rs1262217 | -1.734 | 0.279 | -1.276 | -1.203 | -0.072 | 0.088 | 0.088 | 0 | 0.85 | 0.99 | 0.86 | 1 | 0.82 | 0.89 |
| 10 | 65491733 | rs7095162 | -1.032 | -0.058 | -0.59 | -0.758 | 0 | -0.028 | -0.028 | 0 | 0.33 | 0.34 | 0.18 | 0.07 | 0.24 | 0.15 |
| 10 | 65496107 | rs11599596 | -1.227 | -0.078 | -0.37 | -0.749 | -0.017 | -0.045 | -0.045 | 0.021 | 0.37 | 0.48 | 0.22 | 0.16 | 0.25 | 0.15 |
| 10 | 65497779 | rs4609496 | -2.061 | -0.217 | -0.781 | -0.516 | -0.087 | -0.037 | -0.037 | 0 | 0.56 | 0.91 | 0.42 | 0.38 | 0.5 | 0.42 |
| 10 | 65506596 | rs10822202 | -0.026 | 0.102 | -0.553 | -0.541 | -0.024 | -0.031 | -0.031 | 0 | 0.11 | 0.05 | 0.14 | 0.19 | 0.15 | 0.22 |
| 10 | 65514404 | rs3912164 | -0.948 | -0.076 | -0.021 | -0.92 | -0.002 | -0.006 | -0.006 | 1.726 | 0.26 | 0.35 | 0.18 | 0.24 | 0.27 | 0.17 |
| 10 | 65518790 | rs12241544 | -0.649 | 0.054 | 0.1 | -0.76 | 0.039 | 0.077 | 0.077 | 0 | 0.12 | 0.31 | 0.11 | 0.24 | 0.16 | 0.15 |
| 10 | 65519556 | rs3852415 | -0.072 | -0.174 | -0.239 | -0.913 | 0.032 | 0.095 | 0.095 | 0.882 | 0.21 | 0.06 | 0.12 | 0 | 0.17 | 0.1 |
| 10 | 65519666 | rs4082329 | -0.915 | -0.208 | -0.262 | -0.992 | 0.033 | 0.128 | 0.128 | 0.044 | 0.29 | 0.09 | 0.17 | 0 | 0.24 | 0.15 |
| 10 | 65519754 | rs10740140 | -2.928 | -0.113 | -0.207 | -0.928 | -0.061 | -0.055 | -0.055 | 0.094 | 0.59 | 0.89 | 0.48 | 0.43 | 0.54 | 0.53 |
| 10 | 65523366 | rs10509195 | 0.981 | -0.154 | 0.222 | -0.922 | -0.014 | -0.033 | -0.033 | 0 | 0.58 | 0.54 | 0.66 | 0.8 | 0.65 | 0.66 |
| 10 | 65525237 | rs10995618 | 0.692 | -0.094 | 0.104 | -0.88 | 0.001 | 0.003 | 0.003 | 0.423 | 0.08 | 0.05 | 0.05 | 0 | 0.06 | 0.05 |
| 10 | 65525834 | rs1842189 | 0.382 | -0.09 | 0.128 | -0.923 | 0.048 | 0.122 | 0.122 | 0 | 0.67 | 0.88 | 0.8 | 1 | 0.75 | 0.84 |
| 10 | 65531178 | rs6479920 | -1.859 | -0.126 | 0.364 | -0.765 | -0.001 | -0.003 | -0.003 | 0.026 | 0.28 | 0.73 | 0.3 | 0.41 | 0.29 | 0.37 |
| 10 | 65550686 | rs1904382 | -0.251 | -0.136 | 0.387 | -0.886 | 0.052 | 0.107 | 0.107 | 0 | 0.1 | 0.33 | 0.1 | 0.21 | 0.1 | 0.13 |
| 10 | 65553070 | rs1255509 | 0.078 | -0.144 | -0.049 | -1.097 | 0.043 | 0.123 | 0.123 | 0 | 0.75 | 0.94 | 0.85 | 1 | 0.81 | 0.89 |
| 10 | 65562475 | rs10761796 | 0.964 | -0.085 | 0.462 | -1.074 | 0.012 | 0.06 | 0.06 | 0 | 0.63 | 0.74 | 0.7 | 0.84 | 0.65 | 0.67 |
| 10 | 65563206 | rs17825120 | -0.39 | -0.099 | 0.477 | -1.096 | -0.014 | -0.009 | -0.009 | 0 | 0.12 | 0.07 | 0.15 | 0.15 | 0.18 | 0.25 |
| 10 | 65563647 | rs3999074 | 2.389 | -0.092 | 0.483 | -1.098 | -0.07 | -0.024 | -0.024 | 0.498 | 0.39 | 0.13 | 0.55 | 0.62 | 0.48 | 0.47 |
| 10 | 65568254 | rs7916789 | 0.005 | -0.019 | 0.867 | -1.051 | 0.096 | 0.155 | 0.155 | 0 | 0.09 | 0.33 | 0.09 | 0.2 | 0.12 | 0.12 |
| 10 | 65580158 | rs2619408 | 1.963 | -0.042 | 0.862 | -1.054 | -0.022 | -0.018 | -0.018 | 0 | 0.63 | 0.21 | 0.69 | 0.61 | 0.64 | 0.55 |
| 10 | 65587937 | rs4427464 | 1.133 | -0.017 | 0.867 | -1.046 | 0 | -0.004 | -0.004 | 0.264 | 0.75 | 0.73 | 0.79 | 0.82 | 0.77 | 0.67 |
| 10 | 65592653 | rs10822218 | 0.52 | -0.271 | 0.648 | -1.035 | 0.007 | 0.016 | 0.016 | 0.086 | 0.09 | 0.03 | 0.05 | 0 | 0.05 | 0.07 |
| 10 | 65593561 | rs2140390 | -0.294 | -0.333 | 0.596 | -1.042 | 0.039 | 0.04 | 0.04 | 0 | 0.27 | 0.18 | 0.14 | 0 | 0.17 | 0.08 |
| 10 | 65593781 | rs1880065 | -0.319 | -0.333 | 0.638 | -1.041 | 0.014 | -0.02 | -0.02 | 0 | 0.27 | 0.24 | 0.15 | 0 | 0.17 | 0.08 |
| 10 | 65597702 | rs1880067 | 1.914 | -0.182 | 0.56 | -0.864 | -0.027 | -0.004 | -0.004 | 0.219 | 0.4 | 0.21 | 0.56 | 0.61 | 0.48 | 0.52 |
| 10 | 65617594 | rs10822225 | -0.948 | -0.407 | 0.564 | -1.113 | -0.001 | 0.098 | 0.098 | 0 | 0.37 | 0.14 | 0.27 | 0.16 | 0.4 | 0.35 |
| 10 | 65619267 | rs16918944 | 0.247 | -0.362 | 1.133 | -1.095 | 0.087 | 0.091 | 0.091 | 0.004 | 0.11 | 0.29 | 0.1 | 0.21 | 0.17 | 0.14 |
| 10 | 65621134 | rs11596158 | -0.004 | -0.38 | 0.915 | -1.147 | 0.05 | 0.135 | 0.135 | 1.825 | 0.26 | 0.05 | 0.14 | 0 | 0.19 | 0.13 |
| 10 | 65622494 | rs1916434 | -1.236 | -0.529 | 1.028 | -1.238 | -0.001 | -0.021 | -0.021 | 0 | 0.28 | 0.54 | 0.16 | 0.21 | 0.22 | 0.19 |
| 10 | 65624141 | rs10761805 | 1.024 | -0.522 | 0.995 | -1.254 | 0.008 | 0.04 | 0.04 | 0 | 0.6 | 0.69 | 0.72 | 0.81 | 0.6 | 0.59 |
| 10 | 65624936 | rs6479938 | -0.909 | -0.51 | 0.592 | -1.234 | -0.003 | 0.09 | 0.09 | 0 | 0.37 | 0.14 | 0.26 | 0.18 | 0.39 | 0.35 |
| 10 | 65625814 | rs10509199 | 2.955 | -0.495 | 0.506 | -1.219 | -0.07 | -0.1 | -0.1 | 0 | 0.32 | 0.07 | 0.55 | 0.59 | 0.37 | 0.4 |
| 10 | 65626561 | rs10509200 | -0.636 | -0.394 | 0.627 | -1.116 | -0.039 | -0.026 | -0.026 | 1.627 | 0.1 | 0.04 | 0.12 | 0.14 | 0.2 | 0.21 |
| 10 | 65626926 | rs16918975 | -0.539 | -0.477 | 0.983 | -1.276 | -0.041 | -0.096 | -0.096 | 0.31 | 0.14 | 0.39 | 0.08 | 0 | 0.06 | 0.06 |
| 10 | 65628949 | rs10822233 | -0.496 | -0.47 | 0.985 | -1.275 | -0.035 | -0.094 | -0.094 | 0 | 0.14 | 0.39 | 0.08 | 0 | 0.06 | 0.06 |
| 10 | 65629166 | rs1916440 | -0.315 | -0.325 | 0.755 | -1.2 | 0.023 | 0.022 | 0.022 | 0.138 | 0.15 | 0.23 | 0.1 | 0.21 | 0.18 | 0.14 |
| 10 | 65633037 | rs7074842 | 0.344 | -0.516 | 0.476 | -1.171 | 0 | -0.018 | -0.018 | 0.606 | 0.1 | 0.11 | 0.05 | 0 | 0.05 | 0.06 |
| 10 | 65634031 | rs12774150 | -0.132 | -0.692 | 0.356 | -1.317 | 0.051 | 0.077 | 0.077 | 1.766 | 0.25 | 0.08 | 0.13 | 0 | 0.17 | 0.1 |

| | | | | | | | | | | | | | | | | |
|---|---|---|---|---|---|---|---|---|---|---|---|---|---|---|---|---|
| 10 | 65634112 | rs6479941 | 1.215 | -0.8 | 0.655 | -1.361 | 0.017 | -0.023 | -0.023 | 0 | 0.54 | 0.43 | 0.77 | 0.98 | 0.74 | 0.75 |
| 10 | 65638574 | rs10822238 | 0.96 | -0.603 | 0.503 | -1.32 | 0.004 | 0.076 | 0.076 | 0 | 0.63 | 0.83 | 0.74 | 0.85 | 0.61 | 0.6 |
| 10 | 65641562 | rs12761811 | -0.072 | -0.606 | 0.401 | -1.315 | 0.056 | 0.114 | 0.114 | 1.986 | 0.25 | 0.05 | 0.13 | 0 | 0.17 | 0.1 |
| 10 | 65647308 | rs10822241 | -0.779 | 0.212 | 0.768 | -0.929 | -0.046 | -0.029 | -0.029 | 0 | 0.09 | 0.02 | 0.11 | 0.13 | 0.19 | 0.21 |
| 10 | 65649636 | rs960823 | 2.827 | 0.143 | 0.769 | -0.993 | -0.013 | -0.021 | -0.021 | 0 | 0.39 | 0.14 | 0.57 | 0.61 | 0.39 | 0.42 |
| 10 | 65650779 | rs884242 | 1.307 | 0.14 | 0.84 | -0.913 | -0.015 | -0.012 | -0.012 | 0 | 0.88 | 0.91 | 0.88 | 0.81 | 0.78 | 0.7 |
| 10 | 65654554 | rs1877131 | 1.663 | 0.135 | 0.89 | -0.929 | -0.032 | 0.005 | 0.005 | 0 | 0.58 | 0.25 | 0.69 | 0.61 | 0.55 | 0.51 |
| 10 | 65657716 | rs4745825 | 1.541 | 0.129 | 1.061 | -0.917 | -0.006 | 0.001 | 0.001 | 0 | 0.6 | 0.36 | 0.7 | 0.64 | 0.55 | 0.52 |
| 10 | 65667946 | rs1916418 | -1.387 | 0.133 | 1.045 | -0.916 | 0.015 | 0.002 | 0.002 | 0.451 | 0.3 | 0.37 | 0.27 | 0.34 | 0.42 | 0.39 |
| 10 | 65668164 | rs2177360 | 0.133 | 0.124 | 1.122 | -0.924 | 0.064 | 0.06 | 0.06 | 0 | 0.12 | 0.27 | 0.1 | 0.2 | 0.17 | 0.12 |
| 10 | 65671263 | rs7904940 | 1.014 | -0.065 | 1.143 | -0.868 | 0.001 | -0.002 | -0.002 | 2.602 | 0.58 | 0.43 | 0.69 | 0.64 | 0.54 | 0.5 |
| 10 | 65672482 | rs7085770 | 0.66 | -0.13 | 1.102 | -0.872 | -0.03 | -0.036 | -0.036 | 0 | 0.07 | 0.2 | 0.01 | 0 | 0 | 0 |
| 10 | 65672533 | rs10995689 | 2.167 | -0.162 | 1.035 | -0.909 | 0.002 | -0.008 | -0.008 | 0 | 0.39 | 0.22 | 0.56 | 0.6 | 0.38 | 0.4 |
| 10 | 65674767 | rs10740145 | 1.064 | 0.133 | 1.209 | -0.773 | -0.006 | 0.004 | 0.004 | 0 | 0.59 | 0.38 | 0.68 | 0.64 | 0.54 | 0.5 |
| 10 | 65677634 | rs2244205 | 0.591 | 0.282 | 1.048 | -0.738 | 0.033 | 0.009 | 0.009 | 0.415 | 0.67 | 0.49 | 0.72 | 0.65 | 0.63 | 0.53 |
| 10 | 65680980 | rs2243532 | 0.873 | 0.549 | 1.187 | -0.706 | 0.005 | -0.003 | -0.003 | 0 | 0.6 | 0.38 | 0.67 | 0.63 | 0.54 | 0.51 |
| 10 | 65684472 | rs1916472 | 0.218 | 0.502 | 0.565 | -0.784 | 0.013 | 0.08 | 0.08 | 0 | 0.16 | 0.02 | 0.12 | 0.01 | 0.16 | 0.1 |
| 10 | 65685683 | rs2461892 | 0.218 | 0.502 | 0.565 | -0.784 | 0.013 | 0.08 | 0.08 | 0 | 0.16 | 0.02 | 0.12 | 0.01 | 0.16 | 0.09 |
| 10 | 65686698 | rs16919021 | -0.207 | 0.538 | 0.655 | -0.763 | 0.016 | -0.064 | -0.064 | 0.184 | 0.11 | 0.28 | 0.07 | 0 | 0.1 | 0.03 |
| 10 | 65691928 | rs7096130 | -1.095 | 0.573 | 0.615 | -0.745 | 0.012 | -0.024 | -0.024 | 0 | 0.35 | 0.58 | 0.27 | 0.23 | 0.35 | 0.3 |
| 10 | 65694772 | rs1916463 | -0.268 | 0.57 | 0.629 | -0.745 | 0.037 | 0.054 | 0.054 | 0 | 0.91 | 0.82 | 0.92 | 0.84 | 0.88 | 0.78 |
| 10 | 65700878 | rs7079643 | -0.038 | 0.588 | 0.541 | -0.746 | -0.004 | 0 | 0 | 0.044 | 0.24 | 0.27 | 0.2 | 0.2 | 0.25 | 0.26 |
| 10 | 65702844 | rs2252578 | 0.7 | 0.155 | -0.097 | -0.742 | -0.017 | 0.014 | 0.014 | 0 | 0.56 | 0.22 | 0.65 | 0.61 | 0.52 | 0.48 |
| 10 | 65711686 | rs3858126 | 1.139 | 0.368 | -0.228 | -0.29 | -0.002 | -0.033 | -0.033 | 0 | 0.4 | 0.12 | 0.53 | 0.6 | 0.37 | 0.39 |
| 10 | 65715707 | rs12240813 | -1.013 | 0.241 | 0.615 | -0.2 | 0.016 | -0.054 | -0.054 | 1.196 | 0.09 | 0.23 | 0.06 | 0 | 0.1 | 0.03 |
| 10 | 65718332 | rs12261509 | -1.126 | 0.248 | 0.411 | -0.197 | 0.03 | -0.059 | -0.059 | 0.484 | 0.1 | 0.32 | 0.07 | 0 | 0.1 | 0.03 |
| 10 | 65746001 | rs2394022 | 0.632 | 0.302 | -0.206 | -0.057 | 0 | 0.004 | 0.004 | 0 | 0.91 | 0.92 | 0.93 | 1 | 0.92 | 0.9 |
| 10 | 65762991 | rs2671490 | 1.716 | 0.131 | 0.64 | 0.008 | 0.009 | -0.029 | -0.029 | 0 | 0.87 | 0.58 | 0.87 | 0.93 | 0.9 | 0.89 |
| 10 | 65765874 | rs2578054 | 0.591 | 0.186 | 0.09 | 0.054 | -0.001 | -0.022 | -0.022 | 0 | 0.91 | 0.88 | 0.93 | 1 | 0.92 | 0.91 |
| 10 | 65782691 | rs12220466 | 0.914 | 0.043 | -0.893 | 0.103 | -0.038 | -0.001 | -0.001 | 0 | 0.48 | 0.15 | 0.57 | 0.53 | 0.49 | 0.4 |
| 10 | 65784945 | rs12358989 | -0.943 | -0.274 | -0.452 | 0.26 | -0.003 | -0.006 | -0.006 | 2.357 | 0.1 | 0.11 | 0.09 | 0.07 | 0.09 | 0.11 |
| 10 | 65791943 | rs2671528 | -0.796 | -0.29 | -0.669 | 0.399 | 0.004 | -0.021 | -0.021 | 0.024 | 0.13 | 0.08 | 0.1 | 0.2 | 0.13 | 0.27 |
| 10 | 65794344 | rs7070910 | -0.945 | -0.588 | -0.88 | 0.327 | 0.014 | -0.029 | -0.029 | 0.329 | 0.17 | 0.03 | 0.12 | 0.18 | 0.14 | 0.17 |
| 10 | 65803370 | rs2252686 | 1.413 | -1.23 | 0.1 | -0.002 | -0.041 | -0.008 | -0.008 | 0 | 0.73 | 0.47 | 0.77 | 0.79 | 0.82 | 0.74 |
| 10 | 65810837 | rs2394034 | -1.167 | -1.397 | 0.9 | -0.034 | -0.048 | 0.034 | 0.034 | 1.184 | 0.09 | 0.23 | 0.08 | 0.11 | 0.03 | 0.05 |
| 10 | 65817792 | rs10491072 | -1.682 | -1.412 | 1.225 | -0.058 | 0.003 | 0.003 | 0.003 | 0 | 0.14 | 0.14 | 0.12 | 0.09 | 0.11 | 0.21 |
| 10 | 65817857 | rs10491073 | -1.004 | -1.416 | 1.193 | -0.048 | -0.046 | -0.028 | -0.028 | 0 | 0.09 | 0.23 | 0.06 | 0.03 | 0.02 | 0.04 |
| 10 | 65818459 | rs10822275 | -2.787 | -1.311 | 1.276 | -0.06 | -0.046 | -0.008 | -0.008 | 0.061 | 0.27 | 0.49 | 0.23 | 0.21 | 0.18 | 0.26 |
| 10 | 65820798 | rs2601701 | 2.398 | -1.458 | 1.553 | -0.081 | -0.028 | -0.009 | -0.009 | 0.523 | 0.72 | 0.41 | 0.77 | 0.8 | 0.82 | 0.74 |
| 10 | 65833782 | rs2578017 | 2.535 | -1.727 | 1.178 | -0.338 | 0.082 | 0.038 | 0.038 | 2.779 | 0.65 | 0.1 | 0.7 | 0.79 | 0.79 | 0.72 |
| 10 | 65837750 | rs2671513 | 2.412 | -1.522 | 1.767 | -0.172 | 0.008 | 0.003 | 0.003 | 0.622 | 0.73 | 0.23 | 0.76 | 0.83 | 0.8 | 0.75 |
| 10 | 65844676 | rs10995803 | -1.484 | -1.605 | 1.793 | -0.209 | -0.006 | -0.035 | -0.035 | 0.146 | 0.06 | 0 | 0.07 | 0.09 | 0.06 | 0.16 |
| 10 | 65844922 | rs2578020 | 2.938 | -1.657 | 2.599 | -0.235 | -0.03 | -0.007 | -0.007 | 0.046 | 0.79 | 0.55 | 0.81 | 0.85 | 0.83 | 0.78 |
| 10 | 65845712 | rs10995804 | -3.134 | -1.664 | 2.706 | -0.238 | -0.003 | -0.004 | -0.004 | 0 | 0.19 | 0.24 | 0.15 | 0.1 | 0.17 | 0.22 |
| 10 | 65845866 | rs2578022 | 3.471 | -1.67 | 2.687 | -0.241 | -0.005 | -0.002 | -0.002 | 0 | 0.75 | 0.25 | 0.79 | 0.85 | 0.83 | 0.78 |
| 10 | 65857064 | rs7918323 | -3.332 | -1.83 | 2.712 | -0.291 | -0.065 | -0.013 | -0.013 | 0 | 0.22 | 0.53 | 0.16 | 0.14 | 0.13 | 0.2 |
| 10 | 65870922 | rs2671536 | 3.215 | -1.78 | 2.852 | -0.241 | 0.053 | 0.016 | 0.016 | 2.972 | 0.73 | 0.17 | 0.79 | 0.82 | 0.85 | 0.78 |
| 10 | 65875491 | rs2601705 | 2.829 | -1.604 | 3.029 | -0.271 | 0.059 | 0.083 | 0.083 | 0.006 | 0.93 | 0.57 | 0.92 | 0.95 | 0.96 | 0.98 |
| 10 | 65887214 | rs10740149 | 2.833 | -1.61 | 2.872 | -0.274 | 0.02 | 0.043 | 0.043 | 1.661 | 0.91 | 0.49 | 0.92 | 0.95 | 0.96 | 0.97 |
| 10 | 65898613 | rs2394042 | 2.571 | -1.622 | 3.195 | -0.284 | 0.003 | 0.015 | 0.015 | 6.023 | 0.93 | 0.39 | 0.95 | 0.99 | 1 | 0.99 |
| 10 | 65902028 | rs2394043 | 2.96 | -1.622 | 2.893 | -0.286 | 0.037 | 0.015 | 0.015 | 2.846 | 0.75 | 0.15 | 0.8 | 0.84 | 0.85 | 0.79 |
| 10 | 65902327 | rs2067653 | -2.939 | -1.639 | 2.911 | -0.282 | -0.045 | -0.001 | -0.001 | 0 | 0.21 | 0.44 | 0.17 | 0.17 | 0.15 | 0.21 |
| 10 | 65924250 | rs12415289 | -1.26 | -1.608 | 2.644 | -0.306 | -0.004 | -0.031 | -0.031 | 0.388 | 0.05 | 0.01 | 0.08 | 0.11 | 0.06 | 0.13 |
| 10 | 65929545 | rs7893379 | -2.58 | -1.542 | 2.433 | -0.278 | -0.056 | -0.012 | -0.012 | 0 | 0.21 | 0.53 | 0.18 | 0.16 | 0.15 | 0.2 |
| 10 | 65950993 | rs4746429 | -1.615 | -1.587 | 2.643 | -0.485 | -0.014 | -0.046 | -0.046 | 0 | 0.12 | 0.45 | 0.08 | 0.04 | 0.07 | 0.02 |
| 10 | 65963249 | rs6479970 | 0.655 | -1.224 | 1.699 | -1.381 | -0.014 | 0.182 | 0.182 | 1.735 | 0.42 | 0.12 | 0.51 | 0.87 | 0.37 | 0.7 |
| 10 | 65977811 | rs4271284 | 0.962 | -1.069 | 2.459 | -1.128 | -0.011 | -0.005 | -0.005 | 0 | 0.93 | 0.71 | 0.97 | 0.99 | 0.99 | 1 |
| 10 | 65993415 | rs7919951 | 2.043 | -0.959 | 2.89 | -1.076 | 0.062 | 0.051 | 0.051 | 4.94 | 0.94 | 0.51 | 0.96 | 1 | 1 | 1 |
| 10 | 65998376 | rs4598558 | 2.754 | -0.976 | 2.823 | -1.077 | -0.038 | -0.002 | -0.002 | 0 | 0.91 | 0.32 | 0.95 | 0.94 | 0.99 | 0.99 |
| 10 | 66013548 | rs4402180 | 2.338 | -0.913 | 2.945 | -1.072 | 0.037 | 0.066 | 0.066 | 5.537 | 0.94 | 0.46 | 0.96 | 0.95 | 0.99 | 0.99 |
| 10 | 66013849 | rs10995850 | 0.092 | -0.612 | 2.509 | -1.256 | -0.033 | 0.185 | 0.185 | 0.051 | 0.54 | 0.19 | 0.62 | 0.87 | 0.51 | 0.73 |
| 10 | 66110805 | rs7916532 | -3.547 | 0.052 | 2.535 | -0.727 | 0.071 | 0.117 | 0.117 | 5.17 | 0.05 | 0.45 | 0.04 | 0.06 | 0 | 0.01 |
| 10 | 66134050 | rs2660062 | -1.246 | 0.143 | 2.09 | -0.714 | -0.005 | 0.002 | 0.002 | 0 | 0.05 | 0.13 | 0.01 | 0.06 | 0.02 | 0.01 |
| 10 | 66153612 | rs2660089 | -2.931 | 0.122 | 2.407 | -0.736 | 0.038 | 0.039 | 0.039 | 1.436 | 0.08 | 0.45 | 0.05 | 0.06 | 0.03 | 0.01 |
| 10 | 66163616 | rs2493369 | -3.028 | 0.121 | 2.39 | -0.737 | 0.034 | 0.036 | 0.036 | 1.349 | 0.08 | 0.47 | 0.05 | 0.06 | 0.03 | 0.01 |
| 10 | 66222693 | rs7913086 | -1.64 | 0.168 | 2.343 | -0.739 | 0.051 | 0.008 | 0.008 | 0 | 0.06 | 0.31 | 0.03 | 0 | 0.02 | 0 |
| 10 | 66222994 | rs2493360 | -0.464 | 0.189 | 1.584 | -0.357 | -0.056 | -0.047 | -0.047 | 0 | 0.6 | 0.88 | 0.46 | 0.65 | 0.41 | 0.54 |
| 10 | 66227925 | rs2817991 | -2.068 | 0.352 | 1.712 | -0.213 | 0.044 | 0.056 | 0.056 | 0.253 | 0.07 | 0.39 | 0.04 | 0.06 | 0.02 | 0.01 |
| 10 | 66259625 | rs10995893 | 0.491 | 0.1 | 1.152 | -0.408 | 0.009 | 0.008 | 0.008 | 0 | 0.47 | 0.4 | 0.41 | 0.65 | 0.37 | 0.5 |

| | | | | | | | | | | | | | | | | |
|---|---|---|---|---|---|---|---|---|---|---|---|---|---|---|---|---|
| 10 | 66260362 | rs10822304 | 0.721 | 0.113 | 0.761 | -0.401 | 0.013 | 0.008 | 0.008 | 0 | 0.45 | 0.18 | 0.4 | 0.64 | 0.37 | 0.5 |
| 10 | 66267858 | rs2394072 | 0.101 | 0.04 | 0.944 | -0.31 | 0.016 | 0.006 | 0.006 | 0 | 0.51 | 0.63 | 0.44 | 0.6 | 0.4 | 0.47 |
| 10 | 66269151 | rs4745850 | 1.306 | -0.121 | 0.735 | -0.516 | -0.008 | -0.131 | -0.131 | 0.128 | 0.36 | 0.07 | 0.37 | 0.57 | 0.31 | 0.34 |
| 10 | 66272624 | rs7076459 | -2.176 | 0.253 | 1.013 | -1.038 | -0.019 | -0.085 | -0.085 | 1.475 | 0.16 | 0.56 | 0.07 | 0.04 | 0.09 | 0.13 |
| 10 | 66279382 | rs10822305 | -0.783 | 0.467 | 0.799 | -1.16 | 0.008 | -0.014 | -0.014 | 0 | 0.49 | 0.57 | 0.39 | 0.09 | 0.56 | 0.35 |
| 10 | 66282577 | rs16919911 | -0.04 | 0.593 | 0.503 | -1.118 | 0.001 | -0.027 | -0.027 | 0.274 | 0.06 | 0.11 | 0.03 | 0 | 0.06 | 0.1 |
| 10 | 66286028 | rs7083780 | 0.395 | 0.681 | 0.212 | -1.131 | -0.131 | 0.217 | 0.217 | 0.132 | 0.35 | 0.05 | 0.33 | 0.04 | 0.49 | 0.25 |
| 10 | 66289103 | rs16919914 | -1.677 | 0.92 | 1.368 | -0.341 | 0.002 | -0.07 | -0.07 | 0 | 0.12 | 0.23 | 0.04 | 0 | 0.08 | 0.12 |
| 10 | 66289282 | rs10509208 | -1.996 | 0.949 | 1.352 | -0.113 | 0.011 | -0.044 | -0.044 | 0 | 0.13 | 0.34 | 0.04 | 0.03 | 0.08 | 0.12 |
| 10 | 66290693 | rs4529793 | -2.594 | 1.343 | 1.235 | -0.253 | 0.005 | -0.09 | -0.09 | 0 | 0.28 | 0.51 | 0.15 | 0.1 | 0.25 | 0.16 |
| 10 | 66294394 | rs10822308 | -2.478 | 1.477 | 1.285 | 0.447 | 0.014 | 0.048 | 0.048 | 1.431 | 0.18 | 0.43 | 0.11 | 0.23 | 0.16 | 0.06 |
| 10 | 66299171 | rs12243555 | -1.299 | 1.777 | 0.549 | 0.546 | 0.022 | 0.037 | 0.037 | 0.35 | 0.09 | 0.17 | 0.08 | 0.18 | 0.14 | 0.04 |
| 10 | 66299771 | rs6479987 | 0.483 | 1.312 | 0.589 | 0.448 | 0.009 | -0.026 | -0.026 | 0 | 0.94 | 0.9 | 0.97 | 1 | 0.95 | 0.93 |
| 10 | 66315231 | rs2893964 | 1.041 | 1.752 | 0.11 | 0.658 | 0.012 | 0.012 | 0.012 | 0.935 | 0.83 | 0.77 | 0.88 | 0.79 | 0.81 | 0.93 |
| 10 | 66328453 | rs10995915 | -0.028 | 1.136 | -0.221 | 0.38 | -0.002 | -0.018 | -0.018 | 2.632 | 0.45 | 0.53 | 0.5 | 0.19 | 0.44 | 0.37 |
| 10 | 66333087 | rs10822315 | 0.05 | 1.131 | -0.221 | 0.315 | -0.003 | -0.037 | -0.037 | 0.374 | 0.43 | 0.52 | 0.49 | 0.16 | 0.43 | 0.36 |
| 10 | 66340742 | rs10822318 | 0.461 | 1.411 | -1.141 | 0.959 | -0.106 | -0.099 | -0.099 | 0.251 | 0.33 | 0.04 | 0.23 | 0.38 | 0.36 | 0.3 |
| 10 | 66354252 | rs3999057 | 0.177 | 0.873 | -0.006 | 0.908 | -0.001 | -0.013 | -0.013 | 0.541 | 0.45 | 0.44 | 0.3 | 0.18 | 0.43 | 0.3 |
| 10 | 66364116 | rs10822320 | -0.462 | 0.838 | 0.722 | 1.142 | 0.024 | 0.196 | 0.196 | 0.486 | 0.14 | 0.4 | 0.41 | 0.43 | 0.14 | 0.31 |
| 10 | 66364605 | rs4746454 | 0.079 | 0.836 | 0.711 | 1.157 | 0.025 | 0.087 | 0.087 | 0.107 | 0.13 | 0.28 | 0.4 | 0.31 | 0.14 | 0.29 |
| 10 | 66371729 | rs10822321 | -0.401 | 0.559 | 0.716 | 1.031 | -0.001 | 0 | 0 | 0 | 0.53 | 0.47 | 0.67 | 0.54 | 0.53 | 0.58 |
| 10 | 66373497 | rs2394084 | 0.166 | 0.608 | 0.696 | 1.067 | 0.019 | 0.161 | 0.161 | 0.188 | 0.13 | 0.28 | 0.41 | 0.4 | 0.14 | 0.29 |
| 10 | 66375958 | rs10995933 | 0.097 | 0.55 | 0.855 | 0.962 | -0.005 | -0.008 | -0.008 | 0.549 | 0.08 | 0.13 | 0.04 | 0.04 | 0.06 | 0.02 |
| 10 | 66377000 | rs7921148 | -0.917 | 0.607 | 0.75 | 0.906 | -0.016 | -0.022 | -0.022 | 0.46 | 0.6 | 0.71 | 0.72 | 0.55 | 0.59 | 0.61 |
| 10 | 66377954 | rs2438995 | -0.294 | 0.672 | 0.761 | 0.808 | 0.009 | 0.02 | 0.02 | 0 | 0.5 | 0.41 | 0.31 | 0.07 | 0.46 | 0.32 |
| 10 | 66378372 | rs2683581 | -1.266 | 0.658 | 0.514 | 1.061 | -0.013 | 0.005 | 0.005 | 0 | 0.64 | 0.72 | 0.73 | 0.65 | 0.61 | 0.64 |
| 10 | 66380566 | rs2394081 | -0.288 | 0.706 | 0.064 | 0.603 | -0.008 | 0.064 | 0.064 | 0.353 | 0.09 | 0.2 | 0.04 | 0.16 | 0.06 | 0.06 |
| 10 | 66381526 | rs11812691 | 0.543 | 0.703 | 0.003 | 0.537 | -0.022 | 0.224 | 0.224 | 0.166 | 0.39 | 0.09 | 0.26 | 0.03 | 0.39 | 0.3 |
| 10 | 66381719 | rs10822323 | -0.225 | 0.765 | 0.041 | 0.635 | 0.017 | 0.073 | 0.073 | 0 | 0.49 | 0.33 | 0.31 | 0.19 | 0.45 | 0.35 |
| 10 | 66406859 | rs2394066 | -0.614 | 0.309 | -0.377 | 0.733 | -0.069 | -0.083 | -0.083 | 0 | 0.16 | 0.02 | 0.28 | 0.29 | 0.23 | 0.32 |
| 10 | 66407750 | rs6479995 | 0.42 | -0.619 | -0.146 | 0.387 | 0.012 | -0.016 | -0.016 | 2.993 | 0.27 | 0.22 | 0.11 | 0 | 0.18 | 0.19 |
| 10 | 66413062 | rs7924177 | 0.753 | -0.575 | 0.468 | 0.552 | 0.014 | -0.003 | -0.003 | 0 | 0.46 | 0.4 | 0.47 | 0.49 | 0.59 | 0.43 |
| 10 | 66424808 | rs4746459 | -1.629 | -0.873 | 0.726 | 0.823 | -0.046 | 0.07 | 0.07 | 0.595 | 0.24 | 0.49 | 0.42 | 0.38 | 0.18 | 0.3 |
| 10 | 66430212 | rs2579793 | 0.123 | -0.849 | 0.538 | 0.853 | -0.009 | -0.001 | -0.001 | 0 | 0.07 | 0.06 | 0.03 | 0.04 | 0.05 | 0.02 |
| 10 | 66437606 | rs4466713 | -0.7 | -0.834 | 0.547 | 0.718 | -0.007 | 0.047 | 0.047 | 0 | 0.14 | 0.25 | 0.37 | 0.34 | 0.13 | 0.28 |
| 10 | 66442823 | rs2139911 | -0.541 | -0.817 | 0.486 | 0.706 | -0.006 | -0.001 | -0.001 | 0.05 | 0.93 | 0.95 | 0.97 | 0.96 | 0.95 | 0.98 |
| 10 | 66453375 | rs10995982 | -0.717 | -0.657 | 0.205 | 0.782 | -0.008 | 0.051 | 0.051 | 0.112 | 0.14 | 0.25 | 0.37 | 0.34 | 0.13 | 0.29 |
| 10 | 66459187 | rs2139899 | 0.301 | -0.747 | 0.205 | 0.706 | -0.023 | -0.006 | -0.006 | 0.052 | 0.91 | 0.84 | 0.95 | 0.96 | 0.93 | 0.98 |
| 10 | 66462432 | rs1915604 | 0.92 | -0.578 | 0.058 | 0.768 | 0 | 0 | 0 | 0 | 0.43 | 0.43 | 0.45 | 0.37 | 0.51 | 0.33 |
| 10 | 66462669 | rs12779886 | 0.037 | -0.506 | -0.317 | 0.858 | -0.034 | -0.069 | -0.069 | 0 | 0.12 | 0.02 | 0.17 | 0.27 | 0.15 | 0.33 |
| 10 | 66465095 | rs10822337 | 0.986 | -0.436 | -0.209 | 0.566 | 0.004 | -0.01 | -0.01 | 0 | 0.42 | 0.49 | 0.44 | 0.36 | 0.5 | 0.33 |
| 10 | 66466805 | rs10740162 | -0.12 | -0.446 | -0.171 | 0.582 | -0.02 | 0.099 | 0.099 | 0 | 0.92 | 0.87 | 0.95 | 0.79 | 0.93 | 0.94 |
| 10 | 66468127 | rs1879593 | -0.613 | -0.422 | -0.198 | 0.593 | -0.009 | -0.001 | -0.001 | 0 | 0.93 | 0.93 | 0.96 | 0.96 | 0.93 | 0.98 |
| 10 | 66471147 | rs7908928 | -0.652 | -0.382 | -0.316 | 0.467 | 0.025 | 0.043 | 0.043 | 0 | 0.49 | 0.34 | 0.51 | 0.42 | 0.43 | 0.61 |
| 10 | 66472840 | rs7897869 | 0.125 | -0.376 | -0.649 | -0.046 | -0.001 | 0.177 | 0.177 | 0.876 | 0.32 | 0.06 | 0.13 | 0 | 0.26 | 0.24 |
| 10 | 66480672 | rs7909723 | 0.714 | -0.035 | -0.134 | 0.959 | -0.025 | 0.098 | 0.098 | 0.004 | 0.91 | 0.83 | 0.95 | 0.79 | 0.93 | 0.94 |
| 10 | 66483216 | rs1915613 | 0.256 | -0.137 | -0.558 | 0.253 | 0 | 0 | 0 | 0 | 0.69 | 0.74 | 0.79 | 0.78 | 0.75 | 0.75 |
| 10 | 66484138 | rs1879595 | 0.06 | -0.295 | -0.609 | 0.306 | -0.008 | -0.024 | -0.024 | 0 | 0.9 | 0.96 | 0.96 | 0.88 | 0.88 | 0.89 |
| 10 | 66484990 | rs12359386 | 0.101 | 0.107 | -0.991 | -0.107 | -0.006 | 0.084 | 0.084 | 2.668 | 0.21 | 0.04 | 0.16 | 0.06 | 0.17 | 0.14 |
| 10 | 66486541 | rs1915616 | 0.283 | -0.374 | -0.808 | -0.016 | -0.018 | 0.016 | 0.016 | 2.24 | 0.62 | 0.52 | 0.6 | 0.53 | 0.64 | 0.59 |
| 10 | 66488057 | rs2394109 | 0.676 | -0.67 | -1.092 | -0.121 | -0.004 | 0.017 | 0.017 | 1.04 | 0.17 | 0.22 | 0.29 | 0.36 | 0.17 | 0.26 |
| 10 | 66492674 | rs2177019 | 0.655 | -0.74 | -1.019 | -0.164 | 0.009 | -0.013 | -0.013 | 0 | 0.64 | 0.67 | 0.61 | 0.52 | 0.66 | 0.63 |
| 10 | 66493717 | rs16920271 | 1.426 | -0.756 | -1.024 | -0.173 | -0.004 | -0.023 | -0.023 | 0 | 0.1 | 0.08 | 0.06 | 0 | 0.06 | 0.04 |
| 10 | 66495095 | rs10509211 | 1.191 | -0.589 | -0.954 | -0.139 | 0.006 | 0.006 | 0.006 | 0 | 0.41 | 0.5 | 0.43 | 0.46 | 0.48 | 0.45 |
| 10 | 66501714 | rs1915633 | -0.365 | -0.355 | -1.04 | 0.515 | -0.006 | 0.002 | 0.002 | 0.42 | 0.26 | 0.29 | 0.34 | 0.37 | 0.23 | 0.27 |
| 10 | 66504223 | rs1915637 | 0.144 | 0.242 | -1.291 | 0.562 | -0.008 | -0.024 | -0.024 | 0 | 0.9 | 0.96 | 0.95 | 0.88 | 0.89 | 0.9 |
| 10 | 66509895 | rs1879603 | 0.31 | 0.322 | -1.44 | 0.538 | 0.011 | -0.066 | -0.066 | 0 | 0.68 | 0.83 | 0.54 | 0.5 | 0.64 | 0.7 |
| 10 | 66510544 | rs12253993 | -0.373 | 0.76 | -1.24 | -0.226 | -0.004 | 0.001 | 0.001 | 0.774 | 0.23 | 0.16 | 0.15 | 0.1 | 0.23 | 0.22 |
| 10 | 66515337 | rs2177023 | 0.404 | 0.229 | -1.26 | -0.153 | 0 | 0.004 | 0.004 | 0 | 0.46 | 0.58 | 0.48 | 0.44 | 0.49 | 0.42 |
| 10 | 66523544 | rs10996031 | 0.655 | -0.074 | -0.75 | 0.124 | -0.029 | -0.004 | -0.004 | 0 | 0.42 | 0.21 | 0.45 | 0.44 | 0.49 | 0.42 |
| 10 | 66529552 | rs7091769 | -0.739 | -0.331 | -0.899 | -0.01 | 0.02 | -0.089 | -0.089 | 0.042 | 0.71 | 0.92 | 0.69 | 0.67 | 0.74 | 0.71 |
| 10 | 66551141 | rs16920386 | -0.856 | -0.34 | -0.666 | -0.034 | 0.055 | 0.226 | 0.226 | 2.46 | 0.07 | 0.49 | 0.1 | 0.2 | 0.03 | 0.08 |
| 10 | 66555441 | rs7073506 | 0.266 | -0.131 | -0.977 | -0.211 | 0.005 | -0.049 | -0.049 | 0 | 0.22 | 0.22 | 0.14 | 0.02 | 0.21 | 0.22 |
| 10 | 66561285 | rs10761865 | -0.281 | -0.572 | -1.191 | 0.323 | 0.036 | -0.044 | -0.044 | 0 | 0.69 | 0.84 | 0.69 | 0.65 | 0.75 | 0.71 |
| 10 | 66571457 | rs3912874 | 0.076 | -0.576 | -0.662 | 0.27 | -0.017 | 0.023 | 0.023 | 0.235 | 0.23 | 0.28 | 0.27 | 0.33 | 0.17 | 0.25 |
| 10 | 66572215 | rs1227215 | -0.038 | -0.117 | -0.726 | 0.303 | -0.003 | -0.022 | -0.022 | 0 | 0.64 | 0.84 | 0.72 | 0.59 | 0.61 | 0.63 |
| 10 | 66580852 | rs10996049 | 0.442 | 0.011 | -0.777 | 0.267 | 0.01 | -0.012 | -0.012 | 0 | 0.26 | 0.1 | 0.14 | 0.26 | 0.23 | 0.3 |
| 10 | 66582806 | rs1227230 | 0.604 | -0.011 | -0.552 | -0.357 | 0.006 | 0.024 | 0.024 | 0 | 0.33 | 0.47 | 0.38 | 0.47 | 0.38 | 0.37 |
| 10 | 66586305 | rs1227236 | -0.397 | 0.043 | -0.582 | -0.273 | -0.002 | -0.017 | -0.017 | 0 | 0.64 | 0.84 | 0.71 | 0.58 | 0.61 | 0.62 |

| | | | | | | | | | | | | | | | | |
|---|---|---|---|---|---|---|---|---|---|---|---|---|---|---|---|---|
| 10 | 66592466 | rs10761867 | 0.886 | -0.041 | -0.585 | -0.224 | 0.01 | -0.014 | -0.014 | 0.238 | 0.26 | 0.15 | 0.14 | 0.26 | 0.22 | 0.3 |
| 10 | 66592821 | rs1239017 | 0.536 | -0.01 | -0.62 | -0.463 | 0.005 | -0.085 | -0.085 | 3.597 | 0.13 | 0.38 | 0.09 | 0 | 0.13 | 0.08 |
| 10 | 66594665 | rs3852427 | 0.234 | -0.033 | -0.435 | -0.432 | 0.002 | 0.004 | 0.004 | 0 | 0.51 | 0.51 | 0.63 | 0.59 | 0.5 | 0.55 |
| 10 | 66610911 | rs7087608 | -0.218 | 0.036 | -0.456 | -0.446 | 0.002 | -0.025 | -0.025 | 0 | 0.77 | 0.68 | 0.77 | 0.84 | 0.71 | 0.84 |
| 10 | 66612353 | rs16920484 | 0.993 | -0.154 | -0.596 | -0.549 | -0.03 | 0.04 | 0.04 | 0 | 0.27 | 0.06 | 0.3 | 0.14 | 0.23 | 0.26 |
| 10 | 66642251 | rs2962761 | -0.982 | -0.19 | -0.318 | -0.492 | -0.001 | -0.05 | -0.05 | 0 | 0.88 | 0.76 | 0.92 | 1 | 0.88 | 0.92 |
| 10 | 66643203 | rs2932854 | 1.863 | -0.225 | -0.286 | -0.489 | 0 | 0 | 0 | 0 | 0.11 | 0.06 | 0.15 | 0.15 | 0.16 | 0.08 |
| 10 | 66657497 | rs7922705 | 0.216 | -0.319 | -0.244 | -0.496 | 0.002 | 0.006 | 0.006 | 0 | 0.51 | 0.53 | 0.63 | 0.59 | 0.5 | 0.55 |
| 10 | 66665315 | rs2962755 | -1.05 | -0.254 | -0.23 | -0.538 | -0.003 | -0.05 | -0.05 | 0 | 0.88 | 0.76 | 0.93 | 1 | 0.89 | 0.92 |
| 10 | 66667627 | rs10822370 | 0.136 | -0.339 | -0.261 | -0.244 | 0.002 | -0.001 | -0.001 | 0 | 0.51 | 0.55 | 0.6 | 0.54 | 0.5 | 0.56 |
| 10 | 66680444 | rs2499891 | 1.764 | -0.288 | -0.193 | -0.203 | -0.001 | 0.001 | 0.001 | 0 | 0.11 | 0.09 | 0.18 | 0.2 | 0.16 | 0.08 |
| 10 | 66694248 | rs2452420 | 0.459 | -0.467 | -0.312 | -0.299 | -0.007 | -0.073 | -0.073 | 0 | 0.36 | 0.13 | 0.32 | 0.46 | 0.39 | 0.36 |
| 10 | 66695246 | rs2452421 | -0.795 | -0.214 | -0.099 | -0.219 | -0.006 | -0.062 | -0.062 | 0.006 | 0.86 | 0.71 | 0.92 | 1 | 0.89 | 0.92 |
| 10 | 66702282 | rs4104490 | 1.257 | -0.109 | 0.238 | -0.24 | -0.007 | -0.009 | -0.009 | 0 | 0.21 | 0.21 | 0.29 | 0.39 | 0.27 | 0.3 |
| 10 | 66702789 | rs12357492 | 0.248 | -0.463 | 0.503 | -0.217 | 0.001 | 0 | 0 | 0 | 0.54 | 0.39 | 0.45 | 0.41 | 0.46 | 0.54 |
| 10 | 66705778 | rs2452439 | -0.282 | -0.441 | 0.404 | -0.213 | 0.001 | -0.001 | -0.001 | 0 | 0.66 | 0.54 | 0.63 | 0.61 | 0.63 | 0.61 |
| 10 | 66719093 | rs10822376 | 0.119 | -0.448 | 0.403 | -0.215 | -0.004 | -0.002 | -0.002 | 0 | 0.55 | 0.5 | 0.45 | 0.41 | 0.46 | 0.54 |
| 10 | 66723680 | rs10733810 | -2.083 | -0.452 | 0.167 | -0.164 | -0.037 | -0.062 | -0.062 | 1.373 | 0.89 | 0.99 | 0.82 | 0.8 | 0.84 | 0.92 |
| 10 | 66726865 | rs7911209 | -1.658 | -0.591 | 0.173 | -0.021 | -0.006 | -0.044 | -0.044 | 1.12 | 0.85 | 0.96 | 0.81 | 0.77 | 0.83 | 0.91 |
| 10 | 66727627 | rs12355982 | 1.317 | -0.311 | 0.317 | -0.055 | 0.015 | 0.014 | 0.014 | 0.027 | 0.19 | 0.27 | 0.3 | 0.38 | 0.3 | 0.34 |
| 10 | 66729281 | rs4746478 | 0.826 | -0.569 | 0.326 | -0.183 | 0.002 | -0.007 | -0.007 | 0 | 0.25 | 0.23 | 0.31 | 0.14 | 0.22 | 0.22 |
| 10 | 66733358 | rs7898628 | 0.119 | -0.451 | 0.386 | -0.179 | 0.022 | -0.003 | -0.003 | 0 | 0.32 | 0.57 | 0.36 | 0.38 | 0.39 | 0.41 |
| 10 | 66741449 | rs10128291 | 0.077 | -0.437 | 0.493 | -0.155 | 0.02 | -0.009 | -0.009 | 0 | 0.33 | 0.54 | 0.36 | 0.35 | 0.39 | 0.41 |
| 10 | 66744526 | rs12415141 | 0.33 | -0.478 | 0.607 | -0.164 | 0 | 0 | 0 | 0 | 0.52 | 0.39 | 0.45 | 0.42 | 0.44 | 0.5 |
| 10 | 66757365 | rs11813325 | 1.109 | -0.483 | 0.541 | -0.2 | -0.003 | -0.033 | -0.033 | 0.317 | 0.11 | 0.16 | 0.05 | 0 | 0.09 | 0.07 |
| 10 | 66758101 | rs10822387 | 0.966 | -0.49 | 0.679 | -0.197 | 0.022 | 0.017 | 0.017 | 0.154 | 0.22 | 0.35 | 0.3 | 0.35 | 0.3 | 0.34 |
| 10 | 66760408 | rs10761892 | -0.508 | -0.556 | 0.603 | -0.23 | -0.008 | -0.017 | -0.017 | 0 | 0.61 | 0.77 | 0.51 | 0.6 | 0.61 | 0.68 |
| 10 | 66769196 | rs11814058 | 1.117 | -0.695 | 0.528 | -0.206 | -0.003 | -0.035 | -0.035 | 1.186 | 0.11 | 0.16 | 0.06 | 0 | 0.09 | 0.07 |
| 10 | 66772293 | rs10822396 | 1.314 | -0.693 | 0.419 | -0.156 | 0.033 | -0.006 | -0.006 | 0 | 0.28 | 0.12 | 0.14 | 0.25 | 0.21 | 0.27 |
| 10 | 66788146 | rs4128743 | -0.688 | -0.725 | 0.503 | -0.169 | 0.024 | -0.013 | -0.013 | 0 | 0.56 | 0.73 | 0.66 | 0.53 | 0.61 | 0.64 |
| 10 | 66796908 | rs10740183 | -1.76 | -0.615 | 0.404 | -0.141 | 0 | -0.001 | -0.001 | 1.147 | 0.85 | 0.91 | 0.81 | 0.77 | 0.83 | 0.91 |
| 10 | 66818010 | rs10822424 | -0.523 | -0.591 | 0.535 | -0.154 | -0.014 | -0.007 | -0.007 | 0 | 0.74 | 0.53 | 0.74 | 0.77 | 0.74 | 0.87 |
| 10 | 66827630 | rs10996176 | 1.652 | -0.705 | 0.236 | -0.167 | 0.034 | -0.05 | -0.05 | 0 | 0.25 | 0.04 | 0.2 | 0.25 | 0.21 | 0.28 |
| 10 | 66833436 | rs10822431 | 0.379 | -0.326 | 0.395 | -0.107 | 0 | 0.01 | 0.01 | 0 | 0.48 | 0.39 | 0.44 | 0.6 | 0.49 | 0.6 |
| 10 | 66854292 | rs7097961 | 1.319 | -0.277 | 0.119 | -0.051 | 0.038 | -0.012 | -0.012 | 0 | 0.28 | 0.1 | 0.22 | 0.25 | 0.22 | 0.29 |
| 10 | 66854543 | rs4237299 | 0.959 | -0.156 | 0.457 | -0.103 | -0.028 | 0.003 | 0.003 | 0 | 0.21 | 0.11 | 0.31 | 0.17 | 0.22 | 0.2 |
| 10 | 66877815 | rs7899394 | -0.229 | -0.228 | 0.946 | -0.064 | -0.013 | -0.006 | -0.006 | 0 | 0.74 | 0.58 | 0.75 | 0.76 | 0.74 | 0.89 |
| 10 | 66878139 | rs7917921 | -0.042 | -0.268 | 0.953 | -0.074 | -0.015 | -0.013 | -0.013 | 1.322 | 0.73 | 0.45 | 0.72 | 0.77 | 0.73 | 0.89 |
| 10 | 66880723 | rs4545433 | 0.066 | -0.289 | 0.883 | -0.084 | -0.02 | -0.017 | -0.017 | 0.032 | 0.72 | 0.43 | 0.72 | 0.77 | 0.72 | 0.89 |
| 10 | 66884125 | rs16920977 | 0.549 | -0.199 | 0.698 | -0.32 | 0.005 | -0.005 | -0.005 | 0.077 | 0.23 | 0.26 | 0.32 | 0.18 | 0.21 | 0.21 |
| 10 | 66884210 | rs4501889 | 0.789 | -0.322 | 0.507 | -0.364 | -0.026 | 0.068 | 0.068 | 0 | 0.48 | 0.15 | 0.4 | 0.66 | 0.5 | 0.67 |
| 10 | 66884266 | rs16920987 | 0.838 | -0.338 | 0.545 | -0.387 | -0.043 | 0.018 | 0.018 | 0 | 0.22 | 0.09 | 0.31 | 0.15 | 0.21 | 0.21 |
| 10 | 66885091 | rs10996224 | -0.228 | -0.3 | 0.894 | -0.371 | 0.001 | 0.006 | 0.006 | 0 | 0.59 | 0.41 | 0.47 | 0.73 | 0.61 | 0.73 |
| 10 | 66885915 | rs12248645 | 0.785 | -0.207 | 0.765 | -0.323 | 0.008 | -0.041 | -0.041 | 0 | 0.09 | 0.17 | 0.06 | 0 | 0.1 | 0.04 |
| 10 | 66888615 | rs4133345 | 0.207 | -0.077 | 0.559 | -0.513 | 0.006 | 0.021 | 0.021 | 0 | 0.61 | 0.37 | 0.66 | 0.83 | 0.65 | 0.84 |
| 10 | 66894669 | rs10822465 | -0.314 | -0.023 | 0.373 | -0.488 | 0.005 | -0.005 | -0.005 | 0 | 0.2 | 0.25 | 0.31 | 0.1 | 0.23 | 0.2 |
| 10 | 66896049 | rs10761930 | 0.933 | -0.064 | 0.025 | -0.507 | -0.029 | 0.116 | 0.116 | 0 | 0.48 | 0.16 | 0.44 | 0.73 | 0.52 | 0.68 |
| 10 | 66898841 | rs4570467 | -0.933 | -0.127 | 0.258 | -0.494 | 0.007 | 0.023 | 0.023 | 0 | 0.73 | 0.84 | 0.79 | 0.83 | 0.76 | 0.89 |
| 10 | 66900545 | rs10822467 | 0.555 | 0.109 | -0.059 | -0.161 | 0.012 | 0.004 | 0.004 | 0 | 0.25 | 0.32 | 0.25 | 0.36 | 0.35 | 0.24 |
| 10 | 66901570 | rs10822470 | 0.748 | -0.002 | -0.367 | -0.093 | 0.034 | -0.042 | -0.042 | 0 | 0.33 | 0.14 | 0.26 | 0.37 | 0.28 | 0.49 |
| 10 | 66903171 | rs12262154 | 0.121 | 0.618 | 0.155 | -0.352 | 0.034 | -0.06 | -0.06 | 0.136 | 0.11 | 0.22 | 0.09 | 0 | 0.16 | 0.07 |
| 10 | 66904588 | rs10996232 | 0.939 | 0.786 | 0.514 | 0.052 | 0.023 | 0.049 | 0.049 | 0 | 0.17 | 0.26 | 0.16 | 0.34 | 0.25 | 0.19 |
| 10 | 66909326 | rs10996237 | -0.285 | 0.455 | 0.445 | -0.031 | 0.01 | -0.012 | -0.012 | 0.109 | 0.39 | 0.51 | 0.52 | 0.68 | 0.45 | 0.33 |
| 10 | 66913425 | rs10996243 | 1.28 | 0.424 | 0.543 | 0.051 | 0.009 | 0.021 | 0.021 | 0 | 0.17 | 0.18 | 0.15 | 0.34 | 0.25 | 0.19 |
| 10 | 66917952 | rs17791551 | 1.456 | 0.363 | 0.609 | 0.009 | -0.001 | 0.004 | 0.004 | 0 | 0.16 | 0.13 | 0.15 | 0.34 | 0.25 | 0.16 |
| 10 | 66921411 | rs9787647 | 1.522 | 0.364 | 0.56 | 0.01 | -0.033 | -0.045 | -0.045 | 0 | 0.15 | 0.06 | 0.14 | 0.33 | 0.24 | 0.15 |
| 10 | 66922407 | rs7077159 | -1.464 | 0.03 | 0.535 | -0.293 | 0.036 | 0.01 | 0.01 | 0 | 0.64 | 0.85 | 0.7 | 0.69 | 0.68 | 0.48 |
| 10 | 66925446 | rs4333906 | 0.108 | 0.2 | 0.658 | -0.25 | 0.033 | 0.033 | 0.033 | 0 | 0.09 | 0.23 | 0.26 | 0.11 | 0.11 | 0.11 |
| 10 | 66930074 | rs4345856 | 1.205 | -0.701 | 0.266 | -0.845 | -0.015 | -0.086 | -0.086 | 0 | 0.22 | 0.29 | 0.09 | 0 | 0.18 | 0.07 |
| 10 | 66931651 | rs12258285 | 0.232 | -0.255 | 0.758 | -0.462 | 0.043 | 0.034 | 0.034 | 0 | 0.27 | 0.5 | 0.21 | 0.34 | 0.35 | 0.22 |
| 10 | 66933364 | rs1509939 | 0.717 | -0.271 | 0.784 | -0.458 | 0.022 | 0.024 | 0.024 | 0 | 0.08 | 0.16 | 0.25 | 0.11 | 0.11 | 0.11 |
| 10 | 66935533 | rs10996261 | 1.403 | -0.275 | 0.703 | -0.465 | 0.022 | -0.05 | -0.05 | 0 | 0.09 | 0.23 | 0.05 | 0 | 0.1 | 0.06 |
| 10 | 66937420 | rs10996264 | 2.195 | -0.285 | 0.719 | -0.465 | -0.014 | -0.019 | -0.019 | 0 | 0.16 | 0.1 | 0.14 | 0.34 | 0.25 | 0.16 |
| 10 | 66938581 | rs12243290 | 1.079 | -0.232 | 0.62 | -0.448 | 0.031 | -0.056 | -0.056 | 0.428 | 0.1 | 0.27 | 0.06 | 0 | 0.1 | 0.06 |
| 10 | 66940792 | rs12259213 | 1.387 | -0.227 | 0.484 | -0.444 | 0.022 | -0.05 | -0.05 | 0 | 0.09 | 0.23 | 0.05 | 0 | 0.1 | 0.06 |
| 10 | 66940859 | rs10761935 | -1.604 | -0.728 | 0.251 | -0.574 | 0.016 | 0 | 0 | 0 | 0.68 | 0.82 | 0.69 | 0.69 | 0.69 | 0.49 |
| 10 | 66951748 | rs17202432 | -0.978 | -0.317 | 0.382 | -0.356 | 0.043 | 0.068 | 0.068 | 0 | 0.5 | 0.73 | 0.61 | 0.68 | 0.61 | 0.39 |
| 10 | 66955128 | rs16921046 | 1.779 | -0.154 | 0.159 | -0.804 | -0.025 | -0.065 | -0.065 | 0 | 0.14 | 0.01 | 0.14 | 0.23 | 0.15 | 0.05 |
| 10 | 66956005 | rs1397023 | 0.285 | 0.027 | 0.605 | -0.86 | 0.006 | 0.006 | 0.006 | 0 | 0.08 | 0.08 | 0.25 | 0.11 | 0.11 | 0.11 |

| | | | | | | | | | | | | | | | | |
|---|---|---|---|---|---|---|---|---|---|---|---|---|---|---|---|---|
| 10 | 66957145 | rs10996274 | 1.208 | -0.387 | 0.59 | -0.949 | 0.004 | 0.017 | 0.017 | 0 | 0.14 | 0.07 | 0.08 | 0.01 | 0.08 | 0.09 |
| 10 | 66965168 | rs10996281 | 0.658 | 0.001 | 0.789 | -0.613 | 0.016 | -0.047 | -0.047 | 0 | 0.08 | 0.23 | 0.05 | 0 | 0.08 | 0.06 |
| 10 | 66965617 | rs7090420 | 0.798 | 0.03 | 0.725 | -0.6 | 0.013 | -0.043 | -0.043 | 0.442 | 0.08 | 0.16 | 0.04 | 0 | 0.08 | 0.06 |
| 10 | 66982356 | rs9415798 | -0.467 | -0.186 | 0.175 | -0.789 | 0.006 | 0.011 | 0.011 | 0 | 0.45 | 0.62 | 0.32 | 0.22 | 0.35 | 0.23 |
| 10 | 66982405 | rs9415799 | 0.228 | -0.282 | 0.452 | -0.89 | 0.032 | -0.031 | -0.031 | 0.957 | 0.12 | 0.36 | 0.08 | 0.04 | 0.11 | 0.06 |
| 10 | 66984552 | rs9414881 | -2.106 | -0.39 | -0.077 | -0.787 | 0.002 | -0.105 | -0.105 | 0.138 | 0.75 | 0.92 | 0.78 | 0.45 | 0.73 | 0.63 |
| 10 | 66985436 | rs12780749 | -0.004 | -0.212 | 0.171 | -0.79 | -0.035 | 0 | 0 | 0.206 | 0.35 | 0.24 | 0.58 | 0.37 | 0.51 | 0.27 |
| 10 | 66987629 | rs12411933 | 0.701 | 0.024 | 0.074 | -0.805 | -0.092 | 0.021 | 0.021 | 0.581 | 0.28 | 0.08 | 0.44 | 0.26 | 0.43 | 0.17 |
| 10 | 66990842 | rs1567410 | -1.219 | 0.016 | 0.127 | -0.832 | -0.032 | 0.101 | 0.101 | 0 | 0.52 | 0.82 | 0.55 | 0.33 | 0.43 | 0.34 |
| 10 | 66992498 | rs950998 | 2.117 | 0.022 | 0.107 | -0.826 | -0.033 | -0.026 | -0.026 | 0 | 0.12 | 0.01 | 0.23 | 0.17 | 0.15 | 0.06 |
| 10 | 66996579 | rs6480051 | -0.631 | 0.006 | 0.182 | -0.818 | 0.007 | 0.074 | 0.074 | 0 | 0.4 | 0.76 | 0.32 | 0.16 | 0.28 | 0.28 |
| 10 | 66998577 | rs7912537 | 0.19 | -0.035 | 0.502 | -0.875 | -0.009 | -0.09 | -0.09 | 0.273 | 0.3 | 0.42 | 0.17 | 0.04 | 0.2 | 0.17 |
| 10 | 67003854 | rs7073035 | -1.031 | -0.022 | 0.46 | -0.844 | 0 | 0.001 | 0.001 | 0 | 0.61 | 0.65 | 0.62 | 0.3 | 0.62 | 0.34 |
| 10 | 67007948 | rs16921143 | 0.929 | 0.01 | 0.548 | -0.817 | 0.012 | 0.019 | 0.019 | 0 | 0.06 | 0.07 | 0.13 | 0.11 | 0.08 | 0.11 |
| 10 | 67008048 | rs17204569 | 1.939 | 0.016 | 0.364 | -0.873 | -0.053 | 0.029 | 0.029 | 0 | 0.16 | 0.04 | 0.2 | 0.08 | 0.28 | 0.11 |
| 10 | 67009817 | rs10996305 | -0.019 | -0.041 | 0.42 | -0.914 | -0.018 | -0.121 | -0.121 | 1.215 | 0.32 | 0.49 | 0.17 | 0.04 | 0.2 | 0.17 |
| 10 | 67010005 | rs10822483 | -0.696 | 0.017 | 0.444 | -0.843 | 0.006 | 0.061 | 0.061 | 0 | 0.5 | 0.64 | 0.39 | 0.16 | 0.47 | 0.3 |
| 10 | 67011069 | rs9415809 | 1.168 | 0.019 | 0.544 | -1.038 | 0.015 | -0.046 | -0.046 | 0 | 0.09 | 0.18 | 0.06 | 0 | 0.1 | 0.06 |
| 10 | 67015137 | rs9299479 | -0.537 | 0.067 | 0.278 | -0.945 | 0.004 | 0.057 | 0.057 | 0 | 0.49 | 0.64 | 0.38 | 0.13 | 0.48 | 0.29 |
| 10 | 67019116 | rs9415812 | -1.666 | -0.092 | -0.269 | -0.752 | 0.031 | -0.048 | -0.048 | 0 | 0.69 | 0.88 | 0.76 | 0.41 | 0.71 | 0.45 |
| 10 | 67025244 | rs7898559 | -0.135 | 0.078 | -0.086 | -0.723 | 0.036 | -0.003 | -0.003 | 3.985 | 0.29 | 0.55 | 0.37 | 0.24 | 0.37 | 0.22 |
| 10 | 67029555 | rs12573629 | -1.615 | 0.007 | -0.478 | -0.668 | 0.035 | -0.051 | -0.051 | 0 | 0.68 | 0.87 | 0.76 | 0.41 | 0.71 | 0.45 |
| 10 | 67036162 | rs12413637 | 0.311 | 0.089 | -0.395 | -0.54 | -0.041 | -0.003 | -0.003 | 0 | 0.24 | 0.16 | 0.34 | 0.2 | 0.36 | 0.22 |
| 10 | 67042858 | rs3106131 | -0.687 | -0.133 | -0.19 | -0.715 | -0.005 | -0.063 | -0.063 | 0 | 0.76 | 0.74 | 0.88 | 1 | 0.88 | 0.85 |
| 10 | 67043016 | rs7914535 | -0.166 | 0.057 | -0.506 | -0.342 | -0.058 | -0.007 | -0.007 | 0 | 0.36 | 0.21 | 0.57 | 0.37 | 0.51 | 0.27 |
| 10 | 67055905 | rs1509937 | 0.421 | 0.369 | -0.548 | -0.363 | -0.016 | 0.06 | 0.06 | 0 | 0.3 | 0.16 | 0.27 | 0.15 | 0.33 | 0.18 |
| 10 | 67057853 | rs7904079 | 0.392 | 0.231 | 0.083 | -0.433 | -0.001 | -0.042 | -0.042 | 0 | 0.08 | 0.22 | 0.06 | 0 | 0.05 | 0.05 |
| 10 | 67060612 | rs17205562 | -0.006 | 0.278 | -0.148 | -0.445 | -0.046 | 0.075 | 0.075 | 0.227 | 0.4 | 0.16 | 0.51 | 0.3 | 0.46 | 0.25 |
| 10 | 67060807 | rs10761940 | 1.211 | 0.158 | 0.279 | -0.025 | 0.011 | -0.024 | -0.024 | 4.655 | 0.39 | 0.25 | 0.26 | 0.43 | 0.37 | 0.54 |
| 10 | 67061036 | rs16921201 | 0.292 | 0.158 | 0.45 | -0.113 | 0.008 | -0.036 | -0.036 | 0 | 0.08 | 0.25 | 0.06 | 0 | 0.05 | 0.05 |
| 10 | 67062272 | rs11591785 | 0.465 | 0.171 | 0.38 | -0.113 | 0.005 | -0.039 | -0.039 | 2.868 | 0.07 | 0.21 | 0.06 | 0 | 0.05 | 0.05 |
| 10 | 67066325 | rs17205765 | 1.726 | 0.294 | -0.085 | 0.002 | -0.037 | -0.01 | -0.01 | 0.962 | 0.11 | 0.01 | 0.24 | 0.13 | 0.15 | 0.05 |
| 10 | 67067094 | rs7921022 | 0.035 | 0.253 | 0.364 | 0.168 | 0.003 | 0.01 | 0.01 | 0 | 0.88 | 0.82 | 0.84 | 0.89 | 0.88 | 0.88 |
| 10 | 67072633 | rs10761942 | 0.897 | 0.308 | -0.412 | 0.055 | -0.095 | 0.045 | 0.045 | 0.113 | 0.45 | 0.09 | 0.51 | 0.71 | 0.5 | 0.53 |
| 10 | 67073779 | rs9414897 | 0.704 | 0.201 | -0.282 | -0.022 | -0.002 | 0.005 | 0.005 | 0 | 0.3 | 0.25 | 0.27 | 0.18 | 0.33 | 0.24 |
| 10 | 67075735 | rs4746503 | 0.711 | 0.485 | -1.168 | 0.159 | -0.096 | 0.043 | 0.043 | 0.122 | 0.44 | 0.09 | 0.51 | 0.71 | 0.5 | 0.5 |
| 10 | 67080523 | rs1566851 | 0.764 | 0.371 | -1.173 | 0.29 | -0.111 | -0.002 | -0.002 | 0 | 0.44 | 0.1 | 0.5 | 0.67 | 0.48 | 0.51 |
| 10 | 67082144 | rs10761947 | 1.303 | 0.671 | -0.883 | 0.357 | -0.026 | -0.054 | -0.054 | 0 | 0.31 | 0.14 | 0.25 | 0.57 | 0.31 | 0.41 |
| 10 | 67082879 | rs953458 | -0.231 | 0.566 | -0.66 | -0.032 | -0.002 | -0.011 | -0.011 | 0 | 0.21 | 0.27 | 0.19 | 0.09 | 0.17 | 0.26 |
| 10 | 67083877 | rs10761949 | 0.868 | 0.942 | -1.07 | 0.511 | -0.029 | -0.052 | -0.052 | 0 | 0.3 | 0.14 | 0.24 | 0.4 | 0.31 | 0.4 |
| 10 | 67087439 | rs4746509 | -0.09 | 1.603 | -0.602 | 0.291 | 0.014 | 0.021 | 0.021 | 0 | 0.08 | 0.14 | 0.15 | 0.14 | 0.12 | 0.07 |
| 10 | 67088959 | rs12242604 | -0.39 | 1.744 | -0.793 | -0.061 | 0.002 | -0.022 | -0.022 | 0.115 | 0.16 | 0.16 | 0.14 | 0.01 | 0.25 | 0.2 |
| 10 | 67091080 | rs12708315 | -0.248 | 1.793 | -0.855 | -0.023 | -0.002 | -0.01 | -0.01 | 0 | 0.16 | 0.14 | 0.14 | 0.01 | 0.25 | 0.2 |
| 10 | 67095248 | rs10996337 | 0.633 | 1.782 | -1.103 | 0.521 | -0.039 | -0.045 | -0.045 | 6.03 | 0.08 | 0.02 | 0.24 | 0.24 | 0.14 | 0.08 |
| 10 | 67096368 | rs10509220 | -0.25 | 1.262 | -1.308 | 0.208 | -0.001 | 0.03 | 0.03 | 0 | 0.29 | 0.17 | 0.19 | 0.18 | 0.31 | 0.22 |
| 10 | 67096744 | rs1948853 | 0.974 | 1.027 | -1.389 | 0.018 | -0.027 | -0.071 | -0.071 | 4.62 | 0.25 | 0.14 | 0.34 | 0.49 | 0.31 | 0.37 |
| 10 | 67103240 | rs7909330 | -0.118 | 0.269 | -0.278 | -0.359 | -0.017 | -0.068 | -0.068 | 0 | 0.15 | 0.3 | 0.07 | 0.03 | 0.09 | 0.16 |
| 10 | 67111828 | rs877603 | -0.041 | 0.258 | -0.232 | -0.368 | -0.02 | -0.08 | -0.08 | 0 | 0.15 | 0.35 | 0.08 | 0.03 | 0.09 | 0.16 |
| 10 | 67120713 | rs10996347 | -0.254 | 0.797 | -0.591 | -0.211 | -0.032 | 0.026 | 0.026 | 0.986 | 0.29 | 0.16 | 0.22 | 0.2 | 0.4 | 0.32 |
| 10 | 67129861 | rs7910152 | 1.12 | 0.382 | -0.55 | -0.499 | 0.009 | 0.019 | 0.019 | 0 | 0.16 | 0.12 | 0.09 | 0.05 | 0.11 | 0.17 |
| 10 | 67130644 | rs7071260 | -1.543 | 0.006 | -0.228 | -0.629 | -0.015 | -0.055 | -0.055 | 0 | 0.62 | 0.8 | 0.66 | 0.54 | 0.62 | 0.63 |
| 10 | 67132299 | rs6480063 | 1.068 | 0.018 | 0.382 | -0.606 | -0.018 | -0.049 | -0.049 | 0 | 0.17 | 0.28 | 0.11 | 0.05 | 0.11 | 0.17 |
| 10 | 67135761 | rs1504476 | 0.487 | 0.023 | 0.286 | -0.607 | -0.029 | -0.068 | -0.068 | 2.427 | 0.2 | 0.34 | 0.11 | 0.05 | 0.11 | 0.17 |
| 10 | 67154646 | rs1113930 | -1.264 | 0.08 | 0.38 | -0.576 | -0.004 | 0.009 | 0.009 | 0 | 0.58 | 0.5 | 0.63 | 0.54 | 0.61 | 0.62 |
| 10 | 67159154 | rs7900804 | 0.937 | 0.002 | 0.449 | -0.622 | -0.018 | -0.049 | -0.049 | 0.002 | 0.19 | 0.25 | 0.11 | 0.05 | 0.12 | 0.17 |
| 10 | 67164983 | rs10761972 | -1.033 | 0.157 | 0.352 | -0.591 | 0.003 | -0.012 | -0.012 | 0 | 0.55 | 0.5 | 0.35 | 0.32 | 0.54 | 0.55 |
| 10 | 67168289 | rs6480079 | 0.68 | 0.126 | 0.29 | -0.591 | -0.023 | -0.022 | -0.022 | 0 | 0.2 | 0.28 | 0.12 | 0.12 | 0.12 | 0.21 |
| 10 | 67168401 | rs6480081 | -1.073 | 0.257 | 0.312 | -0.518 | -0.013 | 0.025 | 0.025 | 0 | 0.56 | 0.66 | 0.37 | 0.32 | 0.55 | 0.56 |
| 10 | 67178895 | rs10761977 | 0.045 | 0.214 | 0.342 | -0.575 | 0.006 | -0.01 | -0.01 | 0.152 | 0.3 | 0.34 | 0.23 | 0.2 | 0.4 | 0.32 |
| 10 | 67179126 | rs10996393 | -0.522 | 0.115 | 0.331 | -0.503 | -0.025 | 0 | 0 | 0 | 0.31 | 0.43 | 0.41 | 0.34 | 0.2 | 0.32 |
| 10 | 67183928 | rs10822545 | -0.397 | -0.011 | 0.102 | -0.476 | -0.014 | 0.003 | 0.003 | 0 | 0.29 | 0.31 | 0.38 | 0.34 | 0.16 | 0.31 |
| 10 | 67185099 | rs2655902 | -0.942 | 0.472 | -0.229 | -0.749 | -0.032 | 0.071 | 0.071 | 0 | 0.55 | 0.79 | 0.6 | 0.34 | 0.48 | 0.53 |
| 10 | 67186972 | rs10996405 | -0.266 | 0.321 | 0.564 | -0.168 | -0.003 | 0.155 | 0.155 | 0 | 0.11 | 0.26 | 0.06 | 0.31 | 0.09 | 0.09 |
| 10 | 67192650 | rs7897868 | 0.314 | 0.269 | 0.469 | -0.714 | -0.02 | 0.14 | 0.14 | 0.403 | 0.46 | 0.27 | 0.3 | 0.02 | 0.53 | 0.51 |
| 10 | 67193355 | rs2800210 | -1.942 | 0.167 | 0.265 | -0.471 | -0.086 | -0.141 | -0.141 | 0 | 0.74 | 0.98 | 0.69 | 0.64 | 0.71 | 0.78 |
| 10 | 67193734 | rs10822552 | 0.201 | 0.214 | 0.795 | -0.578 | 0 | -0.074 | -0.074 | 0.375 | 0.5 | 0.53 | 0.32 | 0.02 | 0.54 | 0.51 |
| 10 | 67199769 | rs10509225 | -1.598 | 0.529 | 0.735 | -0.312 | -0.023 | -0.085 | -0.085 | 0.053 | 0.73 | 0.91 | 0.67 | 0.54 | 0.71 | 0.69 |
| 10 | 67203866 | rs7070283 | -0.047 | 0.396 | 0.973 | -0.368 | 0 | 0.019 | 0.019 | 0 | 0.55 | 0.45 | 0.58 | 0.24 | 0.59 | 0.58 |
| 10 | 67204339 | rs7070549 | -1.283 | 0.511 | 0.433 | -0.25 | -0.026 | -0.079 | -0.079 | 0 | 0.69 | 0.87 | 0.66 | 0.54 | 0.69 | 0.67 |

| | | | | | | | | | | | | | | | | |
|---|---|---|---|---|---|---|---|---|---|---|---|---|---|---|---|---|
| 10 | 67212950 | rs10996429 | 0.838 | 0.618 | 0.237 | -0.2 | -0.001 | 0.022 | 0.022 | 0 | 0.08 | 0.06 | 0.04 | 0.3 | 0.07 | 0.08 |
| 10 | 67213589 | rs10996430 | 0.985 | 0.906 | 0.219 | -0.143 | -0.052 | 0.071 | 0.071 | 0 | 0.13 | 0.03 | 0.12 | 0 | 0.25 | 0.19 |
| 10 | 67227685 | rs10996448 | -0.465 | 0.605 | 0.656 | -0.11 | 0.001 | -0.002 | -0.002 | 0.202 | 0.63 | 0.47 | 0.36 | 0.31 | 0.63 | 0.6 |
| 10 | 67244216 | rs11812710 | 0.602 | 0.88 | 0.412 | -0.189 | -0.004 | -0.016 | -0.016 | 0 | 0.14 | 0.15 | 0.12 | 0 | 0.22 | 0.2 |
| 10 | 67251947 | rs10996467 | 0.028 | 0.57 | 0.679 | -0.322 | -0.003 | 0.002 | 0.002 | 0 | 0.58 | 0.57 | 0.28 | 0.32 | 0.42 | 0.44 |
| 10 | 67253807 | rs7087756 | -1.486 | 0.57 | 0.978 | -0.214 | -0.01 | -0.026 | -0.026 | 2.487 | 0.76 | 0.85 | 0.66 | 0.55 | 0.69 | 0.7 |
| 10 | 67255943 | rs10996470 | 0.967 | 0.509 | 1.089 | -0.284 | 0.004 | -0.055 | -0.055 | 0 | 0.15 | 0.21 | 0.13 | 0 | 0.22 | 0.2 |
| 10 | 67257711 | rs1936625 | -1.764 | 0.508 | 1.012 | -0.15 | -0.086 | -0.124 | -0.124 | 0 | 0.78 | 0.98 | 0.68 | 0.62 | 0.7 | 0.78 |
| 10 | 67261194 | rs7908228 | 0.073 | 0.486 | 1.099 | -0.294 | 0.003 | -0.007 | -0.007 | 0.768 | 0.58 | 0.52 | 0.27 | 0.32 | 0.43 | 0.44 |
| 10 | 67263249 | rs2394147 | -1.522 | 0.617 | 0.77 | -0.203 | -0.007 | -0.014 | -0.014 | 0 | 0.77 | 0.84 | 0.66 | 0.51 | 0.69 | 0.69 |
| 10 | 67267265 | rs7094788 | -1.175 | 0.609 | 0.763 | -0.247 | -0.001 | -0.001 | -0.001 | 0 | 0.73 | 0.73 | 0.4 | 0.32 | 0.64 | 0.64 |
| 10 | 67271725 | rs12764106 | 0.94 | 0.352 | 0.561 | -0.422 | 0.007 | 0.03 | 0.03 | 0 | 0.44 | 0.17 | 0.22 | 0.32 | 0.37 | 0.41 |
| 10 | 67291368 | rs10822585 | 0.164 | 0.348 | 0.574 | -0.523 | -0.002 | -0.062 | -0.062 | 0 | 0.22 | 0.25 | 0.13 | 0.01 | 0.24 | 0.21 |
| 10 | 67300028 | rs4271287 | 0.877 | 0.397 | -0.124 | -0.469 | 0.005 | 0.026 | 0.026 | 0 | 0.44 | 0.14 | 0.22 | 0.32 | 0.37 | 0.41 |
| 10 | 67303155 | rs1902367 | 0.616 | 0.784 | 0.141 | -0.178 | -0.008 | -0.008 | -0.008 | 0 | 0.27 | 0.2 | 0.6 | 0.68 | 0.36 | 0.36 |
| 10 | 67318448 | rs7077284 | 0.056 | 0.147 | 0.164 | -0.801 | -0.017 | -0.1 | -0.1 | 0.262 | 0.69 | 0.56 | 0.76 | 0.93 | 0.72 | 0.67 |
| 10 | 67336685 | rs10822593 | -0.468 | -0.137 | 0.94 | -0.706 | -0.028 | -0.074 | -0.074 | 0 | 0.2 | 0.35 | 0.2 | 0.07 | 0.14 | 0.12 |
| 10 | 67344370 | rs6480102 | -1.18 | 0.262 | 0.211 | -0.271 | -0.027 | -0.05 | -0.05 | 0 | 0.7 | 0.84 | 0.67 | 0.59 | 0.57 | 0.61 |
| 10 | 67345326 | rs1902387 | -1.941 | 0.156 | 0.145 | -0.28 | -0.051 | -0.077 | -0.077 | 0 | 0.77 | 0.9 | 0.69 | 0.59 | 0.6 | 0.64 |
| 10 | 67379181 | rs1902365 | 0.722 | 0 | 0.434 | -0.261 | -0.045 | 0.056 | 0.056 | 0 | 0.45 | 0.13 | 0.36 | 0.33 | 0.4 | 0.45 |
| 10 | 67385963 | rs16921640 | 1.232 | -0.013 | 0.147 | -0.578 | -0.003 | -0.005 | -0.005 | 0 | 0.18 | 0.17 | 0.25 | 0.32 | 0.17 | 0.17 |
| 10 | 67395375 | rs12357200 | 0.261 | -0.052 | 0.626 | -0.317 | -0.024 | -0.049 | -0.049 | 5.967 | 0.17 | 0.29 | 0.13 | 0.06 | 0.12 | 0.1 |
| 10 | 67399447 | rs7095596 | -0.228 | -0.041 | 0.59 | -0.298 | -0.018 | 0.003 | 0.003 | 0 | 0.19 | 0.47 | 0.2 | 0.2 | 0.15 | 0.13 |
| 10 | 67418287 | rs10996577 | 2.386 | -0.037 | 0.514 | -0.318 | 0.004 | 0.003 | 0.003 | 0.174 | 0.08 | 0.06 | 0.06 | 0.06 | 0.04 | 0.07 |
| 10 | 67421784 | rs12779021 | -0.545 | -0.406 | 0.479 | -0.337 | 0.019 | -0.038 | -0.038 | 0 | 0.08 | 0.26 | 0.06 | 0 | 0.07 | 0.03 |
| 10 | 67429974 | rs7906316 | -1.536 | -0.76 | 0.455 | -0.652 | -0.03 | -0.013 | -0.013 | 1.758 | 0.29 | 0.54 | 0.3 | 0.26 | 0.2 | 0.19 |
| 10 | 67432125 | rs10509229 | 1.374 | -0.892 | 0.026 | -0.568 | 0.007 | -0.004 | -0.004 | 0.142 | 0.12 | 0.08 | 0.15 | 0.12 | 0.07 | 0.1 |
| 10 | 67455242 | rs1904729 | -0.626 | -0.654 | -0.133 | -0.497 | 0.02 | 0.021 | 0.021 | 0 | 0.67 | 0.46 | 0.61 | 0.59 | 0.53 | 0.61 |
| 10 | 67461627 | rs16921731 | 0.501 | -1.611 | -0.463 | -0.687 | -0.01 | 0.032 | 0.032 | 0 | 0.41 | 0.14 | 0.22 | 0.33 | 0.35 | 0.41 |
| 10 | 67465948 | rs10996604 | -0.007 | -1.631 | -0.223 | -0.685 | 0.015 | 0.043 | 0.043 | 0 | 0.5 | 0.2 | 0.28 | 0.4 | 0.4 | 0.49 |
| 10 | 67470686 | rs4596964 | -1.248 | -1.633 | 0.229 | -0.682 | 0.001 | 0.001 | 0.001 | 0 | 0.7 | 0.72 | 0.54 | 0.58 | 0.5 | 0.58 |
| 10 | 67473745 | rs10509232 | 0.212 | -1.515 | 0.5 | -0.727 | 0.013 | 0.001 | 0.001 | 0 | 0.06 | 0.14 | 0.1 | 0.06 | 0.04 | 0.03 |
| 10 | 67473973 | rs10509234 | -0.611 | -1.598 | 0.487 | -0.769 | -0.008 | -0.016 | -0.016 | 0 | 0.15 | 0.17 | 0.12 | 0.06 | 0.08 | 0.07 |
| 10 | 67474389 | rs4746528 | 0.64 | -1.619 | 0.35 | -0.872 | -0.017 | -0.015 | -0.015 | 0 | 0.39 | 0.29 | 0.51 | 0.51 | 0.53 | 0.48 |
| 10 | 67477049 | rs10996610 | 1.08 | -1.748 | 0.409 | -0.981 | 0.017 | 0.044 | 0.044 | 0 | 0.08 | 0.01 | 0.01 | 0 | 0.04 | 0.05 |
| 10 | 67477964 | rs4333907 | -1.747 | -1.89 | 0.446 | -1.027 | 0.022 | 0.039 | 0.039 | 0 | 0.76 | 0.73 | 0.57 | 0.53 | 0.55 | 0.63 |
| 10 | 67478076 | rs7067834 | -0.677 | -1.864 | 0.378 | -1.018 | 0.014 | 0.011 | 0.011 | 0 | 0.64 | 0.49 | 0.43 | 0.43 | 0.49 | 0.56 |
| 10 | 67478463 | rs10740210 | 0.377 | -1.993 | 0.336 | -1.157 | -0.014 | -0.022 | -0.022 | 0.198 | 0.21 | 0.27 | 0.17 | 0.1 | 0.09 | 0.11 |
| 10 | 67478601 | rs17808647 | 0.581 | -2.021 | 0.125 | -1.113 | -0.029 | 0.037 | 0.037 | 0 | 0.42 | 0.16 | 0.25 | 0.33 | 0.4 | 0.44 |
| 10 | 67481425 | rs17808757 | 0.623 | -1.835 | 0.378 | -0.735 | 0.006 | -0.002 | -0.002 | 0 | 0.05 | 0.08 | 0.09 | 0.04 | 0.04 | 0.03 |
| 10 | 67486746 | rs7921023 | -0.484 | -2.324 | 0.248 | -0.8 | -0.001 | -0.001 | -0.001 | 0 | 0.49 | 0.52 | 0.3 | 0.38 | 0.34 | 0.41 |
| 10 | 67488312 | rs17235168 | -0.027 | -1.84 | -0.28 | -0.621 | 0.01 | 0.008 | 0.008 | 0 | 0.11 | 0.07 | 0.14 | 0.1 | 0.07 | 0.07 |
| 10 | 67495478 | rs11596982 | 1.271 | -1.69 | -0.467 | -0.462 | -0.066 | -0.063 | -0.063 | 2.265 | 0.14 | 0.01 | 0.1 | 0.23 | 0.19 | 0.13 |
| 10 | 67496301 | rs7085632 | 0.608 | -1.666 | 0.204 | -0.452 | -0.02 | -0.02 | -0.02 | 0 | 0.16 | 0.18 | 0.07 | 0.06 | 0.05 | 0.08 |
| 10 | 67498308 | rs4448580 | 0.748 | -1.74 | 0.175 | -0.477 | -0.001 | 0.008 | 0.008 | 0 | 0.35 | 0.24 | 0.27 | 0.3 | 0.34 | 0.35 |
| 10 | 67500183 | rs7097327 | -0.604 | -1.779 | 0.099 | -0.451 | -0.037 | -0.074 | -0.074 | 0.036 | 0.57 | 0.91 | 0.51 | 0.49 | 0.41 | 0.46 |
| 10 | 67516982 | rs7067583 | 0.513 | -1.61 | 0.338 | -0.702 | 0.001 | -0.001 | -0.001 | 0 | 0.38 | 0.31 | 0.35 | 0.37 | 0.36 | 0.37 |
| 10 | 67522622 | rs10822628 | -0.789 | -1.734 | 0.798 | -0.442 | -0.022 | -0.043 | -0.043 | 0 | 0.24 | 0.25 | 0.07 | 0.06 | 0.12 | 0.14 |
| 10 | 67523452 | rs1904741 | 0.159 | -1.794 | 0.902 | -0.481 | -0.007 | -0.015 | -0.015 | 0.058 | 0.81 | 0.8 | 0.92 | 0.94 | 0.91 | 0.88 |
| 10 | 67523912 | rs1904740 | 0.126 | -1.668 | 0.375 | -0.414 | -0.018 | -0.085 | -0.085 | 0 | 0.86 | 0.67 | 0.93 | 1 | 0.92 | 0.94 |
| 10 | 67524893 | rs10822630 | 0.223 | -1.655 | -0.348 | -0.286 | -0.028 | -0.017 | -0.017 | 0 | 0.16 | 0.24 | 0.07 | 0.09 | 0.06 | 0.1 |
| 10 | 67528414 | rs1114148 | 0.228 | -1.19 | -0.857 | -0.07 | 0.069 | 0.061 | 0.061 | 1.012 | 0.66 | 0.2 | 0.73 | 0.8 | 0.85 | 0.81 |
| 10 | 67533460 | rs10762012 | 0.478 | -0.684 | -1.126 | 0.126 | -0.177 | -0.177 | -0.177 | 0.789 | 0.26 | 0.02 | 0.39 | 0.43 | 0.48 | 0.38 |
| 10 | 67539130 | rs1620502 | -0.889 | -1.485 | 0.166 | -0.197 | -0.026 | -0.028 | -0.028 | 0 | 0.31 | 0.63 | 0.25 | 0.17 | 0.14 | 0.18 |
| 10 | 67548278 | rs1670163 | -0.971 | -1.308 | -0.027 | -0.367 | -0.052 | -0.074 | -0.074 | 0 | 0.3 | 0.57 | 0.13 | 0.09 | 0.14 | 0.15 |
| 10 | 67550128 | rs2924301 | -1.157 | -1.79 | 0.053 | -0.746 | -0.046 | -0.106 | -0.106 | 0 | 0.27 | 0.53 | 0.11 | 0.06 | 0.13 | 0.16 |
| 10 | 67553020 | rs7080893 | -1.228 | -1.584 | -0.591 | -0.284 | 0.05 | 0.014 | 0.014 | 0 | 0.41 | 0.79 | 0.34 | 0.36 | 0.31 | 0.25 |
| 10 | 67555074 | rs10762015 | -1.199 | -1.457 | -0.489 | -0.235 | 0.031 | 0.001 | 0.001 | 0 | 0.38 | 0.75 | 0.32 | 0.36 | 0.28 | 0.25 |
| 10 | 67576326 | rs12220499 | -1.054 | -1.417 | -0.476 | -0.205 | -0.003 | 0.002 | 0.002 | 1.173 | 0.31 | 0.62 | 0.28 | 0.34 | 0.24 | 0.22 |
| 10 | 67598906 | rs7090646 | -0.81 | -1.271 | -0.786 | 0.275 | 0.027 | -0.03 | -0.03 | 1.455 | 0.36 | 0.75 | 0.33 | 0.43 | 0.24 | 0.22 |
| 10 | 67606959 | rs4402182 | -1.158 | -1.616 | -0.811 | 0.244 | 0.056 | -0.025 | -0.025 | 0 | 0.43 | 0.79 | 0.35 | 0.52 | 0.28 | 0.29 |
| 10 | 67611745 | rs10996721 | 0.542 | -1.96 | -1.046 | 0.35 | -0.068 | -0.077 | -0.077 | 0 | 0.15 | 0.01 | 0.18 | 0.24 | 0.18 | 0.12 |
| 10 | 67614091 | rs4542295 | -1.01 | -2.121 | -0.307 | 0.27 | -0.068 | -0.068 | -0.068 | 0 | 0.21 | 0.64 | 0.11 | 0.08 | 0.06 | 0.12 |
| 10 | 67617331 | rs7908509 | -0.06 | -1.862 | -0.308 | 0.237 | -0.023 | 0.069 | 0.069 | 0 | 0.07 | 0.14 | 0.06 | 0.2 | 0.03 | 0.05 |
| 10 | 67618247 | rs4379724 | -0.846 | -1.957 | -1.076 | 0.276 | -0.002 | 0.036 | 0.036 | 0.014 | 0.84 | 0.84 | 0.87 | 0.58 | 0.85 | 0.74 |
| 10 | 67619594 | rs1947370 | 1.257 | -1.024 | -1.901 | -0.428 | -0.021 | 0.099 | 0.099 | 0 | 0.23 | 0.08 | 0.2 | 0.04 | 0.26 | 0.14 |
| 10 | 67622739 | rs10509237 | 0.387 | -1.147 | -1.922 | -0.536 | -0.029 | 0.068 | 0.068 | 0.868 | 0.4 | 0.22 | 0.38 | 0.27 | 0.43 | 0.2 |
| 10 | 67624255 | rs10996742 | 1.744 | -1.286 | -1.835 | -0.455 | -0.014 | -0.022 | -0.022 | 3.761 | 0.17 | 0.11 | 0.18 | 0.23 | 0.18 | 0.07 |
| 10 | 67624311 | rs10996743 | 1.782 | -1.285 | -1.839 | -0.454 | -0.02 | -0.029 | -0.029 | 0 | 0.17 | 0.09 | 0.18 | 0.23 | 0.18 | 0.07 |

| | | | | | | | | | | | | | | | | |
|---|---|---|---|---|---|---|---|---|---|---|---|---|---|---|---|---|
| 10 | 67625219 | rs16922054 | 1.386 | -1.352 | -1.78 | -0.493 | -0.015 | -0.016 | -0.016 | 0.133 | 0.06 | 0.1 | 0.06 | 0.03 | 0.01 | 0.01 |
| 10 | 67633950 | rs950054 | 0.66 | -1.34 | -1.697 | -0.449 | -0.001 | -0.006 | -0.006 | 0 | 0.22 | 0.2 | 0.25 | 0.25 | 0.19 | 0.11 |
| 10 | 67634860 | rs10509238 | 0.876 | -1.231 | -1.173 | -0.47 | 0.003 | 0.001 | 0.001 | 0.952 | 0.16 | 0.13 | 0.06 | 0.07 | 0.07 | 0.16 |
| 10 | 67636708 | rs4746533 | 0.271 | -0.702 | -0.643 | -0.544 | 0.035 | 0.039 | 0.039 | 2.099 | 0.45 | 0.36 | 0.41 | 0.32 | 0.36 | 0.25 |
| 10 | 67637104 | rs4399233 | 1.784 | -0.603 | -0.756 | -0.506 | -0.068 | -0.032 | -0.032 | 0.723 | 0.17 | 0.08 | 0.33 | 0.19 | 0.32 | 0.3 |
| 10 | 67637474 | rs7477958 | 1.784 | -0.603 | -0.756 | -0.506 | -0.068 | -0.032 | -0.032 | 0 | 0.17 | 0.08 | 0.32 | 0.19 | 0.32 | 0.3 |
| 10 | 67641201 | rs10996757 | 0.123 | -0.674 | -0.679 | -0.546 | -0.014 | 0.008 | 0.008 | 1.917 | 0.34 | 0.23 | 0.39 | 0.28 | 0.4 | 0.47 |
| 10 | 67643962 | rs10996759 | 0.532 | -0.891 | -0.752 | -0.553 | -0.023 | 0.022 | 0.022 | 0 | 0.1 | 0.18 | 0.06 | 0.16 | 0.05 | 0.07 |
| 10 | 67644475 | rs10822671 | 0.941 | -0.889 | -0.871 | -0.624 | -0.001 | 0 | 0 | 0.013 | 0.23 | 0.25 | 0.26 | 0.25 | 0.19 | 0.11 |
| 10 | 67646861 | rs7350464 | 0.971 | -0.889 | -0.876 | -0.624 | 0 | -0.002 | -0.002 | 0.006 | 0.23 | 0.23 | 0.26 | 0.25 | 0.19 | 0.11 |
| 10 | 67649378 | rs17238902 | 1.837 | -0.916 | -0.976 | -0.663 | 0.039 | 0.114 | 0.114 | 0 | 0.22 | 0.05 | 0.15 | 0.04 | 0.16 | 0.13 |
| 10 | 67653901 | rs1942006 | 1.615 | 0.08 | -0.547 | -0.438 | -0.049 | 0.189 | 0.189 | 0 | 0.33 | 0.03 | 0.25 | 0.07 | 0.35 | 0.22 |
| 10 | 67657144 | rs10822675 | 0.643 | 0.066 | -0.432 | -0.412 | -0.012 | 0.037 | 0.037 | 0.007 | 0.56 | 0.32 | 0.51 | 0.32 | 0.54 | 0.33 |
| 10 | 67666501 | rs17818203 | 1.59 | 0.276 | -0.52 | -0.456 | -0.004 | 0.174 | 0.174 | 0 | 0.33 | 0.07 | 0.25 | 0.07 | 0.33 | 0.21 |
| 10 | 67666582 | rs10740214 | -0.059 | 0.333 | -0.125 | -0.321 | 0.029 | 0.04 | 0.04 | 1.245 | 0.66 | 0.55 | 0.58 | 0.48 | 0.58 | 0.39 |
| 10 | 67667132 | rs1670159 | -0.361 | 0.356 | -0.095 | -0.292 | 0.021 | 0.037 | 0.037 | 0.347 | 0.72 | 0.51 | 0.79 | 0.64 | 0.71 | 0.62 |
| 10 | 67667884 | rs10509245 | 1.088 | 0.282 | 0.16 | -0.254 | -0.01 | 0.004 | 0.004 | 0.162 | 0.12 | 0.26 | 0.06 | 0.09 | 0.06 | 0.14 |
| 10 | 67670747 | rs10996775 | 1.625 | 0.249 | -0.002 | -0.274 | -0.014 | -0.015 | -0.015 | 0 | 0.06 | 0.1 | 0.06 | 0.02 | 0.01 | 0.01 |
| 10 | 67672182 | rs10509244 | 2.077 | 0.032 | -0.305 | -0.43 | -0.016 | 0.016 | 0.016 | 0 | 0.09 | 0.01 | 0.13 | 0.04 | 0.1 | 0.11 |
| 10 | 67677241 | rs1670148 | -0.366 | 0.425 | 0.353 | 0.002 | 0.008 | 0.04 | 0.04 | 0 | 0.34 | 0.47 | 0.47 | 0.48 | 0.37 | 0.47 |
| 10 | 67679048 | rs1786929 | 0.687 | 0.461 | -0.1 | 0.153 | -0.073 | -0.116 | -0.116 | 0 | 0.22 | 0.06 | 0.28 | 0.38 | 0.31 | 0.34 |
| 10 | 67679568 | rs1941996 | 1.355 | 0.446 | -0.054 | 0.043 | -0.012 | -0.012 | -0.012 | 0.94 | 0.06 | 0.06 | 0.21 | 0.16 | 0.13 | 0.23 |
| 10 | 67681086 | rs10996776 | 0.633 | 0.385 | -0.048 | 0.001 | 0.001 | -0.005 | -0.005 | 0 | 0.09 | 0.06 | 0.05 | 0.16 | 0.06 | 0.06 |
| 10 | 67694686 | rs10509240 | -0.308 | 0.399 | 0.221 | -0.009 | 0.001 | -0.005 | -0.005 | 0 | 0.36 | 0.34 | 0.31 | 0.46 | 0.29 | 0.29 |
| 10 | 67697762 | rs1670154 | -0.733 | 0.34 | 0.223 | -0.056 | 0 | 0.005 | 0.005 | 0 | 0.42 | 0.39 | 0.52 | 0.63 | 0.42 | 0.52 |
| 10 | 67700170 | rs1786909 | -1.347 | 0.374 | -0.123 | -0.096 | -0.009 | -0.01 | -0.01 | 2.978 | 0.84 | 0.88 | 0.86 | 0.74 | 0.77 | 0.76 |
| 10 | 67700606 | rs10996784 | 1.859 | 0.29 | -0.207 | -0.123 | -0.002 | -0.003 | -0.003 | 0 | 0.05 | 0.06 | 0.05 | 0.03 | 0.02 | 0.03 |
| 10 | 67701198 | rs1941985 | 1.8 | 0.492 | -0.431 | -0.1 | -0.043 | -0.066 | -0.066 | 0 | 0.17 | 0.01 | 0.2 | 0.24 | 0.17 | 0.11 |
| 10 | 67703119 | rs17819377 | 1.619 | 0.506 | -0.275 | -0.231 | -0.028 | 0.17 | 0.17 | 0 | 0.33 | 0.03 | 0.25 | 0.07 | 0.32 | 0.21 |
| 10 | 67703871 | rs11816737 | 0.881 | 0.541 | -0.077 | -0.175 | 0.019 | 0.09 | 0.09 | 0 | 0.44 | 0.31 | 0.31 | 0.16 | 0.39 | 0.35 |
| 10 | 67709586 | rs1786921 | -0.431 | 0.976 | -0.226 | 0.141 | -0.01 | -0.005 | -0.005 | 0 | 0.82 | 0.87 | 0.83 | 0.76 | 0.71 | 0.74 |
| 10 | 67710200 | rs11599370 | 0.602 | 0.791 | -0.135 | 0.183 | 0.004 | 0.029 | 0.029 | 0.716 | 0.13 | 0.37 | 0.07 | 0.11 | 0.08 | 0.13 |
| 10 | 67719774 | rs1670167 | -1.093 | 0.973 | -0.949 | -0.053 | -0.049 | -0.042 | -0.042 | 0.015 | 0.86 | 0.96 | 0.85 | 0.76 | 0.74 | 0.77 |
| 10 | 67726514 | rs2105702 | -0.426 | 0.408 | -0.606 | -0.732 | -0.002 | 0.003 | 0.003 | 0.241 | 0.3 | 0.31 | 0.47 | 0.39 | 0.26 | 0.38 |
| 10 | 67727223 | rs1670137 | -0.074 | 0.354 | -0.705 | -0.74 | -0.005 | 0.002 | 0.002 | 0.046 | 0.15 | 0.2 | 0.27 | 0.19 | 0.13 | 0.28 |
| 10 | 67735584 | rs12781534 | 0.81 | 0.579 | -0.569 | -1.242 | -0.003 | 0.187 | 0.187 | 0 | 0.35 | 0.06 | 0.25 | 0.07 | 0.31 | 0.21 |
| 10 | 67738700 | rs10996802 | 0.255 | 0.835 | 0.67 | -0.291 | -0.008 | -0.014 | -0.014 | 0 | 0.06 | 0.1 | 0.06 | 0.03 | 0.02 | 0.03 |
| 10 | 67743653 | rs10996809 | 0.129 | 1.05 | 0.385 | -0.255 | 0 | -0.005 | -0.005 | 0 | 0.05 | 0.03 | 0.05 | 0 | 0.12 | 0.03 |
| 10 | 67754225 | rs11599400 | 0.025 | 0.008 | 0.612 | 0.047 | 0.003 | 0.014 | 0.014 | 0.026 | 0.15 | 0.19 | 0.19 | 0.21 | 0.14 | 0.11 |
| 10 | 67755443 | rs7912183 | 0.198 | 0.038 | 0.386 | 0.062 | 0 | 0.004 | 0.004 | 0.332 | 0.14 | 0.14 | 0.19 | 0.21 | 0.14 | 0.11 |
| 10 | 67758920 | rs2893976 | -1.161 | 0.056 | 0.137 | 0.067 | -0.016 | -0.006 | -0.006 | 0 | 0.24 | 0.49 | 0.27 | 0.23 | 0.16 | 0.13 |
| 10 | 67763783 | rs3858135 | -0.069 | 0.694 | -0.884 | 0.421 | -0.004 | -0.011 | -0.011 | 0 | 0.14 | 0.11 | 0.18 | 0.21 | 0.14 | 0.11 |
| 10 | 67766471 | rs1903876 | 0.131 | 0.203 | -0.657 | -0.097 | -0.014 | -0.036 | -0.036 | 0.703 | 0.73 | 0.63 | 0.79 | 0.92 | 0.8 | 0.75 |
| 10 | 67767772 | rs7087052 | -0.632 | 0.873 | -0.93 | 0.081 | -0.036 | -0.004 | -0.004 | 0 | 0.06 | 0.01 | 0.12 | 0.05 | 0.14 | 0.07 |
| 10 | 67770946 | rs7903919 | -0.539 | 0.588 | -0.795 | 0.005 | 0.002 | -0.003 | -0.003 | 0 | 0.45 | 0.52 | 0.57 | 0.43 | 0.55 | 0.5 |
| 10 | 67775907 | rs2394156 | 0.635 | 0.444 | -1.017 | -0.49 | 0.002 | 0.043 | 0.043 | 0.197 | 0.21 | 0.22 | 0.15 | 0.51 | 0.17 | 0.36 |
| 10 | 67776492 | rs10996828 | 0.963 | 0.386 | -0.868 | -0.825 | 0 | 0.002 | 0.002 | 0.124 | 0.13 | 0.06 | 0.13 | 0.06 | 0.14 | 0.1 |
| 10 | 67780727 | rs10822694 | 1.55 | 0.249 | -0.753 | -0.732 | -0.017 | 0.013 | 0.013 | 0 | 0.18 | 0.12 | 0.26 | 0.11 | 0.22 | 0.24 |
| 10 | 67780898 | rs7068134 | -1.412 | 0.174 | -0.712 | -0.761 | 0.01 | 0.03 | 0.03 | 16.749 | 0.66 | 0.73 | 0.71 | 0.94 | 0.71 | 0.86 |
| 10 | 67782742 | rs7074141 | 0.73 | 0.058 | -0.501 | -0.714 | -0.004 | -0.039 | -0.039 | 0 | 0.23 | 0.16 | 0.16 | 0.5 | 0.18 | 0.39 |
| 10 | 67783437 | rs10509258 | -0.992 | 0.143 | -0.409 | -0.756 | -0.002 | -0.003 | -0.003 | 0 | 0.61 | 0.63 | 0.7 | 0.74 | 0.69 | 0.79 |
| 10 | 67784429 | rs10509257 | 0.754 | 0.284 | -0.435 | -0.701 | -0.04 | -0.005 | -0.005 | 0 | 0.07 | 0.01 | 0.13 | 0.06 | 0.15 | 0.07 |
| 10 | 67785037 | rs10996834 | 1.613 | 0.243 | -0.391 | -0.665 | -0.019 | 0.006 | 0.006 | 0 | 0.17 | 0.12 | 0.26 | 0.11 | 0.22 | 0.24 |
| 10 | 67785261 | rs10740218 | -1.285 | -0.002 | -0.232 | -0.728 | 0.001 | 0.001 | 0.001 | 16.6 | 0.65 | 0.68 | 0.72 | 0.76 | 0.71 | 0.84 |
| 10 | 67786048 | rs7076094 | 0.654 | -0.024 | 0.028 | -0.053 | -0.002 | -0.012 | -0.012 | 0 | 0.24 | 0.24 | 0.29 | 0.29 | 0.25 | 0.34 |
| 10 | 67786124 | rs1942007 | -0.702 | -0.083 | 0.434 | -0.078 | -0.003 | -0.007 | -0.007 | 1.11 | 0.52 | 0.53 | 0.57 | 0.68 | 0.57 | 0.73 |
| 10 | 67787837 | rs10996838 | 1.364 | -0.057 | 0.394 | -0.072 | -0.015 | 0.034 | 0.034 | 0 | 0.12 | 0.04 | 0.13 | 0.06 | 0.14 | 0.09 |
| 10 | 67788564 | rs10822695 | -1.925 | -0.692 | 0.178 | -0.416 | -0.001 | 0.009 | 0.009 | 9.938 | 0.77 | 0.83 | 0.88 | 0.8 | 0.78 | 0.91 |
| 10 | 67790146 | rs10740220 | -1.668 | -0.721 | 0.248 | -0.43 | -0.006 | -0.016 | -0.016 | 2.583 | 0.75 | 0.66 | 0.86 | 0.8 | 0.78 | 0.9 |
| 10 | 67792573 | rs4746541 | -1.658 | -0.7 | 0.164 | -0.421 | -0.006 | -0.017 | -0.017 | 0 | 0.75 | 0.67 | 0.86 | 0.8 | 0.78 | 0.9 |
| 10 | 67792731 | rs12355272 | 0.397 | -0.79 | 0.084 | -0.389 | 0.01 | -0.004 | -0.004 | 0 | 0.34 | 0.3 | 0.45 | 0.34 | 0.32 | 0.43 |
| 10 | 67794231 | rs4084961 | 1.157 | -0.476 | 0.573 | -0.196 | -0.002 | -0.014 | -0.014 | 0 | 0.2 | 0.13 | 0.14 | 0.33 | 0.17 | 0.32 |
| 10 | 67799753 | rs7070803 | 0.841 | -0.54 | 0.548 | -0.243 | -0.002 | 0.002 | 0.002 | 0 | 0.27 | 0.22 | 0.15 | 0.33 | 0.17 | 0.33 |
| 10 | 67802101 | rs2199853 | 0.832 | -0.456 | 0.506 | -0.268 | -0.001 | -0.002 | -0.002 | 0 | 0.24 | 0.27 | 0.3 | 0.28 | 0.25 | 0.34 |
| 10 | 67803410 | rs1903863 | 0.031 | -0.463 | 0.485 | -0.298 | 0.005 | 0.016 | 0.016 | 0 | 0.46 | 0.33 | 0.58 | 0.4 | 0.46 | 0.51 |
| 10 | 67803602 | rs10996847 | 1.874 | -0.379 | 0.452 | -0.238 | -0.008 | 0.009 | 0.009 | 0 | 0.17 | 0.14 | 0.26 | 0.11 | 0.22 | 0.24 |
| 10 | 67812810 | rs10996862 | 1.888 | -0.379 | 0.444 | -0.25 | -0.009 | 0.021 | 0.021 | 0 | 0.17 | 0.15 | 0.26 | 0.09 | 0.22 | 0.23 |
| 10 | 67818791 | rs10822701 | -1.939 | -0.175 | 0.51 | -0.259 | 0 | -0.001 | -0.001 | 0 | 0.82 | 0.79 | 0.87 | 0.8 | 0.8 | 0.92 |
| 10 | 67819849 | rs7099996 | 0.241 | -0.189 | 0.383 | -0.23 | -0.041 | -0.063 | -0.063 | 4.406 | 0.33 | 0.12 | 0.27 | 0.4 | 0.34 | 0.41 |

| | | | | | | | | | | | | | | | | |
|---|---|---|---|---|---|---|---|---|---|---|---|---|---|---|---|---|
| 10 | 67821207 | rs10509254 | 1.039 | -0.221 | 0.401 | -0.24 | -0.009 | -0.024 | -0.024 | 0 | 0.23 | 0.25 | 0.29 | 0.12 | 0.21 | 0.17 |
| 10 | 67821720 | rs10509252 | 0.692 | -0.221 | 0.453 | -0.23 | 0.005 | 0.092 | 0.092 | 0 | 0.07 | 0.18 | 0.05 | 0.2 | 0.03 | 0.1 |
| 10 | 67821754 | rs7915266 | 0.692 | -0.221 | 0.453 | -0.23 | 0.005 | 0.092 | 0.092 | 0 | 0.07 | 0.18 | 0.05 | 0.2 | 0.03 | 0.1 |
| 10 | 67828436 | rs10509251 | 0.605 | -0.129 | 0.374 | -0.248 | 0.009 | -0.084 | -0.084 | 0.089 | 0.29 | 0.08 | 0.21 | 0.39 | 0.22 | 0.37 |
| 10 | 67837498 | rs2394189 | -0.982 | -0.208 | 0.402 | -0.256 | 0.004 | -0.017 | -0.017 | 1.714 | 0.63 | 0.54 | 0.61 | 0.71 | 0.59 | 0.68 |
| 10 | 67840879 | rs12572585 | 0.178 | -0.287 | 0.138 | -0.242 | -0.058 | -0.083 | -0.083 | 0.113 | 0.33 | 0.08 | 0.27 | 0.4 | 0.34 | 0.41 |
| 10 | 67847123 | rs10822709 | -0.455 | -0.273 | 0.293 | -0.188 | -0.004 | -0.006 | -0.006 | 0 | 0.56 | 0.34 | 0.57 | 0.51 | 0.56 | 0.59 |
| 10 | 67849668 | rs10822711 | -0.112 | -0.297 | 0.315 | -0.053 | 0.032 | 0.013 | 0.013 | 0 | 0.49 | 0.33 | 0.44 | 0.42 | 0.41 | 0.52 |
| 10 | 67849839 | rs7916078 | -0.468 | -0.296 | 0.431 | -0.044 | -0.001 | -0.005 | -0.005 | 0 | 0.56 | 0.52 | 0.48 | 0.62 | 0.44 | 0.62 |
| 10 | 67850412 | rs7919909 | 0.611 | -0.263 | 0.168 | -0.075 | -0.004 | 0.002 | 0.002 | 0 | 0.27 | 0.27 | 0.16 | 0.31 | 0.19 | 0.34 |
| 10 | 67851530 | rs10996889 | 0.561 | -0.266 | 0.142 | -0.077 | -0.01 | 0.006 | 0.006 | 0 | 0.28 | 0.31 | 0.17 | 0.31 | 0.19 | 0.34 |
| 10 | 67855521 | rs10996892 | 1.256 | -0.234 | -0.296 | -0.1 | -0.01 | 0.045 | 0.045 | 0 | 0.21 | 0.03 | 0.27 | 0.11 | 0.22 | 0.18 |
| 10 | 67859565 | rs10996895 | 2.221 | -0.259 | -0.662 | -0.185 | -0.065 | 0.042 | 0.042 | 0 | 0.15 | 0.01 | 0.24 | 0.06 | 0.19 | 0.21 |
| 10 | 67860320 | rs10430506 | 1.089 | -0.34 | -0.578 | -0.13 | -0.004 | -0.08 | -0.08 | 0.073 | 0.25 | 0.03 | 0.14 | 0.3 | 0.19 | 0.34 |
| 10 | 67863538 | rs4746547 | 1.055 | -0.342 | -0.493 | -0.13 | 0.014 | -0.042 | -0.042 | 1.973 | 0.25 | 0.1 | 0.15 | 0.3 | 0.19 | 0.33 |
| 10 | 67864604 | rs7070570 | 0.158 | -0.193 | -0.213 | -0.039 | 0.012 | -0.004 | -0.004 | 0 | 0.28 | 0.22 | 0.31 | 0.3 | 0.25 | 0.28 |
| 10 | 67871159 | rs2127611 | -0.364 | -0.165 | -0.144 | 0.013 | -0.016 | -0.003 | -0.003 | 0 | 0.34 | 0.75 | 0.46 | 0.36 | 0.37 | 0.41 |
| 10 | 67871774 | rs7903907 | -0.656 | -0.329 | -0.293 | 0.196 | -0.006 | 0.032 | 0.032 | 0.008 | 0.6 | 0.77 | 0.61 | 0.67 | 0.56 | 0.77 |
| 10 | 67879440 | rs10996914 | 0.774 | -0.348 | -0.314 | 0.247 | 0.023 | -0.063 | -0.063 | 0.999 | 0.27 | 0.07 | 0.15 | 0.31 | 0.19 | 0.36 |
| 10 | 67888229 | rs11817346 | 0.607 | -0.322 | 0.102 | 0.247 | -0.008 | 0.021 | 0.021 | 0.004 | 0.11 | 0.29 | 0.06 | 0.15 | 0.07 | 0.09 |
| 10 | 67894481 | rs7895803 | -0.213 | -0.272 | 0.262 | 0.156 | 0.003 | 0.015 | 0.015 | 0 | 0.5 | 0.59 | 0.57 | 0.37 | 0.45 | 0.47 |
| 10 | 67898109 | rs10822729 | 2.108 | -0.202 | 0.332 | 0.147 | -0.014 | 0.009 | 0.009 | 0 | 0.15 | 0.09 | 0.26 | 0.1 | 0.2 | 0.23 |
| 10 | 67899266 | rs16922533 | 1.271 | -0.206 | 0.436 | 0.145 | 0.018 | -0.025 | -0.025 | 4.765 | 0.18 | 0.26 | 0.27 | 0.1 | 0.2 | 0.24 |
| 10 | 67914345 | rs7905595 | -1.701 | -0.297 | -0.073 | 0.231 | 0 | 0.001 | 0.001 | 0.071 | 0.83 | 0.83 | 0.87 | 0.78 | 0.79 | 0.93 |
| 10 | 67929739 | rs10996947 | 0.876 | -0.529 | -0.287 | 0.212 | 0.011 | -0.076 | -0.076 | 2.413 | 0.27 | 0.03 | 0.16 | 0.31 | 0.2 | 0.4 |
| 10 | 67930380 | rs11816309 | 0.302 | -0.547 | -0.061 | 0.21 | -0.002 | 0.067 | 0.067 | 0.054 | 0.14 | 0.5 | 0.07 | 0.16 | 0.07 | 0.1 |
| 10 | 67935471 | rs10822736 | 0.192 | -0.094 | -0.414 | 0.171 | 0.038 | 0.006 | 0.006 | 0 | 0.48 | 0.18 | 0.4 | 0.41 | 0.39 | 0.53 |
| 10 | 67939147 | rs10822739 | 2.079 | -0.018 | -0.389 | 0.149 | -0.003 | 0.003 | 0.003 | 0 | 0.15 | 0.12 | 0.27 | 0.11 | 0.2 | 0.23 |
| 10 | 67940419 | rs4746553 | -0.253 | -0.05 | -0.348 | 0.146 | 0.013 | -0.01 | -0.01 | 0 | 0.43 | 0.64 | 0.26 | 0.46 | 0.32 | 0.49 |
| 10 | 67951369 | rs1462813 | 0.122 | -0.024 | -0.274 | 0.166 | 0.042 | 0.009 | 0.009 | 0 | 0.49 | 0.23 | 0.41 | 0.41 | 0.39 | 0.51 |
| 10 | 67953938 | rs7904019 | 0.586 | -0.03 | -0.199 | 0.15 | -0.001 | 0.048 | 0.048 | 0 | 0.13 | 0.37 | 0.07 | 0.15 | 0.07 | 0.09 |
| 10 | 67954847 | rs10822747 | -0.663 | 0.036 | -0.268 | 0.221 | 0.013 | -0.015 | -0.015 | 0 | 0.64 | 0.5 | 0.58 | 0.68 | 0.55 | 0.67 |
| 10 | 67960463 | rs4746554 | -0.601 | 0.048 | -1.09 | 0.236 | -0.012 | -0.004 | -0.004 | 0 | 0.63 | 0.69 | 0.5 | 0.55 | 0.51 | 0.61 |
| 10 | 67962943 | rs7920585 | 0.949 | -0.081 | -1.336 | 0.21 | 0.016 | -0.083 | -0.083 | 0 | 0.26 | 0.03 | 0.15 | 0.29 | 0.2 | 0.37 |
| 10 | 67963275 | rs12259429 | 1.316 | 0.022 | -1.377 | 0.135 | -0.033 | 0.036 | 0.036 | 0 | 0.21 | 0.03 | 0.25 | 0.11 | 0.19 | 0.13 |
| 10 | 67964514 | rs7910869 | 1.415 | 0.318 | -0.995 | 0.29 | -0.001 | 0.001 | 0.001 | 0 | 0.17 | 0.17 | 0.29 | 0.1 | 0.24 | 0.26 |
| 10 | 67967091 | rs2893985 | -0.983 | 0.315 | -0.94 | 0.35 | 0 | 0 | 0 | 0 | 0.8 | 0.79 | 0.85 | 0.79 | 0.75 | 0.92 |
| 10 | 67969419 | rs7067590 | 0.021 | 0.422 | -0.912 | 0.064 | -0.008 | 0.024 | 0.024 | 0 | 0.34 | 0.29 | 0.44 | 0.26 | 0.4 | 0.3 |
| 10 | 67969845 | rs4745889 | -0.939 | 0.371 | -1.569 | 0.114 | 0.013 | -0.059 | -0.059 | 1.813 | 0.63 | 0.83 | 0.78 | 0.51 | 0.67 | 0.64 |
| 10 | 67969980 | rs10996973 | 0.165 | 0.126 | -1.461 | -0.102 | -0.011 | 0.008 | 0.008 | 0.324 | 0.15 | 0.64 | 0.1 | 0.16 | 0.11 | 0.1 |
| 10 | 67970052 | rs10996975 | 0.382 | 0.458 | -1.729 | -0.198 | -0.028 | 0.078 | 0.078 | 0 | 0.28 | 0.03 | 0.27 | 0.11 | 0.26 | 0.22 |
| 10 | 67971334 | rs12146238 | 0.806 | 0.761 | -0.749 | 0.273 | -0.023 | 0.01 | 0.01 | 0 | 0.15 | 0.06 | 0.27 | 0.1 | 0.2 | 0.25 |
| 10 | 67978859 | rs10822753 | 0.156 | 0.282 | -0.275 | 0.429 | -0.009 | -0.015 | -0.015 | 0 | 0.58 | 0.36 | 0.46 | 0.5 | 0.53 | 0.57 |
| 10 | 67981136 | rs2054384 | 0.694 | 0.353 | 0.259 | 0.579 | 0.005 | 0.043 | 0.043 | 0.292 | 0.12 | 0.18 | 0.1 | 0.24 | 0.17 | 0.04 |
| 10 | 67992622 | rs10822762 | -0.542 | 0.331 | 0.49 | 0.57 | 0.01 | 0.08 | 0.08 | 0.807 | 0.87 | 0.73 | 0.89 | 0.76 | 0.83 | 0.96 |
| 10 | 67995456 | rs1037982 | -0.521 | -0.483 | -0.074 | 0.04 | -0.005 | -0.017 | -0.017 | 0.1 | 0.47 | 0.47 | 0.59 | 0.3 | 0.42 | 0.53 |
| 10 | 67996275 | rs10762033 | 0.999 | -0.483 | -0.091 | 0.039 | -0.034 | -0.038 | -0.038 | 0 | 0.4 | 0.27 | 0.3 | 0.46 | 0.42 | 0.42 |
| 10 | 67997672 | rs7924275 | 0.444 | -0.616 | 0.511 | -0.004 | -0.003 | -0.006 | -0.006 | 0 | 0.21 | 0.22 | 0.29 | 0.19 | 0.19 | 0.21 |
| 10 | 67998336 | rs17247353 | 1.314 | -0.773 | 0.499 | -0.04 | 0.035 | 0.042 | 0.042 | 0 | 0.16 | 0.07 | 0.15 | 0.04 | 0.1 | 0.14 |
| 10 | 67998370 | rs4745895 | 0.254 | -0.363 | 0.731 | 0.197 | -0.004 | 0 | 0 | 1.132 | 0.62 | 0.49 | 0.59 | 0.65 | 0.63 | 0.64 |
| 10 | 68004475 | rs10997006 | 0.415 | -0.336 | 0.633 | 0.202 | -0.003 | -0.015 | -0.015 | 0 | 0.25 | 0.25 | 0.29 | 0.11 | 0.2 | 0.31 |
| 10 | 68005571 | rs1911467 | 1.018 | -0.222 | 0.548 | 0.283 | 0.005 | 0.043 | 0.043 | 0.292 | 0.12 | 0.18 | 0.1 | 0.24 | 0.17 | 0.04 |
| 10 | 68011627 | rs2135731 | 1.255 | 0.034 | 0.569 | 0.356 | -0.02 | 0.021 | 0.021 | 0 | 0.16 | 0.08 | 0.29 | 0.07 | 0.19 | 0.21 |
| 10 | 68012461 | rs4746565 | -0.781 | -0.039 | 0.735 | 0.309 | -0.004 | -0.016 | -0.016 | 0 | 0.34 | 0.68 | 0.41 | 0.38 | 0.37 | 0.37 |
| 10 | 68014395 | rs7097811 | 1.094 | -0.089 | 0.904 | 0.271 | -0.053 | -0.036 | -0.036 | 0 | 0.1 | 0.23 | 0.02 | 0.04 | 0.02 | 0.11 |
| 10 | 68016815 | rs1876334 | 1.294 | -0.019 | 0.786 | 0.275 | -0.007 | 0.008 | 0.008 | 0 | 0.16 | 0.1 | 0.29 | 0.07 | 0.19 | 0.23 |
| 10 | 68018042 | rs10822779 | -0.402 | -0.088 | 0.886 | 0.334 | -0.015 | 0.083 | 0.083 | 0.493 | 0.18 | 0.58 | 0.13 | 0.31 | 0.18 | 0.14 |
| 10 | 68018361 | rs12412285 | 1.164 | 0.156 | 0.673 | 0.259 | 0 | -0.002 | -0.002 | 0.42 | 0.15 | 0.13 | 0.27 | 0.07 | 0.16 | 0.23 |
| 10 | 68024453 | rs6480155 | -0.085 | -0.02 | 0.944 | 0.345 | -0.016 | 0.097 | 0.097 | 0.497 | 0.17 | 0.33 | 0.09 | 0.32 | 0.15 | 0.13 |
| 10 | 68027026 | rs7906924 | 0.076 | -0.035 | 1.165 | 0.345 | 0.06 | 0.19 | 0.19 | 0.002 | 0.08 | 0.2 | 0.08 | 0.27 | 0.14 | 0.04 |
| 10 | 68028720 | rs7912066 | -1.209 | -0.014 | 1.161 | 0.213 | -0.039 | -0.051 | -0.051 | 0 | 0.4 | 0.79 | 0.53 | 0.52 | 0.46 | 0.48 |
| 10 | 68028868 | rs11813599 | 1.57 | -0.042 | 1.177 | 0.234 | -0.021 | -0.006 | -0.006 | 0 | 0.08 | 0.14 | 0.01 | 0.05 | 0.01 | 0.1 |
| 10 | 68040240 | rs10997034 | 1.602 | -0.058 | 1.282 | 0.227 | -0.021 | -0.006 | -0.006 | 0.006 | 0.08 | 0.14 | 0.01 | 0.05 | 0.01 | 0.1 |
| 10 | 68040325 | rs4548513 | -0.809 | 0.031 | 1.261 | 0.236 | -0.002 | -0.021 | -0.021 | 0.021 | 0.35 | 0.73 | 0.41 | 0.42 | 0.37 | 0.39 |
| 10 | 68041535 | rs10822791 | 0.075 | 0.047 | 1.076 | 0.223 | 0 | 0.006 | 0.006 | 0 | 0.25 | 0.27 | 0.31 | 0.12 | 0.22 | 0.33 |
| 10 | 68048196 | rs7907021 | -0.681 | 0.055 | 1.088 | 0.243 | 0.01 | 0.028 | 0.028 | 0 | 0.33 | 0.47 | 0.39 | 0.39 | 0.36 | 0.37 |
| 10 | 68050459 | rs10740229 | -0.853 | 0.101 | 0.883 | 0.237 | 0 | 0.004 | 0.004 | 0 | 0.34 | 0.63 | 0.4 | 0.42 | 0.37 | 0.38 |
| 10 | 68054482 | rs10762051 | -0.165 | 0.087 | 0.938 | 0.22 | -0.014 | -0.05 | -0.05 | 0 | 0.27 | 0.5 | 0.33 | 0.14 | 0.24 | 0.35 |
| 10 | 68057021 | rs1911485 | -0.373 | -0.221 | 0.674 | 0.048 | -0.02 | -0.041 | -0.041 | 0 | 0.37 | 0.51 | 0.45 | 0.18 | 0.27 | 0.4 |

| | | | | | | | | | | | | | | | | |
|---|---|---|---|---|---|---|---|---|---|---|---|---|---|---|---|---|
| 10 | 68057063 | rs1911486 | 0.792 | 0.509 | 0.578 | 0.376 | 0.007 | -0.006 | -0.006 | 0 | 0.18 | 0.24 | 0.31 | 0.1 | 0.22 | 0.25 |
| 10 | 68058454 | rs7920917 | 0.2 | 0.82 | 0.116 | 0.526 | 0.013 | 0.075 | 0.075 | 0 | 0.07 | 0.09 | 0.07 | 0.28 | 0.14 | 0.04 |
| 10 | 68064594 | rs7085547 | -0.575 | 1.029 | -0.797 | 0.18 | 0 | -0.01 | -0.01 | 0 | 0.46 | 0.54 | 0.51 | 0.64 | 0.56 | 0.62 |
| 10 | 68065751 | rs7897609 | -0.118 | 1.293 | -1.048 | 0.762 | -0.021 | -0.037 | -0.037 | 0 | 0.2 | 0.12 | 0.19 | 0.5 | 0.33 | 0.27 |
| 10 | 68072084 | rs4370822 | -0.204 | 0.417 | -1.061 | 0.198 | -0.008 | 0.023 | 0.023 | 0 | 0.87 | 0.92 | 0.89 | 0.93 | 0.84 | 0.87 |
| 10 | 68078481 | rs4297361 | 0.277 | -0.04 | -1.312 | 0.333 | -0.038 | 0.005 | 0.005 | 0.756 | 0.38 | 0.27 | 0.47 | 0.35 | 0.52 | 0.36 |
| 10 | 68079310 | rs7072330 | 0.257 | -1.167 | -1.264 | -0.094 | 0.014 | 0.067 | 0.067 | 0 | 0.47 | 0.3 | 0.53 | 0.26 | 0.57 | 0.41 |
| 10 | 68079505 | rs17511 | -0.327 | -1.656 | -0.885 | 0.215 | 0.013 | 0.007 | 0.007 | 0.026 | 0.56 | 0.43 | 0.63 | 0.62 | 0.65 | 0.55 |
| 10 | 68082391 | rs10822806 | -1.119 | -1.537 | -0.372 | 0.202 | -0.001 | 0.002 | 0.002 | 0 | 0.81 | 0.83 | 0.8 | 0.78 | 0.82 | 0.82 |
| 10 | 68082484 | rs12416656 | 1.762 | -1.394 | -0.415 | 0.242 | -0.044 | 0.032 | 0.032 | 0.044 | 0.24 | 0.18 | 0.41 | 0.11 | 0.4 | 0.17 |
| 10 | 68085353 | rs16922827 | 1.268 | -1.463 | -0.56 | 0.239 | -0.017 | -0.039 | -0.039 | 0.457 | 0.1 | 0.03 | 0.16 | 0.22 | 0.12 | 0.13 |
| 10 | 68085487 | rs16922829 | 0.84 | -1.794 | -0.331 | 0.2 | 0.012 | -0.018 | -0.018 | 1.796 | 0.26 | 0.15 | 0.22 | 0.35 | 0.2 | 0.36 |
| 10 | 68088348 | rs10762057 | 0.109 | -1.92 | 0.019 | 0.119 | -0.019 | -0.005 | -0.005 | 0 | 0.69 | 0.65 | 0.73 | 0.73 | 0.74 | 0.75 |
| 10 | 68089301 | rs7918347 | -1.543 | -1.967 | 0.02 | 0.056 | -0.021 | -0.069 | -0.069 | 0 | 0.88 | 0.81 | 0.94 | 1 | 0.93 | 0.93 |
| 10 | 68096219 | rs4592316 | 0.781 | -2.121 | 0.185 | 0.012 | 0.026 | 0.014 | 0.014 | 0.42 | 0.64 | 0.31 | 0.69 | 0.73 | 0.73 | 0.73 |
| 10 | 68099757 | rs6480160 | -1.336 | -2.403 | 0.517 | 0.275 | -0.021 | -0.069 | -0.069 | 0 | 0.88 | 0.81 | 0.94 | 1 | 0.93 | 0.93 |
| 10 | 68102333 | rs10450284 | 0.79 | -2.531 | 0.433 | 0.317 | 0 | -0.001 | -0.001 | 0.899 | 0.17 | 0.19 | 0.08 | 0.26 | 0.13 | 0.2 |
| 10 | 68107483 | rs4418682 | 0.836 | -2.532 | 0.251 | 0.192 | 0.068 | -0.021 | -0.021 | 0 | 0.58 | 0.27 | 0.66 | 0.46 | 0.69 | 0.57 |
| 10 | 68113595 | rs10997073 | 0.573 | -2.463 | 0.397 | 0.251 | -0.017 | 0.085 | 0.085 | 0 | 0.18 | 0.26 | 0.21 | 0.51 | 0.16 | 0.33 |
| 10 | 68114481 | rs12573282 | 0.902 | -2.515 | 0.049 | 0.195 | 0.063 | 0.034 | 0.034 | 0.991 | 0.55 | 0.26 | 0.61 | 0.35 | 0.63 | 0.46 |
| 10 | 68122473 | rs10997092 | 1.576 | -2.119 | 0.03 | 0.192 | -0.095 | 0.098 | 0.098 | 2.642 | 0.32 | 0.14 | 0.43 | 0.11 | 0.48 | 0.18 |
| 10 | 68126695 | rs4300297 | 0.617 | -2.179 | -0.005 | 0.364 | 0.004 | 0 | 0 | 0 | 0.18 | 0.16 | 0.2 | 0.49 | 0.16 | 0.31 |
| 10 | 68130116 | rs7895266 | 1.594 | -2.157 | -0.22 | 0.239 | -0.112 | 0.107 | 0.107 | 2.054 | 0.32 | 0.11 | 0.43 | 0.11 | 0.48 | 0.2 |
| 10 | 68133906 | rs10997098 | 1.123 | -2.191 | -0.245 | 0.388 | 0.001 | -0.08 | -0.08 | 0 | 0.15 | 0.03 | 0.07 | 0.27 | 0.13 | 0.2 |
| 10 | 68136961 | rs2441718 | 0.09 | -2.426 | -0.015 | 0.309 | 0.017 | 0.028 | 0.028 | 0.781 | 0.39 | 0.3 | 0.22 | 0.16 | 0.3 | 0.34 |
| 10 | 68138358 | rs2939947 | -1.903 | -2.392 | -0.183 | 0.313 | -0.092 | 0.103 | 0.103 | 2.076 | 0.68 | 0.87 | 0.57 | 0.89 | 0.51 | 0.82 |
| 10 | 68139550 | rs16922922 | 0.221 | -2.425 | -0.067 | 0.354 | 0.001 | 0.007 | 0.007 | 0 | 0.08 | 0.08 | 0.11 | 0.24 | 0.06 | 0.14 |
| 10 | 68140020 | rs2007116 | -0.893 | -2.416 | 0.036 | 0.053 | 0.007 | 0.003 | 0.003 | 0.16 | 0.57 | 0.6 | 0.32 | 0.43 | 0.42 | 0.56 |
| 10 | 68150124 | rs2921933 | 0.769 | -2.522 | 0.036 | -0.014 | -0.001 | -0.031 | -0.031 | 0 | 0.16 | 0.37 | 0.13 | 0.11 | 0.14 | 0.18 |
| 10 | 68152745 | rs1911481 | -0.991 | -2.553 | 0.014 | -0.014 | -0.001 | 0.004 | 0.004 | 0 | 0.28 | 0.67 | 0.38 | 0.56 | 0.23 | 0.43 |
| 10 | 68152953 | rs7897082 | -0.475 | -2.493 | -0.004 | 0.093 | -0.001 | 0.222 | 0.222 | 0 | 0.11 | 0.28 | 0.24 | 0.46 | 0.09 | 0.25 |
| 10 | 68153190 | rs12763064 | -0.178 | -2.282 | -0.365 | -0.003 | 0.069 | 0.039 | 0.039 | 0 | 0.41 | 0.23 | 0.2 | 0.33 | 0.28 | 0.39 |
| 10 | 68158254 | rs10509264 | 1.113 | -1.945 | -0.661 | -0.022 | 0.024 | 0.108 | 0.108 | 0.571 | 0.21 | 0.01 | 0.07 | 0.02 | 0.11 | 0.16 |
| 10 | 68165306 | rs2441756 | -0.699 | -1.671 | -0.265 | 0.296 | -0.019 | -0.007 | -0.007 | 0 | 0.61 | 0.67 | 0.38 | 0.46 | 0.46 | 0.62 |
| 10 | 68167292 | rs7079889 | 0.217 | -1.741 | 0.024 | 0.487 | -0.006 | 0.006 | 0.006 | 0 | 0.33 | 0.35 | 0.36 | 0.45 | 0.21 | 0.46 |
| 10 | 68168954 | rs1948946 | -0.079 | -1.987 | 0.04 | 0.381 | -0.018 | 0.015 | 0.015 | 0.189 | 0.23 | 0.3 | 0.13 | 0.3 | 0.19 | 0.24 |
| 10 | 68170551 | rs13376837 | 0.06 | -1.939 | -0.021 | 0.444 | 0.03 | 0.05 | 0.05 | 0.062 | 0.08 | 0.24 | 0.07 | 0.11 | 0.08 | 0.12 |
| 10 | 68171803 | rs1911490 | 0.139 | -1.888 | 0.027 | 0.85 | -0.02 | -0.006 | -0.006 | 0 | 0.38 | 0.49 | 0.4 | 0.37 | 0.27 | 0.49 |
| 10 | 68175450 | rs2394215 | 0.281 | -1.895 | 0.023 | 0.835 | -0.02 | -0.079 | -0.079 | 0.402 | 0.36 | 0.47 | 0.25 | 0.13 | 0.26 | 0.38 |
| 10 | 68179970 | rs10997109 | 0.144 | -1.878 | 0.031 | 0.857 | -0.02 | -0.006 | -0.006 | 0 | 0.38 | 0.49 | 0.4 | 0.37 | 0.27 | 0.49 |
| 10 | 68180999 | rs12774506 | 0.732 | -1.885 | 0.047 | 0.845 | 0.016 | 0.014 | 0.014 | 0.034 | 0.32 | 0.25 | 0.34 | 0.26 | 0.21 | 0.39 |
| 10 | 68183014 | rs10822827 | -0.397 | -1.974 | -0.017 | 0.83 | -0.042 | 0.127 | 0.127 | 0.433 | 0.25 | 0.47 | 0.16 | 0.52 | 0.19 | 0.33 |
| 10 | 68210691 | rs10822833 | 0.713 | -1.861 | -0.121 | 0.62 | 0.055 | 0.095 | 0.095 | 0.394 | 0.05 | 0.24 | 0.06 | 0.11 | 0.07 | 0.09 |
| 10 | 68214970 | rs12777566 | -0.085 | -1.891 | -0.033 | 0.637 | -0.006 | -0.001 | -0.001 | 0 | 0.4 | 0.58 | 0.43 | 0.6 | 0.27 | 0.57 |
| 10 | 68218392 | rs11592249 | -0.224 | -1.643 | -0.749 | 0.222 | 0.035 | 0.051 | 0.051 | 0.542 | 0.07 | 0.22 | 0.07 | 0.11 | 0.08 | 0.11 |
| 10 | 68221549 | rs2921945 | -0.947 | -1.58 | -1.294 | 0.053 | -0.134 | 0.164 | 0.164 | 2.705 | 0.62 | 0.91 | 0.56 | 0.89 | 0.46 | 0.82 |
| 10 | 68223696 | rs7095004 | 0.366 | -1.297 | -0.647 | 0.301 | 0.025 | 0.084 | 0.084 | 0.329 | 0.3 | 0.15 | 0.19 | 0.02 | 0.2 | 0.28 |
| 10 | 68224205 | rs10822834 | -1.165 | -0.234 | -0.098 | 1.293 | -0.006 | 0.082 | 0.082 | 0 | 0.06 | 0.14 | 0.05 | 0.28 | 0.03 | 0.22 |
| 10 | 68237143 | rs11817581 | -0.027 | 0.706 | -0.616 | 0.856 | -0.098 | 0.091 | 0.091 | 0.092 | 0.33 | 0.12 | 0.44 | 0.12 | 0.48 | 0.18 |
| 10 | 68242672 | rs10822837 | 0.046 | 0.738 | -0.85 | 0.871 | -0.156 | 0.078 | 0.078 | 0.151 | 0.33 | 0.03 | 0.43 | 0.12 | 0.49 | 0.17 |
| 10 | 68260306 | rs17179265 | -0.299 | 1.051 | 0.051 | 0.627 | 0.016 | -0.055 | -0.055 | 0 | 0.12 | 0.2 | 0.15 | 0.01 | 0.16 | 0.09 |
| 10 | 68260530 | rs2441749 | -1.132 | 0.822 | 0.111 | 0.558 | 0.024 | 0.057 | 0.057 | 0 | 0.58 | 0.75 | 0.78 | 0.9 | 0.71 | 0.58 |
| 10 | 68274118 | rs10430510 | -0.191 | 1.396 | 0.296 | 1.343 | 0.002 | 0.134 | 0.134 | 0 | 0.05 | 0.07 | 0.04 | 0.31 | 0.05 | 0.12 |
| 10 | 68276333 | rs2660042 | -1.33 | 2.383 | -0.029 | 0.731 | -0.061 | -0.038 | -0.038 | 0.507 | 0.16 | 0.06 | 0.28 | 0.81 | 0.3 | 0.25 |
| 10 | 68282970 | rs997225 | -1.28 | 2.039 | 0.261 | 0.74 | 0.009 | 0.006 | 0.006 | 0.155 | 0.16 | 0.16 | 0.26 | 0.81 | 0.25 | 0.25 |
| 10 | 68283235 | rs2660044 | 0.121 | 1.444 | 0.918 | 0.502 | 0.039 | -0.04 | -0.04 | 0.007 | 0.91 | 0.75 | 0.88 | 0.99 | 0.87 | 0.92 |
| 10 | 68317083 | rs10997232 | -0.284 | 1.488 | 0.497 | 1.079 | 0 | -0.005 | -0.005 | 0 | 0.07 | 0.02 | 0.1 | 0.24 | 0.08 | 0.08 |
| 10 | 68319999 | rs1843666 | 0.199 | 1.51 | 0.694 | 0.486 | 0.003 | -0.007 | -0.007 | 0.162 | 0.27 | 0.26 | 0.47 | 0.11 | 0.42 | 0.2 |
| 10 | 68320326 | rs2631205 | -0.01 | 0.978 | 0.986 | 0.532 | -0.022 | 0.055 | 0.055 | 0 | 0.86 | 0.65 | 0.87 | 0.76 | 0.91 | 0.81 |
| 10 | 68321601 | rs2660020 | 0.481 | 0.995 | 0.996 | 0.523 | -0.02 | 0.026 | 0.026 | 0 | 0.82 | 0.6 | 0.81 | 0.75 | 0.87 | 0.76 |
| 10 | 68324400 | rs991987 | 0.496 | 0.995 | 0.969 | 0.523 | -0.024 | 0.033 | 0.033 | 0 | 0.82 | 0.57 | 0.8 | 0.75 | 0.87 | 0.76 |
| 10 | 68328484 | rs1911294 | 0.426 | 1.135 | 1 | 0.466 | 0.022 | -0.03 | -0.03 | 0 | 0.27 | 0.28 | 0.47 | 0.1 | 0.41 | 0.2 |
| 10 | 68328544 | rs12779355 | -0.072 | 1.054 | 0.922 | 0.57 | 0.027 | 0.008 | 0.008 | 0.399 | 0.65 | 0.54 | 0.42 | 0.65 | 0.51 | 0.71 |
| 10 | 68328936 | rs7919076 | 1.32 | 0.92 | 0.487 | 0.406 | 0.056 | 0.197 | 0.197 | 0 | 0.42 | 0.23 | 0.2 | 0.02 | 0.3 | 0.38 |
| 10 | 68329182 | rs7919336 | 1.006 | 0.9 | 0.521 | 0.433 | 0.053 | 0.112 | 0.112 | 0.002 | 0.44 | 0.27 | 0.27 | 0.26 | 0.32 | 0.51 |
| 10 | 68335254 | rs12256594 | 1.323 | 0.672 | 0.405 | 0.186 | 0.059 | 0.103 | 0.103 | 0 | 0.41 | 0.29 | 0.19 | 0.01 | 0.26 | 0.36 |
| 10 | 68336295 | rs10997242 | 0.004 | 0.639 | 0.871 | 0.29 | -0.016 | 0.024 | 0.024 | 0.163 | 0.19 | 0.3 | 0.16 | 0.25 | 0.16 | 0.13 |
| 10 | 68337401 | rs12268642 | 1.457 | 0.634 | 0.788 | 0.204 | 0.061 | 0.107 | 0.107 | 0 | 0.42 | 0.29 | 0.19 | 0.01 | 0.26 | 0.36 |
| 10 | 68343168 | rs2660016 | 0.716 | 0.989 | 0.96 | 0.354 | -0.032 | -0.072 | -0.072 | 0.012 | 0.71 | 0.54 | 0.54 | 0.13 | 0.73 | 0.67 |

| | | | | | | | | | | | | | | | | |
|---|---|---|---|---|---|---|---|---|---|---|---|---|---|---|---|---|
| 10 | 68346959 | rs10997250 | 0.897 | 1.021 | 0.94 | 0.534 | -0.004 | 0.003 | 0.003 | 0 | 0.07 | 0.03 | 0.05 | 0.35 | 0.09 | 0.06 |
| 10 | 68353739 | rs1911338 | -0.317 | 1.057 | 0.862 | -0.233 | 0.022 | -0.041 | -0.041 | 0 | 0.73 | 0.73 | 0.65 | 0.93 | 0.57 | 0.77 |
| 10 | 68355966 | rs1158119 | -0.957 | 1.047 | 0.898 | -0.191 | -0.024 | -0.059 | -0.059 | 0.588 | 0.28 | 0.42 | 0.45 | 0.87 | 0.27 | 0.33 |
| 10 | 68361408 | rs10822851 | -0.25 | 1.485 | 0.704 | -0.214 | -0.011 | 0.019 | 0.019 | 0.15 | 0.19 | 0.27 | 0.17 | 0.24 | 0.16 | 0.13 |
| 10 | 68370045 | rs2660035 | -0.163 | 2.428 | 0.483 | -0.561 | -0.027 | 0.017 | 0.017 | 3.856 | 0.79 | 0.88 | 0.73 | 0.94 | 0.62 | 0.82 |
| 10 | 68370180 | rs2660034 | 0.065 | 2.217 | 0.485 | -0.526 | -0.007 | -0.006 | -0.006 | 0.003 | 0.75 | 0.83 | 0.66 | 0.93 | 0.57 | 0.77 |
| 10 | 68373403 | rs10997260 | -1.193 | 2.747 | 0.572 | -0.376 | 0.083 | -0.096 | -0.096 | 0 | 0.2 | 0.38 | 0.31 | 0.03 | 0.4 | 0.18 |
| 10 | 68373806 | rs1911328 | -1.782 | 2.774 | 1.123 | 0.169 | 0.082 | 0.06 | 0.06 | 0 | 0.26 | 0.5 | 0.49 | 0.34 | 0.41 | 0.36 |
| 10 | 68374084 | rs7909235 | -0.668 | 2.727 | 0.733 | 0.086 | -0.026 | 0.014 | 0.014 | 0 | 0.16 | 0.14 | 0.25 | 0.03 | 0.32 | 0.18 |
| 10 | 68375667 | rs7090742 | -0.029 | 2.985 | 0.516 | 0.069 | -0.064 | 0.056 | 0.056 | 0.077 | 0.1 | 0.02 | 0.18 | 0.03 | 0.22 | 0.15 |
| 10 | 68376863 | rs1876306 | -0.959 | 2.151 | -0.426 | -0.572 | -0.066 | -0.109 | -0.109 | 2.952 | 0.73 | 0.96 | 0.74 | 0.35 | 0.7 | 0.63 |
| 10 | 68382135 | rs10997263 | 0.828 | 1.676 | -0.208 | -0.835 | 0.048 | 0.042 | 0.042 | 0.054 | 0.47 | 0.23 | 0.43 | 0.32 | 0.36 | 0.4 |
| 10 | 68383827 | rs16923325 | -0.349 | 1.471 | 0.491 | -0.716 | -0.026 | -0.037 | -0.037 | 0.414 | 0.07 | 0.23 | 0.03 | 0 | 0.02 | 0.06 |
| 10 | 68384104 | rs4143861 | -0.662 | 1.814 | 0.621 | -0.546 | 0.111 | -0.12 | -0.12 | 0 | 0.17 | 0.39 | 0.27 | 0.03 | 0.31 | 0.18 |
| 10 | 68393666 | rs4746611 | -1.329 | 0.875 | 0.491 | 0.117 | 0.02 | 0.23 | 0.23 | 0.039 | 0.06 | 0.3 | 0.18 | 0.22 | 0.04 | 0.05 |
| 10 | 68396283 | rs7072222 | -0.662 | 0.837 | 0.045 | -0.743 | -0.003 | -0.093 | -0.093 | 0.018 | 0.6 | 0.46 | 0.65 | 0.94 | 0.63 | 0.75 |
| 10 | 68400418 | rs17274526 | 0.705 | 0.746 | -0.614 | -0.62 | 0.016 | 0.008 | 0.008 | 0.357 | 0.21 | 0.08 | 0.12 | 0.17 | 0.15 | 0.24 |
| 10 | 68401031 | rs745451 | -1.017 | 0.741 | -0.521 | -0.686 | -0.011 | 0.102 | 0.102 | 0 | 0.64 | 0.8 | 0.67 | 0.94 | 0.63 | 0.76 |
| 10 | 68412958 | rs6480192 | -0.99 | 0.775 | -1.318 | -0.671 | -0.009 | 0.1 | 0.1 | 0 | 0.63 | 0.78 | 0.66 | 0.94 | 0.63 | 0.76 |
| 10 | 68415731 | rs2894015 | -0.979 | 0.782 | -1.332 | -0.693 | -0.009 | 0.106 | 0.106 | 0 | 0.63 | 0.78 | 0.66 | 0.94 | 0.63 | 0.76 |
| 10 | 68417589 | rs6480193 | -1.103 | 0.779 | -1.344 | -0.694 | -0.033 | 0.173 | 0.173 | 0 | 0.64 | 0.87 | 0.67 | 0.94 | 0.63 | 0.76 |
| 10 | 68418846 | rs10997287 | 0.81 | 1.032 | -1.332 | -0.528 | -0.079 | -0.155 | -0.155 | 0.319 | 0.27 | 0.03 | 0.27 | 0.52 | 0.32 | 0.4 |
| 10 | 68425467 | rs2394279 | -0.275 | 1.008 | -1.213 | -0.608 | -0.014 | 0.049 | 0.049 | 0.832 | 0.42 | 0.66 | 0.54 | 0.77 | 0.47 | 0.51 |
| 10 | 68428858 | rs10822869 | -0.635 | 0.574 | -0.94 | -0.25 | 0.01 | -0.02 | -0.02 | 0 | 0.35 | 0.74 | 0.39 | 0.43 | 0.31 | 0.36 |
| 10 | 68429372 | rs4746622 | -0.547 | 0.419 | -1.03 | -0.282 | 0.003 | -0.028 | -0.028 | 0.014 | 0.31 | 0.21 | 0.15 | 0.39 | 0.31 | 0.35 |
| 10 | 68429563 | rs7079284 | -0.094 | 0.872 | -0.747 | -0.624 | 0 | -0.018 | -0.018 | 0 | 0.62 | 0.43 | 0.76 | 0.81 | 0.62 | 0.77 |
| 10 | 68435662 | rs2394287 | -0.195 | 0.816 | -1.102 | -0.532 | -0.02 | -0.014 | -0.014 | 0 | 0.56 | 0.69 | 0.38 | 0.44 | 0.49 | 0.48 |
| 10 | 68439358 | rs4746625 | -0.202 | 0.813 | -1.102 | -0.533 | -0.021 | -0.015 | -0.015 | 0 | 0.56 | 0.69 | 0.38 | 0.44 | 0.49 | 0.48 |
| 10 | 68442048 | rs4990414 | 1.276 | 0.334 | -1.371 | -0.999 | 0.012 | 0.083 | 0.083 | 0 | 0.22 | 0.09 | 0.1 | 0.01 | 0.18 | 0.14 |
| 10 | 68443296 | rs713298 | -1.201 | 0.591 | -1.664 | -1.115 | -0.035 | 0.045 | 0.045 | 0 | 0.89 | 0.96 | 0.93 | 1 | 0.85 | 0.91 |
| 10 | 68443699 | rs2394294 | -1.201 | 0.591 | -1.664 | -1.115 | -0.035 | 0.045 | 0.045 | 0 | 0.89 | 0.96 | 0.93 | 1 | 0.85 | 0.91 |
| 10 | 68444734 | rs7915695 | 1.068 | 0.409 | -1.089 | -1.102 | -0.019 | -0.01 | -0.01 | 0.133 | 0.14 | 0.26 | 0.09 | 0.12 | 0.09 | 0.16 |
| 10 | 68448403 | rs6480203 | 0.49 | 0.397 | -0.996 | -1.053 | -0.025 | 0.148 | 0.148 | 0.538 | 0.17 | 0.32 | 0.11 | 0.36 | 0.12 | 0.2 |
| 10 | 68454810 | rs10822887 | 0.561 | 0.396 | -0.968 | -1.053 | -0.023 | 0.12 | 0.12 | 0.494 | 0.17 | 0.27 | 0.1 | 0.36 | 0.12 | 0.2 |
| 10 | 68457608 | rs10997316 | -0.075 | 0.466 | -0.968 | -1.038 | 0.001 | 0.03 | 0.03 | 0 | 0.72 | 0.67 | 0.82 | 0.64 | 0.73 | 0.71 |
| 10 | 68457813 | rs10822890 | 0.848 | 0.503 | -1.286 | -0.882 | -0.024 | -0.075 | -0.075 | 0.269 | 0.31 | 0.17 | 0.54 | 0.56 | 0.36 | 0.43 |
| 10 | 68461866 | rs7893145 | 0.726 | 0.404 | -1.025 | -0.752 | -0.016 | 0.015 | 0.015 | 0 | 0.13 | 0.09 | 0.15 | 0.07 | 0.18 | 0.12 |
| 10 | 68469477 | rs10509276 | 0.151 | 0.033 | -0.998 | -0.626 | -0.001 | 0.005 | 0.005 | 0.041 | 0.16 | 0.15 | 0.07 | 0.27 | 0.11 | 0.13 |
| 10 | 68479424 | rs10997355 | -0.378 | 0.161 | -0.947 | -0.438 | -0.005 | 0.074 | 0.074 | 0 | 0.05 | 0.06 | 0.02 | 0.24 | 0.03 | 0.04 |
| 10 | 68482454 | rs10997361 | 0.708 | 0.433 | -1.295 | -0.649 | -0.047 | -0.014 | -0.014 | 5.74 | 0.2 | 0.07 | 0.5 | 0.79 | 0.3 | 0.38 |
| 10 | 68483603 | rs7908849 | 1.1 | 0.256 | -1.221 | -1.077 | 0.003 | 0.049 | 0.049 | 0 | 0.37 | 0.27 | 0.25 | 0.05 | 0.37 | 0.26 |
| 10 | 68486374 | rs10822904 | 1.48 | 0.325 | -1.193 | -0.873 | -0.073 | -0.123 | -0.123 | 0 | 0.17 | 0.03 | 0.48 | 0.59 | 0.27 | 0.33 |
| 10 | 68486501 | rs12252238 | 1.126 | -0.01 | -0.466 | -0.814 | -0.019 | -0.013 | -0.013 | 0 | 0.14 | 0.21 | 0.09 | 0.12 | 0.1 | 0.15 |
| 10 | 68487347 | rs12254338 | 1.126 | 0.027 | -0.475 | -0.827 | -0.016 | -0.015 | -0.015 | 0 | 0.14 | 0.21 | 0.09 | 0.12 | 0.1 | 0.15 |
| 10 | 68492082 | rs7083773 | -0.777 | 0.103 | -0.683 | -0.842 | 0.008 | 0.03 | 0.03 | 0 | 0.58 | 0.69 | 0.73 | 0.73 | 0.59 | 0.69 |
| 10 | 68499432 | rs7092601 | -0.625 | 0.107 | -0.655 | -0.824 | 0.007 | 0.024 | 0.024 | 0 | 0.57 | 0.63 | 0.71 | 0.73 | 0.58 | 0.66 |
| 10 | 68502854 | rs7094812 | -0.455 | 0.412 | -0.739 | -0.249 | -0.001 | 0.01 | 0.01 | 0 | 0.49 | 0.59 | 0.57 | 0.49 | 0.51 | 0.53 |
| 10 | 68507421 | rs6480209 | -0.874 | 0.428 | -1.014 | -0.257 | 0.014 | -0.035 | -0.035 | 0 | 0.4 | 0.78 | 0.41 | 0.5 | 0.34 | 0.35 |
| 10 | 68522420 | rs7084050 | -0.2 | 0.595 | -0.799 | -0.288 | -0.047 | -0.072 | -0.072 | 0 | 0.4 | 0.15 | 0.54 | 0.48 | 0.47 | 0.48 |
| 10 | 68523326 | rs12763151 | 0.963 | 0.654 | -0.965 | -0.334 | -0.049 | 0.032 | 0.032 | 0.101 | 0.14 | 0.03 | 0.12 | 0.05 | 0.19 | 0.13 |
| 10 | 68524967 | rs4494211 | -1.445 | 0.459 | -0.648 | -0.159 | -0.004 | -0.023 | -0.023 | 0 | 0.76 | 0.86 | 0.82 | 0.75 | 0.77 | 0.81 |
| 10 | 68527738 | rs7899434 | -2.069 | 0.454 | -0.671 | -0.186 | -0.012 | 0.091 | 0.091 | 0 | 0.8 | 0.94 | 0.9 | 0.99 | 0.81 | 0.86 |
| 10 | 68527823 | rs7899834 | -2.069 | 0.454 | -0.671 | -0.186 | -0.012 | 0.091 | 0.091 | 0 | 0.8 | 0.94 | 0.9 | 0.99 | 0.81 | 0.86 |
| 10 | 68535981 | rs4387240 | -1.372 | 0.397 | -0.461 | -0.164 | -0.004 | 0.004 | 0.004 | 0 | 0.75 | 0.64 | 0.8 | 0.75 | 0.77 | 0.81 |
| 10 | 68553995 | rs4272690 | -0.143 | 0.412 | -0.481 | -0.264 | -0.007 | -0.003 | -0.003 | 0.211 | 0.31 | 0.66 | 0.25 | 0.29 | 0.24 | 0.25 |
| 10 | 68556357 | rs4074716 | 0.345 | 0.441 | -0.268 | -0.282 | -0.003 | -0.096 | -0.096 | 0 | 0.26 | 0.39 | 0.15 | 0.05 | 0.21 | 0.21 |
| 10 | 68564566 | rs7076617 | 0.047 | 0.529 | -0.637 | 0.111 | -0.004 | -0.01 | -0.01 | 0 | 0.42 | 0.27 | 0.56 | 0.5 | 0.49 | 0.48 |
| 10 | 68566285 | rs7067838 | -0.357 | 0.443 | -0.608 | -0.078 | -0.001 | -0.003 | -0.003 | 0 | 0.51 | 0.5 | 0.67 | 0.7 | 0.56 | 0.6 |
| 10 | 68570310 | rs10822918 | -0.314 | 0.569 | -0.783 | -0.055 | 0.002 | 0.023 | 0.023 | 0 | 0.51 | 0.4 | 0.67 | 0.75 | 0.56 | 0.59 |
| 10 | 68572526 | rs10822920 | -0.273 | 0.615 | -0.996 | -0.038 | 0.005 | 0.054 | 0.054 | 0 | 0.5 | 0.33 | 0.66 | 0.75 | 0.56 | 0.59 |
| 10 | 68576021 | rs4626963 | -0.172 | 0.999 | -2.036 | -0.068 | -0.057 | 0.004 | 0.004 | 0 | 0.35 | 0.11 | 0.29 | 0.27 | 0.39 | 0.31 |
| 10 | 68576077 | rs12218365 | -0.211 | 1.342 | -2.13 | 0.652 | -0.023 | -0.066 | -0.066 | 0.189 | 0.18 | 0.07 | 0.37 | 0.39 | 0.21 | 0.38 |
| 10 | 68577369 | rs7095347 | -0.516 | 0.585 | -1.802 | -0.021 | 0.013 | 0.02 | 0.02 | 0.902 | 0.48 | 0.6 | 0.62 | 0.51 | 0.44 | 0.5 |
| 10 | 68578970 | rs4746638 | 0.423 | 0.108 | -1.987 | -0.092 | 0 | 0.003 | 0.003 | 0.212 | 0.26 | 0.23 | 0.22 | 0.18 | 0.27 | 0.26 |
| 10 | 68583924 | rs10997397 | 0.814 | -0.297 | -1.493 | -0.437 | -0.041 | -0.103 | -0.103 | 0 | 0.16 | 0.33 | 0.07 | 0.02 | 0.07 | 0.09 |
| 10 | 68585502 | rs6480223 | -2.503 | 0.076 | -1.984 | -0.048 | -0.033 | -0.089 | -0.089 | 0 | 0.83 | 0.99 | 0.92 | 0.78 | 0.85 | 0.9 |
| 10 | 68585916 | rs7919760 | -0.138 | 0.133 | -1.598 | -0.26 | 0.002 | 0.008 | 0.008 | 0.315 | 0.5 | 0.54 | 0.65 | 0.48 | 0.56 | 0.52 |
| 10 | 68586025 | rs6480224 | -1.029 | 0.169 | -1.638 | 0.441 | 0.006 | -0.028 | -0.028 | 3.425 | 0.77 | 0.84 | 0.87 | 0.69 | 0.82 | 0.77 |
| 10 | 68586440 | rs7920397 | 0.542 | 0.781 | -1.505 | 0.512 | -0.029 | -0.062 | -0.062 | 1.296 | 0.15 | 0.04 | 0.37 | 0.41 | 0.21 | 0.31 |

| | | | | | | | | | | | | | | | | |
|---|---|---|---|---|---|---|---|---|---|---|---|---|---|---|---|---|
| 10 | 68587558 | rs4073245 | -0.673 | 0.527 | -1.112 | 0.126 | -0.003 | -0.02 | -0.02 | 0.027 | 0.05 | 0.13 | 0.04 | 0 | 0.04 | 0.02 |
| 10 | 68593916 | rs10733830 | -0.416 | 0.352 | -1.664 | 0.15 | 0.023 | -0.013 | -0.013 | 0.51 | 0.55 | 0.83 | 0.73 | 0.51 | 0.7 | 0.63 |
| 10 | 68593991 | rs10762112 | 1.1 | 0.546 | -0.95 | -0.126 | 0.02 | 0.106 | 0.106 | 2.704 | 0.15 | 0.29 | 0.41 | 0.38 | 0.2 | 0.27 |
| 10 | 68594795 | rs12257423 | 0.6 | 0.278 | -1.588 | -0.091 | 0.028 | 0.052 | 0.052 | 0.032 | 0.29 | 0.15 | 0.18 | 0.14 | 0.15 | 0.23 |
| 10 | 68594908 | rs7904842 | -2.137 | 0.372 | -1.487 | 0.039 | -0.047 | -0.121 | -0.121 | 0.145 | 0.84 | 0.99 | 0.91 | 0.65 | 0.85 | 0.86 |
| 10 | 68597130 | rs11591563 | 1.02 | 0.489 | -1.52 | -0.34 | -0.038 | 0.065 | 0.065 | 0 | 0.21 | 0.09 | 0.22 | 0.09 | 0.33 | 0.19 |
| 10 | 68598292 | rs7902091 | 1.122 | 0.541 | -0.909 | -0.251 | 0.017 | 0.122 | 0.122 | 3.063 | 0.17 | 0.51 | 0.43 | 0.38 | 0.21 | 0.28 |
| 10 | 68598928 | rs12765156 | 0.75 | 0.514 | -0.9 | -0.546 | 0.002 | -0.025 | -0.025 | 0 | 0.08 | 0.13 | 0.04 | 0.02 | 0.1 | 0.1 |
| 10 | 68600498 | rs7919083 | -0.475 | 0.397 | -1.036 | -0.417 | -0.02 | 0.077 | 0.077 | 0 | 0.43 | 0.73 | 0.48 | 0.3 | 0.63 | 0.56 |
| 10 | 68602813 | rs12763021 | 0.514 | 0.638 | -0.943 | -0.515 | 0.001 | -0.029 | -0.029 | 0 | 0.08 | 0.15 | 0.04 | 0.02 | 0.1 | 0.09 |
| 10 | 68603568 | rs12355064 | -0.019 | 0.652 | -0.94 | -0.529 | -0.019 | 0.027 | 0.027 | 0.003 | 0.27 | 0.25 | 0.25 | 0.11 | 0.42 | 0.29 |
| 10 | 68607338 | rs4237311 | -0.509 | 0.752 | -0.944 | -0.343 | -0.008 | -0.01 | -0.01 | 0 | 0.64 | 0.68 | 0.69 | 0.56 | 0.52 | 0.65 |
| 10 | 68613644 | rs4497282 | 0.517 | 0.271 | -0.997 | -0.254 | 0 | 0 | 0 | 0.526 | 0.32 | 0.34 | 0.5 | 0.53 | 0.25 | 0.41 |
| 10 | 68613748 | rs7922094 | -0.521 | 0.409 | -0.728 | -0.777 | 0.011 | 0.091 | 0.091 | 5.303 | 0.52 | 0.65 | 0.42 | 0.12 | 0.6 | 0.45 |
| 10 | 68614840 | rs10997419 | 0.25 | 0.37 | -0.679 | -0.837 | 0.008 | -0.089 | -0.089 | 0 | 0.4 | 0.45 | 0.35 | 0.11 | 0.48 | 0.34 |
| 10 | 68621346 | rs7902209 | -0.685 | 0.326 | -0.519 | -0.065 | -0.001 | -0.045 | -0.045 | 0 | 0.12 | 0.2 | 0.07 | 0.02 | 0.12 | 0.11 |
| 10 | 68626126 | rs9414948 | -0.319 | 0.286 | -0.461 | 0.535 | -0.013 | 0.055 | 0.055 | 0 | 0.24 | 0.39 | 0.18 | 0.33 | 0.16 | 0.15 |
| 10 | 68626963 | rs4746645 | -0.111 | 0.702 | -1.046 | -0.414 | -0.068 | 0.073 | 0.073 | 2.216 | 0.31 | 0.15 | 0.33 | 0.15 | 0.45 | 0.33 |
| 10 | 68627392 | rs11818097 | 0.037 | 0.363 | -0.125 | -0.766 | -0.01 | -0.07 | -0.07 | 0 | 0.17 | 0.3 | 0.1 | 0 | 0.12 | 0.1 |
| 10 | 68630564 | rs7918983 | -1.124 | 0.448 | 0.662 | -0.122 | -0.003 | 0.096 | 0.096 | 0 | 0.26 | 0.58 | 0.23 | 0.35 | 0.2 | 0.19 |
| 10 | 68630940 | rs4746646 | 0.013 | 0.227 | 0.709 | -0.346 | 0.022 | -0.017 | -0.017 | 1.109 | 0.92 | 0.77 | 0.95 | 0.99 | 0.91 | 0.97 |
| 10 | 68635070 | rs10762118 | 1.593 | 0.273 | 0.778 | -0.271 | 0.017 | 0.218 | 0.218 | 0 | 0.82 | 0.51 | 0.82 | 0.64 | 0.82 | 0.85 |
| 10 | 68637811 | rs1880046 | -0.21 | 0.685 | 0.401 | -0.657 | 0.01 | -0.054 | -0.054 | 0 | 0.36 | 0.43 | 0.37 | 0.12 | 0.43 | 0.33 |
| 10 | 68641287 | rs7897872 | -0.993 | 0.731 | 0.511 | -0.695 | -0.003 | -0.068 | -0.068 | 0 | 0.45 | 0.53 | 0.43 | 0.12 | 0.53 | 0.36 |
| 10 | 68645433 | rs1916375 | 0.86 | 0.75 | 0.515 | -0.737 | -0.024 | 0.021 | 0.021 | 0.824 | 0.2 | 0.1 | 0.27 | 0.12 | 0.33 | 0.25 |
| 10 | 68648238 | rs7072319 | -0.248 | 0.552 | 0.611 | -0.474 | 0 | -0.035 | -0.035 | 0.064 | 0.41 | 0.44 | 0.38 | 0.12 | 0.49 | 0.35 |
| 10 | 68649352 | rs10822936 | 1.055 | 0.426 | 0.326 | -0.421 | 0.034 | -0.058 | -0.058 | 6.815 | 0.45 | 0.06 | 0.44 | 0.52 | 0.37 | 0.5 |
| 10 | 68678211 | rs2140380 | -1.214 | 0.777 | 0.73 | -0.304 | 0.002 | -0.032 | -0.032 | 0.219 | 0.09 | 0.15 | 0.05 | 0 | 0.09 | 0.02 |
| 10 | 68680128 | rs16923760 | 0.448 | 0.676 | 0.403 | -0.178 | -0.045 | -0.052 | -0.052 | 0.178 | 0.09 | 0.01 | 0.27 | 0.41 | 0.15 | 0.16 |
| 10 | 68680367 | rs4746648 | 0.953 | 0.367 | 0.951 | -0.215 | 0.018 | 0.366 | 0.366 | 0 | 0.87 | 0.6 | 0.83 | 0.62 | 0.88 | 0.84 |
| 10 | 68684249 | rs1925580 | 0.303 | 0.512 | 0.812 | -0.716 | 0.013 | -0.041 | -0.041 | 0.576 | 0.91 | 0.8 | 0.93 | 0.99 | 0.9 | 0.94 |
| 10 | 68689083 | rs1925576 | 0.192 | 0.416 | 0.512 | -0.868 | -0.064 | 0.093 | 0.093 | 0.391 | 0.33 | 0.17 | 0.35 | 0.13 | 0.44 | 0.36 |
| 10 | 68691199 | rs1925574 | 1.055 | 0.124 | 1.34 | -0.599 | 0.008 | -0.085 | -0.085 | 0 | 0.84 | 0.75 | 0.91 | 1 | 0.84 | 0.92 |
| 10 | 68696578 | rs2140384 | 0.804 | 0.016 | 1.285 | -0.526 | 0.015 | 0.095 | 0.095 | 1.878 | 0.38 | 0.25 | 0.23 | 0.03 | 0.31 | 0.3 |
| 10 | 68698029 | rs7075547 | 1.036 | -0.029 | 1.355 | -0.542 | 0.006 | 0.032 | 0.032 | 2.728 | 0.36 | 0.29 | 0.23 | 0.03 | 0.31 | 0.3 |
| 10 | 68703230 | rs10997449 | -0.671 | -0.109 | 1.449 | -0.175 | 0.03 | -0.017 | -0.017 | 0 | 0.57 | 0.32 | 0.7 | 0.6 | 0.68 | 0.6 |
| 10 | 68715988 | rs1916389 | -0.257 | 0.221 | 1.447 | -0.289 | 0.006 | -0.052 | -0.052 | 0 | 0.15 | 0.19 | 0.11 | 0 | 0.2 | 0.07 |
| 10 | 68716374 | rs10762122 | 0.953 | 0.243 | 0.988 | -0.277 | 0.015 | 0.096 | 0.096 | 2.147 | 0.38 | 0.25 | 0.23 | 0.03 | 0.31 | 0.3 |
| 10 | 68721071 | rs1925564 | -0.47 | 0.406 | 0.638 | -0.155 | 0.01 | 0.114 | 0.114 | 0.05 | 0.18 | 0.31 | 0.16 | 0.37 | 0.22 | 0.17 |
| 10 | 68722086 | rs1925562 | -0.219 | 0.513 | 0.094 | 0.397 | -0.064 | -0.07 | -0.07 | 0 | 0.42 | 0.16 | 0.59 | 0.6 | 0.48 | 0.53 |
| 10 | 68736018 | rs10997462 | -0.338 | 0.993 | -0.848 | 0.638 | -0.015 | -0.02 | -0.02 | 0 | 0.45 | 0.28 | 0.59 | 0.6 | 0.48 | 0.53 |
| 10 | 68741835 | rs942788 | 0.815 | 0.988 | 0.231 | 0.414 | 0.016 | -0.078 | -0.078 | 0 | 0.88 | 0.73 | 0.91 | 1 | 0.87 | 0.95 |
| 10 | 68742030 | rs16923867 | -0.211 | 0.802 | 0.141 | 0.203 | -0.002 | -0.019 | -0.019 | 0 | 0.1 | 0.12 | 0.04 | 0 | 0.07 | 0.06 |
| 10 | 68747691 | rs2840237 | 0.253 | 0.768 | 0.806 | 0.023 | 0.029 | -0.051 | -0.051 | 0 | 0.91 | 0.8 | 0.92 | 1 | 0.86 | 0.97 |
| 10 | 68750412 | rs1925608 | -1.412 | 1.572 | 0.494 | 0.875 | 0.011 | 0.01 | 0.01 | 1.55 | 0.35 | 0.6 | 0.6 | 0.49 | 0.43 | 0.34 |
| 10 | 68750755 | rs7082306 | -0.179 | 1.528 | 0.124 | 0.644 | -0.025 | -0.066 | -0.066 | 0 | 0.12 | 0.23 | 0.05 | 0 | 0.07 | 0.07 |
| 10 | 68754653 | rs716547 | 0.113 | 2.154 | -0.464 | 0.95 | -0.07 | -0.075 | -0.075 | 0 | 0.15 | 0.04 | 0.47 | 0.39 | 0.27 | 0.24 |
| 10 | 68756845 | rs9971243 | -0.531 | 1.58 | -0.018 | 0.551 | 0.032 | -0.031 | -0.031 | 0 | 0.06 | 0.17 | 0.06 | 0 | 0.09 | 0.02 |
| 10 | 68756918 | rs1925611 | 0.464 | 0.861 | -0.803 | 0.121 | 0.015 | 0.108 | 0.108 | 0 | 0.51 | 0.3 | 0.65 | 0.81 | 0.52 | 0.65 |
| 10 | 68757916 | rs10997476 | -0.097 | 0.507 | -1.365 | 0.252 | -0.084 | -0.03 | -0.03 | 4.15 | 0.37 | 0.08 | 0.61 | 0.42 | 0.56 | 0.4 |
| 10 | 68757963 | rs9633546 | -0.7 | -0.275 | -0.261 | 0.409 | -0.011 | 0.096 | 0.096 | 0 | 0.23 | 0.49 | 0.2 | 0.36 | 0.21 | 0.36 |
| 10 | 68764187 | rs10997480 | 0.371 | 0.152 | -0.565 | 0.445 | -0.067 | -0.087 | -0.087 | 0 | 0.19 | 0.11 | 0.49 | 0.53 | 0.34 | 0.33 |
| 10 | 68777013 | rs7900716 | -0.293 | -0.229 | 0.323 | 0.292 | 0.007 | 0.089 | 0.089 | 0 | 0.18 | 0.04 | 0.07 | 0.01 | 0.14 | 0.29 |
| 10 | 68792250 | rs10997487 | -0.286 | -0.055 | 0.381 | 0.551 | -0.108 | -0.118 | -0.118 | 0 | 0.25 | 0.08 | 0.48 | 0.51 | 0.41 | 0.35 |
| 10 | 68796795 | rs10762129 | 1.267 | 0.181 | 0.472 | 0.66 | -0.086 | -0.097 | -0.097 | 0 | 0.13 | 0.01 | 0.39 | 0.44 | 0.23 | 0.29 |
| 10 | 68813983 | rs7908101 | -0.576 | -0.537 | 0.916 | 0.234 | 0.011 | 0.095 | 0.095 | 0 | 0.09 | 0.31 | 0.07 | 0.14 | 0.07 | 0.12 |
| 10 | 68818643 | rs2147886 | -0.189 | -0.491 | 0.669 | 0.17 | -0.101 | -0.11 | -0.11 | 0 | 0.74 | 0.9 | 0.51 | 0.49 | 0.59 | 0.66 |
| 10 | 68822470 | rs1925585 | 0.701 | -0.38 | 0.787 | 0.198 | -0.008 | 0.031 | 0.031 | 0.217 | 0.07 | 0.01 | 0.01 | 0 | 0.07 | 0.03 |
| 10 | 68824626 | rs2764810 | 0.465 | -0.868 | 1.389 | 0.215 | -0.006 | -0.007 | -0.007 | 0 | 0.63 | 0.6 | 0.45 | 0.49 | 0.51 | 0.43 |
| 10 | 68824793 | rs10509279 | 0.081 | -0.872 | 1.499 | 0.207 | -0.004 | -0.051 | -0.051 | 0.015 | 0.11 | 0.17 | 0.06 | 0 | 0.08 | 0.22 |
| 10 | 68825297 | rs2764808 | 0.457 | -0.863 | 1.457 | 0.22 | -0.006 | -0.007 | -0.007 | 2.084 | 0.63 | 0.6 | 0.45 | 0.49 | 0.51 | 0.43 |
| 10 | 68827014 | rs2394323 | 1.467 | -1.223 | 1.327 | -0.068 | 0.046 | 0.119 | 0.119 | 1.734 | 0.45 | 0.21 | 0.3 | 0.19 | 0.4 | 0.23 |
| 10 | 68827794 | rs10822970 | -0.605 | -0.563 | 1.589 | 0.22 | 0.007 | 0.09 | 0.09 | 0 | 0.1 | 0.3 | 0.07 | 0.14 | 0.07 | 0.12 |
| 10 | 68835496 | rs2764813 | -0.151 | -0.532 | 1.517 | 0.21 | -0.01 | -0.012 | -0.012 | 0 | 0.71 | 0.69 | 0.48 | 0.48 | 0.57 | 0.64 |
| 10 | 68842368 | rs2394324 | 0.282 | -0.612 | 1.653 | 0.201 | -0.012 | -0.012 | -0.012 | 0.2 | 0.64 | 0.73 | 0.46 | 0.5 | 0.5 | 0.43 |
| 10 | 68845232 | rs5026468 | 0.643 | -0.603 | 1.658 | 0.2 | -0.001 | 0.003 | 0.003 | 0 | 0.09 | 0.05 | 0.04 | 0 | 0.08 | 0.2 |
| 10 | 68847022 | rs4745927 | 0.135 | -0.608 | 1.682 | 0.228 | -0.022 | -0.022 | -0.022 | 0.053 | 0.65 | 0.77 | 0.46 | 0.5 | 0.5 | 0.43 |
| 10 | 68849988 | rs10822973 | 0.258 | -0.616 | 1.627 | 0.234 | 0.015 | 0.078 | 0.078 | 0 | 0.08 | 0.21 | 0.06 | 0.14 | 0.07 | 0.13 |
| 10 | 68851075 | rs10740266 | -1.379 | -0.651 | 1.496 | 0.251 | -0.015 | 0.164 | 0.164 | 1.105 | 0.13 | 0.51 | 0.16 | 0.26 | 0.08 | 0.15 |

| | | | | | | | | | | | | | | | | |
|---|---|---|---|---|---|---|---|---|---|---|---|---|---|---|---|---|
| 10 | 68855554 | rs7895757 | 0.299 | -0.63 | 1.372 | 0.248 | -0.035 | -0.021 | -0.021 | 0 | 0.63 | 0.78 | 0.41 | 0.39 | 0.5 | 0.41 |
| 10 | 68857184 | rs10822976 | 0.151 | -0.629 | 1.293 | 0.249 | -0.063 | -0.062 | -0.062 | 0.046 | 0.65 | 0.85 | 0.47 | 0.5 | 0.5 | 0.43 |
| 10 | 68869781 | rs10458634 | -1.463 | -0.356 | 1.645 | 0.41 | -0.009 | 0.066 | 0.066 | 0.863 | 0.13 | 0.51 | 0.11 | 0.14 | 0.08 | 0.14 |
| 10 | 68875634 | rs7081413 | -1.388 | -0.317 | 1.577 | 0.392 | -0.015 | 0.065 | 0.065 | 0.121 | 0.19 | 0.44 | 0.18 | 0.26 | 0.14 | 0.36 |
| 10 | 68888483 | rs10822979 | 0.118 | -0.229 | 1.481 | 0.461 | 0.006 | 0.041 | 0.041 | 0 | 0.07 | 0.18 | 0.05 | 0.13 | 0.07 | 0.13 |
| 10 | 68898306 | rs10997527 | -1.106 | -0.19 | 1.121 | 0.446 | 0 | -0.026 | -0.026 | 0 | 0.19 | 0.4 | 0.19 | 0.13 | 0.15 | 0.36 |
| 10 | 68898342 | rs1014329 | -0.637 | -0.151 | 1.016 | 0.438 | -0.029 | -0.033 | -0.033 | 0 | 0.75 | 0.81 | 0.61 | 0.51 | 0.6 | 0.64 |
| 10 | 68901222 | rs10997528 | 1.642 | -0.303 | 0.916 | 0.345 | 0.061 | 0.108 | 0.108 | 3.897 | 0.47 | 0.19 | 0.33 | 0.25 | 0.39 | 0.23 |
| 10 | 68901252 | rs10822982 | 0.373 | -0.204 | 0.75 | 0.424 | 0.013 | 0.003 | 0.003 | 0.204 | 0.65 | 0.58 | 0.52 | 0.38 | 0.54 | 0.59 |
| 10 | 68908982 | rs2394327 | -0.517 | -0.102 | 0.222 | 0.407 | -0.039 | -0.043 | -0.043 | 0 | 0.75 | 0.82 | 0.61 | 0.51 | 0.6 | 0.65 |
| 10 | 68922029 | rs10466008 | -2.303 | -0.73 | 0.972 | 0.278 | 0.034 | -0.017 | -0.017 | 0.473 | 0.32 | 0.87 | 0.31 | 0.28 | 0.22 | 0.42 |
| 10 | 68922242 | rs10822992 | -0.549 | -0.695 | 1.584 | 0.286 | 0.041 | 0.096 | 0.096 | 0 | 0.08 | 0.32 | 0.13 | 0.12 | 0.07 | 0.13 |
| 10 | 68933988 | rs10997551 | -0.363 | -0.676 | 1.325 | 0.257 | -0.005 | -0.045 | -0.045 | 0 | 0.09 | 0.17 | 0.04 | 0 | 0.06 | 0.03 |
| 10 | 68949578 | rs7073762 | -0.747 | -0.68 | 0.928 | 0.313 | -0.01 | 0.031 | 0.031 | 0 | 0.12 | 0.22 | 0.11 | 0.16 | 0.08 | 0.25 |
| 10 | 68954061 | rs10823000 | -1.614 | -0.731 | 0.461 | 0.254 | -0.002 | -0.002 | -0.002 | 0 | 0.21 | 0.71 | 0.2 | 0.12 | 0.14 | 0.17 |
| 10 | 68959645 | rs7905049 | -1.919 | -0.709 | 0.189 | 0.302 | 0.035 | 0.091 | 0.091 | 1.059 | 0.32 | 0.83 | 0.26 | 0.13 | 0.22 | 0.39 |
| 10 | 68961492 | rs1904645 | -1.933 | -0.697 | 0.194 | 0.301 | 0.036 | -0.006 | -0.006 | 0 | 0.32 | 0.83 | 0.31 | 0.28 | 0.22 | 0.42 |
| 10 | 68965526 | rs10823006 | -0.851 | -0.879 | 1.072 | 0.355 | 0.064 | 0.093 | 0.093 | 0 | 0.09 | 0.32 | 0.1 | 0.14 | 0.1 | 0.08 |
| 10 | 68967202 | rs1993992 | 1.431 | -1.117 | 0.433 | -0.093 | -0.033 | 0.084 | 0.084 | 0 | 0.14 | 0.01 | 0.13 | 0 | 0.17 | 0.04 |
| 10 | 68971282 | rs16924233 | 0.135 | -1.008 | 1.12 | 0.356 | -0.011 | -0.021 | -0.021 | 0.032 | 0.05 | 0.08 | 0.02 | 0.01 | 0.03 | 0.08 |
| 10 | 68971354 | rs10733835 | -2.46 | -1.184 | 1.316 | 0.557 | -0.03 | 0.03 | 0.03 | 0.756 | 0.22 | 0.6 | 0.23 | 0.33 | 0.15 | 0.2 |
| 10 | 68979853 | rs9651325 | -0.85 | -1.067 | 1.39 | 0.64 | 0.056 | 0.085 | 0.085 | 0 | 0.1 | 0.34 | 0.11 | 0.14 | 0.11 | 0.08 |
| 10 | 68980275 | rs10823009 | -2.141 | -1.308 | 1.459 | 0.402 | -0.022 | 0.034 | 0.034 | 0 | 0.21 | 0.62 | 0.21 | 0.32 | 0.14 | 0.32 |
| 10 | 69000781 | rs10823013 | -1.901 | -1.309 | 1.28 | 0.435 | -0.024 | 0.13 | 0.13 | 0 | 0.18 | 0.38 | 0.2 | 0.33 | 0.14 | 0.2 |
| 10 | 69003746 | rs7070725 | -2.592 | -1.67 | 0.564 | 0.475 | 0.066 | -0.124 | -0.124 | 0 | 0.41 | 0.93 | 0.33 | 0.49 | 0.29 | 0.48 |
| 10 | 69004247 | rs1904634 | -2.271 | -1.742 | 1.111 | 0.544 | -0.05 | 0.036 | 0.036 | 0.657 | 0.21 | 0.64 | 0.2 | 0.34 | 0.09 | 0.18 |
| 10 | 69005693 | rs10509285 | 0.422 | -1.708 | 1.13 | 0.548 | -0.003 | -0.015 | -0.015 | 0 | 0.07 | 0.05 | 0.01 | 0 | 0.02 | 0.21 |
| 10 | 69011499 | rs10762157 | -1.995 | -1.707 | 1.217 | 0.435 | -0.053 | -0.052 | -0.052 | 0 | 0.18 | 0.62 | 0.14 | 0.07 | 0.09 | 0.15 |
| 10 | 69013466 | rs7076647 | -1.657 | -1.691 | 1.326 | 0.443 | -0.061 | -0.061 | -0.061 | 0 | 0.16 | 0.53 | 0.13 | 0.07 | 0.09 | 0.15 |
| 10 | 69021287 | rs1904608 | -1.688 | -1.78 | 1.418 | 0.419 | -0.071 | -0.075 | -0.075 | 0.491 | 0.19 | 0.47 | 0.11 | 0.06 | 0.08 | 0.29 |
| 10 | 69021594 | rs2170135 | -0.258 | -1.747 | 1.346 | 0.411 | -0.022 | -0.036 | -0.036 | 0 | 0.07 | 0.18 | 0.04 | 0.01 | 0.03 | 0.08 |
| 10 | 69023117 | rs1904610 | -2.626 | -1.81 | 1.213 | 0.577 | 0.101 | -0.043 | -0.043 | 0.624 | 0.3 | 0.84 | 0.23 | 0.34 | 0.12 | 0.4 |
| 10 | 69029763 | rs7085384 | -1.832 | -1.712 | 1.239 | 0.615 | -0.073 | 0.044 | 0.044 | 0 | 0.2 | 0.36 | 0.13 | 0.28 | 0.07 | 0.33 |
| 10 | 69050705 | rs10762159 | -2.247 | -1.688 | 1.389 | 0.596 | -0.057 | 0.035 | 0.035 | 0 | 0.25 | 0.57 | 0.21 | 0.33 | 0.12 | 0.39 |
| 10 | 69054048 | rs4553264 | -2.26 | -1.688 | 1.38 | 0.596 | -0.058 | 0.036 | 0.036 | 0 | 0.25 | 0.57 | 0.21 | 0.33 | 0.12 | 0.39 |
| 10 | 69057549 | rs2894035 | -1.356 | -1.678 | 1.367 | 0.57 | -0.035 | -0.038 | -0.038 | 0.058 | 0.14 | 0.32 | 0.11 | 0.07 | 0.09 | 0.13 |
| 10 | 69063842 | rs10823028 | -1.441 | -1.686 | 1.366 | 0.567 | -0.038 | -0.041 | -0.041 | 0 | 0.14 | 0.32 | 0.11 | 0.07 | 0.09 | 0.13 |
| 10 | 69064448 | rs10762163 | -2.396 | -1.584 | 0.529 | 0.604 | 0.062 | -0.112 | -0.112 | 0.008 | 0.41 | 0.92 | 0.34 | 0.48 | 0.29 | 0.48 |
| 10 | 69072867 | rs2448547 | 0.608 | -1.673 | 1.016 | 0.728 | 0.052 | 0.083 | 0.083 | 0.502 | 0.93 | 0.58 | 0.92 | 0.94 | 0.95 | 0.94 |
| 10 | 69073503 | rs2447659 | 1.397 | -1.623 | 0.704 | 0.734 | 0.058 | -0.035 | -0.035 | 0 | 0.72 | 0.23 | 0.77 | 0.66 | 0.87 | 0.61 |
| 10 | 69088407 | rs2620924 | 0.274 | -1.249 | -0.478 | 0.222 | 0.006 | -0.051 | -0.051 | 0 | 0.44 | 0.16 | 0.56 | 0.52 | 0.54 | 0.51 |
| 10 | 69093086 | rs2616700 | 1.247 | -1.625 | 0.871 | 0.685 | -0.033 | 0.162 | 0.162 | 1.087 | 0.87 | 0.52 | 0.85 | 0.74 | 0.92 | 0.76 |
| 10 | 69093693 | rs1904625 | 1.971 | -1.773 | 0.929 | 0.53 | 0.021 | 0.001 | 0.001 | 0.803 | 0.73 | 0.29 | 0.8 | 0.74 | 0.86 | 0.63 |
| 10 | 69093912 | rs1904627 | 1.812 | -2.116 | 1.049 | 0.547 | -0.032 | 0 | 0 | 0.403 | 0.74 | 0.37 | 0.82 | 0.74 | 0.88 | 0.64 |
| 10 | 69098628 | rs1904630 | 2.356 | -2.24 | 1.039 | 0.502 | 0.074 | 0.007 | 0.007 | 1.968 | 0.7 | 0.2 | 0.8 | 0.73 | 0.88 | 0.63 |
| 10 | 69107773 | rs16924405 | -0.668 | -2.184 | 1.073 | 0.528 | -0.015 | 0.014 | 0.014 | 0 | 0.12 | 0.12 | 0.09 | 0.26 | 0.03 | 0.26 |
| 10 | 69110525 | rs10762166 | 0.825 | -2.186 | 1.494 | 0.48 | 0.071 | 0.029 | 0.029 | 0.706 | 0.93 | 0.58 | 0.95 | 1 | 0.96 | 0.97 |
| 10 | 69111369 | rs9414953 | 0.422 | -2.181 | 1.47 | 0.482 | 0.066 | 0.029 | 0.029 | 0.44 | 0.94 | 0.66 | 0.96 | 1 | 0.96 | 0.97 |
| 10 | 69112493 | rs16924413 | -0.073 | -2.188 | 1.385 | 0.48 | 0.01 | 0 | 0 | 0 | 0.1 | 0.05 | 0.02 | 0 | 0.03 | 0.12 |
| 10 | 69132869 | rs6480258 | 1.818 | -2.269 | 1.403 | 0.53 | -0.066 | 0.038 | 0.038 | 0 | 0.81 | 0.46 | 0.86 | 0.75 | 0.92 | 0.71 |
| 10 | 69140084 | rs10997624 | -2.048 | -2.264 | 1.368 | 0.533 | -0.081 | 0.029 | 0.029 | 0 | 0.21 | 0.4 | 0.15 | 0.26 | 0.08 | 0.33 |
| 10 | 69141544 | rs2394381 | 0.07 | -2.256 | 1.376 | 0.436 | -0.005 | -0.017 | -0.017 | 0 | 0.06 | 0.09 | 0.04 | 0.01 | 0.04 | 0.07 |
| 10 | 69155636 | rs2394375 | -2.094 | -2.276 | 1.385 | 0.511 | -0.079 | 0.051 | 0.051 | 0 | 0.22 | 0.4 | 0.15 | 0.32 | 0.08 | 0.33 |
| 10 | 69161446 | rs10997632 | -0.35 | -2.137 | 1.263 | 0.398 | -0.008 | -0.022 | -0.022 | 0 | 0.07 | 0.09 | 0.04 | 0.01 | 0.04 | 0.07 |
| 10 | 69189889 | rs7920229 | -1.757 | -2.266 | 1.479 | 0.653 | -0.023 | 0.021 | 0.021 | 0 | 0.19 | 0.2 | 0.13 | 0.37 | 0.08 | 0.31 |
| 10 | 69201534 | rs9415880 | 1.34 | -2.331 | 1.577 | 0.609 | 0.012 | 0.409 | 0.409 | 0 | 0.92 | 0.62 | 0.89 | 0.64 | 0.95 | 0.83 |
| 10 | 69201746 | rs9414962 | 2.66 | -2.341 | 1.556 | 0.584 | 0.064 | -0.03 | -0.03 | 0 | 0.7 | 0.2 | 0.8 | 0.63 | 0.87 | 0.66 |
| 10 | 69201760 | rs9414963 | 2.074 | -2.384 | 1.574 | 0.586 | -0.019 | 0.005 | 0.005 | 0.036 | 0.74 | 0.35 | 0.83 | 0.63 | 0.87 | 0.67 |
| 10 | 69222470 | rs10997651 | -2.154 | -2.349 | 1.404 | 0.6 | -0.084 | 0.071 | 0.071 | 0.096 | 0.23 | 0.4 | 0.16 | 0.38 | 0.08 | 0.32 |
| 10 | 69245545 | rs997376 | -3.095 | -2.581 | 1.232 | 0.518 | 0.019 | -0.019 | -0.019 | 0.164 | 0.28 | 0.77 | 0.24 | 0.38 | 0.11 | 0.32 |
| 10 | 69246472 | rs7069964 | -2.302 | -2.578 | 1.445 | 0.52 | -0.106 | 0.094 | 0.094 | 0.021 | 0.21 | 0.39 | 0.18 | 0.38 | 0.06 | 0.28 |
| 10 | 69247842 | rs1875151 | -2.517 | -2.652 | 0.491 | 0.54 | -0.048 | 0.12 | 0.12 | 1.146 | 0.6 | 0.94 | 0.46 | 0.85 | 0.42 | 0.85 |
| 10 | 69254812 | rs7358238 | -1.233 | -2.016 | 1.368 | 0.691 | -0.019 | -0.035 | -0.035 | 0.136 | 0.13 | 0.13 | 0.05 | 0.01 | 0.05 | 0.13 |
| 10 | 69275099 | rs10823054 | -0.406 | -2 | 1.293 | 0.886 | 0.021 | -0.022 | -0.022 | 0 | 0.23 | 0.19 | 0.26 | 0.68 | 0.13 | 0.45 |
| 10 | 69285707 | rs12355429 | -1.324 | -2.073 | 1.411 | 0.827 | 0.041 | 0.007 | 0.007 | 0.228 | 0.08 | 0.43 | 0.05 | 0 | 0.05 | 0.03 |
| 10 | 69286361 | rs10823057 | 0.129 | -2.078 | 1.289 | 0.92 | 0.027 | -0.027 | -0.027 | 0 | 0.2 | 0.08 | 0.15 | 0.3 | 0.12 | 0.3 |
| 10 | 69293282 | rs7918024 | -0.357 | -2.033 | 1.535 | 0.53 | 0.039 | 0.012 | 0.012 | 0 | 0.05 | 0.25 | 0.04 | 0 | 0.05 | 0.03 |
| 10 | 69300672 | rs3920079 | -2.529 | -2.093 | 1.298 | 0.511 | 0.142 | 0.208 | 0.208 | 2.312 | 0.49 | 0.88 | 0.42 | 0.81 | 0.32 | 0.65 |
| 10 | 69304960 | rs10762172 | -2.339 | -2.266 | 1.39 | 0.502 | -0.079 | -0.123 | -0.123 | 0 | 0.21 | 0.57 | 0.1 | 0.01 | 0.1 | 0.15 |

| | | | | | | | | | | | | | | | | |
|---|---|---|---|---|---|---|---|---|---|---|---|---|---|---|---|---|
| 10 | 69327864 | rs7099961 | -2.185 | -2.229 | 1.381 | 0.518 | -0.09 | -0.143 | -0.143 | 0.207 | 0.19 | 0.5 | 0.09 | 0.01 | 0.1 | 0.15 |
| 10 | 69333065 | rs10740278 | -2.051 | -2.099 | 1.15 | 0.616 | 0.109 | 0.039 | 0.039 | 0.657 | 0.44 | 0.76 | 0.35 | 0.69 | 0.23 | 0.6 |
| 10 | 69348758 | rs12251222 | -1.268 | -1.467 | 1.506 | 0.437 | -0.034 | -0.056 | -0.056 | 0.386 | 0.1 | 0.33 | 0.05 | 0.01 | 0.04 | 0.07 |
| 10 | 69359021 | rs10997747 | -0.977 | -1.11 | 1.034 | 0.609 | 0.052 | 0.023 | 0.023 | 0.006 | 0.05 | 0.28 | 0.04 | 0 | 0.05 | 0.03 |
| 10 | 69366602 | rs3750863 | -2.102 | -0.948 | 0.154 | 0.754 | 0.021 | -0.108 | -0.108 | 0 | 0.34 | 0.89 | 0.38 | 0.51 | 0.29 | 0.41 |
| 10 | 69370832 | rs7084028 | -0.41 | -0.991 | 0.493 | 0.719 | -0.011 | -0.028 | -0.028 | 0.803 | 0.07 | 0.08 | 0.09 | 0.01 | 0.05 | 0.07 |
| 10 | 69378278 | rs6480279 | -2.635 | -1.095 | 0.907 | 0.774 | -0.046 | 0.074 | 0.074 | 0.04 | 0.17 | 0.66 | 0.25 | 0.45 | 0.1 | 0.29 |
| 10 | 69380156 | rs16924708 | -1.795 | -1.062 | 1.583 | 0.653 | -0.037 | -0.073 | -0.073 | 0 | 0.12 | 0.38 | 0.12 | 0.01 | 0.06 | 0.07 |
| 10 | 69381155 | rs7901697 | -1.795 | -1.062 | 1.581 | 0.653 | -0.036 | -0.073 | -0.073 | 1.623 | 0.12 | 0.38 | 0.12 | 0.01 | 0.06 | 0.07 |
| 10 | 69410048 | rs10430513 | 0.476 | -1.178 | 1.058 | 0.601 | 0.004 | -0.059 | -0.059 | 2.781 | 0.17 | 0.02 | 0.14 | 0.26 | 0.12 | 0.17 |
| 10 | 69410717 | rs10997770 | 0.141 | -1.054 | 1.219 | 0.543 | 0.051 | 0.003 | 0.003 | 0 | 0.06 | 0.12 | 0.1 | 0.07 | 0.15 | 0.06 |
| 10 | 69411888 | rs3096245 | 0.22 | -1.085 | 1.199 | 0.529 | 0.026 | 0.046 | 0.046 | 0.007 | 0.11 | 0.02 | 0.02 | 0 | 0.05 | 0.07 |
| 10 | 69467040 | rs7091132 | -0.984 | -1.097 | 1.476 | 0.447 | -0.045 | -0.089 | -0.089 | 0.476 | 0.17 | 0.3 | 0.06 | 0 | 0.07 | 0.12 |
| 10 | 69482437 | rs3125309 | -2.82 | -1.409 | 1.247 | 0.423 | 0.004 | 0.129 | 0.129 | 0.519 | 0.53 | 0.93 | 0.52 | 0.78 | 0.4 | 0.55 |
| 10 | 69493240 | rs3125311 | -2.959 | -1.389 | 1.368 | 0.51 | 0.021 | -0.092 | -0.092 | 0 | 0.34 | 0.87 | 0.37 | 0.52 | 0.29 | 0.36 |
| 10 | 69517922 | rs1113976 | 2.479 | -1.417 | 1.332 | 0.51 | 0.019 | -0.11 | -0.11 | 0 | 0.66 | 0.11 | 0.62 | 0.48 | 0.71 | 0.64 |
| 10 | 69523326 | rs7072202 | -0.872 | -1.332 | 1.398 | 0.5 | -0.042 | -0.085 | -0.085 | 0.008 | 0.17 | 0.28 | 0.06 | 0 | 0.07 | 0.12 |
| 10 | 69530678 | rs932656 | 1.236 | -1.262 | 1.531 | 0.59 | -0.009 | -0.029 | -0.029 | 0 | 0.64 | 0.46 | 0.55 | 0.21 | 0.67 | 0.58 |
| 10 | 69543215 | rs7907573 | 0.59 | -1.282 | 1.696 | 0.639 | -0.033 | -0.006 | -0.006 | 0.023 | 0.72 | 0.68 | 0.75 | 0.7 | 0.84 | 0.75 |
| 10 | 69543385 | rs7907719 | 1.111 | -1.236 | 1.36 | 0.374 | -0.009 | -0.026 | -0.026 | 0 | 0.64 | 0.46 | 0.55 | 0.21 | 0.67 | 0.58 |
| 10 | 69546194 | rs12777073 | 1.134 | -1.246 | 1.169 | 0.373 | -0.002 | -0.007 | -0.007 | 2.378 | 0.64 | 0.4 | 0.54 | 0.21 | 0.67 | 0.58 |
| 10 | 69553494 | rs16924881 | -1.334 | -1.042 | 1.374 | 0.426 | -0.061 | -0.108 | -0.108 | 0 | 0.18 | 0.34 | 0.08 | 0 | 0.09 | 0.12 |
| 10 | 69562951 | rs9804180 | -1.023 | -0.421 | 1.099 | 0.703 | 0.01 | -0.006 | -0.006 | 0 | 0.13 | 0.26 | 0.18 | 0.13 | 0.14 | 0.13 |
| 10 | 69563811 | rs10997817 | -1.039 | -0.701 | 0.513 | 0.866 | 0.018 | -0.076 | -0.076 | 0 | 0.18 | 0.04 | 0.16 | 0.32 | 0.14 | 0.2 |
| 10 | 69571336 | rs3740049 | -0.039 | -0.521 | 0.609 | 0.967 | -0.01 | -0.039 | -0.039 | 0 | 0.09 | 0.03 | 0.15 | 0.32 | 0.1 | 0.14 |
| 10 | 69584850 | rs7087595 | 1.261 | -0.879 | 1.001 | 0.852 | -0.024 | -0.008 | -0.008 | 0.045 | 0.65 | 0.62 | 0.68 | 0.28 | 0.79 | 0.55 |
| 10 | 69591262 | rs7918428 | 1.153 | -0.869 | 0.945 | 0.856 | 0.005 | 0.004 | 0.004 | 0 | 0.66 | 0.68 | 0.69 | 0.29 | 0.79 | 0.55 |
| 10 | 69609540 | rs7097008 | 1.754 | -0.825 | 1.076 | 0.851 | -0.015 | -0.033 | -0.033 | 0 | 0.55 | 0.52 | 0.53 | 0.16 | 0.67 | 0.43 |
| 10 | 69622535 | rs11599524 | -0.539 | -0.824 | 0.554 | 0.874 | -0.004 | -0.002 | -0.002 | 2.122 | 0.09 | 0.07 | 0.15 | 0.13 | 0.12 | 0.12 |
| 10 | 69627759 | rs730821 | 1.061 | -0.845 | 0.652 | 0.881 | 0.035 | 0.012 | 0.012 | 0 | 0.67 | 0.73 | 0.69 | 0.29 | 0.79 | 0.55 |
| 10 | 69643342 | rs3758391 | 1.544 | -0.739 | 0.674 | 0.883 | 0.026 | 0.039 | 0.039 | 0 | 0.58 | 0.65 | 0.53 | 0.16 | 0.67 | 0.43 |
| 10 | 69660494 | rs10823108 | 0.156 | -0.768 | 0.609 | 0.917 | 0.012 | -0.036 | -0.036 | 0 | 0.09 | 0.02 | 0.13 | 0.31 | 0.08 | 0.13 |
| 10 | 69664959 | rs10997868 | 2.309 | -0.944 | 0.582 | 0.797 | 0.094 | 0.23 | 0.23 | 0.62 | 0.48 | 0.06 | 0.47 | 0.16 | 0.65 | 0.42 |
| 10 | 69665387 | rs2224573 | 2.207 | -0.953 | 0.897 | 0.793 | 0.056 | 0.081 | 0.081 | 0.021 | 0.51 | 0.34 | 0.5 | 0.16 | 0.65 | 0.42 |
| 10 | 69666598 | rs2273773 | 0.24 | -0.953 | 0.829 | 0.8 | 0.012 | -0.036 | -0.036 | 0.177 | 0.09 | 0.02 | 0.13 | 0.31 | 0.08 | 0.13 |
| 10 | 69668014 | rs10997870 | 2.328 | -0.955 | 0.828 | 0.799 | 0.092 | 0.228 | 0.228 | 0.62 | 0.48 | 0.06 | 0.47 | 0.16 | 0.65 | 0.42 |
| 10 | 69670816 | rs10823112 | 0.276 | -0.919 | 0.823 | 0.818 | 0.011 | -0.033 | -0.033 | 0 | 0.09 | 0.02 | 0.13 | 0.31 | 0.08 | 0.13 |
| 10 | 69679824 | rs10997875 | 1.578 | -0.936 | 0.897 | 0.483 | 0.034 | 0.052 | 0.052 | 0 | 0.58 | 0.66 | 0.53 | 0.16 | 0.67 | 0.43 |
| 10 | 69683445 | rs10823116 | 0.124 | -0.928 | 0.846 | 0.761 | 0.015 | -0.032 | -0.032 | 0.115 | 0.1 | 0.02 | 0.15 | 0.39 | 0.08 | 0.14 |
| 10 | 69713297 | rs10762183 | 2.233 | -0.916 | 0.872 | 0.748 | 0.057 | 0.085 | 0.085 | 0.021 | 0.51 | 0.33 | 0.5 | 0.16 | 0.65 | 0.42 |
| 10 | 69720082 | rs10823120 | 1.697 | -0.918 | 0.901 | 0.747 | 0.012 | 0.013 | 0.013 | 0 | 0.57 | 0.6 | 0.53 | 0.16 | 0.67 | 0.43 |
| 10 | 69725450 | rs16925007 | 0.195 | -0.91 | 0.887 | 0.75 | 0.005 | -0.02 | -0.02 | 0 | 0.07 | 0.26 | 0.02 | 0 | 0.01 | 0.01 |
| 10 | 69726087 | rs10509292 | -0.292 | -0.924 | 0.777 | 0.741 | 0.035 | 0.134 | 0.134 | 0 | 0.2 | 0.03 | 0.07 | 0 | 0.13 | 0.21 |
| 10 | 69727722 | rs10997892 | -0.441 | -0.905 | 0.806 | 0.757 | -0.013 | -0.007 | -0.007 | 0.17 | 0.09 | 0.04 | 0.15 | 0.13 | 0.12 | 0.12 |
| 10 | 69729895 | rs10997893 | 0.019 | -0.904 | 0.812 | 0.757 | 0.005 | 0.028 | 0.028 | 0 | 0.1 | 0.07 | 0.15 | 0.39 | 0.08 | 0.14 |
| 10 | 69753620 | rs497849 | 1.151 | 0.399 | 0.596 | 0.524 | -0.015 | 0.032 | 0.032 | 0 | 0.06 | 0.01 | 0.09 | 0 | 0.21 | 0.02 |
| 10 | 69773841 | rs866255 | -2.239 | 0.165 | 0.933 | 0.804 | 0.013 | 0.013 | 0.013 | 0 | 0.43 | 0.4 | 0.47 | 0.84 | 0.33 | 0.57 |
| 10 | 69805786 | rs10509293 | -0.429 | -1.998 | 0.793 | 0.855 | 0.042 | 0.137 | 0.137 | 0 | 0.21 | 0.03 | 0.07 | 0 | 0.13 | 0.21 |
| 10 | 69816948 | rs6480302 | -0.395 | -1.946 | 0.852 | 0.925 | -0.021 | 0.112 | 0.112 | 0 | 0.12 | 0.2 | 0.17 | 0.39 | 0.08 | 0.14 |
| 10 | 69819865 | rs2485680 | -2.483 | -1.956 | 0.844 | 0.918 | 0.082 | 0.129 | 0.129 | 0.024 | 0.5 | 0.71 | 0.5 | 0.84 | 0.35 | 0.59 |
| 10 | 69820909 | rs4746727 | -0.368 | -1.077 | 0.728 | 1.17 | 0.012 | 0 | 0 | 0 | 0.1 | 0.05 | 0.15 | 0.39 | 0.08 | 0.14 |
| 10 | 69831243 | rs11599081 | -0.872 | -1.036 | 0.725 | 1.111 | -0.013 | -0.007 | -0.007 | 0 | 0.09 | 0.04 | 0.15 | 0.13 | 0.12 | 0.12 |
| 10 | 69850466 | rs8181450 | -0.624 | -1.052 | 0.774 | 1.047 | -0.009 | 0.065 | 0.065 | 0 | 0.11 | 0.13 | 0.17 | 0.39 | 0.08 | 0.14 |
| 10 | 69860245 | rs7086015 | 0.001 | -0.755 | 0.873 | 0.848 | 0.016 | -0.006 | -0.006 | 0 | 0.06 | 0.26 | 0.02 | 0 | 0.01 | 0.01 |
| 10 | 69866173 | rs9919371 | -0.338 | -0.864 | 0.513 | 0.993 | -0.011 | 0.086 | 0.086 | 0 | 0.11 | 0.16 | 0.18 | 0.43 | 0.1 | 0.15 |
| 10 | 69871319 | rs2894058 | -1.681 | -0.967 | -0.097 | 0.952 | 0.017 | -0.001 | -0.001 | 0 | 0.4 | 0.64 | 0.37 | 0.43 | 0.28 | 0.39 |
| 10 | 69879669 | rs6480304 | -1.187 | -0.485 | -0.009 | 1.091 | 0.005 | 0.077 | 0.077 | 0 | 0.24 | 0.42 | 0.31 | 0.45 | 0.26 | 0.24 |
| 10 | 69879709 | rs6480305 | -1.875 | -0.528 | -0.109 | 1.076 | 0.023 | -0.001 | -0.001 | 0 | 0.43 | 0.68 | 0.4 | 0.48 | 0.37 | 0.42 |
| 10 | 69884006 | rs17456902 | -0.234 | -0.509 | -0.701 | 0.92 | 0.02 | 0.025 | 0.025 | 0 | 0.18 | 0.12 | 0.07 | 0.03 | 0.11 | 0.18 |
| 10 | 69888483 | rs12220135 | -1.649 | 0.361 | -0.379 | 1.708 | -0.014 | 0.038 | 0.038 | 0.032 | 0.26 | 0.42 | 0.29 | 0.67 | 0.24 | 0.39 |
| 10 | 69896336 | rs12221035 | -0.869 | 0.59 | -1.126 | 1.781 | -0.003 | -0.036 | -0.036 | 0.214 | 0.09 | 0.02 | 0.15 | 0.39 | 0.11 | 0.11 |
| 10 | 69900240 | rs11598219 | -0.095 | 0.789 | -1.417 | 1.05 | -0.014 | 0.021 | 0.021 | 0 | 0.56 | 0.25 | 0.37 | 0.56 | 0.53 | 0.59 |
| 10 | 69902549 | rs3814180 | -0.365 | 0.532 | -1.402 | 0.751 | 0.026 | 0.028 | 0.028 | 0 | 0.53 | 0.27 | 0.6 | 0.55 | 0.62 | 0.55 |
| 10 | 69903269 | rs2673798 | -0.324 | 0.501 | -0.354 | 0.613 | 0.004 | 0.021 | 0.021 | 0 | 0.51 | 0.6 | 0.62 | 0.61 | 0.56 | 0.55 |
| 10 | 69905300 | rs2817760 | -1.146 | -0.146 | 0.369 | 0.13 | -0.039 | -0.002 | -0.002 | 0.38 | 0.18 | 0.74 | 0.2 | 0.16 | 0.08 | 0.11 |
| 10 | 69908157 | rs11596653 | -1.137 | -0.182 | 1.345 | 0.081 | 0.002 | -0.012 | -0.012 | 0.042 | 0.08 | 0.41 | 0.05 | 0 | 0.02 | 0.02 |
| 10 | 69908241 | rs2200897 | -0.079 | -0.036 | 1.028 | 0.169 | -0.01 | 0.024 | 0.024 | 0 | 0.09 | 0.15 | 0.13 | 0.16 | 0.07 | 0.09 |
| 10 | 69909513 | rs2673801 | -0.894 | 0.253 | 0.969 | -0.386 | 0.014 | -0.038 | -0.038 | 0.412 | 0.34 | 0.58 | 0.5 | 0.17 | 0.44 | 0.28 |
| 10 | 69909802 | rs10997948 | -1.11 | -0.257 | 0.947 | -0.608 | -0.01 | -0.041 | -0.041 | 0 | 0.12 | 0.17 | 0.07 | 0 | 0.07 | 0.08 |

| | | | | | | | | | | | | | | | | |
|---|---|---|---|---|---|---|---|---|---|---|---|---|---|---|---|---|
| 10 | 69910566 | rs7097584 | -0.493 | -0.498 | 0.811 | -0.756 | 0.034 | 0.084 | 0.084 | 0 | 0.49 | 0.34 | 0.62 | 0.31 | 0.63 | 0.59 |
| 10 | 69912278 | rs10997950 | 0.387 | -0.911 | 0.454 | -0.959 | -0.077 | 0.084 | 0.084 | 0.008 | 0.34 | 0.07 | 0.41 | 0.22 | 0.52 | 0.47 |
| 10 | 69913203 | rs7095238 | -0.17 | -1.176 | 0.947 | -1.055 | -0.004 | -0.019 | -0.019 | 0 | 0.43 | 0.44 | 0.47 | 0.22 | 0.57 | 0.47 |
| 10 | 69914009 | rs7081820 | 0.461 | -1.065 | 0.974 | -1.05 | -0.063 | 0.085 | 0.085 | 0.016 | 0.34 | 0.1 | 0.41 | 0.22 | 0.51 | 0.47 |
| 10 | 69914581 | rs1906719 | -1.246 | -1.11 | 0.998 | -1.045 | -0.034 | -0.021 | -0.021 | 0 | 0.59 | 0.5 | 0.74 | 0.31 | 0.74 | 0.6 |
| 10 | 69915035 | rs2673797 | -2.302 | -1.385 | 0.119 | -1.195 | 0.05 | -0.033 | -0.033 | 0 | 0.72 | 0.94 | 0.81 | 0.31 | 0.79 | 0.68 |
| 10 | 69917242 | rs2673796 | -0.811 | -0.973 | 0.201 | -1.01 | 0.057 | 0.05 | 0.05 | 0.082 | 0.55 | 0.28 | 0.69 | 0.32 | 0.7 | 0.61 |
| 10 | 69917811 | rs10997960 | 0.599 | -0.844 | 0.394 | -0.9 | 0 | -0.026 | -0.026 | 0 | 0.06 | 0.09 | 0.04 | 0 | 0.05 | 0.01 |
| 10 | 69921903 | rs10997963 | 0.539 | -0.869 | 0.119 | -0.927 | -0.058 | 0.072 | 0.072 | 0.051 | 0.33 | 0.13 | 0.41 | 0.22 | 0.51 | 0.44 |
| 10 | 69926097 | rs2673794 | -0.64 | -0.729 | 0.243 | -0.844 | 0.074 | 0.068 | 0.068 | 0.021 | 0.53 | 0.28 | 0.68 | 0.31 | 0.69 | 0.58 |
| 10 | 69932878 | rs4518986 | 0.42 | -0.618 | -0.03 | -0.774 | -0.034 | 0.074 | 0.074 | 0 | 0.35 | 0.18 | 0.41 | 0.22 | 0.51 | 0.43 |
| 10 | 69933074 | rs10997974 | 0.544 | -0.617 | -0.022 | -0.739 | 0 | -0.015 | -0.015 | 0.384 | 0.06 | 0.04 | 0.03 | 0 | 0.05 | 0.01 |
| 10 | 69933921 | rs10997975 | 0.212 | -0.478 | -0.161 | -0.5 | -0.055 | 0.046 | 0.046 | 0 | 0.35 | 0.18 | 0.4 | 0.26 | 0.49 | 0.42 |
| 10 | 69933969 | rs7916821 | 0.223 | -0.455 | -0.149 | -0.692 | -0.055 | 0.073 | 0.073 | 0 | 0.35 | 0.18 | 0.4 | 0.22 | 0.49 | 0.42 |
| 10 | 69934467 | rs3814183 | 0.188 | -0.404 | 0.045 | -0.557 | -0.005 | -0.009 | -0.009 | 0 | 0.81 | 0.78 | 0.72 | 0.89 | 0.81 | 0.86 |
| 10 | 69952953 | rs6480308 | -0.694 | -0.596 | 0.053 | -0.454 | 0.051 | 0.041 | 0.041 | 0.06 | 0.52 | 0.29 | 0.66 | 0.36 | 0.67 | 0.55 |
| 10 | 69959097 | rs7079549 | -1.23 | -0.731 | 0.058 | -0.556 | -0.007 | -0.001 | -0.001 | 0 | 0.61 | 0.39 | 0.72 | 0.36 | 0.75 | 0.58 |
| 10 | 69959242 | rs7079481 | 0.404 | -0.462 | -0.212 | -0.374 | -0.026 | 0.021 | 0.021 | 0 | 0.34 | 0.22 | 0.4 | 0.24 | 0.48 | 0.42 |
| 10 | 69962850 | rs7094934 | -0.13 | -0.575 | -0.417 | -0.062 | -0.041 | -0.017 | -0.017 | 0 | 0.15 | 0.01 | 0.27 | 0.12 | 0.2 | 0.14 |
| 10 | 69967447 | rs6480314 | 0.306 | -0.686 | -0.271 | -0.117 | 0 | 0.018 | 0.018 | 0 | 0.82 | 0.9 | 0.73 | 0.87 | 0.81 | 0.83 |
| 10 | 69969986 | rs10998022 | 1.439 | -0.498 | -0.766 | 0.408 | -0.046 | -0.091 | -0.091 | 0 | 0.3 | 0.1 | 0.26 | 0.47 | 0.32 | 0.34 |
| 10 | 69971332 | rs1673 | -0.261 | -0.54 | -0.362 | 0.471 | -0.008 | 0.166 | 0.166 | 0 | 0.5 | 0.74 | 0.43 | 0.21 | 0.46 | 0.43 |
| 10 | 69971567 | rs881976 | -0.114 | -0.538 | -0.362 | 0.472 | -0.008 | 0.166 | 0.166 | 0.025 | 0.5 | 0.74 | 0.43 | 0.21 | 0.46 | 0.43 |
| 10 | 69983127 | rs1867567 | 0.251 | -0.69 | 0.324 | 0.399 | -0.018 | 0.069 | 0.069 | 0 | 0.43 | 0.25 | 0.38 | 0.21 | 0.45 | 0.42 |
| 10 | 69991853 | rs7916697 | 1.575 | -0.848 | 0.769 | 0.566 | 0.048 | -0.045 | -0.045 | 0 | 0.67 | 0.17 | 0.59 | 0.68 | 0.74 | 0.75 |
| 10 | 69993032 | rs2289804 | -1.225 | -0.679 | 1.658 | 0.699 | -0.009 | 0.187 | 0.187 | 1.455 | 0.12 | 0.49 | 0.11 | 0.19 | 0.07 | 0.07 |
| 10 | 70007008 | rs1900020 | -1.677 | -0.649 | 1.71 | 0.75 | -0.001 | 0.283 | 0.283 | 0.002 | 0.09 | 0.43 | 0.08 | 0.19 | 0.03 | 0.1 |
| 10 | 70011838 | rs3858145 | -2.055 | -0.593 | 1.215 | 0.61 | -0.022 | 0.044 | 0.044 | 0 | 0.3 | 0.43 | 0.43 | 0.32 | 0.26 | 0.35 |
| 10 | 70029894 | rs17231637 | -0.091 | -0.523 | 0.853 | 0.597 | -0.029 | -0.047 | -0.047 | 1.322 | 0.13 | 0.01 | 0.26 | 0.14 | 0.16 | 0.23 |
| 10 | 70032522 | rs7100492 | -1.036 | -0.597 | 1.288 | 0.492 | -0.024 | -0.089 | -0.089 | 0 | 0.13 | 0.36 | 0.07 | 0 | 0.05 | 0.02 |
| 10 | 70043190 | rs731063 | -1.336 | -0.49 | 1.316 | 0.817 | -0.001 | 0.081 | 0.081 | 0 | 0.19 | 0.32 | 0.38 | 0.29 | 0.2 | 0.31 |
| 10 | 70043916 | rs736535 | -1.336 | -0.49 | 1.316 | 0.817 | -0.001 | 0.081 | 0.081 | 0 | 0.19 | 0.32 | 0.38 | 0.29 | 0.2 | 0.31 |
| 10 | 70051928 | rs10823171 | 1.167 | -0.607 | 1.651 | 0.729 | -0.026 | -0.024 | -0.024 | 0 | 0.79 | 0.59 | 0.71 | 0.86 | 0.82 | 0.75 |
| 10 | 70061336 | rs10998058 | 1.352 | -0.827 | 0.698 | 0.769 | -0.039 | -0.045 | -0.045 | 0 | 0.35 | 0.1 | 0.28 | 0.41 | 0.38 | 0.25 |
| 10 | 70066553 | rs12359690 | 0.744 | -0.63 | 0.986 | 0.835 | -0.002 | -0.014 | -0.014 | 0 | 0.05 | 0.05 | 0.02 | 0 | 0.03 | 0.01 |
| 10 | 70073669 | rs7895740 | -0.473 | -0.627 | 1.308 | 0.936 | -0.002 | 0.033 | 0.033 | 0 | 0.15 | 0.32 | 0.27 | 0.15 | 0.14 | 0.24 |
| 10 | 70075569 | rs7904473 | -0.915 | -0.638 | 1.41 | 1.041 | -0.007 | 0.178 | 0.178 | 0 | 0.17 | 0.37 | 0.35 | 0.31 | 0.15 | 0.28 |
| 10 | 70082030 | rs1080639 | -0.676 | -0.708 | 1.623 | 1.073 | 0.004 | 0.114 | 0.114 | 0 | 0.1 | 0.25 | 0.13 | 0.17 | 0.06 | 0.06 |
| 10 | 70085529 | rs7893969 | -0.516 | -0.936 | 1.406 | 0.898 | -0.013 | 0.079 | 0.079 | 0 | 0.57 | 0.72 | 0.68 | 0.74 | 0.58 | 0.64 |
| 10 | 70086313 | rs10762224 | -1.502 | -0.94 | 1.554 | 1.137 | -0.021 | 0.064 | 0.064 | 0 | 0.21 | 0.33 | 0.36 | 0.32 | 0.18 | 0.37 |
| 10 | 70091923 | rs10998072 | -1.334 | -0.813 | 1.667 | 1.147 | 0.015 | 0.14 | 0.14 | 0.116 | 0.11 | 0.43 | 0.16 | 0.16 | 0.07 | 0.14 |
| 10 | 70100250 | rs7071140 | 0.098 | -0.884 | 1.143 | 1.167 | -0.001 | -0.01 | -0.01 | 1.857 | 0.1 | 0.07 | 0.21 | 0.14 | 0.11 | 0.28 |
| 10 | 70102749 | rs3199937 | 0.157 | -0.884 | 1.139 | 1.156 | -0.006 | -0.022 | -0.022 | 0 | 0.1 | 0.04 | 0.21 | 0.13 | 0.11 | 0.28 |
| 10 | 70103461 | rs17297439 | 0.108 | -0.886 | 1.138 | 1.156 | -0.004 | -0.022 | -0.022 | 1.13 | 0.1 | 0.04 | 0.21 | 0.13 | 0.11 | 0.28 |
| 10 | 70103958 | rs2281697 | 0.178 | -0.871 | 1.144 | 1.358 | -0.006 | -0.022 | -0.022 | 0.281 | 0.1 | 0.04 | 0.21 | 0.13 | 0.11 | 0.28 |
| 10 | 70105286 | rs3781568 | -0.084 | -0.856 | 1.451 | 1.35 | -0.001 | 0.014 | 0.014 | 0 | 0.11 | 0.21 | 0.23 | 0.13 | 0.11 | 0.28 |
| 10 | 70105560 | rs1162753 | 0.648 | -0.944 | 1.802 | 1.3 | -0.01 | -0.003 | -0.003 | 0 | 0.84 | 0.8 | 0.75 | 0.87 | 0.86 | 0.72 |
| 10 | 70127696 | rs12780135 | 0.135 | -0.933 | 1.8 | 1.292 | -0.002 | -0.025 | -0.025 | 0 | 0.06 | 0.17 | 0.03 | 0 | 0.03 | 0.01 |
| 10 | 70140012 | rs10762227 | 0.432 | -0.742 | 1.337 | 1.025 | 0.03 | 0.015 | 0.015 | 0 | 0.2 | 0.21 | 0.3 | 0.23 | 0.29 | 0.2 |
| 10 | 70143774 | rs7899885 | 1.5 | -1.493 | 1.216 | 0.765 | -0.028 | -0.038 | -0.038 | 0 | 0.73 | 0.39 | 0.67 | 0.86 | 0.8 | 0.69 |
| 10 | 70179746 | rs7897488 | -0.332 | -1.344 | 1.263 | 0.92 | -0.004 | 0.009 | 0.009 | 0.13 | 0.12 | 0.24 | 0.23 | 0.13 | 0.12 | 0.27 |
| 10 | 70189934 | rs10823195 | -0.731 | -1.694 | 1.038 | 0.914 | -0.039 | 0.003 | 0.003 | 0.025 | 0.2 | 0.21 | 0.08 | 0.25 | 0.07 | 0.12 |
| 10 | 70197519 | rs7905556 | -1.68 | -1.661 | 0.961 | 0.925 | -0.055 | 0.009 | 0.009 | 0.791 | 0.33 | 0.53 | 0.33 | 0.37 | 0.18 | 0.4 |
| 10 | 70213168 | rs7897986 | -0.373 | -1.555 | 0.671 | 1.01 | -0.035 | 0.034 | 0.034 | 0 | 0.16 | 0.3 | 0.09 | 0.25 | 0.07 | 0.12 |
| 10 | 70220535 | rs2031098 | 0.105 | -1.672 | 0.502 | 0.967 | -0.001 | -0.037 | -0.037 | 0.928 | 0.11 | 0.02 | 0.23 | 0.13 | 0.11 | 0.28 |
| 10 | 70221194 | rs4388762 | 0.105 | -1.672 | 0.502 | 0.967 | -0.001 | -0.037 | -0.037 | 0.119 | 0.11 | 0.02 | 0.23 | 0.13 | 0.11 | 0.28 |
| 10 | 70221267 | rs7895191 | -1.022 | -1.856 | 0.585 | 0.879 | -0.008 | 0 | 0 | 0 | 0.21 | 0.22 | 0.1 | 0.25 | 0.1 | 0.12 |
| 10 | 70230599 | rs10509303 | -1.257 | -1.864 | 0.564 | 0.903 | -0.041 | 0.011 | 0.011 | 0 | 0.22 | 0.33 | 0.12 | 0.26 | 0.1 | 0.12 |
| 10 | 70230840 | rs12220316 | -1.257 | -1.864 | 0.564 | 0.879 | -0.041 | 0.008 | 0.008 | 0 | 0.22 | 0.33 | 0.12 | 0.25 | 0.1 | 0.12 |
| 10 | 70231701 | rs1535539 | -1.784 | -1.396 | 0.863 | 0.639 | -0.047 | -0.021 | -0.021 | 0 | 0.19 | 0.39 | 0.29 | 0.13 | 0.14 | 0.29 |
| 10 | 70240115 | rs12767790 | -1.788 | -1.389 | 0.94 | 0.641 | -0.052 | -0.022 | -0.022 | 0 | 0.19 | 0.44 | 0.29 | 0.13 | 0.14 | 0.29 |
| 10 | 70242465 | rs10998218 | -0.58 | -1.399 | 0.896 | 0.54 | -0.023 | -0.041 | -0.041 | 0 | 0.07 | 0.19 | 0.04 | 0 | 0.03 | 0.01 |
| 10 | 70242513 | rs1136645 | -1.838 | -1.403 | 0.899 | 0.541 | -0.047 | -0.021 | -0.021 | 0.521 | 0.19 | 0.4 | 0.29 | 0.13 | 0.14 | 0.29 |
| 10 | 70242717 | rs3200066 | -0.547 | -1.387 | 0.888 | 0.538 | -0.022 | -0.04 | -0.04 | 0 | 0.07 | 0.19 | 0.04 | 0 | 0.03 | 0.01 |
| 10 | 70246874 | rs10998222 | -0.965 | -1.389 | 0.894 | 0.537 | -0.02 | -0.041 | -0.041 | 0 | 0.08 | 0.28 | 0.05 | 0 | 0.03 | 0.01 |
| 10 | 70247476 | rs12767599 | -1.588 | -1.371 | 0.86 | 0.558 | -0.047 | -0.021 | -0.021 | 0 | 0.19 | 0.4 | 0.29 | 0.13 | 0.14 | 0.29 |
| 10 | 70255576 | rs10762229 | 1.685 | -1.613 | 0.67 | 0.407 | -0.033 | -0.026 | -0.026 | 0 | 0.73 | 0.65 | 0.66 | 0.86 | 0.82 | 0.61 |
| 10 | 70257238 | rs10159498 | 0.127 | -1.422 | 0.634 | 0.465 | -0.021 | -0.036 | -0.036 | 0 | 0.08 | 0.19 | 0.08 | 0.02 | 0.04 | 0.1 |
| 10 | 70263579 | rs10823215 | -1.164 | -1.403 | 0.377 | 0.487 | -0.004 | -0.001 | -0.001 | 0 | 0.17 | 0.16 | 0.32 | 0.13 | 0.14 | 0.29 |

| | | | | | | | | | | | | | | | | |
|---|---|---|---|---|---|---|---|---|---|---|---|---|---|---|---|---|
| 10 | 70271354 | rs9415909 | 0.336 | -1.011 | 0.206 | 0.572 | -0.003 | -0.01 | -0.01 | 0 | 0.07 | 0.08 | 0.07 | 0.02 | 0.05 | 0.11 |
| 10 | 70319626 | rs1342235 | -0.291 | -0.863 | 0.274 | 0.568 | -0.009 | -0.024 | -0.024 | 0.524 | 0.92 | 0.9 | 0.93 | 0.97 | 0.95 | 0.9 |
| 10 | 70324814 | rs7922602 | -0.593 | -0.548 | -0.137 | 0.691 | -0.039 | -0.04 | -0.04 | 0.062 | 0.12 | 0.01 | 0.14 | 0.13 | 0.15 | 0.26 |
| 10 | 70326994 | rs2795908 | 0.743 | -0.481 | -0.037 | 0.697 | -0.001 | -0.004 | -0.004 | 0 | 0.06 | 0.04 | 0.07 | 0.03 | 0.05 | 0.1 |
| 10 | 70332066 | rs3814177 | 0.592 | -0.188 | -0.54 | 0.663 | -0.003 | -0.01 | -0.01 | 0 | 0.46 | 0.32 | 0.43 | 0.56 | 0.57 | 0.58 |
| 10 | 70332580 | rs10823229 | 1.708 | 0.2 | -0.653 | 0.683 | -0.072 | -0.089 | -0.089 | 0 | 0.33 | 0.02 | 0.26 | 0.39 | 0.39 | 0.3 |
| 10 | 70334316 | rs2795903 | -0.593 | 0.06 | -0.365 | 0.662 | -0.007 | 0.002 | 0.002 | 0.088 | 0.82 | 0.81 | 0.92 | 0.79 | 0.89 | 0.92 |
| 10 | 70334550 | rs7911579 | -0.278 | 0.474 | -0.465 | 0.747 | -0.012 | -0.009 | -0.009 | 0.614 | 0.11 | 0.1 | 0.15 | 0.17 | 0.18 | 0.27 |
| 10 | 70338978 | rs10823231 | 1.463 | 0.423 | -0.456 | 0.741 | 0.005 | -0.002 | -0.002 | 0 | 0.35 | 0.23 | 0.28 | 0.39 | 0.39 | 0.3 |
| 10 | 70341213 | rs1970517 | -0.285 | 0.412 | -0.457 | 0.747 | -0.012 | -0.009 | -0.009 | 0 | 0.11 | 0.07 | 0.14 | 0.17 | 0.18 | 0.27 |
| 10 | 70346506 | rs11815694 | 0.787 | 0.439 | -0.513 | 0.74 | -0.011 | 0.004 | 0.004 | 0 | 0.06 | 0.01 | 0.05 | 0.02 | 0.08 | 0.03 |
| 10 | 70346740 | rs2298117 | 0.412 | 0.387 | -0.479 | 0.734 | -0.002 | -0.009 | -0.009 | 0 | 0.46 | 0.33 | 0.42 | 0.56 | 0.57 | 0.57 |
| 10 | 70349116 | rs7894063 | 1.533 | 0.312 | -0.898 | 0.689 | -0.036 | -0.049 | -0.049 | 0 | 0.33 | 0.06 | 0.26 | 0.39 | 0.39 | 0.31 |
| 10 | 70380498 | rs12764361 | -1.173 | 0.591 | -0.413 | 0.688 | 0.011 | 0.002 | 0.002 | 0 | 0.13 | 0.18 | 0.16 | 0.13 | 0.2 | 0.28 |
| 10 | 70391172 | rs7907322 | 0.651 | 0.285 | -0.461 | 0.655 | 0 | -0.023 | -0.023 | 0 | 0.44 | 0.25 | 0.4 | 0.54 | 0.59 | 0.53 |
| 10 | 70401976 | rs4558056 | -0.089 | 0.714 | -0.645 | 0.659 | -0.07 | -0.04 | -0.04 | 0.556 | 0.18 | 0.02 | 0.3 | 0.17 | 0.24 | 0.21 |
| 10 | 70405237 | rs12241767 | -0.983 | 0.512 | 0.665 | 0.605 | 0.119 | 0.098 | 0.098 | 0.861 | 0.1 | 0.39 | 0.14 | 0.12 | 0.13 | 0.2 |
| 10 | 70405855 | rs3998860 | 1.68 | 0.272 | 0.828 | 0.516 | 0.045 | 0.014 | 0.014 | 0.732 | 0.83 | 0.45 | 0.78 | 0.85 | 0.81 | 0.68 |
| 10 | 70410630 | rs7913568 | 1.803 | 0.275 | 0.836 | 0.517 | 0.046 | 0.011 | 0.011 | 0.557 | 0.82 | 0.44 | 0.79 | 0.85 | 0.81 | 0.68 |
| 10 | 70415515 | rs12266293 | -0.97 | 0.3 | 0.816 | 0.523 | 0.11 | 0.076 | 0.076 | 0.247 | 0.1 | 0.35 | 0.14 | 0.11 | 0.13 | 0.21 |
| 10 | 70423620 | rs1566221 | -1.165 | 0.294 | 0.207 | 0.485 | 0.001 | 0.002 | 0.002 | 0 | 0.94 | 0.95 | 0.93 | 0.97 | 0.94 | 0.92 |
| 10 | 70431420 | rs10762236 | 1.303 | 0.299 | 0.734 | 0.48 | 0.064 | -0.005 | -0.005 | 0 | 0.84 | 0.5 | 0.79 | 0.86 | 0.81 | 0.71 |
| 10 | 70432350 | rs10998379 | 2.052 | 0.224 | 0.179 | 0.577 | -0.043 | -0.028 | -0.028 | 0.056 | 0.32 | 0.09 | 0.23 | 0.38 | 0.39 | 0.28 |
| 10 | 70452471 | rs5030882 | 0.186 | -0.289 | 0.722 | 0.245 | -0.023 | -0.047 | -0.047 | 0 | 0.9 | 0.74 | 0.95 | 0.97 | 0.94 | 0.92 |
| 10 | 70454896 | rs10740308 | 1.018 | -0.573 | 0.684 | 0.113 | 0 | -0.033 | -0.033 | 0 | 0.79 | 0.64 | 0.83 | 0.86 | 0.82 | 0.75 |
| 10 | 70455490 | rs7901888 | 0.997 | -0.568 | 0.396 | 0.116 | -0.001 | -0.018 | -0.018 | 0.208 | 0.79 | 0.7 | 0.83 | 0.86 | 0.82 | 0.75 |
| 10 | 70475978 | rs10998399 | -0.117 | -0.87 | 0.174 | 0.133 | -0.02 | 0.02 | 0.02 | 1.275 | 0.11 | 0.16 | 0.04 | 0.25 | 0.03 | 0.13 |
| 10 | 70477078 | rs1782330 | 1.499 | -0.699 | 0.368 | 0.126 | -0.013 | -0.006 | -0.006 | 0 | 0.66 | 0.52 | 0.76 | 0.64 | 0.74 | 0.63 |
| 10 | 70486325 | rs1782322 | 2.65 | -0.687 | 0.45 | 0.117 | -0.009 | -0.003 | -0.003 | 0 | 0.36 | 0.17 | 0.35 | 0.36 | 0.44 | 0.3 |
| 10 | 70495544 | rs1162966 | -0.11 | -0.658 | 0.254 | 0.143 | 0.01 | -0.04 | -0.04 | 1.356 | 0.85 | 0.93 | 0.92 | 0.74 | 0.86 | 0.85 |
| 10 | 70501910 | rs3740593 | 0.41 | -0.676 | 0.217 | 0.121 | -0.058 | 0.02 | 0.02 | 0.829 | 0.12 | 0.01 | 0.11 | 0.06 | 0.12 | 0.17 |
| 10 | 70502288 | rs1300253 | -0.127 | -0.679 | 0.219 | 0.123 | 0.01 | -0.04 | -0.04 | 0.064 | 0.85 | 0.93 | 0.92 | 0.74 | 0.86 | 0.85 |
| 10 | 70513217 | rs17462632 | 0.723 | -0.638 | 0.18 | 0.117 | -0.042 | 0.009 | 0.009 | 1.911 | 0.1 | 0 | 0.08 | 0.06 | 0.08 | 0.15 |
| 10 | 70516744 | rs7081960 | -0.683 | -0.697 | 0.349 | 0.123 | 0.012 | -0.036 | -0.036 | 3.855 | 0.17 | 0.32 | 0.16 | 0.09 | 0.13 | 0.22 |
| 10 | 70518714 | rs10823256 | 0.353 | -0.72 | 0.075 | -0.008 | -0.09 | -0.111 | -0.111 | 1.888 | 0.19 | 0.01 | 0.31 | 0.29 | 0.22 | 0.25 |
| 10 | 70523932 | rs1162958 | 1.227 | -0.799 | 0.19 | -0.055 | -0.013 | -0.007 | -0.007 | 2.19 | 0.67 | 0.46 | 0.75 | 0.65 | 0.73 | 0.63 |
| 10 | 70549579 | rs11594683 | 0.545 | -0.644 | 0.071 | -0.03 | 0.011 | 0.011 | 0.011 | 0 | 0.08 | 0.04 | 0.06 | 0.01 | 0.06 | 0.04 |
| 10 | 70573834 | rs12771648 | 0.152 | -0.153 | -0.111 | 0.301 | -0.058 | 0.021 | 0.021 | 0.926 | 0.12 | 0.01 | 0.11 | 0.06 | 0.12 | 0.17 |
| 10 | 70577222 | rs2497099 | -1.368 | -0.191 | 0.049 | 0.341 | -0.012 | -0.007 | -0.007 | 0.291 | 0.35 | 0.5 | 0.25 | 0.35 | 0.28 | 0.38 |
| 10 | 70589892 | rs1694505 | -1.163 | -0.077 | -0.27 | 0.328 | -0.008 | -0.01 | -0.01 | 0 | 0.36 | 0.56 | 0.27 | 0.34 | 0.3 | 0.43 |
| 10 | 70592130 | rs10762242 | 0.928 | -0.072 | -0.25 | 0.33 | -0.006 | -0.008 | -0.008 | 0 | 0.64 | 0.44 | 0.73 | 0.66 | 0.7 | 0.58 |
| 10 | 70595426 | rs10762244 | -0.347 | 0.099 | -0.435 | 0.628 | -0.011 | -0.018 | -0.018 | 0.03 | 0.18 | 0.16 | 0.34 | 0.28 | 0.2 | 0.23 |
| 10 | 70613280 | rs10998460 | 0.12 | 0.278 | -0.793 | 0.589 | -0.07 | -0.11 | -0.11 | 0 | 0.17 | 0.01 | 0.27 | 0.3 | 0.18 | 0.21 |
| 10 | 70616517 | rs10998461 | 1.282 | -0.023 | -0.449 | -0.35 | 0.006 | 0.051 | 0.051 | 0.798 | 0.48 | 0.32 | 0.38 | 0.34 | 0.51 | 0.33 |
| 10 | 70616973 | rs1007042 | 1.302 | 0.052 | -0.185 | -0.335 | 0.007 | 0.044 | 0.044 | 0.86 | 0.49 | 0.32 | 0.38 | 0.34 | 0.51 | 0.34 |
| 10 | 70618422 | rs17470053 | -0.423 | 0.214 | -0.25 | -0.433 | -0.045 | 0.001 | 0.001 | 0 | 0.08 | 0.01 | 0.14 | 0.06 | 0.11 | 0.14 |
| 10 | 70618919 | rs1417940 | -1.45 | -0.051 | 0.451 | -0.693 | 0 | -0.093 | -0.093 | 0.017 | 0.25 | 0.42 | 0.29 | 0.09 | 0.21 | 0.28 |
| 10 | 70619276 | rs4746796 | 1.472 | -0.054 | 0.491 | -0.735 | 0.002 | -0.105 | -0.105 | 0 | 0.69 | 0.53 | 0.66 | 0.9 | 0.69 | 0.69 |
| 10 | 70631783 | rs10082399 | -1.518 | -0.054 | 0.709 | -0.724 | -0.001 | -0.072 | -0.072 | 0 | 0.25 | 0.35 | 0.3 | 0.08 | 0.21 | 0.31 |
| 10 | 70636836 | rs1498325 | -0.744 | -0.154 | 0.559 | -0.672 | -0.001 | -0.014 | -0.014 | 0 | 0.93 | 0.9 | 0.95 | 0.99 | 0.9 | 0.93 |
| 10 | 70637445 | rs7095976 | 1.453 | 0.019 | 0.818 | -0.821 | 0.013 | -0.074 | -0.074 | 0 | 0.68 | 0.62 | 0.65 | 0.9 | 0.65 | 0.61 |
| 10 | 70641860 | rs1341667 | 1.734 | -0.502 | 0.893 | -1.02 | 0.003 | 0.012 | 0.012 | 0 | 0.58 | 0.38 | 0.61 | 0.9 | 0.64 | 0.6 |
| 10 | 70645376 | rs10509305 | -0.802 | -0.597 | 0.69 | -1.06 | -0.087 | 0.075 | 0.075 | 0.546 | 0.2 | 0.03 | 0.22 | 0.08 | 0.22 | 0.23 |
| 10 | 70645697 | rs10509306 | 0.976 | -0.495 | 0.741 | -1.007 | -0.039 | -0.004 | -0.004 | 0 | 0.07 | 0.01 | 0.15 | 0.06 | 0.15 | 0.12 |
| 10 | 70655702 | rs2185653 | 0.915 | -0.679 | 0.985 | -1.002 | -0.053 | -0.147 | -0.147 | 0 | 0.79 | 0.59 | 0.85 | 0.98 | 0.86 | 0.84 |
| 10 | 70659942 | rs12263044 | -0.005 | -0.592 | 0.966 | -0.952 | -0.01 | -0.038 | -0.038 | 0.044 | 0.07 | 0.13 | 0.08 | 0.01 | 0.05 | 0.09 |
| 10 | 70661532 | rs10998478 | -0.005 | -0.592 | 0.981 | -0.952 | -0.01 | -0.037 | -0.037 | 0.192 | 0.07 | 0.13 | 0.08 | 0.01 | 0.05 | 0.09 |
| 10 | 70667461 | rs10733848 | 0.439 | -0.609 | 0.971 | -0.97 | -0.003 | -0.037 | -0.037 | 0.06 | 0.87 | 0.86 | 0.88 | 0.98 | 0.86 | 0.84 |
| 10 | 70670696 | rs7918860 | 2.686 | -0.809 | 0.845 | -0.891 | -0.025 | 0.012 | 0.012 | 0.001 | 0.39 | 0.21 | 0.3 | 0.34 | 0.48 | 0.28 |
| 10 | 70680415 | rs10128256 | 0.293 | -0.787 | 1.01 | -0.918 | -0.008 | -0.031 | -0.031 | 0 | 0.07 | 0.11 | 0.08 | 0.01 | 0.05 | 0.09 |
| 10 | 70680767 | rs7084275 | 1.871 | -0.822 | 1.038 | -0.962 | 0.003 | 0.007 | 0.007 | 0 | 0.58 | 0.37 | 0.62 | 0.9 | 0.64 | 0.61 |
| 10 | 70688569 | rs2394497 | -1.385 | -0.74 | 0.877 | -0.897 | -0.008 | 0.023 | 0.023 | 0 | 0.94 | 0.98 | 0.96 | 0.99 | 0.91 | 0.93 |
| 10 | 70697111 | rs10509308 | 0.189 | -0.722 | 0.902 | -0.871 | -0.01 | -0.037 | -0.037 | 0 | 0.07 | 0.14 | 0.08 | 0.01 | 0.05 | 0.09 |
| 10 | 70710358 | rs10509307 | 0.861 | -0.68 | 0.809 | -0.878 | -0.047 | -0.003 | -0.003 | 0 | 0.07 | 0.01 | 0.17 | 0.06 | 0.16 | 0.15 |
| 10 | 70712827 | rs10998499 | -0.668 | -0.744 | 1.005 | -0.894 | -0.058 | -0.108 | -0.108 | 3.852 | 0.15 | 0.38 | 0.1 | 0.01 | 0.05 | 0.09 |
| 10 | 70715155 | rs7073652 | 1.71 | -0.8 | 0.949 | -0.951 | -0.003 | -0.095 | -0.095 | 0 | 0.59 | 0.53 | 0.63 | 0.9 | 0.64 | 0.61 |
| 10 | 70722989 | rs10998504 | 0.281 | -0.698 | 0.956 | -0.921 | -0.013 | -0.036 | -0.036 | 0 | 0.07 | 0.13 | 0.08 | 0.01 | 0.05 | 0.09 |
| 10 | 70725128 | rs3898314 | 0.647 | -0.756 | 0.981 | -0.893 | 0 | 0 | 0 | 0 | 0.18 | 0.13 | 0.31 | 0.56 | 0.16 | 0.33 |
| 10 | 70735135 | rs7908750 | -0.623 | -0.767 | 0.972 | -0.896 | -0.061 | -0.104 | -0.104 | 0 | 0.15 | 0.36 | 0.1 | 0.01 | 0.05 | 0.09 |

| | | | | | | | | | | | | | | | | |
|---|---|---|---|---|---|---|---|---|---|---|---|---|---|---|---|---|
| 10 | 70739630 | rs1539163 | 0.643 | -0.711 | 0.732 | -0.819 | 0.007 | 0.041 | 0.041 | 0 | 0.1 | 0.04 | 0.05 | 0.04 | 0.06 | 0.07 |
| 10 | 70742931 | rs12412103 | 0.112 | -0.673 | 0.79 | -0.815 | -0.014 | -0.04 | -0.04 | 0 | 0.07 | 0.15 | 0.08 | 0.01 | 0.05 | 0.09 |
| 10 | 70749869 | rs17471869 | 2.368 | -0.648 | 0.857 | -0.822 | -0.006 | 0.005 | 0.005 | 0 | 0.4 | 0.33 | 0.32 | 0.36 | 0.48 | 0.28 |
| 10 | 70759289 | rs2487703 | -0.265 | -1.095 | 0.621 | -0.579 | -0.02 | -0.047 | -0.047 | 0.022 | 0.13 | 0.15 | 0.09 | 0.01 | 0.05 | 0.08 |
| 10 | 70762353 | rs2185654 | 0.259 | -0.714 | 0.576 | -0.312 | -0.001 | 0.006 | 0.006 | 0 | 0.74 | 0.74 | 0.64 | 0.43 | 0.74 | 0.58 |
| 10 | 70771895 | rs2256982 | 0.202 | -0.696 | 0.49 | -0.319 | 0 | -0.006 | -0.006 | 0.17 | 0.75 | 0.79 | 0.64 | 0.44 | 0.74 | 0.59 |
| 10 | 70774148 | rs2260294 | -1.141 | -1.033 | 0.258 | -0.005 | 0.003 | -0.003 | -0.003 | 0.107 | 0.39 | 0.58 | 0.51 | 0.55 | 0.25 | 0.48 |
| 10 | 70777713 | rs2246364 | 0.49 | -0.316 | 0.157 | -0.046 | -0.017 | -0.027 | -0.027 | 0.159 | 0.07 | 0.08 | 0.02 | 0 | 0.01 | 0 |
| 10 | 70782126 | rs2487710 | -0.734 | -0.123 | 0.241 | -0.059 | -0.02 | -0.083 | -0.083 | 0 | 0.14 | 0.29 | 0.11 | 0.02 | 0.1 | 0.08 |
| 10 | 70783075 | rs10823277 | -0.349 | 0.186 | -0.337 | 0.446 | -0.001 | 0.019 | 0.019 | 0 | 0.18 | 0.2 | 0.3 | 0.57 | 0.15 | 0.34 |
| 10 | 70799421 | rs2429018 | 0.839 | 0.466 | -0.425 | 0.206 | -0.004 | 0.011 | 0.011 | 0 | 0.46 | 0.32 | 0.33 | 0.36 | 0.58 | 0.41 |
| 10 | 70800623 | rs2998005 | 0.484 | 0.443 | -0.217 | 0.565 | -0.002 | 0.008 | 0.008 | 0 | 0.85 | 0.86 | 0.89 | 0.76 | 0.85 | 0.77 |
| 10 | 70801833 | rs12570170 | 0.317 | 0.09 | -0.288 | 0.323 | 0.04 | -0.054 | -0.054 | 2.206 | 0.21 | 0.07 | 0.29 | 0.37 | 0.09 | 0.25 |
| 10 | 70802309 | rs2491026 | 0.795 | 0.727 | 0.287 | 0.15 | -0.006 | 0.013 | 0.013 | 0 | 0.84 | 0.81 | 0.89 | 0.79 | 0.85 | 0.77 |
| 10 | 70802518 | rs5030931 | -0.449 | 0.849 | 0.332 | 0.241 | 0.064 | -0.108 | -0.108 | 0 | 0.17 | 0.45 | 0.17 | 0 | 0.19 | 0.11 |
| 10 | 70803785 | rs2429014 | 0.204 | 0.3 | -0.126 | 0.645 | -0.01 | 0.031 | 0.031 | 0 | 0.9 | 0.87 | 0.93 | 0.8 | 0.93 | 0.84 |
| 10 | 70812408 | rs2394525 | 0.436 | 1.301 | 0.004 | 0.328 | 0.048 | -0.011 | -0.011 | 2.458 | 0.81 | 0.68 | 0.81 | 0.82 | 0.7 | 0.81 |
| 10 | 70813238 | rs10823283 | 1.466 | 0.676 | -0.29 | 0.278 | 0.006 | 0.009 | 0.009 | 1.13 | 0.37 | 0.23 | 0.28 | 0.26 | 0.38 | 0.21 |
| 10 | 70815810 | rs12415599 | -0.057 | 0.199 | -0.294 | 0.28 | 0.038 | 0.083 | 0.083 | 0 | 0.31 | 0.12 | 0.26 | 0.16 | 0.23 | 0.4 |
| 10 | 70823831 | rs7909789 | -1.046 | 0.451 | 0.611 | 0.349 | 0.001 | -0.012 | -0.012 | 0.635 | 0.3 | 0.32 | 0.3 | 0.13 | 0.41 | 0.34 |
| 10 | 70825489 | rs2394527 | 1.719 | -0.092 | 0.227 | 0.234 | 0.022 | -0.014 | -0.014 | 0.043 | 0.26 | 0.14 | 0.18 | 0.25 | 0.25 | 0.15 |
| 10 | 70827281 | rs10509309 | 0.473 | -0.052 | 0.593 | 0.388 | -0.001 | -0.006 | -0.006 | 0 | 0.38 | 0.4 | 0.32 | 0.24 | 0.32 | 0.34 |
| 10 | 70827582 | rs6480379 | -0.254 | 0.143 | 0.544 | 0.309 | 0.026 | 0.029 | 0.029 | 0 | 0.48 | 0.68 | 0.58 | 0.62 | 0.37 | 0.53 |
| 10 | 70829725 | rs4601652 | 0.735 | 0.467 | 0.505 | 0.221 | 0.03 | 0.034 | 0.034 | 0 | 0.38 | 0.15 | 0.29 | 0.27 | 0.36 | 0.35 |
| 10 | 70830430 | rs4745975 | -0.155 | 0.446 | 0.479 | 0.235 | 0.021 | 0.023 | 0.023 | 0 | 0.48 | 0.66 | 0.57 | 0.62 | 0.37 | 0.53 |
| 10 | 70835612 | rs2001761 | 0.564 | 0.306 | 0.211 | 0.184 | 0.028 | 0.032 | 0.032 | 0 | 0.39 | 0.25 | 0.25 | 0.26 | 0.35 | 0.34 |
| 10 | 70850138 | rs11815117 | 0.635 | 0.528 | -0.422 | 0.273 | -0.081 | 0.046 | 0.046 | 0.033 | 0.11 | 0.01 | 0.11 | 0.02 | 0.27 | 0.12 |
| 10 | 70851106 | rs2855024 | 1.526 | -0.173 | -0.425 | 0.439 | 0.005 | -0.069 | -0.069 | 3.254 | 0.23 | 0.01 | 0.12 | 0.23 | 0.2 | 0.13 |
| 10 | 70864065 | rs12437 | 0.856 | 0.033 | 0.503 | 0.766 | -0.027 | 0.228 | 0.228 | 0.042 | 0.85 | 0.66 | 0.67 | 0.53 | 0.89 | 0.73 |
| 10 | 70871598 | rs2394505 | 0.894 | 0.023 | 0.504 | 0.763 | -0.029 | 0.224 | 0.224 | 0.079 | 0.85 | 0.66 | 0.67 | 0.53 | 0.89 | 0.72 |
| 10 | 70875148 | rs7081677 | -0.074 | 0.744 | 0.462 | 0.387 | 0.103 | -0.059 | -0.059 | 0 | 0.13 | 0.29 | 0.13 | 0.02 | 0.26 | 0.1 |
| 10 | 70882515 | rs4746817 | 0.1 | 0.384 | 0.768 | 0.653 | -0.013 | 0.119 | 0.119 | 0 | 0.89 | 0.84 | 0.7 | 0.61 | 0.89 | 0.8 |
| 10 | 70889053 | rs4572029 | 1.74 | -0.168 | 0.229 | 0.463 | 0.03 | -0.093 | -0.093 | 0 | 0.27 | 0.01 | 0.16 | 0.27 | 0.2 | 0.19 |
| 10 | 70890482 | rs6480383 | 1.057 | -0.108 | 0.785 | 0.584 | -0.046 | 0.223 | 0.223 | 0.673 | 0.82 | 0.49 | 0.65 | 0.49 | 0.88 | 0.72 |
| 10 | 70892791 | rs5030932 | -0.102 | 0.078 | 0.765 | 0.543 | 0.077 | -0.057 | -0.057 | 0 | 0.13 | 0.27 | 0.13 | 0.04 | 0.25 | 0.1 |
| 10 | 70930360 | rs12256853 | 0.229 | 0.288 | 0.743 | 0.441 | 0.087 | -0.047 | -0.047 | 0 | 0.12 | 0.26 | 0.12 | 0.04 | 0.25 | 0.1 |
| 10 | 70931474 | rs1802295 | 1.211 | 0.067 | 0.528 | 0.288 | -0.108 | 0.075 | 0.075 | 0 | 0.28 | 0.02 | 0.25 | 0.11 | 0.29 | 0.26 |
| 10 | 70939542 | rs12257763 | -0.031 | 0.39 | 1.243 | 0.74 | 0.088 | -0.051 | -0.051 | 0 | 0.12 | 0.26 | 0.12 | 0.04 | 0.25 | 0.1 |
| 10 | 70951346 | rs11817569 | 0.63 | 0.509 | 0.939 | 0.794 | -0.049 | 0.029 | 0.029 | 0 | 0.08 | 0.01 | 0.1 | 0.04 | 0.25 | 0.1 |
| 10 | 70955056 | rs5018859 | 0.225 | 0.531 | 1.042 | 0.803 | -0.01 | 0.013 | 0.013 | 0 | 0.1 | 0.09 | 0.11 | 0.04 | 0.25 | 0.1 |
| 10 | 70959710 | rs1506769 | 1.393 | -0.433 | 1.093 | 1.276 | -0.038 | 0.2 | 0.2 | 1.187 | 0.84 | 0.49 | 0.66 | 0.56 | 0.9 | 0.73 |
| 10 | 70960448 | rs7893492 | 0.264 | 0.034 | 0.127 | 0.939 | -0.005 | 0.148 | 0.148 | 0 | 0.91 | 0.84 | 0.71 | 0.63 | 0.9 | 0.8 |
| 10 | 70975239 | rs5030933 | 0.536 | -0.214 | -0.297 | -0.035 | -0.049 | -0.033 | -0.033 | 0 | 0.41 | 0.51 | 0.45 | 0.76 | 0.27 | 0.43 |
| 10 | 70975897 | rs5030937 | -0.359 | -0.245 | -0.849 | -0.195 | 0.162 | 0.162 | 0.162 | 0 | 0.49 | 0.16 | 0.48 | 0.24 | 0.66 | 0.54 |
| 10 | 70975916 | rs5030938 | -0.359 | -0.245 | -0.849 | -0.195 | 0.162 | 0.162 | 0.162 | 0 | 0.49 | 0.16 | 0.48 | 0.24 | 0.66 | 0.53 |
| 10 | 70976702 | rs10998646 | 0.129 | 0.158 | -0.968 | -0.332 | 0.042 | 0.214 | 0.214 | 0 | 0.44 | 0.11 | 0.34 | 0.12 | 0.57 | 0.4 |
| 10 | 70985650 | rs4746826 | -0.174 | 0.109 | -0.678 | -0.275 | 0.05 | 0.103 | 0.103 | 0 | 0.47 | 0.36 | 0.37 | 0.17 | 0.58 | 0.42 |
| 10 | 70986610 | rs9645499 | 0.024 | 1.028 | -0.809 | -0.407 | -0.007 | 0.015 | 0.015 | 0 | 0.42 | 0.41 | 0.42 | 0.25 | 0.42 | 0.47 |
| 10 | 70987319 | rs10762266 | -0.617 | 1.305 | -0.872 | -0.495 | -0.035 | 0.008 | 0.008 | 0.079 | 0.16 | 0.08 | 0.24 | 0.15 | 0.33 | 0.22 |
| 10 | 70989148 | rs4072135 | -0.777 | 1.311 | -0.735 | -0.349 | 0.059 | 0.048 | 0.048 | 0 | 0.08 | 0.26 | 0.16 | 0.12 | 0.09 | 0.13 |
| 10 | 70989157 | rs4075240 | 0.531 | 1.146 | -1.532 | -0.607 | -0.035 | 0.127 | 0.127 | 0 | 0.3 | 0.06 | 0.2 | 0.07 | 0.28 | 0.31 |
| 10 | 70989749 | rs1983128 | -0.985 | 1.809 | -1.008 | -0.138 | -0.086 | 0.089 | 0.089 | 0 | 0.21 | 0.08 | 0.14 | 0.04 | 0.33 | 0.1 |
| 10 | 70992250 | rs4746828 | 0.935 | 1.316 | -0.842 | 0.033 | -0.002 | 0.018 | 0.018 | 1.55 | 0.58 | 0.36 | 0.53 | 0.26 | 0.58 | 0.52 |
| 10 | 70992413 | rs874557 | -1.265 | 1.325 | -0.712 | 0.037 | -0.001 | 0 | 0 | 0 | 0.42 | 0.56 | 0.45 | 0.74 | 0.42 | 0.48 |
| 10 | 70992664 | rs874556 | -1.255 | 1.329 | -0.71 | 0.038 | -0.001 | -0.001 | -0.001 | 0 | 0.42 | 0.55 | 0.45 | 0.74 | 0.41 | 0.48 |
| 10 | 70992716 | rs10998656 | -0.323 | 1.188 | -0.578 | 0.419 | -0.012 | 0.016 | 0.016 | 0 | 0.06 | 0.1 | 0.03 | 0.13 | 0.02 | 0.05 |
| 10 | 70995288 | rs7075720 | -1.268 | 1.361 | -0.979 | 0.121 | -0.004 | 0.052 | 0.052 | 1.627 | 0.43 | 0.73 | 0.47 | 0.74 | 0.41 | 0.48 |
| 10 | 70999492 | rs7085830 | -0.772 | 1.715 | -0.659 | 0.415 | 0.002 | 0.062 | 0.062 | 0 | 0.06 | 0.19 | 0.06 | 0.17 | 0.06 | 0.08 |
| 10 | 71003612 | rs1472814 | -0.946 | 2.881 | -1.179 | -0.066 | -0.09 | 0.052 | 0.052 | 0.607 | 0.12 | 0.01 | 0.16 | 0.01 | 0.26 | 0.13 |
| 10 | 71008656 | rs2279927 | -1.435 | 3.205 | -1.325 | -0.071 | -0.111 | 0.001 | 0.001 | 1.227 | 0.15 | 0.02 | 0.19 | 0.09 | 0.29 | 0.21 |
| 10 | 71008705 | rs7918950 | -0.731 | 2.4 | -1.623 | -1.037 | 0.016 | 0.04 | 0.04 | 1.205 | 0.64 | 0.74 | 0.82 | 0.94 | 0.73 | 0.72 |
| 10 | 71009907 | rs3740600 | 1.023 | 2.065 | -1.48 | -0.665 | -0.013 | 0.068 | 0.068 | 0 | 0.29 | 0.11 | 0.18 | 0.12 | 0.26 | 0.3 |
| 10 | 71010375 | rs5030948 | -0.616 | 2.956 | -1.082 | -0.317 | -0.089 | -0.005 | -0.005 | 0 | 0.13 | 0.01 | 0.19 | 0.08 | 0.27 | 0.19 |
| 10 | 71015389 | rs2102339 | 0.336 | 2.339 | -0.436 | -0.212 | 0.019 | 0.017 | 0.017 | 0 | 0.49 | 0.45 | 0.26 | 0.36 | 0.34 | 0.55 |
| 10 | 71017444 | rs10823325 | -0.569 | 2.642 | -0.893 | -0.333 | -0.102 | 0.007 | 0.007 | 1.105 | 0.14 | 0.02 | 0.21 | 0.08 | 0.3 | 0.19 |
| 10 | 71026510 | rs906219 | 0.225 | 1.205 | -0.521 | -0.23 | 0.048 | -0.074 | -0.074 | 0 | 0.35 | 0.11 | 0.49 | 0.59 | 0.36 | 0.25 |
| 10 | 71029302 | rs2394534 | -0.641 | 1.636 | -0.36 | -0.175 | -0.03 | 0.01 | 0.01 | 0.081 | 0.11 | 0.06 | 0.18 | 0.07 | 0.23 | 0.18 |
| 10 | 71038340 | rs5030949 | 0.532 | 0.944 | -0.358 | -0.127 | 0.059 | -0.086 | -0.086 | 0 | 0.31 | 0.09 | 0.48 | 0.58 | 0.3 | 0.23 |
| 10 | 71040619 | rs10762275 | -0.468 | 1.092 | -0.251 | -0.13 | -0.048 | 0.012 | 0.012 | 0 | 0.42 | 0.15 | 0.67 | 0.65 | 0.58 | 0.4 |

| | | | | | | | | | | | | | | | | |
|---|---|---|---|---|---|---|---|---|---|---|---|---|---|---|---|---|
| 10 | 71041806 | rs4746834 | -0.437 | 1.112 | -0.259 | -0.122 | -0.051 | 0.01 | 0.01 | 0 | 0.42 | 0.13 | 0.67 | 0.65 | 0.58 | 0.4 |
| 10 | 71043781 | rs10998696 | 0.99 | 0.97 | 0.21 | -0.168 | 0.001 | 0.082 | 0.082 | 0 | 0.47 | 0.28 | 0.26 | 0.27 | 0.39 | 0.41 |
| 10 | 71045777 | rs2394540 | -0.992 | 0.936 | 0.756 | -0.164 | -0.006 | 0.042 | 0.042 | 0 | 0.07 | 0.23 | 0.05 | 0.07 | 0.03 | 0.15 |
| 10 | 71049098 | rs5030916 | -1.153 | 0.917 | 0.948 | -0.228 | 0.01 | 0.056 | 0.056 | 0 | 0.52 | 0.62 | 0.74 | 0.72 | 0.62 | 0.59 |
| 10 | 71051643 | rs7907176 | -1.035 | 0.422 | 0.618 | -0.12 | -0.005 | 0.032 | 0.032 | 0 | 0.07 | 0.23 | 0.05 | 0.06 | 0.03 | 0.15 |
| 10 | 71052469 | rs10762276 | 1.36 | 0.435 | 0.752 | -0.161 | -0.046 | -0.044 | -0.044 | 0 | 0.91 | 0.32 | 0.94 | 0.99 | 0.99 | 0.85 |
| 10 | 71052508 | rs10762277 | -0.876 | 1.172 | 0.103 | 0.078 | -0.003 | -0.008 | -0.008 | 0 | 0.44 | 0.47 | 0.67 | 0.66 | 0.49 | 0.54 |
| 10 | 71052888 | rs10823333 | -1.31 | 2.087 | 0.277 | 0.114 | 0.055 | -0.024 | -0.024 | 0.123 | 0.13 | 0.27 | 0.19 | 0.07 | 0.2 | 0.26 |
| 10 | 71054080 | rs10998700 | -1.221 | 1.665 | 0.068 | 0.044 | 0.059 | -0.014 | -0.014 | 0 | 0.18 | 0.27 | 0.25 | 0.1 | 0.37 | 0.13 |
| 10 | 71055232 | rs2894081 | 0.511 | -0.495 | 0.076 | -0.402 | -0.006 | -0.026 | -0.026 | 0 | 0.58 | 0.59 | 0.54 | 0.83 | 0.69 | 0.64 |
| 10 | 71055496 | rs3812689 | -0.015 | -0.554 | 0.47 | 0.473 | -0.002 | 0.313 | 0.313 | 0 | 0.06 | 0.21 | 0.16 | 0.45 | 0.05 | 0.08 |
| 10 | 71056150 | rs7897621 | 0.344 | -1.038 | 0.562 | 0.112 | -0.04 | 0.078 | 0.078 | 0 | 0.79 | 0.7 | 0.75 | 0.47 | 0.87 | 0.81 |
| 10 | 71057289 | rs10998704 | -0.801 | -1.112 | 0.382 | -0.247 | -0.006 | 0.008 | 0.008 | 0.044 | 0.16 | 0.16 | 0.32 | 0.11 | 0.19 | 0.16 |
| 10 | 71058015 | rs10762281 | 1.618 | -1.479 | 0.397 | -0.443 | 0.058 | 0.051 | 0.051 | 0 | 0.54 | 0.28 | 0.38 | 0.29 | 0.67 | 0.63 |
| 10 | 71064512 | rs4746841 | -0.589 | -1.257 | 0.295 | -0.543 | -0.028 | -0.059 | -0.059 | 0 | 0.86 | 0.83 | 0.91 | 0.99 | 0.93 | 0.9 |
| 10 | 71073715 | rs10998710 | -0.658 | -0.594 | 0.084 | 0.049 | -0.009 | 0.105 | 0.105 | 0 | 0.08 | 0.12 | 0.16 | 0.62 | 0.06 | 0.06 |
| 10 | 71079286 | rs7898060 | -0.885 | -0.511 | -0.018 | 0.045 | -0.06 | -0.014 | -0.014 | 0 | 0.14 | 0.01 | 0.3 | 0.11 | 0.2 | 0.15 |
| 10 | 71082214 | rs16926244 | -0.111 | -0.357 | -0.095 | -0.095 | 0.003 | -0.01 | -0.01 | 0 | 0.17 | 0.14 | 0.26 | 0.62 | 0.19 | 0.08 |
| 10 | 71082719 | rs12262904 | -0.068 | -0.358 | 0.06 | -0.192 | -0.034 | -0.019 | -0.019 | 0 | 0.07 | 0.19 | 0.03 | 0.03 | 0.01 | 0.06 |
| 10 | 71084361 | rs7097078 | -1.029 | -0.04 | 0.046 | -0.163 | -0.004 | -0.009 | -0.009 | 0 | 0.4 | 0.48 | 0.61 | 0.75 | 0.4 | 0.29 |
| 10 | 71088538 | rs10998719 | -1.422 | 0.778 | 0.15 | 0.18 | -0.05 | -0.045 | -0.045 | 0.061 | 0.1 | 0.01 | 0.11 | 0.12 | 0.17 | 0.15 |
| 10 | 71091013 | rs10823343 | -2.281 | 0.252 | 0.747 | -0.302 | -0.013 | -0.029 | -0.029 | 0 | 0.35 | 0.62 | 0.27 | 0.13 | 0.29 | 0.36 |
| 10 | 71092803 | rs12220752 | -1.26 | 0.279 | 1.039 | -0.273 | -0.008 | -0.003 | -0.003 | 0 | 0.11 | 0.14 | 0.13 | 0.11 | 0.11 | 0.2 |
| 10 | 71097966 | rs906215 | -1.374 | 0.874 | 0.813 | -0.431 | -0.012 | 0.009 | 0.009 | 0.159 | 0.24 | 0.24 | 0.22 | 0.11 | 0.37 | 0.24 |
| 10 | 71098195 | rs906216 | -2.216 | 0.829 | 0.854 | -0.457 | 0.006 | -0.013 | -0.013 | 0 | 0.35 | 0.66 | 0.27 | 0.14 | 0.42 | 0.34 |
| 10 | 71098376 | rs10998724 | -0.994 | 0.646 | 0.6 | -0.407 | 0.016 | -0.001 | -0.001 | 0 | 0.11 | 0.04 | 0.12 | 0.11 | 0.11 | 0.21 |
| 10 | 71099222 | rs10998726 | -1.214 | 0.642 | 0.632 | -0.408 | 0.011 | 0.003 | 0.003 | 0 | 0.11 | 0.09 | 0.13 | 0.11 | 0.1 | 0.21 |
| 10 | 71099888 | rs10159477 | -0.738 | 0.724 | 0.604 | -0.423 | 0.001 | -0.019 | -0.019 | 0.048 | 0.15 | 0.15 | 0.09 | 0 | 0.15 | 0.13 |
| 10 | 71112394 | rs10823350 | -2.527 | 0.552 | 0.981 | -0.146 | -0.032 | -0.036 | -0.036 | 0.016 | 0.28 | 0.41 | 0.18 | 0.13 | 0.19 | 0.45 |
| 10 | 71113988 | rs11596587 | -0.9 | 0.596 | 0.96 | -0.132 | -0.024 | -0.067 | -0.067 | 0 | 0.12 | 0.24 | 0.03 | 0 | 0.05 | 0.16 |
| 10 | 71119208 | rs7079157 | -0.192 | 0.814 | 0.858 | -0.094 | 0.102 | 0.046 | 0.046 | 0 | 0.11 | 0.24 | 0.14 | 0.12 | 0.22 | 0.13 |
| 10 | 71120168 | rs10823354 | -1.246 | 0.634 | 0.763 | -0.162 | 0.037 | 0.014 | 0.014 | 0.473 | 0.27 | 0.41 | 0.29 | 0.25 | 0.37 | 0.46 |
| 10 | 71128875 | rs2305198 | -0.947 | 0.913 | 0.004 | 0.187 | -0.006 | 0.007 | 0.007 | 1.888 | 0.35 | 0.69 | 0.27 | 0.25 | 0.37 | 0.53 |
| 10 | 71138000 | rs10998751 | -1.228 | 1.176 | -0.939 | 0.494 | -0.055 | -0.037 | -0.037 | 0 | 0.14 | 0.01 | 0.11 | 0.14 | 0.16 | 0.18 |
| 10 | 71138269 | rs10823355 | -1.292 | 1.147 | -0.661 | 0.481 | -0.015 | -0.005 | -0.005 | 0.061 | 0.14 | 0.08 | 0.12 | 0.14 | 0.16 | 0.18 |
| 10 | 71142183 | rs4746846 | -1.678 | 0.957 | -0.547 | 1.268 | -0.003 | -0.023 | -0.023 | 0.129 | 0.22 | 0.17 | 0.41 | 0.5 | 0.19 | 0.25 |
| 10 | 71142420 | rs748235 | 1.089 | 1.699 | -0.853 | 0.999 | -0.025 | -0.042 | -0.042 | 0 | 0.82 | 0.89 | 0.63 | 0.64 | 0.78 | 0.88 |
| 10 | 71143951 | rs2305197 | -0.892 | 2.068 | -0.674 | 1.003 | -0.019 | -0.013 | -0.013 | 0.927 | 0.14 | 0.07 | 0.1 | 0.16 | 0.18 | 0.17 |
| 10 | 71144324 | rs2305196 | -1.601 | 1.748 | -0.449 | 0.934 | -0.001 | -0.035 | -0.035 | 0 | 0.26 | 0.18 | 0.43 | 0.48 | 0.25 | 0.34 |
| 10 | 71144702 | rs749105 | 1.63 | 1.75 | -0.45 | 0.925 | -0.005 | -0.034 | -0.034 | 0 | 0.7 | 0.8 | 0.56 | 0.52 | 0.68 | 0.64 |
| 10 | 71144777 | rs749106 | -0.36 | 1.546 | -0.387 | 1.041 | -0.001 | 0.009 | 0.009 | 0 | 0.1 | 0.16 | 0.07 | 0.15 | 0.11 | 0.25 |
| 10 | 71145486 | rs1867971 | -1.416 | 1.57 | -0.403 | 1.019 | 0.004 | -0.027 | -0.027 | 0 | 0.22 | 0.17 | 0.42 | 0.48 | 0.21 | 0.31 |
| 10 | 71145894 | rs2084274 | -1.861 | 0.293 | -0.141 | 0.173 | -0.006 | 0.003 | 0.003 | 0.952 | 0.55 | 0.62 | 0.61 | 0.65 | 0.48 | 0.6 |
| 10 | 71146874 | rs10823356 | -0.964 | 0.199 | 0.457 | -0.019 | -0.002 | -0.008 | -0.008 | 0.367 | 0.42 | 0.52 | 0.26 | 0.28 | 0.38 | 0.51 |
| 10 | 71149319 | rs3829185 | -1.923 | 0.219 | 0.044 | -0.042 | -0.013 | 0.001 | 0.001 | 0.059 | 0.7 | 0.79 | 0.77 | 0.74 | 0.66 | 0.76 |
| 10 | 71149545 | rs4745987 | -1.003 | -0.067 | 0.437 | -0.466 | 0 | 0.065 | 0.065 | 0.562 | 0.54 | 0.32 | 0.43 | 0.33 | 0.57 | 0.63 |
| 10 | 71149611 | rs4746849 | 0.445 | -0.003 | 0.3 | -0.508 | -0.042 | -0.03 | -0.03 | 0 | 0.13 | 0.05 | 0.18 | 0.17 | 0.2 | 0.16 |
| 10 | 71150635 | rs12414591 | 0.473 | -0.007 | 0.284 | -0.509 | -0.052 | -0.04 | -0.04 | 0 | 0.13 | 0.03 | 0.18 | 0.17 | 0.2 | 0.16 |
| 10 | 71150946 | rs7904932 | -0.902 | -0.088 | 0.662 | -0.541 | 0.058 | 0.048 | 0.048 | 1.614 | 0.06 | 0.41 | 0.04 | 0 | 0 | 0 |
| 10 | 71155633 | rs4745988 | -1.594 | -0.152 | 0.541 | -0.454 | -0.001 | 0 | 0 | 0.02 | 0.6 | 0.66 | 0.45 | 0.35 | 0.58 | 0.7 |
| 10 | 71158268 | rs1227938 | -0.176 | -0.128 | 0.438 | -0.689 | 0.029 | 0.143 | 0.143 | 0 | 0.4 | 0.22 | 0.24 | 0.16 | 0.37 | 0.49 |
| 10 | 71162686 | rs1000251 | 0.674 | -0.294 | 0.227 | -0.754 | -0.049 | 0.067 | 0.067 | 0.416 | 0.14 | 0.01 | 0.13 | 0.01 | 0.19 | 0.17 |
| 10 | 71163727 | rs12244723 | -0.763 | -0.271 | 0.779 | -0.552 | 0.008 | 0.039 | 0.039 | 1.086 | 0.05 | 0.21 | 0.02 | 0.03 | 0 | 0.06 |
| 10 | 71167235 | rs2278746 | -1.484 | -0.181 | 0.694 | -0.489 | -0.003 | 0.003 | 0.003 | 0.069 | 0.6 | 0.62 | 0.45 | 0.3 | 0.56 | 0.72 |
| 10 | 71171968 | rs10733852 | -1.494 | 0.265 | 0.287 | -0.436 | -0.006 | 0.011 | 0.011 | 0.016 | 0.62 | 0.75 | 0.47 | 0.32 | 0.61 | 0.72 |
| 10 | 71175609 | rs10998765 | -0.406 | -0.144 | 0.335 | -0.903 | -0.015 | -0.05 | -0.05 | 1.447 | 0.36 | 0.58 | 0.18 | 0.09 | 0.24 | 0.33 |
| 10 | 71178215 | rs3763737 | -0.886 | 0.074 | -0.327 | -0.906 | -0.008 | 0.043 | 0.043 | 0 | 0.44 | 0.38 | 0.37 | 0.14 | 0.47 | 0.48 |
| 10 | 71180982 | rs10823371 | -0.802 | 0.045 | -0.228 | -0.919 | -0.008 | 0.03 | 0.03 | 0 | 0.43 | 0.39 | 0.37 | 0.13 | 0.47 | 0.48 |
| 10 | 71181770 | rs10509312 | 1.015 | 0.067 | -0.531 | -1.056 | -0.063 | 0.065 | 0.065 | 0.3 | 0.11 | 0.01 | 0.16 | 0 | 0.18 | 0.12 |
| 10 | 71182508 | rs5030923 | -0.753 | -0.158 | -0.282 | -1.087 | -0.025 | 0.071 | 0.071 | 0 | 0.41 | 0.28 | 0.36 | 0.11 | 0.46 | 0.42 |
| 10 | 71191947 | rs1227940 | 0.985 | -0.387 | -0.328 | -0.858 | 0.004 | 0.001 | 0.001 | 0 | 0.13 | 0.12 | 0.1 | 0.13 | 0.14 | 0.19 |
| 10 | 71197722 | rs7098702 | 1.274 | -0.629 | -0.265 | -0.977 | -0.036 | -0.037 | -0.037 | 0 | 0.17 | 0.25 | 0.04 | 0.02 | 0.05 | 0.2 |
| 10 | 71198484 | rs768498 | -0.698 | -0.492 | -0.153 | -1.159 | -0.03 | 0.022 | 0.022 | 1.693 | 0.28 | 0.24 | 0.33 | 0.11 | 0.42 | 0.28 |
| 10 | 71204185 | rs1124367 | -1.893 | -0.494 | -0.079 | -1.487 | -0.004 | -0.013 | -0.013 | 0 | 0.58 | 0.61 | 0.48 | 0.26 | 0.61 | 0.67 |
| 10 | 71206020 | rs1367517 | -0.119 | 0.461 | -0.372 | -1.319 | -0.018 | 0.01 | 0.01 | 0 | 0.19 | 0.15 | 0.24 | 0.11 | 0.29 | 0.22 |
| 10 | 71209768 | rs12414198 | -1.028 | 0.377 | -0.388 | -1.32 | -0.034 | 0.026 | 0.026 | 1.797 | 0.28 | 0.23 | 0.33 | 0.11 | 0.42 | 0.28 |
| 10 | 71213650 | rs7090788 | 1.549 | 0.381 | -0.449 | -0.737 | -0.009 | 0.082 | 0.082 | 0 | 0.11 | 0.14 | 0.3 | 0.44 | 0.08 | 0.06 |
| 10 | 71218059 | rs10998791 | -0.137 | 0.553 | -0.671 | -1.013 | -0.011 | 0.043 | 0.043 | 0 | 0.32 | 0.27 | 0.29 | 0.12 | 0.33 | 0.33 |
| 10 | 71218094 | rs12776158 | 0.833 | 0.736 | -0.71 | -1.039 | -0.011 | -0.01 | -0.01 | 0.172 | 0.07 | 0.07 | 0.08 | 0.11 | 0.09 | 0.09 |

| | | | | | | | | | | | | | | | | |
|---|---|---|---|---|---|---|---|---|---|---|---|---|---|---|---|---|
| 10 | 71218566 | rs10823378 | 0.466 | 0.535 | -0.646 | -1.058 | 0.004 | 0.019 | 0.019 | 0 | 0.26 | 0.2 | 0.21 | 0.04 | 0.23 | 0.28 |
| 10 | 71221128 | rs4565792 | -0.446 | 0.323 | -0.614 | -0.979 | -0.035 | 0.069 | 0.069 | 0 | 0.4 | 0.3 | 0.37 | 0.15 | 0.45 | 0.41 |
| 10 | 71244832 | rs2305201 | -0.606 | 0.313 | -0.256 | -1.01 | 0.009 | -0.036 | -0.036 | 0 | 0.42 | 0.55 | 0.37 | 0.15 | 0.44 | 0.41 |
| 10 | 71245617 | rs1227937 | -1.435 | 0.25 | -0.787 | -0.811 | -0.025 | -0.045 | -0.045 | 0 | 0.7 | 0.93 | 0.78 | 0.71 | 0.68 | 0.67 |
| 10 | 71254765 | rs12414602 | 0.078 | 0.978 | -0.659 | -0.578 | -0.035 | -0.03 | -0.03 | 0 | 0.08 | 0.01 | 0.11 | 0.11 | 0.1 | 0.09 |
| 10 | 71255663 | rs1864588 | -0.175 | 1.104 | -0.683 | -0.716 | -0.003 | 0.009 | 0.009 | 0.244 | 0.12 | 0.13 | 0.12 | 0.04 | 0.13 | 0.14 |
| 10 | 71255747 | rs1864589 | -0.904 | 1.601 | -0.776 | -0.773 | -0.028 | 0.026 | 0.026 | 0.42 | 0.16 | 0.14 | 0.18 | 0.08 | 0.22 | 0.15 |
| 10 | 71256208 | rs4746856 | 1.479 | 1.314 | -0.856 | -0.998 | -0.001 | -0.017 | -0.017 | 0 | 0.07 | 0.05 | 0.04 | 0 | 0.08 | 0.04 |
| 10 | 71262048 | rs1665581 | -0.248 | 1.366 | -0.777 | -1.172 | 0 | -0.026 | -0.026 | 0 | 0.43 | 0.55 | 0.36 | 0.15 | 0.38 | 0.31 |
| 10 | 71262824 | rs12263764 | 1.153 | 1.168 | -0.57 | -1.173 | -0.029 | -0.111 | -0.111 | 0 | 0.27 | 0.3 | 0.14 | 0 | 0.16 | 0.2 |
| 10 | 71267010 | rs1052179 | 0.016 | 1.216 | -0.656 | -1.112 | -0.002 | -0.041 | -0.041 | 0 | 0.4 | 0.54 | 0.33 | 0.16 | 0.31 | 0.31 |
| 10 | 71272018 | rs972570 | -0.662 | 0.672 | -0.685 | -1.07 | -0.001 | 0.044 | 0.044 | 0 | 0.51 | 0.62 | 0.45 | 0.19 | 0.44 | 0.46 |
| 10 | 71273696 | rs4745990 | -1.304 | 0.576 | -0.959 | -0.828 | -0.006 | -0.007 | -0.007 | 0 | 0.64 | 0.75 | 0.69 | 0.63 | 0.51 | 0.55 |
| 10 | 71274737 | rs891562 | 1.295 | 0.526 | -0.911 | -0.892 | -0.013 | -0.081 | -0.081 | 0 | 0.28 | 0.26 | 0.15 | 0 | 0.17 | 0.21 |
| 10 | 71275363 | rs10998840 | 1.238 | 0.463 | -0.843 | -0.251 | 0.006 | -0.026 | -0.026 | 0.466 | 0.1 | 0.05 | 0.24 | 0.43 | 0.08 | 0.06 |
| 10 | 71279367 | rs1311176 | -0.546 | 0.68 | -0.84 | -0.316 | -0.004 | -0.004 | -0.004 | 0 | 0.63 | 0.7 | 0.69 | 0.63 | 0.52 | 0.55 |
| 10 | 71280107 | rs1665585 | -0.546 | 0.68 | -0.84 | -0.316 | -0.004 | -0.004 | -0.004 | 0 | 0.63 | 0.7 | 0.69 | 0.63 | 0.52 | 0.55 |
| 10 | 71293704 | rs12416010 | -0.38 | 0.455 | -0.977 | 0.529 | -0.042 | -0.033 | -0.033 | 0 | 0.11 | 0.01 | 0.13 | 0.15 | 0.11 | 0.18 |
| 10 | 71295253 | rs10998847 | 0.488 | 0.493 | -1.101 | 0.516 | -0.017 | -0.095 | -0.095 | 0.01 | 0.19 | 0.05 | 0.16 | 0.3 | 0.24 | 0.13 |
| 10 | 71297724 | rs10823403 | -0.819 | 0.529 | -0.907 | 0.602 | -0.002 | -0.025 | -0.025 | 0.108 | 0.33 | 0.29 | 0.33 | 0.45 | 0.35 | 0.31 |
| 10 | 71298455 | rs1237105 | -1.517 | 0.447 | -0.842 | 0.512 | 0.001 | -0.032 | -0.032 | 0 | 0.47 | 0.59 | 0.6 | 0.91 | 0.44 | 0.41 |
| 10 | 71300334 | rs754038 | -1.34 | 1.323 | -1.419 | 1.024 | -0.001 | -0.024 | -0.024 | 0 | 0.33 | 0.29 | 0.34 | 0.46 | 0.37 | 0.31 |
| 10 | 71302427 | rs2084273 | -0.428 | 0.96 | -1.71 | 0.99 | 0.003 | -0.065 | -0.065 | 0 | 0.21 | 0.12 | 0.18 | 0.32 | 0.24 | 0.13 |
| 10 | 71303321 | rs12357239 | -0.521 | 1.084 | -1.622 | 1.11 | 0.001 | -0.068 | -0.068 | 0 | 0.23 | 0.12 | 0.21 | 0.32 | 0.27 | 0.15 |
| 10 | 71305267 | rs10998852 | -1.068 | 1.749 | -1.276 | 0.977 | -0.012 | -0.008 | -0.008 | 0 | 0.11 | 0.05 | 0.14 | 0.18 | 0.19 | 0.22 |
| 10 | 71307920 | rs297049 | -0.697 | 1.581 | -1.296 | 1.245 | 0 | -0.003 | -0.003 | 0 | 0.07 | 0.06 | 0.1 | 0.13 | 0.08 | 0.04 |
| 10 | 71310732 | rs2812562 | 0.587 | 1.511 | -1.136 | 1.322 | -0.001 | 0.013 | 0.013 | 0 | 0.92 | 0.91 | 0.87 | 0.85 | 0.92 | 0.96 |
| 10 | 71312023 | rs10823406 | -1.234 | 2.167 | -1.45 | 1.366 | -0.08 | -0.019 | -0.019 | 0.364 | 0.13 | 0.01 | 0.18 | 0.12 | 0.24 | 0.24 |
| 10 | 71312821 | rs297046 | 1.874 | 1.149 | -1.069 | 0.896 | -0.009 | 0.002 | 0.002 | 0 | 0.69 | 0.26 | 0.63 | 0.73 | 0.68 | 0.69 |
| 10 | 71313908 | rs297043 | 0.204 | 1.089 | -1.08 | 1.021 | -0.001 | 0.014 | 0.014 | 0 | 0.92 | 0.91 | 0.87 | 0.85 | 0.92 | 0.96 |
| 10 | 71320358 | rs11592369 | -0.833 | 1.476 | -1.338 | 0.933 | -0.069 | -0.031 | -0.031 | 0 | 0.12 | 0.01 | 0.14 | 0.11 | 0.21 | 0.15 |
| 10 | 71320943 | rs190925 | 0.732 | 0.967 | -1 | 0.571 | -0.006 | -0.001 | -0.001 | 0.16 | 0.53 | 0.46 | 0.34 | 0.6 | 0.33 | 0.58 |
| 10 | 71325721 | rs10823407 | 1.194 | 0.271 | -1.796 | 0.448 | 0.009 | -0.009 | -0.009 | 0 | 0.18 | 0.08 | 0.12 | 0.25 | 0.14 | 0.11 |
| 10 | 71332204 | rs4536103 | -0.704 | 0.54 | -1.39 | 0.718 | 0.038 | 0.037 | 0.037 | 1.224 | 0.43 | 0.26 | 0.61 | 0.31 | 0.67 | 0.41 |
| 10 | 71333783 | rs11592899 | -0.421 | 1.455 | -1.603 | 0.904 | -0.092 | 0.017 | 0.017 | 0.108 | 0.15 | 0.01 | 0.17 | 0.08 | 0.3 | 0.15 |
| 10 | 71334622 | rs3812704 | 0.065 | 0.694 | -1.563 | 1.116 | 0.013 | -0.019 | -0.019 | 0.174 | 0.08 | 0.02 | 0.14 | 0.4 | 0.06 | 0.19 |
| 10 | 71339737 | rs2812545 | 0.675 | 1.412 | -0.963 | 0.538 | 0.003 | -0.003 | -0.003 | 0 | 0.64 | 0.63 | 0.55 | 0.37 | 0.53 | 0.61 |
| 10 | 71341200 | rs4746864 | -0.359 | 1.623 | -1.178 | 0.392 | -0.034 | -0.009 | -0.009 | 0 | 0.24 | 0.16 | 0.42 | 0.28 | 0.35 | 0.23 |
| 10 | 71343400 | rs989485 | 0.111 | 0.438 | -1.422 | -0.757 | 0.001 | 0.08 | 0.08 | 0 | 0.44 | 0.29 | 0.3 | 0.23 | 0.45 | 0.42 |
| 10 | 71346736 | rs7895043 | -0.084 | -0.013 | -0.59 | -0.955 | 0.025 | 0.117 | 0.117 | 0 | 0.5 | 0.29 | 0.65 | 0.77 | 0.55 | 0.55 |
| 10 | 71347276 | rs990394 | 1.379 | -0.56 | -0.909 | -1.225 | 0 | -0.004 | -0.004 | 0 | 0.26 | 0.25 | 0.16 | 0.14 | 0.22 | 0.29 |
| 10 | 71348435 | rs874511 | -0.043 | -0.551 | -0.579 | -1.203 | 0.026 | 0.015 | 0.015 | 0 | 0.09 | 0.25 | 0.08 | 0.11 | 0.08 | 0.07 |
| 10 | 71351750 | rs10998872 | 2.389 | -0.894 | -1.119 | -1.366 | -0.007 | -0.048 | -0.048 | 0 | 0.18 | 0.05 | 0.18 | 0.27 | 0.18 | 0.13 |
| 10 | 71351990 | rs12252820 | 2.431 | -1.015 | -0.987 | -1.477 | 0.001 | -0.022 | -0.022 | 0 | 0.2 | 0.18 | 0.09 | 0.04 | 0.13 | 0.22 |
| 10 | 71352026 | rs12252832 | 0.667 | -0.841 | -0.92 | -1.323 | 0.02 | 0.054 | 0.054 | 0 | 0.39 | 0.27 | 0.25 | 0.13 | 0.36 | 0.38 |
| 10 | 71352551 | rs12779498 | 1.604 | -0.494 | -1.066 | -0.885 | -0.044 | 0.025 | 0.025 | 0 | 0.17 | 0.06 | 0.21 | 0.1 | 0.26 | 0.21 |
| 10 | 71354031 | rs7091934 | 0.414 | -0.554 | -0.884 | -0.849 | -0.039 | 0.001 | 0.001 | 0 | 0.36 | 0.18 | 0.41 | 0.37 | 0.43 | 0.38 |
| 10 | 71357729 | rs7915040 | 2.312 | -0.807 | -0.951 | -0.797 | 0.016 | 0.008 | 0.008 | 0 | 0.11 | 0.06 | 0.03 | 0 | 0.04 | 0.09 |
| 10 | 71357954 | rs731573 | -0.522 | -0.579 | -0.981 | -0.529 | -0.002 | -0.032 | -0.032 | 0.35 | 0.8 | 0.85 | 0.72 | 0.49 | 0.81 | 0.66 |
| 10 | 71360884 | rs1947739 | 0.394 | -0.527 | -0.827 | -1.122 | -0.003 | -0.031 | -0.031 | 0 | 0.23 | 0.25 | 0.16 | 0.11 | 0.22 | 0.22 |
| 10 | 71367698 | rs10998876 | 1.161 | 0.008 | -0.823 | -0.685 | -0.006 | -0.01 | -0.01 | 0 | 0.32 | 0.45 | 0.33 | 0.27 | 0.32 | 0.2 |
| 10 | 71368698 | rs2812555 | -0.026 | 1.31 | -0.663 | -0.015 | -0.006 | -0.032 | -0.032 | 0 | 0.8 | 0.86 | 0.72 | 0.44 | 0.8 | 0.66 |
| 10 | 71372590 | rs2616139 | 0.098 | 0.943 | -0.795 | -0.016 | 0.02 | -0.004 | -0.004 | 0 | 0.64 | 0.62 | 0.52 | 0.38 | 0.58 | 0.52 |
| 10 | 71373767 | rs931028 | 0.239 | 0.821 | -0.85 | 0.026 | 0 | -0.018 | -0.018 | 0.199 | 0.69 | 0.5 | 0.72 | 0.77 | 0.71 | 0.6 |
| 10 | 71374466 | rs7087903 | 0.497 | 0.27 | -1.445 | -0.146 | 0.016 | 0.019 | 0.019 | 0.595 | 0.24 | 0.17 | 0.1 | 0.14 | 0.15 | 0.18 |
| 10 | 71376076 | rs1903791 | 0.656 | 0.603 | -1.48 | 0.24 | -0.007 | 0.012 | 0.012 | 0 | 0.1 | 0.07 | 0.25 | 0.32 | 0.14 | 0.3 |
| 10 | 71376096 | rs1903790 | -0.174 | -0.089 | -1.716 | 0.09 | 0.059 | -0.027 | -0.027 | 0 | 0.41 | 0.19 | 0.38 | 0.51 | 0.33 | 0.37 |
| 10 | 71376535 | rs2927384 | -1.291 | 0.043 | -1.605 | -0.1 | -0.01 | -0.042 | -0.042 | 0.129 | 0.71 | 0.56 | 0.83 | 0.83 | 0.75 | 0.78 |
| 10 | 71377422 | rs2812567 | -0.439 | 0.048 | -1.663 | -0.083 | 0.054 | -0.003 | -0.003 | 0.674 | 0.45 | 0.26 | 0.43 | 0.51 | 0.37 | 0.39 |
| 10 | 71378059 | rs2812568 | 0.336 | 0.754 | -1.411 | 0.444 | 0.013 | -0.031 | -0.031 | 0 | 0.17 | 0.07 | 0.21 | 0.27 | 0.17 | 0.2 |
| 10 | 71378767 | rs4746874 | -0.59 | 1.587 | -0.907 | 0.249 | 0.006 | 0.001 | 0.001 | 0.928 | 0.31 | 0.38 | 0.45 | 0.32 | 0.42 | 0.4 |
| 10 | 71383377 | rs2616104 | -0.545 | 0.978 | -0.951 | -0.381 | 0.001 | 0.005 | 0.005 | 0 | 0.41 | 0.38 | 0.59 | 0.84 | 0.46 | 0.44 |
| 10 | 71389673 | rs10998884 | 0.397 | 0.703 | -1.348 | -0.514 | -0.001 | 0.042 | 0.042 | 0.541 | 0.14 | 0.07 | 0.1 | 0 | 0.17 | 0.03 |
| 10 | 71391621 | rs2306139 | -0.036 | 0.512 | -1.118 | 0.483 | 0.003 | 0.023 | 0.023 | 0 | 0.12 | 0.09 | 0.33 | 0.47 | 0.09 | 0.12 |
| 10 | 71392557 | rs12020 | -0.431 | 0.842 | -0.997 | -0.619 | 0.005 | -0.011 | -0.011 | 0 | 0.6 | 0.6 | 0.74 | 0.89 | 0.67 | 0.69 |
| 10 | 71393025 | rs1132075 | -0.213 | 0.875 | -0.982 | -0.604 | 0.004 | -0.013 | -0.013 | 0 | 0.6 | 0.6 | 0.74 | 0.89 | 0.67 | 0.69 |
| 10 | 71400861 | rs6480413 | 0.084 | 1.119 | -0.887 | -0.189 | 0.017 | -0.025 | -0.025 | 0.1 | 0.62 | 0.53 | 0.67 | 0.64 | 0.57 | 0.48 |
| 10 | 71401017 | rs1870960 | -0.107 | 1.029 | -1.493 | -0.538 | 0 | 0.005 | 0.005 | 0 | 0.55 | 0.58 | 0.37 | 0.25 | 0.44 | 0.39 |
| 10 | 71402902 | rs10509319 | -0.241 | 1.226 | -1.755 | -0.326 | -0.037 | -0.012 | -0.012 | 0.102 | 0.08 | 0.01 | 0.08 | 0.09 | 0.13 | 0.18 |

| | | | | | | | | | | | | | | | | |
|---|---|---|---|---|---|---|---|---|---|---|---|---|---|---|---|---|
| 10 | 71407239 | rs4746882 | 0.346 | 1.707 | -2.243 | 0.322 | -0.089 | -0.122 | -0.122 | 0.037 | 0.17 | 0.01 | 0.39 | 0.4 | 0.23 | 0.28 |
| 10 | 71409200 | rs17578297 | -0.26 | 0.767 | -2.243 | 0.226 | 0.008 | -0.011 | -0.011 | 0.724 | 0.12 | 0.11 | 0.07 | 0.16 | 0.07 | 0.11 |
| 10 | 71409843 | rs12782802 | -1.109 | 0.763 | -2.274 | 0.109 | -0.01 | -0.002 | -0.002 | 0 | 0.23 | 0.16 | 0.17 | 0.25 | 0.22 | 0.29 |
| 10 | 71410314 | rs10998893 | 0.632 | 0.556 | -2.138 | -0.209 | -0.008 | -0.061 | -0.061 | 1.907 | 0.23 | 0.06 | 0.17 | 0.26 | 0.23 | 0.26 |
| 10 | 71415241 | rs17578395 | 0.037 | 1.013 | -1.778 | -0.046 | -0.06 | 0.007 | 0.007 | 0 | 0.1 | 0.01 | 0.1 | 0.08 | 0.17 | 0.19 |
| 10 | 71418213 | rs2616076 | 0.156 | 0.592 | -1.031 | -0.12 | -0.001 | 0 | 0 | 0 | 0.46 | 0.46 | 0.59 | 0.62 | 0.47 | 0.48 |
| 10 | 71419702 | rs2616074 | -0.066 | 0.534 | -0.996 | -0.418 | -0.001 | -0.004 | -0.004 | 0.041 | 0.51 | 0.53 | 0.64 | 0.81 | 0.5 | 0.58 |
| 10 | 71420172 | rs2812500 | -0.603 | 0.475 | -0.924 | -0.404 | -0.015 | -0.056 | -0.056 | 0 | 0.6 | 0.54 | 0.74 | 0.89 | 0.67 | 0.76 |
| 10 | 71421872 | rs2812507 | -0.707 | 0.165 | -0.27 | 0.113 | -0.009 | 0.003 | 0.003 | 0.196 | 0.37 | 0.5 | 0.28 | 0.43 | 0.35 | 0.43 |
| 10 | 71423907 | rs2812510 | -0.685 | 0.074 | -0.301 | 0.06 | -0.002 | 0.001 | 0.001 | 0.237 | 0.37 | 0.56 | 0.44 | 0.48 | 0.35 | 0.43 |
| 10 | 71423976 | rs2812512 | -0.631 | 0.094 | -0.167 | 0.069 | -0.008 | 0.01 | 0.01 | 0 | 0.36 | 0.49 | 0.42 | 0.48 | 0.35 | 0.43 |
| 10 | 71426784 | rs7914685 | 0.982 | 0.102 | -0.027 | -0.044 | -0.024 | -0.037 | -0.037 | 0 | 0.07 | 0.14 | 0.03 | 0 | 0.03 | 0.01 |
| 10 | 71427798 | rs2616069 | 0.601 | 0.034 | 0.361 | -0.086 | 0.004 | -0.073 | -0.073 | 0 | 0.81 | 0.58 | 0.85 | 0.92 | 0.82 | 0.82 |
| 10 | 71432146 | rs2616108 | 1.548 | -0.001 | 0.36 | -0.065 | -0.003 | 0.005 | 0.005 | 0 | 0.14 | 0.13 | 0.08 | 0.21 | 0.14 | 0.15 |
| 10 | 71432279 | rs10509316 | -0.918 | -0.002 | 0.359 | -0.049 | -0.006 | 0.016 | 0.016 | 0 | 0.21 | 0.33 | 0.34 | 0.28 | 0.21 | 0.27 |
| 10 | 71435609 | rs4746885 | 0.872 | 0.026 | -0.076 | -0.046 | 0.009 | -0.007 | -0.007 | 0 | 0.62 | 0.37 | 0.56 | 0.51 | 0.65 | 0.57 |
| 10 | 71439191 | rs10823418 | -0.55 | 0.48 | -0.114 | 0.251 | 0 | -0.004 | -0.004 | 0.019 | 0.28 | 0.27 | 0.39 | 0.49 | 0.3 | 0.4 |
| 10 | 71439391 | rs2616119 | -1.265 | 0.864 | -0.403 | 0.178 | 0.001 | 0 | 0 | 0 | 0.59 | 0.6 | 0.74 | 0.9 | 0.64 | 0.75 |
| 10 | 71442615 | rs12161693 | -0.574 | 1.264 | -0.733 | 0.721 | 0.013 | 0.013 | 0.013 | 0 | 0.23 | 0.35 | 0.19 | 0.29 | 0.27 | 0.31 |
| 10 | 71444076 | rs10998916 | 0.694 | 1.357 | -1.809 | 0.861 | -0.008 | 0.009 | 0.009 | 0 | 0.14 | 0.15 | 0.08 | 0.21 | 0.12 | 0.14 |
| 10 | 71451440 | rs2812530 | -0.334 | 2.223 | -2.211 | 0.894 | -0.007 | -0.055 | -0.055 | 0 | 0.23 | 0.12 | 0.3 | 0.44 | 0.24 | 0.33 |
| 10 | 71451872 | rs2812531 | -1.015 | 2.458 | -2.498 | 0.768 | -0.093 | -0.116 | -0.116 | 0 | 0.33 | 0.11 | 0.4 | 0.46 | 0.43 | 0.52 |
| 10 | 71452285 | rs2812533 | 0.789 | 2.275 | -1.606 | 0.335 | -0.014 | 0.008 | 0.008 | 0 | 0.91 | 0.91 | 0.92 | 0.93 | 0.84 | 0.85 |
| 10 | 71452871 | rs3907199 | -1.159 | 1.816 | -1.795 | 0.382 | -0.053 | 0.009 | 0.009 | 0 | 0.16 | 0.03 | 0.13 | 0.1 | 0.21 | 0.15 |
| 10 | 71456311 | rs2395023 | -0.257 | 1.035 | -2.077 | 0.198 | 0.032 | 0.075 | 0.075 | 0.146 | 0.43 | 0.2 | 0.32 | 0.3 | 0.41 | 0.18 |
| 10 | 71460661 | rs12569387 | -0.834 | 1.651 | -2.322 | 0.175 | -0.005 | 0 | 0 | 0 | 0.16 | 0.15 | 0.16 | 0.03 | 0.27 | 0.07 |
| 10 | 71466578 | rs2642588 | 1.086 | 1.305 | -1.607 | -0.115 | 0.003 | -0.021 | -0.021 | 0.668 | 0.72 | 0.44 | 0.7 | 0.87 | 0.7 | 0.86 |
| 10 | 71466637 | rs2812541 | 1.172 | 0.781 | -2.195 | -0.182 | -0.133 | -0.122 | -0.122 | 0 | 0.35 | 0.1 | 0.41 | 0.47 | 0.48 | 0.61 |
| 10 | 71469137 | rs10823422 | 0.867 | 0.635 | -1.992 | -0.197 | 0.055 | -0.046 | -0.046 | 0 | 0.32 | 0.12 | 0.27 | 0.35 | 0.21 | 0.15 |
| 10 | 71469969 | rs10762307 | 0.699 | 0.701 | -1.942 | -0.179 | -0.039 | -0.039 | -0.039 | 0 | 0.4 | 0.31 | 0.43 | 0.51 | 0.49 | 0.71 |
| 10 | 71470529 | rs2616133 | 1.018 | 0.773 | -1.999 | -0.204 | -0.141 | -0.119 | -0.119 | 0.024 | 0.37 | 0.1 | 0.43 | 0.47 | 0.51 | 0.62 |
| 10 | 71470725 | rs3843584 | -0.132 | 0.566 | -1.396 | -0.185 | 0.006 | 0 | 0 | 6.022 | 0.46 | 0.49 | 0.43 | 0.43 | 0.3 | 0.23 |
| 10 | 71472280 | rs10823423 | 0.453 | 0.472 | -0.768 | -0.304 | 0.032 | 0.001 | 0.001 | 0.787 | 0.38 | 0.23 | 0.28 | 0.34 | 0.24 | 0.16 |
| 10 | 71474493 | rs3847349 | 0.753 | 0.4 | -0.996 | -0.303 | 0.049 | 0.02 | 0.02 | 0.072 | 0.34 | 0.08 | 0.21 | 0.27 | 0.22 | 0.12 |
| 10 | 71476425 | rs3104049 | 0.303 | 0.418 | -0.676 | -0.191 | 0.046 | -0.01 | -0.01 | 0.345 | 0.48 | 0.28 | 0.56 | 0.5 | 0.68 | 0.66 |
| 10 | 71477144 | rs12249608 | -0.38 | 0.323 | -0.425 | -0.197 | 0.001 | 0.027 | 0.027 | 0 | 0.12 | 0.38 | 0.13 | 0.09 | 0.06 | 0.06 |
| 10 | 71482299 | rs2394581 | -1.178 | 0.264 | -0.556 | -0.224 | 0.02 | -0.02 | -0.02 | 0 | 0.67 | 0.97 | 0.79 | 0.73 | 0.78 | 0.87 |
| 10 | 71483516 | rs4746896 | -0.172 | 0.339 | -0.527 | -0.274 | 0.028 | 0.048 | 0.048 | 0.579 | 0.53 | 0.67 | 0.65 | 0.65 | 0.6 | 0.83 |
| 10 | 71484854 | rs10998934 | -1.199 | 0.466 | -0.355 | -0.239 | -0.008 | 0.093 | 0.093 | 0 | 0.16 | 0.45 | 0.21 | 0.22 | 0.09 | 0.17 |
| 10 | 71485458 | rs10740322 | 1.489 | 0.555 | -0.384 | -0.246 | 0 | 0.001 | 0.001 | 0 | 0.69 | 0.24 | 0.67 | 0.73 | 0.72 | 0.79 |
| 10 | 71488223 | rs6480424 | -0.132 | 0.638 | -0.557 | -0.27 | 0 | -0.004 | -0.004 | 0 | 0.87 | 0.88 | 0.89 | 0.94 | 0.81 | 0.96 |
| 10 | 71492543 | rs2011911 | 1.432 | 0.521 | -0.679 | -0.134 | 0.013 | 0.013 | 0.013 | 0 | 0.7 | 0.2 | 0.7 | 0.79 | 0.76 | 0.81 |
| 10 | 71494181 | rs4746900 | 0.89 | 0.51 | -0.875 | -0.232 | -0.044 | -0.022 | -0.022 | 0.033 | 0.38 | 0.23 | 0.56 | 0.59 | 0.54 | 0.66 |
| 10 | 71495645 | rs4746902 | 0.242 | 0.396 | -0.527 | -0.286 | 0.022 | 0.017 | 0.017 | 1.498 | 0.46 | 0.39 | 0.64 | 0.64 | 0.68 | 0.68 |
| 10 | 71495726 | rs10998941 | 1.455 | 0.442 | -0.634 | -0.255 | -0.068 | -0.071 | -0.071 | 0 | 0.32 | 0.14 | 0.46 | 0.49 | 0.49 | 0.56 |
| 10 | 71497404 | rs12357460 | 0.538 | 0.135 | -0.473 | -0.32 | -0.025 | -0.004 | -0.004 | 0 | 0.38 | 0.49 | 0.35 | 0.35 | 0.25 | 0.23 |
| 10 | 71498235 | rs10823427 | 1.142 | 0.132 | -0.628 | -0.315 | 0.043 | 0.005 | 0.005 | 0.882 | 0.32 | 0.03 | 0.17 | 0.26 | 0.21 | 0.16 |
| 10 | 71502192 | rs10740323 | 1.54 | 0.389 | -0.59 | -0.301 | -0.087 | -0.105 | -0.105 | 0 | 0.32 | 0.09 | 0.4 | 0.48 | 0.46 | 0.56 |
| 10 | 71502204 | rs17495523 | 0.459 | 0.322 | -0.547 | -0.342 | -0.003 | -0.019 | -0.019 | 0.14 | 0.08 | 0.08 | 0.02 | 0 | 0.04 | 0.07 |
| 10 | 71502638 | rs10823429 | 0.409 | 0.233 | -0.526 | -0.385 | 0.025 | 0.045 | 0.045 | 0.437 | 0.42 | 0.26 | 0.28 | 0.33 | 0.37 | 0.19 |
| 10 | 71507963 | rs6480426 | -0.61 | 0.217 | -0.512 | -0.388 | 0.003 | 0.008 | 0.008 | 1.748 | 0.55 | 0.44 | 0.38 | 0.38 | 0.44 | 0.27 |
| 10 | 71508424 | rs4617470 | 0.358 | 0.198 | -0.506 | -0.396 | 0.019 | 0.032 | 0.032 | 0.779 | 0.42 | 0.33 | 0.28 | 0.33 | 0.37 | 0.19 |
| 10 | 71508756 | rs4620602 | 1.774 | 0.247 | -0.383 | -0.372 | 0.016 | 0.055 | 0.055 | 0 | 0.82 | 0.52 | 0.71 | 0.81 | 0.87 | 0.81 |
| 10 | 71511081 | rs7894473 | -0.993 | 0.16 | -0.513 | -0.376 | 0.005 | 0.004 | 0.004 | 0 | 0.16 | 0.45 | 0.21 | 0.09 | 0.12 | 0.15 |
| 10 | 71511904 | rs7073220 | 2.02 | 0.233 | -0.534 | -0.391 | -0.011 | -0.006 | -0.006 | 0 | 0.74 | 0.39 | 0.67 | 0.81 | 0.83 | 0.74 |
| 10 | 71513647 | rs2642589 | 1.321 | 0.232 | -0.656 | -0.432 | 0.018 | -0.016 | -0.016 | 0 | 0.81 | 0.58 | 0.77 | 0.83 | 0.79 | 0.85 |
| 10 | 71515143 | rs11814219 | 0.831 | 0.263 | -1.209 | -0.129 | 0.041 | -0.024 | -0.024 | 1.622 | 0.33 | 0.06 | 0.17 | 0.29 | 0.21 | 0.15 |
| 10 | 71515730 | rs4746906 | -0.194 | 0.279 | -1.112 | -0.125 | 0.006 | -0.007 | -0.007 | 0 | 0.09 | 0.14 | 0.1 | 0.07 | 0.15 | 0.04 |
| 10 | 71517044 | rs1077455 | 0.653 | -0.015 | -1.16 | -0.099 | 0.049 | -0.025 | -0.025 | 0 | 0.33 | 0.05 | 0.17 | 0.3 | 0.22 | 0.15 |
| 10 | 71520748 | rs12356090 | -0.627 | -0.412 | -1.031 | -0.111 | 0.01 | -0.02 | -0.02 | 0.193 | 0.2 | 0.15 | 0.12 | 0.23 | 0.15 | 0.18 |
| 10 | 71521878 | rs2704517 | 0.482 | 0.08 | -0.707 | -0.151 | 0.007 | 0.001 | 0.001 | 0 | 0.46 | 0.42 | 0.58 | 0.61 | 0.58 | 0.64 |
| 10 | 71524485 | rs2704463 | -0.947 | 0.338 | -0.829 | -0.146 | -0.001 | -0.011 | -0.011 | 0.169 | 0.25 | 0.23 | 0.25 | 0.25 | 0.23 | 0.25 |
| 10 | 71529084 | rs2252324 | -0.681 | 0.556 | -0.566 | -0.437 | -0.02 | 0.085 | 0.085 | 1.559 | 0.56 | 0.7 | 0.51 | 0.27 | 0.46 | 0.45 |
| 10 | 71529191 | rs2252321 | -0.326 | 0.458 | -0.139 | -0.606 | 0.008 | 0.012 | 0.012 | 0.526 | 0.49 | 0.43 | 0.38 | 0.19 | 0.37 | 0.33 |
| 10 | 71529945 | rs750116 | -0.826 | 0.542 | -0.305 | -0.575 | -0.022 | 0.087 | 0.087 | 1.63 | 0.56 | 0.7 | 0.51 | 0.27 | 0.46 | 0.45 |
| 10 | 71530773 | rs10998959 | 0.517 | 0.193 | -0.211 | -0.699 | 0.013 | 0.003 | 0.003 | 4.082 | 0.38 | 0.26 | 0.21 | 0.13 | 0.25 | 0.14 |
| 10 | 71533042 | rs9919438 | -0.09 | 0.211 | 0.26 | -0.162 | 0.005 | 0.021 | 0.021 | 2.711 | 0.16 | 0.21 | 0.17 | 0.26 | 0.19 | 0.21 |
| 10 | 71534183 | rs2704470 | 0.377 | 0.145 | -0.176 | 0.031 | 0.001 | -0.001 | -0.001 | 0 | 0.47 | 0.29 | 0.53 | 0.56 | 0.5 | 0.5 |
| 10 | 71538292 | rs4746003 | 0.911 | 0.096 | -0.218 | 0.055 | 0.014 | 0.021 | 0.021 | 5.421 | 0.32 | 0.26 | 0.23 | 0.22 | 0.26 | 0.22 |

| | | | | | | | | | | | | | | | | |
|---|---|---|---|---|---|---|---|---|---|---|---|---|---|---|---|---|
| 10 | 71538350 | rs2704474 | -0.907 | 0.102 | -0.207 | 0.06 | 0.012 | -0.009 | -0.009 | 0 | 0.79 | 0.55 | 0.76 | 0.82 | 0.77 | 0.72 |
| 10 | 71539581 | rs2704476 | 0.359 | 0.147 | -0.354 | 0.085 | 0.001 | 0.022 | 0.022 | 0.332 | 0.47 | 0.29 | 0.53 | 0.6 | 0.51 | 0.5 |
| 10 | 71541097 | rs2642576 | -0.662 | -0.001 | -0.024 | -0.034 | -0.026 | 0.038 | 0.038 | 0.117 | 0.85 | 0.57 | 0.88 | 0.77 | 0.94 | 0.79 |
| 10 | 71543053 | rs7903128 | 1.433 | 0.168 | -0.275 | -0.058 | -0.012 | 0.019 | 0.019 | 0 | 0.15 | 0.1 | 0.18 | 0.01 | 0.21 | 0.12 |
| 10 | 71543168 | rs2642575 | 0.221 | -0.324 | -0.103 | 0.525 | 0.021 | -0.002 | -0.002 | 0 | 0.35 | 0.25 | 0.29 | 0.28 | 0.28 | 0.24 |
| 10 | 71546573 | rs2247549 | -0.443 | 0.418 | 0.402 | 0.771 | -0.008 | 0.042 | 0.042 | 0 | 0.88 | 0.76 | 0.91 | 0.76 | 0.95 | 0.82 |
| 10 | 71549168 | rs2642572 | -0.081 | 0.795 | -0.472 | 0.82 | -0.003 | -0.004 | -0.004 | 0.058 | 0.08 | 0.06 | 0.15 | 0.11 | 0.09 | 0.15 |
| 10 | 71552976 | rs3104053 | 0.094 | 0.414 | 0.175 | 0.875 | 0.038 | 0.151 | 0.151 | 5.642 | 0.93 | 0.7 | 0.96 | 0.76 | 0.95 | 0.86 |
| 10 | 71556310 | rs12411657 | 1.056 | 0.654 | -0.568 | 0.947 | -0.055 | -0.019 | -0.019 | 0.905 | 0.27 | 0.15 | 0.33 | 0.26 | 0.35 | 0.27 |
| 10 | 71557944 | rs2704507 | -0.214 | 0.559 | -0.841 | 0.503 | 0.009 | 0.01 | 0.01 | 0.249 | 0.83 | 0.68 | 0.84 | 0.76 | 0.83 | 0.69 |
| 10 | 71559089 | rs2642609 | 0.934 | -0.154 | -1.2 | 0.1 | 0.011 | 0.089 | 0.089 | 0 | 0.29 | 0.13 | 0.16 | 0.05 | 0.25 | 0.18 |
| 10 | 71560367 | rs2642606 | -0.001 | 0.064 | -0.946 | 0.545 | 0.025 | 0.092 | 0.092 | 0 | 0.92 | 0.81 | 0.88 | 0.74 | 0.88 | 0.8 |
| 10 | 71560999 | rs3104051 | -0.806 | 0.122 | -1.008 | 0.571 | -0.009 | -0.009 | -0.009 | 0 | 0.57 | 0.67 | 0.52 | 0.48 | 0.5 | 0.45 |
| 10 | 71564996 | rs4746912 | -0.93 | -0.137 | -0.448 | 0.708 | -0.023 | 0.017 | 0.017 | 4.524 | 0.32 | 0.42 | 0.21 | 0.41 | 0.26 | 0.34 |
| 10 | 71567761 | rs2763336 | 0.283 | -0.05 | -0.48 | 0.532 | -0.006 | 0.019 | 0.019 | 0 | 0.33 | 0.25 | 0.33 | 0.19 | 0.33 | 0.24 |
| 10 | 71568578 | rs2763342 | -0.498 | -0.251 | -0.468 | 0.31 | 0 | 0.001 | 0.001 | 0.225 | 0.29 | 0.26 | 0.18 | 0.24 | 0.29 | 0.29 |
| 10 | 71570363 | rs2763346 | 0.09 | -0.139 | -0.656 | 0.242 | -0.024 | 0.044 | 0.044 | 0 | 0.23 | 0.12 | 0.16 | 0.13 | 0.28 | 0.22 |
| 10 | 71570998 | rs17497293 | 1.375 | -0.033 | -0.678 | 0.068 | -0.01 | 0.014 | 0.014 | 0 | 0.11 | 0.05 | 0.14 | 0.01 | 0.14 | 0.1 |
| 10 | 71571521 | rs2763350 | -0.852 | -0.116 | -0.455 | 0.256 | 0.002 | -0.009 | -0.009 | 0 | 0.87 | 0.9 | 0.89 | 0.76 | 0.88 | 0.79 |
| 10 | 71571792 | rs2683553 | -0.181 | -0.255 | -0.355 | 0.262 | 0.006 | -0.009 | -0.009 | 0 | 0.7 | 0.69 | 0.63 | 0.6 | 0.74 | 0.69 |
| 10 | 71572880 | rs2704495 | 1.302 | -0.114 | -0.442 | 0.19 | -0.01 | 0.017 | 0.017 | 0 | 0.16 | 0.11 | 0.08 | 0.03 | 0.19 | 0.13 |
| 10 | 71573154 | rs7088833 | 0.044 | -0.342 | -0.079 | 0.41 | -0.01 | 0.018 | 0.018 | 0 | 0.25 | 0.36 | 0.32 | 0.29 | 0.21 | 0.17 |
| 10 | 71573945 | rs12768901 | 1.503 | -0.213 | -0.418 | 0.411 | 0.004 | -0.014 | -0.014 | 0.338 | 0.16 | 0.11 | 0.25 | 0.15 | 0.14 | 0.1 |
| 10 | 71575551 | rs2763367 | 0.475 | -0.25 | -0.314 | 0.362 | -0.002 | 0.001 | 0.001 | 0 | 0.59 | 0.53 | 0.56 | 0.46 | 0.64 | 0.67 |
| 10 | 71577319 | rs10998983 | 2.535 | -0.138 | -0.781 | 0.284 | -0.076 | -0.129 | -0.129 | 1.403 | 0.25 | 0.01 | 0.31 | 0.32 | 0.3 | 0.35 |
| 10 | 71577422 | rs2763335 | -1.742 | -0.341 | -0.423 | 0.182 | 0 | 0.003 | 0.003 | 0 | 0.91 | 0.92 | 0.91 | 0.9 | 0.93 | 0.97 |
| 10 | 71578300 | rs1234901 | -0.156 | -0.471 | -0.074 | 0.164 | -0.012 | 0 | 0 | 0.749 | 0.8 | 0.78 | 0.86 | 0.76 | 0.85 | 0.88 |
| 10 | 71578583 | rs10998985 | 1.244 | -0.442 | -0.264 | 0.179 | 0.007 | -0.041 | -0.041 | 0.265 | 0.11 | 0.01 | 0.05 | 0.14 | 0.08 | 0.08 |
| 10 | 71579347 | rs1227773 | 0.787 | -0.485 | -0.112 | 0.242 | 0.028 | 0.001 | 0.001 | 1.334 | 0.57 | 0.37 | 0.57 | 0.58 | 0.67 | 0.73 |
| 10 | 71580120 | rs17497526 | 2.004 | -0.373 | -0.243 | 0.233 | -0.029 | -0.022 | -0.022 | 0 | 0.08 | 0 | 0.08 | 0.11 | 0.1 | 0.03 |
| 10 | 71581783 | rs3847351 | -0.308 | -0.305 | 0.049 | 0.239 | -0.024 | -0.005 | -0.005 | 0.997 | 0.25 | 0.31 | 0.16 | 0.27 | 0.2 | 0.17 |
| 10 | 71582455 | rs2763341 | -1.369 | -0.308 | 0.012 | 0.229 | -0.01 | 0.002 | 0.002 | 0 | 0.82 | 0.7 | 0.72 | 0.85 | 0.85 | 0.9 |
| 10 | 71582996 | rs7076867 | 1.697 | -0.163 | -0.253 | 0.316 | 0.018 | 0.089 | 0.089 | 0 | 0.06 | 0.14 | 0.17 | 0.15 | 0.04 | 0.08 |
| 10 | 71583198 | rs7077164 | -0.329 | -0.039 | -0.394 | 0.271 | -0.016 | -0.004 | -0.004 | 0.287 | 0.42 | 0.49 | 0.42 | 0.41 | 0.32 | 0.26 |
| 10 | 71583757 | rs11598684 | 1.33 | 0.218 | -0.697 | 0.289 | -0.078 | -0.078 | -0.078 | 2.721 | 0.37 | 0.12 | 0.45 | 0.41 | 0.41 | 0.47 |
| 10 | 71588349 | rs7101190 | 1.792 | 0.33 | -0.7 | 0.189 | -0.016 | 0.004 | 0.004 | 0 | 0.25 | 0.13 | 0.29 | 0.26 | 0.34 | 0.3 |
| 10 | 71588504 | rs1227756 | 1.087 | -0.44 | -0.634 | 0.356 | -0.004 | 0.003 | 0.003 | 0.334 | 0.38 | 0.33 | 0.37 | 0.28 | 0.52 | 0.44 |
| 10 | 71588732 | rs10998988 | -0.806 | -0.539 | -0.632 | 0.339 | 0 | -0.002 | -0.002 | 0.157 | 0.28 | 0.26 | 0.31 | 0.33 | 0.22 | 0.18 |
| 10 | 71590887 | rs1227759 | -0.899 | -0.087 | -0.795 | 0.342 | 0.007 | -0.002 | -0.002 | 0 | 0.16 | 0.1 | 0.06 | 0.15 | 0.1 | 0.1 |
| 10 | 71595061 | rs10998990 | 0.056 | 0.214 | -0.998 | 0.529 | -0.023 | -0.047 | -0.047 | 2.038 | 0.08 | 0.01 | 0.14 | 0.19 | 0.11 | 0.06 |
| 10 | 71596040 | rs7079962 | -1.384 | 0.084 | -0.799 | -0.035 | 0.005 | 0.005 | 0.005 | 0 | 0.46 | 0.59 | 0.51 | 0.42 | 0.41 | 0.33 |
| 10 | 71596390 | rs7080527 | -1.354 | -0.008 | -0.533 | -0.175 | -0.006 | 0.01 | 0.01 | 2.856 | 0.29 | 0.46 | 0.41 | 0.28 | 0.27 | 0.16 |
| 10 | 71597997 | rs12357700 | 0.643 | 0.309 | -0.934 | -0.051 | -0.023 | -0.045 | -0.045 | 0 | 0.08 | 0.01 | 0.15 | 0.16 | 0.11 | 0.06 |
| 10 | 71598059 | rs3858154 | -0.987 | 0.207 | -0.499 | -0.116 | -0.015 | -0.004 | -0.004 | 0.184 | 0.26 | 0.34 | 0.18 | 0.25 | 0.19 | 0.24 |
| 10 | 71599042 | rs11593153 | 0.13 | 0.456 | -0.516 | -0.017 | 0.002 | 0.003 | 0.003 | 0 | 0.17 | 0.16 | 0.1 | 0.15 | 0.13 | 0.17 |
| 10 | 71601313 | rs3998144 | 0.768 | 0.262 | -0.655 | -0.258 | -0.001 | 0 | 0 | 0 | 0.49 | 0.51 | 0.39 | 0.54 | 0.46 | 0.62 |
| 10 | 71601764 | rs3886687 | -0.711 | 0.36 | -0.698 | -0.643 | -0.016 | 0.049 | 0.049 | 0.004 | 0.24 | 0.12 | 0.26 | 0.14 | 0.25 | 0.21 |
| 10 | 71603392 | rs11597960 | -0.684 | -0.017 | -0.152 | -0.449 | 0.029 | -0.015 | -0.015 | 0 | 0.06 | 0.29 | 0.05 | 0 | 0.03 | 0.01 |
| 10 | 71604698 | rs10998995 | -0.113 | -0.105 | -1.075 | -0.289 | -0.009 | -0.034 | -0.034 | 0 | 0.15 | 0.01 | 0.13 | 0.14 | 0.13 | 0.12 |
| 10 | 71606416 | rs10998997 | -0.528 | 0.081 | -0.771 | -0.455 | -0.001 | -0.01 | -0.01 | 0 | 0.07 | 0.09 | 0.04 | 0.01 | 0.08 | 0.09 |
| 10 | 71610249 | rs11592327 | 0.324 | -0.856 | -0.368 | -0.832 | -0.052 | -0.067 | -0.067 | 0 | 0.67 | 0.59 | 0.78 | 0.92 | 0.79 | 0.76 |
| 10 | 71613489 | rs2683563 | 0.056 | -0.276 | -0.711 | -0.831 | -0.008 | -0.002 | -0.002 | 0 | 0.39 | 0.33 | 0.46 | 0.6 | 0.45 | 0.54 |
| 10 | 71615124 | rs2683560 | -0.398 | -0.838 | -0.29 | -1.356 | -0.041 | -0.052 | -0.052 | 1.563 | 0.76 | 0.69 | 0.87 | 0.94 | 0.85 | 0.87 |
| 10 | 71617387 | rs3858156 | 0.682 | -0.386 | -1.115 | -1.132 | -0.031 | -0.051 | -0.051 | 0.409 | 0.16 | 0.1 | 0.26 | 0.54 | 0.21 | 0.34 |
| 10 | 71617584 | rs4076957 | 0.712 | -0.388 | -1.116 | -1.133 | -0.03 | -0.049 | -0.049 | 0 | 0.16 | 0.1 | 0.26 | 0.54 | 0.21 | 0.34 |
| 10 | 71618387 | rs2395286 | 0.46 | -0.622 | -1.184 | -1.188 | 0.043 | 0.061 | 0.061 | 0 | 0.23 | 0.1 | 0.12 | 0.07 | 0.17 | 0.14 |
| 10 | 71619166 | rs2395280 | -0.487 | -0.153 | -0.721 | -1.136 | -0.005 | -0.124 | -0.124 | 0 | 0.67 | 0.51 | 0.74 | 0.96 | 0.7 | 0.75 |
| 10 | 71620112 | rs4746922 | -0.383 | 0.158 | -0.905 | -1.002 | -0.006 | -0.077 | -0.077 | 0 | 0.63 | 0.44 | 0.73 | 0.95 | 0.7 | 0.76 |
| 10 | 71620532 | rs12573749 | 0.212 | 0.85 | -1.084 | -0.011 | -0.052 | -0.08 | -0.08 | 0.023 | 0.21 | 0.07 | 0.22 | 0.36 | 0.25 | 0.28 |
| 10 | 71627954 | rs2704526 | 0.627 | 0.149 | 0.03 | -0.609 | 0.007 | -0.042 | -0.042 | 0 | 0.91 | 0.65 | 0.95 | 1 | 0.97 | 0.94 |
| 10 | 71628647 | rs12146165 | -0.187 | -0.264 | -0.6 | -1.155 | 0.017 | 0.05 | 0.05 | 0 | 0.29 | 0.19 | 0.23 | 0.13 | 0.25 | 0.22 |
| 10 | 71632410 | rs4746009 | 0.275 | -0.019 | -1.089 | -1.434 | -0.008 | 0.037 | 0.037 | 0 | 0.22 | 0.12 | 0.17 | 0.07 | 0.22 | 0.08 |
| 10 | 71633256 | rs3998148 | 0.753 | -0.195 | -1.316 | -1.55 | 0.001 | 0.106 | 0.106 | 0 | 0.22 | 0.08 | 0.19 | 0.01 | 0.19 | 0.17 |
| 10 | 71633992 | rs2763359 | 0.633 | 0.005 | -0.38 | -0.907 | 0.004 | -0.001 | -0.001 | 0 | 0.69 | 0.35 | 0.84 | 0.92 | 0.79 | 0.86 |
| 10 | 71634121 | rs2683556 | 0.582 | 0.432 | -0.477 | -0.525 | -0.004 | -0.022 | -0.022 | 0 | 0.71 | 0.61 | 0.78 | 0.8 | 0.72 | 0.86 |
| 10 | 71637986 | rs2683548 | -0.706 | 0.661 | -1.193 | -0.508 | -0.015 | 0.043 | 0.043 | 0 | 0.19 | 0.06 | 0.09 | 0.08 | 0.18 | 0.1 |
| 10 | 71640491 | rs1227783 | -0.858 | 0.59 | -1.255 | -0.819 | -0.026 | 0.057 | 0.057 | 0 | 0.21 | 0.11 | 0.15 | 0.01 | 0.25 | 0.19 |
| 10 | 71640732 | rs999706 | 0.019 | 0.333 | -1.169 | -0.242 | -0.001 | 0.018 | 0.018 | 0 | 0.22 | 0.26 | 0.18 | 0.49 | 0.25 | 0.41 |
| 10 | 71640867 | rs999705 | -0.626 | -0.324 | -0.825 | -0.141 | -0.029 | 0.132 | 0.132 | 6.254 | 0.87 | 0.65 | 0.82 | 0.8 | 0.95 | 0.94 |

| | | | | | | | | | | | | | | | | |
|---|---|---|---|---|---|---|---|---|---|---|---|---|---|---|---|---|
| 10 | 71641240 | rs1227785 | 0.163 | -0.011 | -1.001 | -0.439 | -0.041 | 0.08 | 0.08 | 0 | 0.2 | 0.08 | 0.21 | 0.01 | 0.24 | 0.19 |
| 10 | 71645440 | rs11814083 | 0.476 | -0.013 | -0.861 | 0.007 | -0.001 | 0.075 | 0.075 | 0 | 0.18 | 0.26 | 0.23 | 0.45 | 0.19 | 0.39 |
| 10 | 71647169 | rs11813062 | 0.022 | -0.375 | -1.723 | -0.593 | 0.022 | 0.062 | 0.062 | 0.066 | 0.22 | 0.09 | 0.07 | 0.08 | 0.12 | 0.07 |
| 10 | 71647298 | rs10999011 | 1.909 | -0.132 | -1.723 | -0.573 | -0.012 | 0.012 | 0.012 | 0 | 0.25 | 0.16 | 0.24 | 0.21 | 0.29 | 0.26 |
| 10 | 71647778 | rs12259074 | -0.921 | -0.64 | -1.312 | -0.443 | 0.023 | 0.046 | 0.046 | 0 | 0.29 | 0.21 | 0.11 | 0.14 | 0.21 | 0.12 |
| 10 | 71648316 | rs7100913 | -0.314 | -0.524 | -1.282 | -0.395 | 0.017 | 0.02 | 0.02 | 0 | 0.2 | 0.1 | 0.07 | 0.13 | 0.15 | 0.11 |
| 10 | 71648422 | rs10999012 | -0.512 | -0.193 | -0.904 | -0.245 | 0 | -0.009 | -0.009 | 0 | 0.18 | 0.28 | 0.06 | 0.15 | 0.15 | 0.1 |
| 10 | 71648522 | rs10999014 | -0.389 | -0.335 | -1.183 | -0.333 | 0.021 | 0.028 | 0.028 | 0 | 0.26 | 0.16 | 0.15 | 0.15 | 0.17 | 0.17 |
| 10 | 71650837 | rs2763366 | 0.461 | 0.238 | -0.978 | -0.333 | -0.021 | 0.109 | 0.109 | 0.195 | 0.45 | 0.23 | 0.44 | 0.19 | 0.52 | 0.4 |
| 10 | 71652128 | rs7070915 | 0.285 | -0.267 | -1.111 | -0.312 | -0.024 | 0.098 | 0.098 | 1.567 | 0.48 | 0.25 | 0.3 | 0.32 | 0.52 | 0.39 |
| 10 | 71652501 | rs7071359 | -0.397 | -0.1 | -1.115 | -0.721 | -0.012 | 0.028 | 0.028 | 0 | 0.83 | 0.93 | 0.74 | 0.93 | 0.78 | 0.86 |
| 10 | 71652580 | rs7089663 | 0.357 | -0.319 | -0.707 | -0.205 | -0.008 | 0.036 | 0.036 | 2.286 | 0.31 | 0.54 | 0.4 | 0.4 | 0.22 | 0.32 |
| 10 | 71657040 | rs3812701 | 0.655 | -0.958 | -0.5 | -1.201 | -0.057 | -0.122 | -0.122 | 0 | 0.65 | 0.49 | 0.8 | 0.93 | 0.81 | 0.84 |
| 10 | 71660876 | rs3886895 | 0.038 | -0.483 | -0.553 | -0.895 | -0.03 | -0.091 | -0.091 | 0 | 0.17 | 0.34 | 0.1 | 0 | 0.1 | 0.04 |
| 10 | 71662783 | rs942576 | 0.711 | -0.099 | -1.572 | -0.854 | -0.1 | 0.123 | 0.123 | 0.856 | 0.38 | 0.03 | 0.27 | 0.21 | 0.43 | 0.31 |
| 10 | 71664321 | rs2683576 | 0.766 | -0.29 | -0.037 | -0.185 | -0.047 | -0.127 | -0.127 | 0.286 | 0.78 | 0.53 | 0.88 | 0.98 | 0.87 | 0.89 |
| 10 | 71669953 | rs12412836 | -1.479 | 0.02 | -1.016 | 0.7 | 0.014 | -0.041 | -0.041 | 0.018 | 0.13 | 0.02 | 0.13 | 0.19 | 0.1 | 0.23 |
| 10 | 71670284 | rs3793833 | -1.148 | 0.16 | -1.02 | 0.717 | -0.004 | -0.017 | -0.017 | 0 | 0.05 | 0 | 0.01 | 0.07 | 0.03 | 0.1 |
| 10 | 71671562 | rs2894280 | -1.698 | 0.142 | -0.976 | 1.191 | 0.003 | -0.016 | -0.016 | 0 | 0.07 | 0.02 | 0.06 | 0.16 | 0.05 | 0.1 |
| 10 | 71672879 | rs3793832 | -1.172 | 0.231 | -1.446 | 1.161 | -0.011 | -0.033 | -0.033 | 0.389 | 0.36 | 0.17 | 0.38 | 0.44 | 0.32 | 0.52 |
| 10 | 71673928 | rs7091292 | -0.363 | 0.186 | -1.445 | 0.838 | 0.014 | 0.013 | 0.013 | 0 | 0.16 | 0.11 | 0.06 | 0 | 0.08 | 0.1 |
| 10 | 71675285 | rs16927048 | -1.45 | 0.388 | -1.454 | 0.797 | 0.006 | 0.012 | 0.012 | 0 | 0.22 | 0.16 | 0.13 | 0.17 | 0.13 | 0.18 |
| 10 | 71676400 | rs3793830 | 0.426 | 0.193 | -1.26 | 0.559 | 0 | 0 | 0 | 0 | 0.47 | 0.42 | 0.61 | 0.52 | 0.47 | 0.47 |
| 10 | 71676750 | rs7068393 | -0.335 | 0.104 | -0.921 | 0.505 | -0.003 | -0.026 | -0.026 | 0 | 0.18 | 0.24 | 0.08 | 0.06 | 0.09 | 0.12 |
| 10 | 71679277 | rs3855625 | -0.599 | 0.425 | -0.92 | 0.822 | 0.023 | 0.014 | 0.014 | 0 | 0.67 | 0.48 | 0.72 | 0.7 | 0.58 | 0.66 |
| 10 | 71683024 | rs3793829 | -0.674 | 0.314 | -0.999 | 0.79 | -0.002 | -0.013 | -0.013 | 0 | 0.07 | 0.04 | 0.07 | 0.19 | 0.05 | 0.09 |
| 10 | 71686415 | rs17591282 | -1.258 | 0.521 | -0.925 | 0.6 | 0.002 | -0.009 | -0.009 | 0 | 0.08 | 0.09 | 0.03 | 0.06 | 0.07 | 0.1 |
| 10 | 71687211 | rs1227779 | 0.724 | 0.193 | -0.835 | 0.619 | 0.002 | 0.005 | 0.005 | 0 | 0.6 | 0.72 | 0.64 | 0.51 | 0.69 | 0.47 |
| 10 | 71687492 | rs3793825 | -0.9 | 0.045 | -0.975 | 0.552 | -0.026 | -0.035 | -0.035 | 0 | 0.25 | 0.09 | 0.31 | 0.4 | 0.25 | 0.43 |
| 10 | 71689148 | rs2637229 | 1.488 | 0.178 | -0.714 | 0.473 | -0.003 | 0.005 | 0.005 | 0 | 0.55 | 0.61 | 0.59 | 0.51 | 0.6 | 0.47 |
| 10 | 71689645 | rs2763339 | 0.932 | 0.247 | -0.873 | 0.503 | -0.008 | 0.004 | 0.004 | 0 | 0.64 | 0.76 | 0.63 | 0.58 | 0.69 | 0.57 |
| 10 | 71691569 | rs2683570 | -0.298 | 0.219 | -0.788 | 0.546 | 0 | -0.029 | -0.029 | 0 | 0.85 | 0.81 | 0.92 | 0.91 | 0.9 | 0.9 |
| 10 | 71691585 | rs12241163 | 0.982 | -0.723 | -0.924 | 0.591 | -0.001 | -0.001 | -0.001 | 0 | 0.11 | 0.13 | 0.04 | 0.08 | 0.06 | 0.01 |
| 10 | 71696832 | rs4746933 | -1.2 | -0.53 | -0.748 | 0.361 | 0.006 | 0.004 | 0.004 | 0 | 0.31 | 0.36 | 0.37 | 0.42 | 0.3 | 0.34 |
| 10 | 71698392 | rs3793822 | 0.222 | -0.452 | -0.784 | 0.415 | -0.053 | -0.073 | -0.073 | 0 | 0.12 | 0.01 | 0.26 | 0.33 | 0.17 | 0.24 |
| 10 | 71700200 | rs16927097 | -0.714 | -0.607 | -0.612 | 0.187 | -0.01 | -0.08 | -0.08 | 0.647 | 0.22 | 0.36 | 0.12 | 0.09 | 0.14 | 0.14 |
| 10 | 71701584 | rs3793818 | -0.709 | -0.565 | -0.681 | 0.144 | 0.034 | 0.018 | 0.018 | 0.083 | 0.2 | 0.36 | 0.32 | 0.28 | 0.23 | 0.22 |
| 10 | 71701977 | rs16927103 | -0.595 | -1.216 | -1.298 | -0.141 | 0.05 | 0.118 | 0.118 | 0.534 | 0.27 | 0.09 | 0.1 | 0.07 | 0.11 | 0.23 |
| 10 | 71707415 | rs10999042 | 1.753 | -1.056 | -1.221 | 0.002 | 0 | 0.013 | 0.013 | 0.044 | 0.08 | 0 | 0.02 | 0.06 | 0.03 | 0.12 |
| 10 | 71707687 | rs3793815 | -1.76 | -1.297 | -0.97 | -0.023 | 0.006 | 0.047 | 0.047 | 0.014 | 0.49 | 0.39 | 0.42 | 0.36 | 0.38 | 0.47 |
| 10 | 71709614 | rs10823449 | -0.95 | -1.294 | -0.925 | 0.179 | -0.003 | 0.027 | 0.027 | 0.909 | 0.4 | 0.31 | 0.39 | 0.31 | 0.35 | 0.36 |
| 10 | 71709869 | rs11816683 | 0.198 | -1.239 | -0.957 | 0.173 | -0.026 | -0.029 | -0.029 | 2.979 | 0.17 | 0.08 | 0.26 | 0.22 | 0.21 | 0.21 |
| 10 | 71710262 | rs10762324 | -2.057 | -1.562 | -1.034 | 0.12 | 0.001 | 0.001 | 0.001 | 0 | 0.56 | 0.48 | 0.44 | 0.47 | 0.4 | 0.5 |
| 10 | 71712502 | rs3750779 | -0.238 | -1.07 | -1.394 | 0.243 | -0.045 | 0.02 | 0.02 | 0.17 | 0.22 | 0.06 | 0.25 | 0.19 | 0.24 | 0.29 |
| 10 | 71712896 | rs4746015 | -1.698 | -1.559 | -1.086 | 0.229 | 0.009 | 0.007 | 0.007 | 0.177 | 0.48 | 0.41 | 0.44 | 0.46 | 0.37 | 0.46 |
| 10 | 71713171 | rs10823450 | -0.41 | -1.704 | -1.015 | 0.223 | 0.016 | -0.003 | -0.003 | 0 | 0.27 | 0.23 | 0.15 | 0.25 | 0.15 | 0.1 |
| 10 | 71713924 | rs2394615 | -1.866 | -1.528 | -0.832 | 0.367 | -0.014 | 0.011 | 0.011 | 0 | 0.42 | 0.5 | 0.44 | 0.54 | 0.33 | 0.39 |
| 10 | 71714537 | rs4746937 | -2.394 | -0.245 | -1.206 | 0.395 | 0.084 | 0.091 | 0.091 | 0 | 0.78 | 0.58 | 0.76 | 0.73 | 0.65 | 0.68 |
| 10 | 71714998 | rs4746938 | -0.448 | -0.44 | -1.506 | 0.3 | -0.01 | -0.026 | -0.026 | 0 | 0.25 | 0.19 | 0.37 | 0.39 | 0.29 | 0.23 |
| 10 | 71715359 | rs3750778 | -1.179 | -0.429 | -1.487 | 0.313 | -0.003 | -0.019 | -0.019 | 0 | 0.34 | 0.21 | 0.39 | 0.45 | 0.33 | 0.36 |
| 10 | 71716509 | rs4096395 | 0.34 | -0.08 | -1.569 | 0.509 | -0.049 | -0.04 | -0.04 | 0 | 0.19 | 0.03 | 0.28 | 0.24 | 0.23 | 0.22 |
| 10 | 71720702 | rs10740324 | -1.677 | 0.084 | -0.589 | 0.736 | 0.034 | 0.067 | 0.067 | 0 | 0.78 | 0.66 | 0.7 | 0.64 | 0.68 | 0.67 |
| 10 | 71720900 | rs7072120 | 0.187 | -0.955 | -0.767 | 0.599 | -0.03 | 0 | 0 | 0 | 0.12 | 0.22 | 0.04 | 0.1 | 0.03 | 0.06 |
| 10 | 71725020 | rs3847355 | -1.108 | -0.401 | -0.973 | 0.767 | 0.02 | 0.017 | 0.017 | 0 | 0.62 | 0.47 | 0.56 | 0.54 | 0.47 | 0.48 |
| 10 | 71730948 | rs12570888 | 2.724 | -0.943 | -1.252 | 1.292 | -0.016 | -0.031 | -0.031 | 0 | 0.05 | 0 | 0.05 | 0.12 | 0.04 | 0.06 |
| 10 | 71731851 | rs10509314 | -0.058 | -1.25 | -1.556 | 1.389 | 0.006 | -0.055 | -0.055 | 0 | 0.1 | 0 | 0.14 | 0.22 | 0.09 | 0.15 |
| 10 | 71733321 | rs4109293 | 0.137 | -1.264 | -1.564 | 1.383 | 0 | -0.052 | -0.052 | 0 | 0.09 | 0 | 0.13 | 0.22 | 0.08 | 0.15 |
| 10 | 71736472 | rs10509310 | -1.242 | -1.506 | -1.755 | 1.353 | -0.013 | -0.067 | -0.067 | 0.051 | 0.23 | 0.1 | 0.35 | 0.34 | 0.2 | 0.28 |
| 10 | 71739556 | rs7914288 | -2.219 | -1.832 | -1.666 | 1.098 | 0.015 | 0.01 | 0.01 | 0 | 0.52 | 0.38 | 0.57 | 0.55 | 0.39 | 0.51 |
| 10 | 71740691 | rs3957144 | -0.484 | -1.104 | -1.079 | 1.389 | -0.004 | -0.013 | -0.013 | 0 | 0.16 | 0.1 | 0.12 | 0.14 | 0.13 | 0.2 |
| 10 | 71748962 | rs7897814 | 0.458 | -0.806 | -0.855 | 1.51 | 0 | -0.035 | -0.035 | 0 | 0.09 | 0.04 | 0.23 | 0.2 | 0.08 | 0.09 |
| 10 | 71772019 | rs2394616 | 1.674 | -0.865 | -0.848 | 1.522 | -0.007 | -0.044 | -0.044 | 0 | 0.07 | 0 | 0.21 | 0.22 | 0.04 | 0.07 |
| 10 | 71777660 | rs4746018 | -2.422 | -0.867 | -0.578 | 1.602 | 0.004 | -0.012 | -0.012 | 1.786 | 0.56 | 0.52 | 0.64 | 0.66 | 0.46 | 0.62 |
| 10 | 71790333 | rs7082392 | -0.588 | -0.889 | -0.672 | 1.637 | -0.025 | -0.056 | -0.056 | 0.106 | 0.26 | 0.14 | 0.24 | 0.35 | 0.25 | 0.34 |
| 10 | 71791351 | rs2130791 | -1.208 | -0.908 | -0.646 | 1.624 | -0.022 | -0.047 | -0.047 | 1.925 | 0.33 | 0.17 | 0.46 | 0.57 | 0.29 | 0.41 |
| 10 | 71804649 | rs7899726 | 0.312 | -1.005 | -0.567 | 1.686 | -0.012 | 0.004 | 0.004 | 0 | 0.19 | 0.13 | 0.12 | 0.11 | 0.18 | 0.18 |
| 10 | 71811771 | rs10999108 | 1.304 | -1.086 | -0.497 | 1.524 | 0 | -0.054 | -0.054 | 0 | 0.09 | 0 | 0.22 | 0.22 | 0.04 | 0.07 |
| 10 | 71814832 | rs4746946 | -2.093 | -0.912 | -0.245 | 1.521 | 0 | -0.004 | -0.004 | 0 | 0.53 | 0.53 | 0.64 | 0.66 | 0.46 | 0.64 |
| 10 | 71828021 | rs4746949 | -1.856 | -0.958 | 0.089 | 1.363 | -0.029 | 0.007 | 0.007 | 0.052 | 0.22 | 0.36 | 0.23 | 0.25 | 0.17 | 0.26 |

| | | | | | | | | | | | | | | | | |
|---|---|---|---|---|---|---|---|---|---|---|---|---|---|---|---|---|
| 10 | 71828837 | rs3847356 | -0.685 | -0.917 | 0.218 | 1.383 | 0.006 | 0.073 | 0.073 | 0.184 | 0.1 | 0.2 | 0.16 | 0.25 | 0.12 | 0.19 |
| 10 | 71829100 | rs4385785 | -0.898 | -0.824 | -0.016 | 1.279 | -0.006 | 0 | 0 | 0.968 | 0.1 | 0.1 | 0.22 | 0.12 | 0.05 | 0.08 |
| 10 | 71832430 | rs10999120 | -1.4 | -0.797 | -0.158 | 1.28 | 0.004 | -0.011 | -0.011 | 0 | 0.14 | 0.11 | 0.3 | 0.24 | 0.12 | 0.15 |
| 10 | 71837875 | rs10999122 | -1.454 | -0.672 | 0.259 | 1.341 | -0.007 | -0.026 | -0.026 | 0.812 | 0.37 | 0.5 | 0.34 | 0.29 | 0.33 | 0.39 |
| 10 | 71848434 | rs16927253 | 0.245 | -0.761 | 0.196 | 1.297 | -0.002 | 0.036 | 0.036 | 0.19 | 0.06 | 0.07 | 0.12 | 0.23 | 0.06 | 0.15 |
| 10 | 71849772 | rs16927265 | -0.214 | -0.815 | 0.34 | 1.18 | 0.01 | -0.059 | -0.059 | 0 | 0.21 | 0.31 | 0.15 | 0.06 | 0.2 | 0.18 |
| 10 | 71849936 | rs2271700 | -1.168 | -0.85 | 0.48 | 1.225 | -0.008 | 0.072 | 0.072 | 0.866 | 0.15 | 0.22 | 0.36 | 0.45 | 0.13 | 0.25 |
| 10 | 71850335 | rs7901338 | -1.349 | -0.813 | 0.361 | 1.231 | -0.001 | 0.018 | 0.018 | 1.255 | 0.37 | 0.57 | 0.51 | 0.51 | 0.34 | 0.42 |
| 10 | 71856631 | rs10762337 | 1 | -1.962 | 0.434 | 1.277 | -0.021 | 0.002 | 0.002 | 0 | 0.61 | 0.49 | 0.62 | 0.37 | 0.73 | 0.54 |
| 10 | 71859168 | rs7094103 | -1.532 | -1.946 | 0.394 | 0.944 | -0.004 | 0.072 | 0.072 | 1.715 | 0.16 | 0.26 | 0.37 | 0.43 | 0.17 | 0.25 |
| 10 | 71859747 | rs3750768 | -0.051 | -2.158 | 0.271 | 0.901 | -0.037 | -0.01 | -0.01 | 0.175 | 0.72 | 0.63 | 0.87 | 0.82 | 0.88 | 0.77 |
| 10 | 71859953 | rs3750770 | 0.137 | -2.082 | 0.112 | 0.933 | -0.019 | -0.038 | -0.038 | 0.464 | 0.06 | 0 | 0.14 | 0.18 | 0.08 | 0.14 |
| 10 | 71861626 | rs4746960 | 1.785 | -2.224 | 0.113 | 0.73 | 0.026 | -0.015 | -0.015 | 0.214 | 0.42 | 0.23 | 0.46 | 0.41 | 0.64 | 0.47 |
| 10 | 71862654 | rs7914787 | -0.756 | -2.277 | 0.26 | 1.006 | -0.063 | 0.02 | 0.02 | 0 | 0.32 | 0.48 | 0.32 | 0.37 | 0.14 | 0.29 |
| 10 | 71866161 | rs7917290 | -1.03 | -2.144 | 0.29 | 1.04 | -0.033 | 0.013 | 0.013 | 0 | 0.34 | 0.43 | 0.23 | 0.41 | 0.2 | 0.36 |
| 10 | 71867471 | rs7905431 | -0.366 | -2.13 | 0.273 | 1.003 | -0.052 | -0.019 | -0.019 | 0 | 0.27 | 0.4 | 0.08 | 0.19 | 0.11 | 0.22 |
| 10 | 71868616 | rs2271699 | -0.276 | -1.755 | 0.066 | 1.213 | -0.001 | 0.008 | 0.008 | 1.009 | 0.07 | 0.07 | 0.15 | 0.24 | 0.09 | 0.14 |
| 10 | 71868763 | rs2271698 | 1.492 | -1.798 | -0.003 | 1.178 | 0.058 | -0.006 | -0.006 | 1.055 | 0.44 | 0.14 | 0.5 | 0.37 | 0.7 | 0.51 |
| 10 | 71870114 | rs2394654 | -0.336 | -1.7 | 0.121 | 1.263 | -0.004 | -0.01 | -0.01 | 9.602 | 0.07 | 0.04 | 0.15 | 0.23 | 0.09 | 0.14 |
| 10 | 71872572 | rs10823484 | -0.087 | -1.846 | 0.312 | 1.131 | 0.002 | -0.003 | -0.003 | 0 | 0.28 | 0.3 | 0.24 | 0.39 | 0.11 | 0.29 |
| 10 | 71874048 | rs1053495 | 0.075 | -1.815 | 0.278 | 1.078 | -0.001 | 0.001 | 0.001 | 0.23 | 0.26 | 0.29 | 0.07 | 0.18 | 0.1 | 0.24 |
| 10 | 71874683 | rs2271696 | -0.044 | -1.764 | 0.212 | 1.084 | -0.009 | 0.033 | 0.033 | 0 | 0.09 | 0.21 | 0.23 | 0.17 | 0.06 | 0.04 |
| 10 | 71874734 | rs2271695 | -0.044 | -1.764 | 0.212 | 1.084 | -0.009 | 0.033 | 0.033 | 0 | 0.09 | 0.21 | 0.23 | 0.17 | 0.06 | 0.04 |
| 10 | 71876382 | rs2394656 | 0.377 | -1.124 | -0.558 | 1.007 | 0.037 | 0.026 | 0.026 | 1.93 | 0.56 | 0.37 | 0.66 | 0.64 | 0.77 | 0.65 |
| 10 | 71876819 | rs6480440 | -0.697 | -0.821 | -1.116 | 1.076 | 0.069 | 0.01 | 0.01 | 0.002 | 0.46 | 0.73 | 0.42 | 0.63 | 0.26 | 0.45 |
| 10 | 71881150 | rs3750772 | -1.514 | -0.304 | -0.911 | 1.297 | 0.009 | -0.011 | -0.011 | 0 | 0.09 | 0.08 | 0.2 | 0.31 | 0.05 | 0.12 |
| 10 | 71890630 | rs2394644 | -1.485 | -0.678 | -0.778 | 0.987 | 0.002 | -0.002 | -0.002 | 0.636 | 0.19 | 0.13 | 0.09 | 0.24 | 0.12 | 0.16 |
| 10 | 71895746 | rs4746965 | -0.484 | -0.155 | -0.963 | -0.382 | -0.012 | -0.023 | -0.023 | 0.014 | 0.72 | 0.75 | 0.75 | 1 | 0.62 | 0.67 |
| 10 | 71895781 | rs4746966 | 1.082 | -0.934 | -1.185 | 0.195 | -0.041 | -0.056 | -0.056 | 0 | 0.43 | 0.1 | 0.61 | 0.51 | 0.45 | 0.44 |
| 10 | 71895804 | rs10823485 | -0.891 | -1.033 | -0.474 | 0.276 | -0.01 | 0.016 | 0.016 | 0 | 0.22 | 0.21 | 0.09 | 0.47 | 0.15 | 0.22 |
| 10 | 71896576 | rs10823486 | -0.951 | -1.107 | -0.498 | 0.298 | -0.011 | 0.033 | 0.033 | 0 | 0.12 | 0.08 | 0.03 | 0.39 | 0.06 | 0.1 |
| 10 | 71900189 | rs4746969 | 0.737 | -1.328 | -0.057 | 0.144 | -0.009 | -0.001 | -0.001 | 0.114 | 0.78 | 0.8 | 0.79 | 0.77 | 0.87 | 0.82 |
| 10 | 71900395 | rs10823487 | 1.289 | -1.277 | -0.717 | -0.109 | -0.07 | -0.087 | -0.087 | 0.039 | 0.37 | 0.06 | 0.5 | 0.53 | 0.4 | 0.41 |
| 10 | 71905569 | rs3750774 | 1.188 | -1.253 | -0.32 | -0.126 | -0.044 | -0.045 | -0.045 | 0 | 0.37 | 0.17 | 0.51 | 0.53 | 0.41 | 0.42 |
| 10 | 71910630 | rs1046747 | 0.631 | -1.657 | -0.519 | -0.485 | -0.115 | 0.174 | 0.174 | 0 | 0.29 | 0.09 | 0.24 | 0 | 0.43 | 0.31 |
| 10 | 71910663 | rs15801 | 0.154 | -1.658 | 0.081 | -0.477 | -0.008 | -0.037 | -0.037 | 0 | 0.36 | 0.35 | 0.28 | 0 | 0.44 | 0.33 |
| 10 | 71910888 | rs7919647 | -0.829 | -1.744 | 0.014 | -0.471 | 0.032 | 0.086 | 0.086 | 0 | 0.86 | 0.98 | 0.96 | 1 | 0.93 | 0.93 |
| 10 | 71915507 | rs10762340 | 0.886 | -1.765 | 0.166 | -0.427 | -0.082 | 0.056 | 0.056 | 0 | 0.74 | 0.54 | 0.8 | 0.59 | 0.89 | 0.78 |
| 10 | 71921591 | rs2271690 | 2.267 | -2.145 | -0.581 | -0.42 | -0.075 | -0.141 | -0.141 | 0.005 | 0.24 | 0.03 | 0.4 | 0.42 | 0.31 | 0.19 |
| 10 | 71922900 | rs7094581 | -1.084 | -2.131 | -0.309 | -0.597 | 0.035 | 0.044 | 0.044 | 0 | 0.86 | 0.94 | 0.95 | 0.99 | 0.93 | 0.93 |
| 10 | 71926288 | rs7093644 | -0.12 | -2.086 | -0.047 | -0.565 | -0.045 | 0 | 0 | 0 | 0.79 | 0.71 | 0.83 | 0.76 | 0.91 | 0.85 |
| 10 | 71931012 | rs10999169 | 1.013 | -1.871 | -0.277 | -0.513 | -0.046 | -0.047 | -0.047 | 0 | 0.37 | 0.16 | 0.51 | 0.53 | 0.41 | 0.42 |
| 10 | 71936341 | rs10999170 | 1.489 | -1.836 | -0.279 | -0.56 | -0.038 | -0.056 | -0.056 | 2.056 | 0.3 | 0.16 | 0.48 | 0.53 | 0.38 | 0.38 |
| 10 | 71939374 | rs10762345 | 0.345 | -1.737 | -0.122 | -0.854 | 0.022 | -0.163 | -0.163 | 0 | 0.34 | 0.51 | 0.26 | 0 | 0.4 | 0.3 |
| 10 | 71948787 | rs7905657 | 0.595 | -1.78 | -0.044 | -0.754 | 0.02 | -0.007 | -0.007 | 0 | 0.63 | 0.71 | 0.74 | 0.54 | 0.77 | 0.69 |
| 10 | 71952447 | rs6480445 | 0.972 | -1.833 | -0.079 | -0.823 | -0.006 | 0.002 | 0.002 | 0 | 0.38 | 0.65 | 0.55 | 0.6 | 0.43 | 0.45 |
| 10 | 71959948 | rs12412099 | 1.473 | -1.916 | 0.154 | -0.864 | -0.022 | 0.029 | 0.029 | 0 | 0.33 | 0.15 | 0.49 | 0.71 | 0.38 | 0.39 |
| 10 | 71963608 | rs12355080 | 1.227 | -1.933 | 0.057 | -0.915 | -0.099 | 0.149 | 0.149 | 0 | 0.26 | 0.07 | 0.2 | 0 | 0.37 | 0.3 |
| 10 | 71965793 | rs10999185 | 1.029 | -1.949 | 0.386 | -0.899 | 0.022 | 0.005 | 0.005 | 0.035 | 0.06 | 0.25 | 0.04 | 0 | 0.02 | 0.01 |
| 10 | 71966819 | rs9415004 | 0.691 | -2.026 | 0.45 | -0.928 | -0.045 | -0.048 | -0.048 | 0 | 0.62 | 0.55 | 0.73 | 0.77 | 0.8 | 0.77 |
| 10 | 71978144 | rs7072230 | 1.224 | -1.941 | 0.339 | -0.937 | -0.095 | 0.144 | 0.144 | 0 | 0.26 | 0.07 | 0.19 | 0 | 0.37 | 0.3 |
| 10 | 71979635 | rs10762351 | 0.046 | -2.117 | 0.444 | -0.992 | -0.018 | -0.089 | -0.089 | 0 | 0.76 | 0.62 | 0.88 | 1 | 0.85 | 0.87 |
| 10 | 71983630 | rs12262852 | 1.381 | -2.116 | 0.44 | -0.987 | -0.001 | -0.015 | -0.015 | 0 | 0.05 | 0.18 | 0.04 | 0 | 0.02 | 0.01 |
| 10 | 71984085 | rs9284077 | -0.869 | -2.097 | 0.372 | -0.98 | 0.002 | 0.112 | 0.112 | 0 | 0.82 | 0.94 | 0.93 | 1 | 0.88 | 0.88 |
| 10 | 71986507 | rs7086158 | -0.777 | -2.1 | 0.414 | -0.982 | 0.016 | 0.023 | 0.023 | 0 | 0.82 | 0.87 | 0.93 | 1 | 0.88 | 0.88 |
| 10 | 71989306 | rs7074777 | 1.953 | -1.998 | 0.271 | -0.642 | -0.049 | -0.094 | -0.094 | 0 | 0.29 | 0.07 | 0.48 | 0.49 | 0.37 | 0.37 |
| 10 | 71992518 | rs7073334 | 0.556 | -1.979 | 0.355 | -0.901 | -0.032 | -0.037 | -0.037 | 4.102 | 0.64 | 0.45 | 0.75 | 0.77 | 0.8 | 0.78 |
| 10 | 72004454 | rs7087773 | 0.771 | -1.918 | 0.289 | -1.025 | 0.072 | 0.109 | 0.109 | 0 | 0.58 | 0.24 | 0.82 | 0.89 | 0.79 | 0.77 |
| 10 | 72005190 | rs10823500 | 0.783 | -1.393 | 0.37 | -0.616 | -0.078 | 0.092 | 0.092 | 0 | 0.26 | 0.1 | 0.2 | 0.11 | 0.4 | 0.37 |
| 10 | 72006502 | rs7097067 | 0.693 | -1.489 | 0.34 | -0.699 | -0.012 | 0 | 0 | 0 | 0.1 | 0.13 | 0.03 | 0.1 | 0.05 | 0.08 |
| 10 | 72007149 | rs12246360 | 0.863 | -1.475 | 0.344 | -0.675 | 0.012 | -0.004 | -0.004 | 0.102 | 0.06 | 0.21 | 0.04 | 0 | 0.02 | 0.01 |
| 10 | 72007306 | rs10999204 | -0.088 | -1.352 | 0.031 | -0.63 | -0.003 | -0.011 | -0.011 | 0 | 0.37 | 0.31 | 0.23 | 0.11 | 0.41 | 0.38 |
| 10 | 72012813 | rs7923688 | -0.092 | -0.613 | -0.021 | -0.64 | 0.005 | -0.024 | -0.024 | 0 | 0.29 | 0.3 | 0.2 | 0.16 | 0.35 | 0.39 |
| 10 | 72015573 | rs3812694 | -0.042 | -0.82 | 0.312 | -0.391 | -0.03 | -0.021 | -0.021 | 0.016 | 0.15 | 0.3 | 0.06 | 0.11 | 0.07 | 0.09 |
| 10 | 72018243 | rs6480448 | 1.156 | -0.54 | -0.226 | -0.13 | 0.001 | 0.001 | 0.001 | 0.069 | 0.11 | 0.09 | 0.04 | 0.08 | 0.08 | 0.03 |
| 10 | 72024537 | rs12355480 | 0.413 | -0.061 | -1.111 | -0.103 | -0.094 | -0.018 | -0.018 | 0 | 0.23 | 0.02 | 0.29 | 0.23 | 0.32 | 0.48 |
| 10 | 72025954 | rs3812696 | 1.585 | -0.191 | -0.896 | 0.11 | -0.006 | -0.021 | -0.021 | 0.001 | 0.1 | 0.03 | 0.04 | 0.13 | 0.08 | 0.03 |
| 10 | 72033298 | rs2184331 | -0.626 | -0.209 | -1.022 | -0.143 | 0.024 | 0.015 | 0.015 | 0 | 0.81 | 0.89 | 0.93 | 1 | 0.91 | 0.9 |
| 10 | 72035056 | rs10823505 | 1.417 | -0.1 | -0.734 | 0.063 | 0.005 | 0.005 | 0.005 | 0 | 0.42 | 0.43 | 0.39 | 0.42 | 0.45 | 0.32 |

| | | | | | | | | | | | | | | | | |
|---|---|---|---|---|---|---|---|---|---|---|---|---|---|---|---|---|
| 10 | 72035901 | rs10999231 | 0.585 | 0.2 | -1.181 | 0.089 | -0.093 | -0.053 | -0.053 | 0 | 0.22 | 0.02 | 0.3 | 0.26 | 0.32 | 0.48 |
| 10 | 72038399 | rs868075 | 0.041 | 0.026 | -0.727 | 0.105 | -0.01 | -0.011 | -0.011 | 0 | 0.21 | 0.23 | 0.24 | 0.2 | 0.14 | 0.16 |
| 10 | 72044281 | rs7075666 | 0.459 | 0.43 | -0.784 | 0.284 | 0.002 | 0.002 | 0.002 | 0 | 0.51 | 0.48 | 0.39 | 0.34 | 0.44 | 0.36 |
| 10 | 72045332 | rs10823512 | 0.301 | 0.539 | -1.343 | 0.205 | -0.092 | -0.049 | -0.049 | 0 | 0.22 | 0.02 | 0.31 | 0.26 | 0.33 | 0.49 |
| 10 | 72047013 | rs16913394 | 0.657 | -0.061 | -1.034 | -0.094 | -0.022 | -0.062 | -0.062 | 0.041 | 0.15 | 0.23 | 0.06 | 0 | 0.07 | 0.06 |
| 10 | 72048894 | rs12780221 | 0.58 | 0.575 | -1.586 | 0.155 | -0.109 | 0.012 | 0.012 | 0.018 | 0.19 | 0.02 | 0.16 | 0.12 | 0.33 | 0.33 |
| 10 | 72049765 | rs16927437 | 0.382 | -0.032 | -1.092 | 0.573 | 0.02 | -0.025 | -0.025 | 0 | 0.18 | 0.09 | 0.21 | 0.26 | 0.12 | 0.26 |
| 10 | 72052169 | rs10823513 | -0.524 | -0.132 | -0.595 | 0.488 | -0.002 | -0.001 | -0.001 | 0 | 0.39 | 0.41 | 0.36 | 0.6 | 0.28 | 0.45 |
| 10 | 72052480 | rs10762358 | 0.91 | -0.215 | -0.748 | -0.14 | 0 | 0.007 | 0.007 | 0 | 0.37 | 0.37 | 0.46 | 0.44 | 0.41 | 0.35 |
| 10 | 72055156 | rs4746989 | 1.563 | -0.075 | -1.058 | -0.011 | -0.038 | -0.091 | -0.091 | 0 | 0.27 | 0.09 | 0.41 | 0.41 | 0.35 | 0.27 |
| 10 | 72057163 | rs4746992 | -0.745 | -0.339 | -0.49 | -0.095 | 0.004 | -0.005 | -0.005 | 0 | 0.63 | 0.66 | 0.52 | 0.43 | 0.76 | 0.64 |
| 10 | 72058253 | rs4746996 | 1.475 | -0.169 | -0.914 | 0.182 | -0.04 | -0.06 | -0.06 | 0 | 0.27 | 0.09 | 0.29 | 0.32 | 0.35 | 0.25 |
| 10 | 72058534 | rs7094128 | 0.25 | -0.132 | -0.481 | -0.322 | -0.002 | -0.008 | -0.008 | 0 | 0.11 | 0.17 | 0.06 | 0.1 | 0.1 | 0.16 |
| 10 | 72064976 | rs10823515 | 0.294 | -0.525 | -0.675 | -0.244 | 0.03 | -0.008 | -0.008 | 0.059 | 0.33 | 0.19 | 0.29 | 0.34 | 0.2 | 0.29 |
| 10 | 72071648 | rs3812699 | 0.624 | -0.426 | -0.751 | -0.545 | -0.016 | 0 | 0 | 0 | 0.58 | 0.28 | 0.59 | 0.53 | 0.49 | 0.36 |
| 10 | 72073177 | rs10823520 | -0.828 | -0.774 | -0.134 | -0.763 | 0.003 | -0.006 | -0.006 | 0 | 0.59 | 0.64 | 0.53 | 0.49 | 0.62 | 0.64 |
| 10 | 72074693 | rs7086666 | -0.605 | -0.585 | -0.128 | -0.715 | 0.035 | 0.012 | 0.012 | 0.247 | 0.29 | 0.45 | 0.34 | 0.35 | 0.42 | 0.46 |
| 10 | 72079394 | rs12782697 | 0.352 | -0.175 | -0.178 | -0.816 | 0.005 | 0.008 | 0.008 | 0 | 0.19 | 0.17 | 0.08 | 0.04 | 0.14 | 0.19 |
| 10 | 72081023 | rs10762364 | 1.933 | -0.139 | -0.405 | -0.79 | -0.033 | -0.092 | -0.092 | 0 | 0.27 | 0.1 | 0.41 | 0.47 | 0.33 | 0.22 |
| 10 | 72083835 | rs3750764 | 2.464 | -0.529 | -0.541 | -1.22 | 0.023 | 0.04 | 0.04 | 4.99 | 0.17 | 0.04 | 0.11 | 0.06 | 0.08 | 0.06 |
| 10 | 72084299 | rs3861040 | 0.409 | -0.256 | 0.126 | -1.118 | 0.012 | -0.038 | -0.038 | 0.06 | 0.14 | 0.16 | 0.15 | 0 | 0.27 | 0.12 |
| 10 | 72084377 | rs10999268 | 1.138 | -0.445 | 0.145 | -1.13 | 0.026 | 0.052 | 0.052 | 4.301 | 0.29 | 0.16 | 0.17 | 0.06 | 0.16 | 0.12 |
| 10 | 72088846 | rs4747001 | -0.614 | -0.351 | -0.013 | -1.114 | 0.003 | 0 | 0 | 0 | 0.54 | 0.61 | 0.5 | 0.49 | 0.55 | 0.58 |
| 10 | 72093502 | rs10999275 | 1.656 | -0.318 | -0.014 | -1.1 | -0.009 | -0.049 | -0.049 | 0 | 0.27 | 0.16 | 0.42 | 0.47 | 0.32 | 0.23 |
| 10 | 72102829 | rs4747007 | -0.368 | -0.297 | 0.024 | -1.101 | -0.003 | -0.01 | -0.01 | 3.157 | 0.8 | 0.76 | 0.9 | 0.87 | 0.82 | 0.74 |
| 10 | 72104142 | rs4746028 | -0.128 | -0.389 | 0.235 | -1.165 | 0.008 | 0.027 | 0.027 | 0 | 0.5 | 0.34 | 0.46 | 0.39 | 0.51 | 0.43 |
| 10 | 72107611 | rs7100300 | 0.333 | -0.429 | 0.2 | -1.242 | 0.021 | -0.008 | -0.008 | 0 | 0.43 | 0.25 | 0.58 | 0.55 | 0.65 | 0.42 |
| 10 | 72108175 | rs6480454 | -0.737 | -0.382 | 0.237 | -1.198 | 0.052 | 0.069 | 0.069 | 0 | 0.05 | 0.32 | 0.04 | 0.09 | 0.05 | 0.14 |
| 10 | 72108217 | rs12412698 | 1.334 | -0.436 | -0.051 | -1.255 | 0.028 | 0.071 | 0.071 | 4.572 | 0.28 | 0.12 | 0.18 | 0.06 | 0.16 | 0.12 |
| 10 | 72109482 | rs10762368 | 0.723 | -0.426 | 0.035 | -0.952 | 0.004 | 0.028 | 0.028 | 0 | 0.32 | 0.46 | 0.46 | 0.57 | 0.37 | 0.33 |
| 10 | 72111848 | rs10509322 | 0.649 | -0.563 | -0.036 | -0.712 | -0.021 | 0.015 | 0.015 | 0 | 0.73 | 0.64 | 0.78 | 0.63 | 0.81 | 0.54 |
| 10 | 72112232 | rs10999287 | 0.334 | -0.569 | -0.067 | -0.703 | 0.002 | 0.038 | 0.038 | 0 | 0.07 | 0.05 | 0.13 | 0.31 | 0.06 | 0.19 |
| 10 | 72112825 | rs1078119 | 0.513 | -0.505 | -0.063 | -0.708 | -0.019 | 0.014 | 0.014 | 0 | 0.73 | 0.65 | 0.79 | 0.63 | 0.81 | 0.54 |
| 10 | 72114219 | rs4747010 | 0.559 | -0.224 | 0.242 | -0.991 | -0.012 | -0.048 | -0.048 | 0.293 | 0.69 | 0.55 | 0.84 | 0.87 | 0.76 | 0.57 |
| 10 | 72114245 | rs10999290 | 0.738 | 0.077 | 0.574 | -1.425 | 0 | -0.048 | -0.048 | 0 | 0.36 | 0.35 | 0.31 | 0.07 | 0.37 | 0.14 |
| 10 | 72115839 | rs10823528 | 0.312 | -0.285 | 0.726 | -1.044 | 0 | -0.004 | -0.004 | 0 | 0.1 | 0.1 | 0.05 | 0.06 | 0.07 | 0.17 |
| 10 | 72118655 | rs10823529 | 1.05 | -0.542 | 0.596 | -1.075 | 0.001 | 0 | 0 | 1.281 | 0.42 | 0.32 | 0.57 | 0.54 | 0.61 | 0.35 |
| 10 | 72121024 | rs1904592 | 0.212 | -0.393 | 0.163 | -0.961 | 0.022 | 0.011 | 0.011 | 3.34 | 0.61 | 0.67 | 0.66 | 0.64 | 0.72 | 0.63 |
| 10 | 72131190 | rs7350394 | 0.12 | -0.407 | 0.037 | -0.952 | 0.04 | 0.025 | 0.025 | 4.014 | 0.62 | 0.74 | 0.68 | 0.68 | 0.76 | 0.66 |
| 10 | 72131939 | rs4747019 | -0.072 | -0.402 | -0.198 | -0.916 | 0.016 | -0.006 | -0.006 | 0.262 | 0.19 | 0.1 | 0.2 | 0.24 | 0.15 | 0.32 |
| 10 | 72136000 | rs10999304 | 1.632 | -0.269 | -0.129 | -0.763 | 0.034 | 0.037 | 0.037 | 4.688 | 0.16 | 0.02 | 0.1 | 0.08 | 0.08 | 0.01 |
| 10 | 72137257 | rs7084009 | 0.72 | 0.499 | 0.731 | -0.461 | 0.003 | 0.005 | 0.005 | 0 | 0.76 | 0.64 | 0.75 | 0.71 | 0.76 | 0.64 |
| 10 | 72151884 | rs4607991 | 0.537 | -0.174 | 0.631 | -0.455 | -0.006 | 0.007 | 0.007 | 0 | 0.76 | 0.66 | 0.69 | 0.72 | 0.79 | 0.66 |
| 10 | 72163698 | rs3812706 | 0.75 | 0.037 | 0.302 | -0.701 | 0.002 | 0.053 | 0.053 | 0.621 | 0.48 | 0.3 | 0.44 | 0.68 | 0.46 | 0.48 |
| 10 | 72169625 | rs7918386 | 0.821 | 0.048 | 0.602 | -0.764 | -0.006 | 0.005 | 0.005 | 0.232 | 0.2 | 0.19 | 0.19 | 0.03 | 0.26 | 0.13 |
| 10 | 72172992 | rs7078987 | 1.627 | -0.123 | 0.492 | -0.81 | -0.069 | -0.047 | -0.047 | 0.194 | 0.37 | 0.07 | 0.41 | 0.64 | 0.41 | 0.25 |
| 10 | 72177911 | rs11597457 | 0.745 | -0.097 | 0.548 | -0.758 | -0.021 | -0.018 | -0.018 | 11.089 | 0.1 | 0.15 | 0.04 | 0.05 | 0.05 | 0.23 |
| 10 | 72181460 | rs3793809 | -0.422 | 0.716 | 0.661 | -0.429 | 0.01 | -0.053 | -0.053 | 0 | 0.78 | 0.7 | 0.79 | 0.97 | 0.73 | 0.87 |
| 10 | 72182030 | rs4746035 | 0.318 | 0.746 | 0.674 | -0.417 | -0.001 | 0.024 | 0.024 | 0 | 0.07 | 0.08 | 0.21 | 0.23 | 0.06 | 0.2 |
| 10 | 72190589 | rs4307642 | -0.013 | 0.51 | 0.714 | -0.384 | 0 | 0.015 | 0.015 | 0 | 0.07 | 0.09 | 0.21 | 0.23 | 0.07 | 0.21 |
| 10 | 72191952 | rs2279253 | 1.217 | 0.485 | 0.777 | -0.394 | 0.005 | 0.053 | 0.053 | 0.433 | 0.49 | 0.34 | 0.46 | 0.73 | 0.47 | 0.49 |
| 10 | 72193854 | rs2231947 | 0.249 | 0.586 | 0.782 | -0.361 | 0.001 | -0.042 | -0.042 | 0.02 | 0.21 | 0.19 | 0.11 | 0.01 | 0.2 | 0.18 |
| 10 | 72195439 | rs1904589 | 0.583 | 0.536 | 0.674 | -0.336 | 0.003 | -0.056 | -0.056 | 0 | 0.57 | 0.51 | 0.68 | 0.96 | 0.53 | 0.7 |
| 10 | 72204961 | rs7072552 | -0.271 | 0.641 | 0.238 | -0.243 | -0.011 | 0.088 | 0.088 | 0 | 0.91 | 0.8 | 0.77 | 0.77 | 0.94 | 0.8 |
| 10 | 72206344 | rs10762381 | 0.943 | 0.742 | -0.359 | -0.301 | 0.004 | 0.057 | 0.057 | 0.623 | 0.49 | 0.26 | 0.45 | 0.68 | 0.47 | 0.49 |
| 10 | 72210686 | rs17512976 | -0.713 | 0.29 | -0.342 | 0.041 | 0.001 | -0.043 | -0.043 | 0 | 0.21 | 0.22 | 0.12 | 0.01 | 0.2 | 0.18 |
| 10 | 72216011 | rs7082255 | 0.522 | 0.306 | -0.592 | 0.41 | -0.002 | 0.002 | 0.002 | 0.282 | 0.67 | 0.58 | 0.66 | 0.66 | 0.69 | 0.6 |
| 10 | 72219918 | rs1106887 | -0.116 | -0.139 | -1.626 | 0.394 | -0.047 | -0.046 | -0.046 | 0 | 0.33 | 0.08 | 0.43 | 0.32 | 0.58 | 0.27 |
| 10 | 72225536 | rs7923262 | 0.191 | -1.399 | -1.115 | -0.337 | 0.005 | -0.006 | -0.006 | 0 | 0.46 | 0.43 | 0.28 | 0.15 | 0.35 | 0.54 |
| 10 | 72227693 | rs10823548 | 0.135 | -0.905 | -0.788 | 0.445 | 0.006 | 0.124 | 0.124 | 0 | 0.06 | 0.07 | 0.23 | 0.31 | 0.06 | 0.08 |
| 10 | 72229173 | rs2894123 | -0.388 | -1.328 | -0.524 | 0.043 | -0.038 | -0.054 | -0.054 | 0 | 0.89 | 0.83 | 0.97 | 1 | 0.96 | 0.95 |
| 10 | 72236229 | rs1871598 | 0.263 | -1.73 | -0.744 | -0.205 | -0.078 | -0.102 | -0.102 | 1.224 | 0.74 | 0.57 | 0.85 | 0.92 | 0.88 | 0.65 |
| 10 | 72237693 | rs7922219 | 0.344 | -1.682 | -0.75 | -0.206 | -0.065 | -0.099 | -0.099 | 0 | 0.73 | 0.57 | 0.85 | 0.92 | 0.86 | 0.65 |
| 10 | 72240614 | rs11818000 | 0.342 | -1.667 | -0.743 | -0.215 | -0.065 | -0.102 | -0.102 | 0.012 | 0.73 | 0.58 | 0.85 | 0.92 | 0.86 | 0.65 |
| 10 | 72242374 | rs7920307 | 0.664 | -1.708 | -1.011 | 0.023 | -0.055 | 0.054 | 0.054 | 0 | 0.67 | 0.49 | 0.68 | 0.41 | 0.8 | 0.53 |
| 10 | 72251382 | rs7914811 | -0.007 | -0.912 | -1.085 | 0.186 | 0.008 | -0.003 | -0.003 | 0 | 0.07 | 0.18 | 0.08 | 0.06 | 0.08 | 0.21 |
| 10 | 72252825 | rs10762385 | 0.11 | -1.373 | -1.038 | -0.071 | -0.001 | 0.009 | 0.009 | 0 | 0.17 | 0.24 | 0.29 | 0.32 | 0.15 | 0.22 |
| 10 | 72254808 | rs4747039 | -0.129 | -1.287 | -0.979 | -0.845 | 0.023 | 0.005 | 0.005 | 0 | 0.56 | 0.45 | 0.79 | 0.96 | 0.77 | 0.69 |
| 10 | 72258307 | rs12571825 | -0.118 | 0.062 | -1.164 | -0.661 | -0.002 | 0.065 | 0.065 | 0 | 0.56 | 0.65 | 0.61 | 0.81 | 0.6 | 0.66 |

| | | | | | | | | | | | | | | | | |
|---|---|---|---|---|---|---|---|---|---|---|---|---|---|---|---|---|
| 10 | 72259995 | rs11818284 | -0.149 | 0.405 | -1.04 | -1.077 | 0 | -0.008 | -0.008 | 0 | 0.14 | 0.09 | 0.07 | 0.01 | 0.11 | 0.17 |
| 10 | 72261519 | rs10443943 | 0.37 | -0.012 | -0.321 | -1.466 | -0.014 | -0.067 | -0.067 | 0 | 0.86 | 0.46 | 0.92 | 0.96 | 0.93 | 0.91 |
| 10 | 72263690 | rs4747042 | 0.897 | 0.534 | -0.122 | -0.978 | -0.015 | 0.016 | 0.016 | 1.577 | 0.57 | 0.35 | 0.62 | 0.69 | 0.5 | 0.56 |
| 10 | 72265436 | rs10823556 | 1.127 | 0.748 | -0.743 | -1.245 | 0.011 | -0.055 | -0.055 | 0 | 0.44 | 0.13 | 0.33 | 0.51 | 0.36 | 0.43 |
| 10 | 72272831 | rs7073806 | -1.057 | 0.599 | -0.042 | -1.218 | 0.011 | -0.06 | -0.06 | 0 | 0.9 | 0.72 | 0.95 | 1 | 0.94 | 0.92 |
| 10 | 72273380 | rs4746047 | 0.091 | 0.757 | -0.103 | -1.247 | 0.022 | 0.059 | 0.059 | 0.092 | 0.78 | 0.62 | 0.75 | 0.71 | 0.7 | 0.79 |
| 10 | 72273552 | rs4747046 | 0.462 | 0.704 | -0.517 | -1.148 | -0.006 | 0.024 | 0.024 | 2.261 | 0.58 | 0.17 | 0.63 | 0.7 | 0.58 | 0.57 |
| 10 | 72289244 | rs4746052 | -0.632 | 0.494 | -0.023 | -1.156 | -0.005 | -0.077 | -0.077 | 0 | 0.87 | 0.7 | 0.95 | 1 | 0.94 | 0.92 |
| 10 | 72289701 | rs2275059 | 1.174 | 0.127 | -0.739 | -1.409 | -0.032 | -0.137 | -0.137 | 0 | 0.17 | 0.02 | 0.2 | 0.5 | 0.15 | 0.28 |
| 10 | 72294836 | rs10999379 | 1.431 | 0.316 | -0.83 | -1.559 | -0.051 | -0.036 | -0.036 | 0 | 0.14 | 0.02 | 0.28 | 0.18 | 0.2 | 0.12 |
| 10 | 72297411 | rs871870 | 0.862 | -0.212 | -0.346 | -1.371 | -0.012 | -0.034 | -0.034 | 0 | 0.29 | 0.39 | 0.17 | 0.1 | 0.23 | 0.19 |
| 10 | 72299626 | rs10999382 | 1.371 | -0.307 | -0.682 | -1.385 | 0.007 | 0.041 | 0.041 | 0.27 | 0.24 | 0.12 | 0.13 | 0.06 | 0.2 | 0.17 |
| 10 | 72307101 | rs3740447 | 1.135 | -0.257 | -0.923 | -1.264 | -0.04 | -0.138 | -0.138 | 0 | 0.17 | 0.02 | 0.17 | 0.51 | 0.15 | 0.23 |
| 10 | 72308254 | rs10999385 | -0.798 | -0.261 | -0.634 | -1.324 | -0.027 | -0.127 | -0.127 | 0.18 | 0.65 | 0.56 | 0.83 | 0.98 | 0.76 | 0.78 |
| 10 | 72311394 | rs870457 | -0.083 | 0.013 | -0.769 | -0.649 | 0.016 | -0.056 | -0.056 | 0 | 0.2 | 0.2 | 0.18 | 0.5 | 0.14 | 0.26 |
| 10 | 72312979 | rs10823565 | -0.561 | 0.391 | -0.779 | -1.08 | 0.001 | -0.024 | -0.024 | 0 | 0.37 | 0.57 | 0.43 | 0.86 | 0.4 | 0.45 |
| 10 | 72313642 | rs12415107 | 0.828 | 0.381 | -1.379 | -1.234 | -0.027 | 0.022 | 0.022 | 0 | 0.19 | 0.06 | 0.34 | 0.12 | 0.2 | 0.23 |
| 10 | 72316867 | rs10823571 | 1.403 | 0.225 | -1.253 | -0.565 | -0.011 | -0.006 | -0.006 | 0 | 0.08 | 0.01 | 0.13 | 0.55 | 0.08 | 0.15 |
| 10 | 72318190 | rs12413000 | 0.6 | 0.162 | -0.969 | -0.94 | -0.005 | 0.033 | 0.033 | 0.295 | 0.62 | 0.29 | 0.61 | 0.15 | 0.59 | 0.56 |
| 10 | 72319022 | rs10740354 | -0.051 | 0.009 | -0.768 | -0.288 | 0.003 | -0.019 | -0.019 | 0 | 0.57 | 0.52 | 0.68 | 0.77 | 0.53 | 0.71 |
| 10 | 72319330 | rs10762392 | -0.516 | 0.421 | -0.76 | 0.04 | 0.009 | -0.002 | -0.002 | 0 | 0.77 | 0.71 | 0.76 | 0.79 | 0.68 | 0.87 |
| 10 | 72320504 | rs2394730 | -0.256 | -0.048 | -1.052 | -0.081 | 0.005 | 0.063 | 0.063 | 0 | 0.47 | 0.33 | 0.43 | 0.79 | 0.57 | 0.52 |
| 10 | 72326942 | rs3740442 | 0.445 | 0.344 | -1.429 | 0.796 | 0 | 0 | 0 | 0.158 | 0.12 | 0.11 | 0.07 | 0.41 | 0.11 | 0.2 |
| 10 | 72329470 | rs12768847 | 1.161 | -0.007 | -1.66 | 0.457 | -0.011 | 0.005 | 0.005 | 0 | 0.21 | 0.06 | 0.4 | 0.15 | 0.23 | 0.17 |
| 10 | 72331415 | rs12412518 | 0.826 | -0.053 | -1.7 | 0.442 | -0.019 | 0.016 | 0.016 | 0 | 0.24 | 0.05 | 0.41 | 0.13 | 0.25 | 0.21 |
| 10 | 72331540 | rs10762393 | -0.348 | -0.333 | -1.426 | 0.435 | 0.02 | 0.039 | 0.039 | 0 | 0.48 | 0.31 | 0.4 | 0.72 | 0.57 | 0.52 |
| 10 | 72332440 | rs10999409 | -0.422 | -0.352 | -1.614 | 0.262 | 0.026 | 0.11 | 0.11 | 0.008 | 0.49 | 0.24 | 0.41 | 0.74 | 0.57 | 0.51 |
| 10 | 72333257 | rs10823575 | 0.438 | -0.156 | -1.865 | -0.177 | -0.067 | 0.156 | 0.156 | 0.762 | 0.4 | 0.15 | 0.3 | 0.16 | 0.49 | 0.31 |
| 10 | 72343117 | rs10823577 | 0.152 | -0.172 | -1.688 | -0.062 | -0.006 | 0.025 | 0.025 | 0 | 0.6 | 0.35 | 0.63 | 0.25 | 0.63 | 0.56 |
| 10 | 72344579 | rs2894125 | 0.12 | -0.565 | -1.997 | -0.099 | 0.016 | -0.092 | -0.092 | 0 | 0.27 | 0.09 | 0.31 | 0.55 | 0.2 | 0.24 |
| 10 | 72346060 | rs4747063 | 0.673 | -0.309 | -1.253 | -0.518 | 0.011 | -0.023 | -0.023 | 0.063 | 0.29 | 0.35 | 0.28 | 0.17 | 0.4 | 0.29 |
| 10 | 72346471 | rs1413710 | -0.238 | -0.725 | -0.782 | -0.74 | 0 | 0.003 | 0.003 | 0 | 0.41 | 0.61 | 0.5 | 0.71 | 0.36 | 0.36 |
| 10 | 72348756 | rs4747064 | 0.767 | -0.625 | -0.795 | -0.668 | 0.002 | -0.03 | -0.03 | 0 | 0.29 | 0.19 | 0.41 | 0.67 | 0.28 | 0.28 |
| 10 | 72350779 | rs10509323 | 0.975 | -0.571 | -1.08 | -0.9 | 0.002 | -0.005 | -0.005 | 0 | 0.09 | 0.09 | 0.11 | 0.1 | 0.09 | 0.11 |
| 10 | 72351827 | rs12765134 | 2.407 | -0.449 | -1.24 | -1.13 | -0.036 | 0.061 | 0.061 | 0.013 | 0.1 | 0.02 | 0.07 | 0.01 | 0.14 | 0.18 |
| 10 | 72353577 | rs12358573 | 1.262 | -0.378 | -1.237 | -1.243 | -0.078 | 0.126 | 0.126 | 0 | 0.28 | 0.05 | 0.25 | 0.09 | 0.35 | 0.19 |
| 10 | 72354116 | rs10999423 | 0.267 | -0.394 | -0.834 | -0.984 | -0.001 | -0.004 | -0.004 | 0 | 0.13 | 0.08 | 0.06 | 0 | 0.09 | 0.07 |
| 10 | 72355806 | rs17527619 | 0.911 | -0.311 | -0.804 | -0.763 | -0.013 | -0.016 | -0.016 | 0 | 0.05 | 0.03 | 0.09 | 0.09 | 0.09 | 0.13 |
| 10 | 72356197 | rs12773662 | -0.239 | -0.417 | -0.515 | -0.862 | 0.002 | 0.012 | 0.012 | 0.606 | 0.43 | 0.4 | 0.52 | 0.72 | 0.38 | 0.33 |
| 10 | 72358577 | rs885822 | -0.773 | -0.486 | -0.64 | -0.775 | -0.059 | -0.004 | -0.004 | 0.063 | 0.68 | 0.84 | 0.64 | 0.73 | 0.59 | 0.61 |
| 10 | 72362080 | rs3758562 | -0.97 | -0.526 | -0.228 | -0.443 | 0.001 | 0.001 | 0.001 | 0 | 0.7 | 0.69 | 0.66 | 0.76 | 0.68 | 0.73 |
| 10 | 72365130 | rs1641661 | 0.123 | -0.509 | -0.51 | -1.125 | -0.008 | 0.007 | 0.007 | 0 | 0.33 | 0.3 | 0.33 | 0.26 | 0.38 | 0.28 |
| 10 | 72375801 | rs10999435 | 1.101 | -0.497 | -0.581 | -0.994 | 0.018 | -0.021 | -0.021 | 0 | 0.13 | 0.08 | 0.28 | 0.57 | 0.06 | 0.16 |
| 10 | 72377034 | rs10999436 | 1.247 | -0.387 | -0.826 | -1.211 | -0.068 | 0.139 | 0.139 | 0 | 0.26 | 0.03 | 0.2 | 0.03 | 0.3 | 0.16 |
| 10 | 72387194 | rs2256859 | -0.244 | -0.36 | -0.553 | -1.17 | -0.001 | -0.003 | -0.003 | 0 | 0.56 | 0.47 | 0.45 | 0.27 | 0.62 | 0.64 |
| 10 | 72396646 | rs1761741 | -0.303 | 0.114 | -0.068 | -1 | 0.002 | 0.071 | 0.071 | 0 | 0.49 | 0.37 | 0.61 | 0.81 | 0.45 | 0.46 |
| 10 | 72399179 | rs1641674 | -0.311 | 0.116 | -0.066 | -0.998 | 0.004 | 0.072 | 0.072 | 0 | 0.5 | 0.36 | 0.61 | 0.81 | 0.45 | 0.46 |
| 10 | 72405504 | rs826463 | 0.706 | 0.187 | -0.006 | -0.935 | -0.008 | -0.016 | -0.016 | 0.018 | 0.09 | 0.04 | 0.1 | 0.15 | 0.1 | 0.15 |
| 10 | 72405762 | rs826460 | -0.071 | 0.021 | 0.2 | -1.042 | 0.005 | 0.075 | 0.075 | 0 | 0.5 | 0.37 | 0.61 | 0.81 | 0.46 | 0.46 |
| 10 | 72406967 | rs751644 | -0.986 | 0.052 | 0.311 | -1.002 | 0.026 | -0.036 | -0.036 | 0 | 0.74 | 0.52 | 0.74 | 0.83 | 0.71 | 0.81 |
| 10 | 72412776 | rs12771779 | 1.12 | -0.189 | 0.085 | -0.914 | 0.006 | 0.108 | 0.108 | 0 | 0.23 | 0.07 | 0.13 | 0.02 | 0.23 | 0.35 |
| 10 | 72413827 | rs10999456 | 0.76 | -0.283 | 0.504 | -0.92 | -0.026 | -0.131 | -0.131 | 0 | 0.26 | 0.41 | 0.16 | 0.04 | 0.24 | 0.36 |
| 10 | 72422431 | rs10999459 | -0.067 | -0.298 | 0.071 | -0.877 | -0.002 | 0.006 | 0.006 | 0 | 0.26 | 0.26 | 0.34 | 0.54 | 0.17 | 0.24 |
| 10 | 72424452 | rs17528058 | 1.855 | 0.114 | -0.537 | -0.968 | -0.074 | 0.086 | 0.086 | 0 | 0.18 | 0.04 | 0.17 | 0.02 | 0.27 | 0.13 |
| 10 | 72431602 | rs10999463 | 1.839 | 0.131 | -0.444 | -0.824 | -0.073 | 0.026 | 0.026 | 0 | 0.2 | 0.02 | 0.19 | 0.13 | 0.25 | 0.1 |
| 10 | 72433137 | rs11596281 | 0.171 | 0.037 | 0.17 | -0.674 | -0.041 | 0.141 | 0.141 | 0 | 0.43 | 0.22 | 0.32 | 0.07 | 0.51 | 0.46 |
| 10 | 72434194 | rs614201 | -0.461 | 0.101 | 0.207 | -0.557 | -0.027 | 0.083 | 0.083 | 0 | 0.57 | 0.73 | 0.64 | 0.84 | 0.49 | 0.51 |
| 10 | 72435657 | rs7086591 | 0.009 | -0.045 | 0.596 | -0.364 | 0.002 | 0.009 | 0.009 | 0 | 0.5 | 0.46 | 0.55 | 0.68 | 0.42 | 0.37 |
| 10 | 72436953 | rs1345386 | -0.406 | -0.141 | -0.133 | -0.247 | 0.025 | 0.002 | 0.002 | 0 | 0.67 | 0.75 | 0.73 | 0.71 | 0.69 | 0.51 |
| 10 | 72441228 | rs10999467 | 0.504 | 0.006 | -0.048 | -0.476 | 0.011 | -0.041 | -0.041 | 0 | 0.21 | 0.22 | 0.19 | 0.03 | 0.27 | 0.17 |
| 10 | 72445375 | rs1477069 | 0.621 | -0.3 | -0.478 | -0.204 | 0.013 | -0.002 | -0.002 | 1.442 | 0.13 | 0.08 | 0.27 | 0.55 | 0.07 | 0.21 |
| 10 | 72445630 | rs1477070 | 1.575 | -0.195 | -0.58 | -0.331 | -0.065 | 0.051 | 0.051 | 0 | 0.16 | 0.03 | 0.16 | 0.05 | 0.25 | 0.14 |
| 10 | 72453332 | rs1079351 | 0.554 | -0.382 | -0.783 | -0.544 | -0.011 | 0.064 | 0.064 | 0 | 0.27 | 0.08 | 0.17 | 0.1 | 0.28 | 0.3 |
| 10 | 72456400 | rs4747077 | -1.581 | -0.469 | -0.31 | -0.616 | -0.005 | 0.009 | 0.009 | 0 | 0.68 | 0.72 | 0.72 | 0.76 | 0.67 | 0.78 |
| 10 | 72463045 | rs10762398 | 0.443 | -0.633 | -0.057 | -0.262 | -0.021 | 0.094 | 0.094 | 0 | 0.14 | 0.16 | 0.12 | 0.48 | 0.07 | 0.18 |
| 10 | 72466435 | rs4747079 | 1.341 | -0.616 | -0.394 | -0.262 | 0.002 | 0.049 | 0.049 | 0 | 0.1 | 0.07 | 0.1 | 0.53 | 0.05 | 0.15 |
| 10 | 72467255 | rs4747083 | 1.064 | -0.688 | -0.473 | -0.359 | 0.002 | -0.026 | -0.026 | 0 | 0.07 | 0.02 | 0.1 | 0.16 | 0.06 | 0.14 |
| 10 | 72467944 | rs2024464 | 1.902 | -0.46 | -0.525 | -0.461 | -0.045 | 0.044 | 0.044 | 0 | 0.16 | 0.06 | 0.16 | 0.06 | 0.21 | 0.14 |
| 10 | 72468097 | rs10823600 | -0.059 | -0.485 | -0.181 | -0.492 | 0.009 | 0.002 | 0.002 | 0 | 0.32 | 0.43 | 0.35 | 0.63 | 0.33 | 0.31 |

| | | | | | | | | | | | | | | | | |
|---|---|---|---|---|---|---|---|---|---|---|---|---|---|---|---|---|
| 10 | 72468945 | rs12255288 | -1.888 | -0.618 | -0.104 | -0.543 | -0.006 | 0.032 | 0.032 | 0 | 0.69 | 0.79 | 0.61 | 0.81 | 0.68 | 0.76 |
| 10 | 72469781 | rs10999475 | 1.353 | -0.638 | -0.166 | -0.585 | -0.034 | 0.139 | 0.139 | 0 | 0.22 | 0.01 | 0.12 | 0 | 0.24 | 0.25 |
| 10 | 72471772 | rs12777075 | 1.454 | -0.659 | 0.461 | 0.015 | -0.012 | 0.026 | 0.026 | 0 | 0.27 | 0.19 | 0.23 | 0.16 | 0.31 | 0.17 |
| 10 | 72473079 | rs748343 | -0.72 | -0.473 | 0.479 | 0.067 | -0.003 | -0.039 | -0.039 | 1.227 | 0.63 | 0.49 | 0.59 | 0.79 | 0.66 | 0.72 |
| 10 | 72475476 | rs736189 | -1.36 | -0.361 | -0.122 | 0.119 | -0.005 | 0.026 | 0.026 | 0 | 0.69 | 0.76 | 0.61 | 0.81 | 0.68 | 0.76 |
| 10 | 72475768 | rs10823602 | -0.136 | -0.204 | -0.725 | 0.353 | -0.008 | 0.081 | 0.081 | 0 | 0.15 | 0.24 | 0.17 | 0.56 | 0.11 | 0.19 |
| 10 | 72477186 | rs7093021 | 0.822 | -0.107 | -1.29 | 0.473 | -0.005 | 0.072 | 0.072 | 0 | 0.1 | 0.08 | 0.11 | 0.46 | 0.05 | 0.15 |
| 10 | 72477232 | rs10999479 | 0.269 | 0.348 | -1.448 | 0.006 | -0.098 | 0.221 | 0.221 | 0 | 0.4 | 0.11 | 0.3 | 0.07 | 0.49 | 0.4 |
| 10 | 72477417 | rs10999481 | -0.162 | 0.416 | -1.318 | 0.092 | -0.022 | 0.113 | 0.113 | 0 | 0.47 | 0.21 | 0.39 | 0.25 | 0.55 | 0.53 |
| 10 | 72481720 | rs2894129 | -0.294 | 0.446 | -0.801 | 0.185 | -0.001 | -0.019 | -0.019 | 0 | 0.46 | 0.53 | 0.47 | 0.18 | 0.45 | 0.51 |
| 10 | 72481918 | rs7914626 | -0.038 | 0.287 | -0.766 | 0.979 | 0 | 0.016 | 0.016 | 0 | 0.93 | 0.92 | 0.85 | 0.73 | 0.93 | 0.86 |
| 10 | 72482819 | rs7092269 | -0.113 | 0.141 | -1.144 | 0.283 | -0.003 | 0.048 | 0.048 | 0 | 0.11 | 0.02 | 0.02 | 0 | 0.08 | 0.08 |
| 10 | 72483010 | rs1816002 | -0.77 | 0.149 | -1.11 | 0.197 | -0.002 | 0.017 | 0.017 | 0 | 0.52 | 0.62 | 0.55 | 0.58 | 0.46 | 0.68 |
| 10 | 72483358 | rs7096309 | -0.34 | 0.101 | -0.946 | 0.313 | 0.007 | -0.04 | -0.04 | 0 | 0.4 | 0.48 | 0.29 | 0.14 | 0.38 | 0.31 |
| 10 | 72484943 | rs10762400 | -0.535 | 0.382 | -1.093 | 0.418 | 0.039 | 0.19 | 0.19 | 0.506 | 0.18 | 0.46 | 0.26 | 0.39 | 0.24 | 0.17 |
| 10 | 72485081 | rs10999486 | 1.586 | 0.34 | -1.157 | 0.322 | -0.056 | 0.063 | 0.063 | 0 | 0.18 | 0.03 | 0.15 | 0.06 | 0.2 | 0.2 |
| 10 | 72485801 | rs10159685 | -0.582 | 0.39 | -1.008 | 0.468 | 0.039 | 0.17 | 0.17 | 0.5 | 0.18 | 0.45 | 0.26 | 0.38 | 0.24 | 0.17 |
| 10 | 72486913 | rs4747089 | -0.447 | 0.125 | -0.765 | 0.352 | 0 | -0.005 | -0.005 | 0 | 0.44 | 0.46 | 0.38 | 0.38 | 0.43 | 0.42 |
| 10 | 72493876 | rs10740358 | 0.022 | -0.05 | -0.685 | 0.247 | 0.002 | 0.002 | 0.002 | 0.718 | 0.8 | 0.83 | 0.75 | 0.89 | 0.85 | 0.75 |
| 10 | 72494158 | rs16927852 | -0.773 | -0.124 | 0.203 | 0.861 | 0.017 | 0.276 | 0.276 | 0.099 | 0.1 | 0.4 | 0.19 | 0.34 | 0.13 | 0.15 |
| 10 | 72494778 | rs12765664 | 0.168 | -0.25 | 0.188 | 0.037 | -0.009 | -0.026 | -0.026 | 0 | 0.81 | 0.73 | 0.74 | 0.9 | 0.85 | 0.85 |
| 10 | 72495587 | rs10999499 | 1.175 | -0.233 | -0.44 | -0.36 | -0.089 | 0.089 | 0.089 | 0 | 0.22 | 0.02 | 0.17 | 0.05 | 0.24 | 0.23 |
| 10 | 72496131 | rs1861519 | -0.564 | -0.437 | 0.16 | -0.371 | -0.017 | -0.017 | -0.017 | 1.759 | 0.86 | 0.62 | 0.74 | 0.9 | 0.9 | 0.86 |
| 10 | 72496317 | rs1861518 | 0.614 | -0.302 | -0.183 | -0.335 | 0.019 | 0.058 | 0.058 | 0 | 0.5 | 0.6 | 0.48 | 0.76 | 0.49 | 0.53 |
| 10 | 72498841 | rs4746061 | 0.154 | -0.475 | -0.123 | -0.046 | 0.015 | -0.06 | -0.06 | 0.38 | 0.17 | 0.09 | 0.24 | 0.62 | 0.18 | 0.39 |
| 10 | 72499334 | rs4747094 | -0.766 | -0.272 | 0.045 | -0.243 | -0.002 | -0.044 | -0.044 | 0 | 0.92 | 0.81 | 0.96 | 1 | 0.95 | 0.99 |
| 10 | 72500763 | rs10823607 | -0.132 | -0.272 | -0.017 | -0.224 | -0.011 | -0.048 | -0.048 | 0 | 0.82 | 0.67 | 0.74 | 0.91 | 0.86 | 0.85 |
| 10 | 72502554 | rs7069182 | 1.834 | -0.276 | -0.188 | -0.25 | -0.013 | 0.027 | 0.027 | 0 | 0.26 | 0.15 | 0.19 | 0.15 | 0.28 | 0.13 |
| 10 | 72504556 | rs2541231 | 1.703 | -0.495 | -0.216 | -0.193 | -0.005 | -0.005 | -0.005 | 0 | 0.25 | 0.23 | 0.39 | 0.3 | 0.28 | 0.24 |
| 10 | 72504763 | rs1548906 | -0.579 | -0.643 | -0.331 | -0.369 | -0.013 | 0.058 | 0.058 | 1.019 | 0.63 | 0.78 | 0.7 | 0.84 | 0.63 | 0.69 |
| 10 | 72505315 | rs2791183 | 0.173 | -0.839 | -0.474 | -0.518 | -0.012 | 0.115 | 0.115 | 0.004 | 0.53 | 0.74 | 0.67 | 0.84 | 0.54 | 0.66 |
| 10 | 72505766 | rs12221267 | 0.619 | -0.848 | -0.544 | -0.474 | -0.016 | -0.106 | -0.106 | 1.585 | 0.15 | 0.04 | 0.18 | 0.53 | 0.19 | 0.35 |
| 10 | 72505992 | rs10999507 | 1.502 | -0.898 | -0.35 | -0.549 | -0.048 | 0 | 0 | 0.075 | 0.23 | 0.09 | 0.23 | 0.16 | 0.28 | 0.2 |
| 10 | 72506158 | rs10823610 | -0.548 | -0.882 | -0.01 | -0.608 | 0.001 | 0.039 | 0.039 | 0 | 0.5 | 0.39 | 0.49 | 0.69 | 0.54 | 0.6 |
| 10 | 72508554 | rs7921298 | 0.654 | -0.854 | -0.087 | -0.624 | -0.006 | -0.032 | -0.032 | 0 | 0.1 | 0.11 | 0.04 | 0 | 0.08 | 0.06 |
| 10 | 72508841 | rs10762401 | 0.373 | -0.888 | 0.024 | -0.647 | -0.06 | 0.03 | 0.03 | 0 | 0.38 | 0.15 | 0.41 | 0.72 | 0.47 | 0.56 |
| 10 | 72509080 | rs2994241 | -0.065 | -0.887 | 0.007 | -0.671 | -0.013 | -0.116 | -0.116 | 0 | 0.78 | 0.64 | 0.88 | 1 | 0.85 | 0.92 |
| 10 | 72509321 | rs2541230 | 0.98 | -0.7 | -0.213 | -0.423 | 0 | 0.002 | 0.002 | 0 | 0.36 | 0.35 | 0.43 | 0.29 | 0.36 | 0.3 |
| 10 | 72509809 | rs2541229 | 1.257 | -0.403 | -0.436 | -0.153 | -0.003 | 0 | 0 | 0 | 0.3 | 0.28 | 0.42 | 0.29 | 0.3 | 0.3 |
| 10 | 72511639 | rs12269650 | -0.519 | -0.208 | -0.554 | 0.154 | 0.013 | -0.075 | -0.075 | 0.31 | 0.3 | 0.16 | 0.27 | 0.55 | 0.32 | 0.43 |
| 10 | 72513682 | rs2587475 | 0.945 | -0.104 | -0.11 | -0.072 | -0.001 | 0 | 0 | 0 | 0.55 | 0.52 | 0.66 | 0.44 | 0.54 | 0.51 |
| 10 | 72513957 | rs4746065 | -0.148 | -0.236 | -0.421 | 0.036 | 0.002 | -0.071 | -0.071 | 0.014 | 0.19 | 0.08 | 0.19 | 0.56 | 0.23 | 0.35 |
| 10 | 72514397 | rs2541227 | 0.716 | 0.169 | -0.115 | -0.442 | 0.014 | -0.046 | -0.046 | 0 | 0.74 | 0.44 | 0.79 | 0.91 | 0.75 | 0.75 |
| 10 | 72515691 | rs946742 | 0.346 | -0.15 | -0.383 | -0.153 | -0.002 | 0 | 0 | 0 | 0.53 | 0.46 | 0.47 | 0.48 | 0.58 | 0.61 |
| 10 | 72515775 | rs946743 | 0.476 | -0.238 | -0.448 | -0.213 | -0.001 | -0.001 | -0.001 | 0 | 0.51 | 0.46 | 0.47 | 0.46 | 0.58 | 0.57 |
| 10 | 72515802 | rs946744 | 2.47 | 0.115 | -0.547 | -0.281 | -0.002 | 0 | 0 | 0 | 0.23 | 0.21 | 0.3 | 0.21 | 0.28 | 0.14 |
| 10 | 72515833 | rs2587472 | 0.306 | 0.094 | -0.696 | -0.248 | -0.052 | -0.046 | -0.046 | 1.512 | 0.13 | 0.02 | 0.09 | 0.11 | 0.16 | 0.19 |
| 10 | 72517206 | rs2791189 | 0.133 | -0.026 | -0.339 | -0.328 | -0.006 | 0.018 | 0.018 | 0 | 0.4 | 0.49 | 0.5 | 0.54 | 0.36 | 0.39 |
| 10 | 72517837 | rs6480463 | 0.763 | -0.124 | -0.247 | -0.386 | -0.002 | -0.005 | -0.005 | 0 | 0.58 | 0.55 | 0.61 | 0.4 | 0.55 | 0.48 |
| 10 | 72518009 | rs4747096 | 0.948 | 0.038 | -0.273 | -0.241 | 0.005 | 0.042 | 0.042 | 0 | 0.15 | 0.17 | 0.23 | 0.37 | 0.18 | 0.15 |
| 10 | 72518075 | rs4747097 | -0.175 | 0.046 | -0.314 | -0.269 | -0.011 | -0.02 | -0.02 | 0.602 | 0.23 | 0.15 | 0.27 | 0.39 | 0.28 | 0.26 |
| 10 | 72518536 | rs12413960 | 0.911 | -0.046 | -0.334 | -0.417 | -0.006 | -0.012 | -0.012 | 0 | 0.25 | 0.3 | 0.26 | 0.17 | 0.17 | 0.23 |
| 10 | 72520567 | rs10823616 | 0.665 | -0.038 | -0.322 | -0.406 | -0.005 | -0.014 | -0.014 | 0 | 0.27 | 0.33 | 0.27 | 0.17 | 0.19 | 0.24 |
| 10 | 72521155 | rs2587469 | -1.524 | 0.051 | -0.457 | -0.475 | 0 | -0.004 | -0.004 | 0 | 0.62 | 0.56 | 0.5 | 0.7 | 0.62 | 0.69 |
| 10 | 72521651 | rs3740438 | 0.441 | 0.078 | -0.741 | -0.541 | -0.02 | 0.102 | 0.102 | 0 | 0.28 | 0.1 | 0.15 | 0.1 | 0.26 | 0.18 |
| 10 | 72524341 | rs2688768 | -0.367 | 0.28 | -0.241 | -0.083 | -0.016 | 0.035 | 0.035 | 0 | 0.19 | 0.26 | 0.11 | 0.23 | 0.18 | 0.33 |
| 10 | 72524523 | rs10999521 | -0.159 | 0.285 | -0.182 | -0.08 | -0.002 | 0.004 | 0.004 | 0 | 0.18 | 0.14 | 0.09 | 0.23 | 0.18 | 0.33 |
| 10 | 72525761 | rs10999524 | 0.712 | 0.335 | -0.247 | -0.046 | -0.039 | -0.044 | -0.044 | 0 | 0.08 | 0 | 0.05 | 0.09 | 0.11 | 0.08 |
| 10 | 72529113 | rs4539233 | -0.58 | 0.326 | -0.41 | -0.036 | 0 | -0.006 | -0.006 | 0 | 0.22 | 0.15 | 0.13 | 0.23 | 0.25 | 0.34 |
| 10 | 72532238 | rs2253801 | 0.707 | 0.091 | -0.309 | -0.345 | 0 | 0 | 0 | 0 | 0.44 | 0.46 | 0.55 | 0.39 | 0.45 | 0.42 |
| 10 | 72532443 | rs10823618 | 0.288 | -0.062 | -0.422 | -0.38 | -0.01 | 0.044 | 0.044 | 0 | 0.19 | 0.2 | 0.24 | 0.51 | 0.17 | 0.37 |
| 10 | 72532750 | rs2253901 | -0.762 | 0.008 | -0.226 | -0.703 | -0.002 | -0.006 | -0.006 | 0 | 0.7 | 0.68 | 0.81 | 0.9 | 0.7 | 0.8 |
| 10 | 72533199 | rs12359270 | 1.578 | 0.001 | -0.361 | -0.734 | -0.014 | 0.028 | 0.028 | 0.071 | 0.17 | 0.15 | 0.11 | 0.1 | 0.19 | 0.15 |
| 10 | 72533766 | rs2688752 | 0.761 | -0.069 | -0.181 | -0.638 | 0.004 | 0.013 | 0.013 | 0 | 0.53 | 0.57 | 0.41 | 0.21 | 0.58 | 0.44 |
| 10 | 72533910 | rs7077672 | 0.355 | 0.09 | 0.469 | -0.146 | -0.006 | 0.027 | 0.027 | 0.907 | 0.84 | 0.69 | 0.72 | 0.8 | 0.87 | 0.9 |
| 10 | 72535007 | rs2254174 | 0.583 | -0.012 | 0.185 | 0.092 | -0.014 | 0.037 | 0.037 | 0.326 | 0.81 | 0.79 | 0.69 | 0.63 | 0.88 | 0.85 |
| 10 | 72535492 | rs16927963 | 0.079 | -0.003 | 0.329 | 0.04 | 0.01 | 0.071 | 0.071 | 0.086 | 0.12 | 0.31 | 0.27 | 0.2 | 0.11 | 0.05 |
| 10 | 72535922 | rs12221473 | 0.755 | -0.105 | 0.148 | -0.079 | 0.006 | -0.021 | -0.021 | 0.228 | 0.29 | 0.39 | 0.31 | 0.24 | 0.34 | 0.26 |
| 10 | 72536660 | rs2254415 | -0.119 | -0.274 | 0.156 | 0.595 | 0.006 | 0.049 | 0.049 | 0.102 | 0.12 | 0.25 | 0.26 | 0.18 | 0.11 | 0.05 |

| | | | | | | | | | | | | | | | | |
|---|---|---|---|---|---|---|---|---|---|---|---|---|---|---|---|---|
| 10 | 72537873 | rs2254535 | -0.154 | -0.159 | -0.358 | 0.463 | -0.021 | 0.01 | 0.01 | 0.007 | 0.87 | 0.79 | 0.9 | 0.84 | 0.92 | 0.9 |
| 10 | 72539103 | rs4747100 | -0.134 | 0.25 | -0.641 | 0.411 | 0 | -0.001 | -0.001 | 0 | 0.32 | 0.33 | 0.27 | 0.37 | 0.26 | 0.35 |
| 10 | 72540534 | rs2265549 | 0.121 | 0.889 | -0.488 | 0.366 | -0.001 | 0 | 0 | 0 | 0.83 | 0.84 | 0.89 | 0.92 | 0.83 | 0.89 |
| 10 | 72543579 | rs1538677 | 0.49 | 0.745 | -0.278 | 0.271 | 0.005 | 0.005 | 0.005 | 0 | 0.47 | 0.57 | 0.52 | 0.61 | 0.43 | 0.52 |
| 10 | 72543913 | rs12415051 | 0.403 | 0.75 | -0.282 | 0.206 | -0.028 | 0.015 | 0.015 | 0.019 | 0.33 | 0.21 | 0.35 | 0.31 | 0.39 | 0.33 |
| 10 | 72545587 | rs3824762 | 0 | 0.742 | 0.01 | 0.231 | 0.015 | 0.005 | 0.005 | 1.252 | 0.15 | 0.26 | 0.13 | 0.09 | 0.17 | 0.15 |
| 10 | 72546751 | rs2185349 | 0.237 | 0.726 | 0.076 | 0.224 | -0.009 | -0.005 | -0.005 | 0 | 0.14 | 0.09 | 0.1 | 0.09 | 0.16 | 0.15 |
| 10 | 72550184 | rs12775092 | -0.016 | 0.731 | 0.049 | 0.227 | 0.011 | 0.004 | 0.004 | 0.641 | 0.15 | 0.22 | 0.12 | 0.09 | 0.17 | 0.15 |
| 10 | 72559966 | rs2251370 | 0.733 | 0.809 | -0.017 | 0.146 | 0.009 | 0.002 | 0.002 | 0.613 | 0.73 | 0.62 | 0.77 | 0.79 | 0.72 | 0.81 |
| 10 | 72563607 | rs2251822 | -1.122 | 0.746 | -0.039 | 0.193 | 0.009 | -0.007 | -0.007 | 0.298 | 0.9 | 0.94 | 0.92 | 0.87 | 0.95 | 0.97 |
| 10 | 72565236 | rs7093309 | 0.538 | 0.76 | -0.18 | 0.042 | -0.002 | 0.001 | 0.001 | 0 | 0.29 | 0.28 | 0.31 | 0.3 | 0.33 | 0.31 |
| 10 | 72565359 | rs2791173 | -1.219 | 0.74 | -0.304 | 0.09 | 0.005 | -0.003 | -0.003 | 0 | 0.9 | 0.91 | 0.91 | 0.87 | 0.95 | 0.97 |
| 10 | 72574945 | rs2688760 | -0.244 | 0.64 | -0.255 | 0.116 | -0.006 | -0.001 | -0.001 | 0 | 0.14 | 0.18 | 0.14 | 0.14 | 0.1 | 0.05 |
| 10 | 72592987 | rs3781306 | 1.412 | 0.225 | -0.515 | -0.139 | -0.008 | 0.018 | 0.018 | 0 | 0.19 | 0.21 | 0.3 | 0.41 | 0.12 | 0.19 |
| 10 | 72593893 | rs7087539 | 1.084 | 0.4 | -0.746 | -0.488 | -0.008 | 0.015 | 0.015 | 0.014 | 0.2 | 0.27 | 0.22 | 0.3 | 0.18 | 0.27 |
| 10 | 72594162 | rs10509327 | 1.917 | 0.606 | -1.028 | -0.572 | -0.039 | -0.019 | -0.019 | 0 | 0.11 | 0.03 | 0.1 | 0.08 | 0.18 | 0.1 |
| 10 | 72599497 | rs4746069 | 1.216 | 0.523 | -1.002 | -0.674 | 0.003 | -0.003 | -0.003 | 0 | 0.43 | 0.36 | 0.44 | 0.48 | 0.37 | 0.52 |
| 10 | 72601813 | rs827235 | 0.814 | 0.515 | -1.024 | -0.667 | -0.028 | 0.02 | 0.02 | 2.182 | 0.2 | 0.06 | 0.17 | 0.14 | 0.21 | 0.07 |
| 10 | 72605550 | rs10999565 | 2.311 | 0.298 | -1.179 | -0.669 | -0.017 | -0.066 | -0.066 | 0 | 0.13 | 0.01 | 0.1 | 0.23 | 0.13 | 0.15 |
| 10 | 72629567 | rs827249 | 1.638 | 0.49 | -0.887 | -0.592 | 0.013 | -0.012 | -0.012 | 0 | 0.1 | 0.05 | 0.08 | 0.14 | 0.05 | 0.03 |
| 10 | 72631204 | rs10509326 | 2.582 | 0.507 | -0.95 | -0.635 | -0.065 | 0.064 | 0.064 | 0 | 0.21 | 0.02 | 0.11 | 0.08 | 0.22 | 0.32 |
| 10 | 72631626 | rs865832 | 0.279 | 0.626 | -0.845 | -0.579 | -0.027 | -0.034 | -0.034 | 0 | 0.21 | 0.08 | 0.18 | 0.22 | 0.23 | 0.13 |
| 10 | 72632403 | rs7085273 | 1.489 | 0.602 | -0.815 | -0.584 | 0.015 | -0.025 | -0.025 | 0 | 0.4 | 0.23 | 0.41 | 0.49 | 0.34 | 0.51 |
| 10 | 72636450 | rs923177 | 0.202 | 0.44 | -0.818 | -0.365 | -0.002 | -0.002 | -0.002 | 0 | 0.74 | 0.73 | 0.76 | 0.78 | 0.72 | 0.85 |
| 10 | 72639621 | rs877034 | 0.752 | 0.475 | -1.088 | -0.351 | -0.023 | 0.02 | 0.02 | 0.081 | 0.22 | 0.36 | 0.23 | 0.29 | 0.18 | 0.3 |
| 10 | 72639800 | rs2881 | 1.183 | 0.774 | -1.357 | 0.047 | -0.04 | -0.019 | -0.019 | 0 | 0.11 | 0.03 | 0.1 | 0.08 | 0.18 | 0.1 |
| 10 | 72642702 | rs1767233 | 0.786 | 0.053 | -1.077 | 0.194 | 0.013 | 0.012 | 0.012 | 2.124 | 0.1 | 0.05 | 0.08 | 0.08 | 0.05 | 0.03 |
| 10 | 72646337 | rs12218438 | 1.297 | 0.166 | -1.127 | 0.543 | 0.003 | -0.094 | -0.094 | 0.01 | 0.14 | 0 | 0.12 | 0.35 | 0.07 | 0.12 |
| 10 | 72650574 | rs2691148 | 0.507 | 0.48 | -0.493 | 0.361 | 0.008 | -0.015 | -0.015 | 0.015 | 0.41 | 0.31 | 0.26 | 0.48 | 0.35 | 0.48 |
| 10 | 72651694 | rs1338643 | 0.352 | 0.142 | -0.123 | 0.241 | -0.007 | -0.007 | -0.007 | 0 | 0.82 | 0.8 | 0.68 | 0.86 | 0.85 | 0.89 |
| 10 | 72652518 | rs973982 | 0.378 | 0.139 | -0.08 | 0.239 | -0.018 | -0.018 | -0.018 | 0 | 0.82 | 0.76 | 0.68 | 0.86 | 0.85 | 0.89 |
| 10 | 72653761 | rs7917828 | 0.972 | 0.538 | -0.199 | 0.293 | 0.004 | -0.021 | -0.021 | 0 | 0.66 | 0.53 | 0.53 | 0.77 | 0.64 | 0.75 |
| 10 | 72654381 | rs17533359 | 0.553 | 0.99 | -0.314 | 0.538 | -0.034 | -0.033 | -0.033 | 0 | 0.09 | 0.03 | 0.09 | 0.18 | 0.19 | 0.11 |
| 10 | 72654600 | rs12252993 | -0.268 | 0.284 | 0.038 | 0.183 | -0.012 | -0.035 | -0.035 | 0.374 | 0.07 | 0.2 | 0.07 | 0.01 | 0.06 | 0.05 |
| 10 | 72655450 | rs827349 | -0.571 | 0.295 | -0.288 | 0.093 | -0.021 | -0.058 | -0.058 | 0.574 | 0.58 | 0.79 | 0.45 | 0.55 | 0.53 | 0.58 |
| 10 | 72655798 | rs17533443 | 1.534 | 0.392 | -0.296 | 0.095 | -0.034 | 0.036 | 0.036 | 0 | 0.2 | 0.11 | 0.12 | 0.08 | 0.27 | 0.26 |
| 10 | 72656002 | rs827351 | 0.237 | 0 | 0.357 | 0.21 | -0.001 | -0.001 | -0.001 | 0 | 0.47 | 0.43 | 0.34 | 0.48 | 0.42 | 0.48 |
| 10 | 72656844 | rs7096028 | -0.598 | 0.413 | 0.876 | 0.463 | 0.058 | 0.065 | 0.065 | 0 | 0.06 | 0.26 | 0.05 | 0.06 | 0.04 | 0.03 |
| 10 | 72657973 | rs12253257 | -0.284 | 0.464 | 0.463 | 0.484 | 0.024 | 0.03 | 0.03 | 0 | 0.05 | 0.19 | 0.04 | 0.06 | 0.04 | 0.03 |
| 10 | 72660718 | rs17612029 | -0.227 | 0.474 | -0.423 | 0.57 | -0.011 | 0.008 | 0.008 | 0.458 | 0.07 | 0.04 | 0.17 | 0.05 | 0.14 | 0.12 |
| 10 | 72662237 | rs1624747 | -0.294 | -0.438 | -0.216 | 0.685 | -0.027 | 0.022 | 0.022 | 0 | 0.62 | 0.53 | 0.68 | 0.58 | 0.65 | 0.71 |
| 10 | 72663997 | rs1564961 | 0.784 | -0.145 | -0.913 | 0.239 | -0.074 | 0.105 | 0.105 | 0 | 0.32 | 0.18 | 0.36 | 0.11 | 0.43 | 0.34 |
| 10 | 72665347 | rs10999576 | 0.527 | -0.381 | -0.266 | 1.049 | -0.047 | 0.014 | 0.014 | 0.18 | 0.25 | 0.3 | 0.15 | 0.25 | 0.14 | 0.17 |
| 10 | 72669745 | rs871679 | 0.473 | 0.78 | -0.677 | 1.21 | -0.008 | -0.009 | -0.009 | 0 | 0.05 | 0.03 | 0.07 | 0.06 | 0.11 | 0.06 |
| 10 | 72671684 | rs7095462 | -0.001 | 0.391 | -0.456 | 1.13 | -0.003 | 0.011 | 0.011 | 0 | 0.81 | 0.81 | 0.87 | 0.63 | 0.86 | 0.75 |
| 10 | 72673607 | rs4746070 | -0.402 | -0.063 | -0.338 | 0.694 | -0.02 | 0.016 | 0.016 | 0 | 0.64 | 0.5 | 0.77 | 0.66 | 0.73 | 0.74 |
| 10 | 72674069 | rs827340 | 0.143 | 0.404 | -1.165 | 0.929 | -0.026 | -0.055 | -0.055 | 0 | 0.15 | 0.03 | 0.14 | 0.26 | 0.14 | 0.25 |
| 10 | 72674489 | rs12415816 | 0.371 | 0.282 | -1.037 | 0.617 | -0.043 | 0.013 | 0.013 | 0 | 0.42 | 0.2 | 0.52 | 0.41 | 0.48 | 0.48 |
| 10 | 72680375 | rs7079039 | -0.372 | 0.594 | -0.836 | 0.275 | -0.014 | 0.026 | 0.026 | 0 | 0.39 | 0.34 | 0.55 | 0.32 | 0.48 | 0.47 |
| 10 | 72681365 | rs10999587 | 0.588 | 0.686 | -1.042 | 0.237 | -0.022 | -0.063 | -0.063 | 1.91 | 0.14 | 0.01 | 0.13 | 0.26 | 0.12 | 0.22 |
| 10 | 72683780 | rs4747115 | -0.967 | 0.364 | -0.518 | 0.27 | 0.036 | 0.053 | 0.053 | 0 | 0.48 | 0.75 | 0.69 | 0.41 | 0.56 | 0.59 |
| 10 | 72684682 | rs844128 | -1.686 | -0.373 | -0.919 | 0.267 | 0.047 | -0.037 | -0.037 | 0 | 0.76 | 0.86 | 0.89 | 0.69 | 0.86 | 0.81 |
| 10 | 72688371 | rs1466576 | 0.972 | -0.058 | -0.851 | -0.063 | 0.015 | -0.011 | -0.011 | 0 | 0.23 | 0.26 | 0.43 | 0.1 | 0.38 | 0.23 |
| 10 | 72691651 | rs12764441 | -0.328 | -0.312 | -0.322 | -0.179 | -0.001 | -0.001 | -0.001 | 0 | 0.42 | 0.58 | 0.57 | 0.55 | 0.49 | 0.55 |
| 10 | 72692295 | rs844127 | 1.309 | -0.02 | -0.516 | -0.181 | -0.034 | 0.014 | 0.014 | 0 | 0.06 | 0.01 | 0.08 | 0.01 | 0.14 | 0.03 |
| 10 | 72695261 | rs7915189 | 0.59 | -0.249 | -0.313 | -0.107 | -0.014 | 0.073 | 0.073 | 0.674 | 0.05 | 0.09 | 0.01 | 0.27 | 0.01 | 0.08 |
| 10 | 72701044 | rs1466575 | 1.285 | -0.107 | 0.17 | -0.185 | 0.094 | -0.044 | -0.044 | 0 | 0.21 | 0.34 | 0.42 | 0.12 | 0.37 | 0.23 |
| 10 | 72701888 | rs827294 | -0.943 | -0.467 | 0.203 | -0.418 | 0.005 | 0 | 0 | 0 | 0.6 | 0.6 | 0.73 | 0.42 | 0.75 | 0.55 |
| 10 | 72703816 | rs9962 | -0.283 | -0.402 | 0.115 | -0.386 | -0.006 | 0.016 | 0.016 | 0 | 0.37 | 0.24 | 0.3 | 0.31 | 0.38 | 0.3 |
| 10 | 72705865 | rs827289 | -1.038 | -0.575 | 0.073 | -0.42 | -0.004 | 0.008 | 0.008 | 0 | 0.52 | 0.36 | 0.44 | 0.49 | 0.49 | 0.57 |
| 10 | 72706707 | rs880314 | 0.135 | -0.589 | -0.097 | -0.417 | 0.024 | 0.001 | 0.001 | 0 | 0.31 | 0.2 | 0.22 | 0.31 | 0.24 | 0.26 |
| 10 | 72711156 | rs7083431 | 1.268 | -0.003 | -0.146 | -0.5 | 0.002 | -0.007 | -0.007 | 0 | 0.22 | 0.25 | 0.43 | 0.11 | 0.35 | 0.24 |
| 10 | 72713252 | rs827284 | -2.277 | -1.043 | -0.263 | -0.549 | 0.005 | -0.007 | -0.007 | 0.188 | 0.81 | 0.81 | 0.92 | 0.66 | 0.91 | 0.8 |
| 10 | 72714280 | rs10823644 | 1.102 | -1.01 | -0.351 | -0.601 | 0.009 | 0.033 | 0.033 | 0.261 | 0.24 | 0.16 | 0.2 | 0.04 | 0.22 | 0.18 |
| 10 | 72715367 | rs10823645 | 0.894 | -0.798 | -0.349 | -0.436 | -0.006 | -0.005 | -0.005 | 0.201 | 0.27 | 0.27 | 0.45 | 0.41 | 0.4 | 0.3 |
| 10 | 72717087 | rs10999609 | 1.686 | -0.824 | -0.81 | -0.518 | 0.018 | 0.046 | 0.046 | 0 | 0.2 | 0.07 | 0.12 | 0 | 0.16 | 0.1 |
| 10 | 72718265 | rs1538373 | 1.271 | -0.48 | -0.795 | -0.307 | -0.026 | -0.05 | -0.05 | 0 | 0.1 | 0.02 | 0.11 | 0.2 | 0.1 | 0.26 |
| 10 | 72719407 | rs10999616 | 1.261 | -0.288 | -0.797 | -0.31 | -0.036 | -0.05 | -0.05 | 0 | 0.1 | 0.02 | 0.11 | 0.2 | 0.11 | 0.26 |
| 10 | 72721192 | rs1767201 | -0.229 | -0.269 | -0.733 | -0.423 | 0.008 | 0.016 | 0.016 | 0 | 0.42 | 0.59 | 0.67 | 0.32 | 0.62 | 0.56 |

| | | | | | | | | | | | | | | | | |
|---|---|---|---|---|---|---|---|---|---|---|---|---|---|---|---|---|
| 10 | 72721363 | rs4747121 | 0.704 | -0.196 | -1.047 | -0.236 | -0.013 | 0.04 | 0.04 | 0 | 0.13 | 0.06 | 0.12 | 0.04 | 0.13 | 0.08 |
| 10 | 72721448 | rs12763860 | 1.029 | 0.08 | -0.446 | 0.137 | -0.045 | 0.037 | 0.037 | 0 | 0.21 | 0.13 | 0.42 | 0.11 | 0.37 | 0.25 |
| 10 | 72722497 | rs827273 | -1.412 | -0.027 | -0.576 | 0.268 | -0.012 | 0.011 | 0.011 | 0 | 0.81 | 0.77 | 0.92 | 0.64 | 0.92 | 0.79 |
| 10 | 72723866 | rs827271 | -0.226 | 0.564 | -0.612 | 0.086 | -0.002 | -0.005 | -0.005 | 3.43 | 0.47 | 0.46 | 0.35 | 0.35 | 0.43 | 0.27 |
| 10 | 72729492 | rs11597510 | -0.829 | 0.582 | -0.508 | 0.69 | 0.001 | 0.014 | 0.014 | 0 | 0.22 | 0.24 | 0.24 | 0.45 | 0.23 | 0.25 |
| 10 | 72729729 | rs827266 | -1.258 | 0.04 | 0.011 | 0.629 | 0.009 | 0.187 | 0.187 | 0.367 | 0.17 | 0.44 | 0.34 | 0.44 | 0.13 | 0.19 |
| 10 | 72731151 | rs7078625 | -0.426 | -1.154 | -0.408 | 0.75 | 0.009 | -0.01 | -0.01 | 0.061 | 0.79 | 0.79 | 0.74 | 0.69 | 0.87 | 0.73 |
| 10 | 72733889 | rs827263 | -0.926 | -1.319 | 0.023 | 0.743 | -0.051 | 0.039 | 0.039 | 0 | 0.83 | 0.75 | 0.93 | 0.72 | 0.93 | 0.78 |
| 10 | 72735039 | rs11598947 | 1.142 | -0.745 | -0.97 | 0.04 | -0.091 | -0.01 | -0.01 | 0 | 0.27 | 0.08 | 0.23 | 0.18 | 0.34 | 0.26 |
| 10 | 72735694 | rs1471825 | 0.231 | -1.062 | -0.557 | -0.229 | -0.008 | 0.04 | 0.04 | 0 | 0.44 | 0.33 | 0.36 | 0.23 | 0.49 | 0.45 |
| 10 | 72737390 | rs10999621 | 0.578 | -1.116 | -0.54 | -0.185 | -0.076 | -0.055 | -0.055 | 0 | 0.35 | 0.16 | 0.43 | 0.59 | 0.44 | 0.37 |
| 10 | 72739117 | rs7073396 | 1.599 | -1.437 | -0.177 | -0.431 | -0.036 | -0.053 | -0.053 | 0.56 | 0.11 | 0.21 | 0.05 | 0.01 | 0.04 | 0.08 |
| 10 | 72739317 | rs16928208 | 0.82 | -1.297 | -0.149 | -0.294 | -0.001 | 0.051 | 0.051 | 0 | 0.09 | 0.14 | 0.21 | 0.42 | 0.1 | 0.11 |
| 10 | 72739736 | rs7074431 | 0.29 | -1.228 | -0.237 | -0.442 | -0.002 | -0.012 | -0.012 | 0 | 0.3 | 0.26 | 0.24 | 0.08 | 0.38 | 0.28 |
| 10 | 72739901 | rs4746075 | -1.202 | -1.278 | -0.244 | -0.428 | -0.025 | 0.02 | 0.02 | 0 | 0.77 | 0.69 | 0.73 | 0.68 | 0.87 | 0.73 |
| 10 | 72740115 | rs10823653 | 0.826 | -1.183 | -0.562 | -0.219 | -0.001 | 0.025 | 0.025 | 0 | 0.08 | 0.09 | 0.2 | 0.41 | 0.1 | 0.11 |
| 10 | 72740379 | rs827259 | -0.237 | -1.082 | -0.613 | -0.222 | -0.011 | 0.022 | 0.022 | 0 | 0.36 | 0.27 | 0.44 | 0.14 | 0.43 | 0.33 |
| 10 | 72742933 | rs1767175 | -1.768 | -1.307 | -0.388 | 0.228 | -0.031 | 0.018 | 0.018 | 0 | 0.86 | 0.82 | 0.94 | 0.76 | 0.96 | 0.8 |
| 10 | 72743313 | rs1767180 | 0.268 | -0.935 | -1.103 | -0.001 | 0.018 | 0.051 | 0.051 | 0 | 0.47 | 0.24 | 0.54 | 0.29 | 0.56 | 0.46 |
| 10 | 72744144 | rs1767182 | 0.285 | -1.069 | -1.103 | -0.014 | 0.031 | 0.055 | 0.055 | 0 | 0.46 | 0.23 | 0.53 | 0.29 | 0.57 | 0.47 |
| 10 | 72744630 | rs756322 | -0.384 | -1.453 | -0.991 | 0.046 | -0.017 | 0 | 0 | 0.766 | 0.73 | 0.33 | 0.88 | 0.73 | 0.92 | 0.74 |
| 10 | 72744987 | rs1683863 | 0.125 | 0.142 | -1.141 | -0.224 | -0.038 | 0.037 | 0.037 | 0 | 0.26 | 0.19 | 0.37 | 0.13 | 0.37 | 0.24 |
| 10 | 72746143 | rs1614131 | -0.282 | -0.786 | -0.85 | -0.301 | -0.005 | -0.011 | -0.011 | 0 | 0.16 | 0.16 | 0.08 | 0.04 | 0.08 | 0.13 |
| 10 | 72748410 | rs10999626 | 0.661 | -1.096 | 0.245 | 0.386 | 0.037 | 0.04 | 0.04 | 0.7 | 0.53 | 0.29 | 0.54 | 0.35 | 0.67 | 0.52 |
| 10 | 72749183 | rs1621431 | -0.374 | -1.275 | 0.394 | 0.275 | -0.03 | 0.018 | 0.018 | 0.353 | 0.18 | 0.27 | 0.09 | 0.24 | 0.07 | 0.17 |
| 10 | 72749246 | rs7072890 | 0.502 | -1.362 | 0.457 | 0.25 | -0.066 | 0.026 | 0.026 | 1.815 | 0.79 | 0.53 | 0.89 | 0.76 | 0.93 | 0.83 |
| 10 | 72750636 | rs16928239 | -0.425 | -1.2 | -0.2 | 0.181 | -0.003 | -0.004 | -0.004 | 0 | 0.09 | 0.13 | 0.04 | 0.04 | 0.05 | 0.07 |
| 10 | 72754154 | rs12244234 | 1.195 | -1.243 | -0.334 | 0.396 | 0.012 | -0.014 | -0.014 | 0 | 0.1 | 0.05 | 0.04 | 0.22 | 0.03 | 0.12 |
| 10 | 72755123 | rs1338638 | 0.273 | -1.239 | -0.206 | 0.472 | 0.019 | -0.02 | -0.02 | 0 | 0.66 | 0.69 | 0.69 | 0.56 | 0.72 | 0.57 |
| 10 | 72756088 | rs1111803 | -0.235 | -1.271 | -0.16 | 0.503 | 0.006 | -0.008 | -0.008 | 0.203 | 0.72 | 0.71 | 0.73 | 0.6 | 0.79 | 0.66 |
| 10 | 72756178 | rs10509324 | 0.534 | -1.045 | 0.104 | 0.28 | -0.009 | -0.028 | -0.028 | 0.105 | 0.07 | 0.13 | 0.05 | 0 | 0.07 | 0.04 |
| 10 | 72757521 | rs12771128 | 0.575 | -0.673 | 0.038 | 0.4 | 0.003 | 0.024 | 0.024 | 0 | 0.07 | 0.07 | 0.04 | 0.2 | 0.09 | 0.05 |
| 10 | 72758344 | rs11597098 | 0.901 | -0.76 | -0.188 | 0.314 | -0.024 | -0.008 | -0.008 | 0 | 0.07 | 0 | 0.04 | 0.07 | 0.07 | 0.06 |
| 10 | 72763493 | rs1361239 | -0.178 | -1.256 | 0.123 | 0.289 | -0.047 | 0.084 | 0.084 | 0 | 0.8 | 0.68 | 0.79 | 0.62 | 0.87 | 0.75 |
| 10 | 72764857 | rs11595715 | -0.484 | -0.997 | 0.134 | 0.151 | 0.001 | 0.01 | 0.01 | 0 | 0.16 | 0.17 | 0.1 | 0.26 | 0.16 | 0.1 |
| 10 | 72767409 | rs10509328 | 0.273 | -0.892 | 0.182 | 0.239 | 0.016 | 0.063 | 0.063 | 0 | 0.08 | 0.14 | 0.05 | 0.19 | 0.09 | 0.05 |
| 10 | 72771680 | rs6480474 | 0.152 | -0.589 | -0.306 | -0.3 | -0.004 | 0.028 | 0.028 | 0.019 | 0.75 | 0.44 | 0.71 | 0.4 | 0.72 | 0.68 |
| 10 | 72774514 | rs10999635 | 0.632 | -0.528 | -1.147 | -0.811 | 0.005 | 0.105 | 0.105 | 0 | 0.46 | 0.34 | 0.47 | 0.09 | 0.53 | 0.38 |
| 10 | 72775776 | rs3902977 | 0.023 | -0.196 | -0.951 | -0.251 | -0.007 | -0.043 | -0.043 | 0.871 | 0.22 | 0.14 | 0.27 | 0.6 | 0.28 | 0.31 |
| 10 | 72786497 | rs7072464 | 0.869 | 0.464 | -0.997 | -0.224 | -0.003 | -0.004 | -0.004 | 0 | 0.1 | 0.1 | 0.09 | 0 | 0.12 | 0.04 |
| 10 | 72786849 | rs10762417 | -0.003 | 0.026 | -0.844 | -0.099 | 0.019 | 0.026 | 0.026 | 0.081 | 0.53 | 0.43 | 0.61 | 0.34 | 0.61 | 0.51 |
| 10 | 72788676 | rs7100223 | -0.356 | 0.431 | -0.793 | 0.39 | 0.007 | 0.005 | 0.005 | 0 | 0.49 | 0.6 | 0.4 | 0.45 | 0.45 | 0.54 |
| 10 | 72792488 | rs7076053 | 0.193 | 0.008 | -0.413 | 0.212 | 0.003 | 0.041 | 0.041 | 0 | 0.69 | 0.46 | 0.53 | 0.56 | 0.67 | 0.63 |
| 10 | 72793796 | rs12252440 | 0.395 | -1.107 | -0.161 | -0.131 | -0.005 | 0 | 0 | 0 | 0.65 | 0.64 | 0.54 | 0.64 | 0.7 | 0.65 |
| 10 | 72793852 | rs12261506 | -0.814 | -1.089 | -0.098 | -0.117 | -0.026 | -0.061 | -0.061 | 0 | 0.78 | 0.91 | 0.6 | 0.65 | 0.74 | 0.77 |
| 10 | 72794984 | rs10999645 | 0.524 | -1.196 | 0.092 | -0.159 | -0.025 | -0.004 | -0.004 | 0 | 0.58 | 0.49 | 0.52 | 0.61 | 0.69 | 0.55 |
| 10 | 72809566 | rs7090828 | -0.181 | -0.481 | -0.788 | -0.189 | -0.032 | 0.051 | 0.051 | 0.535 | 0.57 | 0.82 | 0.51 | 0.7 | 0.55 | 0.66 |
| 10 | 72810071 | rs7077129 | -0.51 | -0.578 | -0.859 | -0.246 | -0.085 | -0.002 | -0.002 | 0 | 0.62 | 0.92 | 0.52 | 0.7 | 0.57 | 0.67 |
| 10 | 72810764 | rs7902620 | -0.096 | -0.372 | -0.63 | -0.181 | -0.008 | 0.038 | 0.038 | 0.426 | 0.56 | 0.77 | 0.51 | 0.64 | 0.56 | 0.63 |
| 10 | 72813306 | rs10762420 | 0.116 | -0.373 | -0.41 | -0.193 | 0.002 | 0.006 | 0.006 | 0.507 | 0.52 | 0.55 | 0.48 | 0.61 | 0.54 | 0.52 |
| 10 | 72823084 | rs12416144 | 0.876 | -0.007 | -0.549 | -0.227 | -0.021 | 0.017 | 0.017 | 0.013 | 0.05 | 0 | 0.11 | 0.04 | 0.1 | 0.05 |
| 10 | 72824456 | rs10999657 | -0.76 | 0.209 | -0.658 | 0.109 | -0.013 | -0.049 | -0.049 | 0 | 0.22 | 0.11 | 0.39 | 0.48 | 0.3 | 0.27 |
| 10 | 72824920 | rs4747134 | -0.459 | 0.095 | -0.653 | 0.089 | 0 | 0.078 | 0.078 | 0 | 0.06 | 0.05 | 0.2 | 0.4 | 0.06 | 0.15 |
| 10 | 72832086 | rs7097900 | -0.958 | -0.118 | -0.37 | -1.155 | 0 | -0.031 | -0.031 | 0 | 0.31 | 0.31 | 0.26 | 0.04 | 0.35 | 0.2 |
| 10 | 72847054 | rs9804264 | -0.483 | -0.207 | 0.104 | -0.898 | -0.008 | -0.066 | -0.066 | 0.29 | 0.18 | 0.27 | 0.12 | 0.03 | 0.19 | 0.06 |
| 10 | 72848345 | rs11599200 | -0.507 | -0.208 | 0.103 | -0.898 | -0.008 | -0.066 | -0.066 | 0 | 0.18 | 0.27 | 0.12 | 0.03 | 0.19 | 0.06 |
| 10 | 72849958 | rs16928377 | 0.412 | -0.495 | 0.045 | -1.443 | 0.007 | -0.002 | -0.002 | 0 | 0.27 | 0.24 | 0.15 | 0.02 | 0.2 | 0.21 |
| 10 | 72850385 | rs7075905 | 0.34 | -0.47 | 0.173 | -1.372 | -0.016 | -0.081 | -0.081 | 0.011 | 0.13 | 0.31 | 0.08 | 0 | 0.1 | 0.02 |
| 10 | 72852735 | rs7910164 | -0.923 | -0.518 | 0.077 | -1.409 | 0.01 | -0.003 | -0.003 | 0 | 0.44 | 0.65 | 0.28 | 0.05 | 0.4 | 0.27 |
| 10 | 72855230 | rs7100036 | 0.827 | -0.476 | -0.376 | -1.284 | -0.001 | -0.001 | -0.001 | 0 | 0.06 | 0.09 | 0.05 | 0.03 | 0.1 | 0.04 |
| 10 | 72855411 | rs1935145 | 0.399 | -0.536 | -0.12 | -1.411 | -0.006 | -0.063 | -0.063 | 0.126 | 0.26 | 0.32 | 0.16 | 0.02 | 0.2 | 0.22 |
| 10 | 72857415 | rs10999675 | -0.587 | -0.631 | 0.099 | -1.485 | 0.01 | -0.013 | -0.013 | 0.506 | 0.39 | 0.65 | 0.24 | 0.02 | 0.3 | 0.28 |
| 10 | 72858079 | rs5006438 | -0.795 | -0.577 | 0.038 | -1.351 | 0.029 | 0.094 | 0.094 | 0.101 | 0.45 | 0.74 | 0.29 | 0.05 | 0.4 | 0.31 |
| 10 | 72859876 | rs11497787 | 0.312 | -0.482 | -0.186 | -1.211 | -0.017 | -0.087 | -0.087 | 0.011 | 0.13 | 0.33 | 0.08 | 0.01 | 0.1 | 0.06 |
| 10 | 72880512 | rs7900882 | -0.384 | -0.276 | -1.039 | -1.053 | 0.038 | 0.181 | 0.181 | 0.287 | 0.45 | 0.82 | 0.29 | 0.05 | 0.4 | 0.31 |
| 10 | 72883868 | rs10999688 | 1.089 | 0.463 | -1.137 | -0.015 | -0.067 | -0.102 | -0.102 | 0 | 0.21 | 0.03 | 0.24 | 0.31 | 0.22 | 0.25 |
| 10 | 72893339 | rs947364 | -0.462 | 0.222 | -0.67 | -0.048 | -0.014 | -0.001 | -0.001 | 0 | 0.56 | 0.73 | 0.49 | 0.6 | 0.57 | 0.54 |
| 10 | 72902886 | rs7923921 | -0.139 | 0.121 | -0.221 | 0.393 | -0.005 | 0.115 | 0.115 | 0.469 | 0.1 | 0.23 | 0.15 | 0.32 | 0.16 | 0.14 |
| 10 | 72903531 | rs12246465 | 0.639 | -0.142 | 0.074 | -0.153 | -0.004 | -0.082 | -0.082 | 2.356 | 0.2 | 0.29 | 0.12 | 0 | 0.18 | 0.17 |

| | | | | | | | | | | | | | | | | |
|---|---|---|---|---|---|---|---|---|---|---|---|---|---|---|---|---|
| 10 | 72908675 | rs947366 | 1.869 | -0.055 | -0.421 | -0.224 | -0.053 | -0.117 | -0.117 | 0.329 | 0.22 | 0.04 | 0.27 | 0.37 | 0.23 | 0.28 |
| 10 | 72908983 | rs12220332 | 0.644 | -0.309 | -0.328 | -0.414 | -0.017 | 0.037 | 0.037 | 0.457 | 0.27 | 0.14 | 0.22 | 0.2 | 0.25 | 0.24 |
| 10 | 72913623 | rs7090550 | -1.035 | -0.408 | -0.4 | -0.449 | -0.061 | -0.073 | -0.073 | 0 | 0.6 | 0.91 | 0.51 | 0.52 | 0.59 | 0.54 |
| 10 | 72919002 | rs10509331 | -0.764 | -0.182 | 0.41 | 0.294 | 0 | 0.008 | 0.008 | 0.385 | 0.37 | 0.42 | 0.37 | 0.58 | 0.41 | 0.39 |
| 10 | 72944032 | rs7094373 | -0.444 | -0.102 | 0.603 | 0.639 | -0.002 | -0.013 | -0.013 | 0.013 | 0.29 | 0.27 | 0.27 | 0.49 | 0.29 | 0.33 |
| 10 | 72948577 | rs7080430 | -0.055 | 0.089 | 0.541 | 0.657 | -0.019 | 0.153 | 0.153 | 0.775 | 0.13 | 0.36 | 0.16 | 0.31 | 0.16 | 0.11 |
| 10 | 72958482 | rs10999726 | 0.073 | 0.193 | 0.135 | 0.633 | -0.03 | -0.051 | -0.051 | 0 | 0.15 | 0.01 | 0.1 | 0.2 | 0.14 | 0.17 |
| 10 | 72960558 | rs10999728 | -1.13 | 0.428 | -0.135 | -0.159 | -0.02 | 0.053 | 0.053 | 0 | 0.85 | 0.97 | 0.78 | 0.96 | 0.82 | 0.84 |
| 10 | 72966593 | rs10740369 | -1.663 | 0.346 | 0.641 | 0.596 | 0.017 | 0.019 | 0.019 | 0 | 0.45 | 0.68 | 0.34 | 0.63 | 0.43 | 0.42 |
| 10 | 72974671 | rs16928524 | 0.059 | 0.937 | 0.348 | 0.796 | -0.002 | 0.001 | 0.001 | 0 | 0.15 | 0.14 | 0.08 | 0.04 | 0.12 | 0.09 |
| 10 | 72975434 | rs12264668 | 0.087 | 0.951 | 0.414 | 0.805 | 0.004 | -0.023 | -0.023 | 0 | 0.15 | 0.22 | 0.08 | 0.05 | 0.13 | 0.14 |
| 10 | 72976125 | rs2247491 | -0.848 | 1.028 | 0.187 | 0.864 | -0.001 | 0.009 | 0.009 | 0 | 0.33 | 0.46 | 0.28 | 0.61 | 0.36 | 0.37 |
| 10 | 72982254 | rs2244140 | 0.708 | 1.137 | 0.632 | 0.729 | 0.094 | 0.028 | 0.028 | 1.411 | 0.94 | 0.59 | 0.91 | 1 | 0.94 | 0.99 |
| 10 | 72982985 | rs16928529 | 1.244 | 1.68 | -0.005 | 1.181 | 0.021 | 0.021 | 0.021 | 3.819 | 0.73 | 0.33 | 0.7 | 0.67 | 0.62 | 0.79 |
| 10 | 72984697 | rs875001 | 1.135 | 1.347 | -0.577 | 1.048 | -0.012 | 0.003 | 0.003 | 2.023 | 0.75 | 0.19 | 0.73 | 0.75 | 0.68 | 0.74 |
| 10 | 72991095 | rs12781290 | -0.587 | 1.606 | -0.971 | 1.088 | -0.084 | -0.05 | -0.05 | 1.304 | 0.23 | 0.07 | 0.16 | 0.23 | 0.31 | 0.21 |
| 10 | 72991385 | rs7893421 | -1.239 | 1.471 | -0.055 | 1.009 | 0.029 | -0.003 | -0.003 | 0 | 0.35 | 0.47 | 0.33 | 0.34 | 0.43 | 0.31 |
| 10 | 72998234 | rs10999739 | -0.244 | 1.676 | -0.114 | 1.101 | -0.058 | -0.033 | -0.033 | 3.101 | 0.19 | 0.08 | 0.17 | 0.22 | 0.28 | 0.22 |
| 10 | 73000686 | rs10999742 | -0.8 | 1.493 | -0.302 | 1.002 | -0.023 | -0.023 | -0.023 | 0.267 | 0.09 | 0 | 0.12 | 0.13 | 0.09 | 0.06 |
| 10 | 73000943 | rs2894147 | -1.574 | 1.235 | 0.375 | 1.443 | 0.001 | 0.001 | 0.001 | 0 | 0.34 | 0.35 | 0.34 | 0.48 | 0.42 | 0.36 |
| 10 | 73005037 | rs17545059 | 0.354 | 0.367 | -0.419 | 0.93 | -0.001 | 0.04 | 0.04 | 0 | 0.22 | 0.03 | 0.14 | 0.14 | 0.19 | 0.21 |
| 10 | 73005937 | rs7080150 | -1.354 | 0.31 | 0.294 | 1.559 | 0.008 | 0.005 | 0.005 | 0.008 | 0.38 | 0.43 | 0.55 | 0.5 | 0.44 | 0.4 |
| 10 | 73006620 | rs4746080 | -1.739 | 0.347 | 0.855 | 1.564 | -0.028 | 0.054 | 0.054 | 0 | 0.42 | 0.71 | 0.61 | 0.69 | 0.48 | 0.53 |
| 10 | 73008857 | rs10823706 | 1.611 | 0.228 | 0.9 | 1.44 | 0.015 | 0.13 | 0.13 | 0 | 0.49 | 0.29 | 0.35 | 0.17 | 0.49 | 0.34 |
| 10 | 73010064 | rs7899034 | 0.249 | 0.303 | 0.793 | 1.597 | -0.091 | -0.056 | -0.056 | 0 | 0.2 | 0.06 | 0.16 | 0.2 | 0.27 | 0.21 |
| 10 | 73010322 | rs7915759 | -1.306 | 0.255 | 1.294 | 1.67 | 0 | 0 | 0 | 0.008 | 0.37 | 0.42 | 0.46 | 0.56 | 0.38 | 0.49 |
| 10 | 73011737 | rs7904418 | 2.039 | 0.235 | 1.223 | 1.713 | -0.047 | 0.11 | 0.11 | 0.055 | 0.77 | 0.44 | 0.64 | 0.45 | 0.83 | 0.6 |
| 10 | 73012084 | rs16928560 | -1.319 | 0.301 | 1.306 | 1.726 | -0.003 | 0.029 | 0.029 | 0 | 0.17 | 0.2 | 0.3 | 0.36 | 0.19 | 0.28 |
| 10 | 73012628 | rs4747140 | 0.648 | 0.604 | 1.15 | 1.882 | 0.019 | 0.031 | 0.031 | 0 | 0.7 | 0.59 | 0.75 | 0.61 | 0.68 | 0.65 |
| 10 | 73020768 | rs1538894 | 1.443 | -0.155 | 1.831 | 2.019 | 0.018 | 0.178 | 0.178 | 0 | 0.92 | 0.82 | 0.72 | 0.65 | 0.89 | 0.74 |
| 10 | 73021816 | rs10762431 | 2.088 | -0.152 | 1.724 | 1.935 | 0.003 | 0.157 | 0.157 | 0 | 0.81 | 0.51 | 0.57 | 0.56 | 0.8 | 0.71 |
| 10 | 73027892 | rs4747143 | 1.82 | -0.295 | 1.898 | 1.56 | 0.002 | 0.14 | 0.14 | 0.15 | 0.87 | 0.71 | 0.86 | 0.66 | 0.86 | 0.81 |
| 10 | 73028146 | rs4747144 | 3.092 | -0.483 | 2.195 | 1.544 | 0.013 | -0.019 | -0.019 | 0.398 | 0.77 | 0.22 | 0.72 | 0.58 | 0.83 | 0.75 |
| 10 | 73032455 | rs16928583 | -1.999 | -0.418 | 2.287 | 1.209 | -0.019 | -0.009 | -0.009 | 0.001 | 0.12 | 0.38 | 0.14 | 0.07 | 0.06 | 0.13 |
| 10 | 73034953 | rs11598330 | -1.513 | -0.389 | 2.303 | 1.214 | -0.005 | -0.002 | -0.002 | 0 | 0.09 | 0.11 | 0.12 | 0.07 | 0.06 | 0.13 |
| 10 | 73036899 | rs16928594 | -0.598 | -0.367 | 2.202 | 1.283 | 0 | -0.005 | -0.005 | 6.16 | 0.07 | 0.04 | 0.06 | 0.13 | 0.09 | 0.13 |
| 10 | 73043016 | rs11593534 | -2.479 | -0.373 | 2.331 | 1.264 | -0.038 | 0.007 | 0.007 | 0 | 0.2 | 0.47 | 0.22 | 0.2 | 0.16 | 0.27 |
| 10 | 73045537 | rs4747146 | 2.475 | -0.365 | 2.463 | 1.294 | -0.032 | 0.046 | 0.046 | 0 | 0.76 | 0.44 | 0.71 | 0.64 | 0.78 | 0.66 |
| 10 | 73047690 | rs2277259 | 1.839 | -0.277 | 2.296 | 1.309 | 0 | 0.087 | 0.087 | 0.509 | 0.84 | 0.62 | 0.84 | 0.71 | 0.84 | 0.76 |
| 10 | 73051388 | rs3740460 | -1.559 | -0.053 | 1.471 | 1.086 | 0.002 | 0.031 | 0.031 | 0 | 0.11 | 0.21 | 0.17 | 0.2 | 0.13 | 0.26 |
| 10 | 73055965 | rs1105172 | 1.837 | -0.468 | 0.971 | 0.822 | -0.046 | -0.019 | -0.019 | 0 | 0.81 | 0.42 | 0.82 | 0.84 | 0.87 | 0.77 |
| 10 | 73062048 | rs2275577 | -1.372 | -0.648 | 0.992 | 1.043 | -0.038 | 0.039 | 0.039 | 1.952 | 0.15 | 0.55 | 0.1 | 0.17 | 0.09 | 0.15 |
| 10 | 73063290 | rs7075861 | 0.033 | -0.492 | -0.01 | 1.002 | -0.005 | -0.011 | -0.011 | 0 | 0.3 | 0.21 | 0.28 | 0.37 | 0.34 | 0.32 |
| 10 | 73063438 | rs6480507 | -1.476 | -0.595 | -0.5 | 0.679 | -0.006 | -0.004 | -0.004 | 0 | 0.54 | 0.76 | 0.41 | 0.55 | 0.49 | 0.5 |
| 10 | 73067279 | rs4400714 | 0.555 | -0.425 | -0.967 | 0.758 | -0.028 | -0.048 | -0.048 | 0 | 0.08 | 0.01 | 0.13 | 0.22 | 0.11 | 0.12 |
| 10 | 73069119 | rs17546494 | 0.892 | -0.561 | -0.928 | 0.493 | -0.02 | 0.084 | 0.084 | 0.09 | 0.2 | 0.05 | 0.12 | 0 | 0.21 | 0.12 |
| 10 | 73069492 | rs7079524 | -1.275 | -0.7 | -0.811 | 0.505 | 0.013 | -0.001 | -0.001 | 0.083 | 0.27 | 0.77 | 0.17 | 0.24 | 0.17 | 0.25 |
| 10 | 73076862 | rs869545 | 0.778 | -0.781 | -0.933 | 0.375 | -0.015 | 0.049 | 0.049 | 0.424 | 0.21 | 0.09 | 0.13 | 0.06 | 0.23 | 0.19 |
| 10 | 73078447 | rs4747148 | 1.731 | -0.604 | -1.888 | 0.291 | -0.112 | -0.073 | -0.073 | 0.376 | 0.25 | 0.02 | 0.45 | 0.19 | 0.32 | 0.21 |
| 10 | 73080635 | rs10999769 | 0.225 | -0.545 | -0.718 | 0.121 | 0.017 | 0.136 | 0.136 | 0.105 | 0.12 | 0.44 | 0.09 | 0.2 | 0.08 | 0.09 |
| 10 | 73081911 | rs10999771 | -0.529 | -0.496 | -0.927 | 0.241 | -0.001 | 0.016 | 0.016 | 0 | 0.42 | 0.7 | 0.34 | 0.36 | 0.4 | 0.43 |
| 10 | 73082563 | rs2277257 | -0.529 | -0.496 | -0.937 | 0.241 | -0.001 | 0.016 | 0.016 | 0 | 0.42 | 0.7 | 0.34 | 0.36 | 0.4 | 0.43 |
| 10 | 73087778 | rs10999776 | -0.05 | -0.483 | -0.929 | 0.199 | 0.004 | 0.013 | 0.013 | 0.302 | 0.34 | 0.69 | 0.24 | 0.26 | 0.31 | 0.28 |
| 10 | 73089261 | rs6480511 | 0.91 | -0.463 | -1.315 | 0.169 | -0.009 | -0.021 | -0.021 | 0 | 0.09 | 0.17 | 0.05 | 0.01 | 0.06 | 0.03 |
| 10 | 73093680 | rs6480513 | -1.06 | -0.349 | -1.487 | 0.31 | -0.089 | -0.129 | -0.129 | 0.463 | 0.53 | 0.97 | 0.42 | 0.51 | 0.46 | 0.46 |
| 10 | 73095265 | rs2066211 | 0.803 | -0.076 | -1.22 | 0.01 | 0.017 | -0.043 | -0.043 | 0 | 0.09 | 0.1 | 0.07 | 0 | 0.15 | 0.06 |
| 10 | 73095793 | rs16928705 | 0.061 | -0.259 | -0.967 | 0.048 | -0.008 | 0.034 | 0.034 | 0 | 0.12 | 0.3 | 0.16 | 0.26 | 0.1 | 0.17 |
| 10 | 73096236 | rs12256138 | -0.061 | -0.221 | -1.416 | -0.124 | -0.081 | 0.183 | 0.183 | 0.575 | 0.37 | 0.1 | 0.21 | 0.07 | 0.42 | 0.27 |
| 10 | 73097562 | rs780660 | -1.073 | 0.18 | -0.996 | 0.088 | 0.009 | -0.013 | -0.013 | 0 | 0.25 | 0.36 | 0.23 | 0.26 | 0.29 | 0.24 |
| 10 | 73097953 | rs780662 | -0.06 | 0.248 | -1.606 | 0.166 | -0.009 | 0 | 0 | 0 | 0.13 | 0.12 | 0.13 | 0.18 | 0.14 | 0.18 |
| 10 | 73099240 | rs3781316 | 0.492 | 0.009 | -1.804 | 0.25 | 0.001 | 0.013 | 0.013 | 2.452 | 0.56 | 0.17 | 0.63 | 0.53 | 0.58 | 0.63 |
| 10 | 73100480 | rs3781317 | 0.274 | 1.224 | -1.392 | -0.127 | -0.042 | 0.058 | 0.058 | 0.329 | 0.25 | 0.12 | 0.23 | 0.08 | 0.33 | 0.22 |
| 10 | 73104056 | rs2243540 | -0.112 | 0.915 | -0.835 | 0.722 | 0.009 | 0.114 | 0.114 | 0 | 0.92 | 0.85 | 0.87 | 0.67 | 0.9 | 0.79 |
| 10 | 73104178 | rs883764 | 0.248 | 0.522 | -0.543 | 0.276 | -0.002 | -0.01 | -0.01 | 0 | 0.44 | 0.48 | 0.29 | 0.31 | 0.45 | 0.3 |
| 10 | 73110657 | rs780665 | 0.996 | 0.263 | 0.48 | 0.737 | 0.005 | -0.011 | -0.011 | 0 | 0.69 | 0.32 | 0.5 | 0.44 | 0.77 | 0.49 |
| 10 | 73113790 | rs780675 | 0.975 | 0.236 | 0.734 | 0.76 | -0.008 | 0.014 | 0.014 | 0.133 | 0.69 | 0.39 | 0.51 | 0.43 | 0.76 | 0.48 |
| 10 | 73114486 | rs780678 | 0.846 | 0.197 | 1.069 | 0.724 | -0.019 | 0.083 | 0.083 | 0.81 | 0.73 | 0.53 | 0.54 | 0.43 | 0.78 | 0.49 |
| 10 | 73115942 | rs2252996 | -0.024 | 0.209 | 0.903 | 0.916 | 0.016 | 0.095 | 0.095 | 0 | 0.91 | 0.83 | 0.65 | 0.67 | 0.88 | 0.72 |
| 10 | 73119023 | rs703256 | 1.078 | 0.398 | 0.567 | 0.773 | -0.002 | -0.009 | -0.009 | 0 | 0.67 | 0.64 | 0.5 | 0.44 | 0.68 | 0.43 |

| | | | | | | | | | | | | | | | | |
|---|---|---|---|---|---|---|---|---|---|---|---|---|---|---|---|---|
| 10 | 73121945 | rs1084004 | 0.095 | 0.111 | 0.556 | 0.905 | 0.013 | 0.117 | 0.117 | 0 | 0.9 | 0.75 | 0.66 | 0.72 | 0.9 | 0.72 |
| 10 | 73122468 | rs780680 | 0.174 | 0.171 | 0.038 | 0.484 | 0.009 | 0.005 | 0.005 | 0 | 0.14 | 0.06 | 0.15 | 0.1 | 0.1 | 0.16 |
| 10 | 73124160 | rs780682 | 0.788 | 0.533 | 0.378 | 0.61 | 0 | 0.115 | 0.115 | 0 | 0.8 | 0.57 | 0.59 | 0.66 | 0.83 | 0.62 |
| 10 | 73126031 | rs10999785 | 0.478 | 0.568 | -0.019 | 0.609 | -0.016 | -0.061 | -0.061 | 0 | 0.11 | 0.24 | 0.08 | 0 | 0.06 | 0.08 |
| 10 | 73126727 | rs2185416 | -0.158 | 0.875 | -1.132 | 0.699 | -0.025 | -0.013 | -0.013 | 0 | 0.18 | 0.13 | 0.13 | 0.22 | 0.19 | 0.23 |
| 10 | 73129458 | rs780692 | 0.346 | 0.74 | -1.266 | 0.609 | -0.007 | -0.016 | -0.016 | 0 | 0.59 | 0.29 | 0.4 | 0.44 | 0.63 | 0.37 |
| 10 | 73139333 | rs10762438 | 0.635 | 0.432 | -1.707 | 0.45 | -0.129 | -0.063 | -0.063 | 0 | 0.47 | 0.03 | 0.38 | 0.49 | 0.55 | 0.4 |
| 10 | 73140004 | rs10823725 | -0.492 | -0.375 | -0.451 | -0.173 | -0.005 | -0.027 | -0.027 | 1.046 | 0.27 | 0.38 | 0.21 | 0.12 | 0.23 | 0.25 |
| 10 | 73141020 | rs1417208 | -0.343 | -0.345 | -0.471 | -0.16 | -0.002 | 0.012 | 0.012 | 0 | 0.24 | 0.23 | 0.2 | 0.12 | 0.23 | 0.25 |
| 10 | 73151645 | rs1890886 | -0.3 | -0.582 | -0.389 | -0.166 | 0.006 | 0.01 | 0.01 | 0 | 0.23 | 0.21 | 0.18 | 0.12 | 0.19 | 0.23 |
| 10 | 73153890 | rs16928773 | -0.37 | -0.849 | 0.176 | -0.224 | -0.038 | -0.03 | -0.03 | 0 | 0.15 | 0.25 | 0.07 | 0.07 | 0.06 | 0.12 |
| 10 | 73154073 | rs10823733 | -1.312 | -0.863 | -0.438 | -0.241 | 0.066 | 0.073 | 0.073 | 0 | 0.4 | 0.81 | 0.31 | 0.19 | 0.26 | 0.35 |
| 10 | 73158678 | rs16928777 | 0.405 | -0.903 | -0.41 | -0.235 | 0.011 | 0.008 | 0.008 | 0.022 | 0.11 | 0.09 | 0.03 | 0.07 | 0.04 | 0.1 |
| 10 | 73158707 | rs1417207 | -1.286 | -0.76 | -0.7 | -0.178 | 0.069 | 0.073 | 0.073 | 0 | 0.4 | 0.81 | 0.27 | 0.19 | 0.25 | 0.33 |
| 10 | 73162237 | rs4491141 | -0.364 | -0.25 | -0.964 | 0.023 | 0.007 | 0.017 | 0.017 | 0 | 0.22 | 0.19 | 0.17 | 0.12 | 0.18 | 0.22 |
| 10 | 73163900 | rs7919846 | -0.909 | -0.564 | -1.884 | -0.461 | -0.036 | 0.134 | 0.134 | 2.136 | 0.57 | 0.96 | 0.33 | 0.19 | 0.36 | 0.37 |
| 10 | 73166050 | rs7088571 | -0.245 | -0.908 | -1.156 | -0.207 | 0.002 | 0.003 | 0.003 | 0.743 | 0.37 | 0.36 | 0.27 | 0.23 | 0.33 | 0.3 |
| 10 | 73178389 | rs10823741 | -0.587 | -1.123 | -0.295 | -0.257 | 0.009 | 0.004 | 0.004 | 0.601 | 0.42 | 0.59 | 0.3 | 0.3 | 0.32 | 0.36 |
| 10 | 73180767 | rs10823743 | -0.412 | -1.122 | -0.094 | -0.257 | -0.017 | -0.016 | -0.016 | 0 | 0.39 | 0.52 | 0.29 | 0.23 | 0.31 | 0.3 |
| 10 | 73184175 | rs871118 | -0.276 | -1.167 | -0.242 | -0.267 | 0.005 | 0.005 | 0.005 | 1.228 | 0.38 | 0.36 | 0.28 | 0.22 | 0.31 | 0.3 |
| 10 | 73191336 | rs2394795 | 0.777 | -0.613 | -0.303 | -0.126 | -0.001 | -0.001 | -0.001 | 0 | 0.42 | 0.41 | 0.42 | 0.49 | 0.53 | 0.43 |
| 10 | 73192372 | rs17706726 | 1.422 | -0.765 | -0.271 | -0.429 | -0.049 | -0.055 | -0.055 | 0 | 0.32 | 0.19 | 0.37 | 0.49 | 0.45 | 0.37 |
| 10 | 73197656 | rs10762446 | 0.914 | -0.939 | -0.313 | -0.526 | -0.032 | -0.042 | -0.042 | 0.19 | 0.69 | 0.53 | 0.65 | 0.73 | 0.77 | 0.67 |
| 10 | 73199333 | rs10740381 | -0.47 | -0.953 | -0.436 | -0.514 | -0.058 | 0.001 | 0.001 | 0 | 0.83 | 0.98 | 0.71 | 0.84 | 0.81 | 0.8 |
| 10 | 73200867 | rs7086538 | 0.382 | -0.865 | -0.482 | -0.462 | 0.001 | 0.001 | 0.001 | 0.922 | 0.38 | 0.38 | 0.29 | 0.26 | 0.33 | 0.31 |
| 10 | 73201670 | rs7087853 | 0.096 | -0.898 | -0.486 | -0.09 | -0.023 | 0.01 | 0.01 | 0 | 0.76 | 0.92 | 0.62 | 0.75 | 0.76 | 0.78 |
| 10 | 73201846 | rs10762449 | 0.151 | -0.563 | -0.387 | -0.246 | 0.008 | 0.003 | 0.003 | 0.366 | 0.42 | 0.4 | 0.3 | 0.37 | 0.35 | 0.39 |
| 10 | 73207662 | rs12778279 | 0.086 | -1.026 | -0.365 | -0.333 | 0 | -0.001 | -0.001 | 0 | 0.55 | 0.42 | 0.45 | 0.54 | 0.56 | 0.53 |
| 10 | 73213089 | rs7909560 | 0.396 | -0.72 | -0.185 | -0.032 | 0.006 | 0.004 | 0.004 | 0 | 0.38 | 0.42 | 0.38 | 0.56 | 0.43 | 0.44 |
| 10 | 73220557 | rs7902794 | 1.008 | -0.696 | -0.606 | 0.019 | -0.194 | -0.18 | -0.18 | 2.329 | 0.31 | 0.03 | 0.34 | 0.45 | 0.43 | 0.36 |
| 10 | 73222123 | rs17707219 | 1.539 | 0.425 | -0.792 | 0.488 | -0.026 | -0.033 | -0.033 | 0 | 0.06 | 0.01 | 0.09 | 0.14 | 0.18 | 0.14 |
| 10 | 73225677 | rs2185415 | -0.53 | -0.667 | -0.784 | -0.067 | -0.008 | -0.043 | -0.043 | 0.112 | 0.59 | 0.55 | 0.56 | 0.78 | 0.61 | 0.71 |
| 10 | 73226388 | rs10999828 | 0.034 | -0.544 | -0.264 | -0.003 | -0.046 | 0.039 | 0.039 | 0 | 0.17 | 0.39 | 0.17 | 0.17 | 0.11 | 0.22 |
| 10 | 73227127 | rs12265176 | 0.188 | -0.265 | -0.702 | -0.052 | -0.003 | -0.004 | -0.004 | 0 | 0.08 | 0.12 | 0.05 | 0.06 | 0.08 | 0.05 |
| 10 | 73227739 | rs9783254 | -0.54 | -0.218 | -0.429 | -0.061 | -0.051 | -0.007 | -0.007 | 0 | 0.27 | 0.41 | 0.22 | 0.26 | 0.19 | 0.34 |
| 10 | 73230904 | rs10999834 | -0.078 | 0.812 | -0.782 | 0.337 | -0.006 | 0.01 | 0.01 | 0 | 0.43 | 0.59 | 0.4 | 0.62 | 0.45 | 0.51 |
| 10 | 73235808 | rs10999841 | -0.302 | 0.917 | -0.74 | 0.4 | -0.001 | -0.002 | -0.002 | 0 | 0.45 | 0.52 | 0.52 | 0.35 | 0.47 | 0.49 |
| 10 | 73238973 | rs16928926 | -0.595 | 1.43 | -0.62 | 0.91 | -0.002 | -0.002 | -0.002 | 0 | 0.23 | 0.19 | 0.18 | 0.24 | 0.24 | 0.31 |
| 10 | 73240073 | rs16928929 | 0.029 | 1.588 | -1.256 | 0.789 | -0.038 | -0.015 | -0.015 | 1 | 0.18 | 0.03 | 0.15 | 0.17 | 0.18 | 0.07 |
| 10 | 73240361 | rs10999845 | -0.318 | 1.349 | -1.005 | 0.697 | 0.01 | 0.025 | 0.025 | 0 | 0.23 | 0.12 | 0.18 | 0.17 | 0.2 | 0.07 |
| 10 | 73240527 | rs10999846 | -0.907 | 1.194 | -0.535 | 1.166 | 0.003 | 0.002 | 0.002 | 0 | 0.48 | 0.41 | 0.37 | 0.52 | 0.44 | 0.39 |
| 10 | 73240604 | rs2394800 | 0.878 | 1.52 | -0.609 | 1.183 | -0.004 | -0.005 | -0.005 | 0 | 0.13 | 0.1 | 0.11 | 0.24 | 0.17 | 0.1 |
| 10 | 73240942 | rs2394801 | 0.878 | 1.52 | -0.609 | 1.183 | -0.004 | -0.005 | -0.005 | 0 | 0.13 | 0.1 | 0.11 | 0.23 | 0.17 | 0.1 |
| 10 | 73241440 | rs7901112 | -1.335 | 1.492 | -0.526 | 1.142 | -0.002 | -0.015 | -0.015 | 0 | 0.62 | 0.56 | 0.48 | 0.75 | 0.61 | 0.49 |
| 10 | 73242628 | rs16928934 | 0.655 | 1.391 | -1.179 | 1.313 | 0.002 | 0.008 | 0.008 | 0.01 | 0.13 | 0.12 | 0.11 | 0.23 | 0.17 | 0.12 |
| 10 | 73242975 | rs1538805 | -1.023 | 1.135 | -1.334 | 0.996 | -0.028 | -0.016 | -0.016 | 0.5 | 0.54 | 0.84 | 0.44 | 0.44 | 0.47 | 0.46 |
| 10 | 73245972 | rs7922462 | -0.475 | 1.214 | -0.921 | 1.193 | 0.001 | 0.016 | 0.016 | 1.678 | 0.43 | 0.68 | 0.25 | 0.35 | 0.37 | 0.19 |
| 10 | 73246540 | rs10762451 | -1.068 | 0.501 | -0.641 | 1.185 | -0.019 | -0.025 | -0.025 | 0 | 0.59 | 0.54 | 0.44 | 0.71 | 0.66 | 0.41 |
| 10 | 73260244 | rs1340362 | -0.883 | 0.396 | -0.603 | 1.277 | 0.008 | -0.005 | -0.005 | 4.939 | 0.57 | 0.47 | 0.42 | 0.56 | 0.65 | 0.37 |
| 10 | 73264430 | rs10823772 | 0.452 | 0.65 | -0.769 | 1.316 | -0.002 | -0.001 | -0.001 | 1.455 | 0.2 | 0.2 | 0.21 | 0.26 | 0.26 | 0.17 |
| 10 | 73264879 | rs12778790 | 0.237 | 0.596 | -1.195 | 1.254 | -0.083 | -0.002 | -0.002 | 0.1 | 0.32 | 0.13 | 0.18 | 0.3 | 0.37 | 0.2 |
| 10 | 73266336 | rs10999854 | -0.145 | -0.159 | -0.921 | 1.247 | -0.034 | -0.022 | -0.022 | 0 | 0.41 | 0.69 | 0.36 | 0.51 | 0.45 | 0.27 |
| 10 | 73270906 | rs3802720 | -0.92 | -0.212 | -1.042 | 1.136 | -0.051 | 0.013 | 0.013 | 0.485 | 0.69 | 0.99 | 0.5 | 0.78 | 0.69 | 0.57 |
| 10 | 73270982 | rs3802719 | -0.865 | -0.199 | -1.014 | 1.142 | -0.019 | 0.039 | 0.039 | 0.285 | 0.69 | 0.95 | 0.5 | 0.78 | 0.69 | 0.56 |
| 10 | 73280314 | rs10128409 | -0.498 | -0.167 | -0.9 | 1.118 | -0.004 | 0.001 | 0.001 | 0.001 | 0.66 | 0.71 | 0.47 | 0.65 | 0.69 | 0.58 |
| 10 | 73280607 | rs10999861 | 0.428 | -0.045 | -2.219 | 0.883 | 0.035 | -0.036 | -0.036 | 0 | 0.48 | 0.23 | 0.33 | 0.56 | 0.54 | 0.36 |
| 10 | 73280699 | rs10823774 | -0.124 | 0.638 | -1.978 | 0.513 | 0.01 | 0.044 | 0.044 | 0.101 | 0.68 | 0.92 | 0.5 | 0.76 | 0.7 | 0.59 |
| 10 | 73285009 | rs4747159 | -0.142 | 0.64 | -1.725 | 0.756 | 0.001 | -0.028 | -0.028 | 0.157 | 0.25 | 0.13 | 0.29 | 0.37 | 0.24 | 0.37 |
| 10 | 73285232 | rs4747161 | 0.127 | 0.376 | -1.115 | 0.664 | -0.01 | -0.003 | -0.003 | 0 | 0.81 | 0.7 | 0.78 | 0.8 | 0.81 | 0.92 |
| 10 | 73285391 | rs10999864 | -0.452 | 0.25 | -1.041 | 0.583 | -0.004 | 0.032 | 0.032 | 0 | 0.15 | 0.31 | 0.22 | 0.23 | 0.16 | 0.07 |
| 10 | 73285723 | rs7917660 | -1.104 | 0.208 | -1.125 | 0.444 | 0.004 | -0.022 | -0.022 | 0 | 0.4 | 0.68 | 0.37 | 0.39 | 0.38 | 0.29 |
| 10 | 73286290 | rs17634834 | -0.48 | 0.585 | -0.663 | 0.231 | -0.001 | -0.006 | -0.006 | 0 | 0.16 | 0.13 | 0.11 | 0 | 0.14 | 0.03 |
| 10 | 73288397 | rs9971128 | -0.935 | 0.443 | -0.147 | 0.188 | 0.108 | -0.04 | -0.04 | 0 | 0.1 | 0.46 | 0.08 | 0 | 0.09 | 0.02 |
| 10 | 73290319 | rs2394806 | -1.156 | 0.22 | -0.679 | -0.051 | -0.019 | 0.015 | 0.015 | 0 | 0.8 | 0.83 | 0.7 | 0.91 | 0.67 | 0.72 |
| 10 | 73292293 | rs7893501 | 0.477 | -0.109 | -0.812 | 0.114 | -0.006 | 0.046 | 0.046 | 0 | 0.15 | 0.18 | 0.06 | 0.33 | 0.15 | 0.21 |
| 10 | 73292424 | rs12783388 | 1.032 | -0.275 | -0.481 | 0.063 | 0.043 | 0.005 | 0.005 | 0.034 | 0.46 | 0.3 | 0.38 | 0.49 | 0.35 | 0.46 |
| 10 | 73292788 | rs10999868 | 1.312 | -0.215 | -0.434 | 0.185 | -0.002 | -0.024 | -0.024 | 0.089 | 0.08 | 0.03 | 0.04 | 0.23 | 0.09 | 0.05 |
| 10 | 73294257 | rs3861031 | 0.842 | -0.123 | -0.305 | 0.108 | -0.009 | 0.016 | 0.016 | 0 | 0.16 | 0.2 | 0.07 | 0.16 | 0.14 | 0.2 |
| 10 | 73294743 | rs4423112 | -1.222 | -0.13 | -0.285 | 0.079 | -0.003 | -0.003 | -0.003 | 0 | 0.35 | 0.57 | 0.23 | 0.26 | 0.31 | 0.27 |

| | | | | | | | | | | | | | | | | |
|---|---|---|---|---|---|---|---|---|---|---|---|---|---|---|---|---|
| 10 | 73295121 | rs7922915 | -1.212 | -0.129 | -0.285 | 0.064 | -0.003 | -0.004 | -0.004 | 0 | 0.35 | 0.56 | 0.23 | 0.24 | 0.31 | 0.27 |
| 10 | 73295341 | rs11599279 | -0.496 | -0.079 | -0.347 | 0.06 | 0.011 | -0.037 | -0.037 | 0.004 | 0.19 | 0.36 | 0.16 | 0.08 | 0.17 | 0.06 |
| 10 | 73295727 | rs11594877 | 0.608 | 0.056 | -0.917 | -0.014 | 0 | -0.021 | -0.021 | 0 | 0.12 | 0.13 | 0.09 | 0 | 0.1 | 0.03 |
| 10 | 73297705 | rs10823778 | -1.33 | 0.149 | -1.222 | 0.027 | -0.054 | 0.031 | 0.031 | 0 | 0.81 | 0.89 | 0.61 | 0.9 | 0.67 | 0.72 |
| 10 | 73299302 | rs4747165 | 0.226 | -0.354 | -1.148 | 0.318 | 0.046 | 0.069 | 0.069 | 0.117 | 0.48 | 0.25 | 0.42 | 0.72 | 0.4 | 0.49 |
| 10 | 73300992 | rs10823779 | 1.109 | -0.202 | -1.626 | 0.367 | -0.043 | -0.07 | -0.07 | 0 | 0.24 | 0.03 | 0.13 | 0.34 | 0.25 | 0.38 |
| 10 | 73303755 | rs10762454 | -0.089 | 0.157 | -0.957 | 0.037 | 0.011 | -0.034 | -0.034 | 0.012 | 0.66 | 0.58 | 0.5 | 0.89 | 0.54 | 0.69 |
| 10 | 73306105 | rs10823781 | -0.79 | -0.456 | -1.335 | 0.693 | 0.023 | -0.052 | -0.052 | 0.37 | 0.14 | 0.05 | 0.21 | 0.29 | 0.08 | 0.12 |
| 10 | 73308859 | rs7921151 | 0.302 | -0.418 | -1.417 | 0.654 | 0.009 | -0.036 | -0.036 | 0 | 0.26 | 0.21 | 0.17 | 0.41 | 0.27 | 0.35 |
| 10 | 73309246 | rs875861 | 0.302 | -0.418 | -1.417 | 0.654 | 0.009 | -0.036 | -0.036 | 0 | 0.26 | 0.21 | 0.17 | 0.41 | 0.27 | 0.35 |
| 10 | 73309529 | rs875860 | -0.385 | -0.587 | -0.771 | 0.623 | -0.037 | 0.005 | 0.005 | 0 | 0.18 | 0.28 | 0.26 | 0.16 | 0.12 | 0.14 |
| 10 | 73309890 | rs7910896 | -0.939 | 0.12 | -1.273 | 0.265 | -0.059 | 0.045 | 0.045 | 0 | 0.78 | 0.88 | 0.63 | 0.95 | 0.61 | 0.75 |
| 10 | 73312121 | rs10999876 | -0.133 | -0.44 | -1.139 | 0.905 | -0.012 | 0.009 | 0.009 | 1.217 | 0.09 | 0.06 | 0.06 | 0.4 | 0.15 | 0.18 |
| 10 | 73315940 | rs877783 | -1.443 | -1.192 | -0.405 | 0.28 | -0.024 | 0.01 | 0.01 | 0.532 | 0.33 | 0.44 | 0.33 | 0.61 | 0.27 | 0.31 |
| 10 | 73319089 | rs12247922 | -0.074 | -0.955 | -0.526 | 0.124 | -0.006 | -0.037 | -0.037 | 0.002 | 0.07 | 0.15 | 0.05 | 0 | 0.04 | 0.05 |
| 10 | 73325757 | rs7100757 | -0.198 | -0.634 | -0.58 | 0.142 | -0.002 | -0.001 | -0.001 | 0 | 0.37 | 0.62 | 0.34 | 0.62 | 0.46 | 0.49 |
| 10 | 73325820 | rs4579857 | -0.893 | -1.143 | -0.785 | -0.123 | -0.011 | 0.169 | 0.169 | 0.058 | 0.51 | 0.75 | 0.47 | 0.87 | 0.55 | 0.61 |
| 10 | 73326340 | rs10999882 | -0.217 | -1.241 | -0.626 | -0.055 | 0.004 | -0.013 | -0.013 | 0.431 | 0.4 | 0.57 | 0.45 | 0.74 | 0.47 | 0.55 |
| 10 | 73327405 | rs12264722 | 0.198 | -1.161 | -0.577 | 0.056 | 0 | 0.018 | 0.018 | 0.622 | 0.15 | 0.18 | 0.16 | 0.34 | 0.19 | 0.21 |
| 10 | 73327423 | rs17635709 | 1.257 | -1.13 | -0.841 | -0.039 | 0.023 | 0.019 | 0.019 | 0 | 0.2 | 0.28 | 0.15 | 0.24 | 0.26 | 0.27 |
| 10 | 73327821 | rs885014 | -1.417 | -1.032 | -1.461 | 0.098 | 0.028 | -0.001 | -0.001 | 0 | 0.36 | 0.73 | 0.42 | 0.64 | 0.3 | 0.35 |
| 10 | 73327947 | rs1867996 | -1.615 | -1.229 | -1.822 | -0.257 | -0.118 | 0.126 | 0.126 | 0 | 0.75 | 0.98 | 0.7 | 0.94 | 0.68 | 0.75 |
| 10 | 73328006 | rs719385 | -0.499 | -1.018 | -1.173 | 0.351 | -0.015 | 0.067 | 0.067 | 0.602 | 0.2 | 0.61 | 0.22 | 0.37 | 0.21 | 0.21 |
| 10 | 73328055 | rs719386 | -0.732 | -0.547 | -1.531 | 0.48 | 0.013 | -0.016 | -0.016 | 0 | 0.16 | 0.12 | 0.2 | 0.27 | 0.09 | 0.14 |
| 10 | 73334410 | rs7923245 | -0.814 | 0.123 | -1.834 | -0.165 | -0.095 | 0.144 | 0.144 | 2.207 | 0.7 | 0.94 | 0.62 | 0.92 | 0.6 | 0.75 |
| 10 | 73336877 | rs10999892 | 0.193 | 0.144 | -1.534 | 0.469 | 0.021 | -0.035 | -0.035 | 0 | 0.18 | 0.11 | 0.17 | 0.52 | 0.13 | 0.3 |
| 10 | 73339160 | rs7082486 | -0.824 | 0.225 | -1.924 | 0.328 | -0.051 | 0.103 | 0.103 | 0.027 | 0.61 | 0.94 | 0.6 | 0.81 | 0.61 | 0.7 |
| 10 | 73340767 | rs3998531 | 0.094 | 1.053 | -1.31 | 0.151 | -0.021 | 0.002 | 0.002 | 0 | 0.33 | 0.24 | 0.27 | 0.28 | 0.39 | 0.37 |
| 10 | 73342512 | rs17531870 | -0.804 | 0.501 | -0.751 | 0.832 | -0.001 | 0.005 | 0.005 | 0.01 | 0.3 | 0.32 | 0.28 | 0.48 | 0.32 | 0.36 |
| 10 | 73344838 | rs10509336 | -0.416 | 0.363 | -0.829 | 0.685 | 0.004 | 0.001 | 0.001 | 0.775 | 0.09 | 0.09 | 0.05 | 0.07 | 0.07 | 0.11 |
| 10 | 73345064 | rs7093128 | -0.212 | -0.327 | -0.702 | 0.581 | -0.006 | -0.004 | -0.004 | 0 | 0.23 | 0.21 | 0.12 | 0.26 | 0.18 | 0.38 |
| 10 | 73345380 | rs2394829 | -0.667 | -0.004 | -1.066 | 0.639 | -0.008 | -0.05 | -0.05 | 0 | 0.72 | 0.8 | 0.61 | 0.57 | 0.73 | 0.73 |
| 10 | 73347572 | rs1868001 | 0.04 | -0.25 | -0.558 | 0.241 | -0.022 | -0.036 | -0.036 | 0 | 0.33 | 0.49 | 0.21 | 0.28 | 0.28 | 0.43 |
| 10 | 73348091 | rs10762456 | -1.27 | -0.081 | -1.005 | 0.833 | -0.002 | -0.028 | -0.028 | 0 | 0.8 | 0.9 | 0.72 | 0.8 | 0.84 | 0.78 |
| 10 | 73351149 | rs10999904 | 1.469 | 0.193 | -0.806 | 0.909 | -0.015 | -0.007 | -0.007 | 0 | 0.06 | 0.02 | 0.1 | 0.23 | 0.11 | 0.05 |
| 10 | 73351781 | rs7915479 | -0.271 | -0.092 | -0.718 | 0.834 | 0.009 | 0.023 | 0.023 | 0 | 0.53 | 0.66 | 0.5 | 0.3 | 0.58 | 0.35 |
| 10 | 73352219 | rs6480533 | 1.39 | 0.057 | -0.122 | 1.018 | -0.001 | -0.039 | -0.039 | 0 | 0.1 | 0.2 | 0.07 | 0.02 | 0.1 | 0.06 |
| 10 | 73353426 | rs10999906 | -0.551 | 0.223 | -0.29 | 0.843 | -0.005 | 0.006 | 0.006 | 0.014 | 0.3 | 0.55 | 0.3 | 0.32 | 0.31 | 0.21 |
| 10 | 73355056 | rs6480534 | -0.293 | 0.019 | -0.153 | 0.441 | -0.021 | -0.067 | -0.067 | 0.22 | 0.24 | 0.52 | 0.2 | 0.1 | 0.2 | 0.16 |
| 10 | 73356835 | rs10762458 | 0.581 | 0.128 | -0.763 | 0.252 | -0.113 | 0.175 | 0.175 | 0 | 0.34 | 0.08 | 0.29 | 0.04 | 0.44 | 0.16 |
| 10 | 73370883 | rs7907996 | -0.582 | 0.08 | 0.57 | 0.988 | -0.016 | 0.085 | 0.085 | 0 | 0.25 | 0.56 | 0.3 | 0.51 | 0.22 | 0.22 |
| 10 | 73377330 | rs6480536 | -0.494 | 0.062 | 0.553 | 1.008 | -0.017 | 0.083 | 0.083 | 0 | 0.25 | 0.57 | 0.3 | 0.52 | 0.22 | 0.22 |
| 10 | 73380179 | rs4746087 | -0.315 | 0.185 | 0.392 | 1.103 | 0 | 0.082 | 0.082 | 0.097 | 0.06 | 0.08 | 0.08 | 0.35 | 0.08 | 0.14 |
| 10 | 73386821 | rs10823805 | -0.198 | 0.219 | 0.348 | 0.701 | -0.009 | 0.018 | 0.018 | 0 | 0.36 | 0.46 | 0.34 | 0.6 | 0.33 | 0.45 |
| 10 | 73387685 | rs4746089 | -0.174 | 0.142 | 0.279 | 0.234 | -0.03 | -0.035 | -0.035 | 0 | 0.73 | 0.53 | 0.59 | 0.82 | 0.79 | 0.91 |
| 10 | 73392449 | rs10762462 | -0.178 | 0.264 | 0.274 | 0.23 | -0.017 | -0.035 | -0.035 | 0 | 0.73 | 0.54 | 0.59 | 0.82 | 0.78 | 0.91 |
| 10 | 73392998 | rs2394834 | 0.582 | 0.152 | -0.019 | 0.326 | 0.009 | -0.008 | -0.008 | 0 | 0.53 | 0.35 | 0.42 | 0.49 | 0.57 | 0.49 |
| 10 | 73402788 | rs7913396 | 0.359 | 0.156 | -0.003 | 0.34 | 0.006 | -0.015 | -0.015 | 0.167 | 0.55 | 0.35 | 0.43 | 0.49 | 0.59 | 0.53 |
| 10 | 73405954 | rs2305208 | 0.948 | -0.045 | -0.133 | 0.071 | 0.002 | 0.041 | 0.041 | 0 | 0.17 | 0.08 | 0.11 | 0.05 | 0.18 | 0.17 |
| 10 | 73406504 | rs2305209 | 0.948 | -0.045 | -0.046 | 0.071 | 0.003 | 0.026 | 0.026 | 0 | 0.17 | 0.1 | 0.11 | 0.05 | 0.18 | 0.17 |
| 10 | 73406599 | rs2305210 | 0.348 | 0.124 | -0.124 | -0.38 | 0.043 | -0.1 | -0.1 | 0 | 0.22 | 0.43 | 0.18 | 0.08 | 0.26 | 0.26 |
| 10 | 73406636 | rs11598249 | 1.813 | 0.078 | -0.253 | -0.329 | 0.005 | -0.042 | -0.042 | 0.816 | 0.31 | 0.16 | 0.23 | 0.35 | 0.29 | 0.17 |
| 10 | 73407465 | rs10999929 | 1.529 | 0.077 | -0.123 | -0.296 | -0.011 | -0.039 | -0.039 | 0.093 | 0.18 | 0.08 | 0.16 | 0.33 | 0.19 | 0.38 |
| 10 | 73408326 | rs1900514 | -1.78 | 0.069 | -0.07 | -0.328 | -0.003 | 0.007 | 0.007 | 0 | 0.91 | 0.95 | 0.79 | 1 | 0.92 | 0.98 |
| 10 | 73408462 | rs766200 | 0.515 | 0.057 | -0.095 | -0.349 | -0.009 | 0.029 | 0.029 | 0 | 0.64 | 0.36 | 0.56 | 0.86 | 0.61 | 0.62 |
| 10 | 73409765 | rs12259536 | 0.681 | 0.084 | -0.032 | -0.321 | -0.017 | 0.026 | 0.026 | 0 | 0.2 | 0.13 | 0.16 | 0.08 | 0.26 | 0.26 |
| 10 | 73409978 | rs4746093 | 1.217 | 0.05 | -0.029 | -0.248 | -0.003 | -0.023 | -0.023 | 0 | 0.2 | 0.14 | 0.16 | 0.33 | 0.2 | 0.38 |
| 10 | 73412737 | rs1868007 | 1.754 | 0.064 | -0.082 | -0.202 | 0.01 | -0.029 | -0.029 | 0 | 0.31 | 0.19 | 0.23 | 0.35 | 0.28 | 0.17 |
| 10 | 73413555 | rs1900515 | -1.125 | 0.104 | -0.162 | -0.171 | 0.006 | 0.009 | 0.009 | 0.017 | 0.6 | 0.69 | 0.55 | 0.65 | 0.64 | 0.81 |
| 10 | 73413824 | rs12573587 | -0.476 | 0.149 | -0.164 | -0.174 | -0.001 | -0.001 | -0.001 | 1.375 | 0.28 | 0.28 | 0.29 | 0.31 | 0.31 | 0.36 |
| 10 | 73414010 | rs7896061 | 0.743 | 0.447 | -0.241 | -0.264 | 0.02 | -0.01 | -0.01 | 0 | 0.08 | 0.1 | 0.1 | 0.07 | 0.17 | 0.2 |
| 10 | 73414206 | rs10762465 | 0.069 | 0.256 | -0.018 | -0.214 | -0.001 | 0 | 0 | 0 | 0.37 | 0.38 | 0.29 | 0.38 | 0.38 | 0.54 |
| 10 | 73414502 | rs7087795 | 0.919 | 0.227 | -0.054 | -0.177 | -0.001 | -0.001 | -0.001 | 0.051 | 0.38 | 0.39 | 0.46 | 0.35 | 0.34 | 0.19 |
| 10 | 73414577 | rs7087554 | 0.458 | 0.248 | -0.056 | -0.201 | 0.003 | 0.004 | 0.004 | 0 | 0.52 | 0.42 | 0.63 | 0.53 | 0.48 | 0.26 |
| 10 | 73417236 | rs1665688 | 0.101 | 0.182 | 0.041 | -0.127 | -0.002 | 0.024 | 0.024 | 0 | 0.38 | 0.45 | 0.37 | 0.5 | 0.36 | 0.45 |
| 10 | 73419580 | rs1665692 | 0.182 | 0.145 | -0.102 | -0.043 | -0.04 | -0.002 | -0.002 | 0 | 0.38 | 0.18 | 0.3 | 0.4 | 0.44 | 0.58 |
| 10 | 73421248 | rs1665624 | 0.043 | 0.171 | -0.09 | -0.234 | -0.018 | -0.011 | -0.011 | 0 | 0.41 | 0.28 | 0.4 | 0.57 | 0.46 | 0.6 |
| 10 | 73422284 | rs1625975 | -1.699 | 0.324 | -0.043 | -0.411 | 0 | -0.014 | -0.014 | 0 | 0.93 | 0.95 | 0.8 | 1 | 0.94 | 0.98 |
| 10 | 73426465 | rs11819553 | 0.628 | 0.087 | -0.046 | -0.448 | 0 | -0.006 | -0.006 | 0 | 0.17 | 0.19 | 0.13 | 0.08 | 0.19 | 0.22 |

| | | | | | | | | | | | | | | | | |
|---|---|---|---|---|---|---|---|---|---|---|---|---|---|---|---|---|
| 10 | 73428692 | rs1612048 | 0.354 | -0.327 | -0.113 | -0.506 | -0.012 | -0.042 | -0.042 | 0.556 | 0.66 | 0.45 | 0.63 | 0.86 | 0.73 | 0.74 |
| 10 | 73428758 | rs1625164 | -0.004 | -0.307 | -0.068 | -0.587 | -0.018 | -0.088 | -0.088 | 0 | 0.7 | 0.53 | 0.66 | 0.91 | 0.77 | 0.83 |
| 10 | 73431596 | rs3849958 | 1.112 | 0.013 | -0.301 | -0.457 | -0.004 | 0.006 | 0.006 | 0 | 0.17 | 0.15 | 0.13 | 0.07 | 0.23 | 0.19 |
| 10 | 73432571 | rs1227046 | -0.419 | -0.247 | 0.091 | -0.266 | 0.008 | -0.056 | -0.056 | 0.597 | 0.76 | 0.71 | 0.67 | 0.97 | 0.75 | 0.83 |
| 10 | 73434279 | rs1227048 | 0.271 | -0.397 | 0.04 | 0.063 | -0.005 | -0.013 | -0.013 | 0 | 0.63 | 0.52 | 0.53 | 0.73 | 0.7 | 0.73 |
| 10 | 73434538 | rs10823821 | 1.526 | -0.492 | -0.351 | 0.222 | -0.059 | 0.004 | 0.004 | 0 | 0.37 | 0.15 | 0.33 | 0.38 | 0.48 | 0.37 |
| 10 | 73434906 | rs10999947 | 0.059 | -0.353 | -0.063 | 0.266 | 0.002 | -0.001 | -0.001 | 0 | 0.29 | 0.23 | 0.27 | 0.26 | 0.26 | 0.25 |
| 10 | 73435500 | rs12262033 | 0.238 | -0.221 | -0.302 | 0.025 | -0.001 | -0.008 | -0.008 | 0 | 0.27 | 0.25 | 0.18 | 0.09 | 0.25 | 0.24 |
| 10 | 73439328 | rs3802716 | 0.034 | -0.027 | 0.111 | 0.679 | -0.058 | 0.111 | 0.111 | 0 | 0.11 | 0.24 | 0.14 | 0.28 | 0.04 | 0.04 |
| 10 | 73439975 | rs9415986 | 0.069 | 0.066 | -0.591 | 0.598 | -0.003 | 0.003 | 0.003 | 0 | 0.41 | 0.35 | 0.31 | 0.32 | 0.42 | 0.49 |
| 10 | 73440052 | rs10509334 | 0.384 | 0.401 | -1.543 | 0.624 | -0.015 | 0.021 | 0.021 | 0 | 0.46 | 0.23 | 0.37 | 0.4 | 0.52 | 0.46 |
| 10 | 73445707 | rs1227091 | -0.195 | 0.541 | -1.192 | 0.236 | 0.008 | 0.034 | 0.034 | 1.288 | 0.38 | 0.21 | 0.31 | 0.36 | 0.35 | 0.47 |
| 10 | 73447281 | rs1227087 | -0.125 | -0.073 | -1.012 | 0.3 | 0.029 | 0.019 | 0.019 | 0.674 | 0.36 | 0.24 | 0.26 | 0.35 | 0.27 | 0.41 |
| 10 | 73451573 | rs1227081 | -0.523 | 0.137 | -0.839 | 0.201 | -0.004 | 0.002 | 0.002 | 0 | 0.57 | 0.55 | 0.39 | 0.48 | 0.47 | 0.68 |
| 10 | 73452627 | rs1227079 | 0.025 | 0.028 | -0.605 | 0.025 | -0.01 | -0.024 | -0.024 | 0 | 0.62 | 0.48 | 0.64 | 0.81 | 0.72 | 0.63 |
| 10 | 73454444 | rs1227076 | -0.496 | 0.398 | 0.029 | 0.662 | 0.042 | 0.002 | 0.002 | 0 | 0.7 | 0.5 | 0.56 | 0.8 | 0.58 | 0.8 |
| 10 | 73454637 | rs10823825 | 0.666 | 0.435 | -0.246 | 1.184 | -0.052 | -0.098 | -0.098 | 0.415 | 0.24 | 0.12 | 0.32 | 0.38 | 0.38 | 0.17 |
| 10 | 73455201 | rs3752752 | 0.234 | -0.091 | -0.263 | 0.808 | -0.002 | 0.039 | 0.039 | 0.07 | 0.66 | 0.5 | 0.52 | 0.44 | 0.71 | 0.61 |
| 10 | 73455273 | rs3752751 | 0.234 | -0.091 | -0.263 | 0.8 | -0.002 | 0.04 | 0.04 | 0.07 | 0.66 | 0.5 | 0.52 | 0.43 | 0.71 | 0.61 |
| 10 | 73455821 | rs1227073 | -0.442 | -0.223 | -0.292 | 0.765 | -0.018 | 0.048 | 0.048 | 0.282 | 0.33 | 0.49 | 0.33 | 0.56 | 0.25 | 0.35 |
| 10 | 73456268 | rs10999956 | 1.15 | -0.897 | -0.209 | 0.658 | 0.019 | -0.008 | -0.008 | 2.349 | 0.07 | 0.05 | 0.04 | 0.07 | 0.03 | 0.07 |
| 10 | 73456555 | rs7094761 | 0.179 | -0.848 | -0.056 | 0.58 | 0.02 | 0.028 | 0.028 | 0 | 0.4 | 0.35 | 0.26 | 0.24 | 0.27 | 0.43 |
| 10 | 73458188 | rs1234888 | -0.036 | -0.88 | -0.016 | 0.762 | -0.022 | 0.009 | 0.009 | 0.305 | 0.69 | 0.46 | 0.56 | 0.6 | 0.78 | 0.76 |
| 10 | 73466709 | rs10823829 | -0.586 | -1.445 | 0.87 | 1.011 | -0.065 | 0.186 | 0.186 | 0.264 | 0.13 | 0.41 | 0.15 | 0.27 | 0.03 | 0.09 |
| 10 | 73469442 | rs7072160 | -0.514 | -1.609 | 0.668 | 0.922 | 0.012 | -0.001 | -0.001 | 0 | 0.44 | 0.52 | 0.34 | 0.46 | 0.25 | 0.44 |
| 10 | 73472882 | rs2166631 | -0.473 | -1.588 | 0.771 | 0.928 | 0.038 | -0.003 | -0.003 | 0 | 0.43 | 0.63 | 0.36 | 0.45 | 0.25 | 0.43 |
| 10 | 73481451 | rs7070583 | 0.739 | -1.457 | 0.655 | 1.049 | -0.022 | 0.01 | 0.01 | 0.004 | 0.08 | 0.11 | 0.04 | 0.09 | 0.03 | 0.07 |
| 10 | 73483976 | rs7893748 | -0.529 | -1.479 | 0.689 | 1.009 | -0.063 | 0.176 | 0.176 | 0.122 | 0.13 | 0.44 | 0.16 | 0.3 | 0.03 | 0.09 |
| 10 | 73487127 | rs16929266 | 0.45 | -1.343 | 0.17 | 0.658 | 0.028 | 0.036 | 0.036 | 0 | 0.3 | 0.21 | 0.17 | 0.19 | 0.2 | 0.34 |
| 10 | 73492079 | rs1227065 | -0.567 | -0.743 | 0.692 | 1.206 | 0.002 | -0.011 | -0.011 | 0 | 0.8 | 0.73 | 0.71 | 0.82 | 0.82 | 0.84 |
| 10 | 73493525 | rs1227063 | -0.111 | -0.544 | 0.521 | 1.323 | -0.047 | -0.032 | -0.032 | 0 | 0.09 | 0.01 | 0.1 | 0.15 | 0.14 | 0.19 |
| 10 | 73494229 | rs1227061 | -0.661 | -0.578 | 0.702 | 1.306 | 0.005 | -0.023 | -0.023 | 0 | 0.8 | 0.65 | 0.7 | 0.82 | 0.81 | 0.84 |
| 10 | 73498910 | rs10823837 | 1.303 | -0.61 | 0.483 | 1.426 | 0 | -0.004 | -0.004 | 0 | 0.56 | 0.19 | 0.43 | 0.38 | 0.61 | 0.57 |
| 10 | 73500184 | rs1227054 | 0.235 | -0.563 | 0.357 | 1.462 | -0.002 | 0.004 | 0.004 | 0 | 0.7 | 0.69 | 0.58 | 0.54 | 0.78 | 0.76 |
| 10 | 73501374 | rs1227052 | -0.19 | -0.525 | 0.334 | 1.537 | -0.003 | 0.024 | 0.024 | 0.012 | 0.75 | 0.62 | 0.61 | 0.63 | 0.78 | 0.8 |
| 10 | 73501556 | rs1227051 | -0.111 | -0.447 | 0.409 | 1.013 | -0.001 | -0.066 | -0.066 | 0.118 | 0.75 | 0.56 | 0.69 | 0.83 | 0.79 | 0.82 |
| 10 | 73502992 | rs9415990 | -0.129 | 0.179 | -0.512 | 1.024 | -0.047 | -0.039 | -0.039 | 0 | 0.08 | 0.01 | 0.11 | 0.17 | 0.15 | 0.18 |
| 10 | 73504424 | rs9415037 | -0.077 | 0.278 | -0.401 | 0.892 | 0 | 0.009 | 0.009 | 0 | 0.17 | 0.25 | 0.17 | 0.25 | 0.17 | 0.23 |
| 10 | 73506514 | rs12248205 | -0.027 | 0.626 | -0.66 | 0.303 | 0.007 | 0.052 | 0.052 | 0 | 0.52 | 0.34 | 0.36 | 0.23 | 0.59 | 0.49 |
| 10 | 73507645 | rs1055518 | -0.4 | 1.111 | -0.99 | 0.335 | -0.008 | 0.053 | 0.053 | 0 | 0.6 | 0.68 | 0.57 | 0.8 | 0.36 | 0.61 |
| 10 | 73509465 | rs748503 | -1.156 | 1.761 | -0.765 | 1.348 | -0.004 | 0.014 | 0.014 | 0 | 0.1 | 0.11 | 0.2 | 0.3 | 0.19 | 0.15 |
| 10 | 73514364 | rs7084874 | -0.462 | 2.38 | 0.339 | 0.952 | -0.002 | 0 | 0 | 0 | 0.17 | 0.12 | 0.23 | 0.18 | 0.27 | 0.15 |
| 10 | 73514554 | rs7908979 | 0.691 | 1.908 | -0.486 | 0.401 | -0.002 | 0.085 | 0.085 | 0 | 0.33 | 0.16 | 0.21 | 0.09 | 0.33 | 0.25 |
| 10 | 73515103 | rs3747858 | -0.786 | 2.073 | -0.431 | 1.197 | -0.01 | -0.037 | -0.037 | 0 | 0.63 | 0.79 | 0.57 | 0.58 | 0.62 | 0.65 |
| 10 | 73516463 | rs3747859 | 0.743 | 1.455 | -0.527 | 0.734 | 0.002 | 0.001 | 0.001 | 0.044 | 0.21 | 0.23 | 0.09 | 0.2 | 0.13 | 0.26 |
| 10 | 73517225 | rs9415995 | 0.592 | 0.943 | -0.69 | 0.584 | 0.008 | -0.01 | -0.01 | 0 | 0.39 | 0.28 | 0.35 | 0.41 | 0.39 | 0.38 |
| 10 | 73517520 | rs10823840 | 1.745 | 1.023 | -0.628 | 0.431 | 0 | 0.014 | 0.014 | 0 | 0.25 | 0.17 | 0.18 | 0.1 | 0.28 | 0.23 |
| 10 | 73520391 | rs10999987 | 0.582 | 1.515 | -0.554 | 0.741 | -0.019 | 0.003 | 0.003 | 0 | 0.17 | 0.11 | 0.24 | 0.17 | 0.26 | 0.16 |
| 10 | 73520828 | rs3747868 | 0.753 | 1.645 | -0.547 | 0.871 | -0.024 | -0.008 | -0.008 | 0 | 0.16 | 0.09 | 0.16 | 0.2 | 0.27 | 0.14 |
| 10 | 73521371 | rs3747867 | 0.648 | 1.624 | -0.34 | 0.851 | 0.008 | 0 | 0 | 0 | 0.17 | 0.16 | 0.17 | 0.19 | 0.27 | 0.14 |
| 10 | 73524368 | rs9415996 | 1.062 | 0.964 | -0.691 | 0.327 | -0.003 | 0.216 | 0.216 | 0.009 | 0.46 | 0.11 | 0.27 | 0.19 | 0.41 | 0.48 |
| 10 | 73525525 | rs7919533 | 0.477 | 1.425 | -0.376 | 0.699 | -0.014 | 0.007 | 0.007 | 0.076 | 0.66 | 0.69 | 0.38 | 0.26 | 0.5 | 0.59 |
| 10 | 73526202 | rs2121538 | -0.396 | 1.588 | -0.329 | 0.943 | 0.056 | 0.083 | 0.083 | 0 | 0.79 | 0.64 | 0.61 | 0.55 | 0.63 | 0.79 |
| 10 | 73531069 | rs10762477 | 0.258 | 1.269 | -0.679 | 0.945 | -0.016 | -0.038 | -0.038 | 0 | 0.11 | 0.05 | 0.23 | 0.28 | 0.13 | 0.2 |
| 10 | 73537388 | rs7917781 | -0.499 | 1.312 | -0.302 | 0.739 | 0.004 | 0.004 | 0.004 | 1.708 | 0.33 | 0.31 | 0.43 | 0.49 | 0.45 | 0.4 |
| 10 | 73537614 | rs17712523 | 1.22 | 1.274 | -0.955 | 0.395 | -0.088 | 0.02 | 0.02 | 0.606 | 0.2 | 0.01 | 0.12 | 0.15 | 0.22 | 0.2 |
| 10 | 73538164 | rs10762481 | -0.577 | 1.195 | 0.09 | 0.703 | 0.007 | 0.04 | 0.04 | 0.139 | 0.32 | 0.42 | 0.41 | 0.49 | 0.36 | 0.38 |
| 10 | 73541009 | rs9415047 | 1.311 | 0.944 | 0.212 | 0.398 | -0.006 | 0.153 | 0.153 | 0 | 0.49 | 0.26 | 0.31 | 0.15 | 0.5 | 0.51 |
| 10 | 73544604 | rs7068810 | 0.621 | 1.115 | 0.025 | 0.526 | 0.009 | 0.01 | 0.01 | 0 | 0.16 | 0.25 | 0.17 | 0.24 | 0.2 | 0.13 |
| 10 | 73544638 | rs2394839 | 0.283 | 1.104 | -0.049 | 0.629 | -0.019 | -0.05 | -0.05 | 2.09 | 0.12 | 0.05 | 0.24 | 0.3 | 0.15 | 0.24 |
| 10 | 73545185 | rs7089309 | 0.84 | 1.024 | 0.081 | 0.522 | 0.013 | 0.007 | 0.007 | 0 | 0.11 | 0.09 | 0.04 | 0 | 0.05 | 0.03 |
| 10 | 73550969 | rs10466026 | 0.015 | 1.214 | 0.312 | 0.675 | 0.022 | 0.081 | 0.081 | 0 | 0.19 | 0.26 | 0.35 | 0.53 | 0.28 | 0.28 |
| 10 | 73551258 | rs10823847 | -0.584 | 1.219 | 0.411 | 0.662 | 0.013 | 0.078 | 0.078 | 0 | 0.23 | 0.35 | 0.36 | 0.53 | 0.28 | 0.31 |
| 10 | 73555926 | rs4747192 | -0.898 | 1.355 | 0.354 | 0.764 | 0.03 | 0.066 | 0.066 | 0 | 0.26 | 0.46 | 0.38 | 0.6 | 0.31 | 0.32 |
| 10 | 73558886 | rs4747194 | -0.326 | 1.241 | 0.255 | 0.803 | 0.009 | 0.044 | 0.044 | 0 | 0.21 | 0.27 | 0.35 | 0.53 | 0.28 | 0.31 |
| 10 | 73564453 | rs2121534 | -0.405 | 1.271 | 0.149 | 0.851 | -0.002 | -0.008 | -0.008 | 0 | 0.78 | 0.86 | 0.71 | 0.75 | 0.78 | 0.83 |
| 10 | 73573365 | rs2394840 | -0.483 | 1.34 | 0.175 | 1.092 | 0.002 | 0.188 | 0.188 | 0 | 0.09 | 0.2 | 0.21 | 0.36 | 0.08 | 0.15 |
| 10 | 73575239 | rs2290022 | -0.483 | 1.34 | 0.175 | 1.092 | 0.002 | 0.188 | 0.188 | 0 | 0.09 | 0.2 | 0.21 | 0.36 | 0.08 | 0.15 |
| 10 | 73579217 | rs11000016 | 0.772 | 1.466 | -0.044 | 0.941 | -0.007 | -0.008 | -0.008 | 1.084 | 0.14 | 0.11 | 0.15 | 0.24 | 0.2 | 0.13 |

| | | | | | | | | | | | | | | | | |
|---|---|---|---|---|---|---|---|---|---|---|---|---|---|---|---|---|
| 10 | 73585526 | rs2593163 | -0.586 | 0.92 | 0.102 | 1.07 | 0.016 | 0.182 | 0.182 | 0 | 0.13 | 0.31 | 0.23 | 0.36 | 0.11 | 0.2 |
| 10 | 73586585 | rs2854992 | 0.219 | 0.706 | -0.381 | 0.701 | 0.001 | -0.004 | -0.004 | 0 | 0.76 | 0.87 | 0.7 | 0.75 | 0.76 | 0.81 |
| 10 | 73589418 | rs10740391 | -0.334 | 0.928 | -0.033 | 0.786 | 0.009 | 0.004 | 0.004 | 1.084 | 0.36 | 0.51 | 0.39 | 0.62 | 0.38 | 0.41 |
| 10 | 73591794 | rs11000020 | -0.926 | 0.664 | 0.924 | 1.231 | 0.004 | 0.008 | 0.008 | 0 | 0.06 | 0.16 | 0.13 | 0.04 | 0.05 | 0.09 |
| 10 | 73594499 | rs762571 | 0.782 | 0.617 | 0.706 | 1.197 | 0.001 | -0.007 | -0.007 | 0.307 | 0.64 | 0.63 | 0.55 | 0.24 | 0.6 | 0.6 |
| 10 | 73596364 | rs7905774 | 0.346 | 0.544 | 0.696 | 1.491 | -0.004 | -0.016 | -0.016 | 0 | 0.14 | 0.08 | 0.07 | 0.24 | 0.15 | 0.1 |
| 10 | 73597352 | rs720295 | 2.102 | 0.462 | 0.689 | 1.398 | -0.007 | 0.238 | 0.238 | 0 | 0.51 | 0.16 | 0.37 | 0.13 | 0.53 | 0.48 |
| 10 | 73597960 | rs728289 | 1.716 | 0.628 | 0.696 | 1.454 | -0.001 | 0.064 | 0.064 | 0 | 0.56 | 0.3 | 0.39 | 0.2 | 0.55 | 0.51 |
| 10 | 73598889 | rs4747204 | 0.217 | 0.986 | 0.085 | 1.758 | 0.003 | 0.026 | 0.026 | 0 | 0.81 | 0.78 | 0.72 | 0.68 | 0.79 | 0.83 |
| 10 | 73600155 | rs10762483 | -0.659 | 1.171 | -0.386 | 2.135 | -0.001 | -0.001 | -0.001 | 0 | 0.17 | 0.14 | 0.17 | 0.44 | 0.19 | 0.24 |
| 10 | 73603097 | rs1900498 | -0.313 | 1.317 | -0.707 | 1.774 | -0.013 | 0.027 | 0.027 | 0 | 0.94 | 0.97 | 0.78 | 0.68 | 0.88 | 0.92 |
| 10 | 73610982 | rs2070188 | -0.111 | 0.86 | -0.603 | 1.147 | -0.017 | -0.03 | -0.03 | 0.472 | 0.13 | 0.04 | 0.07 | 0.24 | 0.15 | 0.11 |
| 10 | 73616575 | rs4746097 | -0.778 | 0.603 | -0.138 | 1.108 | 0.001 | 0.022 | 0.022 | 0 | 0.31 | 0.45 | 0.31 | 0.36 | 0.27 | 0.34 |
| 10 | 73619428 | rs10762485 | 0.871 | 0.289 | 0.052 | 1.329 | 0.011 | 0.037 | 0.037 | 0 | 0.6 | 0.24 | 0.46 | 0.31 | 0.66 | 0.55 |
| 10 | 73623277 | rs7908969 | 0.278 | 0.209 | 0.479 | 1.196 | -0.005 | -0.046 | -0.046 | 0 | 0.13 | 0.17 | 0.07 | 0 | 0.09 | 0.07 |
| 10 | 73626223 | rs4747211 | 0.211 | 0.265 | 0.603 | 1.486 | -0.005 | 0.221 | 0.221 | 0 | 0.91 | 0.74 | 0.77 | 0.67 | 0.92 | 0.88 |
| 10 | 73637226 | rs4237333 | -0.275 | 0.406 | 0.306 | 1.272 | 0.012 | 0.073 | 0.073 | 0 | 0.17 | 0.3 | 0.24 | 0.36 | 0.16 | 0.31 |
| 10 | 73641091 | rs7068317 | -0.942 | 0.076 | 0.285 | 0.894 | -0.004 | 0 | 0 | 0.302 | 0.35 | 0.62 | 0.41 | 0.35 | 0.3 | 0.26 |
| 10 | 73642902 | rs4532949 | -0.765 | 0.068 | 0.301 | 0.635 | 0.015 | 0.009 | 0.009 | 0.353 | 0.36 | 0.71 | 0.42 | 0.29 | 0.3 | 0.26 |
| 10 | 73653867 | rs4642997 | -0.365 | 0.055 | 0.847 | 0.834 | -0.02 | 0.061 | 0.061 | 0 | 0.28 | 0.48 | 0.27 | 0.58 | 0.24 | 0.28 |
| 10 | 73656982 | rs10509340 | -0.26 | 0.656 | 0.261 | 0.009 | -0.003 | 0.018 | 0.018 | 0 | 0.26 | 0.19 | 0.17 | 0.08 | 0.28 | 0.19 |
| 10 | 73662181 | rs10823873 | -1.194 | 0.64 | 0.392 | -0.129 | 0.003 | -0.123 | -0.123 | 0 | 0.38 | 0.47 | 0.53 | 0.93 | 0.43 | 0.49 |
| 10 | 73667535 | rs12573684 | 0.321 | 0.509 | 0.081 | 0.43 | 0.012 | -0.019 | -0.019 | 0 | 0.06 | 0.02 | 0.09 | 0.14 | 0.06 | 0.1 |
| 10 | 73667572 | rs1009986 | -1.474 | 0.69 | 0.032 | -0.511 | 0.004 | 0.03 | 0.03 | 0.015 | 0.54 | 0.65 | 0.69 | 0.92 | 0.62 | 0.62 |
| 10 | 73673114 | rs10823878 | 0.356 | 1.193 | -0.106 | -0.441 | 0.004 | -0.004 | -0.004 | 0 | 0.13 | 0.12 | 0.13 | 0.08 | 0.21 | 0.13 |
| 10 | 73680382 | rs7919939 | -0.828 | 0.873 | 0.131 | -0.479 | -0.01 | -0.065 | -0.065 | 0 | 0.37 | 0.48 | 0.54 | 0.83 | 0.39 | 0.48 |
| 10 | 73680393 | rs7919639 | 0.237 | 1.013 | -0.227 | -0.352 | -0.009 | -0.025 | -0.025 | 0 | 0.18 | 0.12 | 0.36 | 0.7 | 0.24 | 0.3 |
| 10 | 73682180 | rs7071860 | 0.804 | 0.77 | -0.219 | -0.534 | 0 | -0.019 | -0.019 | 0 | 0.12 | 0.14 | 0.08 | 0 | 0.09 | 0.08 |
| 10 | 73685039 | rs7086312 | -0.535 | 0.747 | -0.024 | -0.048 | 0 | 0.057 | 0.057 | 0.042 | 0.88 | 0.85 | 0.67 | 0.37 | 0.86 | 0.72 |
| 10 | 73686875 | rs11000090 | -0.121 | 0.857 | -0.181 | -0.027 | 0 | -0.012 | -0.012 | 0.619 | 0.09 | 0.05 | 0.08 | 0.12 | 0.13 | 0.11 |
| 10 | 73687049 | rs7095640 | -0.314 | 0.865 | -0.14 | -0.159 | -0.01 | 0.01 | 0.01 | 0 | 0.17 | 0.12 | 0.25 | 0.13 | 0.22 | 0.2 |
| 10 | 73699145 | rs7086624 | -0.814 | 0.733 | -0.455 | -0.642 | 0.002 | 0.014 | 0.014 | 0 | 0.46 | 0.42 | 0.47 | 0.32 | 0.54 | 0.46 |
| 10 | 73707100 | rs7090957 | -1.128 | 0.617 | -0.386 | -0.376 | 0.009 | -0.005 | -0.005 | 0 | 0.6 | 0.67 | 0.56 | 0.38 | 0.62 | 0.53 |
| 10 | 73708100 | rs11000108 | 1.185 | 0.885 | -0.557 | -0.357 | -0.03 | -0.027 | -0.027 | 0 | 0.1 | 0.05 | 0.09 | 0.14 | 0.15 | 0.13 |
| 10 | 73709990 | rs10762497 | -1.146 | 0.587 | -0.291 | -0.283 | 0.005 | -0.003 | -0.003 | 0 | 0.6 | 0.65 | 0.56 | 0.38 | 0.62 | 0.53 |
| 10 | 73710885 | rs4537681 | -0.931 | 0.533 | -0.156 | -0.32 | -0.011 | -0.002 | -0.002 | 0 | 0.58 | 0.54 | 0.53 | 0.37 | 0.62 | 0.53 |
| 10 | 73717915 | rs11527412 | -0.797 | 0.752 | -0.231 | -0.462 | 0.014 | -0.058 | -0.058 | 0 | 0.29 | 0.45 | 0.26 | 0.13 | 0.34 | 0.19 |
| 10 | 73722023 | rs11000122 | 0.799 | 0.924 | -0.599 | -0.41 | -0.04 | -0.001 | -0.001 | 0 | 0.29 | 0.12 | 0.28 | 0.25 | 0.28 | 0.34 |
| 10 | 73722349 | rs10823892 | -1.041 | 1.075 | 0.054 | -0.155 | 0.015 | -0.033 | -0.033 | 0.75 | 0.28 | 0.37 | 0.26 | 0.15 | 0.34 | 0.19 |
| 10 | 73723370 | rs3808945 | -0.743 | 1.237 | -0.182 | -0.084 | 0.013 | -0.018 | -0.018 | 0 | 0.26 | 0.28 | 0.24 | 0.15 | 0.34 | 0.19 |
| 10 | 73729417 | rs4148911 | 0.437 | 1.409 | -0.696 | -0.031 | -0.04 | 0.014 | 0.014 | 0 | 0.76 | 0.87 | 0.69 | 0.81 | 0.64 | 0.79 |
| 10 | 73729498 | rs4148912 | 1.094 | 0.82 | -0.401 | -0.324 | 0.002 | 0.016 | 0.016 | 1.084 | 0.71 | 0.27 | 0.65 | 0.79 | 0.7 | 0.78 |
| 10 | 73729650 | rs7894516 | 0.848 | 0.497 | -1.097 | -0.227 | -0.06 | 0.095 | 0.095 | 0 | 0.38 | 0.07 | 0.26 | 0.2 | 0.42 | 0.46 |
| 10 | 73729752 | rs4148913 | -0.174 | 0.237 | -0.828 | -0.325 | 0.009 | 0.028 | 0.028 | 0 | 0.52 | 0.58 | 0.4 | 0.25 | 0.41 | 0.42 |
| 10 | 73729806 | rs4148914 | 0.942 | 0.248 | -1.389 | -0.504 | -0.054 | 0.082 | 0.082 | 0 | 0.23 | 0.05 | 0.13 | 0.05 | 0.26 | 0.11 |
| 10 | 73730073 | rs874692 | -0.095 | -0.095 | -0.743 | -0.181 | -0.052 | -0.022 | -0.022 | 0 | 0.87 | 0.99 | 0.8 | 0.84 | 0.83 | 0.79 |
| 10 | 73731442 | rs4148917 | -0.483 | -1.179 | 0.038 | -0.208 | -0.031 | 0.015 | 0.015 | 0 | 0.25 | 0.6 | 0.22 | 0.25 | 0.1 | 0.18 |
| 10 | 73731613 | rs4148919 | 0.445 | -1.29 | 0.366 | -0.216 | 0.007 | 0.015 | 0.015 | 0 | 0.18 | 0.12 | 0.17 | 0.12 | 0.1 | 0.15 |
| 10 | 73731768 | rs4148921 | 0.445 | -1.305 | 0.701 | -0.217 | 0.007 | 0.015 | 0.015 | 0 | 0.18 | 0.12 | 0.17 | 0.12 | 0.1 | 0.15 |
| 10 | 73732419 | rs4148922 | -0.982 | -1.602 | 0.858 | -0.352 | -0.017 | 0.001 | 0.001 | 0 | 0.28 | 0.62 | 0.23 | 0.25 | 0.11 | 0.28 |
| 10 | 73732665 | rs10762498 | -1.074 | -1.581 | 0.65 | -0.491 | 0.017 | -0.001 | -0.001 | 0.199 | 0.58 | 0.75 | 0.61 | 0.35 | 0.57 | 0.42 |
| 10 | 73739261 | rs4148927 | 0.608 | -1.515 | 0.699 | -0.516 | -0.002 | -0.004 | -0.004 | 0 | 0.19 | 0.21 | 0.18 | 0.12 | 0.1 | 0.15 |
| 10 | 73740613 | rs4148928 | -0.36 | -1.429 | 0.824 | -0.424 | 0.036 | 0.077 | 0.077 | 0 | 0.52 | 0.31 | 0.56 | 0.22 | 0.57 | 0.39 |
| 10 | 73740837 | rs4148929 | -0.547 | -1.425 | 1.006 | -0.422 | 0.01 | 0.019 | 0.019 | 0 | 0.54 | 0.39 | 0.57 | 0.22 | 0.57 | 0.39 |
| 10 | 73741706 | rs2091331 | -1.012 | -1.436 | 0.952 | -0.409 | 0 | -0.022 | -0.022 | 0 | 0.61 | 0.84 | 0.62 | 0.37 | 0.58 | 0.42 |
| 10 | 73744470 | rs2394847 | -0.944 | -1.425 | 0.956 | -0.404 | 0.004 | -0.018 | -0.018 | 0.143 | 0.6 | 0.82 | 0.62 | 0.36 | 0.58 | 0.42 |
| 10 | 73744839 | rs16929536 | 0.904 | -1.468 | 0.987 | -0.402 | 0.002 | 0.008 | 0.008 | 0 | 0.16 | 0.11 | 0.17 | 0.12 | 0.09 | 0.12 |
| 10 | 73744987 | rs4746104 | -0.649 | -1.487 | 1.007 | -0.416 | 0.018 | 0.045 | 0.045 | 0 | 0.55 | 0.35 | 0.58 | 0.23 | 0.57 | 0.39 |
| 10 | 73745354 | rs16929537 | 0.834 | -1.479 | 0.96 | -0.41 | 0.001 | 0.003 | 0.003 | 0 | 0.16 | 0.15 | 0.17 | 0.12 | 0.09 | 0.12 |
| 10 | 73745476 | rs1006974 | -0.526 | -1.481 | 1.007 | -0.404 | 0.012 | 0.024 | 0.024 | 0 | 0.54 | 0.4 | 0.57 | 0.22 | 0.57 | 0.39 |
| 10 | 73745599 | rs2127357 | -1.047 | -1.42 | 0.552 | -0.372 | 0.012 | 0.001 | 0.001 | 0 | 0.59 | 0.71 | 0.61 | 0.36 | 0.58 | 0.42 |
| 10 | 73751143 | rs1007242 | 0.694 | -1.193 | 0.063 | -0.488 | -0.136 | 0.161 | 0.161 | 0 | 0.35 | 0.07 | 0.38 | 0.1 | 0.47 | 0.27 |
| 10 | 73755778 | rs7085958 | -0.61 | -1.24 | 0.371 | -0.383 | 0.032 | 0.077 | 0.077 | 0.012 | 0.54 | 0.3 | 0.57 | 0.23 | 0.57 | 0.39 |
| 10 | 73758332 | rs730334 | -0.227 | -1.245 | 0.256 | -0.395 | 0.06 | 0.19 | 0.19 | 0 | 0.49 | 0.13 | 0.54 | 0.21 | 0.56 | 0.39 |
| 10 | 73759794 | rs4148932 | -0.475 | -1.325 | 0.464 | -0.394 | 0.033 | 0.062 | 0.062 | 0 | 0.53 | 0.34 | 0.57 | 0.23 | 0.58 | 0.4 |
| 10 | 73759890 | rs4148933 | -0.492 | -1.325 | 0.451 | -0.394 | 0.029 | 0.054 | 0.054 | 0 | 0.53 | 0.35 | 0.57 | 0.23 | 0.58 | 0.4 |
| 10 | 73761015 | rs751450 | -0.993 | -1.37 | 0.021 | -0.203 | -0.02 | -0.094 | -0.094 | 0 | 0.61 | 0.91 | 0.63 | 0.38 | 0.58 | 0.43 |
| 10 | 73762056 | rs4148936 | 0.139 | -1.177 | 0.533 | -0.474 | -0.008 | 0.005 | 0.005 | 0 | 0.41 | 0.37 | 0.41 | 0.11 | 0.48 | 0.28 |
| 10 | 73762188 | rs4148938 | 0.698 | -1.105 | 0.293 | -0.474 | -0.179 | 0.166 | 0.166 | 0 | 0.34 | 0.03 | 0.37 | 0.06 | 0.45 | 0.25 |

| | | | | | | | | | | | | | | | | |
|---|---|---|---|---|---|---|---|---|---|---|---|---|---|---|---|---|
| 10 | 73762662 | rs4148939 | 0.698 | -1.105 | 0.293 | -0.474 | -0.179 | 0.166 | 0.166 | 0 | 0.34 | 0.03 | 0.37 | 0.06 | 0.45 | 0.24 |
| 10 | 73765995 | rs4148940 | 0.164 | -1.103 | 0.597 | -0.434 | -0.001 | -0.023 | -0.023 | 0 | 0.41 | 0.42 | 0.42 | 0.11 | 0.48 | 0.29 |
| 10 | 73768992 | rs4148941 | -1.203 | -1.333 | 0.241 | -0.246 | -0.026 | -0.098 | -0.098 | 0 | 0.63 | 0.9 | 0.62 | 0.37 | 0.57 | 0.49 |
| 10 | 73769507 | rs4148943 | 0.292 | -1.428 | 0.739 | -0.305 | -0.005 | -0.005 | -0.005 | 3.089 | 0.4 | 0.41 | 0.43 | 0.11 | 0.47 | 0.26 |
| 10 | 73769543 | rs4148944 | 0.532 | -1.483 | 1.032 | -0.313 | -0.008 | 0.119 | 0.119 | 0 | 0.16 | 0.31 | 0.18 | 0.25 | 0.09 | 0.15 |
| 10 | 73770073 | rs4148946 | -1.08 | -1.481 | 0.932 | -0.313 | 0.009 | -0.039 | -0.039 | 0.304 | 0.61 | 0.78 | 0.62 | 0.37 | 0.57 | 0.49 |
| 10 | 73770117 | rs4148947 | 0.612 | -1.498 | 0.945 | -0.329 | -0.04 | 0.076 | 0.076 | 0 | 0.38 | 0.24 | 0.41 | 0.11 | 0.47 | 0.26 |
| 10 | 73770128 | rs17297481 | 1.335 | -1.533 | 0.98 | -0.297 | -0.014 | -0.013 | -0.013 | 0 | 0.13 | 0.02 | 0.14 | 0.11 | 0.08 | 0.12 |
| 10 | 73770651 | rs4148949 | -1.281 | -1.496 | 0.937 | -0.288 | -0.004 | -0.065 | -0.065 | 5.293 | 0.63 | 0.84 | 0.62 | 0.37 | 0.57 | 0.49 |
| 10 | 73771706 | rs4148950 | 0.572 | -1.465 | 0.799 | -0.323 | -0.04 | 0.077 | 0.077 | 0 | 0.38 | 0.25 | 0.41 | 0.1 | 0.47 | 0.26 |
| 10 | 73772014 | rs1871450 | 0.572 | -1.467 | 0.813 | -0.324 | -0.04 | 0.077 | 0.077 | 0 | 0.38 | 0.25 | 0.41 | 0.1 | 0.47 | 0.26 |
| 10 | 73772336 | rs731027 | 0.356 | -1.467 | 0.814 | -0.324 | -0.006 | -0.004 | -0.004 | 0 | 0.4 | 0.41 | 0.43 | 0.11 | 0.47 | 0.26 |
| 10 | 73773014 | rs12418 | 0.579 | -1.433 | 0.429 | -0.315 | -0.04 | 0.075 | 0.075 | 0 | 0.37 | 0.25 | 0.41 | 0.1 | 0.47 | 0.26 |
| 10 | 73775707 | rs896076 | 0.774 | -1.381 | 0.232 | -0.291 | -0.142 | 0.173 | 0.173 | 0 | 0.37 | 0.06 | 0.4 | 0.1 | 0.47 | 0.26 |
| 10 | 73776943 | rs1245584 | 0.774 | -1.381 | 0.232 | -0.291 | -0.142 | 0.173 | 0.173 | 0 | 0.37 | 0.06 | 0.4 | 0.1 | 0.47 | 0.26 |
| 10 | 73778267 | rs1245582 | -1.325 | -1.376 | 0.224 | -0.257 | -0.045 | -0.115 | -0.115 | 0.731 | 0.64 | 0.93 | 0.63 | 0.37 | 0.57 | 0.49 |
| 10 | 73780101 | rs16929563 | 0.696 | -0.94 | 0.84 | -0.072 | -0.002 | -0.002 | -0.002 | 0 | 0.14 | 0.16 | 0.16 | 0.11 | 0.08 | 0.12 |
| 10 | 73784791 | rs1245519 | 0.144 | -0.655 | 0.501 | -0.279 | -0.049 | 0.106 | 0.106 | 0 | 0.38 | 0.19 | 0.42 | 0.1 | 0.47 | 0.26 |
| 10 | 73790819 | rs1245531 | -0.076 | -0.628 | 0.034 | -0.207 | -0.016 | 0.026 | 0.026 | 0 | 0.41 | 0.3 | 0.43 | 0.25 | 0.48 | 0.29 |
| 10 | 73799349 | rs1245507 | 0.035 | -0.38 | -0.167 | -0.494 | -0.029 | 0.208 | 0.208 | 0 | 0.45 | 0.18 | 0.41 | 0.1 | 0.5 | 0.25 |
| 10 | 73806288 | rs1678618 | 0.017 | -0.391 | 0.312 | -0.514 | -0.01 | 0.113 | 0.113 | 0 | 0.46 | 0.24 | 0.41 | 0.1 | 0.49 | 0.25 |
| 10 | 73810402 | rs11000152 | -0.41 | -0.515 | 0.721 | 0.043 | 0.011 | 0.016 | 0.016 | 0 | 0.08 | 0.16 | 0.16 | 0.1 | 0.07 | 0.14 |
| 10 | 73815926 | rs11000157 | -0.391 | -0.531 | 0.702 | 0.036 | -0.001 | 0 | 0 | 0 | 0.08 | 0.07 | 0.15 | 0.1 | 0.07 | 0.12 |
| 10 | 73817299 | rs1678622 | 0.196 | -0.665 | 0.64 | -0.098 | -0.035 | 0.185 | 0.185 | 0 | 0.45 | 0.2 | 0.41 | 0.12 | 0.49 | 0.24 |
| 10 | 73820001 | rs1049269 | -1.057 | -0.781 | 0.453 | -0.109 | -0.106 | -0.145 | -0.145 | 0.206 | 0.63 | 0.94 | 0.6 | 0.42 | 0.54 | 0.42 |
| 10 | 73820622 | rs3180 | -1.057 | -0.781 | 0.453 | -0.083 | -0.106 | -0.148 | -0.148 | 0.369 | 0.63 | 0.93 | 0.59 | 0.42 | 0.54 | 0.42 |
| 10 | 73820911 | rs737080 | -0.865 | -0.799 | 0.934 | -0.09 | -0.013 | 0.259 | 0.259 | 5.349 | 0.08 | 0.45 | 0.05 | 0.19 | 0.01 | 0.05 |
| 10 | 73821633 | rs1678623 | 0.491 | -0.737 | 0.885 | -0.117 | -0.031 | 0.172 | 0.172 | 0 | 0.44 | 0.19 | 0.38 | 0.12 | 0.45 | 0.23 |
| 10 | 73822243 | rs1049245 | 0.764 | -0.893 | 0.653 | -0.193 | -0.118 | 0.218 | 0.218 | 0 | 0.41 | 0.07 | 0.36 | 0.12 | 0.44 | 0.23 |
| 10 | 73822261 | rs11000161 | -1.092 | -0.898 | 0.774 | -0.152 | -0.055 | 0.158 | 0.158 | 4.789 | 0.11 | 0.52 | 0.07 | 0.19 | 0.02 | 0.06 |
| 10 | 73822343 | rs3088251 | -1.439 | -0.834 | 0.448 | -0.217 | -0.007 | 0.015 | 0.015 | 0.52 | 0.19 | 0.68 | 0.22 | 0.29 | 0.1 | 0.18 |
| 10 | 73822507 | rs1530803 | 0.391 | -0.621 | 0.373 | -0.329 | -0.042 | 0.179 | 0.179 | 0 | 0.43 | 0.2 | 0.37 | 0.13 | 0.45 | 0.23 |
| 10 | 73823740 | rs1613186 | 0.531 | -0.565 | 0.26 | -0.299 | -0.062 | 0.192 | 0.192 | 0 | 0.41 | 0.12 | 0.36 | 0.12 | 0.44 | 0.23 |
| 10 | 73828748 | rs1678627 | 0.702 | -0.497 | 0.027 | -0.417 | -0.04 | 0.142 | 0.142 | 0 | 0.36 | 0.16 | 0.33 | 0.08 | 0.36 | 0.18 |
| 10 | 73837420 | rs1245560 | -0.318 | -0.369 | 0.495 | -0.15 | 0.003 | 0.004 | 0.004 | 0 | 0.49 | 0.55 | 0.54 | 0.4 | 0.47 | 0.37 |
| 10 | 73845884 | rs1245548 | -0.378 | -0.004 | 0.115 | -0.65 | -0.059 | 0.056 | 0.056 | 3.864 | 0.59 | 0.87 | 0.52 | 0.27 | 0.4 | 0.49 |
| 10 | 73849472 | rs1245543 | 0.575 | -0.197 | 0.741 | -0.757 | 0.017 | 0.075 | 0.075 | 1.231 | 0.45 | 0.31 | 0.33 | 0.13 | 0.36 | 0.36 |
| 10 | 73856984 | rs3312 | -0.329 | -0.216 | 1.397 | -0.802 | -0.008 | 0.038 | 0.038 | 2.582 | 0.58 | 0.66 | 0.41 | 0.28 | 0.42 | 0.5 |
| 10 | 73858454 | rs3780944 | -0.906 | -0.178 | 1.62 | -0.648 | -0.008 | 0.055 | 0.055 | 0 | 0.08 | 0.19 | 0.05 | 0.14 | 0.04 | 0.13 |
| 10 | 73862032 | rs1668154 | 0.652 | -0.184 | 1.395 | -0.762 | 0.017 | 0.214 | 0.214 | 0.205 | 0.47 | 0.19 | 0.34 | 0.13 | 0.38 | 0.4 |
| 10 | 73865742 | rs7072673 | -0.367 | -0.175 | 1.368 | -0.751 | -0.09 | 0.035 | 0.035 | 2.411 | 0.62 | 0.87 | 0.43 | 0.27 | 0.43 | 0.52 |
| 10 | 73869663 | rs17726002 | -0.681 | -0.025 | 1.374 | -0.717 | -0.014 | -0.012 | -0.012 | 0 | 0.07 | 0.03 | 0.07 | 0.05 | 0.08 | 0.08 |
| 10 | 73875561 | rs16929688 | -1.809 | -0.054 | 1.8 | -0.658 | 0.017 | 0.137 | 0.137 | 2.846 | 0.07 | 0.5 | 0.05 | 0.09 | 0 | 0.05 |
| 10 | 73888394 | rs7098039 | -2.227 | -0.067 | 1.86 | -0.663 | -0.034 | 0.055 | 0.055 | 4.449 | 0.09 | 0.61 | 0.06 | 0.09 | 0 | 0.05 |
| 10 | 73895387 | rs7897444 | -1.509 | -0.06 | 1.91 | -0.66 | 0.013 | 0.121 | 0.121 | 2.084 | 0.07 | 0.41 | 0.04 | 0.09 | 0 | 0.05 |
| 10 | 73903335 | rs10823904 | -0.188 | 0.254 | 1.052 | -0.726 | -0.112 | -0.094 | -0.094 | 0 | 0.83 | 0.96 | 0.55 | 0.34 | 0.61 | 0.63 |
| 10 | 73906150 | rs1668157 | 0.255 | -0.08 | 1.229 | -1.001 | -0.037 | 0.15 | 0.15 | 1.712 | 0.56 | 0.89 | 0.4 | 0.21 | 0.38 | 0.44 |
| 10 | 73917890 | rs7097179 | -1.838 | -0.424 | 1.523 | -0.79 | -0.004 | 0.093 | 0.093 | 3.357 | 0.08 | 0.5 | 0.05 | 0.08 | 0 | 0.04 |
| 10 | 73931564 | rs11000201 | 0.854 | -0.118 | 1.38 | -0.89 | -0.018 | 0.205 | 0.205 | 0 | 0.49 | 0.2 | 0.38 | 0.13 | 0.47 | 0.42 |
| 10 | 73933699 | rs11000202 | 0.895 | -0.119 | 1.254 | -0.89 | -0.041 | 0.25 | 0.25 | 0 | 0.48 | 0.14 | 0.37 | 0.13 | 0.47 | 0.42 |
| 10 | 73936697 | rs7099664 | 0.933 | -0.134 | 1.261 | -0.899 | -0.052 | 0.264 | 0.264 | 0 | 0.48 | 0.1 | 0.37 | 0.13 | 0.47 | 0.42 |
| 10 | 73939338 | rs11000205 | -1.851 | -0.165 | 1.545 | -0.689 | 0.019 | 0.175 | 0.175 | 0.91 | 0.07 | 0.32 | 0.03 | 0.16 | 0.01 | 0.07 |
| 10 | 73940491 | rs12257464 | -2.072 | -0.165 | 1.551 | -0.827 | -0.029 | 0.075 | 0.075 | 2.646 | 0.1 | 0.52 | 0.06 | 0.12 | 0.01 | 0.07 |
| 10 | 73970451 | rs7069703 | 0.884 | -0.095 | 1.277 | -0.923 | -0.009 | 0.183 | 0.183 | 0 | 0.49 | 0.25 | 0.39 | 0.1 | 0.48 | 0.42 |
| 10 | 73988264 | rs4554797 | 0.277 | -0.034 | 1.139 | -0.877 | -0.061 | 0.098 | 0.098 | 0.172 | 0.6 | 0.86 | 0.45 | 0.22 | 0.5 | 0.49 |
| 10 | 73989184 | rs10823909 | -0.394 | 0.276 | 1.942 | -0.029 | -0.001 | 0.039 | 0.039 | 0 | 0.09 | 0.02 | 0.06 | 0 | 0.11 | 0.02 |
| 10 | 73996534 | rs11814574 | -2.612 | 0.349 | 2.32 | 0.143 | -0.017 | 0.093 | 0.093 | 2.078 | 0.1 | 0.52 | 0.05 | 0.12 | 0.01 | 0.05 |
| 10 | 73999284 | rs7921206 | -1.556 | 0.351 | 2.418 | 0.144 | 0.047 | 0.167 | 0.167 | 0.057 | 0.05 | 0.26 | 0.02 | 0.12 | 0.01 | 0.05 |
| 10 | 74003722 | rs7083908 | -2.721 | 0.348 | 2.342 | 0.144 | -0.023 | 0.108 | 0.108 | 2.692 | 0.1 | 0.53 | 0.05 | 0.13 | 0.01 | 0.05 |
| 10 | 74025367 | rs12247568 | -1.873 | 0.543 | 1.759 | 0.051 | 0.062 | 0.169 | 0.169 | 0.011 | 0.06 | 0.35 | 0.02 | 0.1 | 0.01 | 0.05 |
| 10 | 74032366 | rs877767 | -1.944 | 0.555 | 1.757 | 0.069 | 0.055 | 0.159 | 0.159 | 0 | 0.06 | 0.34 | 0.02 | 0.11 | 0.01 | 0.05 |
| 10 | 74090186 | rs11000266 | -0.277 | 0.545 | 0.926 | -0.044 | -0.003 | -0.02 | -0.02 | 0.018 | 0.07 | 0.05 | 0.06 | 0 | 0.11 | 0.02 |
| 10 | 74109459 | rs10400019 | -1.896 | 0.503 | 1.457 | 0.164 | -0.01 | 0.068 | 0.068 | 0.332 | 0.07 | 0.31 | 0.06 | 0.11 | 0.03 | 0.07 |
| 10 | 74181757 | rs7917581 | 0.674 | 0.595 | 1.618 | -0.259 | -0.025 | 0.041 | 0.041 | 0 | 0.69 | 0.83 | 0.53 | 0.15 | 0.62 | 0.46 |
| 10 | 74185638 | rs7079323 | 0.73 | 0.563 | 2.065 | 0.05 | -0.008 | 0.005 | 0.005 | 0.557 | 0.72 | 0.8 | 0.53 | 0.23 | 0.64 | 0.44 |
| 10 | 74262353 | rs16929939 | -1.014 | 0.471 | 2.177 | 0.208 | 0.008 | 0.27 | 0.27 | 0.862 | 0.07 | 0.37 | 0.04 | 0.2 | 0 | 0.04 |
| 10 | 74362355 | rs3009540 | 0.606 | 0.534 | 1.953 | 0.207 | 0.001 | -0.001 | -0.001 | 0.052 | 0.72 | 0.75 | 0.55 | 0.27 | 0.66 | 0.47 |
| 10 | 74490023 | rs2921448 | 0.401 | 1.315 | 2.173 | 0.49 | -0.017 | 0.003 | 0.003 | 0.128 | 0.73 | 0.89 | 0.56 | 0.25 | 0.66 | 0.47 |
| 10 | 74576831 | rs7092031 | 0.466 | 1.398 | 1.743 | 0.415 | -0.025 | 0.053 | 0.053 | 0.178 | 0.73 | 0.88 | 0.54 | 0.07 | 0.65 | 0.43 |

| | | | | | | | | | | | | | | | | |
|---|---|---|---|---|---|---|---|---|---|---|---|---|---|---|---|---|
| 10 | 74611240 | rs6415912 | 0.074 | 1.401 | 1.803 | 0.537 | -0.012 | -0.02 | -0.02 | 0.341 | 0.77 | 0.89 | 0.56 | 0.31 | 0.68 | 0.58 |
| 10 | 74639217 | rs7075988 | -2.16 | 1.341 | 2.077 | 0.561 | 0.031 | 0.285 | 0.285 | 0.972 | 0.1 | 0.52 | 0.07 | 0.3 | 0.05 | 0.19 |
| 10 | 74647824 | rs7081481 | 0.091 | 1.362 | 1.805 | 0.541 | -0.012 | -0.021 | -0.021 | 0.004 | 0.77 | 0.89 | 0.56 | 0.32 | 0.68 | 0.57 |
| 10 | 74652871 | rs11816594 | -1.904 | 1.874 | 2.691 | 1.327 | 0.006 | 0.127 | 0.127 | 0 | 0.07 | 0.16 | 0.04 | 0.29 | 0.05 | 0.18 |
| 10 | 74661041 | rs11818126 | -0.106 | 2.306 | 2.022 | 1.024 | -0.115 | 0.107 | 0.107 | 0 | 0.17 | 0.02 | 0.17 | 0 | 0.31 | 0.05 |
| 10 | 74662104 | rs11000440 | -3.402 | 1.051 | 2.814 | 1.762 | -0.008 | 0.008 | 0.008 | 3.293 | 0.17 | 0.82 | 0.1 | 0.29 | 0.05 | 0.19 |
| 10 | 74666065 | rs2394931 | -3.396 | 1.068 | 2.782 | 1.764 | -0.003 | 0.001 | 0.001 | 2.496 | 0.18 | 0.83 | 0.11 | 0.29 | 0.05 | 0.18 |
| 10 | 74672399 | rs11000445 | -3.372 | 1.067 | 2.789 | 1.762 | -0.01 | 0.011 | 0.011 | 2.253 | 0.18 | 0.82 | 0.11 | 0.32 | 0.05 | 0.18 |
| 10 | 74714177 | rs3829126 | -2.734 | 1.289 | 2.561 | 2.163 | 0.008 | 0.151 | 0.151 | 0 | 0.13 | 0.32 | 0.09 | 0.32 | 0.09 | 0.18 |
| 10 | 74746446 | rs12264150 | -2.694 | 1.112 | 1.702 | 2.327 | -0.002 | 0.072 | 0.072 | 0 | 0.1 | 0.13 | 0.13 | 0.25 | 0.06 | 0.21 |
| 10 | 74757872 | rs4522088 | -1.666 | 1.096 | 1.747 | 2.39 | -0.001 | 0.045 | 0.045 | 0 | 0.06 | 0.04 | 0.09 | 0.3 | 0.06 | 0.26 |
| 10 | 74761503 | rs7897191 | -3.273 | 0.792 | 2.98 | 2.36 | -0.032 | 0.157 | 0.157 | 0.011 | 0.14 | 0.37 | 0.12 | 0.32 | 0.06 | 0.25 |
| 10 | 74767695 | rs16930262 | -2.288 | 0.758 | 3.102 | 2.352 | -0.012 | 0.087 | 0.087 | 0 | 0.1 | 0.17 | 0.1 | 0.3 | 0.05 | 0.24 |
| 10 | 74822666 | rs12243451 | -2.19 | 0.689 | 3.034 | 2.362 | -0.011 | 0.094 | 0.094 | 0 | 0.1 | 0.17 | 0.1 | 0.31 | 0.05 | 0.24 |
| 10 | 74836786 | rs10823977 | -2.19 | 0.689 | 3.034 | 2.362 | -0.011 | 0.09 | 0.09 | 0 | 0.1 | 0.17 | 0.1 | 0.31 | 0.05 | 0.24 |
| 10 | 74874262 | rs4492736 | -3.724 | 0.702 | 3.299 | 2.417 | -0.048 | 0.144 | 0.144 | 0.582 | 0.16 | 0.6 | 0.14 | 0.37 | 0.06 | 0.25 |
| 10 | 74885830 | rs16930346 | -2.197 | 0.711 | 3.248 | 2.406 | -0.012 | 0.081 | 0.081 | 0 | 0.1 | 0.17 | 0.1 | 0.31 | 0.05 | 0.24 |
| 10 | 74886043 | rs12569493 | -2.197 | 0.711 | 3.248 | 2.406 | -0.012 | 0.081 | 0.081 | 0 | 0.1 | 0.17 | 0.1 | 0.31 | 0.05 | 0.24 |
| 10 | 74894258 | rs2271903 | -3.298 | 0.711 | 3.265 | 2.406 | -0.031 | 0.159 | 0.159 | 0.104 | 0.14 | 0.37 | 0.12 | 0.31 | 0.06 | 0.24 |
| 10 | 74894375 | rs2271904 | -2.196 | 0.711 | 3.265 | 2.406 | -0.012 | 0.081 | 0.081 | 0 | 0.1 | 0.17 | 0.1 | 0.31 | 0.05 | 0.24 |
| 10 | 74897816 | rs2271905 | -3.205 | 0.713 | 3.185 | 2.407 | -0.031 | 0.159 | 0.159 | 0.104 | 0.14 | 0.37 | 0.12 | 0.31 | 0.06 | 0.24 |
| 10 | 74912004 | rs7069878 | -3.204 | 0.726 | 3.172 | 2.4 | -0.027 | 0.159 | 0.159 | 0.104 | 0.14 | 0.37 | 0.13 | 0.31 | 0.06 | 0.24 |
| 10 | 74923562 | rs3812619 | -3.202 | 0.691 | 3.181 | 2.404 | -0.031 | 0.161 | 0.161 | 0.104 | 0.14 | 0.37 | 0.12 | 0.31 | 0.06 | 0.24 |
| 10 | 74957896 | rs3998311 | -3.122 | 0.697 | 3.163 | 2.385 | -0.03 | 0.15 | 0.15 | 0.374 | 0.13 | 0.35 | 0.13 | 0.32 | 0.06 | 0.23 |
| 10 | 75000698 | rs11000561 | -2.465 | 0.697 | 3.221 | 2.427 | -0.018 | 0.096 | 0.096 | 0 | 0.11 | 0.22 | 0.1 | 0.32 | 0.06 | 0.22 |
| 10 | 75005420 | rs12258130 | -3.32 | 1.148 | 3.12 | 2.373 | -0.031 | 0.201 | 0.201 | 0.978 | 0.14 | 0.44 | 0.13 | 0.31 | 0.06 | 0.23 |
| 10 | 75006437 | rs16930457 | -2.264 | 1.161 | 2.95 | 2.379 | -0.011 | 0.067 | 0.067 | 0 | 0.11 | 0.18 | 0.1 | 0.31 | 0.05 | 0.22 |
| 10 | 75008561 | rs1064020 | -3.488 | 1.145 | 3.069 | 2.469 | -0.046 | 0.202 | 0.202 | 0.851 | 0.16 | 0.59 | 0.14 | 0.4 | 0.07 | 0.24 |
| 10 | 75056863 | rs6480692 | -2.318 | 1.145 | 2.967 | 2.472 | -0.011 | 0.063 | 0.063 | 0 | 0.11 | 0.18 | 0.1 | 0.31 | 0.05 | 0.22 |
| 10 | 75092686 | rs2183903 | -2.307 | 1.134 | 2.953 | 2.512 | -0.011 | 0.063 | 0.063 | 0 | 0.11 | 0.18 | 0.1 | 0.31 | 0.05 | 0.23 |
| 10 | 75108146 | rs3812622 | -3.449 | 1.118 | 3.02 | 2.507 | -0.047 | 0.229 | 0.229 | 0.993 | 0.16 | 0.52 | 0.14 | 0.37 | 0.07 | 0.24 |
| 10 | 75133893 | rs11000630 | -3.293 | 1.372 | 2.95 | 2.672 | -0.03 | 0.148 | 0.148 | 0.437 | 0.13 | 0.37 | 0.12 | 0.3 | 0.06 | 0.23 |
| 10 | 75146811 | rs12261772 | -2.424 | 0.798 | 3.024 | 2.817 | -0.011 | 0.067 | 0.067 | 0 | 0.11 | 0.18 | 0.1 | 0.3 | 0.06 | 0.22 |
| 10 | 75147579 | rs16930547 | -2.424 | 0.798 | 3.024 | 2.817 | -0.011 | 0.067 | 0.067 | 0 | 0.11 | 0.18 | 0.1 | 0.3 | 0.06 | 0.22 |
| 10 | 75148116 | rs2234965 | -2.424 | 0.798 | 3.025 | 2.817 | -0.011 | 0.063 | 0.063 | 0 | 0.11 | 0.18 | 0.1 | 0.3 | 0.06 | 0.22 |
| 10 | 75160257 | rs12258241 | -3.442 | 0.791 | 3.124 | 2.814 | -0.028 | 0.229 | 0.229 | 1.32 | 0.14 | 0.51 | 0.13 | 0.3 | 0.06 | 0.22 |
| 10 | 75175016 | rs12570678 | -1.175 | 0.797 | 3.056 | 2.817 | -0.007 | 0.035 | 0.035 | 0 | 0.05 | 0.03 | 0.08 | 0.3 | 0.06 | 0.22 |
| 10 | 75197051 | rs12644 | -4.022 | 0.361 | 3.249 | 2.873 | -0.005 | 0.009 | 0.009 | 3.317 | 0.21 | 0.77 | 0.17 | 0.44 | 0.1 | 0.27 |
| 10 | 75207487 | rs1041532 | -2.667 | 0.612 | 2.54 | 2.946 | -0.011 | 0.066 | 0.066 | 0 | 0.11 | 0.18 | 0.1 | 0.3 | 0.06 | 0.22 |
| 10 | 75227789 | rs16930644 | -1.651 | 0.612 | 2.537 | 2.934 | -0.004 | 0.019 | 0.019 | 0 | 0.06 | 0.04 | 0.08 | 0.3 | 0.06 | 0.22 |
| 10 | 75232994 | rs11000666 | -3.181 | 0.612 | 2.539 | 2.934 | -0.027 | 0.145 | 0.145 | 0.525 | 0.13 | 0.35 | 0.12 | 0.3 | 0.06 | 0.22 |
| 10 | 75239977 | rs11000671 | -3.159 | 0.608 | 2.537 | 2.935 | -0.027 | 0.143 | 0.143 | 0.584 | 0.13 | 0.34 | 0.12 | 0.3 | 0.06 | 0.23 |
| 10 | 75241418 | rs11000672 | -3.698 | 0.575 | 2.523 | 2.921 | -0.034 | 0.124 | 0.124 | 0.469 | 0.14 | 0.35 | 0.12 | 0.3 | 0.06 | 0.22 |
| 10 | 75245252 | rs12569942 | -1.231 | 0.714 | 2.579 | 2.982 | -0.007 | 0.033 | 0.033 | 0 | 0.05 | 0.03 | 0.08 | 0.3 | 0.06 | 0.22 |
| 10 | 75285837 | rs12572177 | -2.83 | 0.708 | 2.539 | 2.997 | -0.027 | 0.151 | 0.151 | 0.525 | 0.13 | 0.35 | 0.12 | 0.3 | 0.06 | 0.22 |
| 10 | 75286806 | rs12268338 | -3.385 | 0.716 | 2.552 | 3.126 | -0.028 | 0.271 | 0.271 | 0.94 | 0.14 | 0.51 | 0.14 | 0.33 | 0.07 | 0.22 |
| 10 | 75289359 | rs12255505 | -3.271 | 0.702 | 2.562 | 2.897 | -0.028 | 0.232 | 0.232 | 1.32 | 0.14 | 0.5 | 0.13 | 0.3 | 0.06 | 0.22 |
| 10 | 75300994 | rs4746136 | -1.092 | 0.737 | 1.669 | 2.849 | -0.054 | -0.031 | -0.031 | 0.133 | 0.09 | 0.01 | 0.4 | 0.52 | 0.15 | 0.37 |
| 10 | 75304865 | rs12248024 | -2.753 | 0.227 | 2.017 | 2.515 | -0.021 | 0.16 | 0.16 | 0.324 | 0.12 | 0.34 | 0.12 | 0.3 | 0.06 | 0.22 |
| 10 | 75344761 | rs7899274 | -2.92 | 0.28 | 1.885 | 2.536 | -0.026 | 0.181 | 0.181 | 0.099 | 0.14 | 0.39 | 0.13 | 0.33 | 0.07 | 0.21 |
| 10 | 75384982 | rs11000723 | -2.918 | 0.073 | 1.826 | 2.404 | -0.026 | 0.181 | 0.181 | 0.099 | 0.14 | 0.39 | 0.12 | 0.33 | 0.07 | 0.21 |
| 10 | 75391178 | rs7080436 | -2.311 | 0.328 | 1.047 | 2.497 | -0.007 | 0.111 | 0.111 | 0 | 0.09 | 0.21 | 0.11 | 0.33 | 0.07 | 0.21 |
| 10 | 75391217 | rs7080456 | -1.842 | 0.457 | 1.077 | 2.553 | 0.004 | 0.155 | 0.155 | 0 | 0.08 | 0.2 | 0.1 | 0.33 | 0.07 | 0.21 |
| 10 | 75406141 | rs3740293 | -1.563 | 0.091 | 1.261 | 2.07 | -0.021 | 0.015 | 0.015 | 0.117 | 0.18 | 0.38 | 0.13 | 0.18 | 0.13 | 0.19 |
| 10 | 75407290 | rs3812629 | -1.358 | 0.124 | 1.392 | 2.084 | -0.005 | 0.005 | 0.005 | 0.173 | 0.16 | 0.21 | 0.11 | 0.18 | 0.13 | 0.19 |
| 10 | 75407649 | rs4746139 | -1.299 | 0.112 | 1.422 | 2.079 | 0.001 | -0.003 | -0.003 | 0 | 0.16 | 0.15 | 0.11 | 0.18 | 0.13 | 0.19 |
| 10 | 75409877 | rs2177843 | -1.892 | -0.182 | 2.034 | 2.189 | -0.023 | 0.01 | 0.01 | 0.512 | 0.19 | 0.3 | 0.13 | 0.18 | 0.15 | 0.23 |
| 10 | 75411198 | rs998950 | -1.703 | -0.457 | 2.491 | 2.814 | 0.014 | 0.218 | 0.218 | 3.707 | 0.08 | 0.4 | 0.23 | 0.18 | 0.02 | 0.07 |
| 10 | 75411254 | rs1124351 | -1.703 | -0.457 | 2.492 | 2.814 | 0.013 | 0.215 | 0.215 | 3.707 | 0.08 | 0.4 | 0.23 | 0.18 | 0.02 | 0.07 |
| 10 | 75416789 | rs12570126 | -2.394 | -0.588 | 2.432 | 2.762 | 0 | -0.002 | -0.002 | 0.165 | 0.26 | 0.68 | 0.36 | 0.34 | 0.16 | 0.25 |
| 10 | 75421208 | rs10824026 | 2.189 | -0.705 | 2.375 | 2.694 | 0.026 | -0.018 | -0.018 | 0 | 0.72 | 0.25 | 0.63 | 0.66 | 0.84 | 0.74 |
| 10 | 75424174 | rs3933086 | -2.043 | -0.684 | 2.451 | 2.692 | -0.022 | 0.167 | 0.167 | 3.868 | 0.1 | 0.47 | 0.25 | 0.18 | 0.02 | 0.07 |
| 10 | 75524458 | rs11000771 | -1.707 | -0.284 | 2.513 | 2.682 | 0.008 | 0.005 | 0.005 | 0.103 | 0.19 | 0.17 | 0.12 | 0.16 | 0.13 | 0.2 |
| 10 | 75530962 | rs11541237 | -1.638 | -0.286 | 2.513 | 2.683 | 0.007 | 0.01 | 0.01 | 0.195 | 0.18 | 0.15 | 0.12 | 0.12 | 0.13 | 0.19 |
| 10 | 75534115 | rs11000773 | -1.638 | -0.251 | 2.513 | 3.18 | 0.007 | 0.01 | 0.01 | 0.195 | 0.18 | 0.15 | 0.12 | 0.12 | 0.13 | 0.19 |
| 10 | 75539812 | rs4082339 | -1.583 | -0.253 | 2.533 | 3.179 | 0.009 | 0.014 | 0.014 | 0.138 | 0.18 | 0.12 | 0.1 | 0.12 | 0.13 | 0.19 |
| 10 | 75549916 | rs2292307 | -1.115 | -0.077 | 2.536 | 3.249 | 0.006 | 0.013 | 0.013 | 0.675 | 0.16 | 0.07 | 0.1 | 0.12 | 0.13 | 0.19 |
| 10 | 75559077 | rs11000780 | -1.197 | -0.084 | 2.533 | 3.246 | 0.007 | 0.014 | 0.014 | 0.805 | 0.16 | 0.08 | 0.1 | 0.12 | 0.13 | 0.19 |
| 10 | 75561916 | rs4691 | -0.751 | 0.184 | 2.278 | 3.155 | -0.026 | -0.017 | -0.017 | 0 | 0.13 | 0.08 | 0.26 | 0.54 | 0.22 | 0.46 |

| | | | | | | | | | | | | | | | | |
|---|---|---|---|---|---|---|---|---|---|---|---|---|---|---|---|---|
| 10 | 75565935 | rs4746150 | -0.793 | -0.021 | 2.016 | 2.84 | -0.015 | -0.009 | -0.009 | 0 | 0.14 | 0.01 | 0.08 | 0.12 | 0.11 | 0.18 |
| 10 | 75601688 | rs2254266 | -1.143 | -0.098 | 2.116 | 2.619 | 0.003 | -0.001 | -0.001 | 0 | 0.21 | 0.18 | 0.15 | 0.18 | 0.19 | 0.26 |
| 10 | 75647491 | rs11000801 | -0.969 | -0.09 | 1.629 | 2.64 | 0.006 | -0.02 | -0.02 | 0 | 0.2 | 0.13 | 0.13 | 0.32 | 0.18 | 0.26 |
| 10 | 75648249 | rs2675677 | 1.491 | -0.514 | 1.494 | 2.368 | 0.002 | 0.013 | 0.013 | 0 | 0.77 | 0.78 | 0.51 | 0.48 | 0.77 | 0.54 |
| 10 | 75653800 | rs17741873 | -1.571 | -0.965 | 1.03 | 2.159 | 0.034 | 0.004 | 0.004 | 0 | 0.28 | 0.14 | 0.13 | 0.3 | 0.19 | 0.28 |
| 10 | 75669943 | rs2227553 | -0.431 | -0.675 | 1.258 | 2.166 | -0.004 | 0.028 | 0.028 | 0.034 | 0.05 | 0.22 | 0.03 | 0.05 | 0.01 | 0.05 |
| 10 | 75672961 | rs2227562 | -0.519 | -0.795 | 1.018 | 2.136 | 0.008 | -0.041 | -0.041 | 0.922 | 0.18 | 0.06 | 0.11 | 0.29 | 0.16 | 0.24 |
| 10 | 75673101 | rs2227564 | 0.046 | -0.651 | 0.925 | 2.337 | -0.065 | -0.06 | -0.06 | 0.155 | 0.9 | 0.99 | 0.75 | 0.63 | 0.79 | 0.63 |
| 10 | 75673731 | rs2227566 | 1.798 | -0.652 | 1.233 | 1.337 | 0 | -0.016 | -0.016 | 0 | 0.55 | 0.45 | 0.38 | 0.12 | 0.59 | 0.23 |
| 10 | 75673879 | rs2227568 | -1.037 | -1.075 | -0.023 | 1.67 | 0.005 | -0.048 | -0.048 | 0 | 0.18 | 0.05 | 0.11 | 0.29 | 0.16 | 0.24 |
| 10 | 75692923 | rs16931042 | -0.941 | -1.082 | 0.046 | 1.649 | 0.005 | -0.047 | -0.047 | 0.013 | 0.18 | 0.05 | 0.1 | 0.29 | 0.16 | 0.25 |
| 10 | 75695919 | rs4745724 | -1.271 | -1.039 | 0.181 | 1.689 | 0.001 | 0 | 0 | 0.08 | 0.27 | 0.24 | 0.13 | 0.29 | 0.17 | 0.27 |
| 10 | 75705977 | rs2675661 | -1.309 | -0.794 | 0.323 | 1.942 | -0.02 | 0.072 | 0.072 | 2.013 | 0.09 | 0.16 | 0.29 | 0.35 | 0.06 | 0.07 |
| 10 | 75708906 | rs4746160 | -1.005 | -0.879 | -0.208 | 1.433 | 0.011 | -0.004 | -0.004 | 4.201 | 0.22 | 0.15 | 0.25 | 0.23 | 0.18 | 0.27 |
| 10 | 75733299 | rs11000828 | 0.839 | -1.336 | 0.195 | -0.05 | 0.044 | 0.176 | 0.176 | 0.077 | 0.47 | 0.23 | 0.41 | 0.05 | 0.54 | 0.21 |
| 10 | 75741457 | rs4745727 | -0.981 | -1.75 | 0.542 | 0.124 | -0.013 | -0.005 | -0.005 | 0 | 0.33 | 0.61 | 0.34 | 0.26 | 0.24 | 0.24 |
| 10 | 75743526 | rs7902395 | -0.708 | -1.795 | 1.19 | 0.102 | 0 | 0.001 | 0.001 | 0.057 | 0.28 | 0.23 | 0.26 | 0.2 | 0.22 | 0.25 |
| 10 | 75750764 | rs2675678 | -0.676 | -1.699 | 1.006 | 0.406 | -0.048 | -0.078 | -0.078 | 0 | 0.89 | 0.99 | 0.86 | 0.5 | 0.82 | 0.61 |
| 10 | 75761860 | rs12250729 | -0.853 | -1.71 | 1.174 | 0.012 | -0.04 | -0.012 | -0.012 | 0.781 | 0.3 | 0.42 | 0.31 | 0.26 | 0.24 | 0.28 |
| 10 | 75778687 | rs4746166 | -0.564 | -1.833 | 1.114 | -0.084 | -0.027 | -0.002 | -0.002 | 0 | 0.21 | 0.24 | 0.23 | 0.21 | 0.15 | 0.21 |
| 10 | 75780838 | rs10458640 | 0.977 | -1.063 | 1.013 | -0.242 | 0.001 | -0.01 | -0.01 | 0.053 | 0.68 | 0.65 | 0.66 | 0.25 | 0.64 | 0.41 |
| 10 | 75786395 | rs7914469 | -0.268 | -1.07 | 0.871 | -0.18 | 0.003 | -0.001 | -0.001 | 0 | 0.18 | 0.13 | 0.2 | 0.21 | 0.14 | 0.19 |
| 10 | 75798148 | rs10762573 | 0.806 | -1.47 | 0.944 | -0.385 | -0.006 | -0.004 | -0.004 | 0 | 0.71 | 0.8 | 0.7 | 0.29 | 0.66 | 0.45 |
| 10 | 75808313 | rs4414157 | -0.041 | -1.56 | 0.903 | -0.527 | 0.003 | -0.001 | -0.001 | 0 | 0.18 | 0.13 | 0.2 | 0.21 | 0.14 | 0.19 |
| 10 | 75847804 | rs11596526 | -0.3 | -1.631 | 0.97 | -0.559 | -0.007 | -0.001 | -0.001 | 0.165 | 0.19 | 0.21 | 0.21 | 0.2 | 0.14 | 0.19 |
| 10 | 75847856 | rs11000869 | -1.332 | -1.341 | 1.269 | -0.319 | -0.012 | 0.015 | 0.015 | 0 | 0.29 | 0.18 | 0.32 | 0.76 | 0.35 | 0.53 |
| 10 | 75867193 | rs2279648 | -0.784 | -1.516 | 2.27 | -0.341 | -0.005 | -0.008 | -0.008 | 0 | 0.15 | 0.13 | 0.14 | 0.11 | 0.13 | 0.09 |
| 10 | 75886956 | rs10824076 | -1.202 | -1.538 | 2.419 | -0.297 | -0.025 | -0.037 | -0.037 | 0 | 0.17 | 0.36 | 0.16 | 0.11 | 0.13 | 0.09 |
| 10 | 75900462 | rs7896966 | 0.249 | -1.406 | 2.432 | -0.232 | -0.007 | -0.019 | -0.019 | 0 | 0.81 | 0.79 | 0.68 | 0.38 | 0.76 | 0.5 |
| 10 | 75905996 | rs11000893 | -0.761 | -1.322 | 2.257 | -0.235 | 0.005 | 0.006 | 0.006 | 0.059 | 0.15 | 0.08 | 0.14 | 0.11 | 0.13 | 0.09 |
| 10 | 75911276 | rs11000897 | -0.881 | -1.324 | 2.29 | -0.236 | -0.011 | -0.014 | -0.014 | 0 | 0.16 | 0.19 | 0.14 | 0.11 | 0.13 | 0.09 |
| 10 | 75937554 | rs10824096 | -0.821 | -1.298 | 2.271 | -0.224 | -0.005 | -0.006 | -0.006 | 0 | 0.15 | 0.12 | 0.14 | 0.11 | 0.13 | 0.09 |
| 10 | 75958408 | rs10824106 | -0.852 | -1.307 | 1.933 | -0.235 | 0.004 | 0.005 | 0.005 | 0.321 | 0.16 | 0.1 | 0.14 | 0.12 | 0.13 | 0.09 |
| 10 | 76024496 | rs12217890 | -0.464 | -1.325 | 2.107 | -0.282 | -0.007 | -0.009 | -0.009 | 0 | 0.15 | 0.18 | 0.15 | 0.12 | 0.12 | 0.07 |
| 10 | 76071203 | rs2894234 | 0.15 | -1.338 | 2.04 | -0.261 | 0.008 | 0.029 | 0.029 | 0 | 0.82 | 0.75 | 0.66 | 0.4 | 0.76 | 0.52 |
| 10 | 76214275 | rs10824183 | -0.174 | -1.341 | 1.927 | -0.281 | -0.004 | -0.003 | -0.003 | 0 | 0.83 | 0.88 | 0.68 | 0.4 | 0.76 | 0.51 |
| 10 | 76282793 | rs10824197 | -0.483 | -1.621 | 1.567 | -0.346 | 0 | 0.001 | 0.001 | 0 | 0.15 | 0.12 | 0.14 | 0.12 | 0.12 | 0.07 |
| 10 | 76323150 | rs10762630 | -0.237 | -1.441 | 1.471 | -0.342 | -0.008 | 0.047 | 0.047 | 0 | 0.18 | 0.27 | 0.31 | 0.64 | 0.16 | 0.48 |
| 10 | 76372037 | rs4746209 | -1.865 | -1.148 | -0.184 | -0.791 | 0.099 | 0.198 | 0.198 | 0.182 | 0.48 | 0.9 | 0.52 | 0.79 | 0.33 | 0.6 |
| 10 | 76373904 | rs12360087 | -0.821 | -0.851 | 0.301 | -0.647 | -0.014 | -0.033 | -0.033 | 0 | 0.2 | 0.27 | 0.17 | 0.1 | 0.14 | 0.1 |
| 10 | 76401365 | rs7894466 | -0.824 | -0.791 | 0.291 | -0.629 | -0.019 | -0.038 | -0.038 | 0 | 0.2 | 0.29 | 0.27 | 0.11 | 0.14 | 0.09 |
| 10 | 76403131 | rs10762635 | -0.826 | -0.764 | 0.159 | -0.616 | -0.014 | -0.029 | -0.029 | 0.155 | 0.2 | 0.26 | 0.27 | 0.11 | 0.14 | 0.09 |
| 10 | 76415509 | rs1538311 | 1.495 | -0.211 | -0.429 | -1.125 | -0.033 | 0.134 | 0.134 | 0 | 0.4 | 0.19 | 0.26 | 0.02 | 0.44 | 0.19 |
| 10 | 76437114 | rs1556822 | 0.099 | -0.302 | -0.026 | -0.609 | -0.022 | -0.002 | -0.002 | 0 | 0.83 | 0.91 | 0.86 | 0.83 | 0.76 | 0.79 |
| 10 | 76459669 | rs7086415 | 0.152 | -0.364 | -0.016 | -0.599 | -0.028 | -0.004 | -0.004 | 1.044 | 0.84 | 0.92 | 0.87 | 0.83 | 0.76 | 0.77 |
| 10 | 76461188 | rs2395097 | 0.332 | -0.394 | 0.195 | -0.604 | -0.005 | 0 | 0 | 0 | 0.83 | 0.82 | 0.86 | 0.83 | 0.76 | 0.77 |
| 10 | 76468851 | rs1189553 | 0.041 | -0.403 | 0.154 | -0.606 | -0.066 | -0.032 | -0.032 | 0.648 | 0.85 | 0.97 | 0.89 | 0.83 | 0.76 | 0.78 |
| 10 | 76480450 | rs839698 | -0.327 | -0.639 | 0.727 | -0.576 | -0.027 | 0.045 | 0.045 | 0 | 0.2 | 0.32 | 0.3 | 0.69 | 0.15 | 0.49 |
| 10 | 76481311 | rs10824226 | -0.703 | -0.592 | 1.087 | -0.21 | -0.033 | 0.031 | 0.031 | 0.589 | 0.1 | 0.24 | 0.18 | 0.12 | 0.03 | 0.07 |
| 10 | 76498150 | rs2488089 | 1.366 | -0.567 | 1.088 | -0.264 | 0.002 | 0.002 | 0.002 | 0 | 0.73 | 0.53 | 0.67 | 0.71 | 0.72 | 0.71 |
| 10 | 76498953 | rs2488088 | 0.285 | -0.538 | 1 | -0.252 | -0.033 | -0.004 | -0.004 | 0 | 0.84 | 0.92 | 0.87 | 0.84 | 0.76 | 0.77 |
| 10 | 76505183 | rs12247071 | -0.689 | -0.599 | 1.161 | -0.172 | -0.027 | 0.046 | 0.046 | 1.068 | 0.1 | 0.34 | 0.19 | 0.13 | 0.03 | 0.07 |
| 10 | 76511108 | rs1189555 | 1.297 | -0.155 | 0.885 | 0.179 | 0.014 | 0.02 | 0.02 | 0 | 0.79 | 0.67 | 0.82 | 0.71 | 0.75 | 0.71 |
| 10 | 76515579 | rs839708 | 0.7 | -0.148 | 0.742 | 0.193 | -0.02 | -0.001 | -0.001 | 0 | 0.84 | 0.91 | 0.89 | 0.84 | 0.77 | 0.76 |
| 10 | 76540987 | rs3862556 | -0.837 | -0.55 | 0.222 | 0.062 | -0.001 | 0.003 | 0.003 | 0.065 | 0.08 | 0.01 | 0.14 | 0.07 | 0.06 | 0.34 |
| 10 | 76551502 | rs4746237 | -2.588 | -0.873 | 0.473 | 0.276 | 0.015 | -0.009 | -0.009 | 0 | 0.25 | 0.75 | 0.33 | 0.32 | 0.11 | 0.51 |
| 10 | 76591158 | rs7075439 | -1.343 | -0.848 | 1.687 | 0.264 | 0.049 | 0.101 | 0.101 | 0.279 | 0.07 | 0.35 | 0.04 | 0.09 | 0.03 | 0.05 |
| 10 | 76611753 | rs11001184 | -1.323 | -0.975 | 1.657 | 0.23 | -0.001 | 0.002 | 0.002 | 0 | 0.06 | 0.07 | 0.11 | 0.13 | 0.02 | 0.11 |
| 10 | 76611849 | rs7091085 | -2.599 | -1.032 | 1.706 | 0.205 | -0.002 | -0.001 | -0.001 | 0 | 0.19 | 0.74 | 0.23 | 0.19 | 0.09 | 0.4 |
| 10 | 76617479 | rs951308 | -3.102 | -1.034 | 1.659 | 0.195 | 0.042 | -0.018 | -0.018 | 0 | 0.26 | 0.81 | 0.33 | 0.32 | 0.11 | 0.51 |
| 10 | 76617857 | rs12254441 | 1.254 | -1.01 | 0.647 | -0.719 | -0.081 | 0.163 | 0.163 | 0.692 | 0.28 | 0.06 | 0.23 | 0 | 0.33 | 0.05 |
| 10 | 76642997 | rs10824242 | -2.959 | -1.564 | 1.526 | 0.115 | 0.041 | -0.017 | -0.017 | 0 | 0.26 | 0.81 | 0.24 | 0.32 | 0.11 | 0.5 |
| 10 | 76643827 | rs11001192 | -1.936 | -1.557 | 1.674 | 0.082 | -0.028 | -0.02 | -0.02 | 1.598 | 0.15 | 0.32 | 0.1 | 0.13 | 0.07 | 0.34 |
| 10 | 76658330 | rs4745760 | -3.014 | -1.514 | 1.45 | 0.408 | 0.044 | -0.045 | -0.045 | 0 | 0.26 | 0.82 | 0.25 | 0.38 | 0.11 | 0.5 |
| 10 | 76665108 | rs4307657 | -0.658 | -1.494 | 1.558 | 0.414 | 0 | 0 | 0 | 0 | 0.09 | 0.11 | 0.06 | 0.09 | 0.06 | 0.33 |
| 10 | 76665603 | rs7917746 | -2.424 | -1.501 | 1.67 | 0.415 | -0.034 | 0.073 | 0.073 | 0.053 | 0.21 | 0.56 | 0.2 | 0.34 | 0.1 | 0.5 |
| 10 | 76685101 | rs11001204 | -0.763 | -1.418 | 1.574 | 0.388 | -0.007 | -0.001 | -0.001 | 0 | 0.1 | 0.19 | 0.08 | 0.12 | 0.06 | 0.33 |
| 10 | 76688082 | rs7909358 | -2.63 | -1.442 | 1.549 | 0.388 | 0.017 | 0.006 | 0.006 | 1.335 | 0.22 | 0.8 | 0.15 | 0.22 | 0.1 | 0.39 |
| 10 | 76705881 | rs4746248 | -2.439 | -1.438 | 1.578 | 0.39 | -0.026 | -0.006 | -0.006 | 7.766 | 0.21 | 0.66 | 0.14 | 0.22 | 0.1 | 0.39 |

| | | | | | | | | | | | | | | | | |
|---|---|---|---|---|---|---|---|---|---|---|---|---|---|---|---|---|
| 10 | 76726096 | rs1551068 | -1.037 | -1.471 | 1.507 | 0.372 | -0.012 | 0.001 | 0.001 | 0 | 0.11 | 0.32 | 0.09 | 0.11 | 0.06 | 0.33 |
| 10 | 76729632 | rs3213967 | -3.303 | -1.525 | 1.228 | 0.395 | 0.043 | -0.04 | -0.04 | 0 | 0.3 | 0.82 | 0.3 | 0.38 | 0.17 | 0.54 |
| 10 | 76757407 | rs11001239 | -0.775 | -1.553 | 1.133 | 0.46 | 0.001 | 0.002 | 0.002 | 0 | 0.09 | 0.11 | 0.06 | 0.07 | 0.06 | 0.32 |
| 10 | 76760631 | rs3781252 | -0.866 | -1.554 | 1.193 | 0.459 | -0.001 | -0.001 | -0.001 | 0 | 0.1 | 0.13 | 0.07 | 0.07 | 0.07 | 0.32 |
| 10 | 76784538 | rs10490994 | -1.173 | -1.589 | 1.182 | 0.535 | -0.016 | 0.013 | 0.013 | 0 | 0.05 | 0.09 | 0.06 | 0.09 | 0.02 | 0.03 |
| 10 | 76787128 | rs1551067 | -3.131 | -1.67 | 1.249 | 0.511 | 0.054 | -0.052 | -0.052 | 0 | 0.27 | 0.85 | 0.28 | 0.38 | 0.11 | 0.5 |
| 10 | 76788860 | rs3740322 | -1.354 | -1.398 | 1.388 | 0.617 | -0.016 | 0.013 | 0.013 | 0 | 0.05 | 0.09 | 0.06 | 0.09 | 0.02 | 0.03 |
| 10 | 76789077 | rs3740321 | -1.354 | -1.398 | 1.388 | 0.617 | -0.016 | 0.013 | 0.013 | 0 | 0.05 | 0.09 | 0.06 | 0.09 | 0.02 | 0.03 |
| 10 | 76794788 | rs12255305 | -1.981 | -1.58 | 1.266 | 0.552 | -0.063 | 0.004 | 0.004 | 0.442 | 0.15 | 0.47 | 0.11 | 0.11 | 0.04 | 0.06 |
| 10 | 76803567 | rs16931938 | -2.661 | -1.281 | 1.348 | 0.691 | -0.038 | 0.037 | 0.037 | 0.722 | 0.17 | 0.65 | 0.21 | 0.27 | 0.05 | 0.18 |
| 10 | 76804588 | rs7901279 | -1.659 | -1.245 | 1.463 | 0.623 | -0.022 | 0.049 | 0.049 | 0.199 | 0.12 | 0.42 | 0.1 | 0.1 | 0.04 | 0.05 |
| 10 | 76807788 | rs12218431 | -1.232 | -0.964 | 1.145 | 0.716 | 0.012 | 0.194 | 0.194 | 0 | 0.05 | 0.23 | 0.11 | 0.15 | 0.01 | 0.12 |
| 10 | 76812741 | rs2395129 | -1.463 | -0.828 | 1.156 | 0.515 | 0.007 | 0.115 | 0.115 | 0 | 0.09 | 0.36 | 0.08 | 0.14 | 0.03 | 0.06 |
| 10 | 76817937 | rs1993005 | -0.829 | -1.342 | 0.363 | 0.278 | -0.021 | -0.008 | -0.008 | 0.097 | 0.34 | 0.58 | 0.47 | 0.71 | 0.26 | 0.34 |
| 10 | 76818077 | rs11594934 | -0.328 | -1.423 | 0.343 | 0.053 | -0.003 | -0.01 | -0.01 | 0 | 0.09 | 0.09 | 0.06 | 0.04 | 0.06 | 0.31 |
| 10 | 76821098 | rs11001266 | -0.818 | -1.558 | 0.378 | -0.049 | -0.008 | -0.012 | -0.012 | 0 | 0.11 | 0.21 | 0.08 | 0.08 | 0.07 | 0.33 |
| 10 | 76829374 | rs755228 | -0.307 | -1.981 | 0.358 | 0.008 | -0.036 | 0.01 | 0.01 | 0 | 0.31 | 0.48 | 0.46 | 0.65 | 0.26 | 0.29 |
| 10 | 76834921 | rs4745763 | -0.841 | -1.767 | 0.252 | 0.045 | 0.025 | 0.009 | 0.009 | 0 | 0.37 | 0.67 | 0.49 | 0.69 | 0.28 | 0.32 |
| 10 | 76835788 | rs7905806 | -0.499 | -2.349 | 0.593 | 0.11 | -0.026 | 0.031 | 0.031 | 0 | 0.32 | 0.51 | 0.43 | 0.63 | 0.26 | 0.3 |
| 10 | 76842925 | rs7901680 | -1.746 | -2.176 | 0.85 | -0.138 | -0.011 | -0.066 | -0.066 | 1.173 | 0.21 | 0.35 | 0.13 | 0.1 | 0.15 | 0.43 |
| 10 | 76845632 | rs4746261 | -2.191 | -3.046 | -0.011 | -0.944 | -0.102 | 0.127 | 0.127 | 0 | 0.68 | 0.84 | 0.77 | 0.95 | 0.55 | 0.84 |
| 10 | 76845695 | rs10824268 | -0.327 | -3.016 | 0.061 | -0.895 | -0.006 | -0.028 | -0.028 | 0 | 0.48 | 0.5 | 0.64 | 0.85 | 0.39 | 0.4 |
| 10 | 76852635 | rs1962117 | 0.183 | -3.091 | 0.412 | -0.885 | 0.006 | -0.005 | -0.005 | 0 | 0.25 | 0.21 | 0.24 | 0.27 | 0.2 | 0.17 |
| 10 | 76854564 | rs3088142 | -1.396 | -3.228 | 0.591 | -0.934 | 0.013 | 0.065 | 0.065 | 0 | 0.57 | 0.69 | 0.68 | 0.9 | 0.44 | 0.48 |
| 10 | 76855326 | rs3740318 | 0.159 | -3.278 | 0.715 | -0.948 | -0.025 | 0.028 | 0.028 | 0 | 0.32 | 0.45 | 0.45 | 0.64 | 0.28 | 0.32 |
| 10 | 76855412 | rs3740317 | 0.171 | -3.278 | 0.71 | -0.948 | -0.024 | 0.027 | 0.027 | 0 | 0.32 | 0.43 | 0.44 | 0.64 | 0.28 | 0.32 |
| 10 | 76855421 | rs3740316 | 0.291 | -3.256 | 0.658 | -0.948 | -0.001 | -0.001 | -0.001 | 0 | 0.24 | 0.23 | 0.24 | 0.26 | 0.17 | 0.16 |
| 10 | 76858765 | rs10762657 | 1.315 | -3.228 | 0.721 | -0.926 | -0.005 | 0.031 | 0.031 | 0 | 0.22 | 0.25 | 0.39 | 0.58 | 0.23 | 0.25 |
| 10 | 76867899 | rs7912300 | -0.243 | -3.284 | 0.809 | -0.958 | -0.018 | 0.011 | 0.011 | 0.009 | 0.35 | 0.53 | 0.45 | 0.64 | 0.28 | 0.33 |
| 10 | 76868467 | rs10824274 | 1.32 | -3.218 | 0.723 | -0.926 | -0.005 | 0.032 | 0.032 | 0 | 0.22 | 0.25 | 0.39 | 0.58 | 0.23 | 0.25 |
| 10 | 76873427 | rs11001296 | -0.864 | -3.367 | 0.699 | -0.901 | -0.03 | -0.001 | -0.001 | 0 | 0.34 | 0.41 | 0.29 | 0.34 | 0.22 | 0.24 |
| 10 | 76873606 | rs4745765 | -0.609 | -3.397 | 0.758 | -0.915 | -0.016 | -0.025 | -0.025 | 0 | 0.1 | 0.18 | 0.05 | 0.06 | 0.05 | 0.06 |
| 10 | 76897757 | rs10509354 | 1.537 | -3.601 | 0.685 | -1.02 | 0.013 | 0.012 | 0.012 | 1.335 | 0.15 | 0.1 | 0.1 | 0.02 | 0.09 | 0.05 |
| 10 | 76900187 | rs10509355 | -0.13 | -3.603 | 0.686 | -1.019 | -0.01 | 0.003 | 0.003 | 0.737 | 0.24 | 0.27 | 0.25 | 0.28 | 0.17 | 0.18 |
| 10 | 76909562 | rs4746268 | -0.127 | -3.622 | 0.685 | -1.027 | -0.026 | -0.024 | -0.024 | 0.468 | 0.07 | 0.13 | 0.04 | 0.02 | 0.03 | 0.04 |
| 10 | 76917943 | rs7081862 | -0.15 | -3.53 | 0.48 | -0.994 | -0.01 | 0.003 | 0.003 | 0.737 | 0.24 | 0.27 | 0.25 | 0.28 | 0.17 | 0.18 |
| 10 | 76919273 | rs11001309 | 0.106 | -3.355 | 0.417 | -0.918 | -0.008 | 0.053 | 0.053 | 0 | 0.09 | 0.14 | 0.15 | 0.24 | 0.07 | 0.12 |
| 10 | 76921068 | rs2047951 | -1.336 | -3.29 | 0.242 | -1.035 | 0.019 | 0.149 | 0.149 | 0.007 | 0.57 | 0.78 | 0.69 | 0.9 | 0.45 | 0.49 |
| 10 | 76960310 | rs879155 | -1.301 | -3.301 | -0.099 | -0.944 | -0.026 | 0.029 | 0.029 | 0 | 0.61 | 0.78 | 0.64 | 0.73 | 0.47 | 0.72 |
| 10 | 76961690 | rs11001327 | -0.464 | -3.246 | -0.082 | -0.918 | -0.013 | 0.032 | 0.032 | 0 | 0.11 | 0.18 | 0.16 | 0.22 | 0.09 | 0.13 |
| 10 | 76977117 | rs12269517 | -0.089 | -3.196 | -0.06 | -0.912 | -0.009 | 0.051 | 0.051 | 0 | 0.11 | 0.18 | 0.17 | 0.25 | 0.09 | 0.15 |
| 10 | 76983258 | rs12249405 | -1.923 | -3.436 | 0.099 | -1.002 | -0.041 | 0.026 | 0.026 | 0.237 | 0.24 | 0.64 | 0.25 | 0.31 | 0.14 | 0.24 |
| 10 | 76986741 | rs1259498 | -1.278 | -3.374 | -0.009 | -1.235 | 0.005 | 0.093 | 0.093 | 0 | 0.57 | 0.75 | 0.72 | 0.92 | 0.46 | 0.5 |
| 10 | 76989295 | rs11001334 | 1.641 | -3.483 | -0.133 | -1.142 | 0.011 | 0.01 | 0.01 | 0.298 | 0.16 | 0.1 | 0.11 | 0.08 | 0.1 | 0.04 |
| 10 | 76990185 | rs1259501 | -1.193 | -3.253 | -0.253 | -1.073 | 0.007 | 0.094 | 0.094 | 0 | 0.57 | 0.75 | 0.72 | 0.92 | 0.46 | 0.5 |
| 10 | 76996941 | rs1259505 | -1.273 | -3.254 | -0.254 | -1.074 | 0.004 | 0.087 | 0.087 | 0 | 0.58 | 0.76 | 0.72 | 0.92 | 0.46 | 0.51 |
| 10 | 76998092 | rs1259508 | -1.233 | -3.243 | -0.241 | -1.085 | 0.006 | 0.093 | 0.093 | 0 | 0.57 | 0.75 | 0.72 | 0.92 | 0.46 | 0.51 |
| 10 | 77003479 | rs1259523 | -1.184 | -2.728 | -0.789 | -1.29 | -0.014 | 0.009 | 0.009 | 0 | 0.66 | 0.72 | 0.76 | 0.88 | 0.56 | 0.59 |
| 10 | 77004869 | rs1259525 | 2.264 | -2.625 | -0.989 | -0.976 | -0.024 | -0.071 | -0.071 | 0 | 0.17 | 0.08 | 0.37 | 0.49 | 0.24 | 0.23 |
| 10 | 77011063 | rs11001345 | -0.367 | -2.694 | -0.752 | -0.963 | -0.015 | 0.01 | 0.01 | 0 | 0.25 | 0.3 | 0.25 | 0.27 | 0.18 | 0.27 |
| 10 | 77013701 | rs1113313 | -0.134 | -2.592 | -0.661 | -1.148 | 0.007 | 0.028 | 0.028 | 0 | 0.43 | 0.37 | 0.61 | 0.75 | 0.43 | 0.5 |
| 10 | 77031789 | rs11001349 | -0.676 | -2.488 | -0.792 | -1.241 | 0.031 | 0.035 | 0.035 | 0 | 0.37 | 0.64 | 0.26 | 0.19 | 0.28 | 0.44 |
| 10 | 77032196 | rs11001351 | -0.652 | -2.491 | -0.764 | -1.243 | 0.024 | 0.027 | 0.027 | 0 | 0.37 | 0.63 | 0.26 | 0.19 | 0.29 | 0.44 |
| 10 | 77034063 | rs1777414 | -0.922 | -2.459 | -0.874 | -1.262 | 0.018 | 0.025 | 0.025 | 0 | 0.68 | 0.85 | 0.48 | 0.25 | 0.71 | 0.61 |
| 10 | 77035878 | rs12777214 | 0.276 | -2.653 | -0.528 | -1.364 | -0.021 | -0.029 | -0.029 | 0 | 0.2 | 0.25 | 0.12 | 0.11 | 0.16 | 0.39 |
| 10 | 77036594 | rs11599350 | 0.315 | -2.397 | -0.193 | -1.455 | -0.011 | -0.028 | -0.028 | 0.103 | 0.24 | 0.3 | 0.2 | 0.1 | 0.2 | 0.15 |
| 10 | 77040443 | rs9971134 | -0.164 | -2.109 | 0.126 | -1.129 | -0.049 | 0.086 | 0.086 | 2.131 | 0.14 | 0.24 | 0.15 | 0.21 | 0.04 | 0.12 |
| 10 | 77041169 | rs1149711 | 0.498 | -1.924 | -0.014 | -1.077 | 0.016 | -0.005 | -0.005 | 0 | 0.35 | 0.43 | 0.31 | 0.29 | 0.38 | 0.21 |
| 10 | 77046740 | rs1149713 | -0.416 | -0.575 | -0.018 | -0.63 | -0.007 | 0.047 | 0.047 | 1.939 | 0.51 | 0.29 | 0.37 | 0.29 | 0.51 | 0.57 |
| 10 | 77055853 | rs1966054 | 1.042 | -0.584 | 0.336 | -0.647 | -0.021 | -0.121 | -0.121 | 0 | 0.25 | 0.46 | 0.14 | 0.04 | 0.2 | 0.07 |
| 10 | 77064320 | rs1149718 | 1.378 | -0.514 | 0.23 | -0.705 | -0.025 | 0.009 | 0.009 | 0 | 0.13 | 0.06 | 0.13 | 0.08 | 0.21 | 0.05 |
| 10 | 77067318 | rs11001366 | 1.546 | -0.571 | 0.323 | -0.767 | -0.008 | 0.092 | 0.092 | 2.798 | 0.22 | 0.05 | 0.12 | 0.04 | 0.2 | 0.07 |
| 10 | 77070475 | rs7086353 | 1.091 | -0.523 | 0.435 | -0.746 | -0.014 | -0.089 | -0.089 | 0 | 0.25 | 0.37 | 0.13 | 0.04 | 0.2 | 0.07 |
| 10 | 77072928 | rs6480776 | 0.15 | -0.577 | 0.255 | -0.733 | 0.012 | 0.013 | 0.013 | 0.964 | 0.38 | 0.29 | 0.25 | 0.22 | 0.32 | 0.52 |
| 10 | 77073351 | rs1259511 | -1.788 | -0.578 | 0.367 | -0.301 | -0.019 | -0.046 | -0.046 | 0 | 0.82 | 0.89 | 0.54 | 0.54 | 0.76 | 0.8 |
| 10 | 77081848 | rs2488696 | -0.638 | -0.562 | 0.438 | -0.207 | -0.001 | -0.001 | -0.001 | 0 | 0.56 | 0.43 | 0.4 | 0.5 | 0.56 | 0.71 |
| 10 | 77105739 | rs2435882 | -1.604 | 0.035 | 0.524 | 0.187 | -0.054 | -0.105 | -0.105 | 0.005 | 0.83 | 0.95 | 0.61 | 0.51 | 0.75 | 0.79 |
| 10 | 77108687 | rs1259594 | -0.67 | -0.159 | 0.836 | 0.016 | -0.002 | 0.004 | 0.004 | 1.287 | 0.39 | 0.38 | 0.29 | 0.26 | 0.36 | 0.36 |
| 10 | 77109880 | rs6480779 | 1.006 | -0.079 | 0.24 | 0.059 | 0.024 | 0.039 | 0.039 | 1.638 | 0.41 | 0.22 | 0.27 | 0.25 | 0.38 | 0.4 |

| | | | | | | | | | | | | | | | | |
|---|---|---|---|---|---|---|---|---|---|---|---|---|---|---|---|---|
| 10 | 77111879 | rs1259591 | -1.558 | -0.149 | -0.12 | 0.18 | -0.062 | -0.124 | -0.124 | 1.908 | 0.81 | 0.96 | 0.6 | 0.51 | 0.75 | 0.79 |
| 10 | 77115954 | rs6480780 | -0.537 | -0.213 | -0.202 | -0.211 | 0.002 | -0.037 | -0.037 | 0 | 0.36 | 0.57 | 0.27 | 0.16 | 0.35 | 0.29 |
| 10 | 77117556 | rs17690338 | 0.804 | 0.082 | -1.257 | -0.081 | 0 | 0.05 | 0.05 | 4.438 | 0.4 | 0.12 | 0.27 | 0.25 | 0.39 | 0.4 |
| 10 | 77120456 | rs10509356 | -0.345 | 0.209 | -0.714 | -0.217 | 0.001 | -0.007 | -0.007 | 0 | 0.19 | 0.16 | 0.11 | 0.05 | 0.14 | 0.22 |
| 10 | 77121082 | rs1262398 | -0.825 | 0.636 | -0.723 | 0.12 | 0.001 | -0.001 | -0.001 | 0 | 0.32 | 0.34 | 0.31 | 0.27 | 0.35 | 0.4 |
| 10 | 77123651 | rs1652704 | -1.021 | 0.609 | -1.486 | 0.038 | -0.002 | 0.022 | 0.022 | 0 | 0.37 | 0.69 | 0.34 | 0.22 | 0.35 | 0.42 |
| 10 | 77127233 | rs1259589 | 0.343 | 1.538 | -1.846 | 0.496 | -0.049 | -0.036 | -0.036 | 0 | 0.08 | 0.01 | 0.11 | 0.1 | 0.18 | 0.08 |
| 10 | 77128000 | rs1259590 | -0.695 | 1.006 | -2.073 | 0.43 | 0.027 | -0.014 | -0.014 | 0.776 | 0.54 | 0.26 | 0.42 | 0.5 | 0.61 | 0.62 |
| 10 | 77138156 | rs1259601 | -0.104 | 1.156 | -2.12 | 0.678 | -0.046 | -0.063 | -0.063 | 0 | 0.21 | 0.12 | 0.29 | 0.32 | 0.3 | 0.34 |
| 10 | 77138624 | rs2488704 | -0.342 | 0.606 | -1.846 | 0.577 | 0.053 | -0.007 | -0.007 | 0 | 0.54 | 0.25 | 0.54 | 0.52 | 0.68 | 0.65 |
| 10 | 77140864 | rs7078507 | 0.321 | 0.472 | -2.173 | 0.574 | 0.035 | -0.025 | -0.025 | 0 | 0.55 | 0.16 | 0.48 | 0.57 | 0.61 | 0.42 |
| 10 | 77143467 | rs1897191 | 0.057 | 0.188 | -0.739 | 0.42 | -0.008 | 0.005 | 0.005 | 0 | 0.6 | 0.51 | 0.52 | 0.57 | 0.62 | 0.43 |
| 10 | 77157489 | rs10762666 | -0.14 | -0.038 | -0.186 | 0.696 | -0.001 | -0.002 | -0.002 | 5.405 | 0.75 | 0.76 | 0.73 | 0.87 | 0.76 | 0.72 |
| 10 | 77160682 | rs10824306 | 1.709 | 0.316 | -1.111 | 0.024 | -0.058 | -0.17 | -0.17 | 0.03 | 0.21 | 0.01 | 0.23 | 0.55 | 0.19 | 0.2 |
| 10 | 77170685 | rs12782308 | 0.373 | 0.265 | -0.942 | -0.147 | -0.001 | 0 | 0 | 0 | 0.13 | 0.12 | 0.09 | 0.17 | 0.13 | 0.2 |
| 10 | 77186594 | rs12254607 | -0.667 | 0.292 | -0.08 | -0.122 | 0.088 | 0.053 | 0.053 | 1.83 | 0.06 | 0.44 | 0.05 | 0 | 0.01 | 0.01 |
| 10 | 77213302 | rs7896671 | -0.121 | 0.077 | 0.202 | -0.222 | 0 | -0.005 | -0.005 | 0 | 0.85 | 0.69 | 0.75 | 0.78 | 0.85 | 0.67 |
| 10 | 77213547 | rs7912002 | -0.309 | 0.126 | 0.105 | -0.196 | 0.003 | -0.003 | -0.003 | 0 | 0.31 | 0.31 | 0.29 | 0.23 | 0.32 | 0.26 |
| 10 | 77215176 | rs7096727 | 1.233 | 0.177 | 0.127 | -0.191 | 0.017 | 0.009 | 0.009 | 0 | 0.51 | 0.25 | 0.44 | 0.55 | 0.52 | 0.37 |
| 10 | 77216955 | rs7090265 | 1.167 | 0.11 | 0.123 | -0.024 | 0.024 | 0.092 | 0.092 | 0 | 0.53 | 0.17 | 0.52 | 0.62 | 0.56 | 0.41 |
| 10 | 77219944 | rs7903784 | -0.394 | 0.155 | 0.266 | -0.218 | -0.006 | 0.021 | 0.021 | 0 | 0.2 | 0.33 | 0.25 | 0.23 | 0.18 | 0.25 |
| 10 | 77225753 | rs11001401 | 2.226 | 0.11 | -0.14 | 0.136 | -0.046 | 0.01 | 0.01 | 0.003 | 0.22 | 0.01 | 0.15 | 0.16 | 0.19 | 0.08 |
| 10 | 77227118 | rs4746301 | -0.74 | 0.145 | 1.104 | 0.389 | 0 | -0.007 | -0.007 | 0.484 | 0.22 | 0.29 | 0.14 | 0.2 | 0.23 | 0.25 |
| 10 | 77227763 | rs10824308 | -2.094 | 0.392 | 1.113 | 0.383 | 0.011 | -0.017 | -0.017 | 0 | 0.39 | 0.76 | 0.38 | 0.42 | 0.36 | 0.45 |
| 10 | 77231608 | rs7898528 | -0.793 | 0.405 | 1.243 | 0.389 | 0.019 | 0.075 | 0.075 | 0 | 0.2 | 0.44 | 0.25 | 0.21 | 0.23 | 0.21 |
| 10 | 77249629 | rs2915032 | 0.315 | 0.393 | 1.154 | 0.346 | -0.012 | -0.006 | -0.006 | 0 | 0.83 | 0.72 | 0.88 | 0.8 | 0.87 | 0.76 |
| 10 | 77250488 | rs9416062 | 0.315 | 0.393 | 1.154 | 0.346 | -0.012 | -0.006 | -0.006 | 0.171 | 0.83 | 0.72 | 0.88 | 0.8 | 0.87 | 0.76 |
| 10 | 77255355 | rs10762669 | -0.711 | 0.409 | 1.261 | 0.365 | 0.008 | 0.03 | 0.03 | 0 | 0.19 | 0.35 | 0.24 | 0.2 | 0.23 | 0.21 |
| 10 | 77257376 | rs3012042 | 1.008 | 0.261 | 1.338 | 0.472 | -0.028 | -0.018 | -0.018 | 0.282 | 0.81 | 0.62 | 0.87 | 0.8 | 0.87 | 0.76 |
| 10 | 77257791 | rs4617527 | 0.481 | 0.291 | 1.169 | 0.486 | -0.012 | -0.006 | -0.006 | 0 | 0.83 | 0.72 | 0.88 | 0.8 | 0.87 | 0.76 |
| 10 | 77266467 | rs3012044 | 1.47 | 0.291 | 0.985 | 0.384 | -0.004 | -0.03 | -0.03 | 0.086 | 0.58 | 0.18 | 0.6 | 0.56 | 0.57 | 0.55 |
| 10 | 77273076 | rs2915028 | 1.431 | 0.343 | 0.986 | 0.385 | 0 | -0.031 | -0.031 | 0 | 0.58 | 0.19 | 0.6 | 0.56 | 0.6 | 0.55 |
| 10 | 77274480 | rs9416063 | 1.493 | 0.345 | 0.971 | 0.385 | 0.002 | -0.032 | -0.032 | 0 | 0.58 | 0.18 | 0.6 | 0.57 | 0.6 | 0.55 |
| 10 | 77284374 | rs2395289 | 1.407 | 0.337 | 0.986 | 0.444 | 0.004 | -0.017 | -0.017 | 0 | 0.59 | 0.3 | 0.61 | 0.55 | 0.6 | 0.55 |
| 10 | 77293109 | rs3012053 | 0.19 | 0.264 | 0.817 | 0.405 | -0.021 | 0.006 | 0.006 | 0 | 0.76 | 0.79 | 0.75 | 0.85 | 0.69 | 0.77 |
| 10 | 77294236 | rs4746305 | 0.512 | 0.253 | 1.025 | 0.398 | -0.007 | -0.014 | -0.014 | 0 | 0.06 | 0.22 | 0.03 | 0 | 0 | 0 |
| 10 | 77295843 | rs10824316 | 1.682 | 0.249 | 0.442 | 0.393 | 0.003 | -0.032 | -0.032 | 2.503 | 0.55 | 0.15 | 0.57 | 0.57 | 0.58 | 0.52 |
| 10 | 77297579 | rs2017112 | 0.269 | 0.582 | 0.47 | 0.68 | -0.023 | -0.053 | -0.053 | 0 | 0.08 | 0.01 | 0.08 | 0.29 | 0.1 | 0.25 |
| 10 | 77316235 | rs4746306 | -0.021 | 0.411 | 0.592 | 0.653 | -0.002 | -0.001 | -0.001 | 0 | 0.79 | 0.75 | 0.75 | 0.85 | 0.71 | 0.79 |
| 10 | 77357347 | rs3012037 | -0.046 | 0.37 | 0.296 | 0.627 | -0.003 | -0.001 | -0.001 | 0.32 | 0.79 | 0.75 | 0.75 | 0.85 | 0.71 | 0.79 |
| 10 | 77364207 | rs7095903 | -1.003 | 0.118 | 0.422 | 0.601 | -0.047 | -0.063 | -0.063 | 0.078 | 0.11 | 0.53 | 0.08 | 0 | 0.01 | 0 |
| 10 | 77364570 | rs2059998 | -0.875 | 0.17 | 0.258 | 0.624 | -0.029 | -0.045 | -0.045 | 0 | 0.1 | 0.41 | 0.07 | 0 | 0.01 | 0 |
| 10 | 77372699 | rs4746313 | 0.056 | 0.602 | -0.905 | 0.929 | -0.024 | -0.052 | -0.052 | 0.531 | 0.08 | 0 | 0.08 | 0.29 | 0.11 | 0.25 |
| 10 | 77379512 | rs12778686 | 0.776 | 0.511 | -0.94 | 0.628 | -0.012 | -0.065 | -0.065 | 1.363 | 0.58 | 0.14 | 0.57 | 0.53 | 0.59 | 0.54 |
| 10 | 77380447 | rs7895659 | 1.151 | 0.354 | -1.284 | 0.516 | -0.067 | -0.016 | -0.016 | 0 | 0.41 | 0.14 | 0.38 | 0.41 | 0.45 | 0.33 |
| 10 | 77383569 | rs1430331 | 0.044 | 0.023 | -0.039 | 0.69 | -0.012 | -0.01 | -0.01 | 0 | 0.71 | 0.58 | 0.62 | 0.77 | 0.73 | 0.76 |
| 10 | 77390514 | rs2915026 | 0.546 | -0.039 | -0.028 | 0.624 | -0.016 | -0.017 | -0.017 | 0 | 0.64 | 0.45 | 0.59 | 0.77 | 0.72 | 0.76 |
| 10 | 77390738 | rs12242790 | -0.205 | 0.316 | 0.071 | 0.8 | -0.018 | -0.029 | -0.029 | 0 | 0.06 | 0.09 | 0.02 | 0 | 0.01 | 0 |
| 10 | 77395826 | rs4745785 | -0.706 | 0.802 | -0.18 | 1.092 | 0.004 | 0.024 | 0.024 | 0 | 0.14 | 0.17 | 0.15 | 0.28 | 0.18 | 0.16 |
| 10 | 77400983 | rs2059995 | -0.028 | 0.604 | -0.378 | 0.79 | -0.019 | -0.019 | -0.019 | 0 | 0.07 | 0.01 | 0.05 | 0.09 | 0.07 | 0.12 |
| 10 | 77404582 | rs10160193 | -1.265 | 0.236 | 0.591 | 0.579 | 0.008 | -0.016 | -0.016 | 0 | 0.08 | 0.32 | 0.04 | 0 | 0.02 | 0.04 |
| 10 | 77405436 | rs11001451 | 0.22 | 0.475 | 0.571 | 0.461 | 0.009 | -0.012 | -0.012 | 0 | 0.93 | 0.89 | 0.91 | 0.98 | 0.89 | 0.95 |
| 10 | 77406954 | rs10458641 | -1.768 | 0.098 | 1.615 | 1.533 | -0.013 | 0.33 | 0.33 | 3.285 | 0.1 | 0.51 | 0.07 | 0.36 | 0.02 | 0.17 |
| 10 | 77410361 | rs16932464 | -2.076 | 0.024 | 1.845 | 1.503 | -0.045 | 0.162 | 0.162 | 6.264 | 0.11 | 0.67 | 0.09 | 0.37 | 0.03 | 0.17 |
| 10 | 77415616 | rs10824329 | 1.827 | -0.343 | 1.774 | 1.67 | 0.039 | -0.081 | -0.081 | 0 | 0.65 | 0.14 | 0.74 | 0.44 | 0.74 | 0.54 |
| 10 | 77422702 | rs7068634 | -1.402 | -0.293 | 2.043 | 1.671 | 0.046 | 0.439 | 0.439 | 4.178 | 0.06 | 0.45 | 0.05 | 0.37 | 0.02 | 0.17 |
| 10 | 77436668 | rs2454813 | 1.578 | -0.374 | 1.732 | 1.646 | 0.026 | -0.044 | -0.044 | 0 | 0.69 | 0.17 | 0.75 | 0.54 | 0.75 | 0.64 |
| 10 | 77443038 | rs2463955 | 1.217 | -0.376 | 1.871 | 1.645 | -0.007 | 0.017 | 0.017 | 0.502 | 0.72 | 0.29 | 0.78 | 0.56 | 0.76 | 0.65 |
| 10 | 77448435 | rs7899719 | -1.181 | -0.366 | 1.949 | 1.696 | 0.073 | 0.549 | 0.549 | 1.838 | 0.05 | 0.42 | 0.05 | 0.4 | 0.02 | 0.14 |
| 10 | 77453942 | rs11001471 | 0.108 | -0.364 | 1.597 | 1.636 | -0.001 | 0.007 | 0.007 | 0.329 | 0.14 | 0.12 | 0.11 | 0.02 | 0.14 | 0.07 |
| 10 | 77454739 | rs4745788 | -2.406 | -0.381 | 1.492 | 1.7 | 0.036 | -0.074 | -0.074 | 0 | 0.34 | 0.9 | 0.26 | 0.58 | 0.26 | 0.46 |
| 10 | 77456175 | rs1909677 | -2.083 | -0.296 | 0.238 | 0.863 | -0.057 | 0.149 | 0.149 | 0 | 0.67 | 0.97 | 0.75 | 0.86 | 0.69 | 0.7 |
| 10 | 77461012 | rs16932566 | 0.026 | -0.319 | 0.575 | 1.102 | 0 | 0.003 | 0.003 | 0.57 | 0.09 | 0.07 | 0.06 | 0.17 | 0.1 | 0.21 |
| 10 | 77461214 | rs10824335 | -1.087 | -0.338 | 0.725 | 1.08 | -0.023 | -0.028 | -0.028 | 0.426 | 0.29 | 0.38 | 0.2 | 0.18 | 0.24 | 0.32 |
| 10 | 77464011 | rs16932567 | -0.034 | -0.306 | 0.204 | 1.092 | 0 | 0.002 | 0.002 | 0.731 | 0.09 | 0.07 | 0.06 | 0.17 | 0.1 | 0.22 |
| 10 | 77464314 | rs10740451 | -1.502 | -0.226 | -0.065 | 1.078 | 0.034 | -0.086 | -0.086 | 0.246 | 0.34 | 0.91 | 0.26 | 0.58 | 0.26 | 0.45 |
| 10 | 77489936 | rs11595702 | 0.63 | 0.63 | -0.343 | 1.103 | -0.067 | -0.07 | -0.07 | 0.3 | 0.2 | 0.01 | 0.36 | 0.23 | 0.24 | 0.16 |
| 10 | 77529032 | rs1875160 | -2.462 | 0.216 | -0.442 | 0.473 | -0.028 | 0.148 | 0.148 | 0.407 | 0.78 | 0.97 | 0.89 | 1 | 0.82 | 0.95 |
| 10 | 77529218 | rs12242030 | -1.391 | 0.285 | 0.608 | 0.999 | -0.036 | 0.041 | 0.041 | 0 | 0.1 | 0.27 | 0.05 | 0.1 | 0.04 | 0.14 |

| | | | | | | | | | | | | | | | | |
|---|---|---|---|---|---|---|---|---|---|---|---|---|---|---|---|---|
| 10 | 77548286 | rs12411465 | 0.263 | 0.473 | 0.516 | 1.073 | 0 | 0.029 | 0.029 | 0 | 0.18 | 0.12 | 0.24 | 0.07 | 0.18 | 0.14 |
| 10 | 77550183 | rs4746330 | -0.147 | 0.632 | 0.281 | 0.876 | -0.102 | 0.001 | 0.001 | 3.97 | 0.34 | 0.11 | 0.4 | 0.3 | 0.49 | 0.52 |
| 10 | 77554279 | rs10160008 | -0.126 | 0.676 | 0.448 | 0.897 | -0.056 | 0.021 | 0.021 | 2.443 | 0.36 | 0.19 | 0.42 | 0.3 | 0.49 | 0.52 |
| 10 | 77559168 | rs11001500 | -0.165 | 0.676 | 0.232 | 0.897 | -0.161 | -0.019 | -0.019 | 0.856 | 0.35 | 0.07 | 0.41 | 0.3 | 0.49 | 0.52 |
| 10 | 77560079 | rs4746333 | -0.174 | 0.679 | 0.233 | 0.898 | -0.161 | -0.019 | -0.019 | 3.643 | 0.35 | 0.07 | 0.41 | 0.3 | 0.49 | 0.52 |
| 10 | 77561513 | rs1124372 | -2.325 | 0.614 | 0.425 | 0.834 | -0.012 | 0.169 | 0.169 | 5.821 | 0.65 | 0.87 | 0.78 | 0.99 | 0.7 | 0.89 |
| 10 | 77567728 | rs7072973 | 0.003 | 0.776 | 0.295 | 0.94 | -0.15 | -0.017 | -0.017 | 0 | 0.35 | 0.07 | 0.41 | 0.3 | 0.48 | 0.54 |
| 10 | 77584736 | rs1909694 | 0.688 | 0.99 | 0.149 | 0.893 | -0.06 | -0.009 | -0.009 | 0 | 0.09 | 0.01 | 0.1 | 0.04 | 0.18 | 0.09 |
| 10 | 77586857 | rs2204752 | 0.559 | 0.993 | 0.194 | 1.152 | -0.036 | -0.036 | -0.036 | 0 | 0.1 | 0.07 | 0.11 | 0.19 | 0.18 | 0.18 |
| 10 | 77597792 | rs11595613 | 0.359 | -0.238 | 0.454 | 1.207 | -0.147 | -0.018 | -0.018 | 0 | 0.35 | 0.07 | 0.4 | 0.3 | 0.46 | 0.5 |
| 10 | 77605724 | rs12261071 | -1.074 | -0.3 | 0.742 | 1.208 | -0.007 | -0.012 | -0.012 | 0 | 0.23 | 0.28 | 0.26 | 0.17 | 0.22 | 0.27 |
| 10 | 77606155 | rs17434565 | 0.02 | -0.2 | 0.628 | 1.266 | -0.047 | -0.036 | -0.036 | 4.298 | 0.17 | 0.02 | 0.23 | 0.17 | 0.21 | 0.26 |
| 10 | 77614004 | rs1885822 | -0.992 | -0.274 | 0.766 | 1.231 | 0.008 | 0.007 | 0.007 | 3.401 | 0.56 | 0.41 | 0.67 | 0.38 | 0.65 | 0.68 |
| 10 | 77626164 | rs1885823 | -0.524 | 0.041 | 1.08 | 1.17 | -0.011 | 0.042 | 0.042 | 1.713 | 0.43 | 0.63 | 0.52 | 0.81 | 0.48 | 0.62 |
| 10 | 77631656 | rs7082211 | -1.262 | -0.157 | 1.249 | 1.326 | -0.011 | 0.054 | 0.054 | 0 | 0.24 | 0.48 | 0.28 | 0.27 | 0.22 | 0.33 |
| 10 | 77634338 | rs2181443 | 1.167 | -0.066 | 1.241 | 1.34 | -0.001 | 0.011 | 0.011 | 0.17 | 0.73 | 0.38 | 0.64 | 0.38 | 0.74 | 0.63 |
| 10 | 77636050 | rs11001530 | -0.764 | -0.047 | 0.993 | 1.347 | 0.012 | 0.31 | 0.31 | 0 | 0.06 | 0.16 | 0.08 | 0.45 | 0.06 | 0.14 |
| 10 | 77641999 | rs9415125 | 0.709 | -0.041 | 0.994 | 1.343 | 0.027 | 0.285 | 0.285 | 0 | 0.94 | 0.84 | 0.91 | 0.55 | 0.92 | 0.86 |
| 10 | 77646553 | rs9415126 | 1.478 | -0.127 | 0.94 | 1.162 | 0.023 | 0.058 | 0.058 | 6.451 | 0.36 | 0.21 | 0.21 | 0.02 | 0.25 | 0.13 |
| 10 | 77649224 | rs877531 | -0.547 | -0.226 | 0.109 | 1.217 | -0.003 | 0.008 | 0.008 | 0 | 0.21 | 0.51 | 0.27 | 0.17 | 0.18 | 0.25 |
| 10 | 77656021 | rs10762685 | 0.075 | -0.192 | -1.371 | 0.927 | -0.117 | -0.032 | -0.032 | 0.388 | 0.37 | 0.09 | 0.42 | 0.38 | 0.49 | 0.5 |
| 10 | 77656780 | rs12414570 | 0.029 | 0.003 | -0.657 | 0.425 | -0.01 | 0.042 | 0.042 | 0 | 0.15 | 0.05 | 0.21 | 0.02 | 0.15 | 0.09 |
| 10 | 77660765 | rs2894317 | -0.777 | -0.273 | -0.286 | 0.658 | 0.016 | 0.018 | 0.018 | 6.297 | 0.58 | 0.39 | 0.67 | 0.64 | 0.7 | 0.83 |
| 10 | 77661341 | rs7075958 | 0.636 | -0.96 | 0.016 | 0.422 | -0.015 | -0.016 | -0.016 | 0 | 0.63 | 0.45 | 0.54 | 0.25 | 0.64 | 0.55 |
| 10 | 77662192 | rs11596659 | -0.319 | -1.339 | 0.067 | 0.641 | 0.016 | 0.014 | 0.014 | 0.228 | 0.33 | 0.25 | 0.39 | 0.29 | 0.31 | 0.62 |
| 10 | 77663173 | rs7085352 | -1.107 | -1.476 | -0.446 | 0.285 | -0.125 | 0.143 | 0.143 | 0.068 | 0.68 | 0.95 | 0.72 | 0.91 | 0.5 | 0.76 |
| 10 | 77666528 | rs7082215 | -0.629 | -1.706 | -0.171 | 0.403 | -0.004 | 0.046 | 0.046 | 0.354 | 0.12 | 0.13 | 0.2 | 0.62 | 0.12 | 0.26 |
| 10 | 77670260 | rs7900588 | -0.649 | -1.728 | 0.363 | 0.484 | -0.005 | 0.016 | 0.016 | 0 | 0.6 | 0.69 | 0.52 | 0.3 | 0.45 | 0.57 |
| 10 | 77671322 | rs17435206 | 1.003 | -1.56 | 0.17 | 0.325 | 0.014 | 0.007 | 0.007 | 7.353 | 0.25 | 0.14 | 0.13 | 0 | 0.14 | 0.02 |
| 10 | 77672153 | rs11001545 | -0.308 | -1.264 | 0.074 | 0.662 | -0.005 | 0.053 | 0.053 | 0.637 | 0.09 | 0.14 | 0.08 | 0.24 | 0.1 | 0.2 |
| 10 | 77673629 | rs11816174 | 0.132 | -1.43 | 0.132 | 0.49 | -0.023 | -0.002 | -0.002 | 0 | 0.31 | 0.46 | 0.4 | 0.32 | 0.25 | 0.53 |
| 10 | 77673825 | rs11001547 | -0.48 | -0.222 | -0.031 | 1.014 | -0.01 | -0.054 | -0.054 | 1.282 | 0.09 | 0.18 | 0.05 | 0.01 | 0.08 | 0.12 |
| 10 | 77674635 | rs2057435 | 0.145 | 0.267 | -0.515 | 1.423 | -0.037 | -0.038 | -0.038 | 1.253 | 0.66 | 0.74 | 0.61 | 0.54 | 0.5 | 0.73 |
| 10 | 77677487 | rs2057436 | -0.084 | 0.59 | -1.05 | 0.628 | -0.156 | 0.082 | 0.082 | 0 | 0.71 | 0.95 | 0.73 | 0.88 | 0.52 | 0.79 |
| 10 | 77683306 | rs7903557 | -0.262 | -0.539 | -0.37 | 0.516 | -0.005 | -0.003 | -0.003 | 0 | 0.4 | 0.57 | 0.53 | 0.7 | 0.23 | 0.5 |
| 10 | 77683544 | rs7920276 | -0.972 | 0.26 | -0.194 | -0.416 | 0.01 | -0.101 | -0.101 | 0.003 | 0.28 | 0.36 | 0.23 | 0.04 | 0.35 | 0.31 |
| 10 | 77683563 | rs12572279 | -0.473 | 0.274 | -0.842 | -0.374 | 0.004 | 0.043 | 0.043 | 0 | 0.33 | 0.17 | 0.44 | 0.89 | 0.27 | 0.59 |
| 10 | 77687528 | rs2208948 | 0.802 | 0.602 | -0.361 | 0.365 | 0.002 | 0.04 | 0.04 | 0 | 0.71 | 0.65 | 0.76 | 0.51 | 0.65 | 0.77 |
| 10 | 77698806 | rs4746341 | 0.912 | 0.555 | -0.2 | 0.561 | -0.003 | 0.043 | 0.043 | 0 | 0.7 | 0.62 | 0.72 | 0.47 | 0.66 | 0.78 |
| 10 | 77700235 | rs1323072 | -0.287 | 0.388 | -0.448 | 0.067 | -0.002 | -0.003 | -0.003 | 0 | 0.22 | 0.19 | 0.4 | 0.29 | 0.22 | 0.49 |
| 10 | 77706407 | rs17363658 | -0.14 | 0.359 | -0.688 | 0.408 | -0.017 | -0.028 | -0.028 | 0.11 | 0.12 | 0.01 | 0.06 | 0.13 | 0.13 | 0.1 |
| 10 | 77710913 | rs16932737 | -0.226 | -0.527 | -0.541 | 0.043 | -0.051 | 0.184 | 0.184 | 2.464 | 0.11 | 0.47 | 0.09 | 0.22 | 0.02 | 0.07 |
| 10 | 77719862 | rs9416104 | 0.428 | -0.468 | -0.862 | -0.078 | -0.049 | -0.036 | -0.036 | 0.001 | 0.14 | 0.01 | 0.3 | 0.16 | 0.15 | 0.31 |
| 10 | 77738382 | rs9663814 | 1.597 | -0.697 | -0.5 | -0.237 | -0.001 | 0 | 0 | 0 | 0.58 | 0.39 | 0.49 | 0.51 | 0.6 | 0.41 |
| 10 | 77742856 | rs7077719 | -1.688 | -0.782 | -0.388 | -0.312 | -0.016 | 0.018 | 0.018 | 0 | 0.34 | 0.6 | 0.46 | 0.5 | 0.24 | 0.56 |
| 10 | 77743090 | rs7096129 | -1.121 | -0.769 | -0.409 | -0.306 | -0.001 | -0.005 | -0.005 | 0 | 0.26 | 0.19 | 0.4 | 0.34 | 0.23 | 0.5 |
| 10 | 77750742 | rs7097530 | -2.028 | -0.877 | -0.529 | -0.319 | -0.005 | 0.003 | 0.003 | 0 | 0.38 | 0.63 | 0.48 | 0.52 | 0.27 | 0.56 |
| 10 | 77754199 | rs1323083 | -0.007 | -1.016 | -0.83 | -0.58 | 0.003 | 0.048 | 0.048 | 0 | 0.71 | 0.65 | 0.73 | 0.52 | 0.68 | 0.74 |
| 10 | 77754511 | rs2057438 | -0.442 | -1.114 | -0.572 | -0.315 | -0.023 | -0.034 | -0.034 | 3.993 | 0.22 | 0.08 | 0.36 | 0.29 | 0.21 | 0.49 |
| 10 | 77758013 | rs2057440 | -1.697 | -0.927 | -0.441 | -0.266 | -0.007 | 0.006 | 0.006 | 0 | 0.37 | 0.63 | 0.47 | 0.52 | 0.27 | 0.57 |
| 10 | 77760583 | rs4746343 | 0.209 | 0.151 | -0.603 | -0.061 | 0.002 | -0.007 | -0.007 | 0.589 | 0.09 | 0.07 | 0.09 | 0.15 | 0.06 | 0.21 |
| 10 | 77765857 | rs10824350 | -0.371 | 0.078 | -0.48 | -0.037 | -0.011 | 0.002 | 0.002 | 0 | 0.11 | 0.21 | 0.13 | 0.15 | 0.07 | 0.21 |
| 10 | 77767237 | rs11001574 | -0.279 | 0.345 | -0.486 | 0.116 | -0.01 | -0.047 | -0.047 | 6.018 | 0.21 | 0.09 | 0.32 | 0.35 | 0.21 | 0.33 |
| 10 | 77767412 | rs4745804 | 0.539 | 0.52 | 0.683 | -0.201 | 0 | 0.011 | 0.011 | 0 | 0.48 | 0.5 | 0.39 | 0.3 | 0.47 | 0.44 |
| 10 | 77768678 | rs4746346 | 1.321 | 0.422 | 0.443 | -0.266 | -0.026 | 0.151 | 0.151 | 0 | 0.4 | 0.13 | 0.27 | 0.02 | 0.42 | 0.22 |
| 10 | 77771345 | rs4746348 | 0.482 | 0.535 | 0.521 | -0.153 | 0 | 0.004 | 0.004 | 0 | 0.48 | 0.42 | 0.4 | 0.37 | 0.47 | 0.45 |
| 10 | 77782036 | rs11001589 | -1.529 | 0.549 | 0.683 | -0.608 | -0.014 | 0.083 | 0.083 | 0 | 0.08 | 0.23 | 0.11 | 0.23 | 0.05 | 0.21 |
| 10 | 77795995 | rs4745806 | 0.454 | 0.666 | 0.849 | -0.747 | 0 | 0.044 | 0.044 | 0 | 0.51 | 0.56 | 0.39 | 0.26 | 0.48 | 0.47 |
| 10 | 77800223 | rs12765860 | 1.293 | 0.768 | 0.412 | -0.898 | -0.048 | 0.185 | 0.185 | 0 | 0.41 | 0.14 | 0.27 | 0.01 | 0.43 | 0.25 |
| 10 | 77807027 | rs1898071 | 0.922 | 0.755 | 0.507 | -0.887 | 0.001 | -0.129 | -0.129 | 0 | 0.44 | 0.43 | 0.31 | 0.01 | 0.44 | 0.26 |
| 10 | 77818594 | rs7924288 | 1.077 | 0.798 | 0.562 | -0.868 | -0.007 | -0.005 | -0.005 | 0 | 0.43 | 0.32 | 0.28 | 0.01 | 0.44 | 0.25 |
| 10 | 77850698 | rs4746357 | 1.006 | 0.923 | 0.075 | -0.809 | -0.001 | -0.047 | -0.047 | 0 | 0.43 | 0.42 | 0.35 | 0.09 | 0.42 | 0.24 |
| 10 | 77859740 | rs10824363 | 0.968 | 0.791 | 0.095 | -0.781 | 0.001 | -0.044 | -0.044 | 7.8 | 0.44 | 0.41 | 0.36 | 0.11 | 0.45 | 0.25 |
| 10 | 77863055 | rs10824365 | 0.925 | 0.794 | 0.092 | -0.774 | 0.001 | -0.048 | -0.048 | 0 | 0.44 | 0.43 | 0.36 | 0.11 | 0.45 | 0.25 |
| 10 | 77866150 | rs7919901 | 0.912 | 1.54 | -0.416 | -0.733 | 0.012 | 0.177 | 0.177 | 0 | 0.47 | 0.22 | 0.47 | 0.15 | 0.45 | 0.25 |
| 10 | 77866732 | rs7920628 | -1.041 | 1.319 | -0.315 | -0.563 | -0.006 | 0.094 | 0.094 | 0.018 | 0.48 | 0.72 | 0.54 | 0.78 | 0.49 | 0.56 |
| 10 | 77869761 | rs7910498 | -0.443 | 1.065 | -0.223 | -0.363 | 0.01 | 0.011 | 0.011 | 0.209 | 0.34 | 0.75 | 0.27 | 0.69 | 0.3 | 0.44 |
| 10 | 77870876 | rs7901856 | 0.057 | 0.819 | -0.092 | -0.354 | -0.014 | -0.001 | -0.001 | 0 | 0.31 | 0.65 | 0.22 | 0.65 | 0.24 | 0.4 |
| 10 | 77875070 | rs2164369 | -1.371 | 0.637 | -0.327 | -0.458 | 0.022 | 0.117 | 0.117 | 0 | 0.54 | 0.77 | 0.52 | 0.75 | 0.42 | 0.55 |

| | | | | | | | | | | | | | | | | |
|---|---|---|---|---|---|---|---|---|---|---|---|---|---|---|---|---|
| 10 | 77879106 | rs11001617 | 0.009 | 0.633 | -0.2 | -0.506 | -0.001 | -0.008 | -0.008 | 0 | 0.22 | 0.19 | 0.27 | 0.1 | 0.18 | 0.14 |
| 10 | 77885337 | rs7909516 | 0.329 | 0.635 | -0.167 | -0.466 | -0.025 | 0.008 | 0.008 | 0 | 0.3 | 0.54 | 0.26 | 0.65 | 0.24 | 0.4 |
| 10 | 77887377 | rs11001622 | 0.145 | 0.597 | -0.345 | -0.486 | 0.017 | 0.061 | 0.061 | 0.364 | 0.22 | 0.07 | 0.24 | 0.09 | 0.18 | 0.14 |
| 10 | 77887494 | rs717387 | -1.385 | 0.654 | -0.362 | -0.484 | 0.017 | 0.128 | 0.128 | 0.038 | 0.54 | 0.82 | 0.52 | 0.74 | 0.42 | 0.54 |
| 10 | 77891868 | rs4746364 | -1.698 | 0.906 | -0.461 | -0.468 | -0.019 | 0.165 | 0.165 | 0 | 0.58 | 0.89 | 0.63 | 0.87 | 0.6 | 0.74 |
| 10 | 77893271 | rs5023489 | -0.318 | 0.777 | -0.491 | -0.391 | -0.013 | -0.038 | -0.038 | 1.039 | 0.24 | 0.25 | 0.26 | 0.09 | 0.18 | 0.14 |
| 10 | 77901480 | rs7077442 | -1.42 | 1.218 | -0.866 | -0.301 | 0.002 | 0.104 | 0.104 | 0.247 | 0.57 | 0.8 | 0.62 | 0.87 | 0.6 | 0.74 |
| 10 | 77910502 | rs11001638 | 0.724 | 1.153 | -0.904 | -0.039 | 0.001 | 0.004 | 0.004 | 0.203 | 0.19 | 0.08 | 0.25 | 0.15 | 0.18 | 0.15 |
| 10 | 77911186 | rs12412341 | 0.216 | 1.184 | 0.001 | 1.015 | -0.003 | 0.053 | 0.053 | 0 | 0.12 | 0.19 | 0.16 | 0.24 | 0.1 | 0.09 |
| 10 | 77912416 | rs7895997 | -0.736 | 1.205 | -0.08 | 1.007 | 0.029 | 0.086 | 0.086 | 0 | 0.26 | 0.51 | 0.22 | 0.49 | 0.31 | 0.51 |
| 10 | 77915310 | rs1898102 | 0.01 | 1.434 | -0.095 | 1.113 | -0.005 | 0.059 | 0.059 | 0 | 0.13 | 0.22 | 0.16 | 0.24 | 0.11 | 0.09 |
| 10 | 77916490 | rs12764787 | -0.701 | 1.085 | -0.048 | 1.123 | -0.004 | -0.006 | -0.006 | 0 | 0.07 | 0.06 | 0.11 | 0.13 | 0.18 | 0.21 |
| 10 | 77916959 | rs5007655 | -0.891 | -0.381 | 0.182 | 1.014 | -0.037 | 0.082 | 0.082 | 0 | 0.2 | 0.44 | 0.11 | 0.37 | 0.12 | 0.29 |
| 10 | 77922119 | rs11001649 | -1.017 | -0.672 | -0.048 | 0.548 | -0.025 | -0.001 | -0.001 | 0 | 0.24 | 0.31 | 0.19 | 0.18 | 0.16 | 0.21 |
| 10 | 77923161 | rs11001650 | -0.066 | -0.11 | -0.111 | 0.729 | -0.004 | -0.001 | -0.001 | 1.067 | 0.17 | 0.14 | 0.25 | 0.51 | 0.2 | 0.33 |
| 10 | 77936576 | rs10458660 | -0.123 | -0.147 | -0.037 | 0.712 | -0.005 | -0.008 | -0.008 | 0 | 0.17 | 0.15 | 0.25 | 0.51 | 0.19 | 0.32 |
| 10 | 77943258 | rs1865640 | -1.136 | -0.519 | -0.06 | -0.037 | -0.027 | 0.105 | 0.105 | 0 | 0.58 | 0.8 | 0.6 | 0.98 | 0.51 | 0.75 |
| 10 | 77946340 | rs11816117 | -0.806 | -0.525 | 0.574 | 0.689 | 0.007 | -0.029 | -0.029 | 0.157 | 0.09 | 0.34 | 0.09 | 0.04 | 0.07 | 0.06 |
| 10 | 77949346 | rs1898089 | -1.141 | -0.55 | 0.82 | 0.379 | -0.002 | 0.004 | 0.004 | 4.387 | 0.11 | 0.55 | 0.1 | 0.08 | 0.07 | 0.06 |
| 10 | 77951056 | rs16932997 | -0.023 | -0.388 | 0.194 | 0.619 | -0.06 | -0.11 | -0.11 | 0 | 0.14 | 0.01 | 0.23 | 0.52 | 0.19 | 0.27 |
| 10 | 77954104 | rs1946709 | -0.274 | -0.103 | 0.096 | 0.664 | 0.024 | 0.026 | 0.026 | 0.098 | 0.06 | 0.32 | 0.04 | 0.05 | 0.02 | 0.09 |
| 10 | 77971125 | rs10824385 | -0.992 | -0.032 | -0.455 | 0.705 | 0.003 | 0.126 | 0.126 | 0.208 | 0.22 | 0.45 | 0.28 | 0.61 | 0.2 | 0.34 |
| 10 | 77971599 | rs7094302 | -0.048 | 0.431 | -2.286 | 0.369 | -0.006 | 0.129 | 0.129 | 0.013 | 0.21 | 0.02 | 0.09 | 0.01 | 0.17 | 0.05 |
| 10 | 77983649 | rs16933074 | -0.667 | 0.933 | -1.392 | 1.306 | -0.057 | -0.112 | -0.112 | 0 | 0.15 | 0.01 | 0.23 | 0.49 | 0.18 | 0.26 |
| 10 | 77998443 | rs1247499 | -0.308 | 0.845 | -1.044 | 0.56 | 0 | -0.003 | -0.003 | 1.332 | 0.46 | 0.51 | 0.47 | 0.76 | 0.5 | 0.63 |
| 10 | 78001472 | rs1247494 | -0.229 | 0.81 | -2.059 | 0.723 | -0.068 | -0.06 | -0.06 | 0 | 0.42 | 0.12 | 0.51 | 0.48 | 0.48 | 0.5 |
| 10 | 78005230 | rs1247488 | -0.404 | -0.24 | -1.233 | 0.275 | -0.003 | 0.022 | 0.022 | 0 | 0.67 | 0.63 | 0.64 | 0.58 | 0.62 | 0.61 |
| 10 | 78016427 | rs1247479 | -0.183 | -0.647 | -0.532 | 0.056 | 0.005 | -0.006 | -0.006 | 0 | 0.57 | 0.66 | 0.6 | 0.51 | 0.57 | 0.61 |
| 10 | 78024414 | rs1312911 | 0.042 | -1.119 | -0.237 | 0.064 | 0.011 | 0.021 | 0.021 | 0.059 | 0.63 | 0.35 | 0.6 | 0.75 | 0.66 | 0.76 |
| 10 | 78024856 | rs1247472 | -0.207 | -1.185 | -0.301 | 0.004 | -0.007 | -0.027 | -0.027 | 9.402 | 0.65 | 0.65 | 0.62 | 0.84 | 0.66 | 0.76 |
| 10 | 78028463 | rs10762705 | 0.03 | -0.897 | -0.146 | 0.179 | -0.011 | -0.018 | -0.018 | 0 | 0.57 | 0.79 | 0.56 | 0.56 | 0.49 | 0.58 |
| 10 | 78032988 | rs11819487 | 0.523 | -0.972 | 0.259 | 0.185 | 0.007 | 0.009 | 0.009 | 0 | 0.5 | 0.26 | 0.48 | 0.47 | 0.48 | 0.57 |
| 10 | 78034174 | rs11001715 | 0.598 | -0.895 | 0.07 | 0.232 | -0.043 | -0.07 | -0.07 | 0.128 | 0.1 | 0 | 0.05 | 0.27 | 0.15 | 0.18 |
| 10 | 78046050 | rs1865644 | 0.184 | -0.825 | 0.148 | 0.294 | 0 | -0.002 | -0.002 | 0 | 0.55 | 0.64 | 0.54 | 0.57 | 0.49 | 0.58 |
| 10 | 78049994 | rs6480804 | -0.853 | -0.855 | 0.207 | 0.178 | -0.009 | 0.027 | 0.027 | 2.266 | 0.73 | 0.85 | 0.62 | 0.88 | 0.63 | 0.81 |
| 10 | 78053841 | rs2437424 | 0.561 | -0.856 | 0.026 | -0.089 | -0.016 | 0.056 | 0.056 | 3.46 | 0.27 | 0.13 | 0.35 | 0.04 | 0.35 | 0.2 |
| 10 | 78054402 | rs1992012 | 1.559 | -0.273 | -0.613 | -0.331 | -0.079 | 0.06 | 0.06 | 0.068 | 0.16 | 0.02 | 0.14 | 0.05 | 0.24 | 0.1 |
| 10 | 78055724 | rs2579790 | 0.445 | -1.336 | -0.141 | -0.121 | 0.003 | 0.002 | 0.002 | 0 | 0.44 | 0.37 | 0.51 | 0.11 | 0.59 | 0.29 |
| 10 | 78055891 | rs2573559 | -0.308 | -1.793 | -0.352 | -0.099 | -0.04 | 0.041 | 0.041 | 0 | 0.65 | 0.47 | 0.64 | 0.48 | 0.78 | 0.56 |
| 10 | 78056079 | rs10824389 | 1.539 | -1.928 | -0.552 | -0.087 | -0.041 | -0.056 | -0.056 | 0 | 0.14 | 0.27 | 0.04 | 0.01 | 0.04 | 0.07 |
| 10 | 78057746 | rs11001722 | -0.041 | -1.45 | -1.158 | 0.15 | 0 | -0.06 | -0.06 | 0 | 0.2 | 0.1 | 0.12 | 0.37 | 0.19 | 0.21 |
| 10 | 78058287 | rs1885636 | -0.746 | -1.184 | -0.78 | 0.509 | -0.013 | -0.068 | -0.068 | 0.002 | 0.85 | 0.95 | 0.72 | 0.59 | 0.84 | 0.65 |
| 10 | 78060047 | rs10762709 | 0.465 | -2.029 | -0.065 | 0.398 | -0.08 | -0.037 | -0.037 | 0 | 0.2 | 0.39 | 0.1 | 0.13 | 0.07 | 0.14 |
| 10 | 78063891 | rs2579787 | 0.444 | -2.826 | 0.038 | 0.672 | -0.005 | -0.007 | -0.007 | 0 | 0.66 | 0.33 | 0.64 | 0.44 | 0.88 | 0.49 |
| 10 | 78065708 | rs2758982 | 0.872 | -3.115 | -0.252 | 0.483 | 0.13 | 0.145 | 0.145 | 0.019 | 0.53 | 0.24 | 0.59 | 0.13 | 0.78 | 0.35 |
| 10 | 78066337 | rs2579785 | 0.551 | -3.122 | -0.07 | 0.454 | 0.006 | 0.001 | 0.001 | 0 | 0.57 | 0.38 | 0.6 | 0.14 | 0.78 | 0.34 |
| 10 | 78070752 | rs2579783 | 0.806 | -3.203 | -0.022 | 0.439 | 0.109 | 0.12 | 0.12 | 0 | 0.54 | 0.25 | 0.59 | 0.13 | 0.79 | 0.35 |
| 10 | 78072371 | rs10824393 | 1.179 | -3.114 | -0.118 | 0.562 | -0.001 | -0.052 | -0.052 | 0 | 0.12 | 0.04 | 0.04 | 0.29 | 0.11 | 0.14 |
| 10 | 78078159 | rs967948 | -0.686 | -3.103 | 0.004 | 0.626 | -0.002 | 0.037 | 0.037 | 0 | 0.89 | 0.9 | 0.76 | 0.56 | 0.96 | 0.69 |
| 10 | 78079619 | rs2758964 | 0.5 | -3.1 | -0.004 | 0.609 | -0.034 | 0.083 | 0.083 | 0 | 0.86 | 0.79 | 0.74 | 0.58 | 0.96 | 0.68 |
| 10 | 78084407 | rs1040936 | 1.215 | -3.168 | 0.15 | 0.371 | -0.089 | -0.039 | -0.039 | 0.013 | 0.21 | 0.38 | 0.1 | 0.14 | 0.07 | 0.2 |
| 10 | 78092904 | rs2579776 | 0.667 | -3.171 | 0.112 | 0.42 | 0.007 | 0 | 0 | 0 | 0.74 | 0.76 | 0.7 | 0.26 | 0.85 | 0.55 |
| 10 | 78096526 | rs10824403 | 1.039 | -3.122 | 0.091 | 0.467 | 0.002 | -0.018 | -0.018 | 0 | 0.13 | 0.09 | 0.05 | 0.29 | 0.11 | 0.14 |
| 10 | 78099498 | rs4604815 | -0.227 | -3.212 | 0.147 | 0.43 | -0.07 | 0.037 | 0.037 | 0.459 | 0.33 | 0.49 | 0.14 | 0.41 | 0.17 | 0.34 |
| 10 | 78106937 | rs17450685 | 1.795 | -2.613 | -0.408 | -0.111 | -0.072 | 0.056 | 0.056 | 0 | 0.2 | 0.09 | 0.19 | 0.06 | 0.38 | 0.11 |
| 10 | 78107150 | rs11001742 | 1.385 | -2.625 | -0.361 | -0.078 | 0.002 | -0.016 | -0.016 | 0.105 | 0.13 | 0.1 | 0.04 | 0.29 | 0.11 | 0.15 |
| 10 | 78108946 | rs12416652 | 0.771 | -2.706 | -0.49 | -0.175 | 0.188 | 0.27 | 0.27 | 0 | 0.51 | 0.15 | 0.59 | 0.1 | 0.76 | 0.25 |
| 10 | 78109971 | rs7900522 | -0.381 | -2.717 | -0.413 | -0.152 | -0.025 | 0.074 | 0.074 | 0 | 0.87 | 0.83 | 0.75 | 0.54 | 0.95 | 0.68 |
| 10 | 78115104 | rs10824409 | 1.794 | -2.67 | -0.66 | -0.364 | -0.105 | 0.162 | 0.162 | 0.186 | 0.3 | 0.02 | 0.39 | 0.04 | 0.38 | 0.14 |
| 10 | 78116099 | rs9299535 | -0.129 | -2.592 | -0.534 | -0.189 | 0.021 | 0.001 | 0.001 | 0.678 | 0.37 | 0.67 | 0.15 | 0.37 | 0.18 | 0.29 |
| 10 | 78117093 | rs10509371 | 1.659 | -2.569 | -0.584 | -0.271 | -0.06 | -0.046 | -0.046 | 0.199 | 0.2 | 0.29 | 0.07 | 0.1 | 0.06 | 0.15 |
| 10 | 78122729 | rs12766217 | 0.639 | -2.299 | -1.527 | -0.185 | 0.192 | 0.239 | 0.239 | 0 | 0.51 | 0.15 | 0.6 | 0.14 | 0.76 | 0.38 |
| 10 | 78133074 | rs7083461 | 1.361 | -1.914 | -1.356 | 0.024 | -0.082 | 0.047 | 0.047 | 0 | 0.22 | 0.08 | 0.2 | 0.09 | 0.38 | 0.24 |
| 10 | 78140834 | rs7904646 | -0.414 | -2.784 | -0.717 | 0.781 | 0.023 | -0.003 | -0.003 | 0 | 0.9 | 0.95 | 0.76 | 0.51 | 0.96 | 0.69 |
| 10 | 78143825 | rs6480813 | -0.894 | -2.863 | -0.808 | 0.677 | 0.091 | -0.015 | -0.015 | 0 | 0.39 | 0.77 | 0.18 | 0.41 | 0.18 | 0.44 |
| 10 | 78144393 | rs10824412 | -0.448 | -2.91 | -0.741 | 0.667 | 0.051 | -0.014 | -0.014 | 0.383 | 0.36 | 0.7 | 0.17 | 0.41 | 0.15 | 0.41 |
| 10 | 78145677 | rs12260618 | 0.57 | -2.654 | -0.846 | 0.817 | 0.002 | -0.045 | -0.045 | 0 | 0.13 | 0.05 | 0.04 | 0.28 | 0.1 | 0.15 |
| 10 | 78149043 | rs7909650 | -0.665 | -2.704 | -0.857 | 0.896 | 0 | 0.126 | 0.126 | 0 | 0.94 | 0.93 | 0.77 | 0.5 | 0.97 | 0.69 |
| 10 | 78151687 | rs11001766 | 0.688 | -2.695 | -0.882 | 0.792 | -0.001 | -0.053 | -0.053 | 0 | 0.12 | 0.03 | 0.04 | 0.28 | 0.09 | 0.14 |

| | | | | | | | | | | | | | | | | |
|---|---|---|---|---|---|---|---|---|---|---|---|---|---|---|---|---|
| 10 | 78155142 | rs749128 | 0.524 | -2.868 | -0.648 | 0.002 | -0.085 | -0.061 | -0.061 | 0 | 0.21 | 0.59 | 0.11 | 0.07 | 0.04 | 0.14 |
| 10 | 78155287 | rs749127 | 0.041 | -2.866 | -0.686 | -0.015 | -0.049 | -0.048 | -0.048 | 0 | 0.24 | 0.65 | 0.12 | 0.07 | 0.08 | 0.17 |
| 10 | 78163768 | rs1907342 | 0.143 | -2.554 | -0.901 | 0.073 | 0.099 | 0.1 | 0.1 | 0.027 | 0.59 | 0.26 | 0.62 | 0.14 | 0.83 | 0.41 |
| 10 | 78167388 | rs1907343 | 0.06 | -2.817 | -0.309 | 0.783 | -0.08 | 0.058 | 0.058 | 0 | 0.72 | 0.64 | 0.79 | 0.45 | 0.91 | 0.43 |
| 10 | 78168852 | rs12774333 | 1.441 | -1.978 | -1.333 | 0.295 | -0.104 | 0.098 | 0.098 | 0 | 0.38 | 0.03 | 0.34 | 0.19 | 0.41 | 0.33 |
| 10 | 78171460 | rs1907344 | 0.663 | -2.305 | -1.011 | 0.399 | 0.001 | -0.002 | -0.002 | 0 | 0.17 | 0.16 | 0.1 | 0.22 | 0.09 | 0.12 |
| 10 | 78171477 | rs11598028 | 1.981 | -2.288 | -1.023 | 0.412 | -0.1 | 0.161 | 0.161 | 0 | 0.36 | 0.03 | 0.31 | 0.1 | 0.4 | 0.18 |
| 10 | 78174967 | rs10824415 | 0.384 | -1.726 | -0.198 | 0.584 | 0.006 | 0.011 | 0.011 | 0 | 0.62 | 0.44 | 0.63 | 0.49 | 0.57 | 0.5 |
| 10 | 78183861 | rs12415769 | -0.954 | -1.776 | -0.054 | 0.747 | 0.01 | 0.079 | 0.079 | 0 | 0.08 | 0.23 | 0.2 | 0.12 | 0.07 | 0.11 |
| 10 | 78184816 | rs1907348 | 0.858 | -1.848 | -0.28 | 0.739 | -0.033 | 0.01 | 0.01 | 0 | 0.55 | 0.22 | 0.44 | 0.42 | 0.5 | 0.46 |
| 10 | 78187872 | rs17465450 | 1.775 | -1.762 | -0.649 | 0.53 | -0.006 | 0.013 | 0.013 | 0 | 0.06 | 0.01 | 0.03 | 0 | 0.08 | 0.02 |
| 10 | 78189959 | rs1873465 | -0.353 | -1.584 | 0.264 | 0.999 | -0.055 | 0.037 | 0.037 | 0 | 0.75 | 0.71 | 0.82 | 0.5 | 0.91 | 0.5 |
| 10 | 78190980 | rs4746393 | -0.623 | -1.429 | 0.353 | 0.963 | -0.067 | 0.04 | 0.04 | 0.006 | 0.78 | 0.71 | 0.85 | 0.59 | 0.92 | 0.67 |
| 10 | 78195898 | rs17380780 | 0.227 | 0.215 | 0.505 | 1.119 | -0.01 | 0.007 | 0.007 | 0 | 0.13 | 0.11 | 0.16 | 0 | 0.35 | 0.1 |
| 10 | 78204760 | rs11812165 | -1.14 | -0.473 | 0.871 | 1.303 | 0.019 | 0.024 | 0.024 | 0 | 0.05 | 0.19 | 0.05 | 0.06 | 0.05 | 0.03 |
| 10 | 78205649 | rs7915954 | -0.43 | -0.035 | 1.168 | 1.252 | 0.053 | -0.013 | -0.013 | 0 | 0.15 | 0.2 | 0.2 | 0.09 | 0.34 | 0.23 |
| 10 | 78209331 | rs4746395 | -1.036 | -0.209 | 1.894 | 1.589 | 0.173 | 0 | 0 | 0 | 0.18 | 0.5 | 0.25 | 0.15 | 0.35 | 0.23 |
| 10 | 78216814 | rs4745816 | 1.132 | -0.793 | 1.47 | 1.287 | -0.032 | 0.008 | 0.008 | 0.051 | 0.56 | 0.19 | 0.46 | 0.42 | 0.48 | 0.44 |
| 10 | 78218060 | rs1873468 | 0.362 | -0.833 | 1.68 | 1.261 | -0.029 | 0.026 | 0.026 | 0 | 0.7 | 0.43 | 0.65 | 0.49 | 0.81 | 0.54 |
| 10 | 78218488 | rs3740241 | 0.749 | -0.853 | 1.701 | 1.266 | -0.002 | -0.002 | -0.002 | 0.19 | 0.67 | 0.39 | 0.61 | 0.41 | 0.79 | 0.4 |
| 10 | 78218905 | rs1907356 | 0.889 | -0.829 | 1.647 | 1.246 | -0.007 | 0.009 | 0.009 | 0 | 0.11 | 0.13 | 0.15 | 0 | 0.32 | 0.09 |
| 10 | 78224646 | rs12777728 | 0.171 | -0.913 | 2.062 | 1.323 | 0 | 0.003 | 0.003 | 0 | 0.22 | 0.22 | 0.22 | 0.12 | 0.14 | 0.22 |
| 10 | 78233467 | rs12778475 | 0.435 | -0.942 | 2.043 | 1.265 | 0.015 | -0.005 | -0.005 | 0 | 0.2 | 0.15 | 0.07 | 0 | 0.12 | 0.14 |
| 10 | 78240418 | rs1907333 | 1.37 | -0.883 | 1.921 | 1.304 | -0.02 | 0.005 | 0.005 | 0 | 0.55 | 0.22 | 0.46 | 0.42 | 0.47 | 0.31 |
| 10 | 78243652 | rs2579724 | 0.529 | -0.908 | 1.999 | 1.309 | -0.046 | 0.045 | 0.045 | 0 | 0.71 | 0.59 | 0.64 | 0.48 | 0.81 | 0.53 |
| 10 | 78244948 | rs1907331 | 0.717 | -0.737 | 1.906 | 1.064 | -0.003 | 0.008 | 0.008 | 0 | 0.12 | 0.13 | 0.15 | 0.01 | 0.33 | 0.09 |
| 10 | 78272249 | rs1907322 | -0.355 | -0.775 | 1.964 | 1.079 | 0.073 | -0.018 | -0.018 | 2.802 | 0.17 | 0.35 | 0.23 | 0.11 | 0.34 | 0.23 |
| 10 | 78273721 | rs2637248 | 0.037 | -0.887 | 2.004 | 1.124 | -0.045 | 0.047 | 0.047 | 0 | 0.75 | 0.68 | 0.68 | 0.47 | 0.85 | 0.57 |
| 10 | 78282541 | rs2579739 | 1.578 | -0.529 | 1.849 | 0.883 | 0.027 | 0.058 | 0.058 | 0 | 0.52 | 0.17 | 0.42 | 0.37 | 0.42 | 0.42 |
| 10 | 78282773 | rs2579740 | 2.461 | -0.974 | 1.563 | 0.739 | -0.06 | 0.155 | 0.155 | 0 | 0.37 | 0.03 | 0.31 | 0.13 | 0.36 | 0.18 |
| 10 | 78283023 | rs2451911 | 0.997 | -0.477 | 2.119 | 0.991 | 0.001 | -0.002 | -0.002 | 0 | 0.59 | 0.38 | 0.46 | 0.4 | 0.47 | 0.46 |
| 10 | 78284024 | rs2637251 | 1.925 | -0.437 | 2.125 | 0.996 | 0.021 | 0.071 | 0.071 | 0 | 0.48 | 0.11 | 0.41 | 0.38 | 0.43 | 0.4 |
| 10 | 78289587 | rs11001806 | -1.688 | -0.383 | 2.271 | 0.973 | 0.103 | -0.099 | -0.099 | 0 | 0.25 | 0.53 | 0.26 | 0.07 | 0.39 | 0.28 |
| 10 | 78297974 | rs16933641 | -0.272 | -0.399 | 2.191 | 0.991 | -0.015 | -0.022 | -0.022 | 0.007 | 0.1 | 0.17 | 0.06 | 0.03 | 0.04 | 0.06 |
| 10 | 78299227 | rs1907310 | 0.071 | -0.43 | 2.126 | 1.014 | 0.004 | -0.006 | -0.006 | 0 | 0.75 | 0.75 | 0.69 | 0.45 | 0.81 | 0.67 |
| 10 | 78300542 | rs10824423 | -1.836 | 0.087 | 2.488 | 1.196 | 0.115 | -0.098 | -0.098 | 1.899 | 0.25 | 0.56 | 0.27 | 0.07 | 0.39 | 0.28 |
| 10 | 78300806 | rs2395383 | 0.426 | 0.111 | 2.559 | 1.192 | 0.086 | -0.045 | -0.045 | 0 | 0.11 | 0.2 | 0.17 | 0 | 0.32 | 0.08 |
| 10 | 78301888 | rs11001815 | -1.683 | 0.103 | 2.262 | 1.213 | 0.115 | -0.098 | -0.098 | 0.732 | 0.25 | 0.56 | 0.27 | 0.07 | 0.38 | 0.28 |
| 10 | 78310828 | rs1907302 | 1.615 | 0.085 | 2.005 | 1.344 | -0.012 | 0.018 | 0.018 | 0 | 0.57 | 0.27 | 0.59 | 0.37 | 0.5 | 0.41 |
| 10 | 78312002 | rs2637254 | 1.493 | 0.108 | 1.941 | 1.288 | -0.031 | 0.005 | 0.005 | 0 | 0.58 | 0.2 | 0.6 | 0.41 | 0.52 | 0.55 |
| 10 | 78314705 | rs2637256 | 2.245 | 0.626 | 2.643 | 1.783 | 0.002 | 0.496 | 0.496 | 0.493 | 0.9 | 0.52 | 0.85 | 0.52 | 0.96 | 0.85 |
| 10 | 78315191 | rs16933663 | -1.605 | 0.651 | 2.799 | 1.797 | 0.044 | 0.041 | 0.041 | 0.013 | 0.06 | 0.31 | 0.06 | 0.03 | 0.03 | 0.03 |
| 10 | 78316518 | rs2579766 | 2.286 | 0.653 | 2.688 | 1.791 | 0.006 | 0.504 | 0.504 | 0.418 | 0.9 | 0.52 | 0.85 | 0.52 | 0.96 | 0.85 |
| 10 | 78319770 | rs7091526 | -1.254 | 1.004 | 2.679 | 1.46 | 0.135 | -0.124 | -0.124 | 0 | 0.19 | 0.46 | 0.2 | 0 | 0.35 | 0.12 |
| 10 | 78320346 | rs2637260 | 1.227 | 0.834 | 2.681 | 1.621 | 0.014 | 0.006 | 0.006 | 0 | 0.66 | 0.54 | 0.61 | 0.4 | 0.53 | 0.53 |
| 10 | 78327759 | rs2579747 | 1.631 | 0.799 | 2.607 | 1.627 | -0.056 | -0.009 | -0.009 | 0.083 | 0.59 | 0.17 | 0.59 | 0.41 | 0.51 | 0.51 |
| 10 | 78336015 | rs1907323 | 1.66 | 0.772 | 2.82 | 1.693 | -0.012 | -0.003 | -0.003 | 0 | 0.59 | 0.33 | 0.58 | 0.44 | 0.49 | 0.47 |
| 10 | 78341125 | rs2395385 | 1.87 | 0.765 | 2.888 | 1.677 | -0.014 | -0.003 | -0.003 | 0.26 | 0.61 | 0.31 | 0.59 | 0.41 | 0.51 | 0.51 |
| 10 | 78392318 | rs7909808 | 1.573 | 0.857 | 2.772 | 1.672 | 0.031 | 0.028 | 0.028 | 0 | 0.63 | 0.44 | 0.64 | 0.49 | 0.51 | 0.59 |
| 10 | 78399498 | rs2583056 | 1.502 | 0.861 | 2.743 | 1.674 | 0.032 | 0.029 | 0.029 | 0 | 0.64 | 0.49 | 0.64 | 0.49 | 0.51 | 0.59 |
| 10 | 78415481 | rs2579768 | 1.902 | 0.839 | 2.632 | 1.8 | -0.006 | 0.026 | 0.026 | 0 | 0.56 | 0.23 | 0.55 | 0.38 | 0.46 | 0.38 |
| 10 | 78415841 | rs4979803 | -2.179 | 0.855 | 2.893 | 1.803 | 0.025 | 0.05 | 0.05 | 0 | 0.09 | 0.34 | 0.08 | 0.08 | 0.05 | 0.22 |
| 10 | 78418056 | rs11001846 | -0.257 | 0.865 | 2.96 | 1.756 | 0.052 | -0.053 | -0.053 | 0 | 0.15 | 0.21 | 0.19 | 0 | 0.36 | 0.11 |
| 10 | 78424413 | rs7918668 | -2.164 | 0.855 | 2.938 | 1.775 | 0.025 | 0.093 | 0.093 | 0 | 0.09 | 0.34 | 0.08 | 0.11 | 0.05 | 0.22 |
| 10 | 78427688 | rs2579736 | 2.184 | 0.895 | 2.908 | 1.857 | -0.014 | 0.034 | 0.034 | 0.112 | 0.55 | 0.14 | 0.54 | 0.38 | 0.46 | 0.38 |
| 10 | 78434949 | rs2583042 | 0.125 | 0.912 | 2.955 | 1.88 | 0.01 | -0.009 | -0.009 | 0 | 0.82 | 0.84 | 0.83 | 0.49 | 0.88 | 0.71 |
| 10 | 78444456 | rs241 | 1.382 | 0.953 | 2.831 | 1.882 | 0.025 | 0.024 | 0.024 | 0 | 0.65 | 0.51 | 0.62 | 0.49 | 0.51 | 0.6 |
| 10 | 78457588 | rs4979831 | 1.743 | 1.014 | 2.687 | 1.89 | -0.015 | -0.001 | -0.001 | 3.316 | 0.59 | 0.31 | 0.57 | 0.41 | 0.49 | 0.5 |
| 10 | 78457990 | rs2395423 | 0.405 | 1.011 | 2.719 | 1.884 | 0.001 | 0.006 | 0.006 | 0 | 0.81 | 0.78 | 0.81 | 0.49 | 0.88 | 0.71 |
| 10 | 78459211 | rs1561676 | 2.007 | 0.997 | 2.708 | 1.893 | -0.007 | 0.019 | 0.019 | 0 | 0.57 | 0.29 | 0.54 | 0.38 | 0.46 | 0.38 |
| 10 | 78482228 | rs1992396 | 1.256 | 0.97 | 2.508 | 1.826 | 0.013 | 0.014 | 0.014 | 0 | 0.66 | 0.56 | 0.63 | 0.51 | 0.53 | 0.61 |
| 10 | 78492363 | rs846588 | 0.223 | 0.936 | 2.022 | 1.82 | 0.002 | 0.002 | 0.002 | 0 | 0.82 | 0.82 | 0.82 | 0.52 | 0.9 | 0.72 |
| 10 | 78499053 | rs7090852 | -0.326 | 1.016 | 1.974 | 1.736 | -0.029 | 0.013 | 0.013 | 0 | 0.15 | 0.14 | 0.18 | 0 | 0.36 | 0.12 |
| 10 | 78515883 | rs846646 | 1.893 | 0.952 | 2.11 | 1.822 | -0.001 | 0.001 | 0.001 | 0 | 0.57 | 0.37 | 0.49 | 0.4 | 0.45 | 0.38 |
| 10 | 78522318 | rs846642 | 1.795 | 0.97 | 2.184 | 1.793 | 0.007 | 0.002 | 0.002 | 0 | 0.62 | 0.53 | 0.57 | 0.42 | 0.49 | 0.57 |
| 10 | 78533060 | rs10491078 | 0.043 | 1.064 | 2.067 | 1.588 | 0.03 | 0.02 | 0.02 | 0 | 0.16 | 0.06 | 0.06 | 0.07 | 0.09 | 0.16 |
| 10 | 78545449 | rs846619 | 1.001 | 0.978 | 1.92 | 1.531 | -0.023 | -0.024 | -0.024 | 0.053 | 0.67 | 0.69 | 0.64 | 0.5 | 0.53 | 0.6 |
| 10 | 78587491 | rs2569359 | -0.012 | 0.498 | 1.734 | 1.35 | -0.02 | 0.033 | 0.033 | 0 | 0.82 | 0.76 | 0.84 | 0.53 | 0.9 | 0.73 |
| 10 | 78589532 | rs10824449 | -0.246 | 1.071 | 1.372 | 0.971 | -0.018 | -0.001 | -0.001 | 0 | 0.16 | 0.14 | 0.31 | 0 | 0.41 | 0.12 |

| | | | | | | | | | | | | | | | | |
|---|---|---|---|---|---|---|---|---|---|---|---|---|---|---|---|---|
| 10 | 78593852 | rs7914486 | -1.013 | 1.058 | 1.423 | 1.023 | 0.105 | -0.025 | -0.025 | 0.381 | 0.21 | 0.35 | 0.44 | 0.12 | 0.45 | 0.33 |
| 10 | 78605316 | rs1897584 | -0.462 | 0.884 | 1.174 | 0.786 | -0.072 | 0.081 | 0.081 | 0.028 | 0.2 | 0.39 | 0.17 | 0.39 | 0.1 | 0.24 |
| 10 | 78628331 | rs10824459 | -0.658 | 1.939 | 0.738 | -0.019 | 0.047 | -0.053 | -0.053 | 0 | 0.16 | 0.22 | 0.29 | 0 | 0.38 | 0.13 |
| 10 | 78630781 | rs10824460 | -0.535 | 1.968 | 0.757 | -0.007 | 0.016 | -0.028 | -0.028 | 0.334 | 0.15 | 0.17 | 0.29 | 0 | 0.38 | 0.13 |
| 10 | 78631852 | rs10824462 | -0.586 | 1.975 | 0.671 | -0.004 | 0.049 | -0.054 | -0.054 | 0 | 0.16 | 0.22 | 0.29 | 0 | 0.38 | 0.13 |
| 10 | 78644260 | rs2082415 | 0.328 | 0.71 | -0.975 | 0.38 | -0.014 | 0.101 | 0.101 | 0 | 0.47 | 0.14 | 0.37 | 0.28 | 0.5 | 0.28 |
| 10 | 78645453 | rs10762733 | 0.387 | 0.589 | -0.627 | 0.34 | -0.001 | 0.022 | 0.022 | 0 | 0.57 | 0.35 | 0.36 | 0.3 | 0.58 | 0.32 |
| 10 | 78648972 | rs11001919 | 0.263 | 0.934 | -0.768 | 0.052 | -0.01 | -0.006 | -0.006 | 0.173 | 0.14 | 0.11 | 0.13 | 0 | 0.2 | 0.05 |
| 10 | 78649602 | rs2288839 | -0.087 | 0.222 | -1.193 | -0.029 | -0.01 | -0.111 | -0.111 | 0.102 | 0.72 | 0.89 | 0.71 | 0.46 | 0.77 | 0.61 |
| 10 | 78650261 | rs4639876 | 0.409 | 0.316 | -0.3 | 0.132 | 0 | 0.002 | 0.002 | 0 | 0.67 | 0.36 | 0.49 | 0.41 | 0.71 | 0.5 |
| 10 | 78651796 | rs7087337 | 0.56 | 0.202 | 0.41 | 0.066 | -0.044 | -0.079 | -0.079 | 0 | 0.15 | 0.25 | 0.08 | 0 | 0.08 | 0.11 |
| 10 | 78652814 | rs1897592 | 0.624 | 0.256 | 0.178 | 0.118 | 0.013 | -0.005 | -0.005 | 0 | 0.55 | 0.34 | 0.44 | 0.4 | 0.64 | 0.39 |
| 10 | 78663798 | rs10824465 | 1.463 | 0.435 | -0.405 | 0.314 | -0.011 | 0.025 | 0.025 | 0.097 | 0.4 | 0.06 | 0.25 | 0.27 | 0.41 | 0.32 |
| 10 | 78664629 | rs2116829 | 1.04 | -0.187 | 0.157 | 0.592 | 0.02 | -0.001 | -0.001 | 0 | 0.55 | 0.28 | 0.43 | 0.4 | 0.64 | 0.39 |
| 10 | 78667191 | rs7069982 | 0.768 | 0.269 | 0.086 | 0.99 | -0.003 | -0.032 | -0.032 | 0 | 0.76 | 0.79 | 0.71 | 0.55 | 0.79 | 0.67 |
| 10 | 78670661 | rs1907746 | -1.183 | 0.117 | 0.454 | 0.99 | -0.017 | 0.051 | 0.051 | 0 | 0.24 | 0.44 | 0.22 | 0.41 | 0.18 | 0.34 |
| 10 | 78671776 | rs10762734 | -1.31 | 0.382 | 0.779 | 0.74 | 0.019 | 0.036 | 0.036 | 0 | 0.17 | 0.3 | 0.42 | 0.19 | 0.2 | 0.13 |
| 10 | 78678167 | rs1907729 | 1.287 | 0.041 | 0.933 | 0.953 | 0.028 | 0.02 | 0.02 | 0 | 0.89 | 0.74 | 0.81 | 0.88 | 0.83 | 0.92 |
| 10 | 78685229 | rs12572857 | -0.131 | -0.412 | 0.397 | 0.278 | 0.005 | -0.062 | -0.062 | 0 | 0.29 | 0.09 | 0.2 | 0.61 | 0.26 | 0.55 |
| 10 | 78685477 | rs10740463 | 1.29 | -0.349 | 0.645 | 0.267 | 0.043 | 0.072 | 0.072 | 0 | 0.86 | 0.6 | 0.73 | 0.83 | 0.81 | 0.9 |
| 10 | 78688976 | rs1865020 | -1.18 | -0.81 | 0.826 | 0.078 | 0.007 | 0.007 | 0.007 | 0 | 0.45 | 0.57 | 0.54 | 0.83 | 0.32 | 0.68 |
| 10 | 78694444 | rs1865017 | -0.213 | -0.849 | 0.201 | 0.074 | 0.016 | -0.026 | -0.026 | 0 | 0.68 | 0.91 | 0.79 | 0.32 | 0.72 | 0.41 |
| 10 | 78694965 | rs2131218 | -1.597 | -0.555 | 0.407 | 0.157 | 0.027 | -0.002 | -0.002 | 0 | 0.16 | 0.6 | 0.29 | 0.12 | 0.19 | 0.07 |
| 10 | 78696377 | rs4980117 | 0.925 | -0.518 | 0.259 | -0.006 | 0.008 | 0.085 | 0.085 | 0 | 0.5 | 0.44 | 0.36 | 0.1 | 0.52 | 0.28 |
| 10 | 78697597 | rs12762512 | -0.737 | -0.417 | 0.484 | 0.274 | -0.018 | 0.028 | 0.028 | 0.36 | 0.06 | 0.24 | 0.09 | 0.05 | 0.02 | 0.03 |
| 10 | 78700102 | rs10762739 | 0.062 | -0.207 | 0.121 | 0.299 | -0.025 | -0.018 | -0.018 | 4.541 | 0.69 | 0.61 | 0.62 | 0.78 | 0.77 | 0.71 |
| 10 | 78702871 | rs10762740 | 1.157 | 0.303 | -0.298 | 0.76 | 0 | 0.016 | 0.016 | 0 | 0.81 | 0.74 | 0.75 | 0.32 | 0.78 | 0.76 |
| 10 | 78707603 | rs10762741 | -0.368 | 0.617 | -0.691 | -0.27 | -0.01 | 0.034 | 0.034 | 0 | 0.36 | 0.28 | 0.32 | 0.13 | 0.38 | 0.2 |
| 10 | 78712361 | rs2116833 | -0.063 | 0.038 | -0.693 | -1.044 | 0.001 | -0.01 | -0.01 | 0 | 0.54 | 0.54 | 0.58 | 0.82 | 0.65 | 0.64 |
| 10 | 78715622 | rs4980118 | -0.338 | -0.383 | -0.64 | -0.959 | 0.003 | 0.023 | 0.023 | 0 | 0.68 | 0.73 | 0.57 | 0.89 | 0.69 | 0.69 |
| 10 | 78717892 | rs1907740 | -0.261 | 0.029 | -0.726 | -0.704 | -0.001 | 0.006 | 0.006 | 0 | 0.47 | 0.58 | 0.38 | 0.61 | 0.42 | 0.36 |
| 10 | 78720608 | rs10762742 | -0.246 | -0.066 | -0.694 | -0.623 | -0.004 | -0.005 | -0.005 | 0 | 0.46 | 0.48 | 0.37 | 0.44 | 0.43 | 0.47 |
| 10 | 78727604 | rs7912269 | 0.612 | -0.104 | -0.734 | -0.824 | -0.009 | -0.047 | -0.047 | 0 | 0.11 | 0.26 | 0.06 | 0.03 | 0.07 | 0.04 |
| 10 | 78727967 | rs7915877 | -0.443 | -0.069 | -0.696 | -0.929 | 0.001 | -0.004 | -0.004 | 0 | 0.47 | 0.5 | 0.38 | 0.61 | 0.44 | 0.37 |
| 10 | 78729393 | rs11001950 | -1.409 | -0.175 | -0.881 | -0.871 | -0.032 | 0.037 | 0.037 | 0 | 0.77 | 0.92 | 0.71 | 0.86 | 0.76 | 0.74 |
| 10 | 78731833 | rs10824477 | -1.171 | -0.663 | -0.223 | -0.465 | -0.023 | 0.028 | 0.028 | 0 | 0.33 | 0.42 | 0.4 | 0.51 | 0.24 | 0.31 |
| 10 | 78734666 | rs10824478 | -0.873 | -0.622 | -0.303 | -0.448 | 0 | -0.001 | -0.001 | 0 | 0.31 | 0.3 | 0.39 | 0.51 | 0.23 | 0.31 |
| 10 | 78735498 | rs999995 | -1.448 | -0.536 | -0.282 | -0.528 | -0.011 | -0.02 | -0.02 | 0 | 0.56 | 0.73 | 0.54 | 0.52 | 0.5 | 0.56 |
| 10 | 78735661 | rs999994 | 0.172 | -0.207 | -0.034 | -1.113 | -0.012 | -0.04 | -0.04 | 0 | 0.1 | 0.26 | 0.06 | 0.03 | 0.05 | 0.03 |
| 10 | 78736125 | rs999993 | 0.09 | 0.25 | -0.405 | -0.781 | 0.006 | -0.098 | -0.098 | 0 | 0.25 | 0.42 | 0.16 | 0.03 | 0.29 | 0.25 |
| 10 | 78737595 | rs16934131 | 0.33 | 0.527 | -1.087 | -0.662 | -0.008 | 0.01 | 0.01 | 0 | 0.22 | 0.15 | 0.14 | 0.03 | 0.28 | 0.26 |
| 10 | 78738681 | rs11001957 | -0.022 | 0.288 | -1.029 | -0.129 | -0.001 | -0.047 | -0.047 | 0 | 0.14 | 0.01 | 0.11 | 0.15 | 0.1 | 0.11 |
| 10 | 78739848 | rs1907724 | 0.698 | 0.44 | -0.305 | -0.683 | 0.008 | -0.097 | -0.097 | 0 | 0.77 | 0.65 | 0.82 | 0.99 | 0.75 | 0.81 |
| 10 | 78740285 | rs2163798 | 0.815 | 0.148 | 0.237 | -0.556 | -0.001 | -0.081 | -0.081 | 0 | 0.74 | 0.5 | 0.75 | 0.87 | 0.74 | 0.69 |
| 10 | 78740648 | rs11597111 | -0.996 | 0.388 | 0.006 | -0.362 | 0.04 | 0.034 | 0.034 | 0 | 0.07 | 0.19 | 0.13 | 0.12 | 0.1 | 0.16 |
| 10 | 78745937 | rs17388160 | 0.574 | 0.582 | -0.986 | 0.014 | -0.128 | -0.138 | -0.138 | 0.064 | 0.24 | 0.03 | 0.2 | 0.57 | 0.32 | 0.31 |
| 10 | 78750074 | rs3781153 | -0.075 | 0.278 | -0.405 | -0.154 | -0.006 | 0.005 | 0.005 | 0.065 | 0.64 | 0.68 | 0.67 | 0.24 | 0.58 | 0.57 |
| 10 | 78750810 | rs3824716 | 0.928 | -0.145 | -0.023 | -0.389 | 0.027 | 0.15 | 0.15 | 0.036 | 0.46 | 0.25 | 0.49 | 0.11 | 0.44 | 0.3 |
| 10 | 78752289 | rs10740464 | 0.912 | 0.061 | 0.448 | -0.068 | 0.004 | 0.161 | 0.161 | 0.221 | 0.85 | 0.47 | 0.81 | 0.66 | 0.88 | 0.74 |
| 10 | 78752652 | rs1907720 | 0.364 | 0.118 | 0.573 | -0.009 | 0.009 | 0.119 | 0.119 | 0 | 0.88 | 0.72 | 0.86 | 0.7 | 0.89 | 0.77 |
| 10 | 78756247 | rs3781157 | 0.287 | 0.366 | 0.147 | -0.128 | -0.006 | -0.009 | -0.009 | 0 | 0.32 | 0.31 | 0.26 | 0.54 | 0.36 | 0.4 |
| 10 | 78756815 | rs12243061 | 0.071 | 0.383 | 0.005 | -0.092 | -0.027 | -0.028 | -0.028 | 0.032 | 0.34 | 0.24 | 0.26 | 0.55 | 0.39 | 0.41 |
| 10 | 78756841 | rs7079351 | -0.133 | 0.532 | -0.116 | -0.123 | -0.003 | 0.005 | 0.005 | 0 | 0.08 | 0.03 | 0.04 | 0 | 0.09 | 0.04 |
| 10 | 78758909 | rs3781160 | -1.23 | 0.383 | -0.087 | -0.166 | 0.002 | -0.003 | -0.003 | 0 | 0.15 | 0.18 | 0.15 | 0.16 | 0.15 | 0.22 |
| 10 | 78764793 | rs1873691 | 0.281 | -0.06 | -0.243 | -0.257 | 0.012 | 0.009 | 0.009 | 0 | 0.46 | 0.42 | 0.58 | 0.39 | 0.46 | 0.44 |
| 10 | 78768629 | rs7074008 | -0.765 | 0.119 | 0.111 | -0.443 | 0.033 | 0.012 | 0.012 | 0 | 0.7 | 0.55 | 0.59 | 0.28 | 0.66 | 0.61 |
| 10 | 78770196 | rs17479797 | -0.222 | 0.329 | 0.021 | -0.578 | -0.004 | -0.009 | -0.009 | 0 | 0.11 | 0.1 | 0.08 | 0.01 | 0.17 | 0.07 |
| 10 | 78772456 | rs11001968 | 0.397 | 0.222 | 0.202 | -0.652 | 0.007 | 0.016 | 0.016 | 0 | 0.52 | 0.44 | 0.4 | 0.08 | 0.53 | 0.39 |
| 10 | 78775582 | rs11001971 | 0.454 | 0.027 | 0.118 | -0.704 | 0.039 | 0.127 | 0.127 | 0.078 | 0.5 | 0.34 | 0.39 | 0.07 | 0.47 | 0.38 |
| 10 | 78777704 | rs12219498 | 0.995 | 0.159 | 0.083 | -0.09 | -0.011 | -0.017 | -0.017 | 0.359 | 0.25 | 0.2 | 0.2 | 0.49 | 0.32 | 0.29 |
| 10 | 78786963 | rs11001976 | 0.198 | -0.342 | -0.431 | -0.325 | -0.008 | -0.004 | -0.004 | 0 | 0.16 | 0.01 | 0.09 | 0.06 | 0.1 | 0.14 |
| 10 | 78788660 | rs4980123 | -1.342 | -0.467 | 0.002 | -0.375 | -0.005 | 0.001 | 0.001 | 0 | 0.28 | 0.3 | 0.18 | 0.29 | 0.2 | 0.28 |
| 10 | 78794407 | rs17389253 | 0.541 | -0.515 | 0.056 | -0.363 | -0.001 | -0.013 | -0.013 | 0 | 0.08 | 0.06 | 0.05 | 0 | 0.09 | 0.03 |
| 10 | 78794558 | rs12248514 | -0.111 | -0.583 | 0.03 | -0.353 | -0.007 | 0.001 | 0.001 | 0 | 0.1 | 0.15 | 0.06 | 0.06 | 0.07 | 0.12 |
| 10 | 78794972 | rs3781164 | -0.157 | -0.581 | 0.041 | -0.352 | -0.008 | 0.001 | 0.001 | 0 | 0.1 | 0.16 | 0.06 | 0.06 | 0.07 | 0.12 |
| 10 | 78797205 | rs1247757 | 0.811 | -0.518 | -0.072 | -0.449 | 0.01 | -0.007 | -0.007 | 0 | 0.77 | 0.81 | 0.84 | 0.75 | 0.84 | 0.76 |
| 10 | 78801799 | rs2247557 | 0.59 | -0.498 | 0.19 | -0.443 | 0.035 | 0.099 | 0.099 | 0.964 | 0.93 | 0.75 | 0.92 | 0.81 | 0.94 | 0.89 |
| 10 | 78803716 | rs12248504 | 1.509 | -0.846 | -0.667 | -1.053 | 0.001 | -0.01 | -0.01 | 0 | 0.15 | 0.09 | 0.09 | 0.01 | 0.15 | 0.1 |
| 10 | 78809089 | rs2209666 | 0.531 | -0.804 | -0.792 | -1.103 | -0.037 | 0.112 | 0.112 | 0 | 0.22 | 0.03 | 0.13 | 0.01 | 0.25 | 0.14 |

| | | | | | | | | | | | | | | | | |
|---|---|---|---|---|---|---|---|---|---|---|---|---|---|---|---|---|
| 10 | 78814288 | rs7099467 | 1.225 | -0.683 | -0.44 | -1.038 | -0.009 | 0.209 | 0.209 | 0.726 | 0.47 | 0.21 | 0.43 | 0.01 | 0.55 | 0.31 |
| 10 | 78835356 | rs1247785 | 0.653 | -0.654 | -0.372 | -0.825 | 0.007 | -0.007 | -0.007 | 0 | 0.6 | 0.65 | 0.67 | 0.52 | 0.68 | 0.58 |
| 10 | 78845149 | rs3781167 | 0.806 | -0.724 | -0.51 | -0.791 | 0.008 | -0.025 | -0.025 | 0 | 0.09 | 0 | 0.04 | 0.06 | 0.08 | 0.07 |
| 10 | 78853078 | rs7076995 | 1.124 | -0.728 | 0.073 | -0.672 | 0.005 | 0.174 | 0.174 | 1.4 | 0.48 | 0.23 | 0.44 | 0.01 | 0.56 | 0.34 |
| 10 | 78857975 | rs16934250 | -0.274 | -0.556 | -0.498 | -0.308 | -0.001 | -0.09 | -0.09 | 0 | 0.14 | 0.02 | 0.2 | 0.38 | 0.14 | 0.17 |
| 10 | 78860791 | rs11597359 | 0.972 | -0.019 | -0.611 | -0.445 | -0.012 | 0.051 | 0.051 | 0 | 0.1 | 0.01 | 0.07 | 0.01 | 0.09 | 0.11 |
| 10 | 78866173 | rs10824490 | 1.407 | -0.091 | -0.346 | -0.336 | 0 | -0.034 | -0.034 | 0.199 | 0.16 | 0.14 | 0.1 | 0.01 | 0.16 | 0.1 |
| 10 | 78875386 | rs2616654 | -0.556 | 0.226 | -0.836 | -0.325 | -0.107 | 0.229 | 0.229 | 0.006 | 0.59 | 0.92 | 0.59 | 0.91 | 0.44 | 0.69 |
| 10 | 78875476 | rs2217895 | 2.178 | 0.158 | -1.045 | -0.481 | 0.005 | 0.028 | 0.028 | 0 | 0.08 | 0.03 | 0.04 | 0 | 0.05 | 0.02 |
| 10 | 78875741 | rs2574784 | -0.053 | 0.256 | -0.23 | -0.238 | -0.002 | 0.003 | 0.003 | 0 | 0.54 | 0.47 | 0.47 | 0.44 | 0.42 | 0.47 |
| 10 | 78880943 | rs7907270 | 0.344 | 0.278 | -0.375 | -1.146 | 0.003 | -0.037 | -0.037 | 0.351 | 0.45 | 0.41 | 0.26 | 0.01 | 0.39 | 0.3 |
| 10 | 78891433 | rs7072406 | 0.476 | 0.432 | -0.401 | -0.94 | -0.003 | -0.036 | -0.036 | 2.246 | 0.42 | 0.42 | 0.35 | 0.04 | 0.45 | 0.31 |
| 10 | 78896947 | rs2574793 | 0.427 | 0.476 | -0.729 | -1.038 | 0.011 | 0.039 | 0.039 | 0 | 0.39 | 0.27 | 0.55 | 0.88 | 0.37 | 0.51 |
| 10 | 78896996 | rs2766618 | -1.098 | 0.473 | -0.83 | -0.975 | -0.002 | 0.001 | 0.001 | 0 | 0.89 | 0.91 | 0.93 | 0.92 | 0.85 | 0.87 |
| 10 | 78902454 | rs2574804 | 0.449 | 0.415 | -0.742 | -0.976 | 0 | -0.015 | -0.015 | 0 | 0.38 | 0.32 | 0.55 | 0.88 | 0.37 | 0.51 |
| 10 | 78904528 | rs11002044 | 0.329 | 0.612 | -0.704 | -0.761 | 0.006 | -0.095 | -0.095 | 0 | 0.43 | 0.52 | 0.37 | 0.04 | 0.45 | 0.31 |
| 10 | 78912007 | rs2574807 | 0.413 | 0.471 | -0.965 | -0.497 | 0.006 | -0.016 | -0.016 | 0 | 0.4 | 0.57 | 0.53 | 0.73 | 0.42 | 0.45 |
| 10 | 78912854 | rs2574810 | -0.605 | 0.454 | -1.808 | -1.042 | -0.009 | 0.05 | 0.05 | 1.886 | 0.81 | 0.96 | 0.92 | 0.91 | 0.83 | 0.78 |
| 10 | 78914168 | rs12775320 | 0.538 | -0.008 | -2.097 | -1.367 | -0.003 | 0.239 | 0.239 | 0.276 | 0.36 | 0.05 | 0.35 | 0.01 | 0.38 | 0.22 |
| 10 | 78914597 | rs10824493 | 0.45 | -0.053 | -1.299 | -1.277 | -0.01 | -0.046 | -0.046 | 0 | 0.16 | 0.26 | 0.09 | 0.05 | 0.09 | 0.14 |
| 10 | 78915443 | rs17391134 | 1.725 | -0.219 | -1.967 | -1.529 | 0.025 | 0.067 | 0.067 | 0 | 0.16 | 0.04 | 0.09 | 0.01 | 0.12 | 0.06 |
| 10 | 78919640 | rs573712 | -0.292 | -0.168 | -1.828 | -1.289 | -0.011 | -0.01 | -0.01 | 0 | 0.25 | 0.15 | 0.5 | 0.73 | 0.28 | 0.57 |
| 10 | 78922019 | rs570706 | 0.615 | -0.165 | -1.854 | -1.37 | 0.005 | -0.044 | -0.044 | 0.65 | 0.19 | 0.07 | 0.44 | 0.69 | 0.19 | 0.39 |
| 10 | 78924624 | rs527458 | -0.289 | -0.351 | -1.828 | -1.718 | -0.011 | 0.055 | 0.055 | 0.055 | 0.28 | 0.14 | 0.53 | 0.89 | 0.32 | 0.61 |
| 10 | 78925465 | rs16934331 | 0.5 | -0.266 | -1.504 | -2.092 | -0.013 | -0.063 | -0.063 | 0 | 0.2 | 0.28 | 0.11 | 0 | 0.11 | 0.08 |
| 10 | 78929072 | rs11002057 | 0.207 | -0.271 | -1.64 | -2.099 | 0.014 | -0.007 | -0.007 | 0 | 0.27 | 0.25 | 0.12 | 0 | 0.13 | 0.08 |
| 10 | 78931855 | rs10509382 | 0.604 | 1.036 | -1.659 | -1.767 | -0.075 | 0.106 | 0.106 | 0 | 0.22 | 0.04 | 0.17 | 0.01 | 0.33 | 0.08 |
| 10 | 78933522 | rs17482756 | -0.336 | 0.678 | -1.119 | -1.25 | -0.011 | -0.005 | -0.005 | 0.506 | 0.15 | 0.04 | 0.09 | 0.16 | 0.16 | 0.15 |
| 10 | 78935171 | rs11002065 | -0.811 | 0.48 | -0.431 | -1.463 | 0.003 | -0.002 | -0.002 | 0.873 | 0.38 | 0.42 | 0.26 | 0.13 | 0.34 | 0.22 |
| 10 | 78938880 | rs11002068 | 0.426 | 0.5 | -0.777 | -1.393 | 0.005 | -0.004 | -0.004 | 0 | 0.08 | 0.04 | 0.05 | 0.12 | 0.08 | 0.14 |
| 10 | 78941418 | rs7083476 | -0.311 | 0.499 | -0.724 | -1.446 | 0.026 | 0.088 | 0.088 | 0 | 0.32 | 0.15 | 0.19 | 0.13 | 0.28 | 0.21 |
| 10 | 78944590 | rs1131824 | -0.953 | 0.3 | -0.053 | -1.434 | 0.019 | 0.034 | 0.034 | 0 | 0.42 | 0.71 | 0.27 | 0.25 | 0.34 | 0.33 |
| 10 | 78949728 | rs7914901 | -0.411 | 0.312 | 0.248 | -1.455 | 0.009 | 0.027 | 0.027 | 0 | 0.35 | 0.3 | 0.23 | 0.04 | 0.33 | 0.17 |
| 10 | 78952355 | rs1516509 | 0.989 | 1.035 | 0.486 | -1.143 | -0.011 | 0.069 | 0.069 | 0 | 0.18 | 0.06 | 0.13 | 0.01 | 0.19 | 0.07 |
| 10 | 78960694 | rs16934354 | 0.896 | 1.74 | 0.367 | -0.995 | -0.049 | -0.014 | -0.014 | 0 | 0.08 | 0.01 | 0.11 | 0.09 | 0.18 | 0.11 |
| 10 | 78962815 | rs11002078 | -0.031 | 1.226 | 0.67 | -0.944 | -0.009 | -0.035 | -0.035 | 0 | 0.09 | 0.16 | 0.06 | 0.03 | 0.08 | 0.08 |
| 10 | 78963490 | rs7905358 | -0.217 | 1.19 | 0.808 | -0.924 | 0.037 | 0.24 | 0.24 | 1.461 | 0.06 | 0.34 | 0.04 | 0.21 | 0.01 | 0.17 |
| 10 | 78967952 | rs7070015 | -0.105 | 1.065 | 0.624 | -0.902 | -0.008 | -0.031 | -0.031 | 0 | 0.09 | 0.12 | 0.06 | 0.03 | 0.08 | 0.08 |
| 10 | 78974650 | rs12357970 | 0.225 | 1.477 | -0.268 | -0.874 | 0.001 | -0.006 | -0.006 | 0 | 0.08 | 0.08 | 0.05 | 0.03 | 0.08 | 0.08 |
| 10 | 78982306 | rs11002088 | -0.261 | 1.492 | -0.377 | -0.932 | 0.002 | 0.087 | 0.087 | 0.001 | 0.26 | 0.11 | 0.17 | 0.04 | 0.27 | 0.16 |
| 10 | 78986111 | rs577360 | 0.477 | 1.808 | -0.2 | -0.837 | 0.002 | -0.014 | -0.014 | 0.004 | 0.57 | 0.39 | 0.64 | 0.74 | 0.48 | 0.53 |
| 10 | 78987183 | rs2616612 | 0.462 | 1.752 | -0.035 | -0.692 | 0.029 | -0.009 | -0.009 | 0.081 | 0.76 | 0.49 | 0.76 | 0.75 | 0.71 | 0.61 |
| 10 | 78988848 | rs567857 | -0.305 | 1.835 | -0.405 | -0.779 | -0.028 | 0.038 | 0.038 | 0 | 0.84 | 0.89 | 0.82 | 0.98 | 0.75 | 0.82 |
| 10 | 78993943 | rs499437 | 1.012 | 2.029 | -0.313 | -0.77 | 0.051 | -0.013 | -0.013 | 0 | 0.68 | 0.38 | 0.63 | 0.66 | 0.53 | 0.53 |
| 10 | 78994428 | rs12782077 | -0.407 | 2.008 | -0.375 | -0.926 | 0.004 | -0.018 | -0.018 | 0 | 0.32 | 0.37 | 0.29 | 0.15 | 0.42 | 0.34 |
| 10 | 78994717 | rs527706 | 1.34 | 1.907 | -0.394 | -0.851 | -0.001 | -0.001 | -0.001 | 0 | 0.46 | 0.41 | 0.5 | 0.6 | 0.29 | 0.4 |
| 10 | 78999208 | rs561336 | 1.147 | 1.932 | -0.605 | -0.93 | 0.031 | 0.044 | 0.044 | 1.657 | 0.48 | 0.56 | 0.51 | 0.78 | 0.29 | 0.46 |
| 10 | 78999999 | rs479256 | -1.181 | 1.337 | -0.787 | -0.704 | -0.052 | 0.05 | 0.05 | 0 | 0.88 | 0.99 | 0.87 | 0.97 | 0.83 | 0.85 |
| 10 | 79005918 | rs542007 | 0.763 | 1.238 | -0.573 | -0.702 | 0.002 | -0.016 | -0.016 | 0 | 0.7 | 0.67 | 0.7 | 0.81 | 0.65 | 0.63 |
| 10 | 79006545 | rs1436089 | 0.772 | 1.22 | -0.618 | -1.109 | 0.001 | -0.024 | -0.024 | 0.097 | 0.2 | 0.18 | 0.12 | 0.01 | 0.2 | 0.13 |
| 10 | 79007602 | rs560830 | 0.781 | 1.462 | -1.132 | -0.868 | -0.001 | 0.048 | 0.048 | 0.936 | 0.49 | 0.36 | 0.62 | 0.73 | 0.46 | 0.49 |
| 10 | 79015939 | rs555766 | 0.822 | 2.456 | -1.486 | -0.184 | -0.039 | 0.077 | 0.077 | 0 | 0.78 | 0.9 | 0.82 | 0.96 | 0.71 | 0.81 |
| 10 | 79018276 | rs10762753 | -0.821 | 1.858 | -1.115 | 0.858 | -0.004 | 0.044 | 0.044 | 0 | 0.07 | 0.07 | 0.05 | 0.35 | 0.05 | 0.09 |
| 10 | 79023597 | rs11599090 | -0.571 | 2.917 | -1.312 | 0.054 | -0.052 | 0.035 | 0.035 | 0 | 0.15 | 0.06 | 0.36 | 0.02 | 0.34 | 0.17 |
| 10 | 79028572 | rs4980136 | -0.099 | 2.775 | -1.218 | 0.02 | -0.008 | 0.004 | 0.004 | 0 | 0.11 | 0.07 | 0.21 | 0.01 | 0.22 | 0.13 |
| 10 | 79028810 | rs4980137 | -0.976 | 1.721 | -1.742 | -0.679 | -0.011 | 0.059 | 0.059 | 0.032 | 0.24 | 0.11 | 0.26 | 0.01 | 0.29 | 0.19 |
| 10 | 79033959 | rs1516512 | -0.478 | 0.362 | -2.933 | -0.598 | -0.039 | -0.055 | -0.055 | 0 | 0.62 | 0.89 | 0.62 | 0.52 | 0.51 | 0.64 |
| 10 | 79050161 | rs520887 | -0.322 | 0.424 | -2.494 | -0.977 | 0.009 | 0.043 | 0.043 | 0 | 0.41 | 0.3 | 0.41 | 0.14 | 0.58 | 0.26 |
| 10 | 79051460 | rs1457522 | 0.931 | 0.355 | -2.467 | -0.929 | -0.004 | -0.002 | -0.002 | 0 | 0.42 | 0.61 | 0.38 | 0.51 | 0.26 | 0.49 |
| 10 | 79059483 | rs872794 | 1.072 | 0.178 | -2.196 | -1.001 | -0.049 | 0.008 | 0.008 | 0 | 0.41 | 0.52 | 0.36 | 0.51 | 0.26 | 0.49 |
| 10 | 79065329 | rs583507 | -0.362 | 0.171 | -2.201 | -1.006 | 0.004 | 0.012 | 0.012 | 0 | 0.43 | 0.4 | 0.42 | 0.14 | 0.58 | 0.26 |
| 10 | 79065462 | rs584021 | -0.991 | 0.374 | -2.169 | -0.89 | -0.016 | -0.028 | -0.028 | 0 | 0.83 | 0.92 | 0.79 | 0.65 | 0.84 | 0.75 |
| 10 | 79070902 | rs10824521 | 1.065 | -0.912 | -1.862 | -0.854 | -0.029 | 0.047 | 0.047 | 0 | 0.2 | 0.3 | 0.24 | 0.35 | 0.15 | 0.24 |
| 10 | 79075412 | rs7475710 | 1.295 | -0.931 | -1.777 | -0.862 | 0.008 | -0.009 | -0.009 | 0 | 0.19 | 0.16 | 0.23 | 0.35 | 0.15 | 0.23 |
| 10 | 79077095 | rs611519 | 1.231 | -0.878 | -1.561 | -0.995 | 0.002 | -0.029 | -0.029 | 0 | 0.19 | 0.17 | 0.23 | 0.01 | 0.25 | 0.14 |
| 10 | 79077444 | rs7087190 | 1.696 | -1.344 | -1.931 | -1.253 | -0.073 | -0.042 | -0.042 | 1.11 | 0.18 | 0.43 | 0.14 | 0.13 | 0.09 | 0.25 |
| 10 | 79077847 | rs625148 | -0.647 | -1.325 | -1.967 | -1.248 | -0.008 | 0.005 | 0.005 | 0 | 0.64 | 0.39 | 0.64 | 0.52 | 0.76 | 0.51 |
| 10 | 79080057 | rs655487 | 0.399 | -1.255 | -1.939 | -1.226 | 0.001 | -0.002 | -0.002 | 0 | 0.38 | 0.34 | 0.46 | 0.36 | 0.4 | 0.37 |
| 10 | 79082890 | rs591485 | -1.324 | -1.028 | -1.925 | -1.109 | -0.003 | -0.004 | -0.004 | 0.022 | 0.82 | 0.81 | 0.78 | 0.64 | 0.85 | 0.75 |

| | | | | | | | | | | | | | | | | |
|---|---|---|---|---|---|---|---|---|---|---|---|---|---|---|---|---|
| 10 | 79083609 | rs12359379 | 1.876 | -0.48 | -2.164 | -1.276 | -0.024 | 0.046 | 0.046 | 0 | 0.1 | 0.03 | 0.09 | 0 | 0.15 | 0.05 |
| 10 | 79091222 | rs586182 | -1.852 | -0.832 | -2.059 | -1.149 | -0.02 | -0.033 | -0.033 | 0 | 0.85 | 0.94 | 0.79 | 0.64 | 0.84 | 0.75 |
| 10 | 79123321 | rs250710 | 2.075 | -0.451 | -2.094 | -1.199 | -0.085 | 0.058 | 0.058 | 0 | 0.12 | 0.01 | 0.23 | 0.01 | 0.29 | 0.12 |
| 10 | 79127539 | rs250712 | -1.51 | -0.475 | -2.088 | -1.168 | -0.018 | -0.037 | -0.037 | 0 | 0.79 | 0.91 | 0.77 | 0.65 | 0.8 | 0.73 |
| 10 | 79129757 | rs11002139 | 1.455 | -0.652 | -2.182 | -1.153 | -0.097 | -0.013 | -0.013 | 0 | 0.2 | 0.03 | 0.14 | 0.16 | 0.27 | 0.1 |
| 10 | 79134478 | rs994076 | 0.935 | -0.93 | -1.993 | -1.213 | -0.069 | 0.104 | 0.104 | 0 | 0.22 | 0.43 | 0.25 | 0.34 | 0.15 | 0.24 |
| 10 | 79137838 | rs10824534 | 1.257 | -0.587 | -2.105 | -1.219 | -0.116 | -0.008 | -0.008 | 0 | 0.21 | 0.03 | 0.16 | 0.16 | 0.32 | 0.1 |
| 10 | 79149391 | rs35783 | -0.416 | -0.859 | -1.862 | -1.167 | -0.005 | -0.008 | -0.008 | 0 | 0.57 | 0.63 | 0.53 | 0.3 | 0.64 | 0.49 |
| 10 | 79150700 | rs628948 | 0.757 | -0.75 | -1.831 | -1.173 | 0.009 | -0.039 | -0.039 | 0 | 0.29 | 0.37 | 0.36 | 0.15 | 0.37 | 0.34 |
| 10 | 79152544 | rs12221259 | 2.085 | -0.915 | -1.833 | -0.431 | 0.006 | -0.013 | -0.013 | 0 | 0.07 | 0.03 | 0.08 | 0.2 | 0.07 | 0.17 |
| 10 | 79154151 | rs250704 | -1.283 | -0.995 | -1.928 | -0.442 | -0.025 | -0.042 | -0.042 | 0.258 | 0.78 | 0.93 | 0.73 | 0.63 | 0.79 | 0.72 |
| 10 | 79154537 | rs207675 | 0.143 | -1.223 | -1.908 | -0.995 | 0.068 | 0.048 | 0.048 | 0 | 0.52 | 0.26 | 0.51 | 0.31 | 0.64 | 0.33 |
| 10 | 79158557 | rs10824540 | 1.242 | -0.95 | -1.421 | -0.797 | -0.052 | -0.032 | -0.032 | 0 | 0.17 | 0.3 | 0.1 | 0.11 | 0.08 | 0.24 |
| 10 | 79162342 | rs42455 | 0.073 | -0.981 | -1.712 | -1.029 | 0.017 | 0.101 | 0.101 | 0 | 0.48 | 0.29 | 0.52 | 0.18 | 0.58 | 0.36 |
| 10 | 79164604 | rs593572 | 0.183 | -1.586 | -1.265 | -0.728 | 0.034 | 0.044 | 0.044 | 0 | 0.48 | 0.4 | 0.61 | 0.73 | 0.62 | 0.52 |
| 10 | 79164857 | rs673195 | -0.092 | -0.782 | -1.414 | -0.336 | 0.001 | 0 | 0 | 0 | 0.64 | 0.64 | 0.7 | 0.84 | 0.7 | 0.75 |
| 10 | 79165830 | rs35806 | 0.162 | -0.644 | -1.384 | -0.185 | 0.012 | 0.021 | 0.021 | 0 | 0.55 | 0.63 | 0.46 | 0.33 | 0.6 | 0.48 |
| 10 | 79166345 | rs670898 | -0.151 | -0.929 | -1.099 | -0.297 | 0.022 | 0.021 | 0.021 | 0 | 0.7 | 0.81 | 0.79 | 0.82 | 0.78 | 0.74 |
| 10 | 79167461 | rs686374 | -0.18 | -1.168 | -0.806 | -0.695 | 0.006 | -0.001 | -0.001 | 0.005 | 0.51 | 0.55 | 0.78 | 0.85 | 0.59 | 0.68 |
| 10 | 79167529 | rs7097101 | 0.718 | -1.48 | -0.8 | -0.939 | 0.001 | 0 | 0 | 0 | 0.2 | 0.18 | 0.07 | 0.11 | 0.12 | 0.15 |
| 10 | 79170713 | rs3844118 | -0.474 | -1.839 | -0.517 | -0.663 | -0.067 | 0.029 | 0.029 | 2.633 | 0.23 | 0.58 | 0.3 | 0.29 | 0.1 | 0.19 |
| 10 | 79172944 | rs3844120 | -0.694 | -1.921 | -0.504 | -0.724 | 0.036 | 0.006 | 0.006 | 0.31 | 0.43 | 0.66 | 0.38 | 0.4 | 0.22 | 0.33 |
| 10 | 79173540 | rs12219105 | 0.605 | -1.288 | -0.415 | -0.491 | 0.012 | 0.014 | 0.014 | 0 | 0.23 | 0.18 | 0.09 | 0.11 | 0.13 | 0.15 |
| 10 | 79173697 | rs176286 | 0.078 | -0.112 | -0.421 | -0.098 | -0.031 | -0.035 | -0.035 | 0 | 0.35 | 0.24 | 0.51 | 0.48 | 0.52 | 0.54 |
| 10 | 79175923 | rs287184 | -0.049 | -0.172 | -0.392 | -0.055 | -0.004 | -0.006 | -0.006 | 2.178 | 0.37 | 0.31 | 0.64 | 0.46 | 0.55 | 0.55 |
| 10 | 79176594 | rs10740467 | -0.412 | -0.548 | -0.451 | -1.124 | 0.001 | -0.038 | -0.038 | 0 | 0.47 | 0.45 | 0.59 | 0.08 | 0.45 | 0.24 |
| 10 | 79180099 | rs7898846 | 0.47 | -0.839 | -0.311 | -0.609 | -0.001 | -0.01 | -0.01 | 0.836 | 0.16 | 0.16 | 0.22 | 0.06 | 0.09 | 0.07 |
| 10 | 79181254 | rs35795 | -0.17 | -0.668 | -0.06 | -0.741 | -0.012 | -0.097 | -0.097 | 0.095 | 0.38 | 0.55 | 0.43 | 0.08 | 0.31 | 0.25 |
| 10 | 79181324 | rs35794 | 1.085 | -0.577 | 0.097 | 0.587 | -0.003 | 0.002 | 0.002 | 0 | 0.8 | 0.8 | 0.81 | 0.78 | 0.84 | 0.66 |
| 10 | 79187957 | rs250708 | -0.995 | -0.424 | -0.126 | 0.927 | -0.01 | 0.031 | 0.031 | 0 | 0.14 | 0.2 | 0.15 | 0.21 | 0.09 | 0.29 |
| 10 | 79194348 | rs582449 | 1.177 | -0.545 | -0.134 | 0.953 | -0.012 | 0.006 | 0.006 | 0 | 0.81 | 0.73 | 0.87 | 0.78 | 0.86 | 0.81 |
| 10 | 79195768 | rs7071575 | -1.153 | -0.203 | 0.022 | 1.203 | -0.001 | 0 | 0 | 0 | 0.16 | 0.14 | 0.08 | 0.15 | 0.11 | 0.18 |
| 10 | 79199564 | rs3910606 | -0.539 | -0.434 | 0.5 | 1.104 | 0.01 | 0.018 | 0.018 | 0 | 0.11 | 0.14 | 0.08 | 0.15 | 0.11 | 0.17 |
| 10 | 79200117 | rs10762759 | -1.964 | -0.902 | 0.81 | 1.194 | -0.049 | 0.034 | 0.034 | 0.78 | 0.27 | 0.55 | 0.15 | 0.29 | 0.16 | 0.27 |
| 10 | 79201239 | rs12411508 | -1.003 | -0.951 | 0.849 | 1.183 | -0.021 | 0.031 | 0.031 | 0.367 | 0.08 | 0.18 | 0.04 | 0.14 | 0.04 | 0.09 |
| 10 | 79203228 | rs7922645 | -1.946 | -0.963 | 0.758 | 1.174 | -0.057 | 0.043 | 0.043 | 0.286 | 0.26 | 0.53 | 0.13 | 0.29 | 0.15 | 0.26 |
| 10 | 79204191 | rs7895934 | -1.106 | -0.754 | 0.924 | 1.112 | -0.025 | 0.051 | 0.051 | 0.027 | 0.07 | 0.18 | 0.04 | 0.13 | 0.04 | 0.09 |
| 10 | 79208056 | rs16934799 | 0.2 | 0 | -0.316 | 0.831 | -0.023 | -0.03 | -0.03 | 0 | 0.08 | 0.01 | 0.05 | 0.16 | 0.07 | 0.06 |
| 10 | 79209190 | rs11813909 | 0.147 | -0.521 | -0.191 | 0.565 | -0.022 | -0.021 | -0.021 | 0 | 0.09 | 0.12 | 0.15 | 0.04 | 0.03 | 0.04 |
| 10 | 79212639 | rs16934809 | 0.165 | 0.145 | -0.488 | 0.673 | -0.039 | -0.028 | -0.028 | 0 | 0.08 | 0.01 | 0.09 | 0.16 | 0.1 | 0.07 |
| 10 | 79214413 | rs106242 | 0.252 | 0.439 | -0.831 | 0.442 | -0.018 | 0.037 | 0.037 | 0 | 0.41 | 0.27 | 0.46 | 0.3 | 0.54 | 0.42 |
| 10 | 79215147 | rs193376 | 0.008 | 0.438 | -0.457 | -0.549 | -0.006 | -0.005 | -0.005 | 0 | 0.45 | 0.4 | 0.31 | 0.6 | 0.42 | 0.64 |
| 10 | 79216001 | rs697171 | 0.15 | 0.229 | -0.278 | -0.801 | -0.007 | -0.013 | -0.013 | 0.164 | 0.19 | 0.23 | 0.2 | 0.1 | 0.14 | 0.07 |
| 10 | 79216423 | rs703261 | 0.129 | 0.201 | -0.261 | -0.803 | -0.008 | -0.014 | -0.014 | 0 | 0.19 | 0.24 | 0.21 | 0.1 | 0.14 | 0.07 |
| 10 | 79222539 | rs16934828 | 0.809 | 0.227 | -0.147 | -0.705 | -0.024 | 0.004 | 0.004 | 0 | 0.07 | 0.18 | 0.01 | 0.06 | 0.01 | 0.02 |
| 10 | 79225270 | rs607483 | -0.511 | 0.58 | -0.447 | -0.908 | -0.045 | -0.009 | -0.009 | 0 | 0.81 | 0.92 | 0.69 | 0.86 | 0.77 | 0.8 |
| 10 | 79227405 | rs12775139 | 0.609 | 0.475 | -0.242 | -1.124 | -0.003 | -0.009 | -0.009 | 0 | 0.26 | 0.28 | 0.18 | 0.21 | 0.21 | 0.12 |
| 10 | 79228620 | rs7924171 | 0.842 | 0.47 | -0.173 | -1.15 | -0.002 | -0.023 | -0.023 | 0.097 | 0.25 | 0.26 | 0.18 | 0.14 | 0.21 | 0.1 |
| 10 | 79229608 | rs696211 | 0.225 | 0.385 | 0.071 | -1.278 | -0.004 | -0.029 | -0.029 | 0.436 | 0.35 | 0.44 | 0.36 | 0.27 | 0.32 | 0.2 |
| 10 | 79232531 | rs43204 | 0.414 | 0.625 | 0.297 | -1.042 | -0.015 | -0.009 | -0.009 | 8.846 | 0.16 | 0.32 | 0.19 | 0.12 | 0.11 | 0.11 |
| 10 | 79235196 | rs42311 | 0.107 | 0.737 | 0.174 | -1.15 | 0.003 | -0.012 | -0.012 | 0 | 0.69 | 0.59 | 0.51 | 0.83 | 0.65 | 0.71 |
| 10 | 79238692 | rs612732 | -0.816 | 0.708 | -0.247 | -1.093 | -0.014 | 0.007 | 0.007 | 0 | 0.84 | 0.87 | 0.69 | 0.9 | 0.76 | 0.8 |
| 10 | 79238789 | rs669540 | -0.798 | 0.709 | -0.246 | -1.271 | -0.014 | 0.003 | 0.003 | 0 | 0.83 | 0.87 | 0.69 | 0.89 | 0.76 | 0.8 |
| 10 | 79239107 | rs17412609 | 1.482 | 0.509 | -0.553 | -1.449 | -0.017 | 0.066 | 0.066 | 0 | 0.25 | 0.03 | 0.17 | 0.15 | 0.2 | 0.1 |
| 10 | 79239338 | rs682275 | -0.442 | 0.568 | -0.36 | -1.452 | 0 | -0.002 | -0.002 | 0.043 | 0.54 | 0.52 | 0.5 | 0.68 | 0.56 | 0.64 |
| 10 | 79242092 | rs1866587 | 0.29 | -0.294 | -0.017 | -1.531 | 0 | 0.002 | 0.002 | 0 | 0.22 | 0.2 | 0.2 | 0.55 | 0.21 | 0.43 |
| 10 | 79242574 | rs16934883 | 1.02 | -0.181 | 0.117 | -1.268 | -0.012 | -0.016 | -0.016 | 0.015 | 0.05 | 0.07 | 0.01 | 0 | 0.01 | 0.01 |
| 10 | 79247942 | rs11002179 | 0.218 | -0.2 | 0.17 | -1.394 | 0.014 | -0.018 | -0.018 | 0 | 0.31 | 0.23 | 0.24 | 0.54 | 0.22 | 0.36 |
| 10 | 79249522 | rs1949352 | -0.115 | -0.225 | 0.328 | -1.427 | 0.065 | 0.054 | 0.054 | 1.37 | 0.08 | 0.48 | 0.16 | 0.05 | 0.08 | 0.04 |
| 10 | 79253261 | rs3127447 | -1.133 | -0.249 | 0.188 | -1.552 | -0.083 | 0.052 | 0.052 | 0.71 | 0.68 | 0.83 | 0.57 | 0.8 | 0.52 | 0.58 |
| 10 | 79253986 | rs816852 | -0.563 | -0.398 | 0.212 | -1.547 | 0.011 | -0.024 | -0.024 | 0.055 | 0.62 | 0.55 | 0.54 | 0.8 | 0.49 | 0.57 |
| 10 | 79262018 | rs2119058 | 1.024 | -0.454 | 0.239 | -1.5 | 0.009 | 0.002 | 0.002 | 0.159 | 0.06 | 0.28 | 0.03 | 0 | 0.01 | 0.01 |
| 10 | 79269731 | rs3127449 | -0.687 | -0.48 | 0.161 | -1.531 | 0.064 | 0.065 | 0.065 | 0.11 | 0.43 | 0.77 | 0.41 | 0.66 | 0.32 | 0.46 |
| 10 | 79278832 | rs752372 | -1.211 | -0.43 | 0.028 | -1.479 | -0.056 | 0.034 | 0.034 | 0.621 | 0.68 | 0.78 | 0.55 | 0.8 | 0.51 | 0.58 |
| 10 | 79282798 | rs4979889 | 0.035 | -0.455 | 0.173 | -1.273 | -0.03 | 0.216 | 0.216 | 1.124 | 0.16 | 0.51 | 0.18 | 0.53 | 0.1 | 0.32 |
| 10 | 79298998 | rs11002200 | 0.522 | -0.347 | -0.162 | -1.235 | -0.01 | 0.168 | 0.168 | 0.322 | 0.13 | 0.27 | 0.15 | 0.54 | 0.1 | 0.34 |
| 10 | 79313729 | rs816866 | -0.209 | -0.493 | -0.23 | -1.378 | -0.05 | -0.077 | -0.077 | 0 | 0.3 | 0.49 | 0.25 | 0.14 | 0.22 | 0.13 |
| 10 | 79314590 | rs17413503 | 1.279 | -0.482 | -0.285 | -1.443 | 0.013 | 0.055 | 0.055 | 0 | 0.26 | 0.12 | 0.16 | 0.14 | 0.19 | 0.13 |
| 10 | 79318685 | rs3851040 | 0.019 | -0.446 | -0.302 | -1.32 | -0.007 | -0.022 | -0.022 | 0.949 | 0.27 | 0.28 | 0.12 | 0.08 | 0.15 | 0.06 |

| | | | | | | | | | | | | | | | | |
|---|---|---|---|---|---|---|---|---|---|---|---|---|---|---|---|---|
| 10 | 79320615 | rs2673462 | -0.01 | -0.427 | -0.434 | -1.31 | -0.032 | -0.052 | -0.052 | 0.062 | 0.7 | 0.57 | 0.76 | 0.86 | 0.78 | 0.87 |
| 10 | 79334199 | rs2619607 | -0.148 | -0.3 | -0.336 | -1.28 | -0.044 | -0.093 | -0.093 | 1.079 | 0.72 | 0.52 | 0.76 | 0.91 | 0.78 | 0.89 |
| 10 | 79338239 | rs2673417 | 1.228 | -0.001 | -0.551 | -1.368 | -0.035 | 0.031 | 0.031 | 0.787 | 0.27 | 0.2 | 0.23 | 0.15 | 0.38 | 0.33 |
| 10 | 79340401 | rs2673418 | 0.53 | -0.172 | 0.022 | -0.63 | -0.007 | -0.03 | -0.03 | 0 | 0.68 | 0.62 | 0.55 | 0.88 | 0.69 | 0.79 |
| 10 | 79348735 | rs11002212 | -0.675 | 0.347 | -0.91 | -0.372 | -0.028 | 0.107 | 0.107 | 1.254 | 0.25 | 0.02 | 0.32 | 0.07 | 0.25 | 0.12 |
| 10 | 79352035 | rs1468037 | -0.097 | -0.202 | -0.54 | -0.023 | -0.008 | 0.205 | 0.205 | 0.473 | 0.12 | 0.27 | 0.14 | 0.52 | 0.09 | 0.31 |
| 10 | 79358657 | rs2619634 | -0.437 | -0.012 | -1.964 | -0.826 | -0.006 | 0.105 | 0.105 | 2.306 | 0.19 | 0.01 | 0.09 | 0.02 | 0.14 | 0.03 |
| 10 | 79363388 | rs2673436 | -0.401 | 0.252 | -2.085 | -0.985 | -0.044 | 0.147 | 0.147 | 0.086 | 0.55 | 0.87 | 0.62 | 0.2 | 0.51 | 0.33 |
| 10 | 79363798 | rs1473966 | -0.134 | -0.551 | -1.206 | -1.202 | -0.001 | 0.025 | 0.025 | 0 | 0.45 | 0.63 | 0.55 | 0.19 | 0.5 | 0.32 |
| 10 | 79364564 | rs11598926 | 0.267 | -0.727 | -0.602 | -1.49 | -0.001 | -0.004 | -0.004 | 0 | 0.3 | 0.34 | 0.3 | 0.13 | 0.27 | 0.22 |
| 10 | 79370738 | rs1968013 | 0.369 | -0.858 | -0.497 | -1.552 | -0.002 | 0.05 | 0.05 | 0 | 0.28 | 0.14 | 0.28 | 0.13 | 0.27 | 0.23 |
| 10 | 79371139 | rs2720010 | 0.413 | -0.794 | -0.293 | -1.63 | 0.038 | -0.047 | -0.047 | 2.015 | 0.14 | 0.28 | 0.24 | 0.06 | 0.23 | 0.1 |
| 10 | 79373447 | rs1877993 | -1.011 | -0.917 | -0.147 | -1.668 | 0.007 | 0.044 | 0.044 | 0 | 0.5 | 0.59 | 0.55 | 0.21 | 0.51 | 0.33 |
| 10 | 79374755 | rs2670149 | -1.106 | -1.194 | -0.102 | -1.825 | -0.03 | -0.098 | -0.098 | 0 | 0.28 | 0.48 | 0.29 | 0.09 | 0.25 | 0.11 |
| 10 | 79377940 | rs2673488 | -1.469 | -1.178 | -0.395 | -1.817 | -0.044 | 0.141 | 0.141 | 0.053 | 0.56 | 0.82 | 0.6 | 0.21 | 0.53 | 0.33 |
| 10 | 79378558 | rs1878001 | -0.517 | -1.133 | -0.415 | -1.794 | -0.026 | -0.063 | -0.063 | 0 | 0.38 | 0.48 | 0.35 | 0.14 | 0.3 | 0.23 |
| 10 | 79379937 | rs2280163 | 0.723 | -1.107 | -0.539 | -1.783 | -0.012 | 0.047 | 0.047 | 0 | 0.26 | 0.13 | 0.28 | 0.12 | 0.27 | 0.22 |
| 10 | 79384253 | rs1476318 | 0.292 | -0.845 | -0.453 | -1.612 | 0.01 | -0.046 | -0.046 | 0.264 | 0.16 | 0.24 | 0.24 | 0.06 | 0.23 | 0.1 |
| 10 | 79387112 | rs734949 | -0.821 | -0.515 | -1.201 | -1.297 | -0.018 | 0.129 | 0.129 | 0 | 0.55 | 0.77 | 0.59 | 0.21 | 0.52 | 0.33 |
| 10 | 79393332 | rs2619617 | -0.332 | -0.177 | -1.417 | -1.351 | -0.013 | 0.104 | 0.104 | 0 | 0.55 | 0.73 | 0.58 | 0.21 | 0.58 | 0.4 |
| 10 | 79399370 | rs2673485 | 0.044 | -0.609 | -1.493 | -1.537 | -0.012 | 0.132 | 0.132 | 0 | 0.48 | 0.74 | 0.55 | 0.18 | 0.56 | 0.37 |
| 10 | 79399946 | rs4979902 | 1.168 | -0.645 | -2.016 | -1.341 | -0.01 | -0.08 | -0.08 | 0.419 | 0.18 | 0.1 | 0.17 | 0.53 | 0.16 | 0.35 |
| 10 | 79401316 | rs7071206 | 0.43 | -0.676 | -2.027 | -1.357 | 0.008 | -0.08 | -0.08 | 0.334 | 0.23 | 0.13 | 0.19 | 0.53 | 0.18 | 0.36 |
| 10 | 79405063 | rs2077236 | -0.745 | -0.715 | -2.037 | -1.378 | -0.027 | 0.002 | 0.002 | 0 | 0.75 | 0.91 | 0.76 | 0.75 | 0.74 | 0.74 |
| 10 | 79406780 | rs2244688 | -0.771 | -0.716 | -2.037 | -1.378 | -0.027 | 0.001 | 0.001 | 0 | 0.75 | 0.91 | 0.76 | 0.75 | 0.74 | 0.74 |
| 10 | 79422369 | rs10509391 | 1.107 | -0.667 | -2.017 | -1.359 | -0.007 | -0.083 | -0.083 | 0 | 0.19 | 0.1 | 0.17 | 0.54 | 0.15 | 0.35 |
| 10 | 79425310 | rs2670159 | 0.467 | -0.688 | -1.787 | -1.353 | -0.002 | 0.098 | 0.098 | 0 | 0.48 | 0.67 | 0.55 | 0.18 | 0.55 | 0.37 |
| 10 | 79432167 | rs2619647 | -0.238 | -0.554 | -1.979 | -1.661 | 0.01 | 0.046 | 0.046 | 0 | 0.48 | 0.79 | 0.5 | 0.61 | 0.46 | 0.51 |
| 10 | 79432480 | rs2673467 | -0.851 | -0.623 | -2.173 | -1.766 | -0.003 | -0.013 | -0.013 | 0 | 0.78 | 0.89 | 0.77 | 0.71 | 0.79 | 0.79 |
| 10 | 79433801 | rs17189440 | 1.503 | -0.603 | -2.129 | -1.695 | -0.082 | 0.041 | 0.041 | 0 | 0.23 | 0.05 | 0.25 | 0.11 | 0.3 | 0.26 |
| 10 | 79439753 | rs2619648 | 0.305 | -0.946 | -1.439 | -1.197 | -0.007 | -0.008 | -0.008 | 0 | 0.68 | 0.6 | 0.72 | 0.71 | 0.74 | 0.73 |
| 10 | 79442104 | rs2619649 | 0.708 | -1.113 | -1.884 | -1.069 | -0.096 | -0.029 | -0.029 | 0.593 | 0.19 | 0.02 | 0.19 | 0.2 | 0.25 | 0.22 |
| 10 | 79443520 | rs12254219 | -0.398 | -1.018 | -1.311 | -0.924 | -0.003 | 0.008 | 0.008 | 0 | 0.25 | 0.3 | 0.2 | 0.55 | 0.16 | 0.31 |
| 10 | 79449665 | rs7098023 | -0.816 | -0.761 | -0.942 | -0.822 | -0.043 | 0.074 | 0.074 | 0 | 0.27 | 0.48 | 0.23 | 0.52 | 0.18 | 0.33 |
| 10 | 79459524 | rs7086586 | -0.046 | -0.36 | -1.459 | -0.615 | -0.013 | 0.082 | 0.082 | 0 | 0.19 | 0.35 | 0.21 | 0.52 | 0.14 | 0.3 |
| 10 | 79462793 | rs2048787 | 0.609 | -0.26 | -1.745 | -0.55 | 0.007 | -0.036 | -0.036 | 1.327 | 0.16 | 0.1 | 0.18 | 0.51 | 0.13 | 0.29 |
| 10 | 79464056 | rs2619646 | -0.498 | -0.184 | -1.886 | -0.605 | -0.042 | 0.001 | 0.001 | 0 | 0.71 | 0.92 | 0.74 | 0.73 | 0.7 | 0.69 |
| 10 | 79464078 | rs1609790 | -0.746 | -0.248 | -1.032 | 0.241 | -0.008 | 0.031 | 0.031 | 0 | 0.22 | 0.27 | 0.2 | 0.52 | 0.15 | 0.32 |
| 10 | 79468611 | rs2619645 | -0.905 | -0.037 | -1.425 | 0.417 | -0.015 | -0.001 | -0.001 | 0 | 0.14 | 0.02 | 0.08 | 0.07 | 0.13 | 0.04 |
| 10 | 79469070 | rs6480863 | -1.668 | -0.512 | -1.667 | 0.059 | 0.043 | 0.063 | 0.063 | 2.62 | 0.33 | 0.85 | 0.2 | 0.18 | 0.24 | 0.12 |
| 10 | 79471313 | rs2720008 | 0.473 | -0.169 | -2.162 | 0.604 | -0.001 | -0.05 | -0.05 | 0.006 | 0.22 | 0.08 | 0.16 | 0.29 | 0.21 | 0.15 |
| 10 | 79489015 | rs11002265 | -0.022 | -0.201 | -1.638 | 0.479 | -0.006 | -0.006 | -0.006 | 0 | 0.07 | 0.06 | 0.07 | 0.12 | 0.09 | 0.12 |
| 10 | 79492970 | rs12761718 | -0.555 | -0.216 | -1.173 | 0.297 | -0.002 | -0.015 | -0.015 | 0.271 | 0.13 | 0.15 | 0.07 | 0.05 | 0.14 | 0.05 |
| 10 | 79524387 | rs873740 | -0.69 | -0.22 | -0.503 | 0.518 | -0.029 | -0.03 | -0.03 | 0.035 | 0.19 | 0.52 | 0.11 | 0.12 | 0.11 | 0.1 |
| 10 | 79525505 | rs11002284 | -0.267 | 0.817 | -1.144 | 0.949 | -0.053 | -0.055 | -0.055 | 0 | 0.23 | 0.09 | 0.18 | 0.33 | 0.28 | 0.15 |
| 10 | 79529807 | rs4979920 | -1.12 | 0.12 | -0.003 | 1.222 | -0.023 | 0.09 | 0.09 | 0 | 0.08 | 0.28 | 0.06 | 0.17 | 0.03 | 0.05 |
| 10 | 79530213 | rs2579174 | 0.145 | -0.362 | -0.568 | 1.084 | 0.032 | -0.036 | -0.036 | 0 | 0.48 | 0.13 | 0.69 | 0.47 | 0.62 | 0.59 |
| 10 | 79531931 | rs10824571 | 0.531 | 0.067 | -1.173 | 1.114 | -0.038 | 0.004 | 0.004 | 0 | 0.25 | 0.06 | 0.23 | 0.17 | 0.34 | 0.31 |
| 10 | 79534527 | rs1344966 | 0.441 | -0.141 | -0.203 | 1.024 | -0.019 | 0.009 | 0.009 | 0 | 0.26 | 0.12 | 0.23 | 0.17 | 0.34 | 0.31 |
| 10 | 79538284 | rs2579164 | 0.446 | -0.168 | -0.174 | 1.04 | -0.046 | -0.033 | -0.033 | 0.863 | 0.23 | 0.09 | 0.47 | 0.29 | 0.28 | 0.28 |
| 10 | 79538734 | rs17516020 | 1.211 | -0.152 | -0.337 | 0.92 | -0.069 | 0.097 | 0.097 | 0 | 0.17 | 0.01 | 0.14 | 0 | 0.25 | 0.15 |
| 10 | 79540127 | rs2812440 | 0.495 | -0.144 | -0.192 | 1.017 | -0.045 | -0.032 | -0.032 | 0.076 | 0.23 | 0.09 | 0.47 | 0.29 | 0.28 | 0.28 |
| 10 | 79549242 | rs2812431 | 0.617 | -0.252 | 0.154 | 0.97 | -0.058 | -0.044 | -0.044 | 5.661 | 0.22 | 0.05 | 0.46 | 0.29 | 0.28 | 0.28 |
| 10 | 79550544 | rs2812430 | 0.452 | -0.25 | 0.157 | 0.971 | -0.047 | -0.034 | -0.034 | 0.615 | 0.23 | 0.09 | 0.47 | 0.29 | 0.28 | 0.28 |
| 10 | 79551640 | rs2165047 | 0.426 | -0.253 | 0.319 | 0.969 | -0.029 | -0.019 | -0.019 | 0 | 0.23 | 0.14 | 0.48 | 0.29 | 0.28 | 0.28 |
| 10 | 79552092 | rs2165046 | -0.372 | -0.453 | 0.878 | 1.114 | -0.001 | -0.027 | -0.027 | 0 | 0.33 | 0.65 | 0.29 | 0.44 | 0.37 | 0.39 |
| 10 | 79552107 | rs1058203 | 0.578 | -0.451 | 0.856 | 1.115 | -0.029 | -0.019 | -0.019 | 0 | 0.23 | 0.14 | 0.48 | 0.29 | 0.28 | 0.28 |
| 10 | 79552151 | rs1058202 | -1.477 | -0.545 | 0.876 | 1.074 | -0.025 | 0.048 | 0.048 | 0 | 0.62 | 0.84 | 0.78 | 0.73 | 0.66 | 0.67 |
| 10 | 79552677 | rs2812429 | 0.607 | -0.545 | 0.968 | 1.074 | -0.029 | -0.019 | -0.019 | 0.162 | 0.23 | 0.14 | 0.48 | 0.29 | 0.28 | 0.28 |
| 10 | 79553236 | rs2812427 | -0.373 | -0.536 | 1.125 | 1.087 | -0.003 | 0.029 | 0.029 | 0 | 0.32 | 0.45 | 0.53 | 0.46 | 0.32 | 0.33 |
| 10 | 79553673 | rs2812425 | -1.553 | -0.537 | 1.095 | 1.058 | -0.049 | 0.034 | 0.034 | 0 | 0.63 | 0.9 | 0.79 | 0.73 | 0.66 | 0.67 |
| 10 | 79566632 | rs1058198 | 0.094 | -0.598 | 1.17 | 1.087 | 0.001 | -0.004 | -0.004 | 0 | 0.28 | 0.25 | 0.24 | 0.17 | 0.34 | 0.31 |
| 10 | 79570873 | rs2289310 | -0.411 | -0.706 | 0.807 | 1.066 | 0.019 | -0.02 | -0.02 | 0 | 0.08 | 0.04 | 0.04 | 0.17 | 0.03 | 0.05 |
| 10 | 79571108 | rs2289309 | 0.415 | -0.697 | 0.861 | 0.975 | 0.005 | -0.004 | -0.004 | 0 | 0.26 | 0.24 | 0.24 | 0.17 | 0.34 | 0.31 |
| 10 | 79578292 | rs1268514 | -1.616 | -0.712 | 0.466 | 0.901 | -0.071 | 0.019 | 0.019 | 0 | 0.64 | 0.89 | 0.79 | 0.73 | 0.66 | 0.67 |
| 10 | 79579047 | rs1248635 | -1.239 | -0.709 | 0.519 | 0.902 | -0.004 | 0.061 | 0.061 | 0 | 0.59 | 0.78 | 0.78 | 0.73 | 0.65 | 0.66 |
| 10 | 79593051 | rs1261993 | -1.629 | -0.718 | 0.456 | 0.906 | -0.054 | 0.03 | 0.03 | 0.001 | 0.65 | 0.88 | 0.79 | 0.73 | 0.66 | 0.66 |
| 10 | 79594931 | rs11002309 | 0.495 | -0.68 | 0.642 | 0.931 | -0.026 | 0.01 | 0.01 | 0 | 0.26 | 0.12 | 0.22 | 0.17 | 0.33 | 0.31 |
| 10 | 79595473 | rs1781797 | 0.624 | -0.587 | 0.796 | 0.994 | -0.018 | -0.013 | -0.013 | 1.631 | 0.22 | 0.12 | 0.47 | 0.29 | 0.27 | 0.27 |

| | | | | | | | | | | | | | | | | |
|---|---|---|---|---|---|---|---|---|---|---|---|---|---|---|---|---|
| 10 | 79601822 | rs16935401 | -1.017 | -0.648 | 1.213 | 0.91 | 0.01 | 0.007 | 0.007 | 1.777 | 0.06 | 0.33 | 0.03 | 0 | 0.01 | 0.03 |
| 10 | 79603688 | rs10824582 | 0.333 | -0.689 | 0.774 | 0.921 | -0.026 | 0.011 | 0.011 | 0 | 0.27 | 0.09 | 0.23 | 0.17 | 0.34 | 0.32 |
| 10 | 79603826 | rs1248671 | -1.587 | -0.786 | 0.67 | 1 | -0.057 | 0.053 | 0.053 | 0 | 0.65 | 0.88 | 0.8 | 0.74 | 0.66 | 0.67 |
| 10 | 79604958 | rs1248673 | -0.22 | -0.755 | 0.725 | 1.063 | -0.003 | -0.002 | -0.002 | 0 | 0.3 | 0.65 | 0.51 | 0.3 | 0.28 | 0.3 |
| 10 | 79607372 | rs1248678 | 0.59 | -0.617 | 0.526 | 1.123 | -0.015 | -0.022 | -0.022 | 0 | 0.23 | 0.13 | 0.47 | 0.3 | 0.27 | 0.27 |
| 10 | 79609548 | rs7080321 | -0.078 | -0.544 | 0.387 | 1.135 | 0.003 | 0.014 | 0.014 | 0 | 0.07 | 0.04 | 0.04 | 0.27 | 0.03 | 0.05 |
| 10 | 79649473 | rs7903506 | 1.129 | -0.07 | -0.104 | 0.803 | -0.009 | 0.031 | 0.031 | 0 | 0.11 | 0.02 | 0.17 | 0.04 | 0.11 | 0.13 |
| 10 | 79649653 | rs11002322 | 0.314 | -0.15 | -0.076 | 0.844 | -0.027 | 0.009 | 0.009 | 0.039 | 0.27 | 0.11 | 0.23 | 0.17 | 0.34 | 0.33 |
| 10 | 79652154 | rs1262002 | -1.148 | -0.176 | -0.168 | 0.997 | 0.005 | -0.023 | -0.023 | 1.186 | 0.57 | 0.71 | 0.74 | 0.47 | 0.63 | 0.63 |
| 10 | 79660125 | rs10430531 | 0.688 | 0.42 | -0.881 | 1.183 | -0.053 | -0.041 | -0.041 | 0 | 0.19 | 0.02 | 0.17 | 0.16 | 0.27 | 0.28 |
| 10 | 79662213 | rs1272018 | -0.448 | 0.408 | -0.628 | 1.047 | -0.012 | -0.002 | -0.002 | 0.268 | 0.27 | 0.19 | 0.5 | 0.32 | 0.33 | 0.3 |
| 10 | 79704394 | rs1366047 | -0.614 | 0.253 | -0.568 | 1.135 | -0.006 | 0.064 | 0.064 | 0 | 0.51 | 0.69 | 0.69 | 0.67 | 0.56 | 0.61 |
| 10 | 79713315 | rs1650150 | -1.064 | -0.023 | -0.264 | 1.636 | -0.002 | -0.042 | -0.042 | 0 | 0.11 | 0.03 | 0.14 | 0.26 | 0.09 | 0.2 |
| 10 | 79721317 | rs4979933 | -1.587 | -0.476 | 0.507 | 1.97 | -0.042 | 0.056 | 0.056 | 0.171 | 0.19 | 0.63 | 0.16 | 0.28 | 0.13 | 0.1 |
| 10 | 79722242 | rs2042867 | -1.987 | -0.37 | 0.73 | 1.985 | -0.042 | 0.113 | 0.113 | 0 | 0.26 | 0.47 | 0.28 | 0.55 | 0.21 | 0.29 |
| 10 | 79732721 | rs735638 | -1.61 | 0.025 | 0.763 | 2.187 | -0.002 | 0.089 | 0.089 | 0 | 0.2 | 0.31 | 0.26 | 0.55 | 0.21 | 0.27 |
| 10 | 79735473 | rs4979935 | -1.205 | 0.168 | 0.356 | 1.792 | -0.003 | -0.012 | -0.012 | 0 | 0.15 | 0.11 | 0.14 | 0.29 | 0.18 | 0.12 |
| 10 | 79740262 | rs10824587 | -1.228 | 0.158 | 0.408 | 1.788 | -0.001 | 0 | 0 | 0.27 | 0.15 | 0.12 | 0.14 | 0.3 | 0.18 | 0.12 |
| 10 | 79742617 | rs3781180 | -1.273 | -0.294 | 1.69 | 1.671 | -0.016 | 0.087 | 0.087 | 0 | 0.13 | 0.42 | 0.11 | 0.17 | 0.08 | 0.12 |
| 10 | 79745269 | rs2559658 | -0.62 | -1.031 | 0.412 | 0.545 | 0.013 | 0.175 | 0.175 | 0 | 0.73 | 0.99 | 0.88 | 0.96 | 0.81 | 0.79 |
| 10 | 79745789 | rs3815891 | -1.258 | -0.984 | 1.805 | 1.177 | 0.063 | 0.334 | 0.334 | 0.862 | 0.06 | 0.34 | 0.15 | 0.26 | 0.04 | 0.16 |
| 10 | 79760001 | rs2241551 | -2.304 | -1.186 | 2.267 | 1.089 | -0.014 | 0.186 | 0.186 | 2.419 | 0.09 | 0.6 | 0.17 | 0.27 | 0.04 | 0.16 |
| 10 | 79779150 | rs1863898 | 0.897 | -1.032 | 1.568 | 0.866 | -0.093 | 0.098 | 0.098 | 0 | 0.35 | 0.13 | 0.51 | 0.2 | 0.45 | 0.31 |
| 10 | 79782276 | rs2559825 | -0.546 | -1.508 | 1.569 | 1.019 | 0.02 | 0.038 | 0.038 | 0 | 0.6 | 0.88 | 0.77 | 0.7 | 0.64 | 0.69 |
| 10 | 79795580 | rs16935560 | -1.438 | -1.465 | 1.86 | 1.05 | 0.041 | 0.036 | 0.036 | 2.453 | 0.06 | 0.5 | 0.04 | 0.01 | 0.01 | 0 |
| 10 | 79833352 | rs7913073 | -0.516 | -1.331 | 1.269 | 0.974 | 0.016 | 0.053 | 0.053 | 0 | 0.59 | 0.86 | 0.76 | 0.71 | 0.64 | 0.69 |
| 10 | 79840005 | rs10740469 | 0.418 | -1.391 | 1.505 | 1.04 | 0.064 | 0.006 | 0.006 | 0 | 0.51 | 0.3 | 0.6 | 0.44 | 0.6 | 0.53 |
| 10 | 79844066 | rs4979801 | 0.418 | -1.393 | 1.505 | 1.04 | 0.061 | 0.006 | 0.006 | 0 | 0.51 | 0.29 | 0.6 | 0.44 | 0.59 | 0.53 |
| 10 | 79844239 | rs7100515 | -1.826 | -1.358 | 1.74 | 1.005 | 0.027 | 0.324 | 0.324 | 0.753 | 0.07 | 0.44 | 0.15 | 0.27 | 0.04 | 0.16 |
| 10 | 79871569 | rs2252770 | 0.019 | -1.474 | 1.49 | 0.765 | 0.033 | 0.059 | 0.059 | 0 | 0.43 | 0.3 | 0.24 | 0.24 | 0.35 | 0.31 |
| 10 | 79875209 | rs2256986 | -0.626 | -1.512 | 1.681 | 0.756 | 0.017 | 0.003 | 0.003 | 0 | 0.49 | 0.57 | 0.38 | 0.56 | 0.39 | 0.46 |
| 10 | 79882848 | rs2253184 | -0.603 | -1.495 | 1.701 | 0.741 | 0.009 | 0 | 0 | 0 | 0.49 | 0.62 | 0.38 | 0.55 | 0.39 | 0.46 |
| 10 | 79896196 | rs4979802 | -0.364 | -1.205 | 1.368 | 0.706 | 0.022 | 0.049 | 0.049 | 0 | 0.49 | 0.35 | 0.42 | 0.37 | 0.41 | 0.37 |
| 10 | 79903729 | rs2114907 | -0.828 | -1.121 | 1.306 | 0.75 | -0.001 | 0.022 | 0.022 | 0 | 0.56 | 0.72 | 0.58 | 0.7 | 0.45 | 0.52 |
| 10 | 79904144 | rs7092503 | 0.106 | -1.075 | 0.869 | 0.248 | 0.003 | 0.006 | 0.006 | 0 | 0.06 | 0.12 | 0.19 | 0.13 | 0.05 | 0.05 |
| 10 | 79924544 | rs1249142 | -0.061 | -1.181 | 0.801 | 0.146 | -0.001 | 0.002 | 0.002 | 0 | 0.09 | 0.15 | 0.08 | 0.15 | 0.07 | 0.05 |
| 10 | 79925262 | rs1008019 | -0.061 | -1.181 | 0.801 | 0.146 | -0.001 | 0.002 | 0.002 | 0 | 0.09 | 0.15 | 0.08 | 0.15 | 0.07 | 0.05 |
| 10 | 79929802 | rs1149755 | -0.947 | -1.691 | 0.638 | 0.091 | 0.009 | -0.001 | -0.001 | 1.005 | 0.57 | 0.77 | 0.51 | 0.62 | 0.42 | 0.51 |
| 10 | 79931975 | rs10824592 | 0.019 | -1.754 | 0.836 | 0.098 | -0.007 | -0.024 | -0.024 | 0 | 0.45 | 0.48 | 0.24 | 0.18 | 0.31 | 0.28 |
| 10 | 79931986 | rs1149753 | 0.144 | -1.693 | 0.779 | 0.151 | -0.001 | 0.003 | 0.003 | 0 | 0.09 | 0.15 | 0.08 | 0.15 | 0.07 | 0.05 |
| 10 | 79935837 | rs16935637 | 0.691 | -2.2 | 0.504 | -0.134 | 0.003 | 0.006 | 0.006 | 0 | 0.18 | 0.14 | 0.11 | 0.15 | 0.15 | 0.1 |
| 10 | 79937796 | rs1149750 | 0.2 | -2.226 | 0.479 | -0.11 | -0.001 | 0.002 | 0.002 | 0 | 0.1 | 0.16 | 0.11 | 0.16 | 0.07 | 0.05 |
| 10 | 79937988 | rs10762770 | 0.078 | -2.492 | 0.53 | -0.341 | -0.029 | 0.004 | 0.004 | 0 | 0.31 | 0.35 | 0.27 | 0.32 | 0.2 | 0.34 |
| 10 | 79940471 | rs17522769 | 0.688 | -2.461 | 0.151 | -0.449 | 0.051 | 0.099 | 0.099 | 0 | 0.25 | 0.09 | 0.11 | 0.03 | 0.16 | 0.17 |
| 10 | 79941085 | rs16935649 | 0.751 | -2.328 | 0.211 | -0.431 | -0.002 | -0.004 | -0.004 | 0.047 | 0.16 | 0.16 | 0.13 | 0.14 | 0.15 | 0.09 |
| 10 | 79943595 | rs10824594 | -1.009 | -2.337 | -0.148 | -0.39 | 0.086 | -0.01 | -0.01 | 0.97 | 0.53 | 0.77 | 0.42 | 0.48 | 0.35 | 0.45 |
| 10 | 79944769 | rs765130 | 0.261 | -1.393 | -0.329 | -0.295 | -0.002 | 0.055 | 0.055 | 0 | 0.67 | 0.73 | 0.78 | 0.87 | 0.66 | 0.82 |
| 10 | 79945183 | rs1249136 | -0.988 | -1.208 | -0.285 | -0.199 | -0.014 | 0.043 | 0.043 | 0 | 0.86 | 0.93 | 0.92 | 1 | 0.84 | 0.91 |
| 10 | 79947528 | rs1678632 | 0.714 | -1.188 | -0.169 | -0.112 | 0.004 | -0.003 | -0.003 | 0.316 | 0.58 | 0.39 | 0.61 | 0.56 | 0.61 | 0.62 |
| 10 | 79948848 | rs7089478 | -0.062 | -1.066 | -0.602 | -0.046 | 0.001 | 0.272 | 0.272 | 0 | 0.05 | 0.18 | 0.16 | 0.3 | 0.04 | 0.2 |
| 10 | 79949089 | rs1249123 | 0.406 | -0.982 | -0.772 | -0.388 | 0 | 0.002 | 0.002 | 0 | 0.64 | 0.63 | 0.78 | 0.86 | 0.65 | 0.8 |
| 10 | 79952934 | rs1427827 | 0.468 | -0.61 | -1.014 | -0.272 | -0.008 | 0.001 | 0.001 | 0 | 0.54 | 0.49 | 0.61 | 0.5 | 0.61 | 0.6 |
| 10 | 79955120 | rs1149747 | 0.109 | -0.094 | -1.501 | 0.03 | -0.001 | -0.086 | -0.086 | 0 | 0.22 | 0.08 | 0.27 | 0.5 | 0.23 | 0.3 |
| 10 | 79957503 | rs1720273 | -0.012 | -0.213 | -1.069 | -0.265 | -0.017 | -0.086 | -0.086 | 0 | 0.77 | 0.71 | 0.88 | 1 | 0.83 | 0.92 |
| 10 | 79958794 | rs12217848 | -0.364 | 0.549 | -1.679 | 0.497 | -0.014 | -0.045 | -0.045 | 0 | 0.3 | 0.11 | 0.31 | 0.33 | 0.29 | 0.42 |
| 10 | 79964084 | rs751284 | -0.464 | 0.239 | -1.704 | 0.402 | 0.021 | 0.019 | 0.019 | 0.823 | 0.4 | 0.24 | 0.31 | 0.63 | 0.31 | 0.43 |
| 10 | 79969839 | rs7089561 | -0.483 | 0.441 | -0.622 | 0.135 | 0.058 | 0.002 | 0.002 | 0.013 | 0.06 | 0.29 | 0.05 | 0.01 | 0.06 | 0.07 |
| 10 | 79970100 | rs1249159 | -0.304 | 0.717 | -1.333 | -0.579 | -0.001 | 0.003 | 0.003 | 0 | 0.69 | 0.73 | 0.78 | 0.77 | 0.67 | 0.64 |
| 10 | 79972816 | rs1249187 | 0.748 | 0.225 | -0.092 | -0.887 | -0.021 | 0.03 | 0.03 | 0 | 0.26 | 0.22 | 0.44 | 0.13 | 0.35 | 0.21 |
| 10 | 79973220 | rs7910334 | 0.45 | 0.205 | -0.054 | -0.83 | 0.045 | -0.006 | -0.006 | 0.018 | 0.05 | 0.24 | 0.04 | 0.01 | 0.06 | 0.07 |
| 10 | 79973808 | rs10824596 | -0.465 | -0.219 | -0.326 | -0.062 | -0.019 | 0.104 | 0.104 | 0 | 0.21 | 0.48 | 0.24 | 0.34 | 0.16 | 0.28 |
| 10 | 79975194 | rs11002449 | 1.176 | -0.464 | -0.583 | -0.039 | -0.013 | 0.081 | 0.081 | 0 | 0.22 | 0.07 | 0.11 | 0.07 | 0.2 | 0.16 |
| 10 | 79980507 | rs1249150 | -1.27 | -0.314 | -0.578 | 0.15 | -0.037 | 0.002 | 0.002 | 0.072 | 0.74 | 0.95 | 0.81 | 0.78 | 0.72 | 0.7 |
| 10 | 79984258 | rs11002453 | 0.097 | -0.419 | -0.398 | 0.215 | -0.002 | 0.097 | 0.097 | 0 | 0.17 | 0.32 | 0.23 | 0.35 | 0.15 | 0.27 |
| 10 | 79984355 | rs1720311 | 0.073 | -0.436 | -0.401 | 0.168 | -0.001 | -0.006 | -0.006 | 2.661 | 0.56 | 0.49 | 0.57 | 0.42 | 0.56 | 0.42 |
| 10 | 79984796 | rs2434123 | 0.339 | -0.423 | -0.444 | 0.127 | 0.002 | 0.005 | 0.005 | 0 | 0.53 | 0.45 | 0.57 | 0.19 | 0.56 | 0.38 |
| 10 | 79985673 | rs1149735 | -1.934 | -0.43 | -1.179 | -0.103 | -0.031 | 0.062 | 0.062 | 0.163 | 0.89 | 1 | 0.93 | 1 | 0.86 | 0.91 |
| 10 | 79988956 | rs1249138 | -0.302 | -0.705 | -1.319 | -0.166 | -0.022 | 0.01 | 0.01 | 0 | 0.72 | 0.92 | 0.8 | 0.76 | 0.7 | 0.68 |
| 10 | 79989151 | rs718708 | 1.051 | -0.659 | -1.233 | -0.048 | 0.002 | -0.046 | -0.046 | 0.047 | 0.12 | 0.04 | 0.18 | 0.33 | 0.09 | 0.21 |

| | | | | | | | | | | | | | | | | |
|---|---|---|---|---|---|---|---|---|---|---|---|---|---|---|---|---|
| 10 | 79991063 | rs2461875 | -1.833 | -0.466 | -1.122 | -0.1 | -0.016 | 0.034 | 0.034 | 0.064 | 0.93 | 1 | 0.96 | 1 | 0.92 | 0.93 |
| 10 | 79992965 | rs7097434 | 0.281 | -0.484 | -1.418 | 0.355 | -0.073 | -0.055 | -0.055 | 0 | 0.37 | 0.08 | 0.31 | 0.31 | 0.37 | 0.51 |
| 10 | 79995592 | rs1880051 | 0.428 | -0.542 | -0.92 | 0.373 | 0.001 | -0.002 | -0.002 | 0 | 0.37 | 0.66 | 0.5 | 0.45 | 0.36 | 0.25 |
| 10 | 79997370 | rs1720285 | -0.568 | -0.326 | -1.031 | 0.411 | -0.027 | -0.018 | -0.018 | 0 | 0.76 | 0.95 | 0.83 | 0.76 | 0.74 | 0.72 |
| 10 | 79998769 | rs2448386 | -1.564 | 0.202 | -0.815 | 0.127 | -0.019 | 0.015 | 0.015 | 0 | 0.85 | 0.86 | 0.8 | 0.99 | 0.79 | 0.86 |
| 10 | 80005591 | rs10762774 | 0.481 | 0.261 | -0.826 | 0.079 | -0.031 | -0.003 | -0.003 | 1.795 | 0.28 | 0.15 | 0.27 | 0.26 | 0.39 | 0.33 |
| 10 | 80006747 | rs4575205 | 1.773 | 0.281 | -0.874 | 0.061 | -0.019 | 0.016 | 0.016 | 0 | 0.15 | 0.13 | 0.2 | 0 | 0.22 | 0.14 |
| 10 | 80008668 | rs1880052 | -0.874 | -0.06 | -0.646 | 0.36 | -0.079 | 0.131 | 0.131 | 0.059 | 0.85 | 0.64 | 0.81 | 0.6 | 0.94 | 0.92 |
| 10 | 80012051 | rs11002468 | 0.443 | 0.018 | -0.596 | 0.334 | 0.009 | 0 | 0 | 0.656 | 0.29 | 0.28 | 0.27 | 0.27 | 0.39 | 0.33 |
| 10 | 80016531 | rs11591530 | 1.484 | 0.01 | -0.821 | 0.277 | 0.022 | -0.068 | -0.068 | 0 | 0.17 | 0.3 | 0.2 | 0 | 0.22 | 0.14 |
| 10 | 80017045 | rs4979807 | 0.704 | -0.191 | -1.211 | 0.543 | -0.063 | -0.052 | -0.052 | 0 | 0.38 | 0.04 | 0.32 | 0.34 | 0.35 | 0.45 |
| 10 | 80017446 | rs4979952 | -0.329 | -0.163 | -0.389 | 0.697 | -0.085 | 0.159 | 0.159 | 0.127 | 0.85 | 0.6 | 0.8 | 0.61 | 0.95 | 0.92 |
| 10 | 80019761 | rs1649983 | -0.135 | 0.086 | -0.767 | 0.6 | -0.021 | 0.019 | 0.019 | 0 | 0.83 | 0.7 | 0.77 | 0.73 | 0.86 | 0.9 |
| 10 | 80020907 | rs2140386 | 0.613 | 0.359 | -1.677 | 0.186 | -0.032 | 0.006 | 0.006 | 0 | 0.25 | 0.04 | 0.26 | 0.21 | 0.23 | 0.34 |
| 10 | 80021842 | rs1863833 | 0.619 | 0.352 | -1.041 | 0.144 | 0.054 | 0.041 | 0.041 | 0 | 0.07 | 0.28 | 0.14 | 0.04 | 0.08 | 0.03 |
| 10 | 80021895 | rs1626941 | -1.496 | 0.398 | -1.417 | -0.257 | 0.002 | -0.068 | -0.068 | 0 | 0.76 | 0.75 | 0.8 | 0.98 | 0.76 | 0.81 |
| 10 | 80023759 | rs7921320 | 0.036 | 0.054 | -2.422 | 0.236 | -0.022 | 0.069 | 0.069 | 0.216 | 0.19 | 0.38 | 0.2 | 0.37 | 0.13 | 0.14 |
| 10 | 80023801 | rs7921332 | 0.536 | 0.788 | -3.004 | -0.463 | -0.063 | -0.002 | -0.002 | 0 | 0.28 | 0.05 | 0.21 | 0.22 | 0.27 | 0.3 |
| 10 | 80026039 | rs1720295 | 0.16 | 0.872 | -2.117 | 0.293 | 0.002 | -0.037 | -0.037 | 0.449 | 0.27 | 0.14 | 0.29 | 0.66 | 0.29 | 0.36 |
| 10 | 80027266 | rs1720293 | -0.046 | 0.828 | -2.026 | 0.102 | -0.005 | 0.003 | 0.003 | 0.675 | 0.31 | 0.17 | 0.32 | 0.7 | 0.34 | 0.42 |
| 10 | 80028339 | rs2913115 | -0.429 | 0.663 | -1.872 | -0.211 | 0.009 | 0.056 | 0.056 | 0 | 0.33 | 0.2 | 0.19 | 0.17 | 0.27 | 0.22 |
| 10 | 80029070 | rs1649979 | -0.335 | 0.814 | -2.246 | -0.67 | -0.086 | 0.191 | 0.191 | 0.385 | 0.73 | 0.97 | 0.8 | 1 | 0.73 | 0.84 |
| 10 | 80030941 | rs2913117 | -0.428 | 0.414 | -1.101 | 0.291 | -0.028 | 0.193 | 0.193 | 0.976 | 0.17 | 0.43 | 0.13 | 0.4 | 0.11 | 0.15 |
| 10 | 80035257 | rs11002480 | 0.519 | 0.746 | -1.391 | 0.423 | -0.001 | 0 | 0 | 0 | 0.2 | 0.2 | 0.15 | 0.16 | 0.23 | 0.28 |
| 10 | 80036805 | rs3862505 | 0.037 | 0.932 | -1.147 | 0.387 | -0.002 | -0.007 | -0.007 | 0 | 0.26 | 0.18 | 0.44 | 0.44 | 0.3 | 0.37 |
| 10 | 80041351 | rs1916391 | -0.074 | 0.856 | -1.397 | 0.283 | -0.091 | -0.121 | -0.121 | 1.303 | 0.27 | 0.05 | 0.41 | 0.48 | 0.35 | 0.37 |
| 10 | 80041406 | rs7099487 | 0.808 | 0.23 | -1.572 | -0.084 | 0.013 | 0.005 | 0.005 | 0 | 0.25 | 0.13 | 0.17 | 0.18 | 0.24 | 0.31 |
| 10 | 80041795 | rs11002481 | 0.187 | 0.192 | -1.105 | -0.196 | -0.015 | -0.042 | -0.042 | 0.495 | 0.29 | 0.2 | 0.39 | 0.46 | 0.3 | 0.35 |
| 10 | 80042347 | rs7088190 | -0.289 | 0.036 | -0.542 | -0.421 | -0.008 | 0.019 | 0.019 | 0 | 0.42 | 0.33 | 0.53 | 0.74 | 0.52 | 0.54 |
| 10 | 80043368 | rs1720260 | -1.135 | 0.102 | -0.57 | -0.575 | -0.027 | 0.083 | 0.083 | 0 | 0.77 | 0.93 | 0.74 | 1 | 0.76 | 0.85 |
| 10 | 80043540 | rs10824600 | -0.206 | -0.242 | -0.115 | 0 | 0.003 | 0.02 | 0.02 | 0 | 0.18 | 0.24 | 0.15 | 0.27 | 0.22 | 0.18 |
| 10 | 80044984 | rs1720321 | -0.364 | 0.19 | -0.26 | 0.46 | 0.003 | -0.003 | -0.003 | 0 | 0.43 | 0.39 | 0.49 | 0.51 | 0.47 | 0.45 |
| 10 | 80049593 | rs890685 | -0.242 | 0.448 | -0.963 | 0.209 | 0.009 | -0.004 | -0.004 | 1.196 | 0.68 | 0.79 | 0.65 | 0.63 | 0.72 | 0.79 |
| 10 | 80050769 | rs1649990 | -0.35 | 0.595 | -1.268 | 0.294 | -0.028 | -0.05 | -0.05 | 0 | 0.71 | 0.92 | 0.67 | 0.63 | 0.73 | 0.79 |
| 10 | 80055570 | rs7908220 | 0.307 | 0.287 | -1.118 | -0.098 | -0.001 | -0.003 | -0.003 | 0.015 | 0.3 | 0.28 | 0.29 | 0.16 | 0.29 | 0.27 |
| 10 | 80059372 | rs10762779 | -0.331 | 0.475 | -1.227 | 0.275 | -0.002 | -0.001 | -0.001 | 0 | 0.38 | 0.56 | 0.39 | 0.36 | 0.4 | 0.35 |
| 10 | 80059642 | rs1109073 | -0.648 | 0.818 | -1.107 | 0.042 | -0.059 | 0.021 | 0.021 | 0 | 0.79 | 0.97 | 0.77 | 0.9 | 0.76 | 0.85 |
| 10 | 80066149 | rs7076791 | -0.529 | 0.665 | 0.006 | 0.451 | 0.015 | -0.024 | -0.024 | 0.043 | 0.24 | 0.45 | 0.3 | 0.18 | 0.27 | 0.26 |
| 10 | 80075608 | rs1916395 | -0.978 | -0.966 | 0.224 | 1.068 | -0.06 | 0.064 | 0.064 | 2.212 | 0.27 | 0.54 | 0.24 | 0.47 | 0.17 | 0.28 |
| 10 | 80075655 | rs1649995 | -0.022 | -1.111 | -0.445 | 1.035 | -0.017 | -0.061 | -0.061 | 1.146 | 0.12 | 0.01 | 0.16 | 0.16 | 0.11 | 0.09 |
| 10 | 80080664 | rs16935812 | -0.349 | -1.352 | -0.063 | 0.863 | 0.005 | 0.002 | 0.002 | 0 | 0.11 | 0.1 | 0.08 | 0.02 | 0.09 | 0.03 |
| 10 | 80080866 | rs7920472 | -1.442 | -1.574 | 0.108 | 0.804 | -0.08 | -0.112 | -0.112 | 0 | 0.21 | 0.52 | 0.13 | 0.03 | 0.1 | 0.05 |
| 10 | 80091017 | rs1916399 | -1.453 | -1.466 | 0.705 | 1.132 | -0.013 | 0.286 | 0.286 | 0.023 | 0.1 | 0.23 | 0.16 | 0.45 | 0.08 | 0.25 |
| 10 | 80091833 | rs2395527 | -0.143 | -1.381 | 0.565 | 0.834 | -0.021 | -0.027 | -0.027 | 0.244 | 0.06 | 0.21 | 0.03 | 0 | 0.01 | 0 |
| 10 | 80093253 | rs12773707 | -0.487 | -2.018 | -0.267 | -0.263 | 0.017 | 0.026 | 0.026 | 0 | 0.3 | 0.18 | 0.2 | 0.04 | 0.22 | 0.23 |
| 10 | 80094714 | rs10762784 | 0.824 | -1.66 | -0.474 | -0.487 | -0.037 | 0.068 | 0.068 | 0.347 | 0.42 | 0.22 | 0.51 | 0.21 | 0.52 | 0.32 |
| 10 | 80096527 | rs2894383 | -1.415 | -1.618 | -0.774 | -0.286 | -0.051 | -0.072 | -0.072 | 0 | 0.83 | 0.98 | 0.87 | 0.71 | 0.79 | 0.77 |
| 10 | 80097952 | rs10466038 | 0.012 | -1.79 | -0.525 | 0.073 | -0.006 | 0.01 | 0.01 | 0 | 0.22 | 0.24 | 0.24 | 0.44 | 0.15 | 0.38 |
| 10 | 80099990 | rs11002509 | 1.619 | -1.786 | -0.656 | 0.059 | -0.029 | -0.03 | -0.03 | 0 | 0.05 | 0 | 0.05 | 0.06 | 0.08 | 0.05 |
| 10 | 80101868 | rs4598621 | -1.217 | -1.86 | -0.177 | 0.03 | -0.006 | -0.008 | -0.008 | 0.01 | 0.81 | 0.82 | 0.84 | 0.71 | 0.76 | 0.76 |
| 10 | 80104641 | rs7074180 | 1.461 | -1.864 | -0.019 | 0.356 | -0.028 | 0.046 | 0.046 | 0.217 | 0.42 | 0.25 | 0.53 | 0.26 | 0.53 | 0.33 |
| 10 | 80110906 | rs10490995 | -0.362 | -1.866 | 0.151 | 0.383 | 0.005 | -0.007 | -0.007 | 0 | 0.15 | 0.14 | 0.13 | 0.37 | 0.05 | 0.29 |
| 10 | 80114165 | rs1877591 | -0.604 | -1.699 | 0.232 | 0.503 | 0.005 | -0.021 | -0.021 | 0.9 | 0.17 | 0.15 | 0.18 | 0.43 | 0.12 | 0.34 |
| 10 | 80117445 | rs2670207 | 0.504 | -1.727 | 0.249 | 0.454 | -0.006 | -0.008 | -0.008 | 0.809 | 0.59 | 0.37 | 0.64 | 0.58 | 0.58 | 0.61 |
| 10 | 80122251 | rs2670217 | 0.599 | -1.74 | 0.263 | 0.476 | -0.002 | -0.005 | -0.005 | 0 | 0.61 | 0.71 | 0.63 | 0.33 | 0.62 | 0.39 |
| 10 | 80122477 | rs7915070 | 0.557 | -1.652 | 0.556 | 0.46 | 0.01 | 0.007 | 0.007 | 0.842 | 0.06 | 0.23 | 0.03 | 0 | 0 | 0 |
| 10 | 80125238 | rs1863832 | -0.009 | -1.555 | 0.315 | 0.524 | -0.021 | -0.006 | -0.006 | 0 | 0.28 | 0.2 | 0.25 | 0.3 | 0.34 | 0.31 |
| 10 | 80125781 | rs2670220 | -0.034 | -1.523 | 0.317 | 0.523 | -0.011 | -0.002 | -0.002 | 0.371 | 0.28 | 0.25 | 0.27 | 0.29 | 0.34 | 0.31 |
| 10 | 80127055 | rs2670223 | 0.931 | -1.472 | 0.182 | 0.753 | -0.019 | -0.072 | -0.072 | 0.726 | 0.17 | 0.07 | 0.21 | 0.54 | 0.18 | 0.4 |
| 10 | 80127117 | rs12259980 | 0.816 | -1.446 | 0.377 | 0.75 | 0.015 | 0.026 | 0.026 | 0 | 0.07 | 0.16 | 0.06 | 0.07 | 0.08 | 0.05 |
| 10 | 80127240 | rs2670224 | -0.282 | -1.579 | 0.303 | 0.821 | -0.001 | 0 | 0 | 0 | 0.32 | 0.34 | 0.37 | 0.63 | 0.35 | 0.5 |
| 10 | 80130592 | rs1559990 | -1.476 | -1.684 | 0.513 | 0.705 | -0.006 | -0.003 | -0.003 | 0 | 0.51 | 0.78 | 0.47 | 0.49 | 0.46 | 0.4 |
| 10 | 80131442 | rs10490996 | -0.064 | -1.436 | 0.637 | 0.461 | -0.005 | -0.025 | -0.025 | 0 | 0.13 | 0.11 | 0.12 | 0.03 | 0.08 | 0.09 |
| 10 | 80131550 | rs12257604 | -1.297 | -1.389 | 0.62 | 0.487 | -0.028 | -0.042 | -0.042 | 0 | 0.24 | 0.44 | 0.2 | 0.14 | 0.16 | 0.13 |
| 10 | 80132317 | rs4979967 | -1.4 | -1.395 | 0.579 | 0.485 | -0.028 | -0.037 | -0.037 | 0 | 0.25 | 0.54 | 0.21 | 0.14 | 0.16 | 0.13 |
| 10 | 80140047 | rs12768040 | -0.369 | -1.146 | 0.262 | 0.464 | 0.001 | -0.004 | -0.004 | 0 | 0.16 | 0.12 | 0.13 | 0.08 | 0.1 | 0.09 |
| 10 | 80156597 | rs11002535 | -1.485 | -1.11 | 0.729 | 0.488 | -0.022 | -0.043 | -0.043 | 0 | 0.25 | 0.46 | 0.21 | 0.14 | 0.18 | 0.14 |
| 10 | 80162150 | rs4586094 | -0.814 | -1.097 | 0.708 | 0.479 | -0.033 | -0.067 | -0.067 | 0 | 0.18 | 0.32 | 0.14 | 0.08 | 0.1 | 0.09 |
| 10 | 80168045 | rs2559544 | 0.165 | -1.174 | 0.772 | 0.57 | 0.012 | 0.015 | 0.015 | 0 | 0.26 | 0.36 | 0.26 | 0.29 | 0.29 | 0.28 |

| | | | | | | | | | | | | | | | | |
|---|---|---|---|---|---|---|---|---|---|---|---|---|---|---|---|---|
| 10 | 80171355 | rs7074158 | -0.501 | -1.07 | 0.472 | 0.695 | -0.018 | -0.028 | -0.028 | 0 | 0.16 | 0.25 | 0.13 | 0.08 | 0.1 | 0.09 |
| 10 | 80175240 | rs892392 | -1.133 | -0.454 | -0.425 | 0.593 | -0.015 | -0.1 | -0.1 | 1.308 | 0.75 | 0.92 | 0.69 | 0.52 | 0.76 | 0.61 |
| 10 | 80175782 | rs11002542 | 1.6 | -0.088 | -0.152 | 0.699 | -0.029 | -0.029 | -0.029 | 0 | 0.05 | 0 | 0.05 | 0.07 | 0.1 | 0.05 |
| 10 | 80177764 | rs12261821 | 0.155 | -0.158 | 0.202 | 0.685 | 0.027 | 0.06 | 0.06 | 0 | 0.08 | 0.24 | 0.07 | 0.12 | 0.1 | 0.05 |
| 10 | 80181178 | rs6480890 | -0.325 | -0.206 | 0.259 | 0.663 | 0.031 | 0.069 | 0.069 | 0 | 0.09 | 0.33 | 0.08 | 0.12 | 0.1 | 0.05 |
| 10 | 80184572 | rs2578118 | 0.345 | 0.4 | 0.455 | 0.569 | 0.006 | 0.004 | 0.004 | 0 | 0.25 | 0.27 | 0.28 | 0.28 | 0.27 | 0.3 |
| 10 | 80194620 | rs1347262 | -1.17 | 0.194 | 0.127 | 0.471 | -0.008 | -0.006 | -0.006 | 0.694 | 0.22 | 0.39 | 0.19 | 0.18 | 0.2 | 0.16 |
| 10 | 80196407 | rs11002551 | 1.706 | 0.527 | -0.046 | 0.659 | -0.028 | -0.03 | -0.03 | 0 | 0.05 | 0.01 | 0.06 | 0.1 | 0.1 | 0.05 |
| 10 | 80200607 | rs7894696 | -1.181 | 0.576 | 0.307 | 0.555 | 0.034 | 0.025 | 0.025 | 1.736 | 0.17 | 0.45 | 0.15 | 0.17 | 0.2 | 0.08 |
| 10 | 80202725 | rs2692724 | 0.269 | 0.365 | -0.149 | 0.62 | 0 | 0.01 | 0.01 | 0 | 0.55 | 0.56 | 0.45 | 0.68 | 0.46 | 0.71 |
| 10 | 80203388 | rs989905 | 0.68 | 0.391 | -0.321 | 0.628 | -0.011 | 0.006 | 0.006 | 0 | 0.42 | 0.43 | 0.22 | 0.53 | 0.31 | 0.58 |
| 10 | 80204611 | rs7088476 | -1.401 | 0.429 | -0.372 | 0.017 | -0.078 | 0.086 | 0.086 | 0 | 0.67 | 0.9 | 0.55 | 0.86 | 0.6 | 0.77 |
| 10 | 80206903 | rs10824609 | 1.585 | 0.382 | -0.251 | 0.057 | -0.005 | 0.001 | 0.001 | 0 | 0.13 | 0.07 | 0.22 | 0.15 | 0.16 | 0.12 |
| 10 | 80207497 | rs12784427 | 1.134 | 0.393 | -0.129 | 0.06 | -0.002 | -0.011 | -0.011 | 2.484 | 0.06 | 0.14 | 0.04 | 0.02 | 0.06 | 0.01 |
| 10 | 80208772 | rs2248789 | -0.709 | 0.388 | -0.118 | 0.04 | -0.061 | 0.097 | 0.097 | 0 | 0.62 | 0.83 | 0.5 | 0.86 | 0.52 | 0.72 |
| 10 | 80209510 | rs768395 | -0.123 | 0.425 | -0.107 | 0.073 | -0.003 | 0.049 | 0.049 | 0 | 0.55 | 0.65 | 0.45 | 0.84 | 0.46 | 0.72 |
| 10 | 80210674 | rs2574689 | 0.555 | 0.397 | -0.154 | 0.163 | -0.019 | -0.019 | -0.019 | 0 | 0.42 | 0.48 | 0.23 | 0.69 | 0.31 | 0.6 |
| 10 | 80212856 | rs2578119 | 1.061 | 0.459 | -0.299 | 0.276 | -0.034 | 0.046 | 0.046 | 0 | 0.3 | 0.43 | 0.19 | 0.44 | 0.21 | 0.4 |
| 10 | 80215829 | rs2010029 | 1.507 | 0.71 | -0.619 | 0.371 | 0.001 | -0.046 | -0.046 | 0 | 0.12 | 0.05 | 0.04 | 0.25 | 0.1 | 0.2 |
| 10 | 80217494 | rs12767035 | 2.006 | 0.83 | -0.491 | 0.126 | 0.014 | -0.025 | -0.025 | 0.438 | 0.05 | 0.07 | 0.05 | 0 | 0.08 | 0.05 |
| 10 | 80217968 | rs7918709 | -0.754 | 0.622 | -0.662 | 0.195 | -0.016 | 0.035 | 0.035 | 0 | 0.55 | 0.75 | 0.5 | 0.67 | 0.49 | 0.58 |
| 10 | 80220710 | rs1433794 | -1.379 | 0.817 | -0.607 | 0.257 | 0 | 0.045 | 0.045 | 0 | 0.7 | 0.65 | 0.72 | 0.5 | 0.72 | 0.63 |
| 10 | 80220985 | rs1435961 | 0.387 | 0.812 | -0.442 | 0.02 | 0.004 | 0.027 | 0.027 | 0 | 0.45 | 0.4 | 0.39 | 0.26 | 0.42 | 0.36 |
| 10 | 80221107 | rs7084858 | 0.19 | 1.018 | -0.587 | 0.103 | -0.002 | -0.001 | -0.001 | 0 | 0.23 | 0.23 | 0.32 | 0.2 | 0.3 | 0.27 |
| 10 | 80221806 | rs2670198 | -0.083 | 1.301 | -0.52 | 0.276 | 0.001 | -0.003 | -0.003 | 0 | 0.52 | 0.55 | 0.44 | 0.22 | 0.44 | 0.45 |
| 10 | 80222186 | rs3900506 | -0.498 | 1.249 | -0.377 | 0.315 | 0.003 | 0.002 | 0.002 | 0 | 0.29 | 0.3 | 0.53 | 0.65 | 0.38 | 0.46 |
| 10 | 80222411 | rs2670200 | -0.523 | 1.295 | -0.41 | 0.369 | -0.001 | 0 | 0 | 0 | 0.61 | 0.63 | 0.51 | 0.63 | 0.53 | 0.7 |
| 10 | 80222687 | rs2574692 | 0.976 | 1.18 | -0.457 | 0.296 | 0.019 | 0.02 | 0.02 | 0 | 0.41 | 0.35 | 0.18 | 0.22 | 0.28 | 0.37 |
| 10 | 80222790 | rs2177834 | -1.414 | 1.204 | -0.482 | 0.291 | 0.006 | -0.023 | -0.023 | 0 | 0.72 | 0.68 | 0.72 | 0.87 | 0.67 | 0.82 |
| 10 | 80223615 | rs2578108 | 0.891 | 1.239 | -0.408 | 0.26 | 0.025 | 0.023 | 0.023 | 0 | 0.41 | 0.34 | 0.18 | 0.23 | 0.28 | 0.37 |
| 10 | 80223728 | rs11002569 | -0.678 | 1.762 | -0.519 | 0.83 | 0.024 | 0.052 | 0.052 | 0 | 0.2 | 0.29 | 0.33 | 0.41 | 0.25 | 0.33 |
| 10 | 80224705 | rs10762785 | 0.645 | 1.606 | -0.722 | 0.517 | -0.012 | -0.022 | -0.022 | 0 | 0.1 | 0.05 | 0.15 | 0.21 | 0.14 | 0.13 |
| 10 | 80225164 | rs10762786 | -0.993 | 1.583 | -0.915 | -0.058 | -0.016 | 0.027 | 0.027 | 0 | 0.32 | 0.28 | 0.5 | 0.2 | 0.38 | 0.24 |
| 10 | 80227321 | rs2578109 | 0.252 | 1.411 | -0.714 | 0.377 | 0.001 | 0.012 | 0.012 | 0 | 0.47 | 0.44 | 0.45 | 0.25 | 0.47 | 0.47 |
| 10 | 80230227 | rs10824617 | 0.05 | 1.408 | -0.658 | 1.322 | -0.024 | -0.027 | -0.027 | 0 | 0.08 | 0.01 | 0.18 | 0.13 | 0.09 | 0.13 |
| 10 | 80231861 | rs4979816 | 0.756 | 1.427 | -0.47 | 1.283 | 0.002 | 0.043 | 0.043 | 0 | 0.87 | 0.81 | 0.72 | 0.36 | 0.85 | 0.74 |
| 10 | 80235850 | rs7920318 | -0.79 | 1.189 | -0.701 | 0.935 | -0.005 | -0.018 | -0.018 | 0.123 | 0.13 | 0.09 | 0.12 | 0.21 | 0.12 | 0.15 |
| 10 | 80237449 | rs4979982 | -0.004 | 0.393 | -0.587 | 0.352 | 0.006 | 0.007 | 0.007 | 0 | 0.2 | 0.16 | 0.15 | 0.13 | 0.14 | 0.21 |
| 10 | 80238142 | rs10509398 | 0.131 | 0.17 | 0.288 | 0.44 | 0.009 | 0.042 | 0.042 | 0 | 0.08 | 0.25 | 0.13 | 0.13 | 0.06 | 0.13 |
| 10 | 80244623 | rs4554825 | 0.149 | 0.053 | 0.234 | 0.556 | 0.01 | 0.082 | 0.082 | 0 | 0.08 | 0.25 | 0.13 | 0.17 | 0.06 | 0.14 |
| 10 | 80248755 | rs1369752 | -1.36 | -0.457 | 0.606 | 0.309 | -0.003 | 0 | 0 | 0 | 0.35 | 0.61 | 0.3 | 0.3 | 0.2 | 0.37 |
| 10 | 80253303 | rs1816442 | -0.748 | -0.505 | 0.357 | 0.314 | -0.046 | -0.05 | -0.05 | 1.231 | 0.27 | 0.36 | 0.17 | 0.13 | 0.14 | 0.23 |
| 10 | 80262994 | rs919890 | 0.318 | -0.525 | 0.413 | 0.304 | 0.023 | 0.025 | 0.025 | 0 | 0.18 | 0.07 | 0.14 | 0.13 | 0.14 | 0.21 |
| 10 | 80266239 | rs2099517 | -0.828 | -0.539 | 0.475 | 0.312 | -0.045 | -0.029 | -0.029 | 0 | 0.27 | 0.36 | 0.17 | 0.17 | 0.14 | 0.24 |
| 10 | 80266999 | rs10824620 | 0.367 | -0.355 | 0.187 | 0.369 | 0.018 | 0.013 | 0.013 | 0 | 0.16 | 0.09 | 0.12 | 0.13 | 0.1 | 0.18 |
| 10 | 80271486 | rs12218276 | 0.502 | -0.352 | 0.268 | 0.252 | 0.012 | 0.009 | 0.009 | 0 | 0.15 | 0.08 | 0.11 | 0.12 | 0.1 | 0.17 |
| 10 | 80280541 | rs1811384 | 0.081 | 0.145 | 0.085 | 0.038 | -0.028 | 0.102 | 0.102 | 0 | 0.62 | 0.84 | 0.61 | 0.16 | 0.6 | 0.48 |
| 10 | 80296708 | rs881009 | -0.076 | 0.2 | 0.075 | 0.094 | -0.065 | 0.097 | 0.097 | 0 | 0.64 | 0.91 | 0.62 | 0.2 | 0.6 | 0.49 |
| 10 | 80301056 | rs732187 | -0.139 | 0.196 | 0.041 | 0.503 | -0.085 | -0.025 | -0.025 | 1.535 | 0.34 | 0.07 | 0.32 | 0.34 | 0.4 | 0.39 |
| 10 | 80302281 | rs2177836 | 0.706 | -0.225 | 0.705 | 0.243 | 0 | -0.075 | -0.075 | 0 | 0.52 | 0.51 | 0.49 | 0.04 | 0.55 | 0.32 |
| 10 | 80306807 | rs10762790 | -0.286 | -0.199 | 0.544 | 0.288 | -0.026 | 0.029 | 0.029 | 0 | 0.4 | 0.14 | 0.35 | 0.34 | 0.4 | 0.41 |
| 10 | 80310047 | rs12219399 | -0.653 | -0.281 | 1.545 | 1.463 | -0.016 | 0.287 | 0.287 | 0.102 | 0.1 | 0.33 | 0.13 | 0.52 | 0.05 | 0.26 |
| 10 | 80312407 | rs2019432 | 0.328 | -0.225 | 1.49 | 1.003 | -0.014 | 0.205 | 0.205 | 0.009 | 0.91 | 0.72 | 0.83 | 0.41 | 0.95 | 0.73 |
| 10 | 80316027 | rs16936070 | 0.201 | -0.2 | 1.399 | 0.957 | 0 | 0.036 | 0.036 | 0 | 0.07 | 0.23 | 0.12 | 0.12 | 0.05 | 0.14 |
| 10 | 80318719 | rs1368765 | 1.401 | -0.233 | 1.469 | 0.968 | -0.038 | 0.193 | 0.193 | 0.438 | 0.87 | 0.63 | 0.8 | 0.38 | 0.95 | 0.71 |
| 10 | 80321424 | rs12253312 | -0.646 | 0.031 | 1.134 | 0.333 | -0.025 | 0.009 | 0.009 | 0 | 0.11 | 0.33 | 0.12 | 0.13 | 0.05 | 0.14 |
| 10 | 80325854 | rs164226 | -0.899 | 0.082 | 1.226 | 0.153 | -0.007 | -0.04 | -0.04 | 0.16 | 0.14 | 0.38 | 0.16 | 0.08 | 0.08 | 0.18 |
| 10 | 80328687 | rs164232 | -0.818 | 0.104 | 1.384 | 0.162 | 0.013 | -0.025 | -0.025 | 0 | 0.13 | 0.42 | 0.15 | 0.08 | 0.08 | 0.18 |
| 10 | 80329506 | rs158289 | -0.818 | 0.104 | 1.384 | 0.162 | 0.013 | -0.025 | -0.025 | 0 | 0.13 | 0.42 | 0.15 | 0.09 | 0.08 | 0.18 |
| 10 | 80337464 | rs151551 | -0.949 | 0.048 | 1.478 | 0.242 | 0.002 | -0.007 | -0.007 | 1.265 | 0.14 | 0.48 | 0.16 | 0.12 | 0.08 | 0.18 |
| 10 | 80338173 | rs7071538 | 0.674 | 0.071 | 0.712 | -0.618 | -0.007 | -0.012 | -0.012 | 0 | 0.3 | 0.25 | 0.29 | 0 | 0.32 | 0.17 |
| 10 | 80338908 | rs164247 | 0.17 | 0.107 | 1.33 | -0.034 | 0 | -0.003 | -0.003 | 0 | 0.11 | 0.18 | 0.13 | 0.12 | 0.08 | 0.18 |
| 10 | 80344742 | rs149624 | -1.167 | 0.066 | 0.932 | 0.035 | -0.029 | -0.012 | -0.012 | 0 | 0.17 | 0.59 | 0.18 | 0.16 | 0.08 | 0.2 |
| 10 | 80348121 | rs288444 | 1.189 | 0.307 | 0.843 | 0.678 | 0.018 | 0.338 | 0.338 | 0.031 | 0.87 | 0.59 | 0.78 | 0.55 | 0.91 | 0.86 |
| 10 | 80348890 | rs288442 | -0.354 | 0.404 | 0.424 | 0.698 | -0.008 | 0.07 | 0.07 | 0.042 | 0.93 | 0.93 | 0.83 | 0.59 | 0.91 | 0.87 |
| 10 | 80354027 | rs12766838 | 0.534 | 0.726 | -0.038 | 0.337 | -0.068 | -0.077 | -0.077 | 0 | 0.14 | 0.01 | 0.15 | 0.23 | 0.21 | 0.18 |
| 10 | 80355186 | rs288453 | 0.478 | 0.512 | 0.021 | -0.09 | -0.004 | -0.027 | -0.027 | 0.016 | 0.67 | 0.32 | 0.52 | 0.54 | 0.68 | 0.76 |
| 10 | 80356268 | rs288450 | -0.193 | 0.737 | 0.34 | -0.024 | 0.033 | 0.278 | 0.278 | 0 | 0.91 | 0.76 | 0.82 | 0.56 | 0.91 | 0.86 |
| 10 | 80356443 | rs1816226 | 1.3 | 0.694 | -0.141 | -0.086 | -0.04 | 0.053 | 0.053 | 0 | 0.15 | 0.04 | 0.24 | 0 | 0.22 | 0.09 |

| | | | | | | | | | | | | | | | | |
|---|---|---|---|---|---|---|---|---|---|---|---|---|---|---|---|---|
| 10 | 80359650 | rs288447 | 0.371 | 0.565 | 0.127 | 0.21 | 0.047 | 0.389 | 0.389 | 0.255 | 0.88 | 0.56 | 0.79 | 0.54 | 0.91 | 0.86 |
| 10 | 80371421 | rs11814093 | 0.327 | 0.625 | -1.751 | -0.005 | -0.11 | -0.055 | -0.055 | 0 | 0.39 | 0.01 | 0.3 | 0.38 | 0.42 | 0.47 |
| 10 | 80382636 | rs1684538 | -0.048 | 1.146 | -0.742 | 0.53 | 0.028 | 0.098 | 0.098 | 0 | 0.77 | 0.67 | 0.71 | 0.65 | 0.73 | 0.82 |
| 10 | 80383574 | rs1434437 | 0.563 | 1.19 | -0.566 | 0.154 | 0.007 | -0.079 | -0.079 | 0.997 | 0.16 | 0.23 | 0.13 | 0 | 0.18 | 0.03 |
| 10 | 80386576 | rs1025558 | -0.096 | 0.821 | -0.933 | -0.512 | -0.007 | -0.072 | -0.072 | 0 | 0.4 | 0.37 | 0.46 | 0.14 | 0.4 | 0.33 |
| 10 | 80389499 | rs2593147 | -1.446 | 0.967 | -2.029 | -0.204 | -0.097 | -0.089 | -0.089 | 0.496 | 0.71 | 0.99 | 0.71 | 0.66 | 0.67 | 0.63 |
| 10 | 80397768 | rs2441757 | -0.057 | 0.667 | -1.422 | -0.411 | -0.009 | -0.108 | -0.108 | 0 | 0.29 | 0.53 | 0.26 | 0.12 | 0.23 | 0.18 |
| 10 | 80401458 | rs1434449 | -0.055 | 0.657 | -1.439 | -0.415 | -0.009 | -0.101 | -0.101 | 0 | 0.29 | 0.49 | 0.26 | 0.12 | 0.23 | 0.18 |
| 10 | 80401534 | rs1655630 | 1.065 | 0.414 | -1.317 | -0.55 | 0.018 | -0.084 | -0.084 | 0 | 0.17 | 0.34 | 0.1 | 0.05 | 0.17 | 0.11 |
| 10 | 80404282 | rs1655629 | -1.271 | 0.359 | -1.675 | -0.513 | -0.058 | -0.017 | -0.017 | 0 | 0.53 | 0.94 | 0.52 | 0.53 | 0.48 | 0.37 |
| 10 | 80405937 | rs10762795 | 0.976 | 0.155 | -1.3 | -0.309 | 0.005 | 0.136 | 0.136 | 0.034 | 0.07 | 0.11 | 0.12 | 0.41 | 0.09 | 0.14 |
| 10 | 80410110 | rs1655627 | -0.494 | -0.178 | -1.28 | -0.435 | 0.001 | 0.007 | 0.007 | 0 | 0.36 | 0.66 | 0.38 | 0.52 | 0.32 | 0.31 |
| 10 | 80411629 | rs1665571 | 0.983 | 0.021 | -1.106 | -0.192 | 0.033 | -0.038 | -0.038 | 0 | 0.35 | 0.2 | 0.19 | 0.45 | 0.23 | 0.35 |
| 10 | 80414183 | rs1665568 | 1.481 | 0.375 | -0.32 | -0.474 | -0.021 | 0.012 | 0.012 | 0 | 0.43 | 0.26 | 0.51 | 0.6 | 0.43 | 0.49 |
| 10 | 80415570 | rs1655623 | 0.203 | 0.464 | -0.263 | -0.399 | -0.006 | -0.027 | -0.027 | 0.006 | 0.72 | 0.86 | 0.71 | 0.65 | 0.69 | 0.64 |
| 10 | 80417052 | rs1621754 | 0.652 | 0.678 | -0.09 | -0.201 | -0.007 | 0.019 | 0.019 | 0 | 0.69 | 0.6 | 0.7 | 0.62 | 0.7 | 0.58 |
| 10 | 80421520 | rs35665 | -0.229 | 1.224 | 0.536 | 0.121 | -0.026 | -0.042 | -0.042 | 0 | 0.85 | 0.95 | 0.79 | 0.7 | 0.78 | 0.71 |
| 10 | 80435887 | rs4979997 | -0.268 | 1.226 | 0.576 | 0.116 | -0.02 | -0.016 | -0.016 | 1.083 | 0.1 | 0.24 | 0.07 | 0.06 | 0.06 | 0.02 |
| 10 | 80442471 | rs592242 | 0.789 | 1.428 | 0.403 | -0.043 | 0.002 | 0.034 | 0.034 | 0 | 0.74 | 0.6 | 0.75 | 0.69 | 0.74 | 0.61 |
| 10 | 80447787 | rs11002633 | -0.322 | 1.551 | 0.105 | -0.103 | -0.016 | 0.029 | 0.029 | 0.084 | 0.13 | 0.07 | 0.21 | 0 | 0.18 | 0.05 |
| 10 | 80453448 | rs252822 | 0.376 | 1.358 | 0.566 | 0.511 | 0.001 | 0.002 | 0.002 | 0 | 0.8 | 0.83 | 0.68 | 0.73 | 0.71 | 0.78 |
| 10 | 80458157 | rs3910297 | -1.013 | 0.877 | 0.472 | 0.41 | -0.04 | -0.05 | -0.05 | 0 | 0.09 | 0.2 | 0.03 | 0 | 0.03 | 0.09 |
| 10 | 80459780 | rs4980001 | -1.095 | 0.879 | 0.167 | 0.41 | -0.052 | -0.065 | -0.065 | 0 | 0.11 | 0.36 | 0.04 | 0 | 0.03 | 0.09 |
| 10 | 80462465 | rs190411 | 0.65 | 2.011 | -0.859 | 0.893 | -0.033 | -0.025 | -0.025 | 0.012 | 0.72 | 0.84 | 0.76 | 0.66 | 0.66 | 0.71 |
| 10 | 80462677 | rs3893270 | -1.376 | 1.366 | -0.376 | 0.476 | 0.008 | -0.066 | -0.066 | 0.003 | 0.11 | 0.26 | 0.2 | 0 | 0.13 | 0.08 |
| 10 | 80462841 | rs157994 | 0.714 | 1.794 | -0.934 | 1.143 | -0.001 | -0.001 | -0.001 | 0 | 0.65 | 0.63 | 0.49 | 0.56 | 0.62 | 0.55 |
| 10 | 80466640 | rs650389 | -0.711 | 1.756 | -1.512 | 1.474 | -0.088 | -0.146 | -0.146 | 1.138 | 0.19 | 0.01 | 0.17 | 0.46 | 0.22 | 0.28 |
| 10 | 80469860 | rs153260 | 0.325 | 0.905 | -1.822 | 1.398 | 0.04 | -0.063 | -0.063 | 0 | 0.84 | 0.96 | 0.89 | 0.78 | 0.92 | 0.84 |
| 10 | 80470413 | rs157989 | -0.822 | 1.375 | -1.35 | 1.224 | -0.007 | -0.008 | -0.008 | 0 | 0.24 | 0.21 | 0.22 | 0.45 | 0.29 | 0.3 |
| 10 | 80474579 | rs11002642 | 1.601 | 0.982 | -0.628 | 0.638 | 0 | 0.019 | 0.019 | 0 | 0.49 | 0.42 | 0.47 | 0.33 | 0.5 | 0.49 |
| 10 | 80475376 | rs157983 | 0.409 | 0.488 | 0.19 | 0.512 | -0.029 | 0.018 | 0.018 | 0 | 0.79 | 0.66 | 0.71 | 0.78 | 0.84 | 0.8 |
| 10 | 80478501 | rs11002643 | 1.616 | 0.627 | 0.299 | 0.473 | 0.023 | 0.075 | 0.075 | 0 | 0.49 | 0.25 | 0.48 | 0.33 | 0.55 | 0.49 |
| 10 | 80483345 | rs7100087 | 1.817 | 0.704 | 0.195 | 0.366 | -0.006 | 0.065 | 0.065 | 0 | 0.46 | 0.22 | 0.46 | 0.33 | 0.54 | 0.5 |
| 10 | 80485043 | rs9663663 | 1.233 | 0.723 | 0.33 | 0.35 | 0.006 | 0.011 | 0.011 | 0 | 0.54 | 0.59 | 0.65 | 0.33 | 0.63 | 0.54 |
| 10 | 80487312 | rs6480902 | 1.523 | 0.742 | 0.323 | 0.359 | 0.031 | 0.052 | 0.052 | 0 | 0.52 | 0.33 | 0.64 | 0.33 | 0.63 | 0.54 |
| 10 | 80487897 | rs35660 | -0.346 | 0.729 | 0.268 | 0.452 | -0.001 | -0.002 | -0.002 | 0 | 0.24 | 0.22 | 0.19 | 0.44 | 0.24 | 0.3 |
| 10 | 80488201 | rs35662 | -0.141 | 0.682 | 0.145 | 0.412 | 0.013 | -0.067 | -0.067 | 0 | 0.82 | 0.97 | 0.87 | 0.78 | 0.9 | 0.84 |
| 10 | 80494262 | rs1684543 | 0.612 | 0.67 | 0.111 | 0.535 | -0.077 | 0.064 | 0.064 | 0 | 0.17 | 0.01 | 0.09 | 0.05 | 0.21 | 0.14 |
| 10 | 80496028 | rs35647 | 0.703 | 0.619 | 0.306 | 0.563 | 0.002 | 0 | 0 | 0.32 | 0.73 | 0.76 | 0.83 | 0.74 | 0.84 | 0.8 |
| 10 | 80498032 | rs12356700 | 0.048 | 0.733 | 0.002 | 0.248 | 0.028 | -0.029 | -0.029 | 0 | 0.13 | 0.22 | 0.14 | 0.09 | 0.17 | 0.12 |
| 10 | 80498057 | rs892542 | 2.158 | 0.698 | -0.247 | 0.31 | -0.007 | 0.04 | 0.04 | 0 | 0.38 | 0.2 | 0.49 | 0.25 | 0.44 | 0.42 |
| 10 | 80499572 | rs40429 | 0.558 | 0.579 | -0.856 | 0.382 | -0.009 | -0.001 | -0.001 | 0 | 0.73 | 0.72 | 0.83 | 0.75 | 0.84 | 0.8 |
| 10 | 80502114 | rs10509401 | 2.12 | 0.737 | -1.245 | 0.393 | -0.032 | 0.001 | 0.001 | 0 | 0.34 | 0.09 | 0.32 | 0.25 | 0.37 | 0.38 |
| 10 | 80507797 | rs1881731 | -0.327 | 0.609 | -1.374 | 0.433 | 0.01 | -0.071 | -0.071 | 0 | 0.87 | 0.97 | 0.84 | 0.75 | 0.92 | 0.85 |
| 10 | 80508113 | rs7101045 | -1.481 | 0.546 | -1.904 | 0.499 | -0.021 | 0.027 | 0.027 | 0 | 0.55 | 0.95 | 0.58 | 0.39 | 0.6 | 0.47 |
| 10 | 80508900 | rs727100 | -0.987 | 0.546 | -1.243 | 0.457 | -0.001 | 0.04 | 0.04 | 0 | 0.45 | 0.72 | 0.52 | 0.39 | 0.53 | 0.43 |
| 10 | 80510632 | rs9943334 | 1.075 | 0.562 | -0.729 | 0.844 | 0.018 | 0.005 | 0.005 | 0 | 0.55 | 0.67 | 0.62 | 0.44 | 0.65 | 0.56 |
| 10 | 80511807 | rs4980004 | -0.139 | 0.423 | -0.955 | 0.842 | 0.025 | -0.073 | -0.073 | 0 | 0.84 | 0.97 | 0.84 | 0.75 | 0.91 | 0.84 |
| 10 | 80513859 | rs4547043 | 0.722 | 0.702 | -0.715 | 0.788 | 0.028 | -0.004 | -0.004 | 0 | 0.56 | 0.73 | 0.62 | 0.45 | 0.66 | 0.56 |
| 10 | 80522637 | rs615550 | 0.056 | 1.046 | -0.345 | 1.366 | 0 | 0.024 | 0.024 | 0 | 0.91 | 0.93 | 0.89 | 0.73 | 0.94 | 0.86 |
| 10 | 80527255 | rs12263918 | -0.965 | 1.346 | -0.522 | 1.005 | -0.023 | 0.028 | 0.028 | 0 | 0.07 | 0.02 | 0.06 | 0 | 0.15 | 0.06 |
| 10 | 80529202 | rs946517 | -0.172 | 1.523 | -0.96 | 1.003 | -0.031 | -0.065 | -0.065 | 0 | 0.22 | 0.02 | 0.16 | 0.27 | 0.18 | 0.22 |
| 10 | 80530474 | rs638755 | 1.721 | 1.537 | -0.265 | 0.998 | 0.003 | 0.012 | 0.012 | 0.621 | 0.53 | 0.28 | 0.57 | 0.46 | 0.52 | 0.54 |
| 10 | 80530708 | rs637446 | -0.447 | 1.542 | -0.428 | 1.08 | -0.004 | -0.019 | -0.019 | 0 | 0.92 | 0.98 | 0.9 | 0.73 | 0.94 | 0.86 |
| 10 | 80535287 | rs7910759 | -0.807 | 1.541 | 0.002 | 0.945 | 0 | 0.005 | 0.005 | 0 | 0.27 | 0.21 | 0.22 | 0.27 | 0.27 | 0.25 |
| 10 | 80538509 | rs515372 | 1.538 | 1.31 | 0.281 | 1.059 | 0.011 | 0.01 | 0.01 | 0.429 | 0.55 | 0.29 | 0.59 | 0.45 | 0.55 | 0.54 |
| 10 | 80538860 | rs7904670 | -0.803 | 1.309 | 0.808 | 1.017 | -0.01 | -0.008 | -0.008 | 0 | 0.26 | 0.32 | 0.22 | 0.23 | 0.23 | 0.23 |
| 10 | 80540697 | rs536394 | 0.405 | 1.187 | 1.182 | 1.139 | 0.003 | 0.204 | 0.204 | 0.322 | 0.88 | 0.62 | 0.86 | 0.69 | 0.93 | 0.87 |
| 10 | 80545265 | rs7075511 | -1.243 | 1.226 | 1.035 | 0.909 | 0 | 0 | 0 | 0 | 0.29 | 0.3 | 0.3 | 0.27 | 0.34 | 0.27 |
| 10 | 80552153 | rs11002673 | -0.171 | 1.091 | 0.795 | 0.933 | -0.029 | -0.1 | -0.1 | 0.423 | 0.21 | 0.01 | 0.18 | 0.27 | 0.18 | 0.23 |
| 10 | 80558477 | rs10824643 | -0.778 | 1.349 | 0.847 | 0.83 | 0.01 | -0.027 | -0.027 | 0.676 | 0.09 | 0.13 | 0.13 | 0.02 | 0.16 | 0.09 |
| 10 | 80558801 | rs7913285 | -1.493 | 1.321 | 0.929 | 0.826 | 0 | -0.001 | -0.001 | 0.169 | 0.33 | 0.32 | 0.33 | 0.29 | 0.39 | 0.32 |
| 10 | 80560968 | rs11002681 | -0.54 | 1.18 | 1.024 | 0.837 | -0.012 | 0.012 | 0.012 | 0 | 0.23 | 0.23 | 0.19 | 0.28 | 0.16 | 0.26 |
| 10 | 80563519 | rs11595532 | -0.191 | 1.193 | 0.942 | 0.849 | 0.013 | -0.048 | -0.048 | 0 | 0.21 | 0.05 | 0.18 | 0.28 | 0.16 | 0.28 |
| 10 | 80569876 | rs7477887 | 2.207 | 1.669 | 0.814 | 0.676 | 0.011 | 0.062 | 0.062 | 0.104 | 0.46 | 0.22 | 0.44 | 0.36 | 0.41 | 0.43 |
| 10 | 80575173 | rs7069607 | -1.16 | 1.864 | 0.842 | 0.753 | -0.007 | -0.003 | -0.003 | 0 | 0.11 | 0.1 | 0.13 | 0.02 | 0.2 | 0.07 |
| 10 | 80575870 | rs11002688 | -0.671 | 1.867 | 0.84 | 0.752 | -0.005 | -0.003 | -0.003 | 0 | 0.09 | 0.09 | 0.12 | 0.02 | 0.17 | 0.07 |
| 10 | 80575996 | rs1427818 | -1.586 | 1.892 | 0.904 | 0.772 | 0 | -0.002 | -0.002 | 0 | 0.37 | 0.4 | 0.36 | 0.34 | 0.39 | 0.37 |
| 10 | 80581272 | rs11812282 | -0.522 | 1.898 | 0.512 | 0.802 | -0.016 | -0.109 | -0.109 | 0 | 0.24 | 0.03 | 0.21 | 0.32 | 0.19 | 0.3 |

| | | | | | | | | | | | | | | | | |
|---|---|---|---|---|---|---|---|---|---|---|---|---|---|---|---|---|
| 10 | 80584333 | rs11002693 | -0.939 | 1.928 | 0.7 | 0.723 | -0.005 | -0.006 | -0.006 | 0 | 0.1 | 0.09 | 0.13 | 0.02 | 0.2 | 0.07 |
| 10 | 80591064 | rs753384 | 1.814 | 1.623 | 0.955 | 0.842 | 0.009 | 0.038 | 0.038 | 0.109 | 0.51 | 0.34 | 0.48 | 0.36 | 0.45 | 0.45 |
| 10 | 80591365 | rs1992293 | 0.04 | 1.654 | 0.857 | 0.912 | 0.006 | -0.061 | -0.061 | 0 | 0.19 | 0.08 | 0.16 | 0.28 | 0.17 | 0.23 |
| 10 | 80602061 | rs10740484 | -0.064 | 1.688 | 0.453 | 0.919 | -0.01 | -0.008 | -0.008 | 0 | 0.88 | 0.9 | 0.85 | 0.7 | 0.83 | 0.81 |
| 10 | 80603160 | rs7900915 | -0.564 | 1.688 | 0.479 | 0.909 | 0.021 | -0.045 | -0.045 | 0.158 | 0.15 | 0.24 | 0.19 | 0.06 | 0.22 | 0.13 |
| 10 | 80604614 | rs1439777 | -0.094 | 1.688 | 0.306 | 0.991 | -0.012 | -0.014 | -0.014 | 0 | 0.88 | 0.9 | 0.85 | 0.75 | 0.83 | 0.81 |
| 10 | 80604998 | rs1439778 | 1.918 | 1.743 | -0.065 | 0.876 | 0.021 | 0.069 | 0.069 | 0.279 | 0.49 | 0.27 | 0.47 | 0.36 | 0.44 | 0.45 |
| 10 | 80605477 | rs10824655 | -0.737 | 1.847 | -0.137 | 0.837 | 0.015 | -0.036 | -0.036 | 0 | 0.15 | 0.22 | 0.19 | 0.06 | 0.21 | 0.13 |
| 10 | 80608820 | rs1366057 | 1.522 | 1.874 | -0.191 | 0.89 | 0.008 | 0.004 | 0.004 | 0.018 | 0.71 | 0.4 | 0.64 | 0.65 | 0.61 | 0.73 |
| 10 | 80609135 | rs946516 | 0.699 | 1.109 | -0.372 | 0.792 | 0.001 | 0.017 | 0.017 | 0 | 0.66 | 0.55 | 0.67 | 0.44 | 0.67 | 0.63 |
| 10 | 80612664 | rs1339481 | 0.527 | 0.954 | -1.063 | 0.136 | -0.075 | 0.168 | 0.168 | 0 | 0.7 | 0.94 | 0.81 | 0.97 | 0.63 | 0.76 |
| 10 | 80616150 | rs730324 | -1.411 | 0.123 | 0.263 | 1.097 | -0.003 | 0.085 | 0.085 | 1.211 | 0.16 | 0.48 | 0.12 | 0.27 | 0.15 | 0.12 |
| 10 | 80616723 | rs772830 | -1.643 | -0.477 | 0.44 | 1.469 | -0.066 | 0.17 | 0.17 | 0.045 | 0.23 | 0.47 | 0.24 | 0.42 | 0.12 | 0.43 |
| 10 | 80621021 | rs704018 | -1.03 | -0.39 | 0.307 | 1.852 | -0.009 | 0.135 | 0.135 | 0 | 0.05 | 0.13 | 0.06 | 0.19 | 0.02 | 0.08 |
| 10 | 80622356 | rs570677 | -1.937 | -0.573 | -0.241 | 1.428 | 0.076 | 0.011 | 0.011 | 0.565 | 0.38 | 0.86 | 0.34 | 0.65 | 0.27 | 0.58 |
| 10 | 80627582 | rs2993816 | -1.901 | -0.281 | 0.434 | 1.638 | 0.04 | -0.022 | -0.022 | 0.284 | 0.37 | 0.78 | 0.3 | 0.63 | 0.29 | 0.48 |
| 10 | 80631334 | rs10824657 | -1.36 | -0.407 | 0.803 | 2.12 | -0.005 | -0.037 | -0.037 | 0 | 0.19 | 0.14 | 0.18 | 0.41 | 0.22 | 0.18 |
| 10 | 80635863 | rs522549 | -1.836 | -0.484 | 1.526 | 2.196 | -0.051 | 0.134 | 0.134 | 0.863 | 0.17 | 0.44 | 0.1 | 0.28 | 0.07 | 0.39 |
| 10 | 80636291 | rs497204 | -1.582 | -0.469 | 1.443 | 2.193 | -0.018 | 0.112 | 0.112 | 0.064 | 0.1 | 0.27 | 0.07 | 0.2 | 0.04 | 0.12 |
| 10 | 80640275 | rs10218903 | -0.426 | -0.438 | 0.905 | 2.364 | -0.022 | -0.062 | -0.062 | 0 | 0.11 | 0.02 | 0.06 | 0.33 | 0.14 | 0.12 |
| 10 | 80640638 | rs1981221 | -3.102 | -0.493 | 0.919 | 2.346 | 0.049 | 0.027 | 0.027 | 0.81 | 0.39 | 0.85 | 0.31 | 0.69 | 0.3 | 0.58 |
| 10 | 80641807 | rs10218850 | -0.496 | -0.261 | 0.783 | 2.311 | -0.022 | -0.062 | -0.062 | 0 | 0.11 | 0.03 | 0.06 | 0.32 | 0.14 | 0.12 |
| 10 | 80646221 | rs7078982 | -0.834 | -0.241 | 0.916 | 1.994 | 0 | -0.001 | -0.001 | 0 | 0.08 | 0.09 | 0.12 | 0.08 | 0.08 | 0.06 |
| 10 | 80656930 | rs536030 | -2.236 | -0.045 | 0.787 | 1.783 | -0.005 | 0.005 | 0.005 | 0.78 | 0.32 | 0.4 | 0.27 | 0.34 | 0.27 | 0.51 |
| 10 | 80657477 | rs562757 | -1.204 | 0.052 | 0.699 | 1.843 | -0.016 | -0.008 | -0.008 | 0.553 | 0.05 | 0.02 | 0.05 | 0.2 | 0.07 | 0.09 |
| 10 | 80659151 | rs810083 | -2.576 | 0.158 | 0.867 | 1.722 | -0.001 | -0.003 | -0.003 | 0 | 0.44 | 0.59 | 0.35 | 0.66 | 0.43 | 0.63 |
| 10 | 80663052 | rs2559795 | -2.024 | 0.121 | 0.82 | 1.738 | -0.017 | 0.059 | 0.059 | 0.149 | 0.27 | 0.43 | 0.17 | 0.55 | 0.24 | 0.43 |
| 10 | 80664974 | rs10824662 | -1.281 | 0.287 | 0.524 | 1.659 | 0.006 | -0.017 | -0.017 | 0 | 0.17 | 0.23 | 0.18 | 0.08 | 0.19 | 0.11 |
| 10 | 80666690 | rs2559794 | -2.696 | 0.215 | 0.354 | 1.757 | -0.005 | 0.043 | 0.043 | 1.157 | 0.41 | 0.82 | 0.37 | 0.66 | 0.4 | 0.6 |
| 10 | 80669894 | rs11002726 | -1.28 | 0.239 | 0.322 | 1.721 | -0.014 | 0.015 | 0.015 | 0 | 0.15 | 0.08 | 0.17 | 0.08 | 0.19 | 0.1 |
| 10 | 80672227 | rs531987 | 2.104 | 0.661 | -0.083 | 1.46 | -0.032 | 0.072 | 0.072 | 0.506 | 0.53 | 0.15 | 0.59 | 0.36 | 0.51 | 0.42 |
| 10 | 80673954 | rs12416089 | -1.12 | 0.432 | -0.011 | 1.282 | -0.039 | 0.02 | 0.02 | 0 | 0.16 | 0.03 | 0.15 | 0.12 | 0.19 | 0.26 |
| 10 | 80674984 | rs11819353 | -1.333 | 0.42 | 0.068 | 1.319 | -0.017 | 0.017 | 0.017 | 0 | 0.17 | 0.08 | 0.15 | 0.15 | 0.19 | 0.35 |
| 10 | 80676558 | rs12412915 | -1.719 | 0.38 | 0.434 | 1.313 | 0 | -0.012 | -0.012 | 0 | 0.19 | 0.26 | 0.16 | 0.15 | 0.19 | 0.35 |
| 10 | 80685463 | rs10824671 | -1.981 | 0.503 | 0.543 | 2.236 | 0.024 | 0.041 | 0.041 | 0 | 0.22 | 0.39 | 0.23 | 0.31 | 0.26 | 0.29 |
| 10 | 80687569 | rs7907048 | -2.548 | 0.159 | -0.828 | 1.983 | -0.009 | 0.002 | 0.002 | 1.821 | 0.43 | 0.82 | 0.33 | 0.61 | 0.44 | 0.57 |
| 10 | 80688574 | rs16936637 | 0.233 | 0.249 | -0.89 | 1.397 | -0.002 | -0.035 | -0.035 | 1.504 | 0.06 | 0 | 0.01 | 0.13 | 0.04 | 0.11 |
| 10 | 80689033 | rs7084256 | -1.581 | 0.303 | -0.595 | 0.975 | -0.008 | 0.032 | 0.032 | 0 | 0.22 | 0.13 | 0.19 | 0.14 | 0.26 | 0.26 |
| 10 | 80693615 | rs17463021 | -0.559 | -0.644 | -1.585 | 0.165 | 0.012 | 0.184 | 0.184 | 0 | 0.36 | 0.05 | 0.2 | 0.14 | 0.31 | 0.35 |
| 10 | 80693725 | rs1439037 | -0.976 | -0.88 | -1.06 | 0.338 | 0.062 | 0.108 | 0.108 | 0 | 0.43 | 0.74 | 0.27 | 0.22 | 0.32 | 0.39 |
| 10 | 80694189 | rs6480911 | -1.554 | -1.091 | -1.751 | 0.863 | -0.044 | 0.02 | 0.02 | 0 | 0.57 | 0.88 | 0.42 | 0.65 | 0.46 | 0.57 |
| 10 | 80697529 | rs12763741 | -0.227 | -0.884 | -1.304 | 0.972 | -0.019 | 0.012 | 0.012 | 0 | 0.14 | 0.09 | 0.11 | 0.11 | 0.18 | 0.06 |
| 10 | 80698200 | rs1561439 | -2.049 | -1.311 | -2.025 | 1 | -0.155 | -0.007 | -0.007 | 0 | 0.61 | 0.98 | 0.46 | 0.76 | 0.47 | 0.66 |
| 10 | 80698871 | rs7897711 | -2.229 | -0.061 | 0.283 | 1.695 | -0.003 | -0.003 | -0.003 | 1.736 | 0.37 | 0.63 | 0.33 | 0.64 | 0.37 | 0.56 |
| 10 | 80709254 | rs6480913 | -1.784 | -0.043 | 0.78 | 2.121 | 0.025 | 0.062 | 0.062 | 0 | 0.27 | 0.44 | 0.22 | 0.4 | 0.26 | 0.31 |
| 10 | 80709678 | rs1561442 | -1.101 | -0.037 | 0.831 | 2.139 | 0 | 0.004 | 0.004 | 0 | 0.09 | 0.13 | 0.12 | 0.24 | 0.11 | 0.25 |
| 10 | 80710485 | rs873827 | -2.214 | -0.025 | 0.678 | 2.037 | 0.003 | 0.002 | 0.002 | 0.129 | 0.37 | 0.57 | 0.33 | 0.64 | 0.38 | 0.56 |
| 10 | 80712666 | rs7901632 | -0.395 | 0.058 | 0.498 | 1.835 | 0.004 | 0 | 0 | 0 | 0.07 | 0.16 | 0.03 | 0.06 | 0.04 | 0.15 |
| 10 | 80717362 | rs11002751 | -0.184 | 0.048 | 0.465 | 2.181 | -0.003 | -0.005 | -0.005 | 0 | 0.05 | 0.02 | 0.07 | 0.27 | 0.04 | 0.11 |
| 10 | 80729631 | rs7913392 | -0.578 | 0.345 | 0.612 | 1.845 | -0.013 | 0.009 | 0.009 | 0 | 0.14 | 0.11 | 0.1 | 0.1 | 0.19 | 0.05 |
| 10 | 80739356 | rs11002763 | -2.035 | 0.346 | 1.103 | 2.082 | 0.003 | 0.01 | 0.01 | 0.208 | 0.26 | 0.36 | 0.19 | 0.32 | 0.3 | 0.27 |
| 10 | 80741205 | rs11002764 | -1.208 | 0.255 | 0.955 | 1.894 | 0.001 | -0.043 | -0.043 | 0 | 0.12 | 0.07 | 0.08 | 0.22 | 0.1 | 0.21 |
| 10 | 80748189 | rs11002769 | -0.468 | 0.33 | 1.082 | 1.727 | 0.004 | -0.009 | -0.009 | 0 | 0.13 | 0.14 | 0.12 | 0.06 | 0.22 | 0.06 |
| 10 | 80749115 | rs10824690 | -2.25 | 0.243 | 1.679 | 1.943 | 0.01 | 0.03 | 0.03 | 0 | 0.28 | 0.39 | 0.22 | 0.38 | 0.33 | 0.38 |
| 10 | 80751836 | rs9633597 | -2.466 | 0.231 | 1.906 | 1.948 | 0.009 | 0.041 | 0.041 | 0.112 | 0.3 | 0.52 | 0.22 | 0.39 | 0.33 | 0.38 |
| 10 | 80752333 | rs11002771 | -1.394 | 0.155 | 1.976 | 2.107 | 0.039 | 0.062 | 0.062 | 0 | 0.21 | 0.46 | 0.22 | 0.35 | 0.26 | 0.15 |
| 10 | 80752765 | rs11002772 | -1.865 | 0.305 | 1.709 | 2.037 | -0.003 | 0.041 | 0.041 | 0 | 0.17 | 0.23 | 0.11 | 0.32 | 0.15 | 0.34 |
| 10 | 80756541 | rs10824693 | -2.564 | 0.508 | 1.748 | 2.066 | -0.012 | 0.003 | 0.003 | 0.935 | 0.38 | 0.8 | 0.33 | 0.68 | 0.39 | 0.5 |
| 10 | 80757626 | rs11002773 | -2.593 | 0.659 | 2.097 | 2.115 | -0.016 | 0.13 | 0.13 | 0.246 | 0.23 | 0.5 | 0.2 | 0.38 | 0.18 | 0.42 |
| 10 | 80758155 | rs10824694 | -1.528 | 0.546 | 2.012 | 2.129 | 0.004 | 0.102 | 0.102 | 0.146 | 0.15 | 0.26 | 0.11 | 0.33 | 0.14 | 0.35 |
| 10 | 80763116 | rs7068112 | -2.09 | 0.701 | 2.226 | 2.213 | -0.011 | 0.076 | 0.076 | 0 | 0.2 | 0.43 | 0.14 | 0.27 | 0.17 | 0.25 |
| 10 | 80763534 | rs10762829 | -2.682 | 1.247 | 2.054 | 2.196 | -0.034 | -0.045 | -0.045 | 0 | 0.44 | 0.84 | 0.43 | 0.48 | 0.52 | 0.45 |
| 10 | 80765849 | rs2275283 | -2.179 | 1.439 | 2.225 | 2.142 | 0.011 | -0.001 | -0.001 | 0 | 0.37 | 0.55 | 0.3 | 0.36 | 0.48 | 0.36 |
| 10 | 80766007 | rs1060628 | -2.012 | 1.438 | 2.231 | 2.142 | 0.008 | 0 | 0 | 0 | 0.36 | 0.4 | 0.29 | 0.36 | 0.48 | 0.36 |
| 10 | 80768033 | rs2027260 | -2.825 | 1.387 | 1.95 | 2.196 | 0.029 | 0.117 | 0.117 | 0.352 | 0.5 | 0.86 | 0.5 | 0.71 | 0.56 | 0.6 |
| 10 | 80772541 | rs7068603 | -1.506 | 1.244 | 2.119 | 2.524 | 0.006 | -0.034 | -0.034 | 0 | 0.21 | 0.13 | 0.13 | 0.27 | 0.21 | 0.27 |
| 10 | 80777567 | rs1371104 | -0.454 | 1.334 | 2.222 | 2.493 | 0.072 | -0.042 | -0.042 | 2.03 | 0.14 | 0.37 | 0.14 | 0.1 | 0.21 | 0.07 |
| 10 | 80779846 | rs7921660 | -2.09 | 1.32 | 1.683 | 2.567 | -0.002 | -0.001 | -0.001 | 0 | 0.34 | 0.31 | 0.25 | 0.36 | 0.43 | 0.34 |
| 10 | 80780389 | rs3936399 | -2.459 | 1.346 | 0.96 | 2.553 | -0.046 | 0.017 | 0.017 | 0.489 | 0.45 | 0.81 | 0.48 | 0.65 | 0.51 | 0.52 |

| | | | | | | | | | | | | | | | | |
|---|---|---|---|---|---|---|---|---|---|---|---|---|---|---|---|---|
| 10 | 80783166 | rs3802643 | 2.088 | 0.996 | 1.24 | 2.834 | -0.029 | 0.058 | 0.058 | 0.32 | 0.58 | 0.18 | 0.55 | 0.27 | 0.57 | 0.45 |
| 10 | 80786279 | rs7075882 | 0.771 | 1.352 | 1.162 | 2.69 | -0.008 | -0.082 | -0.082 | 0 | 0.85 | 0.98 | 0.92 | 0.73 | 0.88 | 0.8 |
| 10 | 80790066 | rs10824704 | -1.322 | 1.587 | 1.741 | 2.876 | -0.015 | 0.062 | 0.062 | 0 | 0.14 | 0.26 | 0.08 | 0.27 | 0.08 | 0.26 |
| 10 | 80792301 | rs4980014 | 0.411 | 1.605 | 1.456 | 2.959 | 0 | 0.16 | 0.16 | 0 | 0.94 | 0.78 | 0.85 | 0.71 | 0.96 | 0.84 |
| 10 | 80792528 | rs10824705 | -0.755 | 1.742 | 1.526 | 2.925 | 0.056 | -0.003 | -0.003 | 0 | 0.14 | 0.19 | 0.18 | 0.14 | 0.25 | 0.1 |
| 10 | 80795422 | rs3910579 | -0.665 | 1.751 | 1.362 | 2.924 | 0.024 | -0.005 | -0.005 | 0 | 0.15 | 0.21 | 0.12 | 0.14 | 0.21 | 0.11 |
| 10 | 80797439 | rs1946842 | -1.898 | 1.56 | 0.292 | 3.073 | 0.002 | 0.006 | 0.006 | 1.276 | 0.31 | 0.63 | 0.34 | 0.55 | 0.33 | 0.4 |
| 10 | 80800595 | rs7093140 | -1.785 | 1.901 | -0.082 | 1.556 | 0.019 | 0.024 | 0.024 | 0.682 | 0.3 | 0.55 | 0.26 | 0.4 | 0.33 | 0.39 |
| 10 | 80801482 | rs11002793 | -1.31 | 1.113 | -0.859 | 1.294 | -0.002 | -0.046 | -0.046 | 0 | 0.11 | 0.05 | 0.06 | 0.2 | 0.08 | 0.2 |
| 10 | 80808527 | rs10824709 | 0.717 | 1.838 | -0.091 | 1.413 | 0.027 | 0.034 | 0.034 | 0 | 0.73 | 0.62 | 0.53 | 0.37 | 0.65 | 0.52 |
| 10 | 80808667 | rs10740494 | 0.735 | 1.757 | -0.069 | 2.193 | 0.029 | 0.042 | 0.042 | 0 | 0.77 | 0.63 | 0.68 | 0.65 | 0.69 | 0.7 |
| 10 | 80810343 | rs6480917 | -0.16 | 1.985 | 0.169 | 1.694 | 0.017 | -0.017 | -0.017 | 0 | 0.21 | 0.39 | 0.12 | 0.15 | 0.2 | 0.07 |
| 10 | 80812749 | rs10824710 | 0.465 | 1.265 | -0.679 | 1.266 | 0 | -0.002 | -0.002 | 0 | 0.59 | 0.42 | 0.32 | 0.38 | 0.54 | 0.3 |
| 10 | 80819132 | rs704017 | 0.701 | 1.261 | -0.354 | 1.302 | 0.005 | -0.014 | -0.014 | 0 | 0.6 | 0.56 | 0.38 | 0.28 | 0.56 | 0.42 |
| 10 | 80825449 | rs1370562 | -1.286 | 1.336 | -0.162 | 1.795 | 0.026 | 0.117 | 0.117 | 0 | 0.16 | 0.23 | 0.31 | 0.44 | 0.22 | 0.33 |
| 10 | 80828245 | rs2297922 | -1.199 | 0.676 | -0.294 | 1.766 | -0.008 | 0.108 | 0.108 | 0 | 0.16 | 0.22 | 0.29 | 0.44 | 0.15 | 0.3 |
| 10 | 80832774 | rs704014 | 0.103 | 0.944 | -0.695 | 1.584 | -0.062 | -0.073 | -0.073 | 0.772 | 0.86 | 0.99 | 0.69 | 0.74 | 0.82 | 0.67 |
| 10 | 80834958 | rs7082178 | -1.547 | 0.362 | 0.431 | 1.724 | -0.018 | 0.258 | 0.258 | 0.808 | 0.13 | 0.44 | 0.21 | 0.31 | 0.08 | 0.23 |
| 10 | 80835868 | rs10824713 | -0.205 | 0.509 | -0.358 | 1.083 | -0.006 | -0.011 | -0.011 | 7.791 | 0.58 | 0.66 | 0.49 | 0.48 | 0.56 | 0.55 |
| 10 | 80838651 | rs7920460 | -0.842 | 0.487 | -0.101 | 1.197 | -0.012 | 0.075 | 0.075 | 0 | 0.13 | 0.2 | 0.22 | 0.31 | 0.1 | 0.21 |
| 10 | 80843548 | rs4980027 | 0.125 | 0.61 | -0.048 | 1.482 | 0 | 0.019 | 0.019 | 0 | 0.1 | 0.13 | 0.19 | 0.23 | 0.12 | 0.22 |
| 10 | 80847150 | rs1250002 | -0.76 | 0.029 | 0.17 | 1.501 | -0.039 | 0.102 | 0.102 | 0 | 0.22 | 0.49 | 0.22 | 0.31 | 0.17 | 0.24 |
| 10 | 80847236 | rs4979843 | 1.593 | -0.67 | -0.957 | 0.787 | -0.011 | 0.091 | 0.091 | 0 | 0.32 | 0.11 | 0.18 | 0.13 | 0.3 | 0.22 |
| 10 | 80849153 | rs1269867 | -0.218 | -0.638 | -0.891 | 0.797 | 0 | -0.001 | -0.001 | 0 | 0.55 | 0.59 | 0.4 | 0.44 | 0.47 | 0.46 |
| 10 | 80851257 | rs719338 | -0.941 | -0.491 | -0.88 | 0.637 | -0.02 | 0.007 | 0.007 | 0 | 0.67 | 0.76 | 0.6 | 0.66 | 0.58 | 0.69 |
| 10 | 80852378 | rs1268974 | -0.846 | -0.591 | -0.906 | 0.651 | -0.024 | 0.011 | 0.011 | 0 | 0.66 | 0.77 | 0.59 | 0.67 | 0.58 | 0.69 |
| 10 | 80852602 | rs873823 | -1.159 | -0.632 | -1.029 | 0.692 | -0.144 | -0.025 | -0.025 | 0 | 0.69 | 0.96 | 0.61 | 0.68 | 0.58 | 0.7 |
| 10 | 80853407 | rs2012869 | 1.169 | -0.632 | -0.889 | 0.419 | -0.003 | -0.033 | -0.033 | 0.043 | 0.33 | 0.35 | 0.17 | 0.12 | 0.29 | 0.21 |
| 10 | 80853508 | rs1250013 | 0.633 | -0.828 | -0.998 | 0.706 | -0.044 | -0.033 | -0.033 | 0 | 0.09 | 0.01 | 0.09 | 0.07 | 0.15 | 0.13 |
| 10 | 80854255 | rs1250011 | -0.799 | -0.833 | -0.935 | 0.297 | -0.012 | 0.009 | 0.009 | 0 | 0.37 | 0.57 | 0.44 | 0.53 | 0.32 | 0.5 |
| 10 | 80854413 | rs11002810 | 1.364 | -0.838 | -1.453 | -0.106 | -0.024 | 0.052 | 0.052 | 0 | 0.22 | 0.1 | 0.13 | 0.11 | 0.26 | 0.19 |
| 10 | 80855577 | rs10824714 | 0.216 | -0.487 | -1.406 | -0.182 | -0.005 | 0.041 | 0.041 | 0.146 | 0.47 | 0.37 | 0.42 | 0.21 | 0.52 | 0.28 |
| 10 | 80857533 | rs11002813 | 1.765 | -0.276 | -1.859 | 0.075 | -0.036 | -0.046 | -0.046 | 0 | 0.19 | 0.05 | 0.14 | 0.27 | 0.18 | 0.26 |
| 10 | 80858151 | rs10824715 | 0.94 | -0.237 | -1.837 | -0.15 | 0.004 | -0.022 | -0.022 | 0.015 | 0.25 | 0.3 | 0.16 | 0.1 | 0.29 | 0.18 |
| 10 | 80858505 | rs10762845 | 0.397 | -0.276 | -1.821 | -0.092 | -0.019 | 0.032 | 0.032 | 0 | 0.43 | 0.3 | 0.3 | 0.36 | 0.47 | 0.42 |
| 10 | 80861989 | rs880031 | 0.739 | 0.135 | -1.7 | -0.124 | -0.026 | 0.058 | 0.058 | 0 | 0.24 | 0.12 | 0.14 | 0.12 | 0.28 | 0.18 |
| 10 | 80863226 | rs11597192 | 0.136 | -0.248 | -1.775 | -0.097 | -0.038 | 0.071 | 0.071 | 0 | 0.35 | 0.17 | 0.27 | 0.22 | 0.42 | 0.24 |
| 10 | 80866164 | rs1092116 | -1.667 | -0.339 | -2.09 | 0.162 | -0.02 | -0.07 | -0.07 | 1.113 | 0.88 | 0.99 | 0.75 | 0.75 | 0.87 | 0.73 |
| 10 | 80866709 | rs16936856 | 0.587 | -0.175 | -1.36 | -0.478 | -0.004 | -0.017 | -0.017 | 0 | 0.26 | 0.35 | 0.18 | 0.2 | 0.26 | 0.26 |
| 10 | 80867175 | rs6480919 | -0.484 | -0.313 | -1.732 | -0.537 | 0 | 0.001 | 0.001 | 0 | 0.69 | 0.7 | 0.58 | 0.56 | 0.65 | 0.54 |
| 10 | 80869062 | rs7086204 | -0.161 | 0.129 | -1.422 | -0.477 | -0.002 | 0.003 | 0.003 | 0 | 0.36 | 0.66 | 0.45 | 0.32 | 0.4 | 0.33 |
| 10 | 80872270 | rs1628089 | 1.154 | -0.199 | -1.251 | -0.057 | -0.035 | 0.03 | 0.03 | 0 | 0.21 | 0.11 | 0.19 | 0.01 | 0.37 | 0.15 |
| 10 | 80873515 | rs1658324 | -0.896 | -0.108 | -1.424 | 0.289 | -0.026 | 0.042 | 0.042 | 0 | 0.72 | 0.61 | 0.66 | 0.61 | 0.81 | 0.68 |
| 10 | 80873608 | rs17557872 | -0.19 | 0.174 | -1.791 | 0.48 | -0.021 | -0.042 | -0.042 | 0 | 0.26 | 0.08 | 0.11 | 0.28 | 0.21 | 0.3 |
| 10 | 80874092 | rs11002826 | -0.166 | 0.222 | -1.771 | 0.499 | -0.022 | -0.042 | -0.042 | 0 | 0.26 | 0.07 | 0.11 | 0.28 | 0.21 | 0.3 |
| 10 | 80875213 | rs1545142 | -0.852 | 0.347 | -0.823 | 0.42 | 0.009 | 0.031 | 0.031 | 0 | 0.21 | 0.12 | 0.15 | 0.13 | 0.2 | 0.17 |
| 10 | 80877090 | rs704005 | 1.254 | 0.326 | -0.19 | 0.899 | -0.009 | 0.039 | 0.039 | 0 | 0.89 | 0.84 | 0.91 | 0.82 | 0.87 | 0.87 |
| 10 | 80878219 | rs4336976 | -1.573 | 0.301 | 0.269 | 0.863 | -0.006 | 0.054 | 0.054 | 1.419 | 0.23 | 0.43 | 0.18 | 0.29 | 0.2 | 0.43 |
| 10 | 80879484 | rs10824719 | -0.821 | 0.273 | -0.07 | 0.761 | 0 | -0.004 | -0.004 | 0 | 0.15 | 0.15 | 0.07 | 0.08 | 0.11 | 0.28 |
| 10 | 80881968 | rs704004 | -0.709 | 0.207 | -0.108 | 0.6 | 0.005 | 0.006 | 0.006 | 0.084 | 0.5 | 0.45 | 0.48 | 0.62 | 0.39 | 0.56 |
| 10 | 80882589 | rs1658322 | -0.944 | -0.07 | 0.1 | 0.153 | 0.001 | -0.002 | -0.002 | 3.36 | 0.52 | 0.53 | 0.51 | 0.82 | 0.43 | 0.58 |
| 10 | 80883083 | rs1866159 | -0.911 | -0.109 | -0.05 | 0.89 | -0.012 | 0.043 | 0.043 | 0 | 0.21 | 0.29 | 0.35 | 0.56 | 0.15 | 0.42 |
| 10 | 80888858 | rs7097066 | -1.147 | -0.396 | -0.199 | 1.209 | -0.057 | 0.06 | 0.06 | 0 | 0.29 | 0.49 | 0.34 | 0.44 | 0.14 | 0.39 |
| 10 | 80891631 | rs4980031 | -0.742 | -0.125 | -1.219 | 1.352 | 0.029 | -0.06 | -0.06 | 0.05 | 0.24 | 0.11 | 0.32 | 0.4 | 0.13 | 0.38 |
| 10 | 80892882 | rs704001 | -0.222 | 0.584 | -1.337 | 1.216 | -0.075 | -0.056 | -0.056 | 0 | 0.13 | 0.01 | 0.15 | 0.19 | 0.23 | 0.11 |
| 10 | 80894561 | rs11002846 | -0.985 | 0.381 | -1.189 | 1.274 | -0.004 | -0.079 | -0.079 | 0 | 0.33 | 0.14 | 0.5 | 0.52 | 0.27 | 0.65 |
| 10 | 80903455 | rs4980032 | -0.901 | 1.042 | -0.207 | 1.092 | 0.043 | 0.028 | 0.028 | 0 | 0.2 | 0.49 | 0.4 | 0.19 | 0.23 | 0.39 |
| 10 | 80907147 | rs780159 | -1.231 | 0.156 | -1.179 | 0.485 | -0.056 | 0.083 | 0.083 | 1.326 | 0.66 | 0.85 | 0.67 | 0.86 | 0.57 | 0.73 |
| 10 | 80909508 | rs12359427 | -0.192 | -0.668 | -1.239 | 1.066 | 0.011 | 0.007 | 0.007 | 0 | 0.39 | 0.6 | 0.34 | 0.42 | 0.36 | 0.24 |
| 10 | 80913782 | rs6480922 | -0.048 | -0.786 | -1.294 | 1.36 | -0.013 | -0.077 | -0.077 | 0 | 0.78 | 0.9 | 0.72 | 0.71 | 0.81 | 0.53 |
| 10 | 80919442 | rs749821 | 0.495 | -1.04 | -0.487 | 1.463 | 0.016 | 0.11 | 0.11 | 0 | 0.1 | 0.22 | 0.16 | 0.32 | 0.12 | 0.15 |
| 10 | 80920370 | rs4980038 | 0.466 | -1.042 | -0.488 | 1.462 | 0.015 | 0.108 | 0.108 | 0 | 0.1 | 0.23 | 0.16 | 0.32 | 0.12 | 0.15 |
| 10 | 80921427 | rs697242 | -0.014 | -0.988 | -0.588 | 1.267 | -0.002 | -0.05 | -0.05 | 0.96 | 0.78 | 0.85 | 0.71 | 0.69 | 0.81 | 0.51 |
| 10 | 80924404 | rs1769756 | 1.929 | -1.099 | -0.591 | 1.175 | -0.082 | 0.102 | 0.102 | 1.428 | 0.34 | 0.14 | 0.33 | 0.16 | 0.43 | 0.27 |
| 10 | 80924955 | rs16936991 | 1.266 | -1.02 | -0.774 | 1.492 | 0.027 | -0.022 | -0.022 | 0.049 | 0.1 | 0.05 | 0.04 | 0.14 | 0.04 | 0.09 |
| 10 | 80926036 | rs703994 | 0.548 | -1.1 | -0.673 | 1.409 | 0.016 | -0.019 | -0.019 | 0 | 0.54 | 0.79 | 0.55 | 0.57 | 0.55 | 0.43 |
| 10 | 80929130 | rs703991 | -1.131 | -1.018 | -0.72 | 1.396 | -0.056 | -0.051 | -0.051 | 0 | 0.9 | 0.99 | 0.78 | 0.87 | 0.85 | 0.69 |
| 10 | 80929992 | rs7913964 | -0.472 | -0.934 | -0.465 | 1.241 | -0.011 | 0.012 | 0.012 | 0 | 0.39 | 0.45 | 0.36 | 0.53 | 0.39 | 0.25 |
| 10 | 80930439 | rs703990 | -0.24 | -0.93 | -0.491 | 1.344 | -0.015 | -0.061 | -0.061 | 0.201 | 0.78 | 0.91 | 0.72 | 0.74 | 0.81 | 0.52 |

| | | | | | | | | | | | | | | | | |
|---|---|---|---|---|---|---|---|---|---|---|---|---|---|---|---|---|
| 10 | 80930698 | rs7914926 | 1.293 | -0.789 | -0.399 | 0.67 | 0.01 | -0.035 | -0.035 | 0 | 0.4 | 0.48 | 0.36 | 0.21 | 0.43 | 0.28 |
| 10 | 80932607 | rs10824726 | 0.388 | -0.844 | -0.619 | 0.612 | 0.01 | 0.074 | 0.074 | 0 | 0.23 | 0.09 | 0.17 | 0.12 | 0.25 | 0.09 |
| 10 | 80932877 | rs10762852 | 0.403 | -0.865 | -0.63 | 0.614 | 0.012 | 0.076 | 0.076 | 0.065 | 0.23 | 0.08 | 0.15 | 0.11 | 0.25 | 0.07 |
| 10 | 80936447 | rs3862511 | -0.973 | -2.092 | 0.292 | 1.191 | 0.04 | -0.001 | -0.001 | 0 | 0.36 | 0.74 | 0.36 | 0.65 | 0.16 | 0.48 |
| 10 | 80938499 | rs16937003 | 0.441 | -2.105 | 0.116 | 1.177 | 0.022 | -0.003 | -0.003 | 0.004 | 0.09 | 0.03 | 0.08 | 0.15 | 0.03 | 0.11 |
| 10 | 80940357 | rs4980041 | -1.408 | -2.265 | 0.255 | 1.7 | -0.036 | 0.011 | 0.011 | 0.653 | 0.39 | 0.63 | 0.39 | 0.51 | 0.2 | 0.57 |
| 10 | 80942721 | rs703982 | 1.017 | -1.273 | -0.434 | 1.398 | -0.07 | -0.049 | -0.049 | 0.058 | 0.31 | 0.1 | 0.42 | 0.33 | 0.41 | 0.31 |
| 10 | 80943841 | rs703980 | 0.76 | -1.325 | -0.227 | 1.418 | -0.03 | -0.028 | -0.028 | 0.062 | 0.34 | 0.23 | 0.47 | 0.41 | 0.45 | 0.37 |
| 10 | 80944095 | rs703979 | -0.475 | -1.635 | 0.137 | 1.458 | 0 | 0 | 0 | 0.082 | 0.53 | 0.53 | 0.66 | 0.67 | 0.57 | 0.68 |
| 10 | 80945997 | rs4979848 | 0.059 | -1.609 | 0.081 | 1.31 | -0.017 | 0.009 | 0.009 | 0 | 0.2 | 0.3 | 0.19 | 0.26 | 0.12 | 0.32 |
| 10 | 80953136 | rs703970 | -0.435 | -1.468 | 0.228 | 1.337 | 0 | 0.006 | 0.006 | 0.164 | 0.54 | 0.64 | 0.67 | 0.62 | 0.58 | 0.68 |
| 10 | 80954686 | rs697237 | -0.053 | -1.599 | 0.064 | 1.19 | -0.001 | -0.003 | -0.003 | 0 | 0.48 | 0.42 | 0.37 | 0.42 | 0.46 | 0.38 |
| 10 | 80955067 | rs703965 | 0.22 | -1.601 | 0.019 | 1.115 | 0.001 | -0.006 | -0.006 | 0.359 | 0.45 | 0.42 | 0.33 | 0.4 | 0.46 | 0.3 |
| 10 | 80955507 | rs7894320 | 0.197 | -1.712 | -0.064 | 0.914 | 0.007 | 0.004 | 0.004 | 2.477 | 0.11 | 0.08 | 0.13 | 0.09 | 0.05 | 0.11 |
| 10 | 80957694 | rs4980047 | -1.046 | -1.698 | 0.244 | 0.87 | -0.065 | 0.029 | 0.029 | 0 | 0.24 | 0.47 | 0.25 | 0.26 | 0.14 | 0.38 |
| 10 | 80959517 | rs10509406 | -0.273 | -1.104 | -0.046 | 0.293 | -0.006 | -0.025 | -0.025 | 0.186 | 0.18 | 0.22 | 0.19 | 0.05 | 0.19 | 0.18 |
| 10 | 80961519 | rs2802371 | 0.162 | -1.22 | -0.22 | 0.445 | -0.014 | -0.009 | -0.009 | 0 | 0.41 | 0.38 | 0.37 | 0.42 | 0.47 | 0.43 |
| 10 | 80963010 | rs1250574 | -1.79 | -1.808 | 0.128 | 0.5 | 0.013 | 0 | 0 | 0 | 0.78 | 0.73 | 0.71 | 0.78 | 0.75 | 0.73 |
| 10 | 80963517 | rs16937032 | 0.595 | -1.948 | 0.428 | 0.577 | 0.039 | -0.016 | -0.016 | 1.358 | 0.23 | 0.12 | 0.16 | 0.32 | 0.11 | 0.14 |
| 10 | 80964796 | rs1250576 | 1.438 | -1.735 | 0.416 | 0.657 | -0.055 | 0.054 | 0.054 | 0 | 0.24 | 0.15 | 0.17 | 0.13 | 0.33 | 0.15 |
| 10 | 80967220 | rs942796 | 0.013 | -1.853 | 0.456 | 0.628 | 0.006 | 0.019 | 0.019 | 0 | 0.47 | 0.37 | 0.34 | 0.45 | 0.44 | 0.28 |
| 10 | 80968701 | rs11002868 | 0.605 | -1.884 | 0.388 | 0.617 | 0.009 | -0.002 | -0.002 | 0.199 | 0.23 | 0.21 | 0.12 | 0.3 | 0.11 | 0.12 |
| 10 | 80973447 | rs1250582 | 0.572 | -1.806 | 0.02 | 0.717 | -0.086 | -0.028 | -0.028 | 0 | 0.36 | 0.16 | 0.33 | 0.33 | 0.45 | 0.34 |
| 10 | 80975443 | rs1616493 | -1.416 | -1.22 | 0.217 | 0.515 | 0.002 | -0.002 | -0.002 | 0 | 0.74 | 0.73 | 0.66 | 0.77 | 0.64 | 0.68 |
| 10 | 80985374 | rs1250591 | -0.302 | -1.297 | -0.128 | 0.524 | -0.002 | -0.018 | -0.018 | 1.307 | 0.58 | 0.52 | 0.48 | 0.71 | 0.55 | 0.51 |
| 10 | 80989082 | rs2764818 | -1.076 | -0.994 | 0.199 | 0.617 | 0.035 | -0.027 | -0.027 | 0.013 | 0.73 | 0.57 | 0.62 | 0.77 | 0.63 | 0.68 |
| 10 | 80991900 | rs1749845 | 0.451 | -1.04 | -0.03 | 0.707 | -0.008 | 0.003 | 0.003 | 0 | 0.33 | 0.34 | 0.33 | 0.25 | 0.42 | 0.25 |
| 10 | 80995556 | rs1250593 | -1.327 | -0.996 | 0.019 | 0.627 | 0.011 | -0.029 | -0.029 | 0 | 0.73 | 0.63 | 0.64 | 0.77 | 0.68 | 0.69 |
| 10 | 80996513 | rs10762854 | -0.25 | -1.018 | -0.303 | 0.738 | 0.032 | -0.022 | -0.022 | 0.124 | 0.4 | 0.29 | 0.29 | 0.52 | 0.26 | 0.44 |
| 10 | 80999869 | rs1749849 | -0.188 | -0.947 | -0.363 | 0.834 | -0.006 | -0.041 | -0.041 | 0.498 | 0.42 | 0.24 | 0.44 | 0.44 | 0.54 | 0.62 |
| 10 | 81001175 | rs1782644 | -1.327 | -0.021 | 0.142 | 0.686 | 0.087 | 0.065 | 0.065 | 0.343 | 0.08 | 0.41 | 0.13 | 0.08 | 0.09 | 0.01 |
| 10 | 81010850 | rs1250600 | -0.919 | -0.35 | -0.231 | 0.588 | 0.02 | 0.065 | 0.065 | 0 | 0.58 | 0.69 | 0.59 | 0.8 | 0.64 | 0.7 |
| 10 | 81011589 | rs11002892 | 0.771 | -0.246 | -0.62 | 0.867 | -0.125 | -0.029 | -0.029 | 0 | 0.29 | 0.02 | 0.24 | 0.23 | 0.35 | 0.41 |
| 10 | 81012071 | rs11002893 | 0.293 | -0.139 | -0.196 | 1.099 | 0.018 | 0.093 | 0.093 | 0 | 0.13 | 0.24 | 0.23 | 0.25 | 0.15 | 0.16 |
| 10 | 81013421 | rs4980052 | 0.353 | -0.1 | -1.152 | 0.33 | -0.072 | -0.019 | -0.019 | 0 | 0.36 | 0.09 | 0.3 | 0.32 | 0.38 | 0.4 |
| 10 | 81015896 | rs11593576 | 0.33 | -0.076 | -1.111 | -0.007 | -0.07 | 0.064 | 0.064 | 0 | 0.36 | 0.1 | 0.29 | 0.21 | 0.39 | 0.41 |
| 10 | 81017157 | rs1250532 | -1.264 | -0.016 | -0.844 | -0.38 | -0.002 | 0.081 | 0.081 | 0 | 0.68 | 0.8 | 0.68 | 0.91 | 0.7 | 0.71 |
| 10 | 81018262 | rs11002895 | 0.839 | 0.743 | -0.91 | -0.074 | -0.014 | 0.015 | 0.015 | 0 | 0.19 | 0.14 | 0.14 | 0.1 | 0.25 | 0.09 |
| 10 | 81020402 | rs7908992 | -1.055 | 0.412 | -0.637 | 0.452 | 0.001 | 0.006 | 0.006 | 0 | 0.35 | 0.26 | 0.42 | 0.64 | 0.34 | 0.53 |
| 10 | 81021841 | rs4980054 | -0.943 | 0.801 | -0.223 | 0.589 | -0.013 | 0.005 | 0.005 | 0 | 0.33 | 0.3 | 0.39 | 0.71 | 0.42 | 0.51 |
| 10 | 81030237 | rs4979857 | -1.39 | -0.54 | 0.422 | 0.708 | -0.066 | 0.023 | 0.023 | 0.006 | 0.25 | 0.52 | 0.17 | 0.27 | 0.13 | 0.14 |
| 10 | 81032555 | rs11002902 | -0.894 | -0.54 | 0.48 | 0.703 | 0.005 | 0.147 | 0.147 | 0.018 | 0.11 | 0.48 | 0.14 | 0.22 | 0.09 | 0.04 |
| 10 | 81037229 | rs1250539 | -1.811 | -0.223 | 0.274 | 0.594 | -0.004 | 0.105 | 0.105 | 0.372 | 0.51 | 0.79 | 0.47 | 0.69 | 0.46 | 0.54 |
| 10 | 81037800 | rs1250538 | -0.496 | 0.326 | 0.186 | 0.897 | -0.037 | -0.053 | -0.053 | 0.223 | 0.24 | 0.18 | 0.3 | 0.42 | 0.33 | 0.4 |
| 10 | 81043707 | rs1892497 | 0.319 | 0.457 | -0.042 | 1.034 | -0.1 | -0.103 | -0.103 | 1.435 | 0.14 | 0.02 | 0.24 | 0.34 | 0.27 | 0.32 |
| 10 | 81047575 | rs2802372 | -1.496 | 0.074 | 0.709 | 0.855 | 0.038 | 0.039 | 0.039 | 0 | 0.33 | 0.49 | 0.36 | 0.4 | 0.38 | 0.5 |
| 10 | 81048611 | rs1782645 | -1.379 | 0.104 | 0.999 | 0.766 | 0.035 | -0.006 | -0.006 | 0 | 0.32 | 0.49 | 0.42 | 0.63 | 0.38 | 0.51 |
| 10 | 81049687 | rs12263204 | -0.901 | -0.142 | 1.137 | 0.686 | 0.013 | -0.024 | -0.024 | 0 | 0.07 | 0.24 | 0.05 | 0.01 | 0.05 | 0.04 |
| 10 | 81053475 | rs1250560 | -1.098 | -0.054 | 1.071 | 0.634 | 0.037 | -0.002 | -0.002 | 0 | 0.3 | 0.43 | 0.41 | 0.65 | 0.37 | 0.5 |
| 10 | 81057833 | rs7905321 | -0.07 | -0.08 | 0.978 | 0.873 | -0.034 | 0.082 | 0.082 | 0 | 0.12 | 0.24 | 0.07 | 0.25 | 0.06 | 0.14 |
| 10 | 81058027 | rs1250552 | -1.866 | -0.407 | 0.625 | 0.605 | -0.005 | 0.095 | 0.095 | 0.103 | 0.47 | 0.77 | 0.46 | 0.7 | 0.44 | 0.6 |
| 10 | 81059335 | rs1250551 | 0.108 | -0.314 | 0.422 | 0.701 | -0.045 | -0.05 | -0.05 | 0 | 0.2 | 0.1 | 0.28 | 0.4 | 0.31 | 0.39 |
| 10 | 81060474 | rs1250549 | -1.365 | -0.358 | 0.454 | 0.477 | -0.012 | 0.072 | 0.072 | 0.209 | 0.46 | 0.72 | 0.45 | 0.69 | 0.43 | 0.57 |
| 10 | 81064755 | rs2296424 | 0.55 | -0.443 | -0.301 | 0 | 0.026 | 0.035 | 0.035 | 0 | 0.27 | 0.18 | 0.29 | 0.09 | 0.17 | 0.16 |
| 10 | 81065201 | rs1250558 | 0.744 | -0.128 | -0.973 | 0.164 | -0.121 | -0.128 | -0.128 | 0 | 0.16 | 0.02 | 0.25 | 0.41 | 0.31 | 0.39 |
| 10 | 81065774 | rs1250557 | -1.204 | -0.368 | -0.79 | 0.062 | 0 | 0 | 0 | 0.413 | 0.61 | 0.59 | 0.7 | 0.57 | 0.58 | 0.66 |
| 10 | 81066982 | rs1250555 | -1.209 | 0.652 | -1.156 | 0.442 | -0.072 | -0.063 | -0.063 | 0.124 | 0.32 | 0.2 | 0.36 | 0.44 | 0.41 | 0.5 |
| 10 | 81067040 | rs1250554 | -0.289 | 1.36 | -1.004 | 0.982 | -0.12 | -0.126 | -0.126 | 0.001 | 0.16 | 0.02 | 0.25 | 0.41 | 0.31 | 0.39 |
| 10 | 81074248 | rs11002921 | -0.706 | 0.027 | -0.545 | 0.431 | -0.009 | -0.011 | -0.011 | 0.004 | 0.18 | 0.25 | 0.12 | 0.11 | 0.14 | 0.2 |
| 10 | 81074816 | rs3740259 | 0.273 | 0.097 | -1.003 | 0.295 | -0.007 | 0.014 | 0.014 | 0 | 0.3 | 0.16 | 0.19 | 0.31 | 0.32 | 0.26 |
| 10 | 81077885 | rs884889 | -0.115 | -0.901 | -0.624 | -0.276 | -0.066 | 0.031 | 0.031 | 0.095 | 0.15 | 0.44 | 0.18 | 0.15 | 0.08 | 0.11 |
| 10 | 81089162 | rs11002928 | 1.306 | -0.596 | -0.411 | -0.498 | -0.003 | -0.004 | -0.004 | 0 | 0.22 | 0.21 | 0.15 | 0.15 | 0.19 | 0.18 |
| 10 | 81096071 | rs7077812 | -0.751 | -0.926 | 0.477 | -0.211 | -0.042 | -0.008 | -0.008 | 0 | 0.3 | 0.4 | 0.44 | 0.26 | 0.2 | 0.25 |
| 10 | 81101679 | rs2105461 | -1.18 | -0.202 | 0.336 | -0.309 | -0.069 | 0.026 | 0.026 | 1.426 | 0.57 | 0.85 | 0.67 | 0.64 | 0.53 | 0.47 |
| 10 | 81103862 | rs1892498 | -1.262 | -0.177 | 0.265 | -0.308 | -0.103 | 0.005 | 0.005 | 1.709 | 0.58 | 0.88 | 0.68 | 0.64 | 0.54 | 0.49 |
| 10 | 81104185 | rs4291618 | 0.187 | -0.332 | 0.425 | -0.378 | 0.005 | 0.01 | 0.01 | 0 | 0.13 | 0.08 | 0.18 | 0.01 | 0.1 | 0.11 |
| 10 | 81114060 | rs7332 | -0.781 | -0.252 | 0.334 | -0.089 | -0.083 | 0.017 | 0.017 | 1.802 | 0.56 | 0.87 | 0.67 | 0.65 | 0.53 | 0.46 |
| 10 | 81121960 | rs2181080 | -0.726 | -0.25 | 0.335 | -0.088 | -0.08 | 0.021 | 0.021 | 1.602 | 0.56 | 0.87 | 0.67 | 0.65 | 0.53 | 0.46 |
| 10 | 81132851 | rs1923366 | -0.604 | -0.291 | 0.509 | -0.152 | -0.006 | 0.019 | 0.019 | 0.08 | 0.53 | 0.64 | 0.64 | 0.66 | 0.52 | 0.47 |

| | | | | | | | | | | | | | | | | |
|---|---|---|---|---|---|---|---|---|---|---|---|---|---|---|---|---|
| 10 | 81135045 | rs1923363 | 0.196 | 0.657 | 0.839 | 0.419 | -0.003 | -0.029 | -0.029 | 0 | 0.07 | 0.19 | 0.04 | 0 | 0.04 | 0.02 |
| 10 | 81136015 | rs7082045 | 0.102 | 0.66 | 0.708 | 0.42 | 0.031 | -0.004 | -0.004 | 0 | 0.07 | 0.33 | 0.05 | 0 | 0.04 | 0.02 |
| 10 | 81142365 | rs6858 | -0.69 | 1.274 | -0.919 | 0.53 | -0.017 | 0 | 0 | 0 | 0.39 | 0.35 | 0.52 | 0.62 | 0.48 | 0.37 |
| 10 | 81144697 | rs4979863 | -0.484 | 1.351 | -1.155 | 0.834 | -0.008 | -0.034 | -0.034 | 0 | 0.23 | 0.13 | 0.27 | 0.59 | 0.29 | 0.31 |
| 10 | 81145266 | rs4980069 | 0.886 | 1.412 | -1.956 | 0.709 | -0.049 | -0.041 | -0.041 | 0 | 0.77 | 0.99 | 0.85 | 0.78 | 0.77 | 0.76 |
| 10 | 81145535 | rs1063119 | 0.705 | 2.328 | -1.984 | 0.907 | -0.047 | -0.043 | -0.043 | 2.445 | 0.68 | 0.83 | 0.51 | 0.61 | 0.54 | 0.5 |
| 10 | 81147278 | rs7093996 | 0.634 | 1.95 | -0.876 | 0.105 | 0.037 | 0 | 0 | 0 | 0.81 | 0.58 | 0.82 | 0.8 | 0.72 | 0.77 |
| 10 | 81147948 | rs10740498 | -0.262 | 1.377 | -1.123 | -0.56 | -0.028 | 0.049 | 0.049 | 0 | 0.37 | 0.27 | 0.56 | 0.76 | 0.45 | 0.48 |
| 10 | 81157422 | rs7899991 | -0.057 | 1.271 | -0.939 | -0.35 | -0.002 | -0.001 | -0.001 | 2.568 | 0.85 | 0.77 | 0.83 | 0.9 | 0.84 | 0.89 |
| 10 | 81159038 | rs10824749 | 1.113 | 0.875 | -0.545 | -0.748 | -0.016 | -0.091 | -0.091 | 0.009 | 0.43 | 0.42 | 0.21 | 0.06 | 0.26 | 0.3 |
| 10 | 81160150 | rs4980073 | -0.284 | 1.294 | -1.041 | -0.682 | -0.001 | -0.002 | -0.002 | 2.366 | 0.86 | 0.8 | 0.83 | 0.91 | 0.84 | 0.89 |
| 10 | 81162016 | rs7914731 | -0.284 | 1.294 | -1.041 | -0.534 | -0.001 | 0.002 | 0.002 | 0 | 0.86 | 0.8 | 0.83 | 0.87 | 0.84 | 0.89 |
| 10 | 81164521 | rs2279335 | -0.437 | 1.389 | -1.358 | -0.498 | -0.168 | 0.056 | 0.056 | 0 | 0.34 | 0.04 | 0.54 | 0.79 | 0.49 | 0.52 |
| 10 | 81166079 | rs10740506 | -0.544 | 1.083 | -1.124 | -0.572 | -0.011 | 0.057 | 0.057 | 0 | 0.38 | 0.25 | 0.61 | 0.85 | 0.58 | 0.57 |
| 10 | 81173663 | rs12358729 | 0.839 | 0.644 | -1.502 | -0.29 | -0.011 | 0.033 | 0.033 | 0 | 0.37 | 0.15 | 0.41 | 0.27 | 0.34 | 0.43 |
| 10 | 81175086 | rs2145995 | 0.351 | 0.647 | -1.371 | -0.428 | -0.004 | 0.043 | 0.043 | 0 | 0.45 | 0.27 | 0.46 | 0.26 | 0.4 | 0.44 |
| 10 | 81175110 | rs941871 | -0.631 | 0.56 | -1.524 | -0.388 | -0.001 | -0.01 | -0.01 | 0 | 0.95 | 0.94 | 0.96 | 1 | 0.96 | 0.97 |
| 10 | 81175250 | rs2181079 | 0.903 | 0.739 | -1.558 | -0.25 | -0.01 | 0.035 | 0.035 | 0 | 0.38 | 0.16 | 0.41 | 0.26 | 0.35 | 0.43 |
| 10 | 81180962 | rs11002969 | 0.502 | 0.2 | -1.657 | -0.148 | 0.013 | 0.181 | 0.181 | 0 | 0.49 | 0.09 | 0.43 | 0.21 | 0.48 | 0.49 |
| 10 | 81183884 | rs2395577 | -0.883 | 0.319 | -0.612 | 0.763 | -0.006 | 0.026 | 0.026 | 0 | 0.23 | 0.27 | 0.27 | 0.51 | 0.21 | 0.22 |
| 10 | 81189551 | rs7077340 | -0.726 | 0.415 | -0.548 | 0.255 | 0.004 | 0.004 | 0.004 | 0 | 0.2 | 0.14 | 0.11 | 0.14 | 0.16 | 0.21 |
| 10 | 81190597 | rs2181076 | -1.32 | 0.97 | -0.188 | 0.489 | -0.001 | -0.002 | -0.002 | 0.046 | 0.47 | 0.57 | 0.57 | 0.5 | 0.5 | 0.59 |
| 10 | 81191488 | rs1885433 | 0.478 | 0.173 | 0.224 | 0.595 | -0.004 | 0.006 | 0.006 | 0 | 0.18 | 0.22 | 0.15 | 0.18 | 0.17 | 0.19 |
| 10 | 81192577 | rs7898643 | -1.131 | 0.138 | 0.315 | 0.578 | -0.013 | -0.008 | -0.008 | 0 | 0.4 | 0.52 | 0.26 | 0.32 | 0.33 | 0.39 |
| 10 | 81193126 | rs2395575 | -1.08 | 0.131 | 0.29 | 0.575 | -0.02 | -0.013 | -0.013 | 0 | 0.39 | 0.48 | 0.26 | 0.31 | 0.33 | 0.39 |
| 10 | 81193788 | rs12265906 | -1.096 | -0.018 | 0.51 | 0.51 | 0.027 | 0.019 | 0.019 | 0 | 0.41 | 0.73 | 0.27 | 0.31 | 0.33 | 0.39 |
| 10 | 81194219 | rs4980079 | -1.934 | 0.322 | 0.178 | 0.517 | -0.05 | 0.016 | 0.016 | 0 | 0.61 | 0.86 | 0.67 | 0.66 | 0.58 | 0.75 |
| 10 | 81194236 | rs4980080 | -0.433 | 0.444 | -0.117 | 0.628 | 0.003 | 0.025 | 0.025 | 0.001 | 0.22 | 0.37 | 0.41 | 0.34 | 0.25 | 0.36 |
| 10 | 81195310 | rs2395570 | 0.708 | 0.299 | -0.244 | 0.546 | 0.022 | 0.01 | 0.01 | 0 | 0.13 | 0.02 | 0.04 | 0.13 | 0.08 | 0.18 |
| 10 | 81197177 | rs4980083 | -1.726 | 0.475 | -0.09 | 0.578 | -0.015 | 0.013 | 0.013 | 0 | 0.57 | 0.73 | 0.64 | 0.65 | 0.56 | 0.73 |
| 10 | 81198038 | rs6480941 | 0.046 | 0.165 | 0.145 | 0.619 | -0.017 | 0.017 | 0.017 | 0 | 0.2 | 0.38 | 0.15 | 0.18 | 0.17 | 0.19 |
| 10 | 81208964 | rs16937185 | 0.202 | 0.364 | -0.31 | 0.663 | -0.006 | -0.004 | -0.004 | 0.023 | 0.12 | 0.2 | 0.05 | 0.12 | 0.07 | 0.13 |
| 10 | 81209298 | rs6480943 | 0.433 | 0.431 | -0.329 | 0.697 | -0.007 | 0.017 | 0.017 | 0 | 0.15 | 0.22 | 0.12 | 0.17 | 0.13 | 0.19 |
| 10 | 81212186 | rs12781317 | 0.512 | 0.729 | -0.527 | 0.537 | -0.006 | 0.002 | 0.002 | 0 | 0.16 | 0.14 | 0.16 | 0.16 | 0.22 | 0.15 |
| 10 | 81212511 | rs4979866 | -0.588 | 0.36 | -0.907 | 0.714 | 0.005 | -0.004 | -0.004 | 0.335 | 0.33 | 0.3 | 0.46 | 0.36 | 0.31 | 0.45 |
| 10 | 81212704 | rs6480945 | -1.445 | 0.931 | -0.434 | 0.75 | 0.001 | -0.001 | -0.001 | 0 | 0.45 | 0.51 | 0.36 | 0.44 | 0.49 | 0.48 |
| 10 | 81213070 | rs11002990 | -0.329 | 0.26 | -0.488 | 1.016 | -0.004 | 0.011 | 0.011 | 0.19 | 0.13 | 0.18 | 0.11 | 0.15 | 0.12 | 0.16 |
| 10 | 81215357 | rs11002991 | -0.8 | 0.309 | -0.721 | 0.85 | 0.002 | 0 | 0 | 0 | 0.13 | 0.12 | 0.06 | 0.16 | 0.09 | 0.18 |
| 10 | 81222632 | rs11003003 | -1.246 | 0.47 | -0.921 | 1.266 | 0 | -0.035 | -0.035 | 0.96 | 0.27 | 0.21 | 0.18 | 0.47 | 0.28 | 0.3 |
| 10 | 81223607 | rs10740507 | -1.246 | 0.47 | -0.904 | 1.266 | 0 | -0.029 | -0.029 | 9.924 | 0.27 | 0.22 | 0.18 | 0.47 | 0.28 | 0.3 |
| 10 | 81228041 | rs7909837 | -1.315 | -0.515 | -1.359 | 0.791 | 0.009 | 0.12 | 0.12 | 0.414 | 0.54 | 0.85 | 0.4 | 0.74 | 0.44 | 0.69 |
| 10 | 81232632 | rs12773500 | -0.08 | -0.386 | -1.214 | 0.927 | 0.003 | 0.003 | 0.003 | 0 | 0.16 | 0.13 | 0.07 | 0.16 | 0.16 | 0.19 |
| 10 | 81236210 | rs6480947 | -0.299 | -0.37 | -1.231 | 0.926 | -0.003 | -0.002 | -0.002 | 0 | 0.17 | 0.19 | 0.08 | 0.18 | 0.16 | 0.2 |
| 10 | 81240122 | rs12264210 | -0.424 | -0.548 | -0.18 | 0.998 | -0.003 | 0.1 | 0.1 | 0 | 0.12 | 0.23 | 0.1 | 0.28 | 0.1 | 0.09 |
| 10 | 81244627 | rs11003030 | -1.445 | -0.577 | 0.018 | 0.946 | -0.014 | 0.061 | 0.061 | 0 | 0.3 | 0.46 | 0.19 | 0.46 | 0.28 | 0.29 |
| 10 | 81251344 | rs1351767 | -0.052 | -0.445 | -0.187 | 1.054 | -0.016 | -0.029 | -0.029 | 0.6 | 0.08 | 0.03 | 0.09 | 0.3 | 0.1 | 0.08 |
| 10 | 81266063 | rs1250777 | -0.515 | -0.855 | -0.019 | -0.187 | -0.02 | -0.118 | -0.118 | 0.478 | 0.71 | 0.61 | 0.79 | 0.98 | 0.8 | 0.76 |
| 10 | 81380107 | rs1427815 | 0.154 | -1.664 | -0.449 | 0.154 | 0.051 | 0.021 | 0.021 | 0 | 0.3 | 0.17 | 0.24 | 0.25 | 0.19 | 0.42 |
| 10 | 81388196 | rs1650169 | -0.487 | -1.8 | 0.065 | 0.526 | -0.032 | 0.118 | 0.118 | 0 | 0.2 | 0.36 | 0.23 | 0.49 | 0.14 | 0.17 |
| 10 | 81388222 | rs1713321 | -0.487 | -1.8 | 0.065 | 0.526 | -0.032 | 0.118 | 0.118 | 0 | 0.2 | 0.36 | 0.23 | 0.49 | 0.14 | 0.17 |
| 10 | 81577614 | rs10885307 | 0.363 | -1.322 | -0.08 | 0.166 | 0.017 | 0.03 | 0.03 | 0 | 0.47 | 0.34 | 0.48 | 0.3 | 0.59 | 0.38 |
| 10 | 81652266 | rs927452 | -0.98 | -0.136 | -0.823 | -0.268 | -0.069 | 0.023 | 0.023 | 0 | 0.7 | 0.91 | 0.61 | 0.78 | 0.6 | 0.74 |
| 10 | 81656061 | rs2493719 | -0.331 | -0.352 | -0.749 | -0.538 | -0.053 | 0.064 | 0.064 | 0 | 0.62 | 0.83 | 0.54 | 0.78 | 0.54 | 0.73 |
| 10 | 81656400 | rs7073842 | -1.104 | -0.29 | -0.768 | -0.461 | -0.053 | 0.024 | 0.024 | 0 | 0.7 | 0.89 | 0.62 | 0.78 | 0.6 | 0.74 |
| 10 | 81663575 | rs1983775 | 2.696 | -0.269 | -0.394 | -0.565 | 0.01 | 0.023 | 0.023 | 0 | 0.11 | 0.05 | 0.16 | 0.02 | 0.07 | 0.21 |
| 10 | 81670414 | rs12269573 | -0.169 | 0.106 | 0.371 | -0.459 | 0.045 | -0.027 | -0.027 | 0 | 0.08 | 0.33 | 0.08 | 0 | 0.1 | 0.04 |
| 10 | 81680538 | rs10400059 | -0.086 | -0.023 | 0.195 | -0.409 | -0.004 | -0.061 | -0.061 | 2.909 | 0.1 | 0.25 | 0.09 | 0 | 0.06 | 0.01 |
| 10 | 81680770 | rs11817253 | 0.644 | -0.371 | 0.49 | -0.609 | -0.016 | -0.034 | -0.034 | 0 | 0.06 | 0.17 | 0.02 | 0 | 0.02 | 0 |
| 10 | 81682527 | rs7919775 | -0.159 | -0.349 | 0.426 | -0.6 | -0.004 | -0.066 | -0.066 | 1.238 | 0.11 | 0.28 | 0.1 | 0 | 0.06 | 0.01 |
| 10 | 81684075 | rs7072441 | 0.348 | -0.601 | 0.341 | -0.979 | -0.026 | -0.131 | -0.131 | 0.047 | 0.28 | 0.53 | 0.27 | 0.02 | 0.24 | 0.27 |
| 10 | 81685024 | rs2256581 | 0.038 | -0.7 | -0.172 | -0.406 | 0.001 | -0.006 | -0.006 | 0.941 | 0.37 | 0.35 | 0.31 | 0.48 | 0.33 | 0.38 |
| 10 | 81685062 | rs2256573 | -1.105 | -0.743 | -0.005 | -0.466 | -0.049 | -0.041 | -0.041 | 0 | 0.62 | 0.8 | 0.55 | 0.5 | 0.52 | 0.63 |
| 10 | 81685270 | rs7911085 | -0.387 | -0.839 | 0.026 | -0.357 | 0.009 | 0.106 | 0.106 | 0 | 0.14 | 0.39 | 0.11 | 0.28 | 0.14 | 0.12 |
| 10 | 81690007 | rs1570329 | 1.482 | -0.871 | 0 | -0.383 | -0.015 | -0.029 | -0.029 | 0 | 0.06 | 0.12 | 0.03 | 0 | 0.02 | 0 |
| 10 | 81690295 | rs1923541 | -1.324 | -0.936 | -0.052 | -0.437 | -0.021 | 0.093 | 0.093 | 0 | 0.54 | 0.74 | 0.44 | 0.76 | 0.51 | 0.51 |
| 10 | 81690985 | rs2758556 | -1.973 | -0.916 | -0.103 | -0.432 | -0.05 | 0.021 | 0.021 | 0 | 0.7 | 0.87 | 0.61 | 0.78 | 0.61 | 0.73 |
| 10 | 81691034 | rs2758555 | -1.973 | -0.916 | -0.103 | -0.432 | -0.05 | 0.021 | 0.021 | 0 | 0.7 | 0.87 | 0.61 | 0.78 | 0.61 | 0.73 |
| 10 | 81691080 | rs2758554 | 1.568 | -0.976 | -0.145 | -0.468 | -0.035 | -0.146 | -0.146 | 1.241 | 0.26 | 0.05 | 0.2 | 0.48 | 0.27 | 0.37 |
| 10 | 81694353 | rs12262739 | 1.066 | -0.933 | 0.115 | -0.687 | 0.037 | -0.009 | -0.009 | 0 | 0.07 | 0.27 | 0.06 | 0 | 0.08 | 0.01 |

| | | | | | | | | | | | | | | | | |
|---|---|---|---|---|---|---|---|---|---|---|---|---|---|---|---|---|
| 10 | 81696091 | rs17885031 | 3.695 | -0.893 | -0.027 | -0.65 | 0.003 | 0.052 | 0.052 | 0 | 0.11 | 0.01 | 0.15 | 0.02 | 0.07 | 0.21 |
| 10 | 81696829 | rs17886233 | 1.115 | -0.929 | 0.201 | -0.674 | -0.001 | -0.029 | -0.029 | 0.044 | 0.09 | 0.1 | 0.08 | 0 | 0.06 | 0.01 |
| 10 | 81697818 | rs1051246 | 1.59 | -0.903 | 0.225 | -0.657 | -0.008 | -0.059 | -0.059 | 0 | 0.21 | 0.25 | 0.24 | 0.02 | 0.13 | 0.22 |
| 10 | 81699238 | rs2245121 | -0.704 | -0.873 | 0.187 | -0.595 | 0.011 | 0.03 | 0.03 | 0 | 0.47 | 0.62 | 0.35 | 0.76 | 0.43 | 0.49 |
| 10 | 81701523 | rs911887 | 0.136 | -0.777 | 0.129 | -0.161 | 0 | 0.002 | 0.002 | 0 | 0.36 | 0.39 | 0.3 | 0.36 | 0.41 | 0.41 |
| 10 | 81701722 | rs2243639 | -1.919 | -0.781 | -0.439 | -0.202 | -0.146 | -0.021 | -0.021 | 0.102 | 0.72 | 0.98 | 0.62 | 0.79 | 0.61 | 0.73 |
| 10 | 81702834 | rs10887199 | 1.708 | -0.921 | 0.004 | -0.378 | -0.019 | 0.046 | 0.046 | 0 | 0.2 | 0.32 | 0.21 | 0.43 | 0.11 | 0.3 |
| 10 | 81706135 | rs6413523 | 0.073 | -0.911 | -0.001 | -0.364 | -0.006 | 0.195 | 0.195 | 0.004 | 0.08 | 0.24 | 0.04 | 0.41 | 0.04 | 0.09 |
| 10 | 81706324 | rs721917 | -0.022 | -0.531 | -0.126 | -0.262 | 0.012 | 0.035 | 0.035 | 0 | 0.5 | 0.4 | 0.38 | 0.62 | 0.42 | 0.71 |
| 10 | 81706951 | rs726289 | 3.444 | -0.301 | 0.013 | 0.004 | 0.01 | 0.023 | 0.023 | 0 | 0.11 | 0.06 | 0.16 | 0.02 | 0.07 | 0.21 |
| 10 | 81707613 | rs2819096 | -1.346 | -0.334 | 0.003 | -0.05 | -0.027 | -0.007 | -0.007 | 0.274 | 0.67 | 0.8 | 0.53 | 0.62 | 0.57 | 0.74 |
| 10 | 81708013 | rs723192 | 1.672 | -0.17 | -0.062 | 0.118 | -0.01 | 0.031 | 0.031 | 0 | 0.19 | 0.27 | 0.2 | 0.43 | 0.11 | 0.3 |
| 10 | 81712440 | rs1885550 | 0.786 | -0.14 | -0.325 | 0.252 | -0.005 | 0.027 | 0.027 | 0.09 | 0.24 | 0.31 | 0.25 | 0.56 | 0.18 | 0.41 |
| 10 | 81715738 | rs2146192 | 1.87 | -0.113 | -0.577 | 0.295 | 0.01 | -0.022 | -0.022 | 0.019 | 0.18 | 0.18 | 0.19 | 0.43 | 0.11 | 0.3 |
| 10 | 81733742 | rs4369337 | 0.29 | 0.08 | -0.414 | 0.47 | -0.006 | 0.11 | 0.11 | 0 | 0.07 | 0.16 | 0.04 | 0.41 | 0.03 | 0.09 |
| 10 | 81739920 | rs11201011 | -1.603 | 0.242 | -0.693 | 0.247 | -0.063 | -0.04 | -0.04 | 0 | 0.73 | 0.9 | 0.55 | 0.62 | 0.6 | 0.75 |
| 10 | 81748708 | rs7084667 | 0.727 | 0.228 | -0.308 | -0.095 | 0 | 0.015 | 0.015 | 1.724 | 0.33 | 0.24 | 0.19 | 0.06 | 0.32 | 0.38 |
| 10 | 81754384 | rs4357631 | 0.191 | 0.147 | -0.158 | 0.214 | -0.006 | 0.035 | 0.035 | 0 | 0.13 | 0.17 | 0.06 | 0.41 | 0.06 | 0.09 |
| 10 | 81758718 | rs7908468 | -0.853 | 0.321 | 0.064 | 0.13 | 0.012 | 0.017 | 0.017 | 0.583 | 0.65 | 0.6 | 0.47 | 0.49 | 0.53 | 0.7 |
| 10 | 81758995 | rs7904954 | -1.4 | 0.404 | -0.055 | 0.218 | -0.009 | -0.006 | -0.006 | 0 | 0.71 | 0.74 | 0.53 | 0.64 | 0.6 | 0.81 |
| 10 | 81761401 | rs10887328 | 0.564 | 0.238 | -0.062 | 0.15 | -0.001 | -0.012 | -0.012 | 0 | 0.37 | 0.35 | 0.22 | 0.21 | 0.35 | 0.47 |
| 10 | 81761489 | rs1932571 | 0.348 | 0.073 | -0.24 | 0.375 | 0.017 | -0.014 | -0.014 | 0 | 0.31 | 0.29 | 0.28 | 0.43 | 0.2 | 0.32 |
| 10 | 81764081 | rs10887336 | 0.254 | 0.087 | -0.288 | 0.382 | -0.002 | -0.001 | -0.001 | 0 | 0.32 | 0.36 | 0.29 | 0.43 | 0.2 | 0.32 |
| 10 | 81765502 | rs10788350 | 0.555 | 0.877 | -0.493 | 0.13 | 0.004 | 0.014 | 0.014 | 0 | 0.63 | 0.41 | 0.67 | 0.49 | 0.61 | 0.57 |
| 10 | 81765642 | rs10788351 | -0.541 | 0.425 | -1.072 | -0.624 | 0.008 | -0.041 | -0.041 | 0.899 | 0.38 | 0.47 | 0.3 | 0.12 | 0.48 | 0.22 |
| 10 | 81767388 | rs7070394 | -1.093 | -0.091 | -1.393 | -0.052 | -0.077 | -0.108 | -0.108 | 0.122 | 0.72 | 0.96 | 0.54 | 0.6 | 0.65 | 0.8 |
| 10 | 81788837 | rs7083625 | -0.232 | -0.611 | -0.943 | -0.297 | -0.003 | -0.002 | -0.002 | 0 | 0.46 | 0.43 | 0.28 | 0.51 | 0.47 | 0.53 |
| 10 | 81794163 | rs2342606 | -0.829 | -0.698 | -0.723 | -0.3 | -0.009 | -0.015 | -0.015 | 0.07 | 0.67 | 0.71 | 0.49 | 0.53 | 0.58 | 0.75 |
| 10 | 81807931 | rs11201344 | -0.164 | -0.769 | -1.024 | -0.561 | 0.01 | 0.023 | 0.023 | 0 | 0.45 | 0.3 | 0.32 | 0.65 | 0.38 | 0.45 |
| 10 | 81812036 | rs10887432 | -0.188 | -0.799 | -1.19 | -0.657 | 0.001 | -0.01 | -0.01 | 0 | 0.55 | 0.46 | 0.49 | 0.69 | 0.45 | 0.66 |
| 10 | 81816134 | rs17098169 | 2.849 | -1.06 | -1.124 | -0.573 | 0.004 | 0 | 0 | 0 | 0.12 | 0.11 | 0.16 | 0.02 | 0.08 | 0.21 |
| 10 | 81838656 | rs3824727 | 0.539 | -1.276 | -1.501 | -0.428 | 0.021 | -0.089 | -0.089 | 0.813 | 0.16 | 0.03 | 0.09 | 0.41 | 0.12 | 0.21 |
| 10 | 81841692 | rs1932574 | 0.068 | -0.711 | -1.171 | -0.655 | -0.006 | 0.046 | 0.046 | 0 | 0.44 | 0.23 | 0.34 | 0.69 | 0.41 | 0.47 |
| 10 | 81847914 | rs728616 | 2.87 | -0.798 | -1.072 | -0.642 | -0.009 | -0.04 | -0.04 | 0 | 0.12 | 0.15 | 0.17 | 0.02 | 0.09 | 0.21 |
| 10 | 81852059 | rs3156 | -1.083 | -0.686 | -1.023 | -0.505 | 0.008 | 0.001 | 0.001 | 0.007 | 0.71 | 0.65 | 0.61 | 0.71 | 0.65 | 0.8 |
| 10 | 81859756 | rs2342610 | -1.696 | -0.731 | -1.217 | -0.527 | -0.109 | -0.091 | -0.091 | 0 | 0.77 | 0.98 | 0.63 | 0.71 | 0.66 | 0.8 |
| 10 | 81863021 | rs12259239 | -0.255 | -0.393 | -0.946 | -0.19 | -0.003 | 0.01 | 0.01 | 0 | 0.19 | 0.22 | 0.13 | 0.44 | 0.13 | 0.31 |
| 10 | 81863964 | rs7905482 | 0.262 | -0.343 | -1.05 | -0.35 | -0.001 | 0 | 0 | 0 | 0.35 | 0.37 | 0.29 | 0.25 | 0.33 | 0.25 |
| 10 | 81872073 | rs1078524 | -0.25 | -0.24 | -1.112 | 0 | -0.001 | 0.002 | 0.002 | 0 | 0.57 | 0.6 | 0.52 | 0.28 | 0.54 | 0.53 |
| 10 | 81877845 | rs12249846 | -0.858 | 0.013 | -1.423 | 0.701 | -0.008 | -0.022 | -0.022 | 0.002 | 0.05 | 0.18 | 0.04 | 0.01 | 0.02 | 0.05 |
| 10 | 81881954 | rs11201799 | -0.069 | 0.294 | -2.01 | 1.036 | 0.012 | -0.067 | -0.067 | 0 | 0.13 | 0.03 | 0.09 | 0.41 | 0.11 | 0.27 |
| 10 | 81882370 | rs3851050 | -0.581 | 0.653 | -1.662 | 0.396 | -0.016 | 0.031 | 0.031 | 0 | 0.4 | 0.25 | 0.3 | 0.27 | 0.4 | 0.31 |
| 10 | 81885914 | rs2784761 | 0.81 | 0.701 | -1.519 | 0.485 | -0.023 | 0 | 0 | 0 | 0.85 | 0.94 | 0.87 | 0.9 | 0.84 | 0.89 |
| 10 | 81891206 | rs7078815 | -0.767 | -0.881 | -0.909 | 0.631 | -0.1 | 0.082 | 0.082 | 0.228 | 0.23 | 0.49 | 0.24 | 0.65 | 0.09 | 0.34 |
| 10 | 81891731 | rs2573360 | 0.761 | 0.024 | -0.978 | 0.002 | 0.052 | -0.073 | -0.073 | 0.352 | 0.78 | 0.63 | 0.78 | 0.95 | 0.68 | 0.82 |
| 10 | 81895970 | rs2789698 | 0.026 | -0.515 | -1.639 | 0.17 | -0.022 | -0.058 | -0.058 | 0 | 0.65 | 0.81 | 0.63 | 0.53 | 0.66 | 0.36 |
| 10 | 81900893 | rs2819878 | 0.447 | -0.434 | -1.458 | -0.304 | -0.018 | 0.018 | 0.018 | 0.554 | 0.73 | 0.78 | 0.75 | 0.89 | 0.64 | 0.84 |
| 10 | 81901722 | rs745182 | 0.78 | -1.065 | -0.744 | -0.393 | 0.026 | 0.032 | 0.032 | 0 | 0.66 | 0.48 | 0.71 | 0.56 | 0.61 | 0.61 |
| 10 | 81906119 | rs2819873 | -0.932 | -0.906 | -0.545 | -0.473 | -0.001 | -0.025 | -0.025 | 0.314 | 0.95 | 0.9 | 0.95 | 1 | 0.95 | 0.96 |
| 10 | 81906269 | rs2819872 | 1.025 | -1.324 | -0.372 | -0.331 | -0.001 | 0.039 | 0.039 | 0.05 | 0.48 | 0.36 | 0.44 | 0.34 | 0.51 | 0.3 |
| 10 | 81907884 | rs2789679 | 1.865 | -0.991 | -0.435 | -0.711 | -0.029 | -0.045 | -0.045 | 16.453 | 0.37 | 0.15 | 0.47 | 0.58 | 0.45 | 0.34 |
| 10 | 81908673 | rs10887553 | -0.546 | -1.01 | -0.272 | -0.732 | 0.021 | 0.025 | 0.025 | 0 | 0.27 | 0.32 | 0.24 | 0.4 | 0.34 | 0.37 |
| 10 | 81912179 | rs751740 | -0.883 | -0.989 | -0.363 | -0.725 | -0.001 | 0 | 0 | 0 | 0.46 | 0.55 | 0.42 | 0.43 | 0.44 | 0.62 |
| 10 | 81913561 | rs7091565 | -0.883 | -0.989 | -0.353 | -0.725 | -0.001 | 0 | 0 | 0 | 0.46 | 0.55 | 0.42 | 0.42 | 0.44 | 0.61 |
| 10 | 81915136 | rs2789686 | -0.08 | -1.08 | -0.329 | -0.74 | -0.027 | -0.086 | -0.086 | 0 | 0.83 | 0.67 | 0.89 | 1 | 0.9 | 0.95 |
| 10 | 81915966 | rs2784772 | -1.004 | -1.03 | -0.407 | -0.713 | -0.001 | -0.025 | -0.025 | 1.721 | 0.94 | 0.9 | 0.95 | 1 | 0.95 | 0.96 |
| 10 | 81917708 | rs1079242 | 1.732 | -0.691 | -0.577 | -0.711 | -0.052 | -0.04 | -0.04 | 0.297 | 0.38 | 0.16 | 0.53 | 0.58 | 0.54 | 0.34 |
| 10 | 81917957 | rs3862518 | 2.39 | -1.147 | -0.526 | -1.053 | 0.019 | 0.072 | 0.072 | 0 | 0.15 | 0.03 | 0.17 | 0.02 | 0.1 | 0.24 |
| 10 | 81923205 | rs2236558 | -0.752 | -1.227 | -0.433 | -0.904 | 0.026 | 0.015 | 0.015 | 0 | 0.29 | 0.43 | 0.24 | 0.33 | 0.32 | 0.35 |
| 10 | 81924830 | rs12763624 | 0.743 | -1.113 | -0.665 | -0.882 | -0.039 | 0.01 | 0.01 | 0 | 0.13 | 0.05 | 0.17 | 0.12 | 0.24 | 0.1 |
| 10 | 81926702 | rs1049550 | 1.634 | -0.786 | -0.675 | -0.806 | -0.024 | -0.01 | -0.01 | 0 | 0.37 | 0.16 | 0.47 | 0.64 | 0.45 | 0.34 |
| 10 | 81932680 | rs11201972 | 2 | -0.733 | -0.697 | -0.786 | 0.008 | 0.017 | 0.017 | 0 | 0.17 | 0.09 | 0.18 | 0.02 | 0.13 | 0.24 |
| 10 | 81940701 | rs6585454 | 2 | -0.733 | -0.697 | -0.786 | 0.008 | 0.017 | 0.017 | 0 | 0.17 | 0.09 | 0.19 | 0.02 | 0.13 | 0.24 |
| 10 | 81952956 | rs2819945 | -1.301 | -0.537 | -0.547 | -0.572 | -0.003 | 0.024 | 0.024 | 0 | 0.46 | 0.69 | 0.35 | 0.34 | 0.42 | 0.42 |
| 10 | 81954395 | rs2819944 | 0.213 | -0.526 | -0.419 | -0.306 | -0.003 | -0.024 | -0.024 | 0 | 0.06 | 0.09 | 0.05 | 0 | 0.05 | 0.05 |
| 10 | 81954843 | rs11202059 | -0.254 | -0.57 | -0.368 | -0.206 | -0.025 | 0.06 | 0.06 | 0 | 0.14 | 0.28 | 0.07 | 0.22 | 0.09 | 0.25 |
| 10 | 81961021 | rs2819941 | -1.156 | -0.086 | -1.114 | -0.433 | -0.025 | -0.018 | -0.018 | 0.24 | 0.67 | 0.84 | 0.53 | 0.34 | 0.58 | 0.7 |
| 10 | 81968903 | rs11202112 | 0.489 | -0.129 | -1.258 | -0.712 | -0.023 | 0.055 | 0.055 | 0.248 | 0.1 | 0 | 0.04 | 0 | 0.09 | 0.04 |
| 10 | 81974813 | rs2493948 | -0.231 | -0.623 | -1.177 | -1.052 | -0.001 | -0.029 | -0.029 | 0.13 | 0.19 | 0.23 | 0.11 | 0 | 0.14 | 0.1 |

| | | | | | | | | | | | | | | | | |
|---|---|---|---|---|---|---|---|---|---|---|---|---|---|---|---|---|
| 10 | 81984357 | rs2573377 | -0.168 | -0.155 | -1.245 | -0.746 | -0.002 | 0.036 | 0.036 | 0 | 0.87 | 0.89 | 0.81 | 0.52 | 0.83 | 0.69 |
| 10 | 81984564 | rs11202210 | 0.866 | -0.257 | -1.299 | -0.741 | -0.009 | 0.051 | 0.051 | 0.012 | 0.09 | 0 | 0.04 | 0 | 0.07 | 0.04 |
| 10 | 82012599 | rs12219032 | 1.07 | -0.35 | -1.365 | -1.065 | 0.034 | -0.088 | -0.088 | 0 | 0.32 | 0.14 | 0.3 | 0.48 | 0.29 | 0.22 |
| 10 | 82012748 | rs12217395 | 1.07 | -0.35 | -1.37 | -1.065 | 0.033 | -0.091 | -0.091 | 0 | 0.32 | 0.14 | 0.3 | 0.48 | 0.29 | 0.23 |
| 10 | 82012967 | rs12411782 | 1.128 | -0.363 | -1.379 | -1.081 | 0.03 | -0.094 | -0.094 | 0 | 0.32 | 0.13 | 0.29 | 0.47 | 0.29 | 0.23 |
| 10 | 82013119 | rs12411806 | 1.116 | -0.375 | -1.385 | -1.087 | 0.03 | -0.094 | -0.094 | 0 | 0.32 | 0.14 | 0.29 | 0.47 | 0.29 | 0.23 |
| 10 | 82018594 | rs4120852 | -0.116 | -0.298 | -1.18 | -1.792 | 0.016 | -0.108 | -0.108 | 0 | 0.68 | 0.42 | 0.72 | 0.98 | 0.6 | 0.8 |
| 10 | 82020569 | rs10887701 | 0.239 | -0.434 | -1.249 | -1.864 | 0.022 | 0.054 | 0.054 | 0 | 0.48 | 0.34 | 0.49 | 0.96 | 0.44 | 0.5 |
| 10 | 82024042 | rs1556893 | -0.134 | -0.595 | -1.306 | -1.834 | -0.002 | -0.005 | -0.005 | 0 | 0.19 | 0.13 | 0.21 | 0.46 | 0.2 | 0.29 |
| 10 | 82024062 | rs1538816 | 0.184 | -0.528 | -1.268 | -1.836 | 0.02 | 0.059 | 0.059 | 0 | 0.48 | 0.34 | 0.49 | 0.95 | 0.45 | 0.51 |
| 10 | 82029072 | rs1556897 | -0.222 | -0.554 | -1.237 | -1.872 | 0 | 0.048 | 0.048 | 0 | 0.54 | 0.35 | 0.53 | 0.95 | 0.5 | 0.56 |
| 10 | 82029333 | rs1010245 | -0.222 | -0.554 | -1.237 | -1.872 | 0 | 0.048 | 0.048 | 0 | 0.54 | 0.35 | 0.53 | 0.95 | 0.5 | 0.56 |
| 10 | 82030658 | rs3851059 | -1.011 | -0.57 | -1.272 | -1.826 | 0.005 | -0.035 | -0.035 | 0 | 0.3 | 0.21 | 0.28 | 0.46 | 0.28 | 0.35 |
| 10 | 82031278 | rs3827869 | -0.136 | -0.627 | -1.28 | -1.827 | -0.004 | -0.023 | -0.023 | 0 | 0.19 | 0.12 | 0.21 | 0.46 | 0.21 | 0.3 |
| 10 | 82032240 | rs1985908 | 1.253 | -0.751 | -1.204 | -1.938 | 0.003 | -0.024 | -0.024 | 0 | 0.37 | 0.33 | 0.44 | 0.95 | 0.35 | 0.46 |
| 10 | 82032331 | rs1832683 | 1.657 | -0.864 | -0.605 | -2.125 | 0.002 | -0.026 | -0.026 | 1.183 | 0.25 | 0.25 | 0.23 | 0.02 | 0.17 | 0.29 |
| 10 | 82033594 | rs2993763 | -1.488 | -0.985 | -0.21 | -2.179 | 0.011 | -0.026 | -0.026 | 0 | 0.53 | 0.79 | 0.52 | 0.49 | 0.42 | 0.64 |
| 10 | 82033683 | rs4933327 | -0.02 | -0.991 | -0.203 | -2.183 | -0.003 | -0.013 | -0.013 | 0 | 0.19 | 0.12 | 0.21 | 0.46 | 0.21 | 0.3 |
| 10 | 82034262 | rs2994388 | -1.526 | -1.021 | -0.231 | -2.196 | 0.008 | -0.028 | -0.028 | 0 | 0.54 | 0.79 | 0.52 | 0.49 | 0.42 | 0.64 |
| 10 | 82034854 | rs10788546 | -0.608 | -0.963 | -0.204 | -2.182 | -0.094 | 0.104 | 0.104 | 0 | 0.78 | 0.93 | 0.78 | 0.97 | 0.64 | 0.84 |
| 10 | 82035560 | rs4934027 | -0.059 | -0.921 | -0.146 | -2.134 | -0.003 | -0.013 | -0.013 | 0.534 | 0.19 | 0.12 | 0.21 | 0.46 | 0.21 | 0.3 |
| 10 | 82038779 | rs4934028 | -1.004 | -0.908 | -0.307 | -2.132 | 0.042 | -0.006 | -0.006 | 0 | 0.51 | 0.73 | 0.49 | 0.49 | 0.4 | 0.6 |
| 10 | 82040346 | rs2282367 | -1.209 | -0.722 | -0.443 | -2.243 | -0.12 | 0.106 | 0.106 | 0 | 0.83 | 0.97 | 0.8 | 0.97 | 0.67 | 0.85 |
| 10 | 82041751 | rs756208 | -0.992 | -0.725 | -0.436 | -2.245 | -0.077 | 0.072 | 0.072 | 0 | 0.82 | 0.93 | 0.79 | 0.97 | 0.67 | 0.85 |
| 10 | 82042624 | rs10887718 | 0.683 | -0.887 | -0.404 | -2.313 | 0.022 | -0.116 | -0.116 | 0.189 | 0.64 | 0.41 | 0.67 | 0.97 | 0.54 | 0.76 |
| 10 | 82043226 | rs12242871 | 0.693 | -0.886 | -0.404 | -2.302 | 0.021 | -0.112 | -0.112 | 5.173 | 0.64 | 0.41 | 0.67 | 0.97 | 0.54 | 0.76 |
| 10 | 82043576 | rs2236569 | -1.218 | -0.781 | -0.466 | -2.179 | -0.12 | 0.07 | 0.07 | 0 | 0.83 | 0.97 | 0.8 | 0.94 | 0.67 | 0.85 |
| 10 | 82044920 | rs7081756 | -1.708 | -0.811 | -0.46 | -2.253 | 0.001 | -0.011 | -0.011 | 0 | 0.63 | 0.72 | 0.54 | 0.07 | 0.62 | 0.5 |
| 10 | 82046615 | rs1819684 | 1.672 | -1.028 | -0.266 | -2.128 | -0.002 | -0.025 | -0.025 | 0 | 0.09 | 0.07 | 0.04 | 0 | 0.07 | 0.04 |
| 10 | 82054873 | rs9420359 | -1.348 | -0.861 | -0.362 | -2.124 | -0.073 | 0.066 | 0.066 | 0 | 0.82 | 0.92 | 0.79 | 0.98 | 0.67 | 0.85 |
| 10 | 82057988 | rs10788549 | 0.48 | -1.231 | -0.189 | -2.109 | 0.028 | 0.051 | 0.051 | 0 | 0.5 | 0.36 | 0.51 | 0.95 | 0.44 | 0.48 |
| 10 | 82062661 | rs1591958 | 0.011 | -1.215 | -0.174 | -1.786 | -0.032 | -0.027 | -0.027 | 0.028 | 0.35 | 0.37 | 0.24 | 0.19 | 0.2 | 0.52 |
| 10 | 82077032 | rs10400045 | -1.02 | -1.002 | -0.363 | -1.838 | -0.091 | 0.093 | 0.093 | 0 | 0.8 | 0.92 | 0.78 | 0.98 | 0.67 | 0.86 |
| 10 | 82082340 | rs2066167 | 1.611 | -1.125 | -0.328 | -1.858 | 0.011 | 0.009 | 0.009 | 0 | 0.11 | 0.08 | 0.04 | 0 | 0.06 | 0.04 |
| 10 | 82101849 | rs10788571 | 1.851 | -1.155 | -0.078 | -1.881 | -0.015 | 0.048 | 0.048 | 0.187 | 0.36 | 0.19 | 0.46 | 0.79 | 0.42 | 0.29 |
| 10 | 82103230 | rs12411833 | 1.802 | -1.22 | 0.037 | -1.915 | -0.031 | 0.011 | 0.011 | 0 | 0.2 | 0.29 | 0.19 | 0.19 | 0.11 | 0.47 |
| 10 | 82108610 | rs10788579 | 1.836 | -1.169 | -0.033 | -1.872 | -0.016 | 0.047 | 0.047 | 0.22 | 0.35 | 0.19 | 0.46 | 0.79 | 0.42 | 0.29 |
| 10 | 82112873 | rs1934697 | 0.402 | -1.203 | -0.041 | -1.887 | -0.001 | -0.014 | -0.014 | 0 | 0.53 | 0.54 | 0.54 | 0.79 | 0.51 | 0.34 |
| 10 | 82114425 | rs12570038 | 1.99 | -1.119 | -0.696 | -2.011 | -0.036 | -0.02 | -0.02 | 0.016 | 0.06 | 0.01 | 0.13 | 0.3 | 0.12 | 0.07 |
| 10 | 82118408 | rs17103728 | 0.847 | -1.096 | -0.66 | -2.11 | 0.022 | 0.038 | 0.038 | 0 | 0.15 | 0.08 | 0.06 | 0 | 0.1 | 0.05 |
| 10 | 82118422 | rs7085863 | 1.377 | -0.887 | -0.849 | -2.071 | 0 | -0.005 | -0.005 | 0 | 0.07 | 0.08 | 0.06 | 0 | 0.06 | 0.05 |
| 10 | 82118498 | rs11202666 | 2.427 | -0.784 | -1.058 | -1.926 | -0.041 | -0.15 | -0.15 | 0 | 0.27 | 0.01 | 0.32 | 0.51 | 0.28 | 0.22 |
| 10 | 82122793 | rs1538818 | 2.524 | -0.806 | -0.648 | -2.391 | -0.008 | 0.06 | 0.06 | 0 | 0.14 | 0.02 | 0.17 | 0.03 | 0.11 | 0.3 |
| 10 | 82136664 | rs10749572 | -0.018 | -0.802 | -0.44 | -2.398 | -0.041 | 0.08 | 0.08 | 0 | 0.62 | 0.78 | 0.61 | 0.81 | 0.56 | 0.39 |
| 10 | 82137670 | rs1538815 | -0.67 | -0.468 | -0.426 | -2.172 | -0.049 | 0.044 | 0.044 | 0 | 0.8 | 0.89 | 0.78 | 0.98 | 0.68 | 0.86 |
| 10 | 82143436 | rs2343301 | 1.505 | -0.444 | -0.543 | -2.128 | -0.006 | 0.078 | 0.078 | 0.03 | 0.35 | 0.2 | 0.46 | 0.81 | 0.4 | 0.29 |
| 10 | 82149655 | rs10466235 | 0.229 | 0.108 | -0.188 | -1.791 | 0 | -0.003 | -0.003 | 0.459 | 0.77 | 0.8 | 0.75 | 0.84 | 0.67 | 0.69 |
| 10 | 82165427 | rs7081889 | 0.247 | 0.118 | -0.228 | -1.801 | -0.003 | 0 | 0 | 0 | 0.77 | 0.81 | 0.75 | 0.84 | 0.67 | 0.7 |
| 10 | 82174761 | rs6585989 | -0.471 | -0.515 | -0.051 | -2.087 | 0.008 | 0.005 | 0.005 | 0.24 | 0.39 | 0.6 | 0.32 | 0.16 | 0.25 | 0.58 |
| 10 | 82181434 | rs17679603 | 2.24 | -0.669 | -0.381 | -2.195 | -0.02 | 0.068 | 0.068 | 0 | 0.15 | 0.01 | 0.17 | 0.03 | 0.12 | 0.35 |
| 10 | 82182328 | rs1870143 | 1.673 | -0.586 | -0.44 | -2.357 | -0.006 | 0.151 | 0.151 | 0.717 | 0.4 | 0.11 | 0.45 | 0.82 | 0.43 | 0.3 |
| 10 | 82182558 | rs1870144 | 1.673 | -0.62 | 0.121 | -2.371 | -0.006 | 0.151 | 0.151 | 0.717 | 0.4 | 0.11 | 0.45 | 0.82 | 0.43 | 0.3 |
| 10 | 82189686 | rs7916618 | 1.031 | -0.77 | 0.496 | -2.447 | -0.014 | -0.048 | -0.048 | 0 | 0.1 | 0.18 | 0.04 | 0 | 0.06 | 0.04 |
| 10 | 82189772 | rs17105302 | 0.948 | -0.816 | 0.534 | -2.473 | 0 | -0.036 | -0.036 | 1.568 | 0.08 | 0.14 | 0.05 | 0 | 0.07 | 0.05 |
| 10 | 82191334 | rs7074356 | 1.124 | -0.829 | 0.6 | -2.479 | -0.03 | -0.034 | -0.034 | 0.255 | 0.22 | 0.38 | 0.22 | 0.16 | 0.12 | 0.5 |
| 10 | 82191643 | rs6585994 | -0.03 | -0.813 | 0.432 | -2.483 | -0.004 | -0.001 | -0.001 | 0.088 | 0.6 | 0.61 | 0.56 | 0.82 | 0.56 | 0.39 |
| 10 | 82194330 | rs7907770 | -0.055 | -0.679 | 0.311 | -2.377 | -0.019 | -0.104 | -0.104 | 0.1 | 0.36 | 0.5 | 0.29 | 0.04 | 0.24 | 0.44 |
| 10 | 82196293 | rs3195899 | 1.641 | -0.656 | 0.041 | -2.298 | 0.001 | 0.137 | 0.137 | 0.252 | 0.4 | 0.15 | 0.45 | 0.81 | 0.43 | 0.3 |
| 10 | 82203506 | rs4934167 | 0.817 | -0.439 | 0.456 | -2.199 | -0.001 | -0.036 | -0.036 | 0.174 | 0.59 | 0.61 | 0.6 | 0.95 | 0.51 | 0.59 |
| 10 | 82203663 | rs4934169 | -0.813 | 0 | 0.391 | -2.243 | -0.051 | 0.046 | 0.046 | 0 | 0.85 | 0.91 | 0.84 | 0.98 | 0.73 | 0.91 |
| 10 | 82206348 | rs1870145 | 1.103 | -0.136 | 0.451 | -2.12 | -0.009 | 0.064 | 0.064 | 0.004 | 0.58 | 0.31 | 0.61 | 0.85 | 0.54 | 0.46 |
| 10 | 82208192 | rs2168721 | 0.064 | -0.051 | 0.067 | -2.118 | -0.041 | 0.039 | 0.039 | 0 | 0.73 | 0.8 | 0.74 | 0.95 | 0.64 | 0.66 |
| 10 | 82209807 | rs11202960 | 0.656 | -0.041 | -0.022 | -2.074 | 0.011 | -0.065 | -0.065 | 0 | 0.65 | 0.62 | 0.68 | 0.95 | 0.57 | 0.61 |
| 10 | 82218964 | rs12220642 | 0.675 | 0.402 | -0.133 | -2.028 | 0.023 | -0.041 | -0.041 | 0 | 0.66 | 0.53 | 0.68 | 0.85 | 0.61 | 0.51 |
| 10 | 82219066 | rs12220655 | 0.623 | 0.372 | -0.195 | -2.04 | 0.017 | -0.031 | -0.031 | 0 | 0.66 | 0.57 | 0.68 | 0.85 | 0.61 | 0.51 |
| 10 | 82223767 | rs10887933 | 1.063 | 0.161 | -0.759 | -2.135 | 0.011 | 0.157 | 0.157 | 0 | 0.52 | 0.24 | 0.57 | 0.85 | 0.47 | 0.47 |
| 10 | 82232706 | rs7080169 | -0.112 | 0.18 | -1.117 | -1.738 | 0.006 | 0.034 | 0.034 | 0 | 0.44 | 0.3 | 0.42 | 0.34 | 0.4 | 0.48 |
| 10 | 82234587 | rs878652 | -0.515 | 0.215 | -0.792 | -1.745 | -0.017 | -0.029 | -0.029 | 0 | 0.19 | 0.7 | 0.14 | 0.08 | 0.11 | 0.13 |
| 10 | 82237126 | rs10887949 | 1.913 | 0.086 | -1.391 | -1.513 | -0.052 | -0.148 | -0.148 | 0.088 | 0.22 | 0.01 | 0.29 | 0.53 | 0.25 | 0.22 |

| | | | | | | | | | | | | | | | | |
|---|---|---|---|---|---|---|---|---|---|---|---|---|---|---|---|---|
| 10 | 82241930 | rs7088704 | 0.141 | 0.203 | -0.757 | -1.844 | -0.004 | -0.048 | -0.048 | 0.003 | 0.39 | 0.57 | 0.34 | 0.11 | 0.33 | 0.46 |
| 10 | 82247139 | rs2343308 | -0.345 | 0.509 | -0.71 | -1.927 | 0.007 | -0.056 | -0.056 | 1.515 | 0.61 | 0.52 | 0.61 | 0.83 | 0.59 | 0.52 |
| 10 | 82250831 | rs7097656 | -0.834 | 0.498 | -1.093 | -1.893 | -0.043 | 0.036 | 0.036 | 0 | 0.92 | 0.97 | 0.89 | 0.98 | 0.81 | 0.94 |
| 10 | 82254490 | rs10785884 | 0.268 | 0.459 | -0.73 | -1.871 | 0.003 | 0.148 | 0.148 | 0 | 0.5 | 0.28 | 0.53 | 0.83 | 0.48 | 0.46 |
| 10 | 82256258 | rs10881600 | -0.201 | 0.419 | -0.539 | -1.892 | -0.002 | -0.004 | -0.004 | 0 | 0.57 | 0.37 | 0.59 | 0.83 | 0.55 | 0.52 |
| 10 | 82262826 | rs12260962 | 0.968 | 0.384 | -0.753 | -1.932 | 0.002 | 0.001 | 0.001 | 0 | 0.3 | 0.21 | 0.27 | 0.03 | 0.3 | 0.38 |
| 10 | 82263954 | rs12411484 | 2.311 | 0.588 | -1.696 | -1.941 | -0.049 | -0.069 | -0.069 | 0 | 0.11 | 0.01 | 0.14 | 0.29 | 0.14 | 0.09 |
| 10 | 82264813 | rs9633739 | -0.321 | 0.403 | -1.121 | -2.231 | 0.007 | -0.06 | -0.06 | 0 | 0.59 | 0.53 | 0.59 | 0.83 | 0.55 | 0.51 |
| 10 | 82277341 | rs12264252 | 1.591 | 0.758 | -2.172 | -2.007 | -0.009 | 0.156 | 0.156 | 0.355 | 0.27 | 0.05 | 0.24 | 0.03 | 0.24 | 0.41 |
| 10 | 82279927 | rs873259 | -0.158 | 1.156 | -1.576 | -1.453 | -0.066 | 0.217 | 0.217 | 0 | 0.44 | 0.11 | 0.5 | 0.82 | 0.47 | 0.39 |
| 10 | 82279960 | rs873258 | 0.691 | 1.166 | -1.711 | -1.443 | -0.094 | 0.114 | 0.114 | 0 | 0.34 | 0.04 | 0.44 | 0.82 | 0.39 | 0.3 |
| 10 | 82283538 | rs11599251 | -0.095 | 1.139 | -0.902 | -0.974 | -0.002 | 0.018 | 0.018 | 0.067 | 0.1 | 0.05 | 0.05 | 0 | 0.08 | 0.09 |
| 10 | 82285533 | rs17106495 | -0.634 | 1.327 | -0.238 | -0.898 | 0.037 | -0.041 | -0.041 | 1.691 | 0.08 | 0.21 | 0.09 | 0 | 0.11 | 0.05 |
| 10 | 82295398 | rs11185932 | 0.403 | 1.184 | -0.795 | -0.413 | -0.004 | -0.027 | -0.027 | 0 | 0.35 | 0.11 | 0.33 | 0.55 | 0.32 | 0.3 |
| 10 | 82307117 | rs1870136 | -1.602 | 1.706 | 0.212 | -0.413 | 0.018 | 0.006 | 0.006 | 0.047 | 0.21 | 0.63 | 0.3 | 0.32 | 0.27 | 0.24 |
| 10 | 82313437 | rs7068369 | -1.254 | 1.169 | 0.823 | -0.381 | 0.063 | 0.183 | 0.183 | 0.029 | 0.12 | 0.41 | 0.19 | 0.3 | 0.13 | 0.13 |
| 10 | 82315290 | rs11186029 | 0.763 | 0.896 | 0.199 | -0.26 | -0.005 | -0.106 | -0.106 | 0.244 | 0.31 | 0.08 | 0.26 | 0.5 | 0.29 | 0.27 |
| 10 | 82320993 | rs4038156 | -1.537 | 0.899 | 0.051 | -0.36 | -0.112 | 0.156 | 0.156 | 0 | 0.61 | 0.91 | 0.61 | 0.85 | 0.52 | 0.53 |
| 10 | 82325999 | rs7914650 | -1.777 | 0.864 | 0.775 | -0.211 | -0.008 | 0.003 | 0.003 | 0 | 0.26 | 0.55 | 0.33 | 0.36 | 0.23 | 0.25 |
| 10 | 82331962 | rs7094314 | -0.814 | 0.879 | 1.091 | -0.093 | -0.02 | 0.104 | 0.104 | 0 | 0.18 | 0.36 | 0.31 | 0.4 | 0.12 | 0.16 |
| 10 | 82334411 | rs1573973 | -0.904 | 0.874 | 1.095 | -0.091 | -0.004 | 0.215 | 0.215 | 0 | 0.12 | 0.35 | 0.28 | 0.39 | 0.08 | 0.15 |
| 10 | 82336263 | rs7904557 | -0.924 | 0.872 | 1.112 | -0.096 | -0.023 | 0.123 | 0.123 | 0 | 0.18 | 0.45 | 0.32 | 0.39 | 0.12 | 0.17 |
| 10 | 82338336 | rs7894923 | -1.486 | 1.009 | 0.611 | -0.365 | -0.046 | 0.119 | 0.119 | 0 | 0.62 | 0.87 | 0.61 | 0.84 | 0.55 | 0.47 |
| 10 | 82343793 | rs12777577 | -1.494 | 1 | 1.494 | 0.052 | -0.008 | 0.151 | 0.151 | 0.302 | 0.16 | 0.43 | 0.26 | 0.38 | 0.15 | 0.2 |
| 10 | 82344011 | rs7909966 | -1.537 | 0.99 | 1.508 | 0.025 | -0.022 | 0.074 | 0.074 | 0.262 | 0.22 | 0.49 | 0.3 | 0.39 | 0.17 | 0.21 |
| 10 | 82344455 | rs6586088 | -2.192 | 0.988 | 1.274 | 0.047 | -0.004 | 0.01 | 0.01 | 0.48 | 0.26 | 0.67 | 0.35 | 0.39 | 0.24 | 0.25 |
| 10 | 82347832 | rs2138287 | 1.029 | 0.59 | 0.592 | 0.243 | 0.014 | -0.013 | -0.013 | 0.638 | 0.2 | 0.16 | 0.15 | 0.24 | 0.12 | 0.14 |
| 10 | 82349452 | rs12766852 | 0.168 | 0.594 | 0.751 | 0.351 | -0.004 | 0 | 0 | 0 | 0.08 | 0.13 | 0.18 | 0.07 | 0.08 | 0.11 |
| 10 | 82349461 | rs1878041 | -1.685 | 0.681 | 0.397 | 0.359 | 0.052 | 0.043 | 0.043 | 0.527 | 0.47 | 0.81 | 0.49 | 0.63 | 0.37 | 0.4 |
| 10 | 82355408 | rs17107222 | -0.071 | 1.023 | 0.746 | 0.727 | -0.002 | 0.049 | 0.049 | 0 | 0.12 | 0.2 | 0.09 | 0.31 | 0.1 | 0.1 |
| 10 | 82359534 | rs12569832 | 0.408 | 1.372 | 0.017 | 0.591 | -0.016 | -0.006 | -0.006 | 0 | 0.19 | 0.1 | 0.15 | 0.21 | 0.24 | 0.1 |
| 10 | 82359857 | rs7089872 | 0.961 | 1.094 | 0.118 | 0.785 | 0.012 | -0.08 | -0.08 | 0 | 0.18 | 0.03 | 0.14 | 0.24 | 0.13 | 0.14 |
| 10 | 82362638 | rs10881806 | -0.512 | 1.4 | 0.627 | 0.043 | 0.026 | -0.008 | -0.008 | 0.146 | 0.25 | 0.44 | 0.2 | 0.21 | 0.23 | 0.11 |
| 10 | 82363340 | rs7069048 | -0.518 | 1.093 | 0.429 | 0.371 | -0.019 | 0.103 | 0.103 | 1.045 | 0.28 | 0.52 | 0.25 | 0.51 | 0.24 | 0.26 |
| 10 | 82364513 | rs10785965 | -1.5 | 0.801 | -0.46 | 0.157 | 0.019 | 0.133 | 0.133 | 0 | 0.55 | 0.81 | 0.65 | 0.79 | 0.62 | 0.44 |
| 10 | 82364539 | rs17107333 | -0.48 | 0.822 | 0.465 | 0.752 | 0.043 | 0.08 | 0.08 | 0 | 0.18 | 0.4 | 0.13 | 0.24 | 0.16 | 0.08 |
| 10 | 82366612 | rs7073839 | -0.063 | 0.706 | -0.468 | 0.483 | -0.058 | -0.079 | -0.079 | 0 | 0.28 | 0.1 | 0.33 | 0.55 | 0.34 | 0.2 |
| 10 | 82368221 | rs17693104 | -1.482 | 0.352 | -0.847 | -0.09 | -0.041 | 0.179 | 0.179 | 0 | 0.55 | 0.9 | 0.6 | 0.8 | 0.49 | 0.37 |
| 10 | 82368864 | rs10881815 | -1.481 | 0.495 | -0.623 | 0.051 | -0.053 | 0.21 | 0.21 | 0 | 0.5 | 0.9 | 0.61 | 0.79 | 0.45 | 0.32 |
| 10 | 82369030 | rs10881816 | -0.671 | 0.848 | 0.216 | 0.344 | 0.001 | 0 | 0 | 0 | 0.33 | 0.34 | 0.42 | 0.62 | 0.38 | 0.23 |
| 10 | 82369816 | rs2880655 | -0.317 | 0.761 | -0.13 | 0.443 | -0.065 | -0.067 | -0.067 | 0.108 | 0.29 | 0.11 | 0.37 | 0.54 | 0.33 | 0.19 |
| 10 | 82372300 | rs6586106 | -1.219 | 0.75 | 0.258 | 0.376 | -0.008 | 0.03 | 0.03 | 0 | 0.45 | 0.68 | 0.6 | 0.73 | 0.43 | 0.23 |
| 10 | 82372981 | rs4934372 | -1.225 | 0.789 | -0.187 | 0.408 | -0.03 | 0.094 | 0.094 | 0 | 0.45 | 0.82 | 0.6 | 0.73 | 0.43 | 0.23 |
| 10 | 82374890 | rs12414791 | -1.662 | 0.126 | 0.182 | 0.76 | 0.022 | 0.097 | 0.097 | 0 | 0.08 | 0.32 | 0.19 | 0.19 | 0.08 | 0.03 |
| 10 | 82377997 | rs12414728 | -0.624 | 0.079 | -0.325 | 0.728 | -0.008 | 0.005 | 0.005 | 0 | 0.12 | 0.18 | 0.18 | 0.16 | 0.07 | 0.03 |
| 10 | 82378624 | rs7085490 | -0.912 | 0.207 | -0.871 | 0.537 | 0.013 | 0.05 | 0.05 | 0.023 | 0.42 | 0.84 | 0.56 | 0.73 | 0.35 | 0.27 |
| 10 | 82379403 | rs12220111 | -0.386 | 0.153 | 0.088 | 0.957 | -0.016 | 0.016 | 0.016 | 0 | 0.25 | 0.41 | 0.28 | 0.29 | 0.18 | 0.12 |
| 10 | 82380665 | rs7067934 | -0.183 | 0.557 | -0.54 | 0.949 | -0.034 | -0.041 | -0.041 | 0 | 0.91 | 0.99 | 0.83 | 0.85 | 0.89 | 0.95 |
| 10 | 82382102 | rs17093759 | -0.762 | 0.23 | 0.197 | 0.666 | -0.033 | -0.008 | -0.008 | 0.146 | 0.35 | 0.57 | 0.43 | 0.36 | 0.28 | 0.17 |
| 10 | 82382553 | rs12219619 | -0.687 | 0.309 | 0.414 | 0.655 | -0.006 | -0.005 | -0.005 | 0 | 0.08 | 0.19 | 0.1 | 0.07 | 0.08 | 0.03 |
| 10 | 82385041 | rs9325452 | -0.186 | 0.308 | 0.292 | 0.657 | 0.001 | 0 | 0 | 0 | 0.06 | 0.08 | 0.08 | 0.07 | 0.07 | 0.03 |
| 10 | 82401289 | rs1913752 | -1.102 | 0.245 | 0.157 | 0.464 | 0.051 | 0.044 | 0.044 | 0 | 0.43 | 0.83 | 0.51 | 0.37 | 0.35 | 0.23 |
| 10 | 82405083 | rs11186419 | 0.041 | 0.044 | -0.057 | 0.592 | -0.003 | -0.002 | -0.002 | 0 | 0.06 | 0.11 | 0.09 | 0.06 | 0.05 | 0.02 |
| 10 | 82405254 | rs7087704 | -0.207 | 0.006 | -0.417 | 0.483 | 0.016 | 0.003 | 0.003 | 0 | 0.27 | 0.2 | 0.34 | 0.28 | 0.22 | 0.2 |
| 10 | 82405540 | rs7081754 | -0.276 | 0.005 | -0.445 | 0.483 | 0.019 | 0.005 | 0.005 | 0 | 0.28 | 0.19 | 0.34 | 0.28 | 0.22 | 0.2 |
| 10 | 82406274 | rs12783710 | 0.057 | 0.115 | -1.221 | 0.545 | 0.008 | -0.041 | -0.041 | 0.009 | 0.25 | 0.05 | 0.32 | 0.28 | 0.21 | 0.18 |
| 10 | 82408975 | rs11186429 | 0.377 | 0.578 | -0.815 | 0.402 | -0.028 | 0.043 | 0.043 | 0 | 0.42 | 0.24 | 0.26 | 0.29 | 0.47 | 0.63 |
| 10 | 82411802 | rs1913751 | 1.18 | 0.367 | -0.536 | 0.389 | -0.002 | -0.002 | -0.002 | 0.344 | 0.84 | 0.84 | 0.86 | 0.98 | 0.82 | 0.87 |
| 10 | 82414719 | rs4934395 | -1.326 | -0.257 | -0.325 | 1.174 | 0.001 | 0 | 0 | 3.242 | 0.28 | 0.31 | 0.17 | 0.43 | 0.2 | 0.25 |
| 10 | 82417756 | rs11186460 | -1.461 | -0.356 | 0.232 | 1.462 | -0.028 | 0.062 | 0.062 | 0 | 0.29 | 0.61 | 0.27 | 0.44 | 0.21 | 0.23 |
| 10 | 82419693 | rs11186481 | -1.373 | 0.027 | 0.394 | 1.801 | -0.003 | 0.204 | 0.204 | 0 | 0.11 | 0.23 | 0.18 | 0.38 | 0.07 | 0.16 |
| 10 | 82423260 | rs1157794 | -0.919 | 0.26 | -0.081 | 1.186 | -0.013 | 0.029 | 0.029 | 0 | 0.11 | 0.06 | 0.06 | 0 | 0.09 | 0.06 |
| 10 | 82423687 | rs1913754 | 0.868 | 0.256 | -0.341 | 1.132 | -0.004 | -0.006 | -0.006 | 0 | 0.69 | 0.73 | 0.7 | 0.64 | 0.73 | 0.67 |
| 10 | 82424127 | rs7904536 | 0.595 | 0.494 | -0.703 | 0.557 | -0.002 | -0.001 | -0.001 | 0.131 | 0.45 | 0.46 | 0.45 | 0.56 | 0.4 | 0.54 |
| 10 | 82424161 | rs7908544 | -0.254 | 0.458 | -0.978 | 1.119 | -0.021 | -0.051 | -0.051 | 0 | 0.83 | 0.95 | 0.78 | 0.7 | 0.82 | 0.76 |
| 10 | 82424293 | rs7908701 | -0.254 | 0.458 | -0.991 | 1.114 | -0.023 | -0.055 | -0.055 | 0.056 | 0.83 | 0.95 | 0.78 | 0.7 | 0.82 | 0.76 |
| 10 | 82426023 | rs17637953 | -0.826 | 0.182 | -0.789 | 1 | -0.002 | -0.017 | -0.017 | 0 | 0.09 | 0.03 | 0.14 | 0.1 | 0.08 | 0.04 |
| 10 | 82431413 | rs12255425 | -0.911 | -0.588 | -0.751 | 1.321 | 0.024 | -0.043 | -0.043 | 0 | 0.26 | 0.83 | 0.2 | 0.5 | 0.15 | 0.25 |
| 10 | 82431787 | rs9971172 | -0.638 | -0.43 | 0.163 | 1.432 | -0.02 | 0.155 | 0.155 | 0 | 0.2 | 0.46 | 0.17 | 0.44 | 0.15 | 0.21 |

| | | | | | | | | | | | | | | | |
|---|---|---|---|---|---|---|---|---|---|---|---|---|---|---|---|
| 10 | 82433893 | rs10881912 | 0.462 | -0.306 | -0.053 | 0.987 | -0.01 | -0.015 | -0.015 | 0 | 0.6 | 0.5 | 0.44 | 0.3 | 0.68 | 0.54 |
| 10 | 82445987 | rs17107520 | -0.562 | 0.184 | -0.38 | 0.968 | -0.001 | -0.005 | -0.005 | 0.602 | 0.25 | 0.27 | 0.3 | 0.37 | 0.25 | 0.25 |
| 10 | 82448181 | rs17107524 | -0.929 | -0.136 | -0.758 | 0.378 | 0.004 | 0 | 0 | 0 | 0.36 | 0.32 | 0.34 | 0.37 | 0.34 | 0.35 |
| 10 | 82448841 | rs11186576 | 0.697 | -0.634 | -0.297 | 0.225 | -0.02 | -0.011 | -0.011 | 0 | 0.12 | 0.2 | 0.09 | 0.08 | 0.05 | 0.09 |
| 10 | 82450804 | rs1559554 | 1.526 | -0.777 | -1.133 | -0.165 | -0.114 | 0.045 | 0.045 | 0.006 | 0.25 | 0.06 | 0.3 | 0.15 | 0.39 | 0.29 |
| 10 | 82452030 | rs9420495 | -1.886 | -1.239 | -0.732 | -0.382 | 0.005 | -0.02 | -0.02 | 0.019 | 0.8 | 0.84 | 0.77 | 0.61 | 0.82 | 0.76 |
| 10 | 82452051 | rs2161885 | 0.165 | -1.263 | -0.287 | -0.521 | -0.004 | 0.009 | 0.009 | 0.063 | 0.43 | 0.38 | 0.42 | 0.23 | 0.48 | 0.42 |
| 10 | 82452426 | rs10881916 | 1.589 | -1.208 | -0.299 | -0.509 | -0.019 | 0.016 | 0.016 | 0 | 0.26 | 0.2 | 0.31 | 0.15 | 0.39 | 0.29 |
| 10 | 82454295 | rs4934419 | -0.561 | -1.436 | -0.23 | -0.269 | -0.003 | -0.009 | -0.009 | 0.152 | 0.58 | 0.51 | 0.49 | 0.36 | 0.62 | 0.58 |
| 10 | 82456323 | rs9421683 | -0.29 | -1.504 | -0.256 | -0.486 | -0.003 | 0.002 | 0.002 | 0 | 0.66 | 0.42 | 0.67 | 0.52 | 0.77 | 0.69 |
| 10 | 82456681 | rs11186609 | 0.985 | -1.172 | -0.393 | -0.788 | 0.003 | -0.008 | -0.008 | 0.045 | 0.11 | 0.07 | 0.06 | 0 | 0.11 | 0.1 |
| 10 | 82464478 | rs7900843 | -0.273 | -1.49 | -0.603 | -1.02 | 0 | -0.003 | -0.003 | 0.04 | 0.39 | 0.34 | 0.34 | 0.12 | 0.39 | 0.29 |
| 10 | 82467629 | rs7914246 | 0.393 | -2.37 | -0.431 | -0.806 | 0.005 | 0.003 | 0.003 | 0.537 | 0.48 | 0.44 | 0.43 | 0.42 | 0.65 | 0.51 |
| 10 | 82471832 | rs6586155 | -1.659 | -2.472 | -0.331 | -0.712 | -0.002 | 0.011 | 0.011 | 0 | 0.83 | 0.83 | 0.77 | 0.68 | 0.86 | 0.78 |
| 10 | 82472563 | rs720262 | 0.09 | -2.602 | -0.389 | -1.072 | 0.005 | 0.005 | 0.005 | 0 | 0.54 | 0.44 | 0.55 | 0.19 | 0.58 | 0.46 |
| 10 | 82473065 | rs10509415 | 1.104 | -2.579 | -0.394 | -1.043 | 0.023 | 0.107 | 0.107 | 0 | 0.18 | 0.29 | 0.19 | 0.49 | 0.24 | 0.26 |
| 10 | 82474165 | rs10881952 | 0.717 | -2.587 | -0.351 | -1.505 | -0.043 | 0.187 | 0.187 | 0 | 0.39 | 0.11 | 0.37 | 0.04 | 0.53 | 0.33 |
| 10 | 82476351 | rs9421690 | 0.654 | -2.584 | -0.311 | -1.498 | -0.02 | 0.142 | 0.142 | 0 | 0.4 | 0.14 | 0.37 | 0.04 | 0.54 | 0.34 |
| 10 | 82478836 | rs7896067 | -0.891 | -2.522 | -0.308 | -1.453 | 0 | -0.021 | -0.021 | 0 | 0.66 | 0.63 | 0.58 | 0.19 | 0.62 | 0.52 |
| 10 | 82479714 | rs9421693 | 0.9 | -1.566 | -0.181 | -1.036 | -0.003 | 0.063 | 0.063 | 0.492 | 0.15 | 0.03 | 0.15 | 0.03 | 0.11 | 0.1 |
| 10 | 82480172 | rs7923403 | -0.197 | -1.451 | -0.138 | -0.966 | -0.001 | -0.016 | -0.016 | 0 | 0.46 | 0.54 | 0.52 | 0.19 | 0.55 | 0.4 |
| 10 | 82498194 | rs10882008 | 0.845 | -0.962 | -0.213 | -0.71 | 0.023 | -0.025 | -0.025 | 0.11 | 0.27 | 0.39 | 0.32 | 0.15 | 0.37 | 0.21 |
| 10 | 82502636 | rs1702226 | -0.738 | -2.01 | 0.628 | -0.328 | -0.019 | -0.118 | -0.118 | 0.059 | 0.31 | 0.37 | 0.26 | 0.03 | 0.23 | 0.27 |
| 10 | 82502982 | rs11186793 | 1.839 | -2.226 | 0.266 | -0.067 | -0.046 | 0.065 | 0.065 | 0 | 0.4 | 0.09 | 0.31 | 0.24 | 0.56 | 0.3 |
| 10 | 82506563 | rs1702231 | -0.078 | -2.108 | 0.621 | 0.15 | -0.011 | -0.048 | -0.048 | 1.067 | 0.15 | 0.21 | 0.1 | 0.01 | 0.08 | 0.1 |
| 10 | 82508346 | rs1702235 | -0.234 | -2.021 | 0.615 | 0.694 | -0.009 | 0.024 | 0.024 | 0 | 0.79 | 0.78 | 0.72 | 0.6 | 0.83 | 0.77 |
| 10 | 82510574 | rs11186820 | -0.355 | -1.738 | 0.567 | 0.971 | -0.01 | 0.247 | 0.247 | 0 | 0.06 | 0.27 | 0.14 | 0.34 | 0.01 | 0.14 |
| 10 | 82517519 | rs983808 | -1.51 | -2.002 | 0.022 | -0.148 | 0.124 | 0.164 | 0.164 | 0.178 | 0.5 | 0.85 | 0.53 | 0.79 | 0.3 | 0.63 |
| 10 | 82517845 | rs10882026 | -0.942 | -2.128 | 0.293 | -0.207 | 0.027 | 0.025 | 0.025 | 0.373 | 0.37 | 0.73 | 0.46 | 0.79 | 0.24 | 0.51 |
| 10 | 82532727 | rs2344770 | -1.774 | -2.029 | 0.159 | -0.225 | 0.08 | 0.153 | 0.153 | 0 | 0.53 | 0.91 | 0.55 | 0.81 | 0.32 | 0.63 |
| 10 | 82533854 | rs12772468 | -0.08 | -2.029 | 0.425 | -0.256 | 0.011 | -0.016 | -0.016 | 0.333 | 0.07 | 0.12 | 0.13 | 0.02 | 0.11 | 0.17 |
| 10 | 82541219 | rs7394377 | -0.364 | -2.274 | 0.944 | -0.353 | -0.018 | 0 | 0 | 0.064 | 0.25 | 0.29 | 0.29 | 0.79 | 0.11 | 0.3 |
| 10 | 82543712 | rs11186988 | 0.132 | -2.255 | 0.838 | -0.351 | 0.001 | -0.007 | -0.007 | 0 | 0.06 | 0.09 | 0.11 | 0.02 | 0.09 | 0.16 |
| 10 | 82545386 | rs10882061 | -0.454 | -2.265 | 0.885 | -0.375 | -0.012 | 0 | 0 | 0 | 0.25 | 0.28 | 0.29 | 0.78 | 0.11 | 0.3 |
| 10 | 82547572 | rs11187005 | 0.492 | -2.252 | 0.898 | -0.381 | 0.009 | -0.011 | -0.011 | 0.103 | 0.06 | 0.22 | 0.04 | 0.01 | 0.02 | 0.03 |
| 10 | 82547720 | rs10509416 | 0.131 | -2.115 | 0.549 | -0.312 | 0.007 | -0.015 | -0.015 | 0 | 0.06 | 0.12 | 0.11 | 0.02 | 0.1 | 0.16 |
| 10 | 82553046 | rs7087248 | -1.709 | -2.175 | 0.406 | -0.315 | 0.119 | 0.143 | 0.143 | 0 | 0.48 | 0.82 | 0.51 | 0.81 | 0.28 | 0.55 |
| 10 | 82561214 | rs17107698 | 0.336 | -2.203 | 0.354 | 0.01 | 0.029 | -0.066 | -0.066 | 0 | 0.18 | 0.08 | 0.2 | 0.55 | 0.1 | 0.22 |
| 10 | 82561579 | rs2194539 | -1.109 | -1.994 | 0.329 | -0.672 | -0.022 | -0.113 | -0.113 | 0 | 0.25 | 0.57 | 0.24 | 0.03 | 0.19 | 0.28 |
| 10 | 82563444 | rs1863044 | -0.414 | -1.26 | -0.472 | -0.236 | -0.004 | -0.002 | -0.002 | 0 | 0.4 | 0.58 | 0.39 | 0.79 | 0.19 | 0.41 |
| 10 | 82566702 | rs6583663 | -0.549 | -1.09 | -0.882 | -0.013 | 0.061 | 0.053 | 0.053 | 0 | 0.42 | 0.71 | 0.42 | 0.78 | 0.18 | 0.49 |
| 10 | 82568041 | rs12098864 | -0.574 | -0.319 | -0.855 | -0.035 | 0.037 | -0.014 | -0.014 | 0.699 | 0.06 | 0.16 | 0.11 | 0.03 | 0.09 | 0.14 |
| 10 | 82568684 | rs10882097 | -0.827 | -0.723 | -1.564 | -0.179 | 0.102 | 0.254 | 0.254 | 0.266 | 0.49 | 0.91 | 0.52 | 0.86 | 0.27 | 0.62 |
| 10 | 82570802 | rs1702249 | -0.515 | -0.042 | -1.044 | 0.124 | 0 | -0.001 | -0.001 | 0 | 0.55 | 0.62 | 0.44 | 0.47 | 0.49 | 0.66 |
| 10 | 82572109 | rs12266945 | 0.139 | -0.309 | -1.056 | 0.106 | 0 | 0 | 0 | 0.173 | 0.36 | 0.35 | 0.31 | 0.37 | 0.33 | 0.3 |
| 10 | 82572667 | rs1647196 | -0.827 | -0.39 | -0.898 | 0.036 | 0.002 | 0.003 | 0.003 | 0 | 0.51 | 0.62 | 0.4 | 0.47 | 0.43 | 0.56 |
| 10 | 82573190 | rs1702220 | -0.78 | -0.308 | -0.593 | 0.044 | -0.006 | -0.017 | -0.017 | 0 | 0.18 | 0.43 | 0.12 | 0.12 | 0.14 | 0.29 |
| 10 | 82574780 | rs1647203 | -0.689 | -0.646 | -0.193 | 0.285 | 0 | -0.001 | -0.001 | 0 | 0.44 | 0.4 | 0.32 | 0.42 | 0.38 | 0.55 |
| 10 | 82575289 | rs12253068 | -1.428 | -0.812 | 0.127 | 0.746 | 0.041 | -0.02 | -0.02 | 0 | 0.06 | 0.25 | 0.03 | 0 | 0.04 | 0.08 |
| 10 | 82575778 | rs7075795 | -1.11 | -0.833 | 0.568 | 0.369 | 0.023 | -0.116 | -0.116 | 0.36 | 0.15 | 0.5 | 0.08 | 0 | 0.12 | 0.08 |
| 10 | 82576150 | rs7918269 | -1.325 | -0.861 | 0.627 | 0.357 | 0.015 | -0.124 | -0.124 | 0 | 0.16 | 0.53 | 0.08 | 0 | 0.12 | 0.08 |
| 10 | 82585081 | rs4439456 | -0.87 | -0.875 | 0.821 | 0.35 | 0.015 | -0.077 | -0.077 | 0 | 0.13 | 0.32 | 0.06 | 0 | 0.11 | 0.08 |
| 10 | 82605064 | rs6583698 | -1.256 | -0.759 | 0.057 | 0.214 | -0.008 | 0.012 | 0.012 | 0 | 0.57 | 0.81 | 0.36 | 0.38 | 0.47 | 0.59 |
| 10 | 82613445 | rs7909919 | -0.716 | -0.281 | 0.416 | 0.124 | 0.028 | -0.069 | -0.069 | 0.152 | 0.19 | 0.38 | 0.1 | 0.05 | 0.16 | 0.15 |
| 10 | 82616316 | rs1911456 | -0.681 | -0.281 | 0.438 | 0.124 | 0.036 | -0.075 | -0.075 | 0 | 0.18 | 0.43 | 0.1 | 0.05 | 0.16 | 0.15 |
| 10 | 82616655 | rs7068982 | -1.284 | -0.326 | 0.441 | 0.072 | 0.006 | -0.081 | -0.081 | 0.731 | 0.22 | 0.65 | 0.11 | 0.05 | 0.16 | 0.15 |
| 10 | 82618149 | rs6583704 | -0.577 | -0.259 | 0.305 | 0.104 | 0.034 | -0.067 | -0.067 | 0 | 0.18 | 0.38 | 0.1 | 0.05 | 0.16 | 0.15 |
| 10 | 82621639 | rs17695110 | -0.381 | -0.065 | 0.038 | -0.201 | -0.005 | 0.006 | 0.006 | 0 | 0.33 | 0.27 | 0.19 | 0.09 | 0.28 | 0.33 |
| 10 | 82627787 | rs954405 | -1.337 | 0.595 | -0.279 | -0.409 | -0.08 | 0.064 | 0.064 | 0 | 0.83 | 0.99 | 0.72 | 0.98 | 0.67 | 0.84 |
| 10 | 82634035 | rs7912360 | -0.177 | 0.438 | -0.215 | -0.252 | -0.006 | -0.082 | -0.082 | 0.155 | 0.36 | 0.59 | 0.2 | 0.08 | 0.24 | 0.31 |
| 10 | 82639591 | rs9971180 | 0.76 | 0.389 | -0.32 | 0.823 | 0 | -0.002 | -0.002 | 0 | 0.26 | 0.22 | 0.33 | 0.3 | 0.24 | 0.28 |
| 10 | 82650614 | rs10882244 | -1.009 | 0.547 | 0.363 | 1.446 | -0.018 | 0.098 | 0.098 | 0.091 | 0.25 | 0.47 | 0.33 | 0.39 | 0.23 | 0.29 |
| 10 | 82653860 | rs2345189 | -1.479 | 1.066 | 0.486 | 1.42 | 0.009 | 0.036 | 0.036 | 0.006 | 0.28 | 0.58 | 0.39 | 0.41 | 0.33 | 0.3 |
| 10 | 82655429 | rs1876331 | 1.404 | 0.912 | -0.27 | 1.042 | -0.046 | 0.163 | 0.163 | 0.298 | 0.43 | 0.11 | 0.39 | 0.21 | 0.41 | 0.36 |
| 10 | 82655793 | rs735687 | 0.777 | 0.891 | 0.067 | 1.293 | -0.002 | 0.006 | 0.006 | 1.274 | 0.73 | 0.35 | 0.66 | 0.64 | 0.75 | 0.73 |
| 10 | 82656178 | rs2135728 | -0.657 | 1.038 | 0.11 | 1.448 | -0.001 | 0.037 | 0.037 | 0 | 0.86 | 0.97 | 0.77 | 0.96 | 0.83 | 0.85 |
| 10 | 82656283 | rs1002745 | 0.035 | 1.031 | 0.1 | 1.454 | -0.008 | -0.009 | -0.009 | 0 | 0.29 | 0.24 | 0.26 | 0.4 | 0.35 | 0.37 |
| 10 | 82656896 | rs1911461 | 0.676 | 1.012 | 0.683 | 1.422 | 0.009 | 0.079 | 0.079 | 0 | 0.78 | 0.7 | 0.69 | 0.64 | 0.75 | 0.73 |
| 10 | 82664872 | rs2345193 | 0.97 | 0.487 | 0.283 | 1.304 | 0 | -0.005 | -0.005 | 0 | 0.49 | 0.51 | 0.44 | 0.23 | 0.45 | 0.36 |

| | | | | | | | | | | | | | | | | |
|---|---|---|---|---|---|---|---|---|---|---|---|---|---|---|---|---|
| 10 | 82665743 | rs4933198 | 0.978 | 0.48 | 0.279 | 1.368 | 0 | -0.016 | -0.016 | 1.664 | 0.49 | 0.51 | 0.45 | 0.17 | 0.45 | 0.36 |
| 10 | 82668530 | rs10882295 | -0.367 | 0.467 | 0.029 | 1.293 | -0.015 | 0.018 | 0.018 | 0.149 | 0.35 | 0.24 | 0.33 | 0.74 | 0.4 | 0.5 |
| 10 | 82670626 | rs12245456 | 0.465 | 0.443 | -0.563 | 1.421 | 0 | -0.035 | -0.035 | 0.085 | 0.24 | 0.13 | 0.25 | 0.36 | 0.24 | 0.29 |
| 10 | 82672362 | rs2345196 | -0.507 | 0.455 | -0.59 | 1.366 | 0.002 | 0.036 | 0.036 | 0 | 0.86 | 0.97 | 0.83 | 0.97 | 0.85 | 0.86 |
| 10 | 82672668 | rs2345199 | -0.507 | 0.455 | -0.59 | 1.366 | 0.002 | 0.036 | 0.036 | 0 | 0.86 | 0.97 | 0.83 | 0.97 | 0.85 | 0.86 |
| 10 | 82672817 | rs2345201 | 0.651 | 0.472 | -0.473 | 1.463 | -0.003 | 0.032 | 0.032 | 0 | 0.78 | 0.74 | 0.75 | 0.63 | 0.8 | 0.71 |
| 10 | 82673699 | rs17622815 | 1.14 | 0.408 | -0.651 | 0.849 | -0.066 | 0.148 | 0.148 | 0.201 | 0.45 | 0.14 | 0.42 | 0.23 | 0.44 | 0.36 |
| 10 | 82673974 | rs7922868 | 0.076 | 0.354 | -0.317 | 0.912 | 0.002 | -0.006 | -0.006 | 0 | 0.08 | 0.12 | 0.09 | 0.03 | 0.1 | 0.09 |
| 10 | 82674284 | rs7923313 | -0.688 | 0.137 | -0.29 | 0.932 | -0.007 | 0.005 | 0.005 | 0 | 0.43 | 0.48 | 0.4 | 0.51 | 0.38 | 0.46 |
| 10 | 82674625 | rs10786132 | -0.494 | 0.471 | -0.961 | 0.744 | 0.002 | -0.012 | -0.012 | 0 | 0.39 | 0.27 | 0.33 | 0.43 | 0.36 | 0.42 |
| 10 | 82677160 | rs7893463 | -0.367 | 0.952 | -1.257 | 0.547 | -0.084 | -0.134 | -0.134 | 0.241 | 0.73 | 0.97 | 0.44 | 0.46 | 0.66 | 0.64 |
| 10 | 82678708 | rs4545501 | -0.389 | 0.718 | -1.056 | 0.084 | -0.003 | -0.042 | -0.042 | 0 | 0.14 | 0.19 | 0.07 | 0.02 | 0.13 | 0.22 |
| 10 | 82679542 | rs2881070 | 0.396 | 0.054 | -0.751 | -0.285 | -0.015 | -0.051 | -0.051 | 0 | 0.3 | 0.53 | 0.17 | 0.17 | 0.26 | 0.25 |
| 10 | 82682250 | rs7902968 | 0.103 | 0.021 | -0.751 | -0.279 | 0.005 | 0.004 | 0.004 | 0.126 | 0.36 | 0.71 | 0.19 | 0.16 | 0.24 | 0.35 |
| 10 | 82683220 | rs729724 | -0.809 | 0.062 | -0.752 | -0.261 | -0.09 | -0.101 | -0.101 | 1.966 | 0.69 | 0.93 | 0.41 | 0.44 | 0.59 | 0.58 |
| 10 | 82684928 | rs11187681 | 1.422 | 0.097 | -1.086 | -0.579 | -0.031 | 0.091 | 0.091 | 0 | 0.16 | 0.02 | 0.09 | 0.02 | 0.13 | 0.2 |
| 10 | 82685062 | rs6583746 | -0.962 | 0.141 | -0.94 | -0.555 | -0.041 | -0.047 | -0.047 | 0 | 0.69 | 0.84 | 0.41 | 0.44 | 0.59 | 0.58 |
| 10 | 82696713 | rs2345209 | 0.003 | 0.13 | -0.857 | -0.523 | 0.039 | 0.004 | 0.004 | 0 | 0.5 | 0.66 | 0.3 | 0.43 | 0.46 | 0.36 |
| 10 | 82702830 | rs1484177 | 1.651 | 0.447 | -0.641 | -0.365 | -0.003 | -0.004 | -0.004 | 0 | 0.16 | 0.1 | 0.13 | 0.27 | 0.21 | 0.04 |
| 10 | 82703101 | rs7078742 | 1.13 | 0.2 | -0.992 | -0.548 | -0.002 | -0.008 | -0.008 | 0 | 0.23 | 0.18 | 0.1 | 0.07 | 0.19 | 0.14 |
| 10 | 82704895 | rs7088102 | -0.653 | 1.059 | -0.923 | 0.009 | -0.081 | -0.098 | -0.098 | 1.715 | 0.73 | 0.96 | 0.44 | 0.51 | 0.66 | 0.62 |
| 10 | 82706117 | rs10509420 | -0.173 | 1.2 | -0.103 | 0.736 | 0.057 | 0.155 | 0.155 | 0 | 0.21 | 0.4 | 0.18 | 0.41 | 0.28 | 0.12 |
| 10 | 82707610 | rs11187780 | -0.145 | 0.816 | -1.17 | 0.605 | -0.022 | -0.03 | -0.03 | 0.448 | 0.33 | 0.17 | 0.5 | 0.42 | 0.34 | 0.44 |
| 10 | 82707770 | rs10882390 | -0.18 | 0.345 | -0.578 | 0.488 | -0.016 | 0.006 | 0.006 | 0 | 0.3 | 0.39 | 0.18 | 0.29 | 0.28 | 0.23 |
| 10 | 82708724 | rs12772949 | -0.06 | -0.151 | -0.564 | 0.178 | 0.033 | 0.009 | 0.009 | 0.16 | 0.28 | 0.11 | 0.2 | 0.21 | 0.2 | 0.28 |
| 10 | 82708869 | rs12770949 | -0.039 | -0.021 | -0.574 | 0.208 | 0.024 | 0.006 | 0.006 | 0.056 | 0.28 | 0.1 | 0.2 | 0.21 | 0.2 | 0.28 |
| 10 | 82710334 | rs2881072 | 0.547 | -0.016 | -0.393 | 0.103 | -0.009 | -0.014 | -0.014 | 0 | 0.28 | 0.63 | 0.3 | 0.23 | 0.24 | 0.17 |
| 10 | 82711637 | rs10748620 | -0.1 | 0.359 | -0.798 | -0.24 | 0.012 | 0.039 | 0.039 | 0.978 | 0.53 | 0.65 | 0.64 | 0.24 | 0.59 | 0.35 |
| 10 | 82715404 | rs9988716 | 1.548 | 0.217 | -0.907 | -0.295 | -0.019 | -0.015 | -0.015 | 0 | 0.21 | 0.07 | 0.25 | 0.24 | 0.23 | 0.08 |
| 10 | 82715797 | rs9988765 | -1.032 | 0.337 | -0.889 | -0.276 | 0.007 | 0.009 | 0.009 | 0.147 | 0.7 | 0.78 | 0.76 | 0.79 | 0.74 | 0.72 |
| 10 | 82724076 | rs11187818 | 0.576 | 0.771 | -1.018 | 0.198 | 0 | 0.031 | 0.031 | 0 | 0.11 | 0.11 | 0.07 | 0.47 | 0.09 | 0.29 |
| 10 | 82724799 | rs997312 | 0.197 | 0.85 | -1.14 | -0.31 | -0.089 | 0.163 | 0.163 | 0 | 0.27 | 0.05 | 0.34 | 0.01 | 0.35 | 0.27 |
| 10 | 82727810 | rs1545891 | 0.125 | 0.838 | -1.05 | -0.302 | -0.053 | 0.105 | 0.105 | 0 | 0.28 | 0.09 | 0.34 | 0.01 | 0.36 | 0.27 |
| 10 | 82736647 | rs4511237 | 0.07 | 0.504 | -0.484 | 0.247 | -0.002 | 0.15 | 0.15 | 0 | 0.16 | 0.26 | 0.12 | 0.52 | 0.16 | 0.35 |
| 10 | 82737123 | rs12357777 | 0.2 | 0.859 | -0.667 | -0.604 | -0.032 | 0.075 | 0.075 | 0 | 0.26 | 0.1 | 0.32 | 0.02 | 0.3 | 0.26 |
| 10 | 82738613 | rs10882440 | 0.349 | 0.691 | -0.658 | -0.624 | -0.046 | 0.113 | 0.113 | 0 | 0.23 | 0.06 | 0.32 | 0.01 | 0.28 | 0.26 |
| 10 | 82746127 | rs2345221 | 0.349 | 0.691 | -0.663 | -0.624 | -0.048 | 0.118 | 0.118 | 0 | 0.23 | 0.06 | 0.32 | 0.01 | 0.28 | 0.26 |
| 10 | 82746486 | rs1073077 | -0.322 | 0.38 | -0.648 | -0.854 | 0 | -0.013 | -0.013 | 0 | 0.48 | 0.57 | 0.61 | 0.21 | 0.5 | 0.36 |
| 10 | 82747785 | rs10509423 | 1.129 | 0.44 | -0.601 | -0.753 | -0.028 | -0.002 | -0.002 | 1.593 | 0.2 | 0.09 | 0.25 | 0.2 | 0.21 | 0.08 |
| 10 | 82764044 | rs1484173 | -0.247 | -0.014 | -0.381 | -0.664 | 0 | 0.006 | 0.006 | 0.064 | 0.51 | 0.58 | 0.6 | 0.39 | 0.49 | 0.37 |
| 10 | 82771544 | rs10882491 | 1.362 | -0.399 | -0.415 | -0.549 | 0.01 | -0.008 | -0.008 | 0 | 0.16 | 0.09 | 0.11 | 0.15 | 0.12 | 0.07 |
| 10 | 82776591 | rs3912390 | -0.549 | -0.413 | -0.277 | -0.703 | 0.04 | 0.067 | 0.067 | 0.098 | 0.47 | 0.67 | 0.42 | 0.28 | 0.38 | 0.34 |
| 10 | 82785364 | rs11188088 | 1.593 | -0.142 | -0.326 | -0.577 | -0.017 | -0.01 | -0.01 | 0 | 0.14 | 0.01 | 0.1 | 0.12 | 0.12 | 0.09 |
| 10 | 82795129 | rs900785 | -1.24 | -0.25 | -0.54 | -0.905 | -0.005 | 0.041 | 0.041 | 0 | 0.78 | 0.88 | 0.82 | 0.93 | 0.78 | 0.75 |
| 10 | 82795474 | rs7084611 | -0.295 | -0.086 | 0.309 | -0.509 | -0.019 | -0.031 | -0.031 | 0.437 | 0.27 | 0.45 | 0.3 | 0.21 | 0.25 | 0.2 |
| 10 | 82795855 | rs878824 | 1.413 | -0.083 | 0.315 | -0.475 | -0.001 | 0 | 0 | 1.247 | 0.15 | 0.13 | 0.11 | 0.14 | 0.12 | 0.1 |
| 10 | 82800744 | rs1484166 | 0.069 | -0.095 | 0.535 | -0.494 | -0.001 | -0.003 | -0.003 | 0.296 | 0.24 | 0.29 | 0.32 | 0.21 | 0.25 | 0.19 |
| 10 | 82802046 | rs7897412 | -0.042 | -0.105 | 0.608 | -0.498 | -0.006 | -0.016 | -0.016 | 0 | 0.25 | 0.39 | 0.3 | 0.21 | 0.25 | 0.19 |
| 10 | 82804044 | rs1484167 | 1.461 | -0.111 | 0.613 | -0.501 | -0.001 | 0.001 | 0.001 | 1.4 | 0.15 | 0.13 | 0.11 | 0.14 | 0.12 | 0.09 |
| 10 | 82808595 | rs1001828 | -0.596 | -0.113 | 0.607 | -0.502 | -0.002 | -0.002 | -0.002 | 0.121 | 0.41 | 0.53 | 0.4 | 0.35 | 0.37 | 0.28 |
| 10 | 82813678 | rs10882541 | 1.475 | -0.099 | 0.302 | -0.516 | 0.004 | -0.005 | -0.005 | 0.9 | 0.15 | 0.09 | 0.1 | 0.13 | 0.12 | 0.09 |
| 10 | 82815674 | rs1414909 | -0.706 | -0.091 | 0.34 | -0.515 | 0.007 | 0.006 | 0.006 | 0.131 | 0.42 | 0.6 | 0.42 | 0.35 | 0.37 | 0.29 |
| 10 | 82821606 | rs7922340 | 0.125 | -0.126 | 0.324 | -0.499 | -0.001 | -0.052 | -0.052 | 0 | 0.24 | 0.32 | 0.2 | 0.02 | 0.25 | 0.18 |
| 10 | 82824422 | rs1336442 | -0.391 | -0.104 | 0.328 | -0.482 | -0.016 | -0.026 | -0.026 | 0 | 0.28 | 0.53 | 0.31 | 0.21 | 0.25 | 0.2 |
| 10 | 82832257 | rs1336439 | -0.73 | -0.022 | 0.19 | -0.458 | 0.011 | 0.01 | 0.01 | 0.16 | 0.43 | 0.63 | 0.42 | 0.35 | 0.37 | 0.29 |
| 10 | 82842329 | rs12414042 | -0.323 | 0.403 | 0.213 | -0.224 | 0 | -0.003 | -0.003 | 0 | 0.25 | 0.36 | 0.31 | 0.25 | 0.26 | 0.19 |
| 10 | 82843470 | rs10786205 | -0.293 | 0.415 | 0.01 | -0.204 | -0.001 | -0.002 | -0.002 | 0 | 0.25 | 0.32 | 0.3 | 0.24 | 0.26 | 0.2 |
| 10 | 82854879 | rs7097552 | -0.502 | 0.781 | -0.469 | 0.109 | 0.002 | 0.006 | 0.006 | 0 | 0.63 | 0.64 | 0.56 | 0.47 | 0.56 | 0.56 |
| 10 | 82857977 | rs677210 | -0.052 | 0.684 | -0.454 | 0.19 | -0.016 | -0.023 | -0.023 | 1.102 | 0.32 | 0.43 | 0.3 | 0.26 | 0.25 | 0.29 |
| 10 | 82868149 | rs17097249 | -0.806 | 1.07 | -0.615 | 1.398 | -0.03 | -0.063 | -0.063 | 0.037 | 0.19 | 0.04 | 0.23 | 0.28 | 0.23 | 0.16 |
| 10 | 82887882 | rs11591827 | -1.413 | 1.078 | -0.231 | 1.795 | 0.011 | 0.079 | 0.079 | 0 | 0.06 | 0.12 | 0.07 | 0.16 | 0.07 | 0.11 |
| 10 | 82892288 | rs10509426 | -1.412 | 1.301 | -0.159 | 1.458 | -0.002 | 0.003 | 0.003 | 0 | 0.16 | 0.16 | 0.1 | 0.1 | 0.12 | 0.17 |
| 10 | 82895519 | rs10786248 | -1.968 | 1.118 | 0.534 | 2.62 | 0 | 0.081 | 0.081 | 0 | 0.21 | 0.37 | 0.24 | 0.59 | 0.15 | 0.33 |
| 10 | 82909302 | rs10509427 | 1.901 | 1.706 | 1.238 | 2.578 | 0.038 | 0.003 | 0.003 | 0 | 0.77 | 0.5 | 0.72 | 0.8 | 0.76 | 0.66 |
| 10 | 82917563 | rs7894491 | 0.228 | 0.7 | 1.07 | 1.523 | 0.008 | -0.011 | -0.011 | 0 | 0.22 | 0.16 | 0.1 | 0.02 | 0.15 | 0.11 |
| 10 | 82940303 | rs7915587 | 1.056 | 1.27 | 1.027 | 1.717 | 0.02 | -0.001 | -0.001 | 0 | 0.82 | 0.75 | 0.75 | 0.87 | 0.77 | 0.71 |
| 10 | 82941755 | rs7074341 | -0.121 | 1.087 | 0.952 | 1.776 | 0.021 | 0.031 | 0.031 | 0 | 0.94 | 0.9 | 0.83 | 0.9 | 0.88 | 0.84 |
| 10 | 82974169 | rs7919037 | -1.3 | 1.109 | 0.669 | 1.967 | -0.002 | 0.043 | 0.043 | 0 | 0.12 | 0.15 | 0.12 | 0.43 | 0.1 | 0.19 |
| 10 | 82979066 | rs7908787 | -0.78 | 2.101 | 0.603 | 1.753 | 0.019 | -0.004 | -0.004 | 9.483 | 0.07 | 0.08 | 0.08 | 0.02 | 0.13 | 0.07 |

| | | | | | | | | | | | | | | | | |
|---|---|---|---|---|---|---|---|---|---|---|---|---|---|---|---|---|
| 10 | 82979613 | rs11188912 | -0.317 | 1.884 | 0.553 | 1.318 | -0.002 | -0.04 | -0.04 | 0 | 0.17 | 0.17 | 0.1 | 0.01 | 0.16 | 0.09 |
| 10 | 82982181 | rs4277054 | -2.128 | 1.812 | 0.757 | 1.587 | 0.001 | 0 | 0 | 0 | 0.36 | 0.67 | 0.38 | 0.68 | 0.3 | 0.47 |
| 10 | 82984432 | rs10882846 | -0.315 | 1.812 | 0.741 | 1.566 | -0.006 | -0.067 | -0.067 | 0 | 0.18 | 0.23 | 0.11 | 0.01 | 0.16 | 0.09 |
| 10 | 82986532 | rs10748689 | -2.484 | 1.728 | 0.704 | 1.581 | -0.01 | 0.039 | 0.039 | 0.026 | 0.42 | 0.75 | 0.46 | 0.7 | 0.43 | 0.55 |
| 10 | 83006207 | rs12414453 | -0.843 | 1.565 | 0.817 | 1.882 | -0.007 | 0.099 | 0.099 | 0 | 0.1 | 0.15 | 0.13 | 0.42 | 0.09 | 0.19 |
| 10 | 83012381 | rs12244637 | -0.002 | 1.497 | 0.967 | 1.765 | -0.004 | -0.025 | -0.025 | 0 | 0.07 | 0.09 | 0.04 | 0 | 0.06 | 0.03 |
| 10 | 83019714 | rs10509430 | -1.219 | 1.382 | 0.971 | 1.944 | 0.001 | 0.02 | 0.02 | 0 | 0.15 | 0.3 | 0.21 | 0.14 | 0.16 | 0.21 |
| 10 | 83021731 | rs7919731 | -1.195 | 1.406 | 0.963 | 1.887 | 0.002 | 0.021 | 0.021 | 0 | 0.14 | 0.29 | 0.21 | 0.13 | 0.15 | 0.21 |
| 10 | 83029412 | rs2152745 | -1.931 | 1.59 | 1.069 | 1.909 | -0.016 | 0.062 | 0.062 | 0 | 0.19 | 0.36 | 0.32 | 0.64 | 0.16 | 0.38 |
| 10 | 83045104 | rs3913914 | -1.984 | 1.677 | 1.087 | 1.9 | 0 | 0.089 | 0.089 | 0 | 0.22 | 0.39 | 0.35 | 0.53 | 0.21 | 0.36 |
| 10 | 83059864 | rs11189285 | -0.359 | 1.686 | 0.961 | 1.533 | -0.001 | 0.003 | 0.003 | 0 | 0.07 | 0.05 | 0.04 | 0 | 0.05 | 0.02 |
| 10 | 83060356 | rs17097441 | -0.247 | 1.703 | 0.997 | 1.482 | -0.003 | -0.026 | -0.026 | 0 | 0.05 | 0.11 | 0.03 | 0 | 0.04 | 0.03 |
| 10 | 83070163 | rs11189322 | -2.466 | 1.72 | 0.998 | 1.416 | 0.02 | -0.004 | -0.004 | 0 | 0.14 | 0.27 | 0.12 | 0.12 | 0.16 | 0.16 |
| 10 | 83075488 | rs265454 | 2.219 | 1.283 | 1.188 | 1.295 | -0.005 | 0.009 | 0.009 | 1.702 | 0.62 | 0.29 | 0.54 | 0.35 | 0.64 | 0.46 |
| 10 | 83078959 | rs265477 | 1.207 | 1.415 | 1.114 | 1.218 | 0.011 | -0.055 | -0.055 | 0 | 0.75 | 0.64 | 0.83 | 0.97 | 0.72 | 0.85 |
| 10 | 83090548 | rs10509433 | -1.389 | 1.384 | 0.809 | 1.566 | 0.007 | 0.177 | 0.177 | 0.104 | 0.06 | 0.11 | 0.25 | 0.51 | 0.08 | 0.35 |
| 10 | 83093068 | rs265533 | 1.258 | 0.887 | 0.252 | 0.697 | -0.005 | 0.009 | 0.009 | 0 | 0.49 | 0.27 | 0.51 | 0.55 | 0.51 | 0.56 |
| 10 | 83094375 | rs10748710 | -0.982 | 1.077 | 0.469 | 0.658 | 0.002 | -0.002 | -0.002 | 0.473 | 0.38 | 0.61 | 0.38 | 0.42 | 0.34 | 0.37 |
| 10 | 83096815 | rs265527 | 1.551 | 1.036 | 0.474 | 0.68 | 0.001 | 0.01 | 0.01 | 0 | 0.51 | 0.37 | 0.53 | 0.55 | 0.51 | 0.57 |
| 10 | 83103134 | rs265514 | 1.571 | 1.023 | 0.477 | 0.675 | 0.002 | 0.011 | 0.011 | 0 | 0.51 | 0.37 | 0.53 | 0.55 | 0.51 | 0.57 |
| 10 | 83104421 | rs265511 | 0.312 | 1.041 | 0.375 | 0.682 | -0.042 | 0.028 | 0.028 | 0 | 0.88 | 0.98 | 0.91 | 0.97 | 0.85 | 0.95 |
| 10 | 83118201 | rs7393548 | -0.048 | 0.82 | 0.957 | 0.56 | -0.015 | 0.028 | 0.028 | 0 | 0.19 | 0.28 | 0.25 | 0.3 | 0.16 | 0.21 |
| 10 | 83124784 | rs587973 | 0.253 | 0.916 | 0.829 | 0.54 | -0.043 | 0.013 | 0.013 | 0 | 0.88 | 0.98 | 0.91 | 0.94 | 0.85 | 0.94 |
| 10 | 83125807 | rs12570337 | -0.101 | 0.861 | 0.639 | 0.529 | -0.002 | -0.008 | -0.008 | 0 | 0.06 | 0.04 | 0.05 | 0.09 | 0.06 | 0.06 |
| 10 | 83126887 | rs649207 | 1.599 | 0.653 | 0.845 | 0.415 | 0.016 | 0.028 | 0.028 | 0 | 0.69 | 0.65 | 0.57 | 0.4 | 0.68 | 0.48 |
| 10 | 83128351 | rs632593 | 1.653 | 0.641 | 0.898 | 0.409 | 0.02 | 0.048 | 0.048 | 0 | 0.69 | 0.56 | 0.57 | 0.42 | 0.68 | 0.48 |
| 10 | 83134434 | rs265460 | 2.766 | 0.586 | 0.78 | 0.48 | -0.019 | 0.147 | 0.147 | 0.293 | 0.43 | 0.22 | 0.32 | 0.11 | 0.45 | 0.21 |
| 10 | 83139922 | rs265508 | -0.235 | 0.861 | 0.907 | 0.607 | -0.025 | 0.032 | 0.032 | 0 | 0.89 | 0.98 | 0.93 | 0.97 | 0.9 | 0.95 |
| 10 | 83141004 | rs1329210 | -0.921 | 0.857 | 1.016 | 0.65 | 0.002 | 0.108 | 0.108 | 0 | 0.08 | 0.11 | 0.23 | 0.48 | 0.11 | 0.36 |
| 10 | 83153009 | rs2346230 | -1.291 | 0.851 | 1.128 | 0.64 | 0.02 | 0.141 | 0.141 | 0 | 0.19 | 0.3 | 0.31 | 0.5 | 0.22 | 0.47 |
| 10 | 83153171 | rs496565 | 1.949 | 0.843 | 1.184 | 0.639 | 0.01 | 0.035 | 0.035 | 0 | 0.68 | 0.49 | 0.61 | 0.46 | 0.68 | 0.48 |
| 10 | 83160315 | rs10883138 | 1.624 | 0.847 | 1.039 | 0.906 | 0.016 | 0.034 | 0.034 | 0 | 0.73 | 0.69 | 0.8 | 0.56 | 0.72 | 0.59 |
| 10 | 83161858 | rs10748744 | 0.902 | 0.843 | 1.053 | 0.662 | 0.011 | -0.061 | -0.061 | 0 | 0.78 | 0.73 | 0.85 | 0.94 | 0.79 | 0.84 |
| 10 | 83172937 | rs4933778 | 1.161 | 0.832 | 1.035 | 0.658 | -0.008 | -0.017 | -0.017 | 0.008 | 0.8 | 0.85 | 0.7 | 0.49 | 0.79 | 0.59 |
| 10 | 83175662 | rs975853 | -0.946 | 0.639 | 1.118 | 0.645 | -0.02 | 0.056 | 0.056 | 0 | 0.25 | 0.42 | 0.31 | 0.35 | 0.23 | 0.27 |
| 10 | 83178907 | rs7082300 | 0.489 | 0.646 | 0.891 | 0.647 | 0.004 | -0.018 | -0.018 | 0 | 0.15 | 0.11 | 0.24 | 0.27 | 0.16 | 0.22 |
| 10 | 83184149 | rs11189831 | 0.154 | 0.713 | 0.73 | 0.419 | 0.016 | -0.051 | -0.051 | 0 | 0.11 | 0.19 | 0.08 | 0.03 | 0.11 | 0.1 |
| 10 | 83188768 | rs1329202 | 2.583 | 0.663 | -0.083 | 0.023 | -0.037 | 0.198 | 0.198 | 0.79 | 0.34 | 0.08 | 0.28 | 0.08 | 0.36 | 0.19 |
| 10 | 83188866 | rs4933781 | -1.14 | 0.897 | 0.547 | 0.467 | 0.006 | 0.045 | 0.045 | 0 | 0.28 | 0.39 | 0.36 | 0.55 | 0.26 | 0.44 |
| 10 | 83192498 | rs11189862 | -0.09 | 0.946 | 0.337 | 0.48 | -0.013 | 0.015 | 0.015 | 0 | 0.09 | 0.02 | 0.05 | 0.04 | 0.08 | 0.03 |
| 10 | 83194360 | rs11189865 | -0.078 | 0.861 | 0.51 | 0.502 | 0.008 | 0.098 | 0.098 | 0.167 | 0.17 | 0.21 | 0.3 | 0.51 | 0.18 | 0.41 |
| 10 | 83195321 | rs1329196 | 1.392 | 0.629 | 0.749 | -0.334 | 0.005 | -0.086 | -0.086 | 0 | 0.73 | 0.61 | 0.85 | 0.92 | 0.77 | 0.84 |
| 10 | 83195598 | rs11189882 | -0.699 | 0.889 | 0.348 | -0.041 | -0.009 | -0.025 | -0.025 | 0 | 0.12 | 0.02 | 0.08 | 0.11 | 0.11 | 0.07 |
| 10 | 83199251 | rs1410910 | -0.702 | 0.993 | 0.964 | -0.032 | 0.01 | -0.035 | -0.035 | 0 | 0.16 | 0.31 | 0.09 | 0.07 | 0.14 | 0.12 |
| 10 | 83199854 | rs10883236 | 2.598 | 0.927 | 0.815 | -0.247 | 0.011 | 0.209 | 0.209 | 1.245 | 0.39 | 0.1 | 0.28 | 0.1 | 0.36 | 0.22 |
| 10 | 83203474 | rs9651444 | 2.531 | 0.92 | 0.966 | -0.338 | 0.027 | 0.171 | 0.171 | 0.186 | 0.4 | 0.17 | 0.29 | 0.09 | 0.36 | 0.21 |
| 10 | 83210669 | rs197775 | -1.698 | 0.848 | 0.936 | -0.284 | 0 | -0.022 | -0.022 | 0 | 0.45 | 0.54 | 0.62 | 0.85 | 0.49 | 0.67 |
| 10 | 83212826 | rs4933784 | 0.09 | 0.812 | 1.116 | -0.11 | -0.011 | 0.044 | 0.044 | 0 | 0.21 | 0.33 | 0.23 | 0.64 | 0.19 | 0.33 |
| 10 | 83217694 | rs197781 | -0.226 | 0.871 | 1.042 | -0.091 | 0.004 | 0.004 | 0.004 | 0 | 0.08 | 0.12 | 0.07 | 0.06 | 0.08 | 0.06 |
| 10 | 83223131 | rs4933786 | 0.379 | 0.907 | 0.802 | -0.123 | -0.019 | -0.002 | -0.002 | 0 | 0.15 | 0.07 | 0.32 | 0.14 | 0.23 | 0.28 |
| 10 | 83229149 | rs10458748 | 0.41 | 0.917 | 0.868 | -0.118 | -0.006 | 0 | 0 | 0 | 0.16 | 0.11 | 0.32 | 0.14 | 0.23 | 0.28 |
| 10 | 83236292 | rs10490926 | -0.859 | 0.855 | 0.983 | -0.158 | -0.004 | 0.003 | 0.003 | 0 | 0.28 | 0.37 | 0.29 | 0.7 | 0.27 | 0.39 |
| 10 | 83239566 | rs6584086 | -2.894 | 0.825 | 0.857 | -0.222 | 0.005 | 0.218 | 0.218 | 1.243 | 0.6 | 0.91 | 0.71 | 0.9 | 0.63 | 0.78 |
| 10 | 83245222 | rs11190089 | 0.422 | 0.939 | 0.991 | -0.113 | -0.004 | -0.001 | -0.001 | 1.122 | 0.15 | 0.11 | 0.31 | 0.15 | 0.21 | 0.26 |
| 10 | 83245486 | rs17631622 | 0.886 | 0.935 | 1.06 | -0.114 | 0.001 | -0.001 | -0.001 | 0 | 0.06 | 0.1 | 0.04 | 0.04 | 0.06 | 0.03 |
| 10 | 83253644 | rs1997974 | -1.78 | 0.93 | 1.152 | -0.121 | 0.041 | -0.028 | -0.028 | 0 | 0.3 | 0.45 | 0.4 | 0.21 | 0.35 | 0.38 |
| 10 | 83259758 | rs11190157 | 0.472 | 0.943 | 1.106 | -0.12 | -0.003 | -0.001 | -0.001 | 0 | 0.15 | 0.12 | 0.31 | 0.15 | 0.21 | 0.24 |
| 10 | 83261323 | rs2346520 | 0.614 | 0.901 | 1.017 | -0.106 | 0.027 | -0.033 | -0.033 | 0.17 | 0.09 | 0.24 | 0.04 | 0.03 | 0.08 | 0.09 |
| 10 | 83265201 | rs10786564 | -2.912 | 0.961 | 0.522 | -0.257 | -0.004 | 0.23 | 0.23 | 2.116 | 0.62 | 0.92 | 0.71 | 0.91 | 0.65 | 0.77 |
| 10 | 83273315 | rs9332404 | -1.243 | 1.04 | 0.755 | -0.098 | -0.002 | -0.001 | -0.001 | 0 | 0.3 | 0.36 | 0.29 | 0.69 | 0.3 | 0.39 |
| 10 | 83274383 | rs17550611 | -0.315 | 1.338 | 0.344 | -0.032 | -0.023 | -0.005 | -0.005 | 1.213 | 0.1 | 0.02 | 0.08 | 0.06 | 0.12 | 0.07 |
| 10 | 83287387 | rs7917905 | 0.529 | 1.198 | 0.664 | -0.056 | 0.014 | -0.029 | -0.029 | 0 | 0.09 | 0.15 | 0.04 | 0.03 | 0.08 | 0.09 |
| 10 | 83288438 | rs7907133 | 0.529 | 1.198 | 0.704 | -0.056 | 0.021 | -0.03 | -0.03 | 0 | 0.09 | 0.17 | 0.04 | 0.03 | 0.08 | 0.09 |
| 10 | 83289794 | rs17551124 | -0.624 | 1.215 | 0.359 | 0.006 | -0.02 | 0 | 0 | 0 | 0.22 | 0.17 | 0.38 | 0.19 | 0.31 | 0.31 |
| 10 | 83294798 | rs11190324 | -0.763 | 0.836 | 0.587 | -0.42 | 0.015 | -0.046 | -0.046 | 0.634 | 0.22 | 0.31 | 0.16 | 0.08 | 0.25 | 0.15 |
| 10 | 83295046 | rs10883427 | -1.505 | 0.694 | 0.622 | -0.438 | 0.001 | -0.042 | -0.042 | 0 | 0.27 | 0.58 | 0.19 | 0.13 | 0.26 | 0.19 |
| 10 | 83298806 | rs994789 | -0.97 | 0.904 | 0.2 | -0.496 | 0.028 | -0.077 | -0.077 | 0 | 0.22 | 0.42 | 0.16 | 0.08 | 0.25 | 0.15 |
| 10 | 83301681 | rs2224082 | -1.68 | 0.522 | -0.925 | -0.269 | -0.009 | 0.144 | 0.144 | 0.283 | 0.64 | 0.93 | 0.6 | 0.88 | 0.69 | 0.78 |
| 10 | 83305304 | rs1415918 | -0.853 | 0.452 | 0.376 | 0.03 | 0.005 | -0.02 | -0.02 | 0 | 0.07 | 0.28 | 0.02 | 0 | 0.03 | 0.02 |

| | | | | | | | | | | | | | | | | |
|---|---|---|---|---|---|---|---|---|---|---|---|---|---|---|---|---|
| 10 | 83320898 | rs12253574 | -1.778 | 0.583 | 0.816 | -0.146 | 0.019 | -0.071 | -0.071 | 0.112 | 0.12 | 0.42 | 0.05 | 0.01 | 0.08 | 0.03 |
| 10 | 83325813 | rs10883454 | -2.703 | 0.518 | 1.164 | -0.368 | -0.002 | -0.033 | -0.033 | 0.066 | 0.27 | 0.65 | 0.16 | 0.11 | 0.21 | 0.16 |
| 10 | 83334470 | rs11190474 | -1.857 | 0.468 | 1.366 | 0.413 | -0.002 | 0.003 | 0.003 | 0.409 | 0.33 | 0.69 | 0.25 | 0.41 | 0.31 | 0.23 |
| 10 | 83334512 | rs11599165 | 0.483 | 0.467 | 1.367 | 0.417 | 0.001 | -0.019 | -0.019 | 0 | 0.16 | 0.15 | 0.15 | 0.34 | 0.18 | 0.16 |
| 10 | 83335787 | rs1010468 | 0.147 | 0.478 | 1.462 | 0.408 | 0.011 | -0.023 | -0.023 | 0.373 | 0.05 | 0.14 | 0.03 | 0.01 | 0.05 | 0.01 |
| 10 | 83336273 | rs17098075 | -0.059 | 0.43 | 1.469 | 0.395 | -0.006 | 0.031 | 0.031 | 0 | 0.18 | 0.26 | 0.18 | 0.34 | 0.18 | 0.16 |
| 10 | 83338038 | rs17098082 | -0.802 | 0.444 | 1.403 | 0.675 | -0.001 | 0.011 | 0.011 | 0 | 0.23 | 0.4 | 0.2 | 0.25 | 0.23 | 0.18 |
| 10 | 83339057 | rs7098143 | -0.807 | 0.447 | 1.404 | 0.562 | -0.001 | 0.036 | 0.036 | 0 | 0.23 | 0.4 | 0.21 | 0.35 | 0.23 | 0.18 |
| 10 | 83339105 | rs7067910 | -2.027 | 0.458 | 1.195 | 0.564 | 0.014 | -0.022 | -0.022 | 0 | 0.35 | 0.8 | 0.27 | 0.41 | 0.31 | 0.23 |
| 10 | 83349511 | rs7073315 | -2.206 | 0.46 | 1.53 | 0.596 | 0 | -0.001 | -0.001 | 0.258 | 0.34 | 0.72 | 0.27 | 0.42 | 0.31 | 0.23 |
| 10 | 83373508 | rs1890254 | 0.129 | 0.531 | 1.534 | 0.555 | -0.001 | 0.034 | 0.034 | 0 | 0.17 | 0.24 | 0.17 | 0.37 | 0.19 | 0.2 |
| 10 | 83392668 | rs7918647 | -1.826 | 0.497 | 1.798 | 0.499 | 0.004 | -0.006 | -0.006 | 0 | 0.35 | 0.69 | 0.27 | 0.42 | 0.31 | 0.26 |
| 10 | 83394720 | rs7092099 | -1.545 | 0.524 | 1.913 | 0.42 | -0.009 | 0.008 | 0.008 | 0 | 0.16 | 0.12 | 0.11 | 0.09 | 0.18 | 0.15 |
| 10 | 83414697 | rs7905585 | -1.158 | 0.513 | 2.115 | 0.423 | 0.01 | 0.077 | 0.077 | 0.401 | 0.27 | 0.46 | 0.23 | 0.41 | 0.28 | 0.25 |
| 10 | 83418800 | rs11190855 | 0.153 | 0.529 | 1.992 | 0.418 | -0.016 | 0.016 | 0.016 | 0 | 0.07 | 0.03 | 0.04 | 0.04 | 0.09 | 0.1 |
| 10 | 83423395 | rs1538254 | -2.818 | 0.467 | 2.233 | 0.392 | -0.012 | -0.024 | -0.024 | 0.278 | 0.44 | 0.88 | 0.33 | 0.47 | 0.4 | 0.37 |
| 10 | 83423627 | rs1933286 | -1.781 | 0.461 | 2.646 | 0.13 | 0.041 | -0.08 | -0.08 | 0 | 0.15 | 0.4 | 0.1 | 0.04 | 0.14 | 0.11 |
| 10 | 83433450 | rs12262773 | -2.981 | 0.46 | 2.66 | 0.309 | 0.008 | -0.05 | -0.05 | 0 | 0.25 | 0.53 | 0.14 | 0.11 | 0.21 | 0.16 |
| 10 | 83440938 | rs857919 | -0.477 | 0.485 | 2.667 | 0.42 | 0.001 | 0.001 | 0.001 | 0 | 0.83 | 0.82 | 0.85 | 0.6 | 0.82 | 0.8 |
| 10 | 83449684 | rs506380 | 2.326 | 0.482 | 2.556 | 0.421 | -0.009 | -0.044 | -0.044 | 0 | 0.57 | 0.14 | 0.68 | 0.49 | 0.6 | 0.64 |
| 10 | 83468658 | rs10883672 | -0.924 | 0.5 | 2.565 | 0.127 | 0.026 | -0.033 | -0.033 | 1.193 | 0.09 | 0.23 | 0.07 | 0.04 | 0.1 | 0.11 |
| 10 | 83485025 | rs11191124 | 2.719 | 0.583 | 1.334 | -0.044 | -0.012 | 0.189 | 0.189 | 0 | 0.41 | 0.08 | 0.36 | 0.11 | 0.4 | 0.27 |
| 10 | 83489022 | rs12249692 | -0.756 | 0.64 | 1.944 | 0.196 | 0.015 | -0.018 | -0.018 | 0 | 0.08 | 0.17 | 0.06 | 0.04 | 0.1 | 0.11 |
| 10 | 83494206 | rs1160061 | 1.701 | 0.537 | 1.876 | 0.319 | 0.008 | -0.001 | -0.001 | 0 | 0.66 | 0.21 | 0.72 | 0.6 | 0.69 | 0.66 |
| 10 | 83501262 | rs6584245 | 0.497 | 0.505 | 2.068 | 0.323 | -0.021 | 0.053 | 0.053 | 0.347 | 0.8 | 0.57 | 0.82 | 0.65 | 0.82 | 0.79 |
| 10 | 83509555 | rs7906283 | 2.324 | 0.473 | 1.933 | 0.297 | -0.044 | -0.048 | -0.048 | 0.275 | 0.56 | 0.1 | 0.65 | 0.55 | 0.59 | 0.62 |
| 10 | 83513797 | rs11819459 | -1.004 | 0.605 | 1.838 | 0.332 | 0.011 | 0.005 | 0.005 | 0 | 0.09 | 0.11 | 0.06 | 0.06 | 0.1 | 0.04 |
| 10 | 83515878 | rs7897955 | 1.046 | 0.697 | 1.56 | 0.314 | 0 | 0.002 | 0.002 | 0.521 | 0.73 | 0.69 | 0.77 | 0.63 | 0.72 | 0.68 |
| 10 | 83522580 | rs7905133 | 1.12 | 0.773 | 1.455 | 0.245 | -0.002 | -0.009 | -0.009 | 0 | 0.75 | 0.77 | 0.76 | 0.61 | 0.73 | 0.68 |
| 10 | 83535160 | rs2207653 | -2.796 | 0.798 | 1.582 | -0.434 | 0.034 | -0.052 | -0.052 | 0 | 0.23 | 0.53 | 0.31 | 0.1 | 0.22 | 0.17 |
| 10 | 83542467 | rs11191280 | -3.077 | 0.79 | 1.185 | -0.467 | -0.01 | 0.045 | 0.045 | 0.574 | 0.27 | 0.81 | 0.34 | 0.1 | 0.23 | 0.17 |
| 10 | 83542571 | rs11191281 | -3.052 | 0.79 | 1.188 | -0.467 | -0.01 | 0.042 | 0.042 | 0.584 | 0.27 | 0.8 | 0.33 | 0.1 | 0.23 | 0.17 |
| 10 | 83548750 | rs12247849 | -0.643 | 0.616 | 1.24 | -0.292 | 0.025 | 0.023 | 0.023 | 0.147 | 0.05 | 0.19 | 0.04 | 0.04 | 0.02 | 0.02 |
| 10 | 83552795 | rs1543417 | -1.625 | 0.678 | 0.754 | -0.313 | -0.006 | -0.001 | -0.001 | 0 | 0.1 | 0.11 | 0.19 | 0.05 | 0.06 | 0.11 |
| 10 | 83554248 | rs7086225 | -1.628 | 0.357 | 0.156 | -0.771 | -0.006 | 0.147 | 0.147 | 0 | 0.6 | 0.87 | 0.59 | 0.91 | 0.58 | 0.75 |
| 10 | 83555806 | rs10786676 | -0.809 | 1.032 | 0.981 | 0.131 | -0.002 | 0.009 | 0.009 | 0 | 0.39 | 0.4 | 0.47 | 0.82 | 0.4 | 0.69 |
| 10 | 83556045 | rs9419880 | -0.954 | 1.086 | 1.233 | 0.123 | 0.052 | -0.05 | -0.05 | 0 | 0.09 | 0.15 | 0.06 | 0 | 0.13 | 0.03 |
| 10 | 83557187 | rs11191331 | -0.361 | 1.095 | 1.028 | 0.105 | 0.014 | 0.02 | 0.02 | 0 | 0.33 | 0.23 | 0.39 | 0.82 | 0.29 | 0.64 |
| 10 | 83563647 | rs17712426 | 0.727 | 1.126 | 0.988 | 0.106 | -0.002 | -0.001 | -0.001 | 0.416 | 0.06 | 0.03 | 0.03 | 0.06 | 0.06 | 0.07 |
| 10 | 83566835 | rs11191360 | 0.57 | 1.072 | 1.037 | 0.431 | -0.018 | 0.033 | 0.033 | 0 | 0.21 | 0.23 | 0.22 | 0.57 | 0.09 | 0.45 |
| 10 | 83569687 | rs6584329 | -2.1 | 1.137 | 1.091 | 0.393 | 0.02 | 0.111 | 0.111 | 0 | 0.52 | 0.78 | 0.54 | 0.82 | 0.55 | 0.71 |
| 10 | 83571926 | rs7477565 | -0.998 | 1.154 | 1.07 | 0.407 | 0 | 0.01 | 0.01 | 0 | 0.4 | 0.41 | 0.46 | 0.81 | 0.4 | 0.66 |
| 10 | 83577099 | rs7922516 | -1.624 | 1.31 | 1.251 | 0.458 | 0.059 | -0.003 | -0.003 | 0 | 0.27 | 0.51 | 0.32 | 0.24 | 0.43 | 0.26 |
| 10 | 83578883 | rs11191390 | -2.472 | 1.052 | 1.174 | 0.502 | 0.026 | 0.104 | 0.104 | 0 | 0.58 | 0.82 | 0.58 | 0.85 | 0.61 | 0.71 |
| 10 | 83581933 | rs764866 | -0.163 | 1.267 | 0.344 | 0.627 | -0.004 | -0.004 | -0.004 | 2.376 | 0.15 | 0.15 | 0.19 | 0.24 | 0.21 | 0.19 |
| 10 | 83589705 | rs11191432 | -0.436 | -0.688 | 0.214 | 1.038 | 0.023 | -0.034 | -0.034 | 0.771 | 0.26 | 0.18 | 0.25 | 0.6 | 0.15 | 0.44 |
| 10 | 83592407 | rs11191455 | -1.254 | -0.896 | 0.23 | 0.995 | -0.001 | -0.01 | -0.01 | 0.137 | 0.51 | 0.44 | 0.5 | 0.83 | 0.49 | 0.69 |
| 10 | 83594473 | rs2207654 | -0.655 | -1.685 | 0.018 | 1.684 | 0.02 | -0.03 | -0.03 | 0.061 | 0.19 | 0.14 | 0.22 | 0.54 | 0.09 | 0.38 |
| 10 | 83600968 | rs956429 | -1.728 | -0.816 | 0.707 | 1.169 | -0.013 | -0.062 | -0.062 | 0 | 0.13 | 0.22 | 0.19 | 0.01 | 0.11 | 0.09 |
| 10 | 83601924 | rs10509439 | -1.37 | -0.732 | 0.961 | 1.482 | 0 | 0.001 | 0.001 | 0 | 0.06 | 0.06 | 0.11 | 0.11 | 0.05 | 0.2 |
| 10 | 83602603 | rs2207658 | -1.374 | -0.967 | 1.976 | 2.544 | 0.002 | 0.258 | 0.258 | 0.056 | 0.11 | 0.32 | 0.3 | 0.4 | 0.1 | 0.19 |
| 10 | 83606191 | rs11191528 | -1.264 | -1.18 | 2.814 | 3.324 | 0.024 | 0.138 | 0.138 | 0.132 | 0.07 | 0.25 | 0.27 | 0.18 | 0.09 | 0.11 |
| 10 | 83606306 | rs11191529 | -2.852 | -1.585 | 2.589 | 3.261 | -0.009 | 0.039 | 0.039 | 0.789 | 0.2 | 0.29 | 0.34 | 0.32 | 0.18 | 0.4 |
| 10 | 83608508 | rs10883825 | -2.618 | -1.494 | 3.01 | 3.416 | 0.007 | 0.366 | 0.366 | 0.045 | 0.13 | 0.52 | 0.32 | 0.46 | 0.1 | 0.14 |
| 10 | 83616171 | rs7907714 | -3.008 | -1.622 | 3.24 | 3.112 | -0.048 | 0.091 | 0.091 | 0.966 | 0.21 | 0.61 | 0.46 | 0.34 | 0.16 | 0.26 |
| 10 | 83619940 | rs11191592 | -2.514 | -1.501 | 3.284 | 3.041 | -0.034 | 0.128 | 0.128 | 0.123 | 0.16 | 0.41 | 0.31 | 0.25 | 0.13 | 0.23 |
| 10 | 83625092 | rs2064588 | -1.299 | -1.32 | 3.014 | 3.02 | 0.029 | 0.198 | 0.198 | 0.044 | 0.07 | 0.26 | 0.27 | 0.23 | 0.09 | 0.12 |
| 10 | 83649739 | rs10748842 | -1.947 | -1.464 | 2.969 | 2.387 | -0.001 | -0.025 | -0.025 | 0.42 | 0.11 | 0.18 | 0.09 | 0.06 | 0.1 | 0.22 |
| 10 | 83656526 | rs6584400 | -3.059 | -1.479 | 2.925 | 2.355 | -0.026 | 0.039 | 0.039 | 0.4 | 0.19 | 0.43 | 0.12 | 0.25 | 0.12 | 0.28 |
| 10 | 83664796 | rs2024785 | -2.091 | -1.535 | 2.718 | 2.325 | 0.001 | 0.169 | 0.169 | 0 | 0.07 | 0.21 | 0.03 | 0.22 | 0.01 | 0.05 |
| 10 | 83670383 | rs10883899 | -0.838 | -1.525 | 2.553 | 2.606 | -0.02 | -0.04 | -0.04 | 0 | 0.07 | 0 | 0.07 | 0.14 | 0.09 | 0.23 |
| 10 | 83675990 | rs4933811 | -2.619 | -1.753 | 3.014 | 2.775 | -0.025 | 0.056 | 0.056 | 2.454 | 0.24 | 0.41 | 0.49 | 0.57 | 0.2 | 0.48 |
| 10 | 83677849 | rs7913415 | -2.743 | -1.65 | 2.963 | 2.744 | -0.029 | 0.076 | 0.076 | 0 | 0.11 | 0.24 | 0.05 | 0.22 | 0.04 | 0.07 |
| 10 | 83678141 | rs11191796 | -1.908 | -1.597 | 2.853 | 2.772 | -0.011 | 0.121 | 0.121 | 0 | 0.06 | 0.16 | 0.03 | 0.22 | 0.01 | 0.05 |
| 10 | 83681283 | rs1040674 | -1.608 | -1.461 | 2.541 | 2.437 | -0.02 | 0.1 | 0.1 | 0.435 | 0.15 | 0.29 | 0.41 | 0.44 | 0.12 | 0.25 |
| 10 | 83709652 | rs17098797 | -2.115 | -1.443 | 2.5 | 2.032 | -0.016 | -0.051 | -0.051 | 0 | 0.1 | 0.21 | 0.09 | 0 | 0.06 | 0.04 |
| 10 | 83712877 | rs4933265 | -1.947 | -1.358 | 2.867 | 2.31 | -0.022 | 0.115 | 0.115 | 0 | 0.17 | 0.3 | 0.45 | 0.42 | 0.12 | 0.18 |
| 10 | 83714374 | rs17098803 | -2.355 | -1.339 | 2.83 | 2.257 | -0.014 | -0.053 | -0.053 | 0.008 | 0.11 | 0.21 | 0.12 | 0 | 0.08 | 0.04 |
| 10 | 83717632 | rs10883973 | -1.597 | -1.424 | 2.784 | 2.416 | -0.003 | 0.238 | 0.238 | 0.065 | 0.11 | 0.36 | 0.07 | 0.33 | 0.08 | 0.26 |

| | | | | | | | | | | | | | | | | |
|---|---|---|---|---|---|---|---|---|---|---|---|---|---|---|---|---|
| 10 | 83719507 | rs7099000 | -1.88 | -1.412 | 2.714 | 2.412 | -0.017 | -0.055 | -0.055 | 0 | 0.1 | 0.28 | 0.1 | 0 | 0.05 | 0.03 |
| 10 | 83719888 | rs7100526 | -3.041 | -1.281 | 2.645 | 2.539 | -0.047 | 0.103 | 0.103 | 0.202 | 0.2 | 0.6 | 0.16 | 0.33 | 0.13 | 0.29 |
| 10 | 83720231 | rs10883976 | -1.291 | -1.188 | 2.213 | 2.635 | 0.002 | -0.004 | -0.004 | 1.485 | 0.09 | 0.07 | 0.05 | 0.15 | 0.08 | 0.22 |
| 10 | 83736634 | rs10509440 | -1.284 | -1.152 | 2.156 | 2.291 | -0.012 | 0.005 | 0.005 | 0.811 | 0.07 | 0.21 | 0.08 | 0.11 | 0.05 | 0.04 |
| 10 | 83755315 | rs4933814 | -1.17 | -0.646 | 0.636 | 2.455 | -0.01 | 0.128 | 0.128 | 0.016 | 0.12 | 0.22 | 0.43 | 0.41 | 0.11 | 0.15 |
| 10 | 83755561 | rs12570292 | -1.102 | -0.656 | 0.651 | 2.464 | -0.011 | 0.133 | 0.133 | 0.04 | 0.12 | 0.22 | 0.43 | 0.41 | 0.11 | 0.14 |
| 10 | 83757021 | rs10884019 | -1.488 | -0.239 | -0.103 | 1.948 | -0.003 | 0.036 | 0.036 | 0 | 0.11 | 0.13 | 0.06 | 0.37 | 0.08 | 0.27 |
| 10 | 83764410 | rs11815249 | -0.836 | 0.094 | -0.335 | 0.906 | -0.003 | -0.007 | -0.007 | 0.016 | 0.13 | 0.12 | 0.12 | 0 | 0.14 | 0.04 |
| 10 | 83764742 | rs4933819 | 1.11 | 0.701 | -1.306 | 1.284 | -0.061 | -0.007 | -0.007 | 0 | 0.66 | 0.87 | 0.7 | 0.69 | 0.61 | 0.69 |
| 10 | 83769695 | rs12358407 | -0.603 | 1.117 | -1.3 | 1.157 | -0.006 | -0.014 | -0.014 | 0 | 0.16 | 0.14 | 0.15 | 0 | 0.19 | 0.07 |
| 10 | 83773214 | rs2207778 | -0.409 | 1.141 | -1.447 | 1.245 | -0.045 | -0.034 | -0.034 | 1.963 | 0.15 | 0 | 0.1 | 0.15 | 0.16 | 0.2 |
| 10 | 83775626 | rs7899474 | 1.018 | 0.484 | -1.259 | 1.376 | -0.018 | -0.074 | -0.074 | 0 | 0.77 | 0.87 | 0.83 | 0.64 | 0.76 | 0.66 |
| 10 | 83776457 | rs3903435 | 1.811 | 0.147 | 0.034 | 2.564 | 0.004 | 0.003 | 0.003 | 0.017 | 0.58 | 0.42 | 0.65 | 0.51 | 0.54 | 0.55 |
| 10 | 83777755 | rs9664256 | 1.958 | -0.165 | 0.68 | 2.417 | -0.002 | -0.001 | -0.001 | 1.005 | 0.55 | 0.32 | 0.65 | 0.5 | 0.54 | 0.54 |
| 10 | 83788206 | rs10884056 | -0.786 | -0.508 | 1.092 | 2.826 | -0.001 | 0.039 | 0.039 | 0 | 0.2 | 0.27 | 0.14 | 0.41 | 0.2 | 0.29 |
| 10 | 83789830 | rs10884060 | -0.462 | -0.53 | 1.109 | 2.796 | 0.002 | -0.046 | -0.046 | 0 | 0.17 | 0.26 | 0.17 | 0 | 0.19 | 0.07 |
| 10 | 83790082 | rs17727747 | 2.423 | -0.706 | 1.3 | 2.76 | -0.056 | 0.167 | 0.167 | 1.263 | 0.46 | 0.11 | 0.4 | 0.26 | 0.5 | 0.54 |
| 10 | 83790207 | rs7342133 | 1.882 | -0.709 | 1.38 | 2.737 | -0.007 | -0.004 | -0.004 | 0 | 0.55 | 0.26 | 0.64 | 0.5 | 0.55 | 0.55 |
| 10 | 83790221 | rs10884061 | -0.463 | -0.705 | 1.851 | 2.739 | -0.001 | -0.018 | -0.018 | 2.577 | 0.2 | 0.18 | 0.14 | 0.41 | 0.19 | 0.29 |
| 10 | 83790407 | rs10748871 | -1.456 | -0.703 | 1.809 | 2.742 | 0.004 | 0.007 | 0.007 | 0 | 0.37 | 0.46 | 0.29 | 0.41 | 0.38 | 0.36 |
| 10 | 83794963 | rs10884066 | 0.861 | -0.793 | 1.697 | 2.531 | -0.003 | 0.013 | 0.013 | 0 | 0.74 | 0.62 | 0.71 | 0.65 | 0.73 | 0.85 |
| 10 | 83795839 | rs10509445 | 2.416 | -1.057 | 1.495 | 2.304 | -0.056 | 0.148 | 0.148 | 1.124 | 0.45 | 0.13 | 0.4 | 0.26 | 0.49 | 0.54 |
| 10 | 83798259 | rs11192252 | -1.522 | -0.888 | 1.76 | 2.404 | 0.007 | 0.066 | 0.066 | 0.005 | 0.09 | 0.25 | 0.18 | 0.14 | 0.06 | 0.08 |
| 10 | 83798469 | rs7069727 | 1.516 | -1.422 | 1.173 | 1.808 | 0 | 0.013 | 0.013 | 0.049 | 0.47 | 0.37 | 0.55 | 0.4 | 0.53 | 0.6 |
| 10 | 83802403 | rs2207768 | 0.98 | -1.445 | 1.14 | 1.684 | 0.003 | 0.003 | 0.003 | 0.661 | 0.64 | 0.37 | 0.69 | 0.79 | 0.73 | 0.89 |
| 10 | 83802818 | rs7094235 | 1.26 | -1.385 | 1.187 | 1.703 | 0.048 | 0.123 | 0.123 | 4.117 | 0.92 | 0.55 | 0.94 | 0.91 | 0.96 | 0.99 |
| 10 | 83809837 | rs2881607 | 1.116 | -1.406 | 1.061 | 1.775 | 0.012 | -0.001 | -0.001 | 0 | 0.63 | 0.35 | 0.68 | 0.65 | 0.69 | 0.83 |
| 10 | 83812026 | rs2881608 | 2.595 | -1.163 | 0.758 | 1.782 | -0.086 | 0.087 | 0.087 | 1.135 | 0.37 | 0.09 | 0.37 | 0.26 | 0.45 | 0.5 |
| 10 | 83813700 | rs12767327 | -0.881 | -1.262 | 1.201 | 1.973 | 0.004 | 0.001 | 0.001 | 0.041 | 0.3 | 0.23 | 0.25 | 0.12 | 0.23 | 0.11 |
| 10 | 83819987 | rs7907146 | 1.319 | -1.141 | 1.32 | 2.029 | 0.063 | 0.345 | 0.345 | 2.067 | 0.94 | 0.63 | 0.94 | 0.77 | 0.92 | 0.93 |
| 10 | 83822865 | rs10736173 | 0.707 | -1.042 | 0.529 | 1.84 | -0.001 | 0.026 | 0.026 | 0 | 0.77 | 0.84 | 0.67 | 0.85 | 0.8 | 0.88 |
| 10 | 83822961 | rs10736174 | 0.776 | -0.969 | 0.519 | 1.806 | -0.002 | 0.005 | 0.005 | 0 | 0.77 | 0.84 | 0.64 | 0.81 | 0.79 | 0.87 |
| 10 | 83823289 | rs17099306 | 0.008 | -0.725 | 0.471 | 1.997 | -0.006 | -0.079 | -0.079 | 0 | 0.15 | 0.03 | 0.09 | 0.37 | 0.1 | 0.19 |
| 10 | 83825479 | rs17655972 | 2.177 | -0.611 | 0.251 | 1.763 | -0.109 | 0.148 | 0.148 | 3.007 | 0.44 | 0.07 | 0.41 | 0.21 | 0.54 | 0.51 |
| 10 | 83835232 | rs11192402 | 1.134 | -0.622 | 0.638 | 1.813 | 0.01 | 0.047 | 0.047 | 0 | 0.77 | 0.63 | 0.67 | 0.58 | 0.73 | 0.77 |
| 10 | 83835639 | rs4562759 | 2.621 | -0.902 | 0.436 | 1.641 | -0.15 | 0.141 | 0.141 | 1.87 | 0.42 | 0.03 | 0.41 | 0.19 | 0.53 | 0.56 |
| 10 | 83837703 | rs12411365 | 2.445 | -0.916 | 0.661 | 1.563 | -0.021 | 0.068 | 0.068 | 0.355 | 0.43 | 0.25 | 0.49 | 0.24 | 0.54 | 0.5 |
| 10 | 83847988 | rs926929 | 0.093 | -0.889 | 0.591 | 1.598 | 0.005 | -0.02 | -0.02 | 0.307 | 0.83 | 0.75 | 0.82 | 0.92 | 0.84 | 0.89 |
| 10 | 83851010 | rs2144393 | -0.504 | -0.89 | 0.54 | 1.448 | 0 | -0.023 | -0.023 | 0 | 0.86 | 0.88 | 0.92 | 1 | 0.88 | 0.93 |
| 10 | 83852321 | rs1024193 | 0.589 | -1.001 | 0.484 | 1.476 | 0.005 | 0.042 | 0.042 | 0 | 0.74 | 0.61 | 0.66 | 0.58 | 0.72 | 0.75 |
| 10 | 83865764 | rs7099976 | -1.149 | -0.891 | -0.058 | 1.406 | -0.032 | -0.002 | -0.002 | 0.344 | 0.33 | 0.4 | 0.19 | 0.36 | 0.19 | 0.26 |
| 10 | 83872165 | rs981776 | 1.45 | 0.257 | -0.548 | 1.439 | 0 | 0.002 | 0.002 | 0 | 0.62 | 0.58 | 0.52 | 0.54 | 0.62 | 0.72 |
| 10 | 83873131 | rs4933830 | 0.969 | 0.463 | -0.648 | 1.332 | 0 | 0.006 | 0.006 | 0 | 0.78 | 0.72 | 0.74 | 0.78 | 0.8 | 0.84 |
| 10 | 83874965 | rs1649947 | 0.868 | 0.189 | -1.056 | 0.844 | -0.001 | 0.033 | 0.033 | 0.691 | 0.75 | 0.86 | 0.86 | 0.86 | 0.79 | 0.87 |
| 10 | 83875762 | rs1739766 | 0.356 | 0.353 | -0.941 | 0.879 | -0.026 | 0.071 | 0.071 | 0.275 | 0.45 | 0.18 | 0.41 | 0.36 | 0.54 | 0.49 |
| 10 | 83880290 | rs10884225 | -1.189 | -0.541 | 0.171 | 1.804 | -0.052 | 0.106 | 0.106 | 0 | 0.13 | 0.33 | 0.17 | 0.31 | 0.05 | 0.17 |
| 10 | 83880996 | rs11192604 | -1.263 | -0.629 | 0.675 | 2.327 | -0.037 | 0.032 | 0.032 | 0 | 0.12 | 0.22 | 0.04 | 0.18 | 0.05 | 0.18 |
| 10 | 83881713 | rs1739768 | 0.462 | -0.862 | 0.323 | 2.982 | -0.007 | -0.042 | -0.042 | 0 | 0.93 | 0.98 | 0.72 | 0.79 | 0.92 | 0.87 |
| 10 | 83884894 | rs1649934 | 1.746 | -0.907 | 0.711 | 2.807 | 0.037 | 0.067 | 0.067 | 0 | 0.57 | 0.25 | 0.45 | 0.24 | 0.67 | 0.47 |
| 10 | 83907724 | rs1649967 | 1.732 | -0.947 | 0.807 | 2.952 | 0.041 | 0.04 | 0.04 | 0 | 0.58 | 0.23 | 0.45 | 0.3 | 0.67 | 0.47 |
| 10 | 83908249 | rs10786907 | -0.947 | -0.788 | 1.201 | 3.043 | -0.046 | 0.042 | 0.042 | 0 | 0.13 | 0.3 | 0.05 | 0.18 | 0.05 | 0.18 |
| 10 | 83916424 | rs1649949 | 1.499 | -0.835 | 1.243 | 3.191 | -0.023 | -0.004 | -0.004 | 0.261 | 0.72 | 0.34 | 0.82 | 0.72 | 0.82 | 0.66 |
| 10 | 83920714 | rs12357753 | -1.261 | -0.822 | 1.031 | 3.149 | -0.023 | -0.056 | -0.056 | 2.659 | 0.1 | 0.02 | 0.18 | 0.25 | 0.13 | 0.16 |
| 10 | 83920935 | rs1649952 | 1.673 | -0.743 | 1.275 | 2.89 | -0.003 | 0.004 | 0.004 | 0.074 | 0.7 | 0.28 | 0.69 | 0.55 | 0.82 | 0.63 |
| 10 | 83922529 | rs12249088 | -0.556 | -0.604 | 1.019 | 2.165 | -0.033 | -0.026 | -0.026 | 0 | 0.11 | 0.19 | 0.04 | 0.1 | 0.04 | 0.15 |
| 10 | 83931937 | rs614287 | 1.657 | -0.628 | 0.978 | 2.709 | 0.008 | -0.001 | -0.001 | 0.037 | 0.63 | 0.28 | 0.51 | 0.36 | 0.73 | 0.49 |
| 10 | 83933057 | rs1739779 | 1.201 | -0.627 | 1.472 | 2.807 | 0.005 | -0.01 | -0.01 | 0.026 | 0.85 | 0.8 | 0.83 | 0.87 | 0.82 | 0.88 |
| 10 | 83950540 | rs1739765 | 2.07 | -1.062 | 2.159 | 3.575 | -0.023 | 0.077 | 0.077 | 0 | 0.73 | 0.53 | 0.79 | 0.63 | 0.78 | 0.76 |
| 10 | 83950665 | rs1649943 | 2.567 | -1.062 | 2.561 | 3.457 | 0.035 | 0.065 | 0.065 | 0 | 0.57 | 0.24 | 0.45 | 0.24 | 0.65 | 0.43 |
| 10 | 83951691 | rs1649942 | 1.992 | -0.548 | 2.758 | 3.448 | 0 | -0.012 | -0.012 | 1.009 | 0.75 | 0.69 | 0.79 | 0.79 | 0.74 | 0.84 |
| 10 | 83952605 | rs1336286 | -0.33 | -0.556 | 2.745 | 3.46 | 0.001 | 0.033 | 0.033 | 0 | 0.1 | 0 | 0.04 | 0.02 | 0.06 | 0.16 |
| 10 | 83952753 | rs2152594 | -2.307 | -0.576 | 2.783 | 3.463 | -0.02 | -0.02 | -0.02 | 0 | 0.14 | 0.25 | 0.08 | 0.07 | 0.12 | 0.05 |
| 10 | 83953623 | rs1649940 | 1.946 | -0.486 | 2.77 | 3.463 | -0.002 | -0.008 | -0.008 | 0 | 0.77 | 0.7 | 0.82 | 0.79 | 0.77 | 0.84 |
| 10 | 83953716 | rs1336287 | 2.629 | -0.515 | 2.81 | 3.445 | 0.016 | 0.054 | 0.054 | 0 | 0.57 | 0.26 | 0.43 | 0.24 | 0.6 | 0.42 |
| 10 | 83954965 | rs12772014 | 2.554 | -0.459 | 2.805 | 3.179 | 0.011 | 0.042 | 0.042 | 0 | 0.57 | 0.32 | 0.43 | 0.24 | 0.6 | 0.4 |
| 10 | 83955217 | rs4293073 | -1.103 | -0.506 | 2.74 | 3.142 | -0.02 | 0.008 | 0.008 | 0 | 0.07 | 0.11 | 0.02 | 0.06 | 0.02 | 0.03 |
| 10 | 83958152 | rs12774724 | 2.417 | -0.386 | 2.316 | 3.07 | 0.012 | 0.023 | 0.023 | 0 | 0.57 | 0.28 | 0.43 | 0.39 | 0.6 | 0.42 |
| 10 | 83958312 | rs12769829 | 2.423 | -0.432 | 2.219 | 2.885 | 0.014 | 0.048 | 0.048 | 0 | 0.57 | 0.26 | 0.44 | 0.24 | 0.6 | 0.42 |
| 10 | 83964123 | rs10786961 | 2.196 | -0.823 | 2.085 | 2.915 | 0.02 | 0.023 | 0.023 | 0 | 0.6 | 0.37 | 0.49 | 0.23 | 0.7 | 0.45 |

| | | | | | | | | | | | | | | | | |
|---|---|---|---|---|---|---|---|---|---|---|---|---|---|---|---|---|
| 10 | 83965348 | rs1764073 | 1.678 | -0.862 | 2.29 | 3.115 | -0.022 | 0.06 | 0.06 | 0 | 0.75 | 0.57 | 0.82 | 0.68 | 0.81 | 0.83 |
| 10 | 83966320 | rs7908131 | 2.376 | -1.164 | 1.285 | 2.365 | 0.07 | 0.097 | 0.097 | 0.016 | 0.54 | 0.27 | 0.45 | 0.24 | 0.67 | 0.43 |
| 10 | 83967446 | rs10786964 | 2.317 | -0.859 | 1.22 | 2.332 | 0.039 | 0.071 | 0.071 | 0 | 0.55 | 0.32 | 0.44 | 0.24 | 0.64 | 0.47 |
| 10 | 83968400 | rs1764075 | 0.323 | -0.874 | 0.948 | 2.269 | -0.043 | -0.002 | -0.002 | 0 | 0.88 | 0.96 | 0.86 | 0.88 | 0.83 | 0.88 |
| 10 | 83969034 | rs1764076 | -0.768 | -0.888 | 0.944 | 2.203 | 0.017 | 0.069 | 0.069 | 0 | 0.15 | 0.02 | 0.06 | 0.02 | 0.08 | 0.12 |
| 10 | 83969285 | rs10884340 | 2.021 | -0.773 | 1.122 | 2.261 | 0.012 | 0.011 | 0.011 | 0 | 0.56 | 0.43 | 0.44 | 0.28 | 0.65 | 0.53 |
| 10 | 83973325 | rs1764077 | 0.537 | -0.481 | 0.646 | 1.97 | -0.051 | -0.009 | -0.009 | 0 | 0.88 | 0.97 | 0.87 | 0.88 | 0.83 | 0.88 |
| 10 | 83974087 | rs12255719 | -1.966 | -0.694 | 1.53 | 2.104 | -0.008 | 0.013 | 0.013 | 0 | 0.1 | 0.34 | 0.07 | 0.08 | 0.08 | 0.05 |
| 10 | 83974586 | rs10884342 | -2.649 | -0.644 | 1.863 | 2.384 | -0.011 | 0.01 | 0.01 | 0.22 | 0.31 | 0.58 | 0.47 | 0.45 | 0.3 | 0.41 |
| 10 | 83977442 | rs9419959 | -2.212 | -0.96 | 2.1 | 2.326 | -0.012 | -0.009 | -0.009 | 0 | 0.26 | 0.29 | 0.38 | 0.19 | 0.18 | 0.32 |
| 10 | 83978695 | rs17099627 | -1.822 | -0.992 | 2.173 | 2.356 | -0.001 | 0.011 | 0.011 | 0 | 0.09 | 0.13 | 0.3 | 0.15 | 0.09 | 0.2 |
| 10 | 83981964 | rs10884361 | -1.239 | -1.007 | 2.038 | 2.085 | -0.002 | -0.002 | -0.002 | 0 | 0.12 | 0.13 | 0.09 | 0.06 | 0.12 | 0.12 |
| 10 | 83982090 | rs1649962 | -1.428 | -0.813 | 1.912 | 2.019 | 0.002 | -0.009 | -0.009 | 0 | 0.17 | 0.13 | 0.07 | 0.01 | 0.09 | 0.15 |
| 10 | 83988809 | rs1764094 | -1.486 | -0.81 | 1.882 | 2.014 | -0.006 | -0.025 | -0.025 | 0 | 0.17 | 0.14 | 0.07 | 0.01 | 0.09 | 0.15 |
| 10 | 83993404 | rs7084937 | -2.007 | -1.495 | 1.44 | 2.17 | 0.065 | -0.021 | -0.021 | 0 | 0.39 | 0.78 | 0.35 | 0.55 | 0.26 | 0.25 |
| 10 | 83993952 | rs11598813 | -0.386 | -1.577 | 1.269 | 1.963 | -0.017 | 0.034 | 0.034 | 0 | 0.07 | 0.03 | 0.05 | 0 | 0.12 | 0.05 |
| 10 | 83995904 | rs12249883 | -1.786 | -1.299 | 1.593 | 2.135 | -0.041 | 0.043 | 0.043 | 1.111 | 0.31 | 0.6 | 0.36 | 0.56 | 0.17 | 0.24 |
| 10 | 83996992 | rs7915399 | -0.737 | -1.211 | 1.594 | 2.253 | 0.01 | 0.002 | 0.002 | 2.624 | 0.17 | 0.15 | 0.18 | 0.25 | 0.11 | 0.08 |
| 10 | 84000308 | rs7068961 | -0.451 | -1.189 | 1.604 | 2.23 | 0.005 | 0.005 | 0.005 | 2.882 | 0.15 | 0.12 | 0.16 | 0.22 | 0.1 | 0.07 |
| 10 | 84001018 | rs12265466 | -0.451 | -1.198 | 1.604 | 2.23 | 0.007 | 0.005 | 0.005 | 2.882 | 0.15 | 0.12 | 0.16 | 0.22 | 0.1 | 0.08 |
| 10 | 84005517 | rs17099657 | -2.762 | -1.355 | 2.244 | 2.278 | 0.053 | 0.385 | 0.385 | 0.01 | 0.06 | 0.32 | 0.15 | 0.31 | 0.05 | 0.08 |
| 10 | 84008661 | rs12241441 | -0.449 | -1.379 | 2.263 | 1.818 | 0.007 | 0.005 | 0.005 | 3.121 | 0.16 | 0.12 | 0.17 | 0.22 | 0.1 | 0.07 |
| 10 | 84014053 | rs6584594 | -3.076 | -1.335 | 2.478 | 1.693 | -0.012 | 0.072 | 0.072 | 0 | 0.16 | 0.49 | 0.17 | 0.27 | 0.16 | 0.19 |
| 10 | 84017046 | rs10884421 | -3.109 | -1.319 | 2.137 | 1.567 | 0.052 | 0.011 | 0.011 | 1.058 | 0.4 | 0.77 | 0.38 | 0.51 | 0.31 | 0.27 |
| 10 | 84030522 | rs7358197 | -2.575 | -1.26 | 2.155 | 1.683 | 0.065 | 0.221 | 0.221 | 0.999 | 0.05 | 0.43 | 0.1 | 0.13 | 0.01 | 0.05 |
| 10 | 84030537 | rs7072481 | -2.297 | -1.316 | 2.124 | 1.634 | -0.001 | -0.006 | -0.006 | 0 | 0.29 | 0.34 | 0.32 | 0.26 | 0.31 | 0.18 |
| 10 | 84032154 | rs10509451 | -1.081 | -1.348 | 2.136 | 1.616 | -0.016 | 0.025 | 0.025 | 0 | 0.1 | 0.04 | 0.06 | 0 | 0.17 | 0.05 |
| 10 | 84033986 | rs7096123 | -2.38 | -1.349 | 2.168 | 1.622 | 0.003 | 0.005 | 0.005 | 0.961 | 0.31 | 0.3 | 0.33 | 0.37 | 0.31 | 0.21 |
| 10 | 84035428 | rs4329625 | -3.716 | -1.414 | 2.156 | 1.576 | -0.065 | -0.009 | -0.009 | 0.275 | 0.15 | 0.6 | 0.12 | 0.13 | 0.02 | 0.12 |
| 10 | 84038059 | rs7085170 | -3.466 | -1.39 | 2.129 | 1.593 | 0.058 | 0.055 | 0.055 | 2.032 | 0.41 | 0.86 | 0.43 | 0.39 | 0.33 | 0.25 |
| 10 | 84038093 | rs7085184 | -2.847 | -1.361 | 2.194 | 1.563 | 0 | -0.004 | -0.004 | 0 | 0.34 | 0.38 | 0.33 | 0.25 | 0.32 | 0.19 |
| 10 | 84040680 | rs10884462 | -1.355 | -1.36 | 2.195 | 1.548 | -0.004 | -0.005 | -0.005 | 0 | 0.21 | 0.23 | 0.26 | 0.23 | 0.15 | 0.08 |
| 10 | 84045401 | rs7072670 | -3.428 | -1.35 | 2.054 | 1.59 | 0.066 | 0.02 | 0.02 | 2.361 | 0.41 | 0.85 | 0.44 | 0.47 | 0.32 | 0.27 |
| 10 | 84048346 | rs7070560 | -2.769 | -1.343 | 2.125 | 1.598 | 0.045 | 0.302 | 0.302 | 0.351 | 0.06 | 0.45 | 0.11 | 0.24 | 0.02 | 0.08 |
| 10 | 84048527 | rs4297406 | -0.803 | -1.333 | 1.691 | 1.597 | -0.004 | -0.003 | -0.003 | 0 | 0.18 | 0.19 | 0.25 | 0.23 | 0.13 | 0.13 |
| 10 | 84048765 | rs951204 | -0.803 | -1.333 | 1.691 | 1.597 | -0.004 | -0.003 | -0.003 | 0.008 | 0.18 | 0.19 | 0.25 | 0.23 | 0.13 | 0.13 |
| 10 | 84049409 | rs12244925 | -2.003 | -1.313 | 1.707 | 1.612 | 0.002 | 0.023 | 0.023 | 1.636 | 0.27 | 0.2 | 0.3 | 0.23 | 0.3 | 0.17 |
| 10 | 84051375 | rs2225809 | -3.392 | -1.301 | 1.398 | 1.617 | 0.063 | 0.021 | 0.021 | 2.242 | 0.42 | 0.89 | 0.44 | 0.48 | 0.34 | 0.28 |
| 10 | 84066560 | rs7894092 | -1.69 | -1.12 | 1.465 | 1.459 | -0.013 | 0.043 | 0.043 | 2.561 | 0.24 | 0.11 | 0.29 | 0.23 | 0.29 | 0.11 |
| 10 | 84075777 | rs17099777 | -1.836 | -1.262 | 1.445 | 1.639 | 0.001 | 0.006 | 0.006 | 1.482 | 0.26 | 0.21 | 0.29 | 0.3 | 0.28 | 0.11 |
| 10 | 84076732 | rs11193663 | -0.733 | -1.112 | 1.774 | 2.036 | -0.009 | 0.018 | 0.018 | 1.586 | 0.16 | 0.21 | 0.25 | 0.36 | 0.14 | 0.11 |
| 10 | 84078217 | rs3908834 | -0.585 | -1.098 | 1.702 | 2.001 | -0.006 | -0.002 | -0.002 | 1.457 | 0.16 | 0.19 | 0.24 | 0.26 | 0.14 | 0.04 |
| 10 | 84079184 | rs11193683 | -2.892 | -1.076 | 1.522 | 1.994 | 0.086 | -0.09 | -0.09 | 0 | 0.37 | 0.91 | 0.39 | 0.54 | 0.21 | 0.14 |
| 10 | 84082731 | rs10884546 | -2.854 | -0.947 | 2.472 | 2.026 | -0.031 | 0.081 | 0.081 | 0.558 | 0.15 | 0.37 | 0.11 | 0.26 | 0.07 | 0.06 |
| 10 | 84087761 | rs7476649 | -0.615 | -0.948 | 2.466 | 2.043 | -0.005 | 0.004 | 0.004 | 0.335 | 0.16 | 0.19 | 0.24 | 0.3 | 0.14 | 0.05 |
| 10 | 84088573 | rs17099789 | -1.774 | -0.992 | 2.612 | 2.043 | -0.056 | 0.002 | 0.002 | 1.495 | 0.22 | 0.5 | 0.28 | 0.31 | 0.15 | 0.09 |
| 10 | 84088903 | rs7088954 | -2.973 | -1.006 | 2.77 | 2.074 | 0.055 | -0.032 | -0.032 | 0 | 0.33 | 0.8 | 0.37 | 0.54 | 0.19 | 0.14 |
| 10 | 84089539 | rs7916696 | -1.989 | -1.014 | 2.737 | 2.054 | -0.033 | 0.091 | 0.091 | 0.386 | 0.11 | 0.29 | 0.09 | 0.26 | 0.04 | 0.06 |
| 10 | 84093995 | rs11193797 | -3.567 | -1.02 | 2.875 | 2.045 | -0.053 | 0.044 | 0.044 | 0.509 | 0.18 | 0.62 | 0.13 | 0.26 | 0.07 | 0.1 |
| 10 | 84094771 | rs3862550 | -3.1 | -1.036 | 2.789 | 2.054 | 0.089 | -0.063 | -0.063 | 0 | 0.34 | 0.87 | 0.37 | 0.57 | 0.19 | 0.15 |
| 10 | 84098115 | rs7915217 | -3.56 | -1.031 | 2.87 | 2.019 | -0.057 | 0.022 | 0.022 | 2.57 | 0.18 | 0.7 | 0.13 | 0.26 | 0.05 | 0.1 |
| 10 | 84099660 | rs4143817 | -2.923 | -0.997 | 2.837 | 2.035 | -0.024 | 0.105 | 0.105 | 0.16 | 0.14 | 0.4 | 0.1 | 0.26 | 0.07 | 0.06 |
| 10 | 84100509 | rs7087569 | -2.99 | -1.054 | 2.786 | 2.072 | -0.07 | 0.092 | 0.092 | 1.514 | 0.15 | 0.5 | 0.12 | 0.26 | 0.05 | 0.09 |
| 10 | 84111105 | rs7077048 | -0.661 | -0.945 | 2.319 | 1.994 | 0.001 | -0.001 | -0.001 | 0 | 0.16 | 0.16 | 0.24 | 0.28 | 0.14 | 0.05 |
| 10 | 84121782 | rs17099862 | -1.291 | -0.912 | 2.246 | 1.796 | -0.012 | 0.004 | 0.004 | 0 | 0.19 | 0.29 | 0.26 | 0.29 | 0.16 | 0.1 |
| 10 | 84122467 | rs7895428 | -2.597 | -1.004 | 2.484 | 1.857 | -0.015 | 0.009 | 0.009 | 2.661 | 0.31 | 0.66 | 0.36 | 0.54 | 0.2 | 0.16 |
| 10 | 84126885 | rs4506582 | -2.23 | -0.917 | 2.352 | 1.843 | -0.039 | 0.087 | 0.087 | 0.602 | 0.12 | 0.37 | 0.1 | 0.24 | 0.04 | 0.06 |
| 10 | 84136681 | rs7078253 | -2.249 | -0.187 | 2.348 | 1.839 | 0.055 | 0.133 | 0.133 | 0 | 0.12 | 0.45 | 0.13 | 0.24 | 0.09 | 0.08 |
| 10 | 84151515 | rs3740350 | -2.834 | 0.093 | 1.967 | 1.793 | 0.042 | -0.032 | -0.032 | 0 | 0.34 | 0.82 | 0.41 | 0.54 | 0.26 | 0.18 |
| 10 | 84152990 | rs1937984 | -2.775 | 0.093 | 2.011 | 1.824 | 0.032 | -0.035 | -0.035 | 0.188 | 0.34 | 0.79 | 0.41 | 0.56 | 0.26 | 0.19 |
| 10 | 84173117 | rs11812850 | -1.838 | 0.068 | 2.289 | 1.719 | -0.043 | 0.003 | 0.003 | 0.219 | 0.23 | 0.42 | 0.28 | 0.33 | 0.16 | 0.1 |
| 10 | 84177024 | rs7073820 | -1.567 | 0.058 | 2.248 | 1.716 | -0.027 | 0.004 | 0.004 | 0.07 | 0.22 | 0.32 | 0.27 | 0.33 | 0.16 | 0.1 |
| 10 | 84179887 | rs7069578 | -2.537 | 0.1 | 2.305 | 1.751 | 0.011 | 0.058 | 0.058 | 0.074 | 0.14 | 0.57 | 0.15 | 0.2 | 0.1 | 0.1 |
| 10 | 84206002 | rs7909484 | -2.608 | 0.006 | 1.926 | 1.779 | 0.075 | -0.057 | -0.057 | 0 | 0.35 | 0.9 | 0.4 | 0.54 | 0.23 | 0.19 |
| 10 | 84207574 | rs7902584 | -0.798 | -0.138 | 2.103 | 1.728 | -0.005 | -0.003 | -0.003 | 0 | 0.15 | 0.2 | 0.24 | 0.15 | 0.15 | 0.09 |
| 10 | 84210756 | rs7074029 | -1.864 | 0.185 | 2.272 | 1.669 | 0.152 | 0.192 | 0.192 | 0 | 0.08 | 0.42 | 0.13 | 0.21 | 0.11 | 0.11 |
| 10 | 84213251 | rs17100087 | -2.836 | 0.118 | 2.133 | 1.599 | 0.021 | -0.016 | -0.016 | 0 | 0.36 | 0.8 | 0.5 | 0.5 | 0.28 | 0.24 |
| 10 | 84218664 | rs17100097 | -1.772 | 0.173 | 2.228 | 1.619 | 0.04 | 0.062 | 0.062 | 0 | 0.22 | 0.5 | 0.35 | 0.35 | 0.26 | 0.21 |
| 10 | 84222280 | rs1937968 | -2.838 | 0.22 | 1.973 | 1.89 | 0.021 | -0.056 | -0.056 | 0 | 0.36 | 0.82 | 0.5 | 0.59 | 0.28 | 0.25 |

| | | | | | | | | | | | | | | | | |
|---|---|---|---|---|---|---|---|---|---|---|---|---|---|---|---|---|
| 10 | 84223444 | rs1937969 | -1.879 | 0.252 | 2.073 | 1.937 | 0.015 | 0.085 | 0.085 | 0 | 0.23 | 0.58 | 0.35 | 0.44 | 0.25 | 0.21 |
| 10 | 84223466 | rs1937970 | -2.868 | 0.257 | 1.853 | 1.931 | 0.018 | -0.075 | -0.075 | 1.154 | 0.36 | 0.87 | 0.51 | 0.6 | 0.29 | 0.25 |
| 10 | 84237349 | rs7073972 | 1.794 | 0.031 | 1.986 | 1.874 | -0.009 | 0.05 | 0.05 | 0 | 0.74 | 0.43 | 0.65 | 0.58 | 0.77 | 0.79 |
| 10 | 84237629 | rs10748994 | 2.211 | 0.021 | 1.931 | 1.87 | 0.006 | -0.007 | -0.007 | 0 | 0.68 | 0.28 | 0.62 | 0.58 | 0.75 | 0.79 |
| 10 | 84260043 | rs10509454 | -0.35 | 0.005 | 1.813 | 1.801 | -0.002 | 0.004 | 0.004 | 0 | 0.14 | 0.11 | 0.24 | 0.13 | 0.15 | 0.08 |
| 10 | 84265169 | rs581628 | -2.621 | 0.032 | 1.706 | 1.756 | 0.035 | -0.05 | -0.05 | 0 | 0.36 | 0.78 | 0.49 | 0.57 | 0.26 | 0.23 |
| 10 | 84267826 | rs594612 | -2.932 | 0.116 | 1.469 | 1.704 | 0.046 | -0.082 | -0.082 | 0 | 0.39 | 0.91 | 0.52 | 0.57 | 0.31 | 0.24 |
| 10 | 84279949 | rs999889 | -2.613 | 0.248 | 1.464 | 1.681 | 0.032 | -0.09 | -0.09 | 0 | 0.36 | 0.86 | 0.5 | 0.57 | 0.27 | 0.23 |
| 10 | 84303712 | rs659561 | -1.065 | 0.405 | 1.935 | 1.616 | 0.046 | -0.029 | -0.029 | 0 | 0.06 | 0.15 | 0.05 | 0.02 | 0.1 | 0.04 |
| 10 | 84315531 | rs584702 | -2.017 | 0.384 | 1.839 | 1.605 | 0.1 | -0.068 | -0.068 | 0 | 0.12 | 0.47 | 0.1 | 0.02 | 0.15 | 0.05 |
| 10 | 84315759 | rs585597 | -2.732 | 0.143 | 1.482 | 1.603 | 0.041 | -0.072 | -0.072 | 0.304 | 0.42 | 0.9 | 0.54 | 0.58 | 0.33 | 0.29 |
| 10 | 84322815 | rs617738 | -1.969 | 0.572 | 1.544 | 1.742 | 0.016 | 0.056 | 0.056 | 0 | 0.27 | 0.53 | 0.38 | 0.41 | 0.3 | 0.24 |
| 10 | 84329093 | rs11195057 | -0.088 | 0.95 | 1.001 | 1.552 | -0.015 | -0.013 | -0.013 | 0 | 0.18 | 0.13 | 0.33 | 0.31 | 0.19 | 0.09 |
| 10 | 84333436 | rs2062020 | -0.379 | 0.954 | 0.906 | 1.86 | -0.039 | -0.043 | -0.043 | 0 | 0.08 | 0.02 | 0.05 | 0.16 | 0.1 | 0.11 |
| 10 | 84333628 | rs11195073 | 0.148 | 0.814 | 0.572 | 1.81 | -0.023 | 0.021 | 0.021 | 0 | 0.15 | 0.05 | 0.1 | 0.08 | 0.11 | 0.24 |
| 10 | 84335691 | rs1416851 | -0.624 | 0.702 | 0.798 | 1.91 | 0 | 0.007 | 0.007 | 0 | 0.25 | 0.34 | 0.37 | 0.34 | 0.23 | 0.1 |
| 10 | 84336472 | rs1481127 | -1.816 | 0.641 | 0.416 | 2.166 | 0.006 | 0.016 | 0.016 | 0 | 0.43 | 0.73 | 0.54 | 0.65 | 0.35 | 0.34 |
| 10 | 84350877 | rs660464 | -2.573 | 0.631 | 0.055 | 2.149 | -0.074 | 0.158 | 0.158 | 0 | 0.53 | 0.93 | 0.62 | 0.81 | 0.45 | 0.46 |
| 10 | 84354158 | rs983086 | -1.819 | 0.631 | 0.091 | 2.163 | 0.007 | 0.017 | 0.017 | 0 | 0.44 | 0.73 | 0.54 | 0.65 | 0.35 | 0.34 |
| 10 | 84357748 | rs10884989 | -1.813 | 0.643 | -0.035 | 1.927 | 0.006 | -0.009 | -0.009 | 0 | 0.43 | 0.74 | 0.54 | 0.61 | 0.35 | 0.34 |
| 10 | 84359114 | rs11195188 | -0.234 | 0.657 | -0.577 | 1.926 | -0.006 | -0.003 | -0.003 | 0.354 | 0.22 | 0.13 | 0.34 | 0.31 | 0.2 | 0.09 |
| 10 | 84362174 | rs12413431 | 0.12 | 0.975 | -0.892 | 1.344 | -0.02 | 0.01 | 0.01 | 0 | 0.15 | 0.05 | 0.1 | 0.09 | 0.11 | 0.24 |
| 10 | 84363327 | rs639839 | -0.609 | 0.913 | -0.675 | 1.146 | -0.009 | -0.011 | -0.011 | 0 | 0.08 | 0.07 | 0.06 | 0.16 | 0.1 | 0.11 |
| 10 | 84381535 | rs652183 | -1.446 | 0.913 | -0.843 | 1.124 | -0.075 | 0.104 | 0.104 | 0 | 0.52 | 0.92 | 0.6 | 0.77 | 0.45 | 0.45 |
| 10 | 84383277 | rs2247247 | -1.27 | 0.772 | 0.493 | 1.13 | 0.012 | 0.008 | 0.008 | 0 | 0.17 | 0.35 | 0.1 | 0.16 | 0.15 | 0.12 |
| 10 | 84385896 | rs688413 | 0.915 | 0.752 | 0.645 | 1.388 | 0.024 | 0.098 | 0.098 | 0 | 0.83 | 0.65 | 0.69 | 0.64 | 0.81 | 0.79 |
| 10 | 84386093 | rs661469 | 1.089 | 0.318 | 0.48 | 1.344 | 0.003 | -0.005 | -0.005 | 0 | 0.83 | 0.91 | 0.77 | 0.81 | 0.85 | 0.8 |
| 10 | 84386973 | rs673049 | 0.956 | -0.959 | 0.242 | 1.095 | 0.036 | 0.021 | 0.021 | 0.198 | 0.45 | 0.29 | 0.4 | 0.37 | 0.65 | 0.59 |
| 10 | 84388228 | rs1481129 | -0.016 | -0.738 | -0.02 | 0.458 | -0.015 | 0.023 | 0.023 | 0 | 0.12 | 0.08 | 0.07 | 0 | 0.13 | 0.13 |
| 10 | 84389603 | rs2348332 | -0.657 | -0.992 | -0.027 | 0.483 | 0.006 | -0.029 | -0.029 | 0.439 | 0.12 | 0.06 | 0.11 | 0.23 | 0.09 | 0.15 |
| 10 | 84389839 | rs10885058 | 1.623 | -0.664 | 0.084 | 0.67 | -0.04 | -0.042 | -0.042 | 0 | 0.36 | 0.2 | 0.35 | 0.44 | 0.53 | 0.47 |
| 10 | 84400246 | rs1832586 | 0.784 | -0.764 | 0.93 | 0.524 | 0.02 | -0.047 | -0.047 | 0 | 0.77 | 0.65 | 0.7 | 0.86 | 0.74 | 0.74 |
| 10 | 84401508 | rs10749058 | 1.403 | -0.793 | 1.058 | 0.529 | 0.012 | -0.002 | -0.002 | 0 | 0.7 | 0.62 | 0.6 | 0.64 | 0.67 | 0.64 |
| 10 | 84403376 | rs2255228 | -0.777 | -0.71 | 1.071 | 0.514 | 0.005 | -0.001 | -0.001 | 0.357 | 0.9 | 0.89 | 0.79 | 0.86 | 0.88 | 0.87 |
| 10 | 84404982 | rs1339797 | -0.246 | -0.756 | 1.041 | 0.541 | 0.033 | -0.013 | -0.013 | 0.056 | 0.39 | 0.29 | 0.35 | 0.45 | 0.22 | 0.23 |
| 10 | 84409894 | rs10885087 | 1.626 | -0.69 | 0.621 | 0.551 | -0.03 | -0.033 | -0.033 | 0 | 0.37 | 0.24 | 0.35 | 0.44 | 0.53 | 0.5 |
| 10 | 84412569 | rs604482 | 0.19 | -0.679 | 0.723 | 0.492 | -0.002 | -0.019 | -0.019 | 0 | 0.11 | 0.11 | 0.07 | 0 | 0.13 | 0.13 |
| 10 | 84416129 | rs678139 | -0.756 | -0.756 | 0.788 | 0.697 | 0 | -0.001 | -0.001 | 1.051 | 0.9 | 0.92 | 0.8 | 0.86 | 0.88 | 0.88 |
| 10 | 84423731 | rs17100442 | 0.694 | -0.969 | 0.535 | 0.685 | 0.065 | 0.035 | 0.035 | 1.195 | 0.26 | 0.12 | 0.24 | 0.16 | 0.13 | 0.08 |
| 10 | 84424078 | rs17100444 | 0.038 | -0.888 | 0.551 | 0.702 | 0.038 | 0.026 | 0.026 | 0 | 0.36 | 0.2 | 0.45 | 0.3 | 0.24 | 0.21 |
| 10 | 84430088 | rs4586096 | -0.879 | -0.654 | 0.288 | 0.644 | 0.044 | 0.011 | 0.011 | 0.014 | 0.51 | 0.68 | 0.59 | 0.59 | 0.34 | 0.37 |
| 10 | 84433391 | rs3924461 | -1.409 | 0.346 | 0.838 | 0.879 | 0.008 | -0.003 | -0.003 | 0 | 0.24 | 0.67 | 0.39 | 0.26 | 0.28 | 0.25 |
| 10 | 84440148 | rs1335624 | 1.374 | 0.272 | 0.496 | 0.676 | -0.048 | 0.051 | 0.051 | 0 | 0.66 | 0.14 | 0.54 | 0.74 | 0.62 | 0.65 |
| 10 | 84442373 | rs2494026 | 1.212 | -0.341 | 1.588 | 0.627 | -0.014 | -0.047 | -0.047 | 0 | 0.81 | 0.48 | 0.88 | 0.89 | 0.86 | 0.83 |
| 10 | 84452076 | rs2249964 | 0.156 | -0.803 | 0.881 | 0.469 | 0.104 | 0.033 | 0.033 | 0 | 0.49 | 0.22 | 0.6 | 0.66 | 0.7 | 0.65 |
| 10 | 84457148 | rs2246633 | 0.863 | -0.82 | 0.865 | 0.468 | 0.012 | -0.071 | -0.071 | 0 | 0.41 | 0.17 | 0.58 | 0.56 | 0.65 | 0.54 |
| 10 | 84458075 | rs7068082 | 0.297 | -0.768 | 1.057 | 0.31 | 0.082 | -0.023 | -0.023 | 0 | 0.39 | 0.16 | 0.28 | 0.35 | 0.22 | 0.28 |
| 10 | 84459893 | rs2246386 | -0.359 | -0.76 | 1.267 | 0.015 | -0.002 | -0.032 | -0.032 | 0 | 0.9 | 0.88 | 0.92 | 1 | 0.92 | 0.92 |
| 10 | 84461072 | rs7923865 | 0.221 | -0.761 | 1.268 | 0.096 | 0.071 | 0.029 | 0.029 | 0 | 0.4 | 0.2 | 0.28 | 0.25 | 0.22 | 0.27 |
| 10 | 84461630 | rs2246165 | -0.348 | -0.698 | 1.3 | 0.102 | 0.004 | -0.061 | -0.061 | 0 | 0.9 | 0.78 | 0.92 | 1 | 0.92 | 0.92 |
| 10 | 84462470 | rs1572334 | 0.105 | -0.775 | 1.311 | 0.103 | -0.008 | -0.003 | -0.003 | 0 | 0.42 | 0.56 | 0.31 | 0.25 | 0.22 | 0.28 |
| 10 | 84463662 | rs2245896 | 0.335 | -0.721 | 1.054 | 0.131 | 0.096 | 0.09 | 0.09 | 0 | 0.48 | 0.2 | 0.61 | 0.74 | 0.69 | 0.63 |
| 10 | 84465162 | rs2494028 | -0.099 | -0.228 | 0.221 | -0.095 | -0.002 | -0.034 | -0.034 | 0 | 0.9 | 0.87 | 0.92 | 1 | 0.92 | 0.92 |
| 10 | 84469007 | rs1889709 | 0.017 | -0.375 | -0.121 | 0.019 | 0.044 | 0.051 | 0.051 | 0 | 0.39 | 0.25 | 0.24 | 0.2 | 0.23 | 0.25 |
| 10 | 84470658 | rs10885239 | 0.114 | -0.346 | -0.285 | 0.033 | 0.055 | 0.066 | 0.066 | 0 | 0.38 | 0.2 | 0.24 | 0.2 | 0.23 | 0.24 |
| 10 | 84477898 | rs915349 | 0.344 | -0.04 | -0.2 | 0.14 | 0.015 | 0.057 | 0.057 | 0 | 0.47 | 0.75 | 0.68 | 0.68 | 0.6 | 0.55 |
| 10 | 84479555 | rs1335625 | 0.849 | 0.244 | 0.341 | 0.162 | -0.043 | 0.002 | 0.002 | 0 | 0.4 | 0.24 | 0.59 | 0.63 | 0.54 | 0.52 |
| 10 | 84483977 | rs2249075 | -0.152 | 0.023 | 0.208 | 0.082 | 0.004 | 0.007 | 0.007 | 0 | 0.84 | 0.9 | 0.92 | 0.88 | 0.85 | 0.82 |
| 10 | 84484387 | rs10509459 | 0.852 | -0.679 | -0.112 | -0.276 | 0.003 | -0.003 | -0.003 | 0 | 0.15 | 0.14 | 0.12 | 0.14 | 0.06 | 0.05 |
| 10 | 84484985 | rs2494023 | 1.105 | -0.683 | 0.731 | -0.301 | -0.001 | -0.003 | -0.003 | 0 | 0.44 | 0.46 | 0.65 | 0.69 | 0.6 | 0.54 |
| 10 | 84485323 | rs2248956 | 0.36 | -0.685 | 0.93 | -0.309 | 0.012 | 0.071 | 0.071 | 0 | 0.61 | 0.74 | 0.78 | 0.84 | 0.66 | 0.6 |
| 10 | 84490409 | rs12776248 | 0.738 | -0.692 | 0.937 | -0.357 | -0.015 | 0.11 | 0.11 | 0 | 0.23 | 0.05 | 0.13 | 0.05 | 0.2 | 0.22 |
| 10 | 84491173 | rs2820100 | 0.277 | -0.652 | 0.958 | -0.044 | 0.012 | 0.053 | 0.053 | 0 | 0.61 | 0.74 | 0.78 | 0.8 | 0.66 | 0.6 |
| 10 | 84493564 | rs2644206 | 0.93 | -0.611 | 0.949 | -0.027 | 0.005 | 0.001 | 0.001 | 0 | 0.45 | 0.6 | 0.65 | 0.58 | 0.6 | 0.53 |
| 10 | 84494274 | rs2820107 | 0.999 | -0.602 | 0.925 | -0.023 | -0.002 | 0 | 0 | 0.252 | 0.44 | 0.47 | 0.65 | 0.58 | 0.6 | 0.53 |
| 10 | 84500019 | rs342365 | 0.33 | -0.58 | 0.875 | -0.042 | 0.01 | 0.044 | 0.044 | 0 | 0.61 | 0.71 | 0.78 | 0.79 | 0.66 | 0.6 |
| 10 | 84501091 | rs1010874 | 0.522 | -0.653 | 1.258 | 0.377 | 0.009 | -0.004 | -0.004 | 0 | 0.15 | 0.13 | 0.12 | 0.14 | 0.05 | 0.06 |
| 10 | 84501184 | rs1010875 | 0.415 | -0.657 | 1.264 | 0.375 | -0.001 | -0.001 | -0.001 | 0 | 0.16 | 0.15 | 0.12 | 0.14 | 0.05 | 0.06 |
| 10 | 84501882 | rs578080 | 0.835 | -0.691 | 1.36 | 0.363 | -0.024 | 0.015 | 0.015 | 0 | 0.67 | 0.63 | 0.78 | 0.63 | 0.8 | 0.76 |

| | | | | | | | | | | | | | | | |
|---|---|---|---|---|---|---|---|---|---|---|---|---|---|---|---|
| 10 | 84502161 | rs7903899 | 0.548 | -0.613 | 1.294 | 0.419 | 0.015 | -0.014 | -0.014 | 0 | 0.15 | 0.11 | 0.13 | 0.21 | 0.05 | 0.06 |
| 10 | 84503055 | rs571797 | 0.515 | -0.602 | 1.337 | 0.32 | 0.01 | -0.008 | -0.008 | 0 | 0.7 | 0.74 | 0.8 | 0.7 | 0.82 | 0.78 |
| 10 | 84504702 | rs480220 | 0.92 | -0.556 | 1.324 | 0.314 | -0.019 | 0.011 | 0.011 | 0 | 0.66 | 0.53 | 0.74 | 0.7 | 0.76 | 0.76 |
| 10 | 84505577 | rs514237 | 1.014 | -0.561 | 1.286 | 0.312 | -0.001 | 0 | 0 | 0 | 0.65 | 0.42 | 0.73 | 0.7 | 0.76 | 0.76 |
| 10 | 84515820 | rs342393 | 1.268 | -0.667 | 1.122 | 0.273 | 0.036 | -0.008 | -0.008 | 0 | 0.63 | 0.22 | 0.71 | 0.69 | 0.76 | 0.75 |
| 10 | 84516525 | rs12251206 | 0.4 | -0.536 | 0.805 | 0.335 | 0.003 | -0.002 | -0.002 | 0 | 0.16 | 0.16 | 0.13 | 0.14 | 0.05 | 0.06 |
| 10 | 84519876 | rs2644212 | 0.437 | -0.09 | 0.678 | 0.312 | 0.015 | 0.005 | 0.005 | 0 | 0.82 | 0.73 | 0.87 | 0.84 | 0.83 | 0.81 |
| 10 | 84531521 | rs11195879 | -0.626 | -0.78 | 0.897 | 0.029 | -0.061 | -0.03 | -0.03 | 0.019 | 0.22 | 0.45 | 0.18 | 0.17 | 0.08 | 0.08 |
| 10 | 84537367 | rs2065538 | -0.933 | -0.894 | 0.378 | -0.492 | -0.005 | -0.028 | -0.028 | 12.366 | 0.42 | 0.52 | 0.3 | 0.15 | 0.27 | 0.32 |
| 10 | 84538744 | rs7477796 | -0.933 | -0.894 | 0.378 | -0.469 | -0.005 | -0.026 | -0.026 | 0 | 0.42 | 0.52 | 0.3 | 0.15 | 0.27 | 0.32 |
| 10 | 84551558 | rs11195974 | 0.409 | -0.477 | 0.201 | -0.343 | 0 | 0 | 0 | 0.034 | 0.15 | 0.14 | 0.13 | 0.1 | 0.08 | 0.05 |
| 10 | 84557575 | rs1333214 | -1.333 | -0.741 | -0.259 | -0.48 | -0.005 | 0.045 | 0.045 | 0 | 0.53 | 0.8 | 0.4 | 0.34 | 0.43 | 0.44 |
| 10 | 84559886 | rs12773117 | 0.686 | -0.686 | -0.428 | -0.589 | -0.002 | 0.035 | 0.035 | 0 | 0.17 | 0.12 | 0.09 | 0.07 | 0.17 | 0.24 |
| 10 | 84560528 | rs12415063 | 0.651 | -0.69 | -0.429 | -0.591 | -0.002 | 0.036 | 0.036 | 0 | 0.17 | 0.12 | 0.09 | 0.07 | 0.17 | 0.24 |
| 10 | 84564127 | rs2483307 | -1.562 | -0.787 | -0.642 | -0.182 | -0.052 | 0.002 | 0.002 | 0 | 0.64 | 0.84 | 0.62 | 0.61 | 0.52 | 0.63 |
| 10 | 84565851 | rs342368 | -0.056 | 0.08 | -0.683 | -0.435 | 0.022 | 0.018 | 0.018 | 0 | 0.23 | 0.27 | 0.42 | 0.3 | 0.35 | 0.32 |
| 10 | 84570188 | rs501376 | -1.361 | -0.143 | -0.935 | -0.472 | -0.162 | -0.043 | -0.043 | 0 | 0.63 | 0.96 | 0.66 | 0.62 | 0.51 | 0.63 |
| 10 | 84574375 | rs11196077 | 0.257 | -0.575 | -0.424 | -0.656 | -0.046 | 0.056 | 0.056 | 0 | 0.22 | 0.41 | 0.19 | 0.3 | 0.11 | 0.15 |
| 10 | 84576414 | rs7099749 | 0.475 | -0.348 | -0.47 | -0.672 | -0.001 | -0.043 | -0.043 | 0.22 | 0.11 | 0.16 | 0.08 | 0 | 0.08 | 0.04 |
| 10 | 84583413 | rs193178 | 0.076 | -0.379 | -0.304 | -0.568 | -0.012 | 0.097 | 0.097 | 0 | 0.19 | 0.37 | 0.35 | 0.31 | 0.19 | 0.23 |
| 10 | 84589285 | rs342371 | 0.307 | -0.166 | -0.39 | -0.611 | 0.011 | 0.007 | 0.007 | 0 | 0.22 | 0.22 | 0.41 | 0.31 | 0.35 | 0.32 |
| 10 | 84589336 | rs342372 | -0.877 | -0.362 | -0.481 | -0.702 | -0.001 | -0.001 | -0.001 | 0.202 | 0.47 | 0.53 | 0.52 | 0.34 | 0.54 | 0.55 |
| 10 | 84589789 | rs7069209 | 0.08 | -0.913 | -0.649 | -0.956 | -0.009 | -0.001 | -0.001 | 0 | 0.32 | 0.34 | 0.31 | 0.29 | 0.22 | 0.18 |
| 10 | 84590820 | rs342375 | -0.941 | -0.408 | -0.335 | -0.674 | -0.001 | -0.004 | -0.004 | 0 | 0.46 | 0.49 | 0.52 | 0.33 | 0.54 | 0.55 |
| 10 | 84601325 | rs2348553 | -0.119 | -0.43 | -0.263 | -0.908 | -0.009 | 0.038 | 0.038 | 0 | 0.27 | 0.32 | 0.46 | 0.45 | 0.18 | 0.28 |
| 10 | 84611100 | rs342386 | -0.447 | -0.512 | -0.313 | -1.07 | -0.013 | -0.063 | -0.063 | 0 | 0.34 | 0.44 | 0.25 | 0.14 | 0.33 | 0.34 |
| 10 | 84614844 | rs560083 | -2.58 | -0.432 | -0.531 | -1.176 | 0.024 | -0.03 | -0.03 | 0 | 0.69 | 0.88 | 0.77 | 0.63 | 0.7 | 0.72 |
| 10 | 84618072 | rs7917405 | 0.002 | -0.246 | -0.24 | -0.993 | -0.001 | 0.03 | 0.03 | 0 | 0.22 | 0.24 | 0.42 | 0.48 | 0.18 | 0.28 |
| 10 | 84620220 | rs342398 | 0.731 | -0.336 | -0.092 | -0.975 | 0.012 | 0.034 | 0.034 | 0 | 0.13 | 0.37 | 0.15 | 0.12 | 0.13 | 0.13 |
| 10 | 84621316 | rs17100953 | 0.404 | -0.319 | -0.527 | -1.052 | 0.006 | 0.009 | 0.009 | 0 | 0.2 | 0.14 | 0.09 | 0.02 | 0.17 | 0.21 |
| 10 | 84622010 | rs342395 | 1.054 | -0.245 | -0.527 | -0.879 | 0 | 0.01 | 0.01 | 0 | 0.11 | 0.17 | 0.13 | 0.12 | 0.13 | 0.13 |
| 10 | 84623726 | rs512064 | -1.866 | -0.285 | -0.831 | -0.987 | 0.017 | 0.036 | 0.036 | 0 | 0.57 | 0.82 | 0.72 | 0.6 | 0.56 | 0.57 |
| 10 | 84624369 | rs12254634 | 2.515 | -0.486 | -0.401 | -0.984 | -0.022 | -0.027 | -0.027 | 0.548 | 0.05 | 0.11 | 0.02 | 0 | 0.01 | 0 |
| 10 | 84631585 | rs568271 | -2.045 | -0.321 | -0.672 | -1.051 | -0.083 | 0.031 | 0.031 | 0 | 0.6 | 0.87 | 0.68 | 0.64 | 0.5 | 0.65 |
| 10 | 84634847 | rs11196301 | -0.006 | -0.361 | -0.727 | -1.148 | 0.006 | 0.011 | 0.011 | 0 | 0.25 | 0.18 | 0.1 | 0.03 | 0.2 | 0.23 |
| 10 | 84638905 | rs2132377 | -0.566 | -0.635 | -0.273 | -1.017 | 0.01 | 0.035 | 0.035 | 0.029 | 0.27 | 0.42 | 0.52 | 0.6 | 0.26 | 0.39 |
| 10 | 84647206 | rs581354 | 1.288 | -0.531 | -0.467 | -0.952 | 0.01 | 0.024 | 0.024 | 0 | 0.1 | 0.17 | 0.13 | 0.13 | 0.13 | 0.13 |
| 10 | 84648966 | rs489466 | -0.774 | -1.002 | -0.723 | -1.042 | 0.008 | -0.039 | -0.039 | 0 | 0.41 | 0.51 | 0.32 | 0.18 | 0.48 | 0.47 |
| 10 | 84655185 | rs1609605 | -1.579 | -0.47 | -0.681 | -0.684 | 0.008 | 0.008 | 0.008 | 0 | 0.56 | 0.39 | 0.69 | 0.66 | 0.63 | 0.71 |
| 10 | 84656170 | rs17101029 | 0.616 | -0.867 | -0.456 | -0.268 | -0.002 | -0.013 | -0.013 | 0 | 0.07 | 0.07 | 0.03 | 0.01 | 0.04 | 0.02 |
| 10 | 84661601 | rs4360649 | -1.54 | -0.832 | -0.605 | -0.255 | -0.007 | 0.001 | 0.001 | 0 | 0.43 | 0.46 | 0.49 | 0.5 | 0.29 | 0.42 |
| 10 | 84666081 | rs1333208 | 0.129 | -0.378 | -0.862 | 0.053 | 0.001 | 0.072 | 0.072 | 0 | 0.1 | 0.1 | 0.36 | 0.47 | 0.1 | 0.2 |
| 10 | 84666953 | rs7079963 | -0.902 | -0.287 | -0.873 | 0.128 | 0.002 | -0.005 | -0.005 | 0 | 0.35 | 0.3 | 0.45 | 0.5 | 0.25 | 0.41 |
| 10 | 84675278 | rs12243071 | -0.908 | -0.283 | -0.871 | 0.13 | 0.003 | -0.005 | -0.005 | 0 | 0.35 | 0.3 | 0.45 | 0.5 | 0.25 | 0.41 |
| 10 | 84678397 | rs580298 | -0.702 | -0.134 | -1.174 | 0.455 | 0.012 | -0.045 | -0.045 | 0.232 | 0.22 | 0.06 | 0.21 | 0.28 | 0.16 | 0.36 |
| 10 | 84678512 | rs492203 | -1.698 | 0.486 | -0.85 | -0.192 | 0 | 0.008 | 0.008 | 0 | 0.66 | 0.69 | 0.69 | 0.77 | 0.63 | 0.68 |
| 10 | 84680499 | rs1576987 | -0.628 | 0.823 | -0.337 | -0.184 | 0.005 | -0.004 | -0.004 | 0 | 0.36 | 0.4 | 0.43 | 0.65 | 0.36 | 0.49 |
| 10 | 84681635 | rs10885506 | -1.331 | 0.648 | 0.099 | -0.331 | -0.003 | 0.068 | 0.068 | 0 | 0.46 | 0.65 | 0.6 | 0.69 | 0.42 | 0.56 |
| 10 | 84684932 | rs1475707 | -0.603 | 0.605 | 0.437 | -0.335 | -0.011 | 0.003 | 0.003 | 0 | 0.34 | 0.28 | 0.44 | 0.64 | 0.36 | 0.49 |
| 10 | 84685218 | rs12416404 | 2.73 | 0.762 | 0.207 | -0.575 | 0.033 | 0.076 | 0.076 | 0 | 0.32 | 0.14 | 0.27 | 0.21 | 0.29 | 0.23 |
| 10 | 84685244 | rs1475706 | -0.692 | 0.784 | 0.223 | -0.605 | -0.007 | -0.006 | -0.006 | 0 | 0.92 | 0.94 | 0.94 | 0.89 | 0.87 | 0.86 |
| 10 | 84689060 | rs7078771 | 0.919 | 0.783 | 0.355 | -0.603 | -0.019 | -0.021 | -0.021 | 0 | 0.11 | 0.18 | 0.16 | 0.04 | 0.06 | 0.07 |
| 10 | 84690140 | rs2475797 | -2.496 | 0.784 | 0.258 | -0.683 | 0.003 | 0.107 | 0.107 | 0 | 0.64 | 0.86 | 0.7 | 0.77 | 0.64 | 0.68 |
| 10 | 84698466 | rs17101130 | 0.334 | 0.741 | 0.364 | -0.65 | -0.028 | -0.046 | -0.046 | 0 | 0.12 | 0.29 | 0.18 | 0.03 | 0.05 | 0.07 |
| 10 | 84706653 | rs11196606 | 0.798 | 0.898 | 0.284 | -0.592 | 0.001 | -0.025 | -0.025 | 0 | 0.56 | 0.49 | 0.5 | 0.25 | 0.52 | 0.37 |
| 10 | 84713797 | rs4341481 | 0.802 | 0.892 | 0.313 | -0.596 | 0.002 | -0.032 | -0.032 | 0 | 0.56 | 0.5 | 0.5 | 0.25 | 0.52 | 0.36 |
| 10 | 84715289 | rs2348754 | -0.85 | 0.94 | 0.207 | -0.529 | -0.018 | -0.009 | -0.009 | 0 | 0.93 | 0.96 | 0.95 | 0.92 | 0.87 | 0.84 |
| 10 | 84735241 | rs7092994 | -0.58 | 0.869 | 0.315 | -0.585 | 0.005 | -0.011 | -0.011 | 0 | 0.35 | 0.39 | 0.44 | 0.67 | 0.34 | 0.48 |
| 10 | 84740839 | rs12763193 | 1.684 | 0.838 | 0.209 | -0.707 | 0.012 | 0.133 | 0.133 | 0 | 0.44 | 0.24 | 0.33 | 0.22 | 0.46 | 0.29 |
| 10 | 84745040 | rs17101196 | 0.959 | 0.797 | 0.252 | -0.705 | -0.016 | -0.039 | -0.039 | 0.199 | 0.1 | 0.17 | 0.16 | 0.01 | 0.06 | 0.07 |
| 10 | 84746621 | rs8421 | 1.115 | 0.882 | -0.013 | -0.54 | -0.001 | 0.179 | 0.179 | 0 | 0.49 | 0.24 | 0.37 | 0.23 | 0.54 | 0.39 |
| 10 | 84755074 | rs10787532 | -0.789 | 0.933 | -0.228 | -0.399 | -0.007 | 0.005 | 0.005 | 0 | 0.92 | 0.91 | 0.94 | 0.97 | 0.86 | 0.87 |
| 10 | 84758939 | rs17689242 | 2.799 | 0.424 | -0.656 | -0.6 | -0.003 | 0.134 | 0.134 | 0 | 0.25 | 0.04 | 0.18 | 0.03 | 0.23 | 0.05 |
| 10 | 84764495 | rs11196808 | 0.274 | 0.208 | 0.175 | 0.075 | 0.008 | -0.006 | -0.006 | 0 | 0.13 | 0.05 | 0.07 | 0.15 | 0.08 | 0.19 |
| 10 | 84773077 | rs7906345 | -2.52 | 0.306 | 0.394 | -0.294 | -0.007 | 0.065 | 0.065 | 0 | 0.63 | 0.79 | 0.69 | 0.75 | 0.59 | 0.69 |
| 10 | 84775286 | rs4437972 | 0.618 | 0.319 | 0.472 | -0.348 | -0.007 | 0.014 | 0.014 | 0 | 0.14 | 0.12 | 0.07 | 0.05 | 0.17 | 0.1 |
| 10 | 84775417 | rs10749172 | -1.816 | 0.307 | 0.61 | -0.351 | 0.007 | 0.056 | 0.056 | 0 | 0.48 | 0.6 | 0.62 | 0.7 | 0.42 | 0.59 |
| 10 | 84798552 | rs4244940 | -2.594 | 0.326 | 0.592 | -0.333 | 0 | 0.041 | 0.041 | 0 | 0.63 | 0.73 | 0.69 | 0.75 | 0.59 | 0.69 |
| 10 | 84802738 | rs12784854 | 1.692 | 0.415 | 0.325 | -0.401 | -0.038 | 0.037 | 0.037 | 0 | 0.07 | 0 | 0.06 | 0 | 0.11 | 0.01 |

| | | | | | | | | | | | | | | | | |
|---|---|---|---|---|---|---|---|---|---|---|---|---|---|---|---|---|
| 10 | 84809358 | rs11197049 | 0.197 | 0.121 | 0.629 | 0.048 | 0.011 | 0.017 | 0.017 | 0 | 0.18 | 0.17 | 0.23 | 0.24 | 0.23 | 0.19 |
| 10 | 84813142 | rs11197065 | -0.615 | 0.084 | 0.671 | 0.046 | 0.007 | 0.009 | 0.009 | 0 | 0.25 | 0.22 | 0.39 | 0.25 | 0.28 | 0.27 |
| 10 | 84828676 | rs7087324 | -0.613 | -0.136 | 1.053 | 0.355 | -0.013 | 0.03 | 0.03 | 0 | 0.22 | 0.28 | 0.22 | 0.45 | 0.14 | 0.33 |
| 10 | 84833190 | rs10787560 | -2.695 | -0.06 | 1.024 | 0.274 | -0.005 | 0.045 | 0.045 | 0 | 0.6 | 0.69 | 0.66 | 0.76 | 0.56 | 0.68 |
| 10 | 84836280 | rs12782033 | 0.391 | 0.047 | 0.94 | 0.431 | 0.016 | 0.039 | 0.039 | 0 | 0.15 | 0.17 | 0.21 | 0.24 | 0.2 | 0.19 |
| 10 | 84840318 | rs11197192 | -0.512 | -0.089 | 0.961 | 0.36 | -0.012 | 0.032 | 0.032 | 0 | 0.21 | 0.27 | 0.21 | 0.45 | 0.12 | 0.31 |
| 10 | 84847268 | rs11197219 | -0.676 | -0.012 | 0.863 | 0.334 | 0.006 | 0.017 | 0.017 | 0 | 0.23 | 0.22 | 0.37 | 0.26 | 0.26 | 0.28 |
| 10 | 84855219 | rs10787576 | -2.44 | 0.001 | 0.772 | 0.07 | -0.02 | 0.091 | 0.091 | 0 | 0.6 | 0.78 | 0.66 | 0.76 | 0.55 | 0.69 |
| 10 | 84864276 | rs10885756 | 0.237 | 0.429 | 1.504 | 0.739 | 0.013 | 0.059 | 0.059 | 0 | 0.13 | 0.16 | 0.25 | 0.28 | 0.16 | 0.16 |
| 10 | 84866669 | rs1339844 | -1.64 | 0.277 | 1.738 | 0.65 | 0.001 | -0.023 | -0.023 | 0 | 0.32 | 0.54 | 0.43 | 0.74 | 0.28 | 0.45 |
| 10 | 84870076 | rs7915137 | -1.825 | 0.239 | 2 | 0.583 | -0.006 | 0.007 | 0.007 | 0 | 0.41 | 0.44 | 0.36 | 0.47 | 0.36 | 0.47 |
| 10 | 84879288 | rs11818601 | 0.73 | 0.244 | 1.969 | 0.561 | 0 | -0.011 | -0.011 | 1.556 | 0.08 | 0.05 | 0.12 | 0 | 0.06 | 0.09 |
| 10 | 84887564 | rs10885820 | -0.573 | 0.255 | 1.999 | 0.582 | -0.007 | 0.058 | 0.058 | 0 | 0.18 | 0.24 | 0.16 | 0.45 | 0.12 | 0.29 |
| 10 | 84900150 | rs7911623 | -2.741 | 0.247 | 1.75 | 0.507 | -0.019 | 0.125 | 0.125 | 0 | 0.55 | 0.77 | 0.63 | 0.75 | 0.52 | 0.62 |
| 10 | 84914054 | rs7070132 | -0.85 | 0.298 | 1.806 | 0.517 | -0.002 | -0.079 | -0.079 | 2.402 | 0.23 | 0.3 | 0.21 | 0.01 | 0.25 | 0.18 |
| 10 | 84924164 | rs10749204 | -2.802 | 0.27 | 1.717 | 0.518 | -0.062 | 0.142 | 0.142 | 0 | 0.56 | 0.88 | 0.63 | 0.75 | 0.52 | 0.64 |
| 10 | 84924675 | rs12782058 | -1.697 | 0.271 | 1.768 | 0.515 | -0.001 | -0.014 | -0.014 | 0 | 0.32 | 0.58 | 0.42 | 0.74 | 0.28 | 0.45 |
| 10 | 84925424 | rs1856612 | -0.946 | 0.541 | 1.942 | 0.569 | -0.001 | -0.036 | -0.036 | 0.094 | 0.23 | 0.19 | 0.2 | 0.01 | 0.24 | 0.19 |
| 10 | 84925612 | rs12412876 | 0.122 | 0.577 | 1.868 | 0.595 | 0.001 | -0.025 | -0.025 | 1.128 | 0.14 | 0.15 | 0.09 | 0.01 | 0.18 | 0.09 |
| 10 | 84927666 | rs961435 | -0.184 | 0.496 | 2.229 | 0.678 | 0.024 | 0.099 | 0.099 | 0.049 | 0.14 | 0.26 | 0.25 | 0.28 | 0.16 | 0.16 |
| 10 | 84929207 | rs11197616 | -1.189 | 0.397 | 1.923 | 0.675 | -0.007 | 0.049 | 0.049 | 0.035 | 0.22 | 0.24 | 0.33 | 0.5 | 0.17 | 0.34 |
| 10 | 84932401 | rs2348880 | 2.362 | 0.537 | 0.897 | 0.49 | -0.059 | 0.231 | 0.231 | 0 | 0.46 | 0.09 | 0.3 | 0.19 | 0.43 | 0.38 |
| 10 | 84937146 | rs10787654 | -1.826 | 0.484 | 1.087 | 0.563 | 0.001 | 0.022 | 0.022 | 0 | 0.37 | 0.62 | 0.59 | 0.78 | 0.33 | 0.51 |
| 10 | 84941604 | rs2348886 | -2.393 | 0.524 | 0.677 | 0.546 | -0.018 | 0.229 | 0.229 | 0 | 0.51 | 0.79 | 0.67 | 0.81 | 0.52 | 0.6 |
| 10 | 84944836 | rs6585236 | -2.14 | 0.441 | 1.256 | 0.748 | 0.006 | -0.02 | -0.02 | 1.373 | 0.38 | 0.66 | 0.43 | 0.51 | 0.36 | 0.44 |
| 10 | 84947146 | rs10509470 | -0.031 | 0.466 | 0.986 | 0.699 | -0.025 | 0.05 | 0.05 | 0 | 0.13 | 0.04 | 0.07 | 0.01 | 0.18 | 0.09 |
| 10 | 84951635 | rs17215403 | -0.037 | -0.181 | 1.041 | 0.95 | 0.009 | 0.016 | 0.016 | 0 | 0.09 | 0.02 | 0.11 | 0 | 0.07 | 0.09 |
| 10 | 84969643 | rs1010860 | -1.652 | -0.166 | 1.274 | 1.039 | -0.025 | 0.133 | 0.133 | 0.119 | 0.23 | 0.38 | 0.33 | 0.5 | 0.17 | 0.34 |
| 10 | 84980576 | rs1832591 | -1.926 | -0.417 | 1.058 | 1.046 | -0.015 | 0.013 | 0.013 | 0.248 | 0.33 | 0.59 | 0.34 | 0.45 | 0.25 | 0.39 |
| 10 | 84984421 | rs11197830 | -0.655 | 0.466 | 1.066 | 1.529 | 0.01 | 0.024 | 0.024 | 0 | 0.1 | 0.09 | 0.25 | 0.21 | 0.16 | 0.16 |
| 10 | 85020122 | rs7079919 | -1.443 | 0.295 | 1.62 | 1.464 | -0.021 | 0.071 | 0.071 | 0 | 0.21 | 0.29 | 0.27 | 0.42 | 0.14 | 0.25 |
| 10 | 85029143 | rs4370876 | -2.968 | -0.063 | 1.047 | 1.386 | -0.042 | 0.065 | 0.065 | 0.149 | 0.44 | 0.93 | 0.59 | 0.68 | 0.37 | 0.51 |
| 10 | 85030145 | rs10886053 | -0.453 | -0.057 | 1.175 | 1.441 | 0.008 | 0.027 | 0.027 | 0 | 0.1 | 0.1 | 0.25 | 0.21 | 0.13 | 0.15 |
| 10 | 85032380 | rs7911016 | -0.501 | -0.415 | 1.791 | 1.187 | 0 | -0.008 | -0.008 | 0.895 | 0.1 | 0.1 | 0.06 | 0.01 | 0.1 | 0.07 |
| 10 | 85032996 | rs10886056 | -2.804 | -0.407 | 1.644 | 1.198 | 0.042 | -0.052 | -0.052 | 0.175 | 0.34 | 0.83 | 0.34 | 0.46 | 0.24 | 0.36 |
| 10 | 85043105 | rs11197990 | -0.872 | -0.249 | 1.63 | 1.274 | -0.024 | 0.068 | 0.068 | 0 | 0.13 | 0.18 | 0.13 | 0.42 | 0.07 | 0.21 |
| 10 | 85050535 | rs17101655 | -0.583 | -0.124 | 1.171 | 1.236 | -0.002 | -0.026 | -0.026 | 0.02 | 0.1 | 0.2 | 0.06 | 0.03 | 0.1 | 0.1 |
| 10 | 85060080 | rs6585295 | -0.501 | -0.107 | 1.177 | 1.159 | -0.001 | -0.046 | -0.046 | 0 | 0.1 | 0.2 | 0.06 | 0.01 | 0.1 | 0.07 |
| 10 | 85062296 | rs4362092 | -1.519 | -0.013 | 1.158 | 1.483 | -0.033 | 0.131 | 0.131 | 0 | 0.21 | 0.41 | 0.27 | 0.42 | 0.14 | 0.24 |
| 10 | 85073628 | rs6585303 | -1.399 | 0.582 | 0.92 | 1.381 | 0.018 | 0.01 | 0.01 | 0 | 0.12 | 0.2 | 0.16 | 0.15 | 0.14 | 0.23 |
| 10 | 85074636 | rs4617529 | 0.321 | 0.202 | 1.532 | 1.896 | -0.004 | 0.01 | 0.01 | 0 | 0.83 | 0.83 | 0.85 | 0.63 | 0.88 | 0.8 |
| 10 | 85085640 | rs4933896 | -0.969 | -0.07 | 1.711 | 1.663 | -0.004 | -0.014 | -0.014 | 0 | 0.13 | 0.11 | 0.06 | 0.02 | 0.09 | 0.09 |
| 10 | 85091864 | rs10886170 | -2.332 | -0.065 | 1.722 | 1.67 | 0.003 | -0.014 | -0.014 | 0.037 | 0.25 | 0.31 | 0.22 | 0.17 | 0.24 | 0.31 |
| 10 | 85094893 | rs9888049 | 2.568 | -0.066 | 1.785 | 1.673 | 0.004 | 0.008 | 0.008 | 0 | 0.54 | 0.35 | 0.49 | 0.42 | 0.57 | 0.45 |
| 10 | 85106420 | rs4329627 | 0.416 | -0.091 | 1.905 | 1.668 | -0.023 | 0.037 | 0.037 | 0 | 0.82 | 0.75 | 0.84 | 0.63 | 0.88 | 0.8 |
| 10 | 85108139 | rs11198250 | -0.388 | -0.058 | 1.752 | 1.629 | 0.01 | 0.017 | 0.017 | 0.367 | 0.09 | 0.03 | 0.04 | 0.02 | 0.05 | 0.08 |
| 10 | 85108907 | rs7083654 | -0.897 | -0.043 | 1.756 | 1.64 | 0.018 | 0.01 | 0.01 | 0 | 0.12 | 0.2 | 0.16 | 0.15 | 0.14 | 0.23 |
| 10 | 85127184 | rs12257425 | -0.733 | -0.038 | 1.756 | 1.762 | 0.005 | 0.017 | 0.017 | 0.019 | 0.11 | 0.04 | 0.05 | 0.02 | 0.09 | 0.09 |
| 10 | 85127739 | rs10886214 | -2.168 | -0.05 | 1.861 | 1.763 | 0.005 | -0.009 | -0.009 | 0 | 0.23 | 0.31 | 0.21 | 0.17 | 0.23 | 0.31 |
| 10 | 85133114 | rs4638255 | 2.833 | -0.236 | 1.881 | 1.797 | 0.02 | 0.052 | 0.052 | 0 | 0.51 | 0.21 | 0.48 | 0.39 | 0.57 | 0.4 |
| 10 | 85133956 | rs10886221 | -2.728 | -0.126 | 1.929 | 1.668 | 0.016 | -0.023 | -0.023 | 0.545 | 0.3 | 0.55 | 0.36 | 0.22 | 0.31 | 0.34 |
| 10 | 85139142 | rs4076319 | -1.678 | -0.12 | 1.947 | 1.671 | 0.045 | 0.024 | 0.024 | 0 | 0.17 | 0.38 | 0.3 | 0.19 | 0.21 | 0.26 |
| 10 | 85154286 | rs7913554 | -2.388 | -0.115 | 1.704 | 1.689 | 0.016 | -0.023 | -0.023 | 0 | 0.3 | 0.55 | 0.36 | 0.22 | 0.31 | 0.34 |
| 10 | 85161251 | rs10886274 | 2.112 | -0.675 | 1.396 | 1.609 | 0.021 | 0.005 | 0.005 | 0 | 0.53 | 0.38 | 0.58 | 0.41 | 0.66 | 0.47 |
| 10 | 85163584 | rs4933906 | 0.493 | -0.38 | 1.744 | 1.75 | -0.004 | 0.007 | 0.007 | 0 | 0.82 | 0.82 | 0.84 | 0.7 | 0.86 | 0.81 |
| 10 | 85169558 | rs4933908 | 0.781 | -0.417 | 1.775 | 1.726 | 0.002 | -0.003 | -0.003 | 0 | 0.8 | 0.79 | 0.84 | 0.68 | 0.85 | 0.8 |
| 10 | 85170715 | rs11198560 | -0.461 | -0.453 | 1.841 | 1.812 | -0.002 | 0.009 | 0.009 | 0 | 0.08 | 0.11 | 0.13 | 0.18 | 0.07 | 0.16 |
| 10 | 85179236 | rs2349157 | 0.43 | -0.513 | 1.739 | 1.847 | 0.018 | -0.033 | -0.033 | 0 | 0.82 | 0.93 | 0.91 | 0.75 | 0.88 | 0.82 |
| 10 | 85183390 | rs10886307 | -1.902 | -0.558 | 2.104 | 1.685 | -0.022 | -0.012 | -0.012 | 0.776 | 0.22 | 0.35 | 0.21 | 0.21 | 0.17 | 0.27 |
| 10 | 85187542 | rs1855953 | 2.456 | -0.513 | 2.186 | 1.686 | 0.036 | 0.008 | 0.008 | 0 | 0.55 | 0.27 | 0.55 | 0.46 | 0.64 | 0.44 |
| 10 | 85193946 | rs11198672 | -2.337 | -0.477 | 2.152 | 1.654 | -0.009 | -0.011 | -0.011 | 0 | 0.24 | 0.4 | 0.28 | 0.22 | 0.21 | 0.34 |
| 10 | 85195261 | rs7074730 | -1.452 | -0.502 | 1.857 | 1.659 | -0.005 | 0 | 0 | 0 | 0.17 | 0.2 | 0.17 | 0.2 | 0.11 | 0.24 |
| 10 | 85203737 | rs11198735 | -1.336 | -0.52 | 1.739 | 1.664 | 0.001 | -0.001 | -0.001 | 0.016 | 0.17 | 0.17 | 0.17 | 0.2 | 0.11 | 0.24 |
| 10 | 85205935 | rs2349159 | 1.907 | -0.516 | 1.668 | 1.646 | 0.003 | -0.002 | -0.002 | 0 | 0.59 | 0.39 | 0.63 | 0.47 | 0.69 | 0.51 |
| 10 | 85210888 | rs4523641 | 1.472 | -0.505 | 1.627 | 1.676 | -0.007 | 0.019 | 0.019 | 0 | 0.72 | 0.4 | 0.72 | 0.66 | 0.75 | 0.67 |
| 10 | 85210946 | rs10787907 | 1.632 | -0.619 | 1.915 | 1.619 | -0.03 | 0.06 | 0.06 | 0 | 0.71 | 0.51 | 0.72 | 0.65 | 0.77 | 0.68 |
| 10 | 85210966 | rs10787908 | 1.53 | -0.633 | 1.912 | 1.697 | -0.029 | 0.025 | 0.025 | 0 | 0.72 | 0.51 | 0.78 | 0.72 | 0.8 | 0.7 |
| 10 | 85211865 | rs7913357 | 1.463 | -0.561 | 1.872 | 1.598 | -0.021 | 0.029 | 0.029 | 0 | 0.73 | 0.4 | 0.69 | 0.62 | 0.79 | 0.67 |
| 10 | 85213365 | rs10509475 | 1.84 | -0.592 | 1.81 | 1.584 | -0.002 | 0.001 | 0.001 | 0 | 0.7 | 0.35 | 0.62 | 0.62 | 0.75 | 0.65 |

| | | | | | | | | | | | | | | | | |
|---|---|---|---|---|---|---|---|---|---|---|---|---|---|---|---|---|
| 10 | 85214724 | rs10886391 | 1.658 | -0.124 | 1.783 | 1.545 | 0 | -0.001 | -0.001 | 0 | 0.71 | 0.33 | 0.6 | 0.62 | 0.72 | 0.65 |
| 10 | 85215937 | rs1336400 | 2.039 | -0.155 | 1.796 | 1.579 | 0.01 | -0.044 | -0.044 | 0.082 | 0.65 | 0.22 | 0.53 | 0.56 | 0.66 | 0.57 |
| 10 | 85216947 | rs1984155 | 0.569 | -0.161 | 2.133 | 1.609 | 0.001 | -0.034 | -0.034 | 0 | 0.8 | 0.89 | 0.79 | 0.67 | 0.79 | 0.72 |
| 10 | 85218018 | rs1984156 | -1.685 | -0.124 | 2.222 | 1.52 | 0.046 | -0.031 | -0.031 | 2.529 | 0.07 | 0.24 | 0.11 | 0 | 0.1 | 0.05 |
| 10 | 85229138 | rs10787944 | 1.904 | -0.142 | 2.047 | 1.669 | 0.007 | -0.04 | -0.04 | 0 | 0.65 | 0.25 | 0.53 | 0.51 | 0.66 | 0.57 |
| 10 | 85234050 | rs10787954 | 1.518 | -0.145 | 2.055 | 1.673 | 0.003 | -0.022 | -0.022 | 0 | 0.68 | 0.29 | 0.6 | 0.55 | 0.69 | 0.59 |
| 10 | 85236119 | rs915494 | 1.907 | -0.142 | 2.049 | 1.645 | 0.007 | -0.036 | -0.036 | 0.928 | 0.65 | 0.25 | 0.53 | 0.52 | 0.66 | 0.63 |
| 10 | 85241941 | rs10787962 | 1.121 | -0.147 | 2.386 | 1.641 | -0.018 | 0.074 | 0.074 | 0 | 0.73 | 0.5 | 0.68 | 0.56 | 0.76 | 0.69 |
| 10 | 85247193 | rs17101972 | -2.412 | -0.135 | 2.355 | 1.622 | 0.022 | -0.006 | -0.006 | 0 | 0.08 | 0.25 | 0.15 | 0.04 | 0.1 | 0.06 |
| 10 | 85254470 | rs11198993 | 1.1 | -0.147 | 2.385 | 1.644 | -0.015 | 0.058 | 0.058 | 0 | 0.74 | 0.62 | 0.69 | 0.56 | 0.76 | 0.69 |
| 10 | 85254486 | rs3918861 | -2.421 | -0.142 | 2.401 | 1.621 | 0.063 | 0.021 | 0.021 | 0 | 0.08 | 0.36 | 0.15 | 0.04 | 0.1 | 0.06 |
| 10 | 85259487 | rs10886514 | 2.586 | -0.014 | 1.982 | 1.694 | -0.004 | 0.036 | 0.036 | 0.107 | 0.52 | 0.06 | 0.47 | 0.36 | 0.57 | 0.41 |
| 10 | 85261363 | rs1336405 | 0.218 | -0.135 | 1.124 | 0.686 | 0.027 | -0.045 | -0.045 | 0.028 | 0.36 | 0.22 | 0.25 | 0.52 | 0.29 | 0.54 |
| 10 | 85269860 | rs671918 | 1.396 | 0.953 | 1.95 | 1.889 | -0.004 | 0.012 | 0.012 | 0 | 0.69 | 0.44 | 0.68 | 0.63 | 0.69 | 0.67 |
| 10 | 85271910 | rs7081039 | -2.561 | 1.025 | 2.075 | 1.586 | -0.013 | -0.026 | -0.026 | 0 | 0.25 | 0.63 | 0.3 | 0.15 | 0.23 | 0.2 |
| 10 | 85273305 | rs626939 | 0.754 | 1.233 | 2.133 | 1.812 | -0.001 | -0.001 | -0.001 | 0.024 | 0.86 | 0.83 | 0.86 | 0.73 | 0.82 | 0.81 |
| 10 | 85273952 | rs7915460 | 2.72 | 1.612 | 2.353 | 1.816 | -0.011 | -0.015 | -0.015 | 0 | 0.54 | 0.3 | 0.65 | 0.5 | 0.5 | 0.51 |
| 10 | 85276201 | rs4402239 | 2.07 | 1.622 | 2.859 | 1.974 | 0.012 | 0.008 | 0.008 | 0 | 0.67 | 0.58 | 0.78 | 0.6 | 0.62 | 0.61 |
| 10 | 85296654 | rs7895503 | 2.299 | 1.634 | 2.956 | 1.919 | 0.022 | 0.025 | 0.025 | 0 | 0.65 | 0.51 | 0.75 | 0.53 | 0.6 | 0.6 |
| 10 | 85304788 | rs982345 | 1.99 | 1.604 | 3.052 | 1.918 | 0.015 | 0.024 | 0.024 | 0 | 0.68 | 0.54 | 0.78 | 0.56 | 0.64 | 0.63 |
| 10 | 85312966 | rs10128261 | -1.816 | 1.413 | 3.008 | 1.71 | -0.019 | -0.015 | -0.015 | 0 | 0.1 | 0.23 | 0.09 | 0.04 | 0.07 | 0.1 |
| 10 | 85316258 | rs656127 | -1.341 | 1.442 | 2.988 | 1.688 | -0.005 | -0.003 | -0.003 | 0 | 0.18 | 0.13 | 0.09 | 0.16 | 0.21 | 0.2 |
| 10 | 85317970 | rs602012 | 0.273 | 1.492 | 2.747 | 1.749 | -0.031 | -0.042 | -0.042 | 0.715 | 0.07 | 0 | 0.04 | 0.15 | 0.1 | 0.12 |
| 10 | 85318842 | rs17102157 | -2.42 | 1.431 | 2.915 | 1.685 | -0.03 | -0.029 | -0.029 | 0 | 0.12 | 0.39 | 0.1 | 0.04 | 0.07 | 0.1 |
| 10 | 85330074 | rs1359266 | -2.584 | 1.385 | 2.918 | 1.702 | -0.012 | -0.019 | -0.019 | 0 | 0.13 | 0.34 | 0.12 | 0.07 | 0.09 | 0.12 |
| 10 | 85332848 | rs1408346 | 1.729 | 1.578 | 2.916 | 1.774 | 0.008 | 0.005 | 0.005 | 0.022 | 0.68 | 0.61 | 0.77 | 0.61 | 0.62 | 0.63 |
| 10 | 85358478 | rs7900509 | -1.667 | 1.568 | 2.66 | 1.754 | 0.004 | -0.019 | -0.019 | 0.279 | 0.09 | 0.21 | 0.06 | 0.04 | 0.07 | 0.1 |
| 10 | 85359998 | rs1324250 | -1.222 | 1.623 | 2.258 | 1.748 | -0.019 | 0.027 | 0.027 | 0 | 0.95 | 1 | 0.97 | 1 | 0.93 | 0.99 |
| 10 | 85361271 | rs1535847 | 2.786 | 1.59 | 2.339 | 1.791 | -0.023 | -0.014 | -0.014 | 7.811 | 0.52 | 0.15 | 0.64 | 0.54 | 0.52 | 0.51 |
| 10 | 85385729 | rs12264194 | -1.004 | 1.785 | 2.178 | 1.768 | -0.008 | -0.007 | -0.007 | 0.668 | 0.14 | 0.14 | 0.14 | 0.23 | 0.17 | 0.18 |
| 10 | 85396951 | rs12241673 | -2.652 | 1.384 | 2.352 | 1.517 | -0.005 | 0.006 | 0.006 | 0 | 0.08 | 0.25 | 0.1 | 0.06 | 0.05 | 0.04 |
| 10 | 85397454 | rs7909157 | -2.158 | 1.377 | 2.594 | 1.504 | 0.042 | 0.029 | 0.029 | 0.074 | 0.07 | 0.37 | 0.05 | 0.03 | 0.03 | 0.02 |
| 10 | 85397874 | rs7898756 | 0.915 | 1.428 | 2.365 | 1.641 | -0.006 | -0.012 | -0.012 | 0.112 | 0.72 | 0.76 | 0.82 | 0.59 | 0.68 | 0.64 |
| 10 | 85398017 | rs10749395 | 2.208 | 1.329 | 2.528 | 1.777 | 0.002 | -0.003 | -0.003 | 0 | 0.56 | 0.4 | 0.65 | 0.46 | 0.58 | 0.45 |
| 10 | 85399760 | rs12268063 | -1.175 | 1.324 | 2.52 | 1.625 | -0.027 | -0.001 | -0.001 | 0 | 0.1 | 0.19 | 0.13 | 0.08 | 0.06 | 0.15 |
| 10 | 85407876 | rs944945 | 1.605 | 1.766 | 2.628 | 1.627 | 0.001 | 0.005 | 0.005 | 0 | 0.66 | 0.59 | 0.78 | 0.56 | 0.65 | 0.62 |
| 10 | 85409604 | rs969986 | 1.078 | 1.697 | 2.395 | 1.632 | -0.005 | -0.011 | -0.011 | 0 | 0.72 | 0.76 | 0.85 | 0.59 | 0.68 | 0.65 |
| 10 | 85419023 | rs1408349 | 1.323 | 1.7 | 2.359 | 1.631 | 0.007 | 0.025 | 0.025 | 0 | 0.7 | 0.62 | 0.84 | 0.6 | 0.68 | 0.65 |
| 10 | 85423215 | rs17102381 | -2.219 | 1.437 | 1.948 | 1.507 | 0.083 | -0.017 | -0.017 | 0.19 | 0.11 | 0.5 | 0.18 | 0.03 | 0.1 | 0.11 |
| 10 | 85423893 | rs1029133 | 1.948 | 1.493 | 0.869 | 1.621 | -0.073 | -0.076 | -0.076 | 0.16 | 0.57 | 0.08 | 0.65 | 0.54 | 0.58 | 0.5 |
| 10 | 85423938 | rs1029134 | 1.218 | 0.939 | 1.244 | 1.712 | -0.003 | -0.013 | -0.013 | 0 | 0.67 | 0.71 | 0.84 | 0.58 | 0.68 | 0.63 |
| 10 | 85424102 | rs621681 | 1.378 | 1.027 | 1.063 | 1.957 | 0.005 | -0.008 | -0.008 | 0 | 0.68 | 0.28 | 0.81 | 0.66 | 0.73 | 0.64 |
| 10 | 85424493 | rs7092608 | -2.167 | 0.838 | 1.365 | 2.233 | -0.024 | 0.151 | 0.151 | 0 | 0.17 | 0.32 | 0.16 | 0.46 | 0.12 | 0.42 |
| 10 | 85425691 | rs10886877 | -0.568 | 1.501 | 0.651 | 1.625 | -0.042 | -0.041 | -0.041 | 0 | 0.08 | 0 | 0.11 | 0.13 | 0.13 | 0.13 |
| 10 | 85425986 | rs7921163 | -1.526 | 1.224 | 1.478 | 2.126 | 0 | 0.231 | 0.231 | 0 | 0.07 | 0.23 | 0.13 | 0.26 | 0.04 | 0.15 |
| 10 | 85428066 | rs1333846 | 1.026 | 1.307 | 1.621 | 1.742 | 0.004 | -0.03 | -0.03 | 0.166 | 0.78 | 0.68 | 0.88 | 0.9 | 0.8 | 0.74 |
| 10 | 85431404 | rs10736301 | -2.509 | 0.838 | 2.019 | 2.021 | 0.016 | 0.214 | 0.214 | 0 | 0.2 | 0.47 | 0.27 | 0.46 | 0.23 | 0.31 |
| 10 | 85432838 | rs7072115 | -1.265 | 0.822 | 2.041 | 2.046 | 0.004 | 0.216 | 0.216 | 0 | 0.07 | 0.24 | 0.13 | 0.26 | 0.04 | 0.15 |
| 10 | 85436920 | rs17102402 | -0.896 | 0.642 | 2.081 | 1.832 | -0.02 | -0.048 | -0.048 | 0 | 0.08 | 0.17 | 0.03 | 0 | 0.05 | 0.08 |
| 10 | 85437823 | rs7919387 | -2.495 | 0.733 | 1.882 | 1.916 | -0.006 | 0.027 | 0.027 | 0 | 0.29 | 0.64 | 0.35 | 0.46 | 0.28 | 0.39 |
| 10 | 85438929 | rs4933298 | 0.007 | 0.728 | 1.53 | 1.692 | -0.035 | 0.018 | 0.018 | 0.559 | 0.87 | 0.99 | 0.95 | 0.95 | 0.89 | 0.92 |
| 10 | 85442397 | rs12219420 | -0.701 | 0.813 | 1.666 | 1.781 | 0.002 | 0 | 0 | 1.271 | 0.13 | 0.12 | 0.19 | 0.26 | 0.11 | 0.23 |
| 10 | 85442979 | rs17102416 | -0.953 | 0.83 | 1.543 | 1.788 | -0.036 | 0.07 | 0.07 | 0 | 0.15 | 0.34 | 0.2 | 0.26 | 0.11 | 0.23 |
| 10 | 85444976 | rs581260 | 2.177 | 0.969 | 1.565 | 1.786 | 0.011 | -0.024 | -0.024 | 0.002 | 0.59 | 0.14 | 0.57 | 0.46 | 0.62 | 0.45 |
| 10 | 85445507 | rs3904537 | -1.206 | 1.001 | 1.519 | 1.775 | -0.018 | 0.057 | 0.057 | 0 | 0.17 | 0.37 | 0.21 | 0.26 | 0.15 | 0.25 |
| 10 | 85449661 | rs10788187 | -0.649 | 1 | 1.292 | 1.676 | 0.002 | 0.004 | 0.004 | 0 | 0.09 | 0.13 | 0.09 | 0.16 | 0.15 | 0.14 |
| 10 | 85449791 | rs2065593 | 2.323 | 1.055 | 1.288 | 1.698 | -0.017 | -0.045 | -0.045 | 0.189 | 0.56 | 0.12 | 0.59 | 0.49 | 0.58 | 0.44 |
| 10 | 85451733 | rs10749420 | -2.079 | 0.927 | 1.245 | 1.714 | 0.007 | 0.103 | 0.103 | 0 | 0.28 | 0.54 | 0.33 | 0.46 | 0.31 | 0.46 |
| 10 | 85452103 | rs497022 | -0.279 | 0.916 | 1.13 | 1.613 | -0.006 | -0.006 | -0.006 | 1.319 | 0.89 | 0.87 | 0.97 | 0.95 | 0.94 | 0.91 |
| 10 | 85455159 | rs1572129 | -1.331 | 1.201 | 0.537 | 1.704 | -0.015 | 0.061 | 0.061 | 0 | 0.19 | 0.35 | 0.24 | 0.3 | 0.18 | 0.34 |
| 10 | 85455502 | rs17324239 | 2.003 | 1.19 | 0.26 | 1.753 | -0.026 | 0.003 | 0.003 | 0 | 0.54 | 0.11 | 0.53 | 0.46 | 0.56 | 0.41 |
| 10 | 85455848 | rs10886943 | -1.082 | 1.308 | 0.502 | 2.193 | 0.01 | 0.018 | 0.018 | 0 | 0.09 | 0.17 | 0.15 | 0.2 | 0.17 | 0.14 |
| 10 | 85456184 | rs4933300 | -1.238 | 1.286 | 0.528 | 1.728 | -0.026 | 0.072 | 0.072 | 0 | 0.16 | 0.31 | 0.22 | 0.3 | 0.13 | 0.32 |
| 10 | 85456555 | rs10788189 | -1.952 | 1.271 | 0.424 | 1.558 | 0.008 | 0.085 | 0.085 | 0 | 0.25 | 0.49 | 0.37 | 0.42 | 0.27 | 0.45 |
| 10 | 85456824 | rs11199997 | -0.97 | 1.649 | -0.255 | 0.869 | 0.004 | -0.001 | -0.001 | 0 | 0.08 | 0.14 | 0.1 | 0.07 | 0.15 | 0.12 |
| 10 | 85458234 | rs485220 | -1.028 | 0.942 | -0.361 | 0.781 | -0.005 | 0 | 0 | 0.16 | 0.27 | 0.66 | 0.21 | 0.29 | 0.23 | 0.18 |
| 10 | 85458703 | rs1041626 | -0.937 | 1.004 | -0.584 | 0.029 | -0.003 | -0.027 | -0.027 | 0 | 0.3 | 0.37 | 0.23 | 0.11 | 0.23 | 0.18 |
| 10 | 85461469 | rs609749 | -0.585 | 0.9 | -1.872 | 0.087 | -0.051 | -0.079 | -0.079 | 0 | 0.29 | 0.07 | 0.39 | 0.57 | 0.3 | 0.25 |
| 10 | 85462356 | rs7090131 | -0.121 | 0.134 | -1.178 | -0.51 | -0.004 | -0.088 | -0.088 | 0.01 | 0.14 | 0.27 | 0.14 | 0.02 | 0.13 | 0.08 |

| | | | | | | | | | | | | | | | | |
|---|---|---|---|---|---|---|---|---|---|---|---|---|---|---|---|---|
| 10 | 85463010 | rs524265 | -0.051 | 0.128 | -1.65 | -0.465 | 0.059 | -0.092 | -0.092 | 0 | 0.22 | 0.05 | 0.32 | 0.56 | 0.14 | 0.24 |
| 10 | 85463030 | rs524316 | -0.457 | 0.493 | -1.525 | -0.76 | -0.001 | 0.022 | 0.022 | 0 | 0.94 | 0.93 | 0.85 | 0.69 | 0.89 | 0.66 |
| 10 | 85464025 | rs12573809 | 0.223 | 0.36 | -1.852 | -1.009 | -0.012 | -0.029 | -0.029 | 0 | 0.07 | 0 | 0.03 | 0.08 | 0.04 | 0.08 |
| 10 | 85464038 | rs871955 | 0.272 | -0.072 | -2.081 | -1 | 0.068 | -0.11 | -0.11 | 0 | 0.22 | 0.05 | 0.32 | 0.55 | 0.13 | 0.22 |
| 10 | 85469480 | rs11200070 | -0.246 | -0.307 | -1.371 | -1.758 | -0.025 | -0.116 | -0.116 | 0 | 0.25 | 0.38 | 0.21 | 0.05 | 0.25 | 0.12 |
| 10 | 85471106 | rs1931144 | 0.451 | -0.082 | -1.548 | -2.113 | -0.001 | -0.014 | -0.014 | 0.012 | 0.13 | 0.07 | 0.06 | 0.01 | 0.1 | 0.08 |
| 10 | 85482256 | rs11200166 | 0.719 | -0.112 | -2.019 | -1.809 | -0.011 | -0.103 | -0.103 | 0 | 0.14 | 0.03 | 0.16 | 0.5 | 0.13 | 0.22 |
| 10 | 85482501 | rs4614382 | 2.057 | 0.159 | -1.716 | -2.229 | -0.006 | -0.044 | -0.044 | 0 | 0.24 | 0.31 | 0.14 | 0.06 | 0.22 | 0.11 |
| 10 | 85485473 | rs594467 | 1.206 | 0.885 | -1.94 | -2.205 | -0.029 | -0.044 | -0.044 | 0 | 0.1 | 0.05 | 0.29 | 0.32 | 0.16 | 0.32 |
| 10 | 85496668 | rs474974 | -1.464 | 0.611 | -1.754 | -2.486 | -0.005 | -0.109 | -0.109 | 0 | 0.73 | 0.64 | 0.76 | 0.99 | 0.73 | 0.87 |
| 10 | 85500565 | rs575775 | 0.401 | 0.504 | -1.895 | -2.518 | -0.013 | -0.002 | -0.002 | 0 | 0.43 | 0.39 | 0.49 | 0.48 | 0.48 | 0.56 |
| 10 | 85509615 | rs612715 | -0.333 | 0.562 | -1.768 | -2.471 | 0.002 | -0.009 | -0.009 | 2.683 | 0.58 | 0.4 | 0.65 | 0.98 | 0.6 | 0.79 |
| 10 | 85509861 | rs519076 | 0.823 | 0.412 | -1.792 | -1.987 | -0.026 | -0.041 | -0.041 | 1.08 | 0.1 | 0.05 | 0.29 | 0.32 | 0.16 | 0.32 |
| 10 | 85521252 | rs11200461 | 0.316 | -0.009 | -1.527 | -1.84 | -0.024 | -0.028 | -0.028 | 0 | 0.1 | 0.02 | 0.05 | 0.11 | 0.09 | 0.14 |
| 10 | 85528150 | rs615638 | -0.553 | -0.257 | -1.301 | -1.927 | 0 | -0.044 | -0.044 | 0 | 0.7 | 0.71 | 0.68 | 0.99 | 0.68 | 0.83 |
| 10 | 85540277 | rs594637 | 0.16 | -0.804 | -1.403 | -1.648 | -0.01 | -0.076 | -0.076 | 0 | 0.6 | 0.59 | 0.68 | 0.94 | 0.66 | 0.74 |
| 10 | 85548652 | rs509948 | 1.188 | -0.469 | -2.172 | -1.394 | -0.061 | 0.047 | 0.047 | 0.285 | 0.36 | 0.21 | 0.46 | 0.3 | 0.44 | 0.43 |
| 10 | 85554233 | rs11200657 | 0.349 | -0.355 | -1.657 | -0.289 | -0.046 | -0.122 | -0.122 | 0 | 0.16 | 0 | 0.17 | 0.52 | 0.17 | 0.21 |
| 10 | 85564682 | rs17102623 | -0.504 | -0.462 | -0.882 | -0.658 | -0.027 | -0.022 | -0.022 | 0 | 0.12 | 0.13 | 0.04 | 0.05 | 0.05 | 0.11 |
| 10 | 85567215 | rs12268467 | 0.021 | -0.307 | -1.312 | -0.691 | 0.028 | 0.065 | 0.065 | 0.008 | 0.15 | 0.05 | 0.16 | 0.01 | 0.1 | 0.09 |
| 10 | 85572497 | rs667726 | -0.586 | -0.229 | -2.008 | -0.291 | -0.042 | -0.163 | -0.163 | 0.144 | 0.78 | 0.99 | 0.78 | 0.51 | 0.79 | 0.7 |
| 10 | 85573379 | rs527457 | 0.503 | -0.232 | -2.216 | -0.711 | 0.009 | 0.018 | 0.018 | 0 | 0.14 | 0.05 | 0.04 | 0.06 | 0.1 | 0.18 |
| 10 | 85573752 | rs652142 | -0.639 | -0.023 | -1.476 | -0.952 | -0.01 | -0.02 | -0.02 | 0 | 0.32 | 0.43 | 0.15 | 0.18 | 0.27 | 0.26 |
| 10 | 85576823 | rs607428 | -0.64 | 0.399 | -1.144 | -0.885 | 0 | -0.021 | -0.021 | 0 | 0.7 | 0.66 | 0.49 | 0.2 | 0.69 | 0.47 |
| 10 | 85580449 | rs641286 | 0.411 | 0.552 | -0.829 | -0.392 | 0.015 | 0.027 | 0.027 | 0 | 0.15 | 0.05 | 0.04 | 0.06 | 0.1 | 0.19 |
| 10 | 85582722 | rs7086916 | -0.823 | 0.674 | 0.175 | -0.232 | 0.03 | 0.074 | 0.074 | 0.065 | 0.21 | 0.42 | 0.36 | 0.38 | 0.25 | 0.29 |
| 10 | 85583376 | rs3862561 | 0.162 | 0.685 | 0.238 | -0.236 | -0.006 | 0.009 | 0.009 | 0 | 0.07 | 0.07 | 0.28 | 0.33 | 0.11 | 0.25 |
| 10 | 85583637 | rs4083068 | 0.415 | 0.649 | 0.296 | -0.251 | 0.01 | -0.065 | -0.065 | 0 | 0.11 | 0.27 | 0.07 | 0 | 0.13 | 0.01 |
| 10 | 85586066 | rs662166 | -1.817 | 1.028 | 0.392 | -0.467 | -0.012 | 0.004 | 0.004 | 0.868 | 0.43 | 0.66 | 0.48 | 0.47 | 0.45 | 0.5 |
| 10 | 85592663 | rs1977895 | -1.494 | 1.093 | 0.374 | -0.445 | 0.016 | 0.044 | 0.044 | 0.568 | 0.25 | 0.53 | 0.39 | 0.42 | 0.28 | 0.32 |
| 10 | 85593542 | rs4933938 | -0.149 | 1.193 | 0.132 | -0.393 | 0.017 | 0.07 | 0.07 | 0 | 0.09 | 0.18 | 0.29 | 0.33 | 0.11 | 0.25 |
| 10 | 85599366 | rs532605 | 0.206 | 0.915 | -0.261 | -0.659 | -0.017 | 0.045 | 0.045 | 0 | 0.18 | 0.06 | 0.09 | 0.05 | 0.17 | 0.18 |
| 10 | 85600312 | rs11200683 | -0.025 | 0.906 | 0.042 | -0.235 | 0.005 | -0.014 | -0.014 | 1.158 | 0.11 | 0.19 | 0.07 | 0.06 | 0.14 | 0.03 |
| 10 | 85600447 | rs12355819 | -0.025 | 0.906 | 0.042 | -0.235 | 0.005 | -0.014 | -0.014 | 0 | 0.11 | 0.19 | 0.07 | 0.06 | 0.14 | 0.03 |
| 10 | 85616321 | rs10788287 | -0.623 | 0.915 | 0.01 | -0.225 | 0.007 | -0.018 | -0.018 | 0.133 | 0.14 | 0.24 | 0.09 | 0.1 | 0.17 | 0.07 |
| 10 | 85624169 | rs11200710 | 0.103 | 0.537 | -1.193 | 0.062 | 0.032 | 0.05 | 0.05 | 0 | 0.53 | 0.23 | 0.57 | 0.62 | 0.61 | 0.7 |
| 10 | 85624500 | rs6585624 | 0.158 | 0.218 | -1.068 | 0.172 | -0.001 | -0.005 | -0.005 | 0.016 | 0.25 | 0.22 | 0.35 | 0.33 | 0.24 | 0.31 |
| 10 | 85625165 | rs4426079 | -0.384 | 0.476 | -1.083 | 0.076 | 0.01 | 0.004 | 0.004 | 0.04 | 0.64 | 0.38 | 0.55 | 0.57 | 0.58 | 0.57 |
| 10 | 85626784 | rs7901096 | 1.826 | 0.342 | -1.101 | -0.12 | 0 | -0.01 | -0.01 | 0 | 0.16 | 0.1 | 0.11 | 0 | 0.18 | 0.05 |
| 10 | 85628215 | rs11200713 | 0.267 | -0.048 | -1.465 | -0.075 | 0.018 | 0.013 | 0.013 | 0 | 0.45 | 0.35 | 0.47 | 0.33 | 0.41 | 0.34 |
| 10 | 85632532 | rs1110421 | 0.705 | -0.058 | -1.297 | 0.011 | 0.015 | 0.016 | 0.016 | 0 | 0.26 | 0.47 | 0.38 | 0.33 | 0.28 | 0.32 |
| 10 | 85633383 | rs10887180 | -0.06 | 0.11 | -1.609 | 0.169 | 0.005 | -0.002 | -0.002 | 5.753 | 0.24 | 0.16 | 0.12 | 0.27 | 0.21 | 0.28 |
| 10 | 85634184 | rs1857455 | -1.397 | 0.006 | -1.65 | 0.046 | -0.05 | 0.045 | 0.045 | 0 | 0.74 | 0.86 | 0.8 | 0.87 | 0.67 | 0.77 |
| 10 | 85638143 | rs4933939 | -1.514 | 0.076 | -1.81 | -0.083 | -0.075 | 0.049 | 0.049 | 0 | 0.75 | 0.9 | 0.8 | 0.87 | 0.67 | 0.79 |
| 10 | 85642645 | rs11200718 | -0.697 | -0.017 | -1.209 | 0.604 | 0.007 | -0.016 | -0.016 | 0 | 0.24 | 0.16 | 0.12 | 0.33 | 0.21 | 0.29 |
| 10 | 85645501 | rs7906586 | -0.838 | -0.301 | -0.746 | 0.09 | -0.006 | -0.049 | -0.049 | 0.621 | 0.57 | 0.48 | 0.72 | 0.89 | 0.57 | 0.7 |
| 10 | 85650171 | rs11200721 | 0.795 | 0.102 | -1.544 | 0.172 | -0.104 | -0.098 | -0.098 | 0 | 0.18 | 0.03 | 0.36 | 0.4 | 0.3 | 0.33 |
| 10 | 85651716 | rs7085916 | 0.353 | -0.528 | -1.24 | -0.261 | 0.019 | 0.004 | 0.004 | 0.056 | 0.25 | 0.2 | 0.2 | 0.01 | 0.17 | 0.13 |
| 10 | 85652232 | rs17102772 | -1.022 | -0.307 | -1.051 | 0.446 | 0.001 | -0.001 | -0.001 | 0 | 0.18 | 0.17 | 0.11 | 0.23 | 0.18 | 0.28 |
| 10 | 85658029 | rs4581390 | -0.362 | -0.254 | -1.163 | 0.442 | 0.066 | 0.072 | 0.072 | 0 | 0.49 | 0.21 | 0.62 | 0.68 | 0.58 | 0.67 |
| 10 | 85664031 | rs2349763 | -0.395 | -0.221 | -0.811 | 0.431 | 0.021 | 0.015 | 0.015 | 0 | 0.52 | 0.39 | 0.63 | 0.64 | 0.58 | 0.69 |
| 10 | 85668016 | rs12262107 | 0.065 | -0.382 | -1.18 | 0.251 | 0.021 | 0.039 | 0.039 | 0 | 0.27 | 0.14 | 0.21 | 0.22 | 0.25 | 0.35 |
| 10 | 85672695 | rs7079280 | -0.694 | -1.429 | -0.973 | 0.041 | 0.031 | 0.007 | 0.007 | 0 | 0.7 | 0.87 | 0.78 | 0.74 | 0.77 | 0.78 |
| 10 | 85675348 | rs2349764 | 0.604 | -1.449 | -0.906 | -0.105 | -0.069 | -0.06 | -0.06 | 0.014 | 0.26 | 0.17 | 0.4 | 0.49 | 0.43 | 0.36 |
| 10 | 85677421 | rs6585679 | -0.991 | -1.549 | -0.834 | -0.027 | 0.024 | -0.003 | -0.003 | 0.14 | 0.71 | 0.9 | 0.71 | 0.76 | 0.77 | 0.78 |
| 10 | 85680444 | rs4933943 | -0.996 | -1.727 | -0.632 | 0.029 | 0.019 | -0.011 | -0.011 | 0 | 0.72 | 0.79 | 0.64 | 0.74 | 0.78 | 0.78 |
| 10 | 85682612 | rs12766238 | 0.538 | -2.229 | -0.326 | -0.319 | -0.021 | -0.099 | -0.099 | 0 | 0.23 | 0.32 | 0.17 | 0 | 0.21 | 0.16 |
| 10 | 85684087 | rs10887186 | 0.404 | -2.297 | -0.925 | -0.24 | 0.011 | -0.022 | -0.022 | 0 | 0.13 | 0.06 | 0.08 | 0.24 | 0.08 | 0.22 |
| 10 | 85684595 | rs1857459 | 0.169 | -2.242 | -0.706 | -0.352 | 0.027 | 0.049 | 0.049 | 0 | 0.44 | 0.21 | 0.46 | 0.73 | 0.56 | 0.6 |
| 10 | 85686185 | rs12768886 | 1.022 | -1.96 | -0.804 | -0.337 | 0.01 | 0.01 | 0.01 | 0 | 0.19 | 0.11 | 0.14 | 0 | 0.19 | 0.16 |
| 10 | 85688158 | rs1419271 | -0.723 | -1.872 | -0.699 | 0.098 | 0.002 | -0.002 | -0.002 | 0 | 0.73 | 0.79 | 0.76 | 0.75 | 0.79 | 0.78 |
| 10 | 85689295 | rs10749472 | 0.017 | -1.76 | -1.064 | 0.215 | 0.011 | -0.018 | -0.018 | 2.036 | 0.15 | 0.08 | 0.2 | 0.26 | 0.08 | 0.37 |
| 10 | 85693860 | rs5014028 | -1.002 | -1.937 | -1.366 | 0.172 | -0.09 | -0.074 | -0.074 | 0 | 0.7 | 0.92 | 0.64 | 0.55 | 0.48 | 0.69 |
| 10 | 85700055 | rs7077857 | 0.606 | -2.544 | -0.503 | 0.006 | -0.047 | 0.027 | 0.027 | 0 | 0.26 | 0.44 | 0.22 | 0.24 | 0.18 | 0.12 |
| 10 | 85705980 | rs11200745 | 0.54 | -2.772 | -0.709 | 0.085 | 0.013 | -0.031 | -0.031 | 0.508 | 0.15 | 0.04 | 0.2 | 0.24 | 0.08 | 0.34 |
| 10 | 85708247 | rs7087394 | -1.458 | -2.703 | -1.076 | 0.156 | -0.087 | -0.046 | -0.046 | 1.174 | 0.73 | 0.9 | 0.72 | 0.64 | 0.55 | 0.76 |
| 10 | 85709761 | rs10887188 | -0.624 | -3.001 | -0.019 | 0.072 | 0.009 | 0.064 | 0.064 | 0.446 | 0.1 | 0.46 | 0.15 | 0.12 | 0.08 | 0.14 |
| 10 | 85711308 | rs7067951 | 0.25 | -3.033 | -0.162 | 0.051 | 0.002 | -0.002 | -0.002 | 0.018 | 0.18 | 0.14 | 0.22 | 0.24 | 0.1 | 0.35 |
| 10 | 85715594 | rs7905426 | 0.096 | -2.696 | -0.215 | 0.201 | -0.058 | 0.037 | 0.037 | 0 | 0.32 | 0.53 | 0.34 | 0.36 | 0.26 | 0.24 |

| | | | | | | | | | | | | | | | | |
|---|---|---|---|---|---|---|---|---|---|---|---|---|---|---|---|---|
| 10 | 85715936 | rs7922818 | -1.478 | -1.908 | -0.776 | 0.345 | -0.169 | -0.111 | -0.111 | 0 | 0.79 | 0.98 | 0.74 | 0.66 | 0.57 | 0.78 |
| 10 | 85723816 | rs11200757 | -0.261 | -1.834 | 0.242 | 0.182 | 0.041 | 0.107 | 0.107 | 0 | 0.06 | 0.23 | 0.13 | 0.14 | 0.08 | 0.12 |
| 10 | 85724777 | rs17102865 | -1.327 | -2.036 | 0.569 | 0.045 | 0.019 | 0.029 | 0.029 | 0 | 0.1 | 0.36 | 0.11 | 0.12 | 0.09 | 0.12 |
| 10 | 85725305 | rs1419268 | 0.03 | -2.186 | 0.091 | -0.466 | -0.001 | -0.073 | -0.073 | 0.505 | 0.18 | 0.31 | 0.15 | 0.05 | 0.17 | 0.18 |
| 10 | 85725351 | rs11598552 | -0.232 | -2.979 | -0.638 | -0.446 | 0.086 | 0.03 | 0.03 | 1.03 | 0.34 | 0.13 | 0.2 | 0.22 | 0.21 | 0.17 |
| 10 | 85727995 | rs17102878 | 0.44 | -2.511 | -0.429 | -0.341 | 0.003 | -0.006 | -0.006 | 0.232 | 0.16 | 0.13 | 0.1 | 0.17 | 0.14 | 0.08 |
| 10 | 85737862 | rs4933305 | -0.366 | -2.801 | 0.019 | -0.645 | -0.011 | -0.024 | -0.024 | 0 | 0.35 | 0.45 | 0.26 | 0.18 | 0.31 | 0.23 |
| 10 | 85738811 | rs10887193 | 0.461 | -2.444 | -0.036 | -0.307 | -0.001 | 0.026 | 0.026 | 0 | 0.22 | 0.24 | 0.27 | 0.36 | 0.22 | 0.33 |
| 10 | 85741293 | rs1857454 | -1.992 | -2.537 | -1.096 | -0.372 | -0.117 | -0.088 | -0.088 | 0 | 0.66 | 0.93 | 0.58 | 0.59 | 0.56 | 0.65 |
| 10 | 85741338 | rs4623834 | 0.701 | -2.556 | -1.025 | -0.398 | 0 | 0.001 | 0.001 | 0 | 0.22 | 0.17 | 0.27 | 0.36 | 0.22 | 0.33 |
| 10 | 85741728 | rs6585743 | -1.662 | -2.128 | -1.588 | -0.168 | -0.117 | -0.096 | -0.096 | 1.015 | 0.66 | 0.93 | 0.57 | 0.55 | 0.56 | 0.64 |
| 10 | 85754531 | rs10509484 | 0.572 | -1.591 | -0.903 | 0.342 | 0.006 | 0.058 | 0.058 | 0 | 0.08 | 0.17 | 0.07 | 0.18 | 0.08 | 0.07 |
| 10 | 85755077 | rs17329709 | 0.877 | -1.79 | -0.983 | 0.09 | 0.006 | 0.024 | 0.024 | 0 | 0.18 | 0.12 | 0.16 | 0.02 | 0.18 | 0.2 |
| 10 | 85759155 | rs7909859 | 0.387 | -1.582 | -0.881 | 0.258 | 0.001 | 0.001 | 0.001 | 0 | 0.22 | 0.27 | 0.21 | 0.19 | 0.24 | 0.21 |
| 10 | 85761914 | rs10736313 | -0.896 | -1.108 | -1.501 | 0.361 | -0.033 | -0.09 | -0.09 | 2.001 | 0.79 | 0.97 | 0.75 | 0.63 | 0.79 | 0.67 |
| 10 | 85764135 | rs2350106 | 0.563 | -0.971 | -0.978 | 0.262 | 0.001 | 0.019 | 0.019 | 0 | 0.51 | 0.39 | 0.5 | 0.4 | 0.52 | 0.45 |
| 10 | 85775159 | rs10788306 | -0.956 | -1.561 | -1.13 | 0.646 | 0.091 | -0.019 | -0.019 | 1.836 | 0.37 | 0.78 | 0.4 | 0.41 | 0.18 | 0.36 |
| 10 | 85775538 | rs4145143 | -1.561 | -1.52 | -1.529 | 0.368 | -0.017 | -0.019 | -0.019 | 0 | 0.55 | 0.96 | 0.6 | 0.62 | 0.42 | 0.56 |
| 10 | 85776035 | rs7092579 | -0.351 | -1.706 | -0.807 | 0.62 | -0.054 | 0.007 | 0.007 | 0 | 0.25 | 0.37 | 0.25 | 0.3 | 0.13 | 0.26 |
| 10 | 85778744 | rs4639875 | -1.409 | -1.53 | -1.028 | 0.274 | 0.099 | -0.012 | -0.012 | 0 | 0.4 | 0.85 | 0.45 | 0.6 | 0.24 | 0.42 |
| 10 | 85786443 | rs11593393 | 0.061 | -1.408 | -0.557 | 0.115 | -0.061 | -0.073 | -0.073 | 0 | 0.2 | 0.32 | 0.1 | 0.03 | 0.09 | 0.06 |
| 10 | 85790476 | rs10788309 | 0.365 | -1.388 | -0.8 | -0.004 | 0 | -0.004 | -0.004 | 1.321 | 0.57 | 0.6 | 0.52 | 0.5 | 0.58 | 0.46 |
| 10 | 85796190 | rs6585795 | -1.164 | -1.273 | -0.614 | -0.007 | 0.014 | 0.003 | 0.003 | 0 | 0.88 | 0.87 | 0.85 | 0.92 | 0.86 | 0.89 |
| 10 | 85798580 | rs4615965 | -0.829 | -1.606 | -0.733 | 0.026 | 0.048 | 0.01 | 0.01 | 3.452 | 0.43 | 0.74 | 0.52 | 0.4 | 0.33 | 0.48 |
| 10 | 85800304 | rs11200819 | 2.183 | -0.687 | -1.328 | -0.035 | -0.064 | -0.031 | -0.031 | 0 | 0.12 | 0 | 0.17 | 0.16 | 0.18 | 0.13 |
| 10 | 85802965 | rs7076979 | 0.512 | -1.001 | -0.671 | -0.174 | -0.008 | -0.095 | -0.095 | 0.012 | 0.25 | 0.36 | 0.21 | 0.02 | 0.23 | 0.23 |
| 10 | 85803182 | rs7078319 | 0.998 | -1.049 | -0.432 | -0.022 | -0.023 | -0.064 | -0.064 | 0 | 0.11 | 0.29 | 0.08 | 0.01 | 0.05 | 0.1 |
| 10 | 85804782 | rs2067539 | 1.085 | -0.972 | -0.57 | 0.16 | -0.026 | -0.036 | -0.036 | 0 | 0.07 | 0.01 | 0.17 | 0.27 | 0.08 | 0.19 |
| 10 | 85805384 | rs11814628 | -0.251 | -1.08 | -0.439 | 0.223 | -0.015 | 0.063 | 0.063 | 0 | 0.2 | 0.26 | 0.28 | 0.48 | 0.14 | 0.3 |
| 10 | 85812642 | rs11200822 | 1.321 | -1.141 | -0.876 | -0.046 | -0.02 | -0.047 | -0.047 | 0 | 0.1 | 0.21 | 0.05 | 0 | 0.06 | 0 |
| 10 | 85816113 | rs12247491 | 0.287 | -1.228 | -0.671 | 0.039 | -0.057 | 0.055 | 0.055 | 2.782 | 0.16 | 0.56 | 0.12 | 0.23 | 0.06 | 0.11 |
| 10 | 85821027 | rs4933958 | -0.024 | -1.119 | -0.958 | -0.052 | 0 | -0.008 | -0.008 | 0 | 0.69 | 0.72 | 0.71 | 0.7 | 0.71 | 0.74 |
| 10 | 85824118 | rs11599263 | 1.568 | -1.118 | -1.095 | 0.005 | -0.058 | -0.02 | -0.02 | 0.117 | 0.34 | 0.09 | 0.32 | 0.35 | 0.42 | 0.36 |
| 10 | 85826201 | rs3999209 | 0.071 | -1.158 | -0.976 | 0.02 | -0.054 | 0.064 | 0.064 | 0.212 | 0.21 | 0.51 | 0.26 | 0.34 | 0.11 | 0.25 |
| 10 | 85827858 | rs7923116 | -0.519 | -1.078 | -1.15 | -0.049 | -0.004 | 0.001 | 0.001 | 0 | 0.77 | 0.76 | 0.83 | 0.8 | 0.76 | 0.85 |
| 10 | 85829322 | rs7084533 | -0.598 | -1.07 | -1.146 | -0.019 | -0.004 | 0.002 | 0.002 | 0 | 0.77 | 0.77 | 0.84 | 0.8 | 0.76 | 0.85 |
| 10 | 85833863 | rs12354551 | 0.877 | -0.983 | -1.312 | -0.428 | 0.023 | 0.044 | 0.044 | 0 | 0.48 | 0.14 | 0.45 | 0.37 | 0.6 | 0.5 |
| 10 | 85834685 | rs12774661 | 1.879 | -1.044 | -1.292 | -0.437 | -0.023 | 0.051 | 0.051 | 0.026 | 0.13 | 0.05 | 0.13 | 0.01 | 0.18 | 0.14 |
| 10 | 85838758 | rs7072402 | -0.824 | -1.156 | -0.78 | -0.146 | -0.002 | -0.001 | -0.001 | 0.602 | 0.4 | 0.46 | 0.46 | 0.63 | 0.36 | 0.44 |
| 10 | 85844192 | rs7474895 | -0.452 | -1.216 | -0.822 | -0.549 | 0 | -0.005 | -0.005 | 0 | 0.56 | 0.53 | 0.52 | 0.23 | 0.53 | 0.44 |
| 10 | 85844814 | rs4244944 | 0.802 | -1.343 | -0.714 | -0.8 | -0.001 | 0.017 | 0.017 | 0 | 0.36 | 0.25 | 0.26 | 0.11 | 0.35 | 0.28 |
| 10 | 85846540 | rs11200839 | 0.922 | -1.289 | -0.765 | -0.758 | -0.034 | 0.055 | 0.055 | 0 | 0.13 | 0.03 | 0.12 | 0 | 0.15 | 0.11 |
| 10 | 85853280 | rs10788316 | -0.347 | -1.253 | -0.52 | -0.573 | 0 | -0.03 | -0.03 | 0 | 0.56 | 0.51 | 0.45 | 0.15 | 0.53 | 0.43 |
| 10 | 85853707 | rs11200843 | 0.447 | -0.778 | -0.261 | 0.083 | -0.018 | -0.008 | -0.008 | 0 | 0.07 | 0.2 | 0.06 | 0.03 | 0.03 | 0.05 |
| 10 | 85878070 | rs12412889 | 0.379 | -0.727 | -0.562 | 0.302 | -0.015 | 0.065 | 0.065 | 0 | 0.08 | 0.09 | 0.21 | 0.25 | 0.05 | 0.13 |
| 10 | 85887054 | rs12357826 | 0.153 | -0.938 | -0.816 | -0.154 | -0.007 | 0.041 | 0.041 | 0 | 0.2 | 0.11 | 0.18 | 0.01 | 0.18 | 0.13 |
| 10 | 85887320 | rs11200876 | 1.067 | -1.056 | -0.891 | -0.104 | 0.009 | -0.012 | -0.012 | 3.562 | 0.19 | 0.16 | 0.12 | 0.36 | 0.11 | 0.34 |
| 10 | 85889505 | rs4400739 | -2.188 | -0.96 | -1.348 | -0.34 | -0.107 | -0.031 | -0.031 | 0 | 0.79 | 0.98 | 0.82 | 0.8 | 0.68 | 0.77 |
| 10 | 85891907 | rs10732827 | -0.841 | -1.016 | -1.102 | -0.336 | -0.001 | -0.037 | -0.037 | 0 | 0.64 | 0.77 | 0.59 | 0.55 | 0.62 | 0.62 |
| 10 | 85892211 | rs11200878 | 0.108 | -0.825 | -0.936 | -0.367 | -0.029 | 0.064 | 0.064 | 0 | 0.39 | 0.23 | 0.4 | 0.16 | 0.47 | 0.27 |
| 10 | 85892729 | rs7071034 | 1.047 | -0.77 | -1.225 | -0.335 | -0.066 | 0.04 | 0.04 | 0 | 0.23 | 0.07 | 0.26 | 0.16 | 0.31 | 0.16 |
| 10 | 85898861 | rs3814214 | 1.057 | -0.759 | -1.221 | -0.33 | -0.066 | 0.039 | 0.039 | 0 | 0.23 | 0.07 | 0.26 | 0.16 | 0.31 | 0.16 |
| 10 | 85899713 | rs7087073 | 0.378 | -1.389 | -0.485 | 0.195 | 0.019 | -0.023 | -0.023 | 0 | 0.14 | 0.1 | 0.22 | 0.25 | 0.07 | 0.15 |
| 10 | 85903678 | rs17103143 | -0.118 | -0.121 | -0.027 | -0.143 | 0.019 | -0.003 | -0.003 | 0.03 | 0.06 | 0.31 | 0.05 | 0 | 0.04 | 0 |
| 10 | 85903888 | rs10887241 | -0.185 | 0.166 | -0.827 | -0.002 | -0.005 | -0.007 | -0.007 | 0.676 | 0.15 | 0.1 | 0.14 | 0.01 | 0.16 | 0.11 |
| 10 | 85912606 | rs7576 | -0.185 | 0.166 | -0.827 | -0.002 | -0.005 | -0.007 | -0.007 | 0 | 0.15 | 0.1 | 0.14 | 0.01 | 0.16 | 0.12 |
| 10 | 85915186 | rs7096124 | -0.66 | -0.314 | -0.489 | 0.696 | -0.066 | 0.06 | 0.06 | 0 | 0.25 | 0.52 | 0.21 | 0.39 | 0.15 | 0.35 |
| 10 | 85920085 | rs11200896 | -1.227 | 0.779 | -0.987 | 0.352 | -0.002 | 0.005 | 0.005 | 2.898 | 0.25 | 0.74 | 0.33 | 0.17 | 0.27 | 0.14 |
| 10 | 85923741 | rs4933969 | -0.453 | 0.912 | -1.098 | 0.59 | -0.012 | -0.019 | -0.019 | 0.109 | 0.08 | 0.04 | 0.13 | 0.21 | 0.14 | 0.11 |
| 10 | 85936160 | rs4348833 | -1.235 | 1.119 | -1.635 | 0.127 | -0.061 | -0.013 | -0.013 | 0 | 0.2 | 0.04 | 0.18 | 0.23 | 0.27 | 0.14 |
| 10 | 85936688 | rs11594887 | -0.392 | 1.18 | -1.602 | 0.056 | -0.032 | -0.013 | -0.013 | 0 | 0.09 | 0.02 | 0.1 | 0.14 | 0.12 | 0.12 |
| 10 | 85939031 | rs11200905 | -1.253 | 0.309 | -1.038 | -0.46 | -0.01 | -0.029 | -0.029 | 0 | 0.24 | 0.36 | 0.17 | 0.15 | 0.21 | 0.14 |
| 10 | 85942596 | rs11200909 | -1.652 | 0.244 | -0.764 | -0.492 | -0.011 | -0.044 | -0.044 | 0.334 | 0.3 | 0.41 | 0.21 | 0.15 | 0.26 | 0.15 |
| 10 | 85944141 | rs7070005 | -1.61 | 0.729 | -1.599 | -1.134 | 0.006 | 0.178 | 0.178 | 0 | 0.55 | 0.89 | 0.57 | 0.21 | 0.61 | 0.42 |
| 10 | 85950406 | rs7099098 | -0.214 | 1.006 | -0.858 | -0.863 | 0.073 | -0.109 | -0.109 | 0 | 0.25 | 0.43 | 0.35 | 0.04 | 0.35 | 0.26 |
| 10 | 85958126 | rs11200916 | 0.231 | 0.368 | -1.191 | -0.607 | 0.015 | -0.037 | -0.037 | 0 | 0.1 | 0.03 | 0.12 | 0.2 | 0.08 | 0.14 |
| 10 | 85958218 | rs17103203 | -0.115 | 0.596 | -0.891 | -0.719 | 0.031 | -0.006 | -0.006 | 0 | 0.09 | 0.26 | 0.17 | 0.04 | 0.13 | 0.11 |
| 10 | 85960149 | rs10788333 | -1.252 | 0.759 | -0.985 | -0.628 | -0.012 | 0.008 | 0.008 | 0 | 0.39 | 0.71 | 0.52 | 0.39 | 0.46 | 0.43 |
| 10 | 85960395 | rs4933975 | -1.252 | 0.759 | -0.985 | -0.624 | -0.012 | 0.007 | 0.007 | 0 | 0.39 | 0.71 | 0.52 | 0.39 | 0.46 | 0.43 |

| | | | | | | | | | | | | | | | | |
|---|---|---|---|---|---|---|---|---|---|---|---|---|---|---|---|---|
| 10 | 85965574 | rs4933977 | -1.119 | 0.575 | -0.814 | -0.675 | -0.019 | 0.093 | 0.093 | 0 | 0.06 | 0.2 | 0.14 | 0.19 | 0.03 | 0.08 |
| 10 | 85966873 | rs7092674 | -1.563 | 0.575 | -0.946 | -0.804 | -0.011 | 0.065 | 0.065 | 0 | 0.43 | 0.79 | 0.49 | 0.26 | 0.48 | 0.38 |
| 10 | 85971124 | rs11200924 | -0.358 | 0.444 | -1.114 | -0.772 | -0.039 | 0.034 | 0.034 | 0.446 | 0.2 | 0.01 | 0.17 | 0.15 | 0.23 | 0.12 |
| 10 | 85971347 | rs4933978 | 0.167 | 0.181 | -0.444 | -0.886 | 0.071 | -0.048 | -0.048 | 0 | 0.18 | 0.44 | 0.24 | 0.07 | 0.25 | 0.26 |
| 10 | 85972043 | rs10749482 | -1 | 0.178 | -0.424 | -0.884 | -0.002 | -0.003 | -0.003 | 0 | 0.26 | 0.75 | 0.39 | 0.26 | 0.29 | 0.34 |
| 10 | 85974236 | rs3814213 | -1.863 | 0.165 | -0.413 | -0.867 | -0.012 | 0.04 | 0.04 | 0 | 0.47 | 0.83 | 0.57 | 0.41 | 0.52 | 0.46 |
| 10 | 85974381 | rs3814212 | -2.245 | -0.048 | -0.747 | -1.116 | -0.052 | 0.003 | 0.003 | 0.432 | 0.55 | 0.92 | 0.64 | 0.41 | 0.6 | 0.47 |
| 10 | 85976965 | rs1059341 | -1.547 | -0.037 | -0.582 | -1.09 | -0.006 | 0.02 | 0.02 | 0 | 0.44 | 0.83 | 0.53 | 0.41 | 0.46 | 0.45 |
| 10 | 85978939 | rs4244947 | -1.538 | -0.036 | -0.584 | -1.089 | -0.006 | 0.02 | 0.02 | 0 | 0.43 | 0.82 | 0.53 | 0.41 | 0.46 | 0.45 |
| 10 | 85978991 | rs7895270 | 0.117 | 0.15 | -0.64 | -1.105 | 0.048 | -0.042 | -0.042 | 0.276 | 0.18 | 0.52 | 0.23 | 0.07 | 0.23 | 0.25 |
| 10 | 85980865 | rs4933982 | 0.411 | 0.167 | -0.732 | -1.098 | 0.076 | -0.035 | -0.035 | 0 | 0.17 | 0.44 | 0.23 | 0.07 | 0.23 | 0.25 |
| 10 | 85984323 | rs11200925 | -0.138 | 0.179 | -1.248 | -0.987 | -0.007 | -0.02 | -0.02 | 0 | 0.2 | 0.05 | 0.23 | 0.29 | 0.23 | 0.18 |
| 10 | 85985508 | rs4933984 | -0.138 | 0.181 | -1.243 | -1.004 | -0.007 | -0.05 | -0.05 | 0 | 0.2 | 0.05 | 0.29 | 0.35 | 0.23 | 0.19 |
| 10 | 85990621 | rs10736318 | -1.861 | 0.212 | -1.667 | -0.882 | -0.037 | -0.112 | -0.112 | 0.307 | 0.63 | 0.93 | 0.71 | 0.51 | 0.68 | 0.55 |
| 10 | 85991784 | rs3814211 | 1.876 | -0.785 | -2.012 | -1.205 | 0.04 | 0.062 | 0.062 | 0 | 0.11 | 0 | 0.03 | 0.04 | 0.04 | 0.09 |
| 10 | 85992390 | rs12262099 | -0.037 | -0.545 | -1.411 | -1.084 | 0.044 | 0.075 | 0.075 | 3.257 | 0.07 | 0.55 | 0.11 | 0.04 | 0.04 | 0.09 |
| 10 | 85992478 | rs3814210 | -0.699 | -0.437 | -2.268 | -1.191 | -0.028 | 0.03 | 0.03 | 1.158 | 0.37 | 0.87 | 0.48 | 0.26 | 0.4 | 0.34 |
| 10 | 85992511 | rs3814209 | -0.699 | -0.437 | -2.268 | -1.191 | -0.028 | 0.03 | 0.03 | 1.158 | 0.37 | 0.87 | 0.48 | 0.26 | 0.4 | 0.34 |
| 10 | 85993910 | rs3814206 | -1.154 | -0.019 | -2.369 | -1.234 | -0.037 | 0.007 | 0.007 | 1.035 | 0.43 | 0.89 | 0.54 | 0.41 | 0.47 | 0.43 |
| 10 | 85994719 | rs4933987 | -0.655 | -0.008 | -2.355 | -1.238 | -0.028 | 0.074 | 0.074 | 1.158 | 0.37 | 0.87 | 0.42 | 0.21 | 0.4 | 0.33 |
| 10 | 85997105 | rs11200933 | 1.932 | 0 | -2.282 | -1.182 | -0.001 | -0.015 | -0.015 | 0.082 | 0.06 | 0.03 | 0.06 | 0.15 | 0.07 | 0.1 |
| 10 | 85997250 | rs10887259 | -0.649 | -0.057 | -2.347 | -1.255 | -0.019 | 0.056 | 0.056 | 0.774 | 0.34 | 0.86 | 0.39 | 0.21 | 0.35 | 0.34 |
| 10 | 85999444 | rs4933989 | -1.182 | 0.393 | -2.406 | -1.275 | -0.012 | 0.044 | 0.044 | 0.649 | 0.48 | 0.87 | 0.58 | 0.41 | 0.55 | 0.48 |
| 10 | 86004238 | rs10736319 | -1.525 | -0.079 | -2.77 | -1.349 | -0.075 | -0.075 | -0.075 | 0.773 | 0.57 | 0.95 | 0.59 | 0.5 | 0.49 | 0.48 |
| 10 | 86004269 | rs10887261 | 1.238 | -0.086 | -2.836 | -1.398 | -0.025 | 0.017 | 0.017 | 0 | 0.06 | 0 | 0.1 | 0.02 | 0.08 | 0.04 |
| 10 | 86004311 | rs12217965 | 0.131 | -0.283 | -2.665 | -1.488 | 0.005 | 0.03 | 0.03 | 0.117 | 0.21 | 0.18 | 0.18 | 0.15 | 0.22 | 0.14 |
| 10 | 86004865 | rs11200938 | 1.918 | -0.257 | -2.69 | -1.473 | -0.026 | 0.01 | 0.01 | 0.128 | 0.05 | 0 | 0.1 | 0.02 | 0.07 | 0.03 |
| 10 | 86004873 | rs2279227 | -1.823 | -0.303 | -2.728 | -1.39 | -0.098 | -0.097 | -0.097 | 0.964 | 0.57 | 0.98 | 0.59 | 0.5 | 0.49 | 0.49 |
| 10 | 86005300 | rs2279226 | 0.16 | -0.144 | -2.484 | -1.299 | 0.002 | 0.04 | 0.04 | 0.068 | 0.2 | 0.15 | 0.17 | 0.15 | 0.22 | 0.14 |
| 10 | 86009815 | rs12780894 | -0.541 | -0.109 | -1.799 | -1.112 | 0.01 | 0.054 | 0.054 | 0.708 | 0.15 | 0.57 | 0.31 | 0.2 | 0.15 | 0.2 |
| 10 | 86011743 | rs17103281 | 1.109 | -0.319 | -1.984 | -1.189 | 0.025 | 0.015 | 0.015 | 0 | 0.2 | 0.12 | 0.1 | 0.14 | 0.12 | 0.16 |
| 10 | 86012713 | rs1042454 | -0.635 | -0.24 | -2.486 | -0.939 | 0.06 | 0.047 | 0.047 | 0.033 | 0.45 | 0.85 | 0.48 | 0.33 | 0.36 | 0.37 |
| 10 | 86015195 | rs10788336 | 0.616 | -0.183 | -2.294 | -0.883 | -0.003 | 0.005 | 0.005 | 0 | 0.17 | 0.11 | 0.26 | 0.16 | 0.19 | 0.11 |
| 10 | 86016101 | rs7922232 | -1.912 | -0.127 | -2.715 | -0.8 | -0.113 | -0.188 | -0.188 | 0.061 | 0.74 | 0.99 | 0.72 | 0.59 | 0.69 | 0.76 |
| 10 | 86016166 | rs4933317 | 0.284 | -0.352 | -1.913 | -0.595 | 0.001 | -0.007 | -0.007 | 0.289 | 0.18 | 0.19 | 0.26 | 0.16 | 0.19 | 0.11 |
| 10 | 86016490 | rs736141 | -0.901 | -0.37 | -2.178 | -0.662 | 0.06 | 0.046 | 0.046 | 0.003 | 0.45 | 0.85 | 0.48 | 0.33 | 0.36 | 0.37 |
| 10 | 86016892 | rs10887266 | 0.049 | -0.394 | -1.698 | -0.222 | -0.022 | 0.005 | 0.005 | 0.03 | 0.28 | 0.63 | 0.31 | 0.32 | 0.2 | 0.31 |
| 10 | 86020637 | rs11200947 | 0.573 | -0.269 | -2.118 | -0.306 | 0.026 | 0.029 | 0.029 | 0 | 0.23 | 0.09 | 0.15 | 0.14 | 0.2 | 0.14 |
| 10 | 86020700 | rs4244950 | -0.319 | 0.038 | -2.415 | -0.207 | 0.022 | -0.011 | -0.011 | 0 | 0.37 | 0.77 | 0.43 | 0.36 | 0.35 | 0.32 |
| 10 | 86020898 | rs4481964 | -1.209 | 0.113 | -2.601 | -0.163 | 0.039 | 0.019 | 0.019 | 0.807 | 0.49 | 0.86 | 0.48 | 0.4 | 0.42 | 0.39 |
| 10 | 86021511 | rs10749483 | -0.208 | 0.232 | -1.91 | 0.016 | -0.014 | -0.005 | -0.005 | 0.016 | 0.27 | 0.58 | 0.33 | 0.29 | 0.23 | 0.36 |
| 10 | 86026202 | rs6585852 | -1.575 | 0.438 | -2.264 | -0.148 | -0.04 | -0.094 | -0.094 | 0 | 0.56 | 0.94 | 0.73 | 0.52 | 0.59 | 0.6 |
| 10 | 86030567 | rs12569977 | 0.901 | 0.206 | -2.004 | -0.23 | 0.016 | 0.016 | 0.016 | 0 | 0.19 | 0.13 | 0.1 | 0.1 | 0.12 | 0.19 |
| 10 | 86039278 | rs7894492 | -0.55 | 0.374 | -2.092 | -0.225 | 0.017 | 0.044 | 0.044 | 0 | 0.38 | 0.72 | 0.4 | 0.24 | 0.32 | 0.48 |
| 10 | 86047936 | rs4281435 | 0.257 | 0.864 | -2.079 | -0.211 | -0.02 | 0.085 | 0.085 | 0 | 0.59 | 0.19 | 0.55 | 0.76 | 0.61 | 0.51 |
| 10 | 86048061 | rs4366428 | 0.807 | 0.825 | -2.111 | -0.2 | -0.044 | 0.061 | 0.061 | 0 | 0.54 | 0.18 | 0.44 | 0.62 | 0.52 | 0.48 |
| 10 | 86049040 | rs7914940 | -0.109 | 0.238 | -1.876 | -0.259 | 0.018 | 0.022 | 0.022 | 0 | 0.7 | 0.32 | 0.75 | 0.84 | 0.77 | 0.66 |
| 10 | 86049343 | rs4933997 | -0.743 | 0.381 | -2.047 | -0.193 | 0.017 | 0.036 | 0.036 | 0 | 0.77 | 0.86 | 0.87 | 0.9 | 0.81 | 0.8 |
| 10 | 86058516 | rs10887272 | -0.397 | 0.286 | -2.109 | -0.177 | 0.011 | 0.013 | 0.013 | 0 | 0.72 | 0.76 | 0.77 | 0.84 | 0.78 | 0.67 |
| 10 | 86059232 | rs11200981 | 1.663 | 0.947 | -2.109 | 0.078 | -0.059 | -0.021 | -0.021 | 0 | 0.1 | 0.01 | 0.17 | 0.09 | 0.13 | 0.14 |
| 10 | 86066840 | rs10887273 | 0.114 | 0.483 | -1.634 | 0.001 | -0.037 | -0.018 | -0.018 | 0 | 0.22 | 0.57 | 0.3 | 0.17 | 0.13 | 0.21 |
| 10 | 86070751 | rs1881728 | 0.452 | 0.865 | -1.999 | 0.033 | 0 | 0.016 | 0.016 | 0 | 0.63 | 0.21 | 0.49 | 0.66 | 0.66 | 0.63 |
| 10 | 86093951 | rs11200996 | 1.453 | 0.998 | -2.053 | 0.334 | 0 | 0.009 | 0.009 | 0 | 0.06 | 0.05 | 0.11 | 0.19 | 0.07 | 0.17 |
| 10 | 86122255 | rs11201005 | 1.392 | 1.342 | -2.508 | 0.049 | -0.06 | -0.077 | -0.077 | 0 | 0.48 | 0.06 | 0.29 | 0.47 | 0.48 | 0.44 |
| 10 | 86131059 | rs3814205 | 0.998 | 1.26 | -2.356 | -0.022 | -0.005 | -0.009 | -0.009 | 0 | 0.19 | 0.14 | 0.36 | 0.23 | 0.2 | 0.21 |
| 10 | 86135311 | rs4317927 | 2.431 | 0.949 | -3.033 | -0.011 | -0.044 | -0.125 | -0.125 | 0 | 0.21 | 0.01 | 0.15 | 0.27 | 0.23 | 0.17 |
| 10 | 86147907 | rs11593869 | 2.134 | 1.118 | -2.644 | 0.169 | -0.006 | -0.028 | -0.028 | 3.868 | 0.15 | 0.04 | 0.25 | 0.17 | 0.13 | 0.18 |
| 10 | 86148744 | rs11201017 | 1.718 | 1.021 | -2.859 | 0.024 | -0.056 | -0.115 | -0.115 | 0 | 0.4 | 0.01 | 0.22 | 0.35 | 0.35 | 0.35 |
| 10 | 86152677 | rs11201021 | 1.172 | 1.035 | -2.798 | 0.024 | -0.063 | -0.075 | -0.075 | 2.26 | 0.48 | 0.06 | 0.29 | 0.47 | 0.47 | 0.44 |
| 10 | 86152908 | rs10509494 | 2.298 | 1.056 | -2.653 | 0.007 | -0.008 | 0.006 | 0.006 | 0 | 0.07 | 0.06 | 0.1 | 0 | 0.11 | 0.01 |
| 10 | 86157503 | rs10509497 | 1.178 | 0.985 | -2.778 | -0.011 | -0.052 | -0.059 | -0.059 | 2.843 | 0.48 | 0.07 | 0.29 | 0.47 | 0.48 | 0.44 |
| 10 | 86164877 | rs10887283 | 1.203 | 0.997 | -2.79 | 0.008 | -0.067 | -0.075 | -0.075 | 0 | 0.48 | 0.06 | 0.29 | 0.47 | 0.48 | 0.44 |
| 10 | 86186241 | rs10491033 | 1.733 | 0.9 | -2.484 | -0.015 | 0.081 | 0.076 | 0.076 | 0.219 | 0.05 | 0.44 | 0.05 | 0 | 0.01 | 0.02 |
| 10 | 86194860 | rs868090 | -0.356 | 0.901 | -2.499 | -0.057 | 0.002 | -0.018 | -0.018 | 0.015 | 0.2 | 0.58 | 0.25 | 0.19 | 0.19 | 0.21 |
| 10 | 86199507 | rs1572430 | 3.269 | 0.927 | -2.567 | -0.05 | -0.033 | -0.028 | -0.028 | 0 | 0.08 | 0 | 0.07 | 0.12 | 0.13 | 0.09 |
| 10 | 86206701 | rs10887291 | -0.601 | 0.805 | -2.2 | 0.074 | -0.011 | -0.009 | -0.009 | 0 | 0.34 | 0.28 | 0.57 | 0.42 | 0.38 | 0.39 |
| 10 | 86237184 | rs10887305 | 0.402 | 0.865 | -2.33 | 0.087 | -0.006 | 0.025 | 0.025 | 0 | 0.67 | 0.19 | 0.65 | 0.7 | 0.67 | 0.64 |
| 10 | 86240526 | rs1188793 | -2.474 | 0.905 | -2.298 | 0.078 | 0.009 | -0.008 | -0.008 | 0.671 | 0.87 | 0.8 | 0.9 | 0.89 | 0.86 | 0.85 |
| 10 | 86257447 | rs1763976 | 2.146 | 0.888 | -2.333 | 0.058 | -0.006 | 0.003 | 0.003 | 0 | 0.07 | 0.06 | 0.09 | 0 | 0.11 | 0.01 |

| | | | | | | | | | | | | | | | | |
|---|---|---|---|---|---|---|---|---|---|---|---|---|---|---|---|---|
| 10 | 86267975 | rs10887312 | 1.317 | 0.867 | -2.455 | 0.429 | -0.059 | -0.04 | -0.04 | 0 | 0.48 | 0.05 | 0.29 | 0.4 | 0.47 | 0.43 |
| 10 | 86270394 | rs7905523 | 0.297 | 0.874 | -2.435 | 0.443 | -0.011 | -0.011 | -0.011 | 0.313 | 0.67 | 0.18 | 0.65 | 0.63 | 0.67 | 0.64 |
| 10 | 86274654 | rs10788344 | 0.277 | 0.877 | -2.424 | 0.448 | -0.009 | -0.01 | -0.01 | 0 | 0.67 | 0.21 | 0.65 | 0.63 | 0.67 | 0.64 |
| 10 | 86274871 | rs2784846 | -2.169 | 1.073 | -1.914 | 0.316 | 0.012 | 0.025 | 0.025 | 0 | 0.87 | 0.8 | 0.9 | 0.82 | 0.86 | 0.85 |
| 10 | 86276294 | rs17663697 | 1.77 | 1.024 | -1.981 | 0.294 | -0.033 | 0.005 | 0.005 | 0 | 0.12 | 0.02 | 0.17 | 0.09 | 0.11 | 0.16 |
| 10 | 86276596 | rs2784847 | -0.057 | 1.013 | -1.922 | 0.332 | 0.004 | 0.011 | 0.011 | 0 | 0.68 | 0.64 | 0.53 | 0.59 | 0.67 | 0.65 |
| 10 | 86276956 | rs1050943 | 0.395 | 0.93 | -2.036 | 0.196 | -0.006 | -0.01 | -0.01 | 0 | 0.19 | 0.13 | 0.36 | 0.23 | 0.2 | 0.21 |
| 10 | 86281577 | rs2798070 | 0.131 | 0.802 | -2.367 | -0.133 | -0.011 | -0.005 | -0.005 | 0.244 | 0.51 | 0.23 | 0.46 | 0.52 | 0.45 | 0.45 |
| 10 | 86282208 | rs2784848 | 0.57 | 0.733 | -2.396 | -0.319 | 0.006 | 0.025 | 0.025 | 0 | 0.28 | 0.18 | 0.34 | 0.25 | 0.23 | 0.15 |
| 10 | 86282334 | rs2784849 | -1.482 | 0.957 | -1.681 | -0.26 | 0.02 | -0.024 | -0.024 | 0 | 0.85 | 0.72 | 0.79 | 0.89 | 0.81 | 0.84 |
| 10 | 86297596 | rs1188792 | -2.113 | 0.86 | -1.557 | -0.246 | 0.005 | -0.011 | -0.011 | 5.872 | 0.88 | 0.84 | 0.89 | 0.91 | 0.86 | 0.87 |
| 10 | 86300686 | rs11201093 | 1.5 | 1.045 | -1.816 | -0.266 | -0.025 | -0.038 | -0.038 | 0 | 0.22 | 0.05 | 0.11 | 0.26 | 0.21 | 0.26 |
| 10 | 86308588 | rs11201096 | 1.354 | 1.038 | -1.823 | -0.288 | -0.035 | -0.034 | -0.034 | 0 | 0.24 | 0.04 | 0.13 | 0.26 | 0.25 | 0.28 |
| 10 | 86323214 | rs1188786 | 0.436 | 0.551 | -1.787 | -0.559 | -0.005 | 0.004 | 0.004 | 0 | 0.45 | 0.28 | 0.55 | 0.47 | 0.45 | 0.4 |
| 10 | 86325884 | rs12246215 | 0.915 | 0.496 | -1.218 | -0.651 | 0.07 | 0.034 | 0.034 | 0 | 0.06 | 0.33 | 0.05 | 0 | 0.03 | 0.05 |
| 10 | 86329376 | rs7901807 | 1.122 | 0.504 | -1.153 | -0.648 | 0.08 | 0.043 | 0.043 | 0 | 0.05 | 0.31 | 0.05 | 0 | 0.03 | 0.05 |
| 10 | 86332550 | rs2798074 | 0.408 | 0.53 | -1.692 | -0.603 | -0.003 | 0.01 | 0.01 | 0 | 0.46 | 0.29 | 0.55 | 0.47 | 0.45 | 0.41 |
| 10 | 86337629 | rs12777323 | 0.945 | 0.821 | -1.602 | -0.445 | -0.08 | -0.038 | -0.038 | 0.025 | 0.26 | 0.06 | 0.25 | 0.31 | 0.33 | 0.31 |
| 10 | 86342022 | rs2488624 | 1.649 | 0.659 | -1.498 | -0.315 | -0.003 | -0.013 | -0.013 | 0 | 0.19 | 0.16 | 0.28 | 0.37 | 0.24 | 0.23 |
| 10 | 86342359 | rs7911499 | -0.433 | 0.78 | -1.394 | -0.619 | 0.01 | 0.058 | 0.058 | 0 | 0.52 | 0.63 | 0.45 | 0.38 | 0.55 | 0.57 |
| 10 | 86345863 | rs2350416 | -1.228 | 0.84 | -1.44 | -0.647 | -0.018 | 0.049 | 0.049 | 0 | 0.78 | 0.93 | 0.75 | 0.89 | 0.8 | 0.83 |
| 10 | 86346446 | rs2505765 | -0.276 | 0.93 | -1.38 | -0.641 | -0.026 | 0.065 | 0.065 | 0.013 | 0.51 | 0.84 | 0.52 | 0.59 | 0.53 | 0.53 |
| 10 | 86346642 | rs7902264 | -0.207 | 0.949 | -0.691 | -0.752 | 0.078 | -0.027 | -0.027 | 0 | 0.08 | 0.34 | 0.05 | 0 | 0.09 | 0.05 |
| 10 | 86347955 | rs2488630 | 1.1 | 0.674 | -1.006 | -0.848 | 0.008 | -0.02 | -0.02 | 0 | 0.28 | 0.21 | 0.32 | 0.5 | 0.27 | 0.35 |
| 10 | 86349294 | rs2488631 | 0.823 | 1.023 | -0.779 | -0.796 | 0.026 | 0.036 | 0.036 | 0 | 0.43 | 0.21 | 0.5 | 0.59 | 0.38 | 0.51 |
| 10 | 86349647 | rs9663764 | -0.112 | 0.991 | -0.713 | -1.031 | 0.009 | -0.087 | -0.087 | 0 | 0.24 | 0.62 | 0.19 | 0.07 | 0.25 | 0.2 |
| 10 | 86351066 | rs1539524 | -0.726 | 1.115 | -0.849 | -0.872 | -0.01 | 0.022 | 0.022 | 0 | 0.7 | 0.84 | 0.8 | 0.75 | 0.68 | 0.74 |
| 10 | 86352571 | rs2505758 | -0.411 | 1.071 | -0.832 | -0.949 | -0.014 | 0.022 | 0.022 | 0 | 0.66 | 0.81 | 0.7 | 0.69 | 0.62 | 0.7 |
| 10 | 86353764 | rs1539522 | -0.451 | 1.082 | 0.307 | -0.633 | 0.128 | 0.009 | 0.009 | 0 | 0.07 | 0.41 | 0.06 | 0 | 0.09 | 0.06 |
| 10 | 86355239 | rs11597975 | 1.447 | 0.929 | -0.199 | -0.651 | -0.01 | 0.032 | 0.032 | 0 | 0.15 | 0.1 | 0.13 | 0.08 | 0.16 | 0.13 |
| 10 | 86356555 | rs10887329 | -0.163 | 1.271 | 0.431 | -0.519 | 0.125 | 0.001 | 0.001 | 0 | 0.06 | 0.28 | 0.06 | 0 | 0.1 | 0.06 |
| 10 | 86357285 | rs2350414 | 1.505 | 1.273 | -0.281 | -0.616 | -0.027 | -0.032 | -0.032 | 0 | 0.18 | 0.04 | 0.17 | 0.24 | 0.2 | 0.24 |
| 10 | 86359514 | rs2488638 | -0.645 | 1.259 | -0.242 | -0.546 | -0.021 | -0.017 | -0.017 | 0 | 0.73 | 0.84 | 0.74 | 0.65 | 0.68 | 0.7 |
| 10 | 86363639 | rs2488647 | -0.718 | 0.95 | -0.213 | -0.546 | -0.009 | -0.016 | -0.016 | 0 | 0.74 | 0.85 | 0.75 | 0.65 | 0.71 | 0.71 |
| 10 | 86366489 | rs2488656 | 0.21 | 0.64 | 0.241 | -0.118 | 0.011 | 0.016 | 0.016 | 0.228 | 0.23 | 0.18 | 0.19 | 0.16 | 0.17 | 0.24 |
| 10 | 86370809 | rs11201124 | -0.089 | 0.878 | 0.763 | -0.098 | 0.09 | -0.021 | -0.021 | 0 | 0.06 | 0.25 | 0.06 | 0 | 0.11 | 0.06 |
| 10 | 86375282 | rs2505752 | -1.52 | 0.739 | 0.726 | -0.107 | 0.002 | 0.008 | 0.008 | 0 | 0.4 | 0.67 | 0.45 | 0.35 | 0.38 | 0.37 |
| 10 | 86377269 | rs10887335 | -0.286 | 0.746 | 0.711 | -0.098 | -0.003 | 0.006 | 0.006 | 0.106 | 0.09 | 0.11 | 0.18 | 0.19 | 0.1 | 0.08 |
| 10 | 86380227 | rs11201128 | 1.203 | 0.706 | 0.591 | -0.172 | -0.002 | -0.043 | -0.043 | 0.009 | 0.26 | 0.17 | 0.19 | 0.32 | 0.29 | 0.21 |
| 10 | 86385916 | rs2488649 | 0.539 | 0.786 | 1.088 | -0.013 | 0 | 0.048 | 0.048 | 0 | 0.19 | 0.11 | 0.12 | 0.09 | 0.16 | 0.13 |
| 10 | 86387167 | rs1891655 | -1.127 | 0.919 | 1.136 | 0.019 | 0.073 | 0.001 | 0.001 | 0 | 0.09 | 0.27 | 0.17 | 0.07 | 0.15 | 0.1 |
| 10 | 86392391 | rs2505782 | -1.573 | 0.837 | 1.068 | 0.017 | 0.002 | 0.008 | 0.008 | 0 | 0.4 | 0.67 | 0.45 | 0.35 | 0.39 | 0.37 |
| 10 | 86410057 | rs4388817 | -1.638 | 1.169 | 1.042 | 0.012 | 0.041 | -0.013 | -0.013 | 0.07 | 0.13 | 0.38 | 0.14 | 0.11 | 0.17 | 0.1 |
| 10 | 86412647 | rs4934030 | -0.082 | 1.376 | 0.407 | 0.208 | 0.003 | -0.001 | -0.001 | 0 | 0.88 | 0.88 | 0.87 | 0.89 | 0.79 | 0.92 |
| 10 | 86421486 | rs11201157 | -1.077 | 1.171 | 0.159 | 0.186 | -0.002 | 0.029 | 0.029 | 0 | 0.43 | 0.66 | 0.47 | 0.26 | 0.42 | 0.33 |
| 10 | 86422743 | rs11599612 | -0.554 | 1.421 | 0.072 | 0.452 | 0.011 | 0.019 | 0.019 | 0 | 0.32 | 0.35 | 0.3 | 0.4 | 0.44 | 0.35 |
| 10 | 86422872 | rs11599632 | 1.04 | 1.407 | 0.116 | 0.454 | 0.004 | 0.038 | 0.038 | 0.011 | 0.16 | 0.2 | 0.18 | 0.29 | 0.24 | 0.27 |
| 10 | 86423400 | rs10887346 | -1.034 | 0.663 | 0.386 | 0.394 | 0.006 | 0 | 0 | 0 | 0.4 | 0.53 | 0.53 | 0.51 | 0.55 | 0.47 |
| 10 | 86424325 | rs7068827 | -1.9 | 0.592 | 0.327 | 0.406 | -0.008 | 0.015 | 0.015 | 0.226 | 0.59 | 0.89 | 0.71 | 0.57 | 0.66 | 0.6 |
| 10 | 86426712 | rs17259859 | 0.59 | 0.78 | 0.474 | 0.047 | -0.005 | -0.009 | -0.009 | 0.425 | 0.14 | 0.13 | 0.1 | 0.11 | 0.13 | 0.06 |
| 10 | 86429955 | rs12253879 | -0.963 | 0.752 | 0.595 | 0.089 | -0.015 | -0.008 | -0.008 | 1.802 | 0.25 | 0.36 | 0.33 | 0.22 | 0.23 | 0.19 |
| 10 | 86431626 | rs4400740 | 0.07 | 0.765 | 0.554 | 0.1 | 0.011 | 0.032 | 0.032 | 0 | 0.09 | 0.2 | 0.23 | 0.11 | 0.1 | 0.12 |
| 10 | 86432072 | rs11815535 | 0.439 | 0.765 | 0.262 | 0.092 | -0.005 | -0.008 | -0.008 | 0 | 0.15 | 0.13 | 0.1 | 0.11 | 0.13 | 0.06 |
| 10 | 86438620 | rs7920114 | 1.008 | 0.837 | 0.309 | -0.006 | -0.01 | -0.048 | -0.048 | 0.02 | 0.18 | 0.3 | 0.12 | 0.05 | 0.14 | 0.15 |
| 10 | 86438982 | rs11817609 | 0.667 | 0.857 | 0.23 | 0.006 | 0.001 | 0.001 | 0.001 | 0 | 0.14 | 0.11 | 0.1 | 0.11 | 0.13 | 0.06 |
| 10 | 86439849 | rs4573626 | 0.987 | 1.113 | -0.095 | 0.381 | 0.019 | 0.062 | 0.062 | 0 | 0.16 | 0.21 | 0.19 | 0.44 | 0.29 | 0.26 |
| 10 | 86444162 | rs10887348 | -0.418 | 0.193 | 0.321 | -0.035 | 0.015 | 0.043 | 0.043 | 0.411 | 0.1 | 0.26 | 0.23 | 0.11 | 0.1 | 0.12 |
| 10 | 86448460 | rs6585875 | -0.796 | 0.385 | -0.521 | 0.11 | 0.024 | 0.048 | 0.048 | 0 | 0.43 | 0.68 | 0.43 | 0.27 | 0.37 | 0.3 |
| 10 | 86461085 | rs10509498 | 0.19 | 0.499 | -0.368 | 0.038 | 0.008 | -0.023 | -0.023 | 0 | 0.38 | 0.51 | 0.3 | 0.28 | 0.46 | 0.37 |
| 10 | 86461097 | rs11201179 | 0.429 | 0.055 | -0.528 | -0.509 | -0.017 | -0.086 | -0.086 | 0 | 0.16 | 0.25 | 0.11 | 0 | 0.12 | 0.02 |
| 10 | 86463275 | rs7923246 | 1.246 | 0.204 | -0.171 | -0.332 | 0.004 | 0.001 | 0.001 | 0 | 0.28 | 0.28 | 0.23 | 0.31 | 0.35 | 0.26 |
| 10 | 86464139 | rs1981098 | 0.777 | 0.29 | -0.637 | 0.056 | -0.056 | -0.075 | -0.075 | 0 | 0.12 | 0.01 | 0.19 | 0.27 | 0.16 | 0.2 |
| 10 | 86464587 | rs945146 | -0.959 | -0.351 | -0.407 | -0.674 | 0.017 | -0.011 | -0.011 | 0 | 0.69 | 0.8 | 0.81 | 0.99 | 0.79 | 0.84 |
| 10 | 86466132 | rs1577273 | -0.611 | -0.32 | -0.476 | -0.454 | 0.01 | 0.028 | 0.028 | 2.401 | 0.33 | 0.52 | 0.47 | 0.58 | 0.36 | 0.48 |
| 10 | 86466489 | rs17103959 | 0.677 | -0.333 | -0.502 | -0.441 | -0.011 | -0.039 | -0.039 | 0 | 0.07 | 0 | 0.1 | 0.11 | 0.05 | 0.1 |
| 10 | 86467133 | rs2505777 | -1.038 | -0.336 | -0.457 | -0.465 | 0 | -0.002 | -0.002 | 0 | 0.41 | 0.48 | 0.58 | 0.69 | 0.44 | 0.57 |
| 10 | 86471855 | rs871893 | -0.866 | -0.39 | -0.406 | -0.473 | 0.001 | -0.057 | -0.057 | 0 | 0.68 | 0.72 | 0.81 | 0.98 | 0.79 | 0.84 |
| 10 | 86475581 | rs6585878 | 0.872 | -0.114 | -0.243 | -0.294 | 0.004 | -0.067 | -0.067 | 0.216 | 0.11 | 0.3 | 0.08 | 0.01 | 0.14 | 0.08 |
| 10 | 86481602 | rs2505788 | 0.056 | 0.302 | -0.8 | 0.086 | -0.025 | -0.054 | -0.054 | 0.078 | 0.22 | 0.16 | 0.4 | 0.55 | 0.26 | 0.4 |

| | | | | | | | | | | | | | | | | |
|---|---|---|---|---|---|---|---|---|---|---|---|---|---|---|---|---|
| 10 | 86483892 | rs12261909 | 0.979 | 0.447 | -0.571 | 0.429 | -0.002 | -0.001 | -0.001 | 0 | 0.28 | 0.24 | 0.22 | 0.31 | 0.34 | 0.26 |
| 10 | 86503601 | rs1577286 | -0.801 | 0.156 | -0.536 | 0.572 | 0.001 | -0.007 | -0.007 | 0.957 | 0.45 | 0.75 | 0.53 | 0.43 | 0.45 | 0.41 |
| 10 | 86506192 | rs10887359 | -0.788 | 0.147 | -0.54 | 0.569 | 0 | -0.008 | -0.008 | 0 | 0.44 | 0.75 | 0.53 | 0.43 | 0.45 | 0.41 |
| 10 | 86515816 | rs2960355 | -0.138 | 0.077 | 0.078 | 0.505 | -0.002 | -0.011 | -0.011 | 0 | 0.7 | 0.63 | 0.64 | 0.73 | 0.68 | 0.7 |
| 10 | 86525965 | rs10887363 | 0.813 | 0.202 | 0.027 | 0.735 | 0.018 | 0.062 | 0.062 | 0 | 0.09 | 0.18 | 0.19 | 0.23 | 0.11 | 0.24 |
| 10 | 86528538 | rs10887364 | 0.542 | -0.002 | 0.351 | 0.487 | 0.037 | 0.04 | 0.04 | 0 | 0.12 | 0.34 | 0.1 | 0.12 | 0.12 | 0.1 |
| 10 | 86531527 | rs10788363 | 0.542 | -0.011 | 0.403 | 0.487 | 0.026 | 0.032 | 0.032 | 0 | 0.12 | 0.32 | 0.1 | 0.12 | 0.12 | 0.1 |
| 10 | 86547284 | rs17104125 | 1.034 | 0.01 | 0.35 | 0.364 | 0.021 | -0.061 | -0.061 | 0.103 | 0.11 | 0.25 | 0.1 | 0 | 0.12 | 0.07 |
| 10 | 86552725 | rs2498013 | 1.577 | -0.303 | -0.235 | 0.25 | 0.015 | 0.022 | 0.022 | 0.116 | 0.14 | 0.08 | 0.05 | 0 | 0.07 | 0.05 |
| 10 | 86556250 | rs7916708 | 0.558 | -0.187 | -0.302 | 0.349 | -0.017 | 0.017 | 0.017 | 0 | 0.45 | 0.31 | 0.47 | 0.61 | 0.49 | 0.56 |
| 10 | 86571942 | rs10887382 | 0.386 | -0.376 | -0.634 | 0.28 | 0.028 | 0.018 | 0.018 | 0.1 | 0.48 | 0.11 | 0.51 | 0.58 | 0.6 | 0.62 |
| 10 | 86573350 | rs2960358 | 1.013 | -0.346 | 0.077 | 0.129 | -0.004 | -0.034 | -0.034 | 0 | 0.14 | 0.14 | 0.06 | 0 | 0.07 | 0.05 |
| 10 | 86585840 | rs2498010 | -1.268 | 0.152 | -0.448 | 0.79 | -0.121 | -0.12 | -0.12 | 0.103 | 0.71 | 0.98 | 0.74 | 0.63 | 0.65 | 0.72 |
| 10 | 86587878 | rs7901050 | -0.965 | -0.203 | 0.305 | 0.88 | -0.043 | 0.069 | 0.069 | 0.078 | 0.23 | 0.55 | 0.27 | 0.34 | 0.14 | 0.35 |
| 10 | 86589388 | rs2991922 | -1.326 | 0.248 | 0.469 | 0.712 | -0.021 | -0.019 | -0.019 | 0 | 0.69 | 0.8 | 0.71 | 0.63 | 0.64 | 0.71 |
| 10 | 86592660 | rs11201255 | -0.719 | 0.217 | 0.948 | 0.893 | -0.015 | 0.077 | 0.077 | 0.169 | 0.14 | 0.37 | 0.14 | 0.21 | 0.1 | 0.18 |
| 10 | 86594341 | rs2991926 | 0.431 | 0.075 | 0.625 | 0.786 | -0.028 | 0.023 | 0.023 | 0 | 0.45 | 0.19 | 0.42 | 0.41 | 0.48 | 0.47 |
| 10 | 86599260 | rs11201258 | -0.076 | 0.082 | 0.672 | 0.884 | -0.005 | 0.013 | 0.013 | 0 | 0.1 | 0.16 | 0.09 | 0.16 | 0.06 | 0.14 |
| 10 | 86603486 | rs1577272 | -0.903 | 0.023 | 1.026 | 0.835 | -0.03 | 0.015 | 0.015 | 0 | 0.18 | 0.38 | 0.12 | 0.16 | 0.1 | 0.19 |
| 10 | 86606452 | rs1591315 | 0.013 | 0.274 | 1.127 | 0.838 | -0.001 | 0.001 | 0.001 | 0 | 0.49 | 0.4 | 0.57 | 0.44 | 0.5 | 0.5 |
| 10 | 86619647 | rs12766251 | -1.758 | 0.312 | 1.365 | 1.042 | -0.019 | 0.089 | 0.089 | 0.228 | 0.13 | 0.33 | 0.27 | 0.23 | 0.09 | 0.21 |
| 10 | 86628051 | rs10887386 | 1.286 | 0.63 | 0.901 | 1.116 | -0.039 | -0.029 | -0.029 | 0 | 0.07 | 0.01 | 0.07 | 0.28 | 0.12 | 0.12 |
| 10 | 86631081 | rs11201265 | -2.148 | 0.155 | 1.369 | 0.765 | -0.013 | -0.001 | -0.001 | 0.034 | 0.26 | 0.69 | 0.35 | 0.23 | 0.19 | 0.27 |
| 10 | 86633667 | rs11201272 | 0.317 | 0.149 | 1.446 | 0.771 | 0.002 | -0.01 | -0.01 | 0.024 | 0.06 | 0.14 | 0.04 | 0.05 | 0.06 | 0.05 |
| 10 | 86634344 | rs11201274 | -1.75 | 0.232 | 1.315 | 0.822 | -0.035 | 0.034 | 0.034 | 0.562 | 0.2 | 0.48 | 0.3 | 0.19 | 0.14 | 0.23 |
| 10 | 86634920 | rs7091418 | -0.952 | 0.125 | 1.283 | 0.862 | -0.048 | 0.065 | 0.065 | 0.462 | 0.14 | 0.37 | 0.15 | 0.16 | 0.07 | 0.2 |
| 10 | 86638760 | rs4264108 | -0.626 | 0.363 | 0.841 | 0.814 | -0.005 | -0.003 | -0.003 | 0 | 0.63 | 0.68 | 0.54 | 0.56 | 0.56 | 0.65 |
| 10 | 86639989 | rs4310554 | 0.243 | 0.207 | 1.206 | 0.816 | 0.001 | 0.012 | 0.012 | 0 | 0.51 | 0.39 | 0.44 | 0.39 | 0.51 | 0.47 |
| 10 | 86648725 | rs6585903 | -1.531 | 0.214 | 1.055 | 0.775 | -0.068 | -0.072 | -0.072 | 0 | 0.72 | 0.95 | 0.71 | 0.62 | 0.64 | 0.7 |
| 10 | 86649647 | rs12767486 | 1.015 | 0.707 | 1.206 | 0.756 | 0.003 | -0.015 | -0.015 | 1.076 | 0.3 | 0.19 | 0.31 | 0.38 | 0.3 | 0.29 |
| 10 | 86650206 | rs11201285 | -0.53 | 0.44 | 1.277 | 0.798 | -0.032 | 0.046 | 0.046 | 0 | 0.12 | 0.3 | 0.1 | 0.17 | 0.05 | 0.18 |
| 10 | 86651015 | rs7078668 | -0.547 | 0.441 | 1.294 | 0.798 | -0.032 | 0.047 | 0.047 | 0 | 0.12 | 0.31 | 0.11 | 0.17 | 0.05 | 0.18 |
| 10 | 86655565 | rs6585910 | -0.619 | 0.404 | 1.364 | 0.791 | -0.006 | -0.003 | -0.003 | 0 | 0.63 | 0.68 | 0.54 | 0.58 | 0.56 | 0.63 |
| 10 | 86661963 | rs10887395 | -0.269 | 0.4 | 1.342 | 0.77 | -0.004 | 0.003 | 0.003 | 0.006 | 0.58 | 0.47 | 0.56 | 0.45 | 0.58 | 0.51 |
| 10 | 86662090 | rs7091034 | -1.461 | 0.57 | 1.083 | 0.703 | -0.031 | -0.032 | -0.032 | 0 | 0.72 | 0.85 | 0.71 | 0.62 | 0.64 | 0.69 |
| 10 | 86665044 | rs4244961 | -1.476 | 0.57 | 1.004 | 0.709 | -0.048 | -0.052 | -0.052 | 0 | 0.72 | 0.89 | 0.71 | 0.61 | 0.64 | 0.69 |
| 10 | 86665704 | rs10509500 | 0.105 | 0.658 | 1.02 | 0.703 | 0.003 | -0.021 | -0.021 | 0.268 | 0.05 | 0.09 | 0.11 | 0 | 0.07 | 0.03 |
| 10 | 86669864 | rs10749503 | 0.363 | 0.383 | 1.294 | 0.59 | -0.004 | -0.007 | -0.007 | 0 | 0.47 | 0.3 | 0.45 | 0.53 | 0.45 | 0.55 |
| 10 | 86670623 | rs10749504 | 0.449 | 0.351 | 1.273 | 0.593 | -0.004 | -0.002 | -0.002 | 0 | 0.46 | 0.3 | 0.44 | 0.48 | 0.44 | 0.52 |
| 10 | 86671791 | rs7914715 | 0.377 | 0.35 | 1.274 | 0.587 | -0.005 | -0.008 | -0.008 | 0.516 | 0.47 | 0.3 | 0.45 | 0.53 | 0.45 | 0.55 |
| 10 | 86688193 | rs10788387 | -1.398 | 0.305 | 1.376 | 0.578 | -0.018 | -0.022 | -0.022 | 0 | 0.7 | 0.83 | 0.67 | 0.62 | 0.64 | 0.68 |
| 10 | 86697162 | rs11201313 | -0.269 | 0.096 | 1.67 | 0.454 | 0.001 | -0.001 | -0.001 | 1.105 | 0.09 | 0.11 | 0.04 | 0.08 | 0.05 | 0.04 |
| 10 | 86701332 | rs1933935 | -2.054 | 0.064 | 1.804 | 0.444 | -0.018 | -0.025 | -0.025 | 0 | 0.25 | 0.66 | 0.28 | 0.13 | 0.21 | 0.17 |
| 10 | 86704691 | rs10887406 | -1.368 | 0.046 | 2.039 | 0.394 | -0.012 | -0.017 | -0.017 | 0 | 0.17 | 0.33 | 0.21 | 0.08 | 0.12 | 0.08 |
| 10 | 86712004 | rs10159886 | -2.136 | 0.041 | 2.013 | 0.42 | -0.018 | -0.029 | -0.029 | 0.883 | 0.24 | 0.67 | 0.3 | 0.1 | 0.19 | 0.14 |
| 10 | 86712634 | rs7072257 | -1.45 | 0.178 | 1.529 | 0.498 | -0.011 | -0.015 | -0.015 | 0 | 0.17 | 0.32 | 0.21 | 0.08 | 0.12 | 0.08 |
| 10 | 86721181 | rs12266096 | -1.993 | 0.494 | 1.54 | 0.543 | 0.008 | 0.009 | 0.009 | 0 | 0.19 | 0.59 | 0.28 | 0.18 | 0.19 | 0.12 |
| 10 | 86721948 | rs12266197 | -1.196 | 0.532 | 1.701 | 0.736 | 0.005 | 0.008 | 0.008 | 0 | 0.16 | 0.3 | 0.18 | 0.17 | 0.18 | 0.1 |
| 10 | 86725338 | rs7067845 | -2.108 | 0.517 | 1.73 | 0.727 | 0.003 | 0.023 | 0.023 | 0 | 0.2 | 0.6 | 0.27 | 0.23 | 0.19 | 0.13 |
| 10 | 86742414 | rs1933950 | 0.065 | 0.853 | 1.408 | 0.759 | 0.005 | -0.001 | -0.001 | 0.058 | 0.52 | 0.52 | 0.54 | 0.54 | 0.43 | 0.58 |
| 10 | 86747688 | rs1361264 | 0.133 | 0.803 | 1.325 | 0.675 | 0.013 | -0.001 | -0.001 | 0 | 0.49 | 0.51 | 0.48 | 0.53 | 0.4 | 0.57 |
| 10 | 86751289 | rs1999941 | 0.274 | 0.793 | 1.393 | 0.666 | 0.026 | -0.004 | -0.004 | 0 | 0.48 | 0.32 | 0.46 | 0.49 | 0.38 | 0.49 |
| 10 | 86752528 | rs7918012 | -0.975 | 0.813 | 1.36 | 0.609 | -0.003 | -0.005 | -0.005 | 0 | 0.65 | 0.62 | 0.7 | 0.53 | 0.57 | 0.6 |
| 10 | 86755397 | rs12260827 | -1.521 | 0.816 | 1.346 | 0.679 | -0.042 | -0.025 | -0.025 | 0 | 0.69 | 0.82 | 0.72 | 0.62 | 0.58 | 0.62 |
| 10 | 86758509 | rs1538431 | 0.965 | 0.808 | 0.926 | 0.79 | -0.026 | -0.034 | -0.034 | 0 | 0.1 | 0.06 | 0.12 | 0.22 | 0.13 | 0.18 |
| 10 | 86762257 | rs10788394 | 0.129 | 0.773 | 0.996 | 0.78 | 0.027 | -0.003 | -0.003 | 0 | 0.48 | 0.32 | 0.46 | 0.49 | 0.39 | 0.49 |
| 10 | 86767662 | rs10788395 | 1.01 | 0.453 | 0.463 | 0.577 | -0.017 | -0.017 | -0.017 | 0.37 | 0.32 | 0.1 | 0.3 | 0.34 | 0.28 | 0.31 |
| 10 | 86781967 | rs1933961 | -1.198 | 1.01 | 0.875 | 0.633 | 0.001 | 0 | 0 | 0 | 0.67 | 0.63 | 0.71 | 0.58 | 0.57 | 0.6 |
| 10 | 86798551 | rs1890474 | -0.552 | 1.068 | 0.605 | 0.319 | -0.006 | -0.007 | -0.007 | 0 | 0.13 | 0.12 | 0.16 | 0.01 | 0.16 | 0.07 |
| 10 | 86801029 | rs946324 | -0.508 | 1.069 | 0.591 | 0.319 | -0.013 | 0.009 | 0.009 | 0 | 0.13 | 0.09 | 0.16 | 0.01 | 0.16 | 0.07 |
| 10 | 86804767 | rs17104561 | -0.487 | 1.089 | 0.654 | 0.319 | -0.01 | 0.013 | 0.013 | 0 | 0.09 | 0.07 | 0.14 | 0.01 | 0.11 | 0.07 |
| 10 | 86806057 | rs2211084 | 1.291 | 0.988 | 0.482 | 0.531 | -0.103 | -0.082 | -0.082 | 0 | 0.28 | 0.04 | 0.26 | 0.34 | 0.4 | 0.38 |
| 10 | 86807865 | rs10788413 | 1.206 | 0.972 | 0.62 | 0.523 | -0.084 | -0.065 | -0.065 | 0.008 | 0.29 | 0.09 | 0.27 | 0.34 | 0.4 | 0.38 |
| 10 | 86813756 | rs10509506 | -0.316 | 0.958 | 0.94 | 0.501 | 0.017 | -0.057 | -0.057 | 0 | 0.14 | 0.18 | 0.17 | 0.01 | 0.16 | 0.07 |
| 10 | 86814194 | rs10509507 | 1.295 | 0.964 | 0.908 | 0.523 | -0.103 | -0.082 | -0.082 | 0 | 0.28 | 0.04 | 0.26 | 0.34 | 0.4 | 0.38 |
| 10 | 86815074 | rs10749511 | 1.081 | 0.941 | 1.077 | 0.513 | -0.043 | -0.03 | -0.03 | 0 | 0.3 | 0.14 | 0.26 | 0.34 | 0.4 | 0.38 |
| 10 | 86837430 | rs11816278 | 1.806 | 0.659 | 0.956 | -0.493 | 0.002 | 0.007 | 0.007 | 0 | 0.11 | 0.05 | 0.08 | 0 | 0.1 | 0.03 |
| 10 | 86843226 | rs7067796 | -1.883 | 0.736 | 1.139 | -0.104 | 0.024 | -0.011 | -0.011 | 0 | 0.22 | 0.52 | 0.25 | 0.16 | 0.21 | 0.14 |
| 10 | 86850330 | rs12248262 | -0.611 | 0.847 | 1.05 | -0.543 | 0.007 | -0.037 | -0.037 | 0 | 0.14 | 0.21 | 0.18 | 0.01 | 0.17 | 0.07 |

| | | | | | | | | | | | | | | | | |
|---|---|---|---|---|---|---|---|---|---|---|---|---|---|---|---|---|
| 10 | 86851425 | rs7072549 | 1.148 | 0.938 | 0.91 | -0.706 | -0.003 | -0.032 | -0.032 | 0.012 | 0.25 | 0.22 | 0.16 | 0.01 | 0.32 | 0.15 |
| 10 | 86852648 | rs10509508 | 1.078 | 0.937 | 0.882 | -0.707 | 0.009 | -0.094 | -0.094 | 0.45 | 0.25 | 0.33 | 0.16 | 0.01 | 0.32 | 0.15 |
| 10 | 86885210 | rs17296623 | -0.222 | 0.942 | 0.732 | -0.631 | -0.022 | 0.036 | 0.036 | 0 | 0.13 | 0.07 | 0.16 | 0.01 | 0.16 | 0.07 |
| 10 | 86892095 | rs4934071 | -0.349 | 0.972 | 0.821 | -0.657 | 0.01 | -0.057 | -0.057 | 0 | 0.41 | 0.56 | 0.45 | 0.14 | 0.49 | 0.22 |
| 10 | 86893742 | rs17296807 | 0.221 | 1.007 | 0.711 | -0.908 | -0.066 | 0.133 | 0.133 | 0 | 0.35 | 0.16 | 0.31 | 0.01 | 0.45 | 0.19 |
| 10 | 86895040 | rs11201401 | -0.166 | 0.995 | 0.686 | -0.912 | 0.005 | -0.094 | -0.094 | 1.577 | 0.38 | 0.47 | 0.33 | 0.01 | 0.45 | 0.19 |
| 10 | 86904915 | rs7893958 | 1.492 | 1.092 | 0.409 | -0.842 | -0.043 | 0.092 | 0.092 | 0 | 0.22 | 0.06 | 0.14 | 0 | 0.28 | 0.12 |
| 10 | 86919325 | rs6585933 | -0.221 | 1.074 | 0.515 | -0.852 | 0.01 | -0.103 | -0.103 | 0 | 0.39 | 0.45 | 0.34 | 0.02 | 0.49 | 0.22 |
| 10 | 86924828 | rs17370064 | -0.097 | 1.204 | 0.074 | -1.02 | -0.01 | 0.002 | 0.002 | 0 | 0.12 | 0.1 | 0.17 | 0.01 | 0.17 | 0.07 |
| 10 | 86930900 | rs2350822 | -0.849 | 0.608 | 0.128 | -0.649 | 0.014 | -0.037 | -0.037 | 0 | 0.69 | 0.78 | 0.71 | 0.49 | 0.76 | 0.55 |
| 10 | 86933221 | rs7902392 | 0.094 | 0.515 | 0.453 | -1.287 | 0.01 | -0.109 | -0.109 | 0 | 0.26 | 0.36 | 0.17 | 0.01 | 0.32 | 0.14 |
| 10 | 86953013 | rs11201443 | 0.047 | 0.32 | 0.227 | -0.948 | -0.025 | 0.053 | 0.053 | 0 | 0.23 | 0.12 | 0.18 | 0.04 | 0.29 | 0.12 |
| 10 | 86965265 | rs1416929 | -0.206 | 0.214 | 0.753 | -0.932 | -0.004 | -0.002 | -0.002 | 0.042 | 0.26 | 0.24 | 0.21 | 0.09 | 0.34 | 0.15 |
| 10 | 86972838 | rs7075770 | -0.881 | -0.593 | 0.322 | -1.708 | 0.046 | 0.06 | 0.06 | 0 | 0.77 | 0.89 | 0.85 | 1 | 0.88 | 0.85 |
| 10 | 86973061 | rs7901089 | -0.733 | 0.047 | 1.533 | -1.06 | 0.009 | -0.047 | -0.047 | 0 | 0.3 | 0.54 | 0.35 | 0.18 | 0.34 | 0.16 |
| 10 | 86973767 | rs10788436 | -0.889 | 0.038 | 0.828 | -1.055 | -0.004 | -0.026 | -0.026 | 0 | 0.93 | 0.9 | 0.96 | 1 | 0.94 | 0.97 |
| 10 | 86982746 | rs11201502 | -0.101 | 0.201 | 0.732 | -0.944 | -0.035 | 0.034 | 0.034 | 0 | 0.25 | 0.14 | 0.32 | 0.18 | 0.34 | 0.16 |
| 10 | 86984146 | rs12766108 | -0.085 | 0.224 | 0.707 | -0.802 | -0.056 | 0.053 | 0.053 | 0 | 0.24 | 0.1 | 0.21 | 0.11 | 0.34 | 0.13 |
| 10 | 86992461 | rs2350833 | 0.138 | 0.407 | 0.743 | -0.749 | -0.049 | -0.022 | -0.022 | 0 | 0.21 | 0.12 | 0.31 | 0.2 | 0.31 | 0.14 |
| 10 | 86993448 | rs1105093 | 0.254 | 0.409 | 0.703 | -0.772 | -0.061 | 0.023 | 0.023 | 0 | 0.2 | 0.09 | 0.2 | 0.11 | 0.31 | 0.12 |
| 10 | 87001789 | rs11201519 | -1.854 | 0.377 | 1.205 | -0.633 | 0.029 | 0.118 | 0.118 | 0 | 0.09 | 0.27 | 0.23 | 0.16 | 0.11 | 0.07 |
| 10 | 87008066 | rs11593466 | -1.718 | 0.137 | 1.436 | -0.659 | 0.018 | 0.156 | 0.156 | 0 | 0.09 | 0.33 | 0.21 | 0.16 | 0.07 | 0.07 |
| 10 | 87015549 | rs883800 | -0.506 | 0.129 | 1.725 | -0.924 | 0.004 | -0.055 | -0.055 | 0.166 | 0.25 | 0.4 | 0.2 | 0.07 | 0.27 | 0.12 |
| 10 | 87030407 | rs7895259 | -0.656 | 0.115 | 1.788 | -0.838 | 0.002 | -0.021 | -0.021 | 0 | 0.26 | 0.56 | 0.33 | 0.16 | 0.27 | 0.15 |
| 10 | 87054850 | rs11201540 | -0.769 | 0.055 | 1.963 | -0.863 | -0.002 | -0.015 | -0.015 | 0 | 0.27 | 0.63 | 0.33 | 0.16 | 0.27 | 0.16 |
| 10 | 87066918 | rs7910462 | 0.777 | 0.108 | 1.57 | -0.853 | -0.028 | 0.067 | 0.067 | 0 | 0.17 | 0.08 | 0.1 | 0 | 0.2 | 0.08 |
| 10 | 87071981 | rs7898418 | -0.584 | 0.127 | 1.589 | -0.83 | 0.004 | -0.023 | -0.023 | 0 | 0.26 | 0.53 | 0.33 | 0.16 | 0.27 | 0.16 |
| 10 | 87074877 | rs12146219 | 0.48 | 0 | 1.403 | -0.922 | 0.002 | -0.024 | -0.024 | 0 | 0.19 | 0.27 | 0.23 | 0.1 | 0.18 | 0.16 |
| 10 | 87075046 | rs2037065 | -1.039 | -0.04 | 1.48 | -0.939 | -0.008 | -0.026 | -0.026 | 0 | 0.29 | 0.58 | 0.34 | 0.16 | 0.28 | 0.21 |
| 10 | 87077600 | rs11593225 | -1.05 | -0.052 | 1.325 | -0.903 | -0.004 | 0.007 | 0.007 | 0.164 | 0.11 | 0.3 | 0.12 | 0.07 | 0.09 | 0.05 |
| 10 | 87081507 | rs7915322 | -1.197 | -0.119 | 1.204 | -0.895 | -0.007 | 0.026 | 0.026 | 0.003 | 0.12 | 0.31 | 0.17 | 0.12 | 0.1 | 0.08 |
| 10 | 87081768 | rs2037066 | -1.17 | -0.181 | 1.41 | -0.998 | 0.005 | 0.007 | 0.007 | 0.116 | 0.32 | 0.73 | 0.4 | 0.22 | 0.29 | 0.26 |
| 10 | 87082523 | rs11201563 | 0.573 | -0.154 | 1.385 | -0.995 | 0.005 | -0.12 | -0.12 | 0 | 0.18 | 0.38 | 0.11 | 0 | 0.18 | 0.14 |
| 10 | 87104771 | rs7099093 | -1.147 | -0.186 | 1.164 | -0.999 | 0.003 | 0.003 | 0.003 | 0.054 | 0.32 | 0.7 | 0.4 | 0.22 | 0.29 | 0.26 |
| 10 | 87108770 | rs7068955 | 0.619 | 0.1 | 0.949 | -0.984 | 0.002 | -0.073 | -0.073 | 0 | 0.17 | 0.27 | 0.1 | 0 | 0.18 | 0.13 |
| 10 | 87133028 | rs7902653 | -0.39 | -0.108 | 0.451 | -1.002 | 0 | 0.001 | 0.001 | 0.045 | 0.5 | 0.48 | 0.52 | 0.39 | 0.49 | 0.29 |
| 10 | 87133522 | rs2351335 | 0.376 | -0.167 | 0.557 | -1.11 | -0.018 | -0.011 | -0.011 | 5.249 | 0.42 | 0.15 | 0.42 | 0.39 | 0.41 | 0.26 |
| 10 | 87134919 | rs11201598 | 0.36 | -0.173 | 0.63 | -1.113 | -0.013 | -0.007 | -0.007 | 0 | 0.43 | 0.17 | 0.42 | 0.39 | 0.41 | 0.26 |
| 10 | 87136789 | rs4147059 | -0.393 | -0.218 | 0.66 | -1.113 | -0.001 | 0.006 | 0.006 | 0.048 | 0.49 | 0.53 | 0.53 | 0.39 | 0.49 | 0.29 |
| 10 | 87154121 | rs4934089 | 0.697 | -0.099 | -0.03 | -1.319 | -0.01 | -0.024 | -0.024 | 0 | 0.31 | 0.19 | 0.35 | 0.33 | 0.33 | 0.27 |
| 10 | 87156252 | rs4403749 | -0.37 | -0.039 | 0.115 | -1.148 | -0.005 | 0 | 0 | 2.082 | 0.54 | 0.56 | 0.49 | 0.4 | 0.48 | 0.37 |
| 10 | 87157775 | rs7078897 | -0.373 | -0.03 | 0.115 | -1.148 | -0.005 | 0 | 0 | 1.993 | 0.54 | 0.56 | 0.51 | 0.4 | 0.49 | 0.37 |
| 10 | 87167533 | rs11201609 | 0.636 | -0.508 | -0.276 | -1.744 | 0.022 | 0.031 | 0.031 | 0.178 | 0.16 | 0.11 | 0.12 | 0.05 | 0.13 | 0.11 |
| 10 | 87168136 | rs7091087 | 0.12 | -0.311 | 0.364 | -1.929 | 0.034 | -0.027 | -0.027 | 0 | 0.13 | 0.44 | 0.13 | 0.05 | 0.12 | 0.11 |
| 10 | 87173351 | rs11201614 | 0.054 | 0.246 | 0.13 | -2.16 | 0.004 | 0.105 | 0.105 | 0.969 | 0.33 | 0.19 | 0.22 | 0.05 | 0.32 | 0.19 |
| 10 | 87174520 | rs2351347 | 0.187 | 0.012 | 0.329 | -2.271 | -0.002 | -0.034 | -0.034 | 1.724 | 0.8 | 0.72 | 0.85 | 0.94 | 0.81 | 0.9 |
| 10 | 87186017 | rs11201618 | 0.148 | 0.035 | 0.263 | -2.578 | 0.011 | 0.061 | 0.061 | 0.885 | 0.34 | 0.22 | 0.22 | 0.05 | 0.32 | 0.19 |
| 10 | 87189235 | rs5029676 | 1.02 | -0.029 | 0.277 | -2.541 | -0.01 | 0.08 | 0.08 | 0.17 | 0.45 | 0.3 | 0.61 | 0.89 | 0.49 | 0.68 |
| 10 | 87192280 | rs10788445 | 0.06 | 0 | 0.128 | -2.481 | 0.011 | 0.049 | 0.049 | 0.773 | 0.34 | 0.24 | 0.22 | 0.05 | 0.32 | 0.19 |
| 10 | 87192335 | rs17105579 | 0.719 | -0.076 | 0.295 | -2.249 | 0.016 | -0.017 | -0.017 | 0 | 0.11 | 0.24 | 0.12 | 0.05 | 0.12 | 0.07 |
| 10 | 87205613 | rs1912321 | 0.057 | -0.191 | -0.093 | -2.451 | 0.038 | 0.077 | 0.077 | 0 | 0.58 | 0.71 | 0.76 | 0.93 | 0.63 | 0.82 |
| 10 | 87205723 | rs12354870 | 1.43 | -0.368 | -0.246 | -2.558 | -0.037 | 0.103 | 0.103 | 0 | 0.16 | 0.01 | 0.08 | 0 | 0.18 | 0.07 |
| 10 | 87206781 | rs2351560 | -0.117 | -0.088 | -0.05 | -2.405 | 0.053 | 0.103 | 0.103 | 0 | 0.6 | 0.75 | 0.78 | 0.94 | 0.66 | 0.82 |
| 10 | 87211135 | rs7077501 | 0.122 | -0.037 | -0.031 | -2.366 | 0.018 | 0.013 | 0.013 | 0 | 0.57 | 0.61 | 0.75 | 0.93 | 0.62 | 0.81 |
| 10 | 87214408 | rs1912282 | 0.726 | -0.006 | -0.49 | -2.235 | -0.005 | 0.065 | 0.065 | 0 | 0.43 | 0.29 | 0.63 | 0.84 | 0.49 | 0.71 |
| 10 | 87218772 | rs1912278 | -0.282 | -0.029 | -0.69 | -1.802 | 0.028 | 0.078 | 0.078 | 0.236 | 0.8 | 0.92 | 0.92 | 1 | 0.84 | 0.92 |
| 10 | 87232638 | rs11813392 | -0.168 | 0.278 | -0.283 | -2.106 | 0.034 | -0.073 | -0.073 | 0.006 | 0.26 | 0.43 | 0.19 | 0.09 | 0.29 | 0.15 |
| 10 | 87236564 | rs7898877 | -0.073 | 0.273 | -0.211 | -2.109 | 0.029 | -0.06 | -0.06 | 0.688 | 0.25 | 0.37 | 0.19 | 0.09 | 0.3 | 0.13 |
| 10 | 87251652 | rs1949058 | 0.834 | 0.13 | -0.158 | -2.158 | -0.001 | -0.001 | -0.001 | 0.004 | 0.56 | 0.37 | 0.73 | 0.91 | 0.57 | 0.78 |
| 10 | 87251813 | rs7903512 | -0.269 | 0.315 | -0.06 | -2.039 | 0.043 | -0.091 | -0.091 | 0.234 | 0.26 | 0.49 | 0.19 | 0.09 | 0.3 | 0.13 |
| 10 | 87266072 | rs4543926 | -0.043 | -0.283 | -0.488 | -1.982 | 0.054 | 0.034 | 0.034 | 1.014 | 0.06 | 0.35 | 0.04 | 0.02 | 0.05 | 0.06 |
| 10 | 87268005 | rs4074250 | 0.441 | 0.145 | -0.735 | -2.05 | -0.044 | 0.04 | 0.04 | 0 | 0.21 | 0.12 | 0.17 | 0.08 | 0.31 | 0.1 |
| 10 | 87272113 | rs10887490 | 1.04 | 0.243 | -1.177 | -2.037 | -0.065 | -0.001 | -0.001 | 0 | 0.15 | 0.03 | 0.32 | 0.1 | 0.23 | 0.14 |
| 10 | 87274600 | rs2351563 | 0.085 | 0.332 | -0.606 | -1.683 | -0.031 | -0.121 | -0.121 | 0.351 | 0.78 | 0.56 | 0.89 | 0.98 | 0.83 | 0.87 |
| 10 | 87284920 | rs1912308 | 0.217 | 0.74 | -1.572 | -1.389 | -0.047 | 0.089 | 0.089 | 0 | 0.63 | 0.75 | 0.75 | 0.92 | 0.56 | 0.83 |
| 10 | 87285907 | rs7914555 | 0.154 | 0.62 | -1.793 | -0.814 | -0.023 | 0.168 | 0.168 | 0.417 | 0.52 | 0.17 | 0.49 | 0.19 | 0.53 | 0.38 |
| 10 | 87286630 | rs2202926 | -0.964 | 0.449 | -1.629 | -0.265 | 0.003 | 0.028 | 0.028 | 0.045 | 0.28 | 0.12 | 0.42 | 0.78 | 0.29 | 0.56 |
| 10 | 87287808 | rs1912305 | -0.204 | 0.583 | -1.398 | 0.745 | -0.002 | -0.019 | -0.019 | 0 | 0.12 | 0.06 | 0.21 | 0.19 | 0.14 | 0.05 |
| 10 | 87290629 | rs17105668 | -0.542 | 0.472 | -1.29 | 0.932 | 0 | 0.037 | 0.037 | 0 | 0.15 | 0.11 | 0.26 | 0.57 | 0.17 | 0.38 |

| | | | | | | | | | | | | | | | | |
|---|---|---|---|---|---|---|---|---|---|---|---|---|---|---|---|---|
| 10 | 87293229 | rs10887493 | 0.321 | 0.031 | -2.141 | 0.021 | 0.042 | 0.041 | 0.041 | 2.073 | 0.43 | 0.12 | 0.35 | 0.28 | 0.34 | 0.29 |
| 10 | 87295161 | rs12258553 | 1.214 | -0.034 | -2.122 | 0.013 | 0.051 | 0.134 | 0.134 | 2.491 | 0.31 | 0.12 | 0.15 | 0.09 | 0.21 | 0.23 |
| 10 | 87295210 | rs869963 | -0.016 | 0.343 | -1.61 | 1.018 | -0.004 | -0.05 | -0.05 | 0 | 0.13 | 0.02 | 0.19 | 0.43 | 0.13 | 0.28 |
| 10 | 87295805 | rs869466 | 0.07 | 0.662 | -1.538 | 0.607 | -0.021 | 0.017 | 0.017 | 0.064 | 0.09 | 0.01 | 0.08 | 0.05 | 0.08 | 0.06 |
| 10 | 87297497 | rs1912301 | 0.493 | 0.619 | -0.563 | 0.478 | -0.013 | -0.031 | -0.031 | 0 | 0.84 | 0.68 | 0.87 | 0.88 | 0.9 | 0.86 |
| 10 | 87298662 | rs7092582 | -0.271 | 0.75 | -0.503 | 0.52 | 0.002 | 0.017 | 0.017 | 0 | 0.14 | 0.18 | 0.23 | 0.19 | 0.17 | 0.08 |
| 10 | 87300091 | rs1912299 | 0.71 | 0.369 | -0.289 | 0.4 | 0.002 | 0.003 | 0.003 | 0 | 0.84 | 0.84 | 0.82 | 0.88 | 0.89 | 0.84 |
| 10 | 87301191 | rs1912298 | 0.925 | 0.199 | 0.405 | 0.962 | -0.017 | 0.155 | 0.155 | 0 | 0.91 | 0.71 | 0.87 | 0.76 | 0.96 | 0.92 |
| 10 | 87302253 | rs10887497 | 0.906 | 1.107 | 0.384 | 0.972 | 0.032 | 0.008 | 0.008 | 0 | 0.66 | 0.56 | 0.49 | 0.4 | 0.55 | 0.62 |
| 10 | 87302561 | rs1912296 | 0.211 | 1 | 0.495 | 0.949 | -0.02 | -0.028 | -0.028 | 0 | 0.92 | 0.84 | 0.94 | 0.97 | 0.96 | 0.95 |
| 10 | 87308579 | rs17306423 | 0.517 | 1.551 | -0.039 | 0.789 | -0.033 | 0.021 | 0.021 | 0 | 0.08 | 0.01 | 0.07 | 0.05 | 0.09 | 0.14 |
| 10 | 87309413 | rs2136591 | 0.646 | 1.631 | 0.177 | 0.926 | 0.035 | 0.011 | 0.011 | 0 | 0.66 | 0.54 | 0.5 | 0.4 | 0.55 | 0.62 |
| 10 | 87315581 | rs12243608 | -0.084 | 1.658 | 0.488 | 0.906 | 0.084 | 0.053 | 0.053 | 0.189 | 0.08 | 0.23 | 0.1 | 0.09 | 0.14 | 0.04 |
| 10 | 87316082 | rs10887503 | -0.294 | 1.406 | 0.166 | 0.965 | -0.049 | -0.028 | -0.028 | 0 | 0.33 | 0.16 | 0.45 | 0.35 | 0.49 | 0.38 |
| 10 | 87320674 | rs1876886 | -1.484 | 1.027 | 0.694 | 0.983 | 0.019 | 0.016 | 0.016 | 0.653 | 0.47 | 0.58 | 0.63 | 0.64 | 0.65 | 0.52 |
| 10 | 87330756 | rs2664403 | 1.262 | 0.889 | 0.851 | 0.906 | 0.073 | 0.06 | 0.06 | 0.489 | 0.51 | 0.34 | 0.36 | 0.33 | 0.35 | 0.44 |
| 10 | 87332946 | rs2664402 | -0.054 | 0.932 | 0.693 | 0.964 | -0.071 | -0.04 | -0.04 | 0 | 0.66 | 0.89 | 0.55 | 0.66 | 0.51 | 0.62 |
| 10 | 87333634 | rs10887504 | -0.075 | 0.94 | 0.73 | 0.948 | -0.045 | -0.033 | -0.033 | 0.037 | 0.66 | 0.84 | 0.47 | 0.44 | 0.51 | 0.59 |
| 10 | 87334017 | rs2136590 | 0.554 | 0.747 | 0.909 | 0.888 | 0.017 | -0.019 | -0.019 | 0.265 | 0.08 | 0.11 | 0.09 | 0.02 | 0.13 | 0.04 |
| 10 | 87338403 | rs2616668 | 0.4 | 1.891 | 0.832 | 0.85 | -0.007 | 0.008 | 0.008 | 0.015 | 0.6 | 0.45 | 0.46 | 0.53 | 0.41 | 0.49 |
| 10 | 87341247 | rs1912274 | 1.67 | 0.183 | 0.141 | 0.243 | 0.069 | 0.115 | 0.115 | 0 | 0.36 | 0.15 | 0.25 | 0.14 | 0.25 | 0.27 |
| 10 | 87357738 | rs2175207 | 0.478 | 0.53 | 0.499 | 0.32 | 0.055 | -0.034 | -0.034 | 0.094 | 0.11 | 0.25 | 0.1 | 0.05 | 0.15 | 0.07 |
| 10 | 87358975 | rs1912329 | 0.862 | 0.425 | 0.568 | 0.298 | 0.05 | 0.04 | 0.04 | 0.288 | 0.55 | 0.34 | 0.35 | 0.34 | 0.35 | 0.46 |
| 10 | 87359826 | rs2664405 | -1.153 | 0.473 | 0.543 | 0.353 | -0.064 | -0.043 | -0.043 | 0 | 0.73 | 0.85 | 0.56 | 0.6 | 0.57 | 0.57 |
| 10 | 87360723 | rs2664404 | -1.153 | 0.473 | 0.543 | 0.353 | -0.064 | -0.043 | -0.043 | 0 | 0.73 | 0.85 | 0.56 | 0.6 | 0.57 | 0.57 |
| 10 | 87361291 | rs3812651 | -0.687 | 0.56 | 0.75 | 0.447 | 0.029 | 0.155 | 0.155 | 0 | 0.07 | 0.24 | 0.12 | 0.21 | 0.07 | 0.05 |
| 10 | 87365097 | rs2280011 | -1.262 | 0.564 | 0.7 | 0.419 | 0.055 | 0.089 | 0.089 | 0.138 | 0.18 | 0.5 | 0.22 | 0.27 | 0.22 | 0.12 |
| 10 | 87367366 | rs2616673 | -2.133 | 1.309 | 0.347 | 0.304 | -0.054 | -0.054 | -0.054 | 0 | 0.89 | 1 | 0.87 | 0.85 | 0.86 | 0.8 |
| 10 | 87367395 | rs7907900 | 0.279 | -0.171 | 0.715 | -0.39 | 0.079 | -0.028 | -0.028 | 0 | 0.11 | 0.33 | 0.1 | 0.05 | 0.15 | 0.07 |
| 10 | 87367922 | rs2664400 | 0.548 | -0.375 | 0.328 | -0.458 | -0.032 | -0.036 | -0.036 | 4.142 | 0.08 | 0.01 | 0.13 | 0.12 | 0.08 | 0.1 |
| 10 | 87369264 | rs17105725 | 0.541 | -0.596 | 0.5 | -0.54 | -0.007 | -0.001 | -0.001 | 1.163 | 0.2 | 0.23 | 0.11 | 0.16 | 0.12 | 0.19 |
| 10 | 87379081 | rs1912276 | 0.203 | -0.256 | 0.739 | -0.378 | 0.067 | -0.012 | -0.012 | 0 | 0.09 | 0.26 | 0.09 | 0.04 | 0.14 | 0.03 |
| 10 | 87379811 | rs2292310 | -0.608 | -0.212 | 0.451 | -0.263 | 0.01 | 0.012 | 0.012 | 0.045 | 0.05 | 0.21 | 0.04 | 0.05 | 0.04 | 0.05 |
| 10 | 87406340 | rs7100018 | -0.83 | -0.481 | -0.34 | -0.411 | -0.064 | 0.013 | 0.013 | 0 | 0.64 | 0.85 | 0.45 | 0.35 | 0.48 | 0.49 |
| 10 | 87418456 | rs10788462 | -1.4 | -1.08 | -0.317 | -1.198 | -0.064 | -0.032 | -0.032 | 0 | 0.69 | 0.85 | 0.5 | 0.39 | 0.58 | 0.55 |
| 10 | 87426591 | rs11201725 | 1.693 | -1.809 | -0.663 | -1.639 | 0.062 | 0.083 | 0.083 | 0 | 0.23 | 0.1 | 0.11 | 0 | 0.14 | 0.17 |
| 10 | 87429714 | rs7080323 | 0.11 | -1.81 | -0.197 | -1.657 | 0.04 | -0.013 | -0.013 | 0 | 0.05 | 0.18 | 0.04 | 0.01 | 0.04 | 0.03 |
| 10 | 87430121 | rs7097285 | 0.479 | -2.03 | 0.147 | -1.754 | 0.037 | 0.038 | 0.038 | 0 | 0.39 | 0.34 | 0.29 | 0.2 | 0.22 | 0.32 |
| 10 | 87431548 | rs10887514 | -0.357 | -2.106 | -0.11 | -1.823 | -0.001 | -0.01 | -0.01 | 0 | 0.49 | 0.52 | 0.33 | 0.21 | 0.27 | 0.36 |
| 10 | 87436348 | rs12255557 | 1.53 | -2.053 | -0.233 | -1.739 | 0.067 | 0.083 | 0.083 | 0 | 0.26 | 0.12 | 0.2 | 0.1 | 0.15 | 0.27 |
| 10 | 87448297 | rs2351822 | -0.222 | -1.865 | 0.127 | -1.642 | 0 | -0.006 | -0.006 | 0 | 0.53 | 0.48 | 0.44 | 0.2 | 0.51 | 0.37 |
| 10 | 87455683 | rs4488140 | -0.801 | -1.845 | -0.031 | -1.634 | -0.003 | 0.004 | 0.004 | 0 | 0.59 | 0.64 | 0.47 | 0.21 | 0.55 | 0.4 |
| 10 | 87460984 | rs10749528 | -0.792 | -1.858 | -0.021 | -1.621 | -0.04 | 0.089 | 0.089 | 0 | 0.58 | 0.79 | 0.47 | 0.25 | 0.53 | 0.41 |
| 10 | 87481980 | rs4418740 | -0.375 | -1.803 | 0.12 | -1.627 | -0.001 | -0.012 | -0.012 | 0 | 0.54 | 0.48 | 0.45 | 0.2 | 0.52 | 0.42 |
| 10 | 87485417 | rs10159528 | -1.817 | -1.812 | -0.086 | -1.565 | -0.076 | -0.054 | -0.054 | 0.149 | 0.7 | 0.88 | 0.51 | 0.4 | 0.59 | 0.6 |
| 10 | 87487366 | rs11201754 | 1.627 | -2.586 | 0.374 | -1.583 | 0.055 | 0.097 | 0.097 | 0 | 0.23 | 0.08 | 0.1 | 0.01 | 0.14 | 0.15 |
| 10 | 87490293 | rs7898626 | -2.052 | -2.628 | 0.377 | -1.654 | -0.096 | -0.031 | -0.031 | 0 | 0.7 | 0.87 | 0.48 | 0.39 | 0.52 | 0.59 |
| 10 | 87495534 | rs12773880 | 0.178 | -2.32 | 0.718 | -1.618 | -0.032 | -0.027 | -0.027 | 0 | 0.14 | 0.23 | 0.09 | 0.09 | 0.06 | 0.07 |
| 10 | 87496407 | rs10887523 | -1.603 | -2.253 | 0.787 | -1.584 | -0.069 | 0.075 | 0.075 | 0.02 | 0.63 | 0.83 | 0.47 | 0.25 | 0.51 | 0.48 |
| 10 | 87501765 | rs12098525 | 0.852 | -2.251 | 0.845 | -1.591 | 0.005 | -0.034 | -0.034 | 0 | 0.28 | 0.28 | 0.14 | 0.01 | 0.18 | 0.19 |
| 10 | 87503021 | rs12763937 | 0.796 | -2.215 | 0.822 | -1.608 | 0.025 | -0.026 | -0.026 | 0 | 0.06 | 0.1 | 0.08 | 0.01 | 0.14 | 0.07 |
| 10 | 87503381 | rs10887525 | 0.808 | -2.302 | 0.843 | -1.536 | 0.004 | -0.037 | -0.037 | 1.006 | 0.28 | 0.28 | 0.14 | 0.01 | 0.18 | 0.18 |
| 10 | 87510283 | rs3011698 | 1.274 | -2.288 | 0.784 | -1.528 | -0.059 | 0.068 | 0.068 | 0.004 | 0.37 | 0.21 | 0.53 | 0.75 | 0.49 | 0.51 |
| 10 | 87514045 | rs1917150 | 0.469 | -2.249 | 0.782 | -1.501 | -0.001 | -0.028 | -0.028 | 0 | 0.69 | 0.62 | 0.68 | 0.81 | 0.69 | 0.74 |
| 10 | 87517898 | rs3011701 | 0.444 | -2.177 | 0.813 | -1.764 | 0 | -0.075 | -0.075 | 0 | 0.69 | 0.57 | 0.75 | 0.87 | 0.69 | 0.79 |
| 10 | 87522183 | rs3011704 | 1.302 | -2.171 | 0.69 | -1.762 | -0.067 | 0.075 | 0.075 | 0.059 | 0.37 | 0.17 | 0.53 | 0.75 | 0.49 | 0.51 |
| 10 | 87522371 | rs7091796 | 1.062 | -2.162 | 0.759 | -1.755 | -0.008 | -0.01 | -0.01 | 0 | 0.27 | 0.27 | 0.19 | 0.12 | 0.16 | 0.24 |
| 10 | 87522509 | rs2140661 | 1.012 | -2.121 | 0.603 | -1.735 | -0.055 | 0.081 | 0.081 | 0 | 0.39 | 0.23 | 0.53 | 0.75 | 0.49 | 0.52 |
| 10 | 87536809 | rs1917154 | -0.178 | -1.478 | 1.15 | -1.212 | -0.013 | 0.009 | 0.009 | 0 | 0.82 | 0.88 | 0.85 | 0.96 | 0.75 | 0.85 |
| 10 | 87537219 | rs1880389 | 0.848 | -1.37 | 0.769 | -1.119 | 0.006 | -0.058 | -0.058 | 0.172 | 0.66 | 0.6 | 0.74 | 0.87 | 0.65 | 0.76 |
| 10 | 87553409 | rs1106789 | -0.345 | -2.476 | 0.069 | -0.755 | -0.058 | 0.012 | 0.012 | 0 | 0.3 | 0.48 | 0.26 | 0.33 | 0.18 | 0.34 |
| 10 | 87562775 | rs12245530 | -1.258 | -2.395 | 0.132 | -0.691 | 0.003 | 0.276 | 0.276 | 1.195 | 0.08 | 0.36 | 0.14 | 0.3 | 0.04 | 0.16 |
| 10 | 87592785 | rs947594 | 0.407 | -2.463 | -0.211 | -0.83 | 0.008 | 0.006 | 0.006 | 0 | 0.11 | 0.03 | 0.07 | 0.08 | 0.07 | 0.07 |
| 10 | 87606617 | rs11818522 | -0.822 | -2.441 | -0.048 | -0.837 | -0.034 | 0.001 | 0.001 | 0 | 0.4 | 0.51 | 0.34 | 0.42 | 0.25 | 0.4 |
| 10 | 87611943 | rs1880386 | 0.211 | -2.015 | -0.428 | -0.954 | 0.003 | -0.022 | -0.022 | 2.71 | 0.22 | 0.18 | 0.14 | 0.01 | 0.15 | 0.14 |
| 10 | 87615322 | rs1880385 | 1.725 | -2.18 | -0.762 | -1.078 | -0.076 | -0.096 | -0.096 | 0 | 0.38 | 0.1 | 0.52 | 0.52 | 0.54 | 0.49 |
| 10 | 87625182 | rs7090378 | -1.105 | -2.089 | -0.534 | -1.16 | 0.022 | 0.026 | 0.026 | 0 | 0.35 | 0.29 | 0.21 | 0.19 | 0.23 | 0.24 |
| 10 | 87631521 | rs17401213 | 1.025 | -2.127 | -0.56 | -1.198 | 0.044 | 0.069 | 0.069 | 0 | 0.16 | 0.05 | 0.08 | 0.01 | 0.09 | 0.12 |
| 10 | 87632795 | rs7099032 | -0.543 | -1.709 | -0.341 | -0.999 | 0.009 | 0.01 | 0.01 | 0 | 0.36 | 0.35 | 0.21 | 0.19 | 0.23 | 0.24 |

| | | | | | | | | | | | | | | | | |
|---|---|---|---|---|---|---|---|---|---|---|---|---|---|---|---|---|
| 10 | 87634533 | rs4325261 | 0.531 | -1.625 | -0.383 | -1.12 | -0.003 | -0.004 | -0.004 | 0 | 0.05 | 0.06 | 0.05 | 0 | 0.06 | 0.02 |
| 10 | 87651609 | rs7074854 | -0.552 | -1.643 | -0.381 | -1.106 | 0.022 | 0.028 | 0.028 | 0 | 0.35 | 0.3 | 0.21 | 0.16 | 0.23 | 0.24 |
| 10 | 87660650 | rs11815416 | -0.816 | -1.353 | 0.079 | -1.521 | 0.044 | 0.055 | 0.055 | 0.453 | 0.5 | 0.55 | 0.31 | 0.19 | 0.33 | 0.33 |
| 10 | 87662864 | rs10887536 | -0.699 | -1.283 | -0.377 | -1.492 | 0.038 | 0.045 | 0.045 | 0.233 | 0.5 | 0.55 | 0.31 | 0.19 | 0.33 | 0.34 |
| 10 | 87663255 | rs12246799 | 0.41 | -1.269 | -0.835 | -1.503 | 0.002 | -0.01 | -0.01 | 0.099 | 0.09 | 0.05 | 0.04 | 0 | 0.05 | 0.02 |
| 10 | 87666750 | rs2352097 | 0.23 | -1.152 | -1.155 | -1.436 | 0.072 | 0.146 | 0.146 | 0.013 | 0.34 | 0.12 | 0.19 | 0.01 | 0.26 | 0.23 |
| 10 | 87672771 | rs25434 | -0.06 | 0.047 | -1.303 | -0.756 | 0.05 | 0.066 | 0.066 | 0.185 | 0.52 | 0.6 | 0.31 | 0.19 | 0.33 | 0.36 |
| 10 | 87674655 | rs1997001 | 0.785 | 1.111 | -1.298 | 0.067 | -0.028 | 0.023 | 0.023 | 0 | 0.93 | 0.99 | 0.87 | 0.95 | 0.89 | 0.85 |
| 10 | 87678539 | rs7073759 | -1 | 0.704 | -1.312 | 0.012 | 0.004 | 0.037 | 0.037 | 0 | 0.13 | 0.06 | 0.08 | 0.02 | 0.12 | 0.09 |
| 10 | 87683328 | rs7895100 | -0.875 | 0.578 | -0.821 | 0.431 | -0.004 | 0.036 | 0.036 | 0 | 0.12 | 0.21 | 0.14 | 0.19 | 0.13 | 0.07 |
| 10 | 87685741 | rs7100625 | -0.771 | 0.605 | -1.36 | 0.476 | -0.015 | 0.002 | 0.002 | 0 | 0.53 | 0.74 | 0.47 | 0.62 | 0.53 | 0.51 |
| 10 | 87688537 | rs17105979 | -0.682 | 0.367 | -0.702 | 0.306 | -0.003 | 0.05 | 0.05 | 0 | 0.12 | 0.39 | 0.16 | 0.12 | 0.08 | 0.07 |
| 10 | 87694970 | rs7912519 | -0.812 | 0.729 | -1.509 | 0.241 | -0.05 | 0.014 | 0.014 | 0 | 0.8 | 0.92 | 0.84 | 0.85 | 0.72 | 0.77 |
| 10 | 87699269 | rs11201840 | -0.176 | 0.081 | -1.985 | 0.247 | -0.004 | 0.026 | 0.026 | 0.061 | 0.45 | 0.24 | 0.57 | 0.66 | 0.47 | 0.52 |
| 10 | 87700736 | rs17105990 | 1.139 | -0.314 | -2.201 | 0.313 | 0.013 | -0.029 | -0.029 | 0 | 0.11 | 0.02 | 0.06 | 0.16 | 0.11 | 0.06 |
| 10 | 87700825 | rs1418396 | 0.394 | -1.34 | -2.565 | 0.111 | 0.01 | -0.006 | -0.006 | 0 | 0.32 | 0.28 | 0.39 | 0.33 | 0.2 | 0.38 |
| 10 | 87701479 | rs7899924 | 0.776 | -1.774 | -2.732 | 0.249 | 0.012 | 0.011 | 0.011 | 0 | 0.26 | 0.21 | 0.24 | 0.16 | 0.18 | 0.34 |
| 10 | 87701735 | rs7899222 | 0.751 | -1.774 | -2.731 | 0.249 | 0.012 | 0.01 | 0.01 | 0 | 0.26 | 0.23 | 0.25 | 0.16 | 0.18 | 0.34 |
| 10 | 87701994 | rs7096816 | -0.317 | -1.578 | -2.518 | 0.31 | 0.041 | -0.002 | -0.002 | 0 | 0.43 | 0.33 | 0.46 | 0.49 | 0.31 | 0.43 |
| 10 | 87704350 | rs7069313 | -1.141 | -1.311 | -1.992 | -0.025 | -0.001 | -0.001 | -0.001 | 0 | 0.67 | 0.66 | 0.65 | 0.51 | 0.56 | 0.59 |
| 10 | 87704966 | rs7089160 | -1.071 | -1.316 | -1.977 | -0.038 | -0.01 | -0.009 | -0.009 | 0 | 0.66 | 0.68 | 0.62 | 0.5 | 0.55 | 0.53 |
| 10 | 87708110 | rs11201843 | 2.033 | -1.17 | -1.866 | 0.074 | 0.008 | -0.001 | -0.001 | 0.414 | 0.06 | 0.03 | 0.14 | 0.17 | 0.02 | 0.04 |
| 10 | 87711856 | rs10887538 | -1.628 | -1.116 | -1.413 | -0.042 | 0.032 | 0.021 | 0.021 | 0.248 | 0.74 | 0.74 | 0.68 | 0.53 | 0.62 | 0.6 |
| 10 | 87713420 | rs7090885 | -0.62 | -1.459 | -1.568 | -0.077 | 0 | -0.008 | -0.008 | 0 | 0.61 | 0.75 | 0.62 | 0.5 | 0.45 | 0.58 |
| 10 | 87713604 | rs7094476 | 0.202 | -2.2 | -0.984 | 0.036 | -0.03 | 0.016 | 0.016 | 0.031 | 0.21 | 0.36 | 0.32 | 0.19 | 0.12 | 0.27 |
| 10 | 87715434 | rs11592936 | 0.901 | -2.33 | -0.963 | 0.055 | 0.033 | -0.04 | -0.04 | 7.797 | 0.17 | 0.07 | 0.26 | 0.18 | 0.11 | 0.2 |
| 10 | 87717641 | rs1986508 | 1.077 | -2.474 | -0.877 | 0.027 | -0.045 | 0.047 | 0.047 | 0.082 | 0.09 | 0.32 | 0.11 | 0.12 | 0.02 | 0.04 |
| 10 | 87720085 | rs12765721 | 1.355 | -2.443 | -1.138 | -0.092 | -0.029 | 0.072 | 0.072 | 0 | 0.14 | 0.02 | 0.09 | 0.01 | 0.18 | 0.08 |
| 10 | 87722008 | rs12220682 | 2.668 | -2.692 | -0.784 | 0.194 | 0.026 | -0.028 | -0.028 | 0 | 0.06 | 0.01 | 0.08 | 0.12 | 0.02 | 0.04 |
| 10 | 87722991 | rs11201853 | 0.622 | -2.507 | -0.54 | 0.083 | 0.003 | 0.003 | 0.003 | 0 | 0.27 | 0.28 | 0.16 | 0.18 | 0.24 | 0.25 |
| 10 | 87723254 | rs7914815 | 0.716 | -1.797 | -0.249 | 0.239 | 0.013 | -0.013 | -0.013 | 0 | 0.17 | 0.12 | 0.26 | 0.18 | 0.11 | 0.2 |
| 10 | 87736238 | rs7094074 | 1.966 | -1.639 | -1.238 | 0.268 | -0.003 | -0.017 | -0.017 | 0 | 0.08 | 0.13 | 0.06 | 0.02 | 0.09 | 0.09 |
| 10 | 87737195 | rs2001417 | 0.455 | -1.51 | -1.317 | 0.544 | 0.032 | -0.033 | -0.033 | 0 | 0.17 | 0.09 | 0.26 | 0.18 | 0.11 | 0.2 |
| 10 | 87739205 | rs4448624 | 0.686 | -0.969 | -1.361 | 0.362 | -0.073 | -0.027 | -0.027 | 0.524 | 0.39 | 0.13 | 0.46 | 0.62 | 0.43 | 0.58 |
| 10 | 87742595 | rs4244970 | 1.315 | -1.507 | -0.957 | 0.388 | -0.021 | 0.004 | 0.004 | 0.725 | 0.36 | 0.24 | 0.38 | 0.37 | 0.46 | 0.54 |
| 10 | 87744398 | rs4393254 | 2.185 | -1.667 | -1.07 | 0.147 | -0.003 | 0.017 | 0.017 | 0 | 0.07 | 0.03 | 0.04 | 0 | 0.09 | 0.03 |
| 10 | 87745368 | rs7100188 | -0.813 | -2.085 | -0.421 | 0.026 | 0.011 | -0.001 | -0.001 | 0 | 0.64 | 0.66 | 0.65 | 0.67 | 0.68 | 0.81 |
| 10 | 87747789 | rs2352464 | -0.878 | -2.438 | -0.15 | -0.003 | -0.001 | -0.002 | -0.002 | 0.045 | 0.69 | 0.65 | 0.69 | 0.82 | 0.75 | 0.86 |
| 10 | 87749276 | rs12784735 | 0.692 | -2.425 | -0.184 | -0.053 | 0.003 | 0.016 | 0.016 | 0 | 0.21 | 0.09 | 0.16 | 0.15 | 0.26 | 0.11 |
| 10 | 87750258 | rs17318292 | 1.133 | -2.459 | -0.105 | -0.09 | -0.002 | -0.004 | -0.004 | 0 | 0.47 | 0.46 | 0.51 | 0.67 | 0.49 | 0.75 |
| 10 | 87751705 | rs10887540 | -0.861 | -2.577 | -0.128 | -0.015 | -0.008 | -0.022 | -0.022 | 1.104 | 0.69 | 0.59 | 0.68 | 0.83 | 0.74 | 0.88 |
| 10 | 87754343 | rs12773130 | 1.357 | -2.537 | -0.041 | -0.018 | 0.02 | 0.002 | 0.002 | 0 | 0.17 | 0.09 | 0.09 | 0.15 | 0.17 | 0.08 |
| 10 | 87754821 | rs12779841 | 1.41 | -2.537 | -0.053 | -0.023 | 0.019 | -0.021 | -0.021 | 1.901 | 0.16 | 0.07 | 0.09 | 0.17 | 0.17 | 0.1 |
| 10 | 87755410 | rs4478924 | -1.239 | -2.447 | -0.038 | -0.047 | 0 | -0.002 | -0.002 | 0 | 0.73 | 0.68 | 0.69 | 0.84 | 0.76 | 0.88 |
| 10 | 87761389 | rs4300344 | 2.3 | -2.461 | -0.125 | 0.008 | -0.043 | -0.012 | -0.012 | 0.285 | 0.31 | 0.18 | 0.3 | 0.36 | 0.37 | 0.52 |
| 10 | 87764778 | rs10887544 | -1.185 | -2.006 | -0.055 | -0.275 | 0.004 | 0.014 | 0.014 | 0.201 | 0.74 | 0.75 | 0.7 | 0.84 | 0.76 | 0.88 |
| 10 | 87767044 | rs12776960 | 2.309 | -1.593 | 0.046 | -0.234 | -0.004 | 0 | 0 | 0 | 0.06 | 0.05 | 0.04 | 0 | 0.09 | 0.03 |
| 10 | 87768614 | rs11201870 | -0.138 | -1.66 | 0.242 | -0.227 | 0.033 | -0.036 | -0.036 | 0 | 0.16 | 0.51 | 0.16 | 0.07 | 0.19 | 0.09 |
| 10 | 87769893 | rs7090075 | 0.841 | -1.986 | 0.046 | -0.116 | 0.021 | -0.011 | -0.011 | 0.526 | 0.45 | 0.67 | 0.52 | 0.46 | 0.54 | 0.57 |
| 10 | 87772138 | rs6585992 | -0.337 | -2.103 | -0.083 | -0.14 | 0.017 | 0.028 | 0.028 | 0.009 | 0.6 | 0.81 | 0.59 | 0.62 | 0.64 | 0.65 |
| 10 | 87773471 | rs10749532 | 1.753 | -1.944 | -0.056 | -0.181 | -0.028 | -0.017 | -0.017 | 0.347 | 0.35 | 0.28 | 0.44 | 0.38 | 0.43 | 0.51 |
| 10 | 87774397 | rs10887549 | 2.322 | -1.872 | -0.186 | -0.128 | -0.046 | -0.045 | -0.045 | 0.009 | 0.3 | 0.15 | 0.36 | 0.38 | 0.35 | 0.48 |
| 10 | 87777269 | rs17106160 | 2.631 | -1.588 | -0.053 | -0.29 | -0.019 | -0.027 | -0.027 | 0.57 | 0.06 | 0.09 | 0.01 | 0 | 0.02 | 0 |
| 10 | 87777426 | rs12762291 | 1.461 | -1.501 | -0.095 | -0.201 | 0.007 | -0.031 | -0.031 | 0 | 0.08 | 0.13 | 0.06 | 0 | 0.1 | 0.03 |
| 10 | 87785861 | rs17106172 | 1.954 | -1.557 | -0.339 | -0.061 | 0.008 | -0.009 | -0.009 | 1.34 | 0.08 | 0.05 | 0.05 | 0.16 | 0.07 | 0.07 |
| 10 | 87790138 | rs10788477 | -0.938 | -1.596 | -0.434 | -0.51 | -0.005 | 0.031 | 0.031 | 1.148 | 0.77 | 0.86 | 0.71 | 0.84 | 0.76 | 0.91 |
| 10 | 87795057 | rs17106185 | 0.811 | -1.38 | -0.325 | -0.56 | 0.01 | -0.032 | -0.032 | 1.384 | 0.07 | 0.18 | 0.07 | 0.02 | 0.09 | 0.03 |
| 10 | 87797251 | rs12783158 | 1.518 | -1.499 | -0.424 | -0.544 | -0.001 | -0.009 | -0.009 | 0.035 | 0.07 | 0.07 | 0.06 | 0 | 0.12 | 0.02 |
| 10 | 87799641 | rs1806292 | 1.362 | -1.608 | -0.373 | -0.518 | -0.008 | 0.003 | 0.003 | 0 | 0.08 | 0.12 | 0.07 | 0.13 | 0.06 | 0.07 |
| 10 | 87806564 | rs9421508 | 1.866 | -1.525 | -0.436 | -0.529 | 0.001 | -0.012 | -0.012 | 0.241 | 0.07 | 0.07 | 0.05 | 0 | 0.11 | 0.02 |
| 10 | 87807390 | rs10788478 | 2.442 | -1.716 | -0.725 | -0.466 | -0.066 | -0.051 | -0.051 | 1.267 | 0.3 | 0.12 | 0.35 | 0.41 | 0.37 | 0.49 |
| 10 | 87816720 | rs2003112 | 1.226 | -1.476 | -0.298 | -0.757 | 0.002 | -0.025 | -0.025 | 0 | 0.06 | 0.13 | 0.06 | 0.02 | 0.07 | 0.03 |
| 10 | 87816757 | rs2003111 | -0.04 | -1.602 | -0.742 | -0.797 | 0.013 | 0.034 | 0.034 | 3.639 | 0.7 | 0.77 | 0.69 | 0.87 | 0.74 | 0.85 |
| 10 | 87820706 | rs4934146 | -1.205 | -1.232 | -1.514 | -0.71 | -0.032 | 0.036 | 0.036 | 0 | 0.92 | 0.98 | 0.86 | 1 | 0.85 | 0.92 |
| 10 | 87823099 | rs12266132 | 1.586 | -1.134 | -1.363 | -0.561 | -0.006 | 0.006 | 0.006 | 0 | 0.07 | 0.09 | 0.06 | 0.13 | 0.05 | 0.07 |
| 10 | 87824238 | rs10887554 | 0.993 | -0.95 | -1.589 | -0.473 | 0.015 | 0.081 | 0.081 | 2.542 | 0.47 | 0.22 | 0.5 | 0.71 | 0.51 | 0.72 |
| 10 | 87825498 | rs2001697 | 0.62 | -0.169 | -1.131 | -0.267 | -0.022 | 0.044 | 0.044 | 1.301 | 0.77 | 0.92 | 0.7 | 0.87 | 0.75 | 0.87 |
| 10 | 87840814 | rs4934149 | 0.968 | -0.859 | -0.438 | -0.258 | -0.049 | -0.048 | -0.048 | 0 | 0.69 | 0.55 | 0.72 | 0.86 | 0.74 | 0.68 |
| 10 | 87843398 | rs7909669 | -0.639 | -0.695 | -0.138 | 0.061 | -0.027 | 0.019 | 0.019 | 1.477 | 0.11 | 0.2 | 0.08 | 0.12 | 0.08 | 0.2 |

| | | | | | | | | | | | | | | | | |
|---|---|---|---|---|---|---|---|---|---|---|---|---|---|---|---|---|
| 10 | 87845206 | rs2352178 | -1.411 | -0.65 | -0.514 | 0.117 | -0.014 | 0.021 | 0.021 | 0.334 | 0.24 | 0.31 | 0.21 | 0.35 | 0.19 | 0.38 |
| 10 | 87845548 | rs2352173 | 0.784 | -0.302 | -1.36 | -0.25 | 0 | -0.007 | -0.007 | 0.028 | 0.51 | 0.15 | 0.49 | 0.57 | 0.57 | 0.43 |
| 10 | 87845799 | rs1806590 | -1.606 | -0.296 | -0.078 | -0.411 | 0.004 | 0.003 | 0.003 | 0.57 | 0.33 | 0.61 | 0.25 | 0.25 | 0.24 | 0.36 |
| 10 | 87848346 | rs2352462 | -1.863 | -0.069 | -0.132 | -0.549 | 0.012 | 0.068 | 0.068 | 0 | 0.51 | 0.78 | 0.55 | 0.32 | 0.44 | 0.48 |
| 10 | 87849054 | rs2352461 | -0.678 | -0.531 | 0.521 | -0.606 | -0.007 | 0.009 | 0.009 | 0 | 0.11 | 0.35 | 0.08 | 0.06 | 0.08 | 0.12 |
| 10 | 87857868 | rs7923926 | -1.956 | -0.335 | 0.207 | -0.646 | 0.042 | 0.049 | 0.049 | 0 | 0.47 | 0.74 | 0.51 | 0.33 | 0.36 | 0.47 |
| 10 | 87860250 | rs11201884 | -0.818 | -0.157 | 0.563 | -0.678 | 0.001 | -0.027 | -0.027 | 1.582 | 0.15 | 0.19 | 0.09 | 0.01 | 0.16 | 0.12 |
| 10 | 87861382 | rs7918518 | -1.954 | -0.277 | 0.232 | -0.655 | -0.001 | -0.003 | -0.003 | 0 | 0.33 | 0.69 | 0.19 | 0.18 | 0.25 | 0.34 |
| 10 | 87864802 | rs11593396 | -0.885 | -0.598 | -0.172 | -0.516 | 0.03 | -0.008 | -0.008 | 0 | 0.14 | 0.03 | 0.11 | 0.15 | 0.06 | 0.21 |
| 10 | 87869651 | rs7088100 | 2.194 | -0.421 | -0.178 | -0.643 | 0.03 | 0.092 | 0.092 | 0.025 | 0.54 | 0.16 | 0.65 | 0.69 | 0.65 | 0.53 |
| 10 | 87874877 | rs4933383 | 0.498 | -0.262 | -0.089 | -0.554 | 0.008 | 0.063 | 0.063 | 0.014 | 0.87 | 0.94 | 0.94 | 1 | 0.91 | 0.93 |
| 10 | 87880442 | rs4934154 | 0.952 | -0.254 | 0.314 | -0.534 | -0.02 | -0.022 | -0.022 | 2.972 | 0.83 | 0.69 | 0.91 | 0.91 | 0.9 | 0.89 |
| 10 | 87882916 | rs11595656 | -0.886 | 0.015 | 0.278 | -0.32 | -0.055 | 0.013 | 0.013 | 0 | 0.09 | 0.01 | 0.12 | 0.07 | 0.19 | 0.15 |
| 10 | 87890326 | rs5003944 | -1.851 | -1.555 | 0.531 | -0.469 | -0.042 | -0.045 | -0.045 | 0.335 | 0.38 | 0.49 | 0.27 | 0.2 | 0.21 | 0.42 |
| 10 | 87891912 | rs4132195 | -1.637 | -1.666 | 0.606 | -0.395 | -0.068 | -0.052 | -0.052 | 0 | 0.3 | 0.45 | 0.25 | 0.19 | 0.17 | 0.37 |
| 10 | 87894288 | rs884700 | -1.894 | -1.807 | 1.168 | -0.492 | -0.019 | -0.02 | -0.02 | 0 | 0.33 | 0.6 | 0.33 | 0.23 | 0.23 | 0.32 |
| 10 | 87896157 | rs11816291 | -0.9 | -1.781 | 1.131 | -0.532 | -0.025 | -0.05 | -0.05 | 0.064 | 0.22 | 0.26 | 0.17 | 0.09 | 0.15 | 0.21 |
| 10 | 87897272 | rs17106311 | -1.602 | -1.789 | 1.113 | -0.54 | -0.05 | -0.104 | -0.104 | 0 | 0.31 | 0.4 | 0.2 | 0.09 | 0.2 | 0.26 |
| 10 | 87898729 | rs17106320 | -0.898 | -1.783 | 0.993 | -0.536 | -0.024 | -0.048 | -0.048 | 0 | 0.22 | 0.26 | 0.17 | 0.09 | 0.15 | 0.21 |
| 10 | 87900590 | rs11201896 | -0.16 | -1.768 | 0.924 | -0.52 | 0.012 | 0.02 | 0.02 | 0 | 0.07 | 0.05 | 0.02 | 0 | 0.04 | 0.05 |
| 10 | 87908652 | rs11201909 | -1.248 | -1.827 | 1.008 | -0.524 | -0.014 | -0.012 | -0.012 | 0 | 0.23 | 0.36 | 0.25 | 0.2 | 0.17 | 0.21 |
| 10 | 87909947 | rs7921149 | -2.806 | -1.893 | 0.946 | -0.54 | 0.134 | 0.095 | 0.095 | 0 | 0.49 | 0.85 | 0.44 | 0.33 | 0.28 | 0.54 |
| 10 | 87911573 | rs11599307 | 0.09 | -1.948 | 0.878 | -0.912 | 0.013 | 0.048 | 0.048 | 0 | 0.14 | 0.04 | 0.09 | 0.06 | 0.1 | 0.09 |
| 10 | 87915809 | rs7920781 | -1.677 | -1.885 | 0.967 | -0.83 | -0.082 | -0.017 | -0.017 | 0 | 0.18 | 0.44 | 0.16 | 0.13 | 0.07 | 0.29 |
| 10 | 87919323 | rs10887560 | -2.129 | -1.677 | 0.438 | -0.76 | 0.087 | 0.112 | 0.112 | 0 | 0.45 | 0.77 | 0.31 | 0.19 | 0.24 | 0.46 |
| 10 | 87923404 | rs7912161 | -2.032 | -1.277 | 0.385 | -0.163 | -0.078 | -0.052 | -0.052 | 4.103 | 0.25 | 0.41 | 0.18 | 0.13 | 0.12 | 0.33 |
| 10 | 87924667 | rs4644592 | -1.063 | -1.211 | 0.297 | -0.133 | -0.026 | 0.026 | 0.026 | 0.282 | 0.08 | 0.17 | 0.08 | 0.1 | 0.01 | 0.17 |
| 10 | 87929082 | rs12262754 | -1.936 | -1.148 | -0.075 | -0.107 | -0.064 | -0.042 | -0.042 | 0 | 0.25 | 0.36 | 0.18 | 0.13 | 0.11 | 0.34 |
| 10 | 87932817 | rs11813529 | -1.173 | -0.216 | -0.648 | 0.519 | -0.015 | 0.011 | 0.011 | 0.003 | 0.07 | 0.09 | 0.07 | 0.11 | 0.01 | 0.17 |
| 10 | 87939804 | rs1317833 | -0.751 | -0.286 | -1.353 | 0.611 | 0.059 | -0.045 | -0.045 | 3.163 | 0.43 | 0.86 | 0.39 | 0.57 | 0.36 | 0.49 |
| 10 | 87940997 | rs1870168 | -0.708 | -0.621 | -0.443 | 0.303 | -0.001 | 0.001 | 0.001 | 0 | 0.4 | 0.58 | 0.37 | 0.55 | 0.37 | 0.6 |
| 10 | 87941306 | rs17106392 | -1.443 | -0.757 | 0.205 | 0.351 | -0.006 | 0.004 | 0.004 | 0 | 0.07 | 0.2 | 0.09 | 0.07 | 0.06 | 0.17 |
| 10 | 87944571 | rs11201929 | -1.761 | -1.251 | 0.615 | 0.3 | -0.048 | -0.02 | -0.02 | 0.264 | 0.19 | 0.4 | 0.13 | 0.15 | 0.11 | 0.13 |
| 10 | 87948294 | rs6586000 | -0.883 | -1.233 | 0.662 | 0.31 | -0.02 | 0.025 | 0.025 | 0 | 0.05 | 0.17 | 0.06 | 0.07 | 0.02 | 0.04 |
| 10 | 87951686 | rs2607829 | -0.212 | -1.132 | 1.184 | 0.689 | 0.029 | 0.138 | 0.138 | 0.002 | 0.22 | 0.33 | 0.32 | 0.49 | 0.27 | 0.44 |
| 10 | 87954602 | rs12411971 | -0.926 | -0.836 | 1.051 | 0.552 | 0.003 | -0.027 | -0.027 | 0 | 0.07 | 0.12 | 0.16 | 0 | 0.08 | 0.06 |
| 10 | 87957672 | rs11201937 | 1.756 | -1.293 | 0.005 | 0.086 | -0.129 | -0.025 | -0.025 | 0 | 0.34 | 0.05 | 0.29 | 0.26 | 0.4 | 0.21 |
| 10 | 87958310 | rs1870156 | -0.177 | -1.436 | 0.196 | 0.05 | -0.005 | 0.091 | 0.091 | 0.025 | 0.28 | 0.41 | 0.39 | 0.51 | 0.28 | 0.49 |
| 10 | 87959049 | rs2814315 | -0.207 | -1.425 | 0.198 | 0.044 | 0.001 | -0.004 | -0.004 | 0 | 0.72 | 0.76 | 0.73 | 0.88 | 0.74 | 0.78 |
| 10 | 87960347 | rs7897313 | 0.6 | -1.381 | 0.127 | 0.084 | -0.005 | 0.025 | 0.025 | 0 | 0.48 | 0.41 | 0.4 | 0.4 | 0.48 | 0.32 |
| 10 | 87967102 | rs2607830 | -0.26 | -1.551 | 0.042 | -0.032 | 0.006 | 0.007 | 0.007 | 0 | 0.73 | 0.8 | 0.82 | 0.88 | 0.78 | 0.81 |
| 10 | 87967144 | rs17320881 | 1.195 | -1.606 | -0.112 | -0.328 | -0.041 | 0.012 | 0.012 | 0.731 | 0.37 | 0.23 | 0.38 | 0.27 | 0.42 | 0.24 |
| 10 | 87968625 | rs17320915 | 1.775 | -1.432 | -0.057 | -0.231 | -0.052 | -0.019 | -0.019 | 0 | 0.29 | 0.14 | 0.29 | 0.27 | 0.37 | 0.18 |
| 10 | 87969313 | rs2814322 | -0.487 | -1.387 | -0.225 | -0.187 | 0.006 | 0.004 | 0.004 | 0 | 0.73 | 0.8 | 0.82 | 0.89 | 0.78 | 0.81 |
| 10 | 87978868 | rs2446020 | -0.498 | -1.426 | -0.106 | -0.189 | -0.012 | -0.038 | -0.038 | 0 | 0.73 | 0.75 | 0.82 | 0.9 | 0.78 | 0.81 |
| 10 | 87979532 | rs12355302 | 1.41 | -0.915 | -0.743 | 0.09 | -0.095 | -0.043 | -0.043 | 0 | 0.29 | 0.08 | 0.29 | 0.28 | 0.37 | 0.17 |
| 10 | 87984457 | rs10887562 | -0.402 | -0.616 | -0.687 | 0.216 | 0.009 | 0.129 | 0.129 | 0 | 0.09 | 0.15 | 0.25 | 0.45 | 0.12 | 0.21 |
| 10 | 87985251 | rs10887563 | 1.176 | -0.596 | -0.961 | 0.086 | -0.071 | -0.024 | -0.024 | 0 | 0.3 | 0.15 | 0.29 | 0.28 | 0.37 | 0.16 |
| 10 | 87991224 | rs1870161 | 0.26 | -0.532 | -0.759 | 0.001 | -0.012 | -0.041 | -0.041 | 0.945 | 0.61 | 0.57 | 0.69 | 0.83 | 0.67 | 0.74 |
| 10 | 87991690 | rs2814329 | 0.646 | -0.238 | -2.251 | 0.679 | -0.146 | -0.102 | -0.102 | 0 | 0.3 | 0.05 | 0.42 | 0.4 | 0.49 | 0.27 |
| 10 | 87991916 | rs2607839 | 0.282 | -0.305 | -1.288 | 0.319 | -0.001 | -0.021 | -0.021 | 0.746 | 0.75 | 0.69 | 0.78 | 0.8 | 0.74 | 0.61 |
| 10 | 87996125 | rs1870165 | 0.433 | -0.041 | -1.441 | 1.001 | -0.003 | 0.059 | 0.059 | 0 | 0.92 | 0.93 | 0.91 | 0.44 | 0.9 | 0.85 |
| 10 | 87996451 | rs2607845 | 0.569 | -0.582 | -1.718 | 0.738 | 0.001 | -0.052 | -0.052 | 0 | 0.84 | 0.92 | 0.88 | 0.43 | 0.88 | 0.81 |
| 10 | 87996729 | rs4934164 | -0.064 | -0.652 | -1.8 | 0.323 | 0 | 0.016 | 0.016 | 0.038 | 0.16 | 0.12 | 0.05 | 0.06 | 0.15 | 0.22 |
| 10 | 88004805 | rs2814345 | 1.244 | -1.448 | -2.284 | 0.027 | 0.064 | 0.068 | 0.068 | 0.681 | 0.53 | 0.08 | 0.62 | 0.36 | 0.68 | 0.54 |
| 10 | 88006324 | rs1902681 | -0.182 | -1.514 | -2.176 | 0.217 | -0.005 | -0.057 | -0.057 | 0 | 0.88 | 0.95 | 0.9 | 0.44 | 0.92 | 0.81 |
| 10 | 88006744 | rs2814347 | 0.857 | -1.681 | -2.308 | 0.048 | -0.003 | -0.049 | -0.049 | 0 | 0.67 | 0.11 | 0.79 | 0.38 | 0.74 | 0.64 |
| 10 | 88012223 | rs2255475 | 1.411 | -2.014 | -1.076 | 0.068 | -0.022 | 0.054 | 0.054 | 0 | 0.78 | 0.67 | 0.71 | 0.4 | 0.86 | 0.7 |
| 10 | 88014349 | rs2814352 | 0.86 | -1.899 | -1.104 | -0.069 | 0 | 0.11 | 0.11 | 0 | 0.84 | 0.69 | 0.85 | 0.36 | 0.85 | 0.76 |
| 10 | 88029586 | rs1870167 | 1.246 | -1.969 | -0.704 | -0.107 | -0.016 | 0.156 | 0.156 | 1.814 | 0.8 | 0.52 | 0.75 | 0.4 | 0.87 | 0.79 |
| 10 | 88040764 | rs2168725 | 1.896 | -2.051 | -0.874 | -0.209 | 0.057 | 0.057 | 0.057 | 1.482 | 0.54 | 0.08 | 0.61 | 0.33 | 0.66 | 0.53 |
| 10 | 88048259 | rs867821 | 1.085 | -1.815 | -0.563 | -0.044 | 0.051 | 0.332 | 0.332 | 2.6 | 0.87 | 0.53 | 0.9 | 0.4 | 0.9 | 0.83 |
| 10 | 88066536 | rs1991426 | 0.073 | -1.93 | -0.188 | 0.225 | 0.01 | 0.006 | 0.006 | 0 | 0.89 | 0.91 | 0.83 | 1 | 0.94 | 0.96 |
| 10 | 88067925 | rs11201974 | 1.833 | -2.025 | -1.113 | -0.423 | -0.035 | -0.016 | -0.016 | 0 | 0.21 | 0.01 | 0.25 | 0.22 | 0.22 | 0.31 |
| 10 | 88068120 | rs1896527 | 1.522 | -1.975 | -0.636 | -0.225 | 0.058 | -0.035 | -0.035 | 0 | 0.53 | 0.08 | 0.61 | 0.43 | 0.66 | 0.55 |
| 10 | 88071347 | rs6586002 | 0.206 | -1.952 | -0.538 | -0.302 | -0.047 | -0.09 | -0.09 | 0 | 0.84 | 0.63 | 0.81 | 0.99 | 0.94 | 0.95 |
| 10 | 88072915 | rs7091924 | 0.041 | -1.952 | -0.017 | -0.294 | -0.01 | -0.033 | -0.033 | 1.646 | 0.2 | 0.29 | 0.1 | 0.06 | 0.18 | 0.24 |
| 10 | 88079367 | rs10887577 | 1.471 | -1.844 | -0.188 | -0.215 | 0.062 | -0.003 | -0.003 | 0 | 0.54 | 0.13 | 0.61 | 0.41 | 0.66 | 0.55 |
| 10 | 88081937 | rs7914854 | 0.242 | -1.863 | -0.089 | -0.209 | -0.003 | 0.157 | 0.157 | 0.034 | 0.83 | 0.65 | 0.77 | 0.5 | 0.88 | 0.85 |

| | | | | | | | | | | | | | | | | |
|---|---|---|---|---|---|---|---|---|---|---|---|---|---|---|---|---|
| 10 | 88085715 | rs11201980 | -0.025 | -1.704 | 0.139 | -0.381 | 0.004 | 0.009 | 0.009 | 0.137 | 0.19 | 0.19 | 0.1 | 0.06 | 0.18 | 0.23 |
| 10 | 88088252 | rs7087612 | -1.101 | -1.652 | 0.072 | -0.402 | -0.042 | -0.103 | -0.103 | 0 | 0.28 | 0.5 | 0.16 | 0.06 | 0.22 | 0.3 |
| 10 | 88088456 | rs7903043 | 0.184 | -1.163 | 0.15 | -0.04 | -0.018 | -0.037 | -0.037 | 0 | 0.07 | 0.22 | 0.04 | 0 | 0.02 | 0.02 |
| 10 | 88100089 | rs874758 | 0.865 | -1.29 | -0.196 | 0.11 | 0.002 | -0.004 | -0.004 | 0 | 0.7 | 0.34 | 0.68 | 0.47 | 0.81 | 0.68 |
| 10 | 88104526 | rs10749535 | 1.021 | -0.989 | -1.175 | -0.441 | -0.186 | -0.067 | -0.067 | 0.207 | 0.29 | 0.03 | 0.46 | 0.24 | 0.44 | 0.19 |
| 10 | 88105348 | rs6586006 | -0.311 | -1.165 | -0.504 | -0.417 | -0.013 | 0.191 | 0.191 | 0 | 0.21 | 0.49 | 0.14 | 0.52 | 0.16 | 0.25 |
| 10 | 88112939 | rs1863824 | -0.642 | -1.409 | -0.742 | -0.613 | 0.053 | 0.13 | 0.13 | 0 | 0.56 | 0.84 | 0.65 | 0.77 | 0.63 | 0.47 |
| 10 | 88116479 | rs7904985 | 1.026 | -1.496 | -1.015 | -0.708 | 0.014 | 0.072 | 0.072 | 0 | 0.34 | 0.07 | 0.17 | 0.23 | 0.31 | 0.45 |
| 10 | 88117490 | rs7910309 | 0.174 | -1.552 | -0.576 | -0.706 | 0 | 0.027 | 0.027 | 0 | 0.18 | 0.18 | 0.12 | 0.51 | 0.16 | 0.24 |
| 10 | 88118952 | rs2114826 | -0.426 | -1.484 | -0.972 | -0.693 | -0.01 | 0.062 | 0.062 | 1.748 | 0.23 | 0.64 | 0.15 | 0.52 | 0.17 | 0.26 |
| 10 | 88119015 | rs2114824 | 1.043 | -1.483 | -1.184 | -0.694 | -0.093 | 0.005 | 0.005 | 0.805 | 0.32 | 0.13 | 0.51 | 0.23 | 0.49 | 0.21 |
| 10 | 88122516 | rs7084928 | 0.922 | -1.412 | -1.146 | -0.643 | 0.033 | 0.06 | 0.06 | 0.009 | 0.36 | 0.15 | 0.17 | 0.24 | 0.31 | 0.48 |
| 10 | 88128281 | rs3814614 | -0.605 | -0.981 | -1.148 | -0.49 | -0.086 | 0.03 | 0.03 | 1.242 | 0.62 | 0.86 | 0.35 | 0.77 | 0.52 | 0.76 |
| 10 | 88134257 | rs12220079 | 0.737 | 0.242 | -1.318 | -0.151 | -0.129 | 0.012 | 0.012 | 1.45 | 0.29 | 0.07 | 0.34 | 0.18 | 0.39 | 0.19 |
| 10 | 88141534 | rs1863826 | -0.251 | -0.548 | -0.777 | -0.481 | -0.004 | -0.006 | -0.006 | 0 | 0.61 | 0.61 | 0.64 | 0.77 | 0.72 | 0.55 |
| 10 | 88145602 | rs11202014 | 0.855 | -0.607 | -0.825 | -0.468 | 0.01 | 0.021 | 0.021 | 0 | 0.28 | 0.2 | 0.17 | 0.21 | 0.24 | 0.37 |
| 10 | 88151974 | rs4934199 | 0.464 | -0.657 | -0.36 | -0.456 | -0.016 | -0.031 | -0.031 | 0 | 0.67 | 0.39 | 0.63 | 0.86 | 0.77 | 0.74 |
| 10 | 88158854 | rs4933395 | 0.287 | -0.67 | -0.535 | -0.473 | -0.004 | -0.012 | -0.012 | 0 | 0.69 | 0.66 | 0.65 | 0.86 | 0.77 | 0.76 |
| 10 | 88162238 | rs10458771 | -0.809 | -0.624 | -1.063 | -0.563 | 0.01 | 0.011 | 0.011 | 0.145 | 0.83 | 0.85 | 0.74 | 0.93 | 0.89 | 0.93 |
| 10 | 88162682 | rs731171 | 1.422 | -0.094 | -1.695 | -0.255 | -0.11 | -0.042 | -0.042 | 0.29 | 0.23 | 0.07 | 0.3 | 0.2 | 0.33 | 0.17 |
| 10 | 88163251 | rs2018507 | 0.964 | -0.082 | -1.382 | -0.342 | -0.032 | 0.102 | 0.102 | 1.038 | 0.34 | 0.07 | 0.18 | 0.18 | 0.36 | 0.48 |
| 10 | 88175206 | rs7907234 | 0.235 | -0.127 | -0.97 | -0.132 | 0 | 0.004 | 0.004 | 0 | 0.11 | 0.09 | 0.08 | 0.08 | 0.13 | 0.17 |
| 10 | 88175881 | rs1896514 | 0.28 | -0.644 | -0.606 | -0.323 | -0.007 | 0.016 | 0.016 | 1.731 | 0.45 | 0.43 | 0.42 | 0.32 | 0.46 | 0.29 |
| 10 | 88181002 | rs11202034 | 1.464 | -0.827 | -0.703 | -0.086 | -0.025 | 0.007 | 0.007 | 1.37 | 0.29 | 0.08 | 0.25 | 0.23 | 0.34 | 0.45 |
| 10 | 88193911 | rs10400177 | -0.102 | -1.319 | -0.592 | -0.363 | -0.027 | -0.047 | -0.047 | 0.285 | 0.77 | 0.69 | 0.72 | 0.94 | 0.88 | 0.89 |
| 10 | 88194529 | rs4934207 | 0.488 | -1.347 | 0.065 | -0.289 | -0.041 | 0.027 | 0.027 | 0.003 | 0.72 | 0.45 | 0.67 | 0.57 | 0.85 | 0.84 |
| 10 | 88199488 | rs4934208 | 1.812 | -1.146 | -0.526 | -0.572 | -0.141 | -0.087 | -0.087 | 0 | 0.23 | 0.02 | 0.28 | 0.24 | 0.32 | 0.18 |
| 10 | 88202298 | rs10788507 | -0.383 | -1.281 | -0.072 | -0.457 | -0.011 | 0.101 | 0.101 | 0 | 0.88 | 0.81 | 0.77 | 0.63 | 0.93 | 0.91 |
| 10 | 88203068 | rs7094132 | -0.379 | -1.281 | -0.071 | -0.458 | -0.011 | 0.102 | 0.102 | 0 | 0.88 | 0.81 | 0.77 | 0.63 | 0.93 | 0.91 |
| 10 | 88206702 | rs10887609 | 0.331 | -1.316 | 0.017 | -0.48 | 0 | 0.014 | 0.014 | 0.526 | 0.46 | 0.54 | 0.39 | 0.68 | 0.49 | 0.66 |
| 10 | 88245682 | rs3758521 | 2.136 | -1.192 | 0.007 | -0.492 | 0.003 | -0.008 | -0.008 | 0.136 | 0.11 | 0.09 | 0.08 | 0.05 | 0.08 | 0.02 |
| 10 | 88257602 | rs3858279 | 0.803 | -0.965 | -0.032 | -0.392 | -0.064 | 0.004 | 0.004 | 0 | 0.31 | 0.17 | 0.34 | 0.25 | 0.39 | 0.23 |
| 10 | 88263120 | rs7070233 | 0.558 | -0.868 | 0.472 | -0.189 | 0.005 | -0.003 | -0.003 | 0 | 0.11 | 0.2 | 0.09 | 0.08 | 0.12 | 0.16 |
| 10 | 88276899 | rs7914270 | 0.05 | -0.814 | 0.036 | -0.045 | -0.008 | 0.143 | 0.143 | 0 | 0.88 | 0.73 | 0.76 | 0.64 | 0.93 | 0.91 |
| 10 | 88315181 | rs10887623 | 0.248 | -0.95 | -0.723 | -0.376 | -0.003 | 0.006 | 0.006 | 0 | 0.12 | 0.05 | 0.09 | 0.09 | 0.13 | 0.16 |
| 10 | 88332974 | rs2588279 | 0.093 | -0.885 | -0.414 | -0.389 | 0.024 | 0.052 | 0.052 | 2.725 | 0.23 | 0.14 | 0.16 | 0.09 | 0.2 | 0.09 |
| 10 | 88336279 | rs2588278 | -0.32 | -1.254 | -0.353 | -0.462 | 0 | 0.002 | 0.002 | 0 | 0.57 | 0.43 | 0.63 | 0.66 | 0.6 | 0.63 |
| 10 | 88340471 | rs2353836 | 1.435 | -1.179 | -0.362 | -0.271 | -0.021 | 0.034 | 0.034 | 1 | 0.28 | 0.15 | 0.34 | 0.19 | 0.32 | 0.42 |
| 10 | 88341011 | rs10887625 | -0.547 | -1.126 | -0.477 | -0.196 | -0.001 | 0.006 | 0.006 | 0 | 0.13 | 0.12 | 0.3 | 0.35 | 0.11 | 0.16 |
| 10 | 88343325 | rs6586019 | 0.335 | -1.168 | -0.139 | -0.412 | -0.032 | -0.044 | -0.044 | 0.201 | 0.17 | 0.34 | 0.12 | 0.06 | 0.1 | 0.12 |
| 10 | 88343979 | rs7098517 | 0.36 | -1.026 | -0.141 | -0.415 | -0.007 | -0.046 | -0.046 | 0.033 | 0.17 | 0.34 | 0.12 | 0.06 | 0.13 | 0.13 |
| 10 | 88344497 | rs2353840 | 0.91 | -0.69 | -0.64 | -0.197 | -0.045 | -0.066 | -0.066 | 0 | 0.32 | 0.17 | 0.46 | 0.56 | 0.4 | 0.5 |
| 10 | 88349538 | rs720728 | -0.746 | -0.192 | -0.386 | 0.031 | -0.017 | -0.02 | -0.02 | 0.134 | 0.77 | 0.84 | 0.75 | 0.73 | 0.73 | 0.79 |
| 10 | 88364717 | rs12250999 | -0.673 | -0.114 | 0.409 | 0.183 | -0.004 | 0.113 | 0.113 | 0 | 0.16 | 0.29 | 0.37 | 0.42 | 0.11 | 0.2 |
| 10 | 88372958 | rs2803589 | 1.192 | -0.216 | -0.018 | -0.053 | -0.031 | 0.071 | 0.071 | 0.572 | 0.13 | 0.01 | 0.12 | 0.02 | 0.15 | 0.03 |
| 10 | 88386179 | rs7917872 | -0.333 | -0.229 | 0.604 | 0.002 | 0.02 | 0.178 | 0.178 | 0.025 | 0.16 | 0.29 | 0.27 | 0.48 | 0.14 | 0.25 |
| 10 | 88397886 | rs11202103 | 1.548 | -0.245 | 0.295 | -0.172 | -0.039 | -0.003 | -0.003 | 0.06 | 0.26 | 0.08 | 0.18 | 0.27 | 0.3 | 0.41 |
| 10 | 88397995 | rs10887632 | 1.197 | -0.34 | 0.717 | -0.213 | -0.001 | -0.005 | -0.005 | 0 | 0.29 | 0.27 | 0.2 | 0.27 | 0.31 | 0.42 |
| 10 | 88400760 | rs10887635 | 1.227 | -0.342 | 0.716 | -0.214 | 0 | -0.005 | -0.005 | 0 | 0.29 | 0.27 | 0.2 | 0.27 | 0.31 | 0.42 |
| 10 | 88410251 | rs10887636 | 1.501 | -0.289 | 0.883 | -0.068 | 0.003 | 0.117 | 0.117 | 0.019 | 0.07 | 0.08 | 0.19 | 0.39 | 0.07 | 0.08 |
| 10 | 88413432 | rs2736689 | -0.017 | -0.453 | 0.911 | -0.541 | 0.004 | 0.072 | 0.072 | 0 | 0.49 | 0.36 | 0.47 | 0.33 | 0.5 | 0.56 |
| 10 | 88415619 | rs11202107 | 1.584 | -0.429 | 1.038 | -0.529 | -0.046 | -0.014 | -0.014 | 1.815 | 0.26 | 0.06 | 0.18 | 0.28 | 0.3 | 0.4 |
| 10 | 88419012 | rs3740334 | 0.782 | -0.405 | 0.818 | -0.506 | 0.004 | 0.067 | 0.067 | 0.002 | 0.15 | 0.16 | 0.26 | 0.46 | 0.14 | 0.21 |
| 10 | 88421041 | rs7478092 | -0.072 | -0.463 | 0.667 | -0.508 | -0.024 | -0.001 | -0.001 | 3.203 | 0.06 | 0.18 | 0.03 | 0.04 | 0.02 | 0.07 |
| 10 | 88422116 | rs1079610 | -0.761 | -0.523 | 0.42 | -0.553 | -0.006 | 0.043 | 0.043 | 0 | 0.65 | 0.8 | 0.76 | 0.79 | 0.63 | 0.77 |
| 10 | 88422151 | rs3740337 | 1.664 | -0.454 | 0.242 | -0.501 | -0.061 | -0.027 | -0.027 | 0 | 0.26 | 0.04 | 0.17 | 0.28 | 0.3 | 0.4 |
| 10 | 88426042 | rs3740341 | 0.884 | -0.295 | 0.055 | -0.396 | 0.007 | 0.094 | 0.094 | 0 | 0.15 | 0.2 | 0.26 | 0.43 | 0.13 | 0.18 |
| 10 | 88428076 | rs2675692 | -1.632 | -0.067 | -0.618 | -0.466 | -0.054 | -0.023 | -0.023 | 0 | 0.78 | 0.95 | 0.81 | 0.81 | 0.71 | 0.83 |
| 10 | 88430368 | rs12570315 | 1.434 | 0.079 | -0.596 | -0.337 | -0.057 | -0.023 | -0.023 | 0 | 0.27 | 0.05 | 0.17 | 0.28 | 0.3 | 0.44 |
| 10 | 88432328 | rs11202121 | -0.354 | 0.021 | -0.425 | -0.299 | -0.041 | 0.085 | 0.085 | 1.077 | 0.29 | 0.52 | 0.29 | 0.46 | 0.19 | 0.27 |
| 10 | 88435818 | rs2803566 | -0.649 | 0.298 | -0.853 | -0.107 | 0.027 | -0.008 | -0.008 | 0 | 0.47 | 0.74 | 0.61 | 0.51 | 0.38 | 0.35 |
| 10 | 88436426 | rs2803567 | -1.281 | 0.276 | -0.882 | -0.115 | -0.017 | 0.003 | 0.003 | 0.061 | 0.74 | 0.85 | 0.79 | 0.77 | 0.69 | 0.73 |
| 10 | 88436490 | rs10788525 | -0.934 | 0.451 | -0.691 | 0.625 | 0.001 | 0.017 | 0.017 | 0 | 0.49 | 0.43 | 0.43 | 0.74 | 0.45 | 0.7 |
| 10 | 88437537 | rs4462249 | -0.203 | 0.566 | -1.179 | 0.587 | -0.021 | -0.006 | -0.006 | 0 | 0.27 | 0.11 | 0.18 | 0.31 | 0.31 | 0.49 |
| 10 | 88443060 | rs12244574 | -1.266 | 0.259 | -0.259 | 0.521 | 0.001 | -0.025 | -0.025 | 0.045 | 0.12 | 0.22 | 0.07 | 0.02 | 0.12 | 0.1 |
| 10 | 88446254 | rs1986382 | 0.726 | -0.247 | 0.287 | 0.967 | -0.021 | 0.03 | 0.03 | 0 | 0.73 | 0.5 | 0.5 | 0.3 | 0.74 | 0.65 |
| 10 | 88449377 | rs4934251 | 0.25 | -0.167 | 0.261 | 1.441 | -0.004 | 0.031 | 0.031 | 0 | 0.84 | 0.83 | 0.61 | 0.5 | 0.87 | 0.87 |
| 10 | 88455242 | rs2803572 | 1.05 | -1.156 | 0.477 | 0.924 | 0.044 | 0.009 | 0.009 | 0.123 | 0.64 | 0.21 | 0.5 | 0.32 | 0.75 | 0.64 |
| 10 | 88455879 | rs4933405 | -0.762 | -0.672 | 0.079 | 0.35 | -0.001 | -0.003 | -0.003 | 0 | 0.61 | 0.54 | 0.74 | 0.7 | 0.6 | 0.61 |

| | | | | | | | | | | | | | | | | |
|---|---|---|---|---|---|---|---|---|---|---|---|---|---|---|---|---|
| 10 | 88461699 | rs10887650 | -1.304 | -1.079 | -0.555 | 0.55 | 0.038 | -0.044 | -0.044 | 0.165 | 0.37 | 0.89 | 0.43 | 0.67 | 0.26 | 0.36 |
| 10 | 88468321 | rs10788526 | -1.318 | -1.114 | -0.565 | 0.538 | 0.036 | -0.043 | -0.043 | 0 | 0.37 | 0.89 | 0.43 | 0.68 | 0.27 | 0.36 |
| 10 | 88469587 | rs3740347 | -1.027 | -0.947 | 0.456 | 1.011 | -0.031 | 0.105 | 0.105 | 0 | 0.24 | 0.4 | 0.32 | 0.6 | 0.17 | 0.22 |
| 10 | 88488806 | rs7908765 | -0.834 | -0.851 | -0.017 | 0.937 | -0.044 | -0.101 | -0.101 | 0.064 | 0.24 | 0.11 | 0.29 | 0.6 | 0.32 | 0.35 |
| 10 | 88502670 | rs4244972 | -1.558 | -0.961 | 0.691 | 0.858 | 0 | 0.047 | 0.047 | 0 | 0.31 | 0.45 | 0.33 | 0.6 | 0.33 | 0.38 |
| 10 | 88507643 | rs4934258 | -0.721 | -0.57 | 0.363 | 0.515 | -0.025 | -0.068 | -0.068 | 0.178 | 0.25 | 0.1 | 0.3 | 0.69 | 0.28 | 0.33 |
| 10 | 88515072 | rs3905377 | -0.645 | -0.422 | 0.184 | 0.313 | -0.036 | -0.062 | -0.062 | 0.389 | 0.33 | 0.09 | 0.33 | 0.69 | 0.32 | 0.35 |
| 10 | 88520409 | rs7074582 | -1.292 | -0.724 | 0.444 | 0.15 | -0.001 | -0.003 | -0.003 | 0 | 0.43 | 0.47 | 0.43 | 0.72 | 0.42 | 0.43 |
| 10 | 88520763 | rs11202166 | 0.098 | -0.716 | 0.459 | 0.138 | -0.025 | -0.044 | -0.044 | 0 | 0.09 | 0.21 | 0.05 | 0 | 0.03 | 0.05 |
| 10 | 88530481 | rs3858284 | -1.498 | -0.863 | 0.483 | 0.102 | 0.02 | 0.061 | 0.061 | 0 | 0.47 | 0.7 | 0.45 | 0.73 | 0.42 | 0.43 |
| 10 | 88538911 | rs7096781 | -0.586 | -1.038 | 0.082 | 0.075 | -0.017 | -0.018 | -0.018 | 0 | 0.36 | 0.06 | 0.34 | 0.72 | 0.3 | 0.43 |
| 10 | 88546128 | rs4934262 | -2.176 | -1.031 | 0.172 | 0.07 | 0.009 | 0.123 | 0.123 | 0 | 0.53 | 0.8 | 0.48 | 0.76 | 0.42 | 0.51 |
| 10 | 88555187 | rs6586037 | -1.613 | -1.201 | 0.405 | 0.026 | 0.016 | 0.05 | 0.05 | 0 | 0.5 | 0.63 | 0.46 | 0.77 | 0.42 | 0.51 |
| 10 | 88571339 | rs4934268 | -1.936 | -1.217 | 0.388 | -0.019 | 0.031 | 0.161 | 0.161 | 0 | 0.53 | 0.81 | 0.46 | 0.78 | 0.4 | 0.51 |
| 10 | 88572727 | rs12415784 | 1.394 | -0.011 | -0.204 | -0.141 | -0.029 | -0.029 | -0.029 | 0.047 | 0.05 | 0.01 | 0.25 | 0.1 | 0.16 | 0.13 |
| 10 | 88575498 | rs2354353 | -1.559 | -0.452 | -0.055 | -0.124 | 0.021 | 0.051 | 0.051 | 0 | 0.49 | 0.63 | 0.43 | 0.78 | 0.4 | 0.51 |
| 10 | 88577094 | rs4934272 | -0.045 | -0.392 | -0.483 | 0.081 | -0.011 | -0.102 | -0.102 | 0 | 0.28 | 0.05 | 0.28 | 0.64 | 0.25 | 0.23 |
| 10 | 88585926 | rs722068 | -0.675 | -0.398 | -0.306 | -0.166 | -0.006 | 0.016 | 0.016 | 0 | 0.36 | 0.14 | 0.35 | 0.67 | 0.35 | 0.34 |
| 10 | 88617392 | rs4934275 | 0.531 | -0.467 | -0.21 | -0.761 | 0.005 | -0.078 | -0.078 | 0 | 0.2 | 0.05 | 0.23 | 0.67 | 0.15 | 0.15 |
| 10 | 88618968 | rs11202230 | 0.837 | -0.435 | -0.255 | -0.769 | -0.027 | -0.024 | -0.024 | 0 | 0.05 | 0 | 0.05 | 0.03 | 0.1 | 0.09 |
| 10 | 88639910 | rs4934279 | -2.015 | -0.663 | -0.2 | -0.962 | 0.043 | 0.194 | 0.194 | 0 | 0.52 | 0.8 | 0.43 | 0.78 | 0.39 | 0.5 |
| 10 | 88649176 | rs11597689 | 0.967 | -0.625 | 0.173 | -0.785 | -0.001 | 0 | 0 | 0 | 0.06 | 0.06 | 0.04 | 0.06 | 0.06 | 0.13 |
| 10 | 88654876 | rs7909240 | 0.698 | -0.794 | 0.534 | -0.775 | -0.045 | -0.046 | -0.046 | 0 | 0.09 | 0.18 | 0.03 | 0 | 0.01 | 0.03 |
| 10 | 88662216 | rs10887668 | -0.567 | -0.724 | 0.082 | -0.742 | -0.047 | -0.042 | -0.042 | 0 | 0.2 | 0.56 | 0.12 | 0.04 | 0.12 | 0.16 |
| 10 | 88682555 | rs12765929 | 1.625 | -0.28 | -1.295 | -1.21 | -0.007 | 0.006 | 0.006 | 0 | 0.15 | 0.15 | 0.35 | 0.1 | 0.27 | 0.16 |
| 10 | 88683122 | rs7074064 | -1.258 | -0.699 | -0.638 | -0.828 | 0.06 | 0.072 | 0.072 | 0.999 | 0.42 | 0.73 | 0.34 | 0.7 | 0.27 | 0.31 |
| 10 | 88688336 | rs7094713 | -0.018 | -0.031 | -0.65 | -0.273 | 0.013 | -0.04 | -0.04 | 0.383 | 0.2 | 0.13 | 0.27 | 0.68 | 0.15 | 0.16 |
| 10 | 88697548 | rs7082258 | -0.431 | 0.332 | -0.277 | -0.564 | 0 | -0.004 | -0.004 | 0 | 0.13 | 0.1 | 0.24 | 0.16 | 0.12 | 0.04 |
| 10 | 88700638 | rs746677 | 0.089 | 0.226 | 0.037 | -1.156 | 0.002 | 0.013 | 0.013 | 0 | 0.53 | 0.45 | 0.49 | 0.78 | 0.56 | 0.6 |
| 10 | 88710234 | rs10887679 | -0.57 | 0.39 | -0.129 | -1.064 | 0.001 | -0.007 | -0.007 | 0 | 0.15 | 0.11 | 0.19 | 0.19 | 0.13 | 0.14 |
| 10 | 88716577 | rs10887683 | 0.732 | 0.279 | 0.016 | -1.111 | -0.018 | 0.017 | 0.017 | 0 | 0.42 | 0.27 | 0.42 | 0.69 | 0.49 | 0.44 |
| 10 | 88717154 | rs3750823 | 1.333 | 0.151 | -0.07 | -1.324 | -0.047 | 0.003 | 0.003 | 0.102 | 0.36 | 0.19 | 0.42 | 0.71 | 0.47 | 0.39 |
| 10 | 88718436 | rs1800373 | 0.147 | 0.162 | 0.291 | -1.363 | -0.001 | -0.004 | -0.004 | 0 | 0.49 | 0.52 | 0.48 | 0.71 | 0.55 | 0.42 |
| 10 | 88718833 | rs3793900 | -0.343 | 0.222 | 0.211 | -1.452 | -0.003 | -0.018 | -0.018 | 0 | 0.6 | 0.45 | 0.62 | 0.83 | 0.67 | 0.5 |
| 10 | 88721035 | rs9420407 | 0.12 | 0.515 | 0.219 | -1.402 | -0.006 | -0.063 | -0.063 | 0.021 | 0.37 | 0.41 | 0.29 | 0.07 | 0.3 | 0.39 |
| 10 | 88725984 | rs1240390 | -1.445 | 0.191 | 0.11 | -1.348 | -0.006 | 0.023 | 0.023 | 0.795 | 0.79 | 0.83 | 0.78 | 0.93 | 0.77 | 0.65 |
| 10 | 88726739 | rs1240388 | -1.676 | 0.166 | -0.095 | -1.351 | -0.072 | 0.068 | 0.068 | 1.299 | 0.79 | 0.94 | 0.75 | 0.94 | 0.69 | 0.66 |
| 10 | 88730177 | rs2279601 | 0.041 | -0.025 | -0.127 | -1.315 | 0.002 | 0.035 | 0.035 | 0.301 | 0.55 | 0.36 | 0.44 | 0.82 | 0.6 | 0.5 |
| 10 | 88730312 | rs4869 | 0.19 | -0.014 | -0.076 | -1.307 | 0.006 | 0.054 | 0.054 | 0.171 | 0.53 | 0.36 | 0.43 | 0.81 | 0.6 | 0.46 |
| 10 | 88730374 | rs7960 | 0.837 | 0.041 | -0.183 | -1.261 | -0.028 | 0.022 | 0.022 | 0.316 | 0.41 | 0.21 | 0.29 | 0.69 | 0.48 | 0.38 |
| 10 | 88738864 | rs11202282 | 0.228 | 0.067 | -0.101 | -1.651 | -0.008 | 0.021 | 0.021 | 0 | 0.2 | 0.27 | 0.38 | 0.24 | 0.17 | 0.22 |
| 10 | 88741850 | rs1240402 | -1.735 | -0.07 | -0.164 | -1.692 | -0.071 | 0.075 | 0.075 | 1.59 | 0.79 | 0.93 | 0.75 | 0.94 | 0.71 | 0.67 |
| 10 | 88743691 | rs11202285 | 0.286 | 0.114 | -0.156 | -1.694 | 0.001 | 0.007 | 0.007 | 0 | 0.2 | 0.24 | 0.39 | 0.24 | 0.2 | 0.23 |
| 10 | 88743900 | rs7899599 | 0.822 | 0.02 | -0.348 | -1.665 | -0.001 | 0.014 | 0.014 | 0.094 | 0.4 | 0.26 | 0.28 | 0.69 | 0.44 | 0.36 |
| 10 | 88746547 | rs1240401 | 0.701 | 1.109 | -0.602 | -1.157 | 0 | 0.013 | 0.013 | 0 | 0.09 | 0.04 | 0.03 | 0.01 | 0.08 | 0.05 |
| 10 | 88746717 | rs1240400 | 0.692 | 1.111 | -0.602 | -1.157 | 0 | 0.013 | 0.013 | 0 | 0.09 | 0.04 | 0.03 | 0.01 | 0.08 | 0.05 |
| 10 | 88746826 | rs1240399 | -0.382 | 1.389 | -0.13 | -0.692 | -0.005 | 0.005 | 0.005 | 1.088 | 0.59 | 0.64 | 0.36 | 0.7 | 0.5 | 0.44 |
| 10 | 88768657 | rs1745901 | -0.431 | 1.681 | -0.528 | -0.643 | -0.011 | 0.009 | 0.009 | 1.468 | 0.79 | 0.81 | 0.75 | 0.93 | 0.7 | 0.67 |
| 10 | 88786593 | rs10887686 | -0.501 | 1.363 | -0.273 | -0.077 | 0.002 | 0.019 | 0.019 | 0 | 0.14 | 0.18 | 0.26 | 0.67 | 0.23 | 0.18 |
| 10 | 88794870 | rs9421560 | 0.136 | 0.969 | 0.027 | -1.048 | -0.005 | 0.001 | 0.001 | 0 | 0.42 | 0.35 | 0.32 | 0.07 | 0.4 | 0.33 |
| 10 | 88799072 | rs9421561 | 0.629 | 1.06 | 0.247 | -0.973 | 0.007 | 0.033 | 0.033 | 0 | 0.11 | 0.09 | 0.24 | 0.66 | 0.24 | 0.17 |
| 10 | 88799610 | rs11202308 | 1.106 | 1.022 | -0.266 | -1.401 | -0.039 | 0.046 | 0.046 | 3.035 | 0.2 | 0.01 | 0.19 | 0.06 | 0.16 | 0.15 |
| 10 | 88832502 | rs2181522 | -1.557 | 0.46 | 0.61 | -1.654 | -0.017 | 0.049 | 0.049 | 0 | 0.65 | 0.83 | 0.75 | 0.76 | 0.63 | 0.55 |
| 10 | 88835254 | rs9421580 | 0.777 | 0.606 | 0.742 | -1.529 | 0.036 | 0.098 | 0.098 | 0 | 0.12 | 0.21 | 0.24 | 0.66 | 0.24 | 0.17 |
| 10 | 88847985 | rs7096107 | -1.13 | 0.274 | 1.135 | -1.93 | 0.012 | 0.114 | 0.114 | 0 | 0.58 | 0.76 | 0.73 | 0.84 | 0.63 | 0.52 |
| 10 | 88852942 | rs2296063 | -0.41 | 0.379 | 1.121 | -1.883 | -0.002 | -0.041 | -0.041 | 0 | 0.43 | 0.55 | 0.39 | 0.12 | 0.34 | 0.34 |
| 10 | 88868824 | rs4934297 | -0.969 | 0.077 | 0.627 | -1.825 | 0 | 0.124 | 0.124 | 0 | 0.59 | 0.83 | 0.73 | 0.84 | 0.63 | 0.53 |
| 10 | 88877883 | rs7072149 | -0.893 | 0.258 | 0.109 | -1.746 | 0.012 | 0.096 | 0.096 | 0 | 0.58 | 0.76 | 0.73 | 0.84 | 0.63 | 0.53 |
| 10 | 88945341 | rs7080675 | -0.297 | 0.618 | -0.121 | -1.243 | -0.001 | -0.003 | -0.003 | 1.204 | 0.65 | 0.65 | 0.4 | 0.19 | 0.56 | 0.64 |
| 10 | 89069591 | rs12241520 | -0.89 | 0.442 | -0.349 | -0.758 | 0.001 | -0.023 | -0.023 | 6.185 | 0.24 | 0.23 | 0.16 | 0.02 | 0.2 | 0.18 |
| 10 | 89164194 | rs12251529 | -1.564 | 0.777 | -0.057 | -0.08 | 0.007 | -0.034 | -0.034 | 0 | 0.08 | 0.15 | 0.04 | 0 | 0.06 | 0.05 |
| 10 | 89175636 | rs3128151 | 0.191 | 0.708 | -1.231 | -0.822 | -0.018 | 0.118 | 0.118 | 0 | 0.41 | 0.21 | 0.26 | 0.16 | 0.44 | 0.52 |
| 10 | 89182615 | rs17772401 | -0.445 | 0.471 | -0.826 | -0.706 | 0 | -0.01 | -0.01 | 1.389 | 0.19 | 0.21 | 0.14 | 0.11 | 0.17 | 0.19 |
| 10 | 89260709 | rs11202426 | 0.763 | -0.511 | -1.199 | -0.631 | -0.04 | 0.053 | 0.053 | 0 | 0.37 | 0.21 | 0.27 | 0.22 | 0.47 | 0.45 |
| 10 | 89262717 | rs3758478 | -0.69 | -1.69 | -1.307 | 0.025 | -0.054 | 0.008 | 0.008 | 0.543 | 0.7 | 0.92 | 0.7 | 0.66 | 0.66 | 0.72 |
| 10 | 89262987 | rs10509407 | 1.147 | -1.714 | -1.317 | 0.006 | -0.06 | 0.072 | 0.072 | 0 | 0.39 | 0.17 | 0.28 | 0.22 | 0.48 | 0.46 |
| 10 | 89263372 | rs3758477 | 0.508 | -1.838 | -0.356 | 0.115 | -0.006 | 0.354 | 0.354 | 1.656 | 0.06 | 0.29 | 0.25 | 0.3 | 0.02 | 0.04 |
| 10 | 89267957 | rs3843597 | -0.713 | -1.833 | -0.372 | 0.097 | -0.054 | 0.008 | 0.008 | 0.543 | 0.7 | 0.91 | 0.71 | 0.66 | 0.66 | 0.71 |
| 10 | 89272410 | rs6586073 | -0.713 | -1.833 | -0.372 | 0.097 | -0.054 | 0.008 | 0.008 | 0.543 | 0.7 | 0.91 | 0.71 | 0.66 | 0.66 | 0.71 |

| | | | | | | | | | | | | | | | | |
|---|---|---|---|---|---|---|---|---|---|---|---|---|---|---|---|---|
| 10 | 89273579 | rs2311115 | 0.054 | -1.887 | -0.143 | 0.084 | -0.004 | 0.001 | 0.001 | 0 | 0.62 | 0.5 | 0.44 | 0.36 | 0.64 | 0.67 |
| 10 | 89292637 | rs3847448 | -0.64 | -1.885 | -0.157 | 0.079 | -0.02 | 0.019 | 0.019 | 0.076 | 0.69 | 0.8 | 0.7 | 0.66 | 0.66 | 0.71 |
| 10 | 89294408 | rs11202434 | 0.458 | -1.829 | -0.219 | 0.159 | -0.006 | 0.359 | 0.359 | 1.151 | 0.06 | 0.29 | 0.25 | 0.3 | 0.02 | 0.05 |
| 10 | 89300314 | rs3847452 | 0.862 | -1.819 | -0.472 | -0.044 | 0.017 | 0.006 | 0.006 | 0 | 0.18 | 0.06 | 0.11 | 0.13 | 0.13 | 0.16 |
| 10 | 89309386 | rs3847453 | -1.413 | -1.188 | -0.699 | -0.082 | -0.042 | -0.039 | -0.039 | 0.664 | 0.82 | 0.99 | 0.81 | 0.77 | 0.84 | 0.77 |
| 10 | 89310662 | rs1408377 | -0.547 | -0.877 | -0.173 | 0.026 | 0.02 | -0.04 | -0.04 | 0 | 0.17 | 0.28 | 0.15 | 0.11 | 0.21 | 0.11 |
| 10 | 89315425 | rs2077871 | -0.2 | -0.653 | -0.339 | 0.133 | 0.016 | 0.058 | 0.058 | 0 | 0.53 | 0.72 | 0.61 | 0.69 | 0.59 | 0.54 |
| 10 | 89315881 | rs4934345 | 0.187 | -0.791 | 0.214 | 0.555 | -0.018 | 0.23 | 0.23 | 0.702 | 0.07 | 0.23 | 0.26 | 0.28 | 0.03 | 0.05 |
| 10 | 89318334 | rs2147287 | -0.425 | -0.779 | 0.048 | 0.506 | 0.028 | 0.081 | 0.081 | 0 | 0.54 | 0.82 | 0.66 | 0.7 | 0.64 | 0.55 |
| 10 | 89323177 | rs942423 | 0.536 | -0.862 | -0.42 | 0.324 | -0.136 | -0.066 | -0.066 | 0 | 0.39 | 0.07 | 0.33 | 0.39 | 0.54 | 0.43 |
| 10 | 89324663 | rs1575558 | -0.136 | -0.746 | 0.019 | 0.446 | -0.019 | 0.257 | 0.257 | 0.911 | 0.08 | 0.35 | 0.26 | 0.28 | 0.04 | 0.05 |
| 10 | 89327658 | rs11202449 | 0.822 | -0.489 | -0.194 | 0.226 | -0.033 | -0.035 | -0.035 | 0.245 | 0.1 | 0.01 | 0.08 | 0.15 | 0.13 | 0.05 |
| 10 | 89332144 | rs11202455 | 0.301 | -0.49 | 0.011 | 0.156 | 0.051 | 0.019 | 0.019 | 0.006 | 0.05 | 0.26 | 0.04 | 0.02 | 0.06 | 0.06 |
| 10 | 89335977 | rs7913974 | -0.259 | -0.513 | -0.75 | 0.097 | 0.03 | 0.091 | 0.091 | 0 | 0.52 | 0.77 | 0.64 | 0.7 | 0.62 | 0.51 |
| 10 | 89338633 | rs9664222 | -0.063 | 0.648 | -0.978 | 0.454 | -0.061 | -0.058 | -0.058 | 0 | 0.87 | 0.96 | 0.84 | 0.75 | 0.78 | 0.89 |
| 10 | 89339054 | rs7100988 | -0.544 | 0.1 | -0.796 | 0.3 | 0.001 | -0.002 | -0.002 | 0 | 0.07 | 0.09 | 0.05 | 0.06 | 0.07 | 0.09 |
| 10 | 89339063 | rs6586081 | -0.247 | 0.414 | -0.987 | 0.577 | -0.019 | -0.007 | -0.007 | 0 | 0.63 | 0.73 | 0.56 | 0.6 | 0.5 | 0.66 |
| 10 | 89340206 | rs10749553 | -0.413 | -0.163 | -0.968 | 0.173 | 0.022 | -0.004 | -0.004 | 1.881 | 0.59 | 0.66 | 0.64 | 0.62 | 0.67 | 0.6 |
| 10 | 89343804 | rs4934349 | 0.638 | 0.068 | -1.316 | 0.139 | -0.079 | -0.055 | -0.055 | 0.618 | 0.36 | 0.15 | 0.51 | 0.46 | 0.46 | 0.51 |
| 10 | 89352346 | rs12242864 | 0.771 | -0.348 | -0.831 | 0.003 | -0.039 | -0.04 | -0.04 | 0 | 0.07 | 0.15 | 0.01 | 0 | 0 | 0 |
| 10 | 89355313 | rs12771728 | 0.433 | -0.886 | -1.228 | -0.333 | -0.015 | 0.098 | 0.098 | 0 | 0.36 | 0.13 | 0.24 | 0.19 | 0.36 | 0.31 |
| 10 | 89356933 | rs12778872 | 0.857 | -0.819 | -1.408 | -0.401 | -0.024 | 0.015 | 0.015 | 0 | 0.17 | 0.06 | 0.13 | 0.15 | 0.19 | 0.14 |
| 10 | 89357516 | rs1886280 | -2.043 | -0.7 | -1.726 | -0.259 | -0.134 | -0.098 | -0.098 | 0 | 0.79 | 0.97 | 0.65 | 0.72 | 0.64 | 0.74 |
| 10 | 89360938 | rs12782065 | 1.925 | -0.701 | -1.654 | -0.262 | -0.044 | -0.017 | -0.017 | 0 | 0.11 | 0.01 | 0.09 | 0.12 | 0.14 | 0.07 |
| 10 | 89363781 | rs7894717 | 1.601 | -0.959 | -1.515 | -0.303 | 0.01 | 0.02 | 0.02 | 0 | 0.19 | 0.11 | 0.17 | 0.17 | 0.13 | 0.24 |
| 10 | 89364121 | rs6586085 | 0.462 | -1.196 | -0.684 | -0.32 | -0.078 | 0.142 | 0.142 | 0 | 0.2 | 0.43 | 0.21 | 0.36 | 0.11 | 0.17 |
| 10 | 89366175 | rs17108807 | 1.544 | -1.201 | -0.609 | -0.322 | -0.02 | 0.152 | 0.152 | 0 | 0.13 | 0.24 | 0.2 | 0.36 | 0.11 | 0.17 |
| 10 | 89367086 | rs17344791 | 0.578 | -1.04 | -1.055 | -0.482 | -0.032 | 0.099 | 0.099 | 0 | 0.36 | 0.09 | 0.24 | 0.19 | 0.36 | 0.31 |
| 10 | 89367724 | rs12249606 | 1.148 | -0.778 | -0.553 | -0.183 | -0.036 | -0.038 | -0.038 | 0.087 | 0.07 | 0.15 | 0.01 | 0 | 0 | 0 |
| 10 | 89368767 | rs10788555 | 2.494 | -0.382 | -0.968 | -0.14 | -0.078 | 0.005 | 0.005 | 0 | 0.15 | 0.02 | 0.26 | 0.12 | 0.25 | 0.16 |
| 10 | 89372233 | rs10887732 | -0.721 | -0.936 | -0.732 | -0.127 | -0.02 | -0.034 | -0.034 | 0 | 0.62 | 0.81 | 0.53 | 0.57 | 0.45 | 0.59 |
| 10 | 89374086 | rs17110864 | 0.976 | -1.054 | -0.54 | -0.091 | -0.014 | 0.045 | 0.045 | 0 | 0.15 | 0.15 | 0.2 | 0.36 | 0.11 | 0.18 |
| 10 | 89375896 | rs1324293 | 1.337 | -0.878 | -0.719 | -0.27 | -0.013 | 0 | 0 | 0 | 0.15 | 0.09 | 0.11 | 0.16 | 0.18 | 0.13 |
| 10 | 89376952 | rs11202472 | 0.311 | -1.063 | -0.399 | -0.318 | -0.005 | -0.027 | -0.027 | 0.046 | 0.4 | 0.45 | 0.31 | 0.21 | 0.33 | 0.41 |
| 10 | 89377790 | rs10887733 | 0.643 | -1.064 | -0.775 | -0.318 | 0.022 | 0.096 | 0.096 | 0 | 0.37 | 0.17 | 0.28 | 0.21 | 0.29 | 0.4 |
| 10 | 89379732 | rs7895494 | -0.036 | -1.207 | -0.675 | -0.032 | -0.008 | -0.012 | -0.012 | 0 | 0.4 | 0.57 | 0.3 | 0.28 | 0.32 | 0.49 |
| 10 | 89380317 | rs17111675 | 1.329 | -0.76 | -0.357 | 0.505 | -0.002 | 0.166 | 0.166 | 0 | 0.05 | 0.11 | 0.16 | 0.3 | 0.04 | 0.05 |
| 10 | 89380806 | rs1324295 | -0.169 | -0.948 | -0.742 | 0.362 | 0.003 | -0.006 | -0.006 | 0 | 0.36 | 0.64 | 0.41 | 0.51 | 0.3 | 0.41 |
| 10 | 89381993 | rs11202480 | -0.058 | -0.96 | -1.042 | 0.433 | -0.047 | 0.128 | 0.128 | 1.827 | 0.16 | 0.59 | 0.21 | 0.3 | 0.08 | 0.06 |
| 10 | 89385484 | rs1359277 | 0.884 | -0.821 | -1.678 | 0.449 | -0.029 | -0.03 | -0.03 | 0.095 | 0.31 | 0.08 | 0.29 | 0.32 | 0.3 | 0.29 |
| 10 | 89389578 | rs4934353 | 0.661 | -1.526 | -1.175 | 0.319 | 0.009 | 0.018 | 0.018 | 0.151 | 0.25 | 0.15 | 0.21 | 0.21 | 0.21 | 0.35 |
| 10 | 89393463 | rs1324299 | -0.495 | -1.332 | -1.528 | 0.231 | 0.071 | -0.012 | -0.012 | 0.166 | 0.42 | 0.8 | 0.35 | 0.43 | 0.3 | 0.4 |
| 10 | 89401231 | rs17114050 | 1.258 | -0.792 | -1.382 | 0.03 | -0.008 | 0.044 | 0.044 | 0 | 0.14 | 0.05 | 0.09 | 0.03 | 0.14 | 0.12 |
| 10 | 89406088 | rs791875 | 0.467 | -0.794 | -1.423 | 0.021 | -0.035 | 0.085 | 0.085 | 0 | 0.33 | 0.16 | 0.38 | 0.15 | 0.4 | 0.31 |
| 10 | 89406641 | rs10437508 | -0.085 | -1.386 | -0.914 | 0.163 | -0.016 | 0.009 | 0.009 | 1.332 | 0.28 | 0.66 | 0.26 | 0.34 | 0.18 | 0.31 |
| 10 | 89408246 | rs791881 | -0.755 | -1.418 | -0.91 | 0.156 | -0.058 | -0.068 | -0.068 | 0 | 0.61 | 0.86 | 0.63 | 0.49 | 0.58 | 0.61 |
| 10 | 89411241 | rs7092063 | -0.097 | -1.392 | -0.894 | 0.226 | -0.013 | 0.017 | 0.017 | 1.309 | 0.28 | 0.66 | 0.33 | 0.42 | 0.19 | 0.32 |
| 10 | 89411554 | rs791887 | 1.223 | -1.434 | -1.1 | 0.144 | -0.059 | 0.067 | 0.067 | 0 | 0.28 | 0.1 | 0.35 | 0.15 | 0.36 | 0.26 |
| 10 | 89412575 | rs791888 | 0.738 | -1.318 | -0.98 | 0.203 | -0.01 | 0.044 | 0.044 | 0 | 0.33 | 0.25 | 0.38 | 0.15 | 0.39 | 0.31 |
| 10 | 89415605 | rs2755430 | -0.52 | -0.935 | -1.081 | 0.334 | -0.059 | -0.023 | -0.023 | 0.005 | 0.61 | 0.86 | 0.7 | 0.57 | 0.58 | 0.62 |
| 10 | 89418253 | rs2755425 | -0.566 | -0.844 | -0.903 | 0.406 | -0.027 | -0.056 | -0.056 | 0.061 | 0.58 | 0.85 | 0.58 | 0.5 | 0.59 | 0.59 |
| 10 | 89418887 | rs1555433 | 0.02 | -1.197 | -0.376 | 0.262 | -0.016 | 0.015 | 0.015 | 0.778 | 0.28 | 0.56 | 0.23 | 0.33 | 0.23 | 0.33 |
| 10 | 89418979 | rs2145819 | -0.777 | -1.126 | -0.554 | 0.296 | -0.029 | -0.024 | -0.024 | 0.56 | 0.58 | 0.85 | 0.51 | 0.41 | 0.59 | 0.58 |
| 10 | 89423245 | rs10788561 | -1.063 | -0.954 | 0.298 | 0.208 | 0.027 | -0.089 | -0.089 | 0.241 | 0.73 | 0.59 | 0.73 | 0.91 | 0.66 | 0.79 |
| 10 | 89429897 | rs7085195 | 0.459 | -1.22 | 0.254 | 0.348 | -0.031 | -0.019 | -0.019 | 0 | 0.43 | 0.15 | 0.42 | 0.44 | 0.41 | 0.32 |
| 10 | 89431060 | rs7893652 | -1.986 | -1.309 | 0.33 | 0.288 | -0.025 | 0.028 | 0.028 | 0 | 0.79 | 0.86 | 0.78 | 0.91 | 0.66 | 0.77 |
| 10 | 89432059 | rs17117369 | 0.326 | -1.421 | 0.558 | 0.542 | 0.005 | 0 | 0 | 0.42 | 0.1 | 0.07 | 0.05 | 0.25 | 0.06 | 0.15 |
| 10 | 89432286 | rs9633742 | -0.879 | -1.382 | 0.883 | 0.374 | 0.006 | -0.027 | -0.027 | 0 | 0.59 | 0.45 | 0.59 | 0.78 | 0.51 | 0.49 |
| 10 | 89435501 | rs2077773 | -0.331 | -1.383 | 0.745 | 0.371 | -0.011 | 0.066 | 0.066 | 0.031 | 0.56 | 0.29 | 0.56 | 0.78 | 0.51 | 0.5 |
| 10 | 89440911 | rs7078618 | -1.072 | -1.356 | 0.697 | 0.349 | -0.03 | 0.052 | 0.052 | 0 | 0.63 | 0.73 | 0.61 | 0.78 | 0.53 | 0.51 |
| 10 | 89442697 | rs12570024 | -0.302 | -1.261 | 0.382 | 0.493 | -0.008 | 0.023 | 0.023 | 0.137 | 0.13 | 0.11 | 0.14 | 0.33 | 0.07 | 0.18 |
| 10 | 89444650 | rs7908056 | 0.213 | -1.027 | 0.2 | 0.261 | -0.032 | -0.006 | -0.006 | 0 | 0.45 | 0.2 | 0.44 | 0.45 | 0.46 | 0.33 |
| 10 | 89447687 | rs6586094 | -0.533 | -0.576 | 0.588 | 0.585 | 0.04 | 0.037 | 0.037 | 0.114 | 0.21 | 0.39 | 0.32 | 0.24 | 0.24 | 0.14 |
| 10 | 89450027 | rs7075381 | 1.225 | -0.919 | 0.068 | 0.59 | 0.001 | 0.005 | 0.005 | 0 | 0.17 | 0.15 | 0.16 | 0.13 | 0.14 | 0.25 |
| 10 | 89451830 | rs11202512 | 0.596 | -0.644 | -0.145 | 0.648 | -0.007 | -0.008 | -0.008 | 0 | 0.13 | 0.08 | 0.26 | 0.23 | 0.18 | 0.13 |
| 10 | 89454120 | rs12253888 | -0.093 | -0.892 | -0.042 | 0.611 | -0.031 | 0.01 | 0.01 | 0.619 | 0.27 | 0.47 | 0.29 | 0.33 | 0.18 | 0.33 |
| 10 | 89457071 | rs941829 | -0.804 | -0.91 | -0.937 | 0.639 | 0.016 | 0.028 | 0.028 | 0.794 | 0.48 | 0.62 | 0.61 | 0.64 | 0.43 | 0.61 |
| 10 | 89457161 | rs941830 | -1.275 | -1.058 | -0.977 | 0.561 | 0.008 | 0.04 | 0.04 | 0 | 0.54 | 0.8 | 0.64 | 0.64 | 0.43 | 0.61 |
| 10 | 89457851 | rs4934359 | 0.585 | -0.791 | -0.81 | 0.78 | 0.009 | -0.016 | -0.016 | 0.532 | 0.22 | 0.21 | 0.25 | 0.34 | 0.15 | 0.31 |

| | | | | | | | | | | | | | | | | |
|---|---|---|---|---|---|---|---|---|---|---|---|---|---|---|---|---|
| 10 | 89459858 | rs12267729 | 0.219 | -0.834 | -0.78 | 0.78 | 0.003 | 0.038 | 0.038 | 0 | 0.25 | 0.41 | 0.36 | 0.3 | 0.27 | 0.28 |
| 10 | 89460026 | rs12267849 | 0.435 | -0.529 | -1.406 | 0.948 | 0.011 | -0.018 | -0.018 | 0.732 | 0.22 | 0.19 | 0.25 | 0.34 | 0.15 | 0.32 |
| 10 | 89464125 | rs10887743 | -1.106 | 0.383 | -1.697 | 0.808 | -0.048 | -0.041 | -0.041 | 0 | 0.77 | 0.85 | 0.67 | 0.64 | 0.6 | 0.74 |
| 10 | 89464991 | rs4933458 | -0.333 | -0.064 | -1.495 | 0.66 | 0.004 | -0.013 | -0.013 | 0 | 0.42 | 0.78 | 0.54 | 0.43 | 0.4 | 0.51 |
| 10 | 89473477 | rs7898500 | -0.468 | 0.197 | -1.033 | 0.503 | 0.017 | 0.06 | 0.06 | 0 | 0.5 | 0.62 | 0.56 | 0.73 | 0.45 | 0.66 |
| 10 | 89474072 | rs2302404 | -0.376 | -0.219 | -0.623 | 0.949 | -0.016 | 0.031 | 0.031 | 0.045 | 0.14 | 0.18 | 0.12 | 0.43 | 0.07 | 0.18 |
| 10 | 89474288 | rs3815417 | -1.169 | -0.372 | -0.516 | 0.932 | 0.002 | 0.127 | 0.127 | 0.336 | 0.53 | 0.8 | 0.57 | 0.77 | 0.45 | 0.69 |
| 10 | 89475617 | rs2302405 | 0.361 | -0.066 | -0.348 | 0.856 | 0.003 | -0.002 | -0.002 | 0 | 0.09 | 0.09 | 0.08 | 0.07 | 0.15 | 0.06 |
| 10 | 89476881 | rs3781189 | 0.041 | -0.336 | -0.074 | 0.829 | -0.024 | -0.008 | -0.008 | 0.195 | 0.33 | 0.18 | 0.43 | 0.3 | 0.38 | 0.48 |
| 10 | 89479014 | rs7080768 | 0.035 | -0.284 | -0.356 | 0.862 | 0.008 | -0.002 | -0.002 | 0 | 0.35 | 0.41 | 0.45 | 0.3 | 0.38 | 0.48 |
| 10 | 89495364 | rs1321934 | -0.403 | -0.761 | -0.124 | 0.509 | 0.01 | 0.004 | 0.004 | 2.389 | 0.48 | 0.6 | 0.3 | 0.5 | 0.35 | 0.39 |
| 10 | 89507288 | rs1058798 | 0.836 | -0.87 | 0.152 | 0.652 | 0.004 | 0.012 | 0.012 | 0.62 | 0.32 | 0.3 | 0.24 | 0.09 | 0.29 | 0.24 |
| 10 | 89510229 | rs12765184 | 0.822 | -0.864 | 0.154 | 0.655 | 0.004 | 0.012 | 0.012 | 0.584 | 0.32 | 0.3 | 0.24 | 0.09 | 0.29 | 0.24 |
| 10 | 89512839 | rs12253113 | 0.594 | -0.88 | 0.196 | 0.648 | -0.004 | -0.031 | -0.031 | 0 | 0.33 | 0.39 | 0.25 | 0.09 | 0.29 | 0.24 |
| 10 | 89513118 | rs7481 | 0.774 | -0.875 | 0.207 | 0.66 | 0.003 | 0.005 | 0.005 | 0.467 | 0.32 | 0.32 | 0.25 | 0.09 | 0.29 | 0.24 |
| 10 | 89519376 | rs10509411 | 1.32 | -0.497 | -0.043 | 0.855 | -0.025 | 0.131 | 0.131 | 0.204 | 0.06 | 0.12 | 0.02 | 0.34 | 0.02 | 0.12 |
| 10 | 89534244 | rs2038554 | 0.183 | -0.01 | -0.779 | 0.164 | -0.011 | -0.037 | -0.037 | 0 | 0.38 | 0.46 | 0.27 | 0.15 | 0.3 | 0.25 |
| 10 | 89584284 | rs11202576 | 0.528 | 0.513 | -1.301 | 0.977 | 0.006 | -0.028 | -0.028 | 2.658 | 0.19 | 0.09 | 0.22 | 0.25 | 0.17 | 0.34 |
| 10 | 89585977 | rs17702687 | 0.018 | 0.921 | -1.68 | 0.707 | -0.03 | -0.014 | -0.014 | 0.003 | 0.31 | 0.23 | 0.41 | 0.33 | 0.41 | 0.22 |
| 10 | 89588888 | rs10887755 | 0.889 | 0.824 | -1.523 | 0.733 | -0.001 | 0 | 0 | 1.076 | 0.17 | 0.14 | 0.19 | 0.2 | 0.13 | 0.27 |
| 10 | 89595652 | rs12775504 | 0.426 | 0.569 | -1.791 | 0.002 | 0.026 | 0.078 | 0.078 | 0.02 | 0.28 | 0.15 | 0.21 | 0.04 | 0.21 | 0.15 |
| 10 | 89596558 | rs7078010 | -0.37 | 1.271 | -1.564 | 0.539 | -0.034 | -0.014 | -0.014 | 0 | 0.29 | 0.23 | 0.44 | 0.32 | 0.43 | 0.2 |
| 10 | 89599354 | rs10509412 | -0.171 | 1.789 | -1.918 | 0.866 | -0.092 | -0.102 | -0.102 | 0.017 | 0.24 | 0.09 | 0.36 | 0.52 | 0.39 | 0.18 |
| 10 | 89599576 | rs7076964 | -0.793 | 1.461 | -1.423 | 0.209 | 0 | 0.005 | 0.005 | 0 | 0.49 | 0.57 | 0.53 | 0.55 | 0.49 | 0.33 |
| 10 | 89603315 | rs10887758 | -0.631 | 1.681 | -1.258 | 0.307 | 0 | 0.013 | 0.013 | 0 | 0.14 | 0.12 | 0.24 | 0.35 | 0.2 | 0.22 |
| 10 | 89603424 | rs10887759 | 0.385 | 1.7 | -1.262 | 0.306 | -0.003 | 0.035 | 0.035 | 0 | 0.08 | 0.07 | 0.2 | 0.33 | 0.13 | 0.13 |
| 10 | 89604732 | rs10788569 | -1.077 | 1.24 | -1.383 | 0.029 | -0.009 | -0.016 | -0.016 | 0 | 0.22 | 0.15 | 0.27 | 0.36 | 0.26 | 0.26 |
| 10 | 89616479 | rs1234221 | 1.533 | 0.695 | -1.984 | -0.326 | -0.019 | -0.013 | -0.013 | 5.163 | 0.23 | 0.03 | 0.18 | 0.25 | 0.24 | 0.4 |
| 10 | 89617157 | rs11202589 | -0.466 | 1.111 | -1.524 | -0.275 | 0.003 | 0.017 | 0.017 | 0 | 0.13 | 0.13 | 0.24 | 0.35 | 0.2 | 0.2 |
| 10 | 89630424 | rs35161364 | 0.739 | 0.816 | -0.643 | -0.358 | -0.031 | -0.044 | -0.044 | 1.186 | 0.09 | 0.16 | 0.03 | 0.01 | 0.05 | 0.05 |
| 10 | 89649838 | rs7069904 | 1.636 | 0.65 | -0.671 | -0.336 | 0.008 | 0.006 | 0.006 | 0 | 0.07 | 0.03 | 0.02 | 0 | 0.05 | 0.03 |
| 10 | 89653686 | rs1903858 | -1.425 | 0.338 | -0.723 | -0.533 | -0.005 | 0.01 | 0.01 | 1.928 | 0.41 | 0.49 | 0.43 | 0.51 | 0.35 | 0.29 |
| 10 | 89673569 | rs1234225 | -1.4 | 0.421 | -0.672 | -0.525 | -0.001 | 0.006 | 0.006 | 1.377 | 0.41 | 0.52 | 0.44 | 0.5 | 0.37 | 0.29 |
| 10 | 89674157 | rs12569998 | 1.12 | 0.498 | -0.688 | -0.449 | -0.013 | 0.023 | 0.023 | 0 | 0.07 | 0.06 | 0.2 | 0.33 | 0.13 | 0.13 |
| 10 | 89721412 | rs926091 | 0.844 | 0.497 | -0.493 | -0.251 | 0.034 | 0.113 | 0.113 | 0 | 0.07 | 0.12 | 0.2 | 0.33 | 0.13 | 0.13 |
| 10 | 89727414 | rs11202607 | 2.388 | 0.468 | -0.839 | -0.275 | 0.013 | -0.001 | -0.001 | 0 | 0.1 | 0.05 | 0.13 | 0.14 | 0.08 | 0.03 |
| 10 | 89731870 | rs478839 | 1.604 | 0.416 | -0.909 | -0.387 | -0.019 | -0.06 | -0.06 | 0.499 | 0.34 | 0.12 | 0.36 | 0.43 | 0.39 | 0.66 |
| 10 | 89737554 | rs10509532 | 0.964 | 0.676 | -0.681 | 0.042 | -0.011 | 0.032 | 0.032 | 0 | 0.07 | 0.06 | 0.2 | 0.33 | 0.13 | 0.13 |
| 10 | 89741806 | rs809367 | 0.697 | 0.23 | -0.632 | -0.327 | 0.014 | 0.02 | 0.02 | 3.978 | 0.19 | 0.15 | 0.16 | 0.15 | 0.12 | 0.05 |
| 10 | 89745814 | rs644205 | -0.905 | 0.252 | -0.691 | -0.512 | 0 | -0.018 | -0.018 | 0 | 0.33 | 0.32 | 0.23 | 0.11 | 0.24 | 0.16 |
| 10 | 89746568 | rs647416 | 0.764 | 0.707 | -1.136 | -0.164 | 0.017 | 0.06 | 0.06 | 4.501 | 0.17 | 0.06 | 0.16 | 0.07 | 0.12 | 0.05 |
| 10 | 89754425 | rs11202613 | 1.73 | 0.91 | -1.081 | 0.053 | -0.001 | 0.021 | 0.021 | 0.803 | 0.1 | 0.02 | 0.13 | 0.06 | 0.08 | 0.03 |
| 10 | 89755075 | rs10887766 | 0.039 | 1.331 | -0.974 | 0.38 | 0.017 | 0.077 | 0.077 | 0 | 0.08 | 0.1 | 0.23 | 0.4 | 0.14 | 0.15 |
| 10 | 89756206 | rs10430641 | -0.495 | 1.431 | -1.076 | 0.314 | -0.047 | -0.068 | -0.068 | 1.543 | 0.22 | 0.12 | 0.39 | 0.47 | 0.28 | 0.22 |
| 10 | 89764144 | rs11202620 | 1.796 | 1.227 | -1.007 | 0.354 | -0.001 | 0.016 | 0.016 | 0.929 | 0.1 | 0.02 | 0.13 | 0.06 | 0.08 | 0.03 |
| 10 | 89768584 | rs10788575 | 0.397 | 1.506 | -0.771 | 0.568 | 0.011 | 0.072 | 0.072 | 0 | 0.07 | 0.08 | 0.23 | 0.39 | 0.13 | 0.13 |
| 10 | 89772989 | rs662115 | 0.308 | 0.706 | -0.704 | -0.051 | 0.018 | -0.011 | -0.011 | 0.752 | 0.47 | 0.3 | 0.49 | 0.5 | 0.46 | 0.7 |
| 10 | 89777788 | rs2244092 | -0.757 | 0.832 | -0.427 | -0.068 | 0.044 | 0.058 | 0.058 | 0 | 0.69 | 0.49 | 0.58 | 0.54 | 0.62 | 0.83 |
| 10 | 89778495 | rs2673825 | -0.014 | 0.231 | -0.678 | -0.731 | 0 | -0.013 | -0.013 | 0 | 0.44 | 0.55 | 0.35 | 0.19 | 0.35 | 0.44 |
| 10 | 89779324 | rs2673824 | 0.846 | 0.036 | -0.609 | -0.757 | 0.035 | 0.085 | 0.085 | 0 | 0.37 | 0.22 | 0.31 | 0.16 | 0.28 | 0.4 |
| 10 | 89780851 | rs719426 | -1.061 | 0.732 | -0.343 | -0.114 | 0.013 | 0.041 | 0.041 | 0 | 0.71 | 0.55 | 0.63 | 0.42 | 0.67 | 0.85 |
| 10 | 89782386 | rs2785079 | 0.993 | 0.263 | -0.542 | -0.596 | 0.002 | -0.001 | -0.001 | 0 | 0.23 | 0.2 | 0.15 | 0.09 | 0.19 | 0.3 |
| 10 | 89785190 | rs1414894 | -0.185 | 0.38 | 0.065 | -0.514 | 0.004 | 0.132 | 0.132 | 0 | 0.23 | 0.39 | 0.3 | 0.58 | 0.25 | 0.25 |
| 10 | 89785953 | rs4934374 | 0.967 | 0.256 | -0.262 | -0.719 | -0.044 | -0.038 | -0.038 | 0.454 | 0.24 | 0.04 | 0.26 | 0.21 | 0.28 | 0.37 |
| 10 | 89786353 | rs11202633 | 0.179 | 0.395 | 0.042 | -0.541 | 0.001 | 0.047 | 0.047 | 0 | 0.21 | 0.23 | 0.28 | 0.58 | 0.25 | 0.25 |
| 10 | 89787367 | rs2673823 | -2.245 | 0.381 | -0.073 | -0.667 | 0.005 | -0.042 | -0.042 | 0 | 0.78 | 0.65 | 0.81 | 0.95 | 0.77 | 0.94 |
| 10 | 89788638 | rs7908382 | -0.191 | 0.345 | -0.026 | -0.493 | -0.003 | 0.144 | 0.144 | 0 | 0.23 | 0.45 | 0.3 | 0.6 | 0.24 | 0.26 |
| 10 | 89792371 | rs2248456 | -0.143 | 0.34 | -0.131 | -0.644 | -0.002 | 0.001 | 0.001 | 0 | 0.56 | 0.43 | 0.53 | 0.36 | 0.51 | 0.69 |
| 10 | 89794947 | rs1414895 | 1.189 | 0.233 | -0.271 | -1.074 | -0.002 | 0.072 | 0.072 | 0.005 | 0.16 | 0.02 | 0.09 | 0.02 | 0.15 | 0.15 |
| 10 | 89799613 | rs692788 | 2.854 | 0.156 | -0.458 | -1.24 | -0.025 | 0.041 | 0.041 | 0.186 | 0.12 | 0.04 | 0.11 | 0.07 | 0.15 | 0.21 |
| 10 | 89800100 | rs760177 | -0.23 | 0.167 | -0.286 | -1.137 | -0.008 | 0.019 | 0.019 | 0 | 0.56 | 0.38 | 0.54 | 0.33 | 0.51 | 0.68 |
| 10 | 89801899 | rs2673813 | -0.272 | -0.053 | -0.289 | -1.221 | 0 | -0.002 | -0.002 | 0 | 0.56 | 0.46 | 0.54 | 0.33 | 0.51 | 0.68 |
| 10 | 89802409 | rs2735353 | -2.218 | 0.102 | -0.676 | -1.242 | -0.079 | 0.112 | 0.112 | 0 | 0.82 | 0.99 | 0.83 | 0.99 | 0.76 | 0.94 |
| 10 | 89807339 | rs12240758 | 0.608 | -0.296 | -0.367 | -1.216 | -0.003 | 0.003 | 0.003 | 0.645 | 0.27 | 0.22 | 0.27 | 0.18 | 0.28 | 0.43 |
| 10 | 89807366 | rs10887777 | 0.509 | -0.357 | -0.491 | -1.208 | -0.003 | -0.028 | -0.028 | 0 | 0.21 | 0.14 | 0.28 | 0.63 | 0.24 | 0.29 |
| 10 | 89807852 | rs587039 | 1.885 | -0.448 | -0.595 | -1.232 | 0.032 | 0.047 | 0.047 | 4.766 | 0.17 | 0.02 | 0.14 | 0.07 | 0.08 | 0.05 |
| 10 | 89808124 | rs588307 | -0.045 | -0.267 | -0.271 | -2.065 | 0.017 | 0.053 | 0.053 | 0.966 | 0.45 | 0.3 | 0.42 | 0.25 | 0.36 | 0.48 |
| 10 | 89810463 | rs559783 | -0.051 | -0.262 | -0.269 | -2.063 | 0.017 | 0.053 | 0.053 | 0.955 | 0.45 | 0.3 | 0.42 | 0.25 | 0.36 | 0.47 |
| 10 | 89814565 | rs2785066 | -0.812 | 0.081 | -0.166 | -2.599 | 0.01 | -0.069 | -0.069 | 0.992 | 0.65 | 0.47 | 0.7 | 0.87 | 0.61 | 0.76 |

| | | | | | | | | | | | | | | | | |
|---|---|---|---|---|---|---|---|---|---|---|---|---|---|---|---|---|
| 10 | 89817829 | rs545138 | -0.822 | 0.084 | -0.314 | -2.564 | 0.01 | -0.063 | -0.063 | 1.538 | 0.65 | 0.46 | 0.69 | 0.83 | 0.61 | 0.76 |
| 10 | 89819111 | rs7096511 | 0.428 | 0.227 | -0.586 | -2.187 | 0.01 | -0.039 | -0.039 | 0 | 0.13 | 0.18 | 0.08 | 0.03 | 0.14 | 0.15 |
| 10 | 89819819 | rs4603211 | 1.709 | -0.438 | -1.093 | -2.242 | -0.018 | 0.041 | 0.041 | 0 | 0.15 | 0.07 | 0.13 | 0.08 | 0.17 | 0.07 |
| 10 | 89820152 | rs4344399 | -0.554 | -1.749 | -0.801 | -2.509 | 0.017 | 0.012 | 0.012 | 0 | 0.38 | 0.64 | 0.19 | 0.09 | 0.27 | 0.37 |
| 10 | 89821646 | rs7084813 | 0.034 | -1.223 | -0.331 | -2.213 | -0.044 | -0.1 | -0.1 | 0 | 0.19 | 0.49 | 0.1 | 0.03 | 0.13 | 0.17 |
| 10 | 89822721 | rs1855970 | -1.938 | -1.11 | -0.576 | -2.225 | -0.043 | -0.002 | -0.002 | 0 | 0.9 | 0.98 | 0.89 | 0.9 | 0.85 | 0.83 |
| 10 | 89826130 | rs6586116 | -1.129 | -1.344 | -0.651 | -2.334 | 0.005 | 0.087 | 0.087 | 0 | 0.66 | 0.83 | 0.72 | 0.83 | 0.72 | 0.61 |
| 10 | 89826707 | rs12098338 | 0.363 | -1.403 | -0.584 | -2.338 | -0.04 | -0.091 | -0.091 | 0 | 0.19 | 0.37 | 0.09 | 0.03 | 0.13 | 0.17 |
| 10 | 89826804 | rs17108235 | 1.912 | -1.344 | -0.722 | -2.305 | 0.01 | 0.061 | 0.061 | 2.929 | 0.22 | 0.01 | 0.15 | 0.07 | 0.12 | 0.21 |
| 10 | 89828889 | rs2311740 | 1.145 | -1.094 | -0.616 | -2.167 | -0.052 | -0.019 | -0.019 | 0.051 | 0.29 | 0.18 | 0.42 | 0.68 | 0.43 | 0.2 |
| 10 | 89830746 | rs12247999 | -0.512 | -1.084 | -0.936 | -2.271 | 0.009 | 0.016 | 0.016 | 0.523 | 0.54 | 0.57 | 0.42 | 0.18 | 0.39 | 0.6 |
| 10 | 89833307 | rs7079271 | -0.044 | -1.199 | -0.45 | -2.406 | -0.011 | -0.063 | -0.063 | 0 | 0.7 | 0.46 | 0.78 | 0.87 | 0.72 | 0.65 |
| 10 | 89834396 | rs873418 | -0.447 | -1.022 | -0.996 | -2.328 | -0.006 | 0.029 | 0.029 | 0.55 | 0.77 | 0.82 | 0.8 | 0.88 | 0.73 | 0.65 |
| 10 | 89834899 | rs1889918 | 0.096 | -1.032 | -0.717 | -1.928 | -0.039 | -0.054 | -0.054 | 0 | 0.3 | 0.37 | 0.18 | 0.1 | 0.13 | 0.21 |
| 10 | 89838605 | rs1855969 | -0.361 | 0.259 | -0.453 | -1.005 | -0.05 | -0.147 | -0.147 | 0 | 0.3 | 0.49 | 0.18 | 0.06 | 0.22 | 0.2 |
| 10 | 89849519 | rs4933466 | -0.621 | 1.847 | -0.79 | -0.325 | -0.027 | 0.011 | 0.011 | 0 | 0.23 | 0.15 | 0.45 | 0.78 | 0.39 | 0.44 |
| 10 | 89851815 | rs2184697 | -0.251 | 1.253 | -0.448 | -1.117 | 0 | -0.085 | -0.085 | 0 | 0.34 | 0.35 | 0.21 | 0.03 | 0.31 | 0.19 |
| 10 | 89853442 | rs10887778 | 0.136 | 1.906 | -0.71 | -0.117 | -0.042 | -0.065 | -0.065 | 0 | 0.15 | 0.05 | 0.16 | 0.46 | 0.25 | 0.29 |
| 10 | 89855562 | rs7070956 | -1.159 | 1.763 | -0.261 | -0.137 | 0.034 | 0.02 | 0.02 | 0 | 0.57 | 0.75 | 0.72 | 0.66 | 0.65 | 0.55 |
| 10 | 89857019 | rs4933467 | 0.121 | 1.627 | 0.267 | -0.024 | 0.02 | 0.037 | 0.037 | 0 | 0.08 | 0.22 | 0.35 | 0.17 | 0.11 | 0.06 |
| 10 | 89857817 | rs11202654 | -0.253 | 1.529 | 0.565 | -0.049 | 0.015 | 0.036 | 0.036 | 0.087 | 0.09 | 0.27 | 0.35 | 0.17 | 0.11 | 0.07 |
| 10 | 89865548 | rs10159513 | -0.295 | 1.637 | 0.4 | -0.019 | 0.074 | 0.218 | 0.218 | 0.005 | 0.19 | 0.45 | 0.21 | 0.47 | 0.28 | 0.33 |
| 10 | 89866649 | rs11202656 | 0.2 | 1.655 | 0.229 | 0.011 | 0.009 | 0.044 | 0.044 | 0 | 0.16 | 0.15 | 0.18 | 0.47 | 0.28 | 0.33 |
| 10 | 89868930 | rs1932531 | -0.583 | 1.635 | 0.029 | -0.057 | -0.028 | 0.11 | 0.11 | 0 | 0.45 | 0.8 | 0.56 | 0.27 | 0.43 | 0.23 |
| 10 | 89875683 | rs7079754 | -0.217 | 1.745 | 0.058 | -0.125 | 0.042 | 0.078 | 0.078 | 1.74 | 0.13 | 0.48 | 0.37 | 0.19 | 0.11 | 0.05 |
| 10 | 89876255 | rs17322612 | -0.111 | 1.274 | -0.624 | -0.466 | -0.001 | 0.023 | 0.023 | 0 | 0.36 | 0.25 | 0.21 | 0.09 | 0.35 | 0.21 |
| 10 | 89886204 | rs7069053 | -0.878 | 0.918 | -1.502 | -0.49 | 0.003 | 0.085 | 0.085 | 0.213 | 0.55 | 0.94 | 0.75 | 0.73 | 0.66 | 0.52 |
| 10 | 89887141 | rs17109310 | 0.265 | 0.789 | -0.768 | -0.366 | -0.005 | 0.005 | 0.005 | 0 | 0.06 | 0.11 | 0.05 | 0.06 | 0.04 | 0.05 |
| 10 | 89891695 | rs17420447 | -0.07 | 0.867 | -0.738 | -0.434 | -0.001 | 0.001 | 0.001 | 0 | 0.24 | 0.18 | 0.18 | 0.07 | 0.26 | 0.13 |
| 10 | 89895039 | rs10159855 | -0.758 | 0.64 | -0.569 | -0.257 | -0.008 | -0.005 | -0.005 | 0 | 0.13 | 0.18 | 0.06 | 0.1 | 0.09 | 0.11 |
| 10 | 89895223 | rs11202660 | 0.221 | 1.092 | -1.018 | 0.77 | -0.042 | -0.077 | -0.077 | 0 | 0.11 | 0.01 | 0.09 | 0.31 | 0.18 | 0.18 |
| 10 | 89895807 | rs11202661 | -0.21 | 0.341 | -1.453 | 0.598 | 0.002 | 0 | 0 | 0 | 0.23 | 0.18 | 0.1 | 0.28 | 0.14 | 0.29 |
| 10 | 89897044 | rs10159576 | -0.088 | 0.269 | -1.406 | 0.47 | -0.009 | -0.004 | -0.004 | 0 | 0.13 | 0.15 | 0.04 | 0.12 | 0.06 | 0.12 |
| 10 | 89898756 | rs10887788 | -0.355 | 0.567 | -1.153 | -0.015 | -0.017 | -0.023 | -0.023 | 0 | 0.38 | 0.27 | 0.29 | 0.48 | 0.47 | 0.55 |
| 10 | 89899843 | rs7072702 | -1.624 | 0.364 | -0.884 | 0.037 | -0.001 | 0.011 | 0.011 | 0 | 0.58 | 0.66 | 0.38 | 0.68 | 0.58 | 0.78 |
| 10 | 89903491 | rs10444136 | -0.481 | 0.652 | -1.051 | 0.157 | -0.006 | -0.008 | -0.008 | 0 | 0.4 | 0.37 | 0.3 | 0.48 | 0.47 | 0.57 |
| 10 | 89905498 | rs12782006 | 0.398 | 0.991 | -1.015 | 0.096 | 0.002 | -0.002 | -0.002 | 1.712 | 0.06 | 0.04 | 0.09 | 0 | 0.17 | 0.02 |
| 10 | 89906286 | rs10509536 | -0.452 | -0.092 | -1.232 | 0.53 | -0.061 | -0.023 | -0.023 | 0.039 | 0.37 | 0.13 | 0.29 | 0.4 | 0.43 | 0.59 |
| 10 | 89909218 | rs10887791 | -0.945 | -0.398 | -0.827 | 0.225 | 0.015 | 0.014 | 0.014 | 0 | 0.5 | 0.43 | 0.36 | 0.61 | 0.55 | 0.66 |
| 10 | 89924902 | rs7082696 | -0.046 | -0.579 | -0.461 | -0.177 | -0.001 | 0.001 | 0.001 | 0 | 0.45 | 0.4 | 0.37 | 0.57 | 0.48 | 0.61 |
| 10 | 89931019 | rs531325 | 1.379 | -0.878 | -0.648 | -0.665 | -0.005 | -0.052 | -0.052 | 0.192 | 0.17 | 0.17 | 0.12 | 0 | 0.19 | 0.09 |
| 10 | 89944820 | rs756244 | 1.356 | -0.981 | -0.772 | -0.572 | -0.013 | -0.012 | -0.012 | 0 | 0.27 | 0.2 | 0.25 | 0.4 | 0.38 | 0.28 |
| 10 | 89946829 | rs11598117 | 1.113 | -0.414 | -1.236 | -0.298 | 0.004 | 0.005 | 0.005 | 0.008 | 0.14 | 0.11 | 0.28 | 0.1 | 0.12 | 0.14 |
| 10 | 89950581 | rs868872 | 1.206 | 0.032 | -1.085 | 0.195 | 0 | 0.034 | 0.034 | 0 | 0.16 | 0.04 | 0.11 | 0 | 0.19 | 0.09 |
| 10 | 89953613 | rs590392 | 0.878 | 0.221 | -1.047 | 0.436 | -0.003 | -0.046 | -0.046 | 0 | 0.08 | 0 | 0.09 | 0.13 | 0.06 | 0.16 |
| 10 | 89957523 | rs12250898 | 0.47 | 0.592 | -0.524 | -0.02 | -0.002 | -0.007 | -0.007 | 0 | 0.1 | 0.13 | 0.05 | 0.04 | 0.08 | 0.07 |
| 10 | 89958603 | rs510727 | -0.856 | 0.64 | -0.59 | 0.033 | -0.024 | -0.037 | -0.037 | 0 | 0.91 | 0.93 | 0.86 | 0.74 | 0.91 | 0.89 |
| 10 | 89961657 | rs575510 | 0.68 | 0.611 | -0.454 | -0.012 | 0.007 | 0.039 | 0.039 | 0.043 | 0.85 | 0.72 | 0.75 | 0.68 | 0.85 | 0.74 |
| 10 | 89964299 | rs501751 | -0.862 | 0.65 | -0.555 | -0.009 | -0.01 | -0.008 | -0.008 | 0 | 0.91 | 0.91 | 0.85 | 0.74 | 0.91 | 0.89 |
| 10 | 89965320 | rs545874 | 1.193 | 0.527 | -0.559 | -0.147 | -0.001 | -0.002 | -0.002 | 0 | 0.55 | 0.42 | 0.59 | 0.64 | 0.59 | 0.58 |
| 10 | 89965473 | rs533722 | 1.507 | 1.06 | -0.558 | 0.736 | -0.004 | 0.012 | 0.012 | 0 | 0.73 | 0.59 | 0.7 | 0.64 | 0.77 | 0.68 |
| 10 | 89966471 | rs577880 | -0.012 | 1.464 | -0.619 | 0.885 | 0.003 | 0.026 | 0.026 | 0 | 0.9 | 0.85 | 0.85 | 0.74 | 0.91 | 0.89 |
| 10 | 89968910 | rs629959 | 0.996 | 1.606 | -0.964 | 0.944 | 0 | 0.007 | 0.007 | 0 | 0.79 | 0.73 | 0.73 | 0.69 | 0.79 | 0.68 |
| 10 | 89970531 | rs12266972 | -0.7 | 1.215 | -0.781 | 0.295 | -0.008 | -0.026 | -0.026 | 0.991 | 0.16 | 0.25 | 0.15 | 0.1 | 0.14 | 0.22 |
| 10 | 89976112 | rs12773789 | -1.126 | 1.997 | -1.096 | 0.575 | -0.002 | -0.003 | -0.003 | 2.195 | 0.06 | 0.06 | 0.21 | 0.09 | 0.15 | 0.06 |
| 10 | 89978105 | rs7924020 | 0.179 | 1.574 | -1.713 | 0.486 | -0.013 | -0.008 | -0.008 | 0.013 | 0.63 | 0.8 | 0.68 | 0.65 | 0.57 | 0.64 |
| 10 | 89978800 | rs11202681 | -0.891 | 0.861 | -1.523 | 0.171 | -0.007 | -0.005 | -0.005 | 0 | 0.18 | 0.26 | 0.27 | 0.11 | 0.17 | 0.16 |
| 10 | 89980911 | rs1556382 | 0.319 | 0.716 | -1.484 | -0.254 | 0.015 | 0.005 | 0.005 | 0 | 0.38 | 0.43 | 0.44 | 0.55 | 0.47 | 0.4 |
| 10 | 89982343 | rs4934383 | -0.796 | 0.373 | -1.624 | -0.452 | 0.009 | 0.118 | 0.118 | 0 | 0.58 | 0.82 | 0.75 | 0.78 | 0.65 | 0.59 |
| 10 | 89987311 | rs11202686 | -0.23 | 0.259 | -1.545 | -0.389 | -0.012 | -0.017 | -0.017 | 0.709 | 0.08 | 0.02 | 0.09 | 0.1 | 0.1 | 0.07 |
| 10 | 89990241 | rs472829 | -1.437 | 0.112 | -1.934 | -0.803 | 0.005 | -0.002 | -0.002 | 0.172 | 0.8 | 0.92 | 0.86 | 0.8 | 0.84 | 0.82 |
| 10 | 89990547 | rs12220927 | 0.628 | 0.264 | -2.015 | -0.874 | 0.004 | 0.052 | 0.052 | 0 | 0.19 | 0.09 | 0.1 | 0.01 | 0.16 | 0.22 |
| 10 | 89993044 | rs2871984 | 1.016 | 0.361 | -1.688 | -0.63 | 0.042 | 0.023 | 0.023 | 0 | 0.34 | 0.46 | 0.43 | 0.57 | 0.47 | 0.41 |
| 10 | 89993471 | rs2026914 | -0.055 | 0.315 | -1.61 | -0.736 | 0.022 | 0.053 | 0.053 | 0 | 0.55 | 0.66 | 0.73 | 0.79 | 0.67 | 0.57 |
| 10 | 89997495 | rs7095034 | -0.432 | 1.119 | -1.204 | -1.005 | 0.016 | -0.025 | -0.025 | 0.015 | 0.3 | 0.31 | 0.34 | 0.14 | 0.42 | 0.31 |
| 10 | 89998283 | rs10509539 | 0.618 | -0.126 | -1.183 | -0.831 | 0.002 | 0 | 0 | 0 | 0.35 | 0.31 | 0.4 | 0.58 | 0.3 | 0.43 |
| 10 | 90003768 | rs685578 | -0.249 | 0.444 | -1.066 | -0.578 | -0.002 | 0.031 | 0.031 | 0 | 0.2 | 0.11 | 0.24 | 0.04 | 0.19 | 0.21 |
| 10 | 90006507 | rs17110348 | -0.627 | 0.302 | -0.716 | 0.031 | 0.007 | -0.035 | -0.035 | 0.019 | 0.08 | 0 | 0.07 | 0.13 | 0.05 | 0.04 |
| 10 | 90008568 | rs12776142 | -0.999 | 0.454 | -0.018 | -0.123 | -0.001 | -0.066 | -0.066 | 0.357 | 0.11 | 0.24 | 0.12 | 0 | 0.07 | 0.02 |

| | | | | | | | | | | | | | | | | |
|---|---|---|---|---|---|---|---|---|---|---|---|---|---|---|---|---|
| 10 | 90013189 | rs670472 | 0.912 | 0.524 | -0.753 | 0.301 | 0.003 | 0.044 | 0.044 | 1.299 | 0.6 | 0.17 | 0.61 | 0.66 | 0.65 | 0.53 |
| 10 | 90015274 | rs10887794 | 1.288 | 0.459 | -0.141 | 0.631 | -0.005 | 0.054 | 0.054 | 0.527 | 0.07 | 0.13 | 0.13 | 0.22 | 0.04 | 0.09 |
| 10 | 90015725 | rs17110418 | 0.025 | 0.374 | 0.199 | 0.501 | -0.002 | -0.017 | -0.017 | 0 | 0.09 | 0.1 | 0.09 | 0 | 0.04 | 0.07 |
| 10 | 90015785 | rs11202694 | 1.354 | 0.408 | 0.188 | 0.547 | -0.005 | 0.057 | 0.057 | 0.527 | 0.07 | 0.13 | 0.13 | 0.22 | 0.04 | 0.09 |
| 10 | 90020220 | rs7086615 | 0.488 | 0.359 | 0.491 | 0.547 | -0.012 | 0.054 | 0.054 | 1.95 | 0.09 | 0.18 | 0.15 | 0.24 | 0.05 | 0.09 |
| 10 | 90021251 | rs286493 | -1.931 | 0.19 | 0.334 | 1.16 | 0.01 | 0.04 | 0.04 | 0.504 | 0.45 | 0.81 | 0.49 | 0.67 | 0.43 | 0.55 |
| 10 | 90022647 | rs286492 | -1.697 | 0.073 | 0.886 | 1.318 | -0.052 | 0.071 | 0.071 | 0.421 | 0.23 | 0.58 | 0.25 | 0.45 | 0.17 | 0.21 |
| 10 | 90023033 | rs10509540 | -0.775 | 0.113 | 0.84 | 1.307 | -0.004 | 0 | 0 | 0.011 | 0.23 | 0.22 | 0.24 | 0.19 | 0.26 | 0.34 |
| 10 | 90026566 | rs10788589 | 1.404 | 0.08 | 0.86 | 1.324 | -0.008 | 0.053 | 0.053 | 0.477 | 0.07 | 0.14 | 0.14 | 0.24 | 0.05 | 0.09 |
| 10 | 90028338 | rs286490 | -0.415 | 0.056 | 1.232 | 1.268 | -0.012 | 0.084 | 0.084 | 0.064 | 0.08 | 0.23 | 0.07 | 0.22 | 0.09 | 0.08 |
| 10 | 90028352 | rs286489 | -0.773 | 0.016 | 1.429 | 1.257 | -0.018 | 0.122 | 0.122 | 0.379 | 0.1 | 0.32 | 0.07 | 0.24 | 0.09 | 0.11 |
| 10 | 90029028 | rs286488 | -1.216 | -0.02 | 1.386 | 1.214 | -0.027 | 0.017 | 0.017 | 0 | 0.18 | 0.33 | 0.17 | 0.23 | 0.13 | 0.14 |
| 10 | 90031903 | rs17110523 | 0.448 | 0.084 | 1.285 | 1.268 | -0.005 | -0.004 | -0.004 | 0.026 | 0.13 | 0.11 | 0.14 | 0.18 | 0.21 | 0.26 |
| 10 | 90033593 | rs2114406 | 2.121 | 0.546 | 0.66 | 1.029 | 0.062 | 0.099 | 0.099 | 1.952 | 0.42 | 0.14 | 0.3 | 0.23 | 0.36 | 0.39 |
| 10 | 90034488 | rs10749568 | 0.492 | 0.808 | 0.554 | 0.854 | -0.019 | 0.019 | 0.019 | 0 | 0.84 | 0.7 | 0.84 | 0.78 | 0.88 | 0.86 |
| 10 | 90038033 | rs10749570 | 1.011 | 0.879 | 0.471 | 0.88 | 0.013 | 0.051 | 0.051 | 0.007 | 0.7 | 0.47 | 0.73 | 0.58 | 0.65 | 0.62 |
| 10 | 90044345 | rs10749571 | 0.203 | 0.89 | 0.368 | 0.938 | -0.012 | 0.035 | 0.035 | 0 | 0.89 | 0.82 | 0.89 | 0.78 | 0.87 | 0.91 |
| 10 | 90044789 | rs1048956 | 0.466 | 0.917 | -0.023 | 0.645 | 0 | 0.002 | 0.002 | 0 | 0.63 | 0.37 | 0.65 | 0.56 | 0.61 | 0.55 |
| 10 | 90047709 | rs2162361 | 0.533 | 1.057 | -0.524 | 0.648 | -0.092 | -0.036 | -0.036 | 0 | 0.14 | 0.01 | 0.22 | 0.12 | 0.22 | 0.09 |
| 10 | 90051697 | rs1035796 | 0.799 | 0.594 | -0.021 | 0.62 | 0 | 0.013 | 0.013 | 0 | 0.63 | 0.43 | 0.65 | 0.57 | 0.62 | 0.56 |
| 10 | 90051857 | rs1035797 | 0.668 | 0.759 | -0.338 | 0.678 | 0.004 | 0.022 | 0.022 | 0 | 0.64 | 0.54 | 0.67 | 0.56 | 0.62 | 0.55 |
| 10 | 90053738 | rs1537142 | 0.428 | 0.514 | -1.097 | -0.183 | -0.004 | 0.038 | 0.038 | 0.052 | 0.79 | 0.86 | 0.82 | 1 | 0.82 | 0.91 |
| 10 | 90054268 | rs11202702 | 0.173 | 0.056 | -1.454 | -0.092 | -0.008 | -0.03 | -0.03 | 0.342 | 0.39 | 0.25 | 0.38 | 0.5 | 0.45 | 0.53 |
| 10 | 90069353 | rs12572813 | 0.522 | -0.15 | -1.812 | -0.002 | 0.004 | -0.013 | -0.013 | 0.044 | 0.15 | 0.11 | 0.12 | 0.36 | 0.14 | 0.22 |
| 10 | 90072352 | rs10509543 | 0.854 | -0.307 | -1.941 | -0.047 | -0.04 | 0.015 | 0.015 | 0 | 0.21 | 0.08 | 0.27 | 0.15 | 0.3 | 0.29 |
| 10 | 90075843 | rs10887800 | -0.378 | -0.878 | -2.237 | -0.653 | 0.006 | 0 | 0 | 0 | 0.5 | 0.29 | 0.45 | 0.51 | 0.52 | 0.62 |
| 10 | 90079714 | rs10887801 | -0.064 | -0.744 | -2.205 | -0.585 | -0.005 | -0.02 | -0.02 | 0 | 0.39 | 0.28 | 0.38 | 0.53 | 0.45 | 0.53 |
| 10 | 90093472 | rs7914501 | -0.296 | -0.733 | -2.549 | -0.979 | -0.055 | 0.193 | 0.193 | 0.063 | 0.66 | 0.89 | 0.63 | 0.99 | 0.64 | 0.76 |
| 10 | 90095434 | rs3781201 | 0.259 | -0.86 | -2.62 | -0.921 | 0.031 | -0.016 | -0.016 | 0 | 0.29 | 0.16 | 0.21 | 0.36 | 0.22 | 0.33 |
| 10 | 90095729 | rs1426621 | -0.941 | -0.616 | -2.596 | -0.877 | -0.075 | 0.095 | 0.095 | 0.122 | 0.8 | 0.93 | 0.7 | 0.99 | 0.71 | 0.87 |
| 10 | 90095915 | rs17424286 | 0.402 | -0.614 | -2.385 | -0.723 | 0.009 | 0.09 | 0.09 | 0 | 0.38 | 0.18 | 0.33 | 0.17 | 0.37 | 0.41 |
| 10 | 90119807 | rs1648512 | -0.789 | -0.394 | -2.26 | -0.399 | -0.005 | -0.086 | -0.086 | 0.01 | 0.81 | 0.97 | 0.83 | 0.63 | 0.82 | 0.81 |
| 10 | 90137045 | rs6586129 | 0.663 | -0.705 | -1.765 | -0.523 | -0.007 | 0.041 | 0.041 | 0 | 0.32 | 0.5 | 0.23 | 0.44 | 0.28 | 0.28 |
| 10 | 90139234 | rs792219 | 0.446 | -0.899 | -2.26 | -0.84 | -0.053 | 0.199 | 0.199 | 0 | 0.33 | 0.09 | 0.35 | 0.01 | 0.33 | 0.23 |
| 10 | 90144950 | rs701893 | -0.844 | -0.95 | -1.532 | -0.296 | 0 | -0.002 | -0.002 | 1.161 | 0.67 | 0.69 | 0.6 | 0.71 | 0.62 | 0.57 |
| 10 | 90147344 | rs792232 | -0.327 | -0.952 | -1.618 | 0.011 | 0.006 | -0.002 | -0.002 | 0.027 | 0.87 | 0.9 | 0.93 | 0.62 | 0.9 | 0.8 |
| 10 | 90155863 | rs3891849 | 0.925 | -1.048 | -1.564 | -0.518 | 0.032 | 0.043 | 0.043 | 0 | 0.19 | 0.14 | 0.23 | 0.05 | 0.1 | 0.17 |
| 10 | 90167065 | rs792207 | -0.755 | -1.154 | -1.695 | -0.444 | -0.01 | -0.02 | -0.02 | 0.936 | 0.65 | 0.77 | 0.68 | 0.51 | 0.58 | 0.69 |
| 10 | 90168029 | rs17096268 | 0.679 | -1.23 | -1.714 | -0.482 | -0.015 | -0.013 | -0.013 | 0 | 0.06 | 0.11 | 0.09 | 0.01 | 0.03 | 0.01 |
| 10 | 90168189 | rs813782 | -0.966 | -1.272 | -1.774 | -0.477 | -0.038 | -0.078 | -0.078 | 0 | 0.68 | 0.88 | 0.77 | 0.51 | 0.6 | 0.68 |
| 10 | 90175462 | rs806694 | -0.93 | -1.286 | -1.759 | -0.483 | -0.042 | -0.086 | -0.086 | 0.69 | 0.67 | 0.88 | 0.76 | 0.51 | 0.6 | 0.68 |
| 10 | 90183038 | rs7918921 | -0.306 | -1.413 | -1.628 | -0.539 | 0 | -0.004 | -0.004 | 0 | 0.58 | 0.65 | 0.56 | 0.49 | 0.56 | 0.52 |
| 10 | 90183279 | rs10788597 | 0.895 | -1.176 | -1.749 | -0.645 | -0.01 | 0.065 | 0.065 | 0.225 | 0.29 | 0.49 | 0.22 | 0.43 | 0.24 | 0.29 |
| 10 | 90185847 | rs4934404 | -1.363 | -1.25 | -1.893 | -0.725 | -0.06 | -0.098 | -0.098 | 0 | 0.71 | 0.89 | 0.77 | 0.47 | 0.62 | 0.71 |
| 10 | 90188684 | rs4934405 | 0.838 | -1.259 | -1.805 | -0.726 | -0.015 | -0.013 | -0.013 | 1.11 | 0.06 | 0.1 | 0.1 | 0.01 | 0.03 | 0.01 |
| 10 | 90190619 | rs10788600 | -1.316 | -1.078 | -1.851 | -0.65 | -0.065 | -0.104 | -0.104 | 0 | 0.71 | 0.91 | 0.77 | 0.48 | 0.62 | 0.71 |
| 10 | 90195149 | rs1935581 | -0.57 | -1.181 | -1.851 | -0.664 | -0.009 | -0.066 | -0.066 | 0 | 0.61 | 0.87 | 0.66 | 0.47 | 0.39 | 0.64 |
| 10 | 90198999 | rs10887820 | 0.667 | -0.907 | -1.593 | -0.52 | -0.014 | 0.03 | 0.03 | 0.473 | 0.3 | 0.51 | 0.22 | 0.38 | 0.24 | 0.29 |
| 10 | 90202441 | rs10749586 | -0.064 | -0.813 | -1.892 | -0.453 | -0.019 | 0.016 | 0.016 | 0 | 0.43 | 0.68 | 0.44 | 0.38 | 0.47 | 0.51 |
| 10 | 90211164 | rs10749590 | -1.332 | -1.042 | -2.097 | -0.511 | -0.084 | -0.123 | -0.123 | 0 | 0.72 | 0.94 | 0.77 | 0.49 | 0.62 | 0.73 |
| 10 | 90212999 | rs2153855 | 0.834 | -1.223 | -2.105 | -0.58 | 0.038 | 0.041 | 0.041 | 0 | 0.25 | 0.15 | 0.32 | 0.08 | 0.13 | 0.21 |
| 10 | 90213213 | rs7096710 | 0.775 | -1.15 | -2.061 | -0.643 | -0.01 | 0.006 | 0.006 | 0.236 | 0.32 | 0.59 | 0.33 | 0.38 | 0.24 | 0.44 |
| 10 | 90214051 | rs1342455 | 0.254 | -1.02 | -2.038 | -0.747 | 0.039 | 0.037 | 0.037 | 0 | 0.29 | 0.2 | 0.33 | 0.11 | 0.15 | 0.23 |
| 10 | 90214993 | rs10509549 | 1.314 | -0.945 | -2.069 | -0.735 | -0.024 | 0.019 | 0.019 | 0 | 0.1 | 0.06 | 0.11 | 0 | 0.22 | 0.06 |
| 10 | 90217922 | rs4933483 | 0.352 | -0.985 | -2.067 | -0.742 | 0.021 | 0.02 | 0.02 | 0 | 0.28 | 0.25 | 0.33 | 0.1 | 0.15 | 0.22 |
| 10 | 90219298 | rs10509550 | 1.137 | -0.658 | -1.956 | -0.617 | -0.007 | -0.001 | -0.001 | 4.415 | 0.11 | 0.08 | 0.11 | 0 | 0.22 | 0.06 |
| 10 | 90228162 | rs2488271 | -0.246 | -0.943 | -2.037 | -0.425 | -0.025 | 0.009 | 0.009 | 0.06 | 0.55 | 0.27 | 0.56 | 0.55 | 0.55 | 0.46 |
| 10 | 90237398 | rs1325907 | -0.251 | -0.918 | -2 | -0.437 | -0.013 | 0.007 | 0.007 | 0.007 | 0.55 | 0.32 | 0.56 | 0.55 | 0.52 | 0.47 |
| 10 | 90254417 | rs10509551 | 0.356 | -1.129 | -2.069 | -0.466 | 0.028 | 0.027 | 0.027 | 0 | 0.29 | 0.24 | 0.33 | 0.1 | 0.15 | 0.2 |
| 10 | 90259437 | rs10887829 | 0.675 | -1.055 | -2.08 | -0.476 | -0.004 | 0.004 | 0.004 | 0.478 | 0.35 | 0.61 | 0.34 | 0.45 | 0.27 | 0.47 |
| 10 | 90268918 | rs10887831 | 0.254 | -1.272 | -2.039 | -0.441 | -0.009 | -0.003 | -0.003 | 0 | 0.26 | 0.26 | 0.31 | 0.1 | 0.13 | 0.15 |
| 10 | 90269567 | rs2765440 | 0.006 | -0.426 | -2.352 | -0.657 | 0.072 | -0.071 | -0.071 | 0 | 0.44 | 0.09 | 0.36 | 0.49 | 0.32 | 0.38 |
| 10 | 90278656 | rs2477952 | -0.382 | -0.358 | -2.322 | -0.632 | 0.013 | 0.027 | 0.027 | 0 | 0.51 | 0.3 | 0.53 | 0.57 | 0.46 | 0.44 |
| 10 | 90279372 | rs1952122 | 1.201 | -0.555 | -2.54 | -0.887 | -0.003 | 0.011 | 0.011 | 0.565 | 0.31 | 0.61 | 0.31 | 0.43 | 0.3 | 0.44 |
| 10 | 90280938 | rs1325904 | -1.575 | -0.493 | -2.534 | -0.871 | -0.053 | 0.056 | 0.056 | 0 | 0.85 | 0.94 | 0.85 | 1 | 0.77 | 0.87 |
| 10 | 90286617 | rs2209553 | 0.979 | -0.45 | -2.444 | -0.694 | -0.082 | -0.131 | -0.131 | 6.297 | 0.25 | 0.04 | 0.21 | 0.46 | 0.32 | 0.24 |
| 10 | 90295773 | rs11202766 | 0.053 | -1.042 | -2.001 | -0.567 | 0.002 | 0.003 | 0.003 | 0 | 0.28 | 0.27 | 0.33 | 0.1 | 0.15 | 0.17 |
| 10 | 90297216 | rs11202767 | -0.031 | -0.523 | -1.658 | -0.224 | -0.047 | 0.007 | 0.007 | 0.05 | 0.11 | 0.26 | 0.16 | 0.09 | 0.06 | 0.06 |
| 10 | 90297927 | rs10788606 | -0.016 | -0.034 | -1.452 | -0.216 | 0.017 | 0.008 | 0.008 | 0 | 0.49 | 0.53 | 0.46 | 0.43 | 0.36 | 0.47 |

| | | | | | | | | | | | | | | | |
|---|---|---|---|---|---|---|---|---|---|---|---|---|---|---|---|
| 10 | 90311317 | rs7901991 | -0.018 | 0.125 | -1.558 | -0.197 | 0.033 | 0.013 | 0.013 | 0 | 0.51 | 0.66 | 0.49 | 0.44 | 0.4 | 0.48 |
| 10 | 90334554 | rs7904690 | 0.193 | -0.312 | -1.353 | -0.16 | -0.019 | -0.014 | -0.014 | 0.708 | 0.41 | 0.51 | 0.41 | 0.32 | 0.31 | 0.46 |
| 10 | 90343398 | rs2576178 | 0.637 | -0.22 | -1.753 | 0.011 | 0.005 | -0.025 | -0.025 | 0 | 0.28 | 0.27 | 0.44 | 0.53 | 0.27 | 0.27 |
| 10 | 90345671 | rs2576167 | -1.584 | -0.015 | -1.966 | -0.457 | -0.087 | 0.087 | 0.087 | 0.074 | 0.85 | 0.98 | 0.85 | 0.99 | 0.75 | 0.85 |
| 10 | 90349478 | rs2765452 | 0.541 | -0.03 | -1.97 | -0.46 | 0.008 | -0.019 | -0.019 | 0.164 | 0.31 | 0.32 | 0.45 | 0.55 | 0.27 | 0.37 |
| 10 | 90350151 | rs10887837 | -0.045 | -0.04 | -1.958 | -0.45 | -0.009 | -0.001 | -0.001 | 0.166 | 0.53 | 0.65 | 0.4 | 0.45 | 0.48 | 0.48 |
| 10 | 90358138 | rs17345638 | 1.754 | -0.331 | -2.454 | -0.606 | 0.007 | 0.047 | 0.047 | 0.168 | 0.19 | 0.01 | 0.07 | 0.07 | 0.11 | 0.13 |
| 10 | 90360515 | rs1359581 | -1.349 | 0.051 | -2.024 | -0.329 | -0.087 | 0.091 | 0.091 | 0.143 | 0.84 | 0.98 | 0.85 | 0.99 | 0.75 | 0.85 |
| 10 | 90361836 | rs12777350 | 1.82 | 0.253 | -1.992 | -0.194 | -0.017 | -0.039 | -0.039 | 0 | 0.09 | 0 | 0.05 | 0.13 | 0.05 | 0.04 |
| 10 | 90363908 | rs10887841 | 0.288 | 0.405 | -1.382 | -0.135 | -0.002 | 0.001 | 0.001 | 0 | 0.52 | 0.54 | 0.39 | 0.45 | 0.48 | 0.48 |
| 10 | 90374057 | rs10749593 | -0.865 | 0.643 | -1.755 | -0.053 | -0.062 | 0.085 | 0.085 | 0.149 | 0.81 | 0.93 | 0.84 | 0.97 | 0.74 | 0.75 |
| 10 | 90376462 | rs7912280 | -0.186 | 0.159 | -0.946 | 0.249 | 0.001 | 0.003 | 0.003 | 0 | 0.49 | 0.4 | 0.38 | 0.45 | 0.47 | 0.48 |
| 10 | 90420315 | rs1594601 | -0.229 | -0.138 | -0.672 | 0.409 | -0.006 | 0.049 | 0.049 | 0 | 0.25 | 0.24 | 0.34 | 0.41 | 0.27 | 0.3 |
| 10 | 90422136 | rs814620 | -0.067 | -0.519 | -1.06 | 0.113 | 0.004 | -0.019 | -0.019 | 0 | 0.26 | 0.2 | 0.35 | 0.34 | 0.26 | 0.23 |
| 10 | 90426612 | rs814625 | 0.122 | -0.536 | -1.1 | 0.097 | 0.005 | -0.032 | -0.032 | 0 | 0.23 | 0.11 | 0.25 | 0.31 | 0.22 | 0.21 |
| 10 | 90429652 | rs814628 | 0.789 | -0.574 | -1.228 | 0.075 | -0.038 | -0.106 | -0.106 | 0 | 0.18 | 0.01 | 0.2 | 0.31 | 0.17 | 0.21 |
| 10 | 90434411 | rs3858282 | 0.497 | -1.033 | -1.133 | -0.537 | -0.021 | 0.005 | 0.005 | 0 | 0.43 | 0.5 | 0.4 | 0.51 | 0.34 | 0.47 |
| 10 | 90436201 | rs1432444 | 0.072 | -1.081 | -1.269 | -0.734 | 0.046 | 0.032 | 0.032 | 0 | 0.54 | 0.78 | 0.48 | 0.65 | 0.41 | 0.51 |
| 10 | 90438207 | rs1228187 | 0.922 | -1.015 | -1.172 | -0.583 | 0 | -0.037 | -0.037 | 0 | 0.21 | 0.12 | 0.31 | 0.33 | 0.21 | 0.23 |
| 10 | 90446730 | rs6586147 | 1.477 | -1.123 | -0.948 | -0.481 | -0.012 | 0.031 | 0.031 | 0 | 0.11 | 0.24 | 0.08 | 0.14 | 0.07 | 0.05 |
| 10 | 90452938 | rs1469631 | 1.228 | -1.773 | -1.095 | -0.864 | -0.027 | 0.095 | 0.095 | 0 | 0.26 | 0.43 | 0.33 | 0.48 | 0.23 | 0.29 |
| 10 | 90453691 | rs11202817 | -0.306 | -1.701 | -1.516 | -0.902 | 0.044 | 0.031 | 0.031 | 0 | 0.54 | 0.75 | 0.48 | 0.65 | 0.41 | 0.51 |
| 10 | 90454252 | rs10887848 | -0.69 | -1.706 | -1.661 | -0.975 | 0.002 | 0.019 | 0.019 | 0 | 0.59 | 0.81 | 0.5 | 0.67 | 0.43 | 0.59 |
| 10 | 90461441 | rs12253503 | 0.706 | -1.531 | -1.184 | -0.806 | -0.025 | 0.057 | 0.057 | 0 | 0.29 | 0.43 | 0.35 | 0.48 | 0.24 | 0.3 |
| 10 | 90462716 | rs303511 | -0.843 | -1.273 | -1.412 | -0.975 | -0.019 | 0.014 | 0.014 | 0 | 0.75 | 0.82 | 0.77 | 0.9 | 0.63 | 0.8 |
| 10 | 90465083 | rs192767 | 0.115 | -1.687 | -1.585 | -1.43 | 0.012 | 0.004 | 0.004 | 0.007 | 0.43 | 0.34 | 0.41 | 0.38 | 0.33 | 0.35 |
| 10 | 90465628 | rs303508 | -1.896 | -1.38 | -1.734 | -1.338 | -0.107 | 0.095 | 0.095 | 0 | 0.85 | 0.98 | 0.84 | 1 | 0.7 | 0.84 |
| 10 | 90466296 | rs12267931 | 0.957 | -1.539 | -1.217 | -1.036 | -0.015 | 0.041 | 0.041 | 0 | 0.28 | 0.41 | 0.36 | 0.57 | 0.26 | 0.37 |
| 10 | 90466543 | rs12571524 | 2.04 | -1.507 | -1.287 | -0.919 | -0.038 | -0.145 | -0.145 | 0 | 0.19 | 0.01 | 0.33 | 0.52 | 0.24 | 0.2 |
| 10 | 90477590 | rs12776048 | 0.632 | -0.952 | -0.862 | -0.772 | -0.01 | 0.005 | 0.005 | 0 | 0.37 | 0.45 | 0.39 | 0.6 | 0.33 | 0.47 |
| 10 | 90487519 | rs2163673 | -0.085 | -0.818 | -1.008 | -0.769 | 0.047 | 0.067 | 0.067 | 0 | 0.53 | 0.75 | 0.48 | 0.68 | 0.43 | 0.55 |
| 10 | 90487896 | rs377257 | -1.057 | -0.348 | -1.28 | -0.867 | -0.079 | 0.072 | 0.072 | 0 | 0.83 | 0.94 | 0.83 | 1 | 0.68 | 0.83 |
| 10 | 90489360 | rs390400 | -0.043 | -0.626 | -1.014 | -0.787 | 0.003 | -0.019 | -0.019 | 0 | 0.29 | 0.23 | 0.32 | 0.35 | 0.26 | 0.29 |
| 10 | 90490627 | rs10887850 | 0.838 | -0.654 | -0.845 | -0.775 | -0.008 | 0.001 | 0.001 | 0 | 0.4 | 0.44 | 0.43 | 0.6 | 0.37 | 0.49 |
| 10 | 90490921 | rs412227 | -0.189 | -0.091 | -0.869 | -0.047 | 0.003 | -0.028 | -0.028 | 0 | 0.26 | 0.19 | 0.29 | 0.35 | 0.21 | 0.28 |
| 10 | 90493534 | rs406671 | -0.372 | -0.05 | -0.859 | -0.027 | 0.005 | -0.022 | -0.022 | 0 | 0.27 | 0.22 | 0.3 | 0.35 | 0.21 | 0.26 |
| 10 | 90500521 | rs396394 | -0.167 | 0.03 | -0.904 | 0.143 | 0.003 | -0.041 | -0.041 | 0 | 0.26 | 0.2 | 0.35 | 0.4 | 0.21 | 0.26 |
| 10 | 90506027 | rs400659 | -0.226 | 0.033 | -0.815 | -0.019 | 0 | -0.004 | -0.004 | 0 | 0.26 | 0.25 | 0.3 | 0.35 | 0.22 | 0.26 |
| 10 | 90525792 | rs10509554 | -0.349 | 1.059 | -0.371 | -0.506 | 0.054 | -0.075 | -0.075 | 0 | 0.17 | 0.26 | 0.15 | 0 | 0.31 | 0.12 |
| 10 | 90526379 | rs402781 | 0.067 | 0.878 | -0.411 | -0.718 | -0.001 | -0.014 | -0.014 | 0 | 0.2 | 0.19 | 0.23 | 0.1 | 0.18 | 0.13 |
| 10 | 90527569 | rs17112679 | 1.517 | 0.574 | -0.776 | -0.192 | -0.002 | -0.057 | -0.057 | 3.488 | 0.1 | 0 | 0.15 | 0.27 | 0.05 | 0.2 |
| 10 | 90527823 | rs408011 | 0.865 | 0.536 | -0.736 | -0.235 | -0.003 | -0.003 | -0.003 | 0 | 0.17 | 0.12 | 0.14 | 0.03 | 0.18 | 0.09 |
| 10 | 90532166 | rs11202848 | 1.481 | 0.492 | -0.774 | -0.19 | 0.001 | -0.063 | -0.063 | 0 | 0.1 | 0 | 0.15 | 0.27 | 0.05 | 0.2 |
| 10 | 90532552 | rs12255701 | 0.505 | 0.415 | -0.678 | -0.424 | -0.008 | -0.025 | -0.025 | 0 | 0.11 | 0.27 | 0.05 | 0.04 | 0.1 | 0.05 |
| 10 | 90534698 | rs303471 | 0.124 | 0.451 | -0.351 | -0.526 | 0.046 | -0.038 | -0.038 | 0.148 | 0.75 | 0.61 | 0.79 | 0.92 | 0.59 | 0.81 |
| 10 | 90534759 | rs11202849 | 2.758 | -0.326 | -0.548 | -0.886 | 0.002 | -0.051 | -0.051 | 0.41 | 0.26 | 0.15 | 0.34 | 0.5 | 0.26 | 0.29 |
| 10 | 90536476 | rs303481 | 1.386 | 0.482 | -0.47 | -0.479 | -0.009 | 0.025 | 0.025 | 0 | 0.17 | 0.08 | 0.13 | 0.03 | 0.18 | 0.09 |
| 10 | 90545972 | rs303478 | -1.497 | 0.51 | -0.506 | -0.46 | -0.094 | 0.077 | 0.077 | 0.083 | 0.87 | 0.99 | 0.85 | 1 | 0.69 | 0.87 |
| 10 | 90547576 | rs7081173 | 1.275 | 0.298 | -0.355 | -0.384 | 0.012 | -0.003 | -0.003 | 0 | 0.46 | 0.42 | 0.41 | 0.55 | 0.36 | 0.47 |
| 10 | 90549316 | rs303493 | 0.675 | 0.71 | -0.694 | -0.174 | 0 | -0.013 | -0.013 | 0.016 | 0.18 | 0.18 | 0.23 | 0.1 | 0.18 | 0.13 |
| 10 | 90553051 | rs303499 | -0.462 | 0.756 | -1.06 | -0.15 | -0.086 | -0.056 | -0.056 | 0 | 0.75 | 0.91 | 0.69 | 0.7 | 0.63 | 0.65 |
| 10 | 90559243 | rs1653840 | 0.742 | 0.502 | -0.615 | -0.354 | -0.011 | 0.028 | 0.028 | 0 | 0.16 | 0.07 | 0.13 | 0.02 | 0.18 | 0.09 |
| 10 | 90565399 | rs303536 | 0.095 | 0.479 | -0.54 | -0.289 | 0.001 | -0.007 | -0.007 | 0.187 | 0.19 | 0.2 | 0.24 | 0.14 | 0.19 | 0.13 |
| 10 | 90566907 | rs303528 | -0.28 | 0.533 | -0.532 | -0.225 | 0.004 | -0.003 | -0.003 | 0.84 | 0.22 | 0.31 | 0.32 | 0.19 | 0.23 | 0.14 |
| 10 | 90568182 | rs3899739 | 1.353 | 0.768 | -0.981 | -0.37 | 0.007 | 0.007 | 0.007 | 0.711 | 0.33 | 0.16 | 0.32 | 0.36 | 0.33 | 0.29 |
| 10 | 90578539 | rs1411780 | 0.107 | 0.983 | -0.715 | -0.718 | 0.033 | -0.128 | -0.128 | 0 | 0.78 | 0.66 | 0.86 | 1 | 0.75 | 0.86 |
| 10 | 90579609 | rs3910679 | 0.848 | 0.671 | -0.989 | -0.568 | 0.003 | 0.002 | 0.002 | 0 | 0.52 | 0.56 | 0.49 | 0.62 | 0.46 | 0.54 |
| 10 | 90580144 | rs10788615 | 1.862 | 0.478 | -1.409 | -0.492 | -0.026 | -0.076 | -0.076 | 0.015 | 0.34 | 0.06 | 0.31 | 0.44 | 0.34 | 0.41 |
| 10 | 90580330 | rs1661281 | 0.51 | 0.328 | -1.371 | -0.332 | -0.044 | -0.1 | -0.1 | 0 | 0.27 | 0.07 | 0.37 | 0.37 | 0.29 | 0.32 |
| 10 | 90583092 | rs2304804 | 0.258 | 0.096 | -0.929 | -0.354 | -0.039 | -0.056 | -0.056 | 0 | 0.11 | 0.24 | 0.04 | 0 | 0.03 | 0.03 |
| 10 | 90585566 | rs7894519 | 1.437 | 0.701 | -1.116 | -0.325 | -0.025 | -0.029 | -0.029 | 0 | 0.34 | 0.16 | 0.34 | 0.44 | 0.39 | 0.43 |
| 10 | 90586849 | rs1238449 | 1.161 | 0.807 | -1.191 | -0.416 | -0.041 | 0.009 | 0.009 | 6.478 | 0.42 | 0.17 | 0.42 | 0.44 | 0.53 | 0.42 |
| 10 | 90587303 | rs11202865 | 0.001 | 0.669 | -1.311 | -0.31 | 0.005 | -0.008 | -0.008 | 0.113 | 0.32 | 0.26 | 0.35 | 0.35 | 0.27 | 0.34 |
| 10 | 90588535 | rs1029024 | 0.128 | 1 | -1.161 | -0.333 | 0.017 | -0.01 | -0.01 | 0 | 0.79 | 0.4 | 0.77 | 0.83 | 0.78 | 0.85 |
| 10 | 90589116 | rs10887861 | 0.941 | 1.083 | -1.548 | -0.265 | -0.05 | -0.011 | -0.011 | 0 | 0.46 | 0.14 | 0.51 | 0.6 | 0.51 | 0.49 |
| 10 | 90590746 | rs10736363 | 0.836 | 1.074 | -0.983 | -0.943 | -0.004 | 0.002 | 0.002 | 0.016 | 0.62 | 0.37 | 0.75 | 0.72 | 0.53 | 0.6 |
| 10 | 90591896 | rs1214549 | -0.324 | 0.946 | -0.687 | -1.075 | 0.016 | -0.062 | -0.062 | 0.057 | 0.27 | 0.32 | 0.2 | 0.07 | 0.36 | 0.23 |
| 10 | 90592107 | rs1214550 | -0.645 | 0.94 | -0.64 | -1.009 | 0.028 | -0.08 | -0.08 | 1.255 | 0.29 | 0.5 | 0.21 | 0.1 | 0.36 | 0.25 |
| 10 | 90592929 | rs1214551 | -0.154 | 0.817 | -0.982 | -1.112 | -0.041 | 0.061 | 0.061 | 0 | 0.8 | 0.92 | 0.78 | 0.94 | 0.74 | 0.82 |

| | | | | | | | | | | | | | | | | |
|---|---|---|---|---|---|---|---|---|---|---|---|---|---|---|---|---|
| 10 | 90596859 | rs1214554 | -0.521 | 0.831 | -0.852 | -1.06 | 0.018 | -0.044 | -0.044 | 0 | 0.3 | 0.54 | 0.22 | 0.14 | 0.37 | 0.29 |
| 10 | 90608238 | rs1147601 | 0.713 | 0.889 | -0.857 | -1.135 | -0.059 | 0.065 | 0.065 | 0 | 0.16 | 0.04 | 0.11 | 0.04 | 0.24 | 0.19 |
| 10 | 90609718 | rs4934425 | 1.447 | 0.437 | -0.859 | -1.026 | 0.049 | 0.016 | 0.016 | 0 | 0.45 | 0.15 | 0.54 | 0.66 | 0.37 | 0.51 |
| 10 | 90613517 | rs1214589 | 0.42 | 0.701 | -0.408 | -0.969 | 0.009 | -0.002 | -0.002 | 0 | 0.67 | 0.38 | 0.78 | 0.73 | 0.63 | 0.69 |
| 10 | 90620192 | rs1147604 | 0.62 | 0.366 | -0.43 | -0.966 | -0.007 | 0.009 | 0.009 | 0.062 | 0.6 | 0.39 | 0.71 | 0.76 | 0.51 | 0.61 |
| 10 | 90624386 | rs1214546 | 0.647 | 0.468 | -0.409 | -0.925 | -0.017 | 0.034 | 0.034 | 0 | 0.56 | 0.29 | 0.7 | 0.72 | 0.51 | 0.6 |
| 10 | 90630553 | rs1837836 | 0.536 | 0.471 | -0.392 | -0.931 | -0.019 | 0.037 | 0.037 | 0 | 0.57 | 0.32 | 0.71 | 0.76 | 0.52 | 0.64 |
| 10 | 90633056 | rs7911566 | -0.16 | 0.526 | -0.444 | -0.729 | -0.004 | 0.17 | 0.17 | 0 | 0.09 | 0.33 | 0.03 | 0.17 | 0.02 | 0.11 |
| 10 | 90658059 | rs7912777 | 1.728 | 0.24 | -1.076 | -0.835 | -0.018 | -0.096 | -0.096 | 0.509 | 0.27 | 0.02 | 0.41 | 0.45 | 0.27 | 0.2 |
| 10 | 90662086 | rs1968028 | -0.161 | 0.286 | -0.855 | -0.742 | 0.001 | -0.002 | -0.002 | 1.683 | 0.35 | 0.37 | 0.31 | 0.45 | 0.26 | 0.52 |
| 10 | 90663839 | rs10509557 | -0.097 | 0.323 | -1.039 | -0.857 | 0.011 | -0.026 | -0.026 | 0 | 0.33 | 0.2 | 0.3 | 0.33 | 0.25 | 0.49 |
| 10 | 90671750 | rs10887866 | 1.276 | 0.318 | -0.964 | -0.897 | 0.019 | -0.042 | -0.042 | 0 | 0.18 | 0.08 | 0.2 | 0.25 | 0.09 | 0.38 |
| 10 | 90672057 | rs7077956 | 2.025 | 0.458 | -0.827 | -0.899 | -0.009 | -0.014 | -0.014 | 0 | 0.05 | 0.14 | 0.01 | 0.01 | 0.02 | 0.08 |
| 10 | 90673024 | rs12254856 | 0.964 | 0.491 | -0.675 | -0.965 | 0.011 | -0.043 | -0.043 | 0.358 | 0.09 | 0.14 | 0.09 | 0 | 0.13 | 0.07 |
| 10 | 90675317 | rs1561742 | -0.228 | 0.444 | -0.777 | -1.147 | 0 | -0.013 | -0.013 | 0.088 | 0.66 | 0.69 | 0.74 | 0.9 | 0.53 | 0.82 |
| 10 | 90675836 | rs4431933 | -0.075 | 0.467 | -0.789 | -1.046 | 0.006 | 0.024 | 0.024 | 0 | 0.17 | 0.28 | 0.11 | 0.18 | 0.15 | 0.22 |
| 10 | 90679053 | rs4934429 | -0.101 | 0.437 | -0.955 | -1.104 | -0.002 | -0.011 | -0.011 | 0 | 0.65 | 0.68 | 0.74 | 0.9 | 0.53 | 0.82 |
| 10 | 90683124 | rs1530281 | -0.028 | 0.423 | -0.966 | -0.996 | 0.03 | -0.07 | -0.07 | 0 | 0.63 | 0.57 | 0.69 | 0.9 | 0.49 | 0.78 |
| 10 | 90687557 | rs7098235 | 0.254 | 0.379 | -0.92 | -1.014 | 0.009 | 0.011 | 0.011 | 0 | 0.16 | 0.28 | 0.11 | 0.14 | 0.14 | 0.17 |
| 10 | 90688042 | rs17114186 | 1.319 | 0.385 | -0.998 | -0.998 | -0.011 | 0.033 | 0.033 | 0 | 0.07 | 0.08 | 0.02 | 0.14 | 0.02 | 0.1 |
| 10 | 90690465 | rs11202900 | -0.184 | 0.246 | -1.119 | -0.995 | -0.019 | 0.022 | 0.022 | 0 | 0.37 | 0.56 | 0.3 | 0.39 | 0.23 | 0.54 |
| 10 | 90691412 | rs4934430 | 1.928 | 0.25 | -1.274 | -0.93 | -0.012 | -0.116 | -0.116 | 5.672 | 0.26 | 0.02 | 0.39 | 0.46 | 0.25 | 0.2 |
| 10 | 90693569 | rs1372325 | -1.28 | 1.269 | -1.053 | -0.867 | -0.068 | 0.046 | 0.046 | 0 | 0.86 | 0.99 | 0.9 | 0.96 | 0.8 | 0.89 |
| 10 | 90711743 | rs17114327 | -0.146 | 0.758 | -1.031 | -0.773 | 0.002 | -0.017 | -0.017 | 0.924 | 0.18 | 0.41 | 0.11 | 0.11 | 0.15 | 0.19 |
| 10 | 90713195 | rs17114337 | 1.398 | 1.036 | -1.158 | -0.778 | 0.008 | -0.039 | -0.039 | 0.552 | 0.15 | 0.08 | 0.18 | 0.3 | 0.07 | 0.29 |
| 10 | 90716837 | rs1441734 | 0.142 | 1.799 | -0.801 | -1.165 | 0.002 | -0.02 | -0.02 | 0.162 | 0.64 | 0.67 | 0.7 | 0.9 | 0.49 | 0.79 |
| 10 | 90718677 | rs2902336 | 0.512 | 2.214 | -0.807 | -1.057 | 0.031 | -0.083 | -0.083 | 0.881 | 0.66 | 0.51 | 0.71 | 0.94 | 0.49 | 0.79 |
| 10 | 90719237 | rs7907084 | -0.062 | 2.317 | -1.618 | -0.847 | 0.004 | -0.033 | -0.033 | 0 | 0.24 | 0.22 | 0.21 | 0.03 | 0.38 | 0.11 |
| 10 | 90727334 | rs1926203 | -1.043 | 2.06 | -1.611 | -0.847 | -0.007 | -0.051 | -0.051 | 0.591 | 0.42 | 0.55 | 0.42 | 0.14 | 0.65 | 0.24 |
| 10 | 90728781 | rs7088326 | -0.808 | 2.085 | -2.352 | -0.663 | -0.023 | -0.002 | -0.002 | 0 | 0.32 | 0.24 | 0.33 | 0.03 | 0.52 | 0.16 |
| 10 | 90730211 | rs11202914 | -0.66 | 2.154 | -2.274 | -0.574 | -0.009 | 0.003 | 0.003 | 0 | 0.28 | 0.23 | 0.24 | 0.07 | 0.41 | 0.21 |
| 10 | 90736399 | rs11202916 | -0.341 | 0.458 | -2.604 | 0.175 | 0.02 | -0.117 | -0.117 | 0 | 0.33 | 0.02 | 0.44 | 0.46 | 0.29 | 0.28 |
| 10 | 90739368 | rs1571014 | -0.545 | 1.206 | -2.314 | -0.254 | -0.031 | 0.045 | 0.045 | 0 | 0.31 | 0.15 | 0.37 | 0.78 | 0.41 | 0.54 |
| 10 | 90740501 | rs12765241 | 0.117 | 1.77 | -2.584 | -0.783 | -0.053 | 0.069 | 0.069 | 0.101 | 0.14 | 0.02 | 0.08 | 0.03 | 0.14 | 0.07 |
| 10 | 90740953 | rs4934433 | 0.571 | 1.687 | -1.777 | -0.629 | 0.001 | 0.015 | 0.015 | 0.136 | 0.71 | 0.44 | 0.67 | 0.54 | 0.71 | 0.63 |
| 10 | 90744325 | rs7916294 | 0.531 | 1.365 | -1.214 | -1.206 | -0.008 | -0.102 | -0.102 | 0 | 0.31 | 0.53 | 0.32 | 0.03 | 0.3 | 0.21 |
| 10 | 90745130 | rs10887876 | 0.452 | 1.351 | -1.222 | -1.275 | -0.006 | 0.058 | 0.058 | 0 | 0.13 | 0.18 | 0.21 | 0.43 | 0.12 | 0.21 |
| 10 | 90747169 | rs4934434 | 0.81 | 1.446 | -1.363 | -1.214 | -0.04 | -0.004 | -0.004 | 0 | 0.42 | 0.18 | 0.38 | 0.51 | 0.43 | 0.44 |
| 10 | 90748736 | rs3758483 | -0.808 | 1.49 | -1.376 | -1.229 | -0.052 | 0.059 | 0.059 | 0.783 | 0.9 | 0.99 | 0.92 | 0.99 | 0.86 | 0.9 |
| 10 | 90749963 | rs1800682 | -0.47 | 1.456 | -0.854 | -1.14 | 0.043 | 0.043 | 0.043 | 0 | 0.48 | 0.75 | 0.54 | 0.49 | 0.43 | 0.45 |
| 10 | 90757787 | rs1571011 | 0.802 | 1.357 | -0.788 | -1.174 | 0.01 | 0.023 | 0.023 | 0 | 0.46 | 0.32 | 0.39 | 0.5 | 0.43 | 0.45 |
| 10 | 90759724 | rs4406737 | 0.152 | 1.439 | -0.819 | -1.194 | 0.011 | 0.013 | 0.013 | 0.116 | 0.56 | 0.33 | 0.47 | 0.5 | 0.55 | 0.55 |
| 10 | 90766924 | rs2031613 | -1.191 | 1.362 | -0.756 | -1.308 | -0.003 | -0.016 | -0.016 | 0.013 | 0.74 | 0.77 | 0.8 | 0.96 | 0.7 | 0.82 |
| 10 | 90768965 | rs7911226 | -1.353 | 1.627 | -0.488 | -1.231 | -0.002 | -0.009 | -0.009 | 1.704 | 0.28 | 0.24 | 0.42 | 0.46 | 0.31 | 0.37 |
| 10 | 90771505 | rs9658765 | 0.503 | 1.61 | -0.509 | -1.243 | 0.009 | 0.022 | 0.022 | 0 | 0.05 | 0.01 | 0.12 | 0.02 | 0.04 | 0.05 |
| 10 | 90771829 | rs2234978 | -1.159 | 1.596 | -0.445 | -1.258 | 0.011 | -0.081 | -0.081 | 2.76 | 0.73 | 0.65 | 0.79 | 0.96 | 0.7 | 0.82 |
| 10 | 90773494 | rs1977389 | 0.947 | 1.69 | -0.318 | -1.109 | 0.02 | 0.002 | 0.002 | 0 | 0.45 | 0.33 | 0.37 | 0.5 | 0.4 | 0.46 |
| 10 | 90775291 | rs1468063 | 0.36 | 1.343 | -0.682 | -1.064 | -0.01 | 0.135 | 0.135 | 0 | 0.12 | 0.23 | 0.21 | 0.43 | 0.12 | 0.22 |
| 10 | 90782973 | rs10887883 | -0.394 | 0.868 | -1.061 | -0.483 | 0.004 | 0.001 | 0.001 | 0 | 0.54 | 0.59 | 0.61 | 0.5 | 0.58 | 0.55 |
| 10 | 90787881 | rs11202932 | -0.237 | 0.492 | -1.631 | -0.956 | -0.058 | 0.149 | 0.149 | 0.503 | 0.39 | 0.14 | 0.39 | 0.18 | 0.49 | 0.4 |
| 10 | 90789461 | rs10887887 | -0.551 | -0.49 | -1.396 | -0.573 | 0.002 | -0.003 | -0.003 | 1.177 | 0.22 | 0.24 | 0.28 | 0.49 | 0.16 | 0.4 |
| 10 | 90790476 | rs11202938 | 0.607 | -0.598 | -1.447 | -0.776 | -0.016 | -0.027 | -0.027 | 0 | 0.05 | 0.11 | 0.01 | 0 | 0.02 | 0 |
| 10 | 90790780 | rs10887888 | -0.419 | -0.207 | -1.588 | -0.974 | -0.027 | 0.017 | 0.017 | 0.046 | 0.35 | 0.12 | 0.37 | 0.68 | 0.37 | 0.5 |
| 10 | 90791823 | rs17377783 | 1.164 | -0.778 | -1.693 | -1.42 | 0.004 | 0.063 | 0.063 | 1.223 | 0.34 | 0.13 | 0.31 | 0.23 | 0.34 | 0.19 |
| 10 | 90791968 | rs1926189 | 0.383 | -0.936 | -1.042 | -1.178 | -0.006 | -0.015 | -0.015 | 0.003 | 0.14 | 0.19 | 0.11 | 0.09 | 0.1 | 0.18 |
| 10 | 90793260 | rs7916326 | -0.149 | -0.997 | -1.068 | -1.206 | -0.003 | 0.063 | 0.063 | 0 | 0.38 | 0.26 | 0.49 | 0.88 | 0.38 | 0.54 |
| 10 | 90796151 | rs7898005 | -0.535 | -1.021 | -0.808 | -1.512 | 0.003 | -0.025 | -0.025 | 0 | 0.44 | 0.41 | 0.5 | 0.93 | 0.39 | 0.59 |
| 10 | 90802924 | rs2105436 | -0.535 | -1.021 | -0.808 | -1.512 | 0.003 | -0.025 | -0.025 | 0 | 0.44 | 0.41 | 0.5 | 0.93 | 0.39 | 0.59 |
| 10 | 90818377 | rs7920888 | -0.015 | -0.895 | -0.618 | -1.252 | 0.012 | -0.012 | -0.012 | 0 | 0.38 | 0.53 | 0.48 | 0.3 | 0.46 | 0.4 |
| 10 | 90818777 | rs9633743 | 1.23 | -0.657 | -0.544 | -1.159 | 0.041 | -0.027 | -0.027 | 0 | 0.16 | 0.24 | 0.34 | 0.08 | 0.24 | 0.22 |
| 10 | 90824999 | rs9663252 | 1.177 | -0.717 | -0.621 | -1.682 | -0.016 | -0.024 | -0.024 | 0.637 | 0.05 | 0.1 | 0.01 | 0 | 0.02 | 0 |
| 10 | 90825718 | rs11202952 | -0.249 | -0.95 | -0.646 | -1.846 | 0.002 | 0.005 | 0.005 | 0 | 0.45 | 0.55 | 0.56 | 0.3 | 0.54 | 0.53 |
| 10 | 90830160 | rs7919067 | 1.592 | -0.854 | -0.534 | -1.86 | 0.044 | -0.023 | -0.023 | 0 | 0.15 | 0.21 | 0.33 | 0.08 | 0.22 | 0.2 |
| 10 | 90830468 | rs12244713 | 1.14 | -0.914 | -0.548 | -1.863 | -0.02 | 0.022 | 0.022 | 0 | 0.05 | 0.18 | 0.02 | 0.06 | 0.01 | 0.04 |
| 10 | 90834673 | rs4481939 | -0.466 | -1.429 | -0.923 | -2.283 | -0.015 | 0.064 | 0.064 | 0 | 0.57 | 0.7 | 0.65 | 0.79 | 0.45 | 0.73 |
| 10 | 90835692 | rs2801540 | 0.122 | -1.429 | -0.923 | -2.283 | -0.015 | 0.064 | 0.064 | 0 | 0.43 | 0.3 | 0.35 | 0.21 | 0.55 | 0.27 |
| 10 | 90837919 | rs2841016 | -1.709 | -1.401 | -0.907 | -2.272 | 0.092 | -0.046 | -0.046 | 0.115 | 0.83 | 0.64 | 0.65 | 0.92 | 0.78 | 0.8 |
| 10 | 90843119 | rs7086087 | 0.041 | -1.502 | -0.925 | -2.224 | 0.045 | 0.03 | 0.03 | 0 | 0.4 | 0.31 | 0.3 | 0.7 | 0.23 | 0.53 |
| 10 | 90843886 | rs2211595 | -0.004 | -1.501 | -0.928 | -2.228 | 0.052 | 0.041 | 0.041 | 0 | 0.41 | 0.31 | 0.33 | 0.74 | 0.24 | 0.55 |

| | | | | | | | | | | | | | | | | |
|---|---|---|---|---|---|---|---|---|---|---|---|---|---|---|---|---|
| 10 | 90845035 | rs7094490 | 0.61 | -1.525 | -0.874 | -2.158 | -0.013 | 0.036 | 0.036 | 0 | 0.24 | 0.15 | 0.19 | 0.04 | 0.31 | 0.1 |
| 10 | 90851108 | rs1961812 | 0.494 | -1.582 | -0.859 | -2.139 | -0.017 | -0.006 | -0.006 | 0.311 | 0.15 | 0.18 | 0.07 | 0.11 | 0.08 | 0.19 |
| 10 | 90853352 | rs2801536 | -0.668 | -1.473 | -1.237 | -2.133 | 0.012 | -0.039 | -0.039 | 0 | 0.73 | 0.42 | 0.62 | 0.85 | 0.73 | 0.76 |
| 10 | 90853446 | rs2245548 | -0.032 | -1.453 | -1.292 | -2.153 | -0.002 | 0.022 | 0.022 | 1.054 | 0.63 | 0.34 | 0.56 | 0.81 | 0.66 | 0.61 |
| 10 | 90855419 | rs2801535 | 0.594 | -1.33 | -1.547 | -2.125 | -0.013 | 0.036 | 0.036 | 0 | 0.24 | 0.15 | 0.19 | 0.04 | 0.31 | 0.1 |
| 10 | 90856408 | rs2245863 | 1.546 | -1.269 | -1.589 | -2.173 | -0.007 | -0.002 | -0.002 | 0 | 0.14 | 0.1 | 0.13 | 0.03 | 0.21 | 0.08 |
| 10 | 90856844 | rs10887908 | 0.178 | -1.485 | -1.573 | -2.154 | -0.016 | -0.005 | -0.005 | 0.132 | 0.4 | 0.41 | 0.3 | 0.7 | 0.22 | 0.52 |
| 10 | 90857198 | rs7094057 | 1.887 | -1.216 | -2.338 | -2.057 | -0.043 | -0.005 | -0.005 | 0 | 0.17 | 0.07 | 0.12 | 0.15 | 0.23 | 0.15 |
| 10 | 90871172 | rs2841027 | 0.4 | -1.198 | -2.324 | -2.074 | -0.017 | 0.062 | 0.062 | 0 | 0.25 | 0.12 | 0.19 | 0.04 | 0.31 | 0.1 |
| 10 | 90874029 | rs2253503 | -0.169 | -1.229 | -2.384 | -2.088 | -0.009 | 0.117 | 0.117 | 0 | 0.34 | 0.17 | 0.26 | 0.08 | 0.39 | 0.25 |
| 10 | 90878949 | rs10509564 | 1.948 | -1.186 | -2.373 | -2.071 | -0.03 | 0.001 | 0.001 | 0 | 0.17 | 0.09 | 0.12 | 0.15 | 0.23 | 0.15 |
| 10 | 90879304 | rs1317361 | 0.112 | -1.265 | -2.393 | -2.104 | 0.027 | 0.04 | 0.04 | 0.054 | 0.45 | 0.26 | 0.4 | 0.64 | 0.42 | 0.44 |
| 10 | 90882921 | rs2494647 | 0.549 | -0.508 | -2.44 | -1.713 | -0.04 | 0.106 | 0.106 | 0 | 0.42 | 0.24 | 0.32 | 0.19 | 0.54 | 0.25 |
| 10 | 90889108 | rs10509565 | 1.196 | 0.16 | -2.893 | -1.406 | -0.024 | 0.054 | 0.054 | 0.037 | 0.27 | 0.1 | 0.16 | 0.15 | 0.28 | 0.17 |
| 10 | 90893250 | rs10887915 | 1.205 | 0.169 | -2.889 | -1.472 | -0.024 | 0.054 | 0.054 | 0.041 | 0.27 | 0.1 | 0.16 | 0.15 | 0.28 | 0.17 |
| 10 | 90893825 | rs4934449 | 0.4 | 0.635 | -2.224 | -0.549 | 0.039 | -0.105 | -0.105 | 0.03 | 0.21 | 0.04 | 0.23 | 0.63 | 0.12 | 0.36 |
| 10 | 90895848 | rs9651494 | 0.179 | 1.889 | -1.182 | -0.756 | 0.036 | -0.029 | -0.029 | 0 | 0.83 | 0.79 | 0.65 | 0.93 | 0.76 | 0.8 |
| 10 | 90897052 | rs17380969 | -0.614 | 1.132 | -1.311 | -0.553 | 0 | 0.003 | 0.003 | 0 | 0.09 | 0 | 0.06 | 0.04 | 0.08 | 0.15 |
| 10 | 90903281 | rs2251633 | 0.567 | 2.042 | -0.894 | -0.549 | 0.034 | -0.021 | -0.021 | 0 | 0.81 | 0.78 | 0.6 | 0.9 | 0.68 | 0.75 |
| 10 | 90913029 | rs11203005 | -0.875 | 1.956 | -1.375 | -1.243 | -0.026 | 0.047 | 0.047 | 0 | 0.23 | 0.14 | 0.23 | 0.01 | 0.28 | 0.31 |
| 10 | 90932935 | rs9651424 | 0.092 | 1.572 | -1.969 | -1.176 | 0.005 | -0.037 | -0.037 | 0 | 0.14 | 0.02 | 0.26 | 0.77 | 0.13 | 0.33 |
| 10 | 90940003 | rs2841031 | -1.299 | 1.188 | -2.208 | -2.306 | 0.006 | 0.063 | 0.063 | 0.093 | 0.68 | 0.78 | 0.8 | 0.94 | 0.72 | 0.88 |
| 10 | 90944216 | rs4269843 | -0.315 | 0.217 | -1.112 | -2.422 | -0.043 | -0.037 | -0.037 | 0 | 0.26 | 0.34 | 0.33 | 0.81 | 0.18 | 0.37 |
| 10 | 90944659 | rs10788636 | -1.278 | 0.289 | -0.864 | -2.524 | 0.025 | 0.097 | 0.097 | 0 | 0.49 | 0.63 | 0.49 | 0.82 | 0.44 | 0.71 |
| 10 | 90945756 | rs10509567 | 0.664 | 0.272 | -0.789 | -2.225 | -0.002 | -0.016 | -0.016 | 0.65 | 0.15 | 0.02 | 0.26 | 0.81 | 0.14 | 0.34 |
| 10 | 90957403 | rs7091822 | -0.548 | 0.233 | -0.674 | -2.247 | -0.071 | -0.028 | -0.028 | 0.756 | 0.26 | 0.46 | 0.33 | 0.77 | 0.17 | 0.36 |
| 10 | 90958765 | rs12245538 | 0.439 | 0.262 | -0.695 | -2.209 | -0.056 | -0.068 | -0.068 | 1.087 | 0.11 | 0.29 | 0.05 | 0 | 0.03 | 0.03 |
| 10 | 90964026 | rs17117126 | 0.656 | 0.77 | -1.149 | -1.927 | -0.015 | -0.065 | -0.065 | 0 | 0.14 | 0.01 | 0.26 | 0.74 | 0.13 | 0.33 |
| 10 | 90965691 | rs3802658 | 0.671 | 0.791 | -1.147 | -1.928 | -0.016 | -0.063 | -0.063 | 0.599 | 0.14 | 0.01 | 0.28 | 0.75 | 0.14 | 0.34 |
| 10 | 90966393 | rs17382301 | 0.173 | 1.208 | -0.938 | -1.873 | -0.041 | 0.029 | 0.029 | 0.044 | 0.07 | 0.01 | 0.07 | 0.03 | 0.12 | 0.21 |
| 10 | 90975207 | rs11594137 | 0.054 | 1.309 | -0.751 | -1.888 | -0.001 | -0.004 | -0.004 | 0.539 | 0.07 | 0.07 | 0.08 | 0.03 | 0.12 | 0.09 |
| 10 | 90977996 | rs7908760 | -0.955 | 0.879 | -0.097 | -2.298 | 0.018 | -0.157 | -0.157 | 1.528 | 0.24 | 0.5 | 0.18 | 0.03 | 0.24 | 0.17 |
| 10 | 90978270 | rs6586174 | -1.819 | 0.847 | -0.911 | -2.83 | -0.07 | 0.113 | 0.113 | 0 | 0.82 | 0.99 | 0.87 | 0.96 | 0.79 | 0.92 |
| 10 | 90981876 | rs12780342 | -0.199 | 0.494 | -0.196 | -2.503 | 0.072 | -0.051 | -0.051 | 3.272 | 0.1 | 0.23 | 0.09 | 0.03 | 0.14 | 0.09 |
| 10 | 90981887 | rs7922269 | 0.291 | -0.066 | -0.813 | -2.721 | 0.073 | 0.006 | 0.006 | 0.408 | 0.29 | 0.03 | 0.37 | 0.74 | 0.18 | 0.49 |
| 10 | 90983288 | rs12358054 | 0.839 | 0.292 | -0.663 | -2.428 | 0.001 | 0 | 0 | 0 | 0.13 | 0.11 | 0.1 | 0.03 | 0.11 | 0.17 |
| 10 | 90983629 | rs1556478 | -0.213 | 0.435 | -0.597 | -2.493 | 0.026 | 0.052 | 0.052 | 2.084 | 0.4 | 0.29 | 0.47 | 0.75 | 0.32 | 0.5 |
| 10 | 90984623 | rs2297475 | 0.13 | 0.538 | -0.494 | -2.161 | 0.041 | -0.04 | -0.04 | 0.108 | 0.09 | 0.18 | 0.07 | 0.02 | 0.11 | 0.14 |
| 10 | 90984990 | rs2297472 | 0.281 | 0.585 | -0.916 | -2.05 | 0.023 | -0.005 | -0.005 | 0.071 | 0.26 | 0.1 | 0.37 | 0.7 | 0.2 | 0.31 |
| 10 | 90985948 | rs2254747 | 0.534 | 0.652 | -0.254 | -1.866 | 0.045 | -0.068 | -0.068 | 1.296 | 0.82 | 0.59 | 0.88 | 0.91 | 0.83 | 0.85 |
| 10 | 90987564 | rs2071510 | 0.042 | 0.305 | -1.052 | -0.723 | 0.029 | 0.015 | 0.015 | 0 | 0.48 | 0.25 | 0.43 | 0.42 | 0.56 | 0.41 |
| 10 | 90989109 | rs11203042 | -0.423 | 0.495 | -0.87 | -0.239 | -0.01 | -0.014 | -0.014 | 0 | 0.58 | 0.67 | 0.53 | 0.46 | 0.54 | 0.58 |
| 10 | 90991138 | rs7896502 | 0.081 | 0.092 | -0.423 | -0.575 | -0.035 | -0.051 | -0.051 | 0 | 0.2 | 0.29 | 0.17 | 0.06 | 0.11 | 0.22 |
| 10 | 90991179 | rs1041389 | 0.89 | 0.046 | -0.28 | -0.489 | -0.027 | -0.014 | -0.014 | 0 | 0.36 | 0.21 | 0.36 | 0.36 | 0.44 | 0.36 |
| 10 | 90992104 | rs885561 | -0.544 | 0.199 | -0.264 | -0.48 | -0.003 | -0.007 | -0.007 | 0 | 0.57 | 0.68 | 0.54 | 0.43 | 0.57 | 0.58 |
| 10 | 90993975 | rs6586178 | -1.445 | -0.125 | -0.012 | -0.887 | 0.049 | 0.009 | 0.009 | 1.277 | 0.06 | 0.33 | 0.06 | 0.01 | 0.04 | 0.03 |
| 10 | 90999036 | rs1029074 | -1.274 | 0.032 | -0.532 | -0.895 | -0.017 | -0.048 | -0.048 | 0 | 0.3 | 0.6 | 0.22 | 0.12 | 0.21 | 0.3 |
| 10 | 90999939 | rs2902563 | -0.476 | 0.13 | -0.594 | -0.859 | 0.027 | 0 | 0 | 2.502 | 0.08 | 0.22 | 0.08 | 0.06 | 0.1 | 0.05 |
| 10 | 91001126 | rs11203047 | 0.566 | 0.11 | -0.657 | -0.873 | 0.03 | 0.046 | 0.046 | 0 | 0.15 | 0.05 | 0.05 | 0.04 | 0.08 | 0.15 |
| 10 | 91002927 | rs1412444 | 0.369 | 0.399 | -0.941 | -0.612 | -0.001 | -0.002 | -0.002 | 0 | 0.44 | 0.42 | 0.46 | 0.32 | 0.36 | 0.55 |
| 10 | 91007360 | rs1051338 | 0.768 | 0.663 | -1.85 | -0.464 | -0.025 | -0.009 | -0.009 | 0.235 | 0.35 | 0.11 | 0.35 | 0.31 | 0.33 | 0.41 |
| 10 | 91007470 | rs2250781 | -0.243 | 1.496 | -1.234 | 0.256 | -0.008 | -0.008 | -0.008 | 0.271 | 0.7 | 0.75 | 0.6 | 0.43 | 0.55 | 0.78 |
| 10 | 91017363 | rs1332325 | 0.348 | 1.835 | -1.712 | -0.538 | -0.13 | 0.127 | 0.127 | 0 | 0.79 | 0.96 | 0.79 | 0.97 | 0.66 | 0.75 |
| 10 | 91017996 | rs10887936 | 0.809 | 1.197 | -1.474 | -0.939 | -0.043 | 0.072 | 0.072 | 0 | 0.68 | 0.81 | 0.68 | 0.92 | 0.6 | 0.7 |
| 10 | 91030083 | rs1120668 | 0.086 | 0.431 | -0.599 | -1.258 | -0.003 | -0.004 | -0.004 | 0 | 0.14 | 0.19 | 0.12 | 0.05 | 0.1 | 0.25 |
| 10 | 91033432 | rs1041539 | -0.358 | 0.522 | -0.825 | -1.306 | -0.094 | 0.106 | 0.106 | 0 | 0.78 | 0.94 | 0.79 | 0.97 | 0.66 | 0.74 |
| 10 | 91034358 | rs2151006 | 0.894 | 0.065 | -0.825 | -1.213 | 0.007 | 0.039 | 0.039 | 0 | 0.51 | 0.35 | 0.6 | 0.82 | 0.48 | 0.47 |
| 10 | 91034518 | rs2151004 | 1.334 | 0.007 | -0.845 | -1.225 | 0.006 | 0.033 | 0.033 | 0.823 | 0.44 | 0.32 | 0.5 | 0.79 | 0.44 | 0.42 |
| 10 | 91034967 | rs7068816 | 0.57 | 0.191 | -0.742 | -1.202 | -0.004 | 0.005 | 0.005 | 0.057 | 0.57 | 0.36 | 0.63 | 0.88 | 0.52 | 0.49 |
| 10 | 91041098 | rs12761410 | 0.165 | -0.383 | -1.558 | -1.264 | -0.024 | 0.104 | 0.104 | 0 | 0.43 | 0.13 | 0.31 | 0.22 | 0.4 | 0.37 |
| 10 | 91050991 | rs7087326 | 0.496 | -0.925 | -0.584 | -1.181 | -0.046 | 0.029 | 0.029 | 0 | 0.28 | 0.53 | 0.36 | 0.65 | 0.2 | 0.37 |
| 10 | 91056866 | rs10887941 | 0.452 | -0.26 | -0.575 | -1.135 | -0.008 | -0.031 | -0.031 | 0 | 0.15 | 0.12 | 0.28 | 0.66 | 0.2 | 0.37 |
| 10 | 91059127 | rs12260227 | 0.307 | -0.601 | -0.471 | -1.781 | 0.001 | -0.001 | -0.001 | 1.207 | 0.35 | 0.36 | 0.26 | 0.14 | 0.3 | 0.23 |
| 10 | 91059159 | rs12245107 | 0.315 | -0.579 | -0.469 | -1.778 | 0 | -0.002 | -0.002 | 0 | 0.35 | 0.36 | 0.26 | 0.14 | 0.3 | 0.23 |
| 10 | 91065949 | rs17468739 | 2.26 | -0.387 | -1.215 | -1.928 | -0.046 | 0.098 | 0.098 | 0 | 0.16 | 0.01 | 0.1 | 0 | 0.15 | 0.02 |
| 10 | 91066075 | rs2070845 | 1.082 | -0.441 | -1.036 | -1.68 | -0.003 | 0.009 | 0.009 | 0 | 0.25 | 0.27 | 0.19 | 0.13 | 0.25 | 0.11 |
| 10 | 91071670 | rs4934471 | -0.342 | -0.229 | -1.673 | -1.613 | -0.147 | 0.25 | 0.25 | 0.388 | 0.63 | 0.95 | 0.51 | 0.93 | 0.53 | 0.56 |
| 10 | 91085607 | rs7095398 | 0.101 | -0.397 | -0.991 | -1.689 | -0.009 | 0.002 | 0.002 | 0 | 0.55 | 0.65 | 0.47 | 0.93 | 0.49 | 0.56 |
| 10 | 91090551 | rs7897619 | 1.864 | -0.261 | -0.8 | -1.179 | 0.002 | 0.004 | 0.004 | 0.051 | 0.23 | 0.34 | 0.2 | 0.23 | 0.24 | 0.11 |

| | | | | | | | | | | | | | | | | |
|---|---|---|---|---|---|---|---|---|---|---|---|---|---|---|---|---|
| 10 | 91097554 | rs11203078 | 0.674 | -0.542 | -0.823 | -0.839 | 0 | -0.005 | -0.005 | 0 | 0.25 | 0.23 | 0.27 | 0.6 | 0.25 | 0.45 |
| 10 | 91098008 | rs7916812 | 1.729 | -0.445 | -0.795 | -0.782 | 0 | 0 | 0 | 0 | 0.25 | 0.33 | 0.2 | 0.23 | 0.25 | 0.11 |
| 10 | 91099630 | rs3780878 | 2.02 | -0.446 | -0.796 | -0.779 | -0.002 | -0.003 | -0.003 | 0.815 | 0.23 | 0.28 | 0.19 | 0.23 | 0.24 | 0.11 |
| 10 | 91099657 | rs17119665 | -0.354 | -0.508 | -0.561 | -0.823 | 0.01 | -0.006 | -0.006 | 2.487 | 0.06 | 0.24 | 0.02 | 0.01 | 0.02 | 0 |
| 10 | 91109343 | rs6586188 | 0.699 | -0.649 | -0.667 | -0.845 | -0.016 | 0.023 | 0.023 | 2.197 | 0.34 | 0.55 | 0.22 | 0.34 | 0.28 | 0.12 |
| 10 | 91120287 | rs305374 | -0.591 | -0.673 | -0.854 | -0.823 | 0.025 | -0.026 | -0.026 | 0 | 0.35 | 0.66 | 0.31 | 0.61 | 0.27 | 0.48 |
| 10 | 91129012 | rs17476559 | 1.289 | -0.509 | -1.036 | -1.004 | -0.055 | 0.059 | 0.059 | 0 | 0.1 | 0.01 | 0.09 | 0.01 | 0.19 | 0.12 |
| 10 | 91134786 | rs7913002 | -0.066 | -0.686 | -0.543 | -0.67 | -0.031 | 0.017 | 0.017 | 2.611 | 0.09 | 0.16 | 0.02 | 0.11 | 0.02 | 0.01 |
| 10 | 91135132 | rs304501 | -0.053 | -0.692 | -0.553 | -0.457 | -0.011 | 0.03 | 0.03 | 1.577 | 0.31 | 0.5 | 0.3 | 0.54 | 0.25 | 0.47 |
| 10 | 91136543 | rs11203100 | 0.252 | -0.236 | -0.504 | -0.244 | 0.001 | -0.005 | -0.005 | 0 | 0.08 | 0.09 | 0.02 | 0.11 | 0.02 | 0.01 |
| 10 | 91137521 | rs304502 | 1.468 | -0.331 | -0.725 | -0.098 | -0.064 | -0.13 | -0.13 | 0.331 | 0.18 | 0.03 | 0.24 | 0.46 | 0.2 | 0.39 |
| 10 | 91137826 | rs304504 | -0.592 | -0.381 | -0.852 | -0.286 | -0.146 | 0.181 | 0.181 | 1.662 | 0.63 | 0.95 | 0.51 | 0.87 | 0.53 | 0.56 |
| 10 | 91144909 | rs10887951 | -0.341 | -0.432 | -0.443 | -0.25 | 0.023 | 0.088 | 0.088 | 2.528 | 0.06 | 0.3 | 0.02 | 0.08 | 0.02 | 0.06 |
| 10 | 91145054 | rs10887952 | -1.081 | -0.449 | -0.515 | -0.258 | 0.006 | 0.075 | 0.075 | 3.375 | 0.08 | 0.45 | 0.04 | 0.08 | 0.03 | 0.06 |
| 10 | 91150922 | rs304478 | -0.584 | -0.385 | -1.013 | -0.305 | -0.149 | 0.182 | 0.182 | 1.704 | 0.63 | 0.95 | 0.51 | 0.87 | 0.52 | 0.55 |
| 10 | 91153699 | rs303217 | -0.722 | -0.374 | -1.008 | -0.301 | -0.144 | 0.188 | 0.188 | 1.969 | 0.62 | 0.95 | 0.51 | 0.86 | 0.52 | 0.55 |
| 10 | 91161355 | rs303212 | -0.893 | -0.481 | -1.012 | -0.333 | 0.022 | -0.012 | -0.012 | 0 | 0.37 | 0.69 | 0.33 | 0.62 | 0.28 | 0.48 |
| 10 | 91162677 | rs303210 | 1.646 | -0.681 | -0.943 | -0.211 | -0.067 | -0.129 | -0.129 | 1.808 | 0.18 | 0.03 | 0.25 | 0.44 | 0.21 | 0.36 |
| 10 | 91164170 | rs215493 | -1.002 | -0.682 | -0.928 | -0.239 | -0.131 | 0.155 | 0.155 | 1.28 | 0.64 | 0.95 | 0.53 | 0.86 | 0.55 | 0.57 |
| 10 | 91165833 | rs303208 | -0.324 | -0.657 | -0.784 | -0.213 | -0.001 | 0.004 | 0.004 | 0.02 | 0.31 | 0.29 | 0.3 | 0.62 | 0.28 | 0.48 |
| 10 | 91173397 | rs11203115 | 1.645 | -0.428 | -0.79 | -0.1 | 0.003 | 0.002 | 0.002 | 0.067 | 0.28 | 0.26 | 0.19 | 0.23 | 0.26 | 0.09 |
| 10 | 91178804 | rs304447 | 0.753 | -0.39 | -0.66 | -0.02 | 0.019 | 0.095 | 0.095 | 0.019 | 0.21 | 0.35 | 0.13 | 0.45 | 0.21 | 0.36 |
| 10 | 91179295 | rs184236 | 0.834 | -0.369 | -0.649 | -0.011 | 0.022 | 0.101 | 0.101 | 0.173 | 0.21 | 0.36 | 0.13 | 0.45 | 0.21 | 0.36 |
| 10 | 91179829 | rs303172 | -0.656 | -0.477 | -1.266 | -0.137 | -0.093 | 0.189 | 0.189 | 0.574 | 0.61 | 0.91 | 0.52 | 0.86 | 0.54 | 0.57 |
| 10 | 91181853 | rs10887959 | 1.439 | -0.178 | -1.049 | 0.106 | 0.005 | 0.001 | 0.001 | 0.201 | 0.28 | 0.26 | 0.2 | 0.23 | 0.26 | 0.09 |
| 10 | 91185696 | rs17121248 | 1.249 | -0.143 | -1.038 | 0.122 | -0.006 | -0.003 | -0.003 | 0 | 0.29 | 0.38 | 0.2 | 0.23 | 0.26 | 0.09 |
| 10 | 91186881 | rs304411 | 1.236 | 0.304 | -1.649 | 0.417 | -0.079 | -0.127 | -0.127 | 0 | 0.17 | 0.03 | 0.26 | 0.45 | 0.21 | 0.35 |
| 10 | 91194067 | rs6586196 | 0.606 | 0.162 | -1.134 | 0.157 | 0.005 | 0.001 | 0.001 | 1.172 | 0.37 | 0.7 | 0.24 | 0.34 | 0.3 | 0.2 |
| 10 | 91195080 | rs160052 | -0.612 | 0.599 | -1.49 | -0.193 | -0.049 | 0.069 | 0.069 | 0 | 0.74 | 0.96 | 0.63 | 0.85 | 0.75 | 0.71 |
| 10 | 91195363 | rs303169 | -0.101 | 0.879 | -1.203 | -0.171 | -0.089 | 0.071 | 0.071 | 0.19 | 0.67 | 0.89 | 0.56 | 0.77 | 0.52 | 0.67 |
| 10 | 91199517 | rs7088848 | -1.109 | 0.49 | -0.563 | 0.032 | 0.028 | 0.033 | 0.033 | 0 | 0.2 | 0.1 | 0.09 | 0.11 | 0.13 | 0.33 |
| 10 | 91203214 | rs1556611 | -1.659 | 0.723 | 0.148 | 0.404 | 0.012 | 0.011 | 0.011 | 0.016 | 0.3 | 0.43 | 0.21 | 0.28 | 0.34 | 0.36 |
| 10 | 91212169 | rs10788644 | -1.726 | 0.742 | 0.164 | 0.412 | 0.006 | 0.004 | 0.004 | 0 | 0.31 | 0.48 | 0.21 | 0.28 | 0.35 | 0.36 |
| 10 | 91215809 | rs3740029 | -1.847 | 1.278 | 0.475 | 0.741 | 0.138 | 0.163 | 0.163 | 2.766 | 0.14 | 0.49 | 0.14 | 0.24 | 0.24 | 0.05 |
| 10 | 91218525 | rs11596684 | -1.053 | 1.422 | -0.014 | 0.793 | -0.044 | -0.041 | -0.041 | 0 | 0.09 | 0.03 | 0.11 | 0.23 | 0.23 | 0.05 |
| 10 | 91223352 | rs12359459 | -0.719 | 0.699 | 0.084 | 0.677 | 0.002 | 0.002 | 0.002 | 0 | 0.12 | 0.12 | 0.07 | 0.07 | 0.1 | 0.24 |
| 10 | 91224807 | rs10887965 | -1.566 | 0.705 | 0.141 | 0.672 | -0.002 | -0.012 | -0.012 | 0 | 0.17 | 0.17 | 0.12 | 0.31 | 0.22 | 0.26 |
| 10 | 91226709 | rs304429 | 0.743 | 0.579 | -0.356 | 0.374 | -0.053 | 0.071 | 0.071 | 0.077 | 0.27 | 0.07 | 0.2 | 0.08 | 0.31 | 0.09 |
| 10 | 91233504 | rs992759 | -1.488 | 0.526 | 0.211 | 0.351 | 0.001 | 0.05 | 0.05 | 0 | 0.21 | 0.37 | 0.14 | 0.31 | 0.23 | 0.26 |
| 10 | 91235928 | rs303200 | -1.121 | 0.483 | 0.28 | 0.318 | 0 | 0 | 0 | 0.811 | 0.49 | 0.5 | 0.35 | 0.39 | 0.54 | 0.34 |
| 10 | 91241951 | rs303197 | 0.672 | 0.456 | -0.081 | -0.173 | -0.057 | 0.065 | 0.065 | 0.061 | 0.27 | 0.07 | 0.21 | 0.09 | 0.31 | 0.09 |
| 10 | 91245284 | rs303195 | 0.746 | 0.446 | -0.105 | -0.175 | -0.108 | 0.054 | 0.054 | 0.018 | 0.27 | 0.03 | 0.2 | 0.09 | 0.31 | 0.09 |
| 10 | 91251499 | rs11185728 | -0.302 | 0.527 | -0.368 | -0.09 | 0.002 | 0.003 | 0.003 | 0 | 0.12 | 0.12 | 0.07 | 0.07 | 0.11 | 0.22 |
| 10 | 91251901 | rs11185729 | -0.29 | 0.533 | -0.366 | -0.087 | 0.001 | 0.002 | 0.002 | 0 | 0.12 | 0.12 | 0.07 | 0.07 | 0.11 | 0.22 |
| 10 | 91253508 | rs303175 | 0.601 | 0.633 | -0.439 | -0.079 | -0.061 | 0.066 | 0.066 | 0.033 | 0.27 | 0.07 | 0.21 | 0.09 | 0.31 | 0.09 |
| 10 | 91256166 | rs303178 | 0.383 | 0.64 | -0.439 | -0.07 | -0.058 | 0.068 | 0.068 | 0 | 0.29 | 0.08 | 0.2 | 0.11 | 0.32 | 0.09 |
| 10 | 91263587 | rs188570 | -0.678 | 0.784 | -0.197 | 0.475 | 0.004 | 0.009 | 0.009 | 0 | 0.47 | 0.52 | 0.36 | 0.53 | 0.54 | 0.32 |
| 10 | 91271839 | rs304468 | 0.138 | 0.961 | -1.137 | 0.368 | -0.053 | 0.087 | 0.087 | 0 | 0.37 | 0.16 | 0.26 | 0.18 | 0.4 | 0.27 |
| 10 | 91273754 | rs304466 | 0.841 | 0.756 | -1.876 | -0.065 | -0.018 | 0.118 | 0.118 | 0.145 | 0.25 | 0.04 | 0.28 | 0.07 | 0.24 | 0.12 |
| 10 | 91275580 | rs7087280 | 0.25 | 1.114 | -1.181 | 0.306 | -0.001 | -0.001 | -0.001 | 0 | 0.55 | 0.49 | 0.63 | 0.52 | 0.48 | 0.69 |
| 10 | 91278222 | rs7924208 | -0.206 | 0.78 | -2.322 | -0.551 | -0.088 | 0.128 | 0.128 | 0 | 0.65 | 0.95 | 0.75 | 0.88 | 0.63 | 0.77 |
| 10 | 91279379 | rs4933508 | 0.104 | 0.83 | -2.157 | -0.552 | -0.055 | 0.15 | 0.15 | 0 | 0.61 | 0.89 | 0.74 | 0.84 | 0.6 | 0.73 |
| 10 | 91280569 | rs7080943 | -0.244 | 1.153 | -1.805 | -0.267 | -0.034 | 0.036 | 0.036 | 0 | 0.24 | 0.12 | 0.2 | 0.12 | 0.32 | 0.16 |
| 10 | 91281081 | rs7917083 | -0.518 | 0.412 | -1.516 | 0.252 | 0.012 | -0.008 | -0.008 | 1.284 | 0.35 | 0.71 | 0.35 | 0.33 | 0.28 | 0.19 |
| 10 | 91285135 | rs17123104 | -1.24 | -0.091 | -1.054 | 0.11 | -0.001 | -0.004 | -0.004 | 0.047 | 0.18 | 0.18 | 0.12 | 0.08 | 0.11 | 0.28 |
| 10 | 91286226 | rs7076750 | -0.457 | -0.165 | -0.341 | -0.093 | -0.017 | -0.019 | -0.019 | 2.368 | 0.05 | 0.19 | 0.02 | 0 | 0.01 | 0.02 |
| 10 | 91288049 | rs17123166 | -0.68 | 0.503 | 0.425 | -0.04 | 0.097 | 0.04 | 0.04 | 0 | 0.06 | 0.19 | 0.09 | 0.09 | 0.14 | 0.05 |
| 10 | 91289276 | rs7897886 | -1.626 | 0.363 | 0.763 | 0.435 | 0.007 | 0.237 | 0.237 | 0 | 0.09 | 0.32 | 0.08 | 0.28 | 0.05 | 0.05 |
| 10 | 91290668 | rs7922914 | -1.234 | 0.397 | 0.707 | 0.392 | -0.005 | 0.109 | 0.109 | 0 | 0.07 | 0.21 | 0.14 | 0.26 | 0.05 | 0.05 |
| 10 | 91291468 | rs10509575 | -1.939 | 0.121 | 0.537 | 0.195 | -0.012 | 0.005 | 0.005 | 0 | 0.23 | 0.33 | 0.17 | 0.3 | 0.15 | 0.21 |
| 10 | 91297717 | rs12254170 | -2.152 | 0.077 | -0.066 | 0.183 | 0.003 | -0.029 | -0.029 | 0.126 | 0.37 | 0.79 | 0.3 | 0.44 | 0.32 | 0.42 |
| 10 | 91312337 | rs12784228 | -0.486 | 0.425 | -0.038 | 0.368 | 0.037 | 0.017 | 0.017 | 0 | 0.05 | 0.06 | 0.06 | 0.1 | 0.12 | 0.1 |
| 10 | 91315670 | rs1556610 | -0.355 | 0.409 | -0.055 | 0.369 | 0.03 | 0.015 | 0.015 | 0 | 0.05 | 0.06 | 0.06 | 0.1 | 0.12 | 0.1 |
| 10 | 91315684 | rs12573820 | -0.278 | 0.225 | -0.242 | 0.467 | -0.016 | -0.024 | -0.024 | 0 | 0.06 | 0 | 0.06 | 0.26 | 0.06 | 0.06 |
| 10 | 91318624 | rs10785888 | -1.992 | 0.111 | -0.015 | 0.341 | -0.004 | 0.007 | 0.007 | 0 | 0.33 | 0.63 | 0.27 | 0.42 | 0.28 | 0.39 |
| 10 | 91323777 | rs6586198 | 0.358 | 0.13 | -0.37 | -0.171 | 0.004 | -0.001 | -0.001 | 0 | 0.67 | 0.63 | 0.72 | 0.34 | 0.61 | 0.52 |
| 10 | 91326997 | rs1274044 | 0.714 | -0.113 | -0.346 | -0.458 | 0.001 | -0.063 | -0.063 | 0.01 | 0.59 | 0.43 | 0.55 | 0.1 | 0.6 | 0.48 |
| 10 | 91327719 | rs1147619 | -1.813 | -0.212 | -0.536 | -0.587 | -0.004 | -0.002 | -0.002 | 0 | 0.54 | 0.65 | 0.56 | 0.97 | 0.5 | 0.7 |
| 10 | 91333674 | rs10881603 | -0.171 | 0.235 | -0.632 | -0.178 | 0 | 0.021 | 0.021 | 0 | 0.17 | 0.14 | 0.24 | 0.51 | 0.25 | 0.37 |

| | | | | | | | | | | | | | | | | |
|---|---|---|---|---|---|---|---|---|---|---|---|---|---|---|---|---|
| 10 | 91334162 | rs12354579 | 0.737 | -0.03 | -0.796 | -0.313 | -0.026 | -0.016 | -0.016 | 0.282 | 0.16 | 0.07 | 0.08 | 0.15 | 0.18 | 0.08 |
| 10 | 91336796 | rs11185762 | 0.885 | -0.032 | -0.809 | -0.312 | -0.045 | -0.032 | -0.032 | 0 | 0.16 | 0.04 | 0.08 | 0.15 | 0.18 | 0.08 |
| 10 | 91338675 | rs12773938 | -0.072 | -0.002 | -0.729 | -0.269 | -0.004 | 0.009 | 0.009 | 0 | 0.17 | 0.13 | 0.24 | 0.51 | 0.25 | 0.37 |
| 10 | 91344169 | rs1063943 | 0.523 | -0.325 | -0.35 | -0.537 | 0 | -0.012 | -0.012 | 0.03 | 0.11 | 0.11 | 0.1 | 0.02 | 0.07 | 0.07 |
| 10 | 91346003 | rs7079873 | -0.457 | -0.651 | -0.181 | -0.665 | -0.044 | -0.043 | -0.043 | 0 | 0.15 | 0.26 | 0.07 | 0.07 | 0.09 | 0.22 |
| 10 | 91348352 | rs7913785 | 0.422 | -0.588 | -0.17 | -0.657 | -0.02 | -0.024 | -0.024 | 0.024 | 0.07 | 0.11 | 0.03 | 0 | 0.03 | 0.06 |
| 10 | 91351942 | rs2038921 | -2.282 | -0.343 | -0.417 | -0.796 | -0.014 | 0.079 | 0.079 | 0.189 | 0.59 | 0.71 | 0.63 | 0.94 | 0.56 | 0.64 |
| 10 | 91352850 | rs10160034 | 0.366 | -0.341 | -0.393 | -0.8 | -0.011 | -0.059 | -0.059 | 0 | 0.12 | 0.24 | 0.12 | 0.02 | 0.07 | 0.07 |
| 10 | 91352966 | rs2296616 | -2.17 | -0.293 | -0.841 | -0.775 | -0.013 | 0.077 | 0.077 | 0.129 | 0.59 | 0.71 | 0.63 | 0.94 | 0.56 | 0.64 |
| 10 | 91368638 | rs10881606 | 1.451 | -0.307 | -1.569 | -1.06 | -0.006 | -0.007 | -0.007 | 0.701 | 0.27 | 0.28 | 0.27 | 0.06 | 0.34 | 0.32 |
| 10 | 91369885 | rs6586201 | -1.111 | -0.372 | -1.249 | -0.759 | 0.003 | 0.034 | 0.034 | 1.112 | 0.69 | 0.76 | 0.81 | 0.88 | 0.7 | 0.89 |
| 10 | 91369898 | rs7921294 | 0.276 | -0.303 | -2.099 | -0.612 | 0.043 | -0.003 | -0.003 | 0.142 | 0.48 | 0.35 | 0.55 | 0.55 | 0.59 | 0.62 |
| 10 | 91372533 | rs11185790 | 0.874 | -0.268 | -2.156 | -0.603 | -0.03 | -0.015 | -0.015 | 0 | 0.11 | 0.05 | 0.18 | 0.47 | 0.18 | 0.23 |
| 10 | 91373624 | rs7091626 | 0.257 | -0.278 | -2.17 | -0.612 | 0.045 | -0.002 | -0.002 | 1.694 | 0.48 | 0.35 | 0.55 | 0.55 | 0.59 | 0.62 |
| 10 | 91374967 | rs11185792 | 0.224 | 0.922 | -1.954 | 0.384 | -0.033 | -0.021 | -0.021 | 0 | 0.11 | 0.06 | 0.18 | 0.47 | 0.18 | 0.23 |
| 10 | 91385170 | rs943131 | -0.588 | 0.449 | -1.748 | -0.106 | -0.054 | 0.064 | 0.064 | 2.13 | 0.7 | 0.95 | 0.82 | 0.85 | 0.69 | 0.89 |
| 10 | 91389738 | rs927907 | -0.745 | 0.533 | -1.019 | 0.55 | 0.001 | -0.009 | -0.009 | 0.153 | 0.27 | 0.27 | 0.25 | 0.46 | 0.27 | 0.45 |
| 10 | 91393013 | rs11185822 | -0.231 | 0.383 | -1.007 | 0.648 | -0.015 | 0.006 | 0.006 | 0 | 0.06 | 0.13 | 0.02 | 0.09 | 0.01 | 0.17 |
| 10 | 91395428 | rs1359624 | -1.25 | 0.294 | -0.736 | 1.08 | -0.001 | -0.001 | -0.001 | 1.896 | 0.28 | 0.34 | 0.41 | 0.72 | 0.27 | 0.48 |
| 10 | 91403980 | rs997456 | 0.783 | 0.682 | -1.104 | 0.452 | -0.026 | 0.045 | 0.045 | 0.099 | 0.2 | 0.09 | 0.12 | 0 | 0.23 | 0.17 |
| 10 | 91412602 | rs7076259 | -0.236 | 0.492 | -0.414 | 0.159 | -0.014 | -0.012 | -0.012 | 0 | 0.29 | 0.49 | 0.22 | 0.3 | 0.25 | 0.47 |
| 10 | 91419294 | rs12779880 | 0.699 | 0.475 | -0.249 | 0.152 | 0.008 | -0.015 | -0.015 | 0 | 0.21 | 0.28 | 0.2 | 0.21 | 0.23 | 0.28 |
| 10 | 91422868 | rs7098233 | 0.608 | 0.507 | -0.35 | 0.262 | -0.01 | -0.027 | -0.027 | 0 | 0.2 | 0.13 | 0.23 | 0.37 | 0.27 | 0.29 |
| 10 | 91431298 | rs2420640 | -0.296 | 0.491 | -0.134 | 0.141 | -0.023 | 0.015 | 0.015 | 0.258 | 0.41 | 0.27 | 0.58 | 0.63 | 0.5 | 0.49 |
| 10 | 91440056 | rs11599335 | 1.295 | 0.486 | -0.197 | 0.089 | -0.039 | 0.068 | 0.068 | 1.47 | 0.2 | 0.05 | 0.18 | 0 | 0.23 | 0.17 |
| 10 | 91453196 | rs2104499 | -1.99 | 0.419 | -0.031 | 0.072 | 0.017 | 0.065 | 0.065 | 0.488 | 0.7 | 0.81 | 0.8 | 0.94 | 0.75 | 0.96 |
| 10 | 91457904 | rs7076351 | -0.296 | 0.453 | -0.015 | 0.158 | -0.032 | 0.019 | 0.019 | 0.355 | 0.41 | 0.24 | 0.57 | 0.63 | 0.5 | 0.49 |
| 10 | 91461153 | rs11593892 | 1.21 | 0.503 | -0.136 | 0.113 | -0.039 | 0.068 | 0.068 | 1.47 | 0.2 | 0.05 | 0.18 | 0 | 0.23 | 0.17 |
| 10 | 91470834 | rs1048057 | -0.347 | 0.448 | -0.039 | 0.165 | -0.012 | 0.044 | 0.044 | 2.793 | 0.29 | 0.57 | 0.38 | 0.57 | 0.24 | 0.49 |
| 10 | 91479211 | rs17484219 | 1.216 | 0.469 | -0.051 | -0.161 | -0.027 | 0.045 | 0.045 | 1.172 | 0.2 | 0.09 | 0.19 | 0 | 0.23 | 0.17 |
| 10 | 91484780 | rs8181361 | -0.082 | 0.437 | 0.115 | -0.124 | -0.012 | -0.003 | -0.003 | 1.381 | 0.27 | 0.48 | 0.22 | 0.31 | 0.23 | 0.47 |
| 10 | 91490087 | rs10881645 | -0.672 | 0.44 | 0.11 | -0.131 | -0.007 | 0.043 | 0.043 | 0.11 | 0.47 | 0.57 | 0.4 | 0.31 | 0.46 | 0.64 |
| 10 | 91498127 | rs1886996 | -1.885 | 0.296 | 0.135 | -0.067 | 0.015 | 0.104 | 0.104 | 0.267 | 0.69 | 0.85 | 0.8 | 0.94 | 0.73 | 0.96 |
| 10 | 91498254 | rs1886997 | -0.187 | 0.322 | 0.166 | -0.01 | -0.012 | -0.003 | -0.003 | 0 | 0.27 | 0.48 | 0.22 | 0.31 | 0.23 | 0.47 |
| 10 | 91529457 | rs7096826 | 0.682 | 0.404 | 0.052 | -0.013 | 0.014 | -0.012 | -0.012 | 0 | 0.2 | 0.3 | 0.2 | 0.21 | 0.22 | 0.28 |
| 10 | 91550544 | rs4934499 | -1.832 | 0.341 | 0.027 | -0.064 | 0.016 | 0.074 | 0.074 | 0.28 | 0.69 | 0.82 | 0.81 | 0.94 | 0.73 | 0.96 |
| 10 | 91551019 | rs2419057 | -1.088 | 0.41 | 0.683 | 0.503 | -0.005 | -0.005 | -0.005 | 0 | 0.33 | 0.58 | 0.24 | 0.31 | 0.26 | 0.53 |
| 10 | 91565990 | rs12763178 | 0.481 | 0.481 | 0.645 | 0.489 | -0.003 | 0.001 | 0.001 | 0.051 | 0.18 | 0.13 | 0.18 | 0.21 | 0.21 | 0.27 |
| 10 | 91568130 | rs12776628 | 0.506 | 0.481 | 0.627 | 0.581 | -0.012 | -0.007 | -0.007 | 0 | 0.18 | 0.09 | 0.19 | 0.25 | 0.21 | 0.27 |
| 10 | 91571200 | rs10881679 | -1.287 | 0.447 | 0.514 | 0.735 | 0.001 | -0.003 | -0.003 | 0 | 0.33 | 0.6 | 0.4 | 0.54 | 0.26 | 0.55 |
| 10 | 91573992 | rs10785907 | -0.813 | 0.559 | 0.484 | 0.767 | -0.008 | 0.035 | 0.035 | 0.712 | 0.27 | 0.53 | 0.39 | 0.45 | 0.25 | 0.35 |
| 10 | 91575644 | rs17127588 | 0.27 | 0.989 | 0.08 | 0.889 | -0.035 | -0.062 | -0.062 | 0 | 0.15 | 0.09 | 0.22 | 0.4 | 0.25 | 0.24 |
| 10 | 91575676 | rs11185892 | -0.055 | 0.745 | 0.476 | 0.582 | 0.017 | 0.097 | 0.097 | 0.35 | 0.2 | 0.28 | 0.36 | 0.44 | 0.23 | 0.29 |
| 10 | 91587401 | rs7895851 | -0.825 | 0.521 | 0.416 | 0.451 | -0.032 | 0.031 | 0.031 | 1.706 | 0.23 | 0.49 | 0.28 | 0.29 | 0.18 | 0.39 |
| 10 | 91592198 | rs1125326 | -0.477 | 0.536 | 0.372 | 0.451 | -0.009 | -0.036 | -0.036 | 1.533 | 0.2 | 0.25 | 0.1 | 0.09 | 0.16 | 0.37 |
| 10 | 91593169 | rs10881684 | -0.527 | 0.77 | -0.177 | 0.424 | -0.016 | -0.02 | -0.02 | 0 | 0.09 | 0.09 | 0.15 | 0.19 | 0.13 | 0.07 |
| 10 | 91597426 | rs3740040 | -0.636 | 0.703 | -0.041 | 0.482 | -0.027 | -0.094 | -0.094 | 1.019 | 0.21 | 0.4 | 0.11 | 0.1 | 0.17 | 0.37 |
| 10 | 91598513 | rs3740043 | 0.014 | 1.038 | -0.284 | 0.525 | -0.042 | -0.07 | -0.07 | 0 | 0.14 | 0.05 | 0.22 | 0.51 | 0.24 | 0.24 |
| 10 | 91605122 | rs10881687 | -1.077 | 0.844 | 0.162 | 0.041 | -0.008 | -0.023 | -0.023 | 0 | 0.14 | 0.38 | 0.07 | 0.07 | 0.09 | 0.17 |
| 10 | 91606212 | rs12770938 | 0.264 | 0.875 | 0.218 | 0.003 | 0.02 | -0.031 | -0.031 | 0 | 0.13 | 0.34 | 0.1 | 0.09 | 0.13 | 0.22 |
| 10 | 91611861 | rs7079714 | -1.011 | 0.953 | -0.783 | 0.077 | 0.001 | 0.002 | 0.002 | 3.573 | 0.27 | 0.72 | 0.17 | 0.16 | 0.23 | 0.4 |
| 10 | 91617383 | rs1924715 | -0.482 | 1.342 | -1.24 | 0.596 | -0.033 | -0.07 | -0.07 | 0 | 0.16 | 0.05 | 0.24 | 0.61 | 0.23 | 0.33 |
| 10 | 91618992 | rs1998643 | -0.376 | 1.524 | -1.305 | -0.082 | -0.001 | -0.001 | -0.001 | 0.351 | 0.18 | 0.15 | 0.1 | 0.09 | 0.19 | 0.39 |
| 10 | 91623992 | rs7077710 | -1.065 | 1.286 | -1.172 | -0.119 | -0.032 | 0.08 | 0.08 | 1.018 | 0.34 | 0.16 | 0.49 | 0.85 | 0.44 | 0.73 |
| 10 | 91628519 | rs7085165 | -0.608 | 0.822 | -1.117 | 0.281 | -0.013 | -0.003 | -0.003 | 0 | 0.59 | 0.68 | 0.65 | 0.44 | 0.55 | 0.66 |
| 10 | 91629054 | rs17128448 | -0.178 | 0.558 | -0.623 | 0.171 | 0.015 | 0.043 | 0.043 | 0 | 0.11 | 0.21 | 0.3 | 0.17 | 0.14 | 0.13 |
| 10 | 91630000 | rs10785909 | 0.125 | 0.123 | -0.573 | -0.409 | 0.021 | 0.046 | 0.046 | 0 | 0.35 | 0.26 | 0.29 | 0.18 | 0.31 | 0.37 |
| 10 | 91631922 | rs10785910 | 0.681 | -0.261 | -0.682 | -0.715 | -0.003 | -0.043 | -0.043 | 0 | 0.13 | 0.17 | 0.19 | 0 | 0.17 | 0.06 |
| 10 | 91636235 | rs1886408 | -1.055 | -1.045 | -0.629 | -1.02 | 0.025 | 0.031 | 0.031 | 3.087 | 0.47 | 0.64 | 0.47 | 0.34 | 0.35 | 0.56 |
| 10 | 91637151 | rs1924703 | -1.711 | -0.934 | -0.664 | -1.007 | -0.028 | 0.033 | 0.033 | 3.064 | 0.61 | 0.8 | 0.66 | 0.34 | 0.52 | 0.66 |
| 10 | 91643014 | rs10785914 | -1.239 | -1.22 | -0.145 | -1.069 | 0.002 | -0.002 | -0.002 | 0 | 0.48 | 0.54 | 0.49 | 0.25 | 0.44 | 0.54 |
| 10 | 91643122 | rs10881703 | -0.328 | -1.265 | 0.099 | -1.076 | 0 | 0.001 | 0.001 | 1.755 | 0.3 | 0.28 | 0.3 | 0.25 | 0.26 | 0.41 |
| 10 | 91643343 | rs7084582 | 0.046 | -0.848 | -0.306 | -0.894 | -0.006 | 0.065 | 0.065 | 0 | 0.06 | 0.25 | 0.16 | 0.09 | 0.03 | 0.08 |
| 10 | 91643736 | rs10881704 | 0.096 | -0.748 | -1.048 | -1.108 | -0.045 | -0.018 | -0.018 | 0 | 0.21 | 0.04 | 0.26 | 0.25 | 0.24 | 0.34 |
| 10 | 91644426 | rs10509583 | 0.353 | -0.623 | -0.605 | -1.147 | -0.002 | -0.041 | -0.041 | 0.385 | 0.14 | 0.16 | 0.19 | 0 | 0.17 | 0.1 |
| 10 | 91649717 | rs12782553 | 0.546 | -0.459 | -1 | -1.536 | -0.026 | 0.045 | 0.045 | 0.173 | 0.35 | 0.26 | 0.26 | 0.1 | 0.45 | 0.21 |
| 10 | 91662902 | rs4933165 | -0.667 | -0.797 | -1.027 | -1.267 | 0.005 | -0.009 | -0.009 | 0 | 0.18 | 0.12 | 0.26 | 0.77 | 0.12 | 0.27 |
| 10 | 91673054 | rs10785920 | -0.5 | -1.09 | -0.786 | -1.562 | 0.002 | 0.007 | 0.007 | 0 | 0.31 | 0.3 | 0.48 | 0.82 | 0.26 | 0.48 |
| 10 | 91673756 | rs1111716 | -2.751 | -0.979 | -0.973 | -1.981 | 0.004 | -0.038 | -0.038 | 0 | 0.79 | 0.77 | 0.8 | 1 | 0.62 | 0.91 |

| | | | | | | | | | | | | | | | | |
|---|---|---|---|---|---|---|---|---|---|---|---|---|---|---|---|---|
| 10 | 91680402 | rs11185930 | 1.213 | -1.13 | -0.965 | -1.652 | -0.029 | -0.015 | -0.015 | 0 | 0.07 | 0.01 | 0.22 | 0.66 | 0.07 | 0.18 |
| 10 | 91682281 | rs1274128 | 1.381 | -1.232 | -0.912 | -2.007 | 0.007 | -0.085 | -0.085 | 0.413 | 0.24 | 0.29 | 0.24 | 0.01 | 0.3 | 0.2 |
| 10 | 91682446 | rs7903627 | 0.632 | -1.276 | -0.764 | -2.059 | 0.005 | 0.005 | 0.005 | 0 | 0.24 | 0.23 | 0.25 | 0.13 | 0.16 | 0.3 |
| 10 | 91686710 | rs1274135 | -2.744 | -1.308 | -0.743 | -2.168 | 0.014 | -0.063 | -0.063 | 2.49 | 0.77 | 0.71 | 0.8 | 1 | 0.64 | 0.91 |
| 10 | 91687049 | rs1274136 | 1.421 | -1.302 | -0.745 | -2.15 | 0.007 | -0.085 | -0.085 | 0 | 0.24 | 0.29 | 0.24 | 0.01 | 0.3 | 0.2 |
| 10 | 91700359 | rs1343367 | -0.122 | -1.306 | -0.722 | -2.123 | 0.005 | 0.013 | 0.013 | 0 | 0.31 | 0.3 | 0.48 | 0.84 | 0.24 | 0.48 |
| 10 | 91702027 | rs2798612 | -0.228 | -1.318 | -0.728 | -2.149 | -0.005 | -0.053 | -0.053 | 0 | 0.46 | 0.45 | 0.32 | 0.16 | 0.4 | 0.43 |
| 10 | 91704474 | rs1324698 | 0.297 | -1.202 | -0.784 | -2.016 | 0.008 | 0.007 | 0.007 | 0.275 | 0.22 | 0.15 | 0.08 | 0.12 | 0.1 | 0.23 |
| 10 | 91716028 | rs1936216 | -0.68 | -1.182 | -0.791 | -2.023 | -0.001 | 0 | 0 | 0 | 0.55 | 0.6 | 0.72 | 0.84 | 0.54 | 0.67 |
| 10 | 91723345 | rs17412226 | -0.291 | -1.261 | -1.021 | -2.06 | 0.031 | 0.033 | 0.033 | 0.062 | 0.27 | 0.1 | 0.22 | 0.16 | 0.16 | 0.32 |
| 10 | 91723424 | rs4265510 | 1.591 | -1.374 | -0.757 | -2.158 | -0.001 | -0.028 | -0.028 | 1.681 | 0.23 | 0.2 | 0.23 | 0.04 | 0.3 | 0.21 |
| 10 | 91726635 | rs1936221 | -0.36 | -1.353 | -0.704 | -2.16 | 0.004 | 0.016 | 0.016 | 0 | 0.3 | 0.27 | 0.48 | 0.84 | 0.24 | 0.47 |
| 10 | 91727055 | rs4288675 | -2.662 | -1.13 | -0.585 | -2.014 | 0.011 | -0.063 | -0.063 | 0 | 0.76 | 0.72 | 0.8 | 1 | 0.64 | 0.91 |
| 10 | 91730034 | rs9919479 | -0.749 | -1.093 | -0.584 | -1.701 | 0.008 | 0.006 | 0.006 | 0.447 | 0.29 | 0.23 | 0.29 | 0.82 | 0.17 | 0.4 |
| 10 | 91734888 | rs11185942 | -0.15 | -0.735 | -0.297 | -1.762 | 0.015 | 0.059 | 0.059 | 0 | 0.46 | 0.28 | 0.39 | 0.23 | 0.4 | 0.52 |
| 10 | 91735093 | rs6583667 | -1.556 | -0.825 | -0.127 | -2.014 | -0.004 | -0.042 | -0.042 | 0 | 0.71 | 0.73 | 0.74 | 1 | 0.58 | 0.9 |
| 10 | 91737250 | rs7920725 | -0.545 | -0.744 | -0.094 | -1.836 | -0.001 | 0.024 | 0.024 | 0.235 | 0.32 | 0.26 | 0.53 | 0.84 | 0.28 | 0.51 |
| 10 | 91737711 | rs1891810 | 0.053 | -0.851 | 0.153 | -1.996 | -0.002 | -0.134 | -0.134 | 0 | 0.37 | 0.54 | 0.27 | 0.06 | 0.38 | 0.37 |
| 10 | 91737945 | rs7084035 | -0.662 | -0.709 | 0.143 | -1.753 | -0.001 | -0.052 | -0.052 | 0.903 | 0.1 | 0.23 | 0.09 | 0.01 | 0.1 | 0.04 |
| 10 | 91738263 | rs1547843 | -0.724 | -0.841 | 0.195 | -1.668 | -0.004 | 0 | 0 | 0 | 0.2 | 0.21 | 0.34 | 0.88 | 0.09 | 0.24 |
| 10 | 91738755 | rs10881718 | -1.96 | -0.628 | 0.167 | -1.761 | -0.013 | -0.01 | -0.01 | 0 | 0.74 | 0.82 | 0.8 | 1 | 0.63 | 0.9 |
| 10 | 91743629 | rs7086845 | -0.78 | -0.8 | 0.278 | -1.354 | 0.011 | -0.003 | -0.003 | 0 | 0.64 | 0.59 | 0.6 | 0.38 | 0.54 | 0.76 |
| 10 | 91746100 | rs7097093 | -2.045 | -0.553 | 0.216 | -1.439 | -0.003 | -0.024 | -0.024 | 0 | 0.76 | 0.82 | 0.83 | 1 | 0.7 | 0.9 |
| 10 | 91752552 | rs7070718 | -0.318 | -1.876 | 0.478 | -1.22 | -0.041 | 0.028 | 0.028 | 0 | 0.17 | 0.33 | 0.19 | 0.26 | 0.06 | 0.14 |
| 10 | 91760774 | rs10785927 | -1.801 | -1.38 | 0.211 | -1.234 | -0.028 | 0.012 | 0.012 | 0 | 0.72 | 0.83 | 0.78 | 1 | 0.6 | 0.9 |
| 10 | 91762226 | rs2419344 | -1.012 | -1.447 | 0.262 | -1.171 | -0.007 | 0.001 | 0.001 | 0 | 0.65 | 0.7 | 0.6 | 0.38 | 0.56 | 0.79 |
| 10 | 91763487 | rs10748533 | 0.19 | -1.473 | 0.085 | -1.171 | -0.004 | -0.002 | -0.002 | 0 | 0.35 | 0.39 | 0.37 | 0.67 | 0.34 | 0.33 |
| 10 | 91776003 | rs7914961 | -0.55 | -1.233 | -0.635 | -0.722 | 0.01 | 0.004 | 0.004 | 0 | 0.33 | 0.26 | 0.37 | 0.32 | 0.22 | 0.53 |
| 10 | 91777494 | rs11185967 | -0.185 | -0.978 | -0.848 | -0.22 | 0.023 | -0.004 | -0.004 | 0.015 | 0.07 | 0.04 | 0.16 | 0.6 | 0.04 | 0.1 |
| 10 | 91782728 | rs10881732 | 0.352 | -1.11 | -0.504 | -0.485 | -0.001 | 0.001 | 0.001 | 0 | 0.33 | 0.31 | 0.4 | 0.58 | 0.34 | 0.36 |
| 10 | 91786676 | rs7100106 | -0.939 | -1.113 | -0.204 | -0.24 | 0.007 | 0.006 | 0.006 | 0 | 0.26 | 0.53 | 0.36 | 0.27 | 0.21 | 0.4 |
| 10 | 91787431 | rs11185977 | -0.939 | -1.113 | -0.204 | -0.24 | 0.007 | 0.006 | 0.006 | 0 | 0.26 | 0.53 | 0.36 | 0.27 | 0.21 | 0.4 |
| 10 | 91789627 | rs11185980 | 1.893 | -1.047 | -0.456 | -0.126 | 0.023 | -0.025 | -0.025 | 0.078 | 0.06 | 0.02 | 0.01 | 0.14 | 0.01 | 0.02 |
| 10 | 91798486 | rs10509587 | 1.917 | -1.117 | -0.222 | -0.132 | 0.023 | -0.025 | -0.025 | 0 | 0.06 | 0.02 | 0.01 | 0.14 | 0.01 | 0.02 |
| 10 | 91810719 | rs4288677 | -0.576 | -1.117 | 0.153 | -0.217 | 0.014 | 0.044 | 0.044 | 0 | 0.19 | 0.39 | 0.34 | 0.27 | 0.17 | 0.37 |
| 10 | 91822938 | rs7070137 | -1.06 | -1.587 | 0.342 | -0.468 | -0.045 | 0.044 | 0.044 | 0 | 0.27 | 0.41 | 0.22 | 0.44 | 0.12 | 0.48 |
| 10 | 91824299 | rs7900020 | -1.56 | -1.611 | 0.393 | -0.756 | -0.006 | -0.009 | -0.009 | 0 | 0.29 | 0.59 | 0.4 | 0.3 | 0.22 | 0.48 |
| 10 | 91832066 | rs4933541 | -2.174 | -1.819 | -0.084 | -1.222 | -0.119 | 0.176 | 0.176 | 0 | 0.72 | 0.96 | 0.8 | 0.99 | 0.59 | 0.87 |
| 10 | 91835290 | rs2182239 | 0.063 | -1.78 | -0.015 | -1.164 | 0.014 | 0.023 | 0.023 | 0 | 0.42 | 0.36 | 0.39 | 0.69 | 0.38 | 0.39 |
| 10 | 91846739 | rs4933548 | -0.094 | -1.753 | 0.071 | -1.195 | 0.014 | 0.022 | 0.022 | 0 | 0.44 | 0.34 | 0.4 | 0.75 | 0.38 | 0.41 |
| 10 | 91847663 | rs7070751 | 0.873 | -1.681 | 0.133 | -1.202 | 0.028 | -0.021 | -0.021 | 0.75 | 0.1 | 0.18 | 0.2 | 0.03 | 0.12 | 0.11 |
| 10 | 91855982 | rs7084467 | -2.017 | -1.713 | 0.196 | -1.228 | -0.048 | 0.081 | 0.081 | 0 | 0.7 | 0.86 | 0.81 | 1 | 0.59 | 0.86 |
| 10 | 91859542 | rs6583686 | 0.746 | -1.781 | 0.212 | -0.908 | 0.002 | -0.005 | -0.005 | 0.183 | 0.33 | 0.27 | 0.31 | 0.55 | 0.35 | 0.26 |
| 10 | 91861413 | rs10444056 | 0.225 | -1.712 | 0.347 | -0.865 | -0.008 | 0 | 0 | 0 | 0.15 | 0.24 | 0.2 | 0.22 | 0.09 | 0.34 |
| 10 | 91869383 | rs4933174 | -1.207 | -1.683 | 0.346 | -0.96 | -0.003 | -0.003 | -0.003 | 0 | 0.6 | 0.65 | 0.6 | 0.97 | 0.48 | 0.75 |
| 10 | 91872059 | rs12761190 | 0.88 | -1.605 | 0.385 | -0.885 | 0.028 | -0.02 | -0.02 | 0 | 0.09 | 0.17 | 0.19 | 0.03 | 0.12 | 0.11 |
| 10 | 91881662 | rs11816423 | 0.564 | -1.662 | 0.376 | -0.92 | -0.02 | 0.041 | 0.041 | 0 | 0.07 | 0.13 | 0.19 | 0.22 | 0.03 | 0.23 |
| 10 | 91882323 | rs12265445 | 0.908 | -1.593 | 0.391 | -0.911 | 0.021 | -0.017 | -0.017 | 0 | 0.09 | 0.15 | 0.19 | 0.03 | 0.12 | 0.11 |
| 10 | 91887628 | rs10881756 | 0.289 | -2.184 | 0.203 | -1.094 | -0.074 | 0.215 | 0.215 | 0.102 | 0.15 | 0.36 | 0.15 | 0.55 | 0.1 | 0.17 |
| 10 | 91893394 | rs7894782 | 1.024 | -1.987 | 0.176 | -1.048 | 0.088 | -0.021 | -0.021 | 0 | 0.1 | 0.32 | 0.2 | 0.03 | 0.12 | 0.11 |
| 10 | 91901330 | rs1360116 | 0.171 | -2.333 | -0.05 | -1.051 | 0.006 | -0.039 | -0.039 | 0.91 | 0.34 | 0.17 | 0.41 | 0.41 | 0.31 | 0.48 |
| 10 | 91902898 | rs10785942 | -0.721 | -2.175 | 0.264 | -1.196 | -0.094 | 0.009 | 0.009 | 0.302 | 0.3 | 0.47 | 0.37 | 0.75 | 0.18 | 0.51 |
| 10 | 91907189 | rs750911 | 1.552 | -1.911 | -0.269 | -1.011 | -0.002 | 0.022 | 0.022 | 0 | 0.08 | 0.08 | 0.07 | 0.21 | 0.03 | 0.15 |
| 10 | 91907860 | rs7905491 | -0.922 | -1.648 | -0.514 | -1.225 | 0.023 | 0.063 | 0.063 | 0 | 0.53 | 0.83 | 0.51 | 0.28 | 0.53 | 0.47 |
| 10 | 91911223 | rs7096150 | -0.922 | -1.648 | -0.514 | -1.225 | 0.023 | 0.063 | 0.063 | 0 | 0.53 | 0.83 | 0.51 | 0.28 | 0.53 | 0.47 |
| 10 | 91926859 | rs11186048 | 0.683 | -1.559 | -0.467 | -1.264 | -0.008 | 0.11 | 0.11 | 0 | 0.28 | 0.1 | 0.17 | 0.04 | 0.27 | 0.18 |
| 10 | 91927196 | rs6583692 | 0.315 | -1.604 | -0.325 | -1.223 | 0.001 | 0 | 0 | 0 | 0.31 | 0.38 | 0.25 | 0.25 | 0.31 | 0.19 |
| 10 | 91927669 | rs999901 | 1.103 | -1.54 | -0.292 | -1.175 | -0.001 | 0.127 | 0.127 | 0 | 0.1 | 0.29 | 0.1 | 0.21 | 0.06 | 0.1 |
| 10 | 91930887 | rs986222 | -0.981 | -1.741 | -0.466 | -1.467 | -0.007 | -0.104 | -0.104 | 0 | 0.56 | 0.52 | 0.58 | 0.97 | 0.45 | 0.74 |
| 10 | 91930931 | rs1329180 | 1.218 | -1.302 | -1.04 | -1.379 | -0.028 | 0.023 | 0.023 | 0 | 0.1 | 0.06 | 0.22 | 0.03 | 0.21 | 0.13 |
| 10 | 91933485 | rs1360117 | 1.458 | -1.263 | -1.092 | -1.292 | -0.033 | 0.076 | 0.076 | 0 | 0.2 | 0.08 | 0.16 | 0 | 0.25 | 0.09 |
| 10 | 91936247 | rs3918657 | -1.476 | -1.228 | -0.956 | -1.097 | -0.039 | 0.125 | 0.125 | 0 | 0.7 | 0.93 | 0.83 | 0.96 | 0.67 | 0.88 |
| 10 | 91939348 | rs1329178 | -0.869 | -1.088 | -0.893 | -0.954 | 0.017 | 0.054 | 0.054 | 0 | 0.48 | 0.61 | 0.6 | 0.79 | 0.45 | 0.73 |
| 10 | 91941289 | rs2325 | 0.203 | -1.216 | -1.206 | -0.959 | 0.002 | -0.005 | -0.005 | 3.394 | 0.17 | 0.18 | 0.31 | 0.72 | 0.13 | 0.47 |
| 10 | 91941755 | rs10785943 | -0.579 | -1.366 | -1.23 | -0.38 | 0.007 | 0.005 | 0.005 | 0 | 0.42 | 0.7 | 0.51 | 0.43 | 0.39 | 0.53 |
| 10 | 91947830 | rs10881767 | 0.346 | -0.875 | -1.048 | -0.997 | -0.002 | -0.001 | -0.001 | 0 | 0.18 | 0.2 | 0.31 | 0.73 | 0.12 | 0.46 |
| 10 | 91951170 | rs2148554 | -1.311 | 0.244 | -1.156 | -0.897 | -0.053 | 0.101 | 0.101 | 0.485 | 0.78 | 0.93 | 0.85 | 1 | 0.74 | 0.92 |
| 10 | 91959144 | rs1329169 | -0.685 | -0.136 | -0.743 | -0.452 | 0.006 | 0.069 | 0.069 | 0 | 0.53 | 0.65 | 0.63 | 0.81 | 0.48 | 0.74 |
| 10 | 91960473 | rs1854036 | -0.683 | 0.938 | -0.227 | -0.299 | 0.05 | -0.029 | -0.029 | 0 | 0.15 | 0.23 | 0.17 | 0.05 | 0.26 | 0.19 |

| | | | | | | | | | | | | | | | | |
|---|---|---|---|---|---|---|---|---|---|---|---|---|---|---|---|---|
| 10 | 91962670 | rs2250245 | -0.035 | 0.14 | 0.036 | 0.202 | -0.008 | 0.053 | 0.053 | 0 | 0.27 | 0.53 | 0.26 | 0.47 | 0.25 | 0.32 |
| 10 | 91963463 | rs2250149 | 0.198 | 0.104 | 0.446 | 0.213 | -0.002 | 0.027 | 0.027 | 0 | 0.25 | 0.32 | 0.24 | 0.47 | 0.25 | 0.32 |
| 10 | 91965580 | rs2480268 | -0.614 | -0.447 | 0.029 | -0.026 | 0.041 | -0.031 | -0.031 | 0 | 0.67 | 0.77 | 0.69 | 0.57 | 0.81 | 0.62 |
| 10 | 91965613 | rs2480267 | -1.037 | -0.619 | -0.023 | 0.003 | 0.059 | -0.072 | -0.072 | 0 | 0.72 | 0.9 | 0.7 | 0.66 | 0.81 | 0.72 |
| 10 | 91967241 | rs2766630 | -0.275 | -0.62 | 0.04 | 0 | 0.005 | -0.006 | -0.006 | 0.16 | 0.66 | 0.69 | 0.68 | 0.56 | 0.8 | 0.61 |
| 10 | 91970573 | rs699214 | -0.87 | -0.605 | 0.059 | -0.019 | 0.022 | -0.018 | -0.018 | 0.103 | 0.71 | 0.78 | 0.69 | 0.66 | 0.81 | 0.71 |
| 10 | 91970601 | rs1034075 | 0.233 | -0.499 | -0.038 | -0.054 | 0 | 0.105 | 0.105 | 0.093 | 0.42 | 0.17 | 0.41 | 0.19 | 0.55 | 0.4 |
| 10 | 91977937 | rs699207 | -1.144 | -0.495 | 0.006 | 0.098 | 0.061 | -0.081 | -0.081 | 0 | 0.73 | 0.9 | 0.69 | 0.67 | 0.81 | 0.73 |
| 10 | 91980454 | rs7093513 | -0.645 | -0.402 | 0.117 | 0.078 | 0.012 | 0.042 | 0.042 | 0.465 | 0.06 | 0.29 | 0.03 | 0.1 | 0.02 | 0.12 |
| 10 | 91985192 | rs11186082 | 0.749 | -0.267 | -0.503 | -0.052 | -0.092 | 0.096 | 0.096 | 0.99 | 0.33 | 0.08 | 0.37 | 0.09 | 0.49 | 0.27 |
| 10 | 91996334 | rs2478486 | -0.276 | -0.572 | 0.038 | 0.383 | -0.068 | 0.052 | 0.052 | 0.139 | 0.68 | 0.45 | 0.66 | 0.56 | 0.81 | 0.62 |
| 10 | 91998574 | rs699200 | -0.684 | -0.655 | -0.018 | 0.408 | 0.06 | -0.07 | -0.07 | 0 | 0.72 | 0.86 | 0.69 | 0.62 | 0.82 | 0.69 |
| 10 | 92002112 | rs699191 | -0.478 | -0.459 | -0.276 | 0.403 | -0.069 | 0.045 | 0.045 | 0.143 | 0.69 | 0.45 | 0.67 | 0.57 | 0.82 | 0.63 |
| 10 | 92004794 | rs699190 | -0.427 | -0.408 | -0.634 | 0.371 | -0.062 | 0.042 | 0.042 | 0.059 | 0.67 | 0.49 | 0.66 | 0.53 | 0.81 | 0.6 |
| 10 | 92005165 | rs1410901 | -0.042 | 1.004 | -1.322 | -0.196 | -0.067 | 0.104 | 0.104 | 0 | 0.37 | 0.12 | 0.37 | 0.09 | 0.47 | 0.28 |
| 10 | 92006017 | rs699189 | -0.35 | 0.92 | -0.881 | 0.218 | 0.003 | 0 | 0 | 0 | 0.5 | 0.49 | 0.57 | 0.5 | 0.55 | 0.49 |
| 10 | 92010339 | rs7073768 | -1.059 | 0.206 | 0.026 | -0.257 | -0.024 | -0.037 | -0.037 | 0 | 0.09 | 0.35 | 0.04 | 0 | 0.04 | 0.08 |
| 10 | 92018838 | rs943317 | -0.572 | 1.032 | -0.4 | -0.73 | -0.023 | 0.013 | 0.013 | 0.041 | 0.87 | 0.93 | 0.85 | 1 | 0.76 | 0.91 |
| 10 | 92022480 | rs10219117 | 0.252 | 1.357 | -0.713 | -0.328 | -0.009 | -0.019 | -0.019 | 0 | 0.2 | 0.15 | 0.48 | 0.47 | 0.28 | 0.14 |
| 10 | 92025659 | rs12219014 | 0.417 | 1.117 | -0.698 | -0.291 | 0.012 | 0.011 | 0.011 | 0 | 0.58 | 0.48 | 0.34 | 0.53 | 0.45 | 0.7 |
| 10 | 92027459 | rs17092763 | 0.305 | 1.867 | -1.228 | -0.16 | -0.039 | -0.052 | -0.052 | 0 | 0.11 | 0.03 | 0.1 | 0.37 | 0.15 | 0.19 |
| 10 | 92030619 | rs10509597 | -0.005 | 1.259 | -1.528 | -1.14 | 0.001 | 0.032 | 0.032 | 0 | 0.17 | 0.1 | 0.09 | 0 | 0.17 | 0.06 |
| 10 | 92032294 | rs7069153 | -0.869 | 1.633 | -1.621 | -1.002 | -0.074 | 0.077 | 0.077 | 0.28 | 0.86 | 0.97 | 0.84 | 1 | 0.75 | 0.9 |
| 10 | 92035457 | rs7087199 | -0.847 | 0.916 | -1.297 | -1.159 | 0.011 | 0.028 | 0.028 | 1.196 | 0.34 | 0.71 | 0.2 | 0.05 | 0.31 | 0.37 |
| 10 | 92036154 | rs17092874 | 0.318 | 1.059 | -1.718 | -0.482 | -0.057 | -0.122 | -0.122 | 0 | 0.16 | 0.02 | 0.11 | 0.41 | 0.16 | 0.27 |
| 10 | 92039445 | rs12783212 | -0.013 | 0.777 | -1.165 | -1.28 | -0.025 | 0.003 | 0.003 | 0 | 0.12 | 0.04 | 0.08 | 0.14 | 0.15 | 0.26 |
| 10 | 92053863 | rs12779831 | -0.27 | 0.919 | -1.734 | -1.363 | -0.022 | 0.054 | 0.054 | 0 | 0.18 | 0.06 | 0.11 | 0.05 | 0.2 | 0.28 |
| 10 | 92060639 | rs1926100 | -0.836 | 0.789 | -1.713 | -1.789 | 0 | 0.016 | 0.016 | 0.801 | 0.36 | 0.7 | 0.3 | 0.03 | 0.41 | 0.38 |
| 10 | 92064234 | rs10881780 | -0.867 | 0.732 | -1.696 | -1.814 | 0.002 | 0.039 | 0.039 | 0.819 | 0.37 | 0.72 | 0.29 | 0.03 | 0.41 | 0.38 |
| 10 | 92065512 | rs2900955 | -0.679 | 0.671 | -1.763 | -1.836 | -0.003 | -0.038 | -0.038 | 2.887 | 0.31 | 0.67 | 0.27 | 0.03 | 0.38 | 0.3 |
| 10 | 92076442 | rs12784906 | -0.185 | 0.792 | -1.604 | -1.781 | -0.032 | 0.054 | 0.054 | 0 | 0.21 | 0.08 | 0.16 | 0.03 | 0.31 | 0.3 |
| 10 | 92077274 | rs11186122 | -0.656 | 0.783 | -1.675 | -1.744 | -0.003 | 0.035 | 0.035 | 0.199 | 0.46 | 0.65 | 0.29 | 0.03 | 0.5 | 0.43 |
| 10 | 92078697 | rs2104483 | 0.546 | 0.135 | -2.242 | -1.613 | 0.026 | 0.219 | 0.219 | 0.209 | 0.36 | 0.07 | 0.45 | 0.97 | 0.25 | 0.46 |
| 10 | 92081093 | rs943316 | 0.61 | 0.394 | -1.691 | -0.823 | -0.001 | -0.002 | -0.002 | 0 | 0.61 | 0.6 | 0.51 | 0.72 | 0.65 | 0.65 |
| 10 | 92082250 | rs1326162 | 0.402 | 0.891 | -2.152 | -0.35 | -0.06 | -0.06 | -0.06 | 0 | 0.18 | 0.04 | 0.16 | 0.31 | 0.27 | 0.14 |
| 10 | 92083015 | rs1326163 | 0.185 | 0.868 | -2.046 | -0.317 | -0.003 | -0.005 | -0.005 | 0 | 0.61 | 0.59 | 0.51 | 0.72 | 0.65 | 0.65 |
| 10 | 92090566 | rs1001065 | 0.99 | 0.736 | -1.971 | -0.244 | -0.002 | 0 | 0 | 0 | 0.43 | 0.53 | 0.34 | 0.41 | 0.38 | 0.51 |
| 10 | 92090785 | rs7905472 | -0.062 | 0.654 | -1.933 | -0.215 | -0.003 | -0.007 | -0.007 | 0 | 0.82 | 0.77 | 0.79 | 0.88 | 0.82 | 0.77 |
| 10 | 92095687 | rs11186132 | -0.56 | 0.532 | -2.322 | -0.319 | -0.035 | 0.009 | 0.009 | 0 | 0.31 | 0.19 | 0.33 | 0.31 | 0.38 | 0.23 |
| 10 | 92095819 | rs12571204 | -0.787 | 0.428 | -2.523 | -0.4 | 0.015 | -0.01 | -0.01 | 0 | 0.23 | 0.11 | 0.17 | 0.22 | 0.17 | 0.1 |
| 10 | 92098023 | rs1409298 | -0.228 | 0.447 | -2.536 | -0.406 | -0.014 | 0.016 | 0.016 | 0 | 0.14 | 0.08 | 0.06 | 0.06 | 0.11 | 0.06 |
| 10 | 92098288 | rs12572994 | -0.574 | 0.49 | -2.491 | -0.376 | 0.014 | -0.011 | -0.011 | 0 | 0.23 | 0.11 | 0.17 | 0.22 | 0.17 | 0.1 |
| 10 | 92099644 | rs12571278 | -0.587 | 0.487 | -2.493 | -0.36 | 0.015 | -0.013 | -0.013 | 0 | 0.23 | 0.11 | 0.17 | 0.22 | 0.17 | 0.1 |
| 10 | 92101026 | rs1326167 | 0.73 | 0.668 | -2.505 | -0.333 | -0.069 | -0.082 | -0.082 | 0 | 0.16 | 0.02 | 0.13 | 0.3 | 0.24 | 0.12 |
| 10 | 92102972 | rs1540997 | 1.572 | 0.458 | -2.232 | -0.317 | 0.007 | 0.002 | 0.002 | 0 | 0.43 | 0.69 | 0.37 | 0.37 | 0.39 | 0.51 |
| 10 | 92104855 | rs17518890 | -1.192 | 0.2 | -2.043 | -0.506 | 0.007 | 0.053 | 0.053 | 0 | 0.33 | 0.14 | 0.34 | 0.21 | 0.26 | 0.2 |
| 10 | 92106648 | rs7898091 | -0.644 | -0.013 | -1 | -0.394 | -0.015 | -0.018 | -0.018 | 0.708 | 0.13 | 0.28 | 0.05 | 0.05 | 0.08 | 0.11 |
| 10 | 92106907 | rs7898470 | 0.421 | -0.124 | -1.914 | -0.586 | -0.047 | 0.104 | 0.104 | 0.225 | 0.65 | 0.86 | 0.62 | 0.21 | 0.56 | 0.45 |
| 10 | 92118547 | rs11186140 | 1.539 | -0.056 | -2.484 | -1.197 | -0.109 | 0.134 | 0.134 | 1.048 | 0.21 | 0.01 | 0.29 | 0 | 0.31 | 0.18 |
| 10 | 92124030 | rs17094195 | 2.373 | -0.88 | -1.55 | -1.604 | -0.011 | -0.059 | -0.059 | 0.252 | 0.1 | 0.34 | 0.05 | 0 | 0.05 | 0 |
| 10 | 92132199 | rs10785956 | 0.953 | -1.591 | -1.494 | -1.093 | 0.017 | -0.033 | -0.033 | 0 | 0.31 | 0.23 | 0.31 | 0.5 | 0.23 | 0.52 |
| 10 | 92132871 | rs17094380 | 1.085 | -1.244 | -2.004 | -1.086 | 0.029 | 0.039 | 0.039 | 2.037 | 0.21 | 0.13 | 0.09 | 0.04 | 0.16 | 0.03 |
| 10 | 92137272 | rs10748546 | 0.136 | -1.111 | -1.814 | -0.953 | 0.021 | -0.009 | -0.009 | 0.529 | 0.43 | 0.61 | 0.37 | 0.52 | 0.28 | 0.54 |
| 10 | 92150962 | rs12355470 | -0.061 | -0.718 | -1.805 | -0.64 | 0.039 | 0.043 | 0.043 | 0 | 0.49 | 0.31 | 0.53 | 0.32 | 0.6 | 0.34 |
| 10 | 92159044 | rs10748547 | 0.557 | -0.79 | -1.78 | -0.655 | -0.005 | -0.024 | -0.024 | 0 | 0.62 | 0.7 | 0.65 | 0.51 | 0.56 | 0.71 |
| 10 | 92162586 | rs11186162 | -0.271 | -1.061 | -1.924 | -0.453 | -0.008 | -0.065 | -0.065 | 0 | 0.37 | 0.1 | 0.32 | 0.51 | 0.38 | 0.41 |
| 10 | 92162800 | rs10509601 | 1.529 | -1.106 | -1.807 | -0.512 | -0.017 | 0.007 | 0.007 | 0 | 0.27 | 0.65 | 0.29 | 0.33 | 0.17 | 0.29 |
| 10 | 92167112 | rs10785957 | 0.588 | -0.762 | -1.954 | -0.756 | -0.008 | -0.024 | -0.024 | 0 | 0.57 | 0.73 | 0.63 | 0.51 | 0.51 | 0.72 |
| 10 | 92167737 | rs12762901 | 0.564 | -0.753 | -1.923 | -0.704 | -0.02 | -0.039 | -0.039 | 0 | 0.31 | 0.11 | 0.29 | 0.34 | 0.34 | 0.17 |
| 10 | 92177113 | rs7475087 | 1.023 | -0.822 | -2.062 | -0.69 | 0.023 | -0.03 | -0.03 | 0.021 | 0.33 | 0.71 | 0.32 | 0.5 | 0.21 | 0.53 |
| 10 | 92181166 | rs7091476 | 1.209 | -0.817 | -2.029 | -0.687 | -0.022 | 0.037 | 0.037 | 0 | 0.32 | 0.57 | 0.31 | 0.5 | 0.21 | 0.53 |
| 10 | 92181330 | rs10785962 | 0.922 | -0.53 | -2.089 | -0.537 | 0.023 | -0.015 | -0.015 | 0.026 | 0.41 | 0.66 | 0.35 | 0.5 | 0.26 | 0.53 |
| 10 | 92208256 | rs11186186 | 0.233 | -0.346 | -2.021 | -0.455 | -0.017 | -0.041 | -0.041 | 0 | 0.23 | 0.14 | 0.26 | 0.34 | 0.29 | 0.17 |
| 10 | 92214136 | rs11186188 | -0.611 | -0.369 | -2.087 | -0.498 | 0 | 0 | 0 | 0 | 0.32 | 0.32 | 0.3 | 0.34 | 0.35 | 0.19 |
| 10 | 92214326 | rs2490318 | 0.861 | -0.595 | -1.934 | -0.295 | -0.018 | -0.029 | -0.029 | 0 | 0.06 | 0.01 | 0.05 | 0.18 | 0.09 | 0.1 |
| 10 | 92223676 | rs10509602 | 1.156 | -0.591 | -1.868 | -0.297 | -0.002 | -0.018 | -0.018 | 0 | 0.08 | 0.09 | 0.03 | 0 | 0.06 | 0.02 |
| 10 | 92226605 | rs1782741 | -1.088 | -0.638 | -1.916 | -0.296 | 0.019 | -0.012 | -0.012 | 0.247 | 0.59 | 0.35 | 0.65 | 0.52 | 0.74 | 0.46 |
| 10 | 92230837 | rs2490314 | 0.269 | -0.127 | -1.771 | -0.366 | -0.001 | -0.001 | -0.001 | 3.261 | 0.13 | 0.1 | 0.08 | 0.17 | 0.15 | 0.09 |
| 10 | 92246550 | rs1274352 | 0.361 | -0.06 | -2.015 | -0.721 | -0.034 | 0.086 | 0.086 | 0 | 0.39 | 0.25 | 0.4 | 0.17 | 0.46 | 0.26 |

| | | | | | | | | | | | | | | | | |
|---|---|---|---|---|---|---|---|---|---|---|---|---|---|---|---|---|
| 10 | 92248009 | rs1274351 | 1.357 | 0.078 | -2.181 | -0.789 | -0.031 | -0.029 | -0.029 | 0 | 0.1 | 0.02 | 0.06 | 0.17 | 0.12 | 0.07 |
| 10 | 92252281 | rs7093552 | 1.584 | 0.104 | -2.15 | -0.783 | -0.027 | 0.048 | 0.048 | 0.271 | 0.27 | 0.17 | 0.33 | 0 | 0.33 | 0.19 |
| 10 | 92264124 | rs17521327 | 1.997 | 0.115 | -2.199 | -0.758 | -0.102 | 0.151 | 0.151 | 0.051 | 0.23 | 0.04 | 0.31 | 0 | 0.31 | 0.19 |
| 10 | 92287937 | rs4144884 | 1.152 | 0.44 | -2.149 | -0.252 | -0.059 | -0.052 | -0.052 | 0 | 0.12 | 0.02 | 0.2 | 0.19 | 0.17 | 0.13 |
| 10 | 92292960 | rs1274386 | -1.401 | -0.057 | -2.073 | -0.296 | -0.003 | 0.001 | 0.001 | 0 | 0.68 | 0.39 | 0.68 | 0.41 | 0.74 | 0.43 |
| 10 | 92302869 | rs17443486 | 0.775 | 0.664 | -2.089 | -0.108 | -0.063 | -0.023 | -0.023 | 0 | 0.12 | 0.02 | 0.1 | 0.16 | 0.21 | 0.09 |
| 10 | 92303944 | rs17097104 | 1.028 | 0.083 | -1.352 | 0.138 | 0.014 | 0.199 | 0.199 | 0.599 | 0.07 | 0.34 | 0.12 | 0.19 | 0.03 | 0.25 |
| 10 | 92304910 | rs7094359 | 1.192 | 0.182 | -1.923 | -0.482 | -0.022 | 0.034 | 0.034 | 0 | 0.34 | 0.27 | 0.39 | 0.05 | 0.45 | 0.23 |
| 10 | 92313287 | rs17097279 | 1.28 | 0.264 | -2.012 | -0.159 | -0.024 | -0.051 | -0.051 | 0 | 0.09 | 0.01 | 0.16 | 0.25 | 0.09 | 0.11 |
| 10 | 92345067 | rs1274412 | 0.109 | 0.207 | -2.153 | -0.281 | -0.016 | -0.01 | -0.01 | 0.469 | 0.31 | 0.15 | 0.29 | 0.32 | 0.3 | 0.22 |
| 10 | 92346694 | rs17097510 | 3.07 | 0.024 | -2.002 | -0.488 | -0.009 | -0.03 | -0.03 | 0.073 | 0.09 | 0.11 | 0.05 | 0 | 0.05 | 0.01 |
| 10 | 92346977 | rs1274411 | 0.892 | 0.127 | -1.992 | -0.417 | -0.046 | 0.035 | 0.035 | 0 | 0.13 | 0.02 | 0.08 | 0.05 | 0.15 | 0.1 |
| 10 | 92354474 | rs1274396 | -1.697 | 0.064 | -1.998 | -0.538 | -0.013 | -0.007 | -0.007 | 1.7 | 0.64 | 0.42 | 0.59 | 0.3 | 0.72 | 0.4 |
| 10 | 92355756 | rs2490322 | -0.947 | 0.137 | -2.218 | -0.738 | -0.002 | 0.006 | 0.006 | 1.652 | 0.41 | 0.35 | 0.28 | 0.3 | 0.41 | 0.2 |
| 10 | 92357530 | rs1274399 | -0.666 | -0.061 | -1.699 | -0.331 | 0.001 | 0.003 | 0.003 | 0.106 | 0.29 | 0.22 | 0.18 | 0.27 | 0.26 | 0.17 |
| 10 | 92362797 | rs1274409 | -0.666 | -0.061 | -1.699 | -0.38 | 0.001 | 0.005 | 0.005 | 0.106 | 0.29 | 0.22 | 0.18 | 0.26 | 0.26 | 0.17 |
| 10 | 92371952 | rs2710728 | 0.633 | -0.055 | -1.687 | -0.391 | 0.003 | 0.014 | 0.014 | 0 | 0.46 | 0.58 | 0.5 | 0.74 | 0.43 | 0.63 |
| 10 | 92379924 | rs1419345 | -0.209 | -0.13 | -1.724 | -0.39 | -0.001 | -0.006 | -0.006 | 0 | 0.77 | 0.78 | 0.87 | 0.93 | 0.78 | 0.89 |
| 10 | 92386864 | rs4933596 | 0.199 | -0.258 | -1.682 | -0.344 | -0.012 | 0.047 | 0.047 | 0 | 0.32 | 0.2 | 0.37 | 0.19 | 0.35 | 0.27 |
| 10 | 92401930 | rs2420187 | 2.585 | -0.211 | -1.642 | -0.351 | -0.026 | -0.058 | -0.058 | 0.423 | 0.11 | 0.22 | 0.05 | 0 | 0.06 | 0.01 |
| 10 | 92417767 | rs1592353 | 1.161 | 0.049 | -1.82 | -0.383 | -0.079 | 0.127 | 0.127 | 1.234 | 0.22 | 0.06 | 0.32 | 0 | 0.31 | 0.2 |
| 10 | 92433587 | rs2710766 | 0.584 | -0.035 | -1.755 | -0.161 | -0.009 | -0.018 | -0.018 | 0 | 0.8 | 0.89 | 0.87 | 0.73 | 0.8 | 0.83 |
| 10 | 92434957 | rs1936614 | -0.336 | -0.713 | -1.744 | -0.184 | -0.033 | -0.006 | -0.006 | 0 | 0.23 | 0.65 | 0.2 | 0.2 | 0.13 | 0.3 |
| 10 | 92436384 | rs10509605 | -1.223 | -0.539 | -1.604 | -0.077 | -0.044 | 0.074 | 0.074 | 0.132 | 0.14 | 0.54 | 0.15 | 0.2 | 0.07 | 0.29 |
| 10 | 92437910 | rs2800158 | -0.635 | -0.14 | -1.918 | -0.132 | -0.037 | 0.001 | 0.001 | 0 | 0.89 | 0.98 | 0.93 | 0.92 | 0.84 | 0.9 |
| 10 | 92438659 | rs1892108 | 1.026 | -0.155 | -2.049 | -0.32 | -0.062 | 0.113 | 0.113 | 0.773 | 0.23 | 0.08 | 0.31 | 0 | 0.31 | 0.2 |
| 10 | 92438925 | rs2800157 | -0.708 | 0.116 | -2.028 | -0.081 | -0.051 | 0.01 | 0.01 | 0 | 0.87 | 0.97 | 0.91 | 0.94 | 0.82 | 0.87 |
| 10 | 92439106 | rs2096352 | 1.241 | -0.693 | -2.131 | 0.164 | -0.094 | -0.106 | -0.106 | 0 | 0.35 | 0.08 | 0.49 | 0.56 | 0.4 | 0.37 |
| 10 | 92439690 | rs1936609 | -0.839 | -0.459 | -1.826 | 0.071 | -0.052 | 0.012 | 0.012 | 0.132 | 0.88 | 0.98 | 0.9 | 0.95 | 0.81 | 0.85 |
| 10 | 92440481 | rs12246059 | -0.456 | -0.613 | -1.246 | 0.336 | -0.013 | 0.122 | 0.122 | 0 | 0.1 | 0.28 | 0.14 | 0.23 | 0.07 | 0.22 |
| 10 | 92440843 | rs17098642 | 1.113 | -0.567 | -1.383 | 0.473 | 0.001 | -0.004 | -0.004 | 0 | 0.15 | 0.13 | 0.11 | 0.23 | 0.13 | 0.06 |
| 10 | 92441029 | rs1419351 | -0.419 | -0.589 | -1.516 | 0.395 | 0.035 | 0.117 | 0.117 | 0.649 | 0.48 | 0.76 | 0.43 | 0.23 | 0.41 | 0.28 |
| 10 | 92441337 | rs2800156 | 0.906 | -0.55 | -1.489 | 0.468 | -0.007 | -0.013 | -0.013 | 0 | 0.39 | 0.21 | 0.4 | 0.48 | 0.41 | 0.39 |
| 10 | 92441426 | rs2710771 | 0.238 | -0.109 | -1.307 | 0.551 | -0.012 | -0.018 | -0.018 | 0 | 0.83 | 0.86 | 0.8 | 0.73 | 0.78 | 0.65 |
| 10 | 92442274 | rs1892105 | -0.28 | 0.113 | -1.387 | -0.085 | -0.05 | 0.009 | 0.009 | 0.011 | 0.86 | 0.96 | 0.9 | 0.94 | 0.81 | 0.84 |
| 10 | 92444011 | rs7901850 | 0.117 | -0.336 | -1.027 | -0.287 | -0.012 | -0.113 | -0.113 | 2.029 | 0.33 | 0.59 | 0.31 | 0 | 0.28 | 0.21 |
| 10 | 92444877 | rs2800152 | 0.27 | -0.239 | -1.19 | 0.092 | -0.001 | 0 | 0 | 0 | 0.79 | 0.8 | 0.83 | 0.77 | 0.78 | 0.8 |
| 10 | 92445253 | rs1936602 | -0.303 | -0.528 | -1.575 | 0.153 | 0.039 | 0.013 | 0.013 | 0 | 0.5 | 0.72 | 0.52 | 0.41 | 0.43 | 0.5 |
| 10 | 92456667 | rs17098924 | 0.265 | -0.646 | 0.017 | -0.237 | -0.009 | -0.135 | -0.135 | 0 | 0.19 | 0.41 | 0.13 | 0 | 0.18 | 0.05 |
| 10 | 92457246 | rs1892110 | -0.596 | -0.907 | 0.053 | -0.295 | -0.027 | -0.05 | -0.05 | 0.061 | 0.3 | 0.42 | 0.21 | 0.17 | 0.24 | 0.18 |
| 10 | 92457445 | rs2000322 | -1.086 | -0.973 | 0.078 | -0.178 | 0.01 | -0.008 | -0.008 | 0 | 0.34 | 0.77 | 0.34 | 0.3 | 0.3 | 0.33 |
| 10 | 92457643 | rs1936611 | -0.218 | -1.022 | 0.086 | -0.152 | 0.016 | 0.169 | 0.169 | 0 | 0.14 | 0.37 | 0.21 | 0.3 | 0.11 | 0.28 |
| 10 | 92457733 | rs1892109 | -1.513 | -0.917 | 0.021 | -0.146 | -0.008 | -0.008 | -0.008 | 0 | 0.9 | 0.94 | 0.91 | 0.79 | 0.83 | 0.87 |
| 10 | 92470777 | rs1857586 | 0.661 | -0.924 | 0.056 | -0.153 | -0.001 | -0.005 | -0.005 | 0 | 0.58 | 0.47 | 0.7 | 0.62 | 0.59 | 0.69 |
| 10 | 92473615 | rs2710723 | 0.13 | -0.903 | 0.007 | -0.146 | -0.001 | -0.006 | -0.006 | 1.4 | 0.76 | 0.82 | 0.82 | 0.62 | 0.77 | 0.73 |
| 10 | 92477076 | rs11186286 | 0.342 | -0.865 | 0.034 | -0.145 | -0.01 | -0.118 | -0.118 | 0.053 | 0.2 | 0.38 | 0.13 | 0 | 0.18 | 0.05 |
| 10 | 92479457 | rs1539723 | 0.132 | -0.868 | -0.025 | -0.148 | 0.028 | -0.029 | -0.029 | 1.913 | 0.71 | 0.56 | 0.78 | 0.79 | 0.65 | 0.78 |
| 10 | 92484752 | rs1573253 | 0.9 | -0.796 | 0.145 | 0.051 | 0.012 | 0.041 | 0.041 | 0 | 0.08 | 0.13 | 0.09 | 0.16 | 0.09 | 0.09 |
| 10 | 92486835 | rs2465115 | -1.311 | -0.766 | 0.139 | 0.055 | 0.009 | 0.009 | 0.009 | 0 | 0.89 | 0.87 | 0.9 | 0.79 | 0.83 | 0.83 |
| 10 | 92495336 | rs1107688 | -0.81 | -0.85 | 0.174 | 0.031 | -0.027 | 0.045 | 0.045 | 0 | 0.24 | 0.38 | 0.19 | 0.33 | 0.15 | 0.23 |
| 10 | 92500519 | rs17526697 | 2.487 | -0.93 | 0.097 | -0.089 | 0 | 0.08 | 0.08 | 0.064 | 0.13 | 0 | 0.07 | 0 | 0.09 | 0.08 |
| 10 | 92501468 | rs3740046 | 0.983 | 0.078 | 1.074 | 0.871 | -0.002 | -0.001 | -0.001 | 0 | 0.09 | 0.06 | 0.09 | 0.2 | 0.17 | 0.17 |
| 10 | 92501650 | rs1418213 | -1.328 | 0.064 | 1.074 | 0.876 | -0.027 | 0.045 | 0.045 | 0 | 0.24 | 0.37 | 0.19 | 0.33 | 0.15 | 0.23 |
| 10 | 92505265 | rs11812708 | -0.435 | 0.004 | 1.072 | 0.85 | -0.02 | -0.038 | -0.038 | 0 | 0.23 | 0.35 | 0.22 | 0.14 | 0.21 | 0.2 |
| 10 | 92507863 | rs2420203 | -1.341 | 0.016 | 0.801 | 0.833 | -0.03 | 0.045 | 0.045 | 0 | 0.24 | 0.38 | 0.19 | 0.33 | 0.15 | 0.23 |
| 10 | 92513249 | rs10881830 | -0.459 | 0.022 | 0.627 | 0.734 | -0.05 | -0.023 | -0.023 | 0 | 0.16 | 0.26 | 0.1 | 0.12 | 0.06 | 0.13 |
| 10 | 92514739 | rs11186300 | -2.513 | 0.078 | 0.472 | 0.755 | -0.142 | -0.044 | -0.044 | 0 | 0.58 | 0.93 | 0.51 | 0.63 | 0.52 | 0.61 |
| 10 | 92528523 | rs12762083 | 1.313 | 0.121 | 0.59 | 0.755 | -0.016 | 0.003 | 0.003 | 1.846 | 0.05 | 0.07 | 0.05 | 0.12 | 0.01 | 0.06 |
| 10 | 92542499 | rs955064 | -0.061 | 0.131 | 0.668 | 0.722 | -0.024 | -0.044 | -0.044 | 0.62 | 0.11 | 0.17 | 0.04 | 0 | 0.05 | 0.08 |
| 10 | 92559908 | rs7916403 | -1.937 | 0.133 | 0.732 | 0.741 | 0.022 | 0.019 | 0.019 | 0 | 0.48 | 0.69 | 0.41 | 0.46 | 0.44 | 0.48 |
| 10 | 92561557 | rs11186309 | -1.068 | 0.067 | 0.798 | 0.781 | -0.043 | -0.031 | -0.031 | 1.68 | 0.29 | 0.45 | 0.28 | 0.26 | 0.22 | 0.27 |
| 10 | 92576448 | rs11186320 | -1.499 | 0.085 | 0.772 | 0.803 | -0.003 | 0 | 0 | 0 | 0.38 | 0.56 | 0.38 | 0.42 | 0.31 | 0.38 |
| 10 | 92580807 | rs12261011 | -0.705 | 0.106 | 0.568 | 0.795 | 0.008 | 0.012 | 0.012 | 0 | 0.15 | 0.18 | 0.14 | 0.16 | 0.16 | 0.15 |
| 10 | 92590233 | rs10881838 | -1.671 | 0.088 | 0.63 | 0.79 | 0.013 | -0.007 | -0.007 | 0 | 0.38 | 0.6 | 0.38 | 0.42 | 0.31 | 0.37 |
| 10 | 92595066 | rs2901127 | 0.034 | 0.089 | 0.651 | 0.788 | 0.024 | 0.075 | 0.075 | 0 | 0.09 | 0.21 | 0.1 | 0.16 | 0.09 | 0.1 |
| 10 | 92598171 | rs10785973 | 2.889 | 0.294 | 0.446 | 0.705 | -0.047 | -0.05 | -0.05 | 0 | 0.33 | 0.02 | 0.32 | 0.3 | 0.3 | 0.33 |
| 10 | 92598458 | rs11596991 | 1.575 | -0.723 | 1.104 | 0.858 | 0.003 | 0.001 | 0.001 | 0.029 | 0.08 | 0.03 | 0.04 | 0 | 0.07 | 0.01 |
| 10 | 92598823 | rs11599902 | -1.498 | -0.451 | 0.95 | 0.979 | 0.013 | -0.007 | -0.007 | 0 | 0.38 | 0.6 | 0.38 | 0.42 | 0.31 | 0.37 |
| 10 | 92609306 | rs7074715 | -0.324 | -0.454 | 0.928 | 0.959 | -0.071 | -0.007 | -0.007 | 0 | 0.24 | 0.42 | 0.24 | 0.26 | 0.14 | 0.25 |

| | | | | | | | | | | | | | | | | |
|---|---|---|---|---|---|---|---|---|---|---|---|---|---|---|---|---|
| 10 | 92610045 | rs12259062 | -2.211 | -0.369 | 0.755 | 1.003 | 0.05 | -0.04 | -0.04 | 0 | 0.49 | 0.92 | 0.43 | 0.43 | 0.36 | 0.45 |
| 10 | 92624969 | rs7084125 | -0.19 | -0.479 | 0.792 | 0.947 | -0.054 | -0.005 | -0.005 | 0 | 0.23 | 0.4 | 0.24 | 0.26 | 0.15 | 0.22 |
| 10 | 92631244 | rs4282910 | -0.876 | -0.442 | 0.824 | 0.923 | 0.029 | 0.05 | 0.05 | 0 | 0.15 | 0.32 | 0.15 | 0.16 | 0.16 | 0.14 |
| 10 | 92632899 | rs4586057 | -1.8 | -0.445 | 0.785 | 0.921 | -0.012 | -0.007 | -0.007 | 0.173 | 0.25 | 0.46 | 0.19 | 0.17 | 0.21 | 0.22 |
| 10 | 92637833 | rs11186343 | -0.245 | -0.432 | 0.784 | 0.93 | 0.036 | 0.111 | 0.111 | 0 | 0.09 | 0.27 | 0.11 | 0.16 | 0.08 | 0.11 |
| 10 | 92652850 | rs11186353 | -0.453 | -0.259 | 0.773 | 1.003 | -0.007 | -0.017 | -0.017 | 0 | 0.1 | 0.12 | 0.04 | 0 | 0.05 | 0.08 |
| 10 | 92660702 | rs7917835 | 0.593 | -0.175 | 0.873 | 1.056 | -0.015 | 0.058 | 0.058 | 0.302 | 0.06 | 0.2 | 0.12 | 0.14 | 0.03 | 0.16 |
| 10 | 92667502 | rs11186360 | -0.912 | 0.189 | 0.063 | 1.058 | 0.013 | 0.026 | 0.026 | 0 | 0.15 | 0.25 | 0.15 | 0.16 | 0.16 | 0.14 |
| 10 | 92668029 | rs4933621 | 0.398 | 0.235 | -0.076 | 1.076 | -0.052 | -0.005 | -0.005 | 0 | 0.22 | 0.39 | 0.24 | 0.26 | 0.15 | 0.22 |
| 10 | 92672564 | rs3939 | 1.342 | 0.543 | -0.76 | 1.252 | -0.042 | 0.013 | 0.013 | 0.006 | 0.42 | 0.16 | 0.48 | 0.37 | 0.46 | 0.4 |
| 10 | 92680929 | rs10881855 | 0.859 | 0.605 | -0.511 | 1.003 | 0.019 | 0.026 | 0.026 | 0.124 | 0.7 | 0.61 | 0.76 | 0.66 | 0.68 | 0.65 |
| 10 | 92681475 | rs3824739 | 0.128 | 0.984 | -1.212 | 1.001 | 0.011 | 0.067 | 0.067 | 0 | 0.06 | 0.1 | 0.07 | 0.15 | 0.07 | 0.11 |
| 10 | 92682060 | rs10748555 | 0.721 | 0.917 | -1.256 | 0.962 | -0.052 | 0.026 | 0.026 | 0 | 0.42 | 0.15 | 0.52 | 0.33 | 0.47 | 0.4 |
| 10 | 92682369 | rs11815049 | -1.204 | 0.774 | -0.691 | 1.292 | 0 | 0.023 | 0.023 | 0 | 0.13 | 0.2 | 0.12 | 0.16 | 0.11 | 0.17 |
| 10 | 92685713 | rs10785976 | 0.125 | 0.562 | -0.846 | 0.936 | 0.01 | -0.001 | -0.001 | 0 | 0.55 | 0.42 | 0.65 | 0.48 | 0.62 | 0.57 |
| 10 | 92686421 | rs7905809 | 0.424 | 0.543 | -0.835 | 1.135 | -0.002 | -0.007 | -0.007 | 0.001 | 0.94 | 0.96 | 0.96 | 0.84 | 0.95 | 0.9 |
| 10 | 92686818 | rs12247252 | -1.029 | 0.15 | -0.43 | 1.223 | -0.008 | 0.024 | 0.024 | 0 | 0.17 | 0.22 | 0.11 | 0.28 | 0.15 | 0.17 |
| 10 | 92688076 | rs10881856 | -0.068 | -0.72 | -1.156 | 0.822 | 0.043 | 0.012 | 0.012 | 0 | 0.24 | 0.09 | 0.11 | 0.17 | 0.13 | 0.19 |
| 10 | 92688298 | rs1539479 | -0.401 | 0.591 | -1.37 | 0.683 | 0 | 0.001 | 0.001 | 0 | 0.19 | 0.15 | 0.33 | 0.71 | 0.29 | 0.5 |
| 10 | 92691072 | rs7896199 | -1.335 | 0.514 | -0.518 | 0.053 | -0.001 | -0.031 | -0.031 | 0 | 0.54 | 0.53 | 0.53 | 0.87 | 0.59 | 0.74 |
| 10 | 92699556 | rs11186371 | -0.71 | -0.292 | 0.145 | -0.55 | 0.03 | 0.047 | 0.047 | 0.04 | 0.5 | 0.64 | 0.55 | 0.93 | 0.41 | 0.74 |
| 10 | 92705962 | rs10881863 | 1.428 | -0.289 | -0.096 | 0.14 | 0.062 | -0.091 | -0.091 | 0.51 | 0.19 | 0 | 0.08 | 0.27 | 0.08 | 0.14 |
| 10 | 92706044 | rs10785978 | -0.949 | -0.043 | 0.452 | 0.234 | 0.02 | 0.08 | 0.08 | 0 | 0.51 | 0.39 | 0.51 | 0.82 | 0.56 | 0.6 |
| 10 | 92719584 | rs7099299 | -0.159 | -0.054 | 0.53 | 0.273 | 0.004 | 0.007 | 0.007 | 0.116 | 0.41 | 0.41 | 0.41 | 0.77 | 0.34 | 0.51 |
| 10 | 92720630 | rs1334583 | -0.159 | -0.054 | 0.53 | 0.273 | 0.004 | 0.007 | 0.007 | 0.116 | 0.41 | 0.41 | 0.41 | 0.77 | 0.34 | 0.51 |
| 10 | 92721319 | rs11598058 | -0.571 | 0.114 | 0.453 | 0.319 | -0.054 | 0.032 | 0.032 | 0 | 0.11 | 0.04 | 0.1 | 0.05 | 0.22 | 0.09 |
| 10 | 92732950 | rs1334579 | -1.145 | -0.248 | 0.505 | 0.053 | -0.006 | -0.032 | -0.032 | 0 | 0.55 | 0.56 | 0.53 | 0.82 | 0.59 | 0.61 |
| 10 | 92738106 | rs2118824 | -0.801 | -0.219 | 0.511 | 0.064 | 0.006 | 0.053 | 0.053 | 0 | 0.5 | 0.43 | 0.49 | 0.82 | 0.52 | 0.56 |
| 10 | 92739374 | rs12218436 | 1.382 | -0.23 | 0.595 | 0.24 | 0.023 | -0.015 | -0.015 | 0 | 0.21 | 0.15 | 0.14 | 0.28 | 0.09 | 0.15 |
| 10 | 92745793 | rs7080368 | 0.396 | 0.26 | 0.71 | -0.268 | 0.009 | 0.021 | 0.021 | 0 | 0.4 | 0.36 | 0.4 | 0.77 | 0.34 | 0.5 |
| 10 | 92753721 | rs17535998 | -0.408 | 0.665 | 0.134 | -0.115 | -0.044 | 0.005 | 0.005 | 0 | 0.07 | 0 | 0.07 | 0.05 | 0.15 | 0.04 |
| 10 | 92758267 | rs1318550 | -0.082 | 0.501 | 0.248 | -0.552 | 0.004 | 0.026 | 0.026 | 0 | 0.53 | 0.58 | 0.46 | 0.78 | 0.44 | 0.62 |
| 10 | 92764460 | rs7096430 | -0.643 | 0.423 | 0.347 | -0.584 | -0.003 | 0.052 | 0.052 | 0 | 0.61 | 0.74 | 0.54 | 0.84 | 0.59 | 0.64 |
| 10 | 92769660 | rs934901 | -0.202 | 0.401 | 0.596 | -0.628 | -0.002 | 0.004 | 0.004 | 0 | 0.54 | 0.41 | 0.49 | 0.78 | 0.51 | 0.59 |
| 10 | 92772368 | rs7897249 | -1.422 | 0.331 | 0.548 | -0.879 | -0.092 | 0.23 | 0.23 | 0.351 | 0.67 | 0.97 | 0.67 | 0.97 | 0.64 | 0.84 |
| 10 | 92772522 | rs11596239 | -0.167 | 0.404 | 0.727 | -0.715 | 0.005 | -0.023 | -0.023 | 0.183 | 0.06 | 0.11 | 0.04 | 0 | 0.08 | 0.03 |
| 10 | 92774918 | rs6583745 | -1.289 | 0.401 | 0.765 | -0.711 | 0.067 | 0.147 | 0.147 | 0.475 | 0.09 | 0.46 | 0.14 | 0.15 | 0.08 | 0.04 |
| 10 | 92778715 | rs3864853 | -0.173 | 0.427 | 0.306 | -0.806 | 0.005 | -0.023 | -0.023 | 0 | 0.06 | 0.11 | 0.04 | 0 | 0.08 | 0.03 |
| 10 | 92779333 | rs3895826 | -0.019 | 1.044 | 0.314 | -0.962 | -0.001 | -0.048 | -0.048 | 0.399 | 0.54 | 0.47 | 0.47 | 0.88 | 0.45 | 0.6 |
| 10 | 92779580 | rs4933203 | 0.52 | 1.036 | 0.134 | -0.917 | 0.018 | 0.039 | 0.039 | 0.015 | 0.47 | 0.35 | 0.42 | 0.82 | 0.37 | 0.56 |
| 10 | 92782367 | rs17536636 | -0.274 | 1.176 | -0.215 | -0.777 | -0.039 | 0.018 | 0.018 | 0 | 0.07 | 0 | 0.07 | 0.05 | 0.16 | 0.04 |
| 10 | 92782813 | rs3910157 | -1.411 | 1.031 | 0.015 | -0.683 | 0.046 | 0.051 | 0.051 | 1.305 | 0.09 | 0.37 | 0.05 | 0.07 | 0.08 | 0.03 |
| 10 | 92783513 | rs9787573 | -0.959 | 1.364 | -0.52 | -0.572 | -0.098 | 0.173 | 0.173 | 0 | 0.69 | 0.95 | 0.66 | 0.95 | 0.62 | 0.83 |
| 10 | 92791385 | rs3843023 | -0.611 | 1.474 | -0.466 | -0.412 | 0.002 | -0.01 | -0.01 | 0 | 0.68 | 0.69 | 0.67 | 0.85 | 0.64 | 0.84 |
| 10 | 92791516 | rs10509615 | -1.045 | 1.44 | -0.982 | -0.238 | 0.001 | -0.002 | -0.002 | 0 | 0.1 | 0.12 | 0.1 | 0.05 | 0.17 | 0.05 |
| 10 | 92796244 | rs754313 | -0.763 | 1.599 | -1.498 | -0.326 | -0.108 | 0.049 | 0.049 | 0.532 | 0.74 | 0.97 | 0.82 | 0.85 | 0.69 | 0.88 |
| 10 | 92803788 | rs10881878 | 0.315 | 0.728 | -1.355 | -0.186 | 0.027 | -0.014 | -0.014 | 0 | 0.42 | 0.69 | 0.38 | 0.51 | 0.31 | 0.56 |
| 10 | 92806472 | rs2290715 | -0.728 | 0.545 | -0.717 | -0.287 | 0.047 | -0.01 | -0.01 | 0.797 | 0.15 | 0.42 | 0.21 | 0.13 | 0.18 | 0.14 |
| 10 | 92808264 | rs2066426 | -0.75 | 0.546 | -0.687 | -0.293 | 0.03 | -0.008 | -0.008 | 0.271 | 0.14 | 0.33 | 0.2 | 0.13 | 0.18 | 0.14 |
| 10 | 92811346 | rs12777523 | 0.638 | 0.448 | -0.685 | -0.316 | -0.01 | 0.009 | 0.009 | 0 | 0.38 | 0.42 | 0.36 | 0.51 | 0.31 | 0.51 |
| 10 | 92816196 | rs11594712 | 1.332 | 0.333 | -0.9 | -0.432 | -0.109 | 0.093 | 0.093 | 0.147 | 0.25 | 0.03 | 0.17 | 0.09 | 0.32 | 0.13 |
| 10 | 92820426 | rs7098967 | -0.391 | 0.451 | -0.477 | -0.229 | 0.012 | -0.041 | -0.041 | 0 | 0.09 | 0.29 | 0.04 | 0.01 | 0.07 | 0.05 |
| 10 | 92823539 | rs10785985 | 0.933 | 0.557 | -0.556 | -0.247 | -0.076 | -0.01 | -0.01 | 0.215 | 0.42 | 0.06 | 0.41 | 0.36 | 0.43 | 0.29 |
| 10 | 92824878 | rs7097056 | -0.695 | 0.797 | -0.09 | -0.265 | 0.024 | -0.036 | -0.036 | 0 | 0.09 | 0.34 | 0.04 | 0.01 | 0.07 | 0.05 |
| 10 | 92826020 | rs11595303 | 0.373 | 0.865 | -0.557 | -0.235 | 0.009 | -0.028 | -0.028 | 0 | 0.06 | 0.15 | 0.03 | 0.01 | 0.07 | 0.05 |
| 10 | 92826534 | rs10437464 | 0.222 | 0.354 | -0.899 | -0.064 | -0.009 | -0.011 | -0.011 | 1.317 | 0.3 | 0.61 | 0.24 | 0.17 | 0.27 | 0.31 |
| 10 | 92830260 | rs17093770 | -0.432 | 0.337 | -1.02 | -0.136 | -0.002 | -0.016 | -0.016 | 0.468 | 0.16 | 0.28 | 0.19 | 0.13 | 0.17 | 0.18 |
| 10 | 92835371 | rs7086130 | -1.104 | 0.432 | -1.189 | 0.04 | -0.062 | 0.023 | 0.023 | 1.161 | 0.48 | 0.92 | 0.46 | 0.32 | 0.52 | 0.53 |
| 10 | 92836763 | rs3852502 | -1.052 | 0.64 | -0.918 | 0.097 | -0.029 | 0.011 | 0.011 | 1.241 | 0.43 | 0.8 | 0.43 | 0.32 | 0.5 | 0.47 |
| 10 | 92860281 | rs12267148 | 0.294 | 0.746 | -0.935 | 0.011 | 0.008 | -0.002 | -0.002 | 0.002 | 0.26 | 0.29 | 0.25 | 0.18 | 0.33 | 0.3 |
| 10 | 92860843 | rs893587 | -1.086 | 0.908 | -1.975 | 0.184 | 0 | -0.005 | -0.005 | 0 | 0.65 | 0.8 | 0.7 | 0.55 | 0.69 | 0.67 |
| 10 | 92862341 | rs734627 | 1.088 | 1.178 | -1.62 | 0.495 | 0.021 | -0.01 | -0.01 | 0.077 | 0.22 | 0.14 | 0.26 | 0.24 | 0.17 | 0.2 |
| 10 | 92862763 | rs893586 | -0.867 | 1.761 | -1.653 | 0.845 | -0.001 | -0.001 | -0.001 | 0.013 | 0.75 | 0.8 | 0.77 | 0.56 | 0.76 | 0.71 |
| 10 | 92867818 | rs12572050 | 0.275 | 1.95 | -1.87 | 0.818 | -0.024 | -0.022 | -0.022 | 0 | 0.12 | 0.06 | 0.09 | 0.11 | 0.16 | 0.19 |
| 10 | 92869911 | rs10509619 | -0.798 | 1.391 | -1.488 | 0.561 | -0.003 | -0.003 | -0.003 | 1.239 | 0.07 | 0.06 | 0.15 | 0.14 | 0.1 | 0.12 |
| 10 | 92871894 | rs10509620 | -0.675 | 1.413 | -1.503 | 0.535 | -0.006 | -0.006 | -0.006 | 0 | 0.07 | 0.05 | 0.14 | 0.13 | 0.1 | 0.12 |
| 10 | 92872941 | rs1431864 | -0.379 | 1.087 | -1.762 | 0.333 | -0.015 | 0.009 | 0.009 | 0 | 0.36 | 0.18 | 0.42 | 0.32 | 0.36 | 0.33 |
| 10 | 92880583 | rs10881886 | -0.19 | 1.032 | -1.32 | 0.05 | -0.01 | 0.018 | 0.018 | 0 | 0.59 | 0.55 | 0.56 | 0.72 | 0.53 | 0.64 |
| 10 | 92884288 | rs1560543 | -0.105 | 1.237 | -1.363 | -0.003 | -0.003 | 0.006 | 0.006 | 0 | 0.61 | 0.57 | 0.59 | 0.83 | 0.57 | 0.7 |

| | | | | | | | | | | | | | | | | |
|---|---|---|---|---|---|---|---|---|---|---|---|---|---|---|---|---|
| 10 | 92887250 | rs7083030 | -0.174 | 1.223 | -1.282 | -0.015 | 0 | -0.022 | -0.022 | 0.148 | 0.61 | 0.51 | 0.61 | 0.83 | 0.62 | 0.71 |
| 10 | 92894370 | rs7915101 | -0.012 | 0.886 | -0.874 | -0.034 | 0.002 | -0.01 | -0.01 | 2.491 | 0.58 | 0.39 | 0.56 | 0.71 | 0.53 | 0.67 |
| 10 | 92894740 | rs10881890 | -0.156 | 0.918 | -0.753 | -0.033 | 0.007 | -0.051 | -0.051 | 0.001 | 0.61 | 0.48 | 0.59 | 0.83 | 0.58 | 0.71 |
| 10 | 92897348 | rs7089856 | -0.192 | 0.889 | -0.728 | -0.033 | 0.001 | -0.014 | -0.014 | 0 | 0.61 | 0.53 | 0.6 | 0.82 | 0.59 | 0.71 |
| 10 | 92899400 | rs11186449 | -0.034 | 0.884 | -0.762 | -0.033 | 0.003 | -0.012 | -0.012 | 0 | 0.58 | 0.39 | 0.56 | 0.71 | 0.53 | 0.67 |
| 10 | 92899856 | rs1992245 | -0.15 | 0.81 | -0.793 | -0.1 | -0.002 | 0 | 0 | 0 | 0.61 | 0.56 | 0.6 | 0.82 | 0.59 | 0.71 |
| 10 | 92901247 | rs7090817 | -0.211 | 0.9 | -0.781 | -0.089 | -0.002 | -0.003 | -0.003 | 0.812 | 0.62 | 0.57 | 0.6 | 0.82 | 0.58 | 0.71 |
| 10 | 92903216 | rs7074922 | -0.485 | 0.836 | -1.007 | -0.116 | -0.034 | 0.05 | 0.05 | 0.062 | 0.66 | 0.75 | 0.61 | 0.82 | 0.58 | 0.71 |
| 10 | 92903636 | rs7078922 | 0.471 | 0.762 | -1.47 | 0.335 | -0.021 | 0.008 | 0.008 | 2.081 | 0.25 | 0.02 | 0.22 | 0.14 | 0.22 | 0.24 |
| 10 | 92905682 | rs7088963 | -0.068 | 0.981 | -1.069 | 0.178 | 0.002 | 0.015 | 0.015 | 0 | 0.49 | 0.39 | 0.52 | 0.69 | 0.48 | 0.62 |
| 10 | 92907609 | rs7098281 | 0.531 | 1.282 | -0.69 | 0.602 | 0.004 | 0.014 | 0.014 | 1.64 | 0.8 | 0.75 | 0.87 | 0.77 | 0.82 | 0.73 |
| 10 | 92909779 | rs12242279 | 0.962 | 0.662 | -1.234 | -0.054 | 0.041 | 0.109 | 0.109 | 0 | 0.31 | 0.12 | 0.14 | 0.1 | 0.24 | 0.13 |
| 10 | 92914103 | rs12218128 | 0.84 | 1.477 | -1.136 | 0.238 | -0.005 | -0.019 | -0.019 | 0.32 | 0.05 | 0 | 0.02 | 0.19 | 0.04 | 0.14 |
| 10 | 92914422 | rs2420801 | -0.053 | 1.837 | -1.538 | 0.043 | -0.118 | 0.015 | 0.015 | 0.53 | 0.78 | 0.98 | 0.68 | 0.85 | 0.67 | 0.69 |
| 10 | 92917925 | rs930399 | -0.055 | 1.865 | -1.511 | 0.025 | -0.113 | 0.022 | 0.022 | 0.589 | 0.78 | 0.98 | 0.68 | 0.85 | 0.67 | 0.69 |
| 10 | 92926700 | rs11186478 | 0.471 | 1.732 | -1.271 | 0.114 | -0.019 | 0.01 | 0.01 | 0 | 0.06 | 0.01 | 0.03 | 0.02 | 0.07 | 0.03 |
| 10 | 92927267 | rs10881893 | -0.897 | 1.242 | -1.026 | 0.232 | 0.003 | -0.001 | -0.001 | 0 | 0.22 | 0.67 | 0.26 | 0.19 | 0.19 | 0.12 |
| 10 | 92934593 | rs7908053 | 0.203 | 0.749 | -0.405 | 0.124 | -0.003 | -0.031 | -0.031 | 0 | 0.07 | 0.25 | 0.02 | 0 | 0.03 | 0.03 |
| 10 | 92937921 | rs7098577 | 1.148 | 0.718 | -0.727 | 0.114 | 0.065 | -0.017 | -0.017 | 0 | 0.49 | 0.21 | 0.41 | 0.51 | 0.42 | 0.43 |
| 10 | 92938019 | rs10881895 | -0.027 | 0.833 | -0.795 | -0.218 | -0.158 | 0.041 | 0.041 | 0.01 | 0.26 | 0.03 | 0.38 | 0.14 | 0.41 | 0.34 |
| 10 | 92941679 | rs10881898 | 1.06 | 0.618 | -0.334 | -0.076 | 0.06 | 0.027 | 0.027 | 3.934 | 0.5 | 0.25 | 0.48 | 0.59 | 0.42 | 0.45 |
| 10 | 92946000 | rs7902517 | 0.645 | 0.701 | 0.025 | 0.331 | 0.009 | -0.043 | -0.043 | 0 | 0.2 | 0.14 | 0.33 | 0.48 | 0.17 | 0.3 |
| 10 | 92948478 | rs10881900 | -1.166 | 0.845 | 0.157 | 0.261 | -0.001 | 0.005 | 0.005 | 0 | 0.48 | 0.54 | 0.67 | 0.78 | 0.49 | 0.66 |
| 10 | 92957222 | rs12778694 | -0.11 | 0.3 | 0.701 | 1.012 | 0.004 | -0.008 | -0.008 | 0 | 0.22 | 0.21 | 0.34 | 0.48 | 0.17 | 0.3 |
| 10 | 92960115 | rs11186497 | 1.646 | 0.489 | 0.779 | 0.179 | 0.004 | -0.027 | -0.027 | 0 | 0.83 | 0.32 | 0.88 | 0.91 | 0.87 | 0.92 |
| 10 | 92964477 | rs11186501 | -0.365 | -0.571 | 0.942 | 0.391 | -0.011 | 0.006 | 0.006 | 0.215 | 0.24 | 0.3 | 0.36 | 0.69 | 0.19 | 0.38 |
| 10 | 92967874 | rs12257267 | -0.404 | -0.51 | 0.631 | 0.35 | 0.005 | -0.012 | -0.012 | 0 | 0.07 | 0.26 | 0.02 | 0 | 0.01 | 0.03 |
| 10 | 92971535 | rs17106435 | -0.67 | -0.503 | 0.349 | 0.352 | 0.008 | -0.013 | -0.013 | 0 | 0.08 | 0.41 | 0.03 | 0 | 0.01 | 0.03 |
| 10 | 92972619 | rs2421482 | -0.561 | -0.121 | 0.329 | 0.595 | -0.046 | 0.055 | 0.055 | 0.47 | 0.25 | 0.46 | 0.37 | 0.67 | 0.18 | 0.35 |
| 10 | 92976570 | rs6583759 | -1.322 | -0.21 | 0.626 | 0.092 | -0.053 | -0.096 | -0.096 | 1.296 | 0.14 | 0.49 | 0.06 | 0.01 | 0.06 | 0.11 |
| 10 | 92979176 | rs7894133 | -1.122 | -0.147 | 1.068 | 0.093 | -0.031 | -0.073 | -0.073 | 0 | 0.12 | 0.42 | 0.06 | 0.01 | 0.05 | 0.09 |
| 10 | 92983831 | rs11186507 | -0.046 | 0.975 | 0.443 | -0.15 | -0.072 | -0.016 | -0.016 | 0 | 0.11 | 0.01 | 0.19 | 0.07 | 0.2 | 0.13 |
| 10 | 92987510 | rs1057971 | -0.408 | 0.933 | 0.75 | -0.148 | -0.019 | -0.056 | -0.056 | 0.256 | 0.1 | 0.31 | 0.05 | 0.01 | 0.05 | 0.09 |
| 10 | 93004825 | rs7082510 | -0.784 | 0.933 | 0.783 | -0.149 | -0.023 | -0.067 | -0.067 | 0 | 0.11 | 0.38 | 0.05 | 0 | 0.05 | 0.09 |
| 10 | 93004839 | rs11186517 | -0.064 | 0.273 | 0.733 | 0.645 | -0.022 | 0.148 | 0.148 | 0 | 0.08 | 0.15 | 0.15 | 0.37 | 0.03 | 0.13 |
| 10 | 93032943 | rs2631681 | -1.486 | 0.487 | 0.722 | 0.002 | -0.003 | 0.002 | 0.002 | 0 | 0.36 | 0.7 | 0.31 | 0.07 | 0.34 | 0.21 |
| 10 | 93038530 | rs2450445 | -1.505 | 0.462 | 0.711 | -0.009 | -0.003 | 0.003 | 0.003 | 0 | 0.36 | 0.7 | 0.31 | 0.07 | 0.34 | 0.22 |
| 10 | 93041869 | rs7074612 | 0.065 | 0.555 | 0.656 | 0.003 | -0.01 | -0.033 | -0.033 | 0 | 0.09 | 0.14 | 0.03 | 0 | 0.05 | 0.09 |
| 10 | 93043858 | rs13675 | 0.44 | 0.789 | 0.503 | 0.018 | -0.073 | -0.005 | -0.005 | 0 | 0.11 | 0.01 | 0.2 | 0.07 | 0.21 | 0.13 |
| 10 | 93050333 | rs10509625 | 0.148 | 0.53 | 0.721 | 0.335 | -0.013 | 0.129 | 0.129 | 0 | 0.08 | 0.14 | 0.16 | 0.37 | 0.04 | 0.14 |
| 10 | 93053537 | rs2631686 | -1.698 | 0.772 | 0.281 | 0.253 | -0.032 | -0.026 | -0.026 | 0.108 | 0.44 | 0.85 | 0.43 | 0.44 | 0.39 | 0.32 |
| 10 | 93058075 | rs2245523 | -0.655 | 1.352 | -0.335 | 0.094 | -0.022 | -0.045 | -0.045 | 0.7 | 0.82 | 0.87 | 0.63 | 0.5 | 0.77 | 0.62 |
| 10 | 93059828 | rs10430649 | 0.21 | 0.894 | -0.609 | -0.173 | -0.008 | -0.058 | -0.058 | 0.224 | 0.09 | 0.01 | 0.19 | 0.36 | 0.07 | 0.14 |
| 10 | 93060218 | rs3890159 | 1.222 | 0.225 | -0.362 | -1.179 | 0.018 | 0.025 | 0.025 | 0 | 0.39 | 0.26 | 0.18 | 0.08 | 0.32 | 0.3 |
| 10 | 93060349 | rs2245310 | -0.109 | 0.519 | -0.555 | -0.949 | -0.036 | 0.055 | 0.055 | 0 | 0.27 | 0.16 | 0.19 | 0.07 | 0.32 | 0.13 |
| 10 | 93071262 | rs2255188 | 0.242 | 0.806 | -0.445 | -0.783 | -0.009 | 0.022 | 0.022 | 0 | 0.66 | 0.73 | 0.37 | 0.16 | 0.58 | 0.42 |
| 10 | 93077075 | rs10748559 | -0.155 | 0.6 | -0.594 | -0.462 | -0.097 | -0.045 | -0.045 | 0 | 0.36 | 0.08 | 0.43 | 0.4 | 0.42 | 0.49 |
| 10 | 93136505 | rs7073979 | -1.119 | -0.075 | 1.107 | -0.313 | -0.039 | -0.063 | -0.063 | 0.469 | 0.1 | 0.28 | 0.04 | 0 | 0.04 | 0.02 |
| 10 | 93145440 | rs6583769 | -1.896 | 0.074 | 1.358 | -0.189 | -0.025 | -0.02 | -0.02 | 0 | 0.14 | 0.45 | 0.07 | 0.09 | 0.07 | 0.07 |
| 10 | 93163929 | rs12247672 | -1.867 | 0.182 | 1.624 | -0.095 | -0.027 | -0.004 | -0.004 | 0 | 0.11 | 0.59 | 0.05 | 0.08 | 0.05 | 0.03 |
| 10 | 93177933 | rs12218668 | 0.526 | -0.355 | 1.254 | 0.066 | -0.126 | -0.055 | -0.055 | 0 | 0.36 | 0.09 | 0.51 | 0.62 | 0.5 | 0.55 |
| 10 | 93221077 | rs7920604 | -2.823 | -0.557 | 2.449 | 0.514 | -0.043 | -0.021 | -0.021 | 0 | 0.13 | 0.67 | 0.06 | 0.09 | 0.05 | 0.03 |
| 10 | 93240166 | rs4376822 | -2.138 | -0.539 | 2.449 | 0.444 | 0.002 | 0.029 | 0.029 | 0.629 | 0.1 | 0.55 | 0.05 | 0.08 | 0.04 | 0.02 |
| 10 | 93273792 | rs11186596 | -2.369 | -0.545 | 2.302 | 0.441 | -0.035 | -0.014 | -0.014 | 0.183 | 0.12 | 0.62 | 0.05 | 0.09 | 0.05 | 0.03 |
| 10 | 93284528 | rs17106727 | -0.755 | -0.388 | 1.757 | 0.78 | 0.005 | 0.174 | 0.174 | 0 | 0.05 | 0.25 | 0.14 | 0.18 | 0.02 | 0.01 |
| 10 | 93287580 | rs3904193 | -1.749 | -0.362 | 0.802 | 0.733 | -0.038 | 0.104 | 0.104 | 2.068 | 0.12 | 0.66 | 0.17 | 0.26 | 0.05 | 0.03 |
| 10 | 93296477 | rs10399998 | -1.701 | -0.323 | 0.648 | 0.615 | -0.04 | 0.077 | 0.077 | 3.838 | 0.13 | 0.65 | 0.17 | 0.24 | 0.05 | 0.03 |
| 10 | 93300327 | rs11186608 | -0.482 | -0.579 | 0.421 | -0.404 | -0.003 | -0.009 | -0.009 | 0.1 | 0.44 | 0.56 | 0.28 | 0.15 | 0.36 | 0.26 |
| 10 | 93303294 | rs7068979 | -0.078 | -0.654 | 0.653 | -0.933 | 0.027 | 0.178 | 0.178 | 0.051 | 0.39 | 0.16 | 0.24 | 0.02 | 0.34 | 0.24 |
| 10 | 93303486 | rs4933659 | -0.435 | -0.696 | 0.805 | -0.951 | -0.01 | -0.09 | -0.09 | 3.456 | 0.43 | 0.53 | 0.27 | 0.02 | 0.34 | 0.24 |
| 10 | 93310582 | rs17106748 | -1.959 | -0.551 | 1.025 | -0.726 | 0.06 | 0.056 | 0.056 | 6.43 | 0.05 | 0.41 | 0.03 | 0 | 0 | 0 |
| 10 | 93315271 | rs12268133 | -0.387 | -0.362 | 0.052 | -0.877 | 0.011 | -0.015 | -0.015 | 0.674 | 0.43 | 0.41 | 0.24 | 0.02 | 0.33 | 0.25 |
| 10 | 93338600 | rs4933206 | -0.296 | 0.251 | -0.983 | -0.4 | -0.021 | 0.006 | 0.006 | 0 | 0.31 | 0.06 | 0.4 | 0.76 | 0.28 | 0.62 |
| 10 | 93347297 | rs2149825 | -0.132 | 0.204 | -1.927 | -1.369 | -0.06 | 0.168 | 0.168 | 0.116 | 0.55 | 0.13 | 0.54 | 0.84 | 0.51 | 0.69 |
| 10 | 93348014 | rs1329651 | 0.979 | 0.055 | -2.034 | -1.226 | 0.009 | 0.08 | 0.08 | 1.883 | 0.28 | 0.06 | 0.15 | 0.1 | 0.25 | 0.1 |
| 10 | 93348120 | rs1329650 | 0.1 | 0.049 | -1.913 | -1.074 | -0.024 | -0.034 | -0.034 | 0 | 0.31 | 0.06 | 0.4 | 0.74 | 0.28 | 0.61 |
| 10 | 93349092 | rs1028935 | -0.495 | 0.384 | -1.331 | -1.399 | 0.006 | 0.01 | 0.01 | 0.07 | 0.86 | 0.82 | 0.79 | 0.86 | 0.85 | 0.87 |
| 10 | 93349797 | rs1028936 | 0.563 | 0.536 | -1.643 | -1.213 | 0.021 | -0.076 | -0.076 | 0.682 | 0.23 | 0.06 | 0.24 | 0.65 | 0.19 | 0.5 |
| 10 | 93350014 | rs41403949 | -0.42 | 1.142 | -1.166 | -1.449 | -0.001 | -0.018 | -0.018 | 0 | 0.12 | 0.15 | 0.12 | 0.02 | 0.17 | 0.09 |

| | | | | | | | | | | | | | | | | |
|---|---|---|---|---|---|---|---|---|---|---|---|---|---|---|---|---|
| 10 | 93354689 | rs4933672 | -0.62 | 1.205 | -0.564 | -1.767 | 0.002 | 0.002 | 0.002 | 0 | 0.49 | 0.54 | 0.51 | 0.76 | 0.44 | 0.71 |
| 10 | 93360302 | rs11591635 | 0.649 | 1.237 | 0.193 | -1.884 | 0.031 | -0.003 | -0.003 | 0.082 | 0.09 | 0.22 | 0.19 | 0.08 | 0.08 | 0.1 |
| 10 | 93360330 | rs11597974 | 0.649 | 1.237 | 0.193 | -1.884 | 0.031 | -0.003 | -0.003 | 0 | 0.09 | 0.22 | 0.19 | 0.08 | 0.08 | 0.1 |
| 10 | 93362686 | rs11186623 | 1.642 | 1.475 | 0.16 | -2.084 | -0.054 | 0.16 | 0.16 | 1.409 | 0.31 | 0.11 | 0.19 | 0.01 | 0.34 | 0.1 |
| 10 | 93363594 | rs748401 | 0.897 | 1.484 | 0.42 | -2.071 | 0.008 | -0.007 | -0.007 | 0 | 0.08 | 0.16 | 0.18 | 0.08 | 0.08 | 0.11 |
| 10 | 93364102 | rs7101314 | 0.806 | 1.452 | 0.624 | -2.086 | 0.038 | -0.026 | -0.026 | 0.052 | 0.8 | 0.6 | 0.7 | 0.83 | 0.77 | 0.8 |
| 10 | 93365015 | rs7911956 | 0.133 | 1.437 | 0.632 | -2.113 | 0.002 | -0.017 | -0.017 | 1.307 | 0.07 | 0.09 | 0.05 | 0.02 | 0.08 | 0.06 |
| 10 | 93365431 | rs10509631 | 1.694 | 1.426 | 0.633 | -2.088 | -0.044 | 0.12 | 0.12 | 1.237 | 0.32 | 0.11 | 0.2 | 0.08 | 0.34 | 0.1 |
| 10 | 93366275 | rs4329592 | 0.284 | 1.328 | 0.783 | -1.848 | 0.009 | -0.008 | -0.008 | 0 | 0.88 | 0.84 | 0.89 | 0.92 | 0.85 | 0.91 |
| 10 | 93368163 | rs4933211 | 0.689 | 1.282 | 0.581 | -1.83 | 0.038 | -0.026 | -0.026 | 2.539 | 0.8 | 0.6 | 0.7 | 0.83 | 0.77 | 0.8 |
| 10 | 93368504 | rs4933212 | 1.123 | 0.693 | 0.083 | -1.424 | 0.01 | -0.027 | -0.027 | 0 | 0.19 | 0.11 | 0.17 | 0.67 | 0.18 | 0.44 |
| 10 | 93373287 | rs10881931 | -1.648 | 0.38 | 0.684 | -1.519 | -0.021 | -0.031 | -0.031 | 0 | 0.29 | 0.61 | 0.29 | 0.11 | 0.22 | 0.2 |
| 10 | 93378452 | rs4933679 | -1.239 | 0.375 | 0.987 | -1.529 | -0.026 | -0.125 | -0.125 | 0 | 0.25 | 0.54 | 0.18 | 0.02 | 0.22 | 0.17 |
| 10 | 93387502 | rs11186635 | -0.295 | 0.278 | 1.335 | -1.549 | -0.007 | -0.032 | -0.032 | 0.134 | 0.1 | 0.16 | 0.07 | 0.01 | 0.1 | 0.06 |
| 10 | 93387851 | rs4382814 | 0.448 | 0.271 | 1.533 | -1.572 | -0.003 | -0.052 | -0.052 | 0 | 0.1 | 0.27 | 0.07 | 0.01 | 0.07 | 0.08 |
| 10 | 93389343 | rs12261209 | -0.384 | 0.355 | 1.232 | -1.487 | -0.005 | -0.027 | -0.027 | 0 | 0.1 | 0.13 | 0.07 | 0.01 | 0.1 | 0.06 |
| 10 | 93399130 | rs7475251 | -1.35 | 0.699 | 1.222 | -1.473 | 0 | -0.008 | -0.008 | 0 | 0.55 | 0.48 | 0.46 | 0.72 | 0.57 | 0.75 |
| 10 | 93403068 | rs11186642 | -0.09 | 0.637 | 1.165 | -1.565 | -0.032 | -0.09 | -0.09 | 0 | 0.16 | 0.32 | 0.11 | 0.01 | 0.11 | 0.12 |
| 10 | 93412994 | rs2000177 | -2.168 | 0.822 | 0.886 | -1.626 | -0.033 | 0.11 | 0.11 | 1.408 | 0.67 | 0.9 | 0.79 | 0.92 | 0.65 | 0.89 |
| 10 | 93416361 | rs11186647 | 1.474 | 0.859 | 0.902 | -1.544 | -0.001 | -0.001 | -0.001 | 0.125 | 0.07 | 0.06 | 0.19 | 0.11 | 0.07 | 0.11 |
| 10 | 93430144 | rs11186654 | 0.721 | 0.858 | 0.753 | -1.584 | -0.062 | -0.057 | -0.057 | 0 | 0.29 | 0.1 | 0.29 | 0.7 | 0.36 | 0.57 |
| 10 | 93432632 | rs4933218 | -0.808 | 0.66 | 1.072 | -1.563 | -0.023 | -0.057 | -0.057 | 0 | 0.24 | 0.39 | 0.3 | 0.12 | 0.18 | 0.23 |
| 10 | 93440350 | rs4551673 | -1.557 | 0.704 | 0.545 | -1.558 | 0.025 | 0.05 | 0.05 | 0.019 | 0.36 | 0.76 | 0.38 | 0.13 | 0.29 | 0.29 |
| 10 | 93442223 | rs2421736 | -1.33 | 1.013 | 0.415 | -1.372 | -0.004 | 0.076 | 0.076 | 0.118 | 0.57 | 0.79 | 0.72 | 0.77 | 0.56 | 0.82 |
| 10 | 93442240 | rs1891275 | -1.632 | 0.826 | 0.358 | -1.394 | -0.047 | 0.074 | 0.074 | 1.937 | 0.6 | 0.87 | 0.71 | 0.82 | 0.53 | 0.83 |
| 10 | 93444262 | rs1080496 | -0.525 | 1.157 | 0.849 | -1.36 | 0 | 0.004 | 0.004 | 0 | 0.45 | 0.54 | 0.63 | 0.81 | 0.42 | 0.75 |
| 10 | 93447805 | rs11592233 | 0.873 | 1.102 | 0.192 | -1.193 | 0.001 | -0.001 | -0.001 | 0.148 | 0.12 | 0.11 | 0.19 | 0.15 | 0.07 | 0.14 |
| 10 | 93447863 | rs11186660 | 1.022 | 1.369 | 0.085 | -1.251 | -0.028 | -0.114 | -0.114 | 0 | 0.26 | 0.02 | 0.34 | 0.65 | 0.25 | 0.55 |
| 10 | 93452141 | rs7073486 | -1.76 | 1.189 | 0.013 | -1.38 | -0.054 | 0.107 | 0.107 | 0 | 0.67 | 0.87 | 0.76 | 0.9 | 0.58 | 0.86 |
| 10 | 93462424 | rs2039973 | 0.746 | 1.179 | 0.123 | -1.168 | -0.003 | 0.004 | 0.004 | 0 | 0.13 | 0.17 | 0.19 | 0.16 | 0.07 | 0.14 |
| 10 | 93479949 | rs10786012 | 0.741 | 1.226 | 0.178 | -1.219 | -0.007 | -0.013 | -0.013 | 0 | 0.28 | 0.22 | 0.41 | 0.65 | 0.31 | 0.57 |
| 10 | 93484501 | rs1418160 | 0.238 | 1.185 | 0.172 | -1.255 | -0.003 | -0.005 | -0.005 | 0 | 0.33 | 0.31 | 0.44 | 0.66 | 0.34 | 0.61 |
| 10 | 93501876 | rs10881957 | -0.615 | 0.926 | 0.234 | -1.288 | 0 | 0.013 | 0.013 | 0 | 0.46 | 0.45 | 0.63 | 0.82 | 0.42 | 0.75 |
| 10 | 93526562 | rs17107019 | 0.996 | 0.909 | 0.147 | -1.267 | 0.002 | -0.004 | -0.004 | 0 | 0.12 | 0.11 | 0.19 | 0.16 | 0.07 | 0.13 |
| 10 | 93526988 | rs11594359 | 0.812 | 0.903 | 0.166 | -1.269 | 0.001 | -0.003 | -0.003 | 0 | 0.13 | 0.13 | 0.19 | 0.16 | 0.07 | 0.13 |
| 10 | 93534624 | rs1361552 | -0.607 | 0.913 | 0.249 | -1.272 | -0.001 | 0.017 | 0.017 | 0 | 0.46 | 0.44 | 0.63 | 0.82 | 0.41 | 0.75 |
| 10 | 93541588 | rs6583787 | -1.977 | 0.797 | 0.019 | -1.426 | -0.068 | 0.123 | 0.123 | 0 | 0.67 | 0.88 | 0.76 | 0.91 | 0.58 | 0.86 |
| 10 | 93556855 | rs1341263 | 0.65 | 0.809 | 0.19 | -1.009 | -0.01 | -0.031 | -0.031 | 0.006 | 0.31 | 0.25 | 0.44 | 0.55 | 0.33 | 0.62 |
| 10 | 93557619 | rs2066276 | -2.141 | 0.763 | 0.148 | -1.057 | -0.073 | 0.113 | 0.113 | 0 | 0.68 | 0.88 | 0.76 | 0.91 | 0.58 | 0.89 |
| 10 | 93559751 | rs10786018 | -2.132 | 0.763 | 0.148 | -1.057 | -0.073 | 0.113 | 0.113 | 0 | 0.68 | 0.88 | 0.76 | 0.91 | 0.58 | 0.89 |
| 10 | 93561210 | rs10748568 | -2.132 | 0.763 | 0.148 | -1.057 | -0.073 | 0.113 | 0.113 | 0 | 0.68 | 0.88 | 0.76 | 0.91 | 0.58 | 0.89 |
| 10 | 93564315 | rs2066275 | 1.108 | 0.77 | 0.734 | -1.085 | -0.008 | -0.022 | -0.022 | 0 | 0.27 | 0.25 | 0.41 | 0.54 | 0.3 | 0.58 |
| 10 | 93565641 | rs7072165 | 0.389 | 0.701 | 0.773 | -1.096 | -0.002 | 0.032 | 0.032 | 0 | 0.15 | 0.39 | 0.21 | 0.16 | 0.08 | 0.14 |
| 10 | 93577624 | rs3802650 | -0.873 | 0.761 | 0.778 | -1.101 | 0 | 0.006 | 0.006 | 0 | 0.53 | 0.44 | 0.55 | 0.75 | 0.5 | 0.75 |
| 10 | 93585997 | rs7083959 | -0.309 | 0.738 | 0.733 | -1.099 | -0.007 | -0.005 | -0.005 | 0 | 0.26 | 0.25 | 0.15 | 0.2 | 0.2 | 0.17 |
| 10 | 93586715 | rs2275646 | 0.912 | 0.742 | 0.711 | -1.099 | 0.002 | -0.004 | -0.004 | 1.574 | 0.13 | 0.11 | 0.19 | 0.16 | 0.08 | 0.14 |
| 10 | 93587732 | rs10509637 | 0.399 | 0.749 | 0.765 | -1.079 | 0.012 | 0.005 | 0.005 | 0 | 0.21 | 0.14 | 0.12 | 0.02 | 0.17 | 0.12 |
| 10 | 93600480 | rs17107140 | 1.303 | 0.752 | 0.761 | -1.074 | 0 | -0.017 | -0.017 | 0 | 0.12 | 0.07 | 0.19 | 0.16 | 0.07 | 0.13 |
| 10 | 93608142 | rs3758499 | 1.095 | 0.76 | 0.759 | -1.065 | -0.007 | -0.021 | -0.021 | 0.666 | 0.27 | 0.24 | 0.41 | 0.55 | 0.3 | 0.58 |
| 10 | 93616628 | rs1772188 | 0.386 | 0.746 | 0.757 | -1.073 | 0.012 | 0.005 | 0.005 | 0 | 0.21 | 0.14 | 0.12 | 0.02 | 0.17 | 0.12 |
| 10 | 93624672 | rs2258946 | 0.386 | 0.782 | 0.435 | -1.057 | 0.012 | 0.005 | 0.005 | 0 | 0.21 | 0.14 | 0.12 | 0.02 | 0.17 | 0.12 |
| 10 | 93627158 | rs10881985 | 1.042 | 0.865 | 0.367 | -0.999 | -0.007 | -0.021 | -0.021 | 0 | 0.27 | 0.24 | 0.41 | 0.54 | 0.3 | 0.58 |
| 10 | 93635541 | rs1417731 | 1.086 | 0.888 | 0.245 | -0.955 | 0.002 | -0.01 | -0.01 | 0 | 0.12 | 0.09 | 0.18 | 0.16 | 0.08 | 0.13 |
| 10 | 93643195 | rs7085519 | 1.515 | 0.858 | 0.3 | -1.039 | 0.019 | -0.065 | -0.065 | 0 | 0.35 | 0.43 | 0.26 | 0.09 | 0.43 | 0.11 |
| 10 | 93643371 | rs11186722 | 0.649 | 0.885 | 0.364 | -0.831 | 0.001 | 0.018 | 0.018 | 0 | 0.47 | 0.57 | 0.45 | 0.25 | 0.51 | 0.25 |
| 10 | 93644552 | rs7917710 | 0.706 | 0.939 | 0.134 | -0.769 | -0.009 | -0.025 | -0.025 | 0 | 0.27 | 0.23 | 0.41 | 0.54 | 0.31 | 0.58 |
| 10 | 93648533 | rs7903827 | 0.684 | 0.937 | 0.132 | -0.732 | -0.01 | -0.028 | -0.028 | 0 | 0.27 | 0.23 | 0.4 | 0.54 | 0.31 | 0.58 |
| 10 | 93650847 | rs7089495 | 0.173 | 0.304 | 0.212 | -0.822 | 0.011 | -0.001 | -0.001 | 0 | 0.76 | 0.82 | 0.87 | 0.98 | 0.82 | 0.85 |
| 10 | 93651066 | rs2421705 | 0.162 | 0.314 | 0.104 | -0.818 | 0.012 | 0.002 | 0.002 | 0 | 0.76 | 0.82 | 0.87 | 0.98 | 0.82 | 0.85 |
| 10 | 93657148 | rs12255176 | -0.598 | 0.601 | 0.488 | -0.855 | 0.109 | 0.009 | 0.009 | 0 | 0.06 | 0.31 | 0.09 | 0 | 0.1 | 0.01 |
| 10 | 93657377 | rs7093658 | -0.067 | 0.4 | 0.311 | -0.907 | 0.034 | 0.043 | 0.043 | 0 | 0.77 | 0.88 | 0.89 | 0.98 | 0.85 | 0.87 |
| 10 | 93660727 | rs7919192 | -1.241 | 0.204 | 0.206 | -1.018 | -0.024 | -0.059 | -0.059 | 0 | 0.36 | 0.49 | 0.27 | 0.17 | 0.34 | 0.18 |
| 10 | 93664644 | rs7099658 | -1.237 | 0.14 | 0.205 | -1.016 | -0.034 | -0.068 | -0.068 | 0 | 0.36 | 0.49 | 0.25 | 0.17 | 0.32 | 0.18 |
| 10 | 93665975 | rs7087141 | -0.615 | 0.49 | 0.447 | -0.784 | 0.093 | 0.015 | 0.015 | 0 | 0.06 | 0.33 | 0.07 | 0 | 0.09 | 0.03 |
| 10 | 93673004 | rs7095552 | -0.839 | 0.094 | 0.143 | -1.058 | -0.027 | -0.122 | -0.122 | 0.597 | 0.29 | 0.43 | 0.19 | 0.02 | 0.25 | 0.14 |
| 10 | 93674933 | rs10881996 | -0.067 | 0.554 | -0.457 | -1.242 | 0.009 | -0.015 | -0.015 | 0.056 | 0.64 | 0.57 | 0.42 | 0.28 | 0.64 | 0.27 |
| 10 | 93691840 | rs12416560 | -0.416 | 0.785 | -0.517 | -0.93 | -0.041 | -0.093 | -0.093 | 0 | 0.3 | 0.12 | 0.39 | 0.54 | 0.34 | 0.55 |
| 10 | 93709954 | rs10509641 | 0.944 | 1.063 | -0.683 | -0.937 | -0.032 | 0.065 | 0.065 | 0 | 0.12 | 0 | 0.1 | 0 | 0.11 | 0.08 |
| 10 | 93717432 | rs8181288 | 2.194 | 1.168 | -0.758 | -0.955 | -0.1 | -0.019 | -0.019 | 0 | 0.36 | 0.03 | 0.23 | 0.26 | 0.4 | 0.14 |

| | | | | | | | | | | | | | | | | |
|---|---|---|---|---|---|---|---|---|---|---|---|---|---|---|---|---|
| 10 | 93721025 | rs4642987 | 0.162 | 1.1 | -0.705 | -0.948 | 0.047 | 0.063 | 0.063 | 0.746 | 0.65 | 0.16 | 0.62 | 0.79 | 0.74 | 0.69 |
| 10 | 93729335 | rs7909870 | -0.36 | 1.098 | -0.595 | -0.949 | -0.008 | -0.012 | -0.012 | 0 | 0.33 | 0.68 | 0.36 | 0.21 | 0.26 | 0.31 |
| 10 | 93740429 | rs2792022 | -0.163 | 1.246 | -0.758 | -0.978 | -0.043 | -0.143 | -0.143 | 0 | 0.26 | 0.04 | 0.36 | 0.53 | 0.28 | 0.52 |
| 10 | 93746420 | rs11819184 | 0.928 | 1.038 | -0.575 | -0.955 | 0.015 | 0.048 | 0.048 | 0 | 0.2 | 0.1 | 0.12 | 0.02 | 0.18 | 0.13 |
| 10 | 93778269 | rs2048901 | 1.55 | 0.664 | -0.72 | -0.907 | -0.005 | 0.03 | 0.03 | 0 | 0.11 | 0.23 | 0.21 | 0.19 | 0.08 | 0.18 |
| 10 | 93781926 | rs833368 | 0.215 | 0.713 | -1.207 | -0.902 | 0.044 | 0.063 | 0.063 | 0.746 | 0.65 | 0.16 | 0.62 | 0.79 | 0.74 | 0.69 |
| 10 | 93790505 | rs1059515 | 2.049 | 0.817 | -1.147 | -0.768 | -0.101 | -0.021 | -0.021 | 0.068 | 0.35 | 0.03 | 0.24 | 0.26 | 0.4 | 0.14 |
| 10 | 93801875 | rs1581239 | -0.967 | 0.795 | -0.974 | -0.755 | -0.01 | -0.007 | -0.007 | 0 | 0.5 | 0.21 | 0.51 | 0.55 | 0.52 | 0.69 |
| 10 | 93810872 | rs1776209 | -0.111 | 0.934 | -1.254 | -0.68 | -0.063 | -0.165 | -0.165 | 0.424 | 0.26 | 0.02 | 0.35 | 0.53 | 0.27 | 0.55 |
| 10 | 93812405 | rs10882018 | 1.226 | 0.589 | -1.183 | -0.6 | 0.008 | -0.021 | -0.021 | 0 | 0.36 | 0.24 | 0.36 | 0.34 | 0.36 | 0.21 |
| 10 | 93818504 | rs1099379 | -0.134 | 0.728 | -0.864 | -0.591 | -0.071 | -0.083 | -0.083 | 0.112 | 0.31 | 0.07 | 0.25 | 0.51 | 0.37 | 0.59 |
| 10 | 93828501 | rs1622656 | 1.606 | 0.775 | -0.899 | -0.577 | -0.049 | -0.088 | -0.088 | 0 | 0.14 | 0.01 | 0.11 | 0.48 | 0.18 | 0.47 |
| 10 | 93847722 | rs12256118 | 0.371 | 0.686 | -0.7 | -0.68 | 0.009 | 0.015 | 0.015 | 0 | 0.3 | 0.29 | 0.36 | 0.21 | 0.26 | 0.29 |
| 10 | 93850375 | rs10786028 | 1.686 | 0.755 | -0.71 | -0.64 | -0.074 | -0.083 | -0.083 | 0.371 | 0.34 | 0.03 | 0.35 | 0.29 | 0.36 | 0.12 |
| 10 | 93851714 | rs1229578 | -1.928 | 0.781 | -0.714 | -0.636 | -0.049 | -0.088 | -0.088 | 0 | 0.86 | 0.99 | 0.89 | 0.52 | 0.82 | 0.53 |
| 10 | 93852663 | rs10882023 | 1.512 | 0.759 | -0.509 | -0.625 | -0.004 | 0 | 0 | 2.356 | 0.36 | 0.41 | 0.37 | 0.29 | 0.36 | 0.12 |
| 10 | 93874109 | rs766398 | 1.3 | 0.741 | -0.625 | -0.627 | -0.002 | -0.001 | -0.001 | 0.088 | 0.38 | 0.58 | 0.39 | 0.29 | 0.36 | 0.12 |
| 10 | 93919120 | rs10786029 | -1.899 | 0.883 | -0.853 | -0.503 | -0.051 | -0.088 | -0.088 | 0 | 0.86 | 0.99 | 0.89 | 0.52 | 0.82 | 0.53 |
| 10 | 93930279 | rs7087501 | -0.231 | 0.565 | -0.946 | -0.495 | -0.081 | -0.093 | -0.093 | 0 | 0.67 | 0.95 | 0.75 | 0.5 | 0.62 | 0.4 |
| 10 | 93951793 | rs17107472 | 2.016 | 0.511 | -1.08 | -0.467 | -0.02 | -0.036 | -0.036 | 1.742 | 0.08 | 0.02 | 0.15 | 0.17 | 0.08 | 0.15 |
| 10 | 93977203 | rs10882037 | 1.268 | 0.608 | -1.046 | -0.564 | -0.006 | -0.026 | -0.026 | 0 | 0.37 | 0.57 | 0.21 | 0.12 | 0.32 | 0.1 |
| 10 | 94008580 | rs11186872 | 0.201 | 0.636 | -1.144 | -0.5 | -0.001 | 0.001 | 0.001 | 0 | 0.31 | 0.36 | 0.53 | 0.38 | 0.29 | 0.31 |
| 10 | 94014958 | rs7088903 | 1.629 | 0.624 | -1.158 | -0.535 | 0 | 0.082 | 0.082 | 1.111 | 0.34 | 0.1 | 0.17 | 0.12 | 0.32 | 0.1 |
| 10 | 94027193 | rs9804247 | 0.236 | 0.635 | -1.184 | -0.511 | -0.001 | 0.001 | 0.001 | 0 | 0.31 | 0.36 | 0.53 | 0.38 | 0.29 | 0.31 |
| 10 | 94052164 | rs12411618 | 0.611 | 0.871 | -0.603 | 0.018 | -0.024 | 0.085 | 0.085 | 0 | 0.14 | 0 | 0.11 | 0 | 0.12 | 0.11 |
| 10 | 94090498 | rs10748579 | -0.239 | 0.806 | -0.211 | 0.323 | 0.005 | 0.009 | 0.009 | 0.06 | 0.65 | 0.44 | 0.7 | 0.49 | 0.62 | 0.4 |
| 10 | 94114633 | rs950732 | -0.239 | 0.814 | -0.208 | 0.326 | 0.004 | 0.009 | 0.009 | 0.076 | 0.64 | 0.44 | 0.7 | 0.49 | 0.62 | 0.39 |
| 10 | 94116891 | rs7080813 | 0.504 | 0.802 | -0.344 | -0.452 | 0.002 | 0.02 | 0.02 | 0.266 | 0.52 | 0.44 | 0.33 | 0.14 | 0.5 | 0.21 |
| 10 | 94128012 | rs7090404 | 0.476 | 0.83 | -0.512 | -0.44 | -0.001 | -0.014 | -0.014 | 0.3 | 0.52 | 0.48 | 0.33 | 0.14 | 0.5 | 0.21 |
| 10 | 94147023 | rs2259471 | 0.172 | 0.026 | -0.979 | -0.469 | -0.021 | -0.057 | -0.057 | 1.61 | 0.32 | 0.06 | 0.29 | 0.68 | 0.32 | 0.62 |
| 10 | 94148800 | rs7897267 | 0.137 | 0.02 | -0.611 | -0.526 | 0.006 | 0.054 | 0.054 | 1.283 | 0.15 | 0.4 | 0.3 | 0.21 | 0.12 | 0.19 |
| 10 | 94155499 | rs2259049 | 0.086 | -0.071 | -0.43 | -0.584 | -0.011 | 0.005 | 0.005 | 0.096 | 0.34 | 0.45 | 0.32 | 0.68 | 0.32 | 0.63 |
| 10 | 94162910 | rs11594882 | 0.016 | 0.105 | -0.591 | -0.573 | -0.001 | 0.032 | 0.032 | 1.016 | 0.15 | 0.37 | 0.29 | 0.18 | 0.12 | 0.18 |
| 10 | 94164664 | rs10882058 | -0.126 | 0.093 | -0.583 | -0.703 | -0.001 | 0.013 | 0.013 | 0.892 | 0.15 | 0.37 | 0.24 | 0.18 | 0.12 | 0.19 |
| 10 | 94167087 | rs7084673 | 0.581 | 0.011 | -0.801 | -1.011 | 0.001 | -0.059 | -0.059 | 0 | 0.4 | 0.44 | 0.33 | 0.1 | 0.41 | 0.15 |
| 10 | 94168797 | rs2263638 | -0.524 | 0.639 | -0.885 | -0.801 | 0.001 | -0.005 | -0.005 | 0 | 0.3 | 0.39 | 0.36 | 0.2 | 0.36 | 0.27 |
| 10 | 94170851 | rs2901587 | 0.218 | 0.04 | -0.853 | -0.92 | 0.031 | 0.049 | 0.049 | 0 | 0.35 | 0.28 | 0.45 | 0.81 | 0.23 | 0.55 |
| 10 | 94176088 | rs7086285 | -0.804 | 0.274 | -0.729 | -1.077 | 0.002 | 0.088 | 0.088 | 0 | 0.55 | 0.75 | 0.58 | 0.84 | 0.41 | 0.75 |
| 10 | 94179028 | rs2209972 | 0.178 | 0.231 | -0.768 | -0.991 | 0.047 | 0.091 | 0.091 | 0.409 | 0.41 | 0.21 | 0.47 | 0.76 | 0.31 | 0.6 |
| 10 | 94185670 | rs12257053 | 0.324 | 0.356 | -0.657 | -0.922 | 0.061 | -0.051 | -0.051 | 0 | 0.11 | 0.47 | 0.08 | 0.03 | 0.08 | 0.15 |
| 10 | 94190580 | rs10786044 | -0.079 | 0.319 | -1.438 | -0.751 | 0.053 | 0.099 | 0.099 | 0.229 | 0.43 | 0.23 | 0.48 | 0.76 | 0.31 | 0.6 |
| 10 | 94199337 | rs10882063 | -0.296 | 0.588 | -1.495 | -0.705 | 0.041 | 0.159 | 0.159 | 0.058 | 0.51 | 0.83 | 0.53 | 0.83 | 0.37 | 0.76 |
| 10 | 94209939 | rs6583813 | 0.222 | 0.728 | -2.225 | -0.736 | -0.082 | -0.099 | -0.099 | 1.547 | 0.29 | 0.05 | 0.29 | 0.68 | 0.36 | 0.52 |
| 10 | 94210809 | rs12415807 | 0.389 | 0.659 | -1.134 | -0.86 | 0.011 | 0.031 | 0.031 | 0 | 0.1 | 0.25 | 0.17 | 0.14 | 0.07 | 0.17 |
| 10 | 94222624 | rs551266 | -1.032 | 0.524 | -0.308 | -1.573 | 0.01 | 0.061 | 0.061 | 0 | 0.78 | 0.94 | 0.84 | 0.95 | 0.81 | 0.82 |
| 10 | 94237425 | rs7091270 | 0.571 | 1.023 | -0.088 | -1.528 | 0.043 | 0.092 | 0.092 | 0 | 0.11 | 0.29 | 0.32 | 0.21 | 0.12 | 0.19 |
| 10 | 94239793 | rs7910605 | 0.727 | 1.013 | -0.089 | -1.534 | 0.032 | 0.058 | 0.058 | 0 | 0.11 | 0.28 | 0.26 | 0.18 | 0.11 | 0.18 |
| 10 | 94257976 | rs11187025 | 1.073 | 0.801 | -0.111 | -1.938 | -0.005 | 0.004 | 0.004 | 0.489 | 0.22 | 0.17 | 0.14 | 0.06 | 0.24 | 0.09 |
| 10 | 94290484 | rs7078413 | -0.022 | 0.704 | -0.122 | -2.004 | -0.018 | 0.046 | 0.046 | 0.287 | 0.34 | 0.22 | 0.32 | 0.2 | 0.35 | 0.27 |
| 10 | 94292106 | rs17107709 | 0.652 | 0.585 | -0.108 | -2.064 | 0.009 | 0.007 | 0.007 | 0 | 0.08 | 0.04 | 0.03 | 0 | 0.03 | 0.02 |
| 10 | 94294291 | rs4646955 | 0.789 | 0.643 | -0.082 | -1.999 | -0.045 | 0.092 | 0.092 | 1.992 | 0.25 | 0.08 | 0.15 | 0.06 | 0.28 | 0.1 |
| 10 | 94298331 | rs1832197 | -0.243 | 0.827 | 0.082 | -2.023 | 0.02 | -0.038 | -0.038 | 0.983 | 0.37 | 0.48 | 0.42 | 0.22 | 0.4 | 0.28 |
| 10 | 94304132 | rs11187060 | 0.056 | 0.745 | -0.188 | -1.969 | -0.005 | 0.005 | 0.005 | 0 | 0.32 | 0.33 | 0.39 | 0.22 | 0.35 | 0.27 |
| 10 | 94305409 | rs11187061 | -1.206 | 0.64 | -0.208 | -1.946 | 0.012 | 0.071 | 0.071 | 0 | 0.77 | 0.95 | 0.83 | 0.95 | 0.8 | 0.82 |
| 10 | 94319992 | rs1999763 | 1.336 | 0.703 | -0.201 | -1.871 | -0.002 | 0.01 | 0.01 | 0 | 0.09 | 0.14 | 0.17 | 0.14 | 0.07 | 0.17 |
| 10 | 94331341 | rs2421945 | -0.042 | 0.723 | -0.214 | -1.871 | -0.002 | 0.001 | 0.001 | 0 | 0.33 | 0.34 | 0.45 | 0.27 | 0.36 | 0.28 |
| 10 | 94333827 | rs4646954 | 0.775 | 0.659 | 0.028 | -1.628 | 0.039 | 0.073 | 0.073 | 0 | 0.1 | 0.39 | 0.18 | 0.13 | 0.07 | 0.17 |
| 10 | 94336963 | rs11187076 | -0.463 | 0.604 | -0.062 | -1.704 | 0 | -0.014 | -0.014 | 0.084 | 0.54 | 0.52 | 0.54 | 0.78 | 0.52 | 0.64 |
| 10 | 94347830 | rs6583826 | -0.137 | 0.588 | -0.075 | -1.71 | 0 | -0.003 | -0.003 | 0 | 0.51 | 0.52 | 0.52 | 0.8 | 0.51 | 0.72 |
| 10 | 94353278 | rs12256435 | 0.078 | 0.574 | -0.126 | -1.681 | 0.003 | -0.014 | -0.014 | 0.325 | 0.13 | 0.11 | 0.05 | 0 | 0.08 | 0.03 |
| 10 | 94354204 | rs3824735 | 1.172 | 0.591 | -0.203 | -1.729 | -0.059 | -0.076 | -0.076 | 0 | 0.29 | 0.1 | 0.29 | 0.68 | 0.36 | 0.51 |
| 10 | 94356912 | rs4604791 | 0.51 | 0.544 | -0.025 | -1.759 | 0.005 | 0.079 | 0.079 | 0.313 | 0.1 | 0.45 | 0.12 | 0.09 | 0.03 | 0.09 |
| 10 | 94363993 | rs4933731 | 1.032 | 0.51 | -0.054 | -1.791 | -0.001 | 0.018 | 0.018 | 0 | 0.09 | 0.19 | 0.17 | 0.13 | 0.06 | 0.16 |
| 10 | 94368178 | rs11187094 | 1.822 | 0.627 | -0.184 | -1.706 | -0.037 | -0.137 | -0.137 | 4.536 | 0.23 | 0.03 | 0.22 | 0.6 | 0.22 | 0.47 |
| 10 | 94374377 | rs10882091 | -1.109 | 0.333 | -0.075 | -1.785 | -0.064 | 0.193 | 0.193 | 0 | 0.62 | 0.92 | 0.72 | 0.92 | 0.57 | 0.8 |
| 10 | 94382950 | rs7914814 | -1.109 | 0.333 | -0.075 | -1.785 | -0.064 | 0.193 | 0.193 | 0 | 0.62 | 0.92 | 0.72 | 0.92 | 0.58 | 0.8 |
| 10 | 94388107 | rs12244924 | 0.403 | 0.542 | 0.102 | -1.6 | 0.005 | 0.079 | 0.079 | 0.313 | 0.1 | 0.45 | 0.12 | 0.09 | 0.03 | 0.09 |
| 10 | 94414263 | rs7078243 | -0.825 | 0.54 | -0.09 | -1.648 | -0.023 | 0.127 | 0.127 | 0.01 | 0.54 | 0.78 | 0.56 | 0.79 | 0.51 | 0.64 |
| 10 | 94415078 | rs1044153 | 1.024 | 0.697 | -0.143 | -1.539 | -0.07 | -0.084 | -0.084 | 0.038 | 0.29 | 0.07 | 0.29 | 0.68 | 0.36 | 0.51 |

| | | | | | | | | | | | | | | | | |
|---|---|---|---|---|---|---|---|---|---|---|---|---|---|---|---|---|
| 10 | 94429526 | rs11187126 | 0.009 | 0.501 | 0.041 | -1.451 | 0.004 | -0.01 | -0.01 | 0.664 | 0.12 | 0.1 | 0.05 | 0 | 0.08 | 0.03 |
| 10 | 94446041 | rs2488087 | 0.083 | 0.52 | 0.096 | -1.551 | -0.008 | 0.069 | 0.069 | 0 | 0.54 | 0.29 | 0.47 | 0.24 | 0.48 | 0.4 |
| 10 | 94446635 | rs10786052 | -0.367 | 0.532 | -0.052 | -1.546 | 0.001 | -0.01 | -0.01 | 2.148 | 0.45 | 0.5 | 0.51 | 0.76 | 0.52 | 0.6 |
| 10 | 94452430 | rs2275729 | -0.071 | 0.629 | -0.253 | -1.501 | 0.002 | 0.003 | 0.003 | 0 | 0.13 | 0.13 | 0.12 | 0.05 | 0.09 | 0.03 |
| 10 | 94460650 | rs10882099 | 0.705 | 0.761 | -0.195 | -1.356 | -0.059 | -0.016 | -0.016 | 0 | 0.31 | 0.19 | 0.38 | 0.72 | 0.43 | 0.64 |
| 10 | 94473629 | rs7087591 | 0.645 | 0.501 | -0.155 | -1.745 | 0.005 | -0.007 | -0.007 | 0 | 0.32 | 0.34 | 0.45 | 0.8 | 0.38 | 0.64 |
| 10 | 94480391 | rs2497313 | 0.002 | 0.396 | -0.814 | -1.972 | -0.035 | -0.084 | -0.084 | 0.332 | 0.8 | 0.39 | 0.9 | 0.99 | 0.9 | 0.93 |
| 10 | 94481917 | rs7923837 | 0.825 | 0.446 | -1.333 | -1.944 | -0.124 | -0.008 | -0.008 | 0 | 0.3 | 0.03 | 0.42 | 0.81 | 0.37 | 0.63 |
| 10 | 94485211 | rs2497306 | -0.27 | 0.897 | -0.836 | -1.661 | -0.024 | 0.228 | 0.228 | 0.632 | 0.49 | 0.19 | 0.41 | 0.13 | 0.51 | 0.27 |
| 10 | 94492716 | rs2497304 | 0.262 | 0.897 | -0.89 | -1.676 | -0.106 | 0.188 | 0.188 | 1.406 | 0.41 | 0.11 | 0.38 | 0.11 | 0.5 | 0.21 |
| 10 | 94495753 | rs947591 | 0.133 | 1.425 | -0.554 | -1.479 | 0.005 | 0.143 | 0.143 | 0.017 | 0.46 | 0.24 | 0.4 | 0.14 | 0.47 | 0.26 |
| 10 | 94498436 | rs4933236 | 0.11 | 1.383 | -0.328 | -1.157 | -0.001 | -0.011 | -0.011 | 0 | 0.42 | 0.45 | 0.48 | 0.72 | 0.4 | 0.63 |
| 10 | 94499577 | rs2488071 | 0.019 | 1.541 | -1.019 | -1.112 | -0.001 | 0.062 | 0.062 | 0.1 | 0.52 | 0.34 | 0.48 | 0.2 | 0.52 | 0.3 |
| 10 | 94500111 | rs11187152 | -0.473 | 1.643 | -0.983 | -0.541 | -0.029 | -0.003 | -0.003 | 0.929 | 0.05 | 0.01 | 0.04 | 0.08 | 0.08 | 0.09 |
| 10 | 94502244 | rs11187157 | -0.494 | 1.312 | -0.947 | -0.819 | -0.005 | 0.014 | 0.014 | 0 | 0.39 | 0.36 | 0.41 | 0.27 | 0.43 | 0.28 |
| 10 | 94502738 | rs1539330 | 0.01 | 1.146 | -1.075 | -1.613 | 0 | 0.021 | 0.021 | 0 | 0.13 | 0.08 | 0.08 | 0.02 | 0.13 | 0.05 |
| 10 | 94506628 | rs2488068 | 0.126 | 1.121 | -0.758 | -2.288 | 0.011 | -0.083 | -0.083 | 0.558 | 0.33 | 0.38 | 0.34 | 0.1 | 0.34 | 0.28 |
| 10 | 94507612 | rs2488066 | -0.04 | 0.964 | -0.773 | -2.408 | -0.003 | -0.052 | -0.052 | 0 | 0.78 | 0.73 | 0.87 | 0.97 | 0.8 | 0.92 |
| 10 | 94517064 | rs11187167 | 0.539 | 1.011 | -0.883 | -2.325 | -0.005 | 0.069 | 0.069 | 0 | 0.17 | 0.05 | 0.09 | 0.03 | 0.16 | 0.04 |
| 10 | 94523766 | rs2488062 | 0.658 | 1.069 | -0.551 | -2.652 | 0.004 | -0.03 | -0.03 | 0 | 0.16 | 0.12 | 0.1 | 0 | 0.22 | 0.11 |
| 10 | 94525456 | rs7091916 | 0.184 | 1.105 | -0.44 | -2.658 | -0.008 | 0.033 | 0.033 | 0 | 0.2 | 0.13 | 0.25 | 0.04 | 0.22 | 0.07 |
| 10 | 94525506 | rs1935492 | 0.025 | 0.898 | 0 | -2.602 | 0.001 | -0.025 | -0.025 | 0.8 | 0.31 | 0.28 | 0.29 | 0.11 | 0.3 | 0.27 |
| 10 | 94530832 | rs10882109 | 0.679 | 0.915 | -0.026 | -2.564 | 0 | 0.012 | 0.012 | 0.03 | 0.48 | 0.43 | 0.46 | 0.85 | 0.49 | 0.67 |
| 10 | 94531770 | rs11592067 | 1.301 | 0.823 | -0.271 | -2.973 | -0.004 | 0.071 | 0.071 | 0 | 0.33 | 0.15 | 0.35 | 0.77 | 0.34 | 0.51 |
| 10 | 94535940 | rs2488083 | 0.114 | 0.743 | 0.035 | -2.969 | 0.004 | -0.068 | -0.068 | 0 | 0.31 | 0.47 | 0.44 | 0.13 | 0.32 | 0.28 |
| 10 | 94538960 | rs2497338 | -0.577 | 0.593 | 0 | -2.815 | 0 | -0.022 | -0.022 | 0 | 0.8 | 0.78 | 0.9 | 0.97 | 0.82 | 0.95 |
| 10 | 94542446 | rs7924271 | 0.484 | 0.553 | -0.118 | -2.889 | -0.003 | 0.057 | 0.057 | 0 | 0.45 | 0.28 | 0.43 | 0.87 | 0.43 | 0.67 |
| 10 | 94543870 | rs11596160 | 1.28 | 0.584 | -0.225 | -2.983 | -0.006 | 0.07 | 0.07 | 0 | 0.31 | 0.14 | 0.22 | 0.81 | 0.31 | 0.58 |
| 10 | 94548029 | rs4128214 | 0.871 | 0.432 | 0.238 | -3.077 | -0.005 | -0.042 | -0.042 | 0.469 | 0.36 | 0.38 | 0.37 | 0.83 | 0.34 | 0.6 |
| 10 | 94558538 | rs2490754 | 0.371 | 0.505 | 0.603 | -3.318 | -0.006 | 0.038 | 0.038 | 0 | 0.33 | 0.26 | 0.32 | 0.09 | 0.38 | 0.26 |
| 10 | 94562677 | rs11187182 | 1.438 | 0.523 | 0.381 | -3.283 | 0 | 0.034 | 0.034 | 0 | 0.31 | 0.19 | 0.22 | 0.82 | 0.31 | 0.58 |
| 10 | 94565784 | rs2422067 | 0.446 | 0.474 | 0.316 | -3.287 | -0.004 | -0.063 | -0.063 | 0 | 0.22 | 0.28 | 0.24 | 0.05 | 0.2 | 0.05 |
| 10 | 94568752 | rs2490745 | 0.281 | 0.579 | 0.106 | -3.179 | -0.007 | 0.028 | 0.028 | 0 | 0.32 | 0.26 | 0.32 | 0.08 | 0.37 | 0.26 |
| 10 | 94571660 | rs4933742 | 0.061 | 0.561 | 0.036 | -3.14 | 0 | -0.031 | -0.031 | 2.238 | 0.56 | 0.53 | 0.48 | 0.86 | 0.52 | 0.6 |
| 10 | 94572035 | rs4933743 | 1.258 | 0.287 | -0.657 | -3.259 | -0.015 | 0.05 | 0.05 | 0 | 0.31 | 0.1 | 0.26 | 0.75 | 0.32 | 0.53 |
| 10 | 94576592 | rs10748583 | 0.845 | 0.18 | -0.532 | -3.468 | 0.026 | 0.073 | 0.073 | 0 | 0.36 | 0.21 | 0.38 | 0.88 | 0.31 | 0.54 |
| 10 | 94587755 | rs2497323 | 0.751 | 0.287 | -0.944 | -3.739 | -0.099 | 0.123 | 0.123 | 0 | 0.2 | 0.01 | 0.15 | 0 | 0.3 | 0.12 |
| 10 | 94591607 | rs2497322 | 0.463 | 0.255 | -0.823 | -3.762 | -0.024 | 0.027 | 0.027 | 0 | 0.21 | 0.08 | 0.17 | 0 | 0.32 | 0.13 |
| 10 | 94591841 | rs2490742 | 0.765 | 0.255 | -0.825 | -3.762 | -0.099 | 0.123 | 0.123 | 0 | 0.2 | 0.01 | 0.15 | 0 | 0.31 | 0.12 |
| 10 | 94593407 | rs7916038 | -0.018 | 0.042 | -0.266 | -3.887 | 0.001 | -0.057 | -0.057 | 0 | 0.4 | 0.55 | 0.45 | 0.12 | 0.37 | 0.33 |
| 10 | 94594565 | rs2490741 | -1.558 | -0.01 | -0.248 | -3.849 | 0.021 | 0.044 | 0.044 | 0 | 0.64 | 0.72 | 0.72 | 0.25 | 0.7 | 0.47 |
| 10 | 94596356 | rs2490740 | 0.87 | -0.001 | -0.247 | -3.847 | -0.099 | 0.123 | 0.123 | 0 | 0.2 | 0.01 | 0.15 | 0 | 0.31 | 0.12 |
| 10 | 94597905 | rs2490739 | -1.674 | -0.014 | -0.249 | -3.85 | 0.013 | 0.034 | 0.034 | 0 | 0.66 | 0.72 | 0.72 | 0.25 | 0.7 | 0.48 |
| 10 | 94598259 | rs3740380 | -0.183 | -0.013 | -0.187 | -3.848 | 0 | -0.056 | -0.056 | 0 | 0.42 | 0.54 | 0.45 | 0.12 | 0.37 | 0.33 |
| 10 | 94602822 | rs2490733 | 0.83 | 0.036 | -0.263 | -3.816 | -0.099 | 0.123 | 0.123 | 0 | 0.2 | 0.01 | 0.15 | 0 | 0.31 | 0.12 |
| 10 | 94606337 | rs2497339 | 0.758 | 0.034 | -0.231 | -3.817 | -0.029 | 0.035 | 0.035 | 0 | 0.2 | 0.08 | 0.15 | 0 | 0.31 | 0.12 |
| 10 | 94610368 | rs2250393 | 0.82 | 0.027 | -0.249 | -3.819 | -0.099 | 0.124 | 0.124 | 0 | 0.2 | 0.01 | 0.15 | 0 | 0.31 | 0.12 |
| 10 | 94611055 | rs10882117 | 1.888 | -0.035 | -0.275 | -3.852 | -0.003 | 0.057 | 0.057 | 0 | 0.32 | 0.09 | 0.27 | 0.76 | 0.3 | 0.53 |
| 10 | 94613240 | rs7087174 | -0.333 | 0.021 | -0.215 | -3.849 | 0.002 | 0.026 | 0.026 | 0 | 0.43 | 0.63 | 0.55 | 0.24 | 0.37 | 0.34 |
| 10 | 94618925 | rs11516919 | -0.519 | -0.001 | -0.207 | -3.827 | 0.004 | 0.075 | 0.075 | 0 | 0.45 | 0.67 | 0.57 | 0.24 | 0.39 | 0.35 |
| 10 | 94619980 | rs10748584 | 1.351 | 0.004 | -0.212 | -3.831 | 0.025 | 0.058 | 0.058 | 0 | 0.35 | 0.25 | 0.28 | 0.76 | 0.31 | 0.53 |
| 10 | 94621540 | rs7910006 | -0.297 | 0.008 | -0.206 | -3.828 | 0.002 | 0.02 | 0.02 | 0 | 0.43 | 0.61 | 0.55 | 0.24 | 0.37 | 0.34 |
| 10 | 94622852 | rs10882121 | -0.333 | 0.309 | -0.207 | -3.828 | 0.002 | 0.027 | 0.027 | 0 | 0.43 | 0.63 | 0.55 | 0.24 | 0.37 | 0.34 |
| 10 | 94624759 | rs1987017 | 1.682 | 0.298 | -0.226 | -3.829 | 0.005 | 0.073 | 0.073 | 0.027 | 0.34 | 0.09 | 0.27 | 0.76 | 0.3 | 0.54 |
| 10 | 94628274 | rs7906224 | -0.187 | 0.339 | -0.2 | -3.853 | -0.001 | -0.046 | -0.046 | 0 | 0.42 | 0.55 | 0.45 | 0.12 | 0.36 | 0.32 |
| 10 | 94634842 | rs2422072 | -0.715 | 0.343 | -0.227 | -3.855 | -0.014 | -0.002 | -0.002 | 0 | 0.78 | 0.88 | 0.83 | 1 | 0.67 | 0.87 |
| 10 | 94638491 | rs4508117 | -0.258 | 0.253 | -0.409 | -3.842 | 0.001 | 0.025 | 0.025 | 0 | 0.42 | 0.63 | 0.55 | 0.24 | 0.37 | 0.34 |
| 10 | 94642494 | rs10882123 | 1.736 | 0.254 | -0.455 | -3.825 | 0.002 | 0.066 | 0.066 | 0.022 | 0.33 | 0.09 | 0.27 | 0.76 | 0.3 | 0.53 |
| 10 | 94642659 | rs7068232 | 1.288 | 0.254 | -0.448 | -3.826 | 0.023 | 0.06 | 0.06 | 0 | 0.35 | 0.24 | 0.28 | 0.75 | 0.31 | 0.53 |
| 10 | 94648958 | rs9633694 | 1.89 | 0.283 | -0.462 | -3.812 | -0.006 | 0.056 | 0.056 | 0 | 0.32 | 0.09 | 0.27 | 0.76 | 0.3 | 0.53 |
| 10 | 94656941 | rs7094393 | -0.541 | 0.227 | -0.416 | -3.796 | 0 | 0.138 | 0.138 | 0 | 0.45 | 0.74 | 0.57 | 0.24 | 0.39 | 0.35 |
| 10 | 94660133 | rs17377483 | 2.03 | 0.194 | -0.607 | -3.969 | -0.036 | 0.089 | 0.089 | 0 | 0.16 | 0.03 | 0.09 | 0 | 0.17 | 0.06 |
| 10 | 94660409 | rs7915463 | -0.604 | 0.272 | -0.522 | -3.926 | 0.007 | 0.082 | 0.082 | 0 | 0.44 | 0.68 | 0.54 | 0.24 | 0.37 | 0.34 |
| 10 | 94660689 | rs7918909 | 1.621 | 0.207 | -0.261 | -3.999 | -0.003 | -0.015 | -0.015 | 0 | 0.1 | 0.08 | 0.17 | 0.02 | 0.11 | 0.09 |
| 10 | 94661093 | rs7095772 | -0.69 | 0.206 | -0.265 | -3.997 | 0.006 | 0.081 | 0.081 | 0 | 0.43 | 0.68 | 0.55 | 0.24 | 0.37 | 0.34 |
| 10 | 94663308 | rs7914571 | -0.807 | 0.205 | -0.27 | -3.999 | 0.002 | 0.131 | 0.131 | 0 | 0.45 | 0.73 | 0.57 | 0.24 | 0.39 | 0.35 |
| 10 | 94665431 | rs7923502 | -0.372 | 0.191 | -0.188 | -4.004 | 0 | 0.003 | 0.003 | 0 | 0.4 | 0.66 | 0.44 | 0.11 | 0.36 | 0.33 |
| 10 | 94665458 | rs2148361 | -1.194 | 0.227 | -0.247 | -4.031 | -0.103 | 0.122 | 0.122 | 0 | 0.8 | 0.99 | 0.85 | 1 | 0.68 | 0.88 |
| 10 | 94666581 | rs10786060 | 1.967 | 0.216 | -0.246 | -4.034 | 0.019 | 0.042 | 0.042 | 0 | 0.34 | 0.26 | 0.28 | 0.75 | 0.3 | 0.53 |

| | | | | | | | | | | | | | | | | |
|---|---|---|---|---|---|---|---|---|---|---|---|---|---|---|---|---|
| 10 | 94666983 | rs6583847 | 0.193 | 0.215 | -0.019 | -4.037 | -0.013 | -0.079 | -0.079 | 0 | 0.31 | 0.42 | 0.26 | 0.09 | 0.25 | 0.24 |
| 10 | 94668408 | rs2025730 | -1.177 | 0.223 | -0.058 | -4.017 | -0.098 | 0.119 | 0.119 | 0 | 0.8 | 0.99 | 0.84 | 0.99 | 0.68 | 0.88 |
| 10 | 94668524 | rs10882125 | 1.963 | 0.22 | -0.051 | -4.027 | 0.022 | 0.052 | 0.052 | 0 | 0.33 | 0.25 | 0.28 | 0.75 | 0.3 | 0.53 |
| 10 | 94670849 | rs11187215 | 1.631 | 0.225 | -0.063 | -4.02 | -0.003 | -0.015 | -0.015 | 0 | 0.1 | 0.07 | 0.17 | 0.02 | 0.11 | 0.09 |
| 10 | 94673088 | rs4933753 | 1.891 | 0.214 | -0.076 | -4.013 | 0.018 | 0.037 | 0.037 | 0 | 0.34 | 0.28 | 0.28 | 0.75 | 0.3 | 0.53 |
| 10 | 94679124 | rs10748585 | 1.828 | 0.218 | -0.106 | -4.011 | 0.031 | 0.064 | 0.064 | 0 | 0.35 | 0.25 | 0.28 | 0.76 | 0.3 | 0.53 |
| 10 | 94682835 | rs9633695 | -1.141 | 0.251 | -0.125 | -4.003 | -0.103 | 0.123 | 0.123 | 0 | 0.8 | 0.99 | 0.85 | 0.99 | 0.68 | 0.88 |
| 10 | 94683493 | rs7913340 | -0.939 | 0.251 | -0.11 | -4.004 | -0.061 | 0.084 | 0.084 | 0 | 0.79 | 0.95 | 0.83 | 1 | 0.66 | 0.87 |
| 10 | 94690399 | rs7893270 | -1.157 | 0.25 | -0.127 | -4.004 | -0.103 | 0.123 | 0.123 | 0 | 0.8 | 0.99 | 0.85 | 1 | 0.68 | 0.88 |
| 10 | 94692085 | rs10748586 | -1.174 | 0.25 | -0.127 | -4.004 | -0.103 | 0.122 | 0.122 | 0 | 0.8 | 0.99 | 0.85 | 1 | 0.68 | 0.88 |
| 10 | 94694389 | rs4933755 | -0.789 | 0.199 | -0.132 | -4.034 | 0.004 | 0.125 | 0.125 | 0 | 0.45 | 0.72 | 0.56 | 0.24 | 0.39 | 0.35 |
| 10 | 94695555 | rs1326330 | 0.336 | 0.205 | 0.078 | -4.039 | -0.006 | -0.052 | -0.052 | 0.526 | 0.29 | 0.36 | 0.26 | 0.1 | 0.25 | 0.24 |
| 10 | 94695617 | rs1326331 | -1.172 | 0.25 | -0.187 | -4.018 | -0.103 | 0.123 | 0.123 | 0 | 0.8 | 0.99 | 0.85 | 1 | 0.68 | 0.88 |
| 10 | 94696224 | rs9645558 | 1.831 | 0.235 | -0.127 | -3.975 | 0.016 | 0.075 | 0.075 | 0 | 0.34 | 0.09 | 0.27 | 0.75 | 0.3 | 0.53 |
| 10 | 94703333 | rs17378057 | 1.553 | 0.325 | -0.123 | -3.931 | -0.003 | -0.014 | -0.014 | 0.029 | 0.1 | 0.07 | 0.17 | 0.02 | 0.11 | 0.09 |
| 10 | 94710761 | rs10786063 | 1.821 | 0.268 | 0.227 | -3.955 | 0.018 | 0.036 | 0.036 | 0 | 0.34 | 0.33 | 0.28 | 0.75 | 0.29 | 0.54 |
| 10 | 94711319 | rs10882131 | 1.552 | 0.346 | 0.18 | -3.886 | -0.003 | -0.015 | -0.015 | 0 | 0.1 | 0.07 | 0.17 | 0.02 | 0.11 | 0.09 |
| 10 | 94716313 | rs980204 | -1.115 | 0.338 | 0.08 | -3.919 | -0.108 | 0.123 | 0.123 | 0 | 0.8 | 0.99 | 0.85 | 0.98 | 0.68 | 0.87 |
| 10 | 94718164 | rs11187226 | -0.76 | 0.245 | 0.194 | -3.997 | 0.017 | 0.08 | 0.08 | 0.32 | 0.44 | 0.62 | 0.53 | 0.23 | 0.35 | 0.32 |
| 10 | 94718268 | rs7075749 | 0.278 | 0.27 | 0.32 | -3.999 | -0.005 | -0.043 | -0.043 | 0 | 0.3 | 0.33 | 0.26 | 0.1 | 0.25 | 0.24 |
| 10 | 94719384 | rs7905234 | 1.614 | 0.274 | 0.205 | -3.995 | -0.002 | -0.016 | -0.016 | 2.452 | 0.1 | 0.08 | 0.17 | 0 | 0.1 | 0.08 |
| 10 | 94721905 | rs7069010 | -0.361 | 0.255 | 0.243 | -4.007 | -0.003 | -0.098 | -0.098 | 0 | 0.41 | 0.46 | 0.43 | 0.1 | 0.36 | 0.32 |
| 10 | 94725508 | rs12357515 | 1.553 | 0.273 | -0.139 | -3.99 | -0.002 | -0.016 | -0.016 | 0 | 0.1 | 0.08 | 0.17 | 0 | 0.1 | 0.08 |
| 10 | 94729753 | rs7477130 | -1.083 | 0.303 | -0.244 | -3.899 | -0.108 | 0.124 | 0.124 | 0 | 0.8 | 0.99 | 0.85 | 0.98 | 0.68 | 0.87 |
| 10 | 94736967 | rs2031616 | -0.724 | 0.299 | -0.232 | -3.902 | 0.011 | 0.112 | 0.112 | 0.26 | 0.46 | 0.64 | 0.56 | 0.23 | 0.39 | 0.34 |
| 10 | 94740527 | rs7082214 | -0.655 | 0.271 | -0.246 | -3.913 | 0.004 | 0.098 | 0.098 | 0 | 0.44 | 0.64 | 0.56 | 0.23 | 0.38 | 0.34 |
| 10 | 94744399 | rs1326329 | -1.086 | 0.345 | -0.303 | -3.882 | -0.108 | 0.124 | 0.124 | 0 | 0.8 | 0.99 | 0.85 | 0.98 | 0.68 | 0.87 |
| 10 | 94744904 | rs835247 | -0.739 | 0.347 | -0.3 | -3.881 | 0.012 | 0.113 | 0.113 | 0.236 | 0.46 | 0.64 | 0.56 | 0.23 | 0.38 | 0.34 |
| 10 | 94750745 | rs860668 | -1.087 | 0.396 | -0.291 | -3.839 | -0.108 | 0.118 | 0.118 | 0 | 0.8 | 0.99 | 0.85 | 0.98 | 0.68 | 0.87 |
| 10 | 94755208 | rs835238 | -0.702 | 0.335 | -0.096 | -3.712 | 0.004 | 0.097 | 0.097 | 0 | 0.44 | 0.64 | 0.55 | 0.22 | 0.38 | 0.34 |
| 10 | 94761925 | rs2486690 | -0.64 | -0.052 | -0.206 | -3.657 | -0.126 | 0.22 | 0.22 | 1.409 | 0.64 | 0.94 | 0.75 | 0.98 | 0.51 | 0.82 |
| 10 | 94762056 | rs1409385 | 1.046 | -0.245 | -0.07 | -3.784 | 0.037 | 0.133 | 0.133 | 0 | 0.39 | 0.25 | 0.41 | 0.89 | 0.34 | 0.56 |
| 10 | 94762340 | rs1409386 | 1.251 | -0.247 | -0.066 | -3.785 | 0.026 | 0.112 | 0.112 | 0 | 0.37 | 0.25 | 0.41 | 0.89 | 0.34 | 0.56 |
| 10 | 94764660 | rs7081888 | 1.6 | 0.124 | 0.596 | -3.739 | -0.012 | -0.013 | -0.013 | 0 | 0.1 | 0.17 | 0.14 | 0.07 | 0.07 | 0.17 |
| 10 | 94770488 | rs17108042 | 1.514 | 0.17 | 0.679 | -3.767 | 0.004 | -0.022 | -0.022 | 0 | 0.07 | 0.07 | 0.16 | 0 | 0.09 | 0.07 |
| 10 | 94771496 | rs11187240 | 1.6 | 0.081 | 0.687 | -3.755 | -0.008 | -0.013 | -0.013 | 0 | 0.1 | 0.17 | 0.14 | 0.07 | 0.08 | 0.17 |
| 10 | 94776479 | rs12413181 | 0.193 | -0.147 | 0.764 | -3.79 | -0.03 | -0.001 | -0.001 | 0 | 0.15 | 0.23 | 0.27 | 0.18 | 0.1 | 0.19 |
| 10 | 94778283 | rs2486687 | 0.734 | 0.006 | 0.464 | -3.77 | 0.029 | -0.053 | -0.053 | 0 | 0.1 | 0.21 | 0.18 | 0.01 | 0.11 | 0.08 |
| 10 | 94779550 | rs835256 | 0.761 | 0.007 | 0.464 | -3.77 | 0.03 | -0.053 | -0.053 | 0.74 | 0.1 | 0.21 | 0.18 | 0.01 | 0.11 | 0.08 |
| 10 | 94782735 | rs835259 | -0.666 | -0.143 | 0.308 | -3.926 | -0.119 | 0.237 | 0.237 | 1.659 | 0.62 | 0.94 | 0.74 | 0.98 | 0.52 | 0.81 |
| 10 | 94785856 | rs835264 | -1.486 | 0.042 | 0.211 | -3.976 | -0.035 | 0.048 | 0.048 | 0 | 0.91 | 1 | 0.95 | 1 | 0.88 | 0.94 |
| 10 | 94786259 | rs835265 | 0.73 | -0.299 | 0.443 | -3.802 | 0.029 | -0.053 | -0.053 | 1.938 | 0.1 | 0.21 | 0.18 | 0.01 | 0.11 | 0.08 |
| 10 | 94788835 | rs12416051 | 0.787 | -0.298 | 0.443 | -3.802 | 0.031 | -0.052 | -0.052 | 0.121 | 0.1 | 0.21 | 0.18 | 0.01 | 0.11 | 0.08 |
| 10 | 94788965 | rs9419763 | 0.787 | -0.298 | 0.443 | -3.802 | 0.031 | -0.052 | -0.052 | 0 | 0.1 | 0.21 | 0.18 | 0.01 | 0.11 | 0.08 |
| 10 | 94791099 | rs7895700 | 0.729 | -0.299 | 0.443 | -3.802 | 0.029 | -0.053 | -0.053 | 0 | 0.1 | 0.21 | 0.18 | 0.01 | 0.11 | 0.08 |
| 10 | 94796683 | rs835275 | 0.707 | -0.308 | 0.287 | -3.799 | 0.028 | -0.054 | -0.054 | 0.033 | 0.11 | 0.21 | 0.18 | 0.01 | 0.11 | 0.08 |
| 10 | 94797973 | rs2478236 | -0.255 | -0.398 | 0.213 | -3.887 | -0.019 | 0.262 | 0.262 | 1.25 | 0.54 | 0.89 | 0.61 | 0.91 | 0.44 | 0.64 |
| 10 | 94804530 | rs1750741 | 0.807 | -0.363 | 0.252 | -3.866 | 0.03 | -0.053 | -0.053 | 0.183 | 0.1 | 0.21 | 0.18 | 0.01 | 0.11 | 0.08 |
| 10 | 94805223 | rs10748589 | 1.415 | -0.425 | 0.286 | -3.889 | -0.022 | -0.062 | -0.062 | 0.893 | 0.36 | 0.51 | 0.3 | 0.78 | 0.31 | 0.53 |
| 10 | 94805665 | rs2486681 | -0.187 | -0.399 | 0.174 | -3.877 | -0.016 | 0.269 | 0.269 | 1.009 | 0.54 | 0.89 | 0.6 | 0.91 | 0.44 | 0.64 |
| 10 | 94809609 | rs835276 | -0.196 | -0.391 | 0.174 | -3.877 | -0.024 | 0.268 | 0.268 | 0.968 | 0.54 | 0.89 | 0.61 | 0.91 | 0.44 | 0.64 |
| 10 | 94810665 | rs856534 | 0.401 | -0.393 | 0.144 | -3.876 | 0.014 | 0.158 | 0.158 | 0.441 | 0.48 | 0.75 | 0.48 | 0.79 | 0.42 | 0.62 |
| 10 | 94811786 | rs835280 | 0.79 | -0.348 | 0.123 | -3.858 | 0.032 | -0.054 | -0.054 | 0 | 0.1 | 0.21 | 0.18 | 0.01 | 0.11 | 0.08 |
| 10 | 94813177 | rs11187262 | -0.221 | -0.284 | -0.443 | -3.77 | -0.024 | 0.268 | 0.268 | 0.968 | 0.46 | 0.11 | 0.39 | 0.09 | 0.56 | 0.36 |
| 10 | 94814907 | rs1339821 | -0.221 | -0.284 | -0.443 | -3.77 | -0.024 | 0.268 | 0.268 | 0.968 | 0.46 | 0.11 | 0.39 | 0.09 | 0.56 | 0.36 |
| 10 | 94819053 | rs8211 | -0.89 | -0.292 | -0.404 | -3.768 | 0.011 | 0.15 | 0.15 | 0.471 | 0.53 | 0.26 | 0.52 | 0.21 | 0.58 | 0.39 |
| 10 | 94819606 | rs7922067 | -0.285 | -0.105 | -0.893 | -3.707 | -0.023 | 0.269 | 0.269 | 1.009 | 0.46 | 0.11 | 0.39 | 0.09 | 0.56 | 0.36 |
| 10 | 94820860 | rs1538648 | 0.324 | 0.112 | -0.922 | -3.581 | -0.124 | 0.198 | 0.198 | 0.66 | 0.34 | 0.06 | 0.23 | 0.02 | 0.45 | 0.19 |
| 10 | 94832128 | rs11812460 | 1.049 | -0.056 | -0.569 | -3.254 | -0.007 | 0.056 | 0.056 | 0.346 | 0.3 | 0.2 | 0.27 | 0.78 | 0.29 | 0.52 |
| 10 | 94839642 | rs2068888 | 0.894 | -0.26 | -0.545 | -3.098 | -0.043 | 0.125 | 0.125 | 0 | 0.43 | 0.26 | 0.47 | 0.79 | 0.49 | 0.69 |
| 10 | 94840076 | rs17465590 | 0.942 | -0.361 | -0.56 | -3.131 | -0.052 | 0.088 | 0.088 | 0.757 | 0.14 | 0.01 | 0.07 | 0 | 0.15 | 0.05 |
| 10 | 94841107 | rs7905939 | 0.287 | -0.449 | -0.039 | -3.165 | -0.032 | -0.159 | -0.159 | 0.515 | 0.72 | 0.58 | 0.76 | 0.98 | 0.82 | 0.93 |
| 10 | 94842477 | rs10786068 | -0.491 | -0.221 | -0.178 | -3.125 | 0 | -0.005 | -0.005 | 0.001 | 0.74 | 0.74 | 0.73 | 0.82 | 0.71 | 0.77 |
| 10 | 94842827 | rs6583854 | -0.327 | -0.03 | -0.868 | -3.051 | -0.001 | -0.137 | -0.137 | 0 | 0.71 | 0.54 | 0.71 | 0.91 | 0.7 | 0.76 |
| 10 | 94843535 | rs12359135 | 0.277 | 0.299 | -0.773 | -2.611 | 0.005 | 0.176 | 0.176 | 0.004 | 0.48 | 0.26 | 0.5 | 0.79 | 0.54 | 0.7 |
| 10 | 94845546 | rs10786069 | 0.069 | 0.568 | -1.022 | -2.421 | 0.002 | 0.015 | 0.015 | 0 | 0.52 | 0.47 | 0.63 | 0.81 | 0.55 | 0.7 |
| 10 | 94846093 | rs7080494 | -0.059 | 0.578 | -0.839 | -2.148 | 0.001 | 0.015 | 0.015 | 0 | 0.48 | 0.46 | 0.7 | 0.82 | 0.51 | 0.68 |
| 10 | 94847115 | rs913422 | -0.486 | 0.325 | -0.813 | -2.961 | -0.002 | -0.077 | -0.077 | 0 | 0.41 | 0.56 | 0.49 | 0.92 | 0.43 | 0.62 |
| 10 | 94859247 | rs11187279 | 0.514 | 0.403 | -0.806 | -3.094 | -0.002 | -0.027 | -0.027 | 0 | 0.38 | 0.32 | 0.27 | 0.05 | 0.38 | 0.2 |

| | | | | | | | | | | | | | | | | |
|---|---|---|---|---|---|---|---|---|---|---|---|---|---|---|---|---|
| 10 | 94866678 | rs17108179 | 0.105 | 0.646 | -0.949 | -2.805 | -0.026 | 0.035 | 0.035 | 0 | 0.07 | 0.02 | 0.05 | 0 | 0.11 | 0.05 |
| 10 | 94868454 | rs10882146 | 0.083 | 0.239 | -0.574 | -3.078 | 0.008 | -0.049 | -0.049 | 0.124 | 0.21 | 0.22 | 0.22 | 0.04 | 0.25 | 0.16 |
| 10 | 94879204 | rs17108206 | 0.549 | -0.668 | -0.198 | -3.289 | -0.076 | -0.083 | -0.083 | 0.032 | 0.13 | 0.28 | 0.04 | 0.01 | 0.02 | 0.04 |
| 10 | 94881288 | rs12778515 | 0.678 | -0.156 | -0.009 | -3.761 | 0 | -0.081 | -0.081 | 0 | 0.45 | 0.37 | 0.27 | 0.03 | 0.41 | 0.24 |
| 10 | 94881612 | rs10882148 | -0.378 | -0.381 | 0.211 | -3.937 | -0.001 | -0.031 | -0.031 | 0.683 | 0.32 | 0.32 | 0.21 | 0.07 | 0.36 | 0.2 |
| 10 | 94895653 | rs17466747 | 1.644 | -0.287 | 0.099 | -3.987 | 0.011 | 0.067 | 0.067 | 0 | 0.37 | 0.19 | 0.21 | 0.03 | 0.3 | 0.18 |
| 10 | 94899090 | rs7081913 | 0.563 | -0.415 | -0.066 | -3.958 | 0.005 | -0.088 | -0.088 | 0.015 | 0.59 | 0.52 | 0.74 | 0.94 | 0.52 | 0.74 |
| 10 | 94899803 | rs4918430 | -0.434 | -0.83 | -0.099 | -4.053 | -0.024 | -0.089 | -0.089 | 0.068 | 0.84 | 0.58 | 0.89 | 0.97 | 0.86 | 0.95 |
| 10 | 94900653 | rs4918433 | 1.081 | -0.657 | -0.144 | -4.159 | 0.003 | -0.037 | -0.037 | 4.257 | 0.48 | 0.33 | 0.31 | 0.03 | 0.48 | 0.25 |
| 10 | 94901328 | rs7900846 | 0.603 | -0.667 | -0.088 | -4.177 | -0.002 | 0.016 | 0.016 | 0 | 0.65 | 0.68 | 0.78 | 0.94 | 0.71 | 0.67 |
| 10 | 94903293 | rs932848 | -0.325 | -0.751 | -0.09 | -4.18 | -0.024 | -0.051 | -0.051 | 0 | 0.86 | 0.72 | 0.89 | 0.96 | 0.94 | 0.84 |
| 10 | 94903727 | rs978604 | 0.331 | -0.632 | -0.448 | -4.133 | -0.078 | 0.129 | 0.129 | 0 | 0.21 | 0.03 | 0.11 | 0.03 | 0.24 | 0.18 |
| 10 | 94908144 | rs17382342 | 2.103 | -0.699 | -0.314 | -4.131 | 0 | -0.081 | -0.081 | 0 | 0.38 | 0.32 | 0.25 | 0.03 | 0.37 | 0.18 |
| 10 | 94912188 | rs1927473 | -0.811 | -0.626 | -0.185 | -3.988 | -0.035 | -0.13 | -0.13 | 0 | 0.27 | 0.53 | 0.19 | 0.07 | 0.17 | 0.21 |
| 10 | 94920094 | rs10786074 | 1.304 | -0.512 | -0.241 | -3.954 | -0.002 | 0.002 | 0.002 | 0.047 | 0.48 | 0.37 | 0.65 | 0.9 | 0.5 | 0.59 |
| 10 | 94925825 | rs7085051 | -0.964 | -0.371 | 0.168 | -3.96 | 0 | -0.063 | -0.063 | 0.001 | 0.39 | 0.45 | 0.23 | 0.1 | 0.4 | 0.26 |
| 10 | 94938228 | rs6583867 | 1.553 | -0.312 | 0.127 | -3.909 | 0.002 | 0.014 | 0.014 | 0 | 0.45 | 0.33 | 0.33 | 0.03 | 0.39 | 0.29 |
| 10 | 94946123 | rs7919934 | 1.579 | -0.43 | 0.186 | -3.87 | 0 | -0.004 | -0.004 | 0 | 0.45 | 0.34 | 0.33 | 0.03 | 0.41 | 0.29 |
| 10 | 94948753 | rs4144915 | -1.093 | -0.381 | 0.428 | -3.836 | -0.009 | 0.036 | 0.036 | 0 | 0.44 | 0.39 | 0.63 | 0.92 | 0.56 | 0.64 |
| 10 | 94950273 | rs12778014 | -0.657 | -0.28 | 0.432 | -3.617 | -0.007 | 0.004 | 0.004 | 0 | 0.28 | 0.28 | 0.18 | 0.06 | 0.33 | 0.19 |
| 10 | 94950785 | rs12783719 | 0.219 | 0.045 | 0.094 | -3.364 | -0.078 | 0.071 | 0.071 | 0 | 0.15 | 0.02 | 0.1 | 0.02 | 0.24 | 0.15 |
| 10 | 94953876 | rs12254292 | -0.413 | -0.976 | 0.663 | -2.93 | 0.002 | -0.001 | -0.001 | 2.914 | 0.08 | 0.26 | 0.03 | 0.04 | 0.04 | 0.06 |
| 10 | 94958183 | rs913419 | 0.414 | -0.971 | 0.398 | -2.928 | -0.002 | -0.006 | -0.006 | 0 | 0.91 | 0.65 | 0.96 | 0.96 | 0.96 | 0.94 |
| 10 | 94967076 | rs7070947 | 0.552 | -0.824 | -0.183 | -3.374 | -0.003 | -0.083 | -0.083 | 1.445 | 0.54 | 0.52 | 0.41 | 0.05 | 0.56 | 0.37 |
| 10 | 94968216 | rs11187324 | 0.24 | -0.988 | -0.499 | -3.554 | -0.037 | 0.068 | 0.068 | 0 | 0.14 | 0.01 | 0.06 | 0.02 | 0.15 | 0.12 |
| 10 | 94968665 | rs11596790 | 0.824 | -1.059 | -0.495 | -3.64 | -0.014 | -0.123 | -0.123 | 0.745 | 0.6 | 0.53 | 0.44 | 0.05 | 0.65 | 0.41 |
| 10 | 94973750 | rs1927463 | 1.438 | -1.221 | -0.857 | -3.928 | -0.008 | 0.059 | 0.059 | 0 | 0.44 | 0.3 | 0.38 | 0.03 | 0.5 | 0.31 |
| 10 | 94974129 | rs6583871 | 0.916 | -1.494 | -0.781 | -4.028 | 0.036 | 0.065 | 0.065 | 0.088 | 0.62 | 0.73 | 0.83 | 0.92 | 0.71 | 0.78 |
| 10 | 94974675 | rs7083492 | 0.412 | -1.511 | -0.815 | -4.069 | -0.023 | -0.115 | -0.115 | 0 | 0.73 | 0.63 | 0.87 | 0.99 | 0.8 | 0.9 |
| 10 | 94977719 | rs10882185 | 0.323 | -1.476 | -0.869 | -4.045 | 0 | -0.022 | -0.022 | 3.389 | 0.15 | 0.15 | 0.08 | 0.02 | 0.14 | 0.06 |
| 10 | 94987241 | rs12355604 | 0.576 | -1.41 | -0.898 | -3.942 | -0.034 | 0.011 | 0.011 | 0 | 0.14 | 0.01 | 0.06 | 0.08 | 0.15 | 0.12 |
| 10 | 94988159 | rs10882188 | 1.101 | -1.349 | -0.734 | -3.904 | -0.004 | -0.046 | -0.046 | 0.129 | 0.58 | 0.56 | 0.45 | 0.11 | 0.63 | 0.43 |
| 10 | 94996522 | rs1927465 | -0.379 | -1.267 | -1.15 | -3.746 | 0 | -0.029 | -0.029 | 0.276 | 0.2 | 0.18 | 0.11 | 0.01 | 0.17 | 0.08 |
| 10 | 95006977 | rs11187345 | 0.542 | -1.204 | -1.391 | -3.667 | 0.017 | 0.023 | 0.023 | 0 | 0.13 | 0.09 | 0.07 | 0.01 | 0.08 | 0.04 |
| 10 | 95010777 | rs11596635 | 1.594 | -1.11 | -1.489 | -3.867 | -0.086 | 0.244 | 0.244 | 0.039 | 0.41 | 0.13 | 0.33 | 0.02 | 0.45 | 0.3 |
| 10 | 95018932 | rs12767937 | 1.591 | -1.152 | -1.499 | -3.882 | -0.086 | 0.243 | 0.243 | 0.087 | 0.41 | 0.14 | 0.33 | 0.02 | 0.46 | 0.3 |
| 10 | 95019584 | rs12359829 | -0.502 | -0.932 | -1.671 | -3.023 | -0.021 | 0.018 | 0.018 | 0 | 0.25 | 0.1 | 0.23 | 0.22 | 0.25 | 0.18 |
| 10 | 95023926 | rs11818234 | 0.516 | -1.316 | -1.409 | -3.11 | -0.005 | -0.073 | -0.073 | 0 | 0.12 | 0.34 | 0.11 | 0.02 | 0.06 | 0.15 |
| 10 | 95032661 | rs10882205 | 0.115 | -1.646 | -1.62 | -3.033 | 0.002 | -0.007 | -0.007 | 0 | 0.28 | 0.35 | 0.43 | 0.74 | 0.3 | 0.48 |
| 10 | 95036575 | rs787652 | 1.24 | -1.76 | -1.728 | -3.123 | -0.107 | 0.137 | 0.137 | 0.007 | 0.4 | 0.1 | 0.37 | 0.16 | 0.52 | 0.22 |
| 10 | 95037095 | rs12772206 | 0.462 | -1.731 | -1.642 | -3.084 | 0.004 | 0.011 | 0.011 | 0 | 0.11 | 0.04 | 0.06 | 0.07 | 0.09 | 0.12 |
| 10 | 95039993 | rs12767318 | 0.616 | -1.664 | -1.588 | -3.012 | 0 | -0.001 | -0.001 | 0 | 0.09 | 0.01 | 0.04 | 0.07 | 0.06 | 0.11 |
| 10 | 95041978 | rs1615300 | 0.585 | -1.717 | -1.519 | -2.88 | 0.052 | 0.139 | 0.139 | 0.294 | 0.51 | 0.14 | 0.47 | 0.26 | 0.62 | 0.35 |
| 10 | 95046048 | rs2761304 | 0.744 | -1.348 | -1.203 | -2.864 | 0.021 | 0.072 | 0.072 | 0 | 0.51 | 0.44 | 0.46 | 0.15 | 0.6 | 0.28 |
| 10 | 95046375 | rs2797571 | 0.453 | -1.397 | -1.072 | -2.888 | -0.001 | -0.016 | -0.016 | 0 | 0.54 | 0.62 | 0.48 | 0.15 | 0.61 | 0.28 |
| 10 | 95048686 | rs1776069 | 0.719 | -1.385 | -1.1 | -2.882 | 0.009 | 0.021 | 0.021 | 0 | 0.52 | 0.48 | 0.46 | 0.14 | 0.6 | 0.28 |
| 10 | 95050550 | rs2797573 | 1.499 | -1.331 | -1.269 | -2.957 | -0.109 | 0.119 | 0.119 | 0 | 0.28 | 0.1 | 0.21 | 0.08 | 0.42 | 0.18 |
| 10 | 95050819 | rs2761308 | 0.658 | -1.447 | -1.355 | -3.062 | -0.048 | 0.216 | 0.216 | 0.527 | 0.43 | 0.15 | 0.3 | 0.08 | 0.56 | 0.23 |
| 10 | 95051926 | rs12767227 | 0.754 | -1.272 | -1.267 | -2.934 | -0.007 | -0.05 | -0.05 | 0 | 0.17 | 0.19 | 0.11 | 0 | 0.17 | 0.05 |
| 10 | 95053571 | rs10882212 | -0.426 | -1.314 | -1.392 | -2.953 | 0.006 | -0.025 | -0.025 | 0 | 0.35 | 0.39 | 0.5 | 0.77 | 0.36 | 0.59 |
| 10 | 95063196 | rs7084389 | 0.445 | -0.536 | -1.44 | -2.269 | -0.025 | -0.004 | -0.004 | 0 | 0.08 | 0.14 | 0.03 | 0.07 | 0.01 | 0.1 |
| 10 | 95064137 | rs10882215 | 0.18 | 0.198 | -1.639 | -2.148 | -0.011 | -0.047 | -0.047 | 0 | 0.24 | 0.36 | 0.25 | 0.07 | 0.2 | 0.1 |
| 10 | 95064403 | rs2797577 | -0.111 | 1.217 | -1.935 | -1.289 | -0.087 | 0.107 | 0.107 | 0 | 0.29 | 0.12 | 0.21 | 0.08 | 0.41 | 0.18 |
| 10 | 95069145 | rs2761331 | -0.563 | 1.539 | -1.431 | -0.917 | -0.009 | 0.006 | 0.006 | 0.146 | 0.33 | 0.3 | 0.45 | 0.78 | 0.45 | 0.51 |
| 10 | 95070037 | rs2761286 | 0.26 | 0.922 | -1.594 | -1.475 | 0 | 0.004 | 0.004 | 0 | 0.86 | 0.85 | 0.79 | 0.91 | 0.85 | 0.89 |
| 10 | 95072906 | rs787666 | 0.271 | 0.843 | -1.301 | -1.49 | -0.002 | -0.007 | -0.007 | 0 | 0.85 | 0.79 | 0.78 | 0.91 | 0.85 | 0.89 |
| 10 | 95073046 | rs787667 | 0.59 | 1.251 | -1.901 | -2.099 | -0.047 | 0.094 | 0.094 | 0.238 | 0.62 | 0.76 | 0.6 | 0.9 | 0.51 | 0.78 |
| 10 | 95074412 | rs11596560 | 0.911 | 0.14 | -2.02 | -2.619 | -0.004 | -0.024 | -0.024 | 0 | 0.14 | 0.15 | 0.06 | 0 | 0.08 | 0.03 |
| 10 | 95077553 | rs11187382 | 0.551 | 0.233 | -1.977 | -2.685 | -0.013 | -0.008 | -0.008 | 0 | 0.23 | 0.2 | 0.18 | 0.01 | 0.33 | 0.11 |
| 10 | 95080747 | rs787679 | 0.399 | -0.075 | -1.923 | -2.868 | -0.028 | -0.059 | -0.059 | 0.455 | 0.71 | 0.59 | 0.73 | 0.91 | 0.78 | 0.88 |
| 10 | 95083039 | rs11594445 | 0.485 | -0.21 | -2.513 | -2.891 | 0.009 | -0.006 | -0.006 | 0 | 0.08 | 0.01 | 0.04 | 0.07 | 0.06 | 0.05 |
| 10 | 95083725 | rs2259185 | -0.49 | 0.048 | -2.303 | -2.801 | 0 | 0.003 | 0.003 | 0 | 0.86 | 0.87 | 0.8 | 0.91 | 0.86 | 0.91 |
| 10 | 95089013 | rs1609572 | 0.484 | 0.081 | -2.328 | -2.804 | 0.002 | 0.159 | 0.159 | 0.622 | 0.52 | 0.74 | 0.56 | 0.9 | 0.45 | 0.78 |
| 10 | 95089762 | rs17477816 | 0.655 | 0.365 | -2.368 | -2.612 | -0.098 | 0.118 | 0.118 | 0.252 | 0.21 | 0.03 | 0.18 | 0.01 | 0.33 | 0.11 |
| 10 | 95091023 | rs10509650 | 0.057 | 0.379 | -2.313 | -2.557 | -0.104 | 0.223 | 0.223 | 1.489 | 0.33 | 0.03 | 0.23 | 0.01 | 0.41 | 0.12 |
| 10 | 95097537 | rs1614065 | -0.489 | 0.015 | -1.599 | -2.845 | 0.003 | -0.054 | -0.054 | 0.922 | 0.77 | 0.78 | 0.66 | 0.98 | 0.76 | 0.85 |
| 10 | 95098144 | rs10786089 | -0.048 | -0.514 | -2.283 | -3.426 | 0.047 | 0.174 | 0.174 | 1.627 | 0.38 | 0.11 | 0.41 | 0.88 | 0.33 | 0.64 |
| 10 | 95101480 | rs1408031 | 0.22 | -0.664 | -2.479 | -3.608 | 0.027 | 0.065 | 0.065 | 0 | 0.25 | 0.11 | 0.35 | 0.88 | 0.23 | 0.6 |
| 10 | 95102221 | rs787695 | -0.777 | -0.63 | -1.902 | -3.718 | -0.001 | -0.043 | -0.043 | 0 | 0.72 | 0.38 | 0.63 | 0.98 | 0.75 | 0.85 |

| | | | | | | | | | | | | | | | |
|---|---|---|---|---|---|---|---|---|---|---|---|---|---|---|---|
| 10 | 95102371 | rs812012 | 0.862 | -0.817 | -1.98 | -3.772 | -0.091 | 0.151 | 0.151 | 0.167 | 0.34 | 0.09 | 0.22 | 0.09 | 0.44 | 0.21 |
| 10 | 95110026 | rs878267 | 0.237 | -1.534 | -2.007 | -3.84 | 0.034 | 0.058 | 0.058 | 0 | 0.26 | 0.15 | 0.35 | 0.88 | 0.24 | 0.57 |
| 10 | 95117549 | rs787630 | 1.568 | -1.191 | -1.42 | -3.903 | -0.124 | 0.102 | 0.102 | 0.474 | 0.28 | 0.09 | 0.21 | 0.1 | 0.41 | 0.25 |
| 10 | 95120792 | rs10882220 | -0.217 | -1.415 | -1.018 | -4.023 | 0.014 | 0.006 | 0.006 | 0 | 0.43 | 0.38 | 0.42 | 0.88 | 0.34 | 0.6 |
| 10 | 95122562 | rs4919057 | 2.222 | -1.704 | -1.331 | -3.602 | -0.036 | -0.062 | -0.062 | 0 | 0.1 | 0.01 | 0.22 | 0.73 | 0.11 | 0.47 |
| 10 | 95124531 | rs787639 | 1.601 | -1.302 | -1.207 | -3.601 | -0.14 | 0.096 | 0.096 | 0.584 | 0.27 | 0.05 | 0.21 | 0.1 | 0.41 | 0.25 |
| 10 | 95124673 | rs787640 | -0.084 | -0.564 | -0.608 | -3.229 | 0.008 | -0.112 | -0.112 | 0 | 0.73 | 0.63 | 0.65 | 0.98 | 0.74 | 0.85 |
| 10 | 95132905 | rs787647 | -0.927 | -0.06 | -0.309 | -2.378 | 0.021 | -0.062 | -0.062 | 0 | 0.85 | 0.8 | 0.87 | 0.99 | 0.81 | 0.85 |
| 10 | 95136032 | rs787650 | 0.688 | -0.314 | -0.206 | -2.373 | 0.004 | 0.049 | 0.049 | 0 | 0.52 | 0.39 | 0.3 | 0.12 | 0.54 | 0.24 |
| 10 | 95139526 | rs10882229 | -0.867 | 0.017 | -0.374 | -2.298 | 0.032 | 0.071 | 0.071 | 4.149 | 0.43 | 0.7 | 0.47 | 0.85 | 0.36 | 0.64 |
| 10 | 95148470 | rs696178 | -0.397 | 0.368 | 0.615 | -1.614 | 0.003 | -0.048 | -0.048 | 0.974 | 0.82 | 0.81 | 0.69 | 0.98 | 0.81 | 0.88 |
| 10 | 95148835 | rs17108594 | -0.203 | 0.117 | 0.749 | -1.611 | -0.009 | -0.041 | -0.041 | 0 | 0.07 | 0.14 | 0.05 | 0 | 0.05 | 0.01 |
| 10 | 95150259 | rs696179 | 0.117 | 0.275 | 0.686 | -1.572 | -0.013 | -0.134 | -0.134 | 0.001 | 0.75 | 0.62 | 0.66 | 0.98 | 0.8 | 0.89 |
| 10 | 95151622 | rs720183 | -1.235 | 0.42 | 1.263 | -0.54 | 0.057 | 0.008 | 0.008 | 0.022 | 0.1 | 0.28 | 0.11 | 0.09 | 0.13 | 0.15 |
| 10 | 95158302 | rs788357 | 0.902 | 0.106 | 1.356 | -0.585 | -0.008 | -0.064 | -0.064 | 0 | 0.9 | 0.73 | 0.93 | 1 | 0.93 | 0.98 |
| 10 | 95163809 | rs12262721 | -1.004 | 0.223 | 1.911 | -0.531 | 0.051 | -0.019 | -0.019 | 0.582 | 0.07 | 0.27 | 0.05 | 0 | 0.05 | 0.01 |
| 10 | 95164470 | rs10509651 | -0.709 | 0.194 | 2.032 | -0.544 | -0.006 | -0.045 | -0.045 | 0 | 0.08 | 0.2 | 0.05 | 0 | 0.07 | 0.02 |
| 10 | 95166455 | rs1746489 | -1.557 | 0.453 | 1.959 | 0.184 | 0.134 | -0.03 | -0.03 | 0.229 | 0.1 | 0.38 | 0.1 | 0.03 | 0.13 | 0.09 |
| 10 | 95173966 | rs10882230 | 0.65 | 0.011 | 1.05 | 0.448 | -0.021 | 0.197 | 0.197 | 0 | 0.92 | 0.81 | 0.82 | 0.34 | 0.96 | 0.84 |
| 10 | 95176933 | rs10509657 | -0.757 | 0.185 | 0.554 | 0.349 | 0.002 | -0.021 | -0.021 | 0 | 0.1 | 0.13 | 0.07 | 0.03 | 0.1 | 0.1 |
| 10 | 95189263 | rs10882232 | -0.142 | -0.589 | 0.444 | -0.206 | 0.013 | 0.016 | 0.016 | 0 | 0.54 | 0.4 | 0.55 | 0.96 | 0.38 | 0.55 |
| 10 | 95190568 | rs701873 | 0.515 | -0.628 | 0.046 | -0.38 | -0.1 | 0.165 | 0.165 | 0 | 0.33 | 0.09 | 0.37 | 0.04 | 0.53 | 0.42 |
| 10 | 95194711 | rs7091625 | 0.414 | -1.203 | -0.508 | -0.578 | 0.001 | 0.028 | 0.028 | 0.092 | 0.18 | 0.08 | 0.13 | 0.07 | 0.19 | 0.09 |
| 10 | 95202332 | rs7923523 | 1.133 | -1.26 | -0.596 | -0.481 | -0.005 | 0.034 | 0.034 | 2.187 | 0.3 | 0.04 | 0.28 | 0.21 | 0.21 | 0.2 |
| 10 | 95203670 | rs7075356 | 0.523 | -1.327 | -0.403 | -0.533 | -0.023 | 0.083 | 0.083 | 3.796 | 0.37 | 0.04 | 0.31 | 0.21 | 0.27 | 0.22 |
| 10 | 95212070 | rs2985079 | -1.303 | -1.334 | -0.205 | -0.517 | 0.028 | 0.07 | 0.07 | 1.145 | 0.69 | 0.81 | 0.72 | 0.89 | 0.79 | 0.82 |
| 10 | 95213371 | rs11187433 | -0.064 | -1.422 | 0.341 | -0.523 | -0.013 | -0.045 | -0.045 | 0 | 0.54 | 0.45 | 0.52 | 0.12 | 0.69 | 0.52 |
| 10 | 95216623 | rs701882 | -0.108 | -1.393 | 0.354 | 0.17 | -0.014 | 0.001 | 0.001 | 0.396 | 0.55 | 0.45 | 0.63 | 0.52 | 0.7 | 0.59 |
| 10 | 95217064 | rs701881 | -0.842 | -1.364 | 0.485 | 0.145 | -0.012 | -0.019 | -0.019 | 1.589 | 0.64 | 0.56 | 0.69 | 0.79 | 0.78 | 0.68 |
| 10 | 95228328 | rs1880260 | 0.781 | -1.085 | 0.422 | 0.175 | -0.083 | 0.158 | 0.158 | 0.382 | 0.35 | 0.17 | 0.36 | 0.04 | 0.51 | 0.57 |
| 10 | 95228717 | rs10882240 | 0.982 | -0.469 | 1.456 | 1.263 | -0.04 | -0.009 | -0.009 | 0 | 0.76 | 0.5 | 0.72 | 0.24 | 0.81 | 0.79 |
| 10 | 95233393 | rs871427 | -0.565 | -0.326 | 1.044 | 1.244 | 0.008 | -0.025 | -0.025 | 0 | 0.19 | 0.13 | 0.22 | 0.5 | 0.18 | 0.14 |
| 10 | 95233856 | rs1105340 | 0.503 | -0.324 | 1.075 | 1.242 | -0.023 | 0.133 | 0.133 | 0.153 | 0.79 | 0.65 | 0.77 | 0.49 | 0.82 | 0.86 |
| 10 | 95235408 | rs7922093 | -0.568 | -0.232 | 0.108 | 1.282 | 0.007 | -0.022 | -0.022 | 0 | 0.19 | 0.13 | 0.22 | 0.51 | 0.18 | 0.14 |
| 10 | 95249605 | rs947603 | -1.17 | -0.337 | 0.2 | 0.787 | 0.016 | 0.016 | 0.016 | 0 | 0.3 | 0.24 | 0.31 | 0.76 | 0.25 | 0.28 |
| 10 | 95250157 | rs7917033 | 2.484 | -0.443 | 0.643 | 0.83 | -0.015 | -0.023 | -0.023 | 1.006 | 0.88 | 0.23 | 0.91 | 0.96 | 0.92 | 0.97 |
| 10 | 95252787 | rs954862 | 1.81 | 0.049 | 0.596 | 0.828 | -0.016 | 0.147 | 0.147 | 0.822 | 0.57 | 0.21 | 0.44 | 0.13 | 0.57 | 0.59 |
| 10 | 95253230 | rs11187464 | -1.351 | 0.116 | 1.201 | 0.987 | -0.023 | -0.055 | -0.055 | 0.005 | 0.26 | 0.57 | 0.18 | 0.15 | 0.2 | 0.24 |
| 10 | 95255581 | rs7904542 | 1.052 | 0.162 | 1.175 | 1.389 | 0.013 | 0.155 | 0.155 | 0 | 0.9 | 0.83 | 0.66 | 0.49 | 0.89 | 0.9 |
| 10 | 95256144 | rs1880264 | -1.175 | 0.147 | 1.676 | 1.34 | -0.002 | 0.002 | 0.002 | 0.226 | 0.25 | 0.27 | 0.16 | 0.35 | 0.21 | 0.3 |
| 10 | 95260965 | rs11187476 | -1.302 | 0.133 | 1.72 | 1.299 | -0.008 | 0.007 | 0.007 | 0.355 | 0.26 | 0.37 | 0.18 | 0.29 | 0.22 | 0.29 |
| 10 | 95264507 | rs10509660 | -1.222 | 0.186 | 1.727 | 1.316 | -0.003 | 0.003 | 0.003 | 0.183 | 0.26 | 0.32 | 0.17 | 0.29 | 0.22 | 0.29 |
| 10 | 95266683 | rs947599 | 1.986 | 0.234 | 1.131 | 1.316 | -0.011 | 0.109 | 0.109 | 0.503 | 0.57 | 0.27 | 0.45 | 0.14 | 0.57 | 0.59 |
| 10 | 95270665 | rs10882254 | -1.273 | 0.505 | 1.28 | 1.535 | -0.004 | 0.031 | 0.031 | 0 | 0.1 | 0.08 | 0.32 | 0.57 | 0.12 | 0.1 |
| 10 | 95274972 | rs10882259 | 1.407 | -0.279 | 1.22 | 1.153 | 0 | -0.001 | -0.001 | 0.461 | 0.62 | 0.42 | 0.65 | 0.53 | 0.64 | 0.67 |
| 10 | 95276718 | rs7072484 | -2.16 | 0.231 | 1.661 | 1.255 | 0.059 | 0.034 | 0.034 | 1.274 | 0.05 | 0.32 | 0.04 | 0 | 0.02 | 0 |
| 10 | 95278829 | rs2280605 | 1.616 | 0.232 | 1.384 | 1.482 | 0 | 0.025 | 0.025 | 0.651 | 0.74 | 0.39 | 0.83 | 0.59 | 0.74 | 0.72 |
| 10 | 95294094 | rs7085858 | 1.868 | 0.01 | 1.555 | 1.605 | 0.005 | -0.021 | -0.021 | 0 | 0.7 | 0.28 | 0.74 | 0.54 | 0.76 | 0.69 |
| 10 | 95303052 | rs11187509 | -1.531 | -0.024 | 2.313 | 1.718 | -0.013 | 0.05 | 0.05 | 0 | 0.09 | 0.24 | 0.03 | 0.13 | 0.03 | 0.07 |
| 10 | 95305939 | rs7893412 | 1.34 | -0.032 | 2.315 | 1.708 | -0.008 | 0.101 | 0.101 | 0 | 0.88 | 0.77 | 0.85 | 0.67 | 0.93 | 0.79 |
| 10 | 95309488 | rs7893971 | -1.301 | -0.03 | 2.343 | 1.709 | -0.003 | 0.07 | 0.07 | 0 | 0.11 | 0.15 | 0.14 | 0.33 | 0.07 | 0.21 |
| 10 | 95310539 | rs7091291 | -2.365 | -0.045 | 2.359 | 1.708 | 0.009 | -0.008 | -0.008 | 1.778 | 0.18 | 0.5 | 0.12 | 0.13 | 0.17 | 0.11 |
| 10 | 95310712 | rs4611144 | 2.57 | -0.028 | 1.876 | 1.655 | 0.012 | -0.043 | -0.043 | 0.786 | 0.7 | 0.24 | 0.74 | 0.54 | 0.77 | 0.68 |
| 10 | 95320440 | rs1889931 | -2.884 | -0.382 | 2.118 | 1.487 | -0.007 | 0.03 | 0.03 | 0.146 | 0.28 | 0.62 | 0.35 | 0.48 | 0.22 | 0.35 |
| 10 | 95322044 | rs940390 | -2.161 | -0.474 | 2.42 | 1.687 | -0.018 | 0.166 | 0.166 | 0.066 | 0.19 | 0.39 | 0.34 | 0.45 | 0.15 | 0.22 |
| 10 | 95324503 | rs10882267 | -2.416 | -0.406 | 2.195 | 1.423 | -0.012 | 0.088 | 0.088 | 0 | 0.24 | 0.36 | 0.31 | 0.47 | 0.19 | 0.29 |
| 10 | 95327429 | rs11187515 | -0.513 | -0.224 | 1.583 | 0.705 | 0.01 | -0.047 | -0.047 | 0.47 | 0.09 | 0.16 | 0.11 | 0 | 0.13 | 0.02 |
| 10 | 95332517 | rs12219199 | -1.416 | -0.416 | 1.34 | 1.104 | -0.012 | 0.092 | 0.092 | 0 | 0.1 | 0.17 | 0.13 | 0.33 | 0.05 | 0.25 |
| 10 | 95340811 | rs8181419 | -2.081 | -0.354 | 1.536 | -0.272 | -0.014 | -0.058 | -0.058 | 0 | 0.2 | 0.53 | 0.12 | 0.09 | 0.16 | 0.07 |
| 10 | 95343779 | rs11187529 | -2.284 | -0.356 | 1.612 | -0.39 | -0.01 | -0.033 | -0.033 | 0.227 | 0.07 | 0.21 | 0.05 | 0 | 0.04 | 0.03 |
| 10 | 95346340 | rs11187533 | -1.054 | -0.828 | 1.26 | -0.989 | -0.006 | -0.036 | -0.036 | 0 | 0.34 | 0.4 | 0.2 | 0.1 | 0.28 | 0.37 |
| 10 | 95346386 | rs11187534 | -0.087 | -1.283 | 0.884 | -1.393 | -0.017 | -0.057 | -0.057 | 0 | 0.16 | 0.18 | 0.08 | 0 | 0.1 | 0.05 |
| 10 | 95346555 | rs17108968 | 0.062 | -1.254 | 0.928 | -1.391 | 0.019 | -0.047 | -0.047 | 2.741 | 0.08 | 0.19 | 0.05 | 0 | 0.09 | 0.02 |
| 10 | 95346761 | rs11187536 | -0.797 | -1.248 | 0.818 | -1.385 | -0.006 | -0.036 | -0.036 | 0 | 0.34 | 0.4 | 0.2 | 0.1 | 0.28 | 0.37 |
| 10 | 95348905 | rs10882273 | -1.591 | -1.162 | 0.598 | -1.363 | 0.022 | 0.042 | 0.042 | 0 | 0.51 | 0.62 | 0.28 | 0.11 | 0.39 | 0.42 |
| 10 | 95351310 | rs34571439 | -0.585 | -1.389 | 0.63 | -0.955 | -0.02 | -0.043 | -0.043 | 0.103 | 0.28 | 0.37 | 0.17 | 0.1 | 0.2 | 0.35 |
| 10 | 95352983 | rs17108991 | -0.345 | -1.301 | -0.066 | -0.878 | 0.015 | 0.025 | 0.025 | 0 | 0.26 | 0.2 | 0.15 | 0.1 | 0.2 | 0.35 |
| 10 | 95355617 | rs7094671 | -1.18 | -1.421 | -0.028 | -0.912 | 0.024 | 0.022 | 0.022 | 0 | 0.41 | 0.34 | 0.2 | 0.09 | 0.25 | 0.39 |
| 10 | 95357105 | rs11187545 | -0.716 | -0.561 | 0.071 | -0.36 | -0.005 | -0.033 | -0.033 | 0 | 0.15 | 0.14 | 0.06 | 0 | 0.08 | 0.04 |

| | | | | | | | | | | | | | | | | |
|---|---|---|---|---|---|---|---|---|---|---|---|---|---|---|---|---|
| 10 | 95362050 | rs3758538 | -1.206 | -0.559 | 0.069 | -0.383 | -0.029 | -0.08 | -0.08 | 0 | 0.24 | 0.31 | 0.13 | 0.06 | 0.14 | 0.18 |
| 10 | 95367001 | rs1108197 | -0.584 | 0.048 | -0.47 | -0.067 | 0.002 | 0.009 | 0.009 | 0 | 0.41 | 0.67 | 0.62 | 0.32 | 0.42 | 0.32 |
| 10 | 95368516 | rs12573026 | -0.597 | 0.454 | -0.818 | 0.78 | -0.007 | -0.048 | -0.048 | 0.445 | 0.16 | 0.07 | 0.06 | 0.49 | 0.14 | 0.41 |
| 10 | 95372209 | rs12252137 | -0.941 | 0.291 | -0.41 | 0.591 | -0.003 | -0.006 | -0.006 | 0 | 0.15 | 0.2 | 0.13 | 0.11 | 0.12 | 0.09 |
| 10 | 95376046 | rs6583900 | -0.79 | 0.113 | -0.746 | 1.455 | 0.013 | 0.007 | 0.007 | 0 | 0.1 | 0.07 | 0.03 | 0.32 | 0.04 | 0.26 |
| 10 | 95378547 | rs650058 | -0.141 | 0.281 | -1.002 | 0.75 | -0.014 | 0 | 0 | 0 | 0.73 | 0.81 | 0.5 | 0.67 | 0.69 | 0.74 |
| 10 | 95379543 | rs564626 | 1.254 | -0.022 | -0.611 | 0.216 | -0.015 | 0.062 | 0.062 | 0 | 0.43 | 0.27 | 0.31 | 0.04 | 0.51 | 0.29 |
| 10 | 95381368 | rs2077733 | -0.854 | -0.167 | -0.726 | 1.237 | -0.001 | -0.05 | -0.05 | 0 | 0.09 | 0.01 | 0.25 | 0.31 | 0.06 | 0.21 |
| 10 | 95387857 | rs7080650 | -0.97 | -0.056 | 0.107 | 1.129 | -0.001 | -0.001 | -0.001 | 0 | 0.13 | 0.37 | 0.1 | 0.11 | 0.09 | 0.14 |
| 10 | 95389041 | rs714550 | -1.395 | 0.097 | -0.088 | 1.016 | 0.006 | 0.016 | 0.016 | 0.109 | 0.34 | 0.45 | 0.39 | 0.48 | 0.35 | 0.29 |
| 10 | 95389728 | rs4917871 | -1.48 | 0.081 | -0.052 | 1.019 | 0.007 | 0.02 | 0.02 | 0.063 | 0.34 | 0.47 | 0.39 | 0.47 | 0.35 | 0.29 |
| 10 | 95393621 | rs547759 | 1.278 | 0.23 | 0.082 | 0.982 | -0.006 | -0.013 | -0.013 | 0 | 0.62 | 0.33 | 0.56 | 0.43 | 0.6 | 0.66 |
| 10 | 95395247 | rs616522 | 0.222 | 0.324 | -0.147 | 0.846 | 0.011 | 0.021 | 0.021 | 0 | 0.78 | 0.71 | 0.7 | 0.64 | 0.74 | 0.86 |
| 10 | 95395727 | rs487161 | 0.748 | 0.016 | -0.845 | 0.658 | -0.022 | -0.034 | -0.034 | 0.129 | 0.59 | 0.17 | 0.52 | 0.51 | 0.59 | 0.66 |
| 10 | 95401292 | rs1018634 | -0.754 | -0.305 | -0.541 | 0.575 | 0.002 | -0.011 | -0.011 | 0 | 0.31 | 0.23 | 0.39 | 0.31 | 0.29 | 0.27 |
| 10 | 95402973 | rs3781273 | 1.432 | -0.771 | -0.574 | 0.335 | -0.016 | -0.004 | -0.004 | 0.304 | 0.41 | 0.29 | 0.42 | 0.43 | 0.51 | 0.45 |
| 10 | 95406729 | rs11187569 | 0.096 | -0.591 | -0.842 | 0.333 | -0.024 | 0.002 | 0.002 | 0 | 0.21 | 0.04 | 0.16 | 0.19 | 0.19 | 0.26 |
| 10 | 95406935 | rs503758 | 0.217 | -0.577 | -0.605 | 0.484 | -0.029 | -0.045 | -0.045 | 0 | 0.21 | 0.08 | 0.36 | 0.29 | 0.25 | 0.22 |
| 10 | 95414150 | rs11187575 | 0.019 | -0.702 | -0.508 | 0.396 | -0.023 | -0.001 | -0.001 | 0.481 | 0.19 | 0.04 | 0.16 | 0.19 | 0.19 | 0.26 |
| 10 | 95421249 | rs10748607 | -1.437 | -0.608 | 0.192 | 0.541 | -0.016 | 0.007 | 0.007 | 0.075 | 0.29 | 0.4 | 0.22 | 0.3 | 0.21 | 0.13 |
| 10 | 95426888 | rs11187579 | 0.231 | -0.562 | 0.125 | 0.511 | -0.005 | 0.016 | 0.016 | 0 | 0.06 | 0.05 | 0.1 | 0.26 | 0.08 | 0.06 |
| 10 | 95427281 | rs1547173 | 1.694 | -0.906 | -0.076 | 0.387 | -0.038 | -0.004 | -0.004 | 0 | 0.46 | 0.1 | 0.46 | 0.4 | 0.55 | 0.46 |
| 10 | 95429007 | rs11187580 | 0.747 | -0.924 | -0.056 | 0.358 | 0.004 | 0.028 | 0.028 | 0 | 0.15 | 0.09 | 0.1 | 0.03 | 0.12 | 0.07 |
| 10 | 95431535 | rs913045 | -0.07 | -0.942 | 0.033 | 0.362 | -0.009 | 0.019 | 0.019 | 1.116 | 0.73 | 0.47 | 0.67 | 0.69 | 0.75 | 0.6 |
| 10 | 95440286 | rs10509663 | 0.016 | -0.944 | 0.438 | 0.405 | 0.057 | 0.236 | 0.236 | 0.041 | 0.06 | 0.26 | 0.03 | 0.24 | 0.04 | 0.19 |
| 10 | 95440889 | rs9325465 | 0.293 | -0.938 | 0.302 | 0.377 | -0.002 | 0.014 | 0.014 | 0 | 0.06 | 0.06 | 0.1 | 0.26 | 0.07 | 0.06 |
| 10 | 95441272 | rs11187583 | 0.587 | -0.982 | 0.261 | 0.349 | 0.005 | 0.031 | 0.031 | 0 | 0.16 | 0.09 | 0.1 | 0.03 | 0.12 | 0.07 |
| 10 | 95449292 | rs10748608 | -1.37 | -0.897 | 0.232 | 0.389 | -0.02 | -0.003 | -0.003 | 0 | 0.29 | 0.4 | 0.22 | 0.29 | 0.21 | 0.14 |
| 10 | 95456171 | rs10882306 | 1.808 | -0.868 | 0.034 | 0.383 | -0.067 | -0.034 | -0.034 | 0.017 | 0.44 | 0.07 | 0.45 | 0.4 | 0.55 | 0.46 |
| 10 | 95457992 | rs11187596 | 0.925 | -0.948 | -0.018 | 0.337 | -0.029 | 0.077 | 0.077 | 0 | 0.14 | 0.01 | 0.09 | 0.03 | 0.12 | 0.07 |
| 10 | 95458141 | rs11187597 | 1.86 | -0.894 | 0.004 | 0.358 | -0.071 | -0.034 | -0.034 | 0.014 | 0.44 | 0.07 | 0.45 | 0.4 | 0.54 | 0.46 |
| 10 | 95458872 | rs10882309 | 0.34 | -0.913 | 0.011 | 0.386 | -0.002 | 0.014 | 0.014 | 0 | 0.06 | 0.05 | 0.1 | 0.26 | 0.07 | 0.06 |
| 10 | 95458897 | rs10882310 | 1.842 | -0.884 | 0.011 | 0.41 | -0.092 | -0.053 | -0.053 | 0 | 0.43 | 0.05 | 0.45 | 0.4 | 0.54 | 0.46 |
| 10 | 95459267 | rs1326224 | -0.608 | -0.761 | 0.429 | 0.418 | -0.004 | -0.023 | -0.023 | 0.619 | 0.79 | 0.74 | 0.71 | 0.93 | 0.79 | 0.78 |
| 10 | 95459817 | rs726817 | 0.174 | -0.761 | 0.451 | 0.414 | -0.009 | 0.018 | 0.018 | 1.239 | 0.73 | 0.47 | 0.67 | 0.69 | 0.75 | 0.6 |
| 10 | 95467031 | rs11187609 | 0.367 | -0.748 | 0.223 | 0.219 | -0.037 | -0.088 | -0.088 | 0.546 | 0.71 | 0.55 | 0.69 | 0.93 | 0.79 | 0.78 |
| 10 | 95470199 | rs7902822 | 0.837 | -0.715 | 0.223 | 0.337 | 0.003 | 0.006 | 0.006 | 0.173 | 0.55 | 0.41 | 0.5 | 0.64 | 0.58 | 0.66 |
| 10 | 95472007 | rs12218711 | 0.859 | -0.778 | 1.022 | 0.302 | 0.004 | 0.004 | 0.004 | 0.433 | 0.55 | 0.43 | 0.52 | 0.64 | 0.61 | 0.66 |
| 10 | 95472127 | rs2901694 | 1.666 | -0.821 | 0.803 | 0.466 | -0.081 | -0.051 | -0.051 | 6.082 | 0.44 | 0.06 | 0.46 | 0.4 | 0.55 | 0.46 |
| 10 | 95475336 | rs979117 | 0.505 | -1.02 | 0.93 | 0.377 | -0.005 | 0 | 0 | 0.136 | 0.66 | 0.64 | 0.6 | 0.68 | 0.72 | 0.7 |
| 10 | 95475470 | rs7098209 | -0.341 | -0.621 | 0.945 | 0.372 | 0.01 | 0.016 | 0.016 | 0 | 0.22 | 0.39 | 0.14 | 0.29 | 0.2 | 0.26 |
| 10 | 95476226 | rs7099226 | 0.481 | -0.621 | 0.942 | 0.373 | -0.006 | 0 | 0 | 0.142 | 0.66 | 0.63 | 0.6 | 0.68 | 0.72 | 0.7 |
| 10 | 95480267 | rs863800 | 1.689 | -0.593 | 0.783 | 0.371 | -0.026 | 0.012 | 0.012 | 0.864 | 0.44 | 0.24 | 0.46 | 0.39 | 0.52 | 0.44 |
| 10 | 95484010 | rs7093903 | -0.742 | -0.61 | 0.843 | 0.312 | 0.002 | 0.002 | 0.002 | 0.842 | 0.79 | 0.81 | 0.71 | 0.93 | 0.81 | 0.79 |
| 10 | 95488351 | rs6583904 | -1.041 | -0.515 | 0.792 | 0.371 | -0.01 | 0.08 | 0.08 | 0.527 | 0.13 | 0.28 | 0.12 | 0.25 | 0.09 | 0.08 |
| 10 | 95489806 | rs7088661 | 0.153 | -0.352 | 0.371 | 0.036 | -0.003 | -0.073 | -0.073 | 1.432 | 0.18 | 0.29 | 0.11 | 0.05 | 0.16 | 0.07 |
| 10 | 95500156 | rs10786134 | 1.577 | -0.599 | -0.318 | 0.128 | -0.105 | -0.057 | -0.057 | 0 | 0.42 | 0.11 | 0.44 | 0.42 | 0.52 | 0.45 |
| 10 | 95503759 | rs7071043 | -1.027 | -0.747 | 0.021 | 0.125 | 0.001 | -0.011 | -0.011 | 0 | 0.36 | 0.69 | 0.21 | 0.5 | 0.29 | 0.33 |
| 10 | 95512810 | rs946553 | 0.436 | -0.582 | 0.196 | 0.164 | -0.003 | -0.047 | -0.047 | 2.68 | 0.6 | 0.49 | 0.58 | 0.83 | 0.63 | 0.7 |
| 10 | 95513408 | rs7913442 | 0.034 | -0.302 | 0.333 | 0.274 | 0.013 | -0.051 | -0.051 | 0.049 | 0.21 | 0.37 | 0.13 | 0.1 | 0.18 | 0.09 |
| 10 | 95515515 | rs4919491 | 0.68 | -0.303 | 0.029 | -0.07 | 0.028 | 0.089 | 0.089 | 0.777 | 0.51 | 0.34 | 0.56 | 0.83 | 0.61 | 0.7 |
| 10 | 95519577 | rs1555870 | 0.224 | 0.21 | -0.52 | -0.053 | -0.001 | -0.004 | -0.004 | 0 | 0.28 | 0.22 | 0.34 | 0.36 | 0.33 | 0.43 |
| 10 | 95521230 | rs12769713 | 0.479 | 0.284 | -1.234 | -0.687 | -0.065 | 0.134 | 0.134 | 0 | 0.25 | 0.08 | 0.14 | 0.05 | 0.29 | 0.16 |
| 10 | 95521457 | rs7897180 | 0.204 | 0.45 | -0.452 | -0.434 | 0.001 | 0.012 | 0.012 | 1.416 | 0.29 | 0.34 | 0.35 | 0.36 | 0.33 | 0.43 |
| 10 | 95525331 | rs17109372 | 0.467 | 0.33 | -0.758 | -0.407 | 0.018 | -0.029 | -0.029 | 0.944 | 0.24 | 0.12 | 0.25 | 0.27 | 0.22 | 0.36 |
| 10 | 95526044 | rs11187648 | 0.461 | 0.309 | -0.763 | -0.466 | 0.014 | -0.052 | -0.052 | 0.641 | 0.24 | 0.11 | 0.26 | 0.39 | 0.23 | 0.37 |
| 10 | 95526168 | rs7915994 | 0.486 | 0.472 | -0.873 | -1.063 | -0.018 | 0.022 | 0.022 | 1.263 | 0.32 | 0.17 | 0.39 | 0.03 | 0.39 | 0.22 |
| 10 | 95528442 | rs10509665 | -0.082 | 0.098 | -0.026 | -0.979 | 0.001 | -0.003 | -0.003 | 0.719 | 0.6 | 0.53 | 0.6 | 0.59 | 0.59 | 0.57 |
| 10 | 95531345 | rs10882331 | -1.12 | -0.009 | 0.382 | -1.148 | 0.017 | -0.051 | -0.051 | 0 | 0.84 | 0.81 | 0.88 | 0.99 | 0.81 | 0.94 |
| 10 | 95534265 | rs3781267 | -0.135 | -0.158 | 0.406 | -0.892 | -0.004 | -0.005 | -0.005 | 0 | 0.55 | 0.3 | 0.56 | 0.5 | 0.55 | 0.49 |
| 10 | 95538467 | rs11187664 | 0.586 | -0.338 | 0.105 | -0.847 | 0.019 | -0.044 | -0.044 | 0.633 | 0.13 | 0.06 | 0.13 | 0.47 | 0.09 | 0.24 |
| 10 | 95541735 | rs4531389 | -0.338 | -0.057 | 0.603 | -1.521 | 0.003 | 0.025 | 0.025 | 0 | 0.5 | 0.59 | 0.5 | 0.09 | 0.51 | 0.33 |
| 10 | 95542677 | rs11187665 | -0.216 | -0.266 | 0.651 | -1.425 | -0.001 | -0.037 | -0.037 | 0 | 0.46 | 0.57 | 0.3 | 0.1 | 0.46 | 0.3 |
| 10 | 95543430 | rs17109405 | -1.986 | -0.633 | 1.858 | -0.363 | 0.034 | 0.066 | 0.066 | 0.418 | 0.07 | 0.36 | 0.07 | 0.09 | 0.04 | 0.07 |
| 10 | 95545378 | rs7099034 | 1.942 | -0.602 | 1.242 | -0.368 | -0.044 | -0.027 | -0.027 | 0 | 0.85 | 0.3 | 0.91 | 0.91 | 0.94 | 0.92 |
| 10 | 95548443 | rs10882336 | 1.787 | -0.566 | 1.257 | -0.382 | -0.042 | -0.053 | -0.053 | 0 | 0.87 | 0.36 | 0.93 | 0.97 | 0.96 | 0.93 |
| 10 | 95552925 | rs1108877 | -1.763 | -0.494 | 0.757 | -0.239 | -0.053 | 0.008 | 0.008 | 2.234 | 0.16 | 0.63 | 0.08 | 0.16 | 0.07 | 0.09 |
| 10 | 95554732 | rs12266513 | -1.162 | -0.325 | 0.342 | -0.468 | -0.028 | -0.045 | -0.045 | 5.116 | 0.09 | 0.32 | 0.04 | 0 | 0.03 | 0 |
| 10 | 95559398 | rs12779373 | 0.275 | 0.275 | -0.78 | -0.177 | -0.055 | -0.103 | -0.103 | 0 | 0.21 | 0.05 | 0.32 | 0.34 | 0.25 | 0.34 |

| | | | | | | | | | | | | | | | | |
|---|---|---|---|---|---|---|---|---|---|---|---|---|---|---|---|---|
| 10 | 95561397 | rs1416757 | -0.229 | 0.44 | -0.358 | -0.971 | -0.041 | 0.051 | 0.051 | 0.014 | 0.16 | 0.05 | 0.1 | 0.01 | 0.22 | 0.15 |
| 10 | 95563444 | rs7097242 | -1.052 | 0.287 | -0.014 | -0.988 | 0.003 | -0.125 | -0.125 | 0 | 0.27 | 0.53 | 0.14 | 0.05 | 0.24 | 0.23 |
| 10 | 95579110 | rs4919543 | 0.752 | -0.232 | -0.309 | -0.974 | 0.016 | 0.023 | 0.023 | 0.343 | 0.68 | 0.31 | 0.86 | 0.9 | 0.83 | 0.83 |
| 10 | 95602229 | rs919003 | 0.57 | -0.009 | -1.101 | -1.931 | -0.02 | 0.089 | 0.089 | 0.05 | 0.41 | 0.25 | 0.5 | 0.86 | 0.45 | 0.58 |
| 10 | 95606207 | rs10786144 | -0.064 | -0.345 | -0.24 | -2.182 | 0.023 | 0.064 | 0.064 | 1.381 | 0.43 | 0.3 | 0.4 | 0.04 | 0.42 | 0.28 |
| 10 | 95620060 | rs4917956 | 0.222 | -0.462 | 0.151 | -2.261 | -0.03 | 0.251 | 0.251 | 0.092 | 0.48 | 0.85 | 0.54 | 0.87 | 0.45 | 0.59 |
| 10 | 95640551 | rs10786147 | 0.906 | -0.414 | 0.958 | -2.243 | -0.019 | 0.065 | 0.065 | 0 | 0.39 | 0.28 | 0.5 | 0.87 | 0.45 | 0.6 |
| 10 | 95641536 | rs1477536 | -0.616 | -0.38 | 0.753 | -2.218 | -0.036 | -0.012 | -0.012 | 0 | 0.89 | 0.95 | 0.94 | 0.9 | 0.88 | 0.86 |
| 10 | 95653401 | rs11187723 | 0.085 | -0.504 | 0.786 | -2.32 | 0.011 | 0.142 | 0.142 | 1.723 | 0.41 | 0.19 | 0.41 | 0.03 | 0.43 | 0.27 |
| 10 | 95663285 | rs3088126 | 0.838 | -0.411 | 0.809 | -2.271 | 0.005 | -0.053 | -0.053 | 0 | 0.41 | 0.48 | 0.51 | 0.86 | 0.46 | 0.59 |
| 10 | 95667270 | rs11187729 | 0.933 | -0.379 | 0.432 | -2.258 | -0.005 | 0.005 | 0.005 | 0 | 0.4 | 0.36 | 0.51 | 0.86 | 0.46 | 0.58 |
| 10 | 95669415 | rs1046414 | -0.611 | -0.329 | 0.182 | -2.22 | -0.023 | -0.006 | -0.006 | 0 | 0.89 | 0.94 | 0.94 | 0.89 | 0.88 | 0.86 |
| 10 | 95680019 | rs12359848 | 0.559 | -0.395 | 0.094 | -2.365 | -0.024 | 0.082 | 0.082 | 0.115 | 0.29 | 0.14 | 0.36 | 0.03 | 0.37 | 0.25 |
| 10 | 95683452 | rs1223471 | 0.39 | 0.086 | 0.075 | -1.915 | -0.006 | 0.006 | 0.006 | 0 | 0.41 | 0.33 | 0.5 | 0.95 | 0.45 | 0.59 |
| 10 | 95718037 | rs12768993 | -0.702 | 1.66 | -0.616 | -0.833 | -0.052 | -0.04 | -0.04 | 0 | 0.12 | 0.05 | 0.13 | 0.22 | 0.23 | 0.23 |
| 10 | 95729204 | rs12573786 | -0.656 | 1.842 | -0.659 | -0.549 | -0.057 | -0.058 | -0.058 | 0 | 0.1 | 0.01 | 0.22 | 0.26 | 0.19 | 0.2 |
| 10 | 95732363 | rs978365 | -0.403 | 1.272 | -0.597 | -0.569 | -0.011 | 0.019 | 0.019 | 0 | 0.64 | 0.57 | 0.7 | 0.48 | 0.69 | 0.66 |
| 10 | 95737055 | rs11187749 | 0.515 | 1.134 | -0.211 | -0.639 | 0 | 0.124 | 0.124 | 0 | 0.5 | 0.26 | 0.45 | 0.22 | 0.49 | 0.46 |
| 10 | 95737763 | rs7085672 | -0.47 | 1.061 | -0.12 | -0.593 | -0.006 | 0.011 | 0.011 | 0 | 0.64 | 0.61 | 0.7 | 0.48 | 0.69 | 0.66 |
| 10 | 95743637 | rs9888014 | 0.079 | 0.861 | -0.391 | -0.733 | -0.009 | 0.009 | 0.009 | 0.018 | 0.34 | 0.63 | 0.29 | 0.54 | 0.26 | 0.34 |
| 10 | 95747476 | rs11187757 | -0.487 | 1.242 | -0.517 | -0.708 | -0.06 | -0.05 | -0.05 | 1.237 | 0.15 | 0.05 | 0.26 | 0.26 | 0.25 | 0.19 |
| 10 | 95750827 | rs829232 | -1.02 | 0.791 | -0.827 | -0.918 | -0.022 | 0.06 | 0.06 | 0.285 | 0.43 | 0.73 | 0.53 | 0.77 | 0.46 | 0.53 |
| 10 | 95755058 | rs11187761 | 1.006 | 0.978 | -1.262 | -1.142 | -0.05 | -0.002 | -0.002 | 0 | 0.13 | 0.01 | 0.12 | 0.1 | 0.18 | 0.17 |
| 10 | 95764616 | rs1223585 | -0.797 | 1.144 | -0.702 | -0.902 | -0.004 | 0.017 | 0.017 | 0 | 0.44 | 0.62 | 0.53 | 0.75 | 0.47 | 0.52 |
| 10 | 95769344 | rs829228 | -1.115 | 1.124 | -0.752 | -0.906 | 0.005 | 0.109 | 0.109 | 0 | 0.48 | 0.77 | 0.55 | 0.76 | 0.48 | 0.53 |
| 10 | 95774582 | rs1223613 | -1.147 | 1.109 | -0.768 | -0.163 | 0.007 | 0.115 | 0.115 | 0 | 0.49 | 0.77 | 0.55 | 0.76 | 0.48 | 0.53 |
| 10 | 95777231 | rs1223618 | -1.186 | 1.106 | -0.787 | -0.167 | 0.005 | 0.139 | 0.139 | 0.015 | 0.49 | 0.8 | 0.56 | 0.76 | 0.48 | 0.53 |
| 10 | 95777488 | rs10882381 | -0.12 | 1.032 | -0.519 | 0.102 | -0.021 | 0.081 | 0.081 | 0 | 0.29 | 0.53 | 0.28 | 0.51 | 0.24 | 0.34 |
| 10 | 95782925 | rs1935960 | -0.655 | 1.272 | -0.682 | 0.334 | -0.004 | -0.013 | -0.013 | 0 | 0.15 | 0.08 | 0.24 | 0.21 | 0.15 | 0.15 |
| 10 | 95783242 | rs1935961 | -1.271 | 1.17 | -0.726 | 0.7 | -0.007 | -0.025 | -0.025 | 0 | 0.41 | 0.79 | 0.33 | 0.52 | 0.52 | 0.47 |
| 10 | 95786024 | rs7921117 | -1.749 | 0.646 | -0.807 | 0.389 | -0.007 | -0.006 | -0.006 | 0.036 | 0.91 | 0.9 | 0.95 | 0.95 | 0.95 | 0.98 |
| 10 | 95787977 | rs2182093 | 0.173 | 1.543 | -0.559 | 0.599 | -0.077 | -0.013 | -0.013 | 0 | 0.13 | 0.01 | 0.24 | 0.13 | 0.22 | 0.18 |
| 10 | 95789647 | rs7082530 | -0.903 | 0.931 | 0.027 | 0.837 | -0.008 | -0.019 | -0.019 | 0.061 | 0.3 | 0.73 | 0.26 | 0.48 | 0.3 | 0.31 |
| 10 | 95790063 | rs12254065 | -0.971 | 0.929 | 0.031 | 0.836 | -0.014 | -0.036 | -0.036 | 0 | 0.31 | 0.78 | 0.26 | 0.48 | 0.3 | 0.31 |
| 10 | 95790669 | rs10882386 | -0.676 | 0.939 | 0.258 | 0.877 | 0.057 | 0.133 | 0.133 | 0.04 | 0.25 | 0.5 | 0.21 | 0.44 | 0.29 | 0.3 |
| 10 | 95805955 | rs11187777 | -0.003 | 0.864 | 0.467 | 0.844 | 0.004 | 0.02 | 0.02 | 0.382 | 0.17 | 0.15 | 0.19 | 0.44 | 0.28 | 0.3 |
| 10 | 95826895 | rs3945750 | -0.642 | 0.993 | -0.012 | 0.939 | 0.02 | 0.063 | 0.063 | 0.438 | 0.26 | 0.63 | 0.22 | 0.43 | 0.28 | 0.29 |
| 10 | 95838570 | rs7088723 | -0.611 | 1.278 | -0.235 | 0.971 | 0.025 | 0.078 | 0.078 | 0.064 | 0.26 | 0.57 | 0.23 | 0.43 | 0.28 | 0.3 |
| 10 | 95852129 | rs17109746 | -0.556 | 1.271 | -0.424 | 0.987 | 0.04 | 0.108 | 0.108 | 0.058 | 0.26 | 0.53 | 0.22 | 0.42 | 0.28 | 0.3 |
| 10 | 95859929 | rs4918082 | -0.057 | 0.16 | -0.564 | 0.747 | 0.002 | -0.001 | -0.001 | 1.991 | 0.44 | 0.54 | 0.47 | 0.54 | 0.29 | 0.31 |
| 10 | 95867069 | rs10882393 | 0.934 | 0.432 | 0.254 | 0.825 | 0.073 | 0.273 | 0.273 | 1.317 | 0.93 | 0.6 | 0.92 | 0.83 | 0.97 | 0.97 |
| 10 | 95871655 | rs11187789 | -0.33 | 1.149 | -0.518 | 0.424 | -0.125 | 0.03 | 0.03 | 0.346 | 0.21 | 0.02 | 0.34 | 0.08 | 0.32 | 0.17 |
| 10 | 95872540 | rs4917410 | 1.08 | 0.812 | -0.208 | 0.435 | 0.035 | 0.208 | 0.208 | 2.461 | 0.92 | 0.54 | 0.92 | 0.83 | 0.97 | 0.98 |
| 10 | 95874185 | rs17416616 | 0.881 | 1.042 | -0.002 | 0.343 | 0.018 | -0.002 | -0.002 | 0 | 0.69 | 0.39 | 0.57 | 0.75 | 0.63 | 0.81 |
| 10 | 95885950 | rs10882396 | -0.852 | 0.99 | 0.479 | 0.11 | -0.007 | 0.002 | 0.002 | 0 | 0.26 | 0.27 | 0.36 | 0.08 | 0.35 | 0.15 |
| 10 | 95893514 | rs10786152 | 0.541 | 0.265 | 0.042 | -0.086 | 0 | 0.002 | 0.002 | 0 | 0.53 | 0.51 | 0.61 | 0.69 | 0.47 | 0.51 |
| 10 | 95906315 | rs1223629 | 0.84 | -0.021 | 0.723 | 0.002 | -0.011 | 0.04 | 0.04 | 1.06 | 0.89 | 0.52 | 0.95 | 0.88 | 0.97 | 0.98 |
| 10 | 95928274 | rs932762 | -0.827 | 0.003 | 0.675 | -0.028 | 0.038 | 0.074 | 0.074 | 1.512 | 0.07 | 0.33 | 0.05 | 0.1 | 0.04 | 0.02 |
| 10 | 95933801 | rs1776946 | 0.63 | -0.091 | -0.038 | -0.114 | 0.034 | 0.002 | 0.002 | 1.668 | 0.65 | 0.24 | 0.81 | 0.66 | 0.77 | 0.61 |
| 10 | 95937944 | rs2689694 | 0.266 | 0.085 | -0.553 | -0.478 | -0.056 | 0.091 | 0.091 | 0 | 0.29 | 0.14 | 0.29 | 0.09 | 0.35 | 0.17 |
| 10 | 95946520 | rs2689700 | -0.885 | 0.211 | -0.498 | -0.167 | -0.003 | -0.005 | -0.005 | 0 | 0.79 | 0.78 | 0.89 | 0.87 | 0.82 | 0.64 |
| 10 | 95949432 | rs11187808 | 0.778 | 0.332 | -0.797 | -0.025 | -0.018 | -0.037 | -0.037 | 0.063 | 0.37 | 0.12 | 0.52 | 0.57 | 0.43 | 0.43 |
| 10 | 95951109 | rs2797986 | -0.36 | 0.638 | -0.734 | -0.848 | 0.031 | 0.068 | 0.068 | 0 | 0.51 | 0.41 | 0.55 | 0.21 | 0.6 | 0.58 |
| 10 | 95952934 | rs17093867 | -0.286 | 0.453 | -0.183 | -0.804 | -0.009 | -0.066 | -0.066 | 0 | 0.24 | 0.35 | 0.32 | 0.09 | 0.19 | 0.17 |
| 10 | 95958331 | rs4545470 | -0.504 | 0.394 | 0.06 | -0.655 | -0.008 | -0.032 | -0.032 | 0 | 0.27 | 0.45 | 0.32 | 0.21 | 0.21 | 0.2 |
| 10 | 95959384 | rs2797992 | -1.313 | 0.911 | -0.241 | -0.881 | 0.035 | 0.054 | 0.054 | 0 | 0.52 | 0.63 | 0.59 | 0.29 | 0.58 | 0.38 |
| 10 | 95959620 | rs2689707 | 0.561 | 1.17 | -0.98 | -0.81 | -0.12 | 0.078 | 0.078 | 0.007 | 0.23 | 0.04 | 0.25 | 0.08 | 0.37 | 0.18 |
| 10 | 95966358 | rs7908334 | 0.191 | 1.175 | -0.572 | -0.53 | -0.001 | -0.004 | -0.004 | 0.423 | 0.21 | 0.23 | 0.32 | 0.09 | 0.2 | 0.16 |
| 10 | 95970288 | rs1858608 | 0.563 | 0.19 | -0.497 | -0.495 | 0.039 | 0.046 | 0.046 | 0 | 0.48 | 0.35 | 0.38 | 0.69 | 0.36 | 0.62 |
| 10 | 95971321 | rs11187815 | -0.434 | 0.852 | -0.697 | -1.018 | -0.034 | 0.032 | 0.032 | 0 | 0.68 | 0.74 | 0.61 | 0.96 | 0.58 | 0.77 |
| 10 | 95971466 | rs12769135 | 1.281 | 0.144 | -0.492 | -0.208 | 0.023 | -0.014 | -0.014 | 0 | 0.35 | 0.22 | 0.35 | 0.4 | 0.3 | 0.26 |
| 10 | 95973747 | rs2860750 | -1.133 | 0.043 | -0.66 | -0.368 | -0.016 | 0.01 | 0.01 | 0 | 0.74 | 0.74 | 0.57 | 0.93 | 0.62 | 0.76 |
| 10 | 95976932 | rs10882406 | -0.11 | -0.12 | -0.227 | -0.22 | -0.01 | 0.019 | 0.019 | 0 | 0.34 | 0.4 | 0.23 | 0.54 | 0.34 | 0.49 |
| 10 | 95983053 | rs11187820 | -0.431 | -0.204 | -0.504 | -0.189 | 0.003 | 0 | 0 | 0 | 0.38 | 0.6 | 0.36 | 0.63 | 0.35 | 0.51 |
| 10 | 95987313 | rs11187825 | -0.729 | 0.139 | -0.406 | -0.32 | -0.011 | 0 | 0 | 0 | 0.7 | 0.69 | 0.58 | 0.93 | 0.62 | 0.76 |
| 10 | 95995579 | rs11187828 | 0.118 | 0.039 | -0.083 | -0.171 | 0.039 | -0.011 | -0.011 | 0 | 0.07 | 0.22 | 0.07 | 0.02 | 0.08 | 0.06 |
| 10 | 95997190 | rs12248509 | 0.425 | 0.025 | -0.082 | -0.169 | 0.028 | -0.003 | -0.003 | 0 | 0.06 | 0.22 | 0.06 | 0.02 | 0.05 | 0.06 |
| 10 | 96000162 | rs17109869 | 0.452 | -0.542 | 0.057 | -0.128 | -0.008 | -0.004 | -0.004 | 0.05 | 0.43 | 0.5 | 0.28 | 0.35 | 0.34 | 0.35 |
| 10 | 96000282 | rs7069316 | -1.064 | -0.636 | 0.293 | -0.242 | -0.016 | 0.008 | 0.008 | 0 | 0.7 | 0.71 | 0.57 | 0.93 | 0.61 | 0.75 |

| | | | | | | | | | | | | | | | | |
|---|---|---|---|---|---|---|---|---|---|---|---|---|---|---|---|---|
| 10 | 96001888 | rs17109872 | 0.458 | -0.627 | 0.308 | -0.233 | 0.005 | -0.01 | -0.01 | 0.496 | 0.27 | 0.22 | 0.3 | 0.58 | 0.27 | 0.41 |
| 10 | 96009579 | rs4917450 | 1.313 | -0.721 | 0.129 | -0.255 | 0.017 | 0.002 | 0.002 | 0 | 0.36 | 0.23 | 0.23 | 0.32 | 0.3 | 0.29 |
| 10 | 96011865 | rs1547643 | -1.081 | -0.587 | 0.129 | -0.295 | -0.02 | 0.015 | 0.015 | 0 | 0.7 | 0.72 | 0.58 | 0.93 | 0.61 | 0.76 |
| 10 | 96013824 | rs1408820 | 1.235 | -0.576 | -0.009 | -0.227 | 0.018 | 0.012 | 0.012 | 0 | 0.37 | 0.23 | 0.23 | 0.3 | 0.3 | 0.32 |
| 10 | 96026575 | rs17109875 | 1.959 | -0.671 | -0.188 | 0.031 | -0.009 | -0.023 | -0.023 | 0 | 0.11 | 0.01 | 0.06 | 0.2 | 0.11 | 0.04 |
| 10 | 96033766 | rs4918188 | 0.479 | -0.722 | 0.076 | 0.047 | 0.002 | 0.001 | 0.001 | 0 | 0.44 | 0.54 | 0.29 | 0.34 | 0.34 | 0.38 |
| 10 | 96044734 | rs3736901 | 1.891 | -0.795 | -0.338 | -0.026 | -0.154 | -0.016 | -0.016 | 0 | 0.21 | 0.02 | 0.51 | 0.16 | 0.37 | 0.24 |
| 10 | 96044913 | rs12263737 | 0.31 | -1.399 | -0.223 | -0.304 | -0.011 | -0.023 | -0.023 | 0 | 0.43 | 0.46 | 0.21 | 0.24 | 0.31 | 0.25 |
| 10 | 96045054 | rs10882416 | 0.328 | -1.393 | -0.221 | -0.302 | -0.011 | -0.023 | -0.023 | 0 | 0.42 | 0.46 | 0.21 | 0.24 | 0.31 | 0.25 |
| 10 | 96052511 | rs11187842 | 2.432 | -1.324 | -0.453 | -0.267 | 0.006 | -0.013 | -0.013 | 0 | 0.11 | 0.04 | 0.06 | 0.2 | 0.1 | 0.04 |
| 10 | 96058433 | rs17516758 | 2.573 | -1.315 | -0.242 | -0.316 | 0.037 | 0.063 | 0.063 | 0 | 0.23 | 0.09 | 0.11 | 0 | 0.18 | 0.16 |
| 10 | 96058730 | rs12766693 | 2.199 | -1.362 | -0.168 | -0.313 | 0.02 | 0.035 | 0.035 | 0 | 0.27 | 0.15 | 0.13 | 0.03 | 0.21 | 0.2 |
| 10 | 96061793 | rs4394764 | 2.472 | -1.335 | -0.114 | -0.331 | 0.017 | 0.042 | 0.042 | 0 | 0.25 | 0.12 | 0.12 | 0 | 0.21 | 0.16 |
| 10 | 96066341 | rs2274223 | 0.33 | -1.383 | -0.015 | -0.295 | 0.016 | 0.027 | 0.027 | 0 | 0.43 | 0.38 | 0.21 | 0.24 | 0.34 | 0.27 |
| 10 | 96067947 | rs10509670 | 0.52 | -1.383 | -0.03 | -0.295 | 0.026 | 0.052 | 0.052 | 0 | 0.42 | 0.32 | 0.21 | 0.24 | 0.34 | 0.25 |
| 10 | 96071561 | rs2077218 | -0.694 | -1.579 | -0.19 | -0.201 | -0.004 | -0.028 | -0.028 | 0 | 0.82 | 0.9 | 0.82 | 0.7 | 0.8 | 0.83 |
| 10 | 96081457 | rs11187863 | 1.98 | -1.621 | -0.016 | -0.145 | 0.005 | -0.005 | -0.005 | 0 | 0.11 | 0.08 | 0.06 | 0.21 | 0.09 | 0.04 |
| 10 | 96083920 | rs12781451 | 1.121 | -1.719 | -0.012 | -0.229 | 0.041 | 0.069 | 0.069 | 0.018 | 0.36 | 0.19 | 0.17 | 0.21 | 0.28 | 0.19 |
| 10 | 96086078 | rs10882424 | 0.102 | -1.716 | 0.073 | -0.223 | -0.012 | -0.021 | -0.021 | 0 | 0.43 | 0.46 | 0.2 | 0.25 | 0.31 | 0.24 |
| 10 | 96098093 | rs17109928 | 1.472 | -1.69 | 0.223 | -0.169 | -0.006 | 0.065 | 0.065 | 0 | 0.12 | 0.17 | 0.09 | 0.29 | 0.12 | 0.35 |
| 10 | 96098373 | rs17517578 | 2.065 | -1.691 | 0.175 | -0.25 | 0.026 | 0.041 | 0.041 | 0 | 0.24 | 0.11 | 0.11 | 0 | 0.18 | 0.15 |
| 10 | 96108800 | rs10882436 | 1.287 | -1.309 | 0.22 | -0.018 | 0.02 | 0.031 | 0.031 | 0 | 0.36 | 0.28 | 0.17 | 0.21 | 0.27 | 0.19 |
| 10 | 96114835 | rs12572897 | 1.338 | -1.274 | 0.237 | -0.226 | -0.004 | 0.067 | 0.067 | 0 | 0.12 | 0.16 | 0.09 | 0.29 | 0.12 | 0.34 |
| 10 | 96121022 | rs12780491 | 1.135 | -1.402 | -0.011 | -0.555 | 0.028 | 0.039 | 0.039 | 0 | 0.35 | 0.24 | 0.17 | 0.08 | 0.27 | 0.19 |
| 10 | 96121830 | rs10882437 | 0.461 | -1.377 | -0.079 | -0.508 | -0.024 | -0.07 | -0.07 | 0.041 | 0.42 | 0.48 | 0.2 | 0.12 | 0.3 | 0.24 |
| 10 | 96132106 | rs2764378 | -1.075 | -0.468 | -0.904 | -0.411 | 0.001 | -0.007 | -0.007 | 0 | 0.66 | 0.6 | 0.47 | 0.82 | 0.65 | 0.75 |
| 10 | 96132617 | rs9420636 | 0.166 | -0.402 | -0.947 | 0.014 | -0.001 | -0.053 | -0.053 | 0.203 | 0.17 | 0.07 | 0.15 | 0.41 | 0.18 | 0.17 |
| 10 | 96140822 | rs11598738 | 1.352 | 0.017 | -0.614 | 0.084 | 0.044 | 0.113 | 0.113 | 0 | 0.11 | 0.2 | 0.13 | 0.33 | 0.2 | 0.38 |
| 10 | 96170593 | rs556055 | -0.865 | -0.399 | -0.804 | -0.29 | 0.009 | -0.031 | -0.031 | 1.292 | 0.38 | 0.46 | 0.33 | 0.81 | 0.44 | 0.61 |
| 10 | 96173304 | rs1334596 | 1.851 | -0.815 | -0.984 | -0.641 | -0.015 | -0.064 | -0.064 | 0 | 0.23 | 0.27 | 0.11 | 0.01 | 0.17 | 0.14 |
| 10 | 96176376 | rs11187933 | 0.859 | -0.775 | -1.087 | -0.559 | 0.01 | -0.079 | -0.079 | 0 | 0.19 | 0.05 | 0.16 | 0.41 | 0.19 | 0.18 |
| 10 | 96185590 | rs11187939 | -0.729 | -0.771 | -0.986 | -0.633 | 0.006 | -0.021 | -0.021 | 0.988 | 0.38 | 0.44 | 0.33 | 0.81 | 0.44 | 0.61 |
| 10 | 96204219 | rs17518932 | 3.325 | -0.95 | -1.23 | -0.759 | -0.02 | 0.053 | 0.053 | 0.301 | 0.09 | 0 | 0.05 | 0 | 0.09 | 0.08 |
| 10 | 96229166 | rs1931757 | -1.866 | -0.665 | -1.032 | -0.56 | -0.036 | 0.022 | 0.022 | 0 | 0.72 | 0.85 | 0.48 | 0.8 | 0.66 | 0.75 |
| 10 | 96231169 | rs11187969 | 1.916 | -0.59 | -1.102 | -0.314 | -0.029 | -0.013 | -0.013 | 0 | 0.07 | 0.01 | 0.1 | 0.38 | 0.11 | 0.07 |
| 10 | 96275945 | rs12573638 | 0.167 | -0.475 | -0.852 | -0.325 | -0.012 | -0.002 | -0.002 | 0.128 | 0.3 | 0.22 | 0.29 | 0.73 | 0.39 | 0.55 |
| 10 | 96283928 | rs10882470 | -0.293 | -0.471 | -0.837 | -0.339 | 0.005 | -0.009 | -0.009 | 0.639 | 0.34 | 0.4 | 0.3 | 0.77 | 0.4 | 0.57 |
| 10 | 96305329 | rs10786168 | -0.991 | -0.488 | -0.84 | -0.37 | 0 | -0.01 | -0.01 | 0.162 | 0.41 | 0.58 | 0.33 | 0.81 | 0.44 | 0.61 |
| 10 | 96305729 | rs10882474 | -0.913 | -0.491 | -0.811 | -0.436 | 0.005 | -0.024 | -0.024 | 0.209 | 0.41 | 0.56 | 0.33 | 0.79 | 0.44 | 0.61 |
| 10 | 96307738 | rs1977288 | -1.804 | -0.57 | -0.904 | -0.547 | -0.043 | 0.026 | 0.026 | 0 | 0.72 | 0.87 | 0.48 | 0.83 | 0.66 | 0.77 |
| 10 | 96313960 | rs11188019 | -0.571 | -0.558 | -0.84 | -0.505 | 0.013 | -0.022 | -0.022 | 0 | 0.39 | 0.47 | 0.32 | 0.75 | 0.44 | 0.63 |
| 10 | 96318795 | rs10882476 | 1.121 | -0.554 | -0.891 | -0.418 | 0.004 | -0.066 | -0.066 | 0.001 | 0.2 | 0.05 | 0.17 | 0.39 | 0.19 | 0.19 |
| 10 | 96322189 | rs7093613 | 1.171 | -0.556 | -0.892 | -0.419 | 0.003 | -0.066 | -0.066 | 0.055 | 0.2 | 0.05 | 0.16 | 0.39 | 0.19 | 0.17 |
| 10 | 96324151 | rs1969815 | -0.006 | -0.543 | -0.795 | -0.42 | 0 | 0 | 0 | 0 | 0.31 | 0.29 | 0.29 | 0.71 | 0.4 | 0.56 |
| 10 | 96346949 | rs17519764 | 2.066 | 0.26 | -0.943 | -0.54 | -0.058 | 0.054 | 0.054 | 0 | 0.1 | 0.01 | 0.19 | 0 | 0.15 | 0.13 |
| 10 | 96356181 | rs7078811 | -0.683 | 0.458 | -0.754 | -0.366 | 0.011 | -0.021 | -0.021 | 0 | 0.39 | 0.45 | 0.32 | 0.75 | 0.44 | 0.61 |
| 10 | 96369266 | rs1065026 | 2.123 | 0.365 | -0.743 | -0.422 | -0.01 | -0.051 | -0.051 | 0 | 0.22 | 0.24 | 0.1 | 0.01 | 0.17 | 0.13 |
| 10 | 96370263 | rs2025445 | 1.998 | 0.386 | -0.732 | -0.421 | 0.031 | 0.08 | 0.08 | 0 | 0.11 | 0.17 | 0.12 | 0.32 | 0.21 | 0.38 |
| 10 | 96370853 | rs1047625 | 2.088 | 0.342 | -0.744 | -0.465 | -0.012 | -0.058 | -0.058 | 0 | 0.22 | 0.25 | 0.1 | 0.01 | 0.17 | 0.13 |
| 10 | 96371928 | rs11638 | -0.618 | 0.481 | -0.76 | -0.302 | 0.007 | -0.011 | -0.011 | 0 | 0.38 | 0.42 | 0.32 | 0.75 | 0.44 | 0.61 |
| 10 | 96407978 | rs1998591 | -0.105 | 0.22 | -0.705 | -0.11 | 0.001 | 0.101 | 0.101 | 0 | 0.23 | 0.4 | 0.15 | 0.41 | 0.22 | 0.49 |
| 10 | 96429971 | rs2104543 | -0.049 | 0.222 | -0.701 | -0.113 | -0.001 | 0.048 | 0.048 | 0 | 0.23 | 0.31 | 0.15 | 0.37 | 0.22 | 0.48 |
| 10 | 96453099 | rs2901783 | 0.277 | 0.169 | -0.755 | -0.145 | 0.004 | 0.081 | 0.081 | 0 | 0.21 | 0.32 | 0.15 | 0.41 | 0.22 | 0.48 |
| 10 | 96454156 | rs12243416 | 1.942 | 0.101 | -0.775 | -0.33 | -0.001 | -0.043 | -0.043 | 0 | 0.23 | 0.24 | 0.12 | 0.01 | 0.22 | 0.14 |
| 10 | 96464730 | rs7896133 | 1.432 | 0.093 | -0.774 | -0.312 | -0.001 | -0.001 | -0.001 | 0 | 0.09 | 0.1 | 0.04 | 0.1 | 0.07 | 0.12 |
| 10 | 96466078 | rs1409655 | 1.686 | 0.057 | -0.882 | -0.371 | -0.112 | -0.006 | -0.006 | 0 | 0.26 | 0.04 | 0.52 | 0.17 | 0.34 | 0.23 |
| 10 | 96466342 | rs2296683 | 1.969 | 0.144 | -0.807 | -0.351 | -0.001 | -0.04 | -0.04 | 0 | 0.23 | 0.24 | 0.12 | 0.01 | 0.22 | 0.14 |
| 10 | 96466938 | rs2296680 | 1.824 | 0.107 | -0.762 | -0.344 | 0.011 | 0.1 | 0.1 | 0 | 0.11 | 0.19 | 0.1 | 0.31 | 0.15 | 0.36 |
| 10 | 96468899 | rs11188059 | 2.694 | 0.169 | -0.815 | -0.381 | -0.037 | 0.027 | 0.027 | 0 | 0.06 | 0.01 | 0.17 | 0 | 0.13 | 0.09 |
| 10 | 96469669 | rs7918101 | 1.747 | -0.704 | -0.496 | -0.127 | -0.001 | -0.043 | -0.043 | 0 | 0.23 | 0.24 | 0.12 | 0.01 | 0.22 | 0.14 |
| 10 | 96477854 | rs1926712 | 1.576 | -0.687 | -0.48 | -0.06 | 0.01 | 0.099 | 0.099 | 0 | 0.12 | 0.2 | 0.11 | 0.31 | 0.15 | 0.36 |
| 10 | 96484777 | rs1926711 | 1.275 | -0.688 | -0.473 | -0.059 | 0.031 | 0.184 | 0.184 | 0 | 0.13 | 0.29 | 0.11 | 0.37 | 0.15 | 0.37 |
| 10 | 96485584 | rs1326832 | 1.517 | -0.68 | -0.494 | -0.06 | 0.013 | 0.112 | 0.112 | 0 | 0.12 | 0.21 | 0.11 | 0.31 | 0.15 | 0.36 |
| 10 | 96485680 | rs1326831 | -3.067 | -0.783 | -0.569 | -0.118 | -0.01 | -0.017 | -0.017 | 1.958 | 0.95 | 0.95 | 0.99 | 1 | 0.98 | 0.99 |
| 10 | 96488469 | rs932809 | 1.473 | -0.431 | -0.411 | 0.085 | 0.01 | 0.099 | 0.099 | 0 | 0.12 | 0.19 | 0.11 | 0.31 | 0.15 | 0.36 |
| 10 | 96488888 | rs10509675 | 1.878 | -0.455 | -0.409 | 0.023 | -0.001 | -0.043 | -0.043 | 0.687 | 0.23 | 0.24 | 0.12 | 0.01 | 0.22 | 0.14 |
| 10 | 96493328 | rs2281890 | 1.454 | -0.456 | -0.401 | 0.028 | 0.012 | 0.109 | 0.109 | 0 | 0.12 | 0.21 | 0.11 | 0.31 | 0.15 | 0.36 |
| 10 | 96493358 | rs2281889 | -0.495 | -0.505 | -0.4 | 0.002 | 0.006 | 0.004 | 0.004 | 0 | 0.46 | 0.66 | 0.28 | 0.42 | 0.44 | 0.62 |
| 10 | 96493709 | rs11188067 | 1.911 | -0.504 | -0.4 | 0.001 | -0.001 | -0.048 | -0.048 | 0 | 0.23 | 0.24 | 0.12 | 0.01 | 0.22 | 0.14 |

| | | | | | | | | | | | | | | | | |
|---|---|---|---|---|---|---|---|---|---|---|---|---|---|---|---|---|
| 10 | 96495266 | rs3740367 | 1.65 | -0.504 | -0.416 | 0.005 | 0 | 0.013 | 0.013 | 0 | 0.08 | 0.02 | 0.04 | 0.04 | 0.07 | 0.11 |
| 10 | 96495484 | rs1042194 | 1.478 | -0.487 | -0.414 | 0.007 | 0.015 | 0.112 | 0.112 | 0 | 0.12 | 0.21 | 0.11 | 0.31 | 0.15 | 0.36 |
| 10 | 96499710 | rs1409651 | 1.852 | -0.473 | -0.453 | -0.22 | -0.012 | 0.004 | 0.004 | 0 | 0.07 | 0 | 0.03 | 0.04 | 0.07 | 0.11 |
| 10 | 96499931 | rs10882494 | 0.825 | -0.545 | -0.471 | -0.254 | -0.037 | -0.034 | -0.034 | 3.043 | 0.44 | 0.2 | 0.67 | 0.55 | 0.51 | 0.36 |
| 10 | 96500106 | rs1926707 | -0.475 | -0.52 | -0.476 | -0.242 | 0.006 | 0.004 | 0.004 | 0 | 0.47 | 0.66 | 0.28 | 0.42 | 0.44 | 0.62 |
| 10 | 96519197 | rs4532967 | 1.665 | -0.489 | -0.499 | -0.222 | 0.005 | 0.065 | 0.065 | 0 | 0.11 | 0.17 | 0.1 | 0.31 | 0.15 | 0.36 |
| 10 | 96522365 | rs4986894 | 1.72 | -0.451 | -0.611 | -0.203 | 0.006 | 0.068 | 0.068 | 0 | 0.11 | 0.16 | 0.1 | 0.31 | 0.15 | 0.36 |
| 10 | 96525865 | rs12768009 | 1.632 | -0.45 | -0.611 | -0.217 | 0.005 | 0.079 | 0.079 | 0 | 0.11 | 0.18 | 0.1 | 0.37 | 0.15 | 0.37 |
| 10 | 96534263 | rs6583954 | 1.524 | -0.443 | -0.602 | -0.18 | 0.01 | 0.099 | 0.099 | 0 | 0.12 | 0.2 | 0.11 | 0.31 | 0.15 | 0.36 |
| 10 | 96534584 | rs7916649 | -0.474 | -0.433 | -0.609 | -0.195 | 0.004 | 0.004 | 0.004 | 0 | 0.46 | 0.66 | 0.28 | 0.42 | 0.44 | 0.62 |
| 10 | 96536056 | rs4388808 | 1.5 | -0.378 | -1.043 | -0.128 | 0.004 | -0.016 | -0.016 | 0 | 0.18 | 0.15 | 0.16 | 0.39 | 0.17 | 0.13 |
| 10 | 96536708 | rs7068577 | 1.62 | 0.28 | -1.254 | 0.196 | -0.001 | -0.041 | -0.041 | 0.548 | 0.23 | 0.23 | 0.12 | 0.01 | 0.22 | 0.14 |
| 10 | 96539144 | rs17878673 | 0.713 | 1.038 | -1.516 | 1.005 | 0 | -0.001 | -0.001 | 0 | 0.09 | 0.09 | 0.04 | 0.1 | 0.07 | 0.12 |
| 10 | 96540889 | rs4304697 | 0.899 | 1.106 | -1.592 | 1.012 | 0.001 | -0.001 | -0.001 | 0.127 | 0.08 | 0.07 | 0.04 | 0.1 | 0.07 | 0.12 |
| 10 | 96541616 | rs4244285 | 0.613 | 1.423 | -1.512 | 1.153 | 0.005 | 0.065 | 0.065 | 0 | 0.11 | 0.17 | 0.11 | 0.31 | 0.15 | 0.36 |
| 10 | 96541982 | rs12571421 | 0.698 | 1.422 | -1.513 | 1.153 | 0.006 | 0.069 | 0.069 | 0 | 0.11 | 0.16 | 0.1 | 0.31 | 0.15 | 0.36 |
| 10 | 96543823 | rs35390752 | 0.49 | 1.424 | -1.511 | 1.154 | 0.011 | 0.1 | 0.1 | 0 | 0.11 | 0.19 | 0.1 | 0.31 | 0.15 | 0.36 |
| 10 | 96547463 | rs12767583 | 0.593 | 1.429 | -1.543 | 1.156 | 0.005 | 0.064 | 0.064 | 0 | 0.11 | 0.17 | 0.1 | 0.31 | 0.15 | 0.36 |
| 10 | 96563757 | rs4494250 | 0.735 | 1.443 | -1.586 | 0.784 | -0.133 | -0.025 | -0.025 | 0 | 0.26 | 0.02 | 0.51 | 0.17 | 0.33 | 0.23 |
| 10 | 96566889 | rs12772672 | -0.131 | 0.939 | -0.476 | 1.206 | 0.01 | 0.099 | 0.099 | 0 | 0.12 | 0.2 | 0.11 | 0.31 | 0.15 | 0.36 |
| 10 | 96567386 | rs4641393 | 0.048 | 0.992 | -0.661 | 1.229 | 0.006 | 0.068 | 0.068 | 0 | 0.11 | 0.16 | 0.1 | 0.31 | 0.15 | 0.36 |
| 10 | 96575242 | rs1322179 | 0.069 | 1.002 | -0.658 | 1.232 | 0.006 | 0.069 | 0.069 | 0 | 0.11 | 0.16 | 0.1 | 0.31 | 0.15 | 0.36 |
| 10 | 96576190 | rs10509678 | 0.45 | 0.847 | -0.906 | 1.232 | -0.012 | 0.004 | 0.004 | 0 | 0.07 | 0 | 0.03 | 0.04 | 0.07 | 0.11 |
| 10 | 96581094 | rs10786172 | 0.999 | 1.19 | -1.796 | 0.807 | -0.113 | -0.007 | -0.007 | 0.025 | 0.26 | 0.04 | 0.52 | 0.17 | 0.34 | 0.23 |
| 10 | 96598439 | rs11528090 | 1.155 | 0.565 | -1.868 | 0.201 | 0.001 | -0.014 | -0.014 | 0 | 0.18 | 0.16 | 0.17 | 0.39 | 0.18 | 0.13 |
| 10 | 96602398 | rs28399513 | 1.296 | 0.411 | -1.509 | 0.121 | 0.006 | 0.066 | 0.066 | 0 | 0.11 | 0.17 | 0.1 | 0.31 | 0.15 | 0.36 |
| 10 | 96603414 | rs11592737 | 1.823 | -0.238 | -0.971 | -0.342 | -0.001 | -0.048 | -0.048 | 0.148 | 0.23 | 0.25 | 0.12 | 0.01 | 0.22 | 0.14 |
| 10 | 96609568 | rs4917623 | 0.785 | -0.253 | -0.98 | -0.348 | -0.039 | -0.036 | -0.036 | 0 | 0.44 | 0.2 | 0.67 | 0.55 | 0.51 | 0.36 |
| 10 | 96610631 | rs17878382 | 1.732 | -0.124 | -0.865 | -0.291 | -0.012 | 0.004 | 0.004 | 0 | 0.07 | 0 | 0.03 | 0.04 | 0.07 | 0.11 |
| 10 | 96612040 | rs12779363 | 1.753 | -0.113 | -0.86 | -0.29 | 0 | -0.048 | -0.048 | 0 | 0.23 | 0.24 | 0.12 | 0.01 | 0.23 | 0.14 |
| 10 | 96612371 | rs12268020 | 1.753 | -0.129 | -0.877 | -0.29 | -0.001 | -0.048 | -0.048 | 0 | 0.23 | 0.25 | 0.12 | 0.01 | 0.22 | 0.14 |
| 10 | 96614725 | rs35709381 | 1.198 | -0.056 | -0.6 | -0.126 | 0.013 | 0.112 | 0.112 | 0 | 0.12 | 0.21 | 0.11 | 0.31 | 0.15 | 0.36 |
| 10 | 96619025 | rs3862009 | 1.564 | -0.04 | -0.804 | -0.162 | -0.012 | 0.004 | 0.004 | 0 | 0.08 | 0 | 0.03 | 0.04 | 0.07 | 0.11 |
| 10 | 96632253 | rs4466755 | -0.401 | 0.616 | -1.058 | -0.571 | -0.006 | -0.013 | -0.013 | 0 | 0.59 | 0.7 | 0.38 | 0.42 | 0.57 | 0.65 |
| 10 | 96634310 | rs17521564 | 1.287 | 0.527 | -0.877 | -0.739 | -0.003 | 0.042 | 0.042 | 0 | 0.22 | 0.13 | 0.14 | 0.1 | 0.2 | 0.16 |
| 10 | 96647117 | rs11188105 | -0.407 | 0.517 | -0.897 | -0.648 | -0.005 | -0.007 | -0.007 | 0 | 0.58 | 0.69 | 0.38 | 0.41 | 0.56 | 0.63 |
| 10 | 96648224 | rs9888067 | -1.033 | 0.484 | -0.957 | -0.625 | 0.004 | -0.005 | -0.005 | 0.317 | 0.73 | 0.74 | 0.8 | 0.58 | 0.78 | 0.83 |
| 10 | 96650570 | rs4110518 | 2.094 | 0.502 | -0.954 | -0.628 | -0.023 | -0.025 | -0.025 | 1.051 | 0.15 | 0.1 | 0.42 | 0.23 | 0.21 | 0.21 |
| 10 | 96663819 | rs1853208 | -0.384 | 0.467 | -0.947 | -0.672 | -0.005 | -0.008 | -0.008 | 0.012 | 0.59 | 0.71 | 0.39 | 0.41 | 0.57 | 0.64 |
| 10 | 96666085 | rs2860903 | 1.909 | 0.517 | -0.791 | -0.677 | 0.007 | 0.07 | 0.07 | 0 | 0.11 | 0.16 | 0.1 | 0.3 | 0.15 | 0.34 |
| 10 | 96690371 | rs2475377 | 1.972 | 0.387 | -0.829 | -0.747 | -0.01 | -0.014 | -0.014 | 0 | 0.1 | 0.1 | 0.04 | 0.02 | 0.05 | 0.02 |
| 10 | 96693727 | rs12251688 | 2.181 | 0.352 | -0.728 | -0.827 | 0 | -0.031 | -0.031 | 0 | 0.22 | 0.21 | 0.12 | 0.01 | 0.22 | 0.14 |
| 10 | 96696529 | rs9332092 | 2.081 | 0.338 | -0.758 | -0.833 | -0.012 | 0.008 | 0.008 | 0 | 0.08 | 0 | 0.04 | 0.03 | 0.07 | 0.11 |
| 10 | 96697252 | rs4918758 | 0.359 | 0.547 | -0.693 | -0.833 | -0.004 | -0.006 | -0.006 | 0 | 0.33 | 0.3 | 0.26 | 0.39 | 0.36 | 0.5 |
| 10 | 96697459 | rs9332098 | 1.989 | 0.548 | -0.712 | -0.836 | -0.011 | 0.009 | 0.009 | 0 | 0.08 | 0 | 0.04 | 0.03 | 0.07 | 0.11 |
| 10 | 96698690 | rs9332104 | 2.236 | 0.566 | -0.674 | -0.838 | -0.001 | -0.029 | -0.029 | 0.63 | 0.22 | 0.2 | 0.12 | 0.02 | 0.22 | 0.14 |
| 10 | 96699980 | rs9332108 | 2.164 | 0.592 | -0.717 | -0.807 | -0.013 | 0.007 | 0.007 | 0 | 0.07 | 0 | 0.04 | 0.03 | 0.07 | 0.11 |
| 10 | 96700402 | rs9332113 | 1.892 | 0.534 | -0.631 | -0.815 | -0.043 | -0.055 | -0.055 | 2.247 | 0.1 | 0.01 | 0.1 | 0.31 | 0.16 | 0.34 |
| 10 | 96700537 | rs2253635 | -1.212 | 0.495 | -0.587 | -0.813 | -0.018 | -0.043 | -0.043 | 0 | 0.68 | 0.78 | 0.43 | 0.44 | 0.64 | 0.65 |
| 10 | 96706409 | rs12772675 | 0.321 | 0.507 | -0.591 | -0.778 | -0.003 | -0.006 | -0.006 | 0 | 0.33 | 0.3 | 0.25 | 0.39 | 0.36 | 0.5 |
| 10 | 96707202 | rs4086116 | 1.313 | 0.518 | -0.596 | -0.787 | 0 | 0.004 | 0.004 | 0.157 | 0.22 | 0.21 | 0.15 | 0.09 | 0.2 | 0.16 |
| 10 | 96708226 | rs10509679 | 1.722 | 0.511 | -0.613 | -0.795 | -0.043 | -0.059 | -0.059 | 0.027 | 0.1 | 0.02 | 0.1 | 0.31 | 0.16 | 0.34 |
| 10 | 96711884 | rs4918766 | 0.279 | 0.508 | -0.606 | -0.799 | -0.004 | -0.006 | -0.006 | 0 | 0.33 | 0.3 | 0.25 | 0.39 | 0.36 | 0.5 |
| 10 | 96712400 | rs2475376 | -1.895 | 0.453 | -0.628 | -0.826 | 0.002 | -0.002 | -0.002 | 1.502 | 0.82 | 0.81 | 0.84 | 0.6 | 0.84 | 0.85 |
| 10 | 96720518 | rs7897079 | 2.261 | 0.238 | -0.589 | -0.979 | 0 | -0.031 | -0.031 | 0 | 0.22 | 0.21 | 0.12 | 0.01 | 0.22 | 0.14 |
| 10 | 96723424 | rs2153628 | 2.261 | 0.214 | -0.599 | -0.801 | 0 | -0.031 | -0.031 | 0 | 0.22 | 0.21 | 0.12 | 0.01 | 0.22 | 0.14 |
| 10 | 96725535 | rs4917639 | 1.31 | 0.212 | -0.598 | -0.802 | 0 | 0.004 | 0.004 | 0.157 | 0.22 | 0.21 | 0.15 | 0.09 | 0.2 | 0.16 |
| 10 | 96727160 | rs12569850 | 1.799 | 0.217 | -0.603 | -0.795 | -0.043 | -0.058 | -0.058 | 0 | 0.1 | 0.01 | 0.1 | 0.31 | 0.16 | 0.34 |
| 10 | 96731292 | rs9332168 | 2.247 | 0.27 | -0.595 | -0.797 | 0 | -0.031 | -0.031 | 0 | 0.22 | 0.21 | 0.12 | 0.01 | 0.22 | 0.14 |
| 10 | 96731788 | rs9332172 | 1.201 | 0.261 | -0.585 | -0.799 | -0.001 | -0.026 | -0.026 | 0 | 0.23 | 0.27 | 0.15 | 0.09 | 0.2 | 0.16 |
| 10 | 96732097 | rs9332174 | 2.256 | 0.266 | -0.585 | -0.77 | 0 | -0.029 | -0.029 | 0 | 0.22 | 0.21 | 0.12 | 0.02 | 0.22 | 0.14 |
| 10 | 96732731 | rs1856908 | -0.744 | 0.359 | -0.583 | -0.681 | -0.018 | -0.043 | -0.043 | 0.011 | 0.68 | 0.78 | 0.43 | 0.43 | 0.64 | 0.65 |
| 10 | 96734339 | rs10509680 | 1.807 | 0.618 | -0.566 | -0.496 | -0.012 | 0.008 | 0.008 | 0 | 0.08 | 0.01 | 0.04 | 0.03 | 0.07 | 0.11 |
| 10 | 96734582 | rs9325473 | 1.807 | 0.618 | -0.566 | -0.496 | -0.012 | 0.008 | 0.008 | 0 | 0.08 | 0 | 0.04 | 0.03 | 0.07 | 0.11 |
| 10 | 96741426 | rs1934967 | 2.164 | 0.635 | -0.658 | -0.493 | -0.059 | -0.047 | -0.047 | 0 | 0.13 | 0.04 | 0.41 | 0.17 | 0.21 | 0.2 |
| 10 | 96743108 | rs9332214 | 1.322 | 0.421 | -0.69 | -0.559 | -0.005 | 0.016 | 0.016 | 0 | 0.08 | 0.01 | 0.04 | 0.03 | 0.07 | 0.11 |
| 10 | 96743228 | rs9332217 | 1.665 | 0.422 | -0.69 | -0.559 | -0.012 | 0.008 | 0.008 | 0 | 0.08 | 0 | 0.04 | 0.03 | 0.07 | 0.11 |
| 10 | 96743943 | rs9332220 | 0.845 | 0.42 | -0.695 | -0.578 | 0 | 0.004 | 0.004 | 0.157 | 0.22 | 0.21 | 0.15 | 0.09 | 0.2 | 0.16 |
| 10 | 96745180 | rs9332227 | 1.659 | 0.421 | -0.652 | -0.535 | -0.012 | 0.008 | 0.008 | 0 | 0.08 | 0 | 0.04 | 0.03 | 0.07 | 0.11 |

| | | | | | | | | | | | | | | | | |
|---|---|---|---|---|---|---|---|---|---|---|---|---|---|---|---|---|
| 10 | 96746078 | rs2298037 | 1.341 | 0.505 | -0.785 | -0.496 | -0.041 | -0.058 | -0.058 | 2.603 | 0.1 | 0.01 | 0.1 | 0.31 | 0.16 | 0.34 |
| 10 | 96748492 | rs9332238 | 1.166 | 0.435 | -0.698 | -0.735 | -0.058 | 0.092 | 0.092 | 0 | 0.2 | 0.01 | 0.14 | 0.03 | 0.2 | 0.15 |
| 10 | 96750251 | rs4918797 | 0.717 | 0.546 | -0.85 | -0.67 | 0 | 0.003 | 0.003 | 0.105 | 0.22 | 0.22 | 0.15 | 0.09 | 0.2 | 0.16 |
| 10 | 96766934 | rs2860975 | -0.236 | 0.631 | -1.381 | -0.642 | -0.057 | -0.053 | -0.053 | 0 | 0.71 | 0.81 | 0.44 | 0.42 | 0.61 | 0.67 |
| 10 | 96788561 | rs1934955 | 1.451 | -0.057 | -1.699 | -1.261 | -0.02 | 0.072 | 0.072 | 0 | 0.27 | 0.12 | 0.14 | 0.04 | 0.28 | 0.21 |
| 10 | 96792328 | rs1592037 | 1.451 | -0.057 | -1.699 | -1.261 | -0.02 | 0.072 | 0.072 | 0 | 0.27 | 0.12 | 0.14 | 0.04 | 0.28 | 0.21 |
| 10 | 96793820 | rs11188147 | -0.145 | -0.337 | -1.42 | -1.465 | -0.014 | -0.003 | -0.003 | 0 | 0.42 | 0.49 | 0.29 | 0.38 | 0.36 | 0.41 |
| 10 | 96795524 | rs7087256 | -1.053 | -0.645 | -1.603 | -1.65 | -0.026 | -0.046 | -0.046 | 0 | 0.77 | 0.82 | 0.49 | 0.45 | 0.69 | 0.67 |
| 10 | 96796861 | rs1058932 | 0.197 | -0.83 | -1.253 | -1.744 | 0 | 0.122 | 0.122 | 0 | 0.19 | 0.37 | 0.14 | 0.38 | 0.2 | 0.35 |
| 10 | 96797270 | rs11572177 | 0.927 | -0.831 | -1.271 | -1.721 | 0 | -0.018 | -0.018 | 0 | 0.36 | 0.33 | 0.2 | 0.07 | 0.33 | 0.26 |
| 10 | 96797470 | rs1934953 | -1.135 | -0.839 | -1.168 | -1.742 | -0.026 | -0.045 | -0.045 | 0 | 0.77 | 0.82 | 0.49 | 0.45 | 0.69 | 0.67 |
| 10 | 96797500 | rs1934952 | 0.978 | -0.848 | -1.174 | -1.752 | 0 | -0.018 | -0.018 | 1.17 | 0.35 | 0.33 | 0.19 | 0.07 | 0.33 | 0.26 |
| 10 | 96798548 | rs1934951 | 0.173 | -0.825 | -1.063 | -1.678 | -0.001 | 0.12 | 0.12 | 0 | 0.19 | 0.36 | 0.14 | 0.38 | 0.2 | 0.35 |
| 10 | 96798578 | rs11572170 | 1.471 | -0.832 | -1.12 | -1.691 | -0.013 | -0.023 | -0.023 | 0 | 0.1 | 0.08 | 0.05 | 0 | 0.04 | 0.03 |
| 10 | 96800873 | rs1341164 | 1.387 | -0.794 | -1.139 | -1.729 | 0 | -0.029 | -0.029 | 0 | 0.29 | 0.28 | 0.15 | 0.04 | 0.28 | 0.21 |
| 10 | 96801753 | rs1934985 | -0.553 | -0.826 | -1.145 | -1.729 | -0.022 | -0.039 | -0.039 | 1.042 | 0.7 | 0.77 | 0.45 | 0.42 | 0.64 | 0.62 |
| 10 | 96801929 | rs1934983 | 1.407 | -0.829 | -1.142 | -1.733 | 0 | -0.029 | -0.029 | 2.242 | 0.29 | 0.28 | 0.15 | 0.04 | 0.28 | 0.21 |
| 10 | 96802124 | rs1934982 | 1.607 | -0.832 | -1.127 | -1.735 | -0.018 | 0.077 | 0.077 | 0.294 | 0.27 | 0.12 | 0.14 | 0.04 | 0.28 | 0.21 |
| 10 | 96804451 | rs7910936 | -0.55 | -0.8 | -0.569 | -1.578 | -0.022 | -0.039 | -0.039 | 0 | 0.7 | 0.77 | 0.45 | 0.42 | 0.64 | 0.62 |
| 10 | 96805371 | rs1891071 | -0.55 | -0.792 | -0.661 | -1.574 | -0.022 | -0.039 | -0.039 | 3.145 | 0.7 | 0.77 | 0.45 | 0.42 | 0.64 | 0.62 |
| 10 | 96808973 | rs1934980 | 0.66 | -0.956 | -0.659 | -1.555 | 0 | 0.032 | 0.032 | 0 | 0.16 | 0.17 | 0.13 | 0.38 | 0.19 | 0.34 |
| 10 | 96810552 | rs1341163 | -0.536 | -0.948 | -0.634 | -1.341 | -0.022 | -0.039 | -0.039 | 0 | 0.7 | 0.77 | 0.45 | 0.42 | 0.64 | 0.62 |
| 10 | 96811841 | rs7095531 | -1.074 | -0.919 | -0.637 | -1.326 | -0.026 | -0.046 | -0.046 | 0 | 0.77 | 0.82 | 0.49 | 0.45 | 0.69 | 0.67 |
| 10 | 96814475 | rs6583967 | 1.489 | -0.953 | -0.556 | -1.378 | 0 | -0.032 | -0.032 | 0.728 | 0.28 | 0.28 | 0.16 | 0.04 | 0.28 | 0.21 |
| 10 | 96814915 | rs11572126 | 1.343 | -1.003 | -0.599 | -1.332 | -0.017 | -0.037 | -0.037 | 0.514 | 0.08 | 0 | 0.09 | 0.31 | 0.13 | 0.3 |
| 10 | 96815114 | rs1934957 | 1.483 | -0.986 | -0.573 | -1.348 | 0 | -0.031 | -0.031 | 0 | 0.28 | 0.28 | 0.16 | 0.04 | 0.28 | 0.21 |
| 10 | 96815331 | rs11188156 | -0.318 | -1.03 | -0.581 | -1.366 | -0.013 | -0.003 | -0.003 | 0.166 | 0.42 | 0.49 | 0.29 | 0.38 | 0.36 | 0.41 |
| 10 | 96817297 | rs10882521 | -0.318 | -1.03 | -0.581 | -1.366 | -0.013 | -0.003 | -0.003 | 1.195 | 0.42 | 0.49 | 0.29 | 0.38 | 0.36 | 0.41 |
| 10 | 96818362 | rs11572101 | 0.642 | -0.99 | -0.434 | -1.276 | 0 | 0.032 | 0.032 | 0 | 0.16 | 0.18 | 0.13 | 0.38 | 0.19 | 0.34 |
| 10 | 96818402 | rs11572100 | 1.463 | -0.909 | -0.352 | -1.173 | -0.015 | -0.025 | -0.025 | 0 | 0.1 | 0.09 | 0.04 | 0 | 0.04 | 0.03 |
| 10 | 96824406 | rs11572093 | 1.392 | -0.575 | -0.394 | -0.981 | 0.001 | -0.034 | -0.034 | 0 | 0.28 | 0.28 | 0.16 | 0.04 | 0.28 | 0.21 |
| 10 | 96824883 | rs3752988 | -0.14 | -0.575 | -0.441 | -0.981 | -0.013 | -0.003 | -0.003 | 0.238 | 0.42 | 0.49 | 0.29 | 0.38 | 0.36 | 0.41 |
| 10 | 96824975 | rs2185571 | 1.561 | -0.459 | -0.601 | -0.932 | -0.02 | 0.072 | 0.072 | 0 | 0.27 | 0.12 | 0.14 | 0.04 | 0.28 | 0.21 |
| 10 | 96827178 | rs2275622 | -0.978 | -0.113 | -0.927 | -0.661 | -0.034 | -0.06 | -0.06 | 0.985 | 0.77 | 0.83 | 0.49 | 0.45 | 0.68 | 0.67 |
| 10 | 96828160 | rs1934956 | -0.625 | -0.46 | -1.106 | -0.468 | -0.009 | 0.034 | 0.034 | 0 | 0.82 | 0.77 | 0.72 | 0.52 | 0.86 | 0.82 |
| 10 | 96828323 | rs2071426 | 1.419 | -0.503 | -1.355 | -0.659 | -0.002 | 0.004 | 0.004 | 0 | 0.28 | 0.26 | 0.15 | 0.06 | 0.28 | 0.22 |
| 10 | 96829430 | rs7909236 | 1.981 | 0.057 | -1.482 | -1.12 | -0.077 | -0.017 | -0.017 | 0.313 | 0.12 | 0.02 | 0.28 | 0.08 | 0.22 | 0.2 |
| 10 | 96829529 | rs17110453 | 0.49 | -0.748 | -1.002 | -0.951 | -0.011 | -0.058 | -0.058 | 0.672 | 0.1 | 0 | 0.09 | 0.32 | 0.14 | 0.32 |
| 10 | 96831389 | rs1557044 | -0.147 | -0.911 | -0.542 | -0.896 | 0.001 | 0.006 | 0.006 | 0 | 0.83 | 0.81 | 0.73 | 0.53 | 0.86 | 0.83 |
| 10 | 96854541 | rs7921420 | 0.381 | -0.737 | -0.341 | -1.699 | -0.011 | -0.027 | -0.027 | 0.084 | 0.61 | 0.47 | 0.59 | 0.15 | 0.68 | 0.62 |
| 10 | 96864595 | rs7097340 | 1.443 | -0.705 | -0.225 | -1.682 | 0.017 | 0.003 | 0.003 | 0 | 0.15 | 0.1 | 0.06 | 0 | 0.07 | 0.03 |
| 10 | 96903156 | rs617848 | 0.213 | -0.588 | -0.09 | -1.623 | -0.004 | -0.023 | -0.023 | 0.388 | 0.76 | 0.6 | 0.65 | 0.17 | 0.76 | 0.64 |
| 10 | 96941391 | rs597768 | 0.213 | -0.588 | -0.085 | -1.623 | -0.004 | -0.023 | -0.023 | 0 | 0.76 | 0.6 | 0.65 | 0.17 | 0.76 | 0.64 |
| 10 | 96942283 | rs601746 | 0.537 | -0.797 | -0.489 | -1.976 | 0.053 | 0.11 | 0.11 | 1.303 | 0.3 | 0.12 | 0.17 | 0 | 0.2 | 0.05 |
| 10 | 96965722 | rs12357280 | 1.924 | -0.389 | -0.611 | -2.078 | -0.062 | -0.004 | -0.004 | 0 | 0.12 | 0.02 | 0.25 | 0.08 | 0.18 | 0.17 |
| 10 | 96968285 | rs680829 | 1.71 | -0.479 | -0.601 | -2.116 | 0.011 | -0.006 | -0.006 | 2.137 | 0.16 | 0.12 | 0.07 | 0 | 0.07 | 0.03 |
| 10 | 96970361 | rs16872 | 0.462 | -0.56 | -0.553 | -2.193 | -0.003 | -0.077 | -0.077 | 0 | 0.48 | 0.52 | 0.29 | 0.05 | 0.45 | 0.42 |
| 10 | 96977553 | rs695217 | 1.53 | -0.586 | -0.674 | -2.204 | -0.022 | 0.059 | 0.059 | 0 | 0.3 | 0.18 | 0.2 | 0.05 | 0.38 | 0.39 |
| 10 | 96979504 | rs11188228 | 1.493 | -0.57 | -0.672 | -2.197 | -0.017 | 0.043 | 0.043 | 0 | 0.3 | 0.2 | 0.21 | 0.05 | 0.38 | 0.4 |
| 10 | 96983053 | rs623544 | 0.706 | -0.473 | -0.928 | -2.128 | 0.048 | 0.068 | 0.068 | 0.744 | 0.3 | 0.16 | 0.17 | 0 | 0.19 | 0.05 |
| 10 | 96987103 | rs686016 | -0.736 | 0.043 | -0.795 | -1.842 | 0 | 0.011 | 0.011 | 0 | 0.85 | 0.81 | 0.7 | 0.23 | 0.83 | 0.68 |
| 10 | 96988377 | rs17452714 | 1.829 | 0.263 | -1.581 | -1.776 | -0.02 | 0.079 | 0.079 | 0.156 | 0.14 | 0.02 | 0.1 | 0 | 0.12 | 0.03 |
| 10 | 96988739 | rs11188234 | 1.664 | 0.374 | -1.327 | -1.51 | -0.06 | -0.006 | -0.006 | 0 | 0.11 | 0.02 | 0.25 | 0.08 | 0.18 | 0.17 |
| 10 | 96995971 | rs7909072 | 0.956 | 0.061 | -1.324 | -1.767 | -0.034 | 0.085 | 0.085 | 0.389 | 0.3 | 0.12 | 0.2 | 0.06 | 0.38 | 0.39 |
| 10 | 96996202 | rs9651411 | 0.84 | -0.322 | -1.309 | -1.77 | -0.017 | -0.024 | -0.024 | 0 | 0.21 | 0.23 | 0.09 | 0.04 | 0.09 | 0.06 |
| 10 | 96997820 | rs1049814 | 0.833 | 0.229 | -1.445 | -1.731 | 0.01 | -0.01 | -0.01 | 1.234 | 0.14 | 0.21 | 0.26 | 0.07 | 0.18 | 0.17 |
| 10 | 96998193 | rs12771349 | 0.049 | 0.119 | -1.612 | -1.92 | 0.006 | 0.183 | 0.183 | 1.626 | 0.46 | 0.25 | 0.44 | 0.09 | 0.39 | 0.25 |
| 10 | 97000900 | rs12413028 | 0.82 | 0.962 | -1.327 | -1.508 | -0.038 | 0.08 | 0.08 | 1.739 | 0.29 | 0.11 | 0.3 | 0.06 | 0.39 | 0.41 |
| 10 | 97001263 | rs567229 | -0.749 | 1.228 | -1.304 | -0.792 | -0.029 | 0.005 | 0.005 | 0 | 0.93 | 0.99 | 0.89 | 0.57 | 0.91 | 0.91 |
| 10 | 97009657 | rs10882573 | 1.04 | 1.252 | -1.242 | -1.153 | -0.024 | 0.014 | 0.014 | 0 | 0.12 | 0.07 | 0.25 | 0.07 | 0.17 | 0.16 |
| 10 | 97013497 | rs1887182 | -0.279 | 0.726 | -1.22 | -1.356 | 0.005 | 0.015 | 0.015 | 0.303 | 0.51 | 0.41 | 0.43 | 0.05 | 0.53 | 0.45 |
| 10 | 97020612 | rs7101278 | -0.959 | 0.797 | -1.415 | -1.165 | -0.031 | -0.064 | -0.064 | 0 | 0.65 | 0.55 | 0.68 | 0.13 | 0.71 | 0.61 |
| 10 | 97021389 | rs10509685 | 0.384 | 1.107 | -1.834 | -0.781 | 0.001 | 0.003 | 0.003 | 0 | 0.1 | 0.07 | 0.04 | 0.03 | 0.09 | 0.19 |
| 10 | 97032612 | rs12769370 | -0.38 | 1.275 | -1.815 | -0.757 | 0.003 | 0.034 | 0.034 | 0.036 | 0.27 | 0.18 | 0.17 | 0.03 | 0.23 | 0.22 |
| 10 | 97034610 | rs4917683 | -0.851 | 1.36 | -2.106 | -0.829 | 0.007 | 0.089 | 0.089 | 0.882 | 0.52 | 0.32 | 0.42 | 0.06 | 0.54 | 0.44 |
| 10 | 97035475 | rs11188255 | 0.138 | 1.637 | -2.468 | -0.243 | -0.086 | 0.156 | 0.156 | 0.138 | 0.24 | 0.02 | 0.25 | 0.03 | 0.31 | 0.21 |
| 10 | 97037578 | rs11188256 | -0.237 | 1.436 | -2.174 | 0.353 | -0.03 | -0.093 | -0.093 | 0 | 0.22 | 0.1 | 0.23 | 0.46 | 0.23 | 0.12 |
| 10 | 97039458 | rs10786195 | -0.262 | 1.503 | -1.96 | -0.371 | -0.081 | 0.152 | 0.152 | 0.097 | 0.23 | 0.01 | 0.25 | 0.03 | 0.3 | 0.21 |
| 10 | 97040743 | rs3740508 | -1.19 | 1.657 | -0.929 | 0.392 | 0.045 | 0.064 | 0.064 | 0 | 0.12 | 0.31 | 0.16 | 0.26 | 0.13 | 0.1 |

| | | | | | | | | | | | | | | | | |
|---|---|---|---|---|---|---|---|---|---|---|---|---|---|---|---|---|
| 10 | 97043321 | rs7921287 | -0.199 | 0.998 | -0.718 | 0.545 | -0.017 | 0.094 | 0.094 | 0 | 0.23 | 0.47 | 0.19 | 0.4 | 0.17 | 0.2 |
| 10 | 97043999 | rs4083635 | 0.118 | 2.284 | -1.075 | 0.711 | -0.049 | -0.022 | -0.022 | 0 | 0.14 | 0.02 | 0.28 | 0.15 | 0.21 | 0.2 |
| 10 | 97047200 | rs1471730 | 0.11 | 1.329 | -1.214 | -0.183 | -0.001 | 0.005 | 0.005 | 0 | 0.3 | 0.3 | 0.34 | 0.44 | 0.34 | 0.45 |
| 10 | 97053775 | rs1328599 | -0.209 | 0.671 | -1.081 | -0.352 | 0.005 | 0.003 | 0.003 | 0 | 0.89 | 0.92 | 0.93 | 0.99 | 0.91 | 0.95 |
| 10 | 97057643 | rs7900898 | 0.141 | 0.395 | -0.906 | 0.041 | -0.014 | 0.036 | 0.036 | 0 | 0.14 | 0.22 | 0.2 | 0.29 | 0.12 | 0.2 |
| 10 | 97058246 | rs12761915 | 0.273 | 0.276 | -0.875 | -0.848 | 0.02 | 0.01 | 0.01 | 1.278 | 0.44 | 0.39 | 0.23 | 0.05 | 0.36 | 0.15 |
| 10 | 97070115 | rs7084752 | -0.92 | 0.962 | -0.501 | -0.276 | 0 | -0.001 | -0.001 | 0 | 0.31 | 0.38 | 0.3 | 0.71 | 0.38 | 0.44 |
| 10 | 97070661 | rs1410601 | -0.53 | 1.11 | -0.511 | 0.636 | -0.035 | -0.03 | -0.03 | 0 | 0.19 | 0.14 | 0.17 | 0.27 | 0.27 | 0.16 |
| 10 | 97070709 | rs1410599 | -0.207 | 0.578 | 0.11 | 0.139 | -0.026 | 0.032 | 0.032 | 0 | 0.14 | 0.28 | 0.3 | 0.22 | 0.13 | 0.1 |
| 10 | 97070850 | rs1410598 | 1.961 | 0.366 | 0.202 | -0.079 | 0.003 | 0.2 | 0.2 | 0.567 | 0.5 | 0.21 | 0.4 | 0.06 | 0.47 | 0.45 |
| 10 | 97071334 | rs3814147 | -1.411 | 0.353 | 0.957 | 0.118 | 0.013 | -0.019 | -0.019 | 0 | 0.36 | 0.47 | 0.32 | 0.68 | 0.41 | 0.35 |
| 10 | 97071428 | rs3814149 | -2.252 | 0.172 | 0.54 | -0.329 | 0.015 | 0.164 | 0.164 | 0.733 | 0.49 | 0.73 | 0.48 | 0.94 | 0.43 | 0.59 |
| 10 | 97071872 | rs9313 | 0.348 | 0.097 | 0.54 | 0.085 | 0 | 0.103 | 0.103 | 1.69 | 0.06 | 0.04 | 0.09 | 0.41 | 0.06 | 0.15 |
| 10 | 97074424 | rs2296966 | 0.467 | 0.243 | 0.598 | 0.125 | -0.008 | 0.056 | 0.056 | 0.452 | 0.06 | 0.02 | 0.09 | 0.39 | 0.07 | 0.15 |
| 10 | 97081443 | rs3740510 | 0.304 | 0.233 | 0.57 | 0.126 | -0.021 | 0.024 | 0.024 | 0.66 | 0.06 | 0.01 | 0.09 | 0.41 | 0.07 | 0.15 |
| 10 | 97082300 | rs11188277 | -0.74 | 0.002 | 0.799 | -0.043 | 0.004 | -0.004 | -0.004 | 0 | 0.14 | 0.12 | 0.27 | 0.22 | 0.11 | 0.12 |
| 10 | 97083031 | rs7904853 | 2.146 | -0.018 | 0.831 | -0.295 | 0.016 | 0.191 | 0.191 | 6.6 | 0.49 | 0.26 | 0.4 | 0.05 | 0.47 | 0.42 |
| 10 | 97087059 | rs2078276 | 2.248 | 0.016 | 0.845 | -0.297 | 0.01 | 0.073 | 0.073 | 2.393 | 0.48 | 0.31 | 0.59 | 0.27 | 0.47 | 0.44 |
| 10 | 97087522 | rs7085635 | -1.307 | 0.071 | 0.639 | -0.366 | 0.006 | 0.11 | 0.11 | 0 | 0.18 | 0.04 | 0.1 | 0.01 | 0.14 | 0.14 |
| 10 | 97087852 | rs11188279 | -0.548 | 0.116 | 0.916 | -0.289 | 0.002 | 0.133 | 0.133 | 0.019 | 0.1 | 0.16 | 0.1 | 0.41 | 0.09 | 0.17 |
| 10 | 97095793 | rs10748630 | 2.124 | 0.03 | 1.115 | -0.303 | 0.005 | 0.002 | 0.002 | 0.704 | 0.5 | 0.34 | 0.46 | 0.44 | 0.48 | 0.55 |
| 10 | 97105575 | rs1410602 | -1.985 | -0.013 | 1.009 | -0.317 | 0.012 | -0.008 | -0.008 | 0 | 0.31 | 0.21 | 0.43 | 0.67 | 0.29 | 0.43 |
| 10 | 97106165 | rs726176 | -2.322 | -0.014 | 1.146 | -0.325 | 0.013 | -0.001 | -0.001 | 0.364 | 0.34 | 0.21 | 0.45 | 0.67 | 0.32 | 0.47 |
| 10 | 97107098 | rs11188287 | -1.713 | -0.026 | 1.241 | -0.328 | -0.006 | 0.012 | 0.012 | 0 | 0.25 | 0.32 | 0.41 | 0.68 | 0.24 | 0.39 |
| 10 | 97109019 | rs10882583 | 0.39 | 0.129 | 1.253 | -0.365 | -0.02 | -0.009 | -0.009 | 0 | 0.78 | 0.79 | 0.81 | 0.73 | 0.7 | 0.84 |
| 10 | 97113043 | rs7096152 | -2.172 | 0.102 | 1.323 | -0.36 | 0.002 | 0 | 0 | 0.974 | 0.35 | 0.31 | 0.46 | 0.67 | 0.33 | 0.47 |
| 10 | 97114560 | rs2274632 | 0.383 | 0.115 | 1.238 | -0.307 | 0.001 | 0.131 | 0.131 | 0.502 | 0.07 | 0.11 | 0.1 | 0.4 | 0.07 | 0.15 |
| 10 | 97114941 | rs3758582 | -1.828 | 0.095 | 1.266 | -0.33 | -0.01 | 0.024 | 0.024 | 0 | 0.25 | 0.38 | 0.41 | 0.67 | 0.25 | 0.38 |
| 10 | 97118644 | rs7899506 | 3.169 | 0.075 | 1.287 | -0.38 | 0.004 | 0.128 | 0.128 | 4.166 | 0.4 | 0.19 | 0.34 | 0.05 | 0.4 | 0.36 |
| 10 | 97119049 | rs7900095 | 1.11 | 0.152 | 1.211 | -0.34 | -0.015 | -0.001 | -0.001 | 0 | 0.71 | 0.69 | 0.73 | 0.32 | 0.64 | 0.69 |
| 10 | 97119930 | rs7919009 | -1.737 | 0.146 | 1.226 | -0.323 | -0.001 | -0.001 | -0.001 | 0 | 0.26 | 0.25 | 0.35 | 0.27 | 0.24 | 0.3 |
| 10 | 97122430 | rs10882584 | 0.45 | 0.321 | 1.021 | -0.121 | 0 | 0.106 | 0.106 | 0.522 | 0.07 | 0.09 | 0.1 | 0.41 | 0.07 | 0.15 |
| 10 | 97123885 | rs955760 | 1.573 | 0.313 | 1.019 | -0.2 | -0.031 | 0.022 | 0.022 | 3.367 | 0.63 | 0.69 | 0.68 | 0.32 | 0.58 | 0.64 |
| 10 | 97125874 | rs11188295 | -0.916 | 0.4 | 1.189 | 0.373 | 0.034 | 0.275 | 0.275 | 0.325 | 0.12 | 0.37 | 0.1 | 0.35 | 0.09 | 0.17 |
| 10 | 97127462 | rs6583998 | -0.324 | 0.549 | 0.789 | 0.185 | -0.006 | -0.007 | -0.007 | 0 | 0.86 | 0.87 | 0.86 | 0.76 | 0.75 | 0.88 |
| 10 | 97130450 | rs12269572 | -1.298 | 0.059 | 1.148 | -0.024 | -0.053 | -0.053 | -0.053 | 0.093 | 0.18 | 0.38 | 0.11 | 0.1 | 0.12 | 0.17 |
| 10 | 97141523 | rs2274490 | 1.415 | 0.061 | 1.003 | -0.041 | 0.002 | -0.001 | -0.001 | 0 | 0.65 | 0.43 | 0.72 | 0.65 | 0.6 | 0.68 |
| 10 | 97142245 | rs2025852 | 1.906 | 0.074 | 0.924 | -0.035 | -0.015 | 0.055 | 0.055 | 0 | 0.56 | 0.34 | 0.62 | 0.23 | 0.51 | 0.5 |
| 10 | 97154698 | rs11188311 | 1.871 | 0.114 | 0.875 | 0.012 | -0.011 | 0.049 | 0.049 | 0 | 0.55 | 0.36 | 0.63 | 0.26 | 0.53 | 0.56 |
| 10 | 97155481 | rs10882589 | 0.225 | 0.127 | 0.316 | 0.064 | -0.026 | -0.047 | -0.047 | 0 | 0.08 | 0 | 0.27 | 0.18 | 0.11 | 0.09 |
| 10 | 97156908 | rs17110707 | -0.369 | 0.089 | 0.327 | 0.072 | -0.006 | -0.099 | -0.099 | 0 | 0.15 | 0.05 | 0.14 | 0.42 | 0.14 | 0.18 |
| 10 | 97158713 | rs2274489 | 3.021 | -0.005 | 0.381 | -0.127 | -0.016 | 0.167 | 0.167 | 1.545 | 0.4 | 0.13 | 0.34 | 0.08 | 0.4 | 0.4 |
| 10 | 97160513 | rs1536562 | -2.588 | -0.151 | 0.589 | -0.058 | 0.037 | 0.041 | 0.041 | 0.432 | 0.43 | 0.72 | 0.51 | 0.71 | 0.33 | 0.45 |
| 10 | 97164526 | rs11188312 | -0.683 | 0.087 | 0.616 | 0.063 | -0.001 | -0.008 | -0.008 | 0.422 | 0.16 | 0.06 | 0.32 | 0.19 | 0.18 | 0.15 |
| 10 | 97168140 | rs1327559 | -0.75 | 0.098 | 1.05 | 0.107 | 0.028 | 0.059 | 0.059 | 0.095 | 0.05 | 0.25 | 0.05 | 0.08 | 0.03 | 0.1 |
| 10 | 97169467 | rs4641389 | 1.046 | 0.398 | 0.546 | -0.084 | -0.018 | -0.008 | -0.008 | 0 | 0.67 | 0.68 | 0.48 | 0.57 | 0.53 | 0.69 |
| 10 | 97170181 | rs7907756 | -0.661 | 0.273 | 0.804 | 0.024 | -0.011 | 0.032 | 0.032 | 0 | 0.12 | 0.39 | 0.06 | 0.1 | 0.07 | 0.12 |
| 10 | 97171430 | rs7088546 | 1.812 | 0.289 | 0.21 | 0.087 | 0.032 | -0.001 | -0.001 | 0 | 0.49 | 0.23 | 0.4 | 0.49 | 0.4 | 0.55 |
| 10 | 97173273 | rs7090619 | -0.194 | 1.052 | 0.166 | 0.469 | 0.025 | 0.005 | 0.005 | 0 | 0.87 | 0.84 | 0.86 | 0.8 | 0.72 | 0.88 |
| 10 | 97192452 | rs732101 | -1.241 | 1.102 | 0.402 | 0.262 | 0.024 | 0.019 | 0.019 | 3.544 | 0.31 | 0.5 | 0.43 | 0.66 | 0.33 | 0.47 |
| 10 | 97200044 | rs10882596 | -1.375 | 1.016 | 0.387 | 0.21 | 0.018 | 0.008 | 0.008 | 3.117 | 0.33 | 0.54 | 0.43 | 0.66 | 0.34 | 0.48 |
| 10 | 97210815 | rs17456435 | 0.211 | 1.383 | -0.209 | 0.078 | -0.039 | 0.034 | 0.034 | 0 | 0.09 | 0.02 | 0.09 | 0.02 | 0.19 | 0.03 |
| 10 | 97210983 | rs11592098 | 2.486 | 0.506 | -0.706 | -0.394 | 0.007 | 0.071 | 0.071 | 0.628 | 0.3 | 0.13 | 0.29 | 0.05 | 0.28 | 0.35 |
| 10 | 97212537 | rs11188324 | 0.754 | 0.828 | -0.766 | 0.108 | -0.026 | -0.015 | -0.015 | 0.054 | 0.1 | 0.05 | 0.12 | 0.42 | 0.11 | 0.21 |
| 10 | 97218566 | rs11188326 | 0.352 | 0.746 | -0.81 | -0.025 | -0.012 | -0.018 | -0.018 | 0 | 0.11 | 0.03 | 0.22 | 0.17 | 0.12 | 0.11 |
| 10 | 97219175 | rs7913770 | -0.225 | 0.519 | -0.846 | -0.37 | -0.01 | -0.007 | -0.007 | 0 | 0.69 | 0.85 | 0.75 | 0.69 | 0.64 | 0.81 |
| 10 | 97219419 | rs7899065 | 0.094 | 0.516 | -0.628 | -0.357 | -0.002 | 0.012 | 0.012 | 0 | 0.65 | 0.79 | 0.73 | 0.69 | 0.62 | 0.8 |
| 10 | 97224160 | rs11188327 | -0.86 | 0.684 | -0.61 | -0.072 | -0.012 | -0.03 | -0.03 | 0 | 0.23 | 0.18 | 0.36 | 0.59 | 0.25 | 0.33 |
| 10 | 97228795 | rs12414693 | 1.572 | 0.628 | -0.626 | -0.593 | 0.01 | 0.007 | 0.007 | 0.014 | 0.33 | 0.24 | 0.27 | 0.02 | 0.25 | 0.35 |
| 10 | 97229762 | rs579342 | 0.914 | 0.754 | -0.858 | -0.424 | -0.016 | 0.022 | 0.022 | 0 | 0.08 | 0.02 | 0.05 | 0.02 | 0.1 | 0.08 |
| 10 | 97234991 | rs4917690 | 0.566 | 0.713 | -0.594 | -0.066 | -0.008 | 0.006 | 0.006 | 0 | 0.61 | 0.66 | 0.73 | 0.67 | 0.54 | 0.68 |
| 10 | 97235508 | rs7085831 | -0.493 | 0.682 | -0.312 | 0.233 | 0.002 | 0.02 | 0.02 | 0 | 0.2 | 0.23 | 0.25 | 0.36 | 0.18 | 0.24 |
| 10 | 97236927 | rs10882600 | -0.024 | 0.77 | -0.127 | 0.307 | -0.001 | -0.001 | -0.001 | 3.295 | 0.3 | 0.31 | 0.34 | 0.33 | 0.34 | 0.19 |
| 10 | 97237794 | rs998292 | 0.115 | 0.918 | 0.053 | 0.305 | -0.001 | -0.001 | -0.001 | 0 | 0.62 | 0.58 | 0.61 | 0.63 | 0.57 | 0.71 |
| 10 | 97239708 | rs7071071 | 0.755 | 1.182 | 0.239 | 0.398 | 0.003 | 0.016 | 0.016 | 0 | 0.4 | 0.33 | 0.47 | 0.31 | 0.38 | 0.43 |
| 10 | 97240100 | rs9988729 | -0.355 | 1.163 | 0.094 | 0.355 | -0.002 | 0.003 | 0.003 | 0 | 0.58 | 0.57 | 0.59 | 0.62 | 0.56 | 0.7 |
| 10 | 97246684 | rs725926 | 1.876 | 0.985 | -0.07 | -0.087 | -0.03 | 0.148 | 0.148 | 0 | 0.26 | 0.05 | 0.25 | 0.02 | 0.24 | 0.33 |
| 10 | 97250630 | rs943345 | 0.056 | 1.05 | 0.311 | 0.246 | 0.015 | 0.146 | 0.146 | 0 | 0.1 | 0.22 | 0.23 | 0.36 | 0.14 | 0.14 |
| 10 | 97252314 | rs1571131 | 0.391 | 0.898 | 0.163 | 0.07 | -0.052 | -0.018 | -0.018 | 0 | 0.38 | 0.22 | 0.37 | 0.59 | 0.45 | 0.4 |

| | | | | | | | | | | | | | | | | |
|---|---|---|---|---|---|---|---|---|---|---|---|---|---|---|---|---|
| 10 | 97253512 | rs6584002 | -0.879 | 0.582 | 0.347 | 0.269 | 0.003 | 0.005 | 0.005 | 0 | 0.85 | 0.87 | 0.91 | 0.98 | 0.87 | 0.89 |
| 10 | 97254594 | rs11188334 | 0.632 | 0.541 | 0.335 | 0.338 | 0.03 | 0.186 | 0.186 | 0.023 | 0.08 | 0.26 | 0.23 | 0.27 | 0.12 | 0.07 |
| 10 | 97257034 | rs1571132 | -0.832 | 0.318 | -0.029 | 0.054 | 0.011 | 0.031 | 0.031 | 0 | 0.85 | 0.91 | 0.91 | 0.98 | 0.88 | 0.88 |
| 10 | 97260248 | rs1326935 | 0.001 | 0.477 | 0.188 | 0 | -0.003 | -0.037 | -0.037 | 0 | 0.8 | 0.74 | 0.86 | 0.98 | 0.81 | 0.87 |
| 10 | 97266538 | rs1998911 | 0.424 | 0.549 | 0.543 | 0.226 | -0.012 | 0.018 | 0.018 | 0.309 | 0.71 | 0.41 | 0.64 | 0.67 | 0.71 | 0.79 |
| 10 | 97266574 | rs11188337 | 0.156 | 0.534 | 0.559 | 0.196 | 0.053 | 0.246 | 0.246 | 1.972 | 0.13 | 0.49 | 0.26 | 0.31 | 0.16 | 0.08 |
| 10 | 97268858 | rs6584004 | -0.244 | 0.628 | 0.335 | 0.21 | 0.013 | 0.064 | 0.064 | 0 | 0.52 | 0.61 | 0.62 | 0.78 | 0.56 | 0.7 |
| 10 | 97269652 | rs4918942 | -1.148 | 0.535 | 0.332 | 0.203 | 0.013 | 0.024 | 0.024 | 0 | 0.84 | 0.89 | 0.9 | 0.98 | 0.87 | 0.87 |
| 10 | 97269830 | rs17536344 | 2.077 | 0.586 | 0.195 | 0.117 | -0.022 | 0.097 | 0.097 | 0 | 0.2 | 0.03 | 0.21 | 0.05 | 0.2 | 0.3 |
| 10 | 97269933 | rs12776562 | 1.224 | 0.558 | 0.216 | 0.113 | 0 | 0.019 | 0.019 | 0 | 0.32 | 0.29 | 0.28 | 0.2 | 0.32 | 0.18 |
| 10 | 97270729 | rs12247017 | 1.833 | 0.596 | 0.193 | 0.148 | -0.008 | 0.001 | 0.001 | 0.185 | 0.08 | 0.05 | 0.08 | 0 | 0.12 | 0.02 |
| 10 | 97272774 | rs7912249 | -0.297 | 0.59 | 0.2 | 0.145 | 0.001 | 0.033 | 0.033 | 0 | 0.76 | 0.84 | 0.82 | 0.98 | 0.75 | 0.85 |
| 10 | 97274838 | rs10882607 | 1.165 | 0.625 | 0.278 | 0.297 | 0.002 | 0.03 | 0.03 | 0 | 0.14 | 0.16 | 0.14 | 0.38 | 0.15 | 0.25 |
| 10 | 97276617 | rs7089353 | 1.133 | 0.549 | 0.612 | 0.307 | 0.013 | 0.089 | 0.089 | 0 | 0.1 | 0.15 | 0.23 | 0.31 | 0.16 | 0.08 |
| 10 | 97278123 | rs943347 | 0.176 | 0.556 | 0.526 | 0.303 | 0.044 | 0.148 | 0.148 | 0 | 0.15 | 0.36 | 0.26 | 0.31 | 0.19 | 0.09 |
| 10 | 97283567 | rs17110913 | 1.052 | 0.404 | 0.531 | 0.351 | 0.036 | 0.169 | 0.169 | 0 | 0.1 | 0.26 | 0.24 | 0.31 | 0.16 | 0.08 |
| 10 | 97284081 | rs1326934 | -0.639 | 0.334 | 0.272 | 0.043 | 0.02 | 0.049 | 0.049 | 0 | 0.54 | 0.64 | 0.66 | 0.72 | 0.61 | 0.65 |
| 10 | 97286853 | rs10882610 | 1.288 | 0.396 | 0.181 | -0.057 | 0.004 | 0.01 | 0.01 | 0 | 0.3 | 0.26 | 0.24 | 0.26 | 0.26 | 0.23 |
| 10 | 97289992 | rs11188346 | 1.553 | 0.4 | 0.127 | -0.067 | 0.005 | 0.021 | 0.021 | 0 | 0.27 | 0.2 | 0.22 | 0.2 | 0.25 | 0.17 |
| 10 | 97295922 | rs1536442 | 0.378 | 0.375 | 0.234 | 0.103 | -0.003 | 0.006 | 0.006 | 0 | 0.57 | 0.45 | 0.57 | 0.34 | 0.55 | 0.53 |
| 10 | 97297359 | rs4917696 | 1.113 | 0.329 | 0.398 | 0.414 | 0.005 | 0.008 | 0.008 | 0 | 0.17 | 0.18 | 0.31 | 0.23 | 0.28 | 0.1 |
| 10 | 97299171 | rs10882613 | 0.804 | 0.24 | -0.061 | 0.256 | -0.026 | -0.066 | -0.066 | 0 | 0.4 | 0.16 | 0.29 | 0.57 | 0.4 | 0.37 |
| 10 | 97300128 | rs4357625 | -0.816 | -0.278 | 0.512 | 0.491 | -0.009 | -0.015 | -0.015 | 0 | 0.1 | 0.26 | 0.05 | 0.06 | 0.05 | 0.08 |
| 10 | 97300636 | rs7085647 | 0.148 | -0.283 | 0.514 | 0.48 | -0.015 | -0.022 | -0.022 | 0.016 | 0.26 | 0.39 | 0.29 | 0.18 | 0.19 | 0.42 |
| 10 | 97301004 | rs11188352 | -0.354 | -0.184 | 0.441 | 0.476 | 0.018 | 0.019 | 0.019 | 0 | 0.07 | 0.25 | 0.03 | 0.06 | 0.03 | 0.08 |
| 10 | 97305536 | rs1536444 | 0.51 | -0.107 | 0.3 | 0.535 | -0.003 | -0.009 | -0.009 | 0 | 0.57 | 0.45 | 0.61 | 0.78 | 0.68 | 0.46 |
| 10 | 97308114 | rs1926817 | -0.544 | 0.282 | -0.067 | 0.829 | 0.002 | 0.003 | 0.003 | 0 | 0.87 | 0.9 | 0.91 | 0.99 | 0.88 | 0.89 |
| 10 | 97308571 | rs7098531 | -0.654 | 0.285 | -0.085 | 0.835 | 0.004 | 0.026 | 0.026 | 0 | 0.87 | 0.94 | 0.92 | 0.99 | 0.89 | 0.89 |
| 10 | 97315080 | rs10882614 | 0.774 | 0.288 | -0.069 | 0.919 | 0.001 | 0.001 | 0.001 | 0 | 0.19 | 0.15 | 0.25 | 0.15 | 0.16 | 0.34 |
| 10 | 97315208 | rs10882615 | 0.754 | 0.295 | -0.065 | 0.922 | 0.002 | 0.001 | 0.001 | 0 | 0.19 | 0.15 | 0.25 | 0.15 | 0.16 | 0.34 |
| 10 | 97316040 | rs4918948 | 0.473 | 0.443 | 0.019 | 1.223 | 0 | -0.002 | -0.002 | 0 | 0.54 | 0.44 | 0.57 | 0.46 | 0.57 | 0.34 |
| 10 | 97317158 | rs10786222 | -0.037 | 0.215 | -0.524 | 0.527 | 0.005 | 0.011 | 0.011 | 0 | 0.68 | 0.75 | 0.67 | 0.78 | 0.72 | 0.54 |
| 10 | 97321923 | rs7918235 | -0.808 | -0.149 | 0.308 | 0.654 | -0.024 | 0.074 | 0.074 | 1.523 | 0.08 | 0.29 | 0.07 | 0.15 | 0.02 | 0.24 |
| 10 | 97324127 | rs7070798 | -0.719 | -1.384 | -0.372 | 0.145 | 0.131 | 0.071 | 0.071 | 0 | 0.49 | 0.74 | 0.27 | 0.35 | 0.3 | 0.5 |
| 10 | 97325341 | rs12359410 | 0.643 | -0.963 | -0.313 | 0.083 | 0.004 | -0.004 | -0.004 | 0 | 0.08 | 0.13 | 0.08 | 0.09 | 0.09 | 0.1 |
| 10 | 97325729 | rs7918947 | -1.017 | -0.97 | -0.872 | -0.414 | -0.159 | -0.15 | -0.15 | 3.032 | 0.7 | 0.98 | 0.45 | 0.44 | 0.52 | 0.66 |
| 10 | 97328211 | rs12248661 | 0.71 | -1.629 | -0.627 | -0.678 | -0.006 | 0.006 | 0.006 | 5.165 | 0.08 | 0.17 | 0.04 | 0.08 | 0.05 | 0.04 |
| 10 | 97328227 | rs12255572 | 0.71 | -1.629 | -0.627 | -0.678 | -0.006 | 0.006 | 0.006 | 5.165 | 0.08 | 0.17 | 0.04 | 0.08 | 0.05 | 0.04 |
| 10 | 97331958 | rs7099430 | 0.403 | -1.748 | -0.202 | -0.73 | -0.005 | 0.009 | 0.009 | 4.804 | 0.09 | 0.22 | 0.04 | 0.08 | 0.05 | 0.04 |
| 10 | 97332195 | rs17110984 | 1.3 | -2.618 | -0.001 | -1.153 | 0.021 | 0.019 | 0.019 | 0.607 | 0.25 | 0.21 | 0.13 | 0.13 | 0.17 | 0.18 |
| 10 | 97333879 | rs501603 | -0.416 | -2.731 | 0.116 | -1.164 | -0.049 | -0.015 | -0.015 | 0.003 | 0.86 | 0.72 | 0.9 | 0.86 | 0.93 | 0.71 |
| 10 | 97334479 | rs533383 | 0.918 | -2.368 | -0.127 | -1.258 | -0.055 | -0.018 | -0.018 | 0.12 | 0.4 | 0.09 | 0.63 | 0.62 | 0.6 | 0.39 |
| 10 | 97339438 | rs493193 | 0.809 | -1.548 | -0.142 | -1.251 | -0.061 | -0.026 | -0.026 | 0 | 0.42 | 0.09 | 0.64 | 0.59 | 0.59 | 0.39 |
| 10 | 97339930 | rs7895441 | 0.604 | -1.672 | -0.049 | -1.249 | -0.026 | -0.018 | -0.018 | 0 | 0.34 | 0.42 | 0.17 | 0.2 | 0.22 | 0.21 |
| 10 | 97345469 | rs10509689 | 1.573 | -1.843 | -0.018 | -1.325 | 0.036 | 0.029 | 0.029 | 0.561 | 0.23 | 0.12 | 0.12 | 0.13 | 0.16 | 0.17 |
| 10 | 97348734 | rs524295 | 0.898 | -1.913 | -0.082 | -1.329 | -0.017 | -0.032 | -0.032 | 0 | 0.42 | 0.09 | 0.64 | 0.58 | 0.61 | 0.39 |
| 10 | 97348845 | rs10882635 | 0.371 | -1.904 | 0.156 | -1.318 | -0.008 | 0.003 | 0.003 | 1.411 | 0.11 | 0.24 | 0.04 | 0.08 | 0.06 | 0.05 |
| 10 | 97353426 | rs7906450 | 0.903 | -1.856 | 0.069 | -1.314 | -0.058 | -0.019 | -0.019 | 1.632 | 0.29 | 0.43 | 0.15 | 0.21 | 0.17 | 0.2 |
| 10 | 97353584 | rs1270505 | 1.455 | -1.8 | -0.074 | -1.303 | -0.163 | -0.151 | -0.151 | 0 | 0.29 | 0.02 | 0.55 | 0.53 | 0.48 | 0.34 |
| 10 | 97361407 | rs540963 | 0.078 | -1.831 | 0.089 | -1.294 | -0.032 | -0.041 | -0.041 | 0 | 0.77 | 0.57 | 0.82 | 0.79 | 0.83 | 0.62 |
| 10 | 97362028 | rs476392 | 0.899 | -1.446 | -0.301 | -1.173 | -0.002 | 0.09 | 0.09 | 0 | 0.08 | 0.03 | 0.09 | 0.5 | 0.08 | 0.13 |
| 10 | 97362091 | rs568401 | 0.899 | -1.446 | -0.301 | -1.173 | -0.002 | 0.09 | 0.09 | 0 | 0.08 | 0.03 | 0.09 | 0.5 | 0.08 | 0.13 |
| 10 | 97363344 | rs7905711 | 1.133 | -1.508 | -0.359 | -1.282 | -0.11 | 0.135 | 0.135 | 0 | 0.34 | 0.09 | 0.56 | 0.09 | 0.53 | 0.26 |
| 10 | 97364616 | rs7913204 | 0.607 | -0.549 | -0.291 | -1.304 | -0.001 | -0.003 | -0.003 | 1.872 | 0.61 | 0.5 | 0.64 | 0.31 | 0.56 | 0.47 |
| 10 | 97365820 | rs4037 | 2.105 | -0.619 | -0.403 | -1.472 | -0.056 | 0.055 | 0.055 | 0 | 0.21 | 0.1 | 0.44 | 0.04 | 0.31 | 0.21 |
| 10 | 97368990 | rs10882639 | 1.781 | -0.76 | -0.363 | -1.459 | -0.005 | -0.001 | -0.001 | 1.052 | 0.26 | 0.29 | 0.14 | 0.21 | 0.17 | 0.19 |
| 10 | 97369939 | rs10882640 | 0.813 | -0.824 | -0.414 | -1.498 | -0.062 | 0.138 | 0.138 | 0 | 0.36 | 0.16 | 0.54 | 0.08 | 0.46 | 0.28 |
| 10 | 97370331 | rs497331 | -1.775 | -0.95 | -0.117 | -1.526 | -0.025 | 0.07 | 0.07 | 2.113 | 0.93 | 0.76 | 0.93 | 0.84 | 0.98 | 0.77 |
| 10 | 97372559 | rs1409709 | 0.805 | -0.822 | -0.404 | -1.549 | -0.062 | 0.138 | 0.138 | 0 | 0.36 | 0.16 | 0.54 | 0.08 | 0.46 | 0.28 |
| 10 | 97373966 | rs2296692 | 0.433 | -0.837 | -0.409 | -1.556 | -0.068 | 0.181 | 0.181 | 0.042 | 0.41 | 0.17 | 0.56 | 0.08 | 0.51 | 0.3 |
| 10 | 97378785 | rs535449 | 0.863 | -0.844 | -0.33 | -1.554 | -0.002 | -0.015 | -0.015 | 0.483 | 0.36 | 0.43 | 0.25 | 0.71 | 0.33 | 0.34 |
| 10 | 97383926 | rs501819 | 0.863 | -0.844 | -0.33 | -1.554 | -0.002 | -0.015 | -0.015 | 0 | 0.36 | 0.43 | 0.25 | 0.71 | 0.33 | 0.34 |
| 10 | 97385242 | rs10509688 | 0.698 | -0.808 | -0.303 | -1.491 | -0.008 | 0.039 | 0.039 | 0 | 0.1 | 0.06 | 0.11 | 0.5 | 0.16 | 0.15 |
| 10 | 97385492 | rs10509687 | -1.165 | -0.926 | -0.296 | -1.584 | -0.056 | 0.004 | 0.004 | 0.727 | 0.86 | 0.69 | 0.9 | 0.83 | 0.95 | 0.75 |
| 10 | 97385662 | rs10748634 | -0.086 | -0.814 | -0.291 | -1.537 | -0.04 | -0.038 | -0.038 | 0 | 0.77 | 0.58 | 0.81 | 0.79 | 0.83 | 0.63 |
| 10 | 97388162 | rs2275272 | 1.63 | -0.694 | -0.346 | -1.534 | 0.018 | 0.057 | 0.057 | 0 | 0.13 | 0.04 | 0.09 | 0.04 | 0.14 | 0.06 |
| 10 | 97392969 | rs3750701 | 0.508 | -0.696 | -0.31 | -1.55 | -0.024 | -0.052 | -0.052 | 0 | 0.58 | 0.53 | 0.69 | 0.75 | 0.64 | 0.56 |
| 10 | 97393475 | rs11188409 | -0.228 | -0.654 | -0.282 | -1.533 | 0.014 | 0.046 | 0.046 | 0 | 0.14 | 0.16 | 0.17 | 0.54 | 0.22 | 0.23 |
| 10 | 97397227 | rs2275273 | 0.356 | -0.743 | -0.219 | -1.87 | -0.036 | -0.053 | -0.053 | 0 | 0.64 | 0.54 | 0.71 | 0.75 | 0.69 | 0.57 |

| | | | | | | | | | | | | | | | | |
|---|---|---|---|---|---|---|---|---|---|---|---|---|---|---|---|---|
| 10 | 97408603 | rs4258314 | 1.991 | -0.686 | -0.351 | -1.872 | -0.056 | 0.055 | 0.055 | 0 | 0.21 | 0.1 | 0.44 | 0.04 | 0.31 | 0.21 |
| 10 | 97421345 | rs2282305 | 0.808 | -0.776 | -0.348 | -1.877 | 0.005 | 0.025 | 0.025 | 0 | 0.17 | 0.14 | 0.16 | 0.08 | 0.2 | 0.14 |
| 10 | 97426086 | rs10882649 | 1.977 | -0.401 | -0.373 | -1.903 | -0.052 | 0.058 | 0.058 | 0 | 0.21 | 0.1 | 0.43 | 0.03 | 0.29 | 0.22 |
| 10 | 97438823 | rs10882651 | 0.979 | -0.511 | -0.323 | -1.927 | 0.001 | -0.015 | -0.015 | 1.107 | 0.36 | 0.43 | 0.26 | 0.71 | 0.35 | 0.33 |
| 10 | 97440159 | rs11595248 | 1.557 | -0.544 | -0.305 | -1.883 | 0.023 | 0.056 | 0.056 | 0 | 0.14 | 0.04 | 0.09 | 0.04 | 0.14 | 0.06 |
| 10 | 97440394 | rs12784530 | 0.126 | -0.544 | -0.296 | -1.883 | 0.005 | 0.043 | 0.043 | 0 | 0.23 | 0.16 | 0.19 | 0.08 | 0.25 | 0.15 |
| 10 | 97451518 | rs10748638 | 0.688 | -0.529 | -0.262 | -1.784 | -0.022 | -0.039 | -0.039 | 0.556 | 0.57 | 0.53 | 0.68 | 0.72 | 0.64 | 0.57 |
| 10 | 97455055 | rs10786230 | -0.545 | -0.48 | -0.157 | -1.711 | -0.056 | -0.021 | -0.021 | 5.184 | 0.81 | 0.68 | 0.88 | 0.82 | 0.89 | 0.71 |
| 10 | 97466391 | rs10748641 | -0.545 | -0.415 | 0.086 | -1.72 | -0.056 | -0.021 | -0.021 | 0.468 | 0.81 | 0.68 | 0.88 | 0.82 | 0.89 | 0.71 |
| 10 | 97473214 | rs3949478 | 0.354 | -0.398 | 0.008 | -1.736 | -0.034 | -0.048 | -0.048 | 0 | 0.72 | 0.57 | 0.78 | 0.78 | 0.78 | 0.62 |
| 10 | 97473403 | rs12783226 | 0.816 | -0.771 | -0.374 | -1.98 | -0.032 | -0.049 | -0.049 | 0.491 | 0.16 | 0.07 | 0.13 | 0.57 | 0.21 | 0.21 |
| 10 | 97486825 | rs1933168 | 1.919 | -0.839 | -0.28 | -2.014 | -0.015 | -0.009 | -0.009 | 0 | 0.2 | 0.29 | 0.12 | 0.14 | 0.14 | 0.12 |
| 10 | 97487362 | rs1933166 | 0.368 | -0.684 | -0.282 | -1.917 | -0.056 | -0.021 | -0.021 | 0.084 | 0.19 | 0.32 | 0.12 | 0.18 | 0.11 | 0.29 |
| 10 | 97496989 | rs4501930 | 0.193 | -0.679 | -0.328 | -1.921 | -0.021 | -0.002 | -0.002 | 0 | 0.2 | 0.41 | 0.19 | 0.22 | 0.17 | 0.37 |
| 10 | 97515137 | rs3814159 | -0.075 | -0.605 | -0.499 | -2.021 | 0 | -0.01 | -0.01 | 0.261 | 0.44 | 0.52 | 0.62 | 0.25 | 0.46 | 0.59 |
| 10 | 97515588 | rs3176892 | 2.668 | -0.622 | -0.526 | -2.051 | -0.008 | 0.021 | 0.021 | 7.358 | 0.05 | 0.01 | 0.02 | 0 | 0.05 | 0.01 |
| 10 | 97529318 | rs4582902 | -0.051 | -0.62 | -0.524 | -2.048 | 0 | -0.01 | -0.01 | 0 | 0.44 | 0.51 | 0.62 | 0.25 | 0.46 | 0.59 |
| 10 | 97539737 | rs11188484 | 2.041 | -0.624 | -0.556 | -1.942 | -0.05 | 0.053 | 0.053 | 0 | 0.21 | 0.1 | 0.43 | 0.05 | 0.29 | 0.22 |
| 10 | 97602091 | rs3181131 | -0.443 | -0.64 | -0.475 | -2.021 | 0 | 0 | 0 | 3.419 | 0.49 | 0.52 | 0.65 | 0.25 | 0.51 | 0.6 |
| 10 | 97603442 | rs3181127 | 2.64 | -0.613 | -0.5 | -1.986 | -0.003 | 0.009 | 0.009 | 7.358 | 0.05 | 0.02 | 0.03 | 0 | 0.05 | 0.01 |
| 10 | 97623376 | rs3181118 | 2.435 | -0.628 | -0.481 | -2.007 | -0.002 | 0.007 | 0.007 | 7.22 | 0.06 | 0.02 | 0.03 | 0 | 0.05 | 0.01 |
| 10 | 97624709 | rs2275759 | -0.45 | -0.574 | -0.452 | -2.016 | 0 | -0.001 | -0.001 | 0.355 | 0.49 | 0.52 | 0.65 | 0.25 | 0.51 | 0.6 |
| 10 | 97628170 | rs3181115 | 2.51 | -0.641 | -0.52 | -2.086 | 0.002 | 0.034 | 0.034 | 0 | 0.09 | 0.03 | 0.07 | 0 | 0.12 | 0.03 |
| 10 | 97632082 | rs2051052 | -0.39 | -0.65 | -0.518 | -2.098 | 0 | -0.002 | -0.002 | 0 | 0.49 | 0.52 | 0.65 | 0.25 | 0.51 | 0.6 |
| 10 | 97632808 | rs3192654 | 2.095 | -0.656 | -0.544 | -2.104 | -0.051 | 0.061 | 0.061 | 0 | 0.21 | 0.1 | 0.43 | 0.03 | 0.29 | 0.22 |
| 10 | 97633021 | rs11188513 | -0.835 | -0.677 | -0.514 | -2.11 | 0.002 | 0.028 | 0.028 | 0.94 | 0.61 | 0.67 | 0.73 | 0.26 | 0.64 | 0.64 |
| 10 | 97638224 | rs7897988 | 0.437 | -1.134 | -0.572 | -2.054 | -0.046 | -0.034 | -0.034 | 0.288 | 0.19 | 0.31 | 0.12 | 0.14 | 0.11 | 0.3 |
| 10 | 97649445 | rs11188516 | 2.003 | -0.904 | -0.812 | -1.98 | -0.048 | 0.052 | 0.052 | 0 | 0.19 | 0.09 | 0.43 | 0.03 | 0.28 | 0.17 |
| 10 | 97650509 | rs7074866 | 1.428 | -0.864 | -0.79 | -1.962 | -0.04 | 0.05 | 0.05 | 0 | 0.23 | 0.15 | 0.46 | 0.07 | 0.29 | 0.26 |
| 10 | 97656287 | rs10882684 | -0.915 | -1.028 | -0.776 | -2.027 | 0 | -0.008 | -0.008 | 0.648 | 0.63 | 0.63 | 0.72 | 0.21 | 0.62 | 0.62 |
| 10 | 97657029 | rs7918631 | 0.027 | -1.39 | -0.63 | -2.077 | -0.039 | -0.056 | -0.056 | 0 | 0.3 | 0.39 | 0.21 | 0.14 | 0.24 | 0.37 |
| 10 | 97666798 | rs11188518 | -0.868 | -1.37 | -0.689 | -2.158 | -0.018 | -0.036 | -0.036 | 0 | 0.41 | 0.47 | 0.3 | 0.19 | 0.32 | 0.47 |
| 10 | 97674087 | rs11599779 | 2.423 | -1.359 | -0.86 | -2.141 | -0.001 | 0.034 | 0.034 | 0 | 0.09 | 0.03 | 0.08 | 0 | 0.13 | 0.03 |
| 10 | 97679956 | rs10786244 | -0.018 | -1.351 | -0.785 | -2.138 | -0.053 | -0.083 | -0.083 | 0.129 | 0.31 | 0.46 | 0.22 | 0.15 | 0.25 | 0.36 |
| 10 | 97685352 | rs10882690 | -0.836 | -1.037 | -0.82 | -2.098 | -0.004 | 0.014 | 0.014 | 0.402 | 0.62 | 0.66 | 0.73 | 0.24 | 0.61 | 0.63 |
| 10 | 97695022 | rs10882692 | 1.041 | -1.186 | -0.686 | -1.812 | -0.02 | 0.049 | 0.049 | 0 | 0.11 | 0.33 | 0.1 | 0.16 | 0.05 | 0.25 |
| 10 | 97698222 | rs1336457 | 0.266 | -1.847 | -1.281 | -1.965 | -0.008 | 0.002 | 0.002 | 0 | 0.31 | 0.32 | 0.19 | 0.62 | 0.18 | 0.34 |
| 10 | 97698356 | rs7907476 | 1.551 | -1.909 | -1.403 | -2.01 | 0.006 | 0.006 | 0.006 | 0 | 0.06 | 0.07 | 0.05 | 0.04 | 0.02 | 0.09 |
| 10 | 97698370 | rs4918979 | -1.088 | -1.756 | -1.518 | -2.11 | -0.027 | 0.056 | 0.056 | 0 | 0.67 | 0.83 | 0.77 | 0.86 | 0.55 | 0.81 |
| 10 | 97704739 | rs11188531 | 2.057 | -1.037 | -0.86 | -1.993 | -0.056 | 0.046 | 0.046 | 0 | 0.19 | 0.09 | 0.43 | 0.05 | 0.31 | 0.13 |
| 10 | 97717072 | rs4917718 | -0.234 | -0.719 | -0.892 | -2.073 | 0.002 | 0.026 | 0.026 | 0 | 0.45 | 0.58 | 0.63 | 0.21 | 0.55 | 0.44 |
| 10 | 97728428 | rs7922216 | 0.06 | -0.58 | -0.669 | -1.85 | -0.029 | -0.061 | -0.061 | 0.19 | 0.26 | 0.49 | 0.21 | 0.16 | 0.25 | 0.31 |
| 10 | 97734022 | rs10509696 | 0.087 | 0.005 | -1.133 | -1.789 | -0.008 | 0.036 | 0.036 | 0 | 0.41 | 0.34 | 0.59 | 0.09 | 0.53 | 0.27 |
| 10 | 97735403 | rs17111253 | 1.73 | -0.397 | -0.328 | -1.053 | -0.001 | -0.02 | -0.02 | 0 | 0.09 | 0.13 | 0.06 | 0 | 0.07 | 0.05 |
| 10 | 97740223 | rs11599105 | 0.356 | -0.188 | -0.159 | -0.723 | -0.001 | -0.069 | -0.069 | 1.257 | 0.14 | 0.22 | 0.09 | 0 | 0.17 | 0.04 |
| 10 | 97748040 | rs7900838 | 0.578 | -0.189 | -0.122 | -0.617 | 0.002 | -0.008 | -0.008 | 0.666 | 0.62 | 0.61 | 0.57 | 0.81 | 0.58 | 0.52 |
| 10 | 97753148 | rs11188544 | 0.111 | -0.752 | -0.058 | 0.07 | -0.008 | 0.061 | 0.061 | 0 | 0.12 | 0.11 | 0.06 | 0.55 | 0.08 | 0.2 |
| 10 | 97759203 | rs2039617 | -0.868 | -1.124 | 0.087 | -0.163 | -0.074 | 0.027 | 0.027 | 0.138 | 0.2 | 0.39 | 0.14 | 0.78 | 0.11 | 0.31 |
| 10 | 97764765 | rs7923326 | 0.566 | -0.879 | 0.269 | -0.32 | 0.003 | 0.016 | 0.016 | 0 | 0.53 | 0.44 | 0.6 | 0.82 | 0.44 | 0.54 |
| 10 | 97764915 | rs10882708 | 0.073 | -1.203 | 0.389 | -0.111 | -0.018 | 0.182 | 0.182 | 0.215 | 0.05 | 0.2 | 0.09 | 0.23 | 0.02 | 0.15 |
| 10 | 97766537 | rs1023331 | -1.019 | -1.313 | 0.31 | -0.455 | 0.007 | 0 | 0 | 0 | 0.79 | 0.8 | 0.75 | 0.81 | 0.66 | 0.72 |
| 10 | 97767830 | rs7096743 | 0.249 | -1.493 | 0.124 | -0.589 | -0.005 | 0.005 | 0.005 | 0 | 0.65 | 0.66 | 0.58 | 0.83 | 0.58 | 0.58 |
| 10 | 97772358 | rs17383738 | 1.649 | -1.451 | 0.208 | -0.622 | 0.018 | -0.04 | -0.04 | 0 | 0.09 | 0.17 | 0.09 | 0.01 | 0.16 | 0.04 |
| 10 | 97777315 | rs10748653 | 0.018 | -1.493 | 0.177 | -0.649 | 0.004 | -0.007 | -0.007 | 0 | 0.7 | 0.68 | 0.72 | 0.85 | 0.63 | 0.7 |
| 10 | 97783874 | rs11814327 | 2.531 | -1.559 | -0.032 | -0.412 | 0.016 | -0.02 | -0.02 | 0.653 | 0.36 | 0.12 | 0.41 | 0.34 | 0.32 | 0.28 |
| 10 | 97792352 | rs10509698 | 1.679 | -1.54 | 0.013 | -0.56 | -0.001 | 0.009 | 0.009 | 0 | 0.07 | 0.04 | 0.05 | 0 | 0.08 | 0.04 |
| 10 | 97797422 | rs7916395 | 0.549 | -1.613 | 0.158 | -0.601 | -0.026 | -0.052 | -0.052 | 0 | 0.1 | 0.19 | 0.04 | 0 | 0.04 | 0.02 |
| 10 | 97805074 | rs915506 | 2.246 | -1.673 | 0.146 | -0.556 | 0.027 | -0.056 | -0.056 | 0 | 0.4 | 0.11 | 0.35 | 0.52 | 0.32 | 0.34 |
| 10 | 97814475 | rs1414918 | 0.544 | -1.569 | 0.343 | -0.499 | 0.01 | -0.031 | -0.031 | 0 | 0.61 | 0.5 | 0.57 | 0.8 | 0.46 | 0.62 |
| 10 | 97837013 | rs11188585 | 0.585 | -1.508 | 0.267 | -0.524 | 0.007 | -0.032 | -0.032 | 0 | 0.6 | 0.5 | 0.56 | 0.8 | 0.45 | 0.62 |
| 10 | 97842565 | rs10748659 | 0.016 | -1.407 | 0.29 | -0.534 | 0.005 | -0.005 | -0.005 | 0.028 | 0.7 | 0.66 | 0.65 | 0.81 | 0.62 | 0.67 |
| 10 | 97851432 | rs12263592 | -0.542 | -0.978 | 0.516 | -0.662 | -0.004 | -0.046 | -0.046 | 0.069 | 0.15 | 0.26 | 0.19 | 0.02 | 0.13 | 0.16 |
| 10 | 97868205 | rs7096164 | 1.262 | -0.908 | 0.105 | -0.619 | 0.016 | -0.036 | -0.036 | 2.037 | 0.1 | 0.16 | 0.08 | 0 | 0.16 | 0.05 |
| 10 | 97884731 | rs1172962 | 0.521 | -0.953 | 0.114 | -0.613 | -0.011 | 0.022 | 0.022 | 0.161 | 0.87 | 0.68 | 0.84 | 0.89 | 0.92 | 0.83 |
| 10 | 97884989 | rs7916068 | 1.326 | -0.883 | 0.037 | -0.623 | 0.02 | -0.039 | -0.039 | 0.969 | 0.09 | 0.15 | 0.08 | 0 | 0.16 | 0.05 |
| 10 | 97889423 | rs4917721 | 1.549 | -0.94 | -0.045 | -0.605 | -0.019 | 0.033 | 0.033 | 0 | 0.06 | 0.01 | 0.05 | 0 | 0.08 | 0.03 |
| 10 | 97900024 | rs12255700 | 1.281 | -0.945 | -0.023 | -0.624 | -0.002 | -0.013 | -0.013 | 0.062 | 0.06 | 0.02 | 0.14 | 0.11 | 0.06 | 0.17 |
| 10 | 97919742 | rs7081962 | 0.289 | -1.072 | 0.084 | -0.668 | -0.03 | -0.058 | -0.058 | 0 | 0.1 | 0.25 | 0.04 | 0 | 0.03 | 0.02 |
| 10 | 97920062 | rs3814226 | 1.36 | -1.128 | 0.009 | -0.619 | 0.025 | 0.053 | 0.053 | 0 | 0.05 | 0 | 0.01 | 0.49 | 0.01 | 0.17 |

| | | | | | | | | | | | | | | | | |
|---|---|---|---|---|---|---|---|---|---|---|---|---|---|---|---|---|
| 10 | 97922576 | rs11591374 | 1.384 | -0.98 | 0.127 | -0.553 | 0.02 | -0.037 | -0.037 | 0 | 0.09 | 0.15 | 0.08 | 0.01 | 0.16 | 0.05 |
| 10 | 97927958 | rs10882731 | 0.065 | -0.628 | -0.162 | -0.477 | -0.031 | -0.035 | -0.035 | 0 | 0.21 | 0.12 | 0.32 | 0.68 | 0.34 | 0.43 |
| 10 | 97947041 | rs10748663 | 0.376 | -0.923 | -0.041 | -0.374 | 0.019 | -0.048 | -0.048 | 0 | 0.78 | 0.53 | 0.76 | 0.89 | 0.76 | 0.78 |
| 10 | 97950486 | rs17111391 | -0.218 | -1.005 | -0.538 | 0.121 | -0.029 | -0.052 | -0.052 | 0 | 0.1 | 0.19 | 0.03 | 0 | 0.03 | 0.02 |
| 10 | 97952042 | rs11188657 | 0.171 | -0.407 | -0.888 | 0.433 | 0.029 | 0.031 | 0.031 | 0 | 0.06 | 0 | 0.01 | 0.49 | 0.01 | 0.17 |
| 10 | 97961789 | rs7099132 | -0.352 | -0.049 | -0.708 | -0.077 | 0.002 | -0.003 | -0.003 | 0 | 0.1 | 0.1 | 0.15 | 0.11 | 0.08 | 0.15 |
| 10 | 97962687 | rs12762814 | -1.121 | -0.017 | -0.656 | -0.097 | 0.007 | 0.005 | 0.005 | 0 | 0.45 | 0.45 | 0.6 | 0.3 | 0.67 | 0.57 |
| 10 | 97967269 | rs747259 | -0.557 | 0.563 | -0.834 | 0.022 | -0.02 | 0.006 | 0.006 | 0.013 | 0.27 | 0.24 | 0.38 | 0.19 | 0.5 | 0.35 |
| 10 | 97968148 | rs12772113 | 0.096 | 0.86 | -1.054 | -0.089 | -0.023 | 0.016 | 0.016 | 0 | 0.13 | 0.07 | 0.13 | 0 | 0.22 | 0.11 |
| 10 | 97969694 | rs2290714 | -0.643 | -0.34 | -0.902 | 0.669 | -0.015 | 0.128 | 0.128 | 0 | 0.09 | 0.17 | 0.15 | 0.48 | 0.04 | 0.18 |
| 10 | 97971928 | rs7911684 | 0.792 | -0.589 | -0.871 | 0.292 | 0.001 | -0.007 | -0.007 | 0 | 0.56 | 0.45 | 0.5 | 0.33 | 0.54 | 0.35 |
| 10 | 97980435 | rs7076978 | -0.167 | -0.761 | -1.147 | 0.323 | -0.032 | 0.022 | 0.022 | 0 | 0.22 | 0.29 | 0.18 | 0.45 | 0.08 | 0.24 |
| 10 | 97980721 | rs2125062 | -1.591 | -0.722 | -1.466 | 0.299 | -0.002 | 0.008 | 0.008 | 0 | 0.55 | 0.59 | 0.55 | 0.76 | 0.53 | 0.77 |
| 10 | 97982240 | rs10491067 | 0.951 | -0.29 | -1.469 | 0.471 | -0.074 | -0.022 | -0.022 | 0 | 0.12 | 0.01 | 0.12 | 0.08 | 0.24 | 0.18 |
| 10 | 97982575 | rs4917723 | 0.544 | -1.016 | -1.043 | 0.393 | -0.023 | -0.01 | -0.01 | 0 | 0.3 | 0.24 | 0.36 | 0.3 | 0.42 | 0.49 |
| 10 | 97983028 | rs4917727 | 0.401 | -0.201 | -1.319 | 0.846 | 0.012 | -0.008 | -0.008 | 0 | 0.39 | 0.5 | 0.39 | 0.3 | 0.45 | 0.52 |
| 10 | 97983280 | rs11188666 | -1.125 | -0.843 | -1.517 | 1.247 | 0.014 | -0.018 | -0.018 | 0 | 0.19 | 0.14 | 0.18 | 0.45 | 0.08 | 0.24 |
| 10 | 97988453 | rs12357751 | -0.457 | -0.136 | -1.806 | 0.874 | -0.019 | -0.012 | -0.012 | 2.399 | 0.26 | 0.04 | 0.28 | 0.24 | 0.26 | 0.26 |
| 10 | 97990583 | rs727852 | 0.088 | -1.625 | -2.119 | 1.015 | 0.151 | -0.039 | -0.039 | 0 | 0.55 | 0.16 | 0.66 | 0.45 | 0.71 | 0.49 |
| 10 | 97993942 | rs10882754 | -0.089 | -2.357 | -0.203 | 0.821 | -0.039 | 0.021 | 0.021 | 0 | 0.69 | 0.36 | 0.76 | 0.5 | 0.85 | 0.59 |
| 10 | 97996830 | rs11597989 | -0.221 | -2.35 | -0.053 | 0.651 | -0.027 | 0.025 | 0.025 | 0 | 0.25 | 0.34 | 0.24 | 0.6 | 0.17 | 0.35 |
| 10 | 97997174 | rs12765160 | 0.487 | -0.463 | -0.706 | 0.704 | -0.112 | -0.034 | -0.034 | 0.611 | 0.2 | 0.04 | 0.35 | 0.16 | 0.31 | 0.18 |
| 10 | 98000860 | rs10882759 | 0.703 | -0.339 | -0.585 | 0.257 | -0.047 | 0.078 | 0.078 | 0 | 0.23 | 0.11 | 0.16 | 0.04 | 0.3 | 0.17 |
| 10 | 98002169 | rs10882760 | -0.187 | -0.325 | -0.533 | 0.53 | -0.001 | -0.061 | -0.061 | 1.446 | 0.62 | 0.2 | 0.69 | 0.52 | 0.67 | 0.75 |
| 10 | 98002866 | rs3789927 | -0.294 | -0.555 | -0.234 | 0.537 | 0.021 | 0.028 | 0.028 | 0 | 0.45 | 0.59 | 0.28 | 0.24 | 0.34 | 0.49 |
| 10 | 98004697 | rs3789929 | -1.451 | 0.123 | 0.112 | 1.507 | -0.002 | 0.135 | 0.135 | 0 | 0.09 | 0.09 | 0.19 | 0.55 | 0.06 | 0.2 |
| 10 | 98007545 | rs9794109 | -0.388 | 1.254 | -0.521 | 1.102 | -0.108 | 0.05 | 0.05 | 0 | 0.17 | 0.01 | 0.15 | 0.05 | 0.29 | 0.22 |
| 10 | 98011364 | rs3789930 | -1.927 | 0.362 | 0.872 | 1.322 | -0.012 | 0.285 | 0.285 | 0.424 | 0.15 | 0.39 | 0.23 | 0.57 | 0.08 | 0.21 |
| 10 | 98026182 | rs6584061 | 1.524 | 0.347 | 1.631 | 1.319 | 0.008 | 0.239 | 0.239 | 0.086 | 0.89 | 0.68 | 0.78 | 0.44 | 0.92 | 0.8 |
| 10 | 98041610 | rs941999 | -1.604 | 0.291 | 2.03 | 1.357 | 0.012 | 0.353 | 0.353 | 0.274 | 0.08 | 0.23 | 0.19 | 0.55 | 0.05 | 0.19 |
| 10 | 98055699 | rs7083595 | -2.742 | 0.159 | 2.473 | 1.312 | -0.038 | 0.342 | 0.342 | 1.224 | 0.14 | 0.47 | 0.21 | 0.55 | 0.05 | 0.2 |
| 10 | 98058355 | rs1998409 | 2.173 | 0.457 | 2.352 | 1.292 | -0.005 | 0.027 | 0.027 | 0.178 | 0.74 | 0.45 | 0.67 | 0.34 | 0.79 | 0.63 |
| 10 | 98060917 | rs7073871 | -1.327 | 0.389 | 2.344 | 1.235 | -0.005 | -0.017 | -0.017 | 0 | 0.17 | 0.26 | 0.11 | 0.09 | 0.13 | 0.17 |
| 10 | 98063256 | rs12761715 | -0.158 | 0.479 | 2.119 | 1.274 | -0.002 | 0 | 0 | 0 | 0.11 | 0.07 | 0.09 | 0.09 | 0.12 | 0.16 |
| 10 | 98072541 | rs3789938 | -1.94 | 0.432 | 2.176 | 1.28 | 0.002 | 0.087 | 0.087 | 0 | 0.21 | 0.35 | 0.29 | 0.64 | 0.18 | 0.37 |
| 10 | 98093828 | rs2154183 | 1.985 | 0.207 | 2.459 | 0.899 | 0.002 | -0.002 | -0.002 | 0 | 0.77 | 0.33 | 0.84 | 0.91 | 0.86 | 0.76 |
| 10 | 98095297 | rs12258909 | -1.832 | 0.213 | 2.608 | 0.909 | -0.021 | -0.044 | -0.044 | 0 | 0.09 | 0.53 | 0.05 | 0 | 0.02 | 0.06 |
| 10 | 98103376 | rs12242936 | -0.768 | 0.285 | 2.128 | 1.562 | -0.001 | 0.074 | 0.074 | 0 | 0.06 | 0.04 | 0.15 | 0.44 | 0.04 | 0.19 |
| 10 | 98104703 | rs11188724 | -1.362 | 0.284 | 2.112 | 1.549 | -0.019 | -0.082 | -0.082 | 0 | 0.17 | 0.06 | 0.22 | 0.53 | 0.15 | 0.35 |
| 10 | 98111569 | rs17111736 | -0.626 | 0.293 | 2.103 | 1.566 | -0.014 | 0.04 | 0.04 | 0 | 0.05 | 0.01 | 0.13 | 0.44 | 0.04 | 0.19 |
| 10 | 98115684 | rs1923702 | 1.642 | 0.353 | 2.225 | 1.342 | 0.028 | 0.179 | 0.179 | 0 | 0.87 | 0.74 | 0.8 | 0.54 | 0.86 | 0.76 |
| 10 | 98116030 | rs17479230 | -0.32 | 0.374 | 2.008 | 1.249 | -0.002 | 0.001 | 0.001 | 0 | 0.07 | 0.07 | 0.05 | 0.02 | 0.1 | 0.05 |
| 10 | 98119560 | rs10882777 | 1.849 | 0.289 | 2.465 | 1.187 | -0.001 | -0.052 | -0.052 | 0.2 | 0.73 | 0.58 | 0.83 | 0.9 | 0.77 | 0.71 |
| 10 | 98120109 | rs2901990 | 1.655 | 0.288 | 2.49 | 1.186 | 0.001 | -0.033 | -0.033 | 0 | 0.75 | 0.66 | 0.84 | 0.9 | 0.77 | 0.71 |
| 10 | 98120690 | rs10882779 | 2.029 | 0.292 | 2.486 | 1.343 | 0 | 0.03 | 0.03 | 0 | 0.69 | 0.6 | 0.7 | 0.47 | 0.72 | 0.52 |
| 10 | 98122334 | rs10736104 | 0.053 | 0.346 | 2.208 | 1.149 | 0.016 | -0.018 | -0.018 | 1.144 | 0.93 | 0.89 | 0.95 | 0.99 | 0.9 | 0.95 |
| 10 | 98122808 | rs10786282 | 1.143 | 0.334 | 2.267 | 1.148 | 0.008 | -0.015 | -0.015 | 0.763 | 0.8 | 0.78 | 0.86 | 0.9 | 0.78 | 0.77 |
| 10 | 98124715 | rs7077344 | -0.482 | 0.33 | 2.071 | 1.164 | -0.001 | -0.004 | -0.004 | 0 | 0.13 | 0.14 | 0.1 | 0.09 | 0.12 | 0.19 |
| 10 | 98127590 | rs3814221 | 2.281 | 0.385 | 2.083 | 1.272 | 0 | -0.008 | -0.008 | 0 | 0.66 | 0.35 | 0.66 | 0.43 | 0.7 | 0.52 |
| 10 | 98129058 | rs17388474 | -1.661 | 0.448 | 1.812 | 1.224 | 0.015 | -0.044 | -0.044 | 0.009 | 0.12 | 0.25 | 0.09 | 0.01 | 0.13 | 0.15 |
| 10 | 98130773 | rs17111753 | -0.777 | 0.344 | 1.802 | 1.417 | -0.003 | 0.055 | 0.055 | 0 | 0.06 | 0.03 | 0.13 | 0.44 | 0.04 | 0.2 |
| 10 | 98131446 | rs3789942 | -0.8 | 0.332 | 1.916 | 0.82 | 0 | -0.004 | -0.004 | 7.435 | 0.14 | 0.18 | 0.13 | 0.09 | 0.15 | 0.23 |
| 10 | 98132016 | rs17479671 | 0.644 | -0.079 | 0.626 | 0.025 | -0.109 | 0.121 | 0.121 | 0 | 0.37 | 0.1 | 0.4 | 0.15 | 0.47 | 0.23 |
| 10 | 98134345 | rs11818942 | 0.15 | -0.082 | 1.081 | 0.103 | 0.004 | -0.007 | -0.007 | 0 | 0.13 | 0.21 | 0.1 | 0.09 | 0.13 | 0.14 |
| 10 | 98136250 | rs3762096 | 0.701 | -0.249 | 0.922 | 0.305 | 0.021 | 0.009 | 0.009 | 0 | 0.4 | 0.16 | 0.42 | 0.64 | 0.38 | 0.45 |
| 10 | 98140048 | rs2093558 | -0.645 | -0.342 | 1.56 | 0.126 | 0.001 | -0.011 | -0.011 | 0 | 0.61 | 0.58 | 0.57 | 0.83 | 0.54 | 0.77 |
| 10 | 98145284 | rs2181324 | 0.418 | -0.299 | 1.532 | 0.145 | -0.001 | -0.001 | -0.001 | 0 | 0.12 | 0.08 | 0.1 | 0.08 | 0.12 | 0.15 |
| 10 | 98148857 | rs11188741 | -1.062 | -0.61 | 1.76 | 0 | -0.029 | 0.004 | 0.004 | 0.178 | 0.22 | 0.36 | 0.16 | 0.19 | 0.16 | 0.32 |
| 10 | 98149425 | rs1923700 | 1.425 | -0.611 | 1.668 | -0.075 | -0.021 | -0.033 | -0.033 | 0 | 0.06 | 0.14 | 0.03 | 0 | 0.02 | 0.06 |
| 10 | 98151328 | rs11597420 | 0.171 | -0.334 | 1.49 | -0.138 | -0.003 | 0.003 | 0.003 | 0 | 0.16 | 0.08 | 0.1 | 0.1 | 0.15 | 0.18 |
| 10 | 98151594 | rs11188743 | 1.007 | -0.436 | 1.349 | -0.257 | -0.028 | 0.047 | 0.047 | 0.115 | 0.3 | 0.21 | 0.34 | 0.14 | 0.37 | 0.21 |
| 10 | 98152036 | rs3789945 | -0.889 | -0.496 | 1.353 | -0.248 | -0.01 | 0.024 | 0.024 | 0 | 0.65 | 0.7 | 0.64 | 0.78 | 0.61 | 0.68 |
| 10 | 98158494 | rs7919721 | 0.226 | -0.432 | 1.418 | -0.155 | -0.003 | 0.022 | 0.022 | 0.11 | 0.44 | 0.34 | 0.5 | 0.74 | 0.47 | 0.56 |
| 10 | 98162791 | rs3789950 | 0.6 | -0.453 | 0.957 | -0.132 | -0.009 | 0 | 0 | 0 | 0.38 | 0.18 | 0.48 | 0.67 | 0.43 | 0.45 |
| 10 | 98166356 | rs11188748 | -0.207 | -0.21 | 0.615 | -0.45 | -0.004 | 0.029 | 0.029 | 0 | 0.52 | 0.33 | 0.44 | 0.3 | 0.48 | 0.44 |
| 10 | 98167424 | rs3827866 | 0.986 | -0.326 | 0.228 | -0.584 | -0.013 | 0.083 | 0.083 | 0.803 | 0.31 | 0.14 | 0.29 | 0.11 | 0.32 | 0.1 |
| 10 | 98168154 | rs4917737 | 0.748 | -0.338 | 0.139 | -0.618 | -0.017 | -0.022 | -0.022 | 1.421 | 0.34 | 0.15 | 0.45 | 0.67 | 0.36 | 0.38 |
| 10 | 98171325 | rs730179 | 0.379 | -0.133 | 0.576 | -0.539 | 0.002 | -0.023 | -0.023 | 0 | 0.37 | 0.36 | 0.36 | 0.2 | 0.39 | 0.23 |
| 10 | 98175291 | rs11188751 | 0.357 | -0.214 | 0.169 | -0.422 | 0.005 | 0.018 | 0.018 | 0 | 0.59 | 0.36 | 0.72 | 0.73 | 0.65 | 0.65 |

| | | | | | | | | | | | | | | | | |
|---|---|---|---|---|---|---|---|---|---|---|---|---|---|---|---|---|
| 10 | 98179762 | rs17111841 | -0.477 | -0.485 | 0.854 | -0.538 | 0.001 | 0.014 | 0.014 | 0 | 0.37 | 0.3 | 0.26 | 0.25 | 0.35 | 0.24 |
| 10 | 98180644 | rs3789952 | 0.894 | -0.594 | 0.65 | -0.66 | -0.022 | -0.037 | -0.037 | 0 | 0.15 | 0.02 | 0.11 | 0.17 | 0.14 | 0.13 |
| 10 | 98183070 | rs7078304 | -0.678 | -0.616 | 1.014 | -0.654 | 0 | 0.009 | 0.009 | 0 | 0.36 | 0.3 | 0.25 | 0.25 | 0.34 | 0.24 |
| 10 | 98196029 | rs9988781 | 0.803 | -0.561 | 1.039 | -0.626 | 0.002 | 0.011 | 0.011 | 0 | 0.61 | 0.38 | 0.72 | 0.74 | 0.65 | 0.65 |
| 10 | 98205419 | rs3789955 | 0.691 | -0.137 | 1.039 | -0.38 | -0.03 | 0.019 | 0.019 | 0 | 0.1 | 0.02 | 0.1 | 0.08 | 0.14 | 0.08 |
| 10 | 98207301 | rs2861579 | -0.4 | 0.064 | 0.917 | -0.636 | 0.003 | -0.005 | -0.005 | 0.382 | 0.38 | 0.39 | 0.5 | 0.74 | 0.45 | 0.64 |
| 10 | 98208287 | rs2861580 | -1.426 | 0.004 | 1.074 | -0.7 | 0.01 | 0.048 | 0.048 | 1.589 | 0.54 | 0.64 | 0.64 | 0.91 | 0.58 | 0.84 |
| 10 | 98208544 | rs1316860 | 0.459 | 0.09 | 0.78 | -0.659 | 0.002 | 0.013 | 0.013 | 0 | 0.27 | 0.34 | 0.4 | 0.65 | 0.31 | 0.55 |
| 10 | 98208861 | rs12413925 | 0.971 | 0.346 | 0.449 | -0.535 | -0.008 | -0.02 | -0.02 | 0 | 0.14 | 0.07 | 0.11 | 0.17 | 0.13 | 0.12 |
| 10 | 98208881 | rs17389665 | 0.702 | 0.478 | 0.582 | -0.49 | -0.008 | 0.009 | 0.009 | 0 | 0.1 | 0.06 | 0.1 | 0.08 | 0.14 | 0.09 |
| 10 | 98212575 | rs2154203 | -0.452 | 0.222 | 0.553 | 0.091 | 0.01 | 0.012 | 0.012 | 0 | 0.36 | 0.55 | 0.48 | 0.56 | 0.39 | 0.53 |
| 10 | 98221417 | rs7083245 | 0.473 | 0.227 | 0.826 | 0.135 | 0.015 | 0.024 | 0.024 | 0.016 | 0.65 | 0.41 | 0.64 | 0.47 | 0.6 | 0.55 |
| 10 | 98224776 | rs1937090 | 0.966 | 0.017 | 0.532 | -0.107 | 0.009 | 0.031 | 0.031 | 1.386 | 0.47 | 0.36 | 0.48 | 0.15 | 0.41 | 0.3 |
| 10 | 98226277 | rs12260223 | 0.932 | 0.323 | 0.916 | 0.056 | 0.001 | 0.15 | 0.15 | 0 | 0.14 | 0.23 | 0.17 | 0.43 | 0.14 | 0.2 |
| 10 | 98226497 | rs12261890 | 1.214 | 0.308 | 1.059 | 0.049 | -0.002 | 0.001 | 0.001 | 0 | 0.13 | 0.1 | 0.16 | 0.43 | 0.14 | 0.2 |
| 10 | 98227947 | rs10736109 | 0.705 | 0.291 | 1.136 | 0.036 | 0.024 | 0.034 | 0.034 | 0 | 0.66 | 0.51 | 0.65 | 0.46 | 0.59 | 0.61 |
| 10 | 98230489 | rs7070439 | -0.713 | 0.415 | 0.783 | -0.115 | 0.03 | -0.009 | -0.009 | 0.007 | 0.88 | 0.8 | 0.83 | 0.9 | 0.74 | 0.83 |
| 10 | 98236565 | rs11188782 | 0.283 | 0.466 | 0.817 | -0.129 | -0.009 | 0.008 | 0.008 | 0 | 0.82 | 0.81 | 0.82 | 0.84 | 0.76 | 0.69 |
| 10 | 98239913 | rs2281802 | -0.798 | 0.588 | 0.809 | -0.122 | -0.003 | -0.002 | -0.002 | 0.005 | 0.9 | 0.87 | 0.88 | 0.84 | 0.88 | 0.77 |
| 10 | 98242457 | rs11188784 | 1.373 | 0.215 | 1.178 | -0.386 | 0.01 | 0.042 | 0.042 | 1.492 | 0.48 | 0.39 | 0.41 | 0.11 | 0.4 | 0.23 |
| 10 | 98246740 | rs874370 | 1.367 | 0.196 | 1.201 | -0.394 | 0.004 | -0.003 | -0.003 | 0.486 | 0.48 | 0.42 | 0.42 | 0.11 | 0.4 | 0.24 |
| 10 | 98246777 | rs7097077 | 0.742 | 0.144 | 1.365 | -0.257 | -0.004 | 0.05 | 0.05 | 0 | 0.19 | 0.27 | 0.19 | 0.31 | 0.19 | 0.22 |
| 10 | 98247022 | rs874371 | 1.487 | 0.106 | 1.28 | -0.281 | 0.007 | 0.026 | 0.026 | 1.848 | 0.47 | 0.39 | 0.42 | 0.1 | 0.4 | 0.24 |
| 10 | 98250191 | rs7078184 | 0.823 | 0.127 | 1.285 | -0.184 | -0.003 | 0.026 | 0.026 | 0 | 0.18 | 0.2 | 0.18 | 0.31 | 0.19 | 0.22 |
| 10 | 98252496 | rs6584079 | -0.467 | 0.09 | 1.16 | -0.286 | -0.007 | -0.023 | -0.023 | 0.988 | 0.37 | 0.4 | 0.42 | 0.72 | 0.36 | 0.47 |
| 10 | 98252519 | rs7099756 | 0.874 | 0.253 | 1.093 | -0.205 | 0.002 | -0.009 | -0.009 | 2.066 | 0.07 | 0.07 | 0.06 | 0 | 0.12 | 0.07 |
| 10 | 98254309 | rs11188795 | -0.563 | 0.098 | 0.836 | -0.392 | 0 | -0.014 | -0.014 | 0 | 0.46 | 0.48 | 0.47 | 0.69 | 0.54 | 0.52 |
| 10 | 98255314 | rs734566 | -1.011 | 0.092 | 0.546 | -0.399 | -0.001 | -0.001 | -0.001 | 1.263 | 0.89 | 0.86 | 0.88 | 0.79 | 0.88 | 0.76 |
| 10 | 98255951 | rs6584080 | -0.067 | -0.09 | 0.573 | -0.426 | -0.006 | -0.003 | -0.003 | 1.646 | 0.35 | 0.25 | 0.27 | 0.32 | 0.39 | 0.39 |
| 10 | 98257739 | rs1010719 | 1.367 | 0.324 | 0.18 | -0.264 | 0.02 | 0.137 | 0.137 | 0.465 | 0.4 | 0.19 | 0.37 | 0.11 | 0.34 | 0.17 |
| 10 | 98258871 | rs12569424 | 1.12 | 0.315 | 0.22 | -0.21 | -0.003 | -0.031 | -0.031 | 0 | 0.14 | 0.07 | 0.19 | 0.37 | 0.15 | 0.23 |
| 10 | 98260206 | rs11188799 | -0.296 | 0.353 | 0.369 | -0.299 | 0.002 | 0.02 | 0.02 | 0 | 0.24 | 0.27 | 0.31 | 0.49 | 0.27 | 0.33 |
| 10 | 98263545 | rs879266 | 1.167 | 0.282 | -0.318 | -0.739 | 0.023 | 0.142 | 0.142 | 0.29 | 0.41 | 0.18 | 0.37 | 0.11 | 0.35 | 0.18 |
| 10 | 98266974 | rs4919028 | 0.78 | 0.176 | -0.487 | 0.049 | -0.006 | -0.036 | -0.036 | 0 | 0.13 | 0.06 | 0.14 | 0.28 | 0.14 | 0.21 |
| 10 | 98268119 | rs10882807 | 0.407 | 0.346 | -0.393 | -0.478 | 0.005 | 0.134 | 0.134 | 0 | 0.51 | 0.34 | 0.49 | 0.14 | 0.49 | 0.28 |
| 10 | 98271878 | rs10748681 | 0.33 | -0.021 | -0.435 | -0.161 | 0.004 | -0.002 | -0.002 | 0 | 0.48 | 0.35 | 0.36 | 0.49 | 0.49 | 0.53 |
| 10 | 98274045 | rs4919029 | -0.067 | -0.285 | -0.097 | -0.315 | 0.013 | 0.052 | 0.052 | 1.871 | 0.6 | 0.34 | 0.54 | 0.83 | 0.64 | 0.71 |
| 10 | 98279053 | rs9093 | 0.756 | -0.316 | -0.114 | -0.296 | -0.012 | -0.021 | -0.021 | 0 | 0.46 | 0.23 | 0.35 | 0.49 | 0.48 | 0.53 |
| 10 | 98282986 | rs4534508 | 0.486 | -0.464 | 0.5 | -0.535 | -0.006 | -0.017 | -0.017 | 0 | 0.24 | 0.26 | 0.35 | 0.15 | 0.2 | 0.17 |
| 10 | 98284894 | rs10509711 | 1.125 | -0.474 | 0.464 | -0.518 | -0.013 | -0.03 | -0.03 | 0.324 | 0.46 | 0.23 | 0.35 | 0.48 | 0.48 | 0.53 |
| 10 | 98287792 | rs12779720 | 0.222 | -0.299 | 0.804 | -0.801 | -0.012 | -0.029 | -0.029 | 0 | 0.24 | 0.3 | 0.36 | 0.15 | 0.2 | 0.17 |
| 10 | 98312891 | rs10736113 | -0.718 | -0.164 | 0.806 | -0.731 | -0.006 | 0.01 | 0.01 | 0 | 0.88 | 0.91 | 0.92 | 0.99 | 0.86 | 0.88 |
| 10 | 98328512 | rs11188831 | 0.258 | -0.301 | 1.035 | -0.781 | 0.004 | 0.005 | 0.005 | 3.539 | 0.24 | 0.21 | 0.35 | 0.15 | 0.2 | 0.17 |
| 10 | 98331793 | rs1408367 | 0.317 | -0.258 | 1.061 | -0.78 | 0.013 | 0.034 | 0.034 | 4.02 | 0.23 | 0.15 | 0.36 | 0.15 | 0.21 | 0.17 |
| 10 | 98333169 | rs12761929 | 1.626 | -0.436 | 0.817 | -0.884 | 0.03 | 0.067 | 0.067 | 7.713 | 0.12 | 0.01 | 0.04 | 0 | 0.05 | 0.03 |
| 10 | 98348595 | rs7071044 | -0.273 | -0.398 | 1.025 | -0.866 | -0.007 | -0.018 | -0.018 | 0 | 0.24 | 0.26 | 0.36 | 0.15 | 0.2 | 0.17 |
| 10 | 98353213 | rs7448 | 1.192 | -0.399 | 1.084 | -0.925 | 0.005 | 0.005 | 0.005 | 1.501 | 0.61 | 0.39 | 0.53 | 0.83 | 0.64 | 0.71 |
| 10 | 98353421 | rs8578 | -0.477 | -0.384 | 1.073 | -0.856 | -0.027 | -0.061 | -0.061 | 0.153 | 0.26 | 0.39 | 0.37 | 0.16 | 0.21 | 0.17 |
| 10 | 98353804 | rs1051450 | -0.503 | -0.384 | 1.064 | -0.856 | -0.032 | -0.071 | -0.071 | 0 | 0.26 | 0.44 | 0.37 | 0.16 | 0.21 | 0.17 |
| 10 | 98360072 | rs11188847 | -0.321 | -0.345 | 0.919 | -0.848 | -0.001 | -0.008 | -0.008 | 0 | 0.24 | 0.24 | 0.37 | 0.15 | 0.21 | 0.17 |
| 10 | 98364544 | rs7072865 | 0.089 | -0.384 | 1.135 | -0.886 | 0.006 | -0.089 | -0.089 | 0 | 0.85 | 0.67 | 0.9 | 0.99 | 0.86 | 0.89 |
| 10 | 98366585 | rs10786304 | -0.06 | -0.279 | 0.972 | -0.922 | 0.008 | -0.068 | -0.068 | 0 | 0.86 | 0.77 | 0.9 | 0.99 | 0.84 | 0.87 |
| 10 | 98371084 | rs10882826 | 0.915 | -0.316 | 0.89 | -0.773 | -0.001 | -0.013 | -0.013 | 0 | 0.62 | 0.43 | 0.54 | 0.82 | 0.65 | 0.72 |
| 10 | 98372104 | rs4146727 | -0.618 | -0.321 | 0.732 | -0.773 | 0.007 | 0.004 | 0.004 | 0 | 0.88 | 0.75 | 0.68 | 0.85 | 0.85 | 0.86 |
| 10 | 98374601 | rs10882828 | -0.739 | -0.289 | 0.721 | -0.701 | 0.006 | 0.043 | 0.043 | 1.155 | 0.35 | 0.24 | 0.43 | 0.17 | 0.35 | 0.29 |
| 10 | 98380277 | rs3748234 | 0.759 | -0.19 | 0.569 | -0.722 | 0.013 | 0.065 | 0.065 | 2.711 | 0.58 | 0.32 | 0.51 | 0.82 | 0.6 | 0.68 |
| 10 | 98381191 | rs4917743 | -0.406 | -0.204 | 0.462 | -0.26 | 0.002 | 0.036 | 0.036 | 0.35 | 0.84 | 0.75 | 0.8 | 0.72 | 0.8 | 0.76 |
| 10 | 98381446 | rs880348 | -0.08 | -0.483 | -0.465 | -0.293 | 0.02 | 0.031 | 0.031 | 0 | 0.49 | 0.37 | 0.49 | 0.34 | 0.44 | 0.52 |
| 10 | 98381645 | rs10882829 | 0.941 | -0.423 | -0.443 | -0.021 | -0.052 | -0.033 | -0.033 | 0 | 0.44 | 0.19 | 0.24 | 0.46 | 0.43 | 0.47 |
| 10 | 98382303 | rs9633706 | 0.007 | -0.492 | 0.179 | 0.237 | 0.003 | 0.077 | 0.077 | 1.262 | 0.81 | 0.45 | 0.72 | 0.73 | 0.8 | 0.81 |
| 10 | 98384147 | rs7905024 | -0.086 | -0.779 | -0.002 | -0.243 | -0.035 | -0.068 | -0.068 | 0 | 0.89 | 0.67 | 0.95 | 1 | 0.93 | 0.94 |
| 10 | 98385346 | rs4919041 | 0.912 | -0.86 | -0.702 | -0.382 | 0.017 | 0.161 | 0.161 | 4.071 | 0.51 | 0.13 | 0.44 | 0.76 | 0.54 | 0.63 |
| 10 | 98386558 | rs4344416 | 0.052 | -0.501 | 0.117 | -0.13 | -0.032 | -0.061 | -0.061 | 0 | 0.89 | 0.71 | 0.95 | 1 | 0.93 | 0.94 |
| 10 | 98396739 | rs7906985 | 0.02 | -0.399 | -0.233 | 0.144 | 0.001 | -0.002 | -0.002 | 0 | 0.47 | 0.48 | 0.32 | 0.44 | 0.45 | 0.37 |
| 10 | 98397689 | rs11188863 | 0.412 | -0.65 | -0.934 | 0.149 | 0.031 | -0.047 | -0.047 | 0 | 0.35 | 0.18 | 0.33 | 0.38 | 0.29 | 0.43 |
| 10 | 98403546 | rs11188870 | 0.006 | -0.439 | -0.907 | 0.103 | 0.076 | 0.022 | 0.022 | 0.088 | 0.49 | 0.26 | 0.57 | 0.59 | 0.6 | 0.48 |
| 10 | 98403893 | rs11188871 | 0.825 | -0.227 | -0.943 | 0.122 | -0.049 | -0.055 | -0.055 | 0.347 | 0.35 | 0.2 | 0.26 | 0.5 | 0.41 | 0.39 |
| 10 | 98406194 | rs17392966 | 0.557 | -0.062 | -1.239 | 0.103 | -0.03 | 0.019 | 0.019 | 0 | 0.13 | 0.07 | 0.28 | 0.09 | 0.19 | 0.08 |
| 10 | 98407483 | rs4919048 | 0.181 | -0.532 | -1.135 | 0.177 | 0.054 | 0.04 | 0.04 | 0.123 | 0.41 | 0.18 | 0.3 | 0.25 | 0.33 | 0.34 |

| | | | | | | | | | | | | | | | | |
|---|---|---|---|---|---|---|---|---|---|---|---|---|---|---|---|---|
| 10 | 98410170 | rs10748687 | 0.183 | 0.064 | -0.878 | 0.237 | 0.007 | 0.018 | 0.018 | 0.797 | 0.38 | 0.48 | 0.27 | 0.48 | 0.36 | 0.46 |
| 10 | 98410901 | rs3748232 | 0.52 | -0.362 | -1.723 | 0.441 | -0.127 | -0.08 | -0.08 | 0.439 | 0.33 | 0.09 | 0.33 | 0.46 | 0.43 | 0.34 |
| 10 | 98416324 | rs3827867 | -0.111 | -0.985 | -1.578 | 0.297 | 0.012 | -0.04 | -0.04 | 0 | 0.26 | 0.2 | 0.25 | 0.4 | 0.27 | 0.31 |
| 10 | 98417930 | rs7098785 | -1.305 | -0.902 | -1.94 | 0.017 | -0.101 | -0.049 | -0.049 | 0.014 | 0.68 | 0.92 | 0.64 | 0.61 | 0.54 | 0.65 |
| 10 | 98427049 | rs7083114 | -0.257 | -1.198 | -1.487 | 0.095 | 0.03 | -0.025 | -0.025 | 0.151 | 0.35 | 0.28 | 0.24 | 0.45 | 0.29 | 0.44 |
| 10 | 98429594 | rs7920142 | -0.395 | -1.851 | -0.86 | -0.079 | -0.036 | 0.083 | 0.083 | 0 | 0.27 | 0.43 | 0.23 | 0.33 | 0.24 | 0.3 |
| 10 | 98430592 | rs17112030 | 0.753 | -1.584 | -0.518 | -0.048 | -0.016 | 0.018 | 0.018 | 0 | 0.11 | 0.3 | 0.26 | 0.15 | 0.05 | 0.03 |
| 10 | 98433420 | rs10509712 | -0.603 | -1.851 | -0.602 | -0.209 | -0.04 | -0.031 | -0.031 | 0 | 0.2 | 0.38 | 0.28 | 0.16 | 0.1 | 0.09 |
| 10 | 98435648 | rs519467 | 0.352 | -1.867 | -0.604 | -0.21 | -0.002 | -0.017 | -0.017 | 0 | 0.09 | 0.09 | 0.03 | 0 | 0.05 | 0.06 |
| 10 | 98436548 | rs624029 | 0.082 | -1.741 | -0.61 | -0.137 | -0.04 | -0.028 | -0.028 | 0.328 | 0.19 | 0.36 | 0.27 | 0.16 | 0.09 | 0.09 |
| 10 | 98443405 | rs2282341 | -0.838 | -1.258 | -0.581 | -0.06 | 0.008 | 0.007 | 0.007 | 1.221 | 0.64 | 0.41 | 0.55 | 0.55 | 0.48 | 0.6 |
| 10 | 98454760 | rs1436187 | -0.989 | -0.947 | -1.08 | 0.028 | 0.037 | 0.035 | 0.035 | 0.974 | 0.66 | 0.5 | 0.55 | 0.54 | 0.49 | 0.61 |
| 10 | 98469693 | rs17112076 | 0.167 | -1.18 | -1.379 | 0.696 | -0.042 | -0.084 | -0.084 | 0 | 0.16 | 0.02 | 0.1 | 0.23 | 0.18 | 0.13 |
| 10 | 98477090 | rs4917749 | -0.474 | -1.905 | -0.632 | 0.254 | -0.083 | -0.007 | -0.007 | 0 | 0.27 | 0.48 | 0.41 | 0.27 | 0.14 | 0.23 |
| 10 | 98485105 | rs687562 | -0.463 | -1.995 | -1.248 | 0.166 | 0.046 | -0.014 | -0.014 | 0.122 | 0.29 | 0.23 | 0.42 | 0.67 | 0.13 | 0.36 |
| 10 | 98487678 | rs516321 | -0.809 | -0.747 | -1.643 | -0.586 | -0.151 | 0.105 | 0.105 | 0.028 | 0.74 | 0.97 | 0.75 | 0.89 | 0.54 | 0.75 |
| 10 | 98498310 | rs514408 | 0.035 | -1.132 | -1.053 | -0.707 | -0.005 | -0.014 | -0.014 | 0 | 0.62 | 0.42 | 0.68 | 0.89 | 0.5 | 0.69 |
| 10 | 98502188 | rs12413520 | -0.217 | -1.693 | -0.483 | 0.024 | -0.034 | 0.018 | 0.018 | 0 | 0.3 | 0.36 | 0.38 | 0.48 | 0.18 | 0.34 |
| 10 | 98504184 | rs1172464 | -1 | -1.502 | -1.072 | -0.324 | -0.113 | 0.141 | 0.141 | 0 | 0.72 | 0.95 | 0.75 | 0.92 | 0.58 | 0.78 |
| 10 | 98505437 | rs11188902 | -0.062 | -1.53 | -0.928 | -0.204 | 0.017 | -0.016 | -0.016 | 0 | 0.29 | 0.26 | 0.36 | 0.46 | 0.17 | 0.33 |
| 10 | 98506832 | rs4919054 | 0.392 | -1.004 | -0.675 | -0.025 | -0.033 | -0.037 | -0.037 | 0 | 0.38 | 0.23 | 0.35 | 0.47 | 0.41 | 0.45 |
| 10 | 98513814 | rs7919352 | -1.171 | -0.045 | -0.447 | 0.24 | -0.004 | 0.019 | 0.019 | 0 | 0.1 | 0.12 | 0.33 | 0.17 | 0.11 | 0.05 |
| 10 | 98521516 | rs505010 | -0.164 | 0.037 | -0.734 | -0.206 | -0.112 | 0.152 | 0.152 | 0.013 | 0.7 | 0.94 | 0.74 | 0.96 | 0.56 | 0.76 |
| 10 | 98523061 | rs9420695 | -0.984 | -0.72 | 0.165 | 0.374 | -0.01 | -0.032 | -0.032 | 0 | 0.06 | 0.21 | 0.04 | 0 | 0.04 | 0.05 |
| 10 | 98525185 | rs12412871 | -1.458 | -0.427 | -0.121 | 0.686 | 0 | 0.002 | 0.002 | 0 | 0.17 | 0.13 | 0.41 | 0.19 | 0.21 | 0.16 |
| 10 | 98530682 | rs577981 | -0.288 | -0.047 | -0.218 | 1.199 | -0.083 | -0.032 | -0.032 | 0 | 0.69 | 0.9 | 0.68 | 0.64 | 0.54 | 0.69 |
| 10 | 98532965 | rs17112128 | -2.282 | -0.633 | 1.337 | 2.285 | 0.032 | 0.083 | 0.083 | 1.422 | 0.09 | 0.26 | 0.28 | 0.15 | 0.1 | 0.08 |
| 10 | 98534216 | rs17485349 | -1.582 | -0.628 | 1.315 | 2.296 | 0.016 | 0.083 | 0.083 | 0.305 | 0.06 | 0.16 | 0.27 | 0.15 | 0.08 | 0.07 |
| 10 | 98535269 | rs12771295 | -0.575 | -0.569 | 1.133 | 2.318 | -0.02 | 0.007 | 0.007 | 0 | 0.05 | 0 | 0.1 | 0.33 | 0.1 | 0.06 |
| 10 | 98536104 | rs577715 | 0.836 | -0.75 | 1.59 | 2.306 | -0.016 | 0.09 | 0.09 | 0.385 | 0.91 | 0.59 | 0.93 | 0.89 | 0.98 | 0.82 |
| 10 | 98546460 | rs1172377 | 0.925 | -0.243 | 0.99 | 2.239 | 0.006 | 0.125 | 0.125 | 0 | 0.92 | 0.81 | 0.88 | 0.65 | 0.9 | 0.86 |
| 10 | 98549577 | rs7477274 | -1.612 | -0.261 | 0.89 | 1.723 | 0.05 | 0.154 | 0.154 | 0.096 | 0.05 | 0.18 | 0.27 | 0.16 | 0.08 | 0.04 |
| 10 | 98569570 | rs12264950 | -1.503 | -0.749 | 0.947 | 1.13 | -0.009 | -0.051 | -0.051 | 0.07 | 0.1 | 0.32 | 0.08 | 0 | 0.08 | 0.03 |
| 10 | 98604466 | rs12262784 | -1.599 | -0.784 | 1.013 | 1.099 | -0.012 | -0.054 | -0.054 | 0 | 0.1 | 0.29 | 0.07 | 0.01 | 0.08 | 0.02 |
| 10 | 98624993 | rs7069290 | -1.777 | -2.436 | 0.998 | 1.404 | 0.056 | -0.021 | -0.021 | 0.519 | 0.35 | 0.78 | 0.27 | 0.47 | 0.18 | 0.39 |
| 10 | 98625174 | rs11188952 | -0.804 | -2.604 | 1.604 | 1.63 | -0.033 | 0.085 | 0.085 | 0 | 0.07 | 0.14 | 0.14 | 0.27 | 0.03 | 0.25 |
| 10 | 98639846 | rs17112190 | -0.957 | -2.601 | 1.791 | 1.641 | -0.008 | -0.046 | -0.046 | 0 | 0.1 | 0.31 | 0.07 | 0.01 | 0.08 | 0.02 |
| 10 | 98657516 | rs12265203 | -0.746 | -2.633 | 1.986 | 1.625 | -0.01 | 0.029 | 0.029 | 0 | 0.07 | 0.07 | 0.13 | 0.28 | 0.02 | 0.25 |
| 10 | 98674939 | rs10430660 | -0.884 | -2.633 | 2.01 | 1.625 | -0.02 | 0.044 | 0.044 | 0 | 0.07 | 0.11 | 0.13 | 0.27 | 0.02 | 0.25 |
| 10 | 98684650 | rs10786316 | -0.882 | -2.64 | 1.993 | 1.629 | -0.009 | 0.026 | 0.026 | 0 | 0.07 | 0.07 | 0.13 | 0.28 | 0.02 | 0.25 |
| 10 | 98699136 | rs4919060 | -2.152 | -2.783 | 2.04 | 1.579 | 0.149 | -0.086 | -0.086 | 0.581 | 0.36 | 0.93 | 0.28 | 0.47 | 0.18 | 0.4 |
| 10 | 98709652 | rs10219031 | -0.724 | -2.718 | 2.074 | 1.577 | -0.01 | 0.029 | 0.029 | 0 | 0.07 | 0.07 | 0.13 | 0.28 | 0.02 | 0.25 |
| 10 | 98728921 | rs17112245 | 0.632 | -3.029 | 1.776 | 2.03 | -0.005 | 0.025 | 0.025 | 0.013 | 0.05 | 0 | 0.16 | 0.29 | 0.06 | 0.05 |
| 10 | 98742021 | rs7098255 | -0.515 | -2.987 | 2.339 | 1.485 | 0.003 | 0.012 | 0.012 | 0 | 0.07 | 0.05 | 0.13 | 0.28 | 0.02 | 0.25 |
| 10 | 98742605 | rs3814163 | -1.378 | -2.987 | 2.355 | 1.485 | -0.052 | 0.146 | 0.146 | 0 | 0.1 | 0.28 | 0.14 | 0.28 | 0.02 | 0.26 |
| 10 | 98742750 | rs3829856 | 0.071 | -2.949 | 1.645 | 1.458 | 0.017 | -0.005 | -0.005 | 3.766 | 0.14 | 0.08 | 0.05 | 0.19 | 0.06 | 0.11 |
| 10 | 98747114 | rs4606419 | -1.451 | -2.895 | 1.347 | 1.482 | -0.045 | -0.034 | -0.034 | 0 | 0.27 | 0.63 | 0.14 | 0.19 | 0.15 | 0.15 |
| 10 | 98747753 | rs10399993 | -1.784 | -2.912 | 0.572 | 1.263 | 0.074 | -0.011 | -0.011 | 1.743 | 0.38 | 0.78 | 0.3 | 0.47 | 0.2 | 0.41 |
| 10 | 98762051 | rs3740528 | -1.318 | -2.476 | 0.076 | 0.944 | 0.014 | -0.019 | -0.019 | 0 | 0.07 | 0.03 | 0.04 | 0.11 | 0.06 | 0.1 |
| 10 | 98774102 | rs12254525 | -0.988 | -2.49 | 0.418 | 0.738 | -0.012 | -0.049 | -0.049 | 0 | 0.1 | 0.22 | 0.07 | 0 | 0.09 | 0.02 |
| 10 | 98785776 | rs7922865 | -1.318 | -3.479 | -0.123 | 0.541 | -0.025 | -0.007 | -0.007 | 0.591 | 0.37 | 0.57 | 0.16 | 0.28 | 0.17 | 0.3 |
| 10 | 98786658 | rs17112311 | 0.105 | -3.382 | -1.202 | 0.486 | 0.071 | -0.003 | -0.003 | 0 | 0.19 | 0.04 | 0.04 | 0.18 | 0.06 | 0.21 |
| 10 | 98787354 | rs7896883 | -1.221 | -3.078 | -1.2 | 0.643 | 0.207 | -0.071 | -0.071 | 0 | 0.45 | 0.92 | 0.26 | 0.49 | 0.2 | 0.43 |
| 10 | 98792425 | rs7086186 | -0.872 | -1.385 | -0.294 | -0.086 | 0.043 | -0.08 | -0.08 | 0.36 | 0.23 | 0.39 | 0.19 | 0.1 | 0.32 | 0.22 |
| 10 | 98792520 | rs11188994 | -0.023 | -1.673 | -0.751 | -0.194 | 0.017 | -0.001 | -0.001 | 0.247 | 0.49 | 0.37 | 0.63 | 0.49 | 0.43 | 0.41 |
| 10 | 98794615 | rs12268044 | -1.472 | -2.164 | -0.373 | -1.067 | 0.033 | 0.061 | 0.061 | 0 | 0.54 | 0.65 | 0.32 | 0.29 | 0.41 | 0.41 |
| 10 | 98809229 | rs12098593 | -0.264 | -2.302 | -0.181 | -1.288 | -0.017 | -0.015 | -0.015 | 0 | 0.14 | 0.16 | 0.09 | 0.09 | 0.09 | 0.17 |
| 10 | 98809703 | rs7902871 | -0.691 | -2.258 | 0.888 | -1.525 | -0.008 | -0.021 | -0.021 | 0.019 | 0.28 | 0.35 | 0.13 | 0.11 | 0.21 | 0.21 |
| 10 | 98810881 | rs902470 | 0.957 | -2.306 | 0.823 | -1.552 | -0.004 | -0.021 | -0.021 | 0 | 0.63 | 0.39 | 0.82 | 0.89 | 0.74 | 0.78 |
| 10 | 98812636 | rs10882863 | 1.328 | -2.295 | 0.709 | -1.574 | 0.062 | 0.141 | 0.141 | 0 | 0.55 | 0.21 | 0.72 | 0.83 | 0.67 | 0.68 |
| 10 | 98813371 | rs12269131 | -1.535 | -2.3 | 0.732 | -1.576 | 0.006 | 0.002 | 0.002 | 0 | 0.43 | 0.59 | 0.27 | 0.17 | 0.33 | 0.32 |
| 10 | 98815857 | rs2636790 | 1.34 | -2.269 | 0.611 | -1.59 | 0.056 | 0.121 | 0.121 | 0 | 0.55 | 0.26 | 0.72 | 0.84 | 0.67 | 0.6 |
| 10 | 98816299 | rs2306951 | 0.293 | -2.239 | 0.309 | -1.549 | 0 | -0.018 | -0.018 | 0 | 0.08 | 0.18 | 0.1 | 0.05 | 0.07 | 0.1 |
| 10 | 98817687 | rs7914872 | -1.121 | -2.16 | -0.034 | -1.461 | -0.003 | -0.017 | -0.017 | 0.011 | 0.34 | 0.32 | 0.15 | 0.1 | 0.26 | 0.3 |
| 10 | 98832260 | rs2817682 | 0.966 | -1.709 | 0.057 | -1.421 | -0.004 | -0.031 | -0.031 | 0 | 0.67 | 0.39 | 0.76 | 0.86 | 0.7 | 0.67 |
| 10 | 98832697 | rs12773769 | -0.36 | -1.175 | 0.035 | -1.175 | -0.003 | -0.011 | -0.011 | 2.871 | 0.12 | 0.09 | 0.11 | 0.01 | 0.2 | 0.07 |
| 10 | 98838694 | rs2817683 | -1.66 | -1.08 | -0.527 | -1.545 | 0.007 | 0.218 | 0.218 | 0.463 | 0.49 | 0.85 | 0.39 | 0.16 | 0.44 | 0.39 |
| 10 | 98840679 | rs12268671 | -1.534 | -1.639 | 0.363 | -1.058 | 0.024 | 0.055 | 0.055 | 4.632 | 0.41 | 0.73 | 0.3 | 0.15 | 0.34 | 0.19 |
| 10 | 98840789 | rs11189001 | -1.348 | -1.65 | 0.681 | -1.054 | -0.006 | -0.028 | -0.028 | 0 | 0.38 | 0.62 | 0.28 | 0.15 | 0.34 | 0.18 |

| | | | | | | | | | | | | | | | | |
|---|---|---|---|---|---|---|---|---|---|---|---|---|---|---|---|---|
| 10 | 98844030 | rs2805599 | 1.163 | -1.543 | 0.407 | -1.033 | 0.029 | 0.093 | 0.093 | 2.995 | 0.57 | 0.26 | 0.69 | 0.85 | 0.62 | 0.76 |
| 10 | 98845536 | rs2817685 | 1.186 | -1.541 | 0.389 | -1.032 | 0.031 | 0.101 | 0.101 | 3.358 | 0.57 | 0.25 | 0.69 | 0.84 | 0.62 | 0.76 |
| 10 | 98850800 | rs2817688 | 1.005 | -1.048 | 1.185 | -0.7 | -0.024 | -0.048 | -0.048 | 0 | 0.76 | 0.58 | 0.81 | 0.86 | 0.79 | 0.87 |
| 10 | 98855381 | rs7084418 | -0.716 | -0.727 | 1.297 | -0.705 | 0.02 | -0.075 | -0.075 | 0.143 | 0.16 | 0.33 | 0.1 | 0.02 | 0.16 | 0.05 |
| 10 | 98857923 | rs2817689 | 1.556 | -0.835 | 0.941 | -0.621 | 0.01 | 0.098 | 0.098 | 4.681 | 0.58 | 0.23 | 0.69 | 0.81 | 0.6 | 0.77 |
| 10 | 98862741 | rs2279508 | -1.507 | -0.764 | 1.459 | -0.379 | 0.007 | -0.037 | -0.037 | 0 | 0.25 | 0.48 | 0.28 | 0.15 | 0.23 | 0.12 |
| 10 | 98863189 | rs2279510 | -0.27 | -0.809 | 1.643 | -0.409 | -0.004 | 0.003 | 0.003 | 0.086 | 0.15 | 0.15 | 0.08 | 0.02 | 0.16 | 0.04 |
| 10 | 98864312 | rs10882865 | -1.027 | -0.809 | 1.652 | -0.408 | 0.013 | 0.078 | 0.078 | 0 | 0.1 | 0.32 | 0.2 | 0.13 | 0.07 | 0.09 |
| 10 | 98864927 | rs2636776 | 2.319 | -0.872 | 1.607 | -0.511 | 0.011 | 0.101 | 0.101 | 4.151 | 0.52 | 0.22 | 0.57 | 0.76 | 0.57 | 0.61 |
| 10 | 98865286 | rs2817692 | -0.003 | -0.864 | 1.59 | -0.51 | -0.007 | -0.028 | -0.028 | 0.759 | 0.83 | 0.76 | 0.88 | 0.94 | 0.84 | 0.9 |
| 10 | 98867477 | rs7078655 | -1.522 | -0.834 | 1.492 | -0.531 | 0.006 | -0.028 | -0.028 | 0 | 0.24 | 0.41 | 0.28 | 0.15 | 0.23 | 0.12 |
| 10 | 98871605 | rs2817698 | 1.453 | -0.878 | 1.311 | -0.028 | -0.005 | -0.019 | -0.019 | 0 | 0.59 | 0.44 | 0.75 | 0.69 | 0.63 | 0.68 |
| 10 | 98874113 | rs2817701 | 2.309 | -0.915 | 1.189 | -0.016 | -0.014 | 0.007 | 0.007 | 4.147 | 0.51 | 0.09 | 0.55 | 0.56 | 0.56 | 0.6 |
| 10 | 98874661 | rs2636784 | 2.238 | -0.907 | 1.196 | -0.012 | -0.008 | 0.012 | 0.012 | 3.587 | 0.51 | 0.1 | 0.55 | 0.56 | 0.56 | 0.6 |
| 10 | 98876941 | rs2636788 | 0.358 | -1.215 | 1.404 | 0.12 | -0.012 | 0.011 | 0.011 | 0.381 | 0.81 | 0.72 | 0.86 | 0.72 | 0.82 | 0.85 |
| 10 | 98884402 | rs1336258 | -0.73 | -0.903 | 1.123 | 0.25 | 0.041 | 0.108 | 0.108 | 0 | 0.08 | 0.28 | 0.2 | 0.13 | 0.08 | 0.08 |
| 10 | 98889713 | rs2817708 | 1.191 | -1.003 | 0.839 | 0.062 | -0.003 | -0.004 | -0.004 | 0.972 | 0.59 | 0.38 | 0.64 | 0.65 | 0.63 | 0.63 |
| 10 | 98894444 | rs7097760 | -2.141 | -0.87 | 0.944 | 0.08 | 0.029 | 0.046 | 0.046 | 0 | 0.11 | 0.34 | 0.08 | 0.16 | 0.07 | 0.2 |
| 10 | 98897237 | rs2817647 | 1.666 | -1.077 | 0.148 | 0.131 | 0.051 | 0.033 | 0.033 | 5.752 | 0.51 | 0.11 | 0.57 | 0.62 | 0.59 | 0.67 |
| 10 | 98899114 | rs902469 | -1.259 | -1.186 | 0.505 | 0.05 | 0 | -0.001 | -0.001 | 0 | 0.32 | 0.63 | 0.34 | 0.36 | 0.28 | 0.27 |
| 10 | 98899155 | rs2817649 | 0.393 | -0.696 | 0.797 | -0.182 | -0.002 | -0.02 | -0.02 | 0.705 | 0.85 | 0.8 | 0.92 | 0.98 | 0.87 | 0.95 |
| 10 | 98900882 | rs2817652 | 1.139 | -0.538 | 1.086 | -0.112 | 0.005 | -0.056 | -0.056 | 1.169 | 0.8 | 0.49 | 0.86 | 0.88 | 0.81 | 0.89 |
| 10 | 98903394 | rs17112452 | -2.027 | -1.102 | 1.216 | 0.363 | -0.025 | 0.082 | 0.082 | 0.128 | 0.18 | 0.38 | 0.08 | 0.31 | 0.1 | 0.31 |
| 10 | 98905223 | rs2817656 | 2.024 | -1.408 | 1.121 | 0.516 | 0.014 | -0.01 | -0.01 | 0 | 0.61 | 0.38 | 0.8 | 0.52 | 0.75 | 0.52 |
| 10 | 98907045 | rs2817658 | 2.001 | -1.429 | 1.117 | 0.51 | 0.017 | -0.011 | -0.011 | 0 | 0.62 | 0.37 | 0.81 | 0.58 | 0.75 | 0.53 |
| 10 | 98908733 | rs7072055 | -0.315 | -1.377 | 0.767 | 0.497 | 0.007 | -0.014 | -0.014 | 0 | 0.17 | 0.07 | 0.11 | 0.17 | 0.17 | 0.18 |
| 10 | 98909311 | rs11189013 | -0.845 | -1.31 | 1.134 | 0.572 | -0.021 | -0.004 | -0.004 | 0 | 0.08 | 0.16 | 0.03 | 0.07 | 0.03 | 0.08 |
| 10 | 98909854 | rs2784923 | 2.217 | -1.442 | 1.196 | 0.565 | -0.063 | 0.017 | 0.017 | 0.869 | 0.8 | 0.5 | 0.92 | 0.79 | 0.92 | 0.74 |
| 10 | 98910763 | rs2636809 | 1.724 | -0.838 | 1.111 | 0.677 | -0.007 | 0.039 | 0.039 | 1.989 | 0.7 | 0.5 | 0.8 | 0.62 | 0.72 | 0.57 |
| 10 | 98912176 | rs17410931 | -0.148 | -0.853 | 1.155 | 0.692 | -0.001 | -0.002 | -0.002 | 0 | 0.06 | 0.05 | 0.03 | 0.13 | 0.04 | 0.13 |
| 10 | 98912996 | rs2784921 | 0.117 | -0.868 | 1.234 | 0.685 | -0.001 | -0.009 | -0.009 | 0.541 | 0.84 | 0.9 | 0.89 | 0.83 | 0.82 | 0.79 |
| 10 | 98913628 | rs2784920 | 1.285 | -0.855 | 1.195 | 0.707 | -0.008 | 0.016 | 0.016 | 0.493 | 0.74 | 0.68 | 0.84 | 0.69 | 0.77 | 0.66 |
| 10 | 98915112 | rs11189016 | -0.717 | -0.829 | 1.06 | 0.654 | -0.028 | 0.041 | 0.041 | 0.292 | 0.07 | 0.21 | 0.05 | 0.12 | 0.01 | 0.08 |
| 10 | 98917985 | rs17112469 | -1.186 | -0.765 | 0.847 | 0.602 | -0.038 | 0.094 | 0.094 | 0.058 | 0.14 | 0.37 | 0.09 | 0.25 | 0.06 | 0.21 |
| 10 | 98922759 | rs2817661 | 0.306 | -0.485 | -0.075 | 0.591 | -0.001 | 0.006 | 0.006 | 0.142 | 0.83 | 0.8 | 0.88 | 0.83 | 0.82 | 0.79 |
| 10 | 98923531 | rs902467 | 1.03 | 0.597 | -1.014 | 0.414 | -0.037 | -0.026 | -0.026 | 0 | 0.48 | 0.18 | 0.65 | 0.51 | 0.54 | 0.46 |
| 10 | 98925623 | rs885333 | -0.848 | 1.361 | -0.523 | 0.242 | -0.009 | 0.003 | 0.003 | 0.02 | 0.16 | 0.19 | 0.16 | 0.03 | 0.25 | 0.12 |
| 10 | 98926961 | rs7073919 | -1.478 | 1.792 | -0.203 | 0.446 | 0.037 | -0.003 | -0.003 | 0 | 0.29 | 0.42 | 0.21 | 0.23 | 0.37 | 0.3 |
| 10 | 98930061 | rs2636804 | 0.256 | 1.481 | 0.13 | 0.492 | -0.031 | 0.046 | 0.046 | 0 | 0.85 | 0.69 | 0.84 | 0.79 | 0.9 | 0.85 |
| 10 | 98930348 | rs2636803 | 1.497 | 1.429 | -0.324 | 0.474 | -0.014 | -0.015 | -0.015 | 0.004 | 0.56 | 0.24 | 0.6 | 0.58 | 0.53 | 0.55 |
| 10 | 98930392 | rs2636802 | 0.784 | 1.452 | -0.233 | 0.475 | -0.011 | 0.104 | 0.104 | 0.001 | 0.77 | 0.53 | 0.76 | 0.62 | 0.8 | 0.72 |
| 10 | 98933853 | rs2817666 | 1.674 | 1.324 | -1.114 | -0.033 | 0.034 | 0.001 | 0.001 | 0.078 | 0.45 | 0.24 | 0.55 | 0.59 | 0.39 | 0.53 |
| 10 | 98935272 | rs2784913 | 0.573 | 1.542 | -0.669 | 0.381 | -0.02 | 0.061 | 0.061 | 0.174 | 0.75 | 0.47 | 0.75 | 0.64 | 0.8 | 0.73 |
| 10 | 98936731 | rs2784911 | 1.411 | 1.744 | -0.777 | 0.38 | -0.018 | -0.006 | -0.006 | 0 | 0.57 | 0.28 | 0.62 | 0.59 | 0.54 | 0.58 |
| 10 | 98937054 | rs11189021 | -0.882 | 2.353 | -0.988 | 0.275 | -0.001 | -0.003 | -0.003 | 0 | 0.18 | 0.18 | 0.13 | 0.05 | 0.25 | 0.15 |
| 10 | 98937908 | rs4919079 | -0.85 | 2.337 | -0.984 | 0.278 | 0 | -0.003 | -0.003 | 0 | 0.18 | 0.17 | 0.13 | 0.05 | 0.25 | 0.15 |
| 10 | 98946244 | rs3758587 | -0.711 | 1.693 | -1.411 | 0.636 | -0.049 | -0.041 | -0.041 | 0 | 0.23 | 0.07 | 0.23 | 0.18 | 0.28 | 0.2 |
| 10 | 98950477 | rs7474809 | -0.688 | 1.623 | -0.705 | 0.991 | 0.015 | 0.082 | 0.082 | 0.023 | 0.25 | 0.48 | 0.38 | 0.55 | 0.28 | 0.43 |
| 10 | 98955599 | rs7894232 | -0.265 | 1.667 | -0.992 | 0.5 | -0.027 | 0.018 | 0.018 | 0.302 | 0.23 | 0.1 | 0.23 | 0.16 | 0.26 | 0.21 |
| 10 | 98967359 | rs902466 | -0.687 | 1.128 | -0.292 | 0.097 | 0.003 | -0.073 | -0.073 | 0 | 0.24 | 0.39 | 0.3 | 0.09 | 0.27 | 0.26 |
| 10 | 98971025 | rs1253425 | -0.674 | 0.742 | -0.327 | -0.044 | 0.001 | 0.001 | 0.001 | 0.562 | 0.23 | 0.23 | 0.27 | 0.1 | 0.25 | 0.28 |
| 10 | 98974462 | rs12778761 | 0.46 | 0.96 | -0.744 | 0.598 | -0.112 | -0.119 | -0.119 | 0.167 | 0.22 | 0.03 | 0.28 | 0.34 | 0.29 | 0.2 |
| 10 | 98975893 | rs701815 | -0.652 | 0.808 | -0.089 | 0.261 | 0 | -0.037 | -0.037 | 0 | 0.26 | 0.38 | 0.27 | 0.1 | 0.28 | 0.28 |
| 10 | 98977250 | rs701816 | -0.479 | 0.787 | -0.074 | 0.252 | -0.002 | 0.002 | 0.002 | 2.042 | 0.24 | 0.24 | 0.26 | 0.1 | 0.28 | 0.28 |
| 10 | 98982749 | rs3814164 | 1.222 | 0.459 | 0.411 | 0.48 | 0.01 | 0.036 | 0.036 | 3.453 | 0.62 | 0.27 | 0.6 | 0.71 | 0.64 | 0.54 |
| 10 | 98983955 | rs10748691 | 1.231 | 0.453 | 0.408 | 0.477 | 0.01 | 0.037 | 0.037 | 3.512 | 0.62 | 0.27 | 0.6 | 0.71 | 0.64 | 0.54 |
| 10 | 98988759 | rs1468069 | 0.968 | 0.431 | 0.434 | 0.483 | 0.017 | 0.025 | 0.025 | 3.225 | 0.64 | 0.27 | 0.66 | 0.71 | 0.69 | 0.55 |
| 10 | 98995076 | rs946981 | -0.344 | 0.476 | 0.504 | 0.467 | 0 | 0.002 | 0.002 | 2.318 | 0.25 | 0.25 | 0.25 | 0.1 | 0.27 | 0.28 |
| 10 | 99003375 | rs871988 | 1.876 | 0.239 | 0.395 | 0.648 | 0.011 | 0.031 | 0.031 | 8.659 | 0.35 | 0.11 | 0.27 | 0.27 | 0.34 | 0.25 |
| 10 | 99004171 | rs793529 | 0.446 | 0.4 | 0.54 | 0.593 | -0.002 | 0.108 | 0.108 | 0.08 | 0.91 | 0.63 | 0.93 | 0.81 | 0.97 | 0.83 |
| 10 | 99004220 | rs7905604 | -0.324 | 0.377 | 0.952 | 0.583 | -0.011 | 0.041 | 0.041 | 0 | 0.07 | 0.12 | 0.05 | 0.19 | 0.03 | 0.17 |
| 10 | 99006083 | rs13439 | -0.233 | 0.386 | 0.964 | 0.578 | 0 | -0.028 | -0.028 | 0.161 | 0.25 | 0.33 | 0.27 | 0.1 | 0.27 | 0.28 |
| 10 | 99009969 | rs1253393 | 1.623 | 0.375 | 0.925 | 0.587 | -0.002 | 0.028 | 0.028 | 0 | 0.88 | 0.74 | 0.88 | 0.8 | 0.9 | 0.81 |
| 10 | 99015364 | rs7074678 | 1.645 | 0.412 | 0.907 | 0.628 | 0.032 | 0.015 | 0.015 | 6.763 | 0.38 | 0.14 | 0.3 | 0.36 | 0.35 | 0.34 |
| 10 | 99019177 | rs2253301 | 0.524 | 0.389 | 0.968 | 0.464 | -0.016 | 0.077 | 0.077 | 0 | 0.92 | 0.74 | 0.93 | 0.81 | 0.97 | 0.83 |
| 10 | 99024572 | rs793524 | -0.296 | 0.388 | 0.971 | 0.459 | 0.003 | -0.069 | -0.069 | 0.477 | 0.25 | 0.44 | 0.28 | 0.1 | 0.27 | 0.28 |
| 10 | 99025219 | rs1253383 | 0.52 | 0.388 | 0.936 | 0.469 | -0.016 | 0.077 | 0.077 | 0 | 0.92 | 0.74 | 0.93 | 0.81 | 0.97 | 0.83 |
| 10 | 99026112 | rs10786321 | 0.706 | 0.395 | 0.794 | 0.438 | -0.001 | -0.034 | -0.034 | 0.079 | 0.68 | 0.53 | 0.62 | 0.73 | 0.65 | 0.56 |
| 10 | 99038562 | rs701821 | 0.384 | 0.443 | 0.446 | 0.404 | -0.007 | 0.035 | 0.035 | 2.983 | 0.2 | 0.1 | 0.24 | 0.08 | 0.25 | 0.27 |

| | | | | | | | | | | | | | | | | |
|---|---|---|---|---|---|---|---|---|---|---|---|---|---|---|---|---|
| 10 | 99045876 | rs10882888 | 1.548 | 0.486 | 0.315 | 0.532 | 0.032 | 0.045 | 0.045 | 7.211 | 0.38 | 0.14 | 0.31 | 0.32 | 0.35 | 0.39 |
| 10 | 99059645 | rs10882891 | 1.262 | 0.323 | 0.391 | 0.673 | 0.011 | 0.026 | 0.026 | 2.615 | 0.61 | 0.23 | 0.59 | 0.66 | 0.63 | 0.62 |
| 10 | 99067289 | rs11189119 | -0.933 | 0.33 | 0.472 | 0.676 | 0.002 | -0.002 | -0.002 | 0.859 | 0.06 | 0.09 | 0.07 | 0.02 | 0.08 | 0.03 |
| 10 | 99071408 | rs10786327 | 1.184 | 0.255 | 0.619 | 0.675 | 0.004 | 0.007 | 0.007 | 2.461 | 0.63 | 0.3 | 0.59 | 0.67 | 0.63 | 0.62 |
| 10 | 99072036 | rs11189121 | -0.375 | 0.203 | 0.761 | 0.654 | -0.001 | -0.006 | -0.006 | 0 | 0.26 | 0.3 | 0.26 | 0.14 | 0.27 | 0.23 |
| 10 | 99074959 | rs10786330 | 1.307 | 0.216 | 0.34 | 0.661 | 0.01 | 0.021 | 0.021 | 2.457 | 0.62 | 0.26 | 0.59 | 0.67 | 0.63 | 0.62 |
| 10 | 99078745 | rs3740512 | 1.189 | 0.217 | 0.424 | 0.663 | -0.002 | -0.003 | -0.003 | 0.62 | 0.63 | 0.38 | 0.6 | 0.67 | 0.63 | 0.62 |
| 10 | 99080585 | rs3781373 | 0.835 | 0.219 | 0.406 | 0.663 | -0.008 | -0.017 | -0.017 | 0.931 | 0.67 | 0.47 | 0.62 | 0.7 | 0.65 | 0.63 |
| 10 | 99080871 | rs1334891 | 0.795 | 0.123 | -0.712 | 0.488 | -0.122 | -0.13 | -0.13 | 0.213 | 0.28 | 0.04 | 0.32 | 0.36 | 0.35 | 0.24 |
| 10 | 99083669 | rs7101333 | 0.368 | 0.232 | -0.35 | 0.476 | -0.007 | 0.03 | 0.03 | 1.736 | 0.2 | 0.09 | 0.24 | 0.12 | 0.25 | 0.22 |
| 10 | 99087330 | rs10882898 | -0.653 | 0.213 | -0.264 | 0.482 | -0.014 | 0 | 0 | 0 | 0.32 | 0.19 | 0.35 | 0.32 | 0.36 | 0.37 |
| 10 | 99097191 | rs4917766 | 0.871 | 0.304 | -0.498 | 0.505 | 0.016 | 0.036 | 0.036 | 2.701 | 0.61 | 0.21 | 0.59 | 0.67 | 0.64 | 0.62 |
| 10 | 99107543 | rs11189150 | 0.609 | -0.015 | -1.278 | 0.401 | -0.117 | -0.135 | -0.135 | 0.259 | 0.31 | 0.03 | 0.32 | 0.37 | 0.35 | 0.23 |
| 10 | 99113964 | rs11189159 | 1.075 | -0.534 | -1.231 | 0.026 | 0.021 | 0.006 | 0.006 | 3.558 | 0.3 | 0.18 | 0.27 | 0.31 | 0.26 | 0.38 |
| 10 | 99115683 | rs10882904 | 0.405 | -0.635 | -0.136 | -0.123 | -0.011 | -0.016 | -0.016 | 0.281 | 0.38 | 0.59 | 0.35 | 0.31 | 0.33 | 0.41 |
| 10 | 99116903 | rs1048445 | 0.949 | -0.646 | -0.514 | -0.275 | 0.024 | 0.019 | 0.019 | 1.302 | 0.33 | 0.18 | 0.29 | 0.31 | 0.28 | 0.4 |
| 10 | 99117120 | rs10882905 | 1.085 | -0.549 | -0.52 | -0.191 | -0.126 | -0.14 | -0.14 | 0.301 | 0.28 | 0.03 | 0.32 | 0.37 | 0.35 | 0.23 |
| 10 | 99123006 | rs954148 | -0.268 | -0.115 | -0.189 | -0.178 | 0.036 | 0.002 | 0.002 | 1.54 | 0.47 | 0.74 | 0.42 | 0.47 | 0.38 | 0.5 |
| 10 | 99125949 | rs2275580 | 0.101 | -0.2 | -0.123 | -0.182 | 0.045 | -0.017 | -0.017 | 0.051 | 0.49 | 0.13 | 0.56 | 0.4 | 0.59 | 0.44 |
| 10 | 99126249 | rs2275581 | 1.433 | -0.205 | -0.082 | -0.232 | 0.021 | 0.003 | 0.003 | 3.558 | 0.3 | 0.18 | 0.26 | 0.31 | 0.26 | 0.38 |
| 10 | 99126923 | rs2297990 | 1.685 | -0.244 | -0.037 | -0.176 | -0.007 | -0.011 | -0.011 | 0 | 0.05 | 0.02 | 0.06 | 0.11 | 0.05 | 0.07 |
| 10 | 99129230 | rs6584122 | -0.685 | -0.276 | 0.201 | -0.191 | 0.017 | 0.032 | 0.032 | 0.013 | 0.11 | 0.22 | 0.13 | 0.15 | 0.12 | 0.1 |
| 10 | 99130887 | rs915197 | -0.652 | -0.306 | 0.256 | -0.285 | 0.032 | 0.029 | 0.029 | 2.438 | 0.06 | 0.35 | 0.03 | 0 | 0.01 | 0.01 |
| 10 | 99132083 | rs6584123 | -1.115 | -0.446 | 0.203 | -0.598 | 0.025 | -0.141 | -0.141 | 0 | 0.28 | 0.53 | 0.3 | 0.02 | 0.32 | 0.24 |
| 10 | 99135215 | rs11189172 | 0.778 | -0.594 | 0.283 | -0.337 | 0.005 | 0.02 | 0.02 | 2.527 | 0.34 | 0.2 | 0.31 | 0.31 | 0.33 | 0.4 |
| 10 | 99141205 | rs2275089 | 0.311 | -0.461 | 0.022 | -0.276 | -0.016 | -0.022 | -0.022 | 0 | 0.39 | 0.55 | 0.34 | 0.31 | 0.34 | 0.41 |
| 10 | 99143272 | rs11189178 | 0.311 | -0.461 | 0.017 | -0.276 | -0.015 | -0.021 | -0.021 | 0 | 0.39 | 0.55 | 0.34 | 0.31 | 0.34 | 0.41 |
| 10 | 99155984 | rs3814553 | 0.32 | -0.461 | 0.017 | -0.276 | -0.015 | -0.021 | -0.021 | 0.161 | 0.39 | 0.55 | 0.34 | 0.31 | 0.34 | 0.41 |
| 10 | 99160152 | rs1048442 | 0.503 | -0.412 | 0.025 | -0.254 | -0.013 | -0.02 | -0.02 | 0.121 | 0.38 | 0.54 | 0.34 | 0.31 | 0.34 | 0.41 |
| 10 | 99165975 | rs10786344 | 0.288 | -0.36 | -0.06 | -0.266 | 0.009 | 0.007 | 0.007 | 1.847 | 0.39 | 0.64 | 0.35 | 0.35 | 0.34 | 0.44 |
| 10 | 99167255 | rs7895695 | 0.707 | -0.406 | -0.021 | -0.285 | -0.025 | -0.019 | -0.019 | 0.036 | 0.34 | 0.46 | 0.31 | 0.31 | 0.29 | 0.41 |
| 10 | 99173870 | rs11189211 | 0.426 | -0.28 | -0.294 | -0.387 | -0.019 | 0.017 | 0.017 | 2.427 | 0.37 | 0.55 | 0.3 | 0.52 | 0.27 | 0.44 |
| 10 | 99179206 | rs4418736 | 1.125 | -0.043 | -0.69 | -0.305 | -0.013 | -0.015 | -0.015 | 0 | 0.06 | 0.02 | 0.06 | 0.08 | 0.05 | 0.07 |
| 10 | 99183402 | rs928603 | 1.083 | 0.145 | -1.041 | -0.572 | -0.109 | -0.083 | -0.083 | 0 | 0.31 | 0.04 | 0.34 | 0.33 | 0.36 | 0.23 |
| 10 | 99191935 | rs764223 | 1.145 | 0.12 | -1.062 | -0.565 | -0.115 | -0.089 | -0.089 | 0 | 0.31 | 0.04 | 0.34 | 0.33 | 0.36 | 0.23 |
| 10 | 99204526 | rs7081796 | -0.205 | -0.014 | -0.678 | -0.626 | 0.031 | 0.024 | 0.024 | 0.549 | 0.49 | 0.68 | 0.41 | 0.58 | 0.39 | 0.53 |
| 10 | 99205731 | rs4919098 | -0.103 | -0.27 | -0.577 | -0.592 | -0.007 | 0.004 | 0.004 | 0 | 0.1 | 0.09 | 0.11 | 0.08 | 0.12 | 0.08 |
| 10 | 99206826 | rs11189224 | 1.307 | -0.681 | 0.173 | -0.398 | 0.021 | 0.091 | 0.091 | 0.299 | 0.16 | 0.28 | 0.25 | 0.37 | 0.2 | 0.27 |
| 10 | 99214761 | rs34312169 | 1.043 | -0.432 | 0.141 | -0.944 | 0 | -0.008 | -0.008 | 0 | 0.12 | 0.13 | 0.2 | 0.03 | 0.18 | 0.11 |
| 10 | 99216066 | rs10786349 | -1.265 | -0.876 | 0.611 | -0.988 | -0.002 | 0.028 | 0.028 | 0 | 0.49 | 0.43 | 0.56 | 0.59 | 0.52 | 0.6 |
| 10 | 99216432 | rs10882939 | 0.562 | -0.944 | 0.584 | -0.962 | 0.007 | -0.041 | -0.041 | 0 | 0.3 | 0.23 | 0.32 | 0.49 | 0.27 | 0.39 |
| 10 | 99219885 | rs2152092 | -1.121 | -0.83 | 0.611 | -0.918 | -0.003 | 0.026 | 0.026 | 0 | 0.49 | 0.42 | 0.55 | 0.59 | 0.52 | 0.6 |
| 10 | 99220707 | rs3740526 | 1.275 | -0.757 | 0.473 | -0.927 | -0.036 | 0.022 | 0.022 | 0 | 0.44 | 0.18 | 0.41 | 0.38 | 0.45 | 0.31 |
| 10 | 99243326 | rs10882942 | 1.081 | -0.663 | 0.121 | -0.862 | -0.002 | 0.009 | 0.009 | 0 | 0.46 | 0.38 | 0.42 | 0.38 | 0.45 | 0.33 |
| 10 | 99256557 | rs7068965 | -0.642 | -0.548 | -0.245 | -0.794 | 0.001 | 0.005 | 0.005 | 0 | 0.43 | 0.42 | 0.48 | 0.62 | 0.44 | 0.64 |
| 10 | 99257815 | rs7915501 | 0.845 | -0.558 | -0.264 | -0.696 | -0.002 | 0.016 | 0.016 | 0 | 0.24 | 0.26 | 0.25 | 0.52 | 0.2 | 0.43 |
| 10 | 99259977 | rs7893335 | 1.208 | -0.538 | -0.27 | -0.702 | -0.01 | 0.033 | 0.033 | 0 | 0.22 | 0.24 | 0.21 | 0.52 | 0.15 | 0.43 |
| 10 | 99267639 | rs11189244 | 1.215 | -0.673 | -0.387 | -0.771 | -0.004 | 0.006 | 0.006 | 0 | 0.25 | 0.23 | 0.21 | 0.52 | 0.15 | 0.47 |
| 10 | 99270811 | rs11189247 | 0.346 | -0.219 | -0.131 | -1.415 | -0.016 | 0.017 | 0.017 | 0 | 0.16 | 0.12 | 0.23 | 0.03 | 0.25 | 0.17 |
| 10 | 99271281 | rs10786352 | 0.345 | -0.257 | -0.096 | -1.406 | -0.014 | 0.014 | 0.014 | 0 | 0.16 | 0.12 | 0.14 | 0.03 | 0.24 | 0.16 |
| 10 | 99272669 | rs11189249 | 1.873 | -0.458 | -0.238 | -1.542 | -0.01 | 0.057 | 0.057 | 1.724 | 0.62 | 0.26 | 0.6 | 0.88 | 0.58 | 0.74 |
| 10 | 99274276 | rs10786353 | 1.842 | -0.446 | -0.247 | -1.536 | -0.007 | 0.032 | 0.032 | 1.567 | 0.63 | 0.3 | 0.6 | 0.88 | 0.58 | 0.74 |
| 10 | 99276108 | rs946987 | 1.405 | -0.421 | -0.178 | -1.479 | -0.017 | -0.073 | -0.073 | 0 | 0.79 | 0.43 | 0.84 | 0.92 | 0.83 | 0.92 |
| 10 | 99278269 | rs10882950 | 0.849 | -0.391 | -0.083 | -1.503 | 0.009 | -0.088 | -0.088 | 1.149 | 0.69 | 0.47 | 0.65 | 0.95 | 0.64 | 0.77 |
| 10 | 99278820 | rs7908417 | 1.313 | -0.434 | -0.019 | -1.481 | -0.049 | -0.153 | -0.153 | 0.19 | 0.8 | 0.55 | 0.88 | 0.99 | 0.87 | 0.94 |
| 10 | 99280169 | rs4919104 | 0.679 | -0.766 | -0.063 | -1.173 | 0 | 0.001 | 0.001 | 0.211 | 0.27 | 0.27 | 0.25 | 0.51 | 0.2 | 0.46 |
| 10 | 99280233 | rs4919105 | -0.817 | -0.851 | -0.121 | -1.26 | -0.001 | -0.002 | -0.002 | 0 | 0.48 | 0.55 | 0.53 | 0.62 | 0.45 | 0.67 |
| 10 | 99284409 | rs11189258 | -0.39 | -0.547 | -0.465 | -1.234 | 0.012 | -0.011 | -0.011 | 0 | 0.2 | 0.31 | 0.3 | 0.11 | 0.29 | 0.21 |
| 10 | 99285132 | rs11189259 | 0.399 | -1.029 | -0.103 | -1.254 | -0.053 | 0.06 | 0.06 | 0 | 0.31 | 0.45 | 0.25 | 0.56 | 0.17 | 0.47 |
| 10 | 99286003 | rs7074482 | -0.527 | -0.38 | -0.315 | -1.057 | 0 | -0.001 | -0.001 | 0 | 0.2 | 0.24 | 0.29 | 0.11 | 0.28 | 0.24 |
| 10 | 99294797 | rs12767760 | 0.148 | -1.128 | -0.064 | -1.032 | 0.069 | -0.029 | -0.029 | 0 | 0.63 | 0.2 | 0.69 | 0.4 | 0.8 | 0.55 |
| 10 | 99298476 | rs1417408 | 0.009 | -1.074 | 0.558 | -0.964 | -0.014 | 0.002 | 0.002 | 0 | 0.62 | 0.41 | 0.69 | 0.39 | 0.73 | 0.49 |
| 10 | 99304801 | rs10882959 | 0.875 | -1.083 | 0.716 | -0.953 | -0.002 | 0.011 | 0.011 | 0 | 0.27 | 0.25 | 0.26 | 0.52 | 0.25 | 0.45 |
| 10 | 99313476 | rs7097951 | 0.308 | -0.948 | 0.481 | -0.864 | -0.026 | -0.037 | -0.037 | 0 | 0.92 | 0.75 | 0.98 | 0.99 | 0.98 | 0.96 |
| 10 | 99325062 | rs10748703 | -0.59 | -0.876 | 0.235 | -0.828 | -0.019 | -0.024 | -0.024 | 1.537 | 0.95 | 0.86 | 0.99 | 0.99 | 0.99 | 0.98 |
| 10 | 99329418 | rs11189281 | 0.504 | -0.442 | -0.419 | -0.881 | -0.046 | -0.02 | -0.02 | 0 | 0.46 | 0.15 | 0.44 | 0.43 | 0.51 | 0.3 |
| 10 | 99337572 | rs7094973 | -0.247 | -0.436 | -0.776 | -0.813 | 0.061 | -0.002 | -0.002 | 0 | 0.58 | 0.19 | 0.66 | 0.4 | 0.69 | 0.48 |
| 10 | 99343921 | rs12416267 | -0.284 | 0.085 | -0.767 | -1.111 | -0.018 | 0.02 | 0.02 | 0.818 | 0.31 | 0.16 | 0.25 | 0.22 | 0.36 | 0.41 |
| 10 | 99345406 | rs7070104 | 0.474 | 0.112 | -0.835 | -1.238 | -0.027 | 0.065 | 0.065 | 0.293 | 0.2 | 0.1 | 0.13 | 0.05 | 0.25 | 0.2 |

| | | | | | | | | | | | | | | | | |
|---|---|---|---|---|---|---|---|---|---|---|---|---|---|---|---|---|
| 10 | 99346768 | rs7078759 | -1.718 | -0.142 | -0.491 | -1.257 | 0 | 0.005 | 0.005 | 0 | 0.75 | 0.71 | 0.66 | 0.39 | 0.76 | 0.55 |
| 10 | 99348498 | rs7897232 | 0.198 | -0.13 | -0.873 | -1.163 | -0.001 | -0.002 | -0.002 | 0.019 | 0.06 | 0.06 | 0.12 | 0.09 | 0.02 | 0.06 |
| 10 | 99350128 | rs7093643 | -0.679 | -0.058 | -0.396 | -0.944 | -0.004 | -0.033 | -0.033 | 0 | 0.3 | 0.35 | 0.3 | 0.14 | 0.31 | 0.27 |
| 10 | 99358511 | rs11817730 | -0.266 | 0.233 | -0.908 | -0.697 | 0.007 | -0.002 | -0.002 | 0 | 0.15 | 0.19 | 0.2 | 0.13 | 0.19 | 0.16 |
| 10 | 99359987 | rs2297643 | 0.63 | -0.104 | -0.96 | -0.58 | -0.007 | 0.001 | 0.001 | 0 | 0.18 | 0.22 | 0.09 | 0.22 | 0.13 | 0.22 |
| 10 | 99360533 | rs1105999 | -0.381 | -0.073 | -1.217 | -0.549 | -0.003 | -0.002 | -0.002 | 0.131 | 0.59 | 0.46 | 0.53 | 0.38 | 0.67 | 0.44 |
| 10 | 99360792 | rs1010931 | 0.117 | 0.761 | -1.609 | -0.237 | -0.041 | 0.162 | 0.162 | 0.131 | 0.45 | 0.14 | 0.33 | 0.15 | 0.49 | 0.22 |
| 10 | 99366769 | rs4519024 | 0.071 | 0.631 | -0.717 | 0.31 | -0.005 | -0.003 | -0.003 | 1.425 | 0.67 | 0.57 | 0.56 | 0.31 | 0.73 | 0.49 |
| 10 | 99371729 | rs933057 | -0.719 | 1.866 | -0.71 | 0.033 | -0.016 | 0.011 | 0.011 | 0 | 0.21 | 0.16 | 0.18 | 0.11 | 0.32 | 0.15 |
| 10 | 99375040 | rs9804386 | -1.541 | 1.528 | 0.684 | 0.801 | 0.072 | 0.165 | 0.165 | 1.122 | 0.15 | 0.49 | 0.25 | 0.28 | 0.2 | 0.2 |
| 10 | 99376092 | rs3814555 | -1.579 | 1.531 | 0.723 | 0.801 | 0.071 | 0.154 | 0.154 | 1.175 | 0.16 | 0.49 | 0.25 | 0.28 | 0.2 | 0.2 |
| 10 | 99376152 | rs3814556 | -1.697 | 1.466 | 1.343 | 0.773 | 0.042 | 0.104 | 0.104 | 1.675 | 0.16 | 0.61 | 0.26 | 0.28 | 0.2 | 0.2 |
| 10 | 99376675 | rs3829907 | -1.35 | 1.379 | 1.559 | 0.773 | 0.02 | 0.223 | 0.223 | 0.044 | 0.05 | 0.32 | 0.14 | 0.18 | 0.01 | 0.08 |
| 10 | 99385055 | rs6584138 | -1.239 | 0.988 | 1.255 | 0.575 | -0.042 | 0.012 | 0.012 | 0.593 | 0.53 | 0.85 | 0.52 | 0.56 | 0.48 | 0.56 |
| 10 | 99391286 | rs10786360 | 0.15 | 0.73 | 1.854 | 0.674 | 0.028 | 0.004 | 0.004 | 0.423 | 0.1 | 0.18 | 0.11 | 0.11 | 0.19 | 0.12 |
| 10 | 99400214 | rs3814557 | -1.625 | 0.604 | 1.99 | 0.787 | 0.025 | 0.09 | 0.09 | 2.817 | 0.18 | 0.62 | 0.27 | 0.3 | 0.2 | 0.2 |
| 10 | 99401489 | rs7082459 | -1.536 | 0.595 | 2.108 | 0.791 | 0.009 | 0.213 | 0.213 | 0 | 0.06 | 0.29 | 0.14 | 0.19 | 0.01 | 0.09 |
| 10 | 99410790 | rs7915721 | -1.7 | 0.599 | 1.945 | 0.784 | 0.022 | 0.085 | 0.085 | 2.981 | 0.18 | 0.62 | 0.27 | 0.3 | 0.2 | 0.2 |
| 10 | 99414202 | rs3890727 | -1.018 | 0.506 | 1.805 | 0.701 | 0.003 | -0.001 | -0.001 | 0.021 | 0.19 | 0.21 | 0.18 | 0.19 | 0.28 | 0.22 |
| 10 | 99422648 | rs7905087 | -1.091 | 0.42 | 1.825 | 0.698 | -0.006 | -0.008 | -0.008 | 0 | 0.39 | 0.5 | 0.3 | 0.33 | 0.35 | 0.49 |
| 10 | 99429307 | rs4919128 | -1.322 | 0.415 | 1.564 | 0.708 | 0.013 | -0.018 | -0.018 | 1.177 | 0.42 | 0.7 | 0.44 | 0.48 | 0.36 | 0.49 |
| 10 | 99433297 | rs2065672 | -0.424 | 0.408 | 1.614 | 0.707 | 0.043 | -0.04 | -0.04 | 0 | 0.28 | 0.13 | 0.29 | 0.37 | 0.16 | 0.37 |
| 10 | 99435006 | rs11189321 | -0.123 | 0.414 | 1.544 | 0.704 | 0.042 | 0.006 | 0.006 | 0 | 0.26 | 0.08 | 0.17 | 0.22 | 0.16 | 0.36 |
| 10 | 99435730 | rs1046249 | -1.274 | 0.487 | 1.377 | 0.685 | 0.013 | -0.018 | -0.018 | 0 | 0.42 | 0.7 | 0.44 | 0.48 | 0.36 | 0.49 |
| 10 | 99442836 | rs10444067 | 0.202 | 0.934 | 0.998 | 0.688 | 0.004 | -0.002 | -0.002 | 0 | 0.1 | 0.1 | 0.13 | 0.12 | 0.23 | 0.13 |
| 10 | 99444730 | rs10786364 | -1.304 | 0.65 | 1.155 | 0.579 | -0.001 | 0 | 0 | 0 | 0.43 | 0.58 | 0.45 | 0.49 | 0.39 | 0.53 |
| 10 | 99445122 | rs4919129 | -1.028 | 0.592 | 1.087 | 0.56 | 0.031 | 0.043 | 0.043 | 0.511 | 0.19 | 0.41 | 0.28 | 0.26 | 0.25 | 0.17 |
| 10 | 99446425 | rs4561136 | -0.557 | -0.033 | 0.661 | 0.611 | 0.018 | -0.007 | -0.007 | 0.032 | 0.24 | 0.13 | 0.17 | 0.26 | 0.15 | 0.38 |
| 10 | 99448004 | rs11189328 | -0.408 | -0.048 | 0.662 | 0.512 | 0.003 | 0.003 | 0.003 | 0 | 0.22 | 0.18 | 0.19 | 0.19 | 0.16 | 0.3 |
| 10 | 99459308 | rs1343652 | -1.233 | -0.256 | 0.893 | 0.735 | -0.017 | -0.001 | -0.001 | 0 | 0.34 | 0.54 | 0.27 | 0.37 | 0.22 | 0.48 |
| 10 | 99462277 | rs10786366 | -1.42 | -0.447 | 1.126 | 0.903 | -0.024 | 0.008 | 0.008 | 0.071 | 0.35 | 0.58 | 0.42 | 0.62 | 0.2 | 0.45 |
| 10 | 99468978 | rs7899948 | -1.137 | -0.572 | 1.129 | 0.833 | -0.034 | 0.02 | 0.02 | 0 | 0.31 | 0.6 | 0.41 | 0.63 | 0.18 | 0.42 |
| 10 | 99472710 | rs12415272 | -1.345 | -0.647 | 1.068 | 0.837 | 0.021 | -0.005 | -0.005 | 0 | 0.35 | 0.73 | 0.43 | 0.67 | 0.19 | 0.45 |
| 10 | 99476915 | rs7896853 | -1.662 | -0.527 | 0.748 | 0.717 | 0.066 | 0.012 | 0.012 | 1.882 | 0.41 | 0.79 | 0.46 | 0.68 | 0.25 | 0.51 |
| 10 | 99482850 | rs7914667 | -1.369 | -0.46 | 1.162 | 0.756 | -0.019 | 0.018 | 0.018 | 0 | 0.1 | 0.14 | 0.08 | 0.17 | 0.05 | 0.2 |
| 10 | 99491948 | rs1253027 | 0.548 | -0.722 | 1.425 | 0.573 | 0.013 | -0.018 | -0.018 | 1.767 | 0.67 | 0.65 | 0.58 | 0.55 | 0.76 | 0.68 |
| 10 | 99495578 | rs10786367 | 0.564 | -0.905 | 1.758 | 0.537 | -0.025 | 0.039 | 0.039 | 0 | 0.68 | 0.5 | 0.55 | 0.49 | 0.78 | 0.67 |
| 10 | 99498123 | rs17108375 | -0.991 | -0.411 | 1.653 | 0.495 | -0.001 | -0.001 | -0.001 | 0 | 0.14 | 0.13 | 0.09 | 0.15 | 0.12 | 0.17 |
| 10 | 99498234 | rs3818876 | 1.687 | -0.396 | 1.679 | 0.521 | 0.041 | 0.043 | 0.043 | 0 | 0.53 | 0.29 | 0.45 | 0.33 | 0.66 | 0.51 |
| 10 | 99502430 | rs4917784 | 1.324 | -0.127 | 1.621 | 0.278 | 0.015 | -0.009 | -0.009 | 0.003 | 0.57 | 0.33 | 0.47 | 0.49 | 0.66 | 0.63 |
| 10 | 99504630 | rs10882993 | 0.329 | -0.167 | 1.621 | 0.666 | 0 | 0.003 | 0.003 | 0 | 0.75 | 0.78 | 0.6 | 0.54 | 0.8 | 0.71 |
| 10 | 99509457 | rs946778 | 0.87 | -0.136 | 1.795 | 0.62 | -0.021 | 0.036 | 0.036 | 0 | 0.68 | 0.61 | 0.56 | 0.49 | 0.77 | 0.67 |
| 10 | 99516955 | rs10882995 | 1.15 | -0.448 | 1.645 | 0.634 | -0.004 | 0.001 | 0.001 | 0 | 0.66 | 0.37 | 0.55 | 0.49 | 0.79 | 0.66 |
| 10 | 99517355 | rs3750612 | 0.598 | -0.452 | 1.789 | 0.711 | -0.037 | 0.073 | 0.073 | 0 | 0.72 | 0.5 | 0.59 | 0.53 | 0.83 | 0.71 |
| 10 | 99517848 | rs7080894 | 0.849 | -0.713 | 1.274 | 0.252 | 0.01 | 0 | 0 | 0 | 0.55 | 0.39 | 0.5 | 0.52 | 0.68 | 0.66 |
| 10 | 99518810 | rs1981238 | 1.207 | -0.722 | 2.022 | 0.884 | 0 | -0.002 | -0.002 | 0 | 0.67 | 0.34 | 0.56 | 0.49 | 0.81 | 0.69 |
| 10 | 99519097 | rs1981237 | 0.548 | -0.658 | 1.986 | 0.911 | -0.012 | 0.026 | 0.026 | 0 | 0.73 | 0.68 | 0.61 | 0.53 | 0.83 | 0.73 |
| 10 | 99519424 | rs3814560 | 0.296 | -0.668 | 1.993 | 0.906 | 0.018 | -0.048 | -0.048 | 0 | 0.75 | 0.88 | 0.63 | 0.53 | 0.83 | 0.73 |
| 10 | 99519753 | rs4244329 | 0.555 | -0.656 | 1.664 | 0.912 | -0.012 | 0.025 | 0.025 | 0 | 0.73 | 0.68 | 0.61 | 0.53 | 0.83 | 0.73 |
| 10 | 99526668 | rs2039826 | 1.19 | -0.663 | 1.591 | 0.876 | 0.01 | -0.011 | -0.011 | 0.004 | 0.67 | 0.32 | 0.56 | 0.48 | 0.81 | 0.69 |
| 10 | 99534780 | rs7095551 | -1.505 | -0.204 | 0.676 | 1.073 | 0.03 | 0.086 | 0.086 | 1.607 | 0.06 | 0.32 | 0.04 | 0.03 | 0.02 | 0.04 |
| 10 | 99537490 | rs10748709 | 0.107 | 0.077 | -0.47 | 1.091 | -0.023 | -0.154 | -0.154 | 0 | 0.8 | 0.99 | 0.64 | 0.53 | 0.85 | 0.79 |
| 10 | 99554180 | rs4359141 | 0.538 | 0.054 | 0.042 | 1.031 | -0.014 | 0.054 | 0.054 | 0 | 0.76 | 0.66 | 0.52 | 0.48 | 0.83 | 0.79 |
| 10 | 99557701 | rs12251897 | -1.295 | -0.117 | 0.625 | 0.298 | 0.008 | -0.063 | -0.063 | 0 | 0.11 | 0.49 | 0.1 | 0 | 0.09 | 0.05 |
| 10 | 99558000 | rs1417006 | 0.463 | 0.462 | 0.484 | 0.658 | -0.013 | -0.014 | -0.014 | 0 | 0.8 | 0.85 | 0.65 | 0.74 | 0.76 | 0.79 |
| 10 | 99558310 | rs1112707 | -0.137 | 0.207 | -0.46 | -0.32 | -0.003 | 0.11 | 0.11 | 0.03 | 0.38 | 0.75 | 0.29 | 0.06 | 0.42 | 0.18 |
| 10 | 99558738 | rs10509714 | 0.986 | -0.167 | -0.913 | 0.144 | -0.05 | -0.082 | -0.082 | 0 | 0.12 | 0.01 | 0.08 | 0.32 | 0.14 | 0.28 |
| 10 | 99559595 | rs7078809 | 1.479 | -0.938 | -0.514 | -0.002 | -0.027 | -0.061 | -0.061 | 0 | 0.33 | 0.14 | 0.21 | 0.39 | 0.35 | 0.37 |
| 10 | 99559721 | rs7903338 | 0.832 | -0.58 | 0.174 | -0.256 | -0.016 | -0.106 | -0.106 | 0 | 0.27 | 0.4 | 0.17 | 0.03 | 0.26 | 0.07 |
| 10 | 99560319 | rs11189378 | 1.327 | -0.571 | -0.172 | -0.236 | 0.001 | -0.008 | -0.008 | 0 | 0.23 | 0.22 | 0.14 | 0.03 | 0.22 | 0.07 |
| 10 | 99561644 | rs10882999 | 0.034 | -0.505 | -0.735 | -0.168 | -0.004 | 0.09 | 0.09 | 0 | 0.37 | 0.72 | 0.29 | 0.06 | 0.42 | 0.18 |
| 10 | 99562327 | rs4919144 | 0.215 | -0.411 | -0.772 | -0.108 | 0.002 | -0.05 | -0.05 | 0.18 | 0.34 | 0.57 | 0.27 | 0.06 | 0.41 | 0.14 |
| 10 | 99563198 | rs11189381 | 0.962 | -0.83 | -1.16 | 0.244 | 0.009 | -0.01 | -0.01 | 0.157 | 0.08 | 0.05 | 0.16 | 0.24 | 0.02 | 0.13 |
| 10 | 99563818 | rs4917787 | 0.355 | -0.463 | -0.902 | -0.295 | -0.009 | 0.079 | 0.079 | 0 | 0.49 | 0.24 | 0.46 | 0.7 | 0.52 | 0.67 |
| 10 | 99570333 | rs4919146 | 0.866 | 0.022 | -0.991 | -0.175 | -0.011 | 0.025 | 0.025 | 0 | 0.48 | 0.27 | 0.44 | 0.62 | 0.49 | 0.55 |
| 10 | 99574757 | rs7093131 | -0.516 | 0.504 | -0.761 | 0.089 | -0.059 | -0.072 | -0.072 | 1.542 | 0.83 | 0.93 | 0.68 | 0.73 | 0.81 | 0.77 |
| 10 | 99581256 | rs4919150 | 0.364 | 0.167 | -0.64 | 0.149 | -0.019 | 0.024 | 0.024 | 0.385 | 0.53 | 0.29 | 0.46 | 0.62 | 0.5 | 0.56 |
| 10 | 99600367 | rs872570 | 0.365 | 0.332 | 0.047 | -0.039 | -0.001 | -0.004 | -0.004 | 0 | 0.57 | 0.46 | 0.57 | 0.71 | 0.57 | 0.6 |
| 10 | 99611713 | rs11189412 | -0.35 | 0.252 | 0.216 | -0.196 | -0.012 | 0.016 | 0.016 | 0 | 0.69 | 0.71 | 0.63 | 0.81 | 0.65 | 0.79 |

| | | | | | | | | | | | | | | | | |
|---|---|---|---|---|---|---|---|---|---|---|---|---|---|---|---|---|
| 10 | 99620132 | rs1417001 | 0.396 | 0.209 | 0.649 | -0.326 | 0.001 | 0.004 | 0.004 | 0 | 0.56 | 0.43 | 0.58 | 0.67 | 0.54 | 0.59 |
| 10 | 99624392 | rs7906536 | -0.889 | 0.216 | 0.545 | -0.369 | 0.013 | -0.011 | -0.011 | 0.104 | 0.05 | 0.21 | 0.02 | 0 | 0.05 | 0.02 |
| 10 | 99625818 | rs11189413 | 0.002 | 0.289 | 0.355 | -0.4 | 0.001 | -0.021 | -0.021 | 0 | 0.59 | 0.44 | 0.58 | 0.79 | 0.57 | 0.72 |
| 10 | 99628321 | rs11189414 | 0.134 | 0.081 | 0.326 | -0.046 | -0.006 | 0.09 | 0.09 | 0 | 0.06 | 0.14 | 0.03 | 0.12 | 0.04 | 0.15 |
| 10 | 99630897 | rs553727 | -0.513 | 0.082 | 0.183 | -0.046 | 0.005 | 0.128 | 0.128 | 0.033 | 0.07 | 0.26 | 0.03 | 0.12 | 0.04 | 0.14 |
| 10 | 99633694 | rs11189416 | 0.141 | 0.199 | -1.293 | -0.897 | -0.028 | 0.199 | 0.199 | 2.987 | 0.28 | 0.04 | 0.16 | 0 | 0.28 | 0.07 |
| 10 | 99634541 | rs4919153 | -0.14 | -0.024 | -0.915 | -0.875 | -0.004 | -0.002 | -0.002 | 0 | 0.68 | 0.7 | 0.7 | 0.71 | 0.71 | 0.75 |
| 10 | 99634643 | rs4919154 | -0.027 | 0.135 | -0.608 | -0.925 | 0.006 | 0.024 | 0.024 | 0 | 0.47 | 0.65 | 0.6 | 0.66 | 0.54 | 0.68 |
| 10 | 99635324 | rs484290 | 0.406 | 0.122 | -0.696 | -0.913 | -0.053 | -0.02 | -0.02 | 0 | 0.4 | 0.21 | 0.51 | 0.55 | 0.46 | 0.59 |
| 10 | 99635777 | rs479776 | 0.365 | 0.089 | -0.695 | -0.941 | -0.053 | -0.021 | -0.021 | 0.012 | 0.4 | 0.21 | 0.51 | 0.55 | 0.46 | 0.56 |
| 10 | 99637578 | rs531676 | 0.297 | 0.002 | -0.338 | -0.876 | -0.007 | 0.006 | 0.006 | 0.133 | 0.42 | 0.35 | 0.53 | 0.64 | 0.45 | 0.57 |
| 10 | 99638183 | rs3793695 | 0.343 | -0.074 | -0.428 | -0.985 | 0.005 | 0.194 | 0.194 | 0 | 0.44 | 0.1 | 0.37 | 0.2 | 0.41 | 0.3 |
| 10 | 99638198 | rs3793696 | 0.144 | -0.082 | -0.303 | -0.975 | 0.02 | 0.043 | 0.043 | 0 | 0.46 | 0.33 | 0.41 | 0.28 | 0.42 | 0.31 |
| 10 | 99639573 | rs2297934 | 0.519 | 0.886 | 0.139 | -0.618 | 0.035 | 0.038 | 0.038 | 0 | 0.11 | 0.29 | 0.1 | 0.13 | 0.13 | 0.13 |
| 10 | 99640463 | rs11599481 | 0.644 | 1.09 | -0.22 | -0.453 | -0.063 | 0.004 | 0.004 | 0 | 0.21 | 0.02 | 0.27 | 0.19 | 0.25 | 0.21 |
| 10 | 99640572 | rs10883010 | 0.152 | 1.171 | -0.147 | -0.544 | 0.008 | 0.013 | 0.013 | 0 | 0.24 | 0.43 | 0.31 | 0.28 | 0.25 | 0.22 |
| 10 | 99640651 | rs10786376 | 0.994 | 0.968 | -0.819 | -0.137 | -0.011 | -0.057 | -0.057 | 1.103 | 0.15 | 0.06 | 0.27 | 0.4 | 0.15 | 0.26 |
| 10 | 99640811 | rs10883011 | 0.017 | 1.161 | -0.368 | -0.56 | -0.009 | -0.003 | -0.003 | 0 | 0.24 | 0.18 | 0.32 | 0.28 | 0.31 | 0.21 |
| 10 | 99642393 | rs2297937 | 0.066 | 1.227 | -0.799 | -0.558 | 0.032 | 0.005 | 0.005 | 0 | 0.26 | 0.34 | 0.34 | 0.28 | 0.31 | 0.22 |
| 10 | 99642731 | rs588061 | -0.118 | 0.955 | -1.341 | -0.041 | -0.007 | -0.002 | -0.002 | 0.555 | 0.39 | 0.2 | 0.48 | 0.58 | 0.42 | 0.57 |
| 10 | 99647665 | rs522540 | 0.267 | 1.29 | -0.641 | -0.243 | 0.03 | -0.078 | -0.078 | 0 | 0.66 | 0.53 | 0.57 | 0.85 | 0.55 | 0.71 |
| 10 | 99651450 | rs510393 | -0.561 | 0.802 | -0.891 | 0.526 | -0.006 | -0.041 | -0.041 | 0 | 0.24 | 0.18 | 0.2 | 0.5 | 0.24 | 0.31 |
| 10 | 99654624 | rs11189424 | -0.764 | 0.575 | -0.735 | -0.069 | -0.011 | 0.033 | 0.033 | 0 | 0.2 | 0.16 | 0.13 | 0.1 | 0.19 | 0.19 |
| 10 | 99655648 | rs577537 | -0.718 | 0.696 | -0.739 | -0.231 | -0.002 | -0.002 | -0.002 | 0.483 | 0.07 | 0.03 | 0.04 | 0 | 0.07 | 0.03 |
| 10 | 99657922 | rs611527 | -1.994 | 0.173 | -0.084 | -0.133 | -0.005 | -0.021 | -0.021 | 0 | 0.4 | 0.51 | 0.23 | 0.27 | 0.35 | 0.43 |
| 10 | 99659374 | rs11189425 | -0.866 | 0.036 | 0.067 | -0.229 | -0.003 | 0.002 | 0.002 | 0 | 0.09 | 0.11 | 0.05 | 0.08 | 0.05 | 0.13 |
| 10 | 99666275 | rs7914222 | -0.824 | 0.029 | 0.088 | -0.232 | -0.002 | 0 | 0 | 0 | 0.09 | 0.1 | 0.06 | 0.08 | 0.05 | 0.13 |
| 10 | 99666877 | rs1417004 | 1.885 | -0.194 | 0.477 | -0.426 | -0.008 | -0.022 | -0.022 | 0 | 0.64 | 0.37 | 0.79 | 0.84 | 0.73 | 0.59 |
| 10 | 99674123 | rs894376 | 1.199 | -0.173 | 0.588 | -0.363 | -0.003 | -0.052 | -0.052 | 0 | 0.77 | 0.74 | 0.87 | 0.92 | 0.79 | 0.71 |
| 10 | 99675421 | rs7899412 | 0.948 | -0.297 | 0.585 | -0.358 | -0.015 | -0.037 | -0.037 | 0 | 0.89 | 0.77 | 0.94 | 0.95 | 0.93 | 0.96 |
| 10 | 99678364 | rs1372696 | 1.57 | -0.299 | 0.516 | -0.376 | -0.022 | -0.094 | -0.094 | 0 | 0.73 | 0.58 | 0.85 | 0.92 | 0.79 | 0.69 |
| 10 | 99684366 | rs10883020 | 1.715 | -0.207 | 0.334 | -0.344 | -0.024 | -0.044 | -0.044 | 0 | 0.66 | 0.43 | 0.81 | 0.84 | 0.75 | 0.58 |
| 10 | 99688080 | rs4917793 | 1.708 | -0.21 | 0.207 | -0.333 | -0.026 | -0.049 | -0.049 | 0 | 0.66 | 0.47 | 0.81 | 0.84 | 0.75 | 0.58 |
| 10 | 99689101 | rs11189444 | -0.185 | -0.168 | 0.167 | -0.317 | -0.01 | 0.038 | 0.038 | 0 | 0.13 | 0.07 | 0.07 | 0.03 | 0.14 | 0.28 |
| 10 | 99689883 | rs11189447 | 1.314 | -0.183 | 0.525 | 0.135 | -0.034 | -0.005 | -0.005 | 0 | 0.82 | 0.62 | 0.9 | 0.86 | 0.89 | 0.86 |
| 10 | 99691299 | rs4917794 | 1.22 | -0.06 | 0.231 | 0.189 | -0.045 | -0.018 | -0.018 | 0 | 0.8 | 0.56 | 0.89 | 0.86 | 0.88 | 0.86 |
| 10 | 99695194 | rs474558 | -1.141 | 0.355 | -0.187 | 0.491 | -0.001 | 0.001 | 0.001 | 0 | 0.21 | 0.24 | 0.16 | 0.23 | 0.21 | 0.15 |
| 10 | 99696316 | rs506023 | -1.377 | 0.398 | -0.406 | 0.368 | 0 | -0.003 | -0.003 | 0 | 0.25 | 0.26 | 0.18 | 0.24 | 0.24 | 0.15 |
| 10 | 99703805 | rs556474 | -1.631 | 0.456 | -0.109 | -0.69 | 0.003 | 0.023 | 0.023 | 0 | 0.47 | 0.66 | 0.31 | 0.25 | 0.42 | 0.37 |
| 10 | 99709620 | rs493478 | -1.57 | 0.407 | 0.228 | -0.711 | -0.002 | -0.013 | -0.013 | 0 | 0.47 | 0.53 | 0.3 | 0.25 | 0.42 | 0.37 |
| 10 | 99736603 | rs1925556 | -1.695 | 0.405 | 0.433 | -0.751 | 0.007 | 0.051 | 0.051 | 0 | 0.48 | 0.69 | 0.31 | 0.25 | 0.42 | 0.35 |
| 10 | 99744372 | rs11189457 | -0.729 | 0.321 | 0.539 | -0.825 | 0 | 0 | 0 | 0 | 0.14 | 0.1 | 0.07 | 0.09 | 0.12 | 0.16 |
| 10 | 99754973 | rs508001 | -1.366 | 0.23 | 0.729 | -0.862 | -0.011 | -0.014 | -0.014 | 0 | 0.17 | 0.33 | 0.08 | 0.09 | 0.12 | 0.16 |
| 10 | 99765695 | rs560100 | -1.694 | 0.562 | 0.794 | -0.928 | 0.005 | -0.01 | -0.01 | 0 | 0.35 | 0.49 | 0.22 | 0.24 | 0.33 | 0.23 |
| 10 | 99769125 | rs11189465 | -0.023 | 0.564 | 0.117 | -1.481 | -0.018 | 0.108 | 0.108 | 0.111 | 0.22 | 0.03 | 0.13 | 0.02 | 0.23 | 0.17 |
| 10 | 99770333 | rs11591797 | 2.199 | 0.353 | -0.361 | -0.895 | -0.02 | -0.059 | -0.059 | 0 | 0.26 | 0.07 | 0.3 | 0.36 | 0.26 | 0.37 |
| 10 | 99772373 | rs526748 | -1.282 | 0.351 | -0.245 | -1.07 | 0 | -0.001 | -0.001 | 0 | 0.55 | 0.49 | 0.36 | 0.23 | 0.56 | 0.39 |
| 10 | 99772404 | rs563296 | -1.408 | 0.395 | -0.232 | -1.056 | -0.001 | -0.006 | -0.006 | 0 | 0.57 | 0.61 | 0.37 | 0.23 | 0.56 | 0.39 |
| 10 | 99780908 | rs513564 | -0.26 | 0.22 | -0.739 | -0.995 | -0.021 | 0.039 | 0.039 | 0.222 | 0.3 | 0.12 | 0.15 | 0.16 | 0.25 | 0.19 |
| 10 | 99785636 | rs7922066 | -0.013 | 0.038 | -0.332 | -1.139 | -0.033 | -0.134 | -0.134 | 0.113 | 0.24 | 0.54 | 0.18 | 0.03 | 0.19 | 0.11 |
| 10 | 99790986 | rs514554 | -0.2 | -0.037 | -0.454 | -1.056 | -0.017 | 0.054 | 0.054 | 0 | 0.33 | 0.2 | 0.2 | 0.17 | 0.33 | 0.31 |
| 10 | 99793012 | rs1867073 | 1.696 | 0.042 | -0.806 | -0.476 | -0.076 | -0.124 | -0.124 | 0.133 | 0.29 | 0.03 | 0.37 | 0.4 | 0.3 | 0.35 |
| 10 | 99799824 | rs1867075 | -1.176 | 0.463 | -0.532 | -0.433 | -0.047 | 0.05 | 0.05 | 0 | 0.81 | 0.98 | 0.9 | 0.93 | 0.81 | 0.81 |
| 10 | 99806748 | rs10736121 | 0.245 | 0.397 | 0.17 | -0.341 | 0 | 0.006 | 0.006 | 0 | 0.64 | 0.32 | 0.77 | 0.83 | 0.7 | 0.69 |
| 10 | 99822587 | rs920643 | 0.75 | 0.758 | 0.607 | -0.179 | -0.004 | 0.048 | 0.048 | 0 | 0.51 | 0.25 | 0.54 | 0.41 | 0.51 | 0.47 |
| 10 | 99825756 | rs928577 | 0.208 | 0.823 | 0.523 | -0.15 | -0.024 | 0.013 | 0.013 | 0 | 0.57 | 0.72 | 0.59 | 0.41 | 0.52 | 0.48 |
| 10 | 99835094 | rs4917802 | 1.095 | 0.907 | 0.349 | -0.12 | -0.026 | -0.047 | -0.047 | 0 | 0.12 | 0.06 | 0.23 | 0.43 | 0.18 | 0.22 |
| 10 | 99836031 | rs11189479 | 0.233 | 0.638 | 0.698 | -0.214 | 0 | -0.002 | -0.002 | 0.101 | 0.54 | 0.53 | 0.57 | 0.41 | 0.51 | 0.47 |
| 10 | 99846671 | rs928578 | 1.103 | 0.601 | -0.256 | -0.19 | -0.004 | -0.022 | -0.022 | 0.574 | 0.34 | 0.13 | 0.4 | 0.39 | 0.34 | 0.38 |
| 10 | 99847987 | rs1473795 | 0.071 | 0.612 | -0.069 | 0.211 | -0.001 | 0.032 | 0.032 | 0 | 0.55 | 0.66 | 0.54 | 0.36 | 0.46 | 0.51 |
| 10 | 99849574 | rs10786385 | -0.036 | 0.608 | -0.056 | 0.209 | -0.009 | 0.029 | 0.029 | 0 | 0.56 | 0.66 | 0.55 | 0.35 | 0.49 | 0.51 |
| 10 | 99851976 | rs10883039 | 0.477 | 0.954 | 0.008 | 0.314 | -0.017 | 0.029 | 0.029 | 0 | 0.44 | 0.24 | 0.51 | 0.39 | 0.47 | 0.52 |
| 10 | 99856584 | rs4919173 | -1.288 | 1.078 | -0.27 | 0.216 | -0.003 | 0.036 | 0.036 | 0 | 0.78 | 0.88 | 0.89 | 0.9 | 0.8 | 0.81 |
| 10 | 99862731 | rs12411523 | 1.73 | 1.095 | -0.462 | 0.121 | -0.029 | -0.033 | -0.033 | 0 | 0.31 | 0.05 | 0.38 | 0.34 | 0.31 | 0.42 |
| 10 | 99868267 | rs1334461 | -0.631 | 1.57 | -0.233 | 0.228 | -0.012 | -0.05 | -0.05 | 0 | 0.68 | 0.79 | 0.67 | 0.53 | 0.67 | 0.68 |
| 10 | 99876121 | rs1334465 | -0.662 | 1.573 | -0.086 | 0.23 | -0.011 | -0.045 | -0.045 | 6.84 | 0.69 | 0.79 | 0.67 | 0.53 | 0.67 | 0.68 |
| 10 | 99890470 | rs7911699 | -0.553 | 1.494 | 0.421 | 0.239 | -0.001 | 0 | 0 | 2.252 | 0.68 | 0.68 | 0.65 | 0.53 | 0.67 | 0.68 |
| 10 | 99891010 | rs7088359 | -0.603 | 1.519 | 0.347 | 0.233 | -0.013 | -0.049 | -0.049 | 0 | 0.68 | 0.79 | 0.67 | 0.53 | 0.67 | 0.68 |
| 10 | 99914261 | rs10748715 | -0.642 | 1.492 | 0.367 | 0.245 | -0.011 | -0.051 | -0.051 | 0.02 | 0.68 | 0.79 | 0.67 | 0.53 | 0.68 | 0.68 |

| | | | | | | | | | | | | | | | | |
|---|---|---|---|---|---|---|---|---|---|---|---|---|---|---|---|---|
| 10 | 99917757 | rs4281427 | 0.286 | 1.399 | 0.927 | 0.153 | -0.012 | -0.014 | -0.014 | 0 | 0.21 | 0.32 | 0.12 | 0.14 | 0.14 | 0.16 |
| 10 | 99932320 | rs10509718 | 0.641 | 1.392 | 1.158 | 0.116 | 0.014 | -0.034 | -0.034 | 0.055 | 0.14 | 0.16 | 0.15 | 0.03 | 0.22 | 0.11 |
| 10 | 99962273 | rs11189510 | 0.552 | 1.351 | 1.343 | 0.105 | 0.006 | 0.007 | 0.007 | 0.01 | 0.19 | 0.14 | 0.11 | 0.14 | 0.14 | 0.16 |
| 10 | 99969568 | rs11189513 | 1.962 | 1.34 | 1.398 | 0.104 | 0.002 | -0.017 | -0.017 | 0 | 0.31 | 0.09 | 0.38 | 0.35 | 0.31 | 0.41 |
| 10 | 99990493 | rs10450402 | -0.379 | 1.414 | 1.477 | 0.191 | 0.001 | 0.047 | 0.047 | 0 | 0.66 | 0.47 | 0.64 | 0.53 | 0.67 | 0.68 |
| 10 | 100008436 | rs3793692 | 0.409 | 1.418 | 1.154 | 0.192 | -0.011 | -0.051 | -0.051 | 0.905 | 0.32 | 0.21 | 0.33 | 0.47 | 0.32 | 0.32 |
| 10 | 100013438 | rs17524355 | 0.628 | 1.562 | 0.801 | -0.576 | 0.011 | -0.03 | -0.03 | 0.33 | 0.14 | 0.15 | 0.15 | 0.03 | 0.22 | 0.11 |
| 10 | 100016339 | rs1983865 | 0.394 | 1.345 | 0.582 | -0.157 | 0.013 | 0.111 | 0.111 | 0 | 0.52 | 0.25 | 0.62 | 0.75 | 0.45 | 0.59 |
| 10 | 100017453 | rs1983864 | 1.412 | 1.377 | 0.429 | -0.15 | -0.008 | -0.066 | -0.066 | 0 | 0.36 | 0.06 | 0.52 | 0.61 | 0.34 | 0.43 |
| 10 | 100021983 | rs746257 | 0.735 | 1.49 | -0.013 | -0.227 | 0.001 | -0.001 | -0.001 | 0 | 0.08 | 0.07 | 0.09 | 0.11 | 0.13 | 0.11 |
| 10 | 100025924 | rs10736126 | 0.231 | 0.915 | -0.036 | -0.228 | 0.003 | 0.055 | 0.055 | 0 | 0.53 | 0.32 | 0.61 | 0.75 | 0.45 | 0.6 |
| 10 | 100030768 | rs7899882 | 0.13 | 0.99 | -0.044 | -0.059 | -0.01 | -0.035 | -0.035 | 0 | 0.76 | 0.45 | 0.79 | 0.83 | 0.81 | 0.79 |
| 10 | 100031394 | rs10786408 | 0.407 | 0.938 | -1.161 | -0.104 | -0.045 | -0.016 | -0.016 | 0 | 0.4 | 0.12 | 0.58 | 0.65 | 0.47 | 0.48 |
| 10 | 100032289 | rs878259 | 1.086 | 0.697 | -1.336 | -0.043 | -0.067 | -0.141 | -0.141 | 1.032 | 0.23 | 0.02 | 0.44 | 0.59 | 0.26 | 0.37 |
| 10 | 100034883 | rs12784535 | 0.886 | 0.593 | -0.844 | -0.471 | -0.026 | 0.071 | 0.071 | 0.353 | 0.27 | 0.06 | 0.16 | 0.1 | 0.26 | 0.18 |
| 10 | 100036209 | rs7915743 | 0.941 | 0.258 | 0.578 | -0.086 | -0.008 | -0.035 | -0.035 | 0.038 | 0.91 | 0.69 | 0.94 | 0.99 | 0.95 | 0.91 |
| 10 | 100037683 | rs2094071 | 1.105 | 0.266 | 0.495 | -0.082 | 0.009 | -0.027 | -0.027 | 0 | 0.9 | 0.46 | 0.92 | 0.99 | 0.94 | 0.91 |
| 10 | 100044928 | rs4919201 | 0.455 | 0.312 | -0.147 | 0.254 | -0.018 | 0 | 0 | 0.1 | 0.71 | 0.41 | 0.84 | 0.71 | 0.79 | 0.75 |
| 10 | 100046078 | rs7905091 | 0.673 | 0.806 | -0.351 | 0.212 | 0.038 | -0.02 | -0.02 | 0.032 | 0.91 | 0.8 | 0.91 | 0.96 | 0.84 | 0.84 |
| 10 | 100050787 | rs10736128 | 0.309 | -0.066 | -0.418 | 0.358 | -0.023 | 0.01 | 0.01 | 0 | 0.84 | 0.74 | 0.86 | 0.76 | 0.91 | 0.92 |
| 10 | 100051497 | rs1925637 | 0.16 | -0.144 | -1.16 | -0.084 | 0.071 | 0.192 | 0.192 | 0.261 | 0.5 | 0.19 | 0.37 | 0.2 | 0.57 | 0.52 |
| 10 | 100052901 | rs2182166 | -0.401 | -0.281 | -0.737 | -0.152 | -0.003 | -0.001 | -0.001 | 3.581 | 0.56 | 0.52 | 0.57 | 0.57 | 0.62 | 0.66 |
| 10 | 100055644 | rs6584182 | -0.077 | -0.292 | -0.676 | -0.349 | 0.003 | -0.013 | -0.013 | 0 | 0.44 | 0.63 | 0.52 | 0.54 | 0.44 | 0.49 |
| 10 | 100057461 | rs7917347 | -0.23 | -0.622 | 0.009 | -0.564 | -0.071 | -0.036 | -0.036 | 0 | 0.83 | 0.58 | 0.87 | 0.87 | 0.94 | 0.88 |
| 10 | 100057707 | rs1925636 | 0.461 | -0.411 | -0.522 | -0.459 | -0.073 | 0.167 | 0.167 | 0.302 | 0.39 | 0.14 | 0.32 | 0.09 | 0.49 | 0.37 |
| 10 | 100058711 | rs7098860 | 0.462 | -0.368 | -0.38 | 0.026 | 0 | 0 | 0 | 0 | 0.28 | 0.28 | 0.42 | 0.41 | 0.33 | 0.36 |
| 10 | 100060652 | rs17453066 | 0.714 | 0.294 | -0.695 | -0.464 | -0.009 | 0.006 | 0.006 | 0 | 0.13 | 0.07 | 0.09 | 0.03 | 0.2 | 0.19 |
| 10 | 100066603 | rs11189541 | 1.191 | 0.054 | -1.021 | -0.695 | -0.051 | 0.071 | 0.071 | 0 | 0.25 | 0.08 | 0.23 | 0.02 | 0.32 | 0.21 |
| 10 | 100074813 | rs4917813 | -1.359 | 0.024 | -0.807 | -0.708 | -0.012 | -0.008 | -0.008 | 0.06 | 0.74 | 0.83 | 0.77 | 0.98 | 0.68 | 0.79 |
| 10 | 100076982 | rs10748725 | 0.872 | -0.499 | -0.861 | -0.55 | 0.002 | -0.003 | -0.003 | 0 | 0.31 | 0.3 | 0.5 | 0.6 | 0.37 | 0.39 |
| 10 | 100077152 | rs11189551 | -0.246 | -0.382 | -1.366 | -0.456 | 0.051 | 0.002 | 0.002 | 0.799 | 0.24 | 0.09 | 0.12 | 0.3 | 0.1 | 0.18 |
| 10 | 100077732 | rs927739 | -0.986 | 0.205 | -1.252 | -0.539 | -0.024 | 0.019 | 0.019 | 0 | 0.74 | 0.85 | 0.79 | 0.98 | 0.67 | 0.79 |
| 10 | 100079947 | rs1252895 | 0.025 | 0.184 | -1.635 | -0.102 | -0.083 | 0.172 | 0.172 | 0 | 0.45 | 0.08 | 0.38 | 0.23 | 0.45 | 0.37 |
| 10 | 100080631 | rs11599019 | -0.087 | 0.084 | -0.822 | 0.304 | -0.003 | -0.007 | -0.007 | 0 | 0.41 | 0.42 | 0.55 | 0.57 | 0.44 | 0.59 |
| 10 | 100081443 | rs942800 | 0.181 | 0.202 | -0.747 | 0.217 | 0.003 | 0 | 0 | 0 | 0.49 | 0.57 | 0.59 | 0.47 | 0.45 | 0.5 |
| 10 | 100081757 | rs2862297 | 0.054 | -0.264 | -0.154 | -0.094 | -0.008 | 0.004 | 0.004 | 0 | 0.6 | 0.65 | 0.55 | 0.6 | 0.65 | 0.49 |
| 10 | 100083404 | rs11189552 | 0.42 | -0.446 | 0.283 | -0.137 | 0.039 | 0.013 | 0.013 | 0 | 0.5 | 0.42 | 0.44 | 0.47 | 0.57 | 0.47 |
| 10 | 100085888 | rs4917817 | -0.016 | -0.476 | 0.259 | -0.185 | -0.021 | 0.018 | 0.018 | 0 | 0.64 | 0.53 | 0.49 | 0.47 | 0.68 | 0.54 |
| 10 | 100087228 | rs6584185 | -1.556 | -0.602 | 0.151 | -0.042 | 0.024 | -0.078 | -0.078 | 0.473 | 0.78 | 0.92 | 0.71 | 0.7 | 0.8 | 0.66 |
| 10 | 100088262 | rs4919210 | 0.261 | -0.525 | 0.342 | -0.029 | 0.017 | 0.021 | 0.021 | 0 | 0.08 | 0.2 | 0.11 | 0.14 | 0.08 | 0.02 |
| 10 | 100092279 | rs11189555 | 0.415 | -0.183 | 0.328 | 0.064 | -0.003 | 0.032 | 0.032 | 0 | 0.53 | 0.42 | 0.35 | 0.31 | 0.46 | 0.37 |
| 10 | 100093928 | rs4919213 | 0.438 | -0.104 | 0.165 | 0.087 | 0.041 | 0.045 | 0.045 | 0 | 0.07 | 0.21 | 0.11 | 0.14 | 0.08 | 0.02 |
| 10 | 100096148 | rs1000280 | 0.627 | -0.186 | 0.28 | 0.054 | 0.066 | 0.064 | 0.064 | 0 | 0.76 | 0.47 | 0.71 | 0.67 | 0.69 | 0.73 |
| 10 | 100096859 | rs1536154 | 0.187 | -0.173 | 0.304 | 0.006 | 0.003 | -0.016 | -0.016 | 0 | 0.55 | 0.43 | 0.42 | 0.36 | 0.48 | 0.38 |
| 10 | 100098716 | rs746033 | -0.258 | -0.108 | 0.141 | 0.023 | 0 | -0.002 | -0.002 | 2.3 | 0.59 | 0.53 | 0.45 | 0.44 | 0.54 | 0.39 |
| 10 | 100099508 | rs4917819 | 0.287 | -0.012 | -0.406 | -0.057 | -0.002 | 0.067 | 0.067 | 0.005 | 0.51 | 0.32 | 0.34 | 0.3 | 0.46 | 0.37 |
| 10 | 100100197 | rs10883072 | 0.971 | 0.334 | -1.08 | 0.213 | -0.066 | 0.075 | 0.075 | 0 | 0.35 | 0.08 | 0.27 | 0.27 | 0.35 | 0.21 |
| 10 | 100101261 | rs753299 | 0.515 | 1.1 | -0.733 | 0.796 | -0.069 | -0.068 | -0.068 | 0 | 0.11 | 0.01 | 0.15 | 0.21 | 0.23 | 0.26 |
| 10 | 100102034 | rs750298 | -1.034 | 0.229 | -0.672 | 0.53 | 0.046 | -0.007 | -0.007 | 0 | 0.65 | 0.84 | 0.67 | 0.69 | 0.71 | 0.56 |
| 10 | 100102731 | rs12264514 | 0.132 | 1.185 | -0.439 | 0.776 | 0.004 | 0.001 | 0.001 | 0 | 0.08 | 0.1 | 0.06 | 0.12 | 0.1 | 0.03 |
| 10 | 100102771 | rs11189559 | 0.295 | 1.539 | -0.69 | 0.864 | -0.082 | -0.056 | -0.056 | 0 | 0.12 | 0.02 | 0.2 | 0.14 | 0.29 | 0.24 |
| 10 | 100104006 | rs11189563 | -1.085 | 0.178 | -0.121 | 0.589 | 0.029 | 0.033 | 0.033 | 0 | 0.34 | 0.52 | 0.48 | 0.57 | 0.52 | 0.41 |
| 10 | 100111392 | rs4919218 | -0.294 | -0.171 | -0.004 | 0.146 | -0.002 | -0.005 | -0.005 | 0 | 0.47 | 0.51 | 0.41 | 0.38 | 0.34 | 0.28 |
| 10 | 100114965 | rs765456 | 0.277 | -0.193 | -0.376 | 0.132 | 0.017 | 0.063 | 0.063 | 0 | 0.38 | 0.18 | 0.35 | 0.33 | 0.27 | 0.24 |
| 10 | 100118265 | rs11189572 | -0.073 | -0.42 | 0.25 | 0.497 | -0.01 | 0.005 | 0.005 | 0 | 0.07 | 0.11 | 0.03 | 0.05 | 0.07 | 0.02 |
| 10 | 100119093 | rs11189577 | -1.129 | -0.696 | 1.034 | 0.371 | -0.03 | 0.004 | 0.004 | 1.922 | 0.12 | 0.43 | 0.07 | 0.05 | 0.07 | 0.02 |
| 10 | 100123612 | rs11814584 | -1.347 | -0.714 | 1.054 | 0.442 | -0.03 | 0.042 | 0.042 | 1.119 | 0.14 | 0.48 | 0.1 | 0.13 | 0.08 | 0.03 |
| 10 | 100124075 | rs6584190 | -2.252 | -1.092 | 1.058 | 0.385 | -0.011 | 0 | 0 | 5.687 | 0.21 | 0.77 | 0.27 | 0.27 | 0.12 | 0.12 |
| 10 | 100127551 | rs17109512 | -0.658 | -0.89 | 1.302 | 0.483 | 0.004 | 0.066 | 0.066 | 0 | 0.06 | 0.23 | 0.16 | 0.14 | 0.04 | 0.09 |
| 10 | 100139243 | rs17109576 | -0.947 | -1.008 | 1.666 | 0.447 | -0.001 | 0.015 | 0.015 | 2.779 | 0.06 | 0.35 | 0.07 | 0.02 | 0.01 | 0 |
| 10 | 100140237 | rs7091845 | -0.271 | -1.003 | 1.688 | 0.449 | 0.011 | 0.028 | 0.028 | 2.172 | 0.05 | 0.3 | 0.07 | 0.02 | 0.01 | 0 |
| 10 | 100142154 | rs17109597 | 0.879 | -1.38 | 1.27 | 0.664 | 0.003 | -0.004 | -0.004 | 0.575 | 0.32 | 0.23 | 0.29 | 0.32 | 0.23 | 0.43 |
| 10 | 100144782 | rs2296441 | 0.277 | -1.425 | 1.315 | 0.613 | -0.015 | 0.008 | 0.008 | 0 | 0.38 | 0.37 | 0.41 | 0.44 | 0.26 | 0.55 |
| 10 | 100147060 | rs4345897 | -0.387 | -1.45 | 1.061 | 0.571 | 0.045 | 0.019 | 0.019 | 0 | 0.46 | 0.64 | 0.48 | 0.56 | 0.33 | 0.54 |
| 10 | 100148058 | rs4539242 | -0.641 | -1.459 | 0.93 | 0.566 | 0.152 | 0.073 | 0.073 | 0.204 | 0.51 | 0.88 | 0.54 | 0.58 | 0.35 | 0.54 |
| 10 | 100148176 | rs2147896 | -0.641 | -1.459 | 0.93 | 0.566 | 0.152 | 0.073 | 0.073 | 0.204 | 0.51 | 0.88 | 0.54 | 0.58 | 0.35 | 0.54 |
| 10 | 100155898 | rs6584193 | -1.606 | -0.733 | 0.476 | 0.23 | -0.04 | 0.097 | 0.097 | 0 | 0.7 | 0.93 | 0.77 | 0.86 | 0.7 | 0.81 |
| 10 | 100155963 | rs7924303 | 0.372 | -1.14 | 0.861 | 0.36 | -0.015 | 0.005 | 0.005 | 0.014 | 0.39 | 0.4 | 0.36 | 0.44 | 0.3 | 0.45 |
| 10 | 100157593 | rs4551689 | -0.005 | -1.069 | 0.668 | 0.391 | 0.039 | -0.007 | -0.007 | 0 | 0.44 | 0.65 | 0.39 | 0.44 | 0.3 | 0.45 |

| | | | | | | | | | | | | | | | | |
|---|---|---|---|---|---|---|---|---|---|---|---|---|---|---|---|---|
| 10 | 100159373 | rs17109650 | 0.543 | -0.736 | 0.605 | 0.468 | 0.007 | -0.018 | -0.018 | 0.243 | 0.06 | 0.05 | 0.05 | 0.13 | 0.07 | 0.03 |
| 10 | 100162151 | rs17109653 | -1.098 | -0.862 | 1.098 | 0.422 | -0.028 | 0.064 | 0.064 | 0.627 | 0.12 | 0.42 | 0.12 | 0.15 | 0.09 | 0.03 |
| 10 | 100165967 | rs17109662 | -0.03 | -0.829 | 1.195 | 0.432 | -0.013 | 0.019 | 0.019 | 0 | 0.08 | 0.17 | 0.07 | 0.13 | 0.08 | 0.03 |
| 10 | 100165997 | rs17455577 | 0.373 | -0.317 | 0.879 | 0.37 | -0.093 | -0.044 | -0.044 | 1.422 | 0.2 | 0.05 | 0.21 | 0.28 | 0.35 | 0.28 |
| 10 | 100167396 | rs12763326 | 0.102 | -1.396 | 1.116 | 0.354 | -0.027 | 0.004 | 0.004 | 0 | 0.43 | 0.51 | 0.38 | 0.44 | 0.24 | 0.49 |
| 10 | 100169465 | rs7923729 | -0.03 | -1.455 | 1.044 | 0.328 | 0.002 | -0.002 | -0.002 | 0 | 0.44 | 0.58 | 0.38 | 0.44 | 0.24 | 0.49 |
| 10 | 100170383 | rs6584202 | -0.113 | -1.458 | 1.051 | 0.327 | 0.065 | -0.013 | -0.013 | 0.498 | 0.45 | 0.66 | 0.39 | 0.44 | 0.24 | 0.49 |
| 10 | 100175581 | rs3750604 | 1.134 | -1.524 | 1.006 | 0.348 | 0.064 | -0.058 | -0.058 | 0 | 0.33 | 0.1 | 0.27 | 0.34 | 0.2 | 0.4 |
| 10 | 100176154 | rs1061437 | 0.039 | -1.484 | 0.943 | 0.452 | -0.009 | 0 | 0 | 0.007 | 0.43 | 0.56 | 0.38 | 0.43 | 0.24 | 0.49 |
| 10 | 100176339 | rs1739 | -0.671 | -1.493 | 0.887 | 0.545 | 0.105 | 0.091 | 0.091 | 0.322 | 0.53 | 0.77 | 0.46 | 0.68 | 0.32 | 0.53 |
| 10 | 100176848 | rs3830019 | 0.891 | -1.484 | 0.875 | 0.471 | -0.001 | -0.035 | -0.035 | 0 | 0.06 | 0 | 0.06 | 0.13 | 0.07 | 0.02 |
| 10 | 100176869 | rs701801 | -0.239 | -2.066 | 0.766 | 0.206 | 0.042 | 0 | 0 | 0.719 | 0.43 | 0.65 | 0.41 | 0.49 | 0.28 | 0.39 |
| 10 | 100177049 | rs1061115 | -0.684 | -1.739 | 0.698 | 0.334 | 0.069 | 0.038 | 0.038 | 0 | 0.52 | 0.7 | 0.45 | 0.57 | 0.32 | 0.52 |
| 10 | 100179851 | rs2296436 | -0.219 | -1.394 | 0.32 | 0.431 | -0.004 | 0 | 0 | 0 | 0.12 | 0.17 | 0.12 | 0.15 | 0.11 | 0.04 |
| 10 | 100181847 | rs17109834 | -0.66 | -1.405 | 0.368 | 0.504 | -0.036 | 0.05 | 0.05 | 0 | 0.14 | 0.35 | 0.19 | 0.26 | 0.11 | 0.13 |
| 10 | 100188106 | rs10883094 | 0.521 | -1.019 | -0.125 | 0.639 | 0.029 | -0.009 | -0.009 | 0 | 0.36 | 0.26 | 0.32 | 0.34 | 0.23 | 0.4 |
| 10 | 100189138 | rs1061135 | -0.161 | -0.391 | -0.246 | 0.552 | 0.008 | -0.006 | -0.006 | 0.59 | 0.61 | 0.49 | 0.52 | 0.63 | 0.54 | 0.62 |
| 10 | 100189252 | rs1061134 | -0.643 | -0.642 | -0.395 | 0.448 | 0.005 | 0 | 0 | 0 | 0.16 | 0.12 | 0.09 | 0.15 | 0.13 | 0.06 |
| 10 | 100190264 | rs2296431 | -0.254 | -0.661 | -0.892 | 0.327 | -0.065 | 0.038 | 0.038 | 0 | 0.63 | 0.87 | 0.64 | 0.71 | 0.59 | 0.8 |
| 10 | 100190920 | rs1801287 | 1.461 | -0.564 | -0.803 | 0.851 | 0.03 | -0.031 | -0.031 | 0 | 0.35 | 0.14 | 0.29 | 0.34 | 0.3 | 0.4 |
| 10 | 100194041 | rs10883095 | 0.383 | -0.006 | -0.033 | 0.755 | 0.016 | -0.021 | -0.021 | 1.995 | 0.07 | 0.22 | 0.09 | 0 | 0.04 | 0.01 |
| 10 | 100195996 | rs7082570 | 0.328 | 0.253 | -0.745 | 0.913 | -0.037 | 0.005 | 0.005 | 0 | 0.1 | 0.04 | 0.07 | 0.12 | 0.17 | 0.23 |
| 10 | 100204727 | rs10786422 | 1.327 | -0.301 | -0.588 | 0.927 | 0.028 | -0.036 | -0.036 | 0 | 0.36 | 0.21 | 0.4 | 0.41 | 0.31 | 0.44 |
| 10 | 100212374 | rs7075080 | -0.372 | 0.22 | -0.539 | 0.965 | -0.048 | -0.035 | -0.035 | 0.175 | 0.68 | 0.84 | 0.63 | 0.57 | 0.63 | 0.63 |
| 10 | 100215413 | rs7913009 | -1.33 | -0.114 | 0.164 | 1.1 | -0.01 | -0.004 | -0.004 | 0 | 0.58 | 0.53 | 0.63 | 0.63 | 0.66 | 0.55 |
| 10 | 100217972 | rs4917835 | -1.007 | 0.387 | 0.223 | 1.363 | 0.002 | 0.003 | 0.003 | 0 | 0.49 | 0.57 | 0.53 | 0.49 | 0.51 | 0.44 |
| 10 | 100218766 | rs1060445 | -1.055 | 0.396 | 0.177 | 1.376 | 0.007 | 0.007 | 0.007 | 0 | 0.49 | 0.62 | 0.54 | 0.56 | 0.51 | 0.44 |
| 10 | 100219314 | rs10883099 | -1.055 | 0.396 | 0.177 | 1.252 | 0.007 | -0.003 | -0.003 | 0.012 | 0.49 | 0.62 | 0.53 | 0.51 | 0.51 | 0.44 |
| 10 | 100219519 | rs11189618 | -1.055 | 0.396 | 0.177 | 1.252 | 0.007 | -0.003 | -0.003 | 0 | 0.49 | 0.62 | 0.53 | 0.51 | 0.51 | 0.44 |
| 10 | 100220012 | rs11189619 | -1.056 | 0.397 | 0.178 | 1.253 | 0.007 | -0.003 | -0.003 | 0 | 0.49 | 0.62 | 0.53 | 0.51 | 0.51 | 0.44 |
| 10 | 100229532 | rs1998756 | -1.498 | 0.643 | -0.339 | 1.124 | 0.006 | -0.004 | -0.004 | 0 | 0.34 | 0.72 | 0.34 | 0.39 | 0.35 | 0.3 |
| 10 | 100234276 | rs11189627 | -1.484 | 0.63 | -0.24 | 1.153 | 0 | -0.001 | -0.001 | 0 | 0.34 | 0.66 | 0.34 | 0.41 | 0.35 | 0.3 |
| 10 | 100236682 | rs12265225 | -1.409 | 0.726 | -0.449 | 1.114 | 0.007 | -0.007 | -0.007 | 0 | 0.36 | 0.75 | 0.36 | 0.4 | 0.36 | 0.34 |
| 10 | 100240305 | rs17458353 | -0.825 | 1.498 | 0.044 | 0.949 | 0.099 | 0.034 | 0.034 | 1.158 | 0.07 | 0.25 | 0.12 | 0.08 | 0.16 | 0.08 |
| 10 | 100240334 | rs10883108 | -0.666 | 0.638 | -0.396 | 0.933 | -0.032 | 0.003 | 0.003 | 0 | 0.21 | 0.47 | 0.22 | 0.18 | 0.18 | 0.23 |
| 10 | 100240681 | rs11189628 | -1.045 | 0.154 | -1.689 | 1.216 | 0.057 | -0.097 | -0.097 | 0 | 0.57 | 0.98 | 0.57 | 0.45 | 0.7 | 0.52 |
| 10 | 100242039 | rs10748732 | -1.446 | 0.595 | 0.222 | 1.807 | -0.002 | 0.009 | 0.009 | 0 | 0.26 | 0.3 | 0.23 | 0.62 | 0.26 | 0.35 |
| 10 | 100242195 | rs11189629 | -1.474 | 0.084 | 1.041 | 1.697 | -0.026 | 0.01 | 0.01 | 0 | 0.22 | 0.4 | 0.22 | 0.24 | 0.16 | 0.18 |
| 10 | 100247590 | rs4919231 | -1.494 | 0.237 | 2.01 | 1.634 | 0.002 | 0.009 | 0.009 | 0 | 0.26 | 0.23 | 0.33 | 0.36 | 0.32 | 0.3 |
| 10 | 100249295 | rs4442477 | 1.991 | 0.596 | 1.794 | 1.783 | -0.074 | -0.092 | -0.092 | 0 | 0.67 | 0.06 | 0.62 | 0.62 | 0.66 | 0.6 |
| 10 | 100250186 | rs1925647 | -0.297 | 0.523 | 1.828 | 1.753 | -0.059 | -0.065 | -0.065 | 0.715 | 0.15 | 0.03 | 0.24 | 0.25 | 0.27 | 0.24 |
| 10 | 100268520 | rs1889974 | 2.023 | 0.299 | 1.818 | 1.495 | -0.077 | -0.12 | -0.12 | 1.013 | 0.63 | 0.07 | 0.59 | 0.55 | 0.62 | 0.63 |
| 10 | 100273119 | rs10883112 | 1.904 | 0.345 | 2.082 | 1.45 | -0.02 | -0.075 | -0.075 | 0 | 0.63 | 0.17 | 0.58 | 0.55 | 0.64 | 0.64 |
| 10 | 100278017 | rs10509722 | 2.536 | 0.246 | 2.529 | 1.401 | -0.003 | -0.058 | -0.058 | 0.451 | 0.86 | 0.27 | 0.9 | 0.97 | 0.92 | 0.89 |
| 10 | 100280107 | rs7912749 | -1.566 | 0.226 | 2.522 | 1.494 | 0 | 0.009 | 0.009 | 0.389 | 0.27 | 0.28 | 0.35 | 0.41 | 0.32 | 0.27 |
| 10 | 100283151 | rs4919233 | -1.623 | 0.226 | 2.317 | 1.494 | 0.006 | 0.047 | 0.047 | 0.421 | 0.27 | 0.37 | 0.36 | 0.42 | 0.32 | 0.27 |
| 10 | 100288452 | rs534666 | 1.158 | 0.225 | 1.964 | 1.092 | 0.04 | -0.035 | -0.035 | 0.19 | 0.92 | 0.75 | 0.96 | 1 | 0.94 | 0.95 |
| 10 | 100290770 | rs12358472 | -1.345 | 0.13 | 1.859 | 1.235 | -0.001 | -0.018 | -0.018 | 0.361 | 0.27 | 0.19 | 0.3 | 0.52 | 0.3 | 0.28 |
| 10 | 100292627 | rs922324 | 2.535 | -0.022 | 1.766 | 1.128 | -0.096 | -0.01 | -0.01 | 0 | 0.5 | 0.03 | 0.51 | 0.43 | 0.53 | 0.43 |
| 10 | 100293700 | rs898892 | -1.633 | 0.122 | 1.958 | 1.131 | -0.008 | 0.052 | 0.052 | 0.121 | 0.3 | 0.45 | 0.33 | 0.52 | 0.3 | 0.28 |
| 10 | 100308507 | rs12355571 | 1.857 | 0.134 | 1.413 | 1.14 | -0.075 | -0.03 | -0.03 | 0.209 | 0.58 | 0.06 | 0.59 | 0.44 | 0.59 | 0.56 |
| 10 | 100314339 | rs7907389 | -1.219 | 0.372 | 1.59 | 1.249 | -0.002 | -0.009 | -0.009 | 0.009 | 0.25 | 0.19 | 0.3 | 0.51 | 0.3 | 0.28 |
| 10 | 100321965 | rs471563 | 1.341 | 0.445 | 1.818 | 1.145 | 0.076 | -0.044 | -0.044 | 0 | 0.91 | 0.66 | 0.95 | 1 | 0.92 | 0.94 |
| 10 | 100322658 | rs477950 | 1.401 | 0.443 | 1.876 | 1.144 | 0.091 | -0.04 | -0.04 | 0 | 0.91 | 0.61 | 0.95 | 1 | 0.92 | 0.94 |
| 10 | 100324333 | rs535182 | 1.401 | 0.443 | 1.876 | 1.144 | 0.091 | -0.04 | -0.04 | 0.118 | 0.91 | 0.61 | 0.95 | 1 | 0.92 | 0.94 |
| 10 | 100336680 | rs4919244 | 0.974 | 0.614 | 1.506 | 1.217 | 0.057 | -0.024 | -0.024 | 0 | 0.91 | 0.75 | 0.94 | 0.97 | 0.89 | 0.85 |
| 10 | 100338139 | rs3853516 | 1.706 | 0.558 | 1.542 | 1.277 | 0.028 | 0.013 | 0.013 | 0.237 | 0.65 | 0.5 | 0.63 | 0.46 | 0.59 | 0.54 |
| 10 | 100339741 | rs3936941 | 1.063 | 0.709 | 1.654 | 1.258 | 0.057 | -0.024 | -0.024 | 0 | 0.91 | 0.75 | 0.94 | 0.97 | 0.89 | 0.85 |
| 10 | 100342812 | rs12219601 | -0.117 | 0.532 | 1.531 | 1.212 | 0.026 | 0.038 | 0.038 | 0.296 | 0.1 | 0.02 | 0.08 | 0.03 | 0.06 | 0.13 |
| 10 | 100343534 | rs1111289 | 1.531 | 0.561 | 1.612 | 1.281 | 0.023 | 0.05 | 0.05 | 0.543 | 0.7 | 0.48 | 0.64 | 0.46 | 0.66 | 0.61 |
| 10 | 100344136 | rs11189692 | -0.059 | 0.596 | 1.542 | 1.405 | -0.053 | -0.104 | -0.104 | 0 | 0.14 | 0.02 | 0.2 | 0.4 | 0.22 | 0.23 |
| 10 | 100345376 | rs11189698 | 1.354 | 0.494 | 1.642 | 1.356 | 0.023 | 0.05 | 0.05 | 0.202 | 0.71 | 0.52 | 0.64 | 0.46 | 0.66 | 0.61 |
| 10 | 100347450 | rs6584214 | 0.29 | 0.497 | 1.69 | 1.359 | 0.061 | 0.017 | 0.017 | 1.94 | 0.95 | 0.76 | 0.96 | 0.96 | 0.95 | 0.9 |
| 10 | 100347919 | rs12268248 | 2.013 | 0.485 | 1.64 | 1.557 | -0.003 | 0.001 | 0.001 | 1.233 | 0.6 | 0.48 | 0.62 | 0.6 | 0.61 | 0.49 |
| 10 | 100348138 | rs10786447 | -1.312 | 0.479 | 0.989 | 1.089 | 0.001 | 0.023 | 0.023 | 0 | 0.23 | 0.23 | 0.31 | 0.5 | 0.29 | 0.29 |
| 10 | 100353308 | rs7923554 | -0.758 | 0.416 | 1.017 | 1.099 | 0.003 | 0.051 | 0.051 | 1.999 | 0.05 | 0.21 | 0.07 | 0.06 | 0 | 0 |
| 10 | 100358291 | rs7090780 | -0.781 | 0.637 | 0.788 | 0.956 | -0.001 | -0.001 | -0.001 | 0 | 0.35 | 0.31 | 0.4 | 0.54 | 0.39 | 0.44 |
| 10 | 100368893 | rs7475778 | 0.815 | -0.29 | 0.605 | 0.505 | -0.055 | -0.034 | -0.034 | 0 | 0.78 | 0.51 | 0.82 | 0.85 | 0.91 | 0.88 |
| 10 | 100369397 | rs4554816 | 0.37 | -0.249 | 1.049 | 0.42 | -0.011 | -0.033 | -0.033 | 0 | 0.84 | 0.81 | 0.93 | 0.95 | 0.92 | 0.89 |

| | | | | | | | | | | | | | | | | |
|---|---|---|---|---|---|---|---|---|---|---|---|---|---|---|---|---|
| 10 | 100370002 | rs526465 | -1.541 | -0.1 | 0.975 | 0.449 | -0.013 | -0.024 | -0.024 | 0 | 0.23 | 0.3 | 0.12 | 0.12 | 0.13 | 0.27 |
| 10 | 100376145 | rs10786449 | 0.278 | -0.345 | -0.154 | 0.696 | 0.037 | -0.065 | -0.065 | 0 | 0.65 | 0.15 | 0.5 | 0.51 | 0.71 | 0.77 |
| 10 | 100376798 | rs648003 | 0.287 | -0.653 | 0.06 | 1.014 | -0.009 | 0.017 | 0.017 | 0 | 0.22 | 0.08 | 0.27 | 0.17 | 0.24 | 0.18 |
| 10 | 100378911 | rs688501 | -0.369 | -0.679 | 0.951 | 1.092 | -0.016 | -0.003 | -0.003 | 0.011 | 0.27 | 0.44 | 0.39 | 0.28 | 0.26 | 0.19 |
| 10 | 100380960 | rs10883151 | 1.36 | -0.929 | 1.217 | 1.072 | -0.017 | -0.013 | -0.013 | 0 | 0.53 | 0.09 | 0.43 | 0.4 | 0.6 | 0.51 |
| 10 | 100384100 | rs543289 | -0.655 | -0.902 | 1.349 | 1.115 | -0.012 | -0.011 | -0.011 | 0 | 0.3 | 0.44 | 0.39 | 0.28 | 0.27 | 0.23 |
| 10 | 100384359 | rs656707 | 0.013 | -0.894 | 1.339 | 1.125 | 0.002 | -0.002 | -0.002 | 1.535 | 0.25 | 0.21 | 0.37 | 0.28 | 0.26 | 0.19 |
| 10 | 100401745 | rs693656 | -0.682 | -0.727 | 1.739 | 1.312 | -0.018 | 0.067 | 0.067 | 0 | 0.29 | 0.43 | 0.44 | 0.55 | 0.24 | 0.23 |
| 10 | 100409977 | rs17110576 | 0.278 | -0.666 | 1.626 | 1.308 | -0.021 | -0.023 | -0.023 | 0.049 | 0.05 | 0.01 | 0.04 | 0.08 | 0.07 | 0.2 |
| 10 | 100413929 | rs566626 | 1.36 | -0.734 | 1.784 | 1.287 | 0.036 | 0.093 | 0.093 | 1.946 | 0.92 | 0.55 | 0.87 | 0.89 | 0.96 | 0.98 |
| 10 | 100415158 | rs548079 | -0.009 | -0.754 | 1.757 | 1.286 | 0.006 | -0.039 | -0.039 | 0.122 | 0.24 | 0.13 | 0.35 | 0.35 | 0.22 | 0.18 |
| 10 | 100415279 | rs7086093 | 0.35 | -0.744 | 1.778 | 1.291 | -0.006 | 0.012 | 0.012 | 0.003 | 0.7 | 0.33 | 0.55 | 0.46 | 0.76 | 0.81 |
| 10 | 100420507 | rs11189739 | 0.616 | -1.026 | 0.857 | 0.48 | 0.01 | 0.012 | 0.012 | 0 | 0.53 | 0.41 | 0.52 | 0.67 | 0.56 | 0.56 |
| 10 | 100433762 | rs2484939 | 1.708 | -0.6 | 1.203 | 0.649 | 0.08 | 0.241 | 0.241 | 0.539 | 0.94 | 0.61 | 0.9 | 0.81 | 0.98 | 0.95 |
| 10 | 100442880 | rs766550 | 0.609 | -0.733 | 1.066 | 0.56 | 0.007 | -0.014 | -0.014 | 0.258 | 0.63 | 0.31 | 0.46 | 0.46 | 0.62 | 0.8 |
| 10 | 100455524 | rs10736136 | 0.318 | -0.727 | 1.022 | 0.535 | -0.01 | 0.002 | 0.002 | 0 | 0.66 | 0.47 | 0.52 | 0.65 | 0.63 | 0.81 |
| 10 | 100457056 | rs10883163 | 0.347 | -0.727 | 0.978 | 0.535 | -0.006 | -0.001 | -0.001 | 0 | 0.65 | 0.39 | 0.51 | 0.65 | 0.63 | 0.81 |
| 10 | 100462462 | rs7919776 | 0.764 | -0.723 | 0.979 | 0.523 | 0.014 | -0.02 | -0.02 | 0 | 0.84 | 0.41 | 0.77 | 0.89 | 0.81 | 0.96 |
| 10 | 100476497 | rs11593463 | 0.929 | -0.577 | 0.966 | 0.557 | 0.014 | -0.049 | -0.049 | 0 | 0.6 | 0.2 | 0.44 | 0.46 | 0.61 | 0.76 |
| 10 | 100477385 | rs10883176 | -0.868 | -0.573 | 1.004 | 0.56 | 0.03 | -0.009 | -0.009 | 0 | 0.14 | 0.41 | 0.21 | 0.11 | 0.19 | 0.04 |
| 10 | 100479326 | rs11189770 | 0.704 | -0.603 | 0.868 | 0.558 | 0.011 | -0.043 | -0.043 | 0 | 0.21 | 0.12 | 0.31 | 0.42 | 0.17 | 0.16 |
| 10 | 100487637 | rs4508137 | 1.11 | -0.594 | 0.758 | 0.55 | 0.009 | 0.011 | 0.011 | 0 | 0.54 | 0.18 | 0.4 | 0.38 | 0.54 | 0.56 |
| 10 | 100514409 | rs12775639 | 0.827 | -0.611 | 0.697 | 0.593 | -0.004 | -0.047 | -0.047 | 0.632 | 0.2 | 0.03 | 0.26 | 0.23 | 0.17 | 0.16 |
| 10 | 100538135 | rs861399 | -1.007 | -0.558 | 0.702 | 0.592 | 0.002 | -0.04 | -0.04 | 1.351 | 0.79 | 0.96 | 0.74 | 0.77 | 0.83 | 0.84 |
| 10 | 100552456 | rs10883202 | -0.846 | -0.55 | 1.02 | 0.52 | 0.02 | -0.022 | -0.022 | 0 | 0.15 | 0.29 | 0.2 | 0.11 | 0.2 | 0.08 |
| 10 | 100564408 | rs837745 | -0.985 | -0.517 | 0.929 | 0.528 | -0.005 | -0.026 | -0.026 | 0.619 | 0.8 | 0.96 | 0.74 | 0.79 | 0.83 | 0.85 |
| 10 | 100574251 | rs7088816 | -0.406 | -0.509 | 1.003 | 0.508 | -0.018 | 0.039 | 0.039 | 0 | 0.76 | 0.65 | 0.66 | 0.6 | 0.81 | 0.83 |
| 10 | 100578943 | rs11189833 | -0.105 | -0.51 | 0.91 | 0.472 | 0.018 | -0.062 | -0.062 | 0 | 0.11 | 0.28 | 0.11 | 0 | 0.15 | 0.02 |
| 10 | 100583915 | rs2801405 | -0.6 | -0.496 | 0.567 | 0.534 | 0.007 | -0.017 | -0.017 | 0 | 0.78 | 0.83 | 0.68 | 0.61 | 0.81 | 0.83 |
| 10 | 100603695 | rs2862447 | 1.163 | -0.295 | 0.032 | 0.343 | 0.012 | 0.027 | 0.027 | 0.347 | 0.49 | 0.11 | 0.42 | 0.39 | 0.59 | 0.52 |
| 10 | 100608412 | rs2197417 | -0.919 | -0.321 | -0.053 | 0.456 | -0.021 | 0.006 | 0.006 | 0 | 0.79 | 0.97 | 0.73 | 0.86 | 0.81 | 0.85 |
| 10 | 100612445 | rs7070985 | -0.565 | -0.357 | 0.386 | 0.47 | -0.032 | -0.024 | -0.024 | 0 | 0.17 | 0.32 | 0.16 | 0.12 | 0.09 | 0.06 |
| 10 | 100614789 | rs9988679 | 0.979 | -0.015 | 0.363 | 0.341 | -0.003 | -0.001 | -0.001 | 0 | 0.55 | 0.49 | 0.45 | 0.55 | 0.56 | 0.75 |
| 10 | 100620509 | rs10883221 | 0.647 | -0.212 | 0.455 | 0.44 | -0.013 | 0.003 | 0.003 | 0.436 | 0.63 | 0.45 | 0.54 | 0.6 | 0.74 | 0.79 |
| 10 | 100624132 | rs1444803 | -0.85 | -0.238 | 0.502 | 0.462 | 0.005 | 0.009 | 0.009 | 0 | 0.78 | 0.88 | 0.72 | 0.85 | 0.8 | 0.85 |
| 10 | 100626866 | rs7904255 | -0.319 | -0.254 | 0.5 | 0.45 | -0.002 | 0.002 | 0.002 | 0 | 0.75 | 0.77 | 0.67 | 0.7 | 0.8 | 0.84 |
| 10 | 100637858 | rs895070 | 1.393 | 0.091 | 0.486 | 0.535 | 0.036 | 0.034 | 0.034 | 0.209 | 0.52 | 0.19 | 0.45 | 0.42 | 0.61 | 0.57 |
| 10 | 100640579 | rs7910696 | 0.64 | 0.045 | 0.471 | 0.433 | -0.001 | -0.001 | -0.001 | 0.005 | 0.59 | 0.45 | 0.52 | 0.59 | 0.71 | 0.79 |
| 10 | 100642848 | rs4917849 | -0.75 | 0.05 | 0.472 | 0.423 | 0.001 | -0.005 | -0.005 | 0 | 0.78 | 0.82 | 0.68 | 0.69 | 0.8 | 0.85 |
| 10 | 100648642 | rs7908265 | -0.504 | 0.034 | 0.506 | 0.406 | -0.035 | -0.04 | -0.04 | 0 | 0.17 | 0.31 | 0.16 | 0.1 | 0.09 | 0.06 |
| 10 | 100654000 | rs2018913 | 0.671 | 0.026 | 0.461 | 0.482 | 0 | -0.001 | -0.001 | 0.485 | 0.59 | 0.44 | 0.52 | 0.57 | 0.71 | 0.78 |
| 10 | 100661838 | rs947542 | -0.997 | -0.016 | 0.297 | 0.481 | -0.057 | 0.044 | 0.044 | 0.277 | 0.19 | 0.47 | 0.22 | 0.25 | 0.1 | 0.06 |
| 10 | 100665866 | rs7073636 | -0.589 | -0.011 | 0.295 | 0.482 | -0.041 | -0.045 | -0.045 | 0 | 0.17 | 0.37 | 0.17 | 0.09 | 0.09 | 0.06 |
| 10 | 100683856 | rs11189868 | 0.313 | 0.043 | 0.262 | 0.555 | -0.008 | -0.006 | -0.006 | 0 | 0.05 | 0.04 | 0.05 | 0.14 | 0.09 | 0.22 |
| 10 | 100684443 | rs11189870 | 1.374 | 0.039 | 0.257 | 0.53 | 0.035 | 0.013 | 0.013 | 0.054 | 0.52 | 0.29 | 0.45 | 0.46 | 0.62 | 0.58 |
| 10 | 100685108 | rs34592150 | -1.216 | 0.055 | 0.261 | 0.549 | 0.024 | 0.122 | 0.122 | 0 | 0.09 | 0.23 | 0.11 | 0.3 | 0.1 | 0.23 |
| 10 | 100704597 | rs10786489 | 0.776 | 0.064 | 0.279 | 0.465 | 0.018 | 0.029 | 0.029 | 1.058 | 0.58 | 0.35 | 0.53 | 0.76 | 0.68 | 0.72 |
| 10 | 100710124 | rs947539 | 1.461 | 0.058 | 0.283 | 0.463 | 0.007 | 0.026 | 0.026 | 0.075 | 0.51 | 0.18 | 0.47 | 0.62 | 0.59 | 0.51 |
| 10 | 100713090 | rs4635020 | -0.422 | -0.04 | 0.252 | 0.414 | 0.001 | -0.005 | -0.005 | 0 | 0.76 | 0.7 | 0.7 | 0.85 | 0.78 | 0.79 |
| 10 | 100716236 | rs3897502 | -1.004 | -0.072 | 0.166 | 0.441 | -0.04 | -0.016 | -0.016 | 0.783 | 0.8 | 0.99 | 0.74 | 0.88 | 0.82 | 0.88 |
| 10 | 100730161 | rs10883232 | 0.839 | -0.057 | 0.201 | 0.428 | 0.015 | 0.025 | 0.025 | 0.65 | 0.58 | 0.35 | 0.53 | 0.75 | 0.69 | 0.73 |
| 10 | 100741465 | rs7098469 | -1.256 | -0.107 | 0.198 | 0.418 | -0.009 | 0.002 | 0.002 | 0 | 0.23 | 0.41 | 0.22 | 0.23 | 0.18 | 0.28 |
| 10 | 100742391 | rs10883238 | 1.584 | 0.004 | 0.088 | 0.44 | 0.01 | 0.023 | 0.023 | 0 | 0.5 | 0.17 | 0.43 | 0.45 | 0.59 | 0.5 |
| 10 | 100751620 | rs11189926 | 1.588 | -0.18 | -0.062 | 0.342 | -0.062 | -0.046 | -0.046 | 0 | 0.48 | 0.04 | 0.4 | 0.46 | 0.59 | 0.53 |
| 10 | 100758587 | rs1344214 | -0.061 | -0.039 | 0.047 | 0.375 | 0.011 | 0.02 | 0.02 | 0 | 0.05 | 0.11 | 0.04 | 0.14 | 0.09 | 0.19 |
| 10 | 100762637 | rs2487882 | 0.463 | -0.15 | 0.022 | 0.333 | 0.007 | 0.002 | 0.002 | 0 | 0.22 | 0.09 | 0.27 | 0.17 | 0.19 | 0.15 |
| 10 | 100779825 | rs4579871 | -0.29 | -0.133 | 0.091 | 0.316 | 0.001 | -0.005 | -0.005 | 1.746 | 0.76 | 0.71 | 0.7 | 0.84 | 0.78 | 0.78 |
| 10 | 100792770 | rs4132028 | 1.36 | -0.067 | 0.082 | 0.309 | 0.021 | 0.007 | 0.007 | 0.093 | 0.54 | 0.18 | 0.5 | 0.62 | 0.62 | 0.53 |
| 10 | 100792984 | rs9630091 | -0.658 | -0.076 | 0.135 | 0.296 | -0.049 | -0.052 | -0.052 | 0 | 0.19 | 0.38 | 0.18 | 0.09 | 0.09 | 0.06 |
| 10 | 100794359 | rs10786497 | 0.824 | -0.063 | 0.027 | 0.303 | 0.006 | 0.007 | 0.007 | 0.509 | 0.62 | 0.39 | 0.54 | 0.75 | 0.7 | 0.73 |
| 10 | 100798277 | rs2487889 | 1.018 | -0.064 | -0.058 | 0.303 | 0.008 | -0.005 | -0.005 | 0 | 0.18 | 0.09 | 0.25 | 0.15 | 0.17 | 0.15 |
| 10 | 100799832 | rs2487891 | 1.188 | -0.087 | -0.085 | 0.294 | 0 | 0 | 0 | 0 | 0.74 | 0.29 | 0.74 | 0.76 | 0.79 | 0.71 |
| 10 | 100815322 | rs4491154 | 1.196 | 0.224 | -0.187 | 0.267 | 0.028 | 0.001 | 0.001 | 0.578 | 0.56 | 0.18 | 0.5 | 0.62 | 0.63 | 0.57 |
| 10 | 100823161 | rs1539397 | 0.559 | 0.214 | 0.03 | 0.212 | -0.064 | -0.038 | -0.038 | 0 | 0.82 | 0.48 | 0.79 | 0.9 | 0.92 | 0.93 |
| 10 | 100835731 | rs11818672 | 1.215 | 0.506 | -0.084 | 0.207 | -0.037 | -0.019 | -0.019 | 0 | 0.17 | 0.04 | 0.25 | 0.14 | 0.21 | 0.13 |
| 10 | 100869273 | rs4919276 | 0.164 | 0.492 | 0.039 | 0.203 | -0.056 | -0.032 | -0.032 | 0.258 | 0.84 | 0.62 | 0.82 | 0.9 | 0.92 | 0.98 |
| 10 | 100872973 | rs10883267 | 1.045 | 0.334 | -1.012 | -0.201 | -0.058 | -0.112 | -0.112 | 0 | 0.14 | 0.01 | 0.16 | 0.46 | 0.19 | 0.28 |
| 10 | 100881909 | rs11189982 | 1.126 | 0.643 | -0.443 | 0.002 | 0.008 | 0.014 | 0.014 | 0.015 | 0.57 | 0.23 | 0.5 | 0.65 | 0.63 | 0.6 |
| 10 | 100899295 | rs10786510 | 1.114 | 0.64 | -0.423 | 0.017 | 0 | 0 | 0 | 0.04 | 0.74 | 0.29 | 0.75 | 0.78 | 0.84 | 0.73 |

| | | | | | | | | | | | | | | | | |
|---|---|---|---|---|---|---|---|---|---|---|---|---|---|---|---|---|
| 10 | 100907713 | rs10883282 | 1.103 | 0.652 | -0.442 | 0.02 | 0.009 | 0.015 | 0.015 | 0 | 0.57 | 0.23 | 0.5 | 0.65 | 0.63 | 0.6 |
| 10 | 100920242 | rs4590799 | 1.184 | 0.638 | -0.473 | 0.014 | 0.006 | 0.014 | 0.014 | 0.079 | 0.56 | 0.16 | 0.5 | 0.65 | 0.63 | 0.6 |
| 10 | 100929982 | rs2040023 | 1.24 | 0.626 | -0.475 | 0.01 | 0.004 | 0.009 | 0.009 | 0.172 | 0.56 | 0.14 | 0.49 | 0.65 | 0.63 | 0.6 |
| 10 | 100932793 | rs6584238 | 1.216 | 0.68 | -0.489 | 0.2 | -0.037 | -0.057 | -0.057 | 3.357 | 0.6 | 0.15 | 0.48 | 0.55 | 0.64 | 0.62 |
| 10 | 100942449 | rs2796759 | 0.969 | 0.515 | -0.398 | 0.276 | -0.023 | -0.004 | -0.004 | 0.77 | 0.17 | 0.05 | 0.25 | 0.13 | 0.17 | 0.1 |
| 10 | 100953030 | rs11517066 | 2.071 | 1.129 | -0.936 | 0.023 | -0.015 | 0.05 | 0.05 | 0 | 0.11 | 0.02 | 0.06 | 0 | 0.12 | 0.1 |
| 10 | 100964452 | rs10786518 | 1.305 | 1.128 | -0.823 | 0.15 | 0.011 | -0.068 | -0.068 | 0.54 | 0.57 | 0.11 | 0.53 | 0.5 | 0.66 | 0.61 |
| 10 | 100988002 | rs2862532 | -1.718 | 1.367 | -0.756 | 0.231 | -0.006 | -0.01 | -0.01 | 0 | 0.84 | 0.96 | 0.76 | 0.87 | 0.85 | 0.91 |
| 10 | 101034419 | rs7082580 | -0.519 | 0.938 | -0.615 | 0.283 | -0.016 | -0.046 | -0.046 | 0.399 | 0.77 | 0.85 | 0.67 | 0.6 | 0.75 | 0.69 |
| 10 | 101034921 | rs7910561 | -0.569 | 0.806 | -0.52 | 0.218 | -0.045 | -0.031 | -0.031 | 0 | 0.18 | 0.53 | 0.13 | 0.1 | 0.1 | 0.07 |
| 10 | 101039633 | rs1962213 | -1.557 | 1.012 | -0.969 | 0.325 | 0.001 | -0.046 | -0.046 | 1.342 | 0.84 | 0.94 | 0.72 | 0.72 | 0.84 | 0.89 |
| 10 | 101050978 | rs4144925 | -0.089 | 1.061 | -0.976 | 0.211 | 0.001 | 0.001 | 0.001 | 0 | 0.07 | 0.09 | 0.04 | 0.12 | 0.09 | 0.2 |
| 10 | 101051812 | rs11190037 | 1.487 | 0.931 | -1.251 | -0.028 | -0.024 | 0.082 | 0.082 | 0 | 0.15 | 0.02 | 0.07 | 0 | 0.15 | 0.15 |
| 10 | 101056561 | rs10748771 | -1.012 | 0.93 | -0.316 | 0.918 | -0.006 | -0.01 | -0.01 | 0.049 | 0.84 | 0.96 | 0.76 | 0.88 | 0.85 | 0.91 |
| 10 | 101090428 | rs2862600 | -0.703 | 0.866 | -0.312 | 0.973 | -0.006 | -0.005 | -0.005 | 1.631 | 0.83 | 0.95 | 0.74 | 0.87 | 0.82 | 0.89 |
| 10 | 101102123 | rs2001285 | -2.311 | 0.783 | -0.303 | 1.046 | 0.01 | -0.04 | -0.04 | 0.706 | 0.29 | 0.83 | 0.24 | 0.38 | 0.21 | 0.33 |
| 10 | 101104388 | rs11190061 | 1.815 | 0.797 | -0.374 | 1.021 | -0.014 | -0.048 | -0.048 | 0 | 0.53 | 0.09 | 0.5 | 0.5 | 0.6 | 0.56 |
| 10 | 101104712 | rs10786538 | -0.589 | 0.72 | -0.291 | 1.104 | 0.001 | 0.005 | 0.005 | 0 | 0.82 | 0.92 | 0.74 | 0.87 | 0.81 | 0.89 |
| 10 | 101109264 | rs10883332 | 1.043 | 0.654 | 0.349 | 1.118 | -0.02 | -0.022 | -0.022 | 0 | 0.66 | 0.46 | 0.63 | 0.78 | 0.72 | 0.77 |
| 10 | 101109823 | rs7074815 | 0.879 | 0.646 | 0.63 | 1.114 | -0.02 | -0.022 | -0.022 | 0 | 0.67 | 0.59 | 0.64 | 0.78 | 0.72 | 0.77 |
| 10 | 101115457 | rs11190065 | 1.037 | 0.664 | 0.592 | 1.122 | -0.024 | -0.027 | -0.027 | 0 | 0.66 | 0.48 | 0.63 | 0.78 | 0.72 | 0.77 |
| 10 | 101117557 | rs3750895 | -0.964 | 0.606 | 1.299 | 1.43 | 0.034 | 0.11 | 0.11 | 0.036 | 0.08 | 0.15 | 0.1 | 0.28 | 0.1 | 0.14 |
| 10 | 101117689 | rs12098418 | -1.162 | 0.44 | 1.049 | 0.944 | -0.026 | -0.026 | -0.026 | 0.062 | 0.15 | 0.32 | 0.09 | 0.1 | 0.08 | 0.12 |
| 10 | 101121718 | rs11190074 | -0.979 | 0.804 | 0.489 | 1.121 | 0.013 | 0.021 | 0.021 | 0 | 0.07 | 0.09 | 0.05 | 0.13 | 0.08 | 0.14 |
| 10 | 101129045 | rs12268069 | 2.035 | 0.059 | 0.931 | 1.414 | -0.002 | 0.001 | 0.001 | 0 | 0.66 | 0.35 | 0.56 | 0.43 | 0.7 | 0.57 |
| 10 | 101129296 | rs4919304 | 1.301 | 0.055 | 1.198 | 1.478 | 0.001 | -0.004 | -0.004 | 0.683 | 0.78 | 0.82 | 0.64 | 0.53 | 0.75 | 0.8 |
| 10 | 101130163 | rs1892510 | -2.386 | -0.069 | 1.962 | 1.177 | -0.021 | -0.086 | -0.086 | 1.396 | 0.13 | 0.33 | 0.07 | 0 | 0.07 | 0.14 |
| 10 | 101132282 | rs10509735 | 2.747 | -0.262 | 1.306 | 1.519 | 0.041 | 0.001 | 0.001 | 0.483 | 0.64 | 0.18 | 0.54 | 0.37 | 0.74 | 0.58 |
| 10 | 101134859 | rs7084684 | -2.297 | -0.302 | 1.634 | 1.064 | -0.041 | -0.022 | -0.022 | 0 | 0.16 | 0.65 | 0.1 | 0.1 | 0.06 | 0.16 |
| 10 | 101137063 | rs3829211 | 3.148 | -0.25 | 1.496 | 1.125 | 0.062 | -0.06 | -0.06 | 0 | 0.62 | 0.14 | 0.54 | 0.44 | 0.74 | 0.61 |
| 10 | 101138526 | rs7069535 | 0.737 | -0.104 | 1.831 | 1.463 | 0.006 | -0.01 | -0.01 | 0 | 0.86 | 0.92 | 0.75 | 0.83 | 0.89 | 0.9 |
| 10 | 101138897 | rs3793929 | 1.578 | -0.104 | 1.804 | 1.463 | -0.026 | 0.047 | 0.047 | 0 | 0.83 | 0.73 | 0.71 | 0.73 | 0.87 | 0.76 |
| 10 | 101139262 | rs4917860 | -0.728 | -0.04 | 1.178 | 1.445 | 0 | 0.044 | 0.044 | 0 | 0.08 | 0.09 | 0.11 | 0.28 | 0.1 | 0.06 |
| 10 | 101139381 | rs4919306 | 0.73 | -0.003 | 0.819 | 1.314 | 0.01 | -0.016 | -0.016 | 0 | 0.87 | 0.95 | 0.75 | 0.82 | 0.89 | 0.9 |
| 10 | 101148323 | rs4917862 | -2.648 | -0.21 | 0.92 | 1.338 | 0.014 | -0.018 | -0.018 | 0 | 0.24 | 0.81 | 0.21 | 0.37 | 0.16 | 0.28 |
| 10 | 101149298 | rs4917864 | -2.326 | -0.214 | 1.55 | 1.389 | -0.002 | 0.163 | 0.163 | 1.474 | 0.14 | 0.47 | 0.15 | 0.29 | 0.1 | 0.04 |
| 10 | 101150256 | rs3793931 | 1.76 | -0.99 | 0.397 | 0.698 | 0.155 | 0.005 | 0.005 | 0.56 | 0.51 | 0.1 | 0.51 | 0.44 | 0.71 | 0.57 |
| 10 | 101152191 | rs11592057 | -0.441 | -0.408 | 0.593 | 0.583 | -0.003 | -0.015 | -0.015 | 0.078 | 0.21 | 0.13 | 0.31 | 0.22 | 0.22 | 0.26 |
| 10 | 101155432 | rs12766044 | 0.734 | -0.37 | 0.593 | 0.514 | -0.005 | 0.048 | 0.048 | 0 | 0.14 | 0.03 | 0.05 | 0.04 | 0.11 | 0.2 |
| 10 | 101156759 | rs2641 | -0.509 | -0.563 | 1.312 | 0.555 | -0.038 | 0 | 0 | 0.027 | 0.09 | 0.24 | 0.05 | 0.07 | 0.04 | 0.07 |
| 10 | 101157160 | rs11867 | 0.428 | -0.563 | 1.243 | 0.555 | -0.013 | 0 | 0 | 0.061 | 0.07 | 0.07 | 0.03 | 0.07 | 0.03 | 0.07 |
| 10 | 101163194 | rs2275952 | 0.865 | -0.538 | 1.218 | 0.535 | 0 | 0.046 | 0.046 | 0 | 0.13 | 0.03 | 0.06 | 0.04 | 0.1 | 0.2 |
| 10 | 101165134 | rs10748775 | 0.734 | -0.507 | 1.195 | 0.506 | 0.004 | 0.007 | 0.007 | 0 | 0.87 | 0.91 | 0.95 | 0.93 | 0.92 | 0.85 |
| 10 | 101166544 | rs2234971 | -0.241 | -0.41 | 0.941 | 0.549 | -0.02 | -0.037 | -0.037 | 0 | 0.08 | 0.03 | 0.25 | 0.18 | 0.12 | 0.07 |
| 10 | 101169921 | rs9971274 | 0.595 | -0.455 | 1.044 | 0.449 | 0.003 | 0.001 | 0.001 | 0 | 0.09 | 0.05 | 0.05 | 0 | 0.08 | 0.01 |
| 10 | 101184280 | rs4328160 | 0.246 | -0.507 | 1.391 | 0.428 | -0.019 | -0.06 | -0.06 | 0 | 0.17 | 0.32 | 0.08 | 0.04 | 0.11 | 0.2 |
| 10 | 101185830 | rs11190087 | 1.974 | -0.519 | 1.302 | 0.612 | 0.023 | -0.006 | -0.006 | 0 | 0.48 | 0.11 | 0.46 | 0.44 | 0.6 | 0.54 |
| 10 | 101189494 | rs3793935 | 1.682 | -0.414 | 1.427 | 0.549 | -0.072 | 0.048 | 0.048 | 0.661 | 0.75 | 0.51 | 0.82 | 0.62 | 0.86 | 0.85 |
| 10 | 101193002 | rs17491356 | 0.342 | 0.044 | 0.452 | 0.608 | 0.004 | 0.033 | 0.033 | 0 | 0.09 | 0.03 | 0.05 | 0 | 0.08 | 0.01 |
| 10 | 101205142 | rs2490286 | -0.062 | 0.174 | 0.63 | 0.736 | -0.002 | 0.006 | 0.006 | 0 | 0.79 | 0.78 | 0.85 | 0.65 | 0.85 | 0.69 |
| 10 | 101209006 | rs2494655 | -0.045 | 0.732 | 0.343 | -0.247 | -0.014 | 0.041 | 0.041 | 0.316 | 0.75 | 0.86 | 0.83 | 0.87 | 0.77 | 0.61 |
| 10 | 101213253 | rs2490288 | 0.53 | 0.797 | 0.526 | -0.468 | 0.01 | 0.048 | 0.048 | 0 | 0.6 | 0.7 | 0.78 | 0.9 | 0.68 | 0.59 |
| 10 | 101213280 | rs2494657 | 0.923 | 0.785 | 0.582 | -0.33 | 0.001 | 0.008 | 0.008 | 0 | 0.5 | 0.43 | 0.49 | 0.72 | 0.56 | 0.51 |
| 10 | 101222490 | rs11593108 | -0.746 | 1.301 | 0.964 | 0.406 | 0.005 | 0.012 | 0.012 | 0 | 0.08 | 0.09 | 0.26 | 0.16 | 0.18 | 0.1 |
| 10 | 101224882 | rs10883346 | -0.388 | 0.802 | 0.64 | -0.017 | 0.024 | 0.058 | 0.058 | 0 | 0.22 | 0.12 | 0.16 | 0.08 | 0.16 | 0.32 |
| 10 | 101226028 | rs11190100 | -0.141 | 0.776 | 0.634 | -0.028 | 0.016 | 0.056 | 0.056 | 0 | 0.2 | 0.07 | 0.15 | 0.08 | 0.16 | 0.32 |
| 10 | 101226591 | rs4919320 | -0.49 | 0.756 | 0.938 | -0.037 | 0.011 | 0.017 | 0.017 | 0 | 0.22 | 0.2 | 0.16 | 0.08 | 0.16 | 0.32 |
| 10 | 101227302 | rs10883348 | -1.729 | 0.777 | 1.14 | -0.235 | -0.003 | -0.023 | -0.023 | 0 | 0.42 | 0.55 | 0.49 | 0.24 | 0.43 | 0.44 |
| 10 | 101229527 | rs2862596 | -1.719 | 0.775 | 1.162 | -0.236 | -0.002 | -0.015 | -0.015 | 0 | 0.42 | 0.57 | 0.49 | 0.24 | 0.43 | 0.44 |
| 10 | 101233574 | rs7077930 | -1.271 | 1.018 | 0.793 | -0.127 | 0.048 | 0.014 | 0.014 | 0 | 0.18 | 0.33 | 0.32 | 0.17 | 0.27 | 0.12 |
| 10 | 101237244 | rs4919328 | -1.575 | 0.932 | 0.726 | -0.12 | 0 | -0.021 | -0.021 | 0 | 0.41 | 0.52 | 0.49 | 0.24 | 0.43 | 0.44 |
| 10 | 101250348 | rs9665572 | 1.071 | 0.443 | 0.609 | -0.405 | -0.022 | -0.085 | -0.085 | 3.261 | 0.67 | 0.54 | 0.76 | 0.88 | 0.74 | 0.65 |
| 10 | 101251049 | rs4917868 | 0.92 | 0.431 | 0.626 | -0.416 | 0.025 | -0.077 | -0.077 | 0 | 0.87 | 0.59 | 0.9 | 0.96 | 0.9 | 0.97 |
| 10 | 101257478 | rs4313504 | 0.643 | 0.398 | 0.581 | -0.415 | -0.033 | -0.05 | -0.05 | 1.511 | 0.76 | 0.65 | 0.83 | 0.88 | 0.84 | 0.67 |
| 10 | 101262809 | rs7893268 | 0.495 | 0.578 | 0.013 | -0.259 | 0.002 | -0.003 | -0.003 | 0 | 0.69 | 0.74 | 0.79 | 0.88 | 0.74 | 0.66 |
| 10 | 101264162 | rs4595481 | 0.379 | 0.784 | -0.308 | -0.198 | 0.005 | 0.03 | 0.03 | 0 | 0.72 | 0.82 | 0.8 | 0.92 | 0.75 | 0.67 |
| 10 | 101274365 | rs7078219 | -0.629 | 0.883 | -0.472 | 0.324 | -0.004 | -0.024 | -0.024 | 6.473 | 0.35 | 0.29 | 0.45 | 0.54 | 0.38 | 0.35 |
| 10 | 101277239 | rs7092009 | -1.113 | 0.725 | 0.011 | -0.303 | -0.004 | -0.001 | -0.001 | 0 | 0.53 | 0.69 | 0.62 | 0.56 | 0.53 | 0.63 |
| 10 | 101277816 | rs7096296 | -1.054 | 0.639 | 0.345 | -0.34 | -0.001 | 0 | 0 | 0.916 | 0.53 | 0.43 | 0.6 | 0.56 | 0.52 | 0.63 |

| | | | | | | | | | | | | | | | | |
|---|---|---|---|---|---|---|---|---|---|---|---|---|---|---|---|---|
| 10 | 101280279 | rs17094148 | 0.396 | 0.533 | 0.248 | -0.171 | -0.033 | -0.064 | -0.064 | 0 | 0.21 | 0.11 | 0.33 | 0.44 | 0.3 | 0.2 |
| 10 | 101282200 | rs11190134 | -0.706 | 0.469 | 0.393 | -0.208 | -0.013 | -0.045 | -0.045 | 0.725 | 0.33 | 0.22 | 0.44 | 0.54 | 0.38 | 0.34 |
| 10 | 101284237 | rs4409764 | -1.493 | 0.418 | 0.623 | -0.211 | -0.001 | 0.002 | 0.002 | 1.31 | 0.52 | 0.42 | 0.59 | 0.56 | 0.51 | 0.63 |
| 10 | 101284637 | rs4129132 | -1.493 | 0.418 | 0.623 | -0.211 | -0.001 | 0.002 | 0.002 | 1.31 | 0.52 | 0.42 | 0.59 | 0.56 | 0.51 | 0.63 |
| 10 | 101287764 | rs10883365 | -1.531 | 0.413 | 0.959 | -0.213 | -0.001 | 0.001 | 0.001 | 1.36 | 0.52 | 0.5 | 0.6 | 0.56 | 0.51 | 0.63 |
| 10 | 101289123 | rs7899176 | -1.018 | 0.392 | 1.028 | -0.215 | -0.027 | -0.07 | -0.07 | 0.006 | 0.31 | 0.45 | 0.27 | 0.12 | 0.22 | 0.43 |
| 10 | 101291593 | rs11190140 | -1.493 | 0.446 | 1.085 | -0.186 | -0.005 | 0.004 | 0.004 | 1.267 | 0.52 | 0.63 | 0.61 | 0.56 | 0.51 | 0.63 |
| 10 | 101292484 | rs10883373 | -1.506 | 0.448 | 1.139 | -0.185 | 0 | -0.001 | -0.001 | 0.896 | 0.52 | 0.53 | 0.59 | 0.56 | 0.5 | 0.63 |
| 10 | 101293856 | rs884144 | 0.313 | 0.428 | 1.044 | -0.221 | 0.001 | 0.018 | 0.018 | 0 | 0.17 | 0.11 | 0.15 | 0.02 | 0.14 | 0.29 |
| 10 | 101301346 | rs1332100 | -1.506 | 0.307 | 0.96 | -0.213 | -0.004 | -0.007 | -0.007 | 0.873 | 0.57 | 0.64 | 0.63 | 0.54 | 0.54 | 0.64 |
| 10 | 101312292 | rs7090584 | -1.041 | 0.201 | 0.711 | -0.471 | -0.009 | -0.032 | -0.032 | 0 | 0.4 | 0.54 | 0.32 | 0.12 | 0.28 | 0.45 |
| 10 | 101316142 | rs10883376 | -1.923 | 0.066 | 1.02 | -0.372 | -0.029 | -0.019 | -0.019 | 0 | 0.57 | 0.76 | 0.63 | 0.55 | 0.54 | 0.65 |
| 10 | 101317175 | rs12415804 | 0.468 | -0.01 | 1.06 | -0.092 | 0.006 | 0.021 | 0.021 | 0 | 0.19 | 0.12 | 0.15 | 0.13 | 0.12 | 0.31 |
| 10 | 101318702 | rs12260709 | 0.414 | -0.005 | 1.035 | -0.09 | -0.005 | -0.013 | -0.013 | 0 | 0.19 | 0.21 | 0.16 | 0.13 | 0.13 | 0.31 |
| 10 | 101318875 | rs7913750 | -1.902 | 0.09 | 0.719 | -0.071 | -0.029 | 0.02 | 0.02 | 1.227 | 0.57 | 0.76 | 0.64 | 0.65 | 0.54 | 0.65 |
| 10 | 101320612 | rs10748785 | -1.102 | 0.206 | 1.006 | 0.089 | 0.002 | -0.012 | -0.012 | 0 | 0.4 | 0.57 | 0.48 | 0.56 | 0.38 | 0.59 |
| 10 | 101323976 | rs10458728 | -0.284 | 0.689 | 1.198 | 0.669 | 0.023 | 0.058 | 0.058 | 0 | 0.16 | 0.2 | 0.3 | 0.4 | 0.26 | 0.2 |
| 10 | 101325797 | rs1360522 | -1.148 | 0.245 | 1.228 | 0.477 | 0 | 0.062 | 0.062 | 0 | 0.22 | 0.55 | 0.39 | 0.42 | 0.27 | 0.3 |
| 10 | 101327851 | rs7091572 | -0.358 | 0.754 | 0.366 | 0.681 | 0.018 | 0.058 | 0.058 | 0 | 0.16 | 0.2 | 0.35 | 0.41 | 0.28 | 0.2 |
| 10 | 101329158 | rs12241254 | -1.284 | 0.866 | 0.047 | 0.411 | 0.004 | 0.003 | 0.003 | 0 | 0.13 | 0.18 | 0.21 | 0.07 | 0.17 | 0.07 |
| 10 | 101329967 | rs4919352 | 0.168 | 0.929 | -0.901 | 0.163 | -0.003 | 0.07 | 0.07 | 0.072 | 0.58 | 0.76 | 0.48 | 0.75 | 0.44 | 0.69 |
| 10 | 101334019 | rs4244336 | -0.317 | 1.574 | -0.954 | 0.045 | -0.025 | 0.03 | 0.03 | 0.215 | 0.71 | 0.79 | 0.76 | 0.82 | 0.64 | 0.8 |
| 10 | 101334234 | rs12414093 | -0.083 | 0.424 | -1.162 | -0.3 | 0.001 | 0.047 | 0.047 | 0.063 | 0.17 | 0.08 | 0.36 | 0.02 | 0.16 | 0.27 |
| 10 | 101335003 | rs7906204 | -1.32 | 0.721 | -1.152 | -0.102 | 0.006 | -0.018 | -0.018 | 0 | 0.83 | 0.84 | 0.93 | 1 | 0.93 | 0.97 |
| 10 | 101339828 | rs7896866 | -0.436 | 0.553 | -0.846 | 0.461 | -0.014 | 0.031 | 0.031 | 0 | 0.32 | 0.39 | 0.5 | 0.49 | 0.28 | 0.5 |
| 10 | 101340425 | rs10883379 | 0.738 | 1.271 | -0.718 | 0.452 | 0.001 | -0.001 | -0.001 | 0 | 0.58 | 0.49 | 0.4 | 0.32 | 0.53 | 0.35 |
| 10 | 101342102 | rs4919355 | 0.411 | 1.296 | -0.761 | 0.42 | -0.014 | 0.024 | 0.024 | 0 | 0.64 | 0.54 | 0.48 | 0.49 | 0.7 | 0.47 |
| 10 | 101342941 | rs878939 | 0.628 | 0.962 | -0.699 | 0.266 | -0.02 | 0.041 | 0.041 | 0.044 | 0.24 | 0.15 | 0.15 | 0.16 | 0.26 | 0.16 |
| 10 | 101343726 | rs17112021 | -0.489 | 0.953 | -0.638 | 0.807 | -0.029 | -0.045 | -0.045 | 0 | 0.16 | 0.12 | 0.16 | 0.45 | 0.25 | 0.23 |
| 10 | 101345224 | rs6584294 | 0.706 | 1.181 | -0.638 | 0.568 | 0.005 | -0.001 | -0.001 | 0 | 0.45 | 0.48 | 0.4 | 0.5 | 0.55 | 0.42 |
| 10 | 101345366 | rs1035209 | 1.176 | 1.203 | -0.666 | 0.575 | -0.019 | -0.018 | -0.018 | 0.014 | 0.08 | 0.07 | 0.08 | 0.18 | 0.19 | 0.09 |
| 10 | 101345471 | rs6584295 | 1.356 | 0.911 | -0.628 | 0.518 | -0.001 | -0.002 | -0.002 | 0 | 0.38 | 0.39 | 0.31 | 0.33 | 0.36 | 0.33 |
| 10 | 101345516 | rs1035211 | 0.443 | 0.707 | -0.669 | 0.42 | -0.008 | -0.087 | -0.087 | 0 | 0.22 | 0.29 | 0.38 | 0.02 | 0.19 | 0.3 |
| 10 | 101347202 | rs12573087 | 0.616 | 0.675 | -0.791 | 0.56 | 0.021 | -0.028 | -0.028 | 0 | 0.09 | 0.04 | 0.08 | 0.27 | 0.06 | 0.13 |
| 10 | 101347718 | rs1035213 | -0.243 | 0.871 | -0.637 | 0.384 | -0.005 | -0.012 | -0.012 | 0 | 0.39 | 0.28 | 0.29 | 0.65 | 0.44 | 0.37 |
| 10 | 101350530 | rs10883381 | 0.013 | 0.808 | -0.452 | 0.132 | -0.021 | -0.092 | -0.092 | 0.381 | 0.22 | 0.34 | 0.4 | 0.02 | 0.19 | 0.3 |
| 10 | 101351704 | rs11190164 | 0.546 | 1.717 | -0.641 | 0.781 | -0.044 | -0.048 | -0.048 | 0.48 | 0.08 | 0.03 | 0.17 | 0.23 | 0.28 | 0.14 |
| 10 | 101357635 | rs11190169 | 0.193 | 1.485 | -0.748 | 0.738 | -0.071 | -0.055 | -0.055 | 0 | 0.11 | 0.03 | 0.19 | 0.26 | 0.29 | 0.17 |
| 10 | 101359865 | rs4919360 | 0.308 | 1.291 | -0.493 | 0.671 | -0.011 | 0.004 | 0.004 | 0 | 0.43 | 0.53 | 0.31 | 0.49 | 0.33 | 0.38 |
| 10 | 101362536 | rs2111326 | -0.183 | 1.201 | -0.35 | 0.991 | 0.022 | 0.009 | 0.009 | 0 | 0.48 | 0.65 | 0.4 | 0.47 | 0.53 | 0.42 |
| 10 | 101362751 | rs2111325 | 1.318 | 0.823 | -0.176 | 0.841 | -0.005 | -0.006 | -0.006 | 0 | 0.3 | 0.3 | 0.19 | 0.21 | 0.22 | 0.23 |
| 10 | 101365313 | rs11190179 | 0.711 | 0.882 | -0.313 | 0.943 | 0.001 | 0.003 | 0.003 | 0 | 0.08 | 0.06 | 0.1 | 0.28 | 0.1 | 0.13 |
| 10 | 101366643 | rs4919365 | 0.222 | 0.817 | -0.162 | 0.792 | -0.015 | 0.008 | 0.008 | 0 | 0.42 | 0.5 | 0.31 | 0.49 | 0.33 | 0.39 |
| 10 | 101367315 | rs7068079 | -0.903 | 1.219 | -0.146 | 1.126 | 0.03 | 0.034 | 0.034 | 0.003 | 0.23 | 0.29 | 0.3 | 0.56 | 0.39 | 0.3 |
| 10 | 101367645 | rs7072263 | 0.456 | 1.42 | -0.057 | 0.966 | 0.084 | 0.029 | 0.029 | 0 | 0.13 | 0.2 | 0.21 | 0.23 | 0.28 | 0.16 |
| 10 | 101368917 | rs4917873 | 0.041 | 0.907 | 0.002 | 0.725 | 0 | -0.001 | -0.001 | 0 | 0.46 | 0.45 | 0.38 | 0.48 | 0.36 | 0.42 |
| 10 | 101370015 | rs2297446 | -1.066 | 0.797 | 0.128 | 0.678 | 0.037 | 0.057 | 0.057 | 0 | 0.28 | 0.45 | 0.34 | 0.53 | 0.4 | 0.32 |
| 10 | 101372411 | rs880568 | -0.87 | 0.805 | -0.215 | 0.38 | 0.032 | 0.119 | 0.119 | 0 | 0.92 | 0.79 | 0.88 | 0.78 | 0.88 | 0.89 |
| 10 | 101373729 | rs17494319 | 1.496 | 0.691 | -0.434 | 0.242 | 0.04 | 0.162 | 0.162 | 0 | 0.3 | 0.07 | 0.29 | 0.06 | 0.22 | 0.26 |
| 10 | 101400037 | rs11596134 | 2.294 | 0.612 | -0.522 | 0.301 | -0.052 | 0.022 | 0.022 | 0 | 0.2 | 0.01 | 0.09 | 0.12 | 0.15 | 0.24 |
| 10 | 101412278 | rs1999440 | -0.457 | 0.908 | 0.068 | 0.649 | 0.026 | 0.114 | 0.114 | 0 | 0.91 | 0.75 | 0.88 | 0.78 | 0.88 | 0.89 |
| 10 | 101416647 | rs6584303 | -0.533 | 0.908 | 0.094 | 0.649 | 0.016 | 0.084 | 0.084 | 0 | 0.91 | 0.78 | 0.88 | 0.78 | 0.88 | 0.89 |
| 10 | 101418367 | rs7893702 | -1.052 | 0.913 | 0.047 | 0.651 | -0.01 | -0.013 | -0.013 | 0 | 0.93 | 0.95 | 0.89 | 0.78 | 0.88 | 0.89 |
| 10 | 101433392 | rs911541 | -0.426 | 0.905 | 0.12 | 0.647 | 0.028 | 0.117 | 0.117 | 0 | 0.91 | 0.75 | 0.88 | 0.78 | 0.88 | 0.89 |
| 10 | 101445591 | rs3740078 | 0.186 | 1.067 | 0.257 | 0.669 | 0.024 | 0.015 | 0.015 | 0 | 0.17 | 0.23 | 0.23 | 0.3 | 0.31 | 0.19 |
| 10 | 101451259 | rs11190245 | 0.873 | 1.097 | 0.284 | 0.683 | 0.031 | 0.022 | 0.022 | 0 | 0.13 | 0.18 | 0.21 | 0.3 | 0.3 | 0.17 |
| 10 | 101453066 | rs2013255 | -0.351 | 0.922 | 0.204 | 0.674 | 0.024 | 0.114 | 0.114 | 0 | 0.91 | 0.75 | 0.87 | 0.78 | 0.88 | 0.89 |
| 10 | 101459040 | rs4919384 | -0.802 | 0.931 | 0.181 | 0.678 | 0.014 | 0.075 | 0.075 | 0 | 0.92 | 0.84 | 0.88 | 0.78 | 0.88 | 0.89 |
| 10 | 101463181 | rs1412323 | -0.802 | 0.931 | 0.193 | 0.678 | 0.015 | 0.078 | 0.078 | 0 | 0.92 | 0.84 | 0.88 | 0.78 | 0.88 | 0.89 |
| 10 | 101473218 | rs2231687 | -0.331 | 0.921 | 0.187 | 0.674 | 0.023 | 0.113 | 0.113 | 0 | 0.91 | 0.75 | 0.87 | 0.78 | 0.88 | 0.89 |
| 10 | 101476857 | rs2300984 | -0.738 | 0.952 | -0.002 | 0.6 | 0.023 | 0.021 | 0.021 | 0 | 0.26 | 0.37 | 0.33 | 0.58 | 0.42 | 0.31 |
| 10 | 101485091 | rs2269196 | 0.106 | 0.947 | 0.037 | 0.806 | -0.003 | 0.009 | 0.009 | 0 | 0.84 | 0.87 | 0.8 | 0.49 | 0.78 | 0.77 |
| 10 | 101485770 | rs10786568 | -0.905 | 0.96 | -0.01 | 0.697 | 0.005 | 0.044 | 0.044 | 0 | 0.92 | 0.88 | 0.88 | 0.78 | 0.88 | 0.89 |
| 10 | 101491236 | rs1998290 | -0.664 | 1.165 | -0.021 | 0.687 | 0.005 | 0.031 | 0.031 | 0 | 0.91 | 0.88 | 0.87 | 0.78 | 0.86 | 0.88 |
| 10 | 101492403 | rs2281636 | 0.775 | 1.18 | 0.181 | 0.71 | 0.058 | 0.059 | 0.059 | 0 | 0.72 | 0.45 | 0.65 | 0.4 | 0.6 | 0.67 |
| 10 | 101495847 | rs2295781 | 1.293 | 1.197 | 0.133 | 0.774 | -0.004 | 0.003 | 0.003 | 0 | 0.08 | 0.05 | 0.08 | 0.28 | 0.1 | 0.12 |
| 10 | 101498396 | rs7913190 | -1.015 | 1.163 | 0.153 | 0.746 | 0.034 | 0.034 | 0.034 | 0 | 0.27 | 0.44 | 0.33 | 0.6 | 0.4 | 0.33 |
| 10 | 101500472 | rs767844 | 1.295 | 1.136 | 0.084 | 0.756 | -0.004 | 0.003 | 0.003 | 0 | 0.08 | 0.05 | 0.08 | 0.28 | 0.1 | 0.12 |
| 10 | 101502968 | rs3740076 | 0.607 | 0.967 | -0.256 | 0.485 | -0.005 | 0.036 | 0.036 | 0 | 0.11 | 0.01 | 0.15 | 0.02 | 0.11 | 0.08 |

| | | | | | | | | | | | | | | | | |
|---|---|---|---|---|---|---|---|---|---|---|---|---|---|---|---|---|
| 10 | 101508622 | rs2756115 | 1.168 | 0.735 | -0.214 | 0.38 | -0.002 | 0.047 | 0.047 | 0 | 0.38 | 0.1 | 0.4 | 0.26 | 0.36 | 0.38 |
| 10 | 101509113 | rs2804419 | 0.73 | 0.933 | 0.005 | 0.589 | 0.057 | 0.059 | 0.059 | 0.018 | 0.71 | 0.44 | 0.65 | 0.4 | 0.6 | 0.67 |
| 10 | 101515316 | rs3184991 | 0.302 | 1.325 | -0.249 | 0.567 | 0.021 | 0.018 | 0.018 | 0 | 0.18 | 0.24 | 0.22 | 0.32 | 0.29 | 0.2 |
| 10 | 101530795 | rs7908967 | -0.896 | 1.047 | -0.069 | 0.323 | -0.001 | 0.018 | 0.018 | 0 | 0.59 | 0.63 | 0.59 | 0.75 | 0.57 | 0.65 |
| 10 | 101534006 | rs12268782 | 1.292 | 0.913 | 0.142 | 0.381 | 0.005 | 0.011 | 0.011 | 0 | 0.28 | 0.19 | 0.16 | 0.17 | 0.2 | 0.27 |
| 10 | 101542578 | rs717620 | 1.37 | 0.755 | -0.127 | 0.141 | -0.033 | -0.044 | -0.044 | 0 | 0.12 | 0.03 | 0.17 | 0.22 | 0.21 | 0.1 |
| 10 | 101543047 | rs2756103 | 0.808 | 0.429 | 0.248 | 0.288 | -0.008 | 0.044 | 0.044 | 0 | 0.4 | 0.27 | 0.4 | 0.25 | 0.44 | 0.36 |
| 10 | 101544026 | rs2756104 | 0.79 | 0.425 | 0.322 | 0.286 | -0.005 | 0.034 | 0.034 | 0 | 0.4 | 0.3 | 0.4 | 0.25 | 0.44 | 0.36 |
| 10 | 101545558 | rs4148384 | 1.182 | 0.357 | 0.454 | 0.401 | 0.021 | 0.021 | 0.021 | 0.043 | 0.18 | 0.25 | 0.34 | 0.31 | 0.25 | 0.22 |
| 10 | 101547021 | rs7393105 | -0.921 | 0.35 | 0.409 | 0.175 | -0.001 | 0.001 | 0.001 | 0 | 0.58 | 0.57 | 0.58 | 0.75 | 0.56 | 0.64 |
| 10 | 101547042 | rs2756105 | 0.816 | 0.349 | 0.44 | 0.175 | -0.009 | 0.048 | 0.048 | 0 | 0.4 | 0.27 | 0.4 | 0.25 | 0.44 | 0.36 |
| 10 | 101548177 | rs4148385 | -0.921 | 0.349 | 0.431 | 0.175 | -0.001 | 0.001 | 0.001 | 0 | 0.58 | 0.57 | 0.58 | 0.75 | 0.56 | 0.64 |
| 10 | 101548468 | rs4148386 | -0.872 | 0.344 | 0.453 | 0.173 | -0.001 | -0.005 | -0.005 | 0 | 0.58 | 0.54 | 0.58 | 0.75 | 0.56 | 0.64 |
| 10 | 101548709 | rs2756107 | 0.794 | 0.356 | 0.428 | 0.178 | -0.005 | 0.034 | 0.034 | 0 | 0.4 | 0.3 | 0.4 | 0.25 | 0.44 | 0.36 |
| 10 | 101548795 | rs2145853 | -0.919 | 0.358 | 0.421 | 0.179 | -0.001 | 0.001 | 0.001 | 0 | 0.58 | 0.57 | 0.58 | 0.75 | 0.56 | 0.64 |
| 10 | 101549765 | rs4148388 | -0.919 | 0.358 | 0.421 | 0.179 | -0.001 | 0.001 | 0.001 | 0 | 0.58 | 0.57 | 0.58 | 0.75 | 0.56 | 0.64 |
| 10 | 101553259 | rs2804400 | 0.794 | 0.374 | 0.34 | 0.186 | -0.007 | 0.045 | 0.045 | 0 | 0.4 | 0.27 | 0.4 | 0.25 | 0.44 | 0.36 |
| 10 | 101559415 | rs2804397 | -3.269 | 0.052 | -0.076 | -0.059 | 0.013 | 0.032 | 0.032 | 0.068 | 0.9 | 0.95 | 0.97 | 1 | 0.94 | 0.99 |
| 10 | 101563815 | rs2273697 | 1.611 | -0.054 | -0.005 | -0.113 | 0.004 | 0.012 | 0.012 | 0 | 0.28 | 0.19 | 0.16 | 0.1 | 0.2 | 0.28 |
| 10 | 101566179 | rs2756112 | -3.375 | -0.05 | -0.034 | -0.111 | 0.013 | 0.028 | 0.028 | 0.068 | 0.9 | 0.95 | 0.97 | 1 | 0.94 | 0.99 |
| 10 | 101566866 | rs4148391 | 2.515 | -0.185 | 0.006 | -0.202 | -0.008 | 0.044 | 0.044 | 0.264 | 0.21 | 0.07 | 0.09 | 0.08 | 0.14 | 0.23 |
| 10 | 101567426 | rs2073337 | 0.462 | -0.094 | 0.089 | -0.092 | -0.001 | -0.008 | -0.008 | 0 | 0.35 | 0.41 | 0.39 | 0.25 | 0.39 | 0.38 |
| 10 | 101569371 | rs2756113 | -3.456 | -0.096 | 0.083 | -0.108 | 0.013 | 0.028 | 0.028 | 0.068 | 0.9 | 0.95 | 0.97 | 1 | 0.94 | 0.99 |
| 10 | 101569483 | rs2756114 | 0.428 | -0.031 | 0.114 | -0.056 | -0.001 | -0.009 | -0.009 | 0 | 0.35 | 0.41 | 0.39 | 0.25 | 0.39 | 0.38 |
| 10 | 101571528 | rs3740074 | -0.711 | -0.038 | 0.096 | -0.059 | 0 | -0.006 | -0.006 | 0.115 | 0.65 | 0.61 | 0.61 | 0.75 | 0.61 | 0.62 |
| 10 | 101572343 | rs4148394 | 1.6 | -0.022 | 0.069 | 0.018 | 0.025 | 0.039 | 0.039 | 0.036 | 0.17 | 0.26 | 0.34 | 0.38 | 0.25 | 0.21 |
| 10 | 101576960 | rs3740073 | -1.239 | -0.036 | 0.08 | 0.021 | -0.012 | 0.007 | 0.007 | 0 | 0.69 | 0.78 | 0.64 | 0.75 | 0.62 | 0.64 |
| 10 | 101587569 | rs11595888 | 1.624 | -0.041 | 0.037 | -0.039 | 0.001 | 0.002 | 0.002 | 0 | 0.27 | 0.22 | 0.16 | 0.1 | 0.2 | 0.28 |
| 10 | 101587931 | rs2002042 | 1.436 | 0.091 | 0.114 | 0.109 | 0.023 | 0.039 | 0.039 | 0.028 | 0.18 | 0.25 | 0.35 | 0.37 | 0.25 | 0.21 |
| 10 | 101589925 | rs4148395 | 1.116 | -0.029 | -0.114 | 0.009 | 0.004 | 0.009 | 0.009 | 0 | 0.29 | 0.22 | 0.16 | 0.1 | 0.2 | 0.28 |
| 10 | 101591944 | rs4148396 | -0.659 | 0.212 | -0.002 | 0.315 | -0.013 | 0.009 | 0.009 | 0 | 0.69 | 0.75 | 0.62 | 0.75 | 0.61 | 0.65 |
| 10 | 101592320 | rs4148397 | 0.273 | 0.022 | 0.123 | 0.239 | 0.009 | 0.038 | 0.038 | 0 | 0.53 | 0.6 | 0.57 | 0.73 | 0.54 | 0.63 |
| 10 | 101592622 | rs4148398 | -0.648 | 0.166 | 0.173 | 0.13 | -0.025 | 0.012 | 0.012 | 0 | 0.7 | 0.86 | 0.65 | 0.75 | 0.63 | 0.73 |
| 10 | 101592689 | rs4148399 | 0.303 | 0.016 | 0.432 | 0.558 | -0.001 | 0.064 | 0.064 | 0 | 0.06 | 0.06 | 0.04 | 0.31 | 0.05 | 0.21 |
| 10 | 101594729 | rs7476245 | 0.128 | -0.107 | 0.657 | 0.321 | -0.016 | -0.029 | -0.029 | 0 | 0.09 | 0.2 | 0.07 | 0.02 | 0.06 | 0.04 |
| 10 | 101602004 | rs3758395 | 0.479 | 0.134 | 0.199 | 0.532 | -0.002 | 0.049 | 0.049 | 0 | 0.05 | 0.04 | 0.04 | 0.31 | 0.05 | 0.21 |
| 10 | 101605693 | rs3740065 | 1.041 | -0.015 | 0.251 | 0.42 | -0.005 | 0.012 | 0.012 | 0 | 0.18 | 0.22 | 0.1 | 0.31 | 0.12 | 0.22 |
| 10 | 101610533 | rs8187707 | 2.481 | -0.008 | 0.244 | 0.279 | 0.016 | 0.045 | 0.045 | 0.299 | 0.11 | 0.03 | 0.03 | 0 | 0.07 | 0.02 |
| 10 | 101610723 | rs3740063 | -0.437 | 0.001 | 0.26 | 0.215 | 0.005 | 0.025 | 0.025 | 0 | 0.44 | 0.38 | 0.4 | 0.25 | 0.45 | 0.32 |
| 10 | 101611294 | rs8187710 | 2.218 | -0.003 | 0.238 | 0.214 | -0.005 | -0.032 | -0.032 | 0 | 0.12 | 0.16 | 0.05 | 0 | 0.07 | 0.02 |
| 10 | 101612320 | rs12826 | 0.364 | 0.029 | 0.144 | 0.23 | -0.023 | 0.016 | 0.016 | 0 | 0.31 | 0.15 | 0.34 | 0.25 | 0.38 | 0.31 |
| 10 | 101615015 | rs11816708 | 2.443 | -0.013 | 0.162 | 0.223 | 0.008 | -0.001 | -0.001 | 1.1 | 0.11 | 0.08 | 0.04 | 0 | 0.07 | 0.02 |
| 10 | 101616579 | rs7067971 | 0.314 | 0.038 | 0.185 | 0.237 | -0.007 | 0.014 | 0.014 | 0 | 0.31 | 0.2 | 0.34 | 0.25 | 0.38 | 0.31 |
| 10 | 101619431 | rs7903716 | -0.425 | 0.205 | 0.291 | 0.387 | 0.01 | 0.032 | 0.032 | 0 | 0.46 | 0.41 | 0.4 | 0.25 | 0.46 | 0.37 |
| 10 | 101620948 | rs11190298 | -0.488 | 0.302 | -0.101 | 0.51 | -0.001 | -0.004 | -0.004 | 0 | 0.46 | 0.29 | 0.42 | 0.53 | 0.49 | 0.51 |
| 10 | 101631353 | rs7095052 | -0.045 | 0.268 | -0.229 | 0.255 | -0.006 | 0.065 | 0.065 | 0.077 | 0.41 | 0.23 | 0.37 | 0.25 | 0.46 | 0.35 |
| 10 | 101633447 | rs2256700 | -0.227 | 0.085 | 0.193 | 0.162 | 0.001 | -0.039 | -0.039 | 0.004 | 0.11 | 0.3 | 0.11 | 0.02 | 0.07 | 0.12 |
| 10 | 101636316 | rs1139943 | -0.227 | 0.085 | 0.193 | 0.162 | 0.001 | -0.039 | -0.039 | 0.046 | 0.11 | 0.3 | 0.11 | 0.03 | 0.07 | 0.12 |
| 10 | 101636648 | rs2002109 | -0.908 | 0.047 | 0.128 | 0.233 | 0 | -0.008 | -0.008 | 0.194 | 0.52 | 0.53 | 0.48 | 0.27 | 0.54 | 0.48 |
| 10 | 101645498 | rs2490763 | -0.995 | 0.038 | 0.06 | 0.229 | -0.002 | 0.018 | 0.018 | 0 | 0.52 | 0.59 | 0.48 | 0.27 | 0.54 | 0.48 |
| 10 | 101654696 | rs3758394 | -0.816 | 0.164 | 0.047 | 0.361 | 0 | -0.002 | -0.002 | 0 | 0.51 | 0.47 | 0.48 | 0.28 | 0.57 | 0.49 |
| 10 | 101655327 | rs2252676 | -1.848 | -0.273 | -0.447 | 0.379 | -0.101 | -0.102 | -0.102 | 0 | 0.81 | 0.97 | 0.63 | 0.65 | 0.69 | 0.79 |
| 10 | 101658585 | rs3740057 | -0.455 | -1.107 | 0.008 | 0.259 | 0.008 | -0.021 | -0.021 | 0 | 0.38 | 0.45 | 0.43 | 0.24 | 0.46 | 0.45 |
| 10 | 101664961 | rs4348829 | 0.377 | -1.166 | 0.041 | -0.571 | -0.02 | -0.023 | -0.023 | 0 | 0.09 | 0.25 | 0.08 | 0.02 | 0.04 | 0.02 |
| 10 | 101666475 | rs7898374 | -0.434 | -1.01 | -0.271 | -0.506 | 0.013 | -0.034 | -0.034 | 2.682 | 0.38 | 0.49 | 0.44 | 0.24 | 0.46 | 0.45 |
| 10 | 101668986 | rs6584331 | -0.948 | -0.768 | -0.187 | -0.344 | -0.003 | 0.042 | 0.042 | 0 | 0.52 | 0.6 | 0.48 | 0.24 | 0.54 | 0.48 |
| 10 | 101690050 | rs10786578 | -1.127 | -0.856 | 0.026 | 0.002 | -0.008 | 0.009 | 0.009 | 0 | 0.37 | 0.58 | 0.36 | 0.45 | 0.35 | 0.41 |
| 10 | 101690379 | rs10883428 | -1.127 | -1.429 | 0.208 | -1.009 | -0.01 | -0.022 | -0.022 | 0 | 0.41 | 0.48 | 0.4 | 0.27 | 0.38 | 0.32 |
| 10 | 101691628 | rs7922447 | -1.184 | -1.089 | 0.29 | -1.127 | 0.015 | -0.037 | -0.037 | 0 | 0.34 | 0.51 | 0.36 | 0.23 | 0.38 | 0.35 |
| 10 | 101699304 | rs10883429 | -1.429 | -1.148 | 0.243 | -1.152 | -0.003 | 0.022 | 0.022 | 0.995 | 0.41 | 0.65 | 0.44 | 0.26 | 0.42 | 0.37 |
| 10 | 101703523 | rs4919402 | -0.975 | -1.26 | 0.478 | -1.195 | 0.015 | -0.024 | -0.024 | 0 | 0.32 | 0.46 | 0.36 | 0.23 | 0.37 | 0.33 |
| 10 | 101712951 | rs7906505 | -0.34 | -1.262 | 0.628 | -1.226 | 0.001 | -0.002 | -0.002 | 0.843 | 0.27 | 0.32 | 0.32 | 0.23 | 0.34 | 0.24 |
| 10 | 101715899 | rs2862923 | -1.577 | -1.228 | 0.416 | -1.21 | -0.003 | 0.021 | 0.021 | 0.994 | 0.41 | 0.66 | 0.44 | 0.26 | 0.42 | 0.37 |
| 10 | 101719668 | rs11596987 | 0.551 | -1.195 | 0.067 | -1.184 | -0.094 | -0.061 | -0.061 | 0 | 0.17 | 0.04 | 0.29 | 0.23 | 0.32 | 0.21 |
| 10 | 101735593 | rs11190350 | 0.285 | -1.063 | 0.45 | -1.006 | -0.014 | -0.018 | -0.018 | 0 | 0.07 | 0.14 | 0.07 | 0.03 | 0.04 | 0.02 |
| 10 | 101751700 | rs10159539 | 0.723 | -1.045 | 0.365 | -0.999 | -0.01 | -0.013 | -0.013 | 0 | 0.06 | 0.09 | 0.06 | 0.03 | 0.04 | 0.02 |
| 10 | 101758982 | rs4919409 | -2.045 | -1.445 | -0.153 | -1.31 | 0.043 | 0.069 | 0.069 | 0 | 0.8 | 0.95 | 0.94 | 0.96 | 0.92 | 0.73 |
| 10 | 101762385 | rs11190363 | -1.452 | -0.91 | 0.57 | -1.152 | 0.001 | 0.025 | 0.025 | 0.315 | 0.48 | 0.59 | 0.48 | 0.25 | 0.47 | 0.38 |
| 10 | 101764893 | rs7078766 | -0.505 | -1.131 | 0.081 | -1.052 | -0.009 | -0.015 | -0.015 | 0.272 | 0.17 | 0.27 | 0.08 | 0.05 | 0.07 | 0.21 |

| | | | | | | | | | | | | | | | | |
|---|---|---|---|---|---|---|---|---|---|---|---|---|---|---|---|---|
| 10 | 101775402 | rs11593328 | -1.302 | -1.031 | 0.024 | -1.124 | 0 | 0.024 | 0.024 | 0.559 | 0.44 | 0.63 | 0.42 | 0.22 | 0.4 | 0.36 |
| 10 | 101775611 | rs11591518 | 0.568 | -0.669 | 0.039 | -0.92 | -0.059 | -0.002 | -0.002 | 0 | 0.22 | 0.08 | 0.32 | 0.19 | 0.33 | 0.13 |
| 10 | 101784816 | rs7089847 | -0.644 | -0.941 | 0.232 | -1.004 | -0.073 | -0.095 | -0.095 | 0.069 | 0.21 | 0.36 | 0.13 | 0.03 | 0.06 | 0.06 |
| 10 | 101790389 | rs4919416 | -1.156 | -0.784 | -0.027 | -0.941 | 0.013 | 0.03 | 0.03 | 0.871 | 0.45 | 0.6 | 0.46 | 0.21 | 0.4 | 0.17 |
| 10 | 101798346 | rs2862928 | -1.517 | -1.044 | -0.571 | -0.876 | 0.004 | 0.121 | 0.121 | 0 | 0.53 | 0.78 | 0.51 | 0.24 | 0.46 | 0.22 |
| 10 | 101805442 | rs11599750 | -0.889 | -0.602 | -0.488 | -0.779 | -0.002 | -0.015 | -0.015 | 0 | 0.43 | 0.46 | 0.46 | 0.19 | 0.41 | 0.15 |
| 10 | 101809659 | rs11593916 | 0.079 | 0.256 | -0.735 | -0.385 | -0.072 | 0.005 | 0.005 | 0 | 0.23 | 0.06 | 0.35 | 0.16 | 0.35 | 0.12 |
| 10 | 101817557 | rs2862925 | -0.619 | 0.212 | -0.677 | -0.658 | -0.013 | 0.009 | 0.009 | 0 | 0.31 | 0.36 | 0.39 | 0.08 | 0.43 | 0.23 |
| 10 | 101823543 | rs4462272 | -0.165 | 0.326 | -0.803 | -0.55 | -0.062 | 0.048 | 0.048 | 0 | 0.26 | 0.16 | 0.37 | 0.08 | 0.39 | 0.12 |
| 10 | 101834092 | rs12775433 | 0.009 | 0.299 | 0.027 | 0.085 | -0.037 | 0.041 | 0.041 | 0 | 0.25 | 0.16 | 0.37 | 0.07 | 0.38 | 0.14 |
| 10 | 101835508 | rs3750718 | 0.053 | -0.618 | -0.353 | -0.286 | -0.09 | 0.109 | 0.109 | 0 | 0.15 | 0.34 | 0.04 | 0.28 | 0.03 | 0.07 |
| 10 | 101841153 | rs10883439 | -1.946 | -1.438 | -1.304 | -0.67 | 0.074 | 0.079 | 0.079 | 0 | 0.74 | 0.97 | 0.87 | 0.93 | 0.89 | 0.76 |
| 10 | 101846173 | rs1410079 | -1.779 | -1.097 | -1.04 | -0.347 | -0.111 | -0.095 | -0.095 | 0 | 0.53 | 0.95 | 0.51 | 0.48 | 0.52 | 0.41 |
| 10 | 101848444 | rs7899288 | -0.126 | -0.854 | -0.766 | -0.355 | -0.005 | -0.002 | -0.002 | 0 | 0.29 | 0.27 | 0.4 | 0.07 | 0.41 | 0.23 |
| 10 | 101861435 | rs11597390 | 0.314 | -0.83 | -0.938 | -0.349 | -0.042 | 0.045 | 0.045 | 0 | 0.26 | 0.16 | 0.37 | 0.07 | 0.39 | 0.14 |
| 10 | 101877961 | rs7098331 | -0.569 | -0.815 | -0.827 | -0.238 | 0 | 0.038 | 0.038 | 0 | 0.41 | 0.33 | 0.39 | 0.14 | 0.42 | 0.22 |
| 10 | 101880700 | rs11594333 | -0.67 | -0.855 | -0.815 | -0.267 | 0.001 | 0.028 | 0.028 | 0 | 0.42 | 0.37 | 0.4 | 0.13 | 0.43 | 0.22 |
| 10 | 101888640 | rs11595928 | -0.626 | -0.824 | -0.835 | -0.267 | -0.002 | 0.066 | 0.066 | 0 | 0.4 | 0.26 | 0.39 | 0.13 | 0.43 | 0.22 |
| 10 | 101893418 | rs7893985 | -1.444 | -0.796 | -1.249 | -0.158 | -0.047 | -0.05 | -0.05 | 0 | 0.5 | 0.89 | 0.47 | 0.48 | 0.5 | 0.38 |
| 10 | 101899866 | rs12413812 | 0.51 | -0.348 | -1.297 | -0.259 | -0.097 | 0.016 | 0.016 | 0 | 0.15 | 0.03 | 0.31 | 0.06 | 0.35 | 0.13 |
| 10 | 101900190 | rs12218075 | -0.415 | -0.618 | -0.512 | -0.108 | -0.023 | 0.146 | 0.146 | 2.358 | 0.13 | 0.36 | 0.09 | 0.35 | 0.08 | 0.18 |
| 10 | 101903412 | rs11190401 | -1.328 | -0.96 | -0.825 | -0.338 | -0.009 | -0.018 | -0.018 | 0 | 0.49 | 0.78 | 0.52 | 0.52 | 0.51 | 0.39 |
| 10 | 101906333 | rs975752 | -1.236 | -1.348 | -0.557 | -0.418 | -0.079 | 0.036 | 0.036 | 0.108 | 0.24 | 0.59 | 0.14 | 0.4 | 0.08 | 0.21 |
| 10 | 101908502 | rs17112654 | 0.016 | -1.392 | -0.384 | -0.416 | -0.019 | 0 | 0 | 0 | 0.09 | 0.11 | 0.06 | 0.14 | 0.03 | 0.06 |
| 10 | 101911142 | rs12806 | -1.635 | -1.449 | -0.348 | -0.417 | -0.042 | 0.01 | 0.01 | 0.304 | 0.25 | 0.68 | 0.15 | 0.41 | 0.08 | 0.21 |
| 10 | 101911517 | rs4919428 | -0.208 | -1.441 | -0.335 | -0.462 | -0.03 | -0.005 | -0.005 | 0 | 0.11 | 0.15 | 0.06 | 0.08 | 0.03 | 0.06 |
| 10 | 101912064 | rs2862954 | 0.429 | -0.613 | -0.568 | -0.589 | -0.185 | 0.058 | 0.058 | 0 | 0.23 | 0.03 | 0.3 | 0.07 | 0.48 | 0.15 |
| 10 | 101915629 | rs12769818 | 0.108 | -1.396 | -0.323 | -0.562 | -0.006 | 0 | 0 | 0.193 | 0.05 | 0.12 | 0.05 | 0.06 | 0.04 | 0.09 |
| 10 | 101917520 | rs11190405 | 2.172 | -2.202 | -0.975 | -0.906 | 0.041 | 0.045 | 0.045 | 0 | 0.19 | 0.01 | 0.05 | 0.04 | 0.07 | 0.26 |
| 10 | 101930940 | rs10883452 | 0.07 | -1.838 | -0.725 | -0.81 | -0.031 | 0.074 | 0.074 | 0.307 | 0.12 | 0.2 | 0.07 | 0.36 | 0.06 | 0.15 |
| 10 | 101932337 | rs12262005 | 0.093 | -1.821 | -0.772 | -0.792 | -0.03 | -0.005 | -0.005 | 0 | 0.11 | 0.15 | 0.06 | 0.08 | 0.03 | 0.06 |
| 10 | 101937194 | rs17112705 | 1.792 | -1.903 | -1.07 | -0.923 | -0.16 | -0.145 | -0.145 | 0.432 | 0.23 | 0.02 | 0.44 | 0.42 | 0.38 | 0.37 |
| 10 | 101946432 | rs17881019 | 0.244 | -1.654 | -0.745 | -0.794 | 0.003 | 0.008 | 0.008 | 0.217 | 0.05 | 0.17 | 0.05 | 0.06 | 0.05 | 0.09 |
| 10 | 101953705 | rs11597086 | 0.671 | -1.68 | -0.899 | -0.858 | -0.154 | 0.056 | 0.056 | 0 | 0.23 | 0.03 | 0.27 | 0.07 | 0.43 | 0.14 |
| 10 | 101955612 | rs11190421 | -1.413 | -1.77 | -0.743 | -0.86 | -0.075 | -0.03 | -0.03 | 0 | 0.18 | 0.59 | 0.13 | 0.14 | 0.08 | 0.15 |
| 10 | 101962224 | rs11190423 | 1.801 | -1.737 | -0.76 | -0.824 | -0.013 | 0.057 | 0.057 | 0 | 0.06 | 0.07 | 0.02 | 0.3 | 0.01 | 0.06 |
| 10 | 101977883 | rs2230804 | -1.464 | -1.741 | -0.681 | -0.859 | 0.006 | -0.006 | -0.006 | 0 | 0.49 | 0.8 | 0.43 | 0.51 | 0.52 | 0.35 |
| 10 | 101981274 | rs12570957 | 1.48 | -1.814 | -0.831 | -0.905 | -0.172 | -0.163 | -0.163 | 0.348 | 0.29 | 0.02 | 0.49 | 0.44 | 0.41 | 0.39 |
| 10 | 101985087 | rs12247992 | -1.334 | -1.217 | -0.238 | -0.653 | -0.078 | -0.028 | -0.028 | 1.248 | 0.18 | 0.55 | 0.12 | 0.14 | 0.08 | 0.16 |
| 10 | 101989510 | rs12770784 | 0.108 | -1.185 | -0.28 | -0.644 | 0.002 | 0.007 | 0.007 | 0.217 | 0.05 | 0.17 | 0.05 | 0.06 | 0.05 | 0.09 |
| 10 | 102000701 | rs17668255 | 0.449 | -1.182 | -0.337 | -0.659 | -0.152 | 0.055 | 0.055 | 0 | 0.23 | 0.03 | 0.27 | 0.07 | 0.43 | 0.14 |
| 10 | 102005620 | rs12269345 | -0.229 | -1.211 | -0.289 | -0.65 | -0.033 | -0.008 | -0.008 | 0 | 0.11 | 0.16 | 0.06 | 0.08 | 0.03 | 0.06 |
| 10 | 102006681 | rs17112787 | 1.526 | -1.209 | -0.334 | -0.624 | -0.023 | 0.095 | 0.095 | 0 | 0.06 | 0.12 | 0.02 | 0.3 | 0.01 | 0.06 |
| 10 | 102011211 | rs12265333 | -1.168 | -1.214 | -0.39 | -0.652 | -0.01 | 0.04 | 0.04 | 0 | 0.42 | 0.67 | 0.4 | 0.21 | 0.51 | 0.3 |
| 10 | 102016044 | rs2270962 | -1.679 | -1.221 | -0.384 | -0.651 | -0.075 | -0.031 | -0.031 | 0.115 | 0.18 | 0.6 | 0.13 | 0.14 | 0.08 | 0.16 |
| 10 | 102016268 | rs2270961 | -1.663 | -1.23 | -0.388 | -0.649 | 0.001 | -0.008 | -0.008 | 0.49 | 0.48 | 0.77 | 0.42 | 0.51 | 0.52 | 0.35 |
| 10 | 102018089 | rs7924284 | -1.763 | -0.936 | -0.731 | -0.874 | -0.075 | -0.031 | -0.031 | 0 | 0.18 | 0.6 | 0.13 | 0.14 | 0.08 | 0.16 |
| 10 | 102027365 | rs3750710 | -0.577 | -0.935 | -0.747 | -0.873 | -0.047 | -0.015 | -0.015 | 0 | 0.12 | 0.21 | 0.07 | 0.08 | 0.03 | 0.06 |
| 10 | 102027407 | rs12784396 | 0.324 | -0.918 | -1.041 | -0.868 | -0.136 | 0.066 | 0.066 | 0 | 0.23 | 0.05 | 0.27 | 0.07 | 0.43 | 0.14 |
| 10 | 102028467 | rs12242503 | -1.097 | -0.774 | -0.997 | -0.809 | -0.008 | 0.036 | 0.036 | 0 | 0.42 | 0.66 | 0.4 | 0.21 | 0.51 | 0.3 |
| 10 | 102050242 | rs6584356 | -0.298 | -0.782 | -0.934 | -0.45 | -0.002 | -0.001 | -0.001 | 0 | 0.05 | 0.12 | 0.06 | 0.06 | 0.06 | 0.1 |
| 10 | 102052373 | rs12241379 | -0.911 | -0.868 | -0.965 | -0.484 | -0.002 | 0.001 | 0.001 | 0 | 0.13 | 0.16 | 0.1 | 0.09 | 0.08 | 0.11 |
| 10 | 102054452 | rs11597005 | -0.069 | 0.036 | -1.024 | -0.478 | -0.027 | 0.021 | 0.021 | 0 | 0.1 | 0.04 | 0.12 | 0.03 | 0.22 | 0.08 |
| 10 | 102054602 | rs2242051 | -0.19 | -1.136 | -0.521 | -0.128 | -0.052 | 0.187 | 0.187 | 0.031 | 0.11 | 0.35 | 0.04 | 0.3 | 0.01 | 0.06 |
| 10 | 102056745 | rs2278842 | 0 | -0.828 | -0.963 | -0.58 | 0.006 | 0.003 | 0.003 | 0 | 0.07 | 0.1 | 0.15 | 0.11 | 0.12 | 0.13 |
| 10 | 102062074 | rs3892212 | -0.367 | -1.167 | -1.32 | -0.823 | -0.022 | 0.032 | 0.032 | 0 | 0.45 | 0.75 | 0.66 | 0.62 | 0.52 | 0.51 |
| 10 | 102063560 | rs12360462 | 0.923 | -1.177 | -1.076 | -0.813 | 0.037 | 0.078 | 0.078 | 0 | 0.28 | 0.4 | 0.47 | 0.5 | 0.4 | 0.35 |
| 10 | 102066185 | rs17112901 | -0.964 | -0.888 | -0.811 | -0.614 | 0.028 | 0.021 | 0.021 | 1.548 | 0.12 | 0.32 | 0.18 | 0.12 | 0.12 | 0.15 |
| 10 | 102072092 | rs3844556 | 0.197 | -1.966 | -0.976 | -0.947 | 0.005 | 0.008 | 0.008 | 0 | 0.49 | 0.51 | 0.69 | 0.61 | 0.61 | 0.5 |
| 10 | 102075479 | rs603424 | -0.917 | -3.631 | -0.73 | -1.369 | 0.175 | 0.226 | 0.226 | 0 | 0.51 | 0.76 | 0.2 | 0.07 | 0.2 | 0.46 |
| 10 | 102077925 | rs1993110 | 0.98 | -2.281 | -0.285 | -0.51 | 0.011 | 0.012 | 0.012 | 0 | 0.56 | 0.37 | 0.79 | 0.7 | 0.85 | 0.54 |
| 10 | 102085327 | rs3853519 | 0.748 | -2.116 | -0.621 | -0.384 | 0.025 | 0.017 | 0.017 | 0 | 0.62 | 0.63 | 0.82 | 0.69 | 0.85 | 0.53 |
| 10 | 102087113 | rs6584362 | 0.466 | -1.151 | -0.912 | 0.068 | 0.03 | 0.008 | 0.008 | 0 | 0.7 | 0.79 | 0.86 | 0.69 | 0.86 | 0.55 |
| 10 | 102089436 | rs610745 | -0.119 | -0.913 | -1.131 | -0.533 | 0.025 | 0.051 | 0.051 | 0 | 0.22 | 0.13 | 0.09 | 0.03 | 0.12 | 0.35 |
| 10 | 102090690 | rs7904497 | -1.262 | -0.528 | -0.558 | -0.11 | -0.037 | -0.044 | -0.044 | 1.033 | 0.08 | 0.22 | 0.02 | 0 | 0 | 0.02 |
| 10 | 102097695 | rs2060792 | -0.428 | 0.1 | -1.752 | 0.111 | -0.089 | -0.113 | -0.113 | 0 | 0.28 | 0.03 | 0.42 | 0.36 | 0.32 | 0.14 |
| 10 | 102104521 | rs735877 | 0.414 | -0.237 | -1.817 | -0.041 | -0.003 | -0.002 | -0.002 | 0 | 0.39 | 0.66 | 0.32 | 0.38 | 0.41 | 0.49 |
| 10 | 102106367 | rs670213 | -0.144 | -0.538 | -2.299 | -0.35 | -0.072 | 0.025 | 0.025 | 0 | 0.52 | 0.89 | 0.42 | 0.64 | 0.52 | 0.72 |
| 10 | 102109202 | rs1502593 | 0.6 | -0.99 | -2.225 | -0.488 | -0.07 | -0.015 | -0.015 | 0 | 0.35 | 0.17 | 0.29 | 0.28 | 0.44 | 0.41 |

| | | | | | | | | | | | | | | | | |
|---|---|---|---|---|---|---|---|---|---|---|---|---|---|---|---|---|
| 10 | 102113649 | rs11190483 | 2.027 | -0.613 | -2.407 | -0.653 | -0.092 | -0.103 | -0.103 | 3.28 | 0.16 | 0.02 | 0.22 | 0.32 | 0.33 | 0.33 |
| 10 | 102116311 | rs2234970 | 0.295 | -1.052 | -1.573 | -0.894 | -0.001 | -0.001 | -0.001 | 0.437 | 0.46 | 0.36 | 0.55 | 0.4 | 0.42 | 0.26 |
| 10 | 102122603 | rs7849 | -0.927 | -1.369 | -1.107 | -0.888 | -0.037 | -0.032 | -0.032 | 0.692 | 0.23 | 0.42 | 0.14 | 0.22 | 0.15 | 0.34 |
| 10 | 102124761 | rs508384 | -0.985 | -1.275 | -0.979 | -0.778 | -0.034 | -0.031 | -0.031 | 0.44 | 0.23 | 0.42 | 0.14 | 0.22 | 0.15 | 0.34 |
| 10 | 102127147 | rs1393491 | -0.554 | -1.244 | -0.981 | -0.745 | 0.012 | 0.035 | 0.035 | 0 | 0.19 | 0.12 | 0.1 | 0.11 | 0.14 | 0.24 |
| 10 | 102132914 | rs11190489 | -0.285 | -1.431 | -0.935 | -0.803 | -0.001 | 0.034 | 0.034 | 0 | 0.08 | 0.02 | 0.04 | 0 | 0.05 | 0.11 |
| 10 | 102142421 | rs10883466 | 0.123 | -1.48 | -0.963 | -0.772 | -0.006 | 0.005 | 0.005 | 0.05 | 0.08 | 0.12 | 0.04 | 0.13 | 0.04 | 0.12 |
| 10 | 102143048 | rs685910 | -1.821 | -1.63 | -1.067 | -0.817 | -0.036 | -0.029 | -0.029 | 0 | 0.3 | 0.44 | 0.17 | 0.24 | 0.18 | 0.36 |
| 10 | 102143208 | rs685397 | 0.823 | -1.515 | -1.082 | -0.765 | 0.013 | 0.004 | 0.004 | 0 | 0.1 | 0.05 | 0.05 | 0.11 | 0.07 | 0.13 |
| 10 | 102152297 | rs642370 | -1.553 | -1.684 | -1.135 | -0.829 | -0.039 | -0.031 | -0.031 | 0 | 0.34 | 0.47 | 0.18 | 0.25 | 0.19 | 0.39 |
| 10 | 102154010 | rs7068970 | -0.598 | -0.938 | -1.27 | -0.671 | 0.002 | 0.039 | 0.039 | 0 | 0.09 | 0.02 | 0.05 | 0 | 0.06 | 0.11 |
| 10 | 102165238 | rs11190513 | -0.619 | -0.784 | -1.338 | -0.598 | -0.039 | 0.043 | 0.043 | 0 | 0.24 | 0.04 | 0.19 | 0.13 | 0.21 | 0.21 |
| 10 | 102172752 | rs3829162 | -0.619 | -0.784 | -1.338 | -0.581 | -0.039 | 0.038 | 0.038 | 0 | 0.24 | 0.04 | 0.19 | 0.13 | 0.21 | 0.21 |
| 10 | 102177728 | rs4919458 | 0.6 | -0.96 | -0.933 | -0.666 | 0.001 | 0.001 | 0.001 | 0 | 0.72 | 0.67 | 0.79 | 0.87 | 0.78 | 0.78 |
| 10 | 102190406 | rs10509743 | -0.583 | -0.96 | -0.997 | -0.666 | -0.035 | 0.046 | 0.046 | 0 | 0.24 | 0.04 | 0.19 | 0.13 | 0.21 | 0.2 |
| 10 | 102198879 | rs872290 | -0.634 | -0.955 | -0.924 | -0.664 | 0.005 | 0.041 | 0.041 | 0 | 0.25 | 0.18 | 0.2 | 0.13 | 0.21 | 0.2 |
| 10 | 102203071 | rs11190541 | -0.562 | -1.014 | -0.905 | -0.684 | 0.007 | 0.028 | 0.028 | 0 | 0.24 | 0.21 | 0.19 | 0.13 | 0.2 | 0.2 |
| 10 | 102205292 | rs12251636 | -0.24 | -1.268 | -0.606 | -0.922 | 0.022 | -0.027 | -0.027 | 0 | 0.1 | 0.45 | 0.07 | 0.02 | 0.04 | 0.07 |
| 10 | 102206607 | rs11190552 | -0.519 | -1.279 | -0.655 | -0.782 | 0.007 | 0.028 | 0.028 | 0 | 0.24 | 0.2 | 0.19 | 0.13 | 0.2 | 0.2 |
| 10 | 102209160 | rs11190555 | -0.547 | -1.279 | -0.651 | -0.782 | 0.006 | 0.022 | 0.022 | 0 | 0.24 | 0.23 | 0.19 | 0.13 | 0.2 | 0.2 |
| 10 | 102222825 | rs3793772 | -0.531 | -1.286 | -0.657 | -0.785 | 0.008 | 0.03 | 0.03 | 0 | 0.24 | 0.2 | 0.19 | 0.13 | 0.2 | 0.21 |
| 10 | 102227886 | rs1417823 | -0.707 | -1.29 | -0.65 | -0.785 | 0.001 | 0.001 | 0.001 | 0 | 0.25 | 0.29 | 0.2 | 0.13 | 0.2 | 0.23 |
| 10 | 102243387 | rs11190581 | -0.55 | -1.17 | -1.162 | -0.735 | 0.006 | 0.02 | 0.02 | 0 | 0.24 | 0.21 | 0.19 | 0.13 | 0.2 | 0.21 |
| 10 | 102244911 | rs12355721 | -0.536 | -1.17 | -1.193 | -0.735 | 0.006 | 0.023 | 0.023 | 0 | 0.24 | 0.21 | 0.19 | 0.13 | 0.2 | 0.21 |
| 10 | 102246512 | rs2970 | 0.352 | -0.85 | -1.823 | -0.476 | -0.034 | -0.063 | -0.063 | 1.394 | 0.37 | 0.06 | 0.31 | 0.39 | 0.33 | 0.25 |
| 10 | 102247408 | rs2298075 | -0.327 | -0.431 | -1.54 | -0.787 | 0.007 | 0.024 | 0.024 | 0 | 0.24 | 0.21 | 0.19 | 0.13 | 0.2 | 0.21 |
| 10 | 102259706 | rs2273696 | 0.332 | -0.511 | -1.741 | -0.54 | -0.075 | -0.005 | -0.005 | 0.052 | 0.29 | 0.06 | 0.24 | 0.26 | 0.31 | 0.22 |
| 10 | 102264004 | rs12780796 | 0.792 | -0.371 | -1.76 | -0.448 | -0.099 | -0.06 | -0.06 | 1.964 | 0.23 | 0.02 | 0.21 | 0.26 | 0.27 | 0.21 |
| 10 | 102269085 | rs3793706 | -0.832 | -0.725 | -0.836 | -0.398 | 0.011 | 0.024 | 0.024 | 0 | 0.25 | 0.23 | 0.19 | 0.13 | 0.19 | 0.22 |
| 10 | 102269206 | rs3763695 | -0.743 | -0.729 | -0.83 | -0.4 | 0.013 | 0.034 | 0.034 | 0 | 0.24 | 0.2 | 0.18 | 0.13 | 0.19 | 0.22 |
| 10 | 102275727 | rs3750629 | -0.841 | -0.719 | -0.825 | -0.396 | 0.011 | 0.024 | 0.024 | 0 | 0.25 | 0.23 | 0.19 | 0.13 | 0.19 | 0.22 |
| 10 | 102282126 | rs10786596 | -0.855 | -0.725 | -0.768 | -0.398 | -0.001 | -0.005 | -0.005 | 0.388 | 0.25 | 0.3 | 0.19 | 0.13 | 0.19 | 0.22 |
| 10 | 102291287 | rs10786597 | -0.756 | -0.7 | -0.771 | -0.387 | 0.013 | 0.034 | 0.034 | 0 | 0.24 | 0.19 | 0.18 | 0.13 | 0.19 | 0.22 |
| 10 | 102299239 | rs4919471 | -0.696 | -0.707 | -0.769 | -0.385 | 0.013 | 0.04 | 0.04 | 0 | 0.24 | 0.15 | 0.18 | 0.13 | 0.19 | 0.22 |
| 10 | 102308214 | rs3750633 | -0.566 | -0.701 | -0.768 | -0.541 | 0 | 0.075 | 0.075 | 0 | 0.23 | 0.08 | 0.16 | 0.09 | 0.19 | 0.22 |
| 10 | 102308516 | rs10883512 | -0.767 | -0.698 | -0.771 | -0.54 | 0.013 | 0.044 | 0.044 | 0 | 0.24 | 0.19 | 0.17 | 0.09 | 0.19 | 0.22 |
| 10 | 102313997 | rs11190613 | -0.843 | -0.705 | -0.766 | -0.542 | 0.007 | 0.015 | 0.015 | 0 | 0.25 | 0.24 | 0.17 | 0.09 | 0.19 | 0.21 |
| 10 | 102327260 | rs2495751 | -0.824 | -0.698 | -0.775 | -0.539 | 0.008 | 0.02 | 0.02 | 0 | 0.25 | 0.25 | 0.17 | 0.09 | 0.19 | 0.22 |
| 10 | 102328247 | rs2495747 | -0.911 | -0.735 | -0.742 | -0.556 | 0.002 | -0.003 | -0.003 | 0.268 | 0.26 | 0.32 | 0.18 | 0.09 | 0.19 | 0.22 |
| 10 | 102329409 | rs2495744 | -0.653 | -0.734 | -0.758 | -0.555 | 0.012 | 0.047 | 0.047 | 0 | 0.24 | 0.18 | 0.17 | 0.09 | 0.19 | 0.22 |
| 10 | 102331147 | rs9420797 | -0.931 | -0.716 | -0.802 | -0.547 | -0.005 | -0.021 | -0.021 | 1.487 | 0.26 | 0.35 | 0.19 | 0.09 | 0.19 | 0.22 |
| 10 | 102334327 | rs2489043 | -0.761 | -0.696 | -0.799 | -0.539 | -0.001 | -0.01 | -0.01 | 0.051 | 0.25 | 0.3 | 0.18 | 0.09 | 0.19 | 0.22 |
| 10 | 102338609 | rs2495735 | -0.913 | -0.681 | -0.886 | -0.532 | -0.007 | -0.027 | -0.027 | 0 | 0.26 | 0.35 | 0.19 | 0.09 | 0.19 | 0.22 |
| 10 | 102360855 | rs2489003 | -0.358 | -0.562 | -1.35 | -0.414 | 0.016 | 0.061 | 0.061 | 0 | 0.19 | 0.04 | 0.13 | 0.07 | 0.15 | 0.19 |
| 10 | 102362543 | rs2495726 | -1.027 | -1.376 | -0.741 | -0.759 | -0.067 | -0.095 | -0.095 | 0 | 0.3 | 0.49 | 0.18 | 0.11 | 0.16 | 0.17 |
| 10 | 102375258 | rs7092999 | 0.258 | -0.296 | -1.081 | -0.375 | 0.027 | 0.002 | 0.002 | 0 | 0.34 | 0.4 | 0.52 | 0.67 | 0.44 | 0.61 |
| 10 | 102379091 | rs7902336 | 0.077 | -0.228 | -1.107 | -0.375 | 0.021 | -0.01 | -0.01 | 2.735 | 0.38 | 0.49 | 0.56 | 0.7 | 0.48 | 0.63 |
| 10 | 102383988 | rs1110286 | 0.266 | -0.215 | -1.002 | -0.196 | 0.009 | 0.006 | 0.006 | 0 | 0.35 | 0.43 | 0.55 | 0.61 | 0.43 | 0.48 |
| 10 | 102387274 | rs2495717 | -0.487 | -0.112 | -1.712 | -0.415 | 0.002 | 0.082 | 0.082 | 0.607 | 0.51 | 0.91 | 0.64 | 0.68 | 0.51 | 0.54 |
| 10 | 102390727 | rs2489015 | -0.487 | 0.321 | -1.999 | -0.463 | -0.023 | 0.068 | 0.068 | 0.131 | 0.49 | 0.85 | 0.64 | 0.64 | 0.53 | 0.52 |
| 10 | 102391740 | rs2489013 | 1.179 | 0.379 | -2.085 | -0.185 | -0.056 | -0.091 | -0.091 | 0 | 0.21 | 0.1 | 0.42 | 0.58 | 0.3 | 0.42 |
| 10 | 102396758 | rs17113301 | 1.772 | 0.161 | -2.519 | -0.154 | -0.048 | -0.103 | -0.103 | 0 | 0.17 | 0.03 | 0.23 | 0.29 | 0.2 | 0.11 |
| 10 | 102396789 | rs2489012 | -0.498 | -0.218 | -2.022 | -1.251 | 0.006 | -0.081 | -0.081 | 1.315 | 0.58 | 0.54 | 0.7 | 0.87 | 0.59 | 0.69 |
| 10 | 102407173 | rs7068072 | -0.043 | -0.383 | -1.782 | -0.982 | -0.03 | -0.03 | -0.03 | 0 | 0.18 | 0.26 | 0.1 | 0.08 | 0.11 | 0.16 |
| 10 | 102416555 | rs10883518 | 0.349 | 0.193 | -2.32 | -0.671 | -0.063 | -0.142 | -0.142 | 0 | 0.36 | 0.04 | 0.42 | 0.51 | 0.4 | 0.57 |
| 10 | 102427797 | rs4919478 | 0.025 | 0.175 | -1.622 | -0.726 | -0.001 | 0 | 0 | 0 | 0.33 | 0.33 | 0.32 | 0.36 | 0.27 | 0.24 |
| 10 | 102433046 | rs11190644 | 1.001 | 0.631 | -1.643 | -0.918 | 0.001 | -0.028 | -0.028 | 0 | 0.21 | 0.16 | 0.35 | 0.37 | 0.21 | 0.18 |
| 10 | 102443374 | rs11190658 | 1.06 | 0.63 | -1.638 | -0.919 | 0.002 | -0.052 | -0.052 | 0 | 0.2 | 0.14 | 0.34 | 0.38 | 0.2 | 0.18 |
| 10 | 102444894 | rs1419178 | 1.105 | 0.181 | -1.821 | -1.183 | -0.068 | -0.164 | -0.164 | 0.847 | 0.22 | 0.02 | 0.26 | 0.55 | 0.29 | 0.41 |
| 10 | 102445994 | rs12269237 | 0.567 | 1.022 | -1.224 | -0.811 | -0.055 | -0.094 | -0.094 | 0.476 | 0.24 | 0.07 | 0.39 | 0.32 | 0.3 | 0.17 |
| 10 | 102446578 | rs12248459 | 0.359 | 1.005 | -1.039 | -0.823 | 0.006 | 0.022 | 0.022 | 0 | 0.26 | 0.31 | 0.43 | 0.36 | 0.31 | 0.18 |
| 10 | 102452136 | rs11190661 | 0.627 | 1.125 | -1.614 | -0.936 | -0.065 | -0.101 | -0.101 | 0 | 0.23 | 0.06 | 0.39 | 0.3 | 0.3 | 0.17 |
| 10 | 102460486 | rs12241585 | -0.206 | 0.91 | -1.392 | -1.115 | -0.03 | -0.017 | -0.017 | 1.982 | 0.41 | 0.29 | 0.54 | 0.36 | 0.52 | 0.38 |
| 10 | 102471756 | rs10786603 | 0.269 | 1.143 | -1.047 | -1.985 | -0.019 | 0.02 | 0.02 | 0 | 0.18 | 0.14 | 0.24 | 0.06 | 0.27 | 0.26 |
| 10 | 102478689 | rs10883534 | -0.342 | 0.843 | -0.605 | -2.114 | 0.014 | -0.073 | -0.073 | 0 | 0.28 | 0.39 | 0.28 | 0.06 | 0.32 | 0.26 |
| 10 | 102483131 | rs12254876 | 0.818 | 0.654 | -0.926 | -2.297 | -0.009 | 0.104 | 0.104 | 0 | 0.35 | 0.14 | 0.48 | 0.94 | 0.4 | 0.58 |
| 10 | 102491832 | rs7071427 | 0.396 | 0.353 | -0.234 | -1.596 | -0.001 | -0.056 | -0.056 | 1.446 | 0.78 | 0.74 | 0.83 | 1 | 0.81 | 0.91 |
| 10 | 102496740 | rs4403743 | 0.363 | 0.459 | -0.2 | -1.532 | -0.001 | -0.08 | -0.08 | 0.716 | 0.79 | 0.71 | 0.82 | 1 | 0.8 | 0.94 |
| 10 | 102508577 | rs4244341 | 0.744 | 0.233 | 0.009 | -1.458 | -0.022 | -0.094 | -0.094 | 0.481 | 0.73 | 0.55 | 0.8 | 0.9 | 0.79 | 0.8 |

| | | | | | | | | | | | | | | | | |
|---|---|---|---|---|---|---|---|---|---|---|---|---|---|---|---|---|
| 10 | 102523149 | rs10786607 | -0.48 | 0.058 | 0.429 | -2.223 | -0.012 | -0.102 | -0.102 | 0 | 0.32 | 0.57 | 0.17 | 0.01 | 0.22 | 0.18 |
| 10 | 102524470 | rs11190688 | -0.224 | 0.049 | 0.072 | -2.218 | -0.029 | 0.178 | 0.178 | 0 | 0.27 | 0.03 | 0.11 | 0.01 | 0.2 | 0.17 |
| 10 | 102533297 | rs11190696 | -0.314 | 0.469 | 0.519 | -2.262 | 0.016 | 0.033 | 0.033 | 0 | 0.28 | 0.15 | 0.12 | 0.01 | 0.19 | 0.17 |
| 10 | 102536480 | rs11190699 | 0.275 | 0.423 | 0.446 | -2.317 | -0.03 | 0.135 | 0.135 | 0.019 | 0.22 | 0 | 0.09 | 0.01 | 0.17 | 0.16 |
| 10 | 102536664 | rs12357097 | -0.394 | 0.459 | 0.483 | -2.294 | 0.017 | 0.032 | 0.032 | 0 | 0.28 | 0.15 | 0.12 | 0.01 | 0.19 | 0.17 |
| 10 | 102541820 | rs11190701 | -0.343 | 0.462 | 0.327 | -2.293 | -0.001 | 0.175 | 0.175 | 0 | 0.27 | 0.05 | 0.11 | 0.01 | 0.19 | 0.17 |
| 10 | 102552752 | rs10883543 | -0.489 | 0.523 | 0.471 | -2.326 | -0.008 | 0.005 | 0.005 | 0 | 0.94 | 0.96 | 0.92 | 1 | 0.89 | 0.98 |
| 10 | 102552765 | rs11190710 | -0.705 | 0.612 | 0.471 | -2.427 | -0.001 | 0.002 | 0.002 | 1.358 | 0.05 | 0.11 | 0.05 | 0.09 | 0.04 | 0.13 |
| 10 | 102553647 | rs1006545 | -0.437 | 0.621 | 0.534 | -2.425 | 0 | -0.005 | -0.005 | 0.021 | 0.94 | 0.95 | 0.91 | 1 | 0.89 | 0.98 |
| 10 | 102553995 | rs1006544 | 1.269 | 0.235 | 0.715 | -2.677 | 0.017 | 0.122 | 0.122 | 0.643 | 0.53 | 0.29 | 0.68 | 0.9 | 0.61 | 0.63 |
| 10 | 102554671 | rs10748798 | -0.483 | 0.361 | 0.649 | -2.512 | 0 | -0.005 | -0.005 | 0 | 0.94 | 0.95 | 0.91 | 1 | 0.9 | 0.98 |
| 10 | 102556257 | rs4548562 | -0.077 | 0.293 | 0.638 | -2.542 | -0.001 | -0.007 | -0.007 | 0.122 | 0.93 | 0.94 | 0.92 | 1 | 0.89 | 0.98 |
| 10 | 102559058 | rs12266644 | -1.097 | 0.238 | 0.718 | -2.567 | -0.01 | -0.003 | -0.003 | 0.118 | 0.08 | 0.18 | 0.06 | 0.09 | 0.05 | 0.1 |
| 10 | 102559421 | rs4917911 | -0.605 | 0.269 | 0.613 | -2.557 | -0.016 | 0.015 | 0.015 | 0 | 0.94 | 0.98 | 0.92 | 1 | 0.9 | 0.98 |
| 10 | 102564447 | rs12773509 | 0.106 | 0.017 | 0.559 | -2.598 | -0.019 | 0.174 | 0.174 | 0 | 0.26 | 0.03 | 0.11 | 0 | 0.19 | 0.16 |
| 10 | 102566591 | rs12769518 | -0.139 | 0.02 | 0.669 | -2.595 | 0.002 | 0.153 | 0.153 | 0 | 0.28 | 0.06 | 0.13 | 0 | 0.2 | 0.17 |
| 10 | 102568872 | rs1800897 | -1.343 | -0.199 | 1.012 | -2.681 | -0.025 | 0.008 | 0.008 | 1.818 | 0.1 | 0.39 | 0.06 | 0.09 | 0.04 | 0.1 |
| 10 | 102568983 | rs1800898 | -1.159 | -0.201 | 0.955 | -2.693 | 0.066 | 0.196 | 0.196 | 0.218 | 0.41 | 0.82 | 0.24 | 0.1 | 0.26 | 0.28 |
| 10 | 102575531 | rs10883544 | -0.598 | -0.167 | 1.23 | -2.678 | -0.029 | -0.172 | -0.172 | 0 | 0.33 | 0.46 | 0.16 | 0 | 0.23 | 0.18 |
| 10 | 102581636 | rs11815022 | -0.663 | -0.098 | 0.71 | -2.562 | -0.015 | 0.006 | 0.006 | 0 | 0.07 | 0.17 | 0.05 | 0.1 | 0.03 | 0.1 |
| 10 | 102584597 | rs11190717 | -0.128 | -0.059 | 0.501 | -2.588 | 0.01 | 0.129 | 0.129 | 0 | 0.28 | 0.07 | 0.13 | 0 | 0.2 | 0.17 |
| 10 | 102590753 | rs996359 | 0.933 | 0.568 | -0.033 | -1.798 | -0.074 | -0.129 | -0.129 | 1.188 | 0.17 | 0.02 | 0.21 | 0.42 | 0.22 | 0.3 |
| 10 | 102591454 | rs11816136 | -0.7 | 0.27 | 0.855 | -2.204 | -0.014 | -0.037 | -0.037 | 0.077 | 0.18 | 0.52 | 0.14 | 0.12 | 0.16 | 0.11 |
| 10 | 102592975 | rs2495702 | -0.911 | 0.222 | 0.682 | -2.271 | 0.004 | 0.013 | 0.013 | 2.978 | 0.39 | 0.61 | 0.27 | 0.12 | 0.31 | 0.29 |
| 10 | 102593621 | rs2476964 | -0.17 | 0.354 | 0.058 | -2.256 | -0.054 | 0.164 | 0.164 | 1.13 | 0.29 | 0.06 | 0.18 | 0.01 | 0.27 | 0.19 |
| 10 | 102595920 | rs2476968 | -1.833 | 0.947 | 0.855 | -1.571 | -0.003 | 0.006 | 0.006 | 0.932 | 0.28 | 0.73 | 0.28 | 0.18 | 0.33 | 0.17 |
| 10 | 102602344 | rs12250029 | -0.822 | 0.206 | 1.07 | -1.703 | -0.034 | -0.076 | -0.076 | 0.459 | 0.18 | 0.54 | 0.14 | 0.06 | 0.15 | 0.04 |
| 10 | 102604977 | rs7911849 | -1.581 | 0.218 | 0.445 | -1.743 | 0.007 | 0.013 | 0.013 | 0.23 | 0.29 | 0.71 | 0.25 | 0.06 | 0.27 | 0.08 |
| 10 | 102610360 | rs912478 | -1.407 | 0.265 | 0.246 | -1.652 | -0.001 | -0.018 | -0.018 | 0 | 0.31 | 0.67 | 0.25 | 0.11 | 0.27 | 0.12 |
| 10 | 102618653 | rs12359797 | -0.616 | 0.398 | 0.347 | -1.474 | -0.034 | -0.092 | -0.092 | 0 | 0.29 | 0.4 | 0.15 | 0.06 | 0.18 | 0.13 |
| 10 | 102620497 | rs4919504 | 0.287 | 1.939 | 0.396 | -0.654 | 0.02 | -0.006 | -0.006 | 0 | 0.91 | 0.83 | 0.91 | 0.94 | 0.83 | 0.89 |
| 10 | 102620555 | rs17113518 | -0.895 | 1.462 | 0.365 | -0.519 | 0.071 | -0.029 | -0.029 | 1.901 | 0.07 | 0.24 | 0.1 | 0 | 0.1 | 0.02 |
| 10 | 102627228 | rs10883547 | -0.844 | 1.096 | 0.043 | 0.188 | 0.007 | 0.003 | 0.003 | 0 | 0.4 | 0.49 | 0.38 | 0.56 | 0.45 | 0.48 |
| 10 | 102631263 | rs10883549 | -0.74 | 1.077 | 0.003 | 0.113 | -0.008 | -0.026 | -0.026 | 0 | 0.33 | 0.2 | 0.3 | 0.54 | 0.35 | 0.46 |
| 10 | 102633059 | rs10883551 | -0.302 | 1.04 | -0.359 | 0.528 | -0.081 | -0.115 | -0.115 | 0.37 | 0.19 | 0.03 | 0.25 | 0.35 | 0.28 | 0.25 |
| 10 | 102634940 | rs12248258 | -0.157 | 0.884 | -0.046 | 0.254 | 0.006 | 0.026 | 0.026 | 0 | 0.09 | 0.04 | 0.04 | 0.01 | 0.06 | 0.07 |
| 10 | 102635485 | rs10786609 | 1.174 | 0.849 | -0.006 | 0.142 | -0.009 | -0.021 | -0.021 | 0 | 0.71 | 0.79 | 0.67 | 0.37 | 0.64 | 0.5 |
| 10 | 102638586 | rs912474 | 1.248 | 0.745 | 0.204 | 0.241 | 0.002 | -0.022 | -0.022 | 0 | 0.69 | 0.8 | 0.7 | 0.38 | 0.69 | 0.46 |
| 10 | 102639161 | rs2863073 | 0.889 | 0.933 | 0.178 | 0.394 | -0.007 | -0.046 | -0.046 | 0 | 0.78 | 0.89 | 0.73 | 0.39 | 0.75 | 0.53 |
| 10 | 102639559 | rs912472 | -0.023 | 0.983 | 0.132 | 0.423 | 0.012 | 0.055 | 0.055 | 0.125 | 0.11 | 0.25 | 0.15 | 0.26 | 0.13 | 0.09 |
| 10 | 102643010 | rs11595160 | 0.558 | 0.949 | 0.384 | 0.301 | 0.003 | 0.036 | 0.036 | 0 | 0.74 | 0.63 | 0.71 | 0.34 | 0.74 | 0.56 |
| 10 | 102647686 | rs7899336 | 0.412 | 0.935 | 0.378 | 0.117 | 0.002 | -0.019 | -0.019 | 0 | 0.8 | 0.89 | 0.85 | 0.64 | 0.8 | 0.79 |
| 10 | 102648765 | rs1408343 | -0.942 | 0.857 | 0.662 | 0.175 | 0.009 | 0.097 | 0.097 | 0 | 0.23 | 0.47 | 0.3 | 0.55 | 0.23 | 0.37 |
| 10 | 102653765 | rs10786612 | -0.155 | 0.827 | 0.51 | 0.201 | 0.001 | 0.082 | 0.082 | 0 | 0.18 | 0.27 | 0.23 | 0.55 | 0.17 | 0.35 |
| 10 | 102663197 | rs10883558 | 0.293 | 0.892 | -0.096 | 0.255 | -0.001 | 0.001 | 0.001 | 0 | 0.81 | 0.81 | 0.84 | 0.65 | 0.82 | 0.8 |
| 10 | 102689217 | rs2273654 | 1.92 | 0.975 | -0.411 | -0.03 | -0.055 | 0.23 | 0.23 | 0 | 0.45 | 0.13 | 0.4 | 0.07 | 0.45 | 0.34 |
| 10 | 102691412 | rs10509748 | 1.525 | 0.907 | -0.302 | -0.031 | 0.032 | 0.191 | 0.191 | 0 | 0.52 | 0.23 | 0.48 | 0.1 | 0.58 | 0.4 |
| 10 | 102695193 | rs10748801 | 0.065 | 0.853 | -0.228 | 0.074 | 0.006 | -0.009 | -0.009 | 0 | 0.8 | 0.88 | 0.84 | 0.64 | 0.82 | 0.81 |
| 10 | 102715183 | rs1056295 | -0.211 | 0.671 | -0.133 | 0.04 | -0.004 | 0.149 | 0.149 | 0 | 0.22 | 0.41 | 0.29 | 0.53 | 0.18 | 0.35 |
| 10 | 102725653 | rs2295718 | 0.022 | 0.687 | -0.171 | 0.043 | 0.006 | -0.012 | -0.012 | 0.443 | 0.81 | 0.88 | 0.84 | 0.62 | 0.82 | 0.8 |
| 10 | 102734778 | rs4919510 | -0.21 | 0.683 | -0.084 | 0.048 | -0.007 | 0.158 | 0.158 | 0 | 0.22 | 0.44 | 0.29 | 0.52 | 0.18 | 0.34 |
| 10 | 102744214 | rs2295716 | -0.25 | 0.757 | -0.149 | 0.048 | -0.006 | 0.154 | 0.154 | 0 | 0.22 | 0.43 | 0.29 | 0.53 | 0.18 | 0.34 |
| 10 | 102744376 | rs11190780 | 0.086 | 0.829 | -0.187 | 0.006 | -0.017 | 0.013 | 0.013 | 0 | 0.08 | 0.07 | 0.09 | 0.03 | 0.13 | 0.06 |
| 10 | 102745062 | rs722435 | -0.242 | 0.884 | -0.216 | 0.037 | -0.006 | 0.154 | 0.154 | 0 | 0.22 | 0.43 | 0.29 | 0.53 | 0.18 | 0.34 |
| 10 | 102746231 | rs7184 | 2.048 | 0.853 | -0.287 | -0.025 | -0.079 | 0.253 | 0.253 | 0 | 0.45 | 0.09 | 0.39 | 0.07 | 0.44 | 0.37 |
| 10 | 102746503 | rs2863095 | -0.183 | 0.454 | -0.244 | 0.014 | -0.006 | 0.136 | 0.136 | 0 | 0.22 | 0.39 | 0.29 | 0.53 | 0.18 | 0.34 |
| 10 | 102755779 | rs10883567 | 1.924 | 0.425 | -0.276 | -0.001 | -0.057 | 0.263 | 0.263 | 0 | 0.46 | 0.09 | 0.42 | 0.07 | 0.49 | 0.38 |
| 10 | 102757442 | rs11190786 | -0.245 | 0.414 | -0.256 | 0.017 | -0.011 | 0.138 | 0.138 | 0 | 0.22 | 0.41 | 0.29 | 0.53 | 0.18 | 0.34 |
| 10 | 102762256 | rs752974 | 2.016 | 0.324 | -0.471 | 0.12 | -0.072 | 0.209 | 0.209 | 0 | 0.45 | 0.08 | 0.49 | 0.09 | 0.49 | 0.35 |
| 10 | 102764511 | rs807028 | -0.128 | 0.215 | -0.273 | 0.208 | -0.007 | 0.011 | 0.011 | 0 | 0.83 | 0.83 | 0.87 | 0.62 | 0.87 | 0.79 |
| 10 | 102767117 | rs701836 | -0.522 | 0.285 | -0.228 | 0.15 | -0.005 | 0.112 | 0.112 | 0 | 0.23 | 0.39 | 0.33 | 0.53 | 0.21 | 0.34 |
| 10 | 102770082 | rs807023 | 0.204 | 0.089 | -0.615 | 0.038 | -0.003 | 0 | 0 | 0 | 0.79 | 0.76 | 0.84 | 0.66 | 0.83 | 0.79 |
| 10 | 102771573 | rs807020 | -0.316 | 0.172 | -0.668 | -0.08 | 0.003 | 0.09 | 0.09 | 0 | 0.23 | 0.37 | 0.33 | 0.57 | 0.22 | 0.38 |
| 10 | 102772840 | rs807019 | 0.412 | 0.003 | -0.652 | 0.058 | 0.015 | -0.011 | -0.011 | 0 | 0.79 | 0.85 | 0.84 | 0.67 | 0.82 | 0.79 |
| 10 | 102773011 | rs807018 | 0.654 | 0.135 | -0.989 | 0.065 | -0.027 | -0.006 | -0.006 | 0 | 0.67 | 0.46 | 0.73 | 0.64 | 0.7 | 0.73 |
| 10 | 102774568 | rs10883569 | -0.027 | 0.585 | -2.364 | -0.105 | -0.073 | 0.002 | 0.002 | 2.378 | 0.37 | 0.05 | 0.27 | 0.28 | 0.38 | 0.19 |
| 10 | 102775205 | rs701837 | 0.382 | 0.714 | -1.544 | -0.025 | 0.001 | 0.002 | 0.002 | 0 | 0.65 | 0.7 | 0.6 | 0.24 | 0.65 | 0.42 |
| 10 | 102776870 | rs7094415 | 0.095 | 0.758 | -1.873 | 0.01 | -0.05 | -0.029 | -0.029 | 0.161 | 0.64 | 0.86 | 0.56 | 0.4 | 0.56 | 0.48 |
| 10 | 102780638 | rs1998249 | -0.513 | -0.512 | -0.922 | 0.616 | -0.006 | 0.001 | 0.001 | 0.501 | 0.29 | 0.71 | 0.27 | 0.37 | 0.18 | 0.25 |

| | | | | | | | | | | | | | | | | |
|---|---|---|---|---|---|---|---|---|---|---|---|---|---|---|---|---|
| 10 | 102785771 | rs11190802 | -0.442 | -0.533 | -1.263 | 0.633 | 0.011 | -0.007 | -0.007 | 0.286 | 0.32 | 0.76 | 0.33 | 0.38 | 0.23 | 0.29 |
| 10 | 102787591 | rs967201 | 0.588 | 0.156 | -0.805 | 0.808 | -0.008 | 0.008 | 0.008 | 0 | 0.77 | 0.78 | 0.77 | 0.75 | 0.8 | 0.74 |
| 10 | 102795957 | rs11190804 | -0.957 | 0.934 | -1.198 | 0.097 | -0.031 | 0.057 | 0.057 | 0.038 | 0.15 | 0.06 | 0.15 | 0.01 | 0.2 | 0.25 |
| 10 | 102802394 | rs807052 | -0.456 | 1.039 | -0.86 | -0.772 | 0 | -0.052 | -0.052 | 0 | 0.21 | 0.26 | 0.17 | 0.01 | 0.17 | 0.06 |
| 10 | 102802570 | rs807051 | -0.842 | 0.55 | -0.956 | -0.422 | -0.026 | 0.044 | 0.044 | 0.094 | 0.7 | 0.66 | 0.63 | 0.5 | 0.79 | 0.67 |
| 10 | 102806665 | rs807047 | -0.564 | 0.501 | -0.921 | -0.546 | 0.007 | -0.099 | -0.099 | 0 | 0.33 | 0.49 | 0.3 | 0.02 | 0.31 | 0.31 |
| 10 | 102810715 | rs2863098 | 1.189 | 0.316 | -1.05 | -0.535 | -0.094 | -0.108 | -0.108 | 0 | 0.32 | 0.06 | 0.3 | 0.49 | 0.42 | 0.36 |
| 10 | 102824760 | rs12359843 | 0.714 | 0.543 | -1.067 | -0.568 | -0.039 | 0.032 | 0.032 | 0 | 0.07 | 0.01 | 0.12 | 0.01 | 0.13 | 0.22 |
| 10 | 102827228 | rs2295715 | -0.257 | 0.291 | -1.064 | -0.474 | 0.001 | 0.035 | 0.035 | 0 | 0.14 | 0.33 | 0.27 | 0.2 | 0.18 | 0.07 |
| 10 | 102829164 | rs11190813 | 1.6 | 0.345 | -1.277 | -0.361 | -0.046 | -0.044 | -0.044 | 0 | 0.28 | 0.08 | 0.21 | 0.38 | 0.32 | 0.34 |
| 10 | 102833064 | rs807013 | -0.809 | 0.376 | -1.286 | -0.449 | 0.024 | -0.098 | -0.098 | 0.292 | 0.69 | 0.89 | 0.72 | 0.59 | 0.76 | 0.68 |
| 10 | 102835254 | rs10786617 | 1.399 | 0.422 | -0.958 | -0.497 | 0.001 | 0.012 | 0.012 | 0 | 0.31 | 0.38 | 0.32 | 0.42 | 0.31 | 0.32 |
| 10 | 102841029 | rs10748804 | 0.694 | 0.435 | -0.725 | -0.543 | -0.004 | 0.002 | 0.002 | 0 | 0.39 | 0.54 | 0.45 | 0.43 | 0.44 | 0.53 |
| 10 | 102842961 | rs11190818 | 1.403 | 0.437 | -0.723 | -0.539 | 0.001 | 0.014 | 0.014 | 0 | 0.31 | 0.39 | 0.31 | 0.42 | 0.31 | 0.33 |
| 10 | 102844710 | rs10883579 | 0.329 | 0.389 | -0.762 | -0.488 | -0.008 | -0.01 | -0.01 | 0 | 0.43 | 0.56 | 0.52 | 0.59 | 0.51 | 0.55 |
| 10 | 102845241 | rs17113682 | 0.877 | 0.297 | -1.083 | -0.036 | -0.013 | -0.051 | -0.051 | 0 | 0.27 | 0.03 | 0.26 | 0.23 | 0.21 | 0.26 |
| 10 | 102847237 | rs807043 | -0.966 | 0.301 | -0.924 | -0.14 | -0.046 | 0.08 | 0.08 | 0 | 0.85 | 0.98 | 0.91 | 1 | 0.9 | 0.9 |
| 10 | 102848834 | rs10883580 | 1.219 | 0.204 | -0.576 | 0.131 | 0.003 | 0.014 | 0.014 | 0.013 | 0.3 | 0.35 | 0.32 | 0.42 | 0.32 | 0.33 |
| 10 | 102863097 | rs7085631 | 1.165 | 0.724 | -0.564 | 0.464 | -0.001 | -0.003 | -0.003 | 0 | 0.3 | 0.28 | 0.3 | 0.2 | 0.29 | 0.27 |
| 10 | 102864753 | rs10883581 | 0.279 | 0.692 | -0.57 | 0.491 | -0.002 | 0.013 | 0.013 | 0.365 | 0.41 | 0.44 | 0.44 | 0.59 | 0.48 | 0.36 |
| 10 | 102867868 | rs10786620 | 1.015 | 0.604 | -0.898 | 0.345 | -0.002 | -0.006 | -0.006 | 0 | 0.3 | 0.3 | 0.3 | 0.21 | 0.29 | 0.27 |
| 10 | 102868170 | rs10883586 | 1.003 | 0.605 | -0.899 | 0.345 | -0.003 | -0.008 | -0.008 | 0 | 0.3 | 0.32 | 0.3 | 0.21 | 0.29 | 0.27 |
| 10 | 102873339 | rs11190831 | 0.233 | 0.942 | -1.184 | 0.302 | -0.04 | -0.016 | -0.016 | 0 | 0.14 | 0.04 | 0.17 | 0.17 | 0.24 | 0.06 |
| 10 | 102874668 | rs1361265 | 0.241 | 0.855 | -1.197 | 0.222 | -0.004 | 0.02 | 0.02 | 0.847 | 0.43 | 0.27 | 0.45 | 0.38 | 0.52 | 0.29 |
| 10 | 102879620 | rs11190835 | 0.168 | 1.234 | -1.455 | 0.219 | -0.047 | -0.024 | -0.024 | 0 | 0.14 | 0.04 | 0.17 | 0.19 | 0.23 | 0.06 |
| 10 | 102880793 | rs10786621 | -0.073 | 1.651 | -1.309 | 0.396 | -0.02 | -0.002 | -0.002 | 0 | 0.33 | 0.31 | 0.4 | 0.29 | 0.39 | 0.46 |
| 10 | 102887013 | rs946327 | 0.285 | 1.365 | -1.185 | 0.453 | -0.001 | 0.001 | 0.001 | 0 | 0.58 | 0.4 | 0.64 | 0.6 | 0.58 | 0.75 |
| 10 | 102890421 | rs1111350 | 0.99 | 1.035 | -1.551 | 0.398 | -0.015 | -0.059 | -0.059 | 0.022 | 0.16 | 0.08 | 0.21 | 0.22 | 0.17 | 0.2 |
| 10 | 102893345 | rs946328 | 0.12 | 0.87 | -1.653 | 0.173 | -0.038 | 0.028 | 0.028 | 1.797 | 0.23 | 0.03 | 0.18 | 0.17 | 0.24 | 0.07 |
| 10 | 102893840 | rs12415670 | 0.961 | 0.6 | -1.443 | 0.339 | -0.005 | -0.055 | -0.055 | 0 | 0.16 | 0.08 | 0.21 | 0.22 | 0.16 | 0.2 |
| 10 | 102896801 | rs2235128 | 0.952 | 0.061 | -1.607 | 0.203 | -0.01 | 0.005 | 0.005 | 0 | 0.19 | 0.19 | 0.2 | 0.33 | 0.16 | 0.29 |
| 10 | 102896849 | rs17113735 | 0.49 | 0.179 | -1.476 | -0.085 | 0.011 | -0.028 | -0.028 | 0.228 | 0.33 | 0.46 | 0.27 | 0.24 | 0.32 | 0.25 |
| 10 | 102897057 | rs1051723 | 0.315 | 0.265 | -1.547 | -0.092 | -0.071 | 0.054 | 0.054 | 0.422 | 0.25 | 0.04 | 0.26 | 0.15 | 0.29 | 0.13 |
| 10 | 102897087 | rs2742038 | 1.212 | 0.255 | -1.479 | -0.073 | -0.021 | 0.014 | 0.014 | 0 | 0.19 | 0.21 | 0.2 | 0.33 | 0.16 | 0.29 |
| 10 | 102897311 | rs1051725 | -0.08 | 0.919 | -1.54 | -0.344 | -0.043 | 0.06 | 0.06 | 0.286 | 0.26 | 0.08 | 0.26 | 0.15 | 0.29 | 0.13 |
| 10 | 102907560 | rs792723 | -0.234 | 0.423 | -0.981 | -0.619 | 0.009 | 0.043 | 0.043 | 0 | 0.18 | 0.11 | 0.11 | 0.04 | 0.17 | 0.17 |
| 10 | 102909665 | rs4917928 | 1.052 | 0.001 | -1.34 | -0.873 | -0.023 | -0.005 | -0.005 | 0 | 0.38 | 0.14 | 0.51 | 0.64 | 0.4 | 0.5 |
| 10 | 102915369 | rs4919528 | -0.11 | 0.166 | -1.579 | -1.232 | -0.077 | 0.153 | 0.153 | 0 | 0.66 | 0.87 | 0.7 | 0.95 | 0.57 | 0.62 |
| 10 | 102915753 | rs4919530 | 0.874 | 0.013 | -1.172 | -1.15 | 0.007 | 0.018 | 0.018 | 0 | 0.44 | 0.36 | 0.53 | 0.64 | 0.41 | 0.5 |
| 10 | 102917917 | rs1080358 | 0.453 | -0.107 | -0.886 | -1.118 | 0.013 | 0.028 | 0.028 | 0 | 0.5 | 0.43 | 0.57 | 0.64 | 0.46 | 0.56 |
| 10 | 102924567 | rs12771487 | 0.109 | -0.085 | -0.338 | -1.234 | 0.02 | -0.119 | -0.119 | 0 | 0.68 | 0.49 | 0.75 | 0.95 | 0.66 | 0.65 |
| 10 | 102926766 | rs12240442 | 0.35 | -0.216 | 0.234 | -1.122 | 0 | -0.001 | -0.001 | 0 | 0.24 | 0.24 | 0.19 | 0.3 | 0.22 | 0.12 |
| 10 | 102927396 | rs1618475 | -0.802 | -0.111 | 0.171 | -1.109 | 0.023 | 0.005 | 0.005 | 0.063 | 0.39 | 0.72 | 0.25 | 0.35 | 0.28 | 0.26 |
| 10 | 102930909 | rs2256963 | 0.11 | 0.524 | 0.458 | -1.075 | -0.011 | -0.007 | -0.007 | 0 | 0.09 | 0.23 | 0.04 | 0.04 | 0.06 | 0.12 |
| 10 | 102934720 | rs792715 | -0.381 | 0.841 | 0.02 | -1.044 | 0.006 | -0.045 | -0.045 | 0 | 0.8 | 0.78 | 0.79 | 1 | 0.72 | 0.79 |
| 10 | 102938401 | rs792708 | -0.288 | 0.94 | -0.13 | -0.99 | 0.034 | -0.091 | -0.091 | 3.87 | 0.79 | 0.73 | 0.78 | 1 | 0.72 | 0.83 |
| 10 | 102954897 | rs12357632 | -0.175 | 1.022 | -0.696 | -1.016 | -0.007 | 0.037 | 0.037 | 0 | 0.14 | 0.05 | 0.13 | 0.04 | 0.14 | 0.13 |
| 10 | 102955557 | rs792724 | 0.918 | 0.984 | -0.004 | -1.115 | 0.018 | -0.08 | -0.08 | 0 | 0.6 | 0.51 | 0.62 | 0.82 | 0.56 | 0.67 |
| 10 | 102960061 | rs7908803 | -0.052 | 0.898 | -0.269 | -1.328 | -0.001 | 0.003 | 0.003 | 0 | 0.35 | 0.38 | 0.22 | 0.11 | 0.32 | 0.2 |
| 10 | 102960114 | rs1556862 | -0.907 | 0.572 | -0.606 | -0.63 | 0.001 | -0.017 | -0.017 | 4.911 | 0.73 | 0.81 | 0.53 | 0.62 | 0.76 | 0.51 |
| 10 | 102964512 | rs11598517 | 0.203 | 0.725 | -0.586 | -1.151 | 0.011 | -0.029 | -0.029 | 0 | 0.18 | 0.2 | 0.17 | 0.05 | 0.29 | 0.17 |
| 10 | 102964604 | rs11598564 | -0.495 | 0.405 | -0.748 | -0.806 | 0.001 | 0.004 | 0.004 | 0.587 | 0.4 | 0.41 | 0.33 | 0.48 | 0.46 | 0.44 |
| 10 | 102971968 | rs10786627 | -0.32 | -0.388 | -0.841 | -1.045 | 0.037 | 0.088 | 0.088 | 0 | 0.5 | 0.42 | 0.52 | 0.69 | 0.35 | 0.64 |
| 10 | 102979207 | rs11190870 | -0.537 | -0.496 | -0.491 | -0.458 | -0.002 | -0.007 | -0.007 | 2.359 | 0.35 | 0.4 | 0.32 | 0.46 | 0.41 | 0.3 |
| 10 | 102980156 | rs11598177 | -0.964 | -0.594 | -0.202 | -0.54 | -0.002 | -0.004 | -0.004 | 0 | 0.4 | 0.51 | 0.35 | 0.62 | 0.45 | 0.31 |
| 10 | 102986589 | rs1322331 | -1.007 | -0.638 | -0.126 | -0.575 | 0 | 0.001 | 0.001 | 0.416 | 0.41 | 0.62 | 0.36 | 0.62 | 0.45 | 0.31 |
| 10 | 102993649 | rs625039 | 0.099 | -0.288 | 0.047 | -0.451 | -0.037 | 0.056 | 0.056 | 0.958 | 0.24 | 0.4 | 0.16 | 0.35 | 0.12 | 0.23 |
| 10 | 102995796 | rs594791 | 0.429 | -0.204 | -0.209 | -0.431 | 0.002 | 0.018 | 0.018 | 1.281 | 0.65 | 0.41 | 0.66 | 0.61 | 0.6 | 0.7 |
| 10 | 102996362 | rs11190872 | 0.438 | 0.712 | -0.479 | -0.523 | 0.017 | -0.002 | -0.002 | 0 | 0.08 | 0.08 | 0.15 | 0.06 | 0.24 | 0.06 |
| 10 | 103008802 | rs1885517 | -0.927 | -0.044 | -0.629 | -0.815 | 0.004 | -0.001 | -0.001 | 0.184 | 0.64 | 0.66 | 0.45 | 0.67 | 0.64 | 0.43 |
| 10 | 103018321 | rs17686462 | 0.909 | 0.337 | -1.082 | -0.77 | -0.063 | 0.027 | 0.027 | 0 | 0.11 | 0.03 | 0.15 | 0.06 | 0.31 | 0.07 |
| 10 | 103019884 | rs17113829 | 0.266 | -0.404 | -1.072 | -0.906 | 0.009 | -0.013 | -0.013 | 0 | 0.29 | 0.28 | 0.19 | 0.53 | 0.21 | 0.17 |
| 10 | 103027561 | rs11190888 | 1.065 | -0.369 | -1.26 | -0.91 | -0.055 | 0.026 | 0.026 | 0 | 0.11 | 0.02 | 0.14 | 0.06 | 0.26 | 0.07 |
| 10 | 103029060 | rs10883603 | -0.526 | -0.648 | -1.089 | -1.047 | -0.009 | -0.05 | -0.05 | 0 | 0.4 | 0.46 | 0.25 | 0.14 | 0.39 | 0.28 |
| 10 | 103030046 | rs11190892 | 0.51 | -0.678 | -1.025 | -1.052 | -0.009 | -0.015 | -0.015 | 0 | 0.24 | 0.27 | 0.08 | 0.08 | 0.12 | 0.18 |
| 10 | 103030321 | rs7893126 | 0.548 | -0.522 | -1.029 | -1.028 | 0.003 | -0.002 | -0.002 | 0 | 0.25 | 0.26 | 0.12 | 0.08 | 0.17 | 0.18 |
| 10 | 103031132 | rs17686629 | 0.616 | -0.394 | -1.346 | -0.879 | -0.026 | 0.055 | 0.055 | 0 | 0.2 | 0.06 | 0.16 | 0.07 | 0.29 | 0.16 |
| 10 | 103032109 | rs7924066 | -0.253 | -0.259 | -1.412 | -0.77 | -0.004 | -0.046 | -0.046 | 0 | 0.43 | 0.48 | 0.29 | 0.14 | 0.44 | 0.28 |
| 10 | 103033126 | rs12569599 | 0.682 | 0.108 | -1.499 | -0.575 | -0.016 | -0.09 | -0.09 | 0.039 | 0.22 | 0.11 | 0.18 | 0.53 | 0.23 | 0.14 |

| | | | | | | | | | | | | | | | | |
|---|---|---|---|---|---|---|---|---|---|---|---|---|---|---|---|---|
| 10 | 103037648 | rs7904210 | -0.801 | 0.588 | -1.136 | -0.413 | 0.012 | 0.007 | 0.007 | 0.941 | 0.68 | 0.62 | 0.48 | 0.66 | 0.69 | 0.43 |
| 10 | 103039264 | rs11190901 | -0.374 | 0.107 | -0.885 | -0.057 | -0.002 | -0.001 | -0.001 | 0 | 0.48 | 0.54 | 0.31 | 0.59 | 0.4 | 0.27 |
| 10 | 103050571 | rs644633 | 0.103 | -0.007 | -1.037 | 0.084 | 0.001 | -0.001 | -0.001 | 1.193 | 0.19 | 0.14 | 0.15 | 0.25 | 0.15 | 0.06 |
| 10 | 103054982 | rs614375 | -0.685 | 0.539 | -1.131 | -0.283 | -0.004 | 0.006 | 0.006 | 0 | 0.32 | 0.24 | 0.31 | 0.29 | 0.41 | 0.16 |
| 10 | 103063080 | rs11597176 | -0.199 | 0.527 | -1.773 | -0.833 | -0.086 | 0.09 | 0.09 | 1.306 | 0.31 | 0.12 | 0.29 | 0.18 | 0.41 | 0.42 |
| 10 | 103063437 | rs667707 | 0.046 | 0.133 | -1.942 | -0.7 | 0.021 | 0.013 | 0.013 | 0.254 | 0.35 | 0.2 | 0.29 | 0.65 | 0.32 | 0.25 |
| 10 | 103065599 | rs621106 | -0.269 | -0.214 | -2.31 | -1.075 | -0.012 | 0.107 | 0.107 | 0 | 0.56 | 0.23 | 0.65 | 0.82 | 0.58 | 0.52 |
| 10 | 103066248 | rs589433 | 0.388 | -0.36 | -2.309 | -1.074 | 0.011 | 0.004 | 0.004 | 0 | 0.32 | 0.2 | 0.29 | 0.65 | 0.3 | 0.27 |
| 10 | 103067687 | rs590945 | -0.978 | -0.314 | -1.412 | -1.149 | -0.015 | -0.065 | -0.065 | 0.243 | 0.69 | 0.49 | 0.81 | 0.86 | 0.83 | 0.6 |
| 10 | 103071409 | rs3802727 | -0.905 | -0.165 | -1.358 | -0.761 | 0.02 | 0.047 | 0.047 | 0 | 0.52 | 0.31 | 0.66 | 0.63 | 0.66 | 0.52 |
| 10 | 103071601 | rs752996 | 0.44 | 0.296 | -0.603 | -0.176 | -0.023 | 0.019 | 0.019 | 0 | 0.1 | 0.05 | 0.15 | 0.05 | 0.24 | 0.08 |
| 10 | 103073654 | rs11190921 | -0.536 | 0.265 | -0.649 | -0.128 | 0.003 | -0.004 | -0.004 | 0 | 0.2 | 0.21 | 0.24 | 0.19 | 0.29 | 0.09 |
| 10 | 103079985 | rs633384 | -0.348 | 0.255 | -0.89 | 0.493 | -0.001 | 0.051 | 0.051 | 0.051 | 0.88 | 0.88 | 0.88 | 0.57 | 0.87 | 0.83 |
| 10 | 103083978 | rs11190930 | -0.719 | 0.749 | -1.109 | 0.599 | -0.024 | -0.033 | -0.033 | 0 | 0.1 | 0.03 | 0.15 | 0.52 | 0.16 | 0.14 |
| 10 | 103086222 | rs11591787 | -0.82 | 0.74 | -1.068 | 0.587 | -0.017 | -0.022 | -0.022 | 0 | 0.11 | 0.05 | 0.15 | 0.52 | 0.16 | 0.14 |
| 10 | 103088683 | rs11190934 | -0.052 | 0.734 | -1.741 | -0.145 | -0.021 | 0.085 | 0.085 | 0 | 0.23 | 0.06 | 0.17 | 0.1 | 0.27 | 0.17 |
| 10 | 103095006 | rs7091918 | 1.001 | 0.928 | -0.983 | -0.555 | 0.002 | 0.025 | 0.025 | 0 | 0.53 | 0.43 | 0.54 | 0.26 | 0.46 | 0.53 |
| 10 | 103099577 | rs7081112 | 0.053 | 0.712 | -0.253 | -0.612 | 0 | -0.003 | -0.003 | 0 | 0.26 | 0.25 | 0.19 | 0.13 | 0.27 | 0.17 |
| 10 | 103101504 | rs2025106 | 0.407 | 0.949 | -0.875 | -0.879 | 0 | 0.002 | 0.002 | 0 | 0.41 | 0.41 | 0.34 | 0.29 | 0.33 | 0.5 |
| 10 | 103101769 | rs12781083 | -0.006 | 0.684 | -0.601 | -0.87 | -0.006 | 0.002 | 0.002 | 0 | 0.35 | 0.28 | 0.47 | 0.59 | 0.4 | 0.33 |
| 10 | 103111057 | rs7916830 | -0.62 | 0.996 | -0.73 | -0.961 | -0.012 | -0.021 | -0.021 | 0.013 | 0.36 | 0.47 | 0.41 | 0.69 | 0.33 | 0.38 |
| 10 | 103113035 | rs10883617 | 0.602 | 1.337 | -1.442 | -1.08 | -0.044 | 0.073 | 0.073 | 0 | 0.27 | 0.13 | 0.23 | 0.13 | 0.35 | 0.2 |
| 10 | 103156493 | rs11190990 | 0.099 | 1.233 | -1.481 | -1.121 | -0.066 | 0.168 | 0.168 | 0 | 0.4 | 0.16 | 0.35 | 0.13 | 0.46 | 0.23 |
| 10 | 103164342 | rs10786634 | 0.439 | 1.378 | -1.542 | -1.142 | -0.037 | 0.194 | 0.194 | 0 | 0.37 | 0.07 | 0.3 | 0.13 | 0.38 | 0.19 |
| 10 | 103191124 | rs4919545 | 1.803 | 1.248 | -1.6 | -1.146 | -0.074 | 0.06 | 0.06 | 0 | 0.33 | 0.05 | 0.33 | 0.18 | 0.32 | 0.42 |
| 10 | 103201263 | rs7909404 | 1.186 | 1.265 | -1.431 | -0.916 | 0 | 0.084 | 0.084 | 0 | 0.07 | 0.07 | 0.14 | 0.51 | 0.14 | 0.14 |
| 10 | 103205272 | rs4451650 | -1.773 | 1.284 | -1.352 | -1.085 | 0.002 | 0.066 | 0.066 | 0 | 0.65 | 0.83 | 0.66 | 0.83 | 0.69 | 0.57 |
| 10 | 103247208 | rs9419913 | 0.705 | 2.004 | -1.272 | -1.123 | -0.026 | 0.098 | 0.098 | 0 | 0.25 | 0.02 | 0.18 | 0.11 | 0.19 | 0.12 |
| 10 | 103261414 | rs9420839 | -0.555 | 2.018 | -1.243 | -1.137 | -0.083 | 0.187 | 0.187 | 0 | 0.4 | 0.14 | 0.35 | 0.14 | 0.47 | 0.24 |
| 10 | 103281308 | rs9419923 | 0.722 | 2.099 | -1.563 | -1.102 | -0.015 | 0.105 | 0.105 | 0 | 0.25 | 0.03 | 0.18 | 0.11 | 0.19 | 0.12 |
| 10 | 103315801 | rs1045232 | 0.345 | 1.882 | -1.565 | -1.093 | -0.076 | 0.177 | 0.177 | 0 | 0.6 | 0.84 | 0.65 | 0.86 | 0.53 | 0.76 |
| 10 | 103318303 | rs3127245 | -1.011 | 2.042 | -1.469 | -0.796 | -0.02 | 0.103 | 0.103 | 0 | 0.76 | 0.97 | 0.82 | 0.89 | 0.8 | 0.88 |
| 10 | 103320642 | rs918263 | 0.128 | 1.839 | -1.348 | -0.683 | -0.081 | 0.182 | 0.182 | 0 | 0.6 | 0.85 | 0.65 | 0.86 | 0.53 | 0.76 |
| 10 | 103336402 | rs4919559 | 0.467 | 1.819 | -1.369 | -0.699 | -0.04 | 0.179 | 0.179 | 0 | 0.55 | 0.82 | 0.67 | 0.84 | 0.49 | 0.74 |
| 10 | 103337764 | rs3730484 | -0.208 | 1.765 | -1.153 | -0.696 | -0.014 | 0 | 0 | 0 | 0.2 | 0.16 | 0.18 | 0.04 | 0.3 | 0.14 |
| 10 | 103338008 | rs1549066 | -0.712 | 1.823 | -1.177 | -0.268 | -0.02 | -0.059 | -0.059 | 2.226 | 0.69 | 0.89 | 0.7 | 0.43 | 0.66 | 0.75 |
| 10 | 103338733 | rs1055362 | -0.498 | 2.064 | -0.599 | -0.002 | -0.014 | 0 | 0 | 0 | 0.2 | 0.16 | 0.18 | 0.04 | 0.3 | 0.14 |
| 10 | 103340056 | rs3730477 | -0.049 | 2.214 | -0.608 | -0.017 | -0.024 | 0.078 | 0.078 | 0 | 0.17 | 0.04 | 0.14 | 0.03 | 0.22 | 0.1 |
| 10 | 103340144 | rs3730476 | -0.594 | 2.222 | -0.603 | -0.013 | -0.014 | 0 | 0 | 0 | 0.2 | 0.16 | 0.18 | 0.04 | 0.3 | 0.14 |
| 10 | 103340918 | rs1549067 | -1.063 | 2.166 | -0.318 | -0.037 | 0.031 | -0.099 | -0.099 | 0.084 | 0.23 | 0.35 | 0.2 | 0.04 | 0.3 | 0.14 |
| 10 | 103361088 | rs7874 | -0.115 | 1.841 | -0.654 | 0.103 | -0.029 | 0.073 | 0.073 | 0 | 0.16 | 0.04 | 0.14 | 0.03 | 0.22 | 0.1 |
| 10 | 103367824 | rs11191064 | -1.294 | 1.546 | -0.189 | 0.935 | -0.021 | 0.113 | 0.113 | 0 | 0.14 | 0.26 | 0.21 | 0.28 | 0.11 | 0.14 |
| 10 | 103368654 | rs7006 | -1.483 | 1.874 | -0.044 | 0.391 | 0.01 | -0.03 | -0.03 | 0 | 0.23 | 0.29 | 0.22 | 0.16 | 0.31 | 0.13 |
| 10 | 103370234 | rs3127255 | -1.427 | 1.918 | -0.351 | 0.41 | 0.001 | -0.005 | -0.005 | 0.913 | 0.22 | 0.22 | 0.22 | 0.16 | 0.31 | 0.13 |
| 10 | 103390956 | rs9651449 | -1.328 | 2.136 | -0.314 | 0.087 | -0.005 | -0.012 | -0.012 | 0 | 0.19 | 0.16 | 0.16 | 0.04 | 0.27 | 0.15 |
| 10 | 103392503 | rs3127230 | -1.828 | 2.163 | -0.119 | 0.208 | 0.035 | -0.054 | -0.054 | 0.018 | 0.25 | 0.34 | 0.26 | 0.16 | 0.33 | 0.16 |
| 10 | 103398834 | rs10883668 | -0.465 | 2.161 | -0.234 | 0.183 | -0.014 | 0.056 | 0.056 | 0 | 0.16 | 0.07 | 0.14 | 0.03 | 0.21 | 0.1 |
| 10 | 103445325 | rs10883676 | -0.378 | 2.274 | -0.276 | 0.185 | -0.02 | 0.046 | 0.046 | 0 | 0.14 | 0.05 | 0.13 | 0.03 | 0.2 | 0.1 |
| 10 | 103452645 | rs4244347 | 1.087 | 2.163 | -0.507 | 0.253 | 0.023 | -0.018 | -0.018 | 0 | 0.72 | 0.67 | 0.64 | 0.78 | 0.62 | 0.71 |
| 10 | 103455809 | rs6584444 | 1.11 | 2.176 | -0.396 | 0.427 | 0.044 | -0.026 | -0.026 | 0 | 0.77 | 0.65 | 0.67 | 0.82 | 0.67 | 0.81 |
| 10 | 103488124 | rs6584453 | -0.197 | 1.575 | -0.977 | -0.159 | 0.021 | 0.075 | 0.075 | 0.02 | 0.46 | 0.28 | 0.52 | 0.68 | 0.6 | 0.4 |
| 10 | 103519784 | rs749694 | -0.171 | 1.745 | -0.794 | -0.218 | -0.037 | 0.07 | 0.07 | 0 | 0.41 | 0.23 | 0.45 | 0.74 | 0.53 | 0.34 |
| 10 | 103527782 | rs17697512 | 1.76 | 1.6 | -0.788 | -0.363 | 0.001 | -0.007 | -0.007 | 0 | 0.07 | 0.04 | 0.04 | 0 | 0.06 | 0.02 |
| 10 | 103528516 | rs3218244 | 1.719 | 1.597 | -0.754 | -0.365 | 0 | -0.012 | -0.012 | 0.198 | 0.07 | 0.05 | 0.05 | 0 | 0.06 | 0.02 |
| 10 | 103528987 | rs3218240 | 1.64 | 1.594 | -0.709 | -0.366 | -0.005 | -0.029 | -0.029 | 1.809 | 0.07 | 0.09 | 0.05 | 0 | 0.06 | 0.02 |
| 10 | 103537545 | rs1348870 | -0.726 | 1.526 | -0.476 | -0.439 | 0.009 | 0.023 | 0.023 | 0 | 0.5 | 0.44 | 0.53 | 0.75 | 0.6 | 0.36 |
| 10 | 103550604 | rs1124714 | 1.211 | 1.735 | -0.611 | -0.174 | 0.006 | -0.012 | -0.012 | 0 | 0.25 | 0.21 | 0.2 | 0.51 | 0.19 | 0.27 |
| 10 | 103555611 | rs11191116 | -0.326 | 1.767 | -0.661 | -0.196 | -0.032 | -0.014 | -0.014 | 0.434 | 0.19 | 0.15 | 0.29 | 0.24 | 0.34 | 0.12 |
| 10 | 103558701 | rs17697840 | 1.952 | 1.736 | -0.696 | -0.21 | -0.004 | -0.026 | -0.026 | 0 | 0.07 | 0.08 | 0.05 | 0 | 0.06 | 0.02 |
| 10 | 103573222 | rs11191118 | 1.91 | 1.004 | -1.11 | -0.554 | -0.117 | 0.133 | 0.133 | 0.09 | 0.36 | 0.02 | 0.37 | 0.14 | 0.37 | 0.46 |
| 10 | 103586356 | rs10883689 | 0.406 | 1.724 | -0.843 | -0.304 | -0.073 | -0.059 | -0.059 | 0.128 | 0.15 | 0.03 | 0.25 | 0.22 | 0.28 | 0.09 |
| 10 | 103586429 | rs3209554 | 1.359 | 1.593 | -0.77 | -0.29 | 0.006 | -0.013 | -0.013 | 0.158 | 0.24 | 0.19 | 0.19 | 0.52 | 0.19 | 0.27 |
| 10 | 103587552 | rs2305189 | 1.36 | 1.593 | -0.77 | -0.298 | 0.006 | -0.013 | -0.013 | 0 | 0.24 | 0.19 | 0.19 | 0.51 | 0.19 | 0.27 |
| 10 | 103600517 | rs17094373 | 1.998 | 1.619 | -0.771 | -0.308 | 0.002 | -0.002 | -0.002 | 0 | 0.07 | 0.03 | 0.04 | 0 | 0.06 | 0.02 |
| 10 | 103603162 | rs12413444 | 0.408 | 1.731 | -0.848 | -0.302 | -0.077 | -0.063 | -0.063 | 0 | 0.15 | 0.02 | 0.25 | 0.22 | 0.28 | 0.09 |
| 10 | 103605714 | rs1044258 | -0.369 | 1.89 | -0.859 | -0.301 | -0.094 | -0.051 | -0.051 | 0 | 0.19 | 0.06 | 0.27 | 0.24 | 0.34 | 0.12 |
| 10 | 103618033 | rs1530589 | 0.719 | 1.912 | -0.429 | -0.11 | 0.001 | -0.003 | -0.003 | 0 | 0.32 | 0.28 | 0.25 | 0.47 | 0.25 | 0.29 |
| 10 | 103668286 | rs10786652 | -1.074 | 1.946 | -0.409 | -0.297 | 0.006 | 0.018 | 0.018 | 0 | 0.51 | 0.44 | 0.53 | 0.75 | 0.6 | 0.44 |
| 10 | 103683213 | rs10786655 | 0.18 | 2.172 | -0.362 | -0.245 | -0.001 | 0.003 | 0.003 | 0 | 0.35 | 0.35 | 0.28 | 0.52 | 0.32 | 0.34 |

| | | | | | | | | | | | | | | | | |
|---|---|---|---|---|---|---|---|---|---|---|---|---|---|---|---|---|
| 10 | 103690570 | rs11191150 | 1.42 | 2.153 | -0.389 | -0.264 | 0.008 | -0.018 | -0.018 | 0 | 0.24 | 0.17 | 0.19 | 0.51 | 0.19 | 0.27 |
| 10 | 103690669 | rs10748815 | -0.957 | 2.302 | -0.442 | -0.242 | 0.008 | 0.021 | 0.021 | 0 | 0.51 | 0.44 | 0.53 | 0.75 | 0.6 | 0.44 |
| 10 | 103702763 | rs11191156 | -0.735 | 2.342 | -0.432 | -0.236 | -0.06 | -0.025 | -0.025 | 0 | 0.19 | 0.11 | 0.28 | 0.23 | 0.34 | 0.15 |
| 10 | 103703929 | rs11191157 | 0.458 | 2.338 | -0.321 | -0.238 | -0.048 | -0.036 | -0.036 | 0 | 0.15 | 0.05 | 0.24 | 0.22 | 0.28 | 0.09 |
| 10 | 103721210 | rs3802682 | -1.07 | 2.336 | -0.63 | -0.234 | 0.005 | 0.015 | 0.015 | 0 | 0.52 | 0.43 | 0.53 | 0.75 | 0.6 | 0.44 |
| 10 | 103721697 | rs10509754 | 0.761 | 2.281 | -0.679 | -0.219 | -0.001 | 0.001 | 0.001 | 0 | 0.32 | 0.29 | 0.25 | 0.51 | 0.26 | 0.29 |
| 10 | 103726528 | rs11191161 | 1.07 | 2.416 | -0.764 | -0.144 | -0.053 | 0.009 | 0.009 | 0 | 0.11 | 0.01 | 0.17 | 0.07 | 0.17 | 0.07 |
| 10 | 103747138 | rs10883701 | 0.219 | 2.779 | -0.76 | -0.12 | -0.067 | -0.05 | -0.05 | 0 | 0.16 | 0.06 | 0.25 | 0.22 | 0.28 | 0.09 |
| 10 | 103823135 | rs2786847 | -0.918 | 2.564 | -0.607 | -0.08 | 0.004 | 0.008 | 0.008 | 0 | 0.52 | 0.45 | 0.53 | 0.75 | 0.6 | 0.47 |
| 10 | 103827737 | rs3816 | 0.572 | 1.48 | -0.435 | 0.575 | 0.005 | -0.008 | -0.008 | 0.401 | 0.25 | 0.2 | 0.2 | 0.51 | 0.19 | 0.3 |
| 10 | 103840987 | rs12776515 | -0.889 | 2.232 | -0.585 | 0.528 | -0.013 | -0.021 | -0.021 | 0 | 0.09 | 0.06 | 0.13 | 0.16 | 0.26 | 0.1 |
| 10 | 103862419 | rs2152903 | -0.286 | 1.13 | 0.193 | 0.298 | 0.015 | 0.006 | 0.006 | 0 | 0.06 | 0.35 | 0.04 | 0 | 0.02 | 0.02 |
| 10 | 103880309 | rs12768433 | 0.205 | 1.051 | -0.669 | 0.033 | -0.099 | -0.023 | -0.023 | 0.003 | 0.19 | 0.05 | 0.18 | 0.18 | 0.29 | 0.1 |
| 10 | 103885557 | rs10883710 | 0.809 | 0.324 | -1.566 | -0.523 | -0.106 | 0.136 | 0.136 | 0 | 0.36 | 0.04 | 0.42 | 0.14 | 0.42 | 0.44 |
| 10 | 103891974 | rs2815401 | -2.37 | -0.358 | -1.708 | -0.068 | -0.007 | -0.006 | -0.006 | 0 | 0.84 | 0.98 | 0.92 | 0.86 | 0.85 | 0.85 |
| 10 | 103895979 | rs17114381 | 0.535 | -0.461 | -1.188 | -0.145 | 0.005 | -0.003 | -0.003 | 0 | 0.06 | 0.32 | 0.04 | 0 | 0.02 | 0.02 |
| 10 | 103905779 | rs4919613 | -1.152 | -0.706 | -0.807 | 0.489 | -0.016 | 0.052 | 0.052 | 0 | 0.74 | 0.68 | 0.8 | 0.49 | 0.79 | 0.66 |
| 10 | 103907680 | rs10509755 | 0.76 | -0.428 | -0.432 | 0.574 | -0.019 | -0.053 | -0.053 | 0 | 0.09 | 0.22 | 0.06 | 0 | 0.06 | 0.01 |
| 10 | 103913375 | rs17114406 | 0.728 | -0.424 | -0.431 | 0.295 | -0.02 | -0.055 | -0.055 | 0.928 | 0.1 | 0.23 | 0.06 | 0 | 0.06 | 0.01 |
| 10 | 103920475 | rs1049455 | -0.337 | -0.475 | -0.372 | 0.263 | -0.063 | -0.117 | -0.117 | 0 | 0.16 | 0.55 | 0.11 | 0 | 0.08 | 0.04 |
| 10 | 103922896 | rs7897 | -1.015 | -0.542 | -0.667 | 0.216 | -0.011 | 0.018 | 0.018 | 0 | 0.43 | 0.39 | 0.46 | 0.66 | 0.46 | 0.43 |
| 10 | 103930410 | rs7068663 | 0.601 | -0.281 | -0.861 | -0.596 | -0.022 | -0.073 | -0.073 | 0 | 0.12 | 0.43 | 0.08 | 0 | 0.07 | 0.01 |
| 10 | 103931931 | rs4917962 | 0.426 | -0.143 | -1.7 | -0.411 | 0.004 | -0.013 | -0.013 | 0 | 0.13 | 0.1 | 0.16 | 0.38 | 0.11 | 0.25 |
| 10 | 103931947 | rs4919614 | -1.376 | -0.003 | -2 | -0.642 | -0.087 | 0.139 | 0.139 | 0 | 0.53 | 0.94 | 0.55 | 0.74 | 0.49 | 0.46 |
| 10 | 103940629 | rs11598856 | 0.809 | 0.589 | -1.635 | -0.452 | -0.09 | -0.022 | -0.022 | 0 | 0.18 | 0.03 | 0.16 | 0.16 | 0.25 | 0.08 |
| 10 | 103983087 | rs11191242 | -1.474 | 0.094 | -1.807 | -0.775 | -0.045 | 0.095 | 0.095 | 0 | 0.56 | 0.85 | 0.56 | 0.72 | 0.47 | 0.48 |
| 10 | 103984060 | rs7083450 | -2.622 | 0.157 | -2.037 | -0.938 | -0.037 | 0.013 | 0.013 | 0 | 0.82 | 0.99 | 0.91 | 0.87 | 0.81 | 0.84 |
| 10 | 103987875 | rs1410416 | -1.05 | 0.003 | -1.637 | -0.975 | 0.005 | 0.076 | 0.076 | 0 | 0.52 | 0.66 | 0.52 | 0.71 | 0.46 | 0.45 |
| 10 | 103987986 | rs10748816 | -1.372 | -0.017 | -1.686 | -0.968 | -0.046 | 0.025 | 0.025 | 0 | 0.56 | 0.85 | 0.56 | 0.65 | 0.47 | 0.48 |
| 10 | 103992122 | rs1854291 | -1.609 | 0.27 | -1.695 | -1.27 | -0.014 | 0.09 | 0.09 | 0 | 0.6 | 0.79 | 0.63 | 0.77 | 0.6 | 0.5 |
| 10 | 103997645 | rs1541047 | -1.625 | 0.275 | -1.697 | -1.272 | -0.017 | 0.088 | 0.088 | 0 | 0.6 | 0.79 | 0.62 | 0.77 | 0.6 | 0.5 |
| 10 | 104004195 | rs3758549 | 1.236 | 0.352 | -1.779 | -1.31 | -0.059 | -0.007 | -0.007 | 0.031 | 0.17 | 0.03 | 0.15 | 0.15 | 0.2 | 0.07 |
| 10 | 104005386 | rs1541046 | -1.625 | 0.384 | -1.682 | -1.229 | -0.015 | 0.092 | 0.092 | 0 | 0.59 | 0.79 | 0.62 | 0.77 | 0.6 | 0.5 |
| 10 | 104029307 | rs1628530 | 1.826 | 0.512 | -1.724 | -1.26 | -0.034 | -0.033 | -0.033 | 1.03 | 0.06 | 0.01 | 0.11 | 0.12 | 0.13 | 0.02 |
| 10 | 104047836 | rs17707508 | 1.712 | 0.464 | -1.577 | -1.257 | -0.006 | 0.049 | 0.049 | 0 | 0.22 | 0.13 | 0.28 | 0.1 | 0.21 | 0.34 |
| 10 | 104063367 | rs772012 | 1.376 | 0.487 | -1.485 | -1.162 | -0.015 | 0.092 | 0.092 | 0 | 0.41 | 0.22 | 0.38 | 0.23 | 0.4 | 0.5 |
| 10 | 104084889 | rs10444022 | 1.754 | 0.505 | -1.501 | -0.994 | -0.021 | -0.022 | -0.022 | 0.733 | 0.08 | 0.01 | 0.16 | 0.25 | 0.12 | 0.12 |
| 10 | 104085921 | rs945781 | 0.737 | 0.401 | -1.476 | -1.091 | 0.014 | -0.079 | -0.079 | 1.399 | 0.18 | 0.07 | 0.18 | 0.37 | 0.14 | 0.26 |
| 10 | 104127171 | rs2273555 | -1.272 | 0.323 | -1.321 | -1.187 | 0.004 | 0.027 | 0.027 | 0 | 0.55 | 0.57 | 0.59 | 0.77 | 0.58 | 0.47 |
| 10 | 104157711 | rs7897947 | 0.492 | 0.766 | -1.175 | -1.106 | -0.009 | 0.061 | 0.061 | 0 | 0.22 | 0.38 | 0.21 | 0.37 | 0.19 | 0.26 |
| 10 | 104159696 | rs11574849 | 1.544 | 0.656 | -1.397 | -1.173 | -0.015 | -0.047 | -0.047 | 0.481 | 0.08 | 0.15 | 0.07 | 0 | 0.06 | 0.01 |
| 10 | 104160959 | rs11574851 | 2.025 | 0.704 | -2.301 | -1.084 | -0.002 | 0.017 | 0.017 | 0.215 | 0.09 | 0.01 | 0.02 | 0.06 | 0.05 | 0.03 |
| 10 | 104162770 | rs1056890 | 1.131 | 0.709 | -2.162 | -1.018 | -0.005 | 0.125 | 0.125 | 0 | 0.39 | 0.12 | 0.35 | 0.2 | 0.35 | 0.5 |
| 10 | 104170204 | rs12260430 | 0.299 | 0.743 | -2.088 | -1.077 | -0.017 | 0.073 | 0.073 | 0.003 | 0.23 | 0.43 | 0.22 | 0.37 | 0.19 | 0.3 |
| 10 | 104174895 | rs2282295 | 1.592 | 0.928 | -2.926 | -0.973 | -0.029 | -0.009 | -0.009 | 0.556 | 0.07 | 0.01 | 0.08 | 0.37 | 0.09 | 0.19 |
| 10 | 104184081 | rs1044476 | 2.21 | 0.879 | -2.787 | -0.869 | -0.005 | 0.035 | 0.035 | 0 | 0.17 | 0.01 | 0.08 | 0.09 | 0.15 | 0.11 |
| 10 | 104184981 | rs11593146 | 0.793 | 0.926 | -2.744 | -0.827 | -0.005 | 0.136 | 0.136 | 0 | 0.39 | 0.14 | 0.35 | 0.18 | 0.36 | 0.47 |
| 10 | 104186829 | rs11594302 | 1.19 | 0.878 | -2.282 | -0.387 | -0.05 | -0.039 | -0.039 | 7.067 | 0.16 | 0.02 | 0.13 | 0.18 | 0.18 | 0.05 |
| 10 | 104192567 | rs9665587 | 0.94 | 0.69 | -2.273 | -0.401 | -0.016 | 0.025 | 0.025 | 0 | 0.27 | 0.39 | 0.22 | 0.37 | 0.21 | 0.38 |
| 10 | 104192691 | rs11592718 | 0.934 | 1.505 | -3.319 | 0.02 | -0.012 | 0.038 | 0.038 | 0 | 0.17 | 0.11 | 0.26 | 0.09 | 0.18 | 0.26 |
| 10 | 104195202 | rs1536309 | 0.874 | 1.45 | -2.888 | 0.427 | -0.031 | 0.113 | 0.113 | 0 | 0.38 | 0.2 | 0.38 | 0.18 | 0.39 | 0.33 |
| 10 | 104201995 | rs11597058 | -0.243 | 1.902 | -2.043 | 1.307 | -0.063 | -0.039 | -0.039 | 0 | 0.17 | 0.02 | 0.14 | 0.18 | 0.2 | 0.05 |
| 10 | 104204140 | rs2863707 | -0.424 | 1.369 | -1.935 | 1.423 | -0.015 | -0.034 | -0.034 | 0 | 0.09 | 0.01 | 0.16 | 0.27 | 0.12 | 0.19 |
| 10 | 104207799 | rs927821 | 0.242 | 1.234 | -1.549 | 0.518 | -0.007 | -0.008 | -0.008 | 0 | 0.82 | 0.93 | 0.85 | 0.82 | 0.81 | 0.93 |
| 10 | 104213502 | rs11191291 | -0.64 | 1.832 | -0.986 | 0.241 | 0.005 | -0.012 | -0.012 | 0 | 0.15 | 0.19 | 0.21 | 0.09 | 0.19 | 0.17 |
| 10 | 104217415 | rs4919640 | -0.268 | 1.944 | -1.616 | 0.926 | -0.03 | -0.085 | -0.085 | 0 | 0.16 | 0.07 | 0.25 | 0.41 | 0.2 | 0.29 |
| 10 | 104219132 | rs10786667 | -0.508 | 1.53 | -1.48 | 0.144 | 0 | 0.013 | 0.013 | 1.12 | 0.48 | 0.56 | 0.56 | 0.76 | 0.49 | 0.63 |
| 10 | 104220301 | rs7075281 | 0.452 | 1.307 | -0.777 | 0.323 | 0 | 0.002 | 0.002 | 1.172 | 0.31 | 0.31 | 0.27 | 0.3 | 0.3 | 0.37 |
| 10 | 104220576 | rs4917968 | 0.343 | 1.196 | -0.882 | 0.398 | -0.03 | -0.098 | -0.098 | 0 | 0.14 | 0.04 | 0.26 | 0.41 | 0.18 | 0.25 |
| 10 | 104221539 | rs11191293 | 0.526 | 1.171 | -0.617 | 0.202 | -0.021 | 0.156 | 0.156 | 1.362 | 0.5 | 0.21 | 0.44 | 0.26 | 0.51 | 0.36 |
| 10 | 104225832 | rs10883723 | -0.287 | 1.293 | -0.731 | -0.067 | -0.015 | -0.006 | -0.006 | 0.189 | 0.25 | 0.16 | 0.35 | 0.26 | 0.31 | 0.29 |
| 10 | 104230012 | rs3808937 | 1.407 | 1.534 | -0.95 | -0.434 | -0.014 | 0.054 | 0.054 | 0 | 0.2 | 0.08 | 0.14 | 0.05 | 0.2 | 0.09 |
| 10 | 104230156 | rs2025713 | 0.641 | 1.43 | -0.377 | -0.423 | -0.004 | 0.015 | 0.015 | 0 | 0.58 | 0.37 | 0.37 | 0.31 | 0.56 | 0.44 |
| 10 | 104231054 | rs11191295 | 0.627 | 1.425 | -0.385 | -0.425 | -0.004 | 0.013 | 0.013 | 0 | 0.58 | 0.37 | 0.37 | 0.31 | 0.56 | 0.44 |
| 10 | 104236481 | rs1056738 | -0.101 | 1.431 | -0.328 | -0.422 | -0.009 | -0.003 | -0.003 | 0.149 | 0.25 | 0.19 | 0.36 | 0.26 | 0.31 | 0.3 |
| 10 | 104236542 | rs1056739 | -0.252 | 1.43 | -0.311 | -0.423 | -0.008 | -0.002 | -0.002 | 0 | 0.26 | 0.2 | 0.36 | 0.26 | 0.31 | 0.3 |
| 10 | 104236552 | rs1056740 | -0.265 | 1.43 | -0.311 | -0.423 | -0.008 | -0.002 | -0.002 | 0.061 | 0.26 | 0.2 | 0.36 | 0.26 | 0.31 | 0.3 |
| 10 | 104236687 | rs1056744 | 0.446 | 1.42 | -0.184 | -0.444 | 0.005 | -0.012 | -0.012 | 0 | 0.6 | 0.55 | 0.39 | 0.31 | 0.57 | 0.44 |
| 10 | 104239100 | rs5870 | 0.494 | 1.409 | -0.144 | -0.445 | 0.003 | -0.011 | -0.011 | 0 | 0.59 | 0.49 | 0.38 | 0.31 | 0.57 | 0.46 |

| | | | | | | | | | | | | | | | | |
|---|---|---|---|---|---|---|---|---|---|---|---|---|---|---|---|---|
| 10 | 104240438 | rs13287 | -0.488 | 1.419 | -0.096 | -0.438 | 0.012 | -0.004 | -0.004 | 0 | 0.28 | 0.38 | 0.37 | 0.26 | 0.3 | 0.3 |
| 10 | 104241683 | rs2296580 | 0.526 | 1.371 | -0.127 | -0.483 | -0.028 | 0.049 | 0.049 | 0.946 | 0.27 | 0.12 | 0.18 | 0.13 | 0.3 | 0.33 |
| 10 | 104242886 | rs2296581 | -0.22 | 1.424 | -0.017 | -0.404 | -0.007 | -0.001 | -0.001 | 0.06 | 0.26 | 0.2 | 0.36 | 0.26 | 0.3 | 0.3 |
| 10 | 104245588 | rs2296587 | 0.434 | 1.42 | -0.051 | -0.404 | -0.097 | 0.093 | 0.093 | 0 | 0.21 | 0.02 | 0.26 | 0.08 | 0.28 | 0.27 |
| 10 | 104262628 | rs2296590 | -0.456 | 1.45 | 0.008 | -0.397 | 0.013 | -0.004 | -0.004 | 0.02 | 0.28 | 0.37 | 0.36 | 0.26 | 0.3 | 0.3 |
| 10 | 104264107 | rs2274351 | 0.681 | 1.438 | -0.031 | -0.393 | -0.006 | 0.028 | 0.028 | 0 | 0.58 | 0.3 | 0.36 | 0.31 | 0.57 | 0.46 |
| 10 | 104297791 | rs2031604 | -1.626 | 1.381 | -0.04 | -0.3 | 0.003 | 0.009 | 0.009 | 0 | 0.88 | 0.89 | 0.76 | 0.56 | 0.9 | 0.76 |
| 10 | 104318966 | rs7893954 | -0.732 | 1.673 | -0.088 | -0.463 | -0.023 | -0.002 | -0.002 | 0 | 0.33 | 0.42 | 0.53 | 0.68 | 0.27 | 0.38 |
| 10 | 104343428 | rs7475335 | 0.374 | 1.378 | -0.262 | -0.572 | -0.035 | 0.063 | 0.063 | 0.263 | 0.3 | 0.15 | 0.21 | 0.13 | 0.35 | 0.34 |
| 10 | 104386934 | rs17114803 | 1.159 | 1.217 | -0.042 | -0.441 | -0.013 | 0.109 | 0.109 | 0 | 0.13 | 0.23 | 0.24 | 0.43 | 0.09 | 0.24 |
| 10 | 104391034 | rs11596235 | 0.663 | 1.249 | -0.306 | -0.686 | -0.029 | 0.028 | 0.028 | 0 | 0.24 | 0.15 | 0.19 | 0.13 | 0.32 | 0.2 |
| 10 | 104392497 | rs11191355 | 0.956 | 1.376 | -0.293 | -0.677 | 0.011 | 0.16 | 0.16 | 0 | 0.15 | 0.27 | 0.27 | 0.52 | 0.17 | 0.33 |
| 10 | 104392580 | rs11594179 | 1.356 | 1.36 | -0.678 | -0.742 | -0.028 | 0.054 | 0.054 | 0 | 0.2 | 0.08 | 0.16 | 0.05 | 0.24 | 0.11 |
| 10 | 104395085 | rs2001595 | 0.216 | 1.272 | -0.729 | -0.672 | -0.064 | 0.095 | 0.095 | 0 | 0.22 | 0.04 | 0.26 | 0.08 | 0.26 | 0.26 |
| 10 | 104401632 | rs4146429 | 1.533 | 1.169 | -0.673 | -0.554 | 0.007 | -0.036 | -0.036 | 6.682 | 0.12 | 0.08 | 0.23 | 0.43 | 0.09 | 0.24 |
| 10 | 104412049 | rs10883759 | -0.155 | 1.556 | -0.621 | -0.768 | 0.002 | 0 | 0 | 0 | 0.36 | 0.29 | 0.29 | 0.32 | 0.33 | 0.37 |
| 10 | 104412308 | rs10786701 | 1.699 | 1.524 | -0.643 | -0.755 | 0.006 | -0.035 | -0.035 | 0 | 0.12 | 0.08 | 0.23 | 0.43 | 0.09 | 0.24 |
| 10 | 104414221 | rs3850699 | 1.725 | 1.133 | -0.255 | -0.978 | 0.004 | -0.037 | -0.037 | 0.28 | 0.29 | 0.39 | 0.2 | 0.17 | 0.3 | 0.12 |
| 10 | 104418656 | rs7923415 | -0.62 | 1.062 | -0.196 | -0.854 | 0.001 | -0.003 | -0.003 | 0.035 | 0.47 | 0.48 | 0.52 | 0.75 | 0.4 | 0.61 |
| 10 | 104434194 | rs7088470 | 0.745 | 1.034 | 0.04 | -0.485 | -0.005 | 0.173 | 0.173 | 0 | 0.06 | 0.17 | 0.1 | 0.21 | 0.02 | 0.03 |
| 10 | 104434630 | rs12573077 | 1.355 | 1.076 | -0.094 | -0.675 | 0.005 | -0.037 | -0.037 | 0.745 | 0.3 | 0.4 | 0.21 | 0.16 | 0.32 | 0.12 |
| 10 | 104440050 | rs7907503 | 1.343 | 0.932 | -0.048 | -0.692 | 0.01 | -0.034 | -0.034 | 0.067 | 0.28 | 0.37 | 0.21 | 0.15 | 0.33 | 0.12 |
| 10 | 104442457 | rs3781571 | 0.271 | 0.797 | -0.059 | -0.828 | -0.001 | -0.019 | -0.019 | 0 | 0.3 | 0.17 | 0.26 | 0.33 | 0.31 | 0.25 |
| 10 | 104447629 | rs3977736 | 1.286 | 0.884 | -0.28 | -0.855 | -0.003 | 0.064 | 0.064 | 0 | 0.21 | 0.07 | 0.15 | 0.08 | 0.23 | 0.12 |
| 10 | 104456725 | rs7904252 | 1.431 | 1.054 | -0.174 | -0.762 | 0.01 | -0.038 | -0.038 | 0 | 0.29 | 0.38 | 0.21 | 0.15 | 0.32 | 0.12 |
| 10 | 104456838 | rs7904396 | 1.422 | 1.046 | -0.205 | -0.765 | 0.008 | -0.034 | -0.034 | 0 | 0.29 | 0.37 | 0.21 | 0.15 | 0.32 | 0.12 |
| 10 | 104464763 | rs10883766 | 1.471 | 1.011 | -0.491 | -0.675 | 0.009 | -0.055 | -0.055 | 0 | 0.13 | 0.05 | 0.23 | 0.45 | 0.1 | 0.26 |
| 10 | 104477705 | rs10786705 | 0.481 | 1.036 | -0.459 | -0.663 | -0.002 | -0.016 | -0.016 | 0.327 | 0.29 | 0.18 | 0.26 | 0.33 | 0.31 | 0.25 |
| 10 | 104485301 | rs12775883 | 1.305 | 1.032 | -0.377 | -0.687 | 0.001 | -0.009 | -0.009 | 0.633 | 0.29 | 0.28 | 0.2 | 0.15 | 0.33 | 0.12 |
| 10 | 104487382 | rs2902548 | 0.745 | 1.207 | -0.419 | -0.2 | -0.001 | 0.016 | 0.016 | 0.272 | 0.16 | 0.21 | 0.28 | 0.61 | 0.17 | 0.37 |
| 10 | 104493444 | rs11191381 | 1.648 | 1.053 | -0.952 | -0.471 | -0.089 | 0.063 | 0.063 | 0 | 0.25 | 0.03 | 0.19 | 0.13 | 0.32 | 0.09 |
| 10 | 104500659 | rs10786706 | 0.488 | 0.48 | -0.657 | -0.163 | -0.006 | 0.002 | 0.002 | 0 | 0.42 | 0.43 | 0.48 | 0.65 | 0.47 | 0.46 |
| 10 | 104504564 | rs999867 | 1.522 | 0.057 | -0.781 | 0.179 | 0.005 | -0.085 | -0.085 | 0.218 | 0.12 | 0.02 | 0.22 | 0.46 | 0.09 | 0.26 |
| 10 | 104505884 | rs11191383 | 0.629 | 0.023 | -0.65 | -0.113 | 0.009 | -0.006 | -0.006 | 0 | 0.28 | 0.19 | 0.23 | 0.27 | 0.23 | 0.15 |
| 10 | 104529668 | rs7092340 | 0.188 | 0.02 | -0.666 | -0.13 | 0.008 | 0.001 | 0.001 | 0.03 | 0.33 | 0.24 | 0.24 | 0.26 | 0.25 | 0.28 |
| 10 | 104531050 | rs12266856 | 0.138 | 0.004 | -0.614 | -0.137 | -0.001 | 0 | 0 | 0 | 0.34 | 0.29 | 0.24 | 0.26 | 0.25 | 0.28 |
| 10 | 104532913 | rs549466 | -0.544 | 0.112 | -0.667 | -0.251 | 0.024 | 0.054 | 0.054 | 0 | 0.5 | 0.62 | 0.51 | 0.79 | 0.4 | 0.64 |
| 10 | 104546284 | rs486955 | -0.971 | 0.089 | -0.404 | 0.22 | -0.016 | 0.141 | 0.141 | 0 | 0.86 | 0.68 | 0.77 | 0.52 | 0.89 | 0.68 |
| 10 | 104546467 | rs1475643 | 0.752 | 0.323 | -0.67 | -0.186 | 0.012 | -0.005 | -0.005 | 0 | 0.57 | 0.3 | 0.47 | 0.43 | 0.6 | 0.28 |
| 10 | 104568018 | rs2790914 | -0.087 | 0.413 | -0.607 | -0.199 | 0.005 | -0.029 | -0.029 | 0.021 | 0.22 | 0.22 | 0.28 | 0.09 | 0.27 | 0.27 |
| 10 | 104572276 | rs284861 | -1.179 | 0.204 | -0.627 | 0.044 | -0.021 | 0.134 | 0.134 | 0 | 0.85 | 0.67 | 0.82 | 0.58 | 0.88 | 0.72 |
| 10 | 104572881 | rs17115061 | 1.375 | 0.124 | -0.596 | -0.033 | -0.014 | -0.032 | -0.032 | 0.802 | 0.06 | 0.09 | 0.01 | 0.01 | 0.02 | 0.13 |
| 10 | 104572963 | rs284860 | -0.92 | 0.243 | -1.011 | -0.232 | 0 | 0.001 | 0.001 | 0 | 0.61 | 0.52 | 0.58 | 0.4 | 0.6 | 0.65 |
| 10 | 104573017 | rs284859 | 1.039 | 0.247 | -1.078 | -0.325 | -0.018 | 0.032 | 0.032 | 0 | 0.16 | 0.04 | 0.22 | 0.13 | 0.19 | 0.2 |
| 10 | 104573403 | rs11191401 | 2.142 | 0.289 | -1.065 | -0.292 | -0.037 | 0.034 | 0.034 | 0 | 0.24 | 0.12 | 0.25 | 0.18 | 0.29 | 0.07 |
| 10 | 104574422 | rs284855 | -0.999 | 0.309 | -1.026 | -0.292 | 0 | 0.001 | 0.001 | 0 | 0.61 | 0.52 | 0.58 | 0.4 | 0.59 | 0.65 |
| 10 | 104575870 | rs6892 | 1.165 | 0.335 | -1.238 | -0.272 | -0.043 | 0.016 | 0.016 | 0 | 0.15 | 0.02 | 0.21 | 0.13 | 0.19 | 0.21 |
| 10 | 104581288 | rs619824 | -0.475 | 0.243 | -0.987 | -0.264 | 0.006 | 0.008 | 0.008 | 0 | 0.53 | 0.32 | 0.55 | 0.39 | 0.58 | 0.52 |
| 10 | 104583932 | rs10883782 | 1.465 | 0.138 | -1.891 | -0.205 | 0.016 | 0.018 | 0.018 | 0 | 0.21 | 0.1 | 0.14 | 0.14 | 0.18 | 0.16 |
| 10 | 104584330 | rs4919682 | 2.177 | -0.044 | -2.083 | -0.092 | -0.04 | 0.02 | 0.02 | 0 | 0.21 | 0.1 | 0.25 | 0.2 | 0.29 | 0.07 |
| 10 | 104585125 | rs4919683 | 0.075 | -0.147 | -2.105 | -0.161 | 0.009 | 0.005 | 0.005 | 0 | 0.45 | 0.33 | 0.42 | 0.61 | 0.41 | 0.5 |
| 10 | 104587362 | rs4919685 | 1.746 | -0.034 | -2.388 | -0.143 | -0.04 | 0.052 | 0.052 | 0 | 0.25 | 0.11 | 0.2 | 0.18 | 0.3 | 0.08 |
| 10 | 104591393 | rs17115100 | 1.946 | -0.143 | -2.404 | 0.127 | 0.019 | -0.044 | -0.044 | 0.45 | 0.12 | 0.05 | 0.2 | 0.35 | 0.09 | 0.26 |
| 10 | 104592249 | rs4919686 | 1.785 | -0.038 | -2.391 | 0.023 | -0.085 | 0.055 | 0.055 | 0 | 0.24 | 0.02 | 0.19 | 0.16 | 0.3 | 0.07 |
| 10 | 104595248 | rs4919687 | 1.782 | 0.021 | -2.336 | 0.135 | -0.04 | 0.01 | 0.01 | 0 | 0.25 | 0.11 | 0.2 | 0.25 | 0.3 | 0.1 |
| 10 | 104595420 | rs3781287 | 0.114 | -0.147 | -2.327 | 0.112 | 0.009 | 0.004 | 0.004 | 0 | 0.45 | 0.33 | 0.43 | 0.59 | 0.41 | 0.5 |
| 10 | 104596924 | rs6163 | 0.48 | -0.191 | -2.332 | 0.111 | -0.001 | -0.011 | -0.011 | 0 | 0.39 | 0.33 | 0.42 | 0.59 | 0.39 | 0.35 |
| 10 | 104597152 | rs743572 | 0.464 | -0.207 | -2.34 | 0.104 | -0.001 | -0.011 | -0.011 | 0 | 0.4 | 0.33 | 0.42 | 0.59 | 0.39 | 0.36 |
| 10 | 104597480 | rs2486758 | 0.902 | -0.189 | -2.519 | 0.052 | -0.034 | -0.005 | -0.005 | 0 | 0.19 | 0.04 | 0.24 | 0.19 | 0.23 | 0.25 |
| 10 | 104605328 | rs7089422 | 1.445 | 0.161 | -2.358 | 0.155 | -0.018 | -0.001 | -0.001 | 0 | 0.12 | 0.06 | 0.15 | 0.08 | 0.16 | 0.08 |
| 10 | 104610926 | rs4290163 | 0.12 | 0.115 | -2.239 | 0.268 | 0.004 | 0.001 | 0.001 | 0 | 0.44 | 0.3 | 0.42 | 0.59 | 0.41 | 0.46 |
| 10 | 104616500 | rs4919690 | 1.585 | 0.021 | -2.205 | 0.305 | -0.008 | 0.008 | 0.008 | 0 | 0.25 | 0.12 | 0.19 | 0.26 | 0.26 | 0.09 |
| 10 | 104618695 | rs7096169 | 0.581 | 0.006 | -2.207 | 0.52 | 0.014 | -0.026 | -0.026 | 0.239 | 0.38 | 0.33 | 0.4 | 0.54 | 0.35 | 0.32 |
| 10 | 104623578 | rs9527 | 1.62 | 0.032 | -2.213 | 0.5 | -0.01 | 0.045 | 0.045 | 0 | 0.25 | 0.11 | 0.19 | 0.18 | 0.25 | 0.07 |
| 10 | 104628873 | rs7085104 | 0.599 | 0.007 | -2.208 | 0.528 | 0.014 | -0.027 | -0.027 | 0 | 0.38 | 0.31 | 0.4 | 0.55 | 0.35 | 0.32 |
| 10 | 104629011 | rs12416687 | 1.568 | 0.058 | -2.188 | 0.486 | -0.01 | 0.046 | 0.046 | 0 | 0.25 | 0.11 | 0.19 | 0.18 | 0.25 | 0.07 |
| 10 | 104634107 | rs10509760 | 2.672 | 0.161 | -2.214 | 0.38 | 0.024 | 0.035 | 0.035 | 0 | 0.17 | 0.1 | 0.1 | 0.02 | 0.11 | 0.05 |
| 10 | 104637992 | rs10786719 | -0.093 | 0.254 | -2.284 | 0.407 | 0.019 | 0.002 | 0.002 | 0 | 0.45 | 0.34 | 0.42 | 0.59 | 0.42 | 0.48 |
| 10 | 104638723 | rs11191439 | 2.725 | 0.239 | -2.288 | 0.384 | 0.026 | 0.041 | 0.041 | 0 | 0.17 | 0.09 | 0.1 | 0.02 | 0.11 | 0.06 |

| | | | | | | | | | | | | | | | | |
|---|---|---|---|---|---|---|---|---|---|---|---|---|---|---|---|---|
| 10 | 104640955 | rs10883790 | 0.932 | 0.241 | -2.29 | 0.45 | 0.012 | 0.006 | 0.006 | 0 | 0.31 | 0.16 | 0.19 | 0.3 | 0.27 | 0.25 |
| 10 | 104661484 | rs1046778 | -0.529 | 0.269 | -2.257 | 0.496 | -0.004 | -0.028 | -0.028 | 0 | 0.27 | 0.24 | 0.32 | 0.49 | 0.3 | 0.4 |
| 10 | 104662458 | rs7897654 | -0.529 | 0.269 | -2.257 | 0.496 | -0.004 | -0.028 | -0.028 | 0 | 0.27 | 0.24 | 0.32 | 0.49 | 0.3 | 0.4 |
| 10 | 104671212 | rs10883800 | 0.55 | 0.218 | -2.239 | 0.501 | 0.002 | 0.01 | 0.01 | 0 | 0.34 | 0.29 | 0.22 | 0.3 | 0.33 | 0.26 |
| 10 | 104677126 | rs12221064 | 2.113 | 0.223 | -2.237 | 0.566 | 0.018 | -0.055 | -0.055 | 0 | 0.11 | 0.04 | 0.19 | 0.29 | 0.09 | 0.22 |
| 10 | 104734612 | rs6584535 | -1.411 | 0.362 | -2.487 | 0.36 | -0.062 | -0.025 | -0.025 | 0.008 | 0.81 | 0.98 | 0.75 | 0.82 | 0.77 | 0.72 |
| 10 | 104740568 | rs11191489 | 0.547 | 0.286 | -2.264 | 0.402 | 0.001 | 0.009 | 0.009 | 0 | 0.34 | 0.29 | 0.22 | 0.3 | 0.33 | 0.26 |
| 10 | 104753675 | rs11191494 | 0.312 | 0.393 | -2.214 | 0.492 | -0.011 | -0.013 | -0.013 | 0 | 0.13 | 0.07 | 0.09 | 0.2 | 0.16 | 0.17 |
| 10 | 104773364 | rs11191514 | 2.095 | 0.262 | -2.225 | 0.609 | 0.017 | -0.053 | -0.053 | 0.834 | 0.11 | 0.04 | 0.19 | 0.28 | 0.09 | 0.22 |
| 10 | 104787019 | rs12246739 | 0.703 | 0.235 | -2.282 | 0.49 | -0.002 | 0.019 | 0.019 | 0 | 0.33 | 0.29 | 0.22 | 0.13 | 0.33 | 0.24 |
| 10 | 104814162 | rs2275271 | 0.083 | 0.19 | -1.804 | 0.457 | 0.013 | 0.001 | 0.001 | 0 | 0.44 | 0.33 | 0.41 | 0.59 | 0.41 | 0.44 |
| 10 | 104817709 | rs11191537 | 1.88 | 0.158 | -2.063 | 0.244 | 0.011 | 0.009 | 0.009 | 0 | 0.19 | 0.12 | 0.14 | 0.15 | 0.15 | 0.19 |
| 10 | 104836940 | rs2296568 | 1.102 | 0.644 | -1.978 | 0.424 | -0.034 | -0.029 | -0.029 | 0 | 0.06 | 0.01 | 0.07 | 0.15 | 0.14 | 0.03 |
| 10 | 104843148 | rs11191547 | 0.707 | 0.476 | -1.978 | 0.36 | -0.001 | 0.006 | 0.006 | 0 | 0.32 | 0.3 | 0.22 | 0.22 | 0.32 | 0.2 |
| 10 | 104846797 | rs11191549 | 0.687 | 0.502 | -1.974 | 0.36 | 0 | 0.009 | 0.009 | 0 | 0.33 | 0.28 | 0.22 | 0.22 | 0.32 | 0.24 |
| 10 | 104848123 | rs8139 | 0.133 | 0.411 | -2.035 | 0.428 | 0.01 | -0.016 | -0.016 | 0.018 | 0.43 | 0.34 | 0.41 | 0.51 | 0.41 | 0.46 |
| 10 | 104849468 | rs3740387 | 0.133 | 0.411 | -2.035 | 0.43 | 0.01 | -0.012 | -0.012 | 0.041 | 0.43 | 0.34 | 0.41 | 0.5 | 0.41 | 0.46 |
| 10 | 104851396 | rs11191553 | 0.708 | 0.425 | -2.024 | 0.415 | -0.001 | 0.007 | 0.007 | 0 | 0.33 | 0.3 | 0.22 | 0.22 | 0.33 | 0.25 |
| 10 | 104855656 | rs1926030 | 0.108 | 0.313 | -2.045 | 0.475 | 0.011 | 0.001 | 0.001 | 0 | 0.44 | 0.34 | 0.42 | 0.59 | 0.41 | 0.48 |
| 10 | 104857218 | rs10786737 | 0.492 | 0.294 | -2.106 | 0.44 | -0.006 | -0.006 | -0.006 | 0.277 | 0.12 | 0.08 | 0.08 | 0.2 | 0.16 | 0.17 |
| 10 | 104869038 | rs11191560 | 2.302 | 0.295 | -2.156 | 0.451 | 0.011 | -0.067 | -0.067 | 0 | 0.11 | 0.02 | 0.19 | 0.29 | 0.09 | 0.22 |
| 10 | 104891198 | rs10883836 | 0.372 | 0.478 | -1.709 | 0.186 | -0.001 | 0.003 | 0.003 | 0 | 0.33 | 0.3 | 0.22 | 0.3 | 0.33 | 0.26 |
| 10 | 104897985 | rs11598702 | 0.513 | 0.423 | -1.747 | -0.085 | -0.022 | 0.024 | 0.024 | 0 | 0.33 | 0.18 | 0.31 | 0.23 | 0.35 | 0.24 |
| 10 | 104934709 | rs10883841 | 0.626 | 0.414 | -1.399 | -0.067 | -0.034 | -0.029 | -0.029 | 0 | 0.06 | 0.01 | 0.07 | 0.15 | 0.14 | 0.03 |
| 10 | 104939232 | rs12240508 | 1.438 | 0.308 | -1.377 | -0.11 | 0.013 | 0.038 | 0.038 | 0 | 0.2 | 0.15 | 0.13 | 0.02 | 0.17 | 0.07 |
| 10 | 104954795 | rs7095304 | 0.279 | 0.448 | -1.253 | 0.064 | -0.001 | 0.003 | 0.003 | 0 | 0.32 | 0.3 | 0.22 | 0.22 | 0.32 | 0.24 |
| 10 | 104963721 | rs11191609 | 1.149 | 0.085 | -1.031 | 0.216 | 0.003 | 0.002 | 0.002 | 0 | 0.16 | 0.12 | 0.13 | 0.15 | 0.14 | 0.13 |
| 10 | 105001510 | rs1891292 | -0.964 | 0.061 | -0.338 | 0.135 | -0.001 | 0 | 0 | 0 | 0.44 | 0.51 | 0.55 | 0.52 | 0.53 | 0.32 |
| 10 | 105013147 | rs4918001 | 2.211 | -0.334 | -0.496 | -0.14 | 0.015 | 0.015 | 0.015 | 0 | 0.17 | 0.12 | 0.09 | 0.02 | 0.11 | 0.05 |
| 10 | 105039241 | rs1163085 | -0.202 | -0.057 | -0.279 | -0.103 | 0 | 0.002 | 0.002 | 0 | 0.64 | 0.61 | 0.58 | 0.62 | 0.66 | 0.73 |
| 10 | 105043289 | rs1163087 | 1.116 | -0.273 | -0.4 | -0.232 | -0.027 | -0.047 | -0.047 | 0 | 0.46 | 0.12 | 0.39 | 0.48 | 0.46 | 0.59 |
| 10 | 105044157 | rs4918003 | 0.658 | -0.278 | -0.368 | -0.234 | 0 | 0 | 0 | 0 | 0.53 | 0.41 | 0.43 | 0.48 | 0.46 | 0.68 |
| 10 | 105052624 | rs4293062 | 1.042 | -0.26 | -0.28 | -0.255 | -0.01 | 0.015 | 0.015 | 0 | 0.46 | 0.23 | 0.42 | 0.44 | 0.48 | 0.59 |
| 10 | 105055912 | rs11191642 | 1.604 | -0.321 | -0.398 | -0.189 | -0.08 | -0.064 | -0.064 | 0 | 0.28 | 0.05 | 0.32 | 0.34 | 0.35 | 0.54 |
| 10 | 105065409 | rs6584547 | 0.375 | -0.3 | -0.3 | -0.185 | 0 | -0.001 | -0.001 | 0.386 | 0.54 | 0.42 | 0.46 | 0.44 | 0.52 | 0.7 |
| 10 | 105080575 | rs7093667 | -0.263 | -0.276 | -0.162 | -0.179 | 0.003 | -0.009 | -0.009 | 0 | 0.63 | 0.64 | 0.58 | 0.58 | 0.67 | 0.74 |
| 10 | 105092447 | rs10883855 | -0.263 | -0.276 | -0.162 | -0.179 | 0.003 | -0.009 | -0.009 | 0.31 | 0.63 | 0.64 | 0.58 | 0.58 | 0.67 | 0.74 |
| 10 | 105101701 | rs11191659 | 0.919 | -0.261 | -0.192 | -0.182 | -0.038 | -0.024 | -0.024 | 0 | 0.07 | 0.01 | 0.1 | 0.14 | 0.15 | 0.03 |
| 10 | 105107470 | rs4918007 | -0.52 | -0.431 | -0.098 | -0.271 | -0.042 | 0.001 | 0.001 | 0 | 0.44 | 0.21 | 0.51 | 0.56 | 0.48 | 0.29 |
| 10 | 105118462 | rs11191663 | 0.753 | -0.387 | -0.114 | -0.252 | -0.003 | 0.033 | 0.033 | 0 | 0.51 | 0.19 | 0.42 | 0.43 | 0.49 | 0.69 |
| 10 | 105118476 | rs11191664 | 0.744 | -0.401 | -0.112 | -0.258 | 0 | 0.033 | 0.033 | 0 | 0.51 | 0.21 | 0.42 | 0.43 | 0.49 | 0.69 |
| 10 | 105127278 | rs11191666 | 0.752 | -0.367 | -0.066 | -0.243 | 0 | 0.033 | 0.033 | 0 | 0.51 | 0.21 | 0.42 | 0.44 | 0.49 | 0.69 |
| 10 | 105127708 | rs10883858 | 0.766 | -0.367 | -0.067 | -0.243 | -0.002 | 0.034 | 0.034 | 0 | 0.51 | 0.19 | 0.42 | 0.44 | 0.49 | 0.69 |
| 10 | 105129665 | rs10883861 | 1.371 | -0.592 | -0.244 | -0.228 | 0.04 | -0.068 | -0.068 | 0 | 0.18 | 0.01 | 0.12 | 0.23 | 0.1 | 0.13 |
| 10 | 105138961 | rs11191672 | 0.868 | -0.49 | -0.192 | -0.194 | -0.001 | 0.033 | 0.033 | 0 | 0.51 | 0.2 | 0.42 | 0.43 | 0.49 | 0.69 |
| 10 | 105154089 | rs7911488 | 1.677 | -0.52 | -0.217 | -0.211 | -0.08 | -0.063 | -0.063 | 0 | 0.28 | 0.05 | 0.32 | 0.34 | 0.35 | 0.55 |
| 10 | 105155066 | rs11191676 | 0.867 | -0.487 | -0.2 | -0.191 | -0.003 | 0.033 | 0.033 | 0 | 0.51 | 0.2 | 0.42 | 0.43 | 0.49 | 0.69 |
| 10 | 105155645 | rs2292807 | 0.867 | -0.487 | -0.203 | -0.191 | 0 | 0.033 | 0.033 | 0 | 0.51 | 0.21 | 0.42 | 0.43 | 0.49 | 0.69 |
| 10 | 105157128 | rs10786751 | 0.964 | -0.599 | -0.22 | -0.212 | 0.037 | -0.009 | -0.009 | 0.063 | 0.21 | 0.13 | 0.13 | 0.23 | 0.11 | 0.13 |
| 10 | 105161879 | rs10883863 | -0.474 | -0.057 | -0.265 | -0.427 | 0.001 | -0.003 | -0.003 | 0 | 0.46 | 0.5 | 0.54 | 0.56 | 0.48 | 0.29 |
| 10 | 105174869 | rs2271750 | 1.182 | -0.851 | -0.295 | -0.452 | 0.016 | 0.02 | 0.02 | 0.186 | 0.23 | 0.14 | 0.1 | 0.09 | 0.14 | 0.14 |
| 10 | 105179214 | rs3740382 | 1.086 | -0.3 | -0.326 | -0.21 | -0.034 | -0.011 | -0.011 | 0 | 0.29 | 0.18 | 0.36 | 0.35 | 0.39 | 0.53 |
| 10 | 105194086 | rs2986014 | 0.452 | -0.53 | -0.07 | -0.22 | 0.008 | 0.003 | 0.003 | 0 | 0.37 | 0.48 | 0.4 | 0.42 | 0.42 | 0.64 |
| 10 | 105194476 | rs4917389 | 1.833 | -0.337 | -0.257 | -0.391 | 0.001 | 0.114 | 0.114 | 0 | 0.19 | 0.01 | 0.12 | 0 | 0.17 | 0.07 |
| 10 | 105196814 | rs10883868 | 1.177 | 0.058 | 0.085 | 0 | -0.145 | -0.067 | -0.067 | 0 | 0.27 | 0.04 | 0.35 | 0.32 | 0.4 | 0.52 |
| 10 | 105201577 | rs2281860 | 0.558 | 0.108 | 0.354 | -0.007 | 0.022 | -0.002 | -0.002 | 0 | 0.32 | 0.41 | 0.4 | 0.36 | 0.42 | 0.54 |
| 10 | 105206874 | rs6580 | 0.698 | 0.159 | -0.306 | 0.046 | -0.077 | -0.027 | -0.027 | 0 | 0.3 | 0.15 | 0.36 | 0.36 | 0.41 | 0.54 |
| 10 | 105211432 | rs10883869 | -0.574 | 0.356 | -0.931 | 0.13 | 0.024 | -0.001 | -0.001 | 0 | 0.65 | 0.48 | 0.54 | 0.59 | 0.47 | 0.38 |
| 10 | 105213765 | rs11191692 | 0.765 | 0.705 | -1.953 | 0.127 | -0.12 | -0.055 | -0.055 | 0.06 | 0.28 | 0.06 | 0.31 | 0.31 | 0.39 | 0.51 |
| 10 | 105215016 | rs2986016 | -0.414 | 1.071 | -1.444 | -0.068 | -0.041 | 0.086 | 0.086 | 0 | 0.75 | 0.98 | 0.84 | 0.88 | 0.75 | 0.9 |
| 10 | 105225201 | rs3014192 | 0.726 | 1.027 | -0.974 | 0.144 | 0 | -0.001 | -0.001 | 0 | 0.54 | 0.51 | 0.4 | 0.54 | 0.48 | 0.73 |
| 10 | 105227494 | rs2986029 | -0.019 | 0.993 | -0.47 | -0.208 | -0.009 | 0.066 | 0.066 | 0 | 0.77 | 0.91 | 0.86 | 0.91 | 0.76 | 0.9 |
| 10 | 105232580 | rs2986034 | 0.343 | 0.969 | -0.569 | -0.107 | -0.007 | -0.044 | -0.044 | 0 | 0.73 | 0.57 | 0.79 | 0.77 | 0.76 | 0.89 |
| 10 | 105233110 | rs2986035 | 0.359 | 0.97 | -0.568 | -0.106 | -0.008 | -0.045 | -0.045 | 0 | 0.73 | 0.57 | 0.79 | 0.77 | 0.76 | 0.89 |
| 10 | 105236787 | rs2860194 | -0.512 | 0.835 | -0.978 | -0.476 | -0.013 | 0.104 | 0.104 | 0 | 0.81 | 0.99 | 0.9 | 1 | 0.82 | 0.93 |
| 10 | 105244565 | rs2863733 | 0.299 | 0.898 | -0.686 | -0.498 | 0.015 | -0.1 | -0.1 | 0 | 0.72 | 0.53 | 0.79 | 0.9 | 0.71 | 0.88 |
| 10 | 105245981 | rs3014188 | 0.337 | 0.344 | -1.692 | -0.432 | -0.074 | -0.005 | -0.005 | 0 | 0.62 | 0.92 | 0.5 | 0.75 | 0.47 | 0.74 |
| 10 | 105246283 | rs3014187 | 0.29 | -0.206 | -2.119 | -0.734 | 0.047 | -0.047 | -0.047 | 0 | 0.72 | 0.92 | 0.68 | 0.71 | 0.82 | 0.72 |
| 10 | 105248261 | rs2986037 | -1.289 | -0.616 | -1.216 | -0.98 | -0.004 | 0.01 | 0.01 | 1.124 | 0.52 | 0.7 | 0.56 | 0.59 | 0.44 | 0.52 |

| | | | | | | | | | | | | | | | | |
|---|---|---|---|---|---|---|---|---|---|---|---|---|---|---|---|---|
| 10 | 105248507 | rs2986038 | 0.251 | -0.622 | -0.658 | -1.057 | -0.039 | 0.02 | 0.02 | 0 | 0.22 | 0.47 | 0.19 | 0.34 | 0.11 | 0.26 |
| 10 | 105256773 | rs2986044 | -0.866 | -0.301 | -0.701 | -1.082 | 0.002 | -0.071 | -0.071 | 0 | 0.8 | 0.83 | 0.82 | 1 | 0.75 | 0.92 |
| 10 | 105257786 | rs7071247 | 0.233 | -0.485 | -0.59 | -1.141 | 0.052 | 0.006 | 0.006 | 0.243 | 0.29 | 0.26 | 0.2 | 0.34 | 0.13 | 0.33 |
| 10 | 105259917 | rs10883875 | 0.251 | -0.448 | -0.598 | -1.156 | 0.025 | 0.007 | 0.007 | 0.487 | 0.29 | 0.3 | 0.2 | 0.33 | 0.13 | 0.31 |
| 10 | 105262224 | rs1536222 | -0.626 | -0.146 | -1.047 | -1.153 | 0.012 | -0.083 | -0.083 | 0.007 | 0.79 | 0.79 | 0.8 | 1 | 0.71 | 0.88 |
| 10 | 105262570 | rs10883876 | 1.883 | -0.133 | -1.008 | -1.119 | 0.004 | -0.047 | -0.047 | 0.84 | 0.31 | 0.14 | 0.25 | 0.41 | 0.31 | 0.4 |
| 10 | 105262845 | rs3014206 | -0.555 | -0.036 | -1.158 | -0.995 | 0.003 | -0.074 | -0.074 | 0.003 | 0.8 | 0.83 | 0.82 | 1 | 0.75 | 0.92 |
| 10 | 105269873 | rs17737062 | 0.704 | -0.177 | -1.101 | -0.538 | -0.084 | -0.072 | -0.072 | 0.802 | 0.13 | 0.01 | 0.16 | 0.17 | 0.23 | 0.07 |
| 10 | 105271758 | rs2281859 | -0.949 | -0.263 | -0.974 | -0.486 | -0.01 | -0.004 | -0.004 | 0 | 0.53 | 0.89 | 0.57 | 0.58 | 0.4 | 0.48 |
| 10 | 105271890 | rs2281858 | -0.739 | -0.206 | -0.921 | -0.461 | 0.014 | 0.016 | 0.016 | 1.41 | 0.51 | 0.74 | 0.56 | 0.58 | 0.4 | 0.48 |
| 10 | 105296388 | rs2208322 | -1.005 | 0.846 | -0.868 | -0.429 | -0.006 | 0.014 | 0.014 | 0 | 0.42 | 0.74 | 0.33 | 0.33 | 0.36 | 0.41 |
| 10 | 105299611 | rs6584555 | -0.546 | 0.9 | -0.652 | -0.345 | -0.033 | -0.043 | -0.043 | 0 | 0.28 | 0.53 | 0.18 | 0.16 | 0.15 | 0.31 |
| 10 | 105313039 | rs912889 | -0.656 | 1.416 | -1.698 | -0.184 | -0.057 | 0.059 | 0.059 | 0 | 0.89 | 0.99 | 0.9 | 1 | 0.79 | 0.89 |
| 10 | 105313230 | rs912888 | -0.885 | 0.862 | -1.033 | 0.315 | -0.002 | -0.001 | -0.001 | 0 | 0.33 | 0.29 | 0.28 | 0.32 | 0.32 | 0.39 |
| 10 | 105322373 | rs11191727 | -0.523 | 0.417 | -1.229 | 0.152 | 0.012 | 0.02 | 0.02 | 0 | 0.19 | 0.1 | 0.11 | 0.16 | 0.13 | 0.25 |
| 10 | 105328241 | rs11191730 | 0.445 | 0.53 | -1.731 | 0.311 | -0.008 | -0.003 | -0.003 | 0 | 0.23 | 0.07 | 0.14 | 0.2 | 0.23 | 0.28 |
| 10 | 105331761 | rs11191732 | 0.746 | 1.151 | -1.976 | 0.586 | -0.062 | -0.062 | -0.062 | 0 | 0.1 | 0.01 | 0.13 | 0.18 | 0.18 | 0.08 |
| 10 | 105332452 | rs7067666 | -0.261 | 0.69 | -2.038 | 0.104 | -0.003 | 0.048 | 0.048 | 0 | 0.52 | 0.68 | 0.67 | 0.61 | 0.45 | 0.6 |
| 10 | 105332617 | rs7068081 | -0.345 | 0.7 | -2.055 | 0.105 | -0.012 | 0.051 | 0.051 | 0 | 0.53 | 0.74 | 0.68 | 0.61 | 0.45 | 0.6 |
| 10 | 105340059 | rs7068070 | 1.003 | 0.637 | -2.071 | 0.105 | 0.004 | 0.005 | 0.005 | 0.733 | 0.25 | 0.14 | 0.14 | 0.21 | 0.24 | 0.26 |
| 10 | 105342866 | rs4918025 | 0.255 | 0.534 | -1.291 | 0.019 | -0.01 | -0.019 | -0.019 | 1.587 | 0.24 | 0.18 | 0.36 | 0.45 | 0.28 | 0.29 |
| 10 | 105342900 | rs902991 | -0.9 | 0.49 | -1.145 | -0.072 | -0.041 | 0.028 | 0.028 | 0 | 0.78 | 0.91 | 0.82 | 0.81 | 0.68 | 0.86 |
| 10 | 105349816 | rs2236211 | 0.596 | 0.28 | -0.46 | -0.171 | 0.046 | 0.059 | 0.059 | 0 | 0.47 | 0.3 | 0.43 | 0.36 | 0.4 | 0.56 |
| 10 | 105350447 | rs7918186 | 0.773 | 0.236 | -0.574 | -0.189 | 0.039 | 0.062 | 0.062 | 0 | 0.45 | 0.14 | 0.41 | 0.36 | 0.4 | 0.56 |
| 10 | 105352990 | rs729952 | 0.768 | 0.179 | -0.598 | -0.21 | 0.034 | 0.056 | 0.056 | 0 | 0.44 | 0.14 | 0.41 | 0.36 | 0.4 | 0.55 |
| 10 | 105354556 | rs10879 | 0.39 | 0.263 | -0.479 | -0.195 | -0.014 | -0.028 | -0.028 | 2.316 | 0.24 | 0.17 | 0.35 | 0.45 | 0.28 | 0.28 |
| 10 | 105357181 | rs10883887 | 0.583 | 0.263 | -0.781 | -0.516 | 0.011 | -0.017 | -0.017 | 0.015 | 0.54 | 0.2 | 0.44 | 0.55 | 0.43 | 0.58 |
| 10 | 105358980 | rs2281584 | 0.191 | 0.298 | -0.162 | -0.484 | 0.027 | 0.085 | 0.085 | 0 | 0.27 | 0.51 | 0.4 | 0.45 | 0.29 | 0.3 |
| 10 | 105362104 | rs11191741 | 1.09 | 0.236 | -0.565 | -0.484 | 0.042 | 0.058 | 0.058 | 0 | 0.44 | 0.11 | 0.36 | 0.35 | 0.35 | 0.53 |
| 10 | 105362209 | rs4917396 | -1.395 | 0.531 | -0.509 | -0.466 | -0.055 | 0.061 | 0.061 | 0 | 0.88 | 0.97 | 0.89 | 1 | 0.76 | 0.91 |
| 10 | 105363289 | rs3740471 | 0.794 | 0.604 | -0.488 | -0.165 | -0.047 | -0.029 | -0.029 | 0 | 0.14 | 0.06 | 0.16 | 0.15 | 0.2 | 0.13 |
| 10 | 105376388 | rs3824781 | 1.036 | 0.439 | -0.404 | -0.144 | 0.001 | -0.009 | -0.009 | 0 | 0.1 | 0.06 | 0.09 | 0.2 | 0.1 | 0.03 |
| 10 | 105384262 | rs902997 | -0.088 | 0.537 | -0.561 | -0.304 | -0.108 | 0.111 | 0.111 | 0 | 0.75 | 0.92 | 0.76 | 0.92 | 0.63 | 0.81 |
| 10 | 105385350 | rs3781354 | 0.704 | 0.513 | -0.101 | -0.277 | 0 | -0.001 | -0.001 | 0 | 0.59 | 0.42 | 0.61 | 0.65 | 0.47 | 0.76 |
| 10 | 105388286 | rs11191750 | 0.421 | 0.407 | 0.064 | -0.223 | 0.003 | 0.067 | 0.067 | 0 | 0.14 | 0.26 | 0.14 | 0.26 | 0.16 | 0.05 |
| 10 | 105388691 | rs12415143 | 0.139 | 0.309 | 0.108 | -0.25 | 0.006 | 0.001 | 0.001 | 0 | 0.6 | 0.47 | 0.5 | 0.63 | 0.52 | 0.61 |
| 10 | 105391300 | rs7918134 | -0.022 | 0.498 | 0.164 | -0.239 | 0.014 | 0.127 | 0.127 | 0 | 0.16 | 0.44 | 0.15 | 0.26 | 0.18 | 0.05 |
| 10 | 105401191 | rs3781347 | -0.032 | -0.103 | -0.352 | -0.677 | -0.001 | -0.001 | -0.001 | 0.222 | 0.38 | 0.42 | 0.44 | 0.38 | 0.39 | 0.36 |
| 10 | 105402565 | rs7100298 | 1.365 | -0.239 | -0.392 | -0.789 | -0.015 | -0.018 | -0.018 | 0 | 0.19 | 0.2 | 0.23 | 0.1 | 0.08 | 0.29 |
| 10 | 105405390 | rs880869 | -0.434 | -0.215 | -0.311 | -0.787 | -0.004 | -0.024 | -0.024 | 0 | 0.07 | 0.25 | 0.02 | 0 | 0.01 | 0.01 |
| 10 | 105406129 | rs880549 | 0.691 | -0.102 | -0.448 | -0.733 | -0.039 | -0.036 | -0.036 | 0 | 0.16 | 0.09 | 0.19 | 0.27 | 0.31 | 0.07 |
| 10 | 105407790 | rs7914478 | 0.601 | -0.096 | -0.451 | -0.731 | -0.035 | -0.032 | -0.032 | 0.805 | 0.16 | 0.1 | 0.2 | 0.27 | 0.31 | 0.07 |
| 10 | 105418505 | rs2172459 | 0.035 | -0.191 | -0.547 | -0.769 | 0.003 | 0.033 | 0.033 | 0 | 0.53 | 0.64 | 0.46 | 0.63 | 0.44 | 0.56 |
| 10 | 105423908 | rs4918040 | 0.164 | -0.144 | -0.879 | -0.719 | 0.006 | 0.02 | 0.02 | 0 | 0.5 | 0.46 | 0.45 | 0.61 | 0.44 | 0.56 |
| 10 | 105428152 | rs3781339 | 0.834 | -0.365 | -1.085 | -0.404 | -0.056 | -0.057 | -0.057 | 0 | 0.1 | 0.01 | 0.12 | 0.26 | 0.21 | 0.05 |
| 10 | 105438751 | rs3824774 | 0.606 | -0.833 | -0.938 | -0.319 | 0.014 | -0.01 | -0.01 | 0.107 | 0.34 | 0.29 | 0.31 | 0.51 | 0.25 | 0.47 |
| 10 | 105445588 | rs12413931 | 0.651 | -0.73 | -0.863 | -0.25 | 0.046 | 0.053 | 0.053 | 0 | 0.21 | 0.11 | 0.22 | 0.09 | 0.09 | 0.24 |
| 10 | 105453452 | rs2295590 | 1.045 | -0.284 | -0.637 | 0.248 | -0.005 | 0.045 | 0.045 | 0 | 0.13 | 0.2 | 0.23 | 0.31 | 0.09 | 0.19 |
| 10 | 105460401 | rs11191773 | -0.395 | -0.314 | -0.72 | -0.083 | 0.003 | 0.005 | 0.005 | 0 | 0.45 | 0.54 | 0.47 | 0.43 | 0.44 | 0.39 |
| 10 | 105463885 | rs2145310 | 0.244 | -0.708 | -0.724 | 0.506 | -0.026 | 0.007 | 0.007 | 0 | 0.4 | 0.37 | 0.35 | 0.48 | 0.24 | 0.51 |
| 10 | 105466808 | rs12772784 | 0.165 | 0.563 | -0.309 | 0.506 | -0.003 | -0.004 | -0.004 | 0 | 0.08 | 0.07 | 0.09 | 0 | 0.14 | 0.03 |
| 10 | 105467895 | rs923799 | -1.654 | 0.494 | -0.33 | 0.62 | -0.012 | 0.006 | 0.006 | 0 | 0.39 | 0.62 | 0.43 | 0.6 | 0.45 | 0.34 |
| 10 | 105472989 | rs17115849 | -0.085 | 0.159 | -0.172 | 0.914 | -0.087 | 0.166 | 0.166 | 0.032 | 0.17 | 0.47 | 0.15 | 0.31 | 0.04 | 0.17 |
| 10 | 105473580 | rs11191779 | 0.513 | 0.619 | -0.815 | 0.958 | -0.036 | -0.035 | -0.035 | 0.015 | 0.12 | 0.07 | 0.18 | 0.29 | 0.26 | 0.11 |
| 10 | 105474013 | rs10159470 | 0.567 | 0.634 | -0.833 | 0.966 | -0.053 | -0.052 | -0.052 | 0 | 0.11 | 0.04 | 0.16 | 0.29 | 0.25 | 0.11 |
| 10 | 105474652 | rs10883903 | -1.335 | -0.042 | -0.504 | 0.625 | 0.018 | -0.015 | -0.015 | 0.318 | 0.29 | 0.72 | 0.27 | 0.31 | 0.19 | 0.22 |
| 10 | 105477163 | rs2047036 | 0.224 | 0.417 | -0.387 | 0.345 | -0.005 | 0.001 | 0.001 | 0 | 0.65 | 0.76 | 0.61 | 0.71 | 0.62 | 0.67 |
| 10 | 105483784 | rs12219579 | -1.156 | 0.804 | 0.442 | 0.534 | 0.101 | 0.173 | 0.173 | 0.15 | 0.14 | 0.41 | 0.22 | 0.31 | 0.23 | 0.08 |
| 10 | 105491931 | rs7072004 | -1.283 | 0.625 | 0.426 | 0.293 | 0.018 | 0.005 | 0.005 | 0 | 0.3 | 0.63 | 0.34 | 0.31 | 0.34 | 0.2 |
| 10 | 105500857 | rs879543 | -1.158 | 0.541 | 0.474 | 0.332 | -0.006 | 0.017 | 0.017 | 0.348 | 0.27 | 0.61 | 0.29 | 0.31 | 0.22 | 0.16 |
| 10 | 105510245 | rs17812914 | 0.699 | 0.583 | 0.434 | 0.183 | -0.001 | 0.012 | 0.012 | 0.795 | 0.15 | 0.07 | 0.1 | 0 | 0.11 | 0.14 |
| 10 | 105514473 | rs7920154 | 0.288 | 0.615 | 0.569 | 0.239 | 0.004 | -0.019 | -0.019 | 0 | 0.07 | 0.11 | 0.09 | 0 | 0.11 | 0.01 |
| 10 | 105517583 | rs1076814 | -1.359 | 0.352 | 0.204 | 0.151 | 0.001 | 0.011 | 0.011 | 0.899 | 0.55 | 0.79 | 0.53 | 0.59 | 0.53 | 0.38 |
| 10 | 105518179 | rs10466138 | -0.576 | 0.369 | 0.501 | 0.221 | -0.002 | 0.01 | 0.01 | 1.802 | 0.44 | 0.43 | 0.38 | 0.31 | 0.43 | 0.37 |
| 10 | 105518482 | rs7923100 | -1.332 | 0.394 | 0.455 | 0.196 | 0.004 | 0.017 | 0.017 | 1.189 | 0.54 | 0.73 | 0.53 | 0.6 | 0.53 | 0.38 |
| 10 | 105519725 | rs11191797 | 1.101 | 0.392 | 0.291 | 0.192 | -0.029 | -0.019 | -0.019 | 0 | 0.13 | 0.08 | 0.16 | 0.22 | 0.24 | 0.09 |
| 10 | 105521705 | rs10748848 | -0.485 | 0.328 | 0.371 | 0.062 | -0.003 | 0.023 | 0.023 | 0 | 0.32 | 0.51 | 0.34 | 0.6 | 0.32 | 0.22 |
| 10 | 105524245 | rs9665617 | 0.678 | 0.073 | 0.421 | 0.169 | 0 | -0.029 | -0.029 | 0 | 0.16 | 0.13 | 0.11 | 0 | 0.11 | 0.14 |
| 10 | 105525821 | rs8181413 | 0.121 | 0.054 | 0.673 | 0.831 | -0.029 | -0.042 | -0.042 | 0.057 | 0.08 | 0.02 | 0.14 | 0.16 | 0.13 | 0.12 |

| | | | | | | | | | | | | | | | | |
|---|---|---|---|---|---|---|---|---|---|---|---|---|---|---|---|---|
| 10 | 105527521 | rs12778775 | -1.135 | 0.074 | 1.119 | 0.143 | -0.031 | -0.042 | -0.042 | 0.006 | 0.24 | 0.38 | 0.21 | 0.1 | 0.18 | 0.14 |
| 10 | 105537782 | rs11191804 | -1.155 | 1.075 | 1.383 | 0.251 | 0.033 | 0.158 | 0.158 | 0 | 0.12 | 0.33 | 0.16 | 0.29 | 0.13 | 0.02 |
| 10 | 105543393 | rs2073341 | 1.576 | 1.424 | 1.028 | -0.13 | -0.013 | 0.007 | 0.007 | 0 | 0.67 | 0.27 | 0.59 | 0.7 | 0.6 | 0.79 |
| 10 | 105547517 | rs1015037 | -0.139 | 0.516 | 1.222 | 0.002 | 0.031 | -0.038 | -0.038 | 0 | 0.92 | 0.84 | 0.91 | 1 | 0.89 | 0.99 |
| 10 | 105551219 | rs7092722 | 1.402 | 0.317 | 1.016 | -0.109 | -0.012 | -0.116 | -0.116 | 0 | 0.72 | 0.55 | 0.79 | 0.96 | 0.76 | 0.78 |
| 10 | 105555131 | rs4918052 | 1.563 | 0.273 | 0.863 | 0.007 | -0.004 | -0.002 | -0.002 | 0 | 0.69 | 0.31 | 0.71 | 0.68 | 0.76 | 0.77 |
| 10 | 105556339 | rs6584571 | 1.286 | 0.475 | 0.929 | -0.178 | 0.004 | -0.127 | -0.127 | 0.34 | 0.7 | 0.45 | 0.75 | 0.96 | 0.67 | 0.74 |
| 10 | 105562717 | rs10786769 | 1.782 | 0.109 | 0.726 | -0.109 | 0 | 0.015 | 0.015 | 0 | 0.63 | 0.29 | 0.66 | 0.68 | 0.62 | 0.71 |
| 10 | 105563093 | rs2864000 | 1.838 | 0.121 | 0.728 | -0.104 | 0.001 | 0.018 | 0.018 | 0 | 0.62 | 0.28 | 0.66 | 0.68 | 0.62 | 0.71 |
| 10 | 105568840 | rs7086883 | -1.348 | 0.231 | 0.86 | -0.148 | 0.044 | -0.159 | -0.159 | 0 | 0.24 | 0.46 | 0.2 | 0 | 0.31 | 0.16 |
| 10 | 105574251 | rs7086170 | 0.756 | 0.294 | 0.364 | -0.042 | -0.001 | 0.028 | 0.028 | 0 | 0.94 | 0.89 | 0.89 | 0.71 | 0.96 | 0.95 |
| 10 | 105578730 | rs2902637 | 0.307 | 0.302 | 0.37 | -0.051 | -0.033 | -0.026 | -0.026 | 1.966 | 0.83 | 0.86 | 0.82 | 0.72 | 0.76 | 0.82 |
| 10 | 105580251 | rs11594668 | -0.315 | 0.132 | 0.616 | -0.063 | -0.004 | -0.033 | -0.033 | 0 | 0.09 | 0.12 | 0.09 | 0 | 0.09 | 0.01 |
| 10 | 105581248 | rs725077 | 1.62 | 0.107 | 0.485 | -0.067 | -0.001 | -0.005 | -0.005 | 0.603 | 0.64 | 0.35 | 0.69 | 0.69 | 0.65 | 0.72 |
| 10 | 105581265 | rs725076 | 0.744 | 0.676 | -0.097 | 0.312 | -0.031 | -0.023 | -0.023 | 0 | 0.81 | 0.85 | 0.81 | 0.71 | 0.76 | 0.82 |
| 10 | 105583030 | rs10786770 | 0.105 | 0.295 | 0.044 | 0.151 | -0.004 | -0.016 | -0.016 | 1.164 | 0.63 | 0.46 | 0.55 | 0.35 | 0.64 | 0.69 |
| 10 | 105586509 | rs10883918 | -0.603 | -0.447 | 0.246 | 0.584 | -0.031 | 0.042 | 0.042 | 0 | 0.22 | 0.3 | 0.15 | 0.44 | 0.11 | 0.23 |
| 10 | 105590558 | rs12766594 | -0.008 | -0.091 | 0.204 | 0.103 | -0.012 | -0.009 | -0.009 | 0 | 0.17 | 0.11 | 0.17 | 0 | 0.29 | 0.14 |
| 10 | 105591049 | rs11191817 | 0.853 | -0.33 | 0.316 | 0.286 | -0.02 | 0.062 | 0.062 | 0.244 | 0.13 | 0.18 | 0.11 | 0.42 | 0.05 | 0.14 |
| 10 | 105594855 | rs11191820 | -0.223 | -0.31 | 0.563 | 0.222 | -0.017 | -0.021 | -0.021 | 0 | 0.28 | 0.22 | 0.29 | 0.6 | 0.33 | 0.29 |
| 10 | 105606287 | rs4918060 | 0.216 | -0.401 | 0.973 | -0.091 | -0.017 | -0.062 | -0.062 | 1.583 | 0.89 | 0.64 | 0.94 | 0.98 | 0.94 | 0.92 |
| 10 | 105610326 | rs10786772 | 1.422 | -1.209 | 0.324 | -0.25 | 0.037 | 0.014 | 0.014 | 0.212 | 0.41 | 0.11 | 0.42 | 0.64 | 0.31 | 0.5 |
| 10 | 105623461 | rs11596104 | 1.07 | -1.035 | 0.528 | -0.338 | 0.005 | 0 | 0 | 1.437 | 0.17 | 0.2 | 0.2 | 0.17 | 0.28 | 0.14 |
| 10 | 105623955 | rs10883931 | -0.799 | -1.137 | 0.865 | -0.489 | 0.018 | -0.052 | -0.052 | 0.08 | 0.38 | 0.43 | 0.35 | 0.18 | 0.48 | 0.24 |
| 10 | 105634895 | rs11191839 | 1.622 | -0.999 | 0.703 | -0.346 | 0.021 | 0.024 | 0.024 | 0.412 | 0.4 | 0.11 | 0.38 | 0.65 | 0.31 | 0.5 |
| 10 | 105636989 | rs2902638 | 0.364 | -1.05 | 0.887 | -0.376 | 0.003 | 0.025 | 0.025 | 0 | 0.23 | 0.11 | 0.26 | 0.17 | 0.25 | 0.27 |
| 10 | 105638232 | rs4917405 | -0.488 | -1.121 | 1.045 | -0.36 | -0.006 | -0.025 | -0.025 | 3.008 | 0.91 | 0.91 | 0.96 | 0.98 | 0.94 | 0.92 |
| 10 | 105639421 | rs911547 | 0.89 | -1.154 | 1.159 | -0.361 | -0.017 | -0.11 | -0.11 | 0 | 0.81 | 0.44 | 0.86 | 0.98 | 0.84 | 0.9 |
| 10 | 105639514 | rs10748858 | 1.35 | -1.252 | 1.065 | -0.005 | -0.021 | 0.015 | 0.015 | 0 | 0.38 | 0.14 | 0.38 | 0.29 | 0.35 | 0.41 |
| 10 | 105640978 | rs7100920 | -0.893 | -0.317 | 1.471 | -0.372 | -0.004 | 0.007 | 0.007 | 0.378 | 0.42 | 0.42 | 0.48 | 0.34 | 0.54 | 0.41 |
| 10 | 105643134 | rs2067832 | -0.81 | -0.3 | 1.269 | -0.364 | -0.026 | 0.025 | 0.025 | 0.135 | 0.4 | 0.29 | 0.48 | 0.34 | 0.54 | 0.41 |
| 10 | 105645181 | rs11598840 | -0.619 | -0.195 | 1.459 | -0.333 | 0.033 | -0.04 | -0.04 | 0.026 | 0.93 | 0.84 | 0.93 | 1 | 0.91 | 0.99 |
| 10 | 105647095 | rs3814219 | 0.68 | 0.03 | 0.611 | -0.292 | 0.009 | -0.001 | -0.001 | 0 | 0.16 | 0.21 | 0.21 | 0.16 | 0.25 | 0.12 |
| 10 | 105654391 | rs4918069 | 0.008 | -0.098 | 0.392 | -0.32 | -0.008 | 0.014 | 0.014 | 0 | 0.24 | 0.08 | 0.27 | 0.19 | 0.28 | 0.29 |
| 10 | 105657892 | rs2984132 | 1.215 | -0.199 | 0.559 | -0.34 | 0.044 | 0.061 | 0.061 | 0 | 0.5 | 0.35 | 0.44 | 0.66 | 0.37 | 0.58 |
| 10 | 105659826 | rs2487999 | 0.368 | -0.569 | 1.235 | 0.138 | 0.056 | -0.048 | -0.048 | 0 | 0.9 | 0.64 | 0.92 | 1 | 0.91 | 0.99 |
| 10 | 105671527 | rs1265165 | 0.329 | -0.555 | 1.241 | 0.144 | 0.059 | -0.045 | -0.045 | 0 | 0.9 | 0.64 | 0.92 | 1 | 0.91 | 0.99 |
| 10 | 105674854 | rs1265164 | 0.935 | -0.541 | 0.301 | 0.138 | -0.018 | -0.099 | -0.099 | 0 | 0.81 | 0.41 | 0.86 | 0.98 | 0.87 | 0.91 |
| 10 | 105675946 | rs9419958 | 0.919 | -0.537 | 0.228 | 0.139 | -0.017 | -0.098 | -0.098 | 0 | 0.81 | 0.41 | 0.87 | 0.98 | 0.87 | 0.91 |
| 10 | 105677897 | rs4387287 | 1.066 | -0.383 | -0.242 | 0.197 | -0.001 | -0.002 | -0.002 | 0.568 | 0.78 | 0.28 | 0.78 | 0.82 | 0.85 | 0.81 |
| 10 | 105683494 | rs11814244 | -1.376 | -0.189 | -0.51 | 0.345 | 0.015 | 0.074 | 0.074 | 0 | 0.46 | 0.8 | 0.49 | 0.36 | 0.44 | 0.47 |
| 10 | 105702042 | rs7074532 | -0.584 | -0.29 | -0.288 | 0.336 | 0.001 | 0.039 | 0.039 | 0 | 0.26 | 0.18 | 0.2 | 0.14 | 0.29 | 0.31 |
| 10 | 105708793 | rs7073827 | -1.073 | -0.517 | 0.286 | 0.342 | -0.001 | -0.002 | -0.002 | 0 | 0.32 | 0.35 | 0.32 | 0.22 | 0.32 | 0.37 |
| 10 | 105714925 | rs6584581 | -0.471 | -0.445 | 0.244 | 0.44 | -0.021 | 0.027 | 0.027 | 0 | 0.05 | 0.17 | 0.12 | 0.09 | 0.04 | 0.07 |
| 10 | 105720022 | rs7924310 | -1.354 | -0.549 | 0.21 | 0.833 | 0.043 | -0.05 | -0.05 | 0 | 0.42 | 0.86 | 0.42 | 0.46 | 0.36 | 0.46 |
| 10 | 105724329 | rs10883953 | 1.077 | -0.512 | -0.189 | 0.334 | 0.016 | 0.099 | 0.099 | 0 | 0.56 | 0.21 | 0.52 | 0.71 | 0.57 | 0.47 |
| 10 | 105726338 | rs6584583 | 0.136 | -0.287 | 0.416 | 0.26 | 0.006 | -0.013 | -0.013 | 0 | 0.87 | 0.79 | 0.8 | 0.9 | 0.84 | 0.84 |
| 10 | 105729839 | rs9988667 | -0.081 | -0.353 | 0.358 | 0.231 | -0.016 | 0.014 | 0.014 | 0 | 0.88 | 0.94 | 0.81 | 0.9 | 0.83 | 0.84 |
| 10 | 105729887 | rs9325510 | 0.39 | -0.377 | 0.723 | 0.22 | 0.01 | -0.025 | -0.025 | 0.464 | 0.86 | 0.71 | 0.79 | 0.9 | 0.83 | 0.84 |
| 10 | 105731410 | rs17116203 | 0.955 | -0.513 | -0.078 | 0.722 | 0.005 | -0.058 | -0.058 | 0 | 0.16 | 0.06 | 0.13 | 0.43 | 0.15 | 0.14 |
| 10 | 105739914 | rs2487987 | -0.411 | -0.225 | 0.648 | 0.89 | 0.02 | -0.035 | -0.035 | 0 | 0.15 | 0.36 | 0.21 | 0.1 | 0.17 | 0.16 |
| 10 | 105743943 | rs2487991 | -0.411 | -0.225 | 0.648 | 0.89 | 0.02 | -0.035 | -0.035 | 0 | 0.15 | 0.36 | 0.21 | 0.1 | 0.17 | 0.16 |
| 10 | 105762591 | rs805657 | -0.411 | -0.225 | 0.648 | 0.89 | 0.02 | -0.035 | -0.035 | 0.145 | 0.15 | 0.36 | 0.21 | 0.1 | 0.17 | 0.16 |
| 10 | 105763026 | rs3740469 | -0.014 | -0.331 | 0.753 | 0.848 | -0.011 | 0.009 | 0.009 | 0 | 0.18 | 0.07 | 0.21 | 0.14 | 0.23 | 0.22 |
| 10 | 105788905 | rs3124 | -0.551 | -0.309 | 1.381 | 0.262 | 0.02 | -0.027 | -0.027 | 0 | 0.15 | 0.34 | 0.22 | 0.1 | 0.17 | 0.17 |
| 10 | 105791355 | rs9425 | -0.408 | -0.261 | 1.28 | 0.253 | 0.017 | -0.032 | -0.032 | 1.548 | 0.14 | 0.28 | 0.21 | 0.1 | 0.16 | 0.16 |
| 10 | 105792205 | rs813943 | 0.22 | -0.373 | 1.094 | 0.165 | -0.008 | -0.022 | -0.022 | 0.109 | 0.79 | 0.7 | 0.77 | 0.87 | 0.77 | 0.77 |
| 10 | 105793459 | rs2147268 | 0.962 | -0.305 | 1.203 | 0.171 | -0.009 | -0.062 | -0.062 | 0 | 0.77 | 0.5 | 0.76 | 0.84 | 0.81 | 0.71 |
| 10 | 105797378 | rs2282437 | 1.058 | 0.038 | 1.113 | 0.095 | -0.021 | -0.033 | -0.033 | 0 | 0.74 | 0.42 | 0.77 | 0.82 | 0.82 | 0.71 |
| 10 | 105798872 | rs2274100 | 1.036 | 0.03 | 1.183 | 0.164 | -0.025 | -0.028 | -0.028 | 0 | 0.75 | 0.44 | 0.77 | 0.77 | 0.82 | 0.71 |
| 10 | 105799216 | rs2296219 | 1.133 | -0.032 | 1.093 | 0.069 | -0.021 | -0.033 | -0.033 | 0 | 0.74 | 0.41 | 0.77 | 0.82 | 0.82 | 0.71 |
| 10 | 105800514 | rs10883962 | 0.918 | -0.034 | 1.25 | 0.068 | -0.021 | -0.04 | -0.04 | 5.343 | 0.77 | 0.63 | 0.79 | 0.82 | 0.82 | 0.71 |
| 10 | 105803080 | rs2147269 | 0.843 | 0.207 | 1.102 | 0.069 | -0.015 | -0.1 | -0.1 | 0 | 0.78 | 0.52 | 0.84 | 0.92 | 0.83 | 0.8 |
| 10 | 105805497 | rs813944 | 1.139 | -0.234 | -0.096 | -0.098 | 0.009 | 0.084 | 0.084 | 1.198 | 0.45 | 0.18 | 0.44 | 0.65 | 0.52 | 0.47 |
| 10 | 105806310 | rs805721 | 0.236 | 0.571 | -0.026 | 0.141 | 0.01 | 0.001 | 0.001 | 0 | 0.75 | 0.7 | 0.8 | 0.65 | 0.68 | 0.8 |
| 10 | 105811668 | rs805724 | 0.945 | -0.518 | 0.177 | -0.287 | 0.002 | -0.018 | -0.018 | 0 | 0.62 | 0.47 | 0.69 | 0.69 | 0.64 | 0.61 |
| 10 | 105815116 | rs17821926 | 0.607 | -1.068 | -0.209 | -0.407 | -0.005 | 0.054 | 0.054 | 0.409 | 0.16 | 0.02 | 0.15 | 0.05 | 0.15 | 0.18 |
| 10 | 105815241 | rs1747677 | 1.085 | -0.897 | 0.632 | -0.176 | 0.03 | 0.005 | 0.005 | 1.076 | 0.76 | 0.49 | 0.69 | 0.69 | 0.74 | 0.62 |
| 10 | 105816916 | rs805698 | 0.705 | -1.438 | 0.721 | -0.469 | -0.019 | 0.047 | 0.047 | 7.337 | 0.92 | 0.49 | 0.94 | 0.88 | 0.98 | 0.94 |

| | | | | | | | | | | | | | | | | |
|---|---|---|---|---|---|---|---|---|---|---|---|---|---|---|---|---|
| 10 | 105817891 | rs11191910 | -0.076 | -1.427 | 0.587 | -0.463 | -0.042 | 0.015 | 0.015 | 0 | 0.16 | 0.04 | 0.14 | 0.12 | 0.18 | 0.21 |
| 10 | 105819956 | rs805701 | 1.364 | -1.478 | 0.616 | -0.483 | 0.011 | 0.061 | 0.061 | 1.234 | 0.59 | 0.25 | 0.69 | 0.69 | 0.66 | 0.62 |
| 10 | 105821294 | rs805705 | 0.555 | -1.494 | 0.692 | -0.459 | -0.002 | 0.07 | 0.07 | 6.231 | 0.92 | 0.56 | 0.94 | 0.88 | 0.99 | 0.94 |
| 10 | 105824333 | rs805708 | 1.565 | -1.449 | 0.14 | -0.469 | 0.021 | 0.045 | 0.045 | 1.6 | 0.6 | 0.28 | 0.71 | 0.67 | 0.68 | 0.62 |
| 10 | 105832350 | rs11191916 | 0.979 | -1.19 | 0.154 | -0.342 | 0.008 | 0.002 | 0.002 | 2.946 | 0.14 | 0.12 | 0.08 | 0.11 | 0.12 | 0.1 |
| 10 | 105840003 | rs1570647 | 0.979 | -1.184 | 0.069 | -0.339 | 0.008 | 0.002 | 0.002 | 2.946 | 0.14 | 0.12 | 0.08 | 0.11 | 0.12 | 0.1 |
| 10 | 105870132 | rs10883967 | 0.557 | -0.938 | -0.312 | -0.412 | -0.038 | 0.038 | 0.038 | 0 | 0.11 | 0.04 | 0.14 | 0 | 0.17 | 0.1 |
| 10 | 105875326 | rs471247 | -0.624 | -1.158 | -0.156 | -0.363 | -0.021 | 0.008 | 0.008 | 1.331 | 0.82 | 0.6 | 0.86 | 0.78 | 0.85 | 0.85 |
| 10 | 105876587 | rs809825 | 0.125 | -1.253 | -0.042 | -0.405 | -0.025 | -0.008 | -0.008 | 0 | 0.78 | 0.36 | 0.85 | 0.78 | 0.85 | 0.85 |
| 10 | 105882253 | rs11191927 | 0.111 | -1.242 | -0.041 | -0.426 | -0.032 | 0.001 | 0.001 | 1.143 | 0.2 | 0.45 | 0.14 | 0.23 | 0.15 | 0.17 |
| 10 | 105883395 | rs1338777 | -1.497 | -1.296 | -0.157 | -0.409 | 0.013 | 0.042 | 0.042 | 0.863 | 0.41 | 0.75 | 0.33 | 0.35 | 0.35 | 0.37 |
| 10 | 105888660 | rs4537691 | -0.522 | -1.324 | -0.133 | -0.41 | -0.02 | -0.009 | -0.009 | 0.321 | 0.24 | 0.65 | 0.16 | 0.23 | 0.16 | 0.17 |
| 10 | 105895218 | rs4564255 | -0.496 | -1.325 | -0.134 | -0.416 | -0.02 | -0.01 | -0.01 | 0 | 0.23 | 0.65 | 0.16 | 0.23 | 0.16 | 0.17 |
| 10 | 105905161 | rs2289964 | 0.747 | -1.629 | -0.428 | -0.382 | 0.018 | -0.046 | -0.046 | 0.184 | 0.18 | 0.04 | 0.3 | 0.18 | 0.12 | 0.2 |
| 10 | 105910221 | rs703333 | -0.889 | -0.885 | 0.354 | -0.487 | -0.015 | -0.008 | -0.008 | 0 | 0.24 | 0.66 | 0.16 | 0.23 | 0.16 | 0.17 |
| 10 | 105911963 | rs666156 | -0.123 | -0.873 | 0.332 | -0.482 | -0.026 | -0.006 | -0.006 | 0 | 0.2 | 0.39 | 0.14 | 0.23 | 0.16 | 0.17 |
| 10 | 105911993 | rs11191933 | -0.202 | -0.773 | -0.535 | -0.448 | -0.024 | 0.011 | 0.011 | 0 | 0.16 | 0.08 | 0.17 | 0.13 | 0.19 | 0.2 |
| 10 | 105915340 | rs703336 | -0.914 | -0.918 | -0.175 | -0.398 | -0.026 | -0.013 | -0.013 | 0.342 | 0.24 | 0.64 | 0.16 | 0.23 | 0.16 | 0.17 |
| 10 | 105917031 | rs596229 | -0.821 | -0.903 | -0.177 | -0.392 | -0.044 | -0.02 | -0.02 | 0 | 0.23 | 0.58 | 0.16 | 0.23 | 0.16 | 0.17 |
| 10 | 105926906 | rs10883974 | -1.135 | -0.247 | -1.18 | -0.5 | -0.028 | 0.052 | 0.052 | 0 | 0.27 | 0.13 | 0.19 | 0.14 | 0.3 | 0.26 |
| 10 | 105927104 | rs10748861 | 0.051 | -1.701 | -2.057 | -1.221 | 0.2 | 0.229 | 0.229 | 0.05 | 0.49 | 0.15 | 0.68 | 0.81 | 0.78 | 0.58 |
| 10 | 105927460 | rs17750886 | 0.046 | -0.989 | -1.311 | -0.895 | -0.032 | 0.037 | 0.037 | 0 | 0.1 | 0.03 | 0.09 | 0 | 0.16 | 0.08 |
| 10 | 105928215 | rs2864009 | -0.118 | -1.136 | -1.277 | -0.897 | -0.041 | 0.042 | 0.042 | 0 | 0.12 | 0.04 | 0.1 | 0 | 0.2 | 0.11 |
| 10 | 105933138 | rs581533 | -1.443 | -1.548 | -0.496 | -1.104 | -0.004 | -0.029 | -0.029 | 0 | 0.41 | 0.44 | 0.23 | 0.25 | 0.36 | 0.4 |
| 10 | 105933320 | rs11191937 | 0.739 | -1.624 | -0.365 | -1.142 | 0.024 | 0.021 | 0.021 | 9.076 | 0.17 | 0.12 | 0.08 | 0.12 | 0.11 | 0.14 |
| 10 | 105933417 | rs642743 | -2.126 | -1.801 | 0.069 | -1.132 | 0.022 | 0.077 | 0.077 | 0.253 | 0.49 | 0.69 | 0.36 | 0.39 | 0.43 | 0.52 |
| 10 | 105941240 | rs11191942 | 0.329 | -1.725 | 0.036 | -1.1 | 0.005 | -0.007 | -0.007 | 0 | 0.08 | 0.04 | 0.03 | 0.12 | 0.05 | 0.14 |
| 10 | 105941325 | rs12775992 | -0.587 | -1.742 | 0.045 | -1.117 | -0.017 | 0.074 | 0.074 | 3.152 | 0.29 | 0.16 | 0.18 | 0.12 | 0.31 | 0.25 |
| 10 | 105950516 | rs10883978 | 0.183 | -1.635 | 0.24 | -0.977 | 0.005 | -0.007 | -0.007 | 0 | 0.08 | 0.04 | 0.03 | 0.12 | 0.05 | 0.14 |
| 10 | 105952623 | rs11813171 | -0.702 | -1.649 | 0.319 | -0.962 | -0.002 | 0.04 | 0.04 | 0.702 | 0.08 | 0.23 | 0.13 | 0.15 | 0.07 | 0.13 |
| 10 | 105957714 | rs10883979 | 0.092 | -1.541 | 0.169 | -0.907 | 0.005 | -0.007 | -0.007 | 0 | 0.08 | 0.04 | 0.03 | 0.12 | 0.05 | 0.14 |
| 10 | 105962291 | rs11191946 | -0.522 | -1.612 | 0.177 | -0.937 | 0.001 | 0 | 0 | 0 | 0.1 | 0.06 | 0.04 | 0.12 | 0.05 | 0.14 |
| 10 | 105963553 | rs11191947 | 0.075 | -1.482 | 0.133 | -0.88 | 0.005 | -0.007 | -0.007 | 0 | 0.08 | 0.04 | 0.03 | 0.12 | 0.05 | 0.14 |
| 10 | 105983609 | rs480185 | -0.334 | -1.484 | 0.243 | -0.881 | -0.012 | 0.008 | 0.008 | 0 | 0.1 | 0.18 | 0.04 | 0.12 | 0.05 | 0.14 |
| 10 | 105987542 | rs17752045 | 0.509 | -1.644 | -0.152 | -0.94 | 0.03 | 0.01 | 0.01 | 8.812 | 0.2 | 0.12 | 0.15 | 0.17 | 0.15 | 0.21 |
| 10 | 105989664 | rs703343 | -2.048 | -0.775 | 0.362 | -0.746 | 0.005 | -0.023 | -0.023 | 0.179 | 0.29 | 0.58 | 0.21 | 0.23 | 0.28 | 0.32 |
| 10 | 105997104 | rs12356059 | -1.121 | -0.233 | -0.081 | -0.666 | 0.022 | -0.03 | -0.03 | 0 | 0.18 | 0.26 | 0.16 | 0.11 | 0.23 | 0.17 |
| 10 | 106006181 | rs7896809 | -0.415 | -0.504 | -0.328 | -0.525 | -0.014 | 0.014 | 0.014 | 0.301 | 0.25 | 0.21 | 0.2 | 0.13 | 0.3 | 0.15 |
| 10 | 106006518 | rs12769390 | -0.611 | -0.787 | -0.43 | -0.631 | 0.008 | 0.027 | 0.027 | 0 | 0.08 | 0.02 | 0.03 | 0.01 | 0.06 | 0.16 |
| 10 | 106014467 | rs628480 | -0.42 | -1.184 | -0.161 | -0.786 | 0.018 | 0.023 | 0.023 | 0 | 0.93 | 0.97 | 0.99 | 0.98 | 0.98 | 0.96 |
| 10 | 106022789 | rs4925 | -0.408 | -1.684 | 1.473 | -0.985 | -0.072 | 0.054 | 0.054 | 0.081 | 0.28 | 0.07 | 0.21 | 0.13 | 0.34 | 0.19 |
| 10 | 106024867 | rs11191979 | -0.417 | -1.615 | 1.476 | -0.984 | -0.078 | 0.05 | 0.05 | 0.084 | 0.28 | 0.06 | 0.2 | 0.13 | 0.34 | 0.19 |
| 10 | 106025258 | rs1147611 | -1.515 | -1.718 | 1.583 | -1.04 | 0.011 | 0.109 | 0.109 | 4.833 | 0.45 | 0.81 | 0.29 | 0.25 | 0.4 | 0.35 |
| 10 | 106032213 | rs11191980 | -0.176 | -1.559 | 1.47 | -0.904 | -0.072 | 0.03 | 0.03 | 0.031 | 0.25 | 0.07 | 0.19 | 0.13 | 0.31 | 0.17 |
| 10 | 106037894 | rs157077 | -1.896 | -1.757 | 0.912 | -0.971 | -0.047 | 0 | 0 | 0.082 | 0.57 | 0.81 | 0.34 | 0.41 | 0.48 | 0.48 |
| 10 | 106039185 | rs156697 | -1.617 | -1.002 | 1.678 | -0.854 | -0.001 | 0.083 | 0.083 | 4.476 | 0.42 | 0.8 | 0.28 | 0.26 | 0.37 | 0.33 |
| 10 | 106039303 | rs276203 | -1.617 | -1.002 | 1.678 | -0.854 | -0.001 | 0.083 | 0.083 | 4.476 | 0.42 | 0.8 | 0.28 | 0.26 | 0.37 | 0.33 |
| 10 | 106042502 | rs1865926 | -0.507 | -0.729 | 1.907 | -0.943 | -0.017 | 0.028 | 0.028 | 0.015 | 0.26 | 0.2 | 0.19 | 0.08 | 0.31 | 0.17 |
| 10 | 106053641 | rs156699 | -1.445 | -0.79 | 2.098 | -0.83 | -0.002 | 0.054 | 0.054 | 3.399 | 0.37 | 0.77 | 0.25 | 0.2 | 0.33 | 0.21 |
| 10 | 106055963 | rs157080 | -1.445 | -0.79 | 2.098 | -0.83 | -0.002 | 0.054 | 0.054 | 3.399 | 0.37 | 0.77 | 0.25 | 0.2 | 0.33 | 0.21 |
| 10 | 106061169 | rs3740466 | -0.524 | -0.766 | 2.219 | -0.834 | -0.012 | 0.019 | 0.019 | 0.043 | 0.27 | 0.22 | 0.2 | 0.08 | 0.31 | 0.17 |
| 10 | 106064415 | rs11191994 | -0.843 | -0.776 | 2.203 | -0.806 | -0.023 | 0.027 | 0.027 | 0.029 | 0.07 | 0.14 | 0.03 | 0.12 | 0.02 | 0.03 |
| 10 | 106066863 | rs7085593 | -0.843 | -0.776 | 2.203 | -0.806 | -0.023 | 0.027 | 0.027 | 0.029 | 0.07 | 0.14 | 0.03 | 0.12 | 0.02 | 0.03 |
| 10 | 106067978 | rs11192001 | -0.526 | -0.784 | 2.156 | -0.834 | -0.011 | 0.016 | 0.016 | 0.041 | 0.27 | 0.23 | 0.2 | 0.08 | 0.31 | 0.17 |
| 10 | 106068471 | rs2282092 | -1.32 | -0.792 | 2.223 | -0.832 | -0.009 | 0.074 | 0.074 | 0.693 | 0.08 | 0.3 | 0.04 | 0.12 | 0.02 | 0.03 |
| 10 | 106071001 | rs10491046 | -0.537 | -0.698 | 1.914 | -0.834 | -0.011 | 0.017 | 0.017 | 0.049 | 0.27 | 0.23 | 0.2 | 0.08 | 0.32 | 0.17 |
| 10 | 106073566 | rs505701 | -1.404 | -0.683 | 1.508 | -0.815 | -0.003 | 0.054 | 0.054 | 3.298 | 0.37 | 0.77 | 0.25 | 0.2 | 0.34 | 0.21 |
| 10 | 106081946 | rs276228 | -1.175 | -0.238 | 1.515 | -0.651 | 0.021 | 0.073 | 0.073 | 0 | 0.07 | 0.33 | 0.04 | 0.13 | 0.03 | 0.02 |
| 10 | 106087512 | rs276219 | -1.36 | -0.207 | 1.344 | -0.833 | -0.006 | 0.043 | 0.043 | 4.446 | 0.35 | 0.73 | 0.25 | 0.21 | 0.34 | 0.21 |
| 10 | 106088467 | rs276218 | -0.989 | -0.029 | 1.129 | -0.719 | 0.032 | 0.089 | 0.089 | 0 | 0.07 | 0.34 | 0.04 | 0.13 | 0.03 | 0.03 |
| 10 | 106099408 | rs7095770 | -0.492 | 0.233 | 0.272 | -0.894 | 0.004 | -0.013 | -0.013 | 0.067 | 0.24 | 0.21 | 0.19 | 0.08 | 0.3 | 0.18 |
| 10 | 106125299 | rs17827720 | -0.117 | 0.114 | -0.353 | -0.781 | -0.053 | 0.095 | 0.095 | 0 | 0.26 | 0.06 | 0.17 | 0.07 | 0.27 | 0.18 |
| 10 | 106135117 | rs12267798 | -1.07 | 0.051 | 0.344 | -0.55 | 0.034 | -0.009 | -0.009 | 0 | 0.08 | 0.34 | 0.05 | 0.03 | 0.03 | 0.03 |
| 10 | 106137186 | rs914067 | -1.229 | -0.041 | -1.309 | -0.125 | -0.133 | -0.168 | -0.168 | 0 | 0.72 | 0.98 | 0.56 | 0.58 | 0.6 | 0.62 |
| 10 | 106138539 | rs7083064 | -0.718 | 0.181 | -1.031 | -0.658 | 0.001 | 0.17 | 0.17 | 2.891 | 0.42 | 0.85 | 0.37 | 0.18 | 0.3 | 0.29 |
| 10 | 106139081 | rs7908436 | -0.36 | 0.175 | -0.167 | -0.234 | -0.003 | 0.021 | 0.021 | 0 | 0.47 | 0.4 | 0.28 | 0.29 | 0.47 | 0.3 |
| 10 | 106139294 | rs7908698 | -0.959 | -0.415 | 0.277 | 0.013 | -0.058 | -0.014 | -0.014 | 0.008 | 0.17 | 0.51 | 0.09 | 0.15 | 0.06 | 0.18 |
| 10 | 106145621 | rs12249796 | -0.577 | -0.107 | -0.029 | -0.116 | 0.009 | 0.013 | 0.013 | 3.017 | 0.33 | 0.28 | 0.19 | 0.15 | 0.21 | 0.15 |
| 10 | 106154143 | rs11192037 | -0.151 | -0.278 | 0.41 | -0.193 | 0.005 | 0.073 | 0.073 | 0.268 | 0.28 | 0.06 | 0.17 | 0.13 | 0.2 | 0.11 |

| | | | | | | | | | | | | | | | | |
|---|---|---|---|---|---|---|---|---|---|---|---|---|---|---|---|---|
| 10 | 106159305 | rs10509772 | -1.545 | 0.576 | 0.805 | 0.137 | -0.01 | -0.008 | -0.008 | 0 | 0.54 | 0.79 | 0.42 | 0.51 | 0.49 | 0.45 |
| 10 | 106166570 | rs10786792 | -0.37 | 0.534 | 1.321 | 0.18 | 0.066 | 0.082 | 0.082 | 0 | 0.18 | 0.41 | 0.23 | 0.36 | 0.27 | 0.3 |
| 10 | 106172600 | rs10883998 | 0.072 | 0.507 | 1.32 | 0.175 | 0.015 | 0.005 | 0.005 | 0 | 0.15 | 0.2 | 0.19 | 0.22 | 0.26 | 0.21 |
| 10 | 106173727 | rs12250034 | 0.079 | 0.535 | 1.268 | 0.137 | 0.033 | 0.064 | 0.064 | 0.342 | 0.29 | 0.12 | 0.18 | 0.15 | 0.2 | 0.15 |
| 10 | 106175589 | rs2864011 | -1.296 | 0.472 | 1.297 | 0.171 | -0.005 | 0.006 | 0.006 | 0.398 | 0.51 | 0.72 | 0.41 | 0.51 | 0.48 | 0.45 |
| 10 | 106179063 | rs10884000 | -1.269 | 0.514 | 1.285 | 0.164 | 0 | 0.004 | 0.004 | 0.956 | 0.51 | 0.61 | 0.41 | 0.51 | 0.49 | 0.46 |
| 10 | 106193598 | rs7087214 | -1.981 | 0.52 | 1.067 | 0.25 | 0.032 | 0.159 | 0.159 | 3.117 | 0.07 | 0.43 | 0.04 | 0.14 | 0.01 | 0.14 |
| 10 | 106200600 | rs10786794 | -0.574 | 0.513 | 0.632 | 0.172 | -0.073 | 0.03 | 0.03 | 0 | 0.41 | 0.11 | 0.34 | 0.34 | 0.44 | 0.28 |
| 10 | 106202531 | rs7087113 | -0.546 | 0.352 | 1.102 | 0.35 | -0.052 | 0.018 | 0.018 | 0 | 0.29 | 0.52 | 0.18 | 0.35 | 0.17 | 0.21 |
| 10 | 106207209 | rs9651453 | -0.367 | 1.123 | 0.519 | 0.336 | -0.001 | 0.012 | 0.012 | 0 | 0.28 | 0.23 | 0.23 | 0.23 | 0.26 | 0.16 |
| 10 | 106207933 | rs7095475 | -0.526 | 0.903 | 0.859 | 0.296 | -0.021 | -0.019 | -0.019 | 0 | 0.25 | 0.43 | 0.18 | 0.21 | 0.14 | 0.2 |
| 10 | 106209230 | rs7907074 | -2.637 | 0.937 | 1.034 | 0.384 | -0.016 | 0.068 | 0.068 | 2.998 | 0.08 | 0.35 | 0.06 | 0.14 | 0.03 | 0.17 |
| 10 | 106211449 | rs17761559 | 0.564 | 1.196 | 0.732 | 0.175 | -0.012 | 0.009 | 0.009 | 0 | 0.07 | 0.03 | 0.07 | 0 | 0.13 | 0.08 |
| 10 | 106213055 | rs1334758 | -2.049 | 0.778 | 0.883 | 0.463 | -0.008 | 0.048 | 0.048 | 2.43 | 0.05 | 0.24 | 0.05 | 0.12 | 0.02 | 0.1 |
| 10 | 106214081 | rs17117058 | -1.983 | 0.774 | 0.906 | 0.446 | -0.005 | -0.019 | -0.019 | 0 | 0.05 | 0.24 | 0.04 | 0.01 | 0.03 | 0.07 |
| 10 | 106214937 | rs11192069 | -1.331 | 1.058 | 0.589 | 0.726 | 0.026 | 0.021 | 0.021 | 0 | 0.18 | 0.31 | 0.18 | 0.29 | 0.27 | 0.25 |
| 10 | 106217667 | rs2050363 | -0.165 | 1.359 | -0.596 | 0.657 | -0.051 | -0.021 | -0.021 | 0.358 | 0.12 | 0.04 | 0.13 | 0.15 | 0.24 | 0.15 |
| 10 | 106235467 | rs280839 | 0.451 | 0.974 | -0.682 | 0.993 | -0.058 | -0.008 | -0.008 | 0 | 0.63 | 0.81 | 0.67 | 0.35 | 0.54 | 0.68 |
| 10 | 106237814 | rs3895536 | -0.644 | 0.097 | -0.649 | 1.092 | -0.013 | -0.024 | -0.024 | 0 | 0.25 | 0.18 | 0.15 | 0.27 | 0.27 | 0.29 |
| 10 | 106242515 | rs280845 | 0.731 | -0.243 | -0.705 | 0.826 | 0.005 | -0.062 | -0.062 | 0.083 | 0.37 | 0.13 | 0.54 | 0.47 | 0.37 | 0.37 |
| 10 | 106253104 | rs10884005 | 0.469 | 0.085 | -0.239 | 0.734 | -0.049 | -0.061 | -0.061 | 0 | 0.28 | 0.13 | 0.43 | 0.38 | 0.37 | 0.32 |
| 10 | 106253793 | rs280825 | 0.355 | -0.21 | 0.384 | 0.559 | 0.002 | -0.001 | -0.001 | 0 | 0.54 | 0.46 | 0.45 | 0.55 | 0.42 | 0.56 |
| 10 | 106258039 | rs7072764 | 0.4 | -0.261 | 0.934 | 0.572 | 0.01 | 0.005 | 0.005 | 0 | 0.54 | 0.41 | 0.45 | 0.5 | 0.42 | 0.53 |
| 10 | 106258890 | rs12761070 | -0.159 | -0.232 | 0.675 | 0.549 | -0.005 | 0.003 | 0.003 | 0.162 | 0.12 | 0.1 | 0.09 | 0.06 | 0.19 | 0.11 |
| 10 | 106260146 | rs10884008 | 0.577 | -0.273 | 0.752 | 0.568 | -0.051 | -0.064 | -0.064 | 1.411 | 0.29 | 0.15 | 0.43 | 0.39 | 0.38 | 0.32 |
| 10 | 106265362 | rs1028915 | 0.3 | -0.215 | 0.868 | 0.621 | 0.013 | 0.011 | 0.011 | 0.645 | 0.87 | 0.83 | 0.89 | 0.82 | 0.8 | 0.87 |
| 10 | 106265737 | rs1028917 | 0.299 | -0.214 | 0.846 | 0.622 | 0.025 | 0.022 | 0.022 | 0.672 | 0.87 | 0.81 | 0.89 | 0.82 | 0.8 | 0.87 |
| 10 | 106270406 | rs950129 | 0.45 | -0.414 | 1.004 | 0.331 | -0.004 | -0.005 | -0.005 | 0 | 0.53 | 0.49 | 0.44 | 0.43 | 0.41 | 0.52 |
| 10 | 106272669 | rs11192090 | -0.401 | -0.299 | 0.869 | 0.367 | 0.022 | -0.017 | -0.017 | 1.148 | 0.13 | 0.2 | 0.1 | 0.06 | 0.19 | 0.1 |
| 10 | 106272813 | rs10786801 | -0.683 | -0.247 | 0.965 | 0.393 | -0.021 | -0.021 | -0.021 | 0 | 0.69 | 0.76 | 0.54 | 0.49 | 0.6 | 0.67 |
| 10 | 106273046 | rs4918098 | -0.842 | -0.243 | 0.966 | 0.394 | -0.019 | -0.019 | -0.019 | 1.244 | 0.7 | 0.76 | 0.55 | 0.49 | 0.6 | 0.68 |
| 10 | 106274304 | rs6584604 | 0.144 | -0.176 | 1.013 | 0.323 | 0.013 | 0.009 | 0.009 | 0 | 0.34 | 0.42 | 0.47 | 0.52 | 0.4 | 0.36 |
| 10 | 106275865 | rs962139 | -0.255 | -0.496 | 0.489 | 0.369 | 0.013 | 0.026 | 0.026 | 0 | 0.57 | 0.71 | 0.43 | 0.43 | 0.41 | 0.58 |
| 10 | 106279303 | rs10884013 | 0.558 | -0.486 | 0.106 | 0.1 | 0.028 | 0.068 | 0.068 | 0 | 0.45 | 0.2 | 0.39 | 0.39 | 0.39 | 0.53 |
| 10 | 106281629 | rs6584608 | 1.437 | -0.344 | 0.328 | -0.139 | -0.004 | -0.011 | -0.011 | 0 | 0.76 | 0.32 | 0.86 | 0.85 | 0.79 | 0.7 |
| 10 | 106282595 | rs10884015 | 0.812 | -0.36 | 0.995 | -0.146 | 0.029 | -0.008 | -0.008 | 0 | 0.82 | 0.62 | 0.9 | 0.85 | 0.79 | 0.71 |
| 10 | 106282810 | rs2183167 | -0.225 | -0.336 | 1.036 | -0.137 | 0.007 | 0.002 | 0.002 | 0.231 | 0.88 | 0.86 | 0.92 | 0.86 | 0.81 | 0.76 |
| 10 | 106285308 | rs975145 | -0.643 | -0.713 | 0.987 | -0.398 | -0.007 | -0.06 | -0.06 | 3.521 | 0.29 | 0.46 | 0.17 | 0.14 | 0.27 | 0.33 |
| 10 | 106286419 | rs972243 | 0.432 | -0.71 | 0.499 | -0.397 | -0.035 | 0.022 | 0.022 | 0 | 0.21 | 0.05 | 0.13 | 0.14 | 0.26 | 0.27 |
| 10 | 106287147 | rs11192101 | 0.939 | -0.81 | 0.388 | -0.374 | 0.002 | 0.023 | 0.023 | 0 | 0.33 | 0.18 | 0.37 | 0.33 | 0.33 | 0.41 |
| 10 | 106295606 | rs11192108 | -0.34 | -0.801 | 0.366 | -0.366 | -0.001 | 0.015 | 0.015 | 0 | 0.28 | 0.23 | 0.16 | 0.14 | 0.28 | 0.33 |
| 10 | 106299935 | rs10884017 | 1.002 | -0.735 | -0.258 | -0.262 | -0.051 | -0.059 | -0.059 | 0 | 0.33 | 0.14 | 0.44 | 0.52 | 0.39 | 0.26 |
| 10 | 106315243 | rs2864024 | 0.823 | -0.85 | 0.16 | -0.463 | 0.004 | 0.013 | 0.013 | 0.014 | 0.64 | 0.35 | 0.81 | 0.86 | 0.7 | 0.65 |
| 10 | 106316476 | rs1157171 | -0.079 | -0.624 | 0.008 | -0.366 | -0.02 | 0.027 | 0.027 | 0 | 0.24 | 0.12 | 0.14 | 0.14 | 0.29 | 0.29 |
| 10 | 106318839 | rs1334612 | 0.144 | -0.318 | -0.401 | -0.207 | -0.003 | -0.036 | -0.036 | 0 | 0.67 | 0.47 | 0.82 | 0.86 | 0.69 | 0.65 |
| 10 | 106328791 | rs2902643 | -0.023 | 0.494 | -0.911 | 0.229 | -0.055 | -0.053 | -0.053 | 0 | 0.35 | 0.17 | 0.39 | 0.51 | 0.44 | 0.38 |
| 10 | 106349202 | rs2183164 | 0.801 | -0.388 | 0.146 | -0.02 | -0.061 | 0.016 | 0.016 | 0.235 | 0.85 | 0.52 | 0.85 | 0.86 | 0.93 | 0.9 |
| 10 | 106357532 | rs10786805 | -0.224 | -0.452 | -0.179 | -0.193 | -0.003 | -0.008 | -0.008 | 0 | 0.14 | 0.04 | 0.25 | 0.2 | 0.14 | 0.35 |
| 10 | 106361945 | rs6584618 | 0.445 | -0.632 | 0.3 | -0.198 | -0.038 | 0.072 | 0.072 | 3.017 | 0.9 | 0.65 | 0.92 | 0.87 | 0.99 | 0.93 |
| 10 | 106366594 | rs12244579 | 0.642 | -0.079 | -0.368 | -0.598 | -0.028 | 0.032 | 0.032 | 0 | 0.09 | 0.02 | 0.09 | 0 | 0.13 | 0.03 |
| 10 | 106371784 | rs1855581 | -1.62 | -0.808 | 0.329 | -0.668 | -0.006 | 0.001 | 0.001 | 0.331 | 0.3 | 0.65 | 0.39 | 0.35 | 0.2 | 0.48 |
| 10 | 106377686 | rs1413712 | 1.491 | -0.794 | 0.247 | -0.664 | 0.021 | -0.001 | -0.001 | 0 | 0.67 | 0.26 | 0.6 | 0.65 | 0.8 | 0.52 |
| 10 | 106377705 | rs11192137 | -0.339 | -0.209 | 0.153 | -0.583 | -0.001 | 0.001 | 0.001 | 0 | 0.16 | 0.18 | 0.27 | 0.21 | 0.14 | 0.38 |
| 10 | 106384631 | rs10786809 | 0.717 | -0.373 | 0.262 | -0.687 | -0.014 | -0.001 | -0.001 | 1.867 | 0.8 | 0.64 | 0.73 | 0.77 | 0.84 | 0.62 |
| 10 | 106388689 | rs1577223 | 0.974 | -1.07 | -0.009 | -1.179 | -0.033 | -0.03 | -0.03 | 0 | 0.7 | 0.59 | 0.68 | 0.76 | 0.79 | 0.63 |
| 10 | 106391762 | rs10748866 | 1.334 | 0.145 | 0.346 | -1.245 | -0.004 | -0.008 | -0.008 | 0 | 0.64 | 0.39 | 0.55 | 0.67 | 0.64 | 0.56 |
| 10 | 106392464 | rs4918121 | 0.797 | 0.846 | -0.393 | -1.814 | -0.053 | -0.034 | -0.034 | 0 | 0.22 | 0.09 | 0.2 | 0.65 | 0.33 | 0.33 |
| 10 | 106392549 | rs4918122 | 1.471 | 0.235 | -0.186 | -1.827 | 0.034 | 0.08 | 0.08 | 1.193 | 0.54 | 0.18 | 0.38 | 0.65 | 0.56 | 0.52 |
| 10 | 106403115 | rs7068684 | -0.309 | 0.219 | 0.436 | -1.722 | -0.025 | 0.008 | 0.008 | 0.012 | 0.82 | 0.68 | 0.72 | 0.77 | 0.87 | 0.65 |
| 10 | 106405854 | rs4918123 | 1.601 | 0.75 | 0.041 | -1.879 | -0.013 | 0.114 | 0.114 | 1.178 | 0.51 | 0.16 | 0.32 | 0.66 | 0.47 | 0.51 |
| 10 | 106409545 | rs1467819 | -0.828 | 0.823 | 0.25 | -1.908 | -0.01 | -0.027 | -0.027 | 0 | 0.33 | 0.4 | 0.39 | 0.23 | 0.31 | 0.39 |
| 10 | 106417957 | rs4532962 | 1.859 | 1.115 | 0.234 | -1.781 | -0.037 | 0.045 | 0.045 | 0.044 | 0.6 | 0.16 | 0.36 | 0.66 | 0.53 | 0.52 |
| 10 | 106424921 | rs12412809 | -0.096 | 0.846 | 0.425 | -1.809 | -0.001 | -0.007 | -0.007 | 1.494 | 0.07 | 0.06 | 0.16 | 0.01 | 0.08 | 0.04 |
| 10 | 106439347 | rs7097737 | -0.433 | 0.845 | 0.551 | -1.809 | -0.003 | -0.03 | -0.03 | 0.75 | 0.07 | 0.14 | 0.17 | 0.01 | 0.09 | 0.04 |
| 10 | 106440612 | rs7912178 | -0.31 | 0.818 | 0.361 | -1.621 | 0.011 | 0.006 | 0.006 | 0 | 0.18 | 0.29 | 0.3 | 0.23 | 0.2 | 0.37 |
| 10 | 106450395 | rs9943297 | -0.998 | 0.989 | -0.196 | -1.508 | -0.001 | 0.001 | 0.001 | 0 | 0.13 | 0.12 | 0.24 | 0.25 | 0.15 | 0.31 |
| 10 | 106454672 | rs11599236 | 0.857 | 0.613 | -0.923 | -2.095 | 0.022 | 0.147 | 0.147 | 0 | 0.49 | 0.14 | 0.32 | 0.68 | 0.45 | 0.42 |
| 10 | 106467853 | rs3896224 | 0.838 | 0.568 | -0.896 | -2.081 | 0.028 | 0.125 | 0.125 | 0 | 0.49 | 0.17 | 0.32 | 0.66 | 0.45 | 0.42 |
| 10 | 106469961 | rs11818524 | -0.153 | 0.138 | -0.388 | -2.274 | 0 | -0.001 | -0.001 | 0 | 0.2 | 0.21 | 0.31 | 0.23 | 0.23 | 0.36 |

| | | | | | | | | | | | | | | | | |
|---|---|---|---|---|---|---|---|---|---|---|---|---|---|---|---|---|
| 10 | 106473048 | rs4350297 | 1.16 | -0.429 | -1.173 | -2.638 | 0.007 | 0.136 | 0.136 | 0.265 | 0.24 | 0.02 | 0.13 | 0.03 | 0.2 | 0.15 |
| 10 | 106478366 | rs12256390 | 1.475 | -0.148 | -0.836 | -2.62 | 0.002 | 0.003 | 0.003 | 0 | 0.3 | 0.22 | 0.21 | 0.66 | 0.28 | 0.34 |
| 10 | 106492420 | rs1490183 | -2.049 | -0.174 | -0.171 | -2.708 | 0.008 | -0.014 | -0.014 | 0 | 0.84 | 0.85 | 0.75 | 0.94 | 0.76 | 0.9 |
| 10 | 106498526 | rs999981 | -0.213 | -0.291 | -0.097 | -2.744 | 0.032 | 0.083 | 0.083 | 0 | 0.45 | 0.3 | 0.43 | 0.26 | 0.42 | 0.52 |
| 10 | 106499611 | rs1565415 | -0.467 | -0.282 | 0.038 | -2.716 | 0.035 | 0.092 | 0.092 | 0 | 0.5 | 0.36 | 0.5 | 0.28 | 0.47 | 0.56 |
| 10 | 106512404 | rs2491368 | -2.094 | -0.127 | -0.348 | -2.66 | -0.033 | 0.029 | 0.029 | 0 | 0.85 | 0.93 | 0.76 | 0.94 | 0.76 | 0.9 |
| 10 | 106517759 | rs752053 | -2.174 | -0.127 | -0.346 | -2.66 | -0.05 | 0.042 | 0.042 | 0 | 0.85 | 0.95 | 0.76 | 0.94 | 0.76 | 0.9 |
| 10 | 106520940 | rs4918125 | 1.457 | -0.266 | -0.233 | -2.66 | -0.001 | -0.001 | -0.001 | 0 | 0.31 | 0.27 | 0.22 | 0.62 | 0.28 | 0.34 |
| 10 | 106521003 | rs4918126 | -1.734 | -0.112 | -0.575 | -2.626 | -0.008 | 0.002 | 0.002 | 0 | 0.83 | 0.85 | 0.75 | 0.94 | 0.76 | 0.9 |
| 10 | 106529564 | rs17766570 | 1.725 | -0.36 | -0.953 | -2.687 | -0.009 | 0.112 | 0.112 | 0 | 0.2 | 0.03 | 0.12 | 0 | 0.19 | 0.12 |
| 10 | 106531703 | rs10884043 | 1.66 | -0.516 | -0.958 | -2.66 | -0.021 | -0.017 | -0.017 | 0 | 0.28 | 0.07 | 0.21 | 0.65 | 0.28 | 0.34 |
| 10 | 106534317 | rs970054 | -1.852 | -0.233 | -0.675 | -2.674 | -0.021 | 0.017 | 0.017 | 0 | 0.84 | 0.88 | 0.76 | 0.94 | 0.75 | 0.9 |
| 10 | 106534351 | rs970052 | -1.773 | -0.241 | -0.68 | -2.678 | -0.023 | 0.019 | 0.019 | 0 | 0.83 | 0.88 | 0.76 | 0.94 | 0.75 | 0.9 |
| 10 | 106536976 | rs2491365 | -0.811 | -0.421 | -0.431 | -2.761 | -0.012 | -0.015 | -0.015 | 0.516 | 0.3 | 0.65 | 0.41 | 0.25 | 0.27 | 0.4 |
| 10 | 106539062 | rs2496034 | -2.819 | -0.075 | -0.566 | -2.749 | -0.08 | 0.031 | 0.031 | 0 | 0.88 | 0.99 | 0.91 | 0.94 | 0.82 | 0.9 |
| 10 | 106549900 | rs1490171 | -0.048 | -0.558 | -0.622 | -2.898 | 0.004 | 0.152 | 0.152 | 0 | 0.53 | 0.75 | 0.56 | 0.9 | 0.5 | 0.66 |
| 10 | 106554417 | rs5004408 | -0.778 | -0.51 | -0.679 | -2.885 | -0.024 | 0.024 | 0.024 | 0 | 0.77 | 0.81 | 0.69 | 0.93 | 0.71 | 0.81 |
| 10 | 106555387 | rs2451463 | -0.391 | -0.606 | -0.66 | -2.936 | 0.033 | 0.072 | 0.072 | 0 | 0.47 | 0.66 | 0.47 | 0.27 | 0.43 | 0.47 |
| 10 | 106556804 | rs2496029 | -0.782 | -0.597 | -0.661 | -2.932 | -0.024 | 0.025 | 0.025 | 0 | 0.77 | 0.81 | 0.69 | 0.93 | 0.71 | 0.81 |
| 10 | 106556964 | rs2451462 | 0.772 | -0.644 | -0.192 | -2.935 | -0.004 | -0.002 | -0.002 | 0 | 0.19 | 0.31 | 0.32 | 0.24 | 0.22 | 0.32 |
| 10 | 106558025 | rs4556466 | 1.798 | -0.527 | -0.377 | -2.94 | 0.002 | 0.015 | 0.015 | 0 | 0.29 | 0.15 | 0.32 | 0.67 | 0.27 | 0.34 |
| 10 | 106559787 | rs2491391 | -0.364 | -1.046 | -0.13 | -2.898 | -0.007 | -0.027 | -0.027 | 0.012 | 0.46 | 0.5 | 0.35 | 0.26 | 0.45 | 0.47 |
| 10 | 106562226 | rs2496026 | 0.416 | -1.252 | 0.165 | -2.805 | -0.037 | -0.01 | -0.01 | 0 | 0.18 | 0.27 | 0.13 | 0.2 | 0.12 | 0.28 |
| 10 | 106571525 | rs744823 | 1.548 | -1.019 | -0.243 | -2.693 | -0.02 | 0.011 | 0.011 | 0 | 0.31 | 0.19 | 0.51 | 0.64 | 0.41 | 0.36 |
| 10 | 106582029 | rs1490182 | 0.322 | -0.397 | -0.209 | -2.511 | -0.016 | 0.002 | 0.002 | 0 | 0.86 | 0.91 | 0.86 | 0.83 | 0.79 | 0.85 |
| 10 | 106599009 | rs2496014 | -0.362 | -0.927 | -0.326 | -2.722 | -0.042 | 0.07 | 0.07 | 0.132 | 0.7 | 0.9 | 0.75 | 0.85 | 0.65 | 0.58 |
| 10 | 106600343 | rs7900775 | 0.321 | -1.097 | 0.182 | -2.789 | 0.01 | -0.076 | -0.076 | 0 | 0.65 | 0.56 | 0.72 | 0.83 | 0.65 | 0.55 |
| 10 | 106603824 | rs2451472 | 0.011 | -1.277 | 0.141 | -2.871 | 0.01 | -0.037 | -0.037 | 0 | 0.72 | 0.59 | 0.79 | 0.85 | 0.74 | 0.77 |
| 10 | 106606612 | rs2451470 | -1.72 | -1.225 | -0.149 | -2.901 | -0.031 | 0.031 | 0.031 | 0.221 | 0.83 | 0.93 | 0.88 | 0.99 | 0.79 | 0.85 |
| 10 | 106609900 | rs10786828 | 1.589 | -1.468 | -0.333 | -3.097 | -0.046 | 0.137 | 0.137 | 0 | 0.45 | 0.15 | 0.63 | 0.69 | 0.56 | 0.4 |
| 10 | 106611268 | rs1021362 | -0.552 | -1.381 | -0.319 | -3.059 | -0.059 | 0.09 | 0.09 | 0.189 | 0.69 | 0.93 | 0.76 | 0.85 | 0.66 | 0.58 |
| 10 | 106613417 | rs7895087 | 0.575 | -1.453 | -0.168 | -3.083 | 0.003 | -0.056 | -0.056 | 0 | 0.57 | 0.55 | 0.71 | 0.83 | 0.61 | 0.48 |
| 10 | 106613683 | rs11192206 | 1.175 | -1.424 | -0.215 | -3.019 | 0.026 | -0.016 | -0.016 | 0 | 0.14 | 0.02 | 0.11 | 0.15 | 0.12 | 0.26 |
| 10 | 106614571 | rs10786831 | 1.084 | -1.483 | -0.21 | -3.12 | 0.021 | 0.141 | 0.141 | 0 | 0.54 | 0.16 | 0.66 | 0.69 | 0.62 | 0.47 |
| 10 | 106615387 | rs12259189 | -0.842 | -1.506 | 0.101 | -3.051 | -0.062 | -0.009 | -0.009 | 0.542 | 0.14 | 0.6 | 0.09 | 0.16 | 0.04 | 0.11 |
| 10 | 106616736 | rs3976793 | -1.057 | -1.575 | 0.016 | -3.078 | -0.07 | -0.018 | -0.018 | 0 | 0.15 | 0.6 | 0.09 | 0.16 | 0.04 | 0.11 |
| 10 | 106617755 | rs11192208 | 1.109 | -1.492 | -0.171 | -3.043 | 0.021 | -0.013 | -0.013 | 1.263 | 0.14 | 0.01 | 0.11 | 0.14 | 0.12 | 0.26 |
| 10 | 106622009 | rs7086583 | -0.138 | -1.463 | 0.024 | -3.005 | -0.037 | 0.03 | 0.03 | 1.592 | 0.11 | 0.37 | 0.07 | 0.16 | 0.04 | 0.08 |
| 10 | 106623425 | rs2491380 | 0.4 | -1.317 | 0.104 | -2.98 | 0.002 | -0.077 | -0.077 | 0 | 0.12 | 0.31 | 0.05 | 0.02 | 0.05 | 0.07 |
| 10 | 106629059 | rs1979635 | -1.779 | -0.965 | -0.77 | -2.858 | -0.063 | 0.067 | 0.067 | 0.338 | 0.84 | 0.97 | 0.88 | 0.98 | 0.79 | 0.86 |
| 10 | 106630686 | rs17117815 | 0.997 | -1.022 | -0.814 | -2.877 | 0.026 | 0.005 | 0.005 | 0.014 | 0.14 | 0.01 | 0.11 | 0.11 | 0.11 | 0.27 |
| 10 | 106632655 | rs2451499 | -0.305 | -0.97 | -0.724 | -2.837 | -0.023 | 0.071 | 0.071 | 0.097 | 0.69 | 0.84 | 0.76 | 0.88 | 0.67 | 0.57 |
| 10 | 106645495 | rs12779433 | 2.046 | -1.907 | -0.416 | -2.704 | 0.009 | 0.024 | 0.024 | 0.06 | 0.09 | 0.03 | 0.06 | 0 | 0.06 | 0.01 |
| 10 | 106650949 | rs7073438 | -1.204 | -1.65 | 0.111 | -2.424 | 0.052 | 0.082 | 0.082 | 1.19 | 0.4 | 0.77 | 0.25 | 0.18 | 0.25 | 0.21 |
| 10 | 106672120 | rs11192226 | -0.145 | -1.363 | 0.388 | -2.417 | -0.057 | -0.072 | -0.072 | 0 | 0.26 | 0.54 | 0.21 | 0.16 | 0.19 | 0.13 |
| 10 | 106675585 | rs6584635 | -0.956 | -1.338 | 0.296 | -2.42 | -0.015 | -0.041 | -0.041 | 0 | 0.35 | 0.62 | 0.24 | 0.15 | 0.24 | 0.17 |
| 10 | 106676141 | rs4917432 | -1.907 | -1.345 | -0.003 | -2.419 | 0.089 | 0.182 | 0.182 | 0.246 | 0.53 | 0.93 | 0.37 | 0.28 | 0.36 | 0.47 |
| 10 | 106677384 | rs2864035 | -1.901 | -1.346 | -0.003 | -2.42 | 0.09 | 0.183 | 0.183 | 0.247 | 0.53 | 0.93 | 0.37 | 0.28 | 0.36 | 0.47 |
| 10 | 106689135 | rs7080542 | -0.289 | -1.346 | 0.274 | -2.362 | -0.049 | -0.062 | -0.062 | 0 | 0.28 | 0.6 | 0.21 | 0.15 | 0.19 | 0.13 |
| 10 | 106689597 | rs1525398 | 0.247 | -1.204 | 0.257 | -2.248 | 0.023 | 0.014 | 0.014 | 0.016 | 0.14 | 0.07 | 0.11 | 0.1 | 0.11 | 0.25 |
| 10 | 106694770 | rs1404790 | 0.247 | -1.204 | 0.257 | -2.248 | 0.023 | 0.014 | 0.014 | 0.016 | 0.14 | 0.07 | 0.11 | 0.1 | 0.11 | 0.25 |
| 10 | 106709761 | rs7078084 | 0.088 | -1.131 | 0.374 | -2.275 | 0.015 | -0.065 | -0.065 | 0 | 0.11 | 0.27 | 0.05 | 0.02 | 0.05 | 0.06 |
| 10 | 106713633 | rs2177744 | -1.652 | -1.288 | 0.665 | -2.348 | -0.026 | -0.025 | -0.025 | 0 | 0.31 | 0.51 | 0.21 | 0.28 | 0.2 | 0.42 |
| 10 | 106716775 | rs2140837 | -1.331 | -1.267 | 0.76 | -2.405 | -0.039 | -0.079 | -0.079 | 0 | 0.27 | 0.47 | 0.17 | 0.12 | 0.17 | 0.33 |
| 10 | 106731970 | rs10884071 | -1.104 | -1.19 | 0.282 | -2.432 | 0.059 | 0.126 | 0.126 | 2.001 | 0.34 | 0.86 | 0.22 | 0.02 | 0.22 | 0.1 |
| 10 | 106737007 | rs1340052 | 0.598 | -0.788 | 0.261 | -2.174 | 0 | -0.016 | -0.016 | 0 | 0.08 | 0.07 | 0.03 | 0 | 0.05 | 0.03 |
| 10 | 106752957 | rs7090123 | -0.56 | -0.898 | 0.695 | -2.156 | 0.012 | 0.005 | 0.005 | 0.819 | 0.05 | 0.31 | 0.02 | 0 | 0.01 | 0.02 |
| 10 | 106754277 | rs10884077 | -0.152 | -0.778 | 0.751 | -2.331 | -0.01 | -0.026 | -0.026 | 0 | 0.17 | 0.2 | 0.14 | 0.13 | 0.18 | 0.26 |
| 10 | 106761178 | rs4917436 | 0.842 | -1.278 | 0.717 | -2.291 | -0.031 | 0.017 | 0.017 | 0 | 0.08 | 0.01 | 0.1 | 0.06 | 0.14 | 0.12 |
| 10 | 106761616 | rs11192270 | -0.858 | -1.657 | 0.697 | -2.492 | -0.015 | -0.103 | -0.103 | 0 | 0.36 | 0.65 | 0.2 | 0 | 0.23 | 0.12 |
| 10 | 106763400 | rs17118067 | 0.872 | -1.556 | 0.532 | -2.447 | -0.011 | -0.059 | -0.059 | 0.255 | 0.18 | 0.24 | 0.14 | 0 | 0.16 | 0.03 |
| 10 | 106763703 | rs1997064 | 0.418 | -1.333 | 0.395 | -2.301 | 0.001 | -0.016 | -0.016 | 0.241 | 0.11 | 0.07 | 0.03 | 0 | 0.06 | 0.06 |
| 10 | 106769696 | rs10884078 | 0.125 | -1.438 | 0.607 | -2.364 | -0.002 | -0.02 | -0.02 | 0 | 0.15 | 0.2 | 0.14 | 0.13 | 0.18 | 0.25 |
| 10 | 106771892 | rs813756 | -2.355 | -1.674 | 0.45 | -2.377 | -0.001 | 0.131 | 0.131 | 0.161 | 0.57 | 0.9 | 0.37 | 0.3 | 0.44 | 0.47 |
| 10 | 106773594 | rs11192275 | -0.952 | -1.508 | 0.893 | -2.277 | -0.036 | -0.074 | -0.074 | 0 | 0.22 | 0.45 | 0.16 | 0.13 | 0.19 | 0.28 |
| 10 | 106776484 | rs790647 | -0.878 | -1.464 | 0.935 | -2.184 | -0.011 | 0.009 | 0.009 | 0 | 0.22 | 0.25 | 0.18 | 0.31 | 0.22 | 0.31 |
| 10 | 106780515 | rs790643 | -0.667 | -1.473 | 0.971 | -2.015 | -0.019 | -0.02 | -0.02 | 0 | 0.2 | 0.28 | 0.14 | 0.16 | 0.18 | 0.26 |
| 10 | 106783551 | rs790640 | -0.302 | -1.455 | 0.916 | -2.001 | -0.011 | -0.011 | -0.011 | 1.406 | 0.18 | 0.21 | 0.14 | 0.16 | 0.18 | 0.25 |
| 10 | 106783742 | rs790639 | -2.266 | -1.553 | 0.735 | -2.096 | 0.015 | 0.098 | 0.098 | 0.008 | 0.55 | 0.86 | 0.37 | 0.33 | 0.44 | 0.44 |

| | | | | | | | | | | | | | | | | |
|---|---|---|---|---|---|---|---|---|---|---|---|---|---|---|---|---|
| 10 | 106789291 | rs10509781 | 1.157 | -1.603 | 0.794 | -2.127 | 0.006 | 0.045 | 0.045 | 0.086 | 0.18 | 0.08 | 0.15 | 0 | 0.18 | 0.05 |
| 10 | 106789880 | rs790661 | -2.398 | -1.597 | 0.774 | -2.126 | 0.019 | 0.1 | 0.1 | 0 | 0.55 | 0.85 | 0.37 | 0.33 | 0.44 | 0.44 |
| 10 | 106791365 | rs2791461 | -1.358 | -1.571 | 0.901 | -2.106 | -0.029 | -0.011 | -0.011 | 0 | 0.31 | 0.42 | 0.19 | 0.33 | 0.25 | 0.38 |
| 10 | 106792734 | rs1881224 | -1.142 | -1.525 | 0.956 | -1.929 | 0.005 | 0.001 | 0.001 | 0 | 0.37 | 0.67 | 0.22 | 0.17 | 0.26 | 0.19 |
| 10 | 106832251 | rs790746 | -0.193 | -0.589 | -0.002 | -1.454 | 0 | -0.022 | -0.022 | 0 | 0.9 | 0.9 | 0.97 | 1 | 0.94 | 0.94 |
| 10 | 106838655 | rs790733 | -1.317 | -0.591 | -0.051 | -1.646 | 0.001 | -0.002 | -0.002 | 0 | 0.34 | 0.29 | 0.43 | 0.17 | 0.3 | 0.33 |
| 10 | 106842041 | rs1470804 | 0.04 | -0.575 | -0.078 | -1.661 | -0.001 | -0.001 | -0.001 | 0 | 0.19 | 0.17 | 0.39 | 0.17 | 0.22 | 0.27 |
| 10 | 106845381 | rs17790008 | 0.198 | -0.394 | -0.123 | -1.682 | 0.004 | 0.016 | 0.016 | 0 | 0.15 | 0.05 | 0.05 | 0 | 0.1 | 0.07 |
| 10 | 106849780 | rs703472 | -0.255 | -0.392 | -0.173 | -1.662 | -0.003 | -0.007 | -0.007 | 0 | 0.2 | 0.23 | 0.4 | 0.17 | 0.23 | 0.25 |
| 10 | 106852572 | rs10748873 | 0.273 | -0.267 | -0.148 | -1.627 | 0 | -0.001 | -0.001 | 0 | 0.17 | 0.17 | 0.39 | 0.17 | 0.22 | 0.23 |
| 10 | 106858342 | rs791116 | 0.693 | -0.401 | -0.239 | -1.944 | 0 | -0.012 | -0.012 | 0 | 0.18 | 0.14 | 0.13 | 0 | 0.19 | 0.05 |
| 10 | 106864600 | rs2077677 | -0.207 | -0.412 | -0.285 | -1.75 | 0.005 | 0.003 | 0.003 | 0 | 0.18 | 0.12 | 0.08 | 0.19 | 0.12 | 0.15 |
| 10 | 106865955 | rs703479 | -1.775 | -0.34 | -0.239 | -1.677 | 0.001 | -0.01 | -0.01 | 0.796 | 0.54 | 0.47 | 0.6 | 0.36 | 0.53 | 0.44 |
| 10 | 106872585 | rs11192321 | 1.228 | -0.336 | -0.386 | -1.625 | -0.001 | -0.02 | -0.02 | 0 | 0.05 | 0.06 | 0.2 | 0 | 0.09 | 0.04 |
| 10 | 106882319 | rs1819476 | -1.739 | -0.28 | -0.312 | -1.725 | 0 | -0.005 | -0.005 | 0 | 0.53 | 0.46 | 0.62 | 0.36 | 0.54 | 0.43 |
| 10 | 106883870 | rs6584652 | -0.698 | -0.346 | -0.342 | -1.795 | 0.004 | 0.011 | 0.011 | 0.173 | 0.22 | 0.16 | 0.28 | 0.19 | 0.21 | 0.18 |
| 10 | 106885045 | rs11192323 | 1.216 | -0.374 | -0.411 | -1.833 | -0.042 | -0.044 | -0.044 | 0 | 0.11 | 0.01 | 0.2 | 0.14 | 0.13 | 0.14 |
| 10 | 106887232 | rs10786845 | -1.851 | -0.346 | -0.367 | -1.856 | -0.003 | 0.024 | 0.024 | 0 | 0.52 | 0.37 | 0.61 | 0.36 | 0.54 | 0.43 |
| 10 | 106891264 | rs10884100 | -0.417 | -0.358 | -0.375 | -1.867 | -0.003 | 0 | 0 | 0 | 0.31 | 0.23 | 0.34 | 0.36 | 0.34 | 0.32 |
| 10 | 106895025 | rs4918148 | -0.4 | -0.333 | -0.381 | -1.827 | 0.004 | 0.008 | 0.008 | 0.209 | 0.2 | 0.11 | 0.26 | 0 | 0.19 | 0.11 |
| 10 | 106898766 | rs1377008 | -1.305 | -0.254 | -0.356 | -1.729 | 0.003 | 0.043 | 0.043 | 0 | 0.47 | 0.31 | 0.4 | 0.36 | 0.45 | 0.38 |
| 10 | 106903981 | rs10509783 | -0.361 | -0.394 | -0.612 | -1.8 | -0.002 | 0.138 | 0.138 | 0 | 0.32 | 0.09 | 0.17 | 0 | 0.29 | 0.13 |
| 10 | 106907440 | rs791125 | -1.743 | -0.392 | -0.412 | -1.761 | 0.001 | -0.006 | -0.006 | 0 | 0.55 | 0.48 | 0.63 | 0.35 | 0.55 | 0.49 |
| 10 | 106909528 | rs791140 | -1.447 | -0.393 | -0.414 | -1.753 | -0.001 | 0.022 | 0.022 | 0 | 0.52 | 0.4 | 0.6 | 0.18 | 0.53 | 0.41 |
| 10 | 106917850 | rs7912305 | -0.736 | -0.372 | -0.646 | -1.732 | 0.018 | 0.12 | 0.12 | 0 | 0.37 | 0.16 | 0.2 | 0.17 | 0.32 | 0.22 |
| 10 | 106923973 | rs11192344 | 0.858 | -0.231 | -0.373 | -1.679 | 0.001 | -0.029 | -0.029 | 0.53 | 0.06 | 0.11 | 0.2 | 0 | 0.09 | 0.05 |
| 10 | 106925287 | rs17118671 | 0.499 | -0.197 | -0.182 | -1.724 | 0.009 | 0.004 | 0.004 | 0 | 0.15 | 0.18 | 0.24 | 0.16 | 0.18 | 0.24 |
| 10 | 106929313 | rs11192347 | 1.128 | -0.119 | -0.217 | -1.727 | 0.001 | -0.021 | -0.021 | 0 | 0.05 | 0.06 | 0.21 | 0 | 0.1 | 0.05 |
| 10 | 106930998 | rs7091373 | 0.304 | -0.197 | -0.181 | -1.719 | 0.015 | 0.008 | 0.008 | 0 | 0.16 | 0.26 | 0.25 | 0.16 | 0.18 | 0.25 |
| 10 | 106933181 | rs12250847 | -0.07 | -0.366 | -0.27 | -1.798 | 0.007 | 0.052 | 0.052 | 0 | 0.17 | 0.07 | 0.08 | 0.09 | 0.12 | 0.16 |
| 10 | 106935209 | rs10786848 | -1.697 | -0.507 | -0.208 | -1.444 | 0 | 0.002 | 0.002 | 0 | 0.46 | 0.45 | 0.55 | 0.4 | 0.43 | 0.52 |
| 10 | 106938561 | rs10128431 | -1.72 | -0.743 | -0.257 | -1.438 | -0.003 | -0.007 | -0.007 | 0.147 | 0.46 | 0.49 | 0.55 | 0.39 | 0.42 | 0.51 |
| 10 | 106944379 | rs7069827 | -0.896 | -0.714 | -0.478 | -1.458 | -0.001 | 0.001 | 0.001 | 0 | 0.3 | 0.26 | 0.46 | 0.21 | 0.31 | 0.37 |
| 10 | 106945236 | rs7092104 | -0.858 | -0.724 | -0.465 | -1.438 | -0.001 | 0.001 | 0.001 | 0 | 0.31 | 0.34 | 0.47 | 0.35 | 0.31 | 0.42 |
| 10 | 106955081 | rs1484237 | 0.314 | -0.671 | -0.473 | -1.417 | 0.012 | 0.03 | 0.03 | 0 | 0.19 | 0.23 | 0.26 | 0.34 | 0.21 | 0.31 |
| 10 | 106968669 | rs11192362 | -1.335 | -0.751 | -0.757 | -1.957 | 0.016 | 0.07 | 0.07 | 0 | 0.42 | 0.27 | 0.33 | 0.26 | 0.34 | 0.44 |
| 10 | 106969742 | rs12247132 | 0.303 | -0.763 | -0.789 | -1.967 | 0.024 | 0.086 | 0.086 | 0.111 | 0.16 | 0.03 | 0.05 | 0.03 | 0.1 | 0.1 |
| 10 | 106971524 | rs7894221 | -1.367 | -0.717 | -0.725 | -1.928 | 0.013 | 0.044 | 0.044 | 0 | 0.43 | 0.34 | 0.33 | 0.26 | 0.34 | 0.49 |
| 10 | 106973862 | rs10884119 | -1.765 | -0.739 | -0.704 | -1.907 | 0.01 | 0.032 | 0.032 | 0 | 0.48 | 0.44 | 0.54 | 0.27 | 0.43 | 0.54 |
| 10 | 106981738 | rs2864042 | -2.068 | -0.973 | -0.622 | -2.059 | 0.01 | 0.146 | 0.146 | 0 | 0.53 | 0.82 | 0.59 | 0.26 | 0.44 | 0.54 |
| 10 | 106994756 | rs1484248 | 0.366 | -0.792 | -0.572 | -1.984 | 0.02 | 0.042 | 0.042 | 0 | 0.11 | 0.04 | 0.2 | 0 | 0.1 | 0.06 |
| 10 | 107020914 | rs1484247 | 1.883 | -0.319 | -1.192 | -2.307 | -0.052 | -0.011 | -0.011 | 0.086 | 0.32 | 0.04 | 0.34 | 0.69 | 0.39 | 0.39 |
| 10 | 107021173 | rs6584660 | -0.071 | -0.691 | -0.633 | -1.912 | 0.039 | 0.058 | 0.058 | 0 | 0.22 | 0.12 | 0.08 | 0.05 | 0.1 | 0.15 |
| 10 | 107021332 | rs2220139 | 1.465 | -0.664 | -0.527 | -1.836 | 0.028 | 0.106 | 0.106 | 0 | 0.49 | 0.21 | 0.4 | 0.68 | 0.56 | 0.48 |
| 10 | 107023390 | rs3750261 | 0.341 | -0.259 | -1.048 | -1.694 | -0.001 | 0 | 0 | 0 | 0.18 | 0.17 | 0.27 | 0.25 | 0.22 | 0.32 |
| 10 | 107024459 | rs12264321 | 0.114 | -0.262 | -1.08 | -1.724 | 0.037 | 0.109 | 0.109 | 0 | 0.2 | 0.04 | 0.07 | 0.05 | 0.1 | 0.1 |
| 10 | 107025028 | rs10884126 | 0.503 | -0.243 | -1.053 | -1.7 | -0.009 | -0.007 | -0.007 | 4.552 | 0.17 | 0.13 | 0.27 | 0.25 | 0.22 | 0.32 |
| 10 | 107025800 | rs7099278 | -0.011 | -0.206 | -1.082 | -1.736 | 0.045 | 0.106 | 0.106 | 0 | 0.21 | 0.05 | 0.08 | 0.05 | 0.1 | 0.15 |
| 10 | 107026567 | rs754720 | 1.209 | -0.205 | -1.052 | -1.722 | 0.009 | 0.027 | 0.027 | 0 | 0.54 | 0.59 | 0.45 | 0.68 | 0.58 | 0.48 |
| 10 | 107027076 | rs12248627 | 0.18 | 0.038 | -1.29 | -1.476 | -0.002 | -0.002 | -0.002 | 0.77 | 0.19 | 0.17 | 0.27 | 0.27 | 0.22 | 0.33 |
| 10 | 107031009 | rs11192385 | -0.734 | -0.46 | -1.257 | -1.629 | 0.041 | 0.067 | 0.067 | 0 | 0.21 | 0.11 | 0.08 | 0.05 | 0.1 | 0.15 |
| 10 | 107033965 | rs12244321 | -0.617 | -0.399 | -1.436 | -1.604 | 0.042 | 0.108 | 0.108 | 0 | 0.21 | 0.05 | 0.08 | 0.05 | 0.1 | 0.15 |
| 10 | 107046545 | rs981360 | -0.578 | 0.051 | -0.92 | -1.665 | 0.002 | 0.004 | 0.004 | 0 | 0.25 | 0.26 | 0.12 | 0.06 | 0.19 | 0.35 |
| 10 | 107051639 | rs2930454 | 0.219 | -0.116 | -0.559 | -1.665 | -0.01 | -0.046 | -0.046 | 0.784 | 0.78 | 0.69 | 0.94 | 0.96 | 0.87 | 0.67 |
| 10 | 107069561 | rs10786855 | -0.444 | -0.619 | 0.049 | -1.953 | -0.022 | -0.072 | -0.072 | 0.823 | 0.12 | 0.42 | 0.24 | 0.02 | 0.1 | 0.16 |
| 10 | 107069849 | rs3011682 | 1.069 | -0.757 | 0.017 | -2.087 | 0.022 | 0.022 | 0.022 | 2.468 | 0.65 | 0.26 | 0.7 | 0.94 | 0.78 | 0.51 |
| 10 | 107070536 | rs12248245 | -0.372 | -0.69 | -0.039 | -2.056 | -0.022 | -0.072 | -0.072 | 0.063 | 0.12 | 0.42 | 0.24 | 0.02 | 0.1 | 0.16 |
| 10 | 107070578 | rs11192397 | 0.914 | -0.263 | -0.566 | -1.988 | -0.046 | -0.051 | -0.051 | 4.34 | 0.1 | 0.01 | 0.1 | 0.19 | 0.15 | 0.1 |
| 10 | 107072871 | rs10509789 | -1.178 | -0.359 | -0.326 | -1.951 | -0.025 | -0.093 | -0.093 | 0 | 0.3 | 0.59 | 0.29 | 0.06 | 0.21 | 0.48 |
| 10 | 107085512 | rs6584666 | 0.257 | -0.346 | 0.194 | -1.917 | -0.005 | -0.029 | -0.029 | 0.083 | 0.18 | 0.25 | 0.07 | 0.06 | 0.12 | 0.39 |
| 10 | 107088137 | rs7909673 | 0.503 | -0.345 | 0.187 | -1.917 | 0.002 | 0.021 | 0.021 | 0 | 0.17 | 0.12 | 0.05 | 0.04 | 0.12 | 0.32 |
| 10 | 107089364 | rs11818341 | 0.172 | -0.409 | 0.494 | -1.943 | -0.005 | -0.029 | -0.029 | 0 | 0.18 | 0.26 | 0.07 | 0.06 | 0.12 | 0.39 |
| 10 | 107089929 | rs4528240 | 1.934 | -0.229 | -0.354 | -2.262 | -0.071 | -0.109 | -0.109 | 0.1 | 0.29 | 0.03 | 0.3 | 0.62 | 0.36 | 0.25 |
| 10 | 107092654 | rs4376833 | -1.299 | -0.099 | 0.606 | -2.314 | -0.006 | -0.04 | -0.04 | 0 | 0.31 | 0.67 | 0.3 | 0.06 | 0.22 | 0.48 |
| 10 | 107095157 | rs12265513 | -0.244 | -0.042 | 0.641 | -2.222 | -0.019 | -0.074 | -0.074 | 0 | 0.12 | 0.36 | 0.23 | 0 | 0.09 | 0.09 |
| 10 | 107095370 | rs12359689 | 1.021 | 0.077 | 0.203 | -2.164 | -0.001 | -0.013 | -0.013 | 0 | 0.05 | 0.04 | 0.2 | 0 | 0.09 | 0.05 |
| 10 | 107111730 | rs3011654 | 0.087 | 0.098 | 0.061 | -2.281 | -0.004 | 0.053 | 0.053 | 0 | 0.78 | 0.87 | 0.89 | 0.96 | 0.81 | 0.67 |
| 10 | 107117836 | rs11192417 | -0.407 | 0.096 | 0.118 | -2.341 | -0.007 | -0.035 | -0.035 | 0 | 0.09 | 0.18 | 0.21 | 0.02 | 0.1 | 0.16 |
| 10 | 107126282 | rs7098865 | -0.407 | 0.096 | 0.182 | -2.341 | -0.006 | -0.038 | -0.038 | 0 | 0.09 | 0.21 | 0.21 | 0.02 | 0.1 | 0.16 |

| | | | | | | | | | | | | | | | | |
|---|---|---|---|---|---|---|---|---|---|---|---|---|---|---|---|---|
| 10 | 107126452 | rs7068538 | -0.407 | 0.096 | 0.182 | -2.341 | -0.006 | -0.038 | -0.038 | 0 | 0.09 | 0.21 | 0.21 | 0.02 | 0.1 | 0.16 |
| 10 | 107127507 | rs2999329 | 0.836 | 0.102 | 0.164 | -2.353 | 0.001 | -0.021 | -0.021 | 0.027 | 0.69 | 0.65 | 0.68 | 0.94 | 0.72 | 0.51 |
| 10 | 107127799 | rs11192422 | 0.243 | 0.128 | -0.11 | -2.332 | 0.001 | -0.005 | -0.005 | 0 | 0.08 | 0.04 | 0.2 | 0 | 0.1 | 0.1 |
| 10 | 107136527 | rs4918157 | -0.042 | 0.128 | -0.194 | -2.333 | -0.007 | 0.067 | 0.067 | 0 | 0.79 | 0.89 | 0.89 | 0.96 | 0.82 | 0.67 |
| 10 | 107144262 | rs7478564 | -0.042 | 0.128 | -0.194 | -2.333 | -0.007 | 0.067 | 0.067 | 0 | 0.79 | 0.89 | 0.89 | 0.96 | 0.81 | 0.67 |
| 10 | 107150767 | rs4917442 | 1.328 | -0.365 | -0.067 | -2.673 | 0.039 | 0.082 | 0.082 | 0.156 | 0.58 | 0.25 | 0.64 | 0.94 | 0.71 | 0.51 |
| 10 | 107154018 | rs17118962 | 0.103 | -0.05 | 0.014 | -2.575 | -0.002 | -0.022 | -0.022 | 0 | 0.16 | 0.13 | 0.23 | 0 | 0.16 | 0.17 |
| 10 | 107154256 | rs4468278 | 0.244 | -0.05 | -0.017 | -2.575 | -0.007 | 0.005 | 0.005 | 0 | 0.16 | 0.09 | 0.23 | 0 | 0.16 | 0.17 |
| 10 | 107155625 | rs4333954 | -1.807 | -0.352 | -0.017 | -2.716 | 0.027 | 0.11 | 0.11 | 0.856 | 0.29 | 0.86 | 0.31 | 0.02 | 0.19 | 0.24 |
| 10 | 107157561 | rs4147184 | 2.158 | -0.434 | -0.395 | -2.96 | -0.037 | 0.357 | 0.357 | 0.026 | 0.47 | 0.05 | 0.57 | 0.9 | 0.62 | 0.44 |
| 10 | 107178476 | rs10884145 | 0.292 | -0.208 | 0.007 | -2.635 | -0.013 | 0.031 | 0.031 | 0 | 0.16 | 0.06 | 0.23 | 0 | 0.16 | 0.17 |
| 10 | 107194924 | rs12264818 | -0.221 | -0.223 | 0.496 | -2.644 | -0.005 | -0.075 | -0.075 | 0 | 0.17 | 0.36 | 0.06 | 0.02 | 0.09 | 0.19 |
| 10 | 107196304 | rs1459371 | -1.437 | -0.26 | 0.845 | -2.667 | -0.009 | -0.08 | -0.08 | 0 | 0.26 | 0.67 | 0.29 | 0.02 | 0.19 | 0.25 |
| 10 | 107205083 | rs2813173 | 0.13 | -0.261 | 0.444 | -2.711 | -0.022 | 0.045 | 0.045 | 0 | 0.88 | 0.99 | 0.93 | 0.96 | 0.88 | 0.85 |
| 10 | 107217381 | rs12259354 | -0.184 | -0.231 | 0.503 | -2.689 | -0.005 | -0.076 | -0.076 | 0 | 0.17 | 0.36 | 0.06 | 0.02 | 0.09 | 0.19 |
| 10 | 107241137 | rs2611049 | 0.103 | -0.244 | 0.224 | -2.581 | -0.017 | 0.045 | 0.045 | 0 | 0.88 | 0.99 | 0.93 | 0.96 | 0.88 | 0.85 |
| 10 | 107251107 | rs2936562 | -1.361 | -0.151 | 0.589 | -2.589 | -0.002 | -0.065 | -0.065 | 0 | 0.27 | 0.68 | 0.29 | 0.02 | 0.22 | 0.27 |
| 10 | 107252462 | rs1870461 | 1.324 | -0.146 | 0.78 | -2.841 | -0.004 | -0.075 | -0.075 | 0 | 0.65 | 0.42 | 0.64 | 0.9 | 0.66 | 0.51 |
| 10 | 107255347 | rs2926457 | -0.785 | -0.141 | 0.717 | -2.863 | 0.007 | 0.02 | 0.02 | 0 | 0.91 | 0.95 | 0.93 | 0.95 | 0.92 | 0.75 |
| 10 | 107258947 | rs2936561 | 0.408 | -0.072 | 0.767 | -2.813 | -0.005 | -0.043 | -0.043 | 2.043 | 0.74 | 0.73 | 0.85 | 0.92 | 0.79 | 0.67 |
| 10 | 107260501 | rs2926455 | 0.482 | 0.083 | 0.872 | -2.528 | 0.035 | -0.031 | -0.031 | 1.201 | 0.07 | 0.15 | 0.21 | 0 | 0.12 | 0.07 |
| 10 | 107264645 | rs11192460 | 0.785 | 0.038 | 0.632 | -2.517 | -0.001 | -0.016 | -0.016 | 0 | 0.12 | 0.16 | 0.04 | 0 | 0.08 | 0.1 |
| 10 | 107270764 | rs977980 | 0.677 | 0.036 | 0.677 | -2.518 | -0.001 | -0.048 | -0.048 | 0 | 0.12 | 0.23 | 0.05 | 0 | 0.08 | 0.1 |
| 10 | 107282331 | rs11192469 | 0.862 | 0.173 | 0.548 | -2.491 | 0.006 | -0.012 | -0.012 | 0 | 0.06 | 0.04 | 0.2 | 0 | 0.12 | 0.07 |
| 10 | 107282991 | rs7915785 | -0.553 | -0.026 | 0.738 | -2.541 | -0.011 | -0.105 | -0.105 | 1.149 | 0.18 | 0.33 | 0.06 | 0 | 0.1 | 0.1 |
| 10 | 107295345 | rs2418761 | 0.09 | -0.134 | 0.608 | -2.406 | 0.004 | 0.003 | 0.003 | 0 | 0.86 | 0.88 | 0.93 | 0.91 | 0.9 | 0.74 |
| 10 | 107305108 | rs10159932 | 1.251 | -0.157 | 0.813 | -2.453 | 0.004 | -0.044 | -0.044 | 0.458 | 0.1 | 0.16 | 0.05 | 0 | 0.07 | 0.09 |
| 10 | 107318338 | rs10884172 | 0.805 | -0.187 | 1.067 | -2.465 | 0.013 | -0.07 | -0.07 | 0 | 0.11 | 0.28 | 0.05 | 0 | 0.07 | 0.09 |
| 10 | 107324125 | rs10509792 | 0.267 | -0.205 | 1.068 | -2.858 | -0.002 | -0.055 | -0.055 | 0 | 0.19 | 0.21 | 0.11 | 0.04 | 0.19 | 0.13 |
| 10 | 107328195 | rs7089194 | 0.273 | -0.178 | 0.945 | -2.901 | -0.001 | -0.008 | -0.008 | 0 | 0.8 | 0.8 | 0.72 | 0.92 | 0.78 | 0.67 |
| 10 | 107329126 | rs9703795 | 2.021 | -0.093 | 0.48 | -2.827 | -0.137 | 0.227 | 0.227 | 0.111 | 0.43 | 0.05 | 0.53 | 0.86 | 0.49 | 0.36 |
| 10 | 107337982 | rs10884185 | 1.418 | -0.102 | 0.533 | -2.849 | 0.027 | 0.18 | 0.18 | 0.018 | 0.54 | 0.22 | 0.57 | 0.86 | 0.56 | 0.45 |
| 10 | 107343068 | rs10786877 | 1.404 | -0.054 | 0.573 | -2.83 | 0.018 | 0.143 | 0.143 | 0 | 0.54 | 0.26 | 0.58 | 0.86 | 0.56 | 0.45 |
| 10 | 107351262 | rs11192508 | 1.365 | 0.05 | 0.593 | -2.741 | 0.004 | -0.046 | -0.046 | 0.242 | 0.1 | 0.17 | 0.05 | 0 | 0.07 | 0.09 |
| 10 | 107357563 | rs9633715 | 1.266 | -0.034 | 0.642 | -2.774 | -0.001 | -0.037 | -0.037 | 0.026 | 0.54 | 0.46 | 0.59 | 0.87 | 0.56 | 0.45 |
| 10 | 107366881 | rs17119209 | 0.454 | 0.066 | 0.777 | -2.689 | 0.002 | -0.094 | -0.094 | 0.431 | 0.17 | 0.26 | 0.12 | 0.03 | 0.19 | 0.11 |
| 10 | 107369270 | rs10786886 | 0.382 | -0.201 | 0.69 | -2.8 | 0 | 0.001 | 0.001 | 0 | 0.8 | 0.82 | 0.72 | 0.91 | 0.79 | 0.68 |
| 10 | 107371918 | rs7097382 | 0.898 | -0.316 | 0.556 | -2.943 | -0.005 | -0.051 | -0.051 | 0.258 | 0.12 | 0.21 | 0.05 | 0 | 0.07 | 0.1 |
| 10 | 107373294 | rs9633716 | -1.012 | -0.263 | 0.32 | -2.838 | 0.017 | 0.036 | 0.036 | 0 | 0.89 | 0.97 | 0.93 | 0.95 | 0.91 | 0.74 |
| 10 | 107376457 | rs10884198 | 1.954 | -0.006 | 0.329 | -2.94 | -0.04 | 0.185 | 0.185 | 0.056 | 0.4 | 0.13 | 0.52 | 0.89 | 0.47 | 0.34 |
| 10 | 107376927 | rs7911178 | -0.404 | -0.038 | 0.614 | -2.874 | -0.001 | -0.115 | -0.115 | 0 | 0.32 | 0.33 | 0.17 | 0.03 | 0.3 | 0.23 |
| 10 | 107381646 | rs7894917 | -1.104 | -0.065 | 0.584 | -2.892 | 0.01 | 0.016 | 0.016 | 0.087 | 0.4 | 0.68 | 0.2 | 0.05 | 0.32 | 0.34 |
| 10 | 107389184 | rs2058980 | 1.823 | -0.159 | 0.819 | -3.131 | -0.003 | -0.014 | -0.014 | 0 | 0.43 | 0.31 | 0.53 | 0.92 | 0.47 | 0.34 |
| 10 | 107396147 | rs4537693 | -0.071 | -0.183 | 0.906 | -3.156 | -0.007 | 0.032 | 0.032 | 0 | 0.31 | 0.17 | 0.16 | 0.03 | 0.3 | 0.22 |
| 10 | 107396793 | rs2418770 | 0.243 | -0.18 | 0.884 | -3.149 | 0 | -0.035 | -0.035 | 0 | 0.8 | 0.78 | 0.72 | 0.98 | 0.79 | 0.69 |
| 10 | 107403241 | rs7893754 | -0.449 | -0.193 | 0.784 | -3.152 | 0 | -0.075 | -0.075 | 0.158 | 0.33 | 0.37 | 0.18 | 0.03 | 0.31 | 0.23 |
| 10 | 107409121 | rs11192538 | 0.447 | -0.216 | 0.731 | -3.176 | -0.02 | 0.02 | 0.02 | 0 | 0.1 | 0.05 | 0.22 | 0 | 0.15 | 0.08 |
| 10 | 107409168 | rs12244751 | -0.433 | -0.203 | 0.638 | -3.163 | -0.004 | -0.118 | -0.118 | 0 | 0.35 | 0.47 | 0.18 | 0.05 | 0.31 | 0.33 |
| 10 | 107418068 | rs7081043 | 2.055 | -0.256 | 0.468 | -3.158 | -0.077 | 0.265 | 0.265 | 0.203 | 0.41 | 0.07 | 0.5 | 0.93 | 0.46 | 0.34 |
| 10 | 107419215 | rs11593140 | -0.41 | -0.35 | 0.564 | -3.048 | -0.001 | -0.031 | -0.031 | 0 | 0.33 | 0.29 | 0.17 | 0.05 | 0.3 | 0.32 |
| 10 | 107419892 | rs4917446 | 0.946 | -0.42 | 0.545 | -3.088 | 0.007 | -0.037 | -0.037 | 0.296 | 0.55 | 0.53 | 0.76 | 0.93 | 0.63 | 0.43 |
| 10 | 107420067 | rs7090449 | 0.556 | -0.426 | 0.314 | -3.048 | 0.004 | 0.008 | 0.008 | 0 | 0.14 | 0.08 | 0.05 | 0 | 0.1 | 0.09 |
| 10 | 107420421 | rs7090774 | -1.018 | -0.349 | 0.27 | -2.919 | 0.035 | 0.045 | 0.045 | 0 | 0.89 | 0.95 | 0.94 | 0.98 | 0.93 | 0.75 |
| 10 | 107421250 | rs7919059 | -0.488 | -0.355 | 0.312 | -2.922 | -0.006 | -0.024 | -0.024 | 0 | 0.88 | 0.82 | 0.94 | 0.98 | 0.93 | 0.75 |
| 10 | 107438736 | rs4917447 | 0.512 | 0.268 | 0.045 | -3.205 | -0.027 | 0.044 | 0.044 | 0 | 0.74 | 0.79 | 0.67 | 0.98 | 0.69 | 0.57 |
| 10 | 107441662 | rs4918175 | 0.501 | 0.19 | 0.284 | -3.184 | -0.035 | -0.067 | -0.067 | 0.144 | 0.78 | 0.6 | 0.84 | 0.94 | 0.86 | 0.64 |
| 10 | 107444714 | rs1421488 | 2.288 | 0.114 | 0.335 | -2.807 | -0.001 | -0.004 | -0.004 | 0 | 0.34 | 0.32 | 0.46 | 0.75 | 0.37 | 0.24 |
| 10 | 107451002 | rs12771120 | 0.227 | 0.071 | 0.358 | -2.801 | 0.003 | -0.095 | -0.095 | 0 | 0.21 | 0.42 | 0.13 | 0.07 | 0.2 | 0.21 |
| 10 | 107451498 | rs7073592 | 0.794 | 0.155 | 0.192 | -2.811 | 0.015 | 0.035 | 0.035 | 0 | 0.14 | 0.05 | 0.05 | 0 | 0.07 | 0.1 |
| 10 | 107452683 | rs11192557 | 0.077 | 0.165 | 0.187 | -2.807 | 0.003 | -0.022 | -0.022 | 0 | 0.18 | 0.05 | 0.09 | 0.16 | 0.12 | 0.13 |
| 10 | 107455498 | rs12783109 | 1.069 | -0.007 | 0.457 | -3.085 | 0.003 | -0.09 | -0.09 | 0.146 | 0.17 | 0.3 | 0.12 | 0.03 | 0.19 | 0.11 |
| 10 | 107458408 | rs7895008 | 0.006 | -0.082 | 0.362 | -3.131 | -0.006 | -0.019 | -0.019 | 0.117 | 0.8 | 0.72 | 0.85 | 0.94 | 0.86 | 0.64 |
| 10 | 107470932 | rs11597004 | 1.419 | 0.035 | 0.102 | -3.058 | -0.003 | -0.01 | -0.01 | 0 | 0.15 | 0.13 | 0.11 | 0.03 | 0.19 | 0.11 |
| 10 | 107471078 | rs1362993 | 1.699 | 0.04 | 0.372 | -3.016 | -0.002 | 0.036 | 0.036 | 0 | 0.39 | 0.26 | 0.49 | 0.91 | 0.43 | 0.31 |
| 10 | 107471342 | rs2111996 | 0.756 | 0.144 | 0.382 | -2.516 | 0.006 | -0.095 | -0.095 | 0 | 0.7 | 0.56 | 0.66 | 0.94 | 0.69 | 0.52 |
| 10 | 107479864 | rs7908373 | 0.458 | 0.265 | -0.231 | -2.456 | -0.046 | 0.111 | 0.111 | 0.044 | 0.76 | 0.92 | 0.68 | 0.97 | 0.7 | 0.66 |
| 10 | 107480519 | rs11819719 | 0.673 | -0.671 | 0 | -2.504 | 0.003 | -0.015 | -0.015 | 0 | 0.13 | 0.09 | 0.05 | 0 | 0.07 | 0.14 |
| 10 | 107483655 | rs7917363 | -0.168 | -0.527 | -0.064 | -2.349 | 0.031 | 0.061 | 0.061 | 0 | 0.83 | 0.93 | 0.83 | 0.97 | 0.87 | 0.78 |
| 10 | 107501893 | rs7919079 | -1.183 | -0.647 | 0.961 | -2.3 | -0.014 | 0.014 | 0.014 | 0 | 0.08 | 0.33 | 0.03 | 0.04 | 0.01 | 0.09 |

| | | | | | | | | | | | | | | | | |
|---|---|---|---|---|---|---|---|---|---|---|---|---|---|---|---|---|
| 10 | 107505671 | rs7080489 | -0.664 | -0.692 | 1.145 | -2.318 | -0.005 | 0.023 | 0.023 | 0 | 0.07 | 0.23 | 0.02 | 0.04 | 0.01 | 0.09 |
| 10 | 107507362 | rs2111995 | 1.198 | -0.956 | 0.86 | -2.51 | 0.067 | 0.082 | 0.082 | 0 | 0.6 | 0.28 | 0.73 | 0.93 | 0.78 | 0.49 |
| 10 | 107517345 | rs1362991 | 1.336 | -0.569 | -0.422 | -1.867 | -0.107 | -0.076 | -0.076 | 10.047 | 0.21 | 0.04 | 0.44 | 0.29 | 0.35 | 0.2 |
| 10 | 107526019 | rs7915649 | 0.047 | -1.04 | -0.071 | -1.995 | -0.018 | -0.025 | -0.025 | 0 | 0.78 | 0.75 | 0.83 | 0.93 | 0.86 | 0.64 |
| 10 | 107526521 | rs4437963 | -0.859 | -1.006 | -0.133 | -1.99 | 0.029 | 0.057 | 0.057 | 0 | 0.83 | 0.95 | 0.85 | 0.96 | 0.86 | 0.67 |
| 10 | 107530680 | rs2418774 | 0.977 | -1.007 | -0.42 | -2.327 | 0.024 | 0.236 | 0.236 | 0 | 0.45 | 0.15 | 0.65 | 0.89 | 0.61 | 0.37 |
| 10 | 107543300 | rs11593552 | 1.6 | -1.499 | -0.11 | -2.157 | 0.01 | 0.005 | 0.005 | 0 | 0.12 | 0.06 | 0.04 | 0 | 0.06 | 0.14 |
| 10 | 107544706 | rs17119534 | 0.315 | -1.291 | 0.133 | -2.022 | 0.017 | 0.052 | 0.052 | 0.189 | 0.05 | 0.24 | 0.03 | 0.04 | 0.01 | 0.1 |
| 10 | 107545496 | rs2418776 | 0.262 | -1.344 | 0.286 | -2.048 | -0.026 | -0.057 | -0.057 | 0 | 0.63 | 0.46 | 0.77 | 0.92 | 0.79 | 0.44 |
| 10 | 107559597 | rs10884227 | 0.77 | -0.46 | -0.303 | -1.22 | -0.017 | -0.06 | -0.06 | 0 | 0.23 | 0.12 | 0.26 | 0.62 | 0.26 | 0.16 |
| 10 | 107568998 | rs12571678 | 0.202 | -0.547 | -0.947 | -1.435 | -0.025 | 0.183 | 0.183 | 0 | 0.38 | 0.07 | 0.59 | 0.87 | 0.56 | 0.31 |
| 10 | 107574603 | rs4917452 | -1.077 | -1.555 | -0.22 | -1.839 | 0.032 | 0.035 | 0.035 | 0 | 0.88 | 0.94 | 0.95 | 0.95 | 0.92 | 0.77 |
| 10 | 107577714 | rs11192616 | 0.419 | -1.452 | 0.122 | -1.803 | 0.023 | 0.187 | 0.187 | 0 | 0.38 | 0.11 | 0.26 | 0.04 | 0.34 | 0.29 |
| 10 | 107578214 | rs6584696 | -1.04 | -1.153 | 0.939 | -1.269 | 0.005 | 0.012 | 0.012 | 3.558 | 0.08 | 0.51 | 0.05 | 0.02 | 0.01 | 0.11 |
| 10 | 107582247 | rs11192617 | 0.044 | -1.031 | 0.75 | -0.467 | -0.001 | 0.075 | 0.075 | 0.424 | 0.15 | 0.22 | 0.17 | 0.52 | 0.14 | 0.11 |
| 10 | 107589359 | rs951030 | 0.161 | -1.005 | 1.335 | -0.471 | 0 | 0.045 | 0.045 | 0.489 | 0.14 | 0.19 | 0.17 | 0.52 | 0.15 | 0.11 |
| 10 | 107594356 | rs12259685 | -0.947 | -1.099 | 1.502 | -0.397 | 0.012 | 0.042 | 0.042 | 3.569 | 0.07 | 0.49 | 0.04 | 0.02 | 0 | 0.13 |
| 10 | 107618048 | rs11192637 | -1.002 | -1.099 | 1.142 | -0.104 | 0.014 | 0.21 | 0.21 | 2.453 | 0.07 | 0.49 | 0.17 | 0.14 | 0 | 0.11 |
| 10 | 107625071 | rs2460927 | 0.174 | -1.039 | 1.072 | -0.066 | 0.004 | 0.17 | 0.17 | 0.271 | 0.12 | 0.26 | 0.16 | 0.51 | 0.11 | 0.09 |
| 10 | 107628876 | rs1908643 | 0.174 | -1.039 | 1.072 | -0.066 | 0.004 | 0.17 | 0.17 | 0.271 | 0.12 | 0.26 | 0.16 | 0.51 | 0.11 | 0.09 |
| 10 | 107632783 | rs10884237 | 0.928 | -1.049 | 1.027 | -0.038 | -0.034 | -0.045 | -0.045 | 0 | 0.09 | 0.01 | 0.15 | 0.49 | 0.11 | 0.09 |
| 10 | 107634517 | rs1390491 | 0.117 | -1.055 | 1.07 | -0.127 | 0.003 | 0.158 | 0.158 | 0.352 | 0.12 | 0.27 | 0.16 | 0.53 | 0.11 | 0.09 |
| 10 | 107637740 | rs17236391 | 0.916 | -1.473 | 0.594 | -0.604 | 0 | 0.179 | 0.179 | 0 | 0.32 | 0.08 | 0.17 | 0.04 | 0.28 | 0.26 |
| 10 | 107659545 | rs1496060 | -0.041 | -1.165 | 0.926 | -0.434 | 0.003 | 0.218 | 0.218 | 1.115 | 0.14 | 0.38 | 0.18 | 0.52 | 0.11 | 0.09 |
| 10 | 107661479 | rs2644513 | 0.368 | -1.119 | 0.623 | -0.414 | 0.001 | 0.148 | 0.148 | 0.47 | 0.12 | 0.24 | 0.16 | 0.52 | 0.11 | 0.09 |
| 10 | 107662482 | rs1496063 | 0.368 | -1.119 | 0.623 | -0.391 | 0.001 | 0.15 | 0.15 | 0.47 | 0.12 | 0.24 | 0.17 | 0.51 | 0.11 | 0.09 |
| 10 | 107665118 | rs2683655 | -0.98 | -1.216 | 0.493 | -0.464 | -0.038 | 0.072 | 0.072 | 0.15 | 0.58 | 0.88 | 0.59 | 0.7 | 0.46 | 0.48 |
| 10 | 107665710 | rs7911641 | 0.443 | -0.37 | 1.033 | 0.193 | 0 | 0.068 | 0.068 | 0.051 | 0.11 | 0.12 | 0.16 | 0.51 | 0.11 | 0.09 |
| 10 | 107677186 | rs2644519 | -0.016 | -0.341 | 0.833 | 0.197 | 0.002 | 0.151 | 0.151 | 0.442 | 0.12 | 0.24 | 0.16 | 0.52 | 0.11 | 0.08 |
| 10 | 107680866 | rs10884242 | -0.024 | -0.097 | 0.68 | 0.473 | 0.003 | 0.03 | 0.03 | 0 | 0.15 | 0.18 | 0.2 | 0.42 | 0.23 | 0.13 |
| 10 | 107681935 | rs1496077 | 0.431 | -0.885 | 0.786 | -0.024 | -0.03 | -0.121 | -0.121 | 0.015 | 0.29 | 0.39 | 0.15 | 0 | 0.2 | 0.11 |
| 10 | 107682005 | rs1496078 | 0.306 | -0.912 | 0.818 | 0.237 | -0.041 | -0.003 | -0.003 | 0 | 0.3 | 0.47 | 0.21 | 0.26 | 0.2 | 0.15 |
| 10 | 107699810 | rs574989 | 0.41 | 0.015 | 0.752 | -0.133 | -0.005 | -0.115 | -0.115 | 0 | 0.3 | 0.4 | 0.36 | 0 | 0.28 | 0.12 |
| 10 | 107703278 | rs12259847 | 1.207 | -0.132 | 0.743 | -0.159 | 0.021 | 0.07 | 0.07 | 0 | 0.23 | 0.1 | 0.13 | 0 | 0.2 | 0.11 |
| 10 | 107705511 | rs472548 | 1.206 | -0.061 | 0.479 | -0.14 | -0.018 | 0.106 | 0.106 | 0.611 | 0.46 | 0.15 | 0.35 | 0.33 | 0.46 | 0.64 |
| 10 | 107707091 | rs532243 | -0.855 | 0.128 | 0.625 | -0.174 | -0.057 | -0.179 | -0.179 | 0 | 0.7 | 0.47 | 0.83 | 1 | 0.8 | 0.89 |
| 10 | 107711899 | rs563794 | -0.832 | 0.185 | 0.475 | -0.149 | -0.023 | -0.104 | -0.104 | 2.599 | 0.7 | 0.64 | 0.85 | 1 | 0.81 | 0.88 |
| 10 | 107722094 | rs556646 | 0.205 | 0.454 | 0.551 | 0.068 | -0.001 | -0.003 | -0.003 | 0.181 | 0.59 | 0.38 | 0.39 | 0.29 | 0.5 | 0.68 |
| 10 | 107725810 | rs17119903 | 0.562 | 0.453 | 0.621 | 0.02 | -0.022 | -0.035 | -0.035 | 0.016 | 0.07 | 0.18 | 0.03 | 0 | 0.01 | 0.06 |
| 10 | 107727810 | rs821205 | 0.144 | 0.444 | 0.651 | 0.244 | -0.008 | -0.008 | -0.008 | 0.5 | 0.59 | 0.38 | 0.44 | 0.52 | 0.5 | 0.72 |
| 10 | 107733540 | rs481179 | 0.177 | 0.414 | 0.494 | 0.255 | 0 | 0.014 | 0.014 | 0 | 0.61 | 0.5 | 0.65 | 0.52 | 0.58 | 0.74 |
| 10 | 107735306 | rs10884244 | -0.023 | 0.599 | -0.08 | 0.372 | 0.017 | 0.017 | 0.017 | 0 | 0.14 | 0.16 | 0.24 | 0.71 | 0.24 | 0.17 |
| 10 | 107739028 | rs1032537 | 0.283 | 0.608 | -0.09 | 0.384 | 0.017 | 0.077 | 0.077 | 0 | 0.13 | 0.11 | 0.2 | 0.48 | 0.23 | 0.13 |
| 10 | 107739563 | rs821202 | 0.646 | 0.51 | -0.622 | 0.388 | -0.016 | 0.15 | 0.15 | 1.153 | 0.51 | 0.16 | 0.36 | 0.29 | 0.48 | 0.63 |
| 10 | 107740388 | rs479061 | 1.19 | 0.841 | -1.399 | 0.304 | -0.104 | 0.023 | 0.023 | 0 | 0.25 | 0.02 | 0.18 | 0.18 | 0.29 | 0.23 |
| 10 | 107750766 | rs7089156 | -0.958 | 0.969 | -0.19 | 0.986 | -0.029 | 0.037 | 0.037 | 2.49 | 0.12 | 0.64 | 0.14 | 0.11 | 0.07 | 0.05 |
| 10 | 107752982 | rs517678 | 0.837 | 1.188 | -0.388 | 0.9 | 0.019 | 0.194 | 0.194 | 0 | 0.5 | 0.14 | 0.31 | 0.27 | 0.42 | 0.58 |
| 10 | 107765356 | rs10509810 | -0.211 | 1.619 | -0.649 | 0.974 | -0.019 | -0.01 | -0.01 | 0 | 0.37 | 0.22 | 0.43 | 0.62 | 0.43 | 0.35 |
| 10 | 107770328 | rs2095898 | -1.482 | 1.679 | -0.128 | 0.963 | 0.022 | 0.021 | 0.021 | 2.918 | 0.14 | 0.65 | 0.26 | 0.11 | 0.15 | 0.07 |
| 10 | 107770619 | rs12265647 | -1.102 | 0.959 | -0.226 | 1.034 | -0.022 | 0.069 | 0.069 | 2.139 | 0.12 | 0.57 | 0.14 | 0.11 | 0.07 | 0.05 |
| 10 | 107772704 | rs12571004 | 0.09 | 1.409 | -1.15 | 0.575 | -0.045 | -0.047 | -0.047 | 0 | 0.08 | 0 | 0.17 | 0.3 | 0.13 | 0.1 |
| 10 | 107780090 | rs12265861 | -0.318 | 0.86 | -0.832 | -0.595 | -0.001 | 0.005 | 0.005 | 0 | 0.33 | 0.35 | 0.33 | 0.13 | 0.34 | 0.24 |
| 10 | 107784597 | rs4918192 | -0.188 | 0.46 | -0.622 | -0.928 | 0.005 | 0.024 | 0.024 | 0 | 0.52 | 0.54 | 0.69 | 0.72 | 0.55 | 0.47 |
| 10 | 107786421 | rs955898 | -0.091 | 0.358 | -0.701 | -0.999 | 0.005 | 0.026 | 0.026 | 0 | 0.52 | 0.54 | 0.69 | 0.72 | 0.55 | 0.47 |
| 10 | 107787578 | rs17119996 | 1.079 | 0.427 | -0.817 | -1.091 | -0.035 | -0.043 | -0.043 | 0.026 | 0.13 | 0.05 | 0.12 | 0.23 | 0.16 | 0.31 |
| 10 | 107806595 | rs10884248 | 0.984 | -0.086 | -0.575 | -0.716 | -0.02 | -0.051 | -0.051 | 0 | 0.18 | 0.1 | 0.37 | 0.46 | 0.29 | 0.13 |
| 10 | 107810648 | rs1339634 | -0.415 | 0.087 | -0.275 | -1.167 | 0.01 | 0.035 | 0.035 | 0 | 0.51 | 0.41 | 0.68 | 0.7 | 0.55 | 0.46 |
| 10 | 107814047 | rs10786906 | -0.41 | 0.059 | -0.201 | -1.165 | 0.019 | 0.063 | 0.063 | 0 | 0.51 | 0.37 | 0.68 | 0.7 | 0.55 | 0.46 |
| 10 | 107824423 | rs1339637 | 0.772 | 0.09 | -0.194 | -1.191 | -0.002 | -0.014 | -0.014 | 0 | 0.23 | 0.17 | 0.4 | 0.62 | 0.3 | 0.3 |
| 10 | 107828169 | rs644067 | -0.266 | -0.251 | 0.129 | -0.662 | 0.007 | 0.03 | 0.03 | 0 | 0.46 | 0.44 | 0.31 | 0.3 | 0.44 | 0.54 |
| 10 | 107842158 | rs609295 | -0.475 | -0.251 | 0.131 | -0.66 | 0 | 0.005 | 0.005 | 0 | 0.5 | 0.55 | 0.36 | 0.38 | 0.5 | 0.55 |
| 10 | 107842444 | rs11192711 | 0.532 | -0.032 | 0.555 | -0.36 | 0.002 | -0.022 | -0.022 | 1.27 | 0.19 | 0.15 | 0.07 | 0.02 | 0.09 | 0.1 |
| 10 | 107842529 | rs7100318 | -0.158 | 1.234 | 0.931 | 0.457 | -0.019 | -0.002 | -0.002 | 0 | 0.39 | 0.25 | 0.56 | 0.62 | 0.45 | 0.43 |
| 10 | 107849283 | rs509806 | -0.09 | 0.882 | 0.582 | 0.581 | -0.006 | 0.005 | 0.005 | 0 | 0.57 | 0.61 | 0.41 | 0.36 | 0.49 | 0.57 |
| 10 | 107851952 | rs2486551 | -0.027 | 0.938 | 0.444 | 0.668 | -0.003 | 0.011 | 0.011 | 0 | 0.56 | 0.44 | 0.35 | 0.32 | 0.49 | 0.56 |
| 10 | 107854441 | rs629662 | -0.023 | 0.948 | 0.448 | 0.664 | -0.003 | 0.011 | 0.011 | 0 | 0.55 | 0.44 | 0.35 | 0.32 | 0.49 | 0.56 |
| 10 | 107873989 | rs472664 | -0.035 | 0.747 | -0.609 | 0.221 | -0.004 | 0.008 | 0.008 | 0 | 0.56 | 0.42 | 0.37 | 0.33 | 0.49 | 0.6 |
| 10 | 107886255 | rs1411832 | -0.898 | 0.429 | -0.053 | 0.618 | 0.028 | 0.116 | 0.116 | 1.425 | 0.05 | 0.36 | 0.04 | 0.05 | 0.01 | 0.04 |
| 10 | 107886767 | rs12360303 | -0.319 | 1.024 | -1.561 | 0.064 | -0.092 | 0.13 | 0.13 | 0 | 0.31 | 0.05 | 0.22 | 0.12 | 0.38 | 0.25 |

| | | | | | | | | | | | | | | | |
|---|---|---|---|---|---|---|---|---|---|---|---|---|---|---|---|
| 10 | 107886857 | rs7909165 | -0.329 | 0.381 | -1.054 | 0.855 | -0.002 | 0.102 | 0.102 | 0 | 0.1 | 0.11 | 0.27 | 0.58 | 0.11 | 0.29 |
| 10 | 107888905 | rs11192725 | -0.448 | 0.06 | -1.765 | -0.112 | 0.017 | 0.113 | 0.113 | 0 | 0.27 | 0.03 | 0.18 | 0.11 | 0.24 | 0.19 |
| 10 | 107892872 | rs7070717 | 0.295 | -1.321 | -1.096 | -0.391 | -0.094 | -0.073 | -0.073 | 0.014 | 0.73 | 0.53 | 0.84 | 0.85 | 0.88 | 0.79 |
| 10 | 107908080 | rs10786914 | 0.724 | -0.443 | -1.036 | -0.746 | 0.044 | 0.105 | 0.105 | 0 | 0.65 | 0.16 | 0.67 | 0.85 | 0.76 | 0.76 |
| 10 | 107912176 | rs2150604 | 0.339 | -0.477 | -1.003 | -0.758 | -0.059 | -0.073 | -0.073 | 0.009 | 0.71 | 0.52 | 0.82 | 0.85 | 0.84 | 0.79 |
| 10 | 107916013 | rs17820810 | 0.49 | -0.806 | -0.941 | -0.891 | 0.051 | 0.055 | 0.055 | 0.002 | 0.14 | 0.04 | 0.05 | 0.05 | 0.05 | 0.15 |
| 10 | 107918915 | rs11192739 | 1.096 | -0.744 | -0.935 | -0.891 | 0.058 | 0.083 | 0.083 | 0 | 0.67 | 0.18 | 0.69 | 0.85 | 0.8 | 0.77 |
| 10 | 107928248 | rs4918206 | 0.693 | -1.201 | -1.251 | -0.654 | 0.011 | -0.013 | -0.013 | 0.33 | 0.06 | 0.03 | 0.11 | 0.15 | 0.03 | 0.15 |
| 10 | 107928304 | rs4918207 | 0.889 | -0.587 | -1.944 | -0.45 | -0.085 | -0.068 | -0.068 | 0 | 0.39 | 0.07 | 0.42 | 0.44 | 0.42 | 0.4 |
| 10 | 107934649 | rs12784816 | 0.643 | -0.355 | -1.293 | -0.578 | 0.05 | 0.053 | 0.053 | 0 | 0.14 | 0.04 | 0.05 | 0.05 | 0.05 | 0.15 |
| 10 | 107942299 | rs7918637 | -0.205 | -0.367 | -0.229 | -0.528 | -0.065 | -0.048 | -0.048 | 0 | 0.79 | 0.66 | 0.88 | 0.9 | 0.91 | 0.92 |
| 10 | 107953309 | rs1331149 | 0.728 | -1.185 | -0.378 | -0.472 | 0.067 | 0.07 | 0.07 | 0 | 0.67 | 0.22 | 0.78 | 0.85 | 0.84 | 0.79 |
| 10 | 107955419 | rs7080729 | -0.981 | -1.072 | -0.34 | -0.535 | 0.052 | 0.053 | 0.053 | 0 | 0.31 | 0.79 | 0.2 | 0.05 | 0.17 | 0.21 |
| 10 | 107962771 | rs7068708 | 0.785 | -0.338 | -0.374 | 0.003 | 0.047 | 0.04 | 0.04 | 0.075 | 0.87 | 0.7 | 0.88 | 0.87 | 0.82 | 0.93 |
| 10 | 107968917 | rs6584729 | -0.589 | -0.274 | -1.283 | 0.081 | -0.019 | -0.018 | -0.018 | 0 | 0.93 | 0.98 | 0.93 | 0.86 | 0.87 | 0.94 |
| 10 | 107972720 | rs4917465 | 0.211 | -0.274 | -1.291 | -0.105 | 0.001 | -0.014 | -0.014 | 0.436 | 0.7 | 0.59 | 0.79 | 0.86 | 0.71 | 0.83 |
| 10 | 107972788 | rs4917467 | 0.908 | -0.373 | -2.57 | 0.105 | -0.146 | -0.029 | -0.029 | 1.865 | 0.34 | 0.08 | 0.43 | 0.33 | 0.47 | 0.23 |
| 10 | 107977868 | rs1409403 | -0.709 | -0.575 | -2.549 | 0.174 | -0.021 | -0.029 | -0.029 | 0 | 0.92 | 0.99 | 0.93 | 0.86 | 0.88 | 0.93 |
| 10 | 107979141 | rs2025732 | 0.571 | -1.369 | -2.244 | 0.002 | 0.055 | -0.105 | -0.105 | 0.003 | 0.31 | 0.1 | 0.32 | 0.53 | 0.25 | 0.6 |
| 10 | 107984320 | rs3861988 | -0.3 | -0.151 | -1.934 | 0.656 | -0.032 | -0.106 | -0.106 | 0 | 0.6 | 0.95 | 0.52 | 0.6 | 0.41 | 0.71 |
| 10 | 107990682 | rs7068381 | -0.321 | -0.319 | -1.914 | 0.592 | -0.022 | -0.121 | -0.121 | 0 | 0.61 | 0.95 | 0.53 | 0.57 | 0.4 | 0.71 |
| 10 | 108002270 | rs1160715 | 0.378 | -0.788 | -0.509 | 0.889 | -0.07 | 0.05 | 0.05 | 0 | 0.82 | 0.62 | 0.93 | 0.72 | 0.95 | 0.72 |
| 10 | 108004010 | rs10786928 | 0.137 | -0.153 | -1.12 | 1.016 | -0.047 | -0.051 | -0.051 | 0 | 0.24 | 0.1 | 0.38 | 0.38 | 0.34 | 0.31 |
| 10 | 108005708 | rs11192792 | 0.406 | -0.112 | -1.597 | 0.554 | -0.081 | -0.005 | -0.005 | 1.664 | 0.19 | 0.01 | 0.2 | 0.1 | 0.25 | 0.16 |
| 10 | 108014982 | rs4918219 | 0.483 | 0.073 | -1.321 | 0.474 | -0.009 | 0.054 | 0.054 | 0 | 0.16 | 0.28 | 0.1 | 0.28 | 0.11 | 0.2 |
| 10 | 108015496 | rs4558102 | 0.18 | -0.137 | -0.679 | 0.331 | -0.023 | -0.032 | -0.032 | 0 | 0.92 | 0.65 | 0.96 | 1 | 0.99 | 0.9 |
| 10 | 108027274 | rs10884283 | 0.487 | -0.131 | -1.471 | 0.341 | 0.006 | -0.004 | -0.004 | 0 | 0.56 | 0.13 | 0.65 | 0.62 | 0.65 | 0.55 |
| 10 | 108034320 | rs7096917 | -0.695 | -0.14 | -0.733 | 0.224 | 0.011 | -0.057 | -0.057 | 4.253 | 0.1 | 0.62 | 0.1 | 0 | 0.07 | 0.02 |
| 10 | 108035041 | rs12781996 | 0.32 | -0.377 | -1.113 | 0.424 | -0.064 | 0.008 | 0.008 | 3.663 | 0.26 | 0.02 | 0.21 | 0.16 | 0.25 | 0.19 |
| 10 | 108035356 | rs11192819 | 1.327 | -0.387 | -1.382 | 0.462 | -0.032 | 0.026 | 0.026 | 0 | 0.33 | 0.02 | 0.22 | 0.2 | 0.3 | 0.23 |
| 10 | 108036451 | rs10786936 | 0.353 | -0.278 | -0.831 | 0.661 | 0.008 | -0.012 | -0.012 | 0 | 0.66 | 0.74 | 0.75 | 0.63 | 0.72 | 0.59 |
| 10 | 108045657 | rs1359645 | 0.731 | -0.321 | -0.044 | 0.854 | -0.029 | 0.126 | 0.126 | 0.142 | 0.91 | 0.7 | 0.95 | 0.79 | 0.97 | 0.76 |
| 10 | 108046638 | rs17834272 | 0.742 | -0.553 | -1.091 | 0.437 | 0.004 | 0.089 | 0.089 | 0 | 0.36 | 0.1 | 0.22 | 0.16 | 0.33 | 0.22 |
| 10 | 108046794 | rs11192823 | 0.141 | -0.692 | -1.33 | 0.505 | 0.044 | 0.075 | 0.075 | 0 | 0.49 | 0.16 | 0.38 | 0.33 | 0.41 | 0.32 |
| 10 | 108046865 | rs3934245 | -0.621 | -0.054 | -0.531 | 0.986 | -0.006 | 0.034 | 0.034 | 0 | 0.79 | 0.68 | 0.8 | 0.65 | 0.79 | 0.68 |
| 10 | 108049735 | rs1359646 | -0.015 | 0.18 | -0.367 | 1.328 | 0.002 | 0.058 | 0.058 | 0 | 0.74 | 0.64 | 0.76 | 0.52 | 0.71 | 0.62 |
| 10 | 108051576 | rs10786940 | -0.388 | 0.113 | -0.39 | 1.282 | -0.01 | 0.016 | 0.016 | 0 | 0.78 | 0.68 | 0.83 | 0.69 | 0.79 | 0.69 |
| 10 | 108055302 | rs11192826 | 0.576 | 0.058 | -0.514 | 1.314 | -0.011 | -0.028 | -0.028 | 0.369 | 0.06 | 0.01 | 0.12 | 0.14 | 0.06 | 0.06 |
| 10 | 108058716 | rs10748915 | 1.389 | -0.023 | -0.561 | 1.251 | 0.005 | 0.016 | 0.016 | 0 | 0.44 | 0.18 | 0.4 | 0.36 | 0.4 | 0.34 |
| 10 | 108063765 | rs12766152 | 0.733 | 0.02 | -0.554 | 1.175 | 0.016 | 0.028 | 0.028 | 0 | 0.06 | 0.02 | 0.02 | 0 | 0.02 | 0.02 |
| 10 | 108068283 | rs1887095 | 0.424 | -0.054 | -0.21 | 1.189 | 0.048 | 0.03 | 0.03 | 0 | 0.71 | 0.51 | 0.6 | 0.65 | 0.63 | 0.66 |
| 10 | 108082645 | rs11816843 | 0.271 | 0.516 | 0.351 | 1.003 | 0.09 | -0.063 | -0.063 | 0 | 0.12 | 0.28 | 0.08 | 0 | 0.16 | 0.1 |
| 10 | 108090751 | rs1268565 | 0.697 | 0.496 | 0.307 | 1.051 | 0.035 | -0.038 | -0.038 | 0 | 0.74 | 0.51 | 0.65 | 0.83 | 0.7 | 0.68 |
| 10 | 108096785 | rs10786943 | 1.702 | 0.428 | 0.149 | 1.065 | 0 | 0.042 | 0.042 | 0 | 0.43 | 0.17 | 0.37 | 0.31 | 0.4 | 0.34 |
| 10 | 108103082 | rs2148371 | -1.145 | 0.428 | 0.555 | 1.059 | 0.032 | -0.017 | -0.017 | 0 | 0.23 | 0.37 | 0.33 | 0.17 | 0.27 | 0.26 |
| 10 | 108103104 | rs860273 | -0.651 | 0.466 | 0.215 | 1.464 | 0.001 | -0.046 | -0.046 | 0 | 0.84 | 0.96 | 0.86 | 0.67 | 0.84 | 0.83 |
| 10 | 108116077 | rs2031621 | 2.688 | 0.36 | 0.008 | 1.319 | -0.031 | 0.164 | 0.164 | 0 | 0.41 | 0.13 | 0.28 | 0.12 | 0.42 | 0.28 |
| 10 | 108120190 | rs821693 | -0.579 | 0.398 | 0.114 | 1.381 | 0.001 | -0.036 | -0.036 | 0 | 0.84 | 0.95 | 0.9 | 0.72 | 0.84 | 0.82 |
| 10 | 108131003 | rs12764394 | -0.603 | 0.521 | 0.517 | 1.254 | 0.074 | -0.02 | -0.02 | 0 | 0.15 | 0.37 | 0.12 | 0.07 | 0.16 | 0.24 |
| 10 | 108135176 | rs2248838 | -1.711 | -0.278 | -0.047 | 1.262 | 0.004 | -0.005 | -0.005 | 0 | 0.35 | 0.68 | 0.54 | 0.57 | 0.29 | 0.44 |
| 10 | 108139705 | rs9663603 | -0.501 | -0.369 | 0.406 | 1.591 | 0 | -0.001 | -0.001 | 0.804 | 0.2 | 0.24 | 0.4 | 0.18 | 0.22 | 0.25 |
| 10 | 108141387 | rs821680 | -2.337 | -0.523 | 0.067 | 1.403 | 0.064 | 0.003 | 0.003 | 0 | 0.46 | 0.87 | 0.59 | 0.57 | 0.34 | 0.5 |
| 10 | 108142861 | rs821683 | -1.561 | -0.538 | 0.121 | 1.649 | -0.009 | 0.021 | 0.021 | 0 | 0.28 | 0.63 | 0.46 | 0.42 | 0.23 | 0.39 |
| 10 | 108149165 | rs821688 | -3.172 | -0.256 | -0.293 | 1.378 | -0.105 | 0.116 | 0.116 | 0 | 0.64 | 0.89 | 0.7 | 0.84 | 0.52 | 0.71 |
| 10 | 108155301 | rs7077490 | 0.986 | -0.188 | 0.177 | 1.57 | -0.017 | 0.015 | 0.015 | 0 | 0.75 | 0.46 | 0.55 | 0.75 | 0.77 | 0.63 |
| 10 | 108159560 | rs1568919 | 0.639 | -0.605 | -0.024 | 1.751 | -0.04 | -0.067 | -0.067 | 0 | 0.14 | 0.02 | 0.09 | 0.29 | 0.17 | 0.19 |
| 10 | 108160539 | rs1405044 | 0.639 | -0.605 | -0.039 | 1.751 | -0.044 | -0.071 | -0.071 | 0.565 | 0.14 | 0.02 | 0.09 | 0.29 | 0.17 | 0.2 |
| 10 | 108171388 | rs7089200 | -1.404 | -0.597 | 0.315 | 1.652 | -0.038 | -0.07 | -0.07 | 0 | 0.12 | 0.26 | 0.08 | 0 | 0.04 | 0.07 |
| 10 | 108180183 | rs11192871 | -2.049 | -0.626 | 0.47 | 1.795 | -0.07 | 0.048 | 0.048 | 0 | 0.16 | 0.45 | 0.12 | 0.16 | 0.05 | 0.12 |
| 10 | 108190046 | rs10884317 | 2.683 | -0.008 | 0.14 | 1.632 | -0.027 | 0.011 | 0.011 | 0 | 0.48 | 0.09 | 0.31 | 0.42 | 0.48 | 0.44 |
| 10 | 108192651 | rs7099596 | 1.169 | -0.043 | 0.182 | 1.61 | 0.005 | -0.007 | -0.007 | 0 | 0.11 | 0.08 | 0.09 | 0.17 | 0.06 | 0.03 |
| 10 | 108195087 | rs7075885 | 0.355 | -0.02 | 0.283 | 1.598 | -0.025 | -0.05 | -0.05 | 0 | 0.1 | 0.18 | 0.06 | 0 | 0.04 | 0.07 |
| 10 | 108196592 | rs7894106 | 0.252 | 0.008 | 0.267 | 1.645 | -0.038 | 0.006 | 0.006 | 0 | 0.78 | 0.81 | 0.57 | 0.83 | 0.62 | 0.67 |
| 10 | 108197128 | rs952965 | 2.606 | 0.021 | 0.25 | 1.66 | 0.022 | -0.001 | -0.001 | 0.109 | 0.49 | 0.22 | 0.33 | 0.5 | 0.48 | 0.5 |
| 10 | 108202638 | rs2177828 | 2.695 | 0.027 | 0.187 | 1.784 | -0.042 | 0.037 | 0.037 | 0 | 0.49 | 0.09 | 0.31 | 0.41 | 0.5 | 0.44 |
| 10 | 108208880 | rs7083199 | 1.566 | 0.144 | 0.387 | 1.819 | 0.006 | 0.001 | 0.001 | 0 | 0.66 | 0.6 | 0.47 | 0.67 | 0.57 | 0.55 |
| 10 | 108211643 | rs7071961 | -1.292 | -0.168 | 0.695 | 1.848 | -0.065 | 0.086 | 0.086 | 0.044 | 0.26 | 0.48 | 0.24 | 0.43 | 0.12 | 0.25 |
| 10 | 108216760 | rs7096316 | -1.295 | 0.132 | 0.628 | 2.017 | -0.044 | 0.04 | 0.04 | 0 | 0.24 | 0.54 | 0.2 | 0.35 | 0.12 | 0.18 |
| 10 | 108225769 | rs9630097 | -3.108 | 0.886 | 1.228 | 3.039 | 0.027 | 0.309 | 0.309 | 0.69 | 0.08 | 0.35 | 0.07 | 0.33 | 0.06 | 0.25 |

| | | | | | | | | | | | | | | | | |
|---|---|---|---|---|---|---|---|---|---|---|---|---|---|---|---|---|
| 10 | 108238326 | rs10786950 | 1.123 | 1.048 | 0.999 | 3.036 | 0.024 | 0.012 | 0.012 | 0 | 0.69 | 0.59 | 0.49 | 0.67 | 0.58 | 0.66 |
| 10 | 108242222 | rs10884325 | -3.034 | 1.065 | 0.999 | 2.957 | -0.03 | 0.255 | 0.255 | 0.473 | 0.13 | 0.39 | 0.1 | 0.55 | 0.07 | 0.34 |
| 10 | 108244976 | rs12262411 | -1.805 | 1.223 | 0.745 | 2.207 | -0.031 | 0.122 | 0.122 | 0 | 0.08 | 0.26 | 0.05 | 0.21 | 0.03 | 0.14 |
| 10 | 108256422 | rs12412964 | -1.765 | 2.333 | 0.703 | 1.601 | 0.047 | -0.004 | -0.004 | 0.03 | 0.27 | 0.37 | 0.54 | 0.24 | 0.44 | 0.28 |
| 10 | 108258442 | rs11192926 | -1.838 | 2.389 | 0.699 | 1.572 | 0.046 | -0.012 | -0.012 | 0 | 0.3 | 0.44 | 0.54 | 0.24 | 0.44 | 0.32 |
| 10 | 108260332 | rs2900712 | 2.686 | 2.217 | 0.313 | 1.647 | 0.031 | 0.016 | 0.016 | 0.784 | 0.51 | 0.21 | 0.3 | 0.44 | 0.42 | 0.38 |
| 10 | 108262213 | rs17177040 | -0.763 | 2.356 | 0.679 | 1.29 | -0.004 | 0.003 | 0.003 | 0 | 0.14 | 0.12 | 0.31 | 0.02 | 0.18 | 0.12 |
| 10 | 108262923 | rs4917475 | 1.394 | 2.014 | 0.855 | 1.693 | -0.001 | -0.017 | -0.017 | 0.02 | 0.87 | 0.9 | 0.89 | 0.81 | 0.86 | 0.75 |
| 10 | 108267957 | rs1417949 | -1.752 | 2.07 | 1.638 | 1.548 | 0 | -0.003 | -0.003 | 0 | 0.4 | 0.38 | 0.42 | 0.35 | 0.54 | 0.49 |
| 10 | 108268865 | rs6584754 | -2.403 | 2.265 | 1.725 | 1.624 | 0.078 | 0.004 | 0.004 | 0.181 | 0.49 | 0.78 | 0.49 | 0.45 | 0.6 | 0.54 |
| 10 | 108272226 | rs1361603 | -2.445 | 2.377 | 1.731 | 1.598 | 0.049 | 0 | 0 | 0.163 | 0.49 | 0.78 | 0.52 | 0.46 | 0.57 | 0.59 |
| 10 | 108272675 | rs1891192 | -2.5 | 2.373 | 1.727 | 1.669 | 0.052 | -0.004 | -0.004 | 0 | 0.49 | 0.79 | 0.54 | 0.49 | 0.57 | 0.6 |
| 10 | 108273110 | rs11192931 | -0.473 | 2.717 | 1.963 | 1.631 | 0.006 | -0.06 | -0.06 | 0 | 0.2 | 0.21 | 0.22 | 0.01 | 0.24 | 0.09 |
| 10 | 108273426 | rs10748918 | -2.051 | 2.499 | 1.934 | 1.689 | 0.004 | -0.001 | -0.001 | 0 | 0.4 | 0.39 | 0.46 | 0.39 | 0.51 | 0.55 |
| 10 | 108276397 | rs11599633 | -0.34 | 2.086 | 2.019 | 1.606 | 0 | 0.012 | 0.012 | 0 | 0.18 | 0.09 | 0.18 | 0.01 | 0.18 | 0.09 |
| 10 | 108298091 | rs7088535 | -1.776 | 1.804 | 2.32 | 2.334 | -0.009 | -0.02 | -0.02 | 0.746 | 0.06 | 0.07 | 0.08 | 0.01 | 0.04 | 0.08 |
| 10 | 108302957 | rs7087018 | -2.094 | 1.968 | 2.517 | 2.324 | 0.076 | 0.101 | 0.101 | 3.557 | 0.1 | 0.43 | 0.09 | 0.09 | 0.07 | 0.05 |
| 10 | 108311696 | rs2181019 | -1.544 | 2.162 | 2.592 | 2.603 | 0.064 | 0.144 | 0.144 | 0 | 0.12 | 0.26 | 0.18 | 0.33 | 0.22 | 0.36 |
| 10 | 108314853 | rs7909159 | -2.15 | 1.891 | 2.069 | 2.4 | 0.018 | 0.013 | 0.013 | 1.775 | 0.15 | 0.4 | 0.15 | 0.11 | 0.12 | 0.14 |
| 10 | 108316278 | rs4142097 | -2.185 | 1.803 | 1.327 | 2.394 | 0.009 | 0.014 | 0.014 | 0.617 | 0.3 | 0.66 | 0.38 | 0.46 | 0.4 | 0.51 |
| 10 | 108325470 | rs17120918 | -1.364 | 0.632 | 1.094 | 2.747 | -0.008 | 0.031 | 0.031 | 0 | 0.05 | 0.09 | 0.02 | 0.16 | 0.02 | 0.09 |
| 10 | 108326448 | rs4918237 | -1.75 | 0.739 | 1.411 | 2.999 | 0.003 | 0.005 | 0.005 | 0.016 | 0.15 | 0.16 | 0.2 | 0.25 | 0.18 | 0.3 |
| 10 | 108344559 | rs2418811 | -1.718 | 0.483 | 1.434 | 2.993 | -0.004 | -0.013 | -0.013 | 0 | 0.15 | 0.09 | 0.18 | 0.23 | 0.16 | 0.35 |
| 10 | 108347107 | rs11596520 | -1.811 | 0.094 | 1.501 | 3.002 | 0.004 | -0.014 | -0.014 | 0 | 0.21 | 0.17 | 0.31 | 0.56 | 0.17 | 0.41 |
| 10 | 108350006 | rs4917478 | -2.531 | 0.048 | 2.42 | 2.243 | 0.002 | 0.002 | 0.002 | 0.144 | 0.32 | 0.67 | 0.36 | 0.73 | 0.22 | 0.51 |
| 10 | 108355843 | rs11814145 | -2.414 | 0.136 | 2.243 | 2.254 | -0.009 | -0.011 | -0.011 | 0.109 | 0.32 | 0.59 | 0.36 | 0.73 | 0.23 | 0.53 |
| 10 | 108364803 | rs10884335 | 0.538 | 0.418 | 2.121 | 2.312 | -0.026 | 0.062 | 0.062 | 1.561 | 0.06 | 0.15 | 0.02 | 0.14 | 0.01 | 0.04 |
| 10 | 108369512 | rs6584758 | 1.938 | 0.615 | 2.076 | 2.151 | -0.015 | 0.044 | 0.044 | 1.126 | 0.75 | 0.3 | 0.75 | 0.51 | 0.83 | 0.51 |
| 10 | 108381257 | rs7074484 | -2.381 | 0.989 | 2.222 | 2.212 | 0.022 | 0.173 | 0.173 | 0.021 | 0.2 | 0.47 | 0.25 | 0.51 | 0.17 | 0.42 |
| 10 | 108389111 | rs3802553 | -1.352 | 1.01 | 2.437 | 2.39 | 0.002 | 0.127 | 0.127 | 0 | 0.11 | 0.28 | 0.11 | 0.34 | 0.08 | 0.3 |
| 10 | 108390100 | rs10748921 | 2.159 | 1.5 | 2.264 | 2.056 | 0.016 | 0.016 | 0.016 | 0 | 0.65 | 0.44 | 0.63 | 0.48 | 0.64 | 0.51 |
| 10 | 108394614 | rs10491052 | -2.425 | 1.026 | 2.438 | 2.465 | 0.046 | 0.209 | 0.209 | 0 | 0.16 | 0.4 | 0.18 | 0.5 | 0.16 | 0.35 |
| 10 | 108406800 | rs1322015 | -2.585 | 1.093 | 2.122 | 2.44 | 0.031 | 0.17 | 0.17 | 0.009 | 0.17 | 0.41 | 0.18 | 0.5 | 0.17 | 0.35 |
| 10 | 108418384 | rs11193003 | -2.573 | 1.057 | 2.092 | 1.719 | 0.025 | 0.079 | 0.079 | 0.033 | 0.18 | 0.41 | 0.15 | 0.31 | 0.16 | 0.33 |
| 10 | 108419024 | rs7068978 | -1.995 | 0.54 | 2.214 | 1.887 | -0.006 | 0.09 | 0.09 | 0.006 | 0.13 | 0.3 | 0.1 | 0.31 | 0.09 | 0.31 |
| 10 | 108421337 | rs11193006 | -2.385 | 0.562 | 2.241 | 1.887 | 0.002 | 0.135 | 0.135 | 0.299 | 0.15 | 0.4 | 0.11 | 0.32 | 0.09 | 0.31 |
| 10 | 108424761 | rs11193007 | -1.33 | 0.619 | 1.775 | 1.941 | 0 | 0.021 | 0.021 | 0 | 0.1 | 0.14 | 0.08 | 0.31 | 0.09 | 0.32 |
| 10 | 108426206 | rs7095427 | -2.467 | 0.626 | 1.785 | 1.934 | 0.006 | 0.12 | 0.12 | 0 | 0.16 | 0.38 | 0.18 | 0.31 | 0.13 | 0.35 |
| 10 | 108428709 | rs11598147 | -1.557 | 0.724 | 1.008 | 2.053 | -0.001 | 0.054 | 0.054 | 0 | 0.12 | 0.18 | 0.08 | 0.31 | 0.09 | 0.32 |
| 10 | 108439103 | rs821929 | 1.416 | 0.48 | 1.829 | 2.354 | -0.001 | 0.272 | 0.272 | 0.071 | 0.9 | 0.7 | 0.83 | 0.65 | 0.93 | 0.81 |
| 10 | 108442019 | rs821925 | 2.147 | 0.266 | 2.248 | 2.305 | -0.02 | 0.308 | 0.308 | 1.51 | 0.88 | 0.58 | 0.82 | 0.65 | 0.94 | 0.81 |
| 10 | 108456326 | rs821957 | 2.252 | 0.246 | 2.268 | 2.377 | -0.025 | 0.187 | 0.187 | 1.946 | 0.88 | 0.59 | 0.83 | 0.74 | 0.94 | 0.82 |
| 10 | 108460406 | rs821950 | 1.241 | 0.24 | 2.533 | 2.481 | -0.001 | 0.096 | 0.096 | 0.011 | 0.93 | 0.81 | 0.89 | 0.87 | 0.97 | 0.89 |
| 10 | 108468483 | rs2788677 | 1.745 | 0.509 | 1.47 | 2.197 | -0.07 | 0.041 | 0.041 | 0.216 | 0.51 | 0.04 | 0.51 | 0.32 | 0.57 | 0.33 |
| 10 | 108473395 | rs1252034 | 2.664 | 0.498 | 1.541 | 2.18 | -0.044 | 0.119 | 0.119 | 4.066 | 0.87 | 0.44 | 0.82 | 0.74 | 0.94 | 0.82 |
| 10 | 108479649 | rs10884341 | 0.01 | 0.626 | 1.335 | 1.974 | -0.084 | 0.089 | 0.089 | 0 | 0.14 | 0.01 | 0.13 | 0 | 0.22 | 0.09 |
| 10 | 108482088 | rs7893643 | -1.458 | 0.775 | 1.348 | 1.996 | 0.005 | 0.003 | 0.003 | 0 | 0.36 | 0.41 | 0.32 | 0.42 | 0.39 | 0.48 |
| 10 | 108487180 | rs10748924 | -1.835 | 0.773 | 1.436 | 2.011 | -0.014 | -0.007 | -0.007 | 0 | 0.41 | 0.74 | 0.38 | 0.44 | 0.41 | 0.49 |
| 10 | 108492111 | rs3844062 | -1.622 | 0.763 | 1.472 | 2.048 | 0.016 | 0.01 | 0.01 | 0 | 0.38 | 0.53 | 0.33 | 0.43 | 0.4 | 0.48 |
| 10 | 108495751 | rs2756241 | 2.6 | 0.645 | 1.545 | 2.621 | 0.008 | 0.282 | 0.282 | 1.565 | 0.85 | 0.58 | 0.8 | 0.52 | 0.88 | 0.72 |
| 10 | 108495847 | rs2756242 | 2.6 | 0.645 | 1.547 | 2.621 | 0.008 | 0.286 | 0.286 | 1.565 | 0.85 | 0.56 | 0.8 | 0.52 | 0.88 | 0.72 |
| 10 | 108499902 | rs7903481 | -0.44 | 0.625 | 1.568 | 2.622 | -0.023 | 0.034 | 0.034 | 0.56 | 0.15 | 0.4 | 0.08 | 0.18 | 0.09 | 0.21 |
| 10 | 108517729 | rs10786972 | -2.636 | 0.821 | 1.801 | 2.555 | 0.009 | 0.242 | 0.242 | 0.743 | 0.14 | 0.37 | 0.19 | 0.5 | 0.12 | 0.28 |
| 10 | 108525322 | rs7071222 | -1.694 | 0.833 | 1.78 | 2.532 | 0.022 | 0.092 | 0.092 | 0.646 | 0.09 | 0.23 | 0.15 | 0.24 | 0.11 | 0.19 |
| 10 | 108528687 | rs1890456 | -1.356 | 0.827 | 1.751 | 2.418 | 0.003 | -0.011 | -0.011 | 0 | 0.23 | 0.24 | 0.24 | 0.16 | 0.26 | 0.27 |
| 10 | 108533126 | rs878182 | -0.896 | 0.816 | 1.753 | 2.404 | -0.001 | -0.028 | -0.028 | 0 | 0.21 | 0.18 | 0.22 | 0.04 | 0.25 | 0.21 |
| 10 | 108537499 | rs2184796 | -1.754 | 0.813 | 1.774 | 2.439 | -0.018 | 0.001 | 0.001 | 0 | 0.27 | 0.22 | 0.35 | 0.27 | 0.35 | 0.4 |
| 10 | 108543337 | rs10748927 | -2.317 | 0.787 | 2.007 | 2.407 | 0.002 | 0.007 | 0.007 | 0 | 0.36 | 0.63 | 0.42 | 0.53 | 0.37 | 0.49 |
| 10 | 108543868 | rs12243006 | -1.252 | 0.835 | 1.818 | 2.397 | 0.007 | -0.015 | -0.015 | 0 | 0.23 | 0.25 | 0.24 | 0.16 | 0.27 | 0.28 |
| 10 | 108545071 | rs7100070 | -0.916 | 0.84 | 1.802 | 2.334 | 0.002 | -0.036 | -0.036 | 0.332 | 0.21 | 0.21 | 0.22 | 0.04 | 0.25 | 0.21 |
| 10 | 108553214 | rs2245123 | -1.594 | 0.945 | 1.571 | 2.501 | -0.001 | 0.048 | 0.048 | 0.071 | 0.09 | 0.11 | 0.14 | 0.37 | 0.09 | 0.21 |
| 10 | 108559699 | rs2486154 | -2.328 | 0.757 | 1.655 | 2.241 | -0.006 | -0.004 | -0.004 | 0 | 0.38 | 0.64 | 0.42 | 0.38 | 0.38 | 0.42 |
| 10 | 108562008 | rs12248379 | -1.442 | 1.283 | 1.429 | 2.331 | -0.029 | -0.035 | -0.035 | 0 | 0.1 | 0.05 | 0.14 | 0.28 | 0.17 | 0.21 |
| 10 | 108563116 | rs17121613 | -1.346 | 1.075 | 1.465 | 2.423 | -0.01 | -0.015 | -0.015 | 0 | 0.09 | 0.07 | 0.13 | 0.28 | 0.11 | 0.21 |
| 10 | 108564718 | rs4918255 | -1.573 | 1.404 | 1.788 | 2.221 | 0.026 | -0.048 | -0.048 | 0 | 0.25 | 0.32 | 0.26 | 0.1 | 0.27 | 0.19 |
| 10 | 108566698 | rs17121635 | -1.116 | 1.48 | 1.414 | 2.337 | -0.007 | -0.015 | -0.015 | 0.01 | 0.08 | 0.05 | 0.11 | 0.21 | 0.09 | 0.16 |
| 10 | 108578909 | rs951277 | -1.892 | 1.103 | 1.699 | 2.27 | 0.016 | 0.019 | 0.019 | 0 | 0.37 | 0.54 | 0.42 | 0.44 | 0.38 | 0.43 |
| 10 | 108579956 | rs953591 | -0.852 | 1.235 | 1.197 | 2.351 | -0.104 | -0.039 | -0.039 | 0.255 | 0.23 | 0.04 | 0.23 | 0.2 | 0.29 | 0.23 |
| 10 | 108597250 | rs10786978 | -1.161 | 1.209 | 1.033 | 2.485 | -0.137 | -0.029 | -0.029 | 0 | 0.27 | 0.04 | 0.3 | 0.21 | 0.32 | 0.36 |

| | | | | | | | | | | | | | | | | |
|---|---|---|---|---|---|---|---|---|---|---|---|---|---|---|---|---|
| 10 | 108615370 | rs7904580 | -1.502 | 1.212 | 1.258 | 2.533 | -0.015 | -0.03 | -0.03 | 2.761 | 0.3 | 0.28 | 0.36 | 0.5 | 0.33 | 0.43 |
| 10 | 108621845 | rs7350434 | -1.883 | 1.205 | 1.455 | 2.512 | 0.014 | 0.015 | 0.015 | 0 | 0.33 | 0.44 | 0.45 | 0.37 | 0.36 | 0.42 |
| 10 | 108622962 | rs1107172 | -1.02 | 1.202 | 1.443 | 2.536 | -0.126 | -0.002 | -0.002 | 0 | 0.25 | 0.06 | 0.37 | 0.19 | 0.33 | 0.35 |
| 10 | 108624427 | rs11594752 | -2.033 | 1.169 | 1.443 | 2.541 | 0.012 | 0.027 | 0.027 | 0 | 0.34 | 0.46 | 0.46 | 0.49 | 0.36 | 0.43 |
| 10 | 108624572 | rs11193079 | -1.011 | 1.211 | 1.291 | 2.555 | -0.126 | -0.021 | -0.021 | 0 | 0.25 | 0.06 | 0.37 | 0.2 | 0.33 | 0.35 |
| 10 | 108626609 | rs1336615 | 1.938 | 0.975 | 1.091 | 2.608 | -0.117 | 0.015 | 0.015 | 0.211 | 0.5 | 0.03 | 0.41 | 0.33 | 0.55 | 0.37 |
| 10 | 108628859 | rs12761931 | -2.425 | 0.706 | 1.477 | 2.825 | 0.004 | 0.283 | 0.283 | 0 | 0.06 | 0.25 | 0.08 | 0.31 | 0.03 | 0.08 |
| 10 | 108647761 | rs7916892 | -1.507 | 0.83 | 1.411 | 1.913 | -0.005 | -0.023 | -0.023 | 0.806 | 0.36 | 0.76 | 0.44 | 0.53 | 0.38 | 0.46 |
| 10 | 108650174 | rs4917491 | -0.543 | 0.957 | 0.875 | 1.966 | -0.052 | -0.066 | -0.066 | 0.23 | 0.24 | 0.12 | 0.33 | 0.34 | 0.36 | 0.31 |
| 10 | 108653587 | rs12241863 | -0.644 | 0.973 | 0.855 | 1.973 | -0.035 | -0.047 | -0.047 | 0.402 | 0.24 | 0.18 | 0.33 | 0.34 | 0.36 | 0.3 |
| 10 | 108659048 | rs1004921 | -0.624 | 0.987 | 0.845 | 1.97 | -0.036 | -0.049 | -0.049 | 0.062 | 0.24 | 0.18 | 0.33 | 0.34 | 0.36 | 0.3 |
| 10 | 108663825 | rs12413435 | -0.203 | 1.279 | -0.153 | 1.402 | -0.048 | 0.013 | 0.013 | 0.074 | 0.11 | 0.01 | 0.13 | 0.1 | 0.11 | 0.15 |
| 10 | 108664734 | rs2418821 | -1.221 | 0.99 | 0.318 | 1.761 | 0.002 | 0.01 | 0.01 | 0 | 0.33 | 0.63 | 0.39 | 0.55 | 0.36 | 0.45 |
| 10 | 108679892 | rs7079264 | -0.371 | 0.546 | 0.02 | 0.837 | 0.001 | 0 | 0 | 0 | 0.46 | 0.53 | 0.34 | 0.27 | 0.4 | 0.46 |
| 10 | 108680184 | rs11193101 | -1.071 | 0.392 | -0.041 | 0.718 | -0.001 | -0.002 | -0.002 | 0 | 0.07 | 0.08 | 0.06 | 0.07 | 0.04 | 0.08 |
| 10 | 108681669 | rs7097380 | 0.061 | 0.132 | 0.528 | 0.489 | -0.013 | 0.047 | 0.047 | 0 | 0.35 | 0.24 | 0.27 | 0.14 | 0.36 | 0.24 |
| 10 | 108681712 | rs7097154 | 0.367 | -0.186 | 0.694 | 1.482 | 0.064 | 0.587 | 0.587 | 0.712 | 0.95 | 0.61 | 0.8 | 0.63 | 0.99 | 0.93 |
| 10 | 108688778 | rs11193120 | 0.251 | -0.142 | 0.611 | 1.435 | 0.013 | 0 | 0 | 1.105 | 0.55 | 0.24 | 0.51 | 0.42 | 0.63 | 0.56 |
| 10 | 108692152 | rs10884387 | 0.482 | -0.174 | 0.549 | 1.322 | -0.034 | 0.061 | 0.061 | 0 | 0.32 | 0.16 | 0.23 | 0.17 | 0.34 | 0.23 |
| 10 | 108695409 | rs10128404 | 0.445 | 0.128 | 0.818 | 1.289 | -0.005 | 0.016 | 0.016 | 0 | 0.52 | 0.34 | 0.41 | 0.45 | 0.59 | 0.6 |
| 10 | 108698509 | rs822076 | 0.473 | 0.15 | 0.944 | 1.312 | -0.016 | 0.053 | 0.053 | 0 | 0.33 | 0.16 | 0.21 | 0.17 | 0.32 | 0.23 |
| 10 | 108704547 | rs10786997 | 0.673 | 0.013 | 0.809 | 1.494 | -0.042 | 0.125 | 0.125 | 0.476 | 0.48 | 0.16 | 0.37 | 0.31 | 0.57 | 0.59 |
| 10 | 108709366 | rs10884390 | 0.779 | 0.01 | 0.795 | 1.666 | -0.08 | 0.126 | 0.126 | 0.989 | 0.47 | 0.11 | 0.37 | 0.31 | 0.58 | 0.59 |
| 10 | 108709892 | rs10884391 | 0.943 | -0.041 | 0.786 | 1.526 | -0.144 | 0.026 | 0.026 | 1.484 | 0.45 | 0.04 | 0.37 | 0.29 | 0.58 | 0.46 |
| 10 | 108710127 | rs10786998 | 0.593 | -0.033 | 0.82 | 1.514 | -0.032 | 0.083 | 0.083 | 0.186 | 0.49 | 0.17 | 0.38 | 0.35 | 0.58 | 0.59 |
| 10 | 108711579 | rs822097 | -1.189 | 0.045 | 1.169 | 1.641 | 0.042 | 0.278 | 0.278 | 0.076 | 0.11 | 0.35 | 0.34 | 0.39 | 0.09 | 0.17 |
| 10 | 108718383 | rs822094 | -1.691 | 0.058 | 0.411 | 1.392 | -0.013 | 0.217 | 0.217 | 3.498 | 0.16 | 0.59 | 0.37 | 0.49 | 0.09 | 0.16 |
| 10 | 108720972 | rs10786999 | -0.518 | 0.668 | 0.201 | 0.7 | -0.011 | 0.035 | 0.035 | 0 | 0.67 | 0.84 | 0.79 | 0.8 | 0.69 | 0.77 |
| 10 | 108725864 | rs719965 | 0.119 | 0.651 | 0.148 | 0.537 | -0.003 | 0.009 | 0.009 | 0 | 0.58 | 0.6 | 0.6 | 0.71 | 0.61 | 0.56 |
| 10 | 108726124 | rs1023025 | -0.3 | 0.568 | -0.437 | 0.51 | -0.009 | 0.062 | 0.062 | 0.182 | 0.67 | 0.82 | 0.79 | 0.84 | 0.69 | 0.77 |
| 10 | 108726430 | rs1023024 | -0.132 | 0.456 | -0.496 | 0.463 | 0.007 | 0.027 | 0.027 | 0.38 | 0.66 | 0.74 | 0.8 | 0.85 | 0.73 | 0.8 |
| 10 | 108729960 | rs4918274 | -0.654 | 0.712 | -1.286 | 0.527 | -0.04 | -0.018 | -0.018 | 0.525 | 0.23 | 0.04 | 0.17 | 0.24 | 0.26 | 0.26 |
| 10 | 108738830 | rs11193140 | 0.027 | 0.508 | 0.069 | -0.08 | -0.009 | 0.016 | 0.016 | 0 | 0.09 | 0.03 | 0.09 | 0 | 0.12 | 0.02 |
| 10 | 108740033 | rs10884396 | -1.058 | 0.094 | -0.523 | 0.197 | -0.001 | -0.046 | -0.046 | 0 | 0.74 | 0.9 | 0.59 | 0.66 | 0.78 | 0.59 |
| 10 | 108743550 | rs7897726 | 0.045 | 0.456 | -0.555 | 0.003 | 0 | -0.005 | -0.005 | 0 | 0.27 | 0.28 | 0.21 | 0.23 | 0.29 | 0.24 |
| 10 | 108751846 | rs17121852 | -0.272 | -0.063 | -0.501 | -0.366 | -0.051 | -0.066 | -0.066 | 0 | 0.11 | 0.68 | 0.06 | 0 | 0.02 | 0.01 |
| 10 | 108753379 | rs11193150 | -0.467 | 0.054 | -0.517 | -0.441 | 0.018 | 0.057 | 0.057 | 0.898 | 0.36 | 0.75 | 0.25 | 0.19 | 0.31 | 0.24 |
| 10 | 108754043 | rs7919051 | -0.887 | -0.086 | -0.459 | -0.445 | -0.054 | -0.056 | -0.056 | 0 | 0.1 | 0.71 | 0.07 | 0 | 0.01 | 0 |
| 10 | 108755615 | rs12572392 | 0.493 | -0.064 | -0.529 | -0.437 | -0.02 | 0.028 | 0.028 | 0.346 | 0.26 | 0.08 | 0.18 | 0.19 | 0.29 | 0.23 |
| 10 | 108765792 | rs1887635 | -1.396 | -0.13 | -0.501 | -0.296 | -0.004 | -0.035 | -0.035 | 0 | 0.75 | 0.9 | 0.6 | 0.7 | 0.8 | 0.56 |
| 10 | 108767602 | rs6584778 | -0.991 | -0.22 | -0.161 | -0.382 | -0.021 | -0.024 | -0.024 | 1.085 | 0.08 | 0.62 | 0.06 | 0 | 0 | 0 |
| 10 | 108784789 | rs6584783 | -0.476 | -0.008 | -0.176 | -0.473 | -0.003 | -0.041 | -0.041 | 0 | 0.09 | 0.3 | 0.04 | 0 | 0.03 | 0.01 |
| 10 | 108785115 | rs6584784 | -0.341 | 0.043 | -0.446 | -0.359 | 0.012 | 0.006 | 0.006 | 0.827 | 0.29 | 0.49 | 0.43 | 0.31 | 0.3 | 0.28 |
| 10 | 108789958 | rs10884401 | 0.132 | 0.028 | -0.565 | -0.352 | -0.016 | -0.01 | -0.01 | 0 | 0.26 | 0.16 | 0.39 | 0.3 | 0.3 | 0.27 |
| 10 | 108790758 | rs4918280 | -0.914 | 0.053 | -0.892 | -0.342 | 0.002 | 0.036 | 0.036 | 1.521 | 0.37 | 0.79 | 0.47 | 0.3 | 0.33 | 0.29 |
| 10 | 108791963 | rs12219276 | 0.515 | 0.385 | -1.025 | -0.174 | -0.073 | -0.074 | -0.074 | 0 | 0.21 | 0.04 | 0.4 | 0.31 | 0.29 | 0.29 |
| 10 | 108792942 | rs10884402 | 0.557 | 0.522 | -1.292 | 0.217 | -0.037 | -0.025 | -0.025 | 0 | 0.32 | 0.15 | 0.28 | 0.33 | 0.37 | 0.17 |
| 10 | 108793878 | rs12220840 | -0.347 | 0.95 | -0.742 | 0.944 | 0.001 | 0 | 0 | 0.001 | 0.3 | 0.32 | 0.44 | 0.32 | 0.32 | 0.3 |
| 10 | 108795375 | rs950809 | -0.699 | 0.864 | -0.827 | 0.423 | 0 | -0.003 | -0.003 | 0 | 0.43 | 0.47 | 0.53 | 0.63 | 0.44 | 0.52 |
| 10 | 108798092 | rs17121941 | -0.235 | 0.715 | -0.848 | 0.682 | 0.003 | -0.01 | -0.01 | 2.012 | 0.12 | 0.1 | 0.11 | 0.25 | 0.11 | 0.24 |
| 10 | 108803311 | rs717751 | -0.783 | 1.131 | -0.782 | 0.599 | -0.004 | -0.007 | -0.007 | 0 | 0.23 | 0.18 | 0.46 | 0.74 | 0.27 | 0.35 |
| 10 | 108804840 | rs10509826 | 1.556 | 0.937 | -0.955 | -0.361 | 0.049 | 0.095 | 0.095 | 0 | 0.38 | 0.17 | 0.22 | 0.13 | 0.31 | 0.18 |
| 10 | 108809524 | rs17279997 | 0.924 | 1.448 | -0.905 | -0.591 | -0.034 | 0.043 | 0.043 | 0 | 0.16 | 0.08 | 0.15 | 0.03 | 0.2 | 0.22 |
| 10 | 108811603 | rs10748931 | -0.986 | 1.386 | -0.618 | -0.573 | -0.009 | 0.014 | 0.014 | 0 | 0.84 | 0.9 | 0.89 | 0.91 | 0.81 | 0.78 |
| 10 | 108815475 | rs10748932 | -1.213 | 1.358 | -0.652 | -0.593 | -0.038 | -0.002 | -0.002 | 0 | 0.86 | 0.97 | 0.9 | 0.91 | 0.81 | 0.79 |
| 10 | 108815906 | rs11193188 | 1.253 | 1.19 | -0.703 | -0.228 | -0.042 | -0.117 | -0.117 | 0.02 | 0.15 | 0.01 | 0.15 | 0.38 | 0.17 | 0.2 |
| 10 | 108818287 | rs11193190 | 0.098 | 0.719 | -0.29 | -1.215 | -0.019 | 0.055 | 0.055 | 0 | 0.62 | 0.76 | 0.43 | 0.17 | 0.54 | 0.42 |
| 10 | 108819785 | rs10884409 | 0.513 | 0.635 | 0.445 | -1.288 | 0 | -0.018 | -0.018 | 0.225 | 0.57 | 0.54 | 0.41 | 0.16 | 0.54 | 0.42 |
| 10 | 108821861 | rs6584788 | 0.241 | 0.636 | 0.547 | -1.287 | -0.011 | 0.04 | 0.04 | 0 | 0.61 | 0.71 | 0.42 | 0.16 | 0.54 | 0.42 |
| 10 | 108823457 | rs1328229 | -0.033 | 0.643 | 0.512 | -1.304 | -0.034 | 0.073 | 0.073 | 0 | 0.63 | 0.79 | 0.43 | 0.16 | 0.54 | 0.42 |
| 10 | 108832158 | rs7898742 | 0.496 | 0.543 | 0.983 | -1.383 | -0.005 | 0.044 | 0.044 | 0 | 0.54 | 0.63 | 0.36 | 0.17 | 0.47 | 0.41 |
| 10 | 108835343 | rs2418834 | -0.35 | 0.474 | 0.832 | -1.346 | 0.013 | -0.123 | -0.123 | 0 | 0.21 | 0.57 | 0.19 | 0.03 | 0.19 | 0.23 |
| 10 | 108841019 | rs17209374 | 0.946 | 0.548 | 0.171 | -1.309 | -0.028 | 0.039 | 0.039 | 0 | 0.13 | 0.06 | 0.13 | 0.03 | 0.18 | 0.2 |
| 10 | 108843475 | rs12240854 | 1.713 | 0.576 | 0.142 | -1.302 | 0.055 | 0.102 | 0.102 | 0 | 0.37 | 0.13 | 0.2 | 0.14 | 0.29 | 0.17 |
| 10 | 108847456 | rs7073953 | 0.347 | 0.766 | 0.26 | -1.129 | -0.002 | -0.002 | -0.002 | 0.684 | 0.57 | 0.46 | 0.37 | 0.17 | 0.51 | 0.41 |
| 10 | 108850983 | rs12240947 | 0.632 | 0.265 | 0.029 | -1.181 | 0.01 | 0.093 | 0.093 | 0 | 0.53 | 0.32 | 0.38 | 0.19 | 0.54 | 0.42 |
| 10 | 108851830 | rs1556760 | 0.052 | 0.109 | -0.013 | -1.275 | -0.013 | 0.027 | 0.027 | 0 | 0.59 | 0.66 | 0.48 | 0.17 | 0.52 | 0.42 |
| 10 | 108852936 | rs9783180 | 0.551 | 0.126 | -0.075 | -1.323 | -0.005 | 0.016 | 0.016 | 0 | 0.14 | 0.09 | 0.21 | 0.03 | 0.17 | 0.24 |
| 10 | 108855428 | rs2225176 | -0.475 | 0.09 | -0.434 | -1.389 | -0.01 | 0.01 | 0.01 | 0 | 0.66 | 0.7 | 0.51 | 0.17 | 0.56 | 0.46 |

| | | | | | | | | | | | | | | | | |
|---|---|---|---|---|---|---|---|---|---|---|---|---|---|---|---|---|
| 10 | 108856606 | rs7073924 | 0.506 | 0.21 | -1.522 | -1.329 | -0.002 | 0.008 | 0.008 | 0 | 0.15 | 0.13 | 0.2 | 0.03 | 0.16 | 0.24 |
| 10 | 108859104 | rs1336978 | -0.36 | 0.504 | -1.838 | -1.209 | -0.145 | 0.081 | 0.081 | 0.245 | 0.66 | 0.94 | 0.52 | 0.16 | 0.55 | 0.43 |
| 10 | 108859804 | rs10787009 | 0.445 | 0.707 | -1.154 | -0.457 | 0.012 | 0.114 | 0.114 | 0.054 | 0.41 | 0.19 | 0.25 | 0.13 | 0.36 | 0.19 |
| 10 | 108862741 | rs4918282 | -0.323 | 0.728 | -0.998 | 0.236 | -0.002 | -0.004 | -0.004 | 0 | 0.42 | 0.59 | 0.6 | 0.53 | 0.55 | 0.7 |
| 10 | 108862960 | rs10787010 | 0.156 | 0.365 | -0.743 | 0.296 | 0.001 | 0.001 | 0.001 | 0 | 0.59 | 0.44 | 0.5 | 0.39 | 0.57 | 0.64 |
| 10 | 108876714 | rs822326 | -0.661 | -0.171 | 0.192 | 0.711 | -0.073 | 0.255 | 0.255 | 2.243 | 0.17 | 0.49 | 0.14 | 0.41 | 0.08 | 0.19 |
| 10 | 108877127 | rs669061 | -0.076 | 0.12 | -0.607 | -0.187 | -0.018 | -0.017 | -0.017 | 0 | 0.6 | 0.71 | 0.41 | 0.46 | 0.48 | 0.43 |
| 10 | 108880455 | rs607437 | -0.507 | 0.413 | -0.73 | -0.309 | -0.003 | -0.01 | -0.01 | 0 | 0.67 | 0.72 | 0.49 | 0.47 | 0.61 | 0.44 |
| 10 | 108880521 | rs607523 | -0.442 | 0.04 | 0.069 | 0.048 | -0.001 | 0.324 | 0.324 | 0.208 | 0.06 | 0.29 | 0.09 | 0.36 | 0.02 | 0.15 |
| 10 | 108886293 | rs596577 | -1.342 | 0.048 | -0.08 | -0.148 | -0.02 | 0.036 | 0.036 | 3.69 | 0.24 | 0.7 | 0.22 | 0.42 | 0.15 | 0.23 |
| 10 | 108887899 | rs7089234 | -0.567 | 0.201 | -0.111 | -0.05 | -0.003 | 0.079 | 0.079 | 0 | 0.09 | 0.18 | 0.13 | 0.36 | 0.08 | 0.2 |
| 10 | 108890426 | rs7083707 | 0.783 | 0.218 | -0.105 | -0.032 | -0.025 | -0.003 | -0.003 | 0 | 0.11 | 0.15 | 0.05 | 0.06 | 0.06 | 0.04 |
| 10 | 108893719 | rs685316 | -0.088 | 0.327 | -0.236 | -0.07 | -0.008 | -0.008 | -0.008 | 0 | 0.58 | 0.67 | 0.4 | 0.45 | 0.48 | 0.41 |
| 10 | 108898303 | rs1251753 | -0.507 | -0.178 | 0.291 | 0.553 | -0.001 | -0.002 | -0.002 | 0 | 0.78 | 0.79 | 0.62 | 0.51 | 0.79 | 0.51 |
| 10 | 108906504 | rs613020 | -1.681 | -0.449 | 0.906 | 1.045 | -0.029 | 0.16 | 0.16 | 0 | 0.09 | 0.23 | 0.11 | 0.32 | 0.04 | 0.17 |
| 10 | 108907002 | rs610785 | -1.494 | -0.322 | 0.922 | 1.061 | -0.014 | 0.111 | 0.111 | 0.017 | 0.13 | 0.29 | 0.17 | 0.33 | 0.11 | 0.2 |
| 10 | 108910338 | rs661319 | -1.409 | -0.314 | 0.513 | 0.974 | -0.016 | 0.076 | 0.076 | 0 | 0.13 | 0.28 | 0.17 | 0.26 | 0.11 | 0.2 |
| 10 | 108914383 | rs11815967 | -0.436 | -0.115 | 0.343 | 0.916 | -0.022 | -0.009 | -0.009 | 2.502 | 0.12 | 0.15 | 0.07 | 0.06 | 0.08 | 0.04 |
| 10 | 108919309 | rs615993 | -1.225 | 0.413 | 0.031 | 1.31 | 0.009 | 0.115 | 0.115 | 0 | 0.07 | 0.15 | 0.2 | 0.29 | 0.1 | 0.2 |
| 10 | 108921753 | rs2243581 | -1.174 | 0.353 | 0.21 | 0.856 | 0.003 | -0.016 | -0.016 | 0 | 0.11 | 0.23 | 0.08 | 0.07 | 0.1 | 0.03 |
| 10 | 108929333 | rs7907713 | -1.008 | -0.126 | 0.036 | 0.606 | 0.005 | 0.005 | 0.005 | 0 | 0.26 | 0.26 | 0.18 | 0.1 | 0.2 | 0.14 |
| 10 | 108940147 | rs2485203 | 1.333 | -0.533 | 0.522 | 0.766 | -0.044 | -0.001 | -0.001 | 0.058 | 0.6 | 0.49 | 0.59 | 0.53 | 0.85 | 0.53 |
| 10 | 108950282 | rs665679 | -0.885 | -0.492 | 0.564 | 0.736 | -0.008 | -0.007 | -0.007 | 0 | 0.27 | 0.38 | 0.22 | 0.16 | 0.27 | 0.14 |
| 10 | 108952513 | rs1336979 | 1.16 | -0.513 | 0.518 | 0.777 | 0.033 | -0.002 | -0.002 | 0 | 0.59 | 0.25 | 0.48 | 0.4 | 0.66 | 0.47 |
| 10 | 108953147 | rs12779243 | 1.16 | -0.513 | 0.519 | 0.755 | 0.034 | -0.007 | -0.007 | 3.067 | 0.59 | 0.25 | 0.47 | 0.41 | 0.66 | 0.47 |
| 10 | 108954666 | rs10884432 | 0.377 | -0.538 | 0.408 | 0.649 | 0.001 | -0.001 | -0.001 | 0.004 | 0.13 | 0.09 | 0.08 | 0.09 | 0.11 | 0.08 |
| 10 | 108964544 | rs11193248 | 0.473 | -0.539 | 0.387 | 0.649 | 0.015 | -0.013 | -0.013 | 0.408 | 0.13 | 0.05 | 0.08 | 0.09 | 0.11 | 0.08 |
| 10 | 108965240 | rs10884434 | 0.473 | -0.539 | 0.387 | 0.649 | 0.015 | -0.013 | -0.013 | 0 | 0.13 | 0.05 | 0.08 | 0.09 | 0.11 | 0.08 |
| 10 | 108975575 | rs1415491 | 0.114 | -0.447 | 0.274 | 0.773 | -0.029 | -0.058 | -0.058 | 0 | 0.1 | 0.02 | 0.15 | 0.29 | 0.12 | 0.22 |
| 10 | 108980153 | rs7900362 | -0.779 | -0.581 | 0.336 | 0.852 | 0.024 | 0.005 | 0.005 | 0 | 0.22 | 0.18 | 0.16 | 0.13 | 0.18 | 0.12 |
| 10 | 108982138 | rs10884443 | 0.889 | -0.572 | 0.348 | 0.712 | -0.003 | -0.003 | -0.003 | 0.19 | 0.74 | 0.67 | 0.8 | 0.84 | 0.74 | 0.86 |
| 10 | 108982678 | rs12776541 | -0.589 | -0.596 | 0.219 | 0.695 | 0.026 | 0.019 | 0.019 | 0 | 0.21 | 0.14 | 0.15 | 0.09 | 0.18 | 0.11 |
| 10 | 108986170 | rs10509829 | 1.598 | -0.249 | 0.243 | 0.621 | 0 | 0.002 | 0.002 | 0 | 0.62 | 0.39 | 0.62 | 0.55 | 0.61 | 0.63 |
| 10 | 108987492 | rs4918288 | 0.564 | -0.347 | 0.361 | 0.72 | 0.007 | -0.041 | -0.041 | 0.02 | 0.09 | 0.02 | 0.28 | 0.43 | 0.07 | 0.37 |
| 10 | 108992551 | rs1337429 | -0.22 | -0.346 | 0.364 | 0.713 | -0.022 | 0.195 | 0.195 | 0.044 | 0.14 | 0.36 | 0.32 | 0.43 | 0.09 | 0.39 |
| 10 | 109003441 | rs10509830 | -0.776 | -0.501 | -0.069 | 0.361 | 0.032 | 0.049 | 0.049 | 0 | 0.28 | 0.16 | 0.17 | 0.1 | 0.22 | 0.12 |
| 10 | 109006453 | rs1337430 | -0.785 | -0.283 | 0.297 | 0.536 | -0.03 | 0.116 | 0.116 | 0 | 0.2 | 0.39 | 0.39 | 0.44 | 0.16 | 0.42 |
| 10 | 109013673 | rs1415498 | 0.878 | -0.056 | 0.453 | 0.538 | 0.077 | 0.141 | 0.141 | 0.171 | 0.83 | 0.6 | 0.77 | 0.64 | 0.78 | 0.75 |
| 10 | 109014675 | rs11193284 | -0.762 | -0.055 | 0.464 | 0.559 | 0.023 | 0.069 | 0.069 | 0 | 0.15 | 0.27 | 0.18 | 0.3 | 0.14 | 0.22 |
| 10 | 109028744 | rs1361080 | 0.206 | 0.107 | 0.539 | 0.553 | -0.013 | 0.212 | 0.212 | 0 | 0.13 | 0.36 | 0.33 | 0.43 | 0.1 | 0.37 |
| 10 | 109033022 | rs2210969 | 1.426 | 0.127 | 0.33 | 0.453 | 0 | 0 | 0 | 1.122 | 0.56 | 0.44 | 0.61 | 0.55 | 0.56 | 0.66 |
| 10 | 109043933 | rs11193309 | -0.128 | 0.246 | -0.145 | 0.557 | -0.043 | -0.071 | -0.071 | 0 | 0.12 | 0.03 | 0.15 | 0.29 | 0.14 | 0.17 |
| 10 | 109044467 | rs12416679 | 0.172 | 0.184 | -0.025 | 0.392 | -0.001 | -0.005 | -0.005 | 0 | 0.07 | 0.07 | 0.07 | 0 | 0.07 | 0.04 |
| 10 | 109053481 | rs7090549 | 0.284 | 0.16 | -0.011 | 0.398 | -0.002 | -0.028 | -0.028 | 0 | 0.07 | 0.16 | 0.07 | 0 | 0.07 | 0.03 |
| 10 | 109059174 | rs1338955 | -0.319 | 0.131 | 0.121 | 0.385 | 0.019 | -0.035 | -0.035 | 1.583 | 0.09 | 0.36 | 0.09 | 0 | 0.07 | 0.03 |
| 10 | 109063454 | rs11193325 | -0.097 | 0.201 | 0.008 | 0.511 | -0.032 | -0.052 | -0.052 | 0 | 0.12 | 0.03 | 0.15 | 0.29 | 0.15 | 0.17 |
| 10 | 109063964 | rs7089979 | -0.603 | 0.168 | 0.089 | 0.499 | 0.02 | -0.049 | -0.049 | 0 | 0.1 | 0.37 | 0.1 | 0 | 0.07 | 0.04 |
| 10 | 109066852 | rs1572711 | 0.477 | -0.095 | -0.261 | 0.26 | -0.01 | 0.013 | 0.013 | 2.754 | 0.69 | 0.47 | 0.54 | 0.4 | 0.72 | 0.45 |
| 10 | 109072456 | rs12761509 | 1.071 | -0.064 | -0.431 | 0.392 | -0.028 | -0.041 | -0.041 | 0.273 | 0.11 | 0 | 0.07 | 0.1 | 0.11 | 0.08 |
| 10 | 109078392 | rs1416404 | 1.261 | 0.004 | -0.406 | 0.244 | 0.018 | 0.015 | 0.015 | 1.087 | 0.54 | 0.09 | 0.43 | 0.39 | 0.61 | 0.39 |
| 10 | 109081424 | rs2184916 | -1.138 | 0.168 | -0.188 | 0.351 | 0.009 | 0.022 | 0.022 | 0.637 | 0.2 | 0.27 | 0.23 | 0.29 | 0.22 | 0.21 |
| 10 | 109094097 | rs4917498 | -0.157 | 0.046 | 0.198 | 0.571 | -0.03 | 0.172 | 0.172 | 0.352 | 0.15 | 0.35 | 0.38 | 0.44 | 0.09 | 0.44 |
| 10 | 109097367 | rs11193367 | -0.117 | 0.142 | 0.053 | 0.52 | -0.008 | -0.017 | -0.017 | 0 | 0.13 | 0.07 | 0.11 | 0.29 | 0.15 | 0.17 |
| 10 | 109108123 | rs4439446 | 2.006 | 0.009 | -0.091 | 0.357 | -0.082 | 0.243 | 0.243 | 2.921 | 0.41 | 0.09 | 0.28 | 0.1 | 0.45 | 0.23 |
| 10 | 109109670 | rs7073040 | 0.81 | -0.624 | 0.316 | 0.536 | 0 | -0.001 | -0.001 | 0 | 0.63 | 0.45 | 0.48 | 0.39 | 0.69 | 0.45 |
| 10 | 109114994 | rs1338961 | -0.141 | -0.36 | 0.088 | 0.461 | 0 | 0.001 | 0.001 | 0 | 0.78 | 0.81 | 0.85 | 0.83 | 0.77 | 0.89 |
| 10 | 109115199 | rs12356312 | 0.388 | -0.387 | -0.101 | 0.461 | -0.018 | -0.029 | -0.029 | 0 | 0.18 | 0.01 | 0.1 | 0.1 | 0.16 | 0.1 |
| 10 | 109118904 | rs11193377 | -0.27 | -0.577 | 0.092 | 0.49 | 0.031 | 0.025 | 0.025 | 0 | 0.24 | 0.11 | 0.11 | 0.11 | 0.16 | 0.12 |
| 10 | 109128071 | rs7900177 | -0.823 | -0.371 | 0.168 | 0.724 | -0.028 | 0.074 | 0.074 | 0.167 | 0.13 | 0.26 | 0.14 | 0.33 | 0.07 | 0.12 |
| 10 | 109129874 | rs7084208 | 0.152 | -0.337 | 0.101 | 0.739 | -0.004 | 0.041 | 0.041 | 0 | 0.09 | 0.11 | 0.12 | 0.33 | 0.07 | 0.11 |
| 10 | 109131821 | rs1338967 | 0.152 | -0.337 | 0.101 | 0.739 | -0.004 | 0.041 | 0.041 | 0 | 0.09 | 0.11 | 0.12 | 0.33 | 0.07 | 0.11 |
| 10 | 109132777 | rs1538452 | 0.182 | -0.337 | 0.083 | 0.739 | 0.001 | 0.023 | 0.023 | 0 | 0.09 | 0.08 | 0.12 | 0.33 | 0.07 | 0.11 |
| 10 | 109133534 | rs11193395 | 0.072 | -0.353 | 0.076 | 0.732 | 0.002 | 0.019 | 0.019 | 0 | 0.09 | 0.08 | 0.13 | 0.33 | 0.07 | 0.11 |
| 10 | 109138179 | rs4636589 | 1.291 | -0.319 | 0.35 | 0.756 | 0.015 | 0.184 | 0.184 | 1.084 | 0.83 | 0.55 | 0.8 | 0.61 | 0.83 | 0.84 |
| 10 | 109141973 | rs10748939 | -0.669 | -0.44 | 0.124 | 0.595 | -0.029 | 0.025 | 0.025 | 0 | 0.38 | 0.46 | 0.48 | 0.47 | 0.28 | 0.58 |
| 10 | 109142836 | rs7095625 | 0.098 | -0.425 | 0.024 | 0.598 | -0.003 | 0.034 | 0.034 | 0 | 0.1 | 0.11 | 0.13 | 0.33 | 0.08 | 0.11 |
| 10 | 109144462 | rs1009184 | 1.397 | -0.271 | -0.071 | 0.371 | -0.05 | 0.271 | 0.271 | 2.182 | 0.45 | 0.09 | 0.32 | 0.14 | 0.54 | 0.26 |
| 10 | 109147902 | rs2297855 | 1.436 | -0.311 | -0.111 | 0.371 | -0.062 | 0.27 | 0.27 | 2.526 | 0.44 | 0.06 | 0.32 | 0.14 | 0.55 | 0.26 |
| 10 | 109173829 | rs7901760 | 0.945 | 0.178 | 0.392 | 0.7 | 0.008 | -0.024 | -0.024 | 0 | 0.06 | 0.12 | 0.07 | 0 | 0.07 | 0.03 |

| | | | | | | | | | | | | | | | | |
|---|---|---|---|---|---|---|---|---|---|---|---|---|---|---|---|---|
| 10 | 109183848 | rs11193416 | -0.126 | 0.04 | 0.323 | 0.986 | -0.003 | 0.041 | 0.041 | 0 | 0.1 | 0.11 | 0.13 | 0.35 | 0.08 | 0.11 |
| 10 | 109190042 | rs7917903 | -0.411 | 0.031 | 0.336 | 0.971 | 0.006 | 0.007 | 0.007 | 0 | 0.91 | 0.94 | 0.89 | 0.68 | 0.94 | 0.89 |
| 10 | 109190941 | rs11193420 | 0.122 | -0.379 | -0.19 | 0.401 | -0.013 | 0.101 | 0.101 | 0.274 | 0.31 | 0.04 | 0.2 | 0.08 | 0.25 | 0.18 |
| 10 | 109191459 | rs1338965 | -0.361 | -0.391 | 0.201 | 0.571 | -0.006 | 0.053 | 0.053 | 0 | 0.9 | 0.87 | 0.85 | 0.62 | 0.92 | 0.88 |
| 10 | 109194827 | rs12218576 | 0.293 | -0.383 | 0.208 | 0.579 | -0.002 | 0.052 | 0.052 | 0 | 0.09 | 0.1 | 0.14 | 0.37 | 0.07 | 0.12 |
| 10 | 109198305 | rs2900768 | -0.961 | -0.063 | -0.065 | 0.308 | 0.017 | 0.099 | 0.099 | 0 | 0.18 | 0.31 | 0.23 | 0.38 | 0.22 | 0.16 |
| 10 | 109201632 | rs12571469 | 0.34 | -0.119 | -0.158 | 0.361 | 0.003 | 0.028 | 0.028 | 0 | 0.09 | 0.09 | 0.14 | 0.38 | 0.07 | 0.12 |
| 10 | 109205283 | rs10509835 | 0.429 | -0.458 | 0.056 | 0.357 | -0.027 | 0.232 | 0.232 | 0.009 | 0.11 | 0.35 | 0.3 | 0.38 | 0.06 | 0.42 |
| 10 | 109211106 | rs11592120 | 1.573 | -0.279 | -0.323 | 0.06 | -0.119 | 0.178 | 0.178 | 0.839 | 0.36 | 0.06 | 0.23 | 0.14 | 0.44 | 0.23 |
| 10 | 109215665 | rs10787025 | -0.418 | -0.309 | -0.165 | 0.089 | -0.026 | 0.177 | 0.177 | 0.07 | 0.13 | 0.3 | 0.16 | 0.38 | 0.07 | 0.14 |
| 10 | 109215775 | rs12768572 | 1.404 | -0.192 | -0.315 | 0.05 | -0.024 | 0.021 | 0.021 | 0 | 0.05 | 0.01 | 0.07 | 0 | 0.14 | 0.03 |
| 10 | 109220670 | rs7098468 | -0.306 | -0.349 | -0.317 | 0.032 | -0.003 | 0 | 0 | 0 | 0.44 | 0.44 | 0.53 | 0.48 | 0.34 | 0.61 |
| 10 | 109236658 | rs1676793 | 0.121 | -0.251 | -0.689 | 0.042 | -0.005 | -0.005 | -0.005 | 0 | 0.85 | 0.86 | 0.83 | 0.7 | 0.78 | 0.86 |
| 10 | 109240629 | rs1372184 | -0.031 | -0.416 | -0.728 | -0.004 | 0.034 | -0.039 | -0.039 | 0 | 0.62 | 0.76 | 0.75 | 0.59 | 0.71 | 0.74 |
| 10 | 109253805 | rs1441270 | 0.753 | -0.355 | -0.943 | -0.019 | -0.011 | 0.048 | 0.048 | 0 | 0.26 | 0.05 | 0.16 | 0.09 | 0.22 | 0.14 |
| 10 | 109264291 | rs1992407 | -0.35 | -0.188 | -0.867 | 0.09 | -0.003 | 0.087 | 0.087 | 0 | 0.08 | 0.14 | 0.09 | 0.3 | 0.06 | 0.12 |
| 10 | 109266804 | rs12412473 | 0.999 | -0.358 | -0.605 | 0.141 | -0.005 | 0.099 | 0.099 | 0 | 0.08 | 0.07 | 0.26 | 0.52 | 0.05 | 0.4 |
| 10 | 109269872 | rs1660985 | -0.591 | -0.28 | -0.77 | 0.073 | -0.003 | 0.082 | 0.082 | 0 | 0.93 | 0.9 | 0.92 | 0.67 | 0.94 | 0.89 |
| 10 | 109271386 | rs1903904 | -0.437 | -0.416 | -0.901 | -0.036 | 0.026 | -0.073 | -0.073 | 0 | 0.67 | 0.84 | 0.77 | 0.61 | 0.74 | 0.74 |
| 10 | 109271687 | rs1903905 | 1.62 | -0.375 | -1.054 | -0.226 | -0.137 | 0.249 | 0.249 | 0.91 | 0.37 | 0.05 | 0.25 | 0.05 | 0.44 | 0.2 |
| 10 | 109278432 | rs1676801 | 0.051 | -0.371 | -1.017 | -0.196 | 0.025 | -0.048 | -0.048 | 4.42 | 0.35 | 0.19 | 0.25 | 0.38 | 0.27 | 0.27 |
| 10 | 109280711 | rs7899623 | -0.692 | -0.287 | -1.02 | -0.165 | -0.003 | 0.092 | 0.092 | 0 | 0.93 | 0.89 | 0.91 | 0.67 | 0.93 | 0.88 |
| 10 | 109283156 | rs17291402 | 1.592 | -0.406 | -0.854 | -0.304 | -0.11 | 0.19 | 0.19 | 1.765 | 0.35 | 0.07 | 0.24 | 0.05 | 0.44 | 0.19 |
| 10 | 109285928 | rs11591564 | 1.585 | -0.415 | -0.872 | -0.392 | -0.13 | 0.21 | 0.21 | 1.817 | 0.35 | 0.06 | 0.25 | 0.06 | 0.45 | 0.2 |
| 10 | 109303140 | rs11193535 | 0.741 | -0.752 | -0.59 | -0.037 | 0 | 0.057 | 0.057 | 0 | 0.06 | 0.06 | 0.08 | 0.3 | 0.06 | 0.12 |
| 10 | 109306967 | rs1676784 | -0.636 | -0.753 | -0.351 | -0.071 | 0.007 | 0.01 | 0.01 | 0 | 0.72 | 0.74 | 0.82 | 0.91 | 0.8 | 0.84 |
| 10 | 109314101 | rs1629453 | 0.584 | -0.702 | -0.588 | -0.087 | 0.087 | 0.053 | 0.053 | 0.056 | 0.52 | 0.23 | 0.39 | 0.39 | 0.65 | 0.38 |
| 10 | 109337125 | rs1441279 | -0.498 | -0.392 | -0.599 | -0.089 | 0.019 | 0.041 | 0.041 | 0 | 0.7 | 0.8 | 0.8 | 0.91 | 0.76 | 0.83 |
| 10 | 109340642 | rs12766593 | 0.207 | 0.384 | -0.743 | -0.148 | -0.006 | 0.016 | 0.016 | 0 | 0.45 | 0.4 | 0.32 | 0.36 | 0.47 | 0.4 |
| 10 | 109347602 | rs4917504 | -0.253 | 0.339 | -0.696 | -0.496 | 0.014 | 0.035 | 0.035 | 0 | 0.67 | 0.78 | 0.79 | 0.91 | 0.71 | 0.82 |
| 10 | 109357197 | rs1816759 | -0.401 | 1.62 | -0.262 | -0.608 | 0.092 | -0.075 | -0.075 | 0 | 0.13 | 0.3 | 0.12 | 0.01 | 0.22 | 0.06 |
| 10 | 109361425 | rs7086673 | -0.378 | 1.76 | -0.818 | -1.025 | -0.021 | 0.022 | 0.022 | 0.844 | 0.88 | 0.94 | 0.94 | 1 | 0.82 | 0.94 |
| 10 | 109368043 | rs10748948 | -0.342 | 0.746 | -0.987 | -0.945 | 0.002 | 0.003 | 0.003 | 0 | 0.58 | 0.62 | 0.45 | 0.32 | 0.58 | 0.43 |
| 10 | 109368144 | rs10884481 | 0.919 | -0.314 | -1.734 | -0.688 | 0.047 | -0.03 | -0.03 | 0 | 0.3 | 0.11 | 0.19 | 0.26 | 0.23 | 0.18 |
| 10 | 109372076 | rs11193560 | 0.75 | -0.211 | -1.659 | -0.617 | 0.003 | 0.014 | 0.014 | 0.145 | 0.08 | 0.02 | 0.1 | 0.05 | 0.08 | 0.14 |
| 10 | 109372345 | rs7921765 | -0.608 | -0.046 | -1.422 | -0.515 | 0.033 | -0.005 | -0.005 | 0.069 | 0.65 | 0.85 | 0.79 | 0.74 | 0.7 | 0.81 |
| 10 | 109374384 | rs11193561 | 0.522 | -0.145 | -1.279 | -0.815 | 0.019 | 0.021 | 0.021 | 0.121 | 0.31 | 0.2 | 0.28 | 0.19 | 0.26 | 0.18 |
| 10 | 109375673 | rs2083597 | -0.723 | -0.283 | -0.865 | -1.164 | -0.001 | 0 | 0 | 0 | 0.71 | 0.71 | 0.6 | 0.38 | 0.69 | 0.61 |
| 10 | 109379879 | rs7894314 | 0.612 | 0.27 | -0.128 | -1.383 | 0.026 | -0.071 | -0.071 | 0 | 0.15 | 0.23 | 0.11 | 0 | 0.2 | 0.07 |
| 10 | 109379996 | rs7894460 | 0.814 | 0.104 | 0.021 | -1.347 | 0.001 | 0.009 | 0.009 | 0.624 | 0.58 | 0.38 | 0.49 | 0.2 | 0.58 | 0.3 |
| 10 | 109385893 | rs7897070 | 0.517 | 0.393 | 0.032 | -1.247 | -0.025 | 0.033 | 0.033 | 0 | 0.72 | 0.52 | 0.57 | 0.34 | 0.74 | 0.47 |
| 10 | 109393000 | rs7905111 | 0.303 | 0.4 | 0.04 | -1.244 | -0.015 | 0.029 | 0.029 | 0 | 0.73 | 0.59 | 0.59 | 0.34 | 0.74 | 0.49 |
| 10 | 109393249 | rs17312242 | 1.064 | 0.389 | -0.062 | -1.27 | -0.036 | 0.038 | 0.038 | 0 | 0.09 | 0.02 | 0.06 | 0 | 0.15 | 0.03 |
| 10 | 109393286 | rs2418960 | -0.138 | 0.326 | -0.01 | -1.194 | -0.004 | 0.011 | 0.011 | 0 | 0.76 | 0.71 | 0.6 | 0.34 | 0.74 | 0.49 |
| 10 | 109396661 | rs10748950 | 0.428 | 0.38 | 0.093 | -1.175 | 0.005 | 0.016 | 0.016 | 0.708 | 0.69 | 0.51 | 0.6 | 0.36 | 0.68 | 0.54 |
| 10 | 109397459 | rs2125071 | -0.722 | 0.262 | 0.015 | -1.032 | -0.011 | 0.056 | 0.056 | 0 | 0.81 | 0.71 | 0.69 | 0.36 | 0.83 | 0.59 |
| 10 | 109399558 | rs7903890 | 1.222 | -0.078 | 0.297 | -1.512 | -0.008 | -0.021 | -0.021 | 0.984 | 0.38 | 0.4 | 0.32 | 0.23 | 0.35 | 0.17 |
| 10 | 109401050 | rs1870184 | -0.495 | -0.178 | 0.427 | -1.649 | 0.01 | -0.049 | -0.049 | 0 | 0.31 | 0.43 | 0.2 | 0.11 | 0.35 | 0.32 |
| 10 | 109405754 | rs1902737 | -1.161 | -0.365 | 0.258 | -1.777 | 0.001 | -0.02 | -0.02 | 2.402 | 0.76 | 0.82 | 0.58 | 0.36 | 0.77 | 0.6 |
| 10 | 109405963 | rs1902738 | -0.848 | -0.434 | 0.465 | -1.807 | -0.007 | 0.017 | 0.017 | 0 | 0.74 | 0.69 | 0.56 | 0.36 | 0.77 | 0.6 |
| 10 | 109406793 | rs11814070 | 1.185 | -0.415 | 0.421 | -1.882 | -0.007 | -0.014 | -0.014 | 0 | 0.15 | 0.14 | 0.1 | 0 | 0.2 | 0.12 |
| 10 | 109410486 | rs11193583 | 1.487 | -0.411 | 0.495 | -1.926 | -0.002 | 0.055 | 0.055 | 0.258 | 0.4 | 0.19 | 0.35 | 0.25 | 0.42 | 0.27 |
| 10 | 109411860 | rs12247215 | 0.87 | -0.494 | 0.625 | -1.881 | -0.027 | 0.036 | 0.036 | 0 | 0.05 | 0.16 | 0.03 | 0.11 | 0.02 | 0.16 |
| 10 | 109413483 | rs11193585 | 2.195 | -0.439 | 0.388 | -1.92 | 0.001 | 0.023 | 0.023 | 0.884 | 0.31 | 0.14 | 0.27 | 0.22 | 0.32 | 0.14 |
| 10 | 109416569 | rs12769098 | 0.683 | -0.181 | 0.334 | -1.765 | 0.009 | -0.062 | -0.062 | 1.972 | 0.17 | 0.22 | 0.12 | 0 | 0.2 | 0.12 |
| 10 | 109424604 | rs2168740 | -1.041 | -0.262 | 0.135 | -1.723 | 0 | -0.056 | -0.056 | 0 | 0.76 | 0.9 | 0.58 | 0.35 | 0.78 | 0.6 |
| 10 | 109426605 | rs7098685 | 1.18 | -0.262 | 0.393 | -1.837 | -0.008 | -0.007 | -0.007 | 0 | 0.43 | 0.44 | 0.32 | 0.33 | 0.38 | 0.34 |
| 10 | 109433484 | rs7072807 | -0.132 | -0.233 | 0.305 | -1.828 | 0 | -0.003 | -0.003 | 0 | 0.59 | 0.55 | 0.47 | 0.35 | 0.58 | 0.49 |
| 10 | 109437444 | rs11193608 | 0.477 | 0.077 | 0.472 | -1.888 | 0.031 | -0.124 | -0.124 | 0 | 0.18 | 0.39 | 0.13 | 0 | 0.21 | 0.13 |
| 10 | 109438372 | rs2444215 | -2.062 | 0.028 | 0.424 | -1.961 | 0.023 | 0.005 | 0.005 | 0.046 | 0.64 | 0.77 | 0.71 | 0.67 | 0.66 | 0.66 |
| 10 | 109440246 | rs2249587 | -0.261 | 0.055 | 0.01 | -1.973 | 0.002 | -0.01 | -0.01 | 0 | 0.42 | 0.48 | 0.54 | 0.65 | 0.43 | 0.52 |
| 10 | 109440921 | rs2446026 | -1.877 | 0.034 | -0.062 | -1.945 | 0.003 | 0.001 | 0.001 | 0 | 0.62 | 0.64 | 0.7 | 0.68 | 0.66 | 0.68 |
| 10 | 109446129 | rs2246203 | -0.169 | 0.087 | -0.099 | -1.858 | -0.004 | 0.003 | 0.003 | 0 | 0.41 | 0.36 | 0.53 | 0.65 | 0.43 | 0.52 |
| 10 | 109449483 | rs1870182 | 0.762 | 0.057 | -0.008 | -1.586 | -0.005 | -0.073 | -0.073 | 0 | 0.24 | 0.08 | 0.42 | 0.65 | 0.22 | 0.37 |
| 10 | 109452480 | rs2245437 | 0.97 | -0.003 | 0.499 | -1.403 | -0.053 | -0.038 | -0.038 | 0 | 0.81 | 0.54 | 0.85 | 0.88 | 0.88 | 0.69 |
| 10 | 109453868 | rs12258595 | -0.996 | 0.82 | -0.045 | -1.331 | 0.02 | -0.083 | -0.083 | 0 | 0.21 | 0.28 | 0.18 | 0.02 | 0.23 | 0.16 |
| 10 | 109456317 | rs17314600 | -0.598 | 1.08 | -0.341 | -1.338 | 0.008 | -0.042 | -0.042 | 0 | 0.2 | 0.22 | 0.15 | 0 | 0.32 | 0.13 |
| 10 | 109456956 | rs1902728 | 0.453 | 1.112 | -0.699 | -0.563 | -0.008 | -0.019 | -0.019 | 0 | 0.54 | 0.27 | 0.4 | 0.46 | 0.53 | 0.46 |
| 10 | 109458854 | rs7899718 | 1.381 | 0.385 | -0.693 | -0.517 | 0.002 | 0 | 0 | 0.968 | 0.22 | 0.19 | 0.14 | 0.16 | 0.19 | 0.11 |

| | | | | | | | | | | | | | | | | |
|---|---|---|---|---|---|---|---|---|---|---|---|---|---|---|---|---|
| 10 | 109459645 | rs1902729 | 0.473 | 0.071 | -0.532 | -0.327 | -0.033 | 0.028 | 0.028 | 0 | 0.06 | 0.14 | 0.18 | 0.12 | 0.02 | 0.19 |
| 10 | 109462889 | rs7903669 | 1.603 | 0.149 | -0.768 | -0.344 | 0.013 | 0 | 0 | 0 | 0.21 | 0.14 | 0.14 | 0.16 | 0.19 | 0.11 |
| 10 | 109465153 | rs2252331 | 1.611 | -0.025 | -0.867 | -0.088 | 0 | 0 | 0 | 0.151 | 0.15 | 0.17 | 0.21 | 0.31 | 0.13 | 0.21 |
| 10 | 109466526 | rs7905924 | 1.681 | 0.124 | -1.034 | -0.829 | 0.013 | 0.004 | 0.004 | 0 | 0.23 | 0.11 | 0.16 | 0.16 | 0.22 | 0.12 |
| 10 | 109468251 | rs11193635 | 0.687 | 0.694 | -1.456 | -0.9 | -0.095 | 0.058 | 0.058 | 0 | 0.3 | 0.06 | 0.21 | 0.18 | 0.34 | 0.33 |
| 10 | 109469934 | rs2251717 | -0.955 | 0.607 | -0.348 | -0.522 | 0.004 | -0.043 | -0.043 | 0 | 0.79 | 0.78 | 0.83 | 1 | 0.7 | 0.91 |
| 10 | 109471331 | rs2251521 | 0.697 | 0.413 | -0.148 | -0.28 | -0.002 | 0.043 | 0.043 | 0.552 | 0.18 | 0.21 | 0.26 | 0.51 | 0.16 | 0.28 |
| 10 | 109473579 | rs10884519 | -0.267 | 0.12 | -0.077 | -0.106 | 0.01 | 0.011 | 0.011 | 0 | 0.65 | 0.61 | 0.58 | 0.49 | 0.56 | 0.63 |
| 10 | 109474406 | rs7091009 | 0.133 | -0.718 | -0.002 | -0.054 | 0.009 | -0.033 | -0.033 | 0.896 | 0.06 | 0.15 | 0.05 | 0.01 | 0.06 | 0.02 |
| 10 | 109486011 | rs2244483 | 0.628 | -1.18 | 0.03 | -0.116 | -0.075 | -0.054 | -0.054 | 0 | 0.79 | 0.5 | 0.71 | 0.86 | 0.87 | 0.79 |
| 10 | 109486048 | rs2244482 | 0.424 | -1.675 | -0.148 | 0.123 | -0.014 | 0.049 | 0.049 | 5.756 | 0.19 | 0.25 | 0.21 | 0.52 | 0.11 | 0.23 |
| 10 | 109493215 | rs10884521 | 0.351 | -1.416 | -0.516 | -0.41 | 0.009 | 0.009 | 0.009 | 0.109 | 0.57 | 0.4 | 0.48 | 0.23 | 0.7 | 0.35 |
| 10 | 109503398 | rs10884524 | 0.771 | -1.287 | -0.708 | -0.385 | 0.07 | 0.092 | 0.092 | 0.304 | 0.47 | 0.24 | 0.43 | 0.2 | 0.66 | 0.31 |
| 10 | 109508710 | rs10884527 | -0.485 | -0.804 | -0.049 | -0.071 | 0.007 | 0.017 | 0.017 | 0 | 0.35 | 0.44 | 0.36 | 0.66 | 0.33 | 0.55 |
| 10 | 109509769 | rs6584825 | -0.333 | -0.982 | 0.214 | -0.092 | -0.041 | 0.011 | 0.011 | 0 | 0.91 | 0.8 | 0.83 | 0.88 | 0.96 | 0.84 |
| 10 | 109511008 | rs10884528 | 0.745 | -1.017 | 0.354 | 0.086 | -0.023 | 0.141 | 0.141 | 0.082 | 0.17 | 0.33 | 0.2 | 0.54 | 0.11 | 0.31 |
| 10 | 109514980 | rs11193670 | -0.917 | -0.904 | 0.376 | -0.023 | -0.002 | -0.004 | -0.004 | 0.05 | 0.43 | 0.53 | 0.41 | 0.68 | 0.38 | 0.58 |
| 10 | 109519015 | rs7909332 | 0.538 | -1.05 | -0.139 | -0.466 | -0.022 | -0.008 | -0.008 | 0 | 0.35 | 0.44 | 0.38 | 0.69 | 0.28 | 0.41 |
| 10 | 109519083 | rs17316057 | 1.439 | -0.921 | 0.083 | -0.44 | -0.021 | 0.001 | 0.001 | 0 | 0.14 | 0.07 | 0.14 | 0.11 | 0.17 | 0.21 |
| 10 | 109521588 | rs2418977 | 1.839 | -0.94 | 0.044 | -0.438 | -0.041 | -0.029 | -0.029 | 0 | 0.13 | 0.01 | 0.12 | 0.11 | 0.13 | 0.2 |
| 10 | 109521673 | rs2418978 | -0.626 | -1 | 0.114 | -0.46 | 0.003 | 0.006 | 0.006 | 0 | 0.44 | 0.58 | 0.42 | 0.69 | 0.38 | 0.59 |
| 10 | 109524277 | rs11193676 | -0.739 | -1.099 | 0.01 | -0.5 | -0.031 | 0.054 | 0.054 | 0.021 | 0.92 | 0.78 | 0.82 | 0.86 | 0.97 | 0.86 |
| 10 | 109525067 | rs7912221 | -0.206 | -1.137 | 0.169 | -0.539 | 0.002 | -0.037 | -0.037 | 0.062 | 0.29 | 0.43 | 0.24 | 0.15 | 0.28 | 0.28 |
| 10 | 109525838 | rs10884533 | 0.618 | -1.185 | 0.147 | -0.521 | -0.014 | -0.002 | -0.002 | 0 | 0.44 | 0.2 | 0.4 | 0.49 | 0.5 | 0.45 |
| 10 | 109526561 | rs11193682 | 1.102 | -1.208 | 0.085 | -0.309 | -0.017 | 0.096 | 0.096 | 0.049 | 0.17 | 0.27 | 0.2 | 0.53 | 0.11 | 0.31 |
| 10 | 109527075 | rs2900778 | -0.087 | -1.176 | -0.022 | -0.315 | -0.006 | 0.024 | 0.024 | 0 | 0.34 | 0.4 | 0.35 | 0.6 | 0.28 | 0.55 |
| 10 | 109529803 | rs10884534 | 1.297 | -0.694 | -0.509 | -0.544 | -0.07 | 0.115 | 0.115 | 0 | 0.28 | 0.09 | 0.21 | 0 | 0.36 | 0.14 |
| 10 | 109537148 | rs17123221 | 0.504 | -0.657 | -0.132 | 0.112 | 0 | -0.005 | -0.005 | 0 | 0.1 | 0.13 | 0.07 | 0.04 | 0.1 | 0.04 |
| 10 | 109539279 | rs4431961 | 0.351 | -0.694 | 0.04 | 0.481 | -0.002 | -0.006 | -0.006 | 3.286 | 0.73 | 0.66 | 0.66 | 0.77 | 0.75 | 0.59 |
| 10 | 109543709 | rs10884540 | 0.351 | -0.657 | 0.118 | 0.594 | 0.001 | 0.029 | 0.029 | 0 | 0.75 | 0.66 | 0.66 | 0.66 | 0.77 | 0.71 |
| 10 | 109548708 | rs7908795 | -0.258 | -0.79 | 0.09 | 0.489 | 0.001 | 0.002 | 0.002 | 0 | 0.37 | 0.27 | 0.37 | 0.26 | 0.35 | 0.31 |
| 10 | 109548848 | rs17123241 | 0.181 | -1.147 | 0.191 | 0.502 | -0.002 | -0.001 | -0.001 | 0 | 0.08 | 0.07 | 0.17 | 0.12 | 0.03 | 0.15 |
| 10 | 109549044 | rs4144574 | 0.809 | -1.468 | 0.44 | 0.405 | -0.014 | 0.106 | 0.106 | 0 | 0.11 | 0.19 | 0.18 | 0.55 | 0.09 | 0.34 |
| 10 | 109552580 | rs7902568 | 0.218 | -1.512 | 0.558 | 0.379 | 0 | -0.001 | -0.001 | 0.26 | 0.36 | 0.31 | 0.38 | 0.26 | 0.36 | 0.31 |
| 10 | 109553347 | rs11193693 | -0.01 | -1.778 | 0.982 | 0.273 | -0.034 | 0.125 | 0.125 | 0 | 0.15 | 0.29 | 0.2 | 0.6 | 0.1 | 0.35 |
| 10 | 109554914 | rs11193694 | 0.116 | -1.741 | 0.802 | 0.23 | -0.001 | -0.002 | -0.002 | 0 | 0.37 | 0.37 | 0.38 | 0.25 | 0.36 | 0.31 |
| 10 | 109557818 | rs10884543 | -0.856 | -1.739 | 0.884 | 0.172 | 0.005 | 0.013 | 0.013 | 0 | 0.51 | 0.57 | 0.57 | 0.81 | 0.45 | 0.64 |
| 10 | 109559022 | rs17095155 | 0.283 | -1.665 | 0.687 | 0.221 | -0.003 | -0.009 | -0.009 | 0 | 0.36 | 0.22 | 0.37 | 0.3 | 0.36 | 0.3 |
| 10 | 109561671 | rs10884544 | 0.251 | -1.704 | 0.621 | 0.205 | -0.02 | -0.03 | -0.03 | 0.619 | 0.34 | 0.14 | 0.36 | 0.3 | 0.35 | 0.3 |
| 10 | 109563610 | rs11193702 | 1.019 | -1.726 | 0.354 | 0.154 | -0.071 | -0.063 | -0.063 | 0 | 0.19 | 0.02 | 0.23 | 0.17 | 0.22 | 0.21 |
| 10 | 109565840 | rs10509849 | 1.747 | -1.779 | 0.305 | -0.091 | -0.186 | 0.248 | 0.248 | 0.052 | 0.33 | 0.03 | 0.21 | 0 | 0.46 | 0.16 |
| 10 | 109566211 | rs2243561 | -0.741 | -1.482 | 0.708 | 0.151 | 0.016 | 0.084 | 0.084 | 0 | 0.48 | 0.35 | 0.55 | 0.81 | 0.45 | 0.64 |
| 10 | 109567399 | rs2250938 | -0.638 | -1.603 | 0.468 | 0.165 | 0.021 | 0.1 | 0.1 | 0 | 0.46 | 0.3 | 0.54 | 0.81 | 0.44 | 0.65 |
| 10 | 109567622 | rs17123358 | -1.197 | -1.359 | 0.991 | 0.75 | 0.036 | 0.033 | 0.033 | 1.255 | 0.06 | 0.35 | 0.04 | 0 | 0 | 0 |
| 10 | 109567750 | rs11193706 | 0.807 | -1.046 | 0.025 | 0.934 | -0.026 | -0.047 | -0.047 | 0.717 | 0.13 | 0.01 | 0.11 | 0.12 | 0.13 | 0.09 |
| 10 | 109570518 | rs17123365 | -1.012 | -1.243 | 0.336 | 0.88 | -0.011 | 0.069 | 0.069 | 0 | 0.16 | 0.27 | 0.23 | 0.53 | 0.15 | 0.38 |
| 10 | 109571413 | rs2026777 | -0.686 | -0.963 | 0.167 | 1.008 | -0.005 | 0.106 | 0.106 | 0 | 0.14 | 0.23 | 0.23 | 0.52 | 0.15 | 0.38 |
| 10 | 109573238 | rs11193711 | -0.596 | -0.932 | -0.077 | 0.223 | -0.038 | -0.006 | -0.006 | 0 | 0.1 | 0.18 | 0.18 | 0.08 | 0.03 | 0.14 |
| 10 | 109575094 | rs10509852 | -0.365 | -0.227 | -0.789 | 0.441 | 0.012 | -0.011 | -0.011 | 0.124 | 0.25 | 0.17 | 0.32 | 0.75 | 0.28 | 0.44 |
| 10 | 109575301 | rs10509853 | -0.676 | -0.198 | -0.777 | 0.343 | 0.004 | 0.015 | 0.015 | 0 | 0.28 | 0.21 | 0.34 | 0.77 | 0.32 | 0.49 |
| 10 | 109578567 | rs7916920 | 0.08 | 0.421 | -0.524 | 0.51 | -0.001 | 0 | 0 | 0 | 0.54 | 0.42 | 0.46 | 0.36 | 0.53 | 0.34 |
| 10 | 109578826 | rs1329998 | 0.023 | 0.485 | -0.51 | 0.205 | -0.006 | -0.022 | -0.022 | 0 | 0.58 | 0.48 | 0.52 | 0.29 | 0.58 | 0.42 |
| 10 | 109579127 | rs4918322 | 0.2 | 0.195 | -0.351 | -0.163 | 0.003 | -0.035 | -0.035 | 0.21 | 0.63 | 0.53 | 0.62 | 0.75 | 0.67 | 0.62 |
| 10 | 109579550 | rs12778719 | 0.109 | -0.134 | -0.434 | -0.066 | 0.001 | 0 | 0 | 0.003 | 0.58 | 0.59 | 0.47 | 0.42 | 0.52 | 0.46 |
| 10 | 109579843 | rs12358624 | 0.214 | -0.348 | -0.065 | -0.334 | -0.003 | -0.01 | -0.01 | 0 | 0.65 | 0.7 | 0.72 | 0.89 | 0.66 | 0.69 |
| 10 | 109580141 | rs2246712 | -0.169 | -0.288 | -0.123 | -0.338 | -0.022 | 0.043 | 0.043 | 0 | 0.3 | 0.2 | 0.28 | 0.11 | 0.33 | 0.33 |
| 10 | 109580995 | rs4526707 | 1.104 | -0.61 | -0.607 | -0.912 | -0.011 | 0.013 | 0.013 | 1.364 | 0.27 | 0.25 | 0.2 | 0 | 0.36 | 0.09 |
| 10 | 109581185 | rs17227554 | 1.104 | -0.61 | -0.607 | -0.912 | -0.011 | 0.013 | 0.013 | 1.364 | 0.27 | 0.25 | 0.2 | 0 | 0.36 | 0.09 |
| 10 | 109585709 | rs7924126 | 0.648 | -0.569 | -0.29 | -0.513 | -0.003 | 0.079 | 0.079 | 0 | 0.07 | 0.14 | 0.15 | 0.27 | 0.09 | 0.15 |
| 10 | 109586573 | rs2265217 | 0.978 | -0.683 | -0.354 | -0.586 | -0.027 | 0.023 | 0.023 | 0 | 0.22 | 0.12 | 0.19 | 0.14 | 0.25 | 0.32 |
| 10 | 109587142 | rs11193725 | 0.422 | -0.71 | -0.157 | -0.521 | 0.006 | 0.088 | 0.088 | 0 | 0.08 | 0.22 | 0.16 | 0.2 | 0.09 | 0.15 |
| 10 | 109588382 | rs2551822 | 1.21 | -0.7 | -0.458 | -0.545 | -0.079 | 0.034 | 0.034 | 0 | 0.21 | 0.05 | 0.19 | 0.12 | 0.26 | 0.32 |
| 10 | 109589864 | rs12359040 | 1.487 | -0.712 | -0.683 | -0.449 | -0.009 | -0.022 | -0.022 | 0.57 | 0.11 | 0.01 | 0.04 | 0.08 | 0.05 | 0.17 |
| 10 | 109590355 | rs4090119 | 1.078 | -0.615 | -0.141 | -0.334 | -0.001 | 0.012 | 0.012 | 2.786 | 0.18 | 0.14 | 0.31 | 0.59 | 0.16 | 0.23 |
| 10 | 109590833 | rs12768215 | 1.127 | -0.641 | -0.077 | -0.464 | -0.007 | -0.014 | -0.014 | 0 | 0.28 | 0.28 | 0.2 | 0 | 0.38 | 0.09 |
| 10 | 109593360 | rs1360245 | 0.65 | -0.495 | -0.489 | -0.321 | -0.001 | 0.005 | 0.005 | 0 | 0.38 | 0.31 | 0.25 | 0.2 | 0.3 | 0.51 |
| 10 | 109593556 | rs1360246 | 0.301 | -0.423 | -0.948 | -0.102 | -0.009 | -0.01 | -0.01 | 0 | 0.43 | 0.39 | 0.51 | 0.5 | 0.53 | 0.32 |
| 10 | 109593621 | rs11193730 | -0.217 | -0.209 | -0.815 | 0.938 | -0.008 | 0.058 | 0.058 | 0 | 0.09 | 0.15 | 0.15 | 0.29 | 0.09 | 0.15 |
| 10 | 109599439 | rs1411290 | 0.32 | -0.052 | -0.789 | 0.689 | -0.001 | 0.012 | 0.012 | 2.947 | 0.17 | 0.14 | 0.32 | 0.58 | 0.16 | 0.23 |

| | | | | | | | | | | | | | | | | |
|---|---|---|---|---|---|---|---|---|---|---|---|---|---|---|---|---|
| 10 | 109605065 | rs12768320 | -0.7 | 0.028 | -0.818 | 0.361 | 0.003 | -0.001 | -0.001 | 0 | 0.17 | 0.16 | 0.22 | 0.22 | 0.16 | 0.17 |
| 10 | 109606740 | rs7067538 | -0.691 | 0.708 | -0.614 | 0.238 | 0.009 | 0.02 | 0.02 | 0 | 0.29 | 0.41 | 0.3 | 0.36 | 0.33 | 0.34 |
| 10 | 109609300 | rs765651 | -0.457 | 0.297 | -1.131 | -1.407 | -0.005 | 0.047 | 0.047 | 0 | 0.27 | 0.19 | 0.13 | 0.08 | 0.29 | 0.17 |
| 10 | 109617163 | rs10884552 | -0.431 | 0.225 | -1.158 | -1.471 | -0.009 | 0.061 | 0.061 | 0 | 0.27 | 0.17 | 0.14 | 0.08 | 0.28 | 0.18 |
| 10 | 109618097 | rs2551830 | 0.907 | 0.209 | -1.451 | -2.177 | -0.015 | 0.026 | 0.026 | 0 | 0.25 | 0.17 | 0.21 | 0.07 | 0.33 | 0.22 |
| 10 | 109620271 | rs7087669 | 0.369 | -0.397 | -0.584 | -2.272 | -0.009 | -0.012 | -0.012 | 0 | 0.08 | 0.36 | 0.05 | 0 | 0.01 | 0.01 |
| 10 | 109624234 | rs12217764 | 1.497 | -0.412 | -1.091 | -2.289 | -0.05 | -0.067 | -0.067 | 0 | 0.14 | 0.02 | 0.39 | 0.72 | 0.21 | 0.39 |
| 10 | 109624420 | rs11193752 | -0.09 | -0.817 | -1.087 | -2.492 | 0.002 | -0.032 | -0.032 | 0 | 0.37 | 0.44 | 0.58 | 0.77 | 0.34 | 0.48 |
| 10 | 109625453 | rs2248353 | -0.712 | -0.99 | -1.089 | -2.523 | 0.027 | 0.079 | 0.079 | 0 | 0.47 | 0.64 | 0.63 | 0.85 | 0.37 | 0.61 |
| 10 | 109626008 | rs1424605 | 0.767 | -0.889 | -1.13 | -2.456 | -0.046 | -0.028 | -0.028 | 0 | 0.26 | 0.06 | 0.5 | 0.72 | 0.31 | 0.46 |
| 10 | 109643023 | rs7896459 | -1.662 | -0.734 | -1.1 | -2.667 | -0.052 | 0.087 | 0.087 | 0 | 0.74 | 0.89 | 0.76 | 0.94 | 0.63 | 0.78 |
| 10 | 109649080 | rs1424604 | 0.113 | -0.73 | -0.861 | -2.58 | -0.005 | -0.028 | -0.028 | 0 | 0.36 | 0.43 | 0.55 | 0.78 | 0.32 | 0.49 |
| 10 | 109651849 | rs11193773 | 0.247 | -1.55 | -0.619 | -2.416 | -0.015 | 0.062 | 0.062 | 0 | 0.24 | 0.13 | 0.12 | 0.06 | 0.25 | 0.16 |
| 10 | 109661889 | rs1571553 | -1.642 | -1.646 | -0.564 | -2.491 | -0.014 | 0.116 | 0.116 | 0 | 0.6 | 0.87 | 0.37 | 0.16 | 0.4 | 0.39 |
| 10 | 109662354 | rs1974918 | -0.685 | -1.969 | -0.486 | -2.594 | 0.081 | 0.089 | 0.089 | 0.205 | 0.42 | 0.72 | 0.28 | 0.1 | 0.2 | 0.24 |
| 10 | 109664584 | rs12412457 | -0.588 | -2.015 | -0.399 | -2.633 | -0.047 | -0.068 | -0.068 | 0.1 | 0.41 | 0.47 | 0.26 | 0.1 | 0.21 | 0.24 |
| 10 | 109673127 | rs7905923 | 0.127 | -1.982 | -0.192 | -2.612 | -0.074 | -0.154 | -0.154 | 0 | 0.23 | 0.43 | 0.18 | 0 | 0.11 | 0.1 |
| 10 | 109677767 | rs1834175 | -0.142 | -1.929 | -0.42 | -2.62 | -0.008 | 0.012 | 0.012 | 0 | 0.19 | 0.16 | 0.11 | 0.11 | 0.21 | 0.16 |
| 10 | 109688765 | rs11193802 | -1.84 | -1.899 | -0.518 | -2.751 | -0.005 | 0.129 | 0.129 | 0.825 | 0.59 | 0.88 | 0.38 | 0.19 | 0.4 | 0.33 |
| 10 | 109691871 | rs1863021 | 0.399 | -1.968 | -0.059 | -2.76 | -0.059 | -0.13 | -0.13 | 0 | 0.22 | 0.4 | 0.17 | 0 | 0.11 | 0.1 |
| 10 | 109693841 | rs12775077 | 0.833 | -1.715 | -0.227 | -2.61 | -0.016 | 0.027 | 0.027 | 0 | 0.1 | 0.06 | 0.07 | 0.03 | 0.12 | 0.05 |
| 10 | 109702784 | rs17123614 | 0.922 | -1.736 | -0.259 | -2.601 | -0.023 | -0.025 | -0.025 | 0 | 0.06 | 0.18 | 0.03 | 0 | 0.01 | 0.02 |
| 10 | 109705944 | rs17691916 | 1.314 | -1.767 | -0.326 | -2.633 | -0.007 | -0.003 | -0.003 | 0 | 0.13 | 0.01 | 0.09 | 0.08 | 0.1 | 0.07 |
| 10 | 109707225 | rs2113567 | -1.311 | -1.801 | -0.32 | -2.664 | 0.013 | 0.024 | 0.024 | 0 | 0.37 | 0.34 | 0.2 | 0.19 | 0.28 | 0.23 |
| 10 | 109712692 | rs12785211 | -0.642 | -1.51 | -0.114 | -2.518 | -0.009 | 0.022 | 0.022 | 0.386 | 0.19 | 0.14 | 0.1 | 0.1 | 0.2 | 0.15 |
| 10 | 109728141 | rs11193818 | 0.492 | -1.537 | -0.048 | -2.392 | 0 | 0.001 | 0.001 | 0 | 0.06 | 0.08 | 0.03 | 0.07 | 0.07 | 0.11 |
| 10 | 109730743 | rs7919814 | -1.564 | -1.778 | -0.072 | -2.674 | 0.044 | 0.137 | 0.137 | 0 | 0.48 | 0.75 | 0.33 | 0.1 | 0.37 | 0.28 |
| 10 | 109731584 | rs7915527 | 0.077 | -1.709 | 0.027 | -2.606 | -0.037 | -0.127 | -0.127 | 0 | 0.23 | 0.38 | 0.18 | 0 | 0.14 | 0.11 |
| 10 | 109738180 | rs1330007 | -0.537 | -1.363 | -0.517 | -2.411 | -0.004 | 0.005 | 0.005 | 0.349 | 0.19 | 0.2 | 0.12 | 0.1 | 0.23 | 0.16 |
| 10 | 109743514 | rs10787066 | -0.204 | -2.799 | -0.231 | -2.542 | -0.008 | -0.037 | -0.037 | 0.469 | 0.26 | 0.28 | 0.2 | 0.06 | 0.2 | 0.19 |
| 10 | 109746727 | rs10748962 | -1.45 | -2.829 | -0.463 | -2.57 | 0.026 | 0.093 | 0.093 | 0 | 0.44 | 0.69 | 0.32 | 0.1 | 0.36 | 0.26 |
| 10 | 109755896 | rs7897681 | -2.328 | -3.043 | -0.684 | -2.539 | -0.048 | 0.064 | 0.064 | 0.011 | 0.63 | 0.88 | 0.4 | 0.21 | 0.45 | 0.34 |
| 10 | 109763055 | rs10509856 | 1.317 | -3.105 | -0.705 | -2.559 | 0.025 | -0.007 | -0.007 | 0.2 | 0.13 | 0.06 | 0.07 | 0.1 | 0.08 | 0.06 |
| 10 | 109763752 | rs12774009 | -0.509 | -3.037 | -0.538 | -2.656 | 0.009 | 0.038 | 0.038 | 0 | 0.21 | 0.14 | 0.09 | 0.04 | 0.17 | 0.09 |
| 10 | 109765862 | rs10748965 | -1.868 | -3.057 | -0.573 | -2.654 | 0.073 | 0.229 | 0.229 | 0.041 | 0.49 | 0.82 | 0.33 | 0.1 | 0.36 | 0.28 |
| 10 | 109784849 | rs7094099 | 1.125 | -3.029 | -0.489 | -2.646 | 0.001 | -0.023 | -0.023 | 1.909 | 0.17 | 0.17 | 0.16 | 0 | 0.13 | 0.1 |
| 10 | 109794685 | rs11193857 | 0.958 | -2.939 | -0.735 | -2.596 | -0.027 | 0.055 | 0.055 | 0 | 0.1 | 0.01 | 0.04 | 0 | 0.09 | 0.03 |
| 10 | 109794993 | rs7082958 | -0.489 | -2.961 | -0.592 | -2.586 | -0.006 | -0.036 | -0.036 | 0 | 0.28 | 0.66 | 0.25 | 0.05 | 0.2 | 0.2 |
| 10 | 109800002 | rs1931608 | -1.729 | -2.883 | -0.793 | -2.54 | 0.076 | 0.234 | 0.234 | 0 | 0.5 | 0.82 | 0.34 | 0.11 | 0.36 | 0.29 |
| 10 | 109800376 | rs10509857 | 0.127 | -2.207 | -0.649 | -2.185 | -0.002 | 0.013 | 0.013 | 0 | 0.15 | 0.1 | 0.07 | 0 | 0.14 | 0.05 |
| 10 | 109805032 | rs12764054 | -0.496 | -2.142 | -0.851 | -2.144 | -0.003 | 0.087 | 0.087 | 0 | 0.26 | 0.07 | 0.13 | 0.09 | 0.22 | 0.1 |
| 10 | 109809622 | rs7094944 | -0.22 | -2.137 | -0.395 | -2.258 | -0.004 | -0.057 | -0.057 | 1.606 | 0.25 | 0.35 | 0.22 | 0.07 | 0.22 | 0.2 |
| 10 | 109829770 | rs1556592 | -0.992 | -2.13 | -0.443 | -2.26 | 0.015 | 0.024 | 0.024 | 0 | 0.41 | 0.64 | 0.31 | 0.15 | 0.3 | 0.26 |
| 10 | 109831945 | rs12359408 | 0.102 | -2.182 | -0.458 | -2.309 | -0.001 | 0.017 | 0.017 | 0 | 0.16 | 0.13 | 0.08 | 0 | 0.14 | 0.05 |
| 10 | 109836172 | rs11193880 | -1.826 | -2.199 | -0.418 | -2.32 | -0.005 | 0.081 | 0.081 | 0 | 0.55 | 0.72 | 0.37 | 0.15 | 0.44 | 0.3 |
| 10 | 109836835 | rs10509860 | 0.892 | -2.234 | -0.524 | -2.296 | 0.03 | -0.004 | -0.004 | 0 | 0.14 | 0.01 | 0.05 | 0.08 | 0.08 | 0.06 |
| 10 | 109839002 | rs10509865 | 0.09 | -2.212 | -0.42 | -2.291 | 0.006 | -0.035 | -0.035 | 0 | 0.21 | 0.28 | 0.2 | 0.07 | 0.22 | 0.19 |
| 10 | 109856031 | rs7911003 | -1.114 | -2.059 | -0.629 | -2.209 | -0.002 | -0.008 | -0.008 | 0 | 0.4 | 0.58 | 0.31 | 0.19 | 0.31 | 0.27 |
| 10 | 109861855 | rs1931600 | -0.124 | -2.051 | -0.778 | -2.193 | 0.002 | -0.027 | -0.027 | 0.029 | 0.22 | 0.27 | 0.22 | 0.1 | 0.23 | 0.2 |
| 10 | 109867337 | rs17250227 | -0.066 | -1.618 | -0.348 | -1.734 | -0.016 | 0.052 | 0.052 | 0 | 0.14 | 0.05 | 0.07 | 0 | 0.15 | 0.05 |
| 10 | 109871472 | rs7912486 | -0.572 | -1.263 | -0.818 | -1.236 | -0.008 | -0.004 | -0.004 | 0.249 | 0.21 | 0.24 | 0.11 | 0.15 | 0.17 | 0.16 |
| 10 | 109881982 | rs12217601 | -0.263 | -1.366 | -1.064 | -1.057 | -0.009 | 0.003 | 0.003 | 0 | 0.16 | 0.07 | 0.1 | 0.11 | 0.16 | 0.16 |
| 10 | 109883659 | rs11193900 | -0.918 | -2.525 | -0.949 | -1.952 | -0.016 | -0.09 | -0.09 | 0 | 0.38 | 0.6 | 0.19 | 0.06 | 0.25 | 0.15 |
| 10 | 109884688 | rs7076037 | -1.576 | -2.837 | -1.209 | -1.898 | 0.003 | 0.078 | 0.078 | 0 | 0.59 | 0.81 | 0.39 | 0.27 | 0.42 | 0.38 |
| 10 | 109884711 | rs11193901 | 0.098 | -2.721 | -1.145 | -1.893 | 0.058 | 0.034 | 0.034 | 0 | 0.09 | 0.18 | 0.07 | 0.1 | 0.13 | 0.14 |
| 10 | 109886402 | rs1931603 | -0.059 | -3.132 | -1.116 | -1.913 | 0.006 | 0.111 | 0.111 | 0 | 0.26 | 0.07 | 0.13 | 0 | 0.23 | 0.11 |
| 10 | 109888333 | rs2151875 | -0.114 | -2.602 | -0.89 | -1.617 | 0.08 | 0.085 | 0.085 | 0 | 0.06 | 0.18 | 0.06 | 0.14 | 0.09 | 0.14 |
| 10 | 109894731 | rs6584859 | -1.444 | -3.294 | -1.37 | -1.734 | -0.004 | -0.005 | -0.005 | 2.333 | 0.59 | 0.61 | 0.34 | 0.26 | 0.32 | 0.36 |
| 10 | 109911218 | rs11193921 | 0.454 | -3.182 | -1.393 | -1.879 | 0.005 | 0.001 | 0.001 | 0 | 0.16 | 0.12 | 0.07 | 0.01 | 0.12 | 0.09 |
| 10 | 109911775 | rs17250905 | -0.285 | -3.353 | -1.556 | -1.975 | 0.015 | 0.181 | 0.181 | 1.503 | 0.26 | 0.01 | 0.11 | 0 | 0.17 | 0.17 |
| 10 | 109919044 | rs11193928 | 0.904 | -2.789 | -0.897 | -1.452 | -0.001 | -0.025 | -0.025 | 0 | 0.11 | 0.11 | 0.06 | 0 | 0.11 | 0.05 |
| 10 | 109925206 | rs317455 | -0.792 | -3.138 | -0.718 | -1.385 | -0.059 | -0.016 | -0.016 | 0 | 0.37 | 0.56 | 0.23 | 0.22 | 0.15 | 0.2 |
| 10 | 109934964 | rs11193936 | 0.341 | -2.405 | -0.756 | -1.112 | 0.005 | -0.013 | -0.013 | 0 | 0.15 | 0.12 | 0.07 | 0 | 0.09 | 0.07 |
| 10 | 109949370 | rs317429 | -0.642 | -2.009 | -0.91 | -0.894 | 0.028 | 0.025 | 0.025 | 0 | 0.49 | 0.61 | 0.29 | 0.23 | 0.22 | 0.32 |
| 10 | 109953019 | rs1499247 | 0.098 | -0.04 | -1.272 | -0.812 | -0.026 | 0.013 | 0.013 | 0 | 0.32 | 0.16 | 0.51 | 0.77 | 0.39 | 0.55 |
| 10 | 109956032 | rs12359831 | 0.315 | -0.708 | -1.466 | -1.39 | 0.009 | 0.049 | 0.049 | 0.876 | 0.18 | 0.11 | 0.1 | 0.02 | 0.15 | 0.1 |
| 10 | 109957755 | rs421263 | -0.041 | -0.912 | -0.913 | -1.751 | 0 | -0.017 | -0.017 | 0 | 0.39 | 0.44 | 0.51 | 0.77 | 0.36 | 0.51 |
| 10 | 109959105 | rs1034506 | -1.359 | -1.189 | -0.612 | -1.972 | -0.011 | 0.037 | 0.037 | 0 | 0.62 | 0.74 | 0.66 | 0.79 | 0.56 | 0.69 |
| 10 | 109961825 | rs449442 | -0.982 | -1.048 | -0.419 | -1.952 | -0.001 | 0.049 | 0.049 | 4.183 | 0.55 | 0.65 | 0.62 | 0.79 | 0.51 | 0.58 |

| | | | | | | | | | | | | | | | | |
|---|---|---|---|---|---|---|---|---|---|---|---|---|---|---|---|---|
| 10 | 109962451 | rs317465 | -0.212 | -1.296 | -0.271 | -2.046 | -0.004 | 0.008 | 0.008 | 0.338 | 0.4 | 0.38 | 0.51 | 0.79 | 0.41 | 0.57 |
| 10 | 109963724 | rs317463 | -0.002 | -1.284 | -0.315 | -2.021 | 0.017 | -0.035 | -0.035 | 0 | 0.37 | 0.47 | 0.52 | 0.79 | 0.41 | 0.6 |
| 10 | 109969396 | rs10884600 | 0.85 | -1.226 | -0.475 | -1.976 | -0.004 | 0.052 | 0.052 | 0.018 | 0.16 | 0.09 | 0.09 | 0 | 0.14 | 0.09 |
| 10 | 109988645 | rs10509868 | 0.925 | -1.329 | -0.201 | -2.021 | -0.001 | -0.034 | -0.034 | 0.566 | 0.14 | 0.16 | 0.09 | 0 | 0.14 | 0.06 |
| 10 | 109990024 | rs7081013 | 0.891 | -1.514 | -0.229 | -1.728 | -0.01 | 0.01 | 0.01 | 0 | 0.07 | 0.08 | 0.03 | 0.11 | 0.03 | 0.09 |
| 10 | 109991431 | rs1553146 | 0.344 | -1.429 | -0.314 | -1.742 | -0.001 | -0.024 | -0.024 | 0.455 | 0.2 | 0.21 | 0.12 | 0.04 | 0.2 | 0.16 |
| 10 | 109999601 | rs317506 | -1.099 | -1.279 | 0.037 | -1.685 | -0.013 | -0.023 | -0.023 | 0 | 0.71 | 0.84 | 0.55 | 0.34 | 0.7 | 0.58 |
| 10 | 110005716 | rs317510 | -0.058 | -1.329 | 0.161 | -1.726 | -0.002 | -0.022 | -0.022 | 0 | 0.62 | 0.56 | 0.5 | 0.23 | 0.67 | 0.49 |
| 10 | 110016862 | rs6584864 | 0.69 | -1.508 | -0.221 | -1.775 | -0.013 | -0.038 | -0.038 | 0.571 | 0.1 | 0.16 | 0.06 | 0 | 0.05 | 0.11 |
| 10 | 110022170 | rs317489 | -1.436 | -1.484 | -0.409 | -1.77 | -0.016 | -0.021 | -0.021 | 0.381 | 0.74 | 0.84 | 0.56 | 0.34 | 0.7 | 0.6 |
| 10 | 110023221 | rs12240548 | -0.183 | -1.473 | -0.375 | -1.786 | -0.006 | -0.066 | -0.066 | 0.675 | 0.27 | 0.36 | 0.16 | 0.05 | 0.25 | 0.25 |
| 10 | 110023501 | rs12359500 | 1.544 | -1.532 | -0.998 | -1.745 | 0.014 | -0.044 | -0.044 | 0.636 | 0.07 | 0 | 0.11 | 0.18 | 0.03 | 0.11 |
| 10 | 110052312 | rs161020 | -0.834 | -1.244 | -0.997 | -1.817 | -0.009 | -0.005 | -0.005 | 0 | 0.7 | 0.78 | 0.54 | 0.34 | 0.68 | 0.58 |
| 10 | 110057434 | rs12781557 | 1.995 | -0.794 | -1.275 | -0.937 | -0.029 | -0.092 | -0.092 | 1.111 | 0.16 | 0.02 | 0.27 | 0.36 | 0.19 | 0.31 |
| 10 | 110064014 | rs2799830 | 2.628 | -0.032 | -1.49 | -1.948 | -0.069 | 0.115 | 0.115 | 0.311 | 0.19 | 0.03 | 0.17 | 0 | 0.3 | 0.09 |
| 10 | 110079751 | rs11194016 | 1.796 | 0.158 | -1.286 | -1.795 | -0.046 | 0.066 | 0.066 | 0.281 | 0.13 | 0.04 | 0.09 | 0 | 0.17 | 0.08 |
| 10 | 110083106 | rs319814 | -0.771 | 0.031 | -1.215 | -1.71 | 0 | -0.006 | -0.006 | 0.024 | 0.71 | 0.82 | 0.48 | 0.32 | 0.72 | 0.6 |
| 10 | 110092212 | rs9420040 | -0.806 | -0.667 | -1.002 | -1.635 | -0.003 | -0.004 | -0.004 | 0 | 0.76 | 0.78 | 0.48 | 0.32 | 0.71 | 0.63 |
| 10 | 110093190 | rs10787084 | 1.809 | -0.666 | -1.064 | -1.888 | -0.007 | 0.013 | 0.013 | 0.303 | 0.13 | 0.08 | 0.09 | 0 | 0.14 | 0.08 |
| 10 | 110108512 | rs10732800 | -0.501 | -0.823 | -1.275 | -1.914 | -0.038 | 0.098 | 0.098 | 0.593 | 0.47 | 0.17 | 0.64 | 0.69 | 0.5 | 0.64 |
| 10 | 110111704 | rs10884620 | 2.025 | -0.797 | -1.274 | -1.937 | -0.006 | 0.013 | 0.013 | 0.493 | 0.13 | 0.07 | 0.09 | 0 | 0.14 | 0.08 |
| 10 | 110118673 | rs1411023 | -0.483 | -0.766 | -1.13 | -1.863 | -0.023 | 0.096 | 0.096 | 0.563 | 0.47 | 0.19 | 0.64 | 0.69 | 0.5 | 0.64 |
| 10 | 110126699 | rs10491020 | 1.002 | -0.825 | -0.715 | -1.926 | 0.005 | 0.011 | 0.011 | 0.818 | 0.15 | 0.35 | 0.1 | 0.13 | 0.1 | 0.11 |
| 10 | 110134974 | rs10884639 | -1.499 | -0.924 | -0.925 | -2.056 | -0.001 | -0.06 | -0.06 | 0.402 | 0.66 | 0.35 | 0.74 | 0.99 | 0.6 | 0.84 |
| 10 | 110137844 | rs2183231 | -0.58 | -0.974 | -1.14 | -2.08 | 0 | 0.125 | 0.125 | 0.029 | 0.41 | 0.18 | 0.59 | 0.88 | 0.39 | 0.65 |
| 10 | 110139563 | rs1041670 | -1.663 | -1.07 | -1.041 | -2.11 | -0.008 | 0.043 | 0.043 | 0 | 0.66 | 0.7 | 0.76 | 0.83 | 0.6 | 0.78 |
| 10 | 110140197 | rs2149739 | 0.994 | -1.19 | -1.15 | -1.984 | -0.043 | -0.043 | -0.043 | 0 | 0.22 | 0.08 | 0.52 | 0.68 | 0.3 | 0.37 |
| 10 | 110143610 | rs7476527 | 0.138 | -0.948 | -1.023 | -1.883 | -0.018 | 0.004 | 0.004 | 0.386 | 0.32 | 0.15 | 0.56 | 0.69 | 0.35 | 0.54 |
| 10 | 110146869 | rs1329482 | -0.689 | -0.774 | -0.972 | -1.943 | 0.018 | 0.114 | 0.114 | 0 | 0.51 | 0.61 | 0.66 | 0.86 | 0.43 | 0.64 |
| 10 | 110156433 | rs7083268 | -0.188 | -0.669 | -0.603 | -1.948 | 0.006 | 0.022 | 0.022 | 0 | 0.41 | 0.3 | 0.59 | 0.87 | 0.37 | 0.62 |
| 10 | 110162681 | rs6584866 | -1.495 | -0.171 | -1.011 | -1.756 | -0.014 | 0.026 | 0.026 | 0 | 0.74 | 0.8 | 0.8 | 1 | 0.64 | 0.88 |
| 10 | 110170558 | rs2484105 | 1.602 | 0.039 | -1.097 | -1.501 | 0.023 | 0.008 | 0.008 | 0 | 0.16 | 0.06 | 0.07 | 0.12 | 0.11 | 0.16 |
| 10 | 110171444 | rs1754617 | 0.053 | 0.075 | -0.763 | -1.767 | -0.006 | 0.089 | 0.089 | 0.061 | 0.46 | 0.34 | 0.31 | 0.15 | 0.49 | 0.4 |
| 10 | 110188914 | rs2419006 | -1.556 | 0.431 | -0.594 | -1.62 | 0.005 | -0.062 | -0.062 | 0 | 0.83 | 0.78 | 0.89 | 1 | 0.84 | 0.89 |
| 10 | 110195440 | rs12414634 | 0.95 | 0.392 | -0.754 | -1.605 | -0.026 | -0.038 | -0.038 | 0 | 0.21 | 0.12 | 0.48 | 0.67 | 0.25 | 0.37 |
| 10 | 110201277 | rs17124561 | 1.808 | 0.354 | -1.04 | -1.635 | 0.007 | -0.015 | -0.015 | 0 | 0.2 | 0.05 | 0.11 | 0.22 | 0.17 | 0.12 |
| 10 | 110204286 | rs4918376 | -0.249 | 0.371 | -0.92 | -1.701 | 0.006 | 0.142 | 0.142 | 0 | 0.5 | 0.28 | 0.66 | 0.93 | 0.49 | 0.63 |
| 10 | 110204664 | rs17350394 | 1.967 | 0.313 | -0.932 | -1.725 | -0.062 | 0.11 | 0.11 | 0 | 0.19 | 0.05 | 0.18 | 0 | 0.3 | 0.13 |
| 10 | 110207283 | rs1928449 | 1.505 | 0.393 | -0.921 | -1.703 | -0.028 | 0.023 | 0.023 | 0.85 | 0.07 | 0.02 | 0.05 | 0.03 | 0.08 | 0.13 |
| 10 | 110208539 | rs1329503 | -0.986 | 0.297 | -0.868 | -1.717 | 0 | 0.03 | 0.03 | 0 | 0.63 | 0.74 | 0.78 | 0.77 | 0.67 | 0.75 |
| 10 | 110209221 | rs2183232 | -0.236 | 0.273 | -0.887 | -1.718 | -0.01 | 0.023 | 0.023 | 0 | 0.36 | 0.24 | 0.57 | 0.7 | 0.37 | 0.57 |
| 10 | 110210233 | rs11194100 | 1.638 | 0.548 | -1.253 | -1.271 | -0.047 | -0.01 | -0.01 | 1.186 | 0.07 | 0.02 | 0.31 | 0.35 | 0.18 | 0.19 |
| 10 | 110216429 | rs1329499 | -0.937 | 0.508 | -1.194 | -1.67 | 0 | 0.055 | 0.055 | 0 | 0.63 | 0.73 | 0.78 | 0.86 | 0.67 | 0.79 |
| 10 | 110221750 | rs2419008 | -1.069 | 0.536 | -1.182 | -1.658 | 0.002 | 0.049 | 0.049 | 0 | 0.64 | 0.74 | 0.79 | 0.86 | 0.68 | 0.81 |
| 10 | 110225049 | rs6584869 | -0.949 | 0.555 | -1.191 | -1.658 | -0.004 | 0.054 | 0.054 | 0 | 0.63 | 0.74 | 0.78 | 0.86 | 0.67 | 0.78 |
| 10 | 110225306 | rs9787600 | -0.84 | 0.613 | -1.184 | -1.64 | -0.002 | 0.063 | 0.063 | 0 | 0.62 | 0.73 | 0.78 | 0.86 | 0.66 | 0.75 |
| 10 | 110230050 | rs11194108 | 1.614 | 0.704 | -1.145 | -1.536 | -0.013 | 0.015 | 0.015 | 2.146 | 0.06 | 0.02 | 0.05 | 0.03 | 0.07 | 0.13 |
| 10 | 110238857 | rs7083981 | 0.685 | 0.63 | -1.049 | -1.525 | 0.007 | 0 | 0 | 0 | 0.24 | 0.24 | 0.5 | 0.75 | 0.27 | 0.42 |
| 10 | 110259434 | rs12248785 | 1.482 | 0.6 | -1.081 | -1.501 | -0.015 | 0.017 | 0.017 | 1.393 | 0.07 | 0.03 | 0.05 | 0.03 | 0.07 | 0.13 |
| 10 | 110259662 | rs17124651 | 0.793 | 0.454 | -0.917 | -1.56 | -0.042 | -0.062 | -0.062 | 0.091 | 0.1 | 0.28 | 0.04 | 0 | 0.04 | 0.08 |
| 10 | 110260562 | rs1332270 | 2.023 | 0.554 | -1.509 | -1.705 | -0.053 | 0.101 | 0.101 | 0 | 0.18 | 0.04 | 0.17 | 0 | 0.28 | 0.09 |
| 10 | 110261426 | rs1332272 | 0.433 | 1.022 | -1.073 | -1.287 | 0.002 | 0.01 | 0.01 | 0 | 0.24 | 0.23 | 0.48 | 0.63 | 0.26 | 0.37 |
| 10 | 110262425 | rs4917528 | -0.478 | 1.13 | -1.417 | -1.309 | 0.001 | -0.033 | -0.033 | 0 | 0.77 | 0.75 | 0.84 | 0.97 | 0.77 | 0.76 |
| 10 | 110271335 | rs1332280 | -1.27 | 0.996 | -1.065 | -0.986 | 0.01 | -0.063 | -0.063 | 0 | 0.84 | 0.78 | 0.88 | 1 | 0.82 | 0.88 |
| 10 | 110277915 | rs1929988 | -1.417 | 1.165 | -1.002 | -0.905 | 0.011 | -0.051 | -0.051 | 0 | 0.85 | 0.81 | 0.87 | 1 | 0.81 | 0.88 |
| 10 | 110283067 | rs7901709 | 0.812 | 1.167 | -1.121 | -0.576 | -0.023 | 0.143 | 0.143 | 0 | 0.33 | 0.13 | 0.23 | 0.03 | 0.38 | 0.29 |
| 10 | 110286164 | rs1332283 | -0.506 | 0.939 | -1.001 | -0.009 | 0.005 | 0.024 | 0.024 | 0 | 0.25 | 0.29 | 0.48 | 0.63 | 0.27 | 0.35 |
| 10 | 110300448 | rs11194154 | 0.143 | 1.988 | -1.667 | 0.524 | -0.05 | -0.025 | -0.025 | 0.003 | 0.08 | 0.02 | 0.3 | 0.35 | 0.19 | 0.19 |
| 10 | 110310720 | rs1332266 | 0.085 | 1.027 | -1.27 | -0.466 | -0.017 | 0.03 | 0.03 | 0 | 0.64 | 0.78 | 0.74 | 0.73 | 0.64 | 0.72 |
| 10 | 110316077 | rs6584877 | -0.062 | 0.525 | -0.841 | -0.972 | -0.003 | -0.048 | -0.048 | 0 | 0.54 | 0.38 | 0.42 | 0.06 | 0.53 | 0.43 |
| 10 | 110321696 | rs914600 | 0.54 | -0.138 | -0.55 | 0.079 | -0.027 | 0.07 | 0.07 | 0 | 0.95 | 0.87 | 0.98 | 0.84 | 0.99 | 0.9 |
| 10 | 110330139 | rs7922237 | 0.649 | -0.257 | -0.43 | -0.406 | -0.001 | -0.001 | -0.001 | 0.049 | 0.65 | 0.62 | 0.69 | 0.68 | 0.6 | 0.7 |
| 10 | 110337386 | rs1888902 | 0.408 | 0.059 | -0.531 | -0.641 | 0.017 | -0.038 | -0.038 | 1.051 | 0.11 | 0.2 | 0.09 | 0.05 | 0.14 | 0.05 |
| 10 | 110339682 | rs11194173 | 0.314 | -0.113 | -1.215 | -1.198 | -0.028 | 0.084 | 0.084 | 0 | 0.24 | 0.04 | 0.22 | 0.06 | 0.23 | 0.21 |
| 10 | 110345230 | rs11194176 | 0.158 | -1.228 | -0.853 | -0.759 | 0.007 | 0.01 | 0.01 | 0 | 0.2 | 0.18 | 0.12 | 0.1 | 0.08 | 0.15 |
| 10 | 110348376 | rs7100815 | -0.511 | -0.647 | -0.748 | -0.219 | 0.033 | 0.036 | 0.036 | 0 | 0.51 | 0.69 | 0.42 | 0.36 | 0.44 | 0.41 |
| 10 | 110349412 | rs17760580 | -0.358 | 0.585 | -1.375 | 0.539 | -0.021 | -0.039 | -0.039 | 0 | 0.07 | 0.01 | 0.14 | 0.13 | 0.14 | 0.09 |
| 10 | 110353671 | rs2482594 | 0.365 | 0.895 | -1.437 | 0.73 | -0.041 | -0.035 | -0.035 | 0 | 0.71 | 0.74 | 0.54 | 0.41 | 0.55 | 0.55 |
| 10 | 110354417 | rs2482595 | 0.171 | 1.102 | -1.63 | 0.938 | -0.114 | -0.088 | -0.088 | 0 | 0.78 | 0.93 | 0.5 | 0.4 | 0.51 | 0.58 |

| | | | | | | | | | | | | | | | | |
|---|---|---|---|---|---|---|---|---|---|---|---|---|---|---|---|---|
| 10 | 110354466 | rs11194185 | 0.031 | 0.28 | -1.727 | 0.344 | 0.003 | 0.031 | 0.031 | 0 | 0.09 | 0.03 | 0.03 | 0 | 0.06 | 0.02 |
| 10 | 110356935 | rs2482596 | -0.411 | -0.966 | -1.578 | -0.327 | 0.03 | 0.034 | 0.034 | 0 | 0.42 | 0.64 | 0.19 | 0.17 | 0.21 | 0.24 |
| 10 | 110358531 | rs17124848 | -0.136 | -1.262 | -1.209 | -0.331 | -0.017 | -0.003 | -0.003 | 0 | 0.1 | 0.11 | 0.05 | 0.05 | 0.02 | 0.18 |
| 10 | 110368941 | rs12768159 | -0.731 | -0.631 | -0.38 | -0.468 | 0.018 | -0.055 | -0.055 | 0 | 0.09 | 0.31 | 0.08 | 0 | 0.1 | 0.1 |
| 10 | 110374860 | rs928408 | 0.239 | -1.111 | -0.602 | -0.512 | 0.004 | -0.008 | -0.008 | 0 | 0.22 | 0.11 | 0.16 | 0.19 | 0.23 | 0.12 |
| 10 | 110375178 | rs2482579 | -0.309 | -1.162 | -0.437 | -0.54 | 0.019 | -0.059 | -0.059 | 0 | 0.1 | 0.34 | 0.08 | 0 | 0.1 | 0.1 |
| 10 | 110376820 | rs1332260 | -0.975 | -1.206 | -0.498 | -0.479 | -0.007 | -0.01 | -0.01 | 0.226 | 0.32 | 0.45 | 0.24 | 0.24 | 0.33 | 0.26 |
| 10 | 110377823 | rs10884674 | 1.511 | -0.556 | -0.648 | -1.17 | 0.011 | 0.076 | 0.076 | 0 | 0.36 | 0.2 | 0.23 | 0.09 | 0.32 | 0.19 |
| 10 | 110385967 | rs11194202 | 0.002 | -0.809 | -0.808 | -1.352 | 0.03 | 0.041 | 0.041 | 0.085 | 0.24 | 0.12 | 0.11 | 0.05 | 0.13 | 0.23 |
| 10 | 110387632 | rs4917530 | 0.212 | -0.735 | -0.502 | -0.587 | 0.01 | 0.097 | 0.097 | 0.192 | 0.22 | 0.4 | 0.28 | 0.39 | 0.28 | 0.26 |
| 10 | 110390412 | rs944645 | 1.846 | -0.69 | -0.749 | -0.796 | 0.008 | 0.022 | 0.022 | 0 | 0.34 | 0.21 | 0.21 | 0.01 | 0.28 | 0.18 |
| 10 | 110391261 | rs2482583 | -0.925 | -0.328 | -0.838 | -0.617 | 0.016 | 0.034 | 0.034 | 0.388 | 0.81 | 0.79 | 0.65 | 0.46 | 0.7 | 0.69 |
| 10 | 110391906 | rs12411824 | 1.152 | -0.444 | -0.938 | -0.631 | 0 | -0.005 | -0.005 | 0 | 0.15 | 0.11 | 0.19 | 0.35 | 0.19 | 0.12 |
| 10 | 110397130 | rs2482587 | -0.866 | -0.426 | -0.853 | -0.624 | -0.005 | -0.001 | -0.001 | 0 | 0.81 | 0.84 | 0.61 | 0.45 | 0.69 | 0.67 |
| 10 | 110397614 | rs2026988 | -0.01 | -0.461 | -0.685 | -0.525 | -0.002 | -0.015 | -0.015 | 2.409 | 0.23 | 0.21 | 0.27 | 0.35 | 0.29 | 0.21 |
| 10 | 110404272 | rs12767064 | 1.278 | 0.483 | -1.278 | -0.857 | -0.052 | 0.073 | 0.073 | 0 | 0.12 | 0.01 | 0.09 | 0.01 | 0.14 | 0.04 |
| 10 | 110412755 | rs7899742 | -0.007 | 0.455 | -0.816 | -0.327 | -0.016 | -0.013 | -0.013 | 0 | 0.76 | 0.84 | 0.66 | 0.6 | 0.65 | 0.6 |
| 10 | 110414670 | rs10884679 | -0.694 | 0.674 | -0.759 | -0.222 | 0.024 | 0.031 | 0.031 | 0.2 | 0.82 | 0.78 | 0.73 | 0.61 | 0.71 | 0.68 |
| 10 | 110423549 | rs12253774 | 0.796 | 0.694 | -0.715 | -0.252 | -0.002 | -0.026 | -0.026 | 0 | 0.45 | 0.56 | 0.37 | 0.22 | 0.41 | 0.37 |
| 10 | 110424721 | rs11194219 | 1.109 | 0.727 | -0.812 | -0.373 | -0.003 | -0.049 | -0.049 | 0 | 0.39 | 0.53 | 0.33 | 0.22 | 0.36 | 0.37 |
| 10 | 110428405 | rs10509875 | 1.648 | 0.632 | -0.536 | -0.178 | 0.01 | 0.089 | 0.089 | 0 | 0.37 | 0.2 | 0.26 | 0.17 | 0.33 | 0.29 |
| 10 | 110435038 | rs4244274 | -2.124 | 1.289 | -0.549 | 0.099 | -0.025 | -0.014 | -0.014 | 0 | 0.93 | 0.96 | 0.82 | 0.66 | 0.84 | 0.86 |
| 10 | 110441149 | rs7914735 | 0.251 | 0.634 | -0.404 | -0.023 | 0 | -0.002 | -0.002 | 0.06 | 0.58 | 0.4 | 0.53 | 0.51 | 0.61 | 0.41 |
| 10 | 110452276 | rs4918398 | 0.911 | 0.709 | -0.502 | -0.082 | 0 | -0.013 | -0.013 | 0 | 0.45 | 0.57 | 0.37 | 0.23 | 0.41 | 0.39 |
| 10 | 110452911 | rs11194228 | 2.034 | 0.744 | -1.244 | -0.022 | -0.009 | 0.026 | 0.026 | 0.004 | 0.06 | 0 | 0.02 | 0 | 0.04 | 0.01 |
| 10 | 110453513 | rs7067913 | 0.402 | 0.842 | -1.045 | 0.157 | -0.005 | 0.004 | 0.004 | 0 | 0.54 | 0.38 | 0.48 | 0.56 | 0.53 | 0.48 |
| 10 | 110459956 | rs10884683 | -0.652 | 0.612 | -0.822 | 0.361 | -0.002 | 0 | 0 | 0.235 | 0.44 | 0.55 | 0.47 | 0.46 | 0.41 | 0.39 |
| 10 | 110460471 | rs11194232 | 0.766 | 0.597 | -0.905 | 0.379 | 0.003 | 0 | 0 | 0 | 0.09 | 0.04 | 0.13 | 0.05 | 0.06 | 0.09 |
| 10 | 110463852 | rs12246560 | -0.531 | 0.348 | -0.48 | 0.64 | 0.002 | 0.151 | 0.151 | 0 | 0.19 | 0.33 | 0.22 | 0.38 | 0.19 | 0.1 |
| 10 | 110463874 | rs17124963 | 0.839 | 0.242 | -0.337 | 0.89 | -0.001 | 0.196 | 0.196 | 1.23 | 0.08 | 0.18 | 0.07 | 0.33 | 0.07 | 0.05 |
| 10 | 110465173 | rs17124967 | 0.904 | 0.249 | -0.324 | 1.128 | -0.001 | 0.039 | 0.039 | 3.069 | 0.08 | 0.03 | 0.14 | 0.49 | 0.08 | 0.15 |
| 10 | 110467675 | rs10509876 | 0.128 | 1.014 | -0.347 | 1.058 | 0.018 | -0.004 | -0.004 | 0.393 | 0.1 | 0.15 | 0.16 | 0.06 | 0.15 | 0.05 |
| 10 | 110478002 | rs7093143 | -1.125 | 0.971 | -0.231 | 1.103 | 0.034 | 0.089 | 0.089 | 0 | 0.2 | 0.36 | 0.32 | 0.55 | 0.23 | 0.19 |
| 10 | 110498637 | rs1577858 | 0.095 | 1.035 | -0.296 | 1.046 | 0 | 0.212 | 0.212 | 1.769 | 0.1 | 0.21 | 0.16 | 0.5 | 0.08 | 0.14 |
| 10 | 110502547 | rs1415554 | 0.67 | 0.263 | -0.886 | 0.416 | 0.035 | 0.049 | 0.049 | 0.157 | 0.18 | 0.08 | 0.09 | 0.04 | 0.1 | 0.08 |
| 10 | 110503788 | rs10884686 | 1.029 | 0.889 | -0.98 | 0.461 | -0.045 | -0.009 | -0.009 | 0 | 0.12 | 0.01 | 0.17 | 0.05 | 0.11 | 0.16 |
| 10 | 110504315 | rs6584893 | -0.78 | 0.63 | -0.666 | 0.624 | 0.023 | 0.09 | 0.09 | 0 | 0.19 | 0.29 | 0.25 | 0.49 | 0.23 | 0.16 |
| 10 | 110518598 | rs7893638 | -0.376 | 0.637 | -0.657 | 0.66 | 0.028 | 0.081 | 0.081 | 0 | 0.17 | 0.26 | 0.16 | 0.33 | 0.22 | 0.07 |
| 10 | 110519947 | rs11591528 | 0.611 | 0.649 | -0.66 | 0.631 | -0.003 | -0.011 | -0.011 | 0 | 0.09 | 0.09 | 0.09 | 0 | 0.15 | 0.02 |
| 10 | 110524066 | rs7100848 | -0.481 | 0.636 | -0.602 | 0.687 | 0.028 | 0.107 | 0.107 | 0 | 0.18 | 0.28 | 0.23 | 0.37 | 0.23 | 0.09 |
| 10 | 110527358 | rs17697405 | -0.578 | 0.442 | -0.706 | 0.614 | 0.002 | 0.004 | 0.004 | 0 | 0.13 | 0.15 | 0.25 | 0.2 | 0.16 | 0.16 |
| 10 | 110541074 | rs945824 | -0.578 | 0.442 | -0.706 | 0.614 | 0.002 | 0.004 | 0.004 | 0 | 0.13 | 0.15 | 0.25 | 0.2 | 0.16 | 0.15 |
| 10 | 110552149 | rs3982269 | 0.051 | 0.439 | -0.652 | 0.657 | -0.012 | 0.149 | 0.149 | 1.323 | 0.12 | 0.24 | 0.24 | 0.54 | 0.08 | 0.19 |
| 10 | 110566771 | rs1856165 | -0.925 | -0.962 | -0.553 | -0.196 | 0.031 | 0.036 | 0.036 | 0.448 | 0.54 | 0.69 | 0.44 | 0.58 | 0.36 | 0.36 |
| 10 | 110573160 | rs7100485 | 0.025 | -0.757 | -0.547 | -0.068 | -0.003 | -0.003 | -0.003 | 0.366 | 0.15 | 0.15 | 0.26 | 0.2 | 0.16 | 0.16 |
| 10 | 110576832 | rs9787569 | 0.33 | -0.782 | -0.384 | -0.045 | -0.007 | 0.223 | 0.223 | 1.439 | 0.12 | 0.31 | 0.24 | 0.53 | 0.08 | 0.18 |
| 10 | 110581654 | rs4120665 | 0.127 | -0.752 | -0.417 | -0.06 | 0.021 | 0.068 | 0.068 | 0 | 0.19 | 0.26 | 0.17 | 0.33 | 0.23 | 0.07 |
| 10 | 110582618 | rs17698531 | 1.154 | -0.74 | -0.468 | -0.106 | -0.018 | 0.011 | 0.011 | 0.022 | 0.11 | 0.06 | 0.09 | 0 | 0.16 | 0.02 |
| 10 | 110598065 | rs11813768 | 0.551 | -0.998 | -0.408 | -0.139 | -0.021 | 0.078 | 0.078 | 0 | 0.2 | 0.26 | 0.19 | 0.4 | 0.12 | 0.14 |
| 10 | 110600634 | rs4258310 | 0.69 | -0.946 | -0.451 | -0.164 | -0.005 | -0.003 | -0.003 | 0.144 | 0.13 | 0.11 | 0.17 | 0.16 | 0.16 | 0.1 |
| 10 | 110603382 | rs12780127 | -0.512 | -1.003 | -0.392 | -0.175 | -0.004 | -0.002 | -0.002 | 0.369 | 0.28 | 0.32 | 0.31 | 0.25 | 0.21 | 0.22 |
| 10 | 110609132 | rs1890095 | -0.819 | -0.602 | -0.463 | -0.229 | -0.01 | -0.007 | -0.007 | 0.266 | 0.36 | 0.49 | 0.39 | 0.58 | 0.28 | 0.27 |
| 10 | 110610160 | rs10509879 | 1.338 | -0.57 | -0.508 | -0.421 | -0.053 | 0.062 | 0.062 | 0.033 | 0.11 | 0.02 | 0.09 | 0 | 0.16 | 0.02 |
| 10 | 110612757 | rs2685477 | -1.518 | 0.037 | -0.534 | -0.651 | -0.031 | 0.05 | 0.05 | 0 | 0.65 | 0.82 | 0.62 | 0.81 | 0.55 | 0.57 |
| 10 | 110622053 | rs2685480 | -0.701 | -0.613 | -0.154 | -0.532 | -0.009 | 0.04 | 0.04 | 0 | 0.34 | 0.5 | 0.32 | 0.41 | 0.28 | 0.18 |
| 10 | 110625693 | rs12249791 | 0.558 | 0.21 | 0.149 | -0.354 | -0.043 | 0.06 | 0.06 | 0 | 0.13 | 0.03 | 0.09 | 0 | 0.16 | 0.05 |
| 10 | 110636462 | rs2461320 | -0.618 | 0.263 | 0.14 | -0.372 | -0.002 | 0.055 | 0.055 | 0.324 | 0.56 | 0.81 | 0.48 | 0.59 | 0.39 | 0.39 |
| 10 | 110642737 | rs11194290 | 0.23 | 0.11 | 0.412 | -0.303 | -0.011 | -0.004 | -0.004 | 0 | 0.43 | 0.53 | 0.38 | 0.58 | 0.26 | 0.38 |
| 10 | 110646361 | rs17125303 | 1.13 | 0.07 | 0.45 | -0.269 | 0.053 | 0.024 | 0.024 | 0.139 | 0.29 | 0.18 | 0.18 | 0.22 | 0.13 | 0.23 |
| 10 | 110646659 | rs17125306 | 1.044 | 0.081 | 0.395 | -0.269 | -0.043 | -0.018 | -0.018 | 0 | 0.31 | 0.36 | 0.19 | 0.22 | 0.13 | 0.23 |
| 10 | 110659424 | rs17125402 | 1.05 | 0.251 | 0.323 | -0.197 | -0.002 | 0.033 | 0.033 | 1.038 | 0.1 | 0.09 | 0.1 | 0.32 | 0.12 | 0.08 |
| 10 | 110688807 | rs11194306 | 0.08 | 0.061 | 0.481 | -0.533 | -0.001 | -0.001 | -0.001 | 0 | 0.43 | 0.53 | 0.37 | 0.26 | 0.29 | 0.28 |
| 10 | 110697497 | rs11592721 | 0.717 | 0.137 | 0.509 | -0.531 | 0.002 | -0.024 | -0.024 | 0 | 0.11 | 0.12 | 0.1 | 0 | 0.16 | 0.02 |
| 10 | 110730540 | rs10509882 | -0.303 | 0.075 | 0.625 | -0.555 | 0.064 | 0.072 | 0.072 | 0.656 | 0.47 | 0.69 | 0.38 | 0.25 | 0.29 | 0.31 |
| 10 | 110739497 | rs4918406 | -2.076 | -0.162 | 0.433 | -0.463 | -0.039 | 0.031 | 0.031 | 0 | 0.83 | 0.99 | 0.86 | 0.91 | 0.76 | 0.77 |
| 10 | 110745626 | rs12260848 | 0.718 | -0.393 | 0.864 | -0.388 | -0.072 | -0.03 | -0.03 | 0.022 | 0.32 | 0.39 | 0.19 | 0.22 | 0.12 | 0.25 |
| 10 | 110747704 | rs10509886 | 1.072 | -0.297 | 0.483 | -0.347 | 0.071 | 0.025 | 0.025 | 0.061 | 0.28 | 0.16 | 0.1 | 0.21 | 0.12 | 0.16 |
| 10 | 110751644 | rs11194327 | 0.934 | -0.36 | 0.381 | -0.301 | 0.065 | 0.027 | 0.027 | 0.046 | 0.29 | 0.18 | 0.17 | 0.22 | 0.12 | 0.24 |
| 10 | 110761477 | rs11194336 | 0.868 | -0.278 | 0.469 | -0.174 | -0.089 | 0.001 | 0.001 | 0 | 0.31 | 0.46 | 0.27 | 0.31 | 0.13 | 0.26 |

| | | | | | | | | | | | | | | | | |
|---|---|---|---|---|---|---|---|---|---|---|---|---|---|---|---|---|
| 10 | 110799535 | rs17125615 | 1.022 | 0.567 | 0.221 | -0.059 | -0.069 | -0.078 | -0.078 | 0.055 | 0.17 | 0.03 | 0.11 | 0.3 | 0.22 | 0.12 |
| 10 | 110814528 | rs723483 | -0.623 | 0.323 | 0.442 | -0.215 | -0.036 | 0.005 | 0.005 | 1.755 | 0.59 | 0.77 | 0.49 | 0.59 | 0.44 | 0.43 |
| 10 | 110816967 | rs4918410 | 1.585 | 0.405 | 0.363 | -0.185 | -0.001 | -0.007 | -0.007 | 0 | 0.1 | 0.07 | 0.15 | 0.12 | 0.09 | 0.13 |
| 10 | 110817490 | rs11593905 | 1.782 | 0.11 | 0.423 | -0.106 | -0.053 | -0.073 | -0.073 | 0 | 0.13 | 0.02 | 0.09 | 0.3 | 0.18 | 0.09 |
| 10 | 110818455 | rs35689892 | 1.319 | -0.219 | 0.595 | -0.217 | 0.078 | 0.019 | 0.019 | 0.719 | 0.28 | 0.07 | 0.11 | 0.23 | 0.13 | 0.17 |
| 10 | 110823083 | rs17125651 | 1.118 | -0.206 | 0.579 | -0.212 | 0.048 | 0.022 | 0.022 | 0.304 | 0.3 | 0.17 | 0.12 | 0.23 | 0.13 | 0.17 |
| 10 | 110837359 | rs7098945 | 0.512 | -0.202 | 0.568 | -0.215 | 0.045 | 0.014 | 0.014 | 2.139 | 0.36 | 0.69 | 0.31 | 0.29 | 0.14 | 0.29 |
| 10 | 110850355 | rs17125686 | 0.789 | -0.023 | 0.442 | -0.135 | -0.029 | -0.003 | -0.003 | 1.195 | 0.32 | 0.33 | 0.27 | 0.29 | 0.14 | 0.29 |
| 10 | 110866437 | rs10884698 | 1.244 | 0.545 | 0.042 | -0.112 | -0.021 | -0.005 | -0.005 | 0 | 0.13 | 0.07 | 0.2 | 0.13 | 0.15 | 0.2 |
| 10 | 110868225 | rs7904064 | -1.32 | 0.286 | 0.549 | 0.238 | 0.06 | 0.164 | 0.164 | 2.254 | 0.06 | 0.4 | 0.18 | 0.07 | 0.01 | 0.12 |
| 10 | 110874611 | rs943268 | 0.955 | 0.262 | 0.217 | 0.317 | -0.059 | -0.064 | -0.064 | 0 | 0.1 | 0.01 | 0.23 | 0.27 | 0.18 | 0.17 |
| 10 | 110878391 | rs4406760 | 0.388 | -0.898 | 0.705 | 0.09 | 0.082 | 0.031 | 0.031 | 0.604 | 0.29 | 0.09 | 0.11 | 0.22 | 0.13 | 0.17 |
| 10 | 110905211 | rs11194423 | -1.005 | -0.624 | 0.764 | 0.205 | -0.04 | -0.018 | -0.018 | 0 | 0.6 | 0.77 | 0.41 | 0.56 | 0.44 | 0.44 |
| 10 | 110908214 | rs9783275 | -0.996 | -1.366 | 1.15 | -0.187 | -0.017 | -0.011 | -0.011 | 0.629 | 0.39 | 0.52 | 0.23 | 0.27 | 0.18 | 0.3 |
| 10 | 110916028 | rs2147285 | -0.802 | -1.465 | 1.109 | -0.2 | -0.031 | -0.015 | -0.015 | 0 | 0.38 | 0.52 | 0.22 | 0.27 | 0.17 | 0.32 |
| 10 | 110918774 | rs11194431 | -0.486 | -1.08 | 0.193 | 0.004 | 0.063 | 0.07 | 0.07 | 1.302 | 0.34 | 0.09 | 0.14 | 0.22 | 0.18 | 0.22 |
| 10 | 110922002 | rs10509890 | -0.412 | -1.046 | 0.231 | -0.051 | 0.082 | 0.046 | 0.046 | 1.485 | 0.36 | 0.12 | 0.14 | 0.25 | 0.18 | 0.22 |
| 10 | 110927983 | rs3936867 | -0.581 | -0.997 | 0.142 | -0.005 | 0.023 | 0.017 | 0.017 | 2.202 | 0.38 | 0.32 | 0.15 | 0.25 | 0.18 | 0.22 |
| 10 | 110931761 | rs12249259 | -1.362 | -0.301 | 0.977 | 0.215 | 0.016 | 0.018 | 0.018 | 0.022 | 0.05 | 0.2 | 0.03 | 0.06 | 0.01 | 0.04 |
| 10 | 110934024 | rs504263 | 0.718 | -0.822 | 0.513 | -0.187 | 0.044 | 0.055 | 0.055 | 0.386 | 0.63 | 0.26 | 0.77 | 0.77 | 0.79 | 0.67 |
| 10 | 110936202 | rs511061 | 0.622 | -0.872 | 0.618 | -0.176 | 0.037 | 0.037 | 0.037 | 0.267 | 0.65 | 0.28 | 0.79 | 0.78 | 0.84 | 0.68 |
| 10 | 110938966 | rs578043 | 0.259 | -0.839 | 0.852 | -0.154 | -0.048 | -0.05 | -0.05 | 0 | 0.69 | 0.46 | 0.8 | 0.82 | 0.83 | 0.72 |
| 10 | 110947577 | rs534259 | 0.64 | -1.037 | 1.287 | -0.324 | 0.036 | 0.036 | 0.036 | 0.299 | 0.65 | 0.27 | 0.77 | 0.81 | 0.81 | 0.7 |
| 10 | 110948284 | rs473244 | -0.227 | -1.201 | 1.314 | -0.405 | -0.015 | -0.006 | -0.006 | 0.145 | 0.88 | 0.84 | 0.94 | 0.88 | 0.95 | 0.87 |
| 10 | 110958660 | rs7083413 | -0.677 | -1.2 | 1.596 | -0.408 | -0.005 | -0.011 | -0.011 | 0.049 | 0.19 | 0.24 | 0.12 | 0.05 | 0.13 | 0.15 |
| 10 | 110972191 | rs12249019 | -0.458 | -1.287 | 1.745 | -0.383 | -0.013 | -0.005 | -0.005 | 0 | 0.13 | 0.18 | 0.06 | 0.13 | 0.06 | 0.13 |
| 10 | 110973189 | rs3903858 | -0.243 | -1.285 | 0.701 | -0.355 | 0 | 0.011 | 0.011 | 5.258 | 0.41 | 0.29 | 0.29 | 0.32 | 0.38 | 0.21 |
| 10 | 110980065 | rs10884712 | -0.659 | -1.406 | 0.965 | 0.008 | -0.002 | -0.001 | -0.001 | 0 | 0.45 | 0.54 | 0.32 | 0.4 | 0.38 | 0.29 |
| 10 | 110991026 | rs10787140 | 0.66 | -1.424 | 0.941 | -0.025 | 0 | -0.001 | -0.001 | 0.399 | 0.27 | 0.25 | 0.21 | 0.35 | 0.26 | 0.13 |
| 10 | 110993391 | rs4918415 | -0.125 | -1.454 | 1.044 | -0.047 | -0.003 | -0.016 | -0.016 | 0 | 0.85 | 0.79 | 0.9 | 0.95 | 0.88 | 0.91 |
| 10 | 110994518 | rs4917536 | 0.318 | -1.454 | 1.037 | -0.046 | -0.005 | -0.001 | -0.001 | 0.431 | 0.56 | 0.54 | 0.68 | 0.6 | 0.62 | 0.68 |
| 10 | 110997918 | rs10430676 | -0.278 | -1.431 | 0.944 | -0.062 | -0.004 | -0.01 | -0.01 | 0 | 0.87 | 0.79 | 0.91 | 0.95 | 0.9 | 0.92 |
| 10 | 110998641 | rs1853618 | -0.278 | -1.431 | 0.944 | -0.062 | -0.004 | -0.01 | -0.01 | 0.574 | 0.87 | 0.79 | 0.91 | 0.95 | 0.9 | 0.92 |
| 10 | 110999907 | rs1324288 | 0.139 | -1.472 | 0.931 | -0.072 | 0.003 | -0.004 | -0.004 | 0 | 0.58 | 0.71 | 0.7 | 0.6 | 0.62 | 0.69 |
| 10 | 111000911 | rs4396209 | 0.016 | -1.433 | 0.927 | -0.078 | -0.003 | -0.016 | -0.016 | 0 | 0.85 | 0.79 | 0.9 | 0.95 | 0.88 | 0.91 |
| 10 | 111009243 | rs11194513 | 0.771 | -1.203 | 0.788 | 0.5 | -0.002 | 0.017 | 0.017 | 0 | 0.06 | 0.04 | 0.12 | 0.21 | 0.07 | 0.11 |
| 10 | 111009308 | rs3891910 | 0.36 | -1.506 | 0.563 | 0.256 | 0.009 | -0.013 | -0.013 | 1.818 | 0.14 | 0.01 | 0.06 | 0.15 | 0.08 | 0.18 |
| 10 | 111026156 | rs1853617 | 0.118 | -1.289 | 0.802 | 0.292 | 0.002 | 0.001 | 0.001 | 3.356 | 0.55 | 0.49 | 0.68 | 0.6 | 0.61 | 0.69 |
| 10 | 111028887 | rs830066 | 0.345 | -1.278 | 0.701 | 0.322 | 0.024 | 0.018 | 0.018 | 2.679 | 0.53 | 0.3 | 0.66 | 0.6 | 0.61 | 0.69 |
| 10 | 111035817 | rs7914053 | 1.233 | -1.237 | 0.589 | 0.334 | -0.005 | -0.02 | -0.02 | 0.272 | 0.25 | 0.12 | 0.21 | 0.35 | 0.26 | 0.13 |
| 10 | 111036318 | rs7074781 | -1.002 | -1.209 | 0.62 | 0.313 | -0.018 | -0.048 | -0.048 | 0.063 | 0.2 | 0.4 | 0.12 | 0.06 | 0.13 | 0.18 |
| 10 | 111044737 | rs11194534 | 0.405 | -1.332 | 0.358 | 0.299 | -0.024 | -0.058 | -0.058 | 2.736 | 0.35 | 0.12 | 0.26 | 0.4 | 0.36 | 0.21 |
| 10 | 111050962 | rs10509894 | 0.007 | -1.41 | 0.945 | 0.153 | 0.002 | 0.018 | 0.018 | 0.074 | 0.15 | 0.1 | 0.09 | 0.05 | 0.13 | 0.15 |
| 10 | 111056638 | rs7089953 | -0.992 | -1.451 | 1.075 | 0.135 | -0.033 | -0.061 | -0.061 | 0 | 0.21 | 0.53 | 0.13 | 0.05 | 0.14 | 0.18 |
| 10 | 111062431 | rs4472861 | -0.414 | -1.448 | 1.122 | 0.137 | -0.001 | -0.003 | -0.003 | 1.106 | 0.17 | 0.22 | 0.1 | 0.05 | 0.14 | 0.15 |
| 10 | 111079538 | rs10787147 | -0.815 | -1.147 | 0.613 | -0.519 | 0.008 | 0.082 | 0.082 | 6.629 | 0.67 | 0.81 | 0.71 | 0.81 | 0.68 | 0.6 |
| 10 | 111083188 | rs10450419 | 0.31 | -1.188 | 0.7 | -0.475 | 0.001 | 0.002 | 0.002 | 0 | 0.17 | 0.16 | 0.09 | 0.05 | 0.13 | 0.16 |
| 10 | 111088928 | rs10787149 | -0.673 | -1.219 | 0.787 | -0.488 | -0.032 | -0.046 | -0.046 | 0 | 0.22 | 0.6 | 0.14 | 0.06 | 0.14 | 0.2 |
| 10 | 111089907 | rs7919912 | -0.201 | -1.213 | 0.762 | -0.486 | -0.013 | -0.021 | -0.021 | 0 | 0.19 | 0.28 | 0.1 | 0.05 | 0.13 | 0.16 |
| 10 | 111094155 | rs4918425 | 1.048 | -0.957 | 0.43 | -0.591 | -0.056 | -0.053 | -0.053 | 0.403 | 0.38 | 0.08 | 0.35 | 0.56 | 0.37 | 0.33 |
| 10 | 111094179 | rs7090906 | -0.315 | -0.877 | 0.855 | -0.538 | -0.022 | -0.036 | -0.036 | 0 | 0.19 | 0.35 | 0.11 | 0.05 | 0.14 | 0.16 |
| 10 | 111110455 | rs7475850 | -1.601 | -0.895 | 0.733 | -0.544 | 0.061 | 0.057 | 0.057 | 2.768 | 0.06 | 0.48 | 0.05 | 0 | 0 | 0.04 |
| 10 | 111118596 | rs9645579 | 0.554 | -0.854 | 0.65 | -0.528 | 0.002 | 0.017 | 0.017 | 0 | 0.15 | 0.06 | 0.09 | 0.05 | 0.14 | 0.16 |
| 10 | 111122888 | rs11194592 | 0.043 | -0.842 | 0.644 | -0.523 | -0.009 | -0.016 | -0.016 | 0 | 0.18 | 0.26 | 0.11 | 0.06 | 0.14 | 0.2 |
| 10 | 111162621 | rs7084219 | -0.808 | -0.944 | 0.183 | -0.567 | -0.009 | -0.014 | -0.014 | 0 | 0.23 | 0.71 | 0.14 | 0.05 | 0.13 | 0.16 |
| 10 | 111187116 | rs10884778 | 0.304 | -0.757 | 0.007 | -0.341 | -0.047 | -0.075 | -0.075 | 0 | 0.23 | 0.09 | 0.4 | 0.45 | 0.31 | 0.31 |
| 10 | 111189264 | rs12264293 | -0.407 | -1.014 | 0.278 | -0.541 | -0.023 | -0.055 | -0.055 | 0 | 0.19 | 0.49 | 0.12 | 0.05 | 0.13 | 0.17 |
| 10 | 111195590 | rs10884781 | -0.465 | -1.313 | -0.005 | -0.609 | -0.021 | -0.042 | -0.042 | 0 | 0.25 | 0.34 | 0.15 | 0.06 | 0.19 | 0.14 |
| 10 | 111197730 | rs7092048 | -0.954 | -1.107 | -1.328 | -0.761 | -0.03 | 0.029 | 0.029 | 1.188 | 0.89 | 0.99 | 0.9 | 0.95 | 0.87 | 0.84 |
| 10 | 111202837 | rs7072754 | -0.694 | -1.144 | -1.014 | -0.71 | 0.031 | 0.042 | 0.042 | 0.219 | 0.3 | 0.73 | 0.18 | 0.06 | 0.19 | 0.18 |
| 10 | 111205902 | rs1354298 | -0.501 | -1.165 | -1.322 | -0.624 | -0.08 | 0.03 | 0.03 | 0 | 0.79 | 0.99 | 0.81 | 0.94 | 0.74 | 0.87 |
| 10 | 111214463 | rs7070489 | -1.325 | -1.283 | -0.406 | -0.216 | -0.07 | -0.027 | -0.027 | 0 | 0.12 | 0.65 | 0.05 | 0.05 | 0.01 | 0.08 |
| 10 | 111222113 | rs1503256 | -1.379 | -1.292 | -0.465 | -0.221 | -0.071 | -0.03 | -0.03 | 0.263 | 0.12 | 0.67 | 0.06 | 0.05 | 0.01 | 0.08 |
| 10 | 111224826 | rs1541130 | -0.918 | -0.721 | -1.191 | -0.065 | 0.004 | -0.019 | -0.019 | 0 | 0.49 | 0.84 | 0.56 | 0.57 | 0.43 | 0.57 |
| 10 | 111236512 | rs10884787 | -0.946 | 0.003 | -0.498 | -0.156 | 0.027 | -0.058 | -0.058 | 0 | 0.3 | 0.5 | 0.35 | 0.19 | 0.32 | 0.26 |
| 10 | 111239425 | rs6584940 | -0.799 | -0.007 | -0.547 | -0.16 | 0.012 | -0.035 | -0.035 | 0 | 0.3 | 0.6 | 0.36 | 0.19 | 0.32 | 0.26 |
| 10 | 111244681 | rs12414140 | 0.483 | 0.085 | -0.863 | 0.045 | -0.083 | -0.056 | -0.056 | 0.059 | 0.19 | 0.01 | 0.29 | 0.24 | 0.25 | 0.13 |
| 10 | 111261278 | rs7088966 | -0.029 | -0.033 | -0.279 | 0.039 | 0.015 | -0.003 | -0.003 | 0.461 | 0.45 | 0.41 | 0.56 | 0.5 | 0.56 | 0.32 |
| 10 | 111268887 | rs10884796 | -0.468 | -0.157 | -0.102 | -0.029 | 0.036 | -0.001 | -0.001 | 0.396 | 0.52 | 0.68 | 0.58 | 0.5 | 0.57 | 0.41 |

| | | | | | | | | | | | | | | | | |
|---|---|---|---|---|---|---|---|---|---|---|---|---|---|---|---|---|
| 10 | 111276294 | rs11194697 | -0.448 | -0.118 | -0.112 | -0.002 | 0.029 | 0.001 | 0.001 | 0.238 | 0.52 | 0.65 | 0.57 | 0.49 | 0.56 | 0.41 |
| 10 | 111289799 | rs9787509 | 1.545 | 0.365 | -0.559 | 0.112 | -0.063 | -0.074 | -0.074 | 0 | 0.21 | 0.05 | 0.18 | 0.3 | 0.24 | 0.17 |
| 10 | 111294094 | rs7911732 | -0.256 | 0.163 | -0.263 | -0.007 | 0.017 | 0.003 | 0.003 | 0.167 | 0.5 | 0.46 | 0.55 | 0.49 | 0.55 | 0.41 |
| 10 | 111303358 | rs10884798 | -0.91 | 0.083 | -0.457 | -0.246 | 0.035 | 0 | 0 | 0.293 | 0.57 | 0.69 | 0.61 | 0.55 | 0.66 | 0.45 |
| 10 | 111321939 | rs7078049 | -0.895 | 0.08 | -0.412 | -0.306 | 0.022 | 0.002 | 0.002 | 0.398 | 0.57 | 0.64 | 0.62 | 0.56 | 0.67 | 0.45 |
| 10 | 111322478 | rs12354938 | 0.913 | 0.117 | -0.206 | -0.48 | 0.011 | -0.048 | -0.048 | 0 | 0.1 | 0.15 | 0.1 | 0 | 0.14 | 0.03 |
| 10 | 111330820 | rs11194725 | 0.815 | 0.384 | -0.229 | -0.346 | 0.031 | 0.019 | 0.019 | 0.121 | 0.18 | 0.34 | 0.3 | 0.21 | 0.22 | 0.13 |
| 10 | 111332634 | rs12252704 | 1.525 | 0.44 | -0.727 | -0.287 | -0.045 | -0.058 | -0.058 | 0 | 0.21 | 0.11 | 0.19 | 0.35 | 0.29 | 0.24 |
| 10 | 111335861 | rs11816901 | 0.12 | -0.201 | -0.423 | -0.007 | -0.012 | -0.019 | -0.019 | 3.226 | 0.22 | 0.35 | 0.25 | 0.19 | 0.19 | 0.18 |
| 10 | 111336595 | rs6584950 | -0.063 | -0.472 | -0.812 | 0.154 | 0.03 | -0.004 | -0.004 | 0.22 | 0.57 | 0.69 | 0.56 | 0.54 | 0.66 | 0.45 |
| 10 | 111337260 | rs7914907 | -0.463 | -0.185 | -0.49 | -0.121 | 0.003 | 0.009 | 0.009 | 0.352 | 0.81 | 0.84 | 0.86 | 0.91 | 0.82 | 0.83 |
| 10 | 111346161 | rs2488932 | -0.296 | 0.11 | -0.092 | 0.143 | 0.001 | 0.026 | 0.026 | 0 | 0.51 | 0.67 | 0.38 | 0.41 | 0.51 | 0.41 |
| 10 | 111348202 | rs1936146 | -0.142 | 0.129 | -0.186 | 0.119 | -0.036 | 0.031 | 0.031 | 0 | 0.45 | 0.22 | 0.29 | 0.35 | 0.51 | 0.33 |
| 10 | 111361505 | rs1936126 | 0.078 | 0.036 | 0.059 | 0.062 | -0.041 | 0.001 | 0.001 | 0 | 0.39 | 0.25 | 0.28 | 0.37 | 0.49 | 0.27 |
| 10 | 111375258 | rs10509901 | 0.229 | -0.019 | -0.128 | 0.094 | -0.153 | -0.06 | -0.06 | 0 | 0.37 | 0.07 | 0.27 | 0.36 | 0.49 | 0.27 |
| 10 | 111376954 | rs1936151 | 0.813 | -0.569 | 0.646 | 0.055 | -0.05 | -0.039 | -0.039 | 0 | 0.9 | 0.63 | 0.96 | 0.98 | 0.99 | 0.9 |
| 10 | 111388459 | rs10884815 | 0.047 | -0.088 | 0.064 | 0.051 | -0.005 | 0.019 | 0.019 | 0 | 0.36 | 0.28 | 0.3 | 0.27 | 0.4 | 0.29 |
| 10 | 111405399 | rs7904693 | 0.028 | -0.128 | 0.059 | 0.046 | -0.005 | 0.021 | 0.021 | 0 | 0.36 | 0.28 | 0.3 | 0.27 | 0.4 | 0.29 |
| 10 | 111406295 | rs1936138 | 0.506 | -0.249 | -0.008 | -0.009 | -0.066 | -0.012 | -0.012 | 0 | 0.29 | 0.1 | 0.28 | 0.27 | 0.39 | 0.22 |
| 10 | 111421513 | rs12573817 | 0.008 | -0.46 | 0.833 | -0.129 | 0 | 0.003 | 0.003 | 0 | 0.6 | 0.39 | 0.66 | 0.74 | 0.59 | 0.71 |
| 10 | 111428184 | rs7071813 | 0.496 | -0.411 | 0.684 | -0.085 | -0.01 | 0.008 | 0.008 | 0 | 0.3 | 0.29 | 0.3 | 0.24 | 0.4 | 0.21 |
| 10 | 111429236 | rs7917827 | 0.086 | -0.409 | 0.633 | -0.084 | -0.035 | 0.034 | 0.034 | 0 | 0.35 | 0.13 | 0.29 | 0.26 | 0.4 | 0.29 |
| 10 | 111433627 | rs11194785 | 0.146 | -1.18 | 0.928 | 0.214 | 0.012 | 0.011 | 0.011 | 0 | 0.07 | 0.03 | 0.01 | 0.02 | 0.01 | 0.08 |
| 10 | 111441034 | rs7899245 | -0.071 | -0.872 | 0.768 | 0.021 | -0.005 | 0.059 | 0.059 | 0 | 0.44 | 0.32 | 0.37 | 0.26 | 0.52 | 0.35 |
| 10 | 111465729 | rs10884876 | 0.677 | -0.864 | 0.663 | 0.027 | -0.092 | 0.054 | 0.054 | 0 | 0.36 | 0.1 | 0.33 | 0.24 | 0.52 | 0.26 |
| 10 | 111470354 | rs12241796 | -0.021 | -0.777 | 0.837 | 0.06 | -0.006 | 0.058 | 0.058 | 0 | 0.45 | 0.33 | 0.36 | 0.27 | 0.52 | 0.35 |
| 10 | 111481543 | rs2051107 | 0.779 | -0.792 | 1.245 | 0.007 | 0.079 | 0.075 | 0.075 | 6.065 | 0.95 | 0.49 | 0.95 | 1 | 1 | 1 |
| 10 | 111496271 | rs17736567 | 0.013 | -0.962 | 1.204 | 0.269 | 0.012 | 0.057 | 0.057 | 0 | 0.51 | 0.31 | 0.59 | 0.67 | 0.47 | 0.68 |
| 10 | 111501039 | rs10787213 | 0.23 | -0.976 | 1.233 | 0.309 | -0.025 | 0.079 | 0.079 | 0 | 0.43 | 0.2 | 0.36 | 0.27 | 0.53 | 0.31 |
| 10 | 111505407 | rs4918458 | 0.851 | -1.205 | 1.369 | 0.337 | 0.073 | 0.093 | 0.093 | 6.337 | 0.95 | 0.51 | 0.95 | 0.99 | 1 | 0.98 |
| 10 | 111517109 | rs1245928 | 0.449 | -1.782 | 1.018 | 0.728 | 0.009 | 0.002 | 0.002 | 0.964 | 0.27 | 0.27 | 0.16 | 0.21 | 0.23 | 0.33 |
| 10 | 111523421 | rs754120 | -1.208 | -1.73 | 1.082 | 0.9 | 0.005 | 0.018 | 0.018 | 0 | 0.55 | 0.69 | 0.62 | 0.64 | 0.45 | 0.69 |
| 10 | 111524203 | rs2244680 | 0.165 | -1.989 | 1.171 | 0.846 | -0.04 | -0.009 | -0.009 | 0 | 0.3 | 0.5 | 0.18 | 0.23 | 0.24 | 0.34 |
| 10 | 111528480 | rs1245907 | -0.116 | -1.999 | 1.246 | 1.123 | -0.048 | 0.021 | 0.021 | 0 | 0.33 | 0.53 | 0.18 | 0.34 | 0.25 | 0.47 |
| 10 | 111531409 | rs1245909 | 0.27 | -2.024 | 1.313 | 1.167 | 0.006 | 0.002 | 0.002 | 0.437 | 0.29 | 0.28 | 0.15 | 0.22 | 0.29 | 0.29 |
| 10 | 111534005 | rs11194847 | 0.322 | -2.028 | 1.189 | 1.156 | -0.001 | 0 | 0 | 0 | 0.12 | 0.09 | 0.13 | 0.1 | 0.11 | 0.1 |
| 10 | 111535667 | rs1245916 | -0.876 | -2.236 | 1.181 | 1.166 | -0.009 | -0.004 | -0.004 | 0.015 | 0.46 | 0.56 | 0.48 | 0.62 | 0.34 | 0.62 |
| 10 | 111537595 | rs7091635 | -1.686 | -2.238 | 1.186 | 1.171 | 0.002 | 0.019 | 0.019 | 0 | 0.58 | 0.65 | 0.61 | 0.74 | 0.45 | 0.74 |
| 10 | 111539154 | rs10749021 | -1.686 | -2.238 | 1.157 | 1.171 | 0.001 | 0.015 | 0.015 | 0 | 0.58 | 0.63 | 0.61 | 0.74 | 0.45 | 0.74 |
| 10 | 111542533 | rs2419311 | 0.611 | -2.21 | 1.032 | 1.241 | -0.001 | -0.028 | -0.028 | 0 | 0.07 | 0.04 | 0.29 | 0.28 | 0.04 | 0.1 |
| 10 | 111579787 | rs11194873 | 0.319 | -2.034 | 1.164 | 1.249 | 0 | -0.001 | -0.001 | 0 | 0.11 | 0.09 | 0.13 | 0.12 | 0.11 | 0.1 |
| 10 | 111579895 | rs7917299 | -0.514 | -2.068 | 1.195 | 1.27 | -0.054 | 0.03 | 0.03 | 0 | 0.1 | 0.19 | 0.03 | 0.13 | 0.01 | 0.24 |
| 10 | 111581612 | rs507319 | 0.724 | -2.092 | 1.137 | 1.271 | 0.038 | -0.002 | -0.002 | 0 | 0.5 | 0.41 | 0.69 | 0.54 | 0.59 | 0.38 |
| 10 | 111585360 | rs7901662 | -0.625 | -1.883 | 1.185 | 1.308 | -0.06 | 0.047 | 0.047 | 0 | 0.1 | 0.25 | 0.03 | 0.13 | 0.01 | 0.24 |
| 10 | 111608242 | rs6584960 | -1.08 | -1.993 | 0.921 | 1.269 | -0.029 | 0.048 | 0.048 | 0 | 0.18 | 0.2 | 0.32 | 0.42 | 0.06 | 0.36 |
| 10 | 111609610 | rs11194891 | -1.103 | -1.954 | 0.915 | 1.236 | -0.032 | 0.06 | 0.06 | 0 | 0.18 | 0.21 | 0.32 | 0.42 | 0.06 | 0.38 |
| 10 | 111610382 | rs11194892 | -0.002 | -1.953 | 0.784 | 0.994 | -0.048 | -0.018 | -0.018 | 0.006 | 0.31 | 0.46 | 0.17 | 0.22 | 0.26 | 0.26 |
| 10 | 111612666 | rs11194894 | -0.176 | -1.108 | 0.238 | 0.903 | 0 | 0 | 0 | 0 | 0.08 | 0.08 | 0.13 | 0.1 | 0.13 | 0.08 |
| 10 | 111617509 | rs17811806 | -0.185 | -1.115 | 0.241 | 0.906 | 0 | 0 | 0 | 0 | 0.08 | 0.08 | 0.13 | 0.09 | 0.12 | 0.08 |
| 10 | 111621456 | rs10787218 | -0.985 | -1.592 | 0.039 | 0.923 | -0.015 | -0.036 | -0.036 | 0 | 0.21 | 0.13 | 0.47 | 0.35 | 0.24 | 0.2 |
| 10 | 111622694 | rs10787219 | -0.568 | -1.757 | -0.025 | 0.972 | -0.019 | -0.025 | -0.025 | 0 | 0.18 | 0.12 | 0.47 | 0.34 | 0.23 | 0.16 |
| 10 | 111624769 | rs7834 | -0.767 | -2.329 | 0.721 | 0.599 | -0.062 | -0.06 | -0.06 | 0 | 0.13 | 0.19 | 0.03 | 0 | 0.02 | 0.11 |
| 10 | 111632445 | rs7916801 | -0.06 | -2.856 | 0.386 | 0.748 | -0.017 | 0.014 | 0.014 | 0.005 | 0.24 | 0.26 | 0.11 | 0.35 | 0.18 | 0.43 |
| 10 | 111632879 | rs536343 | 1.604 | -3.036 | 0.739 | 0.734 | 0.004 | 0.008 | 0.008 | 2.438 | 0.45 | 0.45 | 0.4 | 0.26 | 0.57 | 0.28 |
| 10 | 111635212 | rs3818285 | -0.668 | -3.082 | 0.768 | 0.772 | -0.007 | -0.032 | -0.032 | 0.047 | 0.23 | 0.16 | 0.48 | 0.39 | 0.24 | 0.2 |
| 10 | 111637286 | rs550697 | 1.483 | -3.172 | 0.622 | 0.782 | 0.01 | 0.021 | 0.021 | 2.184 | 0.46 | 0.43 | 0.41 | 0.26 | 0.58 | 0.28 |
| 10 | 111641706 | rs3780950 | 0.085 | -3.11 | 0.294 | 0.909 | 0.041 | -0.064 | -0.064 | 0 | 0.22 | 0.11 | 0.09 | 0.35 | 0.18 | 0.44 |
| 10 | 111650150 | rs7100154 | -0.925 | -2.829 | 0.389 | 0.839 | 0.019 | 0.038 | 0.038 | 0 | 0.25 | 0.37 | 0.51 | 0.35 | 0.27 | 0.21 |
| 10 | 111664595 | rs7078789 | -2.017 | -3.508 | -0.367 | 0.623 | -0.011 | 0.099 | 0.099 | 0.176 | 0.59 | 0.85 | 0.63 | 0.76 | 0.44 | 0.74 |
| 10 | 111669239 | rs3758409 | 0.218 | -3.514 | -0.4 | 0.664 | -0.076 | -0.088 | -0.088 | 0 | 0.18 | 0.04 | 0.47 | 0.33 | 0.26 | 0.21 |
| 10 | 111676799 | rs12258440 | -1.286 | -3.661 | -0.254 | 0.593 | 0.012 | 0.026 | 0.026 | 0 | 0.28 | 0.44 | 0.51 | 0.32 | 0.27 | 0.22 |
| 10 | 111705895 | rs975442 | -0.162 | -3.604 | -0.117 | 0.276 | -0.025 | 0.005 | 0.005 | 0 | 0.3 | 0.34 | 0.12 | 0.41 | 0.17 | 0.52 |
| 10 | 111729920 | rs12785152 | 0.685 | -3.505 | -0.395 | 0.327 | -0.084 | -0.006 | -0.006 | 0 | 0.14 | 0.01 | 0.16 | 0.11 | 0.22 | 0.1 |
| 10 | 111732457 | rs11194934 | -0.313 | -3.983 | 0.12 | 0.325 | 0.021 | 0.017 | 0.017 | 8.503 | 0.07 | 0.4 | 0.04 | 0 | 0.01 | 0 |
| 10 | 111738451 | rs9630101 | -0.929 | -3.974 | 0.086 | 0.345 | -0.083 | 0.016 | 0.016 | 0 | 0.35 | 0.46 | 0.13 | 0.42 | 0.17 | 0.53 |
| 10 | 111755416 | rs7079713 | -0.9 | -3.74 | -0.773 | 0.424 | 0.115 | -0.025 | -0.025 | 0 | 0.37 | 0.8 | 0.17 | 0.42 | 0.17 | 0.5 |
| 10 | 111757791 | rs10884919 | -0.706 | -3.637 | -0.336 | 0.454 | -0.061 | 0.046 | 0.046 | 0.124 | 0.26 | 0.56 | 0.16 | 0.43 | 0.15 | 0.4 |
| 10 | 111773423 | rs2060675 | 0.905 | -3.943 | -0.916 | 0.237 | 0.148 | -0.012 | -0.012 | 0 | 0.6 | 0.19 | 0.81 | 0.56 | 0.81 | 0.45 |
| 10 | 111775849 | rs7071509 | -0.769 | -4.106 | -0.601 | 0.188 | -0.045 | 0.025 | 0.025 | 0.015 | 0.28 | 0.6 | 0.16 | 0.43 | 0.15 | 0.43 |

| | | | | | | | | | | | | | | | | |
|---|---|---|---|---|---|---|---|---|---|---|---|---|---|---|---|---|
| 10 | 111781715 | rs10509907 | 0.004 | -4.252 | -0.398 | 0.13 | 0.024 | -0.047 | -0.047 | 0 | 0.18 | 0.1 | 0.1 | 0.42 | 0.14 | 0.41 |
| 10 | 111787716 | rs2225363 | -0.749 | -4.267 | -0.36 | 0.117 | -0.035 | -0.036 | -0.036 | 0.768 | 0.88 | 0.85 | 0.97 | 1 | 0.97 | 0.87 |
| 10 | 111794554 | rs1349348 | 0.463 | -4.257 | -0.354 | 0.139 | 0.018 | -0.011 | -0.011 | 0 | 0.7 | 0.75 | 0.88 | 0.57 | 0.83 | 0.46 |
| 10 | 111812955 | rs17126984 | 0.285 | -5.02 | -1.114 | -0.491 | 0.01 | -0.02 | -0.02 | 0 | 0.14 | 0.1 | 0.08 | 0.42 | 0.11 | 0.38 |
| 10 | 111820311 | rs2076974 | 0.78 | -4.857 | -1.055 | -0.407 | 0.152 | -0.009 | -0.009 | 0.309 | 0.6 | 0.19 | 0.81 | 0.56 | 0.82 | 0.44 |
| 10 | 111830275 | rs11194972 | 0.014 | -4.419 | -0.879 | -0.278 | -0.037 | -0.03 | -0.03 | 0.424 | 0.1 | 0.5 | 0.06 | 0.01 | 0.02 | 0.02 |
| 10 | 111840070 | rs4918478 | 1.091 | -4.102 | -1.734 | -0.138 | -0.055 | -0.11 | -0.11 | 0 | 0.16 | 0.01 | 0.43 | 0.33 | 0.19 | 0.19 |
| 10 | 111850503 | rs2501574 | -2.045 | -4.065 | -1.942 | -0.295 | -0.128 | 0 | 0 | 3.317 | 0.69 | 0.98 | 0.69 | 0.78 | 0.46 | 0.78 |
| 10 | 111886089 | rs3731566 | -2.172 | -3.104 | -1.93 | -0.415 | -0.145 | 0.107 | 0.107 | 0 | 0.82 | 0.98 | 0.76 | 1 | 0.6 | 0.87 |
| 10 | 111893562 | rs1127522 | 0.032 | -3.706 | -0.949 | 0.174 | -0.051 | -0.055 | -0.055 | 0 | 0.11 | 0.53 | 0.06 | 0.01 | 0.02 | 0.02 |
| 10 | 111900010 | rs17127118 | 0.311 | -3.793 | -1.744 | 0.329 | 0.029 | -0.066 | -0.066 | 2.757 | 0.18 | 0.07 | 0.09 | 0.42 | 0.14 | 0.41 |
| 10 | 111901567 | rs6584972 | -0.801 | -3.524 | -1.832 | 0.486 | 0.187 | -0.003 | -0.003 | 0 | 0.41 | 0.88 | 0.19 | 0.42 | 0.19 | 0.54 |
| 10 | 111911687 | rs10884931 | 0.671 | -2.921 | -1.629 | 0.736 | -0.031 | -0.059 | -0.059 | 0 | 0.14 | 0.04 | 0.43 | 0.29 | 0.19 | 0.19 |
| 10 | 111919866 | rs7918758 | -0.968 | -3.511 | -1.393 | 0.45 | 0.166 | -0.024 | -0.024 | 0 | 0.4 | 0.88 | 0.2 | 0.47 | 0.18 | 0.55 |
| 10 | 111920995 | rs12260509 | -0.977 | -3.504 | -1.117 | 0.453 | 0.169 | -0.024 | -0.024 | 0 | 0.4 | 0.87 | 0.2 | 0.46 | 0.18 | 0.55 |
| 10 | 111928766 | rs7918659 | -0.338 | -3.755 | -1.141 | 0.38 | 0.031 | 0.03 | 0.03 | 0.087 | 0.24 | 0.19 | 0.11 | 0.09 | 0.13 | 0.34 |
| 10 | 111928784 | rs6584974 | -1.071 | -3.467 | -1.283 | 0.449 | 0.146 | -0.012 | -0.012 | 0 | 0.4 | 0.83 | 0.2 | 0.46 | 0.18 | 0.55 |
| 10 | 111936671 | rs17127151 | -0.41 | -3.026 | -0.727 | 0.78 | 0.002 | -0.003 | -0.003 | 0 | 0.18 | 0.17 | 0.1 | 0.39 | 0.12 | 0.38 |
| 10 | 111941371 | rs12573269 | -0.337 | -3.51 | -0.908 | 0.782 | -0.017 | 0.004 | 0.004 | 0 | 0.18 | 0.19 | 0.04 | 0.34 | 0.06 | 0.31 |
| 10 | 111943436 | rs10884933 | 0.793 | -3.198 | -1.108 | 0.909 | -0.008 | -0.017 | -0.017 | 0 | 0.16 | 0.12 | 0.39 | 0.25 | 0.2 | 0.18 |
| 10 | 111950733 | rs4918486 | -0.107 | -3.253 | -0.52 | 0.874 | 0.004 | 0.015 | 0.015 | 0 | 0.23 | 0.42 | 0.43 | 0.25 | 0.23 | 0.2 |
| 10 | 111954970 | rs6584977 | -1.631 | -4.054 | -1.422 | 0.522 | 0.033 | 0.032 | 0.032 | 0 | 0.61 | 0.85 | 0.58 | 0.74 | 0.38 | 0.74 |
| 10 | 111956363 | rs6584978 | -1.062 | -4.135 | -1.471 | 0.52 | 0.116 | 0.061 | 0.061 | 0 | 0.48 | 0.83 | 0.51 | 0.64 | 0.31 | 0.6 |
| 10 | 111961868 | rs10884938 | 0.855 | -3.588 | -1.568 | 0.52 | -0.031 | -0.018 | -0.018 | 0 | 0.16 | 0.07 | 0.32 | 0.21 | 0.23 | 0.22 |
| 10 | 111964133 | rs10787225 | -1.306 | -3.711 | -1.904 | 0.34 | 0.089 | 0.05 | 0.05 | 0.227 | 0.48 | 0.94 | 0.6 | 0.74 | 0.33 | 0.6 |
| 10 | 111964276 | rs11195025 | 0.828 | -3.243 | -1.724 | 0.556 | 0.002 | -0.075 | -0.075 | 0 | 0.12 | 0.01 | 0.3 | 0.3 | 0.09 | 0.2 |
| 10 | 111973339 | rs1342273 | 0.167 | -3.48 | -0.986 | 0.357 | -0.017 | -0.099 | -0.099 | 0 | 0.22 | 0.53 | 0.17 | 0.07 | 0.18 | 0.14 |
| 10 | 111992222 | rs17127170 | 1.003 | -3.195 | -1.337 | 0.618 | 0.001 | -0.073 | -0.073 | 1 | 0.11 | 0.01 | 0.28 | 0.28 | 0.07 | 0.2 |
| 10 | 112001009 | rs1891375 | -1.64 | -3.6 | -1.649 | 0.417 | -0.006 | -0.028 | -0.028 | 0 | 0.62 | 0.93 | 0.62 | 0.72 | 0.36 | 0.73 |
| 10 | 112010830 | rs11195047 | 1.51 | -2.366 | -1.544 | -0.088 | -0.105 | -0.053 | -0.053 | 0 | 0.29 | 0.08 | 0.29 | 0.28 | 0.47 | 0.17 |
| 10 | 112011484 | rs2010327 | 1.732 | -2.498 | -1.61 | 0.091 | 0.013 | -0.033 | -0.033 | 0 | 0.07 | 0 | 0.03 | 0.29 | 0.02 | 0.23 |
| 10 | 112029300 | rs1935465 | -0.873 | -2.562 | -1.468 | -0.065 | 0.003 | 0.002 | 0.002 | 0 | 0.55 | 0.6 | 0.55 | 0.7 | 0.36 | 0.71 |
| 10 | 112036975 | rs11195062 | -0.666 | -2.424 | -1.499 | -0.07 | 0.01 | 0.006 | 0.006 | 0 | 0.52 | 0.47 | 0.52 | 0.65 | 0.36 | 0.72 |
| 10 | 112042451 | rs928929 | 0.808 | -2.824 | -1.935 | -0.893 | -0.093 | -0.002 | -0.002 | 0.179 | 0.47 | 0.04 | 0.59 | 0.54 | 0.55 | 0.45 |
| 10 | 112045142 | rs1133195 | 1.282 | -2.936 | -1.968 | -0.964 | 0.045 | 0.089 | 0.089 | 0 | 0.14 | 0 | 0.04 | 0 | 0.07 | 0.1 |
| 10 | 112046150 | rs17658 | 0.407 | -2.982 | -1.82 | -1.003 | -0.006 | -0.051 | -0.051 | 0 | 0.61 | 0.61 | 0.76 | 0.91 | 0.71 | 0.79 |
| 10 | 112046200 | rs1803997 | 1.422 | -2.954 | -1.946 | -0.926 | 0.019 | -0.036 | -0.036 | 0.846 | 0.08 | 0.02 | 0.04 | 0.3 | 0.03 | 0.28 |
| 10 | 112053599 | rs1050755 | 0.396 | -2.756 | -1.835 | -1.192 | -0.063 | 0.228 | 0.228 | 1.458 | 0.55 | 0.06 | 0.62 | 0.84 | 0.58 | 0.7 |
| 10 | 112056098 | rs10509914 | 0.867 | -2.456 | -1.654 | -1.027 | -0.103 | 0.188 | 0.188 | 0.334 | 0.41 | 0.07 | 0.58 | 0.84 | 0.52 | 0.6 |
| 10 | 112067334 | rs7907926 | 0.672 | -1.853 | -0.992 | -0.761 | 0.011 | -0.057 | -0.057 | 0 | 0.2 | 0.47 | 0.17 | 0.11 | 0.19 | 0.17 |
| 10 | 112072649 | rs1673667 | -0.387 | -1.741 | -1.475 | -0.708 | 0 | 0.014 | 0.014 | 0 | 0.77 | 0.87 | 0.88 | 1 | 0.79 | 0.88 |
| 10 | 112079743 | rs10884947 | -0.264 | -1.618 | -0.929 | -0.134 | -0.045 | 0.071 | 0.071 | 0 | 0.18 | 0.52 | 0.13 | 0.31 | 0.08 | 0.39 |
| 10 | 112082992 | rs6584984 | -0.425 | -1.753 | -1.049 | -0.274 | -0.055 | 0.001 | 0.001 | 0 | 0.25 | 0.52 | 0.17 | 0.31 | 0.12 | 0.43 |
| 10 | 112094121 | rs2027126 | 1.342 | -1.297 | -1.665 | -0.217 | -0.062 | -0.071 | -0.071 | 0 | 0.3 | 0.14 | 0.52 | 0.45 | 0.44 | 0.28 |
| 10 | 112097073 | rs963059 | 0.836 | -1.334 | -2.017 | -0.534 | -0.016 | 0.051 | 0.051 | 1.206 | 0.21 | 0.06 | 0.12 | 0 | 0.21 | 0.12 |
| 10 | 112107406 | rs10787229 | 1.394 | -1.044 | -1.982 | -0.03 | -0.15 | -0.16 | -0.16 | 0 | 0.27 | 0.02 | 0.49 | 0.36 | 0.42 | 0.27 |
| 10 | 112109830 | rs1227985 | -0.896 | -0.555 | -1.166 | -0.36 | -0.003 | 0.003 | 0.003 | 0 | 0.82 | 0.79 | 0.86 | 0.9 | 0.78 | 0.9 |
| 10 | 112112032 | rs1023067 | -0.578 | -0.567 | -1.135 | -0.279 | 0.001 | 0.003 | 0.003 | 0.234 | 0.79 | 0.74 | 0.82 | 0.76 | 0.74 | 0.89 |
| 10 | 112112952 | rs1227984 | -0.674 | -0.559 | -1.123 | -0.542 | 0.002 | -0.007 | -0.007 | 0 | 0.8 | 0.74 | 0.82 | 0.9 | 0.76 | 0.9 |
| 10 | 112115492 | rs1342274 | 0.617 | -0.538 | -1.271 | -0.619 | -0.035 | 0.104 | 0.104 | 0 | 0.2 | 0.03 | 0.11 | 0 | 0.2 | 0.12 |
| 10 | 112115666 | rs1774644 | -0.471 | -0.517 | -0.59 | -0.108 | 0.002 | -0.004 | -0.004 | 0 | 0.82 | 0.77 | 0.85 | 0.9 | 0.79 | 0.9 |
| 10 | 112117596 | rs1762279 | 0.656 | -0.532 | -0.019 | -0.284 | -0.001 | -0.008 | -0.008 | 0 | 0.65 | 0.36 | 0.78 | 0.85 | 0.7 | 0.78 |
| 10 | 112120573 | rs2168741 | 0.414 | -0.563 | 0.727 | 0.14 | 0.079 | 0.21 | 0.21 | 0 | 0.95 | 0.71 | 0.85 | 0.8 | 0.94 | 0.96 |
| 10 | 112122958 | rs17127283 | 0.18 | -0.689 | 0.733 | -0.002 | 0.015 | -0.017 | -0.017 | 2.2 | 0.19 | 0.34 | 0.16 | 0.1 | 0.22 | 0.11 |
| 10 | 112127251 | rs12246208 | -0.568 | -0.8 | 0.793 | -0.079 | -0.058 | -0.082 | -0.082 | 0 | 0.15 | 0.35 | 0.03 | 0.05 | 0.04 | 0.09 |
| 10 | 112129539 | rs12252135 | 0.23 | -0.72 | 0.553 | -0.052 | 0.005 | -0.006 | -0.006 | 0 | 0.11 | 0.09 | 0.02 | 0.03 | 0.04 | 0.1 |
| 10 | 112132171 | rs7081449 | -1.136 | -0.776 | 0.428 | 0.335 | -0.002 | -0.001 | -0.001 | 0 | 0.42 | 0.54 | 0.24 | 0.38 | 0.34 | 0.55 |
| 10 | 112133320 | rs11195102 | 0.794 | -0.567 | 0.302 | 0.458 | 0.013 | -0.005 | -0.005 | 1.137 | 0.16 | 0.2 | 0.13 | 0.11 | 0.21 | 0.11 |
| 10 | 112134500 | rs17127304 | 0.136 | -0.358 | -0.193 | 0.563 | 0 | -0.013 | -0.013 | 0 | 0.14 | 0.1 | 0.09 | 0.24 | 0.1 | 0.34 |
| 10 | 112137261 | rs17127310 | 0.515 | 0.045 | -0.734 | 0.353 | -0.008 | 0.006 | 0.006 | 0 | 0.25 | 0.09 | 0.33 | 0.22 | 0.26 | 0.22 |
| 10 | 112143204 | rs17127325 | 0.037 | -0.53 | -0.53 | 0.397 | 0.01 | 0 | 0 | 0 | 0.23 | 0.18 | 0.3 | 0.22 | 0.15 | 0.2 |
| 10 | 112148604 | rs4918502 | -1.344 | -0.175 | -0.59 | 0.405 | -0.049 | 0.049 | 0.049 | 0 | 0.67 | 0.79 | 0.67 | 0.8 | 0.54 | 0.76 |
| 10 | 112148651 | rs4918503 | -1.361 | -0.182 | -0.403 | 0.402 | -0.032 | 0.034 | 0.034 | 0 | 0.67 | 0.76 | 0.67 | 0.8 | 0.54 | 0.76 |
| 10 | 112153284 | rs11195116 | -0.785 | -0.239 | -0.17 | 0.475 | -0.001 | 0 | 0 | 0 | 0.55 | 0.53 | 0.58 | 0.68 | 0.4 | 0.68 |
| 10 | 112154110 | rs4918508 | -1.665 | -0.183 | -0.444 | 0.452 | -0.082 | 0.06 | 0.06 | 0 | 0.69 | 0.85 | 0.68 | 0.8 | 0.55 | 0.76 |
| 10 | 112157994 | rs17824180 | -0.059 | -0.565 | -0.34 | 0.645 | 0.03 | 0.007 | 0.007 | 0 | 0.35 | 0.22 | 0.36 | 0.33 | 0.26 | 0.29 |
| 10 | 112164194 | rs17127354 | 0.076 | -0.338 | -0.239 | 0.588 | 0.001 | -0.007 | -0.007 | 0 | 0.19 | 0.18 | 0.13 | 0.22 | 0.14 | 0.33 |
| 10 | 112164639 | rs4918510 | -1.664 | -0.19 | -0.383 | 0.34 | -0.066 | 0.05 | 0.05 | 0 | 0.69 | 0.83 | 0.69 | 0.79 | 0.57 | 0.76 |
| 10 | 112166708 | rs7098940 | 0.705 | -0.744 | -0.139 | 0.255 | -0.04 | -0.028 | -0.028 | 1.185 | 0.16 | 0.2 | 0.11 | 0.11 | 0.09 | 0.14 |

| | | | | | | | | | | | | | | | | |
|---|---|---|---|---|---|---|---|---|---|---|---|---|---|---|---|---|
| 10 | 112167240 | rs4918511 | 0.761 | -0.726 | -0.298 | 0.325 | -0.001 | -0.008 | -0.008 | 0 | 0.29 | 0.21 | 0.33 | 0.36 | 0.29 | 0.25 |
| 10 | 112167328 | rs4918512 | -0.092 | -0.823 | -0.112 | 0.517 | -0.07 | -0.048 | -0.048 | 0 | 0.18 | 0.34 | 0.12 | 0.11 | 0.09 | 0.13 |
| 10 | 112169511 | rs10884961 | -0.442 | -0.081 | -0.649 | 0.928 | -0.002 | -0.034 | -0.034 | 0.191 | 0.18 | 0.13 | 0.2 | 0.33 | 0.16 | 0.33 |
| 10 | 112171392 | rs763498 | -0.096 | -0.19 | -0.186 | 0.538 | -0.024 | -0.019 | -0.019 | 0 | 0.14 | 0.22 | 0.07 | 0.11 | 0.08 | 0.09 |
| 10 | 112171993 | rs10884962 | -0.33 | -0.415 | -0.122 | 0.944 | -0.008 | 0.015 | 0.015 | 0 | 0.15 | 0.21 | 0.12 | 0.2 | 0.1 | 0.2 |
| 10 | 112175889 | rs7083193 | 1.422 | -0.423 | -0.09 | 0.621 | -0.012 | -0.006 | -0.006 | 1.145 | 0.63 | 0.43 | 0.76 | 0.69 | 0.74 | 0.67 |
| 10 | 112179167 | rs10444086 | 1.242 | -0.197 | -0.285 | 0.569 | -0.001 | -0.001 | -0.001 | 0 | 0.59 | 0.42 | 0.6 | 0.63 | 0.68 | 0.59 |
| 10 | 112179194 | rs17127384 | -0.281 | -0.159 | -0.266 | 0.61 | -0.006 | 0.003 | 0.003 | 0 | 0.18 | 0.21 | 0.12 | 0.2 | 0.12 | 0.26 |
| 10 | 112184984 | rs1541050 | -0.963 | 0.524 | -0.722 | 0.377 | -0.005 | 0.023 | 0.023 | 1.217 | 0.59 | 0.73 | 0.6 | 0.62 | 0.6 | 0.6 |
| 10 | 112185149 | rs12415145 | 1.416 | 0.473 | -0.881 | 0.512 | 0.046 | -0.007 | -0.007 | 0 | 0.24 | 0.09 | 0.23 | 0.29 | 0.15 | 0.27 |
| 10 | 112185596 | rs10884966 | 0.714 | 1.552 | -0.45 | 0.796 | -0.006 | -0.001 | -0.001 | 0 | 0.18 | 0.17 | 0.21 | 0.16 | 0.32 | 0.18 |
| 10 | 112186148 | rs11195128 | 0.725 | 1.538 | -0.451 | 0.685 | -0.008 | 0.004 | 0.004 | 0 | 0.18 | 0.17 | 0.21 | 0.15 | 0.32 | 0.18 |
| 10 | 112191525 | rs4918514 | -0.831 | 0.31 | 0.177 | 0.915 | -0.026 | 0.04 | 0.04 | 0 | 0.32 | 0.48 | 0.24 | 0.4 | 0.26 | 0.38 |
| 10 | 112191884 | rs7915029 | -1.464 | 0.003 | -0.184 | 0.763 | -0.001 | 0 | 0 | 0.114 | 0.58 | 0.43 | 0.63 | 0.71 | 0.74 | 0.67 |
| 10 | 112192112 | rs4918516 | -0.701 | 0.66 | -0.18 | 1.143 | 0.022 | -0.075 | -0.075 | 2.663 | 0.24 | 0.1 | 0.17 | 0.29 | 0.21 | 0.34 |
| 10 | 112194168 | rs10884970 | 1.478 | 1.92 | 0.802 | 1.472 | 0.018 | 0.063 | 0.063 | 0 | 0.77 | 0.67 | 0.77 | 0.55 | 0.74 | 0.68 |
| 10 | 112194290 | rs17127392 | -0.64 | 1.955 | 0.764 | 1.346 | -0.004 | 0.003 | 0.003 | 0 | 0.1 | 0.1 | 0.07 | 0.05 | 0.13 | 0.09 |
| 10 | 112195899 | rs4918517 | -1.614 | 1.987 | 0.825 | 1.467 | 0.004 | 0.027 | 0.027 | 0 | 0.22 | 0.23 | 0.15 | 0.42 | 0.23 | 0.3 |
| 10 | 112197458 | rs1981112 | -1.497 | 1.984 | 0.771 | 1.467 | -0.006 | -0.012 | -0.012 | 0 | 0.21 | 0.16 | 0.15 | 0.42 | 0.23 | 0.3 |
| 10 | 112198864 | rs1000752 | 1.724 | 1.949 | 1.04 | 1.414 | 0.042 | 0.137 | 0.137 | 0 | 0.76 | 0.53 | 0.74 | 0.52 | 0.73 | 0.68 |
| 10 | 112206255 | rs10884971 | 1.608 | 2.021 | 1.018 | 1.413 | 0.034 | 0.1 | 0.1 | 0 | 0.74 | 0.47 | 0.73 | 0.51 | 0.73 | 0.65 |
| 10 | 112208500 | rs11195140 | -1.326 | 1.899 | 0.756 | 1.067 | 0.001 | -0.002 | -0.002 | 1.044 | 0.33 | 0.36 | 0.4 | 0.28 | 0.51 | 0.34 |
| 10 | 112215291 | rs746946 | 0.354 | 2.25 | 0.795 | 1.439 | 0.02 | 0.209 | 0.209 | 0.031 | 0.87 | 0.7 | 0.88 | 0.63 | 0.89 | 0.78 |
| 10 | 112215625 | rs945204 | 1.049 | 2.079 | 0.63 | 1.287 | -0.008 | 0.179 | 0.179 | 0 | 0.79 | 0.6 | 0.76 | 0.59 | 0.82 | 0.75 |
| 10 | 112217707 | rs10884981 | -0.028 | 2.714 | -0.095 | 1.075 | -0.07 | -0.018 | -0.018 | 0.631 | 0.17 | 0.05 | 0.22 | 0.18 | 0.3 | 0.15 |
| 10 | 112225713 | rs12250890 | -0.748 | 2.162 | 0.28 | 1.215 | 0.013 | 0.013 | 0.013 | 0 | 0.1 | 0.24 | 0.1 | 0.06 | 0.09 | 0.11 |
| 10 | 112231446 | rs10509918 | 0.232 | 2.457 | -0.496 | 1.217 | -0.083 | -0.056 | -0.056 | 0 | 0.13 | 0.01 | 0.17 | 0.19 | 0.25 | 0.13 |
| 10 | 112233212 | rs11195144 | -1.423 | 1.501 | -0.202 | 1.018 | -0.01 | 0.013 | 0.013 | 0 | 0.19 | 0.11 | 0.15 | 0.15 | 0.24 | 0.16 |
| 10 | 112235106 | rs5015510 | -1.472 | 1.494 | -0.128 | 1.015 | -0.003 | 0.006 | 0.006 | 0 | 0.19 | 0.14 | 0.16 | 0.15 | 0.24 | 0.17 |
| 10 | 112242851 | rs11195152 | -0.664 | 1.096 | -0.073 | 1.063 | 0.003 | 0.006 | 0.006 | 0 | 0.12 | 0.11 | 0.15 | 0.19 | 0.16 | 0.15 |
| 10 | 112246352 | rs7911973 | -1.355 | 1.019 | 0.082 | 1.077 | 0.008 | 0.03 | 0.03 | 0 | 0.23 | 0.37 | 0.24 | 0.26 | 0.24 | 0.24 |
| 10 | 112253662 | rs11195156 | -0.008 | 0.031 | -0.512 | 1.133 | 0.017 | -0.059 | -0.059 | 0.275 | 0.15 | 0.02 | 0.1 | 0.29 | 0.08 | 0.15 |
| 10 | 112263030 | rs1889568 | 0.91 | 0.472 | -0.234 | 1.02 | 0.005 | 0.035 | 0.035 | 0 | 0.73 | 0.58 | 0.71 | 0.59 | 0.7 | 0.77 |
| 10 | 112266771 | rs1889567 | -1.067 | 0.477 | -0.302 | 1 | 0 | 0.001 | 0.001 | 0 | 0.25 | 0.28 | 0.24 | 0.38 | 0.29 | 0.23 |
| 10 | 112272043 | rs10509919 | -1.185 | 0.488 | -0.313 | 0.933 | 0.002 | 0.006 | 0.006 | 0 | 0.26 | 0.31 | 0.24 | 0.36 | 0.29 | 0.23 |
| 10 | 112279717 | rs11195162 | -0.659 | 0.677 | -0.845 | 1.067 | -0.011 | -0.019 | -0.019 | 1.053 | 0.19 | 0.12 | 0.19 | 0.35 | 0.25 | 0.2 |
| 10 | 112283719 | rs12414450 | 0.971 | -0.531 | -1.157 | 0.641 | 0.039 | 0.012 | 0.012 | 0 | 0.36 | 0.09 | 0.42 | 0.34 | 0.31 | 0.41 |
| 10 | 112294424 | rs882486 | 0.99 | -0.728 | 0 | 0.46 | -0.047 | -0.038 | -0.038 | 4.855 | 0.72 | 0.51 | 0.73 | 0.8 | 0.77 | 0.8 |
| 10 | 112303595 | rs1107442 | 0.684 | -0.856 | 0.403 | 0.413 | -0.029 | -0.01 | -0.01 | 0 | 0.79 | 0.64 | 0.77 | 0.81 | 0.82 | 0.83 |
| 10 | 112305321 | rs10509922 | 0.013 | -0.851 | 0.379 | 0.417 | -0.001 | -0.003 | -0.003 | 0 | 0.1 | 0.07 | 0.12 | 0.16 | 0.12 | 0.14 |
| 10 | 112313903 | rs7901527 | 1.545 | -1.601 | 0.181 | 0.497 | -0.001 | -0.001 | -0.001 | 0 | 0.59 | 0.44 | 0.69 | 0.61 | 0.71 | 0.66 |
| 10 | 112314007 | rs7918064 | 1.195 | -1.505 | 0.153 | 0.571 | -0.018 | 0.008 | 0.008 | 0 | 0.68 | 0.69 | 0.75 | 0.61 | 0.75 | 0.69 |
| 10 | 112314670 | rs6584993 | -0.526 | -1.298 | 0.049 | 0.661 | 0.005 | -0.007 | -0.007 | 0.398 | 0.87 | 0.92 | 0.97 | 0.8 | 0.93 | 0.85 |
| 10 | 112316937 | rs11195182 | 0.14 | -1.017 | -0.025 | 0.554 | 0 | -0.003 | -0.003 | 0.03 | 0.1 | 0.07 | 0.12 | 0.16 | 0.12 | 0.15 |
| 10 | 112316956 | rs7090886 | -0.089 | -1.018 | -0.02 | 0.554 | -0.002 | 0.009 | 0.009 | 0 | 0.11 | 0.13 | 0.13 | 0.16 | 0.12 | 0.15 |
| 10 | 112321969 | rs7913383 | 1.021 | -1.252 | 0.417 | 0.4 | -0.061 | -0.094 | -0.094 | 0 | 0.84 | 0.51 | 0.85 | 0.96 | 0.92 | 0.95 |
| 10 | 112323187 | rs4918540 | 1.311 | -1.46 | 0.312 | 0.593 | -0.048 | 0.027 | 0.027 | 0.606 | 0.67 | 0.44 | 0.65 | 0.59 | 0.79 | 0.71 |
| 10 | 112343591 | rs11195199 | -0.249 | -1.434 | 0.196 | 0.625 | 0.031 | -0.03 | -0.03 | 0.052 | 0.18 | 0.08 | 0.19 | 0.23 | 0.1 | 0.17 |
| 10 | 112349879 | rs3737294 | -1.244 | -1.493 | 0.574 | 0.455 | -0.038 | -0.005 | -0.005 | 0 | 0.28 | 0.39 | 0.26 | 0.23 | 0.15 | 0.2 |
| 10 | 112365598 | rs7898064 | 0.856 | -1.357 | -0.234 | 0.484 | -0.045 | -0.006 | -0.006 | 0.783 | 0.69 | 0.38 | 0.69 | 0.74 | 0.84 | 0.8 |
| 10 | 112375282 | rs11195232 | 0.777 | -1.159 | -1.926 | 0.407 | -0.049 | -0.052 | -0.052 | 0 | 0.35 | 0.02 | 0.33 | 0.35 | 0.31 | 0.41 |
| 10 | 112378718 | rs869966 | -0.8 | -0.869 | -1.117 | 0.478 | 0.019 | -0.013 | -0.013 | 1.633 | 0.2 | 0.12 | 0.27 | 0.24 | 0.12 | 0.21 |
| 10 | 112380329 | rs4350299 | -0.102 | -0.317 | -1.398 | 0.218 | -0.026 | 0.021 | 0.021 | 0 | 0.64 | 0.77 | 0.64 | 0.79 | 0.52 | 0.62 |
| 10 | 112389466 | rs11195242 | -0.132 | -0.446 | -1.592 | 0.312 | 0.008 | -0.044 | -0.044 | 0 | 0.19 | 0.03 | 0.22 | 0.25 | 0.16 | 0.16 |
| 10 | 112392162 | rs1856801 | -0.61 | -0.39 | -1.049 | 0.227 | -0.007 | 0.031 | 0.031 | 0 | 0.11 | 0.18 | 0.1 | 0.17 | 0.11 | 0.17 |
| 10 | 112394034 | rs2419564 | -0.776 | -0.43 | -1.072 | -0.107 | -0.035 | 0.071 | 0.071 | 0 | 0.72 | 0.88 | 0.73 | 0.97 | 0.63 | 0.79 |
| 10 | 112394893 | rs4918546 | -1.371 | -0.779 | -0.145 | 0.042 | -0.031 | 0.038 | 0.038 | 1.876 | 0.16 | 0.64 | 0.13 | 0.17 | 0.12 | 0.19 |
| 10 | 112397144 | rs11195244 | 0.898 | -0.866 | 0.216 | 0.022 | -0.018 | 0.017 | 0.017 | 0.47 | 0.66 | 0.66 | 0.63 | 0.59 | 0.72 | 0.64 |
| 10 | 112398015 | rs7907617 | 0.549 | -0.972 | 0.358 | -0.028 | -0.051 | 0.021 | 0.021 | 0.007 | 0.77 | 0.64 | 0.73 | 0.75 | 0.87 | 0.86 |
| 10 | 112415418 | rs7904854 | -0.625 | -1.195 | 0.404 | -0.027 | -0.068 | 0.041 | 0.041 | 0 | 0.2 | 0.37 | 0.26 | 0.23 | 0.09 | 0.13 |
| 10 | 112418778 | rs10749041 | 0.644 | -1.196 | 0.218 | -0.187 | -0.027 | 0.016 | 0.016 | 0.507 | 0.83 | 0.49 | 0.85 | 0.85 | 0.86 | 0.85 |
| 10 | 112419189 | rs10509923 | 1.256 | -0.881 | -1.029 | -0.622 | -0.097 | 0.174 | 0.174 | 0 | 0.26 | 0.03 | 0.22 | 0 | 0.33 | 0.15 |
| 10 | 112423491 | rs10885008 | 1.012 | -0.708 | -0.944 | -0.353 | -0.124 | -0.067 | -0.067 | 0 | 0.37 | 0.08 | 0.41 | 0.63 | 0.47 | 0.56 |
| 10 | 112425178 | rs12358914 | -0.409 | -0.675 | -0.419 | -0.256 | 0.004 | -0.002 | -0.002 | 0.099 | 0.26 | 0.19 | 0.21 | 0.28 | 0.2 | 0.29 |
| 10 | 112429064 | rs9651418 | 0.296 | -0.267 | -0.013 | -0.169 | -0.004 | 0.007 | 0.007 | 0 | 0.43 | 0.4 | 0.54 | 0.68 | 0.45 | 0.54 |
| 10 | 112431320 | rs12359907 | 0.843 | 0.105 | -0.232 | -0.332 | -0.031 | 0.003 | 0.003 | 0.014 | 0.34 | 0.13 | 0.37 | 0.66 | 0.34 | 0.49 |
| 10 | 112434270 | rs2900925 | 0.828 | -0.279 | -0.049 | -0.469 | 0.004 | 0.002 | 0.002 | 0 | 0.66 | 0.36 | 0.62 | 0.68 | 0.69 | 0.68 |
| 10 | 112435773 | rs12772458 | 1.017 | -0.312 | -0.057 | -0.485 | 0.024 | 0.014 | 0.014 | 0 | 0.65 | 0.21 | 0.61 | 0.68 | 0.68 | 0.68 |
| 10 | 112445662 | rs10885017 | 0.964 | -0.035 | 0.257 | -0.706 | -0.006 | 0.07 | 0.07 | 0 | 0.34 | 0.22 | 0.27 | 0 | 0.36 | 0.18 |

| | | | | | | | | | | | | | | | | |
|---|---|---|---|---|---|---|---|---|---|---|---|---|---|---|---|---|
| 10 | 112449441 | rs11815350 | -0.432 | -0.022 | 0.275 | -0.547 | -0.005 | 0.002 | 0.002 | 0 | 0.24 | 0.22 | 0.18 | 0.31 | 0.2 | 0.28 |
| 10 | 112453917 | rs11195266 | -0.184 | -0.028 | 0.298 | -0.55 | 0.014 | -0.029 | -0.029 | 0 | 0.23 | 0.09 | 0.17 | 0.31 | 0.19 | 0.28 |
| 10 | 112457142 | rs17761619 | 0.334 | 0.022 | 0.319 | -0.578 | 0.016 | -0.038 | -0.038 | 0 | 0.07 | 0.13 | 0.09 | 0 | 0.11 | 0.03 |
| 10 | 112459684 | rs4918565 | -0.89 | 0.009 | 0.571 | 0.348 | 0.003 | -0.02 | -0.02 | 0.449 | 0.3 | 0.25 | 0.35 | 0.52 | 0.31 | 0.32 |
| 10 | 112459918 | rs4918567 | -0.811 | 0.634 | 0.499 | 0.253 | 0.059 | 0.041 | 0.041 | 0 | 0.91 | 0.85 | 0.92 | 0.84 | 0.84 | 0.89 |
| 10 | 112461655 | rs12570388 | 0.997 | 0.628 | -0.097 | 0.161 | -0.116 | -0.065 | -0.065 | 1.51 | 0.31 | 0.03 | 0.34 | 0.67 | 0.33 | 0.51 |
| 10 | 112462244 | rs957306 | 0.307 | 0.583 | 0.274 | 0.204 | 0.021 | -0.045 | -0.045 | 2.003 | 0.08 | 0.18 | 0.1 | 0 | 0.11 | 0.04 |
| 10 | 112464824 | rs11195275 | -0.005 | 0.451 | 0.245 | 0.217 | 0.016 | -0.018 | -0.018 | 0 | 0.23 | 0.11 | 0.16 | 0.31 | 0.19 | 0.27 |
| 10 | 112467694 | rs10885023 | -0.958 | 0.566 | 0.234 | 0.21 | -0.001 | -0.001 | -0.001 | 0 | 0.3 | 0.28 | 0.28 | 0.31 | 0.32 | 0.31 |
| 10 | 112468732 | rs12572685 | 1.646 | 0.488 | -0.135 | 0.262 | -0.021 | -0.172 | -0.172 | 0 | 0.22 | 0.02 | 0.27 | 0.49 | 0.15 | 0.38 |
| 10 | 112471349 | rs11592409 | -0.509 | 0.505 | 0.035 | 0.042 | 0.032 | -0.018 | -0.018 | 0.037 | 0.16 | 0.21 | 0.23 | 0.12 | 0.27 | 0.2 |
| 10 | 112472103 | rs10509928 | 0.172 | 0.493 | -0.033 | 0.047 | -0.006 | -0.004 | -0.004 | 0 | 0.09 | 0.04 | 0.12 | 0.12 | 0.13 | 0.15 |
| 10 | 112473422 | rs7350421 | -1.225 | 0.476 | 0.064 | -0.159 | -0.038 | 0.086 | 0.086 | 0 | 0.59 | 0.77 | 0.63 | 1 | 0.49 | 0.78 |
| 10 | 112474572 | rs1341054 | -0.146 | 0.378 | 0.193 | -0.116 | 0.011 | 0.017 | 0.017 | 0 | 0.48 | 0.45 | 0.4 | 0.86 | 0.36 | 0.63 |
| 10 | 112475768 | rs1557030 | -1.449 | 0.215 | 0.002 | -0.062 | -0.008 | 0.06 | 0.06 | 0 | 0.65 | 0.79 | 0.73 | 1 | 0.63 | 0.82 |
| 10 | 112490002 | rs9971359 | -0.711 | 0.264 | -0.051 | 0.228 | 0.025 | 0.141 | 0.141 | 1.664 | 0.52 | 0.76 | 0.69 | 0.8 | 0.59 | 0.7 |
| 10 | 112490536 | rs7072857 | 0.069 | 0.339 | 0.413 | 0.495 | 0.034 | 0.068 | 0.068 | 0 | 0.12 | 0.42 | 0.09 | 0.15 | 0.13 | 0.12 |
| 10 | 112491620 | rs2419577 | 0.774 | 0.616 | -0.398 | 0.422 | -0.047 | -0.096 | -0.096 | 0 | 0.37 | 0.05 | 0.44 | 0.43 | 0.37 | 0.48 |
| 10 | 112493193 | rs4631816 | -0.045 | 0.615 | 0.104 | 0.438 | -0.003 | 0.069 | 0.069 | 1.255 | 0.5 | 0.68 | 0.4 | 0.34 | 0.48 | 0.38 |
| 10 | 112493216 | rs17127885 | 0.259 | 0.586 | 0.494 | 0.567 | 0.036 | 0.034 | 0.034 | 0 | 0.05 | 0.12 | 0.09 | 0.09 | 0.09 | 0.12 |
| 10 | 112494578 | rs12219040 | 0.731 | 0.921 | 0.317 | 0.371 | -0.014 | 0.032 | 0.032 | 0.577 | 0.16 | 0.31 | 0.07 | 0.25 | 0.1 | 0.1 |
| 10 | 112497651 | rs10787274 | -1.409 | 1.007 | 0.156 | 0.376 | -0.045 | -0.127 | -0.127 | 0 | 0.66 | 0.97 | 0.57 | 0.57 | 0.69 | 0.52 |
| 10 | 112498330 | rs10885031 | -0.486 | 0.986 | 0.317 | 0.358 | -0.002 | 0.005 | 0.005 | 0.304 | 0.57 | 0.56 | 0.46 | 0.49 | 0.59 | 0.4 |
| 10 | 112498657 | rs12782675 | 1.297 | 0.944 | 0.472 | 0.055 | -0.007 | -0.017 | -0.017 | 0 | 0.31 | 0.23 | 0.27 | 0 | 0.35 | 0.16 |
| 10 | 112498838 | rs11195297 | 0.409 | 1.005 | 0.337 | 0.088 | 0.014 | -0.093 | -0.093 | 0 | 0.39 | 0.44 | 0.36 | 0.09 | 0.45 | 0.29 |
| 10 | 112499420 | rs884020 | -0.008 | 1.025 | -0.092 | 0.235 | 0 | 0.027 | 0.027 | 2.682 | 0.48 | 0.42 | 0.32 | 0.25 | 0.45 | 0.28 |
| 10 | 112504147 | rs867331 | -0.511 | 0.922 | -0.683 | 0.107 | -0.021 | -0.018 | -0.018 | 0 | 0.34 | 0.2 | 0.36 | 0.54 | 0.41 | 0.39 |
| 10 | 112505876 | rs2000114 | -0.067 | 1.054 | -0.778 | 0.266 | -0.048 | -0.175 | -0.175 | 1.069 | 0.28 | 0.03 | 0.17 | 0.44 | 0.27 | 0.28 |
| 10 | 112509097 | rs1853372 | 0.64 | 0.742 | -0.619 | -0.709 | -0.001 | -0.112 | -0.112 | 0 | 0.42 | 0.56 | 0.33 | 0 | 0.43 | 0.19 |
| 10 | 112509261 | rs10885039 | -0.183 | 0.762 | -0.605 | -0.622 | -0.04 | -0.044 | -0.044 | 0.015 | 0.33 | 0.14 | 0.35 | 0.51 | 0.4 | 0.41 |
| 10 | 112515735 | rs1885791 | -1.001 | 0.649 | -0.411 | -0.424 | -0.004 | 0.013 | 0.013 | 0 | 0.76 | 0.75 | 0.7 | 0.55 | 0.84 | 0.61 |
| 10 | 112519013 | rs942074 | -1.156 | 0.663 | -0.542 | -0.446 | 0.01 | -0.038 | -0.038 | 0 | 0.77 | 0.88 | 0.72 | 0.58 | 0.84 | 0.62 |
| 10 | 112522967 | rs4522094 | 0.385 | 0.786 | -0.176 | -0.47 | -0.001 | 0.002 | 0.002 | 1.473 | 0.26 | 0.22 | 0.33 | 0.48 | 0.28 | 0.44 |
| 10 | 112531616 | rs10787280 | -0.656 | 0.56 | 0.036 | -0.692 | -0.002 | 0.018 | 0.018 | 1.403 | 0.81 | 0.81 | 0.79 | 0.58 | 0.89 | 0.82 |
| 10 | 112532592 | rs17762956 | 1.288 | 0.56 | -0.046 | -0.593 | -0.002 | 0.009 | 0.009 | 0.471 | 0.2 | 0.11 | 0.21 | 0.18 | 0.16 | 0.37 |
| 10 | 112532796 | rs1322998 | -0.197 | 0.774 | -0.152 | -0.941 | 0.008 | 0.009 | 0.009 | 1.268 | 0.34 | 0.36 | 0.48 | 0.53 | 0.42 | 0.52 |
| 10 | 112535077 | rs2078243 | 0.423 | 0.989 | -0.03 | -1.151 | 0.022 | 0.043 | 0.043 | 0 | 0.48 | 0.35 | 0.3 | 0.04 | 0.46 | 0.32 |
| 10 | 112537734 | rs17763100 | 1.149 | 0.808 | -0.148 | -0.618 | -0.008 | 0.046 | 0.046 | 0 | 0.12 | 0.01 | 0.07 | 0.03 | 0.12 | 0.13 |
| 10 | 112537826 | rs12784388 | 0.219 | 1.388 | 0.523 | -0.021 | -0.045 | -0.002 | -0.002 | 0 | 0.08 | 0.02 | 0.14 | 0.07 | 0.16 | 0.08 |
| 10 | 112542865 | rs12254016 | 0.726 | 1.23 | 0.402 | -0.126 | 0.006 | -0.021 | -0.021 | 0 | 0.48 | 0.2 | 0.52 | 0.5 | 0.43 | 0.63 |
| 10 | 112544063 | rs7898438 | 0.499 | 1.391 | 0.659 | -0.619 | 0.006 | -0.041 | -0.041 | 0 | 0.62 | 0.43 | 0.61 | 0.89 | 0.53 | 0.77 |
| 10 | 112544655 | rs7077757 | 0.083 | 1.506 | 0.318 | -0.594 | 0.128 | 0.013 | 0.013 | 0 | 0.09 | 0.2 | 0.21 | 0.08 | 0.22 | 0.1 |
| 10 | 112547206 | rs10787281 | -0.314 | 1.32 | -0.114 | -0.749 | -0.035 | -0.157 | -0.157 | 0.612 | 0.73 | 0.56 | 0.86 | 1 | 0.78 | 0.91 |
| 10 | 112555409 | rs11195327 | 0.915 | 1.527 | -0.558 | -0.36 | 0.007 | -0.019 | -0.019 | 0.094 | 0.15 | 0.14 | 0.26 | 0.51 | 0.15 | 0.2 |
| 10 | 112555472 | rs10885046 | 0.059 | 1.463 | -0.639 | -0.386 | 0.005 | -0.005 | -0.005 | 0 | 0.25 | 0.22 | 0.23 | 0.22 | 0.32 | 0.16 |
| 10 | 112557773 | rs12356300 | 0.542 | 1.066 | -0.831 | -0.619 | 0.021 | 0.056 | 0.056 | 2.527 | 0.49 | 0.26 | 0.38 | 0.4 | 0.41 | 0.6 |
| 10 | 112559414 | rs7086886 | -0.27 | 0.743 | -0.45 | -0.956 | -0.002 | -0.022 | -0.022 | 0 | 0.21 | 0.18 | 0.17 | 0.07 | 0.23 | 0.15 |
| 10 | 112560012 | rs7090878 | -0.313 | 0.726 | -0.504 | -0.687 | -0.005 | -0.052 | -0.052 | 0 | 0.41 | 0.53 | 0.55 | 0.92 | 0.36 | 0.6 |
| 10 | 112562562 | rs11195332 | -1.378 | 0.828 | -0.222 | -0.026 | 0.011 | -0.006 | -0.006 | 0 | 0.09 | 0.16 | 0.09 | 0.07 | 0.1 | 0.1 |
| 10 | 112563268 | rs6585015 | -0.608 | 1.276 | 0.075 | -0.564 | 0.02 | -0.062 | -0.062 | 0 | 0.22 | 0.31 | 0.16 | 0.1 | 0.28 | 0.12 |
| 10 | 112564104 | rs2146548 | 0.047 | 0.793 | -0.406 | -1.084 | -0.003 | -0.037 | -0.037 | 0 | 0.7 | 0.67 | 0.79 | 0.84 | 0.74 | 0.69 |
| 10 | 112564458 | rs11594321 | -0.077 | 0.349 | -0.366 | -0.153 | -0.016 | -0.024 | -0.024 | 0 | 0.24 | 0.14 | 0.27 | 0.38 | 0.32 | 0.22 |
| 10 | 112565606 | rs7068721 | 1.098 | -0.192 | -0.542 | 0.088 | -0.028 | -0.032 | -0.032 | 1.374 | 0.37 | 0.16 | 0.42 | 0.33 | 0.4 | 0.36 |
| 10 | 112570243 | rs11195338 | -0.174 | -0.621 | -0.646 | 0.174 | -0.043 | -0.087 | -0.087 | 0.673 | 0.3 | 0.1 | 0.29 | 0.55 | 0.38 | 0.32 |
| 10 | 112572873 | rs1832745 | 0.658 | -0.845 | -0.724 | 0.077 | -0.019 | -0.013 | -0.013 | 0 | 0.44 | 0.22 | 0.56 | 0.53 | 0.53 | 0.52 |
| 10 | 112575657 | rs11195342 | 1.258 | -0.957 | -0.699 | 0.163 | -0.032 | -0.006 | -0.006 | 0 | 0.37 | 0.24 | 0.56 | 0.52 | 0.45 | 0.35 |
| 10 | 112576695 | rs10749053 | -0.685 | -0.996 | -0.418 | -0.049 | -0.019 | -0.08 | -0.08 | 0 | 0.8 | 0.68 | 0.86 | 0.96 | 0.86 | 0.81 |
| 10 | 112580880 | rs10787283 | 0.168 | -0.989 | -0.275 | 0.247 | 0 | 0.009 | 0.009 | 0.626 | 0.51 | 0.56 | 0.68 | 0.6 | 0.52 | 0.61 |
| 10 | 112584088 | rs12354578 | 1.445 | -0.972 | -0.91 | 0.323 | -0.084 | -0.012 | -0.012 | 0.395 | 0.14 | 0.01 | 0.26 | 0.09 | 0.22 | 0.18 |
| 10 | 112584451 | rs4918600 | 0.324 | -1.239 | -0.086 | 0.434 | 0.001 | 0.02 | 0.02 | 0.418 | 0.49 | 0.4 | 0.67 | 0.61 | 0.53 | 0.54 |
| 10 | 112586096 | rs726803 | 0.392 | -1.043 | -0.148 | -0.108 | 0 | 0.025 | 0.025 | 1.146 | 0.48 | 0.4 | 0.67 | 0.73 | 0.53 | 0.56 |
| 10 | 112596570 | rs874241 | -0.495 | -0.623 | -0.365 | 0.186 | -0.055 | 0.011 | 0.011 | 2.004 | 0.14 | 0.25 | 0.12 | 0.21 | 0.04 | 0.25 |
| 10 | 112602267 | rs7073243 | -1.215 | -0.554 | -0.764 | 0.132 | 0.006 | -0.009 | -0.009 | 0.079 | 0.58 | 0.43 | 0.5 | 0.5 | 0.54 | 0.57 |
| 10 | 112603026 | rs9633680 | -0.53 | 0.21 | -1.284 | 0.228 | -0.072 | 0.078 | 0.078 | 0.158 | 0.38 | 0.11 | 0.35 | 0.31 | 0.45 | 0.32 |
| 10 | 112604028 | rs11195353 | -0.696 | 0.102 | -0.666 | 0.821 | -0.003 | 0 | 0 | 0.082 | 0.26 | 0.17 | 0.25 | 0.34 | 0.23 | 0.2 |
| 10 | 112606495 | rs7919389 | 0.939 | -0.111 | 0.137 | 0.957 | -0.013 | 0.012 | 0.012 | 2.305 | 0.84 | 0.7 | 0.92 | 0.81 | 0.89 | 0.85 |
| 10 | 112607918 | rs17128067 | -0.167 | -0.309 | 0.226 | 1.478 | -0.001 | 0.014 | 0.014 | 0 | 0.11 | 0.1 | 0.05 | 0.23 | 0.1 | 0.12 |
| 10 | 112609805 | rs4918605 | 1.62 | -1.106 | 0.299 | 1.017 | 0.059 | 0.015 | 0.015 | 0.046 | 0.58 | 0.24 | 0.78 | 0.63 | 0.71 | 0.55 |
| 10 | 112609973 | rs4918606 | 1.123 | -1.228 | 0.949 | 0.973 | -0.012 | 0.002 | 0.002 | 0.326 | 0.78 | 0.67 | 0.88 | 0.79 | 0.81 | 0.7 |

| | | | | | | | | | | | | | | | | |
|---|---|---|---|---|---|---|---|---|---|---|---|---|---|---|---|---|
| 10 | 112610839 | rs10885056 | 1.776 | -1.277 | 1.178 | 0.979 | -0.008 | 0.002 | 0.002 | 0 | 0.67 | 0.39 | 0.82 | 0.62 | 0.76 | 0.55 |
| 10 | 112610872 | rs9943402 | 0.817 | -1.215 | 1.735 | 0.879 | 0.008 | 0.032 | 0.032 | 0 | 0.84 | 0.66 | 0.89 | 0.81 | 0.84 | 0.78 |
| 10 | 112611576 | rs1885794 | -0.152 | -1.194 | 1.683 | 0.902 | -0.002 | 0.038 | 0.038 | 0.01 | 0.89 | 0.83 | 0.94 | 0.81 | 0.91 | 0.78 |
| 10 | 112612471 | rs1885796 | 0.032 | -1.207 | 1.782 | 0.837 | -0.012 | -0.04 | -0.04 | 0.304 | 0.92 | 0.73 | 0.96 | 1 | 0.96 | 0.99 |
| 10 | 112612802 | rs4917589 | 1.679 | -1.312 | 1.829 | 0.781 | -0.041 | -0.137 | -0.137 | 5.499 | 0.8 | 0.54 | 0.9 | 0.97 | 0.85 | 0.81 |
| 10 | 112623274 | rs10787286 | -0.857 | -1.284 | 1.548 | 0.8 | 0.007 | 0.02 | 0.02 | 0 | 0.11 | 0.07 | 0.05 | 0.03 | 0.1 | 0.08 |
| 10 | 112633918 | rs11195360 | 2.283 | -1.231 | 1.615 | 1.473 | 0.036 | 0.029 | 0.029 | 0.077 | 0.57 | 0.26 | 0.61 | 0.3 | 0.7 | 0.38 |
| 10 | 112638028 | rs1407696 | 2.284 | -1.317 | 1.628 | 1.511 | 0.054 | 0.032 | 0.032 | 0.151 | 0.57 | 0.24 | 0.62 | 0.3 | 0.72 | 0.38 |
| 10 | 112638504 | rs17128102 | 0.383 | -1.281 | 1.713 | 1.811 | 0 | 0.071 | 0.071 | 0 | 0.05 | 0.07 | 0.2 | 0.36 | 0.03 | 0.17 |
| 10 | 112641053 | rs7081726 | -0.271 | -1.298 | 1.869 | 1.768 | -0.01 | -0.032 | -0.032 | 1.056 | 0.92 | 0.84 | 0.97 | 0.99 | 0.95 | 0.88 |
| 10 | 112647195 | rs10787287 | 1.999 | -1.38 | 1.87 | 1.905 | -0.023 | -0.049 | -0.049 | 2.327 | 0.73 | 0.63 | 0.88 | 0.87 | 0.79 | 0.71 |
| 10 | 112650055 | rs6585021 | 0.98 | -1.355 | 1.884 | 1.917 | -0.016 | -0.016 | -0.016 | 0 | 0.85 | 0.73 | 0.94 | 0.89 | 0.89 | 0.79 |
| 10 | 112653597 | rs1410052 | 0.236 | -1.352 | 1.883 | 1.971 | 0 | 0.053 | 0.053 | 0 | 0.06 | 0.08 | 0.2 | 0.33 | 0.02 | 0.17 |
| 10 | 112657207 | rs10787289 | 2.229 | -1.365 | 1.891 | 2.006 | -0.016 | 0.012 | 0.012 | 0 | 0.68 | 0.58 | 0.69 | 0.53 | 0.76 | 0.54 |
| 10 | 112658060 | rs12248739 | 0.397 | -1.213 | 1.923 | 2.069 | 0 | 0.066 | 0.066 | 0 | 0.05 | 0.07 | 0.2 | 0.35 | 0.03 | 0.17 |
| 10 | 112658964 | rs2146544 | 0.425 | -1.215 | 1.901 | 1.942 | -0.002 | -0.009 | -0.009 | 0.541 | 0.07 | 0.06 | 0.03 | 0.02 | 0.04 | 0.07 |
| 10 | 112664781 | rs10787291 | -0.569 | -1.192 | 1.913 | 1.953 | 0.005 | 0.009 | 0.009 | 0 | 0.11 | 0.08 | 0.05 | 0.03 | 0.1 | 0.08 |
| 10 | 112668381 | rs6585023 | 0.757 | -1.18 | 1.938 | 1.967 | -0.011 | -0.009 | -0.009 | 0.999 | 0.86 | 0.73 | 0.94 | 0.89 | 0.89 | 0.79 |
| 10 | 112678657 | rs7895872 | 2.085 | -1.189 | 2.001 | 2.007 | -0.023 | 0.023 | 0.023 | 0 | 0.69 | 0.54 | 0.68 | 0.53 | 0.77 | 0.53 |
| 10 | 112679036 | rs3750619 | -0.436 | -1.157 | 1.806 | 1.923 | 0.005 | 0.017 | 0.017 | 0 | 0.11 | 0.06 | 0.05 | 0.03 | 0.1 | 0.08 |
| 10 | 112697241 | rs10787292 | 1.651 | -1.149 | 1.818 | 2.092 | -0.011 | -0.017 | -0.017 | 0 | 0.74 | 0.68 | 0.88 | 0.84 | 0.78 | 0.8 |
| 10 | 112714497 | rs7073594 | 0.072 | -1.177 | 1.829 | 2.162 | -0.01 | 0.058 | 0.058 | 1.49 | 0.11 | 0.18 | 0.06 | 0.23 | 0.05 | 0.15 |
| 10 | 112721408 | rs1327556 | 0.344 | -1.174 | 1.753 | 2.107 | -0.007 | -0.027 | -0.027 | 0 | 0.07 | 0.13 | 0.03 | 0.01 | 0.05 | 0.12 |
| 10 | 112727826 | rs11818771 | 0.537 | -1.173 | 1.753 | 2.108 | -0.005 | -0.024 | -0.024 | 0 | 0.07 | 0.11 | 0.03 | 0 | 0.05 | 0.01 |
| 10 | 112741551 | rs9703807 | 2.088 | -1.25 | 1.842 | 2.164 | 0.023 | -0.014 | -0.014 | 1.335 | 0.68 | 0.27 | 0.64 | 0.38 | 0.81 | 0.48 |
| 10 | 112746735 | rs1927159 | 1.943 | -1.251 | 1.928 | 2.174 | -0.024 | 0.022 | 0.022 | 0 | 0.73 | 0.59 | 0.69 | 0.62 | 0.82 | 0.63 |
| 10 | 112752377 | rs10885076 | 0.962 | -1.25 | 1.912 | 2.155 | -0.013 | -0.027 | -0.027 | 0 | 0.86 | 0.82 | 0.93 | 0.95 | 0.89 | 0.92 |
| 10 | 112753721 | rs6585026 | 0.72 | -1.249 | 1.913 | 2.156 | -0.011 | -0.023 | -0.023 | 1.681 | 0.87 | 0.83 | 0.94 | 0.95 | 0.89 | 0.91 |
| 10 | 112763385 | rs11195400 | 2.552 | -1.249 | 1.881 | 2.193 | 0.065 | 0.088 | 0.088 | 0.095 | 0.56 | 0.16 | 0.6 | 0.28 | 0.71 | 0.37 |
| 10 | 112767049 | rs11816033 | 0.555 | -1.213 | 1.73 | 2.141 | -0.005 | -0.023 | -0.023 | 0 | 0.07 | 0.11 | 0.03 | 0 | 0.05 | 0.01 |
| 10 | 112773079 | rs1327552 | 0.038 | -1.201 | 1.697 | 2.181 | 0 | 0.054 | 0.054 | 0 | 0.06 | 0.09 | 0.21 | 0.33 | 0.02 | 0.17 |
| 10 | 112784182 | rs12412661 | 1.896 | -1.201 | 1.794 | 2.239 | -0.012 | 0.044 | 0.044 | 0 | 0.78 | 0.7 | 0.74 | 0.51 | 0.82 | 0.6 |
| 10 | 112795021 | rs10732804 | -0.11 | -1.205 | 1.812 | 2.239 | -0.001 | 0.019 | 0.019 | 0 | 0.93 | 0.9 | 0.97 | 0.9 | 0.94 | 0.89 |
| 10 | 112799928 | rs7085088 | 1.742 | -1.188 | 1.77 | 2.191 | -0.012 | 0.044 | 0.044 | 0 | 0.78 | 0.7 | 0.74 | 0.51 | 0.82 | 0.6 |
| 10 | 112831994 | rs11195417 | -1.11 | -1.295 | 1.859 | 2.226 | -0.02 | 0.009 | 0.009 | 1.032 | 0.17 | 0.3 | 0.08 | 0.19 | 0.1 | 0.16 |
| 10 | 112832225 | rs17128356 | 0.128 | -1.295 | 1.871 | 2.224 | -0.012 | 0.03 | 0.03 | 1.672 | 0.1 | 0.17 | 0.05 | 0.19 | 0.05 | 0.15 |
| 10 | 112833561 | rs7096359 | -0.447 | -1.323 | 1.826 | 2.213 | 0.008 | 0.012 | 0.012 | 0 | 0.11 | 0.07 | 0.06 | 0.06 | 0.1 | 0.09 |
| 10 | 112835109 | rs12246561 | -0.447 | -1.323 | 1.812 | 2.213 | 0.009 | 0.014 | 0.014 | 0 | 0.11 | 0.07 | 0.06 | 0.06 | 0.1 | 0.09 |
| 10 | 112839368 | rs11195419 | -1.251 | -1.316 | 1.778 | 2.142 | -0.019 | 0.007 | 0.007 | 1.15 | 0.18 | 0.28 | 0.1 | 0.19 | 0.11 | 0.18 |
| 10 | 112839579 | rs553668 | 1.017 | -1.303 | 1.776 | 2.226 | -0.017 | 0.046 | 0.046 | 0 | 0.78 | 0.69 | 0.74 | 0.49 | 0.84 | 0.6 |
| 10 | 112843085 | rs602618 | 1.787 | -1.313 | 1.565 | 2.102 | 0.037 | 0.012 | 0.012 | 0.202 | 0.59 | 0.29 | 0.64 | 0.3 | 0.73 | 0.42 |
| 10 | 112846298 | rs745557 | 1.43 | -0.324 | 1.67 | 2.023 | -0.005 | -0.004 | -0.004 | 0 | 0.67 | 0.44 | 0.61 | 0.28 | 0.7 | 0.43 |
| 10 | 112846502 | rs7905613 | -0.018 | -0.348 | 1.528 | 2.182 | -0.009 | 0.039 | 0.039 | 2.948 | 0.08 | 0.16 | 0.07 | 0.15 | 0.04 | 0.12 |
| 10 | 112856320 | rs7923122 | -0.199 | -0.262 | 0.516 | 1.656 | -0.01 | -0.037 | -0.037 | 0 | 0.08 | 0.14 | 0.03 | 0 | 0.05 | 0.01 |
| 10 | 112856425 | rs7908645 | -1.645 | -0.233 | 0.12 | 1.735 | 0.03 | 0.046 | 0.046 | 0.331 | 0.41 | 0.83 | 0.4 | 0.72 | 0.33 | 0.57 |
| 10 | 112860391 | rs1343451 | -1.391 | -0.237 | -0.182 | 1.677 | 0.019 | -0.009 | -0.009 | 0 | 0.4 | 0.74 | 0.31 | 0.41 | 0.3 | 0.51 |
| 10 | 112862029 | rs945333 | -1.073 | -0.129 | -0.723 | 1.225 | -0.001 | 0.066 | 0.066 | 0.273 | 0.51 | 0.77 | 0.5 | 0.68 | 0.48 | 0.71 |
| 10 | 112863991 | rs10787306 | 0.943 | -0.573 | -0.103 | 0.912 | -0.058 | -0.038 | -0.038 | 3.388 | 0.74 | 0.45 | 0.86 | 0.85 | 0.85 | 0.65 |
| 10 | 112865677 | rs17128407 | -0.666 | -0.17 | -0.601 | 0.926 | -0.012 | -0.01 | -0.01 | 0 | 0.07 | 0 | 0.04 | 0.07 | 0.06 | 0.2 |
| 10 | 112867219 | rs4918625 | 0.765 | -0.026 | -2.087 | 0.521 | -0.115 | -0.067 | -0.067 | 0 | 0.37 | 0.06 | 0.32 | 0.38 | 0.45 | 0.3 |
| 10 | 112867412 | rs1556716 | -0.593 | -0.08 | -1.607 | 0.32 | 0.011 | 0.1 | 0.1 | 0.285 | 0.79 | 0.43 | 0.7 | 0.65 | 0.79 | 0.76 |
| 10 | 112871088 | rs1537769 | 0.244 | -1.044 | -1.294 | 0.212 | -0.002 | 0.011 | 0.011 | 0 | 0.22 | 0.16 | 0.31 | 0.18 | 0.24 | 0.23 |
| 10 | 112872557 | rs7897445 | -0.759 | -1.145 | -1.211 | 0.116 | 0.007 | 0.018 | 0.018 | 0.814 | 0.35 | 0.3 | 0.35 | 0.18 | 0.3 | 0.28 |
| 10 | 112875595 | rs10885089 | 0.324 | -0.979 | -1.268 | 0.207 | -0.005 | 0.013 | 0.013 | 0 | 0.22 | 0.15 | 0.31 | 0.18 | 0.25 | 0.23 |
| 10 | 112877102 | rs1537770 | -2.397 | -0.991 | -1.767 | 0.085 | -0.133 | 0.065 | 0.065 | 1.461 | 0.66 | 0.95 | 0.69 | 0.83 | 0.62 | 0.71 |
| 10 | 112879081 | rs2032023 | -1.913 | -1.239 | -1.53 | 0.37 | -0.023 | -0.025 | -0.025 | 0 | 0.55 | 0.93 | 0.59 | 0.62 | 0.41 | 0.65 |
| 10 | 112882768 | rs7079291 | 1.49 | -1.185 | -1.476 | 0.301 | 0.006 | 0.039 | 0.039 | 0.311 | 0.15 | 0.08 | 0.08 | 0.07 | 0.14 | 0.07 |
| 10 | 112883662 | rs11819365 | 1.13 | -1.086 | -0.662 | 0.246 | -0.014 | -0.045 | -0.045 | 0 | 0.08 | 0.21 | 0.03 | 0 | 0.03 | 0.01 |
| 10 | 112884869 | rs17128431 | 0.845 | -0.91 | -0.976 | 0.378 | 0.004 | 0.006 | 0.006 | 0.047 | 0.06 | 0.08 | 0.22 | 0.11 | 0.11 | 0.15 |
| 10 | 112887549 | rs945335 | -1.324 | -1.2 | -0.991 | 0.23 | 0.097 | -0.034 | -0.034 | 0 | 0.47 | 0.8 | 0.37 | 0.51 | 0.31 | 0.51 |
| 10 | 112897070 | rs7069564 | -0.583 | -0.989 | -0.705 | 0.439 | 0.001 | -0.002 | -0.002 | 0 | 0.17 | 0.13 | 0.26 | 0.38 | 0.15 | 0.3 |
| 10 | 112901012 | rs6585037 | 0.572 | -1.206 | -0.49 | 0.371 | 0.003 | -0.002 | -0.002 | 0.303 | 0.21 | 0.18 | 0.16 | 0.04 | 0.2 | 0.14 |
| 10 | 112903806 | rs954385 | 0.695 | -1.198 | -0.461 | 0.436 | 0.004 | 0.074 | 0.074 | 0 | 0.07 | 0.08 | 0.21 | 0.3 | 0.1 | 0.13 |
| 10 | 112908968 | rs10787308 | -0.482 | -1.223 | -0.47 | 0.381 | 0.014 | 0.026 | 0.026 | 0 | 0.31 | 0.23 | 0.21 | 0.11 | 0.26 | 0.29 |
| 10 | 112909105 | rs869244 | -1.024 | -1.168 | -0.487 | 0.424 | 0.004 | -0.002 | -0.002 | 2.293 | 0.38 | 0.32 | 0.41 | 0.41 | 0.35 | 0.44 |
| 10 | 112909569 | rs11195438 | 0.422 | -0.99 | -0.663 | 0.459 | 0.006 | 0.011 | 0.011 | 0 | 0.11 | 0.03 | 0.08 | 0.08 | 0.09 | 0.21 |
| 10 | 112918176 | rs1335704 | -1.576 | -1.089 | -0.789 | 0.363 | 0.018 | 0.136 | 0.136 | 2.32 | 0.52 | 0.85 | 0.62 | 0.74 | 0.49 | 0.49 |
| 10 | 112918429 | rs7910277 | 0.425 | -0.913 | -0.284 | 0.467 | -0.003 | 0.21 | 0.21 | 0.054 | 0.12 | 0.33 | 0.2 | 0.4 | 0.12 | 0.19 |
| 10 | 112918573 | rs1335703 | -2.045 | -0.883 | -0.455 | 0.416 | -0.112 | 0.106 | 0.106 | 1.279 | 0.62 | 0.93 | 0.66 | 0.81 | 0.54 | 0.66 |

| | | | | | | | | | | | | | | | | |
|---|---|---|---|---|---|---|---|---|---|---|---|---|---|---|---|---|
| 10 | 112922409 | rs4545476 | -1.275 | -0.944 | -0.319 | 0.446 | 0.023 | 0.019 | 0.019 | 0 | 0.5 | 0.59 | 0.46 | 0.41 | 0.42 | 0.47 |
| 10 | 112923191 | rs11195446 | -0.577 | -0.694 | -0.54 | 0.505 | -0.015 | 0.081 | 0.081 | 0.342 | 0.25 | 0.56 | 0.26 | 0.41 | 0.24 | 0.26 |
| 10 | 112923371 | rs7908674 | -1.695 | -0.298 | -1.04 | 0.321 | -0.09 | 0.095 | 0.095 | 0.877 | 0.63 | 0.88 | 0.66 | 0.81 | 0.59 | 0.67 |
| 10 | 112923456 | rs7897248 | -1.742 | -0.298 | -1.216 | 0.321 | -0.14 | 0.089 | 0.089 | 0.845 | 0.63 | 0.94 | 0.67 | 0.81 | 0.59 | 0.67 |
| 10 | 112929753 | rs10885102 | -1.444 | -1.462 | -0.722 | 0.827 | 0.089 | -0.016 | -0.016 | 0 | 0.49 | 0.79 | 0.55 | 0.55 | 0.38 | 0.62 |
| 10 | 112930377 | rs7084236 | -2.109 | -1.129 | -0.993 | 0.879 | -0.162 | -0.079 | -0.079 | 0.009 | 0.67 | 0.94 | 0.66 | 0.7 | 0.54 | 0.76 |
| 10 | 112931645 | rs11195450 | -0.163 | -1.186 | -0.116 | 1.061 | -0.029 | 0.01 | 0.01 | 0 | 0.08 | 0.02 | 0.09 | 0.04 | 0.15 | 0.13 |
| 10 | 112937459 | rs7079973 | -0.772 | -1.49 | 0.29 | 1.09 | -0.042 | 0.086 | 0.086 | 0.802 | 0.19 | 0.56 | 0.23 | 0.34 | 0.14 | 0.23 |
| 10 | 112942838 | rs6585041 | -1.298 | -1.506 | 0.069 | 1.087 | -0.02 | 0.028 | 0.028 | 3.805 | 0.23 | 0.69 | 0.42 | 0.44 | 0.15 | 0.29 |
| 10 | 112944491 | rs4917596 | -0.192 | -1.2 | 0.352 | 1.075 | 0 | 0.01 | 0.01 | 0 | 0.14 | 0.16 | 0.24 | 0.34 | 0.18 | 0.2 |
| 10 | 112947099 | rs17128526 | -1.165 | -1.528 | 0.266 | 0.892 | 0.012 | 0.008 | 0.008 | 0 | 0.3 | 0.2 | 0.28 | 0.24 | 0.22 | 0.47 |
| 10 | 112949684 | rs4417203 | -1.184 | -1.546 | 0.357 | 0.898 | -0.004 | -0.003 | -0.003 | 1.678 | 0.3 | 0.25 | 0.28 | 0.24 | 0.22 | 0.46 |
| 10 | 112954345 | rs11195465 | 0.196 | -1.29 | -0.255 | 1.056 | -0.003 | 0.005 | 0.005 | 0 | 0.13 | 0.13 | 0.24 | 0.34 | 0.18 | 0.11 |
| 10 | 112954412 | rs10787309 | -1.685 | -1.133 | -0.274 | 0.903 | 0.048 | 0.003 | 0.003 | 0.18 | 0.46 | 0.76 | 0.56 | 0.57 | 0.39 | 0.55 |
| 10 | 112962828 | rs12245968 | -0.527 | -0.687 | 0.126 | 0.931 | -0.003 | -0.005 | -0.005 | 0 | 0.07 | 0.11 | 0.03 | 0.06 | 0.07 | 0.18 |
| 10 | 112963152 | rs945332 | -1.273 | -0.804 | 0.199 | 1.056 | -0.002 | 0.001 | 0.001 | 2.501 | 0.34 | 0.63 | 0.47 | 0.38 | 0.34 | 0.42 |
| 10 | 112968165 | rs7073332 | -1.497 | -1.039 | 0.341 | 1.146 | -0.014 | 0.1 | 0.1 | 0 | 0.07 | 0.18 | 0.11 | 0.2 | 0.04 | 0.11 |
| 10 | 112968436 | rs7073296 | -1.495 | -1.034 | 0.344 | 1.221 | -0.014 | 0.126 | 0.126 | 0 | 0.07 | 0.18 | 0.11 | 0.24 | 0.04 | 0.11 |
| 10 | 112968779 | rs12258397 | -0.186 | -0.966 | 0.108 | 0.927 | 0.001 | -0.004 | -0.004 | 4.852 | 0.07 | 0.08 | 0.14 | 0.06 | 0.08 | 0.21 |
| 10 | 112969627 | rs1537768 | -1.639 | -1.587 | 0.281 | 0.777 | -0.014 | 0.012 | 0.012 | 0.99 | 0.39 | 0.63 | 0.42 | 0.48 | 0.34 | 0.47 |
| 10 | 112976584 | rs4307652 | -1.255 | -1.71 | 0.487 | 0.78 | 0.001 | -0.009 | -0.009 | 0 | 0.25 | 0.29 | 0.33 | 0.4 | 0.28 | 0.31 |
| 10 | 112978950 | rs12221264 | 2.041 | -1.818 | 0.257 | 0.835 | -0.019 | -0.046 | -0.046 | 0 | 0.08 | 0.01 | 0.18 | 0.31 | 0.08 | 0.15 |
| 10 | 112979082 | rs17775480 | 0.59 | -1.318 | 0.779 | 0.264 | 0.012 | 0.013 | 0.013 | 0.108 | 0.11 | 0.06 | 0.07 | 0.05 | 0.08 | 0.16 |
| 10 | 112985425 | rs1360864 | -2.105 | -1.412 | 0.827 | 0.173 | 0.002 | -0.004 | -0.004 | 0 | 0.4 | 0.42 | 0.42 | 0.47 | 0.36 | 0.53 |
| 10 | 112986039 | rs1832112 | -2.557 | -1.486 | 1.062 | 0.14 | 0.046 | -0.022 | -0.022 | 0 | 0.44 | 0.79 | 0.44 | 0.48 | 0.36 | 0.56 |
| 10 | 112986661 | rs12244315 | -0.738 | -1.044 | 1.145 | 0.441 | 0 | 0.028 | 0.028 | 0 | 0.09 | 0.11 | 0.12 | 0.33 | 0.1 | 0.21 |
| 10 | 112991416 | rs12778878 | 1.805 | -0.745 | 1.033 | 0.117 | -0.009 | 0.016 | 0.016 | 14.24 | 0.06 | 0.03 | 0.04 | 0 | 0.08 | 0.01 |
| 10 | 112994252 | rs1335714 | -2.707 | -0.917 | 1.051 | 0.333 | 0.063 | -0.099 | -0.099 | 0 | 0.46 | 0.92 | 0.45 | 0.48 | 0.36 | 0.49 |
| 10 | 112994448 | rs17775850 | -1.599 | -0.934 | 1.097 | 0.336 | 0.007 | -0.015 | -0.015 | 0 | 0.36 | 0.32 | 0.4 | 0.47 | 0.35 | 0.47 |
| 10 | 112996845 | rs2256511 | -2.459 | -0.894 | 0.957 | 0.352 | 0.079 | -0.074 | -0.074 | 0.309 | 0.44 | 0.87 | 0.44 | 0.48 | 0.35 | 0.49 |
| 10 | 113011038 | rs4508142 | -1.941 | -0.845 | 0.9 | 0.347 | -0.015 | 0.017 | 0.017 | 0.541 | 0.38 | 0.48 | 0.4 | 0.48 | 0.35 | 0.46 |
| 10 | 113014137 | rs11599394 | 0.398 | -0.85 | 0.574 | 0.224 | 0.015 | 0.015 | 0.015 | 0.005 | 0.11 | 0.06 | 0.07 | 0.05 | 0.08 | 0.16 |
| 10 | 113022555 | rs10885117 | 1.393 | -0.896 | 0.557 | 0.213 | 0.016 | 0.015 | 0.015 | 0.623 | 0.75 | 0.25 | 0.86 | 0.93 | 0.91 | 0.82 |
| 10 | 113032726 | rs10787315 | 1.326 | -0.885 | 0.595 | 0.218 | -0.002 | -0.003 | -0.003 | 0 | 0.76 | 0.27 | 0.87 | 0.93 | 0.91 | 0.82 |
| 10 | 113036693 | rs10509936 | 2.095 | -0.579 | 0.423 | 0.291 | 0.019 | -0.087 | -0.087 | 0.662 | 0.54 | 0.07 | 0.55 | 0.54 | 0.63 | 0.52 |
| 10 | 113040471 | rs10749064 | 2.112 | -0.653 | 1.071 | 0.113 | 0.053 | 0.04 | 0.04 | 0.061 | 0.69 | 0.2 | 0.81 | 0.79 | 0.81 | 0.63 |
| 10 | 113040493 | rs10787318 | 1.798 | -0.663 | 1.175 | 0.109 | -0.002 | 0.041 | 0.041 | 0 | 0.86 | 0.57 | 0.92 | 0.83 | 0.9 | 0.8 |
| 10 | 113040544 | rs10749065 | 1.901 | -0.619 | 0.535 | 0.129 | 0.053 | 0.04 | 0.04 | 0.061 | 0.69 | 0.2 | 0.81 | 0.79 | 0.81 | 0.63 |
| 10 | 113042093 | rs10885122 | 1.535 | -0.733 | 0.56 | 0.146 | 0.034 | 0.038 | 0.038 | 0.539 | 0.75 | 0.22 | 0.85 | 0.93 | 0.88 | 0.8 |
| 10 | 113048223 | rs7084296 | 1.94 | -0.617 | 0.468 | 0.027 | 0.054 | 0.075 | 0.075 | 0 | 0.64 | 0.21 | 0.8 | 0.8 | 0.76 | 0.54 |
| 10 | 113048321 | rs10885123 | -0.875 | -0.344 | 0.129 | 0.185 | -0.014 | -0.036 | -0.036 | 0 | 0.19 | 0.24 | 0.1 | 0.05 | 0.12 | 0.26 |
| 10 | 113048807 | rs12256243 | -0.544 | 0.189 | 0.06 | 0.745 | -0.023 | -0.032 | -0.032 | 0 | 0.06 | 0.01 | 0.03 | 0.14 | 0.07 | 0.17 |
| 10 | 113059322 | rs3885682 | 1.438 | -0.565 | -0.39 | 0.553 | 0.092 | -0.024 | -0.024 | 0.15 | 0.64 | 0.13 | 0.63 | 0.6 | 0.79 | 0.6 |
| 10 | 113060979 | rs10787327 | -0.082 | -0.707 | -0.455 | 0.547 | 0.001 | 0.007 | 0.007 | 0 | 0.11 | 0.01 | 0.17 | 0.1 | 0.1 | 0.13 |
| 10 | 113061132 | rs17128709 | -0.418 | -0.79 | -0.072 | 0.581 | -0.007 | 0.01 | 0.01 | 0 | 0.08 | 0.1 | 0.02 | 0.17 | 0.06 | 0.2 |
| 10 | 113062340 | rs4617520 | 1.183 | -0.811 | 0.855 | 0.545 | -0.033 | 0.018 | 0.018 | 0 | 0.81 | 0.65 | 0.73 | 0.77 | 0.88 | 0.84 |
| 10 | 113068012 | rs10885127 | -1.059 | -0.722 | 0.452 | 0.574 | 0.019 | -0.021 | -0.021 | 2.971 | 0.18 | 0.12 | 0.26 | 0.24 | 0.13 | 0.16 |
| 10 | 113068593 | rs11195514 | -1.226 | -0.66 | 0.185 | 0.376 | 0.002 | -0.04 | -0.04 | 0 | 0.22 | 0.05 | 0.3 | 0.23 | 0.2 | 0.17 |
| 10 | 113077024 | rs7912918 | -0.326 | -0.395 | 0.667 | 0.699 | 0.044 | 0.013 | 0.013 | 0.598 | 0.07 | 0.38 | 0.05 | 0 | 0.01 | 0.02 |
| 10 | 113077472 | rs6585063 | -2.603 | -0.42 | 0.042 | 0.741 | 0.037 | -0.005 | -0.005 | 0.001 | 0.36 | 0.77 | 0.37 | 0.4 | 0.23 | 0.4 |
| 10 | 113079536 | rs4918639 | 0.76 | -0.331 | 0.116 | 0.794 | -0.012 | 0.038 | 0.038 | 0 | 0.89 | 0.7 | 0.94 | 0.83 | 0.91 | 0.78 |
| 10 | 113097146 | rs12268412 | -0.191 | -0.364 | -0.296 | 0.749 | -0.006 | 0 | 0 | 0.008 | 0.11 | 0.01 | 0.17 | 0.11 | 0.11 | 0.13 |
| 10 | 113100630 | rs4348827 | -2.283 | -0.433 | 0.018 | 0.783 | -0.042 | -0.004 | -0.004 | 0 | 0.24 | 0.6 | 0.3 | 0.24 | 0.14 | 0.17 |
| 10 | 113102038 | rs10787335 | 2.149 | 0.203 | -0.218 | 1.048 | -0.015 | 0.02 | 0.02 | 0 | 0.72 | 0.65 | 0.68 | 0.59 | 0.76 | 0.61 |
| 10 | 113104470 | rs7903217 | 2.041 | 0.28 | -0.732 | 1.081 | 0 | -0.001 | -0.001 | 1.565 | 0.73 | 0.75 | 0.68 | 0.59 | 0.76 | 0.61 |
| 10 | 113106287 | rs11195534 | 0.174 | 1.164 | -1.618 | 1.508 | -0.072 | -0.05 | -0.05 | 0 | 0.15 | 0.01 | 0.13 | 0.16 | 0.16 | 0.12 |
| 10 | 113109415 | rs10885140 | -0.492 | 0.858 | -0.947 | 1.506 | 0.003 | 0.005 | 0.005 | 0 | 0.1 | 0.01 | 0.06 | 0.09 | 0.09 | 0.11 |
| 10 | 113111516 | rs4451642 | 0.832 | 0.842 | -0.911 | 1.472 | -0.001 | 0 | 0 | 0 | 0.9 | 0.92 | 0.94 | 0.83 | 0.88 | 0.76 |
| 10 | 113115245 | rs12573790 | -1.066 | 0.619 | -0.876 | 1.316 | 0.005 | -0.058 | -0.058 | 0.385 | 0.13 | 0.02 | 0.24 | 0.24 | 0.12 | 0.15 |
| 10 | 113117827 | rs7082000 | -1.135 | 1.257 | -0.423 | 1.327 | 0.028 | 0.045 | 0.045 | 0 | 0.13 | 0.14 | 0.21 | 0.34 | 0.22 | 0.21 |
| 10 | 113118855 | rs10749075 | -0.22 | 0.318 | -1.178 | 0.571 | -0.078 | 0.087 | 0.087 | 0.036 | 0.43 | 0.16 | 0.66 | 0.74 | 0.49 | 0.54 |
| 10 | 113120504 | rs10509940 | 0.998 | -0.295 | -0.526 | 0.173 | -0.007 | 0.022 | 0.022 | 1.004 | 0.35 | 0.28 | 0.23 | 0.21 | 0.42 | 0.32 |
| 10 | 113121719 | rs7075550 | -0.65 | -0.515 | 0.134 | 0.127 | -0.088 | -0.119 | -0.119 | 0 | 0.16 | 0.42 | 0.05 | 0 | 0.03 | 0.08 |
| 10 | 113124248 | rs7910390 | 0.424 | -0.587 | -0.581 | 0.159 | -0.011 | -0.012 | -0.012 | 0 | 0.21 | 0.11 | 0.18 | 0.26 | 0.21 | 0.24 |
| 10 | 113127257 | rs10509941 | -0.537 | -0.845 | -0.212 | 0 | -0.019 | 0.157 | 0.157 | 0.325 | 0.47 | 0.22 | 0.67 | 0.77 | 0.51 | 0.58 |
| 10 | 113129190 | rs11195546 | -0.418 | -0.813 | 0.121 | 0.003 | -0.013 | 0.109 | 0.109 | 0.595 | 0.46 | 0.27 | 0.67 | 0.77 | 0.49 | 0.53 |
| 10 | 113132884 | rs7895049 | -0.477 | -0.847 | 0.36 | -0.006 | -0.001 | 0.01 | 0.01 | 0.161 | 0.46 | 0.47 | 0.69 | 0.77 | 0.49 | 0.54 |
| 10 | 113135667 | rs7083831 | -0.407 | -0.859 | 0.405 | -0.011 | -0.007 | 0.075 | 0.075 | 0.684 | 0.46 | 0.33 | 0.67 | 0.77 | 0.49 | 0.54 |
| 10 | 113138945 | rs11195550 | 0.274 | -0.863 | 0.679 | -0.056 | 0.001 | 0.023 | 0.023 | 0.188 | 0.52 | 0.37 | 0.3 | 0.23 | 0.5 | 0.46 |

| | | | | | | | | | | | | | | | |
|---|---|---|---|---|---|---|---|---|---|---|---|---|---|---|---|
| 10 | 113139169 | rs7098615 | -1.677 | -1.032 | 0.563 | -0.159 | 0.006 | -0.025 | -0.025 | 0 | 0.65 | 0.6 | 0.74 | 0.79 | 0.57 | 0.7 |
| 10 | 113139813 | rs10787345 | 0.395 | -1.111 | 0.513 | -0.029 | 0.001 | -0.007 | -0.007 | 2.236 | 0.62 | 0.64 | 0.46 | 0.53 | 0.66 | 0.58 |
| 10 | 113140675 | rs10885145 | 0.495 | -1.138 | 0.49 | 0.063 | -0.101 | -0.158 | -0.158 | 0.1 | 0.26 | 0.05 | 0.5 | 0.46 | 0.32 | 0.34 |
| 10 | 113144540 | rs10885147 | -0.675 | -1.188 | 0.56 | 0.024 | 0.003 | -0.007 | -0.007 | 0.426 | 0.4 | 0.38 | 0.55 | 0.47 | 0.35 | 0.42 |
| 10 | 113151980 | rs10885154 | 0.799 | -1.198 | 0.544 | 0.08 | -0.014 | -0.018 | -0.018 | 0 | 0.18 | 0.12 | 0.14 | 0.22 | 0.17 | 0.22 |
| 10 | 113152964 | rs7905487 | 0.361 | -1.029 | 0.3 | 0.152 | 0.005 | -0.012 | -0.012 | 0 | 0.6 | 0.66 | 0.45 | 0.52 | 0.65 | 0.57 |
| 10 | 113155506 | rs7917960 | -0.076 | -0.965 | 0.278 | 0.164 | -0.017 | -0.041 | -0.041 | 0 | 0.26 | 0.5 | 0.18 | 0.2 | 0.19 | 0.24 |
| 10 | 113158514 | rs4128245 | 1.122 | -1.091 | -0.363 | 0.249 | -0.071 | 0.015 | 0.015 | 6.414 | 0.39 | 0.12 | 0.3 | 0.34 | 0.51 | 0.39 |
| 10 | 113166617 | rs9420082 | 0.19 | -0.999 | -0.303 | 0.334 | 0.004 | 0.004 | 0.004 | 1.243 | 0.66 | 0.29 | 0.81 | 0.79 | 0.82 | 0.69 |
| 10 | 113170248 | rs9421093 | -0.307 | -0.4 | -0.005 | 0.666 | -0.028 | -0.015 | -0.015 | 0 | 0.08 | 0.21 | 0.03 | 0.02 | 0.02 | 0.06 |
| 10 | 113173522 | rs10787351 | -0.213 | -0.305 | -0.34 | 0.734 | -0.047 | -0.072 | -0.072 | 0 | 0.26 | 0.12 | 0.51 | 0.45 | 0.32 | 0.31 |
| 10 | 113180191 | rs4269860 | 0.191 | -0.2 | -0.626 | 0.616 | -0.117 | -0.157 | -0.157 | 1.318 | 0.22 | 0.03 | 0.49 | 0.45 | 0.32 | 0.29 |
| 10 | 113185276 | rs4077365 | 0.289 | 0.126 | -0.299 | 0.634 | 0.003 | 0.012 | 0.012 | 0 | 0.61 | 0.6 | 0.42 | 0.52 | 0.62 | 0.57 |
| 10 | 113194673 | rs4342958 | -0.389 | -0.177 | -0.284 | 0.379 | -0.026 | -0.038 | -0.038 | 1.025 | 0.08 | 0.13 | 0.02 | 0 | 0.02 | 0.08 |
| 10 | 113194854 | rs4590798 | 0.139 | -0.272 | -0.208 | 0.431 | 0.002 | 0.013 | 0.013 | 0 | 0.61 | 0.6 | 0.42 | 0.52 | 0.62 | 0.57 |
| 10 | 113198652 | rs7898841 | -0.305 | -0.675 | 0.298 | 0.334 | -0.021 | -0.038 | -0.038 | 0 | 0.69 | 0.81 | 0.45 | 0.53 | 0.64 | 0.64 |
| 10 | 113206374 | rs4593944 | 1.433 | -1.063 | 0.101 | 0.112 | -0.063 | 0.001 | 0.001 | 2.063 | 0.35 | 0.1 | 0.26 | 0.32 | 0.45 | 0.32 |
| 10 | 113221195 | rs9421114 | -0.652 | -1.428 | 0.284 | 0.025 | -0.002 | 0 | 0 | 0 | 0.06 | 0.23 | 0.03 | 0.01 | 0.01 | 0.02 |
| 10 | 113225805 | rs1914140 | -0.947 | -1.473 | 0.286 | 0.092 | 0.018 | 0.016 | 0.016 | 2.305 | 0.32 | 0.74 | 0.18 | 0.23 | 0.17 | 0.27 |
| 10 | 113227145 | rs3125478 | 0.141 | -1.506 | 0.14 | 0.073 | -0.012 | -0.01 | -0.01 | 0 | 0.89 | 0.87 | 0.95 | 0.97 | 0.94 | 0.89 |
| 10 | 113230440 | rs10885194 | 0.862 | -1.532 | 0.518 | 0.056 | -0.004 | -0.001 | -0.001 | 0.742 | 0.17 | 0.26 | 0.12 | 0.2 | 0.15 | 0.17 |
| 10 | 113231124 | rs4372376 | -0.031 | -1.582 | 0.509 | 0.033 | -0.026 | -0.023 | -0.023 | 0 | 0.23 | 0.41 | 0.15 | 0.2 | 0.15 | 0.23 |
| 10 | 113232410 | rs4917610 | 1.065 | -1.6 | 0.481 | 0.055 | 0.051 | 0.05 | 0.05 | 0 | 0.63 | 0.21 | 0.78 | 0.77 | 0.79 | 0.69 |
| 10 | 113240800 | rs11195604 | 0.799 | -1.519 | 0.828 | 0.095 | -0.007 | 0 | 0 | 0.319 | 0.17 | 0.31 | 0.13 | 0.21 | 0.14 | 0.17 |
| 10 | 113245619 | rs3107358 | -0.182 | -0.999 | 0.691 | 0.11 | 0.025 | 0.068 | 0.068 | 0 | 0.29 | 0.46 | 0.52 | 0.43 | 0.32 | 0.31 |
| 10 | 113248666 | rs11527950 | 0.761 | -1.038 | 0.67 | 0.107 | 0 | 0.004 | 0.004 | 0 | 0.6 | 0.46 | 0.4 | 0.52 | 0.63 | 0.56 |
| 10 | 113248711 | rs3107354 | -0.104 | -1.051 | 0.627 | 0.1 | -0.004 | -0.001 | -0.001 | 0.06 | 0.92 | 0.93 | 0.91 | 0.95 | 0.96 | 0.91 |
| 10 | 113261163 | rs3107346 | -0.157 | -1.099 | 0.642 | 0.097 | 0.003 | 0.008 | 0.008 | 0.01 | 0.29 | 0.29 | 0.5 | 0.42 | 0.33 | 0.34 |
| 10 | 113261732 | rs6585082 | 1.154 | -1.075 | 0.663 | 0.086 | -0.041 | -0.028 | -0.028 | 0 | 0.86 | 0.66 | 0.88 | 0.94 | 0.93 | 0.87 |
| 10 | 113267625 | rs10885198 | 1.112 | -1.101 | 0.648 | 0.077 | -0.002 | 0.021 | 0.021 | 0.343 | 0.44 | 0.35 | 0.29 | 0.31 | 0.48 | 0.38 |
| 10 | 113273360 | rs12221261 | 0.996 | -1.112 | 0.454 | 0.791 | -0.098 | -0.027 | -0.027 | 0 | 0.19 | 0.02 | 0.24 | 0.16 | 0.27 | 0.15 |
| 10 | 113280963 | rs7896696 | -0.137 | -1.401 | 0.202 | 0.456 | -0.028 | 0.024 | 0.024 | 0 | 0.15 | 0.26 | 0.05 | 0.21 | 0.07 | 0.22 |
| 10 | 113282741 | rs12412805 | 0.346 | -0.93 | -0.162 | 0.18 | -0.076 | -0.098 | -0.098 | 0.252 | 0.14 | 0.03 | 0.38 | 0.42 | 0.22 | 0.29 |
| 10 | 113282771 | rs17128993 | 0.115 | -1.041 | -0.068 | 0.091 | -0.021 | -0.047 | -0.047 | 0 | 0.16 | 0.21 | 0.15 | 0.04 | 0.1 | 0.08 |
| 10 | 113282918 | rs10509944 | 2.704 | -1.082 | -0.12 | 0.053 | -0.047 | 0.104 | 0.104 | 1.776 | 0.3 | 0.09 | 0.25 | 0.11 | 0.41 | 0.17 |
| 10 | 113283471 | rs10787373 | 0.341 | -1.007 | -0.051 | 0.15 | -0.076 | -0.098 | -0.098 | 0 | 0.14 | 0.03 | 0.38 | 0.42 | 0.22 | 0.29 |
| 10 | 113284609 | rs7921705 | 0.037 | -1.011 | -0.014 | 0.171 | -0.029 | 0.022 | 0.022 | 0 | 0.15 | 0.26 | 0.05 | 0.21 | 0.07 | 0.23 |
| 10 | 113287674 | rs11195620 | 0.346 | -0.903 | -0.191 | 0.199 | -0.08 | -0.101 | -0.101 | 0 | 0.14 | 0.03 | 0.39 | 0.42 | 0.25 | 0.29 |
| 10 | 113290300 | rs953196 | -0.962 | -1.024 | -0.126 | 0.2 | -0.001 | -0.002 | -0.002 | 0.581 | 0.3 | 0.28 | 0.54 | 0.46 | 0.32 | 0.36 |
| 10 | 113295291 | rs7088001 | -0.775 | -0.92 | -0.026 | 0.261 | -0.054 | 0.016 | 0.016 | 0.133 | 0.23 | 0.39 | 0.13 | 0.25 | 0.11 | 0.34 |
| 10 | 113295975 | rs7916268 | 0.165 | -0.616 | -0.358 | 0.403 | 0.004 | 0.009 | 0.009 | 3.953 | 0.28 | 0.24 | 0.19 | 0.2 | 0.22 | 0.19 |
| 10 | 113296233 | rs12573094 | 0.146 | -0.272 | -0.533 | 0.575 | -0.043 | -0.051 | -0.051 | 0 | 0.13 | 0.05 | 0.39 | 0.43 | 0.22 | 0.28 |
| 10 | 113297453 | rs3107338 | -0.769 | -0.601 | -0.558 | 0.188 | 0 | 0.008 | 0.008 | 9.718 | 0.51 | 0.57 | 0.62 | 0.67 | 0.47 | 0.54 |
| 10 | 113305159 | rs10885206 | 1.481 | -0.823 | -1.229 | -0.245 | -0.094 | 0.138 | 0.138 | 0 | 0.34 | 0.07 | 0.3 | 0.09 | 0.51 | 0.18 |
| 10 | 113306185 | rs881103 | -0.836 | -1.412 | -1.559 | -0.006 | 0.109 | -0.015 | -0.015 | 0 | 0.39 | 0.84 | 0.19 | 0.47 | 0.17 | 0.53 |
| 10 | 113309101 | rs10885208 | 1.108 | -1.509 | -0.705 | 0.022 | -0.001 | 0.007 | 0.007 | 0 | 0.53 | 0.42 | 0.36 | 0.28 | 0.57 | 0.47 |
| 10 | 113310787 | rs2138554 | 0.127 | -1.549 | -0.934 | 0.087 | -0.107 | -0.143 | -0.143 | 0 | 0.7 | 0.93 | 0.47 | 0.57 | 0.64 | 0.71 |
| 10 | 113313535 | rs1914143 | 1.019 | -1.518 | -0.814 | 0.072 | 0.01 | -0.01 | -0.01 | 0 | 0.09 | 0.06 | 0.02 | 0.16 | 0.03 | 0.21 |
| 10 | 113314728 | rs2419590 | 1.077 | -1.633 | -1.099 | 0.089 | 0.002 | 0.008 | 0.008 | 0 | 0.5 | 0.42 | 0.33 | 0.28 | 0.52 | 0.47 |
| 10 | 113315445 | rs11195624 | 1.064 | -1.382 | -1.533 | 0.451 | -0.062 | -0.05 | -0.05 | 0 | 0.11 | 0.01 | 0.25 | 0.14 | 0.17 | 0.08 |
| 10 | 113317435 | rs12767680 | 2.426 | -1.376 | -1.441 | 0.02 | -0.074 | 0.185 | 0.185 | 0 | 0.33 | 0.07 | 0.22 | 0.02 | 0.45 | 0.24 |
| 10 | 113327791 | rs1914080 | 0.232 | -1.47 | -1 | 0.259 | -0.003 | -0.004 | -0.004 | 0.274 | 0.22 | 0.2 | 0.15 | 0.31 | 0.21 | 0.26 |
| 10 | 113328934 | rs7090642 | -0.833 | -1.806 | -0.49 | 0.241 | -0.104 | 0.037 | 0.037 | 0 | 0.23 | 0.46 | 0.13 | 0.27 | 0.08 | 0.25 |
| 10 | 113336564 | rs11195640 | 2.178 | -1.788 | -0.502 | 0.319 | 0.004 | -0.033 | -0.033 | 0 | 0.06 | 0 | 0.06 | 0.1 | 0.04 | 0.02 |
| 10 | 113337824 | rs7090533 | 0.769 | -1.771 | -0.582 | 0.225 | -0.001 | 0.005 | 0.005 | 0 | 0.56 | 0.53 | 0.35 | 0.29 | 0.52 | 0.49 |
| 10 | 113337902 | rs10787379 | 0.082 | -1.688 | -0.115 | 0.343 | -0.048 | 0.015 | 0.015 | 0.094 | 0.13 | 0.27 | 0.1 | 0.11 | 0.05 | 0.04 |
| 10 | 113357617 | rs12770980 | 1.656 | -1.667 | -0.029 | 0.088 | 0.001 | -0.003 | -0.003 | 0.943 | 0.45 | 0.57 | 0.28 | 0.18 | 0.45 | 0.4 |
| 10 | 113360081 | rs7077548 | 2.338 | -1.692 | -0.057 | 0.079 | -0.006 | 0.047 | 0.047 | 0.122 | 0.38 | 0.26 | 0.25 | 0.18 | 0.45 | 0.38 |
| 10 | 113366293 | rs7091510 | 1.701 | -1.666 | -0.065 | 0.106 | -0.004 | -0.04 | -0.04 | 0 | 0.45 | 0.49 | 0.28 | 0.18 | 0.45 | 0.4 |
| 10 | 113373981 | rs3120592 | -0.532 | -1.675 | -0.141 | 0.238 | -0.044 | -0.072 | -0.072 | 0 | 0.27 | 0.09 | 0.54 | 0.49 | 0.31 | 0.27 |
| 10 | 113386615 | rs7897542 | 0.817 | -1.694 | 0.007 | 0.166 | 0.004 | 0.001 | 0.001 | 0 | 0.06 | 0.28 | 0.04 | 0.01 | 0.01 | 0.02 |
| 10 | 113389291 | rs4917619 | 1.66 | -1.704 | -0.049 | 0.147 | 0.004 | 0.012 | 0.012 | 0.015 | 0.46 | 0.58 | 0.29 | 0.19 | 0.45 | 0.4 |
| 10 | 113394570 | rs1336414 | -0.274 | -1.694 | -0.097 | 0.155 | -0.003 | -0.006 | -0.006 | 0 | 0.72 | 0.68 | 0.83 | 0.7 | 0.76 | 0.67 |
| 10 | 113406917 | rs7908446 | 1.551 | -1.685 | -0.079 | 0.095 | 0.003 | 0.003 | 0.003 | 0 | 0.46 | 0.57 | 0.29 | 0.18 | 0.45 | 0.4 |
| 10 | 113415997 | rs17129082 | 1.002 | -1.696 | -0.01 | 0.225 | -0.036 | 0.009 | 0.009 | 0 | 0.1 | 0.2 | 0.03 | 0.14 | 0.03 | 0.2 |
| 10 | 113419447 | rs11195666 | 0.453 | -1.818 | -0.312 | 0.211 | -0.064 | -0.017 | -0.017 | 0 | 0.21 | 0.01 | 0.14 | 0.25 | 0.22 | 0.28 |
| 10 | 113422955 | rs17129084 | 0.503 | -1.743 | 0.073 | 0.214 | -0.022 | -0.024 | -0.024 | 0 | 0.07 | 0.22 | 0.03 | 0.01 | 0.01 | 0.02 |
| 10 | 113440271 | rs7922236 | 0.558 | -1.742 | 0.068 | 0.216 | -0.006 | -0.009 | -0.009 | 0 | 0.07 | 0.26 | 0.03 | 0.01 | 0.01 | 0.02 |
| 10 | 113445582 | rs10885243 | -0.601 | -1.533 | -0.121 | 0.582 | -0.029 | -0.07 | -0.07 | 0.039 | 0.26 | 0.1 | 0.52 | 0.5 | 0.29 | 0.25 |

| | | | | | | | | | | | | | | | | |
|---|---|---|---|---|---|---|---|---|---|---|---|---|---|---|---|---|
| 10 | 113449034 | rs11195680 | 1.421 | -1.477 | -0.285 | 0.806 | -0.054 | -0.041 | -0.041 | 1.579 | 0.1 | 0.01 | 0.23 | 0.15 | 0.17 | 0.07 |
| 10 | 113455945 | rs7899581 | 1.082 | -1.542 | -0.163 | 0.655 | -0.026 | 0.008 | 0.008 | 0 | 0.09 | 0.16 | 0.03 | 0.13 | 0.03 | 0.22 |
| 10 | 113460724 | rs7080528 | 2.586 | -1.243 | -0.217 | 0.498 | 0.001 | -0.054 | -0.054 | 0 | 0.35 | 0.32 | 0.26 | 0.06 | 0.43 | 0.2 |
| 10 | 113467206 | rs17129116 | 1.587 | -1.15 | -0.439 | 0.977 | -0.026 | -0.022 | -0.022 | 0.01 | 0.06 | 0.01 | 0.16 | 0.26 | 0.07 | 0.16 |
| 10 | 113476605 | rs9633723 | 0.075 | -1.078 | -0.331 | 0.48 | 0.002 | -0.005 | -0.005 | 0 | 0.21 | 0.17 | 0.14 | 0.24 | 0.17 | 0.32 |
| 10 | 113484881 | rs868258 | -0.494 | -1.172 | -0.331 | 0.475 | -0.002 | 0.006 | 0.006 | 3.245 | 0.28 | 0.23 | 0.18 | 0.28 | 0.26 | 0.33 |
| 10 | 113486148 | rs1414883 | -1.287 | -1.179 | -0.324 | 0.444 | 0.002 | 0.001 | 0.001 | 2.072 | 0.38 | 0.38 | 0.21 | 0.37 | 0.29 | 0.55 |
| 10 | 113498882 | rs7069965 | -0.932 | -1.233 | -0.215 | 0.495 | -0.024 | 0.015 | 0.015 | 0.471 | 0.32 | 0.43 | 0.18 | 0.39 | 0.2 | 0.51 |
| 10 | 113499125 | rs6585109 | 2.454 | -1.095 | -0.246 | 0.113 | 0.001 | -0.047 | -0.047 | 0 | 0.34 | 0.33 | 0.26 | 0.07 | 0.43 | 0.2 |
| 10 | 113501035 | rs11195703 | 1.273 | -1.354 | 0.16 | 0.381 | -0.016 | -0.016 | -0.016 | 0.019 | 0.63 | 0.43 | 0.49 | 0.32 | 0.76 | 0.48 |
| 10 | 113501109 | rs4918686 | 2.196 | -1.233 | 0.47 | 0.121 | -0.001 | -0.014 | -0.014 | 0 | 0.38 | 0.35 | 0.28 | 0.09 | 0.43 | 0.2 |
| 10 | 113501355 | rs4918687 | 2.071 | -1.252 | 0.448 | 0.191 | -0.005 | 0.025 | 0.025 | 0.014 | 0.39 | 0.3 | 0.28 | 0.07 | 0.44 | 0.21 |
| 10 | 113502538 | rs12098793 | 1.619 | -1.197 | 0.449 | 0.268 | -0.024 | 0.015 | 0.015 | 0 | 0.09 | 0.22 | 0.03 | 0.13 | 0.03 | 0.2 |
| 10 | 113503479 | rs10885255 | -0.369 | -1.302 | 0.652 | 0.524 | 0.002 | -0.003 | -0.003 | 0.014 | 0.32 | 0.22 | 0.22 | 0.35 | 0.29 | 0.35 |
| 10 | 113504180 | rs1415848 | 0.908 | -1.282 | 0.511 | 0.548 | 0.02 | -0.014 | -0.014 | 0 | 0.58 | 0.35 | 0.73 | 0.46 | 0.69 | 0.45 |
| 10 | 113505801 | rs12770983 | -0.664 | -1.173 | 0.359 | 0.557 | -0.012 | -0.054 | -0.054 | 0 | 0.22 | 0.14 | 0.44 | 0.42 | 0.22 | 0.25 |
| 10 | 113506422 | rs11195706 | 2.3 | -1.258 | 0.346 | 0.666 | -0.015 | 0.025 | 0.025 | 0 | 0.05 | 0.04 | 0.03 | 0.12 | 0.04 | 0.02 |
| 10 | 113507338 | rs12779426 | 1.481 | -1.163 | -0.576 | 0.558 | 0.068 | 0.006 | 0.006 | 0.502 | 0.52 | 0.24 | 0.71 | 0.44 | 0.67 | 0.43 |
| 10 | 113507744 | rs12357565 | 0.092 | -0.422 | -0.727 | 0.223 | -0.017 | 0.005 | 0.005 | 4.327 | 0.25 | 0.12 | 0.19 | 0.28 | 0.25 | 0.33 |
| 10 | 113511441 | rs7894045 | 1.005 | -0.667 | -0.47 | 0.104 | 0.02 | 0.042 | 0.042 | 0 | 0.58 | 0.27 | 0.44 | 0.3 | 0.66 | 0.48 |
| 10 | 113511642 | rs10885258 | 1.505 | -0.512 | -0.704 | -0.223 | -0.002 | 0.045 | 0.045 | 0 | 0.43 | 0.34 | 0.41 | 0.16 | 0.56 | 0.26 |
| 10 | 113515047 | rs10787403 | 1.635 | -0.766 | -0.927 | 0.076 | -0.031 | 0.045 | 0.045 | 0 | 0.46 | 0.08 | 0.38 | 0.39 | 0.58 | 0.46 |
| 10 | 113516494 | rs10509950 | -0.247 | -1.04 | -0.754 | 0.526 | 0.002 | 0.011 | 0.011 | 0 | 0.24 | 0.26 | 0.4 | 0.37 | 0.27 | 0.25 |
| 10 | 113516620 | rs10509951 | 0.109 | -1.043 | -0.781 | 0.526 | -0.018 | -0.046 | -0.046 | 0 | 0.22 | 0.14 | 0.39 | 0.37 | 0.27 | 0.25 |
| 10 | 113517931 | rs4918688 | -1.109 | -1.214 | -0.375 | 0.502 | -0.036 | 0.01 | 0.01 | 0 | 0.85 | 0.76 | 0.86 | 0.81 | 0.95 | 0.75 |
| 10 | 113518440 | rs12359639 | 1.105 | -1.135 | -0.603 | 0.531 | -0.061 | 0.005 | 0.005 | 1.962 | 0.23 | 0.03 | 0.17 | 0.29 | 0.26 | 0.29 |
| 10 | 113519624 | rs11195715 | 1.411 | -1.111 | 0.309 | 0.929 | -0.016 | -0.004 | -0.004 | 0.268 | 0.58 | 0.47 | 0.46 | 0.41 | 0.68 | 0.49 |
| 10 | 113522754 | rs1923653 | 0.927 | -1.199 | 0.482 | 0.733 | -0.066 | -0.001 | -0.001 | 0 | 0.77 | 0.37 | 0.82 | 0.8 | 0.94 | 0.73 |
| 10 | 113524371 | rs6585111 | 0.634 | -1.195 | 0.838 | 0.739 | -0.077 | -0.002 | -0.002 | 2.403 | 0.79 | 0.61 | 0.84 | 0.78 | 0.94 | 0.73 |
| 10 | 113525960 | rs10787405 | -1.243 | -0.986 | 0.801 | 0.827 | -0.007 | 0.072 | 0.072 | 0 | 0.29 | 0.52 | 0.44 | 0.48 | 0.28 | 0.26 |
| 10 | 113532017 | rs1415852 | -0.798 | -0.994 | 0.899 | 0.852 | -0.001 | -0.01 | -0.01 | 0 | 0.26 | 0.3 | 0.43 | 0.47 | 0.28 | 0.25 |
| 10 | 113532967 | rs11195718 | 1.638 | -0.972 | 0.837 | 0.917 | -0.026 | 0.005 | 0.005 | 0 | 0.11 | 0.06 | 0.15 | 0.11 | 0.18 | 0.08 |
| 10 | 113533305 | rs1415854 | 0.888 | -0.9 | 0.748 | 0.835 | -0.047 | 0.023 | 0.023 | 0 | 0.15 | 0.07 | 0.17 | 0.11 | 0.24 | 0.1 |
| 10 | 113534724 | rs10749099 | -1.675 | -0.904 | 0.775 | 0.844 | 0.004 | 0.046 | 0.046 | 0 | 0.33 | 0.53 | 0.47 | 0.48 | 0.34 | 0.27 |
| 10 | 113536366 | rs12569902 | 0.255 | -0.92 | 0.71 | 0.847 | 0.018 | -0.042 | -0.042 | 0 | 0.13 | 0.06 | 0.26 | 0.36 | 0.1 | 0.17 |
| 10 | 113536606 | rs1415847 | -1.922 | -0.941 | 0.874 | 0.801 | 0.001 | -0.009 | -0.009 | 0 | 0.36 | 0.69 | 0.48 | 0.48 | 0.35 | 0.27 |
| 10 | 113538188 | rs1337987 | -1.103 | -1.123 | 1.092 | 1.059 | -0.001 | -0.003 | -0.003 | 0 | 0.26 | 0.32 | 0.43 | 0.47 | 0.28 | 0.25 |
| 10 | 113538688 | rs11195719 | 2.304 | -1.194 | 0.806 | 1.041 | 0.074 | 0.121 | 0.121 | 0.09 | 0.49 | 0.08 | 0.38 | 0.29 | 0.63 | 0.47 |
| 10 | 113555500 | rs1337997 | -0.901 | -0.679 | 0.686 | 1.081 | -0.025 | -0.075 | -0.075 | 0 | 0.24 | 0.12 | 0.42 | 0.47 | 0.29 | 0.25 |
| 10 | 113563565 | rs12771264 | 1.205 | -0.377 | 0.563 | 0.985 | -0.072 | 0.027 | 0.027 | 0 | 0.15 | 0.06 | 0.23 | 0.1 | 0.33 | 0.11 |
| 10 | 113564630 | rs958249 | -1.723 | -0.626 | 0.801 | 0.951 | -0.021 | -0.025 | -0.025 | 0.115 | 0.41 | 0.78 | 0.56 | 0.49 | 0.45 | 0.36 |
| 10 | 113571480 | rs10787412 | -1.469 | -0.611 | 1.009 | 0.948 | 0.024 | 0.022 | 0.022 | 0 | 0.34 | 0.56 | 0.53 | 0.48 | 0.44 | 0.28 |
| 10 | 113583339 | rs10787413 | -0.856 | -0.577 | 0.762 | 0.936 | -0.018 | -0.02 | -0.02 | 0.025 | 0.3 | 0.32 | 0.51 | 0.51 | 0.44 | 0.27 |
| 10 | 113591240 | rs1923648 | -1.731 | -0.514 | 0.685 | 0.908 | -0.009 | -0.027 | -0.027 | 0.08 | 0.39 | 0.76 | 0.54 | 0.5 | 0.41 | 0.35 |
| 10 | 113593198 | rs10787415 | -1.777 | -0.593 | 0.574 | 0.816 | -0.001 | -0.021 | -0.021 | 2.608 | 0.38 | 0.74 | 0.53 | 0.52 | 0.39 | 0.31 |
| 10 | 113594474 | rs1078288 | -1.86 | -0.312 | 0.748 | 0.799 | -0.001 | 0 | 0 | 0.874 | 0.39 | 0.66 | 0.58 | 0.47 | 0.49 | 0.36 |
| 10 | 113600635 | rs7901004 | -0.062 | -0.216 | 0.642 | 0.421 | 0.011 | -0.009 | -0.009 | 0 | 0.21 | 0.29 | 0.31 | 0.13 | 0.31 | 0.12 |
| 10 | 113610796 | rs12253174 | -1.811 | -0.29 | 0.381 | 0.422 | -0.023 | 0.038 | 0.038 | 0.158 | 0.36 | 0.78 | 0.42 | 0.16 | 0.43 | 0.22 |
| 10 | 113611569 | rs959127 | -1.551 | -0.221 | 0.447 | 0.5 | -0.01 | 0.011 | 0.011 | 0.82 | 0.31 | 0.76 | 0.38 | 0.15 | 0.38 | 0.14 |
| 10 | 113613125 | rs7083899 | 1.205 | -0.25 | 0.053 | 0.669 | -0.081 | 0.001 | 0.001 | 0 | 0.13 | 0.03 | 0.19 | 0.11 | 0.28 | 0.1 |
| 10 | 113619670 | rs7923518 | 0.273 | -0.321 | 0.31 | 0.62 | 0.019 | -0.019 | -0.019 | 0 | 0.17 | 0.25 | 0.21 | 0.09 | 0.29 | 0.11 |
| 10 | 113627886 | rs585070 | 1.762 | -1.517 | 0.244 | 0.587 | 0.007 | 0.047 | 0.047 | 0.867 | 0.47 | 0.25 | 0.34 | 0.33 | 0.54 | 0.42 |
| 10 | 113632807 | rs7917775 | 0.019 | -1.745 | 0.316 | 0.673 | 0.064 | -0.081 | -0.081 | 0 | 0.22 | 0.08 | 0.3 | 0.57 | 0.08 | 0.35 |
| 10 | 113643755 | rs652950 | 0.518 | -1.749 | 0.763 | 0.612 | -0.028 | 0.008 | 0.008 | 0 | 0.7 | 0.68 | 0.59 | 0.4 | 0.87 | 0.6 |
| 10 | 113650316 | rs2193579 | -1.211 | -1.734 | 0.853 | 0.619 | -0.007 | -0.005 | -0.005 | 1.891 | 0.33 | 0.34 | 0.42 | 0.6 | 0.13 | 0.39 |
| 10 | 113650477 | rs1019383 | 1.351 | -1.573 | 0.878 | 0.59 | -0.004 | 0.004 | 0.004 | 0 | 0.14 | 0.14 | 0.2 | 0.06 | 0.29 | 0.07 |
| 10 | 113652440 | rs582128 | -0.576 | -1.596 | 0.935 | 0.622 | 0.087 | -0.051 | -0.051 | 0 | 0.79 | 0.58 | 0.75 | 0.92 | 0.66 | 0.81 |
| 10 | 113656367 | rs7086940 | -1.185 | -1.728 | 0.826 | 0.625 | -0.062 | 0.016 | 0.016 | 0.045 | 0.33 | 0.41 | 0.4 | 0.61 | 0.12 | 0.39 |
| 10 | 113657917 | rs12764680 | 0.37 | -1.439 | 0.593 | 0.544 | -0.016 | 0.015 | 0.015 | 0 | 0.21 | 0.22 | 0.24 | 0.06 | 0.33 | 0.2 |
| 10 | 113686419 | rs2080648 | 1.192 | -1.917 | 0.483 | 0.951 | 0.005 | -0.006 | -0.006 | 0 | 0.12 | 0.07 | 0.11 | 0.14 | 0.02 | 0.2 |
| 10 | 113689700 | rs10885292 | 0.213 | -1.692 | 0.516 | 0.759 | 0.089 | -0.075 | -0.075 | 0 | 0.25 | 0.07 | 0.3 | 0.49 | 0.1 | 0.37 |
| 10 | 113691771 | rs10400070 | 0.39 | -1.573 | 0.664 | 0.714 | -0.002 | -0.003 | -0.003 | 1.197 | 0.21 | 0.24 | 0.24 | 0.06 | 0.33 | 0.21 |
| 10 | 113696042 | rs10787419 | 0.215 | -1.601 | 0.749 | 0.719 | 0.058 | -0.046 | -0.046 | 0 | 0.26 | 0.17 | 0.31 | 0.49 | 0.1 | 0.38 |
| 10 | 113696371 | rs2111639 | -1.447 | -1.568 | 0.751 | 0.633 | 0.023 | 0.001 | 0.001 | 0.252 | 0.47 | 0.36 | 0.54 | 0.58 | 0.43 | 0.59 |
| 10 | 113696815 | rs1421055 | -1.956 | -1.58 | 0.853 | 0.479 | -0.004 | 0.027 | 0.027 | 1.536 | 0.55 | 0.78 | 0.66 | 0.65 | 0.46 | 0.6 |
| 10 | 113697099 | rs953103 | 1.767 | -1.563 | 0.846 | 0.467 | 0.002 | 0.019 | 0.019 | 0 | 0.07 | 0.04 | 0.03 | 0 | 0.04 | 0.13 |
| 10 | 113700265 | rs10787421 | -1.829 | -1.459 | 0.54 | 0.587 | -0.005 | 0.027 | 0.027 | 1.57 | 0.56 | 0.79 | 0.67 | 0.65 | 0.46 | 0.6 |
| 10 | 113713761 | rs2111636 | 0.151 | -1.211 | 0.508 | 0.461 | 0.005 | -0.014 | -0.014 | 0 | 0.24 | 0.24 | 0.28 | 0.06 | 0.4 | 0.22 |
| 10 | 113719764 | rs6585128 | -2.403 | -1.429 | 0.464 | 0.425 | -0.053 | 0.002 | 0.002 | 0.437 | 0.61 | 0.84 | 0.73 | 0.66 | 0.55 | 0.61 |

| | | | | | | | | | | | | | | | | |
|---|---|---|---|---|---|---|---|---|---|---|---|---|---|---|---|---|
| 10 | 113723647 | rs6585131 | -1.891 | -1.41 | 0.625 | 0.663 | 0.003 | 0.024 | 0.024 | 1.111 | 0.55 | 0.71 | 0.66 | 0.64 | 0.46 | 0.59 |
| 10 | 113724245 | rs1019381 | -0.521 | -1.663 | 0.683 | 0.613 | -0.108 | 0.058 | 0.058 | 0.029 | 0.29 | 0.45 | 0.33 | 0.48 | 0.1 | 0.36 |
| 10 | 113731295 | rs3981455 | -1.363 | -1.18 | -0.096 | 0.751 | 0.019 | -0.002 | -0.002 | 0 | 0.51 | 0.67 | 0.56 | 0.57 | 0.44 | 0.58 |
| 10 | 113735645 | rs12766171 | -0.271 | -1.576 | 0.281 | 0.62 | 0.018 | -0.016 | -0.016 | 0 | 0.19 | 0.23 | 0.21 | 0.09 | 0.29 | 0.19 |
| 10 | 113737605 | rs7923493 | -1.04 | -2.934 | 0.651 | 1.197 | -0.036 | -0.003 | -0.003 | 1.794 | 0.18 | 0.54 | 0.18 | 0.15 | 0.12 | 0.03 |
| 10 | 113747007 | rs11195777 | -0.426 | -2.893 | 0.754 | 1.209 | -0.012 | -0.003 | -0.003 | 0 | 0.14 | 0.25 | 0.15 | 0.13 | 0.11 | 0.03 |
| 10 | 113751347 | rs12354545 | -0.113 | -2.829 | 0.697 | 1.338 | -0.007 | 0.013 | 0.013 | 0 | 0.13 | 0.19 | 0.15 | 0.18 | 0.12 | 0.03 |
| 10 | 113753187 | rs11195781 | -1.123 | -3.514 | 0.43 | 1.539 | -0.062 | 0.012 | 0.012 | 0 | 0.36 | 0.56 | 0.45 | 0.39 | 0.18 | 0.37 |
| 10 | 113760817 | rs6585132 | -0.34 | -2.85 | 0.431 | 1.423 | -0.008 | 0.012 | 0.012 | 0 | 0.14 | 0.21 | 0.16 | 0.17 | 0.12 | 0.03 |
| 10 | 113775409 | rs2804576 | -1.687 | -3.297 | 0.007 | 1.755 | 0.009 | -0.011 | -0.011 | 0 | 0.27 | 0.76 | 0.22 | 0.62 | 0.14 | 0.2 |
| 10 | 113783588 | rs2804585 | -0.258 | -3.112 | 0.226 | 1.525 | -0.039 | 0.053 | 0.053 | 0 | 0.16 | 0.38 | 0.16 | 0.17 | 0.09 | 0.04 |
| 10 | 113789598 | rs2792715 | -0.486 | -3.178 | 0.241 | 1.434 | -0.052 | 0.069 | 0.069 | 0 | 0.17 | 0.44 | 0.17 | 0.18 | 0.09 | 0.04 |
| 10 | 113795223 | rs2792709 | -0.587 | -3.254 | 0.114 | 1.436 | -0.071 | 0.108 | 0.108 | 0.386 | 0.14 | 0.57 | 0.14 | 0.2 | 0.03 | 0.03 |
| 10 | 113799201 | rs2792705 | -1.764 | -4.24 | -0.664 | 0.972 | 0.128 | 0.117 | 0.117 | 0.051 | 0.57 | 0.86 | 0.37 | 0.75 | 0.25 | 0.55 |
| 10 | 113806381 | rs2419601 | -0.686 | -3.542 | -0.705 | 1.104 | -0.188 | 0.036 | 0.036 | 0.077 | 0.71 | 0.97 | 0.53 | 0.81 | 0.51 | 0.61 |
| 10 | 113819244 | rs2804602 | -1.393 | -4.648 | 0.383 | 2.084 | 0.071 | -0.014 | -0.014 | 0 | 0.37 | 0.78 | 0.26 | 0.32 | 0.08 | 0.37 |
| 10 | 113831949 | rs2792694 | -0.557 | -3.257 | 0.813 | 2.496 | -0.056 | 0.065 | 0.065 | 0.188 | 0.17 | 0.4 | 0.17 | 0.19 | 0.09 | 0.04 |
| 10 | 113835155 | rs11195809 | -1.46 | -2.306 | 0.608 | 2.315 | 0.004 | 0.043 | 0.043 | 0 | 0.07 | 0.05 | 0.03 | 0.42 | 0.05 | 0.14 |
| 10 | 113847037 | rs2804609 | 1.024 | -2.177 | 0.592 | 2.055 | -0.016 | 0.173 | 0.173 | 0 | 0.92 | 0.79 | 0.95 | 0.59 | 0.96 | 0.84 |
| 10 | 113851841 | rs2804614 | 1.01 | -2.664 | -0.096 | 1.818 | 0.09 | -0.096 | -0.096 | 0 | 0.74 | 0.89 | 0.87 | 0.53 | 0.88 | 0.56 |
| 10 | 113855270 | rs2804616 | -1.728 | -2.572 | 0.26 | 1.297 | 0.021 | -0.003 | -0.003 | 0 | 0.36 | 0.68 | 0.21 | 0.44 | 0.17 | 0.49 |
| 10 | 113855817 | rs17129504 | -0.287 | -2.249 | -0.375 | 1.085 | 0.024 | -0.027 | -0.027 | 0.164 | 0.11 | 0.07 | 0.03 | 0.3 | 0.03 | 0.21 |
| 10 | 113858192 | rs11195813 | -0.471 | -2.172 | -0.351 | 1.163 | 0.02 | 0.001 | 0.001 | 0 | 0.16 | 0.04 | 0.19 | 0.18 | 0.1 | 0.27 |
| 10 | 113859224 | rs11195815 | -0.846 | -1.459 | 0.245 | 1.239 | 0.037 | -0.035 | -0.035 | 0.493 | 0.14 | 0.07 | 0.03 | 0.29 | 0.03 | 0.21 |
| 10 | 113870843 | rs11195817 | -0.908 | -1.426 | 0.118 | 1.166 | 0.033 | -0.031 | -0.031 | 0 | 0.15 | 0.1 | 0.03 | 0.3 | 0.03 | 0.21 |
| 10 | 113874825 | rs9663470 | -1.269 | -0.126 | 0.05 | 0.74 | 0.001 | 0.004 | 0.004 | 0 | 0.07 | 0.14 | 0.07 | 0.1 | 0.08 | 0.07 |
| 10 | 113879794 | rs2792752 | -1.349 | -0.52 | 0.4 | 0.355 | -0.029 | -0.057 | -0.057 | 0 | 0.13 | 0.64 | 0.13 | 0.05 | 0.07 | 0.12 |
| 10 | 113879987 | rs12766562 | -0.055 | 0.848 | -0.713 | 0.376 | -0.152 | -0.061 | -0.061 | 0 | 0.2 | 0.02 | 0.35 | 0.17 | 0.39 | 0.26 |
| 10 | 113884528 | rs2803594 | -1.991 | 0.56 | 0.832 | 0.092 | 0.017 | -0.053 | -0.053 | 0 | 0.1 | 0.53 | 0.07 | 0 | 0.04 | 0.02 |
| 10 | 113885445 | rs2803595 | -1.463 | 0.513 | 0.9 | 0.437 | -0.006 | 0.025 | 0.025 | 4.174 | 0.18 | 0.35 | 0.23 | 0.3 | 0.18 | 0.25 |
| 10 | 113897746 | rs2792747 | -1.864 | 0.505 | 1.046 | 0.39 | -0.008 | -0.003 | -0.003 | 0 | 0.22 | 0.5 | 0.26 | 0.19 | 0.21 | 0.26 |
| 10 | 113902418 | rs2792744 | 0.634 | -0.267 | -0.155 | 0.354 | -0.034 | -0.058 | -0.058 | 0.017 | 0.24 | 0.07 | 0.32 | 0.21 | 0.27 | 0.08 |
| 10 | 113903510 | rs2792743 | -0.672 | -0.287 | 0.065 | 0.427 | -0.009 | 0.011 | 0.011 | 0 | 0.39 | 0.47 | 0.52 | 0.38 | 0.41 | 0.32 |
| 10 | 113903867 | rs2419603 | -0.297 | -0.599 | 0.202 | 0.058 | 0.002 | -0.063 | -0.063 | 0 | 0.13 | 0.2 | 0.08 | 0.01 | 0.12 | 0.1 |
| 10 | 113910721 | rs1129555 | -0.611 | -0.891 | 0.045 | 0.277 | 0 | -0.005 | -0.005 | 0.399 | 0.72 | 0.76 | 0.65 | 0.76 | 0.69 | 0.87 |
| 10 | 113912997 | rs7078775 | -0.099 | -1.079 | -0.006 | 0.274 | 0.004 | -0.005 | -0.005 | 0.353 | 0.13 | 0.1 | 0.18 | 0.16 | 0.15 | 0.25 |
| 10 | 113913193 | rs17129557 | -0.099 | -1.079 | -0.006 | 0.274 | 0.004 | -0.004 | -0.004 | 0 | 0.13 | 0.1 | 0.18 | 0.16 | 0.15 | 0.25 |
| 10 | 113913222 | rs2297991 | -0.52 | -1.07 | -0.046 | 0.26 | -0.001 | 0.001 | 0.001 | 0.461 | 0.71 | 0.69 | 0.65 | 0.76 | 0.68 | 0.87 |
| 10 | 113920465 | rs2277207 | 0.379 | -1.083 | -0.086 | 0.271 | -0.006 | 0.008 | 0.008 | 0.231 | 0.58 | 0.53 | 0.46 | 0.6 | 0.54 | 0.62 |
| 10 | 113926012 | rs10749108 | 0.409 | -1.059 | -0.075 | 0.249 | -0.006 | 0.007 | 0.007 | 0.193 | 0.57 | 0.53 | 0.45 | 0.6 | 0.54 | 0.62 |
| 10 | 113935379 | rs10787428 | 0.403 | -1.03 | -0.275 | 0.261 | -0.007 | 0.007 | 0.007 | 0.23 | 0.57 | 0.53 | 0.46 | 0.6 | 0.54 | 0.62 |
| 10 | 113940329 | rs2792751 | -0.94 | -1.004 | -0.442 | 0.253 | -0.041 | -0.081 | -0.081 | 0 | 0.74 | 0.96 | 0.67 | 0.76 | 0.69 | 0.87 |
| 10 | 113944271 | rs2419604 | -0.504 | -0.868 | -0.408 | 0.308 | -0.005 | 0.008 | 0.008 | 0.195 | 0.7 | 0.66 | 0.64 | 0.76 | 0.69 | 0.87 |
| 10 | 113947510 | rs4918723 | -0.749 | -0.882 | -0.523 | 0.347 | 0.009 | -0.005 | -0.005 | 0 | 0.13 | 0.06 | 0.18 | 0.15 | 0.15 | 0.25 |
| 10 | 113953412 | rs34525937 | -1.213 | -1.106 | -0.143 | 0.453 | -0.017 | 0 | 0 | 0 | 0.22 | 0.25 | 0.07 | 0.25 | 0.1 | 0.24 |
| 10 | 113967744 | rs7913950 | -1.203 | -0.85 | 0.307 | 0.69 | -0.033 | 0.044 | 0.044 | 0 | 0.15 | 0.3 | 0.16 | 0.16 | 0.06 | 0.21 |
| 10 | 113969079 | rs7079286 | -2.238 | -0.995 | 0.104 | 0.532 | 0.029 | -0.01 | -0.01 | 0.044 | 0.27 | 0.85 | 0.28 | 0.27 | 0.13 | 0.25 |
| 10 | 113972869 | rs7088682 | -1.354 | -0.853 | 0.522 | 0.582 | -0.029 | 0.035 | 0.035 | 0 | 0.18 | 0.4 | 0.2 | 0.16 | 0.1 | 0.22 |
| 10 | 113977293 | rs9645581 | -0.577 | -0.893 | 0.116 | 0.304 | -0.002 | -0.003 | -0.003 | 1.608 | 0.12 | 0.09 | 0.07 | 0.09 | 0.08 | 0.04 |
| 10 | 113980294 | rs4306236 | -1.181 | -1.007 | 0.455 | 0.535 | -0.01 | 0.129 | 0.129 | 0.828 | 0.16 | 0.37 | 0.29 | 0.32 | 0.13 | 0.2 |
| 10 | 113982162 | rs4575188 | -2.213 | -1.062 | 0.619 | 0.505 | -0.012 | 0.013 | 0.013 | 0 | 0.29 | 0.58 | 0.38 | 0.42 | 0.22 | 0.24 |
| 10 | 113984760 | rs11195845 | -0.594 | -1.04 | 0.532 | 0.493 | -0.011 | -0.012 | -0.012 | 0 | 0.08 | 0.11 | 0.05 | 0.08 | 0.06 | 0.03 |
| 10 | 113987548 | rs4573621 | -1.245 | -0.651 | 0.635 | 0.561 | -0.001 | 0.005 | 0.005 | 0 | 0.15 | 0.18 | 0.29 | 0.35 | 0.18 | 0.25 |
| 10 | 113990113 | rs12572167 | -0.133 | -0.646 | 0.628 | 0.535 | -0.01 | -0.011 | -0.011 | 0 | 0.07 | 0.09 | 0.05 | 0.06 | 0.05 | 0.03 |
| 10 | 113991205 | rs12572492 | 0.116 | -0.801 | 0.367 | 0.965 | 0.004 | -0.003 | -0.003 | 0 | 0.06 | 0.04 | 0.15 | 0.2 | 0.06 | 0.17 |
| 10 | 113991397 | rs7082582 | -2.371 | -1.088 | 0.618 | 0.891 | -0.005 | 0.026 | 0.026 | 0 | 0.31 | 0.39 | 0.5 | 0.59 | 0.3 | 0.49 |
| 10 | 113993061 | rs12572629 | -0.338 | -1.044 | 0.33 | 0.889 | -0.043 | -0.073 | -0.073 | 0 | 0.13 | 0.03 | 0.27 | 0.33 | 0.18 | 0.26 |
| 10 | 113995809 | rs17129649 | -0.241 | -1.029 | 0.336 | 0.895 | 0.01 | -0.018 | -0.018 | 0 | 0.07 | 0.04 | 0.15 | 0.21 | 0.06 | 0.19 |
| 10 | 114002027 | rs4285809 | -0.623 | -1.011 | 0.366 | 0.874 | -0.018 | -0.043 | -0.043 | 0 | 0.15 | 0.09 | 0.29 | 0.33 | 0.18 | 0.28 |
| 10 | 114002390 | rs4285810 | -1.836 | -1.232 | 0.264 | 0.651 | -0.003 | 0.003 | 0.003 | 0.743 | 0.28 | 0.66 | 0.39 | 0.34 | 0.23 | 0.33 |
| 10 | 114005174 | rs7098802 | -0.147 | -1.127 | 0.012 | 0.711 | -0.009 | 0.057 | 0.057 | 0 | 0.07 | 0.11 | 0.16 | 0.2 | 0.06 | 0.19 |
| 10 | 114007531 | rs6585145 | -1.981 | -1.133 | -0.035 | 0.693 | 0.015 | -0.005 | -0.005 | 0 | 0.35 | 0.74 | 0.43 | 0.4 | 0.28 | 0.36 |
| 10 | 114008568 | rs1926563 | -2.524 | -1.163 | -0.116 | 0.693 | 0.026 | -0.036 | -0.036 | 0 | 0.43 | 0.86 | 0.6 | 0.61 | 0.35 | 0.56 |
| 10 | 114021521 | rs12219777 | -0.393 | -0.958 | 0.017 | 0.777 | 0.01 | -0.02 | -0.02 | 0 | 0.07 | 0.04 | 0.15 | 0.21 | 0.06 | 0.19 |
| 10 | 114036703 | rs11195865 | -1.83 | -1.017 | 0.226 | 0.744 | -0.05 | 0.03 | 0.03 | 0.355 | 0.21 | 0.56 | 0.26 | 0.28 | 0.13 | 0.24 |
| 10 | 114039056 | rs10885324 | -0.928 | -0.94 | 0.258 | 0.718 | -0.038 | -0.051 | -0.051 | 0 | 0.13 | 0.49 | 0.1 | 0.06 | 0.06 | 0.04 |
| 10 | 114045198 | rs943263 | -0.676 | -0.463 | 0.31 | 0.896 | 0.028 | -0.032 | -0.032 | 0.015 | 0.07 | 0.22 | 0.07 | 0 | 0.05 | 0.06 |
| 10 | 114045333 | rs12412743 | -0.489 | -0.45 | 0.205 | 0.97 | 0.001 | 0.019 | 0.019 | 0 | 0.12 | 0.16 | 0.29 | 0.33 | 0.16 | 0.27 |
| 10 | 114047263 | rs943264 | -0.917 | -0.118 | -0.346 | 0.023 | -0.049 | 0.066 | 0.066 | 0 | 0.89 | 0.99 | 0.93 | 1 | 0.88 | 0.92 |

| | | | | | | | | | | | | | | | | |
|---|---|---|---|---|---|---|---|---|---|---|---|---|---|---|---|---|
| 10 | 114047787 | rs11195874 | -1.489 | -0.324 | 0.118 | 0.461 | -0.013 | 0.052 | 0.052 | 0 | 0.15 | 0.35 | 0.23 | 0.21 | 0.11 | 0.24 |
| 10 | 114052917 | rs1926559 | -1.291 | -0.314 | -0.308 | 0.512 | 0.002 | -0.003 | -0.003 | 0 | 0.09 | 0.09 | 0.18 | 0.19 | 0.07 | 0.27 |
| 10 | 114053983 | rs10509957 | -1.17 | 0.177 | -0.469 | 0.431 | 0.004 | -0.016 | -0.016 | 0 | 0.43 | 0.57 | 0.64 | 0.72 | 0.44 | 0.58 |
| 10 | 114054601 | rs10509958 | -0.539 | 0.427 | -0.799 | 0.433 | -0.003 | -0.003 | -0.003 | 4.279 | 0.32 | 0.34 | 0.4 | 0.55 | 0.33 | 0.33 |
| 10 | 114055056 | rs12414050 | -1.098 | 0.252 | -0.867 | 0.481 | -0.001 | -0.014 | -0.014 | 0 | 0.1 | 0.05 | 0.22 | 0.2 | 0.11 | 0.21 |
| 10 | 114056580 | rs12251373 | -0.228 | -0.096 | -1.35 | -0.166 | 0.008 | 0.115 | 0.115 | 2.218 | 0.19 | 0.01 | 0.1 | 0 | 0.15 | 0.06 |
| 10 | 114066230 | rs7904244 | -0.705 | 0.581 | -1.461 | 0.086 | -0.04 | -0.027 | -0.027 | 0 | 0.45 | 0.23 | 0.46 | 0.49 | 0.48 | 0.5 |
| 10 | 114067613 | rs1926562 | -1.452 | 0.134 | -0.486 | -0.107 | -0.001 | 0.019 | 0.019 | 0 | 0.57 | 0.68 | 0.59 | 0.69 | 0.57 | 0.65 |
| 10 | 114072091 | rs10885329 | -1.123 | 0.084 | 0.057 | -0.127 | 0.013 | 0.041 | 0.041 | 0.188 | 0.53 | 0.34 | 0.55 | 0.69 | 0.58 | 0.65 |
| 10 | 114074181 | rs7076400 | -1.297 | 0.065 | 0.526 | -0.135 | -0.002 | -0.007 | -0.007 | 0 | 0.55 | 0.53 | 0.58 | 0.68 | 0.58 | 0.65 |
| 10 | 114075199 | rs1887136 | -1.065 | 0.06 | 0.475 | -0.137 | 0.018 | 0.07 | 0.07 | 0.206 | 0.52 | 0.27 | 0.55 | 0.68 | 0.58 | 0.65 |
| 10 | 114081995 | rs7896275 | -1.38 | 0.041 | 0.702 | -0.123 | -0.011 | 0.039 | 0.039 | 0 | 0.12 | 0.33 | 0.12 | 0.2 | 0.09 | 0.15 |
| 10 | 114082666 | rs10885332 | -0.191 | -0.111 | 0.681 | -0.263 | -0.072 | -0.024 | -0.024 | 0 | 0.33 | 0.13 | 0.44 | 0.68 | 0.44 | 0.53 |
| 10 | 114083272 | rs724548 | 0.697 | 0.022 | 0.748 | -0.219 | -0.052 | -0.081 | -0.081 | 0.559 | 0.25 | 0.11 | 0.35 | 0.47 | 0.35 | 0.43 |
| 10 | 114083768 | rs10787440 | 0.072 | 0.069 | 0.689 | -0.198 | 0.031 | 0.071 | 0.071 | 0 | 0.29 | 0.43 | 0.39 | 0.48 | 0.36 | 0.48 |
| 10 | 114084351 | rs11592670 | 0.409 | 0.044 | 0.546 | -0.455 | 0.024 | 0.058 | 0.058 | 1.716 | 0.2 | 0.13 | 0.11 | 0 | 0.14 | 0.07 |
| 10 | 114094931 | rs7898941 | -0.444 | 0.042 | 0.623 | -0.358 | 0.02 | -0.01 | -0.01 | 0 | 0.34 | 0.42 | 0.45 | 0.69 | 0.43 | 0.57 |
| 10 | 114105294 | rs7097428 | -1.34 | 0.024 | 0.253 | -0.314 | 0 | 0.019 | 0.019 | 0 | 0.57 | 0.67 | 0.57 | 0.69 | 0.58 | 0.63 |
| 10 | 114111202 | rs10885336 | 0.168 | -0.216 | 0.319 | -0.184 | -0.017 | -0.007 | -0.007 | 0 | 0.37 | 0.22 | 0.4 | 0.64 | 0.41 | 0.55 |
| 10 | 114111543 | rs2148493 | -0.049 | -0.583 | 0.812 | -0.429 | -0.002 | -0.049 | -0.049 | 4.292 | 0.69 | 0.55 | 0.65 | 0.84 | 0.67 | 0.78 |
| 10 | 114111660 | rs11195918 | 0.622 | -0.526 | 0.523 | -0.374 | -0.005 | -0.006 | -0.006 | 0 | 0.05 | 0.03 | 0.09 | 0.12 | 0.1 | 0.07 |
| 10 | 114120690 | rs1926565 | 0.419 | -0.599 | 0.604 | -0.395 | -0.011 | -0.002 | -0.002 | 0 | 0.37 | 0.26 | 0.4 | 0.64 | 0.41 | 0.55 |
| 10 | 114121792 | rs11195929 | 0.4 | -0.55 | 0.624 | -0.404 | -0.008 | -0.001 | -0.001 | 0 | 0.37 | 0.27 | 0.4 | 0.65 | 0.41 | 0.55 |
| 10 | 114126734 | rs12268778 | 1.096 | -0.641 | 0.301 | -0.49 | -0.002 | 0.124 | 0.124 | 0.601 | 0.2 | 0.02 | 0.1 | 0.01 | 0.15 | 0.11 |
| 10 | 114135013 | rs2419621 | 1.196 | -0.752 | 0.575 | -0.516 | 0.007 | 0.025 | 0.025 | 0 | 0.29 | 0.2 | 0.23 | 0.2 | 0.26 | 0.23 |
| 10 | 114142399 | rs17129748 | -0.011 | -0.783 | 0.627 | -0.376 | 0.004 | 0.002 | 0.002 | 0 | 0.51 | 0.41 | 0.48 | 0.54 | 0.52 | 0.57 |
| 10 | 114158908 | rs4918748 | -0.105 | -0.836 | 0.14 | -0.488 | 0.011 | 0.017 | 0.017 | 0 | 0.5 | 0.3 | 0.56 | 0.57 | 0.53 | 0.57 |
| 10 | 114159452 | rs10885343 | -0.08 | -0.838 | 0.157 | -0.489 | 0.01 | 0.016 | 0.016 | 0 | 0.5 | 0.29 | 0.56 | 0.57 | 0.53 | 0.57 |
| 10 | 114164146 | rs11195949 | 0.03 | -0.855 | 0.21 | -0.496 | 0.008 | 0.011 | 0.011 | 0 | 0.5 | 0.37 | 0.57 | 0.57 | 0.52 | 0.57 |
| 10 | 114168669 | rs17129769 | 2.09 | -0.92 | 0.139 | -0.503 | 0.011 | -0.038 | -0.038 | 0.289 | 0.08 | 0 | 0.01 | 0.12 | 0.02 | 0.07 |
| 10 | 114169190 | rs3736947 | 1.088 | -0.876 | 0.094 | -0.608 | -0.12 | -0.147 | -0.147 | 2.887 | 0.27 | 0.02 | 0.44 | 0.56 | 0.37 | 0.45 |
| 10 | 114169552 | rs7919710 | 0.643 | -0.883 | 0.209 | -0.622 | -0.02 | -0.01 | -0.01 | 0.043 | 0.41 | 0.51 | 0.29 | 0.33 | 0.3 | 0.31 |
| 10 | 114170750 | rs2419628 | 1.541 | -0.923 | -0.168 | -0.812 | -0.026 | 0.035 | 0.035 | 0 | 0.24 | 0.03 | 0.19 | 0.1 | 0.23 | 0.19 |
| 10 | 114171599 | rs876873 | 0.577 | -0.934 | -0.13 | -0.824 | -0.011 | -0.061 | -0.061 | 0 | 0.23 | 0.27 | 0.12 | 0.01 | 0.15 | 0.11 |
| 10 | 114172768 | rs3740142 | 1.399 | -0.697 | -0.621 | -0.612 | -0.023 | 0.042 | 0.042 | 0 | 0.23 | 0.03 | 0.19 | 0.08 | 0.22 | 0.19 |
| 10 | 114187787 | rs8624 | 0.76 | 0.173 | -0.562 | -0.237 | -0.017 | -0.008 | -0.008 | 0 | 0.4 | 0.54 | 0.3 | 0.32 | 0.3 | 0.31 |
| 10 | 114195645 | rs11598819 | 1.196 | 0.137 | -0.889 | -0.496 | -0.011 | 0.047 | 0.047 | 0 | 0.08 | 0 | 0.06 | 0 | 0.08 | 0.05 |
| 10 | 114197642 | rs7070744 | -0.726 | 0.374 | -0.671 | -0.116 | 0.007 | -0.001 | -0.001 | 0.075 | 0.55 | 0.37 | 0.57 | 0.5 | 0.61 | 0.62 |
| 10 | 114200355 | rs2306159 | -0.726 | 0.374 | -0.671 | -0.112 | 0.007 | 0.001 | 0.001 | 0.011 | 0.55 | 0.37 | 0.57 | 0.48 | 0.6 | 0.62 |
| 10 | 114205389 | rs4918752 | -0.776 | 0.4 | -0.686 | -0.101 | 0.003 | 0 | 0 | 0 | 0.56 | 0.4 | 0.58 | 0.48 | 0.61 | 0.61 |
| 10 | 114206121 | rs6421364 | -0.513 | 0.412 | -0.635 | -0.486 | 0.002 | -0.018 | -0.018 | 0 | 0.26 | 0.24 | 0.15 | 0.02 | 0.23 | 0.14 |
| 10 | 114207480 | rs10885348 | -1.35 | 0.456 | -0.492 | 0.611 | -0.003 | -0.013 | -0.013 | 0 | 0.09 | 0.05 | 0.08 | 0.18 | 0.1 | 0.16 |
| 10 | 114209703 | rs7476679 | 1.191 | 0.261 | -0.629 | -0.151 | -0.001 | 0.149 | 0.149 | 0 | 0.46 | 0.22 | 0.33 | 0.04 | 0.56 | 0.26 |
| 10 | 114211424 | rs7090959 | -0.268 | 0.484 | -0.721 | 0.675 | 0.013 | -0.091 | -0.091 | 0 | 0.85 | 0.98 | 0.7 | 0.53 | 0.93 | 0.76 |
| 10 | 114215769 | rs12098574 | -1.206 | 0.322 | 0.103 | 0.516 | 0.07 | 0.074 | 0.074 | 0 | 0.18 | 0.41 | 0.25 | 0.41 | 0.29 | 0.2 |
| 10 | 114222454 | rs9971308 | 0.571 | 0.244 | 0.282 | 0.41 | -0.001 | -0.023 | -0.023 | 0 | 0.74 | 0.61 | 0.6 | 0.23 | 0.8 | 0.51 |
| 10 | 114223234 | rs11195969 | 1.405 | 0.09 | 0.389 | 0.32 | -0.017 | -0.048 | -0.048 | 0 | 0.63 | 0.46 | 0.52 | 0.23 | 0.75 | 0.44 |
| 10 | 114224639 | rs17129806 | -0.036 | -0.035 | 0.208 | 0.32 | 0.006 | -0.008 | -0.008 | 0.203 | 0.11 | 0.1 | 0.07 | 0 | 0.06 | 0.07 |
| 10 | 114232089 | rs11195970 | -0.003 | 0.357 | -0.104 | 0.518 | -0.034 | -0.038 | -0.038 | 0 | 0.07 | 0 | 0.06 | 0.17 | 0.09 | 0.13 |
| 10 | 114236865 | rs10509963 | 0.612 | 0.446 | 0.145 | 0.121 | 0.021 | 0.042 | 0.042 | 0 | 0.57 | 0.33 | 0.5 | 0.22 | 0.72 | 0.42 |
| 10 | 114238129 | rs7917282 | 0.323 | -0.007 | -0.143 | 0.148 | -0.014 | -0.019 | -0.019 | 0 | 0.63 | 0.56 | 0.55 | 0.26 | 0.77 | 0.46 |
| 10 | 114241570 | rs12269496 | -1.109 | 0.559 | -0.112 | 1.127 | 0.033 | 0.096 | 0.096 | 0.768 | 0.12 | 0.48 | 0.16 | 0.22 | 0.12 | 0.11 |
| 10 | 114258451 | rs10885355 | -0.735 | 0.347 | -1.416 | 1.448 | -0.021 | -0.128 | -0.128 | 0 | 0.18 | 0.01 | 0.28 | 0.4 | 0.16 | 0.26 |
| 10 | 114280702 | rs12241008 | -0.824 | -0.022 | -1.15 | 1.529 | -0.015 | 0.057 | 0.057 | 0 | 0.12 | 0.21 | 0.07 | 0.3 | 0.1 | 0.22 |
| 10 | 114289017 | rs11195986 | -0.996 | -0.089 | -1.185 | 1.496 | -0.014 | 0.048 | 0.048 | 0 | 0.13 | 0.21 | 0.07 | 0.29 | 0.1 | 0.22 |
| 10 | 114296407 | rs12354968 | 0.614 | 0.299 | -2.289 | 0.245 | -0.158 | 0.204 | 0.204 | 0 | 0.3 | 0.02 | 0.23 | 0.03 | 0.43 | 0.23 |
| 10 | 114316652 | rs2028783 | -0.855 | -0.051 | -1.298 | 0.987 | -0.028 | 0.067 | 0.067 | 0 | 0.14 | 0.33 | 0.06 | 0.28 | 0.07 | 0.23 |
| 10 | 114318004 | rs12266418 | -0.719 | 0.018 | -1.258 | 1.018 | -0.02 | 0.066 | 0.066 | 0 | 0.12 | 0.3 | 0.06 | 0.28 | 0.06 | 0.23 |
| 10 | 114342159 | rs11196003 | -0.959 | 0.605 | -1.328 | 0.86 | 0.002 | 0 | 0 | 0 | 0.27 | 0.32 | 0.37 | 0.26 | 0.27 | 0.3 |
| 10 | 114342360 | rs12254159 | -0.633 | 0.572 | -1.158 | 0.832 | 0.04 | 0.008 | 0.008 | 0 | 0.05 | 0.2 | 0.05 | 0.06 | 0.08 | 0.02 |
| 10 | 114345552 | rs11196006 | 0.55 | 0.294 | -2.156 | 0.707 | -0.009 | 0.052 | 0.052 | 0.262 | 0.45 | 0.63 | 0.5 | 0.24 | 0.51 | 0.28 |
| 10 | 114348466 | rs11818906 | -0.285 | 0.386 | -2.14 | 0.667 | 0.003 | 0.004 | 0.004 | 0 | 0.19 | 0.31 | 0.27 | 0.18 | 0.18 | 0.22 |
| 10 | 114352961 | rs7903835 | 0.262 | 0.088 | -2.214 | 0.42 | 0 | -0.019 | -0.019 | 0.059 | 0.11 | 0.16 | 0.06 | 0.05 | 0.11 | 0.06 |
| 10 | 114353465 | rs7099088 | 0.243 | -0.022 | -2.319 | 0.616 | 0.021 | 0.023 | 0.023 | 0.004 | 0.56 | 0.65 | 0.57 | 0.37 | 0.64 | 0.34 |
| 10 | 114358555 | rs1325172 | -0.881 | 0.314 | -2.28 | 0.373 | 0.017 | -0.139 | -0.139 | 0 | 0.75 | 0.96 | 0.84 | 0.54 | 0.81 | 0.54 |
| 10 | 114365647 | rs1536081 | -0.832 | 0.324 | -2.252 | 0.41 | -0.012 | -0.147 | -0.147 | 0 | 0.71 | 0.97 | 0.82 | 0.55 | 0.76 | 0.49 |
| 10 | 114368695 | rs2120932 | 0.136 | 0.245 | -2.089 | 0.353 | -0.002 | -0.029 | -0.029 | 0 | 0.09 | 0.18 | 0.05 | 0 | 0.07 | 0.08 |
| 10 | 114369434 | rs12763623 | 1.069 | 0.94 | -2.464 | -0.058 | -0.047 | 0.035 | 0.035 | 0 | 0.08 | 0.01 | 0.08 | 0 | 0.14 | 0.06 |
| 10 | 114372021 | rs2044367 | 0.69 | 0.471 | -2 | 0.169 | -0.001 | 0 | 0 | 0 | 0.48 | 0.47 | 0.67 | 0.49 | 0.54 | 0.31 |

| | | | | | | | | | | | | | | | | |
|---|---|---|---|---|---|---|---|---|---|---|---|---|---|---|---|---|
| 10 | 114381720 | rs6585163 | 0.515 | 0.493 | -1.569 | 0.189 | 0.006 | 0.002 | 0.002 | 0.474 | 0.5 | 0.57 | 0.67 | 0.5 | 0.54 | 0.31 |
| 10 | 114382062 | rs12267930 | 0.295 | 0.45 | -1.953 | 0.156 | 0 | -0.02 | -0.02 | 0.068 | 0.1 | 0.14 | 0.08 | 0 | 0.11 | 0.1 |
| 10 | 114383724 | rs11196024 | 0.129 | 0.513 | -1.701 | 0.377 | 0 | -0.027 | -0.027 | 0 | 0.11 | 0.16 | 0.06 | 0.05 | 0.11 | 0.06 |
| 10 | 114384057 | rs10787453 | -0.163 | 0.552 | -1.9 | 0.723 | 0.017 | -0.053 | -0.053 | 0 | 0.6 | 0.77 | 0.74 | 0.53 | 0.65 | 0.37 |
| 10 | 114388316 | rs1033922 | 0.012 | 0.764 | -1.419 | 0.811 | 0.029 | 0.02 | 0.02 | 0 | 0.52 | 0.77 | 0.52 | 0.44 | 0.58 | 0.38 |
| 10 | 114388945 | rs10885362 | 0.796 | 0.869 | -1.34 | 0.437 | -0.032 | 0.05 | 0.05 | 1.501 | 0.32 | 0.19 | 0.26 | 0.18 | 0.41 | 0.19 |
| 10 | 114394794 | rs1022907 | 0.019 | 0.761 | -1.005 | 0.939 | -0.011 | -0.014 | -0.014 | 0 | 0.24 | 0.22 | 0.29 | 0.36 | 0.33 | 0.21 |
| 10 | 114400295 | rs2120929 | -0.569 | -0.288 | -0.188 | 1.353 | 0.013 | -0.012 | -0.012 | 0.709 | 0.7 | 0.77 | 0.75 | 0.61 | 0.82 | 0.52 |
| 10 | 114402652 | rs10749124 | 0.683 | -0.525 | 0.136 | 1.327 | -0.009 | 0.001 | 0.001 | 0 | 0.53 | 0.5 | 0.56 | 0.55 | 0.67 | 0.44 |
| 10 | 114407099 | rs11196032 | -0.455 | -0.552 | 0.59 | 1.397 | -0.002 | 0.014 | 0.014 | 0 | 0.19 | 0.19 | 0.32 | 0.31 | 0.17 | 0.19 |
| 10 | 114407316 | rs17585058 | 0.862 | -0.559 | 0.428 | 1.12 | -0.026 | -0.015 | -0.015 | 0.283 | 0.08 | 0 | 0.12 | 0.03 | 0.09 | 0.05 |
| 10 | 114409015 | rs2057577 | 0.429 | -0.71 | 0.603 | 1.06 | -0.024 | -0.084 | -0.084 | 0 | 0.79 | 0.52 | 0.79 | 0.97 | 0.84 | 0.92 |
| 10 | 114409630 | rs10885365 | 0.951 | -0.743 | 0.383 | 1.056 | 0.039 | 0.064 | 0.064 | 0.097 | 0.5 | 0.38 | 0.42 | 0.28 | 0.66 | 0.34 |
| 10 | 114414794 | rs2044366 | 0.065 | -0.588 | 0.444 | 1.244 | 0.034 | -0.053 | -0.053 | 0 | 0.29 | 0.15 | 0.37 | 0.55 | 0.18 | 0.53 |
| 10 | 114428364 | rs2859885 | 0.076 | -0.582 | 0.463 | 0.879 | 0.017 | -0.023 | -0.023 | 0 | 0.29 | 0.16 | 0.26 | 0.42 | 0.18 | 0.45 |
| 10 | 114429148 | rs7898049 | 0.089 | -0.551 | 0.353 | 0.75 | -0.004 | 0.019 | 0.019 | 0 | 0.14 | 0.1 | 0.1 | 0.04 | 0.16 | 0.08 |
| 10 | 114430926 | rs17130000 | 0.442 | -0.57 | 0.354 | 0.74 | -0.001 | -0.001 | -0.001 | 0 | 0.12 | 0.13 | 0.09 | 0.04 | 0.14 | 0.06 |
| 10 | 114435232 | rs7901272 | -1.466 | -0.608 | 0.35 | 0.747 | 0.062 | 0.028 | 0.028 | 0.602 | 0.49 | 0.76 | 0.43 | 0.58 | 0.34 | 0.59 |
| 10 | 114436559 | rs10885372 | -0.626 | -0.554 | 0.399 | 0.788 | 0.017 | -0.02 | -0.02 | 0 | 0.42 | 0.28 | 0.35 | 0.49 | 0.32 | 0.53 |
| 10 | 114440341 | rs7906919 | -1.097 | -0.331 | -0.202 | 0.866 | 0.035 | 0.011 | 0.011 | 0.211 | 0.46 | 0.68 | 0.4 | 0.58 | 0.32 | 0.56 |
| 10 | 114445708 | rs11196056 | -0.157 | -0.829 | -0.732 | 0.677 | 0.043 | -0.052 | -0.052 | 0 | 0.3 | 0.14 | 0.25 | 0.45 | 0.19 | 0.4 |
| 10 | 114448218 | rs7095620 | -1.355 | -0.951 | -0.541 | 0.241 | 0.04 | -0.034 | -0.034 | 0 | 0.42 | 0.73 | 0.38 | 0.53 | 0.3 | 0.52 |
| 10 | 114458929 | rs7088368 | -1.399 | -0.702 | -0.373 | 0.178 | 0.023 | -0.048 | -0.048 | 0 | 0.33 | 0.75 | 0.31 | 0.48 | 0.26 | 0.2 |
| 10 | 114467517 | rs7906312 | -1.049 | -0.865 | 0.899 | 0.177 | -0.065 | 0.166 | 0.166 | 4.163 | 0.15 | 0.61 | 0.16 | 0.38 | 0.03 | 0.09 |
| 10 | 114468438 | rs7907540 | -1.516 | -0.868 | 1.059 | 0.291 | -0.021 | 0.074 | 0.074 | 0.31 | 0.3 | 0.51 | 0.29 | 0.54 | 0.25 | 0.2 |
| 10 | 114468819 | rs7911081 | -1.012 | -0.526 | 1.139 | 0.326 | -0.011 | 0.206 | 0.206 | 1.717 | 0.14 | 0.45 | 0.15 | 0.38 | 0.06 | 0.09 |
| 10 | 114469091 | rs7896810 | -1.08 | -1.062 | 1.142 | 0.318 | -0.055 | 0.204 | 0.204 | 2.19 | 0.14 | 0.47 | 0.15 | 0.37 | 0.03 | 0.09 |
| 10 | 114476466 | rs11196072 | -1.537 | -1.225 | 0.922 | 0.157 | -0.065 | 0.123 | 0.123 | 0.431 | 0.19 | 0.53 | 0.19 | 0.37 | 0.06 | 0.12 |
| 10 | 114482501 | rs6585180 | -1.373 | -1.334 | 0.802 | 0.118 | -0.051 | 0.057 | 0.057 | 2.498 | 0.17 | 0.68 | 0.17 | 0.37 | 0.03 | 0.09 |
| 10 | 114483078 | rs12784726 | 0.002 | -1.148 | 0.624 | 0.103 | -0.022 | -0.008 | -0.008 | 0 | 0.16 | 0.04 | 0.14 | 0.16 | 0.23 | 0.12 |
| 10 | 114492898 | rs1885281 | -1.066 | -1.168 | 0.877 | 0.13 | -0.024 | 0.029 | 0.029 | 0 | 0.14 | 0.23 | 0.15 | 0.29 | 0.06 | 0.09 |
| 10 | 114498145 | rs1408817 | -1.203 | -1.144 | 0.332 | -0.245 | -0.001 | 0.111 | 0.111 | 0.007 | 0.67 | 0.86 | 0.78 | 0.89 | 0.68 | 0.47 |
| 10 | 114500004 | rs11196083 | -0.335 | -1.076 | 0.959 | 0.336 | -0.026 | -0.023 | -0.023 | 0 | 0.17 | 0.08 | 0.15 | 0.21 | 0.23 | 0.15 |
| 10 | 114501685 | rs7075671 | -0.959 | -1.123 | 1.051 | 0.326 | -0.072 | 0.125 | 0.125 | 0.264 | 0.14 | 0.45 | 0.14 | 0.29 | 0.03 | 0.05 |
| 10 | 114505465 | rs7918405 | -1.633 | -1.003 | 0.458 | 0.246 | 0.031 | -0.059 | -0.059 | 0.79 | 0.33 | 0.8 | 0.32 | 0.51 | 0.26 | 0.2 |
| 10 | 114515632 | rs4402225 | -1.271 | -0.359 | 0.356 | 0.38 | -0.004 | 0.026 | 0.026 | 0 | 0.36 | 0.45 | 0.35 | 0.5 | 0.35 | 0.27 |
| 10 | 114518318 | rs4145776 | -0.441 | -0.463 | 0.15 | 0.419 | -0.045 | -0.04 | -0.04 | 0 | 0.19 | 0.05 | 0.22 | 0.23 | 0.25 | 0.12 |
| 10 | 114520879 | rs6585185 | -1.401 | -0.433 | 0.333 | 0.339 | -0.004 | 0.022 | 0.022 | 0 | 0.37 | 0.49 | 0.38 | 0.54 | 0.36 | 0.27 |
| 10 | 114528423 | rs11196098 | -0.298 | -0.283 | 0.531 | 0.314 | -0.008 | 0.03 | 0.03 | 0.959 | 0.13 | 0.38 | 0.16 | 0.14 | 0.11 | 0.12 |
| 10 | 114531094 | rs11196101 | -0.231 | -0.358 | 0.637 | 0.352 | -0.077 | 0.085 | 0.085 | 0.652 | 0.16 | 0.41 | 0.14 | 0.27 | 0.05 | 0.08 |
| 10 | 114531678 | rs6585186 | -1.264 | -0.187 | 0.401 | 0.346 | 0 | 0.058 | 0.058 | 0.117 | 0.24 | 0.35 | 0.25 | 0.42 | 0.22 | 0.15 |
| 10 | 114537131 | rs4918769 | -0.382 | -0.321 | -0.235 | -0.117 | -0.025 | -0.022 | -0.022 | 0.154 | 0.59 | 0.55 | 0.71 | 0.69 | 0.7 | 0.53 |
| 10 | 114541029 | rs7085310 | -0.978 | -0.17 | -0.042 | -0.113 | -0.002 | -0.039 | -0.039 | 0.054 | 0.71 | 0.68 | 0.81 | 0.95 | 0.72 | 0.58 |
| 10 | 114541618 | rs1815239 | -1.338 | -0.164 | -0.059 | -0.107 | 0.008 | 0.031 | 0.031 | 0.321 | 0.73 | 0.82 | 0.84 | 0.95 | 0.75 | 0.61 |
| 10 | 114542699 | rs11196106 | 0.544 | -0.223 | -0.048 | -0.096 | -0.053 | 0.055 | 0.055 | 0 | 0.14 | 0.29 | 0.13 | 0.26 | 0.05 | 0.08 |
| 10 | 114543493 | rs7898648 | 0.853 | 0.73 | -0.743 | -0.661 | -0.001 | -0.013 | -0.013 | 0 | 0.4 | 0.52 | 0.54 | 0.7 | 0.37 | 0.37 |
| 10 | 114546582 | rs4918770 | 0.714 | 0.748 | -0.938 | -0.648 | -0.001 | 0.009 | 0.009 | 0 | 0.42 | 0.63 | 0.55 | 0.7 | 0.4 | 0.37 |
| 10 | 114548220 | rs12243358 | -0.611 | 0.799 | -0.934 | -0.654 | -0.005 | 0.005 | 0.005 | 0.294 | 0.33 | 0.27 | 0.3 | 0.25 | 0.38 | 0.24 |
| 10 | 114548954 | rs11196111 | -0.262 | 0.692 | -1.768 | -0.509 | -0.041 | -0.015 | -0.015 | 0 | 0.3 | 0.06 | 0.26 | 0.25 | 0.32 | 0.22 |
| 10 | 114549196 | rs4918773 | -0.193 | 0.702 | -0.946 | 0.016 | -0.001 | 0.037 | 0.037 | 0 | 0.47 | 0.62 | 0.59 | 0.76 | 0.42 | 0.44 |
| 10 | 114552639 | rs7915497 | 0.126 | 0.646 | -1.062 | 0.032 | -0.009 | -0.015 | -0.015 | 0 | 0.58 | 0.31 | 0.46 | 0.57 | 0.56 | 0.67 |
| 10 | 114554779 | rs10787468 | 0.151 | 0.701 | -0.477 | 0.187 | -0.026 | 0.05 | 0.05 | 0 | 0.8 | 0.54 | 0.71 | 0.78 | 0.86 | 0.85 |
| 10 | 114557647 | rs4554813 | 0.187 | 0.714 | -0.358 | 0.188 | -0.025 | 0.05 | 0.05 | 0 | 0.8 | 0.5 | 0.7 | 0.77 | 0.82 | 0.84 |
| 10 | 114557926 | rs744340 | -0.001 | 0.639 | -0.021 | 0.233 | 0.002 | 0.05 | 0.05 | 0 | 0.86 | 0.64 | 0.74 | 0.83 | 0.9 | 0.85 |
| 10 | 114558887 | rs941827 | 0.165 | 0.396 | -0.246 | 0.155 | -0.056 | -0.045 | -0.045 | 0 | 0.17 | 0.05 | 0.23 | 0.24 | 0.25 | 0.14 |
| 10 | 114571831 | rs1213134 | -0.329 | 0.52 | -0.476 | 0.139 | 0 | 0.096 | 0.096 | 0.078 | 0.15 | 0.45 | 0.28 | 0.2 | 0.15 | 0.16 |
| 10 | 114573002 | rs7903017 | -0.12 | 0.461 | -0.135 | 0.114 | 0.017 | 0.026 | 0.026 | 0 | 0.06 | 0.19 | 0.06 | 0.07 | 0.05 | 0.09 |
| 10 | 114573041 | rs7078407 | 0.804 | -0.091 | -1.374 | -0.916 | 0 | -0.006 | -0.006 | 0 | 0.33 | 0.3 | 0.21 | 0 | 0.38 | 0.2 |
| 10 | 114576753 | rs1047468 | 1.16 | -0.364 | -1.568 | -0.819 | 0 | -0.088 | -0.088 | 0 | 0.25 | 0.06 | 0.26 | 0.51 | 0.23 | 0.49 |
| 10 | 114578875 | rs12766378 | 1.612 | -0.368 | -1.882 | -1.098 | -0.07 | 0.124 | 0.124 | 0.057 | 0.2 | 0.01 | 0.16 | 0 | 0.28 | 0.17 |
| 10 | 114578994 | rs2479124 | -0.144 | 0.3 | -1.661 | -1.053 | -0.001 | -0.006 | -0.006 | 0 | 0.43 | 0.52 | 0.59 | 0.71 | 0.43 | 0.65 |
| 10 | 114581193 | rs12358731 | 0.744 | 0.367 | -1.838 | -1.032 | 0.007 | -0.064 | -0.064 | 0 | 0.3 | 0.17 | 0.31 | 0.57 | 0.32 | 0.53 |
| 10 | 114581913 | rs4083480 | -0.679 | 0.21 | -2.141 | -1.028 | -0.029 | -0.03 | -0.03 | 0 | 0.73 | 0.93 | 0.81 | 0.72 | 0.74 | 0.83 |
| 10 | 114582639 | rs12253794 | 0.723 | -0.268 | -1.929 | -0.866 | -0.002 | -0.027 | -0.027 | 0 | 0.3 | 0.39 | 0.2 | 0.06 | 0.32 | 0.18 |
| 10 | 114585159 | rs2475186 | -0.819 | 0.683 | -2.485 | -0.328 | -0.008 | -0.05 | -0.05 | 0.869 | 0.8 | 0.96 | 0.82 | 0.73 | 0.82 | 0.86 |
| 10 | 114586018 | rs2590 | 0.479 | 0.406 | -2.387 | -0.375 | 0.015 | 0.03 | 0.03 | 0 | 0.43 | 0.69 | 0.46 | 0.23 | 0.42 | 0.31 |
| 10 | 114591305 | rs12569579 | -0.313 | 0.365 | -2.976 | -0.239 | -0.04 | -0.081 | -0.081 | 0 | 0.3 | 0.14 | 0.27 | 0.51 | 0.32 | 0.53 |
| 10 | 114592723 | rs667027 | -0.261 | 0.256 | -2.925 | -0.243 | 0.051 | -0.041 | -0.041 | 0 | 0.73 | 0.92 | 0.84 | 0.66 | 0.84 | 0.85 |
| 10 | 114593006 | rs11196122 | -0.574 | 1.634 | -2.605 | 0.378 | -0.138 | -0.154 | -0.154 | 0 | 0.22 | 0.01 | 0.26 | 0.57 | 0.32 | 0.47 |

| | | | | | | | | | | | | | | | | |
|---|---|---|---|---|---|---|---|---|---|---|---|---|---|---|---|---|
| 10 | 114598202 | rs17746088 | -0.498 | 1.182 | -2.583 | -0.182 | -0.043 | 0.136 | 0.136 | 0 | 0.21 | 0.01 | 0.18 | 0 | 0.23 | 0.14 |
| 10 | 114601712 | rs2003083 | -0.115 | 1.819 | -2.948 | 0.736 | -0.047 | -0.056 | -0.056 | 0 | 0.09 | 0.01 | 0.18 | 0.34 | 0.16 | 0.19 |
| 10 | 114603073 | rs4917637 | -0.07 | 1.031 | -3.362 | -0.163 | 0.011 | 0.099 | 0.099 | 0 | 0.44 | 0.21 | 0.56 | 0.79 | 0.43 | 0.58 |
| 10 | 114604563 | rs12248007 | 0.397 | 0.83 | -2.538 | -0.35 | -0.003 | 0.007 | 0.007 | 0.257 | 0.22 | 0.14 | 0.2 | 0 | 0.26 | 0.14 |
| 10 | 114612972 | rs1547191 | -0.956 | 0.941 | -1.627 | -0.541 | 0.013 | -0.027 | -0.027 | 0 | 0.75 | 0.58 | 0.82 | 0.83 | 0.77 | 0.8 |
| 10 | 114615903 | rs2479118 | -0.011 | 1.037 | -1.328 | -0.676 | -0.002 | -0.051 | -0.051 | 0 | 0.27 | 0.33 | 0.24 | 0.02 | 0.31 | 0.18 |
| 10 | 114620882 | rs2475176 | 0.357 | 1.108 | -1.344 | -0.679 | 0 | -0.046 | -0.046 | 0.964 | 0.23 | 0.27 | 0.21 | 0 | 0.27 | 0.14 |
| 10 | 114635381 | rs1927157 | -0.896 | 0.898 | -1.423 | -0.508 | -0.067 | 0.041 | 0.041 | 0 | 0.83 | 0.96 | 0.89 | 0.94 | 0.79 | 0.85 |
| 10 | 114648198 | rs4918785 | 0.775 | 0.119 | -1.439 | -0.364 | -0.019 | 0.071 | 0.071 | 0 | 0.14 | 0.18 | 0.2 | 0.42 | 0.07 | 0.2 |
| 10 | 114652695 | rs2180964 | 0.088 | -0.005 | -0.814 | -1.182 | -0.003 | -0.081 | -0.081 | 0.232 | 0.31 | 0.44 | 0.26 | 0.02 | 0.35 | 0.2 |
| 10 | 114663997 | rs11196146 | -0.961 | 0.007 | -1.141 | -1.232 | -0.054 | 0.049 | 0.049 | 0 | 0.75 | 0.91 | 0.77 | 0.82 | 0.74 | 0.67 |
| 10 | 114670839 | rs7076475 | 0.198 | -0.076 | -0.62 | -1.027 | 0.003 | -0.042 | -0.042 | 0 | 0.27 | 0.33 | 0.23 | 0.03 | 0.32 | 0.2 |
| 10 | 114676413 | rs7075934 | -0.143 | -0.359 | -0.477 | -1.079 | -0.006 | -0.11 | -0.11 | 0.922 | 0.26 | 0.42 | 0.22 | 0.01 | 0.26 | 0.18 |
| 10 | 114686805 | rs11196152 | -0.741 | -0.192 | -1.059 | -1.384 | 0 | -0.066 | -0.066 | 2.097 | 0.46 | 0.51 | 0.47 | 0.06 | 0.53 | 0.35 |
| 10 | 114687884 | rs2148887 | 0.21 | 0.022 | -1.676 | -1.191 | -0.012 | 0.055 | 0.055 | 0 | 0.23 | 0.12 | 0.27 | 0.05 | 0.29 | 0.23 |
| 10 | 114704781 | rs17746916 | 1.159 | 0.314 | -1.66 | -0.819 | -0.022 | 0.025 | 0.025 | 0.269 | 0.06 | 0 | 0.06 | 0 | 0.07 | 0.03 |
| 10 | 114708510 | rs3814570 | 0.058 | 0.207 | -1.598 | -0.785 | -0.008 | 0.078 | 0.078 | 0.004 | 0.26 | 0.08 | 0.29 | 0.05 | 0.31 | 0.2 |
| 10 | 114711983 | rs7094463 | -0.383 | 0.255 | -1.358 | -0.183 | -0.01 | -0.009 | -0.009 | 0.253 | 0.56 | 0.64 | 0.58 | 0.39 | 0.54 | 0.49 |
| 10 | 114723650 | rs2296783 | -0.302 | -0.364 | -1.397 | -0.612 | 0.04 | -0.028 | -0.028 | 0 | 0.19 | 0.06 | 0.15 | 0.65 | 0.1 | 0.34 |
| 10 | 114723857 | rs2296782 | -1.205 | -0.727 | -1.436 | -1.193 | 0.056 | 0.053 | 0.053 | 0.141 | 0.35 | 0.79 | 0.31 | 0.71 | 0.26 | 0.4 |
| 10 | 114732906 | rs7901275 | -1.035 | -0.535 | -1.889 | -1.886 | -0.014 | 0.059 | 0.059 | 0 | 0.6 | 0.74 | 0.45 | 0.68 | 0.47 | 0.53 |
| 10 | 114734096 | rs11196174 | 0.536 | -0.423 | -2.235 | -2.485 | -0.073 | 0.237 | 0.237 | 0.665 | 0.32 | 0.01 | 0.2 | 0 | 0.31 | 0.2 |
| 10 | 114737050 | rs6585196 | -0.837 | 0.58 | -1.197 | -2.287 | -0.005 | 0.011 | 0.011 | 0 | 0.66 | 0.72 | 0.53 | 0.7 | 0.64 | 0.64 |
| 10 | 114747277 | rs4073288 | -0.292 | 0.423 | -0.267 | -2.531 | 0.012 | 0.138 | 0.138 | 0 | 0.42 | 0.2 | 0.25 | 0.04 | 0.41 | 0.32 |
| 10 | 114748497 | rs4074720 | -1.093 | 0.388 | 0.051 | -2.615 | -0.012 | 0.156 | 0.156 | 0 | 0.53 | 0.78 | 0.34 | 0.05 | 0.5 | 0.38 |
| 10 | 114749018 | rs11196181 | 0.522 | 0.624 | -0.064 | -2.574 | -0.024 | 0.022 | 0.022 | 0 | 0.05 | 0 | 0.03 | 0 | 0.07 | 0.03 |
| 10 | 114754088 | rs7901695 | -0.339 | 0.465 | 0.547 | -2.616 | -0.006 | -0.095 | -0.095 | 1.49 | 0.39 | 0.44 | 0.26 | 0.02 | 0.34 | 0.3 |
| 10 | 114758349 | rs7903146 | -0.122 | 0.48 | 0.744 | -2.606 | 0.015 | 0.047 | 0.047 | 0 | 0.37 | 0.26 | 0.23 | 0.02 | 0.32 | 0.3 |
| 10 | 114766717 | rs7896811 | -0.093 | 0.732 | 0.472 | -2.5 | 0.006 | -0.023 | -0.023 | 0 | 0.11 | 0.14 | 0.06 | 0.02 | 0.14 | 0.08 |
| 10 | 114767771 | rs4132670 | -0.313 | 0.643 | 0.535 | -2.58 | 0.018 | 0.004 | 0.004 | 0 | 0.42 | 0.69 | 0.27 | 0.01 | 0.34 | 0.26 |
| 10 | 114768783 | rs6585201 | -0.884 | 0.557 | 0.505 | -2.532 | -0.005 | 0.214 | 0.214 | 0 | 0.51 | 0.81 | 0.33 | 0.03 | 0.49 | 0.32 |
| 10 | 114777724 | rs10885406 | -1.093 | 0.669 | 0.382 | -2.511 | -0.036 | 0.233 | 0.233 | 0 | 0.54 | 0.85 | 0.34 | 0.04 | 0.49 | 0.38 |
| 10 | 114782287 | rs11196192 | 0.594 | 0.58 | 0.361 | -2.459 | -0.003 | -0.004 | -0.004 | 0.727 | 0.05 | 0.05 | 0.04 | 0.01 | 0.07 | 0.06 |
| 10 | 114788815 | rs12243326 | 0.238 | 0.541 | 0.392 | -2.435 | 0.006 | -0.041 | -0.041 | 0 | 0.34 | 0.32 | 0.21 | 0.01 | 0.29 | 0.22 |
| 10 | 114805860 | rs11196203 | -0.07 | 0.637 | 0.164 | -2.427 | 0 | -0.015 | -0.015 | 0 | 0.14 | 0.14 | 0.07 | 0.02 | 0.16 | 0.1 |
| 10 | 114808902 | rs12255372 | 0.254 | 0.63 | -0.083 | -2.455 | 0.006 | -0.039 | -0.039 | 0.055 | 0.34 | 0.3 | 0.22 | 0.01 | 0.29 | 0.22 |
| 10 | 114810240 | rs12265291 | -0.523 | 0.934 | -0.679 | -2.086 | -0.035 | 0.228 | 0.228 | 0 | 0.54 | 0.86 | 0.35 | 0.03 | 0.49 | 0.37 |
| 10 | 114821249 | rs11196212 | -0.546 | 0.863 | -1.283 | -1.804 | -0.037 | 0.22 | 0.22 | 0 | 0.57 | 0.86 | 0.33 | 0.05 | 0.47 | 0.37 |
| 10 | 114824473 | rs10885410 | -0.391 | 1.66 | -0.663 | -1.039 | 0.005 | -0.012 | -0.012 | 0 | 0.19 | 0.18 | 0.15 | 0.04 | 0.28 | 0.17 |
| 10 | 114824976 | rs7919409 | 0.504 | 0.784 | -0.098 | 0.034 | 0.031 | 0.167 | 0.167 | 0.013 | 0.89 | 0.63 | 0.79 | 0.73 | 0.88 | 0.87 |
| 10 | 114825813 | rs7100388 | -0.841 | 0.746 | -0.997 | -0.146 | -0.003 | -0.012 | -0.012 | 0 | 0.12 | 0.14 | 0.12 | 0.09 | 0.09 | 0.22 |
| 10 | 114833850 | rs3750804 | -0.052 | 0.626 | -1.81 | 0.002 | -0.021 | -0.033 | -0.033 | 0 | 0.31 | 0.05 | 0.2 | 0.25 | 0.27 | 0.3 |
| 10 | 114834498 | rs7908486 | -0.43 | 0.849 | -2.207 | -0.825 | -0.003 | 0.014 | 0.014 | 0 | 0.61 | 0.86 | 0.6 | 0.28 | 0.62 | 0.46 |
| 10 | 114835452 | rs4277044 | 0.476 | 0.83 | -1.418 | -0.892 | 0.01 | -0.069 | -0.069 | 0 | 0.28 | 0.64 | 0.38 | 0.03 | 0.35 | 0.12 |
| 10 | 114840494 | rs11196218 | 0.489 | 0.599 | -1.454 | -0.624 | 0.013 | -0.006 | -0.006 | 0 | 0.32 | 0.17 | 0.21 | 0.28 | 0.27 | 0.34 |
| 10 | 114846649 | rs7082458 | 1.006 | 0.938 | -0.563 | -0.911 | -0.006 | -0.109 | -0.109 | 0 | 0.19 | 0.36 | 0.14 | 0.01 | 0.15 | 0.05 |
| 10 | 114846766 | rs966227 | 0.503 | 0.938 | -0.953 | -0.608 | 0.018 | -0.03 | -0.03 | 0 | 0.33 | 0.15 | 0.31 | 0.34 | 0.28 | 0.34 |
| 10 | 114847723 | rs3814572 | 0.265 | 1.039 | -1.354 | -0.659 | -0.045 | 0.052 | 0.052 | 0 | 0.16 | 0.01 | 0.08 | 0.07 | 0.16 | 0.15 |
| 10 | 114849353 | rs10749127 | 0.908 | 0.443 | -1.189 | -0.892 | 0.001 | -0.006 | -0.006 | 0 | 0.3 | 0.23 | 0.32 | 0.27 | 0.3 | 0.33 |
| 10 | 114849399 | rs4917644 | 0.779 | 0.946 | -1.419 | -1.06 | -0.019 | 0.059 | 0.059 | 0.013 | 0.17 | 0.02 | 0.14 | 0.05 | 0.14 | 0.06 |
| 10 | 114855397 | rs11196224 | 0.971 | 0.429 | -0.607 | -0.852 | 0.003 | 0.001 | 0.001 | 0 | 0.39 | 0.35 | 0.35 | 0.28 | 0.28 | 0.35 |
| 10 | 114859463 | rs7085532 | -0.452 | 1.026 | -0.819 | -0.984 | -0.006 | -0.06 | -0.06 | 0 | 0.7 | 0.83 | 0.72 | 0.44 | 0.68 | 0.57 |
| 10 | 114861304 | rs10885414 | -0.463 | 1.561 | -0.738 | -1.527 | -0.058 | 0.015 | 0.015 | 0 | 0.14 | 0.06 | 0.29 | 0.08 | 0.31 | 0.13 |
| 10 | 114862757 | rs7894421 | 0.536 | 1.013 | -0.417 | -1.439 | 0 | -0.078 | -0.078 | 0.613 | 0.17 | 0.29 | 0.1 | 0.02 | 0.13 | 0.07 |
| 10 | 114865489 | rs10749128 | 0.388 | 1.132 | -1.044 | -1.208 | -0.015 | 0.057 | 0.057 | 0.117 | 0.78 | 0.88 | 0.79 | 0.97 | 0.74 | 0.78 |
| 10 | 114874019 | rs290475 | 0.431 | 1.749 | -1.071 | -0.712 | -0.05 | -0.032 | -0.032 | 1.324 | 0.86 | 0.95 | 0.65 | 0.76 | 0.78 | 0.68 |
| 10 | 114883322 | rs11594681 | -0.64 | 1.826 | -0.924 | -1.218 | 0.011 | -0.035 | -0.035 | 0 | 0.24 | 0.65 | 0.23 | 0.06 | 0.33 | 0.17 |
| 10 | 114887722 | rs11196236 | 1.581 | 1.325 | -1.765 | -1.677 | 0.025 | 0.124 | 0.124 | 0.237 | 0.26 | 0.1 | 0.2 | 0 | 0.22 | 0.1 |
| 10 | 114888608 | rs1362943 | 0.154 | 1.55 | -1.058 | -1.682 | 0.013 | -0.031 | -0.031 | 0.261 | 0.19 | 0.21 | 0.17 | 0.05 | 0.28 | 0.11 |
| 10 | 114893682 | rs10885417 | 0.751 | 1.545 | -1.345 | -1.71 | 0.013 | 0.042 | 0.042 | 0 | 0.34 | 0.26 | 0.47 | 0.23 | 0.29 | 0.33 |
| 10 | 114898093 | rs3814573 | -0.506 | 1.546 | -1.843 | -1.6 | -0.022 | 0.086 | 0.086 | 0 | 0.59 | 0.91 | 0.7 | 0.29 | 0.62 | 0.56 |
| 10 | 114898914 | rs10787476 | 0.69 | 1.242 | -1.439 | -0.963 | -0.023 | -0.03 | -0.03 | 0 | 0.07 | 0.02 | 0.27 | 0.23 | 0.07 | 0.23 |
| 10 | 114904036 | rs7903424 | -0.52 | 1.066 | -1.125 | -1.377 | 0 | -0.015 | -0.015 | 3.271 | 0.25 | 0.23 | 0.17 | 0.06 | 0.25 | 0.14 |
| 10 | 114906714 | rs290490 | 0.426 | 1.983 | -1.126 | -1.237 | 0.087 | -0.067 | -0.067 | 0 | 0.83 | 0.74 | 0.83 | 0.99 | 0.66 | 0.79 |
| 10 | 114907055 | rs290489 | -0.007 | 0.155 | -2.197 | -0.991 | 0.066 | 0.027 | 0.027 | 0 | 0.4 | 0.27 | 0.28 | 0.68 | 0.24 | 0.38 |
| 10 | 114909731 | rs290487 | 0.631 | 0.769 | -2.313 | -0.539 | 0.001 | -0.093 | -0.093 | 0 | 0.21 | 0.05 | 0.19 | 0.62 | 0.2 | 0.25 |
| 10 | 114911079 | rs176632 | -0.703 | 1.57 | -2.045 | -0.699 | -0.008 | 0.002 | 0.002 | 0 | 0.93 | 0.95 | 0.96 | 1 | 0.87 | 0.88 |
| 10 | 114912534 | rs1555485 | 0.66 | 1.845 | -2.332 | -0.687 | -0.042 | -0.023 | -0.023 | 0 | 0.07 | 0.01 | 0.27 | 0.06 | 0.19 | 0.08 |
| 10 | 114912656 | rs1028629 | 0.758 | 0.396 | -2.38 | -0.77 | -0.022 | 0.027 | 0.027 | 0 | 0.18 | 0.07 | 0.17 | 0.14 | 0.16 | 0.24 |

| | | | | | | | | | | | | | | | | |
|---|---|---|---|---|---|---|---|---|---|---|---|---|---|---|---|---|
| 10 | 114915214 | rs290483 | 0.335 | 0.602 | -1.475 | -0.298 | 0 | -0.007 | -0.007 | 0.087 | 0.74 | 0.31 | 0.6 | 0.92 | 0.63 | 0.72 |
| 10 | 114916586 | rs10885421 | 0.515 | 0.502 | -1.637 | -0.299 | -0.046 | 0.064 | 0.064 | 1.503 | 0.7 | 0.22 | 0.59 | 0.89 | 0.63 | 0.68 |
| 10 | 114923825 | rs290481 | -0.496 | -0.777 | -0.995 | 0.636 | 0.023 | -0.025 | -0.025 | 0 | 0.26 | 0.19 | 0.17 | 0.61 | 0.19 | 0.29 |
| 10 | 114925758 | rs1056877 | -1.56 | -0.232 | 0.167 | 0.382 | 0.022 | 0.019 | 0.019 | 4.062 | 0.05 | 0.62 | 0.06 | 0 | 0 | 0 |
| 10 | 114930952 | rs11196253 | -1.543 | -0.308 | -0.286 | 1.026 | -0.015 | 0.037 | 0.037 | 0.144 | 0.07 | 0.12 | 0.06 | 0.22 | 0.03 | 0.18 |
| 10 | 114940502 | rs12243558 | -0.791 | -1.144 | -0.24 | 0.533 | -0.006 | 0.003 | 0.003 | 0 | 0.31 | 0.65 | 0.28 | 0.55 | 0.19 | 0.31 |
| 10 | 114942278 | rs7900784 | -1.332 | -1.079 | -0.245 | 0.489 | 0.083 | 0.085 | 0.085 | 0.468 | 0.39 | 0.84 | 0.33 | 0.78 | 0.23 | 0.58 |
| 10 | 114958159 | rs4918801 | 0.275 | 0.508 | -0.554 | 0.339 | -0.054 | -0.038 | -0.038 | 0 | 0.08 | 0.01 | 0.31 | 0.09 | 0.18 | 0.09 |
| 10 | 114980817 | rs12258181 | -0.433 | -1.039 | 1.178 | 0.906 | -0.034 | 0.209 | 0.209 | 0 | 0.14 | 0.29 | 0.18 | 0.47 | 0.1 | 0.23 |
| 10 | 114992431 | rs6585212 | -1.376 | -1.07 | 1.373 | 0.646 | -0.057 | 0.127 | 0.127 | 0 | 0.22 | 0.48 | 0.23 | 0.57 | 0.14 | 0.3 |
| 10 | 115001714 | rs11819509 | -1.931 | -0.975 | 1.35 | 0.447 | 0.022 | 0.016 | 0.016 | 0.076 | 0.06 | 0.22 | 0.04 | 0.08 | 0.06 | 0.1 |
| 10 | 115005641 | rs1010932 | 0.203 | -1.007 | 1.275 | 0.379 | -0.006 | 0.004 | 0.004 | 0.137 | 0.06 | 0.03 | 0.02 | 0 | 0.05 | 0.02 |
| 10 | 115006062 | rs12098403 | -2.052 | -1.011 | 1.374 | 0.396 | 0.009 | 0.029 | 0.029 | 0.844 | 0.07 | 0.31 | 0.04 | 0.08 | 0.04 | 0.11 |
| 10 | 115010255 | rs7914029 | -2.304 | -1.028 | 1.337 | 0.384 | 0.027 | 0.016 | 0.016 | 0.824 | 0.08 | 0.31 | 0.05 | 0.08 | 0.06 | 0.11 |
| 10 | 115010570 | rs4917647 | -2.13 | -1.278 | 0.8 | 0.184 | 0.043 | 0.078 | 0.078 | 0.751 | 0.37 | 0.77 | 0.33 | 0.88 | 0.23 | 0.49 |
| 10 | 115016030 | rs7075235 | -1.434 | -1.039 | 1.217 | 0.702 | -0.051 | 0.066 | 0.066 | 0 | 0.21 | 0.57 | 0.22 | 0.66 | 0.11 | 0.32 |
| 10 | 115018340 | rs4265536 | -1.298 | -0.963 | 0.933 | 0.539 | -0.003 | 0 | 0 | 0.017 | 0.36 | 0.62 | 0.33 | 0.61 | 0.32 | 0.37 |
| 10 | 115021316 | rs11819326 | -1.296 | -0.988 | 1.347 | 0.771 | -0.05 | 0.233 | 0.233 | 0 | 0.13 | 0.35 | 0.18 | 0.5 | 0.05 | 0.15 |
| 10 | 115021578 | rs7078078 | -1.335 | -0.993 | 1.331 | 0.769 | -0.053 | 0.252 | 0.252 | 0 | 0.13 | 0.39 | 0.17 | 0.49 | 0.05 | 0.16 |
| 10 | 115024801 | rs7092764 | -0.919 | -1.018 | 0.907 | 0.495 | -0.02 | 0.087 | 0.087 | 0 | 0.29 | 0.13 | 0.18 | 0.09 | 0.32 | 0.23 |
| 10 | 115034313 | rs4444012 | -1.735 | -0.863 | 0.647 | 0.59 | 0.047 | 0.192 | 0.192 | 0.067 | 0.48 | 0.79 | 0.43 | 0.83 | 0.41 | 0.53 |
| 10 | 115038675 | rs12219131 | -1.332 | -0.159 | 1.05 | 0.88 | 0.036 | 0.185 | 0.185 | 0.241 | 0.06 | 0.23 | 0.07 | 0.22 | 0.05 | 0.14 |
| 10 | 115049710 | rs4255473 | -0.852 | 0.095 | 0.243 | 0.311 | -0.002 | -0.001 | -0.001 | 0 | 0.21 | 0.21 | 0.15 | 0.09 | 0.26 | 0.16 |
| 10 | 115056097 | rs7906302 | -1.484 | -0.101 | 0.611 | 1.365 | -0.001 | 0.241 | 0.241 | 0.087 | 0.07 | 0.16 | 0.09 | 0.43 | 0.05 | 0.19 |
| 10 | 115057896 | rs2788178 | -1.89 | -0.784 | -0.044 | 0.799 | -0.041 | 0.232 | 0.232 | 0.185 | 0.55 | 0.9 | 0.49 | 0.86 | 0.46 | 0.75 |
| 10 | 115058203 | rs946199 | -0.816 | -1.029 | 0.904 | 0.716 | -0.045 | -0.101 | -0.101 | 0 | 0.21 | 0.47 | 0.1 | 0.05 | 0.11 | 0.12 |
| 10 | 115061448 | rs2788175 | -1.445 | -0.999 | 1.127 | 0.82 | -0.012 | -0.066 | -0.066 | 0 | 0.26 | 0.65 | 0.14 | 0.07 | 0.18 | 0.29 |
| 10 | 115066879 | rs476484 | -1.059 | -1.099 | 1.036 | 0.915 | 0.008 | 0.017 | 0.017 | 0 | 0.27 | 0.19 | 0.33 | 0.79 | 0.25 | 0.43 |
| 10 | 115067809 | rs1538320 | -0.778 | -1.097 | 1.351 | 0.906 | 0.111 | 0.025 | 0.025 | 0 | 0.05 | 0.24 | 0.05 | 0.02 | 0.08 | 0.18 |
| 10 | 115067899 | rs563135 | -1.611 | -1.169 | 1.455 | 0.976 | 0.002 | 0.03 | 0.03 | 0.038 | 0.35 | 0.49 | 0.37 | 0.44 | 0.36 | 0.51 |
| 10 | 115069858 | rs496356 | 1.065 | -1.197 | 1.699 | 0.889 | -0.012 | -0.001 | -0.001 | 0 | 0.88 | 0.68 | 0.92 | 0.9 | 0.91 | 0.83 |
| 10 | 115070140 | rs499832 | 0.334 | -1.195 | 1.72 | 0.884 | -0.005 | -0.031 | -0.031 | 2.353 | 0.91 | 0.77 | 0.94 | 0.95 | 0.92 | 0.9 |
| 10 | 115070465 | rs579902 | -0.458 | -1.165 | 1.493 | 0.916 | -0.007 | -0.06 | -0.06 | 0 | 0.23 | 0.11 | 0.28 | 0.36 | 0.25 | 0.26 |
| 10 | 115071357 | rs550770 | -0.621 | -1.259 | 1.412 | 0.884 | -0.03 | -0.077 | -0.077 | 0 | 0.24 | 0.05 | 0.28 | 0.36 | 0.26 | 0.27 |
| 10 | 115073842 | rs491832 | 1.141 | -1.467 | 1.646 | 1.116 | -0.058 | -0.043 | -0.043 | 0 | 0.78 | 0.61 | 0.9 | 0.9 | 0.89 | 0.82 |
| 10 | 115075143 | rs4918811 | -2.005 | -0.718 | 1.499 | 1.557 | 0.056 | 0.337 | 0.337 | 0.502 | 0.08 | 0.31 | 0.09 | 0.39 | 0.08 | 0.28 |
| 10 | 115077356 | rs1904444 | -0.447 | -0.877 | 0.759 | 1.419 | -0.001 | -0.111 | -0.111 | 0 | 0.16 | 0 | 0.25 | 0.35 | 0.13 | 0.19 |
| 10 | 115080469 | rs490816 | -0.672 | -0.841 | 0.839 | 1.448 | 0.006 | -0.094 | -0.094 | 0 | 0.17 | 0.02 | 0.25 | 0.33 | 0.16 | 0.24 |
| 10 | 115080485 | rs495753 | 1.261 | -0.879 | 1.562 | 1.444 | -0.014 | 0.269 | 0.269 | 0 | 0.88 | 0.65 | 0.88 | 0.51 | 0.91 | 0.72 |
| 10 | 115098417 | rs499050 | 2.066 | -1.142 | 1.924 | 1.264 | -0.036 | 0.135 | 0.135 | 0.186 | 0.8 | 0.38 | 0.83 | 0.49 | 0.88 | 0.72 |
| 10 | 115104798 | rs7080287 | 2.434 | -1.257 | 2.153 | 1.11 | 0.019 | 0.024 | 0.024 | 1.034 | 0.61 | 0.33 | 0.58 | 0.16 | 0.73 | 0.54 |
| 10 | 115105220 | rs2221422 | 2.212 | -1.254 | 2.103 | 1.113 | 0.016 | 0.021 | 0.021 | 1.14 | 0.64 | 0.32 | 0.6 | 0.17 | 0.74 | 0.48 |
| 10 | 115106330 | rs4918814 | 1.956 | -1.205 | 1.923 | 1.13 | -0.014 | -0.038 | -0.038 | 0 | 0.66 | 0.57 | 0.62 | 0.17 | 0.73 | 0.54 |
| 10 | 115107685 | rs536626 | 1.262 | -0.951 | 1.905 | 1.35 | 0 | 0.194 | 0.194 | 0 | 0.88 | 0.71 | 0.9 | 0.55 | 0.91 | 0.8 |
| 10 | 115113013 | rs564782 | 1.751 | -0.824 | 1.455 | 1.223 | -0.014 | 0.162 | 0.162 | 0.313 | 0.78 | 0.46 | 0.84 | 0.53 | 0.84 | 0.62 |
| 10 | 115119640 | rs471447 | -0.844 | -0.947 | 0.97 | 1.593 | 0.009 | -0.016 | -0.016 | 0 | 0.17 | 0.14 | 0.25 | 0.29 | 0.13 | 0.16 |
| 10 | 115119864 | rs7093035 | -1.693 | -0.698 | 0.991 | 1.316 | -0.002 | -0.001 | -0.001 | 0 | 0.08 | 0.08 | 0.04 | 0.11 | 0.09 | 0.1 |
| 10 | 115129261 | rs531790 | 1.205 | -0.869 | 1.864 | 1.357 | -0.008 | -0.015 | -0.015 | 0 | 0.86 | 0.58 | 0.9 | 0.92 | 0.9 | 0.89 |
| 10 | 115132042 | rs491406 | -1.058 | -0.652 | 1.262 | 2.156 | -0.001 | 0.058 | 0.058 | 0 | 0.05 | 0 | 0.19 | 0.5 | 0.03 | 0.21 |
| 10 | 115132537 | rs486759 | -1.242 | -0.668 | 1.445 | 2.159 | -0.005 | 0.157 | 0.157 | 0 | 0.06 | 0.08 | 0.2 | 0.49 | 0.03 | 0.2 |
| 10 | 115140256 | rs17130335 | -1.333 | -0.814 | 0.854 | 1.844 | 0.006 | 0.006 | 0.006 | 0 | 0.07 | 0.07 | 0.04 | 0.05 | 0.05 | 0.19 |
| 10 | 115148569 | rs474616 | -1.585 | -0.831 | 0.79 | 1.874 | 0.008 | 0.003 | 0.003 | 0.013 | 0.12 | 0.13 | 0.23 | 0.57 | 0.08 | 0.41 |
| 10 | 115149323 | rs12763961 | -0.243 | -0.746 | 0.946 | 1.275 | -0.002 | -0.009 | -0.009 | 0 | 0.11 | 0.12 | 0.08 | 0.12 | 0.14 | 0.08 |
| 10 | 115170105 | rs937737 | 1.016 | -0.614 | 0.725 | 0.956 | -0.008 | -0.021 | -0.021 | 0 | 0.75 | 0.37 | 0.81 | 0.82 | 0.77 | 0.82 |
| 10 | 115174263 | rs1566866 | 1.154 | -0.791 | 0.965 | 1.126 | -0.013 | 0.017 | 0.017 | 0.115 | 0.91 | 0.79 | 0.95 | 0.88 | 0.95 | 0.88 |
| 10 | 115175650 | rs11196325 | -0.241 | -0.904 | 0.28 | 0.628 | 0 | 0 | 0 | 0 | 0.42 | 0.57 | 0.64 | 0.56 | 0.53 | 0.68 |
| 10 | 115180454 | rs12414053 | 0.42 | -0.909 | 0.116 | 1.131 | 0.017 | -0.042 | -0.042 | 0.449 | 0.14 | 0.05 | 0.36 | 0.24 | 0.07 | 0.28 |
| 10 | 115180898 | rs10509976 | 1.691 | -0.326 | 0.244 | 1.084 | -0.026 | -0.022 | -0.022 | 0 | 0.17 | 0.11 | 0.2 | 0.29 | 0.36 | 0.26 |
| 10 | 115180995 | rs10787487 | 0.634 | -0.418 | 0.206 | 1.005 | -0.011 | 0.012 | 0.012 | 0.221 | 0.77 | 0.59 | 0.83 | 0.76 | 0.78 | 0.79 |
| 10 | 115192329 | rs6585220 | -0.935 | -0.401 | 0.218 | 0.982 | -0.015 | -0.004 | -0.004 | 0 | 0.94 | 0.87 | 0.98 | 0.95 | 0.98 | 0.93 |
| 10 | 115193523 | rs1416134 | -1.495 | -0.436 | 0.147 | 0.954 | 0.013 | -0.003 | -0.003 | 0.136 | 0.28 | 0.39 | 0.2 | 0.26 | 0.31 | 0.32 |
| 10 | 115193835 | rs7084958 | 0.801 | -0.435 | 0.16 | 0.927 | 0.003 | -0.005 | -0.005 | 0.242 | 0.14 | 0.2 | 0.12 | 0.12 | 0.17 | 0.09 |
| 10 | 115196019 | rs17130385 | 0.608 | -0.524 | 0.011 | 0.953 | 0.021 | -0.043 | -0.043 | 0 | 0.14 | 0.05 | 0.37 | 0.25 | 0.07 | 0.27 |
| 10 | 115198163 | rs7895990 | -0.541 | -0.481 | 0.069 | 0.959 | 0.019 | 0.018 | 0.018 | 0 | 0.38 | 0.47 | 0.37 | 0.54 | 0.58 | 0.45 |
| 10 | 115198443 | rs4917653 | 1.044 | -0.444 | 0.111 | 0.967 | -0.021 | -0.006 | -0.006 | 2.048 | 0.73 | 0.5 | 0.55 | 0.74 | 0.83 | 0.56 |
| 10 | 115205714 | rs10885457 | 2.538 | -1.18 | -0.636 | 0.426 | -0.023 | -0.002 | -0.002 | 1.415 | 0.28 | 0.02 | 0.17 | 0.2 | 0.23 | 0.11 |
| 10 | 115205788 | rs2419820 | 1.324 | -0.783 | -0.28 | 0.614 | -0.027 | -0.027 | -0.027 | 0 | 0.2 | 0.13 | 0.23 | 0.37 | 0.4 | 0.32 |
| 10 | 115209065 | rs1394495 | -1.032 | -1.247 | 0.17 | 0.397 | 0.006 | -0.004 | -0.004 | 0 | 0.37 | 0.62 | 0.56 | 0.59 | 0.26 | 0.45 |
| 10 | 115212692 | rs10736227 | 0.525 | -1.262 | 0.318 | 0.329 | 0.004 | -0.013 | -0.013 | 0 | 0.74 | 0.71 | 0.83 | 0.83 | 0.72 | 0.75 |

| | | | | | | | | | | | | | | | | |
|---|---|---|---|---|---|---|---|---|---|---|---|---|---|---|---|---|
| 10 | 115214154 | rs4532958 | -1.055 | -0.986 | 0.296 | 0.313 | 0.005 | 0.059 | 0.059 | 0.041 | 0.53 | 0.76 | 0.73 | 0.63 | 0.58 | 0.68 |
| 10 | 115219815 | rs6585223 | 0.243 | -0.921 | 0.276 | 0.321 | 0.005 | -0.028 | -0.028 | 0 | 0.76 | 0.74 | 0.84 | 0.89 | 0.73 | 0.79 |
| 10 | 115235656 | rs17769979 | 2.135 | -0.933 | 0.298 | 0.369 | -0.043 | -0.007 | -0.007 | 0.684 | 0.17 | 0.08 | 0.18 | 0.15 | 0.33 | 0.26 |
| 10 | 115236951 | rs7912021 | 0.482 | -0.991 | 0.316 | 0.413 | -0.005 | -0.006 | -0.006 | 1.328 | 0.56 | 0.49 | 0.64 | 0.67 | 0.39 | 0.48 |
| 10 | 115239602 | rs734241 | 0.587 | -1.051 | 0.28 | 0.425 | 0.034 | -0.083 | -0.083 | 1.437 | 0.17 | 0.06 | 0.4 | 0.38 | 0.07 | 0.33 |
| 10 | 115245932 | rs1033772 | 0.515 | -1.029 | 0.321 | 0.355 | -0.007 | -0.008 | -0.008 | 0 | 0.55 | 0.49 | 0.63 | 0.66 | 0.37 | 0.47 |
| 10 | 115248259 | rs1033773 | -0.093 | -0.886 | 0.249 | 0.268 | -0.008 | 0.003 | 0.003 | 0 | 0.9 | 0.92 | 0.95 | 0.99 | 0.9 | 0.88 |
| 10 | 115253601 | rs2900986 | 1.493 | -0.949 | 0.302 | 0.27 | -0.001 | -0.001 | -0.001 | 0.061 | 0.37 | 0.37 | 0.22 | 0.29 | 0.29 | 0.15 |
| 10 | 115257527 | rs1033774 | -0.116 | -0.824 | 0.266 | 0.314 | -0.006 | -0.008 | -0.008 | 0 | 0.38 | 0.34 | 0.61 | 0.53 | 0.42 | 0.61 |
| 10 | 115262306 | rs7899606 | 1.064 | -0.901 | 0.191 | 0.178 | 0.003 | -0.015 | -0.015 | 0 | 0.14 | 0.17 | 0.11 | 0.11 | 0.17 | 0.09 |
| 10 | 115264615 | rs980069 | 1.441 | -0.985 | 0.316 | 0.151 | 0.005 | 0.006 | 0.006 | 1.138 | 0.37 | 0.34 | 0.22 | 0.27 | 0.29 | 0.15 |
| 10 | 115266809 | rs6585225 | 0.691 | -0.91 | 0.226 | 0.231 | 0.003 | 0.01 | 0.01 | 0.935 | 0.51 | 0.52 | 0.34 | 0.38 | 0.47 | 0.24 |
| 10 | 115275327 | rs7077843 | -0.25 | -0.875 | 0.249 | 0.201 | 0.04 | 0.028 | 0.028 | 0 | 0.07 | 0.24 | 0.03 | 0.06 | 0.04 | 0.05 |
| 10 | 115275365 | rs7919248 | 0.295 | -1.184 | 0.103 | 0.31 | 0.02 | -0.036 | -0.036 | 0 | 0.17 | 0.11 | 0.42 | 0.51 | 0.09 | 0.35 |
| 10 | 115276976 | rs2419829 | 1.158 | -1.402 | 0.034 | -0.244 | -0.013 | -0.058 | -0.058 | 0 | 0.33 | 0.49 | 0.19 | 0.17 | 0.26 | 0.14 |
| 10 | 115279811 | rs7100851 | -0.1 | -1.434 | -0.033 | -0.286 | -0.003 | -0.016 | -0.016 | 0 | 0.2 | 0.26 | 0.09 | 0.14 | 0.17 | 0.16 |
| 10 | 115280262 | rs10885470 | 0.418 | -1.43 | 0.002 | -0.278 | -0.001 | -0.006 | -0.006 | 0 | 0.47 | 0.53 | 0.25 | 0.24 | 0.38 | 0.25 |
| 10 | 115285900 | rs11196359 | -0.242 | -1.508 | 0.227 | -0.126 | -0.04 | 0.073 | 0.073 | 0.075 | 0.8 | 0.67 | 0.57 | 0.5 | 0.91 | 0.64 |
| 10 | 115287202 | rs1570269 | 0.531 | -1.506 | 0.247 | -0.166 | -0.05 | 0.078 | 0.078 | 0.793 | 0.73 | 0.44 | 0.55 | 0.44 | 0.87 | 0.58 |
| 10 | 115288501 | rs1570271 | 0.229 | -1.485 | 0.172 | -0.239 | 0.028 | 0.017 | 0.017 | 0.443 | 0.89 | 0.69 | 0.91 | 0.92 | 0.89 | 0.94 |
| 10 | 115292438 | rs11196361 | 2.221 | -1.678 | -0.078 | -0.551 | 0.01 | 0.053 | 0.053 | 1.504 | 0.25 | 0.1 | 0.14 | 0.11 | 0.21 | 0.08 |
| 10 | 115292457 | rs6585227 | -0.189 | -1.554 | 0.277 | -0.39 | -0.014 | 0.022 | 0.022 | 0.051 | 0.78 | 0.75 | 0.58 | 0.5 | 0.9 | 0.62 |
| 10 | 115294705 | rs7088233 | -0.182 | -1.374 | 0.248 | -0.406 | -0.016 | -0.007 | -0.007 | 0.35 | 0.2 | 0.39 | 0.09 | 0.16 | 0.15 | 0.16 |
| 10 | 115298300 | rs11196366 | 0.915 | -1.258 | -0.264 | -0.438 | 0.004 | 0.035 | 0.035 | 0.316 | 0.14 | 0.03 | 0.06 | 0.08 | 0.13 | 0.11 |
| 10 | 115308456 | rs1853204 | 0.789 | -1.329 | -0.082 | -0.437 | -0.01 | -0.005 | -0.005 | 1.18 | 0.58 | 0.45 | 0.49 | 0.33 | 0.74 | 0.45 |
| 10 | 115310098 | rs2000278 | 0.648 | -1.137 | -0.308 | -0.768 | 0.039 | 0.126 | 0.126 | 0.022 | 0.48 | 0.34 | 0.29 | 0.15 | 0.38 | 0.46 |
| 10 | 115310552 | rs3758591 | 0.298 | -1.132 | -0.094 | -0.701 | -0.026 | 0.044 | 0.044 | 0 | 0.68 | 0.57 | 0.49 | 0.41 | 0.73 | 0.75 |
| 10 | 115312606 | rs2240880 | 2.369 | -1.293 | -1.063 | -0.94 | -0.025 | 0.148 | 0.148 | 1.001 | 0.23 | 0.02 | 0.12 | 0.03 | 0.2 | 0.05 |
| 10 | 115314498 | rs10509980 | 0.674 | -0.543 | -0.382 | -0.534 | 0.03 | 0.096 | 0.096 | 0.048 | 0.5 | 0.4 | 0.32 | 0.16 | 0.4 | 0.47 |
| 10 | 115316725 | rs10509981 | 1.946 | -0.472 | -0.525 | -0.446 | -0.055 | -0.03 | -0.03 | 0 | 0.16 | 0.07 | 0.18 | 0.2 | 0.33 | 0.25 |
| 10 | 115316812 | rs4918842 | 0.213 | -0.577 | -0.448 | -0.381 | 0.002 | -0.004 | -0.004 | 0 | 0.14 | 0.08 | 0.4 | 0.47 | 0.12 | 0.14 |
| 10 | 115317262 | rs11575624 | 1.005 | -0.633 | -0.543 | -0.614 | 0 | -0.018 | -0.018 | 0 | 0.12 | 0.11 | 0.09 | 0 | 0.12 | 0.11 |
| 10 | 115319711 | rs4918844 | -0.824 | -0.704 | -0.405 | -0.616 | -0.015 | 0.011 | 0.011 | 0 | 0.32 | 0.47 | 0.5 | 0.65 | 0.27 | 0.27 |
| 10 | 115321129 | rs7094824 | 1.956 | -0.096 | -0.817 | -0.538 | -0.047 | -0.03 | -0.03 | 0 | 0.16 | 0.07 | 0.15 | 0.2 | 0.3 | 0.24 |
| 10 | 115321824 | rs11575634 | 0.886 | -0.22 | -0.613 | -0.496 | -0.002 | -0.017 | -0.017 | 0 | 0.12 | 0.1 | 0.09 | 0 | 0.13 | 0.11 |
| 10 | 115322153 | rs4545483 | 0.703 | -0.11 | -0.689 | -0.366 | -0.059 | -0.024 | -0.024 | 0 | 0.28 | 0.09 | 0.2 | 0.32 | 0.41 | 0.32 |
| 10 | 115322846 | rs3862015 | -0.826 | -0.075 | -0.713 | -0.293 | -0.045 | -0.003 | -0.003 | 1.226 | 0.69 | 0.82 | 0.75 | 0.68 | 0.54 | 0.65 |
| 10 | 115324358 | rs11196379 | -0.875 | -0.281 | -0.5 | -0.242 | -0.015 | 0.065 | 0.065 | 0 | 0.28 | 0.53 | 0.5 | 0.53 | 0.22 | 0.25 |
| 10 | 115325867 | rs3850688 | -0.261 | 0.253 | -0.936 | -0.239 | -0.018 | 0.019 | 0.019 | 0 | 0.62 | 0.74 | 0.75 | 0.71 | 0.56 | 0.62 |
| 10 | 115325999 | rs3911899 | -0.444 | 0.197 | -0.678 | 0.279 | -0.006 | 0.075 | 0.075 | 0 | 0.14 | 0.2 | 0.42 | 0.49 | 0.12 | 0.14 |
| 10 | 115326103 | rs11575663 | -0.127 | 0.364 | -1.014 | -0.021 | -0.004 | -0.041 | -0.041 | 0 | 0.1 | 0.1 | 0.05 | 0 | 0.07 | 0.08 |
| 10 | 115326786 | rs10749134 | -0.872 | 0.726 | -1.109 | 0.301 | 0 | 0.014 | 0.014 | 0.015 | 0.29 | 0.24 | 0.49 | 0.48 | 0.26 | 0.35 |
| 10 | 115327084 | rs11575668 | 0.742 | 0.803 | -1.681 | -0.098 | -0.008 | 0.02 | 0.02 | 0 | 0.1 | 0.03 | 0.06 | 0.04 | 0.09 | 0.11 |
| 10 | 115327399 | rs2419837 | 0.355 | 0.585 | -1.73 | -0.281 | 0.042 | 0.063 | 0.063 | 0.589 | 0.33 | 0.17 | 0.17 | 0.22 | 0.22 | 0.35 |
| 10 | 115327818 | rs4918852 | 0.239 | 1.659 | -1.157 | 0.048 | 0.008 | -0.009 | -0.009 | 0 | 0.19 | 0.22 | 0.21 | 0.09 | 0.34 | 0.23 |
| 10 | 115328860 | rs11575670 | -0.159 | 1.461 | -1.874 | 0.184 | -0.064 | 0.027 | 0.027 | 0 | 0.41 | 0.13 | 0.33 | 0.28 | 0.51 | 0.35 |
| 10 | 115328950 | rs3862019 | -0.247 | 0.658 | -2.189 | -0.038 | 0.018 | 0.064 | 0.064 | 0 | 0.62 | 0.3 | 0.71 | 0.8 | 0.66 | 0.77 |
| 10 | 115329346 | rs3850691 | -0.23 | 0.218 | -1.615 | -0.249 | 0.001 | -0.001 | -0.001 | 0.36 | 0.77 | 0.78 | 0.83 | 0.91 | 0.8 | 0.82 |
| 10 | 115330721 | rs10885476 | 0.797 | 0.12 | -1.317 | -0.275 | -0.001 | 0.013 | 0.013 | 0 | 0.13 | 0.12 | 0.11 | 0.08 | 0.14 | 0.13 |
| 10 | 115331618 | rs911707 | -0.574 | 0.018 | -1.298 | -0.303 | 0.001 | -0.004 | -0.004 | 2.752 | 0.78 | 0.79 | 0.85 | 0.92 | 0.81 | 0.83 |
| 10 | 115332098 | rs2419840 | 1.023 | 0.129 | -2.06 | 0.006 | -0.064 | 0.028 | 0.028 | 0 | 0.38 | 0.12 | 0.46 | 0.3 | 0.43 | 0.5 |
| 10 | 115332240 | rs10749135 | 0.331 | -0.077 | -1.617 | 0.431 | 0.021 | 0.027 | 0.027 | 0 | 0.54 | 0.4 | 0.64 | 0.61 | 0.62 | 0.61 |
| 10 | 115332413 | rs10885477 | 1.026 | -0.054 | -1.541 | 0.445 | 0.001 | -0.012 | -0.012 | 0 | 0.08 | 0.04 | 0.11 | 0.25 | 0.06 | 0.09 |
| 10 | 115334124 | rs3740530 | 0.324 | -0.09 | -1.525 | 0.435 | 0.031 | 0.044 | 0.044 | 0 | 0.54 | 0.3 | 0.64 | 0.6 | 0.62 | 0.63 |
| 10 | 115334636 | rs11196383 | 1.252 | -0.005 | -1.513 | 0.48 | 0.002 | -0.01 | -0.01 | 0.034 | 0.08 | 0.04 | 0.11 | 0.24 | 0.06 | 0.09 |
| 10 | 115336367 | rs7923349 | -0.298 | -0.033 | -1.472 | 0.448 | -0.013 | -0.008 | -0.008 | 0.362 | 0.67 | 0.48 | 0.74 | 0.68 | 0.75 | 0.75 |
| 10 | 115336758 | rs2302374 | 1.014 | 0.13 | -1.222 | 0.836 | -0.005 | -0.009 | -0.009 | 0 | 0.14 | 0.12 | 0.17 | 0.31 | 0.19 | 0.13 |
| 10 | 115337393 | rs1885437 | -0.389 | 0.162 | -1.227 | 0.485 | 0 | -0.001 | -0.001 | 0 | 0.37 | 0.37 | 0.37 | 0.62 | 0.38 | 0.35 |
| 10 | 115340273 | rs2286743 | 1.057 | -0.171 | -1.304 | 0.402 | 0.044 | 0.011 | 0.011 | 1.11 | 0.09 | 0.02 | 0.01 | 0.07 | 0.01 | 0.23 |
| 10 | 115341150 | rs11575687 | -0.249 | 0.404 | -1.057 | 0.485 | 0.004 | -0.008 | -0.008 | 0 | 0.38 | 0.35 | 0.29 | 0.44 | 0.31 | 0.41 |
| 10 | 115341753 | rs2245056 | 0.592 | 0.428 | -1.117 | 0.282 | 0.005 | -0.012 | -0.012 | 15.16 | 0.36 | 0.42 | 0.49 | 0.3 | 0.39 | 0.31 |
| 10 | 115343287 | rs2302373 | -0.377 | 0.596 | -1.175 | 0.238 | -0.018 | 0.01 | 0.01 | 2.428 | 0.66 | 0.73 | 0.74 | 0.7 | 0.6 | 0.66 |
| 10 | 115343700 | rs911704 | 0.634 | 0.591 | -1.189 | 0.253 | 0.001 | -0.003 | -0.003 | 0 | 0.3 | 0.31 | 0.24 | 0.4 | 0.2 | 0.35 |
| 10 | 115344355 | rs11196386 | 2.211 | 0.697 | -0.996 | 0.13 | -0.004 | -0.003 | -0.003 | 1.376 | 0.17 | 0.11 | 0.31 | 0.19 | 0.19 | 0.16 |
| 10 | 115344567 | rs10787491 | 1.173 | 0.696 | -0.973 | 0.085 | -0.009 | 0.001 | 0.001 | 0 | 0.23 | 0.17 | 0.41 | 0.2 | 0.27 | 0.21 |
| 10 | 115345850 | rs3781378 | 0.995 | 0.414 | -1.075 | 0.284 | -0.01 | 0.001 | 0.001 | 0.877 | 0.28 | 0.33 | 0.2 | 0.31 | 0.17 | 0.29 |
| 10 | 115347359 | rs932650 | 0.247 | 0.524 | -0.995 | -0.108 | -0.01 | -0.006 | -0.006 | 0 | 0.33 | 0.38 | 0.47 | 0.3 | 0.3 | 0.32 |
| 10 | 115348916 | rs10885478 | 1.984 | 0.559 | -1.105 | -0.095 | 0.003 | 0.007 | 0.007 | 0 | 0.12 | 0.04 | 0.07 | 0.1 | 0.12 | 0.11 |
| 10 | 115350597 | rs1885434 | 0.224 | 0.507 | -0.869 | -0.124 | -0.015 | -0.008 | -0.008 | 0.008 | 0.33 | 0.42 | 0.47 | 0.3 | 0.3 | 0.32 |

| | | | | | | | | | | | | | | | | |
|---|---|---|---|---|---|---|---|---|---|---|---|---|---|---|---|---|
| 10 | 115350929 | rs10787493 | 1.461 | 0.532 | -1.027 | -0.118 | 0.002 | -0.002 | -0.002 | 0 | 0.19 | 0.12 | 0.38 | 0.19 | 0.19 | 0.21 |
| 10 | 115352005 | rs11196389 | 1.922 | 0.608 | -1.456 | -0.121 | 0.002 | 0.008 | 0.008 | 0 | 0.12 | 0.04 | 0.07 | 0.11 | 0.12 | 0.11 |
| 10 | 115353701 | rs4918859 | 0.772 | 0.538 | -1.503 | -0.166 | -0.015 | 0.025 | 0.025 | 0 | 0.17 | 0.05 | 0.16 | 0.11 | 0.19 | 0.16 |
| 10 | 115354572 | rs1140 | 1.656 | 0.724 | -1.765 | -0.246 | -0.061 | -0.011 | -0.011 | 0 | 0.13 | 0 | 0.09 | 0.07 | 0.2 | 0.17 |
| 10 | 115355328 | rs11196391 | 1.762 | 0.286 | -1.454 | -0.029 | -0.007 | 0.013 | 0.013 | 0 | 0.06 | 0.03 | 0.09 | 0.01 | 0.08 | 0.05 |
| 10 | 115357179 | rs885567 | 1.083 | 0.226 | -1.446 | -0.01 | 0.007 | -0.013 | -0.013 | 0 | 0.3 | 0.29 | 0.26 | 0.4 | 0.25 | 0.33 |
| 10 | 115361317 | rs2419843 | -0.87 | 0.357 | -1.279 | -0.128 | -0.004 | 0.003 | 0.003 | 1.315 | 0.73 | 0.75 | 0.82 | 0.9 | 0.67 | 0.76 |
| 10 | 115363379 | rs2419844 | -0.315 | 0.468 | -1.636 | 0.211 | 0.019 | 0.075 | 0.075 | 0.738 | 0.51 | 0.3 | 0.42 | 0.7 | 0.47 | 0.54 |
| 10 | 115363881 | rs2419846 | -0.703 | 0.559 | -1.512 | 0.317 | -0.011 | 0.024 | 0.024 | 0.283 | 0.57 | 0.4 | 0.51 | 0.71 | 0.55 | 0.6 |
| 10 | 115370274 | rs10749138 | -0.473 | 0.69 | -1.543 | 0.398 | 0.008 | 0.016 | 0.016 | 0 | 0.47 | 0.33 | 0.47 | 0.6 | 0.47 | 0.49 |
| 10 | 115371873 | rs1539588 | 0.591 | 1.232 | -1.556 | 0.724 | -0.057 | -0.089 | -0.089 | 0 | 0.2 | 0.07 | 0.25 | 0.34 | 0.26 | 0.17 |
| 10 | 115373851 | rs2419850 | -1.117 | 0.839 | -0.883 | 0.314 | -0.008 | -0.007 | -0.007 | 0.105 | 0.89 | 0.8 | 0.92 | 0.9 | 0.9 | 0.91 |
| 10 | 115374670 | rs3121453 | -0.003 | 1.31 | -0.881 | -0.055 | 0.003 | -0.009 | -0.009 | 0 | 0.14 | 0.13 | 0.16 | 0.02 | 0.21 | 0.12 |
| 10 | 115376651 | rs3127084 | 0.303 | 0.487 | -0.89 | -0.013 | 0.007 | 0.018 | 0.018 | 0 | 0.53 | 0.33 | 0.56 | 0.35 | 0.53 | 0.64 |
| 10 | 115377503 | rs10509982 | 1.085 | 0.568 | -1.332 | 0.288 | -0.07 | -0.117 | -0.117 | 0 | 0.2 | 0.05 | 0.25 | 0.4 | 0.29 | 0.19 |
| 10 | 115380608 | rs2419851 | -1.198 | 0.375 | -1.028 | 0.107 | -0.009 | -0.014 | -0.014 | 0 | 0.87 | 0.81 | 0.91 | 0.9 | 0.89 | 0.9 |
| 10 | 115381747 | rs868738 | -0.163 | 0.704 | -1.403 | 0.119 | -0.079 | -0.004 | -0.004 | 0 | 0.23 | 0.04 | 0.44 | 0.18 | 0.32 | 0.19 |
| 10 | 115388588 | rs2286741 | -0.201 | 0.798 | -1.258 | -0.017 | -0.014 | -0.008 | -0.008 | 0 | 0.25 | 0.2 | 0.4 | 0.28 | 0.3 | 0.29 |
| 10 | 115389356 | rs2286736 | -0.215 | 0.786 | -1.252 | -0.022 | -0.019 | -0.012 | -0.012 | 0 | 0.25 | 0.18 | 0.39 | 0.28 | 0.3 | 0.29 |
| 10 | 115389366 | rs2286735 | -0.215 | 0.786 | -1.252 | -0.022 | -0.019 | -0.012 | -0.012 | 0 | 0.25 | 0.18 | 0.39 | 0.28 | 0.3 | 0.29 |
| 10 | 115390353 | rs7912899 | -0.543 | -0.094 | -1.27 | -0.273 | 0.014 | 0.02 | 0.02 | 0.224 | 0.56 | 0.43 | 0.68 | 0.71 | 0.62 | 0.64 |
| 10 | 115393386 | rs11592114 | 1.178 | -0.046 | -1.363 | -0.293 | -0.037 | 0.01 | 0.01 | 0 | 0.08 | 0.01 | 0.11 | 0.02 | 0.13 | 0.07 |
| 10 | 115393929 | rs3189030 | 1.048 | -0.083 | -1.333 | -0.305 | -0.027 | -0.075 | -0.075 | 0 | 0.3 | 0.12 | 0.28 | 0.43 | 0.33 | 0.35 |
| 10 | 115393948 | rs11196400 | 1.848 | -0.348 | -1.123 | -0.422 | -0.042 | -0.027 | -0.027 | 1.345 | 0.22 | 0.31 | 0.11 | 0.17 | 0.14 | 0.17 |
| 10 | 115401813 | rs11196402 | 1.228 | -0.283 | -1.111 | -0.376 | -0.028 | -0.077 | -0.077 | 0 | 0.3 | 0.11 | 0.28 | 0.43 | 0.33 | 0.35 |
| 10 | 115405615 | rs3127106 | -0.412 | -0.247 | -1.066 | -0.383 | 0.046 | 0.071 | 0.071 | 0.411 | 0.55 | 0.3 | 0.67 | 0.71 | 0.62 | 0.64 |
| 10 | 115409840 | rs2275799 | 1.024 | -0.187 | -1.093 | -0.26 | -0.019 | -0.026 | -0.026 | 0.741 | 0.3 | 0.34 | 0.22 | 0.22 | 0.27 | 0.25 |
| 10 | 115410234 | rs2185913 | 1.872 | -0.357 | -1.288 | -0.161 | -0.109 | -0.121 | -0.121 | 0 | 0.21 | 0.02 | 0.25 | 0.33 | 0.3 | 0.19 |
| 10 | 115411598 | rs4918861 | 0.8 | -0.033 | -0.827 | -0.204 | -0.018 | -0.024 | -0.024 | 0 | 0.3 | 0.33 | 0.22 | 0.22 | 0.27 | 0.25 |
| 10 | 115412793 | rs3737322 | 0.641 | -0.039 | -0.874 | -0.21 | -0.026 | -0.032 | -0.032 | 0 | 0.31 | 0.49 | 0.24 | 0.22 | 0.27 | 0.25 |
| 10 | 115414911 | rs12775876 | 1.39 | -0.117 | -1.245 | -0.211 | -0.052 | -0.081 | -0.081 | 0 | 0.27 | 0.1 | 0.28 | 0.4 | 0.33 | 0.36 |
| 10 | 115417289 | rs1891762 | -1.594 | -0.174 | -1.053 | -0.244 | -0.006 | -0.011 | -0.011 | 0 | 0.86 | 0.81 | 0.91 | 0.9 | 0.89 | 0.9 |
| 10 | 115418188 | rs10885490 | 0.336 | -0.122 | -0.732 | -0.226 | 0.02 | -0.016 | -0.016 | 0 | 0.11 | 0.27 | 0.13 | 0.05 | 0.13 | 0.08 |
| 10 | 115419049 | rs7907942 | 0.336 | -0.078 | -0.732 | -0.226 | 0.023 | -0.016 | -0.016 | 0 | 0.11 | 0.27 | 0.13 | 0.05 | 0.13 | 0.08 |
| 10 | 115421996 | rs3121491 | -1.01 | -0.203 | -0.908 | -0.265 | -0.01 | -0.025 | -0.025 | 0.093 | 0.58 | 0.7 | 0.63 | 0.5 | 0.56 | 0.54 |
| 10 | 115422474 | rs3127122 | -0.143 | -0.201 | -1.02 | -0.28 | 0.001 | -0.001 | -0.001 | 0 | 0.26 | 0.29 | 0.46 | 0.21 | 0.32 | 0.21 |
| 10 | 115423695 | rs3127124 | 1.569 | -0.119 | -1.19 | -0.244 | -0.053 | -0.035 | -0.035 | 0 | 0.15 | 0.03 | 0.32 | 0.16 | 0.19 | 0.13 |
| 10 | 115423747 | rs1109927 | 0.679 | -0.142 | -1.159 | -0.242 | -0.036 | -0.015 | -0.015 | 0 | 0.33 | 0.41 | 0.18 | 0.29 | 0.24 | 0.33 |
| 10 | 115424373 | rs17773680 | 1.65 | 0.277 | -1.255 | -0.101 | -0.022 | -0.023 | -0.023 | 0.101 | 0.07 | 0.02 | 0.05 | 0.12 | 0.1 | 0.16 |
| 10 | 115426775 | rs3127127 | -0.081 | 0.261 | -1.246 | -0.118 | -0.085 | -0.033 | -0.033 | 0 | 0.24 | 0.04 | 0.43 | 0.22 | 0.32 | 0.21 |
| 10 | 115430933 | rs10885493 | 1.084 | 0.223 | -1.064 | -0.137 | -0.011 | 0.012 | 0.012 | 0 | 0.07 | 0.03 | 0.11 | 0.03 | 0.12 | 0.05 |
| 10 | 115431240 | rs7906704 | 0.681 | -0.007 | -0.876 | -0.116 | -0.015 | -0.002 | -0.002 | 0.524 | 0.29 | 0.32 | 0.17 | 0.29 | 0.24 | 0.33 |
| 10 | 115443090 | rs11196422 | 1.251 | -0.535 | -0.85 | -0.128 | -0.045 | -0.027 | -0.027 | 0 | 0.23 | 0.28 | 0.12 | 0.17 | 0.14 | 0.18 |
| 10 | 115453989 | rs10787495 | -1.466 | -0.285 | -1.224 | -0.116 | -0.07 | -0.059 | -0.059 | 1.119 | 0.86 | 0.99 | 0.68 | 0.82 | 0.82 | 0.88 |
| 10 | 115457597 | rs7100728 | 1.477 | 0.303 | -1.226 | -0.229 | -0.033 | -0.027 | -0.027 | 4.034 | 0.07 | 0.02 | 0.04 | 0.11 | 0.11 | 0.08 |
| 10 | 115461797 | rs12255142 | 1.348 | 0.1 | -1.038 | -0.383 | -0.07 | -0.046 | -0.046 | 1.652 | 0.24 | 0.39 | 0.13 | 0.17 | 0.14 | 0.18 |
| 10 | 115464171 | rs3124743 | -0.572 | 0.192 | -1.218 | -0.478 | 0 | -0.003 | -0.003 | 0.008 | 0.52 | 0.44 | 0.58 | 0.39 | 0.51 | 0.51 |
| 10 | 115469890 | rs11196445 | 2.335 | -0.073 | -1.584 | -0.62 | 0.031 | 0.006 | 0.006 | 0 | 0.17 | 0.07 | 0.09 | 0.17 | 0.13 | 0.17 |
| 10 | 115469992 | rs4457708 | 0.744 | 0.324 | -1.756 | -0.439 | -0.046 | 0.013 | 0.013 | 0.934 | 0.14 | 0.03 | 0.35 | 0.07 | 0.19 | 0.21 |
| 10 | 115471016 | rs11196446 | 1.16 | 0.043 | -1.743 | -0.407 | -0.064 | -0.099 | -0.099 | 2.32 | 0.32 | 0.11 | 0.3 | 0.51 | 0.38 | 0.37 |
| 10 | 115478677 | rs17090907 | 1.563 | 0.192 | -1.652 | -0.438 | 0.008 | 0.011 | 0.011 | 0 | 0.13 | 0.1 | 0.08 | 0.09 | 0.1 | 0.09 |
| 10 | 115479268 | rs11196448 | 1.35 | 0.501 | -1.646 | -0.299 | -0.042 | -0.026 | -0.026 | 0.111 | 0.07 | 0 | 0.04 | 0.09 | 0.11 | 0.08 |
| 10 | 115480582 | rs11196449 | -0.076 | 0.346 | -1.65 | -0.352 | -0.084 | -0.005 | -0.005 | 0 | 0.72 | 0.9 | 0.53 | 0.78 | 0.62 | 0.64 |
| 10 | 115484660 | rs12416109 | 0.465 | 0.204 | -1.285 | -0.313 | -0.002 | 0.004 | 0.004 | 0.11 | 0.46 | 0.49 | 0.4 | 0.63 | 0.49 | 0.44 |
| 10 | 115486610 | rs6585241 | -1.339 | 0.262 | -1.447 | -0.358 | -0.032 | -0.014 | -0.014 | 0.088 | 0.84 | 0.93 | 0.89 | 0.83 | 0.81 | 0.84 |
| 10 | 115489152 | rs2227310 | 1.994 | 0.106 | -1.567 | -0.307 | -0.012 | -0.048 | -0.048 | 0.688 | 0.22 | 0.13 | 0.26 | 0.42 | 0.25 | 0.23 |
| 10 | 115489167 | rs2227309 | 1.981 | 0.101 | -1.57 | -0.283 | -0.011 | -0.048 | -0.048 | 0.245 | 0.22 | 0.13 | 0.26 | 0.41 | 0.25 | 0.23 |
| 10 | 115489589 | rs4353229 | 1.946 | 0.084 | -1.577 | -0.292 | -0.011 | -0.049 | -0.049 | 0 | 0.23 | 0.13 | 0.26 | 0.42 | 0.25 | 0.23 |
| 10 | 115489650 | rs10787498 | 0.015 | -0.026 | -1.578 | -0.373 | -0.021 | -0.049 | -0.049 | 0 | 0.39 | 0.44 | 0.49 | 0.2 | 0.33 | 0.39 |
| 10 | 115490060 | rs12247479 | 1.521 | -0.047 | -1.38 | -0.381 | -0.037 | -0.036 | -0.036 | 0.963 | 0.21 | 0.26 | 0.12 | 0.13 | 0.13 | 0.18 |
| 10 | 115490109 | rs1127687 | 0.127 | 0.558 | -1.356 | -0.15 | 0.001 | -0.004 | -0.004 | 0 | 0.23 | 0.3 | 0.14 | 0.22 | 0.23 | 0.21 |
| 10 | 115491230 | rs17090919 | 1.273 | 0.623 | -1.356 | -0.173 | -0.01 | -0.008 | -0.008 | 2.931 | 0.09 | 0.04 | 0.04 | 0.12 | 0.12 | 0.12 |
| 10 | 115493733 | rs11196454 | 0.001 | 0.581 | -1.377 | -0.11 | 0.001 | -0.002 | -0.002 | 0 | 0.23 | 0.3 | 0.14 | 0.23 | 0.23 | 0.21 |
| 10 | 115497318 | rs12358301 | 1.378 | 0.621 | -1.49 | -0.136 | -0.042 | 0 | 0 | 0 | 0.07 | 0.01 | 0.33 | 0.05 | 0.16 | 0.08 |
| 10 | 115500433 | rs7094186 | 1.016 | 0.361 | -1.192 | 0.027 | 0.008 | 0.006 | 0.006 | 0 | 0.14 | 0.16 | 0.09 | 0.11 | 0.11 | 0.09 |
| 10 | 115505635 | rs3124735 | -0.729 | 0.435 | -1.104 | 0.023 | -0.004 | -0.001 | -0.001 | 0 | 0.4 | 0.41 | 0.25 | 0.41 | 0.41 | 0.38 |
| 10 | 115508382 | rs4394799 | -0.028 | 0.571 | -0.589 | 0.186 | 0.021 | 0.009 | 0.009 | 0 | 0.11 | 0.21 | 0.06 | 0.12 | 0.12 | 0.13 |
| 10 | 115513098 | rs867300 | 0.318 | 0.678 | -0.913 | 0.219 | -0.009 | -0.005 | -0.005 | 0 | 0.1 | 0.07 | 0.06 | 0.12 | 0.12 | 0.13 |
| 10 | 115515120 | rs3981351 | -1.161 | 0.49 | -0.574 | 0.32 | 0.008 | 0.061 | 0.061 | 0.998 | 0.77 | 0.65 | 0.75 | 0.58 | 0.76 | 0.75 |

| | | | | | | | | | | | | | | | | |
|---|---|---|---|---|---|---|---|---|---|---|---|---|---|---|---|---|
| 10 | 115520691 | rs7091079 | 0.191 | 0.609 | -0.907 | 0.058 | 0.019 | -0.03 | -0.03 | 0.064 | 0.2 | 0.18 | 0.17 | 0.32 | 0.16 | 0.26 |
| 10 | 115524913 | rs4528272 | -0.102 | 0.537 | -0.612 | 0.009 | -0.033 | 0.035 | 0.035 | 1.544 | 0.23 | 0.45 | 0.2 | 0.31 | 0.16 | 0.26 |
| 10 | 115525175 | rs4631835 | -1.369 | 0.368 | -0.961 | -0.248 | -0.029 | 0.03 | 0.03 | 4.135 | 0.67 | 0.52 | 0.76 | 0.57 | 0.79 | 0.68 |
| 10 | 115526177 | rs10885500 | 0.672 | 1.032 | -1.518 | -0.398 | -0.128 | 0.081 | 0.081 | 0 | 0.27 | 0.02 | 0.46 | 0.14 | 0.42 | 0.26 |
| 10 | 115532108 | rs3763795 | 0.756 | 1.137 | -1.481 | -0.381 | -0.116 | 0.079 | 0.079 | 0 | 0.26 | 0.02 | 0.44 | 0.14 | 0.4 | 0.27 |
| 10 | 115532291 | rs11196479 | 0.253 | 1.225 | -0.802 | -0.11 | -0.06 | 0.034 | 0.034 | 0 | 0.17 | 0.03 | 0.1 | 0.07 | 0.21 | 0.16 |
| 10 | 115535599 | rs3750901 | 0.415 | 1.156 | -0.793 | -0.075 | 0.016 | -0.06 | -0.06 | 0 | 0.21 | 0.11 | 0.18 | 0.35 | 0.18 | 0.26 |
| 10 | 115540710 | rs3750899 | -1.065 | 1.209 | -0.644 | -0.082 | -0.002 | -0.008 | -0.008 | 0.357 | 0.4 | 0.34 | 0.3 | 0.42 | 0.39 | 0.42 |
| 10 | 115542147 | rs2419874 | -0.985 | 1.167 | -0.64 | -0.079 | 0 | -0.006 | -0.006 | 3.97 | 0.4 | 0.39 | 0.31 | 0.42 | 0.39 | 0.42 |
| 10 | 115542186 | rs2419873 | -0.986 | 1.167 | -0.598 | -0.079 | -0.007 | -0.022 | -0.022 | 2.015 | 0.39 | 0.28 | 0.3 | 0.42 | 0.39 | 0.42 |
| 10 | 115543045 | rs1065209 | 0.441 | 1.2 | -0.586 | -0.027 | 0.013 | -0.062 | -0.062 | 0 | 0.21 | 0.1 | 0.18 | 0.35 | 0.18 | 0.26 |
| 10 | 115544197 | rs2419872 | -0.679 | 1.256 | -0.472 | 0.044 | 0 | -0.006 | -0.006 | 0.169 | 0.4 | 0.39 | 0.3 | 0.42 | 0.38 | 0.42 |
| 10 | 115545724 | rs2419871 | -0.594 | 1.286 | -0.628 | 0.057 | -0.004 | -0.019 | -0.019 | 0.138 | 0.39 | 0.29 | 0.29 | 0.42 | 0.38 | 0.42 |
| 10 | 115547965 | rs7902084 | -0.506 | 1.344 | -0.406 | -0.056 | 0 | 0.005 | 0.005 | 0.534 | 0.4 | 0.6 | 0.32 | 0.38 | 0.36 | 0.36 |
| 10 | 115561171 | rs10509985 | 0.42 | 1.472 | -0.424 | 0.076 | -0.04 | -0.012 | -0.012 | 1.169 | 0.36 | 0.21 | 0.57 | 0.42 | 0.47 | 0.35 |
| 10 | 115561321 | rs10509986 | 0.193 | 1.529 | 0.003 | -0.05 | 0.01 | 0.002 | 0.002 | 0 | 0.14 | 0.15 | 0.09 | 0.11 | 0.17 | 0.11 |
| 10 | 115565998 | rs7071839 | -0.311 | 0.821 | 0.457 | -0.093 | 0.021 | -0.007 | -0.007 | 5.881 | 0.53 | 0.64 | 0.69 | 0.51 | 0.63 | 0.47 |
| 10 | 115583459 | rs2419859 | -0.099 | 1.138 | -0.379 | -0.03 | 0 | -0.001 | -0.001 | 0 | 0.49 | 0.52 | 0.63 | 0.49 | 0.54 | 0.51 |
| 10 | 115587010 | rs7895170 | -0.907 | 0.517 | -0.714 | 0.092 | 0.022 | -0.042 | -0.042 | 0 | 0.69 | 0.8 | 0.81 | 0.56 | 0.79 | 0.66 |
| 10 | 115589161 | rs11196524 | -0.456 | 1.08 | -0.455 | -0.157 | -0.001 | -0.034 | -0.034 | 0.095 | 0.19 | 0.2 | 0.15 | 0.08 | 0.2 | 0.14 |
| 10 | 115590412 | rs7084835 | 0.545 | 1.322 | -0.515 | 0.065 | 0.001 | -0.01 | -0.01 | 0.732 | 0.67 | 0.36 | 0.69 | 0.82 | 0.67 | 0.8 |
| 10 | 115601557 | rs10509990 | -0.311 | 1.571 | -0.86 | 0.667 | -0.018 | -0.023 | -0.023 | 0 | 0.07 | 0.03 | 0.13 | 0.18 | 0.13 | 0.15 |
| 10 | 115622375 | rs11196534 | 0.978 | 1.479 | -1.726 | 0.379 | -0.052 | -0.066 | -0.066 | 0 | 0.53 | 0.09 | 0.57 | 0.43 | 0.5 | 0.51 |
| 10 | 115628218 | rs4373875 | -0.795 | 1.495 | -1.041 | 0.199 | -0.001 | -0.024 | -0.024 | 0.161 | 0.26 | 0.69 | 0.18 | 0.18 | 0.21 | 0.29 |
| 10 | 115631743 | rs6585248 | 0.335 | 1.409 | -0.271 | 0.157 | 0.001 | -0.014 | -0.014 | 0.118 | 0.77 | 0.66 | 0.85 | 0.82 | 0.79 | 0.72 |
| 10 | 115644040 | rs7913176 | 0.036 | 1.409 | -0.549 | 0.242 | -0.032 | -0.055 | -0.055 | 0.445 | 0.19 | 0.12 | 0.23 | 0.31 | 0.29 | 0.19 |
| 10 | 115646295 | rs4323816 | -0.136 | 1.422 | -0.497 | -0.127 | -0.011 | 0.007 | 0.007 | 0 | 0.86 | 0.87 | 0.88 | 0.93 | 0.81 | 0.83 |
| 10 | 115651253 | rs6585251 | 0.005 | 1.597 | -0.754 | -0.031 | -0.003 | -0.001 | -0.001 | 0 | 0.2 | 0.2 | 0.25 | 0.41 | 0.3 | 0.2 |
| 10 | 115660297 | rs10787509 | 1.093 | 1.53 | -1.169 | 0.004 | -0.059 | -0.045 | -0.045 | 0 | 0.52 | 0.08 | 0.57 | 0.4 | 0.49 | 0.51 |
| 10 | 115661165 | rs4345919 | 0.294 | 1.334 | -1.012 | 0.289 | -0.006 | -0.036 | -0.036 | 0.335 | 0.12 | 0.04 | 0.09 | 0.29 | 0.13 | 0.25 |
| 10 | 115679508 | rs7077378 | 0.711 | 0.666 | -0.337 | -0.454 | -0.009 | -0.064 | -0.064 | 0 | 0.66 | 0.53 | 0.81 | 0.8 | 0.73 | 0.61 |
| 10 | 115679965 | rs1576479 | 0.654 | 0.863 | -0.792 | -0.384 | -0.062 | 0.062 | 0.062 | 3.318 | 0.53 | 0.11 | 0.37 | 0.34 | 0.53 | 0.4 |
| 10 | 115679995 | rs1576480 | 0.889 | 0.968 | -0.629 | -0.322 | -0.013 | 0.091 | 0.091 | 0.559 | 0.5 | 0.13 | 0.37 | 0.34 | 0.51 | 0.4 |
| 10 | 115680935 | rs239863 | -0.403 | 0.631 | -0.34 | -0.299 | -0.004 | -0.04 | -0.04 | 0.466 | 0.29 | 0.37 | 0.4 | 0.14 | 0.25 | 0.4 |
| 10 | 115685187 | rs239865 | 0.043 | 0.868 | 0.017 | 0.273 | 0.027 | 0.173 | 0.173 | 0.009 | 0.8 | 0.49 | 0.75 | 0.49 | 0.77 | 0.8 |
| 10 | 115694632 | rs11196543 | -0.236 | 0.808 | -0.435 | 0.271 | -0.006 | -0.03 | -0.03 | 0 | 0.24 | 0.48 | 0.42 | 0.17 | 0.22 | 0.36 |
| 10 | 115697178 | rs151598 | -1.004 | 1.026 | -0.897 | 0.245 | -0.002 | -0.003 | -0.003 | 0.08 | 0.55 | 0.72 | 0.62 | 0.47 | 0.53 | 0.7 |
| 10 | 115698028 | rs114823 | -0.624 | 1.286 | -1.003 | 0.483 | -0.013 | 0.004 | 0.004 | 0 | 0.3 | 0.17 | 0.2 | 0.29 | 0.32 | 0.34 |
| 10 | 115707967 | rs180950 | -0.11 | 0.829 | -1.448 | 0.193 | 0.022 | 0.11 | 0.11 | 0 | 0.46 | 0.2 | 0.59 | 0.26 | 0.6 | 0.53 |
| 10 | 115713566 | rs151591 | -0.48 | -0.086 | -0.842 | 0.786 | 0.016 | -0.02 | -0.02 | 0 | 0.68 | 0.75 | 0.73 | 0.57 | 0.76 | 0.78 |
| 10 | 115728772 | rs7099933 | -0.271 | -0.231 | 0.274 | 0.781 | 0.006 | -0.001 | -0.001 | 0.224 | 0.1 | 0.31 | 0.08 | 0.11 | 0.12 | 0.05 |
| 10 | 115731668 | rs180932 | 0.547 | -0.07 | 0.628 | 1.051 | 0.053 | 0.063 | 0.063 | 0 | 0.52 | 0.19 | 0.63 | 0.34 | 0.61 | 0.6 |
| 10 | 115732630 | rs7077623 | 0.595 | 0.758 | 0.631 | 0.935 | 0.015 | 0.064 | 0.064 | 0 | 0.5 | 0.27 | 0.6 | 0.34 | 0.54 | 0.59 |
| 10 | 115734935 | rs180925 | -0.64 | 0.442 | 0.826 | 1.015 | 0.013 | -0.025 | -0.025 | 0 | 0.66 | 0.79 | 0.73 | 0.58 | 0.73 | 0.74 |
| 10 | 115737849 | rs180915 | 0.215 | 0.453 | 0.717 | 0.965 | 0.03 | -0.012 | -0.012 | 0 | 0.54 | 0.28 | 0.63 | 0.47 | 0.61 | 0.69 |
| 10 | 115746551 | rs6585256 | -0.109 | 0.373 | 0.885 | 0.675 | -0.012 | 0.005 | 0.005 | 0.238 | 0.65 | 0.56 | 0.73 | 0.34 | 0.72 | 0.65 |
| 10 | 115747786 | rs1411407 | -0.637 | 0.895 | 0.871 | 0.77 | 0.001 | 0.002 | 0.002 | 1.737 | 0.4 | 0.55 | 0.54 | 0.55 | 0.53 | 0.56 |
| 10 | 115748333 | rs12767891 | -0.703 | 0.899 | 0.883 | 0.471 | 0.016 | -0.007 | -0.007 | 0.439 | 0.39 | 0.49 | 0.52 | 0.37 | 0.51 | 0.55 |
| 10 | 115748612 | rs6585257 | -0.288 | 0.921 | 1.16 | 0.48 | 0.001 | 0 | 0 | 0.148 | 0.32 | 0.33 | 0.49 | 0.33 | 0.46 | 0.42 |
| 10 | 115749741 | rs10490907 | -0.983 | 0.871 | 1.118 | 0.14 | 0.001 | -0.016 | -0.016 | 0 | 0.09 | 0.19 | 0.05 | 0.04 | 0.07 | 0.14 |
| 10 | 115757207 | rs367502 | 0.987 | 0.834 | 1.692 | -0.107 | 0.12 | 0.039 | 0.039 | 3.088 | 0.94 | 0.53 | 0.93 | 1 | 0.96 | 0.99 |
| 10 | 115763315 | rs426121 | 1.274 | 0.822 | 1.671 | -0.117 | 0.077 | 0.021 | 0.021 | 3.918 | 0.93 | 0.57 | 0.94 | 1 | 0.97 | 0.99 |
| 10 | 115764032 | rs678669 | -0.17 | 0.109 | 0.968 | -0.725 | 0.016 | 0.049 | 0.049 | 1.76 | 0.52 | 0.36 | 0.42 | 0.16 | 0.6 | 0.46 |
| 10 | 115765366 | rs238753 | 0.716 | 0.153 | 0.672 | -0.686 | 0.077 | 0.021 | 0.021 | 3.918 | 0.93 | 0.58 | 0.94 | 1 | 0.97 | 0.99 |
| 10 | 115767540 | rs758586 | 0.528 | 0.059 | 0.141 | -0.651 | 0 | 0.149 | 0.149 | 3.668 | 0.41 | 0.11 | 0.5 | 0.84 | 0.37 | 0.54 |
| 10 | 115777224 | rs10749151 | 0.581 | 0.049 | 0.082 | -0.669 | -0.013 | 0.152 | 0.152 | 4.297 | 0.41 | 0.08 | 0.5 | 0.84 | 0.37 | 0.54 |
| 10 | 115779717 | rs11196589 | -0.166 | 0.413 | -0.167 | -0.799 | -0.012 | 0.008 | 0.008 | 1.724 | 0.24 | 0.26 | 0.28 | 0.13 | 0.35 | 0.25 |
| 10 | 115780369 | rs623499 | -0.778 | 0.283 | -0.897 | -0.848 | -0.066 | 0.101 | 0.101 | 4.919 | 0.37 | 0.83 | 0.37 | 0.16 | 0.41 | 0.38 |
| 10 | 115781367 | rs2782979 | 0.656 | 1.283 | -0.632 | -0.659 | 0 | 0.043 | 0.043 | 1.873 | 0.49 | 0.26 | 0.62 | 0.62 | 0.5 | 0.58 |
| 10 | 115781527 | rs2782980 | -0.362 | 1.376 | -0.064 | -0.921 | -0.019 | -0.063 | -0.063 | 0 | 0.69 | 0.54 | 0.78 | 0.86 | 0.7 | 0.81 |
| 10 | 115782377 | rs11196592 | 0.031 | 1.546 | -0.164 | -0.746 | -0.002 | -0.003 | -0.003 | 0 | 0.13 | 0.14 | 0.07 | 0.22 | 0.1 | 0.18 |
| 10 | 115784493 | rs7090339 | 0.609 | 1.97 | -0.401 | -0.674 | 0.002 | 0.004 | 0.004 | 0 | 0.08 | 0.13 | 0.25 | 0.14 | 0.13 | 0.13 |
| 10 | 115788094 | rs11196597 | 1.483 | 1.365 | -1.152 | -1.002 | 0.034 | -0.144 | -0.144 | 0 | 0.21 | 0.01 | 0.13 | 0.59 | 0.12 | 0.33 |
| 10 | 115788553 | rs9664370 | 0.85 | 1.786 | -0.677 | -0.759 | -0.015 | -0.02 | -0.02 | 1.047 | 0.08 | 0.09 | 0.24 | 0.14 | 0.13 | 0.13 |
| 10 | 115792787 | rs740746 | -0.384 | 1.753 | -0.319 | -0.918 | -0.002 | -0.026 | -0.026 | 0.025 | 0.66 | 0.59 | 0.8 | 0.78 | 0.68 | 0.75 |
| 10 | 115801065 | rs17875474 | 1.09 | 1.76 | -0.298 | -0.814 | -0.014 | -0.014 | -0.014 | 1.768 | 0.05 | 0.03 | 0.04 | 0.17 | 0.05 | 0.05 |
| 10 | 115801253 | rs2429511 | 0.657 | 1.786 | -0.183 | -0.876 | 0.015 | 0.047 | 0.047 | 0.864 | 0.48 | 0.26 | 0.44 | 0.64 | 0.44 | 0.57 |
| 10 | 115801595 | rs2050395 | 0.103 | 1.721 | -0.065 | -0.805 | 0.031 | 0.047 | 0.047 | 0 | 0.1 | 0.3 | 0.26 | 0.14 | 0.13 | 0.13 |
| 10 | 115801750 | rs17875439 | 0.839 | 1.743 | -0.543 | -0.796 | -0.015 | -0.019 | -0.019 | 0 | 0.08 | 0.07 | 0.23 | 0.14 | 0.13 | 0.13 |

| | | | | | | | | | | | | | | | | |
|---|---|---|---|---|---|---|---|---|---|---|---|---|---|---|---|---|
| 10 | 115807016 | rs3813720 | -0.124 | 1.724 | -0.552 | -0.884 | 0.002 | -0.022 | -0.022 | 0 | 0.59 | 0.52 | 0.72 | 0.77 | 0.59 | 0.7 |
| 10 | 115812793 | rs2050394 | 0.932 | 1.904 | -1.252 | -0.647 | -0.03 | -0.032 | -0.032 | 0.572 | 0.08 | 0.04 | 0.23 | 0.14 | 0.13 | 0.13 |
| 10 | 115813924 | rs10787516 | 0.581 | 1.671 | -1.198 | -0.68 | -0.009 | 0.041 | 0.041 | 1.089 | 0.48 | 0.15 | 0.46 | 0.64 | 0.46 | 0.57 |
| 10 | 115826508 | rs4359161 | -0.358 | 1.783 | -0.432 | -0.714 | 0.013 | 0.007 | 0.007 | 0 | 0.11 | 0.17 | 0.29 | 0.16 | 0.17 | 0.15 |
| 10 | 115832294 | rs1034259 | -0.255 | 1.904 | -0.043 | -0.911 | 0.033 | -0.063 | -0.063 | 0 | 0.26 | 0.39 | 0.14 | 0.17 | 0.22 | 0.3 |
| 10 | 115832408 | rs1034258 | -0.255 | 1.904 | -0.061 | -0.911 | 0.033 | -0.063 | -0.063 | 0 | 0.26 | 0.39 | 0.14 | 0.17 | 0.22 | 0.3 |
| 10 | 115839616 | rs10885533 | -1.762 | 1.378 | -0.598 | -1.055 | 0.021 | 0.077 | 0.077 | 0 | 0.49 | 0.86 | 0.55 | 0.33 | 0.54 | 0.46 |
| 10 | 115841641 | rs2419886 | -0.313 | 1.723 | 0.052 | -0.736 | 0.031 | -0.049 | -0.049 | 0 | 0.25 | 0.34 | 0.13 | 0.14 | 0.22 | 0.29 |
| 10 | 115844949 | rs686239 | -0.632 | 1.788 | -0.167 | -0.413 | 0.037 | 0.064 | 0.064 | 0 | 0.07 | 0.19 | 0.27 | 0.18 | 0.14 | 0.13 |
| 10 | 115848372 | rs17776203 | -1.156 | 1.6 | -0.578 | -0.604 | -0.051 | 0.005 | 0.005 | 0.335 | 0.36 | 0.74 | 0.43 | 0.35 | 0.42 | 0.44 |
| 10 | 115850176 | rs7905846 | -1.592 | 1.923 | -0.412 | -0.48 | -0.023 | -0.01 | -0.01 | 0 | 0.28 | 0.71 | 0.4 | 0.35 | 0.36 | 0.43 |
| 10 | 115856833 | rs12778917 | -0.84 | 1.359 | -0.665 | -1.043 | 0.007 | 0.011 | 0.011 | 0 | 0.28 | 0.23 | 0.22 | 0.04 | 0.25 | 0.11 |
| 10 | 115873670 | rs582413 | -0.344 | 1.509 | 0.018 | -1.03 | 0.044 | 0.041 | 0.041 | 0.025 | 0.1 | 0.24 | 0.21 | 0.15 | 0.15 | 0.18 |
| 10 | 115893434 | rs682836 | -2.024 | 1.569 | 0.084 | -1.049 | -0.048 | 0.093 | 0.093 | 0 | 0.46 | 0.85 | 0.51 | 0.29 | 0.51 | 0.37 |
| 10 | 115894242 | rs643148 | -0.286 | 1.593 | 0.187 | -1.013 | 0.048 | 0.041 | 0.041 | 0.05 | 0.1 | 0.26 | 0.21 | 0.15 | 0.16 | 0.17 |
| 10 | 115899581 | rs659280 | -0.979 | 1.479 | 0.511 | -0.908 | 0.04 | 0.039 | 0.039 | 1.178 | 0.11 | 0.33 | 0.23 | 0.12 | 0.15 | 0.21 |
| 10 | 115899972 | rs657497 | -1.34 | 1.543 | 0.214 | -1.185 | -0.007 | -0.064 | -0.064 | 0 | 0.24 | 0.4 | 0.25 | 0.09 | 0.18 | 0.21 |
| 10 | 115910928 | rs7095762 | -1.327 | 1.245 | 0.141 | -1.127 | -0.008 | -0.048 | -0.048 | 0 | 0.29 | 0.34 | 0.23 | 0.06 | 0.25 | 0.11 |
| 10 | 115915674 | rs11598654 | 0.795 | 1.244 | 0.159 | -1.137 | 0.001 | 0.019 | 0.019 | 0 | 0.1 | 0.04 | 0.09 | 0 | 0.12 | 0.02 |
| 10 | 115922774 | rs1061159 | -1.325 | 1.237 | 0.139 | -1.146 | -0.008 | -0.048 | -0.048 | 0 | 0.29 | 0.34 | 0.23 | 0.06 | 0.25 | 0.11 |
| 10 | 115936626 | rs12242939 | -1.438 | 1.135 | 0.11 | -1.179 | -0.009 | -0.045 | -0.045 | 0 | 0.3 | 0.35 | 0.24 | 0.06 | 0.26 | 0.12 |
| 10 | 115941162 | rs12251656 | -0.293 | 1.146 | 0.087 | -1.158 | 0.002 | 0.003 | 0.003 | 0 | 0.18 | 0.18 | 0.14 | 0.06 | 0.13 | 0.1 |
| 10 | 115941330 | rs12774590 | 0.672 | 1.15 | 0.067 | -1.171 | 0.008 | 0.027 | 0.027 | 0 | 0.11 | 0.05 | 0.09 | 0 | 0.13 | 0.02 |
| 10 | 115949262 | rs11196652 | -1.01 | 1.291 | 0.171 | -1.09 | -0.01 | -0.049 | -0.049 | 0 | 0.3 | 0.35 | 0.24 | 0.06 | 0.26 | 0.11 |
| 10 | 115962623 | rs12268539 | 1.109 | 1.522 | 0.177 | -1.025 | -0.006 | -0.005 | -0.005 | 0.714 | 0.05 | 0.04 | 0.1 | 0.06 | 0.08 | 0.02 |
| 10 | 115971995 | rs10510000 | -0.229 | 1.469 | 0.2 | -1.042 | 0.011 | -0.007 | -0.007 | 1.771 | 0.23 | 0.2 | 0.13 | 0 | 0.17 | 0.09 |
| 10 | 115975065 | rs10160173 | 1.119 | 1.469 | 0.151 | -1.043 | 0.001 | 0.002 | 0.002 | 0 | 0.05 | 0.09 | 0.1 | 0.06 | 0.08 | 0.03 |
| 10 | 115983112 | rs7078513 | -0.322 | 1.468 | 0.185 | -1.044 | -0.012 | -0.078 | -0.078 | 0.189 | 0.23 | 0.27 | 0.14 | 0 | 0.17 | 0.09 |
| 10 | 115986065 | rs12265237 | -0.922 | 1.471 | 0.185 | -1.044 | -0.009 | -0.052 | -0.052 | 0.707 | 0.29 | 0.35 | 0.24 | 0.06 | 0.25 | 0.11 |
| 10 | 115991745 | rs12266538 | 0.173 | 1.489 | 0.147 | -1.073 | -0.002 | 0.002 | 0.002 | 0 | 0.08 | 0.08 | 0.19 | 0.06 | 0.09 | 0.03 |
| 10 | 115993813 | rs1978224 | -0.008 | 1.545 | 0.232 | -0.994 | 0.003 | -0.034 | -0.034 | 0 | 0.23 | 0.23 | 0.13 | 0 | 0.17 | 0.09 |
| 10 | 116000453 | rs12219192 | 1.721 | 1.209 | -0.323 | -0.888 | -0.005 | -0.151 | -0.151 | 1.455 | 0.18 | 0.01 | 0.13 | 0.54 | 0.13 | 0.21 |
| 10 | 116003053 | rs672785 | -1.949 | 1.312 | 0.89 | -1 | 0.001 | 0.03 | 0.03 | 0 | 0.4 | 0.68 | 0.46 | 0.19 | 0.41 | 0.32 |
| 10 | 116008497 | rs9664945 | -0.977 | 1.253 | 1.039 | -0.807 | -0.009 | -0.051 | -0.051 | 1.965 | 0.27 | 0.34 | 0.22 | 0.06 | 0.25 | 0.11 |
| 10 | 116014827 | rs12262953 | 0.498 | 1.29 | 0.855 | -0.783 | 0.013 | 0.017 | 0.017 | 0 | 0.06 | 0.15 | 0.09 | 0.06 | 0.08 | 0.02 |
| 10 | 116019598 | rs572208 | -0.043 | 1.278 | 0.51 | -0.765 | 0.025 | 0.013 | 0.013 | 0.178 | 0.13 | 0.3 | 0.26 | 0.12 | 0.15 | 0.18 |
| 10 | 116020148 | rs12773168 | -0.694 | 1.318 | -0.205 | -0.751 | -0.015 | -0.083 | -0.083 | 0 | 0.28 | 0.42 | 0.22 | 0.07 | 0.25 | 0.11 |
| 10 | 116027958 | rs17091481 | -0.54 | 0.537 | 0.248 | -0.448 | -0.039 | -0.085 | -0.085 | 0 | 0.24 | 0.42 | 0.15 | 0.07 | 0.16 | 0.1 |
| 10 | 116031663 | rs17091488 | -0.433 | 0.558 | -0.17 | -0.433 | -0.019 | -0.051 | -0.051 | 0 | 0.24 | 0.36 | 0.15 | 0.07 | 0.16 | 0.1 |
| 10 | 116032519 | rs597371 | -1.071 | 0.509 | -1.229 | -0.13 | -0.062 | 0.102 | 0.102 | 2.55 | 0.49 | 0.88 | 0.47 | 0.34 | 0.43 | 0.54 |
| 10 | 116040001 | rs540609 | -1.021 | 0.243 | 0.135 | 0.135 | 0.039 | 0 | 0 | 0.001 | 0.16 | 0.29 | 0.3 | 0.15 | 0.21 | 0.23 |
| 10 | 116044587 | rs10787521 | -1.271 | -0.847 | 0.537 | 0.039 | -0.013 | 0.013 | 0.013 | 0 | 0.37 | 0.5 | 0.33 | 0.45 | 0.29 | 0.46 |
| 10 | 116045577 | rs1889446 | 1.393 | -1.525 | 0.643 | -0.285 | -0.021 | 0.003 | 0.003 | 0.177 | 0.73 | 0.42 | 0.84 | 0.72 | 0.83 | 0.7 |
| 10 | 116046579 | rs539946 | -0.157 | -1.558 | 1.345 | -0.307 | -0.006 | -0.043 | -0.043 | 0.104 | 0.91 | 0.82 | 0.95 | 0.98 | 0.93 | 0.89 |
| 10 | 116047921 | rs11196682 | -0.683 | -1.595 | 1.155 | -0.304 | -0.006 | -0.012 | -0.012 | 0 | 0.25 | 0.34 | 0.26 | 0.18 | 0.21 | 0.28 |
| 10 | 116048058 | rs1889447 | 2.178 | -1.786 | 1.374 | -0.364 | 0.032 | -0.01 | -0.01 | 0 | 0.47 | 0.2 | 0.56 | 0.51 | 0.61 | 0.36 |
| 10 | 116051568 | rs1889449 | -0.536 | -1.756 | 1.509 | -0.362 | 0.003 | 0.015 | 0.015 | 0 | 0.23 | 0.15 | 0.24 | 0.18 | 0.21 | 0.27 |
| 10 | 116053000 | rs12572135 | -0.271 | -1.814 | 2.018 | -0.688 | 0.001 | -0.011 | -0.011 | 0 | 0.1 | 0.07 | 0.06 | 0.26 | 0.08 | 0.18 |
| 10 | 116055417 | rs3750821 | 2.085 | -1.799 | 1.808 | -0.717 | 0.048 | -0.002 | -0.002 | 0 | 0.69 | 0.2 | 0.82 | 0.69 | 0.82 | 0.64 |
| 10 | 116058933 | rs1889450 | 1.505 | -1.872 | 1.893 | -0.719 | -0.057 | -0.125 | -0.125 | 0 | 0.81 | 0.56 | 0.9 | 0.98 | 0.91 | 0.82 |
| 10 | 116060387 | rs2286396 | 1.806 | -1.701 | 1.232 | -0.588 | 0.008 | -0.002 | -0.002 | 0 | 0.72 | 0.3 | 0.84 | 0.71 | 0.83 | 0.7 |
| 10 | 116062510 | rs507098 | -1.608 | -1.151 | 0.949 | -0.256 | -0.042 | 0.027 | 0.027 | 0.067 | 0.26 | 0.42 | 0.12 | 0.29 | 0.16 | 0.29 |
| 10 | 116073345 | rs1005109 | -1.248 | -0.875 | 0.662 | -0.59 | -0.003 | -0.005 | -0.005 | 0 | 0.1 | 0.2 | 0.05 | 0.08 | 0.06 | 0.16 |
| 10 | 116073801 | rs621375 | -1.667 | -1.261 | 0.337 | -1.054 | 0.022 | 0.025 | 0.025 | 0 | 0.33 | 0.73 | 0.17 | 0.11 | 0.21 | 0.38 |
| 10 | 116079188 | rs11196693 | 0.474 | -1.186 | 0.077 | -1.325 | -0.047 | -0.14 | -0.14 | 0.68 | 0.74 | 0.49 | 0.79 | 0.94 | 0.82 | 0.73 |
| 10 | 116079535 | rs17586071 | -0.414 | -1.133 | -0.541 | -1.345 | 0.006 | 0.008 | 0.008 | 0 | 0.12 | 0.05 | 0.06 | 0.02 | 0.08 | 0.15 |
| 10 | 116090503 | rs4918900 | 0.81 | -0.936 | -0.542 | -1.2 | 0.096 | 0.267 | 0.267 | 1.076 | 0.58 | 0.12 | 0.7 | 0.92 | 0.73 | 0.58 |
| 10 | 116109658 | rs4751933 | 0.25 | -0.008 | -0.777 | -1.128 | 0.066 | 0.226 | 0.226 | 0.658 | 0.53 | 0.18 | 0.65 | 0.87 | 0.66 | 0.63 |
| 10 | 116111028 | rs1334469 | 0.851 | 0.268 | -1.105 | -0.916 | -0.108 | 0.019 | 0.019 | 0.135 | 0.37 | 0.09 | 0.52 | 0.62 | 0.46 | 0.39 |
| 10 | 116114636 | rs10885557 | 0.962 | 0.437 | -0.113 | -0.727 | -0.001 | 0.006 | 0.006 | 0.424 | 0.66 | 0.37 | 0.73 | 0.7 | 0.64 | 0.69 |
| 10 | 116118908 | rs4144104 | 0.685 | 0.29 | 0.984 | -0.275 | 0.052 | 0.042 | 0.042 | 0.907 | 0.94 | 0.6 | 0.94 | 1 | 0.98 | 0.98 |
| 10 | 116119983 | rs7076281 | -0.788 | 0.299 | 1.094 | -0.272 | 0.055 | 0.046 | 0.046 | 0.24 | 0.05 | 0.33 | 0.05 | 0 | 0.02 | 0.02 |
| 10 | 116122453 | rs7907030 | 1.138 | 0.364 | 0.462 | -0.265 | 0.001 | 0.024 | 0.024 | 1.642 | 0.83 | 0.46 | 0.81 | 0.75 | 0.83 | 0.78 |
| 10 | 116127780 | rs10787526 | 0.718 | 0.427 | -0.131 | -0.365 | -0.004 | 0.081 | 0.081 | 0 | 0.49 | 0.22 | 0.59 | 0.7 | 0.57 | 0.48 |
| 10 | 116131214 | rs2244178 | -0.036 | 0.498 | -0.353 | -0.499 | -0.079 | -0.046 | -0.046 | 0 | 0.14 | 0.03 | 0.11 | 0.16 | 0.23 | 0.19 |
| 10 | 116131841 | rs10437496 | -0.08 | -0.062 | 0.425 | -0.681 | -0.014 | -0.017 | -0.017 | 3.082 | 0.2 | 0.28 | 0.22 | 0.13 | 0.14 | 0.27 |
| 10 | 116133734 | rs493347 | 0.367 | -0.107 | 0.986 | -0.707 | 0.049 | 0.026 | 0.026 | 0.929 | 0.95 | 0.69 | 0.96 | 1 | 0.99 | 0.99 |
| 10 | 116135410 | rs2475245 | 0.503 | -0.149 | 1.017 | -0.726 | -0.013 | -0.019 | -0.019 | 0.161 | 0.74 | 0.34 | 0.74 | 0.86 | 0.85 | 0.72 |
| 10 | 116136525 | rs11196717 | -0.121 | -0.124 | 0.839 | -0.679 | -0.015 | -0.022 | -0.022 | 0.238 | 0.21 | 0.28 | 0.21 | 0.14 | 0.15 | 0.27 |

| | | | | | | | | | | | | | | | | |
|---|---|---|---|---|---|---|---|---|---|---|---|---|---|---|---|---|
| 10 | 116137961 | rs496783 | -1.47 | 0.465 | 0.462 | -0.471 | -0.003 | 0.065 | 0.065 | 0 | 0.5 | 0.69 | 0.41 | 0.31 | 0.45 | 0.52 |
| 10 | 116139029 | rs4751674 | 0.762 | 1.471 | 0.662 | 0.123 | 0.065 | -0.002 | -0.002 | 0 | 0.8 | 0.61 | 0.81 | 0.83 | 0.73 | 0.81 |
| 10 | 116150161 | rs4751756 | -0.108 | 0.715 | -0.015 | 0.312 | 0.005 | 0.046 | 0.046 | 0 | 0.5 | 0.36 | 0.56 | 0.37 | 0.46 | 0.57 |
| 10 | 116152354 | rs537432 | -0.288 | 0.557 | 0.602 | 0.817 | -0.001 | -0.001 | -0.001 | 0 | 0.44 | 0.59 | 0.47 | 0.57 | 0.53 | 0.52 |
| 10 | 116156803 | rs475150 | 0.682 | 0.404 | 1.387 | 0.817 | 0.027 | 0.005 | 0.005 | 0 | 0.9 | 0.74 | 0.9 | 0.91 | 0.9 | 0.91 |
| 10 | 116168638 | rs11196730 | 1.692 | 0.348 | 1.349 | 0.761 | 0.003 | 0.101 | 0.101 | 0 | 0.78 | 0.41 | 0.77 | 0.5 | 0.79 | 0.77 |
| 10 | 116170504 | rs10736236 | 1.736 | 0.088 | 1.579 | 0.691 | -0.006 | 0.103 | 0.103 | 0 | 0.76 | 0.49 | 0.77 | 0.49 | 0.79 | 0.76 |
| 10 | 116171638 | rs7081297 | -1.757 | 0.066 | 1.658 | 0.957 | -0.018 | 0.078 | 0.078 | 0 | 0.21 | 0.43 | 0.2 | 0.56 | 0.15 | 0.24 |
| 10 | 116172486 | rs7910088 | -0.69 | 0.041 | 1.772 | 0.976 | -0.003 | -0.009 | -0.009 | 0.361 | 0.13 | 0.16 | 0.09 | 0.09 | 0.1 | 0.09 |
| 10 | 116175677 | rs4752697 | 0.931 | -0.071 | 1.121 | 0.811 | -0.13 | -0.077 | -0.077 | 1.124 | 0.3 | 0.04 | 0.29 | 0.37 | 0.39 | 0.36 |
| 10 | 116177609 | rs1570224 | -0.825 | 0.109 | 1.298 | 0.747 | 0.001 | 0.131 | 0.131 | 0 | 0.1 | 0.13 | 0.12 | 0.45 | 0.09 | 0.16 |
| 10 | 116177976 | rs4751902 | 1.021 | -0.151 | 1.693 | 0.611 | 0.028 | 0.184 | 0.184 | 0 | 0.43 | 0.18 | 0.44 | 0.02 | 0.37 | 0.35 |
| 10 | 116179616 | rs11196736 | -0.936 | -0.229 | 1.788 | 0.905 | 0.001 | 0.131 | 0.131 | 0 | 0.1 | 0.13 | 0.12 | 0.45 | 0.09 | 0.16 |
| 10 | 116183068 | rs7075375 | 1.855 | -0.213 | 1.815 | 0.904 | 0.003 | 0.057 | 0.057 | 0 | 0.74 | 0.48 | 0.74 | 0.45 | 0.76 | 0.74 |
| 10 | 116189070 | rs4752727 | 1.477 | -0.153 | 1.648 | 1.081 | 0.028 | 0.187 | 0.187 | 0 | 0.84 | 0.61 | 0.77 | 0.5 | 0.82 | 0.83 |
| 10 | 116190741 | rs4751905 | 1.234 | -0.167 | 1.598 | 1.076 | 0.023 | 0.148 | 0.148 | 0 | 0.85 | 0.75 | 0.78 | 0.5 | 0.82 | 0.83 |
| 10 | 116191174 | rs6646 | 1.269 | -0.308 | 1.431 | 1.113 | 0.007 | 0.118 | 0.118 | 0 | 0.84 | 0.77 | 0.79 | 0.56 | 0.82 | 0.84 |
| 10 | 116191799 | rs7916374 | -0.544 | -0.324 | 1.613 | 1.143 | -0.005 | -0.014 | -0.014 | 0.07 | 0.13 | 0.17 | 0.09 | 0.09 | 0.1 | 0.09 |
| 10 | 116192626 | rs3750820 | -1.868 | -0.214 | 1.748 | 1.383 | 0.015 | 0.251 | 0.251 | 0.226 | 0.14 | 0.41 | 0.19 | 0.51 | 0.12 | 0.17 |
| 10 | 116193167 | rs17091846 | -1.21 | -0.088 | 1.136 | 1.458 | 0.004 | 0.087 | 0.087 | 0 | 0.12 | 0.14 | 0.17 | 0.52 | 0.12 | 0.17 |
| 10 | 116193917 | rs10787530 | -0.96 | -0.071 | 0.904 | 1.329 | -0.032 | 0.014 | 0.014 | 0.224 | 0.7 | 0.79 | 0.67 | 0.72 | 0.59 | 0.63 |
| 10 | 116194210 | rs3802734 | -1.088 | -0.177 | 1.357 | 1.888 | -0.019 | 0.172 | 0.172 | 0.567 | 0.17 | 0.5 | 0.18 | 0.3 | 0.14 | 0.14 |
| 10 | 116198370 | rs4751908 | 0.116 | -0.815 | 1.154 | 1.43 | -0.008 | 0.032 | 0.032 | 0 | 0.45 | 0.27 | 0.37 | 0.39 | 0.43 | 0.41 |
| 10 | 116199175 | rs941853 | -0.597 | -0.798 | 1.232 | 1.611 | -0.002 | 0.128 | 0.128 | 0 | 0.15 | 0.35 | 0.17 | 0.26 | 0.14 | 0.15 |
| 10 | 116202474 | rs1885335 | -1.202 | -0.799 | 0.73 | 1.502 | -0.051 | 0.002 | 0.002 | 0 | 0.63 | 0.85 | 0.55 | 0.64 | 0.56 | 0.55 |
| 10 | 116218728 | rs10749166 | -0.218 | -0.607 | 0.864 | 0.954 | -0.003 | 0.028 | 0.028 | 0 | 0.45 | 0.38 | 0.4 | 0.31 | 0.42 | 0.45 |
| 10 | 116222167 | rs10885570 | 0.546 | -0.575 | 0.715 | 0.847 | -0.013 | 0.072 | 0.072 | 0 | 0.34 | 0.21 | 0.36 | 0.15 | 0.34 | 0.33 |
| 10 | 116222749 | rs7920878 | -0.115 | -0.477 | 0.849 | 0.977 | 0.003 | 0.021 | 0.021 | 0 | 0.42 | 0.66 | 0.44 | 0.35 | 0.4 | 0.39 |
| 10 | 116224579 | rs10510004 | 0.127 | -0.2 | -0.126 | 1.058 | -0.004 | 0.019 | 0.019 | 0 | 0.39 | 0.36 | 0.42 | 0.35 | 0.39 | 0.38 |
| 10 | 116227321 | rs6585283 | -0.421 | -0.085 | -0.139 | 1.063 | 0 | 0.018 | 0.018 | 0 | 0.1 | 0.13 | 0.06 | 0.2 | 0.09 | 0.11 |
| 10 | 116233234 | rs1535386 | 0.642 | 0.247 | -0.639 | 0.621 | -0.023 | -0.054 | -0.054 | 0 | 0.44 | 0.13 | 0.39 | 0.39 | 0.46 | 0.47 |
| 10 | 116233824 | rs11814528 | -0.121 | -0.014 | -0.121 | 0.077 | -0.006 | -0.108 | -0.108 | 0 | 0.3 | 0.55 | 0.24 | 0.03 | 0.28 | 0.17 |
| 10 | 116252408 | rs11196764 | 0.28 | -0.104 | -0.371 | 0.265 | 0.012 | -0.09 | -0.09 | 0 | 0.17 | 0.04 | 0.29 | 0.52 | 0.14 | 0.31 |
| 10 | 116252865 | rs2475228 | -1.525 | -0.188 | -0.297 | 0.162 | -0.009 | -0.013 | -0.013 | 0 | 0.53 | 0.79 | 0.59 | 0.62 | 0.48 | 0.51 |
| 10 | 116257523 | rs7910989 | -0.284 | 0.029 | -0.006 | 0.064 | 0.003 | -0.042 | -0.042 | 0 | 0.3 | 0.63 | 0.25 | 0.08 | 0.32 | 0.19 |
| 10 | 116259930 | rs2475230 | -1.273 | -0.114 | -0.199 | 0.333 | -0.013 | 0.03 | 0.03 | 0.045 | 0.3 | 0.41 | 0.42 | 0.67 | 0.26 | 0.45 |
| 10 | 116263488 | rs17091953 | 0.68 | -0.083 | -0.654 | 0.361 | -0.012 | 0.012 | 0.012 | 0.548 | 0.14 | 0.08 | 0.11 | 0 | 0.17 | 0.06 |
| 10 | 116275097 | rs941852 | -0.766 | -0.837 | -0.507 | 0.931 | 0.018 | -0.05 | -0.05 | 0 | 0.22 | 0.1 | 0.31 | 0.6 | 0.16 | 0.35 |
| 10 | 116279023 | rs11196775 | 0.855 | -0.55 | -0.248 | 0.289 | -0.013 | 0.017 | 0.017 | 0 | 0.38 | 0.29 | 0.34 | 0.24 | 0.42 | 0.49 |
| 10 | 116281550 | rs2497708 | 0.214 | -1.012 | -0.249 | 0.562 | -0.022 | 0.099 | 0.099 | 0 | 0.53 | 0.84 | 0.61 | 0.67 | 0.49 | 0.61 |
| 10 | 116284210 | rs2475240 | 0.562 | -0.568 | 0.399 | 0.736 | -0.011 | -0.008 | -0.008 | 0 | 0.73 | 0.79 | 0.71 | 0.68 | 0.7 | 0.71 |
| 10 | 116285341 | rs2145894 | 0.208 | -0.218 | 0.494 | 0.884 | -0.062 | -0.07 | -0.07 | 0 | 0.81 | 0.92 | 0.75 | 0.67 | 0.74 | 0.74 |
| 10 | 116292561 | rs2497714 | -0.905 | -0.178 | 0.449 | 0.849 | -0.012 | 0.034 | 0.034 | 0 | 0.94 | 0.98 | 0.97 | 1 | 0.94 | 0.98 |
| 10 | 116294497 | rs7913199 | -0.34 | -0.508 | 0.849 | 0.923 | -0.046 | 0.053 | 0.053 | 0.022 | 0.71 | 0.49 | 0.63 | 0.71 | 0.79 | 0.79 |
| 10 | 116294749 | rs12569428 | -0.548 | -0.449 | 0.763 | 0.94 | -0.075 | -0.081 | -0.081 | 0 | 0.19 | 0.05 | 0.26 | 0.33 | 0.26 | 0.26 |
| 10 | 116296654 | rs11196780 | -0.548 | -0.45 | 0.773 | 0.94 | -0.073 | -0.081 | -0.081 | 0.108 | 0.19 | 0.05 | 0.26 | 0.33 | 0.26 | 0.26 |
| 10 | 116311531 | rs986212 | -0.579 | -0.472 | 0.856 | 1.07 | 0.001 | 0.003 | 0.003 | 0.006 | 0.18 | 0.21 | 0.26 | 0.35 | 0.24 | 0.28 |
| 10 | 116315109 | rs2497730 | -0.569 | -0.602 | 0.248 | 0.968 | 0.051 | -0.031 | -0.031 | 1.363 | 0.35 | 0.8 | 0.4 | 0.36 | 0.25 | 0.27 |
| 10 | 116319266 | rs7894443 | 0.466 | -0.512 | 0.357 | 0.464 | 0.006 | 0.004 | 0.004 | 0 | 0.06 | 0.2 | 0.02 | 0.06 | 0.04 | 0.07 |
| 10 | 116323720 | rs7923449 | 1.144 | -0.372 | -0.243 | 0.464 | -0.027 | 0.099 | 0.099 | 0.038 | 0.46 | 0.19 | 0.35 | 0.31 | 0.49 | 0.46 |
| 10 | 116327449 | rs2901044 | 0.326 | -0.173 | -0.547 | 0.151 | -0.073 | -0.091 | -0.091 | 0 | 0.12 | 0.01 | 0.23 | 0.56 | 0.2 | 0.23 |
| 10 | 116332770 | rs2483567 | 0.946 | -0.303 | -0.329 | 0.096 | 0.001 | 0.003 | 0.003 | 0 | 0.74 | 0.74 | 0.71 | 0.35 | 0.7 | 0.66 |
| 10 | 116333974 | rs11196791 | 0.361 | -0.272 | -0.656 | 0.118 | -0.074 | -0.092 | -0.092 | 1.609 | 0.12 | 0.01 | 0.22 | 0.56 | 0.2 | 0.23 |
| 10 | 116335246 | rs985273 | 0.219 | -0.525 | -0.466 | 0.007 | -0.035 | 0.005 | 0.005 | 0 | 0.27 | 0.48 | 0.36 | 0.23 | 0.21 | 0.21 |
| 10 | 116346929 | rs11196793 | 1.091 | -0.485 | -0.888 | -0.006 | -0.047 | 0.257 | 0.257 | 0.074 | 0.46 | 0.1 | 0.34 | 0.12 | 0.52 | 0.45 |
| 10 | 116347338 | rs2483580 | -0.064 | 0.177 | -0.773 | 0.405 | -0.09 | -0.132 | -0.132 | 0 | 0.81 | 0.96 | 0.75 | 0.42 | 0.75 | 0.72 |
| 10 | 116354196 | rs2483583 | -0.557 | -0.392 | 0.082 | 0.48 | -0.037 | 0.004 | 0.004 | 0 | 0.27 | 0.56 | 0.38 | 0.23 | 0.21 | 0.21 |
| 10 | 116362320 | rs7914325 | -0.717 | -0.017 | 0.129 | 0.187 | -0.009 | -0.061 | -0.061 | 0 | 0.13 | 0.23 | 0.06 | 0 | 0.1 | 0.03 |
| 10 | 116362835 | rs7094673 | 0.716 | -0.225 | 0.434 | 0.135 | -0.007 | -0.015 | -0.015 | 0 | 0.7 | 0.65 | 0.75 | 0.84 | 0.69 | 0.78 |
| 10 | 116364311 | rs10787539 | 0.885 | -0.342 | 1.03 | 0.019 | -0.023 | -0.098 | -0.098 | 0.665 | 0.68 | 0.47 | 0.72 | 0.88 | 0.69 | 0.78 |
| 10 | 116364424 | rs10787540 | 0.937 | -0.357 | 0.964 | 0.013 | -0.019 | -0.079 | -0.079 | 0.264 | 0.68 | 0.43 | 0.72 | 0.88 | 0.69 | 0.78 |
| 10 | 116365213 | rs11196799 | 1.233 | -0.159 | 0.836 | 0.684 | 0.021 | -0.058 | -0.058 | 1.012 | 0.56 | 0.16 | 0.56 | 0.5 | 0.58 | 0.58 |
| 10 | 116365316 | rs2483589 | 0.5 | -0.317 | 1.224 | 0.366 | -0.023 | -0.116 | -0.116 | 0 | 0.84 | 0.56 | 0.89 | 1 | 0.88 | 0.96 |
| 10 | 116377962 | rs2483603 | -1.494 | -1.002 | 1.308 | 0.7 | -0.037 | 0.022 | 0.022 | 0 | 0.33 | 0.58 | 0.26 | 0.51 | 0.24 | 0.34 |
| 10 | 116380703 | rs11814078 | 0.121 | -1.097 | 1.559 | 0.645 | -0.022 | -0.044 | -0.044 | 0.118 | 0.09 | 0.32 | 0.04 | 0 | 0.04 | 0.01 |
| 10 | 116383343 | rs17719089 | 1.422 | -0.601 | 0.578 | 0.374 | -0.111 | 0.044 | 0.044 | 0.185 | 0.37 | 0.1 | 0.39 | 0.2 | 0.43 | 0.43 |
| 10 | 116383619 | rs11196810 | 0.214 | -0.731 | 0.509 | 0.504 | 0.024 | 0.019 | 0.019 | 0 | 0.14 | 0.01 | 0.06 | 0.11 | 0.11 | 0.13 |
| 10 | 116384062 | rs11196811 | 0.317 | -0.645 | 0.6 | 0.576 | -0.021 | 0.005 | 0.005 | 0.032 | 0.06 | 0.01 | 0.13 | 0.4 | 0.08 | 0.19 |
| 10 | 116390475 | rs1333689 | -0.78 | -0.665 | 0.769 | 0.567 | -0.032 | 0.178 | 0.178 | 0 | 0.17 | 0.44 | 0.18 | 0.41 | 0.12 | 0.21 |

| | | | | | | | | | | | | | | | | |
|---|---|---|---|---|---|---|---|---|---|---|---|---|---|---|---|---|
| 10 | 116402108 | rs2483519 | -0.655 | -0.768 | 0.421 | 0.587 | -0.063 | -0.024 | -0.024 | 0 | 0.7 | 0.88 | 0.67 | 0.7 | 0.64 | 0.75 |
| 10 | 116407406 | rs11196826 | 0.08 | -0.857 | 0.202 | 0.603 | 0.03 | 0.007 | 0.007 | 0 | 0.13 | 0.01 | 0.06 | 0.11 | 0.09 | 0.13 |
| 10 | 116409913 | rs10885591 | 0.399 | -0.815 | 0.206 | 0.887 | -0.019 | 0.043 | 0.043 | 0.147 | 0.05 | 0.01 | 0.13 | 0.43 | 0.09 | 0.17 |
| 10 | 116410970 | rs10787544 | 1.914 | -0.755 | 0.156 | 0.742 | -0.073 | 0.026 | 0.026 | 1.374 | 0.32 | 0.09 | 0.33 | 0.18 | 0.35 | 0.41 |
| 10 | 116416959 | rs12570718 | 1.914 | -0.767 | 0.427 | 0.735 | -0.085 | 0.019 | 0.019 | 1.374 | 0.32 | 0.08 | 0.33 | 0.19 | 0.35 | 0.41 |
| 10 | 116417012 | rs17092237 | -0.586 | -0.658 | 0.697 | 0.846 | 0 | 0.113 | 0.113 | 0 | 0.07 | 0.12 | 0.15 | 0.43 | 0.09 | 0.18 |
| 10 | 116419321 | rs7893317 | 1.465 | -0.71 | 0.945 | 0.516 | -0.003 | -0.021 | -0.021 | 0.917 | 0.05 | 0.08 | 0.04 | 0 | 0.06 | 0.02 |
| 10 | 116421842 | rs11196845 | -0.555 | -0.836 | 0.884 | 0.472 | 0.005 | 0.004 | 0.004 | 0.234 | 0.19 | 0.16 | 0.12 | 0.11 | 0.16 | 0.13 |
| 10 | 116430566 | rs11196855 | 0.768 | -0.823 | 1.593 | 0.479 | 0.001 | -0.005 | -0.005 | 0 | 0.45 | 0.52 | 0.59 | 0.63 | 0.46 | 0.61 |
| 10 | 116433347 | rs11597188 | 1.74 | -0.429 | 1.534 | 0.665 | -0.057 | 0.134 | 0.134 | 2.715 | 0.34 | 0.09 | 0.32 | 0.08 | 0.34 | 0.42 |
| 10 | 116434810 | rs4751639 | 0.763 | -0.379 | 1.629 | 0.65 | -0.001 | 0.001 | 0.001 | 1.761 | 0.45 | 0.56 | 0.59 | 0.63 | 0.46 | 0.63 |
| 10 | 116437174 | rs11196865 | 0.417 | -0.323 | 1.491 | 1.011 | -0.028 | 0.067 | 0.067 | 0.015 | 0.05 | 0.01 | 0.26 | 0.39 | 0.13 | 0.17 |
| 10 | 116438345 | rs10885595 | 0.827 | -0.342 | 2.045 | 0.913 | -0.001 | -0.001 | -0.001 | 0.234 | 0.45 | 0.4 | 0.5 | 0.44 | 0.51 | 0.62 |
| 10 | 116442618 | rs11196869 | -1.132 | -0.564 | 2.058 | 0.76 | -0.084 | -0.138 | -0.138 | 0 | 0.27 | 0.58 | 0.16 | 0 | 0.14 | 0.12 |
| 10 | 116453733 | rs882870 | 1.069 | -0.53 | 2.027 | 0.793 | -0.011 | -0.013 | -0.013 | 1.521 | 0.43 | 0.27 | 0.49 | 0.43 | 0.5 | 0.6 |
| 10 | 116461422 | rs217190 | -1.109 | -0.563 | 2.026 | 0.629 | -0.1 | -0.16 | -0.16 | 0 | 0.27 | 0.52 | 0.16 | 0 | 0.14 | 0.09 |
| 10 | 116463434 | rs1005968 | 1.757 | -0.475 | 1.938 | 0.693 | -0.028 | 0.105 | 0.105 | 1.645 | 0.36 | 0.17 | 0.22 | 0.04 | 0.37 | 0.43 |
| 10 | 116469048 | rs808260 | 0.292 | -0.654 | 1.784 | 0.794 | -0.032 | -0.057 | -0.057 | 0 | 0.12 | 0.2 | 0.07 | 0 | 0.07 | 0.06 |
| 10 | 116469381 | rs808259 | -0.467 | -0.225 | 1.349 | 0.854 | 0 | 0.003 | 0.003 | 0.008 | 0.71 | 0.71 | 0.63 | 0.44 | 0.68 | 0.71 |
| 10 | 116479396 | rs808263 | -0.634 | -0.137 | 1 | 0.849 | -0.021 | -0.054 | -0.054 | 1.926 | 0.73 | 0.86 | 0.67 | 0.44 | 0.69 | 0.7 |
| 10 | 116489493 | rs808355 | -0.452 | -0.148 | 1.087 | 0.837 | -0.007 | -0.027 | -0.027 | 0 | 0.72 | 0.8 | 0.66 | 0.44 | 0.69 | 0.7 |
| 10 | 116493083 | rs7093476 | 1.451 | -0.245 | 1.331 | 0.783 | -0.017 | -0.026 | -0.026 | 0.53 | 0.05 | 0.15 | 0.03 | 0 | 0.03 | 0.02 |
| 10 | 116493516 | rs808340 | 0.047 | -0.257 | 1.488 | 0.78 | -0.004 | 0.017 | 0.017 | 0.079 | 0.66 | 0.6 | 0.63 | 0.44 | 0.66 | 0.69 |
| 10 | 116498910 | rs808332 | -0.401 | -0.266 | 1.421 | 0.788 | 0 | 0.001 | 0.001 | 0 | 0.72 | 0.74 | 0.66 | 0.44 | 0.69 | 0.7 |
| 10 | 116507307 | rs808316 | -1.603 | -0.173 | 1.229 | 0.648 | -0.036 | 0.039 | 0.039 | 0.676 | 0.93 | 1 | 0.96 | 1 | 0.89 | 0.89 |
| 10 | 116508147 | rs808314 | -0.355 | -0.102 | 1.297 | 0.545 | 0.001 | 0.014 | 0.014 | 0 | 0.71 | 0.67 | 0.65 | 0.44 | 0.69 | 0.7 |
| 10 | 116508805 | rs11196898 | 0.632 | -0.029 | 1.183 | 0.563 | 0.005 | 0.139 | 0.139 | 0.021 | 0.05 | 0.04 | 0.26 | 0.4 | 0.12 | 0.15 |
| 10 | 116509129 | rs808312 | -0.09 | -0.061 | 1.163 | 0.45 | 0.011 | -0.008 | -0.008 | 0 | 0.16 | 0.13 | 0.08 | 0 | 0.1 | 0.1 |
| 10 | 116511189 | rs7070890 | 1.35 | 0.032 | 0.937 | 0.521 | 0.02 | 0.019 | 0.019 | 1.099 | 0.08 | 0.02 | 0.02 | 0.01 | 0.04 | 0.02 |
| 10 | 116515430 | rs7080890 | 1.218 | 0.081 | 1.05 | 0.438 | 0.003 | -0.07 | -0.07 | 0.014 | 0.45 | 0.43 | 0.3 | 0.04 | 0.46 | 0.46 |
| 10 | 116525603 | rs808294 | -0.85 | 0.161 | 0.422 | 0.596 | -0.032 | -0.086 | -0.086 | 0 | 0.75 | 0.89 | 0.66 | 0.44 | 0.69 | 0.69 |
| 10 | 116528294 | rs11196905 | -0.157 | 0.416 | 0.577 | 0.71 | -0.001 | -0.001 | -0.001 | 3.797 | 0.14 | 0.11 | 0.28 | 0.4 | 0.16 | 0.18 |
| 10 | 116557080 | rs11196917 | 0.049 | 0.284 | 0.436 | 0.756 | 0.012 | -0.013 | -0.013 | 0 | 0.6 | 0.74 | 0.59 | 0.48 | 0.61 | 0.64 |
| 10 | 116566550 | rs808275 | 0.038 | 0.236 | 0.056 | 0.467 | 0.018 | -0.01 | -0.01 | 1.978 | 0.55 | 0.64 | 0.54 | 0.36 | 0.56 | 0.62 |
| 10 | 116571796 | rs11196924 | -0.128 | 0.153 | 0.208 | -0.447 | -0.024 | 0.019 | 0.019 | 0 | 0.35 | 0.21 | 0.35 | 0.23 | 0.47 | 0.48 |
| 10 | 116576790 | rs4752255 | -0.009 | 0.172 | 0.861 | -0.386 | 0.01 | -0.007 | -0.007 | 0 | 0.36 | 0.36 | 0.49 | 0.25 | 0.48 | 0.48 |
| 10 | 116582560 | rs758585 | -0.375 | -0.012 | 1.37 | -0.471 | -0.003 | 0.004 | 0.004 | 0.043 | 0.39 | 0.62 | 0.51 | 0.26 | 0.49 | 0.51 |
| 10 | 116593175 | rs11196939 | -0.093 | 0.015 | 1.393 | -0.459 | -0.016 | 0.012 | 0.012 | 0 | 0.35 | 0.25 | 0.35 | 0.23 | 0.47 | 0.48 |
| 10 | 116596071 | rs17095662 | -0.14 | -0.004 | 1.417 | -0.466 | 0.009 | -0.012 | -0.012 | 0 | 0.37 | 0.39 | 0.36 | 0.24 | 0.47 | 0.48 |
| 10 | 116604344 | rs2181563 | -0.462 | -0.067 | 1.618 | -0.492 | -0.014 | 0.026 | 0.026 | 0.251 | 0.39 | 0.72 | 0.53 | 0.25 | 0.49 | 0.53 |
| 10 | 116619651 | rs11196954 | -0.873 | 0.354 | 1.12 | -0.495 | 0.025 | 0.066 | 0.066 | 0 | 0.07 | 0.16 | 0.16 | 0.2 | 0.09 | 0.08 |
| 10 | 116625077 | rs2016966 | -0.873 | 0.354 | 1.12 | -0.495 | 0.025 | 0.066 | 0.066 | 0 | 0.07 | 0.16 | 0.16 | 0.2 | 0.09 | 0.08 |
| 10 | 116627910 | rs1966384 | 0.441 | 0.485 | 1.041 | -0.521 | -0.007 | -0.007 | -0.007 | 0 | 0.3 | 0.23 | 0.2 | 0.04 | 0.37 | 0.4 |
| 10 | 116630289 | rs17092542 | 0.432 | 0.473 | 1.042 | -0.52 | -0.008 | -0.007 | -0.007 | 0 | 0.3 | 0.23 | 0.2 | 0.04 | 0.38 | 0.43 |
| 10 | 116636721 | rs10510007 | 0.437 | 0.367 | 0.929 | -0.432 | 0.004 | -0.009 | -0.009 | 0 | 0.28 | 0.25 | 0.22 | 0.11 | 0.38 | 0.42 |
| 10 | 116643844 | rs10885622 | -0.413 | 0.404 | 0.899 | -0.615 | 0.004 | -0.002 | -0.002 | 0 | 0.33 | 0.34 | 0.33 | 0.24 | 0.44 | 0.51 |
| 10 | 116644141 | rs17721283 | 0.834 | 0.191 | 0.619 | -0.702 | -0.031 | 0.026 | 0.026 | 0.091 | 0.23 | 0.13 | 0.18 | 0.04 | 0.34 | 0.37 |
| 10 | 116647977 | rs11196963 | 0.124 | -0.346 | 0.307 | -1.023 | -0.015 | 0.121 | 0.121 | 0 | 0.44 | 0.22 | 0.27 | 0.05 | 0.45 | 0.37 |
| 10 | 116652502 | rs10885625 | 0.475 | -0.396 | -0.064 | -0.786 | -0.013 | -0.028 | -0.028 | 0 | 0.09 | 0.03 | 0.16 | 0.2 | 0.12 | 0.11 |
| 10 | 116653044 | rs10885626 | -1.179 | -0.313 | -0.04 | -0.866 | -0.035 | 0.039 | 0.039 | 0 | 0.57 | 0.81 | 0.58 | 0.29 | 0.54 | 0.5 |
| 10 | 116654560 | rs17794472 | 0.624 | -0.396 | -0.168 | -0.937 | -0.05 | 0.197 | 0.197 | 0 | 0.39 | 0.1 | 0.22 | 0.05 | 0.39 | 0.33 |
| 10 | 116655660 | rs10885628 | 0.624 | -0.395 | -0.219 | -0.937 | -0.05 | 0.197 | 0.197 | 0 | 0.39 | 0.1 | 0.22 | 0.05 | 0.39 | 0.34 |
| 10 | 116658106 | rs10787548 | -1.105 | -0.263 | -0.28 | -0.828 | -0.044 | 0.041 | 0.041 | 0 | 0.57 | 0.83 | 0.58 | 0.28 | 0.54 | 0.5 |
| 10 | 116676281 | rs12265048 | 0.011 | -0.448 | 0.237 | -1.405 | 0.003 | 0.119 | 0.119 | 0.009 | 0.44 | 0.24 | 0.25 | 0.03 | 0.43 | 0.36 |
| 10 | 116685463 | rs10885632 | 0.542 | -0.373 | 0.149 | -1.437 | -0.031 | 0.195 | 0.195 | 0.013 | 0.39 | 0.11 | 0.22 | 0.03 | 0.39 | 0.33 |
| 10 | 116687214 | rs7085964 | -0.627 | -0.535 | 0.602 | -1.545 | 0.016 | 0.173 | 0.173 | 0.049 | 0.5 | 0.74 | 0.32 | 0.04 | 0.47 | 0.38 |
| 10 | 116689936 | rs7100159 | -0.522 | -0.558 | 0.621 | -1.553 | 0.022 | 0.101 | 0.101 | 0 | 0.49 | 0.68 | 0.29 | 0.04 | 0.44 | 0.37 |
| 10 | 116696928 | rs6585304 | 0.122 | -0.539 | 0.672 | -1.539 | 0.007 | -0.031 | -0.031 | 0 | 0.09 | 0.53 | 0.07 | 0 | 0.05 | 0.04 |
| 10 | 116697194 | rs10885633 | 0.484 | -0.539 | 0.593 | -1.548 | -0.013 | 0.17 | 0.17 | 0 | 0.4 | 0.15 | 0.22 | 0.03 | 0.39 | 0.33 |
| 10 | 116700436 | rs10885634 | -0.895 | -0.544 | 0.606 | -1.55 | -0.006 | 0.196 | 0.196 | 0.442 | 0.53 | 0.82 | 0.33 | 0.04 | 0.48 | 0.4 |
| 10 | 116708190 | rs12269261 | 0.5 | -0.528 | 0.505 | -1.523 | -0.018 | 0.183 | 0.183 | 0 | 0.4 | 0.14 | 0.22 | 0.03 | 0.39 | 0.33 |
| 10 | 116719543 | rs7099565 | -0.722 | -0.516 | 0.514 | -1.533 | 0.01 | 0.189 | 0.189 | 0.177 | 0.52 | 0.78 | 0.3 | 0.04 | 0.45 | 0.39 |
| 10 | 116722834 | rs12764662 | 0.751 | 0.386 | 1.147 | -0.901 | -0.035 | 0.043 | 0.043 | 0.239 | 0.2 | 0.1 | 0.13 | 0.03 | 0.29 | 0.29 |
| 10 | 116730538 | rs7092748 | 1.123 | 0.329 | 1.914 | -0.792 | 0.043 | 0.01 | 0.01 | 0.173 | 0.06 | 0.29 | 0.05 | 0 | 0.04 | 0.01 |
| 10 | 116735341 | rs7071789 | -1.052 | 0.195 | 1.589 | -0.845 | -0.014 | 0.08 | 0.08 | 0 | 0.36 | 0.78 | 0.24 | 0.03 | 0.37 | 0.36 |
| 10 | 116750911 | rs6585314 | 1.054 | 0.415 | 1.604 | -0.658 | 0.06 | 0.024 | 0.024 | 0.232 | 0.06 | 0.33 | 0.05 | 0 | 0.04 | 0.01 |
| 10 | 116757856 | rs968848 | 0.158 | 0.54 | 0.896 | -0.64 | -0.015 | 0.01 | 0.01 | 0 | 0.24 | 0.2 | 0.16 | 0.03 | 0.31 | 0.32 |
| 10 | 116779483 | rs17092671 | 0.067 | 0.644 | 0.745 | -0.515 | -0.014 | 0.016 | 0.016 | 0 | 0.25 | 0.2 | 0.16 | 0.06 | 0.31 | 0.28 |
| 10 | 116802563 | rs808421 | 0.274 | 0.498 | 1.606 | -0.417 | -0.002 | -0.014 | -0.014 | 0 | 0.7 | 0.31 | 0.79 | 0.86 | 0.73 | 0.74 |

| | | | | | | | | | | | | | | | | |
|---|---|---|---|---|---|---|---|---|---|---|---|---|---|---|---|---|
| 10 | 116815448 | rs1147626 | 0.301 | 0.883 | 1.049 | -0.062 | -0.047 | -0.046 | -0.046 | 0 | 0.08 | 0.01 | 0.13 | 0.14 | 0.13 | 0.12 |
| 10 | 116828592 | rs11197045 | -0.05 | 0.372 | 1.132 | -0.379 | 0 | -0.001 | -0.001 | 0.02 | 0.27 | 0.27 | 0.17 | 0.15 | 0.27 | 0.26 |
| 10 | 116830383 | rs808427 | 0.416 | 0.389 | 0.599 | -0.361 | 0.007 | 0.038 | 0.038 | 0 | 0.67 | 0.23 | 0.78 | 0.85 | 0.7 | 0.7 |
| 10 | 116834272 | rs11197050 | 0.412 | 0.258 | -0.924 | -0.08 | -0.013 | 0.007 | 0.007 | 0.614 | 0.16 | 0 | 0.16 | 0.11 | 0.14 | 0.17 |
| 10 | 116843474 | rs10885649 | 0.017 | 0.934 | -1.153 | 0.13 | -0.121 | 0.031 | 0.031 | 0 | 0.28 | 0.01 | 0.15 | 0.17 | 0.28 | 0.27 |
| 10 | 116848454 | rs1889610 | -0.306 | 0.581 | -0.551 | 0.374 | -0.008 | 0.024 | 0.024 | 0.016 | 0.29 | 0.35 | 0.49 | 0.62 | 0.28 | 0.38 |
| 10 | 116875810 | rs1272383 | 1.553 | 0.451 | -0.736 | 0.842 | 0.008 | -0.074 | -0.074 | 0 | 0.13 | 0 | 0.3 | 0.27 | 0.12 | 0.18 |
| 10 | 116884793 | rs1264782 | -0.384 | 0.493 | 0.144 | 0.085 | 0.01 | -0.038 | -0.038 | 0 | 0.66 | 0.57 | 0.78 | 0.82 | 0.69 | 0.67 |
| 10 | 116902943 | rs1264794 | -0.727 | 0.457 | 0.69 | 0.169 | -0.006 | 0.008 | 0.008 | 0 | 0.69 | 0.73 | 0.81 | 0.78 | 0.72 | 0.72 |
| 10 | 116947148 | rs1264764 | 0.293 | 0.41 | 1.022 | -0.004 | -0.003 | -0.038 | -0.038 | 0 | 0.26 | 0.14 | 0.49 | 0.69 | 0.28 | 0.4 |
| 10 | 116965037 | rs1899721 | 0.426 | 0.462 | 1.057 | -0.095 | 0.006 | -0.021 | -0.021 | 0 | 0.31 | 0.35 | 0.2 | 0.18 | 0.28 | 0.3 |
| 10 | 117033561 | rs11498596 | -0.117 | 0.452 | 1.199 | -0.048 | -0.009 | 0.007 | 0.007 | 0.017 | 0.29 | 0.36 | 0.52 | 0.69 | 0.28 | 0.38 |
| 10 | 117061793 | rs2291669 | 1.418 | 0.411 | 1.12 | -0.157 | -0.035 | 0.112 | 0.112 | 0.135 | 0.35 | 0.1 | 0.26 | 0.13 | 0.41 | 0.28 |
| 10 | 117147589 | rs10885701 | 1.441 | 0.424 | 1.119 | -0.13 | -0.055 | 0.115 | 0.115 | 0.099 | 0.35 | 0.07 | 0.26 | 0.12 | 0.41 | 0.26 |
| 10 | 117171076 | rs589508 | -0.061 | 0.484 | 1.392 | -0.111 | -0.002 | -0.001 | -0.001 | 0 | 0.35 | 0.63 | 0.23 | 0.18 | 0.3 | 0.3 |
| 10 | 117173622 | rs1170884 | -0.021 | 0.53 | 1.41 | -0.11 | 0.007 | -0.04 | -0.04 | 0 | 0.34 | 0.56 | 0.22 | 0.18 | 0.3 | 0.3 |
| 10 | 117190413 | rs653295 | -0.137 | 0.631 | 1.379 | -0.185 | 0 | -0.011 | -0.011 | 0.036 | 0.34 | 0.62 | 0.21 | 0.18 | 0.29 | 0.28 |
| 10 | 117205246 | rs10787573 | -0.581 | 0.313 | 1.794 | -0.104 | 0.001 | -0.01 | -0.01 | 0.372 | 0.72 | 0.81 | 0.48 | 0.29 | 0.71 | 0.59 |
| 10 | 117223605 | rs1414636 | -0.481 | 0.443 | 1.356 | -0.03 | -0.006 | -0.005 | -0.005 | 0 | 0.32 | 0.6 | 0.32 | 0.29 | 0.24 | 0.28 |
| 10 | 117225906 | rs6585333 | 0.664 | 0.611 | 1.119 | -0.172 | -0.005 | -0.016 | -0.016 | 0 | 0.4 | 0.42 | 0.32 | 0.04 | 0.47 | 0.31 |
| 10 | 117228794 | rs10885721 | 0.5 | 0.636 | 0.937 | -0.131 | -0.02 | 0.088 | 0.088 | 0.055 | 0.43 | 0.29 | 0.47 | 0.17 | 0.49 | 0.36 |
| 10 | 117244968 | rs1414632 | 0.321 | 0.651 | 0.909 | -0.063 | -0.005 | 0.023 | 0.023 | 0.062 | 0.25 | 0.13 | 0.13 | 0.16 | 0.22 | 0.22 |
| 10 | 117268817 | rs7899493 | -0.8 | 0.606 | 1.374 | -0.093 | -0.012 | 0.002 | 0.002 | 0.107 | 0.68 | 0.62 | 0.62 | 0.33 | 0.71 | 0.58 |
| 10 | 117273552 | rs17724010 | 0.323 | 0.582 | 1.444 | -0.222 | -0.001 | 0.004 | 0.004 | 0 | 0.25 | 0.23 | 0.14 | 0.16 | 0.22 | 0.23 |
| 10 | 117276069 | rs1005000 | 0.525 | 0.583 | 1.492 | -0.222 | -0.008 | 0.043 | 0.043 | 0 | 0.43 | 0.35 | 0.33 | 0.14 | 0.49 | 0.35 |
| 10 | 117277009 | rs11595630 | 0.593 | 0.619 | 1.416 | -0.238 | -0.089 | 0.01 | 0.01 | 0 | 0.23 | 0.02 | 0.13 | 0.16 | 0.24 | 0.23 |
| 10 | 117282016 | rs7895853 | -0.708 | 0.627 | 1.463 | -0.223 | -0.016 | -0.009 | -0.009 | 0 | 0.67 | 0.52 | 0.46 | 0.31 | 0.71 | 0.57 |
| 10 | 117299863 | rs11197282 | 0.65 | 0.632 | 1.327 | -0.252 | -0.008 | 0.064 | 0.064 | 0.038 | 0.42 | 0.32 | 0.32 | 0.14 | 0.46 | 0.34 |
| 10 | 117302717 | rs12781613 | 0.627 | 0.653 | 1.197 | -0.279 | -0.09 | 0.008 | 0.008 | 0 | 0.23 | 0.01 | 0.13 | 0.16 | 0.24 | 0.22 |
| 10 | 117303444 | rs11197286 | 1.261 | 0.568 | 1.301 | -0.331 | -0.072 | 0.191 | 0.191 | 0.059 | 0.36 | 0.12 | 0.27 | 0.04 | 0.44 | 0.3 |
| 10 | 117324239 | rs1572400 | -0.399 | 0.492 | 1.446 | -0.195 | -0.014 | -0.035 | -0.035 | 0 | 0.64 | 0.57 | 0.46 | 0.2 | 0.68 | 0.52 |
| 10 | 117327420 | rs1932582 | 0.735 | 0.499 | 1.377 | -0.156 | 0.005 | 0.001 | 0.001 | 0.227 | 0.3 | 0.18 | 0.51 | 0.69 | 0.29 | 0.42 |
| 10 | 117328426 | rs1336115 | 0.735 | 0.499 | 1.377 | -0.156 | 0.005 | 0.001 | 0.001 | 0.227 | 0.3 | 0.18 | 0.5 | 0.69 | 0.29 | 0.42 |
| 10 | 117331064 | rs17093295 | -1.126 | 0.406 | 1.722 | -0.177 | 0.001 | 0.067 | 0.067 | 0 | 0.07 | 0.28 | 0.05 | 0.11 | 0.02 | 0.06 |
| 10 | 117331998 | rs1328008 | 0.069 | 0.5 | 1.455 | 0.046 | -0.002 | 0.02 | 0.02 | 0.152 | 0.46 | 0.61 | 0.39 | 0.22 | 0.49 | 0.36 |
| 10 | 117339523 | rs2804192 | -0.281 | 0.501 | 1.134 | -0.147 | -0.004 | 0.074 | 0.074 | 0.158 | 0.53 | 0.3 | 0.63 | 0.85 | 0.51 | 0.64 |
| 10 | 117372854 | rs2769378 | -0.264 | 0.533 | 1.08 | -0.147 | -0.006 | 0.133 | 0.133 | 0.323 | 0.53 | 0.22 | 0.62 | 0.85 | 0.51 | 0.64 |
| 10 | 117390995 | rs2769423 | 0.296 | 0.376 | 0.538 | -0.682 | 0.004 | 0.004 | 0.004 | 1.336 | 0.72 | 0.25 | 0.78 | 0.85 | 0.78 | 0.7 |
| 10 | 117395006 | rs2769419 | -0.693 | 0.244 | 0.805 | -0.6 | 0.005 | -0.064 | -0.064 | 0.208 | 0.63 | 0.56 | 0.67 | 0.96 | 0.53 | 0.69 |
| 10 | 117395349 | rs2769418 | 0.069 | 0.247 | 0.831 | -0.597 | -0.003 | 0.042 | 0.042 | 0.084 | 0.54 | 0.35 | 0.63 | 0.85 | 0.5 | 0.64 |
| 10 | 117420115 | rs7089599 | -0.59 | 0.451 | 0.546 | -0.598 | 0.005 | -0.065 | -0.065 | 0 | 0.63 | 0.57 | 0.68 | 0.96 | 0.53 | 0.69 |
| 10 | 117427484 | rs12219110 | 0.884 | 0.456 | -0.054 | 0.18 | -0.076 | -0.083 | -0.083 | 0.024 | 0.19 | 0.01 | 0.1 | 0.26 | 0.2 | 0.22 |
| 10 | 117432639 | rs2532725 | 0.567 | 0.199 | -0.109 | -0.156 | -0.071 | 0.041 | 0.041 | 0.015 | 0.2 | 0.01 | 0.1 | 0.13 | 0.19 | 0.2 |
| 10 | 117433011 | rs1981663 | -1.044 | 0.235 | 0.306 | -0.483 | -0.001 | -0.033 | -0.033 | 0 | 0.69 | 0.64 | 0.73 | 0.96 | 0.61 | 0.74 |
| 10 | 117433657 | rs714849 | 0.974 | 0.118 | 0.545 | -0.069 | -0.014 | -0.045 | -0.045 | 0.274 | 0.36 | 0.16 | 0.54 | 0.59 | 0.35 | 0.48 |
| 10 | 117439034 | rs2804147 | 0.079 | 0.329 | 1.037 | 0.081 | -0.017 | -0.036 | -0.036 | 0 | 0.26 | 0.17 | 0.18 | 0.38 | 0.28 | 0.26 |
| 10 | 117441174 | rs4751923 | 0.692 | 0.174 | 1.07 | 0.217 | 0.011 | 0.008 | 0.008 | 0 | 0.41 | 0.24 | 0.54 | 0.61 | 0.35 | 0.49 |
| 10 | 117442572 | rs2052067 | 0.608 | 0.203 | 0.996 | 0.209 | 0.008 | 0.016 | 0.016 | 0.019 | 0.43 | 0.23 | 0.56 | 0.61 | 0.38 | 0.52 |
| 10 | 117443212 | rs2804148 | 0.613 | 0.234 | 0.987 | 0.198 | -0.066 | -0.101 | -0.101 | 0.02 | 0.21 | 0.05 | 0.13 | 0.35 | 0.22 | 0.22 |
| 10 | 117443822 | rs10885769 | 0.598 | 0.259 | 1.229 | 0.31 | -0.014 | -0.072 | -0.072 | 0 | 0.42 | 0.12 | 0.55 | 0.54 | 0.37 | 0.51 |
| 10 | 117444741 | rs2420102 | 0.593 | 0.296 | 1.148 | 0.298 | 0.007 | -0.03 | -0.03 | 0.61 | 0.42 | 0.23 | 0.55 | 0.54 | 0.35 | 0.49 |
| 10 | 117449079 | rs7908611 | -0.079 | 0.32 | 1.144 | 0.33 | 0.006 | 0.013 | 0.013 | 0 | 0.48 | 0.52 | 0.62 | 0.72 | 0.4 | 0.55 |
| 10 | 117469964 | rs7076927 | 0.484 | 0.339 | 0.908 | 0.347 | 0.01 | 0.027 | 0.027 | 0.04 | 0.43 | 0.24 | 0.56 | 0.65 | 0.38 | 0.52 |
| 10 | 117473636 | rs7069993 | 0.485 | 0.358 | 0.914 | 0.308 | 0.01 | 0.014 | 0.014 | 0.015 | 0.43 | 0.25 | 0.56 | 0.61 | 0.37 | 0.52 |
| 10 | 117481431 | rs4751927 | -0.153 | 0.386 | 0.766 | -0.215 | 0.024 | 0.045 | 0.045 | 0 | 0.49 | 0.64 | 0.63 | 0.71 | 0.39 | 0.55 |
| 10 | 117484461 | rs17093513 | 1.136 | 0.555 | -0.092 | -0.395 | -0.079 | -0.051 | -0.051 | 0 | 0.15 | 0.01 | 0.09 | 0.18 | 0.19 | 0.19 |
| 10 | 117486910 | rs2286631 | 0.428 | 0.546 | 0.151 | -0.454 | 0.003 | 0.114 | 0.114 | 0.422 | 0.44 | 0.16 | 0.57 | 0.78 | 0.4 | 0.53 |
| 10 | 117489278 | rs2804161 | -0.028 | 0.077 | 0.339 | -0.439 | -0.066 | 0.071 | 0.071 | 0 | 0.4 | 0.12 | 0.33 | 0.25 | 0.48 | 0.47 |
| 10 | 117490762 | rs2804164 | 0.422 | 0.056 | 0.347 | -0.472 | -0.015 | 0.062 | 0.062 | 0.07 | 0.18 | 0.07 | 0.15 | 0.05 | 0.23 | 0.22 |
| 10 | 117492507 | rs2804165 | -0.454 | -0.128 | 1.146 | -0.514 | -0.028 | -0.086 | -0.086 | 0.547 | 0.23 | 0.55 | 0.19 | 0.05 | 0.22 | 0.28 |
| 10 | 117497003 | rs2532709 | -0.174 | -0.043 | 1.667 | -0.46 | -0.011 | 0.038 | 0.038 | 0 | 0.41 | 0.28 | 0.32 | 0.25 | 0.46 | 0.47 |
| 10 | 117504086 | rs2907543 | 0.63 | -0.053 | 1.547 | -0.485 | -0.001 | -0.009 | -0.009 | 0.051 | 0.17 | 0.15 | 0.15 | 0.04 | 0.22 | 0.2 |
| 10 | 117505905 | rs11197430 | 1.316 | -0.012 | 0.92 | -0.281 | -0.044 | -0.063 | -0.063 | 0 | 0.22 | 0.05 | 0.17 | 0.26 | 0.24 | 0.27 |
| 10 | 117522784 | rs11197439 | 1.473 | 0.004 | 0.78 | -0.32 | -0.08 | -0.041 | -0.041 | 0 | 0.21 | 0.01 | 0.16 | 0.2 | 0.24 | 0.27 |
| 10 | 117531155 | rs10490918 | 1.087 | 0.154 | 0.613 | -0.169 | -0.061 | 0.043 | 0.043 | 0 | 0.22 | 0.03 | 0.31 | 0.08 | 0.27 | 0.11 |
| 10 | 117533271 | rs10885810 | 0.809 | 0.025 | 1.619 | 0.078 | 0 | 0.039 | 0.039 | 0 | 0.52 | 0.18 | 0.62 | 0.68 | 0.54 | 0.44 |
| 10 | 117534146 | rs10885811 | 0.941 | 0.131 | 1.42 | -0.579 | -0.013 | 0.003 | 0.003 | 0 | 0.22 | 0.12 | 0.16 | 0.19 | 0.23 | 0.25 |
| 10 | 117536249 | rs2901061 | -0.279 | 0.126 | 1.488 | -0.511 | -0.001 | 0.012 | 0.012 | 0 | 0.83 | 0.85 | 0.85 | 0.96 | 0.78 | 0.78 |
| 10 | 117543550 | rs11593928 | 0.792 | 0.181 | 1.34 | -0.47 | -0.023 | 0.007 | 0.007 | 0 | 0.24 | 0.11 | 0.17 | 0.19 | 0.25 | 0.26 |

| | | | | | | | | | | | | | | | | |
|---|---|---|---|---|---|---|---|---|---|---|---|---|---|---|---|---|
| 10 | 117548240 | rs10490916 | 0.673 | 0.147 | 1.598 | -0.512 | 0.004 | 0.104 | 0.104 | 0 | 0.51 | 0.24 | 0.62 | 0.76 | 0.53 | 0.43 |
| 10 | 117567760 | rs12777802 | 0.642 | 0.133 | 2.018 | -0.498 | 0.007 | 0.074 | 0.074 | 0 | 0.52 | 0.27 | 0.62 | 0.76 | 0.53 | 0.47 |
| 10 | 117568955 | rs4531378 | 0.538 | 0.125 | 2.104 | -0.492 | 0.003 | 0.046 | 0.046 | 0 | 0.53 | 0.3 | 0.63 | 0.75 | 0.53 | 0.52 |
| 10 | 117574391 | rs2420121 | -1.081 | 0.01 | 2.212 | -0.57 | -0.022 | 0.012 | 0.012 | 0 | 0.76 | 0.85 | 0.81 | 0.81 | 0.75 | 0.75 |
| 10 | 117575619 | rs2960662 | -0.599 | -0.017 | 2.704 | -0.574 | 0.004 | -0.017 | -0.017 | 0 | 0.71 | 0.6 | 0.79 | 0.81 | 0.74 | 0.73 |
| 10 | 117578449 | rs2960665 | -1.089 | -0.011 | 2.621 | -0.589 | -0.022 | 0.012 | 0.012 | 0 | 0.76 | 0.85 | 0.81 | 0.81 | 0.75 | 0.75 |
| 10 | 117591081 | rs2024179 | 0.507 | -0.022 | 2.721 | -0.615 | -0.001 | 0.022 | 0.022 | 0 | 0.16 | 0.07 | 0.13 | 0.04 | 0.2 | 0.2 |
| 10 | 117592731 | rs979416 | 0.746 | -0.052 | 2.845 | -0.632 | 0.005 | 0.111 | 0.111 | 0 | 0.52 | 0.22 | 0.62 | 0.76 | 0.53 | 0.47 |
| 10 | 117594744 | rs1541466 | 0.49 | -0.05 | 2.836 | -0.631 | -0.006 | 0.043 | 0.043 | 0 | 0.16 | 0.04 | 0.13 | 0.04 | 0.2 | 0.2 |
| 10 | 117595544 | rs2420124 | 0.658 | -0.049 | 2.835 | -0.63 | 0.003 | 0.086 | 0.086 | 0 | 0.52 | 0.26 | 0.63 | 0.76 | 0.53 | 0.53 |
| 10 | 117597417 | rs725556 | -0.183 | 0.013 | 2.821 | -0.604 | 0.003 | 0.023 | 0.023 | 0 | 0.67 | 0.23 | 0.75 | 0.8 | 0.74 | 0.66 |
| 10 | 117598178 | rs878709 | -1.935 | -0.004 | 2.965 | -0.58 | 0.045 | 0.058 | 0.058 | 6.787 | 0.06 | 0.4 | 0.05 | 0.01 | 0.01 | 0.02 |
| 10 | 117605972 | rs2960679 | -1.004 | 0.035 | 2.496 | -0.573 | 0.001 | -0.001 | -0.001 | 2.339 | 0.75 | 0.77 | 0.81 | 0.8 | 0.75 | 0.75 |
| 10 | 117610700 | rs2901062 | 0.695 | -0.022 | 2.209 | -0.626 | 0.002 | 0.121 | 0.121 | 0 | 0.52 | 0.18 | 0.62 | 0.75 | 0.54 | 0.46 |
| 10 | 117618612 | rs2907567 | -1.132 | -0.32 | 1.89 | -0.543 | -0.003 | 0.006 | 0.006 | 0 | 0.76 | 0.83 | 0.82 | 0.8 | 0.79 | 0.75 |
| 10 | 117634434 | rs2927372 | 1.449 | -0.315 | 1.924 | -0.558 | 0.063 | 0.058 | 0.058 | 1.536 | 0.94 | 0.54 | 0.96 | 1 | 1 | 0.93 |
| 10 | 117635798 | rs17093700 | 0.038 | -0.137 | 1.596 | -0.533 | 0 | -0.022 | -0.022 | 0.16 | 0.17 | 0.21 | 0.14 | 0.04 | 0.22 | 0.2 |
| 10 | 117638490 | rs2615886 | 0.049 | 0.024 | 1.305 | -0.328 | 0.001 | 0.007 | 0.007 | 0 | 0.56 | 0.38 | 0.65 | 0.75 | 0.58 | 0.49 |
| 10 | 117649124 | rs7904803 | -1.884 | -0.592 | 2.118 | -0.269 | -0.058 | -0.052 | -0.052 | 0 | 0.13 | 0.76 | 0.1 | 0.01 | 0.01 | 0.02 |
| 10 | 117655028 | rs10490985 | -0.065 | -0.9 | 1.772 | -0.234 | 0.004 | -0.036 | -0.036 | 0 | 0.13 | 0.01 | 0.07 | 0.14 | 0.11 | 0.12 |
| 10 | 117661802 | rs17095696 | -0.661 | -0.778 | 3.026 | -0.228 | -0.032 | -0.02 | -0.02 | 0 | 0.08 | 0.33 | 0.04 | 0.01 | 0.01 | 0.02 |
| 10 | 117662179 | rs7086294 | -2.004 | -0.779 | 2.693 | -0.229 | -0.062 | -0.056 | -0.056 | 0.004 | 0.13 | 0.74 | 0.1 | 0.01 | 0.01 | 0.02 |
| 10 | 117664648 | rs2615874 | 1.185 | -0.615 | 2.328 | -0.377 | 0.018 | 0.095 | 0.095 | 7.484 | 0.69 | 0.13 | 0.75 | 0.94 | 0.74 | 0.78 |
| 10 | 117674306 | rs10787612 | 1.216 | -0.533 | 2.458 | -0.168 | 0.001 | -0.006 | -0.006 | 0 | 0.73 | 0.27 | 0.78 | 0.92 | 0.78 | 0.8 |
| 10 | 117679971 | rs17093786 | -0.342 | -0.527 | 2.421 | -0.116 | 0.006 | -0.016 | -0.016 | 0 | 0.17 | 0.22 | 0.16 | 0.07 | 0.22 | 0.17 |
| 10 | 117680914 | rs17093794 | -0.351 | -0.554 | 2.799 | -0.125 | -0.037 | -0.026 | -0.026 | 0.699 | 0.08 | 0.24 | 0.04 | 0.01 | 0.01 | 0.02 |
| 10 | 117682532 | rs2249295 | 1.31 | -0.559 | 2.427 | -0.129 | -0.015 | -0.083 | -0.083 | 1.099 | 0.75 | 0.4 | 0.81 | 0.95 | 0.77 | 0.81 |
| 10 | 117682626 | rs7083759 | -0.745 | -0.471 | 2.217 | -0.088 | -0.028 | -0.015 | -0.015 | 0 | 0.09 | 0.38 | 0.05 | 0.01 | 0.01 | 0.02 |
| 10 | 117687402 | rs10490986 | -0.21 | -0.16 | 1.526 | 0.019 | -0.001 | -0.002 | -0.002 | 0.151 | 0.16 | 0.12 | 0.15 | 0.07 | 0.21 | 0.18 |
| 10 | 117695712 | rs17270580 | -0.324 | -0.183 | 1.523 | 0.006 | -0.002 | -0.002 | -0.002 | 0 | 0.16 | 0.13 | 0.14 | 0.06 | 0.22 | 0.18 |
| 10 | 117701577 | rs17270699 | -0.062 | 0.004 | 0.839 | 0.064 | -0.025 | 0.03 | 0.03 | 0.999 | 0.16 | 0.07 | 0.13 | 0.05 | 0.22 | 0.18 |
| 10 | 117705146 | rs3740571 | -1.033 | -0.836 | 1.434 | 0.417 | -0.055 | 0.013 | 0.013 | 0.003 | 0.2 | 0.34 | 0.1 | 0.23 | 0.11 | 0.12 |
| 10 | 117707733 | rs10885833 | -1.065 | -0.681 | 1.144 | 0.489 | -0.031 | 0.013 | 0.013 | 0 | 0.2 | 0.29 | 0.1 | 0.23 | 0.11 | 0.11 |
| 10 | 117708868 | rs4751939 | -0.468 | 0.098 | 0.835 | 0.395 | 0.005 | 0.005 | 0.005 | 0.742 | 0.4 | 0.44 | 0.47 | 0.58 | 0.49 | 0.36 |
| 10 | 117713221 | rs4751940 | -0.683 | 0.109 | 0.532 | 0.422 | 0.022 | -0.011 | -0.011 | 0.567 | 0.47 | 0.7 | 0.55 | 0.52 | 0.58 | 0.35 |
| 10 | 117720208 | rs917382 | -0.476 | 0.284 | -0.435 | 0.502 | 0.006 | -0.016 | -0.016 | 0 | 0.43 | 0.29 | 0.51 | 0.51 | 0.58 | 0.35 |
| 10 | 117725755 | rs10787618 | -0.798 | 0.42 | -0.42 | 0.54 | 0.032 | 0.055 | 0.055 | 1.593 | 0.44 | 0.72 | 0.39 | 0.26 | 0.37 | 0.26 |
| 10 | 117727166 | rs180727 | 0.139 | 0.725 | -0.691 | 0.843 | -0.058 | -0.026 | -0.026 | 0 | 0.21 | 0.03 | 0.12 | 0.22 | 0.25 | 0.26 |
| 10 | 117728684 | rs180723 | 0.966 | 0.629 | 0.525 | 1.33 | -0.009 | 0.009 | 0.009 | 0.137 | 0.61 | 0.24 | 0.48 | 0.39 | 0.62 | 0.61 |
| 10 | 117728741 | rs180722 | 0.761 | 0.627 | 1.166 | 1.22 | 0.011 | 0.019 | 0.019 | 0 | 0.64 | 0.41 | 0.58 | 0.48 | 0.61 | 0.64 |
| 10 | 117730587 | rs11197487 | -0.642 | 0.632 | 1.673 | 1.513 | -0.001 | -0.001 | -0.001 | 0 | 0.3 | 0.3 | 0.24 | 0.28 | 0.31 | 0.23 |
| 10 | 117732413 | rs180714 | 0.849 | 0.673 | 1.301 | 0.97 | 0.01 | 0.022 | 0.022 | 0 | 0.63 | 0.45 | 0.59 | 0.49 | 0.62 | 0.64 |
| 10 | 117735419 | rs180707 | 1.455 | -0.4 | 2.073 | 1.659 | -0.02 | 0.145 | 0.145 | 0 | 0.77 | 0.47 | 0.74 | 0.49 | 0.85 | 0.78 |
| 10 | 117735450 | rs180706 | 1.581 | -0.474 | 2.481 | 1.65 | -0.009 | 0.013 | 0.013 | 0.354 | 0.75 | 0.27 | 0.73 | 0.44 | 0.85 | 0.77 |
| 10 | 117736585 | rs180703 | 1.312 | -0.455 | 2.529 | 1.64 | -0.031 | 0.125 | 0.125 | 0 | 0.78 | 0.53 | 0.76 | 0.5 | 0.87 | 0.79 |
| 10 | 117741578 | rs180696 | 1.643 | -0.446 | 2.602 | 1.521 | -0.024 | 0.131 | 0.131 | 0 | 0.75 | 0.51 | 0.75 | 0.48 | 0.85 | 0.76 |
| 10 | 117747653 | rs180680 | 1.739 | -0.438 | 2.707 | 1.518 | -0.004 | 0.01 | 0.01 | 0 | 0.73 | 0.3 | 0.73 | 0.49 | 0.85 | 0.74 |
| 10 | 117753912 | rs180671 | 1.515 | -0.46 | 2.736 | 1.5 | -0.019 | 0.112 | 0.112 | 0 | 0.74 | 0.46 | 0.74 | 0.48 | 0.84 | 0.73 |
| 10 | 117757648 | rs180665 | 1.683 | -0.465 | 2.802 | 1.492 | -0.016 | 0.075 | 0.075 | 0 | 0.73 | 0.4 | 0.73 | 0.48 | 0.84 | 0.71 |
| 10 | 117759983 | rs180660 | 1.512 | -0.523 | 2.741 | 1.429 | -0.008 | 0.014 | 0.014 | 0 | 0.64 | 0.38 | 0.69 | 0.48 | 0.74 | 0.65 |
| 10 | 117761676 | rs11197496 | -0.572 | -0.571 | 2.863 | 1.556 | 0 | 0.057 | 0.057 | 0 | 0.09 | 0.13 | 0.18 | 0.41 | 0.09 | 0.16 |
| 10 | 117763084 | rs180654 | 1.688 | -0.606 | 2.8 | 1.543 | 0.013 | -0.022 | -0.022 | 0 | 0.64 | 0.27 | 0.68 | 0.48 | 0.74 | 0.66 |
| 10 | 117773116 | rs180627 | 1.746 | -0.541 | 2.812 | 1.42 | 0.02 | -0.018 | -0.018 | 0.054 | 0.63 | 0.23 | 0.68 | 0.44 | 0.74 | 0.66 |
| 10 | 117784322 | rs180623 | 1.577 | -0.54 | 2.833 | 1.42 | 0.002 | -0.004 | -0.004 | 0 | 0.65 | 0.32 | 0.67 | 0.45 | 0.74 | 0.67 |
| 10 | 117786756 | rs180620 | 0.786 | -0.492 | 2.993 | 1.393 | -0.014 | -0.035 | -0.035 | 0.103 | 0.75 | 0.65 | 0.88 | 0.85 | 0.81 | 0.85 |
| 10 | 117786867 | rs180619 | 1.779 | -0.481 | 2.623 | 1.208 | 0.024 | -0.016 | -0.016 | 0 | 0.62 | 0.18 | 0.66 | 0.4 | 0.71 | 0.64 |
| 10 | 117790223 | rs1679542 | 1.542 | -0.36 | 2.67 | 1.26 | -0.005 | 0.005 | 0.005 | 0 | 0.64 | 0.38 | 0.67 | 0.4 | 0.72 | 0.64 |
| 10 | 117807835 | rs7907680 | -0.692 | -0.329 | 2.353 | 1.463 | 0.003 | 0.097 | 0.097 | 0 | 0.1 | 0.17 | 0.21 | 0.45 | 0.09 | 0.19 |
| 10 | 117809158 | rs180580 | -0.795 | -0.425 | 2.646 | 1.32 | -0.023 | -0.052 | -0.052 | 0 | 0.22 | 0.39 | 0.12 | 0.15 | 0.18 | 0.16 |
| 10 | 117809866 | rs180571 | -0.795 | -0.425 | 2.644 | 1.32 | -0.022 | -0.048 | -0.048 | 0 | 0.22 | 0.37 | 0.12 | 0.15 | 0.18 | 0.16 |
| 10 | 117813312 | rs180561 | -0.716 | -0.399 | 2.564 | 1.306 | -0.017 | -0.039 | -0.039 | 0.475 | 0.22 | 0.32 | 0.13 | 0.15 | 0.19 | 0.18 |
| 10 | 117818560 | rs180554 | -1.698 | -0.397 | 2.122 | 1.337 | 0.024 | -0.017 | -0.017 | 0 | 0.36 | 0.78 | 0.36 | 0.6 | 0.28 | 0.36 |
| 10 | 117818842 | rs180552 | -1.849 | -0.402 | 2.141 | 1.336 | 0.013 | -0.01 | -0.01 | 0 | 0.35 | 0.74 | 0.35 | 0.6 | 0.28 | 0.35 |
| 10 | 117821350 | rs7071452 | -1.918 | -0.558 | 2.165 | 1.343 | 0.018 | -0.011 | -0.011 | 0.986 | 0.37 | 0.77 | 0.38 | 0.61 | 0.3 | 0.35 |
| 10 | 117823204 | rs2072277 | -0.627 | -0.511 | 2.247 | 1.389 | 0.006 | 0.202 | 0.202 | 0 | 0.1 | 0.23 | 0.21 | 0.47 | 0.07 | 0.17 |
| 10 | 117823739 | rs1078080 | -1.948 | -0.515 | 2.143 | 1.388 | 0.026 | -0.018 | -0.018 | 0 | 0.38 | 0.82 | 0.39 | 0.61 | 0.3 | 0.37 |
| 10 | 117826992 | rs11598215 | -1.205 | -0.662 | 2.102 | 0.487 | -0.038 | -0.076 | -0.076 | 0 | 0.25 | 0.53 | 0.15 | 0.14 | 0.21 | 0.19 |
| 10 | 117828486 | rs3781504 | -1.156 | -0.743 | 1.969 | 0.656 | -0.038 | -0.05 | -0.05 | 0 | 0.27 | 0.53 | 0.15 | 0.2 | 0.2 | 0.2 |
| 10 | 117830114 | rs7070763 | -1.86 | -0.25 | 2.126 | 0.978 | 0.076 | 0.071 | 0.071 | 6.276 | 0.05 | 0.52 | 0.04 | 0 | 0 | 0.03 |

| | | | | | | | | | | | | | | | | |
|---|---|---|---|---|---|---|---|---|---|---|---|---|---|---|---|---|
| 10 | 117842360 | rs4751949 | -1.298 | 0.049 | 0.6 | 1.731 | 0.009 | 0.17 | 0.17 | 0 | 0.13 | 0.24 | 0.21 | 0.48 | 0.12 | 0.2 |
| 10 | 117843015 | rs10490904 | -1.191 | -0.181 | 0.575 | 1.426 | -0.016 | -0.012 | -0.012 | 0 | 0.16 | 0.27 | 0.07 | 0.13 | 0.11 | 0.22 |
| 10 | 117843954 | rs11197522 | -0.639 | -0.01 | 0.381 | 1.79 | -0.038 | -0.015 | -0.015 | 0 | 0.07 | 0.01 | 0.19 | 0.41 | 0.1 | 0.17 |
| 10 | 117844044 | rs7087152 | -1.433 | -0.726 | 0.451 | 1.177 | -0.008 | 0.067 | 0.067 | 0 | 0.24 | 0.58 | 0.29 | 0.43 | 0.21 | 0.29 |
| 10 | 117844216 | rs7087213 | -1.686 | -1.313 | -0.538 | 0.879 | 0.074 | -0.093 | -0.093 | 0.968 | 0.46 | 0.93 | 0.38 | 0.57 | 0.32 | 0.5 |
| 10 | 117844652 | rs2694773 | -1.952 | -0.978 | 0.298 | 1.596 | 0.059 | 0.368 | 0.368 | 0 | 0.08 | 0.25 | 0.21 | 0.42 | 0.1 | 0.25 |
| 10 | 117845895 | rs11197523 | 0.124 | -1.415 | -1.156 | 0.702 | -0.004 | -0.012 | -0.012 | 1.836 | 0.34 | 0.09 | 0.45 | 0.7 | 0.29 | 0.37 |
| 10 | 117846736 | rs3901216 | 0.814 | -1.45 | -1.711 | 0.211 | -0.147 | 0.186 | 0.186 | 0 | 0.36 | 0.02 | 0.29 | 0.11 | 0.53 | 0.38 |
| 10 | 117851318 | rs2530343 | -0.147 | -1.221 | -0.515 | 0.709 | -0.021 | 0.013 | 0.013 | 0 | 0.15 | 0.21 | 0.14 | 0.16 | 0.09 | 0.14 |
| 10 | 117852470 | rs7070180 | -0.238 | -1.274 | -0.518 | 0.935 | 0.008 | 0.01 | 0.01 | 0 | 0.32 | 0.2 | 0.45 | 0.72 | 0.27 | 0.36 |
| 10 | 117852569 | rs11197525 | -1.118 | -0.905 | 0.455 | 1.671 | -0.023 | 0.098 | 0.098 | 0 | 0.07 | 0.18 | 0.23 | 0.19 | 0.03 | 0.06 |
| 10 | 117861297 | rs3781514 | -1.321 | -0.252 | 0.681 | 1.814 | -0.047 | -0.022 | -0.022 | 0 | 0.11 | 0.01 | 0.1 | 0.17 | 0.17 | 0.12 |
| 10 | 117862573 | rs2694785 | -0.856 | -0.191 | 0.7 | 1.543 | 0.003 | 0.006 | 0.006 | 0 | 0.25 | 0.22 | 0.14 | 0.2 | 0.19 | 0.19 |
| 10 | 117862850 | rs2694787 | -0.761 | 0.03 | 0.863 | 1.645 | -0.007 | 0.001 | 0.001 | 0 | 0.2 | 0.14 | 0.13 | 0.2 | 0.18 | 0.18 |
| 10 | 117863334 | rs7070267 | 1.725 | -0.323 | 1.394 | 1.41 | -0.001 | -0.001 | -0.001 | 0 | 0.56 | 0.4 | 0.54 | 0.39 | 0.56 | 0.56 |
| 10 | 117864680 | rs9787473 | 1.722 | -0.128 | 1.464 | 1.456 | -0.014 | -0.014 | -0.014 | 3.697 | 0.57 | 0.26 | 0.54 | 0.42 | 0.57 | 0.57 |
| 10 | 117869850 | rs10787619 | 1.477 | -0.094 | 1.487 | 0.839 | -0.005 | 0.006 | 0.006 | 0 | 0.58 | 0.35 | 0.53 | 0.56 | 0.57 | 0.59 |
| 10 | 117877793 | rs2530348 | 1.655 | -0.105 | 2.459 | 0.835 | 0.066 | 0.044 | 0.044 | 0.885 | 0.93 | 0.52 | 0.9 | 0.98 | 0.97 | 0.93 |
| 10 | 117878601 | rs2530347 | 1.655 | -0.106 | 2.459 | 0.835 | 0.068 | 0.046 | 0.046 | 0.968 | 0.93 | 0.53 | 0.9 | 0.98 | 0.97 | 0.94 |
| 10 | 117881996 | rs3843600 | 2.086 | -0.208 | 2.178 | 0.793 | -0.006 | 0.018 | 0.018 | 0 | 0.55 | 0.25 | 0.64 | 0.55 | 0.55 | 0.56 |
| 10 | 117884950 | rs2245020 | 2.067 | -0.213 | 2.176 | 0.791 | -0.007 | 0.016 | 0.016 | 0 | 0.56 | 0.25 | 0.65 | 0.55 | 0.55 | 0.56 |
| 10 | 117887344 | rs12781677 | -1.805 | -0.147 | 2.246 | 0.788 | -0.009 | 0.001 | 0.001 | 0 | 0.38 | 0.28 | 0.25 | 0.42 | 0.42 | 0.37 |
| 10 | 117888908 | rs12771220 | -1.833 | -0.151 | 2.244 | 0.801 | -0.009 | 0 | 0 | 0 | 0.38 | 0.28 | 0.25 | 0.43 | 0.42 | 0.37 |
| 10 | 117892248 | rs3781531 | -1.338 | -0.082 | 1.942 | 0.829 | -0.034 | -0.033 | -0.033 | 0 | 0.13 | 0.04 | 0.08 | 0.2 | 0.18 | 0.18 |
| 10 | 117894381 | rs4237493 | 1.193 | -0.122 | 2.271 | 0.818 | -0.003 | -0.022 | -0.022 | 0 | 0.73 | 0.48 | 0.81 | 0.77 | 0.76 | 0.81 |
| 10 | 117895604 | rs3781532 | 2.132 | -0.117 | 1.995 | 0.821 | -0.007 | 0.017 | 0.017 | 0 | 0.56 | 0.25 | 0.53 | 0.55 | 0.55 | 0.57 |
| 10 | 117900691 | rs2286732 | -2.568 | 0.411 | 2.207 | 1.043 | 0.032 | 0.027 | 0.027 | 0 | 0.29 | 0.45 | 0.41 | 0.41 | 0.38 | 0.39 |
| 10 | 117902564 | rs10787628 | 2.368 | 0.417 | 2.23 | 1.011 | 0.023 | 0.013 | 0.013 | 0 | 0.65 | 0.46 | 0.56 | 0.59 | 0.6 | 0.59 |
| 10 | 117915108 | rs7090606 | -1.677 | 0.735 | 2.07 | 0.963 | 0.035 | 0.015 | 0.015 | 0 | 0.16 | 0.32 | 0.12 | 0.21 | 0.18 | 0.21 |
| 10 | 117918067 | rs11197557 | -1.381 | 0.375 | 1.909 | 1.14 | -0.07 | -0.08 | -0.08 | 0 | 0.13 | 0.01 | 0.32 | 0.26 | 0.21 | 0.25 |
| 10 | 117918311 | rs10749189 | -2.706 | 0.388 | 2.244 | 1.071 | 0.022 | 0.033 | 0.033 | 0 | 0.28 | 0.37 | 0.43 | 0.43 | 0.36 | 0.4 |
| 10 | 117920341 | rs10510015 | -2.506 | 0.349 | 2.309 | 1.045 | 0.062 | 0.04 | 0.04 | 0.628 | 0.06 | 0.42 | 0.05 | 0 | 0.01 | 0.03 |
| 10 | 117922201 | rs11197560 | -1.233 | 0.382 | 1.715 | 1.126 | -0.048 | -0.062 | -0.062 | 2.876 | 0.13 | 0.02 | 0.32 | 0.26 | 0.2 | 0.24 |
| 10 | 117924062 | rs4553287 | -2.585 | -0.533 | 1.935 | 0.948 | -0.008 | -0.001 | -0.001 | 0 | 0.43 | 0.54 | 0.46 | 0.44 | 0.4 | 0.47 |
| 10 | 117929704 | rs10749194 | 1.987 | -0.572 | 1.898 | 0.93 | 0.005 | -0.003 | -0.003 | 0.144 | 0.6 | 0.6 | 0.55 | 0.56 | 0.62 | 0.54 |
| 10 | 117932193 | rs11197567 | -1.486 | -0.544 | 1.881 | 0.88 | -0.004 | 0.016 | 0.016 | 0 | 0.24 | 0.16 | 0.14 | 0.17 | 0.23 | 0.21 |
| 10 | 117936356 | rs10749199 | 0.331 | -0.438 | 1.863 | 0.961 | -0.001 | 0.002 | 0.002 | 0 | 0.94 | 0.92 | 0.97 | 0.9 | 0.95 | 0.9 |
| 10 | 117941797 | rs7917174 | 1.395 | -0.486 | 1.845 | 0.892 | 0.001 | 0.001 | 0.001 | 0 | 0.87 | 0.84 | 0.93 | 0.9 | 0.92 | 0.87 |
| 10 | 117942634 | rs11197571 | -0.355 | -0.473 | 1.481 | 0.876 | -0.002 | 0.011 | 0.011 | 0 | 0.15 | 0.13 | 0.07 | 0.06 | 0.15 | 0.07 |
| 10 | 117944165 | rs7909875 | 2.156 | -0.458 | 1.482 | 0.894 | -0.019 | 0.036 | 0.036 | 0 | 0.73 | 0.44 | 0.62 | 0.65 | 0.76 | 0.62 |
| 10 | 117946337 | rs7904143 | -1.869 | -0.303 | 1.308 | 1.016 | -0.008 | 0.036 | 0.036 | 0 | 0.29 | 0.48 | 0.38 | 0.37 | 0.31 | 0.27 |
| 10 | 117947143 | rs953920 | 0.038 | -0.266 | 1.243 | 1.043 | 0.002 | -0.02 | -0.02 | 0 | 0.14 | 0.04 | 0.06 | 0.17 | 0.13 | 0.07 |
| 10 | 117947419 | rs11197576 | -1.887 | -0.014 | 1.309 | 1.23 | 0.024 | 0.107 | 0.107 | 0 | 0.2 | 0.47 | 0.39 | 0.3 | 0.23 | 0.34 |
| 10 | 117949032 | rs4751581 | 1.299 | 0.167 | -0.336 | 1.26 | 0.026 | -0.009 | -0.009 | 0 | 0.54 | 0.23 | 0.48 | 0.49 | 0.59 | 0.54 |
| 10 | 117949460 | rs3781545 | -1.404 | 0.932 | -0.335 | 1.505 | -0.023 | -0.035 | -0.035 | 0.019 | 0.1 | 0.04 | 0.29 | 0.19 | 0.18 | 0.2 |
| 10 | 117957130 | rs3824842 | -1.697 | 0.686 | 0.109 | 1.323 | 0.015 | 0.065 | 0.065 | 0 | 0.14 | 0.32 | 0.29 | 0.26 | 0.16 | 0.32 |
| 10 | 117961273 | rs3781550 | -0.671 | -0.448 | -0.394 | 0.747 | 0.012 | 0.022 | 0.022 | 0.029 | 0.26 | 0.2 | 0.14 | 0.15 | 0.22 | 0.27 |
| 10 | 117963418 | rs12269391 | -0.904 | -0.632 | 1.003 | 0.663 | -0.035 | -0.064 | -0.064 | 0 | 0.28 | 0.46 | 0.17 | 0.16 | 0.22 | 0.27 |
| 10 | 117966778 | rs2577383 | -0.698 | -0.808 | 1.325 | 0.681 | -0.004 | -0.007 | -0.007 | 0.978 | 0.25 | 0.26 | 0.15 | 0.17 | 0.22 | 0.28 |
| 10 | 117967763 | rs2694765 | -1.121 | -1.447 | 0.323 | 0.536 | -0.082 | -0.077 | -0.077 | 0 | 0.67 | 0.89 | 0.47 | 0.46 | 0.53 | 0.71 |
| 10 | 117968330 | rs2694767 | -1.097 | -1.477 | 0.56 | 0.509 | -0.051 | -0.05 | -0.05 | 3.262 | 0.67 | 0.81 | 0.46 | 0.46 | 0.53 | 0.71 |
| 10 | 117968826 | rs2694769 | -1.233 | -1.311 | 0.547 | 0.482 | -0.115 | -0.099 | -0.099 | 0 | 0.69 | 0.93 | 0.48 | 0.46 | 0.54 | 0.72 |
| 10 | 117979318 | rs4611119 | 0.199 | -1.449 | 0.68 | 0.443 | -0.001 | -0.001 | -0.001 | 0 | 0.43 | 0.54 | 0.31 | 0.29 | 0.32 | 0.43 |
| 10 | 117987546 | rs7894329 | 0.753 | -1.403 | 0.585 | 0.457 | -0.047 | -0.044 | -0.044 | 0 | 0.33 | 0.17 | 0.53 | 0.54 | 0.46 | 0.29 |
| 10 | 117987805 | rs7911126 | 0.775 | -1.4 | 0.515 | 0.453 | -0.08 | -0.073 | -0.073 | 0 | 0.32 | 0.12 | 0.53 | 0.54 | 0.46 | 0.29 |
| 10 | 117988864 | rs881726 | 0.533 | -1.82 | 0.137 | 0.251 | 0.032 | 0.159 | 0.159 | 0 | 0.33 | 0.01 | 0.22 | 0.06 | 0.27 | 0.37 |
| 10 | 117991693 | rs10787637 | 0.01 | -1.869 | 0.198 | 0.215 | 0.063 | 0.051 | 0.051 | 0.005 | 0.4 | 0.16 | 0.29 | 0.3 | 0.34 | 0.44 |
| 10 | 117992418 | rs7097998 | 0.213 | -1.71 | 0.595 | 0.046 | -0.001 | -0.044 | -0.044 | 0.393 | 0.1 | 0.18 | 0.06 | 0 | 0.1 | 0.07 |
| 10 | 117993920 | rs11812459 | 0.009 | -2.212 | 0.942 | 0.206 | 0.008 | 0.006 | 0.006 | 0 | 0.49 | 0.54 | 0.57 | 0.56 | 0.55 | 0.44 |
| 10 | 117995793 | rs3781554 | -0.093 | -2.189 | 1.204 | 0.126 | 0.015 | -0.022 | -0.022 | 0.171 | 0.12 | 0.27 | 0.09 | 0.06 | 0.12 | 0.1 |
| 10 | 117996150 | rs7907314 | 0.204 | -2.534 | 1.028 | 0.091 | -0.001 | 0 | 0 | 0.513 | 0.55 | 0.59 | 0.63 | 0.57 | 0.59 | 0.51 |
| 10 | 118000519 | rs10787638 | 1.002 | -2.824 | 0.512 | 0.352 | -0.068 | 0.062 | 0.062 | 0 | 0.34 | 0.11 | 0.33 | 0.23 | 0.48 | 0.3 |
| 10 | 118000945 | rs2420255 | -0.026 | -2.988 | 0.864 | 0.574 | -0.023 | 0.008 | 0.008 | 0 | 0.29 | 0.35 | 0.32 | 0.34 | 0.2 | 0.3 |
| 10 | 118002927 | rs3740572 | 0.764 | -2.635 | 0.8 | 0.657 | -0.034 | 0.07 | 0.07 | 0 | 0.37 | 0.16 | 0.35 | 0.23 | 0.5 | 0.31 |
| 10 | 118014205 | rs1077114 | -0.425 | -3.147 | 0.557 | 0.437 | 0.024 | 0.001 | 0.001 | 0 | 0.42 | 0.62 | 0.38 | 0.42 | 0.24 | 0.46 |
| 10 | 118016664 | rs12413585 | 0.265 | -2.68 | 1.114 | 0.924 | 0 | 0 | 0 | 0 | 0.44 | 0.53 | 0.5 | 0.53 | 0.46 | 0.36 |
| 10 | 118018805 | rs11816383 | -0.183 | -2.396 | 1.317 | 0.965 | -0.021 | 0.056 | 0.056 | 0 | 0.13 | 0.25 | 0.19 | 0.27 | 0.07 | 0.11 |
| 10 | 118020066 | rs12774375 | 1.479 | -2.432 | 1.314 | 0.95 | -0.008 | 0.002 | 0.002 | 0 | 0.31 | 0.24 | 0.31 | 0.28 | 0.39 | 0.25 |
| 10 | 118022076 | rs7920266 | 0.378 | -2.556 | 1.351 | 0.938 | -0.021 | 0.017 | 0.017 | 0 | 0.66 | 0.5 | 0.54 | 0.4 | 0.68 | 0.67 |

| | | | | | | | | | | | | | | | |
|---|---|---|---|---|---|---|---|---|---|---|---|---|---|---|---|
| 10 | 118024544 | rs2577355 | -0.624 | -2.505 | 1.295 | 0.942 | 0.005 | 0.011 | 0.011 | 0 | 0.66 | 0.72 | 0.75 | 0.88 | 0.71 | 0.59 |
| 10 | 118028614 | rs2074986 | -0.707 | -2.281 | 1.524 | 1.173 | -0.028 | 0.195 | 0.195 | 0.375 | 0.15 | 0.43 | 0.2 | 0.34 | 0.07 | 0.11 |
| 10 | 118035740 | rs2532689 | -0.481 | -2.172 | 1.211 | 0.843 | -0.001 | -0.002 | -0.002 | 0 | 0.54 | 0.5 | 0.51 | 0.5 | 0.52 | 0.64 |
| 10 | 118046534 | rs881786 | -0.403 | -1.973 | 1.338 | 1.155 | 0 | 0 | 0 | 0 | 0.46 | 0.52 | 0.45 | 0.38 | 0.35 | 0.55 |
| 10 | 118047298 | rs758577 | -0.517 | -1.971 | 1.398 | 1.196 | -0.004 | 0.066 | 0.066 | 0 | 0.8 | 0.7 | 0.76 | 0.63 | 0.77 | 0.82 |
| 10 | 118047671 | rs758575 | 0.194 | -1.668 | 1.413 | 1.131 | 0.008 | 0.007 | 0.007 | 0 | 0.54 | 0.45 | 0.55 | 0.62 | 0.64 | 0.47 |
| 10 | 118049105 | rs12767703 | 0.219 | -1.6 | 1.398 | 1.167 | 0.022 | 0.019 | 0.019 | 0 | 0.53 | 0.37 | 0.54 | 0.62 | 0.65 | 0.46 |
| 10 | 118052051 | rs11812522 | 0.082 | -1.802 | 1.379 | 0.995 | -0.008 | 0.022 | 0.022 | 0 | 0.1 | 0.15 | 0.17 | 0.22 | 0.06 | 0.11 |
| 10 | 118056943 | rs988977 | 0.189 | -1.922 | 1.36 | 0.921 | -0.006 | -0.007 | -0.007 | 0 | 0.54 | 0.47 | 0.55 | 0.67 | 0.66 | 0.45 |
| 10 | 118059344 | rs2577346 | 0.257 | -1.884 | 1.126 | 0.94 | 0.048 | 0.046 | 0.046 | 0 | 0.53 | 0.31 | 0.56 | 0.66 | 0.67 | 0.45 |
| 10 | 118075798 | rs1819810 | 0.26 | -1.779 | 0.322 | 0.974 | 0.048 | 0.054 | 0.054 | 0 | 0.53 | 0.33 | 0.54 | 0.67 | 0.66 | 0.47 |
| 10 | 118076455 | rs10490910 | -0.367 | -1.754 | 0.059 | 1.039 | -0.03 | 0.052 | 0.052 | 0 | 0.12 | 0.31 | 0.18 | 0.2 | 0.05 | 0.11 |
| 10 | 118087298 | rs10430687 | 0.894 | -0.792 | -0.189 | 1.111 | -0.09 | 0.025 | 0.025 | 0.119 | 0.35 | 0.14 | 0.34 | 0.26 | 0.44 | 0.3 |
| 10 | 118093608 | rs12217215 | 0.867 | -0.848 | -0.487 | 1.138 | -0.137 | 0.011 | 0.011 | 0.223 | 0.36 | 0.07 | 0.33 | 0.26 | 0.44 | 0.31 |
| 10 | 118102986 | rs2577373 | -0.559 | -1.082 | -0.39 | 1.052 | 0.002 | -0.009 | -0.009 | 0 | 0.63 | 0.71 | 0.73 | 0.87 | 0.72 | 0.59 |
| 10 | 118106468 | rs10885912 | 0.021 | 0.096 | -0.842 | 1.481 | -0.141 | 0.03 | 0.03 | 0.155 | 0.35 | 0.07 | 0.32 | 0.24 | 0.43 | 0.28 |
| 10 | 118123862 | rs6585391 | -1.576 | -0.054 | 0.665 | 2.924 | 0 | 0.24 | 0.24 | 0 | 0.05 | 0.19 | 0.22 | 0.29 | 0.03 | 0.02 |
| 10 | 118131382 | rs11197640 | -1.757 | 0.054 | 1.824 | 2.988 | 0.004 | 0.291 | 0.291 | 0 | 0.05 | 0.23 | 0.23 | 0.31 | 0.03 | 0.02 |
| 10 | 118133552 | rs7093477 | -2.861 | 0.055 | 1.98 | 2.988 | 0.06 | 0.413 | 0.413 | 0.298 | 0.09 | 0.48 | 0.31 | 0.35 | 0.11 | 0.05 |
| 10 | 118142066 | rs11197647 | -3.126 | 0.022 | 2.661 | 2.87 | 0.067 | 0.27 | 0.27 | 0.477 | 0.11 | 0.51 | 0.27 | 0.26 | 0.12 | 0.06 |
| 10 | 118147645 | rs17094643 | -1.854 | -0.128 | 3.088 | 3.186 | 0.065 | 0.184 | 0.184 | 0.979 | 0.07 | 0.51 | 0.06 | 0.13 | 0.02 | 0.02 |
| 10 | 118147665 | rs17094644 | -3.211 | -0.187 | 2.738 | 3.168 | -0.003 | 0.053 | 0.053 | 2.571 | 0.13 | 0.71 | 0.24 | 0.26 | 0.09 | 0.05 |
| 10 | 118156983 | rs17734465 | 0.241 | -0.33 | 1.969 | 2.853 | -0.036 | 0.112 | 0.112 | 0.598 | 0.34 | 0.09 | 0.18 | 0.16 | 0.26 | 0.43 |
| 10 | 118167864 | rs11197666 | -2.449 | -0.374 | 3.62 | 3.82 | -0.005 | 0.289 | 0.289 | 0.004 | 0.08 | 0.29 | 0.16 | 0.31 | 0.03 | 0.06 |
| 10 | 118185623 | rs17094717 | -1.735 | -0.316 | 4.036 | 4.344 | 0.006 | 0.056 | 0.056 | 0.108 | 0.06 | 0.13 | 0.16 | 0.17 | 0.04 | 0.06 |
| 10 | 118186269 | rs1897511 | 2.19 | -0.281 | 4.182 | 4.4 | 0.041 | 0.138 | 0.138 | 0.022 | 0.95 | 0.73 | 0.77 | 0.88 | 0.93 | 0.97 |
| 10 | 118193216 | rs1431485 | 2.757 | -0.234 | 3.387 | 4.022 | 0.014 | 0.222 | 0.222 | 0.895 | 0.89 | 0.61 | 0.62 | 0.71 | 0.9 | 0.92 |
| 10 | 118196141 | rs11197681 | -2.615 | -0.189 | 3.051 | 3.84 | 0.008 | 0.068 | 0.068 | 0.077 | 0.09 | 0.2 | 0.29 | 0.19 | 0.07 | 0.07 |
| 10 | 118198276 | rs10885935 | -2.16 | -0.433 | 3.059 | 3.861 | -0.004 | 0.075 | 0.075 | 0.06 | 0.08 | 0.18 | 0.28 | 0.19 | 0.05 | 0.06 |
| 10 | 118199796 | rs2116287 | 2.747 | -0.484 | 3.6 | 4.178 | 0.079 | 0.719 | 0.719 | 2.48 | 0.94 | 0.56 | 0.77 | 0.62 | 0.96 | 0.97 |
| 10 | 118209910 | rs7071213 | 3.32 | -0.431 | 3.602 | 3.939 | 0.03 | 0.381 | 0.381 | 1.907 | 0.93 | 0.45 | 0.77 | 0.73 | 0.96 | 0.96 |
| 10 | 118211224 | rs4751967 | 2.64 | -0.415 | 3.668 | 3.905 | 0.049 | 0.344 | 0.344 | 0.68 | 0.95 | 0.69 | 0.87 | 0.77 | 0.98 | 0.96 |
| 10 | 118218904 | rs891665 | 2.56 | -0.407 | 3.557 | 4.015 | -0.016 | 0.091 | 0.091 | 0 | 0.8 | 0.44 | 0.63 | 0.38 | 0.82 | 0.82 |
| 10 | 118221256 | rs1897515 | 2.588 | -0.408 | 3.962 | 3.944 | -0.017 | 0.089 | 0.089 | 0 | 0.8 | 0.44 | 0.63 | 0.38 | 0.82 | 0.82 |
| 10 | 118228763 | rs7077408 | -2.529 | -0.378 | 3.987 | 4.01 | 0.099 | 0.334 | 0.334 | 0.064 | 0.05 | 0.33 | 0.14 | 0.23 | 0.06 | 0.08 |
| 10 | 118229239 | rs4751586 | 1.363 | -0.395 | 4.203 | 3.984 | -0.005 | -0.039 | -0.039 | 0 | 0.89 | 0.85 | 0.94 | 1 | 0.9 | 0.92 |
| 10 | 118231305 | rs10510020 | -2.184 | -0.5 | 4.13 | 3.971 | 0.05 | 0.252 | 0.252 | 0 | 0.07 | 0.32 | 0.15 | 0.24 | 0.06 | 0.08 |
| 10 | 118237356 | rs7898331 | 2.19 | -0.464 | 3.285 | 3.663 | 0.041 | 0.238 | 0.238 | 0.43 | 0.92 | 0.62 | 0.84 | 0.76 | 0.94 | 0.92 |
| 10 | 118238417 | rs10787677 | 2.259 | -0.464 | 3.28 | 3.663 | 0.049 | 0.248 | 0.248 | 0.543 | 0.92 | 0.6 | 0.84 | 0.76 | 0.94 | 0.92 |
| 10 | 118238635 | rs10787678 | 2.539 | -0.466 | 3.271 | 3.65 | -0.004 | 0.067 | 0.067 | 0 | 0.82 | 0.6 | 0.78 | 0.76 | 0.84 | 0.85 |
| 10 | 118255985 | rs11197705 | 1.844 | -0.365 | 2.728 | 3.6 | 0.014 | 0.158 | 0.158 | 0.079 | 0.92 | 0.74 | 0.83 | 0.68 | 0.94 | 0.93 |
| 10 | 118264236 | rs10736256 | 1.844 | -0.365 | 2.728 | 3.6 | 0.014 | 0.158 | 0.158 | 0.079 | 0.92 | 0.74 | 0.83 | 0.68 | 0.94 | 0.93 |
| 10 | 118266623 | rs11592646 | 1.844 | -0.365 | 2.728 | 3.6 | 0.014 | 0.158 | 0.158 | 0.079 | 0.92 | 0.74 | 0.83 | 0.68 | 0.94 | 0.93 |
| 10 | 118272580 | rs10885978 | 2.077 | -0.357 | 2.618 | 3.681 | 0.006 | 0.192 | 0.192 | 0.034 | 0.92 | 0.75 | 0.76 | 0.62 | 0.91 | 0.91 |
| 10 | 118278528 | rs10787704 | 1.594 | -0.341 | 2.577 | 3.492 | 0.016 | 0.165 | 0.165 | 0 | 0.93 | 0.78 | 0.84 | 0.67 | 0.94 | 0.93 |
| 10 | 118285633 | rs10510022 | -2.336 | -0.344 | 2.598 | 3.466 | 0.017 | 0.176 | 0.176 | 0.1 | 0.08 | 0.24 | 0.17 | 0.33 | 0.06 | 0.07 |
| 10 | 118287900 | rs10885983 | 1.037 | -0.571 | 2.531 | 2.808 | -0.051 | 0.249 | 0.249 | 1.95 | 0.89 | 0.59 | 0.8 | 0.75 | 0.96 | 0.95 |
| 10 | 118297033 | rs12245521 | -1.35 | 0.032 | 1.655 | 2.437 | 0.007 | 0.024 | 0.024 | 0 | 0.09 | 0.03 | 0.04 | 0 | 0.09 | 0.08 |
| 10 | 118298706 | rs12249434 | -1.412 | 0.024 | 1.7 | 2.431 | -0.001 | -0.021 | -0.021 | 0 | 0.09 | 0.09 | 0.04 | 0 | 0.09 | 0.08 |
| 10 | 118313265 | rs2915748 | 1.329 | 0.039 | 1.647 | 2.59 | 0.003 | 0.121 | 0.121 | 0 | 0.93 | 0.86 | 0.84 | 0.67 | 0.92 | 0.92 |
| 10 | 118315835 | rs2915775 | 2.63 | 0.017 | 1.706 | 2.576 | -0.01 | 0.071 | 0.071 | 0 | 0.82 | 0.63 | 0.79 | 0.66 | 0.84 | 0.85 |
| 10 | 118322269 | rs2915772 | 1.39 | 0.022 | 1.614 | 2.58 | 0.003 | 0.133 | 0.133 | 0 | 0.93 | 0.83 | 0.84 | 0.66 | 0.92 | 0.92 |
| 10 | 118328664 | rs3010494 | 2.698 | 0.011 | 1.364 | 2.576 | -0.01 | 0.163 | 0.163 | 0.021 | 0.82 | 0.49 | 0.79 | 0.66 | 0.84 | 0.85 |
| 10 | 118345097 | rs7908775 | -1.474 | 0.184 | 0.049 | 1.604 | 0.006 | 0 | 0 | 0 | 0.1 | 0.08 | 0.04 | 0 | 0.08 | 0.07 |
| 10 | 118347262 | rs11197742 | -1.198 | 0.398 | 0.034 | 1.664 | -0.02 | -0.015 | -0.015 | 1.545 | 0.06 | 0.03 | 0.06 | 0.19 | 0.1 | 0.08 |
| 10 | 118347886 | rs7906829 | -1.496 | -0.03 | 0.179 | 1.008 | -0.007 | -0.034 | -0.034 | 0.405 | 0.1 | 0.14 | 0.04 | 0 | 0.08 | 0.07 |
| 10 | 118351182 | rs7904117 | -1.739 | 0.054 | 0.082 | 0.955 | -0.007 | -0.024 | -0.024 | 1.23 | 0.09 | 0.18 | 0.07 | 0.03 | 0.06 | 0.06 |
| 10 | 118351905 | rs1867991 | 0.161 | 0.007 | 0.118 | 1.118 | -0.002 | 0.014 | 0.014 | 0 | 0.66 | 0.58 | 0.72 | 0.6 | 0.68 | 0.64 |
| 10 | 118355530 | rs2305206 | 0.65 | -0.364 | -0.667 | 0.881 | 0 | -0.021 | -0.021 | 0 | 0.47 | 0.42 | 0.59 | 0.2 | 0.51 | 0.5 |
| 10 | 118357139 | rs7906926 | -0.033 | -0.557 | -0.753 | 0.933 | -0.035 | -0.069 | -0.069 | 0.522 | 0.3 | 0.14 | 0.26 | 0.4 | 0.35 | 0.34 |
| 10 | 118359548 | rs3816355 | -1.496 | -0.368 | -0.77 | 0.291 | 0.001 | -0.013 | -0.013 | 0 | 0.64 | 0.43 | 0.61 | 0.94 | 0.64 | 0.66 |
| 10 | 118359989 | rs17094995 | -0.106 | -0.436 | -0.832 | 0.161 | 0.006 | -0.003 | -0.003 | 0 | 0.12 | 0.15 | 0.06 | 0 | 0.08 | 0.06 |
| 10 | 118360899 | rs3781565 | 0.651 | -0.297 | -1.118 | 0.209 | 0 | 0 | 0 | 0.529 | 0.14 | 0.14 | 0.16 | 0.1 | 0.21 | 0.21 |
| 10 | 118360994 | rs3781564 | 1.314 | -0.326 | -1.145 | 0.366 | 0.002 | -0.048 | -0.048 | 0.037 | 0.09 | 0.01 | 0.06 | 0.41 | 0.06 | 0.11 |
| 10 | 118361370 | rs12414627 | 0.509 | -0.868 | -0.597 | 0.104 | 0 | 0.008 | 0.008 | 0 | 0.24 | 0.22 | 0.35 | 0.46 | 0.28 | 0.29 |
| 10 | 118363642 | rs3010501 | 0.583 | -1.062 | -0.474 | 0.068 | 0 | -0.029 | -0.029 | 0.368 | 0.69 | 0.54 | 0.76 | 0.9 | 0.69 | 0.71 |
| 10 | 118364842 | rs2915753 | 1.108 | -1.049 | -0.466 | 0.057 | 0.002 | -0.007 | -0.007 | 0 | 0.6 | 0.53 | 0.71 | 0.49 | 0.64 | 0.6 |
| 10 | 118367168 | rs7905887 | -2.191 | -1.044 | -0.673 | -0.289 | 0.019 | -0.112 | -0.112 | 0.384 | 0.72 | 0.68 | 0.65 | 0.97 | 0.66 | 0.71 |
| 10 | 118367744 | rs7909787 | 0.24 | -1.03 | -0.668 | -0.245 | -0.013 | -0.047 | -0.047 | 0 | 0.14 | 0.25 | 0.08 | 0 | 0.09 | 0.06 |

| | | | | | | | | | | | | | | | | |
|---|---|---|---|---|---|---|---|---|---|---|---|---|---|---|---|---|
| 10 | 118368606 | rs1049125 | 0.987 | -0.943 | -0.841 | -0.14 | -0.064 | -0.035 | -0.035 | 0 | 0.14 | 0.01 | 0.18 | 0.15 | 0.18 | 0.12 |
| 10 | 118370202 | rs12251289 | 1.322 | -0.968 | -0.84 | -0.102 | -0.01 | 0.021 | 0.021 | 0 | 0.19 | 0.2 | 0.22 | 0.31 | 0.14 | 0.18 |
| 10 | 118373329 | rs7358159 | 0.164 | -0.961 | -0.844 | -0.116 | -0.003 | -0.028 | -0.028 | 0 | 0.32 | 0.2 | 0.4 | 0.46 | 0.32 | 0.28 |
| 10 | 118380345 | rs12415178 | -0.194 | -1.038 | -0.84 | -0.701 | -0.002 | 0.06 | 0.06 | 0.562 | 0.38 | 0.23 | 0.42 | 0.87 | 0.35 | 0.45 |
| 10 | 118381945 | rs12783584 | -0.725 | -0.818 | -0.84 | -0.47 | -0.032 | 0.01 | 0.01 | 0 | 0.35 | 0.25 | 0.34 | 0.28 | 0.42 | 0.37 |
| 10 | 118382857 | rs11197762 | 0.532 | -0.876 | -0.834 | -0.505 | -0.006 | 0 | 0 | 0 | 0.19 | 0.16 | 0.15 | 0.15 | 0.23 | 0.25 |
| 10 | 118383703 | rs2286776 | -0.229 | -0.804 | -0.719 | -0.164 | -0.006 | -0.022 | -0.022 | 5.117 | 0.34 | 0.24 | 0.42 | 0.58 | 0.34 | 0.3 |
| 10 | 118386706 | rs9787661 | 0.57 | -0.794 | -0.736 | -0.197 | -0.014 | 0.009 | 0.009 | 0 | 0.16 | 0.11 | 0.2 | 0.13 | 0.2 | 0.11 |
| 10 | 118392545 | rs10047377 | 1.274 | -0.8 | -0.735 | -0.16 | 0.005 | -0.009 | -0.009 | 0 | 0.19 | 0.12 | 0.22 | 0.45 | 0.14 | 0.18 |
| 10 | 118396138 | rs11197775 | -0.788 | -0.361 | -0.617 | -0.659 | -0.045 | 0.011 | 0.011 | 0 | 0.33 | 0.18 | 0.33 | 0.27 | 0.42 | 0.37 |
| 10 | 118397884 | rs4751995 | -1.822 | -0.191 | -0.476 | -0.786 | 0.002 | 0.01 | 0.01 | 0.007 | 0.5 | 0.63 | 0.46 | 0.27 | 0.54 | 0.44 |
| 10 | 118398658 | rs2420314 | -0.28 | -0.109 | -0.486 | -0.742 | -0.018 | -0.075 | -0.075 | 0.002 | 0.14 | 0.34 | 0.11 | 0 | 0.11 | 0.07 |
| 10 | 118402207 | rs12774180 | 0.624 | -0.06 | -0.513 | -0.734 | -0.043 | 0.002 | 0.002 | 0 | 0.13 | 0.04 | 0.2 | 0.13 | 0.19 | 0.11 |
| 10 | 118404620 | rs2301179 | -1.535 | -0.028 | -0.326 | -0.599 | 0 | 0 | 0 | 0 | 0.49 | 0.62 | 0.45 | 0.27 | 0.52 | 0.44 |
| 10 | 118404700 | rs2301178 | -0.486 | 0.067 | -0.307 | -0.443 | 0.001 | -0.02 | -0.02 | 0 | 0.27 | 0.38 | 0.3 | 0.13 | 0.29 | 0.18 |
| 10 | 118412310 | rs12571521 | -0.284 | 0.019 | 0.071 | -0.148 | 0.027 | 0.073 | 0.073 | 0 | 0.51 | 0.38 | 0.45 | 0.27 | 0.57 | 0.49 |
| 10 | 118414855 | rs10787712 | -0.598 | 0.089 | -0.084 | -0.136 | 0.004 | 0.013 | 0.013 | 0 | 0.55 | 0.46 | 0.45 | 0.26 | 0.58 | 0.49 |
| 10 | 118416020 | rs7087620 | -0.436 | 0.324 | -0.689 | 0.062 | 0.009 | 0.059 | 0.059 | 0 | 0.49 | 0.41 | 0.42 | 0.26 | 0.52 | 0.46 |
| 10 | 118417453 | rs1867987 | -0.29 | 0.505 | -1.38 | 0.168 | -0.009 | 0.115 | 0.115 | 0 | 0.47 | 0.2 | 0.4 | 0.26 | 0.53 | 0.49 |
| 10 | 118418781 | rs2420320 | 0.866 | 0.907 | -0.283 | 0.817 | 0.008 | 0.068 | 0.068 | 0 | 0.81 | 0.49 | 0.69 | 0.33 | 0.81 | 0.82 |
| 10 | 118419231 | rs880977 | -0.294 | 1.109 | -0.198 | 0.656 | -0.001 | -0.001 | -0.001 | 0 | 0.45 | 0.6 | 0.39 | 0.47 | 0.5 | 0.36 |
| 10 | 118421376 | rs1955164 | -0.447 | 1.297 | 0.263 | 0.813 | -0.015 | 0.02 | 0.02 | 0.151 | 0.47 | 0.74 | 0.46 | 0.58 | 0.5 | 0.38 |
| 10 | 118426048 | rs3010455 | 0.018 | 1.202 | 0.649 | 0.307 | -0.025 | 0.138 | 0.138 | 0 | 0.43 | 0.21 | 0.34 | 0.03 | 0.45 | 0.3 |
| 10 | 118426363 | rs9633727 | 0.175 | 1.215 | 1.139 | 1.234 | 0 | 0.016 | 0.016 | 0 | 0.5 | 0.27 | 0.5 | 0.52 | 0.48 | 0.6 |
| 10 | 118429123 | rs877396 | -0.229 | 0.796 | 0.528 | 0.849 | -0.007 | 0.027 | 0.027 | 0 | 0.29 | 0.19 | 0.26 | 0.24 | 0.28 | 0.31 |
| 10 | 118436068 | rs2166634 | -0.577 | 0.71 | 0.372 | 0.882 | 0.007 | 0.003 | 0.003 | 0 | 0.63 | 0.39 | 0.62 | 0.73 | 0.67 | 0.82 |
| 10 | 118444339 | rs12771818 | 0.83 | 0.724 | 0.099 | 0.481 | 0.036 | 0.105 | 0.105 | 0 | 0.3 | 0.14 | 0.27 | 0.07 | 0.25 | 0.14 |
| 10 | 118448078 | rs1900500 | -0.17 | 0.969 | 0.173 | 0.593 | -0.014 | 0.02 | 0.02 | 0 | 0.6 | 0.64 | 0.52 | 0.32 | 0.51 | 0.5 |
| 10 | 118448956 | rs12268494 | 0.132 | 0.956 | 0.013 | 0.492 | -0.037 | 0.04 | 0.04 | 0 | 0.18 | 0.06 | 0.17 | 0.13 | 0.21 | 0.27 |
| 10 | 118452921 | rs12263233 | -0.397 | 0.975 | 0.209 | 0.502 | -0.022 | 0.023 | 0.023 | 0 | 0.24 | 0.06 | 0.23 | 0.21 | 0.22 | 0.37 |
| 10 | 118454499 | rs11197797 | 0.986 | 0.944 | 0.736 | 0.835 | -0.006 | 0.14 | 0.14 | 0 | 0.32 | 0.08 | 0.24 | 0.12 | 0.29 | 0.16 |
| 10 | 118457481 | rs2420330 | 0.759 | 0.961 | 0.833 | 0.9 | -0.019 | -0.082 | -0.082 | 1.074 | 0.35 | 0.48 | 0.29 | 0.13 | 0.29 | 0.16 |
| 10 | 118466042 | rs1014246 | 0.227 | 0.955 | 0.522 | 0.892 | 0.001 | -0.007 | -0.007 | 0.056 | 0.4 | 0.58 | 0.35 | 0.2 | 0.29 | 0.21 |
| 10 | 118470750 | rs1900508 | -0.454 | 0.906 | 0.603 | 1.022 | 0.008 | -0.003 | -0.003 | 0 | 0.57 | 0.38 | 0.55 | 0.54 | 0.62 | 0.72 |
| 10 | 118472715 | rs4752003 | -0.182 | 1.313 | 0.737 | 1.2 | 0.051 | 0.178 | 0.178 | 0 | 0.15 | 0.32 | 0.27 | 0.43 | 0.22 | 0.16 |
| 10 | 118474186 | rs1628985 | 0.506 | 1.092 | 0.663 | 0.746 | 0.003 | 0.027 | 0.027 | 0.765 | 0.93 | 0.72 | 0.91 | 0.92 | 0.98 | 0.98 |
| 10 | 118485201 | rs758367 | -0.86 | 1.433 | 0.087 | 0.886 | -0.014 | 0.01 | 0.01 | 0 | 0.56 | 0.73 | 0.57 | 0.61 | 0.51 | 0.73 |
| 10 | 118492612 | rs7093835 | -0.217 | 1.527 | 0.818 | 1.038 | 0.055 | 0.085 | 0.085 | 0 | 0.14 | 0.27 | 0.16 | 0.28 | 0.24 | 0.17 |
| 10 | 118493063 | rs1638419 | 0.721 | 1.47 | 0.313 | 0.998 | 0.009 | 0.026 | 0.026 | 0.505 | 0.78 | 0.32 | 0.74 | 0.66 | 0.74 | 0.77 |
| 10 | 118506755 | rs740599 | -0.669 | 0.808 | 0.116 | 0.766 | 0.013 | -0.002 | -0.002 | 0 | 0.53 | 0.56 | 0.6 | 0.56 | 0.6 | 0.75 |
| 10 | 118507976 | rs887195 | -0.304 | 0.91 | 0.037 | 0.822 | 0.033 | 0.001 | 0.001 | 0.275 | 0.46 | 0.29 | 0.5 | 0.48 | 0.58 | 0.73 |
| 10 | 118519724 | rs1638412 | 0.635 | 1.236 | 0.398 | 0.746 | 0.005 | -0.042 | -0.042 | 0 | 0.84 | 0.76 | 0.88 | 0.92 | 0.84 | 0.88 |
| 10 | 118520282 | rs1665676 | -0.466 | 1.075 | 0.207 | 1.431 | 0.004 | 0.14 | 0.14 | 0 | 0.07 | 0.12 | 0.11 | 0.28 | 0.07 | 0.07 |
| 10 | 118522665 | rs1665672 | -0.175 | 1.356 | 0.217 | 0.862 | 0.004 | -0.034 | -0.034 | 0.258 | 0.7 | 0.63 | 0.7 | 0.77 | 0.7 | 0.61 |
| 10 | 118523964 | rs1990275 | -0.829 | 1.388 | 0.089 | 0.884 | 0.013 | -0.006 | -0.006 | 0 | 0.47 | 0.68 | 0.51 | 0.47 | 0.54 | 0.51 |
| 10 | 118528593 | rs1630816 | 0.088 | 1.589 | 0.099 | 1.059 | -0.046 | -0.048 | -0.048 | 0.21 | 0.15 | 0.05 | 0.21 | 0.22 | 0.26 | 0.14 |
| 10 | 118535961 | rs1638407 | -0.328 | 1.426 | 0.678 | 1.349 | 0.087 | 0.086 | 0.086 | 0 | 0.17 | 0.39 | 0.27 | 0.23 | 0.25 | 0.15 |
| 10 | 118536492 | rs1900501 | 0.78 | 0.686 | 0.531 | 1.617 | 0.031 | 0.083 | 0.083 | 0 | 0.47 | 0.2 | 0.42 | 0.35 | 0.44 | 0.4 |
| 10 | 118537768 | rs1622217 | -0.585 | 0.834 | 0.437 | 1.405 | -0.001 | -0.001 | -0.001 | 0 | 0.46 | 0.47 | 0.53 | 0.39 | 0.53 | 0.54 |
| 10 | 118543807 | rs2420333 | 0.272 | 1.013 | 0.727 | 1.532 | 0.104 | 0.041 | 0.041 | 0 | 0.83 | 0.56 | 0.73 | 0.82 | 0.76 | 0.84 |
| 10 | 118544640 | rs1860261 | -0.07 | 1.028 | 0.776 | 1.563 | 0.111 | 0.033 | 0.033 | 0 | 0.84 | 0.58 | 0.77 | 0.86 | 0.77 | 0.85 |
| 10 | 118546743 | rs3125617 | 0.62 | 0.355 | 0.417 | 1.386 | -0.038 | 0.138 | 0.138 | 0 | 0.76 | 0.47 | 0.82 | 0.57 | 0.82 | 0.74 |
| 10 | 118549007 | rs3125619 | 0.447 | 0.196 | -0.284 | 0.999 | -0.002 | 0.102 | 0.102 | 0 | 0.77 | 0.45 | 0.76 | 0.66 | 0.79 | 0.76 |
| 10 | 118550763 | rs3095122 | 1.226 | -0.159 | -0.635 | 0.509 | -0.086 | 0.122 | 0.122 | 0 | 0.32 | 0.05 | 0.37 | 0.12 | 0.4 | 0.35 |
| 10 | 118553139 | rs1637557 | -0.182 | -0.192 | -0.477 | 0.625 | -0.009 | 0.273 | 0.273 | 0.242 | 0.89 | 0.46 | 0.91 | 0.73 | 0.95 | 0.85 |
| 10 | 118553863 | rs17095255 | 1.115 | -0.226 | -0.627 | 0.646 | 0.018 | -0.018 | -0.018 | 0 | 0.14 | 0.03 | 0.08 | 0.16 | 0.11 | 0.15 |
| 10 | 118554760 | rs1681731 | 0.037 | 0.03 | -0.139 | 0.591 | -0.02 | 0.215 | 0.215 | 0.255 | 0.88 | 0.44 | 0.9 | 0.73 | 0.95 | 0.85 |
| 10 | 118560477 | rs7097645 | -0.366 | 0.102 | -0.296 | 0.368 | -0.001 | -0.001 | -0.001 | 0.014 | 0.6 | 0.45 | 0.56 | 0.61 | 0.57 | 0.53 |
| 10 | 118563329 | rs1681739 | 0.855 | 0.308 | -0.785 | -0.07 | -0.09 | 0.117 | 0.117 | 0 | 0.31 | 0.07 | 0.36 | 0.12 | 0.41 | 0.32 |
| 10 | 118565501 | rs1681743 | 0.744 | 0.314 | -0.543 | -0.099 | -0.017 | 0.019 | 0.019 | 0 | 0.3 | 0.18 | 0.3 | 0.12 | 0.39 | 0.28 |
| 10 | 118568352 | rs1637556 | 0.908 | 0.116 | -0.731 | -0.36 | -0.083 | 0.138 | 0.138 | 0.129 | 0.35 | 0.08 | 0.38 | 0.12 | 0.42 | 0.25 |
| 10 | 118575606 | rs740363 | 0.398 | -0.27 | -0.598 | -0.542 | -0.009 | 0.045 | 0.045 | 0.01 | 0.43 | 0.28 | 0.44 | 0.12 | 0.46 | 0.29 |
| 10 | 118576146 | rs1626678 | 0.651 | -0.226 | -0.596 | -0.547 | -0.012 | 0.039 | 0.039 | 0 | 0.41 | 0.27 | 0.43 | 0.12 | 0.46 | 0.3 |
| 10 | 118589179 | rs1614092 | -0.211 | -0.371 | -0.343 | -0.586 | -0.001 | -0.027 | -0.027 | 0 | 0.51 | 0.5 | 0.53 | 0.87 | 0.52 | 0.64 |
| 10 | 118589566 | rs17095303 | 1.228 | -0.425 | -0.737 | -0.139 | -0.003 | -0.072 | -0.072 | 1.92 | 0.16 | 0.01 | 0.08 | 0.27 | 0.13 | 0.24 |
| 10 | 118594392 | rs1681720 | 0.788 | -0.59 | -0.176 | -0.015 | -0.031 | -0.043 | -0.043 | 0 | 0.31 | 0.17 | 0.41 | 0.46 | 0.39 | 0.4 |
| 10 | 118596756 | rs1637565 | -1.459 | 0.009 | -0.323 | -0.505 | -0.003 | -0.122 | -0.122 | 0 | 0.78 | 0.55 | 0.73 | 0.96 | 0.84 | 0.78 |
| 10 | 118603228 | rs10886014 | 1.159 | 0.109 | -0.424 | 0.14 | -0.047 | 0.147 | 0.147 | 0.081 | 0.4 | 0.17 | 0.41 | 0.13 | 0.46 | 0.29 |
| 10 | 118608102 | rs9421246 | -0.446 | 0.525 | 0.155 | 0.721 | 0.005 | 0.017 | 0.017 | 0 | 0.31 | 0.34 | 0.39 | 0.44 | 0.3 | 0.35 |

| | | | | | | | | | | | | | | | | |
|---|---|---|---|---|---|---|---|---|---|---|---|---|---|---|---|---|
| 10 | 118611403 | rs1681752 | 0.61 | 0.443 | 0.173 | 0.676 | -0.001 | -0.011 | -0.011 | 0.424 | 0.65 | 0.36 | 0.57 | 0.56 | 0.69 | 0.63 |
| 10 | 118620459 | rs11197835 | 1.657 | 0.372 | -0.231 | 0.444 | -0.101 | 0.203 | 0.203 | 0.903 | 0.4 | 0.06 | 0.34 | 0.1 | 0.5 | 0.29 |
| 10 | 118620996 | rs2170862 | -0.771 | 0.267 | 0.261 | 0.735 | 0.01 | 0.041 | 0.041 | 1.285 | 0.31 | 0.43 | 0.36 | 0.45 | 0.27 | 0.33 |
| 10 | 118637809 | rs2531687 | 2.186 | 0.403 | -0.086 | 0.379 | -0.034 | 0.147 | 0.147 | 0 | 0.32 | 0.06 | 0.25 | 0.07 | 0.35 | 0.25 |
| 10 | 118641396 | rs11197843 | -0.42 | 0.874 | 1.323 | 1.825 | -0.036 | -0.029 | -0.029 | 0.086 | 0.15 | 0.05 | 0.21 | 0.2 | 0.18 | 0.11 |
| 10 | 118643670 | rs2257791 | 1.45 | 0.852 | 1.337 | 1.992 | -0.002 | 0.015 | 0.015 | 2.024 | 0.71 | 0.6 | 0.72 | 0.7 | 0.76 | 0.75 |
| 10 | 118648043 | rs879255 | -0.716 | 0.956 | 1.755 | 2.228 | 0.028 | 0.143 | 0.143 | 0.085 | 0.16 | 0.38 | 0.32 | 0.46 | 0.19 | 0.19 |
| 10 | 118650996 | rs10510025 | -1.263 | 1.023 | 1.888 | 1.763 | 0 | 0.033 | 0.033 | 0.075 | 0.27 | 0.36 | 0.35 | 0.55 | 0.26 | 0.31 |
| 10 | 118651882 | rs740600 | 1.372 | 1.023 | 1.667 | 1.765 | 0.012 | -0.046 | -0.046 | 0 | 0.68 | 0.21 | 0.61 | 0.45 | 0.74 | 0.65 |
| 10 | 118672531 | rs10886017 | -1.744 | 1.03 | 1.582 | 1.794 | -0.009 | 0.062 | 0.062 | 0.075 | 0.3 | 0.5 | 0.29 | 0.42 | 0.26 | 0.34 |
| 10 | 118689195 | rs719012 | 1.499 | 1.007 | 1.381 | 2 | 0.015 | -0.074 | -0.074 | 0.724 | 0.67 | 0.18 | 0.6 | 0.45 | 0.74 | 0.65 |
| 10 | 118715399 | rs7902527 | -0.97 | 0.438 | 1.817 | 2.211 | -0.001 | -0.005 | -0.005 | 0 | 0.25 | 0.22 | 0.25 | 0.29 | 0.24 | 0.24 |
| 10 | 118750248 | rs10787735 | 1.352 | 0.424 | 1.78 | 1.908 | 0.015 | -0.048 | -0.048 | 0 | 0.68 | 0.19 | 0.6 | 0.44 | 0.73 | 0.65 |
| 10 | 118757139 | rs10218905 | -1.442 | 0.463 | 1.597 | 1.931 | -0.003 | 0.029 | 0.029 | 0.045 | 0.28 | 0.36 | 0.35 | 0.54 | 0.27 | 0.32 |
| 10 | 118776306 | rs12769006 | 1.305 | 0.5 | 0.74 | 1.943 | 0.012 | -0.084 | -0.084 | 0 | 0.67 | 0.12 | 0.59 | 0.44 | 0.73 | 0.64 |
| 10 | 118801980 | rs12766760 | -1.268 | 1.144 | 1.058 | 2.105 | -0.06 | 0.013 | 0.013 | 0 | 0.19 | 0.01 | 0.14 | 0.14 | 0.23 | 0.21 |
| 10 | 118826441 | rs10787751 | 1.608 | 0.861 | 2.12 | 2.81 | 0.024 | 0.217 | 0.217 | 2.583 | 0.84 | 0.39 | 0.75 | 0.56 | 0.82 | 0.85 |
| 10 | 118827560 | rs7078160 | -1.01 | 0.879 | 2.157 | 2.822 | 0.025 | 0.138 | 0.138 | 0.263 | 0.12 | 0.25 | 0.21 | 0.41 | 0.18 | 0.12 |
| 10 | 118834991 | rs4752028 | 1.665 | 0.873 | 1.903 | 2.887 | 0.043 | 0.199 | 0.199 | 1.932 | 0.85 | 0.45 | 0.75 | 0.68 | 0.83 | 0.86 |
| 10 | 118890573 | rs7086344 | 1.285 | 0.688 | 1.106 | 2.291 | -0.013 | 0.157 | 0.157 | 0.088 | 0.83 | 0.57 | 0.8 | 0.57 | 0.89 | 0.7 |
| 10 | 118895368 | rs1871345 | -1.767 | 0.943 | 1.013 | 2.311 | 0.042 | 0.27 | 0.27 | 0.628 | 0.17 | 0.42 | 0.33 | 0.42 | 0.19 | 0.22 |
| 10 | 118904495 | rs716183 | 0.646 | 0.53 | 0.306 | 2.022 | 0.017 | -0.043 | -0.043 | 1.616 | 0.54 | 0.2 | 0.53 | 0.58 | 0.57 | 0.4 |
| 10 | 118906143 | rs11593856 | 0.643 | 0.542 | 0.311 | 2.17 | 0.016 | -0.043 | -0.043 | 0.222 | 0.54 | 0.2 | 0.53 | 0.58 | 0.57 | 0.4 |
| 10 | 118909263 | rs2420309 | 0.682 | 0.379 | 0.149 | 2.196 | -0.063 | -0.07 | -0.07 | 4.397 | 0.21 | 0.01 | 0.25 | 0.2 | 0.23 | 0.19 |
| 10 | 118916215 | rs10787762 | 1.404 | 0.27 | 0.883 | 1.989 | -0.021 | -0.047 | -0.047 | 0 | 0.92 | 0.31 | 0.9 | 1 | 0.97 | 0.98 |
| 10 | 118916227 | rs17095748 | 0.214 | 0.543 | 0.808 | 2.152 | 0.011 | -0.025 | -0.025 | 0 | 0.56 | 0.28 | 0.57 | 0.59 | 0.58 | 0.42 |
| 10 | 118920455 | rs17095763 | -1.277 | 0.66 | 1.345 | 2.626 | 0.011 | 0.139 | 0.139 | 0 | 0.13 | 0.25 | 0.26 | 0.45 | 0.12 | 0.15 |
| 10 | 118934887 | rs1638440 | 1.307 | 0.558 | 1.568 | 1.876 | -0.004 | -0.031 | -0.031 | 0 | 0.87 | 0.67 | 0.89 | 0.94 | 0.89 | 0.8 |
| 10 | 118944413 | rs1678046 | 1.116 | 0.577 | 1.466 | 1.878 | -0.003 | -0.027 | -0.027 | 0 | 0.88 | 0.76 | 0.9 | 0.95 | 0.89 | 0.81 |
| 10 | 118949255 | rs11197921 | -1.791 | 0.464 | 1.121 | 2.331 | -0.02 | -0.02 | -0.02 | 0 | 0.23 | 0.15 | 0.29 | 0.74 | 0.25 | 0.5 |
| 10 | 118952876 | rs10886049 | -2.388 | 0 | 1.883 | 2.783 | -0.013 | 0.069 | 0.069 | 0 | 0.13 | 0.29 | 0.21 | 0.33 | 0.09 | 0.2 |
| 10 | 118961742 | rs363308 | -2.585 | -0.046 | 2.268 | 2.821 | -0.022 | 0.107 | 0.107 | 0 | 0.15 | 0.38 | 0.32 | 0.33 | 0.09 | 0.2 |
| 10 | 118961849 | rs11197926 | -1.242 | -0.046 | 2.241 | 2.845 | -0.013 | -0.04 | -0.04 | 0 | 0.07 | 0.01 | 0.14 | 0.22 | 0.05 | 0.14 |
| 10 | 118962081 | rs363311 | -2.252 | -0.041 | 2.281 | 2.841 | -0.015 | 0.065 | 0.065 | 0 | 0.12 | 0.22 | 0.17 | 0.24 | 0.06 | 0.14 |
| 10 | 118968222 | rs2803827 | 1.694 | -0.249 | 2.275 | 2.992 | 0 | 0.091 | 0.091 | 0 | 0.78 | 0.59 | 0.81 | 0.52 | 0.8 | 0.71 |
| 10 | 118977123 | rs2619105 | 2.449 | -0.558 | 2.193 | 2.735 | 0.015 | 0.026 | 0.026 | 0 | 0.62 | 0.28 | 0.62 | 0.2 | 0.76 | 0.53 |
| 10 | 118980138 | rs2803825 | 1.023 | -0.395 | 2.174 | 3.106 | 0.004 | -0.018 | -0.018 | 3.687 | 0.92 | 0.95 | 0.86 | 0.78 | 0.95 | 0.87 |
| 10 | 118981071 | rs2803823 | 1.939 | -0.107 | 3.005 | 3.437 | 0.027 | 0.191 | 0.191 | 0.451 | 0.9 | 0.66 | 0.93 | 0.74 | 0.92 | 0.88 |
| 10 | 118982402 | rs363366 | 2.254 | -0.147 | 3.032 | 3.263 | -0.034 | 0.073 | 0.073 | 2.008 | 0.85 | 0.45 | 0.76 | 0.79 | 0.95 | 0.93 |
| 10 | 118986396 | rs363371 | -1.444 | -0.109 | 2.935 | 3.376 | -0.01 | -0.027 | -0.027 | 0 | 0.17 | 0.09 | 0.15 | 0.57 | 0.17 | 0.36 |
| 10 | 118987670 | rs2532798 | 2.598 | -0.203 | 2.947 | 3.303 | 0.015 | 0.047 | 0.047 | 0.084 | 0.65 | 0.27 | 0.6 | 0.22 | 0.77 | 0.57 |
| 10 | 118989162 | rs363324 | 2.589 | -0.199 | 2.949 | 3.305 | 0.014 | 0.047 | 0.047 | 0.088 | 0.65 | 0.27 | 0.6 | 0.22 | 0.77 | 0.57 |
| 10 | 118990516 | rs363326 | -2.121 | -0.161 | 2.934 | 3.288 | -0.032 | 0.079 | 0.079 | 0.99 | 0.15 | 0.5 | 0.23 | 0.21 | 0.06 | 0.07 |
| 10 | 118994479 | rs10886051 | -1.278 | 0.002 | 2.336 | 3.319 | -0.047 | -0.093 | -0.093 | 1.154 | 0.17 | 0.03 | 0.15 | 0.58 | 0.18 | 0.34 |
| 10 | 119004095 | rs363333 | 2.293 | -0.169 | 2.382 | 2.672 | -0.035 | 0.024 | 0.024 | 1.61 | 0.82 | 0.32 | 0.75 | 0.73 | 0.94 | 0.9 |
| 10 | 119007980 | rs363336 | -0.749 | -0.047 | 2.022 | 2.546 | 0.015 | -0.007 | -0.007 | 0 | 0.09 | 0.06 | 0.18 | 0.12 | 0.04 | 0.03 |
| 10 | 119009389 | rs363338 | 2.147 | -0.059 | 2.026 | 2.545 | 0.008 | 0.027 | 0.027 | 0.072 | 0.64 | 0.3 | 0.59 | 0.2 | 0.75 | 0.54 |
| 10 | 119009966 | rs2532805 | 0.858 | -0.01 | 1.615 | 2.119 | -0.028 | -0.002 | -0.002 | 0 | 0.89 | 0.72 | 0.81 | 0.88 | 0.96 | 0.97 |
| 10 | 119010465 | rs363341 | -1.912 | 0.519 | 1.369 | 2.393 | 0.022 | 0.033 | 0.033 | 0.361 | 0.24 | 0.41 | 0.21 | 0.68 | 0.2 | 0.42 |
| 10 | 119014948 | rs363343 | 1.678 | 0.275 | 1.103 | 2.609 | -0.011 | 0.163 | 0.163 | 0.258 | 0.81 | 0.63 | 0.74 | 0.42 | 0.82 | 0.61 |
| 10 | 119018679 | rs2283138 | 1.595 | 0.202 | 1.114 | 2.73 | -0.016 | 0.105 | 0.105 | 0.261 | 0.87 | 0.7 | 0.9 | 0.71 | 0.9 | 0.74 |
| 10 | 119019467 | rs363251 | -0.233 | -0.134 | 0.178 | 1.846 | 0.05 | 0.053 | 0.053 | 0 | 0.5 | 0.37 | 0.43 | 0.66 | 0.64 | 0.58 |
| 10 | 119019658 | rs11197936 | -0.103 | -0.087 | -0.282 | 0.227 | 0.017 | 0.067 | 0.067 | 0 | 0.46 | 0.32 | 0.42 | 0.23 | 0.39 | 0.35 |
| 10 | 119021407 | rs10082463 | 0.307 | -0.532 | 0.162 | 0.057 | 0.007 | -0.01 | -0.01 | 0 | 0.17 | 0.12 | 0.2 | 0.39 | 0.09 | 0.21 |
| 10 | 119021737 | rs2015586 | 0.216 | -0.171 | 0.525 | -0.374 | -0.001 | 0 | 0 | 0 | 0.46 | 0.54 | 0.59 | 0.5 | 0.57 | 0.53 |
| 10 | 119032926 | rs2255321 | -0.493 | -0.097 | 0.616 | -0.688 | -0.001 | 0.005 | 0.005 | 0.035 | 0.82 | 0.81 | 0.8 | 0.6 | 0.88 | 0.74 |
| 10 | 119042408 | rs12337 | 1.107 | -0.207 | 0.535 | -1.004 | -0.068 | 0.117 | 0.117 | 0 | 0.25 | 0.09 | 0.16 | 0.01 | 0.31 | 0.17 |
| 10 | 119043554 | rs363294 | 0.661 | -0.277 | 0.488 | -1.043 | -0.004 | 0.032 | 0.032 | 0 | 0.08 | 0 | 0.04 | 0.01 | 0.05 | 0.07 |
| 10 | 119051788 | rs2532812 | 0.273 | -0.277 | 0.74 | -1.042 | -0.009 | -0.028 | -0.028 | 1.782 | 0.09 | 0.11 | 0.05 | 0.01 | 0.05 | 0.07 |
| 10 | 119061632 | rs17627160 | 1.156 | -0.248 | 0.681 | -1.038 | -0.065 | 0.116 | 0.116 | 0 | 0.24 | 0.09 | 0.15 | 0.01 | 0.3 | 0.16 |
| 10 | 119069842 | rs2532841 | 0.243 | -0.261 | 1.165 | -1.089 | 0.002 | 0.027 | 0.027 | 0.377 | 0.5 | 0.3 | 0.58 | 0.55 | 0.55 | 0.54 |
| 10 | 119075731 | rs2532784 | -0.04 | -0.229 | 1.302 | -1.061 | -0.023 | 0.046 | 0.046 | 0 | 0.77 | 0.63 | 0.76 | 0.56 | 0.85 | 0.71 |
| 10 | 119075741 | rs2532838 | -0.007 | -0.229 | 1.307 | -1.061 | -0.024 | 0.047 | 0.047 | 0 | 0.77 | 0.62 | 0.76 | 0.56 | 0.85 | 0.71 |
| 10 | 119098162 | rs181438 | -0.301 | -0.087 | 1.187 | -1.149 | 0.007 | 0.055 | 0.055 | 0.254 | 0.21 | 0.33 | 0.41 | 0.48 | 0.33 | 0.25 |
| 10 | 119102636 | rs4752052 | 1.18 | -0.008 | 0.726 | -1.295 | 0.025 | -0.039 | -0.039 | 0 | 0.16 | 0.06 | 0.18 | 0.34 | 0.08 | 0.2 |
| 10 | 119103471 | rs181440 | 0.518 | 0.033 | 0.648 | -1.305 | -0.012 | -0.005 | -0.005 | 1.218 | 0.42 | 0.29 | 0.55 | 0.53 | 0.5 | 0.47 |
| 10 | 119107982 | rs11815112 | 0.988 | 0.382 | 0.228 | -1.2 | -0.041 | 0.074 | 0.074 | 0 | 0.25 | 0.14 | 0.16 | 0.01 | 0.3 | 0.16 |
| 10 | 119111723 | rs17545387 | 1.021 | 0.424 | -0.209 | -1.186 | -0.065 | 0.115 | 0.115 | 0 | 0.24 | 0.09 | 0.15 | 0.01 | 0.3 | 0.16 |
| 10 | 119140895 | rs11197981 | -0.634 | 0.094 | 0.382 | -0.388 | -0.034 | 0.073 | 0.073 | 0 | 0.24 | 0.42 | 0.24 | 0.44 | 0.15 | 0.31 |

| | | | | | | | | | | | | | | | | |
|---|---|---|---|---|---|---|---|---|---|---|---|---|---|---|---|---|
| 10 | 119142449 | rs10787771 | -0.014 | 0.305 | -0.472 | -0.42 | 0.011 | -0.018 | -0.018 | 1.417 | 0.19 | 0.15 | 0.2 | 0.4 | 0.12 | 0.28 |
| 10 | 119157485 | rs1772810 | -0.428 | 1.229 | -0.837 | -0.986 | -0.001 | -0.01 | -0.01 | 2.015 | 0.28 | 0.25 | 0.19 | 0.07 | 0.25 | 0.29 |
| 10 | 119170681 | rs1341626 | -0.008 | 1.517 | -0.829 | -0.541 | -0.004 | 0.009 | 0.009 | 0 | 0.56 | 0.57 | 0.51 | 0.12 | 0.48 | 0.39 |
| 10 | 119177760 | rs1638662 | -1.13 | 1.597 | -0.405 | 0.983 | -0.001 | 0.076 | 0.076 | 0 | 0.17 | 0.27 | 0.19 | 0.4 | 0.19 | 0.28 |
| 10 | 119177940 | rs6585435 | -0.145 | 1.008 | -0.426 | 1.049 | -0.027 | 0.014 | 0.014 | 0 | 0.33 | 0.42 | 0.41 | 0.55 | 0.23 | 0.48 |
| 10 | 119178574 | rs17546598 | -0.619 | 1.437 | -0.398 | 0.937 | -0.001 | -0.001 | -0.001 | 0 | 0.06 | 0.08 | 0.06 | 0.09 | 0.1 | 0.06 |
| 10 | 119179082 | rs181502 | -0.864 | 1.442 | -0.739 | 0.759 | -0.011 | 0.029 | 0.029 | 0 | 0.43 | 0.32 | 0.26 | 0.3 | 0.45 | 0.56 |
| 10 | 119180751 | rs730080 | 0.576 | 1.332 | -0.201 | 1.374 | 0.015 | 0.085 | 0.085 | 0 | 0.08 | 0.14 | 0.15 | 0.29 | 0.1 | 0.16 |
| 10 | 119183424 | rs181499 | -0.718 | 1.13 | -0.078 | 1.167 | 0.013 | 0.017 | 0.017 | 0 | 0.43 | 0.32 | 0.44 | 0.32 | 0.36 | 0.54 |
| 10 | 119185451 | rs10886073 | 0.027 | 1.25 | -0.743 | 0.795 | -0.055 | 0.033 | 0.033 | 0.792 | 0.27 | 0.05 | 0.2 | 0.16 | 0.31 | 0.44 |
| 10 | 119193153 | rs758569 | -1.352 | 1.172 | -0.816 | 0.799 | 0.011 | 0.018 | 0.018 | 0 | 0.58 | 0.36 | 0.7 | 0.74 | 0.64 | 0.8 |
| 10 | 119195756 | rs2768335 | -1.772 | 1.117 | -0.587 | 0.757 | -0.006 | -0.009 | -0.009 | 0 | 0.65 | 0.48 | 0.73 | 0.75 | 0.69 | 0.82 |
| 10 | 119196241 | rs2794423 | -0.146 | 1.17 | -0.959 | 0.709 | -0.003 | -0.008 | -0.008 | 0 | 0.12 | 0.14 | 0.05 | 0.14 | 0.08 | 0.08 |
| 10 | 119197179 | rs12783276 | -0.247 | 1.355 | -1.09 | 0.784 | -0.081 | -0.086 | -0.086 | 0 | 0.41 | 0.15 | 0.5 | 0.5 | 0.49 | 0.67 |
| 10 | 119197977 | rs887336 | 0.624 | 0.978 | -0.583 | 1.037 | -0.002 | 0.033 | 0.033 | 0 | 0.08 | 0.07 | 0.26 | 0.34 | 0.11 | 0.22 |
| 10 | 119199967 | rs2794427 | -0.498 | 0.726 | -0.557 | 0.886 | -0.003 | -0.006 | -0.006 | 0.354 | 0.12 | 0.14 | 0.05 | 0.14 | 0.08 | 0.08 |
| 10 | 119200948 | rs2768312 | -0.862 | 0.527 | -0.809 | 0.694 | 0.002 | 0.006 | 0.006 | 0.043 | 0.39 | 0.26 | 0.35 | 0.46 | 0.31 | 0.46 |
| 10 | 119202762 | rs385872 | -0.319 | 0.47 | -0.847 | 0.685 | -0.008 | -0.002 | -0.002 | 0 | 0.31 | 0.2 | 0.33 | 0.36 | 0.28 | 0.42 |
| 10 | 119203902 | rs366755 | 0.932 | 0.469 | -0.888 | 0.648 | -0.002 | 0.017 | 0.017 | 0.998 | 0.2 | 0.11 | 0.13 | 0.03 | 0.15 | 0.2 |
| 10 | 119204344 | rs452035 | -1.655 | 0.428 | -0.777 | 0.758 | -0.001 | -0.001 | -0.001 | 0 | 0.51 | 0.5 | 0.57 | 0.6 | 0.44 | 0.68 |
| 10 | 119208169 | rs7095357 | 0.081 | 0.64 | -0.801 | 1.053 | 0.008 | 0.004 | 0.004 | 0 | 0.08 | 0.05 | 0.01 | 0.1 | 0.03 | 0.03 |
| 10 | 119210375 | rs6585436 | 0.081 | 0.64 | -0.802 | 1.053 | -0.002 | -0.001 | -0.001 | 0 | 0.08 | 0.08 | 0.01 | 0.1 | 0.03 | 0.03 |
| 10 | 119211641 | rs17096263 | -0.388 | 0.982 | -0.881 | 1.15 | -0.004 | -0.006 | -0.006 | 0 | 0.11 | 0.08 | 0.2 | 0.32 | 0.13 | 0.22 |
| 10 | 119216600 | rs12258862 | 0.006 | 0.554 | -0.88 | 0.658 | 0 | -0.004 | -0.004 | 0.606 | 0.05 | 0.04 | 0.04 | 0.01 | 0.04 | 0.03 |
| 10 | 119227342 | rs4752066 | -1.109 | 0.481 | -1.138 | 0.548 | 0.004 | 0.011 | 0.011 | 0 | 0.1 | 0.07 | 0.05 | 0.04 | 0.07 | 0.08 |
| 10 | 119233375 | rs242967 | -1.49 | 0.202 | -1.6 | 0.408 | 0.009 | -0.025 | -0.025 | 0 | 0.58 | 0.83 | 0.55 | 0.62 | 0.6 | 0.83 |
| 10 | 119234786 | rs242966 | -1.561 | 0.372 | -1.581 | 0.455 | 0.006 | 0.008 | 0.008 | 0 | 0.14 | 0.12 | 0.05 | 0.14 | 0.11 | 0.17 |
| 10 | 119237397 | rs417348 | -0.505 | 0.367 | -1.481 | 0.502 | -0.001 | -0.001 | -0.001 | 0 | 0.09 | 0.11 | 0.04 | 0.1 | 0.08 | 0.11 |
| 10 | 119237988 | rs242971 | -0.567 | 0.507 | -1.34 | 0.539 | 0.005 | -0.007 | -0.007 | 0.562 | 0.4 | 0.46 | 0.39 | 0.4 | 0.39 | 0.64 |
| 10 | 119243566 | rs2532664 | -0.806 | 0.46 | -2.197 | 0.577 | -0.06 | -0.144 | -0.144 | 0 | 0.64 | 0.95 | 0.6 | 0.58 | 0.65 | 0.87 |
| 10 | 119247667 | rs2768327 | -0.594 | 0.247 | -1.772 | 0.583 | 0.006 | -0.017 | -0.017 | 0 | 0.34 | 0.72 | 0.4 | 0.5 | 0.28 | 0.36 |
| 10 | 119250211 | rs242965 | -0.647 | 0.194 | -1.749 | 0.167 | 0.011 | 0.008 | 0.008 | 0.965 | 0.34 | 0.74 | 0.3 | 0.32 | 0.26 | 0.33 |
| 10 | 119251232 | rs10886079 | 0.688 | -0.116 | -2.24 | -0.104 | -0.002 | 0.001 | 0.001 | 0 | 0.15 | 0.16 | 0.11 | 0.17 | 0.15 | 0.3 |
| 10 | 119251745 | rs183607 | -1.587 | -0.337 | -2.694 | -0.383 | -0.115 | -0.098 | -0.098 | 0 | 0.59 | 0.94 | 0.47 | 0.51 | 0.51 | 0.69 |
| 10 | 119251763 | rs11597619 | 1.103 | -0.339 | -2.313 | -0.404 | 0 | -0.018 | -0.018 | 0 | 0.1 | 0.1 | 0.06 | 0 | 0.09 | 0.04 |
| 10 | 119254301 | rs451123 | -1.462 | -0.201 | -2.426 | -0.295 | -0.116 | -0.095 | -0.095 | 0 | 0.59 | 0.94 | 0.47 | 0.51 | 0.51 | 0.69 |
| 10 | 119257454 | rs2420459 | -0.079 | 0.343 | -1.674 | -0.613 | -0.021 | 0.097 | 0.097 | 1.209 | 0.56 | 0.74 | 0.69 | 0.81 | 0.5 | 0.39 |
| 10 | 119261549 | rs7900490 | 0.321 | 0.226 | -1.57 | -0.57 | -0.012 | -0.018 | -0.018 | 1.206 | 0.18 | 0.25 | 0.22 | 0.15 | 0.17 | 0.37 |
| 10 | 119261945 | rs758573 | -1.114 | 0.273 | -1.911 | -0.777 | -0.086 | 0.165 | 0.165 | 0.577 | 0.75 | 0.97 | 0.82 | 0.97 | 0.68 | 0.73 |
| 10 | 119262365 | rs2901157 | 0.813 | 0.3 | -2.118 | -0.57 | -0.034 | -0.066 | -0.066 | 0 | 0.1 | 0.01 | 0.06 | 0.23 | 0.13 | 0.1 |
| 10 | 119262900 | rs10787774 | -0.586 | 0.001 | -2.004 | -0.981 | -0.036 | 0.19 | 0.19 | 0.115 | 0.66 | 0.93 | 0.8 | 0.95 | 0.66 | 0.71 |
| 10 | 119264020 | rs7909884 | 1.44 | -0.128 | -1.296 | -0.993 | -0.002 | 0.005 | 0.005 | 0 | 0.3 | 0.35 | 0.45 | 0.57 | 0.29 | 0.14 |
| 10 | 119264592 | rs242975 | 0.6 | -0.152 | -0.484 | -1.091 | -0.008 | 0.001 | 0.001 | 0 | 0.77 | 0.67 | 0.86 | 0.79 | 0.83 | 0.88 |
| 10 | 119268208 | rs181586 | 0.451 | -0.131 | -0.331 | -1.092 | -0.013 | 0.07 | 0.07 | 0.476 | 0.44 | 0.27 | 0.4 | 0.18 | 0.51 | 0.63 |
| 10 | 119271013 | rs11198020 | 1.461 | -0.064 | -0.209 | -0.984 | -0.003 | 0.013 | 0.013 | 0 | 0.3 | 0.38 | 0.45 | 0.57 | 0.29 | 0.14 |
| 10 | 119272851 | rs242985 | -0.93 | 0.038 | -0.269 | -0.946 | 0.005 | 0.023 | 0.023 | 0 | 0.94 | 0.97 | 0.97 | 0.99 | 0.96 | 0.97 |
| 10 | 119283708 | rs10886094 | 1.604 | -0.071 | -0.075 | -0.971 | 0 | 0.016 | 0.016 | 0 | 0.27 | 0.32 | 0.43 | 0.54 | 0.29 | 0.11 |
| 10 | 119284408 | rs12570134 | 0.756 | 0.005 | -0.041 | -0.893 | -0.003 | 0.051 | 0.051 | 0.742 | 0.11 | 0.23 | 0.09 | 0.19 | 0.1 | 0.07 |
| 10 | 119290006 | rs10510030 | 1.927 | 0.214 | -0.185 | -0.875 | -0.06 | -0.123 | -0.123 | 0.002 | 0.23 | 0.04 | 0.4 | 0.54 | 0.28 | 0.09 |
| 10 | 119295929 | rs82612 | 1.235 | 0.45 | 0.789 | -0.329 | -0.007 | -0.035 | -0.035 | 1.559 | 0.87 | 0.77 | 0.93 | 0.94 | 0.92 | 0.93 |
| 10 | 119301454 | rs1638626 | 0.402 | 0.378 | 0.835 | -0.203 | 0.016 | 0.116 | 0.116 | 0 | 0.08 | 0.23 | 0.12 | 0.23 | 0.09 | 0.06 |
| 10 | 119313030 | rs242962 | 0.39 | 0.391 | 0.941 | -0.355 | 0.012 | 0.025 | 0.025 | 0 | 0.72 | 0.59 | 0.64 | 0.57 | 0.67 | 0.5 |
| 10 | 119315324 | rs8181280 | 1.352 | 0.344 | 0.952 | -0.4 | 0.001 | 0.009 | 0.009 | 0.013 | 0.57 | 0.23 | 0.56 | 0.54 | 0.62 | 0.43 |
| 10 | 119316080 | rs855767 | -1.166 | 0.375 | 0.947 | -0.381 | 0.004 | 0.01 | 0.01 | 0 | 0.34 | 0.55 | 0.39 | 0.44 | 0.33 | 0.54 |
| 10 | 119321449 | rs2255424 | 0.827 | 0.684 | 0.536 | -0.225 | 0 | -0.001 | -0.001 | 0 | 0.59 | 0.44 | 0.58 | 0.55 | 0.61 | 0.42 |
| 10 | 119324960 | rs703417 | -0.652 | 0.746 | 0.65 | 0.155 | 0.009 | 0.094 | 0.094 | 0 | 0.11 | 0.29 | 0.13 | 0.23 | 0.11 | 0.08 |
| 10 | 119327321 | rs242959 | -1.142 | 0.624 | 0.4 | -0.042 | -0.007 | -0.033 | -0.033 | 0.016 | 0.06 | 0.12 | 0.02 | 0.02 | 0.02 | 0.04 |
| 10 | 119327897 | rs181585 | 0.174 | 0.832 | 0.306 | -0.072 | -0.003 | 0.002 | 0.002 | 0 | 0.91 | 0.94 | 0.94 | 0.96 | 0.9 | 0.97 |
| 10 | 119329341 | rs855775 | 0.108 | 0.811 | 0.485 | 0.163 | -0.015 | -0.002 | -0.002 | 0 | 0.22 | 0.41 | 0.19 | 0.24 | 0.18 | 0.43 |
| 10 | 119331597 | rs430152 | 0.629 | 1.246 | 0.212 | 0.196 | -0.01 | -0.024 | -0.024 | 0 | 0.66 | 0.26 | 0.53 | 0.53 | 0.67 | 0.5 |
| 10 | 119336992 | rs7088735 | 1.182 | 1.223 | -0.086 | 0.042 | -0.032 | -0.039 | -0.039 | 0.321 | 0.55 | 0.13 | 0.5 | 0.51 | 0.61 | 0.42 |
| 10 | 119340830 | rs2768313 | -0.876 | 1.23 | 0.067 | 0.049 | -0.013 | -0.027 | -0.027 | 0 | 0.36 | 0.76 | 0.47 | 0.47 | 0.34 | 0.49 |
| 10 | 119341820 | rs703361 | -0.632 | 1.261 | 0.435 | 0.086 | -0.014 | -0.007 | -0.007 | 0 | 0.78 | 0.61 | 0.81 | 0.76 | 0.82 | 0.58 |
| 10 | 119345838 | rs181601 | 0.269 | 1.31 | 0.405 | 0.223 | 0.087 | 0.176 | 0.176 | 0 | 0.09 | 0.27 | 0.25 | 0.25 | 0.12 | 0.07 |
| 10 | 119347241 | rs7077608 | 0.024 | 1.507 | -0.117 | -0.191 | -0.005 | 0.005 | 0.005 | 0 | 0.12 | 0.07 | 0.09 | 0.02 | 0.16 | 0.11 |
| 10 | 119347918 | rs703364 | -0.113 | 1.391 | -0.103 | -0.029 | -0.002 | -0.03 | -0.03 | 0.608 | 0.71 | 0.58 | 0.74 | 0.77 | 0.7 | 0.53 |
| 10 | 119348009 | rs4752084 | 0.716 | 0.837 | -1.197 | 0.337 | -0.009 | -0.08 | -0.08 | 0 | 0.14 | 0.04 | 0.23 | 0.38 | 0.15 | 0.1 |
| 10 | 119348243 | rs703365 | -0.945 | 1.12 | -0.979 | 0.409 | 0.026 | 0.052 | 0.052 | 0.456 | 0.55 | 0.8 | 0.62 | 0.64 | 0.56 | 0.67 |
| 10 | 119348596 | rs181604 | -0.544 | 1.127 | -0.134 | 0.803 | 0.002 | 0.02 | 0.02 | 0 | 0.24 | 0.48 | 0.2 | 0.29 | 0.23 | 0.36 |

| | | | | | | | | | | | | | | | | |
|---|---|---|---|---|---|---|---|---|---|---|---|---|---|---|---|---|
| 10 | 119348693 | rs181605 | -0.441 | 1.292 | -0.566 | 0.53 | -0.031 | 0.068 | 0.068 | 0 | 0.17 | 0.05 | 0.07 | 0.02 | 0.16 | 0.16 |
| 10 | 119348729 | rs181606 | -1.289 | 1.099 | 0.474 | 1.51 | 0.089 | 0.49 | 0.49 | 0.137 | 0.06 | 0.34 | 0.16 | 0.27 | 0.06 | 0.03 |
| 10 | 119349761 | rs71003 | 0.032 | 0.961 | -0.159 | -0.175 | 0.007 | 0.01 | 0.01 | 0 | 0.64 | 0.62 | 0.63 | 0.54 | 0.59 | 0.68 |
| 10 | 119351325 | rs703370 | 0.803 | 0.274 | -0.018 | -0.595 | -0.052 | 0.025 | 0.025 | 0.017 | 0.81 | 0.45 | 0.87 | 0.72 | 0.9 | 0.88 |
| 10 | 119351904 | rs181610 | 0.255 | 0.299 | -0.179 | -0.783 | -0.022 | -0.036 | -0.036 | 0.717 | 0.33 | 0.22 | 0.38 | 0.4 | 0.44 | 0.37 |
| 10 | 119354829 | rs703377 | 0.604 | 0.266 | -0.156 | -0.724 | -0.052 | 0.025 | 0.025 | 0.017 | 0.81 | 0.45 | 0.87 | 0.72 | 0.9 | 0.88 |
| 10 | 119355131 | rs12780073 | 1.321 | 0.426 | -0.346 | -0.891 | 0.006 | -0.026 | -0.026 | 1.157 | 0.16 | 0.09 | 0.17 | 0.39 | 0.13 | 0.32 |
| 10 | 119360043 | rs181615 | -0.878 | 0.519 | -0.577 | -1.411 | -0.011 | 0.033 | 0.033 | 0 | 0.69 | 0.74 | 0.61 | 0.81 | 0.65 | 0.71 |
| 10 | 119362455 | rs7358112 | -0.249 | 0.491 | -0.595 | -1.426 | -0.023 | -0.103 | -0.103 | 0 | 0.14 | 0.42 | 0.05 | 0.02 | 0.08 | 0.05 |
| 10 | 119363648 | rs181622 | -0.992 | 0.953 | -0.46 | -1.321 | -0.001 | -0.001 | -0.001 | 0 | 0.6 | 0.6 | 0.6 | 0.46 | 0.68 | 0.48 |
| 10 | 119363679 | rs181623 | -1.199 | 1.027 | -0.505 | -1.288 | 0.002 | 0.001 | 0.001 | 0 | 0.62 | 0.65 | 0.61 | 0.47 | 0.68 | 0.49 |
| 10 | 119364995 | rs181632 | 0.298 | 2.378 | -0.353 | -0.553 | -0.031 | 0.019 | 0.019 | 0 | 0.19 | 0.07 | 0.32 | 0.18 | 0.27 | 0.17 |
| 10 | 119366745 | rs4751631 | 0.122 | 2.216 | -0.408 | -0.52 | -0.045 | -0.029 | -0.029 | 2.077 | 0.17 | 0.06 | 0.17 | 0.26 | 0.25 | 0.15 |
| 10 | 119375835 | rs181646 | -0.558 | 1.876 | 0.107 | -1.251 | 0.019 | -0.059 | -0.059 | 0 | 0.24 | 0.48 | 0.38 | 0.19 | 0.26 | 0.16 |
| 10 | 119376627 | rs181649 | 0.703 | 1.925 | 0.087 | -1.291 | 0.01 | -0.041 | -0.041 | 0.304 | 0.72 | 0.4 | 0.6 | 0.79 | 0.69 | 0.79 |
| 10 | 119381386 | rs703393 | 0.55 | 2.408 | -0.018 | -0.787 | 0.01 | -0.026 | -0.026 | 0.683 | 0.95 | 0.85 | 0.96 | 0.98 | 0.95 | 0.95 |
| 10 | 119386122 | rs181669 | 0.988 | 2.593 | 0.26 | -0.75 | 0.022 | -0.047 | -0.047 | 0 | 0.67 | 0.6 | 0.56 | 0.81 | 0.56 | 0.78 |
| 10 | 119388494 | rs181672 | 0.499 | 2.62 | 0.187 | -0.792 | -0.001 | -0.038 | -0.038 | 1.051 | 0.79 | 0.76 | 0.86 | 0.99 | 0.73 | 0.9 |
| 10 | 119391213 | rs181676 | 1.305 | 2.601 | 0.149 | -0.758 | 0.023 | 0.016 | 0.016 | 0 | 0.48 | 0.4 | 0.33 | 0.6 | 0.31 | 0.54 |
| 10 | 119400061 | rs181679 | 1.31 | 2.826 | 0.524 | -0.386 | 0.015 | 0.024 | 0.024 | 0 | 0.54 | 0.36 | 0.38 | 0.64 | 0.4 | 0.57 |
| 10 | 119406434 | rs703404 | 1.191 | 3.07 | 0.39 | -0.296 | 0.042 | -0.05 | -0.05 | 0 | 0.73 | 0.46 | 0.6 | 0.81 | 0.65 | 0.78 |
| 10 | 119407615 | rs855994 | 0.219 | 2.597 | 0.559 | -0.111 | 0.044 | 0.005 | 0.005 | 0 | 0.84 | 0.62 | 0.68 | 0.81 | 0.81 | 0.82 |
| 10 | 119407988 | rs855993 | -0.291 | 2.605 | 0.493 | -0.108 | 0.014 | 0.003 | 0.003 | 0 | 0.87 | 0.8 | 0.7 | 0.82 | 0.81 | 0.86 |
| 10 | 119408093 | rs855992 | 1.083 | 2.6 | 0.387 | -0.106 | 0.001 | 0.002 | 0.002 | 0 | 0.56 | 0.41 | 0.4 | 0.64 | 0.42 | 0.57 |
| 10 | 119411797 | rs855986 | -0.178 | 2.883 | -0.341 | 0.086 | 0.029 | 0.011 | 0.011 | 0 | 0.87 | 0.74 | 0.7 | 0.82 | 0.82 | 0.86 |
| 10 | 119418104 | rs2184898 | -0.968 | 2.37 | -0.84 | 0.043 | -0.016 | 0.013 | 0.013 | 0 | 0.28 | 0.19 | 0.27 | 0.17 | 0.39 | 0.25 |
| 10 | 119420722 | rs12770749 | -1.385 | 2.095 | -0.781 | 0.238 | -0.028 | 0.043 | 0.043 | 0 | 0.13 | 0.09 | 0.12 | 0 | 0.2 | 0.05 |
| 10 | 119421127 | rs703405 | -1.358 | 1.661 | 0.121 | 1.192 | 0.01 | 0.167 | 0.167 | 0 | 0.11 | 0.36 | 0.19 | 0.31 | 0.1 | 0.13 |
| 10 | 119422893 | rs12773399 | -1.315 | 2.57 | -0.19 | 0.751 | -0.008 | 0.003 | 0.003 | 0 | 0.06 | 0.05 | 0.08 | 0 | 0.17 | 0.03 |
| 10 | 119424467 | rs10787781 | -0.913 | 2.034 | 0.447 | 1.235 | -0.023 | 0.055 | 0.055 | 0 | 0.27 | 0.42 | 0.2 | 0.51 | 0.26 | 0.13 |
| 10 | 119425873 | rs7093450 | -1.214 | 2.404 | 0.54 | 1.216 | 0.001 | 0.032 | 0.032 | 0 | 0.31 | 0.44 | 0.27 | 0.51 | 0.41 | 0.14 |
| 10 | 119425908 | rs7093567 | -1.033 | 2.303 | 0.457 | 1.233 | 0.029 | 0.106 | 0.106 | 0 | 0.21 | 0.43 | 0.24 | 0.47 | 0.34 | 0.1 |
| 10 | 119426280 | rs7078168 | -0.662 | 2.187 | 0.568 | 1.179 | -0.008 | 0.168 | 0.168 | 0.068 | 0.18 | 0.45 | 0.16 | 0.51 | 0.19 | 0.09 |
| 10 | 119426958 | rs855974 | -0.704 | 2.229 | 0.434 | 1.381 | -0.022 | -0.063 | -0.063 | 0 | 0.21 | 0.08 | 0.35 | 0.66 | 0.26 | 0.16 |
| 10 | 119427941 | rs7894825 | -1.902 | 2.436 | 1.01 | 1.421 | 0.024 | -0.007 | -0.007 | 3.9 | 0.22 | 0.58 | 0.3 | 0.22 | 0.29 | 0.27 |
| 10 | 119433496 | rs12572932 | -0.917 | 2.052 | 1.029 | 1.528 | -0.031 | 0.044 | 0.044 | 0.063 | 0.16 | 0.36 | 0.2 | 0.23 | 0.11 | 0.27 |
| 10 | 119442975 | rs855966 | -1.477 | 2.144 | 0.896 | 1.529 | -0.005 | -0.036 | -0.036 | 0.042 | 0.35 | 0.48 | 0.56 | 0.85 | 0.39 | 0.4 |
| 10 | 119446868 | rs855964 | 0.161 | 2.202 | 0.57 | 1.876 | -0.029 | -0.019 | -0.019 | 0 | 0.05 | 0.01 | 0.24 | 0.2 | 0.08 | 0.08 |
| 10 | 119447247 | rs4752091 | 1.269 | 2.073 | 1.281 | 1.703 | -0.003 | -0.033 | -0.033 | 0 | 0.65 | 0.52 | 0.4 | 0.15 | 0.61 | 0.61 |
| 10 | 119449952 | rs11198069 | 1.236 | 1.673 | 0.031 | 0.894 | -0.051 | -0.123 | -0.123 | 0 | 0.34 | 0.01 | 0.19 | 0.36 | 0.24 | 0.41 |
| 10 | 119457780 | rs7894337 | -0.051 | 2.21 | 1.384 | 0.939 | 0.055 | -0.027 | -0.027 | 0 | 0.05 | 0.14 | 0.09 | 0 | 0.16 | 0.02 |
| 10 | 119460525 | rs12218538 | 1.617 | 2.171 | 1.691 | 0.875 | -0.014 | -0.006 | -0.006 | 0 | 0.55 | 0.32 | 0.32 | 0.43 | 0.43 | 0.62 |
| 10 | 119462812 | rs1538080 | 0.648 | 2.131 | 1.732 | 0.82 | 0.002 | 0.013 | 0.013 | 0 | 0.73 | 0.69 | 0.49 | 0.48 | 0.68 | 0.72 |
| 10 | 119468305 | rs4752095 | 0.614 | 2.129 | 1.611 | 0.841 | -0.008 | -0.025 | -0.025 | 1.192 | 0.73 | 0.74 | 0.5 | 0.49 | 0.68 | 0.72 |
| 10 | 119471333 | rs11597064 | 0.568 | 2.147 | 1.54 | 0.89 | -0.027 | -0.018 | -0.018 | 0.341 | 0.14 | 0.01 | 0.09 | 0.16 | 0.18 | 0.04 |
| 10 | 119471827 | rs4752098 | 1.783 | 2.145 | 1.574 | 0.875 | -0.028 | -0.009 | -0.009 | 0 | 0.55 | 0.25 | 0.32 | 0.42 | 0.43 | 0.62 |
| 10 | 119474930 | rs10787783 | 0.45 | 2.087 | 1.176 | 0.848 | -0.046 | -0.107 | -0.107 | 0 | 0.74 | 0.89 | 0.51 | 0.48 | 0.68 | 0.72 |
| 10 | 119477150 | rs1925272 | 1.543 | 2.118 | 1.151 | 0.766 | -0.001 | 0.009 | 0.009 | 0 | 0.61 | 0.47 | 0.42 | 0.43 | 0.59 | 0.65 |
| 10 | 119478641 | rs4752101 | 1.524 | 2.167 | 1.151 | 0.547 | -0.001 | 0.011 | 0.011 | 0 | 0.61 | 0.46 | 0.42 | 0.5 | 0.61 | 0.66 |
| 10 | 119480499 | rs1925274 | 1.061 | 2.203 | 1.121 | 0.556 | -0.001 | 0.003 | 0.003 | 0 | 0.65 | 0.58 | 0.47 | 0.51 | 0.66 | 0.68 |
| 10 | 119481889 | rs12358696 | 1.484 | 2.173 | 0.615 | 0.579 | -0.009 | -0.006 | -0.006 | 0 | 0.58 | 0.42 | 0.35 | 0.51 | 0.46 | 0.62 |
| 10 | 119482303 | rs1886603 | -0.802 | 1.851 | -0.063 | 0.699 | -0.052 | -0.062 | -0.062 | 0 | 0.1 | 0.01 | 0.37 | 0.34 | 0.13 | 0.25 |
| 10 | 119485262 | rs7099192 | -0.292 | 1.289 | 0.398 | -0.081 | -0.035 | -0.024 | -0.024 | 0 | 0.1 | 0.22 | 0.06 | 0.03 | 0.05 | 0.03 |
| 10 | 119487704 | rs1024265 | -0.672 | 1.337 | 0.168 | -0.067 | -0.035 | -0.045 | -0.045 | 0 | 0.13 | 0.34 | 0.09 | 0.03 | 0.08 | 0.06 |
| 10 | 119490038 | rs11198078 | -0.144 | 2.082 | -0.912 | 0.027 | -0.054 | 0.021 | 0.021 | 0 | 0.14 | 0.01 | 0.1 | 0.11 | 0.16 | 0.27 |
| 10 | 119490410 | rs17552671 | -1.038 | 1.772 | -0.313 | 0.257 | 0.033 | -0.026 | -0.026 | 0 | 0.05 | 0.08 | 0.1 | 0 | 0.16 | 0.04 |
| 10 | 119493253 | rs6585456 | 0.296 | 1.107 | -0.637 | 0.043 | 0.005 | -0.015 | -0.015 | 0 | 0.58 | 0.54 | 0.53 | 0.76 | 0.51 | 0.57 |
| 10 | 119493806 | rs10510032 | 0.341 | 0.55 | -0.265 | 0.127 | -0.008 | -0.013 | -0.013 | 1.11 | 0.82 | 0.54 | 0.84 | 0.88 | 0.86 | 0.89 |
| 10 | 119498160 | rs912745 | -0.264 | -0.177 | -0.94 | 0.024 | 0.03 | 0.024 | 0.024 | 0 | 0.16 | 0.07 | 0.09 | 0.08 | 0.09 | 0.14 |
| 10 | 119509897 | rs1570787 | 0.934 | -0.107 | -0.915 | 0.412 | -0.006 | -0.001 | -0.001 | 0 | 0.66 | 0.38 | 0.72 | 0.64 | 0.78 | 0.74 |
| 10 | 119519010 | rs1006286 | -0.667 | -0.447 | -1.648 | 0.672 | 0.045 | -0.016 | -0.016 | 0.003 | 0.2 | 0.12 | 0.1 | 0.28 | 0.1 | 0.15 |
| 10 | 119520842 | rs10886126 | -0.326 | 0.692 | -1.407 | 0.662 | -0.023 | -0.029 | -0.029 | 0 | 0.18 | 0.08 | 0.33 | 0.26 | 0.19 | 0.3 |
| 10 | 119522728 | rs6585459 | -0.867 | -0.023 | -1.363 | 0.05 | 0.005 | -0.018 | -0.018 | 0.031 | 0.16 | 0.15 | 0.08 | 0 | 0.11 | 0.07 |
| 10 | 119522856 | rs1886606 | -0.914 | 1.271 | -1.741 | 0.56 | -0.028 | -0.029 | -0.029 | 0 | 0.11 | 0.07 | 0.32 | 0.2 | 0.24 | 0.22 |
| 10 | 119522943 | rs7077625 | -0.745 | 0.778 | -1.698 | 0.558 | 0.001 | 0 | 0 | 0 | 0.54 | 0.48 | 0.48 | 0.38 | 0.38 | 0.59 |
| 10 | 119522964 | rs878747 | 0.548 | 0.528 | -2.574 | 0.533 | -0.108 | -0.045 | -0.045 | 0 | 0.28 | 0.09 | 0.29 | 0.37 | 0.49 | 0.28 |
| 10 | 119523207 | rs1925279 | -0.373 | -0.406 | -2.575 | -0.06 | 0.07 | 0.132 | 0.132 | 0 | 0.4 | 0.17 | 0.26 | 0.15 | 0.3 | 0.27 |
| 10 | 119523717 | rs10430688 | -0.326 | 0.164 | -2.038 | 0.633 | 0.007 | 0.016 | 0.016 | 0 | 0.49 | 0.37 | 0.5 | 0.61 | 0.44 | 0.55 |
| 10 | 119524240 | rs7083178 | -0.345 | -0.551 | -2.105 | 1.087 | 0.04 | -0.058 | -0.058 | 0 | 0.24 | 0.04 | 0.13 | 0.31 | 0.12 | 0.34 |

| | | | | | | | | | | | | | | | | |
|---|---|---|---|---|---|---|---|---|---|---|---|---|---|---|---|---|
| 10 | 119526414 | rs11198092 | -0.236 | -0.457 | -2.184 | 1.525 | -0.005 | -0.086 | -0.086 | 0 | 0.13 | 0.03 | 0.24 | 0.4 | 0.14 | 0.15 |
| 10 | 119528591 | rs7897056 | 0.513 | -0.127 | -2.504 | 0.536 | -0.041 | 0.054 | 0.054 | 0.801 | 0.28 | 0.17 | 0.37 | 0.08 | 0.38 | 0.17 |
| 10 | 119529361 | rs1925280 | -0.032 | -0.887 | -2.643 | 0.53 | 0.018 | 0.022 | 0.022 | 0.051 | 0.56 | 0.3 | 0.54 | 0.39 | 0.55 | 0.51 |
| 10 | 119531719 | rs7918427 | 0.743 | -1.244 | -2.611 | 0.712 | 0.026 | -0.037 | -0.037 | 0 | 0.26 | 0.13 | 0.17 | 0.33 | 0.17 | 0.31 |
| 10 | 119533041 | rs12358707 | 1.006 | -1.396 | -2.314 | 0.692 | 0.004 | -0.003 | -0.003 | 0 | 0.13 | 0.06 | 0.09 | 0.13 | 0.1 | 0.12 |
| 10 | 119533345 | rs1925283 | 0.551 | -1.376 | -1.671 | 0.598 | -0.004 | -0.001 | -0.001 | 0 | 0.6 | 0.41 | 0.58 | 0.41 | 0.68 | 0.6 |
| 10 | 119533757 | rs7088969 | 0.6 | -1.395 | -1.643 | 0.573 | -0.006 | -0.001 | -0.001 | 0 | 0.59 | 0.43 | 0.58 | 0.41 | 0.67 | 0.6 |
| 10 | 119536279 | rs10510033 | 0.669 | -1.226 | -1.655 | 0.418 | 0.005 | -0.002 | -0.002 | 0 | 0.13 | 0.06 | 0.09 | 0.13 | 0.11 | 0.14 |
| 10 | 119539294 | rs912748 | -0.323 | -1.648 | -1.366 | 0.568 | 0.02 | 0.002 | 0.002 | 0 | 0.62 | 0.67 | 0.76 | 0.62 | 0.77 | 0.59 |
| 10 | 119542045 | rs7476605 | 0.149 | -2.076 | -1.298 | 0.145 | 0.062 | -0.018 | -0.018 | 3.155 | 0.37 | 0.22 | 0.44 | 0.64 | 0.3 | 0.42 |
| 10 | 119542582 | rs17096821 | 1.118 | -1.661 | -0.749 | -0.169 | -0.005 | -0.005 | -0.005 | 0 | 0.12 | 0.12 | 0.03 | 0.09 | 0.09 | 0.17 |
| 10 | 119544669 | rs1925284 | 0.622 | -1.708 | -0.382 | -0.137 | -0.017 | -0.036 | -0.036 | 0 | 0.81 | 0.66 | 0.79 | 0.85 | 0.81 | 0.87 |
| 10 | 119547678 | rs4752112 | 0.087 | -1.705 | -0.384 | -0.146 | -0.002 | -0.046 | -0.046 | 0 | 0.91 | 0.82 | 0.93 | 1 | 0.91 | 0.96 |
| 10 | 119548382 | rs7904075 | 0.668 | -1.708 | -0.409 | -0.137 | -0.021 | -0.045 | -0.045 | 0 | 0.81 | 0.6 | 0.73 | 0.85 | 0.8 | 0.87 |
| 10 | 119553310 | rs10787795 | 0.664 | -1.801 | -0.376 | -0.159 | -0.021 | -0.038 | -0.038 | 0 | 0.65 | 0.43 | 0.69 | 0.77 | 0.71 | 0.73 |
| 10 | 119559170 | rs10886139 | 0.113 | -0.924 | -0.733 | 0.093 | -0.046 | 0.073 | 0.073 | 1.266 | 0.38 | 0.21 | 0.45 | 0.25 | 0.49 | 0.26 |
| 10 | 119564143 | rs11198112 | -0.669 | -0.28 | 0.282 | 0.724 | 0.014 | 0.007 | 0.007 | 0 | 0.11 | 0.19 | 0.15 | 0.1 | 0.15 | 0.21 |
| 10 | 119565254 | rs1925251 | 0.391 | -0.707 | 0.673 | 0.863 | -0.017 | 0.079 | 0.079 | 0 | 0.83 | 0.57 | 0.84 | 0.79 | 0.84 | 0.83 |
| 10 | 119568133 | rs17096846 | -0.353 | -0.724 | 0.248 | 0.463 | -0.01 | -0.005 | -0.005 | 0 | 0.24 | 0.28 | 0.17 | 0.24 | 0.18 | 0.18 |
| 10 | 119568331 | rs11198117 | 1.336 | -0.539 | -0.072 | 0.533 | -0.012 | -0.052 | -0.052 | 0.832 | 0.51 | 0.09 | 0.55 | 0.53 | 0.54 | 0.52 |
| 10 | 119576277 | rs729465 | 1.121 | -0.415 | 0.53 | 0.695 | -0.001 | 0 | 0 | 0 | 0.62 | 0.43 | 0.6 | 0.61 | 0.66 | 0.75 |
| 10 | 119582073 | rs11198125 | -1.097 | 0.483 | 1.012 | 0.786 | 0.013 | 0.032 | 0.032 | 0.146 | 0.07 | 0.25 | 0.05 | 0.05 | 0.05 | 0.01 |
| 10 | 119584861 | rs6585463 | 0.649 | 0.408 | 0.914 | 0.817 | -0.007 | 0.014 | 0.014 | 0.012 | 0.79 | 0.59 | 0.83 | 0.76 | 0.8 | 0.65 |
| 10 | 119587413 | rs4752120 | 0.852 | 0.426 | 0.166 | 0.801 | -0.002 | 0.001 | 0.001 | 0 | 0.76 | 0.32 | 0.79 | 0.71 | 0.75 | 0.64 |
| 10 | 119587425 | rs4751641 | 1.379 | 0.625 | -0.218 | 0.911 | -0.007 | -0.031 | -0.031 | 0 | 0.67 | 0.22 | 0.63 | 0.63 | 0.66 | 0.62 |
| 10 | 119590330 | rs11198133 | -0.202 | 1.253 | -0.13 | 0.947 | -0.032 | -0.032 | -0.032 | 0 | 0.06 | 0.01 | 0.15 | 0.08 | 0.09 | 0.02 |
| 10 | 119601743 | rs12250372 | -1.493 | 0.902 | 0.307 | 0.632 | 0.027 | 0.061 | 0.061 | 0.475 | 0.1 | 0.34 | 0.09 | 0.12 | 0.09 | 0.23 |
| 10 | 119604938 | rs1925265 | -0.657 | 1.444 | -0.351 | 0.147 | -0.006 | 0.004 | 0.004 | 0 | 0.3 | 0.32 | 0.34 | 0.17 | 0.43 | 0.39 |
| 10 | 119604976 | rs1925266 | 0.085 | 1.334 | -0.235 | 0.386 | 0.002 | -0.009 | -0.009 | 0 | 0.66 | 0.64 | 0.67 | 0.8 | 0.62 | 0.77 |
| 10 | 119606709 | rs7906809 | 0.79 | 0.818 | -0.047 | 0.708 | -0.018 | 0.175 | 0.175 | 0 | 0.83 | 0.61 | 0.69 | 0.4 | 0.85 | 0.87 |
| 10 | 119608564 | rs12718340 | -0.311 | 0.807 | -0.684 | 0.48 | -0.013 | 0.003 | 0.003 | 0.172 | 0.08 | 0.16 | 0.05 | 0.12 | 0.04 | 0.07 |
| 10 | 119609983 | rs11198153 | 0.095 | 0.501 | -1.83 | -0.106 | 0.001 | -0.001 | -0.001 | 0 | 0.2 | 0.01 | 0.17 | 0.15 | 0.16 | 0.24 |
| 10 | 119611043 | rs17096923 | 0.218 | 0.609 | -1.825 | -0.046 | -0.016 | -0.058 | -0.058 | 0.037 | 0.19 | 0.02 | 0.19 | 0.23 | 0.16 | 0.25 |
| 10 | 119611130 | rs17641472 | 0.27 | 1.214 | -1.846 | -0.932 | -0.056 | 0.118 | 0.118 | 0 | 0.21 | 0.03 | 0.13 | 0.01 | 0.22 | 0.12 |
| 10 | 119611204 | rs12784400 | 0.047 | 1.097 | -1.012 | -0.303 | 0 | 0.026 | 0.026 | 0 | 0.36 | 0.27 | 0.24 | 0.14 | 0.33 | 0.37 |
| 10 | 119613089 | rs2147741 | -1.045 | 1.767 | -0.817 | 0.128 | -0.002 | -0.002 | -0.002 | 0 | 0.15 | 0.12 | 0.2 | 0.03 | 0.25 | 0.17 |
| 10 | 119613610 | rs10886159 | -0.735 | 0.711 | -0.631 | 0.97 | 0.002 | -0.003 | -0.003 | 0 | 0.26 | 0.24 | 0.35 | 0.55 | 0.24 | 0.12 |
| 10 | 119613633 | rs4752123 | 0.169 | 0.563 | -0.265 | 1.055 | -0.026 | -0.011 | -0.011 | 0 | 0.7 | 0.43 | 0.69 | 0.74 | 0.78 | 0.87 |
| 10 | 119615257 | rs11198159 | -1.179 | 1.303 | -0.349 | 1.681 | 0 | 0.004 | 0.004 | 0 | 0.23 | 0.26 | 0.27 | 0.39 | 0.24 | 0.3 |
| 10 | 119619382 | rs4612716 | -1.211 | 1.79 | -0.274 | 1.737 | -0.003 | -0.006 | -0.006 | 0 | 0.36 | 0.7 | 0.56 | 0.55 | 0.39 | 0.51 |
| 10 | 119620559 | rs6585465 | -1.271 | 1.769 | -0.27 | 1.781 | -0.003 | -0.004 | -0.004 | 0 | 0.37 | 0.71 | 0.56 | 0.58 | 0.4 | 0.52 |
| 10 | 119621697 | rs6585466 | -1.262 | 1.744 | -0.165 | 1.774 | -0.005 | -0.007 | -0.007 | 0 | 0.36 | 0.69 | 0.55 | 0.59 | 0.39 | 0.52 |
| 10 | 119623656 | rs7904164 | -1.212 | 2.405 | 0.461 | 1.901 | 0.036 | 0.057 | 0.057 | 0 | 0.23 | 0.38 | 0.46 | 0.6 | 0.32 | 0.36 |
| 10 | 119625944 | rs10510036 | -0.625 | 2.091 | 0.16 | 1.666 | -0.028 | 0.027 | 0.027 | 0 | 0.05 | 0.02 | 0.04 | 0 | 0.1 | 0.06 |
| 10 | 119627204 | rs4751642 | 1.221 | 2.492 | 0.616 | 1.679 | 0.014 | 0.025 | 0.025 | 0 | 0.67 | 0.48 | 0.54 | 0.39 | 0.62 | 0.51 |
| 10 | 119628241 | rs17641954 | -0.604 | 2.383 | 0.331 | 1.735 | -0.028 | 0.027 | 0.027 | 0 | 0.05 | 0.02 | 0.04 | 0 | 0.1 | 0.06 |
| 10 | 119628641 | rs12414011 | 0.077 | 2.429 | 0.321 | 2.006 | -0.042 | -0.033 | -0.033 | 3.267 | 0.09 | 0.01 | 0.16 | 0.13 | 0.15 | 0.1 |
| 10 | 119632192 | rs11198164 | -2.13 | 2.124 | 0.756 | 1.557 | 0.012 | 0.014 | 0.014 | 0 | 0.49 | 0.66 | 0.56 | 0.59 | 0.54 | 0.62 |
| 10 | 119632453 | rs11198165 | -1.582 | 1.961 | 1.165 | 1.79 | 0 | -0.009 | -0.009 | 0.193 | 0.42 | 0.35 | 0.44 | 0.48 | 0.4 | 0.44 |
| 10 | 119633278 | rs6585469 | -1.012 | 1.971 | 1.287 | 1.76 | 0.045 | -0.038 | -0.038 | 0 | 0.07 | 0.2 | 0.06 | 0 | 0.1 | 0.06 |
| 10 | 119635950 | rs1343418 | 1.827 | 1.905 | 0.726 | 1.931 | -0.041 | 0.078 | 0.078 | 0.035 | 0.49 | 0.09 | 0.35 | 0.3 | 0.46 | 0.37 |
| 10 | 119638197 | rs1419140 | 0.499 | 1.831 | -0.026 | 2.11 | -0.067 | -0.086 | -0.086 | 2.393 | 0.69 | 0.92 | 0.63 | 0.63 | 0.65 | 0.57 |
| 10 | 119641986 | rs1936298 | 1.291 | 1.607 | -0.206 | 1.922 | -0.02 | 0.147 | 0.147 | 0.012 | 0.51 | 0.17 | 0.3 | 0.22 | 0.5 | 0.46 |
| 10 | 119644686 | rs12218761 | -0.26 | 0.943 | 0.378 | 1.694 | 0.012 | 0.028 | 0.028 | 0 | 0.07 | 0.02 | 0.03 | 0.02 | 0.04 | 0.05 |
| 10 | 119646217 | rs2255624 | 1.656 | 0.989 | 1.233 | 1.997 | 0.006 | 0.155 | 0.155 | 2.891 | 0.9 | 0.6 | 0.89 | 0.8 | 0.97 | 0.88 |
| 10 | 119648854 | rs10510038 | 0.216 | 1.375 | 0.262 | 1.855 | -0.048 | 0.006 | 0.006 | 0.049 | 0.31 | 0.08 | 0.37 | 0.26 | 0.37 | 0.42 |
| 10 | 119649298 | rs2243640 | 0.197 | 1.383 | 1.177 | 2.306 | -0.005 | -0.009 | -0.009 | 0.01 | 0.59 | 0.4 | 0.36 | 0.43 | 0.51 | 0.46 |
| 10 | 119652158 | rs1104941 | 1.337 | 1.323 | 2.034 | 2.275 | 0.048 | 0.168 | 0.168 | 1.247 | 0.95 | 0.71 | 0.85 | 0.86 | 0.97 | 0.96 |
| 10 | 119662092 | rs1936295 | -0.371 | 0.971 | 1.164 | 1.48 | 0.001 | -0.007 | -0.007 | 0.22 | 0.8 | 0.81 | 0.87 | 0.94 | 0.82 | 0.87 |
| 10 | 119662153 | rs2265987 | 1.014 | 0.947 | 0.94 | 1.85 | 0.01 | -0.015 | -0.015 | 0 | 0.66 | 0.18 | 0.56 | 0.37 | 0.7 | 0.68 |
| 10 | 119663968 | rs2246493 | 1.111 | 0.536 | 0.63 | 1.48 | 0.009 | 0.15 | 0.15 | 0 | 0.36 | 0.13 | 0.2 | 0.14 | 0.31 | 0.29 |
| 10 | 119670998 | rs8181443 | 0.99 | 0.533 | 0.874 | 1.467 | -0.167 | -0.052 | -0.052 | 0.066 | 0.3 | 0.02 | 0.37 | 0.26 | 0.41 | 0.41 |
| 10 | 119671282 | rs2250044 | 1.133 | 0.452 | 0.93 | 1.4 | 0.01 | 0.147 | 0.147 | 0 | 0.36 | 0.14 | 0.2 | 0.14 | 0.31 | 0.29 |
| 10 | 119679259 | rs2253812 | -0.829 | 0.493 | 1.013 | 1.509 | 0.026 | 0.058 | 0.058 | 0 | 0.81 | 0.93 | 0.9 | 0.91 | 0.88 | 0.89 |
| 10 | 119683609 | rs2254426 | 0.29 | 0.516 | 1.239 | 1.537 | 0.001 | 0 | 0 | 0 | 0.44 | 0.43 | 0.33 | 0.41 | 0.42 | 0.43 |
| 10 | 119690695 | rs10886179 | 0.947 | 0.598 | 0.474 | 1.305 | 0.023 | 0.116 | 0.116 | 0 | 0.37 | 0.15 | 0.24 | 0.17 | 0.31 | 0.29 |
| 10 | 119692335 | rs7069291 | -0.191 | 0.536 | 0.38 | 1.427 | 0.004 | -0.016 | -0.016 | 0.257 | 0.42 | 0.49 | 0.58 | 0.69 | 0.52 | 0.62 |
| 10 | 119694504 | rs1419141 | -1.286 | 0.584 | 0.376 | 1.414 | 0.002 | 0.031 | 0.031 | 0 | 0.63 | 0.73 | 0.68 | 0.75 | 0.64 | 0.72 |
| 10 | 119694680 | rs12412410 | -0.849 | 0.8 | 0.179 | 1.36 | 0.11 | 0.423 | 0.423 | 0.032 | 0.05 | 0.36 | 0.16 | 0.24 | 0.07 | 0.14 |

| | | | | | | | | | | | | | | | | |
|---|---|---|---|---|---|---|---|---|---|---|---|---|---|---|---|---|
| 10 | 119695314 | rs4752135 | 0.382 | 0.716 | 0.259 | 0.684 | 0.019 | 0.01 | 0.01 | 0 | 0.35 | 0.38 | 0.53 | 0.52 | 0.49 | 0.53 |
| 10 | 119697225 | rs1361863 | -0.619 | 0.183 | 0.038 | 0.239 | -0.019 | -0.049 | -0.049 | 0 | 0.23 | 0.33 | 0.12 | 0.08 | 0.15 | 0.18 |
| 10 | 119702642 | rs11198190 | -0.133 | 0.916 | -0.247 | 0.639 | 0.005 | -0.01 | -0.01 | 0.185 | 0.63 | 0.54 | 0.48 | 0.29 | 0.7 | 0.65 |
| 10 | 119705438 | rs1797 | 0.012 | 0.998 | -1.217 | 0.684 | -0.031 | 0.052 | 0.052 | 0 | 0.56 | 0.21 | 0.38 | 0.29 | 0.61 | 0.61 |
| 10 | 119714635 | rs12415995 | -0.456 | 0.178 | -1.341 | 0.584 | 0.024 | 0.029 | 0.029 | 0 | 0.33 | 0.15 | 0.42 | 0.25 | 0.24 | 0.28 |
| 10 | 119723015 | rs7073514 | 0.49 | -0.921 | -0.44 | 1.228 | -0.073 | 0.103 | 0.103 | 0.62 | 0.83 | 0.4 | 0.76 | 0.41 | 0.95 | 0.88 |
| 10 | 119729762 | rs1936296 | 0.412 | -0.295 | 0.511 | 1.013 | -0.072 | 0.203 | 0.203 | 0.181 | 0.86 | 0.64 | 0.79 | 0.42 | 0.95 | 0.88 |
| 10 | 119749103 | rs12218935 | 0.783 | -0.648 | 0.069 | 0.9 | 0.032 | 0.225 | 0.225 | 0.008 | 0.54 | 0.13 | 0.37 | 0.16 | 0.61 | 0.58 |
| 10 | 119752955 | rs11198218 | 0.772 | -0.644 | 0.071 | 0.912 | 0.031 | 0.227 | 0.227 | 0.034 | 0.54 | 0.13 | 0.37 | 0.15 | 0.61 | 0.58 |
| 10 | 119765851 | rs11198226 | 0.97 | -0.619 | -0.137 | 0.763 | -0.111 | 0.007 | 0.007 | 0 | 0.22 | 0.02 | 0.18 | 0.15 | 0.36 | 0.28 |
| 10 | 119768551 | rs7923321 | -0.645 | -1.038 | 0.849 | 0.53 | 0.039 | 0.493 | 0.493 | 0 | 0.95 | 0.7 | 0.88 | 0.58 | 0.96 | 0.88 |
| 10 | 119769667 | rs10430689 | 0.116 | -1.042 | 0.755 | -0.316 | 0.096 | 0.16 | 0.16 | 0.192 | 0.59 | 0.17 | 0.46 | 0.16 | 0.74 | 0.6 |
| 10 | 119788183 | rs1016112 | 0.562 | -1.039 | 0.761 | -0.318 | 0.033 | 0.217 | 0.217 | 0.023 | 0.55 | 0.14 | 0.37 | 0.16 | 0.61 | 0.58 |
| 10 | 119796664 | rs10886196 | 0.121 | -1.053 | 0.798 | -0.315 | 0.097 | 0.162 | 0.162 | 0.192 | 0.59 | 0.17 | 0.46 | 0.16 | 0.74 | 0.61 |
| 10 | 119820073 | rs853576 | -0.417 | -1.056 | 1.158 | -0.189 | -0.037 | 0.227 | 0.227 | 0 | 0.9 | 0.71 | 0.81 | 0.39 | 0.95 | 0.87 |
| 10 | 119840532 | rs7906694 | -0.273 | -1.02 | 1.015 | -0.377 | -0.003 | -0.015 | -0.015 | 0.836 | 0.37 | 0.58 | 0.45 | 0.23 | 0.34 | 0.29 |
| 10 | 119849144 | rs1932644 | -0.214 | -1.278 | 0.952 | -0.668 | -0.005 | 0.108 | 0.108 | 0 | 0.33 | 0.17 | 0.25 | 0.05 | 0.34 | 0.21 |
| 10 | 119854863 | rs853585 | -0.325 | -0.802 | 1.096 | -0.422 | 0.02 | -0.002 | -0.002 | 0 | 0.06 | 0.31 | 0.03 | 0 | 0.02 | 0.01 |
| 10 | 119904921 | rs853604 | -0.958 | -0.849 | 0.551 | -0.505 | 0.057 | 0.188 | 0.188 | 0.141 | 0.48 | 0.8 | 0.64 | 0.86 | 0.41 | 0.41 |
| 10 | 119928781 | rs1336326 | -0.413 | -0.86 | 0.624 | -0.43 | 0.027 | 0.055 | 0.055 | 0 | 0.41 | 0.29 | 0.41 | 0.7 | 0.37 | 0.34 |
| 10 | 119949748 | rs703424 | -1.299 | -0.847 | -0.545 | -0.766 | -0.167 | 0.074 | 0.074 | 0.362 | 0.73 | 0.98 | 0.79 | 0.88 | 0.57 | 0.61 |
| 10 | 119963569 | rs10510042 | 0.303 | -0.87 | 0.209 | -0.032 | 0.002 | 0.139 | 0.139 | 1.098 | 0.28 | 0.02 | 0.14 | 0.04 | 0.21 | 0.16 |
| 10 | 119967243 | rs1369 | -0.939 | -0.918 | 0.503 | 0.1 | 0.058 | 0.162 | 0.162 | 0.05 | 0.47 | 0.77 | 0.64 | 0.87 | 0.4 | 0.41 |
| 10 | 119973974 | rs853571 | -0.394 | -0.358 | 0.244 | -0.002 | 0.01 | 0.026 | 0.026 | 0 | 0.38 | 0.3 | 0.42 | 0.22 | 0.36 | 0.25 |
| 10 | 119983056 | rs1577074 | -1.841 | 0.033 | 0.836 | 0.442 | 0.019 | 0.047 | 0.047 | 0.56 | 0.09 | 0.46 | 0.08 | 0.06 | 0.06 | 0.08 |
| 10 | 119994465 | rs11198317 | 1.134 | 0.266 | 0.85 | 0.447 | 0.05 | 0.067 | 0.067 | 0 | 0.93 | 0.62 | 0.94 | 0.94 | 0.96 | 0.92 |
| 10 | 119994707 | rs10787813 | 2.099 | 0.188 | 1.038 | 0.404 | 0.017 | 0.012 | 0.012 | 0.644 | 0.89 | 0.44 | 0.9 | 0.93 | 0.91 | 0.91 |
| 10 | 119996257 | rs2184169 | -0.499 | 0.755 | -0.134 | 0.458 | -0.037 | -0.012 | -0.012 | 6.443 | 0.24 | 0.1 | 0.21 | 0.26 | 0.27 | 0.23 |
| 10 | 120001191 | rs2210604 | 0.503 | 0.984 | -0.003 | 0.617 | -0.005 | -0.006 | -0.006 | 0 | 0.64 | 0.74 | 0.47 | 0.59 | 0.6 | 0.68 |
| 10 | 120008097 | rs11198323 | 0.856 | 0.852 | -1.039 | 0.007 | -0.014 | 0.092 | 0.092 | 0 | 0.23 | 0.01 | 0.09 | 0.06 | 0.17 | 0.2 |
| 10 | 120014511 | rs11198331 | 1.034 | 0.87 | -0.228 | 0.635 | 0.004 | -0.001 | -0.001 | 0 | 0.33 | 0.39 | 0.3 | 0.31 | 0.37 | 0.35 |
| 10 | 120016868 | rs7922488 | 0.821 | 0.843 | 0.309 | 0.728 | 0.005 | 0.019 | 0.019 | 0 | 0.36 | 0.6 | 0.34 | 0.5 | 0.41 | 0.44 |
| 10 | 120022278 | rs10886215 | -0.2 | 0.916 | 0.332 | 0.695 | -0.001 | -0.006 | -0.006 | 0 | 0.61 | 0.71 | 0.45 | 0.57 | 0.6 | 0.65 |
| 10 | 120024591 | rs9919453 | 0.949 | 0.878 | 0.076 | 0.69 | -0.02 | -0.034 | -0.034 | 0 | 0.34 | 0.27 | 0.32 | 0.52 | 0.42 | 0.45 |
| 10 | 120025435 | rs11198338 | 0.846 | 0.567 | -0.024 | 0.575 | 0.029 | 0.106 | 0.106 | 0 | 0.25 | 0.05 | 0.1 | 0.06 | 0.17 | 0.2 |
| 10 | 120031952 | rs11198344 | 0.434 | 0.868 | 0.1 | 0.838 | -0.028 | -0.023 | -0.023 | 0.982 | 0.39 | 0.31 | 0.35 | 0.43 | 0.5 | 0.43 |
| 10 | 120032904 | rs12249758 | 0.251 | 0.91 | -0.382 | 0.454 | 0.022 | -0.011 | -0.011 | 0 | 0.22 | 0.29 | 0.18 | 0.12 | 0.33 | 0.23 |
| 10 | 120033929 | rs1776649 | -0.017 | 1.023 | -0.269 | 0.508 | 0.008 | 0.009 | 0.009 | 0 | 0.54 | 0.56 | 0.58 | 0.4 | 0.41 | 0.48 |
| 10 | 120035787 | rs11198351 | 1.265 | 1.044 | -0.698 | 0.595 | -0.014 | -0.088 | -0.088 | 0 | 0.23 | 0.1 | 0.23 | 0.48 | 0.26 | 0.28 |
| 10 | 120038770 | rs984428 | 0.008 | 0.664 | -0.601 | 0.563 | -0.004 | -0.058 | -0.058 | 0.49 | 0.43 | 0.2 | 0.38 | 0.46 | 0.56 | 0.47 |
| 10 | 120044002 | rs830763 | -0.081 | 0.519 | -0.201 | 0.568 | -0.013 | 0.012 | 0.012 | 0 | 0.33 | 0.36 | 0.52 | 0.51 | 0.33 | 0.32 |
| 10 | 120044395 | rs830762 | -1.023 | 0.293 | -0.808 | 0.125 | 0.028 | 0.054 | 0.054 | 0 | 0.64 | 0.86 | 0.73 | 0.81 | 0.7 | 0.69 |
| 10 | 120059997 | rs7068621 | -0.137 | 0.044 | -0.341 | -0.139 | -0.02 | -0.063 | -0.063 | 0.766 | 0.75 | 0.49 | 0.67 | 0.79 | 0.78 | 0.81 |
| 10 | 120063661 | rs17097460 | 1.282 | 0.027 | -0.208 | -0.126 | -0.006 | 0.007 | 0.007 | 0 | 0.1 | 0.17 | 0.23 | 0.2 | 0.1 | 0.04 |
| 10 | 120067450 | rs17618810 | 1.815 | -0.076 | -0.817 | -0.263 | -0.064 | -0.022 | -0.022 | 0 | 0.28 | 0.05 | 0.28 | 0.18 | 0.34 | 0.26 |
| 10 | 120070427 | rs4751651 | 0.805 | -0.051 | -0.709 | -0.098 | -0.02 | 0.005 | 0.005 | 0.001 | 0.51 | 0.2 | 0.53 | 0.45 | 0.54 | 0.48 |
| 10 | 120070709 | rs3213691 | 0.723 | -0.1 | -0.724 | -0.201 | -0.019 | -0.019 | -0.019 | 0 | 0.5 | 0.16 | 0.53 | 0.41 | 0.54 | 0.47 |
| 10 | 120071546 | rs853907 | 0.533 | -0.447 | -0.813 | -0.369 | -0.016 | -0.032 | -0.032 | 0.374 | 0.56 | 0.23 | 0.47 | 0.4 | 0.47 | 0.5 |
| 10 | 120073905 | rs6585481 | 0.623 | -0.437 | -0.533 | -0.248 | -0.009 | -0.01 | -0.01 | 0 | 0.53 | 0.22 | 0.56 | 0.46 | 0.59 | 0.5 |
| 10 | 120079195 | rs4752151 | 0.904 | -0.057 | -0.794 | -0.03 | 0.015 | -0.038 | -0.038 | 0.345 | 0.21 | 0.09 | 0.26 | 0.29 | 0.2 | 0.24 |
| 10 | 120082538 | rs10886222 | 1.138 | 0.427 | -0.128 | 0.115 | -0.001 | 0 | 0 | 0 | 0.63 | 0.34 | 0.79 | 0.65 | 0.69 | 0.55 |
| 10 | 120096866 | rs1935917 | 0.466 | 1.475 | -1.011 | 0.627 | 0.006 | 0.014 | 0.014 | 0 | 0.16 | 0.16 | 0.12 | 0.35 | 0.23 | 0.26 |
| 10 | 120099005 | rs236195 | 0.237 | 0.492 | -0.716 | 0.076 | -0.008 | -0.037 | -0.037 | 0.063 | 0.12 | 0.16 | 0.05 | 0.01 | 0.07 | 0.12 |
| 10 | 120106540 | rs236217 | -0.395 | 0.901 | -0.877 | 0.508 | 0.005 | 0.005 | 0.005 | 0 | 0.17 | 0.2 | 0.26 | 0.26 | 0.24 | 0.16 |
| 10 | 120109482 | rs11198373 | 0.587 | 0.782 | -1.828 | 0.342 | -0.068 | -0.17 | -0.17 | 0 | 0.22 | 0.02 | 0.2 | 0.41 | 0.22 | 0.24 |
| 10 | 120117595 | rs703447 | -0.787 | 0.049 | -1.241 | 0.052 | 0.008 | -0.008 | -0.008 | 0 | 0.51 | 0.77 | 0.66 | 0.53 | 0.47 | 0.48 |
| 10 | 120118598 | rs236212 | 0.273 | 0 | -1.444 | 0.037 | -0.028 | -0.042 | -0.042 | 0.428 | 0.16 | 0.08 | 0.28 | 0.26 | 0.21 | 0.17 |
| 10 | 120119237 | rs1930971 | 0.641 | -0.017 | -1.391 | 0.025 | -0.014 | -0.032 | -0.032 | 0 | 0.48 | 0.14 | 0.34 | 0.47 | 0.53 | 0.52 |
| 10 | 120122016 | rs236213 | 0.333 | 0.169 | -1.431 | 0.029 | -0.005 | -0.038 | -0.038 | 0 | 0.14 | 0.07 | 0.24 | 0.26 | 0.15 | 0.14 |
| 10 | 120122033 | rs236214 | 0.48 | -0.053 | -1.148 | -0.118 | -0.004 | 0.009 | 0.009 | 0 | 0.24 | 0.21 | 0.36 | 0.23 | 0.28 | 0.21 |
| 10 | 120140042 | rs10886227 | 0.716 | -0.16 | -1.32 | -0.062 | -0.005 | -0.018 | -0.018 | 0 | 0.48 | 0.19 | 0.35 | 0.47 | 0.52 | 0.51 |
| 10 | 120140127 | rs78756 | 0.368 | -0.094 | -0.897 | 0.054 | 0.004 | 0.013 | 0.013 | 0 | 0.21 | 0.08 | 0.31 | 0.23 | 0.22 | 0.19 |
| 10 | 120142022 | rs180481 | -0.034 | 0.031 | -0.247 | 0.168 | -0.008 | 0.042 | 0.042 | 0 | 0.26 | 0.31 | 0.41 | 0.43 | 0.25 | 0.3 |
| 10 | 120144787 | rs236205 | 0.65 | 0.048 | -0.063 | -0.47 | 0 | -0.017 | -0.017 | 0.123 | 0.2 | 0.18 | 0.18 | 0.01 | 0.2 | 0.13 |
| 10 | 120146612 | rs10787819 | 0.419 | 0.146 | 0.189 | -0.571 | -0.001 | 0.003 | 0.003 | 0.303 | 0.43 | 0.4 | 0.32 | 0.58 | 0.41 | 0.42 |
| 10 | 120152304 | rs853922 | -0.584 | 0.187 | 0.223 | -0.649 | 0.01 | -0.028 | -0.028 | 0 | 0.92 | 0.92 | 0.92 | 1 | 0.86 | 0.92 |
| 10 | 120153201 | rs9325554 | 0.652 | 0.298 | -0.027 | -0.967 | -0.03 | -0.131 | -0.131 | 2.143 | 0.22 | 0.04 | 0.19 | 0.51 | 0.24 | 0.17 |
| 10 | 120154436 | rs10490913 | 0.377 | 0.313 | 0.187 | -0.824 | -0.011 | -0.019 | -0.019 | 2.785 | 0.43 | 0.31 | 0.29 | 0.54 | 0.41 | 0.37 |
| 10 | 120154866 | rs853925 | 1.386 | 0.26 | 0.177 | -0.847 | -0.007 | -0.085 | -0.085 | 0 | 0.2 | 0.04 | 0.35 | 0.4 | 0.2 | 0.29 |

| | | | | | | | | | | | | | | | | |
|---|---|---|---|---|---|---|---|---|---|---|---|---|---|---|---|---|
| 10 | 120155733 | rs853926 | 0.359 | 0.457 | 0.434 | -0.394 | 0.002 | 0.002 | 0.002 | 0 | 0.47 | 0.47 | 0.58 | 0.41 | 0.45 | 0.5 |
| 10 | 120157792 | rs10128523 | -0.082 | 0.535 | -0.024 | -0.461 | -0.005 | 0.005 | 0.005 | 0.321 | 0.43 | 0.34 | 0.29 | 0.56 | 0.41 | 0.43 |
| 10 | 120159880 | rs2153953 | 0.006 | 0.883 | 0.002 | -0.21 | 0.001 | 0.001 | 0.001 | 0 | 0.38 | 0.4 | 0.28 | 0.57 | 0.37 | 0.42 |
| 10 | 120162568 | rs853937 | -0.084 | 0.823 | 0.028 | -0.248 | 0.021 | -0.044 | -0.044 | 0 | 0.91 | 0.85 | 0.91 | 1 | 0.86 | 0.92 |
| 10 | 120164251 | rs853943 | 0.454 | 0.811 | 0.044 | -0.253 | 0 | 0.006 | 0.006 | 0 | 0.52 | 0.48 | 0.6 | 0.38 | 0.49 | 0.46 |
| 10 | 120164906 | rs766478 | -0.208 | 0.591 | -0.133 | 0.206 | -0.002 | -0.006 | -0.006 | 0 | 0.38 | 0.34 | 0.28 | 0.54 | 0.36 | 0.42 |
| 10 | 120168873 | rs11198393 | -0.306 | 0.652 | -0.215 | 0.091 | -0.001 | -0.005 | -0.005 | 0 | 0.39 | 0.43 | 0.32 | 0.62 | 0.37 | 0.47 |
| 10 | 120172638 | rs1857269 | -0.294 | 0.688 | -0.507 | 0.219 | -0.001 | 0.001 | 0.001 | 0 | 0.39 | 0.36 | 0.31 | 0.62 | 0.37 | 0.48 |
| 10 | 120175019 | rs4752158 | -0.049 | 0.66 | -1.013 | 0.701 | -0.045 | -0.126 | -0.126 | 0 | 0.15 | 0.02 | 0.18 | 0.5 | 0.19 | 0.23 |
| 10 | 120175805 | rs1418895 | -0.073 | 0.911 | -0.524 | 0.949 | -0.002 | 0.023 | 0.023 | 0 | 0.12 | 0.11 | 0.15 | 0.39 | 0.16 | 0.18 |
| 10 | 120176423 | rs853956 | 0.297 | 1.11 | -0.12 | 0.206 | 0.01 | -0.004 | -0.004 | 0 | 0.27 | 0.34 | 0.39 | 0.31 | 0.31 | 0.24 |
| 10 | 120180717 | rs11198403 | 1.155 | 1.003 | -0.673 | -0.177 | -0.044 | 0.134 | 0.134 | 0 | 0.21 | 0.02 | 0.16 | 0.01 | 0.21 | 0.13 |
| 10 | 120181561 | rs2153952 | 0.025 | 1.113 | -0.475 | 0.056 | -0.014 | -0.026 | -0.026 | 0.204 | 0.14 | 0.11 | 0.25 | 0.58 | 0.2 | 0.26 |
| 10 | 120185902 | rs1981304 | 0.484 | 1.311 | -0.306 | -0.072 | 0.03 | -0.033 | -0.033 | 0.033 | 0.08 | 0.13 | 0.1 | 0 | 0.13 | 0.07 |
| 10 | 120187722 | rs1013620 | 0.171 | 1.275 | -0.326 | -0.052 | -0.015 | -0.026 | -0.026 | 1.06 | 0.14 | 0.09 | 0.25 | 0.58 | 0.2 | 0.25 |
| 10 | 120192326 | rs10886244 | -1.202 | 1.216 | -0.041 | -0.128 | 0.024 | 0.005 | 0.005 | 0.059 | 0.25 | 0.32 | 0.38 | 0.66 | 0.35 | 0.38 |
| 10 | 120197496 | rs1004421 | 0.331 | 1.246 | -0.048 | -0.121 | 0.038 | -0.038 | -0.038 | 0 | 0.08 | 0.17 | 0.1 | 0 | 0.12 | 0.06 |
| 10 | 120198913 | rs12413353 | 1.317 | 1.191 | 0.166 | -0.213 | -0.002 | 0.003 | 0.003 | 0 | 0.19 | 0.21 | 0.29 | 0.3 | 0.18 | 0.18 |
| 10 | 120204872 | rs729700 | -0.102 | 1.264 | 0.135 | -0.215 | 0.02 | 0.007 | 0.007 | 0 | 0.1 | 0.16 | 0.13 | 0.08 | 0.14 | 0.13 |
| 10 | 120206869 | rs7091683 | -1.067 | 1.117 | 0.262 | -0.206 | 0.034 | 0.02 | 0.02 | 0.071 | 0.25 | 0.35 | 0.31 | 0.62 | 0.34 | 0.38 |
| 10 | 120209516 | rs874231 | -0.426 | 0.871 | 0.436 | -0.348 | -0.003 | 0.028 | 0.028 | 0.245 | 0.46 | 0.34 | 0.43 | 0.64 | 0.51 | 0.51 |
| 10 | 120210112 | rs11198424 | 0.288 | 1.361 | 0.997 | 0.385 | 0.015 | 0.076 | 0.076 | 0 | 0.13 | 0.18 | 0.16 | 0.44 | 0.19 | 0.16 |
| 10 | 120210168 | rs7088823 | 0.653 | 1.357 | 0.977 | 0.26 | 0.001 | -0.002 | -0.002 | 0.036 | 0.6 | 0.4 | 0.38 | 0.24 | 0.53 | 0.6 |
| 10 | 120210221 | rs7088624 | 0.486 | 0.97 | 0.859 | 0.527 | 0.003 | -0.009 | -0.009 | 0 | 0.64 | 0.55 | 0.45 | 0.24 | 0.59 | 0.61 |
| 10 | 120212789 | rs1932542 | -0.269 | 1.227 | 0.27 | 0.722 | -0.015 | -0.019 | -0.019 | 0 | 0.15 | 0.11 | 0.19 | 0.55 | 0.23 | 0.24 |
| 10 | 120214924 | rs12220340 | 0.372 | 1.245 | 0.217 | 0.666 | -0.024 | -0.029 | -0.029 | 0.008 | 0.12 | 0.04 | 0.15 | 0.39 | 0.19 | 0.16 |
| 10 | 120215198 | rs1932543 | -0.902 | 1.053 | 0.369 | -0.02 | 0 | -0.061 | -0.061 | 0 | 0.38 | 0.41 | 0.59 | 0.96 | 0.43 | 0.5 |
| 10 | 120215827 | rs11198431 | 0.231 | 0.662 | 0.102 | 0.506 | 0.003 | -0.008 | -0.008 | 0 | 0.19 | 0.17 | 0.35 | 0.33 | 0.18 | 0.18 |
| 10 | 120216655 | rs11198434 | 0.294 | 0.731 | 0.129 | 0.533 | 0.001 | -0.004 | -0.004 | 3.925 | 0.18 | 0.17 | 0.35 | 0.33 | 0.18 | 0.16 |
| 10 | 120216783 | rs1932544 | 0.336 | 0.741 | 0.169 | 0.51 | -0.001 | 0.001 | 0.001 | 0 | 0.17 | 0.18 | 0.36 | 0.31 | 0.18 | 0.16 |
| 10 | 120217225 | rs11198435 | -0.773 | 0.99 | 0.761 | 0.205 | 0.03 | -0.038 | -0.038 | 0 | 0.1 | 0.28 | 0.06 | 0.03 | 0.11 | 0.15 |
| 10 | 120218162 | rs11819557 | -1.163 | 0.666 | 1.564 | 0.062 | -0.003 | -0.045 | -0.045 | 0 | 0.1 | 0.3 | 0.05 | 0.03 | 0.08 | 0.17 |
| 10 | 120218268 | rs2184699 | 1.637 | 0.63 | 1.692 | 0.046 | 0.004 | -0.068 | -0.068 | 0 | 0.87 | 0.45 | 0.92 | 0.97 | 0.92 | 0.82 |
| 10 | 120220709 | rs6585487 | -0.445 | 0.442 | 0.954 | -0.435 | -0.011 | -0.04 | -0.04 | 0 | 0.66 | 0.62 | 0.72 | 0.92 | 0.71 | 0.73 |
| 10 | 120221323 | rs11198436 | 0.859 | 1.031 | 0.025 | 0.012 | -0.071 | -0.104 | -0.104 | 0 | 0.15 | 0.02 | 0.17 | 0.36 | 0.22 | 0.17 |
| 10 | 120222457 | rs4411217 | 0.066 | 0.771 | 0.459 | -0.105 | 0.013 | 0.028 | 0.028 | 0.559 | 0.57 | 0.34 | 0.67 | 0.89 | 0.65 | 0.61 |
| 10 | 120229090 | rs11198439 | 0.482 | 0.973 | -0.687 | 0.141 | -0.027 | -0.014 | -0.014 | 0 | 0.49 | 0.08 | 0.47 | 0.62 | 0.55 | 0.44 |
| 10 | 120237076 | rs10510044 | 0.148 | 0.818 | -0.368 | -0.149 | 0.014 | 0.064 | 0.064 | 0 | 0.32 | 0.18 | 0.27 | 0.08 | 0.3 | 0.29 |
| 10 | 120239987 | rs12263039 | -0.499 | 0.863 | 0.223 | -0.266 | -0.001 | -0.055 | -0.055 | 0.142 | 0.23 | 0.37 | 0.2 | 0.07 | 0.21 | 0.19 |
| 10 | 120240041 | rs7088160 | 0.099 | 0.591 | 0.597 | -0.386 | -0.025 | -0.019 | -0.019 | 0 | 0.93 | 0.78 | 0.97 | 0.97 | 0.97 | 0.88 |
| 10 | 120242187 | rs2420477 | -1.021 | 0.904 | 0.369 | -0.174 | -0.001 | -0.001 | -0.001 | 0 | 0.51 | 0.47 | 0.41 | 0.53 | 0.52 | 0.56 |
| 10 | 120242212 | rs11198446 | 0.032 | 0.588 | 0.228 | -0.103 | -0.024 | -0.018 | -0.018 | 0.137 | 0.07 | 0.23 | 0.03 | 0.03 | 0.03 | 0.12 |
| 10 | 120242929 | rs885545 | 0.164 | 0.599 | 0.084 | -0.098 | -0.025 | -0.019 | -0.019 | 0 | 0.06 | 0.18 | 0.02 | 0.03 | 0.03 | 0.12 |
| 10 | 120245474 | rs4751662 | -0.346 | 0.857 | -0.095 | -0.091 | 0.007 | -0.062 | -0.062 | 1.363 | 0.38 | 0.15 | 0.32 | 0.49 | 0.37 | 0.37 |
| 10 | 120248460 | rs7071466 | -0.846 | 1.414 | 0.049 | -0.075 | -0.001 | -0.007 | -0.007 | 0 | 0.47 | 0.53 | 0.48 | 0.68 | 0.56 | 0.56 |
| 10 | 120251996 | rs945488 | -0.568 | 2.086 | 0.425 | 0.06 | 0.02 | 0.008 | 0.008 | 0 | 0.36 | 0.55 | 0.38 | 0.59 | 0.51 | 0.37 |
| 10 | 120252855 | rs945491 | -0.269 | 2.597 | 0.658 | -0.007 | 0.021 | 0.017 | 0.017 | 0 | 0.29 | 0.44 | 0.32 | 0.59 | 0.4 | 0.32 |
| 10 | 120253399 | rs12242910 | 0.511 | 2.598 | 0.648 | -0.011 | 0.015 | -0.018 | -0.018 | 0.362 | 0.06 | 0.08 | 0.06 | 0 | 0.11 | 0.05 |
| 10 | 120254372 | rs10886259 | -0.77 | 2.458 | 0.637 | 0.004 | 0.02 | 0.008 | 0.008 | 0 | 0.36 | 0.55 | 0.38 | 0.59 | 0.51 | 0.38 |
| 10 | 120254734 | rs11198458 | 1.427 | 1.716 | 0.346 | -0.474 | 0.066 | 0.053 | 0.053 | 0.073 | 0.41 | 0.17 | 0.37 | 0.33 | 0.28 | 0.41 |
| 10 | 120254909 | rs10787841 | -0.681 | 1.616 | 0.412 | -0.507 | -0.002 | -0.01 | -0.01 | 0 | 0.77 | 0.81 | 0.76 | 0.92 | 0.78 | 0.79 |
| 10 | 120260837 | rs2420480 | -0.438 | 2.003 | 0.487 | -0.315 | 0.016 | 0.012 | 0.012 | 0 | 0.36 | 0.53 | 0.37 | 0.56 | 0.5 | 0.38 |
| 10 | 120277577 | rs667335 | -1.751 | 1.308 | 0.485 | -0.502 | -0.019 | -0.006 | -0.006 | 0 | 0.63 | 0.63 | 0.72 | 0.63 | 0.75 | 0.6 |
| 10 | 120279177 | rs483523 | 0.417 | 1.321 | 0.48 | -0.559 | -0.004 | -0.021 | -0.021 | 0 | 0.22 | 0.23 | 0.25 | 0.08 | 0.23 | 0.18 |
| 10 | 120283292 | rs4523612 | 0.337 | 1.56 | 0.326 | -0.186 | -0.093 | -0.048 | -0.048 | 0 | 0.33 | 0.1 | 0.42 | 0.37 | 0.43 | 0.31 |
| 10 | 120292312 | rs645662 | -1.854 | 1.429 | 0.441 | -0.192 | -0.017 | -0.006 | -0.006 | 0 | 0.63 | 0.64 | 0.72 | 0.63 | 0.75 | 0.61 |
| 10 | 120301859 | rs17097790 | 1.681 | 1.207 | 0.62 | -0.207 | -0.008 | 0.005 | 0.005 | 0 | 0.13 | 0.05 | 0.16 | 0.08 | 0.17 | 0.13 |
| 10 | 120304062 | rs1002525 | 0.279 | 1.23 | 0.57 | -0.223 | -0.112 | -0.045 | -0.045 | 0 | 0.34 | 0.1 | 0.45 | 0.37 | 0.48 | 0.29 |
| 10 | 120309813 | rs626114 | 1.573 | 1.032 | 0.762 | -0.291 | -0.024 | -0.025 | -0.025 | 0 | 0.05 | 0.12 | 0.02 | 0 | 0.01 | 0.03 |
| 10 | 120311534 | rs477874 | -1.958 | 1.047 | 0.692 | -0.242 | -0.047 | -0.011 | -0.011 | 0 | 0.64 | 0.57 | 0.75 | 0.63 | 0.8 | 0.58 |
| 10 | 120326513 | rs10886271 | 1.944 | 1.018 | 0.634 | -0.239 | 0.044 | 0.005 | 0.005 | 0.159 | 0.34 | 0.19 | 0.21 | 0.36 | 0.2 | 0.41 |
| 10 | 120327457 | rs2676830 | -0.5 | 1.202 | 0.45 | -0.293 | 0.001 | -0.018 | -0.018 | 0 | 0.27 | 0.27 | 0.28 | 0.08 | 0.29 | 0.24 |
| 10 | 120328074 | rs1105500 | -0.787 | 1.475 | 0.333 | -0.317 | 0.017 | 0.002 | 0.002 | 0.598 | 0.51 | 0.42 | 0.6 | 0.45 | 0.63 | 0.51 |
| 10 | 120330259 | rs1711882 | 0.305 | 0.896 | 0.326 | -0.267 | -0.012 | -0.038 | -0.038 | 0 | 0.2 | 0.33 | 0.2 | 0.08 | 0.19 | 0.17 |
| 10 | 120332025 | rs1711879 | -1.546 | 1.126 | 0.26 | 0.081 | -0.011 | -0.003 | -0.003 | 0 | 0.59 | 0.58 | 0.66 | 0.6 | 0.72 | 0.52 |
| 10 | 120333733 | rs17097816 | 1.788 | 1.362 | -0.273 | 0.176 | -0.032 | -0.013 | -0.013 | 0 | 0.11 | 0.02 | 0.16 | 0.08 | 0.16 | 0.13 |
| 10 | 120334125 | rs12569768 | 0.262 | 1.311 | -0.158 | 0.212 | -0.005 | -0.005 | -0.005 | 0.263 | 0.19 | 0.22 | 0.2 | 0.27 | 0.25 | 0.23 |
| 10 | 120335636 | rs10886273 | 0.273 | 1.275 | -0.233 | 0.167 | -0.095 | -0.002 | -0.002 | 0 | 0.33 | 0.09 | 0.42 | 0.31 | 0.44 | 0.24 |
| 10 | 120342865 | rs1711860 | 0.441 | 1.053 | 0.142 | 0.235 | -0.013 | -0.043 | -0.043 | 2.216 | 0.08 | 0.15 | 0.11 | 0 | 0.06 | 0.05 |

| | | | | | | | | | | | | | | | | |
|---|---|---|---|---|---|---|---|---|---|---|---|---|---|---|---|---|
| 10 | 120343777 | rs1766249 | 0.411 | 1.054 | 0.16 | 0.254 | -0.009 | -0.043 | -0.043 | 0.045 | 0.08 | 0.21 | 0.11 | 0.01 | 0.06 | 0.06 |
| 10 | 120348226 | rs1766253 | -0.657 | 0.873 | -0.327 | 0.055 | 0.001 | 0.007 | 0.007 | 0 | 0.47 | 0.45 | 0.57 | 0.28 | 0.56 | 0.42 |
| 10 | 120348368 | rs2901164 | 0.444 | 0.852 | -0.951 | 0.12 | -0.135 | 0.061 | 0.061 | 0.289 | 0.37 | 0.08 | 0.45 | 0.26 | 0.5 | 0.28 |
| 10 | 120350036 | rs2292767 | 1.333 | 1.507 | -0.893 | 0.28 | -0.12 | -0.024 | -0.024 | 0 | 0.25 | 0.05 | 0.27 | 0.25 | 0.37 | 0.23 |
| 10 | 120353422 | rs1711869 | -0.386 | 1.123 | -0.265 | 0.245 | 0.028 | -0.032 | -0.032 | 0 | 0.11 | 0.22 | 0.25 | 0.02 | 0.13 | 0.09 |
| 10 | 120354472 | rs938700 | 1.15 | 1.11 | -0.37 | 0.249 | -0.002 | -0.017 | -0.017 | 0.032 | 0.06 | 0.07 | 0.07 | 0 | 0.05 | 0.05 |
| 10 | 120354818 | rs3740551 | 0.687 | 0.95 | -0.159 | 0.365 | 0.025 | 0.019 | 0.019 | 1.21 | 0.42 | 0.26 | 0.27 | 0.38 | 0.29 | 0.48 |
| 10 | 120357856 | rs4752177 | 0.461 | 1.155 | 0.017 | -0.033 | 0.043 | -0.08 | -0.08 | 0 | 0.82 | 0.6 | 0.71 | 0.98 | 0.82 | 0.86 |
| 10 | 120363944 | rs4751667 | 0.595 | 0.327 | -0.348 | 0.402 | 0.02 | 0.005 | 0.005 | 0.96 | 0.41 | 0.28 | 0.27 | 0.64 | 0.26 | 0.49 |
| 10 | 120366341 | rs11198499 | -0.198 | 0.493 | -0.313 | 0.362 | -0.003 | 0.008 | 0.008 | 0 | 0.53 | 0.32 | 0.46 | 0.4 | 0.5 | 0.63 |
| 10 | 120368581 | rs881902 | 1.068 | 0.648 | -0.176 | 0.523 | -0.015 | -0.03 | -0.03 | 0.294 | 0.29 | 0.22 | 0.26 | 0.43 | 0.33 | 0.25 |
| 10 | 120371322 | rs11198510 | 0.677 | 0.687 | -0.059 | 0.508 | 0.006 | 0.008 | 0.008 | 0 | 0.33 | 0.39 | 0.31 | 0.43 | 0.37 | 0.28 |
| 10 | 120371617 | rs1247109 | -0.793 | 0.661 | -0.39 | 0.18 | -0.019 | 0.054 | 0.054 | 0 | 0.72 | 0.86 | 0.75 | 0.87 | 0.68 | 0.67 |
| 10 | 120382875 | rs1247158 | -0.65 | 0.565 | -0.145 | 0.16 | 0.001 | 0.03 | 0.03 | 0 | 0.7 | 0.79 | 0.77 | 0.9 | 0.7 | 0.69 |
| 10 | 120383625 | rs1247157 | 0.574 | 0.231 | -0.193 | 0.197 | -0.003 | 0.009 | 0.009 | 0.174 | 0.31 | 0.33 | 0.43 | 0.48 | 0.28 | 0.37 |
| 10 | 120392479 | rs2765473 | 0.015 | 0.535 | -0.148 | -0.179 | -0.001 | 0.007 | 0.007 | 0 | 0.65 | 0.71 | 0.72 | 0.9 | 0.65 | 0.65 |
| 10 | 120398369 | rs4752183 | 0.602 | 0.57 | -0.278 | -0.165 | 0.005 | -0.04 | -0.04 | 0 | 0.58 | 0.54 | 0.54 | 0.86 | 0.56 | 0.57 |
| 10 | 120400613 | rs2765471 | -0.113 | 0.763 | -0.349 | 0.314 | -0.002 | -0.001 | -0.001 | 0 | 0.07 | 0.06 | 0.18 | 0.01 | 0.09 | 0.06 |
| 10 | 120423184 | rs873598 | -0.039 | 0.691 | -0.425 | 0.395 | -0.002 | 0 | 0 | 0 | 0.39 | 0.41 | 0.5 | 0.62 | 0.4 | 0.42 |
| 10 | 120426849 | rs1247122 | -0.214 | 0.581 | -0.081 | 0.346 | 0.002 | -0.002 | -0.002 | 0.022 | 0.59 | 0.39 | 0.47 | 0.38 | 0.6 | 0.57 |
| 10 | 120428781 | rs17097926 | 1.042 | 0.621 | -0.074 | 0.666 | 0.001 | 0.006 | 0.006 | 0 | 0.26 | 0.29 | 0.22 | 0.4 | 0.29 | 0.3 |
| 10 | 120432485 | rs3887073 | -1.087 | 0.69 | -0.203 | 0.619 | -0.003 | -0.004 | -0.004 | 2.742 | 0.12 | 0.13 | 0.27 | 0.22 | 0.1 | 0.12 |
| 10 | 120436448 | rs10510045 | 1.246 | 0.152 | -0.728 | 0.192 | 0.03 | 0.1 | 0.1 | 4.281 | 0.2 | 0.02 | 0.08 | 0.1 | 0.16 | 0.17 |
| 10 | 120442882 | rs17662895 | -0.344 | 0.267 | -0.667 | 0.24 | -0.008 | 0.006 | 0.006 | 0 | 0.09 | 0.09 | 0.15 | 0.02 | 0.09 | 0.08 |
| 10 | 120447437 | rs10886285 | 1.474 | 0.134 | -0.268 | 0.136 | -0.002 | -0.005 | -0.005 | 2.939 | 0.26 | 0.24 | 0.21 | 0.4 | 0.29 | 0.3 |
| 10 | 120448570 | rs3824821 | -1.117 | 0.15 | -0.236 | 0.204 | 0.029 | 0.091 | 0.091 | 0 | 0.11 | 0.32 | 0.29 | 0.22 | 0.09 | 0.11 |
| 10 | 120453168 | rs10510046 | -1.181 | 0.15 | -0.245 | 0.204 | 0.031 | 0.095 | 0.095 | 0 | 0.12 | 0.33 | 0.29 | 0.22 | 0.09 | 0.11 |
| 10 | 120463720 | rs10886289 | -1.115 | -0.675 | -0.361 | 0.218 | -0.01 | 0.003 | 0.003 | 2.582 | 0.77 | 0.75 | 0.86 | 0.71 | 0.8 | 0.78 |
| 10 | 120480163 | rs17586536 | 0.83 | -0.644 | -0.59 | 0.101 | -0.093 | 0.115 | 0.115 | 0 | 0.32 | 0.02 | 0.26 | 0.08 | 0.37 | 0.34 |
| 10 | 120480688 | rs1925943 | 1.293 | -0.537 | -0.405 | 0.177 | 0.006 | 0.013 | 0.013 | 0 | 0.28 | 0.32 | 0.23 | 0.4 | 0.3 | 0.3 |
| 10 | 120493700 | rs7099523 | 0.595 | -0.594 | -0.101 | 0.36 | 0.011 | -0.016 | -0.016 | 1.217 | 0.61 | 0.35 | 0.49 | 0.48 | 0.67 | 0.64 |
| 10 | 120508692 | rs17587238 | -0.164 | -0.547 | -0.273 | 0.202 | 0.004 | -0.007 | -0.007 | 1.637 | 0.06 | 0.08 | 0.15 | 0.02 | 0.08 | 0.08 |
| 10 | 120515182 | rs4319431 | -0.572 | -0.515 | -0.182 | 0.41 | -0.019 | -0.007 | -0.007 | 0 | 0.72 | 0.68 | 0.79 | 0.7 | 0.77 | 0.77 |
| 10 | 120522232 | rs10886296 | 0.862 | -0.301 | 0.253 | 0.552 | -0.012 | -0.015 | -0.015 | 0 | 0.29 | 0.23 | 0.33 | 0.59 | 0.32 | 0.34 |
| 10 | 120523463 | rs4752200 | -1.013 | -0.268 | 0.282 | 0.459 | -0.001 | -0.001 | -0.001 | 0 | 0.74 | 0.77 | 0.83 | 0.73 | 0.76 | 0.77 |
| 10 | 120523997 | rs12358672 | -0.661 | -0.242 | 0.341 | 0.369 | 0.058 | -0.003 | -0.003 | 0 | 0.06 | 0.3 | 0.06 | 0.02 | 0.06 | 0.04 |
| 10 | 120526300 | rs10886297 | -1.011 | -0.246 | 0.129 | 0.492 | -0.001 | 0.001 | 0.001 | 1.183 | 0.74 | 0.76 | 0.75 | 0.69 | 0.76 | 0.77 |
| 10 | 120527394 | rs7080934 | -0.207 | -0.236 | 0.031 | 0.474 | -0.002 | -0.008 | -0.008 | 0 | 0.67 | 0.44 | 0.53 | 0.67 | 0.7 | 0.71 |
| 10 | 120531816 | rs11198602 | 1.161 | -0.3 | -0.196 | 0.502 | -0.075 | -0.085 | -0.085 | 0.777 | 0.27 | 0.07 | 0.2 | 0.39 | 0.31 | 0.31 |
| 10 | 120535370 | rs7917438 | 0.759 | -0.511 | -0.244 | 0.345 | 0.038 | 0.094 | 0.094 | 0 | 0.33 | 0.09 | 0.22 | 0.1 | 0.3 | 0.34 |
| 10 | 120539995 | rs10886304 | -0.294 | -0.088 | 0.002 | 0.787 | -0.009 | 0.025 | 0.025 | 0.193 | 0.7 | 0.54 | 0.65 | 0.62 | 0.72 | 0.75 |
| 10 | 120542084 | rs10886305 | 0.942 | -0.022 | -0.526 | 0.728 | -0.075 | -0.078 | -0.078 | 0 | 0.26 | 0.07 | 0.17 | 0.35 | 0.31 | 0.31 |
| 10 | 120542167 | rs10749270 | -0.388 | -0.14 | 0.045 | 0.943 | 0 | 0.02 | 0.02 | 0.292 | 0.72 | 0.72 | 0.6 | 0.51 | 0.73 | 0.69 |
| 10 | 120545165 | rs11813458 | -0.944 | -0.434 | 0.171 | 0.595 | -0.005 | -0.039 | -0.039 | 0 | 0.12 | 0.18 | 0.15 | 0.02 | 0.1 | 0.07 |
| 10 | 120548139 | rs7096525 | 0.385 | -0.629 | 0.167 | 0.775 | -0.001 | 0 | 0 | 0 | 0.3 | 0.24 | 0.28 | 0.3 | 0.33 | 0.32 |
| 10 | 120561875 | rs10787862 | -0.186 | -0.746 | 0.27 | 0.803 | 0.022 | -0.01 | -0.01 | 0 | 0.54 | 0.75 | 0.47 | 0.49 | 0.49 | 0.47 |
| 10 | 120566921 | rs7902478 | 0.453 | -0.799 | 0.396 | 0.981 | -0.011 | 0.023 | 0.023 | 0 | 0.35 | 0.52 | 0.45 | 0.38 | 0.39 | 0.39 |
| 10 | 120568512 | rs10749271 | -1.016 | -0.787 | 0.374 | 0.93 | 0.008 | 0.005 | 0.005 | 0 | 0.66 | 0.66 | 0.66 | 0.57 | 0.61 | 0.61 |
| 10 | 120571732 | rs7917673 | -0.985 | -0.978 | 0.434 | 0.806 | 0.002 | 0.002 | 0.002 | 0 | 0.65 | 0.69 | 0.57 | 0.51 | 0.58 | 0.52 |
| 10 | 120572246 | rs11596456 | -0.039 | -1.035 | 0.064 | 0.555 | 0.044 | 0.044 | 0.044 | 1.464 | 0.24 | 0.09 | 0.11 | 0.16 | 0.14 | 0.21 |
| 10 | 120573988 | rs10886311 | -0.156 | -0.893 | 0.002 | 0.678 | 0.027 | 0.023 | 0.023 | 3.824 | 0.26 | 0.12 | 0.21 | 0.23 | 0.21 | 0.18 |
| 10 | 120575178 | rs17098065 | -0.723 | -0.682 | 0.481 | 0.575 | 0.011 | 0.054 | 0.054 | 0 | 0.1 | 0.27 | 0.18 | 0.17 | 0.12 | 0.05 |
| 10 | 120577228 | rs12573699 | 0.79 | -0.762 | 0.039 | 0.357 | 0.002 | -0.008 | -0.008 | 1.29 | 0.3 | 0.18 | 0.36 | 0.31 | 0.35 | 0.33 |
| 10 | 120577700 | rs876414 | 0.85 | -0.668 | 0.043 | 0.397 | 0 | -0.001 | -0.001 | 0 | 0.32 | 0.34 | 0.38 | 0.25 | 0.34 | 0.33 |
| 10 | 120577833 | rs876413 | -0.632 | -0.447 | -0.385 | 0.521 | -0.006 | -0.018 | -0.018 | 0.142 | 0.59 | 0.24 | 0.52 | 0.59 | 0.57 | 0.5 |
| 10 | 120577900 | rs12763514 | 0.772 | -1.23 | -0.203 | -0.102 | 0.015 | 0.01 | 0.01 | 3.841 | 0.19 | 0.14 | 0.11 | 0.12 | 0.12 | 0.12 |
| 10 | 120582664 | rs10886317 | -0.007 | -1.053 | -0.393 | 0.022 | 0.024 | 0.012 | 0.012 | 4.78 | 0.27 | 0.26 | 0.22 | 0.22 | 0.2 | 0.19 |
| 10 | 120584211 | rs10787868 | -0.441 | -1.063 | -0.434 | 0.012 | 0.015 | 0.013 | 0.013 | 2.278 | 0.33 | 0.32 | 0.24 | 0.22 | 0.23 | 0.23 |
| 10 | 120585915 | rs4752208 | 0.841 | -0.912 | -0.356 | 0.03 | -0.002 | -0.025 | -0.025 | 0 | 0.33 | 0.43 | 0.39 | 0.24 | 0.35 | 0.34 |
| 10 | 120586332 | rs4752212 | -0.424 | -0.575 | -0.534 | 0.224 | 0.009 | 0.001 | 0.001 | 0 | 0.63 | 0.6 | 0.6 | 0.41 | 0.56 | 0.48 |
| 10 | 120587265 | rs751973 | -0.852 | -0.208 | -0.783 | 0.252 | -0.02 | -0.016 | -0.016 | 0 | 0.69 | 0.77 | 0.64 | 0.48 | 0.61 | 0.64 |
| 10 | 120588918 | rs7098657 | -0.202 | 0.229 | -0.724 | 0.032 | -0.007 | -0.026 | -0.026 | 0 | 0.37 | 0.45 | 0.4 | 0.25 | 0.38 | 0.36 |
| 10 | 120592015 | rs10787871 | -0.019 | 0.575 | -1.153 | -0.071 | -0.011 | 0.044 | 0.044 | 0 | 0.59 | 0.34 | 0.66 | 0.72 | 0.57 | 0.71 |
| 10 | 120592826 | rs7092363 | 0.562 | -0.169 | -1.742 | 0.088 | 0.035 | -0.026 | -0.026 | 0 | 0.59 | 0.11 | 0.67 | 0.58 | 0.64 | 0.61 |
| 10 | 120594481 | rs4752213 | -0.295 | -0.343 | -0.983 | -0.082 | -0.005 | -0.023 | -0.023 | 0 | 0.18 | 0.17 | 0.12 | 0.06 | 0.15 | 0.08 |
| 10 | 120596386 | rs6585514 | 0.337 | -0.074 | -0.76 | -0.048 | -0.006 | -0.007 | -0.007 | 0 | 0.64 | 0.47 | 0.67 | 0.62 | 0.68 | 0.64 |
| 10 | 120603342 | rs4642993 | 0.561 | -0.123 | -0.434 | 0.066 | -0.005 | -0.018 | -0.018 | 0 | 0.59 | 0.28 | 0.6 | 0.46 | 0.56 | 0.48 |
| 10 | 120604209 | rs10886323 | -0.043 | -0.179 | -0.363 | 0.16 | -0.033 | 0.021 | 0.021 | 0 | 0.22 | 0.06 | 0.16 | 0.18 | 0.24 | 0.22 |
| 10 | 120606890 | rs7923177 | -1.984 | -0.092 | 0.286 | 0.163 | 0.046 | 0.128 | 0.128 | 0.123 | 0.05 | 0.24 | 0.07 | 0.15 | 0.07 | 0.12 |

| | | | | | | | | | | | | | | | | |
|---|---|---|---|---|---|---|---|---|---|---|---|---|---|---|---|---|
| 10 | 120613996 | rs4752217 | 0.964 | -0.451 | 0.206 | -0.125 | -0.003 | 0.04 | 0.04 | 0 | 0.46 | 0.28 | 0.57 | 0.57 | 0.54 | 0.52 |
| 10 | 120615752 | rs1409314 | 0.744 | -0.454 | 0.254 | -0.144 | 0.004 | 0.026 | 0.026 | 0 | 0.48 | 0.36 | 0.58 | 0.58 | 0.53 | 0.52 |
| 10 | 120626562 | rs10510050 | 0.654 | -0.604 | 0.171 | -0.207 | 0.022 | 0.043 | 0.043 | 0 | 0.52 | 0.34 | 0.6 | 0.62 | 0.57 | 0.55 |
| 10 | 120645820 | rs7908018 | 0.382 | -0.632 | 0.447 | -0.312 | -0.005 | -0.018 | -0.018 | 0.711 | 0.55 | 0.52 | 0.57 | 0.66 | 0.57 | 0.59 |
| 10 | 120647146 | rs1071895 | 0.644 | -0.747 | 0.231 | -0.361 | 0.015 | 0.034 | 0.034 | 0 | 0.51 | 0.46 | 0.57 | 0.66 | 0.57 | 0.53 |
| 10 | 120648726 | rs4751680 | 1.342 | -0.737 | -0.053 | 0.025 | -0.061 | -0.026 | -0.026 | 0 | 0.42 | 0.15 | 0.47 | 0.39 | 0.48 | 0.37 |
| 10 | 120650662 | rs10886332 | 0.768 | -0.719 | -0.041 | 0.066 | 0.015 | 0.008 | 0.008 | 0 | 0.48 | 0.33 | 0.51 | 0.42 | 0.51 | 0.42 |
| 10 | 120663911 | rs611959 | -0.368 | -0.578 | -0.02 | -0.072 | 0.001 | 0.006 | 0.006 | 0.265 | 0.8 | 0.87 | 0.87 | 0.93 | 0.79 | 0.81 |
| 10 | 120668089 | rs11198669 | -0.789 | -0.586 | -0.194 | -0.049 | -0.03 | -0.057 | -0.057 | 0.035 | 0.18 | 0.36 | 0.1 | 0.04 | 0.11 | 0.1 |
| 10 | 120668290 | rs12415021 | 0.084 | 0.248 | -1.213 | 0.182 | -0.04 | -0.053 | -0.053 | 0 | 0.11 | 0.03 | 0.19 | 0.24 | 0.2 | 0.21 |
| 10 | 120670912 | rs11198670 | 0.852 | -0.023 | -1.442 | -0.02 | -0.131 | -0.072 | -0.072 | 0 | 0.38 | 0.07 | 0.47 | 0.38 | 0.46 | 0.32 |
| 10 | 120672502 | rs10749275 | 0.856 | -0.021 | -1.453 | -0.039 | -0.132 | -0.079 | -0.079 | 1.124 | 0.38 | 0.07 | 0.47 | 0.39 | 0.46 | 0.31 |
| 10 | 120685596 | rs10430633 | 0.253 | -0.478 | -1.167 | -0.375 | 0.091 | 0.119 | 0.119 | 0.626 | 0.57 | 0.22 | 0.67 | 0.77 | 0.7 | 0.62 |
| 10 | 120687570 | rs10787879 | -0.408 | -0.803 | -0.472 | -0.558 | -0.044 | -0.057 | -0.057 | 0.054 | 0.7 | 0.49 | 0.75 | 0.81 | 0.8 | 0.71 |
| 10 | 120690639 | rs11198678 | 0.67 | -0.618 | -0.664 | -0.229 | 0.06 | 0.005 | 0.005 | 0.912 | 0.55 | 0.18 | 0.57 | 0.6 | 0.63 | 0.51 |
| 10 | 120690832 | rs10886342 | 1.714 | -0.069 | -0.999 | -0.362 | -0.063 | -0.088 | -0.088 | 1.264 | 0.29 | 0.09 | 0.38 | 0.45 | 0.3 | 0.25 |
| 10 | 120692771 | rs7899785 | -0.599 | -0.079 | -0.935 | -0.456 | -0.023 | -0.028 | -0.028 | 0 | 0.69 | 0.42 | 0.75 | 0.82 | 0.8 | 0.72 |
| 10 | 120697555 | rs17098212 | -0.394 | 0.055 | -1.268 | -0.427 | 0.005 | 0.027 | 0.027 | 0 | 0.14 | 0.2 | 0.18 | 0.22 | 0.17 | 0.21 |
| 10 | 120703685 | rs943271 | -0.611 | 0.041 | -1.369 | -0.441 | -0.034 | -0.01 | -0.01 | 0 | 0.4 | 0.23 | 0.35 | 0.35 | 0.5 | 0.46 |
| 10 | 120706575 | rs11198686 | 0.519 | -0.127 | -1.362 | -0.844 | -0.08 | 0.056 | 0.056 | 0.462 | 0.26 | 0.02 | 0.18 | 0.14 | 0.32 | 0.25 |
| 10 | 120706950 | rs1247278 | -0.049 | -0.164 | -1.056 | -0.837 | -0.011 | -0.028 | -0.028 | 0 | 0.87 | 0.78 | 0.91 | 0.95 | 0.9 | 0.9 |
| 10 | 120708167 | rs7069995 | -0.612 | -0.268 | -0.71 | -0.724 | -0.045 | -0.058 | -0.058 | 0 | 0.74 | 0.63 | 0.76 | 0.83 | 0.8 | 0.73 |
| 10 | 120709474 | rs4384316 | -0.15 | -0.102 | -1.344 | -0.453 | -0.001 | 0.074 | 0.074 | 0.818 | 0.36 | 0.22 | 0.24 | 0.2 | 0.4 | 0.33 |
| 10 | 120711337 | rs2148496 | 0.068 | 0.572 | -1.77 | -0.7 | 0.008 | 0.067 | 0.067 | 0.013 | 0.61 | 0.79 | 0.68 | 0.84 | 0.65 | 0.65 |
| 10 | 120712433 | rs4752234 | 0.711 | 0.451 | -2.121 | -0.094 | -0.03 | -0.074 | -0.074 | 0.012 | 0.33 | 0.11 | 0.41 | 0.46 | 0.36 | 0.34 |
| 10 | 120715118 | rs943269 | -0.475 | 0.118 | -1.537 | -0.503 | -0.001 | 0.07 | 0.07 | 0.155 | 0.32 | 0.11 | 0.3 | 0.17 | 0.33 | 0.26 |
| 10 | 120726228 | rs4752236 | 0.447 | -0.554 | -1.217 | -0.282 | 0.019 | 0.018 | 0.018 | 2.828 | 0.74 | 0.25 | 0.69 | 0.83 | 0.84 | 0.82 |
| 10 | 120726847 | rs7900920 | 0.538 | -0.622 | -1.246 | -0.311 | 0.025 | 0.024 | 0.024 | 3.107 | 0.72 | 0.25 | 0.69 | 0.83 | 0.84 | 0.79 |
| 10 | 120730977 | rs4751686 | 0.418 | -0.427 | -0.505 | -0.475 | -0.01 | -0.062 | -0.062 | 0.917 | 0.89 | 0.72 | 0.89 | 1 | 0.87 | 0.98 |
| 10 | 120731229 | rs11198700 | 0.002 | -0.547 | -0.557 | 0.18 | -0.007 | 0.037 | 0.037 | 0 | 0.17 | 0.23 | 0.07 | 0.36 | 0.1 | 0.38 |
| 10 | 120732488 | rs11198701 | 0.192 | 0.258 | -0.986 | 0.263 | -0.02 | -0.027 | -0.027 | 0.062 | 0.27 | 0.16 | 0.15 | 0.35 | 0.32 | 0.22 |
| 10 | 120737256 | rs4752238 | -0.186 | -0.083 | -0.647 | -0.144 | -0.002 | -0.006 | -0.006 | 0 | 0.34 | 0.42 | 0.39 | 0.25 | 0.33 | 0.38 |
| 10 | 120738653 | rs683817 | -0.67 | 0.19 | -0.688 | 0.087 | -0.013 | -0.022 | -0.022 | 0 | 0.76 | 0.79 | 0.81 | 0.55 | 0.75 | 0.82 |
| 10 | 120739449 | rs12242686 | -0.32 | -0.224 | -0.418 | -0.297 | -0.015 | -0.015 | -0.015 | 0.562 | 0.41 | 0.3 | 0.43 | 0.52 | 0.42 | 0.45 |
| 10 | 120742200 | rs10886356 | 0.347 | -0.306 | -0.199 | -0.329 | -0.001 | -0.014 | -0.014 | 0 | 0.16 | 0.14 | 0.25 | 0.06 | 0.16 | 0.24 |
| 10 | 120743172 | rs691222 | -0.304 | -0.363 | 0.025 | -0.314 | -0.003 | 0.001 | 0.001 | 0.732 | 0.54 | 0.59 | 0.63 | 0.47 | 0.54 | 0.67 |
| 10 | 120753787 | rs10886360 | 0.793 | -0.389 | -0.445 | -0.174 | 0.006 | 0.003 | 0.003 | 0.322 | 0.35 | 0.24 | 0.34 | 0.33 | 0.36 | 0.34 |
| 10 | 120753933 | rs690870 | -0.357 | -0.397 | -0.086 | -0.29 | 0.003 | 0.016 | 0.016 | 0.4 | 0.34 | 0.27 | 0.39 | 0.13 | 0.33 | 0.35 |
| 10 | 120754331 | rs17667035 | 0.965 | -0.436 | -0.635 | -0.363 | 0.004 | 0.019 | 0.019 | 1.317 | 0.1 | 0.01 | 0.06 | 0.05 | 0.04 | 0.12 |
| 10 | 120758093 | rs12413092 | -0.156 | -0.442 | -0.445 | -0.288 | -0.01 | 0.073 | 0.073 | 1.744 | 0.31 | 0.12 | 0.37 | 0.13 | 0.33 | 0.35 |
| 10 | 120758823 | rs10787889 | 0.63 | -0.27 | -0.239 | -0.098 | 0.001 | 0.001 | 0.001 | 0.045 | 0.37 | 0.33 | 0.38 | 0.33 | 0.38 | 0.35 |
| 10 | 120762125 | rs10787892 | -0.536 | -0.22 | -0.185 | 0.075 | 0 | 0.003 | 0.003 | 0 | 0.3 | 0.29 | 0.39 | 0.31 | 0.33 | 0.34 |
| 10 | 120762715 | rs7089236 | -0.478 | 0.068 | -0.12 | 0.064 | 0.002 | 0 | 0 | 0 | 0.31 | 0.26 | 0.4 | 0.32 | 0.37 | 0.34 |
| 10 | 120764967 | rs10886361 | 1.253 | -0.123 | -0.648 | -0.221 | 0.039 | 0.087 | 0.087 | 0 | 0.32 | 0.08 | 0.29 | 0.16 | 0.26 | 0.35 |
| 10 | 120767097 | rs11198727 | 0.726 | -0.003 | -0.346 | 0.099 | -0.053 | -0.042 | -0.042 | 0 | 0.31 | 0.16 | 0.24 | 0.45 | 0.37 | 0.27 |
| 10 | 120769168 | rs7393514 | 1.45 | -0.184 | -0.258 | -0.001 | 0.015 | 0.024 | 0.024 | 0 | 0.31 | 0.22 | 0.29 | 0.17 | 0.26 | 0.34 |
| 10 | 120770637 | rs7093507 | -0.166 | -0.17 | -0.175 | -0.007 | -0.003 | -0.004 | -0.004 | 0.261 | 0.33 | 0.38 | 0.39 | 0.31 | 0.35 | 0.33 |
| 10 | 120774821 | rs10749283 | -1.24 | -0.11 | -0.217 | 0.002 | -0.021 | -0.012 | -0.012 | 0.002 | 0.66 | 0.55 | 0.66 | 0.75 | 0.74 | 0.61 |
| 10 | 120778784 | rs12411582 | 0.815 | 0.335 | -0.44 | 0.005 | -0.04 | 0.019 | 0.019 | 0 | 0.11 | 0.01 | 0.19 | 0.06 | 0.13 | 0.13 |
| 10 | 120782186 | rs10886368 | -0.493 | 0.384 | -0.722 | 0.288 | -0.109 | -0.066 | -0.066 | 0.149 | 0.37 | 0.07 | 0.4 | 0.43 | 0.41 | 0.3 |
| 10 | 120784977 | rs494469 | -1.063 | 0.47 | 0.246 | 0.571 | 0.02 | 0.03 | 0.03 | 0.044 | 0.36 | 0.57 | 0.44 | 0.53 | 0.43 | 0.35 |
| 10 | 120792703 | rs546808 | 1.887 | 0.774 | 1.045 | 0.744 | 0.014 | 0.148 | 0.148 | 3.816 | 0.92 | 0.42 | 0.92 | 0.88 | 0.97 | 0.96 |
| 10 | 120817450 | rs10886375 | 1.79 | 0.809 | 1.322 | 0.759 | 0.042 | 0.184 | 0.184 | 2.548 | 0.92 | 0.48 | 0.93 | 0.88 | 0.97 | 0.96 |
| 10 | 120821625 | rs9325559 | 0.227 | 1.296 | 1.173 | 0.709 | -0.011 | 0.006 | 0.006 | 0 | 0.13 | 0.11 | 0.21 | 0.01 | 0.16 | 0.11 |
| 10 | 120828969 | rs7908387 | 2.071 | 0.602 | 1.377 | 0.86 | 0.01 | 0.141 | 0.141 | 4.168 | 0.92 | 0.39 | 0.92 | 0.88 | 0.97 | 0.96 |
| 10 | 120850164 | rs4751691 | 0.74 | 0.691 | 1.161 | 0.855 | 0.033 | 0.039 | 0.039 | 0.13 | 0.67 | 0.35 | 0.56 | 0.46 | 0.57 | 0.69 |
| 10 | 120852654 | rs10886381 | -0.289 | 0.51 | 1.031 | 0.817 | -0.065 | -0.049 | -0.049 | 0 | 0.26 | 0.14 | 0.35 | 0.42 | 0.4 | 0.27 |
| 10 | 120862541 | rs7911636 | -0.534 | 0.525 | 1.063 | 0.823 | -0.022 | -0.016 | -0.016 | 0 | 0.27 | 0.27 | 0.36 | 0.42 | 0.4 | 0.27 |
| 10 | 120863232 | rs11198774 | 0.373 | 0.51 | 1.109 | 0.827 | 0.036 | 0.041 | 0.041 | 0 | 0.7 | 0.53 | 0.61 | 0.53 | 0.59 | 0.73 |
| 10 | 120864781 | rs1511034 | 0.376 | 0.516 | 1.111 | 0.832 | 0.036 | 0.041 | 0.041 | 0 | 0.7 | 0.53 | 0.61 | 0.53 | 0.59 | 0.73 |
| 10 | 120868099 | rs7078261 | 1.332 | 0.507 | 1.073 | 0.926 | 0.024 | 0.221 | 0.221 | 0 | 0.95 | 0.66 | 0.96 | 0.86 | 1 | 0.96 |
| 10 | 120874259 | rs11198781 | 0.673 | 0.546 | 0.881 | 0.938 | 0.026 | 0.016 | 0.016 | 0.329 | 0.67 | 0.41 | 0.61 | 0.6 | 0.59 | 0.72 |
| 10 | 120874802 | rs17098456 | -0.7 | 0.565 | 0.817 | 0.944 | 0.011 | 0.002 | 0.002 | 0.025 | 0.28 | 0.35 | 0.35 | 0.36 | 0.4 | 0.27 |
| 10 | 120880910 | rs12355492 | 0.586 | 0.529 | 0.504 | 0.825 | 0.044 | 0.041 | 0.041 | 0 | 0.67 | 0.48 | 0.62 | 0.56 | 0.57 | 0.71 |
| 10 | 120889022 | rs3802739 | -0.288 | 0.723 | 0.11 | 0.88 | -0.106 | -0.072 | -0.072 | 0 | 0.26 | 0.08 | 0.3 | 0.39 | 0.42 | 0.28 |
| 10 | 120892542 | rs915272 | 0.225 | 0.79 | 0.136 | 0.909 | -0.029 | -0.016 | -0.016 | 2.12 | 0.71 | 0.74 | 0.69 | 0.61 | 0.58 | 0.72 |
| 10 | 120895131 | rs7920891 | 0.214 | 0.794 | 0.138 | 0.843 | -0.029 | -0.016 | -0.016 | 0 | 0.71 | 0.74 | 0.69 | 0.61 | 0.58 | 0.72 |
| 10 | 120897479 | rs34179455 | 0.254 | 0.789 | 0.123 | 0.828 | -0.024 | -0.014 | -0.014 | 0 | 0.71 | 0.71 | 0.69 | 0.61 | 0.58 | 0.72 |
| 10 | 120900154 | rs7912115 | 0.314 | -0.272 | -0.278 | 0.325 | 0.026 | 0.03 | 0.03 | 0 | 0.3 | 0.21 | 0.2 | 0.06 | 0.19 | 0.27 |

| | | | | | | | | | | | | | | | | |
|---|---|---|---|---|---|---|---|---|---|---|---|---|---|---|---|---|
| 10 | 120904013 | rs6585536 | -0.229 | -0.238 | 0.405 | 0.704 | -0.004 | 0.018 | 0.018 | 0 | 0.61 | 0.48 | 0.51 | 0.46 | 0.63 | 0.53 |
| 10 | 120905214 | rs7086828 | -0.249 | -0.654 | 0.303 | 0.221 | -0.001 | -0.015 | -0.015 | 0 | 0.21 | 0.25 | 0.19 | 0.1 | 0.17 | 0.18 |
| 10 | 120907828 | rs10510053 | -0.096 | -0.948 | 0.244 | 0.274 | 0.001 | 0.083 | 0.083 | 0 | 0.25 | 0.32 | 0.25 | 0.5 | 0.28 | 0.22 |
| 10 | 120916467 | rs2297696 | 1.481 | -1.103 | 0.014 | 0.098 | -0.066 | 0.079 | 0.079 | 0 | 0.34 | 0.13 | 0.31 | 0.18 | 0.39 | 0.39 |
| 10 | 120917445 | rs2275111 | -0.773 | -0.851 | 0.03 | 0.392 | -0.005 | -0.014 | -0.014 | 0.19 | 0.44 | 0.45 | 0.49 | 0.71 | 0.46 | 0.44 |
| 10 | 120920588 | rs10749291 | -0.733 | -1.389 | 0.101 | 0.442 | -0.009 | -0.014 | -0.014 | 0 | 0.42 | 0.43 | 0.45 | 0.72 | 0.44 | 0.44 |
| 10 | 120922304 | rs10886406 | -0.332 | -0.861 | 0.417 | 0.796 | -0.006 | 0.096 | 0.096 | 0 | 0.23 | 0.3 | 0.21 | 0.52 | 0.27 | 0.23 |
| 10 | 120927544 | rs3377 | 0.86 | -1.143 | -0.364 | 0.233 | 0.037 | 0.164 | 0.164 | 1.585 | 0.49 | 0.12 | 0.5 | 0.31 | 0.56 | 0.57 |
| 10 | 120936701 | rs3740562 | -0.591 | -1.195 | 0.596 | 0.442 | -0.005 | 0.061 | 0.061 | 0 | 0.27 | 0.52 | 0.29 | 0.55 | 0.3 | 0.23 |
| 10 | 120943664 | rs10886412 | -0.521 | -1.337 | 0.865 | 0.426 | -0.006 | 0.077 | 0.077 | 0 | 0.27 | 0.47 | 0.35 | 0.56 | 0.29 | 0.23 |
| 10 | 120949557 | rs1889742 | -0.607 | -1.278 | 0.86 | 0.428 | -0.008 | 0.038 | 0.038 | 0 | 0.28 | 0.55 | 0.29 | 0.56 | 0.29 | 0.25 |
| 10 | 120951095 | rs1107942 | -0.651 | -1.292 | 0.854 | 0.422 | -0.008 | 0.035 | 0.035 | 0 | 0.28 | 0.57 | 0.29 | 0.56 | 0.29 | 0.25 |
| 10 | 120952674 | rs10886416 | 0.027 | -1.391 | 0.796 | 0.321 | -0.01 | -0.04 | -0.04 | 2.485 | 0.14 | 0.25 | 0.1 | 0.03 | 0.1 | 0.15 |
| 10 | 120954350 | rs1108472 | -1.117 | -1.462 | 0.683 | 0.328 | 0.014 | 0.01 | 0.01 | 0 | 0.41 | 0.75 | 0.38 | 0.59 | 0.39 | 0.39 |
| 10 | 120957680 | rs10510054 | 0.963 | -1.449 | 0.433 | 0.328 | 0.025 | 0.082 | 0.082 | 0.971 | 0.58 | 0.13 | 0.56 | 0.32 | 0.62 | 0.58 |
| 10 | 120959450 | rs10749299 | -1.14 | -1.332 | 0.748 | 0.329 | 0.006 | 0.001 | 0.001 | 0.023 | 0.38 | 0.69 | 0.35 | 0.6 | 0.38 | 0.39 |
| 10 | 120960702 | rs11198819 | -0.489 | -1.117 | 0.732 | 0.6 | 0.003 | 0.128 | 0.128 | 0 | 0.24 | 0.46 | 0.25 | 0.52 | 0.28 | 0.25 |
| 10 | 120970543 | rs9325562 | -0.667 | -1.033 | 0.447 | 0.364 | 0.003 | 0.058 | 0.058 | 0 | 0.28 | 0.44 | 0.28 | 0.58 | 0.33 | 0.23 |
| 10 | 120976553 | rs4336954 | 1.275 | -1.143 | -0.312 | 0.012 | -0.087 | 0.078 | 0.078 | 0.012 | 0.34 | 0.1 | 0.37 | 0.2 | 0.42 | 0.38 |
| 10 | 120978183 | rs17098556 | -0.812 | -1.497 | 0.292 | -0.21 | -0.002 | 0.051 | 0.051 | 0.038 | 0.06 | 0.23 | 0.09 | 0.06 | 0.02 | 0.09 |
| 10 | 120979749 | rs4751702 | -0.131 | -1.486 | -0.21 | -0.551 | 0.016 | 0.014 | 0.014 | 0.929 | 0.85 | 0.91 | 0.92 | 0.93 | 0.92 | 0.88 |
| 10 | 120982356 | rs7093673 | -0.407 | -1.636 | -0.002 | -0.533 | -0.006 | -0.016 | -0.016 | 0.06 | 0.3 | 0.38 | 0.31 | 0.18 | 0.28 | 0.29 |
| 10 | 120986693 | rs17606354 | -0.161 | -1.577 | -0.603 | -0.404 | 0.04 | 0.021 | 0.021 | 0.958 | 0.15 | 0.04 | 0.08 | 0.07 | 0.07 | 0.12 |
| 10 | 120993661 | rs4237510 | 0.04 | -1.058 | -0.153 | -0.378 | -0.018 | -0.034 | -0.034 | 0 | 0.79 | 0.75 | 0.82 | 0.9 | 0.86 | 0.78 |
| 10 | 121001183 | rs7914808 | 1.411 | -1.1 | -0.303 | -0.184 | -0.002 | -0.001 | -0.001 | 0 | 0.81 | 0.21 | 0.66 | 0.78 | 0.9 | 0.77 |
| 10 | 121010817 | rs17098589 | -1.075 | -0.916 | 0.084 | -0.184 | -0.001 | 0.021 | 0.021 | 0 | 0.06 | 0.13 | 0.17 | 0.12 | 0.07 | 0.09 |
| 10 | 121010851 | rs11198845 | -1.881 | -0.58 | 0.804 | -0.711 | 0.013 | -0.038 | -0.038 | 0 | 0.23 | 0.46 | 0.38 | 0.17 | 0.25 | 0.21 |
| 10 | 121013417 | rs11198846 | -0.77 | -0.562 | 0.808 | -0.741 | -0.001 | -0.002 | -0.002 | 0.25 | 0.15 | 0.12 | 0.19 | 0.05 | 0.18 | 0.12 |
| 10 | 121025097 | rs7095989 | -1.749 | -1.205 | 0.52 | -1.248 | -0.004 | -0.016 | -0.016 | 0.761 | 0.38 | 0.42 | 0.46 | 0.24 | 0.36 | 0.34 |
| 10 | 121031659 | rs928670 | 1.556 | -0.95 | 0.278 | -1.601 | 0.001 | 0.006 | 0.006 | 0 | 0.52 | 0.47 | 0.45 | 0.73 | 0.53 | 0.51 |
| 10 | 121034914 | rs12358835 | 0.803 | -0.912 | -0.149 | -1.598 | 0.013 | -0.042 | -0.042 | 3.308 | 0.11 | 0.12 | 0.13 | 0.01 | 0.16 | 0.11 |
| 10 | 121037952 | rs4752269 | -0.902 | -1.013 | -0.368 | -1.605 | -0.011 | 0.034 | 0.034 | 0.059 | 0.31 | 0.1 | 0.3 | 0.21 | 0.28 | 0.27 |
| 10 | 121041698 | rs11198856 | 1.058 | -0.151 | -0.566 | -1.719 | -0.028 | -0.04 | -0.04 | 0 | 0.23 | 0.11 | 0.25 | 0.67 | 0.3 | 0.31 |
| 10 | 121049079 | rs4752275 | 0.876 | -0.286 | 0.08 | -1.738 | 0.006 | -0.074 | -0.074 | 0.031 | 0.73 | 0.54 | 0.67 | 0.88 | 0.73 | 0.71 |
| 10 | 121049565 | rs10886439 | -1.028 | -0.164 | -0.188 | -1.716 | 0.012 | -0.054 | -0.054 | 0 | 0.88 | 0.82 | 0.87 | 0.99 | 0.84 | 0.89 |
| 10 | 121050308 | rs7097630 | 0.184 | -0.099 | -0.144 | -1.685 | 0.005 | -0.042 | -0.042 | 0.241 | 0.79 | 0.73 | 0.78 | 0.94 | 0.77 | 0.82 |
| 10 | 121052908 | rs10128498 | 0.816 | -0.681 | -0.232 | -1.737 | -0.009 | -0.019 | -0.019 | 0.14 | 0.59 | 0.45 | 0.52 | 0.78 | 0.66 | 0.65 |
| 10 | 121054142 | rs4752281 | -0.057 | -0.17 | -0.452 | -1.366 | -0.002 | -0.072 | -0.072 | 0 | 0.83 | 0.76 | 0.86 | 0.99 | 0.84 | 0.88 |
| 10 | 121054378 | rs10886442 | 0.261 | -0.211 | -0.962 | -1.004 | -0.038 | -0.048 | -0.048 | 0 | 0.25 | 0.13 | 0.29 | 0.67 | 0.32 | 0.35 |
| 10 | 121062068 | rs10886445 | 0.531 | -0.173 | -0.809 | -1.179 | 0.032 | 0.138 | 0.138 | 0 | 0.5 | 0.31 | 0.44 | 0.18 | 0.55 | 0.39 |
| 10 | 121064720 | rs12416019 | -0.253 | -0.639 | -0.735 | -0.943 | 0.005 | -0.015 | -0.015 | 0 | 0.31 | 0.21 | 0.33 | 0.65 | 0.3 | 0.38 |
| 10 | 121065079 | rs2420614 | -0.138 | -0.743 | -0.197 | -0.984 | 0.003 | 0.003 | 0.003 | 0 | 0.32 | 0.21 | 0.35 | 0.75 | 0.31 | 0.43 |
| 10 | 121069074 | rs871196 | -0.417 | -0.395 | -0.412 | -0.945 | 0.016 | 0.008 | 0.008 | 0.014 | 0.37 | 0.2 | 0.29 | 0.67 | 0.35 | 0.41 |
| 10 | 121069546 | rs10886447 | 1.023 | -0.389 | -0.557 | -0.929 | -0.003 | 0.166 | 0.166 | 0.007 | 0.44 | 0.19 | 0.49 | 0.19 | 0.53 | 0.36 |
| 10 | 121074420 | rs2420615 | 1.005 | -0.399 | 0.672 | 0.38 | -0.027 | 0.022 | 0.022 | 0 | 0.75 | 0.43 | 0.72 | 0.7 | 0.79 | 0.73 |
| 10 | 121084909 | rs17098705 | -0.698 | -0.433 | 0.689 | 0.259 | -0.031 | -0.029 | -0.029 | 0 | 0.17 | 0.36 | 0.23 | 0.08 | 0.14 | 0.2 |
| 10 | 121098290 | rs11198886 | 1.099 | 0.071 | 0.234 | 0.598 | -0.015 | -0.001 | -0.001 | 0 | 0.05 | 0.01 | 0.02 | 0.21 | 0.08 | 0.1 |
| 10 | 121100153 | rs4752292 | 1.601 | -0.234 | 0.598 | 0.454 | -0.022 | 0.071 | 0.071 | 0 | 0.82 | 0.43 | 0.84 | 0.71 | 0.87 | 0.76 |
| 10 | 121105311 | rs10886462 | 0.858 | 0.135 | -0.849 | -0.352 | -0.085 | 0.13 | 0.13 | 0.023 | 0.26 | 0.01 | 0.17 | 0.05 | 0.32 | 0.17 |
| 10 | 121106620 | rs7092272 | 0.191 | -0.271 | -0.228 | 0.274 | -0.004 | 0.001 | 0.001 | 0 | 0.33 | 0.33 | 0.3 | 0.38 | 0.33 | 0.25 |
| 10 | 121107900 | rs11198893 | 1.468 | -0.085 | -0.393 | 0.683 | -0.025 | -0.012 | -0.012 | 0 | 0.05 | 0 | 0.02 | 0.23 | 0.08 | 0.15 |
| 10 | 121114292 | rs10886464 | 1.045 | -0.509 | 0.015 | 0.626 | -0.045 | 0.172 | 0.172 | 0 | 0.79 | 0.54 | 0.69 | 0.59 | 0.87 | 0.64 |
| 10 | 121122182 | rs17098773 | 0.287 | -0.641 | -0.278 | 0.445 | -0.003 | 0.013 | 0.013 | 0 | 0.22 | 0.26 | 0.17 | 0.38 | 0.2 | 0.22 |
| 10 | 121123633 | rs291970 | 0.639 | -0.694 | -0.566 | 0.376 | -0.007 | -0.01 | -0.01 | 0.389 | 0.19 | 0.07 | 0.2 | 0.22 | 0.22 | 0.22 |
| 10 | 121129075 | rs17608274 | 1.444 | -0.689 | -0.554 | 0.224 | 0.009 | 0.032 | 0.032 | 0 | 0.14 | 0.06 | 0.18 | 0 | 0.14 | 0.07 |
| 10 | 121129113 | rs2901211 | -0.143 | -0.882 | 0.086 | 0.287 | -0.026 | 0.048 | 0.048 | 0.012 | 0.26 | 0.47 | 0.2 | 0.4 | 0.21 | 0.25 |
| 10 | 121129167 | rs17608302 | 1.958 | -0.535 | -0.514 | 0.059 | -0.088 | -0.039 | -0.039 | 0 | 0.16 | 0.01 | 0.21 | 0.13 | 0.23 | 0.18 |
| 10 | 121132192 | rs1413582 | 1.278 | -1.52 | 0.764 | 0.605 | 0.044 | 0.1 | 0.1 | 0.185 | 0.46 | 0.28 | 0.48 | 0.24 | 0.59 | 0.35 |
| 10 | 121137182 | rs506657 | -0.992 | -1.682 | 1.098 | 0.741 | -0.024 | 0.021 | 0.021 | 0 | 0.33 | 0.4 | 0.15 | 0.63 | 0.23 | 0.47 |
| 10 | 121139045 | rs1248077 | -0.112 | -1.626 | 1.209 | 0.791 | 0.003 | 0.118 | 0.118 | 0 | 0.93 | 0.79 | 0.8 | 0.86 | 0.96 | 0.9 |
| 10 | 121139331 | rs6585546 | 1.103 | -1.702 | 1.303 | 0.708 | 0.027 | 0.042 | 0.042 | 1.392 | 0.49 | 0.4 | 0.54 | 0.23 | 0.6 | 0.42 |
| 10 | 121140321 | rs2275036 | 0.173 | -1.566 | 1.4 | 0.955 | 0.005 | -0.008 | -0.008 | 0.232 | 0.2 | 0.21 | 0.09 | 0.37 | 0.13 | 0.19 |
| 10 | 121140671 | rs3818779 | 0.116 | -1.375 | 1.341 | 1.063 | 0.006 | 0.127 | 0.127 | 0 | 0.93 | 0.78 | 0.8 | 0.86 | 0.96 | 0.9 |
| 10 | 121146290 | rs7077176 | -0.011 | -1.331 | 1.255 | 0.985 | 0.008 | 0.107 | 0.107 | 0 | 0.09 | 0.15 | 0.11 | 0.23 | 0.06 | 0.24 |
| 10 | 121149403 | rs10886471 | 1.686 | -1.523 | 0.902 | 0.729 | -0.067 | 0.12 | 0.12 | 0.234 | 0.41 | 0.13 | 0.45 | 0.22 | 0.48 | 0.38 |
| 10 | 121150893 | rs12780837 | 0.907 | -1.526 | 1.294 | 0.795 | -0.001 | 0.012 | 0.012 | 0 | 0.66 | 0.49 | 0.72 | 0.4 | 0.66 | 0.56 |
| 10 | 121151332 | rs12781823 | 1.268 | -1.514 | 0.94 | 0.799 | -0.049 | -0.044 | -0.044 | 0.475 | 0.08 | 0 | 0.06 | 0.3 | 0.18 | 0.08 |
| 10 | 121153108 | rs11198905 | -0.364 | -1.451 | 1.188 | 0.736 | -0.001 | 0.005 | 0.005 | 0 | 0.24 | 0.29 | 0.15 | 0.37 | 0.25 | 0.19 |
| 10 | 121154736 | rs17608873 | -0.805 | -1.06 | 0.465 | 0.328 | -0.002 | -0.005 | -0.005 | 0.045 | 0.08 | 0.08 | 0.12 | 0.03 | 0.1 | 0.06 |

| | | | | | | | | | | | | | | | | |
|---|---|---|---|---|---|---|---|---|---|---|---|---|---|---|---|---|
| 10 | 121157897 | rs915110 | -1.084 | -1.123 | 0.609 | 0.393 | -0.009 | -0.056 | -0.056 | 0.095 | 0.84 | 0.74 | 0.85 | 1 | 0.85 | 0.93 |
| 10 | 121161798 | rs4623810 | -1.031 | -1.04 | 0.615 | 0.631 | 0.001 | -0.002 | -0.002 | 0 | 0.4 | 0.58 | 0.3 | 0.6 | 0.38 | 0.51 |
| 10 | 121165616 | rs4752300 | 0.94 | -1.059 | 0.136 | 0.52 | 0.018 | 0.132 | 0.132 | 0 | 0.55 | 0.18 | 0.57 | 0.23 | 0.61 | 0.44 |
| 10 | 121175674 | rs17098857 | -0.256 | -1.41 | 0.466 | 0.449 | -0.004 | -0.001 | -0.001 | 0 | 0.23 | 0.3 | 0.28 | 0.16 | 0.18 | 0.15 |
| 10 | 121176601 | rs4752305 | -1.844 | -1.388 | 0.29 | 0.452 | -0.062 | 0.123 | 0.123 | 0.283 | 0.59 | 0.87 | 0.53 | 0.8 | 0.54 | 0.62 |
| 10 | 121180765 | rs7076555 | -0.723 | -1.498 | 0.821 | 0.812 | -0.024 | 0.044 | 0.044 | 0 | 0.31 | 0.51 | 0.28 | 0.56 | 0.28 | 0.33 |
| 10 | 121184828 | rs4752308 | 0.841 | -1.439 | 0.986 | 0.826 | 0.003 | 0.015 | 0.015 | 0 | 0.51 | 0.55 | 0.63 | 0.36 | 0.49 | 0.47 |
| 10 | 121185522 | rs11198919 | 1.117 | -1.367 | 0.351 | 0.844 | -0.029 | -0.032 | -0.032 | 0 | 0.07 | 0 | 0.04 | 0.25 | 0.08 | 0.2 |
| 10 | 121186001 | rs1889432 | 1.618 | -1.386 | 0.386 | 0.762 | -0.082 | 0.128 | 0.128 | 0.34 | 0.41 | 0.12 | 0.47 | 0.21 | 0.46 | 0.38 |
| 10 | 121189480 | rs915121 | 0.19 | -1.525 | 0.55 | 0.688 | 0 | -0.003 | -0.003 | 0 | 0.27 | 0.2 | 0.15 | 0.4 | 0.27 | 0.25 |
| 10 | 121190846 | rs2275038 | -0.592 | -1.571 | 0.855 | 0.723 | -0.036 | 0.049 | 0.049 | 0 | 0.09 | 0.21 | 0.15 | 0.15 | 0.03 | 0.08 |
| 10 | 121191186 | rs7099478 | 0.485 | -1.567 | 0.669 | 0.747 | 0.009 | 0.053 | 0.053 | 0 | 0.52 | 0.73 | 0.63 | 0.31 | 0.51 | 0.45 |
| 10 | 121191551 | rs915119 | -0.29 | -1.561 | 0.603 | 0.721 | 0.014 | 0.004 | 0.004 | 0 | 0.41 | 0.26 | 0.28 | 0.4 | 0.41 | 0.34 |
| 10 | 121196062 | rs2275040 | 1.54 | -1.481 | 0.494 | 0.574 | -0.075 | 0.136 | 0.136 | 0 | 0.38 | 0.15 | 0.41 | 0.16 | 0.47 | 0.37 |
| 10 | 121197057 | rs11198925 | 0.504 | -1.524 | 0.333 | 0.628 | -0.004 | -0.025 | -0.025 | 0.095 | 0.59 | 0.62 | 0.61 | 0.22 | 0.62 | 0.49 |
| 10 | 121197945 | rs1475753 | -0.027 | -1.369 | -0.42 | 0.735 | 0.018 | -0.054 | -0.054 | 0 | 0.3 | 0.13 | 0.23 | 0.49 | 0.27 | 0.29 |
| 10 | 121200954 | rs10886484 | 0.852 | -0.784 | -0.701 | 0.974 | -0.068 | 0.129 | 0.129 | 0 | 0.38 | 0.18 | 0.42 | 0.17 | 0.47 | 0.37 |
| 10 | 121202677 | rs11198927 | 0.634 | -0.421 | -1.048 | 1.143 | -0.031 | -0.016 | -0.016 | 0 | 0.07 | 0 | 0.1 | 0.28 | 0.08 | 0.2 |
| 10 | 121207272 | rs11198929 | 0.259 | -0.632 | -1.037 | 1.175 | 0.015 | 0.018 | 0.018 | 0 | 0.4 | 0.3 | 0.31 | 0.36 | 0.4 | 0.28 |
| 10 | 121207961 | rs10749321 | 0.364 | 0.175 | -1.55 | 0.707 | -0.139 | 0.203 | 0.203 | 0 | 0.39 | 0.08 | 0.33 | 0.1 | 0.44 | 0.33 |
| 10 | 121212405 | rs2297641 | -0.596 | 0.265 | -0.305 | 1.656 | -0.028 | -0.013 | -0.013 | 0 | 0.07 | 0.02 | 0.12 | 0.32 | 0.09 | 0.2 |
| 10 | 121218572 | rs10886486 | -1.111 | 0.291 | 0.091 | 1.319 | 0.036 | 0.128 | 0.128 | 0 | 0.08 | 0.19 | 0.12 | 0.27 | 0.09 | 0.17 |
| 10 | 121219038 | rs10886487 | -1.252 | 0.449 | -0.25 | 1.187 | 0.003 | 0.009 | 0.009 | 0 | 0.1 | 0.09 | 0.13 | 0.24 | 0.12 | 0.19 |
| 10 | 121225352 | rs3009874 | 0.665 | -0.354 | -0.46 | 0.453 | 0.02 | 0.041 | 0.041 | 0 | 0.56 | 0.33 | 0.55 | 0.7 | 0.61 | 0.66 |
| 10 | 121225476 | rs10886492 | -0.824 | -0.326 | -0.864 | 0.428 | -0.012 | 0.072 | 0.072 | 0 | 0.38 | 0.11 | 0.33 | 0.26 | 0.38 | 0.32 |
| 10 | 121226447 | rs2901212 | -0.624 | -0.426 | -0.913 | 0.284 | 0.001 | 0.005 | 0.005 | 0.081 | 0.22 | 0.17 | 0.27 | 0.21 | 0.21 | 0.18 |
| 10 | 121232454 | rs12783252 | 0.322 | -0.357 | -1.217 | 0.185 | -0.027 | 0.062 | 0.062 | 1.44 | 0.11 | 0 | 0.06 | 0 | 0.12 | 0.03 |
| 10 | 121234063 | rs2991770 | -1.117 | -0.693 | -1.42 | 0.859 | 0.006 | -0.044 | -0.044 | 0.02 | 0.33 | 0.1 | 0.37 | 0.29 | 0.32 | 0.21 |
| 10 | 121238902 | rs11198957 | -1.196 | -0.543 | -0.829 | 0.897 | -0.004 | -0.008 | -0.008 | 0.514 | 0.08 | 0.03 | 0.21 | 0.15 | 0.08 | 0.14 |
| 10 | 121241482 | rs11198959 | -1.131 | -0.539 | -0.874 | 0.899 | -0.028 | -0.038 | -0.038 | 0 | 0.07 | 0.01 | 0.2 | 0.15 | 0.08 | 0.14 |
| 10 | 121244739 | rs2039488 | -1.208 | -0.685 | -0.435 | 1.076 | 0.003 | 0.065 | 0.065 | 0 | 0.08 | 0.18 | 0.15 | 0.25 | 0.09 | 0.14 |
| 10 | 121249171 | rs3009892 | -1.692 | -0.824 | -0.302 | 0.807 | 0.002 | 0.003 | 0.003 | 0 | 0.27 | 0.32 | 0.23 | 0.24 | 0.23 | 0.42 |
| 10 | 121257200 | rs10787973 | 1.4 | -1.049 | 0.37 | 1.119 | -0.025 | 0.085 | 0.085 | 0 | 0.8 | 0.62 | 0.64 | 0.62 | 0.84 | 0.85 |
| 10 | 121258718 | rs12259923 | 0.034 | -1.019 | 0.346 | 1.141 | 0.019 | 0.014 | 0.014 | 0 | 0.1 | 0.03 | 0.08 | 0 | 0.05 | 0.02 |
| 10 | 121259879 | rs756279 | 0.934 | -0.901 | 0.446 | 1.254 | 0.019 | 0.203 | 0.203 | 0 | 0.91 | 0.77 | 0.73 | 0.62 | 0.89 | 0.87 |
| 10 | 121260017 | rs756278 | -0.456 | -0.889 | 0.434 | 1.253 | 0.001 | 0.099 | 0.099 | 0 | 0.06 | 0.17 | 0.19 | 0.19 | 0.07 | 0.07 |
| 10 | 121262454 | rs3009907 | -1.425 | -0.884 | 0.441 | 1.22 | 0.014 | 0.08 | 0.08 | 0 | 0.08 | 0.22 | 0.17 | 0.23 | 0.09 | 0.1 |
| 10 | 121266607 | rs7100441 | 0.304 | -0.901 | 0.438 | 1.204 | 0.019 | 0.171 | 0.171 | 0 | 0.93 | 0.75 | 0.8 | 0.81 | 0.94 | 0.93 |
| 10 | 121272968 | rs7088764 | -0.621 | -1.517 | -0.216 | 0.648 | 0.031 | 0.091 | 0.091 | 0 | 0.24 | 0.09 | 0.08 | 0.05 | 0.17 | 0.34 |
| 10 | 121274163 | rs2420628 | 1.447 | -2.231 | 0.473 | 0.387 | 0.038 | 0.007 | 0.007 | 0 | 0.53 | 0.33 | 0.62 | 0.6 | 0.69 | 0.54 |
| 10 | 121275903 | rs1556591 | 0.334 | -1.94 | 0.082 | 0.178 | 0.015 | 0.017 | 0.017 | 0 | 0.73 | 0.87 | 0.85 | 0.76 | 0.78 | 0.61 |
| 10 | 121278072 | rs12262842 | -0.707 | -1.531 | 0.085 | 0.158 | 0.044 | 0.124 | 0.124 | 0 | 0.31 | 0.11 | 0.17 | 0.04 | 0.23 | 0.16 |
| 10 | 121280443 | rs11198992 | -1.063 | -1.492 | 0.591 | 0.252 | 0.011 | 0.015 | 0.015 | 0 | 0.35 | 0.29 | 0.24 | 0.25 | 0.27 | 0.18 |
| 10 | 121281607 | rs1467813 | 0.62 | -1.423 | 0.883 | 0.228 | 0.014 | 0.021 | 0.021 | 0 | 0.65 | 0.68 | 0.74 | 0.76 | 0.73 | 0.8 |
| 10 | 121284536 | rs1537577 | -0.039 | -0.402 | 0.826 | 0.649 | 0 | -0.01 | -0.01 | 0 | 0.32 | 0.31 | 0.36 | 0.53 | 0.28 | 0.41 |
| 10 | 121288501 | rs17098973 | 0.062 | -0.361 | 1.129 | 0.625 | 0 | 0.034 | 0.034 | 0 | 0.28 | 0.35 | 0.32 | 0.59 | 0.23 | 0.37 |
| 10 | 121290571 | rs4752324 | 0.849 | 0.019 | 1.344 | 0.606 | 0.005 | 0.034 | 0.034 | 0 | 0.8 | 0.7 | 0.68 | 0.75 | 0.8 | 0.78 |
| 10 | 121291941 | rs7076994 | -0.493 | -0.236 | 1.359 | 0.528 | -0.003 | 0.065 | 0.065 | 0 | 0.12 | 0.37 | 0.21 | 0.13 | 0.08 | 0.04 |
| 10 | 121296029 | rs11199005 | 0.277 | -0.051 | 0.435 | 0.564 | -0.004 | 0.037 | 0.037 | 0 | 0.09 | 0.02 | 0.09 | 0 | 0.07 | 0.02 |
| 10 | 121296366 | rs2277266 | 0.387 | 0.145 | 0.785 | 0.832 | 0.036 | 0.168 | 0.168 | 0 | 0.89 | 0.69 | 0.79 | 0.76 | 0.88 | 0.8 |
| 10 | 121296970 | rs6585551 | 0.362 | 0.151 | 0.775 | 0.85 | 0.032 | 0.16 | 0.16 | 0 | 0.89 | 0.71 | 0.79 | 0.76 | 0.88 | 0.8 |
| 10 | 121297851 | rs3808954 | -0.156 | 0.269 | -0.195 | 0.475 | -0.002 | -0.002 | -0.002 | 0 | 0.91 | 0.93 | 0.9 | 0.89 | 0.9 | 0.82 |
| 10 | 121309199 | rs10444117 | -0.003 | 0.227 | -0.139 | 0.309 | 0 | -0.018 | -0.018 | 0 | 0.08 | 0.08 | 0.12 | 0 | 0.06 | 0.03 |
| 10 | 121311308 | rs11528321 | 0.367 | 0.475 | -0.245 | 0.796 | -0.001 | 0.013 | 0.013 | 0.53 | 0.18 | 0.17 | 0.21 | 0.49 | 0.19 | 0.11 |
| 10 | 121311606 | rs7071853 | 0.082 | 0.366 | -0.072 | 0.773 | 0.003 | 0.035 | 0.035 | 0.481 | 0.19 | 0.22 | 0.22 | 0.49 | 0.19 | 0.09 |
| 10 | 121313621 | rs2917939 | -1.374 | -0.404 | 0.466 | 0.594 | 0.002 | 0.001 | 0.001 | 0.763 | 0.39 | 0.62 | 0.37 | 0.69 | 0.28 | 0.47 |
| 10 | 121328009 | rs12246119 | -0.98 | -0.68 | 0.533 | 0.476 | -0.038 | -0.04 | -0.04 | 0 | 0.2 | 0.31 | 0.1 | 0.11 | 0.09 | 0.29 |
| 10 | 121336248 | rs12356865 | 1.095 | -0.599 | 0.307 | 0.466 | 0.001 | -0.004 | -0.004 | 0.017 | 0.07 | 0.06 | 0.05 | 0 | 0.05 | 0.02 |
| 10 | 121343589 | rs10886515 | -1.591 | -0.472 | 0.36 | 0.373 | -0.012 | 0.007 | 0.007 | 3.073 | 0.38 | 0.5 | 0.31 | 0.59 | 0.25 | 0.38 |
| 10 | 121356382 | rs7919868 | -0.736 | -0.365 | 0.247 | 0.611 | -0.003 | 0.011 | 0.011 | 1.384 | 0.28 | 0.32 | 0.23 | 0.59 | 0.2 | 0.39 |
| 10 | 121357825 | rs11199037 | -0.748 | -0.366 | 0.247 | 0.61 | -0.003 | 0.01 | 0.01 | 1.402 | 0.28 | 0.32 | 0.23 | 0.59 | 0.2 | 0.41 |
| 10 | 121362660 | rs12569630 | 1.699 | -0.556 | -0.251 | 0.278 | -0.083 | 0.246 | 0.246 | 0.882 | 0.48 | 0.03 | 0.43 | 0.12 | 0.61 | 0.37 |
| 10 | 121365398 | rs11199041 | -1.384 | -0.578 | 0.567 | 0.421 | -0.022 | -0.023 | -0.023 | 0.053 | 0.32 | 0.38 | 0.24 | 0.26 | 0.22 | 0.45 |
| 10 | 121367763 | rs17099080 | 0.449 | -0.895 | 0.153 | 0.165 | -0.003 | 0.04 | 0.04 | 1.887 | 0.23 | 0.31 | 0.25 | 0.48 | 0.19 | 0.19 |
| 10 | 121369394 | rs2420636 | -1.902 | -0.465 | -0.294 | -0.055 | -0.031 | 0.05 | 0.05 | 2.583 | 0.58 | 0.76 | 0.52 | 0.75 | 0.47 | 0.67 |
| 10 | 121375181 | rs2577335 | -1.241 | -0.505 | 0.029 | 0.058 | -0.008 | 0.003 | 0.003 | 0 | 0.37 | 0.56 | 0.39 | 0.41 | 0.28 | 0.49 |
| 10 | 121375761 | rs17099083 | 0.736 | -0.611 | -0.137 | 0.03 | 0.026 | -0.022 | -0.022 | 0 | 0.08 | 0.01 | 0.02 | 0.12 | 0.02 | 0.2 |
| 10 | 121377286 | rs10886523 | 0.081 | -0.646 | 0.211 | -0.044 | -0.008 | -0.02 | -0.02 | 0 | 0.13 | 0.23 | 0.13 | 0.08 | 0.1 | 0.16 |
| 10 | 121378250 | rs84086 | -2.094 | -0.419 | -0.105 | -0.101 | -0.031 | 0.129 | 0.129 | 1.205 | 0.58 | 0.84 | 0.65 | 0.88 | 0.51 | 0.7 |

| | | | | | | | | | | | | | | | | |
|---|---|---|---|---|---|---|---|---|---|---|---|---|---|---|---|---|
| 10 | 121378493 | rs3847487 | -0.012 | -0.453 | -0.023 | -0.065 | 0.003 | 0.023 | 0.023 | 2.265 | 0.29 | 0.38 | 0.31 | 0.47 | 0.28 | 0.23 |
| 10 | 121379389 | rs196307 | 0.282 | -0.391 | 0.082 | -0.05 | 0.018 | 0.051 | 0.051 | 0 | 0.15 | 0.27 | 0.25 | 0.22 | 0.16 | 0.12 |
| 10 | 121386251 | rs196260 | -1.278 | -0.365 | 0.089 | -0.022 | -0.015 | -0.013 | -0.013 | 0 | 0.28 | 0.35 | 0.23 | 0.27 | 0.22 | 0.46 |
| 10 | 121387549 | rs7099126 | 0.091 | -0.122 | 0.129 | 0.02 | 0.002 | 0.014 | 0.014 | 2.032 | 0.28 | 0.34 | 0.3 | 0.47 | 0.27 | 0.24 |
| 10 | 121394687 | rs196272 | -0.525 | -0.028 | -0.079 | 0.042 | 0.006 | -0.024 | -0.024 | 0 | 0.19 | 0.25 | 0.2 | 0.15 | 0.2 | 0.29 |
| 10 | 121397276 | rs196276 | -0.621 | -0.024 | -0.08 | 0.004 | 0.007 | 0.029 | 0.029 | 0 | 0.2 | 0.27 | 0.31 | 0.3 | 0.21 | 0.3 |
| 10 | 121401490 | rs196279 | -1.694 | 0.536 | -0.953 | -0.621 | -0.039 | 0.059 | 0.059 | 0.348 | 0.85 | 0.98 | 0.87 | 1 | 0.8 | 0.85 |
| 10 | 121408071 | rs196290 | 0.101 | 0.281 | -0.811 | -0.648 | 0.016 | 0.076 | 0.076 | 0 | 0.49 | 0.29 | 0.4 | 0.3 | 0.48 | 0.54 |
| 10 | 121410026 | rs2420641 | 0.331 | 0.16 | -0.308 | -0.422 | 0.001 | 0.035 | 0.035 | 2.448 | 0.27 | 0.37 | 0.29 | 0.48 | 0.26 | 0.22 |
| 10 | 121411861 | rs196335 | 0.723 | 0.176 | -0.497 | -0.703 | 0.009 | -0.034 | -0.034 | 0.019 | 0.07 | 0.11 | 0.07 | 0 | 0.09 | 0.02 |
| 10 | 121416240 | rs196340 | 0.211 | -0.168 | -0.87 | -0.795 | -0.009 | 0.046 | 0.046 | 0 | 0.13 | 0.04 | 0.09 | 0.05 | 0.1 | 0.14 |
| 10 | 121416611 | rs196341 | -0.43 | -0.337 | -0.706 | -0.338 | -0.004 | 0.034 | 0.034 | 0 | 0.75 | 0.7 | 0.69 | 0.46 | 0.75 | 0.53 |
| 10 | 121417338 | rs3847489 | 0.978 | -0.068 | -0.861 | -0.748 | 0.012 | 0.014 | 0.014 | 0 | 0.3 | 0.35 | 0.4 | 0.36 | 0.37 | 0.21 |
| 10 | 121418228 | rs7906577 | 1.284 | 0.052 | -0.64 | -0.694 | 0.017 | 0.109 | 0.109 | 0 | 0.08 | 0.17 | 0.17 | 0.26 | 0.1 | 0.09 |
| 10 | 121420922 | rs196345 | -0.083 | 0.002 | -0.645 | -0.789 | 0.006 | 0.07 | 0.07 | 0.013 | 0.45 | 0.33 | 0.29 | 0.1 | 0.37 | 0.33 |
| 10 | 121423364 | rs17099142 | 1.312 | 0.12 | -0.564 | -0.653 | 0.019 | 0.116 | 0.116 | 0 | 0.08 | 0.17 | 0.17 | 0.26 | 0.1 | 0.09 |
| 10 | 121423961 | rs7909225 | 1.348 | 0.126 | -0.561 | -0.651 | 0.02 | 0.118 | 0.118 | 0 | 0.08 | 0.17 | 0.17 | 0.26 | 0.1 | 0.09 |
| 10 | 121427330 | rs7099687 | 1.757 | 0.149 | -0.726 | -0.74 | -0.011 | 0.016 | 0.016 | 0 | 0.23 | 0.17 | 0.23 | 0.1 | 0.28 | 0.12 |
| 10 | 121428828 | rs196325 | 0.025 | -0.365 | -1.169 | -0.901 | 0.001 | 0.004 | 0.004 | 1.854 | 0.59 | 0.41 | 0.66 | 0.66 | 0.6 | 0.45 |
| 10 | 121429633 | rs2234962 | 1.324 | 0.085 | -1.206 | -0.809 | -0.074 | 0.123 | 0.123 | 0.447 | 0.2 | 0.01 | 0.1 | 0 | 0.22 | 0.18 |
| 10 | 121436362 | rs196295 | -1.09 | -0.241 | -0.578 | -0.287 | -0.002 | 0.052 | 0.052 | 0 | 0.8 | 0.75 | 0.72 | 0.49 | 0.8 | 0.76 |
| 10 | 121440528 | rs112029 | -0.534 | -0.466 | -0.395 | -0.217 | -0.007 | 0.02 | 0.02 | 0 | 0.24 | 0.43 | 0.29 | 0.29 | 0.19 | 0.24 |
| 10 | 121443168 | rs7091662 | 0.749 | -0.108 | -0.541 | -0.304 | -0.018 | 0 | 0 | 0.248 | 0.11 | 0.28 | 0.09 | 0.05 | 0.05 | 0.01 |
| 10 | 121450146 | rs196239 | -0.29 | -0.072 | -0.982 | -0.501 | 0.004 | -0.01 | -0.01 | 0.069 | 0.29 | 0.27 | 0.19 | 0.07 | 0.18 | 0.25 |
| 10 | 121462047 | rs11199075 | 0.853 | 0.815 | -1.419 | -0.104 | -0.071 | 0.005 | 0.005 | 0 | 0.18 | 0.03 | 0.16 | 0.1 | 0.28 | 0.09 |
| 10 | 121462638 | rs2223082 | -0.017 | 0.62 | -1.222 | 0.223 | -0.008 | -0.01 | -0.01 | 0 | 0.42 | 0.31 | 0.53 | 0.55 | 0.54 | 0.29 |
| 10 | 121468629 | rs714095 | 1.568 | 0.242 | -1.014 | 0.291 | 0.01 | -0.02 | -0.02 | 1.629 | 0.06 | 0.02 | 0.13 | 0.1 | 0.06 | 0.02 |
| 10 | 121475180 | rs12217213 | 1.568 | 0.242 | -1.014 | 0.291 | 0.01 | -0.02 | -0.02 | 0 | 0.06 | 0.02 | 0.13 | 0.1 | 0.06 | 0.02 |
| 10 | 121480161 | rs196255 | 0.486 | 0.085 | -0.559 | 0.048 | -0.002 | 0.008 | 0.008 | 0.33 | 0.42 | 0.33 | 0.38 | 0.29 | 0.4 | 0.45 |
| 10 | 121484123 | rs196212 | 0.389 | 0.058 | -0.362 | 0.037 | 0 | -0.02 | -0.02 | 0.801 | 0.43 | 0.39 | 0.39 | 0.29 | 0.4 | 0.46 |
| 10 | 121484999 | rs196213 | 0.485 | 0.082 | -0.387 | 0.047 | -0.002 | 0.008 | 0.008 | 0.294 | 0.42 | 0.33 | 0.38 | 0.29 | 0.4 | 0.46 |
| 10 | 121487640 | rs196214 | -2.805 | 0.067 | -0.483 | -0.056 | -0.004 | -0.015 | -0.015 | 0 | 0.95 | 0.93 | 0.98 | 1 | 0.97 | 1 |
| 10 | 121513063 | rs2420642 | 0.737 | 0.144 | -0.342 | 0.203 | 0 | 0.006 | 0.006 | 0.026 | 0.25 | 0.25 | 0.29 | 0.5 | 0.26 | 0.19 |
| 10 | 121551581 | rs2289306 | -0.159 | 0.179 | -0.74 | -0.271 | 0.008 | 0.104 | 0.104 | 0.003 | 0.49 | 0.33 | 0.35 | 0.23 | 0.45 | 0.72 |
| 10 | 121555971 | rs12774619 | 0.392 | 0.127 | -0.821 | -0.317 | -0.012 | 0.065 | 0.065 | 0.129 | 0.4 | 0.23 | 0.32 | 0.21 | 0.39 | 0.42 |
| 10 | 121571507 | rs3781503 | -0.24 | 0.13 | -0.712 | -0.302 | 0.006 | 0.047 | 0.047 | 0 | 0.49 | 0.44 | 0.36 | 0.23 | 0.45 | 0.72 |
| 10 | 121580883 | rs10787993 | 1.228 | 0.084 | -1.03 | 0.438 | -0.056 | -0.126 | -0.126 | 0 | 0.19 | 0.03 | 0.27 | 0.39 | 0.19 | 0.16 |
| 10 | 121582556 | rs2289307 | 0.734 | 0.101 | -1.334 | -0.198 | -0.003 | 0.028 | 0.028 | 0 | 0.34 | 0.27 | 0.28 | 0.18 | 0.34 | 0.28 |
| 10 | 121585598 | rs3847492 | 2.35 | 0.022 | -1.315 | -0.157 | -0.026 | 0.012 | 0.012 | 0 | 0.15 | 0.06 | 0.23 | 0.09 | 0.25 | 0.07 |
| 10 | 121586882 | rs3188055 | 0.869 | 0.034 | -1.295 | -0.175 | -0.004 | 0.029 | 0.029 | 0 | 0.33 | 0.27 | 0.28 | 0.18 | 0.34 | 0.28 |
| 10 | 121588537 | rs1048120 | 0.94 | -0.065 | -1.135 | -0.074 | 0.011 | 0.018 | 0.018 | 0 | 0.21 | 0.13 | 0.13 | 0.19 | 0.17 | 0.46 |
| 10 | 121589913 | rs12777827 | 2 | -0.043 | -1.105 | -0.065 | 0.017 | -0.04 | -0.04 | 0 | 0.08 | 0 | 0.03 | 0.19 | 0.05 | 0.32 |
| 10 | 121603681 | rs2475321 | -0.033 | -0.18 | -0.896 | 0.032 | 0.004 | -0.016 | -0.016 | 0 | 0.44 | 0.46 | 0.57 | 0.63 | 0.5 | 0.26 |
| 10 | 121605530 | rs7072901 | 1.662 | -0.172 | -0.9 | 0.045 | 0 | -0.026 | -0.026 | 0 | 0.13 | 0.12 | 0.1 | 0 | 0.11 | 0.14 |
| 10 | 121607024 | rs3740568 | 1.859 | -0.13 | -0.8 | 0.199 | 0.017 | -0.04 | -0.04 | 0 | 0.08 | 0 | 0.03 | 0.19 | 0.05 | 0.32 |
| 10 | 121616262 | rs2456709 | -0.278 | -0.11 | -0.739 | 0.17 | -0.001 | 0.002 | 0.002 | 0 | 0.45 | 0.53 | 0.58 | 0.63 | 0.5 | 0.26 |
| 10 | 121631543 | rs2252156 | 0.193 | -0.123 | -0.642 | 0.169 | -0.002 | 0.027 | 0.027 | 0 | 0.28 | 0.33 | 0.34 | 0.55 | 0.25 | 0.15 |
| 10 | 121635427 | rs6585563 | 1.881 | -0.027 | -0.623 | 0.515 | -0.013 | 0.052 | 0.052 | 1.738 | 0.12 | 0.25 | 0.07 | 0.15 | 0.06 | 0.04 |
| 10 | 121636406 | rs2939931 | -0.19 | -0.077 | -0.927 | -0.261 | -0.005 | 0.023 | 0.023 | 0 | 0.46 | 0.63 | 0.59 | 0.63 | 0.51 | 0.26 |
| 10 | 121648868 | rs11199112 | 1.68 | -0.118 | -0.906 | -0.158 | -0.001 | 0.002 | 0.002 | 0 | 0.07 | 0.05 | 0.05 | 0.11 | 0.06 | 0.02 |
| 10 | 121652180 | rs3808959 | 1.729 | -0.118 | -0.943 | -0.189 | 0.016 | -0.039 | -0.039 | 0 | 0.08 | 0.01 | 0.03 | 0.2 | 0.05 | 0.32 |
| 10 | 121667020 | rs12241478 | 0.75 | -0.035 | -1.047 | -0.168 | -0.024 | -0.026 | -0.026 | 1.439 | 0.23 | 0.08 | 0.25 | 0.28 | 0.3 | 0.4 |
| 10 | 121685559 | rs12771873 | 1.746 | -0.041 | -1.041 | -0.153 | 0.018 | -0.045 | -0.045 | 0 | 0.08 | 0 | 0.03 | 0.21 | 0.05 | 0.31 |
| 10 | 121686097 | rs10788000 | 0.417 | 0.23 | -0.8 | -0.051 | -0.005 | 0.027 | 0.027 | 0 | 0.3 | 0.55 | 0.37 | 0.56 | 0.26 | 0.16 |
| 10 | 121690312 | rs17099385 | 1.629 | 0.237 | -0.816 | -0.029 | 0.001 | 0 | 0 | 0 | 0.07 | 0.05 | 0.05 | 0.12 | 0.06 | 0.02 |
| 10 | 121692908 | rs2901218 | 1.804 | 0.492 | -0.874 | -0.528 | -0.027 | 0.019 | 0.019 | 0 | 0.15 | 0.08 | 0.23 | 0.08 | 0.25 | 0.1 |
| 10 | 121700255 | rs1052289 | -0.091 | 0.523 | -0.907 | -0.504 | -0.006 | 0.022 | 0.022 | 0 | 0.46 | 0.64 | 0.6 | 0.63 | 0.51 | 0.26 |
| 10 | 121701774 | rs3981102 | 0.468 | 0.398 | -0.724 | -0.17 | -0.005 | 0.029 | 0.029 | 0 | 0.3 | 0.42 | 0.35 | 0.56 | 0.26 | 0.16 |
| 10 | 121711232 | rs10749330 | -0.206 | 0.621 | -1.071 | -0.8 | -0.013 | 0.052 | 0.052 | 0 | 0.66 | 0.8 | 0.72 | 0.84 | 0.66 | 0.71 |
| 10 | 121714108 | rs10749331 | 0.019 | 0.087 | -0.336 | -1.121 | -0.004 | -0.109 | -0.109 | 0.061 | 0.81 | 0.64 | 0.86 | 1 | 0.85 | 0.87 |
| 10 | 121715908 | rs10736277 | 0.252 | -0.166 | -0.302 | -0.832 | -0.009 | 0.036 | 0.036 | 0 | 0.83 | 0.73 | 0.88 | 0.73 | 0.87 | 0.76 |
| 10 | 121718112 | rs11199134 | 0.814 | -0.224 | -0.385 | -1.24 | -0.001 | 0.001 | 0.001 | 0.493 | 0.46 | 0.43 | 0.48 | 0.61 | 0.4 | 0.32 |
| 10 | 121721342 | rs10749335 | -0.265 | -0.084 | -0.279 | -0.747 | -0.016 | 0.042 | 0.042 | 0 | 0.86 | 0.77 | 0.91 | 0.72 | 0.9 | 0.74 |
| 10 | 121727655 | rs10886551 | 0.575 | -0.202 | -0.354 | -1.062 | 0.007 | 0.007 | 0.007 | 0 | 0.68 | 0.57 | 0.6 | 0.63 | 0.59 | 0.59 |
| 10 | 121733572 | rs2420654 | -0.428 | -0.011 | -0.417 | -1.283 | -0.008 | -0.009 | -0.009 | 0.031 | 0.86 | 0.79 | 0.88 | 0.88 | 0.87 | 0.75 |
| 10 | 121733993 | rs10788008 | -0.267 | -0.049 | -0.397 | -1.298 | -0.015 | -0.017 | -0.017 | 0 | 0.84 | 0.69 | 0.87 | 0.88 | 0.87 | 0.75 |
| 10 | 121734542 | rs4751759 | 0.775 | -0.038 | -0.451 | -1.338 | -0.008 | -0.009 | -0.009 | 0 | 0.17 | 0.14 | 0.3 | 0.26 | 0.27 | 0.13 |
| 10 | 121736699 | rs3999564 | 0.824 | -0.025 | -0.446 | -1.333 | -0.007 | -0.008 | -0.008 | 0.018 | 0.17 | 0.14 | 0.3 | 0.26 | 0.27 | 0.13 |
| 10 | 121736770 | rs10886553 | 0.282 | -0.059 | -0.492 | -1.304 | 0.004 | 0.007 | 0.007 | 0 | 0.67 | 0.54 | 0.57 | 0.61 | 0.6 | 0.52 |

| | | | | | | | | | | | | | | | | |
|---|---|---|---|---|---|---|---|---|---|---|---|---|---|---|---|---|
| 10 | 121742431 | rs4752360 | 1.232 | -0.955 | -0.691 | -1.782 | -0.011 | 0.013 | 0.013 | 1.003 | 0.37 | 0.21 | 0.42 | 0.67 | 0.34 | 0.22 |
| 10 | 121748543 | rs2050306 | 0.487 | -2.186 | 0.298 | -1.31 | -0.001 | -0.035 | -0.035 | 0 | 0.82 | 0.7 | 0.84 | 0.89 | 0.85 | 0.74 |
| 10 | 121759747 | rs10886561 | 0.993 | -2.887 | 0.013 | -1.67 | 0.067 | 0.071 | 0.071 | 4.005 | 0.53 | 0.22 | 0.56 | 0.68 | 0.67 | 0.51 |
| 10 | 121762190 | rs954639 | 0.795 | -3.049 | 0.319 | -1.749 | -0.018 | 0.011 | 0.011 | 0.015 | 0.72 | 0.52 | 0.65 | 0.68 | 0.77 | 0.6 |
| 10 | 121764195 | rs2065660 | -0.29 | -3.062 | 0.483 | -1.766 | -0.008 | -0.01 | -0.01 | 0 | 0.86 | 0.8 | 0.88 | 0.89 | 0.9 | 0.78 |
| 10 | 121768863 | rs10788013 | 1.146 | -3.085 | 0.534 | -1.769 | 0.046 | 0.049 | 0.049 | 3.427 | 0.54 | 0.32 | 0.57 | 0.68 | 0.66 | 0.52 |
| 10 | 121769013 | rs7096102 | 0.232 | -2.939 | 0.673 | -1.761 | 0.006 | 0.075 | 0.075 | 0.124 | 0.84 | 0.55 | 0.75 | 0.79 | 0.87 | 0.82 |
| 10 | 121772452 | rs2184352 | 0.104 | -2.957 | 0.706 | -1.72 | -0.027 | 0.094 | 0.094 | 0 | 0.12 | 0.28 | 0.2 | 0.21 | 0.08 | 0.17 |
| 10 | 121773070 | rs950367 | 1.145 | -3.084 | 0.696 | -1.798 | 0.082 | 0.091 | 0.091 | 4.973 | 0.53 | 0.16 | 0.56 | 0.68 | 0.67 | 0.52 |
| 10 | 121773850 | rs6585571 | 0.992 | -3.107 | 0.771 | -1.793 | -0.007 | 0 | 0 | 0.087 | 0.72 | 0.37 | 0.68 | 0.68 | 0.82 | 0.61 |
| 10 | 121781774 | rs12768228 | 0.862 | -3.418 | 0.451 | -1.987 | 0.002 | 0.054 | 0.054 | 0.167 | 0.16 | 0.01 | 0.06 | 0.03 | 0.12 | 0.24 |
| 10 | 121785471 | rs2225692 | 0.635 | -3.344 | 0.542 | -1.94 | -0.045 | 0.02 | 0.02 | 0 | 0.74 | 0.51 | 0.69 | 0.68 | 0.82 | 0.61 |
| 10 | 121790576 | rs11199152 | -0.888 | -3.226 | 0.562 | -1.906 | -0.01 | -0.011 | -0.011 | 0 | 0.43 | 0.39 | 0.4 | 0.32 | 0.33 | 0.48 |
| 10 | 121792095 | rs4752371 | 1.778 | -3.195 | 0.479 | -1.904 | -0.033 | 0.002 | 0.002 | 0.48 | 0.37 | 0.17 | 0.5 | 0.65 | 0.55 | 0.27 |
| 10 | 121792404 | rs11594689 | 0 | -3.004 | 0.522 | -1.836 | -0.014 | 0.119 | 0.119 | 4.757 | 0.29 | 0.05 | 0.19 | 0.11 | 0.25 | 0.3 |
| 10 | 121794372 | rs10886576 | -0.111 | -2.966 | 0.664 | -1.82 | 0 | -0.015 | -0.015 | 0 | 0.68 | 0.68 | 0.78 | 0.89 | 0.75 | 0.7 |
| 10 | 121798022 | rs11598346 | 0.863 | -2.973 | 0.621 | -2.088 | 0.002 | 0.071 | 0.071 | 1.152 | 0.21 | 0.08 | 0.14 | 0.02 | 0.17 | 0.15 |
| 10 | 121814553 | rs4288683 | 1.525 | -2.845 | -0.182 | -1.965 | -0.014 | 0.048 | 0.048 | 1.847 | 0.4 | 0.23 | 0.53 | 0.72 | 0.55 | 0.32 |
| 10 | 121819920 | rs10886586 | -0.101 | -2.777 | 0.008 | -1.979 | -0.032 | 0.109 | 0.109 | 0.84 | 0.87 | 0.54 | 0.8 | 0.81 | 0.92 | 0.82 |
| 10 | 121821502 | rs1889441 | 0.52 | -2.703 | -0.207 | -1.984 | 0.051 | 0.082 | 0.082 | 3.707 | 0.55 | 0.33 | 0.6 | 0.75 | 0.67 | 0.56 |
| 10 | 121825959 | rs4415675 | 0.53 | -2.315 | 0.098 | -2.046 | -0.057 | 0.027 | 0.027 | 0 | 0.74 | 0.56 | 0.73 | 0.71 | 0.81 | 0.56 |
| 10 | 121836959 | rs11596124 | 0 | -2.477 | -0.049 | -2.188 | 0.008 | 0.003 | 0.003 | 2.801 | 0.31 | 0.26 | 0.19 | 0.08 | 0.25 | 0.23 |
| 10 | 121837285 | rs10886590 | 1.554 | -2.558 | 0.038 | -2.214 | -0.004 | 0.018 | 0.018 | 0.52 | 0.41 | 0.22 | 0.52 | 0.64 | 0.56 | 0.33 |
| 10 | 121838912 | rs12355817 | 0.528 | -2.553 | 0.06 | -2.203 | 0.006 | 0.202 | 0.202 | 0.327 | 0.1 | 0.29 | 0.21 | 0.25 | 0.08 | 0.15 |
| 10 | 121839738 | rs11593681 | 0.062 | -2.552 | -0.542 | -2.223 | 0.016 | 0.046 | 0.046 | 2.874 | 0.31 | 0.18 | 0.18 | 0.08 | 0.25 | 0.24 |
| 10 | 121841944 | rs11593974 | -0.838 | -2.541 | -0.547 | -2.217 | 0.054 | 0.161 | 0.161 | 0 | 0.46 | 0.23 | 0.24 | 0.11 | 0.36 | 0.52 |
| 10 | 121845376 | rs4752377 | 0.143 | -2.285 | -0.651 | -2.156 | 0.02 | 0.02 | 0.02 | 5.878 | 0.59 | 0.38 | 0.59 | 0.67 | 0.67 | 0.6 |
| 10 | 121852519 | rs11594353 | 0.758 | -2.449 | -0.712 | -2.201 | 0.005 | -0.02 | -0.02 | 3.451 | 0.23 | 0.2 | 0.14 | 0.02 | 0.17 | 0.16 |
| 10 | 121852646 | rs4751765 | 1.563 | -2.386 | -0.774 | -2.225 | -0.036 | -0.001 | -0.001 | 0.002 | 0.4 | 0.18 | 0.51 | 0.63 | 0.54 | 0.28 |
| 10 | 121855554 | rs10788034 | 0.943 | -2.429 | -0.598 | -2.23 | 0 | 0.019 | 0.019 | 0.002 | 0.08 | 0.08 | 0.2 | 0.25 | 0.08 | 0.16 |
| 10 | 121856417 | rs7906272 | -0.764 | -2.147 | -0.955 | -2.161 | 0.008 | -0.031 | -0.031 | 0 | 0.42 | 0.39 | 0.22 | 0.05 | 0.28 | 0.46 |
| 10 | 121859814 | rs1467785 | -0.03 | -1.873 | -0.856 | -1.875 | 0.025 | 0.067 | 0.067 | 0.029 | 0.16 | 0.04 | 0.06 | 0.03 | 0.11 | 0.33 |
| 10 | 121860349 | rs7923882 | -0.374 | -1.187 | -0.305 | -1.427 | 0.005 | 0.098 | 0.098 | 0 | 0.81 | 0.6 | 0.7 | 0.68 | 0.82 | 0.76 |
| 10 | 121862405 | rs1929677 | -0.35 | -1.137 | -0.858 | -1.612 | -0.012 | 0.049 | 0.049 | 0 | 0.79 | 0.44 | 0.78 | 0.69 | 0.82 | 0.75 |
| 10 | 121865347 | rs10886596 | 0.854 | -1.051 | -1.275 | -1.615 | -0.009 | 0.039 | 0.039 | 1.42 | 0.44 | 0.12 | 0.6 | 0.64 | 0.57 | 0.28 |
| 10 | 121868586 | rs1929670 | -0.468 | -0.952 | -1.143 | -1.563 | -0.01 | 0.049 | 0.049 | 0 | 0.79 | 0.42 | 0.79 | 0.7 | 0.83 | 0.76 |
| 10 | 121871353 | rs4752384 | 0.078 | 0.189 | -1.608 | -0.9 | 0.014 | 0.049 | 0.049 | 0.247 | 0.46 | 0.25 | 0.62 | 0.64 | 0.57 | 0.27 |
| 10 | 121872112 | rs7084094 | 0.077 | 0.161 | -1.596 | -0.864 | 0.02 | 0.051 | 0.051 | 0.356 | 0.46 | 0.25 | 0.62 | 0.64 | 0.58 | 0.27 |
| 10 | 121874756 | rs4237517 | 0.203 | 0.575 | -1.357 | -0.591 | -0.03 | 0.011 | 0.011 | 0 | 0.39 | 0.26 | 0.51 | 0.62 | 0.46 | 0.27 |
| 10 | 121875612 | rs17622645 | -1.15 | 0.794 | -1.06 | -0.561 | -0.005 | 0.031 | 0.031 | 0 | 0.1 | 0.01 | 0.06 | 0.03 | 0.08 | 0.19 |
| 10 | 121875721 | rs7894011 | -1.78 | 1.227 | -0.624 | -0.287 | 0.013 | 0.013 | 0.013 | 0 | 0.06 | 0.13 | 0.02 | 0.09 | 0.07 | 0.17 |
| 10 | 121876227 | rs10788037 | -0.873 | 1.37 | -0.36 | -1.181 | 0.009 | -0.124 | -0.124 | 2.822 | 0.34 | 0.42 | 0.22 | 0 | 0.36 | 0.19 |
| 10 | 121876520 | rs10788038 | 0.53 | 1.005 | 0.003 | -0.865 | -0.003 | -0.085 | -0.085 | 0 | 0.66 | 0.46 | 0.65 | 0.91 | 0.67 | 0.69 |
| 10 | 121901799 | rs11199183 | -0.871 | 0.859 | 0.137 | 0.232 | -0.047 | -0.098 | -0.098 | 0 | 0.16 | 0.01 | 0.2 | 0.26 | 0.16 | 0.15 |
| 10 | 121902150 | rs928381 | 1.035 | 0.555 | 0.919 | -0.352 | -0.029 | -0.1 | -0.1 | 0 | 0.86 | 0.46 | 0.89 | 0.99 | 0.92 | 0.85 |
| 10 | 121903974 | rs1041517 | 1.202 | 0.69 | 1.014 | -0.045 | 0.002 | -0.004 | -0.004 | 0 | 0.73 | 0.67 | 0.81 | 0.77 | 0.75 | 0.62 |
| 10 | 121905620 | rs10749351 | 0.853 | 0.528 | 1.076 | 0.114 | -0.005 | 0.001 | 0.001 | 0.188 | 0.8 | 0.78 | 0.87 | 0.76 | 0.84 | 0.69 |
| 10 | 121907055 | rs11199185 | 2.099 | 0.538 | 1.285 | 0.067 | -0.006 | 0.032 | 0.032 | 0.289 | 0.47 | 0.22 | 0.58 | 0.6 | 0.52 | 0.38 |
| 10 | 121916062 | rs7915072 | 1.716 | 0.486 | 1.487 | 0.05 | -0.002 | -0.018 | -0.018 | 0 | 0.56 | 0.49 | 0.65 | 0.71 | 0.59 | 0.49 |
| 10 | 121920122 | rs6585586 | 1.391 | 0.498 | 1.479 | 0.016 | -0.002 | -0.015 | -0.015 | 0 | 0.66 | 0.58 | 0.77 | 0.72 | 0.7 | 0.57 |
| 10 | 121922699 | rs7907490 | -2.582 | 0.426 | 1.804 | 0.007 | -0.006 | -0.037 | -0.037 | 0 | 0.35 | 0.62 | 0.27 | 0.19 | 0.3 | 0.43 |
| 10 | 121923674 | rs10886614 | -0.002 | 0.37 | 1.499 | 0.033 | -0.036 | -0.06 | -0.06 | 0 | 0.16 | 0.01 | 0.1 | 0.22 | 0.15 | 0.19 |
| 10 | 121926720 | rs4390268 | 1.491 | 0.331 | 2.003 | 0.015 | -0.047 | 0.009 | 0.009 | 0.928 | 0.7 | 0.48 | 0.81 | 0.65 | 0.79 | 0.71 |
| 10 | 121930730 | rs10749352 | 0.971 | 0.421 | 2.211 | 0.061 | -0.038 | 0.031 | 0.031 | 1.629 | 0.75 | 0.46 | 0.82 | 0.65 | 0.81 | 0.73 |
| 10 | 121932750 | rs4529826 | 0.094 | 0.419 | 2.243 | 0.069 | 0.001 | -0.004 | -0.004 | 0.799 | 0.83 | 0.85 | 0.89 | 0.77 | 0.85 | 0.81 |
| 10 | 121932833 | rs4427485 | 2.187 | 0.463 | 2.207 | 0.037 | 0.006 | 0.043 | 0.043 | 0.185 | 0.49 | 0.24 | 0.6 | 0.6 | 0.52 | 0.51 |
| 10 | 121936401 | rs10886617 | 1.141 | 0.414 | 2.09 | 0.049 | -0.044 | 0.029 | 0.029 | 1.544 | 0.73 | 0.46 | 0.81 | 0.64 | 0.81 | 0.59 |
| 10 | 121938619 | rs2420658 | 2.212 | 0.46 | 1.865 | 0.044 | -0.006 | 0.046 | 0.046 | 0.294 | 0.48 | 0.19 | 0.61 | 0.59 | 0.54 | 0.38 |
| 10 | 121940717 | rs7915835 | -0.309 | 0.484 | 1.95 | -0.064 | 0 | -0.046 | -0.046 | 0.02 | 0.1 | 0.16 | 0.1 | 0 | 0.1 | 0.08 |
| 10 | 121945631 | rs928385 | 2.28 | 0.473 | 1.584 | -0.006 | -0.033 | 0.066 | 0.066 | 0.375 | 0.47 | 0.11 | 0.6 | 0.63 | 0.54 | 0.38 |
| 10 | 121947114 | rs1007771 | 2.094 | 0.463 | 1.631 | -0.012 | -0.002 | 0.078 | 0.078 | 0.095 | 0.49 | 0.19 | 0.61 | 0.63 | 0.54 | 0.51 |
| 10 | 121947698 | rs10886624 | 2.235 | 0.47 | 1.534 | -0.006 | -0.031 | 0.061 | 0.061 | 0.265 | 0.47 | 0.11 | 0.6 | 0.62 | 0.54 | 0.39 |
| 10 | 121950989 | rs11199206 | -1.312 | 0.533 | 1.693 | 0.048 | -0.001 | 0.013 | 0.013 | 0 | 0.26 | 0.16 | 0.21 | 0.2 | 0.27 | 0.26 |
| 10 | 121952250 | rs4376826 | -1.951 | 0.522 | 1.983 | 0.026 | 0 | -0.006 | -0.006 | 0 | 0.4 | 0.38 | 0.33 | 0.26 | 0.41 | 0.38 |
| 10 | 121952638 | rs12770480 | -0.252 | 0.496 | 1.975 | 0.059 | 0 | -0.016 | -0.016 | 0 | 0.09 | 0.06 | 0.1 | 0 | 0.1 | 0.08 |
| 10 | 121954927 | rs7092944 | 0.677 | 0.456 | 2.215 | 0.207 | -0.027 | 0.048 | 0.048 | 0.109 | 0.93 | 0.7 | 0.94 | 0.87 | 0.99 | 0.92 |
| 10 | 121955348 | rs10886626 | -0.233 | 0.46 | 2.215 | 0.237 | -0.002 | -0.008 | -0.008 | 0 | 0.16 | 0.09 | 0.11 | 0.21 | 0.16 | 0.18 |
| 10 | 121955896 | rs1929698 | 0.748 | 0.458 | 2.49 | 0.24 | -0.024 | 0.053 | 0.053 | 0 | 0.93 | 0.7 | 0.95 | 0.87 | 0.99 | 0.91 |
| 10 | 121962068 | rs1929690 | -0.705 | 0.49 | 2.023 | 0.235 | 0.001 | -0.007 | -0.007 | 0 | 0.16 | 0.17 | 0.11 | 0.05 | 0.17 | 0.13 |

| | | | | | | | | | | | | | | | | |
|---|---|---|---|---|---|---|---|---|---|---|---|---|---|---|---|---|
| 10 | 121963461 | rs10736286 | 1.852 | 0.47 | 2.104 | 0.229 | -0.011 | -0.021 | -0.021 | 0 | 0.73 | 0.28 | 0.8 | 0.82 | 0.81 | 0.67 |
| 10 | 121964330 | rs7069652 | -0.835 | 0.971 | 1.171 | 0.231 | 0.002 | -0.008 | -0.008 | 0 | 0.16 | 0.17 | 0.11 | 0.05 | 0.17 | 0.13 |
| 10 | 121965565 | rs7078050 | -0.894 | 0.991 | 0.986 | 0.24 | 0.004 | -0.015 | -0.015 | 0.11 | 0.16 | 0.19 | 0.13 | 0.05 | 0.17 | 0.13 |
| 10 | 121969111 | rs7093218 | -1.752 | 0.787 | 1.331 | 0.724 | -0.015 | 0.054 | 0.054 | 0 | 0.19 | 0.42 | 0.15 | 0.23 | 0.13 | 0.25 |
| 10 | 121973334 | rs7896655 | 1.539 | 0.614 | 1.999 | 0.584 | -0.03 | -0.05 | -0.05 | 0.068 | 0.83 | 0.6 | 0.87 | 0.91 | 0.89 | 0.78 |
| 10 | 121981248 | rs7075789 | 0.947 | 0.581 | 2.05 | 0.571 | 0.008 | 0.074 | 0.074 | 0 | 0.94 | 0.72 | 0.97 | 0.91 | 0.98 | 0.92 |
| 10 | 121983126 | rs1929666 | 2.041 | 0.562 | 2.072 | 0.562 | -0.032 | -0.049 | -0.049 | 0 | 0.83 | 0.59 | 0.87 | 0.91 | 0.89 | 0.78 |
| 10 | 122003502 | rs1929648 | 0.52 | 0.664 | 1.775 | 0.576 | 0 | -0.028 | -0.028 | 0 | 0.91 | 0.89 | 0.91 | 1 | 0.91 | 0.94 |
| 10 | 122007448 | rs1999414 | 1.404 | 0.987 | 1.476 | 0.473 | 0 | -0.002 | -0.002 | 0 | 0.6 | 0.38 | 0.67 | 0.76 | 0.57 | 0.71 |
| 10 | 122008026 | rs7394051 | -0.538 | 1.275 | 1.754 | 0.457 | -0.003 | 0.003 | 0.003 | 0 | 0.24 | 0.24 | 0.22 | 0.14 | 0.31 | 0.12 |
| 10 | 122008230 | rs12243526 | 2.33 | 1.581 | 0.976 | 0.515 | -0.007 | 0.023 | 0.023 | 0.007 | 0.37 | 0.03 | 0.22 | 0.31 | 0.31 | 0.33 |
| 10 | 122008352 | rs2026959 | -1.394 | 1.717 | 0.936 | 0.421 | 0.002 | 0.023 | 0.023 | 0 | 0.47 | 0.59 | 0.65 | 0.68 | 0.57 | 0.43 |
| 10 | 122013624 | rs7079943 | 0.123 | 2.058 | 1.207 | 1.076 | 0.001 | 0.035 | 0.035 | 0 | 0.12 | 0.15 | 0.23 | 0.31 | 0.14 | 0.2 |
| 10 | 122013947 | rs11199245 | -0.949 | 2.075 | 1.03 | 0.932 | 0.025 | 0.107 | 0.107 | 0 | 0.19 | 0.33 | 0.42 | 0.54 | 0.22 | 0.31 |
| 10 | 122020271 | rs10788063 | 1.133 | 2.072 | 1.269 | 0.72 | -0.002 | -0.002 | -0.002 | 0 | 0.6 | 0.37 | 0.65 | 0.86 | 0.57 | 0.64 |
| 10 | 122020617 | rs10510063 | 0.084 | 2.028 | 0.874 | 0.84 | -0.009 | 0.01 | 0.01 | 0 | 0.23 | 0.17 | 0.2 | 0.14 | 0.3 | 0.11 |
| 10 | 122030053 | rs10886652 | 1.108 | 1.928 | 1.249 | 0.972 | 0.004 | -0.001 | -0.001 | 0 | 0.53 | 0.39 | 0.43 | 0.6 | 0.49 | 0.57 |
| 10 | 122033526 | rs10510066 | -0.436 | 2.198 | 0.929 | 1.219 | 0.015 | 0.028 | 0.028 | 0 | 0.13 | 0.19 | 0.31 | 0.24 | 0.17 | 0.14 |
| 10 | 122035486 | rs2420680 | 0.92 | 2.504 | 0.977 | 1.297 | -0.01 | -0.007 | -0.007 | 0 | 0.6 | 0.59 | 0.37 | 0.53 | 0.59 | 0.56 |
| 10 | 122035596 | rs2420679 | -0.001 | 2.454 | 0.621 | 1.189 | -0.007 | -0.009 | -0.009 | 0.008 | 0.75 | 0.81 | 0.69 | 0.77 | 0.77 | 0.89 |
| 10 | 122041934 | rs4339955 | -1.609 | 2.801 | 1.35 | 1.522 | 0.02 | 0.049 | 0.049 | 0.035 | 0.19 | 0.3 | 0.17 | 0.36 | 0.26 | 0.37 |
| 10 | 122046508 | rs10886654 | -0.683 | 2.558 | 1.253 | 1.42 | 0.024 | 0.013 | 0.013 | 0 | 0.16 | 0.22 | 0.34 | 0.21 | 0.21 | 0.22 |
| 10 | 122049925 | rs10430700 | -1.274 | 2.369 | 1.119 | 1.414 | 0.043 | 0.031 | 0.031 | 0 | 0.2 | 0.45 | 0.37 | 0.23 | 0.19 | 0.39 |
| 10 | 122057157 | rs11199264 | -0.855 | 2.385 | 1.133 | 1.403 | 0.039 | 0.028 | 0.028 | 0 | 0.18 | 0.33 | 0.35 | 0.2 | 0.18 | 0.22 |
| 10 | 122065032 | rs2103332 | -1.494 | 2.259 | 1.276 | 1.348 | 0.022 | 0.007 | 0.007 | 0 | 0.23 | 0.48 | 0.37 | 0.2 | 0.18 | 0.37 |
| 10 | 122066549 | rs11199268 | -0.707 | 2.276 | 1.251 | 1.355 | 0.042 | 0.03 | 0.03 | 0 | 0.17 | 0.33 | 0.35 | 0.2 | 0.18 | 0.22 |
| 10 | 122067677 | rs11199269 | -1.255 | 2.269 | 1.22 | 1.352 | 0.03 | 0.014 | 0.014 | 0 | 0.21 | 0.46 | 0.37 | 0.2 | 0.18 | 0.22 |
| 10 | 122068766 | rs10788066 | -2.334 | 2.279 | 1.236 | 1.477 | -0.001 | -0.004 | -0.004 | 0.159 | 0.33 | 0.69 | 0.43 | 0.42 | 0.25 | 0.45 |
| 10 | 122069164 | rs10886662 | -1.1 | 2.24 | 1.253 | 1.46 | 0.05 | 0.034 | 0.034 | 0 | 0.2 | 0.43 | 0.36 | 0.21 | 0.18 | 0.37 |
| 10 | 122070436 | rs4300322 | -1.373 | 1.719 | 1.15 | 1.503 | -0.022 | 0.036 | 0.036 | 0 | 0.14 | 0.31 | 0.06 | 0.21 | 0.07 | 0.12 |
| 10 | 122072802 | rs2420684 | -0.971 | 2.185 | 1.02 | 1.257 | 0.012 | 0.009 | 0.009 | 0 | 0.23 | 0.38 | 0.35 | 0.21 | 0.2 | 0.42 |
| 10 | 122073670 | rs11199272 | -2.071 | 2.144 | 0.779 | 1.332 | 0.005 | -0.014 | -0.014 | 0.384 | 0.35 | 0.78 | 0.43 | 0.33 | 0.27 | 0.52 |
| 10 | 122079594 | rs9732028 | -0.643 | 1.998 | 0.91 | 1.231 | 0.017 | 0.022 | 0.022 | 0 | 0.21 | 0.37 | 0.35 | 0.21 | 0.2 | 0.22 |
| 10 | 122087833 | rs3847499 | -1.87 | 2.701 | 0.111 | 1.356 | -0.046 | -0.05 | -0.05 | 0 | 0.4 | 0.79 | 0.53 | 0.47 | 0.48 | 0.6 |
| 10 | 122088351 | rs3847501 | -1.138 | 2.574 | 0.172 | 1.39 | 0.01 | 0.007 | 0.007 | 0 | 0.15 | 0.18 | 0.16 | 0.26 | 0.28 | 0.16 |
| 10 | 122094271 | rs7067775 | -1.111 | 1.511 | 0.202 | 1.289 | 0.002 | 0.01 | 0.01 | 0 | 0.25 | 0.47 | 0.25 | 0.22 | 0.22 | 0.44 |
| 10 | 122099219 | rs7075775 | -1.764 | 1.509 | 0.215 | 1.498 | 0.006 | -0.017 | -0.017 | 0.375 | 0.34 | 0.67 | 0.3 | 0.46 | 0.27 | 0.57 |
| 10 | 122101832 | rs11813317 | -0.806 | 1.503 | 0.285 | 1.592 | -0.009 | 0.157 | 0.157 | 0 | 0.1 | 0.29 | 0.06 | 0.26 | 0.05 | 0.15 |
| 10 | 122105511 | rs2420699 | -0.357 | 1.471 | -0.891 | 1.073 | -0.01 | 0.045 | 0.045 | 0 | 0.76 | 0.88 | 0.59 | 0.83 | 0.77 | 0.86 |
| 10 | 122113893 | rs997856 | 0.8 | 1.476 | -1.18 | 1.168 | 0.002 | 0.029 | 0.029 | 0 | 0.53 | 0.33 | 0.35 | 0.62 | 0.57 | 0.43 |
| 10 | 122119574 | rs12765856 | 1.096 | 1.551 | -1.8 | 1.108 | -0.105 | -0.03 | -0.03 | 0 | 0.47 | 0.05 | 0.31 | 0.39 | 0.55 | 0.38 |
| 10 | 122121602 | rs10886670 | -0.26 | 1.474 | -1.473 | 0.721 | -0.035 | 0.067 | 0.067 | 0 | 0.24 | 0.05 | 0.42 | 0.18 | 0.27 | 0.14 |
| 10 | 122121788 | rs10886671 | 0.587 | 1.397 | -1.168 | 1.046 | -0.003 | 0.002 | 0.002 | 0 | 0.49 | 0.24 | 0.33 | 0.52 | 0.52 | 0.47 |
| 10 | 122124595 | rs11199297 | 0.114 | 0.869 | -0.987 | 1.426 | -0.004 | -0.042 | -0.042 | 1.098 | 0.09 | 0.01 | 0.06 | 0.21 | 0.07 | 0.04 |
| 10 | 122128960 | rs2420703 | 0.938 | 0.582 | 0.244 | 1.838 | -0.007 | 0.23 | 0.23 | 0.839 | 0.9 | 0.43 | 0.84 | 0.72 | 0.94 | 0.81 |
| 10 | 122133913 | rs2420706 | 0.82 | 0.592 | 0.479 | 1.842 | 0.009 | 0.279 | 0.279 | 0.762 | 0.9 | 0.48 | 0.84 | 0.73 | 0.94 | 0.82 |
| 10 | 122137715 | rs11199302 | 0.022 | 0.59 | 0.351 | 2.159 | -0.003 | 0.089 | 0.089 | 0.506 | 0.05 | 0.07 | 0.11 | 0.3 | 0.06 | 0.18 |
| 10 | 122141486 | rs2420707 | -1.497 | 0.469 | 0.805 | 1.746 | -0.006 | 0.134 | 0.134 | 0.499 | 0.09 | 0.3 | 0.05 | 0.2 | 0.06 | 0.21 |
| 10 | 122141876 | rs7069042 | 1.59 | 0.24 | 0.71 | 1.625 | -0.026 | 0.029 | 0.029 | 1.011 | 0.83 | 0.27 | 0.81 | 0.66 | 0.92 | 0.69 |
| 10 | 122150076 | rs10788079 | 0.3 | 0.7 | -0.209 | 1.189 | -0.154 | 0.026 | 0.026 | 0 | 0.43 | 0.03 | 0.43 | 0.31 | 0.52 | 0.37 |
| 10 | 122150121 | rs7073577 | -0.388 | 0.842 | 0.981 | 1.123 | 0.002 | 0.116 | 0.116 | 0.265 | 0.06 | 0.17 | 0.03 | 0.19 | 0.06 | 0.17 |
| 10 | 122155151 | rs11595096 | -0.059 | 0.848 | 0.683 | 0.919 | -0.061 | 0.111 | 0.111 | 0 | 0.18 | 0.02 | 0.24 | 0 | 0.21 | 0.15 |
| 10 | 122159833 | rs12571802 | -0.476 | 0.761 | 1.635 | 1.768 | -0.003 | 0.068 | 0.068 | 0.499 | 0.05 | 0.1 | 0.03 | 0.19 | 0.06 | 0.18 |
| 10 | 122161794 | rs7913921 | -0.681 | 0.761 | 1.62 | 1.768 | -0.003 | 0.087 | 0.087 | 0.312 | 0.06 | 0.14 | 0.04 | 0.19 | 0.06 | 0.18 |
| 10 | 122170293 | rs17100130 | -0.743 | 0.989 | 0.06 | 1.66 | -0.005 | 0.07 | 0.07 | 0.588 | 0.06 | 0.11 | 0.03 | 0.19 | 0.06 | 0.18 |
| 10 | 122174562 | rs7916482 | -1.659 | 1.363 | -0.051 | 1.549 | -0.001 | 0 | 0 | 0.21 | 0.18 | 0.74 | 0.18 | 0.31 | 0.19 | 0.28 |
| 10 | 122175979 | rs2420717 | -1.952 | 1.162 | 0.641 | 1.765 | 0.02 | 0.094 | 0.094 | 0.376 | 0.17 | 0.61 | 0.26 | 0.32 | 0.2 | 0.33 |
| 10 | 122176608 | rs12254609 | -1.952 | 1.213 | 0.641 | 1.768 | 0.023 | 0.084 | 0.084 | 0.376 | 0.17 | 0.61 | 0.26 | 0.3 | 0.2 | 0.33 |
| 10 | 122178140 | rs10788085 | -0.693 | 1.271 | 0.672 | 1.818 | -0.006 | 0.01 | 0.01 | 0 | 0.06 | 0.06 | 0.15 | 0.17 | 0.13 | 0.25 |
| 10 | 122181385 | rs10749375 | -0.74 | 1.244 | 0.687 | 1.747 | -0.004 | 0.025 | 0.025 | 0 | 0.06 | 0.07 | 0.15 | 0.21 | 0.12 | 0.29 |
| 10 | 122185868 | rs1326647 | 0.286 | 1.201 | 0.284 | 1.458 | 0 | 0 | 0 | 0 | 0.54 | 0.52 | 0.57 | 0.55 | 0.53 | 0.48 |
| 10 | 122189275 | rs7068028 | 0.407 | 1.061 | 0.26 | 1.124 | -0.009 | -0.03 | -0.03 | 0 | 0.06 | 0.14 | 0.03 | 0 | 0.03 | 0 |
| 10 | 122191973 | rs10886685 | 1.272 | 1.06 | -0.145 | 1.229 | -0.06 | 0.128 | 0.128 | 2.386 | 0.4 | 0.03 | 0.46 | 0.18 | 0.41 | 0.33 |
| 10 | 122192911 | rs2478408 | 1.695 | 1.085 | 1.104 | 1.511 | 0.016 | 0.144 | 0.144 | 0 | 0.8 | 0.55 | 0.76 | 0.59 | 0.8 | 0.55 |
| 10 | 122199895 | rs6585625 | -0.613 | 1.106 | 1.367 | 1.002 | 0.001 | -0.025 | -0.025 | 0 | 0.05 | 0.15 | 0.04 | 0.01 | 0.03 | 0.1 |
| 10 | 122213468 | rs11199345 | -0.648 | 1.241 | 1.11 | 1.245 | 0.037 | 0.063 | 0.063 | 0 | 0.05 | 0.22 | 0.04 | 0.09 | 0.05 | 0.04 |
| 10 | 122213646 | rs10886690 | -1.664 | 1.575 | 1.149 | 1.48 | 0.067 | 0.132 | 0.132 | 0 | 0.15 | 0.37 | 0.23 | 0.32 | 0.24 | 0.37 |
| 10 | 122215377 | rs2463147 | 0.426 | 0.914 | 0.777 | 0.816 | -0.006 | -0.01 | -0.01 | 0 | 0.54 | 0.59 | 0.7 | 0.62 | 0.64 | 0.79 |
| 10 | 122217471 | rs1571103 | -0.712 | 0.769 | 0.869 | 0.855 | -0.009 | -0.014 | -0.014 | 0 | 0.4 | 0.52 | 0.25 | 0.28 | 0.25 | 0.15 |

| | | | | | | | | | | | | | | | | |
|---|---|---|---|---|---|---|---|---|---|---|---|---|---|---|---|---|
| 10 | 122218655 | rs1041168 | -0.725 | 0.865 | 0.778 | 0.833 | -0.005 | 0.04 | 0.04 | 0.059 | 0.27 | 0.45 | 0.24 | 0.38 | 0.2 | 0.2 |
| 10 | 122221495 | rs2182514 | -1.381 | 0.423 | 1.069 | 0.777 | -0.018 | -0.005 | -0.005 | 1.498 | 0.2 | 0.67 | 0.14 | 0.22 | 0.14 | 0.11 |
| 10 | 122225978 | rs2478432 | 1.126 | -0.117 | 1.537 | 0.649 | -0.044 | -0.059 | -0.059 | 0.005 | 0.76 | 0.55 | 0.88 | 0.91 | 0.88 | 0.82 |
| 10 | 122231245 | rs10886697 | 1.161 | -0.358 | 1.659 | 0.573 | -0.018 | -0.029 | -0.029 | 0 | 0.65 | 0.42 | 0.85 | 0.91 | 0.84 | 0.8 |
| 10 | 122238850 | rs4752423 | 1.134 | -0.061 | 1.676 | 0.54 | -0.016 | -0.031 | -0.031 | 0.002 | 0.65 | 0.42 | 0.84 | 0.91 | 0.81 | 0.8 |
| 10 | 122239999 | rs4237520 | 1.087 | -0.081 | 1.964 | 0.535 | -0.04 | -0.065 | -0.065 | 0 | 0.66 | 0.56 | 0.85 | 0.91 | 0.81 | 0.8 |
| 10 | 122240609 | rs2254865 | 0.134 | -0.143 | 1.991 | 0.529 | -0.023 | 0.025 | 0.025 | 0 | 0.89 | 0.73 | 0.92 | 0.92 | 0.95 | 0.96 |
| 10 | 122246287 | rs4595466 | 0.422 | -0.029 | 0.919 | 0.401 | -0.014 | -0.018 | -0.018 | 0 | 0.33 | 0.25 | 0.45 | 0.41 | 0.36 | 0.37 |
| 10 | 122246966 | rs2244544 | -1.204 | -0.017 | 1.017 | 0.286 | -0.01 | 0.005 | 0.005 | 0.18 | 0.61 | 0.77 | 0.61 | 0.7 | 0.57 | 0.59 |
| 10 | 122248600 | rs1326662 | 0.137 | 0.071 | 0.699 | 0.324 | 0.004 | 0 | 0 | 0.036 | 0.37 | 0.43 | 0.52 | 0.69 | 0.41 | 0.45 |
| 10 | 122249485 | rs2478425 | -0.787 | -0.159 | 0.582 | 0.222 | -0.013 | -0.024 | -0.024 | 0 | 0.26 | 0.35 | 0.13 | 0.24 | 0.2 | 0.23 |
| 10 | 122256857 | rs2478422 | -0.692 | -0.465 | 0.608 | 0.395 | -0.038 | -0.038 | -0.038 | 0 | 0.2 | 0.35 | 0.17 | 0.11 | 0.1 | 0.15 |
| 10 | 122264200 | rs1536389 | -0.234 | -0.351 | -0.124 | 0.454 | -0.031 | 0.031 | 0.031 | 0.578 | 0.65 | 0.86 | 0.68 | 0.74 | 0.59 | 0.56 |
| 10 | 122265668 | rs10510070 | -0.443 | -0.477 | -0.111 | 0.385 | 0.002 | -0.002 | -0.002 | 2.396 | 0.1 | 0.08 | 0.13 | 0.11 | 0.08 | 0.17 |
| 10 | 122270126 | rs12769643 | 1.531 | -0.685 | 0.633 | 0.698 | -0.003 | -0.003 | -0.003 | 0 | 0.39 | 0.36 | 0.46 | 0.5 | 0.45 | 0.33 |
| 10 | 122270209 | rs12770951 | 1.127 | -0.696 | 0.972 | 0.498 | 0.006 | -0.009 | -0.009 | 0.176 | 0.43 | 0.43 | 0.49 | 0.66 | 0.48 | 0.35 |
| 10 | 122270417 | rs1571101 | -0.707 | -0.694 | 1.004 | 0.477 | 0.001 | 0.024 | 0.024 | 0.017 | 0.45 | 0.34 | 0.46 | 0.33 | 0.47 | 0.61 |
| 10 | 122271671 | rs7077111 | -0.715 | -0.693 | 0.951 | 0.477 | -0.002 | 0.046 | 0.046 | 0.023 | 0.45 | 0.28 | 0.45 | 0.33 | 0.47 | 0.61 |
| 10 | 122275933 | rs4752425 | -0.747 | -0.696 | 1.335 | 0.684 | -0.049 | -0.01 | -0.01 | 0 | 0.86 | 0.54 | 0.92 | 0.92 | 0.95 | 0.89 |
| 10 | 122277211 | rs4237521 | -0.948 | -0.67 | 1.352 | 0.674 | -0.043 | -0.028 | -0.028 | 0.025 | 0.87 | 0.55 | 0.93 | 0.95 | 0.95 | 0.9 |
| 10 | 122285398 | rs11199378 | 1.068 | -0.424 | 0.457 | 0.771 | -0.03 | -0.006 | -0.006 | 0 | 0.42 | 0.22 | 0.46 | 0.59 | 0.5 | 0.31 |
| 10 | 122291305 | rs12768425 | 1.142 | -0.304 | 0.213 | 0.625 | -0.042 | 0.015 | 0.015 | 0.065 | 0.41 | 0.23 | 0.48 | 0.62 | 0.5 | 0.28 |
| 10 | 122292066 | rs11597798 | 0.004 | 0.315 | -0.345 | 0.678 | -0.05 | -0.018 | -0.018 | 0 | 0.08 | 0.01 | 0.12 | 0.08 | 0.17 | 0.2 |
| 10 | 122292377 | rs10886706 | 0.41 | -0.474 | -0.246 | 0.521 | 0.016 | 0.005 | 0.005 | 2.318 | 0.2 | 0.15 | 0.27 | 0.15 | 0.14 | 0.14 |
| 10 | 122294000 | rs10788099 | 1.171 | -0.002 | -0.011 | 0.719 | 0 | -0.001 | -0.001 | 0 | 0.41 | 0.41 | 0.45 | 0.6 | 0.43 | 0.28 |
| 10 | 122294034 | rs10886707 | 0.486 | 0.072 | -0.172 | 0.541 | -0.002 | -0.002 | -0.002 | 0 | 0.55 | 0.55 | 0.41 | 0.48 | 0.5 | 0.36 |
| 10 | 122295036 | rs10788100 | -1.37 | -0.722 | 0.173 | 0.29 | -0.02 | -0.061 | -0.061 | 0.399 | 0.89 | 0.79 | 0.96 | 1 | 0.95 | 0.98 |
| 10 | 122295262 | rs7069941 | -0.256 | -0.875 | 0.202 | 0.17 | -0.004 | -0.009 | -0.009 | 0.575 | 0.11 | 0.13 | 0.15 | 0.04 | 0.07 | 0.11 |
| 10 | 122298030 | rs2420724 | 1.178 | -1.247 | 0.129 | 0.246 | 0.038 | 0.002 | 0.002 | 0.071 | 0.46 | 0.32 | 0.47 | 0.57 | 0.62 | 0.36 |
| 10 | 122298366 | rs2901245 | 0.678 | -1.371 | 0.209 | 0.156 | -0.011 | -0.015 | -0.015 | 0 | 0.82 | 0.77 | 0.82 | 0.86 | 0.88 | 0.68 |
| 10 | 122301705 | rs978855 | 2.005 | -1.321 | 0.28 | 0.103 | -0.016 | -0.046 | -0.046 | 0 | 0.1 | 0.12 | 0.04 | 0 | 0.05 | 0.02 |
| 10 | 122302038 | rs978854 | 0.298 | -1.411 | 0.394 | 0.168 | 0.007 | 0.009 | 0.009 | 0 | 0.65 | 0.68 | 0.64 | 0.86 | 0.73 | 0.73 |
| 10 | 122303500 | rs7910507 | 0.684 | -1.379 | 0.339 | 0.181 | -0.017 | -0.031 | -0.031 | 0 | 0.16 | 0.19 | 0.06 | 0.1 | 0.1 | 0.25 |
| 10 | 122307101 | rs4752432 | -0.131 | -1.395 | 0.348 | 0.172 | -0.03 | -0.089 | -0.089 | 0 | 0.79 | 0.69 | 0.81 | 0.97 | 0.88 | 0.87 |
| 10 | 122307518 | rs11199393 | 1.161 | -1.374 | 0.255 | 0.192 | 0.008 | 0.002 | 0.002 | 0 | 0.07 | 0.03 | 0.02 | 0.1 | 0.05 | 0.22 |
| 10 | 122319776 | rs7070520 | -0.131 | -1.413 | 0.326 | 0.218 | -0.006 | -0.01 | -0.01 | 0 | 0.71 | 0.68 | 0.76 | 0.86 | 0.76 | 0.8 |
| 10 | 122320621 | rs4752438 | 1.85 | -0.862 | 0.054 | 0.32 | -0.003 | -0.003 | -0.003 | 0.833 | 0.34 | 0.36 | 0.37 | 0.54 | 0.39 | 0.26 |
| 10 | 122321096 | rs11199404 | 1.289 | -0.83 | -0.108 | 0.317 | 0.007 | 0.001 | 0.001 | 0 | 0.06 | 0.03 | 0.03 | 0.1 | 0.05 | 0.24 |
| 10 | 122322285 | rs10886711 | 2.006 | -0.795 | -0.133 | 0.331 | -0.013 | -0.013 | -0.013 | 0 | 0.33 | 0.29 | 0.28 | 0.39 | 0.38 | 0.15 |
| 10 | 122327563 | rs4752440 | 0.6 | -0.71 | -0.243 | 0.367 | 0.01 | 0.013 | 0.013 | 0.677 | 0.24 | 0.21 | 0.21 | 0.11 | 0.2 | 0.16 |
| 10 | 122330886 | rs10886714 | 1.502 | -0.71 | -0.199 | 0.347 | -0.022 | 0.034 | 0.034 | 0.963 | 0.12 | 0.2 | 0.12 | 0.15 | 0.06 | 0.14 |
| 10 | 122332952 | rs11596919 | -0.668 | -0.675 | -0.77 | 0.299 | -0.011 | 0.037 | 0.037 | 0 | 0.43 | 0.29 | 0.43 | 0.33 | 0.48 | 0.42 |
| 10 | 122332991 | rs11592655 | -1.503 | -0.639 | -0.731 | 0.325 | -0.005 | -0.006 | -0.006 | 0.128 | 0.6 | 0.62 | 0.69 | 0.51 | 0.57 | 0.62 |
| 10 | 122335333 | rs7915990 | -0.538 | -0.57 | -0.647 | 0.399 | 0.007 | 0.003 | 0.003 | 1.05 | 0.43 | 0.55 | 0.48 | 0.32 | 0.29 | 0.36 |
| 10 | 122336310 | rs11199417 | 0.31 | -0.056 | -1.18 | 0.218 | 0.016 | 0.018 | 0.018 | 0.404 | 0.24 | 0.18 | 0.21 | 0.13 | 0.2 | 0.16 |
| 10 | 122340062 | rs12776883 | -0.064 | 0.541 | -1.043 | 0.321 | -0.022 | -0.003 | -0.003 | 0.396 | 0.17 | 0.13 | 0.2 | 0.19 | 0.26 | 0.25 |
| 10 | 122345459 | rs10444120 | 1.055 | 0.416 | -1.015 | 0.437 | -0.028 | -0.007 | -0.007 | 0.017 | 0.3 | 0.22 | 0.27 | 0.34 | 0.39 | 0.15 |
| 10 | 122346065 | rs11597338 | -0.302 | 0.109 | -0.808 | 0.031 | 0.004 | -0.023 | -0.023 | 0 | 0.28 | 0.35 | 0.25 | 0.19 | 0.3 | 0.28 |
| 10 | 122346343 | rs3816785 | 0.007 | -0.122 | -1.105 | 0.155 | 0.022 | -0.009 | -0.009 | 0.199 | 0.24 | 0.13 | 0.21 | 0.22 | 0.22 | 0.16 |
| 10 | 122348969 | rs1047369 | -0.314 | 0.342 | -0.939 | -0.167 | -0.004 | 0.006 | 0.006 | 0.792 | 0.16 | 0.15 | 0.16 | 0.07 | 0.21 | 0.25 |
| 10 | 122351696 | rs12260045 | -0.44 | -0.107 | -0.776 | -0.593 | 0.007 | -0.004 | -0.004 | 0.008 | 0.49 | 0.43 | 0.41 | 0.11 | 0.41 | 0.5 |
| 10 | 122354955 | rs11199430 | 0.141 | -0.148 | -0.634 | -0.61 | -0.002 | -0.04 | -0.04 | 0 | 0.51 | 0.52 | 0.59 | 0.89 | 0.59 | 0.53 |
| 10 | 122355441 | rs7894815 | -0.114 | -0.398 | -1.199 | -0.238 | 0.024 | -0.013 | -0.013 | 0 | 0.36 | 0.24 | 0.43 | 0.53 | 0.25 | 0.55 |
| 10 | 122358463 | rs2901257 | 0.14 | 0.604 | -0.97 | -0.178 | 0.035 | 0.032 | 0.032 | 0 | 0.68 | 0.57 | 0.56 | 0.49 | 0.56 | 0.59 |
| 10 | 122360844 | rs11199434 | 0.535 | 0.485 | -1.423 | 0.334 | 0.006 | -0.04 | -0.04 | 0 | 0.11 | 0.03 | 0.12 | 0.25 | 0.07 | 0.05 |
| 10 | 122361313 | rs11199436 | 0.062 | 0.553 | -1.415 | -0.268 | 0.004 | 0.082 | 0.082 | 0 | 0.18 | 0.04 | 0.14 | 0 | 0.16 | 0.11 |
| 10 | 122365049 | rs11199437 | -0.614 | 1.921 | -0.909 | 0.204 | 0.055 | 0.009 | 0.009 | 0 | 0.19 | 0.29 | 0.33 | 0.21 | 0.4 | 0.4 |
| 10 | 122365160 | rs4751793 | 0.488 | 0.755 | -0.64 | -0.101 | -0.033 | 0.021 | 0.021 | 0 | 0.34 | 0.46 | 0.26 | 0.46 | 0.21 | 0.22 |
| 10 | 122367961 | rs2420783 | -0.509 | 2.383 | -0.399 | -0.611 | 0 | -0.028 | -0.028 | 0 | 0.28 | 0.33 | 0.3 | 0.04 | 0.37 | 0.2 |
| 10 | 122374644 | rs4506553 | 0.474 | 2.58 | 0.225 | -0.136 | 0.086 | 0.046 | 0.046 | 0 | 0.92 | 0.72 | 0.89 | 0.9 | 0.86 | 0.86 |
| 10 | 122374787 | rs2420779 | 0.528 | 2.608 | -0.122 | -0.086 | -0.002 | -0.014 | -0.014 | 0.093 | 0.57 | 0.44 | 0.48 | 0.74 | 0.47 | 0.44 |
| 10 | 122383804 | rs2901256 | 0.457 | 2.248 | 0.039 | -0.07 | 0.009 | 0.032 | 0.032 | 0 | 0.55 | 0.33 | 0.45 | 0.73 | 0.42 | 0.43 |
| 10 | 122386059 | rs10886725 | -0.295 | 2.099 | 0.04 | 0.212 | -0.003 | -0.013 | -0.013 | 0 | 0.13 | 0.08 | 0.12 | 0.28 | 0.17 | 0.17 |
| 10 | 122388196 | rs10749379 | -0.185 | 2.119 | 0.335 | 0.127 | -0.003 | 0 | 0 | 0.605 | 0.34 | 0.3 | 0.49 | 0.31 | 0.43 | 0.36 |
| 10 | 122392318 | rs10788106 | 0.205 | 2.163 | 0.281 | 0.127 | 0.033 | -0.043 | -0.043 | 0.227 | 0.79 | 0.55 | 0.83 | 0.87 | 0.72 | 0.75 |
| 10 | 122394214 | rs10886730 | 0.672 | 2.234 | 0.135 | 0.14 | -0.01 | 0.007 | 0.007 | 0 | 0.24 | 0.18 | 0.39 | 0.14 | 0.31 | 0.28 |
| 10 | 122395310 | rs7068008 | 0.911 | 2.133 | 0.42 | 0.151 | 0.003 | 0 | 0 | 0.251 | 0.68 | 0.63 | 0.73 | 0.59 | 0.55 | 0.57 |
| 10 | 122396106 | rs7072655 | -0.897 | 2.19 | 0.393 | 0.137 | 0 | -0.001 | -0.001 | 0.097 | 0.34 | 0.36 | 0.24 | 0.35 | 0.38 | 0.31 |
| 10 | 122397020 | rs7475720 | -0.775 | 2.156 | 0.081 | 0.187 | 0.009 | -0.01 | -0.01 | 0 | 0.75 | 0.75 | 0.61 | 0.86 | 0.69 | 0.72 |

| | | | | | | | | | | | | | | | | |
|---|---|---|---|---|---|---|---|---|---|---|---|---|---|---|---|---|
| 10 | 122397531 | rs11199452 | 0.798 | 2.149 | 0.046 | 0.194 | -0.029 | -0.039 | -0.039 | 0 | 0.09 | 0.03 | 0.09 | 0.17 | 0.12 | 0.08 |
| 10 | 122397634 | rs10886732 | 0.583 | 1.895 | 0.423 | 0.333 | 0.003 | 0 | 0 | 0 | 0.43 | 0.39 | 0.34 | 0.56 | 0.28 | 0.37 |
| 10 | 122398063 | rs4752444 | 0.404 | 2.057 | -0.052 | 0.355 | -0.006 | 0.003 | 0.003 | 0 | 0.23 | 0.17 | 0.38 | 0.14 | 0.31 | 0.28 |
| 10 | 122399277 | rs7087588 | -0.095 | 1.974 | -0.159 | 0.375 | 0.008 | -0.013 | -0.013 | 0 | 0.39 | 0.54 | 0.3 | 0.4 | 0.47 | 0.44 |
| 10 | 122405421 | rs7914572 | 0.433 | 0.685 | -1.103 | 0.648 | 0.043 | -0.045 | -0.045 | 0 | 0.37 | 0.21 | 0.31 | 0.46 | 0.22 | 0.28 |
| 10 | 122407186 | rs10749381 | 0.092 | 1.536 | -0.652 | 0.475 | 0.024 | -0.004 | -0.004 | 0 | 0.69 | 0.62 | 0.54 | 0.66 | 0.59 | 0.54 |
| 10 | 122409257 | rs4287269 | -0.32 | 1.795 | -1.469 | 0.491 | -0.133 | -0.131 | -0.131 | 0 | 0.25 | 0.03 | 0.37 | 0.39 | 0.35 | 0.34 |
| 10 | 122418864 | rs7921049 | -0.035 | 0.988 | -0.739 | 0.222 | -0.01 | 0.003 | 0.003 | 0 | 0.07 | 0.14 | 0.05 | 0.06 | 0.04 | 0.17 |
| 10 | 122430075 | rs10458766 | -0.191 | 0.947 | -0.293 | 0.169 | -0.046 | -0.046 | -0.046 | 0 | 0.27 | 0.13 | 0.37 | 0.39 | 0.35 | 0.33 |
| 10 | 122441466 | rs11199483 | -0.007 | 0.621 | -0.695 | -0.674 | -0.001 | -0.011 | -0.011 | 2.352 | 0.25 | 0.24 | 0.18 | 0.07 | 0.3 | 0.25 |
| 10 | 122443197 | rs12164772 | 0.799 | -0.204 | -0.356 | -0.851 | 0.001 | 0.013 | 0.013 | 0 | 0.28 | 0.28 | 0.3 | 0.23 | 0.27 | 0.26 |
| 10 | 122446453 | rs4751798 | -0.153 | 0.024 | -0.158 | -0.705 | -0.001 | 0.005 | 0.005 | 0 | 0.53 | 0.47 | 0.45 | 0.29 | 0.51 | 0.51 |
| 10 | 122448893 | rs12570141 | 1.241 | -0.082 | 0.133 | -0.667 | 0.013 | -0.012 | -0.012 | 0.662 | 0.13 | 0.09 | 0.12 | 0.32 | 0.09 | 0.14 |
| 10 | 122449733 | rs7893835 | 1.018 | -0.11 | 0.287 | -1.089 | 0.011 | 0.006 | 0.006 | 0 | 0.12 | 0.22 | 0.22 | 0.23 | 0.18 | 0.21 |
| 10 | 122455229 | rs2420730 | -0.499 | -0.307 | 0.274 | -1.001 | -0.027 | 0.001 | 0.001 | 0 | 0.9 | 0.73 | 0.94 | 0.94 | 0.97 | 0.83 |
| 10 | 122460186 | rs10886746 | 0.046 | -0.435 | 0.033 | -1.099 | -0.001 | 0.001 | 0.001 | 0 | 0.49 | 0.44 | 0.55 | 0.7 | 0.5 | 0.5 |
| 10 | 122461483 | rs6585658 | 0.553 | -0.207 | 0.705 | -0.359 | 0.004 | -0.005 | -0.005 | 0.001 | 0.09 | 0.08 | 0.04 | 0 | 0.05 | 0.02 |
| 10 | 122463594 | rs7078946 | 0.139 | 0.245 | 0.867 | -0.026 | -0.028 | -0.003 | -0.003 | 0 | 0.91 | 0.79 | 0.94 | 0.93 | 0.97 | 0.84 |
| 10 | 122471683 | rs6585659 | -0.38 | 0.277 | 0.652 | -0.139 | -0.016 | -0.017 | -0.017 | 0 | 0.93 | 0.82 | 0.95 | 0.97 | 0.97 | 0.84 |
| 10 | 122472364 | rs7916658 | 0.406 | 0.364 | 0.566 | 0.003 | 0.015 | 0.043 | 0.043 | 0 | 0.1 | 0.24 | 0.2 | 0.23 | 0.12 | 0.16 |
| 10 | 122473979 | rs11199509 | 0.475 | 0.39 | 0.054 | 0.007 | -0.016 | 0.003 | 0.003 | 0.228 | 0.79 | 0.4 | 0.72 | 0.75 | 0.84 | 0.66 |
| 10 | 122480687 | rs4751799 | 0.61 | 0.57 | 0.025 | -0.009 | 0 | -0.004 | -0.004 | 0 | 0.77 | 0.39 | 0.7 | 0.74 | 0.8 | 0.64 |
| 10 | 122482531 | rs10886747 | 0.559 | 0.787 | -0.574 | -0.151 | 0.04 | 0.005 | 0.005 | 0 | 0.44 | 0.31 | 0.39 | 0.39 | 0.33 | 0.33 |
| 10 | 122482705 | rs12780849 | -0.418 | 1.25 | -0.529 | -0.147 | -0.023 | -0.001 | -0.001 | 0 | 0.39 | 0.3 | 0.34 | 0.38 | 0.48 | 0.46 |
| 10 | 122483407 | rs10788116 | -0.402 | 1.871 | -0.412 | -0.509 | 0.025 | -0.072 | -0.072 | 0.319 | 0.21 | 0.24 | 0.17 | 0 | 0.32 | 0.19 |
| 10 | 122483828 | rs2997262 | -0.539 | 1.663 | -0.455 | -0.71 | 0.004 | -0.024 | -0.024 | 0 | 0.32 | 0.38 | 0.31 | 0.17 | 0.35 | 0.23 |
| 10 | 122485809 | rs11199515 | 0.025 | 0.869 | -0.621 | -0.084 | -0.011 | 0.03 | 0.03 | 0 | 0.52 | 0.24 | 0.38 | 0.42 | 0.54 | 0.52 |
| 10 | 122489318 | rs2997228 | -1.615 | 0.436 | -0.6 | 0.233 | 0 | -0.001 | -0.001 | 0 | 0.88 | 0.89 | 0.8 | 0.88 | 0.94 | 0.94 |
| 10 | 122494719 | rs2997238 | 0.177 | 0.826 | 0.022 | 0.117 | 0.005 | 0.012 | 0.012 | 0 | 0.31 | 0.63 | 0.37 | 0.46 | 0.34 | 0.33 |
| 10 | 122499700 | rs9421389 | -0.692 | 0.65 | -0.245 | 0.067 | -0.042 | -0.051 | -0.051 | 0 | 0.44 | 0.78 | 0.48 | 0.55 | 0.45 | 0.49 |
| 10 | 122500375 | rs4752460 | 0.7 | 0.682 | 0.044 | 0.147 | 0 | 0.044 | 0.044 | 0 | 0.23 | 0.15 | 0.11 | 0.13 | 0.21 | 0.22 |
| 10 | 122501487 | rs3011399 | 0.389 | 0.746 | 0.319 | 0.315 | 0.035 | 0.02 | 0.02 | 0.572 | 0.14 | 0.42 | 0.15 | 0.12 | 0.12 | 0.16 |
| 10 | 122502586 | rs1937826 | -0.924 | 0.815 | -0.383 | 0.481 | -0.027 | -0.027 | -0.027 | 0 | 0.44 | 0.75 | 0.46 | 0.5 | 0.43 | 0.48 |
| 10 | 122503390 | rs1937825 | 1.46 | 1.043 | -0.216 | 0.812 | -0.029 | 0.003 | 0.003 | 0 | 0.05 | 0 | 0.18 | 0.25 | 0.09 | 0.11 |
| 10 | 122504952 | rs3011408 | 0.364 | 0.586 | 0.184 | 0.42 | 0.014 | 0.005 | 0.005 | 0.356 | 0.13 | 0.27 | 0.13 | 0.12 | 0.12 | 0.15 |
| 10 | 122508325 | rs3011415 | -0.57 | 0.496 | -0.581 | 0.48 | 0.005 | -0.01 | -0.01 | 0.185 | 0.76 | 0.78 | 0.74 | 0.65 | 0.82 | 0.74 |
| 10 | 122508540 | rs3011417 | 0.088 | 0.669 | -0.703 | 0.355 | 0.017 | 0.017 | 0.017 | 0 | 0.52 | 0.61 | 0.63 | 0.53 | 0.6 | 0.54 |
| 10 | 122509251 | rs17100738 | 0.536 | 1.124 | -1.296 | 0.785 | -0.055 | -0.087 | -0.087 | 0.319 | 0.14 | 0.03 | 0.23 | 0.26 | 0.16 | 0.19 |
| 10 | 122509311 | rs1937819 | 0.023 | 0.529 | -1.007 | 0.467 | 0.008 | -0.006 | -0.006 | 0 | 0.32 | 0.28 | 0.31 | 0.39 | 0.23 | 0.33 |
| 10 | 122510352 | rs7097321 | 0.341 | 0.72 | -1.512 | 0.269 | -0.039 | 0.003 | 0.003 | 0.416 | 0.43 | 0.14 | 0.36 | 0.39 | 0.48 | 0.38 |
| 10 | 122513248 | rs10886755 | 0.354 | 0.7 | -1.521 | 0.251 | -0.04 | 0.006 | 0.006 | 0.515 | 0.42 | 0.14 | 0.35 | 0.39 | 0.49 | 0.38 |
| 10 | 122513414 | rs2997247 | 0.726 | 0.726 | -0.558 | -0.063 | -0.004 | -0.09 | -0.09 | 0 | 0.2 | 0.42 | 0.13 | 0.1 | 0.17 | 0.2 |
| 10 | 122515278 | rs2997248 | 1.219 | 0.732 | -0.366 | -0.141 | -0.006 | -0.011 | -0.011 | 0 | 0.1 | 0.16 | 0.06 | 0.11 | 0.07 | 0.17 |
| 10 | 122517640 | rs1937821 | 0.685 | 1.036 | -1.125 | -0.139 | -0.099 | -0.14 | -0.14 | 1.24 | 0.27 | 0.05 | 0.42 | 0.42 | 0.36 | 0.31 |
| 10 | 122521706 | rs2997254 | 0.938 | 0.905 | -0.154 | -0.329 | 0 | -0.015 | -0.015 | 0.342 | 0.13 | 0.44 | 0.08 | 0.11 | 0.07 | 0.17 |
| 10 | 122526305 | rs1937832 | 0.1 | 0.967 | -0.372 | -0.361 | 0.041 | 0.083 | 0.083 | 0 | 0.56 | 0.22 | 0.63 | 0.7 | 0.67 | 0.56 |
| 10 | 122526973 | rs7093648 | 1.809 | 0.957 | -0.353 | -0.369 | 0.002 | -0.047 | -0.047 | 0 | 0.1 | 0.12 | 0.06 | 0 | 0.11 | 0.03 |
| 10 | 122530138 | rs2186171 | -1.453 | 0.824 | -0.108 | -0.308 | -0.012 | 0.004 | 0.004 | 0 | 0.78 | 0.72 | 0.78 | 0.8 | 0.85 | 0.77 |
| 10 | 122530485 | rs2997256 | 0.765 | 0.684 | -0.249 | -0.52 | -0.033 | -0.104 | -0.104 | 0.116 | 0.31 | 0.48 | 0.2 | 0.13 | 0.23 | 0.3 |
| 10 | 122534587 | rs11199546 | 1.544 | 1.344 | -0.383 | -0.067 | -0.06 | -0.08 | -0.08 | 2.143 | 0.12 | 0.02 | 0.3 | 0.32 | 0.18 | 0.17 |
| 10 | 122536612 | rs7097481 | -0.505 | 1.188 | -0.17 | -0.184 | -0.024 | -0.008 | -0.008 | 0 | 0.69 | 0.57 | 0.71 | 0.7 | 0.79 | 0.57 |
| 10 | 122537487 | rs12221320 | 1.554 | 1.471 | -0.484 | 0.02 | -0.017 | -0.038 | -0.038 | 1.056 | 0.11 | 0.03 | 0.05 | 0.16 | 0.12 | 0.12 |
| 10 | 122541906 | rs2997264 | 0.012 | 1.366 | -0.179 | -0.011 | 0.044 | 0.096 | 0.096 | 0.023 | 0.23 | 0.38 | 0.4 | 0.45 | 0.27 | 0.36 |
| 10 | 122542027 | rs11199558 | 0.514 | 1.551 | -0.666 | -0.103 | -0.034 | -0.067 | -0.067 | 0.29 | 0.17 | 0.07 | 0.15 | 0.31 | 0.22 | 0.22 |
| 10 | 122542659 | rs11199560 | 1.178 | 1.559 | -0.667 | -0.077 | -0.013 | -0.016 | -0.016 | 0 | 0.06 | 0.03 | 0.1 | 0.15 | 0.1 | 0.09 |
| 10 | 122542898 | rs12268319 | 0.825 | 1.436 | -0.605 | -0.219 | -0.034 | -0.067 | -0.067 | 0.37 | 0.17 | 0.07 | 0.15 | 0.31 | 0.22 | 0.22 |
| 10 | 122543587 | rs12255639 | 1.461 | 1.176 | -0.408 | -0.304 | 0.004 | -0.024 | -0.024 | 0 | 0.3 | 0.3 | 0.18 | 0.03 | 0.25 | 0.14 |
| 10 | 122544355 | rs6585662 | -0.165 | 1.219 | -0.373 | -0.117 | 0.027 | 0.096 | 0.096 | 0.077 | 0.76 | 0.53 | 0.61 | 0.59 | 0.77 | 0.65 |
| 10 | 122545237 | rs2997267 | 0.539 | 1.09 | -0.212 | -0.362 | 0.064 | 0.032 | 0.032 | 0 | 0.11 | 0.41 | 0.1 | 0.14 | 0.1 | 0.19 |
| 10 | 122545247 | rs17591472 | 1.589 | 1.054 | -0.52 | -0.481 | 0.017 | 0.145 | 0.145 | 1.926 | 0.27 | 0.06 | 0.16 | 0.03 | 0.25 | 0.14 |
| 10 | 122546404 | rs2997275 | -1.261 | 1.412 | -0.038 | 0.006 | -0.01 | 0.011 | 0.011 | 0.427 | 0.43 | 0.59 | 0.33 | 0.45 | 0.48 | 0.47 |
| 10 | 122547756 | rs2997279 | 0.119 | 1.382 | 0.576 | 0.235 | 0.032 | 0.01 | 0.01 | 0 | 0.1 | 0.27 | 0.08 | 0.14 | 0.1 | 0.2 |
| 10 | 122553175 | rs2997288 | -0.386 | 1.216 | 0.539 | 0.359 | 0.101 | 0.134 | 0.134 | 0 | 0.07 | 0.41 | 0.06 | 0.14 | 0.05 | 0.13 |
| 10 | 122553260 | rs2997289 | -0.386 | 1.242 | 0.539 | 0.359 | 0.107 | 0.134 | 0.134 | 0 | 0.07 | 0.41 | 0.06 | 0.14 | 0.05 | 0.13 |
| 10 | 122554218 | rs2997291 | 0.482 | 1.187 | 0.615 | 0.347 | 0.017 | 0.043 | 0.043 | 0 | 0.05 | 0.22 | 0.03 | 0.1 | 0.01 | 0.12 |
| 10 | 122555846 | rs3011369 | -0.375 | 1.179 | 0.199 | 0.341 | 0.093 | 0.118 | 0.118 | 0 | 0.07 | 0.41 | 0.06 | 0.14 | 0.05 | 0.13 |
| 10 | 122557286 | rs3011373 | -0.375 | 1.179 | 0.199 | 0.341 | 0.093 | 0.118 | 0.118 | 0 | 0.07 | 0.41 | 0.06 | 0.14 | 0.05 | 0.13 |
| 10 | 122560915 | rs2096420 | 0.067 | 1.599 | -0.825 | 0.482 | -0.06 | -0.078 | -0.078 | 0 | 0.19 | 0.07 | 0.15 | 0.32 | 0.26 | 0.27 |
| 10 | 122561704 | rs4631806 | -0.319 | 1.688 | -0.911 | 0.42 | -0.076 | -0.081 | -0.081 | 0.145 | 0.22 | 0.07 | 0.2 | 0.33 | 0.32 | 0.3 |

| | | | | | | | | | | | | | | | | |
|---|---|---|---|---|---|---|---|---|---|---|---|---|---|---|---|---|
| 10 | 122564081 | rs7093135 | 0.198 | 1.933 | -1.091 | 0.427 | -0.028 | -0.045 | -0.045 | 0.006 | 0.17 | 0.08 | 0.36 | 0.32 | 0.24 | 0.19 |
| 10 | 122564882 | rs11199574 | 0.339 | 1.374 | -0.857 | -0.171 | -0.003 | 0.002 | 0.002 | 0 | 0.46 | 0.41 | 0.54 | 0.39 | 0.49 | 0.34 |
| 10 | 122565481 | rs11199575 | 1.065 | 1.347 | -0.842 | -0.164 | -0.016 | -0.027 | -0.027 | 0 | 0.15 | 0.08 | 0.31 | 0.32 | 0.19 | 0.16 |
| 10 | 122566328 | rs2997310 | 1.289 | 1.202 | -0.465 | -0.185 | 0.019 | 0.05 | 0.05 | 0 | 0.05 | 0.23 | 0.03 | 0.09 | 0.01 | 0.12 |
| 10 | 122567432 | rs10886772 | 0.66 | 1.265 | -0.563 | -0.187 | -0.001 | -0.001 | -0.001 | 0.723 | 0.43 | 0.41 | 0.49 | 0.39 | 0.44 | 0.31 |
| 10 | 122567702 | rs2997312 | 0.123 | 1.039 | -0.598 | -0.411 | -0.021 | -0.082 | -0.082 | 0 | 0.18 | 0.37 | 0.1 | 0.09 | 0.14 | 0.15 |
| 10 | 122568882 | rs10788123 | -0.079 | 1.078 | -0.686 | -0.381 | -0.023 | -0.022 | -0.022 | 0.233 | 0.64 | 0.48 | 0.7 | 0.72 | 0.76 | 0.61 |
| 10 | 122569984 | rs7071661 | 1.845 | 1.125 | -0.735 | -0.408 | -0.008 | -0.047 | -0.047 | 0 | 0.28 | 0.34 | 0.18 | 0.07 | 0.25 | 0.15 |
| 10 | 122571422 | rs7901805 | 0.421 | 1.118 | -0.735 | -0.395 | 0.009 | 0.026 | 0.026 | 0 | 0.49 | 0.41 | 0.38 | 0.4 | 0.57 | 0.46 |
| 10 | 122572373 | rs9919349 | 0.08 | 1.119 | -0.623 | -0.37 | 0.02 | 0.024 | 0.024 | 0 | 0.24 | 0.36 | 0.25 | 0.38 | 0.34 | 0.32 |
| 10 | 122573258 | rs7094965 | -0.124 | 1.084 | -0.51 | -0.311 | -0.005 | 0.004 | 0.004 | 0 | 0.62 | 0.5 | 0.46 | 0.41 | 0.69 | 0.49 |
| 10 | 122578338 | rs10510081 | 1.319 | 1.137 | -0.557 | -0.294 | 0.013 | -0.013 | -0.013 | 0 | 0.17 | 0.11 | 0.21 | 0.19 | 0.1 | 0.23 |
| 10 | 122581611 | rs10886776 | -0.144 | 1.285 | -0.501 | -0.234 | -0.026 | -0.035 | -0.035 | 0.053 | 0.38 | 0.31 | 0.39 | 0.53 | 0.42 | 0.52 |
| 10 | 122585280 | rs1254150 | 0.185 | 1.295 | -0.295 | -0.259 | 0.004 | 0.001 | 0.001 | 0.02 | 0.57 | 0.49 | 0.55 | 0.46 | 0.53 | 0.44 |
| 10 | 122585600 | rs1254148 | -0.625 | 1.696 | -0.25 | -0.132 | 0.009 | 0.031 | 0.031 | 0.573 | 0.82 | 0.57 | 0.84 | 0.79 | 0.8 | 0.79 |
| 10 | 122589158 | rs1254185 | -0.383 | 1.807 | -0.205 | -0.143 | 0.067 | 0.038 | 0.038 | 0 | 0.92 | 0.79 | 0.94 | 0.89 | 0.89 | 0.92 |
| 10 | 122591373 | rs2919009 | 1.526 | 1.711 | -0.202 | -0.294 | 0 | -0.02 | -0.02 | 0 | 0.29 | 0.23 | 0.15 | 0.1 | 0.22 | 0.24 |
| 10 | 122593020 | rs11199588 | 1.324 | 2.254 | -0.338 | 0.055 | -0.028 | -0.017 | -0.017 | 0 | 0.08 | 0.01 | 0.25 | 0.35 | 0.11 | 0.16 |
| 10 | 122595790 | rs1254178 | 0.951 | 2.168 | -0.331 | -0.109 | -0.002 | 0.003 | 0.003 | 0 | 0.08 | 0.04 | 0.05 | 0 | 0.11 | 0.03 |
| 10 | 122598339 | rs1271526 | 0.427 | 2.176 | -0.141 | 0.084 | 0.069 | 0.015 | 0.015 | 0 | 0.89 | 0.77 | 0.88 | 0.88 | 0.84 | 0.9 |
| 10 | 122601003 | rs10886780 | 0.56 | 1.746 | -0.502 | 0.19 | -0.048 | -0.056 | -0.056 | 0.681 | 0.1 | 0.01 | 0.3 | 0.34 | 0.17 | 0.19 |
| 10 | 122604051 | rs11199591 | 0.703 | 1.743 | -0.497 | 0.189 | -0.044 | -0.053 | -0.053 | 0 | 0.1 | 0.02 | 0.3 | 0.34 | 0.17 | 0.19 |
| 10 | 122610639 | rs12778011 | 0.567 | 1.356 | -0.689 | -0.06 | -0.011 | 0.047 | 0.047 | 0 | 0.17 | 0.08 | 0.1 | 0.03 | 0.17 | 0.18 |
| 10 | 122610657 | rs11199598 | -0.549 | 1.366 | -0.691 | 0.086 | -0.053 | -0.082 | -0.082 | 0 | 0.28 | 0.12 | 0.4 | 0.38 | 0.33 | 0.37 |
| 10 | 122620616 | rs11199604 | -0.657 | 1.393 | -0.571 | -0.006 | -0.04 | -0.07 | -0.07 | 0 | 0.29 | 0.17 | 0.41 | 0.42 | 0.33 | 0.36 |
| 10 | 122624679 | rs10886789 | -0.657 | 1.393 | -0.571 | -0.006 | -0.04 | -0.07 | -0.07 | 0 | 0.29 | 0.17 | 0.41 | 0.42 | 0.33 | 0.36 |
| 10 | 122636730 | rs4751800 | 0.55 | 1.471 | -0.609 | -0.07 | -0.021 | 0.056 | 0.056 | 0 | 0.18 | 0.08 | 0.1 | 0.03 | 0.18 | 0.18 |
| 10 | 122639170 | rs3758510 | -0.621 | 1.287 | -0.012 | -0.066 | 0.016 | 0.043 | 0.043 | 0 | 0.31 | 0.43 | 0.44 | 0.42 | 0.33 | 0.36 |
| 10 | 122649482 | rs2289337 | 1.879 | 0.992 | -0.037 | -0.402 | -0.001 | -0.015 | -0.015 | 0.511 | 0.27 | 0.26 | 0.16 | 0.1 | 0.22 | 0.22 |
| 10 | 122656619 | rs12415091 | -0.109 | 1.019 | -0.101 | -0.374 | -0.062 | -0.099 | -0.099 | 0 | 0.28 | 0.11 | 0.4 | 0.42 | 0.33 | 0.36 |
| 10 | 122658017 | rs6585674 | -0.344 | 1.068 | -0.111 | -0.341 | 0.016 | 0.045 | 0.045 | 0 | 0.31 | 0.43 | 0.44 | 0.42 | 0.33 | 0.36 |
| 10 | 122668296 | rs7077126 | -0.354 | 1.064 | -0.242 | -0.358 | 0.021 | 0.055 | 0.055 | 0 | 0.31 | 0.45 | 0.44 | 0.42 | 0.33 | 0.36 |
| 10 | 122668604 | rs1045154 | -0.226 | 1.061 | -0.203 | -0.359 | -0.036 | -0.065 | -0.065 | 0 | 0.29 | 0.19 | 0.41 | 0.42 | 0.33 | 0.36 |
| 10 | 122668801 | rs1045179 | -0.329 | 1.062 | -0.204 | -0.359 | 0.021 | 0.055 | 0.055 | 0 | 0.31 | 0.45 | 0.44 | 0.42 | 0.33 | 0.36 |
| 10 | 122677062 | rs4751808 | -0.408 | 1.044 | -0.467 | -0.357 | -0.026 | -0.068 | -0.068 | 0 | 0.61 | 0.84 | 0.63 | 0.52 | 0.59 | 0.6 |
| 10 | 122684525 | rs10510083 | 1.694 | 1.011 | -0.485 | -0.432 | -0.008 | -0.037 | -0.037 | 1.273 | 0.26 | 0.32 | 0.17 | 0.1 | 0.22 | 0.22 |
| 10 | 122686546 | rs7906521 | 1.694 | 0.983 | -0.584 | -0.401 | -0.01 | -0.041 | -0.041 | 0 | 0.26 | 0.34 | 0.17 | 0.1 | 0.22 | 0.22 |
| 10 | 122687144 | rs6585677 | -0.118 | 1.127 | -0.566 | -0.192 | -0.001 | 0 | 0 | 0 | 0.57 | 0.53 | 0.58 | 0.51 | 0.57 | 0.56 |
| 10 | 122689964 | rs1652728 | -0.128 | 1.02 | -0.959 | -0.682 | -0.098 | 0.208 | 0.208 | 0 | 0.43 | 0.11 | 0.64 | 0.85 | 0.52 | 0.57 |
| 10 | 122693564 | rs1659592 | -0.233 | 0.784 | -0.425 | -0.932 | -0.004 | -0.062 | -0.062 | 0.135 | 0.75 | 0.65 | 0.87 | 0.95 | 0.81 | 0.8 |
| 10 | 122697234 | rs2118962 | 1.933 | 0.829 | -0.441 | -0.927 | 0.009 | 0.034 | 0.034 | 1.702 | 0.25 | 0.16 | 0.16 | 0.02 | 0.23 | 0.14 |
| 10 | 122698414 | rs12356763 | 0.418 | 0.852 | -0.309 | -0.832 | 0.009 | 0.064 | 0.064 | 0.108 | 0.47 | 0.35 | 0.29 | 0.07 | 0.42 | 0.34 |
| 10 | 122700499 | rs7899002 | -0.05 | 1.109 | -0.892 | -0.679 | -0.01 | 0.126 | 0.126 | 0 | 0.55 | 0.72 | 0.35 | 0.15 | 0.46 | 0.42 |
| 10 | 122704013 | rs934320 | 0.12 | 1.312 | -1.209 | -0.416 | -0.077 | -0.041 | -0.041 | 1.523 | 0.31 | 0.1 | 0.47 | 0.64 | 0.41 | 0.48 |
| 10 | 122710328 | rs10886816 | 0.07 | 1.148 | -1.184 | -0.478 | 0.005 | 0.01 | 0.01 | 0 | 0.64 | 0.65 | 0.71 | 0.72 | 0.72 | 0.69 |
| 10 | 122712110 | rs11199675 | -1.442 | 1.069 | -1.202 | -0.524 | -0.002 | 0 | 0 | 0 | 0.84 | 0.85 | 0.83 | 0.77 | 0.9 | 0.88 |
| 10 | 122717301 | rs11199680 | 1.481 | 1.124 | -1.543 | -0.662 | -0.006 | -0.065 | -0.065 | 0 | 0.33 | 0.56 | 0.23 | 0.01 | 0.27 | 0.22 |
| 10 | 122717705 | rs11199681 | 0.229 | 1.18 | -1.601 | -0.577 | -0.139 | -0.064 | -0.064 | 0 | 0.31 | 0.09 | 0.49 | 0.63 | 0.45 | 0.47 |
| 10 | 122720309 | rs7906032 | 0.606 | 0.963 | -1.361 | -0.537 | 0.005 | 0.031 | 0.031 | 0 | 0.19 | 0.14 | 0.11 | 0.05 | 0.16 | 0.18 |
| 10 | 122720568 | rs12782586 | 1.133 | 1.326 | -1.374 | 0.076 | -0.041 | -0.05 | -0.05 | 0 | 0.11 | 0.03 | 0.29 | 0.36 | 0.18 | 0.17 |
| 10 | 122720974 | rs4752485 | -0.155 | 0.726 | -1.181 | -0.455 | 0.006 | 0.014 | 0.014 | 0 | 0.62 | 0.65 | 0.7 | 0.71 | 0.7 | 0.7 |
| 10 | 122721684 | rs7080781 | 1.163 | 0.395 | -0.829 | -0.458 | -0.004 | -0.049 | -0.049 | 0.666 | 0.24 | 0.19 | 0.17 | 0.01 | 0.22 | 0.13 |
| 10 | 122724230 | rs7914796 | -0.563 | 0.53 | -1.01 | 0.237 | 0.005 | 0.01 | 0.01 | 0 | 0.22 | 0.17 | 0.19 | 0.14 | 0.21 | 0.38 |
| 10 | 122727732 | rs12570961 | -0.677 | 0.323 | -0.388 | 0.582 | -0.008 | 0.008 | 0.008 | 0 | 0.16 | 0.3 | 0.11 | 0.18 | 0.13 | 0.27 |
| 10 | 122728687 | rs12412237 | 0.217 | 0.362 | -0.624 | 0.549 | -0.029 | 0.026 | 0.026 | 0 | 0.11 | 0.01 | 0.06 | 0.05 | 0.11 | 0.17 |
| 10 | 122731074 | rs10510085 | 0.613 | 0.357 | -0.64 | 0.564 | -0.025 | -0.001 | -0.001 | 0.014 | 0.07 | 0.01 | 0.05 | 0.05 | 0.09 | 0.12 |
| 10 | 122731511 | rs4751819 | 0.336 | 0.16 | -0.803 | 0.649 | 0.002 | -0.054 | -0.054 | 0.292 | 0.09 | 0.01 | 0.1 | 0.16 | 0.09 | 0.18 |
| 10 | 122732356 | rs10886825 | 0.18 | 0.77 | -1.284 | 0.692 | -0.025 | -0.023 | -0.023 | 1.241 | 0.28 | 0.15 | 0.26 | 0.36 | 0.28 | 0.28 |
| 10 | 122734544 | rs1897195 | 1.78 | 0.731 | -1.057 | 0.461 | -0.021 | 0.025 | 0.025 | 0.191 | 0.38 | 0.2 | 0.47 | 0.28 | 0.44 | 0.27 |
| 10 | 122734927 | rs1897196 | -1.235 | 0.595 | -0.644 | 0.493 | 0.01 | -0.035 | -0.035 | 2.796 | 0.93 | 0.89 | 0.94 | 1 | 0.91 | 0.9 |
| 10 | 122736421 | rs4752487 | 1.285 | 0.492 | -0.784 | 0.529 | 0 | 0.001 | 0.001 | 0 | 0.07 | 0.13 | 0.06 | 0.11 | 0.1 | 0.08 |
| 10 | 122746804 | rs10886827 | 1.302 | 0.214 | -0.595 | 0.378 | -0.017 | 0.01 | 0.01 | 0 | 0.63 | 0.37 | 0.71 | 0.58 | 0.69 | 0.51 |
| 10 | 122747777 | rs2435892 | -0.136 | 0.298 | -0.514 | 0.105 | -0.016 | 0.025 | 0.025 | 0 | 0.08 | 0.38 | 0.07 | 0.04 | 0.01 | 0.09 |
| 10 | 122747879 | rs11199689 | 1.017 | 0.961 | -1.614 | -0.166 | -0.103 | -0.059 | -0.059 | 0 | 0.17 | 0.02 | 0.31 | 0.19 | 0.28 | 0.13 |
| 10 | 122749548 | rs11199690 | -0.266 | -0.334 | -1.902 | -0.957 | 0.009 | 0.039 | 0.039 | 0 | 0.2 | 0.02 | 0.12 | 0.12 | 0.14 | 0.25 |
| 10 | 122749930 | rs2258739 | -0.021 | 0.22 | -1.303 | -0.939 | 0.006 | 0.034 | 0.034 | 0.078 | 0.55 | 0.69 | 0.52 | 0.37 | 0.61 | 0.56 |
| 10 | 122750096 | rs10886829 | 1.637 | 0.404 | -1.35 | -1.261 | -0.076 | 0.103 | 0.103 | 0 | 0.22 | 0.04 | 0.2 | 0.03 | 0.34 | 0.15 |
| 10 | 122752212 | rs12768523 | 1.247 | 0.434 | -1.289 | -1.158 | -0.013 | 0.057 | 0.057 | 0 | 0.22 | 0.05 | 0.14 | 0.14 | 0.2 | 0.29 |
| 10 | 122754609 | rs6585682 | -0.855 | 0.527 | -0.371 | -1.067 | 0.001 | -0.034 | -0.034 | 0 | 0.54 | 0.45 | 0.6 | 0.82 | 0.48 | 0.6 |

| | | | | | | | | | | | | | | | | |
|---|---|---|---|---|---|---|---|---|---|---|---|---|---|---|---|---|
| 10 | 122756578 | rs2257619 | -0.923 | 0.535 | -0.236 | -1.062 | -0.006 | 0.029 | 0.029 | 0 | 0.54 | 0.56 | 0.62 | 0.82 | 0.48 | 0.6 |
| 10 | 122756727 | rs12414987 | 0.795 | 0.395 | -0.39 | -0.995 | 0.014 | -0.035 | -0.035 | 0 | 0.25 | 0.12 | 0.33 | 0.6 | 0.19 | 0.2 |
| 10 | 122758389 | rs11199697 | 0.106 | 0.508 | -0.21 | -0.968 | -0.004 | -0.032 | -0.032 | 0 | 0.25 | 0.31 | 0.15 | 0.11 | 0.22 | 0.29 |
| 10 | 122759891 | rs2260801 | -0.236 | 0.397 | -0.245 | -1.057 | 0.034 | 0.09 | 0.09 | 0 | 0.51 | 0.34 | 0.47 | 0.25 | 0.61 | 0.56 |
| 10 | 122760542 | rs10886834 | 0.853 | 0.343 | -0.103 | -1.076 | -0.001 | -0.001 | -0.001 | 0 | 0.27 | 0.21 | 0.34 | 0.6 | 0.21 | 0.21 |
| 10 | 122761091 | rs10510088 | 1.735 | 0.385 | -0.382 | -1.097 | -0.005 | 0.031 | 0.031 | 0 | 0.19 | 0.1 | 0.14 | 0.15 | 0.18 | 0.21 |
| 10 | 122764181 | rs2263920 | 1.223 | 0.366 | -0.477 | -1.241 | -0.001 | -0.023 | -0.023 | 0 | 0.32 | 0.32 | 0.25 | 0.1 | 0.42 | 0.2 |
| 10 | 122765401 | rs3962130 | -0.662 | 0.352 | -0.413 | -1.186 | 0.002 | -0.006 | -0.006 | 0.378 | 0.48 | 0.46 | 0.6 | 0.76 | 0.4 | 0.6 |
| 10 | 122780131 | rs10886838 | 1.663 | 0.008 | -1.252 | -1.092 | -0.002 | 0.027 | 0.027 | 0 | 0.2 | 0.12 | 0.14 | 0.15 | 0.18 | 0.22 |
| 10 | 122782626 | rs12413407 | 1.157 | -0.116 | -1.342 | -1.149 | 0.01 | -0.02 | -0.02 | 1.162 | 0.24 | 0.12 | 0.33 | 0.6 | 0.19 | 0.19 |
| 10 | 122787762 | rs11199717 | 1.211 | 0.146 | -1.375 | -1.373 | 0.002 | 0.044 | 0.044 | 0 | 0.23 | 0.11 | 0.16 | 0.14 | 0.19 | 0.27 |
| 10 | 122789818 | rs11199718 | 1.088 | 0.5 | -1.119 | -0.85 | -0.033 | 0.018 | 0.018 | 0 | 0.13 | 0.01 | 0.11 | 0.05 | 0.13 | 0.25 |
| 10 | 122797096 | rs11199725 | 1.271 | 0.784 | -0.827 | -0.692 | -0.001 | -0.013 | -0.013 | 0.129 | 0.07 | 0.09 | 0.03 | 0 | 0.09 | 0.02 |
| 10 | 122799724 | rs6585687 | -0.22 | 0.826 | -0.93 | -0.732 | -0.005 | 0.005 | 0.005 | 0.145 | 0.38 | 0.36 | 0.33 | 0.22 | 0.46 | 0.33 |
| 10 | 122801652 | rs11199729 | 1.367 | 0.885 | -1.957 | -0.688 | -0.084 | 0.094 | 0.094 | 0 | 0.21 | 0.03 | 0.19 | 0.04 | 0.34 | 0.11 |
| 10 | 122801759 | rs10886845 | 0.214 | 0.935 | -1.867 | -0.619 | -0.034 | 0.139 | 0.139 | 0.014 | 0.58 | 0.15 | 0.62 | 0.77 | 0.53 | 0.66 |
| 10 | 122802462 | rs748476 | -0.012 | 0.745 | -1.279 | -0.336 | -0.01 | -0.093 | -0.093 | 0 | 0.25 | 0.51 | 0.16 | 0.06 | 0.21 | 0.23 |
| 10 | 122802506 | rs2420903 | -0.211 | 1.242 | -1.986 | 0.47 | -0.022 | -0.005 | -0.005 | 0 | 0.35 | 0.18 | 0.39 | 0.4 | 0.42 | 0.16 |
| 10 | 122805351 | rs12781367 | 0.297 | 0.664 | -1.795 | 0.742 | 0.009 | -0.022 | -0.022 | 0.01 | 0.48 | 0.17 | 0.5 | 0.44 | 0.42 | 0.44 |
| 10 | 122805547 | rs7911360 | -0.358 | 1.153 | -1.059 | 1.119 | -0.047 | -0.028 | -0.028 | 1.617 | 0.27 | 0.12 | 0.29 | 0.28 | 0.36 | 0.21 |
| 10 | 122808053 | rs7920826 | 0.809 | 0.239 | 0.093 | 1.344 | -0.001 | 0.001 | 0.001 | 0 | 0.82 | 0.72 | 0.88 | 0.82 | 0.81 | 0.64 |
| 10 | 122815724 | rs17597155 | -0.413 | 0.227 | -0.17 | 1.225 | -0.005 | 0.067 | 0.067 | 0 | 0.12 | 0.01 | 0.04 | 0.01 | 0.09 | 0.23 |
| 10 | 122821238 | rs4752496 | 1.549 | -0.147 | 0.255 | 1.54 | 0.017 | -0.005 | -0.005 | 0 | 0.56 | 0.34 | 0.56 | 0.4 | 0.64 | 0.52 |
| 10 | 122827577 | rs11199742 | -0.801 | -0.458 | 0.199 | 1.777 | 0.005 | -0.06 | -0.06 | 0 | 0.19 | 0.05 | 0.15 | 0.38 | 0.16 | 0.11 |
| 10 | 122834630 | rs7906864 | -1.01 | -0.688 | 0.964 | 1.826 | -0.036 | -0.012 | -0.012 | 1.133 | 0.18 | 0.28 | 0.27 | 0.18 | 0.08 | 0.35 |
| 10 | 122834737 | rs7907003 | -1.539 | -0.775 | 1.624 | 1.982 | -0.059 | 0.05 | 0.05 | 0 | 0.22 | 0.6 | 0.3 | 0.42 | 0.09 | 0.38 |
| 10 | 122848522 | rs2248774 | -1.522 | -0.777 | 1.722 | 1.991 | -0.047 | 0.019 | 0.019 | 0.002 | 0.22 | 0.64 | 0.31 | 0.37 | 0.09 | 0.37 |
| 10 | 122854917 | rs2248039 | -1.363 | -0.789 | 2.084 | 2.012 | -0.066 | 0.052 | 0.052 | 0 | 0.21 | 0.55 | 0.3 | 0.38 | 0.09 | 0.37 |
| 10 | 122857963 | rs10886849 | -1.144 | -0.91 | 1.85 | 2.008 | -0.013 | 0.025 | 0.025 | 0 | 0.2 | 0.25 | 0.15 | 0.38 | 0.13 | 0.1 |
| 10 | 122859177 | rs7079858 | -0.801 | -0.902 | 1.763 | 1.872 | -0.039 | -0.033 | -0.033 | 0 | 0.17 | 0.32 | 0.14 | 0.13 | 0.08 | 0.32 |
| 10 | 122859623 | rs7083597 | -0.413 | -0.901 | 1.773 | 1.872 | -0.001 | 0 | 0 | 0 | 0.15 | 0.13 | 0.12 | 0.13 | 0.08 | 0.32 |
| 10 | 122863793 | rs7087772 | -0.915 | -0.894 | 1.711 | 1.907 | -0.041 | -0.004 | -0.004 | 0.033 | 0.18 | 0.31 | 0.12 | 0.18 | 0.08 | 0.35 |
| 10 | 122868170 | rs11199770 | -0.581 | -0.681 | 1.575 | 1.923 | 0.006 | -0.016 | -0.016 | 0.211 | 0.17 | 0.1 | 0.15 | 0.18 | 0.17 | 0.08 |
| 10 | 122870316 | rs6585692 | -0.463 | -0.725 | 1.702 | 1.919 | -0.01 | -0.009 | -0.009 | 0 | 0.15 | 0.19 | 0.12 | 0.13 | 0.09 | 0.32 |
| 10 | 122871763 | rs11199775 | -0.621 | -0.706 | 1.632 | 1.911 | 0.005 | -0.015 | -0.015 | 0 | 0.18 | 0.1 | 0.15 | 0.18 | 0.17 | 0.08 |
| 10 | 122872046 | rs2438673 | -1.818 | -0.764 | 1.799 | 2.022 | -0.017 | 0.016 | 0.016 | 0 | 0.36 | 0.6 | 0.44 | 0.57 | 0.26 | 0.44 |
| 10 | 122873026 | rs7072244 | -0.518 | -0.706 | 1.768 | 2.05 | -0.017 | -0.013 | -0.013 | 0 | 0.15 | 0.22 | 0.13 | 0.13 | 0.09 | 0.31 |
| 10 | 122874348 | rs2243609 | -1.495 | -0.626 | 1.783 | 2.037 | -0.033 | 0.067 | 0.067 | 0 | 0.21 | 0.58 | 0.32 | 0.44 | 0.12 | 0.38 |
| 10 | 122875356 | rs2254079 | -1.169 | -0.614 | 1.754 | 2.009 | -0.023 | 0 | 0 | 0 | 0.19 | 0.38 | 0.29 | 0.2 | 0.12 | 0.35 |
| 10 | 122875446 | rs10444045 | -0.724 | -1.05 | 1.38 | 1.865 | -0.017 | -0.053 | -0.053 | 0.484 | 0.19 | 0.01 | 0.14 | 0.18 | 0.17 | 0.08 |
| 10 | 122878281 | rs2250942 | -1.987 | -0.809 | 2.011 | 1.86 | 0.07 | 0.352 | 0.352 | 0.175 | 0.05 | 0.42 | 0.17 | 0.26 | 0.01 | 0.04 |
| 10 | 122879732 | rs2130779 | -0.508 | -0.845 | 1.995 | 1.856 | -0.004 | -0.002 | -0.002 | 0 | 0.14 | 0.14 | 0.09 | 0.13 | 0.07 | 0.31 |
| 10 | 122879869 | rs2250810 | -2.219 | -0.322 | 2.121 | 2.069 | 0.07 | 0.35 | 0.35 | 0.142 | 0.05 | 0.42 | 0.17 | 0.26 | 0.01 | 0.04 |
| 10 | 122888555 | rs1907235 | 0.819 | -0.438 | 1.352 | 1.987 | -0.009 | 0.008 | 0.008 | 0.585 | 0.76 | 0.73 | 0.8 | 0.72 | 0.78 | 0.86 |
| 10 | 122889466 | rs2246868 | 1.201 | -0.605 | 2.529 | 2.371 | 0.059 | 0.334 | 0.334 | 0.105 | 0.94 | 0.53 | 0.89 | 0.77 | 0.98 | 0.98 |
| 10 | 122892234 | rs12259125 | -0.488 | -0.479 | 2.354 | 2.146 | -0.009 | 0.016 | 0.016 | 0 | 0.07 | 0.04 | 0.06 | 0.01 | 0.09 | 0.04 |
| 10 | 122892846 | rs1907238 | -0.846 | -0.235 | 2.324 | 2.201 | -0.001 | -0.031 | -0.031 | 0 | 0.23 | 0.13 | 0.3 | 0.28 | 0.28 | 0.13 |
| 10 | 122897913 | rs2250266 | 1.773 | -0.259 | 2.538 | 2.427 | 0.007 | 0.025 | 0.025 | 0 | 0.62 | 0.17 | 0.43 | 0.34 | 0.59 | 0.77 |
| 10 | 122902140 | rs2172072 | 1.002 | -0.307 | 3.09 | 2.531 | 0.059 | 0.403 | 0.403 | 0 | 0.94 | 0.55 | 0.9 | 0.74 | 0.98 | 0.97 |
| 10 | 122911702 | rs1907260 | 0.96 | -0.307 | 3.038 | 2.526 | 0.061 | 0.412 | 0.412 | 0 | 0.94 | 0.56 | 0.9 | 0.74 | 0.98 | 0.97 |
| 10 | 122912390 | rs11199800 | -1.262 | -0.306 | 2.88 | 2.499 | -0.011 | -0.006 | -0.006 | 0 | 0.3 | 0.16 | 0.34 | 0.26 | 0.37 | 0.17 |
| 10 | 122924854 | rs10788141 | 1.459 | -0.306 | 2.61 | 2.543 | 0.019 | 0.416 | 0.416 | 0.132 | 0.93 | 0.43 | 0.78 | 0.61 | 0.97 | 0.95 |
| 10 | 122939374 | rs7085142 | 1.387 | -0.077 | 1.463 | 2.007 | 0.006 | -0.012 | -0.012 | 0 | 0.65 | 0.28 | 0.48 | 0.39 | 0.63 | 0.78 |
| 10 | 122941364 | rs6585709 | -1.817 | 0.057 | 2.043 | 2.395 | 0.006 | 0.048 | 0.048 | 0 | 0.23 | 0.61 | 0.38 | 0.38 | 0.25 | 0.18 |
| 10 | 122949506 | rs11199820 | -1.546 | 0.224 | 2.217 | 2.106 | 0.022 | 0.076 | 0.076 | 0 | 0.19 | 0.3 | 0.35 | 0.38 | 0.25 | 0.17 |
| 10 | 122956997 | rs11199823 | -1.3 | 0.368 | 1.34 | 2.161 | -0.005 | -0.002 | -0.002 | 0 | 0.17 | 0.16 | 0.32 | 0.38 | 0.25 | 0.17 |
| 10 | 122966573 | rs12571414 | -0.842 | 0.21 | 1.055 | 2.147 | -0.028 | -0.03 | -0.03 | 0.626 | 0.05 | 0.01 | 0.16 | 0.11 | 0.09 | 0.13 |
| 10 | 122970923 | rs1907220 | -0.397 | -0.134 | 1.322 | 1.734 | 0.003 | 0 | 0 | 0 | 0.69 | 0.71 | 0.78 | 0.71 | 0.69 | 0.61 |
| 10 | 122975633 | rs1907225 | -0.284 | -0.331 | 1.438 | 1.964 | 0.007 | 0.002 | 0.002 | 0 | 0.67 | 0.71 | 0.77 | 0.71 | 0.69 | 0.61 |
| 10 | 122977030 | rs10510092 | -0.413 | -0.441 | 1.209 | 1.958 | 0.006 | 0.005 | 0.005 | 0 | 0.13 | 0.18 | 0.22 | 0.16 | 0.18 | 0.21 |
| 10 | 122981248 | rs7093348 | 0.48 | -0.928 | 0.765 | 1.338 | -0.009 | -0.018 | -0.018 | 0 | 0.38 | 0.31 | 0.5 | 0.55 | 0.5 | 0.24 |
| 10 | 122982711 | rs10886872 | 0.189 | -0.943 | 0.732 | 1.415 | 0.007 | 0.029 | 0.029 | 0 | 0.31 | 0.23 | 0.3 | 0.17 | 0.3 | 0.38 |
| 10 | 122985118 | rs10886873 | 0.982 | -0.944 | 0.236 | 1.415 | -0.079 | -0.004 | -0.004 | 0 | 0.31 | 0.1 | 0.36 | 0.36 | 0.49 | 0.22 |
| 10 | 122987126 | rs1907232 | 0.546 | -0.963 | 0.281 | 1.401 | -0.008 | -0.013 | -0.013 | 0 | 0.36 | 0.32 | 0.5 | 0.54 | 0.49 | 0.23 |
| 10 | 122995141 | rs12220647 | 0.944 | -0.926 | -0.707 | 1.395 | -0.058 | 0.006 | 0.006 | 0 | 0.32 | 0.12 | 0.36 | 0.35 | 0.49 | 0.22 |
| 10 | 122995312 | rs11199849 | 0.371 | -0.893 | -0.898 | 1.022 | 0.005 | 0.034 | 0.034 | 0 | 0.31 | 0.19 | 0.3 | 0.17 | 0.3 | 0.39 |
| 10 | 122996499 | rs11599530 | 1.121 | -0.51 | -1.663 | 0.712 | -0.072 | 0.054 | 0.054 | 0 | 0.24 | 0.1 | 0.23 | 0.14 | 0.34 | 0.13 |
| 10 | 123000544 | rs4751828 | 0.063 | -1.285 | -1.343 | 1.125 | 0.072 | -0.01 | -0.01 | 0 | 0.31 | 0.1 | 0.16 | 0.26 | 0.2 | 0.38 |
| 10 | 123002806 | rs729014 | -1.111 | -0.827 | -0.994 | 1.582 | 0.004 | -0.031 | -0.031 | 0.831 | 0.14 | 0.09 | 0.21 | 0.28 | 0.15 | 0.1 |

| | | | | | | | | | | | | | | | | |
|---|---|---|---|---|---|---|---|---|---|---|---|---|---|---|---|---|
| 10 | 123003689 | rs7899611 | -0.632 | -1.208 | -0.54 | 1.48 | -0.018 | 0.02 | 0.02 | 0 | 0.36 | 0.54 | 0.35 | 0.47 | 0.26 | 0.53 |
| 10 | 123004639 | rs731009 | -0.657 | -1.329 | 0.251 | 1.655 | -0.01 | 0.007 | 0.007 | 0 | 0.34 | 0.37 | 0.34 | 0.42 | 0.25 | 0.5 |
| 10 | 123009133 | rs4237529 | -0.391 | -1.276 | 0.089 | 1.395 | -0.002 | 0.001 | 0.001 | 0.747 | 0.68 | 0.66 | 0.63 | 0.68 | 0.65 | 0.85 |
| 10 | 123019551 | rs11199859 | 0.844 | -1.236 | -0.045 | 1.525 | -0.033 | -0.033 | -0.033 | 0.191 | 0.06 | 0 | 0.14 | 0.13 | 0.11 | 0.2 |
| 10 | 123019616 | rs1125527 | -0.021 | -1.97 | 0.183 | 1.445 | -0.011 | 0.017 | 0.017 | 0.387 | 0.64 | 0.46 | 0.54 | 0.58 | 0.64 | 0.83 |
| 10 | 123021268 | rs7081844 | -0.257 | -1.914 | 0.214 | 1.39 | -0.009 | 0.028 | 0.028 | 0.406 | 0.67 | 0.51 | 0.62 | 0.63 | 0.64 | 0.81 |
| 10 | 123021731 | rs7076500 | -0.072 | -1.919 | 0.301 | 1.399 | -0.005 | 0.01 | 0.01 | 0.351 | 0.33 | 0.39 | 0.36 | 0.37 | 0.36 | 0.18 |
| 10 | 123022956 | rs11199862 | 0.569 | -2.024 | 0.11 | 1.43 | -0.012 | -0.025 | -0.025 | 0 | 0.24 | 0.05 | 0.24 | 0.25 | 0.28 | 0.11 |
| 10 | 123023243 | rs4752526 | -0.05 | -2.293 | 0.345 | 1.338 | -0.067 | 0.133 | 0.133 | 0 | 0.85 | 0.64 | 0.93 | 0.71 | 0.96 | 0.7 |
| 10 | 123024757 | rs7091083 | -0.257 | -1.669 | 0.385 | 1.099 | 0.002 | 0.003 | 0.003 | 2.101 | 0.33 | 0.29 | 0.41 | 0.29 | 0.32 | 0.17 |
| 10 | 123029744 | rs7080775 | -0.542 | -1.479 | 0.428 | 1.149 | 0.029 | 0.049 | 0.049 | 0 | 0.06 | 0.22 | 0.19 | 0.11 | 0.07 | 0.05 |
| 10 | 123031410 | rs12761612 | 0.575 | -1.561 | 0.245 | 1.143 | -0.014 | 0.006 | 0.006 | 4.836 | 0.25 | 0.04 | 0.21 | 0.19 | 0.24 | 0.15 |
| 10 | 123032168 | rs4575197 | 1.701 | -1.674 | -0.052 | 0.952 | -0.166 | 0.07 | 0.07 | 0.804 | 0.29 | 0.06 | 0.33 | 0.14 | 0.47 | 0.22 |
| 10 | 123034391 | rs4468286 | 0.473 | -1.441 | 0.237 | 1.076 | 0.004 | 0.007 | 0.007 | 2.872 | 0.27 | 0.24 | 0.29 | 0.23 | 0.26 | 0.15 |
| 10 | 123037309 | rs10788162 | 0.186 | -1.438 | 0.118 | 1.085 | -0.014 | -0.008 | -0.008 | 1.196 | 0.31 | 0.15 | 0.39 | 0.3 | 0.34 | 0.2 |
| 10 | 123039881 | rs11199877 | 0.813 | -1.484 | -0.514 | 1.279 | -0.008 | -0.031 | -0.031 | 0.254 | 0.05 | 0 | 0.12 | 0.2 | 0.04 | 0.2 |
| 10 | 123044383 | rs9630106 | 0.435 | -1.529 | -0.284 | 1.129 | 0.017 | -0.006 | -0.006 | 0.127 | 0.56 | 0.33 | 0.46 | 0.45 | 0.64 | 0.58 |
| 10 | 123047313 | rs10886895 | 0.41 | -1.385 | -0.776 | 1.047 | 0.016 | 0.014 | 0.014 | 9.175 | 0.29 | 0.12 | 0.26 | 0.23 | 0.24 | 0.16 |
| 10 | 123049264 | rs10886902 | 0.481 | -1.358 | -1.177 | 1.058 | -0.032 | -0.033 | -0.033 | 0 | 0.28 | 0.02 | 0.25 | 0.23 | 0.24 | 0.16 |
| 10 | 123052410 | rs12220387 | 0.341 | -1.315 | -0.591 | 1.017 | 0.018 | -0.008 | -0.008 | 0.024 | 0.26 | 0.16 | 0.13 | 0.31 | 0.17 | 0.35 |
| 10 | 123057781 | rs7071101 | 0.892 | -0.601 | -0.82 | 0.899 | -0.089 | 0.061 | 0.061 | 1.531 | 0.3 | 0.17 | 0.33 | 0.17 | 0.47 | 0.25 |
| 10 | 123057965 | rs7896863 | 0.172 | -1.095 | -0.651 | 1.012 | 0.059 | 0.08 | 0.08 | 0 | 0.51 | 0.26 | 0.51 | 0.37 | 0.62 | 0.42 |
| 10 | 123062540 | rs7097525 | 0.19 | -1.164 | -0.281 | 0.987 | 0.061 | 0.107 | 0.107 | 0 | 0.51 | 0.24 | 0.47 | 0.32 | 0.62 | 0.42 |
| 10 | 123067805 | rs12411339 | -0.201 | -1.187 | -0.246 | 1.321 | -0.001 | 0.021 | 0.021 | 0.021 | 0.1 | 0.09 | 0.26 | 0.27 | 0.09 | 0.2 |
| 10 | 123069717 | rs10736302 | -0.929 | -1.201 | -0.322 | 1.056 | -0.001 | -0.001 | -0.001 | 0 | 0.78 | 0.75 | 0.86 | 0.85 | 0.85 | 0.85 |
| 10 | 123076181 | rs11199892 | 0.732 | -1.134 | -0.233 | 1.12 | -0.002 | -0.001 | -0.001 | 0 | 0.22 | 0.23 | 0.14 | 0.2 | 0.15 | 0.17 |
| 10 | 123081449 | rs4752530 | 0.043 | -0.803 | -0.32 | 0.984 | 0.005 | 0.003 | 0.003 | 0 | 0.35 | 0.27 | 0.29 | 0.3 | 0.26 | 0.34 |
| 10 | 123088199 | rs4752534 | 0.416 | 0.103 | -0.648 | 0.958 | -0.047 | 0.019 | 0.019 | 0.42 | 0.31 | 0.18 | 0.41 | 0.23 | 0.44 | 0.34 |
| 10 | 123089079 | rs1896419 | 0.488 | 0.15 | -0.915 | 0.81 | -0.122 | 0.094 | 0.094 | 0.98 | 0.29 | 0.08 | 0.33 | 0.12 | 0.46 | 0.31 |
| 10 | 123089808 | rs9420317 | -0.142 | -0.214 | -0.779 | 1.156 | 0.004 | -0.011 | -0.011 | 2.504 | 0.26 | 0.19 | 0.33 | 0.31 | 0.23 | 0.28 |
| 10 | 123092807 | rs4752536 | -0.032 | 0.486 | -0.952 | 1.321 | -0.037 | -0.063 | -0.063 | 0 | 0.72 | 0.9 | 0.8 | 0.59 | 0.7 | 0.69 |
| 10 | 123094861 | rs11199916 | -0.124 | 0.47 | -0.946 | 1.436 | -0.1 | -0.095 | -0.095 | 0.196 | 0.27 | 0.03 | 0.19 | 0.38 | 0.3 | 0.23 |
| 10 | 123094884 | rs7089943 | -0.019 | 0.471 | -0.838 | 1.407 | -0.026 | -0.044 | -0.044 | 0 | 0.72 | 0.87 | 0.79 | 0.59 | 0.7 | 0.69 |
| 10 | 123095186 | rs7913694 | 0.453 | 0.711 | -0.45 | 1.024 | -0.017 | 0.006 | 0.006 | 4.179 | 0.41 | 0.54 | 0.41 | 0.46 | 0.33 | 0.31 |
| 10 | 123095366 | rs7914065 | 0.883 | 0.526 | -0.364 | 1.039 | 0.012 | -0.003 | -0.003 | 0 | 0.35 | 0.29 | 0.33 | 0.32 | 0.24 | 0.23 |
| 10 | 123095376 | rs7914068 | -0.28 | 0.774 | -0.145 | 0.528 | 0.018 | -0.037 | -0.037 | 0.392 | 0.68 | 0.58 | 0.62 | 0.8 | 0.62 | 0.54 |
| 10 | 123096559 | rs9421411 | 0.292 | 0.932 | -0.191 | 0.675 | -0.005 | -0.005 | -0.005 | 0 | 0.43 | 0.57 | 0.44 | 0.38 | 0.4 | 0.26 |
| 10 | 123096719 | rs10466214 | 0.002 | 1.625 | -0.339 | 0.306 | 0.002 | -0.017 | -0.017 | 1.216 | 0.2 | 0.2 | 0.16 | 0.1 | 0.24 | 0.31 |
| 10 | 123096747 | rs10466215 | 0.026 | 1.667 | -0.359 | 0.301 | 0.001 | -0.015 | -0.015 | 0.172 | 0.2 | 0.18 | 0.16 | 0.09 | 0.25 | 0.33 |
| 10 | 123098249 | rs7895870 | -0.302 | 1.504 | -0.106 | 0.751 | 0.008 | -0.005 | -0.005 | 0.255 | 0.38 | 0.48 | 0.45 | 0.6 | 0.41 | 0.37 |
| 10 | 123105123 | rs7903175 | -0.028 | 1.379 | 0.098 | 1.082 | 0.01 | 0.075 | 0.075 | 0.016 | 0.27 | 0.44 | 0.31 | 0.51 | 0.29 | 0.27 |
| 10 | 123115687 | rs7071430 | 0.089 | 1.671 | 0.138 | 0.97 | 0.069 | -0.028 | -0.028 | 0.008 | 0.82 | 0.64 | 0.7 | 0.84 | 0.73 | 0.9 |
| 10 | 123117653 | rs10886918 | -0.659 | 1 | -0.769 | 0.714 | 0.023 | -0.009 | -0.009 | 2.162 | 0.66 | 0.51 | 0.52 | 0.68 | 0.61 | 0.81 |
| 10 | 123119045 | rs7070463 | -1.034 | 1.443 | -0.389 | 1.292 | 0.004 | 0.009 | 0.009 | 0.218 | 0.35 | 0.38 | 0.33 | 0.52 | 0.38 | 0.3 |
| 10 | 123124214 | rs2420931 | -0.114 | 1.743 | -1.198 | 1.46 | -0.042 | -0.063 | -0.063 | 0.166 | 0.23 | 0.08 | 0.21 | 0.36 | 0.28 | 0.23 |
| 10 | 123130495 | rs12220114 | -0.447 | 1.406 | -1.15 | 1.535 | -0.05 | -0.031 | -0.031 | 0 | 0.32 | 0.16 | 0.24 | 0.39 | 0.39 | 0.29 |
| 10 | 123132045 | rs10886927 | -0.413 | 1.435 | -1.137 | 1.266 | -0.05 | -0.001 | -0.001 | 0.15 | 0.31 | 0.16 | 0.24 | 0.36 | 0.39 | 0.25 |
| 10 | 123133203 | rs10510094 | -0.536 | 1.481 | -1.12 | 1.022 | -0.033 | -0.036 | -0.036 | 0 | 0.33 | 0.23 | 0.3 | 0.45 | 0.39 | 0.29 |
| 10 | 123135200 | rs10788173 | 0.073 | 1.235 | -0.746 | 0.787 | -0.001 | -0.002 | -0.002 | 0 | 0.63 | 0.36 | 0.61 | 0.63 | 0.57 | 0.66 |
| 10 | 123135345 | rs12244218 | 0.033 | 1.071 | -0.843 | 0.887 | -0.001 | 0.003 | 0.003 | 0 | 0.26 | 0.29 | 0.24 | 0.31 | 0.21 | 0.19 |
| 10 | 123135724 | rs10886929 | -0.233 | 1.585 | -0.596 | 0.937 | -0.025 | -0.013 | -0.013 | 0 | 0.25 | 0.19 | 0.34 | 0.32 | 0.37 | 0.15 |
| 10 | 123135952 | rs12570261 | -0.788 | 0.673 | 0.031 | 0.749 | 0.024 | 0.065 | 0.065 | 0 | 0.05 | 0.16 | 0.03 | 0.07 | 0.05 | 0.12 |
| 10 | 123138257 | rs4752541 | 0.04 | 0.235 | -0.241 | -0.101 | 0.001 | -0.019 | -0.019 | 0 | 0.6 | 0.62 | 0.46 | 0.09 | 0.58 | 0.28 |
| 10 | 123138537 | rs4752542 | 0.213 | 0.072 | -0.137 | 0.472 | 0.003 | 0.019 | 0.019 | 0.144 | 0.61 | 0.69 | 0.59 | 0.22 | 0.6 | 0.29 |
| 10 | 123140794 | rs9421417 | 0.453 | -0.74 | 0.18 | -0.2 | 0.014 | 0.095 | 0.095 | 0 | 0.48 | 0.36 | 0.36 | 0.08 | 0.53 | 0.24 |
| 10 | 123146405 | rs4752545 | -0.466 | -1.253 | 0.444 | 0.351 | 0.009 | 0.003 | 0.003 | 0 | 0.45 | 0.35 | 0.4 | 0.59 | 0.41 | 0.64 |
| 10 | 123148756 | rs4752546 | 0.84 | -1.372 | 0.644 | 0.308 | -0.007 | 0.229 | 0.229 | 0 | 0.46 | 0.19 | 0.36 | 0.08 | 0.53 | 0.24 |
| 10 | 123150301 | rs4752549 | -0.041 | -1.339 | 0.631 | 0.37 | -0.012 | 0.083 | 0.083 | 0 | 0.91 | 0.74 | 0.9 | 0.82 | 0.94 | 0.88 |
| 10 | 123151488 | rs4752550 | -0.082 | -1.361 | 0.417 | 0.363 | 0.007 | -0.005 | -0.005 | 0 | 0.41 | 0.29 | 0.39 | 0.58 | 0.38 | 0.63 |
| 10 | 123153176 | rs2420929 | 0.799 | -1.362 | 0.435 | 0.395 | -0.05 | 0.166 | 0.166 | 0.512 | 0.85 | 0.56 | 0.75 | 0.65 | 0.9 | 0.86 |
| 10 | 123153472 | rs2420928 | -0.375 | -1.002 | 0.164 | 0.536 | 0.009 | -0.002 | -0.002 | 0 | 0.42 | 0.29 | 0.39 | 0.59 | 0.38 | 0.63 |
| 10 | 123155416 | rs4751835 | -0.367 | -0.996 | 0.122 | 1.158 | -0.001 | 0.009 | 0.009 | 0 | 0.34 | 0.25 | 0.3 | 0.3 | 0.33 | 0.5 |
| 10 | 123174477 | rs11199952 | -1.754 | -1.601 | 0.592 | 1.98 | -0.04 | 0.155 | 0.155 | 0.074 | 0.12 | 0.22 | 0.28 | 0.61 | 0.06 | 0.25 |
| 10 | 123177160 | rs10510095 | -1.058 | -1.496 | 0.437 | 1.748 | -0.005 | -0.003 | -0.003 | 0 | 0.07 | 0.06 | 0.19 | 0.04 | 0.04 | 0.09 |
| 10 | 123184875 | rs10788178 | 0.548 | -1.324 | 1.091 | 1.771 | -0.015 | 0.048 | 0.048 | 0.205 | 0.75 | 0.57 | 0.67 | 0.7 | 0.76 | 0.48 |
| 10 | 123185303 | rs10749415 | 0.768 | -1.316 | 1.229 | 1.752 | -0.004 | -0.011 | -0.011 | 0 | 0.92 | 0.74 | 0.78 | 0.98 | 0.96 | 0.91 |
| 10 | 123188167 | rs11199957 | -1.425 | -0.443 | 1.195 | 2.475 | 0.026 | 0.242 | 0.242 | 0 | 0.07 | 0.16 | 0.13 | 0.57 | 0.1 | 0.17 |
| 10 | 123189052 | rs17101977 | -1.516 | -0.44 | 1.21 | 2.522 | 0.057 | 0.363 | 0.363 | 0 | 0.07 | 0.27 | 0.14 | 0.53 | 0.1 | 0.17 |
| 10 | 123190112 | rs17101979 | -0.046 | -1.496 | 1.003 | 1.92 | -0.023 | -0.005 | -0.005 | 0.016 | 0.1 | 0.16 | 0.07 | 0.08 | 0.04 | 0.04 |

| | | | | | | | | | | | | | | | | |
|---|---|---|---|---|---|---|---|---|---|---|---|---|---|---|---|---|
| 10 | 123191391 | rs10788181 | -0.318 | -1.323 | 0.913 | 1.995 | -0.001 | 0.005 | 0.005 | 0 | 0.79 | 0.75 | 0.9 | 0.79 | 0.78 | 0.59 |
| 10 | 123193998 | rs11199964 | -0.377 | -2.091 | 0.136 | 1.125 | -0.026 | 0.008 | 0.008 | 0 | 0.39 | 0.42 | 0.29 | 0.43 | 0.25 | 0.25 |
| 10 | 123194255 | rs7084101 | -0.209 | -2.136 | 0.082 | 1.053 | -0.039 | 0.012 | 0.012 | 0 | 0.37 | 0.42 | 0.27 | 0.43 | 0.2 | 0.24 |
| 10 | 123194321 | rs12781987 | 1.198 | -1.908 | -0.574 | 0.942 | -0.064 | -0.06 | -0.06 | 0 | 0.17 | 0.02 | 0.07 | 0.21 | 0.24 | 0.43 |
| 10 | 123197382 | rs7098583 | -1.406 | -2.075 | -0.23 | 0.628 | 0.015 | -0.027 | -0.027 | 0 | 0.63 | 0.56 | 0.51 | 0.71 | 0.5 | 0.76 |
| 10 | 123198136 | rs7894494 | 1.122 | -2.182 | -0.319 | 0.44 | 0.009 | 0.064 | 0.064 | 0.024 | 0.41 | 0.26 | 0.5 | 0.28 | 0.55 | 0.29 |
| 10 | 123198286 | rs7911028 | -0.959 | -2.081 | -0.185 | 0.547 | -0.012 | -0.007 | -0.007 | 2.97 | 0.81 | 0.87 | 0.9 | 0.79 | 0.76 | 0.56 |
| 10 | 123198332 | rs2162543 | -0.837 | -1.935 | 0.109 | 0.818 | -0.015 | 0.18 | 0.18 | 0 | 0.08 | 0.26 | 0.22 | 0.2 | 0.03 | 0.12 |
| 10 | 123203286 | rs10788182 | 0.141 | -1.821 | 0.036 | 0.809 | 0.105 | 0.319 | 0.319 | 0 | 0.94 | 0.7 | 0.9 | 0.74 | 0.93 | 0.88 |
| 10 | 123203921 | rs6585735 | -0.499 | -1.979 | -0.075 | 0.671 | 0.034 | 0.005 | 0.005 | 0 | 0.49 | 0.37 | 0.25 | 0.51 | 0.42 | 0.6 |
| 10 | 123205203 | rs7918550 | 0.641 | -1.972 | -0.258 | 0.662 | -0.009 | 0.117 | 0.117 | 0 | 0.44 | 0.29 | 0.57 | 0.05 | 0.5 | 0.25 |
| 10 | 123208881 | rs2420936 | 0.635 | -1.847 | -0.714 | 0.133 | -0.009 | 0.092 | 0.092 | 0 | 0.44 | 0.27 | 0.63 | 0.22 | 0.51 | 0.28 |
| 10 | 123215089 | rs10788184 | -0.387 | -2.192 | -0.55 | 0.523 | 0.005 | -0.007 | -0.007 | 0 | 0.3 | 0.29 | 0.22 | 0.57 | 0.19 | 0.3 |
| 10 | 123217600 | rs4644589 | -0.486 | -1.323 | -1.191 | 0.145 | 0.006 | 0 | 0 | 0.35 | 0.49 | 0.26 | 0.45 | 0.54 | 0.58 | 0.71 |
| 10 | 123219430 | rs2420938 | 1.198 | -0.772 | -1.21 | -0.022 | -0.002 | 0.002 | 0.002 | 0 | 0.31 | 0.31 | 0.41 | 0.18 | 0.3 | 0.17 |
| 10 | 123221890 | rs2114684 | 0.482 | -0.2 | -2.057 | 0.195 | -0.162 | 0.02 | 0.02 | 0 | 0.33 | 0.03 | 0.21 | 0.23 | 0.49 | 0.51 |
| 10 | 123224879 | rs7911727 | -0.993 | -0.109 | -1.96 | -0.198 | -0.001 | -0.015 | -0.015 | 0.08 | 0.6 | 0.39 | 0.54 | 0.84 | 0.69 | 0.76 |
| 10 | 123228046 | rs6585739 | -0.438 | -0.504 | -1.393 | 0.245 | -0.001 | 0 | 0 | 0 | 0.27 | 0.26 | 0.23 | 0.4 | 0.2 | 0.31 |
| 10 | 123228660 | rs7072877 | -0.535 | -0.403 | -1.421 | 0.255 | -0.006 | 0.005 | 0.005 | 0 | 0.3 | 0.3 | 0.33 | 0.42 | 0.21 | 0.33 |
| 10 | 123229626 | rs2420941 | -0.181 | 0.741 | -1.982 | 0.07 | -0.067 | -0.023 | -0.023 | 0 | 0.42 | 0.08 | 0.28 | 0.39 | 0.51 | 0.46 |
| 10 | 123230953 | rs2250218 | 0.812 | 0.935 | -0.737 | -0.353 | 0.034 | -0.086 | -0.086 | 0.552 | 0.84 | 0.71 | 0.88 | 0.99 | 0.74 | 0.96 |
| 10 | 123235872 | rs1649161 | -0.142 | 0.198 | -1.123 | -0.262 | -0.002 | 0 | 0 | 0 | 0.36 | 0.6 | 0.19 | 0.64 | 0.26 | 0.34 |
| 10 | 123236125 | rs1649204 | -0.021 | 0.203 | -1.122 | -0.268 | -0.006 | 0.001 | 0.001 | 0.014 | 0.35 | 0.6 | 0.19 | 0.64 | 0.24 | 0.34 |
| 10 | 123236920 | rs3135831 | 1.219 | 0.161 | -1.29 | -0.544 | -0.139 | -0.012 | -0.012 | 0.723 | 0.36 | 0.05 | 0.5 | 0.25 | 0.4 | 0.44 |
| 10 | 123239112 | rs1047057 | 0.69 | 0.128 | -0.894 | -0.373 | -0.035 | -0.003 | -0.003 | 0.609 | 0.49 | 0.09 | 0.69 | 0.35 | 0.51 | 0.62 |
| 10 | 123241794 | rs2556537 | -0.525 | 0.02 | -0.387 | -0.277 | 0.015 | -0.01 | -0.01 | 0 | 0.35 | 0.65 | 0.2 | 0.65 | 0.25 | 0.32 |
| 10 | 123243197 | rs2278202 | 0.922 | 0.136 | -0.516 | -0.332 | 0.001 | 0.012 | 0.012 | 0.707 | 0.54 | 0.11 | 0.69 | 0.33 | 0.54 | 0.63 |
| 10 | 123243730 | rs1649200 | 0.858 | 0.43 | -0.129 | -0.484 | 0.053 | -0.083 | -0.083 | 0 | 0.87 | 0.65 | 0.88 | 1 | 0.81 | 0.95 |
| 10 | 123244834 | rs1613776 | -0.557 | 0.507 | -0.617 | -0.376 | 0.016 | -0.024 | -0.024 | 0 | 0.05 | 0.21 | 0.02 | 0 | 0.06 | 0.01 |
| 10 | 123247037 | rs2981461 | 0.681 | 0.491 | -0.934 | -0.246 | 0.006 | 0.044 | 0.044 | 1.087 | 0.55 | 0.15 | 0.66 | 0.3 | 0.55 | 0.62 |
| 10 | 123249057 | rs3135797 | -0.968 | 1.163 | -0.284 | -0.256 | -0.005 | -0.004 | -0.004 | 0.041 | 0.08 | 0.05 | 0.09 | 0 | 0.18 | 0.03 |
| 10 | 123257402 | rs2912753 | 1.008 | 0.619 | 0.205 | -0.144 | 0.046 | -0.008 | -0.008 | 0 | 0.94 | 0.74 | 0.95 | 1 | 0.95 | 0.96 |
| 10 | 123262708 | rs2912755 | 1.534 | 0.147 | 0.788 | -0.391 | -0.014 | -0.053 | -0.053 | 0 | 0.89 | 0.46 | 0.88 | 0.99 | 0.97 | 0.96 |
| 10 | 123263616 | rs3135772 | -0.224 | 0.004 | -0.062 | 0.405 | -0.011 | -0.104 | -0.104 | 0 | 0.13 | 0 | 0.11 | 0.51 | 0.13 | 0.15 |
| 10 | 123267631 | rs4752566 | -0.046 | 0.092 | -0.908 | -0.852 | -0.028 | 0.231 | 0.231 | 0 | 0.44 | 0.11 | 0.41 | 0.91 | 0.46 | 0.66 |
| 10 | 123268889 | rs7090018 | -0.279 | 0.217 | -0.39 | -0.723 | 0.015 | -0.052 | -0.052 | 0.092 | 0.23 | 0.31 | 0.39 | 0.1 | 0.28 | 0.19 |
| 10 | 123272296 | rs3135766 | 0.309 | 0.042 | -0.799 | -0.869 | -0.008 | 0.124 | 0.124 | 0.664 | 0.57 | 0.17 | 0.47 | 0.8 | 0.6 | 0.58 |
| 10 | 123276091 | rs3135761 | 0.432 | 0.28 | -0.95 | -0.461 | -0.011 | -0.155 | -0.155 | 0 | 0.18 | 0.01 | 0.11 | 0.52 | 0.16 | 0.22 |
| 10 | 123276290 | rs2912762 | -0.049 | 0.354 | -0.146 | -0.719 | -0.007 | -0.004 | -0.004 | 0 | 0.24 | 0.21 | 0.25 | 0.06 | 0.31 | 0.27 |
| 10 | 123279745 | rs2981448 | 0.213 | -0.032 | 0.806 | -0.545 | 0.047 | 0.043 | 0.043 | 3.69 | 0.94 | 0.6 | 0.95 | 1 | 1 | 1 |
| 10 | 123296319 | rs2912770 | -0.276 | 0.267 | 0.697 | -0.553 | 0.017 | -0.048 | -0.048 | 0 | 0.79 | 0.73 | 0.78 | 0.94 | 0.76 | 0.73 |
| 10 | 123298158 | rs1047100 | -0.406 | 0.324 | 0.096 | -0.479 | 0.018 | -0.048 | -0.048 | 0 | 0.8 | 0.73 | 0.8 | 0.94 | 0.79 | 0.74 |
| 10 | 123317731 | rs2981427 | 0.24 | 0.119 | 0.195 | 0.021 | 0.054 | -0.002 | -0.002 | 0 | 0.5 | 0.25 | 0.55 | 0.4 | 0.6 | 0.47 |
| 10 | 123318961 | rs2912791 | -0.206 | 0.2 | 0.244 | 0.106 | 0.02 | -0.007 | -0.007 | 0 | 0.56 | 0.74 | 0.68 | 0.47 | 0.61 | 0.47 |
| 10 | 123325538 | rs2912787 | -0.189 | 0.822 | 0.641 | -0.238 | 0.04 | -0.105 | -0.105 | 0.073 | 0.18 | 0.45 | 0.14 | 0.03 | 0.18 | 0.08 |
| 10 | 123325794 | rs3135730 | 0.2 | 0.605 | 0.762 | -0.087 | 0.038 | 0.013 | 0.013 | 0 | 0.12 | 0.44 | 0.18 | 0.07 | 0.09 | 0.01 |
| 10 | 123326646 | rs2981432 | -0.572 | 0.517 | 0.034 | 0.149 | -0.003 | -0.014 | -0.014 | 0 | 0.57 | 0.65 | 0.6 | 0.4 | 0.55 | 0.49 |
| 10 | 123327876 | rs10510097 | -0.144 | 0.405 | 0.117 | 0.605 | -0.002 | 0.037 | 0.037 | 0 | 0.15 | 0.39 | 0.17 | 0.19 | 0.12 | 0.13 |
| 10 | 123329429 | rs2981428 | 0.132 | 0.508 | -0.941 | 0.713 | -0.116 | -0.094 | -0.094 | 0.112 | 0.46 | 0.05 | 0.42 | 0.56 | 0.55 | 0.55 |
| 10 | 123332577 | rs3750817 | 0.492 | 0.587 | -1.052 | 0.622 | -0.089 | -0.147 | -0.147 | 0 | 0.34 | 0.05 | 0.37 | 0.52 | 0.38 | 0.47 |
| 10 | 123334930 | rs11200014 | 0.306 | -0.017 | -0.721 | 0.458 | -0.02 | 0.059 | 0.059 | 0 | 0.42 | 0.19 | 0.38 | 0.29 | 0.44 | 0.33 |
| 10 | 123337335 | rs2981579 | -0.172 | -0.055 | 0.123 | 0.438 | 0.004 | 0.014 | 0.014 | 0 | 0.52 | 0.34 | 0.57 | 0.55 | 0.55 | 0.61 |
| 10 | 123340311 | rs2981578 | 0.837 | -0.009 | -0.082 | 0.809 | -0.108 | -0.078 | -0.078 | 1.493 | 0.38 | 0.08 | 0.49 | 0.44 | 0.48 | 0.51 |
| 10 | 123346190 | rs1219648 | -0.012 | -0.031 | 0.538 | 0.617 | 0.001 | -0.004 | -0.004 | 1.138 | 0.43 | 0.44 | 0.41 | 0.38 | 0.43 | 0.38 |
| 10 | 123348355 | rs1219643 | 0.127 | -0.141 | 0.516 | 0.379 | 0.014 | -0.05 | -0.05 | 0 | 0.21 | 0.31 | 0.13 | 0.09 | 0.2 | 0.16 |
| 10 | 123350191 | rs17102287 | 1.26 | -0.183 | 0.282 | 0.421 | -0.003 | -0.027 | -0.027 | 0 | 0.22 | 0.13 | 0.28 | 0.29 | 0.23 | 0.22 |
| 10 | 123351302 | rs2860197 | -0.187 | -0.245 | 0.524 | 0.594 | 0 | 0.006 | 0.006 | 0 | 0.55 | 0.43 | 0.57 | 0.62 | 0.58 | 0.61 |
| 10 | 123351324 | rs2420946 | -0.13 | -0.248 | 0.533 | 0.593 | -0.001 | 0.005 | 0.005 | 0 | 0.55 | 0.44 | 0.57 | 0.62 | 0.58 | 0.61 |
| 10 | 123352317 | rs2981582 | -0.186 | -0.181 | 0.581 | 0.461 | 0 | -0.01 | -0.01 | 0.519 | 0.56 | 0.51 | 0.58 | 0.68 | 0.58 | 0.66 |
| 10 | 123355688 | rs7899765 | -0.581 | -0.727 | 1.167 | 0.467 | -0.043 | -0.008 | -0.008 | 0 | 0.14 | 0.32 | 0.08 | 0.09 | 0.07 | 0.11 |
| 10 | 123358312 | rs1219639 | -0.238 | -0.401 | 1.126 | 0.429 | 0.014 | -0.05 | -0.05 | 0 | 0.88 | 0.7 | 0.92 | 0.97 | 0.92 | 0.96 |
| 10 | 123380393 | rs1649187 | 0.52 | -1.403 | 1.214 | 0.484 | -0.052 | 0.003 | 0.003 | 1.1 | 0.71 | 0.54 | 0.8 | 0.72 | 0.82 | 0.68 |
| 10 | 123386254 | rs1909669 | 1.339 | -1.254 | 0.832 | 0.893 | 0.078 | -0.053 | -0.053 | 0.851 | 0.56 | 0.12 | 0.6 | 0.49 | 0.71 | 0.61 |
| 10 | 123387029 | rs10788190 | -0.72 | -1.253 | 1.16 | 0.875 | -0.035 | -0.002 | -0.002 | 0 | 0.3 | 0.41 | 0.19 | 0.29 | 0.18 | 0.32 |
| 10 | 123390785 | rs1631281 | 0.235 | -1.025 | 0.62 | 0.939 | 0.001 | 0.016 | 0.016 | 0 | 0.11 | 0.17 | 0.16 | 0.21 | 0.1 | 0.07 |
| 10 | 123392378 | rs7896565 | 1.595 | -1.181 | -0.009 | 0.555 | 0.044 | -0.01 | -0.01 | 0.029 | 0.49 | 0.19 | 0.52 | 0.55 | 0.59 | 0.49 |
| 10 | 123396728 | rs1909670 | 0.913 | -0.989 | -0.067 | 0.593 | -0.023 | -0.044 | -0.044 | 0 | 0.09 | 0.03 | 0.13 | 0.22 | 0.1 | 0.07 |
| 10 | 123396806 | rs1614303 | 0.663 | -1.312 | 0.275 | 0.329 | -0.05 | -0.022 | -0.022 | 0 | 0.75 | 0.58 | 0.81 | 0.82 | 0.81 | 0.74 |
| 10 | 123399847 | rs10466158 | -1.19 | -1.243 | 0.322 | 0.36 | -0.033 | -0.044 | -0.044 | 0 | 0.32 | 0.41 | 0.23 | 0.18 | 0.25 | 0.27 |

| | | | | | | | | | | | | | | | | |
|---|---|---|---|---|---|---|---|---|---|---|---|---|---|---|---|---|
| 10 | 123402175 | rs11200051 | 0.221 | -0.85 | 0.076 | 0.627 | -0.005 | 0.014 | 0.014 | 0 | 0.12 | 0.09 | 0.07 | 0.06 | 0.11 | 0.17 |
| 10 | 123410746 | rs12413814 | 1.725 | -1.29 | -0.264 | 0.542 | 0.039 | -0.065 | -0.065 | 0 | 0.49 | 0.09 | 0.56 | 0.53 | 0.61 | 0.54 |
| 10 | 123410940 | rs7068354 | 1.417 | -1.436 | 0.483 | 0.575 | 0.012 | 0.006 | 0.006 | 0 | 0.56 | 0.39 | 0.68 | 0.65 | 0.67 | 0.66 |
| 10 | 123412399 | rs7906816 | 0.225 | -1.411 | 0.518 | 0.589 | 0.021 | 0.084 | 0.084 | 0.021 | 0.07 | 0.3 | 0.12 | 0.09 | 0.06 | 0.11 |
| 10 | 123415590 | rs11200059 | 1.891 | -1.433 | 0.18 | 0.714 | 0.05 | -0.028 | -0.028 | 0 | 0.49 | 0.1 | 0.54 | 0.47 | 0.62 | 0.55 |
| 10 | 123417635 | rs10788196 | -0.314 | -1.424 | 0.524 | 0.639 | -0.001 | -0.003 | -0.003 | 0.122 | 0.84 | 0.87 | 0.84 | 0.78 | 0.85 | 0.9 |
| 10 | 123419652 | rs7910243 | -1.066 | -1.422 | 0.857 | 0.583 | 0.005 | 0.007 | 0.007 | 0 | 0.23 | 0.19 | 0.16 | 0.14 | 0.17 | 0.22 |
| 10 | 123421024 | rs11200063 | -0.645 | -1.487 | 0.814 | 0.589 | 0.009 | 0.021 | 0.021 | 0 | 0.2 | 0.08 | 0.14 | 0.13 | 0.16 | 0.21 |
| 10 | 123422434 | rs7071275 | -1.114 | -1.501 | 0.884 | 0.559 | 0.002 | 0.002 | 0.002 | 0 | 0.24 | 0.22 | 0.15 | 0.14 | 0.17 | 0.22 |
| 10 | 123422766 | rs2935715 | -0.352 | -1.349 | 1.08 | 0.617 | 0.024 | 0.126 | 0.126 | 0 | 0.92 | 0.7 | 0.85 | 0.85 | 0.93 | 0.9 |
| 10 | 123423254 | rs2936854 | -1.767 | -1.433 | 1.094 | 0.73 | -0.008 | 0.003 | 0.003 | 0 | 0.38 | 0.42 | 0.39 | 0.41 | 0.3 | 0.3 |
| 10 | 123431515 | rs9971131 | 2.192 | -1.574 | 0.956 | 0.615 | 0.067 | 0.02 | 0.02 | 0.007 | 0.51 | 0.23 | 0.44 | 0.43 | 0.6 | 0.6 |
| 10 | 123431846 | rs10510099 | -0.839 | -1.698 | 0.986 | 0.557 | 0.017 | 0.02 | 0.02 | 0 | 0.23 | 0.16 | 0.15 | 0.15 | 0.18 | 0.22 |
| 10 | 123432188 | rs2935713 | 1.424 | -1.656 | 0.976 | 0.54 | 0.001 | -0.011 | -0.011 | 0 | 0.1 | 0.05 | 0.06 | 0 | 0.07 | 0.02 |
| 10 | 123432652 | rs12356902 | 0.609 | -1.707 | 0.924 | 0.532 | -0.038 | 0.023 | 0.023 | 0 | 0.12 | 0.01 | 0.1 | 0.07 | 0.13 | 0.13 |
| 10 | 123436246 | rs2935717 | 0.019 | -1.597 | 1.103 | 0.546 | -0.016 | -0.024 | -0.024 | 0 | 0.82 | 0.64 | 0.77 | 0.86 | 0.85 | 0.88 |
| 10 | 123436465 | rs3104688 | -0.587 | -1.554 | 1.104 | 0.577 | 0.014 | 0.084 | 0.084 | 0 | 0.91 | 0.68 | 0.83 | 0.86 | 0.92 | 0.9 |
| 10 | 123436696 | rs1693688 | 1.249 | -1.359 | 0.621 | 0.567 | 0.048 | 0.055 | 0.055 | 0.042 | 0.55 | 0.32 | 0.63 | 0.72 | 0.67 | 0.66 |
| 10 | 123437453 | rs2935718 | 1.314 | -1.336 | 0.512 | 0.56 | 0.027 | 0.003 | 0.003 | 0 | 0.54 | 0.37 | 0.49 | 0.45 | 0.62 | 0.6 |
| 10 | 123438113 | rs1693687 | 1.523 | -1.311 | 0.27 | 0.581 | 0.061 | 0.013 | 0.013 | 0.002 | 0.52 | 0.26 | 0.48 | 0.45 | 0.61 | 0.6 |
| 10 | 123441384 | rs4752580 | 1.133 | -1.449 | 0.326 | 0.51 | 0.024 | -0.024 | -0.024 | 0 | 0.63 | 0.27 | 0.53 | 0.47 | 0.68 | 0.62 |
| 10 | 123443605 | rs4752582 | -1.281 | -1.315 | -0.114 | 0.618 | -0.01 | -0.009 | -0.009 | 0.311 | 0.08 | 0.04 | 0.17 | 0.26 | 0.09 | 0.06 |
| 10 | 123443636 | rs6585751 | 1.24 | -1.481 | -0.336 | 0.45 | 0.076 | -0.022 | -0.022 | 0.019 | 0.55 | 0.12 | 0.48 | 0.43 | 0.64 | 0.58 |
| 10 | 123450233 | rs2132902 | 0.898 | -1.437 | -0.439 | 0.447 | 0.109 | 0.106 | 0.106 | 2.801 | 0.52 | 0.19 | 0.58 | 0.67 | 0.66 | 0.62 |
| 10 | 123452085 | rs10788197 | 1.45 | -1.403 | -0.499 | 0.447 | -0.009 | -0.04 | -0.04 | 0.535 | 0.45 | 0.09 | 0.41 | 0.41 | 0.58 | 0.55 |
| 10 | 123455602 | rs10749425 | -0.171 | -1.672 | 0.126 | 0.053 | -0.06 | 0.004 | 0.004 | 0.202 | 0.79 | 0.48 | 0.77 | 0.78 | 0.86 | 0.76 |
| 10 | 123456270 | rs4751848 | 1.092 | -1.621 | -0.601 | 0.042 | 0.118 | 0.127 | 0.127 | 3.033 | 0.52 | 0.18 | 0.58 | 0.68 | 0.66 | 0.62 |
| 10 | 123458644 | rs11200083 | -0.23 | -1.449 | -0.5 | 0.064 | 0.036 | 0.009 | 0.009 | 0.152 | 0.22 | 0.1 | 0.19 | 0.19 | 0.16 | 0.27 |
| 10 | 123461066 | rs1219506 | -1.358 | -1.413 | 0.033 | -0.051 | 0.039 | 0.07 | 0.07 | 0.117 | 0.48 | 0.65 | 0.42 | 0.24 | 0.36 | 0.41 |
| 10 | 123461654 | rs1219508 | -0.414 | -1.325 | 0.537 | 0.043 | -0.009 | -0.078 | -0.078 | 0 | 0.25 | 0.31 | 0.19 | 0.05 | 0.2 | 0.13 |
| 10 | 123463320 | rs1693681 | -0.488 | -1.689 | 0.324 | 0.119 | 0.018 | 0.017 | 0.017 | 0.206 | 0.35 | 0.29 | 0.31 | 0.27 | 0.25 | 0.32 |
| 10 | 123463359 | rs17544933 | 0.978 | -1.764 | 0.036 | 0.008 | 0.007 | 0.043 | 0.043 | 0.269 | 0.12 | 0.04 | 0.08 | 0 | 0.07 | 0.02 |
| 10 | 123463733 | rs1693682 | -1.415 | -2.088 | -0.027 | -0.025 | 0.04 | 0.042 | 0.042 | 1.586 | 0.53 | 0.72 | 0.49 | 0.4 | 0.41 | 0.48 |
| 10 | 123465033 | rs10886968 | -0.249 | -1.868 | -1.053 | -0.001 | 0.01 | 0.039 | 0.039 | 2.318 | 0.22 | 0.12 | 0.14 | 0.06 | 0.17 | 0.14 |
| 10 | 123467204 | rs11200087 | 1.335 | -1.723 | -1.561 | 0.172 | -0.097 | -0.08 | -0.08 | 0 | 0.37 | 0.06 | 0.4 | 0.43 | 0.53 | 0.46 |
| 10 | 123468677 | rs4752586 | 1.06 | -1.488 | -1.202 | 0.033 | 0.005 | 0.03 | 0.03 | 0.062 | 0.11 | 0.07 | 0.08 | 0 | 0.07 | 0.02 |
| 10 | 123470963 | rs12358208 | -0.226 | -1.604 | -1.015 | 0.233 | 0.006 | 0.012 | 0.012 | 0 | 0.15 | 0.09 | 0.07 | 0.06 | 0.14 | 0.15 |
| 10 | 123473022 | rs2901295 | 0.481 | -1.439 | -1.061 | 0.547 | -0.008 | -0.034 | -0.034 | 0.31 | 0.26 | 0.02 | 0.22 | 0.27 | 0.22 | 0.37 |
| 10 | 123481200 | rs2935695 | 0.583 | -1.17 | -0.376 | 0.573 | -0.017 | 0.019 | 0.019 | 0 | 0.77 | 0.68 | 0.8 | 0.71 | 0.83 | 0.76 |
| 10 | 123483828 | rs12263297 | 1.09 | -1.023 | -0.784 | 0.179 | -0.084 | 0.146 | 0.146 | 0.045 | 0.34 | 0.11 | 0.23 | 0.11 | 0.42 | 0.18 |
| 10 | 123483879 | rs2420953 | 1.097 | -0.907 | -0.345 | 0.579 | 0.004 | 0.031 | 0.031 | 0 | 0.16 | 0.19 | 0.34 | 0.25 | 0.17 | 0.13 |
| 10 | 123484737 | rs2936862 | 0.08 | -0.961 | -0.411 | 0.572 | 0.015 | 0.021 | 0.021 | 0 | 0.54 | 0.59 | 0.6 | 0.35 | 0.59 | 0.32 |
| 10 | 123486219 | rs2936863 | 0.052 | -0.726 | -0.736 | 0.648 | 0 | -0.02 | -0.02 | 0 | 0.39 | 0.3 | 0.54 | 0.55 | 0.41 | 0.58 |
| 10 | 123486626 | rs2935689 | 0.258 | -0.621 | 0.14 | 0.775 | 0.002 | 0.029 | 0.029 | 0 | 0.17 | 0.26 | 0.35 | 0.24 | 0.17 | 0.14 |
| 10 | 123488067 | rs11200108 | -0.073 | -0.694 | -0.029 | 0.648 | -0.001 | -0.005 | -0.005 | 2.654 | 0.13 | 0.14 | 0.05 | 0.07 | 0.07 | 0.13 |
| 10 | 123490869 | rs2936874 | -1.303 | -0.728 | 0.272 | 0.706 | -0.01 | -0.004 | -0.004 | 0 | 0.38 | 0.53 | 0.46 | 0.38 | 0.33 | 0.33 |
| 10 | 123491643 | rs2936875 | -0.748 | -0.371 | 0.51 | 0.944 | 0.007 | 0.037 | 0.037 | 0 | 0.25 | 0.39 | 0.41 | 0.3 | 0.25 | 0.2 |
| 10 | 123494644 | rs753455 | 0.618 | -0.656 | 0.459 | 1.248 | -0.016 | 0.058 | 0.058 | 0 | 0.79 | 0.59 | 0.61 | 0.66 | 0.83 | 0.79 |
| 10 | 123501732 | rs1874664 | -0.451 | -0.616 | 0.365 | 1.074 | -0.005 | -0.015 | -0.015 | 0 | 0.17 | 0.21 | 0.31 | 0.09 | 0.13 | 0.2 |
| 10 | 123501836 | rs3763755 | -0.832 | -0.965 | -0.31 | 1.398 | 0.008 | -0.012 | -0.012 | 0.086 | 0.21 | 0.21 | 0.15 | 0.5 | 0.12 | 0.16 |
| 10 | 123506701 | rs17628732 | -0.146 | -0.966 | -0.471 | 0.804 | 0.004 | -0.011 | -0.011 | 0 | 0.12 | 0.1 | 0.05 | 0 | 0.07 | 0.02 |
| 10 | 123509687 | rs1696839 | 1.046 | 0.231 | -1.238 | 0.364 | -0.12 | 0.083 | 0.083 | 0 | 0.18 | 0.01 | 0.17 | 0.03 | 0.32 | 0.15 |
| 10 | 123518349 | rs6585758 | 0.201 | -0.031 | -0.415 | 0.497 | 0 | 0 | 0 | 0 | 0.14 | 0.1 | 0.21 | 0.08 | 0.1 | 0.16 |
| 10 | 123521484 | rs2901297 | 0.032 | -0.033 | -0.465 | 0.839 | -0.003 | 0.064 | 0.064 | 0 | 0.84 | 0.75 | 0.75 | 0.69 | 0.87 | 0.82 |
| 10 | 123540904 | rs2420963 | 0.394 | 0.561 | -0.557 | 0.752 | -0.001 | -0.001 | -0.001 | 0 | 0.61 | 0.6 | 0.58 | 0.56 | 0.6 | 0.7 |
| 10 | 123541866 | rs11200146 | 0.697 | 0.561 | -0.58 | 0.764 | -0.002 | -0.006 | -0.006 | 0 | 0.1 | 0.06 | 0.21 | 0.09 | 0.11 | 0.11 |
| 10 | 123543042 | rs7085499 | 1.241 | 0.331 | -0.687 | 0.547 | -0.007 | 0.003 | 0.003 | 0 | 0.44 | 0.48 | 0.48 | 0.53 | 0.42 | 0.58 |
| 10 | 123552966 | rs2132911 | 1.861 | 0.175 | -1.394 | 0.754 | -0.075 | -0.008 | -0.008 | 0.052 | 0.35 | 0.04 | 0.43 | 0.29 | 0.4 | 0.52 |
| 10 | 123555466 | rs4752595 | 0.032 | 0.29 | -1.047 | 0.777 | -0.012 | -0.017 | -0.017 | 0.011 | 0.63 | 0.73 | 0.61 | 0.56 | 0.61 | 0.7 |
| 10 | 123566720 | rs1909009 | 0.6 | 0.386 | -0.739 | 0.493 | -0.004 | 0.049 | 0.049 | 0 | 0.77 | 0.69 | 0.66 | 0.61 | 0.79 | 0.71 |
| 10 | 123569487 | rs9651480 | 1.181 | 0.515 | -1.033 | -0.052 | -0.055 | 0.142 | 0.142 | 0 | 0.44 | 0.14 | 0.45 | 0.2 | 0.49 | 0.5 |
| 10 | 123570191 | rs7913185 | 0.67 | 0.558 | -0.712 | 0.158 | -0.002 | 0.017 | 0.017 | 0 | 0.76 | 0.7 | 0.63 | 0.4 | 0.78 | 0.7 |
| 10 | 123576479 | rs10788212 | 1.139 | 0.811 | -1.03 | 0.015 | -0.044 | 0.053 | 0.053 | 0 | 0.39 | 0.13 | 0.43 | 0.29 | 0.46 | 0.55 |
| 10 | 123578337 | rs11200176 | 0.729 | 0.703 | -0.634 | -0.193 | -0.008 | 0.002 | 0.002 | 0 | 0.63 | 0.66 | 0.55 | 0.41 | 0.61 | 0.62 |
| 10 | 123584192 | rs7476932 | -0.702 | 0.75 | -0.762 | -0.056 | -0.061 | -0.109 | -0.109 | 0 | 0.3 | 0.09 | 0.22 | 0.52 | 0.31 | 0.27 |
| 10 | 123587732 | rs4752606 | -0.891 | 0.847 | -0.448 | 0.07 | 0.021 | 0.019 | 0.019 | 0 | 0.94 | 0.88 | 0.79 | 0.93 | 0.92 | 0.9 |
| 10 | 123591176 | rs12255859 | 1.312 | 0.66 | -0.257 | 0.179 | -0.021 | -0.013 | -0.013 | 0 | 0.13 | 0.04 | 0.14 | 0.18 | 0.17 | 0.08 |
| 10 | 123594941 | rs4752607 | 0.857 | 0.554 | -0.272 | 0.114 | -0.003 | 0.013 | 0.013 | 1.291 | 0.72 | 0.18 | 0.58 | 0.73 | 0.74 | 0.8 |
| 10 | 123596254 | rs4237536 | -1.001 | 0.631 | -0.269 | 0.15 | 0.021 | 0.019 | 0.019 | 0 | 0.94 | 0.88 | 0.79 | 0.93 | 0.92 | 0.9 |

| | | | | | | | | | | | | | | | | |
|---|---|---|---|---|---|---|---|---|---|---|---|---|---|---|---|---|
| 10 | 123600590 | rs10886994 | 1.384 | 0.644 | -0.281 | 0.148 | -0.022 | -0.013 | -0.013 | 0 | 0.13 | 0.04 | 0.14 | 0.18 | 0.17 | 0.08 |
| 10 | 123606502 | rs6585765 | -0.719 | 0.628 | -0.101 | 0.134 | 0.041 | 0.038 | 0.038 | 0 | 0.93 | 0.8 | 0.78 | 0.93 | 0.92 | 0.9 |
| 10 | 123615028 | rs7092052 | -1 | 0.638 | -0.082 | 0.139 | 0.011 | 0.01 | 0.01 | 0 | 0.94 | 0.91 | 0.79 | 0.93 | 0.92 | 0.92 |
| 10 | 123624305 | rs11200201 | 1.311 | 0.632 | -0.115 | 0.126 | -0.009 | -0.003 | -0.003 | 0 | 0.13 | 0.07 | 0.14 | 0.18 | 0.17 | 0.08 |
| 10 | 123624510 | rs10510103 | 1.613 | 0.585 | -0.333 | -0.085 | 0.01 | 0.019 | 0.019 | 0 | 0.34 | 0.2 | 0.38 | 0.31 | 0.32 | 0.44 |
| 10 | 123625190 | rs10510102 | 1.092 | 0.587 | -0.546 | -0.088 | 0.001 | -0.002 | -0.002 | 0 | 0.15 | 0.16 | 0.15 | 0.18 | 0.18 | 0.09 |
| 10 | 123627512 | rs4752614 | 0.02 | 0.585 | -0.988 | -0.081 | -0.117 | -0.146 | -0.146 | 0 | 0.4 | 0.06 | 0.21 | 0.4 | 0.42 | 0.38 |
| 10 | 123633118 | rs7917343 | -1.105 | 0.742 | -0.891 | -0.028 | 0.019 | 0.017 | 0.017 | 0 | 0.94 | 0.89 | 0.79 | 0.93 | 0.92 | 0.92 |
| 10 | 123635883 | rs11200206 | 1.497 | 0.704 | -1.035 | -0.046 | 0.011 | 0.027 | 0.027 | 0 | 0.34 | 0.18 | 0.37 | 0.3 | 0.32 | 0.44 |
| 10 | 123642539 | rs7081719 | 0.786 | 0.693 | -1.045 | -0.055 | -0.007 | 0.022 | 0.022 | 0 | 0.46 | 0.26 | 0.51 | 0.48 | 0.49 | 0.53 |
| 10 | 123643822 | rs11200215 | 1.581 | 0.687 | -1.034 | -0.052 | 0.008 | 0.021 | 0.021 | 0 | 0.33 | 0.22 | 0.38 | 0.3 | 0.32 | 0.44 |
| 10 | 123661722 | rs2420970 | 0.72 | 0.732 | -1.235 | -0.047 | -0.008 | 0.027 | 0.027 | 0 | 0.47 | 0.2 | 0.5 | 0.48 | 0.48 | 0.53 |
| 10 | 123669255 | rs12253283 | 1.595 | 0.769 | -1.331 | -0.053 | -0.021 | -0.007 | -0.007 | 0 | 0.13 | 0.04 | 0.14 | 0.18 | 0.17 | 0.08 |
| 10 | 123673339 | rs10749435 | -1.037 | 0.785 | -1.227 | -0.039 | 0.014 | 0.013 | 0.013 | 0 | 0.94 | 0.9 | 0.78 | 0.93 | 0.92 | 0.92 |
| 10 | 123681962 | rs10887023 | 1.495 | 0.701 | -1.331 | -0.071 | -0.005 | 0.014 | 0.014 | 0 | 0.33 | 0.09 | 0.36 | 0.3 | 0.32 | 0.45 |
| 10 | 123685227 | rs7914615 | -0.631 | 0.579 | -1.244 | -0.199 | 0.017 | 0.099 | 0.099 | 0.329 | 0.48 | 0.69 | 0.29 | 0.14 | 0.45 | 0.21 |
| 10 | 123702736 | rs11594070 | 0.686 | 0.707 | -1.305 | 0.044 | -0.117 | 0.08 | 0.08 | 0 | 0.28 | 0.02 | 0.19 | 0.16 | 0.37 | 0.18 |
| 10 | 123715402 | rs10887032 | 1.34 | 1.054 | -1.054 | 0.284 | -0.015 | -0.004 | -0.004 | 0.218 | 0.13 | 0.04 | 0.14 | 0.15 | 0.17 | 0.08 |
| 10 | 123715870 | rs7095093 | 0.239 | 1.065 | -0.975 | 0.338 | 0.024 | 0.022 | 0.022 | 0 | 0.07 | 0.15 | 0.23 | 0.08 | 0.09 | 0.09 |
| 10 | 123722906 | rs12240306 | 1.202 | 0.794 | -1.362 | 0.437 | -0.016 | -0.004 | -0.004 | 0 | 0.12 | 0.04 | 0.14 | 0.15 | 0.17 | 0.08 |
| 10 | 123724389 | rs10788226 | 0.943 | 0.492 | -0.964 | 0.492 | 0.042 | 0.048 | 0.048 | 0 | 0.39 | 0.2 | 0.35 | 0.29 | 0.28 | 0.43 |
| 10 | 123728523 | rs17102809 | -0.666 | 1.02 | -0.604 | 1.215 | 0.002 | 0.096 | 0.096 | 0.228 | 0.06 | 0.06 | 0.05 | 0.32 | 0.08 | 0.2 |
| 10 | 123740377 | rs11200306 | 0.527 | 0.902 | -0.603 | 0.807 | -0.002 | -0.002 | -0.002 | 0 | 0.13 | 0.08 | 0.15 | 0.15 | 0.15 | 0.09 |
| 10 | 123744366 | rs11200308 | 0.547 | 0.904 | -0.614 | 0.808 | -0.002 | -0.003 | -0.003 | 0 | 0.13 | 0.07 | 0.15 | 0.15 | 0.15 | 0.09 |
| 10 | 123744571 | rs7913538 | -1.076 | 0.499 | -0.555 | 0.733 | 0.038 | 0.023 | 0.023 | 0.154 | 0.6 | 0.74 | 0.77 | 0.71 | 0.74 | 0.56 |
| 10 | 123747807 | rs10159856 | 0.655 | 0.666 | -0.561 | 0.356 | -0.001 | -0.001 | -0.001 | 0 | 0.14 | 0.07 | 0.15 | 0.15 | 0.16 | 0.09 |
| 10 | 123749116 | rs12257711 | 0.429 | 0.652 | -0.181 | 0.338 | -0.001 | -0.001 | -0.001 | 4.035 | 0.16 | 0.16 | 0.27 | 0.14 | 0.18 | 0.09 |
| 10 | 123751997 | rs6585775 | -0.926 | 0.669 | -0.325 | 0.319 | 0.024 | 0.028 | 0.028 | 0 | 0.53 | 0.64 | 0.7 | 0.42 | 0.64 | 0.39 |
| 10 | 123754074 | rs10160002 | 0.556 | 0.968 | -0.734 | 0.394 | -0.016 | 0 | 0 | 0 | 0.14 | 0.04 | 0.26 | 0.14 | 0.18 | 0.09 |
| 10 | 123766854 | rs12762668 | 0.075 | 1.177 | -0.201 | 0.13 | 0.019 | -0.048 | -0.048 | 3.777 | 0.27 | 0.34 | 0.2 | 0.14 | 0.37 | 0.15 |
| 10 | 123767296 | rs10887040 | 0.579 | 0.758 | -0.501 | 0.305 | 0.044 | 0.082 | 0.082 | 0 | 0.47 | 0.24 | 0.31 | 0.6 | 0.36 | 0.62 |
| 10 | 123770094 | rs6585776 | 0.82 | 0.901 | -0.644 | 0.513 | 0.003 | -0.028 | -0.028 | 0.32 | 0.42 | 0.23 | 0.32 | 0.47 | 0.39 | 0.6 |
| 10 | 123772907 | rs3750839 | 0.892 | 1.339 | -0.611 | 0.364 | -0.012 | 0.042 | 0.042 | 0 | 0.19 | 0.09 | 0.11 | 0.14 | 0.22 | 0.14 |
| 10 | 123773023 | rs3750842 | 0.467 | 1.812 | -0.429 | 0.946 | -0.025 | -0.025 | -0.025 | 0.409 | 0.09 | 0.03 | 0.09 | 0.19 | 0.14 | 0.15 |
| 10 | 123780283 | rs17631911 | 0.907 | 1.624 | -0.459 | 0.96 | -0.034 | -0.043 | -0.043 | 0.002 | 0.07 | 0.01 | 0.08 | 0.14 | 0.12 | 0.06 |
| 10 | 123780378 | rs17102867 | 0.326 | 1.33 | -0.58 | 1.091 | -0.006 | 0.023 | 0.023 | 0 | 0.09 | 0.12 | 0.08 | 0.14 | 0.05 | 0.17 |
| 10 | 123780784 | rs10887046 | -0.702 | 1.206 | -0.155 | 0.965 | -0.014 | -0.005 | -0.005 | 0 | 0.17 | 0.24 | 0.1 | 0.14 | 0.09 | 0.2 |
| 10 | 123781339 | rs2461221 | -1.255 | 1.22 | -0.113 | 0.837 | 0.002 | 0.059 | 0.059 | 0 | 0.48 | 0.65 | 0.73 | 0.68 | 0.53 | 0.52 |
| 10 | 123782114 | rs17102880 | 0.198 | 1.377 | 0.465 | 1.009 | 0.019 | 0.064 | 0.064 | 0 | 0.1 | 0.19 | 0.14 | 0.27 | 0.14 | 0.16 |
| 10 | 123782894 | rs4752635 | 0.561 | 1.022 | 0.378 | 0.907 | -0.004 | -0.037 | -0.037 | 0 | 0.07 | 0.12 | 0.03 | 0 | 0.04 | 0.03 |
| 10 | 123790901 | rs17102882 | -0.155 | 1.029 | 0.491 | 1 | -0.012 | 0.073 | 0.073 | 0 | 0.14 | 0.21 | 0.13 | 0.28 | 0.08 | 0.22 |
| 10 | 123793394 | rs5013487 | 0.442 | 1.173 | 0.522 | 0.901 | 0.018 | 0.027 | 0.027 | 0 | 0.15 | 0.21 | 0.17 | 0.26 | 0.22 | 0.18 |
| 10 | 123793875 | rs2461211 | -1.034 | 1.066 | 1.291 | 0.724 | -0.019 | -0.086 | -0.086 | 0.045 | 0.16 | 0.35 | 0.21 | 0.02 | 0.09 | 0.05 |
| 10 | 123794115 | rs7898269 | -0.086 | 1.064 | 1.181 | 0.724 | 0 | -0.046 | -0.046 | 0 | 0.08 | 0.23 | 0.04 | 0 | 0.04 | 0.03 |
| 10 | 123794568 | rs10887054 | 1.543 | 0.92 | 0.307 | 0.692 | 0.017 | 0.005 | 0.005 | 0 | 0.44 | 0.23 | 0.32 | 0.39 | 0.45 | 0.47 |
| 10 | 123795016 | rs17102893 | 0.876 | 0.969 | 0.263 | 0.679 | -0.002 | 0.031 | 0.031 | 0 | 0.07 | 0.04 | 0.16 | 0.02 | 0.04 | 0.03 |
| 10 | 123795293 | rs2459068 | -1.265 | 1.129 | 0.338 | 0.75 | -0.011 | -0.003 | -0.003 | 0 | 0.86 | 0.87 | 0.91 | 0.82 | 0.79 | 0.83 |
| 10 | 123795655 | rs10788231 | -1.265 | 1.129 | 0.338 | 0.75 | -0.011 | -0.003 | -0.003 | 0 | 0.86 | 0.87 | 0.91 | 0.82 | 0.79 | 0.83 |
| 10 | 123796743 | rs2461204 | -1.139 | 1.035 | 0.423 | 0.734 | 0.011 | -0.003 | -0.003 | 0.988 | 0.42 | 0.63 | 0.59 | 0.43 | 0.35 | 0.37 |
| 10 | 123799687 | rs10887055 | 1.375 | 1.066 | 0.195 | 0.695 | 0.02 | 0.001 | 0.001 | 0 | 0.45 | 0.24 | 0.31 | 0.4 | 0.45 | 0.47 |
| 10 | 123800550 | rs10887056 | -0.577 | 0.34 | 0.335 | 0.786 | 0 | 0.013 | 0.013 | 0 | 0.28 | 0.31 | 0.41 | 0.41 | 0.28 | 0.32 |
| 10 | 123801422 | rs2253762 | -0.832 | 0.393 | 0.248 | 0.569 | -0.006 | -0.062 | -0.062 | 0 | 0.3 | 0.39 | 0.49 | 0.1 | 0.27 | 0.18 |
| 10 | 123802646 | rs2459075 | -0.587 | 0.351 | 0.474 | 1.056 | 0 | 0.001 | 0.001 | 0 | 0.86 | 0.86 | 0.89 | 0.76 | 0.79 | 0.83 |
| 10 | 123803584 | rs10788232 | 1.584 | 0.205 | 0.254 | 0.921 | 0.006 | -0.043 | -0.043 | 0.054 | 0.6 | 0.24 | 0.6 | 0.5 | 0.64 | 0.55 |
| 10 | 123804553 | rs2943764 | -1.598 | -0.037 | 0.424 | 0.534 | -0.029 | -0.026 | -0.026 | 0 | 0.2 | 0.44 | 0.24 | 0.14 | 0.1 | 0.22 |
| 10 | 123807240 | rs3981027 | -0.927 | 0.279 | 0.149 | 0.657 | -0.005 | -0.005 | -0.005 | 0.479 | 0.15 | 0.29 | 0.22 | 0.14 | 0.1 | 0.22 |
| 10 | 123810060 | rs11200367 | -0.72 | 0.293 | 0.087 | 0.782 | 0 | 0.007 | 0.007 | 0 | 0.1 | 0.2 | 0.26 | 0.13 | 0.08 | 0.11 |
| 10 | 123815371 | rs10788233 | -0.524 | -1.314 | -0.133 | -0.237 | -0.046 | -0.105 | -0.105 | 0 | 0.71 | 0.66 | 0.87 | 0.95 | 0.83 | 0.8 |
| 10 | 123815394 | rs11591752 | 0.176 | -0.699 | -0.578 | -0.017 | -0.032 | -0.021 | -0.021 | 0 | 0.12 | 0.08 | 0.13 | 0.15 | 0.18 | 0.11 |
| 10 | 123817654 | rs7894692 | 0.478 | -1.454 | -0.568 | 0.273 | -0.001 | 0.008 | 0.008 | 0 | 0.17 | 0.17 | 0.27 | 0.32 | 0.17 | 0.16 |
| 10 | 123818827 | rs2459093 | 0.112 | -1.285 | -0.644 | 0.349 | 0.066 | 0.043 | 0.043 | 0.146 | 0.52 | 0.37 | 0.58 | 0.63 | 0.65 | 0.63 |
| 10 | 123828066 | rs2461230 | 0.186 | -0.79 | -0.016 | 0.312 | 0.017 | -0.011 | -0.011 | 0 | 0.93 | 0.71 | 0.96 | 1 | 0.98 | 0.95 |
| 10 | 123833081 | rs2461227 | 1.026 | -0.836 | -1.319 | 0.416 | -0.075 | 0.07 | 0.07 | 0 | 0.45 | 0.09 | 0.37 | 0.34 | 0.51 | 0.36 |
| 10 | 123835093 | rs11200381 | 0.25 | -0.93 | -0.514 | 0.73 | 0.029 | 0.028 | 0.028 | 0 | 0.1 | 0.2 | 0.16 | 0.12 | 0.18 | 0.1 |
| 10 | 123838026 | rs2461225 | 0.494 | -1.978 | -0.265 | 0.68 | -0.032 | 0.003 | 0.003 | 0 | 0.72 | 0.56 | 0.68 | 0.7 | 0.8 | 0.75 |
| 10 | 123839549 | rs2461224 | 1.809 | -2.134 | -0.52 | 0.473 | -0.043 | -0.042 | -0.042 | 0.098 | 0.38 | 0.2 | 0.44 | 0.48 | 0.51 | 0.43 |
| 10 | 123842523 | rs11200385 | -0.508 | -1.728 | 0.162 | 0.744 | 0.014 | -0.024 | -0.024 | 0 | 0.18 | 0.31 | 0.19 | 0.1 | 0.21 | 0.18 |
| 10 | 123844503 | rs10887063 | 0.007 | -1.709 | 0.098 | 0.744 | 0.034 | -0.001 | -0.001 | 0.048 | 0.13 | 0.29 | 0.17 | 0.1 | 0.18 | 0.14 |
| 10 | 123846288 | rs4752642 | -0.702 | -1.749 | 0.059 | 0.725 | 0.011 | -0.022 | -0.022 | 0 | 0.18 | 0.31 | 0.18 | 0.1 | 0.21 | 0.18 |

| | | | | | | | | | | | | | | | | |
|---|---|---|---|---|---|---|---|---|---|---|---|---|---|---|---|---|
| 10 | 123846485 | rs4751871 | -0.702 | -1.749 | 0.059 | 0.736 | 0.011 | -0.02 | -0.02 | 0 | 0.18 | 0.31 | 0.18 | 0.1 | 0.21 | 0.18 |
| 10 | 123846490 | rs7920896 | 0.511 | -1.325 | -0.43 | 0.729 | -0.006 | -0.002 | -0.002 | 0 | 0.06 | 0.11 | 0.14 | 0.05 | 0.04 | 0.07 |
| 10 | 123849465 | rs12242677 | 0.108 | -1.202 | -0.762 | 0.706 | 0.003 | 0.003 | 0.003 | 0 | 0.08 | 0.05 | 0.02 | 0.04 | 0.04 | 0.15 |
| 10 | 123849605 | rs12258830 | -0.499 | -1.215 | -0.815 | 0.854 | 0.024 | -0.048 | -0.048 | 0 | 0.17 | 0.06 | 0.04 | 0.32 | 0.13 | 0.21 |
| 10 | 123851369 | rs10788238 | 0.545 | -0.428 | -0.295 | 0.87 | 0.032 | -0.018 | -0.018 | 0 | 0.47 | 0.2 | 0.34 | 0.53 | 0.61 | 0.34 |
| 10 | 123853004 | rs2420995 | 0.441 | -0.243 | -0.971 | 0.888 | 0.042 | -0.013 | -0.013 | 0 | 0.48 | 0.27 | 0.35 | 0.54 | 0.62 | 0.34 |
| 10 | 123855878 | rs11200397 | -0.796 | -0.682 | -0.606 | 0.713 | 0.005 | 0.003 | 0.003 | 0.068 | 0.46 | 0.52 | 0.42 | 0.66 | 0.28 | 0.55 |
| 10 | 123857433 | rs2420992 | 0.171 | 0.1 | -0.524 | 0.738 | -0.001 | -0.007 | -0.007 | 1.107 | 0.6 | 0.59 | 0.53 | 0.7 | 0.57 | 0.57 |
| 10 | 123858724 | rs884147 | -0.482 | 0.014 | 0.257 | 0.799 | 0.002 | -0.012 | -0.012 | 0 | 0.38 | 0.32 | 0.47 | 0.33 | 0.45 | 0.43 |
| 10 | 123862706 | rs11595073 | -0.235 | 0.305 | -0.001 | 1.141 | -0.007 | -0.006 | -0.006 | 0 | 0.2 | 0.2 | 0.28 | 0.47 | 0.23 | 0.38 |
| 10 | 123862952 | rs2420994 | -0.888 | 0.158 | 0.381 | 1.094 | 0.002 | 0.001 | 0.001 | 0.543 | 0.56 | 0.42 | 0.73 | 0.79 | 0.67 | 0.81 |
| 10 | 123864291 | rs12764791 | -0.101 | 0.176 | 0.432 | 1.529 | -0.004 | -0.022 | -0.022 | 0 | 0.11 | 0.08 | 0.05 | 0.26 | 0.12 | 0.18 |
| 10 | 123864745 | rs7076095 | 1.204 | 0.199 | 0.607 | 1.726 | -0.015 | -0.005 | -0.005 | 0 | 0.39 | 0.25 | 0.38 | 0.37 | 0.39 | 0.25 |
| 10 | 123869825 | rs4462273 | 1.124 | 0.4 | 0.815 | 1.628 | 0.002 | -0.067 | -0.067 | 0 | 0.87 | 0.58 | 0.9 | 0.98 | 0.9 | 0.96 |
| 10 | 123875554 | rs11200406 | 0.03 | 0.457 | 0.768 | 1.709 | 0.001 | 0.011 | 0.011 | 0.068 | 0.1 | 0.11 | 0.05 | 0.24 | 0.12 | 0.18 |
| 10 | 123878029 | rs7097450 | 0.443 | 0.145 | 0.87 | 1.563 | -0.051 | -0.083 | -0.083 | 0 | 0.79 | 0.58 | 0.81 | 0.96 | 0.89 | 0.91 |
| 10 | 123878679 | rs2421005 | 1.623 | 0.231 | 0.479 | 1.692 | 0.009 | -0.057 | -0.057 | 0 | 0.62 | 0.2 | 0.7 | 0.54 | 0.69 | 0.66 |
| 10 | 123880823 | rs4751873 | 1.008 | 0.2 | 0.142 | 1.694 | 0.015 | 0.006 | 0.006 | 0.147 | 0.72 | 0.24 | 0.74 | 0.78 | 0.82 | 0.84 |
| 10 | 123882245 | rs11200413 | 0.057 | 0.286 | -0.024 | 1.714 | -0.006 | -0.017 | -0.017 | 0 | 0.1 | 0.07 | 0.05 | 0.24 | 0.12 | 0.18 |
| 10 | 123882956 | rs10887082 | 0.621 | -0.029 | -0.024 | 1.609 | -0.084 | -0.056 | -0.056 | 1.552 | 0.26 | 0.04 | 0.35 | 0.33 | 0.36 | 0.46 |
| 10 | 123885789 | rs2421001 | 1.865 | -0.452 | 1.14 | 1.723 | 0.03 | -0.017 | -0.017 | 0 | 0.66 | 0.12 | 0.72 | 0.64 | 0.77 | 0.74 |
| 10 | 123887925 | rs17103093 | -1.529 | -0.34 | 1.275 | 1.712 | 0.113 | 0.17 | 0.17 | 0 | 0.07 | 0.23 | 0.07 | 0.18 | 0.09 | 0.06 |
| 10 | 123892874 | rs7908768 | -1.11 | -0.359 | 0.327 | 1.552 | -0.015 | 0.026 | 0.026 | 0 | 0.22 | 0.59 | 0.19 | 0.35 | 0.13 | 0.23 |
| 10 | 123893386 | rs4237537 | 0.935 | -0.113 | -0.465 | 1.574 | 0.019 | -0.029 | -0.029 | 0 | 0.66 | 0.11 | 0.7 | 0.62 | 0.76 | 0.72 |
| 10 | 123894412 | rs2420998 | 1.079 | 0.017 | 0.633 | 1.729 | -0.049 | 0.294 | 0.294 | 0 | 0.88 | 0.68 | 0.78 | 0.54 | 0.93 | 0.82 |
| 10 | 123894522 | rs2420999 | -0.68 | 0.937 | -0.041 | 0.537 | 0.006 | 0.001 | 0.001 | 0 | 0.3 | 0.47 | 0.35 | 0.32 | 0.31 | 0.28 |
| 10 | 123901964 | rs10788251 | -1.288 | 0.931 | 0.259 | -0.006 | 0.009 | -0.134 | -0.134 | 0 | 0.19 | 0.46 | 0.12 | 0.02 | 0.14 | 0.06 |
| 10 | 123902771 | rs10788252 | -1.644 | 1.194 | 0.394 | 0.738 | 0.004 | 0.02 | 0.02 | 0 | 0.22 | 0.28 | 0.34 | 0.39 | 0.23 | 0.19 |
| 10 | 123903133 | rs12765679 | -0.114 | 2.482 | -0.002 | 0.143 | -0.018 | 0.007 | 0.007 | 0 | 0.11 | 0.07 | 0.11 | 0 | 0.23 | 0.05 |
| 10 | 123903694 | rs7917144 | 1.186 | 1.867 | 0.764 | 1.224 | -0.04 | 0.158 | 0.158 | 0 | 0.84 | 0.6 | 0.86 | 0.54 | 0.91 | 0.82 |
| 10 | 123904213 | rs7918107 | -0.602 | 1.868 | 0.685 | 1.269 | -0.015 | -0.014 | -0.014 | 0.256 | 0.11 | 0.25 | 0.06 | 0.04 | 0.06 | 0.02 |
| 10 | 123908325 | rs7097751 | -0.963 | 1.315 | 0.197 | 0.254 | 0 | 0.003 | 0.003 | 1.024 | 0.51 | 0.5 | 0.55 | 0.4 | 0.47 | 0.39 |
| 10 | 123911345 | rs7078373 | -2.076 | 0.717 | 0.982 | 0.503 | -0.035 | 0.003 | 0.003 | 0 | 0.28 | 0.41 | 0.18 | 0.33 | 0.19 | 0.15 |
| 10 | 123911886 | rs4752659 | 0.803 | 0.534 | 0.884 | 0.64 | 0.003 | 0.059 | 0.059 | 0 | 0.89 | 0.8 | 0.74 | 0.7 | 0.89 | 0.79 |
| 10 | 123912338 | rs4752662 | 1.806 | 0.54 | 0.98 | 0.686 | -0.031 | 0.07 | 0.07 | 0.305 | 0.7 | 0.5 | 0.62 | 0.41 | 0.78 | 0.71 |
| 10 | 123912848 | rs10887092 | -0.937 | 0.707 | 0.426 | 0.729 | -0.001 | 0.004 | 0.004 | 0 | 0.05 | 0.03 | 0.06 | 0.21 | 0.04 | 0.04 |
| 10 | 123913330 | rs17103138 | -1.206 | 0.398 | 0.42 | 0.7 | -0.034 | -0.003 | -0.003 | 0 | 0.13 | 0.23 | 0.08 | 0.09 | 0.07 | 0.07 |
| 10 | 123913826 | rs7100333 | 1.055 | 0.79 | -0.206 | 0.873 | -0.004 | 0.069 | 0.069 | 0 | 0.82 | 0.72 | 0.7 | 0.51 | 0.85 | 0.76 |
| 10 | 123914242 | rs10788253 | 0.302 | 0.73 | -0.221 | 0.71 | 0.002 | -0.008 | -0.008 | 1.207 | 0.89 | 0.91 | 0.79 | 0.79 | 0.91 | 0.81 |
| 10 | 123918935 | rs12241688 | -0.768 | 0.78 | 0.04 | 0.592 | 0.002 | 0.02 | 0.02 | 0 | 0.07 | 0.19 | 0.04 | 0.07 | 0.04 | 0.01 |
| 10 | 123926310 | rs10887096 | -0.504 | 0.878 | -0.435 | 0.999 | -0.001 | -0.009 | -0.009 | 0 | 0.11 | 0.07 | 0.15 | 0.29 | 0.11 | 0.09 |
| 10 | 123933923 | rs10788257 | -0.91 | 0.701 | -0.814 | 1.025 | -0.007 | -0.01 | -0.01 | 0 | 0.39 | 0.64 | 0.35 | 0.55 | 0.38 | 0.23 |
| 10 | 123937461 | rs7078370 | -0.846 | 0.557 | -1.286 | 1.011 | -0.007 | -0.019 | -0.019 | 0 | 0.31 | 0.73 | 0.24 | 0.37 | 0.31 | 0.22 |
| 10 | 123944489 | rs4752665 | 0.452 | 0.973 | -1.387 | 1.106 | -0.008 | -0.011 | -0.011 | 0.502 | 0.89 | 0.94 | 0.95 | 0.87 | 0.88 | 0.92 |
| 10 | 123948200 | rs4752666 | 0.404 | 0.89 | -2.147 | 0.167 | -0.036 | 0.086 | 0.086 | 0 | 0.63 | 0.84 | 0.78 | 0.83 | 0.57 | 0.66 |
| 10 | 123950002 | rs4751878 | 1.05 | 0.631 | -1.99 | 0.817 | -0.026 | 0.098 | 0.098 | 0 | 0.47 | 0.18 | 0.36 | 0.3 | 0.48 | 0.45 |
| 10 | 123965678 | rs10887104 | -0.432 | 0.321 | -0.587 | 1.55 | -0.021 | 0.137 | 0.137 | 0 | 0.15 | 0.42 | 0.32 | 0.3 | 0.09 | 0.16 |
| 10 | 123967911 | rs11200480 | 0.817 | 0.258 | -0.854 | 1.526 | -0.023 | -0.036 | -0.036 | 0 | 0.08 | 0.01 | 0.26 | 0.29 | 0.08 | 0.15 |
| 10 | 123970530 | rs2295873 | -1.008 | -0.054 | -0.808 | 1.134 | -0.024 | 0.017 | 0.017 | 0.697 | 0.21 | 0.71 | 0.28 | 0.35 | 0.09 | 0.19 |
| 10 | 123970569 | rs2295874 | 0.573 | 0.243 | -1.057 | 1.176 | 0.005 | -0.008 | -0.008 | 0 | 0.12 | 0.06 | 0.14 | 0.12 | 0.08 | 0.15 |
| 10 | 123973846 | rs7916162 | 0.442 | 0.21 | -0.975 | 1.181 | 0.004 | -0.001 | -0.001 | 3.689 | 0.12 | 0.08 | 0.14 | 0.11 | 0.08 | 0.15 |
| 10 | 123976029 | rs2281670 | -0.266 | 0.194 | -1.056 | 1.359 | -0.001 | -0.034 | -0.034 | 0 | 0.13 | 0.07 | 0.3 | 0.41 | 0.12 | 0.24 |
| 10 | 123980108 | rs6585804 | -0.273 | 0.484 | -0.843 | 0.849 | -0.003 | 0.003 | 0.003 | 0 | 0.59 | 0.35 | 0.69 | 0.86 | 0.51 | 0.7 |
| 10 | 123980945 | rs11592039 | 0.421 | 0.372 | -1.14 | 0.64 | 0.031 | 0.112 | 0.112 | 0.021 | 0.49 | 0.22 | 0.36 | 0.24 | 0.41 | 0.45 |
| 10 | 123981619 | rs1010897 | 0.368 | 0.362 | -1.205 | 0.66 | 0.032 | 0.019 | 0.019 | 0.016 | 0.49 | 0.24 | 0.39 | 0.47 | 0.41 | 0.48 |
| 10 | 123983959 | rs12416376 | 0.64 | 0.329 | -1.247 | 0.668 | 0.038 | 0 | 0 | 0.108 | 0.46 | 0.18 | 0.37 | 0.47 | 0.39 | 0.45 |
| 10 | 123991774 | rs10788262 | -0.763 | 0.6 | -1.578 | 0.566 | -0.069 | -0.089 | -0.089 | 0.057 | 0.23 | 0.08 | 0.43 | 0.54 | 0.32 | 0.36 |
| 10 | 123996503 | rs4752673 | -1.312 | 0.321 | -1.644 | 0.319 | -0.016 | 0.056 | 0.056 | 0.179 | 0.77 | 0.89 | 0.85 | 1 | 0.71 | 0.82 |
| 10 | 123996976 | rs2295879 | 1.496 | 0.427 | -1.787 | 0.4 | 0.014 | 0.056 | 0.056 | 0.649 | 0.39 | 0.13 | 0.29 | 0.24 | 0.34 | 0.41 |
| 10 | 123997717 | rs1010657 | 0.512 | 0.525 | -1.729 | 0.54 | 0.02 | 0.041 | 0.041 | 0.141 | 0.5 | 0.14 | 0.42 | 0.35 | 0.42 | 0.55 |
| 10 | 123999005 | rs1535460 | 1.115 | 0.107 | -2.264 | 0.204 | 0.021 | -0.053 | -0.053 | 0.252 | 0.37 | 0.12 | 0.3 | 0.41 | 0.35 | 0.45 |
| 10 | 123999656 | rs12777747 | 0.785 | -0.037 | -2.576 | -0.004 | -0.007 | 0.046 | 0.046 | 0 | 0.21 | 0.02 | 0.14 | 0.07 | 0.21 | 0.26 |
| 10 | 124001205 | rs746070 | 0.621 | 0.248 | -2.175 | 0.051 | 0.015 | -0.022 | -0.022 | 0 | 0.39 | 0.2 | 0.33 | 0.4 | 0.4 | 0.46 |
| 10 | 124007573 | rs10887110 | 0.416 | 0.342 | -2.58 | 0.064 | -0.139 | -0.106 | -0.106 | 0 | 0.31 | 0.08 | 0.45 | 0.39 | 0.37 | 0.38 |
| 10 | 124008927 | rs2273794 | 0.179 | -0.099 | -1.925 | 0.472 | 0.012 | -0.043 | -0.043 | 0 | 0.39 | 0.14 | 0.33 | 0.4 | 0.39 | 0.46 |
| 10 | 124012858 | rs911784 | -0.589 | -0.1 | -1.165 | 0.497 | 0.035 | -0.007 | -0.007 | 0.543 | 0.47 | 0.63 | 0.55 | 0.49 | 0.41 | 0.49 |
| 10 | 124013604 | rs3763763 | 0.102 | -0.417 | -1.237 | 0.126 | 0.01 | -0.001 | -0.001 | 0 | 0.28 | 0.22 | 0.26 | 0.25 | 0.26 | 0.33 |
| 10 | 124014939 | rs12262185 | -0.621 | -0.233 | -1.529 | 0.192 | 0.083 | 0.091 | 0.091 | 0 | 0.46 | 0.8 | 0.36 | 0.25 | 0.36 | 0.44 |
| 10 | 124015462 | rs4751880 | -0.62 | -0.095 | -0.948 | 0.282 | -0.012 | -0.001 | -0.001 | 0.659 | 0.39 | 0.52 | 0.53 | 0.6 | 0.34 | 0.43 |

| | | | | | | | | | | | | | | | | |
|---|---|---|---|---|---|---|---|---|---|---|---|---|---|---|---|---|
| 10 | 124015709 | rs10887112 | -0.423 | 0.087 | -1.39 | -0.417 | 0.036 | 0.103 | 0.103 | 0 | 0.49 | 0.62 | 0.36 | 0.18 | 0.4 | 0.38 |
| 10 | 124015986 | rs10788264 | -0.307 | 0.448 | -1.538 | -0.823 | 0.019 | 0.186 | 0.186 | 0 | 0.49 | 0.3 | 0.61 | 0.93 | 0.48 | 0.64 |
| 10 | 124017404 | rs10887113 | -0.601 | 0.325 | -0.858 | -0.539 | -0.005 | 0.025 | 0.025 | 0 | 0.55 | 0.61 | 0.45 | 0.72 | 0.52 | 0.63 |
| 10 | 124017423 | rs9630083 | -0.796 | 0.145 | -0.76 | -0.659 | -0.026 | 0.154 | 0.154 | 0 | 0.58 | 0.83 | 0.69 | 0.93 | 0.55 | 0.67 |
| 10 | 124021200 | rs7081032 | 0.026 | -0.184 | -0.558 | -0.377 | -0.011 | 0.025 | 0.025 | 0.553 | 0.35 | 0.56 | 0.24 | 0.44 | 0.32 | 0.35 |
| 10 | 124022694 | rs7098767 | 1.02 | -0.229 | -0.65 | -0.404 | -0.034 | -0.148 | -0.148 | 0.19 | 0.25 | 0.04 | 0.16 | 0.44 | 0.27 | 0.33 |
| 10 | 124023434 | rs10736311 | -1.094 | -0.171 | -0.614 | -0.507 | -0.03 | 0.086 | 0.086 | 0 | 0.57 | 0.8 | 0.47 | 0.72 | 0.51 | 0.63 |
| 10 | 124027433 | rs3901289 | -0.567 | -0.259 | -0.448 | -0.54 | 0.002 | 0.006 | 0.006 | 0 | 0.51 | 0.5 | 0.43 | 0.72 | 0.47 | 0.61 |
| 10 | 124029436 | rs1885512 | -0.769 | -0.211 | -0.36 | -0.44 | 0.002 | -0.014 | -0.014 | 0.067 | 0.32 | 0.75 | 0.51 | 0.48 | 0.28 | 0.34 |
| 10 | 124034528 | rs7096744 | -0.256 | -0.27 | -0.05 | -0.246 | 0.001 | 0.001 | 0.001 | 0.328 | 0.36 | 0.54 | 0.47 | 0.62 | 0.37 | 0.36 |
| 10 | 124035694 | rs7893461 | 0.526 | -0.1 | 0.058 | -0.221 | 0.05 | -0.024 | -0.024 | 0 | 0.07 | 0.32 | 0.06 | 0 | 0.08 | 0.02 |
| 10 | 124036619 | rs10788271 | 0.427 | -0.18 | 0.01 | -0.267 | 0.007 | -0.022 | -0.022 | 0 | 0.28 | 0.26 | 0.19 | 0.41 | 0.25 | 0.27 |
| 10 | 124039455 | rs7086791 | 0.593 | -0.427 | -0.163 | -0.406 | -0.002 | 0.006 | 0.006 | 0 | 0.22 | 0.24 | 0.23 | 0.26 | 0.19 | 0.28 |
| 10 | 124041103 | rs7076232 | 1.043 | -0.41 | -0.209 | -0.422 | 0.005 | -0.031 | -0.031 | 0 | 0.06 | 0.16 | 0.05 | 0 | 0.07 | 0.02 |
| 10 | 124041447 | rs6585810 | -1.348 | -0.44 | -0.277 | -0.446 | -0.047 | 0.189 | 0.189 | 0 | 0.59 | 0.87 | 0.71 | 0.89 | 0.54 | 0.61 |
| 10 | 124041701 | rs17103336 | 0.931 | -0.46 | -0.351 | -0.379 | -0.012 | -0.117 | -0.117 | 0.024 | 0.26 | 0.05 | 0.17 | 0.41 | 0.24 | 0.27 |
| 10 | 124042438 | rs11200527 | 1.394 | -0.149 | -0.726 | -0.627 | -0.039 | 0.08 | 0.08 | 0 | 0.24 | 0 | 0.16 | 0.11 | 0.2 | 0.32 |
| 10 | 124043184 | rs2273796 | 0.302 | -0.509 | -0.033 | -0.103 | 0 | 0.011 | 0.011 | 0 | 0.23 | 0.23 | 0.43 | 0.48 | 0.23 | 0.32 |
| 10 | 124048905 | rs7896211 | 0.025 | -0.416 | -0.255 | 0.253 | -0.026 | 0.007 | 0.007 | 0 | 0.38 | 0.46 | 0.33 | 0.41 | 0.3 | 0.39 |
| 10 | 124050130 | rs1007865 | -1.239 | -0.303 | -0.827 | 0.315 | -0.072 | 0.041 | 0.041 | 0 | 0.58 | 0.86 | 0.5 | 0.63 | 0.51 | 0.57 |
| 10 | 124050575 | rs11592545 | 0.865 | -0.032 | -0.436 | 0.744 | -0.003 | 0.003 | 0.003 | 0 | 0.12 | 0.07 | 0.1 | 0.26 | 0.14 | 0.17 |
| 10 | 124061501 | rs10788275 | 0.092 | -0.319 | -0.425 | 0.526 | 0.048 | -0.009 | -0.009 | 0 | 0.39 | 0.24 | 0.31 | 0.4 | 0.29 | 0.38 |
| 10 | 124062106 | rs17103365 | 0.099 | -0.324 | -0.427 | 0.524 | 0.048 | -0.01 | -0.01 | 0 | 0.39 | 0.24 | 0.31 | 0.4 | 0.29 | 0.38 |
| 10 | 124063140 | rs4237538 | -0.718 | -0.186 | -0.484 | 0.042 | -0.01 | 0.148 | 0.148 | 0 | 0.56 | 0.73 | 0.66 | 0.9 | 0.48 | 0.57 |
| 10 | 124066652 | rs7082865 | -0.702 | -0.178 | -0.404 | 0.134 | -0.003 | 0.134 | 0.134 | 0 | 0.57 | 0.74 | 0.59 | 0.88 | 0.46 | 0.54 |
| 10 | 124074891 | rs3905303 | -0.918 | -0.24 | -0.342 | 0.189 | -0.044 | 0.104 | 0.104 | 0 | 0.62 | 0.83 | 0.65 | 0.87 | 0.51 | 0.56 |
| 10 | 124086757 | rs6585812 | -0.483 | -0.329 | 0.431 | 0.316 | 0.001 | 0.006 | 0.006 | 0 | 0.55 | 0.43 | 0.38 | 0.64 | 0.48 | 0.45 |
| 10 | 124087092 | rs6585813 | -0.785 | -0.312 | 0.193 | 0.194 | -0.007 | 0.032 | 0.032 | 0 | 0.58 | 0.69 | 0.62 | 0.87 | 0.51 | 0.5 |
| 10 | 124088367 | rs10736312 | -0.551 | -0.421 | 0.225 | 0.149 | 0 | -0.005 | -0.005 | 0 | 0.55 | 0.59 | 0.6 | 0.87 | 0.5 | 0.47 |
| 10 | 124089036 | rs2421013 | -0.918 | -0.344 | 0.242 | 0.206 | 0.002 | -0.023 | -0.023 | 4.475 | 0.59 | 0.6 | 0.6 | 0.87 | 0.51 | 0.5 |
| 10 | 124091921 | rs4627016 | -0.742 | 0.057 | -0.014 | 0.111 | 0.001 | -0.004 | -0.004 | 0 | 0.58 | 0.44 | 0.51 | 0.73 | 0.51 | 0.6 |
| 10 | 124095982 | rs3817280 | -0.672 | -0.398 | -0.299 | 0.559 | -0.001 | 0.002 | 0.002 | 0 | 0.33 | 0.4 | 0.49 | 0.69 | 0.33 | 0.42 |
| 10 | 124096035 | rs3817281 | -0.672 | -0.398 | -0.299 | 0.559 | -0.001 | 0.002 | 0.002 | 0 | 0.33 | 0.4 | 0.49 | 0.69 | 0.33 | 0.42 |
| 10 | 124096061 | rs1048347 | -0.668 | -0.395 | -0.297 | 0.561 | 0 | 0.002 | 0.002 | 0 | 0.33 | 0.4 | 0.49 | 0.69 | 0.33 | 0.42 |
| 10 | 124096160 | rs10510108 | 1.177 | -0.778 | -0.813 | 0.908 | 0.016 | -0.013 | -0.013 | 0.332 | 0.08 | 0 | 0.06 | 0.08 | 0.03 | 0.08 |
| 10 | 124096859 | rs11200558 | 0.397 | -0.342 | -0.526 | 1.123 | 0.011 | -0.036 | -0.036 | 0 | 0.12 | 0.07 | 0.09 | 0.3 | 0.09 | 0.13 |
| 10 | 124097427 | rs11200559 | 0.76 | -0.341 | -0.778 | 1.073 | -0.018 | -0.094 | -0.094 | 0 | 0.22 | 0.02 | 0.2 | 0.26 | 0.15 | 0.26 |
| 10 | 124097755 | rs2271597 | -0.429 | -0.305 | -0.3 | 0.778 | -0.001 | 0 | 0 | 0 | 0.37 | 0.33 | 0.34 | 0.57 | 0.29 | 0.39 |
| 10 | 124099344 | rs17103452 | -0.515 | -0.167 | -0.624 | 0.762 | 0.02 | -0.022 | -0.022 | 0 | 0.34 | 0.18 | 0.29 | 0.6 | 0.26 | 0.37 |
| 10 | 124100849 | rs7085944 | 0.148 | -0.228 | -0.373 | 0.889 | 0.011 | -0.022 | -0.022 | 0 | 0.24 | 0.26 | 0.21 | 0.48 | 0.13 | 0.28 |
| 10 | 124102664 | rs7085821 | -0.304 | 0.657 | -0.432 | 0.582 | 0.023 | -0.002 | -0.002 | 0 | 0.82 | 0.79 | 0.81 | 0.81 | 0.72 | 0.84 |
| 10 | 124103876 | rs7091160 | 0.479 | 0.265 | -0.837 | 0.194 | 0.054 | 0.016 | 0.016 | 0 | 0.45 | 0.23 | 0.34 | 0.35 | 0.36 | 0.43 |
| 10 | 124104396 | rs6585817 | -1.508 | 0.436 | -0.935 | 0.14 | -0.028 | 0.023 | 0.023 | 0 | 0.88 | 0.93 | 0.89 | 1 | 0.79 | 0.95 |
| 10 | 124104476 | rs10788283 | -0.107 | 0.251 | -0.749 | 0.146 | 0.014 | -0.003 | -0.003 | 5.481 | 0.63 | 0.54 | 0.61 | 0.35 | 0.61 | 0.57 |
| 10 | 124104793 | rs7095410 | 0.426 | 0.191 | -0.815 | 0.127 | -0.021 | 0.018 | 0.018 | 0 | 0.33 | 0.55 | 0.31 | 0.41 | 0.31 | 0.33 |
| 10 | 124106759 | rs11595694 | 0.481 | 0.264 | -0.802 | 0.135 | 0.001 | -0.01 | -0.01 | 0 | 0.36 | 0.35 | 0.42 | 0.3 | 0.37 | 0.38 |
| 10 | 124107045 | rs11598094 | 0.835 | 0.027 | -0.68 | 0.313 | 0.01 | 0.02 | 0.02 | 0 | 0.29 | 0.13 | 0.23 | 0.23 | 0.3 | 0.27 |
| 10 | 124108629 | rs6585819 | -0.114 | 0.063 | -0.675 | 0.243 | 0.001 | 0.001 | 0.001 | 0 | 0.52 | 0.58 | 0.47 | 0.7 | 0.42 | 0.58 |
| 10 | 124114356 | rs7904674 | 0.993 | 0.125 | -1.153 | -0.146 | 0.016 | 0.092 | 0.092 | 0 | 0.25 | 0.09 | 0.16 | 0.06 | 0.24 | 0.16 |
| 10 | 124115576 | rs7088780 | -0.652 | 0.498 | -0.965 | -0.365 | 0.003 | -0.045 | -0.045 | 0 | 0.8 | 0.78 | 0.86 | 1 | 0.74 | 0.93 |
| 10 | 124122430 | rs7916855 | 0.27 | 0.23 | -0.777 | -0.099 | 0.01 | 0.038 | 0.038 | 0 | 0.31 | 0.35 | 0.46 | 0.46 | 0.37 | 0.44 |
| 10 | 124122632 | rs7080960 | -0.548 | 0.244 | -1.373 | -0.482 | -0.035 | 0.195 | 0.195 | 0 | 0.54 | 0.78 | 0.7 | 0.95 | 0.5 | 0.75 |
| 10 | 124123470 | rs11200583 | 1.16 | 0.428 | -1.124 | 0.041 | 0.014 | -0.056 | -0.056 | 0 | 0.19 | 0.08 | 0.18 | 0.29 | 0.12 | 0.23 |
| 10 | 124124306 | rs1998345 | -0.189 | 0.753 | -1.219 | -0.039 | -0.064 | -0.013 | -0.013 | 0.263 | 0.65 | 0.9 | 0.72 | 0.71 | 0.65 | 0.62 |
| 10 | 124126761 | rs12246005 | 1.081 | 0.443 | -0.751 | -0.507 | 0.001 | -0.003 | -0.003 | 0 | 0.22 | 0.21 | 0.13 | 0.05 | 0.15 | 0.14 |
| 10 | 124132188 | rs4751889 | 0.026 | 0.757 | -0.378 | -0.535 | 0.008 | -0.034 | -0.034 | 0 | 0.36 | 0.5 | 0.29 | 0.19 | 0.41 | 0.27 |
| 10 | 124155381 | rs4146894 | 0.311 | 0.787 | -0.618 | -0.563 | 0.015 | 0.049 | 0.049 | 0 | 0.47 | 0.35 | 0.62 | 0.6 | 0.47 | 0.53 |
| 10 | 124156852 | rs4752692 | 1.478 | 0.773 | -0.745 | -0.631 | -0.004 | 0.055 | 0.055 | 0 | 0.16 | 0.06 | 0.07 | 0 | 0.12 | 0.14 |
| 10 | 124161546 | rs11592338 | 1.624 | 0.819 | -0.704 | -0.577 | 0.007 | 0.018 | 0.018 | 0 | 0.21 | 0.13 | 0.12 | 0.06 | 0.16 | 0.14 |
| 10 | 124164654 | rs10082476 | 1.103 | 0.838 | -0.713 | -0.566 | 0.003 | 0.017 | 0.017 | 0 | 0.25 | 0.15 | 0.17 | 0.18 | 0.23 | 0.24 |
| 10 | 124167004 | rs10887149 | 0.682 | 0.891 | -0.732 | -0.609 | -0.024 | 0.069 | 0.069 | 0 | 0.3 | 0.15 | 0.2 | 0.06 | 0.34 | 0.18 |
| 10 | 124178978 | rs11200621 | 1.258 | 1.08 | -0.985 | -0.577 | -0.028 | -0.01 | -0.01 | 0 | 0.09 | 0.01 | 0.28 | 0.08 | 0.14 | 0.09 |
| 10 | 124183732 | rs4405249 | -1.832 | 1.001 | -0.774 | -0.47 | -0.044 | 0.042 | 0.042 | 0 | 0.92 | 0.98 | 0.92 | 1 | 0.82 | 0.96 |
| 10 | 124186127 | rs2292623 | 0.712 | 0.834 | -0.618 | -0.417 | -0.007 | -0.016 | -0.016 | 0 | 0.19 | 0.48 | 0.13 | 0.21 | 0.12 | 0.19 |
| 10 | 124193181 | rs2280141 | -0.169 | 0.904 | -0.975 | -0.286 | 0.025 | 0.066 | 0.066 | 0 | 0.49 | 0.68 | 0.65 | 0.61 | 0.47 | 0.54 |
| 10 | 124201665 | rs2292627 | 1.3 | 1.231 | -0.916 | 0.437 | -0.009 | -0.12 | -0.12 | 0 | 0.16 | 0.01 | 0.18 | 0.36 | 0.11 | 0.24 |
| 10 | 124214448 | rs10490924 | 0.566 | 1.635 | -0.358 | 0.305 | 0 | 0.002 | 0.002 | 0 | 0.23 | 0.25 | 0.25 | 0.4 | 0.19 | 0.34 |
| 10 | 124214921 | rs17623531 | 0.896 | 1.53 | -0.564 | 0.077 | -0.007 | 0.087 | 0.087 | 0 | 0.17 | 0.03 | 0.07 | 0.01 | 0.13 | 0.15 |
| 10 | 124215421 | rs3750847 | 0.732 | 1.491 | -0.542 | 0.108 | 0 | 0.002 | 0.002 | 0 | 0.23 | 0.25 | 0.25 | 0.4 | 0.19 | 0.34 |

| | | | | | | | | | | | | | | | | |
|---|---|---|---|---|---|---|---|---|---|---|---|---|---|---|---|---|
| 10 | 124215594 | rs2736912 | 1.024 | 1.621 | -0.539 | 0.165 | -0.01 | -0.007 | -0.007 | 1.912 | 0.09 | 0.06 | 0.09 | 0.19 | 0.17 | 0.07 |
| 10 | 124217632 | rs2014307 | -1.652 | 1.458 | 0.016 | 0.112 | 0.017 | 0.015 | 0.015 | 0 | 0.59 | 0.74 | 0.5 | 0.68 | 0.6 | 0.64 |
| 10 | 124223944 | rs2248799 | -0.848 | 1.643 | 0.248 | 0.281 | 0.007 | -0.018 | -0.018 | 0 | 0.41 | 0.58 | 0.42 | 0.75 | 0.49 | 0.51 |
| 10 | 124226215 | rs2249049 | -0.395 | 1.801 | 0.095 | 0.35 | 0.013 | 0.001 | 0.001 | 0 | 0.15 | 0.19 | 0.15 | 0.25 | 0.27 | 0.15 |
| 10 | 124226630 | rs2284665 | 0.503 | 1.311 | 0.126 | 0.503 | -0.001 | 0.009 | 0.009 | 0 | 0.23 | 0.24 | 0.25 | 0.43 | 0.2 | 0.34 |
| 10 | 124230513 | rs12252027 | -1.029 | 1.5 | 0.077 | 0.496 | 0.034 | 0.059 | 0.059 | 0 | 0.12 | 0.34 | 0.09 | 0.13 | 0.13 | 0.1 |
| 10 | 124230612 | rs12571363 | -0.217 | 1.738 | -0.435 | 0.265 | -0.021 | -0.011 | -0.011 | 0 | 0.06 | 0.03 | 0.06 | 0.06 | 0.11 | 0.07 |
| 10 | 124234880 | rs7093894 | -0.408 | 1.339 | -0.126 | 0.201 | -0.002 | 0.018 | 0.018 | 0.04 | 0.19 | 0.4 | 0.15 | 0.26 | 0.15 | 0.23 |
| 10 | 124234988 | rs2672589 | -1.276 | 1.348 | -0.904 | -0.164 | 0.037 | 0.066 | 0.066 | 0 | 0.57 | 0.74 | 0.5 | 0.77 | 0.63 | 0.64 |
| 10 | 124235059 | rs17696741 | 0.195 | 1.774 | -1.035 | -0.28 | -0.025 | 0.077 | 0.077 | 0 | 0.14 | 0.01 | 0.07 | 0.02 | 0.13 | 0.09 |
| 10 | 124235296 | rs2672588 | -0.539 | 2.449 | -0.577 | -0.07 | 0.13 | -0.036 | -0.036 | 0 | 0.15 | 0.34 | 0.18 | 0.07 | 0.31 | 0.09 |
| 10 | 124237612 | rs4752700 | 0.159 | 1.587 | -1.762 | 0.138 | -0.016 | -0.019 | -0.019 | 0 | 0.65 | 0.78 | 0.4 | 0.43 | 0.46 | 0.54 |
| 10 | 124239437 | rs11200647 | -0.545 | 1.647 | -2.078 | 0.077 | -0.032 | 0.081 | 0.081 | 0 | 0.4 | 0.17 | 0.65 | 0.74 | 0.49 | 0.59 |
| 10 | 124241557 | rs2253755 | 0.413 | 1.698 | -2.143 | 0.278 | -0.071 | -0.073 | -0.073 | 0.079 | 0.2 | 0.09 | 0.45 | 0.37 | 0.31 | 0.31 |
| 10 | 124242817 | rs2300431 | -0.674 | 0.121 | -2.094 | 0.059 | 0.007 | -0.013 | -0.013 | 0 | 0.31 | 0.3 | 0.25 | 0.43 | 0.27 | 0.45 |
| 10 | 124243189 | rs2300432 | -0.407 | 0.043 | -2.128 | 0.048 | 0.042 | -0.048 | -0.048 | 0 | 0.27 | 0.13 | 0.19 | 0.36 | 0.18 | 0.43 |
| 10 | 124245483 | rs2268345 | -0.587 | -0.01 | -2.126 | 0.067 | 0.026 | -0.015 | -0.015 | 0.16 | 0.28 | 0.2 | 0.19 | 0.34 | 0.19 | 0.43 |
| 10 | 124249235 | rs2239586 | 0.32 | -0.038 | -1.749 | 0.237 | -0.007 | 0.025 | 0.025 | 0 | 0.17 | 0.19 | 0.16 | 0.36 | 0.12 | 0.36 |
| 10 | 124255340 | rs10887154 | 0.47 | -0.036 | -1.738 | 0.341 | 0.006 | -0.005 | -0.005 | 1 | 0.17 | 0.15 | 0.16 | 0.18 | 0.1 | 0.34 |
| 10 | 124256345 | rs714816 | 0.386 | 0.173 | -1.885 | 0.279 | -0.079 | -0.089 | -0.089 | 0 | 0.26 | 0.09 | 0.49 | 0.38 | 0.35 | 0.31 |
| 10 | 124259062 | rs2736923 | 0.22 | 0.607 | -1.583 | 0.807 | -0.018 | 0.023 | 0.023 | 0 | 0.79 | 0.72 | 0.82 | 0.64 | 0.88 | 0.64 |
| 10 | 124262444 | rs763720 | 1.007 | 0.948 | -1.837 | 0.986 | -0.084 | -0.09 | -0.09 | 1.898 | 0.14 | 0.02 | 0.13 | 0.3 | 0.23 | 0.14 |
| 10 | 124277017 | rs2736928 | 0.937 | 0.3 | -1.106 | 0.972 | 0.022 | -0.03 | -0.03 | 0 | 0.69 | 0.77 | 0.78 | 0.6 | 0.76 | 0.58 |
| 10 | 124277713 | rs17103667 | -0.138 | 0.192 | -1.095 | 0.953 | 0.018 | -0.019 | -0.019 | 0 | 0.15 | 0.07 | 0.13 | 0.18 | 0.1 | 0.35 |
| 10 | 124281928 | rs2975746 | -0.427 | 0.398 | -1.016 | 0.891 | 0.016 | 0.011 | 0.011 | 0 | 0.9 | 0.91 | 0.97 | 1 | 0.94 | 0.96 |
| 10 | 124285188 | rs7909454 | 0.651 | 0.343 | -0.855 | 1.009 | -0.017 | 0.01 | 0.01 | 0.026 | 0.85 | 0.74 | 0.93 | 0.78 | 0.91 | 0.93 |
| 10 | 124288724 | rs7902878 | -0.083 | 0.998 | -0.603 | 1.316 | -0.015 | 0.007 | 0.007 | 0 | 0.22 | 0.1 | 0.19 | 0.21 | 0.29 | 0.16 |
| 10 | 124290957 | rs3013211 | -0.995 | 1.132 | -0.406 | 1.294 | -0.002 | 0.002 | 0.002 | 0 | 0.45 | 0.42 | 0.4 | 0.61 | 0.49 | 0.55 |
| 10 | 124291502 | rs11528742 | -0.701 | 0.707 | -0.707 | 1.064 | 0.005 | -0.005 | -0.005 | 1.44 | 0.38 | 0.21 | 0.32 | 0.39 | 0.39 | 0.51 |
| 10 | 124308252 | rs3013223 | 0.508 | 0.713 | -0.239 | 1.054 | -0.016 | 0.029 | 0.029 | 0 | 0.69 | 0.81 | 0.77 | 0.81 | 0.58 | 0.7 |
| 10 | 124322595 | rs2981756 | 0.633 | 0.445 | 0.261 | 1.519 | 0.003 | -0.016 | -0.016 | 0 | 0.71 | 0.56 | 0.78 | 0.77 | 0.68 | 0.73 |
| 10 | 124329217 | rs2981786 | 0.57 | 0.441 | 0.281 | 1.517 | 0.004 | -0.012 | -0.012 | 0 | 0.71 | 0.61 | 0.79 | 0.77 | 0.68 | 0.73 |
| 10 | 124334810 | rs2005952 | 0.548 | 0.41 | 0.181 | 1.827 | -0.002 | 0.048 | 0.048 | 0.014 | 0.69 | 0.5 | 0.77 | 0.59 | 0.68 | 0.49 |
| 10 | 124390982 | rs17103755 | -1.516 | 0.003 | 0.331 | 2.341 | -0.016 | 0.24 | 0.24 | 0 | 0.08 | 0.26 | 0.08 | 0.39 | 0.07 | 0.21 |
| 10 | 124393339 | rs4752712 | -1.481 | -0.023 | 0.318 | 2.316 | -0.015 | 0.215 | 0.215 | 0 | 0.08 | 0.21 | 0.08 | 0.38 | 0.07 | 0.23 |
| 10 | 124398289 | rs4752713 | -2.297 | -0.148 | 0.696 | 2.142 | -0.022 | 0.207 | 0.207 | 1.089 | 0.13 | 0.38 | 0.15 | 0.38 | 0.09 | 0.29 |
| 10 | 124398834 | rs2684742 | -0.56 | -1.012 | -0.409 | 1.278 | 0.057 | 0.012 | 0.012 | 0 | 0.42 | 0.79 | 0.42 | 0.66 | 0.34 | 0.55 |
| 10 | 124399467 | rs2261655 | -1.044 | -0.945 | -0.866 | 0.977 | 0.034 | 0.047 | 0.047 | 0 | 0.53 | 0.89 | 0.5 | 0.67 | 0.41 | 0.65 |
| 10 | 124404122 | rs2742226 | -0.853 | -1.371 | -0.838 | 0.822 | 0.052 | -0.028 | -0.028 | 0 | 0.44 | 0.81 | 0.38 | 0.53 | 0.37 | 0.41 |
| 10 | 124406547 | rs7091947 | 0.034 | -1.518 | -0.402 | 0.806 | -0.014 | -0.001 | -0.001 | 0 | 0.1 | 0.1 | 0.03 | 0.13 | 0.05 | 0.05 |
| 10 | 124412538 | rs9731015 | -0.314 | -1.49 | -0.393 | 0.708 | 0 | -0.016 | -0.016 | 1.843 | 0.1 | 0.1 | 0.05 | 0 | 0.05 | 0.16 |
| 10 | 124417654 | rs2947586 | 0.075 | -1.259 | 0.062 | 0.949 | -0.003 | 0 | 0 | 0 | 0.25 | 0.3 | 0.16 | 0.29 | 0.25 | 0.21 |
| 10 | 124417900 | rs2421030 | -0.027 | -0.787 | -0.794 | 0.555 | 0.004 | -0.001 | -0.001 | 1.592 | 0.4 | 0.33 | 0.47 | 0.42 | 0.37 | 0.6 |
| 10 | 124436530 | rs3019500 | -0.317 | -0.198 | -0.614 | 0.381 | 0.026 | 0.029 | 0.029 | 0 | 0.45 | 0.71 | 0.42 | 0.4 | 0.39 | 0.4 |
| 10 | 124456033 | rs7913531 | 0.31 | 0.323 | -0.593 | 0.646 | 0 | -0.001 | -0.001 | 0 | 0.36 | 0.29 | 0.32 | 0.4 | 0.36 | 0.48 |
| 10 | 124457452 | rs2947594 | -0.853 | 0.379 | -0.547 | 0.571 | -0.018 | -0.014 | -0.014 | 0 | 0.79 | 0.84 | 0.73 | 0.81 | 0.75 | 0.88 |
| 10 | 124459139 | rs11813597 | 0.326 | 0.158 | -0.277 | 0.592 | -0.003 | 0.005 | 0.005 | 0 | 0.4 | 0.41 | 0.33 | 0.53 | 0.41 | 0.52 |
| 10 | 124460069 | rs10902929 | -0.709 | -0.183 | -0.239 | 0.36 | -0.019 | 0.014 | 0.014 | 0 | 0.17 | 0.34 | 0.17 | 0.23 | 0.16 | 0.21 |
| 10 | 124462596 | rs12266077 | -0.487 | -0.174 | -0.153 | 0.396 | 0.018 | 0.022 | 0.022 | 0 | 0.45 | 0.67 | 0.43 | 0.39 | 0.39 | 0.4 |
| 10 | 124465874 | rs10902833 | -1.089 | -0.054 | 0.099 | 0.477 | -0.002 | 0.061 | 0.061 | 0 | 0.07 | 0.3 | 0.04 | 0.12 | 0.04 | 0.05 |
| 10 | 124470089 | rs1917847 | -0.368 | 0.106 | 0.121 | 0.591 | -0.017 | 0.1 | 0.1 | 0 | 0.2 | 0.27 | 0.34 | 0.41 | 0.18 | 0.22 |
| 10 | 124476677 | rs884644 | 0.961 | 0.253 | 0.248 | 0.623 | -0.004 | -0.023 | -0.023 | 0.032 | 0.12 | 0.04 | 0.22 | 0.2 | 0.14 | 0.1 |
| 10 | 124481043 | rs10902851 | 1.032 | 0.683 | 0.133 | 0.613 | -0.009 | 0.001 | 0.001 | 0 | 0.22 | 0.11 | 0.17 | 0.29 | 0.27 | 0.24 |
| 10 | 124488143 | rs2901344 | 0.439 | 0.748 | 0.446 | 0.492 | 0.006 | 0.008 | 0.008 | 0 | 0.1 | 0.16 | 0.16 | 0.12 | 0.1 | 0.14 |
| 10 | 124490368 | rs2901345 | 0.97 | 0.741 | 0.441 | 0.198 | 0.007 | -0.009 | -0.009 | 0.213 | 0.29 | 0.23 | 0.27 | 0.39 | 0.22 | 0.31 |
| 10 | 124491040 | rs17104017 | -0.124 | 0.729 | 0.37 | 0.217 | 0.013 | 0.016 | 0.016 | 0 | 0.13 | 0.35 | 0.18 | 0.12 | 0.1 | 0.14 |
| 10 | 124503149 | rs10902895 | 0.681 | 0.669 | 0.17 | 0.29 | -0.01 | 0.007 | 0.007 | 0 | 0.3 | 0.34 | 0.27 | 0.39 | 0.22 | 0.31 |
| 10 | 124504595 | rs13376724 | -0.841 | 1.272 | 0.683 | 0.053 | 0.013 | -0.025 | -0.025 | 2.013 | 0.2 | 0.25 | 0.35 | 0.12 | 0.25 | 0.28 |
| 10 | 124505217 | rs12411856 | 0.022 | 1.22 | 0.882 | 0.384 | -0.007 | 0.058 | 0.058 | 0 | 0.12 | 0.26 | 0.19 | 0.22 | 0.08 | 0.24 |
| 10 | 124506084 | rs7894294 | -0.246 | 1.576 | 0.827 | -0.122 | -0.012 | 0.011 | 0.011 | 0.22 | 0.63 | 0.68 | 0.55 | 0.73 | 0.47 | 0.71 |
| 10 | 124510966 | rs10794610 | 0.576 | 1.19 | 0.396 | -0.128 | 0.009 | 0.079 | 0.079 | 1.306 | 0.49 | 0.12 | 0.39 | 0.57 | 0.39 | 0.46 |
| 10 | 124511073 | rs11248479 | 1.254 | 1.821 | 0.449 | 0.065 | -0.011 | -0.006 | -0.006 | 0.522 | 0.2 | 0.12 | 0.18 | 0.29 | 0.28 | 0.19 |
| 10 | 124512091 | rs11248487 | -0.288 | 1.809 | 0.522 | 0.095 | -0.014 | 0.011 | 0.011 | 1.175 | 0.41 | 0.31 | 0.47 | 0.43 | 0.54 | 0.5 |
| 10 | 124515295 | rs953355 | 0.187 | 1.711 | 0.328 | 0.007 | 0.009 | -0.069 | -0.069 | 2.376 | 0.16 | 0.28 | 0.12 | 0 | 0.19 | 0.24 |
| 10 | 124515520 | rs10751758 | 0.276 | 1.422 | 0.214 | 0.477 | 0.002 | 0.032 | 0.032 | 0 | 0.08 | 0.07 | 0.16 | 0.31 | 0.08 | 0.22 |
| 10 | 124516699 | rs10736935 | 0.619 | 1.379 | 0.03 | 0.232 | 0.025 | 0.005 | 0.005 | 2.447 | 0.4 | 0.3 | 0.31 | 0.5 | 0.27 | 0.43 |
| 10 | 124520349 | rs4980284 | -1.418 | 1.44 | -0.045 | -0.167 | -0.005 | 0.001 | 0.001 | 0.006 | 0.75 | 0.84 | 0.63 | 0.7 | 0.66 | 0.78 |
| 10 | 124520493 | rs11595572 | 1.604 | 1.468 | -0.174 | -0.143 | 0.02 | 0.027 | 0.027 | 2.652 | 0.31 | 0.1 | 0.18 | 0.2 | 0.19 | 0.21 |
| 10 | 124521655 | rs7906712 | 0.067 | 1.891 | 0.13 | -0.63 | 0 | -0.014 | -0.014 | 7.488 | 0.56 | 0.48 | 0.47 | 0.69 | 0.41 | 0.53 |

| | | | | | | | | | | | | | | | | |
|---|---|---|---|---|---|---|---|---|---|---|---|---|---|---|---|---|
| 10 | 124523514 | rs17104093 | 0.348 | 1.589 | -0.299 | -0.265 | 0.001 | -0.022 | -0.022 | 0.001 | 0.15 | 0.13 | 0.09 | 0 | 0.14 | 0.04 |
| 10 | 124524241 | rs1341391 | -0.412 | 1.727 | -0.305 | 0.348 | 0.004 | -0.004 | -0.004 | 0 | 0.74 | 0.77 | 0.66 | 0.71 | 0.66 | 0.76 |
| 10 | 124528930 | rs947260 | -0.465 | 1.591 | -0.461 | 0.534 | -0.001 | -0.003 | -0.003 | 0 | 0.48 | 0.52 | 0.5 | 0.45 | 0.51 | 0.6 |
| 10 | 124530915 | rs7100288 | -0.804 | 2.666 | -0.293 | 1.338 | 0.026 | 0.04 | 0.04 | 0 | 0.16 | 0.22 | 0.32 | 0.42 | 0.3 | 0.18 |
| 10 | 124532133 | rs1578474 | -0.767 | 2.591 | -0.308 | 1.526 | -0.021 | -0.02 | -0.02 | 0 | 0.16 | 0.14 | 0.3 | 0.29 | 0.29 | 0.17 |
| 10 | 124536352 | rs869742 | -0.801 | 2.624 | -0.312 | 1.52 | -0.058 | -0.054 | -0.054 | 0 | 0.15 | 0.07 | 0.29 | 0.29 | 0.3 | 0.17 |
| 10 | 124539105 | rs1539065 | 1.222 | 2.029 | -0.032 | 0.967 | 0.035 | 0.034 | 0.034 | 0.029 | 0.66 | 0.53 | 0.58 | 0.53 | 0.6 | 0.74 |
| 10 | 124539718 | rs1970473 | 1.17 | 2.528 | -0.032 | 0.938 | -0.01 | 0.037 | 0.037 | 0.153 | 0.67 | 0.51 | 0.6 | 0.53 | 0.68 | 0.75 |
| 10 | 124543343 | rs7096042 | 1.19 | 2.584 | 0.005 | 0.938 | -0.002 | 0.037 | 0.037 | 0.101 | 0.66 | 0.53 | 0.59 | 0.53 | 0.66 | 0.75 |
| 10 | 124544283 | rs7923560 | -0.706 | 2.586 | 0.039 | 0.92 | 0.005 | 0.003 | 0.003 | 0 | 0.17 | 0.2 | 0.31 | 0.29 | 0.31 | 0.18 |
| 10 | 124544336 | rs10510115 | 0.628 | 2.747 | -0.407 | 0.789 | -0.032 | -0.001 | -0.001 | 0.395 | 0.09 | 0 | 0.03 | 0.07 | 0.09 | 0.17 |
| 10 | 124545656 | rs7911308 | 0.857 | 2.737 | -0.058 | 0.652 | -0.001 | -0.001 | -0.001 | 0 | 0.67 | 0.63 | 0.6 | 0.67 | 0.6 | 0.75 |
| 10 | 124547332 | rs6599711 | -0.701 | 2.515 | -0.018 | 0.642 | 0.011 | 0.022 | 0.022 | 0.105 | 0.84 | 0.89 | 0.92 | 0.96 | 0.91 | 0.94 |
| 10 | 124548859 | rs7896663 | 0.075 | 1.835 | 0.401 | 0.881 | -0.016 | 0.074 | 0.074 | 0 | 0.18 | 0.33 | 0.23 | 0.38 | 0.12 | 0.33 |
| 10 | 124551486 | rs10751763 | -0.52 | 1.953 | 0.653 | 0.81 | -0.025 | -0.068 | -0.068 | 0.669 | 0.83 | 0.7 | 0.9 | 0.96 | 0.91 | 0.94 |
| 10 | 124553691 | rs2421130 | -0.569 | 2.07 | 0.531 | 0.825 | 0.001 | 0.02 | 0.02 | 0 | 0.18 | 0.2 | 0.31 | 0.43 | 0.24 | 0.19 |
| 10 | 124554089 | rs11248243 | 0.946 | 2.089 | 0.513 | 0.832 | -0.005 | 0.027 | 0.027 | 0.186 | 0.66 | 0.54 | 0.59 | 0.53 | 0.67 | 0.74 |
| 10 | 124565743 | rs7915633 | 0.095 | 2.158 | 0.384 | 0.811 | -0.016 | -0.03 | -0.03 | 0 | 0.15 | 0.1 | 0.28 | 0.29 | 0.23 | 0.17 |
| 10 | 124577515 | rs4980167 | -0.402 | 2.125 | 0.485 | 0.788 | -0.019 | -0.024 | -0.024 | 0 | 0.83 | 0.73 | 0.88 | 0.86 | 0.9 | 0.92 |
| 10 | 124579796 | rs1557064 | -0.332 | 2.415 | 0.31 | 0.945 | 0.002 | 0.013 | 0.013 | 0 | 0.15 | 0.13 | 0.28 | 0.29 | 0.23 | 0.14 |
| 10 | 124586574 | rs10510116 | -0.249 | 2.416 | 0.227 | 0.946 | -0.034 | -0.056 | -0.056 | 0.257 | 0.14 | 0.07 | 0.28 | 0.3 | 0.23 | 0.14 |
| 10 | 124589041 | rs1417794 | -0.444 | 2.366 | 0.284 | 0.936 | 0.019 | 0.013 | 0.013 | 0 | 0.16 | 0.16 | 0.28 | 0.3 | 0.31 | 0.15 |
| 10 | 124606165 | rs1891111 | -0.154 | 1.8 | 0.11 | 0.414 | 0.001 | -0.001 | -0.001 | 0 | 0.47 | 0.44 | 0.44 | 0.51 | 0.57 | 0.49 |
| 10 | 124609283 | rs2153669 | -0.052 | 1.658 | 0.2 | 0.321 | 0 | -0.001 | -0.001 | 0 | 0.45 | 0.35 | 0.44 | 0.48 | 0.58 | 0.49 |
| 10 | 124610027 | rs1891110 | -0.284 | 1.59 | 0.875 | 0.295 | -0.001 | -0.001 | -0.001 | 0.082 | 0.48 | 0.48 | 0.46 | 0.49 | 0.58 | 0.49 |
| 10 | 124624436 | rs10794575 | 0.022 | 1.626 | 0.495 | 0.334 | -0.007 | 0.005 | 0.005 | 0.081 | 0.44 | 0.3 | 0.41 | 0.42 | 0.56 | 0.47 |
| 10 | 124638827 | rs4638251 | 0.381 | 2.002 | 0.31 | 0.349 | -0.05 | -0.011 | -0.011 | 0.079 | 0.13 | 0.04 | 0.13 | 0.16 | 0.21 | 0.18 |
| 10 | 124648285 | rs10902839 | -0.059 | 1.726 | 0.386 | 0.336 | -0.008 | 0.006 | 0.006 | 0.013 | 0.44 | 0.34 | 0.42 | 0.42 | 0.56 | 0.48 |
| 10 | 124660458 | rs7915165 | 0.016 | 1.735 | 0.305 | 0.359 | -0.01 | 0.001 | 0.001 | 0.058 | 0.43 | 0.33 | 0.41 | 0.42 | 0.56 | 0.48 |
| 10 | 124671655 | rs7921647 | -0.243 | 1.733 | 0.382 | 0.384 | -0.001 | -0.001 | -0.001 | 0.428 | 0.46 | 0.46 | 0.45 | 0.49 | 0.58 | 0.49 |
| 10 | 124676335 | rs11248332 | -0.07 | 1.673 | 0.348 | 0.366 | -0.006 | 0.004 | 0.004 | 0 | 0.44 | 0.36 | 0.41 | 0.42 | 0.56 | 0.47 |
| 10 | 124680929 | rs2421162 | -0.097 | 1.674 | 0.348 | 0.366 | -0.006 | 0.005 | 0.005 | 0 | 0.44 | 0.36 | 0.41 | 0.42 | 0.56 | 0.47 |
| 10 | 124686656 | rs10794579 | -0.401 | 1.584 | 0.32 | 0.235 | 0.01 | 0.003 | 0.003 | 0 | 0.49 | 0.42 | 0.46 | 0.51 | 0.6 | 0.51 |
| 10 | 124692586 | rs4980169 | -0.414 | 1.574 | 0.292 | 0.223 | 0.004 | 0.002 | 0.002 | 0 | 0.51 | 0.52 | 0.47 | 0.53 | 0.6 | 0.5 |
| 10 | 124697198 | rs10794580 | 0.156 | 1.57 | 0.047 | 0.216 | -0.017 | -0.018 | -0.018 | 0 | 0.43 | 0.3 | 0.35 | 0.45 | 0.49 | 0.46 |
| 10 | 124704149 | rs6599638 | 0.008 | 1.592 | -0.011 | 0.224 | -0.01 | -0.007 | -0.007 | 0 | 0.44 | 0.36 | 0.38 | 0.51 | 0.51 | 0.49 |
| 10 | 124710659 | rs10902847 | 0.068 | 1.593 | -0.006 | 0.191 | -0.009 | -0.01 | -0.01 | 0.156 | 0.44 | 0.36 | 0.36 | 0.45 | 0.49 | 0.48 |
| 10 | 124712961 | rs1106056 | -0.412 | 1.807 | -0.049 | -0.699 | 0.01 | -0.022 | -0.022 | 0.364 | 0.06 | 0.08 | 0.09 | 0 | 0.1 | 0.02 |
| 10 | 124716092 | rs9423292 | -0.477 | 0.978 | -0.301 | -1.189 | 0.004 | 0.002 | 0.002 | 0.063 | 0.61 | 0.66 | 0.55 | 0.39 | 0.72 | 0.54 |
| 10 | 124720263 | rs12268329 | 1.069 | 1.155 | -0.775 | -1.467 | -0.051 | 0.069 | 0.069 | 0 | 0.25 | 0.07 | 0.24 | 0.11 | 0.31 | 0.29 |
| 10 | 124720884 | rs7085513 | 0.241 | 0.947 | -0.387 | -1.44 | -0.009 | 0.016 | 0.016 | 0.706 | 0.41 | 0.35 | 0.39 | 0.26 | 0.5 | 0.47 |
| 10 | 124721136 | rs2901358 | 0.859 | 0.973 | -0.347 | -1.419 | 0.006 | -0.073 | -0.073 | 0 | 0.19 | 0.32 | 0.14 | 0.01 | 0.22 | 0.06 |
| 10 | 124721761 | rs2421157 | 1.199 | 1.031 | -0.417 | -1.457 | 0.007 | -0.054 | -0.054 | 0.152 | 0.18 | 0.25 | 0.13 | 0.01 | 0.22 | 0.06 |
| 10 | 124733038 | rs2901354 | -0.545 | 0.758 | -0.621 | -1.292 | 0.008 | 0.004 | 0.004 | 0.12 | 0.61 | 0.66 | 0.56 | 0.39 | 0.73 | 0.54 |
| 10 | 124740621 | rs10902853 | -0.545 | 0.758 | -0.658 | -1.289 | 0.004 | 0.002 | 0.002 | 0.12 | 0.61 | 0.66 | 0.56 | 0.39 | 0.73 | 0.54 |
| 10 | 124742792 | rs3736583 | -0.545 | 0.758 | -0.658 | -1.305 | 0.004 | 0.002 | 0.002 | 0.12 | 0.61 | 0.66 | 0.56 | 0.39 | 0.73 | 0.54 |
| 10 | 124749740 | rs11190 | 0.917 | 1.095 | -1.101 | -1.545 | -0.102 | 0.091 | 0.091 | 0 | 0.27 | 0.04 | 0.28 | 0.1 | 0.39 | 0.22 |
| 10 | 124750087 | rs9423240 | -0.234 | 0.482 | -1.046 | -2.132 | 0.001 | -0.002 | -0.002 | 0 | 0.61 | 0.4 | 0.49 | 0.22 | 0.7 | 0.36 |
| 10 | 124753060 | rs9423302 | 1.518 | 0.405 | -1.001 | -2.161 | 0.005 | 0.006 | 0.006 | 0 | 0.16 | 0.12 | 0.08 | 0.11 | 0.09 | 0.09 |
| 10 | 124768583 | rs12263012 | 1.431 | 0.527 | -0.952 | -2.096 | -0.058 | 0.08 | 0.08 | 0 | 0.28 | 0.09 | 0.28 | 0.1 | 0.39 | 0.21 |
| 10 | 124774439 | rs10794583 | -1.281 | 0.323 | -0.649 | -1.92 | -0.028 | 0.032 | 0.032 | 0 | 0.77 | 0.68 | 0.69 | 0.38 | 0.86 | 0.63 |
| 10 | 124779296 | rs10902861 | 1.466 | 0.704 | -1.034 | -1.752 | -0.007 | -0.003 | -0.003 | 0.12 | 0.14 | 0.1 | 0.17 | 0.16 | 0.15 | 0.27 |
| 10 | 124789186 | rs7070793 | -0.216 | 0.785 | -0.805 | -1.74 | 0.017 | 0.078 | 0.078 | 0 | 0.47 | 0.31 | 0.44 | 0.22 | 0.63 | 0.29 |
| 10 | 124793597 | rs4980245 | 1.16 | 1.062 | -0.639 | -1.325 | -0.012 | 0.023 | 0.023 | 0 | 0.14 | 0.1 | 0.12 | 0.01 | 0.18 | 0.05 |
| 10 | 124794977 | rs17104496 | -0.088 | 1.113 | -0.394 | -1.07 | 0 | -0.001 | -0.001 | 0.589 | 0.09 | 0.06 | 0.1 | 0.12 | 0.16 | 0.25 |
| 10 | 124796215 | rs2421139 | -1.325 | 1.004 | -0.173 | -1.317 | 0.004 | 0.008 | 0.008 | 0 | 0.51 | 0.6 | 0.47 | 0.4 | 0.56 | 0.64 |
| 10 | 124805633 | rs4128783 | 0.044 | 1.259 | -0.334 | -1.034 | -0.001 | 0.001 | 0.001 | 0 | 0.91 | 0.89 | 0.9 | 0.86 | 0.88 | 0.96 |
| 10 | 124818438 | rs7100031 | 0.088 | 0.788 | -0.374 | -1.178 | -0.02 | 0.077 | 0.077 | 0 | 0.38 | 0.28 | 0.37 | 0.11 | 0.39 | 0.46 |
| 10 | 124822417 | rs10902869 | 0.062 | 0.577 | 0.254 | -1.189 | -0.003 | 0.013 | 0.013 | 0.012 | 0.57 | 0.39 | 0.58 | 0.75 | 0.6 | 0.53 |
| 10 | 124824340 | rs7919930 | 0.823 | 0.468 | -0.039 | -1.15 | 0.001 | 0.008 | 0.008 | 0 | 0.12 | 0.11 | 0.19 | 0.6 | 0.09 | 0.19 |
| 10 | 124824630 | rs11248373 | 0.49 | 0.471 | 0.166 | -1.262 | 0 | -0.005 | -0.005 | 0.784 | 0.48 | 0.5 | 0.4 | 0.15 | 0.51 | 0.34 |
| 10 | 124827165 | rs4980248 | 0.018 | 0.646 | 0.052 | -1.431 | -0.005 | -0.004 | -0.004 | 0 | 0.39 | 0.38 | 0.38 | 0.1 | 0.39 | 0.46 |
| 10 | 124843497 | rs11248386 | -0.503 | 0.548 | 0.703 | -0.677 | -0.013 | 0.042 | 0.042 | 0 | 0.21 | 0.41 | 0.31 | 0.67 | 0.19 | 0.31 |
| 10 | 124844537 | rs10902878 | -0.414 | 0.56 | 0.696 | -0.648 | -0.003 | 0.007 | 0.007 | 0 | 0.2 | 0.22 | 0.3 | 0.67 | 0.19 | 0.31 |
| 10 | 124851234 | rs2421170 | -0.313 | 1.142 | 0.577 | -0.681 | 0.041 | -0.032 | -0.032 | 0 | 0.08 | 0.17 | 0.1 | 0.04 | 0.1 | 0.04 |
| 10 | 124852915 | rs7074044 | 0.215 | 1.157 | 0.617 | -0.635 | 0.06 | 0.044 | 0.044 | 0.088 | 0.92 | 0.83 | 0.88 | 0.75 | 0.88 | 0.95 |
| 10 | 124859974 | rs12262149 | 0.614 | 0.812 | -0.453 | -1.295 | -0.014 | 0.078 | 0.078 | 0 | 0.32 | 0.19 | 0.3 | 0.07 | 0.31 | 0.39 |
| 10 | 124865051 | rs12416137 | 0.8 | 0.591 | -0.382 | -1.015 | 0.002 | 0.104 | 0.104 | 0 | 0.51 | 0.32 | 0.41 | 0.09 | 0.49 | 0.39 |
| 10 | 124866446 | rs11248393 | -0.746 | 0.575 | -0.422 | -0.241 | -0.007 | 0.001 | 0.001 | 0 | 0.23 | 0.15 | 0.37 | 0.73 | 0.27 | 0.37 |

| | | | | | | | | | | | | | | | | |
|---|---|---|---|---|---|---|---|---|---|---|---|---|---|---|---|---|
| 10 | 124871148 | rs11248395 | -0.627 | -0.065 | 0.021 | 0.894 | 0.013 | 0.327 | 0.327 | 0.09 | 0.05 | 0.29 | 0.04 | 0.23 | 0.01 | 0.09 |
| 10 | 124872514 | rs4980259 | -0.529 | -0.226 | -0.306 | 0.688 | -0.004 | 0.014 | 0.014 | 0 | 0.69 | 0.63 | 0.78 | 0.48 | 0.7 | 0.87 |
| 10 | 124872687 | rs12764514 | 0.68 | -0.04 | -1.054 | 0.337 | -0.03 | 0.074 | 0.074 | 0 | 0.25 | 0.06 | 0.15 | 0.13 | 0.27 | 0.03 |
| 10 | 124877448 | rs12411348 | -0.233 | -0.02 | -0.571 | 0.802 | 0 | 0.002 | 0.002 | 0.545 | 0.24 | 0.17 | 0.37 | 0.34 | 0.25 | 0.49 |
| 10 | 124879121 | rs7094343 | 0.574 | 0.065 | -0.526 | 0.75 | 0 | 0.006 | 0.006 | 0 | 0.2 | 0.12 | 0.33 | 0.4 | 0.17 | 0.4 |
| 10 | 124879183 | rs11594133 | 0.488 | -0.297 | -0.484 | 0.272 | 0.002 | -0.027 | -0.027 | 1.061 | 0.18 | 0.13 | 0.1 | 0 | 0.14 | 0.1 |
| 10 | 124881502 | rs10794595 | -0.21 | 0.145 | -0.458 | 0.468 | 0.07 | -0.079 | -0.079 | 0 | 0.76 | 0.48 | 0.7 | 0.88 | 0.69 | 0.8 |
| 10 | 124887244 | rs10751752 | -0.241 | 0.208 | -0.54 | 0.728 | 0.008 | 0.013 | 0.013 | 2.069 | 0.45 | 0.31 | 0.46 | 0.63 | 0.39 | 0.69 |
| 10 | 124887778 | rs11248405 | 0.399 | 0.191 | -0.531 | 0.796 | 0.005 | 0 | 0 | 0.66 | 0.37 | 0.21 | 0.41 | 0.36 | 0.33 | 0.52 |
| 10 | 124887825 | rs4980180 | 0.355 | -0.274 | 0.02 | 1.517 | 0.016 | 0.028 | 0.028 | 0 | 0.33 | 0.27 | 0.26 | 0.24 | 0.3 | 0.1 |
| 10 | 124891137 | rs17104692 | -0.06 | -0.176 | -0.49 | 1.363 | 0.001 | 0.026 | 0.026 | 0 | 0.09 | 0.1 | 0.04 | 0.27 | 0.05 | 0.19 |
| 10 | 124891395 | rs4980264 | -0.124 | 0.002 | -0.728 | 0.628 | -0.011 | -0.006 | -0.006 | 0 | 0.43 | 0.54 | 0.37 | 0.63 | 0.35 | 0.29 |
| 10 | 124898260 | rs12243192 | 0.286 | 0.812 | -0.624 | 0.716 | 0.006 | -0.008 | -0.008 | 0 | 0.65 | 0.36 | 0.48 | 0.33 | 0.57 | 0.59 |
| 10 | 124904611 | rs10902882 | -0.954 | 0.151 | -0.647 | 1.228 | -0.001 | 0.002 | 0.002 | 0 | 0.22 | 0.26 | 0.29 | 0.33 | 0.24 | 0.29 |
| 10 | 124907389 | rs4980181 | -1.484 | -0.333 | -0.149 | 1.84 | -0.025 | 0.152 | 0.152 | 0 | 0.17 | 0.36 | 0.14 | 0.4 | 0.12 | 0.18 |
| 10 | 124913576 | rs3808960 | -1.46 | -0.366 | -0.084 | 2.071 | -0.01 | 0.175 | 0.175 | 0 | 0.15 | 0.34 | 0.14 | 0.38 | 0.12 | 0.18 |
| 10 | 124913996 | rs7897156 | -1.479 | -0.366 | -0.084 | 2.071 | -0.011 | 0.173 | 0.173 | 0 | 0.15 | 0.34 | 0.14 | 0.38 | 0.12 | 0.18 |
| 10 | 124917238 | rs2304550 | -1.507 | -0.361 | -0.064 | 2.162 | -0.018 | 0.128 | 0.128 | 0 | 0.16 | 0.33 | 0.14 | 0.37 | 0.12 | 0.18 |
| 10 | 124928952 | rs7918913 | -1.789 | -1.114 | 0.012 | 1.605 | -0.018 | 0.029 | 0.029 | 0.151 | 0.31 | 0.62 | 0.32 | 0.41 | 0.24 | 0.31 |
| 10 | 124929763 | rs11248424 | -0.474 | -0.843 | -0.046 | 1.709 | 0.003 | 0.002 | 0.002 | 0 | 0.07 | 0.07 | 0.06 | 0.1 | 0.1 | 0.12 |
| 10 | 124931515 | rs11248426 | -1.711 | -0.885 | 0.107 | 1.713 | -0.002 | 0.03 | 0.03 | 0 | 0.15 | 0.19 | 0.13 | 0.38 | 0.11 | 0.15 |
| 10 | 124932086 | rs2495774 | -0.968 | -1.082 | -0.82 | 1.055 | -0.099 | 0.116 | 0.116 | 0 | 0.68 | 0.95 | 0.67 | 0.82 | 0.47 | 0.63 |
| 10 | 124941951 | rs4980268 | -0.234 | -0.859 | -0.132 | 1.327 | -0.028 | -0.023 | -0.023 | 0 | 0.07 | 0.01 | 0.04 | 0.1 | 0.1 | 0.12 |
| 10 | 124949389 | rs10902892 | -0.78 | -0.966 | 0.158 | 1.359 | 0.016 | 0.275 | 0.275 | 3.055 | 0.06 | 0.59 | 0.08 | 0.19 | 0 | 0.06 |
| 10 | 124951379 | rs4980270 | -1.252 | -0.684 | -0.908 | 0.649 | 0.002 | -0.003 | -0.003 | 0 | 0.14 | 0.11 | 0.1 | 0.18 | 0.12 | 0.14 |
| 10 | 124957828 | rs2495769 | -0.218 | -0.586 | -1.626 | 0.145 | 0.04 | 0.049 | 0.049 | 0 | 0.41 | 0.79 | 0.31 | 0.28 | 0.33 | 0.25 |
| 10 | 124958574 | rs2495767 | -0.952 | 0.61 | -2.19 | -0.125 | 0.016 | -0.025 | -0.025 | 0 | 0.63 | 0.86 | 0.54 | 0.38 | 0.66 | 0.46 |
| 10 | 124959614 | rs11248442 | 0.4 | 1.101 | -2.199 | -0.208 | -0.052 | 0.04 | 0.04 | 0 | 0.08 | 0.01 | 0.1 | 0.03 | 0.2 | 0.05 |
| 10 | 124962758 | rs2099586 | 0.008 | 0.74 | -1.298 | -0.292 | 0.04 | 0.062 | 0.062 | 0 | 0.11 | 0.37 | 0.08 | 0.14 | 0.12 | 0.13 |
| 10 | 124966517 | rs9423256 | 1.156 | 0.716 | -1.307 | -0.383 | -0.004 | -0.006 | -0.006 | 0 | 0.28 | 0.29 | 0.19 | 0.14 | 0.2 | 0.13 |
| 10 | 124967722 | rs10902901 | -0.636 | 1.279 | -0.931 | -0.291 | -0.08 | 0.065 | 0.065 | 0 | 0.21 | 0.09 | 0.24 | 0.09 | 0.34 | 0.21 |
| 10 | 124974126 | rs6599665 | -0.602 | 1.181 | -0.824 | -0.257 | 0.032 | 0.056 | 0.056 | 0.125 | 0.13 | 0.47 | 0.1 | 0.14 | 0.11 | 0.13 |
| 10 | 124979929 | rs7083581 | -0.971 | 1.16 | -1.047 | -0.352 | -0.069 | 0.073 | 0.073 | 0 | 0.23 | 0.13 | 0.25 | 0.09 | 0.34 | 0.21 |
| 10 | 124980600 | rs17104841 | 0.095 | 1.004 | -0.996 | -0.307 | 0 | 0.023 | 0.023 | 0 | 0.15 | 0.1 | 0.14 | 0.06 | 0.14 | 0.16 |
| 10 | 124981862 | rs6599666 | -1.309 | 0.917 | -0.987 | -0.255 | 0.005 | -0.009 | -0.009 | 0 | 0.37 | 0.67 | 0.35 | 0.23 | 0.46 | 0.34 |
| 10 | 124987484 | rs11595616 | -0.631 | 1.562 | -0.998 | -0.179 | -0.016 | 0.003 | 0.003 | 0.89 | 0.08 | 0.04 | 0.13 | 0.06 | 0.2 | 0.05 |
| 10 | 124988153 | rs11597068 | 0.147 | 1.313 | -1.197 | -0.682 | 0.009 | 0.024 | 0.024 | 0 | 0.19 | 0.12 | 0.09 | 0.06 | 0.12 | 0.14 |
| 10 | 124988589 | rs766295 | -0.859 | 1.461 | -1.077 | -0.711 | -0.02 | 0.014 | 0.014 | 1.897 | 0.1 | 0.12 | 0.17 | 0.05 | 0.24 | 0.08 |
| 10 | 124995307 | rs6599671 | -1.339 | 0.802 | -0.915 | -1.27 | -0.057 | -0.094 | -0.094 | 0 | 0.63 | 0.57 | 0.85 | 0.91 | 0.8 | 0.89 |
| 10 | 124996706 | rs11248466 | -0.618 | 1.108 | -0.953 | -1.235 | -0.03 | 0.016 | 0.016 | 0.106 | 0.11 | 0.12 | 0.18 | 0.06 | 0.27 | 0.11 |
| 10 | 124998593 | rs10794605 | 0.554 | 0.93 | -0.995 | -1.343 | -0.005 | 0.007 | 0.007 | 0 | 0.32 | 0.28 | 0.55 | 0.8 | 0.4 | 0.65 |
| 10 | 124999294 | rs11248472 | 1.22 | 1.033 | -1.298 | -0.639 | -0.109 | -0.071 | -0.071 | 5.425 | 0.24 | 0.01 | 0.34 | 0.24 | 0.34 | 0.48 |
| 10 | 125002485 | rs10510126 | 0.596 | 0.653 | -1.108 | -1.128 | 0.011 | 0.031 | 0.031 | 0 | 0.2 | 0.13 | 0.1 | 0.06 | 0.13 | 0.14 |
| 10 | 125005193 | rs17663978 | 0.838 | 0.625 | -1.129 | -1.142 | 0.021 | 0.062 | 0.062 | 0 | 0.19 | 0.08 | 0.07 | 0.05 | 0.1 | 0.11 |
| 10 | 125008718 | rs6599673 | -1.356 | 0.712 | -0.977 | -1.092 | -0.053 | -0.094 | -0.094 | 0 | 0.63 | 0.57 | 0.85 | 0.91 | 0.8 | 0.88 |
| 10 | 125020203 | rs17587364 | -0.387 | 1.374 | -0.655 | -0.635 | -0.046 | 0.019 | 0.019 | 0 | 0.08 | 0.02 | 0.14 | 0.04 | 0.24 | 0.07 |
| 10 | 125030886 | rs7894928 | -0.744 | 1.202 | -0.581 | -0.64 | -0.017 | 0.017 | 0.017 | 0 | 0.29 | 0.26 | 0.26 | 0.18 | 0.34 | 0.21 |
| 10 | 125031183 | rs7916600 | 0.504 | 0.959 | -0.581 | -0.595 | 0.018 | 0.034 | 0.034 | 0 | 0.19 | 0.14 | 0.09 | 0.07 | 0.1 | 0.11 |
| 10 | 125039776 | rs7917062 | -1.369 | 1.201 | -0.38 | -0.543 | 0.037 | -0.026 | -0.026 | 0 | 0.07 | 0.19 | 0.11 | 0.02 | 0.13 | 0.1 |
| 10 | 125043537 | rs10902920 | -1.129 | 1.322 | -0.234 | -0.473 | -0.004 | 0.001 | 0.001 | 0 | 0.33 | 0.69 | 0.32 | 0.25 | 0.38 | 0.25 |
| 10 | 125050134 | rs7099339 | -1.205 | 1.297 | -0.746 | -0.6 | -0.127 | -0.08 | -0.08 | 0 | 0.71 | 0.98 | 0.52 | 0.3 | 0.6 | 0.41 |
| 10 | 125057140 | rs11596786 | -1.975 | 1.6 | 0.183 | -0.019 | 0.171 | 0.104 | 0.104 | 0.856 | 0.15 | 0.59 | 0.25 | 0.22 | 0.29 | 0.15 |
| 10 | 125078855 | rs9328844 | -0.743 | 1.486 | 0.154 | -0.114 | -0.013 | -0.039 | -0.039 | 0 | 0.27 | 0.62 | 0.21 | 0.13 | 0.22 | 0.22 |
| 10 | 125079287 | rs9328845 | -1.134 | 1.473 | 0.137 | -0.113 | 0.009 | -0.003 | -0.003 | 0 | 0.31 | 0.68 | 0.31 | 0.24 | 0.37 | 0.25 |
| 10 | 125084639 | rs913524 | -1.179 | 1.469 | 0.139 | -0.119 | 0.006 | -0.002 | -0.002 | 0 | 0.31 | 0.69 | 0.31 | 0.26 | 0.37 | 0.25 |
| 10 | 125092615 | rs2149407 | -1.239 | 1.368 | 0.231 | 0.113 | 0.006 | -0.002 | -0.002 | 0 | 0.32 | 0.69 | 0.32 | 0.25 | 0.38 | 0.25 |
| 10 | 125093310 | rs1023045 | -0.627 | 1.495 | 0.233 | 0.13 | -0.012 | 0.008 | 0.008 | 0 | 0.26 | 0.27 | 0.26 | 0.14 | 0.37 | 0.23 |
| 10 | 125095478 | rs6599685 | -0.216 | 1.465 | 0.245 | 0.08 | 0 | -0.02 | -0.02 | 0 | 0.23 | 0.26 | 0.17 | 0.05 | 0.22 | 0.2 |
| 10 | 125099422 | rs7906793 | -0.194 | 1.464 | 0.238 | 0.08 | 0 | -0.021 | -0.021 | 0 | 0.22 | 0.27 | 0.17 | 0.05 | 0.22 | 0.2 |
| 10 | 125106194 | rs6599689 | -1.163 | 1.003 | -0.158 | -0.145 | -0.01 | 0.057 | 0.057 | 0 | 0.53 | 0.73 | 0.39 | 0.28 | 0.47 | 0.33 |
| 10 | 125109478 | rs7906403 | 0.304 | 0.64 | 0.101 | -0.268 | -0.083 | -0.14 | -0.14 | 2.046 | 0.27 | 0.52 | 0.13 | 0.03 | 0.1 | 0.07 |
| 10 | 125111052 | rs6599691 | -0.237 | 1.483 | -0.072 | 0.146 | -0.022 | -0.016 | -0.016 | 0.01 | 0.06 | 0.04 | 0.1 | 0.2 | 0.16 | 0.07 |
| 10 | 125114051 | rs1887821 | -0.761 | 1.464 | 0.451 | 0.054 | 0.045 | 0.01 | 0.01 | 0 | 0.3 | 0.42 | 0.29 | 0.31 | 0.38 | 0.24 |
| 10 | 125114667 | rs11593654 | -1.125 | 1.505 | 0.483 | 0.029 | 0.05 | -0.002 | -0.002 | 0 | 0.05 | 0.14 | 0.09 | 0.05 | 0.11 | 0.05 |
| 10 | 125115722 | rs12776663 | -0.051 | 1.478 | 0.169 | 0.066 | 0.005 | 0 | 0 | 0 | 0.22 | 0.23 | 0.19 | 0.22 | 0.26 | 0.16 |
| 10 | 125124500 | rs9423335 | 0.055 | 1.459 | 0.087 | 0.028 | -0.025 | -0.009 | -0.009 | 0 | 0.23 | 0.11 | 0.17 | 0.22 | 0.25 | 0.15 |
| 10 | 125124884 | rs6599694 | -1.319 | 1.322 | 0.336 | -0.004 | -0.014 | 0.002 | 0.002 | 0.387 | 0.35 | 0.73 | 0.34 | 0.32 | 0.4 | 0.25 |
| 10 | 125125151 | rs7894765 | -0.866 | 1.334 | 0.486 | -0.002 | 0.03 | -0.008 | -0.008 | 0.048 | 0.32 | 0.56 | 0.3 | 0.28 | 0.38 | 0.23 |
| 10 | 125127837 | rs7083401 | 0.259 | 1.114 | 0.03 | -0.184 | -0.012 | 0.039 | 0.039 | 0.021 | 0.29 | 0.18 | 0.2 | 0.15 | 0.3 | 0.13 |

| | | | | | | | | | | | | | | | | |
|---|---|---|---|---|---|---|---|---|---|---|---|---|---|---|---|---|
| 10 | 125130306 | rs12264884 | 1.225 | 1.123 | -0.016 | -0.174 | 0.033 | 0.144 | 0.144 | 0.354 | 0.22 | 0.02 | 0.08 | 0 | 0.13 | 0.1 |
| 10 | 125134743 | rs4980285 | 1.309 | 1.532 | -0.435 | -0.242 | -0.072 | -0.053 | -0.053 | 0.094 | 0.37 | 0.03 | 0.47 | 0.64 | 0.38 | 0.54 |
| 10 | 125143250 | rs1467692 | -1.219 | 1.347 | -0.285 | -0.161 | -0.033 | 0.037 | 0.037 | 0 | 0.58 | 0.8 | 0.48 | 0.35 | 0.5 | 0.41 |
| 10 | 125143334 | rs7896499 | -0.122 | 0.905 | -0.028 | -0.127 | 0.054 | 0.046 | 0.046 | 0.046 | 0.05 | 0.32 | 0.1 | 0.03 | 0.02 | 0.08 |
| 10 | 125143668 | rs17665876 | 0.405 | 1.108 | -0.926 | -0.029 | 0.021 | 0.046 | 0.046 | 0 | 0.19 | 0.1 | 0.11 | 0.09 | 0.12 | 0.12 |
| 10 | 125151935 | rs9423263 | 0.436 | 1.239 | -0.928 | 0.01 | 0.023 | 0.056 | 0.056 | 0 | 0.2 | 0.06 | 0.11 | 0.09 | 0.12 | 0.14 |
| 10 | 125160933 | rs1328600 | 0.502 | 2.211 | -0.41 | 0.812 | -0.01 | 0.023 | 0.023 | 0.379 | 0.54 | 0.23 | 0.47 | 0.4 | 0.45 | 0.63 |
| 10 | 125160967 | rs9423343 | -0.434 | 1.6 | -0.094 | 0.896 | 0.012 | 0.057 | 0.057 | 0.527 | 0.38 | 0.69 | 0.37 | 0.23 | 0.33 | 0.22 |
| 10 | 125162193 | rs7916586 | -0.545 | 1.667 | -0.253 | 0.966 | -0.02 | -0.037 | -0.037 | 0 | 0.41 | 0.8 | 0.48 | 0.59 | 0.39 | 0.32 |
| 10 | 125164858 | rs17105163 | -1.32 | 2.009 | 0.934 | 0.794 | -0.001 | 0.005 | 0.005 | 0.855 | 0.06 | 0.19 | 0.1 | 0.07 | 0.04 | 0.03 |
| 10 | 125185442 | rs845102 | 0.594 | 1.952 | -1.361 | 0.474 | -0.046 | -0.186 | -0.186 | 0 | 0.31 | 0.02 | 0.32 | 0.41 | 0.28 | 0.41 |
| 10 | 125190029 | rs7073471 | -0.055 | 2.151 | -0.713 | 0.103 | -0.002 | -0.021 | -0.021 | 0 | 0.44 | 0.48 | 0.31 | 0.22 | 0.42 | 0.38 |
| 10 | 125192081 | rs17105212 | 0.15 | 2.512 | -0.329 | 0.005 | 0.001 | -0.028 | -0.028 | 0 | 0.12 | 0.12 | 0.13 | 0.04 | 0.13 | 0.07 |
| 10 | 125192325 | rs7916709 | 0.076 | 1.888 | -0.574 | -0.257 | 0.028 | -0.024 | -0.024 | 0 | 0.27 | 0.38 | 0.26 | 0.21 | 0.33 | 0.3 |
| 10 | 125195639 | rs9423349 | -0.126 | 1.319 | -0.602 | -0.156 | -0.023 | -0.042 | -0.042 | 0.2 | 0.37 | 0.42 | 0.17 | 0.14 | 0.22 | 0.32 |
| 10 | 125206633 | rs11248533 | 0.098 | 1.404 | -0.486 | 0.032 | -0.004 | 0.006 | 0.006 | 0 | 0.16 | 0.19 | 0.18 | 0.35 | 0.1 | 0.25 |
| 10 | 125213711 | rs845087 | 0.603 | 1.711 | 0.127 | 0.014 | 0.053 | -0.023 | -0.023 | 0 | 0.79 | 0.54 | 0.78 | 0.85 | 0.77 | 0.79 |
| 10 | 125214607 | rs6599697 | -0.219 | 1.663 | 0.333 | -0.015 | 0.024 | 0.002 | 0.002 | 1.014 | 0.14 | 0.33 | 0.16 | 0.11 | 0.12 | 0.17 |
| 10 | 125214716 | rs768733 | 0.992 | 1.533 | 0.134 | -0.084 | 0.024 | 0.048 | 0.048 | 0 | 0.13 | 0.05 | 0.05 | 0.04 | 0.07 | 0.15 |
| 10 | 125215290 | rs705137 | 1.223 | 1.644 | 0.661 | -0.025 | 0.01 | 0.034 | 0.034 | 0.59 | 0.75 | 0.34 | 0.64 | 0.58 | 0.74 | 0.69 |
| 10 | 125217896 | rs845085 | 0.47 | 2.184 | 0.422 | -0.449 | 0.003 | -0.004 | -0.004 | 0 | 0.59 | 0.46 | 0.59 | 0.59 | 0.56 | 0.58 |
| 10 | 125219941 | rs12572414 | 0.074 | 2.028 | 0.384 | -0.56 | -0.012 | -0.024 | -0.024 | 0 | 0.31 | 0.35 | 0.19 | 0.1 | 0.18 | 0.26 |
| 10 | 125220036 | rs845084 | 2.322 | 2.237 | 0.116 | -0.574 | 0.007 | -0.025 | -0.025 | 0.864 | 0.29 | 0.22 | 0.27 | 0.4 | 0.25 | 0.35 |
| 10 | 125220256 | rs11597044 | -0.134 | 2.464 | 0.158 | -0.517 | -0.002 | -0.002 | -0.002 | 0.676 | 0.09 | 0.08 | 0.11 | 0.15 | 0.13 | 0.04 |
| 10 | 125221734 | rs845083 | -0.412 | 2.571 | 0.321 | -0.65 | 0 | -0.01 | -0.01 | 0 | 0.84 | 0.84 | 0.81 | 0.96 | 0.78 | 0.83 |
| 10 | 125222037 | rs2282015 | -0.781 | 2.544 | 0.194 | -0.651 | -0.001 | -0.008 | -0.008 | 0 | 0.39 | 0.37 | 0.46 | 0.52 | 0.43 | 0.3 |
| 10 | 125223672 | rs705141 | 0.897 | 2.599 | -0.08 | -0.669 | -0.038 | 0.215 | 0.215 | 0 | 0.54 | 0.17 | 0.59 | 0.9 | 0.5 | 0.65 |
| 10 | 125224260 | rs705142 | 0.926 | 2.62 | 0.145 | -0.698 | 0.053 | -0.102 | -0.102 | 0.077 | 0.75 | 0.62 | 0.69 | 0.96 | 0.63 | 0.8 |
| 10 | 125225057 | rs845080 | 1.436 | 2.721 | 0.173 | -0.587 | 0.05 | 0.011 | 0.011 | 0 | 0.46 | 0.15 | 0.32 | 0.55 | 0.37 | 0.53 |
| 10 | 125226178 | rs705145 | 1.748 | 2.729 | 0.111 | -0.598 | 0.044 | -0.093 | -0.093 | 0 | 0.43 | 0.08 | 0.3 | 0.44 | 0.34 | 0.52 |
| 10 | 125226200 | rs705146 | 0.925 | 2.76 | 0.139 | -0.696 | -0.001 | -0.069 | -0.069 | 0.668 | 0.54 | 0.46 | 0.55 | 0.92 | 0.48 | 0.73 |
| 10 | 125229021 | rs2479903 | 1.194 | 2.759 | 0.237 | -0.494 | 0.019 | -0.033 | -0.033 | 0 | 0.07 | 0.14 | 0.09 | 0 | 0.11 | 0.01 |
| 10 | 125230186 | rs705150 | 0.783 | 2.688 | 0.142 | -0.573 | 0.003 | -0.013 | -0.013 | 0 | 0.56 | 0.46 | 0.46 | 0.66 | 0.51 | 0.66 |
| 10 | 125230352 | rs705151 | 0.085 | 2.738 | 0.165 | -0.556 | -0.002 | 0.011 | 0.011 | 0 | 0.65 | 0.74 | 0.58 | 0.66 | 0.64 | 0.71 |
| 10 | 125233053 | rs705159 | 0.848 | 2.879 | 0.46 | -0.56 | -0.001 | 0.009 | 0.009 | 0 | 0.55 | 0.4 | 0.44 | 0.66 | 0.49 | 0.62 |
| 10 | 125233868 | rs7922033 | 0.29 | 2.513 | 0.662 | -0.372 | 0.117 | 0.093 | 0.093 | 0.222 | 0.06 | 0.24 | 0.13 | 0.09 | 0.12 | 0.07 |
| 10 | 125234492 | rs7090367 | -0.689 | 2.585 | 0.63 | -0.356 | -0.007 | -0.013 | -0.013 | 0 | 0.38 | 0.51 | 0.48 | 0.36 | 0.33 | 0.33 |
| 10 | 125236844 | rs11248545 | 0.427 | 2.435 | 0.527 | -0.391 | 0.035 | 0.06 | 0.06 | 0 | 0.25 | 0.14 | 0.16 | 0.06 | 0.13 | 0.17 |
| 10 | 125242283 | rs11248546 | -1.396 | 2.592 | 0.617 | -0.357 | -0.021 | 0.006 | 0.006 | 0 | 0.45 | 0.74 | 0.6 | 0.45 | 0.46 | 0.4 |
| 10 | 125245215 | rs6599699 | -1.386 | 2.586 | 0.599 | -0.36 | -0.061 | -0.01 | -0.01 | 0 | 0.46 | 0.88 | 0.62 | 0.45 | 0.46 | 0.4 |
| 10 | 125247462 | rs1557614 | -0.73 | 2.504 | 0.226 | -0.387 | 0.002 | 0.001 | 0.001 | 0 | 0.35 | 0.61 | 0.47 | 0.39 | 0.39 | 0.23 |
| 10 | 125251675 | rs705162 | 2.267 | 2.585 | -0.022 | -0.351 | -0.002 | -0.166 | -0.166 | 3.358 | 0.31 | 0.02 | 0.23 | 0.39 | 0.24 | 0.4 |
| 10 | 125252093 | rs7916049 | -0.042 | 2.682 | 0.727 | -0.082 | 0.005 | -0.005 | -0.005 | 0 | 0.27 | 0.23 | 0.2 | 0.06 | 0.17 | 0.2 |
| 10 | 125252316 | rs860912 | -1.173 | 2.736 | 0.777 | -0.117 | 0.073 | -0.018 | -0.018 | 0.101 | 0.15 | 0.61 | 0.23 | 0.09 | 0.24 | 0.08 |
| 10 | 125254416 | rs12250242 | -0.296 | 2.516 | 0.585 | 0.044 | -0.001 | -0.001 | -0.001 | 0.66 | 0.36 | 0.35 | 0.44 | 0.36 | 0.3 | 0.27 |
| 10 | 125256216 | rs10794619 | -1.149 | 2.555 | 0.327 | 0.208 | -0.021 | -0.018 | -0.018 | 0 | 0.44 | 0.71 | 0.6 | 0.49 | 0.46 | 0.38 |
| 10 | 125257608 | rs7916003 | -1.695 | 2.591 | 0.815 | 0.275 | 0.13 | 0.07 | 0.07 | 0.059 | 0.09 | 0.38 | 0.16 | 0.1 | 0.16 | 0.11 |
| 10 | 125259616 | rs7091074 | -0.287 | 2.252 | 0.014 | 0.341 | -0.001 | 0.002 | 0.002 | 0 | 0.22 | 0.21 | 0.29 | 0.41 | 0.23 | 0.14 |
| 10 | 125259780 | rs10902937 | 0.562 | 2.413 | -0.127 | 0.351 | -0.037 | 0.043 | 0.043 | 0 | 0.16 | 0.02 | 0.07 | 0.08 | 0.13 | 0.05 |
| 10 | 125260923 | rs7897598 | 1.073 | 2.224 | -0.187 | -0.329 | 0.029 | -0.009 | -0.009 | 0 | 0.43 | 0.31 | 0.31 | 0.51 | 0.37 | 0.59 |
| 10 | 125261923 | rs10751760 | -0.009 | 2.019 | 0.09 | -0.429 | 0.042 | -0.083 | -0.083 | 1.159 | 0.74 | 0.59 | 0.6 | 0.88 | 0.63 | 0.78 |
| 10 | 125262764 | rs7903114 | -0.711 | 2.273 | 0.482 | -0.471 | 0.006 | 0.002 | 0.002 | 0 | 0.36 | 0.48 | 0.34 | 0.37 | 0.33 | 0.22 |
| 10 | 125265804 | rs1627852 | -0.312 | 2.229 | 0.125 | -0.538 | 0.004 | -0.014 | -0.014 | 0.029 | 0.8 | 0.77 | 0.66 | 0.89 | 0.71 | 0.81 |
| 10 | 125266207 | rs6599700 | -0.457 | 2.318 | -0.402 | -0.39 | 0.001 | -0.006 | -0.006 | 0 | 0.31 | 0.28 | 0.29 | 0.37 | 0.25 | 0.2 |
| 10 | 125274134 | rs1767123 | 0.219 | 1.717 | -1.392 | -0.153 | -0.04 | 0.046 | 0.046 | 0 | 0.3 | 0.15 | 0.33 | 0.2 | 0.36 | 0.26 |
| 10 | 125274226 | rs10902940 | 1.232 | 1.541 | -1.953 | -0.198 | 0.01 | -0.046 | -0.046 | 0.119 | 0.25 | 0.15 | 0.22 | 0.45 | 0.24 | 0.45 |
| 10 | 125275363 | rs4980297 | -0.028 | 2.285 | -1.44 | -0.005 | -0.045 | -0.044 | -0.044 | 0.005 | 0.31 | 0.16 | 0.48 | 0.49 | 0.42 | 0.24 |
| 10 | 125275458 | rs11248566 | -0.411 | 2.17 | -1.177 | -0.238 | -0.033 | 0.093 | 0.093 | 0 | 0.2 | 0.05 | 0.13 | 0.01 | 0.22 | 0.12 |
| 10 | 125275978 | rs10732908 | 1.081 | 2.433 | -0.391 | 0.579 | 0.035 | 0.026 | 0.026 | 0 | 0.89 | 0.73 | 0.82 | 0.85 | 0.85 | 0.87 |
| 10 | 125278806 | rs1474016 | -0.139 | 1.375 | -0.958 | 0.399 | 0.011 | 0.013 | 0.013 | 1.143 | 0.52 | 0.54 | 0.49 | 0.47 | 0.55 | 0.47 |
| 10 | 125284832 | rs841018 | 0.82 | 0.85 | -0.689 | -0.085 | -0.012 | -0.035 | -0.035 | 0.618 | 0.89 | 0.8 | 0.94 | 0.98 | 0.94 | 0.83 |
| 10 | 125285443 | rs705169 | 0.286 | 1.041 | -1.027 | -0.558 | -0.002 | 0.024 | 0.024 | 0 | 0.52 | 0.58 | 0.64 | 0.71 | 0.45 | 0.53 |
| 10 | 125285875 | rs951631 | 0.478 | 1.109 | -1.019 | -0.59 | -0.013 | -0.019 | -0.019 | 0 | 0.16 | 0.1 | 0.37 | 0.61 | 0.19 | 0.32 |
| 10 | 125288304 | rs7909799 | 0.75 | 1.992 | -0.574 | -0.949 | 0.023 | 0.04 | 0.04 | 0.61 | 0.38 | 0.24 | 0.25 | 0.16 | 0.35 | 0.29 |
| 10 | 125289785 | rs2486032 | 0.09 | 1.835 | -1.114 | -1.071 | -0.033 | -0.031 | -0.031 | 0.054 | 0.61 | 0.78 | 0.55 | 0.39 | 0.64 | 0.39 |
| 10 | 125290738 | rs4556475 | 1.485 | 1.728 | -1.011 | -1.271 | 0.044 | 0.074 | 0.074 | 0 | 0.35 | 0.17 | 0.22 | 0.15 | 0.31 | 0.24 |
| 10 | 125294649 | rs841825 | -0.115 | 2.11 | -0.605 | -1.329 | 0 | -0.003 | -0.003 | 0 | 0.72 | 0.71 | 0.58 | 0.37 | 0.77 | 0.52 |
| 10 | 125294814 | rs4980195 | 1.599 | 1.973 | -0.451 | -1.34 | 0.044 | 0.104 | 0.104 | 0.156 | 0.33 | 0.09 | 0.22 | 0.18 | 0.31 | 0.25 |
| 10 | 125295750 | rs12415721 | -0.734 | 1.588 | -0.633 | -1.734 | -0.003 | -0.008 | -0.008 | 0 | 0.52 | 0.48 | 0.67 | 0.76 | 0.62 | 0.61 |

| | | | | | | | | | | | | | | | | |
|---|---|---|---|---|---|---|---|---|---|---|---|---|---|---|---|---|
| 10 | 125296372 | rs10902947 | 0.841 | 1.87 | -0.619 | -1.432 | 0.009 | -0.005 | -0.005 | 0 | 0.06 | 0.05 | 0.1 | 0.12 | 0.21 | 0.04 |
| 10 | 125298585 | rs705177 | -0.898 | 1.05 | -0.511 | -1.485 | 0.014 | 0.027 | 0.027 | 0 | 0.61 | 0.68 | 0.7 | 0.82 | 0.63 | 0.71 |
| 10 | 125303383 | rs7898111 | -1.114 | 1.156 | -0.988 | -1.455 | 0.02 | 0.128 | 0.128 | 0.109 | 0.63 | 0.85 | 0.73 | 0.82 | 0.63 | 0.71 |
| 10 | 125307696 | rs932147 | 1.251 | 1.239 | -0.886 | -1.237 | -0.016 | 0.013 | 0.013 | 0 | 0.11 | 0.01 | 0.06 | 0.05 | 0.08 | 0.08 |
| 10 | 125310002 | rs705179 | 0.137 | 0.912 | -0.579 | -1.086 | -0.001 | -0.007 | -0.007 | 0 | 0.21 | 0.24 | 0.32 | 0.61 | 0.17 | 0.41 |
| 10 | 125310499 | rs11592288 | 1.284 | 1.201 | -1.052 | -1.481 | -0.052 | 0.058 | 0.058 | 0.096 | 0.22 | 0.06 | 0.19 | 0.12 | 0.28 | 0.09 |
| 10 | 125310660 | rs7908279 | 0.1 | 0.932 | -0.594 | -1.475 | -0.005 | -0.017 | -0.017 | 0.013 | 0.11 | 0.2 | 0.11 | 0.05 | 0.09 | 0.05 |
| 10 | 125312671 | rs9783185 | 0.26 | 0.927 | -0.614 | -1.504 | -0.002 | -0.039 | -0.039 | 0.305 | 0.09 | 0.12 | 0.07 | 0 | 0.07 | 0.02 |
| 10 | 125318231 | rs10902952 | 1.001 | 1.015 | -1.045 | -1.554 | -0.031 | 0.063 | 0.063 | 0.138 | 0.23 | 0.07 | 0.18 | 0.11 | 0.26 | 0.12 |
| 10 | 125319543 | rs7893296 | -0.188 | 0.725 | -0.397 | -1.751 | 0.01 | -0.092 | -0.092 | 0.45 | 0.14 | 0.36 | 0.14 | 0 | 0.15 | 0.07 |
| 10 | 125320027 | rs2486048 | 0.406 | 0.531 | -1.045 | -1.545 | -0.001 | 0.001 | 0.001 | 0.056 | 0.2 | 0.21 | 0.3 | 0.58 | 0.17 | 0.38 |
| 10 | 125320905 | rs1878165 | 1.353 | 0.484 | -1.37 | -1.735 | 0.004 | 0.073 | 0.073 | 0 | 0.31 | 0.02 | 0.2 | 0.18 | 0.27 | 0.21 |
| 10 | 125322297 | rs1767103 | 0.436 | 0.655 | -0.673 | -1.429 | -0.001 | 0.001 | 0.001 | 0.056 | 0.2 | 0.21 | 0.3 | 0.59 | 0.17 | 0.38 |
| 10 | 125333494 | rs1710958 | 0.741 | 0.578 | -0.962 | -1.355 | -0.017 | -0.082 | -0.082 | 0 | 0.2 | 0.11 | 0.32 | 0.58 | 0.2 | 0.38 |
| 10 | 125334949 | rs1317737 | 1.558 | 0.682 | -1.153 | -1.721 | -0.031 | 0.047 | 0.047 | 0.071 | 0.27 | 0.12 | 0.24 | 0.14 | 0.33 | 0.16 |
| 10 | 125336789 | rs10902956 | -0.127 | 0.314 | -0.376 | -1.726 | -0.013 | -0.001 | -0.001 | 0 | 0.71 | 0.55 | 0.64 | 0.33 | 0.79 | 0.53 |
| 10 | 125340640 | rs841042 | -0.023 | 0.318 | -0.235 | -1.671 | -0.014 | 0.05 | 0.05 | 0 | 0.92 | 0.64 | 0.96 | 0.91 | 0.98 | 0.9 |
| 10 | 125342395 | rs12254057 | 0.133 | 0.143 | 0.055 | -1.781 | -0.008 | -0.08 | -0.08 | 0 | 0.13 | 0.25 | 0.11 | 0 | 0.13 | 0.06 |
| 10 | 125348154 | rs10902957 | -0.343 | 0.224 | 0.084 | -1.814 | 0.036 | -0.056 | -0.056 | 0 | 0.16 | 0.4 | 0.16 | 0.05 | 0.17 | 0.13 |
| 10 | 125349956 | rs1588960 | 2.012 | 0.261 | -0.341 | -1.991 | -0.095 | 0.107 | 0.107 | 0.001 | 0.26 | 0.02 | 0.2 | 0.09 | 0.33 | 0.16 |
| 10 | 125356716 | rs953557 | 0.138 | 0.36 | -0.054 | -1.905 | 0.006 | -0.065 | -0.065 | 0.067 | 0.14 | 0.2 | 0.11 | 0 | 0.17 | 0.11 |
| 10 | 125360806 | rs12253624 | 1.434 | 0.273 | -0.294 | -1.987 | -0.036 | 0.068 | 0.068 | 0 | 0.28 | 0.1 | 0.22 | 0.16 | 0.31 | 0.22 |
| 10 | 125360989 | rs2486042 | 0.226 | 0.23 | -0.22 | -1.98 | -0.022 | 0.155 | 0.155 | 1.198 | 0.49 | 0.21 | 0.52 | 0.74 | 0.48 | 0.55 |
| 10 | 125361059 | rs2486041 | 0.226 | 0.23 | -0.22 | -1.98 | -0.022 | 0.155 | 0.155 | 1.198 | 0.49 | 0.21 | 0.52 | 0.74 | 0.48 | 0.55 |
| 10 | 125361269 | rs2486040 | -1.585 | 0.404 | 0.068 | -1.9 | -0.001 | 0.001 | 0.001 | 0 | 0.68 | 0.68 | 0.7 | 0.89 | 0.66 | 0.74 |
| 10 | 125368406 | rs1009350 | -0.35 | 0.414 | 0.459 | -1.844 | 0.078 | -0.001 | -0.001 | 0 | 0.11 | 0.35 | 0.15 | 0.05 | 0.16 | 0.11 |
| 10 | 125370973 | rs1011056 | 0.444 | 0.34 | 0.066 | -1.897 | 0 | -0.001 | -0.001 | 0.139 | 0.64 | 0.39 | 0.52 | 0.32 | 0.66 | 0.47 |
| 10 | 125372707 | rs6599708 | 1.601 | 0.343 | -0.026 | -1.968 | -0.003 | 0.007 | 0.007 | 0 | 0.31 | 0.29 | 0.29 | 0.11 | 0.34 | 0.25 |
| 10 | 125377212 | rs841015 | 1.737 | 0.053 | -0.698 | -2.001 | -0.004 | -0.092 | -0.092 | 0 | 0.19 | 0.07 | 0.3 | 0.58 | 0.18 | 0.32 |
| 10 | 125379406 | rs12241392 | 0.757 | 0.241 | -0.396 | -1.962 | -0.031 | 0.06 | 0.06 | 0.17 | 0.33 | 0.1 | 0.22 | 0.21 | 0.32 | 0.23 |
| 10 | 125380508 | rs2486037 | 0.565 | -0.022 | 0.153 | -1.739 | 0.025 | -0.103 | -0.103 | 0.825 | 0.86 | 0.59 | 0.88 | 1 | 0.85 | 0.85 |
| 10 | 125383033 | rs2459344 | 0.307 | -0.262 | -0.53 | -1.833 | 0.054 | 0.006 | 0.006 | 0 | 0.39 | 0.12 | 0.41 | 0.63 | 0.28 | 0.5 |
| 10 | 125383328 | rs11248604 | 1.033 | 0.13 | -0.37 | -2.11 | -0.006 | 0.01 | 0.01 | 0 | 0.37 | 0.26 | 0.26 | 0.09 | 0.41 | 0.29 |
| 10 | 125385066 | rs11248605 | -0.075 | 0.004 | -0.162 | -2.173 | -0.004 | 0.004 | 0.004 | 0.044 | 0.53 | 0.58 | 0.33 | 0.09 | 0.48 | 0.38 |
| 10 | 125388653 | rs12358123 | 0.134 | 0.238 | -0.083 | -2.089 | 0.008 | -0.041 | -0.041 | 0 | 0.14 | 0.18 | 0.12 | 0 | 0.2 | 0.13 |
| 10 | 125396161 | rs10902967 | 1.259 | 0.241 | -0.075 | -2.057 | -0.032 | 0.001 | 0.001 | 0 | 0.19 | 0.11 | 0.23 | 0.19 | 0.31 | 0.18 |
| 10 | 125399080 | rs10794626 | -0.924 | 0.022 | 0.141 | -2.214 | 0 | -0.034 | -0.034 | 0 | 0.6 | 0.53 | 0.7 | 0.91 | 0.57 | 0.7 |
| 10 | 125399966 | rs12762474 | -0.106 | 0.063 | -0.253 | -2.116 | 0.001 | -0.052 | -0.052 | 0 | 0.54 | 0.43 | 0.37 | 0.15 | 0.49 | 0.43 |
| 10 | 125400162 | rs12243970 | -0.047 | 0.095 | -0.526 | -2.104 | -0.001 | 0.015 | 0.015 | 0 | 0.54 | 0.39 | 0.32 | 0.12 | 0.51 | 0.41 |
| 10 | 125400554 | rs12763734 | 2.297 | 0.106 | -1.228 | -1.834 | -0.058 | 0.061 | 0.061 | 0 | 0.23 | 0.01 | 0.15 | 0.09 | 0.23 | 0.15 |
| 10 | 125402461 | rs1228412 | 0.805 | -0.085 | -0.823 | -1.384 | 0.024 | -0.058 | -0.058 | 0 | 0.21 | 0.12 | 0.38 | 0.63 | 0.19 | 0.4 |
| 10 | 125407194 | rs705185 | -0.27 | -0.156 | -0.442 | -1.626 | -0.008 | -0.058 | -0.058 | 0 | 0.76 | 0.47 | 0.76 | 0.87 | 0.81 | 0.73 |
| 10 | 125415905 | rs3850766 | -1.187 | -0.237 | -0.781 | -1.483 | 0.013 | -0.072 | -0.072 | 0.028 | 0.79 | 0.9 | 0.69 | 0.41 | 0.81 | 0.77 |
| 10 | 125421436 | rs11248631 | 1.723 | -0.086 | -0.813 | -1.705 | 0.004 | 0.047 | 0.047 | 0 | 0.3 | 0.16 | 0.21 | 0.12 | 0.28 | 0.29 |
| 10 | 125421696 | rs1219826 | -0.427 | -0.032 | -0.898 | -1.674 | 0.002 | 0.032 | 0.032 | 0 | 0.45 | 0.2 | 0.44 | 0.62 | 0.42 | 0.42 |
| 10 | 125422121 | rs1122370 | 0.111 | -0.048 | -0.81 | -1.678 | 0.019 | 0.055 | 0.055 | 0 | 0.54 | 0.71 | 0.55 | 0.35 | 0.58 | 0.55 |
| 10 | 125423919 | rs734000 | 0.073 | -0.021 | -0.997 | -1.695 | 0.006 | 0.034 | 0.034 | 0 | 0.55 | 0.78 | 0.56 | 0.38 | 0.58 | 0.58 |
| 10 | 125424306 | rs17676342 | 0.956 | -0.735 | -0.946 | -2.002 | 0.017 | 0.068 | 0.068 | 0.756 | 0.19 | 0.1 | 0.07 | 0.03 | 0.12 | 0.15 |
| 10 | 125427898 | rs11596507 | 0.476 | -0.213 | -0.42 | -1.544 | -0.036 | 0.044 | 0.044 | 0 | 0.21 | 0.1 | 0.2 | 0.12 | 0.3 | 0.14 |
| 10 | 125429706 | rs12261064 | -0.278 | -0.403 | -0.25 | -1.522 | -0.012 | -0.048 | -0.048 | 0.097 | 0.09 | 0.17 | 0.04 | 0 | 0.05 | 0.08 |
| 10 | 125436555 | rs1219812 | 1.092 | -0.37 | -0.061 | -1.316 | 0 | 0.008 | 0.008 | 0.005 | 0.18 | 0.27 | 0.36 | 0.68 | 0.19 | 0.25 |
| 10 | 125438459 | rs748219 | -0.398 | -0.237 | -0.208 | -1.414 | 0.005 | -0.03 | -0.03 | 0 | 0.33 | 0.34 | 0.28 | 0.16 | 0.4 | 0.25 |
| 10 | 125439038 | rs908471 | -0.58 | -0.275 | -0.246 | -1.43 | 0.005 | -0.072 | -0.072 | 0 | 0.37 | 0.46 | 0.36 | 0.18 | 0.41 | 0.34 |
| 10 | 125444002 | rs10902975 | 1.379 | -0.106 | -0.603 | -1.386 | -0.03 | 0.165 | 0.165 | 0 | 0.39 | 0.1 | 0.24 | 0.13 | 0.39 | 0.33 |
| 10 | 125444104 | rs7098285 | -0.618 | 0.422 | -0.016 | -0.994 | 0.019 | -0.049 | -0.049 | 0.058 | 0.27 | 0.32 | 0.23 | 0.13 | 0.34 | 0.22 |
| 10 | 125447705 | rs1878166 | 0.149 | 0.587 | 0.178 | -0.895 | 0.077 | -0.077 | -0.077 | 0 | 0.15 | 0.47 | 0.16 | 0.05 | 0.22 | 0.11 |
| 10 | 125448745 | rs12414023 | 0.479 | 0.412 | -0.266 | -1.057 | -0.016 | 0.03 | 0.03 | 0.565 | 0.1 | 0 | 0.13 | 0.03 | 0.09 | 0.21 |
| 10 | 125451208 | rs7084727 | 2.131 | -0.023 | -1.084 | -1.105 | -0.067 | -0.02 | -0.02 | 0 | 0.27 | 0.05 | 0.22 | 0.25 | 0.31 | 0.29 |
| 10 | 125451452 | rs12265793 | 0.956 | 0.025 | -1.003 | -1.086 | 0.08 | -0.075 | -0.075 | 0 | 0.15 | 0.45 | 0.16 | 0.05 | 0.22 | 0.12 |
| 10 | 125452085 | rs1356232 | 1.568 | -0.296 | -1.296 | -0.761 | -0.016 | -0.031 | -0.031 | 0.09 | 0.14 | 0.05 | 0.21 | 0.22 | 0.16 | 0.14 |
| 10 | 125452232 | rs7900707 | 0.629 | -0.634 | -1.399 | -0.877 | 0.011 | -0.13 | -0.13 | 0 | 0.26 | 0.06 | 0.22 | 0.43 | 0.2 | 0.19 |
| 10 | 125457622 | rs4980200 | -1.231 | -0.447 | -1.062 | -0.853 | 0.002 | 0.013 | 0.013 | 0 | 0.84 | 0.74 | 0.76 | 0.78 | 0.84 | 0.85 |
| 10 | 125457952 | rs7098190 | -0.358 | -0.75 | -1.022 | -0.898 | 0.029 | -0.1 | -0.1 | 0.115 | 0.25 | 0.47 | 0.29 | 0.09 | 0.31 | 0.33 |
| 10 | 125462847 | rs7097661 | 2.554 | -0.689 | -1.227 | -0.852 | -0.065 | -0.008 | -0.008 | 0 | 0.27 | 0.08 | 0.24 | 0.25 | 0.34 | 0.32 |
| 10 | 125463604 | rs4980202 | 0.09 | -0.601 | -0.898 | -0.844 | 0.008 | -0.087 | -0.087 | 0 | 0.74 | 0.42 | 0.71 | 0.94 | 0.69 | 0.66 |
| 10 | 125464431 | rs2362963 | -0.732 | -0.613 | -0.971 | -0.832 | 0.027 | 0.045 | 0.045 | 0 | 0.43 | 0.28 | 0.46 | 0.67 | 0.34 | 0.34 |
| 10 | 125465305 | rs4980203 | -0.224 | -0.604 | -0.866 | -0.825 | -0.02 | -0.045 | -0.045 | 0.009 | 0.85 | 0.76 | 0.85 | 0.97 | 0.9 | 0.79 |
| 10 | 125466049 | rs12779404 | 2.328 | -0.276 | -0.887 | -0.666 | -0.062 | -0.004 | -0.004 | 0.012 | 0.28 | 0.08 | 0.24 | 0.25 | 0.34 | 0.32 |
| 10 | 125466915 | rs732712 | 1.107 | -0.192 | -0.9 | -0.469 | 0.001 | -0.01 | -0.01 | 0 | 0.14 | 0.14 | 0.21 | 0.21 | 0.13 | 0.14 |

| | | | | | | | | | | | | | | | | |
|---|---|---|---|---|---|---|---|---|---|---|---|---|---|---|---|---|
| 10 | 125468066 | rs6599618 | 0.863 | 0.016 | -0.735 | -0.523 | 0 | 0.006 | 0.006 | 0.136 | 0.44 | 0.55 | 0.4 | 0.3 | 0.56 | 0.45 |
| 10 | 125469725 | rs4980204 | 0.349 | 0.147 | -0.779 | -0.074 | 0 | -0.002 | -0.002 | 1.773 | 0.58 | 0.34 | 0.67 | 0.59 | 0.59 | 0.7 |
| 10 | 125476325 | rs10902797 | -0.645 | 0.063 | -0.623 | -0.656 | 0.002 | 0.022 | 0.022 | 0.027 | 0.53 | 0.57 | 0.73 | 0.74 | 0.54 | 0.59 |
| 10 | 125480831 | rs10902800 | 1.802 | 0.185 | -0.905 | -0.951 | 0.018 | 0.11 | 0.11 | 0 | 0.27 | 0.09 | 0.14 | 0.01 | 0.21 | 0.16 |
| 10 | 125481863 | rs4980206 | -0.028 | 0.271 | -0.703 | -0.913 | 0.006 | 0.005 | 0.005 | 0 | 0.81 | 0.75 | 0.87 | 0.74 | 0.75 | 0.75 |
| 10 | 125485091 | rs10902803 | -0.287 | -0.153 | -0.195 | -0.856 | -0.007 | -0.012 | -0.012 | 0.976 | 0.11 | 0.24 | 0.16 | 0.07 | 0.07 | 0.19 |
| 10 | 125486119 | rs1896384 | 0.109 | -0.319 | -0.278 | -0.888 | -0.037 | 0.03 | 0.03 | 0.341 | 0.39 | 0.21 | 0.56 | 0.66 | 0.47 | 0.4 |
| 10 | 125487915 | rs12358957 | 0.088 | -0.315 | -0.27 | -0.886 | -0.034 | 0.03 | 0.03 | 0.232 | 0.39 | 0.23 | 0.55 | 0.66 | 0.46 | 0.4 |
| 10 | 125504915 | rs7918257 | -1.563 | -0.293 | 0.715 | -0.552 | -0.066 | -0.103 | -0.103 | 1.092 | 0.24 | 0.53 | 0.18 | 0.06 | 0.1 | 0.23 |
| 10 | 125506086 | rs12414446 | -1.321 | -0.167 | 0.046 | -0.552 | 0.007 | 0.004 | 0.004 | 0 | 0.23 | 0.14 | 0.14 | 0.05 | 0.13 | 0.25 |
| 10 | 125510357 | rs10736932 | 2.043 | -0.374 | -0.024 | -0.184 | -0.039 | 0.022 | 0.022 | 0 | 0.62 | 0.52 | 0.71 | 0.57 | 0.8 | 0.62 |
| 10 | 125511450 | rs7078710 | -0.423 | -0.417 | -0.025 | -0.147 | -0.009 | 0.158 | 0.158 | 0 | 0.91 | 0.83 | 0.88 | 0.72 | 0.95 | 0.92 |
| 10 | 125512142 | rs17105920 | -1.954 | -0.325 | -0.236 | -0.428 | 0.008 | 0.005 | 0.005 | 0 | 0.28 | 0.27 | 0.15 | 0.12 | 0.14 | 0.27 |
| 10 | 125512721 | rs12240880 | -0.929 | 0.178 | -0.137 | -0.186 | 0 | 0.001 | 0.001 | 0 | 0.08 | 0.12 | 0.12 | 0.07 | 0.07 | 0.16 |
| 10 | 125515829 | rs1896376 | -0.672 | 0.163 | -0.732 | -0.17 | 0.01 | -0.016 | -0.016 | 0.03 | 0.08 | 0 | 0.1 | 0.06 | 0.04 | 0.15 |
| 10 | 125517080 | rs4246949 | 0.951 | 0.233 | -0.735 | -0.127 | 0.01 | -0.019 | -0.019 | 2.968 | 0.68 | 0.76 | 0.76 | 0.63 | 0.77 | 0.69 |
| 10 | 125521533 | rs10794566 | 1.346 | 0.397 | -0.526 | -0.231 | -0.021 | 0.027 | 0.027 | 0 | 0.64 | 0.46 | 0.73 | 0.57 | 0.77 | 0.59 |
| 10 | 125521721 | rs10751743 | 0.036 | 0.434 | -0.938 | -0.088 | 0.002 | -0.01 | -0.01 | 0 | 0.83 | 0.87 | 0.89 | 0.68 | 0.87 | 0.86 |
| 10 | 125522409 | rs7909042 | -0.468 | 0.511 | -0.877 | -0.267 | 0.043 | -0.009 | -0.009 | 1.521 | 0.05 | 0.21 | 0.06 | 0 | 0.05 | 0.01 |
| 10 | 125525978 | rs1219738 | 0.193 | -0.052 | -1.268 | -0.11 | 0.014 | -0.022 | -0.022 | 0.655 | 0.14 | 0.08 | 0.05 | 0.27 | 0.06 | 0.1 |
| 10 | 125526689 | rs1896392 | 0.956 | -0.05 | -1.258 | -0.226 | -0.043 | 0.04 | 0.04 | 0 | 0.66 | 0.5 | 0.74 | 0.58 | 0.79 | 0.69 |
| 10 | 125527460 | rs1219732 | 1.362 | 1.599 | -0.815 | -0.667 | 0.081 | 0.068 | 0.068 | 0.34 | 0.5 | 0.12 | 0.46 | 0.6 | 0.35 | 0.38 |
| 10 | 125528048 | rs1219725 | 1.536 | 1.479 | -0.812 | -0.643 | 0.075 | 0.075 | 0.075 | 0.325 | 0.5 | 0.15 | 0.48 | 0.6 | 0.37 | 0.39 |
| 10 | 125530746 | rs727416 | 0.704 | 1.582 | -1.08 | -0.655 | 0.032 | -0.041 | -0.041 | 0 | 0.64 | 0.45 | 0.65 | 0.74 | 0.48 | 0.55 |
| 10 | 125534912 | rs1649151 | 0.646 | 0.79 | -1.562 | -0.689 | 0.064 | 0.043 | 0.043 | 1.604 | 0.68 | 0.22 | 0.76 | 0.76 | 0.83 | 0.64 |
| 10 | 125538630 | rs1237731 | -0.858 | 1.583 | -1.131 | -0.333 | -0.007 | 0.024 | 0.024 | 0 | 0.12 | 0 | 0.03 | 0.04 | 0.07 | 0.1 |
| 10 | 125540177 | rs17678853 | -1.089 | 2.72 | -0.307 | -0.353 | 0.111 | -0.086 | -0.086 | 0.263 | 0.13 | 0.34 | 0.2 | 0 | 0.27 | 0.06 |
| 10 | 125543163 | rs765335 | -1.078 | 2.165 | -0.808 | -0.216 | -0.007 | 0.006 | 0.006 | 0 | 0.07 | 0.04 | 0.06 | 0.03 | 0.1 | 0.12 |
| 10 | 125548464 | rs7922528 | 0.289 | 1.999 | -1.065 | -0.614 | -0.004 | 0.034 | 0.034 | 0.962 | 0.59 | 0.71 | 0.68 | 0.81 | 0.55 | 0.49 |
| 10 | 125551436 | rs1219719 | 0.617 | 2.716 | -1.332 | -0.155 | -0.026 | -0.011 | -0.011 | 0 | 0.85 | 0.9 | 0.8 | 0.82 | 0.77 | 0.81 |
| 10 | 125551931 | rs2162536 | 0.472 | 1.525 | -1.581 | -0.676 | 0.01 | 0.01 | 0.01 | 0 | 0.58 | 0.72 | 0.65 | 0.62 | 0.61 | 0.59 |
| 10 | 125556374 | rs9651511 | 0.489 | 1.882 | -2.012 | -0.674 | -0.121 | 0.002 | 0.002 | 1.168 | 0.76 | 0.94 | 0.76 | 0.88 | 0.58 | 0.74 |
| 10 | 125556955 | rs17106063 | -0.125 | 1.008 | -0.837 | -0.698 | 0.016 | -0.047 | -0.047 | 0 | 0.17 | 0.49 | 0.13 | 0.08 | 0.16 | 0.11 |
| 10 | 125561547 | rs1991398 | 0.585 | 0.172 | -1.541 | -0.142 | 0.021 | -0.04 | -0.04 | 0 | 0.19 | 0.12 | 0.17 | 0.31 | 0.13 | 0.17 |
| 10 | 125567936 | rs11248286 | -0.128 | -0.084 | -1.307 | 0.025 | -0.003 | -0.033 | -0.033 | 0.022 | 0.21 | 0.13 | 0.43 | 0.44 | 0.2 | 0.32 |
| 10 | 125571743 | rs7909601 | -1.099 | -0.565 | -1.326 | -0.177 | 0.011 | 0.251 | 0.251 | 0 | 0.52 | 0.82 | 0.69 | 0.85 | 0.42 | 0.58 |
| 10 | 125576927 | rs12218449 | -0.146 | 0.019 | -1.083 | 0.121 | -0.003 | -0.033 | -0.033 | 0 | 0.21 | 0.13 | 0.43 | 0.44 | 0.2 | 0.3 |
| 10 | 125584893 | rs1219707 | 0.406 | -0.451 | -0.973 | 0.236 | 0.021 | -0.013 | -0.013 | 0 | 0.66 | 0.27 | 0.73 | 0.57 | 0.77 | 0.69 |
| 10 | 125585750 | rs1219703 | 0.406 | -0.451 | -0.973 | 0.236 | 0.021 | -0.013 | -0.013 | 1.847 | 0.66 | 0.27 | 0.73 | 0.57 | 0.78 | 0.69 |
| 10 | 125594966 | rs7071884 | 0.186 | -0.849 | -0.744 | -0.248 | -0.008 | -0.028 | -0.028 | 0 | 0.17 | 0.2 | 0.1 | 0.03 | 0.11 | 0.11 |
| 10 | 125600308 | rs2176487 | 0.732 | -1.279 | -0.363 | -0.213 | 0.01 | -0.012 | -0.012 | 0 | 0.19 | 0.19 | 0.15 | 0.33 | 0.1 | 0.15 |
| 10 | 125600481 | rs2138919 | -0.943 | -1.43 | 0.512 | -0.307 | -0.032 | -0.003 | -0.003 | 0 | 0.4 | 0.53 | 0.26 | 0.41 | 0.22 | 0.33 |
| 10 | 125601307 | rs17106110 | 1.575 | -1.142 | -0.085 | -0.397 | -0.067 | -0.119 | -0.119 | 0 | 0.15 | 0.02 | 0.41 | 0.4 | 0.2 | 0.24 |
| 10 | 125601457 | rs7924189 | 0.329 | -1.586 | 0.187 | -0.644 | 0.002 | 0.03 | 0.03 | 0 | 0.16 | 0.07 | 0.08 | 0.03 | 0.12 | 0.12 |
| 10 | 125601681 | rs7923509 | -1.566 | -1.602 | 0.456 | -0.646 | -0.003 | 0 | 0 | 0 | 0.4 | 0.44 | 0.27 | 0.41 | 0.24 | 0.33 |
| 10 | 125602021 | rs10794571 | -0.88 | -1.652 | 0.433 | -0.9 | 0.023 | -0.001 | -0.001 | 0 | 0.35 | 0.32 | 0.22 | 0.37 | 0.19 | 0.25 |
| 10 | 125602943 | rs6599625 | -1.418 | -1.331 | 0.325 | -0.755 | -0.017 | 0 | 0 | 0 | 0.38 | 0.59 | 0.27 | 0.41 | 0.23 | 0.33 |
| 10 | 125615869 | rs876814 | -0.891 | -0.521 | 0.543 | -1.316 | -0.019 | -0.103 | -0.103 | 0 | 0.23 | 0.45 | 0.16 | 0.08 | 0.18 | 0.2 |
| 10 | 125625215 | rs28634188 | -0.348 | 0.207 | 0.05 | -1.539 | 0 | -0.007 | -0.007 | 0.091 | 0.64 | 0.62 | 0.44 | 0.11 | 0.66 | 0.45 |
| 10 | 125637766 | rs7070153 | -0.118 | 0.555 | 0.144 | -1.156 | 0.002 | -0.065 | -0.065 | 0.903 | 0.13 | 0.22 | 0.12 | 0 | 0.13 | 0.1 |
| 10 | 125640380 | rs7909120 | -0.536 | 0.752 | -0.252 | -1.426 | -0.013 | -0.079 | -0.079 | 0.003 | 0.38 | 0.47 | 0.26 | 0.1 | 0.3 | 0.29 |
| 10 | 125649549 | rs28627761 | -0.723 | 1.762 | -0.588 | -0.973 | 0.006 | -0.05 | -0.05 | 0 | 0.2 | 0.18 | 0.2 | 0.01 | 0.27 | 0.19 |
| 10 | 125655550 | rs28687195 | -1.155 | 1.525 | -0.72 | 0.04 | -0.01 | -0.011 | -0.011 | 2.332 | 0.1 | 0.05 | 0.29 | 0.17 | 0.15 | 0.16 |
| 10 | 125673547 | rs4329614 | 0.153 | 0.922 | -0.61 | -0.197 | 0.009 | -0.026 | -0.026 | 0 | 0.65 | 0.59 | 0.55 | 0.24 | 0.65 | 0.46 |
| 10 | 125681274 | rs1914533 | -0.147 | 0.254 | -0.975 | 0.227 | 0.022 | -0.014 | -0.014 | 0 | 0.31 | 0.18 | 0.33 | 0.58 | 0.24 | 0.31 |
| 10 | 125683133 | rs2362924 | -0.177 | 0.61 | -0.461 | -0.521 | 0.003 | -0.088 | -0.088 | 0 | 0.73 | 0.58 | 0.74 | 0.89 | 0.75 | 0.75 |
| 10 | 125684849 | rs9794150 | -0.109 | 0.604 | -0.601 | -0.664 | 0.002 | -0.03 | -0.03 | 0 | 0.25 | 0.27 | 0.21 | 0.06 | 0.26 | 0.17 |
| 10 | 125690164 | rs9702289 | -0.586 | 0.992 | -1.352 | -0.242 | -0.041 | 0.002 | 0.002 | 0 | 0.83 | 0.94 | 0.85 | 0.89 | 0.74 | 0.89 |
| 10 | 125698169 | rs11511761 | 0.466 | 0.166 | -1.568 | 0.391 | 0.041 | -0.071 | -0.071 | 0.493 | 0.37 | 0.14 | 0.4 | 0.57 | 0.3 | 0.53 |
| 10 | 125703102 | rs7071108 | 0.055 | 0.108 | -0.531 | 0.242 | 0.012 | 0.1 | 0.1 | 0.966 | 0.12 | 0.43 | 0.1 | 0.12 | 0.08 | 0.02 |
| 10 | 125703263 | rs7070936 | 0.105 | -1.024 | -1.044 | 0.025 | -0.086 | -0.017 | -0.017 | 0 | 0.3 | 0.45 | 0.13 | 0.21 | 0.14 | 0.12 |
| 10 | 125709109 | rs7096722 | -0.363 | -0.118 | -1.336 | -0.262 | 0 | -0.013 | -0.013 | 0.264 | 0.47 | 0.56 | 0.61 | 0.77 | 0.49 | 0.72 |
| 10 | 125711882 | rs7087529 | -0.27 | 0.027 | -1.24 | 0.374 | -0.002 | -0.003 | -0.003 | 0 | 0.35 | 0.33 | 0.44 | 0.59 | 0.4 | 0.46 |
| 10 | 125719813 | rs6588743 | 0.419 | 0.152 | -1.358 | 0.912 | -0.032 | -0.094 | -0.094 | 0 | 0.14 | 0.01 | 0.13 | 0.3 | 0.17 | 0.26 |
| 10 | 125725603 | rs4075911 | -0.767 | -0.14 | -0.755 | 1.041 | -0.017 | 0.082 | 0.082 | 0 | 0.2 | 0.32 | 0.21 | 0.47 | 0.14 | 0.39 |
| 10 | 125725809 | rs4460752 | -0.039 | -0.175 | -0.627 | 0.604 | -0.004 | 0.062 | 0.062 | 1.442 | 0.62 | 0.78 | 0.65 | 0.85 | 0.64 | 0.7 |
| 10 | 125726028 | rs6588741 | -0.303 | -0.144 | -0.272 | 0.617 | -0.001 | 0.009 | 0.009 | 0 | 0.67 | 0.41 | 0.73 | 0.66 | 0.67 | 0.65 |
| 10 | 125727175 | rs7074389 | 0.055 | 0.101 | -0.314 | 0.713 | 0 | 0 | 0 | 1.959 | 0.56 | 0.59 | 0.6 | 0.46 | 0.59 | 0.5 |
| 10 | 125734152 | rs28759196 | -0.082 | -0.026 | -0.481 | 0.729 | 0.012 | -0.02 | -0.02 | 2.199 | 0.57 | 0.74 | 0.62 | 0.56 | 0.62 | 0.51 |

| | | | | | | | | | | | | | | | | |
|---|---|---|---|---|---|---|---|---|---|---|---|---|---|---|---|---|
| 10 | 125741845 | rs7394262 | 0.09 | 0.018 | -0.624 | 0.579 | 0.012 | 0.006 | 0.006 | 0.326 | 0.56 | 0.63 | 0.61 | 0.46 | 0.62 | 0.51 |
| 10 | 125750120 | rs4929810 | -0.168 | -0.595 | -0.722 | 0.439 | 0.002 | -0.009 | -0.009 | 0 | 0.7 | 0.72 | 0.82 | 0.61 | 0.76 | 0.59 |
| 10 | 125750525 | rs9422277 | -0.101 | -0.263 | -1.22 | -0.001 | -0.018 | -0.01 | -0.01 | 0 | 0.56 | 0.71 | 0.65 | 0.57 | 0.54 | 0.56 |
| 10 | 125757321 | rs28619954 | 0.12 | -0.648 | -0.933 | -0.147 | -0.015 | 0.056 | 0.056 | 0 | 0.19 | 0.28 | 0.25 | 0.51 | 0.15 | 0.27 |
| 10 | 125761439 | rs4590807 | 0.269 | -0.634 | -1.012 | -0.386 | -0.002 | 0.009 | 0.009 | 0 | 0.52 | 0.4 | 0.45 | 0.4 | 0.5 | 0.55 |
| 10 | 125764026 | rs4929801 | 0.677 | -0.586 | -0.775 | -0.214 | -0.012 | -0.044 | -0.044 | 8.256 | 0.31 | 0.38 | 0.21 | 0.12 | 0.27 | 0.2 |
| 10 | 125765973 | rs4590806 | -1.269 | -1.14 | -1.029 | -0.194 | -0.003 | -0.015 | -0.015 | 0 | 0.69 | 0.74 | 0.68 | 0.92 | 0.62 | 0.79 |
| 10 | 125768220 | rs10899 | -0.006 | -0.843 | -0.901 | -0.026 | -0.004 | -0.013 | -0.013 | 0 | 0.38 | 0.31 | 0.39 | 0.44 | 0.43 | 0.43 |
| 10 | 125771433 | rs7897629 | 0.441 | -0.615 | -0.655 | 0.007 | 0.003 | 0.011 | 0.011 | 0 | 0.25 | 0.29 | 0.27 | 0.33 | 0.27 | 0.25 |
| 10 | 125771517 | rs6588759 | 0.409 | -0.591 | -0.627 | 0.017 | 0.004 | 0.014 | 0.014 | 0 | 0.25 | 0.31 | 0.28 | 0.33 | 0.27 | 0.25 |
| 10 | 125774483 | rs7915736 | 0.204 | 0.565 | -0.555 | 0.536 | 0.064 | 0.092 | 0.092 | 0 | 0.12 | 0.23 | 0.25 | 0.33 | 0.27 | 0.24 |
| 10 | 125778225 | rs7901504 | 0.251 | 0.575 | -0.566 | 0.393 | 0.073 | 0.066 | 0.066 | 0 | 0.12 | 0.22 | 0.25 | 0.25 | 0.27 | 0.24 |
| 10 | 125780360 | rs4072197 | -0.766 | 0.511 | -1.388 | -0.104 | 0.011 | 0.02 | 0.02 | 0 | 0.52 | 0.67 | 0.64 | 0.62 | 0.59 | 0.65 |
| 10 | 125786695 | rs9422267 | 0.449 | -0.054 | -1.452 | -0.154 | -0.014 | 0.05 | 0.05 | 0.15 | 0.43 | 0.26 | 0.37 | 0.01 | 0.52 | 0.33 |
| 10 | 125788094 | rs4472872 | -0.85 | 0.273 | -1.126 | 1.474 | -0.001 | -0.036 | -0.036 | 0.049 | 0.22 | 0.19 | 0.33 | 0.44 | 0.24 | 0.29 |
| 10 | 125803303 | rs7085454 | -0.704 | -0.091 | -1.181 | 1.161 | 0.051 | -0.037 | -0.037 | 0 | 0.35 | 0.83 | 0.42 | 0.53 | 0.26 | 0.32 |
| 10 | 125803672 | rs12778619 | -0.539 | -0.191 | -0.609 | 1.654 | 0.013 | -0.003 | -0.003 | 0 | 0.24 | 0.13 | 0.2 | 0.25 | 0.22 | 0.28 |
| 10 | 125824058 | rs6588752 | 0.604 | 0.265 | -0.709 | 0.747 | 0.001 | 0.049 | 0.049 | 0 | 0.33 | 0.18 | 0.22 | 0.08 | 0.3 | 0.31 |
| 10 | 125824450 | rs4929821 | -0.859 | 0.657 | -0.473 | 0.964 | -0.022 | 0.04 | 0.04 | 0 | 0.65 | 0.82 | 0.7 | 0.73 | 0.61 | 0.82 |
| 10 | 125827502 | rs4073218 | -0.75 | 0.649 | 0.185 | 0.949 | 0.002 | -0.012 | -0.012 | 0 | 0.63 | 0.58 | 0.68 | 0.73 | 0.6 | 0.82 |
| 10 | 125828819 | rs4929820 | -0.889 | 0.719 | 0.084 | 0.948 | -0.005 | 0.029 | 0.029 | 0 | 0.65 | 0.75 | 0.7 | 0.73 | 0.62 | 0.81 |
| 10 | 125840222 | rs4397784 | 0.589 | 0.782 | -0.301 | 1.202 | -0.054 | -0.074 | -0.074 | 0 | 0.17 | 0.01 | 0.15 | 0.2 | 0.17 | 0.27 |
| 10 | 125842025 | rs4929816 | -0.235 | 0.619 | -0.075 | 1.16 | -0.036 | -0.038 | -0.038 | 0 | 0.25 | 0.09 | 0.26 | 0.33 | 0.3 | 0.41 |
| 10 | 125849425 | rs4397783 | -0.679 | 0.724 | 0.19 | 1.085 | 0.003 | 0.004 | 0.004 | 0 | 0.63 | 0.66 | 0.64 | 0.63 | 0.64 | 0.71 |
| 10 | 125856308 | rs7909900 | -0.317 | 0.898 | -0.15 | 1.014 | -0.004 | -0.008 | -0.008 | 0.105 | 0.27 | 0.26 | 0.31 | 0.41 | 0.3 | 0.5 |
| 10 | 125863117 | rs7078763 | -0.344 | 0.891 | -0.345 | 0.714 | 0.005 | 0.011 | 0.011 | 0 | 0.67 | 0.74 | 0.71 | 0.66 | 0.69 | 0.82 |
| 10 | 125868602 | rs7078438 | -0.231 | 0.934 | -0.746 | 0.677 | -0.01 | -0.04 | -0.04 | 0 | 0.66 | 0.81 | 0.68 | 0.54 | 0.65 | 0.74 |
| 10 | 125868860 | rs28689800 | -1.036 | 1.311 | -0.153 | 0.149 | 0.062 | -0.104 | -0.104 | 1.06 | 0.21 | 0.39 | 0.26 | 0.04 | 0.32 | 0.27 |
| 10 | 125921423 | rs12266204 | -0.049 | 0.329 | -1.373 | 0.402 | 0.044 | 0.034 | 0.034 | 2.063 | 0.47 | 0.16 | 0.29 | 0.39 | 0.39 | 0.37 |
| 10 | 125921909 | rs10901642 | -0.374 | 0.426 | -0.492 | 0.399 | 0.006 | -0.004 | -0.004 | 0 | 0.64 | 0.57 | 0.52 | 0.68 | 0.59 | 0.7 |
| 10 | 125921963 | rs11245116 | -0.054 | -0.321 | -0.775 | 0.304 | 0.005 | 0.008 | 0.008 | 0 | 0.31 | 0.31 | 0.34 | 0.46 | 0.4 | 0.26 |
| 10 | 125922133 | rs11592850 | 0.899 | -0.554 | -0.783 | 0.34 | 0.091 | 0.071 | 0.071 | 0 | 0.28 | 0.09 | 0.1 | 0.14 | 0.13 | 0.3 |
| 10 | 125923400 | rs11593516 | -0.507 | -0.575 | -0.523 | 0.258 | 0.001 | -0.017 | -0.017 | 0.001 | 0.07 | 0.11 | 0.08 | 0.01 | 0.07 | 0.14 |
| 10 | 125923664 | rs4578341 | -0.093 | -0.873 | -0.211 | 0.507 | -0.027 | 0.002 | 0.002 | 0 | 0.4 | 0.46 | 0.22 | 0.42 | 0.24 | 0.49 |
| 10 | 125924051 | rs3884528 | -1.027 | -0.693 | -0.206 | 0.393 | -0.004 | -0.001 | -0.001 | 0 | 0.63 | 0.66 | 0.51 | 0.86 | 0.55 | 0.71 |
| 10 | 125924595 | rs7905355 | -0.22 | -0.525 | -0.205 | 0.709 | -0.007 | -0.01 | -0.01 | 2.4 | 0.28 | 0.25 | 0.33 | 0.63 | 0.36 | 0.36 |
| 10 | 125925273 | rs11245373 | 0.356 | -0.836 | -0.135 | 0.76 | -0.003 | 0.016 | 0.016 | 0 | 0.05 | 0.06 | 0.02 | 0.18 | 0.04 | 0.08 |
| 10 | 125932457 | rs7907696 | 0.39 | -1.219 | 0.215 | 0.604 | -0.028 | -0.002 | -0.002 | 0 | 0.34 | 0.38 | 0.11 | 0.32 | 0.13 | 0.41 |
| 10 | 125933075 | rs7922403 | 1.255 | -1.241 | 0.142 | 0.551 | 0.076 | 0.049 | 0.049 | 0.14 | 0.27 | 0.14 | 0.05 | 0.12 | 0.08 | 0.24 |
| 10 | 125934405 | rs3824767 | -0.967 | -1.073 | 0.411 | 0.559 | -0.01 | 0.015 | 0.015 | 0 | 0.64 | 0.69 | 0.51 | 0.85 | 0.55 | 0.68 |
| 10 | 125934816 | rs3824766 | 0.901 | -1.024 | 0.361 | 0.576 | 0.046 | 0.021 | 0.021 | 0 | 0.31 | 0.24 | 0.12 | 0.21 | 0.14 | 0.28 |
| 10 | 125936152 | rs11244691 | 0.491 | -0.918 | 0.197 | 0.694 | -0.057 | -0.057 | -0.057 | 0.039 | 0.25 | 0.1 | 0.33 | 0.39 | 0.35 | 0.23 |
| 10 | 125936341 | rs10901467 | -0.072 | -0.805 | 0.494 | 0.73 | -0.055 | 0.004 | 0.004 | 0 | 0.38 | 0.52 | 0.17 | 0.41 | 0.18 | 0.46 |
| 10 | 125937820 | rs11244794 | 0.637 | -1.046 | 0.223 | 0.253 | -0.041 | -0.037 | -0.037 | 1.502 | 0.29 | 0.33 | 0.09 | 0.1 | 0.09 | 0.21 |
| 10 | 125944772 | rs6597799 | 0.293 | -0.56 | -0.119 | 0.517 | 0.032 | 0.027 | 0.027 | 0.341 | 0.41 | 0.35 | 0.15 | 0.24 | 0.2 | 0.48 |
| 10 | 125946185 | rs7082886 | -0.514 | 0.448 | -0.373 | 0.933 | -0.011 | -0.021 | -0.021 | 0 | 0.29 | 0.23 | 0.36 | 0.46 | 0.4 | 0.26 |
| 10 | 125954944 | rs7100068 | -0.607 | 0.367 | -0.327 | 0.703 | 0.01 | 0.002 | 0.002 | 0 | 0.36 | 0.6 | 0.46 | 0.61 | 0.44 | 0.42 |
| 10 | 125956348 | rs11245310 | -0.958 | 0.115 | -1.122 | 0.749 | -0.002 | -0.096 | -0.096 | 0 | 0.24 | 0.09 | 0.23 | 0.57 | 0.26 | 0.31 |
| 10 | 125956605 | rs4411244 | 0.786 | -0.657 | -2.207 | -0.042 | 0.017 | 0.049 | 0.049 | 0 | 0.32 | 0.05 | 0.33 | 0.19 | 0.22 | 0.24 |
| 10 | 125959231 | rs7900187 | -0.487 | -0.165 | -1.113 | -0.221 | 0.022 | 0.009 | 0.009 | 0.429 | 0.78 | 0.65 | 0.76 | 0.76 | 0.71 | 0.74 |
| 10 | 125961498 | rs4962394 | -0.405 | -0.393 | -1.507 | -0.444 | -0.011 | -0.067 | -0.067 | 0.322 | 0.33 | 0.09 | 0.45 | 0.62 | 0.33 | 0.38 |
| 10 | 125967161 | rs7921749 | -0.415 | -0.329 | -1.495 | -0.459 | 0 | -0.058 | -0.058 | 0 | 0.33 | 0.13 | 0.45 | 0.63 | 0.32 | 0.38 |
| 10 | 125967560 | rs7088629 | -1.152 | -0.525 | -1.569 | -0.523 | -0.087 | -0.009 | -0.009 | 2.294 | 0.82 | 0.97 | 0.8 | 0.86 | 0.71 | 0.74 |
| 10 | 125970511 | rs11591371 | -0.267 | -0.639 | -0.926 | -0.557 | 0.003 | 0.001 | 0.001 | 1.292 | 0.32 | 0.21 | 0.44 | 0.72 | 0.32 | 0.39 |
| 10 | 125973009 | rs7900964 | -1.056 | -0.657 | -0.904 | -0.471 | -0.087 | 0.061 | 0.061 | 0.296 | 0.82 | 0.96 | 0.81 | 0.9 | 0.71 | 0.84 |
| 10 | 125975658 | rs4336987 | 1.468 | -1.173 | -0.562 | -0.734 | -0.004 | 0.059 | 0.059 | 0 | 0.34 | 0.16 | 0.24 | 0.14 | 0.33 | 0.23 |
| 10 | 125977968 | rs10901826 | 1.36 | -1.342 | -0.038 | -0.773 | -0.001 | -0.006 | -0.006 | 0.01 | 0.35 | 0.32 | 0.27 | 0.17 | 0.33 | 0.32 |
| 10 | 125980099 | rs7919469 | 1.066 | -1.261 | 0.374 | -0.795 | -0.005 | -0.038 | -0.038 | 0 | 0.38 | 0.5 | 0.29 | 0.14 | 0.33 | 0.23 |
| 10 | 125980308 | rs10794182 | 0.674 | -1.247 | 0.396 | -0.676 | -0.004 | -0.018 | -0.018 | 0 | 0.43 | 0.5 | 0.3 | 0.22 | 0.37 | 0.3 |
| 10 | 125992472 | rs11598366 | 0.252 | -1.519 | 0.131 | 0.012 | -0.011 | -0.1 | -0.1 | 0.002 | 0.21 | 0.03 | 0.16 | 0.37 | 0.19 | 0.26 |
| 10 | 125992726 | rs7098566 | -0.587 | -1.42 | 0.221 | -0.091 | -0.023 | 0.035 | 0.035 | 0 | 0.74 | 0.83 | 0.76 | 0.87 | 0.66 | 0.76 |
| 10 | 125995771 | rs7082638 | -0.613 | -1.486 | 0.23 | 0.113 | -0.015 | 0.016 | 0.016 | 0.044 | 0.34 | 0.39 | 0.43 | 0.64 | 0.28 | 0.39 |
| 10 | 125996619 | rs4293082 | 1.654 | -1.431 | 0.045 | -0.08 | -0.009 | 0.052 | 0.052 | 0 | 0.34 | 0.18 | 0.26 | 0.13 | 0.34 | 0.23 |
| 10 | 125996970 | rs12411939 | 1.781 | -1.429 | 0.03 | -0.076 | -0.013 | 0.058 | 0.058 | 0 | 0.34 | 0.16 | 0.26 | 0.13 | 0.34 | 0.23 |
| 10 | 125997171 | rs4505015 | -0.825 | -1.289 | 0.021 | -0.025 | -0.047 | 0.041 | 0.041 | 0 | 0.76 | 0.88 | 0.76 | 0.87 | 0.66 | 0.76 |
| 10 | 126003837 | rs11245493 | -1.161 | -1.324 | 0.219 | 0.121 | 0.03 | 0.035 | 0.035 | 0 | 0.43 | 0.7 | 0.53 | 0.74 | 0.36 | 0.49 |
| 10 | 126007164 | rs6597881 | -0.838 | -1.306 | 0.034 | 0.081 | -0.033 | 0.033 | 0.033 | 0 | 0.76 | 0.87 | 0.78 | 0.86 | 0.7 | 0.75 |
| 10 | 126007929 | rs7081107 | 0.395 | -1.149 | 0.146 | 0.169 | 0.004 | 0.013 | 0.013 | 0 | 0.06 | 0.22 | 0.07 | 0.04 | 0.03 | 0.06 |
| 10 | 126008767 | rs11245507 | 1.931 | -1.053 | -0.282 | 0.182 | -0.103 | 0.057 | 0.057 | 0 | 0.31 | 0.02 | 0.22 | 0.13 | 0.34 | 0.25 |

| | | | | | | | | | | | | | | | | |
|---|---|---|---|---|---|---|---|---|---|---|---|---|---|---|---|---|
| 10 | 126010920 | rs11594231 | 0.163 | -1.071 | 0.017 | 0.362 | 0 | -0.004 | -0.004 | 0 | 0.23 | 0.23 | 0.21 | 0.4 | 0.23 | 0.21 |
| 10 | 126012384 | rs7917787 | -0.767 | -0.915 | 0.235 | 0.353 | 0.002 | 0.004 | 0.004 | 0 | 0.44 | 0.72 | 0.56 | 0.47 | 0.43 | 0.49 |
| 10 | 126014762 | rs6597885 | -1.844 | -1.098 | 0.24 | 0.137 | -0.007 | 0.041 | 0.041 | 0 | 0.65 | 0.75 | 0.72 | 0.89 | 0.63 | 0.72 |
| 10 | 126014869 | rs6597886 | -0.271 | -1.134 | 0.448 | 0.157 | -0.003 | -0.021 | -0.021 | 0.971 | 0.38 | 0.44 | 0.3 | 0.2 | 0.34 | 0.39 |
| 10 | 126016549 | rs4962732 | -0.777 | -1.143 | 0.225 | 0.207 | 0.006 | 0.016 | 0.016 | 0 | 0.43 | 0.72 | 0.53 | 0.43 | 0.4 | 0.49 |
| 10 | 126018115 | rs2807068 | -2.067 | -1.121 | 0.197 | 0.211 | -0.02 | 0.068 | 0.068 | 0 | 0.66 | 0.82 | 0.74 | 0.89 | 0.64 | 0.74 |
| 10 | 126020788 | rs994844 | -0.227 | -1.087 | 0.567 | 0.738 | 0.035 | 0.14 | 0.14 | 0 | 0.06 | 0.23 | 0.22 | 0.23 | 0.06 | 0.1 |
| 10 | 126025035 | rs2674353 | -0.025 | -1.101 | -0.19 | 0.736 | 0.004 | -0.073 | -0.073 | 0 | 0.76 | 0.87 | 0.79 | 0.54 | 0.76 | 0.74 |
| 10 | 126031474 | rs7911910 | -1.13 | -0.875 | 0.237 | 0.567 | -0.027 | 0.007 | 0.007 | 0.11 | 0.21 | 0.68 | 0.16 | 0.28 | 0.12 | 0.22 |
| 10 | 126031511 | rs7915136 | -1.11 | -0.863 | 0.247 | 0.604 | -0.028 | 0.011 | 0.011 | 0.09 | 0.2 | 0.68 | 0.16 | 0.3 | 0.12 | 0.22 |
| 10 | 126036778 | rs11244450 | -0.43 | -0.913 | 0.318 | 0.424 | -0.022 | -0.01 | -0.01 | 0 | 0.16 | 0.21 | 0.18 | 0.08 | 0.1 | 0.18 |
| 10 | 126037221 | rs885042 | -0.186 | -0.931 | 0.054 | 0.319 | 0.001 | 0.004 | 0.004 | 0 | 0.48 | 0.56 | 0.53 | 0.4 | 0.48 | 0.47 |
| 10 | 126037442 | rs2998194 | 0.837 | -0.995 | -0.728 | 0.155 | -0.119 | -0.017 | -0.017 | 0 | 0.4 | 0.1 | 0.39 | 0.33 | 0.45 | 0.33 |
| 10 | 126037889 | rs7910801 | 0.23 | -0.711 | 0.027 | 0.218 | 0.004 | -0.023 | -0.023 | 2.101 | 0.07 | 0.19 | 0.05 | 0.02 | 0.05 | 0.07 |
| 10 | 126046479 | rs10794004 | -0.875 | -0.508 | 0.209 | 0.689 | -0.004 | 0.1 | 0.1 | 0.015 | 0.18 | 0.45 | 0.25 | 0.26 | 0.15 | 0.17 |
| 10 | 126047795 | rs2674326 | -0.771 | -0.764 | -0.17 | 0.609 | -0.099 | -0.088 | -0.088 | 0 | 0.6 | 0.92 | 0.62 | 0.52 | 0.53 | 0.56 |
| 10 | 126047857 | rs7068637 | 1.09 | -0.672 | 0.117 | 0.698 | -0.02 | 0.064 | 0.064 | 0.626 | 0.1 | 0.31 | 0.16 | 0.15 | 0.05 | 0.14 |
| 10 | 126049426 | rs2674322 | -0.376 | -0.639 | -0.125 | 0.682 | -0.024 | 0.004 | 0.004 | 0 | 0.53 | 0.77 | 0.58 | 0.51 | 0.48 | 0.5 |
| 10 | 126051366 | rs867907 | -0.337 | -0.63 | -0.151 | 0.589 | -0.023 | -0.018 | -0.018 | 3.023 | 0.53 | 0.76 | 0.56 | 0.44 | 0.48 | 0.49 |
| 10 | 126052440 | rs867997 | 1.35 | -0.226 | -0.482 | 0.354 | 0.002 | 0.022 | 0.022 | 0 | 0.3 | 0.18 | 0.29 | 0.15 | 0.28 | 0.29 |
| 10 | 126053047 | rs1105778 | 0.899 | 0.863 | -0.241 | 0.541 | -0.081 | 0.004 | 0.004 | 0.08 | 0.13 | 0.01 | 0.16 | 0.09 | 0.24 | 0.14 |
| 10 | 126053705 | rs17151852 | -1.163 | 0.64 | 0.033 | 0.581 | 0.041 | 0.003 | 0.003 | 0 | 0.1 | 0.27 | 0.09 | 0.04 | 0.1 | 0.04 |
| 10 | 126054225 | rs2674328 | 0.692 | 0.577 | -0.489 | 0.715 | 0.017 | 0.004 | 0.004 | 0 | 0.61 | 0.5 | 0.62 | 0.61 | 0.49 | 0.67 |
| 10 | 126057541 | rs7902179 | -0.838 | 0.423 | -0.656 | 0.79 | 0.002 | 0.019 | 0.019 | 0 | 0.09 | 0.16 | 0.07 | 0.09 | 0.08 | 0.05 |
| 10 | 126057825 | rs2459219 | -0.42 | 0.113 | -1.481 | 0.425 | 0.029 | -0.023 | -0.023 | 0.345 | 0.41 | 0.72 | 0.53 | 0.56 | 0.39 | 0.44 |
| 10 | 126058904 | rs17151594 | -0.392 | 0.442 | -0.595 | -0.587 | 0.001 | -0.063 | -0.063 | 0 | 0.14 | 0.16 | 0.07 | 0 | 0.13 | 0.12 |
| 10 | 126059884 | rs11244591 | -0.381 | 0.675 | -0.831 | -0.337 | -0.036 | -0.016 | -0.016 | 0 | 0.07 | 0 | 0.04 | 0.03 | 0.09 | 0.06 |
| 10 | 126062722 | rs10901414 | 0.549 | 0.549 | -0.909 | -0.353 | 0.004 | -0.004 | -0.004 | 0.392 | 0.13 | 0.11 | 0.05 | 0.21 | 0.08 | 0.12 |
| 10 | 126062763 | rs1040151 | -1.133 | 0.224 | -0.447 | -0.581 | -0.007 | -0.032 | -0.032 | 0 | 0.22 | 0.33 | 0.12 | 0.09 | 0.21 | 0.17 |
| 10 | 126064956 | rs11244602 | 0.461 | 0.562 | -0.395 | -0.281 | -0.002 | 0 | 0 | 0 | 0.14 | 0.14 | 0.05 | 0.2 | 0.08 | 0.12 |
| 10 | 126065943 | rs10794015 | -0.483 | 0.563 | -0.349 | -0.334 | 0.005 | 0.024 | 0.024 | 0.087 | 0.19 | 0.36 | 0.11 | 0.23 | 0.17 | 0.17 |
| 10 | 126074233 | rs870210 | -0.4 | 1.175 | -0.805 | -0.428 | 0.013 | -0.01 | -0.01 | 1.481 | 0.35 | 0.39 | 0.24 | 0.25 | 0.39 | 0.32 |
| 10 | 126078126 | rs2674346 | -1.29 | 1.403 | -1.555 | -0.592 | -0.015 | 0.026 | 0.026 | 0 | 0.87 | 0.93 | 0.83 | 0.97 | 0.81 | 0.92 |
| 10 | 126085115 | rs10430741 | -0.047 | 1.719 | -1.313 | -0.39 | -0.008 | 0.021 | 0.021 | 0 | 0.31 | 0.26 | 0.2 | 0.24 | 0.3 | 0.3 |
| 10 | 126086326 | rs2459216 | 0.431 | 1.465 | -1.261 | -0.445 | -0.029 | -0.017 | -0.017 | 0 | 0.11 | 0.22 | 0.06 | 0.05 | 0.07 | 0.11 |
| 10 | 126090430 | rs17619406 | 0.57 | 1.875 | -2.037 | -0.323 | -0.062 | 0.028 | 0.028 | 0 | 0.11 | 0.01 | 0.09 | 0.05 | 0.2 | 0.08 |
| 10 | 126100523 | rs10794058 | -0.383 | 1.067 | -1.486 | 0.053 | -0.008 | 0.02 | 0.02 | 0 | 0.31 | 0.25 | 0.2 | 0.22 | 0.29 | 0.3 |
| 10 | 126128552 | rs1710563 | -0.632 | 0.551 | -1.303 | 0.455 | -0.003 | 0 | 0 | 0 | 0.55 | 0.67 | 0.38 | 0.53 | 0.46 | 0.6 |
| 10 | 126135262 | rs1399928 | -1.136 | 0.967 | -1.206 | 0.433 | -0.042 | -0.019 | -0.019 | 0 | 0.12 | 0.02 | 0.08 | 0.1 | 0.17 | 0.16 |
| 10 | 126144231 | rs11244995 | -0.963 | -0.084 | -0.456 | 0.097 | -0.065 | -0.069 | -0.069 | 0 | 0.21 | 0.36 | 0.09 | 0.11 | 0.08 | 0.17 |
| 10 | 126144839 | rs908366 | -0.045 | 0.71 | -1.584 | 0.525 | -0.082 | -0.107 | -0.107 | 0 | 0.25 | 0.04 | 0.28 | 0.39 | 0.29 | 0.23 |
| 10 | 126150148 | rs2459214 | -0.292 | 0.528 | -1.348 | -0.159 | -0.002 | -0.053 | -0.053 | 0.071 | 0.71 | 0.36 | 0.67 | 0.86 | 0.7 | 0.66 |
| 10 | 126152978 | rs10794092 | 0.457 | 0.337 | -1.594 | 0.026 | -0.093 | -0.129 | -0.129 | 0 | 0.28 | 0.05 | 0.28 | 0.45 | 0.32 | 0.28 |
| 10 | 126153362 | rs2807045 | -0.112 | 0.377 | -1.165 | -0.162 | 0 | -0.022 | -0.022 | 0.123 | 0.71 | 0.31 | 0.67 | 0.86 | 0.7 | 0.64 |
| 10 | 126157910 | rs11245061 | -0.547 | 1.356 | -1.04 | 0.636 | -0.004 | 0.005 | 0.005 | 0 | 0.14 | 0.13 | 0.21 | 0.27 | 0.19 | 0.05 |
| 10 | 126163164 | rs11245086 | -0.142 | 1.198 | -1.004 | -0.076 | 0.011 | 0.031 | 0.031 | 0 | 0.33 | 0.43 | 0.36 | 0.62 | 0.35 | 0.33 |
| 10 | 126169373 | rs7070581 | -0.086 | 1.486 | -1.041 | 0.336 | -0.063 | -0.042 | -0.042 | 0 | 0.12 | 0.02 | 0.18 | 0.15 | 0.19 | 0.05 |
| 10 | 126173339 | rs12785110 | -0.767 | 1.53 | -0.302 | 0.196 | 0.089 | 0.014 | 0.014 | 0 | 0.07 | 0.24 | 0.11 | 0.05 | 0.14 | 0.11 |
| 10 | 126173556 | rs2001515 | -0.19 | 1.444 | -0.761 | 0.157 | -0.024 | -0.032 | -0.032 | 0.019 | 0.16 | 0.01 | 0.18 | 0.14 | 0.16 | 0.07 |
| 10 | 126177925 | rs9422989 | -0.874 | 1.366 | -0.115 | -0.288 | 0.032 | -0.034 | -0.034 | 0 | 0.85 | 0.68 | 0.78 | 0.92 | 0.83 | 0.77 |
| 10 | 126178925 | rs11245177 | 0.039 | 1.331 | -0.137 | -0.341 | -0.004 | -0.003 | -0.003 | 0.785 | 0.17 | 0.04 | 0.19 | 0.14 | 0.19 | 0.07 |
| 10 | 126180644 | rs4246200 | 0.221 | 1.2 | 0.075 | 0.122 | -0.007 | 0.003 | 0.003 | 0.322 | 0.73 | 0.45 | 0.71 | 0.78 | 0.76 | 0.59 |
| 10 | 126181006 | rs4962632 | 0.222 | 1.197 | 0.071 | 0.12 | -0.007 | 0.003 | 0.003 | 0.378 | 0.73 | 0.44 | 0.71 | 0.78 | 0.76 | 0.59 |
| 10 | 126181395 | rs4962633 | 1.287 | 1.291 | 0.043 | 0.076 | -0.011 | 0.02 | 0.02 | 0.889 | 0.52 | 0.28 | 0.42 | 0.64 | 0.43 | 0.49 |
| 10 | 126181762 | rs4962635 | -0.894 | 1.567 | 0.505 | 0.342 | 0.026 | -0.031 | -0.031 | 0 | 0.85 | 0.72 | 0.78 | 0.92 | 0.83 | 0.77 |
| 10 | 126182680 | rs2154394 | -1.267 | 1.161 | 1.082 | 0.565 | 0.126 | 0.234 | 0.234 | 0.184 | 0.07 | 0.38 | 0.1 | 0.17 | 0.11 | 0.13 |
| 10 | 126184988 | rs12569498 | 0.304 | 0.707 | 0.422 | 0.387 | 0.011 | 0.008 | 0.008 | 0 | 0.62 | 0.45 | 0.56 | 0.38 | 0.55 | 0.6 |
| 10 | 126185451 | rs10794134 | -0.735 | 0.767 | 0.362 | 0.686 | -0.028 | -0.028 | -0.028 | 0.071 | 0.76 | 0.83 | 0.69 | 0.59 | 0.69 | 0.81 |
| 10 | 126185618 | rs3824810 | -1.107 | 0.924 | 0.456 | 0.742 | -0.013 | -0.013 | -0.013 | 2.427 | 0.78 | 0.82 | 0.68 | 0.59 | 0.69 | 0.82 |
| 10 | 126186771 | rs17620824 | -1.298 | -0.392 | 0.551 | 0.367 | -0.02 | -0.052 | -0.052 | 0 | 0.3 | 0.49 | 0.21 | 0.22 | 0.3 | 0.32 |
| 10 | 126188580 | rs9423014 | 0.134 | -0.404 | 0.746 | 0.326 | -0.018 | -0.056 | -0.056 | 0 | 0.1 | 0.26 | 0.06 | 0 | 0.05 | 0.06 |
| 10 | 126189866 | rs7904991 | -1.369 | -0.599 | 0.77 | 0.894 | -0.038 | 0.111 | 0.111 | 0 | 0.2 | 0.58 | 0.15 | 0.39 | 0.14 | 0.28 |
| 10 | 126194249 | rs11593207 | 0.389 | -0.491 | 0.626 | 0.782 | 0.019 | 0.158 | 0.158 | 0 | 0.09 | 0.28 | 0.08 | 0.3 | 0.08 | 0.16 |
| 10 | 126195145 | rs2604155 | -1.177 | -0.535 | 0.18 | 0.694 | -0.022 | -0.006 | -0.006 | 0 | 0.76 | 0.81 | 0.63 | 0.68 | 0.6 | 0.79 |
| 10 | 126202890 | rs12413427 | 1.925 | -1.246 | -0.204 | 0.34 | 0.034 | 0.066 | 0.066 | 3.796 | 0.34 | 0.17 | 0.26 | 0.14 | 0.24 | 0.15 |
| 10 | 126203813 | rs4501944 | 0.753 | -1.042 | -0.005 | 0.453 | -0.008 | -0.03 | -0.03 | 0 | 0.08 | 0.11 | 0.04 | 0 | 0.05 | 0.05 |
| 10 | 126205537 | rs11245253 | -0.215 | -0.945 | -0.217 | 0.686 | 0.028 | -0.059 | -0.059 | 0.02 | 0.11 | 0.05 | 0.16 | 0.28 | 0.08 | 0.35 |
| 10 | 126207206 | rs10901745 | -0.873 | -0.95 | 0.023 | 0.678 | -0.048 | 0.131 | 0.131 | 0.245 | 0.15 | 0.36 | 0.19 | 0.3 | 0.08 | 0.35 |
| 10 | 126215607 | rs2015483 | -0.597 | -0.648 | -0.627 | 0.292 | -0.001 | -0.038 | -0.038 | 0 | 0.85 | 0.82 | 0.91 | 1 | 0.84 | 0.9 |

| | | | | | | | | | | | | | | | | |
|---|---|---|---|---|---|---|---|---|---|---|---|---|---|---|---|---|
| 10 | 126215674 | rs2045183 | 0.894 | -0.781 | -0.831 | 0.25 | -0.074 | -0.019 | -0.019 | 0 | 0.26 | 0.13 | 0.36 | 0.33 | 0.4 | 0.22 |
| 10 | 126217563 | rs17151975 | -0.101 | -0.808 | -0.828 | 0.254 | 0.001 | 0.03 | 0.03 | 0 | 0.59 | 0.7 | 0.55 | 0.68 | 0.43 | 0.68 |
| 10 | 126219694 | rs11245267 | -0.546 | -0.607 | -0.56 | 0.478 | 0.005 | 0 | 0 | 4.147 | 0.72 | 0.69 | 0.62 | 0.68 | 0.59 | 0.79 |
| 10 | 126221855 | rs4962645 | 1.416 | -0.667 | -0.77 | 0.666 | -0.012 | -0.04 | -0.04 | 0 | 0.07 | 0 | 0.06 | 0.21 | 0.07 | 0.08 |
| 10 | 126222735 | rs4143052 | -1.041 | -0.656 | -0.717 | 0.697 | -0.003 | -0.002 | -0.002 | 0.008 | 0.74 | 0.76 | 0.62 | 0.66 | 0.6 | 0.79 |
| 10 | 126224704 | rs12265012 | 0.794 | -0.64 | -0.747 | 0.703 | -0.066 | -0.02 | -0.02 | 2.714 | 0.25 | 0.12 | 0.36 | 0.32 | 0.4 | 0.22 |
| 10 | 126227537 | rs1869660 | 0.658 | -1.004 | -0.451 | 0.717 | -0.047 | 0.05 | 0.05 | 0 | 0.79 | 0.48 | 0.86 | 0.69 | 0.86 | 0.74 |
| 10 | 126228742 | rs4962379 | 0.223 | -1.05 | -0.736 | 0.845 | 0.009 | -0.016 | -0.016 | 0 | 0.84 | 0.89 | 0.77 | 0.57 | 0.85 | 0.64 |
| 10 | 126229235 | rs12570000 | 0.216 | -0.922 | -0.616 | 0.885 | 0.002 | 0.001 | 0.001 | 0 | 0.09 | 0.07 | 0.17 | 0.22 | 0.07 | 0.29 |
| 10 | 126229876 | rs7916782 | -0.07 | -0.78 | -0.56 | 0.511 | -0.026 | 0.017 | 0.017 | 0.205 | 0.13 | 0.36 | 0.06 | 0.1 | 0.06 | 0.18 |
| 10 | 126230878 | rs7077436 | 0.001 | -0.296 | -1.428 | 0.595 | -0.094 | -0.082 | -0.082 | 0 | 0.33 | 0.12 | 0.52 | 0.56 | 0.47 | 0.51 |
| 10 | 126235677 | rs11245281 | 0.997 | -0.925 | -1.068 | 0.237 | 0.018 | 0.023 | 0.023 | 2.051 | 0.28 | 0.14 | 0.13 | 0.14 | 0.21 | 0.16 |
| 10 | 126236581 | rs12413892 | -0.52 | -1.643 | -0.881 | 0.076 | 0.003 | -0.002 | -0.002 | 0 | 0.39 | 0.59 | 0.33 | 0.41 | 0.3 | 0.42 |
| 10 | 126240819 | rs11245286 | 1.035 | -1.187 | -1.467 | 0.129 | -0.067 | -0.07 | -0.07 | 0 | 0.28 | 0.07 | 0.3 | 0.36 | 0.31 | 0.37 |
| 10 | 126241577 | rs17622092 | 1.205 | -1.285 | -1.401 | 0 | -0.065 | 0.008 | 0.008 | 0 | 0.24 | 0.1 | 0.31 | 0.26 | 0.35 | 0.16 |
| 10 | 126245041 | rs7915771 | -0.131 | -1.539 | -0.702 | 0.081 | 0.017 | 0.008 | 0.008 | 0 | 0.49 | 0.6 | 0.39 | 0.48 | 0.41 | 0.51 |
| 10 | 126245297 | rs12258489 | 0.391 | -1.523 | -0.761 | 0.19 | -0.018 | 0.059 | 0.059 | 0 | 0.09 | 0.18 | 0.18 | 0.28 | 0.07 | 0.19 |
| 10 | 126248200 | rs11245290 | 0.138 | -1.601 | -0.751 | 0.145 | -0.022 | 0.028 | 0.028 | 0 | 0.23 | 0.27 | 0.21 | 0.37 | 0.13 | 0.31 |
| 10 | 126249847 | rs7918137 | -0.346 | -1.409 | -0.754 | 0.081 | 0.003 | 0.009 | 0.009 | 0 | 0.57 | 0.33 | 0.46 | 0.37 | 0.59 | 0.39 |
| 10 | 126253625 | rs10901760 | 0.939 | -1.082 | -0.938 | 0.067 | -0.026 | -0.047 | -0.047 | 0 | 0.14 | 0.07 | 0.24 | 0.35 | 0.17 | 0.27 |
| 10 | 126254468 | rs10901761 | 1.381 | -1.14 | -1.107 | 0.041 | -0.069 | -0.048 | -0.048 | 1.495 | 0.23 | 0.08 | 0.31 | 0.31 | 0.34 | 0.17 |
| 10 | 126254695 | rs12264955 | 0.587 | -1.208 | -1.045 | 0.016 | -0.003 | 0.003 | 0.003 | 0 | 0.09 | 0.13 | 0.17 | 0.2 | 0.07 | 0.22 |
| 10 | 126258162 | rs10794152 | 1.336 | -1.087 | -1.042 | 0.032 | -0.057 | -0.033 | -0.033 | 0 | 0.23 | 0.08 | 0.3 | 0.3 | 0.34 | 0.17 |
| 10 | 126258711 | rs10751594 | -1.322 | -1.437 | -0.663 | -0.185 | -0.02 | -0.021 | -0.021 | 0.243 | 0.68 | 0.42 | 0.84 | 0.86 | 0.81 | 0.86 |
| 10 | 126258772 | rs10751595 | -1.322 | -1.437 | -0.663 | -0.185 | -0.02 | -0.021 | -0.021 | 0 | 0.68 | 0.42 | 0.81 | 0.86 | 0.81 | 0.86 |
| 10 | 126259339 | rs3891517 | 0.897 | -1.303 | -0.376 | -0.028 | 0.017 | 0.075 | 0.075 | 0.255 | 0.15 | 0.24 | 0.25 | 0.35 | 0.18 | 0.29 |
| 10 | 126260223 | rs12269470 | -0.132 | -1.343 | -0.409 | -0.069 | 0.013 | 0.038 | 0.038 | 0 | 0.31 | 0.23 | 0.3 | 0.2 | 0.3 | 0.4 |
| 10 | 126260246 | rs12264692 | -0.793 | -1.123 | -0.517 | -0.061 | 0 | 0 | 0 | 0.077 | 0.47 | 0.48 | 0.55 | 0.55 | 0.48 | 0.7 |
| 10 | 126263119 | rs4962656 | -0.848 | -1.274 | -0.834 | -0.184 | 0.074 | 0.095 | 0.095 | 0 | 0.5 | 0.79 | 0.52 | 0.67 | 0.37 | 0.64 |
| 10 | 126265489 | rs17622597 | 0.724 | -0.698 | -0.297 | -0.137 | 0.007 | -0.031 | -0.031 | 0 | 0.27 | 0.48 | 0.24 | 0.21 | 0.27 | 0.3 |
| 10 | 126265529 | rs10901770 | 0.537 | -1.045 | -0.561 | -0.008 | -0.017 | -0.041 | -0.041 | 1.025 | 0.19 | 0.08 | 0.25 | 0.32 | 0.22 | 0.37 |
| 10 | 126265591 | rs4962660 | -0.306 | -1.198 | -0.438 | 0.062 | 0.006 | -0.004 | -0.004 | 0.104 | 0.48 | 0.68 | 0.55 | 0.45 | 0.45 | 0.44 |
| 10 | 126266790 | rs4962661 | -0.241 | -1.365 | -0.141 | 0.169 | 0 | 0 | 0 | 0 | 0.55 | 0.58 | 0.65 | 0.4 | 0.65 | 0.53 |
| 10 | 126267759 | rs17152064 | 0.644 | -1.223 | -0.304 | 0.19 | 0.007 | -0.002 | -0.002 | 0.265 | 0.09 | 0.04 | 0.04 | 0.11 | 0.05 | 0.25 |
| 10 | 126268260 | rs7077413 | 0.892 | -1.34 | 0.051 | 0.194 | -0.014 | 0.009 | 0.009 | 0 | 0.27 | 0.29 | 0.15 | 0.34 | 0.21 | 0.33 |
| 10 | 126269695 | rs10901773 | -0.54 | -1.404 | 0.062 | 0.117 | 0 | 0 | 0 | 0 | 0.49 | 0.43 | 0.44 | 0.47 | 0.47 | 0.51 |
| 10 | 126270056 | rs4962390 | 1.227 | -1.364 | 0.01 | 0.13 | -0.002 | 0.016 | 0.016 | 0 | 0.26 | 0.31 | 0.3 | 0.36 | 0.25 | 0.16 |
| 10 | 126273079 | rs10751597 | -0.543 | -1.381 | -0.146 | 0.095 | 0.004 | 0.017 | 0.017 | 0 | 0.52 | 0.63 | 0.46 | 0.47 | 0.48 | 0.52 |
| 10 | 126275630 | rs12768101 | 0.261 | -1.415 | -0.013 | 0.111 | 0.009 | 0.017 | 0.017 | 0 | 0.25 | 0.14 | 0.12 | 0.12 | 0.21 | 0.16 |
| 10 | 126275914 | rs12773846 | -0.798 | -1.463 | 0.021 | 0.15 | -0.016 | -0.012 | -0.012 | 0 | 0.38 | 0.43 | 0.19 | 0.24 | 0.27 | 0.44 |
| 10 | 126276587 | rs12572285 | 0.041 | -1.03 | -0.191 | 0.393 | 0.008 | -0.003 | -0.003 | 0 | 0.09 | 0.06 | 0.05 | 0.13 | 0.06 | 0.27 |
| 10 | 126278156 | rs17152066 | 1.522 | -1.024 | 0.162 | 0.354 | -0.017 | 0 | 0 | 0.367 | 0.2 | 0.34 | 0.12 | 0.23 | 0.15 | 0.08 |
| 10 | 126278458 | rs11595762 | -0.43 | -1.274 | 0.022 | 0.212 | 0 | 0 | 0 | 0 | 0.54 | 0.5 | 0.43 | 0.46 | 0.5 | 0.39 |
| 10 | 126278648 | rs10430737 | -0.673 | -1.266 | -0.283 | 0.155 | 0.005 | 0.013 | 0.013 | 0 | 0.56 | 0.62 | 0.64 | 0.68 | 0.58 | 0.61 |
| 10 | 126279181 | rs1124734 | 0.481 | -0.911 | -0.532 | 0.216 | -0.051 | 0.03 | 0.03 | 0 | 0.19 | 0.08 | 0.27 | 0.11 | 0.26 | 0.2 |
| 10 | 126280593 | rs4962664 | -1.155 | -1.476 | -0.356 | 0.102 | -0.005 | 0.002 | 0.002 | 0 | 0.58 | 0.39 | 0.59 | 0.56 | 0.55 | 0.63 |
| 10 | 126284935 | rs7898793 | 1.145 | -1.522 | -0.335 | 0.168 | -0.029 | 0.031 | 0.031 | 0.082 | 0.3 | 0.38 | 0.13 | 0.38 | 0.19 | 0.25 |
| 10 | 126290050 | rs897293 | -0.082 | -1.162 | -0.352 | 0.376 | -0.004 | 0.003 | 0.003 | 0 | 0.63 | 0.63 | 0.49 | 0.68 | 0.56 | 0.54 |
| 10 | 126294167 | rs3740540 | 1.454 | -1.088 | -0.728 | 0.369 | -0.068 | 0.013 | 0.013 | 0 | 0.2 | 0.05 | 0.25 | 0.2 | 0.3 | 0.28 |
| 10 | 126294239 | rs10751600 | -1.051 | -0.969 | -0.667 | 0.125 | -0.006 | -0.005 | -0.005 | 0 | 0.79 | 0.8 | 0.84 | 0.98 | 0.74 | 0.82 |
| 10 | 126294359 | rs10901782 | 0.031 | -0.737 | -0.45 | 0.621 | 0.001 | -0.001 | -0.001 | 0 | 0.72 | 0.71 | 0.71 | 0.71 | 0.76 | 0.67 |
| 10 | 126294480 | rs12777136 | 0.623 | -0.709 | -0.344 | 0.659 | -0.013 | 0.013 | 0.013 | 0 | 0.63 | 0.5 | 0.52 | 0.59 | 0.67 | 0.54 |
| 10 | 126294873 | rs4962392 | 0.385 | -0.144 | -0.376 | 1.063 | -0.001 | -0.002 | -0.002 | 0.245 | 0.08 | 0.08 | 0.18 | 0.11 | 0.09 | 0.07 |
| 10 | 126296383 | rs2061069 | 0.145 | -0.156 | -0.388 | 1.08 | 0 | 0.004 | 0.004 | 0 | 0.08 | 0.1 | 0.19 | 0.16 | 0.09 | 0.15 |
| 10 | 126297670 | rs11245315 | 0.428 | -0.298 | -0.378 | 0.76 | 0.001 | -0.004 | -0.004 | 0 | 0.27 | 0.29 | 0.18 | 0.25 | 0.3 | 0.14 |
| 10 | 126303580 | rs11245318 | -0.584 | -0.129 | -0.487 | 1.074 | 0 | -0.002 | -0.002 | 0 | 0.13 | 0.15 | 0.2 | 0.18 | 0.13 | 0.21 |
| 10 | 126308683 | rs3802703 | -0.304 | -0.062 | -0.676 | 1.352 | -0.009 | -0.039 | -0.039 | 0 | 0.07 | 0 | 0.03 | 0.13 | 0.05 | 0.11 |
| 10 | 126310042 | rs3740537 | -1.478 | -0.183 | -0.436 | 1.378 | -0.004 | 0.018 | 0.018 | 0 | 0.09 | 0.09 | 0.04 | 0.2 | 0.06 | 0.2 |
| 10 | 126310781 | rs3802701 | -0.298 | -0.032 | -0.238 | 1.269 | -0.008 | 0.003 | 0.003 | 0 | 0.37 | 0.5 | 0.21 | 0.42 | 0.34 | 0.26 |
| 10 | 126312289 | rs4962672 | 0.092 | 0.014 | -0.409 | 1.241 | -0.001 | -0.001 | -0.001 | 0 | 0.12 | 0.16 | 0.19 | 0.11 | 0.11 | 0.12 |
| 10 | 126313386 | rs2061071 | -1.257 | 0.038 | -0.44 | 1.179 | 0.006 | 0.027 | 0.027 | 0 | 0.51 | 0.79 | 0.43 | 0.6 | 0.48 | 0.44 |
| 10 | 126314475 | rs3781476 | 1.803 | 0.003 | -0.883 | 0.872 | -0.068 | 0.033 | 0.033 | 0 | 0.25 | 0.01 | 0.28 | 0.13 | 0.3 | 0.3 |
| 10 | 126317857 | rs10901791 | -1.296 | -0.184 | -0.363 | 0.805 | 0.006 | -0.002 | -0.002 | 0.464 | 0.78 | 0.8 | 0.92 | 0.73 | 0.88 | 0.91 |
| 10 | 126319153 | rs11599339 | -0.856 | -0.121 | -0.033 | 0.941 | -0.002 | 0.035 | 0.035 | 0 | 0.17 | 0.24 | 0.22 | 0.28 | 0.17 | 0.21 |
| 10 | 126319487 | rs897297 | -1.021 | 0.049 | 0.114 | 0.926 | 0.013 | 0 | 0 | 0 | 0.5 | 0.41 | 0.58 | 0.44 | 0.56 | 0.51 |
| 10 | 126319600 | rs748129 | 0.475 | 0.111 | 0.071 | 0.834 | -0.046 | 0.052 | 0.052 | 0 | 0.32 | 0.16 | 0.35 | 0.16 | 0.38 | 0.27 |
| 10 | 126319992 | rs3781470 | 0.677 | 0.311 | 0.526 | 0.917 | -0.007 | 0.013 | 0.013 | 0 | 0.34 | 0.44 | 0.34 | 0.4 | 0.38 | 0.48 |
| 10 | 126330565 | rs10794173 | -0.017 | 0.368 | -0.018 | 0.924 | 0.002 | 0.064 | 0.064 | 0.842 | 0.49 | 0.11 | 0.39 | 0.4 | 0.49 | 0.34 |
| 10 | 126331555 | rs1470996 | -0.914 | 0.44 | 0.208 | 1.074 | -0.003 | 0.037 | 0.037 | 0 | 0.16 | 0.22 | 0.22 | 0.28 | 0.17 | 0.21 |

| | | | | | | | | | | | | | | | | |
|---|---|---|---|---|---|---|---|---|---|---|---|---|---|---|---|---|
| 10 | 126333874 | rs7096880 | -1.254 | 0.437 | 0.161 | 0.965 | -0.001 | 0.026 | 0.026 | 0.081 | 0.18 | 0.27 | 0.24 | 0.26 | 0.19 | 0.23 |
| 10 | 126335697 | rs12411439 | -0.858 | 0.481 | 0.322 | 1.034 | 0.005 | -0.012 | -0.012 | 0 | 0.07 | 0.03 | 0.04 | 0.12 | 0.07 | 0.1 |
| 10 | 126343931 | rs3781458 | -0.798 | 0.461 | 0.424 | 1.01 | 0.011 | -0.002 | -0.002 | 0 | 0.58 | 0.38 | 0.59 | 0.56 | 0.6 | 0.45 |
| 10 | 126348274 | rs10901799 | -0.019 | 0.446 | 0.37 | 1.01 | 0.01 | 0.038 | 0.038 | 0.254 | 0.49 | 0.16 | 0.39 | 0.42 | 0.49 | 0.35 |
| 10 | 126348761 | rs3781453 | 1.747 | 0.426 | 0.403 | 1.008 | -0.005 | -0.007 | -0.007 | 0.619 | 0.29 | 0.09 | 0.32 | 0.3 | 0.34 | 0.41 |
| 10 | 126350341 | rs17152175 | -0.529 | 0.496 | 0.488 | 1.045 | 0.001 | -0.003 | -0.003 | 0 | 0.06 | 0.03 | 0.04 | 0.12 | 0.06 | 0.06 |
| 10 | 126358094 | rs7073663 | -1.062 | 0.497 | 0.626 | 1.057 | 0.009 | 0.112 | 0.112 | 0.077 | 0.17 | 0.49 | 0.25 | 0.26 | 0.16 | 0.2 |
| 10 | 126370067 | rs7093768 | 0.411 | 0.523 | 0.42 | 1.025 | 0.009 | 0.031 | 0.031 | 0 | 0.1 | 0.2 | 0.19 | 0.14 | 0.11 | 0.11 |
| 10 | 126380338 | rs11245330 | 0.376 | 0.521 | 0.422 | 1.024 | 0.007 | 0.027 | 0.027 | 0 | 0.1 | 0.2 | 0.19 | 0.14 | 0.11 | 0.11 |
| 10 | 126383088 | rs11245333 | 0.884 | 0.47 | 0.368 | 0.996 | -0.003 | -0.013 | -0.013 | 0.186 | 0.38 | 0.31 | 0.5 | 0.44 | 0.43 | 0.52 |
| 10 | 126383363 | rs4450131 | -0.035 | 0.449 | 0.352 | 0.94 | 0.012 | 0.041 | 0.041 | 0.217 | 0.49 | 0.16 | 0.4 | 0.42 | 0.49 | 0.34 |
| 10 | 126383857 | rs11597613 | 0.871 | 0.747 | 0.225 | 0.886 | -0.044 | 0.01 | 0.01 | 0 | 0.24 | 0.06 | 0.27 | 0.19 | 0.26 | 0.26 |
| 10 | 126399159 | rs10901808 | 0.597 | 0.465 | 0.362 | 0.852 | -0.034 | 0.016 | 0.016 | 0.555 | 0.27 | 0.1 | 0.15 | 0.24 | 0.29 | 0.11 |
| 10 | 126404204 | rs4962688 | 0.029 | 0.544 | 0.467 | 0.889 | 0.073 | 0.093 | 0.093 | 1.158 | 0.95 | 0.52 | 0.94 | 0.98 | 1 | 0.96 |
| 10 | 126410804 | rs10901809 | 0.018 | 0.552 | 0.368 | 0.914 | -0.026 | -0.015 | -0.015 | 0 | 0.33 | 0.13 | 0.19 | 0.36 | 0.34 | 0.17 |
| 10 | 126411924 | rs10794178 | 0.208 | 0.629 | 0.285 | 0.871 | -0.006 | -0.006 | -0.006 | 0 | 0.32 | 0.28 | 0.44 | 0.32 | 0.31 | 0.34 |
| 10 | 126414452 | rs869303 | 1.213 | 0.577 | 0.192 | 0.874 | -0.023 | 0.003 | 0.003 | 0 | 0.22 | 0.06 | 0.25 | 0.18 | 0.21 | 0.23 |
| 10 | 126418782 | rs11245344 | 1.689 | 0.41 | 1.17 | 1.06 | -0.008 | -0.021 | -0.021 | 0 | 0.51 | 0.15 | 0.57 | 0.53 | 0.55 | 0.68 |
| 10 | 126419153 | rs7894089 | -1.514 | 0.335 | 1.036 | 1.054 | -0.004 | 0.033 | 0.033 | 0 | 0.17 | 0.27 | 0.22 | 0.23 | 0.16 | 0.19 |
| 10 | 126438019 | rs10510138 | -0.537 | 0.384 | 0.905 | 1.064 | 0.001 | -0.023 | -0.023 | 0 | 0.07 | 0 | 0.03 | 0.08 | 0.05 | 0.09 |
| 10 | 126445630 | rs4962696 | 0.128 | 0.377 | 1.006 | 1.063 | 0.001 | -0.003 | -0.003 | 0 | 0.71 | 0.73 | 0.82 | 0.73 | 0.72 | 0.89 |
| 10 | 126446473 | rs2306702 | -0.51 | 0.397 | 0.785 | 1.069 | 0.001 | -0.023 | -0.023 | 0.926 | 0.07 | 0 | 0.03 | 0.08 | 0.05 | 0.09 |
| 10 | 126446592 | rs1055256 | 1.397 | 0.424 | 0.872 | 1.127 | -0.001 | -0.002 | -0.002 | 0 | 0.54 | 0.42 | 0.59 | 0.51 | 0.55 | 0.68 |
| 10 | 126449428 | rs10901818 | 1.36 | 0.426 | 0.823 | 1.127 | -0.001 | -0.002 | -0.002 | 0 | 0.54 | 0.43 | 0.59 | 0.51 | 0.55 | 0.68 |
| 10 | 126482849 | rs11245366 | 1.568 | 0.495 | -0.005 | 0.67 | -0.009 | 0.018 | 0.018 | 0 | 0.51 | 0.17 | 0.56 | 0.59 | 0.55 | 0.68 |
| 10 | 126488660 | rs10901823 | 1.574 | 0.408 | 0.003 | 0.677 | -0.009 | 0.019 | 0.019 | 0 | 0.51 | 0.17 | 0.56 | 0.59 | 0.55 | 0.68 |
| 10 | 126505969 | rs10901827 | -0.025 | 0.502 | 0.071 | 0.286 | 0.006 | 0.025 | 0.025 | 0 | 0.1 | 0.21 | 0.19 | 0.14 | 0.11 | 0.1 |
| 10 | 126512248 | rs7097254 | 0.939 | 0.497 | 0.069 | 0.281 | 0.001 | 0.008 | 0.008 | 0 | 0.61 | 0.38 | 0.76 | 0.73 | 0.66 | 0.78 |
| 10 | 126517989 | rs2303611 | 1.501 | 0.47 | -0.079 | 0.262 | -0.009 | 0.015 | 0.015 | 0 | 0.51 | 0.17 | 0.57 | 0.59 | 0.55 | 0.68 |
| 10 | 126553531 | rs2015176 | -0.615 | -0.665 | -0.698 | 0.367 | 0.007 | -0.024 | -0.024 | 0 | 0.26 | 0.23 | 0.23 | 0.37 | 0.22 | 0.17 |
| 10 | 126553571 | rs2015178 | 1.055 | -0.463 | -1.485 | 0.436 | 0.067 | 0 | 0 | 1.278 | 0.55 | 0.19 | 0.64 | 0.6 | 0.68 | 0.68 |
| 10 | 126554570 | rs2629544 | 1.092 | -0.399 | -1.451 | 0.468 | 0.076 | 0.011 | 0.011 | 0 | 0.54 | 0.19 | 0.64 | 0.59 | 0.67 | 0.66 |
| 10 | 126557206 | rs17152324 | -0.184 | 0.582 | -1.131 | 0.534 | -0.005 | -0.054 | -0.054 | 0.022 | 0.17 | 0.01 | 0.06 | 0.23 | 0.12 | 0.06 |
| 10 | 126569203 | rs11245401 | 0.648 | 1.41 | -0.292 | 0.294 | 0.005 | 0 | 0 | 0 | 0.69 | 0.67 | 0.54 | 0.64 | 0.59 | 0.68 |
| 10 | 126569279 | rs11595972 | -1.051 | 1.679 | 0.267 | -0.159 | -0.002 | -0.032 | -0.032 | 0 | 0.22 | 0.42 | 0.17 | 0.14 | 0.19 | 0.21 |
| 10 | 126569993 | rs7914368 | -1.608 | 2.149 | 0.242 | -0.07 | 0.058 | -0.05 | -0.05 | 0 | 0.09 | 0.24 | 0.12 | 0 | 0.2 | 0.09 |
| 10 | 126575210 | rs12247671 | -1.404 | 2.081 | -0.305 | -0.05 | -0.032 | 0.027 | 0.027 | 0 | 0.09 | 0.05 | 0.1 | 0 | 0.24 | 0.05 |
| 10 | 126581984 | rs1896529 | 1.569 | 1.058 | 0.779 | -0.091 | 0.001 | -0.014 | -0.014 | 0.682 | 0.9 | 0.52 | 0.71 | 0.95 | 0.93 | 0.9 |
| 10 | 126590820 | rs7079199 | -0.573 | 1.014 | 0.347 | 0.056 | -0.012 | -0.02 | -0.02 | 0 | 0.14 | 0.07 | 0.19 | 0.17 | 0.14 | 0.15 |
| 10 | 126629204 | rs7074163 | 1.801 | 0.656 | 1.304 | -0.228 | -0.024 | -0.071 | -0.071 | 2.523 | 0.85 | 0.54 | 0.68 | 0.95 | 0.89 | 0.89 |
| 10 | 126629701 | rs11245438 | 1.944 | 0.427 | 2.047 | -0.307 | 0.021 | 0.026 | 0.026 | 0.252 | 0.68 | 0.17 | 0.46 | 0.78 | 0.74 | 0.74 |
| 10 | 126630557 | rs2114835 | 1.114 | 0.5 | 2.083 | -0.269 | 0.011 | -0.026 | -0.026 | 0 | 0.9 | 0.69 | 0.71 | 0.95 | 0.89 | 0.89 |
| 10 | 126635262 | rs10901842 | 1.711 | 0.501 | 2.304 | -0.267 | -0.004 | -0.005 | -0.005 | 0 | 0.73 | 0.31 | 0.48 | 0.78 | 0.75 | 0.74 |
| 10 | 126650696 | rs11245446 | 1.135 | 0.217 | 0.982 | -1.612 | -0.038 | 0.238 | 0.238 | 0 | 0.35 | 0.04 | 0.19 | 0.01 | 0.33 | 0.32 |
| 10 | 126658075 | rs11245450 | 0.771 | 0.157 | 1.073 | -1.599 | -0.043 | 0.06 | 0.06 | 0.247 | 0.37 | 0.14 | 0.27 | 0.77 | 0.42 | 0.42 |
| 10 | 126670356 | rs17152433 | -0.397 | 0.182 | 1.436 | -1.338 | 0.008 | 0.04 | 0.04 | 0 | 0.18 | 0.5 | 0.23 | 0.19 | 0.14 | 0.16 |
| 10 | 126671673 | rs2241541 | 0.265 | 0.181 | 1.435 | -1.337 | 0.001 | 0.004 | 0.004 | 0 | 0.16 | 0.19 | 0.2 | 0.17 | 0.14 | 0.15 |
| 10 | 126672779 | rs6597869 | 0.676 | 0.182 | 1.335 | -1.346 | -0.002 | -0.031 | -0.031 | 0 | 0.91 | 0.83 | 0.71 | 0.96 | 0.89 | 0.9 |
| 10 | 126673998 | rs3906796 | 1.348 | 0.181 | 1.212 | -1.346 | 0.011 | -0.035 | -0.035 | 0.912 | 0.88 | 0.53 | 0.68 | 0.95 | 0.89 | 0.89 |
| 10 | 126676686 | rs3740535 | 1.16 | 0.208 | 0.667 | -1.337 | 0.006 | 0.009 | 0.009 | 0.916 | 0.72 | 0.21 | 0.47 | 0.78 | 0.75 | 0.74 |
| 10 | 126681688 | rs7097802 | -0.069 | 0.077 | 0.475 | -1.417 | -0.006 | -0.015 | -0.015 | 0.841 | 0.9 | 0.89 | 0.97 | 0.96 | 0.95 | 0.82 |
| 10 | 126683485 | rs10901849 | 0.396 | -0.452 | 0.188 | -2.119 | -0.002 | -0.046 | -0.046 | 0 | 0.31 | 0.35 | 0.38 | 0.01 | 0.37 | 0.31 |
| 10 | 126683766 | rs11245453 | 1.098 | -0.465 | -0.097 | -2.143 | -0.08 | 0.171 | 0.171 | 0 | 0.25 | 0.04 | 0.16 | 0.01 | 0.26 | 0.27 |
| 10 | 126684134 | rs3781446 | -0.811 | -0.578 | -0.048 | -1.16 | 0.008 | 0.002 | 0.002 | 1.058 | 0.52 | 0.8 | 0.62 | 0.42 | 0.55 | 0.49 |
| 10 | 126687330 | rs4962717 | 1.287 | -0.501 | 0.664 | -0.951 | 0.026 | -0.035 | -0.035 | 0 | 0.89 | 0.58 | 0.67 | 0.94 | 0.87 | 0.88 |
| 10 | 126687528 | rs4962718 | 0.801 | -0.846 | -0.194 | -1.299 | 0.034 | -0.034 | -0.034 | 0 | 0.61 | 0.19 | 0.43 | 0.55 | 0.68 | 0.69 |
| 10 | 126688370 | rs3781445 | 1.012 | -1.041 | -0.434 | -1.174 | -0.076 | 0.07 | 0.07 | 0 | 0.28 | 0.1 | 0.24 | 0.14 | 0.38 | 0.3 |
| 10 | 126688968 | rs3781444 | -1.093 | -0.894 | 0.06 | -0.774 | -0.022 | 0.039 | 0.039 | 0 | 0.54 | 0.88 | 0.69 | 0.35 | 0.5 | 0.54 |
| 10 | 126690383 | rs7923776 | 0.746 | -0.844 | 0.049 | -0.939 | -0.032 | 0.062 | 0.062 | 0 | 0.46 | 0.13 | 0.33 | 0.7 | 0.51 | 0.49 |
| 10 | 126691180 | rs12771627 | 0.638 | -0.855 | 0.041 | -1.053 | -0.001 | 0.177 | 0.177 | 0 | 0.33 | 0.06 | 0.17 | 0.01 | 0.27 | 0.3 |
| 10 | 126691471 | rs4348846 | -1.262 | -0.813 | 0.616 | -0.879 | 0.004 | 0.093 | 0.093 | 0.545 | 0.17 | 0.58 | 0.36 | 0.25 | 0.19 | 0.12 |
| 10 | 126695269 | rs3781436 | -0.324 | -0.808 | 1.184 | -0.868 | 0.027 | 0.171 | 0.171 | 0.088 | 0.1 | 0.35 | 0.19 | 0.22 | 0.09 | 0.09 |
| 10 | 126695673 | rs1561589 | -0.351 | -0.931 | 1.11 | -1.026 | 0.009 | 0.026 | 0.026 | 0 | 0.44 | 0.61 | 0.39 | 0.23 | 0.37 | 0.39 |
| 10 | 126696061 | rs11593921 | 0.87 | -0.777 | 0.496 | -1.055 | -0.017 | 0.02 | 0.02 | 0 | 0.05 | 0 | 0.15 | 0 | 0.09 | 0.02 |
| 10 | 126696872 | rs4962416 | 0.433 | -0.811 | 0.727 | -1.088 | 0.015 | 0.047 | 0.047 | 0 | 0.34 | 0.17 | 0.19 | 0.01 | 0.28 | 0.3 |
| 10 | 126697086 | rs4962418 | -0.047 | -0.812 | 0.732 | -1.082 | 0.013 | 0.098 | 0.098 | 0 | 0.39 | 0.17 | 0.33 | 0.01 | 0.37 | 0.33 |
| 10 | 126704345 | rs12776353 | 0.566 | -1.166 | 0.424 | -1.23 | 0.035 | 0.019 | 0.019 | 0 | 0.22 | 0.1 | 0.06 | 0.01 | 0.09 | 0.22 |
| 10 | 126706676 | rs11595908 | 0.245 | -0.943 | 0.324 | -0.991 | 0.008 | 0.029 | 0.029 | 0 | 0.09 | 0.03 | 0.17 | 0 | 0.11 | 0.03 |
| 10 | 126707881 | rs10901850 | -0.703 | -0.967 | 0.887 | -0.965 | 0.018 | -0.002 | -0.002 | 0 | 0.31 | 0.43 | 0.46 | 0.23 | 0.38 | 0.2 |

| | | | | | | | | | | | | | | | | |
|---|---|---|---|---|---|---|---|---|---|---|---|---|---|---|---|---|
| 10 | 126711107 | rs3781418 | -0.649 | -0.967 | 1.038 | -0.987 | -0.013 | -0.066 | -0.066 | 0.105 | 0.12 | 0.32 | 0.19 | 0.01 | 0.11 | 0.03 |
| 10 | 126711633 | rs7073257 | 0.541 | -0.935 | 1.044 | -0.978 | -0.005 | -0.005 | -0.005 | 0 | 0.19 | 0.14 | 0.26 | 0.22 | 0.29 | 0.17 |
| 10 | 126711691 | rs7086797 | -0.112 | -1.014 | 1.201 | -0.933 | 0.012 | 0.012 | 0.012 | 0 | 0.23 | 0.23 | 0.38 | 0.27 | 0.32 | 0.22 |
| 10 | 126714200 | rs2363893 | 1.259 | -1.324 | 1.046 | -1.043 | -0.019 | 0.024 | 0.024 | 0 | 0.45 | 0.26 | 0.33 | 0.66 | 0.48 | 0.48 |
| 10 | 126714966 | rs3012075 | 1.181 | -1.455 | 1.283 | -1.097 | -0.012 | 0.019 | 0.019 | 0 | 0.46 | 0.3 | 0.33 | 0.66 | 0.48 | 0.48 |
| 10 | 126715436 | rs3781411 | -0.6 | -1.439 | 1.321 | -1.097 | 0.001 | 0.002 | 0.002 | 0.054 | 0.13 | 0.11 | 0.29 | 0.05 | 0.14 | 0.09 |
| 10 | 126715629 | rs3781409 | 0.874 | -1.42 | 1.291 | -1.089 | -0.026 | -0.028 | -0.028 | 0.273 | 0.18 | 0.1 | 0.27 | 0.24 | 0.28 | 0.17 |
| 10 | 126716346 | rs2938006 | 1.533 | -1.718 | 0.976 | -1.312 | -0.012 | 0.004 | 0.004 | 0 | 0.42 | 0.32 | 0.33 | 0.66 | 0.48 | 0.48 |
| 10 | 126716870 | rs3012067 | 1.694 | -1.728 | 0.808 | -1.308 | -0.045 | -0.006 | -0.006 | 0 | 0.39 | 0.18 | 0.3 | 0.66 | 0.47 | 0.47 |
| 10 | 126721702 | rs3781399 | -0.178 | -1.584 | 0.624 | -1.23 | -0.005 | -0.006 | -0.006 | 0 | 0.06 | 0.11 | 0.14 | 0.04 | 0.04 | 0.06 |
| 10 | 126723133 | rs1077497 | 1.283 | -1.681 | 0.664 | -1.295 | 0.024 | 0.024 | 0.024 | 0 | 0.17 | 0.13 | 0.08 | 0.03 | 0.08 | 0.2 |
| 10 | 126723567 | rs893856 | -0.805 | -1.692 | 0.762 | -1.312 | -0.012 | -0.04 | -0.04 | 1.083 | 0.15 | 0.26 | 0.32 | 0.05 | 0.15 | 0.09 |
| 10 | 126724294 | rs11818446 | -0.252 | -1.68 | 0.771 | -1.298 | 0.002 | -0.003 | -0.003 | 0 | 0.06 | 0.16 | 0.14 | 0.04 | 0.05 | 0.07 |
| 10 | 126724927 | rs4962421 | 1.199 | -1.616 | 0.569 | -1.286 | -0.104 | -0.107 | -0.107 | 0 | 0.18 | 0.01 | 0.23 | 0.24 | 0.28 | 0.17 |
| 10 | 126726885 | rs3781394 | 1.266 | -1.62 | 0.626 | -1.311 | 0.024 | 0.024 | 0.024 | 0 | 0.17 | 0.13 | 0.08 | 0.03 | 0.08 | 0.2 |
| 10 | 126727268 | rs11599762 | 0.733 | -1.545 | 0.584 | -1.264 | -0.01 | 0.044 | 0.044 | 0 | 0.09 | 0.01 | 0.17 | 0.01 | 0.11 | 0.03 |
| 10 | 126730491 | rs2935653 | -1.821 | -1.509 | 0.863 | -1.132 | 0.01 | -0.001 | -0.001 | 0.225 | 0.59 | 0.57 | 0.66 | 0.34 | 0.54 | 0.51 |
| 10 | 126731736 | rs2936546 | -1.792 | -1.454 | 0.762 | -1.138 | 0.007 | -0.001 | -0.001 | 0 | 0.59 | 0.5 | 0.66 | 0.34 | 0.54 | 0.52 |
| 10 | 126733607 | rs3012064 | -1.248 | -1.491 | 0.438 | -1.185 | 0.075 | 0.103 | 0.103 | 0 | 0.43 | 0.77 | 0.45 | 0.07 | 0.25 | 0.35 |
| 10 | 126733643 | rs10901854 | 0.955 | -1.268 | -0.258 | -1.056 | -0.04 | -0.048 | -0.048 | 0.064 | 0.19 | 0.09 | 0.26 | 0.27 | 0.29 | 0.17 |
| 10 | 126734419 | rs12243317 | -0.722 | -1.569 | -0.333 | -1.326 | 0.01 | -0.002 | -0.002 | 0 | 0.33 | 0.34 | 0.38 | 0.07 | 0.24 | 0.31 |
| 10 | 126734779 | rs7075394 | -0.711 | -1.567 | -0.332 | -1.335 | 0.01 | -0.008 | -0.008 | 0.006 | 0.33 | 0.34 | 0.39 | 0.05 | 0.24 | 0.29 |
| 10 | 126737997 | rs3012066 | -0.406 | -1.182 | -0.473 | -0.847 | 0.008 | 0.019 | 0.019 | 0 | 0.38 | 0.33 | 0.31 | 0.25 | 0.3 | 0.41 |
| 10 | 126739029 | rs718949 | -0.605 | -1.209 | -0.246 | -0.796 | -0.047 | -0.104 | -0.104 | 0 | 0.34 | 0.53 | 0.25 | 0.04 | 0.21 | 0.26 |
| 10 | 126739149 | rs718947 | 0.033 | -0.092 | -0.817 | -0.313 | -0.055 | 0.011 | 0.011 | 0 | 0.32 | 0.13 | 0.42 | 0.26 | 0.36 | 0.28 |
| 10 | 126739338 | rs718948 | 0.115 | -0.004 | -0.682 | -0.226 | -0.039 | 0.033 | 0.033 | 0 | 0.33 | 0.19 | 0.42 | 0.24 | 0.34 | 0.27 |
| 10 | 126740797 | rs2938010 | -1.011 | -0.69 | 0.3 | 0.431 | -0.018 | 0.066 | 0.066 | 0 | 0.29 | 0.52 | 0.37 | 0.49 | 0.26 | 0.38 |
| 10 | 126741541 | rs998836 | -0.042 | -0.843 | 0.353 | 0.327 | -0.01 | -0.035 | -0.035 | 0 | 0.15 | 0.25 | 0.07 | 0.02 | 0.12 | 0.18 |
| 10 | 126743096 | rs7893862 | 0.322 | -0.807 | 0.322 | 0.342 | -0.006 | -0.029 | -0.029 | 0 | 0.13 | 0.2 | 0.06 | 0.02 | 0.12 | 0.18 |
| 10 | 126743195 | rs10901856 | -0.51 | -0.745 | 0.286 | 0.591 | 0.002 | 0.092 | 0.092 | 0 | 0.13 | 0.17 | 0.29 | 0.5 | 0.14 | 0.21 |
| 10 | 126747542 | rs2936540 | -1.577 | -0.836 | -0.773 | 0.328 | 0.048 | 0.049 | 0.049 | 0.179 | 0.48 | 0.93 | 0.62 | 0.61 | 0.4 | 0.6 |
| 10 | 126747809 | rs3012069 | -1.584 | -0.84 | -0.775 | 0.336 | 0.049 | 0 | 0 | 0.19 | 0.48 | 0.93 | 0.62 | 0.59 | 0.4 | 0.6 |
| 10 | 126749162 | rs2946998 | -0.98 | -0.703 | -0.282 | 0.248 | -0.019 | 0.015 | 0.015 | 0 | 0.32 | 0.45 | 0.51 | 0.62 | 0.28 | 0.43 |
| 10 | 126750945 | rs11245486 | -0.657 | -0.455 | -0.439 | -0.089 | -0.007 | -0.034 | -0.034 | 0 | 0.3 | 0.7 | 0.24 | 0.08 | 0.27 | 0.29 |
| 10 | 126753504 | rs2936541 | -0.484 | 0.435 | -1.119 | 0.38 | -0.079 | 0.048 | 0.048 | 0 | 0.63 | 0.88 | 0.62 | 0.78 | 0.59 | 0.71 |
| 10 | 126754834 | rs2919289 | -0.858 | 0.519 | -0.06 | 0.791 | 0.039 | 0.061 | 0.061 | 0 | 0.1 | 0.25 | 0.1 | 0.21 | 0.12 | 0.12 |
| 10 | 126755536 | rs2936542 | -1.227 | -0.191 | 0.398 | 1.45 | -0.021 | 0.186 | 0.186 | 0 | 0.15 | 0.33 | 0.34 | 0.38 | 0.11 | 0.22 |
| 10 | 126755887 | rs2935652 | 0.243 | -0.365 | -0.116 | 0.474 | 0.005 | 0.016 | 0.016 | 0 | 0.42 | 0.4 | 0.6 | 0.65 | 0.42 | 0.49 |
| 10 | 126755930 | rs2935651 | 0.032 | 0.157 | -0.413 | 0.331 | -0.001 | 0.082 | 0.082 | 0 | 0.53 | 0.4 | 0.61 | 0.73 | 0.5 | 0.55 |
| 10 | 126756553 | rs2935650 | -0.952 | 0.408 | -1.343 | 0.411 | -0.062 | 0.018 | 0.018 | 0 | 0.74 | 0.88 | 0.73 | 0.74 | 0.59 | 0.75 |
| 10 | 126757251 | rs7087286 | 0.011 | 0.318 | -1.393 | 0.582 | 0.033 | 0.062 | 0.062 | 0 | 0.47 | 0.18 | 0.42 | 0.35 | 0.42 | 0.4 |
| 10 | 126764413 | rs3012070 | -0.658 | 0.856 | -1.431 | 0.627 | 0.005 | 0.027 | 0.027 | 0 | 0.51 | 0.83 | 0.49 | 0.51 | 0.43 | 0.38 |
| 10 | 126773746 | rs11245496 | -0.48 | 0.771 | -0.981 | 0.376 | 0 | 0.003 | 0.003 | 0 | 0.18 | 0.17 | 0.09 | 0.11 | 0.13 | 0.08 |
| 10 | 126775236 | rs10794197 | 1.095 | 0.712 | -1.69 | -0.105 | 0.027 | 0.059 | 0.059 | 0 | 0.27 | 0.13 | 0.16 | 0.1 | 0.25 | 0.35 |
| 10 | 126776488 | rs1152659 | -0.314 | 1.01 | -0.821 | -0.611 | 0.002 | -0.127 | -0.127 | 0 | 0.28 | 0.39 | 0.2 | 0 | 0.28 | 0.11 |
| 10 | 126776717 | rs17152617 | 1.077 | 0.931 | -0.433 | -0.659 | 0.024 | -0.074 | -0.074 | 0.212 | 0.14 | 0.23 | 0.13 | 0 | 0.18 | 0.06 |
| 10 | 126777042 | rs1152661 | -1.031 | 1.3 | -0.384 | -0.041 | -0.002 | -0.024 | -0.024 | 0 | 0.65 | 0.68 | 0.58 | 0.48 | 0.66 | 0.57 |
| 10 | 126778078 | rs1152663 | -0.87 | 1.982 | 0.293 | 0.282 | -0.002 | -0.171 | -0.171 | 0.544 | 0.3 | 0.5 | 0.22 | 0 | 0.28 | 0.11 |
| 10 | 126781929 | rs1152676 | -0.255 | 1.905 | 0.063 | 0.7 | -0.097 | -0.04 | -0.04 | 0 | 0.77 | 0.94 | 0.77 | 0.83 | 0.66 | 0.77 |
| 10 | 126782331 | rs1152677 | -1.597 | 1.227 | 0.903 | 0.785 | -0.033 | 0.025 | 0.025 | 0 | 0.23 | 0.56 | 0.26 | 0.27 | 0.13 | 0.14 |
| 10 | 126788191 | rs1152686 | -1.555 | 1.33 | 1.036 | 0.821 | -0.038 | 0.068 | 0.068 | 0 | 0.24 | 0.52 | 0.28 | 0.34 | 0.14 | 0.27 |
| 10 | 126793086 | rs913165 | -1.513 | 1.893 | 1.1 | 0.758 | 0.023 | -0.091 | -0.091 | 10.248 | 0.23 | 0.46 | 0.15 | 0.09 | 0.19 | 0.22 |
| 10 | 126796382 | rs1152690 | -1.05 | 1.646 | 1.074 | 0.972 | -0.014 | 0.152 | 0.152 | 0 | 0.15 | 0.39 | 0.25 | 0.34 | 0.09 | 0.25 |
| 10 | 126798920 | rs11814706 | -0.28 | 1.6 | 1.068 | 0.749 | -0.016 | -0.042 | -0.042 | 0.331 | 0.12 | 0.28 | 0.07 | 0.04 | 0.06 | 0.17 |
| 10 | 126802394 | rs7908461 | -0.4 | 1.518 | 0.637 | 0.833 | -0.027 | -0.046 | -0.046 | 0 | 0.12 | 0.3 | 0.07 | 0.04 | 0.06 | 0.17 |
| 10 | 126802740 | rs1152695 | -1.432 | 1.81 | -0.142 | 0.876 | 0 | -0.003 | -0.003 | 0.057 | 0.29 | 0.71 | 0.35 | 0.31 | 0.24 | 0.3 |
| 10 | 126808877 | rs1254702 | -0.691 | 2.35 | -0.563 | 1.037 | -0.045 | -0.042 | -0.042 | 0 | 0.93 | 1 | 0.89 | 0.92 | 0.86 | 0.91 |
| 10 | 126811170 | rs1254699 | -1.319 | 1.872 | -0.111 | 0.966 | -0.003 | -0.002 | -0.002 | 0 | 0.28 | 0.69 | 0.31 | 0.31 | 0.21 | 0.3 |
| 10 | 126812270 | rs10901863 | 0.385 | 1.187 | -1.202 | 0.029 | 0.003 | 0.077 | 0.077 | 1.562 | 0.29 | 0.07 | 0.31 | 0.17 | 0.31 | 0.29 |
| 10 | 126814138 | rs12264453 | 0.736 | 0.657 | -1.005 | -0.448 | -0.011 | 0.056 | 0.056 | 0 | 0.52 | 0.23 | 0.53 | 0.71 | 0.53 | 0.57 |
| 10 | 126815036 | rs2225079 | 0.226 | 0.62 | -0.333 | -0.579 | -0.001 | -0.002 | -0.002 | 0.089 | 0.2 | 0.26 | 0.16 | 0.16 | 0.23 | 0.24 |
| 10 | 126815404 | rs2209684 | -0.711 | 0.533 | -0.405 | -0.666 | 0 | -0.001 | -0.001 | 0 | 0.34 | 0.31 | 0.28 | 0.19 | 0.31 | 0.29 |
| 10 | 126818270 | rs10510141 | 0.878 | 0.541 | -0.359 | -0.656 | 0.006 | 0.039 | 0.039 | 0 | 0.13 | 0.03 | 0.12 | 0.03 | 0.09 | 0.05 |
| 10 | 126820152 | rs10901865 | -0.988 | 0.509 | 0.086 | -0.674 | -0.013 | -0.027 | -0.027 | 0 | 0.38 | 0.53 | 0.32 | 0.2 | 0.32 | 0.33 |
| 10 | 126822134 | rs10901867 | 0.024 | 0.734 | -0.087 | -0.477 | -0.002 | -0.002 | -0.002 | 0 | 0.21 | 0.23 | 0.16 | 0.16 | 0.23 | 0.24 |
| 10 | 126823911 | rs1475384 | -0.607 | 1.059 | -0.553 | -0.336 | 0.005 | -0.001 | -0.001 | 0.634 | 0.9 | 0.87 | 0.84 | 0.9 | 0.83 | 0.85 |
| 10 | 126828084 | rs7090538 | 0.211 | 0.5 | -0.908 | -0.835 | 0.017 | 0.052 | 0.052 | 0 | 0.46 | 0.59 | 0.54 | 0.2 | 0.37 | 0.44 |
| 10 | 126833066 | rs11245518 | 0.89 | 0.346 | -0.982 | -0.563 | -0.029 | -0.032 | -0.032 | 0 | 0.15 | 0.03 | 0.12 | 0.17 | 0.2 | 0.24 |
| 10 | 126837944 | rs6597883 | -0.014 | 0.248 | -0.677 | -1.064 | -0.044 | -0.125 | -0.125 | 0 | 0.23 | 0.48 | 0.2 | 0.04 | 0.16 | 0.07 |

| | | | | | | | | | | | | | | | | |
|---|---|---|---|---|---|---|---|---|---|---|---|---|---|---|---|---|
| 10 | 126837986 | rs1693628 | -1.578 | 0.673 | -1.299 | -0.827 | -0.046 | -0.042 | -0.042 | 0 | 0.93 | 1 | 0.88 | 0.9 | 0.85 | 0.9 |
| 10 | 126843661 | rs1693655 | -0.478 | 0.594 | -1.349 | -0.912 | -0.082 | -0.02 | -0.02 | 0 | 0.77 | 0.98 | 0.74 | 0.86 | 0.75 | 0.82 |
| 10 | 126846214 | rs1915146 | 0.462 | 0.562 | -0.375 | -0.807 | 0.008 | 0.052 | 0.052 | 0 | 0.44 | 0.58 | 0.54 | 0.21 | 0.41 | 0.45 |
| 10 | 126856255 | rs1715862 | -0.055 | 0.584 | -0.846 | -0.375 | -0.067 | 0.084 | 0.084 | 0 | 0.59 | 0.85 | 0.65 | 0.31 | 0.5 | 0.48 |
| 10 | 126863721 | rs11245533 | -0.278 | 0.621 | -0.545 | 0.188 | -0.048 | -0.03 | -0.03 | 0.161 | 0.15 | 0.01 | 0.12 | 0.17 | 0.19 | 0.2 |
| 10 | 126875396 | rs1693663 | -1.202 | 0.051 | 0.221 | 0.559 | -0.042 | 0.035 | 0.035 | 0.313 | 0.21 | 0.61 | 0.14 | 0.31 | 0.09 | 0.21 |
| 10 | 126875920 | rs1715869 | -1.748 | 0.447 | 0.16 | 0.784 | -0.001 | 0.013 | 0.013 | 0.866 | 0.14 | 0.16 | 0.07 | 0.22 | 0.1 | 0.18 |
| 10 | 126879881 | rs7074501 | -1.434 | 0.297 | 0.32 | 0.414 | -0.018 | 0.021 | 0.021 | 2.517 | 0.13 | 0.45 | 0.09 | 0.1 | 0.07 | 0.1 |
| 10 | 126885159 | rs7909594 | -0.178 | 0.612 | -0.873 | 0.241 | -0.025 | 0.034 | 0.034 | 0 | 0.3 | 0.12 | 0.34 | 0.21 | 0.31 | 0.28 |
| 10 | 126885852 | rs11245535 | -0.755 | 1.051 | -0.52 | 0.207 | 0.014 | 0.019 | 0.019 | 0 | 0.47 | 0.39 | 0.6 | 0.27 | 0.59 | 0.35 |
| 10 | 126890983 | rs4393270 | -0.66 | 0.478 | -1.112 | 0.218 | 0.02 | 0.03 | 0.03 | 0 | 0.68 | 0.85 | 0.7 | 0.76 | 0.72 | 0.59 |
| 10 | 126895917 | rs10901874 | -1.073 | 0.776 | -0.235 | 0.633 | 0 | 0.012 | 0.012 | 0.313 | 0.26 | 0.69 | 0.26 | 0.32 | 0.27 | 0.16 |
| 10 | 126896927 | rs7096790 | -0.243 | 1.038 | 0.258 | 0.267 | 0.039 | 0.005 | 0.005 | 0 | 0.11 | 0.19 | 0.14 | 0.08 | 0.18 | 0.05 |
| 10 | 126899646 | rs7919596 | -1.597 | 0.891 | 0.641 | 0.127 | 0.018 | -0.065 | -0.065 | 0.212 | 0.28 | 0.52 | 0.24 | 0.13 | 0.28 | 0.14 |
| 10 | 126900741 | rs12242774 | -0.067 | 1.072 | 0.939 | -0.056 | 0.033 | -0.052 | -0.052 | 0 | 0.09 | 0.22 | 0.11 | 0 | 0.1 | 0.01 |
| 10 | 126901596 | rs10901876 | 0.709 | 0.503 | 0.877 | 0.067 | -0.002 | 0.003 | 0.003 | 0 | 0.67 | 0.6 | 0.65 | 0.62 | 0.65 | 0.78 |
| 10 | 126901930 | rs10128157 | 0.39 | -0.301 | 0.939 | -0.449 | 0.022 | -0.046 | -0.046 | 1.652 | 0.08 | 0.16 | 0.09 | 0 | 0.1 | 0.01 |
| 10 | 126903068 | rs11245545 | -0.495 | -0.458 | 0.791 | -0.536 | 0.035 | 0.043 | 0.043 | 0 | 0.19 | 0.11 | 0.1 | 0.02 | 0.14 | 0.06 |
| 10 | 126905011 | rs7086919 | -0.417 | 0.056 | 1.451 | -0.164 | 0.031 | -0.069 | -0.069 | 0.085 | 0.11 | 0.27 | 0.1 | 0 | 0.11 | 0.01 |
| 10 | 126906820 | rs11596972 | -0.292 | 0.047 | 1.362 | -0.16 | 0.016 | -0.063 | -0.063 | 0 | 0.11 | 0.21 | 0.09 | 0 | 0.11 | 0.01 |
| 10 | 126910750 | rs10794206 | 1.044 | -0.257 | 0.624 | -0.51 | 0.023 | 0.226 | 0.226 | 0 | 0.55 | 0.12 | 0.64 | 0.9 | 0.6 | 0.68 |
| 10 | 126911910 | rs4962739 | 0.876 | -0.313 | 1.51 | -0.565 | 0.032 | -0.086 | -0.086 | 0.785 | 0.83 | 0.54 | 0.83 | 0.96 | 0.8 | 0.86 |
| 10 | 126912186 | rs12414556 | 0.816 | -0.468 | 0.796 | -0.181 | -0.164 | -0.159 | -0.159 | 0 | 0.22 | 0.01 | 0.36 | 0.49 | 0.36 | 0.33 |
| 10 | 126912670 | rs10901880 | 1.455 | -0.364 | 1.234 | -0.67 | 0 | 0.249 | 0.249 | 0 | 0.56 | 0.07 | 0.64 | 0.9 | 0.6 | 0.7 |
| 10 | 126914896 | rs17152738 | 0 | -0.214 | 1.347 | -0.76 | -0.003 | -0.023 | -0.023 | 0 | 0.25 | 0.31 | 0.15 | 0.05 | 0.19 | 0.16 |
| 10 | 126916123 | rs9422860 | 0.301 | 0.078 | 1.324 | -0.473 | 0.006 | -0.035 | -0.035 | 1.384 | 0.09 | 0.11 | 0.08 | 0 | 0.11 | 0.01 |
| 10 | 126937369 | rs9422750 | 0.296 | 0.086 | 1.177 | -0.469 | 0 | -0.019 | -0.019 | 2.227 | 0.09 | 0.08 | 0.08 | 0 | 0.11 | 0.01 |
| 10 | 126944539 | rs9422864 | 0.646 | 0.036 | 1.433 | -0.333 | 0.022 | -0.021 | -0.021 | 0.016 | 0.92 | 0.75 | 0.91 | 0.97 | 0.9 | 0.86 |
| 10 | 126958611 | rs11592111 | 0.241 | -0.076 | 1.036 | 0.02 | -0.004 | -0.008 | -0.008 | 0 | 0.09 | 0.07 | 0.06 | 0.16 | 0.1 | 0.02 |
| 10 | 126962284 | rs9422868 | -0.098 | -0.338 | 1.764 | -0.43 | -0.022 | -0.133 | -0.133 | 1.489 | 0.23 | 0.43 | 0.14 | 0.01 | 0.2 | 0.19 |
| 10 | 126963368 | rs9422871 | -0.51 | -0.488 | 2.103 | -0.459 | -0.012 | -0.055 | -0.055 | 0 | 0.27 | 0.67 | 0.19 | 0.04 | 0.21 | 0.24 |
| 10 | 126969961 | rs6597689 | 0.524 | -0.569 | 1.922 | -0.541 | 0.014 | -0.001 | -0.001 | 0 | 0.68 | 0.28 | 0.79 | 0.95 | 0.77 | 0.74 |
| 10 | 126976810 | rs4962447 | -0.729 | -0.232 | 2.048 | -0.382 | -0.003 | -0.03 | -0.03 | 2.092 | 0.28 | 0.7 | 0.2 | 0.05 | 0.22 | 0.23 |
| 10 | 126981173 | rs3851566 | 0.259 | -0.129 | 1.458 | -0.235 | -0.179 | -0.175 | -0.175 | 0.064 | 0.25 | 0.02 | 0.41 | 0.58 | 0.42 | 0.35 |
| 10 | 126995050 | rs2169036 | -0.352 | -0.127 | 2.101 | -0.012 | -0.02 | -0.041 | -0.041 | 0.097 | 0.25 | 0.55 | 0.2 | 0.1 | 0.22 | 0.22 |
| 10 | 127010681 | rs1459377 | -0.154 | -0.138 | 2.353 | -0.017 | -0.02 | -0.044 | -0.044 | 0.039 | 0.23 | 0.48 | 0.19 | 0.1 | 0.21 | 0.22 |
| 10 | 127014001 | rs12784803 | 0.43 | -0.149 | 2.119 | 0.009 | -0.006 | -0.017 | -0.017 | 0 | 0.09 | 0.05 | 0.06 | 0.17 | 0.11 | 0.02 |
| 10 | 127016670 | rs4020030 | -0.573 | -0.187 | 2.166 | -0.027 | -0.017 | -0.027 | -0.027 | 0 | 0.27 | 0.59 | 0.22 | 0.11 | 0.21 | 0.22 |
| 10 | 127040155 | rs3863288 | 0.369 | -0.223 | 1.758 | -0.09 | 0.002 | 0.007 | 0.007 | 0.224 | 0.2 | 0.21 | 0.16 | 0.09 | 0.2 | 0.19 |
| 10 | 127041985 | rs11244426 | 1.989 | -0.336 | 1.461 | -0.244 | 0.045 | 0.171 | 0.171 | 0 | 0.36 | 0.02 | 0.15 | 0.11 | 0.27 | 0.36 |
| 10 | 127048474 | rs2365577 | -1.383 | -0.066 | 3.006 | 0.291 | -0.008 | 0.018 | 0.018 | 0 | 0.33 | 0.52 | 0.27 | 0.32 | 0.31 | 0.25 |
| 10 | 127049407 | rs12217399 | -1.331 | -0.061 | 2.953 | 0.293 | -0.004 | 0.011 | 0.011 | 0 | 0.32 | 0.46 | 0.26 | 0.32 | 0.31 | 0.25 |
| 10 | 127053583 | rs12782166 | -0.581 | -0.013 | 2.688 | 0.324 | 0.01 | 0.06 | 0.06 | 0 | 0.11 | 0.21 | 0.1 | 0.23 | 0.11 | 0.06 |
| 10 | 127057080 | rs12762641 | -0.494 | 0.015 | 2.688 | 0.14 | 0.013 | 0.044 | 0.044 | 0 | 0.11 | 0.2 | 0.09 | 0.19 | 0.11 | 0.05 |
| 10 | 127058586 | rs2045566 | -0.946 | -0.06 | 2.704 | -0.055 | -0.031 | -0.048 | -0.048 | 4.307 | 0.26 | 0.48 | 0.2 | 0.1 | 0.2 | 0.22 |
| 10 | 127063501 | rs1380319 | -1.128 | 0.403 | 2.368 | -0.115 | -0.003 | 0.001 | 0.001 | 0 | 0.32 | 0.28 | 0.19 | 0.06 | 0.33 | 0.22 |
| 10 | 127064523 | rs4597024 | -1.745 | -0.074 | 2.407 | 0.187 | -0.011 | -0.006 | -0.006 | 0 | 0.35 | 0.56 | 0.24 | 0.3 | 0.29 | 0.29 |
| 10 | 127066971 | rs7089140 | -1.661 | 0.308 | 2.21 | 0.187 | -0.003 | 0.008 | 0.008 | 0 | 0.36 | 0.7 | 0.29 | 0.3 | 0.33 | 0.3 |
| 10 | 127080951 | rs17646925 | -0.226 | 0.27 | 2.123 | 0.083 | 0.008 | 0.073 | 0.073 | 0 | 0.21 | 0.07 | 0.08 | 0.06 | 0.18 | 0.21 |
| 10 | 127081079 | rs11812204 | -0.765 | 0.259 | 2.381 | 0.073 | 0.039 | -0.077 | -0.077 | 1.391 | 0.12 | 0.43 | 0.11 | 0.02 | 0.1 | 0.05 |
| 10 | 127081226 | rs4962295 | -0.406 | 0.279 | 1.889 | 0.103 | -0.009 | 0.039 | 0.039 | 0 | 0.22 | 0.14 | 0.12 | 0.06 | 0.22 | 0.21 |
| 10 | 127097031 | rs6597703 | -1.008 | 0.4 | 1.854 | -0.65 | -0.001 | -0.041 | -0.041 | 0.501 | 0.26 | 0.32 | 0.15 | 0.1 | 0.23 | 0.24 |
| 10 | 127099425 | rs11244455 | -1.315 | 0.489 | 1.489 | -0.603 | 0.085 | 0.009 | 0.009 | 0.558 | 0.06 | 0.25 | 0.07 | 0.03 | 0.05 | 0.03 |
| 10 | 127100408 | rs9422878 | -1.496 | 0.466 | 0.786 | -0.55 | -0.004 | 0.099 | 0.099 | 0 | 0.38 | 0.75 | 0.25 | 0.11 | 0.33 | 0.29 |
| 10 | 127101883 | rs9422879 | -0.74 | 0.85 | 0.802 | -0.107 | 0.045 | -0.055 | -0.055 | 0 | 0.1 | 0.31 | 0.1 | 0 | 0.1 | 0.02 |
| 10 | 127104136 | rs2125669 | 0.763 | 0.502 | 0.208 | 0.17 | 0.001 | 0.015 | 0.015 | 0 | 0.68 | 0.22 | 0.63 | 0.76 | 0.71 | 0.72 |
| 10 | 127104258 | rs2125668 | -0.577 | 0.368 | 1.095 | -0.101 | 0.004 | -0.041 | -0.041 | 0 | 0.18 | 0.26 | 0.12 | 0.06 | 0.18 | 0.2 |
| 10 | 127107129 | rs2169038 | -1.375 | -0.07 | 1.52 | -0.218 | -0.042 | -0.08 | -0.08 | 0.067 | 0.22 | 0.43 | 0.15 | 0.08 | 0.16 | 0.21 |
| 10 | 127113394 | rs4534531 | -2.16 | -0.006 | 1.92 | -0.359 | 0 | -0.012 | -0.012 | 0 | 0.29 | 0.67 | 0.24 | 0.12 | 0.24 | 0.25 |
| 10 | 127113626 | rs898262 | -1.243 | -0.007 | 2.122 | -0.357 | -0.049 | -0.086 | -0.086 | 0 | 0.22 | 0.46 | 0.14 | 0.09 | 0.16 | 0.2 |
| 10 | 127121196 | rs7895894 | -1.66 | 0.094 | 2.046 | -0.26 | 0.063 | -0.015 | -0.015 | 0 | 0.05 | 0.16 | 0.1 | 0.03 | 0.08 | 0.04 |
| 10 | 127132024 | rs11244490 | -2.275 | 0.099 | 2.08 | -0.277 | 0.064 | -0.024 | -0.024 | 0 | 0.07 | 0.22 | 0.11 | 0.03 | 0.09 | 0.05 |
| 10 | 127133053 | rs957416 | 0.735 | -0.148 | 1.885 | -0.633 | -0.03 | -0.073 | -0.073 | 0.605 | 0.73 | 0.61 | 0.85 | 0.91 | 0.81 | 0.78 |
| 10 | 127133684 | rs9422762 | 1.487 | -0.234 | 1.67 | -0.639 | 0.002 | -0.003 | -0.003 | 0 | 0.65 | 0.36 | 0.73 | 0.88 | 0.71 | 0.73 |
| 10 | 127135838 | rs989507 | 0.676 | -0.278 | 1.824 | -0.653 | -0.017 | -0.078 | -0.078 | 0 | 0.74 | 0.58 | 0.83 | 0.91 | 0.77 | 0.77 |
| 10 | 127136511 | rs3886847 | 1.4 | -0.183 | 1.781 | -0.652 | -0.001 | -0.068 | -0.068 | 0.318 | 0.67 | 0.43 | 0.75 | 0.87 | 0.7 | 0.74 |
| 10 | 127139427 | rs11244498 | -1.891 | -0.167 | 1.697 | -0.634 | 0.016 | -0.02 | -0.02 | 0.922 | 0.07 | 0.11 | 0.07 | 0.04 | 0.07 | 0.03 |
| 10 | 127140233 | rs3929230 | 1.063 | -0.455 | 0.489 | -0.55 | 0.007 | -0.05 | -0.05 | 0 | 0.4 | 0.12 | 0.25 | 0.45 | 0.37 | 0.34 |
| 10 | 127142806 | rs1350606 | -2.037 | -0.266 | 0.772 | -0.559 | 0.015 | -0.087 | -0.087 | 0 | 0.16 | 0.45 | 0.15 | 0.06 | 0.13 | 0.1 |

| | | | | | | | | | | | | | | | | |
|---|---|---|---|---|---|---|---|---|---|---|---|---|---|---|---|---|
| 10 | 127143515 | rs898260 | -1.187 | -0.201 | 0.677 | -0.518 | -0.002 | -0.073 | -0.073 | 0 | 0.33 | 0.56 | 0.25 | 0.13 | 0.3 | 0.26 |
| 10 | 127146175 | rs10793997 | 0.665 | -0.454 | -0.031 | -0.259 | 0.023 | -0.043 | -0.043 | 0.013 | 0.39 | 0.15 | 0.22 | 0.52 | 0.36 | 0.42 |
| 10 | 127148124 | rs6597706 | 0.188 | -0.496 | 0.283 | -0.355 | -0.001 | -0.018 | -0.018 | 0 | 0.21 | 0.3 | 0.13 | 0.06 | 0.19 | 0.17 |
| 10 | 127152944 | rs2886405 | 0.736 | -0.533 | 0.335 | -0.34 | 0.022 | -0.032 | -0.032 | 0 | 0.39 | 0.17 | 0.22 | 0.52 | 0.36 | 0.42 |
| 10 | 127153752 | rs17153010 | -0.546 | -0.578 | 0.566 | -0.398 | 0.007 | -0.075 | -0.075 | 0.014 | 0.27 | 0.49 | 0.2 | 0.11 | 0.26 | 0.28 |
| 10 | 127154372 | rs2045565 | 0.201 | -0.575 | 0.197 | -0.391 | -0.004 | -0.036 | -0.036 | 0.016 | 0.21 | 0.32 | 0.13 | 0.06 | 0.2 | 0.17 |
| 10 | 127163681 | rs3858289 | 1.065 | -0.63 | 0.085 | -0.254 | 0.048 | -0.07 | -0.07 | 0 | 0.35 | 0.08 | 0.24 | 0.5 | 0.3 | 0.31 |
| 10 | 127165054 | rs11244513 | -0.308 | -0.591 | 0.195 | -0.282 | -0.02 | -0.009 | -0.009 | 0 | 0.58 | 0.25 | 0.41 | 0.6 | 0.55 | 0.58 |
| 10 | 127168157 | rs3847459 | 1.298 | -0.587 | 0.155 | -0.25 | 0.036 | -0.105 | -0.105 | 0 | 0.33 | 0.03 | 0.22 | 0.44 | 0.27 | 0.3 |
| 10 | 127174440 | rs4962300 | -0.211 | -0.521 | -0.337 | -0.196 | -0.024 | -0.013 | -0.013 | 0.298 | 0.58 | 0.23 | 0.41 | 0.59 | 0.55 | 0.58 |
| 10 | 127175202 | rs4962461 | -0.075 | -0.567 | -0.528 | -0.197 | -0.037 | -0.023 | -0.023 | 0 | 0.57 | 0.18 | 0.38 | 0.54 | 0.53 | 0.57 |
| 10 | 127184196 | rs4962463 | 0.397 | -0.39 | -0.843 | -0.272 | -0.04 | 0.078 | 0.078 | 0 | 0.19 | 0.02 | 0.1 | 0.05 | 0.2 | 0.17 |
| 10 | 127190028 | rs1545045 | 0.936 | -1.207 | -0.918 | 0.059 | 0.035 | -0.133 | -0.133 | 0 | 0.33 | 0.02 | 0.22 | 0.46 | 0.27 | 0.3 |
| 10 | 127199963 | rs7088247 | 1.248 | -0.659 | -0.798 | 0.308 | -0.011 | -0.11 | -0.11 | 0.597 | 0.66 | 0.1 | 0.51 | 0.52 | 0.73 | 0.55 |
| 10 | 127201663 | rs4962465 | 0.568 | -0.779 | 0.32 | 0.251 | -0.006 | 0.072 | 0.072 | 0.003 | 0.74 | 0.53 | 0.64 | 0.53 | 0.78 | 0.59 |
| 10 | 127203938 | rs17153091 | -1.877 | -0.752 | 0.569 | 0.241 | 0.06 | 0.056 | 0.056 | 0.366 | 0.06 | 0.35 | 0.05 | 0 | 0.01 | 0.02 |
| 10 | 127209126 | rs10901399 | 1.167 | -0.785 | 0.403 | 0.244 | 0.004 | -0.03 | -0.03 | 0 | 0.68 | 0.19 | 0.54 | 0.53 | 0.73 | 0.57 |
| 10 | 127209521 | rs7077246 | 0.935 | -0.76 | 0.577 | 0.292 | -0.003 | 0.029 | 0.029 | 0.47 | 0.73 | 0.33 | 0.59 | 0.57 | 0.77 | 0.62 |
| 10 | 127213302 | rs4962309 | 0.567 | -0.738 | 0.616 | 0.284 | -0.02 | 0.085 | 0.085 | 0.377 | 0.78 | 0.43 | 0.7 | 0.64 | 0.84 | 0.65 |
| 10 | 127219324 | rs9630114 | 1.164 | -0.878 | 0.116 | 0.088 | 0.011 | -0.041 | -0.041 | 0.179 | 0.68 | 0.19 | 0.54 | 0.57 | 0.72 | 0.59 |
| 10 | 127221190 | rs755606 | -1.513 | -0.911 | 0.441 | 0.091 | 0.002 | -0.01 | -0.01 | 0 | 0.3 | 0.64 | 0.45 | 0.43 | 0.28 | 0.41 |
| 10 | 127229571 | rs17153141 | -0.863 | -1.033 | 0.718 | 0.14 | 0.001 | 0.059 | 0.059 | 0 | 0.24 | 0.39 | 0.38 | 0.37 | 0.24 | 0.37 |
| 10 | 127236712 | rs4310531 | 0.405 | -1.006 | 0.238 | 0.199 | 0 | -0.077 | -0.077 | 0.117 | 0.14 | 0.01 | 0.14 | 0.23 | 0.11 | 0.15 |
| 10 | 127237449 | rs1025512 | 0.089 | -0.894 | 0.193 | -0.14 | -0.001 | -0.003 | -0.003 | 0 | 0.07 | 0.11 | 0.06 | 0.05 | 0.08 | 0.04 |
| 10 | 127237918 | rs10794008 | 1.652 | -0.971 | 0.059 | -0.159 | -0.036 | -0.113 | -0.113 | 0.138 | 0.58 | 0.04 | 0.44 | 0.52 | 0.62 | 0.53 |
| 10 | 127239628 | rs4962471 | -1.571 | -0.62 | 0.568 | -0.131 | 0.011 | -0.033 | -0.033 | 0 | 0.13 | 0.28 | 0.16 | 0.05 | 0.13 | 0.09 |
| 10 | 127243392 | rs7068053 | -0.426 | -0.868 | 0.26 | -0.272 | 0.001 | -0.002 | -0.002 | 0.22 | 0.4 | 0.35 | 0.44 | 0.46 | 0.4 | 0.58 |
| 10 | 127243627 | rs4962473 | 0.339 | -0.743 | 0.35 | -0.292 | 0.046 | -0.002 | -0.002 | 0 | 0.95 | 0.76 | 0.91 | 1 | 0.95 | 0.97 |
| 10 | 127244526 | rs4962311 | -0.84 | -0.83 | 0.253 | -0.281 | -0.002 | 0 | 0 | 0 | 0.44 | 0.5 | 0.47 | 0.47 | 0.43 | 0.6 |
| 10 | 127246319 | rs10794009 | 0.11 | -0.65 | -0.671 | -0.216 | -0.014 | 0.011 | 0.011 | 0 | 0.61 | 0.53 | 0.56 | 0.54 | 0.62 | 0.44 |
| 10 | 127247815 | rs10901406 | 0.749 | -0.135 | -1.663 | 0.022 | -0.115 | -0.129 | -0.129 | 2.032 | 0.43 | 0.03 | 0.37 | 0.49 | 0.46 | 0.34 |
| 10 | 127255373 | rs7084723 | 0.209 | 0.24 | -0.25 | -0.089 | -0.003 | -0.015 | -0.015 | 0 | 0.78 | 0.72 | 0.75 | 0.82 | 0.8 | 0.76 |
| 10 | 127255951 | rs12242997 | 0.533 | 0.325 | -1.036 | -0.136 | -0.034 | -0.018 | -0.018 | 0 | 0.3 | 0.06 | 0.23 | 0.29 | 0.3 | 0.48 |
| 10 | 127256423 | rs7896345 | 0.173 | 0.267 | -0.839 | -0.346 | -0.002 | -0.016 | -0.016 | 3.284 | 0.71 | 0.68 | 0.71 | 0.8 | 0.71 | 0.74 |
| 10 | 127262447 | rs17153200 | 0.784 | 0.121 | -0.932 | 0.175 | -0.096 | -0.141 | -0.141 | 0.278 | 0.31 | 0.04 | 0.36 | 0.45 | 0.4 | 0.23 |
| 10 | 127264189 | rs7083854 | -0.016 | 0.146 | -0.257 | 0.206 | 0.004 | -0.013 | -0.013 | 0 | 0.41 | 0.56 | 0.65 | 0.81 | 0.52 | 0.38 |
| 10 | 127265374 | rs11244568 | 0.776 | 0.141 | -0.722 | 0.241 | -0.087 | -0.14 | -0.14 | 1.051 | 0.32 | 0.04 | 0.36 | 0.45 | 0.4 | 0.23 |
| 10 | 127269015 | rs1560749 | -1.572 | 0.125 | 0.084 | 0.344 | 0.118 | 0.228 | 0.228 | 1.039 | 0.07 | 0.41 | 0.24 | 0.24 | 0.11 | 0.04 |
| 10 | 127270143 | rs2043356 | -0.707 | 0.047 | -1.09 | 0.111 | 0 | 0.146 | 0.146 | 1.41 | 0.54 | 0.9 | 0.76 | 0.83 | 0.61 | 0.42 |
| 10 | 127279571 | rs17153228 | 0.253 | 0.072 | -0.372 | 0.339 | -0.011 | -0.014 | -0.014 | 3.442 | 0.06 | 0.02 | 0.07 | 0.13 | 0.06 | 0.12 |
| 10 | 127280109 | rs2365818 | -0.001 | -0.095 | -0.172 | 0.339 | -0.01 | 0.008 | 0.008 | 1.553 | 0.4 | 0.8 | 0.5 | 0.6 | 0.41 | 0.32 |
| 10 | 127295915 | rs297223 | -1.11 | -0.011 | 0.89 | 0.372 | -0.029 | -0.054 | -0.054 | 0 | 0.21 | 0.53 | 0.19 | 0.11 | 0.13 | 0.27 |
| 10 | 127297038 | rs12254965 | -1.286 | 0.286 | 1.061 | 0.867 | 0.044 | 0.048 | 0.048 | 0 | 0.05 | 0.18 | 0.09 | 0.17 | 0.11 | 0.02 |
| 10 | 127299121 | rs297226 | -1.654 | -0.201 | 1.235 | 0.676 | -0.003 | 0.013 | 0.013 | 0.273 | 0.35 | 0.75 | 0.4 | 0.29 | 0.32 | 0.34 |
| 10 | 127300265 | rs297227 | -0.436 | -0.226 | 2.04 | 0.63 | 0.03 | -0.068 | -0.068 | 0 | 0.19 | 0.36 | 0.15 | 0.1 | 0.21 | 0.13 |
| 10 | 127308741 | rs297256 | -1.938 | -0.298 | 1.847 | 0.607 | -0.029 | 0.045 | 0.045 | 1.05 | 0.37 | 0.9 | 0.42 | 0.29 | 0.32 | 0.34 |
| 10 | 127308873 | rs297255 | -1.938 | -0.298 | 1.847 | 0.607 | -0.029 | 0.045 | 0.045 | 1.05 | 0.37 | 0.9 | 0.42 | 0.29 | 0.32 | 0.34 |
| 10 | 127314030 | rs11244585 | -0.401 | -0.669 | 1.944 | 1.044 | -0.002 | -0.037 | -0.037 | 0 | 0.07 | 0.02 | 0.07 | 0.12 | 0.04 | 0.18 |
| 10 | 127316262 | rs297238 | -2.553 | -0.716 | 2.552 | 1.025 | -0.025 | 0.047 | 0.047 | 0.089 | 0.17 | 0.52 | 0.26 | 0.19 | 0.1 | 0.22 |
| 10 | 127317447 | rs11244587 | -0.436 | -0.685 | 2.3 | 1.027 | -0.002 | -0.037 | -0.037 | 0 | 0.07 | 0.02 | 0.07 | 0.12 | 0.04 | 0.19 |
| 10 | 127320259 | rs297236 | -2.053 | -0.708 | 2.455 | 0.997 | 0.023 | 0.043 | 0.043 | 0.095 | 0.1 | 0.46 | 0.19 | 0.08 | 0.06 | 0.03 |
| 10 | 127321822 | rs9422902 | -0.597 | -0.573 | 1.884 | 0.998 | 0 | 0 | 0 | 0.299 | 0.07 | 0.06 | 0.06 | 0.12 | 0.04 | 0.21 |
| 10 | 127325378 | rs9422904 | -0.748 | -0.602 | 1.851 | 0.988 | 0.009 | -0.01 | -0.01 | 0 | 0.07 | 0.05 | 0.05 | 0.1 | 0.04 | 0.21 |
| 10 | 127325453 | rs9422778 | -2.227 | -0.644 | 2.173 | 1.766 | -0.028 | 0.262 | 0.262 | 0.029 | 0.17 | 0.49 | 0.38 | 0.46 | 0.11 | 0.28 |
| 10 | 127329391 | rs9422906 | -1.133 | -0.875 | 1.317 | 1.68 | -0.123 | -0.016 | -0.016 | 0 | 0.6 | 0.98 | 0.69 | 0.71 | 0.53 | 0.64 |
| 10 | 127347690 | rs9422914 | -2.613 | -1.121 | 3.156 | 2.16 | -0.072 | -0.001 | -0.001 | 0 | 0.16 | 0.51 | 0.14 | 0.14 | 0.07 | 0.26 |
| 10 | 127368542 | rs12414594 | -1.566 | -0.89 | 2.32 | 2.12 | 0 | 0.096 | 0.096 | 0 | 0.09 | 0.24 | 0.28 | 0.24 | 0.07 | 0.04 |
| 10 | 127370912 | rs1483017 | -0.757 | -0.926 | 1.516 | 1.91 | 0.001 | 0.001 | 0.001 | 0 | 0.07 | 0.05 | 0.03 | 0.03 | 0.05 | 0.18 |
| 10 | 127371995 | rs3814625 | -2.264 | -0.899 | 1.327 | 1.942 | -0.015 | 0.021 | 0.021 | 0.009 | 0.24 | 0.7 | 0.44 | 0.43 | 0.15 | 0.28 |
| 10 | 127372751 | rs9422941 | -2.133 | -0.714 | 0.871 | 1.989 | -0.023 | 0.04 | 0.04 | 0.07 | 0.23 | 0.69 | 0.46 | 0.43 | 0.15 | 0.28 |
| 10 | 127375368 | rs9422804 | -1.285 | -0.34 | 0.328 | 0.777 | -0.037 | -0.045 | -0.045 | 0 | 0.12 | 0.33 | 0.09 | 0.04 | 0.07 | 0.18 |
| 10 | 127380451 | rs11244609 | -1.054 | 0.113 | -0.373 | 1.327 | -0.001 | 0.019 | 0.019 | 0 | 0.05 | 0.05 | 0.04 | 0.12 | 0.05 | 0.03 |
| 10 | 127387433 | rs11244615 | -0.889 | -0.644 | 0.03 | 1.678 | -0.05 | 0.165 | 0.165 | 2.18 | 0.23 | 0.48 | 0.33 | 0.53 | 0.12 | 0.44 |
| 10 | 127387775 | rs10901418 | -0.664 | -0.735 | 0 | 2.373 | -0.039 | 0.134 | 0.134 | 3.303 | 0.15 | 0.52 | 0.27 | 0.3 | 0.06 | 0.35 |
| 10 | 127388282 | rs4962480 | -0.749 | -0.145 | -1.654 | 1.097 | -0.089 | -0.019 | -0.019 | 0 | 0.67 | 0.96 | 0.76 | 0.72 | 0.61 | 0.65 |
| 10 | 127389009 | rs1872006 | -0.759 | -0.095 | -1.388 | 1.063 | -0.022 | 0.039 | 0.039 | 0.273 | 0.68 | 0.86 | 0.75 | 0.76 | 0.61 | 0.7 |
| 10 | 127394745 | rs10794017 | 0.284 | 0.101 | -1.751 | 0.709 | -0.074 | 0.017 | 0.017 | 0.271 | 0.21 | 0.05 | 0.22 | 0.14 | 0.34 | 0.13 |
| 10 | 127394788 | rs10794018 | -0.746 | -0.931 | -1.396 | 0.214 | 0.004 | 0.003 | 0.003 | 0 | 0.65 | 0.67 | 0.51 | 0.9 | 0.73 | 0.75 |
| 10 | 127400433 | rs7909932 | -0.095 | -1.37 | -1.399 | -0.068 | -0.003 | 0.001 | 0.001 | 0 | 0.87 | 0.84 | 0.86 | 0.92 | 0.89 | 0.92 |

| | | | | | | | | | | | | | | | | |
|---|---|---|---|---|---|---|---|---|---|---|---|---|---|---|---|---|
| 10 | 127425059 | rs11244626 | 0.207 | -2.024 | 0.518 | 0.312 | 0.057 | 0.034 | 0.034 | 0.087 | 0.61 | 0.25 | 0.48 | 0.77 | 0.75 | 0.76 |
| 10 | 127434834 | rs4962487 | -0.319 | -2.131 | 0.599 | 0.263 | 0 | 0.007 | 0.007 | 0 | 0.89 | 0.93 | 0.88 | 0.95 | 0.88 | 0.91 |
| 10 | 127436048 | rs2296833 | 0.775 | -2.686 | 0.4 | 0.852 | -0.024 | -0.061 | -0.061 | 0 | 0.28 | 0.1 | 0.21 | 0.41 | 0.32 | 0.48 |
| 10 | 127436411 | rs2296832 | -0.509 | -2.744 | 0.68 | 0.846 | -0.027 | -0.04 | -0.04 | 0.408 | 0.64 | 0.55 | 0.52 | 0.89 | 0.75 | 0.79 |
| 10 | 127436774 | rs11244632 | 0.654 | -2.726 | 0.469 | 1.073 | -0.006 | -0.045 | -0.045 | 0 | 0.29 | 0.2 | 0.22 | 0.45 | 0.32 | 0.48 |
| 10 | 127441439 | rs17153510 | 0.585 | -2.502 | 0.334 | 1.14 | -0.013 | -0.054 | -0.054 | 0 | 0.1 | 0 | 0.04 | 0.21 | 0.08 | 0.1 |
| 10 | 127444650 | rs10901423 | 1.546 | -2.295 | 0.414 | 1.087 | -0.083 | 0.025 | 0.025 | 0.023 | 0.22 | 0.05 | 0.23 | 0.14 | 0.35 | 0.18 |
| 10 | 127460753 | rs7922546 | -0.108 | -2.284 | 0.628 | 1.269 | -0.012 | 0.032 | 0.032 | 0.043 | 0.34 | 0.18 | 0.28 | 0.17 | 0.37 | 0.31 |
| 10 | 127463239 | rs12775804 | -0.069 | -2.164 | 0.608 | 1.481 | -0.002 | -0.01 | -0.01 | 0 | 0.33 | 0.21 | 0.29 | 0.3 | 0.34 | 0.3 |
| 10 | 127465348 | rs2170643 | -0.593 | -2.358 | 0.12 | 1.314 | 0.037 | 0.049 | 0.049 | 0 | 0.64 | 0.84 | 0.58 | 0.68 | 0.75 | 0.56 |
| 10 | 127469087 | rs10901430 | -1.037 | -2.25 | -0.164 | 1.28 | 0.024 | 0.022 | 0.022 | 0.157 | 0.69 | 0.83 | 0.62 | 0.72 | 0.78 | 0.79 |
| 10 | 127474375 | rs10901431 | 0.473 | -2.239 | 0.213 | 1.384 | 0.024 | -0.004 | -0.004 | 0.306 | 0.34 | 0.53 | 0.35 | 0.33 | 0.44 | 0.48 |
| 10 | 127476323 | rs10901432 | 0.561 | -2.25 | 0.569 | 1.41 | 0.038 | -0.011 | -0.011 | 0 | 0.34 | 0.44 | 0.34 | 0.31 | 0.44 | 0.45 |
| 10 | 127483573 | rs3740179 | 0.575 | -2.252 | 0.577 | 1.408 | 0.039 | -0.009 | -0.009 | 1.27 | 0.34 | 0.44 | 0.34 | 0.31 | 0.44 | 0.45 |
| 10 | 127484957 | rs2281953 | -0.48 | -2.295 | 0.464 | 1.467 | -0.017 | -0.027 | -0.027 | 0 | 0.93 | 1 | 0.92 | 0.86 | 0.94 | 0.76 |
| 10 | 127486429 | rs10901436 | 0.572 | -2.293 | 0.755 | 1.413 | 0.041 | -0.004 | -0.004 | 0 | 0.34 | 0.48 | 0.34 | 0.33 | 0.44 | 0.48 |
| 10 | 127486885 | rs10901437 | 0.581 | -2.642 | 0.587 | 0.74 | 0.051 | -0.001 | -0.001 | 0 | 0.3 | 0.4 | 0.33 | 0.3 | 0.43 | 0.43 |
| 10 | 127493414 | rs10794023 | -1.339 | -2.798 | 0.568 | 0.701 | 0.022 | 0.033 | 0.033 | 0.426 | 0.7 | 0.8 | 0.62 | 0.83 | 0.78 | 0.82 |
| 10 | 127495197 | rs12251135 | 0.121 | -2.828 | 0.56 | 0.743 | 0.004 | -0.05 | -0.05 | 0 | 0.34 | 0.21 | 0.26 | 0.4 | 0.34 | 0.33 |
| 10 | 127495807 | rs10510149 | 0.45 | -2.776 | 0.582 | 0.749 | 0.031 | -0.005 | -0.005 | 0 | 0.33 | 0.4 | 0.34 | 0.31 | 0.44 | 0.45 |
| 10 | 127520723 | rs4962491 | -1.115 | -2.822 | 0.467 | 0.666 | 0.002 | -0.005 | -0.005 | 0.05 | 0.92 | 0.97 | 0.97 | 0.86 | 0.94 | 0.92 |
| 10 | 127524612 | rs4075326 | 0.234 | -2.62 | 0.4 | 0.664 | 0.012 | -0.004 | -0.004 | 0 | 0.35 | 0.57 | 0.35 | 0.31 | 0.44 | 0.48 |
| 10 | 127528267 | rs3781058 | 0.404 | -0.262 | 0.68 | 0.589 | -0.007 | 0.002 | 0.002 | 0.009 | 0.32 | 0.31 | 0.31 | 0.29 | 0.41 | 0.43 |
| 10 | 127529135 | rs2282439 | 1.014 | -0.612 | 0.281 | 0.604 | -0.078 | -0.123 | -0.123 | 0 | 0.24 | 0.03 | 0.21 | 0.33 | 0.29 | 0.29 |
| 10 | 127532514 | rs17737550 | 1.549 | -0.468 | 0.399 | 0.651 | 0.006 | -0.029 | -0.029 | 0 | 0.07 | 0.01 | 0.03 | 0.07 | 0.05 | 0.02 |
| 10 | 127540321 | rs11244672 | 2.554 | -0.453 | 0.493 | 0.474 | 0.039 | 0.114 | 0.114 | 1.903 | 0.23 | 0.04 | 0.34 | 0.03 | 0.17 | 0.1 |
| 10 | 127561568 | rs10794029 | -1.616 | -0.558 | 0.499 | 0.4 | 0.028 | 0.038 | 0.038 | 0.385 | 0.67 | 0.78 | 0.6 | 0.83 | 0.77 | 0.81 |
| 10 | 127563509 | rs10901457 | 0.718 | -0.43 | 0.679 | 0.357 | -0.039 | -0.001 | -0.001 | 0.641 | 0.29 | 0.17 | 0.29 | 0.29 | 0.4 | 0.42 |
| 10 | 127565320 | rs10794031 | 0.274 | -0.388 | 0.502 | 0.378 | 0.025 | -0.001 | -0.001 | 0 | 0.33 | 0.56 | 0.34 | 0.32 | 0.43 | 0.47 |
| 10 | 127626702 | rs7070013 | 0.194 | -0.478 | 0.447 | 0.374 | -0.002 | -0.068 | -0.068 | 0.013 | 0.33 | 0.21 | 0.25 | 0.43 | 0.33 | 0.32 |
| 10 | 127630671 | rs2176101 | -1.431 | -0.558 | 0.556 | 0.34 | 0.014 | 0.013 | 0.013 | 0.346 | 0.66 | 0.76 | 0.59 | 0.72 | 0.76 | 0.79 |
| 10 | 127640999 | rs1878029 | 2.533 | -0.562 | 0.243 | 0.317 | 0.035 | 0.119 | 0.119 | 1.903 | 0.23 | 0.04 | 0.35 | 0.01 | 0.17 | 0.1 |
| 10 | 127641275 | rs10794041 | -1.536 | -0.332 | 0.411 | 0.471 | -0.032 | -0.029 | -0.029 | 0 | 0.65 | 0.62 | 0.57 | 0.73 | 0.76 | 0.8 |
| 10 | 127641756 | rs10794042 | 0.368 | -0.118 | 0.105 | 0.502 | 0.003 | -0.002 | -0.002 | 0.418 | 0.3 | 0.31 | 0.3 | 0.28 | 0.4 | 0.44 |
| 10 | 127642617 | rs10901476 | 0.461 | -0.096 | -0.015 | 0.498 | -0.01 | 0.004 | 0.004 | 0 | 0.29 | 0.27 | 0.29 | 0.27 | 0.4 | 0.43 |
| 10 | 127656067 | rs7907300 | -0.214 | 0.107 | -0.498 | 0.647 | -0.026 | -0.063 | -0.063 | 0.623 | 0.34 | 0.21 | 0.28 | 0.42 | 0.39 | 0.36 |
| 10 | 127662935 | rs7918092 | 0.708 | 0.773 | -0.333 | 0.911 | -0.006 | -0.005 | -0.005 | 0 | 0.16 | 0.13 | 0.21 | 0.29 | 0.25 | 0.19 |
| 10 | 127668487 | rs2176097 | 0.237 | 0.609 | -0.017 | 0.641 | 0.008 | -0.021 | -0.021 | 0.159 | 0.36 | 0.42 | 0.33 | 0.31 | 0.39 | 0.47 |
| 10 | 127671377 | rs10794049 | 0.193 | 0.596 | 0.111 | 0.662 | 0.008 | -0.018 | -0.018 | 0 | 0.37 | 0.46 | 0.33 | 0.32 | 0.39 | 0.48 |
| 10 | 127671666 | rs2366343 | 0.453 | 0.58 | 0.187 | 0.651 | -0.013 | 0.01 | 0.01 | 0 | 0.33 | 0.2 | 0.29 | 0.31 | 0.38 | 0.46 |
| 10 | 127675130 | rs11244752 | 0.156 | 0.577 | 0.193 | 0.604 | 0.006 | -0.007 | -0.007 | 0 | 0.37 | 0.47 | 0.33 | 0.35 | 0.39 | 0.47 |
| 10 | 127677374 | rs2280173 | 0.435 | 0.604 | 0.096 | 0.613 | -0.013 | 0.002 | 0.002 | 0 | 0.33 | 0.2 | 0.29 | 0.33 | 0.39 | 0.47 |
| 10 | 127680508 | rs10901498 | 1.692 | 0.264 | -0.024 | 0.441 | 0.004 | 0.003 | 0.003 | 1.034 | 0.31 | 0.33 | 0.41 | 0.12 | 0.24 | 0.15 |
| 10 | 127683083 | rs11244754 | 1.378 | 0.408 | -0.088 | 0.567 | -0.028 | -0.03 | -0.03 | 0 | 0.15 | 0.09 | 0.21 | 0.3 | 0.25 | 0.2 |
| 10 | 127685045 | rs3736478 | 1.066 | 0.051 | 0.038 | 0.483 | -0.03 | 0.058 | 0.058 | 0 | 0.06 | 0.16 | 0.03 | 0.17 | 0.01 | 0.05 |
| 10 | 127689040 | rs10901507 | -0.419 | 0.018 | -0.008 | 0.419 | 0.026 | 0.021 | 0.021 | 0 | 0.48 | 0.34 | 0.5 | 0.64 | 0.64 | 0.66 |
| 10 | 127693203 | rs7897850 | -0.82 | 0.078 | -0.019 | 0.351 | 0.004 | 0.002 | 0.002 | 0 | 0.54 | 0.5 | 0.53 | 0.8 | 0.65 | 0.7 |
| 10 | 127698643 | rs1878027 | -1.263 | -0.082 | 0.026 | 0.334 | 0.044 | 0.09 | 0.09 | 0 | 0.58 | 0.79 | 0.56 | 0.79 | 0.65 | 0.71 |
| 10 | 127703110 | rs1044169 | -1.126 | 0.001 | 0.019 | 0.372 | 0.044 | 0.079 | 0.079 | 0 | 0.56 | 0.73 | 0.55 | 0.79 | 0.65 | 0.71 |
| 10 | 127705404 | rs7913591 | -0.851 | -0.01 | 0.011 | 0.363 | -0.007 | -0.015 | -0.015 | 0.033 | 0.55 | 0.49 | 0.54 | 0.79 | 0.65 | 0.71 |
| 10 | 127710314 | rs1878026 | -1.368 | 0.084 | -0.642 | 0.382 | 0.008 | 0.019 | 0.019 | 0.612 | 0.86 | 0.94 | 0.94 | 0.89 | 0.89 | 0.86 |
| 10 | 127711296 | rs11244772 | 1.515 | 0.788 | -0.63 | 0.723 | -0.093 | 0.008 | 0.008 | 0 | 0.24 | 0.01 | 0.21 | 0.17 | 0.3 | 0.18 |
| 10 | 127716551 | rs9971063 | -1.266 | 0.786 | -0.196 | 0.574 | 0.049 | 0.086 | 0.086 | 0 | 0.56 | 0.74 | 0.56 | 0.79 | 0.65 | 0.7 |
| 10 | 127720923 | rs12251249 | 0.14 | 1.618 | -0.072 | 0.897 | 0.015 | 0.026 | 0.026 | 0 | 0.2 | 0.24 | 0.27 | 0.45 | 0.31 | 0.24 |
| 10 | 127722122 | rs11244778 | 0.313 | 1.642 | -0.472 | 0.874 | -0.013 | -0.012 | -0.012 | 0.435 | 0.19 | 0.16 | 0.25 | 0.45 | 0.32 | 0.24 |
| 10 | 127722422 | rs7920091 | 0.4 | 0.822 | -0.447 | 0.9 | -0.009 | -0.008 | -0.008 | 0.397 | 0.19 | 0.16 | 0.23 | 0.45 | 0.27 | 0.24 |
| 10 | 127729222 | rs7099772 | -0.703 | 0.112 | -0.656 | 0.242 | 0.038 | 0.128 | 0.128 | 0.087 | 0.51 | 0.73 | 0.52 | 0.79 | 0.58 | 0.69 |
| 10 | 127729734 | rs10794057 | 0.365 | 0.173 | -0.861 | 0.547 | -0.079 | -0.087 | -0.087 | 0 | 0.37 | 0.08 | 0.39 | 0.58 | 0.54 | 0.39 |
| 10 | 127729754 | rs1290149 | -0.703 | -0.269 | -0.618 | 0.329 | 0.049 | 0.045 | 0.045 | 0 | 0.59 | 0.82 | 0.56 | 0.7 | 0.69 | 0.47 |
| 10 | 127730360 | rs6693 | 0.394 | 0.396 | -0.205 | 0.388 | -0.007 | 0.033 | 0.033 | 0 | 0.52 | 0.31 | 0.56 | 0.56 | 0.52 | 0.46 |
| 10 | 127731197 | rs1278260 | 0.214 | 0.517 | 0.037 | 0.405 | 0.033 | 0.055 | 0.055 | 0 | 0.11 | 0.18 | 0.2 | 0.28 | 0.19 | 0.21 |
| 10 | 127731309 | rs7920207 | -0.085 | 0.219 | 0.273 | 0.08 | -0.007 | -0.043 | -0.043 | 0 | 0.36 | 0.44 | 0.41 | 0.16 | 0.3 | 0.37 |
| 10 | 127733872 | rs17154105 | 0.921 | 0.054 | 0.108 | -0.134 | 0.007 | 0.024 | 0.024 | 0.262 | 0.2 | 0.12 | 0.15 | 0.03 | 0.16 | 0.16 |
| 10 | 127734513 | rs1710313 | -1.205 | -0.593 | -0.372 | 0.117 | 0.071 | 0.039 | 0.039 | 1.65 | 0.51 | 0.92 | 0.51 | 0.62 | 0.34 | 0.61 |
| 10 | 127736134 | rs1878032 | -0.624 | -0.592 | 0.573 | 0.264 | -0.033 | 0.053 | 0.053 | 0 | 0.27 | 0.4 | 0.33 | 0.55 | 0.18 | 0.42 |
| 10 | 127738918 | rs1710278 | -1.157 | -0.323 | 0.453 | -0.106 | 0 | 0.027 | 0.027 | 0 | 0.45 | 0.6 | 0.6 | 0.9 | 0.46 | 0.63 |
| 10 | 127739633 | rs1710281 | -0.73 | -0.405 | 0.65 | -0.02 | 0.003 | 0.005 | 0.005 | 0 | 0.35 | 0.35 | 0.52 | 0.48 | 0.41 | 0.45 |
| 10 | 127740389 | rs3824637 | 0.83 | -0.387 | 0.493 | 0.013 | -0.064 | -0.077 | -0.077 | 0 | 0.17 | 0.04 | 0.27 | 0.32 | 0.28 | 0.19 |
| 10 | 127742255 | rs10901528 | 0.497 | -0.411 | 0.67 | 0.025 | -0.036 | 0.282 | 0.282 | 0 | 0.11 | 0.3 | 0.09 | 0.42 | 0.05 | 0.2 |

| | | | | | | | | | | | | | | | | |
|---|---|---|---|---|---|---|---|---|---|---|---|---|---|---|---|---|
| 10 | 127742664 | rs2138278 | -2.007 | -0.481 | 0.586 | -0.128 | -0.028 | 0.105 | 0.105 | 0 | 0.66 | 0.84 | 0.76 | 0.93 | 0.62 | 0.81 |
| 10 | 127742994 | rs11813508 | 0.056 | -0.233 | 0.437 | -0.008 | 0.007 | 0.018 | 0.018 | 0 | 0.26 | 0.16 | 0.32 | 0.2 | 0.23 | 0.25 |
| 10 | 127743048 | rs1674888 | -1.2 | -0.373 | 0.48 | -0.107 | -0.003 | 0.001 | 0.001 | 0 | 0.54 | 0.48 | 0.67 | 0.51 | 0.57 | 0.59 |
| 10 | 127744299 | rs17154161 | 1.164 | -0.233 | 0.197 | 0.165 | -0.016 | 0.018 | 0.018 | 0 | 0.13 | 0.01 | 0.06 | 0.1 | 0.08 | 0.2 |
| 10 | 127746893 | rs1278264 | -0.762 | -0.155 | 0.361 | 0.198 | -0.009 | -0.014 | -0.014 | 0 | 0.34 | 0.3 | 0.52 | 0.48 | 0.41 | 0.43 |
| 10 | 127747228 | rs1278266 | -0.744 | -0.197 | 0.295 | 0.2 | -0.013 | -0.023 | -0.023 | 0 | 0.34 | 0.28 | 0.52 | 0.48 | 0.41 | 0.43 |
| 10 | 127747972 | rs1278271 | -0.672 | -0.149 | -0.086 | 0.189 | -0.035 | -0.039 | -0.039 | 0.297 | 0.33 | 0.2 | 0.51 | 0.44 | 0.41 | 0.41 |
| 10 | 127748839 | rs7076519 | -0.347 | -0.286 | 0.05 | 0.078 | 0.002 | -0.001 | -0.001 | 0 | 0.31 | 0.33 | 0.37 | 0.68 | 0.35 | 0.41 |
| 10 | 127748908 | rs1278272 | 0.969 | -0.684 | 0.017 | -0.158 | -0.013 | -0.015 | -0.015 | 0 | 0.22 | 0.25 | 0.15 | 0.13 | 0.15 | 0.17 |
| 10 | 127749598 | rs1278274 | -1.712 | -0.718 | -0.498 | -0.34 | -0.1 | 0.156 | 0.156 | 0 | 0.67 | 0.97 | 0.77 | 0.89 | 0.6 | 0.82 |
| 10 | 127753209 | rs2279089 | 0.768 | -0.953 | 0.049 | -0.172 | -0.003 | -0.004 | -0.004 | 0 | 0.14 | 0.18 | 0.28 | 0.13 | 0.1 | 0.18 |
| 10 | 127753274 | rs1278278 | -0.595 | -1.203 | 0.414 | 0.211 | -0.01 | 0.001 | 0.001 | 0.076 | 0.42 | 0.56 | 0.5 | 0.56 | 0.27 | 0.47 |
| 10 | 127753478 | rs1278279 | -0.08 | -1.083 | 0.086 | -0.04 | -0.009 | -0.034 | -0.034 | 0 | 0.33 | 0.38 | 0.43 | 0.16 | 0.24 | 0.32 |
| 10 | 127753985 | rs12098279 | 1.551 | -0.623 | -0.588 | 0.178 | 0 | 0.005 | 0.005 | 0 | 0.12 | 0.07 | 0.07 | 0.06 | 0.12 | 0.06 |
| 10 | 127755544 | rs1278280 | -2.021 | -0.476 | -1.248 | -0.132 | -0.109 | 0.075 | 0.075 | 0 | 0.75 | 0.98 | 0.81 | 0.89 | 0.66 | 0.84 |
| 10 | 127755642 | rs1278281 | 0.265 | -0.823 | -0.931 | -0.213 | 0 | 0 | 0 | 1.861 | 0.35 | 0.33 | 0.25 | 0.15 | 0.3 | 0.24 |
| 10 | 127755954 | rs1466356 | 0.703 | -0.608 | -1.487 | 0.06 | 0.001 | 0.003 | 0.003 | 0 | 0.12 | 0.1 | 0.17 | 0.26 | 0.2 | 0.27 |
| 10 | 127756280 | rs11244796 | 1.585 | -0.753 | -1.708 | -0.122 | 0.012 | 0.027 | 0.027 | 0.115 | 0.19 | 0.04 | 0.14 | 0.09 | 0.14 | 0.1 |
| 10 | 127759571 | rs1152655 | -1.796 | -0.47 | -1.749 | 0.13 | -0.082 | 0.006 | 0.006 | 0.173 | 0.73 | 0.97 | 0.81 | 0.77 | 0.65 | 0.82 |
| 10 | 127760309 | rs2290841 | 0.896 | -0.146 | -1.56 | 0.465 | -0.051 | -0.052 | -0.052 | 0 | 0.1 | 0.02 | 0.11 | 0.23 | 0.18 | 0.3 |
| 10 | 127764148 | rs3781013 | 0.728 | -0.565 | -1.369 | 0.349 | -0.008 | -0.008 | -0.008 | 0 | 0.16 | 0.2 | 0.12 | 0.12 | 0.09 | 0.13 |
| 10 | 127768110 | rs1380439 | -0.336 | -0.553 | -1.119 | 0.394 | 0.001 | 0.004 | 0.004 | 0 | 0.27 | 0.32 | 0.24 | 0.35 | 0.28 | 0.43 |
| 10 | 127769545 | rs985461 | -1.913 | -0.677 | -1.127 | 0.256 | -0.011 | 0.041 | 0.041 | 0 | 0.69 | 0.88 | 0.79 | 0.77 | 0.64 | 0.8 |
| 10 | 127769598 | rs985462 | 0.23 | -0.722 | -1.057 | 0.287 | -0.004 | 0.003 | 0.003 | 0 | 0.42 | 0.54 | 0.55 | 0.4 | 0.37 | 0.36 |
| 10 | 127772919 | rs11813475 | 0.706 | -0.901 | -1.237 | 0.217 | -0.021 | -0.012 | -0.012 | 0 | 0.11 | 0.19 | 0.11 | 0.12 | 0.07 | 0.09 |
| 10 | 127773376 | rs1278352 | -0.699 | -0.755 | -1.274 | 0.234 | 0.012 | 0.028 | 0.028 | 0 | 0.58 | 0.68 | 0.68 | 0.65 | 0.58 | 0.71 |
| 10 | 127780811 | rs10794059 | 0.148 | -1.123 | -1.14 | 0.583 | 0.003 | -0.007 | -0.007 | 0.051 | 0.7 | 0.24 | 0.5 | 0.61 | 0.75 | 0.7 |
| 10 | 127782061 | rs11244810 | 1.934 | -1.052 | -1.652 | 0.295 | -0.122 | 0.127 | 0.127 | 0.415 | 0.31 | 0.02 | 0.21 | 0.1 | 0.38 | 0.16 |
| 10 | 127782409 | rs1871054 | -0.141 | -1.276 | -1.378 | 0.517 | 0.009 | -0.003 | -0.003 | 0 | 0.59 | 0.51 | 0.43 | 0.44 | 0.51 | 0.43 |
| 10 | 127783034 | rs3781031 | -0.492 | -1.426 | -1.314 | 0.474 | -0.048 | -0.009 | -0.009 | 0 | 0.25 | 0.39 | 0.32 | 0.23 | 0.11 | 0.24 |
| 10 | 127784936 | rs3758420 | 1.109 | -1.19 | -1.714 | 0.373 | -0.057 | -0.053 | -0.053 | 0.53 | 0.19 | 0.03 | 0.16 | 0.28 | 0.25 | 0.34 |
| 10 | 127785087 | rs3758422 | 0.485 | -1.93 | -0.97 | 0.614 | 0.036 | -0.05 | -0.05 | 0 | 0.66 | 0.15 | 0.49 | 0.48 | 0.73 | 0.66 |
| 10 | 127786969 | rs2290842 | -0.291 | -1.814 | -0.854 | 0.695 | -0.001 | 0.088 | 0.088 | 0 | 0.74 | 0.52 | 0.52 | 0.64 | 0.76 | 0.68 |
| 10 | 127787142 | rs2290844 | -0.47 | -1.411 | -0.644 | 0.362 | 0.054 | 0.02 | 0.02 | 0.027 | 0.21 | 0.08 | 0.12 | 0.21 | 0.11 | 0.2 |
| 10 | 127789598 | rs2290845 | -0.561 | -0.79 | -0.499 | 0.368 | -0.057 | -0.01 | -0.01 | 0.018 | 0.2 | 0.4 | 0.14 | 0.17 | 0.07 | 0.17 |
| 10 | 127791184 | rs1278300 | -0.701 | 0.368 | -0.82 | 0.688 | -0.013 | 0.01 | 0.01 | 0 | 0.88 | 0.98 | 0.92 | 0.96 | 0.87 | 0.92 |
| 10 | 127798951 | rs1278302 | -0.492 | 0.689 | -0.273 | 0.536 | -0.015 | 0.037 | 0.037 | 0 | 0.44 | 0.73 | 0.49 | 0.65 | 0.46 | 0.59 |
| 10 | 127802660 | rs1278384 | -0.328 | 0.383 | -0.426 | 0.13 | 0.034 | 0.053 | 0.053 | 0 | 0.52 | 0.8 | 0.56 | 0.61 | 0.58 | 0.53 |
| 10 | 127804531 | rs1278387 | 0.825 | 0.476 | -0.474 | -0.012 | -0.082 | 0.116 | 0.116 | 0.321 | 0.27 | 0.1 | 0.22 | 0.01 | 0.39 | 0.2 |
| 10 | 127807171 | rs1278394 | 0.191 | 0.154 | 0.022 | 0.668 | 0.003 | 0.002 | 0.002 | 0 | 0.49 | 0.54 | 0.52 | 0.54 | 0.57 | 0.54 |
| 10 | 127808710 | rs1278396 | 0.075 | 0.246 | -0.252 | 0.363 | 0.02 | 0.196 | 0.196 | 0.052 | 0.53 | 0.86 | 0.65 | 0.76 | 0.58 | 0.59 |
| 10 | 127808724 | rs1278397 | 0.084 | 0.246 | -0.245 | 0.363 | 0.025 | 0.193 | 0.193 | 0.026 | 0.53 | 0.82 | 0.64 | 0.76 | 0.58 | 0.59 |
| 10 | 127814489 | rs1278313 | -0.214 | 0.043 | -0.425 | 0.235 | 0.008 | -0.015 | -0.015 | 1.319 | 0.76 | 0.84 | 0.7 | 0.69 | 0.77 | 0.63 |
| 10 | 127815531 | rs1278315 | 0.463 | 0.033 | -0.461 | 0.2 | 0.03 | 0.135 | 0.135 | 0 | 0.53 | 0.77 | 0.65 | 0.77 | 0.58 | 0.59 |
| 10 | 127816662 | rs1278318 | -0.187 | 0.167 | -1.263 | -0.429 | 0.002 | 0.003 | 0.003 | 0.434 | 0.76 | 0.77 | 0.78 | 0.83 | 0.78 | 0.66 |
| 10 | 127819788 | rs12248384 | 0.347 | 0.325 | -1.207 | -0.373 | 0 | -0.022 | -0.022 | 0 | 0.26 | 0.22 | 0.15 | 0.06 | 0.2 | 0.07 |
| 10 | 127820978 | rs1278325 | -0.002 | 0.578 | -1.091 | -0.096 | 0.023 | 0.079 | 0.079 | 0.003 | 0.52 | 0.7 | 0.64 | 0.73 | 0.58 | 0.6 |
| 10 | 127824919 | rs10794060 | -0.073 | 0.688 | -1.069 | 0.348 | -0.008 | -0.009 | -0.009 | 0 | 0.42 | 0.29 | 0.44 | 0.48 | 0.44 | 0.35 |
| 10 | 127825898 | rs12762349 | 0.266 | 0.571 | -1.366 | 0.338 | 0.004 | -0.021 | -0.021 | 0 | 0.17 | 0.1 | 0.11 | 0.15 | 0.15 | 0.19 |
| 10 | 127826762 | rs1278333 | -0.775 | 0.409 | -0.648 | 0.111 | 0.034 | -0.05 | -0.05 | 0 | 0.8 | 0.69 | 0.8 | 0.91 | 0.74 | 0.83 |
| 10 | 127827684 | rs7896870 | 0.369 | -0.113 | -0.795 | -0.079 | 0.003 | 0.003 | 0.003 | 0.839 | 0.55 | 0.37 | 0.53 | 0.36 | 0.55 | 0.36 |
| 10 | 127829304 | rs17154306 | 0.266 | -0.19 | -0.981 | -0.331 | -0.01 | 0.014 | 0.014 | 0 | 0.25 | 0.14 | 0.16 | 0.15 | 0.26 | 0.29 |
| 10 | 127829788 | rs1278336 | -1.422 | -0.157 | -0.657 | -0.487 | -0.011 | 0.023 | 0.023 | 0 | 0.8 | 0.87 | 0.86 | 0.94 | 0.75 | 0.81 |
| 10 | 127831501 | rs12761170 | 1.179 | -0.104 | -0.746 | -0.355 | -0.01 | 0.025 | 0.025 | 0.128 | 0.33 | 0.22 | 0.39 | 0.21 | 0.34 | 0.19 |
| 10 | 127834566 | rs11244841 | 1.872 | -0.079 | -1.112 | -0.537 | -0.039 | -0.008 | -0.008 | 0.167 | 0.21 | 0.06 | 0.26 | 0.18 | 0.24 | 0.13 |
| 10 | 127842432 | rs4272714 | 0.763 | -0.217 | -0.971 | -0.615 | 0.019 | 0.126 | 0.126 | 0.015 | 0.5 | 0.21 | 0.48 | 0.26 | 0.56 | 0.38 |
| 10 | 127843126 | rs12244973 | 1.429 | -0.175 | -1.033 | -0.693 | 0.006 | 0.017 | 0.017 | 0 | 0.16 | 0.12 | 0.14 | 0.05 | 0.14 | 0.21 |
| 10 | 127843169 | rs1676719 | -0.506 | 0.106 | -0.456 | -0.521 | 0.003 | -0.067 | -0.067 | 4.159 | 0.81 | 0.55 | 0.79 | 0.9 | 0.85 | 0.79 |
| 10 | 127843740 | rs1466359 | -0.289 | 0.111 | -0.435 | -0.502 | 0.002 | 0.006 | 0.006 | 0 | 0.47 | 0.47 | 0.46 | 0.72 | 0.43 | 0.62 |
| 10 | 127845310 | rs12412988 | 1.467 | 0.176 | -0.393 | -0.528 | 0 | -0.002 | -0.002 | 0.046 | 0.11 | 0.09 | 0.13 | 0.03 | 0.1 | 0.07 |
| 10 | 127845432 | rs11244849 | 0.821 | 0.144 | -0.615 | -0.302 | -0.005 | 0.006 | 0.006 | 0 | 0.32 | 0.33 | 0.34 | 0.5 | 0.22 | 0.52 |
| 10 | 127845775 | rs10901541 | 1.194 | 0.116 | -0.626 | -0.414 | -0.109 | 0.077 | 0.077 | 0 | 0.36 | 0.06 | 0.34 | 0.19 | 0.43 | 0.18 |
| 10 | 127846128 | rs1674919 | -0.357 | -0.106 | -0.315 | -0.248 | 0.001 | 0.014 | 0.014 | 0.5 | 0.58 | 0.35 | 0.64 | 0.72 | 0.61 | 0.66 |
| 10 | 127846669 | rs12778749 | -1.1 | 0.055 | -0.251 | -0.297 | -0.003 | -0.016 | -0.016 | 0 | 0.69 | 0.41 | 0.77 | 0.77 | 0.71 | 0.74 |
| 10 | 127848239 | rs1676731 | -0.113 | 0.092 | -0.153 | -0.249 | -0.001 | -0.001 | -0.001 | 2.154 | 0.6 | 0.41 | 0.66 | 0.73 | 0.62 | 0.69 |
| 10 | 127848468 | rs10794063 | -0.113 | 0.093 | -0.153 | -0.249 | -0.001 | -0.001 | -0.001 | 0 | 0.6 | 0.41 | 0.66 | 0.73 | 0.62 | 0.68 |
| 10 | 127848780 | rs12767127 | 1.51 | 0.112 | -0.305 | -0.281 | -0.005 | 0.031 | 0.031 | 0 | 0.11 | 0.05 | 0.12 | 0.03 | 0.1 | 0.06 |
| 10 | 127849127 | rs2169076 | 0.386 | 0.35 | -0.44 | -0.041 | -0.025 | 0.005 | 0.005 | 0 | 0.43 | 0.25 | 0.47 | 0.58 | 0.45 | 0.62 |
| 10 | 127849546 | rs1676733 | 1.857 | 0.35 | -0.461 | -0.047 | 0.003 | 0.012 | 0.012 | 0 | 0.16 | 0.07 | 0.18 | 0.09 | 0.17 | 0.07 |

| | | | | | | | | | | | | | | | | |
|---|---|---|---|---|---|---|---|---|---|---|---|---|---|---|---|---|
| 10 | 127850507 | rs1674924 | -0.425 | 0.4 | -0.376 | -0.028 | -0.002 | -0.027 | -0.027 | 0.007 | 0.61 | 0.48 | 0.66 | 0.75 | 0.62 | 0.69 |
| 10 | 127850629 | rs11244854 | 0.634 | 0.553 | -0.714 | 0.684 | -0.002 | -0.045 | -0.045 | 0.223 | 0.32 | 0.16 | 0.35 | 0.41 | 0.29 | 0.54 |
| 10 | 127851530 | rs1676741 | -0.971 | 0.722 | -0.893 | 0.787 | 0.006 | 0.067 | 0.067 | 0 | 0.95 | 0.88 | 0.96 | 0.85 | 0.95 | 0.87 |
| 10 | 127851943 | rs11244857 | 0.839 | 0.735 | -0.996 | 0.62 | -0.03 | 0.026 | 0.026 | 0.046 | 0.3 | 0.15 | 0.25 | 0.22 | 0.39 | 0.19 |
| 10 | 127851977 | rs10794064 | 0.358 | 0.702 | -0.852 | 1.203 | 0.001 | -0.001 | -0.001 | 0 | 0.62 | 0.8 | 0.61 | 0.37 | 0.66 | 0.32 |
| 10 | 127852395 | rs1674927 | -0.209 | 0.706 | -1.441 | 0.989 | -0.05 | -0.12 | -0.12 | 0.169 | 0.3 | 0.02 | 0.35 | 0.4 | 0.27 | 0.54 |
| 10 | 127855523 | rs898325 | 0.973 | 0.804 | -0.091 | 1.246 | 0.031 | 0.05 | 0.05 | 0 | 0.86 | 0.77 | 0.75 | 0.74 | 0.82 | 0.91 |
| 10 | 127856213 | rs1676744 | -0.934 | 0.323 | 0.021 | 0.954 | -0.026 | 0.033 | 0.033 | 0.055 | 0.1 | 0.37 | 0.09 | 0.12 | 0.03 | 0.1 |
| 10 | 127859150 | rs11592681 | -0.864 | 0.24 | -0.309 | 1.177 | -0.005 | 0.011 | 0.011 | 0 | 0.13 | 0.07 | 0.07 | 0.1 | 0.14 | 0.06 |
| 10 | 127859846 | rs1380444 | 0.053 | -0.455 | -0.338 | 0.89 | -0.003 | 0.004 | 0.004 | 0 | 0.45 | 0.48 | 0.49 | 0.29 | 0.42 | 0.46 |
| 10 | 127862926 | rs10901549 | -0.602 | -0.439 | -0.259 | 1.698 | 0.017 | -0.037 | -0.037 | 0 | 0.15 | 0.12 | 0.08 | 0.33 | 0.15 | 0.21 |
| 10 | 127868584 | rs12256460 | -0.661 | -0.471 | -0.195 | 1.702 | 0.015 | -0.028 | -0.028 | 0 | 0.16 | 0.15 | 0.09 | 0.34 | 0.15 | 0.21 |
| 10 | 127868744 | rs10901553 | -1.72 | -0.823 | 0.306 | 1.752 | -0.021 | 0.04 | 0.04 | 1.814 | 0.23 | 0.71 | 0.27 | 0.46 | 0.17 | 0.25 |
| 10 | 127869478 | rs1278377 | -1.549 | -1.06 | -0.225 | 1.248 | -0.144 | 0.176 | 0.176 | 0.011 | 0.64 | 0.97 | 0.68 | 0.91 | 0.5 | 0.57 |
| 10 | 127872166 | rs11244871 | 0.974 | -1.188 | -0.011 | 1.124 | 0.005 | 0.118 | 0.118 | 0 | 0.28 | 0.09 | 0.19 | 0.02 | 0.25 | 0.15 |
| 10 | 127874740 | rs2930121 | -2.364 | -0.813 | 1.295 | 2.454 | -0.018 | -0.003 | -0.003 | 0.098 | 0.15 | 0.72 | 0.22 | 0.12 | 0.07 | 0.08 |
| 10 | 127874820 | rs2927510 | -1.4 | -0.737 | 1.415 | 2.488 | -0.013 | 0.031 | 0.031 | 0 | 0.11 | 0.26 | 0.17 | 0.12 | 0.06 | 0.07 |
| 10 | 127879242 | rs7912236 | -2.333 | -0.458 | 0.91 | 3.198 | -0.054 | 0.134 | 0.134 | 1.055 | 0.23 | 0.56 | 0.26 | 0.51 | 0.17 | 0.24 |
| 10 | 127881422 | rs4962514 | -1.96 | -0.223 | 0.988 | 2.356 | -0.011 | 0.448 | 0.448 | 1.04 | 0.1 | 0.48 | 0.27 | 0.43 | 0.04 | 0.15 |
| 10 | 127890323 | rs10901555 | -1.343 | 0.38 | 0.217 | 1.578 | -0.002 | 0.007 | 0.007 | 0 | 0.16 | 0.17 | 0.16 | 0.25 | 0.18 | 0.2 |
| 10 | 127890797 | rs12416625 | -1.8 | 0.341 | 0.405 | 1.79 | -0.012 | 0.044 | 0.044 | 0 | 0.08 | 0.15 | 0.04 | 0.19 | 0.04 | 0.13 |
| 10 | 127898686 | rs10901559 | -1.283 | 0.964 | 1.108 | 1.296 | 0.072 | -0.049 | -0.049 | 0 | 0.16 | 0.36 | 0.24 | 0.1 | 0.24 | 0.07 |
| 10 | 127904710 | rs7089454 | -1.621 | 1.157 | 1.311 | 1.25 | 0.094 | -0.057 | -0.057 | 0 | 0.17 | 0.42 | 0.26 | 0.12 | 0.26 | 0.08 |
| 10 | 127911533 | rs10794069 | -1.458 | 1.097 | 1.517 | 1.968 | 0.05 | 0.038 | 0.038 | 0 | 0.16 | 0.26 | 0.33 | 0.25 | 0.27 | 0.13 |
| 10 | 127911584 | rs7097356 | -1.786 | 1.066 | 1.656 | 1.954 | 0.093 | 0.071 | 0.071 | 0 | 0.17 | 0.42 | 0.34 | 0.25 | 0.27 | 0.13 |
| 10 | 127915891 | rs10466250 | -0.169 | 0.52 | 1.305 | 1.759 | 0.022 | 0.047 | 0.047 | 0 | 0.51 | 0.69 | 0.5 | 0.58 | 0.53 | 0.36 |
| 10 | 127927283 | rs3858316 | -0.537 | 0.052 | 1.97 | 1.84 | -0.023 | 0.014 | 0.014 | 0.02 | 0.32 | 0.54 | 0.45 | 0.36 | 0.27 | 0.34 |
| 10 | 127929715 | rs4962528 | -1.159 | -0.051 | 2.137 | 2.021 | 0.001 | 0.029 | 0.029 | 0 | 0.54 | 0.73 | 0.62 | 0.65 | 0.49 | 0.68 |
| 10 | 127930820 | rs6597749 | -0.18 | -0.168 | 2.376 | 1.952 | -0.013 | 0.005 | 0.005 | 0 | 0.22 | 0.34 | 0.33 | 0.23 | 0.18 | 0.24 |
| 10 | 127936662 | rs17155554 | 0.681 | -0.14 | 2.396 | 1.861 | -0.001 | -0.005 | -0.005 | 0 | 0.22 | 0.15 | 0.15 | 0.3 | 0.22 | 0.34 |
| 10 | 127943457 | rs12771688 | 0.228 | -0.272 | 2.538 | 1.821 | -0.008 | -0.01 | -0.01 | 0.003 | 0.19 | 0.28 | 0.26 | 0.16 | 0.17 | 0.2 |
| 10 | 127944593 | rs11244909 | 0.532 | -0.338 | 2.512 | 1.782 | 0.036 | -0.006 | -0.006 | 0 | 0.54 | 0.23 | 0.51 | 0.5 | 0.6 | 0.42 |
| 10 | 127945261 | rs10794073 | 0.705 | -0.335 | 2.607 | 1.786 | -0.026 | 0.014 | 0.014 | 0.043 | 0.88 | 0.71 | 0.88 | 0.87 | 0.91 | 0.89 |
| 10 | 127946213 | rs11598454 | -0.256 | -0.325 | 2.348 | 1.845 | 0.011 | -0.001 | -0.001 | 0 | 0.32 | 0.21 | 0.31 | 0.35 | 0.31 | 0.45 |
| 10 | 127946858 | rs11244911 | -0.29 | -0.309 | 2.352 | 1.772 | 0.007 | -0.025 | -0.025 | 0 | 0.33 | 0.22 | 0.33 | 0.47 | 0.31 | 0.48 |
| 10 | 127948079 | rs6597753 | 0.779 | -0.35 | 2.484 | 1.761 | 0.003 | 0 | 0 | 0 | 0.49 | 0.46 | 0.47 | 0.42 | 0.53 | 0.35 |
| 10 | 127948238 | rs6597754 | -0.105 | -0.296 | 2.51 | 1.716 | -0.021 | -0.025 | -0.025 | 0 | 0.79 | 0.58 | 0.74 | 0.84 | 0.83 | 0.81 |
| 10 | 127951861 | rs3952692 | -1.142 | -0.397 | 2.442 | 1.701 | 0.008 | 0.001 | 0.001 | 0 | 0.52 | 0.37 | 0.52 | 0.59 | 0.48 | 0.66 |
| 10 | 127952874 | rs17684713 | 1.073 | -0.092 | 2.163 | 1.769 | -0.043 | -0.02 | -0.02 | 4.646 | 0.07 | 0.01 | 0.12 | 0.06 | 0.2 | 0.06 |
| 10 | 127952901 | rs7902734 | -0.388 | -0.579 | 2.341 | 1.693 | 0.017 | -0.015 | -0.015 | 0.079 | 0.34 | 0.19 | 0.25 | 0.42 | 0.31 | 0.45 |
| 10 | 127953097 | rs11244922 | 1.092 | -0.508 | 2.369 | 1.695 | -0.015 | 0.004 | 0.004 | 0 | 0.43 | 0.2 | 0.4 | 0.37 | 0.51 | 0.31 |
| 10 | 127956996 | rs4962331 | -0.295 | -0.603 | 2.247 | 1.542 | -0.001 | 0.011 | 0.011 | 0 | 0.4 | 0.5 | 0.46 | 0.6 | 0.4 | 0.54 |
| 10 | 127957104 | rs17684825 | 1.198 | -0.716 | 2.001 | 1.482 | -0.044 | -0.047 | -0.047 | 0 | 0.43 | 0.09 | 0.41 | 0.38 | 0.55 | 0.36 |
| 10 | 127958529 | rs12251014 | -0.312 | -0.702 | 2.353 | 1.509 | 0.003 | 0 | 0 | 0 | 0.34 | 0.27 | 0.28 | 0.4 | 0.33 | 0.42 |
| 10 | 127959634 | rs7070441 | 1.325 | -0.681 | 2.171 | 1.525 | -0.034 | -0.037 | -0.037 | 0.415 | 0.43 | 0.12 | 0.41 | 0.39 | 0.53 | 0.39 |
| 10 | 127961602 | rs7095359 | 0.841 | -0.746 | 2.467 | 1.539 | -0.034 | -0.005 | -0.005 | 0 | 0.69 | 0.54 | 0.67 | 0.71 | 0.8 | 0.75 |
| 10 | 127962222 | rs10901580 | 1.472 | -0.727 | 2.16 | 1.577 | -0.075 | -0.06 | -0.06 | 0.595 | 0.42 | 0.08 | 0.39 | 0.39 | 0.52 | 0.38 |
| 10 | 127962700 | rs4412676 | 1.132 | -0.722 | 2.269 | 1.589 | 0.005 | -0.002 | -0.002 | 0 | 0.67 | 0.3 | 0.58 | 0.65 | 0.79 | 0.72 |
| 10 | 127964149 | rs7072808 | 0.716 | -0.664 | 2.085 | 1.604 | 0.001 | 0.007 | 0.007 | 0 | 0.14 | 0.22 | 0.2 | 0.16 | 0.12 | 0.14 |
| 10 | 127964859 | rs17154643 | 0.132 | -0.648 | 1.811 | 1.567 | 0.003 | -0.036 | -0.036 | 0.226 | 0.11 | 0.01 | 0.1 | 0.12 | 0.07 | 0.08 |
| 10 | 127967660 | rs17154652 | 0.356 | -0.554 | 2.001 | 1.681 | 0.001 | 0.003 | 0.003 | 0 | 0.25 | 0.2 | 0.18 | 0.27 | 0.27 | 0.34 |
| 10 | 127973248 | rs11816765 | 0.453 | -0.5 | 1.932 | 1.535 | 0.002 | 0.014 | 0.014 | 0 | 0.14 | 0.29 | 0.22 | 0.16 | 0.13 | 0.11 |
| 10 | 127974470 | rs12220042 | 0.415 | -0.575 | 1.9 | 1.551 | -0.009 | 0.003 | 0.003 | 0 | 0.25 | 0.07 | 0.19 | 0.27 | 0.26 | 0.35 |
| 10 | 127975022 | rs7083626 | 0.261 | -0.487 | 2.117 | 1.605 | 0.015 | 0.07 | 0.07 | 0 | 0.16 | 0.4 | 0.29 | 0.23 | 0.14 | 0.17 |
| 10 | 127975335 | rs2366698 | 0.424 | -0.481 | 2.153 | 1.588 | 0.002 | 0.021 | 0.021 | 0 | 0.15 | 0.38 | 0.23 | 0.17 | 0.13 | 0.12 |
| 10 | 127976339 | rs2366699 | -0.772 | -0.467 | 1.713 | 1.581 | 0.017 | -0.019 | -0.019 | 0 | 0.44 | 0.74 | 0.51 | 0.5 | 0.41 | 0.51 |
| 10 | 127978216 | rs11244932 | 0.36 | -0.44 | 1.616 | 1.575 | -0.003 | -0.003 | -0.003 | 0 | 0.27 | 0.25 | 0.19 | 0.28 | 0.27 | 0.36 |
| 10 | 127978792 | rs3858320 | -0.578 | -0.218 | 0.8 | 1.624 | 0.016 | -0.011 | -0.011 | 0 | 0.44 | 0.66 | 0.48 | 0.5 | 0.41 | 0.52 |
| 10 | 127980603 | rs728104 | 0.314 | -0.097 | 0.369 | 1.424 | -0.021 | 0.002 | 0.002 | 0 | 0.76 | 0.62 | 0.69 | 0.76 | 0.82 | 0.85 |
| 10 | 127982469 | rs11244938 | 0.899 | 0.691 | -0.799 | 1.263 | -0.094 | 0.033 | 0.033 | 0.259 | 0.31 | 0.03 | 0.24 | 0.19 | 0.3 | 0.31 |
| 10 | 127996741 | rs4420178 | -0.467 | 1.416 | -0.041 | 1.364 | 0.016 | 0.014 | 0.014 | 0 | 0.12 | 0.2 | 0.16 | 0.15 | 0.21 | 0.07 |
| 10 | 127997303 | rs9422969 | -0.297 | 1.194 | -0.53 | 1.001 | -0.008 | 0.007 | 0.007 | 0 | 0.77 | 0.83 | 0.82 | 0.75 | 0.75 | 0.74 |
| 10 | 127998826 | rs11244948 | -1.021 | 1.38 | -0.089 | 1.329 | 0.037 | 0.044 | 0.044 | 0 | 0.28 | 0.45 | 0.37 | 0.37 | 0.35 | 0.26 |
| 10 | 127999424 | rs4962548 | -0.715 | 1.948 | -0.68 | 1.43 | -0.041 | -0.044 | -0.044 | 0 | 0.08 | 0.04 | 0.23 | 0.25 | 0.13 | 0.17 |
| 10 | 128001759 | rs10901587 | 0.171 | 1.626 | -1.087 | 1.126 | -0.002 | -0.003 | -0.003 | 0 | 0.24 | 0.19 | 0.23 | 0.22 | 0.23 | 0.23 |
| 10 | 128002008 | rs868589 | 0.393 | 1.697 | -1.879 | 0.383 | -0.029 | 0.082 | 0.082 | 0 | 0.23 | 0.06 | 0.18 | 0.13 | 0.26 | 0.14 |
| 10 | 128002338 | rs17154713 | 0.893 | 1.55 | -1.71 | 0.564 | 0.024 | 0.02 | 0.02 | 0 | 0.31 | 0.16 | 0.17 | 0.22 | 0.27 | 0.25 |
| 10 | 128003149 | rs873957 | 0.137 | 1.589 | -0.628 | 0.694 | 0.003 | -0.008 | -0.008 | 5.714 | 0.65 | 0.57 | 0.51 | 0.59 | 0.66 | 0.68 |
| 10 | 128005524 | rs10901589 | -0.388 | 0.89 | -0.582 | 0.842 | 0.006 | 0.027 | 0.027 | 0.115 | 0.51 | 0.34 | 0.53 | 0.54 | 0.48 | 0.43 |

| | | | | | | | | | | | | | | | | |
|---|---|---|---|---|---|---|---|---|---|---|---|---|---|---|---|---|
| 10 | 128007683 | rs7097346 | -1.192 | 0.993 | -0.355 | 0.567 | -0.001 | -0.004 | -0.004 | 1.525 | 0.64 | 0.6 | 0.75 | 0.82 | 0.65 | 0.65 |
| 10 | 128008557 | rs10901590 | 0.695 | 0.841 | -0.905 | 0.974 | -0.038 | -0.069 | -0.069 | 0.697 | 0.24 | 0.02 | 0.21 | 0.25 | 0.2 | 0.22 |
| 10 | 128021575 | rs10901596 | -0.743 | 0.06 | -0.102 | 0.746 | 0.001 | 0.01 | 0.01 | 0 | 0.54 | 0.54 | 0.45 | 0.25 | 0.49 | 0.39 |
| 10 | 128025384 | rs10901597 | 0.362 | 0.135 | -0.261 | 0.932 | -0.028 | -0.045 | -0.045 | 0 | 0.07 | 0.01 | 0.14 | 0.16 | 0.09 | 0.16 |
| 10 | 128027301 | rs10901598 | -0.818 | 0.198 | -0.03 | 1.169 | -0.012 | -0.009 | -0.009 | 0.18 | 0.56 | 0.61 | 0.6 | 0.5 | 0.55 | 0.46 |
| 10 | 128030262 | rs7910333 | 0.23 | -0.085 | 0.181 | 0.97 | 0.018 | 0.075 | 0.075 | 0 | 0.23 | 0.4 | 0.3 | 0.3 | 0.27 | 0.17 |
| 10 | 128039386 | rs10901600 | -0.397 | -0.21 | 0.236 | 0.866 | -0.006 | 0.006 | 0.006 | 0 | 0.28 | 0.21 | 0.29 | 0.24 | 0.3 | 0.23 |
| 10 | 128041169 | rs3892767 | 0.86 | -0.226 | 0.159 | 0.829 | -0.017 | 0.063 | 0.063 | 0 | 0.19 | 0.04 | 0.13 | 0.07 | 0.21 | 0.09 |
| 10 | 128043536 | rs10751545 | -2.057 | -0.295 | 0.22 | 0.817 | -0.02 | 0.014 | 0.014 | 0 | 0.65 | 0.74 | 0.76 | 0.66 | 0.64 | 0.62 |
| 10 | 128045741 | rs10901601 | -0.497 | -0.304 | 0.311 | 0.845 | 0.016 | 0.017 | 0.017 | 0 | 0.41 | 0.26 | 0.39 | 0.36 | 0.37 | 0.43 |
| 10 | 128053597 | rs10751548 | -1.987 | -0.286 | 0.265 | 0.851 | -0.014 | 0.015 | 0.015 | 0 | 0.64 | 0.72 | 0.76 | 0.67 | 0.64 | 0.65 |
| 10 | 128053754 | rs10794080 | -1.106 | -0.29 | 0.365 | 0.801 | -0.01 | -0.011 | -0.011 | 4.094 | 0.55 | 0.61 | 0.59 | 0.48 | 0.55 | 0.48 |
| 10 | 128058851 | rs2366705 | 0.083 | -0.252 | -0.345 | 0.714 | 0.022 | 0.06 | 0.06 | 0.055 | 0.33 | 0.18 | 0.23 | 0.18 | 0.28 | 0.28 |
| 10 | 128063396 | rs4357606 | 0.839 | -0.364 | -0.418 | 0.692 | -0.015 | 0.061 | 0.061 | 0 | 0.19 | 0.04 | 0.13 | 0.07 | 0.21 | 0.08 |
| 10 | 128066982 | rs17759651 | 0.458 | -0.641 | -0.779 | 0.548 | 0.015 | 0.048 | 0.048 | 0 | 0.25 | 0.15 | 0.13 | 0.03 | 0.22 | 0.2 |
| 10 | 128068677 | rs11244970 | -0.67 | -0.496 | -0.498 | 0.479 | -0.007 | 0.107 | 0.107 | 0 | 0.12 | 0.15 | 0.32 | 0.41 | 0.11 | 0.17 |
| 10 | 128074094 | rs12355374 | 1.024 | -0.499 | -0.807 | 0.082 | -0.007 | -0.038 | -0.038 | 0 | 0.06 | 0 | 0.1 | 0.16 | 0.05 | 0.09 |
| 10 | 128074225 | rs7081193 | 0.37 | -0.551 | -1.024 | -0.191 | 0.005 | 0.007 | 0.007 | 0 | 0.12 | 0.06 | 0.07 | 0.06 | 0.09 | 0.06 |
| 10 | 128074661 | rs7081505 | -1.45 | -0.287 | -0.951 | 0.012 | 0.008 | -0.009 | -0.009 | 0.056 | 0.65 | 0.57 | 0.75 | 0.67 | 0.65 | 0.61 |
| 10 | 128075119 | rs11244975 | 0.374 | -0.736 | -1.213 | 0.521 | -0.006 | -0.039 | -0.039 | 0 | 0.07 | 0.01 | 0.11 | 0.16 | 0.05 | 0.09 |
| 10 | 128080672 | rs11244976 | -0.883 | -1.278 | -1.182 | 0.367 | 0.005 | -0.004 | -0.004 | 0.145 | 0.31 | 0.32 | 0.25 | 0.35 | 0.25 | 0.3 |
| 10 | 128082813 | rs12783662 | 1.085 | -0.778 | -1.87 | 0.545 | -0.031 | -0.031 | -0.031 | 0 | 0.08 | 0.01 | 0.14 | 0.12 | 0.12 | 0.1 |
| 10 | 128083065 | rs17154787 | 0.975 | -0.593 | -1.641 | 0.498 | 0.009 | 0.01 | 0.01 | 0 | 0.1 | 0.08 | 0.05 | 0.06 | 0.07 | 0.05 |
| 10 | 128085524 | rs4370848 | -1.18 | 0.227 | -1.901 | 0.641 | -0.186 | -0.026 | -0.026 | 0 | 0.71 | 0.98 | 0.59 | 0.71 | 0.58 | 0.74 |
| 10 | 128085765 | rs10794082 | -1.362 | 0.413 | -0.482 | 1.368 | -0.002 | 0.015 | 0.015 | 0 | 0.28 | 0.36 | 0.27 | 0.34 | 0.27 | 0.32 |
| 10 | 128086622 | rs4962553 | -1.113 | 0.706 | -0.928 | 1.664 | 0 | -0.02 | -0.02 | 0 | 0.16 | 0.12 | 0.18 | 0.23 | 0.19 | 0.2 |
| 10 | 128089672 | rs12569571 | -0.704 | 0.624 | -0.737 | 1.345 | 0 | 0.099 | 0.099 | 0 | 0.14 | 0.23 | 0.26 | 0.48 | 0.16 | 0.26 |
| 10 | 128094493 | rs7919486 | -0.593 | 0.477 | -1.309 | 0.225 | 0.06 | 0.162 | 0.162 | 0 | 0.5 | 0.78 | 0.51 | 0.79 | 0.39 | 0.63 |
| 10 | 128097413 | rs7084957 | 0.362 | 0.909 | -1.145 | 0.12 | -0.008 | 0.009 | 0.009 | 0 | 0.6 | 0.66 | 0.6 | 0.34 | 0.49 | 0.46 |
| 10 | 128099361 | rs7095414 | -0.109 | 1.496 | -1.025 | 0.013 | 0.02 | -0.027 | -0.027 | 0 | 0.94 | 0.89 | 0.94 | 0.99 | 0.89 | 0.92 |
| 10 | 128103811 | rs4246191 | -0.109 | 1.496 | -1.025 | 0.013 | 0.02 | -0.027 | -0.027 | 0.037 | 0.94 | 0.89 | 0.94 | 0.99 | 0.89 | 0.92 |
| 10 | 128107981 | rs4077952 | -0.421 | 1.594 | -1.409 | -0.474 | 0 | 0.036 | 0.036 | 0 | 0.58 | 0.32 | 0.58 | 0.73 | 0.59 | 0.69 |
| 10 | 128113320 | rs10901609 | 1.032 | 1.165 | -1.601 | -0.207 | -0.009 | 0.071 | 0.071 | 0 | 0.26 | 0.08 | 0.25 | 0.1 | 0.21 | 0.22 |
| 10 | 128114325 | rs3812676 | 0.387 | 0.601 | -0.991 | 0.025 | 0.011 | 0.023 | 0.023 | 0 | 0.31 | 0.19 | 0.3 | 0.27 | 0.25 | 0.21 |
| 10 | 128117100 | rs12260659 | 0.366 | 0.584 | -0.995 | 0.091 | 0.01 | 0.018 | 0.018 | 0 | 0.31 | 0.18 | 0.3 | 0.27 | 0.26 | 0.22 |
| 10 | 128117765 | rs7099858 | -0.266 | 0.677 | -0.75 | 1.427 | 0.006 | -0.009 | -0.009 | 1.288 | 0.2 | 0.13 | 0.11 | 0.32 | 0.16 | 0.19 |
| 10 | 128126229 | rs10794083 | 0.272 | 0.649 | -0.536 | 0.735 | -0.007 | -0.01 | -0.01 | 0 | 0.86 | 0.85 | 0.93 | 0.93 | 0.9 | 0.93 |
| 10 | 128138653 | rs7076654 | 0.828 | 0.887 | -0.542 | 0.639 | 0 | 0.006 | 0.006 | 0 | 0.48 | 0.52 | 0.47 | 0.36 | 0.43 | 0.5 |
| 10 | 128139066 | rs6597770 | -0.514 | 1.387 | -1.243 | -0.089 | -0.009 | 0.006 | 0.006 | 0.077 | 0.83 | 0.9 | 0.85 | 0.98 | 0.77 | 0.93 |
| 10 | 128141955 | rs4962558 | -0.337 | 1 | -0.808 | 0.805 | 0.01 | 0.052 | 0.052 | 0 | 0.18 | 0.35 | 0.3 | 0.28 | 0.19 | 0.29 |
| 10 | 128143338 | rs10751556 | 0.64 | 0.88 | -1.396 | 0.412 | -0.007 | 0.007 | 0.007 | 0 | 0.83 | 0.97 | 0.9 | 0.89 | 0.85 | 0.91 |
| 10 | 128144078 | rs7091466 | 0.787 | 0.992 | -1.315 | 0.34 | -0.001 | 0.026 | 0.026 | 0 | 0.79 | 0.87 | 0.89 | 0.87 | 0.81 | 0.87 |
| 10 | 128145099 | rs10901612 | -0.322 | 1.101 | -0.893 | 0.348 | 0.004 | -0.018 | -0.018 | 1.632 | 0.57 | 0.46 | 0.47 | 0.25 | 0.48 | 0.6 |
| 10 | 128148941 | rs7082278 | -0.478 | 1.354 | -0.946 | 0.7 | -0.024 | -0.004 | -0.004 | 0 | 0.13 | 0.06 | 0.09 | 0.13 | 0.14 | 0.14 |
| 10 | 128153623 | rs7079392 | -1.021 | 1.07 | -0.903 | 0.953 | 0.014 | -0.001 | -0.001 | 0 | 0.52 | 0.32 | 0.47 | 0.52 | 0.54 | 0.66 |
| 10 | 128156254 | rs13328738 | -1.021 | 0.775 | -0.021 | 1.22 | 0.006 | -0.005 | -0.005 | 0 | 0.32 | 0.34 | 0.19 | 0.25 | 0.31 | 0.18 |
| 10 | 128160275 | rs17154831 | -0.047 | 0.736 | -0.008 | 1.185 | 0 | -0.005 | -0.005 | 0 | 0.18 | 0.2 | 0.08 | 0.07 | 0.17 | 0.05 |
| 10 | 128165332 | rs7919269 | -1.26 | 0.688 | 0.353 | 1.172 | 0.001 | -0.003 | -0.003 | 0 | 0.33 | 0.58 | 0.21 | 0.25 | 0.31 | 0.19 |
| 10 | 128167728 | rs11244996 | -0.478 | 0.758 | 0.593 | 1.275 | 0.108 | 0.104 | 0.104 | 0 | 0.15 | 0.43 | 0.13 | 0.18 | 0.14 | 0.14 |
| 10 | 128172651 | rs7086835 | -0.191 | 0.585 | -0.6 | 1.109 | 0.003 | -0.062 | -0.062 | 0.113 | 0.23 | 0.04 | 0.28 | 0.31 | 0.23 | 0.47 |
| 10 | 128174706 | rs10901616 | -0.009 | 0.883 | -0.283 | 1.023 | -0.029 | -0.009 | -0.009 | 0.053 | 0.12 | 0.06 | 0.1 | 0.12 | 0.14 | 0.12 |
| 10 | 128187245 | rs7918878 | -1.27 | -0.003 | -0.381 | 1.012 | 0.041 | 0.018 | 0.018 | 0 | 0.52 | 0.76 | 0.4 | 0.57 | 0.42 | 0.6 |
| 10 | 128187301 | rs7907159 | -1.312 | -0.023 | -0.509 | 1.002 | 0.033 | 0.009 | 0.009 | 0 | 0.52 | 0.8 | 0.41 | 0.57 | 0.42 | 0.6 |
| 10 | 128190918 | rs10901621 | 0.109 | -0.018 | -0.64 | 1.024 | -0.044 | -0.055 | -0.055 | 0 | 0.09 | 0.01 | 0.18 | 0.19 | 0.1 | 0.11 |
| 10 | 128191368 | rs10510159 | -0.695 | -0.69 | 0.204 | 0.646 | -0.012 | -0.041 | -0.041 | 0.138 | 0.35 | 0.37 | 0.17 | 0.09 | 0.24 | 0.23 |
| 10 | 128192049 | rs4246194 | -1.668 | -0.026 | 0.743 | 1.346 | 0.029 | 0.231 | 0.231 | 0 | 0.14 | 0.44 | 0.1 | 0.32 | 0.12 | 0.25 |
| 10 | 128192985 | rs11245007 | -1.794 | -0.769 | 0.473 | 0.913 | 0.042 | -0.017 | -0.017 | 0 | 0.41 | 0.69 | 0.41 | 0.51 | 0.31 | 0.5 |
| 10 | 128193368 | rs11245008 | -1.735 | -0.54 | 1.239 | 0.733 | -0.032 | -0.037 | -0.037 | 0 | 0.18 | 0.46 | 0.16 | 0.13 | 0.12 | 0.12 |
| 10 | 128193754 | rs10901623 | -0.988 | -0.474 | 0.666 | 0.778 | 0.004 | 0.003 | 0.003 | 0.287 | 0.13 | 0.08 | 0.13 | 0.14 | 0.11 | 0.12 |
| 10 | 128200697 | rs11245011 | 1.278 | -0.108 | 0.656 | 0.789 | -0.003 | 0.004 | 0.004 | 0 | 0.86 | 0.9 | 0.86 | 0.85 | 0.86 | 0.87 |
| 10 | 128202435 | rs1436803 | 1.278 | -0.108 | 0.657 | 0.789 | -0.004 | 0.004 | 0.004 | 0 | 0.86 | 0.91 | 0.86 | 0.86 | 0.86 | 0.87 |
| 10 | 128203522 | rs4962565 | 1.836 | -0.191 | 1.048 | 0.752 | -0.003 | -0.025 | -0.025 | 0 | 0.82 | 0.59 | 0.84 | 0.83 | 0.85 | 0.87 |
| 10 | 128203545 | rs4962566 | 2.258 | -0.05 | 1.039 | 0.801 | -0.007 | 0.002 | 0.002 | 0.01 | 0.66 | 0.43 | 0.68 | 0.62 | 0.71 | 0.56 |
| 10 | 128208436 | rs11819200 | -0.859 | -0.015 | 0.985 | 0.8 | 0.001 | -0.011 | -0.011 | 1.183 | 0.16 | 0.12 | 0.16 | 0.21 | 0.14 | 0.31 |
| 10 | 128209491 | rs1865904 | 1.157 | 0.094 | 0.732 | 0.826 | -0.025 | -0.006 | -0.006 | 0 | 0.86 | 0.97 | 0.86 | 0.86 | 0.86 | 0.87 |
| 10 | 128221817 | rs7895050 | 0.875 | 0.285 | -0.205 | 0.909 | -0.045 | -0.022 | -0.022 | 0 | 0.88 | 0.99 | 0.87 | 0.86 | 0.85 | 0.88 |
| 10 | 128226017 | rs11245023 | -1.274 | -0.026 | -0.033 | 0.87 | -0.025 | 0 | 0 | 0.152 | 0.24 | 0.57 | 0.16 | 0.22 | 0.18 | 0.32 |
| 10 | 128226557 | rs7350443 | -1.122 | 0.71 | -0.418 | 1.206 | -0.023 | -0.023 | -0.023 | 0 | 0.08 | 0.04 | 0.1 | 0.16 | 0.12 | 0.08 |
| 10 | 128227080 | rs7916680 | -1.195 | 0.71 | -0.392 | 1.359 | -0.021 | -0.021 | -0.021 | 0 | 0.07 | 0.03 | 0.1 | 0.23 | 0.13 | 0.05 |

| | | | | | | | | | | | | | | | | |
|---|---|---|---|---|---|---|---|---|---|---|---|---|---|---|---|---|
| 10 | 128227329 | rs12259670 | 0.335 | 0.157 | -0.544 | 0.992 | -0.001 | 0.002 | 0.002 | 0 | 0.43 | 0.39 | 0.64 | 0.38 | 0.51 | 0.46 |
| 10 | 128229313 | rs10751560 | 0.086 | 0.126 | -1.036 | 0.995 | 0.037 | -0.037 | -0.037 | 0 | 0.51 | 0.87 | 0.7 | 0.47 | 0.56 | 0.48 |
| 10 | 128231556 | rs12780232 | 1.125 | -0.126 | -1.282 | 0.814 | 0.01 | 0.044 | 0.044 | 0.052 | 0.27 | 0.01 | 0.1 | 0.14 | 0.22 | 0.2 |
| 10 | 128232460 | rs7904837 | -0.027 | 0.171 | -0.744 | 0.882 | 0.039 | 0.026 | 0.026 | 0 | 0.5 | 0.75 | 0.62 | 0.44 | 0.55 | 0.47 |
| 10 | 128240277 | rs7916733 | 0.536 | 0.743 | -0.488 | 0.747 | -0.006 | -0.006 | -0.006 | 0.022 | 0.83 | 0.84 | 0.67 | 0.76 | 0.76 | 0.7 |
| 10 | 128244955 | rs7917910 | 0.12 | 0.298 | -0.75 | 0.721 | 0.057 | 0.003 | 0.003 | 0 | 0.39 | 0.15 | 0.24 | 0.38 | 0.29 | 0.33 |
| 10 | 128249486 | rs2118111 | 0.661 | 0.051 | -0.052 | 0.755 | 0.008 | -0.003 | -0.003 | 0 | 0.4 | 0.48 | 0.36 | 0.37 | 0.44 | 0.36 |
| 10 | 128249738 | rs7093782 | -0.253 | 0.231 | -0.53 | 0.776 | -0.001 | 0.032 | 0.032 | 0 | 0.93 | 0.92 | 0.81 | 0.79 | 0.92 | 0.76 |
| 10 | 128251618 | rs10794091 | -0.704 | 0.145 | -0.061 | 0.57 | -0.005 | -0.002 | -0.002 | 0.5 | 0.54 | 0.57 | 0.45 | 0.51 | 0.51 | 0.4 |
| 10 | 128252212 | rs11245044 | -0.975 | 0.053 | -0.763 | 0.755 | 0.004 | -0.014 | -0.014 | 0 | 0.18 | 0.12 | 0.18 | 0.35 | 0.16 | 0.23 |
| 10 | 128255032 | rs7067739 | -0.297 | 0.074 | -0.534 | 0.553 | 0 | 0.014 | 0.014 | 0 | 0.43 | 0.23 | 0.33 | 0.4 | 0.42 | 0.38 |
| 10 | 128257882 | rs4531372 | -0.131 | 0.127 | -0.86 | 0.587 | -0.043 | -0.027 | -0.027 | 0 | 0.4 | 0.09 | 0.32 | 0.4 | 0.42 | 0.35 |
| 10 | 128264561 | rs2082988 | 0.41 | 0.053 | -0.713 | 0.618 | -0.034 | -0.032 | -0.032 | 0 | 0.31 | 0.13 | 0.31 | 0.35 | 0.37 | 0.24 |
| 10 | 128266996 | rs11245051 | -0.015 | -0.162 | -0.453 | 0.854 | -0.003 | -0.005 | -0.005 | 0 | 0.61 | 0.7 | 0.53 | 0.43 | 0.58 | 0.49 |
| 10 | 128267560 | rs7898721 | 0.795 | -0.307 | -0.05 | 0.709 | -0.015 | 0.034 | 0.034 | 0 | 0.44 | 0.3 | 0.36 | 0.34 | 0.46 | 0.39 |
| 10 | 128271632 | rs10736874 | 1.606 | -0.362 | 0.492 | 0.935 | -0.039 | 0.031 | 0.031 | 0 | 0.77 | 0.65 | 0.69 | 0.7 | 0.85 | 0.61 |
| 10 | 128273149 | rs10901639 | -0.786 | 0.107 | 0.048 | 0.447 | -0.007 | -0.012 | -0.012 | 0 | 0.14 | 0.14 | 0.15 | 0.09 | 0.11 | 0.1 |
| 10 | 128283448 | rs2576027 | -1.42 | 0.496 | 0.653 | 1.015 | 0.015 | 0.137 | 0.137 | 0 | 0.09 | 0.27 | 0.14 | 0.2 | 0.04 | 0.26 |
| 10 | 128287185 | rs2576022 | -1.428 | 0.486 | 0.732 | 1 | 0.017 | 0.128 | 0.128 | 0 | 0.09 | 0.29 | 0.15 | 0.19 | 0.04 | 0.27 |
| 10 | 128291437 | rs2842164 | -1.093 | 0.497 | 0.488 | 0.976 | 0.001 | -0.005 | -0.005 | 0 | 0.07 | 0.06 | 0.13 | 0.13 | 0.04 | 0.26 |
| 10 | 128292604 | rs11597150 | 0.106 | 0.043 | 0.429 | 0.523 | 0.003 | -0.005 | -0.005 | 0 | 0.09 | 0.08 | 0.05 | 0 | 0.07 | 0.02 |
| 10 | 128294934 | rs12359545 | -0.753 | -0.22 | 0.507 | 0.47 | 0 | 0.004 | 0.004 | 0 | 0.19 | 0.18 | 0.25 | 0.25 | 0.18 | 0.38 |
| 10 | 128298104 | rs4962347 | -1.945 | -0.171 | 1.152 | 0.181 | 0.006 | 0.005 | 0.005 | 0 | 0.37 | 0.65 | 0.38 | 0.28 | 0.31 | 0.49 |
| 10 | 128299831 | rs10901640 | 0.992 | -0.306 | 0.045 | 0.11 | -0.146 | -0.082 | -0.082 | 0.38 | 0.3 | 0.07 | 0.37 | 0.4 | 0.47 | 0.34 |
| 10 | 128300327 | rs5023821 | -0.162 | -0.236 | 0.135 | -0.002 | 0.001 | -0.003 | -0.003 | 0 | 0.21 | 0.17 | 0.25 | 0.25 | 0.19 | 0.39 |
| 10 | 128307825 | rs2164583 | -0.458 | -0.239 | 0.232 | -0.026 | 0 | 0.003 | 0.003 | 0 | 0.23 | 0.25 | 0.28 | 0.25 | 0.24 | 0.39 |
| 10 | 128310426 | rs17155051 | 1.083 | 0.05 | 0.439 | -0.116 | 0.006 | 0.005 | 0.005 | 0 | 0.09 | 0.07 | 0.04 | 0 | 0.07 | 0.02 |
| 10 | 128311456 | rs2804432 | -0.93 | -0.003 | 0.571 | -0.11 | 0.007 | 0.009 | 0.009 | 0 | 0.08 | 0.35 | 0.08 | 0.03 | 0.05 | 0.08 |
| 10 | 128312030 | rs2804433 | -0.613 | -0.188 | 0.494 | -0.165 | 0.002 | 0.001 | 0.001 | 0.083 | 0.65 | 0.36 | 0.59 | 0.53 | 0.57 | 0.66 |
| 10 | 128316496 | rs2804439 | -0.834 | -0.218 | 0.231 | -0.181 | -0.008 | -0.005 | -0.005 | 0 | 0.66 | 0.66 | 0.62 | 0.53 | 0.53 | 0.63 |
| 10 | 128323682 | rs1865898 | -0.073 | -0.452 | 0.312 | 0.023 | -0.001 | 0 | 0 | 0 | 0.5 | 0.44 | 0.49 | 0.5 | 0.37 | 0.52 |
| 10 | 128333011 | rs1436789 | 0.623 | 0.347 | -0.495 | 0.068 | -0.024 | -0.059 | -0.059 | 0 | 0.41 | 0.14 | 0.51 | 0.48 | 0.43 | 0.4 |
| 10 | 128335225 | rs2842149 | -0.412 | 0.633 | -0.503 | 0.208 | -0.006 | -0.004 | -0.004 | 0 | 0.61 | 0.62 | 0.63 | 0.53 | 0.54 | 0.57 |
| 10 | 128338095 | rs7069588 | -0.358 | 0.672 | -0.8 | 0.096 | 0.004 | 0.013 | 0.013 | 1.614 | 0.13 | 0.1 | 0.07 | 0.05 | 0.09 | 0.17 |
| 10 | 128339929 | rs12258667 | 0.4 | 1.089 | -1.098 | 0.53 | -0.031 | -0.047 | -0.047 | 0 | 0.27 | 0.14 | 0.3 | 0.39 | 0.34 | 0.33 |
| 10 | 128340976 | rs11245078 | 0.254 | 1.039 | -1.2 | 0.526 | -0.073 | -0.079 | -0.079 | 0.048 | 0.29 | 0.08 | 0.31 | 0.38 | 0.38 | 0.35 |
| 10 | 128341404 | rs2002950 | 1.032 | 1.028 | -1.679 | 0.383 | -0.028 | -0.032 | -0.032 | 0 | 0.13 | 0.03 | 0.17 | 0.16 | 0.16 | 0.07 |
| 10 | 128343567 | rs2366867 | 1.454 | 0.545 | -1.346 | 0.176 | -0.003 | -0.004 | -0.004 | 0 | 0.26 | 0.27 | 0.2 | 0.2 | 0.16 | 0.15 |
| 10 | 128348107 | rs17155120 | 0.299 | 0.617 | -1.806 | 0.257 | -0.011 | -0.018 | -0.018 | 0 | 0.16 | 0.07 | 0.18 | 0.18 | 0.17 | 0.24 |
| 10 | 128353684 | rs12412594 | 0.728 | 0.814 | -2.022 | 0.361 | -0.026 | -0.046 | -0.046 | 0.284 | 0.11 | 0.01 | 0.11 | 0.14 | 0.12 | 0.18 |
| 10 | 128354226 | rs10736875 | 0.021 | 1.272 | -1.664 | 0.279 | -0.061 | -0.003 | -0.003 | 0 | 0.42 | 0.17 | 0.47 | 0.59 | 0.53 | 0.43 |
| 10 | 128355114 | rs11245085 | 1.137 | 1.35 | -1.901 | 0.369 | -0.028 | -0.033 | -0.033 | 0 | 0.13 | 0.04 | 0.16 | 0.16 | 0.16 | 0.07 |
| 10 | 128356095 | rs12354898 | 0.875 | 1.416 | -1.79 | 0.378 | -0.105 | -0.156 | -0.156 | 0.124 | 0.27 | 0.02 | 0.3 | 0.41 | 0.35 | 0.36 |
| 10 | 128362622 | rs7095083 | 0.286 | 1.278 | -1.119 | 0.283 | 0.001 | -0.017 | -0.017 | 0.168 | 0.12 | 0.17 | 0.1 | 0.03 | 0.12 | 0.17 |
| 10 | 128372802 | rs10751572 | 0.219 | 1.377 | -1.633 | 0.42 | -0.014 | -0.035 | -0.035 | 0 | 0.4 | 0.2 | 0.42 | 0.52 | 0.42 | 0.42 |
| 10 | 128372997 | rs4962587 | 0.096 | 1.145 | -1.347 | 0.328 | 0.035 | 0.055 | 0.055 | 0 | 0.47 | 0.7 | 0.46 | 0.38 | 0.4 | 0.39 |
| 10 | 128373226 | rs6597790 | -0.689 | 1.031 | -0.733 | 0.557 | -0.001 | 0.003 | 0.003 | 0 | 0.73 | 0.73 | 0.72 | 0.63 | 0.73 | 0.65 |
| 10 | 128373707 | rs10736877 | 0.734 | 0.595 | -0.28 | 0.335 | -0.023 | -0.015 | -0.015 | 0 | 0.77 | 0.44 | 0.77 | 0.83 | 0.82 | 0.76 |
| 10 | 128378598 | rs2366870 | -0.054 | 0.809 | -0.683 | -0.063 | -0.016 | 0.064 | 0.064 | 0 | 0.42 | 0.27 | 0.4 | 0.14 | 0.51 | 0.39 |
| 10 | 128386150 | rs10901656 | -0.408 | 0.697 | -0.282 | -0.123 | 0.002 | 0.017 | 0.017 | 0 | 0.51 | 0.45 | 0.43 | 0.22 | 0.53 | 0.53 |
| 10 | 128387149 | rs11245103 | 0.797 | 0.728 | -0.337 | -0.093 | 0.008 | 0 | 0 | 0.271 | 0.11 | 0.09 | 0.05 | 0.09 | 0.07 | 0.15 |
| 10 | 128387376 | rs10751575 | 0.576 | 0.53 | -0.371 | 0.565 | -0.011 | -0.021 | -0.021 | 0 | 0.15 | 0.11 | 0.36 | 0.52 | 0.16 | 0.23 |
| 10 | 128390001 | rs7921680 | 0.541 | 0.521 | -0.375 | 0.655 | -0.011 | -0.023 | -0.023 | 0 | 0.15 | 0.11 | 0.36 | 0.57 | 0.17 | 0.23 |
| 10 | 128390546 | rs11245105 | 0.846 | 0.44 | 0.097 | 0.461 | -0.009 | -0.014 | -0.014 | 4.932 | 0.32 | 0.31 | 0.18 | 0.2 | 0.26 | 0.23 |
| 10 | 128390803 | rs7908838 | 0.729 | 0.226 | 0.183 | 0.44 | -0.03 | 0.093 | 0.093 | 0 | 0.4 | 0.22 | 0.35 | 0.13 | 0.46 | 0.35 |
| 10 | 128390819 | rs10901663 | 0.57 | 0.115 | -0.1 | 0.243 | 0.001 | -0.003 | -0.003 | 0 | 0.35 | 0.31 | 0.22 | 0.2 | 0.32 | 0.28 |
| 10 | 128391896 | rs4962355 | 1.061 | 0.642 | -0.769 | -0.011 | -0.052 | 0.192 | 0.192 | 0 | 0.28 | 0.01 | 0.26 | 0 | 0.27 | 0.14 |
| 10 | 128393767 | rs2366875 | 0.083 | 0.729 | 0.201 | 0.694 | -0.008 | -0.005 | -0.005 | 0.233 | 0.87 | 0.8 | 0.93 | 0.88 | 0.9 | 0.79 |
| 10 | 128394130 | rs10736878 | -1.414 | 0.403 | 0.587 | 1.001 | -0.003 | -0.007 | -0.007 | 0 | 0.33 | 0.45 | 0.5 | 0.68 | 0.33 | 0.58 |
| 10 | 128400379 | rs10901666 | 0.909 | 0.643 | 0.439 | 1.399 | 0.003 | 0.001 | 0.001 | 0.091 | 0.6 | 0.52 | 0.67 | 0.52 | 0.53 | 0.5 |
| 10 | 128400660 | rs12251322 | 1.143 | 0.977 | -0.024 | 1.059 | -0.039 | -0.019 | -0.019 | 0.212 | 0.09 | 0.04 | 0.12 | 0.11 | 0.16 | 0.04 |
| 10 | 128401270 | rs7074840 | -0.721 | 1.019 | -0.003 | 1.126 | -0.018 | -0.014 | -0.014 | 0 | 0.22 | 0.2 | 0.21 | 0.24 | 0.29 | 0.17 |
| 10 | 128404888 | rs7905277 | -0.646 | -0.07 | 0.486 | 1.168 | -0.02 | 0.066 | 0.066 | 0 | 0.13 | 0.36 | 0.1 | 0.17 | 0.08 | 0.12 |
| 10 | 128405292 | rs7902474 | -0.136 | 0.004 | 0.021 | 1.272 | -0.013 | 0.032 | 0.032 | 0 | 0.11 | 0.18 | 0.07 | 0.19 | 0.08 | 0.11 |
| 10 | 128405932 | rs2366878 | -0.769 | 0.133 | -0.091 | 0.95 | -0.007 | 0.051 | 0.051 | 2 | 0.22 | 0.27 | 0.3 | 0.48 | 0.2 | 0.41 |
| 10 | 128406096 | rs2366879 | 0.58 | -0.093 | -0.492 | -0.052 | -0.051 | 0.055 | 0.055 | 0.987 | 0.43 | 0.19 | 0.34 | 0.36 | 0.47 | 0.44 |
| 10 | 128407765 | rs4962360 | -0.23 | 0.412 | -0.201 | -0.49 | -0.021 | 0.064 | 0.064 | 0.044 | 0.39 | 0.24 | 0.35 | 0.02 | 0.44 | 0.17 |
| 10 | 128408688 | rs17155177 | -0.66 | 0.116 | -0.204 | -0.045 | 0.007 | 0.003 | 0.003 | 0 | 0.47 | 0.44 | 0.58 | 0.48 | 0.46 | 0.36 |
| 10 | 128410393 | rs4628596 | -0.562 | 0.288 | -0.176 | -0.016 | -0.001 | 0.005 | 0.005 | 0 | 0.39 | 0.44 | 0.53 | 0.47 | 0.4 | 0.31 |

| | | | | | | | | | | | | | | | | |
|---|---|---|---|---|---|---|---|---|---|---|---|---|---|---|---|---|
| 10 | 128413223 | rs12771596 | 0.772 | -0.603 | -0.107 | -1.118 | 0.01 | 0.002 | 0.002 | 0 | 0.56 | 0.6 | 0.63 | 0.88 | 0.58 | 0.73 |
| 10 | 128413520 | rs2366886 | 0.733 | -0.669 | 0.001 | -1.044 | 0.002 | -0.019 | -0.019 | 1.652 | 0.59 | 0.56 | 0.65 | 0.87 | 0.64 | 0.74 |
| 10 | 128422770 | rs4962598 | 0.88 | -0.552 | -0.055 | -0.923 | 0.019 | 0.033 | 0.033 | 0.591 | 0.63 | 0.68 | 0.68 | 0.86 | 0.67 | 0.76 |
| 10 | 128423873 | rs10794108 | -0.726 | -0.687 | -0.14 | -1.003 | -0.001 | 0.114 | 0.114 | 0 | 0.28 | 0.05 | 0.23 | 0.11 | 0.29 | 0.17 |
| 10 | 128425046 | rs11245122 | -0.117 | -0.703 | -0.221 | -1.03 | -0.03 | 0.126 | 0.126 | 0 | 0.19 | 0.01 | 0.15 | 0 | 0.24 | 0.08 |
| 10 | 128429139 | rs7911765 | -1.186 | -0.469 | -0.056 | -0.926 | 0.02 | 0.052 | 0.052 | 0.271 | 0.38 | 0.27 | 0.33 | 0.14 | 0.35 | 0.24 |
| 10 | 128430469 | rs11245126 | 1.057 | -0.467 | -0.082 | -0.926 | 0.007 | 0.001 | 0.001 | 0.213 | 0.61 | 0.63 | 0.66 | 0.86 | 0.64 | 0.76 |
| 10 | 128431056 | rs10901683 | -1.062 | -0.311 | -0.272 | -0.824 | 0.021 | 0.057 | 0.057 | 0 | 0.34 | 0.23 | 0.27 | 0.11 | 0.32 | 0.16 |
| 10 | 128431443 | rs4494239 | -0.992 | -0.295 | -0.378 | -0.817 | 0.023 | 0.078 | 0.078 | 0 | 0.33 | 0.2 | 0.26 | 0.11 | 0.32 | 0.16 |
| 10 | 128438264 | rs10219075 | 0.422 | 0.196 | -0.073 | -0.316 | -0.013 | -0.063 | -0.063 | 0 | 0.78 | 0.67 | 0.73 | 0.94 | 0.81 | 0.82 |
| 10 | 128443813 | rs11245140 | -0.476 | 0.995 | -0.363 | 0.389 | 0.029 | 0.024 | 0.024 | 1.329 | 0.27 | 0.31 | 0.59 | 0.61 | 0.42 | 0.46 |
| 10 | 128445385 | rs2005556 | -0.519 | 0.034 | -0.705 | 0.21 | -0.016 | -0.011 | -0.011 | 0 | 0.36 | 0.29 | 0.49 | 0.43 | 0.44 | 0.59 |
| 10 | 128449111 | rs6597797 | 0.447 | -0.146 | -1.153 | -0.515 | 0.013 | 0.009 | 0.009 | 0 | 0.74 | 0.78 | 0.69 | 0.96 | 0.85 | 0.72 |
| 10 | 128458075 | rs4962610 | -0.085 | 0.468 | -0.956 | -0.122 | -0.002 | -0.014 | -0.014 | 0 | 0.89 | 0.84 | 0.76 | 0.96 | 0.91 | 0.8 |
| 10 | 128458218 | rs4962611 | -0.085 | 0.468 | -0.956 | -0.122 | -0.002 | -0.014 | -0.014 | 0 | 0.89 | 0.84 | 0.76 | 0.96 | 0.91 | 0.8 |
| 10 | 128459850 | rs4962613 | 0.035 | 0.435 | -0.981 | 0.201 | -0.002 | 0.031 | 0.031 | 0 | 0.47 | 0.34 | 0.43 | 0.64 | 0.51 | 0.36 |
| 10 | 128466412 | rs17690810 | -0.51 | 0.339 | -1.22 | 0.263 | -0.012 | 0.013 | 0.013 | 0 | 0.08 | 0 | 0.03 | 0.06 | 0.08 | 0.06 |
| 10 | 128466845 | rs10901701 | -0.714 | -0.35 | -0.853 | -0.084 | -0.019 | -0.04 | -0.04 | 0 | 0.36 | 0.48 | 0.31 | 0.23 | 0.3 | 0.33 |
| 10 | 128467258 | rs7100708 | 0.68 | -1.186 | -0.694 | -0.465 | 0.066 | 0.12 | 0.12 | 2.217 | 0.57 | 0.2 | 0.65 | 0.81 | 0.69 | 0.67 |
| 10 | 128469703 | rs4403722 | 0.915 | -0.869 | -1.619 | 0.057 | -0.075 | -0.046 | -0.046 | 0 | 0.29 | 0.11 | 0.27 | 0.36 | 0.42 | 0.33 |
| 10 | 128470148 | rs10901703 | -0.397 | -1.3 | -1.073 | 0.313 | -0.002 | 0 | 0 | 0 | 0.56 | 0.4 | 0.49 | 0.43 | 0.5 | 0.51 |
| 10 | 128479589 | rs10901705 | -0.779 | -1.249 | -0.633 | 0.369 | 0.006 | 0 | 0 | 0.022 | 0.6 | 0.52 | 0.5 | 0.44 | 0.5 | 0.51 |
| 10 | 128481434 | rs11597250 | 1.484 | -2.058 | -0.885 | 0.115 | -0.078 | -0.009 | -0.009 | 0 | 0.15 | 0.31 | 0.06 | 0.12 | 0.04 | 0.04 |
| 10 | 128485923 | rs7909809 | 0.366 | -1.912 | -0.961 | 0.024 | -0.056 | -0.034 | -0.034 | 0 | 0.3 | 0.01 | 0.21 | 0.27 | 0.27 | 0.23 |
| 10 | 128487203 | rs7090934 | -0.127 | -2.087 | -0.774 | 0.003 | -0.006 | -0.018 | -0.018 | 0.004 | 0.65 | 0.61 | 0.75 | 0.78 | 0.72 | 0.77 |
| 10 | 128488397 | rs4962623 | -0.086 | -2.337 | -0.222 | -0.127 | 0.013 | 0.012 | 0.012 | 3.078 | 0.67 | 0.35 | 0.71 | 0.91 | 0.78 | 0.73 |
| 10 | 128491473 | rs4635001 | -0.123 | -2.495 | -0.51 | -0.042 | -0.049 | 0.032 | 0.032 | 0.36 | 0.83 | 0.59 | 0.91 | 0.82 | 0.87 | 0.84 |
| 10 | 128494765 | rs10751582 | -0.765 | -2.502 | -0.548 | -0.069 | -0.018 | -0.067 | -0.067 | 0 | 0.7 | 0.45 | 0.72 | 0.91 | 0.78 | 0.73 |
| 10 | 128496291 | rs10901713 | 0.116 | -2.558 | -0.403 | -0.081 | 0.006 | 0.008 | 0.008 | 1.413 | 0.59 | 0.38 | 0.65 | 0.73 | 0.66 | 0.57 |
| 10 | 128507234 | rs4492724 | -0.691 | -2.513 | -0.399 | -0.056 | -0.02 | 0.058 | 0.058 | 0.389 | 0.87 | 0.67 | 0.91 | 0.82 | 0.88 | 0.86 |
| 10 | 128516156 | rs7084855 | 0.46 | -2.551 | -0.489 | -0.081 | 0.057 | 0.075 | 0.075 | 2.015 | 0.57 | 0.26 | 0.64 | 0.73 | 0.65 | 0.59 |
| 10 | 128527330 | rs6597817 | 0.725 | -2.586 | -0.493 | -0.228 | -0.016 | -0.08 | -0.08 | 0.063 | 0.27 | 0.37 | 0.27 | 0.07 | 0.21 | 0.27 |
| 10 | 128529658 | rs12267639 | 1.446 | -2.454 | -0.474 | -0.084 | 0.004 | -0.011 | -0.011 | 0 | 0.09 | 0.03 | 0.06 | 0.19 | 0.12 | 0.14 |
| 10 | 128530796 | rs4615932 | 0.205 | -2.42 | -0.337 | -0.077 | 0.04 | 0.037 | 0.037 | 5.539 | 0.95 | 0.63 | 0.97 | 1 | 1 | 1 |
| 10 | 128531680 | rs7911459 | -0.019 | -2.456 | -0.419 | -0.106 | -0.021 | -0.029 | -0.029 | 0 | 0.36 | 0.37 | 0.32 | 0.25 | 0.31 | 0.39 |
| 10 | 128532675 | rs4293044 | -0.123 | -2.463 | -0.661 | -0.115 | -0.006 | -0.008 | -0.008 | 0.136 | 0.38 | 0.37 | 0.31 | 0.26 | 0.32 | 0.41 |
| 10 | 128535769 | rs10901724 | -0.703 | -3.028 | -1.089 | -0.434 | 0.127 | 0.158 | 0.158 | 0.172 | 0.5 | 0.79 | 0.37 | 0.27 | 0.35 | 0.42 |
| 10 | 128537708 | rs961347 | 0.236 | -3.155 | -0.276 | -1.151 | -0.052 | -0.093 | -0.093 | 0.006 | 0.23 | 0.29 | 0.12 | 0.03 | 0.1 | 0.06 |
| 10 | 128538253 | rs17155407 | 1.28 | -3.068 | -0.116 | -1.119 | 0.031 | -0.065 | -0.065 | 0 | 0.17 | 0.47 | 0.2 | 0.04 | 0.18 | 0.24 |
| 10 | 128541675 | rs10901726 | -0.497 | -2.995 | -0.357 | -0.763 | 0.002 | 0.002 | 0.002 | 0 | 0.46 | 0.39 | 0.39 | 0.31 | 0.39 | 0.4 |
| 10 | 128546740 | rs6597823 | -0.174 | -2.765 | -0.607 | -0.744 | 0.027 | -0.01 | -0.01 | 0.14 | 0.32 | 0.21 | 0.22 | 0.34 | 0.21 | 0.18 |
| 10 | 128549120 | rs4474358 | -0.088 | -2.313 | -0.369 | -0.636 | 0.003 | 0.001 | 0.001 | 0 | 0.46 | 0.39 | 0.4 | 0.39 | 0.39 | 0.41 |
| 10 | 128553795 | rs7074496 | -0.535 | -1.675 | -0.55 | -0.678 | 0.015 | 0.017 | 0.017 | 0 | 0.64 | 0.38 | 0.49 | 0.46 | 0.55 | 0.56 |
| 10 | 128554642 | rs4962373 | -0.473 | -1.254 | -0.957 | -0.986 | -0.032 | -0.072 | -0.072 | 0 | 0.84 | 0.73 | 0.89 | 0.97 | 0.9 | 0.97 |
| 10 | 128556341 | rs11814321 | 0.031 | 0.243 | -1.797 | -0.874 | 0.006 | 0.006 | 0.006 | 0 | 0.38 | 0.4 | 0.23 | 0.24 | 0.29 | 0.38 |
| 10 | 128556410 | rs11814342 | 0.039 | 0.243 | -1.797 | -0.874 | 0.006 | 0.006 | 0.006 | 0 | 0.38 | 0.4 | 0.23 | 0.24 | 0.29 | 0.38 |
| 10 | 128556741 | rs4562717 | -1.07 | 0.729 | -1.873 | -0.785 | -0.007 | 0.11 | 0.11 | 0 | 0.75 | 0.59 | 0.63 | 0.4 | 0.8 | 0.67 |
| 10 | 128556923 | rs4570501 | -0.468 | 0.839 | -2.022 | -0.807 | -0.001 | -0.01 | -0.01 | 0 | 0.55 | 0.53 | 0.5 | 0.28 | 0.58 | 0.45 |
| 10 | 128558061 | rs11596809 | 1.576 | 1.099 | -1.92 | -0.666 | -0.038 | 0.022 | 0.022 | 0 | 0.2 | 0.05 | 0.13 | 0.13 | 0.21 | 0.21 |
| 10 | 128562826 | rs11245203 | -0.36 | 1.024 | -2.011 | -0.601 | 0.004 | 0.031 | 0.031 | 0 | 0.52 | 0.41 | 0.38 | 0.29 | 0.54 | 0.42 |
| 10 | 128565076 | rs11245204 | 0.915 | 0.953 | -2.076 | 0.355 | -0.074 | 0.092 | 0.092 | 0 | 0.36 | 0.1 | 0.31 | 0.17 | 0.45 | 0.26 |
| 10 | 128565859 | rs11245205 | 0.588 | 0.63 | -1.673 | 0.523 | -0.007 | 0.1 | 0.1 | 0 | 0.23 | 0.07 | 0.12 | 0.04 | 0.2 | 0.26 |
| 10 | 128569545 | rs9422843 | 0.71 | 0.844 | -0.512 | 1.537 | -0.018 | 0.048 | 0.048 | 0.25 | 0.72 | 0.45 | 0.65 | 0.42 | 0.81 | 0.65 |
| 10 | 128581315 | rs11245214 | -1.37 | 2.222 | -0.999 | 1.73 | -0.063 | -0.064 | -0.064 | 0 | 0.11 | 0.01 | 0.15 | 0.29 | 0.2 | 0.25 |
| 10 | 128581683 | rs12783987 | -1.68 | 2.466 | -0.817 | 2.347 | -0.022 | -0.01 | -0.01 | 0 | 0.11 | 0.04 | 0.2 | 0.43 | 0.22 | 0.23 |
| 10 | 128582657 | rs4553283 | -1.378 | 2.651 | -0.792 | 2.129 | -0.014 | -0.007 | -0.007 | 0 | 0.13 | 0.07 | 0.26 | 0.41 | 0.23 | 0.3 |
| 10 | 128584997 | rs4448606 | -1.815 | 2.605 | -0.187 | 1.709 | 0.042 | 0.041 | 0.041 | 0.055 | 0.23 | 0.32 | 0.36 | 0.41 | 0.34 | 0.34 |
| 10 | 128589414 | rs7910439 | -0.379 | 1.958 | 0.154 | 1.263 | 0.009 | -0.061 | -0.061 | 0 | 0.1 | 0.25 | 0.1 | 0 | 0.11 | 0.02 |
| 10 | 128592872 | rs7092254 | 0.63 | 1.362 | 0.322 | 1.783 | -0.032 | 0.083 | 0.083 | 0.706 | 0.88 | 0.76 | 0.85 | 0.75 | 0.95 | 0.89 |
| 10 | 128597733 | rs9422853 | -0.12 | 1.478 | -0.449 | 1.674 | -0.015 | -0.039 | -0.039 | 0 | 0.2 | 0.12 | 0.34 | 0.5 | 0.25 | 0.39 |
| 10 | 128598629 | rs7474861 | 0.167 | 1.465 | -0.394 | 1.669 | -0.006 | -0.024 | -0.024 | 0 | 0.77 | 0.83 | 0.66 | 0.5 | 0.75 | 0.61 |
| 10 | 128602101 | rs8181424 | 0.878 | 1.063 | -0.651 | 1 | -0.028 | 0.116 | 0.116 | 0.808 | 0.46 | 0.2 | 0.35 | 0.25 | 0.52 | 0.42 |
| 10 | 128606124 | rs10901740 | -0.206 | 1.024 | -0.137 | 1.46 | -0.001 | 0.091 | 0.091 | 0 | 0.06 | 0.17 | 0.15 | 0.15 | 0.04 | 0.1 |
| 10 | 128607787 | rs10794142 | -0.14 | 1.945 | -1.243 | 0.95 | -0.15 | -0.043 | -0.043 | 0 | 0.23 | 0.02 | 0.21 | 0.21 | 0.37 | 0.2 |
| 10 | 128608215 | rs10901741 | -0.502 | 2.038 | -0.458 | 1.423 | 0.033 | 0.057 | 0.057 | 0.48 | 0.24 | 0.3 | 0.3 | 0.48 | 0.37 | 0.29 |
| 10 | 128608985 | rs4255455 | 0.367 | 0.896 | -0.559 | 0.857 | 0.004 | -0.003 | -0.003 | 0 | 0.58 | 0.63 | 0.58 | 0.45 | 0.38 | 0.6 |
| 10 | 128613618 | rs7915184 | 0.393 | -0.244 | -0.859 | 0.102 | 0.005 | 0.005 | 0.005 | 0.97 | 0.18 | 0.21 | 0.1 | 0.11 | 0.11 | 0.12 |
| 10 | 128766533 | rs11018080 | -0.363 | 0.323 | -1.118 | 0.195 | -0.053 | -0.017 | -0.017 | 0 | 0.69 | 0.85 | 0.7 | 0.65 | 0.56 | 0.65 |
| 10 | 128768736 | rs11016850 | 1.086 | -0.186 | -0.969 | 0.349 | 0.013 | -0.036 | -0.036 | 0 | 0.19 | 0.11 | 0.22 | 0.56 | 0.13 | 0.29 |

| | | | | | | | | | | | | | | | | |
|---|---|---|---|---|---|---|---|---|---|---|---|---|---|---|---|---|
| 10 | 128768902 | rs11016943 | 1.254 | -0.368 | -1.021 | 0.102 | 0.004 | -0.002 | -0.002 | 0 | 0.09 | 0.08 | 0.04 | 0 | 0.07 | 0.02 |
| 10 | 128771878 | rs4075924 | 0.642 | -0.32 | -0.61 | 0.089 | 0.007 | 0.028 | 0.028 | 0 | 0.05 | 0.12 | 0.06 | 0.07 | 0.09 | 0.04 |
| 10 | 128774872 | rs10830183 | 1.367 | -0.347 | -0.6 | 0.286 | 0.01 | -0.031 | -0.031 | 1.115 | 0.18 | 0.1 | 0.21 | 0.48 | 0.13 | 0.28 |
| 10 | 128776941 | rs9732211 | -0.72 | -0.42 | -0.463 | -0.234 | -0.013 | 0.014 | 0.014 | 0 | 0.59 | 0.72 | 0.53 | 0.64 | 0.47 | 0.57 |
| 10 | 128778271 | rs10732439 | -0.561 | -0.434 | -0.47 | -0.24 | -0.003 | 0.006 | 0.006 | 0 | 0.57 | 0.6 | 0.52 | 0.64 | 0.47 | 0.57 |
| 10 | 128779958 | rs10741150 | 0.181 | -0.35 | -0.5 | -0.186 | 0.008 | -0.003 | -0.003 | 0.016 | 0.45 | 0.34 | 0.46 | 0.56 | 0.4 | 0.54 |
| 10 | 128786366 | rs2066158 | 0.774 | -0.158 | -0.699 | -0.355 | -0.007 | 0.038 | 0.038 | 0 | 0.29 | 0.16 | 0.39 | 0.15 | 0.26 | 0.4 |
| 10 | 128788311 | rs4500401 | -1.006 | 0.189 | -0.811 | -0.238 | -0.005 | 0.003 | 0.003 | 0 | 0.7 | 0.75 | 0.76 | 0.76 | 0.59 | 0.77 |
| 10 | 128788633 | rs3750560 | 0.141 | 0.425 | -1.064 | 0.447 | -0.005 | 0.012 | 0.012 | 0.139 | 0.28 | 0.3 | 0.32 | 0.59 | 0.26 | 0.34 |
| 10 | 128789070 | rs3829124 | 0.141 | 0.425 | -1.064 | 0.447 | -0.005 | 0.012 | 0.012 | 0.139 | 0.28 | 0.3 | 0.32 | 0.59 | 0.26 | 0.34 |
| 10 | 128791805 | rs4750951 | -0.037 | 0.703 | -1.518 | 0.327 | -0.027 | 0.026 | 0.026 | 0 | 0.28 | 0.11 | 0.37 | 0.18 | 0.26 | 0.41 |
| 10 | 128797717 | rs2275540 | -0.445 | 0.849 | -1.173 | 0.904 | -0.002 | -0.019 | -0.019 | 0 | 0.22 | 0.2 | 0.26 | 0.32 | 0.21 | 0.18 |
| 10 | 128799999 | rs9418753 | 0.357 | 0.654 | -0.909 | 0.541 | -0.011 | 0.014 | 0.014 | 0.281 | 0.41 | 0.26 | 0.36 | 0.33 | 0.47 | 0.55 |
| 10 | 128805407 | rs9418677 | -0.492 | 1.027 | -0.68 | 0.387 | 0.002 | 0.001 | 0.001 | 0 | 0.38 | 0.46 | 0.39 | 0.51 | 0.42 | 0.33 |
| 10 | 128808908 | rs7912111 | -0.034 | 0.946 | -0.255 | 0.311 | -0.011 | 0.087 | 0.087 | 0 | 0.27 | 0.48 | 0.29 | 0.41 | 0.21 | 0.2 |
| 10 | 128809541 | rs4751175 | -0.454 | 0.765 | -0.076 | 0.196 | 0.005 | 0.006 | 0.006 | 0 | 0.77 | 0.77 | 0.81 | 0.73 | 0.69 | 0.79 |
| 10 | 128810820 | rs1340242 | -0.159 | 0.729 | -0.187 | 0.41 | 0 | 0 | 0 | 1.066 | 0.68 | 0.7 | 0.59 | 0.65 | 0.66 | 0.63 |
| 10 | 128813524 | rs9418776 | 1.878 | 0.767 | -0.315 | 0.363 | -0.04 | 0.04 | 0.04 | 0 | 0.16 | 0.04 | 0.2 | 0.05 | 0.19 | 0.25 |
| 10 | 128815530 | rs11018116 | 0.836 | 0.68 | -0.15 | 0.385 | 0.001 | 0 | 0 | 0.907 | 0.09 | 0.07 | 0.04 | 0.09 | 0.07 | 0.03 |
| 10 | 128820544 | rs9418790 | 2.216 | 0.675 | -0.085 | 0.322 | 0.023 | 0.019 | 0.019 | 0 | 0.26 | 0.07 | 0.27 | 0.27 | 0.22 | 0.22 |
| 10 | 128820978 | rs7094555 | -0.326 | 0.905 | 0.165 | 0.437 | 0.006 | 0.061 | 0.061 | 1.407 | 0.84 | 0.75 | 0.79 | 0.7 | 0.84 | 0.88 |
| 10 | 128821415 | rs2050812 | 1.163 | 0.929 | 0.135 | 0.482 | 0.006 | 0.064 | 0.064 | 0.788 | 0.07 | 0.12 | 0.14 | 0.22 | 0.07 | 0.06 |
| 10 | 128821616 | rs9418791 | 0.778 | 0.872 | 0.346 | 0.334 | -0.005 | -0.008 | -0.008 | 0 | 0.45 | 0.5 | 0.42 | 0.37 | 0.37 | 0.34 |
| 10 | 128822886 | rs3818880 | 0.989 | 0.906 | 0.162 | 0.349 | 0.018 | 0.03 | 0.03 | 0 | 0.43 | 0.29 | 0.41 | 0.37 | 0.37 | 0.34 |
| 10 | 128825311 | rs7092134 | 0.595 | 0.966 | 0.19 | 0.337 | 0.004 | 0.015 | 0.015 | 1.429 | 0.49 | 0.38 | 0.44 | 0.46 | 0.51 | 0.38 |
| 10 | 128826012 | rs4615933 | -0.102 | 0.604 | -0.018 | -0.03 | -0.004 | 0.025 | 0.025 | 0 | 0.38 | 0.32 | 0.36 | 0.14 | 0.36 | 0.49 |
| 10 | 128827346 | rs9418800 | 1.463 | 0.578 | -0.06 | 0.203 | -0.004 | 0.008 | 0.008 | 0 | 0.18 | 0.12 | 0.21 | 0.06 | 0.19 | 0.37 |
| 10 | 128828137 | rs7084307 | 1.892 | 0.466 | 0.128 | 0.286 | 0.003 | 0.003 | 0.003 | 0.092 | 0.28 | 0.29 | 0.24 | 0.24 | 0.22 | 0.22 |
| 10 | 128829337 | rs9418715 | -0.196 | 0.697 | 0.032 | 0.457 | -0.055 | -0.032 | -0.032 | 0.022 | 0.36 | 0.17 | 0.42 | 0.37 | 0.42 | 0.29 |
| 10 | 128830409 | rs2229597 | 0.677 | 0.749 | -0.037 | 0.409 | -0.033 | 0.056 | 0.056 | 0 | 0.21 | 0.05 | 0.2 | 0.09 | 0.21 | 0.13 |
| 10 | 128831248 | rs9418803 | 1.24 | 0.701 | 0.132 | 0.432 | 0.008 | 0.026 | 0.026 | 0 | 0.46 | 0.39 | 0.45 | 0.3 | 0.41 | 0.59 |
| 10 | 128831500 | rs9418804 | 0.183 | 0.681 | 0.148 | 0.469 | -0.01 | 0.12 | 0.12 | 0 | 0.83 | 0.56 | 0.86 | 0.67 | 0.83 | 0.88 |
| 10 | 128832151 | rs10830146 | 0.701 | 0.729 | -0.162 | 0.384 | -0.057 | 0.038 | 0.038 | 0 | 0.21 | 0.02 | 0.2 | 0.09 | 0.21 | 0.13 |
| 10 | 128833433 | rs2185404 | -1.337 | 0.679 | -0.063 | 0.43 | 0 | 0.001 | 0.001 | 0 | 0.92 | 0.95 | 0.94 | 0.92 | 0.91 | 0.94 |
| 10 | 128835320 | rs17771462 | 0.064 | 0.708 | -0.088 | 0.426 | -0.056 | -0.067 | -0.067 | 0 | 0.3 | 0.12 | 0.38 | 0.38 | 0.36 | 0.28 |
| 10 | 128837068 | rs4750895 | 0.131 | 0.513 | 0.371 | 0.486 | -0.009 | 0.083 | 0.083 | 0 | 0.83 | 0.73 | 0.86 | 0.65 | 0.83 | 0.88 |
| 10 | 128837927 | rs9418809 | 2.168 | 0.468 | 0.243 | 0.351 | 0.013 | 0.02 | 0.02 | 0.014 | 0.27 | 0.22 | 0.23 | 0.21 | 0.22 | 0.22 |
| 10 | 128840842 | rs880272 | -1.279 | 0.28 | 0.344 | 0.553 | -0.001 | -0.002 | -0.002 | 0 | 0.92 | 0.95 | 0.94 | 0.92 | 0.91 | 0.94 |
| 10 | 128845623 | rs2890087 | 0.824 | 0.298 | 0.364 | 0.542 | -0.002 | 0.01 | 0.01 | 0 | 0.48 | 0.35 | 0.48 | 0.5 | 0.48 | 0.4 |
| 10 | 128845857 | rs1538786 | 0.576 | 0.377 | 0.463 | 0.601 | -0.007 | 0.029 | 0.029 | 0 | 0.66 | 0.46 | 0.68 | 0.56 | 0.67 | 0.75 |
| 10 | 128847220 | rs7092964 | -0.118 | 0.304 | 0.487 | 0.683 | -0.016 | 0.037 | 0.037 | 0 | 0.87 | 0.78 | 0.91 | 0.79 | 0.91 | 0.92 |
| 10 | 128852144 | rs9418726 | 0.466 | 0.254 | 0.485 | 0.654 | -0.012 | 0.03 | 0.03 | 0 | 0.67 | 0.45 | 0.69 | 0.56 | 0.68 | 0.75 |
| 10 | 128853568 | rs7100020 | 0.488 | 0.272 | 0.513 | 0.664 | 0 | 0 | 0 | 0 | 0.2 | 0.24 | 0.22 | 0.23 | 0.22 | 0.17 |
| 10 | 128857810 | rs12411416 | 1.094 | 0.372 | 0.332 | 0.765 | -0.026 | -0.039 | -0.039 | 0.073 | 0.15 | 0.09 | 0.21 | 0.27 | 0.21 | 0.16 |
| 10 | 128858176 | rs9418730 | 1.196 | 0.286 | 0.259 | 0.417 | -0.014 | 0.025 | 0.025 | 0 | 0.19 | 0.09 | 0.21 | 0.06 | 0.19 | 0.37 |
| 10 | 128858508 | rs6482980 | 1.773 | 0.218 | 0.281 | 0.405 | 0.012 | 0.03 | 0.03 | 0 | 0.32 | 0.24 | 0.26 | 0.22 | 0.27 | 0.23 |
| 10 | 128860971 | rs9418731 | 0.738 | 0.591 | 0.252 | 0.625 | -0.009 | 0.017 | 0.017 | 0 | 0.2 | 0.11 | 0.22 | 0.06 | 0.19 | 0.36 |
| 10 | 128863969 | rs6482989 | 0.012 | 0.587 | 0.211 | 0.643 | -0.004 | 0.018 | 0.018 | 0 | 0.71 | 0.8 | 0.72 | 0.86 | 0.71 | 0.57 |
| 10 | 128864601 | rs9418733 | -0.025 | 0.778 | -0.057 | 0.678 | 0.001 | -0.002 | -0.002 | 0 | 0.24 | 0.27 | 0.25 | 0.19 | 0.21 | 0.38 |
| 10 | 128869673 | rs9418740 | 0.668 | 0.661 | -0.509 | 0.651 | -0.038 | -0.02 | -0.02 | 0.052 | 0.12 | 0.01 | 0.16 | 0.1 | 0.14 | 0.12 |
| 10 | 128871301 | rs9418834 | 0.007 | 0.483 | -0.177 | 0.755 | 0 | 0 | 0 | 0 | 0.69 | 0.66 | 0.68 | 0.32 | 0.64 | 0.75 |
| 10 | 128880582 | rs7068057 | 1.177 | 0.058 | -0.486 | 0.516 | -0.013 | 0.012 | 0.012 | 0 | 0.29 | 0.21 | 0.3 | 0.23 | 0.33 | 0.27 |
| 10 | 128886284 | rs7895398 | 0.212 | -0.574 | -0.177 | 0.594 | 0.009 | 0 | 0 | 0 | 0.48 | 0.74 | 0.51 | 0.39 | 0.46 | 0.38 |
| 10 | 128887890 | rs9783262 | -0.013 | -0.441 | 0.118 | 0.475 | -0.046 | 0.037 | 0.037 | 0.106 | 0.19 | 0.53 | 0.21 | 0.16 | 0.13 | 0.11 |
| 10 | 128889858 | rs2000062 | 0.215 | -0.421 | -0.206 | 0.207 | -0.066 | 0.131 | 0.131 | 0 | 0.23 | 0.03 | 0.21 | 0.03 | 0.25 | 0.39 |
| 10 | 128892927 | rs9988689 | 0.885 | -0.72 | 0.703 | 0.501 | -0.021 | -0.026 | -0.026 | 0.05 | 0.14 | 0.26 | 0.18 | 0.05 | 0.1 | 0.09 |
| 10 | 128894539 | rs10829363 | 0.531 | -0.805 | 0.623 | 0.744 | -0.048 | -0.017 | -0.017 | 0.002 | 0.19 | 0.31 | 0.25 | 0.12 | 0.11 | 0.1 |
| 10 | 128901414 | rs7923977 | 0.623 | -0.74 | 0.626 | 0.448 | -0.047 | -0.052 | -0.052 | 0 | 0.19 | 0.3 | 0.18 | 0.05 | 0.11 | 0.08 |
| 10 | 128902948 | rs6482668 | -0.082 | -0.766 | 0.725 | 0.47 | -0.067 | 0.01 | 0.01 | 0.227 | 0.22 | 0.5 | 0.2 | 0.16 | 0.12 | 0.1 |
| 10 | 128911984 | rs2489401 | -0.094 | -0.76 | 0.671 | 0.471 | -0.06 | 0 | 0 | 0.384 | 0.77 | 0.45 | 0.79 | 0.84 | 0.87 | 0.9 |
| 10 | 128918330 | rs2991794 | -0.157 | -0.824 | 0.693 | 0.465 | -0.07 | 0.002 | 0.002 | 0.565 | 0.77 | 0.46 | 0.8 | 0.85 | 0.88 | 0.91 |
| 10 | 128920546 | rs2255863 | -0.714 | -0.808 | 0.606 | 0.437 | -0.055 | -0.062 | -0.062 | 0.098 | 0.81 | 0.61 | 0.81 | 0.95 | 0.89 | 0.92 |
| 10 | 128926013 | rs1060561 | 0.045 | -0.849 | 0.601 | 0.409 | -0.056 | -0.012 | -0.012 | 0 | 0.75 | 0.44 | 0.79 | 0.84 | 0.86 | 0.9 |
| 10 | 128930240 | rs9299865 | -1.165 | -0.771 | 0.435 | 0.418 | 0.002 | 0.081 | 0.081 | 0.644 | 0.06 | 0.17 | 0.03 | 0.11 | 0.03 | 0.02 |
| 10 | 128933992 | rs2483855 | 0.3 | -0.919 | 0.593 | 0.358 | 0.027 | 0.008 | 0.008 | 0.608 | 0.73 | 0.25 | 0.78 | 0.84 | 0.86 | 0.9 |
| 10 | 128934510 | rs1334419 | -0.905 | -0.737 | 0.568 | 0.363 | 0.002 | 0.088 | 0.088 | 0.472 | 0.06 | 0.17 | 0.02 | 0.11 | 0.02 | 0.02 |
| 10 | 128935483 | rs11595374 | -0.471 | -0.595 | 0.254 | 0.087 | 0.011 | -0.018 | -0.018 | 0 | 0.25 | 0.34 | 0.24 | 0.15 | 0.27 | 0.39 |
| 10 | 128940306 | rs2483853 | 1.255 | -0.839 | -0.026 | 0.358 | 0.021 | 0.039 | 0.039 | 0.028 | 0.5 | 0.13 | 0.53 | 0.58 | 0.59 | 0.43 |
| 10 | 128942752 | rs2489394 | -0.993 | -0.633 | 0.27 | 0.105 | -0.055 | -0.032 | -0.032 | 0 | 0.82 | 0.57 | 0.82 | 0.88 | 0.9 | 0.92 |

| | | | | | | | | | | | | | | | | |
|---|---|---|---|---|---|---|---|---|---|---|---|---|---|---|---|---|
| 10 | 128944080 | rs2489392 | 0.038 | -0.672 | -0.105 | 0.093 | 0.031 | -0.005 | -0.005 | 0.698 | 0.74 | 0.22 | 0.78 | 0.77 | 0.88 | 0.9 |
| 10 | 128953049 | rs2890091 | 0.059 | -0.207 | -0.177 | 0.081 | -0.048 | 0.061 | 0.061 | 0.199 | 0.2 | 0.06 | 0.21 | 0.03 | 0.28 | 0.31 |
| 10 | 128959164 | rs7077003 | -0.871 | -0.223 | -0.003 | 0.106 | 0.022 | -0.044 | -0.044 | 0 | 0.27 | 0.42 | 0.24 | 0.14 | 0.28 | 0.33 |
| 10 | 128972799 | rs1892135 | 0.4 | -0.171 | -0.001 | 0.155 | 0.018 | -0.021 | -0.021 | 0 | 0.76 | 0.65 | 0.77 | 0.86 | 0.71 | 0.67 |
| 10 | 128974220 | rs11594560 | 0.149 | -0.151 | -0.325 | 0.132 | -0.052 | 0.06 | 0.06 | 0 | 0.19 | 0.04 | 0.21 | 0.03 | 0.28 | 0.31 |
| 10 | 128974621 | rs2489386 | 0.799 | -0.216 | -0.572 | 0.188 | -0.001 | 0.072 | 0.072 | 0 | 0.55 | 0.24 | 0.56 | 0.74 | 0.59 | 0.61 |
| 10 | 128975256 | rs1182885 | 0.798 | -0.208 | -0.725 | 0.158 | -0.011 | 0.088 | 0.088 | 0 | 0.55 | 0.21 | 0.56 | 0.74 | 0.59 | 0.61 |
| 10 | 128986854 | rs2489382 | 0.006 | -0.245 | -0.503 | 0.623 | -0.01 | -0.056 | -0.056 | 0 | 0.24 | 0.17 | 0.28 | 0.54 | 0.29 | 0.29 |
| 10 | 128989670 | rs1167522 | -0.095 | -0.296 | 0.053 | 0.611 | -0.039 | -0.034 | -0.034 | 0.385 | 0.8 | 0.66 | 0.81 | 0.88 | 0.87 | 0.94 |
| 10 | 128990711 | rs12269277 | 0.368 | -0.26 | 0.052 | 0.624 | -0.039 | -0.019 | -0.019 | 0 | 0.17 | 0.26 | 0.18 | 0.12 | 0.1 | 0.05 |
| 10 | 128991629 | rs2483869 | -0.099 | -0.256 | 0.031 | 0.75 | -0.035 | -0.005 | -0.005 | 0.314 | 0.8 | 0.68 | 0.81 | 0.85 | 0.87 | 0.94 |
| 10 | 128996576 | rs2489379 | -0.07 | -0.232 | 0.015 | 0.76 | -0.036 | -0.006 | -0.006 | 0 | 0.8 | 0.69 | 0.82 | 0.85 | 0.87 | 0.94 |
| 10 | 129001986 | rs12246104 | 0.433 | -0.283 | -0.088 | 0.736 | -0.024 | -0.009 | -0.009 | 0 | 0.17 | 0.19 | 0.17 | 0.12 | 0.1 | 0.05 |
| 10 | 129005367 | rs2483865 | 0.139 | -0.037 | 0.009 | 0.726 | -0.001 | 0.01 | 0.01 | 0 | 0.84 | 0.72 | 0.89 | 0.84 | 0.86 | 0.77 |
| 10 | 129005720 | rs928571 | 0.096 | 0.034 | -0.472 | 0.855 | -0.011 | -0.01 | -0.01 | 0 | 0.3 | 0.24 | 0.37 | 0.41 | 0.34 | 0.36 |
| 10 | 129007159 | rs12255713 | 0.593 | -0.151 | -0.402 | 0.752 | -0.037 | 0.011 | 0.011 | 1.424 | 0.17 | 0.21 | 0.17 | 0.16 | 0.1 | 0.06 |
| 10 | 129012678 | rs4522083 | 1.037 | 0.696 | -1.628 | 0.684 | -0.085 | -0.051 | -0.051 | 0 | 0.3 | 0.03 | 0.28 | 0.22 | 0.33 | 0.3 |
| 10 | 129015005 | rs10829597 | -0.397 | 1.255 | -0.889 | 0.732 | -0.014 | -0.013 | -0.013 | 0 | 0.41 | 0.32 | 0.4 | 0.59 | 0.41 | 0.49 |
| 10 | 129016861 | rs4751079 | -0.124 | 1.27 | -0.903 | 1.025 | -0.038 | -0.023 | -0.023 | 0 | 0.29 | 0.19 | 0.36 | 0.43 | 0.37 | 0.3 |
| 10 | 129017883 | rs11016808 | 1.872 | 1.245 | -0.78 | 0.976 | 0.06 | 0.033 | 0.033 | 0 | 0.36 | 0.12 | 0.26 | 0.24 | 0.27 | 0.36 |
| 10 | 129019025 | rs1334420 | 1.127 | 1.399 | -0.526 | 0.983 | 0.017 | -0.027 | -0.027 | 0 | 0.16 | 0.08 | 0.18 | 0.22 | 0.12 | 0.1 |
| 10 | 129019678 | rs2050305 | -0.047 | 1.397 | -0.557 | 0.964 | -0.039 | -0.005 | -0.005 | 0 | 0.32 | 0.23 | 0.38 | 0.42 | 0.38 | 0.41 |
| 10 | 129023096 | rs11599788 | 0.866 | 1.669 | -0.932 | 1.041 | -0.113 | -0.077 | -0.077 | 0 | 0.17 | 0.01 | 0.29 | 0.25 | 0.29 | 0.18 |
| 10 | 129025576 | rs1186361 | -1.487 | 1.617 | -0.13 | 0.653 | -0.005 | 0.037 | 0.037 | 0 | 0.57 | 0.67 | 0.61 | 0.74 | 0.52 | 0.58 |
| 10 | 129030343 | rs10829607 | 1.579 | 1.629 | -0.372 | 0.405 | 0.03 | 0.073 | 0.073 | 0 | 0.39 | 0.17 | 0.32 | 0.23 | 0.35 | 0.39 |
| 10 | 129031127 | rs1334417 | -0.53 | 1.632 | -0.25 | 0.391 | -0.009 | -0.01 | -0.01 | 0 | 0.38 | 0.34 | 0.4 | 0.6 | 0.4 | 0.47 |
| 10 | 129031243 | rs10829608 | 1.536 | 1.563 | -0.612 | 0.431 | 0.031 | 0.077 | 0.077 | 0 | 0.39 | 0.17 | 0.32 | 0.22 | 0.35 | 0.39 |
| 10 | 129034159 | rs2489426 | 0.893 | 1.641 | -0.39 | 0.681 | 0.014 | -0.016 | -0.016 | 0 | 0.16 | 0.08 | 0.18 | 0.18 | 0.12 | 0.1 |
| 10 | 129034251 | rs2489425 | 0.893 | 1.647 | -0.403 | 0.684 | 0.014 | -0.017 | -0.017 | 1.323 | 0.16 | 0.08 | 0.18 | 0.18 | 0.12 | 0.1 |
| 10 | 129037070 | rs12777771 | 1.298 | 2.152 | -0.563 | 0.506 | 0.048 | 0.122 | 0.122 | 0.012 | 0.43 | 0.1 | 0.38 | 0.21 | 0.35 | 0.51 |
| 10 | 129041541 | rs7077776 | -0.421 | 2.014 | -0.055 | 0.823 | -0.001 | -0.001 | -0.001 | 0 | 0.28 | 0.28 | 0.33 | 0.44 | 0.26 | 0.24 |
| 10 | 129046407 | rs3740013 | -0.62 | 1.868 | 0.267 | 0.824 | 0.003 | 0.003 | 0.003 | 0 | 0.38 | 0.57 | 0.45 | 0.57 | 0.42 | 0.28 |
| 10 | 129048450 | rs2498935 | 0.299 | 1.713 | 0.116 | 0.768 | 0.001 | -0.002 | -0.002 | 0.065 | 0.16 | 0.14 | 0.12 | 0.2 | 0.12 | 0.11 |
| 10 | 129048746 | rs11016936 | 1.311 | 1.778 | -0.181 | 0.707 | 0.006 | 0.112 | 0.112 | 0.05 | 0.43 | 0.09 | 0.37 | 0.23 | 0.38 | 0.58 |
| 10 | 129058024 | rs2489414 | 0.059 | 1.576 | 0.261 | 0.873 | -0.014 | 0.015 | 0.015 | 0 | 0.18 | 0.26 | 0.14 | 0.21 | 0.13 | 0.11 |
| 10 | 129060759 | rs11017012 | 0.006 | 2.068 | 0.343 | 0.752 | -0.003 | -0.005 | -0.005 | 0 | 0.32 | 0.31 | 0.4 | 0.43 | 0.35 | 0.24 |
| 10 | 129064193 | rs7895120 | -0.121 | 1.935 | 0.342 | 0.74 | -0.025 | 0.062 | 0.062 | 0 | 0.81 | 0.61 | 0.85 | 0.75 | 0.87 | 0.88 |
| 10 | 129064394 | rs10458718 | 1.162 | 1.976 | -0.238 | 0.594 | -0.061 | 0.123 | 0.123 | 0.157 | 0.44 | 0.07 | 0.38 | 0.22 | 0.45 | 0.59 |
| 10 | 129066154 | rs10430605 | 1.34 | 1.883 | -0.219 | 0.62 | -0.04 | 0.108 | 0.108 | 0.093 | 0.42 | 0.07 | 0.35 | 0.22 | 0.39 | 0.55 |
| 10 | 129070313 | rs2498946 | 1.16 | 2.265 | 0.377 | 0.915 | 0.069 | 0.171 | 0.171 | 0.165 | 0.5 | 0.15 | 0.4 | 0.22 | 0.38 | 0.56 |
| 10 | 129075674 | rs10741209 | -0.59 | 2.392 | 0.765 | 1.091 | -0.009 | -0.007 | -0.007 | 0 | 0.34 | 0.3 | 0.44 | 0.43 | 0.41 | 0.27 |
| 10 | 129089317 | rs10829711 | -0.563 | 2.379 | 0.773 | 1.127 | -0.009 | -0.007 | -0.007 | 0.386 | 0.34 | 0.31 | 0.44 | 0.43 | 0.42 | 0.27 |
| 10 | 129090007 | rs6482774 | -1.185 | 2.207 | 1.149 | 1.215 | 0.087 | 0.127 | 0.127 | 0.423 | 0.13 | 0.37 | 0.15 | 0.25 | 0.2 | 0.14 |
| 10 | 129096856 | rs12767483 | -0.653 | 2.142 | 1.506 | 1.111 | 0.035 | 0.117 | 0.117 | 0.039 | 0.11 | 0.29 | 0.1 | 0.23 | 0.12 | 0.11 |
| 10 | 129096956 | rs2766032 | -1.937 | 2.139 | 1.439 | 1.103 | 0.052 | 0.154 | 0.154 | 0.599 | 0.17 | 0.53 | 0.17 | 0.33 | 0.2 | 0.18 |
| 10 | 129100245 | rs7072458 | -1.793 | 1.959 | 1.18 | 0.951 | 0.074 | -0.004 | -0.004 | 0 | 0.05 | 0.22 | 0.07 | 0 | 0.08 | 0.04 |
| 10 | 129117721 | rs10829749 | -0.4 | 2.032 | 0.728 | 1.07 | -0.025 | -0.042 | -0.042 | 0 | 0.32 | 0.18 | 0.4 | 0.45 | 0.37 | 0.28 |
| 10 | 129118884 | rs2720972 | -1.767 | 2.155 | 0.61 | 0.91 | 0.002 | 0.161 | 0.161 | 0.404 | 0.51 | 0.95 | 0.61 | 0.78 | 0.59 | 0.44 |
| 10 | 129119981 | rs7083352 | -0.8 | 2.274 | 0.778 | 1.086 | -0.01 | 0.014 | 0.014 | 0.371 | 0.18 | 0.4 | 0.12 | 0.19 | 0.13 | 0.17 |
| 10 | 129121087 | rs2766047 | -0.906 | 2.389 | 0.411 | 0.697 | -0.006 | -0.011 | -0.011 | 0 | 0.4 | 0.64 | 0.47 | 0.55 | 0.44 | 0.32 |
| 10 | 129121861 | rs2766050 | -1.56 | 2.215 | 0.174 | 0.697 | -0.026 | 0.165 | 0.165 | 0 | 0.48 | 0.94 | 0.58 | 0.78 | 0.57 | 0.41 |
| 10 | 129123033 | rs2766053 | -0.505 | 2.146 | 0.308 | 0.712 | 0.003 | -0.011 | -0.011 | 0 | 0.37 | 0.66 | 0.42 | 0.54 | 0.36 | 0.31 |
| 10 | 129127446 | rs2766059 | -1.532 | 2.158 | -0.566 | 0.635 | -0.025 | 0.206 | 0.206 | 0 | 0.49 | 0.95 | 0.58 | 0.79 | 0.57 | 0.42 |
| 10 | 129128317 | rs9663318 | -0.867 | 1.677 | -0.078 | 0.849 | 0.056 | 0.125 | 0.125 | 0.265 | 0.13 | 0.62 | 0.15 | 0.24 | 0.15 | 0.08 |
| 10 | 129132502 | rs10764972 | -0.682 | 1.174 | -0.652 | 0.785 | -0.002 | -0.041 | -0.041 | 0 | 0.34 | 0.2 | 0.37 | 0.56 | 0.38 | 0.26 |
| 10 | 129136409 | rs4751240 | -1.418 | 0.919 | -0.027 | 0.384 | -0.01 | -0.038 | -0.038 | 0 | 0.08 | 0.15 | 0.05 | 0 | 0.08 | 0.05 |
| 10 | 129139130 | rs10829775 | -0.412 | 1.213 | -0.281 | 0.937 | -0.001 | 0 | 0 | 0 | 0.2 | 0.2 | 0.27 | 0.4 | 0.22 | 0.11 |
| 10 | 129142265 | rs943207 | 0.572 | 1.05 | -0.385 | 0.138 | -0.004 | 0.016 | 0.016 | 0 | 0.6 | 0.35 | 0.48 | 0.32 | 0.59 | 0.56 |
| 10 | 129142417 | rs751008 | 0.586 | 0.88 | -0.806 | -0.067 | 0.012 | 0.109 | 0.109 | 0 | 0.5 | 0.24 | 0.39 | 0.32 | 0.47 | 0.5 |
| 10 | 129149464 | rs1006217 | 0.035 | 0.543 | -1.205 | -0.255 | -0.037 | 0.082 | 0.082 | 0 | 0.17 | 0.01 | 0.09 | 0.04 | 0.17 | 0.3 |
| 10 | 129152153 | rs10741228 | 0.216 | 0.665 | -0.35 | -0.004 | 0 | 0 | 0 | 0.065 | 0.53 | 0.43 | 0.35 | 0.38 | 0.43 | 0.55 |
| 10 | 129152608 | rs10764990 | -0.402 | 1.017 | -0.471 | 0.522 | -0.001 | 0.002 | 0.002 | 0 | 0.74 | 0.74 | 0.51 | 0.51 | 0.62 | 0.7 |
| 10 | 129156987 | rs11017537 | 0.083 | -0.042 | -0.374 | 0.498 | -0.024 | -0.019 | -0.019 | 1.855 | 0.36 | 0.46 | 0.28 | 0.24 | 0.21 | 0.25 |
| 10 | 129158853 | rs10829838 | 0.234 | 0.136 | -0.804 | 0.668 | 0.028 | -0.001 | -0.001 | 0 | 0.32 | 0.27 | 0.25 | 0.26 | 0.2 | 0.24 |
| 10 | 129159888 | rs12251654 | -0.951 | 0.923 | -0.532 | 0.607 | 0.001 | -0.004 | -0.004 | 0 | 0.24 | 0.28 | 0.15 | 0.18 | 0.25 | 0.36 |
| 10 | 129160101 | rs2296637 | -0.64 | 1.014 | -0.581 | 0.7 | -0.011 | 0.011 | 0.011 | 0 | 0.15 | 0.21 | 0.07 | 0.16 | 0.1 | 0.24 |
| 10 | 129160313 | rs2296636 | -0.924 | 1.36 | -0.641 | 0.979 | -0.008 | -0.009 | -0.009 | 0.971 | 0.19 | 0.19 | 0.13 | 0.24 | 0.26 | 0.31 |
| 10 | 129160458 | rs2296635 | -1.128 | 1.664 | -0.765 | 0.732 | -0.024 | 0.018 | 0.018 | 0 | 0.21 | 0.17 | 0.16 | 0.14 | 0.27 | 0.28 |
| 10 | 129160691 | rs2296634 | -0.913 | 1.578 | -0.614 | 0.699 | 0.006 | -0.001 | -0.001 | 2.174 | 0.21 | 0.26 | 0.18 | 0.17 | 0.26 | 0.28 |

| | | | | | | | | | | | | | | | | |
|---|---|---|---|---|---|---|---|---|---|---|---|---|---|---|---|---|
| 10 | 129164681 | rs4751282 | 0.316 | 1.115 | -0.5 | 0.548 | -0.002 | -0.002 | -0.002 | 0.422 | 0.57 | 0.42 | 0.59 | 0.51 | 0.48 | 0.44 |
| 10 | 129165419 | rs11017614 | 0.645 | 0.477 | -0.984 | 0.55 | 0.018 | -0.03 | -0.03 | 0 | 0.14 | 0.08 | 0.23 | 0.29 | 0.11 | 0.1 |
| 10 | 129165510 | rs11594969 | 0.751 | 0.457 | -0.808 | 0.432 | -0.001 | 0.037 | 0.037 | 0 | 0.41 | 0.3 | 0.35 | 0.22 | 0.37 | 0.31 |
| 10 | 129168150 | rs12570609 | 0.252 | 0.433 | -0.268 | -0.081 | 0.018 | 0 | 0 | 0 | 0.45 | 0.37 | 0.51 | 0.43 | 0.36 | 0.38 |
| 10 | 129168894 | rs10765017 | 0.91 | 0.198 | -0.253 | -0.152 | 0.021 | -0.039 | -0.039 | 0.163 | 0.33 | 0.19 | 0.41 | 0.43 | 0.25 | 0.32 |
| 10 | 129169498 | rs7895951 | -1.226 | 0.538 | -0.5 | -0.077 | -0.031 | -0.008 | -0.008 | 0 | 0.82 | 0.9 | 0.76 | 0.77 | 0.73 | 0.74 |
| 10 | 129179426 | rs7099958 | 0.228 | 0.606 | -0.125 | -0.15 | 0.013 | 0.004 | 0.004 | 0 | 0.27 | 0.36 | 0.22 | 0.26 | 0.33 | 0.32 |
| 10 | 129182308 | rs877748 | -0.992 | 0.618 | -0.119 | -0.052 | -0.024 | -0.003 | -0.003 | 0.17 | 0.81 | 0.85 | 0.75 | 0.77 | 0.72 | 0.72 |
| 10 | 129184040 | rs7078619 | 0.795 | 0.38 | 0.145 | 0.08 | -0.025 | -0.019 | -0.019 | 0 | 0.12 | 0.24 | 0.06 | 0.08 | 0.06 | 0.04 |
| 10 | 129186195 | rs11017780 | 0.863 | 0.901 | -0.116 | 0.172 | 0.021 | -0.026 | -0.026 | 0 | 0.38 | 0.21 | 0.48 | 0.42 | 0.31 | 0.37 |
| 10 | 129186924 | rs2148379 | -1.205 | 1.068 | -0.039 | 0.226 | 0 | -0.001 | -0.001 | 0.009 | 0.87 | 0.86 | 0.85 | 0.77 | 0.79 | 0.78 |
| 10 | 129189646 | rs2104511 | 0.017 | 0.508 | -0.048 | -0.116 | 0.016 | 0.014 | 0.014 | 0 | 0.18 | 0.25 | 0.18 | 0.26 | 0.28 | 0.3 |
| 10 | 129190678 | rs6482855 | -0.464 | 0.772 | -0.749 | 0.206 | -0.001 | -0.001 | -0.001 | 0 | 0.47 | 0.51 | 0.34 | 0.34 | 0.48 | 0.41 |
| 10 | 129192143 | rs4751323 | 0.788 | 1.571 | -0.834 | 1.071 | 0.006 | 0.044 | 0.044 | 0.19 | 0.9 | 0.9 | 0.9 | 0.72 | 0.86 | 0.94 |
| 10 | 129193311 | rs4750853 | 0.216 | 1.316 | -1.055 | 0.773 | -0.003 | -0.003 | -0.003 | 0 | 0.81 | 0.82 | 0.73 | 0.72 | 0.72 | 0.76 |
| 10 | 129193390 | rs4750855 | 0.276 | 0.325 | -1.137 | 0.238 | -0.002 | -0.027 | -0.027 | 0.757 | 0.15 | 0.16 | 0.07 | 0 | 0.08 | 0.06 |
| 10 | 129197384 | rs2296633 | -0.108 | 1.314 | -1.377 | 0.89 | -0.045 | -0.031 | -0.031 | 0 | 0.16 | 0.03 | 0.1 | 0.15 | 0.2 | 0.23 |
| 10 | 129200463 | rs10829972 | -0.618 | 1.343 | -1.086 | 0.825 | 0.02 | 0.082 | 0.082 | 0.765 | 0.51 | 0.63 | 0.61 | 0.78 | 0.5 | 0.61 |
| 10 | 129200964 | rs11017876 | -0.067 | 1.656 | -1.206 | 1.095 | -0.015 | 0.003 | 0.003 | 0 | 0.26 | 0.16 | 0.22 | 0.28 | 0.24 | 0.26 |
| 10 | 129201649 | rs2249656 | -0.692 | 1.127 | -0.709 | 0.634 | 0.008 | 0.032 | 0.032 | 0.811 | 0.48 | 0.39 | 0.58 | 0.77 | 0.5 | 0.61 |
| 10 | 129206913 | rs3740547 | 0.21 | 1.194 | -0.702 | 0.812 | -0.001 | 0.002 | 0.002 | 0 | 0.52 | 0.55 | 0.56 | 0.62 | 0.48 | 0.45 |
| 10 | 129207429 | rs2229599 | 0.569 | 1.344 | -0.921 | 0.646 | -0.014 | 0.027 | 0.027 | 0 | 0.05 | 0.01 | 0.03 | 0 | 0.07 | 0.02 |
| 10 | 129208021 | rs11017927 | 0.382 | 1.238 | -0.619 | 0.524 | 0.025 | 0 | 0 | 0 | 0.19 | 0.35 | 0.17 | 0.18 | 0.23 | 0.13 |
| 10 | 129208050 | rs11017928 | 1.234 | 1.158 | -1.148 | 0.338 | -0.008 | 0.012 | 0.012 | 0 | 0.15 | 0.01 | 0.14 | 0.09 | 0.12 | 0.15 |
| 10 | 129208186 | rs7895671 | 0.814 | 1.161 | -1.127 | 0.382 | 0.027 | 0.001 | 0.001 | 0 | 0.18 | 0.35 | 0.17 | 0.18 | 0.23 | 0.13 |
| 10 | 129209084 | rs2296628 | 0.525 | 0.981 | -1.093 | 0.487 | 0.064 | -0.01 | -0.01 | 0 | 0.49 | 0.16 | 0.56 | 0.57 | 0.37 | 0.47 |
| 10 | 129213586 | rs3736940 | 1.367 | 0.922 | -0.978 | 0.498 | 0.023 | -0.012 | -0.012 | 0.645 | 0.32 | 0.19 | 0.39 | 0.44 | 0.25 | 0.32 |
| 10 | 129213780 | rs3736938 | 0.713 | 1.174 | -1.061 | 0.678 | -0.001 | -0.012 | -0.012 | 1.176 | 0.09 | 0.04 | 0.09 | 0.18 | 0.11 | 0.19 |
| 10 | 129218901 | rs7097399 | 0.987 | 1.164 | -0.797 | 0.625 | 0.022 | -0.002 | -0.002 | 0 | 0.19 | 0.35 | 0.17 | 0.19 | 0.23 | 0.13 |
| 10 | 129219857 | rs7092530 | -0.566 | 1.374 | -1.032 | 0.883 | -0.052 | -0.078 | -0.078 | 0 | 0.75 | 0.95 | 0.76 | 0.72 | 0.75 | 0.69 |
| 10 | 129220931 | rs7920574 | -1.011 | 1.467 | -0.944 | 0.886 | -0.026 | -0.031 | -0.031 | 5.642 | 0.9 | 0.97 | 0.9 | 0.89 | 0.87 | 0.84 |
| 10 | 129221607 | rs7097526 | 0.855 | 1.329 | -0.836 | 0.713 | 0.026 | -0.002 | -0.002 | 0 | 0.2 | 0.39 | 0.18 | 0.19 | 0.23 | 0.13 |
| 10 | 129222061 | rs1078336 | 0.249 | 1.257 | -0.637 | 0.643 | -0.001 | -0.046 | -0.046 | 0.336 | 0.11 | 0.26 | 0.09 | 0.02 | 0.11 | 0.03 |
| 10 | 129225796 | rs10765093 | -0.626 | 1.321 | -0.7 | 0.771 | -0.019 | -0.041 | -0.041 | 0.101 | 0.83 | 0.93 | 0.82 | 0.77 | 0.79 | 0.66 |
| 10 | 129225930 | rs10830045 | 0.637 | 1.067 | -0.727 | 0.71 | 0.029 | -0.006 | -0.006 | 0 | 0.2 | 0.51 | 0.19 | 0.18 | 0.23 | 0.12 |
| 10 | 129227488 | rs11018077 | 0.531 | 1.001 | -0.762 | 0.724 | 0.031 | 0.005 | 0.005 | 0 | 0.21 | 0.42 | 0.3 | 0.19 | 0.24 | 0.12 |
| 10 | 129228268 | rs1573822 | 2.161 | 0.297 | -1.511 | 0.432 | 0.016 | 0.069 | 0.069 | 1.311 | 0.21 | 0.07 | 0.11 | 0.08 | 0.15 | 0.06 |
| 10 | 129229880 | rs2386491 | -1.273 | 0.593 | -1.341 | 0.751 | -0.019 | -0.045 | -0.045 | 0.275 | 0.89 | 0.97 | 0.89 | 0.81 | 0.86 | 0.72 |
| 10 | 129229919 | rs12354451 | 1.366 | 0.644 | -1.48 | 0.642 | -0.038 | -0.03 | -0.03 | 0 | 0.15 | 0.01 | 0.15 | 0.16 | 0.14 | 0.15 |
| 10 | 129230728 | rs7095368 | -0.292 | 0.747 | -0.867 | 0.896 | -0.008 | 0.099 | 0.099 | 0 | 0.83 | 0.67 | 0.82 | 0.75 | 0.85 | 0.71 |
| 10 | 129230909 | rs7095670 | -0.156 | 0.666 | -0.855 | 0.26 | 0.028 | -0.03 | -0.03 | 0 | 0.25 | 0.41 | 0.23 | 0.17 | 0.34 | 0.19 |
| 10 | 129233816 | rs11816982 | 0.143 | 0.44 | -0.989 | 0.514 | 0.002 | -0.016 | -0.016 | 0 | 0.15 | 0.14 | 0.35 | 0.4 | 0.13 | 0.34 |
| 10 | 129234393 | rs7923936 | -0.177 | 0.323 | -1.238 | 0.19 | -0.042 | -0.066 | -0.066 | 0 | 0.73 | 0.88 | 0.75 | 0.59 | 0.63 | 0.6 |
| 10 | 129235680 | rs4553284 | -0.024 | 0.307 | -1.294 | 0.064 | -0.085 | -0.097 | -0.097 | 0 | 0.67 | 0.88 | 0.71 | 0.59 | 0.57 | 0.59 |
| 10 | 129236512 | rs1887097 | 0.047 | -0.298 | -1.18 | -0.339 | -0.002 | 0.002 | 0.002 | 0 | 0.36 | 0.62 | 0.48 | 0.39 | 0.4 | 0.42 |
| 10 | 129237572 | rs1926427 | -0.067 | -0.134 | -1.238 | -0.209 | -0.065 | -0.05 | -0.05 | 3.125 | 0.33 | 0.18 | 0.34 | 0.5 | 0.44 | 0.42 |
| 10 | 129238058 | rs11815834 | 0.657 | -0.168 | -0.867 | -0.288 | 0.004 | 0.006 | 0.006 | 0 | 0.24 | 0.29 | 0.39 | 0.38 | 0.29 | 0.39 |
| 10 | 129238206 | rs9418709 | -0.23 | 0.146 | -1.278 | 0.011 | -0.065 | -0.05 | -0.05 | 0.615 | 0.33 | 0.18 | 0.34 | 0.5 | 0.44 | 0.42 |
| 10 | 129239267 | rs4750886 | -1.6 | -0.407 | -1.42 | -0.349 | -0.021 | 0.054 | 0.054 | 0 | 0.86 | 0.96 | 0.87 | 0.97 | 0.88 | 0.89 |
| 10 | 129240182 | rs11018089 | -0.001 | -0.114 | -1.264 | -0.125 | -0.049 | -0.086 | -0.086 | 0.255 | 0.28 | 0.12 | 0.27 | 0.46 | 0.36 | 0.39 |
| 10 | 129240618 | rs10765097 | -0.116 | -0.096 | -1.188 | -0.117 | -0.031 | -0.06 | -0.06 | 1 | 0.29 | 0.16 | 0.28 | 0.46 | 0.36 | 0.38 |
| 10 | 129243551 | rs10765100 | -0.232 | -0.103 | -1.137 | -0.12 | -0.018 | -0.038 | -0.038 | 0.878 | 0.3 | 0.22 | 0.28 | 0.47 | 0.36 | 0.38 |
| 10 | 129245244 | rs1931702 | -0.366 | -0.308 | -1.036 | -0.586 | -0.007 | -0.007 | -0.007 | 0.092 | 0.44 | 0.62 | 0.56 | 0.57 | 0.55 | 0.51 |
| 10 | 129245267 | rs1931703 | -0.367 | -0.707 | -1.328 | -0.685 | -0.035 | -0.029 | -0.029 | 0 | 0.58 | 0.74 | 0.62 | 0.57 | 0.55 | 0.56 |
| 10 | 129245426 | rs869799 | -0.073 | -0.712 | -1.11 | -0.938 | 0.011 | 0.021 | 0.021 | 0 | 0.53 | 0.71 | 0.58 | 0.59 | 0.47 | 0.48 |
| 10 | 129245684 | rs869801 | -0.634 | -1.224 | -0.974 | -1.54 | 0.001 | 0.081 | 0.081 | 0 | 0.29 | 0.21 | 0.22 | 0.12 | 0.26 | 0.21 |
| 10 | 129245967 | rs873185 | -0.271 | -1.169 | -0.906 | -1.49 | 0.008 | 0.039 | 0.039 | 0 | 0.76 | 0.78 | 0.82 | 0.88 | 0.83 | 0.83 |
| 10 | 129246555 | rs7915648 | -0.094 | -1.329 | -0.489 | -1.798 | -0.023 | -0.129 | -0.129 | 0 | 0.65 | 0.51 | 0.85 | 0.99 | 0.74 | 0.9 |
| 10 | 129246566 | rs10765101 | 0.041 | -1.353 | -0.574 | -1.783 | 0 | -0.021 | -0.021 | 0 | 0.63 | 0.38 | 0.83 | 0.91 | 0.74 | 0.89 |
| 10 | 129246759 | rs11018102 | -0.049 | -1.691 | -0.404 | -1.608 | 0.01 | -0.033 | -0.033 | 0 | 0.33 | 0.28 | 0.18 | 0.03 | 0.16 | 0.14 |
| 10 | 129247444 | rs7079945 | -0.468 | -1.354 | -0.072 | -1.06 | -0.055 | -0.103 | -0.103 | 1.183 | 0.29 | 0.47 | 0.15 | 0.08 | 0.19 | 0.12 |
| 10 | 129248480 | rs7904011 | -0.149 | -1.317 | 0.038 | -0.826 | -0.096 | -0.1 | -0.1 | 0 | 0.27 | 0.5 | 0.12 | 0.1 | 0.1 | 0.06 |
| 10 | 129248817 | rs6482916 | -0.173 | -0.098 | -0.674 | -0.183 | 0.001 | 0.001 | 0.001 | 0 | 0.48 | 0.45 | 0.56 | 0.53 | 0.48 | 0.55 |
| 10 | 129249511 | rs3740546 | -0.083 | 0.722 | -1.172 | 0.185 | -0.038 | -0.075 | -0.075 | 0.966 | 0.21 | 0.12 | 0.28 | 0.52 | 0.35 | 0.35 |
| 10 | 129249693 | rs2229605 | 0.722 | 0.063 | -1.337 | -0.294 | 0.01 | -0.004 | -0.004 | 0 | 0.5 | 0.21 | 0.58 | 0.46 | 0.52 | 0.52 |
| 10 | 129250075 | rs3740543 | 0.71 | -0.019 | -1.309 | -0.405 | 0.01 | 0.017 | 0.017 | 0 | 0.5 | 0.22 | 0.58 | 0.39 | 0.52 | 0.52 |
| 10 | 129256715 | rs7087658 | -2.054 | -0.457 | -0.402 | -0.535 | -0.053 | -0.043 | -0.043 | 2.242 | 0.85 | 0.68 | 0.94 | 0.92 | 0.98 | 0.94 |
| 10 | 129256933 | rs10830072 | 0.842 | -0.507 | -0.477 | -0.927 | -0.02 | 0.002 | 0.002 | 0 | 0.47 | 0.2 | 0.59 | 0.39 | 0.51 | 0.49 |
| 10 | 129258578 | rs915144 | 0.511 | -0.463 | -0.458 | -0.932 | -0.027 | -0.069 | -0.069 | 0 | 0.26 | 0.15 | 0.27 | 0.53 | 0.37 | 0.38 |

| | | | | | | | | | | | | | | | | |
|---|---|---|---|---|---|---|---|---|---|---|---|---|---|---|---|---|
| 10 | 129261583 | rs10830078 | 0.479 | -0.481 | -0.2 | -0.908 | 0.001 | 0.011 | 0.011 | 0 | 0.5 | 0.35 | 0.6 | 0.39 | 0.51 | 0.52 |
| 10 | 129261904 | rs7090762 | 0.522 | -0.48 | -0.178 | -0.908 | 0.001 | 0.011 | 0.011 | 0 | 0.5 | 0.35 | 0.6 | 0.39 | 0.51 | 0.52 |
| 10 | 129262490 | rs2151935 | -2.141 | -0.483 | -0.144 | -0.921 | -0.054 | -0.044 | -0.044 | 0.493 | 0.85 | 0.67 | 0.94 | 0.92 | 0.98 | 0.95 |
| 10 | 129263045 | rs10830080 | 0.28 | 0.066 | -1.121 | -1.349 | -0.002 | 0.032 | 0.032 | 0 | 0.47 | 0.32 | 0.31 | 0.01 | 0.53 | 0.36 |
| 10 | 129263278 | rs2184378 | -0.047 | 0.064 | -1.742 | -1.009 | 0.009 | 0.032 | 0.032 | 0 | 0.47 | 0.19 | 0.4 | 0.47 | 0.6 | 0.43 |
| 10 | 129263589 | rs11018119 | -0.609 | -0.44 | -1.4 | -0.899 | 0.017 | -0.009 | -0.009 | 0 | 0.54 | 0.32 | 0.62 | 0.41 | 0.61 | 0.55 |
| 10 | 129264247 | rs915146 | 1.171 | -0.29 | -1.433 | -0.745 | -0.044 | -0.078 | -0.078 | 0.01 | 0.27 | 0.14 | 0.31 | 0.51 | 0.37 | 0.39 |
| 10 | 129265653 | rs885587 | 0.116 | -0.64 | -1.602 | -0.942 | -0.019 | -0.021 | -0.021 | 0 | 0.42 | 0.29 | 0.55 | 0.41 | 0.49 | 0.46 |
| 10 | 129265798 | rs10741271 | -1.054 | -0.724 | -1.427 | -0.936 | -0.085 | -0.119 | -0.119 | 0.265 | 0.74 | 0.49 | 0.88 | 0.91 | 0.89 | 0.89 |
| 10 | 129266910 | rs868567 | 1.144 | -0.662 | -1.476 | -0.798 | -0.036 | -0.071 | -0.071 | 0 | 0.3 | 0.13 | 0.31 | 0.5 | 0.37 | 0.42 |
| 10 | 129267207 | rs10741272 | -1.754 | -0.756 | -1.173 | -1.255 | -0.079 | -0.074 | -0.074 | 0 | 0.82 | 0.57 | 0.93 | 0.91 | 0.96 | 0.93 |
| 10 | 129267236 | rs11018121 | 1.992 | -0.506 | -1.44 | -1.273 | -0.002 | -0.032 | -0.032 | 0 | 0.09 | 0.12 | 0.04 | 0 | 0.09 | 0.06 |
| 10 | 129267595 | rs11018123 | 0.874 | -0.699 | -1.538 | -1.385 | 0.031 | 0.03 | 0.03 | 0.572 | 0.23 | 0.16 | 0.06 | 0 | 0.13 | 0.1 |
| 10 | 129267643 | rs11018124 | 1.55 | -0.506 | -1.436 | -1.286 | -0.001 | -0.027 | -0.027 | 0 | 0.12 | 0.16 | 0.05 | 0 | 0.11 | 0.07 |
| 10 | 129267759 | rs11018125 | 0.098 | -0.318 | -1.519 | -1.379 | -0.004 | -0.002 | -0.002 | 0 | 0.21 | 0.17 | 0.14 | 0 | 0.24 | 0.11 |
| 10 | 129268527 | rs1931725 | -0.706 | -0.32 | -1.288 | -0.763 | -0.023 | 0.085 | 0.085 | 0 | 0.75 | 0.38 | 0.62 | 0.45 | 0.88 | 0.76 |
| 10 | 129269059 | rs11018127 | 1.004 | 0.125 | -1.513 | -0.946 | -0.047 | 0.042 | 0.042 | 1.128 | 0.3 | 0.17 | 0.56 | 0.82 | 0.39 | 0.53 |
| 10 | 129270400 | rs12250382 | 0.824 | 0.823 | -1.268 | -0.495 | -0.005 | 0.026 | 0.026 | 0 | 0.21 | 0.08 | 0.16 | 0.09 | 0.23 | 0.24 |
| 10 | 129270746 | rs10830085 | -1.459 | 0.674 | -0.92 | -0.473 | -0.006 | 0.019 | 0.019 | 0.394 | 0.9 | 0.6 | 0.94 | 0.91 | 0.98 | 0.96 |
| 10 | 129271850 | rs10765110 | -0.243 | 0.719 | -1.056 | -0.572 | 0.004 | 0.019 | 0.019 | 0.066 | 0.34 | 0.3 | 0.21 | 0.09 | 0.33 | 0.3 |
| 10 | 129272635 | rs11018132 | 0.569 | 0.988 | -1.073 | -0.422 | -0.01 | 0.037 | 0.037 | 0 | 0.09 | 0.07 | 0.31 | 0.51 | 0.1 | 0.18 |
| 10 | 129273003 | rs10830088 | -0.213 | 0.972 | -1.075 | -0.45 | 0.004 | 0.019 | 0.019 | 0.066 | 0.34 | 0.3 | 0.21 | 0.09 | 0.33 | 0.3 |
| 10 | 129273257 | rs11018134 | -1.082 | 0.976 | -1.083 | -0.465 | -0.007 | -0.013 | -0.013 | 0 | 0.3 | 0.31 | 0.46 | 0.51 | 0.34 | 0.29 |
| 10 | 129273673 | rs10830090 | -0.739 | 0.965 | -1.053 | -0.444 | -0.004 | 0.004 | 0.004 | 0 | 0.43 | 0.37 | 0.52 | 0.6 | 0.43 | 0.48 |
| 10 | 129274503 | rs4363506 | -0.722 | 0.971 | -1.056 | -0.447 | -0.004 | 0.004 | 0.004 | 0 | 0.43 | 0.37 | 0.52 | 0.59 | 0.43 | 0.48 |
| 10 | 129274752 | rs10765114 | -0.733 | 0.979 | -1.052 | -0.434 | -0.004 | 0.004 | 0.004 | 0 | 0.43 | 0.37 | 0.52 | 0.59 | 0.43 | 0.48 |
| 10 | 129276920 | rs1931729 | -1.493 | 0.979 | -0.934 | -0.378 | -0.007 | 0.018 | 0.018 | 0.394 | 0.9 | 0.6 | 0.95 | 0.9 | 0.98 | 0.96 |
| 10 | 129277551 | rs11018141 | -0.418 | 1.317 | -1.418 | -0.233 | -0.038 | -0.061 | -0.061 | 0 | 0.18 | 0.1 | 0.4 | 0.51 | 0.24 | 0.23 |
| 10 | 129280178 | rs884436 | -0.515 | 1.16 | -1.331 | -0.506 | -0.06 | -0.016 | -0.016 | 0.558 | 0.39 | 0.18 | 0.55 | 0.59 | 0.47 | 0.47 |
| 10 | 129281900 | rs1931737 | 0.031 | 1.18 | -1.244 | -0.491 | -0.034 | -0.04 | -0.04 | 0 | 0.3 | 0.14 | 0.46 | 0.59 | 0.34 | 0.46 |
| 10 | 129282460 | rs10830096 | 0.573 | 0.849 | -1.067 | -0.424 | -0.011 | -0.06 | -0.06 | 0 | 0.13 | 0.23 | 0.06 | 0 | 0.1 | 0.04 |
| 10 | 129284365 | rs10830099 | -1.07 | 0.892 | -1.139 | -0.421 | 0.028 | 0.027 | 0.027 | 0 | 0.51 | 0.36 | 0.61 | 0.59 | 0.58 | 0.53 |
| 10 | 129284413 | rs10830100 | -0.398 | 0.891 | -1.123 | -0.426 | -0.031 | -0.001 | -0.001 | 0 | 0.4 | 0.22 | 0.56 | 0.59 | 0.47 | 0.5 |
| 10 | 129285183 | rs10765118 | -0.982 | 0.884 | -1.16 | -0.335 | 0.026 | 0.025 | 0.025 | 0 | 0.52 | 0.36 | 0.61 | 0.59 | 0.58 | 0.53 |
| 10 | 129288183 | rs4751427 | 0.643 | 0.868 | -1.194 | -0.401 | -0.02 | 0.051 | 0.051 | 0.427 | 0.12 | 0.07 | 0.06 | 0 | 0.13 | 0.04 |
| 10 | 129288498 | rs4751428 | 1.166 | 0.682 | -1.018 | -0.428 | 0.011 | 0.002 | 0.002 | 0 | 0.41 | 0.26 | 0.34 | 0.33 | 0.4 | 0.43 |
| 10 | 129289451 | rs10830103 | 0.372 | 0.69 | -0.922 | -0.254 | -0.015 | 0.027 | 0.027 | 0 | 0.16 | 0.12 | 0.15 | 0.08 | 0.18 | 0.1 |
| 10 | 129289558 | rs7090712 | 1.196 | 0.613 | -0.875 | -0.239 | 0.014 | -0.002 | -0.002 | 0 | 0.38 | 0.26 | 0.32 | 0.33 | 0.34 | 0.4 |
| 10 | 129291366 | rs2065674 | 1.135 | 0.523 | -0.783 | -0.247 | 0.009 | -0.008 | -0.008 | 0 | 0.39 | 0.29 | 0.33 | 0.43 | 0.34 | 0.44 |
| 10 | 129292364 | rs10741275 | -1.218 | 0.911 | -0.4 | -0.551 | -0.012 | -0.051 | -0.051 | 0 | 0.91 | 0.66 | 0.95 | 1 | 0.98 | 0.99 |
| 10 | 129294641 | rs2065675 | 0.313 | 1.118 | -0.489 | -0.64 | 0.017 | -0.087 | -0.087 | 0 | 0.76 | 0.56 | 0.82 | 0.9 | 0.75 | 0.87 |
| 10 | 129296719 | rs4751430 | 0.204 | 0.903 | -0.833 | -0.6 | -0.008 | -0.004 | -0.004 | 0 | 0.15 | 0.13 | 0.15 | 0.18 | 0.15 | 0.11 |
| 10 | 129297103 | rs6482919 | 0.925 | 0.919 | -0.758 | -0.474 | 0.003 | 0.002 | 0.002 | 0 | 0.58 | 0.5 | 0.49 | 0.47 | 0.56 | 0.71 |
| 10 | 129298233 | rs2151937 | 0.472 | 0.398 | -0.948 | -0.703 | 0 | -0.003 | -0.003 | 0 | 0.45 | 0.44 | 0.46 | 0.42 | 0.56 | 0.52 |
| 10 | 129300066 | rs9665721 | -1.401 | 0.34 | -0.748 | -0.66 | -0.017 | -0.052 | -0.052 | 0.144 | 0.9 | 0.64 | 0.95 | 1 | 0.98 | 0.98 |
| 10 | 129300464 | rs7069740 | 1.739 | 0.371 | -1.356 | -0.714 | -0.023 | -0.027 | -0.027 | 7.498 | 0.28 | 0.12 | 0.31 | 0.32 | 0.34 | 0.35 |
| 10 | 129304075 | rs10830107 | 0.508 | 0.152 | -0.937 | -0.506 | 0.037 | 0.102 | 0.102 | 0 | 0.28 | 0.08 | 0.13 | 0.05 | 0.22 | 0.29 |
| 10 | 129304998 | rs4751432 | 0.071 | 0.261 | -0.867 | -0.444 | 0.053 | 0.072 | 0.072 | 0 | 0.6 | 0.33 | 0.8 | 0.94 | 0.76 | 0.66 |
| 10 | 129307068 | rs11018168 | -0.439 | 0.85 | -1.418 | -0.574 | -0.049 | 0.065 | 0.065 | 0 | 0.19 | 0.08 | 0.12 | 0 | 0.26 | 0.14 |
| 10 | 129309600 | rs7920899 | 0.704 | 0.344 | -1.337 | -0.412 | 0.031 | 0.024 | 0.024 | 0 | 0.21 | 0.09 | 0.08 | 0.15 | 0.14 | 0.29 |
| 10 | 129311165 | rs7907561 | -0.723 | 0.545 | -0.985 | -0.426 | -0.004 | -0.034 | -0.034 | 0.279 | 0.93 | 0.87 | 0.96 | 1 | 0.94 | 0.98 |
| 10 | 129312691 | rs1572237 | -0.174 | 0.42 | -0.984 | -0.488 | -0.021 | -0.038 | -0.038 | 0.081 | 0.76 | 0.72 | 0.9 | 0.89 | 0.83 | 0.77 |
| 10 | 129312952 | rs999069 | 0.998 | 0.57 | -2.081 | -0.26 | -0.115 | -0.045 | -0.045 | 0 | 0.46 | 0.07 | 0.52 | 0.39 | 0.52 | 0.46 |
| 10 | 129313964 | rs12257984 | 0.728 | 0.749 | -2.127 | -0.305 | -0.032 | -0.017 | -0.017 | 0.123 | 0.51 | 0.07 | 0.56 | 0.39 | 0.64 | 0.54 |
| 10 | 129315562 | rs7085844 | 0.474 | 0.763 | -1.534 | -0.067 | -0.016 | -0.023 | -0.023 | 0 | 0.63 | 0.29 | 0.57 | 0.5 | 0.63 | 0.67 |
| 10 | 129315699 | rs10830112 | -0.247 | 0.982 | -1.974 | 0.46 | -0.002 | 0.102 | 0.102 | 0 | 0.06 | 0.04 | 0.23 | 0.5 | 0.06 | 0.14 |
| 10 | 129316419 | rs1009983 | 0.553 | 0.985 | -2.051 | 0.19 | -0.095 | -0.021 | -0.021 | 0 | 0.36 | 0.05 | 0.36 | 0.28 | 0.41 | 0.39 |
| 10 | 129321009 | rs4582883 | 0.5 | 1.179 | -0.748 | 0.691 | 0.009 | 0.04 | 0.04 | 0 | 0.76 | 0.62 | 0.66 | 0.62 | 0.73 | 0.73 |
| 10 | 129321875 | rs4751436 | 0.535 | 1.022 | -0.309 | 0.172 | 0.02 | -0.024 | -0.024 | 0.97 | 0.95 | 0.84 | 0.87 | 1 | 0.96 | 0.97 |
| 10 | 129322065 | rs1999637 | -0.552 | 0.284 | -0.931 | -0.65 | 0.012 | 0.146 | 0.146 | 0.367 | 0.44 | 0.77 | 0.27 | 0.13 | 0.36 | 0.36 |
| 10 | 129328254 | rs11599062 | 0.659 | -0.213 | 0.006 | -0.881 | -0.032 | -0.03 | -0.03 | 0 | 0.21 | 0.29 | 0.11 | 0.11 | 0.12 | 0.26 |
| 10 | 129330043 | rs12245703 | -0.465 | -0.123 | 0.589 | -1.003 | -0.019 | -0.092 | -0.092 | 0 | 0.33 | 0.5 | 0.19 | 0.11 | 0.25 | 0.3 |
| 10 | 129333397 | rs7094852 | -0.618 | -0.1 | 0.587 | -1.038 | -0.001 | -0.025 | -0.025 | 0 | 0.34 | 0.64 | 0.2 | 0.11 | 0.25 | 0.3 |
| 10 | 129334448 | rs4751439 | 0.436 | -0.625 | 0.202 | -1.109 | 0.001 | -0.005 | -0.005 | 5.043 | 0.87 | 0.91 | 0.9 | 1 | 0.9 | 0.9 |
| 10 | 129335512 | rs11594607 | 0.423 | -0.607 | 0.583 | -0.965 | -0.052 | -0.057 | -0.057 | 0.045 | 0.23 | 0.39 | 0.12 | 0.1 | 0.12 | 0.26 |
| 10 | 129336988 | rs7912885 | -0.479 | -1.231 | -0.061 | -0.518 | -0.035 | -0.001 | -0.001 | 0 | 0.6 | 0.52 | 0.56 | 0.57 | 0.69 | 0.63 |
| 10 | 129337138 | rs11018181 | -0.624 | -1.755 | -0.129 | -0.64 | -0.02 | -0.019 | -0.019 | 0 | 0.67 | 0.58 | 0.82 | 0.77 | 0.84 | 0.71 |
| 10 | 129337420 | rs12771712 | 0.732 | -1.94 | 0.018 | -0.275 | -0.036 | 0.096 | 0.096 | 1.555 | 0.75 | 0.53 | 0.63 | 0.57 | 0.82 | 0.69 |
| 10 | 129339304 | rs2050334 | 0.033 | -2.018 | -0.847 | -0.461 | 0 | 0.143 | 0.143 | 1.909 | 0.41 | 0.15 | 0.69 | 0.79 | 0.6 | 0.53 |

| | | | | | | | | | | | | | | | | |
|---|---|---|---|---|---|---|---|---|---|---|---|---|---|---|---|---|
| 10 | 129339695 | rs11018185 | 0.991 | -1.963 | -0.918 | -0.347 | -0.118 | -0.102 | -0.102 | 2.639 | 0.26 | 0.08 | 0.38 | 0.43 | 0.48 | 0.35 |
| 10 | 129346199 | rs1931706 | 0.115 | -2.446 | -0.581 | -0.653 | 0.023 | 0.034 | 0.034 | 0 | 0.44 | 0.64 | 0.25 | 0.21 | 0.3 | 0.38 |
| 10 | 129347785 | rs990310 | 0.93 | -2.568 | -0.166 | -0.685 | -0.013 | -0.021 | -0.021 | 0 | 0.09 | 0.32 | 0.05 | 0.05 | 0.05 | 0.12 |
| 10 | 129348918 | rs2184375 | 1.095 | -2.609 | -0.304 | -0.684 | 0.016 | -0.063 | -0.063 | 0 | 0.15 | 0.07 | 0.31 | 0.36 | 0.12 | 0.18 |
| 10 | 129349748 | rs11591259 | 1.733 | -2.515 | -0.17 | -0.739 | 0.016 | 0.014 | 0.014 | 0.018 | 0.26 | 0.26 | 0.15 | 0.15 | 0.13 | 0.21 |
| 10 | 129349788 | rs11018194 | 0.83 | -2.505 | -0.277 | -0.722 | 0.051 | -0.063 | -0.063 | 1.848 | 0.31 | 0.1 | 0.37 | 0.37 | 0.23 | 0.28 |
| 10 | 129349833 | rs1931708 | 2.119 | -1.053 | -0.497 | -0.761 | -0.028 | -0.05 | -0.05 | 0.032 | 0.15 | 0.07 | 0.16 | 0.34 | 0.2 | 0.23 |
| 10 | 129349946 | rs11018195 | 1.557 | -1.185 | -0.413 | -0.778 | -0.023 | -0.029 | -0.029 | 0.048 | 0.09 | 0.23 | 0.03 | 0.05 | 0.05 | 0.12 |
| 10 | 129351543 | rs11018198 | 0.942 | -1.014 | -0.904 | -0.694 | 0.05 | -0.063 | -0.063 | 0 | 0.3 | 0.1 | 0.37 | 0.37 | 0.23 | 0.27 |
| 10 | 129352101 | rs3740549 | 0.105 | -0.943 | -0.754 | -0.784 | 0.024 | 0.035 | 0.035 | 0 | 0.44 | 0.64 | 0.25 | 0.19 | 0.3 | 0.38 |
| 10 | 129352866 | rs4750892 | 1.813 | -0.366 | -1.01 | -0.631 | -0.011 | 0.002 | 0.002 | 0 | 0.08 | 0.06 | 0.05 | 0 | 0.12 | 0.05 |
| 10 | 129353080 | rs10830126 | 2.037 | -0.146 | -1.079 | -0.577 | -0.061 | -0.009 | -0.009 | 0 | 0.09 | 0.01 | 0.2 | 0.1 | 0.27 | 0.12 |
| 10 | 129353132 | rs10734107 | -0.648 | -0.677 | -0.918 | -0.597 | 0.002 | -0.003 | -0.003 | 0 | 0.49 | 0.38 | 0.62 | 0.53 | 0.55 | 0.51 |
| 10 | 129353489 | rs10830127 | -0.038 | -0.501 | -1.213 | -0.581 | -0.097 | -0.071 | -0.071 | 0 | 0.4 | 0.11 | 0.57 | 0.44 | 0.51 | 0.39 |
| 10 | 129354555 | rs4390263 | -0.083 | -0.497 | -1.1 | -0.554 | -0.02 | -0.017 | -0.017 | 0 | 0.4 | 0.23 | 0.58 | 0.42 | 0.48 | 0.35 |
| 10 | 129357315 | rs11018200 | -0.091 | -0.98 | -1.488 | -0.638 | 0.015 | -0.057 | -0.057 | 0 | 0.63 | 0.21 | 0.67 | 0.48 | 0.71 | 0.55 |
| 10 | 129359425 | rs4751452 | 0.943 | -1.219 | -1.625 | -0.65 | 0.02 | -0.062 | -0.062 | 0 | 0.15 | 0.07 | 0.31 | 0.31 | 0.11 | 0.17 |
| 10 | 129359499 | rs4751453 | 0.423 | -0.926 | -1.488 | -0.606 | -0.045 | -0.048 | -0.048 | 0 | 0.55 | 0.15 | 0.62 | 0.48 | 0.58 | 0.5 |
| 10 | 129366558 | rs1547321 | -0.182 | -0.664 | -0.851 | -0.429 | 0.001 | 0.004 | 0.004 | 0 | 0.44 | 0.35 | 0.52 | 0.38 | 0.56 | 0.43 |
| 10 | 129368590 | rs1931713 | 1.481 | -0.69 | -0.98 | -0.247 | -0.037 | -0.045 | -0.045 | 0 | 0.14 | 0.07 | 0.22 | 0.3 | 0.19 | 0.2 |
| 10 | 129372024 | rs12412993 | 0.738 | -0.727 | -0.426 | -0.261 | 0.044 | 0.037 | 0.037 | 0.032 | 0.34 | 0.26 | 0.18 | 0.19 | 0.2 | 0.27 |
| 10 | 129373991 | rs971175 | 0.881 | -0.625 | -0.12 | -0.055 | -0.013 | 0.038 | 0.038 | 0 | 0.12 | 0.14 | 0.19 | 0.27 | 0.07 | 0.14 |
| 10 | 129376084 | rs11018209 | 1.819 | -0.422 | -0.1 | -0.509 | 0.05 | -0.015 | -0.015 | 0 | 0.1 | 0.1 | 0.23 | 0.06 | 0.28 | 0.12 |
| 10 | 129380861 | rs4363508 | -1.279 | -0.539 | 0.027 | -0.503 | -0.022 | 0.014 | 0.014 | 0 | 0.92 | 0.82 | 0.94 | 0.87 | 0.96 | 0.89 |
| 10 | 129381096 | rs12252756 | 0.571 | 0.057 | -0.058 | -0.622 | 0.028 | 0.083 | 0.083 | 0 | 0.41 | 0.28 | 0.23 | 0.19 | 0.34 | 0.3 |
| 10 | 129384094 | rs10830137 | 0.685 | 0.165 | -0.024 | -0.446 | -0.016 | 0.027 | 0.027 | 0 | 0.13 | 0.16 | 0.11 | 0.23 | 0.08 | 0.13 |
| 10 | 129384543 | rs12774813 | 1.524 | 0.316 | -0.492 | -0.505 | 0.07 | -0.021 | -0.021 | 0.599 | 0.1 | 0.13 | 0.23 | 0.05 | 0.28 | 0.12 |
| 10 | 129385588 | rs12267188 | -0.116 | 0.344 | -0.486 | -0.458 | 0.006 | 0.116 | 0.116 | 0 | 0.46 | 0.3 | 0.26 | 0.19 | 0.41 | 0.34 |
| 10 | 129385698 | rs10430603 | -0.525 | 2.285 | 0.141 | 0.457 | -0.002 | -0.003 | -0.003 | 0 | 0.2 | 0.17 | 0.3 | 0.27 | 0.33 | 0.25 |
| 10 | 129388646 | rs10765137 | -1.172 | 2.251 | 0.134 | 0.419 | 0.009 | 0.004 | 0.004 | 0 | 0.25 | 0.22 | 0.33 | 0.31 | 0.39 | 0.28 |
| 10 | 129389743 | rs4751459 | -0.552 | 2.309 | 0.068 | 0.452 | -0.007 | -0.008 | -0.008 | 0 | 0.2 | 0.16 | 0.3 | 0.3 | 0.31 | 0.25 |
| 10 | 129394566 | rs10443996 | 1.076 | 2.374 | -0.016 | 0.414 | -0.048 | -0.011 | -0.011 | 0.101 | 0.08 | 0 | 0.19 | 0.09 | 0.26 | 0.11 |
| 10 | 129394587 | rs12777985 | 0.036 | 2.222 | 0.097 | 0.356 | 0.005 | -0.001 | -0.001 | 0 | 0.2 | 0.26 | 0.17 | 0.2 | 0.2 | 0.24 |
| 10 | 129400407 | rs6482940 | 0.114 | 1.569 | -0.648 | 0.503 | -0.038 | 0.048 | 0.048 | 0 | 0.12 | 0.18 | 0.11 | 0.21 | 0.06 | 0.13 |
| 10 | 129400615 | rs7082785 | 1.014 | 2.043 | -1.291 | 0.227 | 0.095 | -0.002 | -0.002 | 0 | 0.51 | 0.19 | 0.47 | 0.49 | 0.4 | 0.48 |
| 10 | 129401906 | rs7901816 | 0.265 | 3.06 | -0.911 | 0.264 | -0.043 | -0.015 | -0.015 | 0 | 0.08 | 0.01 | 0.19 | 0.09 | 0.26 | 0.11 |
| 10 | 129405259 | rs11018222 | 0.234 | 2.393 | -0.882 | 0.261 | -0.065 | -0.016 | -0.016 | 0 | 0.1 | 0.01 | 0.18 | 0.07 | 0.19 | 0.1 |
| 10 | 129405859 | rs10830139 | -0.918 | 0.588 | -0.406 | -0.092 | 0 | -0.034 | -0.034 | 0 | 0.43 | 0.46 | 0.7 | 0.89 | 0.54 | 0.56 |
| 10 | 129406109 | rs1113103 | -0.907 | 0.614 | -0.455 | -0.079 | 0.002 | -0.022 | -0.022 | 0 | 0.43 | 0.44 | 0.7 | 0.89 | 0.54 | 0.56 |
| 10 | 129418956 | rs11018230 | -0.201 | 0.941 | -0.392 | 0.198 | 0.029 | -0.056 | -0.056 | 0 | 0.3 | 0.39 | 0.27 | 0.06 | 0.42 | 0.24 |
| 10 | 129420087 | rs11597076 | -0.937 | 0.082 | -0.544 | 1.571 | -0.001 | 0.041 | 0.041 | 0 | 0.15 | 0.16 | 0.38 | 0.49 | 0.12 | 0.25 |
| 10 | 129443941 | rs1936845 | 0.197 | 0.043 | -0.57 | 1.307 | 0 | -0.002 | -0.002 | 2.381 | 0.77 | 0.78 | 0.78 | 0.74 | 0.8 | 0.72 |
| 10 | 129445709 | rs7071097 | 0.139 | 0.02 | -0.576 | 1.372 | -0.001 | -0.001 | -0.001 | 0 | 0.78 | 0.8 | 0.78 | 0.78 | 0.8 | 0.75 |
| 10 | 129445779 | rs7071119 | -0.502 | 0.332 | -0.476 | 2.06 | 0.002 | 0.036 | 0.036 | 0 | 0.1 | 0.12 | 0.23 | 0.38 | 0.14 | 0.2 |
| 10 | 129447502 | rs11596988 | 0.333 | -0.503 | -0.953 | 1.389 | 0.026 | 0.029 | 0.029 | 0 | 0.25 | 0.08 | 0.16 | 0.13 | 0.19 | 0.17 |
| 10 | 129451365 | rs10830150 | 1.002 | -0.969 | -1.304 | 1.237 | -0.049 | 0.104 | 0.104 | 0.267 | 0.37 | 0.16 | 0.33 | 0.2 | 0.42 | 0.35 |
| 10 | 129453002 | rs7079699 | 1.174 | -1.014 | -1.011 | 1.264 | -0.008 | 0.04 | 0.04 | 0 | 0.19 | 0.05 | 0.09 | 0.07 | 0.16 | 0.2 |
| 10 | 129454892 | rs12245508 | -0.127 | -1.06 | 0.161 | 1.362 | 0.019 | 0.049 | 0.049 | 0 | 0.69 | 0.56 | 0.53 | 0.53 | 0.65 | 0.61 |
| 10 | 129455935 | rs12765922 | 1.18 | -0.919 | 0.107 | 1.389 | -0.125 | 0.23 | 0.23 | 0.303 | 0.33 | 0.04 | 0.31 | 0.03 | 0.41 | 0.26 |
| 10 | 129461659 | rs12248827 | -0.264 | -0.822 | 0.663 | 1.194 | 0.003 | 0.002 | 0.002 | 0 | 0.48 | 0.49 | 0.29 | 0.54 | 0.37 | 0.5 |
| 10 | 129476525 | rs7912681 | 0.218 | -0.228 | 0.789 | 1.627 | 0.031 | 0.117 | 0.117 | 0 | 0.88 | 0.77 | 0.64 | 0.58 | 0.82 | 0.81 |
| 10 | 129477828 | rs2186047 | -0.172 | -0.6 | 0.408 | 1.612 | -0.004 | -0.004 | -0.004 | 0 | 0.15 | 0.13 | 0.25 | 0.45 | 0.19 | 0.22 |
| 10 | 129478145 | rs10454472 | -0.788 | -0.951 | 0.993 | 1.417 | 0.005 | 0.109 | 0.109 | 0 | 0.16 | 0.3 | 0.25 | 0.46 | 0.17 | 0.29 |
| 10 | 129478197 | rs10454473 | -0.864 | -1.236 | 0.782 | 0.997 | 0.008 | -0.069 | -0.069 | 0.39 | 0.42 | 0.53 | 0.57 | 0.9 | 0.48 | 0.58 |
| 10 | 129479279 | rs1892183 | -0.215 | -1.366 | 0.858 | 1.083 | 0.004 | 0.034 | 0.034 | 0.616 | 0.73 | 0.46 | 0.71 | 0.62 | 0.75 | 0.64 |
| 10 | 129479305 | rs1892184 | 0.834 | -1.475 | 0.578 | 1.036 | -0.009 | -0.049 | -0.049 | 0 | 0.12 | 0.06 | 0.13 | 0.32 | 0.16 | 0.17 |
| 10 | 129479727 | rs10765143 | 0.245 | -1.356 | 0.716 | 0.472 | 0.001 | -0.005 | -0.005 | 0 | 0.68 | 0.37 | 0.69 | 0.63 | 0.74 | 0.61 |
| 10 | 129480209 | rs10830157 | 0.06 | -1.412 | 0.906 | 0.402 | -0.003 | 0 | 0 | 0 | 0.6 | 0.44 | 0.59 | 0.4 | 0.65 | 0.61 |
| 10 | 129482187 | rs7921601 | -0.416 | -0.63 | 0.945 | 0.626 | 0.052 | 0.015 | 0.015 | 0.156 | 0.95 | 0.77 | 0.95 | 1 | 0.97 | 0.96 |
| 10 | 129482294 | rs7921731 | 0.26 | -0.5 | 0.113 | 0.785 | -0.014 | -0.021 | -0.021 | 0 | 0.2 | 0.12 | 0.22 | 0.22 | 0.21 | 0.21 |
| 10 | 129485809 | rs10741281 | 0.445 | -1.718 | 0.769 | 0.429 | 0.005 | 0.001 | 0.001 | 0 | 0.45 | 0.42 | 0.61 | 0.42 | 0.62 | 0.49 |
| 10 | 129486233 | rs10830159 | -0.526 | -1.587 | 0.673 | 0.486 | 0.035 | 0.02 | 0.02 | 0 | 0.51 | 0.33 | 0.36 | 0.58 | 0.38 | 0.5 |
| 10 | 129488347 | rs10765148 | -0.092 | -2.064 | 0.086 | 0.354 | 0.038 | -0.029 | -0.029 | 1.266 | 0.33 | 0.21 | 0.27 | 0.57 | 0.21 | 0.38 |
| 10 | 129488763 | rs11018266 | -0.007 | -1.959 | 0.092 | 0.408 | 0.024 | -0.037 | -0.037 | 0.274 | 0.29 | 0.2 | 0.17 | 0.54 | 0.18 | 0.35 |
| 10 | 129490094 | rs4751468 | 0.044 | -1.972 | 0.217 | 0.412 | 0.014 | -0.018 | -0.018 | 0.826 | 0.3 | 0.24 | 0.18 | 0.54 | 0.18 | 0.35 |
| 10 | 129502398 | rs7919990 | 0.864 | -1.479 | 1.258 | -0.101 | -0.001 | -0.074 | -0.074 | 0.943 | 0.77 | 0.7 | 0.88 | 1 | 0.81 | 0.83 |
| 10 | 129513191 | rs7904713 | 1.111 | -1.522 | 1.533 | -0.119 | -0.006 | -0.171 | -0.171 | 0 | 0.75 | 0.53 | 0.87 | 1 | 0.81 | 0.83 |
| 10 | 129519132 | rs7083606 | 1.014 | -1.573 | 1.451 | 0.158 | -0.01 | -0.02 | -0.02 | 0.773 | 0.74 | 0.4 | 0.86 | 0.81 | 0.81 | 0.71 |
| 10 | 129526128 | rs11018288 | 1.222 | -1.533 | 1.444 | 0.212 | 0.001 | -0.004 | -0.004 | 1.128 | 0.8 | 0.55 | 0.89 | 0.83 | 0.86 | 0.75 |

| | | | | | | | | | | | | | | | | |
|---|---|---|---|---|---|---|---|---|---|---|---|---|---|---|---|---|
| 10 | 129526605 | rs5002823 | 1.564 | -1.545 | 1.583 | 0.199 | 0.025 | 0.166 | 0.166 | 0 | 0.94 | 0.74 | 0.96 | 0.83 | 0.98 | 0.85 |
| 10 | 129528865 | rs4751499 | 1.528 | -1.559 | 1.534 | 0.189 | 0.003 | -0.005 | -0.005 | 0 | 0.8 | 0.56 | 0.89 | 0.83 | 0.85 | 0.74 |
| 10 | 129532117 | rs1926156 | 1.922 | -1.553 | 1.643 | 0.245 | 0.018 | 0.161 | 0.161 | 0 | 0.93 | 0.68 | 0.94 | 0.83 | 0.97 | 0.85 |
| 10 | 129536575 | rs7075733 | 1.352 | -1.578 | 1.29 | -0.026 | 0.001 | -0.117 | -0.117 | 0 | 0.82 | 0.66 | 0.89 | 1 | 0.84 | 0.88 |
| 10 | 129542526 | rs12264783 | -0.969 | -1.066 | 1.049 | -0.006 | 0.015 | -0.063 | -0.063 | 0.017 | 0.15 | 0.21 | 0.12 | 0 | 0.18 | 0.11 |
| 10 | 129551672 | rs1926150 | 0.353 | -1.651 | 0.615 | -0.224 | -0.015 | 0.037 | 0.037 | 0 | 0.87 | 0.96 | 0.92 | 1 | 0.87 | 0.94 |
| 10 | 129551736 | rs7916596 | -0.974 | -2.54 | 1.173 | -0.423 | -0.073 | 0.082 | 0.082 | 0 | 0.14 | 0.47 | 0.07 | 0.15 | 0.04 | 0.21 |
| 10 | 129553772 | rs12784644 | 1.253 | -2.181 | 0.389 | -0.662 | 0.029 | 0.016 | 0.016 | 0 | 0.19 | 0.08 | 0.1 | 0.18 | 0.12 | 0.15 |
| 10 | 129566665 | rs2182406 | 1.214 | -2.057 | 0.818 | -0.748 | -0.003 | -0.004 | -0.004 | 0 | 0.69 | 0.36 | 0.83 | 0.85 | 0.8 | 0.7 |
| 10 | 129571206 | rs1926145 | -1.457 | -1.957 | 1.015 | -0.698 | 0.011 | -0.017 | -0.017 | 0 | 0.05 | 0.2 | 0.03 | 0 | 0.03 | 0.02 |
| 10 | 129574877 | rs1926143 | 1.048 | -2.576 | 0.928 | -0.892 | 0.035 | 0.033 | 0.033 | 0 | 0.5 | 0.59 | 0.64 | 0.68 | 0.68 | 0.52 |
| 10 | 129581662 | rs4751515 | 0.935 | -2.52 | 0.937 | -0.933 | 0 | 0.001 | 0.001 | 0 | 0.49 | 0.51 | 0.77 | 0.68 | 0.71 | 0.53 |
| 10 | 129585004 | rs2094209 | 1.099 | -2.64 | 0.938 | -0.992 | 0.021 | 0.017 | 0.017 | 0.002 | 0.21 | 0.15 | 0.11 | 0.16 | 0.12 | 0.17 |
| 10 | 129586689 | rs1926181 | 0.683 | -2.65 | 0.932 | -0.999 | -0.023 | -0.029 | -0.029 | 0 | 0.7 | 0.62 | 0.87 | 0.85 | 0.8 | 0.69 |
| 10 | 129594596 | rs10830179 | 1.209 | -1.517 | 0.303 | -0.619 | 0.028 | 0.073 | 0.073 | 0 | 0.2 | 0.26 | 0.33 | 0.4 | 0.31 | 0.19 |
| 10 | 129596555 | rs12780271 | 1.19 | -1.706 | 0.311 | -0.781 | 0.019 | 0.02 | 0.02 | 0 | 0.22 | 0.15 | 0.1 | 0.16 | 0.15 | 0.17 |
| 10 | 129597382 | rs12762583 | 1.148 | -1.707 | 0.334 | -0.781 | 0.011 | 0.012 | 0.012 | 0 | 0.22 | 0.19 | 0.11 | 0.16 | 0.15 | 0.17 |
| 10 | 129602871 | rs1887018 | 0.74 | -1.633 | 0.359 | -0.729 | 0.035 | 0.036 | 0.036 | 0 | 0.48 | 0.39 | 0.76 | 0.69 | 0.69 | 0.53 |
| 10 | 129604375 | rs11816297 | -0.492 | -1.761 | 0.334 | -0.828 | -0.021 | -0.052 | -0.052 | 0 | 0.42 | 0.45 | 0.2 | 0.16 | 0.25 | 0.27 |
| 10 | 129614354 | rs7093392 | 0.015 | -1.613 | -0.108 | -0.85 | 0.011 | 0.002 | 0.002 | 0 | 0.59 | 0.63 | 0.8 | 0.88 | 0.75 | 0.74 |
| 10 | 129615722 | rs4620636 | 0.288 | -1.263 | -0.253 | -0.766 | -0.007 | 0.033 | 0.033 | 0 | 0.66 | 0.73 | 0.8 | 0.9 | 0.64 | 0.78 |
| 10 | 129617888 | rs7901629 | 0.675 | -1.764 | 0.215 | -0.511 | 0.001 | 0.001 | 0.001 | 0 | 0.76 | 0.73 | 0.88 | 0.85 | 0.84 | 0.75 |
| 10 | 129621741 | rs11018330 | 0.565 | -1.673 | 0.146 | -0.507 | -0.015 | 0.041 | 0.041 | 0 | 0.13 | 0.08 | 0.08 | 0 | 0.13 | 0.07 |
| 10 | 129623851 | rs7905689 | 0.216 | -1.814 | 0.167 | -0.53 | -0.009 | -0.009 | -0.009 | 0 | 0.28 | 0.25 | 0.12 | 0.12 | 0.13 | 0.2 |
| 10 | 129626293 | rs649611 | 1.277 | -1.17 | 0.356 | -0.384 | 0.004 | 0.008 | 0.008 | 0 | 0.46 | 0.29 | 0.58 | 0.63 | 0.57 | 0.46 |
| 10 | 129628377 | rs7089263 | 0.026 | -1.342 | 0.48 | -0.317 | -0.018 | 0.006 | 0.006 | 0 | 0.12 | 0.14 | 0.03 | 0.14 | 0.04 | 0.19 |
| 10 | 129629493 | rs7896329 | -0.032 | -1.192 | 0.504 | -0.39 | 0.004 | -0.11 | -0.11 | 0.025 | 0.17 | 0.39 | 0.1 | 0 | 0.13 | 0.07 |
| 10 | 129630873 | rs518525 | 0.803 | -1.058 | 0.374 | -0.325 | 0.004 | 0.014 | 0.014 | 0.121 | 0.55 | 0.4 | 0.63 | 0.77 | 0.64 | 0.63 |
| 10 | 129631750 | rs12412686 | -0.103 | -0.902 | -0.148 | -0.144 | 0.001 | 0.001 | 0.001 | 1.157 | 0.29 | 0.26 | 0.28 | 0.23 | 0.22 | 0.3 |
| 10 | 129632141 | rs486238 | 0.252 | -0.615 | -0.08 | 0.104 | -0.004 | -0.001 | -0.001 | 0 | 0.54 | 0.52 | 0.62 | 0.61 | 0.63 | 0.53 |
| 10 | 129633649 | rs540729 | 0.602 | -1.503 | -0.027 | 0.471 | 0.115 | -0.009 | -0.009 | 0.047 | 0.5 | 0.15 | 0.56 | 0.59 | 0.68 | 0.54 |
| 10 | 129634584 | rs11816315 | 0.408 | -1.522 | -0.005 | 0.263 | 0.024 | 0.007 | 0.007 | 4.968 | 0.21 | 0.12 | 0.12 | 0.18 | 0.12 | 0.15 |
| 10 | 129641502 | rs11018336 | -0.107 | -1.444 | 0.439 | 0.284 | -0.017 | 0.027 | 0.027 | 0.197 | 0.12 | 0.31 | 0.22 | 0.12 | 0.07 | 0.11 |
| 10 | 129642300 | rs7920167 | -0.603 | -1.433 | 0.178 | 0.281 | -0.037 | -0.028 | -0.028 | 0 | 0.2 | 0.47 | 0.28 | 0.11 | 0.15 | 0.12 |
| 10 | 129643446 | rs749791 | -0.727 | -1.107 | -0.011 | 0.352 | 0.01 | 0.007 | 0.007 | 3.325 | 0.43 | 0.64 | 0.4 | 0.27 | 0.27 | 0.24 |
| 10 | 129647972 | rs516465 | -0.491 | -0.617 | -0.027 | 0.49 | 0.018 | 0.017 | 0.017 | 0.466 | 0.39 | 0.28 | 0.28 | 0.27 | 0.31 | 0.17 |
| 10 | 129649793 | rs11018352 | 0.533 | -0.201 | 0.216 | 0.621 | -0.003 | 0 | 0 | 0 | 0.17 | 0.13 | 0.22 | 0.21 | 0.22 | 0.1 |
| 10 | 129650739 | rs582667 | -0.377 | -0.385 | 0.295 | 0.5 | -0.015 | -0.046 | -0.046 | 0 | 0.08 | 0.19 | 0.05 | 0 | 0.05 | 0.02 |
| 10 | 129652787 | rs1000797 | 0.417 | -0.421 | 0.081 | 0.516 | 0.002 | 0.008 | 0.008 | 0 | 0.11 | 0.14 | 0.08 | 0.14 | 0.15 | 0.07 |
| 10 | 129653425 | rs4750921 | 0.748 | -0.515 | 0.539 | 0.761 | 0.014 | 0.109 | 0.109 | 0.109 | 0.1 | 0.29 | 0.2 | 0.29 | 0.1 | 0.22 |
| 10 | 129653725 | rs11018361 | 1.215 | -0.524 | 0.539 | 0.743 | -0.017 | -0.01 | -0.01 | 0 | 0.15 | 0.05 | 0.2 | 0.21 | 0.21 | 0.09 |
| 10 | 129654224 | rs633941 | 0.711 | -0.518 | 0.542 | 0.722 | -0.003 | -0.019 | -0.019 | 0 | 0.06 | 0.09 | 0.04 | 0 | 0.05 | 0.02 |
| 10 | 129655008 | rs10741296 | -0.788 | -0.502 | 0.557 | 0.736 | 0.011 | 0.018 | 0.018 | 0 | 0.37 | 0.47 | 0.49 | 0.64 | 0.46 | 0.39 |
| 10 | 129657368 | rs680168 | -1.554 | -0.445 | 0.385 | 0.781 | 0 | -0.001 | -0.001 | 0 | 0.64 | 0.64 | 0.67 | 0.77 | 0.63 | 0.47 |
| 10 | 129660060 | rs4590782 | 0.58 | -0.309 | 0.06 | 0.863 | 0.033 | 0.108 | 0.108 | 0.1 | 0.1 | 0.29 | 0.21 | 0.29 | 0.12 | 0.23 |
| 10 | 129664418 | rs12357845 | 0.781 | -0.315 | -1.176 | 0.566 | -0.006 | -0.002 | -0.002 | 0 | 0.16 | 0.11 | 0.22 | 0.21 | 0.22 | 0.09 |
| 10 | 129664877 | rs12254032 | 0.388 | -0.109 | -0.873 | 0.605 | 0.01 | 0.018 | 0.018 | 0 | 0.38 | 0.18 | 0.37 | 0.34 | 0.33 | 0.16 |
| 10 | 129665073 | rs11018370 | 0.422 | -0.118 | -0.877 | 0.601 | 0.008 | 0.017 | 0.017 | 0 | 0.37 | 0.18 | 0.36 | 0.34 | 0.33 | 0.16 |
| 10 | 129667338 | rs2182407 | 0.313 | 0.078 | -0.923 | 0.68 | 0.006 | 0.008 | 0.008 | 0 | 0.39 | 0.35 | 0.39 | 0.34 | 0.34 | 0.16 |
| 10 | 129670197 | rs7898699 | 0.365 | 0.365 | -1.489 | 0.718 | -0.124 | -0.049 | -0.049 | 0.348 | 0.32 | 0.04 | 0.22 | 0.23 | 0.37 | 0.54 |
| 10 | 129670910 | rs4751545 | -0.182 | 0.136 | -0.762 | 0.695 | 0.015 | 0.021 | 0.021 | 0 | 0.5 | 0.36 | 0.52 | 0.45 | 0.47 | 0.22 |
| 10 | 129671751 | rs11018376 | -0.765 | -0.151 | 0.022 | 0.574 | 0.027 | 0.04 | 0.04 | 0 | 0.06 | 0.16 | 0.06 | 0.08 | 0.06 | 0.11 |
| 10 | 129671905 | rs575552 | -0.432 | -0.567 | -0.214 | 0.181 | 0.005 | -0.005 | -0.005 | 0 | 0.11 | 0.06 | 0.06 | 0 | 0.11 | 0.03 |
| 10 | 129672427 | rs7069800 | -0.153 | -0.739 | -0.218 | -0.058 | 0.005 | 0.003 | 0.003 | 0.249 | 0.47 | 0.57 | 0.22 | 0.28 | 0.31 | 0.28 |
| 10 | 129673226 | rs4750922 | 0.71 | -1.057 | -0.286 | 0.079 | 0.014 | 0.049 | 0.049 | 0 | 0.41 | 0.18 | 0.71 | 0.67 | 0.63 | 0.61 |
| 10 | 129673530 | rs7896581 | 0.713 | -1.625 | 0.087 | -0.199 | 0.002 | 0.002 | 0.002 | 0 | 0.27 | 0.24 | 0.11 | 0.16 | 0.16 | 0.17 |
| 10 | 129676169 | rs2296785 | 0.469 | -1.582 | -0.675 | 0.351 | -0.003 | 0.005 | 0.005 | 0 | 0.1 | 0.16 | 0.07 | 0.11 | 0.05 | 0.11 |
| 10 | 129677202 | rs11591839 | 0.044 | -1.578 | -1.286 | 0.189 | 0.033 | 0.033 | 0.033 | 0 | 0.36 | 0.28 | 0.13 | 0.22 | 0.2 | 0.26 |
| 10 | 129677328 | rs11018382 | -0.031 | -1.489 | -1.318 | 0.164 | -0.008 | 0.007 | 0.007 | 0 | 0.74 | 0.36 | 0.69 | 0.68 | 0.81 | 0.85 |
| 10 | 129677797 | rs11018383 | 0.728 | -0.824 | -1.742 | 0.349 | -0.079 | -0.122 | -0.122 | 0 | 0.38 | 0.07 | 0.55 | 0.43 | 0.59 | 0.59 |
| 10 | 129681029 | rs10830191 | 0.543 | -0.786 | -0.975 | 0.424 | 0.043 | -0.01 | -0.01 | 0 | 0.65 | 0.25 | 0.66 | 0.58 | 0.75 | 0.77 |
| 10 | 129682249 | rs10765177 | -0.722 | -0.494 | -0.483 | -0.511 | -0.009 | -0.107 | -0.107 | 0.197 | 0.3 | 0.39 | 0.12 | 0.03 | 0.24 | 0.23 |
| 10 | 129684456 | rs6482993 | -1.171 | -0.909 | -0.179 | -0.446 | -0.001 | -0.007 | -0.007 | 0.005 | 0.37 | 0.38 | 0.23 | 0.15 | 0.32 | 0.3 |
| 10 | 129686235 | rs4751546 | -0.713 | -0.959 | -0.331 | -0.615 | -0.002 | 0.003 | 0.003 | 0 | 0.3 | 0.35 | 0.38 | 0.26 | 0.34 | 0.21 |
| 10 | 129686579 | rs10741297 | -2.031 | -0.997 | -0.206 | -0.626 | -0.041 | 0.004 | 0.004 | 0.682 | 0.62 | 0.76 | 0.54 | 0.29 | 0.57 | 0.47 |
| 10 | 129688467 | rs12413939 | 1.123 | -0.946 | -0.178 | -0.593 | -0.007 | -0.02 | -0.02 | 0.092 | 0.06 | 0.02 | 0.07 | 0.09 | 0.09 | 0.05 |
| 10 | 129692293 | rs2148895 | -1.386 | -1.117 | -0.14 | -0.29 | -0.023 | 0.004 | 0.004 | 0 | 0.39 | 0.5 | 0.43 | 0.37 | 0.36 | 0.36 |
| 10 | 129695410 | rs4751548 | -1.157 | -1.061 | -0.243 | -0.473 | 0.02 | 0.022 | 0.022 | 0 | 0.47 | 0.55 | 0.38 | 0.31 | 0.35 | 0.5 |
| 10 | 129698493 | rs7074497 | 0.688 | -0.686 | -0.208 | -0.353 | 0.002 | -0.032 | -0.032 | 0.152 | 0.08 | 0.15 | 0.07 | 0 | 0.11 | 0.02 |

| | | | | | | | | | | | | | | | | |
|---|---|---|---|---|---|---|---|---|---|---|---|---|---|---|---|---|
| 10 | 129708049 | rs12248686 | 0.557 | -0.849 | -0.247 | -0.383 | 0.009 | 0.043 | 0.043 | 2.678 | 0.17 | 0.06 | 0.07 | 0 | 0.15 | 0.08 |
| 10 | 129713541 | rs11018401 | -0.022 | -0.567 | 0.28 | 0.278 | 0.004 | -0.011 | -0.011 | 0 | 0.11 | 0.06 | 0.17 | 0.14 | 0.1 | 0.18 |
| 10 | 129718678 | rs913287 | -0.332 | -0.235 | 0.682 | 0.23 | 0.083 | 0.023 | 0.023 | 0 | 0.1 | 0.38 | 0.09 | 0.04 | 0.07 | 0.06 |
| 10 | 129723712 | rs10830196 | -1.899 | -0.468 | 0.322 | 0.292 | -0.008 | 0.014 | 0.014 | 0.105 | 0.3 | 0.61 | 0.34 | 0.32 | 0.23 | 0.4 |
| 10 | 129730027 | rs3824761 | -1.702 | -0.77 | 0.351 | 0.535 | -0.024 | 0.021 | 0.021 | 0 | 0.38 | 0.57 | 0.46 | 0.42 | 0.32 | 0.35 |
| 10 | 129733149 | rs2275805 | -2.227 | -0.845 | 0.172 | 0.501 | 0.046 | -0.013 | -0.013 | 0 | 0.48 | 0.86 | 0.53 | 0.51 | 0.4 | 0.45 |
| 10 | 129736477 | rs932837 | -0.435 | -0.889 | 0.245 | 0.958 | 0.02 | -0.037 | -0.037 | 0 | 0.17 | 0.08 | 0.21 | 0.24 | 0.15 | 0.19 |
| 10 | 129737478 | rs4751551 | -1.49 | -0.925 | 0.461 | 0.943 | -0.011 | 0.039 | 0.039 | 0.196 | 0.24 | 0.32 | 0.39 | 0.42 | 0.2 | 0.3 |
| 10 | 129740668 | rs12769331 | -0.66 | -0.741 | 0.019 | 0.603 | 0.018 | -0.048 | -0.048 | 0.274 | 0.18 | 0.08 | 0.24 | 0.27 | 0.16 | 0.19 |
| 10 | 129747902 | rs11018412 | -1.612 | -0.746 | 0.17 | 0.513 | -0.017 | -0.001 | -0.001 | 0 | 0.31 | 0.4 | 0.3 | 0.26 | 0.26 | 0.24 |
| 10 | 129748903 | rs7900735 | -0.694 | -0.562 | -0.502 | 0.773 | 0.015 | -0.042 | -0.042 | 0 | 0.17 | 0.07 | 0.21 | 0.24 | 0.16 | 0.19 |
| 10 | 129751331 | rs7099752 | 0.024 | -0.665 | -0.256 | 0.593 | -0.007 | -0.004 | -0.004 | 0.632 | 0.2 | 0.25 | 0.15 | 0.22 | 0.16 | 0.16 |
| 10 | 129753712 | rs7922467 | -0.056 | -0.523 | -0.671 | 0.241 | -0.003 | -0.041 | -0.041 | 0 | 0.14 | 0.16 | 0.05 | 0 | 0.1 | 0.06 |
| 10 | 129757850 | rs985521 | -0.443 | -0.155 | -1.189 | 0.782 | -0.007 | -0.041 | -0.041 | 0 | 0.18 | 0.15 | 0.23 | 0.39 | 0.22 | 0.36 |
| 10 | 129758271 | rs4347296 | -1.565 | 0.454 | -1.674 | 0.554 | -0.037 | -0.028 | -0.028 | 0 | 0.76 | 0.95 | 0.83 | 0.76 | 0.71 | 0.85 |
| 10 | 129759892 | rs747716 | 0.878 | -0.02 | -1.447 | 0.341 | 0.005 | 0.063 | 0.063 | 0.048 | 0.19 | 0.04 | 0.21 | 0.05 | 0.16 | 0.14 |
| 10 | 129760613 | rs747715 | 0.754 | 0.081 | -1.136 | 0.629 | 0.007 | 0.029 | 0.029 | 0 | 0.07 | 0.16 | 0.05 | 0.15 | 0.08 | 0.12 |
| 10 | 129761287 | rs12412502 | -0.228 | -0.086 | -0.727 | 0.856 | -0.001 | 0.066 | 0.066 | 0 | 0.16 | 0.24 | 0.11 | 0.32 | 0.16 | 0.28 |
| 10 | 129761686 | rs11018424 | -0.449 | -0.377 | -0.91 | 0.039 | 0.02 | 0.036 | 0.036 | 0 | 0.41 | 0.65 | 0.35 | 0.19 | 0.32 | 0.33 |
| 10 | 129762754 | rs1812886 | -1.228 | -0.369 | -0.852 | 0.163 | 0.014 | 0.013 | 0.013 | 0 | 0.53 | 0.68 | 0.44 | 0.4 | 0.44 | 0.6 |
| 10 | 129763277 | rs4751554 | 0.091 | 0.313 | -0.581 | 0.497 | 0.002 | 0.006 | 0.006 | 0 | 0.26 | 0.22 | 0.25 | 0.19 | 0.24 | 0.26 |
| 10 | 129767097 | rs12266909 | -0.503 | 0.572 | -0.701 | 0.539 | 0 | 0.018 | 0.018 | 0 | 0.12 | 0.16 | 0.11 | 0.19 | 0.13 | 0.28 |
| 10 | 129767555 | rs10764730 | -0.175 | 1.168 | -1.216 | -0.067 | -0.071 | 0.101 | 0.101 | 0.451 | 0.39 | 0.09 | 0.5 | 0.26 | 0.38 | 0.49 |
| 10 | 129769546 | rs7084584 | -1.528 | 1.761 | 0.16 | 0.358 | 0.034 | -0.017 | -0.017 | 0 | 0.08 | 0.28 | 0.18 | 0.02 | 0.09 | 0.06 |
| 10 | 129772642 | rs11015984 | -1.354 | 2.262 | -0.121 | 0.523 | -0.013 | -0.01 | -0.01 | 0.976 | 0.13 | 0.08 | 0.13 | 0.22 | 0.21 | 0.22 |
| 10 | 129774063 | rs2148894 | -1.639 | 2.116 | -0.67 | 0.509 | -0.033 | -0.015 | -0.015 | 0.565 | 0.16 | 0.06 | 0.26 | 0.22 | 0.25 | 0.26 |
| 10 | 129775360 | rs7475525 | -0.662 | 1.965 | -0.653 | 0.63 | -0.022 | -0.023 | -0.023 | 0 | 0.08 | 0.05 | 0.08 | 0.2 | 0.15 | 0.16 |
| 10 | 129782299 | rs11015994 | -1.219 | 2.144 | -0.632 | 0.589 | -0.024 | -0.014 | -0.014 | 0.16 | 0.14 | 0.08 | 0.14 | 0.22 | 0.23 | 0.22 |
| 10 | 129784473 | rs11015999 | -0.971 | 1.465 | 0.216 | 0.594 | -0.012 | 0.081 | 0.081 | 0 | 0.11 | 0.43 | 0.11 | 0.2 | 0.05 | 0.2 |
| 10 | 129789784 | rs4531377 | -0.583 | 1.993 | -0.234 | 0.099 | -0.043 | 0.044 | 0.044 | 0 | 0.09 | 0.04 | 0.11 | 0 | 0.16 | 0.05 |
| 10 | 129790840 | rs7911506 | -0.594 | 2.268 | -0.361 | 0.055 | -0.029 | 0.033 | 0.033 | 0 | 0.29 | 0.26 | 0.32 | 0.17 | 0.38 | 0.23 |
| 10 | 129792021 | rs7904367 | -1.554 | 1.845 | -0.762 | -0.11 | 0.031 | -0.056 | -0.056 | 0.423 | 0.61 | 0.9 | 0.66 | 0.63 | 0.67 | 0.65 |
| 10 | 129793485 | rs11016003 | 0.294 | 1.904 | -0.801 | -0.092 | 0.005 | 0.021 | 0.021 | 0.816 | 0.25 | 0.23 | 0.23 | 0.17 | 0.24 | 0.19 |
| 10 | 129795715 | rs11593952 | -0.469 | 1.958 | 0.036 | -0.041 | 0.04 | 0.043 | 0.043 | 0 | 0.18 | 0.27 | 0.22 | 0.42 | 0.3 | 0.34 |
| 10 | 129796246 | rs7922288 | 0.093 | 1.921 | 0.119 | -0.032 | -0.001 | 0.013 | 0.013 | 0 | 0.14 | 0.16 | 0.13 | 0.42 | 0.17 | 0.31 |
| 10 | 129796530 | rs7922406 | 0.1 | 1.787 | -0.178 | -0.175 | 0.008 | 0.042 | 0.042 | 0.376 | 0.29 | 0.23 | 0.27 | 0.15 | 0.27 | 0.31 |
| 10 | 129798995 | rs7088062 | -1.179 | 2.107 | 0.146 | 0.266 | 0.013 | -0.022 | -0.022 | 0 | 0.64 | 0.79 | 0.63 | 0.62 | 0.66 | 0.77 |
| 10 | 129803120 | rs7085243 | -0.672 | 2.143 | -0.096 | 0.263 | -0.001 | -0.001 | -0.001 | 0 | 0.57 | 0.49 | 0.52 | 0.55 | 0.51 | 0.73 |
| 10 | 129804052 | rs7089981 | -0.479 | 1.208 | -0.503 | -0.207 | 0 | -0.001 | -0.001 | 0 | 0.49 | 0.56 | 0.51 | 0.22 | 0.48 | 0.43 |
| 10 | 129816246 | rs7089539 | -0.632 | 1.638 | -0.425 | 0.24 | 0.014 | 0.008 | 0.008 | 0 | 0.32 | 0.39 | 0.25 | 0.43 | 0.37 | 0.38 |
| 10 | 129818448 | rs7915092 | -0.949 | 0.937 | -0.858 | 0.076 | 0.009 | -0.023 | -0.023 | 0.416 | 0.66 | 0.79 | 0.53 | 0.62 | 0.68 | 0.61 |
| 10 | 129822049 | rs7900939 | -0.907 | 0.924 | -0.72 | 0.09 | 0.002 | -0.002 | -0.002 | 0 | 0.64 | 0.7 | 0.53 | 0.63 | 0.68 | 0.61 |
| 10 | 129825454 | rs10764735 | 0.384 | 1.597 | -0.886 | 0.109 | -0.015 | -0.013 | -0.013 | 0 | 0.13 | 0.08 | 0.13 | 0.38 | 0.21 | 0.27 |
| 10 | 129826373 | rs6482647 | -0.49 | 1.06 | -0.85 | -0.475 | 0 | 0.002 | 0.002 | 0 | 0.49 | 0.59 | 0.43 | 0.58 | 0.51 | 0.52 |
| 10 | 129831030 | rs4547012 | -0.453 | 1.061 | -1.038 | -0.308 | -0.021 | 0.008 | 0.008 | 0.151 | 0.45 | 0.3 | 0.44 | 0.63 | 0.51 | 0.5 |
| 10 | 129834886 | rs10764739 | 0.06 | 0.752 | -1.076 | -0.255 | 0.002 | 0.011 | 0.011 | 0 | 0.37 | 0.27 | 0.48 | 0.28 | 0.43 | 0.36 |
| 10 | 129835928 | rs7083801 | 0.241 | 0.621 | -0.694 | -0.546 | -0.003 | -0.051 | -0.051 | 0 | 0.16 | 0.2 | 0.07 | 0 | 0.13 | 0.07 |
| 10 | 129843858 | rs10764740 | 0.588 | 0.858 | -1.207 | -0.076 | -0.038 | -0.057 | -0.057 | 0 | 0.3 | 0.11 | 0.55 | 0.38 | 0.3 | 0.33 |
| 10 | 129845547 | rs4262642 | -0.364 | 0.614 | -0.995 | 0.139 | 0.002 | 0.011 | 0.011 | 0.094 | 0.73 | 0.74 | 0.85 | 0.83 | 0.74 | 0.8 |
| 10 | 129851621 | rs10829321 | 0.236 | 0.473 | -0.288 | 0.193 | -0.007 | 0.014 | 0.014 | 0 | 0.63 | 0.44 | 0.76 | 0.57 | 0.68 | 0.53 |
| 10 | 129854145 | rs11016039 | 0.453 | 0.507 | 0.438 | 0.599 | -0.004 | 0.092 | 0.092 | 0 | 0.11 | 0.21 | 0.08 | 0.25 | 0.08 | 0.26 |
| 10 | 129855795 | rs12249657 | -0.098 | 0.567 | 0.547 | 0.551 | 0.016 | 0.119 | 0.119 | 0 | 0.13 | 0.31 | 0.11 | 0.25 | 0.13 | 0.27 |
| 10 | 129858897 | rs10764742 | -0.985 | 0.665 | -0.279 | 0.461 | -0.024 | 0.009 | 0.009 | 0 | 0.82 | 0.96 | 0.9 | 0.82 | 0.79 | 0.86 |
| 10 | 129859992 | rs4462251 | 0.744 | 0.681 | -0.065 | 0.345 | -0.004 | 0.002 | 0.002 | 0 | 0.58 | 0.42 | 0.7 | 0.57 | 0.53 | 0.55 |
| 10 | 129860813 | rs7093446 | -0.918 | 0.799 | -0.586 | 0.341 | -0.02 | 0.013 | 0.013 | 0 | 0.82 | 0.93 | 0.9 | 0.82 | 0.78 | 0.87 |
| 10 | 129864515 | rs10764743 | 0.635 | 0.588 | -0.323 | 0.503 | -0.004 | 0.003 | 0.003 | 0 | 0.55 | 0.27 | 0.51 | 0.41 | 0.51 | 0.47 |
| 10 | 129871690 | rs2275803 | -0.787 | 0.586 | -0.141 | 0.657 | 0 | -0.001 | -0.001 | 0 | 0.23 | 0.28 | 0.24 | 0.26 | 0.2 | 0.29 |
| 10 | 129871795 | rs11016053 | 0.04 | 0.696 | -0.08 | 0.717 | -0.003 | 0.017 | 0.017 | 0 | 0.13 | 0.2 | 0.17 | 0.26 | 0.1 | 0.25 |
| 10 | 129874447 | rs12708321 | -0.768 | 0.699 | 0.109 | 0.777 | 0 | 0.059 | 0.059 | 0.361 | 0.2 | 0.33 | 0.23 | 0.42 | 0.2 | 0.28 |
| 10 | 129875382 | rs1539598 | 0.782 | 0.943 | -0.388 | 0.334 | 0.007 | 0.013 | 0.013 | 0 | 0.41 | 0.32 | 0.4 | 0.35 | 0.35 | 0.32 |
| 10 | 129875714 | rs2298197 | -0.035 | 0.891 | -0.396 | 0.414 | 0.006 | -0.004 | -0.004 | 0 | 0.53 | 0.43 | 0.53 | 0.51 | 0.54 | 0.53 |
| 10 | 129877567 | rs3763791 | 0.775 | 0.785 | -1.155 | 0.415 | -0.001 | 0.043 | 0.043 | 0 | 0.25 | 0.04 | 0.27 | 0.16 | 0.22 | 0.14 |
| 10 | 129878602 | rs7911438 | 0.115 | 0.69 | -0.547 | 0.633 | -0.01 | 0.021 | 0.021 | 0 | 0.11 | 0.17 | 0.08 | 0.15 | 0.09 | 0.13 |
| 10 | 129879700 | rs2096285 | 0.86 | 0.796 | -1.414 | 0.442 | -0.057 | -0.038 | -0.038 | 0.195 | 0.35 | 0.09 | 0.37 | 0.35 | 0.32 | 0.33 |
| 10 | 129882119 | rs1127299 | -0.004 | 0.895 | -0.508 | 0.406 | 0.002 | -0.002 | -0.002 | 0 | 0.54 | 0.59 | 0.66 | 0.48 | 0.53 | 0.48 |
| 10 | 129882640 | rs4568912 | 0.001 | 0.897 | -0.515 | 0.407 | 0.003 | -0.002 | -0.002 | 1.134 | 0.54 | 0.59 | 0.66 | 0.48 | 0.53 | 0.48 |
| 10 | 129882699 | rs4367863 | -0.006 | 0.929 | -0.523 | 0.547 | 0.003 | -0.001 | -0.001 | 0.165 | 0.54 | 0.59 | 0.74 | 0.52 | 0.54 | 0.5 |
| 10 | 129883110 | rs4237449 | 0.013 | 0.924 | -0.518 | 0.495 | 0.003 | -0.002 | -0.002 | 0.036 | 0.54 | 0.59 | 0.66 | 0.48 | 0.53 | 0.48 |
| 10 | 129883121 | rs3186827 | 0.013 | 0.924 | -0.518 | 0.495 | 0.003 | -0.002 | -0.002 | 1.512 | 0.54 | 0.59 | 0.66 | 0.48 | 0.53 | 0.48 |

| | | | | | | | | | | | | | | | | |
|---|---|---|---|---|---|---|---|---|---|---|---|---|---|---|---|---|
| 10 | 129883933 | rs6911 | 0.013 | 0.916 | -0.518 | 0.495 | 0.003 | -0.002 | -0.002 | 0.694 | 0.54 | 0.59 | 0.66 | 0.48 | 0.53 | 0.48 |
| 10 | 129885668 | rs12261926 | 0.944 | 1.005 | -0.529 | 0.431 | 0 | 0.007 | 0.007 | 0 | 0.09 | 0.15 | 0.16 | 0.12 | 0.09 | 0.12 |
| 10 | 129888727 | rs10764747 | -0.85 | 0.959 | -0.899 | 0.355 | -0.021 | -0.005 | -0.005 | 0 | 0.74 | 0.97 | 0.8 | 0.79 | 0.72 | 0.77 |
| 10 | 129894034 | rs10732438 | 0.217 | 1.055 | -0.296 | 0.531 | 0.006 | 0.005 | 0.005 | 0.542 | 0.66 | 0.58 | 0.69 | 0.62 | 0.65 | 0.6 |
| 10 | 129897565 | rs3781301 | 1.099 | 1.151 | -0.345 | 0.668 | -0.001 | 0.001 | 0.001 | 0 | 0.07 | 0.07 | 0.15 | 0.15 | 0.08 | 0.11 |
| 10 | 129897791 | rs3781303 | 0.987 | 1.019 | -0.515 | 0.605 | 0.004 | 0.005 | 0.005 | 0 | 0.32 | 0.31 | 0.35 | 0.25 | 0.29 | 0.22 |
| 10 | 129897935 | rs2857037 | -0.427 | 1.045 | -0.342 | 0.527 | -0.001 | -0.001 | -0.001 | 0 | 0.51 | 0.49 | 0.61 | 0.42 | 0.53 | 0.39 |
| 10 | 129898890 | rs3781305 | -0.009 | 1.105 | -0.42 | 0.5 | -0.001 | 0 | 0 | 0 | 0.44 | 0.45 | 0.54 | 0.39 | 0.43 | 0.33 |
| 10 | 129899578 | rs8473 | -0.436 | 1.099 | -0.414 | 0.517 | 0.003 | 0.011 | 0.011 | 0 | 0.51 | 0.46 | 0.61 | 0.39 | 0.52 | 0.39 |
| 10 | 129901393 | rs11016071 | -0.569 | 1.104 | -0.377 | 0.533 | 0.024 | 0.049 | 0.049 | 0 | 0.17 | 0.34 | 0.23 | 0.26 | 0.16 | 0.14 |
| 10 | 129901722 | rs10082432 | -0.569 | 1.1 | -0.37 | 0.526 | 0.024 | 0.049 | 0.049 | 0 | 0.17 | 0.34 | 0.23 | 0.26 | 0.16 | 0.14 |
| 10 | 129901825 | rs10082391 | -0.569 | 1.1 | -0.37 | 0.526 | 0.024 | 0.049 | 0.049 | 0 | 0.17 | 0.34 | 0.23 | 0.26 | 0.16 | 0.14 |
| 10 | 129901946 | rs1050767 | -0.497 | 1.099 | -0.367 | 0.525 | 0.02 | 0.042 | 0.042 | 0 | 0.16 | 0.31 | 0.22 | 0.26 | 0.16 | 0.14 |
| 10 | 129902158 | rs12777740 | -0.497 | 1.099 | -0.367 | 0.525 | 0.02 | 0.042 | 0.042 | 0 | 0.16 | 0.31 | 0.22 | 0.26 | 0.16 | 0.14 |
| 10 | 129902281 | rs1063535 | -0.443 | 1.075 | -0.435 | 0.497 | 0.004 | 0.015 | 0.015 | 0 | 0.5 | 0.46 | 0.61 | 0.38 | 0.52 | 0.39 |
| 10 | 129903016 | rs7071768 | -0.443 | 1.075 | -0.435 | 0.497 | 0.004 | 0.015 | 0.015 | 0 | 0.5 | 0.46 | 0.61 | 0.38 | 0.52 | 0.39 |
| 10 | 129905429 | rs7918199 | 1.351 | 1.139 | -0.254 | 0.626 | 0 | 0.001 | 0.001 | 0 | 0.07 | 0.07 | 0.16 | 0.14 | 0.08 | 0.11 |
| 10 | 129906364 | rs4750685 | 1.351 | 1.139 | -0.254 | 0.626 | 0 | 0.001 | 0.001 | 0 | 0.07 | 0.07 | 0.16 | 0.14 | 0.08 | 0.11 |
| 10 | 129906468 | rs4750936 | -0.747 | 1.14 | -0.256 | 0.626 | 0.024 | 0.049 | 0.049 | 0 | 0.17 | 0.34 | 0.23 | 0.26 | 0.16 | 0.14 |
| 10 | 129907741 | rs2065718 | 1.16 | 1.187 | -0.266 | 0.615 | 0.021 | 0.001 | 0.001 | 0.069 | 0.76 | 0.66 | 0.7 | 0.74 | 0.73 | 0.8 |
| 10 | 129910475 | rs997983 | 0.794 | 1.124 | -0.46 | 0.712 | 0.001 | 0.002 | 0.002 | 0 | 0.08 | 0.09 | 0.16 | 0.14 | 0.08 | 0.11 |
| 10 | 129917560 | rs2071498 | 0.082 | 1.195 | -0.262 | 0.689 | 0.011 | 0.018 | 0.018 | 0 | 0.68 | 0.58 | 0.72 | 0.64 | 0.67 | 0.61 |
| 10 | 129920241 | rs769005 | 1.32 | 0.926 | -0.62 | 0.354 | -0.001 | 0.053 | 0.053 | 0 | 0.27 | 0.13 | 0.33 | 0.12 | 0.28 | 0.2 |
| 10 | 129921652 | rs1335009 | -1.547 | 0.993 | -0.615 | 0.483 | -0.034 | 0.009 | 0.009 | 0 | 0.84 | 0.93 | 0.84 | 0.92 | 0.83 | 0.85 |
| 10 | 129930349 | rs2387327 | -0.212 | 0.953 | -0.329 | 0.655 | 0.015 | 0.005 | 0.005 | 0 | 0.31 | 0.43 | 0.36 | 0.35 | 0.32 | 0.29 |
| 10 | 129933456 | rs2387326 | 0.974 | 0.761 | -0.331 | 0.443 | 0.006 | 0.014 | 0.014 | 0 | 0.28 | 0.27 | 0.34 | 0.13 | 0.26 | 0.2 |
| 10 | 129941513 | rs954650 | 1.091 | 0.834 | -0.35 | 0.373 | -0.005 | -0.014 | -0.014 | 0 | 0.27 | 0.29 | 0.17 | 0.12 | 0.23 | 0.18 |
| 10 | 129949720 | rs4750687 | -0.508 | 1.09 | -0.42 | 0.757 | 0 | 0 | 0 | 0.004 | 0.35 | 0.39 | 0.52 | 0.4 | 0.35 | 0.3 |
| 10 | 129954773 | rs7912288 | 0.138 | 0.93 | -0.515 | 0.613 | 0.006 | 0.009 | 0.009 | 0.203 | 0.51 | 0.39 | 0.37 | 0.41 | 0.45 | 0.56 |
| 10 | 129954972 | rs12572759 | -0.388 | 0.682 | -0.45 | 0.802 | 0.013 | 0.012 | 0.012 | 0.537 | 0.35 | 0.45 | 0.3 | 0.38 | 0.38 | 0.38 |
| 10 | 129958296 | rs7906241 | -0.82 | 0.152 | -0.671 | 0.711 | 0.012 | -0.008 | -0.008 | 3.998 | 0.21 | 0.16 | 0.34 | 0.27 | 0.14 | 0.23 |
| 10 | 129961901 | rs1467702 | 0.018 | 0.57 | -0.61 | 0.913 | -0.001 | 0.008 | 0.008 | 0.626 | 0.09 | 0.1 | 0.29 | 0.19 | 0.08 | 0.15 |
| 10 | 129962286 | rs6482661 | -0.864 | 0.927 | -0.659 | 0.791 | 0 | 0.002 | 0.002 | 0 | 0.76 | 0.72 | 0.79 | 0.7 | 0.72 | 0.8 |
| 10 | 129962572 | rs12571194 | 0.109 | 0.947 | -0.524 | 0.474 | 0.002 | -0.026 | -0.026 | 0 | 0.12 | 0.23 | 0.11 | 0.05 | 0.13 | 0.07 |
| 10 | 129962672 | rs7101063 | -0.107 | 0.995 | -0.528 | 0.445 | 0.005 | -0.035 | -0.035 | 0.23 | 0.13 | 0.23 | 0.19 | 0.05 | 0.14 | 0.09 |
| 10 | 129963871 | rs10764759 | -0.074 | 0.985 | -1.499 | 0.661 | -0.01 | -0.046 | -0.046 | 0 | 0.57 | 0.81 | 0.52 | 0.48 | 0.5 | 0.49 |
| 10 | 129967700 | rs2183171 | -0.872 | 1.143 | -1.55 | 0.733 | -0.043 | -0.06 | -0.06 | 9.558 | 0.87 | 0.99 | 0.77 | 0.82 | 0.84 | 0.8 |
| 10 | 129969368 | rs10741148 | -1.343 | 0.666 | -1.574 | 0.048 | 0.006 | 0.022 | 0.022 | 0 | 0.79 | 0.9 | 0.89 | 0.86 | 0.82 | 0.84 |
| 10 | 129975342 | rs10829351 | -0.064 | 0.119 | -1.948 | 0.002 | 0.035 | -0.076 | -0.076 | 0 | 0.27 | 0.17 | 0.39 | 0.53 | 0.22 | 0.43 |
| 10 | 129977477 | rs10764760 | 1.223 | 0.608 | -1.741 | -0.153 | -0.003 | -0.002 | -0.002 | 0 | 0.21 | 0.18 | 0.19 | 0.28 | 0.29 | 0.17 |
| 10 | 129979010 | rs11016118 | 1.371 | 0.9 | -1.341 | -0.313 | -0.005 | -0.019 | -0.019 | 0.179 | 0.15 | 0.13 | 0.12 | 0.01 | 0.17 | 0.13 |
| 10 | 129984460 | rs7076143 | 0.612 | 0.523 | -1.311 | 0 | -0.004 | -0.072 | -0.072 | 0 | 0.17 | 0.06 | 0.36 | 0.52 | 0.19 | 0.37 |
| 10 | 129987575 | rs10764761 | -1.468 | 0.394 | -1.12 | -0.305 | 0.01 | 0.018 | 0.018 | 0 | 0.79 | 0.87 | 0.89 | 0.86 | 0.83 | 0.84 |
| 10 | 129998372 | rs1329189 | -0.877 | 0.925 | -0.788 | -0.023 | 0.037 | -0.029 | -0.029 | 0 | 0.63 | 0.83 | 0.64 | 0.58 | 0.69 | 0.51 |
| 10 | 130008259 | rs12413650 | 0.043 | 1.407 | -0.441 | -0.103 | 0.005 | -0.031 | -0.031 | 0 | 0.13 | 0.17 | 0.18 | 0.03 | 0.17 | 0.1 |
| 10 | 130018768 | rs10829360 | -0.759 | 1.804 | -0.363 | 0.166 | 0.003 | 0.007 | 0.007 | 0 | 0.48 | 0.76 | 0.43 | 0.44 | 0.49 | 0.41 |
| 10 | 130031585 | rs1335012 | 0.561 | 1.654 | -0.089 | -0.876 | 0.004 | -0.001 | -0.001 | 0 | 0.18 | 0.23 | 0.12 | 0.18 | 0.22 | 0.14 |
| 10 | 130033858 | rs7086416 | -0.621 | 1.472 | -0.169 | -0.951 | -0.038 | 0.111 | 0.111 | 0 | 0.39 | 0.85 | 0.29 | 0.18 | 0.39 | 0.29 |
| 10 | 130035410 | rs1335010 | -0.901 | 1.274 | 0.014 | -0.724 | -0.044 | 0.166 | 0.166 | 1.015 | 0.47 | 0.84 | 0.33 | 0.2 | 0.46 | 0.34 |
| 10 | 130044240 | rs12358598 | 0.421 | 1.501 | -0.034 | -0.84 | 0.002 | -0.001 | -0.001 | 0 | 0.2 | 0.22 | 0.14 | 0.18 | 0.26 | 0.15 |
| 10 | 130044668 | rs1153018 | -0.604 | 1.278 | -0.65 | -0.684 | -0.013 | -0.038 | -0.038 | 0 | 0.93 | 0.99 | 0.89 | 0.85 | 0.93 | 0.96 |
| 10 | 130048165 | rs12262774 | 0.033 | 1.298 | -0.749 | -0.742 | -0.001 | -0.069 | -0.069 | 1.67 | 0.27 | 0.66 | 0.19 | 0.01 | 0.21 | 0.19 |
| 10 | 130054651 | rs11591431 | -0.072 | 1.558 | -0.857 | 0.421 | -0.045 | -0.02 | -0.02 | 0 | 0.18 | 0.07 | 0.12 | 0.18 | 0.23 | 0.12 |
| 10 | 130057875 | rs10829364 | -0.097 | 1.774 | -1.453 | 0.514 | -0.044 | -0.018 | -0.018 | 0 | 0.19 | 0.07 | 0.12 | 0.18 | 0.23 | 0.14 |
| 10 | 130059276 | rs988520 | -1.051 | 1.137 | -1.017 | 0.549 | -0.001 | 0.002 | 0.002 | 0 | 0.09 | 0.09 | 0.13 | 0.15 | 0.08 | 0.06 |
| 10 | 130063360 | rs978462 | -0.476 | 1.504 | -0.982 | 0.247 | -0.024 | 0.023 | 0.023 | 0 | 0.37 | 0.31 | 0.41 | 0.23 | 0.46 | 0.46 |
| 10 | 130070134 | rs11016151 | -1.024 | 1.364 | -0.489 | 0.629 | 0.008 | 0.037 | 0.037 | 0 | 0.11 | 0.17 | 0.22 | 0.14 | 0.14 | 0.24 |
| 10 | 130074063 | rs7900125 | -0.736 | 1.456 | -0.786 | -0.283 | 0.01 | 0.06 | 0.06 | 0 | 0.55 | 0.75 | 0.57 | 0.28 | 0.54 | 0.57 |
| 10 | 130084909 | rs2387608 | -0.75 | 2.211 | -0.069 | -0.444 | 0.06 | 0.01 | 0.01 | 0 | 0.17 | 0.34 | 0.28 | 0.14 | 0.23 | 0.29 |
| 10 | 130092362 | rs11595866 | 0.385 | 2.107 | -0.878 | -0.632 | -0.024 | 0.034 | 0.034 | 0.215 | 0.28 | 0.09 | 0.29 | 0.22 | 0.29 | 0.22 |
| 10 | 130093238 | rs11016162 | 0.718 | 1.89 | -0.368 | -0.748 | -0.009 | -0.006 | -0.006 | 0 | 0.09 | 0.17 | 0.06 | 0.08 | 0.06 | 0.09 |
| 10 | 130097446 | rs7922731 | -1.585 | 1.925 | -0.203 | -0.721 | 0.004 | -0.004 | -0.004 | 0 | 0.61 | 0.66 | 0.68 | 0.47 | 0.65 | 0.64 |
| 10 | 130098969 | rs2244545 | 0.208 | 1.975 | -0.316 | -0.578 | -0.009 | 0.001 | 0.001 | 0.004 | 0.56 | 0.68 | 0.4 | 0.57 | 0.48 | 0.5 |
| 10 | 130102051 | rs1262207 | -0.751 | 1.879 | -0.582 | -0.553 | -0.075 | 0 | 0 | 1.055 | 0.69 | 0.97 | 0.69 | 0.73 | 0.67 | 0.72 |
| 10 | 130105852 | rs10829379 | -0.123 | 1.689 | -0.442 | -0.578 | 0.029 | 0.031 | 0.031 | 0 | 0.17 | 0.3 | 0.3 | 0.19 | 0.2 | 0.27 |
| 10 | 130107905 | rs2254606 | -0.914 | 1.675 | -0.699 | -0.529 | -0.08 | -0.019 | -0.019 | 0 | 0.7 | 0.98 | 0.7 | 0.75 | 0.68 | 0.75 |
| 10 | 130108418 | rs7921212 | -0.102 | 1.661 | -0.5 | -0.452 | 0.007 | 0.021 | 0.021 | 0 | 0.17 | 0.27 | 0.29 | 0.2 | 0.17 | 0.28 |
| 10 | 130111702 | rs1273180 | 0.695 | 1.517 | 0.262 | -0.543 | 0.003 | -0.005 | -0.005 | 0 | 0.45 | 0.41 | 0.35 | 0.54 | 0.42 | 0.32 |
| 10 | 130113107 | rs2799400 | 1.783 | 1.283 | -0.316 | -0.066 | -0.022 | -0.037 | -0.037 | 0 | 0.17 | 0.04 | 0.12 | 0.17 | 0.19 | 0.12 |

| | | | | | | | | | | | | | | | | |
|---|---|---|---|---|---|---|---|---|---|---|---|---|---|---|---|---|
| 10 | 130113462 | rs2799399 | 0.124 | 1.121 | -0.23 | -0.161 | 0.004 | -0.005 | -0.005 | 0 | 0.45 | 0.41 | 0.35 | 0.54 | 0.42 | 0.32 |
| 10 | 130113999 | rs2251103 | 0.926 | 1.11 | -0.706 | -0.173 | -0.011 | -0.106 | -0.106 | 0 | 0.3 | 0.08 | 0.26 | 0.5 | 0.31 | 0.2 |
| 10 | 130114766 | rs10829382 | 0.095 | 1.24 | -0.446 | -0.123 | 0.002 | 0.013 | 0.013 | 0 | 0.17 | 0.24 | 0.29 | 0.21 | 0.17 | 0.28 |
| 10 | 130116817 | rs10829383 | 0.794 | 0.998 | -0.565 | -0.593 | -0.002 | -0.044 | -0.044 | 0 | 0.27 | 0.26 | 0.25 | 0.07 | 0.23 | 0.15 |
| 10 | 130121584 | rs731092 | -0.519 | 0.481 | 0.11 | -0.119 | 0.002 | 0.103 | 0.103 | 0.345 | 0.16 | 0.44 | 0.29 | 0.24 | 0.11 | 0.28 |
| 10 | 130122697 | rs12772143 | 1.83 | 0.468 | -0.145 | -0.166 | -0.014 | 0.019 | 0.019 | 0.104 | 0.16 | 0.07 | 0.16 | 0.1 | 0.19 | 0.11 |
| 10 | 130123291 | rs7080260 | 0.073 | 0.433 | -0.124 | 0.137 | -0.001 | 0.006 | 0.006 | 0 | 0.4 | 0.31 | 0.32 | 0.36 | 0.4 | 0.35 |
| 10 | 130128297 | rs1754233 | -0.111 | 0.407 | -0.049 | 0.085 | -0.001 | 0.004 | 0.004 | 0 | 0.44 | 0.39 | 0.36 | 0.36 | 0.44 | 0.36 |
| 10 | 130144139 | rs10829385 | -0.855 | 0.391 | -0.202 | 0.134 | -0.04 | 0.024 | 0.024 | 0 | 0.56 | 0.78 | 0.63 | 0.61 | 0.52 | 0.62 |
| 10 | 130150129 | rs6482672 | -2.117 | 0.408 | -0.461 | 0.159 | -0.095 | -0.03 | -0.03 | 0.391 | 0.74 | 0.98 | 0.72 | 0.76 | 0.67 | 0.78 |
| 10 | 130150319 | rs1936423 | 0.649 | -0.175 | 0.424 | 0.429 | -0.008 | 0.164 | 0.164 | 0 | 0.08 | 0.23 | 0.23 | 0.24 | 0.06 | 0.25 |
| 10 | 130151744 | rs11595317 | 1.185 | -0.245 | 0.411 | 0.34 | -0.006 | 0.014 | 0.014 | 0.492 | 0.19 | 0.16 | 0.2 | 0.11 | 0.22 | 0.13 |
| 10 | 130152006 | rs12263749 | 0.718 | -0.253 | 0.431 | 0.362 | -0.008 | 0.164 | 0.164 | 0 | 0.08 | 0.23 | 0.23 | 0.24 | 0.06 | 0.25 |
| 10 | 130152724 | rs1891977 | 1.537 | -0.331 | 0.083 | 0.278 | -0.027 | -0.077 | -0.077 | 0 | 0.13 | 0.01 | 0.12 | 0.25 | 0.13 | 0.07 |
| 10 | 130153428 | rs1419240 | -0.849 | -0.307 | 0.327 | 0.031 | 0.001 | 0.03 | 0.03 | 0 | 0.5 | 0.58 | 0.59 | 0.61 | 0.5 | 0.6 |
| 10 | 130154533 | rs2387611 | -1.271 | -0.29 | 0.309 | 0.033 | -0.022 | 0.028 | 0.028 | 0 | 0.55 | 0.72 | 0.62 | 0.61 | 0.52 | 0.62 |
| 10 | 130154648 | rs7909327 | 0.368 | -0.223 | 0.309 | 0.137 | 0.015 | -0.048 | -0.048 | 0 | 0.09 | 0.17 | 0.05 | 0.01 | 0.08 | 0.15 |
| 10 | 130160426 | rs12246905 | 0.249 | -0.211 | 0.107 | 0.246 | -0.007 | 0.038 | 0.038 | 0 | 0.22 | 0.25 | 0.36 | 0.49 | 0.19 | 0.32 |
| 10 | 130163010 | rs10829388 | -0.247 | -0.155 | 0.341 | 0.122 | 0.023 | -0.071 | -0.071 | 0 | 0.12 | 0.26 | 0.06 | 0.01 | 0.1 | 0.15 |
| 10 | 130163026 | rs11594406 | 0.56 | -0.102 | -0.048 | 0.106 | -0.025 | 0.047 | 0.047 | 0.362 | 0.25 | 0.12 | 0.22 | 0.11 | 0.3 | 0.19 |
| 10 | 130163490 | rs7922283 | 0.386 | -0.275 | 0.103 | 0.371 | 0.007 | -0.074 | -0.074 | 0.498 | 0.2 | 0.09 | 0.35 | 0.49 | 0.19 | 0.32 |
| 10 | 130163871 | rs7898000 | -0.468 | -0.218 | 0.296 | 0.36 | -0.019 | 0.098 | 0.098 | 0 | 0.26 | 0.41 | 0.39 | 0.5 | 0.21 | 0.33 |
| 10 | 130169134 | rs1857389 | -0.56 | -0.481 | -0.035 | 0.014 | 0.027 | 0.088 | 0.088 | 1.462 | 0.39 | 0.75 | 0.29 | 0.13 | 0.32 | 0.3 |
| 10 | 130169857 | rs2154081 | -0.274 | -0.977 | 0.232 | 0.109 | -0.006 | -0.006 | -0.006 | 0 | 0.41 | 0.43 | 0.54 | 0.64 | 0.39 | 0.56 |
| 10 | 130170270 | rs2032031 | -2.303 | -1.175 | -0.302 | 0.043 | -0.188 | 0.045 | 0.045 | 2.388 | 0.66 | 0.97 | 0.67 | 0.78 | 0.53 | 0.75 |
| 10 | 130171171 | rs7917861 | -1.247 | -1.251 | 0.13 | 0.246 | 0.006 | 0.051 | 0.051 | 1.126 | 0.53 | 0.74 | 0.62 | 0.64 | 0.45 | 0.59 |
| 10 | 130173498 | rs11016199 | 1.797 | -1.162 | 0.161 | 0.545 | -0.004 | 0.086 | 0.086 | 0 | 0.06 | 0.09 | 0.22 | 0.24 | 0.06 | 0.22 |
| 10 | 130176586 | rs10741151 | -1.763 | -1.14 | 0.052 | -0.408 | -0.024 | 0.063 | 0.063 | 2.782 | 0.59 | 0.72 | 0.63 | 0.75 | 0.48 | 0.67 |
| 10 | 130178143 | rs11599135 | 1.386 | -1.274 | -0.362 | -0.436 | -0.006 | 0.019 | 0.019 | 0.844 | 0.19 | 0.01 | 0.19 | 0.11 | 0.17 | 0.15 |
| 10 | 130185535 | rs11016216 | 0.661 | -1.129 | 0.074 | -0.202 | -0.01 | 0 | 0 | 1.386 | 0.1 | 0.15 | 0.03 | 0.11 | 0.02 | 0.04 |
| 10 | 130185899 | rs11016217 | -0.758 | -1.125 | 0.031 | -0.199 | -0.009 | 0.003 | 0.003 | 0 | 0.31 | 0.61 | 0.38 | 0.61 | 0.23 | 0.42 |
| 10 | 130187293 | rs1857383 | 0.54 | -1.133 | -0.193 | -0.276 | 0.01 | 0.036 | 0.036 | 3.649 | 0.28 | 0.2 | 0.27 | 0.13 | 0.25 | 0.26 |
| 10 | 130190217 | rs4750950 | 0.956 | -0.488 | -0.471 | 0.045 | -0.095 | 0.06 | 0.06 | 2.136 | 0.39 | 0.1 | 0.34 | 0.26 | 0.52 | 0.33 |
| 10 | 130191499 | rs11016224 | 0.764 | -0.372 | -0.296 | 0.134 | 0 | 0.013 | 0.013 | 0 | 0.17 | 0.15 | 0.29 | 0.5 | 0.15 | 0.33 |
| 10 | 130193141 | rs1934261 | -1.13 | -0.349 | -0.38 | 0.099 | -0.067 | 0.087 | 0.087 | 2.651 | 0.59 | 0.87 | 0.65 | 0.74 | 0.48 | 0.67 |
| 10 | 130197453 | rs870493 | 0.721 | -0.128 | -0.427 | 0.24 | -0.001 | 0.04 | 0.04 | 0 | 0.16 | 0.19 | 0.29 | 0.5 | 0.16 | 0.34 |
| 10 | 130198861 | rs11016232 | -0.526 | 0.336 | -0.405 | -0.019 | -0.005 | 0.028 | 0.028 | 2.123 | 0.57 | 0.66 | 0.63 | 0.74 | 0.48 | 0.67 |
| 10 | 130205063 | rs950047 | 0.213 | 0.618 | -0.372 | -0.199 | 0.024 | -0.071 | -0.071 | 0 | 0.33 | 0.42 | 0.33 | 0.12 | 0.41 | 0.36 |
| 10 | 130205150 | rs950045 | -0.29 | 0.061 | -0.564 | 0.032 | 0.001 | -0.003 | -0.003 | 0 | 0.37 | 0.32 | 0.39 | 0.57 | 0.34 | 0.44 |
| 10 | 130205245 | rs2387858 | 0.415 | -0.275 | -0.363 | -0.132 | -0.001 | -0.018 | -0.018 | 0 | 0.57 | 0.43 | 0.55 | 0.26 | 0.59 | 0.52 |
| 10 | 130205385 | rs1416720 | -0.088 | -0.824 | -0.795 | 0.23 | 0.026 | -0.028 | -0.028 | 0.176 | 0.76 | 0.87 | 0.88 | 0.74 | 0.85 | 0.89 |
| 10 | 130206013 | rs2211132 | 0.194 | -0.516 | -0.568 | 0.631 | -0.001 | 0.04 | 0.04 | 0 | 0.15 | 0.13 | 0.29 | 0.48 | 0.16 | 0.34 |
| 10 | 130210370 | rs946536 | -0.95 | -0.738 | -0.546 | 0.137 | -0.009 | -0.009 | -0.009 | 0 | 0.44 | 0.49 | 0.44 | 0.62 | 0.33 | 0.51 |
| 10 | 130212570 | rs1339520 | -0.562 | -0.668 | -0.155 | 0.464 | -0.016 | 0.212 | 0.212 | 0 | 0.19 | 0.45 | 0.33 | 0.48 | 0.17 | 0.36 |
| 10 | 130214365 | rs7915042 | -0.179 | -0.564 | -0.432 | 0.51 | -0.006 | 0.15 | 0.15 | 0 | 0.17 | 0.29 | 0.32 | 0.48 | 0.17 | 0.36 |
| 10 | 130217130 | rs11016235 | 0.968 | -0.496 | -0.815 | 1.156 | -0.009 | -0.041 | -0.041 | 0.956 | 0.07 | 0 | 0.15 | 0.17 | 0.06 | 0.07 |
| 10 | 130217404 | rs11016236 | -0.121 | -0.511 | -0.71 | 1.044 | -0.001 | 0.061 | 0.061 | 0 | 0.15 | 0.15 | 0.3 | 0.48 | 0.16 | 0.35 |
| 10 | 130220305 | rs7902013 | 0.274 | -0.649 | -1.377 | 0.395 | -0.013 | 0.04 | 0.04 | 0 | 0.21 | 0.05 | 0.21 | 0.17 | 0.18 | 0.15 |
| 10 | 130221524 | rs4750695 | 0.115 | -0.59 | -1.04 | 0.749 | -0.001 | 0.009 | 0.009 | 0.427 | 0.25 | 0.22 | 0.3 | 0.47 | 0.18 | 0.3 |
| 10 | 130222045 | rs4750952 | 0.07 | -0.404 | -0.81 | 0.95 | 0.001 | 0.143 | 0.143 | 0 | 0.15 | 0.25 | 0.19 | 0.47 | 0.15 | 0.28 |
| 10 | 130222673 | rs1316687 | -1.033 | -0.43 | -0.765 | 0.393 | 0 | -0.003 | -0.003 | 0 | 0.46 | 0.5 | 0.55 | 0.78 | 0.4 | 0.47 |
| 10 | 130223257 | rs1339521 | 1.633 | 0 | -1.265 | 0.069 | -0.057 | 0.066 | 0.066 | 0 | 0.29 | 0.13 | 0.31 | 0.15 | 0.45 | 0.37 |
| 10 | 130224595 | rs1890629 | -0.395 | -1.109 | -0.713 | 0.391 | -0.013 | -0.003 | -0.003 | 3.592 | 0.37 | 0.39 | 0.28 | 0.4 | 0.3 | 0.45 |
| 10 | 130225540 | rs1541178 | -0.848 | -1.183 | -0.769 | 0.433 | -0.008 | 0.003 | 0.003 | 0 | 0.45 | 0.44 | 0.4 | 0.52 | 0.35 | 0.53 |
| 10 | 130226011 | rs1541179 | -0.497 | -1.353 | -0.612 | 0.37 | -0.001 | 0.008 | 0.008 | 0 | 0.5 | 0.53 | 0.53 | 0.56 | 0.5 | 0.53 |
| 10 | 130226289 | rs2387859 | -1.909 | -1.167 | -0.533 | 0.423 | 0.003 | -0.009 | -0.009 | 0 | 0.69 | 0.61 | 0.71 | 0.84 | 0.62 | 0.66 |
| 10 | 130228363 | rs12768121 | 1.422 | -1.125 | -0.38 | 0.46 | -0.01 | -0.027 | -0.027 | 0 | 0.07 | 0.1 | 0.03 | 0.03 | 0.06 | 0.1 |
| 10 | 130229843 | rs12782052 | -0.838 | -1.109 | -0.416 | 0.383 | -0.001 | 0.001 | 0.001 | 0 | 0.43 | 0.41 | 0.48 | 0.47 | 0.4 | 0.33 |
| 10 | 130231857 | rs11016239 | 1.622 | -1.057 | -0.742 | 0.087 | 0.009 | -0.031 | -0.031 | 0 | 0.1 | 0.03 | 0.05 | 0.15 | 0.06 | 0.05 |
| 10 | 130232922 | rs12773241 | 0.986 | -0.969 | -0.767 | -0.046 | -0.07 | 0.048 | 0.048 | 1.157 | 0.2 | 0.02 | 0.27 | 0.16 | 0.26 | 0.11 |
| 10 | 130233808 | rs10764775 | -0.741 | -1.118 | -0.558 | -0.009 | 0.024 | 0.031 | 0.031 | 0.295 | 0.45 | 0.59 | 0.42 | 0.69 | 0.31 | 0.51 |
| 10 | 130242160 | rs7096748 | 0.709 | -0.674 | -0.48 | 0.381 | -0.054 | 0.097 | 0.097 | 0 | 0.37 | 0.08 | 0.44 | 0.26 | 0.34 | 0.19 |
| 10 | 130247380 | rs1416711 | 1.153 | -0.662 | -0.382 | 0.372 | -0.036 | 0.118 | 0.118 | 0 | 0.32 | 0.11 | 0.35 | 0.15 | 0.33 | 0.14 |
| 10 | 130248300 | rs7912500 | -1.48 | -0.682 | -0.153 | 0.444 | -0.005 | 0.008 | 0.008 | 0 | 0.69 | 0.67 | 0.71 | 0.84 | 0.62 | 0.67 |
| 10 | 130248440 | rs9804350 | -0.617 | -0.465 | -0.167 | 0.604 | -0.026 | 0.066 | 0.066 | 0 | 0.25 | 0.4 | 0.23 | 0.58 | 0.2 | 0.42 |
| 10 | 130248926 | rs9804317 | 0.268 | -0.439 | -0.028 | 0.644 | -0.008 | 0.192 | 0.192 | 0 | 0.11 | 0.23 | 0.15 | 0.4 | 0.09 | 0.27 |
| 10 | 130251523 | rs7475176 | 0.419 | -0.598 | -1.403 | 0.598 | -0.022 | -0.031 | -0.031 | 0.116 | 0.37 | 0.19 | 0.45 | 0.5 | 0.38 | 0.33 |
| 10 | 130256859 | rs2387861 | 0.098 | -0.614 | -1.429 | 0.667 | -0.023 | -0.011 | -0.011 | 0 | 0.44 | 0.21 | 0.47 | 0.53 | 0.44 | 0.38 |
| 10 | 130261435 | rs4489659 | -0.168 | -0.9 | -1.282 | 0.864 | 0.016 | -0.023 | -0.023 | 0 | 0.85 | 0.9 | 0.74 | 0.74 | 0.9 | 0.89 |

| | | | | | | | | | | | | | | | | |
|---|---|---|---|---|---|---|---|---|---|---|---|---|---|---|---|---|
| 10 | 130263348 | rs11016257 | 0.654 | -0.762 | -1.239 | 0.856 | 0.016 | -0.031 | -0.031 | 0 | 0.11 | 0.02 | 0.16 | 0.16 | 0.08 | 0.08 |
| 10 | 130263843 | rs11016258 | 0.061 | -0.561 | -0.741 | 0.642 | -0.027 | -0.044 | -0.044 | 0 | 0.16 | 0.53 | 0.13 | 0.07 | 0.14 | 0.14 |
| 10 | 130264979 | rs11016259 | -0.441 | -0.362 | -1.219 | 0.646 | -0.019 | 0.063 | 0.063 | 0 | 0.46 | 0.73 | 0.35 | 0.25 | 0.49 | 0.55 |
| 10 | 130271740 | rs1572760 | -0.056 | -0.297 | -1.52 | 0.702 | -0.05 | 0.014 | 0.014 | 0 | 0.41 | 0.21 | 0.44 | 0.67 | 0.47 | 0.61 |
| 10 | 130271977 | rs10829414 | 1.099 | -0.445 | -1.381 | 0.594 | 0.001 | 0.052 | 0.052 | 0 | 0.4 | 0.19 | 0.42 | 0.26 | 0.36 | 0.23 |
| 10 | 130272033 | rs7096455 | 0.234 | -0.529 | -1.366 | 0.676 | -0.001 | 0.026 | 0.026 | 0 | 0.86 | 0.82 | 0.77 | 0.53 | 0.87 | 0.8 |
| 10 | 130272477 | rs7097015 | -0.019 | -0.484 | -1.36 | 0.615 | -0.03 | -0.06 | -0.06 | 0 | 0.19 | 0.57 | 0.14 | 0.07 | 0.17 | 0.16 |
| 10 | 130273179 | rs4750961 | -0.447 | -0.496 | -1.376 | 0.829 | 0 | -0.005 | -0.005 | 0 | 0.89 | 0.93 | 0.88 | 0.64 | 0.89 | 0.83 |
| 10 | 130275947 | rs4750705 | 1.085 | -0.473 | -1.472 | 0.773 | -0.099 | -0.029 | -0.029 | 1.014 | 0.26 | 0.04 | 0.21 | 0.2 | 0.35 | 0.41 |
| 10 | 130276566 | rs1572759 | 0.705 | -0.606 | -1.14 | 0.881 | -0.027 | 0.049 | 0.049 | 0.766 | 0.71 | 0.41 | 0.66 | 0.46 | 0.74 | 0.65 |
| 10 | 130276925 | rs11016267 | 0.719 | -0.479 | -0.966 | 0.221 | -0.005 | 0.034 | 0.034 | 0 | 0.36 | 0.19 | 0.39 | 0.26 | 0.32 | 0.22 |
| 10 | 130278058 | rs951827 | 0.335 | -0.092 | -0.732 | 0.427 | -0.062 | 0.009 | 0.009 | 0.209 | 0.8 | 0.46 | 0.78 | 0.82 | 0.84 | 0.83 |
| 10 | 130278179 | rs7901182 | -0.192 | 0.09 | -0.706 | 0.207 | -0.053 | -0.061 | -0.061 | 0 | 0.17 | 0.45 | 0.11 | 0.07 | 0.14 | 0.14 |
| 10 | 130279912 | rs4750963 | -0.239 | 1.036 | -1.005 | 0.593 | -0.017 | -0.015 | -0.015 | 0.02 | 0.36 | 0.25 | 0.35 | 0.56 | 0.45 | 0.55 |
| 10 | 130284223 | rs11818187 | -1.151 | 1.545 | 0.152 | 0.372 | 0.1 | 0.008 | 0.008 | 0 | 0.12 | 0.37 | 0.17 | 0.12 | 0.19 | 0.11 |
| 10 | 130285936 | rs7905909 | -0.334 | 0.35 | -0.816 | -0.205 | 0.015 | 0.122 | 0.122 | 0.194 | 0.4 | 0.14 | 0.21 | 0.18 | 0.4 | 0.23 |
| 10 | 130288721 | rs2185261 | 0.027 | 0.527 | -0.083 | -0.538 | 0.033 | -0.034 | -0.034 | 0 | 0.12 | 0.36 | 0.13 | 0.05 | 0.12 | 0.09 |
| 10 | 130289519 | rs7915794 | 0.677 | 0.337 | -0.29 | -0.665 | 0.053 | 0.109 | 0.109 | 0.381 | 0.29 | 0.09 | 0.12 | 0.08 | 0.24 | 0.14 |
| 10 | 130290359 | rs7900768 | 1.234 | -0.232 | -0.222 | -0.223 | -0.045 | -0.007 | -0.007 | 0 | 0.35 | 0.19 | 0.56 | 0.41 | 0.43 | 0.5 |
| 10 | 130290476 | rs2050766 | 0.941 | -0.176 | 0.098 | -0.189 | 0.072 | 0.003 | 0.003 | 0 | 0.07 | 0.25 | 0.11 | 0.03 | 0.1 | 0.06 |
| 10 | 130291738 | rs7893667 | -0.028 | -0.17 | -0.511 | -0.164 | 0.006 | 0.046 | 0.046 | 0 | 0.69 | 0.41 | 0.7 | 0.51 | 0.68 | 0.67 |
| 10 | 130292525 | rs10764780 | 0.359 | -0.046 | -0.97 | -0.099 | 0.004 | -0.014 | -0.014 | 0 | 0.64 | 0.28 | 0.68 | 0.48 | 0.66 | 0.64 |
| 10 | 130294844 | rs1541175 | 0.529 | 0.449 | -0.879 | 0.082 | -0.059 | -0.012 | -0.012 | 0.198 | 0.35 | 0.17 | 0.56 | 0.41 | 0.43 | 0.5 |
| 10 | 130299910 | rs11016281 | 0.664 | 0.103 | -1.05 | -0.133 | 0.044 | 0.109 | 0.109 | 0.503 | 0.29 | 0.08 | 0.12 | 0.06 | 0.2 | 0.19 |
| 10 | 130300253 | rs11016282 | -0.441 | 0.721 | -0.744 | 0.271 | -0.003 | 0.026 | 0.026 | 0 | 0.27 | 0.44 | 0.23 | 0.39 | 0.27 | 0.24 |
| 10 | 130305654 | rs7904038 | -0.167 | 1.369 | -1.06 | 0.787 | 0.001 | 0.001 | 0.001 | 0 | 0.15 | 0.13 | 0.17 | 0.19 | 0.2 | 0.12 |
| 10 | 130307593 | rs7093850 | -0.597 | 1.061 | -1.005 | 0.608 | 0.025 | 0.014 | 0.014 | 0 | 0.49 | 0.33 | 0.66 | 0.57 | 0.59 | 0.61 |
| 10 | 130309299 | rs1890627 | -0.111 | 0.823 | -1.206 | 0.509 | -0.025 | -0.004 | -0.004 | 0 | 0.33 | 0.14 | 0.48 | 0.36 | 0.38 | 0.48 |
| 10 | 130318217 | rs10829419 | 0.263 | 0.224 | -0.811 | 0.403 | 0.005 | -0.005 | -0.005 | 0 | 0.67 | 0.64 | 0.4 | 0.25 | 0.58 | 0.58 |
| 10 | 130320343 | rs11016300 | 0.118 | -0.026 | -1.502 | -0.062 | -0.022 | 0.005 | 0.005 | 0 | 0.21 | 0.05 | 0.13 | 0.15 | 0.21 | 0.33 |
| 10 | 130322451 | rs10764786 | -0.541 | 0.377 | -0.727 | 0.677 | 0.051 | 0.105 | 0.105 | 0.035 | 0.85 | 0.67 | 0.54 | 0.65 | 0.73 | 0.91 |
| 10 | 130322483 | rs10764787 | -0.606 | 0.373 | -0.728 | 0.836 | 0.055 | 0.072 | 0.072 | 0.011 | 0.85 | 0.67 | 0.54 | 0.73 | 0.73 | 0.92 |
| 10 | 130325971 | rs10764788 | 0.326 | -0.069 | -0.854 | 0.708 | 0.015 | 0.036 | 0.036 | 0 | 0.72 | 0.6 | 0.5 | 0.6 | 0.66 | 0.69 |
| 10 | 130328926 | rs1544087 | -0.766 | -0.824 | -0.966 | 0.274 | -0.006 | 0.042 | 0.042 | 0.013 | 0.59 | 0.67 | 0.74 | 0.85 | 0.61 | 0.65 |
| 10 | 130344003 | rs7913051 | 0.037 | -0.068 | -0.506 | 0.849 | 0.06 | 0.067 | 0.067 | 0.061 | 0.28 | 0.5 | 0.55 | 0.4 | 0.41 | 0.34 |
| 10 | 130345804 | rs11016315 | 0.002 | -0.112 | -1.136 | 0.913 | 0.008 | -0.024 | -0.024 | 0.025 | 0.17 | 0.09 | 0.14 | 0.35 | 0.13 | 0.22 |
| 10 | 130345929 | rs11598841 | -0.429 | -0.07 | -0.712 | 0.836 | -0.005 | -0.01 | -0.01 | 0 | 0.41 | 0.6 | 0.59 | 0.47 | 0.48 | 0.57 |
| 10 | 130347772 | rs4750710 | 0.603 | -0.81 | -0.112 | 1.183 | -0.062 | 0.044 | 0.044 | 0 | 0.73 | 0.55 | 0.83 | 0.59 | 0.83 | 0.63 |
| 10 | 130347966 | rs7086546 | -0.775 | -0.613 | 0.19 | 1.117 | -0.014 | 0.005 | 0.005 | 0.332 | 0.13 | 0.24 | 0.08 | 0.14 | 0.08 | 0.15 |
| 10 | 130349382 | rs7912195 | 0.288 | 0.019 | 0.464 | 1.111 | 0.008 | -0.001 | -0.001 | 0.56 | 0.79 | 0.7 | 0.82 | 0.79 | 0.78 | 0.89 |
| 10 | 130353231 | rs7898393 | -1.325 | -0.102 | 1.117 | 1.775 | 0.023 | 0.465 | 0.465 | 0 | 0.08 | 0.38 | 0.12 | 0.46 | 0.06 | 0.23 |
| 10 | 130354186 | rs7078053 | 0.385 | 0.068 | 1.171 | 1.419 | 0.014 | -0.012 | -0.012 | 0 | 0.7 | 0.59 | 0.77 | 0.75 | 0.7 | 0.76 |
| 10 | 130355180 | rs943810 | 1.248 | 0.268 | 1.2 | 1.393 | 0.003 | -0.001 | -0.001 | 0.541 | 0.61 | 0.46 | 0.67 | 0.38 | 0.61 | 0.58 |
| 10 | 130355794 | rs752658 | 1.349 | 0.234 | 1.152 | 1.406 | -0.048 | 0.066 | 0.066 | 0.632 | 0.57 | 0.12 | 0.63 | 0.3 | 0.59 | 0.5 |
| 10 | 130356576 | rs7899042 | 1.362 | 0.246 | 1.157 | 1.409 | -0.047 | 0.068 | 0.068 | 0.668 | 0.57 | 0.12 | 0.63 | 0.3 | 0.59 | 0.5 |
| 10 | 130359012 | rs10764792 | 0.772 | 0.29 | 1.175 | 1.425 | 0.015 | 0.013 | 0.013 | 0.007 | 0.66 | 0.53 | 0.7 | 0.38 | 0.66 | 0.62 |
| 10 | 130363238 | rs1004344 | 1.238 | 0.336 | 1.053 | 1.383 | -0.02 | 0.011 | 0.011 | 0.043 | 0.61 | 0.27 | 0.65 | 0.35 | 0.6 | 0.55 |
| 10 | 130370900 | rs2031854 | 0.574 | 0.665 | 1.367 | 1.314 | 0 | -0.016 | -0.016 | 0 | 0.64 | 0.57 | 0.71 | 0.72 | 0.64 | 0.59 |
| 10 | 130371881 | rs7082932 | 0.589 | 0.638 | 1.374 | 1.31 | -0.001 | -0.015 | -0.015 | 0.377 | 0.64 | 0.58 | 0.71 | 0.72 | 0.64 | 0.59 |
| 10 | 130372412 | rs1571370 | 1.495 | 0.67 | 1.35 | 1.416 | -0.027 | 0.124 | 0.124 | 0.675 | 0.52 | 0.16 | 0.61 | 0.28 | 0.53 | 0.28 |
| 10 | 130372799 | rs7087654 | -2.399 | 0.526 | 1.754 | 1.764 | 0.039 | 0.336 | 0.336 | 0 | 0.08 | 0.3 | 0.08 | 0.42 | 0.08 | 0.22 |
| 10 | 130374161 | rs11016342 | 0.974 | 0.658 | 1.201 | 1.266 | -0.022 | 0.072 | 0.072 | 0 | 0.42 | 0.13 | 0.5 | 0.27 | 0.4 | 0.23 |
| 10 | 130374405 | rs7089798 | -1.794 | 0.449 | 1.365 | 1.323 | 0.051 | 0.417 | 0.417 | 0 | 0.07 | 0.3 | 0.07 | 0.42 | 0.07 | 0.21 |
| 10 | 130375341 | rs12569501 | -2.04 | 0.437 | 1.519 | 1.302 | 0.066 | 0.494 | 0.494 | 0 | 0.08 | 0.38 | 0.08 | 0.42 | 0.06 | 0.21 |
| 10 | 130382101 | rs7093970 | 0.6 | 0.506 | 1.338 | 1.185 | -0.016 | 0.078 | 0.078 | 0.02 | 0.46 | 0.26 | 0.52 | 0.29 | 0.43 | 0.32 |
| 10 | 130384010 | rs12357464 | 0.6 | 0.506 | 1.338 | 1.185 | -0.016 | 0.078 | 0.078 | 0.02 | 0.46 | 0.26 | 0.52 | 0.29 | 0.43 | 0.32 |
| 10 | 130392132 | rs10829443 | 0.912 | 0.503 | 1.088 | 1.279 | -0.04 | 0.077 | 0.077 | 0 | 0.43 | 0.13 | 0.5 | 0.28 | 0.42 | 0.24 |
| 10 | 130394371 | rs1359974 | 0.189 | 0.369 | 0.97 | 1.094 | -0.006 | 0.043 | 0.043 | 0 | 0.52 | 0.66 | 0.59 | 0.72 | 0.48 | 0.53 |
| 10 | 130396140 | rs7474694 | 0.241 | 0.438 | 0.916 | 1.091 | 0 | 0.005 | 0.005 | 0 | 0.5 | 0.54 | 0.58 | 0.68 | 0.47 | 0.45 |
| 10 | 130401197 | rs4460733 | 1.122 | 0.244 | 1.35 | 1.509 | -0.014 | 0.075 | 0.075 | 0 | 0.85 | 0.79 | 0.9 | 0.61 | 0.86 | 0.75 |
| 10 | 130403699 | rs11016379 | -2.401 | 0.041 | 1.752 | 1.466 | 0.027 | 0.181 | 0.181 | 1.232 | 0.07 | 0.41 | 0.09 | 0.14 | 0.01 | 0.13 |
| 10 | 130407679 | rs943808 | 0.922 | 0.107 | 1.556 | 1.612 | -0.01 | 0.035 | 0.035 | 0 | 0.85 | 0.81 | 0.9 | 0.62 | 0.87 | 0.75 |
| 10 | 130408126 | rs943809 | 1.755 | 0.39 | 1.581 | 1.401 | 0.003 | -0.017 | -0.017 | 0.366 | 0.76 | 0.64 | 0.83 | 0.81 | 0.7 | 0.72 |
| 10 | 130408854 | rs1001990 | -1.888 | 0.317 | 1.526 | 1.47 | 0.007 | 0.044 | 0.044 | 0.361 | 0.18 | 0.38 | 0.2 | 0.27 | 0.22 | 0.28 |
| 10 | 130409727 | rs10829446 | -1.072 | 0.308 | 1.682 | 1.565 | 0.018 | 0.046 | 0.046 | 0 | 0.08 | 0.18 | 0.08 | 0.14 | 0.13 | 0.08 |
| 10 | 130416520 | rs7083081 | -0.903 | 0.359 | 1.484 | 1.493 | 0.017 | 0.027 | 0.027 | 0 | 0.08 | 0.16 | 0.08 | 0.12 | 0.13 | 0.1 |
| 10 | 130416893 | rs943817 | 1.984 | 0.081 | 1.715 | 1.558 | -0.051 | 0.15 | 0.15 | 0 | 0.79 | 0.43 | 0.82 | 0.48 | 0.86 | 0.61 |
| 10 | 130417506 | rs998717 | -1.462 | 0.159 | 1.59 | 1.639 | -0.009 | 0.145 | 0.145 | 0.005 | 0.12 | 0.25 | 0.13 | 0.47 | 0.09 | 0.27 |
| 10 | 130418334 | rs6482693 | 2.072 | 0.032 | 1.563 | 1.438 | 0.02 | -0.018 | -0.018 | 0 | 0.68 | 0.24 | 0.71 | 0.36 | 0.72 | 0.52 |

| | | | | | | | | | | | | | | | | |
|---|---|---|---|---|---|---|---|---|---|---|---|---|---|---|---|---|
| 10 | 130418455 | rs6482695 | 1.529 | 0.027 | 1.847 | 1.292 | -0.03 | 0.036 | 0.036 | 0 | 0.78 | 0.66 | 0.87 | 0.62 | 0.86 | 0.69 |
| 10 | 130419070 | rs7894394 | 1.742 | 0.443 | 1.644 | 1.197 | 0 | 0.064 | 0.064 | 0 | 0.72 | 0.5 | 0.77 | 0.53 | 0.68 | 0.69 |
| 10 | 130419737 | rs4750983 | -0.017 | 0.374 | 1.491 | 1.254 | -0.007 | -0.001 | -0.001 | 0.141 | 0.94 | 0.88 | 0.96 | 0.96 | 0.98 | 0.92 |
| 10 | 130420221 | rs7074022 | 2.251 | 0.528 | 1.713 | 1.25 | 0.003 | 0.075 | 0.075 | 0 | 0.77 | 0.45 | 0.81 | 0.61 | 0.77 | 0.69 |
| 10 | 130420678 | rs12359499 | -1.593 | 0.564 | 1.737 | 1.261 | 0.003 | 0.028 | 0.028 | 0.446 | 0.16 | 0.22 | 0.13 | 0.36 | 0.2 | 0.22 |
| 10 | 130423301 | rs7912469 | 0.209 | 0.64 | 1.589 | 1.205 | 0.006 | 0.04 | 0.04 | 0 | 0.87 | 0.85 | 0.91 | 0.62 | 0.86 | 0.73 |
| 10 | 130424011 | rs2182965 | 0.673 | 0.4 | 0.855 | 0.464 | -0.003 | 0.164 | 0.164 | 0 | 0.26 | 0.02 | 0.14 | 0.01 | 0.2 | 0.18 |
| 10 | 130427652 | rs11016394 | -1.222 | 0.975 | 2.099 | 1.389 | 0.037 | 0.006 | 0.006 | 0.297 | 0.06 | 0.25 | 0.05 | 0 | 0.04 | 0.03 |
| 10 | 130430401 | rs10829450 | -2.013 | 0.967 | 1.918 | 1.617 | 0.001 | 0.067 | 0.067 | 0 | 0.24 | 0.58 | 0.24 | 0.49 | 0.24 | 0.38 |
| 10 | 130439482 | rs7098205 | -0.897 | 1.013 | 1.48 | 1.549 | 0.008 | 0.039 | 0.039 | 0.044 | 0.09 | 0.24 | 0.12 | 0.1 | 0.1 | 0.12 |
| 10 | 130445346 | rs10829462 | -0.915 | 1.073 | 0.746 | 1.501 | 0.027 | 0.089 | 0.089 | 0.66 | 0.17 | 0.47 | 0.21 | 0.22 | 0.24 | 0.17 |
| 10 | 130450189 | rs992250 | -1.967 | 1.07 | 0.106 | 1.518 | -0.01 | -0.017 | -0.017 | 0 | 0.33 | 0.76 | 0.31 | 0.62 | 0.38 | 0.4 |
| 10 | 130452507 | rs12360327 | -1.381 | 0.898 | 0.329 | 1.033 | 0.011 | 0.054 | 0.054 | 1.519 | 0.18 | 0.54 | 0.22 | 0.22 | 0.24 | 0.17 |
| 10 | 130455063 | rs710930 | -0.327 | 0.933 | -0.339 | 0.733 | 0.002 | -0.009 | -0.009 | 0 | 0.38 | 0.51 | 0.34 | 0.67 | 0.36 | 0.48 |
| 10 | 130456798 | rs782401 | -0.764 | 0.452 | -0.705 | 0.641 | 0.004 | 0.006 | 0.006 | 0 | 0.28 | 0.26 | 0.29 | 0.21 | 0.25 | 0.16 |
| 10 | 130457732 | rs3847457 | -0.412 | 1.341 | -1.014 | 1.065 | -0.038 | -0.045 | -0.045 | 0 | 0.1 | 0.03 | 0.13 | 0.19 | 0.14 | 0.13 |
| 10 | 130458099 | rs782396 | -0.526 | 1.453 | -0.811 | 1.033 | 0.011 | 0.015 | 0.015 | 0 | 0.26 | 0.19 | 0.28 | 0.21 | 0.22 | 0.16 |
| 10 | 130470565 | rs2767290 | -1.239 | 1.078 | -0.628 | 0.768 | -0.009 | -0.019 | -0.019 | 0 | 0.39 | 0.43 | 0.45 | 0.24 | 0.36 | 0.39 |
| 10 | 130472020 | rs12269586 | -0.766 | 1.106 | -0.591 | 0.808 | -0.024 | -0.058 | -0.058 | 0 | 0.1 | 0.2 | 0.06 | 0 | 0.08 | 0.08 |
| 10 | 130477668 | rs1409757 | -1.417 | 1.304 | -0.845 | 1.046 | -0.005 | -0.014 | -0.014 | 0 | 0.36 | 0.3 | 0.48 | 0.56 | 0.39 | 0.48 |
| 10 | 130477824 | rs1409756 | -1.535 | 1.404 | -0.779 | 0.939 | 0.007 | 0.03 | 0.03 | 0 | 0.18 | 0.23 | 0.27 | 0.38 | 0.24 | 0.35 |
| 10 | 130478722 | rs11594084 | -0.149 | 1.57 | -0.84 | 0.951 | -0.034 | -0.032 | -0.032 | 0 | 0.06 | 0.01 | 0.09 | 0.17 | 0.12 | 0.09 |
| 10 | 130479587 | rs11016427 | -0.712 | 1.374 | -0.719 | 0.601 | 0.006 | -0.019 | -0.019 | 1.168 | 0.05 | 0.08 | 0.05 | 0 | 0.08 | 0.01 |
| 10 | 130479642 | rs11599262 | -0.897 | 1.091 | -0.755 | 0.782 | -0.007 | -0.019 | -0.019 | 0 | 0.35 | 0.28 | 0.47 | 0.47 | 0.38 | 0.45 |
| 10 | 130480719 | rs7088130 | -1.042 | 0.421 | -0.006 | 0.456 | -0.029 | -0.025 | -0.025 | 0 | 0.16 | 0.47 | 0.09 | 0.13 | 0.12 | 0.18 |
| 10 | 130481274 | rs10764803 | -0.205 | 0.692 | -0.517 | 0.501 | 0.006 | -0.002 | -0.002 | 4.252 | 0.48 | 0.39 | 0.49 | 0.51 | 0.57 | 0.51 |
| 10 | 130482188 | rs913172 | -0.168 | 0.923 | -0.976 | -0.123 | -0.015 | 0.06 | 0.06 | 0.059 | 0.59 | 0.68 | 0.62 | 0.79 | 0.53 | 0.55 |
| 10 | 130483163 | rs12412378 | 0.089 | 1.411 | -0.957 | -0.005 | -0.016 | 0.023 | 0.023 | 0 | 0.16 | 0.07 | 0.18 | 0.09 | 0.15 | 0.16 |
| 10 | 130484409 | rs1107247 | 0.109 | 1.273 | -0.441 | 0.355 | 0 | 0.058 | 0.058 | 0 | 0.3 | 0.53 | 0.27 | 0.48 | 0.27 | 0.26 |
| 10 | 130487055 | rs2038968 | -0.628 | 1.24 | -0.29 | -0.407 | 0.004 | 0.064 | 0.064 | 0 | 0.5 | 0.58 | 0.57 | 0.83 | 0.45 | 0.54 |
| 10 | 130487671 | rs4750989 | 0.007 | 1.647 | -0.2 | -0.568 | 0.005 | -0.036 | -0.036 | 0 | 0.33 | 0.38 | 0.38 | 0.79 | 0.3 | 0.36 |
| 10 | 130488026 | rs4750990 | 0.148 | 1.225 | -0.678 | -0.154 | -0.005 | -0.005 | -0.005 | 0 | 0.33 | 0.3 | 0.35 | 0.58 | 0.39 | 0.42 |
| 10 | 130488204 | rs2038967 | -1.423 | 1.241 | -1.41 | -0.738 | 0.002 | 0.102 | 0.102 | 0 | 0.6 | 0.86 | 0.62 | 0.79 | 0.57 | 0.7 |
| 10 | 130490759 | rs11016432 | 0.892 | 1.53 | -1.119 | -0.416 | -0.04 | 0.043 | 0.043 | 0 | 0.12 | 0.02 | 0.11 | 0.03 | 0.1 | 0.17 |
| 10 | 130497635 | rs1555930 | 0.076 | 1.564 | -1.057 | -0.392 | -0.035 | -0.012 | -0.012 | 0.727 | 0.32 | 0.2 | 0.38 | 0.65 | 0.38 | 0.29 |
| 10 | 130499303 | rs10829475 | -1.035 | 1.119 | -0.573 | -0.885 | 0.002 | 0.018 | 0.018 | 0 | 0.7 | 0.78 | 0.8 | 0.91 | 0.71 | 0.71 |
| 10 | 130499829 | rs1543591 | 0.289 | 1.142 | -0.481 | -0.83 | 0.004 | 0.045 | 0.045 | 0 | 0.38 | 0.29 | 0.37 | 0.17 | 0.42 | 0.32 |
| 10 | 130500410 | rs11016441 | -1.01 | 0.949 | 0.077 | -0.681 | -0.001 | -0.056 | -0.056 | 0 | 0.13 | 0.23 | 0.15 | 0.03 | 0.14 | 0.08 |
| 10 | 130504906 | rs1887244 | -0.147 | 0.989 | -0.264 | -0.603 | 0 | -0.022 | -0.022 | 0.062 | 0.89 | 0.9 | 0.91 | 1 | 0.88 | 0.96 |
| 10 | 130509103 | rs7078934 | -0.262 | 0.773 | -0.191 | -0.523 | 0.001 | -0.015 | -0.015 | 0.556 | 0.25 | 0.29 | 0.28 | 0.2 | 0.24 | 0.28 |
| 10 | 130511895 | rs1327012 | 0.066 | 0.754 | -0.486 | -0.262 | 0 | 0.016 | 0.016 | 0 | 0.85 | 0.81 | 0.71 | 0.38 | 0.83 | 0.71 |
| 10 | 130512229 | rs11016451 | 1.301 | 0.761 | -0.953 | -0.8 | -0.046 | 0.055 | 0.055 | 0 | 0.18 | 0.02 | 0.07 | 0.06 | 0.13 | 0.24 |
| 10 | 130512503 | rs12258312 | 0.343 | 0.292 | -0.135 | -0.174 | -0.03 | 0.025 | 0.025 | 0 | 0.07 | 0.02 | 0.09 | 0.03 | 0.15 | 0.09 |
| 10 | 130518039 | rs4750993 | 1.504 | 0.212 | 0.374 | 0.16 | 0.006 | 0.107 | 0.107 | 0 | 0.75 | 0.54 | 0.61 | 0.45 | 0.71 | 0.63 |
| 10 | 130518417 | rs913170 | 0.649 | 0.245 | 0.447 | 0.48 | -0.001 | 0.055 | 0.055 | 0 | 0.86 | 0.79 | 0.71 | 0.49 | 0.86 | 0.72 |
| 10 | 130521354 | rs12571096 | 0.338 | 0.315 | 0.823 | -0.439 | -0.009 | -0.003 | -0.003 | 0 | 0.16 | 0.19 | 0.12 | 0.14 | 0.08 | 0.18 |
| 10 | 130525985 | rs10829482 | 0.175 | 0.4 | 0.84 | -0.285 | 0.021 | 0.009 | 0.009 | 0 | 0.9 | 0.79 | 0.8 | 0.92 | 0.89 | 0.85 |
| 10 | 130526241 | rs7100181 | 1.034 | 0.303 | 0.526 | -0.351 | 0.028 | -0.049 | -0.049 | 0 | 0.8 | 0.55 | 0.7 | 0.91 | 0.77 | 0.8 |
| 10 | 130527944 | rs881403 | -0.003 | -0.489 | -0.099 | -0.347 | -0.004 | 0 | 0 | 0 | 0.17 | 0.17 | 0.19 | 0.19 | 0.08 | 0.18 |
| 10 | 130529771 | rs11598555 | -0.474 | -0.086 | -0.222 | -0.528 | -0.003 | -0.045 | -0.045 | 1.143 | 0.13 | 0.19 | 0.11 | 0 | 0.15 | 0.08 |
| 10 | 130530328 | rs4750717 | -0.819 | -0.253 | -0.612 | -0.63 | 0.039 | 0.079 | 0.079 | 0.667 | 0.45 | 0.68 | 0.37 | 0.13 | 0.37 | 0.27 |
| 10 | 130535073 | rs2387866 | 0.236 | -0.157 | -0.3 | -0.12 | 0.015 | 0.024 | 0.024 | 0 | 0.95 | 0.82 | 0.96 | 0.95 | 0.97 | 0.95 |
| 10 | 130538787 | rs11016457 | -0.131 | -0.6 | -0.987 | -0.521 | 0.022 | 0.113 | 0.113 | 1.257 | 0.29 | 0.02 | 0.16 | 0.1 | 0.23 | 0.19 |
| 10 | 130539111 | rs7078785 | 1.289 | -0.369 | 0.415 | -0.12 | -0.024 | -0.067 | -0.067 | 0.302 | 0.81 | 0.54 | 0.79 | 0.93 | 0.84 | 0.86 |
| 10 | 130541923 | rs10764807 | 1.688 | -0.479 | 0.837 | 0.074 | -0.027 | -0.009 | -0.009 | 0 | 0.87 | 0.42 | 0.83 | 0.93 | 0.93 | 0.93 |
| 10 | 130542341 | rs10764808 | 0.889 | -0.242 | 0.486 | 0.228 | -0.001 | 0.015 | 0.015 | 0 | 0.93 | 0.8 | 0.95 | 0.95 | 0.97 | 0.94 |
| 10 | 130543960 | rs7910094 | -0.571 | 0.575 | 0.104 | 1.224 | 0.057 | 0.125 | 0.125 | 0 | 0.06 | 0.11 | 0.26 | 0.26 | 0.15 | 0.16 |
| 10 | 130549015 | rs11594990 | -0.03 | -0.288 | -0.535 | 0.478 | 0.02 | 0.071 | 0.071 | 0 | 0.31 | 0.13 | 0.13 | 0.12 | 0.26 | 0.17 |
| 10 | 130549553 | rs10829485 | 0.053 | -0.326 | -0.523 | 0.467 | 0.028 | 0.084 | 0.084 | 0 | 0.3 | 0.12 | 0.13 | 0.09 | 0.25 | 0.17 |
| 10 | 130550033 | rs11016463 | 0.077 | -0.323 | -0.522 | 0.468 | 0.02 | 0.073 | 0.073 | 0 | 0.29 | 0.12 | 0.12 | 0.09 | 0.24 | 0.17 |
| 10 | 130552334 | rs1536452 | -0.744 | -0.206 | -0.664 | 1.262 | 0.048 | 0.031 | 0.031 | 0.369 | 0.47 | 0.85 | 0.35 | 0.6 | 0.38 | 0.39 |
| 10 | 130553741 | rs7090877 | -0.944 | 0.074 | -0.272 | 1.604 | 0.023 | 0.14 | 0.14 | 0.046 | 0.06 | 0.14 | 0.05 | 0.33 | 0.06 | 0.09 |
| 10 | 130555220 | rs10829489 | -0.326 | -0.257 | 0.123 | 1.46 | -0.007 | 0.017 | 0.017 | 0.016 | 0.4 | 0.46 | 0.29 | 0.49 | 0.34 | 0.3 |
| 10 | 130557238 | rs11016469 | 0.107 | -0.053 | -0.25 | 1.492 | -0.009 | -0.038 | -0.038 | 0 | 0.4 | 0.56 | 0.26 | 0.13 | 0.28 | 0.28 |
| 10 | 130564068 | rs2254816 | -0.662 | -0.152 | -0.368 | 1.744 | 0.021 | -0.014 | -0.014 | 0 | 0.48 | 0.86 | 0.37 | 0.59 | 0.42 | 0.43 |
| 10 | 130566038 | rs17136 | -1.568 | 0.052 | -0.152 | 1.839 | 0.052 | 0.283 | 0.283 | 0.639 | 0.09 | 0.27 | 0.1 | 0.47 | 0.15 | 0.14 |
| 10 | 130566749 | rs561020 | -0.649 | -0.045 | -0.255 | 1.624 | -0.001 | 0.068 | 0.068 | 0 | 0.05 | 0.07 | 0.06 | 0.35 | 0.08 | 0.08 |
| 10 | 130567929 | rs638395 | -0.09 | -0.213 | -0.252 | 1.339 | -0.018 | -0.061 | -0.061 | 0.196 | 0.38 | 0.54 | 0.27 | 0.12 | 0.27 | 0.28 |
| 10 | 130571475 | rs12412363 | -0.7 | -0.191 | -0.629 | 2.06 | 0.002 | -0.002 | -0.002 | 0.578 | 0.09 | 0.07 | 0.08 | 0.1 | 0.03 | 0.11 |

| | | | | | | | | | | | | | | | | |
|---|---|---|---|---|---|---|---|---|---|---|---|---|---|---|---|---|
| 10 | 130577585 | rs10829492 | -1.429 | 0.063 | -0.73 | 2.053 | -0.001 | 0.015 | 0.015 | 0 | 0.13 | 0.14 | 0.32 | 0.28 | 0.12 | 0.2 |
| 10 | 130580251 | rs12219067 | -1.341 | 0.002 | -0.7 | 1.862 | -0.009 | 0.002 | 0.002 | 0 | 0.09 | 0.13 | 0.18 | 0.1 | 0.06 | 0.15 |
| 10 | 130580494 | rs646066 | -0.134 | 0.401 | -1.286 | 1.222 | -0.043 | 0.083 | 0.083 | 0 | 0.61 | 0.83 | 0.62 | 0.81 | 0.52 | 0.62 |
| 10 | 130581094 | rs579655 | -0.273 | 0.511 | -1.588 | 1.189 | -0.114 | 0.093 | 0.093 | 0 | 0.63 | 0.95 | 0.66 | 0.82 | 0.56 | 0.66 |
| 10 | 130587452 | rs497414 | -0.234 | 1.409 | -1.493 | 1.369 | -0.144 | 0.015 | 0.015 | 0 | 0.77 | 0.97 | 0.71 | 0.85 | 0.62 | 0.82 |
| 10 | 130587803 | rs11596125 | 1.078 | 0.407 | -2.414 | 0.154 | -0.023 | 0.181 | 0.181 | 0.648 | 0.3 | 0.05 | 0.13 | 0.01 | 0.25 | 0.15 |
| 10 | 130588415 | rs10829495 | 0.088 | 0.102 | -2.154 | 0.466 | 0.004 | 0.003 | 0.003 | 0 | 0.11 | 0.06 | 0.06 | 0.07 | 0.11 | 0.16 |
| 10 | 130589876 | rs472752 | -0.961 | 0.159 | -2.144 | 0.469 | -0.149 | 0.089 | 0.089 | 0 | 0.77 | 0.97 | 0.71 | 0.9 | 0.62 | 0.82 |
| 10 | 130595233 | rs7082333 | 0.219 | -0.303 | -1.118 | 1.025 | -0.049 | 0.141 | 0.141 | 0.062 | 0.19 | 0.48 | 0.28 | 0.4 | 0.11 | 0.37 |
| 10 | 130595286 | rs11016487 | 0.246 | -0.384 | -0.773 | 1.022 | -0.06 | 0.154 | 0.154 | 0.112 | 0.19 | 0.43 | 0.28 | 0.4 | 0.1 | 0.37 |
| 10 | 130598982 | rs12783941 | 0.84 | -0.234 | -0.794 | 0.677 | 0.023 | 0.022 | 0.022 | 0 | 0.42 | 0.26 | 0.33 | 0.33 | 0.36 | 0.26 |
| 10 | 130607361 | rs4751007 | 1.268 | -0.102 | -1.151 | 0.665 | -0.006 | -0.017 | -0.017 | 0 | 0.37 | 0.09 | 0.29 | 0.33 | 0.3 | 0.27 |
| 10 | 130611022 | rs10741165 | -0.973 | 0.363 | -0.999 | -0.214 | -0.088 | 0.085 | 0.085 | 0 | 0.76 | 0.9 | 0.71 | 0.9 | 0.62 | 0.81 |
| 10 | 130619367 | rs5026308 | -0.823 | 0.328 | -0.914 | -0.231 | -0.069 | 0.075 | 0.075 | 0 | 0.75 | 0.87 | 0.7 | 0.9 | 0.62 | 0.81 |
| 10 | 130622498 | rs1555702 | 0.595 | 0.046 | -0.826 | -0.212 | 0.057 | 0.11 | 0.11 | 1.641 | 0.49 | 0.76 | 0.51 | 0.66 | 0.37 | 0.36 |
| 10 | 130623157 | rs4266990 | -0.724 | 0.141 | -0.301 | -0.201 | -0.001 | -0.013 | -0.013 | 5.491 | 0.74 | 0.76 | 0.7 | 0.9 | 0.62 | 0.81 |
| 10 | 130624838 | rs942525 | -0.249 | 0.051 | -0.616 | -0.148 | 0.002 | 0.002 | 0.002 | 0.295 | 0.73 | 0.76 | 0.78 | 0.75 | 0.74 | 0.55 |
| 10 | 130627887 | rs7895842 | 0.213 | 0.342 | -1.033 | -0.068 | -0.073 | 0.061 | 0.061 | 0 | 0.27 | 0.13 | 0.36 | 0.16 | 0.4 | 0.27 |
| 10 | 130632009 | rs7893468 | 1.102 | 0.802 | -1.46 | -0.038 | -0.105 | 0.048 | 0.048 | 0 | 0.19 | 0.01 | 0.16 | 0.11 | 0.28 | 0.12 |
| 10 | 130632974 | rs7091108 | 0.048 | 0.28 | -1.006 | -0.116 | -0.003 | -0.002 | -0.002 | 0 | 0.34 | 0.37 | 0.38 | 0.19 | 0.44 | 0.29 |
| 10 | 130633376 | rs10764818 | 0.17 | 0.104 | -1.766 | 0.111 | 0.019 | 0.007 | 0.007 | 0 | 0.41 | 0.25 | 0.42 | 0.57 | 0.34 | 0.28 |
| 10 | 130635586 | rs11016509 | -0.605 | -0.104 | -1.996 | -0.366 | 0.037 | 0.082 | 0.082 | 0.04 | 0.24 | 0.09 | 0.09 | 0.08 | 0.17 | 0.14 |
| 10 | 130635614 | rs3940168 | 0.019 | 0.885 | -1.935 | -0.203 | -0.059 | 0.089 | 0.089 | 0 | 0.39 | 0.15 | 0.69 | 0.77 | 0.52 | 0.54 |
| 10 | 130639298 | rs4751013 | -0.05 | 0.635 | -1.144 | 0.378 | -0.022 | 0.03 | 0.03 | 0 | 0.54 | 0.78 | 0.48 | 0.37 | 0.51 | 0.42 |
| 10 | 130641688 | rs12250515 | 0.229 | 0.527 | -0.584 | 0.347 | 0 | 0 | 0 | 0 | 0.51 | 0.48 | 0.45 | 0.31 | 0.51 | 0.38 |
| 10 | 130652909 | rs753712 | 0.139 | 0.594 | -0.532 | 0.361 | 0 | 0.011 | 0.011 | 0 | 0.5 | 0.42 | 0.4 | 0.3 | 0.5 | 0.36 |
| 10 | 130656497 | rs12770017 | -0.247 | 0.967 | -1.37 | 0.391 | -0.032 | -0.012 | -0.012 | 0 | 0.09 | 0.03 | 0.24 | 0.08 | 0.13 | 0.13 |
| 10 | 130659224 | rs10741174 | 0.131 | 0.961 | -1.282 | 0.389 | 0.005 | -0.006 | -0.006 | 1.216 | 0.87 | 0.82 | 0.77 | 0.89 | 0.87 | 0.82 |
| 10 | 130662740 | rs1359361 | -0.112 | 0.833 | -1.656 | 0.596 | -0.045 | -0.11 | -0.11 | 0.97 | 0.34 | 0.09 | 0.3 | 0.46 | 0.36 | 0.44 |
| 10 | 130669638 | rs2027140 | 0.11 | 0.415 | -0.639 | 0.101 | -0.016 | -0.07 | -0.07 | 2.824 | 0.8 | 0.71 | 0.88 | 0.95 | 0.82 | 0.94 |
| 10 | 130673783 | rs10741175 | 0.72 | 0.016 | -0.179 | 0.442 | -0.028 | 0.03 | 0.03 | 0.447 | 0.9 | 0.73 | 0.86 | 0.86 | 0.97 | 0.91 |
| 10 | 130673804 | rs12249496 | -0.377 | 1.156 | -0.721 | 0.378 | -0.043 | -0.009 | -0.009 | 0 | 0.07 | 0.01 | 0.06 | 0.04 | 0.14 | 0.07 |
| 10 | 130675517 | rs942528 | 1.405 | 0.881 | -1.116 | 1.161 | -0.039 | -0.127 | -0.127 | 0.037 | 0.68 | 0.12 | 0.58 | 0.51 | 0.73 | 0.66 |
| 10 | 130677548 | rs11016531 | -0.097 | 0.537 | -0.652 | 1.536 | 0.001 | 0 | 0 | 0 | 0.08 | 0.08 | 0.12 | 0.11 | 0.08 | 0.13 |
| 10 | 130678789 | rs10829511 | -0.64 | 0.503 | -0.603 | 1.724 | 0 | 0.017 | 0.017 | 0 | 0.1 | 0.08 | 0.16 | 0.28 | 0.09 | 0.17 |
| 10 | 130678890 | rs10829512 | -0.613 | 0.508 | -0.611 | 1.733 | 0 | 0.014 | 0.014 | 0 | 0.1 | 0.08 | 0.16 | 0.27 | 0.09 | 0.17 |
| 10 | 130684713 | rs11016536 | 0.629 | 0.685 | -1.466 | 1.494 | -0.072 | -0.085 | -0.085 | 0.009 | 0.21 | 0.02 | 0.15 | 0.25 | 0.24 | 0.28 |
| 10 | 130685641 | rs12572126 | 1.035 | 0.495 | -0.614 | 1.416 | -0.008 | 0.041 | 0.041 | 1.875 | 0.82 | 0.72 | 0.73 | 0.72 | 0.88 | 0.78 |
| 10 | 130686476 | rs10764826 | 0.152 | 0.292 | -0.583 | 1.347 | -0.025 | 0.051 | 0.051 | 0.566 | 0.91 | 0.8 | 0.86 | 0.82 | 0.97 | 0.91 |
| 10 | 130696681 | rs10764829 | 0.481 | 0.346 | -0.458 | 0.97 | -0.004 | -0.013 | -0.013 | 0.191 | 0.67 | 0.39 | 0.76 | 0.73 | 0.7 | 0.66 |
| 10 | 130704327 | rs4750722 | -0.149 | 0.286 | -0.225 | 1.092 | 0.004 | 0.005 | 0.005 | 0 | 0.82 | 0.87 | 0.89 | 0.86 | 0.86 | 0.85 |
| 10 | 130706844 | rs1886380 | 1.052 | 0.217 | -0.114 | 1.17 | -0.004 | -0.009 | -0.009 | 0.015 | 0.67 | 0.43 | 0.77 | 0.7 | 0.7 | 0.71 |
| 10 | 130710167 | rs12570459 | 0.533 | 0.487 | -0.842 | 1.119 | -0.043 | -0.109 | -0.109 | 0 | 0.28 | 0.07 | 0.47 | 0.45 | 0.34 | 0.34 |
| 10 | 130710923 | rs7085203 | -0.053 | 0.345 | 0.142 | 1.117 | 0.006 | 0 | 0 | 0 | 0.12 | 0.16 | 0.1 | 0.15 | 0.16 | 0.14 |
| 10 | 130712316 | rs3824759 | 0.485 | 0.586 | -0.311 | 1.349 | -0.03 | -0.063 | -0.063 | 0 | 0.1 | 0 | 0.32 | 0.21 | 0.13 | 0.16 |
| 10 | 130713288 | rs3740427 | -0.087 | 0.489 | 0.448 | 0.642 | -0.003 | -0.003 | -0.003 | 0.151 | 0.82 | 0.78 | 0.88 | 0.87 | 0.86 | 0.85 |
| 10 | 130715389 | rs10829518 | 0.326 | 0.416 | 0.657 | 0.7 | 0 | -0.012 | -0.012 | 0 | 0.68 | 0.52 | 0.78 | 0.72 | 0.7 | 0.71 |
| 10 | 130729771 | rs7074804 | -0.501 | 0.298 | 0.508 | 1.087 | -0.001 | 0.105 | 0.105 | 0 | 0.14 | 0.37 | 0.12 | 0.22 | 0.13 | 0.04 |
| 10 | 130738463 | rs7092525 | -0.493 | 0.31 | 0.552 | 1.098 | 0.002 | 0.11 | 0.11 | 0 | 0.13 | 0.38 | 0.11 | 0.2 | 0.13 | 0.05 |
| 10 | 130744348 | rs765934 | 0.1 | 0.396 | 0.104 | 0.693 | -0.001 | -0.001 | -0.001 | 0.588 | 0.66 | 0.73 | 0.64 | 0.72 | 0.67 | 0.61 |
| 10 | 130749975 | rs1466196 | 0.886 | 0.655 | -0.187 | 0.7 | -0.002 | 0.003 | 0.003 | 0 | 0.52 | 0.43 | 0.42 | 0.51 | 0.51 | 0.48 |
| 10 | 130750248 | rs10829520 | 0.229 | 0.626 | -0.202 | 0.693 | -0.001 | -0.001 | -0.001 | 0 | 0.61 | 0.46 | 0.57 | 0.65 | 0.64 | 0.57 |
| 10 | 130750456 | rs17295100 | -0.289 | 0.993 | -0.693 | 0.839 | -0.039 | -0.039 | -0.039 | 0 | 0.1 | 0.04 | 0.15 | 0.15 | 0.13 | 0.11 |
| 10 | 130752570 | rs11016569 | 0.233 | 0.736 | -0.909 | 0.65 | -0.086 | -0.085 | -0.085 | 0 | 0.21 | 0.03 | 0.23 | 0.29 | 0.27 | 0.33 |
| 10 | 130753520 | rs4751035 | 0.42 | 0.833 | -0.931 | 0.629 | -0.069 | -0.086 | -0.086 | 0 | 0.19 | 0.03 | 0.2 | 0.29 | 0.24 | 0.31 |
| 10 | 130755548 | rs4619065 | -0.003 | 0.15 | -0.382 | 0.442 | -0.064 | -0.074 | -0.074 | 0 | 0.86 | 0.53 | 0.81 | 0.98 | 0.94 | 0.96 |
| 10 | 130755913 | rs11016570 | -0.965 | 0.902 | -1.118 | 0.768 | -0.022 | -0.02 | -0.02 | 0 | 0.34 | 0.24 | 0.43 | 0.47 | 0.4 | 0.47 |
| 10 | 130757816 | rs2060894 | -0.729 | 0.747 | -0.891 | 0.735 | 0.011 | 0.048 | 0.048 | 0 | 0.06 | 0.11 | 0.12 | 0.15 | 0.06 | 0.05 |
| 10 | 130761949 | rs10829531 | 0.194 | 0.566 | -1.617 | 0.162 | -0.018 | 0.012 | 0.012 | 0 | 0.21 | 0.02 | 0.17 | 0.13 | 0.16 | 0.2 |
| 10 | 130767319 | rs7091722 | -1.152 | 0.967 | -1.244 | 0.032 | -0.007 | 0.012 | 0.012 | 0 | 0.14 | 0.12 | 0.11 | 0.06 | 0.13 | 0.17 |
| 10 | 130767361 | rs10764836 | 0.276 | 1.201 | -1.303 | 0.329 | -0.01 | -0.038 | -0.038 | 0 | 0.36 | 0.18 | 0.33 | 0.51 | 0.35 | 0.48 |
| 10 | 130768215 | rs10764837 | 0.276 | 1.187 | -1.231 | 0.323 | -0.012 | -0.042 | -0.042 | 0 | 0.36 | 0.18 | 0.34 | 0.51 | 0.35 | 0.48 |
| 10 | 130769047 | rs7919827 | -0.026 | 0.745 | -1.501 | -0.18 | 0.004 | -0.008 | -0.008 | 0 | 0.37 | 0.33 | 0.3 | 0.1 | 0.28 | 0.33 |
| 10 | 130769170 | rs4751039 | 0.732 | 0.766 | -1.567 | -0.031 | -0.062 | -0.118 | -0.118 | 0 | 0.16 | 0.02 | 0.19 | 0.41 | 0.18 | 0.27 |
| 10 | 130770501 | rs7896687 | 0.035 | 0.607 | -1.294 | -0.123 | -0.021 | -0.038 | -0.038 | 0 | 0.07 | 0.24 | 0.04 | 0 | 0.02 | 0.01 |
| 10 | 130772391 | rs10829534 | -0.987 | 0.559 | -0.671 | -0.161 | -0.006 | -0.007 | -0.007 | 1.19 | 0.55 | 0.63 | 0.53 | 0.51 | 0.46 | 0.6 |
| 10 | 130772488 | rs11016580 | 1.359 | 0.548 | -0.579 | -0.124 | 0.004 | 0.018 | 0.018 | 0 | 0.23 | 0.13 | 0.14 | 0.08 | 0.2 | 0.23 |
| 10 | 130777415 | rs4750725 | -0.302 | 0.396 | -0.387 | -0.1 | 0.006 | 0.009 | 0.009 | 0 | 0.82 | 0.78 | 0.73 | 0.69 | 0.78 | 0.71 |
| 10 | 130778452 | rs7086428 | -0.233 | 0.247 | -0.344 | -0.231 | 0 | 0.001 | 0.001 | 0 | 0.66 | 0.63 | 0.49 | 0.58 | 0.66 | 0.57 |

| | | | | | | | | | | | | | | | | |
|---|---|---|---|---|---|---|---|---|---|---|---|---|---|---|---|---|
| 10 | 130782526 | rs7900076 | 1.721 | 0.446 | -0.625 | 0.09 | -0.007 | 0.013 | 0.013 | 0 | 0.22 | 0.1 | 0.14 | 0.17 | 0.21 | 0.22 |
| 10 | 130783543 | rs7904490 | 1.721 | 0.446 | -0.636 | 0.09 | -0.004 | 0.011 | 0.011 | 0 | 0.22 | 0.11 | 0.14 | 0.17 | 0.21 | 0.22 |
| 10 | 130784324 | rs10491032 | 0.906 | 0.41 | -0.852 | 0.131 | -0.059 | -0.098 | -0.098 | 3.388 | 0.14 | 0.02 | 0.18 | 0.32 | 0.18 | 0.26 |
| 10 | 130786548 | rs7094322 | 1.224 | 0.383 | -0.798 | 0.111 | 0.006 | 0.028 | 0.028 | 0.038 | 0.25 | 0.22 | 0.13 | 0.11 | 0.28 | 0.07 |
| 10 | 130786576 | rs11016590 | 0.29 | 0.399 | -0.581 | 0.424 | 0.003 | 0.022 | 0.022 | 0 | 0.16 | 0.19 | 0.2 | 0.32 | 0.18 | 0.26 |
| 10 | 130793666 | rs12357147 | -0.011 | 0.321 | -0.567 | 0.024 | -0.009 | -0.052 | -0.052 | 0.438 | 0.31 | 0.47 | 0.21 | 0.17 | 0.24 | 0.26 |
| 10 | 130794916 | rs7914150 | 0.705 | 0.83 | -1.374 | 0.339 | -0.031 | 0.004 | 0.004 | 0 | 0.23 | 0.09 | 0.15 | 0.17 | 0.22 | 0.25 |
| 10 | 130800090 | rs10734080 | -0.079 | 0.685 | -1.386 | 0.431 | -0.015 | 0.052 | 0.052 | 0.525 | 0.52 | 0.83 | 0.34 | 0.37 | 0.47 | 0.41 |
| 10 | 130801822 | rs4751041 | 0.227 | 0.707 | -1.473 | 0.588 | -0.006 | -0.009 | -0.009 | 0 | 0.17 | 0.18 | 0.35 | 0.32 | 0.23 | 0.23 |
| 10 | 130803158 | rs1810899 | -0.082 | 0.282 | -0.819 | 0.449 | -0.043 | -0.043 | -0.043 | 0 | 0.58 | 0.78 | 0.51 | 0.51 | 0.56 | 0.48 |
| 10 | 130804669 | rs992574 | 1.851 | 0.076 | -0.459 | -0.164 | 0 | -0.05 | -0.05 | 0.384 | 0.21 | 0.2 | 0.09 | 0.01 | 0.22 | 0.07 |
| 10 | 130807895 | rs10764838 | -0.562 | -0.31 | -0.124 | -0.159 | 0.004 | -0.028 | -0.028 | 0 | 0.21 | 0.37 | 0.25 | 0.19 | 0.22 | 0.19 |
| 10 | 130810465 | rs11016609 | -1.156 | -0.295 | -0.62 | -0.075 | -0.007 | 0.005 | 0.005 | 0.457 | 0.38 | 0.59 | 0.42 | 0.48 | 0.33 | 0.4 |
| 10 | 130811855 | rs16908946 | 0.349 | -0.342 | -0.629 | -0.096 | 0.025 | -0.03 | -0.03 | 1.766 | 0.22 | 0.15 | 0.21 | 0.3 | 0.19 | 0.25 |
| 10 | 130812555 | rs1869251 | 0.804 | -0.293 | -0.673 | -0.096 | -0.004 | 0.001 | 0.001 | 0 | 0.38 | 0.31 | 0.43 | 0.34 | 0.46 | 0.3 |
| 10 | 130817733 | rs914572 | -1.914 | 0.058 | -0.627 | -0.085 | -0.032 | -0.003 | -0.003 | 0 | 0.78 | 0.92 | 0.85 | 0.83 | 0.79 | 0.7 |
| 10 | 130817977 | rs914573 | -0.923 | 0.617 | 0.033 | 0.398 | 0.027 | 0.008 | 0.008 | 1.299 | 0.16 | 0.37 | 0.2 | 0.19 | 0.13 | 0.15 |
| 10 | 130833157 | rs11016630 | -1.183 | -0.111 | 0.202 | 0.705 | 0.016 | 0.116 | 0.116 | 0.565 | 0.06 | 0.29 | 0.07 | 0.11 | 0.02 | 0.01 |
| 10 | 130834698 | rs1556659 | -0.959 | 0.06 | -0.044 | 0.873 | -0.003 | -0.005 | -0.005 | 0.609 | 0.27 | 0.21 | 0.43 | 0.29 | 0.34 | 0.25 |
| 10 | 130835311 | rs12572420 | -0.631 | -0.172 | -0.024 | 0.936 | -0.034 | 0.111 | 0.111 | 0 | 0.23 | 0.4 | 0.22 | 0.4 | 0.16 | 0.3 |
| 10 | 130837490 | rs10491026 | -0.086 | -0.192 | -0.023 | 0.849 | -0.001 | 0.004 | 0.004 | 0 | 0.2 | 0.19 | 0.15 | 0.34 | 0.13 | 0.26 |
| 10 | 130837802 | rs16909010 | -1.579 | 0.262 | -0.204 | 0.861 | 0.027 | 0.054 | 0.054 | 0.02 | 0.1 | 0.28 | 0.14 | 0.2 | 0.09 | 0.16 |
| 10 | 130839012 | rs10829553 | -2.138 | 0.302 | -0.216 | 0.861 | -0.013 | 0.055 | 0.055 | 0 | 0.57 | 0.69 | 0.72 | 0.79 | 0.56 | 0.64 |
| 10 | 130839557 | rs10829555 | -1.692 | 0.42 | 0.147 | 1.169 | 0.013 | 0.036 | 0.036 | 0 | 0.11 | 0.28 | 0.14 | 0.19 | 0.1 | 0.13 |
| 10 | 130839812 | rs11016638 | -0.532 | 0.603 | -0.56 | 1.1 | -0.004 | 0.003 | 0.003 | 0 | 0.06 | 0.07 | 0.07 | 0.09 | 0.08 | 0.16 |
| 10 | 130840798 | rs10764840 | -0.798 | 0.575 | -0.706 | 1.141 | 0.008 | -0.011 | -0.011 | 0 | 0.36 | 0.65 | 0.31 | 0.54 | 0.31 | 0.46 |
| 10 | 130842093 | rs11598539 | -0.198 | 1.189 | -1.165 | 1.308 | -0.057 | -0.073 | -0.073 | 0 | 0.13 | 0.02 | 0.38 | 0.22 | 0.21 | 0.15 |
| 10 | 130842899 | rs10829557 | -0.656 | 0.687 | -0.476 | 1.178 | -0.001 | 0.005 | 0.005 | 1.412 | 0.22 | 0.25 | 0.18 | 0.41 | 0.18 | 0.42 |
| 10 | 130844717 | rs12355487 | -1.103 | 1.068 | -0.289 | 0.676 | -0.003 | -0.011 | -0.011 | 0 | 0.11 | 0.06 | 0.3 | 0.25 | 0.15 | 0.15 |
| 10 | 130844805 | rs7101205 | 0.89 | 1.16 | 0.43 | -0.038 | 0.003 | -0.07 | -0.07 | 0.123 | 0.8 | 0.64 | 0.85 | 0.93 | 0.77 | 0.8 |
| 10 | 130849424 | rs11016642 | -1.653 | 1.5 | 1.248 | -0.165 | 0.059 | -0.044 | -0.044 | 0.677 | 0.1 | 0.34 | 0.09 | 0.02 | 0.14 | 0.06 |
| 10 | 130851955 | rs10829558 | -3.16 | 1.48 | 1.567 | -0.194 | 0.022 | -0.026 | -0.026 | 0 | 0.25 | 0.44 | 0.39 | 0.24 | 0.28 | 0.28 |
| 10 | 130852170 | rs12265750 | -1.979 | 1.482 | 1.516 | -0.195 | 0.066 | -0.045 | -0.045 | 0.189 | 0.1 | 0.34 | 0.09 | 0.03 | 0.15 | 0.1 |
| 10 | 130852739 | rs11016646 | -3.037 | 1.478 | 1.521 | -0.178 | 0.014 | -0.027 | -0.027 | 0 | 0.24 | 0.35 | 0.22 | 0.2 | 0.28 | 0.28 |
| 10 | 130852876 | rs7079442 | -3.463 | 1.364 | 2.152 | -0.244 | 0.001 | -0.008 | -0.008 | 0 | 0.27 | 0.73 | 0.41 | 0.24 | 0.29 | 0.28 |
| 10 | 130854095 | rs4751050 | -2.116 | 1.388 | 2.343 | -0.211 | 0.019 | -0.071 | -0.071 | 0 | 0.16 | 0.27 | 0.11 | 0.02 | 0.2 | 0.11 |
| 10 | 130854243 | rs4751051 | -2.802 | 1.375 | 2.386 | -0.243 | 0.01 | -0.11 | -0.11 | 0.145 | 0.2 | 0.63 | 0.15 | 0.02 | 0.2 | 0.11 |
| 10 | 130856644 | rs7099230 | -3.459 | 1.533 | 2.571 | -0.013 | 0.018 | 0 | 0 | 0.255 | 0.19 | 0.69 | 0.36 | 0.24 | 0.21 | 0.2 |
| 10 | 130859801 | rs10829565 | -2.478 | 1.561 | 2.685 | -0.041 | 0.044 | 0.118 | 0.118 | 0.648 | 0.1 | 0.45 | 0.29 | 0.23 | 0.08 | 0.17 |
| 10 | 130860965 | rs1377273 | -3.52 | 1.504 | 2.652 | -0.073 | 0.008 | -0.007 | -0.007 | 0 | 0.22 | 0.76 | 0.37 | 0.23 | 0.23 | 0.24 |
| 10 | 130863266 | rs1888874 | -2.042 | 1.373 | 2.6 | -0.193 | 0.096 | -0.038 | -0.038 | 0 | 0.1 | 0.37 | 0.07 | 0.01 | 0.13 | 0.05 |
| 10 | 130863854 | rs10829566 | -3.065 | 1.374 | 2.436 | -0.191 | 0.015 | -0.088 | -0.088 | 0 | 0.17 | 0.69 | 0.12 | 0.01 | 0.18 | 0.12 |
| 10 | 130874125 | rs4751058 | -1.759 | 1.33 | 2.389 | -0.084 | 0.025 | 0.018 | 0.018 | 0 | 0.11 | 0.18 | 0.28 | 0.22 | 0.18 | 0.15 |
| 10 | 130878393 | rs12412608 | -1.269 | 0.948 | 2.545 | 0.023 | 0.014 | 0.076 | 0.076 | 0 | 0.06 | 0.18 | 0.24 | 0.22 | 0.06 | 0.14 |
| 10 | 130882099 | rs12269683 | -2.857 | 0.897 | 2.74 | -0.012 | 0.025 | 0.004 | 0.004 | 0.422 | 0.05 | 0.31 | 0.04 | 0.02 | 0.02 | 0.05 |
| 10 | 130883170 | rs2123425 | -3.014 | 0.905 | 2.918 | -0.009 | 0.062 | 0.037 | 0.037 | 1.229 | 0.05 | 0.38 | 0.04 | 0.01 | 0.02 | 0.04 |
| 10 | 130889679 | rs2167743 | -2.863 | 0.891 | 2.944 | -0.015 | 0.051 | 0.027 | 0.027 | 0.875 | 0.06 | 0.38 | 0.04 | 0.01 | 0.02 | 0.05 |
| 10 | 130890818 | rs7069581 | -2.509 | 1.689 | 2.205 | -0.269 | -0.011 | 0.023 | 0.023 | 1.315 | 0.27 | 0.8 | 0.4 | 0.24 | 0.28 | 0.25 |
| 10 | 130894033 | rs7084245 | -2.451 | 1.556 | 2.183 | -0.423 | 0.094 | 0.055 | 0.055 | 0.977 | 0.05 | 0.41 | 0.04 | 0 | 0.02 | 0.03 |
| 10 | 130900389 | rs2039372 | -1.194 | 1.629 | 1.368 | -0.432 | 0.017 | -0.084 | -0.084 | 0 | 0.17 | 0.25 | 0.11 | 0 | 0.19 | 0.09 |
| 10 | 130902337 | rs12778834 | -1.194 | 1.629 | 1.368 | -0.432 | 0.017 | -0.084 | -0.084 | 0.185 | 0.17 | 0.25 | 0.11 | 0 | 0.19 | 0.09 |
| 10 | 130904542 | rs11598274 | -0.925 | 1.699 | 0.72 | -0.427 | -0.032 | 0.042 | 0.042 | 0.149 | 0.15 | 0.07 | 0.1 | 0 | 0.21 | 0.11 |
| 10 | 130905792 | rs12774599 | -1.052 | 1.624 | 0.979 | -0.427 | -0.002 | -0.024 | -0.024 | 0 | 0.14 | 0.12 | 0.1 | 0 | 0.19 | 0.11 |
| 10 | 130910213 | rs2218983 | 0.331 | 1.555 | 1.168 | -0.459 | -0.003 | -0.017 | -0.017 | 0 | 0.87 | 0.91 | 0.9 | 1 | 0.82 | 0.94 |
| 10 | 130913600 | rs12775933 | -1.287 | 1.258 | 0.875 | -0.584 | 0.032 | 0.035 | 0.035 | 1.259 | 0.51 | 0.45 | 0.72 | 0.75 | 0.67 | 0.56 |
| 10 | 130915346 | rs11016681 | 1.502 | 0.945 | 0.706 | -0.732 | 0.093 | 0.081 | 0.081 | 0.838 | 0.42 | 0.07 | 0.21 | 0.25 | 0.31 | 0.38 |
| 10 | 130917824 | rs10764851 | -1.182 | 1.046 | 0.799 | -0.68 | 0.073 | 0.072 | 0.072 | 1.403 | 0.5 | 0.41 | 0.72 | 0.75 | 0.69 | 0.61 |
| 10 | 130919703 | rs11016684 | -0.585 | 1.322 | 1.207 | -0.695 | 0.009 | -0.014 | -0.014 | 0.516 | 0.39 | 0.45 | 0.65 | 0.75 | 0.49 | 0.5 |
| 10 | 130919942 | rs10764855 | 0.613 | 1.337 | 1.038 | -0.692 | 0.029 | -0.122 | -0.122 | 0 | 0.82 | 0.47 | 0.85 | 1 | 0.8 | 0.89 |
| 10 | 130925103 | rs1452880 | -0.328 | 1.343 | 0.29 | -0.686 | -0.035 | 0.026 | 0.026 | 0 | 0.36 | 0.3 | 0.64 | 0.75 | 0.49 | 0.49 |
| 10 | 130926027 | rs7093724 | 0.967 | 0.951 | -0.252 | -0.502 | 0.094 | 0.098 | 0.098 | 2.026 | 0.45 | 0.16 | 0.21 | 0.25 | 0.31 | 0.39 |
| 10 | 130931337 | rs2123420 | -0.351 | 1.68 | -0.752 | 0.083 | -0.067 | -0.013 | -0.013 | 0 | 0.23 | 0.11 | 0.43 | 0.3 | 0.41 | 0.23 |
| 10 | 130931432 | rs1563040 | 0.933 | 0.575 | -0.86 | 0.15 | 0.045 | 0.066 | 0.066 | 2.594 | 0.42 | 0.04 | 0.21 | 0.25 | 0.31 | 0.38 |
| 10 | 130932314 | rs10741184 | 0.587 | -0.18 | 0.388 | 0.107 | -0.046 | -0.05 | -0.05 | 0 | 0.75 | 0.54 | 0.83 | 0.87 | 0.85 | 0.92 |
| 10 | 130933744 | rs914570 | 0.511 | -0.166 | 0.304 | 0.333 | 0.023 | 0.012 | 0.012 | 0 | 0.52 | 0.4 | 0.51 | 0.61 | 0.63 | 0.63 |
| 10 | 130934779 | rs1998824 | 1.306 | -0.376 | 1.086 | 0.366 | -0.011 | 0.174 | 0.174 | 0.68 | 0.88 | 0.63 | 0.68 | 0.73 | 0.91 | 0.81 |
| 10 | 130935586 | rs4750741 | 0.739 | -0.376 | 0.861 | 0.471 | -0.002 | 0.053 | 0.053 | 0 | 0.93 | 0.79 | 0.81 | 0.9 | 0.94 | 0.87 |
| 10 | 130937457 | rs7074836 | 0.299 | -0.114 | 0.603 | 0.415 | -0.001 | 0.015 | 0.015 | 0 | 0.92 | 0.88 | 0.73 | 0.82 | 0.94 | 0.91 |
| 10 | 130938873 | rs7912442 | 0.839 | -0.149 | 0.627 | 0.329 | -0.002 | 0.142 | 0.142 | 0.957 | 0.84 | 0.53 | 0.63 | 0.71 | 0.88 | 0.79 |

| | | | | | | | | | | | | | | | | |
|---|---|---|---|---|---|---|---|---|---|---|---|---|---|---|---|---|
| 10 | 130939105 | rs7916199 | 1.062 | -0.351 | 0.456 | 0.151 | 0.018 | 0.018 | 0.018 | 0 | 0.46 | 0.28 | 0.46 | 0.6 | 0.58 | 0.61 |
| 10 | 130939211 | rs7912924 | -0.176 | -0.347 | 0.375 | 0.075 | 0.027 | 0.066 | 0.066 | 0 | 0.38 | 0.25 | 0.18 | 0.11 | 0.3 | 0.18 |
| 10 | 130941163 | rs10764860 | 0.599 | 0.018 | -0.214 | 0.258 | 0.018 | 0.024 | 0.024 | 0 | 0.46 | 0.3 | 0.46 | 0.61 | 0.58 | 0.61 |
| 10 | 130947216 | rs7899979 | -0.832 | 0.349 | -0.286 | 0.253 | 0.004 | 0 | 0 | 0 | 0.54 | 0.65 | 0.58 | 0.46 | 0.53 | 0.44 |
| 10 | 130947838 | rs10128422 | 0.508 | 0.315 | -0.147 | 0.239 | 0.006 | -0.002 | -0.002 | 0 | 0.46 | 0.3 | 0.42 | 0.54 | 0.47 | 0.55 |
| 10 | 130951498 | rs11016694 | 0.545 | 0.285 | -0.071 | 0.612 | 0.001 | -0.001 | -0.001 | 0 | 0.45 | 0.25 | 0.35 | 0.42 | 0.47 | 0.54 |
| 10 | 130952994 | rs10734083 | 0.424 | 0.093 | 0.319 | -0.228 | -0.001 | 0.003 | 0.003 | 0 | 0.54 | 0.37 | 0.5 | 0.55 | 0.52 | 0.56 |
| 10 | 130954227 | rs7475003 | 0.53 | 0.103 | 0.318 | -0.241 | -0.006 | 0.016 | 0.016 | 0 | 0.53 | 0.29 | 0.49 | 0.55 | 0.52 | 0.56 |
| 10 | 130954407 | rs6482722 | 0.043 | 0.108 | 0.487 | -0.553 | 0.018 | 0.031 | 0.031 | 0 | 0.7 | 0.53 | 0.6 | 0.55 | 0.67 | 0.61 |
| 10 | 130957352 | rs11016695 | 0.31 | 0.115 | 0.811 | -0.577 | 0.008 | 0.016 | 0.016 | 0 | 0.71 | 0.6 | 0.6 | 0.56 | 0.67 | 0.59 |
| 10 | 130957528 | rs10829585 | 1.096 | 0.01 | 0.9 | -1.112 | -0.007 | 0.059 | 0.059 | 0 | 0.55 | 0.28 | 0.56 | 0.74 | 0.55 | 0.58 |
| 10 | 130957661 | rs7901035 | 0.561 | -0.234 | 1.432 | -0.822 | 0.007 | -0.01 | -0.01 | 1.464 | 0.7 | 0.62 | 0.67 | 0.75 | 0.67 | 0.61 |
| 10 | 130963500 | rs2209592 | 0.556 | -0.28 | 1.491 | -0.834 | 0.014 | -0.016 | -0.016 | 0 | 0.71 | 0.57 | 0.66 | 0.74 | 0.67 | 0.61 |
| 10 | 130975999 | rs716909 | 0.989 | -0.32 | 1.126 | -0.858 | 0.002 | 0.027 | 0.027 | 0 | 0.64 | 0.27 | 0.62 | 0.74 | 0.67 | 0.58 |
| 10 | 130984128 | rs7905824 | 0.409 | -0.659 | 0.84 | -1.501 | -0.006 | 0.007 | 0.007 | 0 | 0.74 | 0.76 | 0.72 | 0.83 | 0.71 | 0.73 |
| 10 | 130984835 | rs1452859 | 1.527 | -0.858 | 1.304 | -1.604 | 0.004 | 0.012 | 0.012 | 0 | 0.6 | 0.36 | 0.64 | 0.84 | 0.66 | 0.64 |
| 10 | 130986137 | rs997161 | 0.457 | -0.794 | 1.263 | -1.573 | -0.012 | 0.015 | 0.015 | 0 | 0.75 | 0.78 | 0.72 | 0.84 | 0.71 | 0.73 |
| 10 | 130998435 | rs4264079 | 1.35 | -0.826 | 0.914 | -1.54 | -0.001 | -0.015 | -0.015 | 0.259 | 0.6 | 0.41 | 0.64 | 0.81 | 0.66 | 0.64 |
| 10 | 131003181 | rs11016706 | -0.663 | -0.681 | 0.507 | -1.439 | -0.04 | 0.02 | 0.02 | 0 | 0.26 | 0.12 | 0.28 | 0.2 | 0.29 | 0.27 |
| 10 | 131009253 | rs4751072 | 1.215 | -0.791 | 0.51 | -1.827 | 0.027 | 0.163 | 0.163 | 0 | 0.52 | 0.25 | 0.62 | 0.73 | 0.62 | 0.59 |
| 10 | 131013164 | rs9888101 | -0.957 | -0.658 | 0.575 | -1.751 | -0.012 | -0.05 | -0.05 | 0 | 0.31 | 0.41 | 0.31 | 0.21 | 0.24 | 0.33 |
| 10 | 131023108 | rs543469 | 0.42 | -0.463 | 0.326 | -1.56 | -0.036 | -0.143 | -0.143 | 0 | 0.77 | 0.54 | 0.9 | 0.94 | 0.84 | 0.9 |
| 10 | 131028858 | rs1990783 | 0.077 | -0.519 | -0.143 | -1.291 | 0.001 | -0.028 | -0.028 | 0 | 0.1 | 0.01 | 0.07 | 0.08 | 0.1 | 0.13 |
| 10 | 131032716 | rs562950 | 0.509 | -0.447 | 0.048 | -1.364 | -0.002 | -0.031 | -0.031 | 0 | 0.61 | 0.48 | 0.65 | 0.68 | 0.67 | 0.64 |
| 10 | 131033938 | rs555788 | 0.89 | -0.269 | -0.405 | -1.273 | -0.005 | 0.069 | 0.069 | 0 | 0.46 | 0.26 | 0.54 | 0.61 | 0.54 | 0.48 |
| 10 | 131035372 | rs12258833 | -0.816 | 0.073 | 0.065 | -1.558 | -0.001 | -0.017 | -0.017 | 0 | 0.38 | 0.42 | 0.36 | 0.26 | 0.31 | 0.39 |
| 10 | 131037021 | rs570306 | 0.459 | -0.303 | 0.184 | -1.332 | -0.003 | -0.013 | -0.013 | 0 | 0.82 | 0.78 | 0.93 | 0.91 | 0.88 | 0.85 |
| 10 | 131040813 | rs507033 | 0.934 | -0.4 | 0.043 | -1.534 | -0.028 | -0.113 | -0.113 | 0.207 | 0.68 | 0.53 | 0.85 | 0.9 | 0.76 | 0.77 |
| 10 | 131041657 | rs476628 | 0.995 | -0.401 | 0 | -1.535 | -0.031 | -0.115 | -0.115 | 0 | 0.67 | 0.51 | 0.84 | 0.9 | 0.76 | 0.77 |
| 10 | 131051873 | rs534451 | 0.536 | 0.15 | -0.605 | -1.056 | 0.006 | 0.012 | 0.012 | 0 | 0.83 | 0.86 | 0.93 | 0.98 | 0.88 | 0.86 |
| 10 | 131054677 | rs7895517 | -0.772 | 0.508 | -0.3 | -1.023 | 0.003 | -0.071 | -0.071 | 0.019 | 0.15 | 0.2 | 0.11 | 0 | 0.19 | 0.06 |
| 10 | 131054701 | rs6482726 | -0.859 | 0.561 | -0.319 | -1.098 | 0.004 | -0.074 | -0.074 | 0.395 | 0.16 | 0.2 | 0.11 | 0 | 0.2 | 0.08 |
| 10 | 131054861 | rs7085947 | -0.47 | 0.507 | -0.517 | -1.395 | -0.002 | 0.005 | 0.005 | 1.475 | 0.13 | 0.08 | 0.08 | 0.01 | 0.16 | 0.06 |
| 10 | 131055789 | rs10741187 | 0.537 | 0.618 | -0.699 | -1.268 | -0.009 | 0.024 | 0.024 | 0.149 | 0.42 | 0.38 | 0.33 | 0.7 | 0.44 | 0.65 |
| 10 | 131056071 | rs580556 | -0.12 | 0.286 | -0.679 | -1.493 | 0 | 0.001 | 0.001 | 0 | 0.44 | 0.55 | 0.61 | 0.3 | 0.46 | 0.27 |
| 10 | 131056779 | rs11016723 | 1.088 | 0.46 | -1.273 | -0.778 | -0.06 | -0.103 | -0.103 | 0 | 0.32 | 0.08 | 0.26 | 0.46 | 0.38 | 0.59 |
| 10 | 131060844 | rs2803458 | 0.913 | 0.263 | -0.273 | -0.23 | -0.039 | -0.124 | -0.124 | 1.536 | 0.8 | 0.63 | 0.9 | 0.99 | 0.89 | 0.92 |
| 10 | 131073967 | rs12358748 | -0.301 | 0.858 | -0.781 | 0.016 | -0.024 | -0.018 | -0.018 | 0 | 0.1 | 0.03 | 0.09 | 0.12 | 0.12 | 0.12 |
| 10 | 131083245 | rs4750744 | -0.944 | 0.934 | -0.834 | -0.036 | -0.035 | 0.047 | 0.047 | 0 | 0.11 | 0.03 | 0.07 | 0.02 | 0.14 | 0.07 |
| 10 | 131092082 | rs2246538 | 1.894 | 1.4 | -0.015 | 0.691 | 0.016 | 0.035 | 0.035 | 6.36 | 0.72 | 0.41 | 0.77 | 0.61 | 0.71 | 0.84 |
| 10 | 131092913 | rs7917272 | -0.793 | 1.412 | 0.485 | 0.821 | 0.057 | 0.083 | 0.083 | 0.161 | 0.08 | 0.42 | 0.06 | 0.05 | 0.04 | 0 |
| 10 | 131093567 | rs2246405 | 0.468 | 1.478 | 0.295 | 0.858 | 0.006 | 0.042 | 0.042 | 4.786 | 0.9 | 0.87 | 0.9 | 0.68 | 0.89 | 0.92 |
| 10 | 131101856 | rs12356849 | -0.836 | 1.439 | 0.591 | 0.798 | -0.002 | 0.003 | 0.003 | 0 | 0.12 | 0.13 | 0.08 | 0.06 | 0.15 | 0.07 |
| 10 | 131107275 | rs2032704 | -2.324 | 1.681 | 0.848 | 1.01 | -0.014 | -0.018 | -0.018 | 1.313 | 0.25 | 0.23 | 0.22 | 0.4 | 0.32 | 0.16 |
| 10 | 131107692 | rs11016754 | -0.665 | 1.707 | 0.805 | 0.982 | -0.038 | 0.044 | 0.044 | 0 | 0.11 | 0.04 | 0.07 | 0.02 | 0.16 | 0.07 |
| 10 | 131114514 | rs2622441 | -1.273 | 1.369 | 0.103 | 0.756 | -0.004 | 0.011 | 0.011 | 0.003 | 0.64 | 0.76 | 0.73 | 0.63 | 0.65 | 0.34 |
| 10 | 131119642 | rs12267072 | -0.706 | 1.487 | 0.113 | 0.984 | 0.006 | 0.048 | 0.048 | 4.183 | 0.13 | 0.13 | 0.15 | 0.4 | 0.16 | 0.09 |
| 10 | 131128952 | rs2542615 | -1.152 | 1.383 | 0.022 | 0.967 | 0 | 0.012 | 0.012 | 0.019 | 0.63 | 0.7 | 0.72 | 0.63 | 0.64 | 0.33 |
| 10 | 131129656 | rs2622436 | 0.657 | 1.244 | 0.191 | 0.854 | 0.032 | 0.055 | 0.055 | 0.411 | 0.37 | 0.2 | 0.49 | 0.2 | 0.33 | 0.18 |
| 10 | 131132672 | rs6482731 | 0.441 | 1.063 | 0.707 | 1.128 | -0.001 | -0.001 | -0.001 | 1.247 | 0.39 | 0.4 | 0.28 | 0.43 | 0.41 | 0.66 |
| 10 | 131140513 | rs7900988 | 0.918 | 1.033 | 0.4 | 1.419 | -0.022 | 0.043 | 0.043 | 0.786 | 0.34 | 0.19 | 0.23 | 0.26 | 0.36 | 0.65 |
| 10 | 131144113 | rs11598707 | -0.631 | 0.702 | 0.513 | 1.42 | 0.001 | 0.016 | 0.016 | 6.539 | 0.09 | 0.14 | 0.16 | 0.15 | 0.07 | 0.04 |
| 10 | 131146611 | rs2542612 | 0.041 | 0.395 | 0.555 | 1.206 | 0.025 | -0.02 | -0.02 | 0 | 0.43 | 0.23 | 0.47 | 0.42 | 0.36 | 0.26 |
| 10 | 131147692 | rs2676441 | 0.091 | 0.369 | 0.223 | 1.169 | 0.024 | -0.032 | -0.032 | 0.746 | 0.42 | 0.16 | 0.47 | 0.41 | 0.36 | 0.26 |
| 10 | 131151105 | rs2676446 | 0.672 | 0.491 | 0.186 | 0.693 | 0.007 | 0.067 | 0.067 | 0 | 0.51 | 0.24 | 0.48 | 0.68 | 0.49 | 0.6 |
| 10 | 131157446 | rs1554224 | -0.367 | 0.164 | 0.193 | -0.13 | 0.007 | 0.041 | 0.041 | 0 | 0.42 | 0.26 | 0.6 | 0.37 | 0.37 | 0.29 |
| 10 | 131160866 | rs2111627 | 1.048 | -0.076 | 0.224 | -0.756 | 0.024 | 0.211 | 0.211 | 1.665 | 0.54 | 0.09 | 0.4 | 0.8 | 0.62 | 0.72 |
| 10 | 131172666 | rs12253191 | -1.446 | 0.011 | 1.322 | -0.932 | -0.012 | -0.069 | -0.069 | 0.683 | 0.17 | 0.49 | 0.21 | 0.15 | 0.14 | 0.17 |
| 10 | 131179027 | rs2676445 | -0.09 | -0.548 | 1.498 | -1.058 | -0.012 | -0.09 | -0.09 | 0.924 | 0.73 | 0.63 | 0.64 | 0.98 | 0.77 | 0.88 |
| 10 | 131188383 | rs2542642 | -0.198 | -0.455 | 1.426 | -1.018 | -0.005 | -0.055 | -0.055 | 0.042 | 0.73 | 0.7 | 0.66 | 0.98 | 0.77 | 0.88 |
| 10 | 131192152 | rs7070281 | -1.36 | 0.443 | 1.11 | -0.256 | -0.02 | -0.013 | -0.013 | 0 | 0.1 | 0.12 | 0.15 | 0.12 | 0.06 | 0.11 |
| 10 | 131196160 | rs1711677 | 0.353 | 0.781 | 1.18 | -0.694 | 0.01 | -0.112 | -0.112 | 0.908 | 0.74 | 0.66 | 0.66 | 0.99 | 0.73 | 0.8 |
| 10 | 131196447 | rs1891008 | -0.414 | 0.659 | 1.49 | -0.611 | 0.001 | -0.051 | -0.051 | 0 | 0.22 | 0.28 | 0.3 | 0.01 | 0.24 | 0.21 |
| 10 | 131200443 | rs2792980 | 2.379 | 0.606 | 2.221 | 0.303 | 0.005 | -0.007 | -0.007 | 0.034 | 0.84 | 0.17 | 0.8 | 0.76 | 0.89 | 0.81 |
| 10 | 131203176 | rs1711644 | 2.405 | 0.541 | 2.853 | 0.287 | -0.034 | -0.052 | -0.052 | 0.718 | 0.89 | 0.36 | 0.92 | 0.99 | 0.96 | 0.9 |
| 10 | 131203805 | rs1771445 | 2.893 | 0.51 | 2.529 | 0.292 | 0.006 | -0.008 | -0.008 | 0.391 | 0.84 | 0.19 | 0.8 | 0.76 | 0.9 | 0.81 |
| 10 | 131205707 | rs7076371 | -2.709 | 0.645 | 3.015 | 0.298 | 0.063 | 0.188 | 0.188 | 1.474 | 0.1 | 0.48 | 0.16 | 0.23 | 0.09 | 0.16 |
| 10 | 131210677 | rs2005492 | -2.993 | 0.658 | 2.732 | 0.279 | -0.002 | 0.052 | 0.052 | 1.632 | 0.12 | 0.66 | 0.19 | 0.25 | 0.1 | 0.17 |
| 10 | 131212448 | rs7099267 | -2.412 | 0.828 | 2.541 | 0.348 | 0.048 | 0.137 | 0.137 | 0 | 0.09 | 0.25 | 0.16 | 0.24 | 0.1 | 0.15 |

| | | | | | | | | | | | | | | | | |
|---|---|---|---|---|---|---|---|---|---|---|---|---|---|---|---|---|
| 10 | 131218373 | rs1711639 | -1.139 | 1.037 | 1.727 | 0.101 | 0.003 | -0.001 | -0.001 | 0 | 0.07 | 0.08 | 0.05 | 0 | 0.06 | 0.03 |
| 10 | 131225551 | rs6482736 | 2.839 | 1.085 | 2.736 | 0.265 | -0.001 | 0.018 | 0.018 | 1.601 | 0.83 | 0.29 | 0.77 | 0.78 | 0.84 | 0.75 |
| 10 | 131233033 | rs7084480 | 2.773 | 1.329 | 2.674 | -0.025 | 0 | -0.001 | -0.001 | 0.005 | 0.75 | 0.27 | 0.68 | 0.77 | 0.73 | 0.73 |
| 10 | 131236879 | rs10734086 | 2.671 | 1.067 | 2.721 | 0.213 | 0.01 | 0.107 | 0.107 | 0.664 | 0.86 | 0.48 | 0.79 | 0.77 | 0.88 | 0.8 |
| 10 | 131237179 | rs3864859 | 1.784 | 1.111 | 2.589 | 0.188 | 0.022 | -0.051 | -0.051 | 0 | 0.9 | 0.67 | 0.91 | 1 | 0.91 | 0.89 |
| 10 | 131239375 | rs1626983 | -1.441 | 1.245 | 1.688 | 0.227 | 0.008 | -0.037 | -0.037 | 0 | 0.08 | 0.16 | 0.07 | 0 | 0.09 | 0.09 |
| 10 | 131241249 | rs1711656 | -2.015 | 1.205 | 1.556 | 0.268 | 0.033 | 0.055 | 0.055 | 0 | 0.13 | 0.31 | 0.12 | 0.2 | 0.16 | 0.25 |
| 10 | 131250665 | rs1762444 | -0.899 | 1.314 | 1.009 | 0.359 | 0.004 | 0.006 | 0.006 | 0 | 0.33 | 0.38 | 0.37 | 0.53 | 0.38 | 0.4 |
| 10 | 131253796 | rs1771449 | -1.158 | 1.502 | 1.411 | 0.14 | -0.058 | -0.036 | -0.036 | 0 | 0.39 | 0.78 | 0.5 | 0.54 | 0.46 | 0.44 |
| 10 | 131256585 | rs11016798 | 0.816 | 1.312 | 1.68 | 0.061 | -0.125 | -0.086 | -0.086 | 0.826 | 0.59 | 0.12 | 0.5 | 0.45 | 0.54 | 0.55 |
| 10 | 131260478 | rs7070624 | -0.815 | 1.288 | 2.183 | 0.088 | 0.024 | 0.02 | 0.02 | 0 | 0.31 | 0.55 | 0.38 | 0.35 | 0.32 | 0.39 |
| 10 | 131262486 | rs1762440 | 1.036 | 0.74 | 1.703 | -0.434 | -0.006 | 0.009 | 0.009 | 0 | 0.55 | 0.38 | 0.5 | 0.62 | 0.56 | 0.55 |
| 10 | 131265852 | rs12356501 | 0.795 | 0.57 | 1.471 | -0.517 | -0.061 | -0.015 | -0.015 | 1.078 | 0.63 | 0.15 | 0.55 | 0.64 | 0.61 | 0.56 |
| 10 | 131267193 | rs7069786 | -0.935 | 0.239 | 2.088 | -0.467 | -0.008 | 0.216 | 0.216 | 0.552 | 0.13 | 0.51 | 0.18 | 0.33 | 0.09 | 0.2 |
| 10 | 131269508 | rs4751086 | 0.064 | 0.192 | 2.112 | -0.489 | -0.001 | 0.043 | 0.043 | 0 | 0.93 | 0.92 | 0.88 | 0.72 | 0.92 | 0.87 |
| 10 | 131269840 | rs1762433 | 1.566 | 0.541 | 1.663 | -0.756 | -0.103 | -0.005 | -0.005 | 0 | 0.61 | 0.15 | 0.52 | 0.66 | 0.56 | 0.58 |
| 10 | 131271838 | rs1762431 | 1.877 | 0.535 | 1.836 | -0.722 | -0.018 | 0.044 | 0.044 | 1.201 | 0.85 | 0.33 | 0.81 | 0.67 | 0.9 | 0.81 |
| 10 | 131271870 | rs7079768 | -1.359 | 0.542 | 1.927 | -0.718 | 0.056 | 0.057 | 0.057 | 0.451 | 0.06 | 0.45 | 0.06 | 0.04 | 0.02 | 0.05 |
| 10 | 131273707 | rs1762429 | 1.257 | 0.616 | 1.532 | -0.736 | -0.084 | -0.028 | -0.028 | 0 | 0.64 | 0.14 | 0.55 | 0.64 | 0.61 | 0.62 |
| 10 | 131278410 | rs1711667 | 1.152 | 0.413 | 1.761 | -1.008 | -0.007 | -0.024 | -0.024 | 0.004 | 0.6 | 0.63 | 0.61 | 0.92 | 0.58 | 0.67 |
| 10 | 131280140 | rs1762427 | -0.281 | 0.634 | 1.692 | -0.842 | 0.002 | 0.079 | 0.079 | 0 | 0.92 | 0.89 | 0.87 | 0.71 | 0.91 | 0.86 |
| 10 | 131295828 | rs10741190 | 1.349 | 0.606 | 2.053 | -0.88 | -0.02 | 0.009 | 0.009 | 0 | 0.69 | 0.43 | 0.7 | 0.65 | 0.77 | 0.73 |
| 10 | 131299647 | rs12251275 | -0.291 | 0.605 | 2.014 | -0.974 | 0.059 | -0.091 | -0.091 | 0 | 0.22 | 0.32 | 0.28 | 0.03 | 0.29 | 0.18 |
| 10 | 131299704 | rs12246239 | 0.021 | 0.607 | 1.917 | -0.973 | -0.075 | 0.079 | 0.079 | 0.2 | 0.2 | 0.08 | 0.25 | 0.03 | 0.29 | 0.19 |
| 10 | 131307521 | rs7080949 | 0.661 | 0.541 | 1.831 | -0.991 | 0.006 | 0.066 | 0.066 | 0.785 | 0.12 | 0.01 | 0.09 | 0.02 | 0.08 | 0.03 |
| 10 | 131308055 | rs11016823 | 0.049 | 0.579 | 1.842 | -0.992 | -0.075 | 0.078 | 0.078 | 0.21 | 0.2 | 0.08 | 0.25 | 0.03 | 0.29 | 0.19 |
| 10 | 131309444 | rs7922772 | -0.753 | 0.561 | 1.903 | -1 | -0.002 | -0.028 | -0.028 | 0 | 0.2 | 0.2 | 0.14 | 0.05 | 0.13 | 0.12 |
| 10 | 131309979 | rs7087541 | 1.729 | 0.542 | 1.971 | -0.967 | 0.043 | 0.063 | 0.063 | 0 | 0.5 | 0.24 | 0.44 | 0.64 | 0.48 | 0.56 |
| 10 | 131310420 | rs12248682 | -0.497 | 0.53 | 1.868 | -0.842 | 0.052 | 0.303 | 0.303 | 0 | 0.09 | 0.33 | 0.16 | 0.28 | 0.09 | 0.13 |
| 10 | 131319562 | rs12256491 | -1.331 | 0.5 | 1.654 | -0.888 | -0.031 | 0.043 | 0.043 | 0.268 | 0.24 | 0.49 | 0.26 | 0.33 | 0.17 | 0.2 |
| 10 | 131320026 | rs12264367 | -1.331 | 0.5 | 1.654 | -0.888 | -0.031 | 0.043 | 0.043 | 0.268 | 0.24 | 0.49 | 0.26 | 0.33 | 0.17 | 0.2 |
| 10 | 131323689 | rs10741194 | 1.429 | 0.507 | 1.682 | -0.912 | -0.01 | 0.005 | 0.005 | 0 | 0.71 | 0.41 | 0.7 | 0.67 | 0.78 | 0.75 |
| 10 | 131324278 | rs12266768 | -0.347 | 0.52 | 1.811 | -0.906 | -0.046 | -0.114 | -0.114 | 0.057 | 0.17 | 0.38 | 0.12 | 0.02 | 0.08 | 0.07 |
| 10 | 131325138 | rs12268726 | -1.237 | 0.535 | 1.563 | -0.834 | -0.031 | 0.027 | 0.027 | 0.268 | 0.24 | 0.49 | 0.26 | 0.3 | 0.17 | 0.2 |
| 10 | 131325299 | rs12268840 | -0.018 | 0.721 | 1.255 | -0.93 | -0.058 | 0.06 | 0.06 | 0.21 | 0.2 | 0.09 | 0.25 | 0.03 | 0.29 | 0.19 |
| 10 | 131331612 | rs11016833 | -0.004 | 0.722 | 1.256 | -0.93 | -0.058 | 0.059 | 0.059 | 0.034 | 0.2 | 0.09 | 0.25 | 0.03 | 0.29 | 0.19 |
| 10 | 131338714 | rs7909964 | 2.145 | 0.501 | 1.312 | -1.043 | 0.026 | 0.024 | 0.024 | 0 | 0.44 | 0.23 | 0.42 | 0.64 | 0.44 | 0.55 |
| 10 | 131355130 | rs496298 | 0.531 | 0.597 | 1.267 | -1.087 | -0.036 | 0.037 | 0.037 | 0 | 0.72 | 0.81 | 0.72 | 0.97 | 0.66 | 0.81 |
| 10 | 131367368 | rs10466114 | -0.017 | 0.678 | 1.105 | -1.03 | -0.039 | 0.035 | 0.035 | 0 | 0.21 | 0.12 | 0.27 | 0.05 | 0.29 | 0.18 |
| 10 | 131370724 | rs578932 | 1.965 | 0.301 | 0.88 | -1.002 | 0.028 | 0.017 | 0.017 | 0 | 0.44 | 0.2 | 0.42 | 0.62 | 0.42 | 0.55 |
| 10 | 131386731 | rs12768004 | -0.011 | 0.209 | 0.928 | -0.978 | 0.012 | 0.001 | 0.001 | 0.868 | 0.14 | 0.13 | 0.1 | 0.01 | 0.09 | 0.03 |
| 10 | 131397821 | rs551491 | 1.64 | 0.295 | 0.483 | -0.974 | -0.005 | 0.038 | 0.038 | 0.142 | 0.46 | 0.2 | 0.45 | 0.63 | 0.48 | 0.59 |
| 10 | 131399616 | rs3852507 | -1.674 | 0.235 | 0.697 | -1.002 | 0.024 | 0.051 | 0.051 | 0.013 | 0.51 | 0.63 | 0.53 | 0.37 | 0.51 | 0.41 |
| 10 | 131403041 | rs12775171 | 0.714 | -0.091 | 0.529 | -1.071 | 0.01 | 0.071 | 0.071 | 0.257 | 0.12 | 0.01 | 0.09 | 0.01 | 0.07 | 0.03 |
| 10 | 131406489 | rs513736 | 1.913 | 0.134 | 0.577 | -1.014 | -0.036 | 0.018 | 0.018 | 0 | 0.42 | 0.18 | 0.44 | 0.64 | 0.45 | 0.56 |
| 10 | 131413676 | rs553371 | 1.83 | 0.157 | 0.805 | -1.015 | -0.011 | 0.029 | 0.029 | 0 | 0.43 | 0.23 | 0.45 | 0.63 | 0.45 | 0.57 |
| 10 | 131420584 | rs544217 | 1.209 | 0.231 | 0.857 | -1.02 | 0.007 | 0.051 | 0.051 | 0.182 | 0.49 | 0.35 | 0.48 | 0.63 | 0.5 | 0.61 |
| 10 | 131426022 | rs477692 | 1.581 | 0.214 | 0.604 | -1.016 | 0.007 | 0.036 | 0.036 | 0 | 0.46 | 0.33 | 0.46 | 0.63 | 0.46 | 0.57 |
| 10 | 131445731 | rs1008982 | -1.011 | 0.225 | 0.671 | -1.155 | 0.009 | -0.067 | -0.067 | 0.007 | 0.33 | 0.56 | 0.37 | 0.04 | 0.38 | 0.25 |
| 10 | 131448143 | rs7905095 | -1.589 | 0.407 | 0.366 | -1.003 | -0.004 | 0.2 | 0.2 | 0 | 0.48 | 0.72 | 0.46 | 0.05 | 0.44 | 0.27 |
| 10 | 131452917 | rs4750761 | -0.62 | 0.563 | 0.609 | -0.831 | 0.014 | -0.115 | -0.115 | 0 | 0.3 | 0.49 | 0.33 | 0.04 | 0.3 | 0.17 |
| 10 | 131453616 | rs10829613 | -0.527 | 0.746 | 0.105 | -0.811 | 0.003 | -0.081 | -0.081 | 1.357 | 0.29 | 0.36 | 0.3 | 0.04 | 0.3 | 0.17 |
| 10 | 131455235 | rs7080419 | -0.987 | 0.874 | -1.269 | -0.519 | -0.026 | 0.031 | 0.031 | 0 | 0.56 | 0.74 | 0.53 | 0.37 | 0.47 | 0.44 |
| 10 | 131457179 | rs6482743 | -0.623 | 0.802 | -1.226 | -0.439 | 0.002 | 0.059 | 0.059 | 0.342 | 0.52 | 0.7 | 0.35 | 0.38 | 0.44 | 0.39 |
| 10 | 131466410 | rs1537257 | 0.414 | 0.875 | -1.754 | -0.377 | -0.063 | -0.023 | -0.023 | 0.205 | 0.34 | 0.19 | 0.47 | 0.61 | 0.46 | 0.51 |
| 10 | 131470024 | rs7068301 | -0.311 | 0.86 | -2.362 | -0.379 | -0.097 | 0.043 | 0.043 | 0 | 0.69 | 0.92 | 0.73 | 0.86 | 0.66 | 0.76 |
| 10 | 131471179 | rs10829615 | -0.182 | 0.325 | -2.33 | -0.637 | -0.004 | -0.009 | -0.009 | 0.028 | 0.17 | 0.26 | 0.23 | 0.08 | 0.15 | 0.2 |
| 10 | 131471398 | rs11016864 | 1.248 | -0.131 | -1.904 | -0.042 | -0.004 | 0.037 | 0.037 | 0 | 0.17 | 0.2 | 0.1 | 0.28 | 0.15 | 0.16 |
| 10 | 131474247 | rs7071424 | 0.183 | -0.081 | -1.697 | -0.079 | -0.008 | -0.021 | -0.021 | 0.007 | 0.17 | 0.28 | 0.23 | 0.08 | 0.16 | 0.13 |
| 10 | 131474474 | rs7087131 | 1.254 | -0.062 | -1.874 | 0.179 | 0.001 | 0.002 | 0.002 | 0 | 0.11 | 0.11 | 0.11 | 0.21 | 0.18 | 0.18 |
| 10 | 131475845 | rs7095819 | -0.556 | -0.358 | -1.955 | 0.072 | 0.039 | 0.001 | 0.001 | 1.01 | 0.39 | 0.8 | 0.4 | 0.36 | 0.34 | 0.38 |
| 10 | 131483282 | rs2039374 | 0.505 | -0.211 | -1.885 | 0.06 | 0.018 | -0.034 | -0.034 | 0 | 0.36 | 0.21 | 0.33 | 0.57 | 0.31 | 0.37 |
| 10 | 131484410 | rs11016875 | 1.009 | -0.33 | -1.774 | 0.027 | -0.018 | -0.031 | -0.031 | 1.066 | 0.08 | 0.16 | 0.05 | 0.01 | 0.04 | 0.1 |
| 10 | 131485306 | rs12219606 | 0.05 | -0.576 | -2.061 | -0.072 | -0.004 | -0.024 | -0.024 | 0.043 | 0.39 | 0.23 | 0.53 | 0.55 | 0.44 | 0.44 |
| 10 | 131485345 | rs11016877 | -1.946 | -0.437 | -2.362 | -0.139 | -0.118 | -0.003 | -0.003 | 0 | 0.76 | 0.97 | 0.75 | 0.87 | 0.71 | 0.79 |
| 10 | 131489782 | rs11016879 | -0.378 | -0.243 | -2.179 | -0.07 | -0.009 | -0.009 | -0.009 | 1.147 | 0.65 | 0.85 | 0.56 | 0.57 | 0.62 | 0.59 |
| 10 | 131490594 | rs7917453 | -1.352 | -0.172 | -2.024 | -0.05 | -0.016 | -0.049 | -0.049 | 2.005 | 0.79 | 0.92 | 0.67 | 0.7 | 0.79 | 0.8 |
| 10 | 131493946 | rs7073892 | 0.535 | -0.164 | -2.151 | -0.129 | -0.059 | 0.005 | 0.005 | 0 | 0.39 | 0.17 | 0.28 | 0.33 | 0.44 | 0.4 |
| 10 | 131496490 | rs12771882 | 1.18 | -0.338 | -1.834 | -0.195 | 0.006 | -0.009 | -0.009 | 0 | 0.24 | 0.16 | 0.13 | 0.31 | 0.21 | 0.25 |

| | | | | | | | | | | | | | | | | |
|---|---|---|---|---|---|---|---|---|---|---|---|---|---|---|---|---|
| 10 | 131496629 | rs4751110 | 0.227 | -0.189 | -1.669 | -0.129 | 0.002 | 0.001 | 0.001 | 0 | 0.63 | 0.33 | 0.41 | 0.64 | 0.65 | 0.66 |
| 10 | 131501938 | rs2026975 | 1.461 | 0.182 | -2.024 | 0.058 | -0.083 | -0.019 | -0.019 | 0.306 | 0.22 | 0.03 | 0.16 | 0.14 | 0.23 | 0.21 |
| 10 | 131502448 | rs1978759 | -0.729 | 0.053 | -1.314 | 0.332 | 0.004 | -0.011 | -0.011 | 0 | 0.78 | 0.77 | 0.65 | 0.71 | 0.79 | 0.8 |
| 10 | 131504495 | rs11016884 | -0.489 | -0.215 | -0.611 | 0.511 | -0.004 | -0.001 | -0.001 | 0 | 0.33 | 0.32 | 0.36 | 0.32 | 0.33 | 0.33 |
| 10 | 131506192 | rs1803965 | -0.572 | -0.335 | -0.145 | 0.568 | -0.006 | -0.004 | -0.004 | 1.468 | 0.11 | 0.18 | 0.2 | 0.1 | 0.12 | 0.14 |
| 10 | 131506283 | rs12917 | -0.524 | -0.304 | -0.161 | 0.573 | -0.003 | -0.001 | -0.001 | 0 | 0.11 | 0.16 | 0.2 | 0.11 | 0.14 | 0.15 |
| 10 | 131507505 | rs4751113 | -1.079 | -0.415 | 0.018 | 0.203 | -0.022 | 0.07 | 0.07 | 0 | 0.53 | 0.86 | 0.54 | 0.65 | 0.5 | 0.56 |
| 10 | 131509132 | rs2053139 | -0.015 | -0.103 | 0.204 | 0.162 | 0.001 | 0.002 | 0.002 | 0 | 0.33 | 0.34 | 0.28 | 0.54 | 0.36 | 0.38 |
| 10 | 131509617 | rs10764900 | -0.043 | -0.092 | 0.181 | 0.166 | 0.008 | 0.019 | 0.019 | 0 | 0.33 | 0.44 | 0.29 | 0.54 | 0.36 | 0.38 |
| 10 | 131510293 | rs4751115 | -0.991 | -0.091 | 0.095 | 0.158 | -0.01 | 0.048 | 0.048 | 2.292 | 0.46 | 0.75 | 0.53 | 0.65 | 0.5 | 0.56 |
| 10 | 131512933 | rs7081359 | -0.361 | 0.232 | 0.147 | 0.446 | 0.013 | 0.033 | 0.033 | 0 | 0.33 | 0.48 | 0.3 | 0.54 | 0.36 | 0.38 |
| 10 | 131514391 | rs4750768 | -0.171 | 0.102 | -0.213 | 0.174 | -0.008 | -0.028 | -0.028 | 0 | 0.38 | 0.42 | 0.34 | 0.7 | 0.34 | 0.51 |
| 10 | 131514862 | rs10764901 | 0.144 | -0.493 | -0.064 | 0.335 | -0.022 | -0.002 | -0.002 | 0 | 0.67 | 0.45 | 0.49 | 0.63 | 0.71 | 0.66 |
| 10 | 131519274 | rs748190 | -1.006 | -0.465 | -0.381 | 0.251 | 0.013 | 0.005 | 0.005 | 0 | 0.72 | 0.74 | 0.72 | 0.64 | 0.54 | 0.69 |
| 10 | 131522055 | rs2183878 | 0.665 | -0.716 | -0.84 | 0.192 | -0.021 | -0.076 | -0.076 | 0 | 0.2 | 0.08 | 0.38 | 0.44 | 0.23 | 0.21 |
| 10 | 131525017 | rs7078706 | -0.077 | -0.931 | -0.468 | 0.089 | 0.038 | 0.057 | 0.057 | 0 | 0.47 | 0.33 | 0.3 | 0.21 | 0.3 | 0.46 |
| 10 | 131526661 | rs11592158 | 0.593 | -0.527 | -0.95 | -0.234 | 0.024 | 0.104 | 0.104 | 0.008 | 0.32 | 0.11 | 0.19 | 0.06 | 0.23 | 0.2 |
| 10 | 131529254 | rs7897876 | -0.393 | 0.263 | -0.305 | 0.473 | 0.034 | 0.01 | 0.01 | 0 | 0.68 | 0.43 | 0.69 | 0.64 | 0.53 | 0.66 |
| 10 | 131540063 | rs11596935 | 0.453 | -0.101 | -0.492 | 0.434 | 0.013 | 0.021 | 0.021 | 0 | 0.19 | 0.1 | 0.15 | 0.12 | 0.13 | 0.15 |
| 10 | 131548336 | rs11511217 | -0.346 | 0.411 | -0.352 | 0.222 | -0.004 | 0.019 | 0.019 | 0 | 0.27 | 0.22 | 0.21 | 0.14 | 0.27 | 0.21 |
| 10 | 131558651 | rs7071825 | -1.31 | -0.149 | -0.413 | 0.152 | -0.048 | 0.005 | 0.005 | 0 | 0.65 | 0.84 | 0.73 | 0.65 | 0.55 | 0.65 |
| 10 | 131559213 | rs3793906 | -0.322 | -0.524 | -0.069 | 0.685 | 0.065 | 0.023 | 0.023 | 0.112 | 0.5 | 0.25 | 0.56 | 0.42 | 0.38 | 0.52 |
| 10 | 131560452 | rs3829194 | -0.468 | -0.498 | 0.628 | 0.701 | 0.076 | 0.374 | 0.374 | 0.523 | 0.05 | 0.33 | 0.1 | 0.21 | 0.03 | 0.06 |
| 10 | 131562744 | rs3793908 | 0.361 | -0.575 | 0.563 | 0.649 | -0.036 | 0.053 | 0.053 | 0 | 0.25 | 0.46 | 0.25 | 0.34 | 0.17 | 0.22 |
| 10 | 131562993 | rs3793910 | 1.558 | -0.572 | 0.862 | 0.631 | -0.006 | 0.024 | 0.024 | 0 | 0.18 | 0.05 | 0.14 | 0.13 | 0.14 | 0.16 |
| 10 | 131563151 | rs3793912 | -1.338 | -0.258 | 1.027 | 0.756 | -0.07 | 0.002 | 0.002 | 0 | 0.66 | 0.88 | 0.74 | 0.66 | 0.55 | 0.65 |
| 10 | 131565064 | rs2308321 | 0.423 | -0.364 | 0.973 | 0.761 | -0.046 | 0.042 | 0.042 | 0 | 0.08 | 0.01 | 0.05 | 0.01 | 0.14 | 0.06 |
| 10 | 131565170 | rs2308327 | 0.423 | -0.364 | 0.973 | 0.761 | -0.046 | 0.042 | 0.042 | 0 | 0.08 | 0.01 | 0.05 | 0.01 | 0.14 | 0.06 |
| 10 | 131570133 | rs10764906 | -0.787 | -0.301 | 1.047 | 0.795 | 0.053 | 0.156 | 0.156 | 0 | 0.07 | 0.19 | 0.12 | 0.22 | 0.11 | 0.08 |
| 10 | 131572285 | rs10829630 | -0.837 | -0.289 | 0.957 | 0.611 | -0.009 | -0.008 | -0.008 | 0 | 0.59 | 0.43 | 0.62 | 0.44 | 0.52 | 0.58 |
| 10 | 131574729 | rs1547177 | -0.869 | -0.343 | 0.979 | 0.9 | -0.006 | 0.261 | 0.261 | 0.898 | 0.12 | 0.44 | 0.13 | 0.29 | 0.07 | 0.1 |
| 10 | 131577319 | rs11814529 | 0.875 | -0.26 | 0.897 | 0.823 | -0.014 | 0.018 | 0.018 | 0 | 0.36 | 0.35 | 0.28 | 0.28 | 0.46 | 0.35 |
| 10 | 131577449 | rs11016925 | -0.577 | -0.508 | 0.712 | 0.769 | -0.023 | -0.01 | -0.01 | 0.654 | 0.39 | 0.58 | 0.51 | 0.59 | 0.3 | 0.45 |
| 10 | 131578789 | rs4750772 | -0.644 | -0.018 | 0.386 | 0.792 | -0.001 | 0 | 0 | 0 | 0.71 | 0.75 | 0.73 | 0.6 | 0.73 | 0.74 |
| 10 | 131582096 | rs7075383 | -0.357 | -0.048 | 0.005 | 0.74 | 0.072 | 0.129 | 0.129 | 0 | 0.55 | 0.29 | 0.73 | 0.82 | 0.72 | 0.62 |
| 10 | 131583388 | rs4750773 | 0.469 | 0.241 | -0.14 | 1.034 | 0.021 | 0.036 | 0.036 | 0 | 0.34 | 0.46 | 0.42 | 0.49 | 0.41 | 0.34 |
| 10 | 131585151 | rs2094290 | -0.813 | 0.484 | -0.378 | 1.093 | -0.026 | -0.07 | -0.07 | 0.751 | 0.61 | 0.53 | 0.75 | 0.86 | 0.72 | 0.61 |
| 10 | 131587140 | rs3750832 | 0.014 | 0.652 | -0.456 | 1.197 | -0.006 | -0.005 | -0.005 | 0.884 | 0.41 | 0.4 | 0.43 | 0.5 | 0.47 | 0.38 |
| 10 | 131592692 | rs3750829 | -0.335 | 0.847 | -0.276 | 1.837 | -0.007 | 0.034 | 0.034 | 0 | 0.06 | 0 | 0.01 | 0.01 | 0.04 | 0.04 |
| 10 | 131594850 | rs1887181 | -0.762 | 0.779 | -0.058 | 1.976 | 0.014 | 0.135 | 0.135 | 0.3 | 0.11 | 0.45 | 0.08 | 0.13 | 0.05 | 0.05 |
| 10 | 131597962 | rs11016942 | -0.878 | 1.382 | -1.245 | 1.802 | -0.059 | -0.099 | -0.099 | 0.374 | 0.19 | 0.04 | 0.3 | 0.35 | 0.24 | 0.21 |
| 10 | 131598860 | rs6482756 | -0.816 | 0.892 | -1.112 | 1.687 | 0.003 | -0.048 | -0.048 | 0.005 | 0.18 | 0.06 | 0.21 | 0.32 | 0.15 | 0.14 |
| 10 | 131600411 | rs10829637 | -1.009 | 1.205 | -0.931 | 1.765 | -0.002 | -0.009 | -0.009 | 0.018 | 0.11 | 0.08 | 0.18 | 0.32 | 0.14 | 0.11 |
| 10 | 131601154 | rs11016948 | -0.921 | 1.725 | -0.091 | 1.844 | 0.103 | 0.06 | 0.06 | 0.409 | 0.08 | 0.31 | 0.1 | 0.08 | 0.11 | 0.05 |
| 10 | 131601957 | rs11016949 | -1.173 | 1.43 | -0.122 | 1.718 | 0.002 | -0.005 | -0.005 | 0 | 0.15 | 0.14 | 0.18 | 0.32 | 0.16 | 0.16 |
| 10 | 131602046 | rs10829638 | -0.699 | 1.438 | -0.147 | 1.711 | 0.005 | -0.055 | -0.055 | 0.096 | 0.12 | 0.01 | 0.12 | 0.28 | 0.1 | 0.15 |
| 10 | 131602484 | rs10734090 | -2.366 | 1.561 | -0.135 | 2.079 | 0.031 | 0.1 | 0.1 | 0 | 0.24 | 0.48 | 0.33 | 0.46 | 0.29 | 0.25 |
| 10 | 131603126 | rs7899958 | 0.302 | 1.798 | -1.597 | 1.975 | -0.073 | 0.025 | 0.025 | 0.05 | 0.31 | 0.04 | 0.41 | 0.23 | 0.34 | 0.3 |
| 10 | 131606426 | rs11016954 | -0.765 | 1.008 | -1.227 | 2.14 | 0.01 | -0.022 | -0.022 | 0 | 0.14 | 0.11 | 0.13 | 0.52 | 0.11 | 0.19 |
| 10 | 131622170 | rs10829645 | -1.09 | 0.936 | -1.074 | 1.82 | -0.005 | 0.033 | 0.033 | 0 | 0.19 | 0.31 | 0.26 | 0.62 | 0.19 | 0.26 |
| 10 | 131626650 | rs7911165 | -0.615 | 0.588 | -1.091 | 1.179 | 0.008 | 0.005 | 0.005 | 0 | 0.54 | 0.47 | 0.74 | 0.68 | 0.63 | 0.54 |
| 10 | 131633720 | rs8217 | -0.84 | 0.947 | -1.66 | 1.282 | -0.092 | -0.141 | -0.141 | 0.392 | 0.25 | 0.02 | 0.45 | 0.44 | 0.29 | 0.39 |
| 10 | 131641242 | rs7895783 | -1.303 | 0.003 | -0.414 | 0.682 | -0.045 | -0.067 | -0.067 | 0.091 | 0.17 | 0.5 | 0.19 | 0.09 | 0.07 | 0.21 |
| 10 | 131648687 | rs7100028 | -1.292 | -0.647 | -0.843 | 0.855 | 0.021 | 0.002 | 0.002 | 0.646 | 0.22 | 0.78 | 0.26 | 0.2 | 0.11 | 0.17 |
| 10 | 131648947 | rs12783441 | -0.339 | -0.494 | -1.33 | 1.628 | -0.02 | -0.024 | -0.024 | 0 | 0.07 | 0 | 0.12 | 0.29 | 0.08 | 0.19 |
| 10 | 131650401 | rs11016976 | -1.013 | -1.455 | -1.471 | 1.003 | 0.019 | 0.026 | 0.026 | 0.057 | 0.24 | 0.13 | 0.22 | 0.11 | 0.17 | 0.16 |
| 10 | 131651838 | rs1028830 | -1.586 | -1.29 | -1.719 | 1.005 | 0.058 | -0.012 | -0.012 | 0 | 0.52 | 0.73 | 0.54 | 0.46 | 0.36 | 0.47 |
| 10 | 131656162 | rs4751139 | -0.316 | -0.86 | -1.681 | 0.896 | 0.012 | 0.017 | 0.017 | 0 | 0.18 | 0.05 | 0.2 | 0.1 | 0.14 | 0.13 |
| 10 | 131658530 | rs10829654 | -0.105 | -0.742 | -1.425 | 1.146 | -0.002 | 0.001 | 0.001 | 0.056 | 0.1 | 0.13 | 0.16 | 0.21 | 0.1 | 0.14 |
| 10 | 131660498 | rs10734091 | -1.21 | -1.045 | -0.876 | 1.059 | -0.007 | 0.002 | 0.002 | 0.618 | 0.33 | 0.38 | 0.37 | 0.33 | 0.26 | 0.28 |
| 10 | 131663651 | rs10764919 | -1.001 | -1.074 | -2.154 | 0.393 | -0.102 | 0.096 | 0.096 | 0 | 0.61 | 0.95 | 0.71 | 0.77 | 0.41 | 0.83 |
| 10 | 131664108 | rs4751140 | 0.818 | -0.314 | -2.481 | 0.376 | -0.098 | 0.065 | 0.065 | 0 | 0.35 | 0.06 | 0.44 | 0.15 | 0.54 | 0.28 |
| 10 | 131669792 | rs4750779 | 1.06 | -0.406 | -1.444 | 0.636 | -0.001 | 0.014 | 0.014 | 0 | 0.44 | 0.37 | 0.49 | 0.37 | 0.55 | 0.28 |
| 10 | 131671299 | rs7895769 | -0.586 | -1.141 | -0.978 | 0.352 | -0.036 | -0.061 | -0.061 | 0 | 0.28 | 0.38 | 0.13 | 0.09 | 0.13 | 0.24 |
| 10 | 131672269 | rs7074160 | -0.157 | -1.222 | -0.822 | 0.31 | -0.015 | -0.057 | -0.057 | 1.556 | 0.22 | 0.26 | 0.11 | 0.04 | 0.12 | 0.22 |
| 10 | 131673986 | rs9804200 | 0.83 | -0.94 | -0.63 | 0.581 | -0.016 | 0.014 | 0.014 | 0.244 | 0.65 | 0.59 | 0.61 | 0.45 | 0.71 | 0.49 |
| 10 | 131675252 | rs7076708 | -0.284 | -1.033 | -0.59 | 0.779 | 0.024 | 0.06 | 0.06 | 2.981 | 0.15 | 0.3 | 0.23 | 0.19 | 0.15 | 0.19 |
| 10 | 131675444 | rs6482761 | -0.619 | -0.969 | -1.064 | 0.83 | -0.005 | -0.007 | -0.007 | 0 | 0.36 | 0.51 | 0.32 | 0.24 | 0.25 | 0.39 |
| 10 | 131679393 | rs4142058 | 0.527 | -0.058 | -1.177 | 0.945 | -0.007 | 0 | 0 | 0 | 0.55 | 0.58 | 0.58 | 0.46 | 0.67 | 0.37 |

| | | | | | | | | | | | | | | | | |
|---|---|---|---|---|---|---|---|---|---|---|---|---|---|---|---|---|
| 10 | 131684101 | rs7082533 | 0.53 | 0.522 | -1.668 | 1.648 | -0.034 | -0.109 | -0.109 | 0 | 0.86 | 0.98 | 0.78 | 0.54 | 0.85 | 0.79 |
| 10 | 131688008 | rs944963 | 0.309 | 0.702 | -1.906 | 1.102 | 0.058 | 0.044 | 0.044 | 0.933 | 0.48 | 0.75 | 0.44 | 0.58 | 0.43 | 0.44 |
| 10 | 131693411 | rs7100586 | 0.164 | 1.575 | -1.961 | 1.373 | -0.066 | -0.084 | -0.084 | 0.123 | 0.86 | 0.99 | 0.86 | 0.77 | 0.83 | 0.82 |
| 10 | 131694085 | rs6482764 | 0.339 | 0.583 | -1.284 | 1.385 | -0.007 | 0.028 | 0.028 | 0 | 0.35 | 0.5 | 0.35 | 0.41 | 0.35 | 0.27 |
| 10 | 131699296 | rs944962 | -0.284 | 0.747 | -1.795 | 1.291 | -0.025 | -0.097 | -0.097 | 0.232 | 0.59 | 0.91 | 0.64 | 0.49 | 0.6 | 0.62 |
| 10 | 131700735 | rs11017007 | 0.108 | 1.073 | -1.405 | 0.814 | 0.002 | -0.014 | -0.014 | 0 | 0.55 | 0.49 | 0.61 | 0.69 | 0.53 | 0.61 |
| 10 | 131701925 | rs4751143 | 0.491 | 0.926 | -1.525 | 0.881 | -0.006 | -0.001 | -0.001 | 0 | 0.45 | 0.26 | 0.51 | 0.42 | 0.45 | 0.42 |
| 10 | 131702075 | rs4751144 | -0.464 | 0.55 | -1.341 | 0.507 | -0.001 | -0.002 | -0.002 | 0.814 | 0.67 | 0.62 | 0.71 | 0.68 | 0.67 | 0.63 |
| 10 | 131704434 | rs4751145 | -1.254 | 0.627 | -1.913 | -0.232 | -0.01 | 0.058 | 0.058 | 0 | 0.8 | 0.92 | 0.86 | 0.94 | 0.77 | 0.8 |
| 10 | 131711365 | rs11597635 | 0.038 | 0.856 | -1.806 | 0.059 | 0.022 | 0.083 | 0.083 | 0 | 0.36 | 0.17 | 0.18 | 0.19 | 0.32 | 0.32 |
| 10 | 131721056 | rs11598130 | -0.285 | 1.121 | -1.81 | 0.323 | -0.148 | -0.123 | -0.123 | 0 | 0.25 | 0.06 | 0.52 | 0.38 | 0.43 | 0.25 |
| 10 | 131722110 | rs7095306 | -0.486 | 0.214 | -0.559 | -0.38 | -0.008 | 0.05 | 0.05 | 0.539 | 0.14 | 0.42 | 0.12 | 0.16 | 0.07 | 0.12 |
| 10 | 131726740 | rs10829661 | 0.315 | -0.251 | -0.653 | -1.105 | 0.012 | 0.02 | 0.02 | 0 | 0.55 | 0.43 | 0.72 | 0.8 | 0.66 | 0.63 |
| 10 | 131727813 | rs4751149 | -0.662 | -0.153 | -0.986 | -1.182 | -0.031 | 0.097 | 0.097 | 0 | 0.77 | 0.94 | 0.88 | 0.95 | 0.74 | 0.8 |
| 10 | 131733077 | rs1984103 | -0.248 | 0.116 | -0.787 | -1.001 | -0.005 | -0.064 | -0.064 | 0 | 0.65 | 0.63 | 0.82 | 0.92 | 0.7 | 0.74 |
| 10 | 131739733 | rs11017022 | -0.398 | 1.071 | -1.134 | -0.114 | -0.012 | 0.02 | 0.02 | 0 | 0.33 | 0.33 | 0.53 | 0.75 | 0.39 | 0.34 |
| 10 | 131741604 | rs4751153 | -0.347 | 0.747 | -1.377 | 0.354 | 0.011 | -0.009 | -0.009 | 0 | 0.38 | 0.26 | 0.24 | 0.37 | 0.36 | 0.25 |
| 10 | 131741695 | rs744680 | -0.81 | 0.759 | -2.047 | 0.21 | -0.032 | 0.009 | 0.009 | 0 | 0.23 | 0.03 | 0.18 | 0.17 | 0.19 | 0.43 |
| 10 | 131744131 | rs10829663 | -0.625 | 0.732 | -1.948 | 0.7 | 0.003 | -0.044 | -0.044 | 0 | 0.29 | 0.2 | 0.28 | 0.42 | 0.23 | 0.26 |
| 10 | 131747705 | rs556327 | 0.341 | 0.767 | -1.466 | 0.69 | -0.004 | 0 | 0 | 0 | 0.82 | 0.76 | 0.82 | 0.74 | 0.85 | 0.83 |
| 10 | 131753010 | rs10829665 | 0.2 | 0.92 | -1.719 | 0.797 | -0.019 | -0.033 | -0.033 | 0 | 0.19 | 0.13 | 0.22 | 0.24 | 0.2 | 0.16 |
| 10 | 131760352 | rs573370 | 0.956 | 0.166 | -1.239 | 0.418 | 0.015 | -0.02 | -0.02 | 0 | 0.55 | 0.27 | 0.57 | 0.44 | 0.64 | 0.63 |
| 10 | 131766660 | rs10829669 | 0.36 | 0.44 | -1.315 | 0.358 | -0.024 | -0.039 | -0.039 | 0 | 0.18 | 0.12 | 0.21 | 0.25 | 0.2 | 0.16 |
| 10 | 131772971 | rs7099076 | 0.3 | 0.718 | -1.388 | 0.089 | -0.059 | -0.014 | -0.014 | 0 | 0.23 | 0.12 | 0.28 | 0.18 | 0.33 | 0.18 |
| 10 | 131777935 | rs670243 | 0.472 | 0.671 | -0.706 | 0.052 | 0.003 | -0.001 | -0.001 | 0.731 | 0.6 | 0.57 | 0.54 | 0.56 | 0.53 | 0.67 |
| 10 | 131798192 | rs10741205 | 0.067 | 0.234 | -0.585 | -0.224 | -0.001 | -0.005 | -0.005 | 0 | 0.82 | 0.72 | 0.82 | 0.74 | 0.85 | 0.84 |
| 10 | 131801337 | rs11017045 | 1.55 | -0.078 | -1.13 | -0.424 | -0.02 | -0.085 | -0.085 | 4.803 | 0.3 | 0.04 | 0.29 | 0.32 | 0.29 | 0.41 |
| 10 | 131802533 | rs7910342 | 1.18 | -0.601 | 0.013 | -1.065 | -0.002 | -0.008 | -0.008 | 0 | 0.68 | 0.31 | 0.71 | 0.74 | 0.73 | 0.75 |
| 10 | 131807625 | rs10741206 | -0.299 | -0.116 | -0.165 | -0.778 | 0.005 | -0.046 | -0.046 | 1.31 | 0.92 | 0.8 | 0.92 | 1 | 0.91 | 0.97 |
| 10 | 131811356 | rs12572664 | -0.697 | 0.063 | -0.61 | -0.497 | 0.004 | -0.003 | -0.003 | 0.012 | 0.17 | 0.3 | 0.18 | 0.27 | 0.15 | 0.17 |
| 10 | 131813841 | rs11017049 | -0.804 | 0.026 | -0.772 | -0.613 | -0.009 | -0.013 | -0.013 | 2.576 | 0.2 | 0.24 | 0.17 | 0.22 | 0.14 | 0.24 |
| 10 | 131815454 | rs7069452 | -1.1 | -0.111 | -1.436 | -1.245 | 0.033 | 0.126 | 0.126 | 0 | 0.47 | 0.73 | 0.38 | 0.18 | 0.36 | 0.27 |
| 10 | 131815674 | rs942950 | 0.745 | 0.261 | -1.622 | -1.149 | 0.009 | 0.039 | 0.039 | 0.525 | 0.71 | 0.21 | 0.7 | 0.83 | 0.77 | 0.81 |
| 10 | 131815779 | rs942951 | -0.423 | 0.874 | -1.837 | -0.501 | -0.049 | 0.015 | 0.015 | 0 | 0.45 | 0.18 | 0.34 | 0.41 | 0.46 | 0.37 |
| 10 | 131819496 | rs942952 | 0.182 | 0.966 | -0.973 | -0.253 | 0.003 | 0.011 | 0.011 | 0 | 0.59 | 0.38 | 0.64 | 0.7 | 0.62 | 0.71 |
| 10 | 131825297 | rs1171754 | 0.233 | 1.853 | -1.033 | 0.184 | -0.019 | 0.002 | 0.002 | 0 | 0.92 | 0.96 | 0.79 | 0.91 | 0.82 | 0.96 |
| 10 | 131830640 | rs11017056 | -0.057 | 0.834 | -1.111 | 0 | 0.017 | 0.012 | 0.012 | 0 | 0.28 | 0.24 | 0.16 | 0.23 | 0.22 | 0.23 |
| 10 | 131830899 | rs2477970 | 0.518 | 0.878 | -0.624 | 0.188 | 0 | -0.001 | -0.001 | 0 | 0.55 | 0.56 | 0.54 | 0.5 | 0.57 | 0.67 |
| 10 | 131833065 | rs7899076 | -0.27 | 1.251 | -0.797 | -0.048 | 0.004 | 0.05 | 0.05 | 0 | 0.39 | 0.28 | 0.36 | 0.25 | 0.4 | 0.35 |
| 10 | 131839522 | rs611788 | 0.443 | 1.306 | -0.148 | 0.287 | 0.008 | 0.003 | 0.003 | 0 | 0.52 | 0.64 | 0.37 | 0.42 | 0.42 | 0.52 |
| 10 | 131841169 | rs1536090 | 0.505 | 1.283 | -0.379 | 0.169 | -0.001 | -0.001 | -0.001 | 0 | 0.46 | 0.52 | 0.51 | 0.46 | 0.4 | 0.52 |
| 10 | 131844235 | rs2765480 | -0.044 | 1.46 | -0.273 | 0.27 | 0.021 | 0.024 | 0.024 | 0 | 0.86 | 0.83 | 0.84 | 0.77 | 0.8 | 0.9 |
| 10 | 131845331 | rs2765478 | -0.429 | 1.102 | -0.56 | 0.188 | -0.01 | 0.008 | 0.008 | 0 | 0.59 | 0.65 | 0.62 | 0.71 | 0.53 | 0.58 |
| 10 | 131850891 | rs647216 | 0.115 | 0.551 | -0.561 | -0.285 | 0.025 | 0.093 | 0.093 | 0 | 0.5 | 0.67 | 0.75 | 0.75 | 0.57 | 0.5 |
| 10 | 131852222 | rs1016314 | 0.375 | 0.033 | -0.55 | -0.274 | -0.004 | -0.006 | -0.006 | 0 | 0.24 | 0.21 | 0.08 | 0.13 | 0.16 | 0.2 |
| 10 | 131856651 | rs10829685 | 0.064 | -0.073 | -0.494 | 0.032 | 0.027 | -0.014 | -0.014 | 0.367 | 0.27 | 0.18 | 0.14 | 0.37 | 0.19 | 0.4 |
| 10 | 131857008 | rs671799 | 0.595 | 0.091 | -0.545 | -0.163 | 0.004 | 0.006 | 0.006 | 0 | 0.47 | 0.58 | 0.73 | 0.54 | 0.58 | 0.35 |
| 10 | 131859987 | rs1180610 | 1.1 | 0.449 | 0.115 | 0.359 | 0.006 | -0.008 | -0.008 | 0 | 0.61 | 0.25 | 0.73 | 0.54 | 0.67 | 0.48 |
| 10 | 131863069 | rs2244309 | 0.744 | 0.42 | 0.653 | 0.37 | 0 | -0.001 | -0.001 | 0 | 0.66 | 0.63 | 0.77 | 0.64 | 0.67 | 0.51 |
| 10 | 131871869 | rs662041 | -0.233 | 0.382 | 0.378 | 0.286 | -0.006 | 0.005 | 0.005 | 0.273 | 0.9 | 0.95 | 0.96 | 0.92 | 0.9 | 0.86 |
| 10 | 131875743 | rs2764369 | 1.418 | 0.37 | 0.375 | 0.285 | -0.001 | 0.004 | 0.004 | 0.022 | 0.63 | 0.46 | 0.75 | 0.54 | 0.67 | 0.48 |
| 10 | 131884171 | rs654367 | 1.227 | 0.46 | 0.747 | 0.61 | 0.005 | 0.097 | 0.097 | 0.681 | 0.83 | 0.61 | 0.87 | 0.65 | 0.89 | 0.64 |
| 10 | 131887319 | rs942675 | 0.025 | 0.802 | 0.132 | 0.712 | -0.002 | 0.003 | 0.003 | 0.256 | 0.92 | 0.98 | 0.97 | 0.92 | 0.93 | 0.9 |
| 10 | 131907568 | rs1171728 | 0.109 | 1.146 | -0.393 | 0.041 | -0.006 | 0.007 | 0.007 | 0 | 0.92 | 0.97 | 0.96 | 0.97 | 0.9 | 0.91 |
| 10 | 131908016 | rs1171727 | 0.209 | 1.185 | -0.828 | -0.211 | -0.003 | 0.039 | 0.039 | 0 | 0.29 | 0.17 | 0.31 | 0.14 | 0.29 | 0.24 |
| 10 | 131922572 | rs942674 | 0.49 | 1.259 | -0.634 | -0.333 | 0.04 | 0.053 | 0.053 | 0.011 | 0.47 | 0.37 | 0.5 | 0.68 | 0.39 | 0.5 |
| 10 | 131923726 | rs3949927 | -0.085 | 1.558 | -0.301 | -0.299 | 0.039 | 0.005 | 0.005 | 0 | 0.14 | 0.4 | 0.11 | 0.12 | 0.14 | 0.13 |
| 10 | 131925912 | rs2147753 | 0.059 | 1.592 | -0.259 | -0.284 | 0.039 | 0.008 | 0.008 | 0 | 0.13 | 0.35 | 0.11 | 0.12 | 0.14 | 0.13 |
| 10 | 131926121 | rs11017093 | -0.501 | 1.887 | -0.956 | -0.293 | -0.029 | 0.052 | 0.052 | 0 | 0.69 | 0.81 | 0.65 | 0.83 | 0.64 | 0.7 |
| 10 | 131928643 | rs12414821 | 0.021 | 2.181 | -0.886 | -0.046 | -0.038 | 0.014 | 0.014 | 0.297 | 0.06 | 0 | 0.1 | 0.04 | 0.13 | 0.09 |
| 10 | 131933599 | rs3763759 | -0.206 | 2.498 | -1.416 | 0.794 | -0.042 | -0.075 | -0.075 | 0 | 0.85 | 0.94 | 0.78 | 0.66 | 0.8 | 0.73 |
| 10 | 131939898 | rs2388757 | 0.114 | 2.214 | -1.366 | 0.186 | 0 | 0.015 | 0.015 | 0.741 | 0.42 | 0.4 | 0.28 | 0.23 | 0.39 | 0.42 |
| 10 | 131941265 | rs4750790 | -0.824 | 2.161 | -1.447 | 0.445 | -0.062 | -0.108 | -0.108 | 0.652 | 0.84 | 0.97 | 0.79 | 0.54 | 0.79 | 0.59 |
| 10 | 131943423 | rs7073590 | -0.591 | 2.126 | -0.83 | 0.367 | 0.009 | 0.064 | 0.064 | 4.792 | 0.53 | 0.7 | 0.46 | 0.31 | 0.56 | 0.52 |
| 10 | 131945179 | rs12269149 | 0.977 | 2.125 | -0.916 | 0.085 | -0.004 | 0.025 | 0.025 | 0 | 0.31 | 0.23 | 0.21 | 0.12 | 0.29 | 0.24 |
| 10 | 131945252 | rs12248841 | -0.306 | 2.114 | -0.507 | 0.158 | -0.001 | -0.01 | -0.01 | 0.155 | 0.45 | 0.56 | 0.36 | 0.23 | 0.45 | 0.43 |
| 10 | 131949021 | rs12415867 | 1.348 | 2.079 | -0.52 | 0.191 | -0.031 | -0.022 | -0.022 | 0.004 | 0.07 | 0 | 0.1 | 0.08 | 0.12 | 0.09 |
| 10 | 131959150 | rs2274217 | 0.242 | 2.305 | -1.256 | -0.11 | 0.002 | 0.106 | 0.106 | 0 | 0.28 | 0.12 | 0.13 | 0.11 | 0.29 | 0.31 |
| 10 | 131961891 | rs7904722 | -0.57 | 2.293 | -1.137 | -0.053 | 0.003 | 0.045 | 0.045 | 0.948 | 0.38 | 0.28 | 0.3 | 0.29 | 0.41 | 0.45 |

| | | | | | | | | | | | | | | | | |
|---|---|---|---|---|---|---|---|---|---|---|---|---|---|---|---|---|
| 10 | 131966636 | rs2182098 | 1.545 | 2.127 | -1.046 | -0.052 | 0 | -0.001 | -0.001 | 0 | 0.36 | 0.62 | 0.47 | 0.33 | 0.29 | 0.13 |
| 10 | 131970877 | rs1016376 | -1.648 | 2.215 | -1.119 | 0.074 | -0.059 | -0.1 | -0.1 | 0 | 0.85 | 0.98 | 0.79 | 0.65 | 0.8 | 0.72 |
| 10 | 131978358 | rs3750834 | 1.315 | 1.97 | -0.936 | -0.075 | 0.008 | 0.009 | 0.009 | 5.629 | 0.36 | 0.67 | 0.48 | 0.33 | 0.29 | 0.13 |
| 10 | 131979761 | rs2281842 | 0.291 | 1.996 | -1.279 | -0.075 | -0.033 | 0.128 | 0.128 | 0 | 0.54 | 0.76 | 0.59 | 0.33 | 0.5 | 0.23 |
| 10 | 131984960 | rs7906617 | -0.87 | 2.142 | -0.906 | 0.133 | 0 | -0.002 | -0.002 | 0 | 0.27 | 0.7 | 0.36 | 0.2 | 0.28 | 0.21 |
| 10 | 131989576 | rs10764938 | 0.141 | 2.35 | -1.544 | 0.809 | -0.083 | -0.087 | -0.087 | 0 | 0.73 | 0.9 | 0.71 | 0.68 | 0.62 | 0.62 |
| 10 | 131990036 | rs11017128 | 0.607 | 2.165 | -1.029 | 0.99 | 0.036 | -0.01 | -0.01 | 0 | 0.13 | 0.25 | 0.15 | 0.08 | 0.13 | 0.06 |
| 10 | 131991181 | rs16910188 | -0.085 | 1.723 | -1.099 | 1.023 | 0.002 | 0.081 | 0.081 | 0.007 | 0.17 | 0.52 | 0.14 | 0.27 | 0.14 | 0.35 |
| 10 | 131992245 | rs4751178 | -0.177 | 1.937 | -1.253 | 1.078 | -0.01 | -0.013 | -0.013 | 0 | 0.75 | 0.81 | 0.71 | 0.69 | 0.72 | 0.71 |
| 10 | 131992367 | rs10764940 | 0.834 | 1.855 | -1.713 | 1.126 | 0.044 | 0.118 | 0.118 | 1.084 | 0.5 | 0.25 | 0.48 | 0.33 | 0.46 | 0.27 |
| 10 | 131993713 | rs6482776 | -1.407 | 2.125 | -1.507 | 1.757 | -0.03 | -0.001 | -0.001 | 1.324 | 0.95 | 0.99 | 0.85 | 0.8 | 0.88 | 0.93 |
| 10 | 131996932 | rs11017131 | -0.277 | 1.543 | -1.588 | 0.947 | -0.019 | -0.007 | -0.007 | 0 | 0.22 | 0.04 | 0.08 | 0.23 | 0.22 | 0.31 |
| 10 | 132003803 | rs10764942 | -0.658 | 1.639 | -1.177 | 1.116 | -0.03 | -0.03 | -0.03 | 0 | 0.93 | 1 | 0.9 | 0.9 | 0.88 | 0.91 |
| 10 | 132007282 | rs1410906 | -0.291 | 1.567 | -0.347 | 0.995 | 0.058 | 0.044 | 0.044 | 0.272 | 0.09 | 0.34 | 0.11 | 0.17 | 0.09 | 0.11 |
| 10 | 132009221 | rs7086678 | 0.809 | 1.352 | -0.657 | 0.789 | 0.001 | 0.023 | 0.023 | 1.075 | 0.55 | 0.38 | 0.46 | 0.32 | 0.44 | 0.44 |
| 10 | 132012962 | rs11017144 | 0.842 | 1.052 | -0.665 | 0.594 | 0.031 | 0.136 | 0.136 | 2.038 | 0.5 | 0.25 | 0.46 | 0.24 | 0.41 | 0.27 |
| 10 | 132012970 | rs11017145 | 0.844 | 0.995 | -0.856 | 1.503 | 0.016 | -0.065 | -0.065 | 0 | 0.11 | 0.02 | 0.11 | 0.34 | 0.07 | 0.33 |
| 10 | 132016610 | rs11017147 | 0.7 | 1.124 | -0.71 | 1.85 | -0.014 | -0.006 | -0.006 | 0 | 0.58 | 0.32 | 0.56 | 0.58 | 0.48 | 0.61 |
| 10 | 132018853 | rs748788 | 0.138 | 1.193 | -0.823 | 1.919 | -0.008 | -0.006 | -0.006 | 0 | 0.08 | 0.02 | 0.09 | 0.33 | 0.07 | 0.31 |
| 10 | 132021145 | rs2182097 | -0.743 | 1.538 | -0.912 | 1.787 | -0.024 | -0.024 | -0.024 | 1.194 | 0.91 | 0.96 | 0.82 | 0.75 | 0.85 | 0.85 |
| 10 | 132031664 | rs7077478 | 0.065 | 1.711 | -0.441 | 1.456 | 0.083 | 0.068 | 0.068 | 0.193 | 0.1 | 0.39 | 0.11 | 0.17 | 0.1 | 0.12 |
| 10 | 132032454 | rs11597007 | 0.119 | 1.585 | -0.493 | 1.307 | -0.011 | 0.057 | 0.057 | 0.395 | 0.22 | 0.14 | 0.15 | 0 | 0.27 | 0.13 |
| 10 | 132034281 | rs7088014 | -1.131 | 1.243 | -0.019 | 1.867 | -0.036 | -0.02 | -0.02 | 0 | 0.93 | 1 | 0.94 | 0.94 | 0.88 | 0.91 |
| 10 | 132043419 | rs7094199 | -1.251 | 1.144 | 0.155 | 2.107 | 0.04 | 0.179 | 0.179 | 0.579 | 0.17 | 0.4 | 0.2 | 0.46 | 0.17 | 0.44 |
| 10 | 132044710 | rs12570092 | 0.879 | 1.02 | -0.083 | 2.209 | -0.014 | -0.01 | -0.01 | 0.045 | 0.05 | 0.01 | 0.09 | 0.17 | 0.07 | 0.17 |
| 10 | 132045911 | rs11017158 | -0.092 | 1.021 | -0.083 | 2.198 | -0.011 | -0.017 | -0.017 | 0 | 0.08 | 0.02 | 0.09 | 0.29 | 0.07 | 0.29 |
| 10 | 132046689 | rs10829720 | 0.381 | 0.392 | -0.235 | 1.378 | -0.011 | -0.015 | -0.015 | 0 | 0.2 | 0.21 | 0.13 | 0.15 | 0.14 | 0.16 |
| 10 | 132046816 | rs4750795 | 1.393 | 0.443 | 0.211 | 1.466 | 0.002 | 0.004 | 0.004 | 0.424 | 0.48 | 0.42 | 0.49 | 0.46 | 0.51 | 0.37 |
| 10 | 132048851 | rs11017162 | -0.092 | 0.082 | 0.057 | 1.032 | 0.015 | 0.02 | 0.02 | 0.23 | 0.25 | 0.14 | 0.1 | 0.08 | 0.17 | 0.17 |
| 10 | 132052655 | rs11017165 | -1.051 | 0.612 | 0.88 | 1.745 | -0.003 | 0.003 | 0.003 | 0.165 | 0.32 | 0.31 | 0.31 | 0.38 | 0.26 | 0.47 |
| 10 | 132054599 | rs2892592 | 0.52 | 0.652 | 0.807 | 1.804 | 0.002 | -0.005 | -0.005 | 0.211 | 0.09 | 0.09 | 0.05 | 0.14 | 0.05 | 0.11 |
| 10 | 132059592 | rs1028920 | -0.641 | 0.539 | 0.849 | 1.837 | -0.002 | 0 | 0 | 0 | 0.28 | 0.31 | 0.31 | 0.29 | 0.24 | 0.44 |
| 10 | 132059763 | rs1028919 | 0.036 | 0.514 | 0.919 | 1.927 | 0.006 | -0.011 | -0.011 | 0 | 0.13 | 0.09 | 0.05 | 0.22 | 0.07 | 0.16 |
| 10 | 132061197 | rs2480271 | 1.444 | 0.553 | 0.963 | 1.691 | 0 | 0.043 | 0.043 | 0.292 | 0.72 | 0.43 | 0.75 | 0.44 | 0.74 | 0.66 |
| 10 | 132061518 | rs2771227 | -0.258 | 0.655 | 0.936 | 1.241 | 0.029 | 0.025 | 0.025 | 0.197 | 0.89 | 0.63 | 0.85 | 0.91 | 0.88 | 0.92 |
| 10 | 132061768 | rs2771226 | -1.31 | 0.709 | 0.686 | 1.265 | 0.013 | 0.012 | 0.012 | 0.176 | 0.93 | 0.84 | 0.86 | 0.91 | 0.88 | 0.92 |
| 10 | 132063889 | rs2628152 | 2.248 | 0.59 | 0.416 | 1.079 | -0.036 | 0.234 | 0.234 | 3.222 | 0.44 | 0.12 | 0.43 | 0.14 | 0.5 | 0.21 |
| 10 | 132063909 | rs2628153 | 2.271 | 0.615 | 0.417 | 1.08 | -0.045 | 0.231 | 0.231 | 2.739 | 0.44 | 0.12 | 0.43 | 0.14 | 0.5 | 0.21 |
| 10 | 132064890 | rs2771223 | 0.587 | 0.43 | 0.815 | 1.286 | 0.011 | 0.113 | 0.113 | 0.032 | 0.81 | 0.6 | 0.64 | 0.66 | 0.8 | 0.72 |
| 10 | 132072271 | rs7909507 | -0.354 | 0.726 | 0.568 | 1.341 | -0.014 | -0.012 | -0.012 | 0 | 0.22 | 0.07 | 0.11 | 0.22 | 0.21 | 0.21 |
| 10 | 132073879 | rs2628148 | 1.958 | 0.818 | 0.599 | 1.325 | 0.002 | 0.099 | 0.099 | 0.361 | 0.49 | 0.29 | 0.48 | 0.32 | 0.55 | 0.3 |
| 10 | 132079770 | rs2449882 | -1.724 | 0.921 | 0.52 | 1.434 | -0.036 | -0.034 | -0.034 | 0.826 | 0.94 | 1 | 0.87 | 0.91 | 0.88 | 0.93 |
| 10 | 132080749 | rs12265437 | 1.324 | 0.886 | 0.632 | 1.379 | 0.038 | 0.065 | 0.065 | 0 | 0.05 | 0.13 | 0.04 | 0.18 | 0.06 | 0.09 |
| 10 | 132083611 | rs184846 | -2.655 | 0.822 | 0.448 | 1.271 | -0.035 | 0.25 | 0.25 | 3.613 | 0.55 | 0.88 | 0.56 | 0.86 | 0.5 | 0.79 |
| 10 | 132096167 | rs11017185 | 0.162 | 0.814 | 0.624 | 1.646 | 0.004 | -0.025 | -0.025 | 0 | 0.1 | 0.04 | 0.02 | 0.21 | 0.06 | 0.24 |
| 10 | 132102683 | rs11017192 | 1.059 | 0.92 | 0.835 | 1.648 | 0.046 | 0.076 | 0.076 | 0 | 0.05 | 0.17 | 0.04 | 0.18 | 0.06 | 0.09 |
| 10 | 132108832 | rs1030782 | 0.021 | 0.912 | 0.856 | 1.738 | -0.001 | 0.071 | 0.071 | 0 | 0.1 | 0.15 | 0.21 | 0.26 | 0.1 | 0.21 |
| 10 | 132118122 | rs359053 | 0.247 | 1.046 | 0.779 | 1.423 | 0.027 | 0.038 | 0.038 | 0.945 | 0.11 | 0.13 | 0.17 | 0.27 | 0.18 | 0.17 |
| 10 | 132121961 | rs12247973 | 0.242 | 0.721 | 1.309 | 1.281 | 0.027 | 0.075 | 0.075 | 0 | 0.08 | 0.34 | 0.09 | 0.11 | 0.08 | 0.07 |
| 10 | 132124481 | rs2117650 | 2.578 | 0.564 | 0.727 | 0.954 | -0.082 | 0.23 | 0.23 | 1.316 | 0.42 | 0.06 | 0.42 | 0.14 | 0.44 | 0.2 |
| 10 | 132124523 | rs1469808 | 1.902 | 0.539 | 0.912 | 1.11 | -0.027 | 0.135 | 0.135 | 1.704 | 0.5 | 0.16 | 0.45 | 0.34 | 0.5 | 0.31 |
| 10 | 132126118 | rs12413734 | 0.296 | 0.52 | 0.875 | 1.143 | -0.007 | 0.092 | 0.092 | 0 | 0.1 | 0.16 | 0.2 | 0.28 | 0.09 | 0.19 |
| 10 | 132126298 | rs10764948 | 1.433 | 0.553 | 0.885 | 1.061 | -0.001 | -0.001 | -0.001 | 0 | 0.58 | 0.45 | 0.54 | 0.45 | 0.58 | 0.38 |
| 10 | 132134695 | rs435701 | -0.967 | 0.832 | 0.836 | 1.107 | 0.041 | 0.063 | 0.063 | 2.255 | 0.17 | 0.44 | 0.19 | 0.25 | 0.2 | 0.32 |
| 10 | 132138397 | rs6482785 | 1.817 | 0.798 | -0.002 | 0.572 | 0.001 | 0.009 | 0.009 | 0 | 0.53 | 0.52 | 0.45 | 0.16 | 0.46 | 0.26 |
| 10 | 132139508 | rs447363 | -1.871 | 0.789 | 0.032 | 0.621 | -0.016 | 0.07 | 0.07 | 0 | 0.43 | 0.31 | 0.46 | 0.69 | 0.47 | 0.63 |
| 10 | 132140841 | rs366346 | -0.979 | 0.884 | -0.021 | 0.674 | 0.013 | 0.019 | 0.019 | 0 | 0.93 | 0.83 | 0.91 | 0.84 | 0.93 | 0.89 |
| 10 | 132141392 | rs7903921 | 1.513 | 0.859 | -0.197 | 0.855 | 0.01 | 0.025 | 0.025 | 1.193 | 0.68 | 0.55 | 0.71 | 0.57 | 0.65 | 0.61 |
| 10 | 132144195 | rs1385259 | 2.948 | 1.012 | -0.212 | 0.795 | 0.005 | 0.009 | 0.009 | 1.136 | 0.42 | 0.34 | 0.4 | 0.15 | 0.38 | 0.19 |
| 10 | 132144804 | rs11017211 | 0.168 | 0.749 | -0.891 | 1.263 | -0.001 | 0.001 | 0.001 | 0 | 0.16 | 0.17 | 0.12 | 0.14 | 0.17 | 0.1 |
| 10 | 132144959 | rs654789 | 0.901 | 0.77 | -0.918 | 1.309 | -0.046 | -0.034 | -0.034 | 0 | 0.08 | 0.01 | 0.12 | 0.07 | 0.16 | 0.1 |
| 10 | 132145331 | rs12360159 | 0.18 | 0.767 | -0.917 | 1.308 | -0.001 | 0.001 | 0.001 | 0 | 0.16 | 0.17 | 0.12 | 0.14 | 0.17 | 0.1 |
| 10 | 132145351 | rs359080 | -0.594 | 0.578 | -0.838 | 1.306 | -0.023 | -0.008 | -0.008 | 0.014 | 0.24 | 0.19 | 0.24 | 0.21 | 0.33 | 0.2 |
| 10 | 132145664 | rs168596 | -1.106 | 0.486 | -0.772 | 1.13 | -0.028 | -0.028 | -0.028 | 0 | 0.33 | 0.23 | 0.43 | 0.49 | 0.42 | 0.52 |
| 10 | 132147366 | rs7914348 | 1.082 | -0.018 | -0.654 | 1.152 | -0.006 | -0.004 | -0.004 | 0 | 0.05 | 0.02 | 0.02 | 0.2 | 0.05 | 0.11 |
| 10 | 132150299 | rs359069 | -0.032 | 0.035 | -0.599 | 1.013 | -0.002 | -0.001 | -0.001 | 0 | 0.88 | 0.88 | 0.89 | 0.85 | 0.87 | 0.85 |
| 10 | 132150822 | rs7083725 | 1.192 | -0.302 | -0.635 | 0.942 | -0.002 | 0.031 | 0.031 | 0.001 | 0.56 | 0.66 | 0.48 | 0.39 | 0.48 | 0.34 |
| 10 | 132151982 | rs10829728 | 1.19 | -0.304 | -0.615 | 0.941 | -0.001 | 0.024 | 0.024 | 0 | 0.56 | 0.64 | 0.48 | 0.39 | 0.48 | 0.34 |
| 10 | 132152178 | rs12569655 | 0.523 | -0.749 | -0.373 | 1.103 | -0.008 | 0.029 | 0.029 | 0 | 0.06 | 0.02 | 0.18 | 0.27 | 0.08 | 0.18 |

| | | | | | | | | | | | | | | | | |
|---|---|---|---|---|---|---|---|---|---|---|---|---|---|---|---|---|
| 10 | 132153491 | rs4751197 | 1.465 | -2.089 | -0.264 | 0.192 | 0.048 | -0.026 | -0.026 | 0 | 0.45 | 0.22 | 0.55 | 0.42 | 0.63 | 0.4 |
| 10 | 132154475 | rs359066 | 0.895 | -1.659 | 0.336 | 0.142 | -0.008 | -0.022 | -0.022 | 0 | 0.68 | 0.58 | 0.77 | 0.83 | 0.76 | 0.8 |
| 10 | 132157137 | rs452678 | 1.303 | -1.835 | 0.577 | 0.123 | -0.016 | -0.055 | -0.055 | 0.888 | 0.63 | 0.47 | 0.65 | 0.8 | 0.68 | 0.75 |
| 10 | 132159651 | rs10829730 | 0.032 | -1.842 | 0.581 | 0.537 | -0.021 | -0.022 | -0.022 | 0 | 0.05 | 0.01 | 0.1 | 0.14 | 0.09 | 0.1 |
| 10 | 132160456 | rs7913323 | 1.485 | -2.258 | 1.091 | 0.509 | -0.024 | 0.009 | 0.009 | 0 | 0.67 | 0.55 | 0.74 | 0.56 | 0.82 | 0.58 |
| 10 | 132163841 | rs243267 | -0.013 | -2.086 | 1.047 | 0.462 | 0.02 | 0.02 | 0.02 | 0 | 0.86 | 0.89 | 0.95 | 0.96 | 0.95 | 0.95 |
| 10 | 132166896 | rs173379 | 0.926 | -2.132 | 1.208 | 0.433 | -0.023 | -0.044 | -0.044 | 0.361 | 0.79 | 0.7 | 0.82 | 0.94 | 0.86 | 0.89 |
| 10 | 132167165 | rs652182 | -0.277 | -2.221 | 1.278 | 0.555 | -0.001 | 0.02 | 0.02 | 3.123 | 0.13 | 0.13 | 0.09 | 0.38 | 0.06 | 0.32 |
| 10 | 132168065 | rs11017221 | 0.856 | -2.161 | 1.205 | 0.527 | -0.006 | -0.002 | -0.002 | 0.036 | 0.13 | 0.14 | 0.14 | 0.16 | 0.15 | 0.11 |
| 10 | 132168379 | rs359114 | -0.221 | -2.384 | 1.363 | 0.478 | -0.002 | -0.003 | -0.003 | 0 | 0.22 | 0.19 | 0.1 | 0.12 | 0.14 | 0.21 |
| 10 | 132168849 | rs1001694 | 2.274 | -2.182 | 1.176 | 0.568 | -0.044 | -0.035 | -0.035 | 0.113 | 0.39 | 0.14 | 0.52 | 0.39 | 0.56 | 0.42 |
| 10 | 132170008 | rs877023 | -0.692 | -1.204 | 2.36 | 1.129 | 0.016 | 0.013 | 0.013 | 0 | 0.18 | 0.27 | 0.25 | 0.24 | 0.24 | 0.17 |
| 10 | 132172598 | rs7915054 | -0.093 | -1.18 | 1.952 | 0.812 | -0.011 | 0.005 | 0.005 | 0 | 0.05 | 0.02 | 0.14 | 0.32 | 0.09 | 0.09 |
| 10 | 132173596 | rs359116 | 1.195 | -1.255 | 1.945 | 0.425 | -0.074 | -0.142 | -0.142 | 0 | 0.71 | 0.52 | 0.86 | 0.96 | 0.85 | 0.8 |
| 10 | 132174900 | rs10764951 | -0.931 | -1.079 | 1.972 | 0.528 | 0.037 | 0.125 | 0.125 | 0 | 0.21 | 0.49 | 0.32 | 0.48 | 0.24 | 0.2 |
| 10 | 132176907 | rs427721 | 0.702 | -1.148 | 1.99 | 0.489 | -0.021 | -0.091 | -0.091 | 2.385 | 0.8 | 0.59 | 0.81 | 0.96 | 0.82 | 0.9 |
| 10 | 132177303 | rs612028 | 1.25 | -1.211 | 2.071 | 0.411 | 0.001 | -0.004 | -0.004 | 0 | 0.64 | 0.3 | 0.62 | 0.52 | 0.67 | 0.75 |
| 10 | 132179421 | rs7084067 | -0.448 | -1.204 | 2.076 | 0.416 | 0.022 | 0.076 | 0.076 | 0 | 0.21 | 0.39 | 0.31 | 0.46 | 0.24 | 0.2 |
| 10 | 132179610 | rs435880 | -0.109 | -1.2 | 2.097 | 0.414 | -0.031 | -0.096 | -0.096 | 0.664 | 0.85 | 0.66 | 0.92 | 1 | 0.91 | 0.96 |
| 10 | 132190059 | rs440075 | 1.076 | -1.139 | 1.578 | 0.45 | 0.004 | -0.01 | -0.01 | 0 | 0.64 | 0.23 | 0.61 | 0.52 | 0.67 | 0.75 |
| 10 | 132191520 | rs12253830 | 1.949 | -1.155 | 1.439 | 0.413 | 0.047 | 0.022 | 0.022 | 0.001 | 0.51 | 0.15 | 0.56 | 0.5 | 0.62 | 0.6 |
| 10 | 132193733 | rs415219 | 1.498 | -1.14 | 1.351 | 0.42 | 0.012 | -0.004 | -0.004 | 0 | 0.6 | 0.22 | 0.61 | 0.54 | 0.66 | 0.64 |
| 10 | 132194315 | rs10764954 | -2.018 | -0.392 | 1.96 | 1.227 | -0.001 | 0.044 | 0.044 | 0 | 0.34 | 0.49 | 0.43 | 0.54 | 0.3 | 0.36 |
| 10 | 132194414 | rs10764955 | -0.963 | -0.341 | 1.662 | 1.28 | 0.041 | 0.141 | 0.141 | 0 | 0.21 | 0.43 | 0.38 | 0.5 | 0.25 | 0.2 |
| 10 | 132194847 | rs7923478 | -1.95 | -0.295 | 1.011 | 1.164 | 0.002 | 0.024 | 0.024 | 0 | 0.35 | 0.55 | 0.38 | 0.51 | 0.31 | 0.37 |
| 10 | 132194943 | rs10764956 | -1.366 | 0.601 | 1.153 | 1.485 | 0.06 | 0.102 | 0.102 | 0 | 0.23 | 0.43 | 0.32 | 0.48 | 0.27 | 0.21 |
| 10 | 132200602 | rs12571833 | -0.715 | 0.027 | 1.027 | 1.532 | 0 | 0.009 | 0.009 | 0 | 0.06 | 0.05 | 0.04 | 0.15 | 0.09 | 0.05 |
| 10 | 132202054 | rs10430606 | -1.042 | 0.232 | 1.559 | 1.838 | 0.072 | 0.145 | 0.145 | 0 | 0.13 | 0.35 | 0.19 | 0.32 | 0.15 | 0.22 |
| 10 | 132205196 | rs10829742 | -1.019 | 0.233 | 1.488 | 1.838 | 0.049 | 0.105 | 0.105 | 0 | 0.13 | 0.3 | 0.19 | 0.32 | 0.15 | 0.22 |
| 10 | 132208261 | rs11017239 | 1.04 | -0.043 | -0.194 | 1.293 | -0.057 | -0.01 | -0.01 | 0 | 0.57 | 0.08 | 0.55 | 0.37 | 0.6 | 0.5 |
| 10 | 132212976 | rs672996 | 0.354 | -0.392 | 0.603 | 0.902 | 0 | -0.014 | -0.014 | 0.005 | 0.8 | 0.8 | 0.85 | 0.83 | 0.81 | 0.75 |
| 10 | 132220631 | rs165517 | 0.104 | -0.507 | 1.564 | 0.931 | 0 | -0.002 | -0.002 | 0 | 0.83 | 0.86 | 0.87 | 0.83 | 0.82 | 0.73 |
| 10 | 132221435 | rs7082179 | -1.191 | -1.065 | 1.678 | 1.284 | -0.008 | 0.045 | 0.045 | 0 | 0.14 | 0.16 | 0.32 | 0.51 | 0.07 | 0.28 |
| 10 | 132223115 | rs11017245 | -1.899 | -1.64 | 1.19 | 0.932 | -0.073 | 0.089 | 0.089 | 0 | 0.24 | 0.46 | 0.38 | 0.56 | 0.14 | 0.32 |
| 10 | 132224296 | rs165516 | 1.874 | -1.637 | 1.086 | 0.926 | 0.052 | 0.097 | 0.097 | 0 | 0.52 | 0.33 | 0.42 | 0.07 | 0.63 | 0.34 |
| 10 | 132224722 | rs4750806 | -1.163 | -1.484 | 1.223 | 1.089 | -0.048 | 0.213 | 0.213 | 0 | 0.15 | 0.38 | 0.35 | 0.51 | 0.07 | 0.28 |
| 10 | 132225095 | rs7912913 | -0.577 | -1.451 | 1.271 | 1.292 | 0.004 | 0.395 | 0.395 | 0 | 0.08 | 0.32 | 0.32 | 0.45 | 0.05 | 0.13 |
| 10 | 132233642 | rs11017255 | -0.716 | -1.454 | 1.094 | 1.374 | 0.001 | 0.356 | 0.356 | 0 | 0.09 | 0.34 | 0.32 | 0.43 | 0.05 | 0.13 |
| 10 | 132238741 | rs165507 | 1.103 | -1.276 | 0.141 | 0.97 | 0.014 | 0.005 | 0.005 | 0 | 0.67 | 0.74 | 0.8 | 0.7 | 0.72 | 0.54 |
| 10 | 132250590 | rs359063 | 0.481 | -0.538 | 0.148 | 0.9 | -0.001 | -0.002 | -0.002 | 2.74 | 0.79 | 0.81 | 0.84 | 0.76 | 0.75 | 0.68 |
| 10 | 132254561 | rs10829748 | -0.934 | -0.397 | 0.2 | 0.443 | 0.004 | -0.038 | -0.038 | 0.067 | 0.34 | 0.44 | 0.51 | 0.74 | 0.34 | 0.48 |
| 10 | 132262631 | rs416737 | -0.703 | -0.377 | 0.791 | 1.158 | 0.04 | 0.231 | 0.231 | 0 | 0.14 | 0.3 | 0.29 | 0.39 | 0.12 | 0.19 |
| 10 | 132267750 | rs88722 | -1.535 | -0.898 | 0.802 | 0.661 | -0.001 | 0.013 | 0.013 | 0 | 0.3 | 0.56 | 0.46 | 0.53 | 0.26 | 0.33 |
| 10 | 132268039 | rs3858306 | -0.429 | -1.133 | 0.603 | 0.524 | -0.017 | -0.069 | -0.069 | 0 | 0.12 | 0.26 | 0.15 | 0 | 0.08 | 0.08 |
| 10 | 132273788 | rs11597160 | -1.146 | -1.236 | 0.782 | 0.548 | -0.004 | 0.005 | 0.005 | 0.219 | 0.13 | 0.22 | 0.15 | 0.14 | 0.12 | 0.09 |
| 10 | 132279008 | rs410411 | -1.092 | -1.167 | 1.059 | 0.677 | 0.005 | 0.086 | 0.086 | 1.172 | 0.2 | 0.3 | 0.4 | 0.39 | 0.17 | 0.23 |
| 10 | 132281238 | rs430345 | 0.226 | -1.323 | 0.775 | 0.482 | -0.003 | -0.002 | -0.002 | 0 | 0.94 | 0.94 | 0.94 | 0.86 | 0.94 | 0.9 |
| 10 | 132289763 | rs7075908 | 0.047 | -1.409 | 0.758 | 0.423 | 0.004 | 0.005 | 0.005 | 0.365 | 0.07 | 0.1 | 0.12 | 0 | 0.07 | 0.03 |
| 10 | 132292622 | rs12411435 | -0.962 | -1.584 | 1.502 | 0.407 | -0.009 | 0.081 | 0.081 | 2.827 | 0.21 | 0.36 | 0.39 | 0.39 | 0.16 | 0.22 |
| 10 | 132296407 | rs11017288 | 1.828 | -1.634 | 1.545 | 0.386 | 0.006 | -0.007 | -0.007 | 0.011 | 0.67 | 0.31 | 0.5 | 0.47 | 0.77 | 0.64 |
| 10 | 132296750 | rs4506548 | -0.835 | -1.578 | 1.582 | 0.412 | -0.004 | 0.035 | 0.035 | 2.371 | 0.2 | 0.28 | 0.38 | 0.39 | 0.16 | 0.22 |
| 10 | 132297773 | rs10764960 | 1.606 | -2.851 | 0.715 | -0.262 | 0.082 | 0.058 | 0.058 | 0 | 0.5 | 0.31 | 0.46 | 0.38 | 0.7 | 0.55 |
| 10 | 132300535 | rs1475779 | -0.877 | -3.059 | 0.688 | -0.348 | -0.01 | 0.009 | 0.009 | 1.766 | 0.32 | 0.38 | 0.42 | 0.47 | 0.21 | 0.29 |
| 10 | 132300883 | rs10450337 | -1.608 | -3.018 | 0.508 | -0.326 | 0.101 | -0.022 | -0.022 | 0 | 0.45 | 0.75 | 0.49 | 0.48 | 0.24 | 0.36 |
| 10 | 132311372 | rs1334913 | 0.043 | -2.86 | 0.546 | -0.273 | 0.009 | 0.009 | 0.009 | 0 | 0.18 | 0.22 | 0.15 | 0.09 | 0.13 | 0.1 |
| 10 | 132312927 | rs933077 | 1.149 | -2.901 | 0.731 | -0.254 | -0.011 | -0.001 | -0.001 | 0 | 0.63 | 0.52 | 0.51 | 0.39 | 0.73 | 0.62 |
| 10 | 132313026 | rs10829762 | 0.226 | -2.789 | 0.643 | -0.236 | -0.002 | 0.1 | 0.1 | 0 | 0.14 | 0.2 | 0.27 | 0.39 | 0.09 | 0.19 |
| 10 | 132319854 | rs7899888 | -0.183 | -2.861 | 0.078 | -0.721 | 0.015 | 0.01 | 0.01 | 0 | 0.43 | 0.44 | 0.36 | 0.36 | 0.35 | 0.41 |
| 10 | 132323265 | rs10829764 | 0.602 | -2.713 | 0.161 | -0.667 | -0.002 | 0.148 | 0.148 | 0 | 0.14 | 0.25 | 0.29 | 0.44 | 0.09 | 0.18 |
| 10 | 132328300 | rs11017310 | -0.112 | -2.806 | 0.01 | -0.785 | -0.002 | -0.001 | -0.001 | 0 | 0.42 | 0.46 | 0.35 | 0.31 | 0.35 | 0.41 |
| 10 | 132328529 | rs11017313 | 1.234 | -2.88 | -0.037 | -0.836 | -0.17 | 0.272 | 0.272 | 0.105 | 0.38 | 0.05 | 0.27 | 0.05 | 0.5 | 0.32 |
| 10 | 132329272 | rs7094321 | 0.905 | -2.885 | 0.134 | -0.771 | 0.001 | 0.009 | 0.009 | 0 | 0.23 | 0.3 | 0.19 | 0.23 | 0.22 | 0.28 |
| 10 | 132333323 | rs12260158 | 0.555 | -2.407 | -0.186 | -0.823 | 0.026 | 0.048 | 0.048 | 0 | 0.2 | 0.15 | 0.17 | 0.08 | 0.15 | 0.14 |
| 10 | 132339630 | rs7074043 | 0.855 | -2.236 | -0.122 | -0.712 | -0.003 | 0.003 | 0.003 | 0 | 0.23 | 0.29 | 0.17 | 0.22 | 0.21 | 0.28 |
| 10 | 132341125 | rs10829769 | 0.147 | -2.231 | -0.428 | -0.787 | 0.005 | 0.074 | 0.074 | 0 | 0.58 | 0.34 | 0.45 | 0.14 | 0.64 | 0.44 |
| 10 | 132342120 | rs11592555 | 2.043 | -1.945 | -0.828 | -0.903 | -0.16 | 0.153 | 0.153 | 0.06 | 0.23 | 0.01 | 0.22 | 0.01 | 0.42 | 0.14 |
| 10 | 132342752 | rs915206 | -0.181 | -1.316 | -0.198 | -0.287 | -0.002 | 0.057 | 0.057 | 0 | 0.71 | 0.52 | 0.74 | 0.59 | 0.73 | 0.62 |
| 10 | 132345497 | rs4750814 | -0.899 | -1.286 | 0.503 | 0.528 | -0.03 | 0.027 | 0.027 | 0.219 | 0.37 | 0.49 | 0.48 | 0.51 | 0.26 | 0.32 |
| 10 | 132346727 | rs10829770 | 0.767 | -1.217 | 0.364 | -0.197 | -0.005 | -0.008 | -0.008 | 0 | 0.58 | 0.44 | 0.43 | 0.26 | 0.69 | 0.58 |

| | | | | | | | | | | | | | | | | |
|---|---|---|---|---|---|---|---|---|---|---|---|---|---|---|---|---|
| 10 | 132348289 | rs7099645 | 0.634 | -0.492 | 0.586 | -0.146 | -0.002 | -0.003 | -0.003 | 0 | 0.84 | 0.78 | 0.91 | 0.89 | 0.89 | 0.74 |
| 10 | 132348416 | rs7088267 | -1.847 | -0.16 | 1.237 | -0.224 | -0.003 | -0.061 | -0.061 | 0 | 0.21 | 0.66 | 0.29 | 0.05 | 0.21 | 0.1 |
| 10 | 132357304 | rs4750818 | -1.752 | 0.202 | 1.212 | 0.025 | 0.045 | -0.052 | -0.052 | 0 | 0.14 | 0.45 | 0.25 | 0.03 | 0.16 | 0.09 |
| 10 | 132360110 | rs7910053 | -1.494 | 0.258 | 1.05 | 0.033 | 0.044 | -0.044 | -0.044 | 0 | 0.12 | 0.35 | 0.23 | 0.03 | 0.16 | 0.09 |
| 10 | 132366530 | rs4751239 | -1.456 | 0.172 | 0.784 | -0.129 | -0.014 | -0.138 | -0.138 | 0 | 0.23 | 0.49 | 0.25 | 0.02 | 0.18 | 0.15 |
| 10 | 132369266 | rs11017338 | -0.252 | 0.417 | 1.041 | 1.329 | 0.042 | 0.28 | 0.28 | 0 | 0.06 | 0.17 | 0.2 | 0.37 | 0.08 | 0.07 |
| 10 | 132373205 | rs11597303 | 0.134 | 0.729 | 0.952 | 0.845 | 0.013 | -0.02 | -0.02 | 0 | 0.87 | 0.75 | 0.88 | 0.91 | 0.85 | 0.81 |
| 10 | 132374743 | rs7076942 | 0.982 | 0.649 | 0.823 | 1.114 | 0.002 | 0.005 | 0.005 | 0.038 | 0.71 | 0.54 | 0.59 | 0.33 | 0.71 | 0.51 |
| 10 | 132374877 | rs3858307 | -0.105 | 0.488 | 0.685 | 1.237 | 0.001 | -0.048 | -0.048 | 0.716 | 0.14 | 0.06 | 0.27 | 0.58 | 0.14 | 0.29 |
| 10 | 132376251 | rs4751241 | -0.444 | 0.798 | 0.853 | 1.49 | -0.007 | -0.022 | -0.022 | 0 | 0.17 | 0.11 | 0.24 | 0.31 | 0.18 | 0.29 |
| 10 | 132377231 | rs10764969 | -1.69 | 0.244 | 1.32 | 1.135 | -0.009 | -0.003 | -0.003 | 0 | 0.36 | 0.45 | 0.5 | 0.6 | 0.34 | 0.45 |
| 10 | 132384335 | rs7070951 | 0.018 | 0.343 | 0.935 | 1.178 | -0.002 | -0.05 | -0.05 | 0 | 0.16 | 0.06 | 0.21 | 0.3 | 0.14 | 0.28 |
| 10 | 132386550 | rs9299876 | -0.03 | 0.333 | 1.088 | 1.176 | 0 | -0.018 | -0.018 | 0 | 0.16 | 0.1 | 0.21 | 0.31 | 0.15 | 0.28 |
| 10 | 132386874 | rs11017357 | -0.422 | 0.111 | 1.656 | 0.857 | -0.016 | -0.076 | -0.076 | 0 | 0.17 | 0.26 | 0.09 | 0 | 0.14 | 0.05 |
| 10 | 132387676 | rs10829774 | -1.594 | 0.188 | 1.611 | 0.956 | -0.014 | 0.007 | 0.007 | 0 | 0.34 | 0.54 | 0.47 | 0.55 | 0.29 | 0.36 |
| 10 | 132387898 | rs11017359 | -1.595 | 0.204 | 1.627 | 0.944 | -0.008 | -0.003 | -0.003 | 0 | 0.36 | 0.54 | 0.52 | 0.6 | 0.34 | 0.44 |
| 10 | 132406880 | rs4751252 | -1.225 | 0.088 | 1.807 | 0.672 | 0.032 | 0.115 | 0.115 | 0 | 0.15 | 0.45 | 0.24 | 0.22 | 0.16 | 0.11 |
| 10 | 132407513 | rs1177847 | 1.629 | -0.115 | 1.709 | 0.618 | 0.003 | 0.088 | 0.088 | 0 | 0.82 | 0.52 | 0.77 | 0.73 | 0.82 | 0.81 |
| 10 | 132408125 | rs12251302 | 0.511 | 0.191 | 0.882 | 0.046 | -0.003 | 0.028 | 0.028 | 0 | 0.51 | 0.59 | 0.43 | 0.34 | 0.52 | 0.35 |
| 10 | 132409262 | rs12360112 | 0.734 | -0.305 | 0.345 | -0.479 | -0.024 | 0.007 | 0.007 | 0 | 0.17 | 0.3 | 0.23 | 0.16 | 0.11 | 0.2 |
| 10 | 132412890 | rs11017373 | -0.713 | -0.363 | 0.05 | -0.803 | -0.005 | -0.038 | -0.038 | 0 | 0.1 | 0.15 | 0.07 | 0.02 | 0.09 | 0.13 |
| 10 | 132416075 | rs11591632 | 1.61 | -0.301 | -0.454 | -0.994 | -0.032 | 0.06 | 0.06 | 0 | 0.22 | 0.11 | 0.14 | 0 | 0.3 | 0.09 |
| 10 | 132418881 | rs1184394 | -0.183 | -0.651 | -0.59 | -1.111 | -0.022 | 0.031 | 0.031 | 0 | 0.41 | 0.21 | 0.42 | 0.66 | 0.41 | 0.57 |
| 10 | 132419819 | rs1184393 | 0.218 | -0.535 | -1.124 | -1.52 | -0.075 | 0.005 | 0.005 | 0 | 0.27 | 0.08 | 0.46 | 0.71 | 0.31 | 0.47 |
| 10 | 132423915 | rs1177844 | -1.066 | -1.194 | 0.132 | -1.668 | 0.017 | 0.091 | 0.091 | 0 | 0.54 | 0.65 | 0.68 | 0.83 | 0.55 | 0.7 |
| 10 | 132429184 | rs12251170 | -1.177 | -1.199 | 0.624 | -1.616 | -0.002 | -0.011 | -0.011 | 0 | 0.14 | 0.42 | 0.13 | 0.11 | 0.1 | 0.19 |
| 10 | 132431119 | rs16911018 | -1.164 | -0.971 | 1.202 | -1.498 | 0.063 | 0.096 | 0.096 | 0 | 0.05 | 0.24 | 0.09 | 0.11 | 0.04 | 0.12 |
| 10 | 132434981 | rs1176471 | -0.284 | -1.042 | 1.135 | -1.593 | -0.001 | -0.001 | -0.001 | 0 | 0.46 | 0.47 | 0.43 | 0.74 | 0.47 | 0.63 |
| 10 | 132439825 | rs1176445 | -1.449 | -1.204 | 0.794 | -1.741 | -0.002 | 0.002 | 0.002 | 0 | 0.66 | 0.68 | 0.75 | 0.89 | 0.63 | 0.91 |
| 10 | 132448750 | rs1176502 | 0.256 | -1.075 | 1.01 | -1.522 | -0.003 | 0.005 | 0.005 | 0 | 0.39 | 0.33 | 0.33 | 0.63 | 0.4 | 0.5 |
| 10 | 132456892 | rs1702200 | -0.381 | -1.262 | 1.138 | -1.619 | 0 | -0.009 | -0.009 | 0.306 | 0.48 | 0.47 | 0.43 | 0.73 | 0.47 | 0.65 |
| 10 | 132459738 | rs1665539 | -0.413 | -1.251 | 1.254 | -1.601 | 0.006 | 0.056 | 0.056 | 0 | 0.5 | 0.56 | 0.45 | 0.73 | 0.48 | 0.69 |
| 10 | 132460015 | rs1665537 | -0.413 | -1.251 | 1.254 | -1.601 | 0.006 | 0.056 | 0.056 | 0 | 0.5 | 0.56 | 0.45 | 0.73 | 0.48 | 0.69 |
| 10 | 132461165 | rs1702206 | -0.29 | -1.305 | 1.286 | -1.614 | -0.009 | 0.061 | 0.061 | 0 | 0.45 | 0.61 | 0.45 | 0.73 | 0.48 | 0.68 |
| 10 | 132462185 | rs7898050 | 0.338 | -0.999 | 1.656 | -1.241 | 0.009 | 0.023 | 0.023 | 0 | 0.05 | 0.1 | 0.21 | 0.09 | 0.07 | 0.05 |
| 10 | 132468412 | rs7917753 | -2.127 | -1.522 | 1.973 | -1.318 | -0.008 | -0.016 | -0.016 | 0.002 | 0.15 | 0.39 | 0.13 | 0.11 | 0.11 | 0.17 |
| 10 | 132471605 | rs1176493 | 1.589 | -1.517 | 1.842 | -1.328 | -0.035 | 0.013 | 0.013 | 0 | 0.87 | 0.56 | 0.88 | 0.9 | 0.93 | 0.89 |
| 10 | 132477287 | rs2921907 | 0.173 | -1.741 | 1.52 | -1.516 | 0.001 | 0.078 | 0.078 | 0 | 0.46 | 0.27 | 0.48 | 0.83 | 0.49 | 0.66 |
| 10 | 132483929 | rs1176461 | 1.346 | -1.735 | 1.422 | -1.521 | -0.085 | -0.017 | -0.017 | 0 | 0.34 | 0.09 | 0.31 | 0.64 | 0.4 | 0.52 |
| 10 | 132489040 | rs1185023 | 1.346 | -1.735 | 1.422 | -1.521 | -0.085 | -0.017 | -0.017 | 0 | 0.34 | 0.09 | 0.31 | 0.64 | 0.4 | 0.52 |
| 10 | 132491701 | rs2939924 | 0.285 | -1.669 | 1.491 | -1.53 | -0.006 | 0.114 | 0.114 | 0 | 0.46 | 0.18 | 0.47 | 0.74 | 0.49 | 0.74 |
| 10 | 132492920 | rs16911059 | 0.06 | -1.382 | 1.39 | -1.411 | 0.014 | -0.004 | -0.004 | 0 | 0.08 | 0.14 | 0.12 | 0.1 | 0.06 | 0.06 |
| 10 | 132500555 | rs12245254 | 0.275 | -1.45 | 1.238 | -1.469 | -0.003 | 0.123 | 0.123 | 0 | 0.42 | 0.29 | 0.33 | 0.08 | 0.43 | 0.2 |
| 10 | 132504534 | rs10829777 | -0.08 | -1.462 | 1.157 | -1.458 | 0.003 | 0.02 | 0.02 | 0 | 0.45 | 0.41 | 0.47 | 0.16 | 0.48 | 0.22 |
| 10 | 132515601 | rs1876314 | -0.93 | -1.364 | 0.875 | -1.456 | -0.009 | -0.016 | -0.016 | 0 | 0.67 | 0.71 | 0.76 | 1 | 0.61 | 0.9 |
| 10 | 132515808 | rs1911370 | 0.551 | -1.258 | 0.223 | -1.341 | 0.018 | 0.014 | 0.014 | 0 | 0.12 | 0.09 | 0.09 | 0.09 | 0.06 | 0.15 |
| 10 | 132516104 | rs2174715 | 0.66 | -0.487 | 0.738 | -1.062 | -0.014 | 0.111 | 0.111 | 0 | 0.49 | 0.27 | 0.43 | 0.72 | 0.47 | 0.74 |
| 10 | 132522563 | rs10829783 | -0.612 | -0.173 | 0.84 | -0.742 | -0.001 | 0.004 | 0.004 | 0.021 | 0.14 | 0.16 | 0.2 | 0.24 | 0.15 | 0.29 |
| 10 | 132525181 | rs4751263 | 1.185 | -0.327 | 0.779 | -0.701 | -0.041 | 0.009 | 0.009 | 0 | 0.44 | 0.13 | 0.34 | 0.6 | 0.41 | 0.53 |
| 10 | 132526482 | rs7089879 | -1.436 | -0.092 | 1.096 | -0.945 | -0.004 | 0.037 | 0.037 | 0.536 | 0.22 | 0.48 | 0.36 | 0.29 | 0.2 | 0.32 |
| 10 | 132529833 | rs1014637 | -0.451 | -0.131 | 0.924 | -1.031 | -0.005 | 0.014 | 0.014 | 0 | 0.67 | 0.73 | 0.78 | 0.98 | 0.63 | 0.88 |
| 10 | 132531025 | rs12219907 | -1.159 | -0.326 | 1.326 | 0.096 | 0.022 | 0.077 | 0.077 | 0 | 0.05 | 0.28 | 0.12 | 0.05 | 0.02 | 0.04 |
| 10 | 132538195 | rs1876319 | 1.361 | -0.172 | 0.166 | 0.075 | 0.008 | -0.01 | -0.01 | 0 | 0.72 | 0.27 | 0.58 | 0.64 | 0.76 | 0.61 |
| 10 | 132543170 | rs12146195 | -0.785 | -0.327 | 0.075 | -0.109 | -0.005 | 0 | 0 | 0 | 0.14 | 0.17 | 0.1 | 0.17 | 0.12 | 0.29 |
| 10 | 132547516 | rs11017438 | 0.009 | -0.799 | 0.902 | -0.708 | -0.026 | -0.09 | -0.09 | 0 | 0.75 | 0.48 | 0.73 | 0.93 | 0.81 | 0.88 |
| 10 | 132547998 | rs9733640 | -0.416 | -0.866 | 0.894 | -0.738 | -0.021 | -0.073 | -0.073 | 0 | 0.25 | 0.37 | 0.18 | 0.07 | 0.18 | 0.12 |
| 10 | 132555804 | rs7100411 | -1.075 | -1.166 | 0.901 | -0.709 | -0.019 | 0.011 | 0.011 | 0.022 | 0.25 | 0.36 | 0.24 | 0.29 | 0.22 | 0.39 |
| 10 | 132555904 | rs7893312 | 0.982 | -1.141 | 0.854 | -0.7 | -0.007 | 0.013 | 0.013 | 0.062 | 0.72 | 0.62 | 0.71 | 0.65 | 0.73 | 0.55 |
| 10 | 132561458 | rs10829791 | -0.865 | -1.39 | 0.78 | -0.709 | 0.006 | -0.016 | -0.016 | 0 | 0.21 | 0.12 | 0.17 | 0.24 | 0.21 | 0.36 |
| 10 | 132561953 | rs10829792 | 1.263 | -1.245 | 0.943 | -0.761 | 0.06 | 0.063 | 0.063 | 0.033 | 0.5 | 0.28 | 0.6 | 0.64 | 0.57 | 0.49 |
| 10 | 132563074 | rs2660054 | 0.534 | -1.492 | 0.511 | -0.765 | 0.002 | -0.031 | -0.031 | 6.592 | 0.2 | 0.2 | 0.12 | 0 | 0.17 | 0.06 |
| 10 | 132568315 | rs11017458 | -0.203 | -1.533 | 0.407 | -0.747 | 0.009 | -0.006 | -0.006 | 0 | 0.15 | 0.11 | 0.11 | 0.17 | 0.14 | 0.3 |
| 10 | 132572369 | rs10741226 | 1.378 | -1.612 | 0.525 | -0.73 | 0.088 | 0.055 | 0.055 | 0.659 | 0.49 | 0.19 | 0.56 | 0.64 | 0.57 | 0.48 |
| 10 | 132572375 | rs7093355 | -0.443 | -1.387 | 0.298 | -0.97 | 0.01 | -0.007 | -0.007 | 1.439 | 0.15 | 0.12 | 0.11 | 0.17 | 0.14 | 0.3 |
| 10 | 132574025 | rs1176413 | -0.197 | -0.965 | 0.443 | -0.996 | -0.032 | -0.121 | -0.121 | 0.009 | 0.21 | 0.35 | 0.14 | 0 | 0.18 | 0.06 |
| 10 | 132574696 | rs1912001 | -0.091 | -0.928 | -0.302 | -0.98 | 0.008 | -0.017 | -0.017 | 0 | 0.21 | 0.2 | 0.13 | 0 | 0.18 | 0.06 |
| 10 | 132577431 | rs12763340 | -0.926 | -0.876 | -0.189 | -0.774 | -0.018 | -0.068 | -0.068 | 0 | 0.18 | 0.22 | 0.07 | 0 | 0.15 | 0.11 |
| 10 | 132578847 | rs2663014 | 0.75 | -0.468 | -0.602 | -0.862 | 0.002 | 0.004 | 0.004 | 0 | 0.51 | 0.4 | 0.6 | 0.75 | 0.58 | 0.53 |
| 10 | 132579195 | rs2663013 | -0.159 | -0.324 | -0.86 | -0.456 | -0.003 | 0.003 | 0.003 | 0 | 0.94 | 0.94 | 0.95 | 0.95 | 0.95 | 0.93 |

| | | | | | | | | | | | | | | | | |
|---|---|---|---|---|---|---|---|---|---|---|---|---|---|---|---|---|
| 10 | 132580167 | rs1912000 | -0.341 | -0.358 | -0.777 | -0.528 | 0.001 | -0.023 | -0.023 | 0 | 0.11 | 0.12 | 0.04 | 0 | 0.13 | 0.09 |
| 10 | 132581334 | rs1911999 | 1.085 | -0.919 | -1.598 | -0.404 | 0.012 | 0.03 | 0.03 | 2.405 | 0.6 | 0.05 | 0.6 | 0.75 | 0.67 | 0.51 |
| 10 | 132583456 | rs4397751 | -1.317 | -0.961 | -1.443 | -0.351 | 0.037 | 0.069 | 0.069 | 2.189 | 0.4 | 0.9 | 0.31 | 0.25 | 0.34 | 0.52 |
| 10 | 132585831 | rs7077726 | 1.093 | -0.654 | -0.331 | 0.377 | -0.008 | 0.03 | 0.03 | 0 | 0.75 | 0.6 | 0.71 | 0.73 | 0.74 | 0.65 |
| 10 | 132590821 | rs10764980 | 0.868 | -1.317 | 0.116 | -0.035 | -0.047 | -0.025 | -0.025 | 0.248 | 0.78 | 0.67 | 0.74 | 0.81 | 0.83 | 0.78 |
| 10 | 132591791 | rs4553286 | -1.431 | -1.675 | 0.641 | 0.134 | -0.027 | -0.005 | -0.005 | 0 | 0.17 | 0.29 | 0.19 | 0.16 | 0.12 | 0.31 |
| 10 | 132595663 | rs7907842 | 0.339 | -1.942 | -0.172 | -0.582 | 0.039 | 0.051 | 0.051 | 0.017 | 0.51 | 0.7 | 0.55 | 0.66 | 0.57 | 0.46 |
| 10 | 132595971 | rs11017475 | 1.207 | -1.983 | 0.279 | -0.347 | -0.011 | -0.005 | -0.005 | 0 | 0.61 | 0.43 | 0.56 | 0.66 | 0.69 | 0.5 |
| 10 | 132596342 | rs12265537 | 0.633 | -1.425 | -0.145 | -0.128 | 0.004 | -0.008 | -0.008 | 0 | 0.84 | 0.83 | 0.69 | 0.72 | 0.81 | 0.65 |
| 10 | 132596470 | rs12245420 | 0.579 | -1.409 | -0.138 | -0.121 | 0.004 | -0.007 | -0.007 | 0 | 0.84 | 0.83 | 0.69 | 0.72 | 0.81 | 0.65 |
| 10 | 132597423 | rs10764985 | 0.508 | -1.202 | -0.133 | -0.161 | -0.001 | 0.001 | 0.001 | 0 | 0.86 | 0.85 | 0.82 | 0.77 | 0.82 | 0.71 |
| 10 | 132598275 | rs7085125 | 0.697 | -1.009 | -0.223 | 0.016 | 0.033 | 0.018 | 0.018 | 0 | 0.51 | 0.39 | 0.61 | 0.59 | 0.64 | 0.36 |
| 10 | 132602033 | rs11813883 | -0.73 | -0.077 | 0.214 | -0.27 | 0.051 | -0.073 | -0.073 | 0 | 0.16 | 0.34 | 0.18 | 0.06 | 0.21 | 0.1 |
| 10 | 132602111 | rs1655162 | -1.445 | -1.107 | 0.271 | -0.982 | -0.004 | -0.133 | -0.133 | 0 | 0.32 | 0.52 | 0.26 | 0.06 | 0.33 | 0.2 |
| 10 | 132602659 | rs12570660 | 0.425 | -0.159 | -0.59 | -0.86 | -0.073 | -0.067 | -0.067 | 0.068 | 0.16 | 0.01 | 0.09 | 0.23 | 0.19 | 0.22 |
| 10 | 132603060 | rs1655164 | -1.276 | -0.154 | -0.126 | -0.444 | 0.014 | 0.06 | 0.06 | 0.37 | 0.4 | 0.72 | 0.39 | 0.23 | 0.37 | 0.32 |
| 10 | 132604292 | rs10829802 | 1.899 | -0.585 | -0.046 | -0.337 | -0.049 | -0.079 | -0.079 | 0 | 0.28 | 0.14 | 0.38 | 0.44 | 0.38 | 0.24 |
| 10 | 132605497 | rs1703660 | 0.097 | -0.655 | 0.06 | -0.372 | 0.054 | 0.069 | 0.069 | 0.062 | 0.52 | 0.79 | 0.7 | 0.67 | 0.64 | 0.47 |
| 10 | 132606517 | rs1703663 | -0.877 | -0.647 | 0.586 | -0.396 | -0.004 | -0.039 | -0.039 | 0 | 0.38 | 0.44 | 0.36 | 0.24 | 0.37 | 0.32 |
| 10 | 132607399 | rs11017491 | 1.757 | -0.627 | 0.592 | -0.316 | -0.018 | -0.04 | -0.04 | 0.008 | 0.29 | 0.26 | 0.38 | 0.46 | 0.39 | 0.25 |
| 10 | 132608211 | rs2136286 | 0.626 | -0.515 | 0.616 | -0.429 | -0.004 | 0.002 | 0.002 | 0 | 0.43 | 0.48 | 0.58 | 0.64 | 0.51 | 0.44 |
| 10 | 132609125 | rs1631311 | -0.536 | -0.483 | 0.743 | -0.548 | 0.01 | 0.015 | 0.015 | 0 | 0.16 | 0.34 | 0.22 | 0.16 | 0.13 | 0.21 |
| 10 | 132615026 | rs7897190 | -0.73 | -0.591 | 0.68 | -0.642 | -0.002 | -0.008 | -0.008 | 0.211 | 0.32 | 0.29 | 0.29 | 0.18 | 0.28 | 0.25 |
| 10 | 132615497 | rs1655178 | -0.853 | -0.623 | 0.638 | -0.479 | 0.017 | -0.044 | -0.044 | 0 | 0.68 | 0.85 | 0.77 | 0.57 | 0.76 | 0.57 |
| 10 | 132618471 | rs4750832 | 0.221 | -0.537 | 0.818 | -0.273 | -0.011 | 0.037 | 0.037 | 0.007 | 0.1 | 0.22 | 0.13 | 0.15 | 0.04 | 0.15 |
| 10 | 132620849 | rs12242470 | -0.58 | -0.594 | 0.862 | -0.339 | -0.037 | -0.032 | -0.032 | 0 | 0.25 | 0.35 | 0.21 | 0.15 | 0.17 | 0.23 |
| 10 | 132623093 | rs10829813 | 0.282 | -0.548 | 0.858 | -0.331 | -0.009 | 0.021 | 0.021 | 0.008 | 0.1 | 0.19 | 0.13 | 0.15 | 0.04 | 0.15 |
| 10 | 132626327 | rs7088120 | 0.708 | -0.252 | 0.741 | -0.475 | -0.017 | -0.057 | -0.057 | 0 | 0.15 | 0.17 | 0.08 | 0 | 0.12 | 0.08 |
| 10 | 132627994 | rs7893625 | 1.156 | 0.231 | 0.563 | -0.16 | -0.013 | -0.019 | -0.019 | 0.065 | 0.29 | 0.27 | 0.39 | 0.38 | 0.39 | 0.24 |
| 10 | 132632688 | rs892330 | 0.724 | 0.265 | 0.434 | -0.114 | 0.029 | -0.02 | -0.02 | 0 | 0.05 | 0.15 | 0.08 | 0.01 | 0.09 | 0.06 |
| 10 | 132633081 | rs1655182 | 0.753 | 0.407 | 0.394 | -0.155 | -0.022 | -0.019 | -0.019 | 0 | 0.32 | 0.3 | 0.46 | 0.39 | 0.48 | 0.3 |
| 10 | 132634610 | rs1703720 | 0.52 | 0.686 | 0.332 | -0.266 | 0.002 | 0.004 | 0.004 | 0.027 | 0.3 | 0.33 | 0.39 | 0.38 | 0.35 | 0.3 |
| 10 | 132635031 | rs10829817 | 1.4 | 0.74 | 0.074 | -0.208 | -0.027 | -0.058 | -0.058 | 0 | 0.23 | 0.14 | 0.31 | 0.37 | 0.26 | 0.23 |
| 10 | 132645033 | rs1655097 | 0.806 | 0.653 | 0.437 | -0.168 | 0.005 | 0.008 | 0.008 | 0 | 0.28 | 0.35 | 0.38 | 0.36 | 0.37 | 0.24 |
| 10 | 132646047 | rs7076678 | 0.032 | 0.827 | 0.475 | -0.146 | 0.013 | -0.029 | -0.029 | 0 | 0.07 | 0.16 | 0.08 | 0.01 | 0.08 | 0.06 |
| 10 | 132647568 | rs919857 | 0.236 | 0.238 | 0.38 | -0.156 | 0.001 | -0.019 | -0.019 | 1.526 | 0.43 | 0.5 | 0.43 | 0.23 | 0.47 | 0.28 |
| 10 | 132648767 | rs1655120 | -0.541 | 0.211 | -0.055 | -0.862 | -0.032 | 0.082 | 0.082 | 0.096 | 0.69 | 0.84 | 0.86 | 0.97 | 0.7 | 0.75 |
| 10 | 132652504 | rs12360242 | -0.162 | 0.002 | 0.483 | 0.041 | 0.002 | 0.009 | 0.009 | 0 | 0.07 | 0.12 | 0.13 | 0.12 | 0.09 | 0.2 |
| 10 | 132662898 | rs1703702 | 0.181 | -0.21 | 0.078 | 0.12 | -0.018 | -0.012 | -0.012 | 0 | 0.72 | 0.66 | 0.8 | 0.74 | 0.81 | 0.85 |
| 10 | 132669618 | rs4751287 | 0.565 | -0.037 | -0.735 | -0.303 | -0.028 | 0.074 | 0.074 | 0.004 | 0.4 | 0.25 | 0.57 | 0.77 | 0.47 | 0.46 |
| 10 | 132673234 | rs1655152 | -0.863 | -0.506 | -0.822 | -0.645 | -0.01 | 0.027 | 0.027 | 0 | 0.78 | 0.85 | 0.87 | 0.98 | 0.77 | 0.8 |
| 10 | 132674944 | rs1655155 | -0.782 | -0.468 | -0.968 | -0.347 | -0.001 | -0.001 | -0.001 | 2.058 | 0.42 | 0.6 | 0.39 | 0.45 | 0.4 | 0.49 |
| 10 | 132675510 | rs1655156 | -0.591 | -1.095 | -1.083 | -0.742 | 0.002 | -0.012 | -0.012 | 0 | 0.32 | 0.58 | 0.29 | 0.28 | 0.35 | 0.29 |
| 10 | 132676113 | rs7902465 | 1.571 | -1.357 | -1.093 | -0.75 | -0.083 | -0.032 | -0.032 | 0 | 0.4 | 0.11 | 0.53 | 0.55 | 0.51 | 0.53 |
| 10 | 132676283 | rs4750838 | 0.599 | -1.367 | -0.937 | -0.795 | 0.008 | 0.019 | 0.019 | 0.58 | 0.57 | 0.35 | 0.7 | 0.67 | 0.64 | 0.66 |
| 10 | 132677088 | rs1703688 | -0.505 | -1.431 | -0.493 | -0.845 | 0.015 | -0.105 | -0.105 | 0.32 | 0.74 | 0.61 | 0.8 | 0.98 | 0.73 | 0.78 |
| 10 | 132677376 | rs1703687 | -0.045 | -1.081 | -0.338 | -0.635 | -0.003 | -0.011 | -0.011 | 0 | 0.62 | 0.54 | 0.72 | 0.67 | 0.67 | 0.71 |
| 10 | 132677593 | rs10829833 | 0.754 | -0.948 | -0.294 | -0.538 | -0.015 | -0.01 | -0.01 | 0 | 0.17 | 0.24 | 0.17 | 0.12 | 0.14 | 0.13 |
| 10 | 132677907 | rs1560797 | 0.603 | -0.636 | -0.042 | -0.545 | 0.008 | 0.021 | 0.021 | 0.58 | 0.57 | 0.35 | 0.7 | 0.67 | 0.64 | 0.66 |
| 10 | 132678584 | rs4237463 | -0.04 | -0.482 | -0.14 | -0.253 | -0.026 | -0.022 | -0.022 | 0 | 0.7 | 0.49 | 0.79 | 0.7 | 0.77 | 0.76 |
| 10 | 132679344 | rs1655159 | 0.231 | -0.176 | -0.328 | -0.433 | -0.004 | 0.037 | 0.037 | 0.007 | 0.44 | 0.38 | 0.7 | 0.78 | 0.53 | 0.53 |
| 10 | 132681679 | rs10741231 | 0.061 | 0.149 | -0.183 | -1.062 | 0.005 | -0.019 | -0.019 | 0 | 0.33 | 0.48 | 0.63 | 0.78 | 0.38 | 0.43 |
| 10 | 132682127 | rs12769679 | 0.176 | -0.244 | -0.156 | -0.92 | -0.005 | -0.001 | -0.001 | 0 | 0.45 | 0.36 | 0.56 | 0.44 | 0.47 | 0.46 |
| 10 | 132682436 | rs12768898 | 1.184 | -0.295 | -0.268 | -0.975 | 0.003 | 0.011 | 0.011 | 0.526 | 0.22 | 0.16 | 0.13 | 0.09 | 0.19 | 0.11 |
| 10 | 132682729 | rs4751295 | -0.739 | -0.1 | -0.39 | -1.044 | -0.01 | -0.049 | -0.049 | 0 | 0.74 | 0.65 | 0.87 | 0.9 | 0.78 | 0.71 |
| 10 | 132683814 | rs10829834 | 0.222 | 0.14 | -1.194 | -0.913 | -0.09 | -0.009 | -0.009 | 0 | 0.4 | 0.09 | 0.58 | 0.39 | 0.49 | 0.51 |
| 10 | 132684470 | rs12217313 | 0.929 | 0.147 | -0.721 | -0.692 | -0.034 | -0.05 | -0.05 | 0.02 | 0.21 | 0.08 | 0.47 | 0.36 | 0.28 | 0.34 |
| 10 | 132688084 | rs4751296 | -0.094 | 0.092 | -1.323 | -1.477 | 0 | 0.085 | 0.085 | 0 | 0.55 | 0.72 | 0.76 | 0.87 | 0.57 | 0.55 |
| 10 | 132689538 | rs892326 | -0.639 | 0.33 | -1.407 | -0.987 | -0.015 | -0.015 | -0.015 | 0 | 0.22 | 0.1 | 0.28 | 0.27 | 0.27 | 0.27 |
| 10 | 132690451 | rs1036268 | 0.504 | 0.166 | -1.084 | -1.404 | 0 | -0.006 | -0.006 | 0 | 0.35 | 0.44 | 0.51 | 0.65 | 0.31 | 0.47 |
| 10 | 132690651 | rs1036270 | 0.548 | 0.204 | -0.992 | -1.371 | -0.004 | -0.007 | -0.007 | 0 | 0.37 | 0.44 | 0.53 | 0.66 | 0.33 | 0.5 |
| 10 | 132690778 | rs1655122 | -0.287 | 0.174 | -1.359 | -1.644 | 0.006 | -0.019 | -0.019 | 0 | 0.41 | 0.47 | 0.34 | 0.33 | 0.43 | 0.45 |
| 10 | 132690980 | rs1703682 | -0.089 | 0.069 | -1.375 | -1.731 | -0.043 | 0.028 | 0.028 | 0 | 0.25 | 0.45 | 0.22 | 0.31 | 0.22 | 0.32 |
| 10 | 132692121 | rs1655125 | -0.922 | 0.359 | -2.256 | -1.639 | 0.004 | 0.039 | 0.039 | 0.825 | 0.78 | 0.87 | 0.86 | 0.98 | 0.76 | 0.96 |
| 10 | 132693122 | rs1655127 | -1.08 | 0.438 | -2.421 | -1.606 | -0.011 | 0.093 | 0.093 | 0.934 | 0.79 | 0.92 | 0.86 | 0.98 | 0.76 | 0.96 |
| 10 | 132694631 | rs11017550 | 0.736 | 1.162 | -1.902 | -0.897 | -0.096 | 0.091 | 0.091 | 0 | 0.16 | 0.01 | 0.11 | 0.01 | 0.22 | 0.13 |
| 10 | 132697051 | rs1655132 | -0.261 | 0.943 | -1.722 | -0.814 | 0.002 | -0.004 | -0.004 | 0 | 0.22 | 0.22 | 0.21 | 0.29 | 0.22 | 0.31 |
| 10 | 132698419 | rs964681 | -0.282 | 1.079 | -1.494 | -0.537 | -0.028 | 0.027 | 0.027 | 0.046 | 0.38 | 0.24 | 0.33 | 0.3 | 0.43 | 0.44 |
| 10 | 132704645 | rs1655089 | -0.465 | 0.997 | -1.012 | -0.224 | 0 | -0.001 | -0.001 | 0 | 0.4 | 0.39 | 0.5 | 0.81 | 0.36 | 0.63 |

| | | | | | | | | | | | | | | | | |
|---|---|---|---|---|---|---|---|---|---|---|---|---|---|---|---|---|
| 10 | 132708309 | rs7072329 | -0.769 | 1.351 | -1.091 | 0.564 | -0.001 | 0.009 | 0.009 | 0 | 0.15 | 0.2 | 0.15 | 0.31 | 0.16 | 0.13 |
| 10 | 132710560 | rs7085596 | -1.14 | 1.803 | -1.563 | 0.29 | -0.017 | -0.033 | -0.033 | 0.404 | 0.41 | 0.29 | 0.4 | 0.48 | 0.54 | 0.4 |
| 10 | 132710574 | rs7072753 | -1.136 | 1.805 | -1.562 | 0.291 | -0.018 | -0.033 | -0.033 | 0 | 0.41 | 0.29 | 0.4 | 0.48 | 0.54 | 0.4 |
| 10 | 132712986 | rs11017563 | -0.495 | 1.91 | -1.492 | -0.032 | -0.061 | 0.06 | 0.06 | 2.782 | 0.26 | 0.12 | 0.23 | 0.16 | 0.36 | 0.27 |
| 10 | 132714218 | rs7099843 | -0.452 | 1.937 | -1.53 | -0.021 | -0.073 | 0.061 | 0.061 | 3.055 | 0.26 | 0.08 | 0.23 | 0.16 | 0.36 | 0.27 |
| 10 | 132720292 | rs1615656 | -0.408 | 1.987 | -0.756 | -0.035 | 0.027 | -0.03 | -0.03 | 0.043 | 0.16 | 0.22 | 0.11 | 0.06 | 0.16 | 0.09 |
| 10 | 132720780 | rs11017578 | -1.019 | 2.434 | -0.592 | -0.199 | 0.011 | -0.027 | -0.027 | 0.018 | 0.34 | 0.36 | 0.37 | 0.26 | 0.37 | 0.32 |
| 10 | 132721599 | rs10829844 | -0.074 | 2.431 | -0.632 | -0.207 | -0.012 | 0.009 | 0.009 | 3.844 | 0.22 | 0.12 | 0.27 | 0.26 | 0.26 | 0.28 |
| 10 | 132723003 | rs1655098 | -1.317 | 2.278 | -1.281 | -0.091 | -0.028 | -0.047 | -0.047 | 0 | 0.87 | 1 | 0.92 | 0.86 | 0.87 | 0.86 |
| 10 | 132723132 | rs7893744 | -0.741 | 2.358 | -0.547 | -0.123 | 0.001 | -0.008 | -0.008 | 0 | 0.33 | 0.3 | 0.35 | 0.26 | 0.35 | 0.32 |
| 10 | 132727601 | rs2397395 | -0.265 | 2.53 | -0.471 | -0.239 | -0.015 | 0.017 | 0.017 | 2.744 | 0.27 | 0.19 | 0.31 | 0.26 | 0.36 | 0.3 |
| 10 | 132733095 | rs1368658 | -1.105 | 2.573 | -0.381 | -0.235 | 0.005 | -0.028 | -0.028 | 0 | 0.41 | 0.44 | 0.41 | 0.31 | 0.51 | 0.38 |
| 10 | 132733753 | rs10829846 | -0.552 | 2.521 | -0.358 | -0.262 | -0.001 | 0 | 0 | 4.095 | 0.31 | 0.3 | 0.34 | 0.31 | 0.42 | 0.34 |
| 10 | 132734162 | rs10765004 | -1.128 | 2.499 | -0.409 | -0.253 | 0.004 | -0.026 | -0.026 | 0 | 0.42 | 0.47 | 0.42 | 0.31 | 0.51 | 0.38 |
| 10 | 132736239 | rs1655107 | 0.806 | 1.895 | -0.274 | -0.15 | 0.003 | -0.005 | -0.005 | 0.89 | 0.18 | 0.2 | 0.14 | 0.29 | 0.14 | 0.14 |
| 10 | 132740034 | rs11017590 | 0.923 | 1.874 | -0.217 | -0.149 | 0.008 | -0.026 | -0.026 | 0 | 0.18 | 0.13 | 0.13 | 0.29 | 0.14 | 0.14 |
| 10 | 132741287 | rs892323 | 0.098 | 1.985 | -0.208 | -0.26 | -0.011 | 0.019 | 0.019 | 2.776 | 0.28 | 0.22 | 0.31 | 0.19 | 0.37 | 0.29 |
| 10 | 132742585 | rs1655118 | 1.067 | 1.908 | -0.1 | -0.324 | 0 | -0.022 | -0.022 | 0 | 0.1 | 0.12 | 0.08 | 0 | 0.09 | 0.04 |
| 10 | 132748661 | rs3097956 | -1.556 | 1.841 | -0.059 | -0.283 | 0.018 | -0.003 | -0.003 | 0.537 | 0.59 | 0.67 | 0.57 | 0.61 | 0.65 | 0.52 |
| 10 | 132749766 | rs7069620 | 0.896 | 1.753 | -0.043 | -0.238 | 0.007 | -0.029 | -0.029 | 0.408 | 0.17 | 0.13 | 0.14 | 0.37 | 0.14 | 0.14 |
| 10 | 132753083 | rs3125602 | -1.348 | 1.896 | -0.01 | -0.271 | -0.003 | -0.001 | -0.001 | 0 | 0.57 | 0.48 | 0.54 | 0.61 | 0.63 | 0.51 |
| 10 | 132755235 | rs10765007 | -0.672 | 2.017 | -0.223 | -0.32 | 0.011 | -0.041 | -0.041 | 1.995 | 0.4 | 0.47 | 0.41 | 0.24 | 0.49 | 0.38 |
| 10 | 132768365 | rs7916787 | 0.388 | 1.811 | -0.042 | -0.251 | 0.042 | -0.068 | -0.068 | 0 | 0.11 | 0.27 | 0.09 | 0 | 0.09 | 0.04 |
| 10 | 132768632 | rs7916919 | -1.057 | 1.736 | -0.097 | -0.179 | 0.02 | 0.028 | 0.028 | 0 | 0.31 | 0.53 | 0.28 | 0.39 | 0.33 | 0.22 |
| 10 | 132768738 | rs12260998 | -0.587 | 1.783 | -0.028 | -0.126 | 0.071 | -0.103 | -0.103 | 0 | 0.15 | 0.39 | 0.14 | 0 | 0.18 | 0.06 |
| 10 | 132772441 | rs10829862 | -0.673 | 1.659 | -0.125 | -0.165 | 0 | 0.013 | 0.013 | 0 | 0.33 | 0.41 | 0.25 | 0.38 | 0.27 | 0.25 |
| 10 | 132781042 | rs10829866 | -0.904 | 1.616 | -0.256 | -0.159 | 0.008 | 0.005 | 0.005 | 0 | 0.36 | 0.54 | 0.29 | 0.38 | 0.36 | 0.26 |
| 10 | 132795062 | rs4622176 | -0.04 | 1.449 | -0.59 | 0.057 | -0.017 | -0.062 | -0.062 | 0.443 | 0.23 | 0.12 | 0.19 | 0.37 | 0.22 | 0.21 |
| 10 | 132802525 | rs4488114 | -0.357 | 1.457 | -0.271 | 0.074 | 0.004 | 0.015 | 0.015 | 0 | 0.27 | 0.33 | 0.25 | 0.38 | 0.3 | 0.23 |
| 10 | 132803392 | rs7092194 | 0.212 | 1.263 | -0.226 | -0.158 | -0.003 | -0.052 | -0.052 | 2.18 | 0.09 | 0.21 | 0.05 | 0 | 0.04 | 0.03 |
| 10 | 132811254 | rs6482834 | 0.041 | 1.072 | -1.131 | -0.559 | -0.015 | 0.07 | 0.07 | 0 | 0.8 | 0.96 | 0.78 | 0.91 | 0.83 | 0.83 |
| 10 | 132811555 | rs7896393 | -0.371 | 0.566 | -1.03 | -0.581 | 0.016 | 0.021 | 0.021 | 0 | 0.51 | 0.58 | 0.33 | 0.44 | 0.42 | 0.41 |
| 10 | 132812391 | rs11017656 | 0.394 | 1.066 | -1.138 | -0.816 | -0.009 | -0.031 | -0.031 | 5.191 | 0.3 | 0.24 | 0.2 | 0.36 | 0.33 | 0.36 |
| 10 | 132814391 | rs4132180 | 0.174 | 0.677 | -0.491 | -0.936 | -0.001 | -0.011 | -0.011 | 0 | 0.33 | 0.25 | 0.43 | 0.58 | 0.32 | 0.42 |
| 10 | 132814853 | rs7924276 | -0.609 | 0.476 | -0.338 | -1.038 | -0.002 | 0 | 0 | 0 | 0.43 | 0.43 | 0.49 | 0.59 | 0.39 | 0.46 |
| 10 | 132815843 | rs7073330 | -0.215 | 0.408 | -0.139 | -0.969 | 0 | 0 | 0 | 0 | 0.42 | 0.37 | 0.49 | 0.58 | 0.4 | 0.46 |
| 10 | 132817082 | rs10765021 | 0.291 | 0.407 | -0.138 | -0.969 | -0.01 | -0.022 | -0.022 | 0 | 0.37 | 0.21 | 0.46 | 0.58 | 0.38 | 0.44 |
| 10 | 132817276 | rs12255502 | -0.444 | 0.419 | -0.119 | -0.948 | -0.018 | -0.032 | -0.032 | 0 | 0.06 | 0.16 | 0.03 | 0 | 0.02 | 0.05 |
| 10 | 132822263 | rs2051113 | 0.187 | 0.605 | -0.208 | -0.866 | -0.005 | -0.009 | -0.009 | 0.553 | 0.37 | 0.27 | 0.41 | 0.59 | 0.38 | 0.44 |
| 10 | 132826089 | rs7904315 | -0.662 | 0.75 | -0.252 | -0.765 | -0.002 | 0.003 | 0.003 | 0 | 0.48 | 0.56 | 0.48 | 0.59 | 0.46 | 0.49 |
| 10 | 132827852 | rs7904651 | -0.659 | 0.764 | -0.324 | -0.77 | -0.001 | 0.001 | 0.001 | 0 | 0.48 | 0.55 | 0.48 | 0.59 | 0.46 | 0.48 |
| 10 | 132831819 | rs4539224 | 0.106 | 0.839 | -0.3 | -0.628 | -0.02 | -0.014 | -0.014 | 0 | 0.36 | 0.2 | 0.43 | 0.62 | 0.39 | 0.45 |
| 10 | 132837998 | rs9664920 | 0.124 | 0.798 | -0.255 | -0.652 | -0.025 | -0.013 | -0.013 | 0.794 | 0.36 | 0.18 | 0.42 | 0.62 | 0.38 | 0.44 |
| 10 | 132839543 | rs1419107 | -0.997 | 0.986 | 0.085 | -0.955 | 0.005 | 0.008 | 0.008 | 0 | 0.58 | 0.58 | 0.76 | 0.71 | 0.64 | 0.58 |
| 10 | 132846150 | rs12416478 | -0.301 | 0.756 | 0.473 | -0.84 | 0.007 | -0.032 | -0.032 | 0 | 0.21 | 0.29 | 0.32 | 0.08 | 0.25 | 0.16 |
| 10 | 132851437 | rs10741236 | -1.469 | 0.822 | -0.111 | -0.851 | -0.035 | -0.007 | -0.007 | 0 | 0.61 | 0.92 | 0.71 | 0.64 | 0.63 | 0.59 |
| 10 | 132852826 | rs7905804 | -0.021 | 0.723 | 0.115 | -0.801 | -0.011 | -0.043 | -0.043 | 0 | 0.34 | 0.18 | 0.32 | 0.54 | 0.33 | 0.41 |
| 10 | 132854084 | rs7910464 | -0.666 | 0.717 | 0.366 | -0.826 | -0.002 | -0.002 | -0.002 | 0.112 | 0.45 | 0.36 | 0.44 | 0.54 | 0.44 | 0.46 |
| 10 | 132859258 | rs1419108 | -1.907 | 0.43 | 0.149 | -0.907 | -0.049 | -0.022 | -0.022 | 0 | 0.63 | 0.93 | 0.69 | 0.64 | 0.61 | 0.6 |
| 10 | 132860024 | rs10829891 | -2.036 | 0.364 | 0.111 | -0.935 | -0.063 | -0.037 | -0.037 | 0 | 0.65 | 0.93 | 0.69 | 0.64 | 0.62 | 0.62 |
| 10 | 132864476 | rs10765031 | -0.012 | 0.579 | 0.468 | -0.721 | 0.008 | -0.011 | -0.011 | 0 | 0.32 | 0.26 | 0.25 | 0.54 | 0.24 | 0.34 |
| 10 | 132866405 | rs10765032 | -0.723 | 0.698 | 0.523 | -0.634 | 0.006 | -0.001 | -0.001 | 0.304 | 0.44 | 0.59 | 0.33 | 0.54 | 0.36 | 0.42 |
| 10 | 132869514 | rs4751314 | -1.097 | 0.97 | 0.577 | -0.601 | 0.062 | -0.059 | -0.059 | 2.174 | 0.23 | 0.47 | 0.39 | 0.1 | 0.27 | 0.18 |
| 10 | 132870415 | rs10765034 | -0.466 | 0.632 | 0.196 | -0.457 | -0.031 | 0.07 | 0.07 | 0 | 0.28 | 0.43 | 0.2 | 0.42 | 0.2 | 0.24 |
| 10 | 132872967 | rs10829895 | -2.018 | 0.703 | -0.747 | -0.491 | -0.043 | -0.02 | -0.02 | 0 | 0.64 | 0.92 | 0.7 | 0.65 | 0.62 | 0.6 |
| 10 | 132874665 | rs7067955 | 0.323 | 0.991 | -0.478 | -0.686 | -0.032 | 0.038 | 0.038 | 0 | 0.09 | 0.02 | 0.15 | 0.01 | 0.11 | 0.09 |
| 10 | 132878541 | rs11017701 | 0.936 | 0.904 | -0.361 | -0.75 | 0 | -0.012 | -0.012 | 1.792 | 0.1 | 0.13 | 0.14 | 0.01 | 0.08 | 0.05 |
| 10 | 132879852 | rs7098146 | -1.506 | 0.905 | -0.172 | -0.804 | -0.005 | 0.001 | 0.001 | 0 | 0.34 | 0.65 | 0.55 | 0.11 | 0.38 | 0.28 |
| 10 | 132881469 | rs7082854 | -0.863 | 0.897 | -0.278 | -0.811 | -0.026 | 0.055 | 0.055 | 0 | 0.29 | 0.14 | 0.34 | 0.2 | 0.29 | 0.31 |
| 10 | 132882247 | rs1891792 | -1.603 | 0.897 | -0.104 | -0.816 | -0.008 | 0.019 | 0.019 | 0 | 0.43 | 0.6 | 0.61 | 0.28 | 0.48 | 0.43 |
| 10 | 132884337 | rs12245418 | -1.312 | 0.944 | -0.142 | -0.732 | 0.021 | -0.025 | -0.025 | 0 | 0.33 | 0.4 | 0.47 | 0.27 | 0.4 | 0.37 |
| 10 | 132884578 | rs4751319 | -0.036 | 0.963 | -0.125 | -0.729 | 0.072 | -0.022 | -0.022 | 0 | 0.14 | 0.32 | 0.26 | 0.08 | 0.19 | 0.12 |
| 10 | 132885066 | rs11017706 | -0.291 | 0.608 | -0.754 | -0.695 | -0.029 | 0.002 | 0.002 | 0 | 0.19 | 0.08 | 0.21 | 0.19 | 0.21 | 0.24 |
| 10 | 132889970 | rs7098775 | -1.369 | 0.56 | -0.604 | -0.785 | 0.004 | 0.116 | 0.116 | 0 | 0.5 | 0.71 | 0.53 | 0.26 | 0.48 | 0.45 |
| 10 | 132890603 | rs7070294 | -0.604 | 0.754 | -0.576 | -0.885 | 0.022 | -0.068 | -0.068 | 0 | 0.2 | 0.38 | 0.19 | 0.06 | 0.23 | 0.18 |
| 10 | 132891026 | rs7088377 | -1.499 | 0.73 | -1.438 | -0.593 | -0.131 | -0.06 | -0.06 | 0 | 0.68 | 0.98 | 0.7 | 0.68 | 0.6 | 0.62 |
| 10 | 132891202 | rs7071145 | -0.728 | 0.742 | -0.332 | -0.808 | 0.004 | -0.073 | -0.073 | 0 | 0.24 | 0.36 | 0.22 | 0.05 | 0.26 | 0.15 |
| 10 | 132894844 | rs7073264 | -0.076 | 0.881 | -0.368 | -0.435 | 0.002 | -0.036 | -0.036 | 0 | 0.09 | 0.11 | 0.05 | 0 | 0.09 | 0.06 |
| 10 | 132897429 | rs4751321 | -0.549 | 0.346 | -0.258 | -0.017 | -0.026 | 0.104 | 0.104 | 0 | 0.22 | 0.42 | 0.27 | 0.48 | 0.22 | 0.21 |

| | | | | | | | | | | | | | | | | |
|---|---|---|---|---|---|---|---|---|---|---|---|---|---|---|---|---|
| 10 | 132901331 | rs11017718 | 0.063 | 0.437 | -0.478 | -0.172 | -0.002 | 0 | 0 | 0 | 0.12 | 0.14 | 0.08 | 0.18 | 0.09 | 0.14 |
| 10 | 132903839 | rs6482850 | -1.696 | 0.802 | -1.096 | -0.702 | -0.001 | 0.001 | 0.001 | 0 | 0.68 | 0.67 | 0.63 | 0.72 | 0.68 | 0.66 |
| 10 | 132904116 | rs11017719 | 0.361 | 0.67 | -1.371 | -0.435 | 0 | -0.003 | -0.003 | 0.537 | 0.09 | 0.07 | 0.05 | 0 | 0.09 | 0.06 |
| 10 | 132907255 | rs1419109 | 0.081 | 0.783 | -1.361 | -0.089 | 0.001 | -0.008 | -0.008 | 0 | 0.11 | 0.06 | 0.14 | 0.19 | 0.12 | 0.15 |
| 10 | 132908210 | rs4418720 | -1.206 | 0.867 | -1.387 | -0.13 | -0.003 | -0.002 | -0.002 | 0 | 0.29 | 0.19 | 0.18 | 0.36 | 0.33 | 0.47 |
| 10 | 132908468 | rs1891787 | -0.607 | 0.554 | -1.64 | 0.077 | -0.002 | -0.038 | -0.038 | 0 | 0.22 | 0.09 | 0.14 | 0.35 | 0.22 | 0.4 |
| 10 | 132914909 | rs11017735 | -0.338 | 0.428 | -1.531 | -0.204 | 0.01 | 0.065 | 0.065 | 0 | 0.24 | 0.14 | 0.24 | 0.01 | 0.19 | 0.15 |
| 10 | 132915202 | rs2298193 | -0.434 | 0.534 | -1.485 | 0.531 | 0.002 | -0.047 | -0.047 | 0 | 0.27 | 0.2 | 0.26 | 0.42 | 0.24 | 0.25 |
| 10 | 132924946 | rs1106524 | -0.486 | 0.355 | -1.254 | -0.317 | -0.01 | -0.093 | -0.093 | 0 | 0.68 | 0.49 | 0.71 | 0.89 | 0.7 | 0.81 |
| 10 | 132926683 | rs10765036 | -0.032 | 0.256 | -1.501 | 0.482 | -0.006 | 0.001 | 0.001 | 0 | 0.61 | 0.49 | 0.62 | 0.55 | 0.62 | 0.64 |
| 10 | 132927646 | rs12250538 | 0.868 | 0.22 | -1.65 | 0.519 | -0.009 | -0.009 | -0.009 | 0 | 0.29 | 0.27 | 0.35 | 0.35 | 0.39 | 0.27 |
| 10 | 132928393 | rs7923325 | 1.131 | 0.334 | -1.776 | 0.251 | -0.042 | 0.061 | 0.061 | 0 | 0.27 | 0.13 | 0.31 | 0.13 | 0.36 | 0.18 |
| 10 | 132928907 | rs10829909 | 0.502 | -0.117 | -1.885 | 0.012 | 0.008 | 0.015 | 0.015 | 0 | 0.19 | 0.14 | 0.16 | 0.12 | 0.13 | 0.3 |
| 10 | 132931236 | rs10829912 | 1.353 | -0.244 | -1.962 | 0.093 | -0.061 | 0.065 | 0.065 | 0 | 0.28 | 0.09 | 0.33 | 0.19 | 0.37 | 0.22 |
| 10 | 132931252 | rs10829913 | 1.204 | -0.216 | -1.956 | 0.105 | -0.073 | 0.078 | 0.078 | 0 | 0.29 | 0.07 | 0.33 | 0.19 | 0.37 | 0.23 |
| 10 | 132931753 | rs7919857 | -0.899 | -0.482 | -2.058 | -0.19 | -0.036 | -0.076 | -0.076 | 0.47 | 0.73 | 0.45 | 0.74 | 0.89 | 0.82 | 0.89 |
| 10 | 132931959 | rs6482862 | 0.651 | -0.269 | -2.224 | -0.064 | -0.029 | 0.054 | 0.054 | 0 | 0.38 | 0.19 | 0.38 | 0.31 | 0.42 | 0.47 |
| 10 | 132934019 | rs10829918 | 0.267 | 0.56 | -1.724 | 0.609 | -0.023 | -0.032 | -0.032 | 0 | 0.19 | 0.11 | 0.21 | 0.35 | 0.3 | 0.26 |
| 10 | 132935492 | rs4751326 | 0.004 | 0.602 | -1.013 | 0.468 | -0.002 | 0.017 | 0.017 | 0 | 0.69 | 0.51 | 0.72 | 0.67 | 0.72 | 0.74 |
| 10 | 132939307 | rs12415758 | -1.399 | 0.535 | -1.112 | 0.316 | -0.003 | 0.012 | 0.012 | 0 | 0.88 | 0.58 | 0.86 | 0.89 | 0.93 | 0.9 |
| 10 | 132942160 | rs11017747 | 0.639 | 0.328 | -1.444 | 0.384 | -0.004 | 0.005 | 0.005 | 0 | 0.05 | 0.06 | 0.03 | 0.1 | 0.03 | 0.22 |
| 10 | 132943161 | rs4751329 | 1.149 | 0.252 | -1.983 | 0.197 | 0.009 | -0.057 | -0.057 | 0 | 0.38 | 0.16 | 0.43 | 0.44 | 0.32 | 0.32 |
| 10 | 132944167 | rs11017755 | 1.355 | 0.239 | -2.049 | 0.121 | -0.041 | 0.11 | 0.11 | 0 | 0.36 | 0.03 | 0.42 | 0.2 | 0.32 | 0.25 |
| 10 | 132945676 | rs10741242 | -0.654 | 0.281 | -1.825 | 0.239 | -0.054 | 0.019 | 0.019 | 0 | 0.82 | 0.47 | 0.83 | 0.8 | 0.9 | 0.72 |
| 10 | 132947578 | rs12411368 | 0.592 | 0.533 | -1.987 | 0.256 | -0.06 | -0.091 | -0.091 | 0 | 0.21 | 0.05 | 0.22 | 0.58 | 0.27 | 0.3 |
| 10 | 132948840 | rs7081792 | -1.653 | 0.371 | -1.618 | 0.344 | -0.017 | 0.013 | 0.013 | 0.026 | 0.87 | 0.55 | 0.86 | 0.89 | 0.93 | 0.93 |
| 10 | 132948977 | rs10765042 | 0.177 | 0.384 | -1.703 | 0.313 | -0.002 | 0.043 | 0.043 | 0 | 0.69 | 0.42 | 0.71 | 0.55 | 0.7 | 0.59 |
| 10 | 132952277 | rs7074563 | -0.187 | 0.422 | -1.721 | 0.333 | -0.017 | -0.05 | -0.05 | 0 | 0.19 | 0.13 | 0.15 | 0.34 | 0.23 | 0.37 |
| 10 | 132953008 | rs2918145 | 0.13 | 0.432 | -1.679 | -0.157 | -0.003 | -0.031 | -0.031 | 2.839 | 0.69 | 0.48 | 0.73 | 0.78 | 0.69 | 0.63 |
| 10 | 132954749 | rs2944531 | -0.496 | 0.102 | -1.415 | -0.095 | -0.004 | -0.051 | -0.051 | 0 | 0.89 | 0.83 | 0.94 | 1 | 0.9 | 0.92 |
| 10 | 132955745 | rs2918147 | 0.969 | 0.231 | -1.415 | 0.201 | 0.001 | 0.011 | 0.011 | 0.018 | 0.13 | 0.45 | 0.14 | 0.11 | 0.09 | 0.08 |
| 10 | 132957703 | rs11017774 | 0.425 | 0.474 | -2.615 | 0.427 | -0.003 | -0.049 | -0.049 | 0 | 0.25 | 0.12 | 0.37 | 0.33 | 0.24 | 0.16 |
| 10 | 132957895 | rs10765045 | 0.544 | 0.925 | -2.62 | 0.376 | -0.004 | 0.127 | 0.127 | 0 | 0.42 | 0.11 | 0.42 | 0.23 | 0.39 | 0.24 |
| 10 | 132958245 | rs2944530 | 0.548 | 1.169 | -2.044 | 0.973 | 0.003 | 0.076 | 0.076 | 0 | 0.94 | 0.88 | 0.87 | 0.83 | 0.94 | 0.89 |
| 10 | 132958926 | rs2944529 | 0.177 | 1.305 | -2.192 | 0.735 | -0.06 | 0 | 0 | 0 | 0.71 | 0.84 | 0.66 | 0.67 | 0.61 | 0.67 |
| 10 | 132962430 | rs2944527 | 0.126 | 0.39 | -1.834 | 0.748 | 0.006 | -0.069 | -0.069 | 0.791 | 0.75 | 0.88 | 0.72 | 0.49 | 0.76 | 0.54 |
| 10 | 132963432 | rs4751335 | 0.485 | 0.375 | -1.731 | 0.106 | -0.001 | 0.103 | 0.103 | 0.014 | 0.46 | 0.22 | 0.48 | 0.27 | 0.52 | 0.31 |
| 10 | 132966465 | rs2996069 | -0.286 | 0.123 | -0.968 | -0.105 | -0.039 | -0.101 | -0.101 | 0.858 | 0.24 | 0.52 | 0.12 | 0.05 | 0.16 | 0.16 |
| 10 | 132967242 | rs10829939 | 0.333 | 0.132 | -1.485 | 0.232 | 0.017 | 0.022 | 0.022 | 0 | 0.59 | 0.28 | 0.64 | 0.76 | 0.62 | 0.74 |
| 10 | 132968374 | rs11017781 | 0.356 | 0.563 | -2.365 | -0.018 | -0.145 | -0.025 | -0.025 | 0 | 0.25 | 0.04 | 0.31 | 0.26 | 0.37 | 0.17 |
| 10 | 132968393 | rs2918152 | -0.394 | 0.007 | -1.934 | 0.106 | -0.001 | 0.001 | 0.001 | 0 | 0.14 | 0.15 | 0.16 | 0.2 | 0.13 | 0.05 |
| 10 | 132969094 | rs893504 | 0.804 | -0.313 | -1.069 | 0.049 | -0.011 | 0.087 | 0.087 | 0 | 0.82 | 0.67 | 0.7 | 0.71 | 0.87 | 0.85 |
| 10 | 132970636 | rs2996081 | 0.073 | -0.24 | -1.31 | 0.055 | -0.005 | -0.017 | -0.017 | 0 | 0.12 | 0.09 | 0.15 | 0.21 | 0.12 | 0.04 |
| 10 | 132972901 | rs7903016 | 0.403 | -0.905 | -0.937 | -0.311 | -0.038 | -0.068 | -0.068 | 0 | 0.11 | 0.29 | 0.04 | 0.01 | 0.05 | 0.03 |
| 10 | 132974253 | rs10829941 | 0.656 | -0.948 | -2.053 | -0.552 | -0.017 | 0.012 | 0.012 | 0.535 | 0.49 | 0.08 | 0.54 | 0.7 | 0.56 | 0.7 |
| 10 | 132976517 | rs11017797 | 0.51 | -0.973 | -2.002 | -0.561 | 0.016 | 0.082 | 0.082 | 0.808 | 0.53 | 0.16 | 0.59 | 0.77 | 0.57 | 0.79 |
| 10 | 132980357 | rs11017802 | -0.856 | -0.61 | -1.33 | -0.292 | 0.017 | 0.02 | 0.02 | 0 | 0.13 | 0.07 | 0.1 | 0.08 | 0.06 | 0.16 |
| 10 | 132980435 | rs11017804 | -1.013 | -0.656 | -1.401 | -0.362 | 0.022 | 0.032 | 0.032 | 0 | 0.15 | 0.07 | 0.1 | 0.08 | 0.07 | 0.15 |
| 10 | 132980932 | rs999191 | -0.134 | -0.349 | -1.251 | -0.273 | -0.002 | 0.014 | 0.014 | 0.324 | 0.62 | 0.73 | 0.66 | 0.66 | 0.6 | 0.61 |
| 10 | 132982293 | rs10829942 | -1.017 | 0.084 | -0.657 | 0.41 | 0.006 | 0.117 | 0.117 | 0 | 0.11 | 0.18 | 0.23 | 0.26 | 0.12 | 0.12 |
| 10 | 132982494 | rs12255029 | -0.391 | 0.21 | -0.685 | 0.389 | 0.018 | 0.113 | 0.113 | 0 | 0.08 | 0.12 | 0.18 | 0.19 | 0.12 | 0.04 |
| 10 | 132983821 | rs2397752 | -1.105 | -0.401 | -0.658 | 0.168 | -0.01 | 0 | 0 | 0 | 0.33 | 0.59 | 0.36 | 0.24 | 0.27 | 0.27 |
| 10 | 132985051 | rs2944462 | 0.54 | -0.164 | -0.312 | 0.586 | -0.022 | 0.058 | 0.058 | 0 | 0.82 | 0.58 | 0.81 | 0.75 | 0.88 | 0.95 |
| 10 | 132987665 | rs2397753 | -0.309 | -0.086 | -0.625 | 0.071 | -0.002 | -0.043 | -0.043 | 0.339 | 0.2 | 0.23 | 0.16 | 0.01 | 0.19 | 0.06 |
| 10 | 132992902 | rs2918142 | 1.134 | 0.11 | -0.463 | 0.367 | -0.001 | 0 | 0 | 0 | 0.67 | 0.34 | 0.49 | 0.54 | 0.63 | 0.84 |
| 10 | 132993796 | rs11594799 | 0.298 | 0.366 | -0.949 | 0.512 | 0.042 | 0.155 | 0.155 | 0 | 0.09 | 0.22 | 0.25 | 0.2 | 0.14 | 0.09 |
| 10 | 132995046 | rs11597694 | 0.52 | 0.336 | -1.059 | 0.033 | -0.002 | 0.038 | 0.038 | 0 | 0.13 | 0.07 | 0.11 | 0.01 | 0.15 | 0.05 |
| 10 | 132995300 | rs10765051 | -0.886 | 0.258 | -0.998 | 0.133 | 0 | 0.005 | 0.005 | 0 | 0.34 | 0.4 | 0.49 | 0.5 | 0.38 | 0.28 |
| 10 | 132999333 | rs4751345 | 0.503 | 0.811 | -1.738 | -0.142 | -0.059 | 0.025 | 0.025 | 0.194 | 0.2 | 0.05 | 0.17 | 0.14 | 0.19 | 0.41 |
| 10 | 133000984 | rs10829945 | 0.259 | -0.203 | -2.024 | -0.151 | -0.108 | 0.043 | 0.043 | 0 | 0.25 | 0.04 | 0.17 | 0.19 | 0.32 | 0.18 |
| 10 | 133001451 | rs10829946 | -0.739 | -0.815 | -1.535 | 0.037 | 0.011 | 0.013 | 0.013 | 0 | 0.55 | 0.61 | 0.67 | 0.64 | 0.64 | 0.42 |
| 10 | 133001706 | rs10829947 | -0.442 | -0.891 | -1.486 | 0.046 | -0.002 | 0.013 | 0.013 | 0 | 0.46 | 0.58 | 0.57 | 0.63 | 0.5 | 0.36 |
| 10 | 133003604 | rs2918128 | 0.255 | -1.55 | -0.608 | 0.088 | 0.038 | -0.005 | -0.005 | 0 | 0.69 | 0.81 | 0.83 | 0.78 | 0.79 | 0.85 |
| 10 | 133004468 | rs2918125 | -0.214 | -1.086 | -0.039 | -0.177 | -0.006 | -0.037 | -0.037 | 1.105 | 0.57 | 0.53 | 0.76 | 0.75 | 0.63 | 0.79 |
| 10 | 133005521 | rs893500 | -0.781 | -1.847 | 0.967 | 0.084 | 0.012 | -0.074 | -0.074 | 0 | 0.13 | 0.33 | 0.09 | 0.01 | 0.15 | 0.15 |
| 10 | 133006725 | rs11017818 | -0.901 | -2.087 | 0.911 | 0.343 | -0.003 | 0.007 | 0.007 | 0.01 | 0.2 | 0.2 | 0.16 | 0.24 | 0.15 | 0.12 |
| 10 | 133007958 | rs11596938 | 0.528 | -2.108 | 0.632 | 0.155 | -0.014 | 0.047 | 0.047 | 0 | 0.11 | 0.02 | 0.09 | 0.01 | 0.11 | 0.08 |
| 10 | 133009203 | rs10741246 | 1.869 | -2.653 | 0.948 | 0.094 | 0.06 | 0.066 | 0.066 | 1.951 | 0.49 | 0.24 | 0.64 | 0.73 | 0.59 | 0.65 |
| 10 | 133009690 | rs2918123 | -0.849 | -2.502 | 1.01 | 0.213 | -0.026 | 0.017 | 0.017 | 0 | 0.94 | 0.85 | 0.94 | 0.93 | 0.99 | 0.96 |

| | | | | | | | | | | | | | | | | |
|---|---|---|---|---|---|---|---|---|---|---|---|---|---|---|---|---|
| 10 | 133011726 | rs2944490 | 0.61 | -2.566 | 1.169 | 0.179 | -0.017 | 0.078 | 0.078 | 0 | 0.85 | 0.73 | 0.86 | 0.76 | 0.9 | 0.9 |
| 10 | 133012146 | rs10765056 | 1.577 | -2.556 | 1.315 | 0.159 | -0.006 | -0.006 | -0.006 | 0.431 | 0.54 | 0.5 | 0.67 | 0.72 | 0.6 | 0.66 |
| 10 | 133013187 | rs893507 | -0.879 | -2.336 | 1.064 | 0.085 | -0.028 | -0.103 | -0.103 | 0 | 0.19 | 0.31 | 0.14 | 0.01 | 0.15 | 0.1 |
| 10 | 133013442 | rs893509 | -2.184 | -2.267 | 0.506 | 0.476 | 0.014 | 0.013 | 0.013 | 0 | 0.57 | 0.76 | 0.53 | 0.54 | 0.45 | 0.38 |
| 10 | 133014759 | rs10829950 | -0.799 | -1.636 | 0.412 | 0.677 | 0.001 | 0.006 | 0.006 | 0 | 0.26 | 0.23 | 0.2 | 0.1 | 0.27 | 0.26 |
| 10 | 133016385 | rs2944493 | -1.416 | -1.052 | 0.58 | 1.171 | -0.044 | 0.093 | 0.093 | 0 | 0.28 | 0.43 | 0.33 | 0.43 | 0.22 | 0.13 |
| 10 | 133025748 | rs10765057 | -1.394 | -0.033 | 0.321 | 1.449 | -0.011 | -0.004 | -0.004 | 0.035 | 0.27 | 0.2 | 0.23 | 0.22 | 0.37 | 0.25 |
| 10 | 133028266 | rs10765059 | -1.271 | -0.038 | 0.437 | 1.756 | 0.001 | 0.107 | 0.107 | 0 | 0.17 | 0.23 | 0.3 | 0.51 | 0.17 | 0.14 |
| 10 | 133033315 | rs2944499 | -2.402 | -0.283 | -0.102 | 1.627 | 0.004 | 0.015 | 0.015 | 0 | 0.44 | 0.6 | 0.5 | 0.62 | 0.46 | 0.37 |
| 10 | 133036058 | rs754575 | -1.882 | 0.162 | 0.103 | 1.668 | -0.017 | 0.018 | 0.018 | 0 | 0.29 | 0.52 | 0.33 | 0.37 | 0.27 | 0.25 |
| 10 | 133036574 | rs10829956 | -2.287 | 0.206 | 0.051 | 1.13 | -0.001 | 0 | 0 | 0 | 0.38 | 0.57 | 0.42 | 0.55 | 0.42 | 0.31 |
| 10 | 133036876 | rs11599122 | -0.209 | 0.361 | -0.36 | 1.323 | -0.004 | 0.054 | 0.054 | 0 | 0.1 | 0.01 | 0.06 | 0.01 | 0.08 | 0.07 |
| 10 | 133039080 | rs11598030 | -0.666 | 0.644 | 0.132 | 1.793 | -0.005 | 0.028 | 0.028 | 0 | 0.13 | 0.04 | 0.08 | 0.03 | 0.17 | 0.14 |
| 10 | 133040379 | rs1020105 | -1.166 | 0.655 | 0.147 | 1.735 | 0.023 | 0.118 | 0.118 | 0 | 0.22 | 0.39 | 0.33 | 0.54 | 0.29 | 0.22 |
| 10 | 133048749 | rs1968517 | -0.891 | 0.781 | 0.202 | 2.071 | -0.006 | 0.056 | 0.056 | 0 | 0.27 | 0.62 | 0.45 | 0.42 | 0.26 | 0.24 |
| 10 | 133048792 | rs1530064 | 0.515 | 0.145 | -0.945 | 0.984 | 0.068 | -0.055 | -0.055 | 0 | 0.41 | 0.15 | 0.3 | 0.54 | 0.3 | 0.55 |
| 10 | 133050831 | rs2918087 | -0.827 | -0.428 | -0.403 | 0.372 | -0.006 | -0.016 | -0.016 | 0 | 0.2 | 0.3 | 0.27 | 0.1 | 0.18 | 0.16 |
| 10 | 133056128 | rs2918091 | -0.417 | -0.728 | 0.182 | 0.257 | -0.048 | -0.033 | -0.033 | 0 | 0.19 | 0.44 | 0.23 | 0.11 | 0.14 | 0.18 |
| 10 | 133056853 | rs2944507 | -1.173 | -0.722 | -0.324 | -0.017 | -0.038 | 0.06 | 0.06 | 0 | 0.68 | 0.81 | 0.74 | 0.88 | 0.63 | 0.77 |
| 10 | 133057246 | rs2280199 | -0.216 | -1.597 | -0.67 | -0.078 | -0.003 | 0.025 | 0.025 | 0.196 | 0.43 | 0.41 | 0.6 | 0.69 | 0.46 | 0.51 |
| 10 | 133057358 | rs1055043 | 0.732 | -1.508 | -1.124 | 0.074 | -0.102 | -0.157 | -0.157 | 0 | 0.29 | 0.08 | 0.38 | 0.52 | 0.35 | 0.38 |
| 10 | 133060988 | rs12573222 | 1.443 | -1.58 | -1.279 | -0.329 | -0.169 | 0.161 | 0.161 | 0 | 0.27 | 0.02 | 0.18 | 0.04 | 0.4 | 0.24 |
| 10 | 133063347 | rs7909338 | 1.387 | -0.985 | -1.24 | -0.483 | -0.079 | 0.137 | 0.137 | 0 | 0.27 | 0.06 | 0.16 | 0.04 | 0.33 | 0.25 |
| 10 | 133068269 | rs12265476 | 0.371 | -1.346 | -0.863 | -0.256 | -0.029 | 0.08 | 0.08 | 0 | 0.43 | 0.25 | 0.27 | 0.22 | 0.48 | 0.43 |
| 10 | 133075148 | rs2918108 | -0.424 | -1.744 | -0.555 | -0.142 | -0.06 | 0.08 | 0.08 | 0.75 | 0.87 | 0.58 | 0.75 | 0.84 | 0.96 | 0.87 |
| 10 | 133078257 | rs4751353 | 1.396 | -1.488 | -0.899 | -0.101 | -0.055 | -0.082 | -0.082 | 3.622 | 0.22 | 0.07 | 0.19 | 0.28 | 0.25 | 0.34 |
| 10 | 133079937 | rs4751354 | -0.779 | -1.712 | -0.731 | -0.026 | -0.053 | -0.045 | -0.045 | 0.164 | 0.83 | 0.62 | 0.84 | 0.92 | 0.92 | 0.92 |
| 10 | 133081756 | rs4751357 | 0.903 | -1.538 | -0.752 | 0.216 | -0.036 | -0.119 | -0.119 | 0 | 0.25 | 0.08 | 0.21 | 0.46 | 0.3 | 0.47 |
| 10 | 133083887 | rs7093292 | -0.9 | -1.347 | -0.065 | -0.045 | -0.01 | 0.008 | 0.008 | 0 | 0.08 | 0.1 | 0.06 | 0.08 | 0.06 | 0.02 |
| 10 | 133083894 | rs10741249 | 0.157 | -0.978 | -0.088 | -0.232 | -0.039 | -0.014 | -0.014 | 0.288 | 0.73 | 0.54 | 0.78 | 0.78 | 0.82 | 0.75 |
| 10 | 133084992 | rs2918117 | -0.559 | -0.977 | -0.314 | -0.21 | -0.026 | -0.031 | -0.031 | 0 | 0.8 | 0.65 | 0.84 | 0.87 | 0.88 | 0.77 |
| 10 | 133086735 | rs10128158 | 0.432 | -0.929 | -0.262 | -0.242 | -0.009 | 0.043 | 0.043 | 0 | 0.09 | 0.19 | 0.07 | 0.18 | 0.08 | 0.15 |
| 10 | 133086814 | rs2918118 | -0.404 | -0.906 | -0.433 | -0.229 | -0.021 | 0.004 | 0.004 | 0 | 0.78 | 0.66 | 0.81 | 0.78 | 0.86 | 0.76 |
| 10 | 133099271 | rs6482875 | 0.983 | -0.746 | -1.155 | -0.274 | -0.001 | 0.021 | 0.021 | 0.703 | 0.46 | 0.26 | 0.56 | 0.6 | 0.47 | 0.53 |
| 10 | 133102434 | rs2033790 | 0.359 | 0.059 | -1.181 | 0.035 | -0.102 | 0.12 | 0.12 | 4.734 | 0.18 | 0.03 | 0.15 | 0 | 0.28 | 0.06 |
| 10 | 133107775 | rs10829970 | 0.293 | 0.336 | -0.924 | 0.303 | 0 | 0.005 | 0.005 | 0 | 0.08 | 0.08 | 0.04 | 0.18 | 0.07 | 0.16 |
| 10 | 133108761 | rs1438507 | -0.34 | 0.425 | -0.824 | 0.39 | -0.013 | 0.024 | 0.024 | 0 | 0.31 | 0.25 | 0.24 | 0.18 | 0.39 | 0.22 |
| 10 | 133114983 | rs10829973 | 0.611 | -0.501 | -0.112 | 1.08 | -0.071 | 0.028 | 0.028 | 0.219 | 0.77 | 0.47 | 0.69 | 0.74 | 0.9 | 0.86 |
| 10 | 133117157 | rs11017878 | -0.941 | 0.62 | -0.042 | 1.215 | 0.019 | 0.049 | 0.049 | 0 | 0.18 | 0.28 | 0.14 | 0.25 | 0.22 | 0.16 |
| 10 | 133120674 | rs868316 | -0.701 | 1.066 | -0.007 | 1.725 | 0.04 | 0.112 | 0.112 | 0 | 0.08 | 0.14 | 0.11 | 0.29 | 0.16 | 0.13 |
| 10 | 133120700 | rs2397769 | 0.13 | -0.052 | -0.705 | 0.821 | 0.035 | 0.039 | 0.039 | 0.801 | 0.73 | 0.83 | 0.74 | 0.88 | 0.82 | 0.74 |
| 10 | 133125924 | rs7078641 | -0.982 | 0.203 | -0.451 | 1.166 | -0.001 | 0.054 | 0.054 | 0 | 0.06 | 0.11 | 0.04 | 0.21 | 0.06 | 0.19 |
| 10 | 133130078 | rs9733876 | 0.01 | 0.987 | -1.106 | 0.923 | -0.082 | -0.05 | -0.05 | 0.423 | 0.18 | 0.05 | 0.26 | 0.26 | 0.34 | 0.11 |
| 10 | 133132032 | rs7093992 | -1.381 | 0.192 | -1.455 | -0.021 | -0.002 | -0.039 | -0.039 | 0.356 | 0.73 | 0.8 | 0.58 | 0.48 | 0.75 | 0.61 |
| 10 | 133132701 | rs11017887 | 0.623 | -0.261 | -0.729 | -0.501 | -0.006 | -0.017 | -0.017 | 0 | 0.08 | 0.21 | 0.06 | 0.04 | 0.05 | 0.12 |
| 10 | 133136539 | rs2397763 | -0.611 | -1.347 | -0.928 | -1.111 | -0.014 | -0.127 | -0.127 | 0 | 0.76 | 0.59 | 0.86 | 1 | 0.79 | 0.93 |
| 10 | 133137016 | rs11591724 | 1.472 | -1.175 | -1.086 | -0.921 | -0.081 | -0.035 | -0.035 | 1.153 | 0.18 | 0.05 | 0.26 | 0.25 | 0.34 | 0.11 |
| 10 | 133139043 | rs7090413 | 1.018 | -1.493 | -0.832 | -0.752 | -0.034 | -0.02 | -0.02 | 0 | 0.17 | 0.22 | 0.08 | 0.09 | 0.08 | 0.21 |
| 10 | 133139767 | rs7897686 | 1.346 | -0.81 | -1.01 | -0.38 | -0.05 | -0.122 | -0.122 | 0 | 0.23 | 0.06 | 0.41 | 0.53 | 0.25 | 0.4 |
| 10 | 133141257 | rs10734104 | -0.272 | -0.516 | -0.631 | -0.517 | -0.014 | -0.076 | -0.076 | 0 | 0.83 | 0.75 | 0.92 | 1 | 0.87 | 0.95 |
| 10 | 133142938 | rs4259751 | -0.016 | -0.054 | -0.919 | -0.118 | -0.011 | 0.004 | 0.004 | 0 | 0.3 | 0.18 | 0.44 | 0.67 | 0.3 | 0.52 |
| 10 | 133143956 | rs11594412 | -0.466 | 0.151 | -0.891 | 0.081 | -0.022 | -0.025 | -0.025 | 1.543 | 0.08 | 0.01 | 0.06 | 0.12 | 0.09 | 0.16 |
| 10 | 133146480 | rs10829977 | -0.197 | 0.079 | -0.642 | -0.366 | -0.002 | -0.019 | -0.019 | 0 | 0.54 | 0.5 | 0.63 | 0.71 | 0.45 | 0.72 |
| 10 | 133146488 | rs11017901 | 0.975 | -0.341 | -0.867 | -0.011 | 0.004 | -0.017 | -0.017 | 0 | 0.2 | 0.1 | 0.26 | 0.23 | 0.19 | 0.26 |
| 10 | 133146930 | rs7898539 | -0.673 | -0.165 | -0.68 | -0.296 | -0.019 | 0.063 | 0.063 | 0.004 | 0.62 | 0.77 | 0.73 | 0.79 | 0.56 | 0.79 |
| 10 | 133149493 | rs3935531 | -0.031 | -0.139 | -0.346 | -0.136 | -0.003 | -0.001 | -0.001 | 0 | 0.28 | 0.31 | 0.31 | 0.28 | 0.22 | 0.35 |
| 10 | 133150638 | rs7099724 | 0.453 | -0.06 | -0.278 | -0.105 | -0.006 | -0.001 | -0.001 | 0 | 0.23 | 0.3 | 0.3 | 0.24 | 0.2 | 0.3 |
| 10 | 133151778 | rs6482876 | -1.218 | -0.131 | -0.372 | -0.032 | 0.002 | -0.013 | -0.013 | 10.109 | 0.74 | 0.79 | 0.68 | 0.59 | 0.77 | 0.64 |
| 10 | 133153053 | rs11017904 | -0.009 | -0.571 | 0.105 | -0.09 | -0.002 | -0.014 | -0.014 | 0 | 0.17 | 0.35 | 0.19 | 0.13 | 0.17 | 0.1 |
| 10 | 133154069 | rs10829981 | -0.73 | -0.238 | -0.048 | -0.163 | -0.005 | -0.002 | -0.002 | 0 | 0.47 | 0.54 | 0.48 | 0.41 | 0.42 | 0.44 |
| 10 | 133154905 | rs4751365 | -0.113 | 0.252 | 0.12 | -0.057 | 0.016 | 0.04 | 0.04 | 0 | 0.73 | 0.5 | 0.67 | 0.64 | 0.69 | 0.78 |
| 10 | 133156268 | rs6482877 | 0.744 | 0.197 | -0.34 | -0.285 | 0.024 | 0.027 | 0.027 | 0.201 | 0.32 | 0.14 | 0.31 | 0.26 | 0.27 | 0.33 |
| 10 | 133158421 | rs7900159 | 1.466 | 0.186 | -0.1 | -0.313 | 0 | 0.001 | 0.001 | 0 | 0.29 | 0.24 | 0.3 | 0.37 | 0.3 | 0.37 |
| 10 | 133159554 | rs7896909 | 1.583 | 0.219 | -0.17 | -0.295 | -0.015 | -0.027 | -0.027 | 0 | 0.12 | 0.05 | 0.14 | 0.23 | 0.18 | 0.15 |
| 10 | 133162311 | rs10829987 | 0.012 | -0.004 | -0.231 | -0.336 | 0.003 | -0.027 | -0.027 | 3.913 | 0.44 | 0.2 | 0.45 | 0.48 | 0.44 | 0.47 |
| 10 | 133162862 | rs7098312 | 0.611 | 0.092 | 0.03 | -0.244 | 0.005 | 0.004 | 0.004 | 0 | 0.39 | 0.43 | 0.35 | 0.39 | 0.37 | 0.44 |
| 10 | 133165646 | rs7078529 | 0.903 | -0.171 | 0.099 | -0.274 | -0.004 | 0.001 | 0.001 | 0 | 0.41 | 0.32 | 0.45 | 0.6 | 0.49 | 0.52 |
| 10 | 133165887 | rs7100844 | 0.362 | -0.015 | -0.022 | -0.089 | 0.002 | 0.001 | 0.001 | 0 | 0.51 | 0.47 | 0.5 | 0.59 | 0.55 | 0.58 |
| 10 | 133169986 | rs12573492 | 0.162 | 0.244 | 0.081 | -0.11 | -0.003 | 0 | 0 | 0.315 | 0.13 | 0.26 | 0.15 | 0.13 | 0.14 | 0.06 |

| | | | | | | | | | | | | | | | | |
|---|---|---|---|---|---|---|---|---|---|---|---|---|---|---|---|---|
| 10 | 133172248 | rs4295961 | 0.035 | 0.167 | 0.128 | -0.309 | -0.006 | -0.028 | -0.028 | 0.809 | 0.75 | 0.67 | 0.74 | 0.87 | 0.75 | 0.9 |
| 10 | 133172455 | rs7083148 | 0.71 | 0.642 | 0.009 | 0.017 | -0.022 | -0.026 | -0.026 | 0 | 0.14 | 0.06 | 0.12 | 0.22 | 0.17 | 0.17 |
| 10 | 133175059 | rs11017922 | -0.083 | -0.915 | 0.068 | -0.727 | -0.015 | -0.056 | -0.056 | 2.722 | 0.56 | 0.47 | 0.39 | 0.13 | 0.64 | 0.49 |
| 10 | 133178201 | rs10829998 | 1.326 | -0.571 | -0.296 | -0.78 | -0.086 | 0.045 | 0.045 | 0 | 0.19 | 0.06 | 0.19 | 0.11 | 0.34 | 0.28 |
| 10 | 133178622 | rs3123221 | -0.271 | -0.912 | 0.056 | -0.409 | 0.04 | 0.019 | 0.019 | 0 | 0.35 | 0.51 | 0.4 | 0.4 | 0.43 | 0.47 |
| 10 | 133180530 | rs3123217 | 0.211 | -0.693 | -0.116 | -0.143 | 0 | 0.069 | 0.069 | 0 | 0.07 | 0.11 | 0.12 | 0.29 | 0.09 | 0.1 |
| 10 | 133183611 | rs7091428 | -0.164 | -0.795 | -0.002 | -0.48 | -0.008 | -0.037 | -0.037 | 2.924 | 0.46 | 0.47 | 0.36 | 0.14 | 0.38 | 0.34 |
| 10 | 133187958 | rs4750868 | 0.88 | -0.394 | 0.15 | -0.36 | 0.06 | -0.04 | -0.04 | 0 | 0.21 | 0.29 | 0.21 | 0.12 | 0.34 | 0.3 |
| 10 | 133189077 | rs3123212 | -0.3 | -0.636 | -0.193 | -0.356 | -0.018 | 0.014 | 0.014 | 0.052 | 0.34 | 0.24 | 0.27 | 0.27 | 0.47 | 0.39 |
| 10 | 133189191 | rs3123209 | -0.772 | -0.856 | -0.064 | -0.293 | 0.006 | 0 | 0 | 1.3 | 0.43 | 0.63 | 0.41 | 0.42 | 0.54 | 0.44 |
| 10 | 133189399 | rs3123208 | -0.831 | -0.813 | 0.052 | -0.268 | 0.029 | 0.007 | 0.007 | 0 | 0.48 | 0.65 | 0.54 | 0.54 | 0.56 | 0.55 |
| 10 | 133189478 | rs3123207 | 0.491 | -0.772 | -0.003 | -0.232 | -0.001 | 0.036 | 0.036 | 0 | 0.1 | 0.17 | 0.05 | 0.15 | 0.07 | 0.08 |
| 10 | 133189605 | rs3123205 | -0.983 | -0.957 | -0.302 | -0.38 | 0.02 | -0.029 | -0.029 | 2.296 | 0.69 | 0.76 | 0.64 | 0.54 | 0.74 | 0.69 |
| 10 | 133189864 | rs11812842 | -0.19 | -0.953 | -0.244 | -0.393 | -0.004 | -0.001 | -0.001 | 0.068 | 0.58 | 0.57 | 0.59 | 0.39 | 0.67 | 0.61 |
| 10 | 133189982 | rs6482878 | 0.198 | -0.927 | -0.245 | -0.397 | 0.04 | 0.016 | 0.016 | 0 | 0.34 | 0.53 | 0.47 | 0.39 | 0.42 | 0.44 |
| 10 | 133190744 | rs7922656 | 0.337 | -0.912 | -0.256 | -0.391 | 0.014 | 0.006 | 0.006 | 0 | 0.33 | 0.38 | 0.46 | 0.39 | 0.42 | 0.44 |
| 10 | 133191575 | rs3123202 | -1.291 | -1.105 | -0.302 | -0.089 | 0.02 | -0.032 | -0.032 | 0 | 0.73 | 0.8 | 0.68 | 0.55 | 0.8 | 0.72 |
| 10 | 133192781 | rs3123201 | 0.29 | -0.916 | -0.403 | -0.001 | -0.002 | 0.057 | 0.057 | 0 | 0.1 | 0.19 | 0.05 | 0.17 | 0.07 | 0.08 |
| 10 | 133193798 | rs4751379 | -0.473 | -1.036 | -0.545 | -0.143 | -0.024 | 0.019 | 0.019 | 0 | 0.67 | 0.55 | 0.63 | 0.39 | 0.74 | 0.65 |
| 10 | 133194410 | rs7893895 | -0.219 | -0.488 | -0.938 | -0.351 | 0.052 | 0.088 | 0.088 | 0.016 | 0.55 | 0.3 | 0.49 | 0.24 | 0.66 | 0.6 |
| 10 | 133195255 | rs11017948 | 0.208 | -0.11 | -0.976 | -0.167 | -0.008 | 0.147 | 0.147 | 0 | 0.25 | 0.04 | 0.13 | 0 | 0.24 | 0.17 |
| 10 | 133196163 | rs3123194 | -0.522 | 0.151 | -0.204 | 0.67 | 0.008 | 0.068 | 0.068 | 0 | 0.06 | 0.15 | 0.03 | 0.13 | 0.05 | 0.06 |
| 10 | 133200104 | rs4076152 | -0.322 | -0.475 | -0.732 | 0.22 | 0.013 | 0.014 | 0.014 | 0 | 0.39 | 0.36 | 0.29 | 0.19 | 0.28 | 0.29 |
| 10 | 133200384 | rs4075537 | 0.183 | -0.115 | -0.894 | 0.413 | 0.001 | -0.001 | -0.001 | 2.6 | 0.68 | 0.67 | 0.7 | 0.66 | 0.73 | 0.8 |
| 10 | 133204121 | rs4601678 | 0.313 | 0.454 | -1.296 | 0.053 | -0.015 | 0.001 | 0.001 | 0 | 0.17 | 0.05 | 0.11 | 0.18 | 0.17 | 0.2 |
| 10 | 133204350 | rs3125510 | 0.202 | 0.189 | -1.765 | -0.493 | 0.02 | 0.079 | 0.079 | 0.234 | 0.18 | 0.08 | 0.09 | 0 | 0.16 | 0.13 |
| 10 | 133204527 | rs12263081 | 0.208 | 0.986 | -1.685 | -0.26 | 0.011 | 0.016 | 0.016 | 0 | 0.15 | 0.19 | 0.35 | 0.27 | 0.26 | 0.19 |
| 10 | 133206629 | rs10765069 | 1.835 | 0.3 | -2.023 | -0.721 | -0.042 | -0.109 | -0.109 | 0 | 0.25 | 0.08 | 0.3 | 0.52 | 0.26 | 0.33 |
| 10 | 133212223 | rs10830004 | 0.101 | 0.303 | -1.789 | -0.822 | -0.015 | -0.014 | -0.014 | 0 | 0.35 | 0.25 | 0.45 | 0.46 | 0.43 | 0.42 |
| 10 | 133214941 | rs3125524 | -0.226 | -0.247 | -1.063 | -1.174 | 0.008 | -0.026 | -0.026 | 0 | 0.26 | 0.28 | 0.14 | 0 | 0.19 | 0.19 |
| 10 | 133215131 | rs3123183 | 1.298 | -0.36 | -1.144 | -1.251 | 0.021 | 0.069 | 0.069 | 0.89 | 0.14 | 0.02 | 0.07 | 0 | 0.11 | 0.04 |
| 10 | 133216922 | rs3123180 | -0.156 | -0.199 | -1.045 | -1.149 | 0.008 | -0.026 | -0.026 | 0.309 | 0.26 | 0.28 | 0.14 | 0 | 0.19 | 0.19 |
| 10 | 133217630 | rs4751388 | -0.961 | 0.162 | -0.824 | -0.81 | 0.017 | -0.082 | -0.082 | 0 | 0.85 | 0.62 | 0.89 | 0.98 | 0.87 | 0.93 |
| 10 | 133225447 | rs10830006 | 1.508 | 0.385 | -1.479 | -0.608 | -0.038 | -0.105 | -0.105 | 0 | 0.24 | 0.12 | 0.3 | 0.49 | 0.27 | 0.34 |
| 10 | 133227554 | rs4751390 | 0.738 | 0.224 | -1.49 | -0.623 | -0.035 | -0.009 | -0.009 | 0 | 0.23 | 0.05 | 0.17 | 0.22 | 0.27 | 0.2 |
| 10 | 133231622 | rs3922590 | 0.439 | 0.316 | -0.471 | -0.544 | 0.046 | 0.165 | 0.165 | 0.097 | 0.11 | 0.36 | 0.29 | 0.28 | 0.14 | 0.25 |
| 10 | 133235583 | rs3125535 | 0.386 | -0.004 | -0.824 | -0.784 | 0.02 | 0.1 | 0.1 | 0 | 0.2 | 0.04 | 0.09 | 0 | 0.15 | 0.1 |
| 10 | 133237375 | rs3125537 | -1.409 | -0.103 | -0.631 | -0.971 | -0.002 | -0.012 | -0.012 | 0.488 | 0.64 | 0.63 | 0.63 | 0.49 | 0.68 | 0.6 |
| 10 | 133237565 | rs3125538 | -0.249 | -0.114 | -0.835 | -1.02 | 0.008 | 0.097 | 0.097 | 0 | 0.46 | 0.22 | 0.28 | 0.21 | 0.45 | 0.3 |
| 10 | 133238622 | rs4077607 | -0.823 | -0.025 | -0.673 | -1.035 | 0.005 | -0.001 | -0.001 | 0.648 | 0.49 | 0.56 | 0.57 | 0.5 | 0.55 | 0.55 |
| 10 | 133238930 | rs12778086 | 0.55 | 0.043 | -0.627 | -0.919 | 0.004 | 0.08 | 0.08 | 0.131 | 0.15 | 0.32 | 0.32 | 0.27 | 0.18 | 0.29 |
| 10 | 133239056 | rs3123155 | 0.218 | 0.03 | -1.149 | -1.04 | 0.014 | 0.083 | 0.083 | 0 | 0.23 | 0.07 | 0.12 | 0.01 | 0.2 | 0.11 |
| 10 | 133239763 | rs4593913 | 0.855 | 0.312 | -0.799 | -0.642 | -0.031 | -0.003 | -0.003 | 0 | 0.23 | 0.07 | 0.18 | 0.21 | 0.28 | 0.2 |
| 10 | 133246492 | rs4751392 | 0.832 | 0.326 | -0.814 | -0.678 | -0.036 | -0.013 | -0.013 | 0.273 | 0.24 | 0.06 | 0.19 | 0.22 | 0.28 | 0.21 |
| 10 | 133247995 | rs12780934 | -1.016 | 0.14 | -0.355 | -0.581 | -0.002 | 0.029 | 0.029 | 0 | 0.66 | 0.52 | 0.65 | 0.5 | 0.7 | 0.65 |
| 10 | 133251838 | rs3125551 | -0.854 | 0.167 | -0.678 | -0.799 | -0.052 | 0.022 | 0.022 | 0 | 0.68 | 0.87 | 0.74 | 0.75 | 0.63 | 0.79 |
| 10 | 133253673 | rs4384310 | 0.255 | -0.171 | -0.071 | -0.55 | -0.008 | 0.043 | 0.043 | 0 | 0.22 | 0.32 | 0.32 | 0.34 | 0.23 | 0.27 |
| 10 | 133256404 | rs3123154 | 0.782 | -0.304 | -0.038 | -0.543 | -0.004 | -0.008 | -0.008 | 2.762 | 0.33 | 0.32 | 0.33 | 0.39 | 0.31 | 0.41 |
| 10 | 133257254 | rs3123150 | 1.262 | -0.379 | 0.021 | -0.499 | -0.002 | -0.009 | -0.009 | 1.692 | 0.27 | 0.26 | 0.29 | 0.39 | 0.25 | 0.37 |
| 10 | 133262270 | rs7077171 | 0.052 | -0.474 | 0.063 | -0.626 | -0.005 | -0.028 | -0.028 | 4.837 | 0.18 | 0.16 | 0.09 | 0.01 | 0.14 | 0.11 |
| 10 | 133262798 | rs7081384 | -1.635 | -0.367 | -0.14 | -0.393 | -0.011 | -0.032 | -0.032 | 0 | 0.76 | 0.86 | 0.71 | 0.62 | 0.75 | 0.63 |
| 10 | 133262919 | rs4462254 | 0.94 | -0.202 | 0.036 | -0.425 | -0.001 | 0.059 | 0.059 | 0 | 0.16 | 0.24 | 0.3 | 0.34 | 0.17 | 0.25 |
| 10 | 133273781 | rs7091184 | -0.96 | -0.483 | -0.315 | -0.066 | -0.006 | -0.025 | -0.025 | 0 | 0.73 | 0.88 | 0.76 | 0.68 | 0.75 | 0.58 |
| 10 | 133275948 | rs4750869 | -0.733 | -0.478 | 0.012 | -0.038 | -0.008 | 0.015 | 0.015 | 0 | 0.65 | 0.81 | 0.71 | 0.71 | 0.64 | 0.53 |
| 10 | 133276361 | rs10830011 | 1.071 | -0.265 | -0.288 | -0.097 | -0.004 | -0.006 | -0.006 | 0 | 0.25 | 0.26 | 0.41 | 0.37 | 0.33 | 0.28 |
| 10 | 133280976 | rs7899857 | 0.519 | -0.422 | -0.581 | -0.178 | -0.067 | -0.096 | -0.096 | 1.173 | 0.26 | 0.02 | 0.22 | 0.33 | 0.27 | 0.23 |
| 10 | 133288281 | rs4456177 | -0.179 | -0.472 | -0.079 | -0.254 | 0.008 | 0.022 | 0.022 | 0 | 0.56 | 0.45 | 0.65 | 0.7 | 0.62 | 0.52 |
| 10 | 133296653 | rs10765074 | -0.58 | -0.295 | -0.491 | -0.158 | -0.001 | 0.03 | 0.03 | 0 | 0.64 | 0.84 | 0.71 | 0.71 | 0.63 | 0.53 |
| 10 | 133298563 | rs4074862 | 0.949 | -0.316 | 0.075 | -0.127 | 0.001 | 0.003 | 0.003 | 0.222 | 0.29 | 0.33 | 0.42 | 0.38 | 0.34 | 0.27 |
| 10 | 133300058 | rs10830019 | -0.406 | -0.42 | 0.37 | -0.296 | 0.003 | 0.034 | 0.034 | 0 | 0.56 | 0.66 | 0.67 | 0.71 | 0.59 | 0.46 |
| 10 | 133300258 | rs10765075 | -0.953 | -0.409 | 0.317 | -0.29 | 0.001 | 0.01 | 0.01 | 0 | 0.65 | 0.71 | 0.71 | 0.71 | 0.65 | 0.52 |
| 10 | 133309117 | rs12049746 | 0.597 | -0.308 | 0.353 | -0.266 | -0.006 | -0.025 | -0.025 | 0.9 | 0.21 | 0.1 | 0.2 | 0.33 | 0.21 | 0.18 |
| 10 | 133309942 | rs7912381 | -0.707 | -0.133 | 0.118 | -0.141 | -0.005 | -0.005 | -0.005 | 0.06 | 0.38 | 0.52 | 0.3 | 0.33 | 0.31 | 0.25 |
| 10 | 133316452 | rs7922424 | -0.287 | -0.273 | 0.061 | -0.527 | -0.001 | -0.044 | -0.044 | 0 | 0.09 | 0.13 | 0.06 | 0 | 0.05 | 0.05 |
| 10 | 133317940 | rs10741254 | -0.407 | -0.446 | -0.299 | -0.724 | 0.008 | 0.044 | 0.044 | 0 | 0.49 | 0.4 | 0.43 | 0.73 | 0.54 | 0.52 |
| 10 | 133320805 | rs7920351 | -0.222 | -0.283 | -0.207 | -0.531 | -0.003 | 0.021 | 0.021 | 0 | 0.64 | 0.72 | 0.74 | 0.77 | 0.6 | 0.54 |
| 10 | 133322295 | rs11018014 | 0.995 | -0.169 | -0.19 | -0.424 | -0.022 | -0.023 | -0.023 | 0.342 | 0.21 | 0.08 | 0.18 | 0.23 | 0.28 | 0.36 |
| 10 | 133322701 | rs12164734 | 0.52 | -0.133 | 0.388 | -0.355 | 0.031 | -0.023 | -0.023 | 0 | 0.07 | 0.25 | 0.06 | 0 | 0.06 | 0.05 |
| 10 | 133323095 | rs12414552 | 0.611 | -0.03 | -0.34 | -0.183 | -0.008 | 0.007 | 0.007 | 0 | 0.28 | 0.25 | 0.47 | 0.27 | 0.31 | 0.3 |

| | | | | | | | | | | | | | | | | |
|---|---|---|---|---|---|---|---|---|---|---|---|---|---|---|---|---|
| 10 | 133325470 | rs12219278 | 0.337 | 0.477 | -0.686 | 0.512 | -0.027 | -0.072 | -0.072 | 0 | 0.18 | 0.09 | 0.18 | 0.49 | 0.2 | 0.15 |
| 10 | 133325496 | rs4130923 | 0.095 | 0.064 | -0.798 | -0.019 | -0.013 | 0.072 | 0.072 | 0.215 | 0.61 | 0.88 | 0.76 | 0.79 | 0.59 | 0.48 |
| 10 | 133326536 | rs11018019 | 0.348 | -0.153 | -0.659 | -0.025 | -0.003 | -0.003 | -0.003 | 0 | 0.75 | 0.84 | 0.81 | 0.76 | 0.73 | 0.57 |
| 10 | 133328670 | rs10741259 | 0.36 | -0.279 | -0.306 | -0.073 | 0 | 0 | 0 | 0 | 0.73 | 0.76 | 0.8 | 0.76 | 0.72 | 0.57 |
| 10 | 133329049 | rs4442457 | -0.83 | -0.306 | -0.374 | -0.073 | -0.003 | -0.022 | -0.022 | 0 | 0.87 | 0.93 | 0.85 | 0.77 | 0.84 | 0.66 |
| 10 | 133330352 | rs7895295 | 0.411 | -0.514 | 0.025 | -0.126 | 0.001 | 0.01 | 0.01 | 0 | 0.5 | 0.5 | 0.52 | 0.32 | 0.5 | 0.49 |
| 10 | 133336844 | rs10765087 | -0.153 | -0.605 | -0.005 | -0.101 | -0.001 | 0.001 | 0.001 | 0.066 | 0.37 | 0.38 | 0.3 | 0.44 | 0.34 | 0.18 |
| 10 | 133339122 | rs11018027 | 0.951 | -0.777 | -0.149 | -0.216 | 0.011 | 0.003 | 0.003 | 0 | 0.15 | 0.09 | 0.08 | 0 | 0.13 | 0.04 |
| 10 | 133344752 | rs4578293 | -0.926 | -0.654 | 0.017 | -0.124 | -0.002 | 0.003 | 0.003 | 0 | 0.86 | 0.81 | 0.84 | 0.81 | 0.84 | 0.66 |
| 10 | 133347995 | rs11018031 | 1.523 | -0.596 | -0.016 | -0.157 | 0.004 | -0.004 | -0.004 | 0 | 0.37 | 0.39 | 0.5 | 0.31 | 0.38 | 0.39 |
| 10 | 133349223 | rs4301710 | 0.939 | -0.647 | -0.03 | -0.274 | -0.016 | -0.055 | -0.055 | 0 | 0.18 | 0.11 | 0.18 | 0.44 | 0.19 | 0.13 |
| 10 | 133350658 | rs4620634 | -0.88 | -0.559 | -0.241 | -0.286 | -0.002 | 0.001 | 0.001 | 0 | 0.85 | 0.81 | 0.84 | 0.81 | 0.84 | 0.65 |
| 10 | 133352612 | rs7898450 | 0.712 | 0.06 | -0.011 | -0.162 | 0.003 | -0.002 | -0.002 | 0 | 0.36 | 0.39 | 0.5 | 0.36 | 0.38 | 0.39 |
| 10 | 133353416 | rs7092114 | -0.34 | 0.155 | -0.342 | -0.104 | 0.001 | 0.014 | 0.014 | 0.084 | 0.54 | 0.57 | 0.6 | 0.36 | 0.53 | 0.44 |
| 10 | 133355589 | rs7071307 | -0.341 | 0.882 | -0.689 | 0.57 | -0.022 | 0.008 | 0.008 | 0.068 | 0.64 | 0.67 | 0.52 | 0.65 | 0.61 | 0.63 |
| 10 | 133357401 | rs11018045 | -0.853 | 0.543 | -0.103 | 1.206 | -0.01 | 0.123 | 0.123 | 0.128 | 0.21 | 0.43 | 0.18 | 0.43 | 0.19 | 0.26 |
| 10 | 133357513 | rs10830037 | -1.105 | 0.938 | 0.044 | 0.842 | 0.003 | -0.002 | -0.002 | 0 | 0.39 | 0.53 | 0.44 | 0.61 | 0.39 | 0.4 |
| 10 | 133357613 | rs4388790 | -0.951 | 0.915 | 0.211 | 1.201 | 0.002 | 0.035 | 0.035 | 0 | 0.32 | 0.48 | 0.27 | 0.48 | 0.28 | 0.26 |
| 10 | 133357766 | rs4486534 | 0.194 | 1.152 | -0.335 | 0.703 | -0.003 | 0.01 | 0.01 | 0.006 | 0.57 | 0.68 | 0.45 | 0.39 | 0.59 | 0.58 |
| 10 | 133358014 | rs12356074 | -0.483 | 1.268 | -0.09 | 0.94 | 0 | 0.004 | 0.004 | 0 | 0.05 | 0.06 | 0.12 | 0.1 | 0.08 | 0.04 |
| 10 | 133360399 | rs10765092 | -1.486 | 1.223 | -0.342 | 0.752 | -0.037 | 0.047 | 0.047 | 0.321 | 0.6 | 0.78 | 0.73 | 0.79 | 0.57 | 0.74 |
| 10 | 133362116 | rs12265483 | -0.341 | 0.975 | 0.302 | 0.786 | -0.001 | 0.005 | 0.005 | 0.808 | 0.16 | 0.29 | 0.24 | 0.23 | 0.13 | 0.3 |
| 10 | 133366939 | rs7099134 | -0.365 | 0.612 | 0.576 | 0.708 | 0.001 | 0.028 | 0.028 | 0.034 | 0.34 | 0.46 | 0.39 | 0.44 | 0.37 | 0.35 |
| 10 | 133367723 | rs6482910 | -0.294 | 0.593 | 0.788 | 0.701 | 0.001 | 0.013 | 0.013 | 0.01 | 0.34 | 0.37 | 0.38 | 0.44 | 0.37 | 0.35 |
| 10 | 133368824 | rs11018057 | -1.219 | 0.481 | 0.79 | 0.71 | -0.031 | 0.001 | 0.001 | 3.558 | 0.27 | 0.45 | 0.34 | 0.34 | 0.2 | 0.4 |
| 10 | 133370057 | rs6482912 | -0.116 | 0.622 | 0.921 | 0.688 | -0.005 | 0.009 | 0.009 | 0 | 0.32 | 0.32 | 0.36 | 0.44 | 0.31 | 0.34 |
| 10 | 133373941 | rs11018064 | -0.158 | 0.637 | 0.844 | 0.695 | -0.001 | 0.003 | 0.003 | 0 | 0.33 | 0.33 | 0.36 | 0.45 | 0.33 | 0.34 |
| 10 | 133374747 | rs7906001 | -2.19 | 0.571 | 0.574 | 0.689 | -0.041 | 0.032 | 0.032 | 0.391 | 0.62 | 0.85 | 0.73 | 0.78 | 0.59 | 0.75 |
| 10 | 133375319 | rs4405224 | -1.413 | 0.736 | 0.639 | 0.71 | 0 | 0.003 | 0.003 | 0.681 | 0.49 | 0.53 | 0.6 | 0.68 | 0.52 | 0.66 |
| 10 | 133377136 | rs10830053 | -0.308 | 0.777 | 0.041 | 0.593 | -0.002 | -0.004 | -0.004 | 0.142 | 0.14 | 0.1 | 0.21 | 0.23 | 0.13 | 0.3 |
| 10 | 133437506 | rs4129966 | 1.41 | 0.698 | 0.232 | 0.398 | 0 | -0.043 | -0.043 | 0.204 | 0.08 | 0.14 | 0.05 | 0 | 0.09 | 0.07 |
| 10 | 133437939 | rs4631813 | -0.474 | 0.6 | 0.298 | 0.827 | -0.004 | -0.011 | -0.011 | 4.026 | 0.79 | 0.83 | 0.7 | 0.63 | 0.77 | 0.74 |
| 10 | 133441851 | rs11156442 | 1.608 | 0.524 | 0.148 | 0.651 | -0.007 | 0.051 | 0.051 | 0 | 0.48 | 0.3 | 0.32 | 0.27 | 0.5 | 0.32 |
| 10 | 133441876 | rs11816792 | -0.013 | 0.464 | 0.184 | 0.688 | -0.001 | -0.002 | -0.002 | 0 | 0.14 | 0.12 | 0.17 | 0.19 | 0.13 | 0.33 |
| 10 | 133443512 | rs10872807 | 1.485 | 0.608 | 0.013 | 0.545 | 0.021 | 0.074 | 0.074 | 0 | 0.49 | 0.23 | 0.36 | 0.27 | 0.56 | 0.33 |
| 10 | 133445033 | rs11156453 | -1.438 | 0.429 | 0.188 | 0.449 | -0.014 | -0.009 | -0.009 | 0 | 0.32 | 0.6 | 0.37 | 0.3 | 0.27 | 0.43 |
| 10 | 133446983 | rs7071489 | -0.533 | -0.059 | 0.272 | 0.075 | -0.01 | 0.008 | 0.008 | 0 | 0.27 | 0.38 | 0.32 | 0.31 | 0.22 | 0.29 |
| 10 | 133448637 | rs9419596 | 1.628 | -0.01 | 0.123 | 0.117 | 0.044 | 0.068 | 0.068 | 0 | 0.48 | 0.16 | 0.36 | 0.28 | 0.6 | 0.27 |
| 10 | 133453734 | rs9419665 | 0.251 | 0.243 | -0.121 | 0.239 | -0.008 | 0.004 | 0.004 | 0.178 | 0.09 | 0.07 | 0.27 | 0.38 | 0.12 | 0.37 |
| 10 | 133458722 | rs4993019 | -0.705 | 0.021 | -0.256 | 0.2 | 0.031 | -0.046 | -0.046 | 0.066 | 0.79 | 0.93 | 0.75 | 0.68 | 0.85 | 0.77 |
| 10 | 133461663 | rs10782374 | 1.074 | 0.051 | 0.101 | 0.058 | 0.002 | 0.004 | 0.004 | 0 | 0.56 | 0.36 | 0.38 | 0.29 | 0.62 | 0.34 |
| 10 | 133466045 | rs6571245 | -1.093 | -0.188 | 0.478 | -0.035 | -0.012 | 0.14 | 0.14 | 0 | 0.2 | 0.52 | 0.32 | 0.43 | 0.15 | 0.41 |
| 10 | 133467924 | rs9419608 | 1.783 | 0.024 | 0.159 | -0.331 | -0.05 | 0.013 | 0.013 | 0 | 0.38 | 0.16 | 0.3 | 0.27 | 0.5 | 0.22 |
| 10 | 133470557 | rs4076579 | -0.802 | -0.021 | 0.152 | -0.384 | 0.026 | -0.034 | -0.034 | 0 | 0.78 | 0.9 | 0.73 | 0.69 | 0.84 | 0.74 |
| 10 | 133471652 | rs4611134 | 1.634 | 0.174 | 0.111 | -0.225 | -0.05 | 0.014 | 0.014 | 0 | 0.38 | 0.16 | 0.3 | 0.27 | 0.5 | 0.22 |
| 10 | 133471888 | rs3924169 | -0.231 | 0.144 | 0.463 | 0.279 | -0.002 | 0.083 | 0.083 | 0.058 | 0.11 | 0.21 | 0.27 | 0.43 | 0.1 | 0.4 |
| 10 | 133475164 | rs11156539 | 0.097 | 0.163 | 0.245 | 0.308 | -0.003 | 0.054 | 0.054 | 0.048 | 0.09 | 0.12 | 0.25 | 0.43 | 0.07 | 0.39 |
| 10 | 133479621 | rs12250241 | -0.416 | 0.374 | 0.392 | -0.092 | 0.072 | -0.027 | -0.027 | 0 | 0.08 | 0.33 | 0.06 | 0 | 0.07 | 0.01 |
| 10 | 133481888 | rs7069785 | -0.976 | 0.314 | 0.041 | -0.185 | 0 | -0.007 | -0.007 | 0 | 0.4 | 0.48 | 0.63 | 0.73 | 0.38 | 0.66 |
| 10 | 133482605 | rs7096432 | -0.833 | 0.445 | -0.229 | -0.624 | -0.023 | 0.088 | 0.088 | 0.098 | 0.82 | 0.96 | 0.93 | 1 | 0.85 | 0.94 |
| 10 | 133482633 | rs11156544 | 0.029 | 0.527 | 0.315 | -0.11 | 0.058 | 0.004 | 0.004 | 0.024 | 0.06 | 0.29 | 0.05 | 0 | 0.04 | 0 |
| 10 | 133484666 | rs11156545 | -0.629 | 0.47 | -0.136 | 0.141 | -0.003 | -0.003 | -0.003 | 0.493 | 0.21 | 0.21 | 0.31 | 0.32 | 0.23 | 0.21 |
| 10 | 133490397 | rs11156546 | 0.921 | 0.267 | -0.472 | -0.118 | -0.008 | -0.021 | -0.021 | 0 | 0.26 | 0.16 | 0.2 | 0.29 | 0.3 | 0.23 |
| 10 | 133491775 | rs9419615 | -0.905 | 0.16 | -0.157 | -0.68 | 0.002 | -0.022 | -0.022 | 0 | 0.63 | 0.66 | 0.79 | 1 | 0.64 | 0.75 |
| 10 | 133493141 | rs4545466 | 0.263 | 0.118 | 0.141 | -0.414 | 0 | 0.005 | 0.005 | 3.026 | 0.43 | 0.36 | 0.49 | 0.62 | 0.44 | 0.55 |
| 10 | 133499307 | rs12765115 | 0.385 | 0.035 | 0.183 | -0.448 | 0.004 | 0.023 | 0.023 | 0 | 0.15 | 0.16 | 0.28 | 0.36 | 0.2 | 0.2 |
| 10 | 133499753 | rs4897813 | -1.178 | 0.023 | 0.23 | -0.496 | 0.008 | -0.004 | -0.004 | 0.04 | 0.64 | 0.7 | 0.81 | 1 | 0.67 | 0.75 |
| 10 | 133501535 | rs9419689 | 1.165 | -0.487 | 1.251 | 0.37 | 0.004 | 0.007 | 0.007 | 1.12 | 0.68 | 0.37 | 0.6 | 0.62 | 0.66 | 0.71 |
| 10 | 133506848 | rs10872824 | -0.275 | -0.183 | 0.416 | 0.558 | 0 | 0.014 | 0.014 | 0 | 0.15 | 0.13 | 0.28 | 0.36 | 0.2 | 0.2 |
| 10 | 133507268 | rs9419617 | 1.15 | -0.195 | -0.216 | 0.595 | -0.04 | 0.101 | 0.101 | 1.145 | 0.49 | 0.1 | 0.28 | 0.27 | 0.51 | 0.38 |
| 10 | 133509442 | rs4897759 | 0.528 | 0.502 | 0.787 | 0.786 | 0.068 | -0.071 | -0.071 | 0 | 0.82 | 0.56 | 0.84 | 0.92 | 0.79 | 0.83 |
| 10 | 133511176 | rs9664915 | -1.272 | 0.205 | 0.359 | 1.366 | -0.008 | 0.024 | 0.024 | 0.355 | 0.16 | 0.2 | 0.23 | 0.32 | 0.14 | 0.15 |
| 10 | 133513897 | rs4387283 | 1.275 | 0.266 | 0.409 | 1.36 | -0.01 | 0.052 | 0.052 | 0.243 | 0.81 | 0.69 | 0.74 | 0.66 | 0.83 | 0.8 |
| 10 | 133514796 | rs10872825 | -1.822 | 0.355 | 0.138 | 1.348 | 0.001 | -0.003 | -0.003 | 0 | 0.18 | 0.17 | 0.15 | 0.23 | 0.2 | 0.17 |
| 10 | 133517373 | rs9419618 | -1.59 | 0.557 | 0.445 | 1.839 | -0.005 | 0.06 | 0.06 | 0 | 0.19 | 0.29 | 0.26 | 0.48 | 0.23 | 0.14 |
| 10 | 133518041 | rs9419698 | 0.813 | 0.804 | 0.967 | 1.663 | 0.005 | 0.122 | 0.122 | 0.011 | 0.75 | 0.55 | 0.71 | 0.52 | 0.74 | 0.51 |
| 10 | 133521382 | rs11156567 | -0.77 | 0.738 | 1.338 | 1.898 | 0.009 | 0.1 | 0.1 | 0 | 0.09 | 0.15 | 0.21 | 0.33 | 0.17 | 0.07 |
| 10 | 133525348 | rs10872828 | -0.243 | 0.924 | 1.507 | 1.147 | 0.057 | 0.092 | 0.092 | 0 | 0.07 | 0.28 | 0.08 | 0.12 | 0.14 | 0.04 |
| 10 | 133531153 | rs9419702 | -2.34 | 0.657 | 1.585 | 0.877 | 0.017 | -0.006 | -0.006 | 1.15 | 0.2 | 0.45 | 0.16 | 0.19 | 0.24 | 0.16 |

| | | | | | | | | | | | | | | | | |
|---|---|---|---|---|---|---|---|---|---|---|---|---|---|---|---|---|
| 10 | 133532269 | rs7083395 | -2.097 | 0.648 | 1.562 | 0.872 | -0.004 | -0.032 | -0.032 | 0 | 0.12 | 0.35 | 0.09 | 0.11 | 0.08 | 0.11 |
| 10 | 133533485 | rs4367873 | 2.446 | 0.393 | 1.777 | 0.81 | 0.004 | 0.007 | 0.007 | 0.13 | 0.7 | 0.23 | 0.76 | 0.78 | 0.66 | 0.79 |
| 10 | 133534822 | rs9419624 | 2.268 | 0.374 | 1.75 | 0.784 | 0.005 | 0.011 | 0.011 | 0.089 | 0.69 | 0.22 | 0.75 | 0.78 | 0.65 | 0.77 |
| 10 | 133536902 | rs7906770 | 0.533 | 0.118 | 1.834 | 0.83 | -0.007 | -0.003 | -0.003 | 0 | 0.89 | 0.69 | 0.9 | 0.96 | 0.88 | 0.92 |
| 10 | 133540749 | rs9419568 | -1.084 | 0.124 | 1.497 | 0.815 | -0.018 | 0 | 0 | 0 | 0.06 | 0.15 | 0.03 | 0.1 | 0.03 | 0.09 |
| 10 | 133541881 | rs9419569 | 1.428 | 0.128 | 1.465 | 0.817 | -0.009 | -0.003 | -0.003 | 0 | 0.74 | 0.33 | 0.79 | 0.78 | 0.71 | 0.79 |
| 10 | 133555393 | rs9419646 | -1.752 | 0.134 | 1.255 | 0.846 | -0.025 | -0.011 | -0.011 | 0 | 0.1 | 0.24 | 0.06 | 0.13 | 0.06 | 0.1 |
| 10 | 133555654 | rs4243514 | 0.809 | 0.335 | 0.965 | 0.881 | 0.017 | 0.01 | 0.01 | 0 | 0.81 | 0.56 | 0.84 | 0.88 | 0.73 | 0.88 |
| 10 | 133557922 | rs9419583 | -2.125 | 0.252 | 0.425 | 0.799 | -0.018 | -0.008 | -0.008 | 0 | 0.1 | 0.23 | 0.07 | 0.13 | 0.06 | 0.1 |
| 10 | 133559274 | rs9419647 | -1.762 | 0.262 | 0.403 | 0.795 | -0.019 | -0.01 | -0.01 | 0.002 | 0.07 | 0.2 | 0.05 | 0.1 | 0.04 | 0.11 |
| 10 | 133561666 | rs9419651 | 0.055 | 0.6 | -0.337 | 0.854 | -0.033 | -0.037 | -0.037 | 0 | 0.63 | 0.81 | 0.59 | 0.48 | 0.59 | 0.39 |
| 10 | 133562475 | rs9419652 | 0.053 | 0.58 | -0.206 | 0.496 | 0.002 | 0 | 0 | 0 | 0.2 | 0.22 | 0.22 | 0.21 | 0.3 | 0.12 |
| 10 | 133567096 | rs7898015 | -0.012 | 0.464 | 0.423 | 0.468 | 0.007 | 0.001 | 0.001 | 0 | 0.21 | 0.25 | 0.22 | 0.21 | 0.3 | 0.12 |
| 10 | 133569448 | rs4307646 | -0.367 | 0.238 | 1.004 | 0.443 | 0.005 | -0.001 | -0.001 | 0 | 0.59 | 0.57 | 0.57 | 0.41 | 0.57 | 0.34 |
| 10 | 133575396 | rs7095978 | 0.561 | -0.337 | 0.731 | -0.228 | 0.015 | 0.029 | 0.029 | 0 | 0.34 | 0.22 | 0.28 | 0.22 | 0.26 | 0.25 |
| 10 | 133578066 | rs4243517 | -0.269 | -0.306 | 1.264 | -0.384 | 0.043 | -0.127 | -0.127 | 0 | 0.79 | 0.66 | 0.83 | 0.99 | 0.69 | 0.9 |
| 10 | 133579416 | rs4897770 | -0.104 | -0.172 | 0.852 | 0.354 | -0.003 | -0.005 | -0.005 | 1.693 | 0.66 | 0.71 | 0.74 | 0.48 | 0.55 | 0.48 |
| 10 | 133590090 | rs11156470 | 0.388 | -0.173 | 0.539 | 0.186 | -0.011 | -0.007 | -0.007 | 0 | 0.19 | 0.14 | 0.29 | 0.19 | 0.24 | 0.17 |
| 10 | 133592726 | rs10782370 | -0.053 | -0.107 | 0.663 | 0.354 | 0.018 | 0.027 | 0.027 | 0 | 0.64 | 0.52 | 0.72 | 0.51 | 0.55 | 0.48 |
| 10 | 133606145 | rs4897779 | -0.083 | -0.025 | 0.491 | 0.313 | -0.004 | -0.004 | -0.004 | 0.738 | 0.63 | 0.63 | 0.71 | 0.52 | 0.55 | 0.47 |
| 10 | 133607892 | rs4897739 | 0.142 | -0.229 | 0.348 | 0.188 | 0 | 0 | 0 | 0.064 | 0.52 | 0.7 | 0.69 | 0.48 | 0.46 | 0.42 |
| 10 | 133608210 | rs4897741 | -0.397 | -0.288 | 0.319 | 0.132 | -0.022 | -0.027 | -0.027 | 2.639 | 0.61 | 0.75 | 0.71 | 0.48 | 0.53 | 0.44 |
| 10 | 133608237 | rs4897783 | 0.217 | -0.266 | 0.578 | 0.165 | 0 | 0 | 0 | 0 | 0.51 | 0.54 | 0.67 | 0.48 | 0.46 | 0.42 |
| 10 | 133624539 | rs11156478 | 0.139 | -0.258 | 0.515 | 0.317 | 0 | 0 | 0 | 0 | 0.49 | 0.5 | 0.57 | 0.41 | 0.45 | 0.4 |
| 10 | 133629469 | rs11156479 | 1.142 | -0.598 | -0.453 | 0.086 | 0.017 | -0.015 | -0.015 | 0.249 | 0.22 | 0.07 | 0.2 | 0.24 | 0.13 | 0.22 |
| 10 | 133632779 | rs9419604 | -0.645 | -0.459 | -0.328 | 0.348 | -0.027 | -0.02 | -0.02 | 0.829 | 0.6 | 0.78 | 0.72 | 0.52 | 0.53 | 0.44 |
| 10 | 133637385 | rs7893330 | -0.138 | -0.548 | 0.036 | 0.311 | -0.032 | -0.006 | -0.006 | 0 | 0.13 | 0.32 | 0.16 | 0.1 | 0.08 | 0.05 |
| 10 | 133639202 | rs4897796 | -0.659 | 0.014 | -0.715 | 0.312 | -0.01 | 0.016 | 0.016 | 0 | 0.63 | 0.84 | 0.67 | 0.69 | 0.67 | 0.74 |
| 10 | 133647554 | rs11156481 | 0.089 | 0.703 | -0.413 | 0.501 | 0.005 | 0.01 | 0.01 | 0 | 0.45 | 0.33 | 0.31 | 0.45 | 0.42 | 0.59 |
| 10 | 133649010 | rs7099404 | -0.643 | 0.791 | -0.451 | 0.434 | -0.015 | 0.051 | 0.051 | 0 | 0.59 | 0.8 | 0.66 | 0.79 | 0.55 | 0.72 |
| 10 | 133649089 | rs6571241 | -0.613 | 0.743 | -0.414 | 0.391 | -0.021 | 0.06 | 0.06 | 0 | 0.59 | 0.78 | 0.66 | 0.82 | 0.55 | 0.72 |
| 10 | 133649265 | rs6571242 | 0.046 | 0.506 | -0.031 | 0.566 | -0.01 | -0.003 | -0.003 | 3.439 | 0.37 | 0.62 | 0.39 | 0.73 | 0.32 | 0.53 |
| 10 | 133649738 | rs4897752 | -0.276 | 0.796 | 0.025 | 0.578 | 0.002 | 0.006 | 0.006 | 0 | 0.43 | 0.64 | 0.42 | 0.73 | 0.4 | 0.56 |
| 10 | 133655020 | rs11814195 | 0.065 | 0.082 | 0.448 | 0.946 | 0.003 | -0.004 | -0.004 | 0.5 | 0.36 | 0.35 | 0.26 | 0.5 | 0.33 | 0.45 |
| 10 | 133657703 | rs7900951 | -0.416 | 0.068 | 0.434 | 0.964 | 0 | 0.001 | 0.001 | 0 | 0.4 | 0.63 | 0.37 | 0.73 | 0.34 | 0.52 |
| 10 | 133664823 | rs4078375 | 0.193 | -0.007 | 0.188 | 0.935 | 0.005 | -0.019 | -0.019 | 0 | 0.27 | 0.21 | 0.21 | 0.49 | 0.28 | 0.39 |
| 10 | 133666812 | rs4078150 | 0.213 | 0.007 | 0.195 | 0.941 | 0.004 | -0.018 | -0.018 | 0 | 0.27 | 0.21 | 0.21 | 0.49 | 0.28 | 0.38 |
| 10 | 133667886 | rs10430621 | -1.237 | 0.209 | 0.316 | 0.919 | -0.003 | 0.17 | 0.17 | 0.337 | 0.05 | 0.26 | 0.02 | 0.19 | 0.01 | 0.08 |
| 10 | 133676255 | rs10747291 | -0.642 | 0.033 | -0.347 | 0.169 | 0.039 | 0.077 | 0.077 | 0 | 0.49 | 0.69 | 0.56 | 0.77 | 0.45 | 0.57 |
| 10 | 133676869 | rs9322841 | -0.177 | -0.189 | -0.272 | 0.007 | 0.02 | 0.02 | 0.02 | 0 | 0.4 | 0.7 | 0.52 | 0.78 | 0.32 | 0.55 |
| 10 | 133729749 | rs9419374 | -0.91 | -0.328 | -0.893 | 0.011 | -0.002 | -0.021 | -0.021 | 0.014 | 0.57 | 0.86 | 0.67 | 0.65 | 0.61 | 0.65 |
| 10 | 133742677 | rs10159746 | -0.12 | -0.275 | -0.421 | -0.012 | 0 | -0.002 | -0.002 | 2.351 | 0.28 | 0.29 | 0.26 | 0.45 | 0.26 | 0.3 |
| 10 | 133743166 | rs9419338 | 1.217 | -0.33 | -0.204 | 0.065 | -0.006 | 0.035 | 0.035 | 0 | 0.11 | 0.18 | 0.14 | 0.36 | 0.08 | 0.21 |
| 10 | 133745628 | rs7072006 | 0.711 | -0.144 | 0.139 | -0.065 | 0.026 | -0.043 | -0.043 | 0.099 | 0.1 | 0.25 | 0.12 | 0.02 | 0.12 | 0.05 |
| 10 | 133750141 | rs7085239 | 0.944 | -0.195 | 0.322 | -0.084 | 0.03 | 0.006 | 0.006 | 0.181 | 0.18 | 0.26 | 0.27 | 0.17 | 0.22 | 0.3 |
| 10 | 133757922 | rs10781564 | -0.382 | -0.18 | 0.029 | -0.071 | 0 | -0.002 | -0.002 | 1.119 | 0.29 | 0.3 | 0.27 | 0.45 | 0.27 | 0.31 |
| 10 | 133758358 | rs9419387 | -0.453 | -0.191 | -0.001 | -0.082 | 0.017 | -0.021 | -0.021 | 0.614 | 0.29 | 0.58 | 0.4 | 0.19 | 0.34 | 0.35 |
| 10 | 133758379 | rs9419217 | -1.216 | -0.116 | 0 | -0.03 | -0.006 | 0 | 0 | 0 | 0.39 | 0.54 | 0.39 | 0.47 | 0.39 | 0.35 |
| 10 | 133777122 | rs7898755 | -0.489 | -0.32 | -0.441 | 0.122 | 0.001 | -0.004 | -0.004 | 0 | 0.28 | 0.29 | 0.26 | 0.45 | 0.26 | 0.3 |
| 10 | 133784881 | rs3737058 | -1.299 | -0.132 | -0.823 | 0.129 | 0 | -0.02 | -0.02 | 3.01 | 0.59 | 0.81 | 0.65 | 0.64 | 0.61 | 0.66 |
| 10 | 133792672 | rs3793740 | -0.712 | -0.218 | -1.026 | -0.095 | -0.003 | -0.016 | -0.016 | 0 | 0.21 | 0.29 | 0.13 | 0.11 | 0.2 | 0.11 |
| 10 | 133798428 | rs7098827 | 0.023 | 0.148 | -1.269 | 0.059 | -0.002 | 0.026 | 0.026 | 0 | 0.16 | 0.12 | 0.11 | 0.09 | 0.19 | 0.08 |
| 10 | 133803156 | rs7915610 | 0.919 | 0.13 | -1.486 | 0.278 | -0.027 | 0.055 | 0.055 | 0 | 0.28 | 0.05 | 0.19 | 0.14 | 0.28 | 0.26 |
| 10 | 133806576 | rs9419235 | -1.018 | 0.105 | -0.806 | 0.719 | -0.006 | 0.024 | 0.024 | 0 | 0.51 | 0.75 | 0.6 | 0.63 | 0.47 | 0.59 |
| 10 | 133806618 | rs11146471 | 0.392 | 0.29 | -0.733 | 0.801 | 0.029 | 0.008 | 0.008 | 0.429 | 0.13 | 0.16 | 0.24 | 0.14 | 0.18 | 0.24 |
| 10 | 133808341 | rs7914539 | -0.514 | 0.141 | -0.798 | 0.793 | -0.024 | 0.01 | 0.01 | 0 | 0.37 | 0.51 | 0.35 | 0.48 | 0.29 | 0.34 |
| 10 | 133809325 | rs11146481 | -0.222 | -0.003 | -1.291 | 0.585 | 0.017 | 0.009 | 0.009 | 0 | 0.11 | 0.11 | 0.05 | 0.09 | 0.04 | 0.06 |
| 10 | 133811201 | rs9419406 | -0.519 | 0.443 | -1.351 | 0.694 | -0.002 | 0.02 | 0.02 | 0.053 | 0.48 | 0.7 | 0.59 | 0.65 | 0.47 | 0.58 |
| 10 | 133816967 | rs11146526 | -0.605 | 0.563 | -1.367 | 0.981 | 0.008 | -0.021 | -0.021 | 0 | 0.25 | 0.16 | 0.29 | 0.35 | 0.18 | 0.12 |
| 10 | 133829719 | rs4880224 | -0.857 | 0.334 | -1.529 | 0.828 | 0.016 | -0.033 | -0.033 | 0 | 0.37 | 0.26 | 0.34 | 0.48 | 0.27 | 0.4 |
| 10 | 133831989 | rs9419257 | -0.544 | 0.572 | -1.636 | 1.028 | 0.004 | -0.02 | -0.02 | 2.011 | 0.3 | 0.26 | 0.33 | 0.5 | 0.23 | 0.3 |
| 10 | 133833317 | rs10870215 | -1.164 | 0.61 | -1.452 | 0.94 | -0.002 | 0.048 | 0.048 | 2.832 | 0.45 | 0.39 | 0.6 | 0.82 | 0.43 | 0.58 |
| 10 | 133833615 | rs2149945 | -1.164 | 0.61 | -1.452 | 0.94 | -0.002 | 0.048 | 0.048 | 2.832 | 0.45 | 0.39 | 0.6 | 0.82 | 0.43 | 0.58 |
| 10 | 133834146 | rs9419138 | -0.06 | 1.115 | -1.124 | 0.975 | 0.009 | 0.043 | 0.043 | 0.109 | 0.15 | 0.25 | 0.28 | 0.3 | 0.2 | 0.27 |
| 10 | 133836143 | rs11146100 | -0.381 | 0.964 | -1.314 | 0.433 | -0.014 | -0.016 | -0.016 | 0 | 0.27 | 0.13 | 0.26 | 0.33 | 0.29 | 0.26 |
| 10 | 133844484 | rs9419313 | 0.255 | 0.63 | -0.924 | 0.215 | 0.004 | -0.014 | -0.014 | 2.419 | 0.72 | 0.76 | 0.78 | 0.63 | 0.74 | 0.7 |
| 10 | 133848760 | rs6560669 | 0.059 | 0.412 | -0.436 | 0.109 | 0.001 | -0.045 | -0.045 | 0.042 | 0.1 | 0.18 | 0.06 | 0 | 0.08 | 0.04 |
| 10 | 133853288 | rs7084312 | 0.725 | 0.528 | -0.475 | -0.13 | 0 | -0.006 | -0.006 | 0.005 | 0.12 | 0.1 | 0.07 | 0.04 | 0.15 | 0.12 |
| 10 | 133855259 | rs2026121 | 0.854 | 0.043 | -0.421 | -0.538 | -0.017 | 0.039 | 0.039 | 0 | 0.23 | 0.3 | 0.25 | 0.58 | 0.16 | 0.39 |

| | | | | | | | | | | | | | | | | |
|---|---|---|---|---|---|---|---|---|---|---|---|---|---|---|---|---|
| 10 | 133858154 | rs28539701 | 0.774 | 0.154 | -0.293 | -0.71 | -0.01 | 0.017 | 0.017 | 0 | 0.26 | 0.32 | 0.4 | 0.61 | 0.18 | 0.4 |
| 10 | 133858248 | rs7919591 | 0.602 | 0.964 | -0.689 | -0.853 | -0.02 | -0.03 | -0.03 | 0.061 | 0.2 | 0.1 | 0.2 | 0.36 | 0.26 | 0.27 |
| 10 | 133858767 | rs9419165 | -1.035 | 0.818 | -0.504 | -1.303 | 0 | -0.026 | -0.026 | 0 | 0.88 | 0.86 | 0.84 | 1 | 0.85 | 0.94 |
| 10 | 133859537 | rs2379214 | -0.286 | 0.739 | -0.614 | -1.368 | -0.003 | 0.042 | 0.042 | 2 | 0.72 | 0.85 | 0.75 | 1 | 0.71 | 0.85 |
| 10 | 133862007 | rs9419167 | 0.006 | 0.883 | -0.26 | -1.168 | -0.001 | -0.155 | -0.155 | 0 | 0.63 | 0.48 | 0.67 | 1 | 0.63 | 0.79 |
| 10 | 133862022 | rs9419168 | 0.402 | 0.845 | -0.36 | -1.176 | -0.004 | 0.068 | 0.068 | 0.561 | 0.49 | 0.38 | 0.59 | 0.97 | 0.48 | 0.66 |
| 10 | 133862452 | rs9419326 | 0.248 | 0.76 | -0.342 | -0.252 | -0.006 | -0.033 | -0.033 | 2.252 | 0.63 | 0.43 | 0.67 | 0.79 | 0.64 | 0.78 |
| 10 | 133862943 | rs7897589 | -0.129 | 0.717 | -0.388 | 0.036 | -0.001 | 0 | 0 | 0.075 | 0.21 | 0.26 | 0.14 | 0.22 | 0.21 | 0.15 |
| 10 | 133863091 | rs1970005 | 0.181 | 0.794 | -0.798 | 0.338 | -0.005 | -0.018 | -0.018 | 0 | 0.72 | 0.78 | 0.67 | 0.63 | 0.72 | 0.6 |
| 10 | 133863637 | rs9419328 | 0.007 | 0.712 | -0.847 | 0.52 | -0.012 | -0.034 | -0.034 | 0 | 0.76 | 0.86 | 0.86 | 0.57 | 0.76 | 0.83 |
| 10 | 133863719 | rs9419331 | 0.104 | 0.075 | -0.487 | -0.395 | -0.021 | -0.146 | -0.146 | 0 | 0.23 | 0.49 | 0.15 | 0.03 | 0.22 | 0.12 |
| 10 | 133864146 | rs1888120 | 0.397 | -0.258 | -0.089 | -0.268 | 0.002 | 0.018 | 0.018 | 0 | 0.51 | 0.57 | 0.72 | 0.75 | 0.53 | 0.71 |
| 10 | 133868904 | rs913868 | 0.899 | -0.424 | -0.249 | -0.55 | 0.004 | -0.001 | -0.001 | 0 | 0.16 | 0.08 | 0.09 | 0.01 | 0.14 | 0.08 |
| 10 | 133869023 | rs11813033 | 1.881 | -0.527 | -0.192 | -0.615 | 0.004 | -0.001 | -0.001 | 0 | 0.11 | 0.07 | 0.04 | 0.03 | 0.07 | 0.06 |
| 10 | 133871019 | rs9419332 | 0.604 | -0.509 | 0.042 | -0.674 | -0.004 | 0.011 | 0.011 | 0 | 0.45 | 0.58 | 0.7 | 0.75 | 0.51 | 0.69 |
| 10 | 133873308 | rs4880231 | -0.817 | -0.52 | -0.106 | -0.755 | -0.024 | 0.036 | 0.036 | 0.211 | 0.76 | 0.82 | 0.85 | 1 | 0.73 | 0.85 |
| 10 | 133875481 | rs9419334 | 0.725 | -0.38 | -0.418 | -0.916 | 0.014 | -0.001 | -0.001 | 0 | 0.29 | 0.2 | 0.3 | 0.31 | 0.26 | 0.15 |
| 10 | 133875524 | rs9419335 | 2.15 | -0.32 | -0.51 | -0.837 | 0.007 | -0.005 | -0.005 | 0.741 | 0.11 | 0.07 | 0.04 | 0.3 | 0.07 | 0.07 |
| 10 | 133875557 | rs12572115 | 1.089 | -0.081 | -0.052 | -0.87 | 0.024 | 0.099 | 0.099 | 0 | 0.2 | 0.32 | 0.39 | 0.45 | 0.25 | 0.43 |
| 10 | 133878345 | rs11146142 | -0.228 | 0.02 | -0.224 | -0.883 | -0.003 | 0.064 | 0.064 | 0 | 0.48 | 0.66 | 0.71 | 0.68 | 0.49 | 0.71 |
| 10 | 133880163 | rs4880317 | -1.13 | 0.36 | -0.748 | -1.05 | -0.045 | 0.087 | 0.087 | 0.422 | 0.75 | 0.86 | 0.83 | 1 | 0.69 | 0.86 |
| 10 | 133884354 | rs11146145 | 0.581 | 0.447 | -0.636 | -0.811 | -0.005 | -0.008 | -0.008 | 0 | 0.33 | 0.31 | 0.59 | 0.44 | 0.4 | 0.43 |
| 10 | 133889541 | rs9419346 | -0.98 | 0.406 | -1.097 | -0.908 | -0.12 | 0.199 | 0.199 | 0.87 | 0.72 | 0.98 | 0.84 | 1 | 0.67 | 0.83 |
| 10 | 133889814 | rs9419348 | 0.716 | 0.364 | -0.363 | -0.443 | -0.012 | 0.011 | 0.011 | 0 | 0.19 | 0.25 | 0.1 | 0.31 | 0.1 | 0.26 |
| 10 | 133892178 | rs11146148 | -0.16 | 0.355 | -0.287 | -0.708 | -0.001 | -0.005 | -0.005 | 0.138 | 0.37 | 0.58 | 0.47 | 0.74 | 0.36 | 0.63 |
| 10 | 133909650 | rs2026782 | -1.346 | 0.193 | -0.774 | -0.718 | 0.028 | 0.079 | 0.079 | 0 | 0.77 | 0.94 | 0.87 | 0.89 | 0.83 | 0.84 |
| 10 | 133909957 | rs2172130 | 0.885 | 1.191 | 0.139 | 0.14 | 0.007 | 0.014 | 0.014 | 0 | 0.14 | 0.23 | 0.27 | 0.13 | 0.15 | 0.2 |
| 10 | 133911868 | rs9419184 | -1.258 | 1.319 | -0.035 | -0.032 | 0 | -0.003 | -0.003 | 0 | 0.57 | 0.51 | 0.61 | 0.73 | 0.65 | 0.72 |
| 10 | 133912388 | rs11598779 | -0.251 | 1.771 | -0.137 | 0.071 | -0.003 | 0.033 | 0.033 | 0 | 0.35 | 0.41 | 0.4 | 0.42 | 0.34 | 0.44 |
| 10 | 133912838 | rs2172129 | 0.323 | 1.809 | -0.191 | 0.098 | -0.051 | -0.035 | -0.035 | 0 | 0.23 | 0.11 | 0.2 | 0.31 | 0.3 | 0.28 |
| 10 | 133913725 | rs7924197 | -1.221 | 1.812 | -0.088 | 0.126 | -0.001 | -0.006 | -0.006 | 4.256 | 0.58 | 0.51 | 0.59 | 0.73 | 0.62 | 0.72 |
| 10 | 133913882 | rs9419188 | -1.391 | 1.842 | -0.084 | 0.129 | 0.018 | 0.065 | 0.065 | 0 | 0.61 | 0.7 | 0.65 | 0.89 | 0.66 | 0.77 |
| 10 | 133914617 | rs913871 | -0.667 | 1.855 | 0.182 | 0.424 | -0.005 | 0.001 | 0.001 | 0 | 0.4 | 0.36 | 0.47 | 0.62 | 0.43 | 0.67 |
| 10 | 133915225 | rs9419364 | 0.373 | 1.588 | 0.37 | 0.569 | 0.024 | 0.133 | 0.133 | 0 | 0.18 | 0.49 | 0.16 | 0.29 | 0.18 | 0.09 |
| 10 | 133918372 | rs10870252 | 0.547 | 1.63 | 0.291 | 0.584 | 0.018 | 0.11 | 0.11 | 0 | 0.17 | 0.39 | 0.14 | 0.29 | 0.18 | 0.09 |
| 10 | 133923348 | rs7897877 | 0.766 | 1.653 | 0.082 | 0.605 | 0.017 | 0.078 | 0.078 | 0 | 0.15 | 0.23 | 0.12 | 0.29 | 0.18 | 0.09 |
| 10 | 133924823 | rs7897094 | -1.102 | 1.793 | -0.146 | 0.585 | -0.005 | 0.046 | 0.046 | 0 | 0.44 | 0.67 | 0.37 | 0.79 | 0.48 | 0.58 |
| 10 | 133926108 | rs7923067 | 0.478 | 1.901 | -0.268 | 1.041 | 0.02 | 0.091 | 0.091 | 0 | 0.16 | 0.27 | 0.13 | 0.29 | 0.18 | 0.09 |
| 10 | 133926165 | rs4880335 | -0.642 | 2.013 | -0.899 | 1.042 | 0.001 | 0.02 | 0.02 | 0 | 0.19 | 0.21 | 0.33 | 0.42 | 0.19 | 0.35 |
| 10 | 133932946 | rs7084508 | -0.44 | 2.221 | -1.378 | 0.836 | -0.084 | -0.044 | -0.044 | 0.397 | 0.21 | 0.05 | 0.18 | 0.29 | 0.33 | 0.32 |
| 10 | 133933380 | rs10870261 | -0.231 | 2.12 | -1.014 | 0.588 | 0.002 | -0.003 | -0.003 | 0 | 0.19 | 0.23 | 0.31 | 0.17 | 0.23 | 0.28 |
| 10 | 133938279 | rs7083603 | -1.036 | 1.508 | -1.013 | 0.707 | -0.017 | 0.033 | 0.033 | 0.478 | 0.47 | 0.78 | 0.53 | 0.64 | 0.45 | 0.71 |
| 10 | 133942779 | rs1491192 | 0.363 | 1.629 | -0.985 | 1.175 | -0.005 | -0.014 | -0.014 | 0.039 | 0.15 | 0.12 | 0.26 | 0.33 | 0.16 | 0.13 |
| 10 | 133943800 | rs902633 | -1.716 | 0.744 | -1.853 | 0.294 | 0.008 | 0.078 | 0.078 | 0.285 | 0.86 | 0.99 | 0.95 | 0.99 | 0.88 | 0.91 |
| 10 | 133943951 | rs902634 | -0.935 | 1.323 | -2.088 | 0.071 | -0.092 | 0.175 | 0.175 | 1.153 | 0.73 | 0.98 | 0.85 | 0.99 | 0.69 | 0.75 |
| 10 | 133947657 | rs2637642 | 0.74 | 0.903 | -1.576 | 0.405 | -0.027 | 0.086 | 0.086 | 1.255 | 0.45 | 0.19 | 0.37 | 0.29 | 0.48 | 0.19 |
| 10 | 133948347 | rs2814183 | 0.866 | 0.875 | -1.45 | 0.353 | -0.046 | 0.11 | 0.11 | 1.48 | 0.43 | 0.16 | 0.37 | 0.25 | 0.48 | 0.19 |
| 10 | 133948634 | rs2814182 | -0.905 | 1.035 | -1.37 | 0.467 | -0.092 | 0.175 | 0.175 | 1.078 | 0.73 | 0.98 | 0.85 | 0.99 | 0.7 | 0.74 |
| 10 | 133953469 | rs7096456 | 0.381 | 0.981 | -1.707 | 0.212 | -0.071 | -0.042 | -0.042 | 0.141 | 0.17 | 0.05 | 0.23 | 0.23 | 0.34 | 0.14 |
| 10 | 133954011 | rs2818387 | -0.439 | 0.179 | -1.104 | 0.135 | -0.001 | 0 | 0 | 0.079 | 0.46 | 0.46 | 0.58 | 0.59 | 0.48 | 0.5 |
| 10 | 133954831 | rs10870265 | 0.302 | 0.277 | -0.657 | 0.245 | -0.025 | 0.026 | 0.026 | 0.568 | 0.3 | 0.54 | 0.4 | 0.62 | 0.23 | 0.44 |
| 10 | 133955427 | rs11592585 | 0.412 | 0.685 | -1.271 | 0.21 | -0.098 | -0.062 | -0.062 | 0.104 | 0.17 | 0.04 | 0.23 | 0.24 | 0.34 | 0.14 |
| 10 | 133955901 | rs12778246 | -0.228 | 0.731 | -0.987 | -0.11 | 0.01 | 0.001 | 0.001 | 0 | 0.39 | 0.45 | 0.39 | 0.36 | 0.46 | 0.25 |
| 10 | 133956675 | rs2637624 | -0.372 | 0.388 | -0.849 | 0.113 | 0.009 | 0.067 | 0.067 | 0 | 0.49 | 0.71 | 0.62 | 0.69 | 0.48 | 0.57 |
| 10 | 133960422 | rs4534498 | -0.035 | -0.223 | -0.418 | -0.06 | 0.008 | 0.001 | 0.001 | 0 | 0.67 | 0.73 | 0.81 | 0.94 | 0.7 | 0.68 |
| 10 | 133960517 | rs7914962 | 0.425 | -0.06 | -0.617 | -0.197 | 0 | 0.101 | 0.101 | 0 | 0.54 | 0.7 | 0.74 | 0.91 | 0.54 | 0.52 |
| 10 | 133961494 | rs2818384 | 0.553 | -0.365 | -0.45 | 0.433 | 0.008 | 0.04 | 0.04 | 1.475 | 0.55 | 0.28 | 0.72 | 0.31 | 0.59 | 0.36 |
| 10 | 133966434 | rs11146215 | -0.35 | -0.396 | -0.318 | 1.031 | -0.009 | 0.226 | 0.226 | 0 | 0.05 | 0.09 | 0.02 | 0.49 | 0.03 | 0.1 |
| 10 | 133970588 | rs870552 | 0.611 | -0.457 | -0.273 | 0.366 | 0.031 | 0.031 | 0.031 | 0 | 0.53 | 0.31 | 0.57 | 0.38 | 0.57 | 0.39 |
| 10 | 133977531 | rs7915260 | 1.156 | -0.482 | -0.646 | 0.221 | -0.095 | 0.029 | 0.029 | 0 | 0.41 | 0.11 | 0.53 | 0.32 | 0.49 | 0.33 |
| 10 | 133992803 | rs2637658 | 1.092 | -0.602 | -0.317 | 0.028 | 0 | -0.012 | -0.012 | 0 | 0.13 | 0.1 | 0.11 | 0.02 | 0.11 | 0.05 |
| 10 | 133996225 | rs2637648 | 1.247 | -0.606 | -0.342 | 0.026 | -0.016 | 0.056 | 0.056 | 4.636 | 0.12 | 0.03 | 0.09 | 0.02 | 0.11 | 0.04 |
| 10 | 133997031 | rs2637656 | 0.977 | -0.647 | -0.338 | 0.008 | 0.003 | 0.012 | 0.012 | 3.25 | 0.14 | 0.09 | 0.1 | 0.02 | 0.12 | 0.05 |
| 10 | 134000146 | rs902632 | -1.047 | -0.719 | -0.132 | -0.053 | -0.001 | -0.024 | -0.024 | 0 | 0.81 | 0.78 | 0.88 | 0.96 | 0.81 | 0.85 |
| 10 | 134002767 | rs2101613 | 1.629 | -0.749 | -0.214 | 0.06 | -0.112 | 0.013 | 0.013 | 0 | 0.4 | 0.09 | 0.53 | 0.32 | 0.49 | 0.32 |
| 10 | 134007114 | rs2818421 | -1.063 | -0.797 | -0.185 | 0.012 | -0.002 | -0.026 | -0.026 | 0 | 0.81 | 0.78 | 0.88 | 0.96 | 0.8 | 0.85 |
| 10 | 134008812 | rs10781563 | 0.91 | -0.9 | 0.26 | 0.006 | -0.006 | 0.116 | 0.116 | 0 | 0.47 | 0.29 | 0.56 | 0.91 | 0.51 | 0.44 |
| 10 | 134013856 | rs871883 | 1.636 | -0.786 | -0.438 | 0.431 | -0.141 | 0.024 | 0.024 | 0 | 0.41 | 0.05 | 0.51 | 0.68 | 0.48 | 0.39 |
| 10 | 134013990 | rs2247705 | -0.858 | -0.825 | -0.229 | 0.452 | -0.004 | -0.016 | -0.016 | 0 | 0.83 | 0.8 | 0.89 | 0.93 | 0.81 | 0.87 |

| | | | | | | | | | | | | | | | | |
|---|---|---|---|---|---|---|---|---|---|---|---|---|---|---|---|---|
| 10 | 134016396 | rs3736258 | 1.599 | -0.757 | -0.659 | 0.599 | -0.122 | 0.011 | 0.011 | 0 | 0.4 | 0.09 | 0.52 | 0.34 | 0.49 | 0.32 |
| 10 | 134020606 | rs2818415 | -0.555 | -0.115 | -0.075 | 1.221 | -0.004 | -0.024 | -0.024 | 0 | 0.81 | 0.77 | 0.88 | 0.92 | 0.81 | 0.85 |
| 10 | 134020697 | rs11146249 | -0.503 | -0.139 | -0.244 | 1.116 | -0.014 | -0.019 | -0.019 | 0 | 0.19 | 0.09 | 0.09 | 0.22 | 0.18 | 0.26 |
| 10 | 134021870 | rs923661 | 1.106 | -0.09 | -0.053 | 0.626 | -0.002 | -0.003 | -0.003 | 0 | 0.57 | 0.43 | 0.67 | 0.36 | 0.59 | 0.43 |
| 10 | 134026212 | rs11146253 | -1.702 | -0.244 | -0.516 | 1.274 | -0.019 | 0.189 | 0.189 | 0 | 0.07 | 0.13 | 0.04 | 0.56 | 0.03 | 0.1 |
| 10 | 134034379 | rs2035804 | 0.239 | 0.071 | -0.754 | 0.203 | -0.009 | -0.025 | -0.025 | 0.26 | 0.77 | 0.69 | 0.86 | 0.84 | 0.79 | 0.75 |
| 10 | 134039891 | rs870427 | 0.01 | 0.546 | -1.566 | 0.358 | -0.029 | 0.058 | 0.058 | 2.804 | 0.48 | 0.15 | 0.51 | 0.33 | 0.5 | 0.24 |
| 10 | 134041065 | rs2637637 | -0.011 | 0.346 | -1.35 | -0.379 | -0.003 | -0.032 | -0.032 | 0 | 0.62 | 0.43 | 0.74 | 0.92 | 0.68 | 0.8 |
| 10 | 134045675 | rs4880352 | -0.398 | 0.435 | -0.963 | 0.361 | -0.006 | -0.003 | -0.003 | 0 | 0.66 | 0.38 | 0.7 | 0.7 | 0.73 | 0.54 |
| 10 | 134046708 | rs7076997 | 0.318 | 0.512 | -1.088 | 0.464 | -0.006 | 0.025 | 0.025 | 3.987 | 0.36 | 0.17 | 0.46 | 0.32 | 0.37 | 0.21 |
| 10 | 134052164 | rs7913492 | -0.448 | 0.438 | -0.529 | 0.797 | 0.002 | 0.019 | 0.019 | 0 | 0.11 | 0.12 | 0.09 | 0.25 | 0.14 | 0.2 |
| 10 | 134069131 | rs2818400 | -0.821 | 0.329 | -0.57 | 0.134 | -0.013 | 0.028 | 0.028 | 0.021 | 0.69 | 0.73 | 0.51 | 0.77 | 0.64 | 0.64 |
| 10 | 134069351 | rs12764265 | 0.847 | -0.117 | -0.701 | -0.303 | -0.033 | 0.098 | 0.098 | 0 | 0.3 | 0.12 | 0.17 | 0.07 | 0.29 | 0.13 |
| 10 | 134069610 | rs2814193 | -1.199 | -0.155 | -0.805 | -0.315 | -0.011 | 0.036 | 0.036 | 0.004 | 0.67 | 0.72 | 0.5 | 0.76 | 0.63 | 0.61 |
| 10 | 134069930 | rs11146268 | 0.84 | 0.067 | -0.591 | -0.283 | -0.015 | 0.059 | 0.059 | 0 | 0.3 | 0.2 | 0.17 | 0.07 | 0.29 | 0.13 |
| 10 | 134073393 | rs11146273 | 0.438 | -0.018 | -0.175 | -0.302 | -0.002 | -0.08 | -0.08 | 0 | 0.4 | 0.43 | 0.23 | 0.06 | 0.34 | 0.31 |
| 10 | 134073886 | rs11146274 | 0.835 | -0.095 | -0.158 | -0.148 | -0.003 | 0.017 | 0.017 | 0.475 | 0.35 | 0.31 | 0.5 | 0.25 | 0.37 | 0.38 |
| 10 | 134079092 | rs10781565 | 0.036 | -0.097 | -0.688 | 0.205 | -0.005 | -0.031 | -0.031 | 0 | 0.81 | 0.88 | 0.78 | 0.4 | 0.76 | 0.69 |
| 10 | 134087990 | rs11146301 | -0.07 | -0.001 | -0.296 | -0.137 | 0.004 | 0.024 | 0.024 | 0.006 | 0.44 | 0.64 | 0.3 | 0.07 | 0.37 | 0.33 |
| 10 | 134091509 | rs7094372 | -0.688 | 0.099 | -1.115 | 0.463 | -0.073 | -0.053 | -0.053 | 0 | 0.56 | 0.91 | 0.43 | 0.43 | 0.52 | 0.54 |
| 10 | 134096205 | rs12247439 | 0.207 | 0.516 | -0.142 | 0.808 | 0.045 | 0.17 | 0.17 | 0 | 0.06 | 0.19 | 0.07 | 0.27 | 0.1 | 0.14 |
| 10 | 134102055 | rs7911172 | 0.033 | -0.225 | 0.006 | -0.246 | -0.001 | 0.04 | 0.04 | 0.502 | 0.4 | 0.33 | 0.27 | 0.06 | 0.35 | 0.29 |
| 10 | 134104861 | rs10781566 | -0.113 | -0.279 | 0.19 | -0.219 | -0.002 | -0.056 | -0.056 | 0 | 0.42 | 0.5 | 0.28 | 0.09 | 0.35 | 0.29 |
| 10 | 134105957 | rs10870288 | 0.836 | -0.387 | -0.145 | -0.454 | -0.065 | 0.106 | 0.106 | 0 | 0.25 | 0.08 | 0.17 | 0 | 0.28 | 0.1 |
| 10 | 134107216 | rs4462257 | -0.115 | -0.256 | -0.005 | -0.316 | 0 | -0.033 | -0.033 | 0 | 0.4 | 0.43 | 0.27 | 0.06 | 0.35 | 0.21 |
| 10 | 134114295 | rs2492643 | -0.181 | -0.394 | -0.126 | 0.343 | -0.006 | 0.054 | 0.054 | 0 | 0.13 | 0.15 | 0.11 | 0.56 | 0.11 | 0.22 |
| 10 | 134114599 | rs7075670 | -0.068 | -0.663 | 0.245 | -0.687 | -0.006 | -0.136 | -0.136 | 0.42 | 0.42 | 0.5 | 0.3 | 0.01 | 0.36 | 0.23 |
| 10 | 134116354 | rs6560688 | 0.839 | -0.678 | 0.252 | -0.908 | 0.008 | -0.087 | -0.087 | 0 | 0.21 | 0.29 | 0.15 | 0 | 0.19 | 0.07 |
| 10 | 134116771 | rs11146323 | 1.555 | -0.546 | 0.02 | -0.91 | -0.044 | 0.06 | 0.06 | 0 | 0.17 | 0.04 | 0.12 | 0 | 0.2 | 0.07 |
| 10 | 134118000 | rs2492651 | -1.671 | -0.099 | -0.834 | -1.218 | -0.072 | -0.007 | -0.007 | 0.001 | 0.85 | 0.99 | 0.88 | 0.9 | 0.76 | 0.84 |
| 10 | 134124871 | rs7894429 | 0.076 | -0.419 | -0.395 | -1.383 | -0.006 | -0.092 | -0.092 | 0.66 | 0.38 | 0.42 | 0.24 | 0.02 | 0.28 | 0.26 |
| 10 | 134129509 | rs7895073 | 0.283 | -0.409 | -0.524 | -1.368 | -0.003 | -0.084 | -0.084 | 0.334 | 0.34 | 0.37 | 0.21 | 0 | 0.26 | 0.2 |
| 10 | 134134703 | rs12570033 | 0.847 | 0.297 | -0.755 | -0.5 | -0.017 | -0.024 | -0.024 | 0 | 0.24 | 0.18 | 0.2 | 0.37 | 0.29 | 0.16 |
| 10 | 134137638 | rs11146329 | 1.057 | 0.256 | -1.106 | -0.625 | -0.043 | -0.044 | -0.044 | 0 | 0.12 | 0.01 | 0.1 | 0.15 | 0.13 | 0.2 |
| 10 | 134142321 | rs7922627 | -0.078 | 0.045 | -0.486 | -0.791 | -0.001 | 0.001 | 0.001 | 0.012 | 0.39 | 0.36 | 0.27 | 0.35 | 0.37 | 0.27 |
| 10 | 134143942 | rs1052281 | -0.463 | -0.07 | -0.214 | -0.871 | -0.002 | -0.007 | -0.007 | 0.014 | 0.45 | 0.47 | 0.29 | 0.35 | 0.39 | 0.33 |
| 10 | 134143985 | rs1132165 | -0.077 | -0.066 | -0.228 | -0.869 | 0 | 0 | 0 | 0 | 0.4 | 0.37 | 0.27 | 0.35 | 0.37 | 0.27 |
| 10 | 134146811 | rs7099831 | 2.116 | 0.074 | -0.268 | -0.987 | -0.025 | 0.036 | 0.036 | 0 | 0.18 | 0.08 | 0.12 | 0 | 0.19 | 0.05 |
| 10 | 134147172 | rs12414718 | 1.128 | 0.019 | -0.144 | -1.019 | -0.022 | -0.049 | -0.049 | 0 | 0.14 | 0.16 | 0.07 | 0 | 0.07 | 0.12 |
| 10 | 134149946 | rs4554798 | -0.443 | 0.068 | -0.023 | -0.918 | -0.002 | -0.009 | -0.009 | 0 | 0.43 | 0.48 | 0.29 | 0.36 | 0.39 | 0.32 |
| 10 | 134150920 | rs2295273 | 0.319 | 0.318 | -0.235 | -0.672 | 0.025 | 0.141 | 0.141 | 0 | 0.06 | 0.13 | 0.08 | 0.35 | 0.12 | 0.09 |
| 10 | 134161517 | rs2474329 | 1.845 | -0.068 | -0.612 | -0.84 | -0.035 | 0.057 | 0.057 | 0 | 0.17 | 0.05 | 0.11 | 0 | 0.19 | 0.05 |
| 10 | 134173354 | rs2474339 | -0.799 | -0.236 | -1.16 | -0.905 | -0.081 | 0.018 | 0.018 | 0 | 0.79 | 0.98 | 0.79 | 0.86 | 0.75 | 0.77 |
| 10 | 134175578 | rs10870295 | 1.301 | -0.582 | -1.155 | -0.772 | -0.017 | -0.182 | -0.182 | 0.064 | 0.23 | 0.01 | 0.23 | 0.49 | 0.2 | 0.37 |
| 10 | 134178456 | rs12358251 | -0.817 | -0.478 | 0.695 | -1.194 | -0.003 | -0.042 | -0.042 | 0 | 0.15 | 0.31 | 0.17 | 0.06 | 0.14 | 0.14 |
| 10 | 134186772 | rs10870301 | -1.223 | -0.421 | 0.354 | -1.304 | -0.014 | 0.188 | 0.188 | 0.239 | 0.52 | 0.79 | 0.5 | 0.15 | 0.46 | 0.32 |
| 10 | 134188874 | rs1054038 | 0.304 | -0.488 | 0.538 | -1.337 | 0.006 | 0.011 | 0.011 | 0.017 | 0.32 | 0.19 | 0.18 | 0 | 0.26 | 0.17 |
| 10 | 134190734 | rs4481949 | 1.786 | -0.417 | -0.037 | -1.321 | -0.046 | 0.083 | 0.083 | 0 | 0.18 | 0.03 | 0.11 | 0 | 0.19 | 0.05 |
| 10 | 134191005 | rs4443996 | -0.918 | -0.014 | 0.164 | -1.047 | -0.024 | 0.218 | 0.218 | 0.405 | 0.52 | 0.85 | 0.5 | 0.14 | 0.46 | 0.31 |
| 10 | 134191738 | rs10870302 | -0.894 | -0.002 | 0.142 | -1.045 | -0.011 | 0.189 | 0.189 | 0.284 | 0.52 | 0.79 | 0.5 | 0.14 | 0.46 | 0.31 |
| 10 | 134191811 | rs10870303 | -0.894 | -0.002 | 0.13 | -1.045 | -0.011 | 0.192 | 0.192 | 0.284 | 0.52 | 0.79 | 0.49 | 0.14 | 0.46 | 0.31 |
| 10 | 134195971 | rs11146352 | 0.181 | 0.282 | 0.364 | -0.89 | -0.009 | -0.118 | -0.118 | 0 | 0.34 | 0.34 | 0.2 | 0 | 0.26 | 0.18 |
| 10 | 134199109 | rs11814053 | -1.223 | 1.145 | 0.551 | 0.242 | 0.022 | 0.02 | 0.02 | 0 | 0.09 | 0.17 | 0.22 | 0.13 | 0.14 | 0.13 |
| 10 | 134213897 | rs11146360 | -1.067 | 0.626 | 0.748 | -0.05 | -0.02 | -0.083 | -0.083 | 0 | 0.16 | 0.59 | 0.23 | 0.05 | 0.1 | 0.11 |
| 10 | 134214937 | rs7893611 | -0.115 | 1.318 | 0.418 | 0.541 | 0.031 | 0.078 | 0.078 | 0 | 0.14 | 0.21 | 0.36 | 0.36 | 0.22 | 0.15 |
| 10 | 134216982 | rs4880251 | -1.211 | 1.2 | 0.589 | 0.461 | 0.046 | 0.121 | 0.121 | 0 | 0.25 | 0.51 | 0.46 | 0.51 | 0.34 | 0.39 |
| 10 | 134218269 | rs11146363 | 1.297 | 1.197 | -0.105 | -0.159 | 0 | -0.001 | -0.001 | 0 | 0.61 | 0.41 | 0.47 | 0.63 | 0.59 | 0.51 |
| 10 | 134225058 | rs10781572 | -0.904 | 0.777 | -0.688 | -0.473 | 0.03 | 0.043 | 0.043 | 0 | 0.53 | 0.91 | 0.75 | 0.36 | 0.58 | 0.47 |
| 10 | 134226240 | rs4075187 | -0.889 | 0.721 | 0.497 | -0.714 | 0.01 | 0.011 | 0.011 | 0 | 0.08 | 0.19 | 0.21 | 0.09 | 0.09 | 0.13 |
| 10 | 134228278 | rs7920199 | -1.35 | 0.742 | 0.498 | -0.743 | -0.001 | -0.019 | -0.019 | 0 | 0.16 | 0.21 | 0.26 | 0.1 | 0.15 | 0.18 |
| 10 | 134232065 | rs10870304 | 1.022 | 0.735 | 0.741 | -0.781 | 0.004 | -0.068 | -0.068 | 0 | 0.8 | 0.58 | 0.79 | 0.94 | 0.84 | 0.84 |
| 10 | 134233608 | rs10781573 | 0.88 | 0.808 | 0.701 | -0.753 | 0.01 | -0.051 | -0.051 | 0 | 0.74 | 0.48 | 0.6 | 0.85 | 0.76 | 0.63 |
| 10 | 134246697 | rs4880373 | 0.258 | 0.487 | 0.399 | -1.597 | -0.001 | -0.006 | -0.006 | 0 | 0.52 | 0.57 | 0.41 | 0.15 | 0.5 | 0.42 |
| 10 | 134258047 | rs12269409 | -0.752 | -0.233 | 0.841 | -1.461 | -0.023 | -0.033 | -0.033 | 3.193 | 0.09 | 0.45 | 0.06 | 0 | 0.02 | 0.02 |
| 10 | 134261810 | rs11146378 | 0.307 | -0.155 | 0.444 | -1.427 | -0.019 | -0.026 | -0.026 | 0 | 0.07 | 0.26 | 0.04 | 0 | 0.02 | 0.01 |
| 10 | 134267284 | rs7922380 | 0.716 | -0.299 | -0.642 | -1.548 | 0.01 | 0.034 | 0.034 | 1.078 | 0.72 | 0.11 | 0.7 | 0.78 | 0.77 | 0.72 |
| 10 | 134272808 | rs6560698 | -0.476 | -0.182 | -0.377 | -2.294 | -0.002 | 0.136 | 0.136 | 0 | 0.46 | 0.24 | 0.52 | 0.82 | 0.49 | 0.48 |
| 10 | 134277154 | rs7918074 | -1.236 | -0.337 | -0.519 | -2.509 | -0.074 | 0.145 | 0.145 | 0 | 0.61 | 0.85 | 0.75 | 0.85 | 0.57 | 0.71 |
| 10 | 134281537 | rs4593926 | -1.428 | -0.339 | -0.607 | -2.592 | -0.143 | 0.212 | 0.212 | 0 | 0.64 | 0.96 | 0.78 | 0.88 | 0.59 | 0.71 |

| | | | | | | | | | | | | | | | | |
|---|---|---|---|---|---|---|---|---|---|---|---|---|---|---|---|---|
| 10 | 134283875 | rs7897741 | -1.842 | -0.577 | -0.343 | -2.648 | 0.003 | 0.005 | 0.005 | 2.89 | 0.24 | 0.76 | 0.28 | 0.08 | 0.12 | 0.14 |
| 10 | 134285403 | rs10747061 | 0.063 | -0.416 | -0.353 | -2.51 | -0.057 | 0.137 | 0.137 | 1.61 | 0.4 | 0.11 | 0.49 | 0.81 | 0.48 | 0.57 |
| 10 | 134290526 | rs10870311 | 0.049 | -0.361 | 0.445 | -2.427 | -0.002 | 0.004 | 0.004 | 0 | 0.61 | 0.65 | 0.38 | 0.17 | 0.54 | 0.44 |
| 10 | 134294859 | rs12772482 | -1.338 | -0.402 | 1.036 | -2.176 | -0.048 | 0.015 | 0.015 | 0.857 | 0.14 | 0.52 | 0.12 | 0.08 | 0.08 | 0.11 |
| 10 | 134295644 | rs7904728 | 0.378 | -0.278 | 0.84 | -2.287 | 0 | -0.006 | -0.006 | 0 | 0.53 | 0.53 | 0.34 | 0.19 | 0.48 | 0.34 |
| 10 | 134296241 | rs4880385 | -0.066 | -0.166 | 0.74 | -2.199 | -0.008 | 0.031 | 0.031 | 2.59 | 0.62 | 0.69 | 0.41 | 0.21 | 0.55 | 0.42 |
| 10 | 134300136 | rs4492738 | -0.162 | -0.372 | 0.6 | -2.135 | -0.011 | 0.028 | 0.028 | 1.13 | 0.59 | 0.65 | 0.38 | 0.22 | 0.51 | 0.43 |
| 10 | 134305663 | rs7894289 | 0.365 | -0.351 | 0.633 | -2.285 | 0 | 0.037 | 0.037 | 0 | 0.51 | 0.44 | 0.33 | 0.19 | 0.49 | 0.36 |
| 10 | 134309194 | rs4469802 | 0.599 | 0.112 | 0.815 | -2.166 | 0.005 | -0.012 | -0.012 | 0 | 0.5 | 0.46 | 0.3 | 0.17 | 0.44 | 0.35 |
| 10 | 134315903 | rs4291594 | 0.056 | 0.052 | 0.716 | -2.213 | -0.003 | -0.015 | -0.015 | 0 | 0.91 | 0.86 | 0.93 | 0.97 | 0.91 | 0.95 |
| 10 | 134317884 | rs10747063 | 0.09 | 0.232 | 0.428 | -2.247 | -0.029 | 0.055 | 0.055 | 0.035 | 0.36 | 0.29 | 0.59 | 0.86 | 0.46 | 0.55 |
| 10 | 134320913 | rs5025116 | 0.133 | 0.272 | 0.351 | -2.229 | -0.035 | 0.064 | 0.064 | 0.307 | 0.36 | 0.26 | 0.59 | 0.83 | 0.46 | 0.54 |
| 10 | 134325083 | rs10781577 | -0.649 | 0.265 | 0.283 | -2.099 | -0.001 | -0.058 | -0.058 | 0 | 0.48 | 0.47 | 0.66 | 0.86 | 0.53 | 0.62 |
| 10 | 134335157 | rs10747068 | -0.286 | 0.53 | -0.338 | -1.976 | -0.015 | 0.074 | 0.074 | 0 | 0.42 | 0.3 | 0.63 | 0.84 | 0.53 | 0.57 |
| 10 | 134336503 | rs7923119 | 0.459 | 0.658 | -0.503 | -2.022 | -0.001 | -0.075 | -0.075 | 0 | 0.66 | 0.4 | 0.64 | 0.97 | 0.68 | 0.77 |
| 10 | 134341296 | rs7903076 | 0.469 | 0.378 | -0.576 | -1.983 | -0.012 | -0.075 | -0.075 | 0 | 0.64 | 0.56 | 0.75 | 0.92 | 0.62 | 0.61 |
| 10 | 134343062 | rs4394754 | 0.212 | 0.29 | -0.546 | -1.993 | -0.013 | -0.075 | -0.075 | 0 | 0.73 | 0.64 | 0.81 | 0.93 | 0.73 | 0.71 |
| 10 | 134352620 | rs11818345 | -0.127 | 0.097 | -0.176 | -2.096 | -0.018 | -0.088 | -0.088 | 0 | 0.19 | 0.39 | 0.14 | 0.06 | 0.18 | 0.22 |
| 10 | 134359062 | rs12241389 | -0.189 | 0.189 | -0.662 | -1.966 | 0 | 0 | 0 | 0 | 0.08 | 0.09 | 0.07 | 0.02 | 0.09 | 0.07 |
| 10 | 134393338 | rs12412216 | 0.854 | 0.684 | -1.666 | -2.114 | -0.059 | 0.023 | 0.023 | 0 | 0.2 | 0.03 | 0.27 | 0.15 | 0.19 | 0.13 |
| 10 | 134397125 | rs7084105 | -0.804 | 0.615 | -2.052 | -1.321 | -0.006 | -0.034 | -0.034 | 0 | 0.61 | 0.8 | 0.7 | 0.47 | 0.61 | 0.54 |
| 10 | 134404571 | rs2819718 | 1.624 | 0.557 | -2.232 | -1.356 | -0.05 | -0.136 | -0.136 | 1.823 | 0.27 | 0.05 | 0.2 | 0.5 | 0.28 | 0.31 |
| 10 | 134405041 | rs7907513 | 1.204 | 0.425 | -2.475 | -1.244 | -0.008 | -0.028 | -0.028 | 0 | 0.09 | 0.01 | 0.05 | 0.07 | 0.08 | 0.11 |
| 10 | 134406401 | rs2814452 | 1.315 | 0.376 | -2.417 | -1.287 | -0.048 | -0.118 | -0.118 | 3.928 | 0.3 | 0.06 | 0.23 | 0.52 | 0.3 | 0.36 |
| 10 | 134443032 | rs12767762 | 1.393 | 0.466 | -2.398 | -1.248 | -0.071 | 0.017 | 0.017 | 0 | 0.2 | 0.02 | 0.27 | 0.13 | 0.19 | 0.14 |
| 10 | 134453920 | rs945944 | -1.75 | 0.426 | -2.291 | -1.023 | 0 | -0.006 | -0.006 | 0.007 | 0.93 | 0.91 | 0.94 | 1 | 0.9 | 0.9 |
| 10 | 134456109 | rs4880419 | 0.703 | 0.371 | -2.517 | -1.224 | -0.096 | 0.09 | 0.09 | 0 | 0.39 | 0.1 | 0.42 | 0.79 | 0.44 | 0.49 |
| 10 | 134459388 | rs1133400 | 0.14 | 0.978 | -2.032 | -0.993 | -0.028 | -0.032 | -0.032 | 0 | 0.14 | 0.1 | 0.26 | 0.3 | 0.21 | 0.24 |
| 10 | 134466163 | rs10781583 | -1.475 | 0.91 | -1.873 | -1.289 | 0 | -0.011 | -0.011 | 0.238 | 0.91 | 0.9 | 0.94 | 0.99 | 0.9 | 0.9 |
| 10 | 134491348 | rs1890177 | -1.071 | 0.077 | -1.374 | -1.266 | -0.002 | -0.028 | -0.028 | 0 | 0.44 | 0.45 | 0.47 | 0.16 | 0.38 | 0.32 |
| 10 | 134499718 | rs12770172 | 1.075 | 0.116 | -1.454 | -1.239 | -0.065 | 0.029 | 0.029 | 0.007 | 0.2 | 0.01 | 0.27 | 0.13 | 0.19 | 0.14 |
| 10 | 134504693 | rs2786894 | 0.501 | 0.124 | -1.291 | -1.199 | -0.007 | -0.013 | -0.013 | 0 | 0.13 | 0.16 | 0.1 | 0.03 | 0.1 | 0.14 |
| 10 | 134512521 | rs2767419 | -0.863 | 0.062 | -1.356 | -1.218 | -0.003 | -0.032 | -0.032 | 0.289 | 0.43 | 0.46 | 0.45 | 0.15 | 0.37 | 0.31 |
| 10 | 134522995 | rs913196 | 0.374 | 0.308 | -1.807 | -1.029 | -0.048 | 0.033 | 0.033 | 0 | 0.21 | 0.02 | 0.14 | 0.1 | 0.18 | 0.14 |
| 10 | 134525314 | rs2803997 | 0.938 | 0.425 | -1.378 | -0.855 | -0.003 | -0.033 | -0.033 | 0.01 | 0.1 | 0.15 | 0.05 | 0 | 0.09 | 0.06 |
| 10 | 134529972 | rs873946 | 1.102 | 0.171 | -1.551 | -0.866 | -0.043 | 0.004 | 0.004 | 0.038 | 0.16 | 0.01 | 0.11 | 0.11 | 0.15 | 0.13 |
| 10 | 134540400 | rs945943 | 0.925 | 0.13 | -1.38 | -0.762 | -0.003 | -0.019 | -0.019 | 0 | 0.09 | 0.13 | 0.04 | 0 | 0.08 | 0.06 |
| 10 | 134547732 | rs3793673 | 0.835 | 0.29 | -1.05 | -0.603 | 0.004 | 0.044 | 0.044 | 0 | 0.25 | 0.13 | 0.15 | 0.11 | 0.22 | 0.21 |
| 10 | 134550882 | rs2803992 | 0.258 | 0.204 | -1.039 | -0.678 | 0.004 | 0.011 | 0.011 | 0 | 0.3 | 0.27 | 0.18 | 0.12 | 0.25 | 0.26 |
| 10 | 134551731 | rs11146487 | 1.02 | 0.146 | -1.648 | -0.396 | -0.152 | -0.148 | -0.148 | 2.966 | 0.34 | 0.02 | 0.24 | 0.48 | 0.39 | 0.3 |
| 10 | 134553369 | rs10781585 | 0.491 | 0.406 | -0.648 | -0.873 | -0.002 | -0.03 | -0.03 | 0.094 | 0.55 | 0.46 | 0.72 | 0.88 | 0.64 | 0.65 |
| 10 | 134553785 | rs2148666 | 0.466 | 0.403 | -0.649 | -0.875 | -0.003 | -0.031 | -0.031 | 0 | 0.56 | 0.46 | 0.72 | 0.88 | 0.64 | 0.65 |
| 10 | 134553909 | rs12780199 | -0.068 | 0.624 | -0.817 | -0.624 | -0.01 | 0.098 | 0.098 | 0.097 | 0.33 | 0.12 | 0.19 | 0.11 | 0.31 | 0.23 |
| 10 | 134557068 | rs2803991 | -0.184 | 0.305 | -0.265 | -0.258 | 0.004 | 0.011 | 0.011 | 0 | 0.3 | 0.27 | 0.19 | 0.12 | 0.25 | 0.26 |
| 10 | 134564831 | rs12776770 | 0.307 | 0.341 | -0.7 | -0.195 | 0.007 | 0.042 | 0.042 | 0 | 0.25 | 0.12 | 0.15 | 0.11 | 0.21 | 0.21 |
| 10 | 134579026 | rs12769959 | -0.064 | 0.342 | -0.9 | -0.156 | 0.01 | 0.059 | 0.059 | 0 | 0.29 | 0.13 | 0.17 | 0.12 | 0.24 | 0.25 |
| 10 | 134582521 | rs2767428 | -0.232 | 0.376 | -1.16 | -0.213 | 0.003 | -0.051 | -0.051 | 0 | 0.91 | 0.84 | 0.94 | 1 | 0.94 | 0.96 |
| 10 | 134589631 | rs12784407 | -0.359 | 0.382 | -1.726 | 0.25 | -0.008 | 0.041 | 0.041 | 0 | 0.29 | 0.04 | 0.16 | 0.15 | 0.25 | 0.25 |
| 10 | 134594316 | rs3793687 | 0.062 | 0.383 | -1.853 | 0.261 | -0.027 | 0.013 | 0.013 | 0 | 0.27 | 0.03 | 0.14 | 0.14 | 0.22 | 0.22 |
| 10 | 134597211 | rs3750579 | -0.039 | 0.534 | -1.737 | 0.455 | -0.022 | 0.029 | 0.029 | 0 | 0.3 | 0.03 | 0.15 | 0.15 | 0.25 | 0.25 |
| 10 | 134607938 | rs7096226 | -0.789 | 0.83 | -0.746 | 1.248 | 0.001 | 0.096 | 0.096 | 0 | 0.12 | 0.25 | 0.39 | 0.3 | 0.17 | 0.12 |
| 10 | 134609699 | rs7092049 | 0.311 | 0.771 | -1.03 | 0.891 | 0.014 | -0.029 | -0.029 | 0 | 0.6 | 0.25 | 0.37 | 0.54 | 0.61 | 0.64 |
| 10 | 134615651 | rs913191 | -0.182 | 0.961 | -0.644 | 0.641 | -0.006 | -0.004 | -0.004 | 0 | 0.65 | 0.73 | 0.47 | 0.64 | 0.67 | 0.7 |
| 10 | 134616641 | rs2767438 | 0.496 | 1.209 | -1.206 | 0.484 | 0.016 | 0.018 | 0.018 | 0 | 0.49 | 0.25 | 0.33 | 0.59 | 0.49 | 0.59 |
| 10 | 134623959 | rs2254419 | -0.447 | 1.15 | -1.642 | 0.272 | -0.102 | -0.013 | -0.013 | 0 | 0.68 | 0.97 | 0.54 | 0.74 | 0.7 | 0.7 |
| 10 | 134646988 | rs4880433 | -0.062 | 1.06 | -1.668 | -0.073 | -0.053 | 0.062 | 0.062 | 0.183 | 0.38 | 0.14 | 0.51 | 0.28 | 0.4 | 0.39 |
| 10 | 134650440 | rs3750580 | -0.055 | 1.041 | -1.609 | -0.074 | -0.078 | 0.062 | 0.062 | 0.385 | 0.38 | 0.1 | 0.51 | 0.27 | 0.39 | 0.39 |
| 10 | 134659626 | rs10870341 | -0.819 | 0.566 | 0.135 | -0.35 | 0.005 | -0.015 | -0.015 | 0 | 0.12 | 0.29 | 0.12 | 0.07 | 0.14 | 0.18 |
| 10 | 134664230 | rs10747072 | -1.109 | 0.831 | 0.318 | -0.315 | 0.003 | -0.006 | -0.006 | 1.103 | 0.6 | 0.73 | 0.72 | 0.4 | 0.61 | 0.65 |
| 10 | 134666424 | rs7095353 | -1.193 | 0.705 | 0.252 | -0.359 | -0.019 | -0.033 | -0.033 | 0 | 0.61 | 0.87 | 0.73 | 0.4 | 0.61 | 0.65 |
| 10 | 134669463 | rs9419242 | 0.184 | 0.717 | 0.387 | -0.371 | -0.003 | 0.006 | 0.006 | 0 | 0.07 | 0.05 | 0.05 | 0.06 | 0.1 | 0.1 |
| 10 | 134681932 | rs10870347 | -0.655 | 0.746 | 0.424 | -0.396 | 0.019 | 0.003 | 0.003 | 0 | 0.07 | 0.14 | 0.09 | 0.05 | 0.13 | 0.09 |
| 10 | 134684769 | rs11146557 | 0.391 | 0.704 | 0.336 | -0.423 | -0.068 | 0.068 | 0.068 | 0.466 | 0.38 | 0.12 | 0.52 | 0.26 | 0.39 | 0.39 |
| 10 | 134692110 | rs4880459 | -0.464 | 0.682 | 0.364 | -0.411 | 0.048 | 0.118 | 0.118 | 0 | 0.49 | 0.73 | 0.59 | 0.29 | 0.39 | 0.46 |
| 10 | 134693532 | rs2387074 | -0.96 | 0.649 | 0.302 | -0.41 | -0.036 | 0.028 | 0.028 | 0 | 0.56 | 0.84 | 0.68 | 0.35 | 0.52 | 0.56 |
| 10 | 134705373 | rs7907613 | -0.898 | 0.698 | 0.097 | -0.433 | -0.015 | 0.055 | 0.055 | 0 | 0.56 | 0.79 | 0.67 | 0.29 | 0.52 | 0.55 |
| 10 | 134706234 | rs7097501 | 0.633 | 0.806 | -0.161 | -0.358 | -0.101 | 0.022 | 0.022 | 0.597 | 0.33 | 0.03 | 0.48 | 0.21 | 0.33 | 0.34 |
| 10 | 134714100 | rs7913509 | 0.645 | 0.851 | -0.165 | -0.352 | -0.083 | 0.025 | 0.025 | 0.611 | 0.33 | 0.03 | 0.46 | 0.21 | 0.31 | 0.33 |
| 10 | 134715685 | rs4268437 | 0.553 | 0.861 | -0.107 | -0.357 | -0.034 | 0.04 | 0.04 | 0.355 | 0.33 | 0.06 | 0.47 | 0.22 | 0.3 | 0.4 |

| | | | | | | | | | | | | | | | | |
|---|---|---|---|---|---|---|---|---|---|---|---|---|---|---|---|---|
| 10 | 134724151 | rs12416523 | 0.455 | 0.97 | 0.338 | -0.391 | 0 | 0.028 | 0.028 | 0.02 | 0.37 | 0.27 | 0.5 | 0.21 | 0.35 | 0.37 |
| 10 | 134731079 | rs11597534 | 0.098 | 0.3 | -0.083 | -0.467 | -0.031 | -0.024 | -0.024 | 0 | 0.39 | 0.27 | 0.34 | 0.61 | 0.49 | 0.42 |
| 10 | 134738232 | rs4075501 | -0.267 | 0.179 | -0.057 | -0.485 | -0.02 | 0.009 | 0.009 | 0 | 0.57 | 0.7 | 0.62 | 0.35 | 0.47 | 0.49 |
| 10 | 134747190 | rs12357485 | -0.033 | 0.155 | 0.288 | -0.629 | -0.002 | 0.039 | 0.039 | 0 | 0.13 | 0.02 | 0.05 | 0.02 | 0.1 | 0.08 |
| 10 | 134748331 | rs12781609 | 0.827 | 0.126 | 0.725 | -0.644 | 0.001 | 0 | 0 | 0 | 0.39 | 0.32 | 0.5 | 0.34 | 0.35 | 0.41 |
| 10 | 134751492 | rs10870212 | -0.491 | 0.126 | 0.654 | -0.647 | -0.012 | -0.029 | -0.029 | 0 | 0.16 | 0.26 | 0.09 | 0.02 | 0.11 | 0.08 |
| 10 | 134751911 | rs12269281 | -0.382 | 0.118 | 0.66 | -0.649 | -0.02 | -0.029 | -0.029 | 0 | 0.14 | 0.25 | 0.06 | 0.02 | 0.08 | 0.08 |
| 10 | 134754297 | rs4242712 | -0.436 | 0.121 | 0.697 | -0.645 | -0.02 | 0.009 | 0.009 | 0 | 0.56 | 0.7 | 0.61 | 0.36 | 0.47 | 0.49 |
| 10 | 134760584 | rs7083897 | 0.746 | 0.002 | 0.702 | -0.698 | -0.006 | -0.002 | -0.002 | 0 | 0.4 | 0.43 | 0.51 | 0.34 | 0.34 | 0.4 |
| 10 | 134773713 | rs7074212 | -0.297 | 0.037 | 0.485 | -0.738 | -0.003 | 0.025 | 0.025 | 0 | 0.55 | 0.66 | 0.59 | 0.33 | 0.44 | 0.53 |
| 10 | 134777671 | rs12780014 | 0.685 | 0.072 | 0.542 | -0.69 | -0.003 | 0.007 | 0.007 | 0 | 0.4 | 0.26 | 0.51 | 0.33 | 0.39 | 0.41 |
| 10 | 134782824 | rs10781545 | -0.644 | -0.117 | 0.438 | -0.839 | 0.047 | 0.064 | 0.064 | 0 | 0.49 | 0.83 | 0.46 | 0.67 | 0.59 | 0.58 |
| 10 | 134783181 | rs10870214 | -0.002 | -0.339 | 0.663 | -0.969 | -0.008 | 0.013 | 0.013 | 0 | 0.41 | 0.29 | 0.37 | 0.67 | 0.54 | 0.49 |
| 10 | 134783778 | rs4880300 | -0.723 | -0.327 | 0.297 | -0.906 | 0.047 | 0.063 | 0.063 | 0 | 0.49 | 0.82 | 0.46 | 0.67 | 0.59 | 0.58 |
| 10 | 134784486 | rs10781546 | -0.333 | -0.319 | 0.497 | -0.899 | 0 | 0 | 0 | 0 | 0.45 | 0.57 | 0.42 | 0.67 | 0.56 | 0.49 |
| 10 | 134803058 | rs7081698 | -0.592 | -0.359 | -0.114 | -1.117 | 0.039 | 0.02 | 0.02 | 0 | 0.49 | 0.61 | 0.44 | 0.66 | 0.61 | 0.5 |
| 10 | 134803789 | rs9419142 | -1.298 | -0.578 | -0.383 | -1.19 | 0.032 | -0.008 | -0.008 | 0 | 0.62 | 0.79 | 0.48 | 0.66 | 0.66 | 0.66 |
| 10 | 134810401 | rs12413155 | -0.596 | -0.065 | -0.574 | -1.371 | 0.013 | 0.013 | 0.013 | 0 | 0.54 | 0.35 | 0.42 | 0.65 | 0.6 | 0.51 |
| 10 | 134810519 | rs9419288 | -1.823 | -0.106 | -0.707 | -1.359 | 0.021 | -0.028 | -0.028 | 0.261 | 0.68 | 0.83 | 0.52 | 0.67 | 0.7 | 0.66 |
| 10 | 134811475 | rs11146095 | -0.118 | 0.806 | -0.912 | -1.488 | -0.013 | 0.01 | 0.01 | 0 | 0.23 | 0.18 | 0.16 | 0.01 | 0.26 | 0.15 |
| 10 | 134821422 | rs10870228 | -0.691 | 0.974 | -1.006 | -1.004 | -0.008 | 0.004 | 0.004 | 0 | 0.43 | 0.33 | 0.35 | 0.65 | 0.47 | 0.43 |
| 10 | 134832400 | rs7071996 | -0.285 | 1.159 | -0.97 | -0.784 | 0.024 | 0.038 | 0.038 | 0 | 0.3 | 0.35 | 0.26 | 0.6 | 0.37 | 0.32 |
| 10 | 134832782 | rs12220781 | -0.094 | 0.92 | -0.969 | -0.675 | 0.001 | 0.003 | 0.003 | 0 | 0.35 | 0.35 | 0.28 | 0.59 | 0.37 | 0.39 |
| 10 | 134834077 | rs34130117 | -0.996 | 0.923 | -1.996 | -0.757 | 0.019 | 0.029 | 0.029 | 0 | 0.49 | 0.73 | 0.38 | 0.63 | 0.44 | 0.48 |
| 10 | 134846982 | rs7073479 | 0.078 | 0.857 | -2.007 | -0.376 | -0.001 | -0.001 | -0.001 | 2.213 | 0.35 | 0.59 | 0.31 | 0.62 | 0.38 | 0.41 |
| 10 | 134850350 | rs11101908 | 0.425 | 0.541 | -1.999 | -0.392 | -0.032 | -0.015 | -0.015 | 0.007 | 0.08 | 0.1 | 0.03 | 0.03 | 0.01 | 0.08 |
| 10 | 134852702 | rs12265931 | -1.159 | 1.037 | -2.202 | -0.348 | -0.083 | 0.103 | 0.103 | 0 | 0.64 | 0.98 | 0.66 | 0.85 | 0.66 | 0.66 |
| 10 | 134858046 | rs4626998 | -0.796 | 0.695 | -0.895 | 0.209 | -0.007 | 0.002 | 0.002 | 0 | 0.26 | 0.36 | 0.31 | 0.24 | 0.22 | 0.26 |
| 10 | 134868127 | rs10745292 | -0.413 | 1.036 | -0.9 | -0.351 | 0.002 | -0.028 | -0.028 | 0.108 | 0.43 | 0.53 | 0.68 | 0.87 | 0.46 | 0.54 |
| 10 | 134872169 | rs11591580 | -0.473 | 1.019 | -0.902 | -0.36 | 0 | -0.013 | -0.013 | 0 | 0.44 | 0.51 | 0.7 | 0.87 | 0.49 | 0.58 |
| 10 | 134884595 | rs4271313 | -0.23 | 1.057 | -1.141 | -0.283 | -0.037 | 0.032 | 0.032 | 0 | 0.62 | 0.66 | 0.54 | 0.66 | 0.57 | 0.65 |
| 10 | 134886879 | rs10857718 | -0.131 | 0.777 | -0.531 | 0.005 | -0.003 | 0 | 0 | 0 | 0.12 | 0.21 | 0.13 | 0.09 | 0.18 | 0.08 |
| 10 | 134889866 | rs7912475 | -0.099 | 0.545 | -0.656 | -0.534 | 0.001 | 0 | 0 | 1.438 | 0.39 | 0.55 | 0.37 | 0.56 | 0.3 | 0.47 |
| 10 | 134897512 | rs12569550 | -0.065 | 0.347 | -0.074 | -0.607 | 0.003 | 0.001 | 0.001 | 0 | 0.35 | 0.25 | 0.36 | 0.64 | 0.29 | 0.52 |
| 10 | 134898842 | rs3827691 | -1.043 | 0.366 | -0.036 | -0.898 | -0.036 | 0.096 | 0.096 | 1.516 | 0.57 | 0.75 | 0.56 | 0.72 | 0.56 | 0.64 |
| 10 | 134900807 | rs10776691 | -1.114 | 0.364 | -0.274 | -0.789 | -0.135 | 0.106 | 0.106 | 2.052 | 0.57 | 0.92 | 0.58 | 0.71 | 0.56 | 0.64 |
| 10 | 134902795 | rs11101896 | -1.095 | 0.377 | -0.268 | -0.724 | -0.143 | 0.086 | 0.086 | 1.963 | 0.57 | 0.92 | 0.58 | 0.71 | 0.55 | 0.64 |
| 10 | 134907239 | rs11101913 | 0.388 | 0.286 | 0.014 | -0.722 | 0.002 | -0.01 | -0.01 | 0 | 0.31 | 0.22 | 0.35 | 0.62 | 0.27 | 0.5 |
| 10 | 134913358 | rs11101925 | -0.835 | 0.29 | 0.29 | -0.761 | 0.007 | 0.082 | 0.082 | 0.298 | 0.52 | 0.69 | 0.54 | 0.69 | 0.51 | 0.62 |
| 10 | 134915832 | rs9419103 | -0.548 | 0.484 | 0.034 | -1.015 | -0.007 | 0.005 | 0.005 | 0 | 0.11 | 0.11 | 0.09 | 0.05 | 0.19 | 0.19 |
| 10 | 134916045 | rs7918741 | 0.206 | 0.446 | 0.384 | -0.988 | 0 | -0.032 | -0.032 | 0 | 0.16 | 0.23 | 0.14 | 0.06 | 0.19 | 0.13 |
| 10 | 134917297 | rs736017 | 0.315 | 0.477 | 0.025 | -0.975 | -0.001 | 0.006 | 0.006 | 0 | 0.16 | 0.13 | 0.11 | 0.05 | 0.19 | 0.13 |
| 10 | 134928139 | rs761774 | 0.288 | 0.186 | 0.213 | -0.228 | -0.004 | 0.006 | 0.006 | 0 | 0.22 | 0.22 | 0.31 | 0.56 | 0.16 | 0.31 |
| 10 | 134931784 | rs9419011 | 0.184 | 0.096 | 0.225 | -0.234 | -0.021 | 0.028 | 0.028 | 0 | 0.23 | 0.26 | 0.3 | 0.56 | 0.14 | 0.33 |
| 10 | 134935916 | rs7904992 | -0.692 | 0.45 | 0.238 | -0.463 | 0.001 | 0.01 | 0.01 | 0 | 0.51 | 0.48 | 0.54 | 0.62 | 0.58 | 0.6 |
| 10 | 134937394 | rs10857782 | 0.232 | -0.028 | 0.577 | 0.103 | -0.025 | 0.116 | 0.116 | 0 | 0.17 | 0.27 | 0.29 | 0.42 | 0.12 | 0.24 |
| 10 | 134947643 | rs4838698 | -0.961 | -0.047 | 0.223 | -0.534 | 0.046 | -0.038 | -0.038 | 0 | 0.43 | 0.72 | 0.41 | 0.46 | 0.33 | 0.5 |
| 10 | 134948776 | rs10857665 | 0.283 | 0.126 | 0.562 | -0.276 | -0.035 | 0.219 | 0.219 | 0 | 0.19 | 0.38 | 0.3 | 0.5 | 0.14 | 0.32 |
| 10 | 134959064 | rs10857669 | 0.52 | 0.181 | 0.355 | -0.44 | -0.02 | 0.114 | 0.114 | 0 | 0.18 | 0.27 | 0.28 | 0.46 | 0.15 | 0.27 |
| 10 | 134967702 | rs7095632 | 0.127 | 0.179 | 0.101 | -0.657 | -0.005 | -0.074 | -0.074 | 0 | 0.21 | 0.32 | 0.1 | 0 | 0.18 | 0.13 |
| 10 | 134968464 | rs7923021 | 0.253 | 0.2 | -0.04 | -0.667 | 0 | -0.032 | -0.032 | 0.727 | 0.2 | 0.24 | 0.09 | 0 | 0.18 | 0.13 |
| 10 | 134971602 | rs4326710 | -1.036 | 0.872 | -0.658 | -0.5 | -0.005 | -0.011 | -0.011 | 0 | 0.65 | 0.73 | 0.73 | 0.55 | 0.61 | 0.63 |
| 10 | 134972196 | rs12776037 | 0.033 | 1.042 | -0.619 | -0.364 | 0.001 | -0.012 | -0.012 | 0 | 0.27 | 0.23 | 0.35 | 0.09 | 0.28 | 0.25 |
| 10 | 134979963 | rs2998153 | -0.461 | 0.974 | -0.953 | 0.157 | -0.003 | -0.015 | -0.015 | 0 | 0.72 | 0.74 | 0.76 | 0.55 | 0.66 | 0.67 |
| 10 | 134987146 | rs4240518 | 0.273 | 1.102 | -1.345 | -0.018 | 0 | -0.059 | -0.059 | 0 | 0.43 | 0.1 | 0.36 | 0.57 | 0.38 | 0.62 |
| 10 | 134993107 | rs11101615 | -0.006 | 1.176 | -1.497 | -0.197 | -0.011 | 0.002 | 0.002 | 2.415 | 0.5 | 0.12 | 0.39 | 0.58 | 0.44 | 0.66 |
| 10 | 134994032 | rs3810967 | -0.118 | 1.059 | -0.613 | -0.248 | 0.013 | 0.017 | 0.017 | 1.619 | 0.52 | 0.34 | 0.42 | 0.58 | 0.44 | 0.66 |
| 10 | 135000148 | rs3810965 | -0.023 | 1.038 | -0.692 | -0.257 | -0.007 | 0.001 | 0.001 | 2.267 | 0.5 | 0.12 | 0.38 | 0.58 | 0.44 | 0.66 |
| 10 | 135000159 | rs3810964 | -0.023 | 1.038 | -0.692 | -0.257 | -0.007 | 0.001 | 0.001 | 2.267 | 0.5 | 0.12 | 0.38 | 0.58 | 0.44 | 0.66 |
| 10 | 135002837 | rs2998144 | -0.713 | 1.319 | -0.886 | -0.459 | -0.003 | -0.009 | -0.009 | 0 | 0.87 | 0.9 | 0.83 | 0.74 | 0.85 | 0.85 |
| 10 | 135003551 | rs12359206 | 0.972 | 1.229 | -0.328 | -0.582 | 0.003 | -0.007 | -0.007 | 1.726 | 0.25 | 0.23 | 0.33 | 0.11 | 0.37 | 0.14 |
| 10 | 135003724 | rs11101622 | 0.581 | 1.276 | -0.401 | -0.553 | -0.002 | 0.055 | 0.055 | 2.545 | 0.5 | 0.12 | 0.4 | 0.61 | 0.44 | 0.67 |
| 10 | 135012162 | rs2998139 | -0.005 | 1.114 | 0.194 | -0.49 | 0.014 | -0.005 | -0.005 | 0 | 0.6 | 0.52 | 0.49 | 0.61 | 0.53 | 0.63 |
| 10 | 135023586 | rs9418943 | 0.466 | 1.192 | -0.261 | -0.475 | 0.025 | 0.06 | 0.06 | 0.24 | 0.49 | 0.21 | 0.43 | 0.62 | 0.46 | 0.57 |
| 10 | 135027248 | rs7070941 | -0.245 | 1.055 | -0.265 | -0.316 | -0.001 | -0.007 | -0.007 | 0.018 | 0.05 | 0.04 | 0.03 | 0 | 0.06 | 0.03 |
| 10 | 135034542 | rs9419030 | -0.422 | 1.06 | 0.005 | -0.114 | -0.006 | 0.115 | 0.115 | 0 | 0.84 | 0.61 | 0.8 | 0.66 | 0.89 | 0.78 |
| 10 | 135038088 | rs11597218 | 0.438 | 1.157 | -0.382 | -0.362 | -0.056 | 0.112 | 0.112 | 0.017 | 0.35 | 0.17 | 0.37 | 0.01 | 0.45 | 0.16 |
| 10 | 135040283 | rs3008363 | 0.039 | 1.179 | -0.235 | -0.314 | 0.001 | 0.001 | 0.001 | 0.244 | 0.5 | 0.49 | 0.44 | 0.65 | 0.44 | 0.62 |
| 10 | 135045750 | rs3008360 | 1.422 | 0.62 | 0.54 | 0.507 | -0.007 | 0.02 | 0.02 | 0 | 0.76 | 0.34 | 0.75 | 0.62 | 0.83 | 0.76 |

| | | | | | | | | | | | | | | | | |
|---|---|---|---|---|---|---|---|---|---|---|---|---|---|---|---|---|
| 10 | 135045989 | rs2998110 | 0.77 | 0.601 | 0.055 | 0.453 | -0.014 | 0.053 | 0.053 | 0 | 0.8 | 0.62 | 0.79 | 0.66 | 0.86 | 0.79 |
| 10 | 135046903 | rs2980728 | 0.135 | 0.689 | 0.037 | 0.639 | -0.005 | 0.092 | 0.092 | 0 | 0.85 | 0.65 | 0.81 | 0.66 | 0.89 | 0.82 |
| 10 | 135053123 | rs880340 | -0.27 | 0.685 | -0.71 | 0.21 | -0.016 | 0.025 | 0.025 | 0 | 0.58 | 0.78 | 0.56 | 0.37 | 0.53 | 0.31 |
| 10 | 135055251 | rs3750849 | 0.008 | -0.38 | -0.49 | -0.447 | 0.009 | 0.009 | 0.009 | 1.339 | 0.15 | 0.11 | 0.16 | 0.08 | 0.1 | 0.14 |
| 10 | 135055349 | rs1054776 | 0.008 | -0.38 | -0.49 | -0.447 | 0.009 | 0.009 | 0.009 | 1.339 | 0.15 | 0.12 | 0.16 | 0.08 | 0.1 | 0.14 |
| 10 | 135056496 | rs7393606 | -1.083 | -0.851 | -0.198 | -0.652 | -0.028 | -0.029 | -0.029 | 0 | 0.23 | 0.7 | 0.22 | 0.09 | 0.11 | 0.15 |
| 10 | 135060864 | rs2185743 | -1.723 | -1.239 | 0.532 | -0.929 | -0.038 | -0.081 | -0.081 | 0.029 | 0.13 | 0.3 | 0.08 | 0.01 | 0.07 | 0.09 |
| 10 | 135061804 | rs2995337 | 1.147 | -1.362 | 0.397 | -1.217 | -0.026 | -0.063 | -0.063 | 0 | 0.76 | 0.61 | 0.77 | 0.92 | 0.83 | 0.83 |
| 10 | 135063027 | rs4838710 | 2.032 | -1.519 | 0.281 | -1.342 | 0.03 | 0.042 | 0.042 | 0 | 0.61 | 0.3 | 0.69 | 0.85 | 0.73 | 0.71 |
| 10 | 135063652 | rs11101659 | -1.07 | -1.33 | 0.667 | -1.293 | -0.007 | -0.064 | -0.064 | 0 | 0.11 | 0.29 | 0.07 | 0 | 0.08 | 0.05 |
| 10 | 135068257 | rs3008307 | -0.491 | -1.271 | 0.501 | -1.183 | 0.008 | 0.029 | 0.029 | 0 | 0.88 | 0.95 | 0.85 | 0.94 | 0.92 | 0.87 |
| 10 | 135068724 | rs3008309 | -0.339 | -1.46 | 0.49 | -1.259 | 0.011 | 0.016 | 0.016 | 0 | 0.87 | 0.91 | 0.85 | 0.94 | 0.92 | 0.87 |
| 10 | 135071004 | rs7912335 | 1.293 | -1.721 | 0.985 | -1.389 | -0.042 | -0.071 | -0.071 | 1.214 | 0.78 | 0.34 | 0.75 | 0.92 | 0.86 | 0.84 |
| 10 | 135076523 | rs2995300 | 0.217 | -1.686 | 1.448 | -1.363 | -0.022 | -0.038 | -0.038 | 0 | 0.86 | 0.69 | 0.81 | 0.92 | 0.89 | 0.87 |
| 10 | 135082346 | rs2275720 | 0.386 | -1.266 | 1.688 | -1.354 | -0.008 | -0.055 | -0.055 | 0 | 0.84 | 0.69 | 0.8 | 0.93 | 0.88 | 0.85 |
| 10 | 135085754 | rs1131718 | 0.098 | -1.246 | 1.387 | -1.424 | 0.007 | 0.018 | 0.018 | 0.027 | 0.86 | 0.9 | 0.82 | 0.93 | 0.88 | 0.86 |
| 10 | 135086380 | rs3008324 | -0.25 | -1.286 | 1.415 | -1.397 | 0.007 | 0.012 | 0.012 | 0.12 | 0.87 | 0.91 | 0.84 | 0.93 | 0.9 | 0.88 |
| 10 | 135088741 | rs2275724 | 1.752 | -1.401 | 1.438 | -1.547 | 0.018 | 0.021 | 0.021 | 0.082 | 0.71 | 0.24 | 0.73 | 0.91 | 0.81 | 0.79 |
| 10 | 135089035 | rs2275725 | -0.678 | -1.25 | 1.273 | -1.448 | 0.008 | 0.016 | 0.016 | 0.631 | 0.88 | 0.94 | 0.84 | 0.93 | 0.9 | 0.88 |
| 10 | 135091802 | rs1573041 | 0.487 | -1.429 | 0.689 | -1.184 | 0.008 | -0.061 | -0.061 | 0 | 0.28 | 0.1 | 0.18 | 0.37 | 0.22 | 0.39 |
| 10 | 135098830 | rs3008335 | -0.514 | -0.994 | 1.289 | -0.961 | 0.001 | 0 | 0 | 0.483 | 0.88 | 0.92 | 0.84 | 0.93 | 0.9 | 0.88 |
| 10 | 135112940 | rs2298120 | -0.506 | -0.767 | 1.734 | -1.037 | -0.004 | -0.025 | -0.025 | 0 | 0.15 | 0.19 | 0.19 | 0.05 | 0.14 | 0.14 |
| 10 | 135121732 | rs11101690 | -0.992 | -0.787 | 1.667 | -1.042 | -0.019 | -0.066 | -0.066 | 0 | 0.17 | 0.38 | 0.2 | 0.06 | 0.14 | 0.14 |
| 10 | 135139635 | rs2275723 | 0.481 | -0.821 | 1.343 | -1.022 | -0.009 | 0.031 | 0.031 | 1.486 | 0.11 | 0.03 | 0.16 | 0.05 | 0.11 | 0.11 |
| 10 | 135141572 | rs2298122 | 1.399 | -0.919 | 1.817 | -1.092 | -0.03 | -0.095 | -0.095 | 0 | 0.74 | 0.44 | 0.75 | 0.93 | 0.81 | 0.8 |
| 10 | 135146714 | rs11101694 | -0.671 | -0.825 | 1.699 | -1.006 | -0.012 | -0.043 | -0.043 | 0 | 0.16 | 0.28 | 0.2 | 0.07 | 0.14 | 0.15 |
| 10 | 135157955 | rs10776672 | 2.337 | -0.9 | 2.422 | -0.944 | -0.011 | -0.05 | -0.05 | 0 | 0.82 | 0.31 | 0.87 | 0.99 | 0.9 | 0.9 |
| 10 | 135167121 | rs1891371 | 2.075 | -0.77 | 2.531 | -0.878 | 0.026 | -0.021 | -0.021 | 0 | 0.92 | 0.42 | 0.92 | 0.99 | 0.96 | 0.96 |
| 10 | 135175844 | rs11101707 | 0.992 | -0.826 | 3.329 | -1.136 | 0.006 | -0.049 | -0.049 | 0 | 0.9 | 0.67 | 0.93 | 1 | 0.95 | 0.95 |
| 10 | 135183094 | rs10776675 | 1.958 | -0.602 | 2.764 | -0.45 | -0.032 | -0.073 | -0.073 | 0 | 0.87 | 0.47 | 0.92 | 0.99 | 0.94 | 0.94 |
| 10 | 135184126 | rs1049951 | 2.026 | -0.608 | 2.735 | -0.453 | -0.035 | -0.076 | -0.076 | 0.006 | 0.87 | 0.47 | 0.91 | 0.99 | 0.95 | 0.93 |
| 10 | 135192153 | rs11101724 | -1.555 | -0.357 | 2.342 | -0.305 | -0.012 | -0.031 | -0.031 | 0.02 | 0.1 | 0.27 | 0.06 | 0.02 | 0.04 | 0.06 |
| 10 | 135195332 | rs2185746 | 1.666 | -0.346 | 2.183 | -0.283 | -0.001 | -0.017 | -0.017 | 4.446 | 0.89 | 0.6 | 0.92 | 0.98 | 0.96 | 0.94 |
| 11 | 285920 | rs3817637 | 0.266 | 1.225 | -0.279 | 1.111 | -0.004 | 0.164 | 0.164 | 5.767 | 0.83 | 0.68 | 0.73 | 0.49 | 0.88 | 0.8 |
| 11 | 286621 | rs9737419 | -0.959 | 1.576 | -0.592 | 0.894 | -0.001 | 0.008 | 0.008 | 0 | 0.14 | 0.16 | 0.3 | 0.29 | 0.16 | 0.23 |
| 11 | 290158 | rs3809107 | -0.321 | 1.302 | -0.562 | 0.881 | -0.001 | 0.027 | 0.027 | 0 | 0.08 | 0.13 | 0.19 | 0.26 | 0.05 | 0.12 |
| 11 | 293188 | rs12801980 | -0.05 | 1.235 | -0.688 | 0.101 | 0.006 | -0.005 | -0.005 | 0.175 | 0.39 | 0.36 | 0.37 | 0.4 | 0.37 | 0.51 |
| 11 | 295343 | rs2242565 | 0.026 | 1.319 | 0.132 | 0.193 | 0.045 | 0.105 | 0.105 | 6.137 | 0.77 | 0.64 | 0.65 | 0.46 | 0.74 | 0.7 |
| 11 | 295876 | rs2242566 | -0.03 | 0.909 | 0.007 | 0.12 | -0.002 | 0.001 | 0.001 | 0.514 | 0.38 | 0.42 | 0.35 | 0.4 | 0.34 | 0.5 |
| 11 | 298316 | rs2293744 | -0.202 | 0.707 | -0.406 | 0.491 | -0.005 | 0.017 | 0.017 | 0 | 0.12 | 0.13 | 0.21 | 0.28 | 0.05 | 0.12 |
| 11 | 298758 | rs2293745 | -0.605 | 1.256 | -0.513 | 0.579 | 0.004 | 0.029 | 0.029 | 0 | 0.22 | 0.26 | 0.2 | 0.51 | 0.22 | 0.3 |
| 11 | 300364 | rs7111803 | -0.372 | 0.565 | -0.81 | 0.544 | -0.011 | -0.007 | -0.007 | 1.603 | 0.52 | 0.71 | 0.46 | 0.43 | 0.5 | 0.55 |
| 11 | 305961 | rs11246059 | -0.058 | -0.259 | -1.457 | 0.158 | -0.006 | 0.006 | 0.006 | 0 | 0.21 | 0.28 | 0.13 | 0.27 | 0.15 | 0.33 |
| 11 | 307036 | rs3809112 | -0.391 | -0.827 | -1.85 | -0.929 | 0 | -0.001 | -0.001 | 0 | 0.52 | 0.52 | 0.47 | 0.69 | 0.34 | 0.55 |
| 11 | 308180 | rs10398 | -0.124 | 0.001 | -1.777 | -0.415 | -0.015 | 0.027 | 0.027 | 0 | 0.26 | 0.38 | 0.36 | 0.63 | 0.19 | 0.41 |
| 11 | 316299 | rs7947900 | -0.225 | 1.546 | -2.452 | -1.207 | -0.125 | 0.119 | 0.119 | 1.102 | 0.79 | 0.96 | 0.78 | 1 | 0.53 | 0.87 |
| 11 | 323649 | rs7944394 | 0.255 | 1.112 | -2.375 | -1.107 | -0.097 | 0.082 | 0.082 | 2.273 | 0.2 | 0.08 | 0.23 | 0 | 0.43 | 0.18 |
| 11 | 325800 | rs7944917 | 0.384 | 0.86 | -2.253 | -0.852 | 0.003 | 0.001 | 0.001 | 0 | 0.06 | 0.05 | 0.03 | 0 | 0.03 | 0.08 |
| 11 | 327334 | rs11246068 | 0.349 | 0.485 | -2.566 | -0.494 | 0.017 | -0.075 | -0.075 | 0.726 | 0.32 | 0.08 | 0.3 | 0.56 | 0.26 | 0.22 |
| 11 | 331326 | rs7395319 | 0.204 | 0.165 | -1.629 | -0.718 | 0.001 | -0.024 | -0.024 | 0.173 | 0.86 | 0.84 | 0.92 | 0.98 | 0.82 | 0.83 |
| 11 | 335017 | rs10160565 | -0.415 | -0.624 | -1.327 | -0.337 | -0.033 | -0.008 | -0.008 | 0 | 0.28 | 0.65 | 0.21 | 0.24 | 0.11 | 0.4 |
| 11 | 336332 | rs11246074 | -1.114 | 1.159 | -1.458 | -0.263 | -0.02 | 0.017 | 0.017 | 5.585 | 0.11 | 0.1 | 0.09 | 0.03 | 0.18 | 0.06 |
| 11 | 343420 | rs11246083 | -1.157 | 0.436 | -1.549 | -0.098 | -0.008 | 0 | 0 | 0 | 0.1 | 0.03 | 0.05 | 0.13 | 0.11 | 0.08 |
| 11 | 345595 | rs6598035 | -0.278 | -0.514 | -1.395 | -0.499 | -0.005 | -0.006 | -0.006 | 0 | 0.52 | 0.55 | 0.39 | 0.75 | 0.32 | 0.66 |
| 11 | 350917 | rs11246108 | -0.27 | -0.976 | -1.431 | -0.081 | -0.019 | 0.013 | 0.013 | 2.028 | 0.39 | 0.55 | 0.35 | 0.48 | 0.28 | 0.54 |
| 11 | 353376 | rs11246114 | -0.511 | -0.134 | -1.137 | -1.187 | -0.001 | -0.013 | -0.013 | 0.095 | 0.45 | 0.48 | 0.31 | 0.19 | 0.52 | 0.43 |
| 11 | 353757 | rs7947784 | -0.311 | 0.293 | -1.112 | -1.124 | -0.027 | 0.023 | 0.023 | 0.57 | 0.28 | 0.3 | 0.26 | 0.09 | 0.42 | 0.32 |
| 11 | 362668 | rs11246126 | -0.134 | 0.217 | -0.89 | -1.19 | 0.022 | 0.041 | 0.041 | 0 | 0.52 | 0.61 | 0.54 | 0.82 | 0.41 | 0.55 |
| 11 | 365161 | rs10902140 | 0.05 | 0.125 | -0.473 | -1.19 | 0.001 | -0.024 | -0.024 | 0 | 0.23 | 0.24 | 0.16 | 0.08 | 0.26 | 0.14 |
| 11 | 366710 | rs7394810 | 0.183 | 0.447 | -0.095 | -1.349 | 0.011 | -0.05 | -0.05 | 0 | 0.73 | 0.38 | 0.76 | 0.92 | 0.7 | 0.86 |
| 11 | 367139 | rs11246131 | -0.18 | 0.324 | -0.579 | -1.217 | -0.006 | -0.003 | -0.003 | 0 | 0.08 | 0 | 0.04 | 0.06 | 0.08 | 0.06 |
| 11 | 368778 | rs11246132 | 0.196 | 0.676 | -0.464 | -0.55 | -0.014 | -0.004 | -0.004 | 0 | 0.58 | 0.33 | 0.61 | 0.46 | 0.52 | 0.65 |
| 11 | 370831 | rs11246135 | -0.438 | 0.903 | -0.577 | -0.366 | 0.035 | 0.025 | 0.025 | 0.06 | 0.5 | 0.64 | 0.49 | 0.38 | 0.4 | 0.45 |
| 11 | 371265 | rs11246136 | -0.003 | 0.975 | -0.783 | -0.566 | -0.033 | 0.046 | 0.046 | 0 | 0.08 | 0.01 | 0.16 | 0 | 0.1 | 0.07 |
| 11 | 373404 | rs7396812 | -0.175 | 0.851 | -0.182 | -0.271 | 0.002 | -0.002 | -0.002 | 0 | 0.41 | 0.55 | 0.33 | 0.41 | 0.3 | 0.37 |
| 11 | 377795 | rs10751657 | -0.408 | 1.171 | -0.069 | -0.295 | 0.022 | -0.018 | -0.018 | 0.415 | 0.41 | 0.63 | 0.34 | 0.49 | 0.33 | 0.38 |
| 11 | 378188 | rs7481525 | -0.844 | 1.201 | -0.084 | -0.307 | 0.016 | -0.011 | -0.011 | 0 | 0.45 | 0.66 | 0.47 | 0.49 | 0.4 | 0.41 |
| 11 | 382912 | rs11819881 | 0.678 | 1.425 | 0.039 | -0.257 | -0.023 | 0.005 | 0.005 | 0 | 0.55 | 0.35 | 0.48 | 0.42 | 0.51 | 0.54 |
| 11 | 383855 | rs7481739 | -0.289 | 1.023 | 0.394 | -0.371 | -0.034 | 0.026 | 0.026 | 0 | 0.29 | 0.57 | 0.27 | 0.58 | 0.17 | 0.42 |

| | | | | | | | | | | | | | | | | |
|---|---|---|---|---|---|---|---|---|---|---|---|---|---|---|---|---|
| 11 | 392953 | rs12788610 | 0.027 | 1.372 | 0.046 | -0.421 | -0.093 | 0.084 | 0.084 | 0 | 0.86 | 0.98 | 0.76 | 1 | 0.72 | 0.96 |
| 11 | 394005 | rs10902157 | 1.097 | 1.277 | 0.235 | -0.411 | -0.003 | -0.002 | -0.002 | 0 | 0.56 | 0.32 | 0.49 | 0.43 | 0.56 | 0.55 |
| 11 | 396308 | rs10902158 | 0.083 | 1.133 | 0.816 | -0.215 | -0.053 | 0.076 | 0.076 | 0 | 0.26 | 0.39 | 0.25 | 0.57 | 0.17 | 0.41 |
| 11 | 403545 | rs35828848 | -0.255 | 1.237 | 0.766 | -0.256 | -0.054 | 0.031 | 0.031 | 0 | 0.31 | 0.45 | 0.3 | 0.64 | 0.22 | 0.42 |
| 11 | 403980 | rs11748 | 0.645 | 1.461 | 0.761 | -0.274 | 0.009 | -0.002 | -0.002 | 0 | 0.59 | 0.52 | 0.61 | 0.37 | 0.56 | 0.53 |
| 11 | 404317 | rs1134578 | 0.998 | 1.46 | 0.45 | -0.347 | -0.017 | 0.022 | 0.022 | 0 | 0.56 | 0.31 | 0.59 | 0.35 | 0.56 | 0.53 |
| 11 | 406043 | rs7947 | -0.102 | 1.716 | 0.047 | -0.179 | 0.025 | -0.018 | -0.018 | 0 | 0.91 | 0.91 | 0.89 | 1 | 0.78 | 0.95 |
| 11 | 406473 | rs3087588 | -0.154 | 1.742 | 0.036 | -0.17 | 0.021 | -0.013 | -0.013 | 0.96 | 0.91 | 0.91 | 0.89 | 1 | 0.78 | 0.95 |
| 11 | 406483 | rs3210908 | -0.154 | 1.726 | -0.192 | -0.17 | -0.055 | 0.051 | 0.051 | 0 | 0.91 | 0.98 | 0.9 | 1 | 0.78 | 0.95 |
| 11 | 408352 | rs7396562 | -0.012 | 0.237 | 0.264 | 0.236 | -0.014 | 0.016 | 0.016 | 0 | 0.25 | 0.28 | 0.26 | 0.58 | 0.18 | 0.41 |
| 11 | 419906 | rs7484182 | 0.971 | 1.074 | 0.327 | -0.331 | 0.036 | -0.058 | -0.058 | 0 | 0.89 | 0.75 | 0.76 | 1 | 0.83 | 0.97 |
| 11 | 424291 | rs6598023 | -0.08 | 1.19 | 0.036 | -0.264 | -0.03 | 0.025 | 0.025 | 0 | 0.95 | 0.99 | 0.75 | 1 | 0.82 | 0.97 |
| 11 | 429659 | rs7111432 | 0.384 | 0.717 | 0.316 | -0.299 | -0.007 | -0.003 | -0.003 | 0.185 | 0.61 | 0.55 | 0.51 | 0.37 | 0.62 | 0.53 |
| 11 | 436461 | rs11246159 | 0.032 | 0.665 | 0.48 | -0.3 | -0.011 | 0.005 | 0.005 | 0.183 | 0.31 | 0.35 | 0.24 | 0.63 | 0.21 | 0.45 |
| 11 | 443731 | rs6421975 | 0.65 | 0.992 | 0.722 | -0.431 | 0.012 | -0.03 | -0.03 | 0.273 | 0.69 | 0.49 | 0.59 | 0.79 | 0.63 | 0.77 |
| 11 | 444628 | rs6598020 | -0.042 | 0.883 | 0.851 | -0.39 | -0.003 | -0.003 | -0.003 | 0.111 | 0.2 | 0.35 | 0.14 | 0.18 | 0.21 | 0.19 |
| 11 | 460303 | rs11821341 | -0.876 | 0.876 | 0.895 | -0.427 | 0.015 | -0.028 | -0.028 | 0.445 | 0.27 | 0.53 | 0.36 | 0.18 | 0.3 | 0.22 |
| 11 | 470331 | rs2613996 | -1.433 | 0.788 | 0.789 | -0.617 | 0.006 | 0 | 0 | 0 | 0.35 | 0.53 | 0.51 | 0.33 | 0.36 | 0.31 |
| 11 | 502013 | rs7930593 | -0.758 | 1.273 | 0.433 | -0.604 | 0.01 | -0.008 | -0.008 | 0 | 0.27 | 0.38 | 0.4 | 0.19 | 0.34 | 0.23 |
| 11 | 502217 | rs7944548 | -0.664 | 0.931 | 0.47 | -0.601 | 0.007 | -0.009 | -0.009 | 0 | 0.26 | 0.39 | 0.34 | 0.18 | 0.3 | 0.22 |
| 11 | 511805 | rs12420868 | -0.459 | 0.82 | 0.367 | -0.67 | -0.004 | -0.003 | -0.003 | 0.102 | 0.23 | 0.32 | 0.14 | 0.18 | 0.21 | 0.2 |
| 11 | 538940 | rs12420647 | -1.244 | 1.025 | 0.55 | -0.583 | 0.005 | -0.013 | -0.013 | 5.858 | 0.28 | 0.37 | 0.37 | 0.16 | 0.33 | 0.28 |
| 11 | 542844 | rs2894649 | -0.254 | 0.963 | 0.544 | -0.468 | -0.003 | -0.006 | -0.006 | 0 | 0.12 | 0.08 | 0.08 | 0.12 | 0.12 | 0.2 |
| 11 | 545087 | rs35579818 | -1.831 | 0.769 | 0.436 | -0.518 | 0.007 | 0.003 | 0.003 | 0 | 0.32 | 0.4 | 0.51 | 0.33 | 0.38 | 0.34 |
| 11 | 549119 | rs4963136 | 0.67 | 0.396 | 0.704 | -0.936 | -0.004 | -0.036 | -0.036 | 0 | 0.82 | 0.67 | 0.86 | 0.92 | 0.83 | 0.83 |
| 11 | 549959 | rs7942030 | 0.727 | 0.327 | 0.783 | -1.103 | -0.016 | -0.055 | -0.055 | 1.213 | 0.79 | 0.62 | 0.84 | 0.92 | 0.82 | 0.77 |
| 11 | 554798 | rs12277611 | 1.636 | 0.424 | 1.065 | -0.775 | -0.005 | 0.021 | 0.021 | 0 | 0.63 | 0.32 | 0.59 | 0.77 | 0.62 | 0.71 |
| 11 | 557342 | rs936466 | -0.005 | 0.208 | 1.112 | -0.873 | 0.044 | 0.08 | 0.08 | 0.789 | 0.94 | 0.77 | 0.96 | 0.91 | 0.94 | 0.88 |
| 11 | 558884 | rs2061586 | 1.141 | 0.287 | 1.068 | -0.92 | 0 | -0.035 | -0.035 | 0 | 0.76 | 0.56 | 0.85 | 0.86 | 0.82 | 0.79 |
| 11 | 565840 | rs1056812 | 0.372 | 0.201 | 0.704 | -0.811 | -0.004 | -0.028 | -0.028 | 0 | 0.08 | 0.06 | 0.16 | 0.16 | 0.06 | 0.1 |
| 11 | 566300 | rs3740651 | -1.615 | -0.021 | 1.095 | -1.152 | -0.012 | -0.098 | -0.098 | 1.263 | 0.31 | 0.51 | 0.31 | 0.08 | 0.28 | 0.18 |
| 11 | 566936 | rs12802200 | -0.044 | 0.007 | 1.219 | -1.246 | 0.041 | -0.058 | -0.058 | 0 | 0.12 | 0.25 | 0.24 | 0.02 | 0.19 | 0.06 |
| 11 | 567391 | rs7927267 | 2.229 | -0.012 | 1.002 | -1.288 | -0.001 | 0.088 | 0.088 | 0 | 0.54 | 0.16 | 0.48 | 0.71 | 0.6 | 0.6 |
| 11 | 573554 | rs12286521 | -2.031 | -0.001 | 1.03 | -1.176 | -0.006 | -0.024 | -0.024 | 0.888 | 0.4 | 0.53 | 0.47 | 0.22 | 0.34 | 0.29 |
| 11 | 577534 | rs7936397 | -1.239 | 0.054 | 1.083 | -1.421 | 0.005 | -0.16 | -0.16 | 1.372 | 0.25 | 0.48 | 0.29 | 0.02 | 0.26 | 0.11 |
| 11 | 589564 | rs4963128 | 1.406 | -0.449 | 1.093 | -1.754 | -0.006 | -0.112 | -0.112 | 1.379 | 0.61 | 0.53 | 0.67 | 0.94 | 0.67 | 0.76 |
| 11 | 606294 | rs11246211 | 0.093 | -0.476 | 0.456 | -1.683 | 0.019 | 0.006 | 0.006 | 4.881 | 0.15 | 0.14 | 0.06 | 0 | 0.06 | 0.02 |
| 11 | 610277 | rs11246212 | -0.204 | -0.235 | 0.497 | -1.53 | 0.013 | 0.041 | 0.041 | 0 | 0.87 | 0.78 | 0.79 | 0.77 | 0.87 | 0.81 |
| 11 | 612355 | rs12805435 | -1.142 | -0.293 | 0.518 | -1.657 | -0.009 | -0.1 | -0.1 | 0.024 | 0.32 | 0.58 | 0.3 | 0.02 | 0.27 | 0.11 |
| 11 | 612843 | rs10902178 | -1.153 | -0.294 | 0.517 | -1.657 | -0.009 | -0.1 | -0.1 | 0 | 0.32 | 0.58 | 0.3 | 0.02 | 0.27 | 0.11 |
| 11 | 612967 | rs11246213 | -1.142 | -0.293 | 0.518 | -1.657 | -0.009 | -0.1 | -0.1 | 0.283 | 0.32 | 0.58 | 0.3 | 0.02 | 0.27 | 0.11 |
| 11 | 618172 | rs6598008 | 2.275 | -0.365 | 0.341 | -1.562 | -0.021 | 0.068 | 0.068 | 0.328 | 0.46 | 0.22 | 0.46 | 0.7 | 0.56 | 0.6 |
| 11 | 619671 | rs1124816 | -0.136 | -0.359 | 0.593 | -1.586 | 0.059 | -0.11 | -0.11 | 0.154 | 0.16 | 0.4 | 0.26 | 0.02 | 0.22 | 0.09 |
| 11 | 619789 | rs2246614 | 1.095 | -0.41 | 0.602 | -1.615 | 0.017 | 0.002 | 0.002 | 0 | 0.71 | 0.45 | 0.54 | 0.73 | 0.66 | 0.7 |
| 11 | 620599 | rs7932167 | 1.304 | -0.445 | 0.249 | -1.564 | 0.009 | 0.007 | 0.007 | 1.801 | 0.62 | 0.59 | 0.71 | 0.72 | 0.78 | 0.69 |
| 11 | 620927 | rs12360820 | 0.704 | -0.218 | 0.167 | -1.408 | 0 | -0.001 | -0.001 | 0 | 0.77 | 0.75 | 0.76 | 0.73 | 0.83 | 0.71 |
| 11 | 623765 | rs11246217 | -0.86 | -0.222 | 0.496 | -1.532 | -0.011 | -0.116 | -0.116 | 2.337 | 0.32 | 0.43 | 0.3 | 0.02 | 0.27 | 0.11 |
| 11 | 630124 | rs11246221 | -0.973 | -0.214 | 0.208 | -1.531 | -0.002 | -0.104 | -0.104 | 0.719 | 0.36 | 0.63 | 0.34 | 0.02 | 0.3 | 0.13 |
| 11 | 636399 | rs3758653 | -0.812 | -0.807 | -0.361 | -1.301 | 0.018 | 0.001 | 0.001 | 0 | 0.28 | 0.18 | 0.26 | 0.26 | 0.2 | 0.37 |
| 11 | 637622 | rs752306 | -0.137 | -0.341 | -0.443 | -0.859 | -0.013 | -0.025 | -0.025 | 0 | 0.06 | 0.01 | 0.04 | 0.19 | 0.07 | 0.09 |
| 11 | 643348 | rs7395429 | 0.462 | -0.214 | -0.526 | -1.312 | -0.001 | 0.005 | 0.005 | 0 | 0.49 | 0.43 | 0.49 | 0.72 | 0.52 | 0.61 |
| 11 | 654043 | rs10902188 | 1.008 | -0.159 | -0.779 | -0.94 | 0.005 | -0.018 | -0.018 | 0 | 0.11 | 0.06 | 0.17 | 0.15 | 0.08 | 0.09 |
| 11 | 656845 | rs4963126 | 1.544 | -0.452 | -0.938 | -1.128 | -0.044 | -0.033 | -0.033 | 0 | 0.38 | 0.15 | 0.35 | 0.53 | 0.48 | 0.5 |
| 11 | 662407 | rs6598004 | -0.354 | -0.302 | -0.543 | -1.34 | -0.028 | 0.01 | 0.01 | 0 | 0.17 | 0.52 | 0.13 | 0.22 | 0.13 | 0.14 |
| 11 | 674499 | rs6597996 | 0.602 | -0.336 | -0.828 | -1.419 | -0.004 | 0.122 | 0.122 | 0 | 0.53 | 0.24 | 0.52 | 0.72 | 0.57 | 0.59 |
| 11 | 678973 | rs7935419 | 0.125 | -0.331 | -0.73 | -1.4 | -0.027 | -0.006 | -0.006 | 0 | 0.82 | 0.4 | 0.86 | 0.78 | 0.87 | 0.85 |
| 11 | 680647 | rs11246262 | 1.307 | -0.337 | -0.868 | -1.415 | -0.025 | 0.016 | 0.016 | 0 | 0.42 | 0.19 | 0.36 | 0.57 | 0.49 | 0.51 |
| 11 | 680866 | rs7483337 | 0.442 | -0.165 | -0.876 | -1.432 | -0.058 | 0.124 | 0.124 | 0.653 | 0.28 | 0.1 | 0.33 | 0.06 | 0.3 | 0.25 |
| 11 | 684056 | rs9734266 | -0.444 | -0.218 | -0.368 | -1.38 | -0.028 | 0 | 0 | 0 | 0.17 | 0.37 | 0.12 | 0.21 | 0.13 | 0.14 |
| 11 | 684697 | rs12790950 | 0.503 | -0.165 | -0.432 | -1.473 | -0.025 | 0.049 | 0.049 | 0.841 | 0.27 | 0.15 | 0.33 | 0.06 | 0.3 | 0.26 |
| 11 | 692221 | rs7128243 | -0.677 | -0.376 | -0.339 | -1.101 | -0.031 | -0.006 | -0.006 | 0 | 0.17 | 0.41 | 0.12 | 0.21 | 0.13 | 0.15 |
| 11 | 697761 | rs6597990 | -0.462 | -0.233 | -0.537 | -1.542 | 0.003 | -0.069 | -0.069 | 0 | 0.68 | 0.61 | 0.62 | 0.94 | 0.7 | 0.72 |
| 11 | 698807 | rs4963166 | 1.234 | -0.418 | -0.844 | -1.002 | -0.052 | -0.054 | -0.054 | 0 | 0.38 | 0.14 | 0.34 | 0.5 | 0.49 | 0.46 |
| 11 | 699852 | rs7480436 | -0.308 | -0.171 | -0.618 | -0.813 | 0.012 | -0.024 | -0.024 | 0 | 0.67 | 0.59 | 0.49 | 0.78 | 0.67 | 0.63 |
| 11 | 702250 | rs7479861 | 0.799 | -0.027 | -1.216 | -0.223 | -0.075 | -0.083 | -0.083 | 0 | 0.4 | 0.08 | 0.35 | 0.48 | 0.5 | 0.47 |
| 11 | 705404 | rs12293143 | -1.244 | 0.361 | -0.402 | 0.224 | 0.033 | 0.159 | 0.159 | 0.134 | 0.05 | 0.22 | 0.17 | 0.19 | 0.06 | 0.11 |
| 11 | 709861 | rs11246281 | -0.455 | 0.368 | -0.399 | -0.129 | 0.001 | -0.002 | -0.002 | 0.458 | 0.37 | 0.61 | 0.51 | 0.28 | 0.36 | 0.42 |
| 11 | 710139 | rs7942569 | 0.209 | 0.22 | -0.362 | -0.519 | 0 | -0.03 | -0.03 | 0 | 0.3 | 0.3 | 0.34 | 0.09 | 0.28 | 0.29 |
| 11 | 710558 | rs7395822 | -0.723 | 0.102 | -1.193 | -0.297 | 0.03 | 0.097 | 0.097 | 0 | 0.51 | 0.77 | 0.48 | 0.33 | 0.43 | 0.47 |

| | | | | | | | | | | | | | | | | |
|---|---|---|---|---|---|---|---|---|---|---|---|---|---|---|---|---|
| 11 | 715420 | rs11246286 | -0.019 | 0.133 | -1.259 | -0.451 | 0.009 | 0.068 | 0.068 | 1.819 | 0.17 | 0.06 | 0.12 | 0.03 | 0.18 | 0.05 |
| 11 | 716765 | rs12222447 | -0.365 | 0.582 | -1.536 | -0.311 | -0.066 | 0 | 0 | 0 | 0.71 | 0.9 | 0.7 | 0.26 | 0.58 | 0.55 |
| 11 | 720197 | rs3087546 | 0.12 | 0.264 | -0.828 | -0.535 | 0.017 | -0.027 | -0.027 | 0 | 0.63 | 0.57 | 0.65 | 0.22 | 0.55 | 0.51 |
| 11 | 721570 | rs7635 | 0.72 | -0.071 | -0.977 | -0.713 | -0.011 | 0.079 | 0.079 | 3.288 | 0.38 | 0.19 | 0.56 | 0.17 | 0.42 | 0.17 |
| 11 | 723429 | rs4078520 | 0.492 | 0.026 | -0.663 | -0.649 | 0.007 | 0.143 | 0.143 | 5.658 | 0.5 | 0.28 | 0.62 | 0.17 | 0.48 | 0.26 |
| 11 | 729570 | rs10902208 | 1.239 | 0.703 | -0.665 | -0.315 | -0.05 | 0.057 | 0.057 | 0.146 | 0.19 | 0.04 | 0.1 | 0.12 | 0.21 | 0.3 |
| 11 | 733639 | rs10902210 | -1.11 | 1.011 | 0.282 | 0.609 | 0.026 | 0.212 | 0.212 | 2.947 | 0.1 | 0.39 | 0.08 | 0.31 | 0.09 | 0.21 |
| 11 | 745884 | rs7936322 | 0.353 | 0.892 | -0.194 | 0.667 | 0.003 | 0.021 | 0.021 | 3.509 | 0.45 | 0.31 | 0.61 | 0.57 | 0.45 | 0.34 |
| 11 | 752059 | rs7478765 | -1.051 | 1.048 | -0.231 | 0.717 | 0.013 | 0.03 | 0.03 | 1.031 | 0.6 | 0.89 | 0.77 | 0.64 | 0.67 | 0.48 |
| 11 | 766479 | rs7937869 | -0.925 | 1.142 | 0.283 | 0.657 | 0.104 | -0.011 | -0.011 | 0.046 | 0.14 | 0.47 | 0.17 | 0.08 | 0.22 | 0.15 |
| 11 | 771034 | rs7930569 | 0.365 | 1.068 | 0.053 | 0.655 | 0.006 | 0.019 | 0.019 | 3.404 | 0.45 | 0.31 | 0.6 | 0.56 | 0.44 | 0.34 |
| 11 | 772490 | rs12224894 | 0.577 | 0.999 | 0.055 | 0.882 | -0.023 | 0.044 | 0.044 | 6.095 | 0.42 | 0.17 | 0.59 | 0.28 | 0.44 | 0.19 |
| 11 | 772701 | rs12223324 | 0.294 | 1.09 | 0.04 | 0.544 | 0.005 | 0.011 | 0.011 | 3.206 | 0.46 | 0.4 | 0.61 | 0.56 | 0.45 | 0.34 |
| 11 | 772982 | rs11823172 | -0.797 | 1.4 | 0.317 | 0.485 | 0.096 | -0.009 | -0.009 | 0 | 0.12 | 0.35 | 0.15 | 0.07 | 0.22 | 0.15 |
| 11 | 787679 | rs7946354 | -0.21 | 0.82 | -0.153 | 0.505 | 0.002 | 0.011 | 0.011 | 0.182 | 0.5 | 0.46 | 0.66 | 0.58 | 0.53 | 0.4 |
| 11 | 788007 | rs6597982 | -1.02 | 0.876 | -0.069 | 0.427 | 0.006 | 0.007 | 0.007 | 0.057 | 0.58 | 0.68 | 0.75 | 0.62 | 0.65 | 0.48 |
| 11 | 791462 | rs4963153 | 0.541 | 0.748 | -0.183 | 0.626 | -0.056 | 0.06 | 0.06 | 2.244 | 0.4 | 0.14 | 0.59 | 0.28 | 0.46 | 0.17 |
| 11 | 805712 | rs11246316 | 0.381 | 0.62 | -0.169 | 0.619 | -0.051 | 0.107 | 0.107 | 0.798 | 0.44 | 0.19 | 0.62 | 0.27 | 0.49 | 0.16 |
| 11 | 810882 | rs10902222 | -0.87 | 0.619 | 0.002 | 0.893 | 0.011 | 0.006 | 0.006 | 0.054 | 0.58 | 0.7 | 0.75 | 0.6 | 0.64 | 0.48 |
| 11 | 813524 | rs11246319 | 0.458 | 0.493 | -0.266 | 0.903 | -0.071 | 0.102 | 0.102 | 1.676 | 0.43 | 0.17 | 0.61 | 0.25 | 0.5 | 0.17 |
| 11 | 816017 | rs11246321 | -0.895 | 0.131 | 0.081 | 1.156 | -0.018 | 0.159 | 0.159 | 0 | 0.07 | 0.18 | 0.04 | 0.27 | 0.02 | 0.17 |
| 11 | 822622 | rs7942159 | 0.078 | 0.454 | -0.149 | 0.583 | 0.009 | 0.029 | 0.029 | 0.246 | 0.5 | 0.49 | 0.68 | 0.28 | 0.57 | 0.24 |
| 11 | 825110 | rs1138714 | 0.364 | 0.522 | -0.635 | 0.549 | -0.006 | 0.117 | 0.117 | 0.488 | 0.46 | 0.23 | 0.64 | 0.24 | 0.55 | 0.18 |
| 11 | 827993 | rs28407517 | -0.283 | 0.12 | -0.977 | 0.82 | -0.007 | -0.062 | -0.062 | 0.089 | 0.84 | 0.98 | 0.91 | 0.76 | 0.89 | 0.72 |
| 11 | 828916 | rs7126805 | -0.403 | -0.073 | -0.3 | 0.564 | -0.017 | -0.013 | -0.013 | 0 | 0.55 | 0.46 | 0.77 | 0.66 | 0.69 | 0.35 |
| 11 | 829539 | rs28558789 | -0.396 | 0.091 | -0.194 | 0.095 | 0.004 | -0.019 | -0.019 | 0 | 0.56 | 0.58 | 0.78 | 0.89 | 0.67 | 0.45 |
| 11 | 830670 | rs4075289 | 0.149 | -0.288 | -0.372 | -0.042 | 0.015 | -0.004 | -0.004 | 0 | 0.5 | 0.54 | 0.78 | 0.89 | 0.67 | 0.45 |
| 11 | 831809 | rs1130276 | -0.793 | -0.024 | -0.004 | 0.101 | 0.061 | 0.129 | 0.129 | 0 | 0.61 | 0.82 | 0.82 | 0.89 | 0.72 | 0.47 |
| 11 | 833668 | rs7924806 | -0.241 | 0.023 | 0.01 | 0.174 | 0.003 | -0.021 | -0.021 | 0 | 0.55 | 0.56 | 0.79 | 0.88 | 0.66 | 0.44 |
| 11 | 837582 | rs1130663 | -0.058 | 0.108 | 0.056 | 0.269 | -0.005 | -0.032 | -0.032 | 0 | 0.54 | 0.53 | 0.79 | 0.86 | 0.66 | 0.44 |
| 11 | 837772 | rs5030779 | 1.224 | -0.09 | -0.02 | 0.114 | 0.049 | 0.077 | 0.077 | 0 | 0.31 | 0.14 | 0.13 | 0.14 | 0.23 | 0.47 |
| 11 | 838424 | rs1130680 | -0.2 | 0.032 | -0.052 | 0.2 | -0.003 | -0.025 | -0.025 | 0 | 0.56 | 0.56 | 0.8 | 0.85 | 0.66 | 0.44 |
| 11 | 838542 | rs1130698 | -0.105 | 0.027 | -0.052 | 0.2 | -0.002 | -0.035 | -0.035 | 0 | 0.56 | 0.56 | 0.8 | 0.88 | 0.66 | 0.45 |
| 11 | 838672 | rs1803956 | -0.956 | -0.11 | 0.214 | 0.711 | -0.006 | 0.071 | 0.071 | 0.079 | 0.12 | 0.16 | 0.09 | 0.65 | 0.08 | 0.27 |
| 11 | 838722 | rs6762 | 0.002 | -0.254 | 0.242 | 0.659 | 0.011 | 0.004 | 0.004 | 0 | 0.43 | 0.4 | 0.18 | 0.14 | 0.29 | 0.55 |
| 11 | 838760 | rs1130719 | -0.026 | -0.255 | 0.044 | 0.669 | 0 | -0.02 | -0.02 | 0 | 0.52 | 0.53 | 0.79 | 0.85 | 0.66 | 0.44 |
| 11 | 840034 | rs11246327 | 1.093 | 0.272 | -0.04 | 1.415 | -0.046 | -0.045 | -0.045 | 0.877 | 0.08 | 0.01 | 0.17 | 0.24 | 0.13 | 0.15 |
| 11 | 840319 | rs3059 | -0.25 | 0.54 | 0.149 | 0.719 | -0.006 | -0.035 | -0.035 | 0 | 0.54 | 0.53 | 0.79 | 0.86 | 0.66 | 0.44 |
| 11 | 840363 | rs6591 | 0.468 | 0.673 | 0.258 | 0.928 | -0.011 | 0.022 | 0.022 | 0.255 | 0.4 | 0.38 | 0.6 | 0.2 | 0.53 | 0.15 |
| 11 | 843927 | rs12577104 | 0.657 | 0.297 | 0.196 | 1.147 | -0.042 | 0.045 | 0.045 | 0 | 0.71 | 0.53 | 0.86 | 0.57 | 0.8 | 0.68 |
| 11 | 845789 | rs10902227 | 1.486 | 0.68 | -0.14 | 1.273 | -0.017 | -0.088 | -0.088 | 0 | 0.28 | 0.06 | 0.22 | 0.35 | 0.23 | 0.47 |
| 11 | 858424 | rs7946539 | 0.376 | 1.14 | 0.032 | 0.927 | -0.011 | 0.027 | 0.027 | 0.568 | 0.42 | 0.39 | 0.58 | 0.23 | 0.52 | 0.18 |
| 11 | 867824 | rs4963206 | 1.807 | 1.419 | -0.059 | 0.759 | -0.003 | 0.002 | 0.002 | 0 | 0.13 | 0.04 | 0.09 | 0.12 | 0.11 | 0.21 |
| 11 | 870446 | rs6682 | -0.519 | 1.465 | 0.124 | 0.446 | 0 | -0.005 | -0.005 | 0 | 0.56 | 0.41 | 0.41 | 0.76 | 0.49 | 0.79 |
| 11 | 876624 | rs12422106 | -0.665 | 1.528 | -0.448 | 0.456 | -0.016 | 0.044 | 0.044 | 0.686 | 0.57 | 0.64 | 0.42 | 0.76 | 0.48 | 0.81 |
| 11 | 876755 | rs7941920 | -0.665 | 1.528 | -0.448 | 0.462 | -0.016 | 0.043 | 0.043 | 0.686 | 0.57 | 0.64 | 0.42 | 0.76 | 0.48 | 0.82 |
| 11 | 884543 | rs7479983 | 1.768 | 1.536 | -0.734 | 0.572 | -0.021 | -0.096 | -0.096 | 0 | 0.28 | 0.07 | 0.23 | 0.43 | 0.24 | 0.5 |
| 11 | 892760 | rs7121173 | 0.128 | 1.522 | -0.467 | -0.118 | -0.058 | 0.076 | 0.076 | 0.472 | 0.42 | 0.21 | 0.57 | 0.24 | 0.52 | 0.19 |
| 11 | 898155 | rs4963183 | -0.466 | 1.476 | -0.478 | -0.118 | -0.036 | 0.065 | 0.065 | 0.415 | 0.58 | 0.75 | 0.43 | 0.76 | 0.48 | 0.81 |
| 11 | 898344 | rs7946095 | 1.134 | 1.567 | -0.655 | -0.104 | 0.006 | 0.039 | 0.039 | 0 | 0.14 | 0.04 | 0.09 | 0.05 | 0.11 | 0.18 |
| 11 | 906708 | rs11246342 | -0.085 | 1.305 | -0.55 | -0.079 | 0.032 | 0.021 | 0.021 | 0 | 0.38 | 0.27 | 0.44 | 0.27 | 0.35 | 0.22 |
| 11 | 918219 | rs10902230 | -0.438 | 1.154 | -0.435 | -0.691 | -0.001 | -0.021 | -0.021 | 0 | 0.44 | 0.52 | 0.59 | 0.18 | 0.46 | 0.18 |
| 11 | 918793 | rs4963159 | 2.181 | 1.222 | -0.63 | -0.827 | 0.004 | 0.028 | 0.028 | 0 | 0.13 | 0.05 | 0.11 | 0.03 | 0.11 | 0.1 |
| 11 | 924904 | rs10794342 | -0.282 | 1.066 | -0.414 | -0.766 | 0.015 | 0.074 | 0.074 | 0.647 | 0.51 | 0.35 | 0.67 | 0.27 | 0.56 | 0.27 |
| 11 | 927020 | rs7482484 | -0.898 | 1.211 | -0.454 | -0.825 | 0.001 | 0.003 | 0.003 | 0.115 | 0.59 | 0.38 | 0.72 | 0.28 | 0.66 | 0.39 |
| 11 | 934594 | rs11605562 | 0.944 | 1.229 | -0.497 | -0.787 | -0.023 | 0.024 | 0.024 | 0 | 0.07 | 0.01 | 0.05 | 0.02 | 0.1 | 0.12 |
| 11 | 944466 | rs10751670 | -0.869 | 1.347 | -0.37 | -0.439 | 0.02 | 0.12 | 0.12 | 0 | 0.92 | 0.83 | 0.95 | 0.69 | 0.89 | 0.92 |
| 11 | 955638 | rs7950967 | -0.347 | 1.308 | -0.929 | -0.812 | 0.005 | 0.024 | 0.024 | 0.448 | 0.52 | 0.46 | 0.69 | 0.27 | 0.56 | 0.27 |
| 11 | 976019 | rs7483870 | 2.105 | 1.308 | -1.412 | -0.912 | -0.062 | -0.045 | -0.045 | 0.614 | 0.14 | 0.01 | 0.31 | 0.16 | 0.23 | 0.1 |
| 11 | 985547 | rs10794358 | -0.517 | 0.819 | -0.879 | -0.576 | 0.006 | 0.017 | 0.017 | 0.249 | 0.58 | 0.36 | 0.71 | 0.28 | 0.66 | 0.39 |
| 11 | 996522 | rs4077293 | 1.244 | 0.953 | -0.946 | -0.582 | -0.008 | 0.016 | 0.016 | 0 | 0.32 | 0.25 | 0.53 | 0.27 | 0.37 | 0.23 |
| 11 | 1005308 | rs6597948 | -0.117 | 0.813 | -1.058 | -0.668 | 0.024 | 0.096 | 0.096 | 1.738 | 0.5 | 0.32 | 0.66 | 0.3 | 0.57 | 0.36 |
| 11 | 1011490 | rs7950955 | -0.125 | 0.811 | -1.024 | -0.668 | 0.02 | 0.077 | 0.077 | 1.732 | 0.5 | 0.36 | 0.66 | 0.29 | 0.57 | 0.36 |
| 11 | 1015021 | rs4076950 | 2.037 | 0.841 | -1.143 | -0.678 | -0.009 | 0.007 | 0.007 | 0 | 0.28 | 0.22 | 0.51 | 0.27 | 0.35 | 0.19 |
| 11 | 1019194 | rs11246381 | 0.095 | 0.771 | -0.955 | -0.45 | -0.062 | 0.084 | 0.084 | 0 | 0.15 | 0.02 | 0.13 | 0.02 | 0.19 | 0.12 |
| 11 | 1027811 | rs7481521 | 0.349 | 0.647 | -0.847 | -0.345 | 0.009 | 0.106 | 0.106 | 0.86 | 0.47 | 0.26 | 0.65 | 0.3 | 0.57 | 0.32 |
| 11 | 1037029 | rs6597947 | 0.59 | 0.329 | -0.875 | -0.283 | -0.032 | -0.037 | -0.037 | 0.059 | 0.07 | 0 | 0.04 | 0.29 | 0.1 | 0.12 |
| 11 | 1038110 | rs11605303 | 1.759 | 0.149 | -0.744 | -0.278 | -0.001 | -0.015 | -0.015 | 0 | 0.09 | 0.08 | 0.06 | 0 | 0.08 | 0.01 |
| 11 | 1039867 | rs9794921 | -0.088 | 0.323 | -0.732 | -0.175 | 0.059 | 0.097 | 0.097 | 0.126 | 0.83 | 0.63 | 0.79 | 0.65 | 0.78 | 0.73 |

| | | | | | | | | | | | | | | | | |
|---|---|---|---|---|---|---|---|---|---|---|---|---|---|---|---|---|
| 11 | 1040790 | rs12296021 | 0.324 | 0.457 | -0.974 | 0.072 | 0.014 | 0.035 | 0.035 | 0.387 | 0.48 | 0.35 | 0.66 | 0.38 | 0.58 | 0.34 |
| 11 | 1044938 | rs11246386 | 0.088 | 0.835 | -0.772 | 0.499 | 0.042 | 0.076 | 0.076 | 0 | 0.82 | 0.7 | 0.8 | 0.66 | 0.78 | 0.73 |
| 11 | 1051715 | rs10794359 | -0.884 | 0.047 | -1.421 | 0.64 | -0.032 | -0.01 | -0.01 | 0.081 | 0.62 | 0.71 | 0.36 | 0.59 | 0.51 | 0.68 |
| 11 | 1054625 | rs11826309 | 0.268 | -0.736 | -1.274 | 0.76 | 0.028 | -0.017 | -0.017 | 5.343 | 0.23 | 0.19 | 0.08 | 0.28 | 0.12 | 0.39 |
| 11 | 1060934 | rs10902073 | 1.053 | -0.197 | -1.421 | 0.079 | 0.055 | 0.142 | 0.142 | 0.222 | 0.4 | 0.21 | 0.27 | 0.17 | 0.34 | 0.41 |
| 11 | 1061781 | rs10794279 | 1.059 | -0.29 | -1.244 | 0.029 | 0.05 | 0.116 | 0.116 | 0 | 0.4 | 0.24 | 0.27 | 0.16 | 0.34 | 0.41 |
| 11 | 1062519 | rs11245920 | -0.52 | -0.199 | -1.477 | 0.788 | -0.008 | -0.029 | -0.029 | 0 | 0.08 | 0.02 | 0.02 | 0.25 | 0.08 | 0.08 |
| 11 | 1062867 | rs10794280 | 1.469 | -0.217 | -1.466 | 0.777 | 0.049 | 0.16 | 0.16 | 0 | 0.39 | 0.16 | 0.26 | 0.15 | 0.34 | 0.41 |
| 11 | 1062990 | rs10751635 | 0.48 | -0.225 | -0.973 | 0.759 | 0.043 | 0.038 | 0.038 | 0 | 0.49 | 0.36 | 0.29 | 0.41 | 0.41 | 0.49 |
| 11 | 1068233 | rs12146505 | -0.861 | 0.117 | -0.162 | 1.29 | -0.032 | 0.065 | 0.065 | 0 | 0.11 | 0.2 | 0.15 | 0.22 | 0.04 | 0.13 |
| 11 | 1068900 | rs7942850 | 0.956 | 0.461 | -0.23 | 0.481 | 0.044 | 0.105 | 0.105 | 0 | 0.46 | 0.31 | 0.29 | 0.16 | 0.38 | 0.43 |
| 11 | 1073378 | rs2071173 | -1.721 | 0.687 | 1.409 | 0.525 | 0.022 | 0.252 | 0.252 | 0 | 0.14 | 0.31 | 0.09 | 0.43 | 0.14 | 0.2 |
| 11 | 1073712 | rs2071174 | -1.265 | 1.083 | 1.539 | 0.375 | 0.024 | 0.061 | 0.061 | 0.534 | 0.26 | 0.54 | 0.18 | 0.44 | 0.34 | 0.2 |
| 11 | 1075747 | rs2856111 | -1.985 | 0.413 | 1.61 | 0.639 | 0.025 | 0.311 | 0.311 | 0.211 | 0.15 | 0.41 | 0.11 | 0.44 | 0.14 | 0.17 |
| 11 | 1083364 | rs7396030 | -1.711 | 0.636 | 1.147 | -0.655 | -0.007 | 0.085 | 0.085 | 0 | 0.23 | 0.45 | 0.37 | 0.39 | 0.22 | 0.42 |
| 11 | 1087972 | rs10902088 | -1.464 | 0.834 | 0.769 | -0.705 | 0.002 | 0.076 | 0.076 | 0 | 0.21 | 0.38 | 0.36 | 0.39 | 0.22 | 0.29 |
| 11 | 1094357 | rs10902089 | 0.325 | 0.728 | 0.198 | -1.809 | -0.033 | 0.097 | 0.097 | 0 | 0.61 | 0.31 | 0.69 | 0.93 | 0.51 | 0.7 |
| 11 | 1101078 | rs11245954 | -0.403 | 1.046 | 0.861 | -0.692 | 0.022 | 0.002 | 0.002 | 0 | 0.06 | 0.16 | 0.04 | 0.05 | 0.08 | 0.07 |
| 11 | 1105976 | rs4077759 | 1.254 | 1.062 | 0.03 | -0.726 | -0.009 | 0.026 | 0.026 | 0 | 0.62 | 0.24 | 0.53 | 0.66 | 0.62 | 0.63 |
| 11 | 1111474 | rs6597976 | 0.414 | 0.271 | 0.027 | -0.299 | 0.01 | -0.011 | -0.011 | 0 | 0.85 | 0.91 | 0.65 | 0.81 | 0.91 | 0.73 |
| 11 | 1126979 | rs6421966 | 0.58 | 0.052 | -0.048 | -0.203 | 0.001 | 0 | 0 | 0.008 | 0.69 | 0.33 | 0.54 | 0.71 | 0.8 | 0.59 |
| 11 | 1129721 | rs7130988 | 0.587 | 0.054 | -0.044 | -0.199 | 0.001 | 0 | 0 | 0.005 | 0.69 | 0.32 | 0.54 | 0.71 | 0.8 | 0.59 |
| 11 | 1131998 | rs11602160 | 0.562 | -0.067 | -0.061 | -0.181 | 0.002 | 0 | 0 | 0.082 | 0.7 | 0.31 | 0.55 | 0.71 | 0.8 | 0.59 |
| 11 | 1138100 | rs10902101 | 0.717 | 0.047 | -0.246 | -0.125 | -0.013 | -0.002 | -0.002 | 0 | 0.7 | 0.35 | 0.56 | 0.71 | 0.81 | 0.59 |
| 11 | 1150353 | rs17859811 | -1.03 | 0.149 | -0.28 | -0.065 | -0.034 | -0.003 | -0.003 | 0.059 | 0.3 | 0.56 | 0.43 | 0.31 | 0.18 | 0.39 |
| 11 | 1151695 | rs28737416 | -0.775 | 0.119 | -0.226 | 0.153 | -0.02 | -0.003 | -0.003 | 0 | 0.31 | 0.61 | 0.35 | 0.32 | 0.19 | 0.39 |
| 11 | 1214349 | rs35705491 | -0.704 | 1.302 | -0.29 | -0.035 | 0.038 | -0.05 | -0.05 | 0 | 0.1 | 0.16 | 0.1 | 0 | 0.2 | 0.05 |
| 11 | 1220760 | rs28514396 | -1.12 | 0.667 | -0.398 | -0.152 | 0.012 | 0.044 | 0.044 | 1.109 | 0.43 | 0.7 | 0.45 | 0.35 | 0.41 | 0.55 |
| 11 | 1229005 | rs28678421 | -0.971 | 0.329 | 0.355 | -0.209 | -0.006 | -0.048 | -0.048 | 0 | 0.22 | 0.42 | 0.14 | 0.11 | 0.24 | 0.29 |
| 11 | 1238828 | rs7120886 | -1.383 | 0.434 | 0.777 | -0.273 | -0.009 | -0.056 | -0.056 | 0 | 0.25 | 0.51 | 0.2 | 0.11 | 0.26 | 0.29 |
| 11 | 1241005 | rs2672794 | -1.026 | 0.725 | -0.667 | -1.077 | -0.08 | -0.069 | -0.069 | 0 | 0.69 | 0.95 | 0.73 | 0.41 | 0.61 | 0.67 |
| 11 | 1243391 | rs885454 | -1.051 | 0.541 | -0.443 | 0.086 | -0.005 | 0.038 | 0.038 | 0 | 0.1 | 0.14 | 0.24 | 0.28 | 0.07 | 0.22 |
| 11 | 1246941 | rs2672785 | -1.739 | 0.569 | 0.213 | 0.896 | -0.005 | 0.024 | 0.024 | 0 | 0.26 | 0.34 | 0.28 | 0.44 | 0.24 | 0.32 |
| 11 | 1247458 | rs2075853 | -0.878 | 0.552 | 0.448 | 0.882 | -0.001 | 0.187 | 0.187 | 0 | 0.08 | 0.19 | 0.11 | 0.38 | 0.09 | 0.16 |
| 11 | 1250488 | rs2075859 | -0.018 | 1.215 | -0.044 | 0.238 | 0.001 | -0.008 | -0.008 | 0 | 0.32 | 0.28 | 0.44 | 0.61 | 0.34 | 0.35 |
| 11 | 1258803 | rs2014486 | -1.339 | 1.065 | 0.058 | 0.122 | -0.001 | 0.004 | 0.004 | 0 | 0.52 | 0.49 | 0.6 | 0.65 | 0.5 | 0.52 |
| 11 | 1263523 | rs1541314 | 0.206 | 1.082 | 0.04 | 0.104 | -0.011 | 0.019 | 0.019 | 0 | 0.09 | 0.02 | 0.11 | 0.05 | 0.08 | 0.09 |
| 11 | 1272858 | rs4963059 | 0.267 | 1.166 | 0.028 | 0.192 | 0.007 | -0.022 | -0.022 | 0 | 0.28 | 0.21 | 0.33 | 0.61 | 0.29 | 0.29 |
| 11 | 1281134 | rs2857476 | -1.5 | 1.223 | 0.42 | 0.188 | -0.002 | 0.01 | 0.01 | 0 | 0.52 | 0.58 | 0.61 | 0.67 | 0.5 | 0.53 |
| 11 | 1292831 | rs3750919 | 1.208 | 1.194 | 0.381 | 0.258 | -0.044 | -0.104 | -0.104 | 0 | 0.21 | 0.02 | 0.21 | 0.26 | 0.2 | 0.15 |
| 11 | 1296649 | rs3168046 | 0.533 | 1.598 | -0.583 | -0.203 | 0 | 0.016 | 0.016 | 0 | 0.48 | 0.39 | 0.37 | 0.33 | 0.46 | 0.47 |
| 11 | 1300406 | rs3829223 | 0.304 | 1.925 | -0.617 | -0.31 | 0 | 0.003 | 0.003 | 0 | 0.51 | 0.46 | 0.47 | 0.33 | 0.51 | 0.51 |
| 11 | 1301982 | rs3793964 | -1.101 | 1.943 | -0.599 | -0.32 | -0.001 | -0.004 | -0.004 | 0.111 | 0.7 | 0.73 | 0.64 | 0.39 | 0.65 | 0.67 |
| 11 | 1302707 | rs3793965 | 0.521 | 1.923 | -0.662 | -0.349 | 0 | 0.014 | 0.014 | 0 | 0.47 | 0.39 | 0.37 | 0.33 | 0.46 | 0.47 |
| 11 | 1309956 | rs3750920 | 0.838 | 1.49 | -0.138 | -0.495 | -0.01 | 0.021 | 0.021 | 0 | 0.4 | 0.28 | 0.34 | 0.32 | 0.46 | 0.43 |
| 11 | 1314028 | rs5743942 | -0.323 | 1.311 | -0.287 | -0.728 | 0 | -0.028 | -0.028 | 0.077 | 0.53 | 0.45 | 0.45 | 0.09 | 0.51 | 0.44 |
| 11 | 1318312 | rs11042484 | 1.186 | 1.348 | -0.406 | -0.555 | -0.013 | 0.027 | 0.027 | 0 | 0.09 | 0.03 | 0.11 | 0.06 | 0.08 | 0.09 |
| 11 | 1323564 | rs5743899 | 0.953 | 1.294 | -0.13 | -0.118 | 0.004 | 0.098 | 0.098 | 0 | 0.79 | 0.54 | 0.72 | 0.59 | 0.79 | 0.71 |
| 11 | 1327752 | rs5743869 | 1.805 | 0.847 | -0.456 | -0.218 | -0.002 | -0.018 | -0.018 | 0.23 | 0.08 | 0.07 | 0.04 | 0 | 0.04 | 0.07 |
| 11 | 1328351 | rs5743867 | -1.324 | 0.867 | -0.433 | -0.183 | 0.005 | 0.245 | 0.245 | 0 | 0.93 | 0.76 | 0.88 | 0.6 | 0.91 | 0.84 |
| 11 | 1333402 | rs7481967 | 0.075 | 0.878 | -0.483 | -0.182 | -0.021 | 0.12 | 0.12 | 0 | 0.81 | 0.55 | 0.74 | 0.59 | 0.82 | 0.73 |
| 11 | 1365531 | rs4963039 | -1.594 | 0.618 | -0.977 | 0.064 | -0.011 | 0.037 | 0.037 | 0.239 | 0.9 | 0.89 | 0.96 | 0.71 | 0.95 | 0.89 |
| 11 | 1369555 | rs2056083 | 2.676 | 0.687 | -1.249 | -0.145 | -0.041 | 0.077 | 0.077 | 0 | 0.13 | 0.01 | 0.09 | 0 | 0.15 | 0.06 |
| 11 | 1369851 | rs11023369 | -2.065 | 0.869 | -0.632 | -0.201 | -0.012 | 0.023 | 0.023 | 0 | 0.68 | 0.49 | 0.74 | 0.59 | 0.69 | 0.69 |
| 11 | 1374456 | rs7122936 | -1.898 | 0.984 | -0.487 | -0.274 | -0.004 | -0.005 | -0.005 | 0 | 0.66 | 0.73 | 0.64 | 0.59 | 0.61 | 0.67 |
| 11 | 1375531 | rs10832442 | -1.287 | 0.995 | -0.517 | -0.269 | 0 | 0.001 | 0.001 | 0 | 0.64 | 0.39 | 0.61 | 0.59 | 0.61 | 0.64 |
| 11 | 1375960 | rs11023590 | -1.268 | 1.005 | -0.513 | -0.27 | 0 | 0.002 | 0.002 | 0 | 0.63 | 0.39 | 0.61 | 0.59 | 0.61 | 0.63 |
| 11 | 1377230 | rs10832483 | -2.758 | 1.107 | -0.804 | -0.231 | 0.006 | -0.013 | -0.013 | 0 | 0.75 | 0.83 | 0.85 | 0.7 | 0.77 | 0.8 |
| 11 | 1381530 | rs7109736 | -2.751 | 1.105 | -0.91 | -0.232 | 0.007 | -0.017 | -0.017 | 1.446 | 0.75 | 0.86 | 0.85 | 0.7 | 0.77 | 0.8 |
| 11 | 1383289 | rs908228 | -2.225 | 1.128 | -0.887 | -0.239 | -0.006 | -0.005 | -0.005 | 0.902 | 0.71 | 0.79 | 0.73 | 0.69 | 0.69 | 0.75 |
| 11 | 1384207 | rs6421029 | -2.727 | 0.973 | -0.949 | -0.192 | 0.007 | -0.017 | -0.017 | 0 | 0.75 | 0.85 | 0.85 | 0.69 | 0.77 | 0.8 |
| 11 | 1384920 | rs7395465 | -2.727 | 0.973 | -0.949 | -0.192 | 0.007 | -0.017 | -0.017 | 0 | 0.75 | 0.85 | 0.85 | 0.7 | 0.77 | 0.8 |
| 11 | 1391359 | rs7481546 | 0.884 | 0.941 | -1.304 | -0.429 | 0.002 | 0.008 | 0.008 | 0 | 0.38 | 0.31 | 0.32 | 0.21 | 0.36 | 0.29 |
| 11 | 1393384 | rs7125627 | -2.82 | 0.886 | -1.543 | -0.311 | 0.006 | -0.013 | -0.013 | 0 | 0.75 | 0.84 | 0.85 | 0.7 | 0.77 | 0.8 |
| 11 | 1395321 | rs11024281 | 2.465 | 0.911 | -1.799 | -0.524 | -0.071 | 0.172 | 0.172 | 0 | 0.27 | 0.02 | 0.17 | 0.02 | 0.31 | 0.21 |
| 11 | 1398019 | rs10832843 | 1.044 | 0.798 | -1.279 | 0.001 | 0.007 | -0.097 | -0.097 | 0 | 0.17 | 0.04 | 0.3 | 0.44 | 0.14 | 0.3 |
| 11 | 1398316 | rs4881745 | 0.868 | 0.891 | -1.191 | -0.005 | 0.001 | -0.05 | -0.05 | 0 | 0.17 | 0.11 | 0.31 | 0.44 | 0.18 | 0.31 |
| 11 | 1401890 | rs2090192 | 0.687 | 0.731 | -0.945 | -0.024 | -0.005 | 0.113 | 0.113 | 0 | 0.19 | 0.44 | 0.37 | 0.4 | 0.15 | 0.31 |
| 11 | 1404838 | rs7944761 | -1.418 | 1.046 | -1.37 | -0.258 | -0.002 | 0.004 | 0.004 | 0 | 0.55 | 0.77 | 0.55 | 0.54 | 0.54 | 0.57 |

| | | | | | | | | | | | | | | | | |
|---|---|---|---|---|---|---|---|---|---|---|---|---|---|---|---|---|
| 11 | 1408590 | rs4881729 | 1.059 | 0.826 | -1.192 | -0.73 | -0.002 | 0.089 | 0.089 | 0.103 | 0.52 | 0.31 | 0.44 | 0.19 | 0.56 | 0.37 |
| 11 | 1418877 | rs11822087 | 1.505 | 0.945 | -0.904 | -1.053 | 0.007 | -0.08 | -0.08 | 0 | 0.22 | 0.27 | 0.15 | 0.03 | 0.25 | 0.12 |
| 11 | 1423176 | rs4963076 | 0.283 | 0.793 | -1.141 | -0.917 | -0.004 | 0.033 | 0.033 | 0 | 0.51 | 0.28 | 0.46 | 0.39 | 0.51 | 0.5 |
| 11 | 1429616 | rs7941517 | 1.075 | 0.941 | -1.402 | -1.448 | 0.004 | -0.078 | -0.078 | 0 | 0.35 | 0.36 | 0.3 | 0.03 | 0.46 | 0.22 |
| 11 | 1435379 | rs4963079 | -1.255 | 0.681 | -1.329 | -1.346 | -0.011 | 0.053 | 0.053 | 0 | 0.73 | 0.61 | 0.61 | 0.41 | 0.77 | 0.62 |
| 11 | 1439894 | rs10833655 | 1.061 | 0.001 | -1.457 | -1.081 | 0.009 | -0.033 | -0.033 | 0 | 0.23 | 0.14 | 0.25 | 0.33 | 0.21 | 0.28 |
| 11 | 1442226 | rs10833703 | 0.092 | 0.157 | -1.348 | -1.082 | 0.031 | -0.021 | -0.021 | 0 | 0.35 | 0.23 | 0.3 | 0.38 | 0.27 | 0.4 |
| 11 | 1443398 | rs7114346 | 2.337 | 0.871 | -1.097 | -1.651 | -0.004 | -0.021 | -0.021 | 0 | 0.19 | 0.13 | 0.14 | 0.02 | 0.25 | 0.1 |
| 11 | 1462217 | rs9651643 | -0.137 | 0.54 | -1.082 | -1.409 | 0 | -0.027 | -0.027 | 0 | 0.47 | 0.42 | 0.3 | 0.06 | 0.46 | 0.24 |
| 11 | 1473931 | rs11602013 | -0.151 | 0.395 | -1.528 | -2.016 | 0 | 0.162 | 0.162 | 0 | 0.46 | 0.25 | 0.32 | 0.05 | 0.51 | 0.3 |
| 11 | 1479601 | rs12221580 | -0.12 | 0.231 | -1.084 | -1.709 | -0.004 | -0.02 | -0.02 | 0 | 0.37 | 0.52 | 0.35 | 0.29 | 0.34 | 0.38 |
| 11 | 1485629 | rs1106245 | -0.709 | 0.041 | -1.128 | -1.397 | -0.013 | -0.001 | -0.001 | 0 | 0.61 | 0.55 | 0.6 | 0.39 | 0.68 | 0.58 |
| 11 | 1491096 | rs9440 | -0.226 | 0.193 | -0.664 | -1.581 | 0.03 | 0.036 | 0.036 | 0 | 0.53 | 0.7 | 0.52 | 0.41 | 0.57 | 0.49 |
| 11 | 1492445 | rs4881724 | -1.275 | 0.403 | -0.646 | -1.531 | -0.032 | 0.036 | 0.036 | 0 | 0.93 | 0.99 | 0.88 | 1 | 0.88 | 0.91 |
| 11 | 1507512 | rs1881502 | 0.134 | 0.2 | -0.805 | -1.15 | -0.038 | 0.016 | 0.016 | 0 | 0.86 | 0.93 | 0.84 | 0.92 | 0.81 | 0.86 |
| 11 | 1515271 | rs4881736 | 0.224 | -0.041 | -0.867 | -1.235 | 0.052 | 0.008 | 0.008 | 0 | 0.39 | 0.24 | 0.36 | 0.61 | 0.31 | 0.38 |
| 11 | 1516110 | rs10835531 | 1.703 | -0.46 | -1.023 | -1.466 | 0.016 | -0.037 | -0.037 | 0 | 0.23 | 0.15 | 0.3 | 0.53 | 0.2 | 0.25 |
| 11 | 1520063 | rs11599981 | 1.424 | -0.181 | -1.01 | -1.651 | -0.053 | 0.065 | 0.065 | 0.077 | 0.2 | 0.08 | 0.14 | 0.06 | 0.27 | 0.24 |
| 11 | 1521917 | rs12278822 | 0.91 | -0.544 | -0.523 | -1.111 | -0.001 | -0.002 | -0.002 | 0 | 0.5 | 0.47 | 0.49 | 0.39 | 0.57 | 0.54 |
| 11 | 1544922 | rs11031756 | 0.927 | -0.384 | -0.555 | -1.102 | -0.006 | -0.011 | -0.011 | 0.05 | 0.06 | 0.05 | 0.04 | 0.08 | 0.07 | 0.05 |
| 11 | 1561484 | rs7127282 | -1.4 | -0.475 | -0.617 | -0.863 | 0.013 | -0.031 | -0.031 | 0 | 0.77 | 0.84 | 0.7 | 0.51 | 0.8 | 0.75 |
| 11 | 1562713 | rs1038723 | 0.714 | -0.466 | -0.557 | -0.961 | 0.003 | 0.002 | 0.002 | 0 | 0.52 | 0.51 | 0.47 | 0.43 | 0.57 | 0.52 |
| 11 | 1575068 | rs7951820 | -1.383 | -0.442 | -0.526 | -0.908 | 0.018 | -0.049 | -0.049 | 0.76 | 0.75 | 0.85 | 0.69 | 0.51 | 0.79 | 0.71 |
| 11 | 1576079 | rs1982827 | 0.636 | -0.301 | -0.539 | -0.907 | 0.004 | 0.007 | 0.007 | 0 | 0.49 | 0.54 | 0.45 | 0.43 | 0.53 | 0.48 |
| 11 | 1581028 | rs10836478 | 0.66 | -0.418 | -0.536 | -0.903 | 0.004 | 0.006 | 0.006 | 0 | 0.49 | 0.54 | 0.46 | 0.42 | 0.53 | 0.48 |
| 11 | 1607346 | rs10734456 | -0.368 | -0.486 | -0.501 | -0.924 | 0.017 | -0.014 | -0.014 | 0 | 0.62 | 0.63 | 0.63 | 0.5 | 0.72 | 0.62 |
| 11 | 1608229 | rs11034930 | 0.611 | -0.425 | -0.5 | -0.936 | 0.018 | 0.03 | 0.03 | 0 | 0.49 | 0.6 | 0.46 | 0.42 | 0.53 | 0.48 |
| 11 | 1608295 | rs11606312 | 1.214 | -0.435 | -0.461 | -0.961 | 0.009 | -0.001 | -0.001 | 0.044 | 0.12 | 0.06 | 0.04 | 0 | 0.06 | 0.09 |
| 11 | 1619859 | rs1809667 | 0.083 | -0.409 | -0.642 | -0.839 | -0.013 | -0.019 | -0.019 | 0 | 0.2 | 0.28 | 0.14 | 0.11 | 0.14 | 0.15 |
| 11 | 1619907 | rs1809668 | 0.81 | -0.513 | -0.695 | -0.781 | 0.003 | 0.004 | 0.004 | 0 | 0.47 | 0.5 | 0.45 | 0.42 | 0.53 | 0.45 |
| 11 | 1620784 | rs2334501 | -0.736 | -0.446 | -0.759 | -1.11 | -0.001 | -0.012 | -0.012 | 3.299 | 0.27 | 0.3 | 0.25 | 0.11 | 0.26 | 0.24 |
| 11 | 1625363 | rs10837407 | -0.52 | -0.599 | -1.196 | -0.925 | 0.006 | -0.116 | -0.116 | 0.023 | 0.67 | 0.91 | 0.59 | 0.52 | 0.66 | 0.59 |
| 11 | 1627296 | rs7117114 | 0.161 | -0.942 | -0.583 | -1.028 | 0.024 | 0.017 | 0.017 | 0 | 0.54 | 0.58 | 0.57 | 0.36 | 0.65 | 0.53 |
| 11 | 1627926 | rs2004828 | 1.428 | -0.414 | -0.69 | -1.469 | -0.077 | 0.131 | 0.131 | 0 | 0.34 | 0.1 | 0.23 | 0.11 | 0.39 | 0.33 |
| 11 | 1628844 | rs1989592 | -1.389 | -0.809 | -0.807 | -1.464 | 0.044 | -0.114 | -0.114 | 0 | 0.74 | 0.92 | 0.7 | 0.52 | 0.79 | 0.67 |
| 11 | 1632180 | rs7947734 | 0.492 | -0.564 | -0.53 | -1.31 | -0.002 | -0.004 | -0.004 | 0 | 0.43 | 0.5 | 0.45 | 0.39 | 0.52 | 0.44 |
| 11 | 1633661 | rs10768675 | 0.181 | -0.255 | -0.566 | -1.051 | 0.002 | -0.007 | -0.007 | 0 | 0.07 | 0.09 | 0.07 | 0.11 | 0.07 | 0.07 |
| 11 | 1638347 | rs12284916 | 0.717 | -0.23 | -0.591 | -1.037 | -0.007 | -0.02 | -0.02 | 0.129 | 0.06 | 0.04 | 0.06 | 0.11 | 0.07 | 0.06 |
| 11 | 1640024 | rs9651654 | -0.504 | -0.895 | -0.82 | -1.506 | 0.005 | -0.007 | -0.007 | 0 | 0.66 | 0.65 | 0.63 | 0.51 | 0.73 | 0.61 |
| 11 | 1641649 | rs7116239 | -0.461 | -0.992 | -1.074 | -1.532 | 0.02 | -0.068 | -0.068 | 0 | 0.73 | 0.85 | 0.63 | 0.4 | 0.74 | 0.66 |
| 11 | 1643728 | rs10768791 | 0.179 | -0.332 | -0.671 | -1.991 | 0.005 | -0.019 | -0.019 | 0.773 | 0.57 | 0.68 | 0.47 | 0.5 | 0.54 | 0.55 |
| 11 | 1644905 | rs11605284 | 1.718 | -0.318 | -1.057 | -2.261 | 0.015 | 0.032 | 0.032 | 0 | 0.16 | 0.14 | 0.21 | 0.37 | 0.25 | 0.19 |
| 11 | 1644916 | rs10837841 | -0.448 | -0.607 | -1.255 | -1.86 | 0.022 | 0.052 | 0.052 | 0 | 0.26 | 0.17 | 0.21 | 0.12 | 0.24 | 0.21 |
| 11 | 1645885 | rs10837857 | -0.457 | -0.389 | -1.116 | -1.705 | -0.001 | 0 | 0 | 3.205 | 0.13 | 0.15 | 0.16 | 0.12 | 0.16 | 0.1 |
| 11 | 1647110 | rs7937618 | -1.339 | -0.412 | -1.119 | -1.668 | -0.001 | 0.017 | 0.017 | 0 | 0.81 | 0.8 | 0.69 | 0.54 | 0.81 | 0.73 |
| 11 | 1649822 | rs12366207 | 0.123 | -0.524 | -1.335 | -1.967 | -0.008 | 0.072 | 0.072 | 0 | 0.26 | 0.1 | 0.15 | 0.12 | 0.23 | 0.15 |
| 11 | 1650047 | rs1021345 | -0.785 | -0.185 | -0.513 | -1.262 | -0.004 | -0.014 | -0.014 | 0.731 | 0.26 | 0.28 | 0.21 | 0.14 | 0.24 | 0.22 |
| 11 | 1650264 | rs1021342 | 0.521 | -0.084 | -0.386 | -1.231 | 0.03 | -0.027 | -0.027 | 0.036 | 0.06 | 0.18 | 0.08 | 0 | 0.09 | 0.05 |
| 11 | 1650805 | rs17832809 | 0.227 | -0.029 | -0.697 | -1.202 | 0.002 | -0.026 | -0.026 | 0 | 0.19 | 0.13 | 0.14 | 0.02 | 0.19 | 0.16 |
| 11 | 1651830 | rs4752770 | 0.608 | 0.152 | -0.573 | -1.039 | 0.038 | 0.026 | 0.026 | 0 | 0.54 | 0.71 | 0.48 | 0.39 | 0.55 | 0.49 |
| 11 | 1651917 | rs4752769 | 0.666 | 0.169 | -0.608 | -1.055 | 0.038 | 0.018 | 0.018 | 0 | 0.55 | 0.75 | 0.49 | 0.4 | 0.55 | 0.52 |
| 11 | 1652422 | rs10838027 | -0.263 | 0.273 | -0.271 | -0.788 | 0.048 | 0.084 | 0.084 | 0 | 0.07 | 0.23 | 0.09 | 0.12 | 0.09 | 0.06 |
| 11 | 1653445 | rs902219 | 0.648 | 0.429 | -0.548 | -1.043 | 0.003 | 0.008 | 0.008 | 0.398 | 0.56 | 0.63 | 0.53 | 0.41 | 0.52 | 0.46 |
| 11 | 1657308 | rs10082701 | 0.398 | 0.637 | -0.495 | -1.043 | -0.003 | -0.071 | -0.071 | 0 | 0.2 | 0.3 | 0.18 | 0 | 0.22 | 0.16 |
| 11 | 1659308 | rs10768964 | 0.241 | 0.681 | -0.51 | -0.947 | -0.005 | -0.005 | -0.005 | 0.437 | 0.17 | 0.13 | 0.35 | 0.58 | 0.22 | 0.3 |
| 11 | 1659810 | rs7927995 | -0.714 | 0.554 | -0.552 | -0.942 | -0.001 | 0 | 0 | 0 | 0.39 | 0.43 | 0.53 | 0.58 | 0.43 | 0.47 |
| 11 | 1667746 | rs16927442 | -0.077 | 0.93 | -0.652 | -0.675 | 0.011 | 0.001 | 0.001 | 0 | 0.07 | 0.07 | 0.2 | 0.1 | 0.16 | 0.07 |
| 11 | 1667948 | rs11821612 | 1.16 | 1 | -0.731 | -0.615 | -0.015 | -0.033 | -0.033 | 0.009 | 0.11 | 0.07 | 0.21 | 0.48 | 0.13 | 0.24 |
| 11 | 1672567 | rs12364282 | -0.066 | 1.079 | -0.651 | -0.601 | 0.011 | 0.001 | 0.001 | 0 | 0.07 | 0.08 | 0.2 | 0.1 | 0.16 | 0.07 |
| 11 | 1673636 | rs7108440 | -0.945 | 1.837 | -0.338 | -0.757 | 0.187 | 0.007 | 0.007 | 0 | 0.13 | 0.47 | 0.36 | 0.12 | 0.31 | 0.1 |
| 11 | 1674938 | rs11604009 | 1.056 | 1.244 | -0.734 | -1.115 | 0.007 | 0.011 | 0.011 | 0.007 | 0.46 | 0.34 | 0.28 | 0.4 | 0.35 | 0.43 |
| 11 | 1675559 | rs11038344 | -0.003 | 1.474 | -0.747 | -0.936 | -0.009 | -0.101 | -0.101 | 0 | 0.29 | 0.11 | 0.33 | 0.49 | 0.27 | 0.38 |
| 11 | 1675607 | rs7950561 | 0.607 | 1.583 | -0.841 | -0.994 | -0.007 | -0.012 | -0.012 | 0 | 0.11 | 0.04 | 0.17 | 0.23 | 0.14 | 0.11 |
| 11 | 1676922 | rs16927458 | -0.785 | 1.507 | -0.711 | -1.132 | 0.008 | -0.013 | -0.013 | 5.747 | 0.4 | 0.57 | 0.5 | 0.28 | 0.51 | 0.32 |
| 11 | 1677351 | rs10400297 | 0.627 | 1.699 | -0.495 | -1.022 | 0.187 | 0.027 | 0.027 | 0 | 0.05 | 0.3 | 0.15 | 0.01 | 0.14 | 0.02 |
| 11 | 1678014 | rs10769146 | -0.314 | 1.404 | -1.025 | -1.394 | -0.014 | -0.003 | -0.003 | 4.157 | 0.72 | 0.79 | 0.66 | 0.3 | 0.68 | 0.6 |
| 11 | 1679464 | rs12146638 | 0.089 | 1.627 | -0.68 | -0.867 | 0.109 | 0.094 | 0.094 | 0 | 0.16 | 0.39 | 0.32 | 0.27 | 0.27 | 0.12 |
| 11 | 1679711 | rs17173843 | 0.818 | 1.135 | -0.807 | -1.016 | 0.028 | 0.011 | 0.011 | 0 | 0.33 | 0.2 | 0.16 | 0.03 | 0.17 | 0.27 |
| 11 | 1680161 | rs7119110 | -0.465 | 2.314 | -0.36 | 0.067 | 0.086 | 0.1 | 0.1 | 0 | 0.16 | 0.4 | 0.31 | 0.28 | 0.25 | 0.12 |

| | | | | | | | | | | | | | | | | |
|---|---|---|---|---|---|---|---|---|---|---|---|---|---|---|---|---|
| 11 | 1681001 | rs1038727 | 0.399 | 1.799 | -1.024 | 0.155 | -0.017 | -0.015 | -0.015 | 0 | 0.11 | 0.07 | 0.12 | 0.19 | 0.19 | 0.14 |
| 11 | 1689325 | rs7131362 | 0.112 | 0.872 | -1.045 | -0.196 | 0.017 | 0.003 | 0.003 | 0 | 0.48 | 0.35 | 0.47 | 0.54 | 0.44 | 0.64 |
| 11 | 1690969 | rs7112918 | -0.137 | 0.444 | -1.243 | -1.427 | 0 | 0.026 | 0.026 | 0 | 0.5 | 0.59 | 0.53 | 0.76 | 0.49 | 0.5 |
| 11 | 1691621 | rs1489945 | -0.139 | 0.29 | -1.355 | -1.806 | -0.006 | 0.01 | 0.01 | 0 | 0.57 | 0.65 | 0.42 | 0.38 | 0.52 | 0.61 |
| 11 | 1693052 | rs12806058 | 0.972 | -0.067 | -1.451 | -2.16 | -0.017 | 0.005 | 0.005 | 0.016 | 0.13 | 0.01 | 0.08 | 0.08 | 0.08 | 0.09 |
| 11 | 1696849 | rs2334499 | 0.688 | -0.007 | -1.556 | -2.366 | -0.13 | 0.082 | 0.082 | 0 | 0.31 | 0.07 | 0.46 | 0.82 | 0.43 | 0.28 |
| 11 | 1701470 | rs4752779 | -1.311 | -0.245 | -1.256 | -2.514 | -0.012 | -0.094 | -0.094 | 0.014 | 0.67 | 0.56 | 0.67 | 0.91 | 0.67 | 0.64 |
| 11 | 1702321 | rs11039428 | 0.52 | -0.318 | -2.109 | -2.388 | 0.036 | 0.108 | 0.108 | 0 | 0.21 | 0.04 | 0.1 | 0.01 | 0.15 | 0.26 |
| 11 | 1703564 | rs4417225 | 0.36 | 0.131 | -1.796 | -1.982 | -0.128 | 0.084 | 0.084 | 0 | 0.31 | 0.06 | 0.45 | 0.83 | 0.42 | 0.27 |
| 11 | 1713618 | rs10839203 | -0.342 | -0.4 | -1.833 | -2.067 | 0.047 | 0.161 | 0.161 | 0 | 0.33 | 0.06 | 0.17 | 0.1 | 0.21 | 0.34 |
| 11 | 1714061 | rs10769560 | -1.029 | -0.261 | -1.114 | -2.246 | 0.004 | -0.001 | -0.001 | 0.1 | 0.64 | 0.32 | 0.65 | 0.93 | 0.64 | 0.63 |
| 11 | 1714688 | rs11607954 | -0.52 | -0.518 | -0.871 | -2.371 | 0.004 | 0.076 | 0.076 | 0 | 0.54 | 0.66 | 0.61 | 0.84 | 0.55 | 0.53 |
| 11 | 1716153 | rs4752741 | -1.029 | -1.452 | -0.51 | -2.055 | 0.042 | 0.062 | 0.062 | 0 | 0.34 | 0.23 | 0.18 | 0.11 | 0.22 | 0.34 |
| 11 | 1716749 | rs16927507 | 0.272 | -1.053 | -0.639 | -1.701 | -0.006 | 0.012 | 0.012 | 0 | 0.13 | 0.02 | 0.08 | 0.09 | 0.08 | 0.09 |
| 11 | 1718266 | rs4752744 | -0.704 | -1.228 | -0.28 | -1.681 | -0.015 | -0.011 | -0.011 | 2.214 | 0.88 | 0.87 | 0.87 | 0.93 | 0.92 | 0.89 |
| 11 | 1721089 | rs11040489 | -0.772 | -1.408 | -0.411 | -2.28 | 0.005 | -0.035 | -0.035 | 0 | 0.33 | 0.37 | 0.13 | 0.03 | 0.2 | 0.25 |
| 11 | 1729250 | rs7925551 | -0.217 | -1.453 | -0.464 | -2.26 | -0.003 | -0.023 | -0.023 | 0 | 0.15 | 0.21 | 0.04 | 0.04 | 0.07 | 0.14 |
| 11 | 1729593 | rs7937045 | -1.793 | -1.494 | -0.406 | -2.535 | 0.021 | 0.037 | 0.037 | 0.817 | 0.49 | 0.65 | 0.35 | 0.14 | 0.33 | 0.43 |
| 11 | 1736664 | rs7102751 | -0.662 | -1.461 | -0.757 | -1.966 | -0.003 | -0.015 | -0.015 | 0 | 0.26 | 0.27 | 0.26 | 0.1 | 0.18 | 0.28 |
| 11 | 1736750 | rs7115056 | -0.197 | -0.835 | -0.599 | -1.661 | -0.001 | -0.041 | -0.041 | 0 | 0.14 | 0.21 | 0.16 | 0.04 | 0.13 | 0.16 |
| 11 | 1751986 | rs17834326 | -0.071 | -0.618 | -1.438 | -1.326 | -0.01 | 0.031 | 0.031 | 0 | 0.11 | 0.03 | 0.06 | 0.04 | 0.09 | 0.1 |
| 11 | 1754400 | rs3740627 | 0.719 | -1.293 | -1.38 | -1.517 | -0.046 | -0.045 | -0.045 | 0 | 0.26 | 0.55 | 0.22 | 0.15 | 0.14 | 0.17 |
| 11 | 1756291 | rs2292958 | -0.522 | -1.74 | -1.965 | -1.745 | 0.017 | 0.028 | 0.028 | 0 | 0.56 | 0.63 | 0.33 | 0.25 | 0.33 | 0.47 |
| 11 | 1756848 | rs4752755 | 0.929 | -0.836 | -1.983 | -0.613 | -0.157 | -0.138 | -0.138 | 0 | 0.24 | 0.05 | 0.51 | 0.61 | 0.49 | 0.34 |
| 11 | 1771797 | rs3740621 | 0.035 | -1.151 | -1.848 | -0.683 | -0.002 | -0.006 | -0.006 | 0.023 | 0.38 | 0.6 | 0.28 | 0.19 | 0.24 | 0.28 |
| 11 | 1773477 | rs2334411 | -0.012 | -1.133 | -2.037 | -0.674 | 0.023 | 0.032 | 0.032 | 0 | 0.39 | 0.64 | 0.28 | 0.19 | 0.24 | 0.28 |
| 11 | 1779138 | rs1317356 | 0.224 | -0.17 | -2.173 | -0.043 | -0.17 | -0.138 | -0.138 | 0 | 0.26 | 0.04 | 0.51 | 0.61 | 0.51 | 0.34 |
| 11 | 1782594 | rs17571 | -0.293 | -0.097 | -1.415 | -0.297 | -0.005 | 0.025 | 0.025 | 0 | 0.1 | 0.02 | 0.05 | 0.04 | 0.07 | 0.1 |
| 11 | 1803095 | rs10839641 | -0.577 | 0.466 | -1.317 | -0.248 | -0.02 | 0.003 | 0.003 | 0 | 0.26 | 0.22 | 0.38 | 0.21 | 0.44 | 0.33 |
| 11 | 1807114 | rs11041105 | -0.203 | -0.138 | -1.141 | -0.651 | 0.004 | -0.001 | -0.001 | 0 | 0.28 | 0.24 | 0.27 | 0.23 | 0.42 | 0.37 |
| 11 | 1812367 | rs10839670 | 0.111 | -0.799 | -1.104 | -1.182 | 0.005 | -0.035 | -0.035 | 0.158 | 0.23 | 0.29 | 0.24 | 0.07 | 0.25 | 0.22 |
| 11 | 1818095 | rs4752759 | -1.017 | -1.227 | -1.109 | -0.708 | 0.018 | -0.002 | -0.002 | 0 | 0.61 | 0.63 | 0.68 | 0.37 | 0.76 | 0.72 |
| 11 | 1824253 | rs10743009 | -0.312 | -0.662 | -1.719 | -0.58 | 0.004 | 0.009 | 0.009 | 0.608 | 0.38 | 0.22 | 0.45 | 0.31 | 0.6 | 0.37 |
| 11 | 1829564 | rs7101654 | -0.429 | -0.987 | -2.487 | -0.413 | 0.069 | 0.024 | 0.024 | 0.008 | 0.53 | 0.84 | 0.6 | 0.63 | 0.42 | 0.56 |
| 11 | 1830754 | rs7482841 | 0.244 | -0.864 | -2.82 | -0.565 | -0.009 | 0.018 | 0.018 | 0 | 0.13 | 0.04 | 0.23 | 0.11 | 0.16 | 0.22 |
| 11 | 1834254 | rs939685 | 0.658 | -0.676 | -3.198 | -0.762 | -0.007 | 0.103 | 0.103 | 0 | 0.33 | 0.02 | 0.1 | 0.12 | 0.22 | 0.28 |
| 11 | 1837050 | rs17835310 | -0.188 | -0.703 | -3.06 | -0.768 | 0.039 | 0.172 | 0.172 | 0.261 | 0.47 | 0.17 | 0.35 | 0.22 | 0.38 | 0.5 |
| 11 | 1840954 | rs7481661 | -0.292 | -0.58 | -3.026 | -0.738 | 0.077 | 0.2 | 0.2 | 0.417 | 0.5 | 0.18 | 0.35 | 0.22 | 0.38 | 0.51 |
| 11 | 1849333 | rs907603 | 0.178 | -0.088 | -2.684 | -0.517 | 0.002 | 0.013 | 0.013 | 0 | 0.14 | 0.14 | 0.25 | 0.1 | 0.17 | 0.21 |
| 11 | 1849354 | rs907604 | -0.154 | 0.342 | -2.092 | -0.207 | 0.056 | 0.007 | 0.007 | 0.287 | 0.78 | 0.4 | 0.7 | 0.81 | 0.63 | 0.76 |
| 11 | 1854571 | rs498337 | -0.162 | -0.215 | -2.537 | 0.106 | 0.004 | -0.084 | -0.084 | 1.829 | 0.31 | 0.14 | 0.35 | 0.53 | 0.3 | 0.27 |
| 11 | 1857042 | rs907605 | -0.163 | -0.043 | -2.818 | -0.033 | -0.023 | 0.053 | 0.053 | 0 | 0.49 | 0.16 | 0.39 | 0.34 | 0.55 | 0.57 |
| 11 | 1857270 | rs907609 | -0.578 | -0.444 | -2.216 | 0.16 | 0.035 | 0.047 | 0.047 | 0 | 0.18 | 0.1 | 0.19 | 0.09 | 0.15 | 0.2 |
| 11 | 1858262 | rs484955 | -0.379 | 0.756 | -1.749 | 1.125 | -0.014 | -0.011 | -0.011 | 0 | 0.09 | 0.05 | 0.23 | 0.41 | 0.13 | 0.16 |
| 11 | 1865091 | rs626738 | 0.206 | 0.528 | -1.109 | -0.259 | 0 | -0.099 | -0.099 | 0.296 | 0.73 | 0.52 | 0.69 | 0.92 | 0.73 | 0.72 |
| 11 | 1865560 | rs11822805 | -0.52 | 0.339 | -1.672 | -0.354 | 0.002 | 0.046 | 0.046 | 0.36 | 0.13 | 0.02 | 0.11 | 0.07 | 0.12 | 0.07 |
| 11 | 1874072 | rs907611 | -0.514 | 0.926 | -2.022 | -0.3 | -0.035 | 0.01 | 0.01 | 0 | 0.27 | 0.09 | 0.35 | 0.24 | 0.33 | 0.24 |
| 11 | 1874404 | rs2089910 | 0.345 | 0.906 | -1.763 | -0.14 | 0 | 0.002 | 0.002 | 0 | 0.26 | 0.28 | 0.28 | 0.46 | 0.27 | 0.17 |
| 11 | 1878636 | rs599774 | -0.255 | 0.923 | -0.874 | -0.42 | 0 | 0.026 | 0.026 | 0 | 0.34 | 0.47 | 0.35 | 0.54 | 0.32 | 0.25 |
| 11 | 1881256 | rs587961 | -1.066 | 0.417 | -0.633 | -0.866 | 0.003 | 0.039 | 0.039 | 0 | 0.66 | 0.76 | 0.78 | 0.85 | 0.68 | 0.6 |
| 11 | 1884342 | rs2137320 | 0.084 | 0.683 | -0.641 | -0.926 | -0.035 | -0.006 | -0.006 | 0 | 0.33 | 0.16 | 0.54 | 0.63 | 0.42 | 0.3 |
| 11 | 1886147 | rs622173 | -0.779 | 0.8 | -0.766 | -1.103 | 0.049 | 0.041 | 0.041 | 0 | 0.92 | 0.69 | 0.89 | 0.89 | 0.88 | 0.9 |
| 11 | 1888614 | rs4980379 | 0.454 | 1.264 | -1.282 | -1.126 | -0.043 | -0.022 | -0.022 | 0 | 0.32 | 0.09 | 0.54 | 0.63 | 0.39 | 0.31 |
| 11 | 1890990 | rs592373 | -0.57 | 0.761 | -1.289 | -1.504 | -0.001 | 0.059 | 0.059 | 0 | 0.64 | 0.25 | 0.77 | 0.78 | 0.68 | 0.5 |
| 11 | 1901467 | rs2271439 | -1.161 | 0.734 | -0.718 | -0.802 | -0.01 | 0.042 | 0.042 | 0 | 0.78 | 0.74 | 0.75 | 0.45 | 0.79 | 0.79 |
| 11 | 1901847 | rs4980384 | 0.069 | 0.636 | -0.898 | -0.958 | 0.004 | 0.007 | 0.007 | 1.063 | 0.11 | 0.05 | 0.08 | 0.08 | 0.06 | 0.13 |
| 11 | 1905292 | rs661348 | 0.585 | 0.982 | -1.047 | -0.551 | -0.063 | -0.036 | -0.036 | 0 | 0.32 | 0.14 | 0.58 | 0.59 | 0.43 | 0.31 |
| 11 | 1906174 | rs3817197 | -0.234 | 0.865 | -0.552 | -0.967 | 0.002 | 0.038 | 0.038 | 0.02 | 0.53 | 0.37 | 0.66 | 0.8 | 0.49 | 0.54 |
| 11 | 1909006 | rs3817198 | 1.419 | 0.938 | -0.691 | -1.166 | 0.008 | 0.112 | 0.112 | 0.638 | 0.34 | 0.14 | 0.18 | 0.1 | 0.31 | 0.36 |
| 11 | 1911439 | rs17173834 | 1.035 | 1.105 | -0.18 | -1.035 | 0.024 | 0.007 | 0.007 | 0 | 0.08 | 0.23 | 0.07 | 0.09 | 0.08 | 0.1 |
| 11 | 1912726 | rs542605 | 0.155 | 1.066 | -0.128 | -1.19 | 0.013 | 0.126 | 0.126 | 0.074 | 0.52 | 0.25 | 0.65 | 0.81 | 0.49 | 0.54 |
| 11 | 1920255 | rs7483477 | 1.083 | 0.731 | -0.27 | -1.084 | 0.006 | 0.013 | 0.013 | 0 | 0.25 | 0.22 | 0.37 | 0.2 | 0.25 | 0.2 |
| 11 | 1920285 | rs4264135 | 0.497 | 0.746 | -0.009 | -1.062 | -0.002 | -0.03 | -0.03 | 0 | 0.3 | 0.44 | 0.45 | 0.24 | 0.29 | 0.27 |
| 11 | 1928217 | rs7115847 | -0.942 | 0.391 | -0.33 | -0.405 | 0.003 | -0.001 | -0.001 | 0 | 0.6 | 0.33 | 0.72 | 0.47 | 0.61 | 0.61 |
| 11 | 1941946 | rs909116 | 0.198 | 0.68 | -0.443 | -0.525 | -0.024 | 0.005 | 0.005 | 0 | 0.35 | 0.26 | 0.61 | 0.64 | 0.46 | 0.35 |
| 11 | 1942170 | rs2668870 | -0.775 | 0.918 | -0.285 | -0.53 | 0.108 | 0.003 | 0.003 | 0.289 | 0.95 | 0.78 | 0.93 | 1 | 0.9 | 0.99 |
| 11 | 1943214 | rs2734498 | -0.581 | 0.736 | -1.132 | -0.73 | 0.033 | -0.062 | -0.062 | 0.046 | 0.69 | 0.43 | 0.73 | 0.88 | 0.61 | 0.82 |
| 11 | 1943605 | rs1398256 | 0.092 | -0.177 | -0.492 | 0 | 0.149 | 0.005 | 0.005 | 1.005 | 0.88 | 0.52 | 0.84 | 0.92 | 0.81 | 0.92 |
| 11 | 1943708 | rs2334385 | 0.211 | -0.188 | -0.516 | 0.02 | 0.04 | -0.011 | -0.011 | 0 | 0.8 | 0.49 | 0.78 | 0.82 | 0.79 | 0.79 |

| | | | | | | | | | | | | | | | | |
|---|---|---|---|---|---|---|---|---|---|---|---|---|---|---|---|---|
| 11 | 1945884 | rs3781956 | -0.541 | -0.721 | -0.253 | 1.258 | -0.025 | -0.023 | -0.023 | 0 | 0.29 | 0.18 | 0.54 | 0.61 | 0.36 | 0.28 |
| 11 | 1961869 | rs11041941 | 1.508 | -1.088 | -0.065 | 1.648 | 0.003 | -0.05 | -0.05 | 3.295 | 0.4 | 0.18 | 0.31 | 0.49 | 0.55 | 0.31 |
| 11 | 1969888 | rs1618613 | -2.075 | -0.842 | 0.317 | 1.987 | -0.051 | 0.06 | 0.06 | 0 | 0.62 | 0.78 | 0.72 | 0.74 | 0.53 | 0.75 |
| 11 | 1972205 | rs2240197 | -0.385 | -1.082 | 0.493 | 2.074 | -0.007 | 0.007 | 0.007 | 0 | 0.07 | 0.08 | 0.03 | 0.14 | 0.02 | 0.17 |
| 11 | 1972919 | rs1635155 | -1.158 | -1.095 | 1.048 | 2.184 | -0.001 | -0.002 | -0.002 | 0.352 | 0.5 | 0.49 | 0.64 | 0.38 | 0.42 | 0.52 |
| 11 | 1974542 | rs217206 | -1.63 | -1.089 | 1.145 | 2.154 | 0.005 | -0.004 | -0.004 | 0 | 0.56 | 0.56 | 0.66 | 0.53 | 0.44 | 0.69 |
| 11 | 1976306 | rs11600982 | 1.601 | -0.945 | 0.82 | 2.16 | -0.082 | 0.041 | 0.041 | 0 | 0.42 | 0.13 | 0.27 | 0.35 | 0.48 | 0.36 |
| 11 | 1977552 | rs12812 | -0.387 | -1.276 | 0.628 | 1.883 | 0.004 | 0.008 | 0.008 | 0 | 0.19 | 0.01 | 0.2 | 0.14 | 0.17 | 0.25 |
| 11 | 1984911 | rs217242 | -0.474 | -1.289 | 0.581 | 1.871 | 0.009 | 0.018 | 0.018 | 0 | 0.21 | 0.01 | 0.2 | 0.14 | 0.17 | 0.25 |
| 11 | 1988576 | rs184267 | -1.373 | -1.4 | 1.036 | 1.935 | -0.012 | -0.015 | -0.015 | 0 | 0.42 | 0.45 | 0.57 | 0.3 | 0.38 | 0.44 |
| 11 | 1993975 | rs190063 | 0.768 | -1.297 | 0.982 | 1.729 | 0.017 | 0.009 | 0.009 | 0 | 0.73 | 0.79 | 0.89 | 0.79 | 0.82 | 0.64 |
| 11 | 2001895 | rs10840138 | -0.165 | -1.29 | 0.94 | 1.812 | 0 | 0 | 0 | 0.292 | 0.25 | 0.29 | 0.38 | 0.22 | 0.23 | 0.29 |
| 11 | 2003541 | rs217704 | 1.907 | -1.164 | 1.059 | 1.258 | -0.013 | 0 | 0 | 0 | 0.43 | 0.33 | 0.42 | 0.48 | 0.49 | 0.34 |
| 11 | 2009895 | rs1706879 | -1.585 | -1.308 | 0.548 | 1.396 | 0.037 | 0.013 | 0.013 | 0 | 0.47 | 0.37 | 0.45 | 0.37 | 0.37 | 0.55 |
| 11 | 2014646 | rs2285935 | 2.222 | -1.041 | 0.087 | 1.456 | -0.133 | 0.048 | 0.048 | 0.15 | 0.39 | 0.06 | 0.42 | 0.33 | 0.52 | 0.27 |
| 11 | 2014709 | rs217228 | 0.798 | -0.739 | 0.309 | 1.386 | -0.002 | -0.078 | -0.078 | 0.065 | 0.64 | 0.15 | 0.5 | 0.54 | 0.69 | 0.64 |
| 11 | 2016908 | rs217727 | 0.523 | -0.568 | 0.285 | 1.313 | -0.007 | -0.022 | -0.022 | 0 | 0.13 | 0.07 | 0.27 | 0.32 | 0.18 | 0.23 |
| 11 | 2021075 | rs2107425 | -0.948 | -0.342 | 0.489 | 1.406 | -0.013 | 0.003 | 0.003 | 0.005 | 0.39 | 0.55 | 0.47 | 0.4 | 0.3 | 0.49 |
| 11 | 2022899 | rs17658052 | -0.332 | -0.011 | -0.266 | 1.269 | -0.024 | 0.04 | 0.04 | 0 | 0.1 | 0 | 0.05 | 0.03 | 0.08 | 0.16 |
| 11 | 2024544 | rs4930103 | 0.888 | -0.079 | -0.183 | 1.349 | -0.006 | 0.013 | 0.013 | 0 | 0.47 | 0.29 | 0.45 | 0.48 | 0.53 | 0.36 |
| 11 | 2029278 | rs7950932 | -2.037 | 0.79 | 0.258 | 1.773 | -0.034 | 0.035 | 0.035 | 0 | 0.18 | 0.07 | 0.14 | 0.12 | 0.22 | 0.25 |
| 11 | 2034099 | rs7116438 | -2.195 | 1.123 | 0.597 | 2.228 | 0.015 | 0.045 | 0.045 | 0.469 | 0.2 | 0.68 | 0.17 | 0.38 | 0.22 | 0.21 |
| 11 | 2037163 | rs4384367 | -1.01 | 1 | 0.019 | 1.757 | -0.001 | 0.011 | 0.011 | 0 | 0.07 | 0.13 | 0.04 | 0.12 | 0.08 | 0.07 |
| 11 | 2040997 | rs7124169 | -1.409 | 0.976 | 0.183 | 1.748 | 0.038 | 0.201 | 0.201 | 0 | 0.17 | 0.55 | 0.18 | 0.39 | 0.17 | 0.19 |
| 11 | 2043867 | rs4930001 | -0.416 | 0.975 | -0.748 | 0.82 | 0.001 | 0.007 | 0.007 | 0 | 0.37 | 0.35 | 0.32 | 0.24 | 0.32 | 0.36 |
| 11 | 2044150 | rs7115466 | -0.858 | 1.797 | -1.135 | 0.521 | -0.038 | 0.03 | 0.03 | 0 | 0.19 | 0.09 | 0.27 | 0.16 | 0.29 | 0.23 |
| 11 | 2047640 | rs12282459 | -0.174 | 0.957 | -0.687 | 0.677 | -0.011 | -0.01 | -0.01 | 0.956 | 0.7 | 0.62 | 0.74 | 0.71 | 0.76 | 0.61 |
| 11 | 2048488 | rs4930144 | -0.806 | 1.356 | -1.04 | 1.038 | -0.011 | -0.019 | -0.019 | 0 | 0.23 | 0.18 | 0.47 | 0.72 | 0.31 | 0.43 |
| 11 | 2049317 | rs11042301 | -0.396 | 0.704 | -0.511 | 1.316 | -0.015 | 0.095 | 0.095 | 0 | 0.24 | 0.49 | 0.12 | 0.38 | 0.21 | 0.19 |
| 11 | 2054045 | rs12293815 | -0.676 | 0.258 | -0.058 | 0.889 | -0.008 | 0.001 | 0.001 | 0 | 0.23 | 0.45 | 0.18 | 0.2 | 0.24 | 0.16 |
| 11 | 2057647 | rs3858522 | 0.231 | 0.058 | -0.674 | 0.164 | 0.001 | -0.001 | -0.001 | 0 | 0.31 | 0.32 | 0.28 | 0.14 | 0.23 | 0.26 |
| 11 | 2066031 | rs7125325 | -0.383 | -0.112 | -0.649 | 0.067 | 0.012 | 0.025 | 0.025 | 0 | 0.41 | 0.35 | 0.33 | 0.24 | 0.3 | 0.34 |
| 11 | 2068717 | rs7479065 | -1.688 | -0.16 | -1.513 | 0.195 | -0.077 | -0.157 | -0.157 | 0 | 0.8 | 0.98 | 0.79 | 0.58 | 0.74 | 0.74 |
| 11 | 2068776 | rs12806495 | 0.973 | -0.212 | -1.232 | -0.268 | -0.014 | 0.06 | 0.06 | 0 | 0.26 | 0.06 | 0.23 | 0.14 | 0.19 | 0.24 |
| 11 | 2070570 | rs7932964 | 0.085 | -0.541 | -0.153 | -0.106 | 0.016 | -0.022 | -0.022 | 0.009 | 0.25 | 0.4 | 0.36 | 0.25 | 0.28 | 0.34 |
| 11 | 2073476 | rs10743110 | 0.145 | -0.977 | -0.143 | -0.42 | -0.006 | 0.051 | 0.051 | 0 | 0.48 | 0.32 | 0.59 | 0.32 | 0.46 | 0.42 |
| 11 | 2073913 | rs7925515 | -1.096 | -1.109 | -0.233 | -0.215 | 0 | -0.106 | -0.106 | 0.06 | 0.71 | 0.92 | 0.85 | 0.61 | 0.73 | 0.68 |
| 11 | 2074576 | rs10743111 | -1.124 | -1.1 | -0.182 | -0.211 | 0.002 | -0.096 | -0.096 | 0 | 0.72 | 0.92 | 0.85 | 0.6 | 0.73 | 0.68 |
| 11 | 2076043 | rs11825579 | -0.758 | -1.052 | -0.071 | -0.443 | 0 | -0.059 | -0.059 | 0 | 0.13 | 0.24 | 0.1 | 0 | 0.13 | 0.07 |
| 11 | 2087220 | rs4930030 | 0.319 | -1.059 | -0.113 | -0.398 | -0.003 | 0.05 | 0.05 | 0 | 0.49 | 0.34 | 0.61 | 0.33 | 0.47 | 0.42 |
| 11 | 2090456 | rs7130222 | -1.418 | -0.53 | 0.488 | -0.092 | 0.002 | -0.004 | -0.004 | 0 | 0.28 | 0.34 | 0.26 | 0.28 | 0.32 | 0.22 |
| 11 | 2095733 | rs6578974 | -0.62 | -0.382 | 1.001 | 0.234 | 0.049 | -0.02 | -0.02 | 0.076 | 0.08 | 0.45 | 0.07 | 0 | 0.08 | 0.03 |
| 11 | 2095799 | rs6578975 | -0.776 | -0.34 | 0.656 | 0.262 | 0.022 | 0.032 | 0.032 | 0 | 0.21 | 0.51 | 0.23 | 0.27 | 0.25 | 0.18 |
| 11 | 2095968 | rs6578976 | -0.776 | -0.34 | 0.617 | 0.262 | 0.022 | 0.032 | 0.032 | 0 | 0.21 | 0.5 | 0.23 | 0.26 | 0.25 | 0.18 |
| 11 | 2101692 | rs7949013 | -0.215 | -0.32 | 0.007 | -0.109 | 0.01 | -0.04 | -0.04 | 0 | 0.07 | 0.23 | 0.06 | 0 | 0.08 | 0.04 |
| 11 | 2107655 | rs12575480 | 0.743 | -0.287 | -0.729 | 0.025 | -0.041 | -0.052 | -0.052 | 0 | 0.13 | 0.03 | 0.15 | 0.25 | 0.18 | 0.15 |
| 11 | 2108072 | rs7924768 | -0.937 | -0.578 | -0.714 | -0.113 | -0.011 | 0.027 | 0.027 | 0 | 0.71 | 0.68 | 0.83 | 0.57 | 0.77 | 0.66 |
| 11 | 2110067 | rs7939857 | -0.04 | -0.122 | -1.576 | -0.368 | 0.002 | 0.121 | 0.121 | 0 | 0.49 | 0.17 | 0.61 | 0.32 | 0.52 | 0.48 |
| 11 | 2117403 | rs11042594 | -0.671 | -0.294 | -1.908 | -0.404 | -0.013 | -0.131 | -0.131 | 0 | 0.59 | 0.94 | 0.8 | 0.52 | 0.67 | 0.5 |
| 11 | 2126229 | rs4601751 | -0.808 | -0.061 | -0.747 | -0.378 | 0.003 | -0.02 | -0.02 | 0 | 0.65 | 0.77 | 0.8 | 0.51 | 0.68 | 0.48 |
| 11 | 2141603 | rs4341514 | 0.054 | -0.131 | -0.39 | -0.74 | 0.029 | 0.001 | 0.001 | 0 | 0.5 | 0.21 | 0.66 | 0.51 | 0.58 | 0.45 |
| 11 | 2149995 | rs11042753 | 0.228 | -0.07 | 0.043 | -0.756 | 0.007 | -0.045 | -0.045 | 0 | 0.08 | 0.14 | 0.09 | 0 | 0.1 | 0.03 |
| 11 | 2150444 | rs2585 | -1.076 | -0.176 | -0.116 | -0.802 | -0.048 | -0.136 | -0.136 | 6.372 | 0.65 | 0.92 | 0.81 | 0.51 | 0.68 | 0.47 |
| 11 | 2150697 | rs7873 | -0.303 | -0.066 | 0.497 | -0.769 | 0.048 | -0.045 | -0.045 | 0 | 0.1 | 0.29 | 0.11 | 0 | 0.1 | 0.03 |
| 11 | 2152052 | rs3802971 | 0.549 | -0.095 | -0.264 | -0.796 | 0.008 | -0.022 | -0.022 | 0.056 | 0.12 | 0.1 | 0.06 | 0.21 | 0.09 | 0.07 |
| 11 | 2154188 | rs2230949 | 0.483 | 0.014 | -0.15 | -0.957 | -0.001 | -0.009 | -0.009 | 1.226 | 0.08 | 0.03 | 0.08 | 0 | 0.09 | 0.03 |
| 11 | 2154964 | rs3213232 | -0.121 | -0.032 | 0.288 | -0.979 | 0.007 | -0.056 | -0.056 | 4.912 | 0.1 | 0.3 | 0.1 | 0 | 0.09 | 0.03 |
| 11 | 2156213 | rs734351 | -0.717 | -0.118 | -0.204 | -1.093 | 0 | 0.16 | 0.16 | 0 | 0.51 | 0.81 | 0.73 | 0.29 | 0.58 | 0.37 |
| 11 | 2156930 | rs3213223 | 0.136 | 1.046 | -0.311 | -1.07 | -0.048 | -0.012 | -0.012 | 0 | 0.08 | 0.01 | 0.15 | 0.09 | 0.21 | 0.09 |
| 11 | 2158179 | rs3213216 | -0.569 | 0.503 | -0.248 | -1.081 | -0.005 | 0.039 | 0.039 | 0 | 0.62 | 0.89 | 0.6 | 0.69 | 0.66 | 0.69 |
| 11 | 2161858 | rs3741212 | 0.835 | 1.395 | 0.385 | -1.141 | 0.066 | -0.046 | -0.046 | 0 | 0.8 | 0.39 | 0.71 | 0.89 | 0.68 | 0.84 |
| 11 | 2165173 | rs11603378 | -0.529 | 1.504 | 0.698 | -1.101 | 0.097 | -0.009 | -0.009 | 0 | 0.13 | 0.3 | 0.19 | 0.11 | 0.25 | 0.13 |
| 11 | 2167543 | rs1003483 | 0.12 | 1.066 | -0.095 | -1.216 | -0.064 | 0.001 | 0.001 | 0 | 0.45 | 0.11 | 0.57 | 0.4 | 0.46 | 0.33 |
| 11 | 2167618 | rs1003484 | -1.097 | 0.924 | 0.206 | -1.349 | -0.007 | 0.008 | 0.008 | 0 | 0.63 | 0.58 | 0.78 | 0.5 | 0.7 | 0.48 |
| 11 | 2167641 | rs17880206 | 2.108 | 0.734 | -0.194 | -1.588 | -0.012 | 0.039 | 0.039 | 0 | 0.07 | 0.01 | 0.04 | 0 | 0.05 | 0.02 |
| 11 | 2167850 | rs17885785 | 0.603 | 1.178 | 0.152 | -1.274 | -0.057 | 0.025 | 0.025 | 0 | 0.1 | 0.01 | 0.14 | 0.04 | 0.19 | 0.11 |
| 11 | 2167981 | rs2239681 | 1.034 | 1.094 | 0.384 | -1.249 | 0.021 | -0.045 | -0.045 | 0.64 | 0.34 | 0.19 | 0.19 | 0.5 | 0.3 | 0.51 |
| 11 | 2169014 | rs10770125 | -0.051 | 1.125 | 0.298 | -1.28 | -0.047 | 0.019 | 0.019 | 0 | 0.47 | 0.12 | 0.58 | 0.4 | 0.46 | 0.33 |
| 11 | 2169110 | rs3741211 | -0.557 | 1.168 | 0.46 | -1.39 | 0.054 | 0.023 | 0.023 | 0.185 | 0.26 | 0.41 | 0.26 | 0.3 | 0.35 | 0.23 |

| | | | | | | | | | | | | | | | | |
|---|---|---|---|---|---|---|---|---|---|---|---|---|---|---|---|---|
| 11 | 2169774 | rs3741208 | -0.4 | 1.144 | 0.49 | -1.405 | 0.011 | 0.042 | 0.042 | 0.075 | 0.61 | 0.65 | 0.45 | 0.8 | 0.66 | 0.75 |
| 11 | 2169864 | rs3741206 | 0.352 | 1.188 | 0.505 | -1.407 | 0.006 | 0.063 | 0.063 | 0 | 0.37 | 0.28 | 0.54 | 0.2 | 0.34 | 0.25 |
| 11 | 2169884 | rs3741205 | -1.23 | 1.199 | 0.523 | -1.403 | 0.008 | -0.01 | -0.01 | 0 | 0.63 | 0.68 | 0.8 | 0.5 | 0.7 | 0.49 |
| 11 | 2170143 | rs1004446 | -0.591 | 1.253 | 0.267 | -1.225 | 0.063 | 0.042 | 0.042 | 0.305 | 0.26 | 0.43 | 0.27 | 0.31 | 0.35 | 0.23 |
| 11 | 2171601 | rs4320932 | 1.655 | 0.958 | -0.228 | -1.561 | 0.02 | -0.097 | -0.097 | 0 | 0.27 | 0.12 | 0.13 | 0.49 | 0.22 | 0.4 |
| 11 | 2171922 | rs10840442 | -0.091 | 1.245 | -0.571 | -1.551 | 0.006 | -0.032 | -0.032 | 0 | 0.12 | 0.15 | 0.1 | 0.01 | 0.18 | 0.15 |
| 11 | 2173447 | rs7924316 | -0.339 | 1.295 | -0.548 | -1.304 | 0.015 | 0.022 | 0.022 | 0 | 0.5 | 0.33 | 0.63 | 0.45 | 0.52 | 0.47 |
| 11 | 2176042 | rs10840447 | -0.288 | 1.311 | -1.32 | -1.376 | 0.013 | 0.16 | 0.16 | 0 | 0.39 | 0.15 | 0.37 | 0.14 | 0.36 | 0.31 |
| 11 | 2179313 | rs3842762 | -0.303 | 1.403 | -0.541 | -0.644 | -0.001 | -0.011 | -0.011 | 0.354 | 0.07 | 0.11 | 0.05 | 0.01 | 0.05 | 0.02 |
| 11 | 2181073 | rs3842752 | -0.154 | 1.951 | -0.417 | -0.425 | 0.032 | -0.006 | -0.006 | 0 | 0.09 | 0.09 | 0.17 | 0.05 | 0.21 | 0.13 |
| 11 | 2186335 | rs2070762 | -0.327 | 1.08 | -0.68 | -0.366 | -0.038 | -0.022 | -0.022 | 0 | 0.57 | 0.2 | 0.63 | 0.47 | 0.48 | 0.48 |
| 11 | 2190519 | rs7483056 | -0.262 | 0.934 | -0.142 | -0.362 | 0 | -0.002 | -0.002 | 0.07 | 0.55 | 0.45 | 0.62 | 0.51 | 0.44 | 0.47 |
| 11 | 2192798 | rs10840489 | -0.378 | 0.856 | 0.3 | -0.472 | 0.002 | -0.009 | -0.009 | 0 | 0.12 | 0.16 | 0.22 | 0.11 | 0.12 | 0.17 |
| 11 | 2193840 | rs10770141 | 0.738 | 1.232 | 0.601 | -0.926 | 0.053 | -0.087 | -0.087 | 0.05 | 0.73 | 0.37 | 0.62 | 0.92 | 0.64 | 0.83 |
| 11 | 2194390 | rs10840491 | -0.525 | 1.031 | 0.425 | -0.949 | -0.002 | -0.008 | -0.008 | 0 | 0.14 | 0.13 | 0.22 | 0.12 | 0.12 | 0.18 |
| 11 | 2198418 | rs11042978 | 1.514 | 0.896 | 0.763 | -1.011 | -0.025 | 0.064 | 0.064 | 0 | 0.59 | 0.29 | 0.41 | 0.79 | 0.52 | 0.65 |
| 11 | 2200329 | rs11564710 | 1.748 | 0.594 | 0.537 | -0.819 | -0.073 | 0.089 | 0.089 | 0.864 | 0.61 | 0.09 | 0.43 | 0.78 | 0.6 | 0.68 |
| 11 | 2201163 | rs6578993 | 0.278 | 0.423 | 0.658 | -0.786 | -0.001 | 0.001 | 0.001 | 0 | 0.87 | 0.93 | 0.77 | 0.88 | 0.85 | 0.83 |
| 11 | 2206027 | rs11043016 | -0.876 | 0.578 | 1.187 | -0.742 | 0.032 | -0.058 | -0.058 | 0.358 | 0.22 | 0.55 | 0.3 | 0.1 | 0.23 | 0.14 |
| 11 | 2207209 | rs11564703 | 1.636 | 0.472 | 0.43 | -0.735 | -0.062 | 0.068 | 0.068 | 1.34 | 0.62 | 0.09 | 0.44 | 0.76 | 0.61 | 0.68 |
| 11 | 2213166 | rs7111341 | -1.008 | 0.603 | 1.281 | -0.622 | 0.053 | -0.052 | -0.052 | 0.075 | 0.21 | 0.41 | 0.21 | 0.1 | 0.26 | 0.14 |
| 11 | 2219962 | rs3922756 | 1.461 | 0.48 | 0.674 | -0.536 | -0.066 | 0.065 | 0.065 | 0.587 | 0.67 | 0.09 | 0.37 | 0.8 | 0.67 | 0.72 |
| 11 | 2220558 | rs7946543 | -1.022 | 0.505 | 1.627 | -0.43 | 0.07 | -0.022 | -0.022 | 0.565 | 0.1 | 0.41 | 0.06 | 0.03 | 0.08 | 0.01 |
| 11 | 2220754 | rs10840546 | -1.745 | 0.498 | 1.709 | -0.438 | 0.049 | -0.019 | -0.019 | 0 | 0.13 | 0.49 | 0.25 | 0.09 | 0.11 | 0.05 |
| 11 | 2222913 | rs12292212 | 0.258 | 0.589 | 1.294 | -0.412 | -0.016 | 0.019 | 0.019 | 0 | 0.06 | 0.04 | 0.03 | 0.03 | 0.08 | 0.01 |
| 11 | 2229759 | rs10743181 | 0.42 | 0.522 | 1.518 | -0.462 | 0.055 | -0.002 | -0.002 | 0 | 0.85 | 0.61 | 0.66 | 0.9 | 0.83 | 0.77 |
| 11 | 2229782 | rs10743182 | 0.837 | 0.489 | 1.644 | -0.478 | 0.021 | -0.021 | -0.021 | 0 | 0.83 | 0.42 | 0.64 | 0.89 | 0.83 | 0.77 |
| 11 | 2231273 | rs10840595 | 0.17 | -0.518 | 0.863 | -0.523 | 0 | -0.004 | -0.004 | 0.35 | 0.36 | 0.35 | 0.3 | 0.45 | 0.16 | 0.3 |
| 11 | 2232229 | rs6579002 | 0.513 | 0.208 | 0.978 | -0.595 | 0.028 | -0.02 | -0.02 | 1.212 | 0.83 | 0.67 | 0.66 | 0.9 | 0.82 | 0.75 |
| 11 | 2233574 | rs7127900 | 0.588 | 0.206 | 0.831 | -0.596 | 0.038 | -0.025 | -0.025 | 0 | 0.82 | 0.59 | 0.65 | 0.89 | 0.82 | 0.75 |
| 11 | 2233951 | rs11043140 | 1.007 | 0.185 | 0.429 | -0.572 | 0.041 | -0.053 | -0.053 | 0.003 | 0.76 | 0.48 | 0.61 | 0.86 | 0.74 | 0.75 |
| 11 | 2236689 | rs11021705 | -0.619 | -0.291 | -0.101 | -0.798 | 0.001 | -0.08 | -0.08 | 3.744 | 0.28 | 0.45 | 0.19 | 0.11 | 0.24 | 0.35 |
| 11 | 2238288 | rs10831551 | 0.523 | -1.095 | -0.651 | -0.555 | 0.075 | -0.127 | -0.127 | 0 | 0.2 | 0.03 | 0.05 | 0.34 | 0.07 | 0.14 |
| 11 | 2240416 | rs7480235 | 1.294 | -1.149 | -0.578 | -0.625 | 0.021 | -0.024 | -0.024 | 0.187 | 0.45 | 0.22 | 0.39 | 0.48 | 0.65 | 0.4 |
| 11 | 2241272 | rs11021732 | -1.09 | -1.16 | -0.383 | -0.629 | 0.038 | -0.002 | -0.002 | 0 | 0.48 | 0.56 | 0.24 | 0.45 | 0.31 | 0.48 |
| 11 | 2242612 | rs10831570 | 0.42 | -1.149 | -0.305 | -0.593 | 0.011 | -0.016 | -0.016 | 0 | 0.21 | 0.19 | 0.07 | 0.34 | 0.07 | 0.14 |
| 11 | 2250149 | rs4644645 | 0.402 | -1.173 | -0.382 | -0.701 | -0.002 | 0.001 | 0.001 | 0 | 0.13 | 0.12 | 0.22 | 0.08 | 0.16 | 0.24 |
| 11 | 2250672 | rs12365075 | 0.496 | -1.01 | -0.569 | -0.814 | 0.007 | -0.019 | -0.019 | 0 | 0.08 | 0.17 | 0.07 | 0.04 | 0.13 | 0.01 |
| 11 | 2250713 | rs4595544 | 1.896 | -1.043 | -0.605 | -0.802 | -0.162 | -0.116 | -0.116 | 0 | 0.38 | 0.04 | 0.32 | 0.45 | 0.53 | 0.4 |
| 11 | 2253079 | rs7949253 | -0.682 | -1.222 | -0.454 | -0.718 | 0.045 | -0.009 | -0.009 | 0.487 | 0.39 | 0.62 | 0.15 | 0.38 | 0.16 | 0.33 |
| 11 | 2260854 | rs11021859 | 1.126 | -1.101 | -0.712 | -0.6 | 0.114 | -0.013 | -0.013 | 0.786 | 0.51 | 0.16 | 0.54 | 0.55 | 0.69 | 0.64 |
| 11 | 2261338 | rs11021861 | 0.413 | -1.567 | -0.557 | -0.333 | 0.069 | -0.126 | -0.126 | 1.547 | 0.2 | 0.01 | 0.05 | 0.32 | 0.07 | 0.14 |
| 11 | 2281501 | rs7941585 | -0.069 | -1.205 | 0.179 | -0.657 | -0.042 | -0.121 | -0.121 | 0 | 0.18 | 0.35 | 0.09 | 0.03 | 0.11 | 0.19 |
| 11 | 2282360 | rs11021984 | 0.524 | -1.112 | -0.485 | -0.628 | 0.006 | 0.062 | 0.062 | 0.152 | 0.13 | 0.01 | 0.05 | 0.03 | 0.08 | 0.18 |
| 11 | 2283459 | rs17737713 | 0.216 | -0.867 | -0.556 | -0.568 | -0.015 | 0.014 | 0.014 | 0 | 0.12 | 0.12 | 0.2 | 0.09 | 0.15 | 0.23 |
| 11 | 2291715 | rs7126721 | -0.701 | -1.715 | -0.35 | -0.126 | -0.027 | 0.012 | 0.012 | 0 | 0.31 | 0.71 | 0.33 | 0.46 | 0.12 | 0.24 |
| 11 | 2298259 | rs4929971 | -0.001 | -1.737 | -0.757 | -0.052 | 0.049 | -0.068 | -0.068 | 0.189 | 0.24 | 0.13 | 0.27 | 0.44 | 0.09 | 0.18 |
| 11 | 2299035 | rs1011476 | -0.039 | -1.778 | -1.082 | -0.346 | -0.051 | 0.093 | 0.093 | 0 | 0.24 | 0.04 | 0.26 | 0.11 | 0.25 | 0.34 |
| 11 | 2304502 | rs10765903 | -1.648 | -1.435 | -0.949 | -0.203 | -0.063 | -0.08 | -0.08 | 0.111 | 0.58 | 0.96 | 0.62 | 0.58 | 0.37 | 0.6 |
| 11 | 2305638 | rs4930074 | -0.935 | -1.192 | -0.615 | -0.008 | 0.048 | 0.025 | 0.025 | 0 | 0.48 | 0.28 | 0.54 | 0.58 | 0.32 | 0.52 |
| 11 | 2306026 | rs10831706 | -0.469 | -1.723 | 0.171 | 0.396 | -0.076 | 0.092 | 0.092 | 0 | 0.27 | 0.51 | 0.33 | 0.53 | 0.12 | 0.22 |
| 11 | 2306168 | rs7104786 | 0.007 | -0.727 | 0.009 | 0.269 | 0.007 | 0.021 | 0.021 | 0 | 0.35 | 0.5 | 0.3 | 0.44 | 0.37 | 0.58 |
| 11 | 2306339 | rs2521259 | -0.055 | -0.854 | -0.06 | 0.757 | -0.001 | -0.002 | -0.002 | 0.02 | 0.54 | 0.49 | 0.47 | 0.37 | 0.53 | 0.42 |
| 11 | 2308762 | rs2521263 | 0.192 | -1.405 | 0.667 | 1.234 | -0.01 | 0.319 | 0.319 | 0 | 0.88 | 0.67 | 0.71 | 0.51 | 0.92 | 0.89 |
| 11 | 2309898 | rs1029409 | 0.061 | -1.408 | 0.451 | 0.575 | 0.001 | 0.047 | 0.047 | 0 | 0.49 | 0.42 | 0.42 | 0.19 | 0.53 | 0.54 |
| 11 | 2310605 | rs756919 | -0.041 | -1.42 | 0.454 | 0.795 | -0.044 | 0.351 | 0.351 | 0 | 0.87 | 0.64 | 0.81 | 0.52 | 0.93 | 0.89 |
| 11 | 2311152 | rs7944004 | 0.149 | -1.233 | 0.383 | -0.335 | -0.004 | 0.003 | 0.003 | 0 | 0.45 | 0.45 | 0.41 | 0.18 | 0.5 | 0.49 |
| 11 | 2315404 | rs7941928 | 0.754 | -1.426 | 0.618 | -0.179 | 0.008 | -0.028 | -0.028 | 0 | 0.35 | 0.38 | 0.3 | 0.19 | 0.36 | 0.42 |
| 11 | 2316720 | rs2651836 | 0.655 | -1.505 | 0.876 | 0.315 | -0.015 | 0.071 | 0.071 | 0 | 0.79 | 0.69 | 0.71 | 0.4 | 0.79 | 0.81 |
| 11 | 2318867 | rs2074019 | 0.426 | -1.701 | 0.839 | 0.15 | 0.002 | 0.021 | 0.021 | 0 | 0.45 | 0.6 | 0.34 | 0.19 | 0.41 | 0.45 |
| 11 | 2320884 | rs1501466 | -1.236 | -1.667 | 1.006 | 0.203 | 0.071 | -0.009 | -0.009 | 0 | 0.11 | 0.39 | 0.23 | 0.07 | 0.16 | 0.16 |
| 11 | 2322136 | rs2077078 | -0.316 | -1.662 | 0.712 | 0.197 | 0.043 | -0.03 | -0.03 | 0 | 0.08 | 0.26 | 0.11 | 0.01 | 0.13 | 0.07 |
| 11 | 2322662 | rs756923 | -0.095 | -1.692 | -0.03 | 0.306 | 0.007 | -0.068 | -0.068 | 2.879 | 0.85 | 0.95 | 0.71 | 0.34 | 0.89 | 0.84 |
| 11 | 2322829 | rs11022157 | 1.153 | -1.762 | -0.071 | 0.238 | -0.08 | 0.084 | 0.084 | 0.023 | 0.32 | 0.08 | 0.26 | 0.14 | 0.36 | 0.27 |
| 11 | 2323029 | rs756924 | -0.297 | -1.717 | 0.144 | 0.601 | 0.029 | -0.056 | -0.056 | 0.068 | 0.87 | 0.96 | 0.83 | 0.41 | 0.93 | 0.88 |
| 11 | 2324041 | rs2074022 | -0.297 | -1.717 | 0.128 | 0.598 | 0.029 | -0.056 | -0.056 | 0 | 0.87 | 0.96 | 0.83 | 0.41 | 0.93 | 0.88 |
| 11 | 2324440 | rs10831733 | 0.481 | -1.681 | 0.369 | 0.261 | -0.008 | 0.051 | 0.051 | 0 | 0.39 | 0.23 | 0.31 | 0.19 | 0.38 | 0.4 |
| 11 | 2329078 | rs756926 | 0.708 | -2.382 | 1.194 | 1.562 | 0.017 | -0.02 | -0.02 | 0 | 0.85 | 0.93 | 0.81 | 0.38 | 0.93 | 0.83 |
| 11 | 2332037 | rs808963 | 1.435 | -2.403 | 2.059 | 1.489 | -0.045 | 0.085 | 0.085 | 0 | 0.81 | 0.58 | 0.67 | 0.32 | 0.9 | 0.78 |

| | | | | | | | | | | | | | | | | |
|---|---|---|---|---|---|---|---|---|---|---|---|---|---|---|---|---|
| 11 | 2339737 | rs2521282 | 1.214 | -2.341 | 2.152 | 1.64 | -0.019 | 0.044 | 0.044 | 0 | 0.8 | 0.75 | 0.78 | 0.42 | 0.83 | 0.79 |
| 11 | 2339822 | rs4930081 | -2.019 | -2.644 | 2.703 | 1.511 | -0.011 | 0.07 | 0.07 | 0.022 | 0.12 | 0.35 | 0.19 | 0.16 | 0.08 | 0.12 |
| 11 | 2341732 | rs2651843 | 1.881 | -2.657 | 2.633 | 1.509 | -0.07 | 0.051 | 0.051 | 0 | 0.7 | 0.51 | 0.72 | 0.42 | 0.78 | 0.72 |
| 11 | 2348778 | rs2521294 | 0.836 | -2.656 | 2.539 | 1.528 | 0.006 | -0.008 | -0.008 | 0 | 0.82 | 0.9 | 0.78 | 0.44 | 0.83 | 0.81 |
| 11 | 2349462 | rs2651803 | 1.873 | -2.6 | 2.381 | 1.284 | -0.027 | -0.015 | -0.015 | 0 | 0.68 | 0.42 | 0.59 | 0.26 | 0.75 | 0.67 |
| 11 | 2364234 | rs800123 | 1.915 | -2.62 | 2.431 | 1.277 | -0.023 | -0.01 | -0.01 | 0 | 0.67 | 0.38 | 0.59 | 0.28 | 0.75 | 0.67 |
| 11 | 2373515 | rs800153 | 1.112 | -2.232 | 2.305 | 1.647 | -0.003 | 0.014 | 0.014 | 0 | 0.82 | 0.79 | 0.68 | 0.39 | 0.83 | 0.81 |
| 11 | 2386600 | rs808964 | 2.361 | -2.304 | 2.481 | 1.552 | -0.003 | -0.001 | -0.001 | 0 | 0.7 | 0.3 | 0.6 | 0.3 | 0.76 | 0.69 |
| 11 | 2386636 | rs7116047 | -2.584 | -1.868 | 2.717 | 1.627 | -0.004 | 0.039 | 0.039 | 1.148 | 0.13 | 0.52 | 0.09 | 0.12 | 0.07 | 0.12 |
| 11 | 2396666 | rs708154 | 0.414 | -2.082 | 1.928 | 1.898 | 0.04 | -0.133 | -0.133 | 0.038 | 0.79 | 0.96 | 0.66 | 0.52 | 0.85 | 0.76 |
| 11 | 2396995 | rs708155 | 0.602 | -2.067 | 2.097 | 1.887 | 0.015 | -0.054 | -0.054 | 0.482 | 0.77 | 0.85 | 0.64 | 0.51 | 0.83 | 0.72 |
| 11 | 2402747 | rs2019938 | 1.204 | -2.184 | 2.202 | 1.867 | -0.034 | 0.035 | 0.035 | 0 | 0.69 | 0.41 | 0.61 | 0.45 | 0.81 | 0.66 |
| 11 | 2408523 | rs11022565 | -1.1 | -2.067 | 2.337 | 1.825 | 0.011 | 0.008 | 0.008 | 0 | 0.05 | 0.26 | 0.03 | 0.03 | 0.01 | 0.04 |
| 11 | 2410099 | rs708564 | 0.989 | -2.021 | 1.601 | 1.883 | 0.039 | -0.044 | -0.044 | 0 | 0.68 | 0.24 | 0.58 | 0.47 | 0.8 | 0.65 |
| 11 | 2420500 | rs739673 | -1.223 | -1.448 | 1.777 | 1.494 | 0.031 | 0.163 | 0.163 | 0 | 0.06 | 0.29 | 0.07 | 0.11 | 0.02 | 0.02 |
| 11 | 2429733 | rs800345 | 0.71 | -1.53 | -0.17 | 1.687 | 0 | -0.002 | -0.002 | 0 | 0.77 | 0.81 | 0.69 | 0.52 | 0.92 | 0.75 |
| 11 | 2432964 | rs2074234 | 0.092 | -0.848 | -0.58 | 1.163 | 0.034 | -0.043 | -0.043 | 0 | 0.39 | 0.2 | 0.37 | 0.46 | 0.35 | 0.32 |
| 11 | 2437425 | rs2301698 | -0.891 | -0.245 | 0.105 | 0.812 | 0 | 0.003 | 0.003 | 0 | 0.48 | 0.47 | 0.58 | 0.66 | 0.48 | 0.48 |
| 11 | 2439767 | rs886277 | -0.781 | -0.292 | 0.235 | 1.014 | 0.006 | 0.009 | 0.009 | 0.061 | 0.4 | 0.64 | 0.54 | 0.66 | 0.37 | 0.44 |
| 11 | 2440989 | rs757091 | -0.689 | 0.099 | 0.469 | 1.221 | 0 | 0.001 | 0.001 | 0 | 0.36 | 0.51 | 0.52 | 0.61 | 0.36 | 0.44 |
| 11 | 2447855 | rs4930105 | -0.732 | -0.068 | 0.183 | 1.188 | -0.006 | 0.01 | 0.01 | 0 | 0.27 | 0.29 | 0.42 | 0.34 | 0.26 | 0.25 |
| 11 | 2469915 | rs12287029 | -1.766 | 0.104 | 0.886 | 0.588 | 0.044 | -0.055 | -0.055 | 5.616 | 0.11 | 0.56 | 0.09 | 0 | 0.12 | 0.08 |
| 11 | 2474956 | rs12274567 | -2.146 | -0.088 | 0.868 | 0.417 | 0.001 | -0.033 | -0.033 | 0.952 | 0.09 | 0.13 | 0.06 | 0 | 0.11 | 0.08 |
| 11 | 2479195 | rs11022922 | -1.633 | -0.879 | 0.478 | 0.985 | 0.037 | -0.031 | -0.031 | 0 | 0.31 | 0.81 | 0.36 | 0.37 | 0.22 | 0.43 |
| 11 | 2482092 | rs3864884 | -1.47 | -1.098 | 0.856 | 0.991 | -0.037 | 0.086 | 0.086 | 2.005 | 0.26 | 0.53 | 0.32 | 0.38 | 0.21 | 0.34 |
| 11 | 2482555 | rs2075873 | -1.149 | -1.197 | 0.984 | 1.25 | -0.001 | 0.057 | 0.057 | 0.19 | 0.07 | 0.09 | 0.13 | 0.28 | 0.07 | 0.22 |
| 11 | 2485486 | rs7939542 | -1.912 | -1.228 | 1.138 | 1.278 | -0.026 | 0.098 | 0.098 | 2.111 | 0.24 | 0.43 | 0.32 | 0.36 | 0.22 | 0.3 |
| 11 | 2486120 | rs16928297 | -2.419 | -1.206 | 1.092 | 1.245 | -0.038 | 0.034 | 0.034 | 1.691 | 0.31 | 0.57 | 0.35 | 0.37 | 0.24 | 0.42 |
| 11 | 2492629 | rs11601717 | 1.348 | -0.86 | 0.045 | 1.379 | -0.014 | -0.042 | -0.042 | 0 | 0.59 | 0.22 | 0.43 | 0.56 | 0.6 | 0.58 |
| 11 | 2495515 | rs3815064 | 1.085 | -0.738 | 0.194 | 1.316 | -0.01 | 0.003 | 0.003 | 0 | 0.65 | 0.33 | 0.56 | 0.65 | 0.63 | 0.67 |
| 11 | 2502319 | rs12576239 | -0.873 | -0.728 | 0.269 | 1.201 | -0.002 | -0.011 | -0.011 | 1.717 | 0.14 | 0.18 | 0.18 | 0.1 | 0.15 | 0.08 |
| 11 | 2505733 | rs4578370 | -1.058 | -0.782 | 0.016 | 1.503 | -0.023 | 0.049 | 0.049 | 3.731 | 0.11 | 0.22 | 0.14 | 0.23 | 0.07 | 0.22 |
| 11 | 2511527 | rs1080015 | 1.203 | -0.761 | -0.115 | 1.248 | -0.005 | 0.004 | 0.004 | 0 | 0.61 | 0.28 | 0.54 | 0.61 | 0.61 | 0.57 |
| 11 | 2526898 | rs179407 | -1.147 | -0.99 | 0.308 | 0.958 | -0.01 | -0.023 | -0.023 | 0 | 0.13 | 0.16 | 0.09 | 0.02 | 0.08 | 0.05 |
| 11 | 2527361 | rs11023094 | -1.844 | -1.123 | 1.098 | 1.231 | -0.006 | 0.067 | 0.067 | 2.192 | 0.23 | 0.61 | 0.32 | 0.44 | 0.2 | 0.33 |
| 11 | 2528003 | rs11023096 | -1.275 | -0.962 | 1.816 | 1.264 | 0.008 | 0.036 | 0.036 | 0.568 | 0.17 | 0.21 | 0.3 | 0.39 | 0.22 | 0.3 |
| 11 | 2528233 | rs4929992 | 1.954 | -1.549 | 1.223 | 0.604 | -0.028 | 0.049 | 0.049 | 0.015 | 0.44 | 0.18 | 0.37 | 0.38 | 0.51 | 0.3 |
| 11 | 2531256 | rs2012618 | 0.297 | -1.327 | 0.76 | 0.418 | -0.032 | 0.007 | 0.007 | 0 | 0.7 | 0.86 | 0.71 | 0.79 | 0.74 | 0.65 |
| 11 | 2534483 | rs2011750 | -2.121 | -1.823 | 1.086 | 0.288 | -0.028 | 0.037 | 0.037 | 1.543 | 0.3 | 0.4 | 0.32 | 0.43 | 0.21 | 0.35 |
| 11 | 2536131 | rs7924915 | -0.881 | -1.915 | 1.286 | 0.316 | -0.021 | -0.013 | -0.013 | 0 | 0.16 | 0.19 | 0.08 | 0.04 | 0.07 | 0.11 |
| 11 | 2537751 | rs2237869 | -1.197 | -1.896 | 1.299 | 0.339 | 0.011 | 0.065 | 0.065 | 0 | 0.17 | 0.26 | 0.26 | 0.38 | 0.17 | 0.25 |
| 11 | 2538580 | rs736609 | 2.328 | -1.845 | 1.335 | 0.38 | -0.005 | 0.004 | 0.004 | 0 | 0.43 | 0.4 | 0.4 | 0.39 | 0.56 | 0.32 |
| 11 | 2539107 | rs729979 | 0.228 | -1.485 | 1.096 | 0.484 | 0 | 0.011 | 0.011 | 0 | 0.72 | 0.76 | 0.72 | 0.82 | 0.77 | 0.67 |
| 11 | 2541165 | rs11826873 | -1.303 | -1.413 | 0.868 | 0.53 | 0.01 | 0.064 | 0.064 | 0 | 0.17 | 0.25 | 0.26 | 0.38 | 0.17 | 0.25 |
| 11 | 2544871 | rs2283152 | 0.626 | -1.368 | 0.644 | 0.505 | -0.011 | -0.019 | -0.019 | 0 | 0.64 | 0.75 | 0.5 | 0.48 | 0.65 | 0.46 |
| 11 | 2548175 | rs2283155 | 0.691 | -1.465 | -0.343 | 0.292 | -0.02 | -0.007 | -0.007 | 0 | 0.58 | 0.73 | 0.64 | 0.53 | 0.55 | 0.46 |
| 11 | 2550730 | rs179429 | 0.211 | -1.915 | -0.173 | -0.136 | 0.001 | -0.02 | -0.02 | 0 | 0.78 | 0.79 | 0.9 | 0.97 | 0.81 | 0.81 |
| 11 | 2552148 | rs179431 | 1.45 | -2.095 | -0.04 | 0.049 | 0.003 | 0.002 | 0.002 | 0.696 | 0.58 | 0.39 | 0.68 | 0.7 | 0.7 | 0.6 |
| 11 | 2553703 | rs179435 | 0.904 | -1.81 | -0.193 | 0.184 | -0.006 | -0.003 | -0.003 | 0.036 | 0.67 | 0.58 | 0.74 | 0.72 | 0.77 | 0.61 |
| 11 | 2560335 | rs179441 | -0.212 | -0.858 | -0.555 | 0.434 | -0.008 | -0.005 | -0.005 | 0.079 | 0.9 | 0.92 | 0.92 | 0.77 | 0.86 | 0.83 |
| 11 | 2562231 | rs1997619 | -0.411 | -1.603 | -0.562 | 0.499 | -0.008 | 0.002 | 0.002 | 0 | 0.2 | 0.28 | 0.22 | 0.3 | 0.12 | 0.22 |
| 11 | 2562501 | rs9299929 | -2.326 | -1.883 | -0.874 | 0.173 | -0.038 | -0.012 | -0.012 | 0.023 | 0.56 | 0.81 | 0.5 | 0.58 | 0.45 | 0.58 |
| 11 | 2563137 | rs3944148 | 0.052 | -1.985 | -0.719 | 0.272 | -0.002 | -0.001 | -0.001 | 0 | 0.19 | 0.2 | 0.21 | 0.31 | 0.12 | 0.21 |
| 11 | 2566400 | rs7130209 | -2.013 | -1.994 | -0.968 | 0.187 | 0.065 | 0.015 | 0.015 | 0 | 0.48 | 0.79 | 0.42 | 0.59 | 0.32 | 0.56 |
| 11 | 2569989 | rs2283161 | -0.055 | -1.331 | -0.885 | 0.391 | -0.008 | 0.004 | 0.004 | 0 | 0.2 | 0.22 | 0.21 | 0.3 | 0.12 | 0.25 |
| 11 | 2570845 | rs12292841 | -1.331 | -0.542 | -0.713 | 0.113 | 0.025 | 0 | 0 | 0 | 0.21 | 0.43 | 0.16 | 0.2 | 0.18 | 0.29 |
| 11 | 2573121 | rs4539307 | -1.505 | 0.065 | -1.593 | 0.447 | -0.029 | -0.018 | -0.018 | 0.168 | 0.21 | 0.13 | 0.15 | 0.24 | 0.25 | 0.21 |
| 11 | 2574708 | rs10430833 | -0.836 | -0.157 | -1.21 | 0.866 | -0.004 | -0.003 | -0.003 | 0 | 0.13 | 0.13 | 0.1 | 0.24 | 0.14 | 0.19 |
| 11 | 2579889 | rs2283166 | -0.43 | -1.502 | -1.484 | 0.419 | 0.062 | 0.031 | 0.031 | 0 | 0.29 | 0.16 | 0.14 | 0.18 | 0.14 | 0.36 |
| 11 | 2580299 | rs2283169 | -0.72 | -1.48 | -1.359 | 0.496 | 0.125 | -0.002 | -0.002 | 1.726 | 0.49 | 0.76 | 0.48 | 0.48 | 0.27 | 0.56 |
| 11 | 2580737 | rs7936703 | -0.694 | -0.826 | -1.512 | 0.507 | 0.102 | 0.006 | 0.006 | 0 | 0.49 | 0.81 | 0.52 | 0.46 | 0.34 | 0.55 |
| 11 | 2583141 | rs2283170 | -0.553 | -0.834 | -0.838 | 0.483 | 0.029 | 0 | 0 | 0 | 0.47 | 0.63 | 0.5 | 0.47 | 0.33 | 0.54 |
| 11 | 2598552 | rs12282937 | -0.715 | -0.304 | -0.785 | -0.096 | 0 | -0.002 | -0.002 | 0.009 | 0.43 | 0.56 | 0.61 | 0.82 | 0.43 | 0.55 |
| 11 | 2603026 | rs2283174 | -0.989 | -0.441 | -0.456 | 0.599 | -0.004 | -0.001 | -0.001 | 2.562 | 0.27 | 0.73 | 0.34 | 0.26 | 0.18 | 0.31 |
| 11 | 2604437 | rs2012323 | -0.937 | 0.026 | -0.891 | 0.077 | -0.052 | 0.061 | 0.061 | 0.603 | 0.7 | 0.84 | 0.76 | 0.92 | 0.55 | 0.82 |
| 11 | 2607850 | rs12808620 | -0.421 | -0.304 | -0.129 | 0.614 | 0.009 | 0.011 | 0.011 | 0 | 0.14 | 0.11 | 0.17 | 0.06 | 0.11 | 0.13 |
| 11 | 2610466 | rs12804487 | -0.629 | -0.184 | -0.02 | 0.712 | -0.001 | 0.003 | 0.003 | 0 | 0.18 | 0.22 | 0.27 | 0.21 | 0.16 | 0.24 |
| 11 | 2612561 | rs2283175 | -0.663 | -0.673 | -0.043 | 1.479 | -0.004 | 0.026 | 0.026 | 0 | 0.18 | 0.21 | 0.4 | 0.5 | 0.13 | 0.3 |
| 11 | 2617782 | rs1116714 | -1.245 | -0.701 | 0.507 | 0.523 | -0.038 | 0.023 | 0.023 | 0.057 | 0.22 | 0.62 | 0.32 | 0.28 | 0.13 | 0.3 |

| | | | | | | | | | | | | | | | | |
|---|---|---|---|---|---|---|---|---|---|---|---|---|---|---|---|---|
| 11 | 2619390 | rs4930129 | -0.814 | -0.662 | 0.695 | 0.511 | -0.03 | 0.035 | 0.035 | 0 | 0.19 | 0.36 | 0.3 | 0.26 | 0.13 | 0.24 |
| 11 | 2623009 | rs4930000 | -0.861 | -0.527 | 0.479 | -0.036 | -0.035 | -0.004 | -0.004 | 0.023 | 0.2 | 0.4 | 0.23 | 0.2 | 0.14 | 0.28 |
| 11 | 2630594 | rs4930137 | -1.363 | -0.588 | -0.458 | -0.155 | 0.008 | 0.001 | 0.001 | 0.417 | 0.28 | 0.73 | 0.4 | 0.25 | 0.21 | 0.32 |
| 11 | 2631303 | rs4930139 | -0.757 | -0.298 | -0.403 | -0.03 | 0.004 | 0.004 | 0.004 | 0 | 0.22 | 0.22 | 0.24 | 0.16 | 0.19 | 0.32 |
| 11 | 2633152 | rs10766212 | -0.165 | 0.324 | -1.232 | -0.463 | 0.041 | 0.07 | 0.07 | 1.3 | 0.46 | 0.3 | 0.53 | 0.78 | 0.61 | 0.63 |
| 11 | 2639712 | rs11023485 | 0.312 | 1.031 | -1.683 | -0.156 | -0.075 | -0.102 | -0.102 | 0 | 0.23 | 0.05 | 0.3 | 0.67 | 0.27 | 0.46 |
| 11 | 2640824 | rs7929804 | -0.436 | 0.853 | -1.451 | -0.679 | 0.015 | 0.08 | 0.08 | 1.098 | 0.45 | 0.28 | 0.6 | 0.78 | 0.56 | 0.61 |
| 11 | 2641129 | rs2412058 | -0.397 | 0.2 | -1.127 | -0.925 | -0.029 | -0.003 | -0.003 | 0 | 0.23 | 0.57 | 0.29 | 0.2 | 0.2 | 0.26 |
| 11 | 2643480 | rs10741669 | 0.033 | 0.107 | -1.254 | -1.382 | -0.009 | -0.039 | -0.039 | 1.377 | 0.88 | 0.87 | 0.92 | 0.99 | 0.88 | 0.89 |
| 11 | 2654673 | rs10832430 | 0.35 | 0.714 | -1.926 | -1.375 | -0.111 | -0.103 | -0.103 | 0 | 0.24 | 0.03 | 0.46 | 0.72 | 0.27 | 0.42 |
| 11 | 2673575 | rs6578283 | 1.001 | 0.626 | -1.013 | -1.439 | -0.014 | -0.034 | -0.034 | 0 | 0.68 | 0.48 | 0.69 | 0.86 | 0.68 | 0.65 |
| 11 | 2673681 | rs231348 | -0.323 | 0.62 | -1.42 | -1.102 | -0.006 | 0 | 0 | 0.861 | 0.12 | 0.12 | 0.14 | 0.09 | 0.13 | 0.11 |
| 11 | 2680815 | rs151216 | -0.232 | 0.596 | -1.36 | -1.08 | -0.013 | 0.004 | 0.004 | 0.449 | 0.11 | 0.13 | 0.13 | 0.09 | 0.12 | 0.1 |
| 11 | 2683357 | rs760419 | -0.629 | 0.956 | -2.118 | -1.599 | -0.083 | -0.024 | -0.024 | 0 | 0.32 | 0.86 | 0.37 | 0.33 | 0.45 | 0.44 |
| 11 | 2685864 | rs9666537 | 0.867 | 0.729 | -1.623 | -1.97 | -0.001 | 0.001 | 0.001 | 0 | 0.72 | 0.71 | 0.82 | 0.89 | 0.69 | 0.71 |
| 11 | 2688526 | rs10832514 | -0.605 | 0.391 | -1.467 | -1.989 | -0.005 | 0.067 | 0.067 | 0 | 0.3 | 0.16 | 0.46 | 0.91 | 0.31 | 0.48 |
| 11 | 2689100 | rs12418076 | -0.531 | 0.096 | -1.063 | -1.886 | -0.006 | -0.039 | -0.039 | 0 | 0.13 | 0.13 | 0.06 | 0 | 0.1 | 0.06 |
| 11 | 2689546 | rs1459825 | -1.526 | -0.32 | -0.453 | -1.849 | -0.021 | -0.091 | -0.091 | 0 | 0.26 | 0.49 | 0.23 | 0.11 | 0.2 | 0.26 |
| 11 | 2690653 | rs4930006 | 0.231 | -0.178 | -0.837 | -1.666 | -0.012 | -0.01 | -0.01 | 0.88 | 0.06 | 0.02 | 0.06 | 0.07 | 0.07 | 0.07 |
| 11 | 2691471 | rs231362 | -1.125 | -0.37 | -1.011 | -2.048 | -0.039 | 0.143 | 0.143 | 2.985 | 0.57 | 0.8 | 0.69 | 0.88 | 0.5 | 0.75 |
| 11 | 2696744 | rs7128926 | -1.56 | -0.58 | 0.008 | -1.268 | -0.027 | -0.058 | -0.058 | 0 | 0.28 | 0.34 | 0.22 | 0.1 | 0.21 | 0.26 |
| 11 | 2697595 | rs9666604 | -0.824 | -0.608 | -0.004 | -1.251 | -0.032 | -0.079 | -0.079 | 0 | 0.2 | 0.27 | 0.14 | 0.03 | 0.13 | 0.18 |
| 11 | 2712286 | rs7939976 | -1.998 | 0.214 | -0.173 | -0.633 | -0.019 | -0.029 | -0.029 | 0 | 0.19 | 0.32 | 0.18 | 0.11 | 0.17 | 0.32 |
| 11 | 2729340 | rs2283200 | -0.43 | 0.638 | -0.401 | -0.181 | -0.003 | 0.052 | 0.052 | 0 | 0.06 | 0.12 | 0.13 | 0.15 | 0.06 | 0.06 |
| 11 | 2736683 | rs231845 | -0.888 | 1.035 | -0.509 | -0.209 | 0.004 | -0.006 | -0.006 | 0.161 | 0.08 | 0.13 | 0.07 | 0.05 | 0.13 | 0.18 |
| 11 | 2737905 | rs2283202 | 0.143 | 0.767 | -0.665 | -0.019 | -0.001 | 0 | 0 | 0 | 0.57 | 0.55 | 0.6 | 0.69 | 0.42 | 0.69 |
| 11 | 2739110 | rs231888 | -0.747 | 1.022 | -0.954 | 0.282 | -0.036 | 0.035 | 0.035 | 0 | 0.34 | 0.15 | 0.45 | 0.75 | 0.4 | 0.52 |
| 11 | 2739180 | rs231887 | -1.474 | 0.654 | 0.243 | 0.519 | 0.054 | 0.069 | 0.069 | 0 | 0.1 | 0.52 | 0.12 | 0.11 | 0.1 | 0.12 |
| 11 | 2741327 | rs2283205 | -0.708 | 0.491 | -0.336 | 0.649 | -0.002 | 0.002 | 0.002 | 0 | 0.21 | 0.2 | 0.13 | 0.19 | 0.16 | 0.2 |
| 11 | 2743859 | rs2283208 | -1.007 | 0.452 | -1.077 | -0.435 | 0.005 | 0.102 | 0.102 | 0.159 | 0.5 | 0.27 | 0.31 | 0.22 | 0.48 | 0.36 |
| 11 | 2748826 | rs2283212 | -0.244 | -0.206 | -0.232 | -0.507 | -0.051 | -0.075 | -0.075 | 0 | 0.3 | 0.44 | 0.26 | 0.11 | 0.21 | 0.17 |
| 11 | 2749015 | rs231915 | -0.33 | 0.376 | -0.639 | -0.245 | 0.035 | -0.009 | -0.009 | 0 | 0.72 | 0.65 | 0.54 | 0.7 | 0.56 | 0.58 |
| 11 | 2750703 | rs170786 | 1.041 | 0.154 | -0.474 | 0.491 | -0.007 | -0.011 | -0.011 | 0 | 0.34 | 0.49 | 0.21 | 0.32 | 0.27 | 0.31 |
| 11 | 2750720 | rs2283214 | -0.112 | 0.256 | -0.347 | 0.666 | -0.027 | -0.009 | -0.009 | 0 | 0.34 | 0.43 | 0.33 | 0.25 | 0.29 | 0.22 |
| 11 | 2752593 | rs17743926 | 1.109 | -0.177 | -0.759 | 0.296 | 0.037 | 0.064 | 0.064 | 1.652 | 0.22 | 0.06 | 0.19 | 0.07 | 0.12 | 0.2 |
| 11 | 2755334 | rs422316 | -0.056 | -0.096 | -0.563 | 0.508 | 0 | 0 | 0 | 1.017 | 0.52 | 0.48 | 0.39 | 0.48 | 0.49 | 0.51 |
| 11 | 2755346 | rs422314 | -0.266 | -0.104 | -0.684 | 0.509 | -0.01 | 0.032 | 0.032 | 0.123 | 0.56 | 0.73 | 0.44 | 0.62 | 0.51 | 0.51 |
| 11 | 2755601 | rs2283222 | 0.401 | -0.044 | -0.762 | 0.337 | 0.003 | 0.037 | 0.037 | 1.321 | 0.37 | 0.22 | 0.26 | 0.21 | 0.32 | 0.26 |
| 11 | 2757985 | rs108961 | 0.407 | -0.064 | -0.623 | 0.37 | 0.002 | -0.024 | -0.024 | 0 | 0.55 | 0.51 | 0.31 | 0.23 | 0.45 | 0.37 |
| 11 | 2759090 | rs231880 | 0.276 | -0.044 | -1.016 | 0.732 | 0.026 | 0.047 | 0.047 | 0 | 0.41 | 0.21 | 0.23 | 0.31 | 0.34 | 0.33 |
| 11 | 2760741 | rs1079714 | -0.301 | 0.075 | -1.023 | 1.498 | -0.003 | 0.123 | 0.123 | 0 | 0.07 | 0.13 | 0.14 | 0.29 | 0.09 | 0.11 |
| 11 | 2761382 | rs231879 | 0.764 | 0.033 | -1.528 | 0.917 | -0.019 | -0.071 | -0.071 | 0 | 0.11 | 0.01 | 0.1 | 0.27 | 0.1 | 0.13 |
| 11 | 2763228 | rs7114479 | 0.099 | -0.141 | -1.671 | 0.273 | 0.03 | 0.14 | 0.14 | 0.177 | 0.52 | 0.73 | 0.37 | 0.17 | 0.39 | 0.51 |
| 11 | 2764118 | rs2237875 | -0.831 | -0.381 | -1.383 | 1.673 | 0.005 | 0.009 | 0.009 | 0 | 0.14 | 0.11 | 0.15 | 0.52 | 0.08 | 0.25 |
| 11 | 2766234 | rs2237877 | -0.296 | -0.276 | -1.463 | 1.327 | 0.013 | -0.028 | -0.028 | 0 | 0.12 | 0.07 | 0.15 | 0.24 | 0.11 | 0.18 |
| 11 | 2766282 | rs2237878 | 0.257 | 0.04 | -1.325 | 0.55 | 0 | 0.027 | 0.027 | 0 | 0.09 | 0.09 | 0.09 | 0.28 | 0.1 | 0.12 |
| 11 | 2767967 | rs231899 | -0.443 | 0.05 | -0.972 | -0.225 | 0.002 | 0.005 | 0.005 | 0 | 0.73 | 0.67 | 0.77 | 0.69 | 0.69 | 0.78 |
| 11 | 2773471 | rs2056892 | 0.549 | -0.016 | -0.506 | -0.61 | 0.006 | -0.002 | -0.002 | 0 | 0.57 | 0.52 | 0.41 | 0.39 | 0.45 | 0.39 |
| 11 | 2774839 | rs151291 | 0.755 | -0.357 | -0.695 | -1.011 | -0.002 | 0.157 | 0.157 | 0 | 0.37 | 0.12 | 0.2 | 0.04 | 0.34 | 0.3 |
| 11 | 2776059 | rs231890 | -1.786 | -0.356 | -0.59 | -0.84 | -0.03 | 0.036 | 0.036 | 0 | 0.88 | 0.95 | 0.89 | 0.94 | 0.82 | 0.74 |
| 11 | 2776448 | rs11023996 | 0.83 | -0.362 | -0.483 | -0.653 | 0.009 | 0.048 | 0.048 | 0 | 0.09 | 0.12 | 0.12 | 0.29 | 0.16 | 0.11 |
| 11 | 2778083 | rs11023999 | 0.385 | -0.793 | -0.171 | -0.746 | 0.003 | 0 | 0 | 0 | 0.4 | 0.37 | 0.34 | 0.37 | 0.28 | 0.31 |
| 11 | 2779970 | rs6578296 | -1.579 | -0.526 | -0.288 | -0.665 | 0 | -0.007 | -0.007 | 0 | 0.87 | 0.87 | 0.89 | 0.93 | 0.82 | 0.74 |
| 11 | 2781519 | rs179785 | 0.08 | -0.609 | -0.391 | -0.689 | 0.004 | -0.001 | -0.001 | 0 | 0.51 | 0.63 | 0.37 | 0.52 | 0.48 | 0.44 |
| 11 | 2782648 | rs548566 | -1.878 | -0.501 | -0.566 | -0.63 | -0.011 | -0.023 | -0.023 | 0 | 0.88 | 0.92 | 0.81 | 0.79 | 0.82 | 0.75 |
| 11 | 2785623 | rs4255520 | 1.273 | -0.546 | -0.765 | -0.729 | 0.002 | -0.012 | -0.012 | 0 | 0.2 | 0.15 | 0.15 | 0.03 | 0.16 | 0.2 |
| 11 | 2786302 | rs151288 | 1.233 | -0.452 | -1.068 | -0.682 | -0.001 | -0.001 | -0.001 | 0 | 0.26 | 0.27 | 0.16 | 0.29 | 0.28 | 0.15 |
| 11 | 2787061 | rs231872 | -0.18 | -0.688 | -1.02 | -0.459 | -0.013 | 0.015 | 0.015 | 0 | 0.62 | 0.51 | 0.68 | 0.55 | 0.65 | 0.44 |
| 11 | 2791208 | rs2237882 | -0.263 | 0.142 | -1.726 | -0.287 | -0.016 | -0.034 | -0.034 | 0 | 0.27 | 0.01 | 0.41 | 0.26 | 0.25 | 0.23 |
| 11 | 2794114 | rs4930007 | -0.547 | 0.828 | -1.092 | -0.066 | 0.001 | 0.006 | 0.006 | 0 | 0.48 | 0.43 | 0.63 | 0.77 | 0.49 | 0.44 |
| 11 | 2797237 | rs1057128 | -0.62 | 1.344 | -1.127 | 1.122 | -0.046 | -0.089 | -0.089 | 0 | 0.16 | 0.01 | 0.33 | 0.29 | 0.2 | 0.17 |
| 11 | 2798305 | rs81204 | -0.882 | 0.674 | -0.631 | 0.906 | -0.001 | -0.017 | -0.017 | 0 | 0.18 | 0.14 | 0.23 | 0.27 | 0.25 | 0.26 |
| 11 | 2807522 | rs234873 | 0.548 | 0.903 | -0.557 | 1.258 | -0.001 | -0.002 | -0.002 | 0 | 0.57 | 0.36 | 0.51 | 0.47 | 0.56 | 0.43 |
| 11 | 2810731 | rs2237886 | -1.136 | 0.962 | -0.593 | 1.535 | -0.003 | 0.003 | 0.003 | 0 | 0.09 | 0.07 | 0.1 | 0.19 | 0.1 | 0.11 |
| 11 | 2819620 | rs151316 | 0.657 | 0.618 | -0.161 | 0.857 | -0.017 | -0.029 | -0.029 | 0 | 0.74 | 0.56 | 0.75 | 0.79 | 0.82 | 0.59 |
| 11 | 2821065 | rs163171 | 0.226 | 0.557 | 0.255 | 1.2 | -0.009 | 0.09 | 0.09 | 0.094 | 0.76 | 0.41 | 0.64 | 0.59 | 0.79 | 0.88 |
| 11 | 2825228 | rs163166 | -0.322 | 0.58 | 0.412 | 1.016 | 0 | 0.024 | 0.024 | 0 | 0.82 | 0.92 | 0.92 | 1 | 0.84 | 0.91 |
| 11 | 2832905 | rs2237888 | -0.704 | 0.055 | 1.474 | 0.801 | -0.023 | 0.202 | 0.202 | 0 | 0.17 | 0.41 | 0.38 | 0.42 | 0.14 | 0.16 |
| 11 | 2837625 | rs233446 | -0.232 | 0.727 | 1.265 | 0.529 | -0.002 | 0.007 | 0.007 | 0 | 0.27 | 0.19 | 0.16 | 0.17 | 0.26 | 0.3 |

| | | | | | | | | | | | | | | | | |
|---|---|---|---|---|---|---|---|---|---|---|---|---|---|---|---|---|
| 11 | 2838413 | rs163177 | 1.001 | 1.658 | 1.462 | 0.44 | -0.001 | 0.006 | 0.006 | 0 | 0.49 | 0.31 | 0.43 | 0.43 | 0.51 | 0.48 |
| 11 | 2839188 | rs12284372 | -0.116 | 1.62 | 1.436 | 0.426 | -0.003 | 0.017 | 0.017 | 0 | 0.2 | 0.13 | 0.11 | 0.04 | 0.2 | 0.25 |
| 11 | 2839899 | rs2237893 | -0.048 | 1.633 | 1.469 | 0.346 | -0.006 | 0.046 | 0.046 | 0 | 0.19 | 0.11 | 0.11 | 0.04 | 0.2 | 0.21 |
| 11 | 2840424 | rs233448 | 0.097 | 1.905 | 1.322 | 0.307 | -0.015 | 0.033 | 0.033 | 0 | 0.26 | 0.11 | 0.17 | 0.16 | 0.26 | 0.26 |
| 11 | 2843676 | rs11606985 | 1.085 | 1.607 | 0.416 | -0.659 | -0.007 | 0.12 | 0.12 | 0 | 0.22 | 0.05 | 0.15 | 0 | 0.24 | 0.13 |
| 11 | 2843803 | rs233449 | 0.678 | 1.69 | 0.667 | -0.066 | -0.014 | 0.006 | 0.006 | 0 | 0.25 | 0.1 | 0.15 | 0.24 | 0.26 | 0.28 |
| 11 | 2847069 | rs163184 | 0.446 | 1.735 | 0.191 | 0.04 | -0.003 | -0.021 | -0.021 | 2.148 | 0.48 | 0.14 | 0.4 | 0.42 | 0.49 | 0.49 |
| 11 | 2850782 | rs234852 | -0.052 | 1.548 | 0.202 | -0.05 | 0.004 | 0.034 | 0.034 | 0 | 0.73 | 0.44 | 0.57 | 0.61 | 0.74 | 0.77 |
| 11 | 2852529 | rs234857 | 0.198 | 1.698 | 0.018 | -0.036 | -0.006 | 0.013 | 0.013 | 0 | 0.28 | 0.24 | 0.19 | 0.18 | 0.28 | 0.3 |
| 11 | 2857194 | rs2237895 | 0.053 | 1.625 | -0.34 | 0.427 | 0 | 0.024 | 0.024 | 0 | 0.41 | 0.15 | 0.4 | 0.33 | 0.42 | 0.4 |
| 11 | 2859055 | rs234868 | 0.115 | 1.409 | -0.37 | 0.844 | -0.001 | -0.002 | -0.002 | 0 | 0.63 | 0.61 | 0.46 | 0.56 | 0.58 | 0.77 |
| 11 | 2862159 | rs4930013 | -0.161 | 1.533 | -0.17 | 0.228 | 0.014 | -0.046 | -0.046 | 0 | 0.23 | 0.32 | 0.32 | 0.06 | 0.3 | 0.14 |
| 11 | 2862244 | rs2283229 | 0.058 | 1.595 | -0.746 | 1.1 | -0.031 | -0.112 | -0.112 | 0.953 | 0.18 | 0.03 | 0.37 | 0.56 | 0.2 | 0.29 |
| 11 | 2862945 | rs11024304 | 0.444 | 1.399 | -0.313 | 1.063 | -0.003 | 0.061 | 0.061 | 0 | 0.07 | 0.16 | 0.03 | 0.2 | 0.05 | 0.07 |
| 11 | 2863056 | rs433052 | -0.551 | 1.373 | -0.196 | 1.037 | -0.002 | 0.076 | 0.076 | 0 | 0.92 | 0.81 | 0.97 | 0.8 | 0.95 | 0.92 |
| 11 | 2863820 | rs2237898 | -0.713 | 1.256 | -0.272 | 0.893 | -0.009 | 0.07 | 0.07 | 0 | 0.12 | 0.25 | 0.07 | 0.26 | 0.08 | 0.15 |
| 11 | 2868259 | rs12419347 | -1.003 | 1.694 | -0.172 | 0.04 | 0.006 | -0.034 | -0.034 | 0 | 0.34 | 0.41 | 0.5 | 0.83 | 0.38 | 0.51 |
| 11 | 2870108 | rs8234 | -0.905 | 1.72 | -0.103 | 0.01 | 0.007 | -0.043 | -0.043 | 0 | 0.33 | 0.43 | 0.48 | 0.84 | 0.36 | 0.52 |
| 11 | 2870165 | rs10798 | -0.905 | 1.72 | -0.145 | 0.01 | 0.006 | -0.037 | -0.037 | 0.285 | 0.33 | 0.41 | 0.48 | 0.84 | 0.36 | 0.52 |
| 11 | 2875163 | rs7108037 | -0.44 | 1.239 | -0.528 | -0.095 | -0.003 | 0.012 | 0.012 | 0.047 | 0.46 | 0.6 | 0.57 | 0.81 | 0.51 | 0.52 |
| 11 | 2880009 | rs10832835 | -0.703 | 1.219 | -0.541 | -0.149 | -0.004 | 0.014 | 0.014 | 0 | 0.46 | 0.6 | 0.57 | 0.81 | 0.5 | 0.52 |
| 11 | 2885728 | rs10766424 | -1.104 | 1.142 | -0.776 | -0.224 | 0.009 | 0.142 | 0.142 | 0 | 0.53 | 0.78 | 0.7 | 0.89 | 0.59 | 0.59 |
| 11 | 2891965 | rs2239897 | 0.114 | 0.806 | -0.783 | 0.995 | 0.018 | -0.044 | -0.044 | 0.609 | 0.11 | 0.02 | 0.18 | 0.54 | 0.08 | 0.29 |
| 11 | 2895800 | rs3987740 | -0.124 | 0.521 | -0.661 | -0.059 | -0.001 | 0.004 | 0.004 | 0 | 0.29 | 0.22 | 0.32 | 0.61 | 0.26 | 0.39 |
| 11 | 2896128 | rs2078786 | -1.405 | 0.818 | -0.854 | -1.05 | 0.005 | 0.041 | 0.041 | 0 | 0.67 | 0.79 | 0.8 | 0.99 | 0.63 | 0.63 |
| 11 | 2896313 | rs12786533 | 1.429 | 0.811 | -1.15 | -0.961 | -0.009 | -0.044 | -0.044 | 0 | 0.1 | 0.02 | 0.17 | 0.2 | 0.09 | 0.09 |
| 11 | 2896783 | rs10488671 | 0.797 | 1.051 | -1.042 | -0.82 | -0.024 | -0.044 | -0.044 | 0 | 0.07 | 0.02 | 0.13 | 0.16 | 0.06 | 0.11 |
| 11 | 2897026 | rs10766441 | -0.432 | 0.967 | -0.993 | -0.941 | 0.012 | 0.038 | 0.038 | 0 | 0.43 | 0.27 | 0.48 | 0.78 | 0.35 | 0.51 |
| 11 | 2900418 | rs4930026 | -1.181 | 1.298 | -0.519 | -0.887 | 0.001 | -0.003 | -0.003 | 0 | 0.24 | 0.28 | 0.26 | 0.16 | 0.23 | 0.14 |
| 11 | 2902105 | rs3864883 | 0.717 | 1.564 | -0.548 | -0.593 | 0.045 | -0.025 | -0.025 | 0 | 0.05 | 0.1 | 0.09 | 0 | 0.12 | 0.02 |
| 11 | 2908195 | rs452338 | 0.608 | 0.778 | -0.871 | -0.88 | -0.013 | 0.025 | 0.025 | 0 | 0.25 | 0.17 | 0.28 | 0.16 | 0.29 | 0.16 |
| 11 | 2908656 | rs11024437 | 1.037 | 0.795 | -1.02 | -0.554 | -0.006 | -0.004 | -0.004 | 0 | 0.09 | 0.02 | 0.14 | 0.55 | 0.09 | 0.24 |
| 11 | 2908965 | rs450563 | 0.173 | 0.659 | -0.829 | -0.902 | 0.001 | 0.041 | 0.041 | 0 | 0.3 | 0.21 | 0.3 | 0.16 | 0.32 | 0.16 |
| 11 | 2910621 | rs10832869 | 0.333 | 0.488 | -0.561 | -1.006 | -0.001 | -0.001 | -0.001 | 0 | 0.41 | 0.37 | 0.4 | 0.7 | 0.33 | 0.42 |
| 11 | 2912553 | rs384037 | 0.129 | 0.956 | 0.075 | -0.562 | 0.011 | -0.03 | -0.03 | 0 | 0.61 | 0.42 | 0.62 | 0.8 | 0.54 | 0.53 |
| 11 | 2912946 | rs400643 | 0.009 | 1.009 | -0.506 | -0.535 | 0.001 | -0.009 | -0.009 | 0.14 | 0.63 | 0.59 | 0.64 | 0.81 | 0.55 | 0.53 |
| 11 | 2919469 | rs390266 | -0.717 | 0.856 | -0.453 | -0.196 | 0.006 | -0.024 | -0.024 | 0 | 0.28 | 0.6 | 0.42 | 0.17 | 0.3 | 0.13 |
| 11 | 2920283 | rs10741735 | 0.375 | 0.096 | -1.174 | 0.064 | -0.003 | 0.052 | 0.052 | 0.091 | 0.51 | 0.25 | 0.43 | 0.56 | 0.54 | 0.73 |
| 11 | 2920666 | rs365605 | -0.654 | 0.334 | -1.098 | 0.146 | -0.001 | -0.015 | -0.015 | 0 | 0.64 | 0.73 | 0.66 | 0.4 | 0.66 | 0.59 |
| 11 | 2922493 | rs4930027 | 0.249 | 0.615 | -0.78 | -0.318 | -0.013 | 0.031 | 0.031 | 0 | 0.4 | 0.36 | 0.4 | 0.27 | 0.44 | 0.47 |
| 11 | 2923091 | rs7934793 | -0.419 | 1.009 | -0.549 | -0.676 | 0.019 | -0.042 | -0.042 | 0 | 0.35 | 0.53 | 0.34 | 0.19 | 0.44 | 0.45 |
| 11 | 2923826 | rs367035 | 0.939 | 0.953 | -1.188 | -0.791 | 0.027 | 0.152 | 0.152 | 0 | 0.5 | 0.18 | 0.51 | 0.75 | 0.43 | 0.49 |
| 11 | 2924610 | rs1048047 | 0.989 | 0.918 | -0.704 | -0.225 | 0.001 | -0.003 | -0.003 | 0 | 0.64 | 0.45 | 0.59 | 0.76 | 0.6 | 0.68 |
| 11 | 2925761 | rs389646 | 0.141 | 0.986 | -1.235 | 0.001 | 0.015 | -0.043 | -0.043 | 1.024 | 0.89 | 0.81 | 0.86 | 1 | 0.86 | 0.94 |
| 11 | 2925915 | rs451443 | 0.141 | 0.986 | -1.235 | 0.001 | 0.015 | -0.043 | -0.043 | 0 | 0.89 | 0.81 | 0.86 | 1 | 0.86 | 0.94 |
| 11 | 2931981 | rs450208 | -0.059 | 0.96 | -1.373 | 0.11 | -0.001 | 0 | 0 | 0 | 0.84 | 0.84 | 0.71 | 0.91 | 0.8 | 0.86 |
| 11 | 2935303 | rs3782067 | -0.179 | 0.392 | -2.307 | 0.271 | -0.006 | -0.113 | -0.113 | 1.687 | 0.34 | 0.04 | 0.26 | 0.61 | 0.31 | 0.31 |
| 11 | 2936434 | rs450876 | 0.262 | 0.3 | -2.113 | -0.213 | -0.021 | 0.003 | 0.003 | 0 | 0.36 | 0.26 | 0.4 | 0.34 | 0.49 | 0.43 |
| 11 | 2936952 | rs16928809 | 1.508 | 0.21 | -2.364 | -0.241 | -0.022 | -0.033 | -0.033 | 0 | 0.08 | 0.01 | 0.15 | 0.08 | 0.09 | 0.08 |
| 11 | 2937631 | rs367480 | -0.258 | 0.136 | -2.385 | -0.272 | 0.013 | -0.003 | -0.003 | 0 | 0.44 | 0.26 | 0.54 | 0.42 | 0.57 | 0.49 |
| 11 | 2939050 | rs427398 | 0.351 | -0.447 | -1.886 | -1.113 | 0.021 | 0.045 | 0.045 | 0 | 0.48 | 0.35 | 0.47 | 0.27 | 0.57 | 0.45 |
| 11 | 2939126 | rs11024580 | 0.318 | -1.043 | -2.287 | -1.547 | 0.007 | -0.089 | -0.089 | 0 | 0.21 | 0.08 | 0.24 | 0.63 | 0.2 | 0.26 |
| 11 | 2939705 | rs11024581 | 1.252 | -1.075 | -2.254 | -1.407 | 0.035 | -0.09 | -0.09 | 0 | 0.12 | 0.01 | 0.17 | 0.63 | 0.07 | 0.22 |
| 11 | 2940236 | rs444198 | 0.535 | -0.882 | -2.125 | -1.525 | 0.045 | 0.105 | 0.105 | 0 | 0.49 | 0.27 | 0.47 | 0.25 | 0.57 | 0.46 |
| 11 | 2943115 | rs3847644 | -0.965 | -0.186 | -1.225 | -2.455 | 0 | 0 | 0 | 0 | 0.5 | 0.55 | 0.46 | 0.23 | 0.53 | 0.53 |
| 11 | 2945945 | rs3764895 | 0.568 | 0.234 | -1.4 | -1.967 | -0.036 | -0.013 | -0.013 | 0 | 0.15 | 0.05 | 0.13 | 0.12 | 0.19 | 0.21 |
| 11 | 2949861 | rs1056819 | 0.509 | 0.253 | -1.39 | -1.948 | -0.039 | -0.011 | -0.011 | 0.682 | 0.16 | 0.02 | 0.13 | 0.13 | 0.19 | 0.21 |
| 11 | 2956166 | rs2583442 | 1.407 | 0.385 | -1.393 | -2.122 | -0.064 | -0.073 | -0.073 | 0.246 | 0.34 | 0.03 | 0.51 | 0.7 | 0.3 | 0.42 |
| 11 | 2969242 | rs9971558 | 0.577 | 0.583 | -1.352 | -2.165 | -0.028 | 0.002 | 0.002 | 0 | 0.2 | 0.03 | 0.16 | 0.18 | 0.22 | 0.37 |
| 11 | 2973740 | rs3213614 | -0.477 | 0.706 | -0.4 | -2.37 | -0.035 | -0.001 | -0.001 | 0 | 0.87 | 0.96 | 0.9 | 0.91 | 0.79 | 0.91 |
| 11 | 2974912 | rs2001006 | 1.877 | 0.649 | -0.498 | -2.437 | -0.07 | -0.064 | -0.064 | 0.457 | 0.33 | 0.02 | 0.51 | 0.72 | 0.3 | 0.42 |
| 11 | 2986700 | rs10832974 | 0.591 | 0.644 | 1.009 | -2.43 | -0.002 | 0.014 | 0.014 | 0 | 0.22 | 0.14 | 0.16 | 0.18 | 0.22 | 0.37 |
| 11 | 2987303 | rs10766497 | -0.606 | 0.719 | 0.96 | -2.407 | -0.036 | 0.007 | 0.007 | 0 | 0.88 | 0.96 | 0.9 | 0.93 | 0.79 | 0.91 |
| 11 | 2992160 | rs16928945 | 0.92 | 0.674 | 1.077 | -2.515 | -0.004 | -0.04 | -0.04 | 0.205 | 0.11 | 0.18 | 0.06 | 0.01 | 0.1 | 0.06 |
| 11 | 2997698 | rs11024814 | 0.731 | 0.647 | 1.097 | -2.535 | 0.001 | 0.026 | 0.026 | 0 | 0.17 | 0.11 | 0.12 | 0.02 | 0.17 | 0.07 |
| 11 | 2999507 | rs3814964 | 2.066 | 0.645 | 1.256 | -2.553 | -0.006 | -0.015 | -0.015 | 0.11 | 0.34 | 0.12 | 0.52 | 0.7 | 0.3 | 0.42 |
| 11 | 2999919 | rs11024843 | 0.471 | 0.6 | 1.346 | -2.527 | -0.004 | -0.057 | -0.057 | 0 | 0.19 | 0.22 | 0.14 | 0.02 | 0.17 | 0.07 |
| 11 | 3003801 | rs2071106 | 0.348 | 0.692 | 1.36 | -2.46 | -0.007 | -0.025 | -0.025 | 1.888 | 0.23 | 0.44 | 0.19 | 0.18 | 0.22 | 0.37 |
| 11 | 3009438 | rs7127415 | -0.526 | 0.989 | 1.142 | -2.454 | 0.001 | -0.001 | -0.001 | 0.792 | 0.35 | 0.66 | 0.25 | 0.19 | 0.33 | 0.43 |

| | | | | | | | | | | | | | | | | |
|---|---|---|---|---|---|---|---|---|---|---|---|---|---|---|---|---|
| 11 | 3020789 | rs10219272 | -0.584 | 1.06 | 0.943 | -2.403 | -0.002 | -0.012 | -0.012 | 0 | 0.41 | 0.57 | 0.34 | 0.21 | 0.39 | 0.42 |
| 11 | 3024624 | rs2239899 | -0.596 | 1.054 | 1.05 | -2.423 | 0.003 | 0.03 | 0.03 | 0 | 0.4 | 0.35 | 0.32 | 0.2 | 0.39 | 0.42 |
| 11 | 3028140 | rs729662 | 2.298 | 1.099 | 0.774 | -2.408 | -0.062 | -0.1 | -0.1 | 0.356 | 0.33 | 0.03 | 0.5 | 0.66 | 0.29 | 0.42 |
| 11 | 3030104 | rs12363575 | 0.074 | 0.542 | 1.53 | -1.943 | -0.006 | 0.006 | 0.006 | 0 | 0.2 | 0.12 | 0.15 | 0.13 | 0.21 | 0.08 |
| 11 | 3030992 | rs2239900 | -0.758 | 0.663 | 1.582 | -1.857 | 0 | -0.005 | -0.005 | 0.07 | 0.41 | 0.59 | 0.34 | 0.2 | 0.39 | 0.42 |
| 11 | 3036613 | rs739402 | 0.843 | 0.711 | 1.47 | -1.824 | 0.001 | -0.043 | -0.043 | 0.05 | 0.11 | 0.15 | 0.05 | 0.01 | 0.1 | 0.06 |
| 11 | 3046201 | rs428726 | 1.961 | 0.76 | 1.175 | -1.873 | -0.11 | -0.065 | -0.065 | 0.118 | 0.39 | 0.05 | 0.54 | 0.66 | 0.39 | 0.44 |
| 11 | 3046231 | rs417957 | -0.797 | 0.795 | 1.241 | -1.864 | 0.01 | 0.053 | 0.053 | 0 | 0.42 | 0.7 | 0.35 | 0.2 | 0.4 | 0.42 |
| 11 | 3054873 | rs4758510 | 0.263 | 0.811 | 1.137 | -2.01 | 0.002 | -0.003 | -0.003 | 0 | 0.82 | 0.82 | 0.89 | 0.86 | 0.79 | 0.86 |
| 11 | 3060725 | rs451041 | 1.049 | 0.822 | 1.031 | -1.927 | -0.001 | 0.016 | 0.016 | 0 | 0.52 | 0.53 | 0.61 | 0.71 | 0.49 | 0.5 |
| 11 | 3067687 | rs3782077 | 0.914 | 0.931 | 0.452 | -1.868 | -0.026 | 0.005 | 0.005 | 0 | 0.17 | 0.04 | 0.16 | 0.16 | 0.22 | 0.32 |
| 11 | 3076257 | rs7945933 | -0.795 | 0.918 | 0.484 | -1.825 | -0.004 | -0.002 | -0.002 | 0 | 0.94 | 0.97 | 0.96 | 0.87 | 0.91 | 0.93 |
| 11 | 3078885 | rs384490 | -0.965 | 0.773 | 0.745 | -1.976 | 0.001 | 0 | 0 | 3.586 | 0.42 | 0.61 | 0.36 | 0.18 | 0.4 | 0.4 |
| 11 | 3090062 | rs11025240 | -1.136 | 1.084 | 0.263 | -1.693 | 0.008 | -0.041 | -0.041 | 0 | 0.35 | 0.49 | 0.3 | 0.18 | 0.4 | 0.39 |
| 11 | 3098800 | rs35134285 | -0.088 | 1.113 | -0.481 | -1.182 | 0.007 | -0.029 | -0.029 | 0 | 0.06 | 0.14 | 0.06 | 0 | 0.06 | 0.01 |
| 11 | 3105508 | rs4758517 | -0.005 | 1.066 | -0.575 | -1.112 | 0 | -0.01 | -0.01 | 0.694 | 0.19 | 0.25 | 0.19 | 0.16 | 0.2 | 0.31 |
| 11 | 3106659 | rs2084239 | 0.685 | 1.024 | -1.53 | -1.063 | -0.104 | -0.056 | -0.056 | 0 | 0.38 | 0.04 | 0.51 | 0.67 | 0.38 | 0.47 |
| 11 | 3110956 | rs895564 | 0.674 | 1.407 | -0.803 | -1.982 | -0.007 | 0.157 | 0.157 | 0 | 0.48 | 0.2 | 0.56 | 0.78 | 0.44 | 0.55 |
| 11 | 3110976 | rs895563 | -0.077 | 1.587 | -0.295 | -2.227 | 0 | 0 | 0 | 0.154 | 0.77 | 0.73 | 0.81 | 0.81 | 0.76 | 0.87 |
| 11 | 3115354 | rs9795009 | 0.08 | 1.705 | -0.44 | -2.373 | -0.018 | 0.004 | 0.004 | 0 | 0.79 | 0.89 | 0.88 | 0.83 | 0.78 | 0.87 |
| 11 | 3115943 | rs10833220 | 1.963 | 2.085 | -0.845 | -2.729 | -0.029 | 0.084 | 0.084 | 0 | 0.38 | 0.02 | 0.48 | 0.78 | 0.3 | 0.45 |
| 11 | 3116073 | rs4758519 | 0.304 | 2.11 | 0.133 | -2.695 | 0.016 | -0.135 | -0.135 | 0 | 0.2 | 0.37 | 0.12 | 0.03 | 0.23 | 0.29 |
| 11 | 3116632 | rs10833221 | -0.761 | 1.847 | 0.308 | -2.152 | 0.006 | -0.044 | -0.044 | 0 | 0.19 | 0.25 | 0.25 | 0.02 | 0.25 | 0.14 |
| 11 | 3121785 | rs12270081 | -0.025 | 1.717 | 0.354 | -2.133 | 0.041 | -0.078 | -0.078 | 2.341 | 0.12 | 0.32 | 0.09 | 0 | 0.14 | 0.1 |
| 11 | 3121853 | rs12282905 | -0.011 | 1.72 | 0.355 | -2.132 | 0.043 | -0.077 | -0.077 | 0.282 | 0.12 | 0.32 | 0.09 | 0 | 0.14 | 0.1 |
| 11 | 3123829 | rs4758468 | 0.17 | 1.281 | -0.156 | -2.14 | 0.024 | 0.005 | 0.005 | 0 | 0.8 | 0.86 | 0.91 | 0.82 | 0.85 | 0.88 |
| 11 | 3124435 | rs1542188 | -0.212 | 1.308 | -0.27 | -2.063 | -0.002 | -0.003 | -0.003 | 0 | 0.1 | 0.13 | 0.21 | 0.02 | 0.18 | 0.06 |
| 11 | 3128380 | rs3741355 | -0.477 | 1.185 | -0.37 | -2.039 | 0.007 | -0.008 | -0.008 | 0 | 0.86 | 0.92 | 0.93 | 0.82 | 0.88 | 0.91 |
| 11 | 3128774 | rs3741353 | 1.743 | 1.388 | -0.74 | -2.344 | -0.105 | 0.147 | 0.147 | 0.331 | 0.45 | 0.03 | 0.55 | 0.78 | 0.44 | 0.4 |
| 11 | 3129395 | rs17178796 | 0.711 | 1.69 | -0.479 | -2.198 | -0.02 | 0.016 | 0.016 | 0 | 0.11 | 0.06 | 0.08 | 0 | 0.14 | 0.1 |
| 11 | 3130708 | rs1900372 | -0.101 | 1.511 | -0.372 | -2.03 | 0.033 | -0.036 | -0.036 | 0 | 0.08 | 0.18 | 0.2 | 0.01 | 0.17 | 0.05 |
| 11 | 3131418 | rs7925089 | -0.736 | 0.986 | -0.727 | -1.927 | -0.034 | 0.078 | 0.078 | 1.282 | 0.39 | 0.77 | 0.37 | 0.04 | 0.45 | 0.51 |
| 11 | 3131741 | rs4237782 | 0.465 | 0.491 | -0.027 | -0.749 | -0.006 | 0.008 | 0.008 | 0 | 0.83 | 0.74 | 0.91 | 0.82 | 0.83 | 0.9 |
| 11 | 3140196 | rs4758528 | -0.761 | 0.03 | -0.24 | -0.467 | -0.04 | -0.005 | -0.005 | 0.178 | 0.27 | 0.44 | 0.16 | 0.22 | 0.19 | 0.24 |
| 11 | 3141109 | rs4758529 | -0.81 | -0.29 | -1.465 | -0.278 | -0.001 | -0.173 | -0.173 | 2.002 | 0.7 | 0.99 | 0.75 | 0.47 | 0.77 | 0.64 |
| 11 | 3143030 | rs6578322 | -1.005 | -0.697 | -1.449 | -0.233 | 0.057 | -0.161 | -0.161 | 0 | 0.73 | 0.98 | 0.76 | 0.49 | 0.86 | 0.65 |
| 11 | 3148037 | rs734458 | 0.685 | -0.358 | -1.685 | -0.557 | -0.017 | -0.007 | -0.007 | 0 | 0.15 | 0.05 | 0.09 | 0.13 | 0.15 | 0.12 |
| 11 | 3149397 | rs1466045 | -0.004 | -0.506 | -0.845 | -0.782 | -0.001 | -0.004 | -0.004 | 0 | 0.44 | 0.44 | 0.51 | 0.34 | 0.62 | 0.44 |
| 11 | 3150086 | rs4758533 | -1.148 | -0.454 | -0.964 | -0.362 | 0.076 | -0.085 | -0.085 | 0 | 0.67 | 0.91 | 0.67 | 0.55 | 0.82 | 0.61 |
| 11 | 3153423 | rs4758469 | -1.849 | -0.559 | -1.106 | -0.419 | 0.058 | -0.14 | -0.14 | 0 | 0.75 | 0.98 | 0.77 | 0.56 | 0.87 | 0.67 |
| 11 | 3154105 | rs11025546 | 0.093 | -0.403 | -0.713 | -0.518 | -0.002 | -0.018 | -0.018 | 1.433 | 0.08 | 0.09 | 0.1 | 0 | 0.05 | 0.04 |
| 11 | 3163653 | rs1562353 | -0.54 | -0.651 | -0.292 | -0.577 | -0.008 | -0.011 | -0.011 | 0.053 | 0.31 | 0.61 | 0.25 | 0.2 | 0.22 | 0.22 |
| 11 | 3164791 | rs4758470 | -1.531 | -0.615 | -0.566 | -0.431 | 0.055 | -0.106 | -0.106 | 0.364 | 0.7 | 0.91 | 0.74 | 0.55 | 0.82 | 0.6 |
| 11 | 3167177 | rs11025603 | 1.022 | -0.39 | -0.097 | -0.349 | -0.007 | 0.057 | 0.057 | 0 | 0.14 | 0.31 | 0.12 | 0.15 | 0.14 | 0.09 |
| 11 | 3168538 | rs11820744 | -0.427 | -0.068 | -0.383 | -0.597 | -0.002 | 0.016 | 0.016 | 0 | 0.3 | 0.24 | 0.22 | 0.15 | 0.26 | 0.24 |
| 11 | 3168755 | rs11025610 | -0.197 | 0.5 | 0.157 | -0.361 | -0.025 | 0.051 | 0.051 | 0 | 0.21 | 0.45 | 0.14 | 0.24 | 0.17 | 0.14 |
| 11 | 3170174 | rs10466385 | 0.037 | 0.869 | -0.278 | -0.614 | 0.034 | -0.007 | -0.007 | 0 | 0.19 | 0.25 | 0.34 | 0.18 | 0.43 | 0.19 |
| 11 | 3170866 | rs4758471 | -0.246 | 1.092 | -0.304 | -0.575 | 0.122 | -0.027 | -0.027 | 7.376 | 0.21 | 0.41 | 0.34 | 0.18 | 0.43 | 0.19 |
| 11 | 3178136 | rs4370914 | -0.997 | 0.32 | -1.323 | -0.674 | 0.068 | 0.072 | 0.072 | 0 | 0.49 | 0.74 | 0.5 | 0.37 | 0.64 | 0.39 |
| 11 | 3178235 | rs4261264 | 1.096 | 0.09 | -1.405 | -0.534 | 0.003 | -0.014 | -0.014 | 0 | 0.14 | 0.11 | 0.17 | 0.16 | 0.15 | 0.11 |
| 11 | 3179253 | rs7942908 | -1.069 | 0.049 | -1.401 | -0.585 | 0.087 | 0.072 | 0.072 | 0 | 0.53 | 0.83 | 0.52 | 0.38 | 0.67 | 0.42 |
| 11 | 3179460 | rs7928630 | -0.39 | 0.803 | -1.441 | -0.711 | -0.011 | 0.037 | 0.037 | 0 | 0.41 | 0.65 | 0.34 | 0.28 | 0.51 | 0.25 |
| 11 | 3179942 | rs7102375 | 0.576 | 0.918 | -1.306 | -0.447 | 0 | -0.001 | -0.001 | 0 | 0.16 | 0.16 | 0.13 | 0.13 | 0.14 | 0.1 |
| 11 | 3180959 | rs4758544 | -0.079 | 0.797 | -1.242 | -0.34 | 0.001 | -0.003 | -0.003 | 0 | 0.11 | 0.17 | 0.24 | 0.1 | 0.16 | 0.17 |
| 11 | 3181085 | rs4237786 | -1.14 | 0.202 | -1.426 | -0.174 | 0.061 | -0.087 | -0.087 | 0.237 | 0.68 | 0.91 | 0.75 | 0.54 | 0.83 | 0.54 |
| 11 | 3181629 | rs7935956 | -0.588 | 0.673 | -1.41 | -0.255 | 0.052 | 0.045 | 0.045 | 0 | 0.53 | 0.75 | 0.64 | 0.42 | 0.72 | 0.43 |
| 11 | 3188629 | rs4758548 | -0.702 | 1.584 | -1.429 | -0.26 | -0.058 | 0.043 | 0.043 | 0 | 0.19 | 0.06 | 0.26 | 0.13 | 0.34 | 0.17 |
| 11 | 3197290 | rs11025773 | -1.077 | 0.871 | -1.686 | -0.103 | 0.044 | -0.018 | -0.018 | 0.074 | 0.65 | 0.79 | 0.78 | 0.4 | 0.78 | 0.54 |
| 11 | 3199846 | rs2412143 | -1.256 | 1.057 | -1.714 | -0.594 | 0.041 | 0.075 | 0.075 | 0.016 | 0.5 | 0.71 | 0.6 | 0.33 | 0.59 | 0.4 |
| 11 | 3202576 | rs16929180 | 0.819 | 0.821 | -1.597 | -0.641 | 0.003 | -0.004 | -0.004 | 0 | 0.07 | 0.05 | 0.11 | 0.11 | 0.03 | 0.06 |
| 11 | 3203057 | rs4758478 | -0.743 | 0.758 | -0.715 | -0.479 | 0.019 | 0.031 | 0.031 | 0 | 0.28 | 0.42 | 0.4 | 0.38 | 0.37 | 0.26 |
| 11 | 3204655 | rs895558 | -1.856 | 0.586 | -0.76 | 0.334 | 0.038 | -0.016 | -0.016 | 0.625 | 0.7 | 0.79 | 0.83 | 0.62 | 0.86 | 0.58 |
| 11 | 3205005 | rs7103638 | -0.635 | 1.301 | 0.24 | 0.485 | 0.03 | 0.003 | 0.003 | 0 | 0.21 | 0.29 | 0.32 | 0.26 | 0.36 | 0.2 |
| 11 | 3208101 | rs2121438 | 0.047 | 1.071 | 0.385 | 0.551 | 0.002 | -0.011 | -0.011 | 0 | 0.26 | 0.35 | 0.27 | 0.18 | 0.27 | 0.2 |
| 11 | 3211112 | rs9971622 | 0.242 | 1.079 | 0.135 | 0.388 | 0 | -0.009 | -0.009 | 0.17 | 0.24 | 0.32 | 0.26 | 0.14 | 0.25 | 0.22 |
| 11 | 3211468 | rs4237787 | -1.258 | 1.187 | -0.548 | 0.615 | 0.094 | -0.013 | -0.013 | 0 | 0.5 | 0.85 | 0.66 | 0.52 | 0.67 | 0.44 |
| 11 | 3212259 | rs11025844 | -0.484 | 1.942 | 0.156 | 1.008 | 0.049 | 0.051 | 0.051 | 0 | 0.15 | 0.18 | 0.34 | 0.37 | 0.35 | 0.2 |
| 11 | 3214158 | rs7926105 | -0.805 | 1.322 | 0.359 | 0.517 | 0.017 | 0.01 | 0.01 | 0 | 0.28 | 0.49 | 0.47 | 0.31 | 0.35 | 0.18 |
| 11 | 3215218 | rs7930262 | -0.542 | 1.307 | 0.497 | 0.545 | 0.001 | 0.001 | 0.001 | 0 | 0.25 | 0.3 | 0.44 | 0.3 | 0.34 | 0.19 |

| | | | | | | | | | | | | | | | | |
|---|---|---|---|---|---|---|---|---|---|---|---|---|---|---|---|---|
| 11 | 3215318 | rs7930392 | -0.371 | 1.106 | 0.289 | 0.024 | 0.014 | -0.039 | -0.039 | 0.161 | 0.66 | 0.56 | 0.67 | 0.8 | 0.61 | 0.67 |
| 11 | 3216129 | rs12802907 | 0.552 | 1.241 | -0.21 | -0.13 | -0.017 | 0.065 | 0.065 | 0.132 | 0.28 | 0.12 | 0.28 | 0.1 | 0.31 | 0.32 |
| 11 | 3218709 | rs4758480 | 0.032 | 1.068 | -0.089 | 0.085 | 0.025 | 0.03 | 0.03 | 0.128 | 0.7 | 0.52 | 0.53 | 0.58 | 0.61 | 0.82 |
| 11 | 3218934 | rs4758481 | 0.089 | 0.886 | -0.501 | 0.424 | -0.006 | 0.015 | 0.015 | 0 | 0.07 | 0.12 | 0.19 | 0.12 | 0.05 | 0.06 |
| 11 | 3220364 | rs12577685 | 1.057 | 0.559 | -1.154 | 0.274 | -0.056 | -0.179 | -0.179 | 0 | 0.26 | 0.01 | 0.18 | 0.48 | 0.23 | 0.41 |
| 11 | 3222869 | rs4758483 | -0.742 | 0.382 | -0.359 | 0.247 | -0.001 | 0.008 | 0.008 | 0 | 0.65 | 0.64 | 0.7 | 0.41 | 0.65 | 0.53 |
| 11 | 3223297 | rs11025909 | 0.89 | 0.42 | -0.361 | 0.274 | 0 | -0.066 | -0.066 | 0 | 0.31 | 0.11 | 0.24 | 0.51 | 0.3 | 0.47 |
| 11 | 3225964 | rs10833460 | 0.225 | 0.271 | -0.407 | -0.315 | 0.004 | 0.008 | 0.008 | 0 | 0.35 | 0.36 | 0.37 | 0.11 | 0.26 | 0.24 |
| 11 | 3226221 | rs10833462 | -1.078 | 0.426 | -0.856 | -0.184 | -0.009 | 0.017 | 0.017 | 0 | 0.63 | 0.78 | 0.69 | 0.31 | 0.59 | 0.45 |
| 11 | 3226934 | rs11025930 | 0.186 | 0.502 | -1.02 | -0.125 | -0.002 | 0 | 0 | 0 | 0.2 | 0.19 | 0.24 | 0.2 | 0.27 | 0.12 |
| 11 | 3230469 | rs4367927 | 0.106 | 0.209 | -0.944 | 0.269 | -0.007 | 0.014 | 0.014 | 0.06 | 0.65 | 0.4 | 0.74 | 0.56 | 0.69 | 0.71 |
| 11 | 3247346 | rs4556520 | 0.128 | 0.259 | -1.733 | 0.154 | -0.028 | -0.05 | -0.05 | 0 | 0.39 | 0.12 | 0.33 | 0.56 | 0.43 | 0.51 |
| 11 | 3248615 | rs4998520 | -0.342 | 0.173 | -1.684 | -0.352 | 0.018 | -0.041 | -0.041 | 0.328 | 0.13 | 0.04 | 0.09 | 0.17 | 0.08 | 0.11 |
| 11 | 3248942 | rs10833512 | 0.045 | 0.403 | -1.299 | -1.069 | -0.006 | 0.013 | 0.013 | 0.597 | 0.31 | 0.22 | 0.39 | 0.07 | 0.34 | 0.23 |
| 11 | 3249829 | rs11026040 | -0.155 | -0.938 | -0.891 | -0.788 | -0.016 | -0.019 | -0.019 | 2.826 | 0.27 | 0.28 | 0.32 | 0.13 | 0.14 | 0.27 |
| 11 | 3252320 | rs11026055 | 0.001 | -0.802 | -0.793 | -0.805 | -0.025 | -0.053 | -0.053 | 0 | 0.28 | 0.32 | 0.29 | 0.12 | 0.17 | 0.22 |
| 11 | 3253856 | rs4758558 | -0.465 | -0.668 | -1.185 | -0.442 | -0.004 | -0.04 | -0.04 | 1.006 | 0.09 | 0.02 | 0.21 | 0.2 | 0.1 | 0.14 |
| 11 | 3257511 | rs10741878 | -0.196 | -0.919 | -0.776 | -0.588 | 0.02 | 0.019 | 0.019 | 0 | 0.69 | 0.76 | 0.59 | 0.79 | 0.81 | 0.7 |
| 11 | 3258705 | rs10766807 | 0.689 | -0.839 | -0.74 | -1.065 | -0.002 | 0.012 | 0.012 | 0 | 0.41 | 0.65 | 0.42 | 0.75 | 0.57 | 0.47 |
| 11 | 3259083 | rs11026086 | 0.552 | -0.809 | -0.975 | -1.047 | 0.046 | 0.088 | 0.088 | 0.051 | 0.28 | 0.13 | 0.28 | 0.12 | 0.18 | 0.23 |
| 11 | 3259535 | rs4545536 | -0.875 | -0.643 | -0.869 | -0.9 | -0.016 | 0.061 | 0.061 | 0 | 0.8 | 0.94 | 0.89 | 0.95 | 0.82 | 0.83 |
| 11 | 3259809 | rs11026091 | 1.035 | -0.51 | -0.455 | -0.859 | 0.005 | 0 | 0 | 0 | 0.33 | 0.39 | 0.32 | 0.63 | 0.47 | 0.39 |
| 11 | 3261658 | rs11026102 | -0.417 | -0.347 | -1.278 | -0.739 | 0.054 | 0.066 | 0.066 | 0 | 0.45 | 0.26 | 0.53 | 0.33 | 0.36 | 0.4 |
| 11 | 3263298 | rs7103275 | -0.842 | -0.209 | -1.552 | -1.123 | -0.023 | 0.067 | 0.067 | 0 | 0.8 | 0.95 | 0.89 | 0.96 | 0.82 | 0.82 |
| 11 | 3263621 | rs11026107 | -0.369 | -0.726 | -1.135 | -0.596 | 0.003 | -0.001 | -0.001 | 0.667 | 0.13 | 0.14 | 0.13 | 0.11 | 0.19 | 0.12 |
| 11 | 3263822 | rs11026108 | -0.892 | -0.748 | -1.139 | -0.551 | 0.056 | 0.081 | 0.081 | 0 | 0.45 | 0.28 | 0.45 | 0.3 | 0.36 | 0.39 |
| 11 | 3265951 | rs7940835 | -1.127 | 0.708 | -0.358 | 0.162 | 0.003 | -0.001 | -0.001 | 0 | 0.13 | 0.14 | 0.13 | 0.11 | 0.19 | 0.12 |
| 11 | 3361087 | rs2073805 | 0.235 | 0.414 | -0.217 | 0.332 | 0.004 | 0.002 | 0.002 | 0 | 0.56 | 0.39 | 0.47 | 0.56 | 0.58 | 0.7 |
| 11 | 3364358 | rs2412166 | 1.329 | 0.811 | 0.761 | 0.815 | -0.005 | 0.062 | 0.062 | 0 | 0.72 | 0.58 | 0.7 | 0.52 | 0.72 | 0.6 |
| 11 | 3364517 | rs7109838 | 1.285 | 0.881 | 0.792 | 0.849 | -0.002 | 0.07 | 0.07 | 0 | 0.73 | 0.58 | 0.7 | 0.52 | 0.72 | 0.62 |
| 11 | 3367976 | rs969725 | 1.004 | 0.995 | 0.525 | 0.725 | -0.004 | 0.01 | 0.01 | 0 | 0.87 | 0.9 | 0.89 | 1 | 0.83 | 0.91 |
| 11 | 3371687 | rs11026661 | -1.727 | 0.899 | 0.675 | 0.879 | -0.001 | 0.039 | 0.039 | 0 | 0.13 | 0.16 | 0.19 | 0.47 | 0.11 | 0.29 |
| 11 | 3371853 | rs764477 | 1.365 | 0.839 | 1.428 | 0.41 | 0.005 | -0.114 | -0.114 | 0 | 0.83 | 0.59 | 0.85 | 0.99 | 0.82 | 0.89 |
| 11 | 3371863 | rs17761375 | -1.129 | 0.981 | 1.128 | 0.491 | 0.002 | -0.042 | -0.042 | 0.445 | 0.08 | 0.14 | 0.04 | 0 | 0.08 | 0.07 |
| 11 | 3374745 | rs11026691 | -0.933 | 1.23 | 0.84 | 0.831 | -0.019 | 0.021 | 0.021 | 0 | 0.05 | 0.01 | 0.08 | 0.35 | 0.05 | 0.1 |
| 11 | 3377046 | rs7395353 | -2.439 | 1.158 | 1.12 | 0.584 | -0.001 | 0.047 | 0.047 | 0 | 0.23 | 0.38 | 0.18 | 0.41 | 0.2 | 0.37 |
| 11 | 3377660 | rs11827296 | 1.315 | 1.181 | 1.02 | 0.474 | -0.001 | -0.022 | -0.022 | 0 | 0.84 | 0.81 | 0.86 | 0.99 | 0.82 | 0.89 |
| 11 | 3377794 | rs10766959 | 1.315 | 1.181 | 1.012 | 0.474 | -0.001 | -0.017 | -0.017 | 0 | 0.84 | 0.82 | 0.86 | 0.99 | 0.82 | 0.89 |
| 11 | 3378163 | rs4758573 | 0.091 | 1.164 | 1.003 | 0.539 | -0.001 | -0.003 | -0.003 | 0 | 0.92 | 0.9 | 0.91 | 0.99 | 0.9 | 0.95 |
| 11 | 3386251 | rs4758574 | 0.173 | 1.218 | 1.008 | 0.538 | -0.001 | -0.004 | -0.004 | 0 | 0.92 | 0.9 | 0.91 | 0.99 | 0.9 | 0.96 |
| 11 | 3392745 | rs2269733 | 1.799 | 1.156 | 1.151 | 0.585 | -0.001 | 0.029 | 0.029 | 0 | 0.78 | 0.68 | 0.79 | 0.58 | 0.79 | 0.66 |
| 11 | 3394438 | rs12417350 | -1.358 | 1.157 | 1.086 | 0.59 | -0.004 | -0.02 | -0.02 | 0 | 0.13 | 0.11 | 0.11 | 0.4 | 0.11 | 0.29 |
| 11 | 3397231 | rs7103207 | 1.875 | 1.146 | 1.122 | 0.589 | -0.002 | 0.049 | 0.049 | 0 | 0.78 | 0.63 | 0.78 | 0.58 | 0.79 | 0.65 |
| 11 | 3398067 | rs4758578 | 0.151 | 1.151 | 1.061 | 0.584 | 0 | -0.007 | -0.007 | 0 | 0.92 | 0.89 | 0.91 | 0.99 | 0.9 | 0.96 |
| 11 | 3398662 | rs10833833 | -1.452 | 1.15 | 1.079 | 0.601 | -0.001 | -0.007 | -0.007 | 0.314 | 0.14 | 0.13 | 0.11 | 0.41 | 0.11 | 0.3 |
| 11 | 3400334 | rs7931660 | 0.62 | 1.183 | 1.231 | 0.593 | 0.005 | -0.048 | -0.048 | 0.153 | 0.9 | 0.7 | 0.88 | 0.99 | 0.9 | 0.96 |
| 11 | 3401374 | rs7116856 | 0.899 | 1.757 | 1.085 | 0.548 | 0.053 | -0.06 | -0.06 | 0.717 | 0.89 | 0.66 | 0.86 | 0.99 | 0.83 | 0.93 |
| 11 | 3402094 | rs10741955 | 0.647 | 1.305 | 1.086 | 0.587 | 0.012 | -0.049 | -0.049 | 0 | 0.9 | 0.66 | 0.87 | 0.99 | 0.89 | 0.96 |
| 11 | 3403162 | rs3928927 | 1.19 | 1.255 | 0.935 | 0.659 | 0.002 | 0.041 | 0.041 | 0.312 | 0.69 | 0.46 | 0.73 | 0.58 | 0.7 | 0.6 |
| 11 | 3407529 | rs956453 | -1.487 | 1.978 | 0.147 | -0.361 | 0.024 | -0.109 | -0.109 | 0 | 0.3 | 0.64 | 0.23 | 0.03 | 0.35 | 0.28 |
| 11 | 3460621 | rs7105826 | -0.313 | 1.018 | -0.724 | 0.235 | -0.022 | 0.005 | 0.005 | 0 | 0.42 | 0.26 | 0.47 | 0.43 | 0.47 | 0.42 |
| 11 | 3568044 | rs10430923 | -0.918 | -0.617 | -0.894 | 0.065 | 0.006 | 0.001 | 0.001 | 0.13 | 0.37 | 0.35 | 0.23 | 0.3 | 0.26 | 0.29 |
| 11 | 3568166 | rs10430924 | -0.918 | -0.617 | -0.894 | 0.065 | 0.006 | 0.001 | 0.001 | 0.13 | 0.37 | 0.34 | 0.23 | 0.29 | 0.25 | 0.29 |
| 11 | 3583135 | rs12417683 | -0.387 | -0.909 | -1.272 | -0.355 | 0.03 | 0.023 | 0.023 | 0.886 | 0.17 | 0.08 | 0.05 | 0.11 | 0.07 | 0.11 |
| 11 | 3583753 | rs7127562 | -1.456 | -0.419 | -2.069 | -0.695 | -0.074 | 0.079 | 0.079 | 0 | 0.55 | 0.85 | 0.39 | 0.3 | 0.54 | 0.45 |
| 11 | 3613163 | rs11028320 | -0.374 | -0.89 | -1.565 | -0.558 | -0.05 | -0.009 | -0.009 | 0 | 0.1 | 0.02 | 0.13 | 0.08 | 0.16 | 0.13 |
| 11 | 3624237 | rs4910716 | -0.67 | -1.726 | -1.812 | -1.099 | 0.074 | 0.11 | 0.11 | 8.178 | 0.25 | 0.07 | 0.12 | 0.03 | 0.13 | 0.15 |
| 11 | 3625720 | rs7111164 | -2.424 | -2.296 | -1.401 | -1.176 | 0.085 | 0.105 | 0.105 | 1.719 | 0.52 | 0.84 | 0.35 | 0.29 | 0.36 | 0.33 |
| 11 | 3626177 | rs10834605 | -1.171 | -2.33 | -0.278 | -1.165 | 0.01 | 0.04 | 0.04 | 0 | 0.22 | 0.5 | 0.21 | 0.25 | 0.23 | 0.17 |
| 11 | 3629024 | rs4432051 | -2.148 | -2.627 | 0.299 | -1.159 | 0.023 | 0.022 | 0.022 | 2.569 | 0.48 | 0.61 | 0.31 | 0.29 | 0.32 | 0.31 |
| 11 | 3629578 | rs11028462 | -0.201 | -2.575 | 0.509 | -1.149 | 0.001 | -0.007 | -0.007 | 2.237 | 0.12 | 0.2 | 0.14 | 0.1 | 0.15 | 0.14 |
| 11 | 3631058 | rs10767317 | -0.761 | -2.757 | 0.505 | -1.259 | 0.012 | 0.025 | 0.025 | 2.695 | 0.26 | 0.23 | 0.22 | 0.1 | 0.23 | 0.15 |
| 11 | 3631121 | rs2203402 | -1.946 | -2.531 | 0.97 | -1.095 | 0.052 | 0.049 | 0.049 | 2.164 | 0.48 | 0.66 | 0.32 | 0.29 | 0.33 | 0.31 |
| 11 | 3631189 | rs10834615 | -0.357 | -1.765 | 1.147 | -0.718 | -0.001 | -0.013 | -0.013 | 0.222 | 0.11 | 0.2 | 0.12 | 0.08 | 0.14 | 0.12 |
| 11 | 3633789 | rs7107053 | -0.152 | -1.68 | 1.181 | -0.608 | -0.002 | 0.016 | 0.016 | 0 | 0.08 | 0.21 | 0.04 | 0.06 | 0.06 | 0.04 |
| 11 | 3638061 | rs6578398 | -1.358 | -1.902 | 0.501 | -0.922 | 0 | -0.005 | -0.005 | 0 | 0.39 | 0.65 | 0.27 | 0.15 | 0.29 | 0.28 |
| 11 | 3638280 | rs6578399 | -0.086 | -1.756 | 0.191 | -0.769 | 0.02 | 0.009 | 0.009 | 5.312 | 0.21 | 0.14 | 0.12 | 0.03 | 0.12 | 0.11 |
| 11 | 3639339 | rs11028536 | 0.788 | -0.573 | -0.145 | 0.061 | -0.124 | -0.084 | -0.084 | 0.18 | 0.34 | 0.1 | 0.5 | 0.62 | 0.43 | 0.46 |
| 11 | 3643778 | rs879114 | 0.114 | -1.125 | 0.886 | -0.116 | 0.018 | 0.013 | 0.013 | 0.814 | 0.35 | 0.27 | 0.19 | 0.18 | 0.19 | 0.27 |

| | | | | | | | | | | | | | | | | |
|---|---|---|---|---|---|---|---|---|---|---|---|---|---|---|---|---|
| 11 | 3645057 | rs2898934 | -0.765 | -0.964 | 1.5 | -0.083 | 0.033 | 0.017 | 0.017 | 0 | 0.11 | 0.33 | 0.12 | 0.06 | 0.12 | 0.05 |
| 11 | 3645133 | rs6578401 | 0.965 | -0.977 | 1.757 | 0.118 | 0.032 | 0.113 | 0.113 | 0 | 0.87 | 0.45 | 0.84 | 0.79 | 0.88 | 0.94 |
| 11 | 3645345 | rs10834648 | 0.578 | -0.996 | 1.008 | 0.124 | 0.13 | 0.074 | 0.074 | 0.16 | 0.49 | 0.05 | 0.63 | 0.61 | 0.68 | 0.67 |
| 11 | 3646127 | rs7115284 | -1.131 | -0.804 | 1.256 | 0.333 | 0.036 | 0.112 | 0.112 | 0 | 0.12 | 0.34 | 0.15 | 0.21 | 0.12 | 0.06 |
| 11 | 3647528 | rs17173897 | -0.969 | -0.806 | 1.261 | 0.332 | 0.018 | 0.066 | 0.066 | 0 | 0.11 | 0.25 | 0.14 | 0.2 | 0.12 | 0.05 |
| 11 | 3648067 | rs2271581 | -0.966 | -0.832 | 1.247 | 0.255 | 0.005 | -0.004 | -0.004 | 0 | 0.11 | 0.24 | 0.11 | 0.07 | 0.12 | 0.05 |
| 11 | 3648731 | rs11028621 | 0.536 | -0.643 | 0.833 | 0.245 | 0.007 | 0.041 | 0.041 | 0.198 | 0.44 | 0.05 | 0.6 | 0.62 | 0.63 | 0.65 |
| 11 | 3656521 | rs2271584 | -2.168 | 0.033 | 2.026 | 0.894 | 0.009 | 0.07 | 0.07 | 0.081 | 0.15 | 0.48 | 0.16 | 0.17 | 0.11 | 0.06 |
| 11 | 3657047 | rs1514690 | -1.824 | 0.075 | 1.91 | 0.889 | 0.011 | 0.063 | 0.063 | 0 | 0.13 | 0.35 | 0.15 | 0.16 | 0.11 | 0.06 |
| 11 | 3657436 | rs1514691 | -1.602 | 0.096 | 1.569 | 0.899 | -0.002 | 0.021 | 0.021 | 0 | 0.12 | 0.22 | 0.14 | 0.15 | 0.11 | 0.06 |
| 11 | 3659768 | rs2271585 | -1.663 | 0.155 | 1.236 | 0.865 | 0.003 | -0.004 | -0.004 | 0.453 | 0.14 | 0.12 | 0.11 | 0.14 | 0.11 | 0.06 |
| 11 | 3659993 | rs2271586 | -2.244 | 0.39 | 1.384 | 0.968 | 0.013 | 0.037 | 0.037 | 0.166 | 0.22 | 0.59 | 0.2 | 0.23 | 0.19 | 0.17 |
| 11 | 3660295 | rs1996208 | -1.011 | 0.026 | 1.306 | 1.088 | 0.013 | 0.118 | 0.118 | 0 | 0.11 | 0.33 | 0.27 | 0.18 | 0.08 | 0.04 |
| 11 | 3660914 | rs2271589 | -1.45 | 0.218 | 0.654 | 0.785 | -0.004 | 0.042 | 0.042 | 0 | 0.07 | 0.27 | 0.08 | 0.08 | 0.04 | 0.02 |
| 11 | 3663201 | rs7128137 | -0.042 | 0.481 | -0.164 | 0.582 | 0.023 | 0.019 | 0.019 | 0 | 0.47 | 0.32 | 0.36 | 0.57 | 0.58 | 0.62 |
| 11 | 3664427 | rs6578410 | -1.505 | 1.442 | 0.587 | 0.454 | 0.125 | -0.016 | -0.016 | 2.143 | 0.14 | 0.4 | 0.15 | 0.06 | 0.18 | 0.12 |
| 11 | 3676824 | rs11028806 | 0.326 | 0.509 | 0.106 | 0.555 | -0.012 | -0.007 | -0.007 | 3.979 | 0.62 | 0.52 | 0.82 | 0.29 | 0.68 | 0.55 |
| 11 | 3677075 | rs16925377 | 0.073 | 0.371 | 0.137 | 0.01 | 0.001 | -0.045 | -0.045 | 0 | 0.09 | 0.14 | 0.08 | 0 | 0.11 | 0.02 |
| 11 | 3679811 | rs925697 | 0.031 | -0.314 | -0.176 | -0.344 | -0.029 | -0.016 | -0.016 | 0 | 0.09 | 0.21 | 0.03 | 0.04 | 0.02 | 0.15 |
| 11 | 3681519 | rs2280134 | -0.405 | -0.421 | 0.311 | -0.39 | -0.002 | -0.001 | -0.001 | 0.056 | 0.64 | 0.45 | 0.81 | 0.29 | 0.7 | 0.63 |
| 11 | 3681850 | rs10834744 | -0.102 | -0.755 | 0.665 | 0.324 | -0.008 | 0.026 | 0.026 | 0 | 0.25 | 0.27 | 0.12 | 0.48 | 0.15 | 0.27 |
| 11 | 3685655 | rs2271583 | -0.899 | -0.684 | 1.066 | 0.16 | 0.004 | 0.029 | 0.029 | 0.265 | 0.86 | 0.77 | 0.93 | 0.72 | 0.82 | 0.82 |
| 11 | 3686221 | rs10834761 | 0.78 | -0.64 | 0.944 | -0.082 | 0.006 | 0.008 | 0.008 | 1.506 | 0.67 | 0.28 | 0.79 | 0.68 | 0.66 | 0.65 |
| 11 | 3691417 | rs2672213 | 1.961 | -0.953 | -0.061 | -0.985 | 0.025 | 0.093 | 0.093 | 3.669 | 0.22 | 0.07 | 0.18 | 0.03 | 0.15 | 0.1 |
| 11 | 3691649 | rs2672215 | 0.694 | -1.228 | 0.419 | -0.634 | 0.014 | 0.017 | 0.017 | 0 | 0.5 | 0.45 | 0.74 | 0.54 | 0.56 | 0.48 |
| 11 | 3693120 | rs2231532 | 0.718 | -1.323 | 0.237 | -0.714 | 0.034 | 0.088 | 0.088 | 0 | 0.5 | 0.28 | 0.73 | 0.64 | 0.56 | 0.47 |
| 11 | 3693253 | rs2231529 | 0.679 | -1.332 | 0.093 | -0.701 | 0.023 | 0.056 | 0.056 | 0 | 0.19 | 0.05 | 0.07 | 0.04 | 0.1 | 0.16 |
| 11 | 3695972 | rs6578411 | 0.539 | -1.343 | 0.064 | -0.701 | 0.027 | 0.063 | 0.063 | 0 | 0.19 | 0.05 | 0.07 | 0.04 | 0.1 | 0.17 |
| 11 | 3698843 | rs2741863 | 0.163 | -1.547 | 0.253 | -0.738 | -0.022 | -0.015 | -0.015 | 0 | 0.6 | 0.52 | 0.76 | 0.64 | 0.69 | 0.5 |
| 11 | 3703371 | rs12793302 | 0.574 | -1.513 | 0.154 | -0.72 | 0.023 | 0.062 | 0.062 | 0 | 0.19 | 0.05 | 0.07 | 0.04 | 0.1 | 0.17 |
| 11 | 3704770 | rs12293627 | 0.877 | -1.437 | 0.14 | -0.72 | -0.04 | 0.051 | 0.051 | 0.158 | 0.1 | 0 | 0.08 | 0 | 0.14 | 0.03 |
| 11 | 3706446 | rs4575303 | -0.684 | -1.467 | 0.309 | -0.772 | -0.02 | 0.037 | 0.037 | 1.304 | 0.79 | 0.62 | 0.9 | 0.68 | 0.81 | 0.65 |
| 11 | 3714489 | rs1875 | 0.459 | -1.568 | 0.268 | -0.384 | -0.018 | 0.034 | 0.034 | 1.215 | 0.21 | 0.36 | 0.1 | 0.32 | 0.19 | 0.33 |
| 11 | 3715036 | rs683142 | -1.511 | -1.64 | 0.185 | -0.402 | -0.036 | -0.001 | -0.001 | 0 | 0.56 | 0.84 | 0.33 | 0.64 | 0.46 | 0.62 |
| 11 | 3717120 | rs2957865 | 0.393 | -1.693 | 0.372 | -1.139 | -0.018 | 0.032 | 0.032 | 1.193 | 0.21 | 0.38 | 0.1 | 0.31 | 0.19 | 0.31 |
| 11 | 3718129 | rs12362694 | 0.269 | -1.765 | -0.037 | -1.268 | 0.012 | 0.013 | 0.013 | 0 | 0.15 | 0.08 | 0.04 | 0.06 | 0.07 | 0.12 |
| 11 | 3722304 | rs685782 | 1.336 | -1.643 | 0.104 | -1.395 | 0.005 | 0.014 | 0.014 | 0.056 | 0.14 | 0.19 | 0.08 | 0.27 | 0.2 | 0.18 |
| 11 | 3729676 | rs602879 | -0.14 | -1.493 | 0.237 | -1.65 | -0.029 | -0.017 | -0.017 | 0 | 0.3 | 0.47 | 0.13 | 0.32 | 0.22 | 0.4 |
| 11 | 3732960 | rs658731 | -1.668 | -1.708 | 0.002 | -2.058 | -0.004 | 0.005 | 0.005 | 0 | 0.59 | 0.61 | 0.3 | 0.43 | 0.51 | 0.5 |
| 11 | 3742758 | rs7941153 | 0.746 | -1.735 | 0.071 | -2.067 | -0.011 | -0.011 | -0.011 | 0 | 0.13 | 0.19 | 0.03 | 0.05 | 0.03 | 0.21 |
| 11 | 3743046 | rs2957866 | -1.758 | -1.742 | -0.383 | -2.077 | 0.006 | -0.009 | -0.009 | 0.355 | 0.59 | 0.57 | 0.3 | 0.43 | 0.52 | 0.56 |
| 11 | 3744079 | rs7124090 | 0.607 | -1.712 | -0.331 | -2.05 | -0.035 | -0.038 | -0.038 | 0 | 0.14 | 0.31 | 0.06 | 0.05 | 0.03 | 0.21 |
| 11 | 3745165 | rs7931173 | 0.155 | -1.725 | -0.861 | -2.127 | -0.016 | 0.029 | 0.029 | 0 | 0.16 | 0.07 | 0.11 | 0.11 | 0.2 | 0.13 |
| 11 | 3763168 | rs2672221 | -2.648 | -1.73 | -1.061 | -2.262 | 0.004 | 0.123 | 0.123 | 1.872 | 0.77 | 0.97 | 0.81 | 0.97 | 0.83 | 0.9 |
| 11 | 3768719 | rs2957869 | 0.596 | -1.949 | -0.27 | -1.769 | 0.005 | 0 | 0 | 0 | 0.22 | 0.11 | 0.22 | 0.27 | 0.22 | 0.21 |
| 11 | 3776371 | rs276892 | -2.318 | -1.868 | -0.181 | -1.759 | -0.011 | 0.011 | 0.011 | 0.019 | 0.6 | 0.66 | 0.44 | 0.43 | 0.52 | 0.56 |
| 11 | 3778432 | rs7933512 | -0.343 | -1.685 | -0.135 | -1.631 | -0.007 | 0.029 | 0.029 | 0 | 0.2 | 0.14 | 0.13 | 0.11 | 0.23 | 0.15 |
| 11 | 3792657 | rs7931756 | 0.203 | -1.697 | -0.096 | -1.608 | -0.027 | -0.034 | -0.034 | 0 | 0.14 | 0.28 | 0.06 | 0.05 | 0.04 | 0.16 |
| 11 | 3792691 | rs169346 | -0.543 | -1.435 | -0.623 | -1.479 | -0.024 | -0.023 | -0.023 | 0.059 | 0.41 | 0.48 | 0.31 | 0.32 | 0.3 | 0.36 |
| 11 | 3799985 | rs188044 | -1.921 | -1.229 | -1.141 | -1.411 | -0.038 | 0.028 | 0.028 | 0 | 0.6 | 0.77 | 0.32 | 0.42 | 0.52 | 0.5 |
| 11 | 3808138 | rs10767587 | -1.021 | -0.69 | -0.725 | -1.25 | -0.006 | 0.022 | 0.022 | 0 | 0.2 | 0.15 | 0.13 | 0.11 | 0.23 | 0.15 |
| 11 | 3813961 | rs4910810 | -0.552 | -0.741 | -0.336 | -0.558 | -0.024 | -0.044 | -0.044 | 3.385 | 0.1 | 0.17 | 0.06 | 0 | 0.08 | 0.01 |
| 11 | 3820785 | rs4910818 | 0.502 | -0.094 | -0.613 | -0.152 | 0.006 | 0 | 0 | 0 | 0.9 | 0.87 | 0.79 | 0.87 | 0.89 | 0.9 |
| 11 | 3827301 | rs10767612 | 1.154 | -0.707 | -1.329 | -0.744 | 0.064 | 0.114 | 0.114 | 0.368 | 0.6 | 0.2 | 0.55 | 0.81 | 0.74 | 0.68 |
| 11 | 3831304 | rs3781615 | 1.11 | -0.763 | -1.35 | -0.617 | 0.057 | 0.083 | 0.083 | 0.045 | 0.61 | 0.24 | 0.57 | 0.79 | 0.75 | 0.68 |
| 11 | 3838953 | rs10835147 | 0.664 | -0.962 | -1.226 | -0.689 | -0.006 | -0.005 | -0.005 | 0 | 0.75 | 0.34 | 0.68 | 0.81 | 0.88 | 0.71 |
| 11 | 3841463 | rs7120836 | 0.954 | -0.692 | -1.383 | -0.505 | 0.069 | 0.075 | 0.075 | 0.478 | 0.63 | 0.16 | 0.56 | 0.77 | 0.78 | 0.68 |
| 11 | 3847261 | rs1055639 | -0.302 | -0.013 | -1.341 | -0.314 | -0.011 | 0.055 | 0.055 | 0 | 0.45 | 0.24 | 0.43 | 0.29 | 0.46 | 0.43 |
| 11 | 3847293 | rs1055640 | -0.175 | -0.003 | -1.385 | -0.049 | 0 | -0.001 | -0.001 | 0 | 0.33 | 0.67 | 0.38 | 0.32 | 0.34 | 0.24 |
| 11 | 3847430 | rs15820 | 0.349 | 0.078 | -0.198 | -0.404 | 0.027 | -0.058 | -0.058 | 3.773 | 0.15 | 0.4 | 0.11 | 0.02 | 0.15 | 0.07 |
| 11 | 3848086 | rs4910852 | 0.405 | -0.083 | 0.011 | -0.613 | -0.008 | -0.056 | -0.056 | 0 | 0.11 | 0.14 | 0.09 | 0 | 0.1 | 0.04 |
| 11 | 3849711 | rs12272393 | -0.503 | -0.144 | 0.264 | -0.575 | 0 | -0.084 | -0.084 | 0 | 0.23 | 0.4 | 0.19 | 0.05 | 0.23 | 0.08 |
| 11 | 3851383 | rs7933771 | 1.143 | -0.067 | 0.369 | -0.499 | 0.015 | -0.01 | -0.01 | 0 | 0.12 | 0.15 | 0.1 | 0.05 | 0.14 | 0.06 |
| 11 | 3854765 | rs962199 | -1.054 | -0.209 | -0.182 | -0.4 | 0 | -0.001 | -0.001 | 0.987 | 0.64 | 0.62 | 0.63 | 0.59 | 0.66 | 0.62 |
| 11 | 3856553 | rs1451722 | -0.302 | -0.085 | -0.728 | -0.401 | -0.003 | -0.014 | -0.014 | 0 | 0.5 | 0.18 | 0.52 | 0.53 | 0.57 | 0.57 |
| 11 | 3856668 | rs1451724 | 0.192 | -0.007 | -0.884 | -0.345 | -0.074 | -0.092 | -0.092 | 0 | 0.39 | 0.12 | 0.45 | 0.53 | 0.47 | 0.53 |
| 11 | 3859386 | rs1349542 | -0.856 | -0.071 | -0.468 | -0.564 | -0.006 | -0.037 | -0.037 | 0.263 | 0.33 | 0.63 | 0.25 | 0.06 | 0.28 | 0.11 |
| 11 | 3863718 | rs7934650 | 0.459 | 0.304 | -0.918 | 0.431 | 0.003 | 0.068 | 0.068 | 0 | 0.92 | 0.85 | 0.84 | 0.79 | 0.91 | 0.88 |
| 11 | 3870463 | rs1869002 | 0.999 | 0.849 | -1.772 | 0.199 | -0.085 | -0.08 | -0.08 | 0 | 0.18 | 0.02 | 0.3 | 0.34 | 0.29 | 0.21 |

| | | | | | | | | | | | | | | | | |
|---|---|---|---|---|---|---|---|---|---|---|---|---|---|---|---|---|
| 11 | 3876433 | rs4367942 | 0.047 | 0.367 | -0.949 | -0.144 | -0.012 | -0.073 | -0.073 | 0 | 0.84 | 0.77 | 0.89 | 1 | 0.89 | 0.97 |
| 11 | 3881730 | rs11030106 | 0.488 | 0.551 | -1.883 | 0.134 | -0.062 | -0.06 | -0.06 | 0 | 0.23 | 0.06 | 0.25 | 0.36 | 0.31 | 0.42 |
| 11 | 3885511 | rs4414209 | 0.251 | -0.091 | -2.239 | -0.128 | 0.006 | -0.01 | -0.01 | 0 | 0.33 | 0.12 | 0.29 | 0.34 | 0.34 | 0.4 |
| 11 | 3886070 | rs4243965 | -1.322 | -0.467 | -2.544 | -0.251 | 0.013 | 0.005 | 0.005 | 0.058 | 0.91 | 0.98 | 0.94 | 0.97 | 0.95 | 0.97 |
| 11 | 3888234 | rs7932404 | 0.109 | -0.683 | -2.135 | -0.371 | 0.047 | 0.029 | 0.029 | 0 | 0.54 | 0.73 | 0.63 | 0.62 | 0.61 | 0.56 |
| 11 | 3889383 | rs4910582 | 0.568 | -0.849 | -2.086 | -0.631 | -0.007 | 0.007 | 0.007 | 0 | 0.41 | 0.68 | 0.54 | 0.62 | 0.5 | 0.51 |
| 11 | 3890059 | rs7126870 | 0.731 | -0.871 | -2.205 | -0.693 | -0.019 | -0.002 | -0.002 | 0 | 0.38 | 0.67 | 0.54 | 0.62 | 0.48 | 0.5 |
| 11 | 3890853 | rs7951076 | 1.765 | -0.381 | -1.957 | -0.34 | -0.026 | -0.004 | -0.004 | 0 | 0.23 | 0.11 | 0.26 | 0.28 | 0.3 | 0.26 |
| 11 | 3890960 | rs7952083 | -0.055 | -0.443 | -1.999 | -0.307 | -0.019 | 0.016 | 0.016 | 0 | 0.42 | 0.82 | 0.56 | 0.63 | 0.5 | 0.52 |
| 11 | 3891380 | rs11825543 | -0.108 | -0.354 | -1.871 | -0.209 | 0.082 | 0.167 | 0.167 | 0 | 0.18 | 0.51 | 0.28 | 0.35 | 0.21 | 0.28 |
| 11 | 3892053 | rs16929712 | 0.374 | -0.392 | -1.37 | -0.298 | 0.047 | -0.013 | -0.013 | 0.918 | 0.07 | 0.18 | 0.04 | 0.04 | 0.13 | 0.16 |
| 11 | 3893041 | rs16929706 | 0.299 | -0.483 | -1.397 | -0.283 | 0.031 | -0.014 | -0.014 | 0 | 0.08 | 0.16 | 0.04 | 0.04 | 0.13 | 0.15 |
| 11 | 3900133 | rs10835232 | 0.612 | -0.923 | -1.935 | -0.357 | 0.041 | 0.072 | 0.072 | 0.442 | 0.46 | 0.2 | 0.38 | 0.35 | 0.45 | 0.45 |
| 11 | 3900915 | rs7120828 | -0.09 | -0.85 | -1.789 | -0.261 | 0.069 | 0.142 | 0.142 | 0 | 0.19 | 0.52 | 0.28 | 0.34 | 0.21 | 0.3 |
| 11 | 3909286 | rs1869084 | 1.325 | -0.809 | -2.41 | -0.13 | 0.003 | 0.004 | 0.004 | 0 | 0.32 | 0.13 | 0.29 | 0.28 | 0.33 | 0.4 |
| 11 | 3910169 | rs11030173 | 1.37 | -0.791 | -2.333 | -0.224 | 0.005 | -0.008 | -0.008 | 0.102 | 0.13 | 0.11 | 0.09 | 0 | 0.11 | 0.05 |
| 11 | 3915880 | rs12792666 | 1.799 | -0.663 | -2.196 | 0.036 | -0.101 | -0.057 | -0.057 | 0 | 0.22 | 0.02 | 0.25 | 0.28 | 0.29 | 0.21 |
| 11 | 3916506 | rs10835262 | 0.487 | -0.808 | -1.609 | 0.061 | 0.012 | -0.001 | -0.001 | 0 | 0.33 | 0.25 | 0.31 | 0.34 | 0.33 | 0.4 |
| 11 | 3918601 | rs4622250 | 0.547 | -0.685 | -1.735 | 0.152 | -0.001 | -0.01 | -0.01 | 0 | 0.3 | 0.11 | 0.28 | 0.3 | 0.32 | 0.34 |
| 11 | 3924712 | rs11030210 | -0.955 | -0.494 | -1.434 | 0.205 | 0.011 | 0.01 | 0.01 | 0.059 | 0.91 | 0.98 | 0.95 | 0.96 | 0.95 | 0.95 |
| 11 | 3925776 | rs7929653 | 0.278 | -0.454 | -1.088 | 0.349 | 0.006 | 0.015 | 0.015 | 0 | 0.53 | 0.31 | 0.53 | 0.57 | 0.58 | 0.53 |
| 11 | 3928752 | rs10742189 | 0.371 | -0.116 | -0.399 | 1.024 | -0.008 | 0.005 | 0.005 | 0 | 0.45 | 0.35 | 0.59 | 0.57 | 0.52 | 0.46 |
| 11 | 3929164 | rs11030224 | 0.617 | 0.1 | -0.415 | 1.114 | -0.005 | -0.001 | -0.001 | 0 | 0.41 | 0.34 | 0.45 | 0.4 | 0.5 | 0.43 |
| 11 | 3931152 | rs2060822 | -0.397 | 0.122 | -0.384 | 1.069 | 0.003 | -0.001 | -0.001 | 0 | 0.92 | 0.98 | 0.96 | 0.95 | 0.94 | 0.95 |
| 11 | 3931619 | rs4365044 | 0.956 | 0.105 | -0.335 | 1.036 | 0 | -0.003 | -0.003 | 0 | 0.87 | 0.89 | 0.8 | 0.77 | 0.9 | 0.91 |
| 11 | 3933564 | rs10430939 | -1.188 | 0.242 | -0.215 | 1.062 | 0.02 | 0.087 | 0.087 | 0.042 | 0.07 | 0.14 | 0.15 | 0.18 | 0.06 | 0.04 |
| 11 | 3934132 | rs10767702 | 0.553 | 0.04 | -0.607 | 0.908 | -0.041 | 0.017 | 0.017 | 0 | 0.38 | 0.02 | 0.28 | 0.31 | 0.31 | 0.45 |
| 11 | 3938345 | rs7120938 | -0.733 | 0.078 | -0.403 | 0.915 | -0.002 | -0.012 | -0.012 | 0.596 | 0.52 | 0.78 | 0.45 | 0.54 | 0.44 | 0.53 |
| 11 | 3938385 | rs10160649 | -0.136 | 0.084 | 0.341 | 0.853 | 0.119 | 0.061 | 0.061 | 0.627 | 0.07 | 0.48 | 0.08 | 0.05 | 0.08 | 0.05 |
| 11 | 3948894 | rs750525 | -0.07 | 0.102 | 0.244 | 0.919 | 0.023 | -0.006 | -0.006 | 0 | 0.45 | 0.28 | 0.38 | 0.5 | 0.36 | 0.49 |
| 11 | 3957766 | rs10835333 | -0.153 | 0.086 | 0.252 | 0.912 | 0.004 | -0.001 | -0.001 | 0 | 0.46 | 0.37 | 0.38 | 0.5 | 0.36 | 0.49 |
| 11 | 3993231 | rs10835407 | 0.555 | -0.06 | 0.102 | 0.838 | -0.038 | 0.001 | 0.001 | 0 | 0.38 | 0.03 | 0.28 | 0.32 | 0.31 | 0.44 |
| 11 | 4011806 | rs2923946 | 0.536 | -0.022 | 0.318 | 0.836 | 0.032 | -0.029 | -0.029 | 0.217 | 0.83 | 0.52 | 0.88 | 0.91 | 0.84 | 0.9 |
| 11 | 4019445 | rs2923945 | 1.153 | -0.026 | -0.078 | 1.007 | -0.145 | -0.093 | -0.093 | 0.423 | 0.38 | 0.02 | 0.44 | 0.37 | 0.49 | 0.39 |
| 11 | 4027040 | rs2959081 | 1.054 | -0.015 | -0.018 | 0.948 | -0.055 | -0.037 | -0.037 | 0.012 | 0.39 | 0.15 | 0.46 | 0.4 | 0.5 | 0.43 |
| 11 | 4039056 | rs11030639 | 0.101 | -0.511 | 0.028 | 0.869 | 0.023 | 0.034 | 0.034 | 0 | 0.42 | 0.26 | 0.3 | 0.34 | 0.32 | 0.44 |
| 11 | 4081504 | rs894457 | 0.087 | -0.397 | 0.25 | 0.905 | -0.026 | -0.03 | -0.03 | 0.761 | 0.87 | 0.51 | 0.92 | 0.96 | 0.95 | 0.94 |
| 11 | 4083258 | rs11030781 | 0.589 | -0.427 | -0.219 | 0.767 | -0.041 | 0.014 | 0.014 | 0.181 | 0.36 | 0.06 | 0.29 | 0.33 | 0.34 | 0.44 |
| 11 | 4088356 | rs2920142 | 0.666 | -0.629 | -0.024 | 0.83 | -0.032 | -0.059 | -0.059 | 0 | 0.84 | 0.39 | 0.9 | 0.96 | 0.92 | 0.93 |
| 11 | 4093950 | rs7933164 | 1.4 | -0.652 | -0.269 | 0.92 | 0.026 | -0.019 | -0.019 | 0.047 | 0.78 | 0.18 | 0.7 | 0.75 | 0.87 | 0.9 |
| 11 | 4099013 | rs11030841 | 1.419 | -0.679 | -0.264 | 0.883 | 0.028 | -0.011 | -0.011 | 0 | 0.77 | 0.19 | 0.77 | 0.78 | 0.88 | 0.9 |
| 11 | 4103524 | rs2304891 | 0.71 | -0.517 | -0.305 | 0.951 | -0.054 | -0.053 | -0.053 | 0 | 0.44 | 0.1 | 0.48 | 0.44 | 0.57 | 0.48 |
| 11 | 4114539 | rs1465952 | 0.598 | -0.405 | 0.31 | 0.922 | -0.016 | -0.038 | -0.038 | 0 | 0.85 | 0.48 | 0.91 | 0.95 | 0.92 | 0.93 |
| 11 | 4115264 | rs1662162 | 0.67 | -0.477 | 0.073 | 1.075 | -0.044 | 0.065 | 0.065 | 0.466 | 0.83 | 0.43 | 0.8 | 0.77 | 0.92 | 0.91 |
| 11 | 4115974 | rs12806698 | 0.577 | -0.316 | -0.583 | 0.861 | -0.057 | -0.01 | -0.01 | 0.182 | 0.3 | 0.03 | 0.22 | 0.3 | 0.28 | 0.36 |
| 11 | 4117155 | rs1662172 | 0.349 | -0.166 | -0.017 | 1.089 | 0.002 | -0.006 | -0.006 | 0 | 0.45 | 0.39 | 0.51 | 0.48 | 0.59 | 0.49 |
| 11 | 4129859 | rs11030976 | 0.43 | 0.039 | 0.014 | 1.239 | -0.007 | -0.025 | -0.025 | 0 | 0.43 | 0.26 | 0.48 | 0.47 | 0.57 | 0.48 |
| 11 | 4142592 | rs3817657 | 0.309 | -0.169 | 0.527 | 1.192 | -0.005 | -0.018 | -0.018 | 0 | 0.41 | 0.43 | 0.33 | 0.3 | 0.35 | 0.44 |
| 11 | 4148948 | rs1980412 | 0.697 | -0.223 | 0.26 | 1.21 | -0.177 | -0.072 | -0.072 | 0.488 | 0.4 | 0.02 | 0.44 | 0.38 | 0.52 | 0.39 |
| 11 | 4159929 | rs1042927 | 0.362 | -0.172 | 0.33 | 1.258 | -0.015 | 0.02 | 0.02 | 1.015 | 0.85 | 0.8 | 0.83 | 0.76 | 0.93 | 0.93 |
| 11 | 4164967 | rs2898949 | 0.362 | -0.172 | 0.33 | 1.258 | -0.015 | 0.02 | 0.02 | 1.015 | 0.85 | 0.8 | 0.83 | 0.76 | 0.93 | 0.93 |
| 11 | 4165529 | rs4910904 | 0.365 | -0.174 | 0.643 | 1.255 | 0.003 | 0.009 | 0.009 | 0.005 | 0.41 | 0.38 | 0.34 | 0.29 | 0.35 | 0.44 |
| 11 | 4167416 | rs10835678 | 0.014 | -0.131 | 0.478 | 1.312 | 0.002 | 0.003 | 0.003 | 0.646 | 0.88 | 0.87 | 0.95 | 0.95 | 0.95 | 0.95 |
| 11 | 4168343 | rs4593998 | 1.041 | -0.125 | 0.485 | 1.3 | -0.016 | 0.084 | 0.084 | 0.444 | 0.83 | 0.65 | 0.79 | 0.68 | 0.88 | 0.84 |
| 11 | 4170922 | rs2412341 | 0.1 | -0.181 | -0.016 | 0.515 | 0.002 | 0.003 | 0.003 | 0.646 | 0.88 | 0.87 | 0.95 | 0.95 | 0.95 | 0.95 |
| 11 | 4173173 | rs2099875 | 0.205 | -0.153 | 0.247 | 0.565 | 0 | -0.001 | -0.001 | 0 | 0.46 | 0.49 | 0.59 | 0.47 | 0.55 | 0.47 |
| 11 | 4173253 | rs2099876 | 0.923 | 0.375 | 0.09 | 0.255 | -0.01 | -0.061 | -0.061 | 0.009 | 0.28 | 0.11 | 0.43 | 0.43 | 0.34 | 0.32 |
| 11 | 4176502 | rs9919651 | 0.059 | -0.021 | 0.088 | 0.357 | -0.055 | 0.06 | 0.06 | 2.584 | 0.82 | 0.59 | 0.84 | 0.76 | 0.92 | 0.93 |
| 11 | 4185595 | rs979942 | 0.063 | 0.267 | 0.11 | 0.182 | 0.011 | 0.022 | 0.022 | 1.539 | 0.51 | 0.43 | 0.38 | 0.31 | 0.55 | 0.6 |
| 11 | 4187233 | rs2044135 | 0.988 | 0.295 | -0.331 | 0.191 | -0.118 | -0.003 | -0.003 | 0 | 0.35 | 0.02 | 0.24 | 0.27 | 0.37 | 0.46 |
| 11 | 4188188 | rs1442727 | 0.832 | 0.332 | -0.14 | 0.209 | -0.044 | 0.049 | 0.049 | 0.971 | 0.37 | 0.14 | 0.26 | 0.27 | 0.37 | 0.46 |
| 11 | 4200169 | rs11031226 | 0.312 | 0.403 | 0.004 | 0.21 | 0.017 | -0.018 | -0.018 | 0 | 0.06 | 0.09 | 0.06 | 0 | 0.13 | 0.06 |
| 11 | 4213590 | rs894459 | 0.727 | 0.359 | -0.134 | 0.263 | -0.026 | 0.055 | 0.055 | 1.708 | 0.39 | 0.19 | 0.32 | 0.27 | 0.39 | 0.42 |
| 11 | 4213853 | rs894460 | -0.207 | 0.199 | 0.34 | 0.299 | -0.038 | 0.062 | 0.062 | 3.161 | 0.84 | 0.76 | 0.87 | 0.74 | 0.93 | 0.93 |
| 11 | 4215172 | rs894461 | 1.078 | 0.266 | 0.243 | 0.266 | -0.022 | -0.049 | -0.049 | 2.083 | 0.3 | 0.15 | 0.45 | 0.43 | 0.37 | 0.38 |
| 11 | 4215597 | rs11031262 | 0.824 | 0.42 | 0.195 | 0.215 | -0.035 | 0.029 | 0.029 | 0 | 0.06 | 0.01 | 0.06 | 0 | 0.14 | 0.08 |
| 11 | 4218391 | rs10767886 | 0.839 | 0.382 | 0.224 | 0.26 | -0.033 | 0.057 | 0.057 | 1.615 | 0.38 | 0.16 | 0.28 | 0.27 | 0.39 | 0.42 |
| 11 | 4222978 | rs7104032 | 1.363 | 0.096 | 0.655 | 0.398 | 0.012 | 0.001 | 0.001 | 3.908 | 0.67 | 0.28 | 0.72 | 0.7 | 0.75 | 0.8 |
| 11 | 4227908 | rs6578461 | -0.037 | 0.044 | 0.704 | 0.404 | 0.015 | -0.022 | -0.022 | 0 | 0.85 | 0.9 | 0.87 | 0.75 | 0.93 | 0.93 |

| | | | | | | | | | | | | | | | | |
|---|---|---|---|---|---|---|---|---|---|---|---|---|---|---|---|---|
| 11 | 4231825 | rs4910604 | 0.977 | 0.112 | 0.646 | 0.449 | -0.004 | 0 | 0 | 0.252 | 0.68 | 0.35 | 0.72 | 0.71 | 0.73 | 0.8 |
| 11 | 4387760 | rs12293167 | -0.134 | -0.176 | -0.279 | 0.05 | -0.001 | 0.06 | 0.06 | 0 | 0.52 | 0.29 | 0.43 | 0.3 | 0.5 | 0.37 |
| 11 | 4389639 | rs4910623 | -0.139 | -0.19 | -0.285 | 0.044 | -0.001 | 0.059 | 0.059 | 0 | 0.52 | 0.29 | 0.43 | 0.3 | 0.5 | 0.37 |
| 11 | 4391231 | rs17214070 | 1.352 | -0.381 | -0.949 | -0.186 | -0.015 | 0.009 | 0.009 | 0 | 0.15 | 0.04 | 0.14 | 0.13 | 0.14 | 0.22 |
| 11 | 4391529 | rs6578490 | -0.789 | -0.444 | -0.965 | 0.096 | 0.002 | 0.013 | 0.013 | 0 | 0.65 | 0.56 | 0.55 | 0.6 | 0.64 | 0.63 |
| 11 | 4396185 | rs10835979 | -1.651 | -0.199 | -1.395 | 0.158 | -0.012 | 0.009 | 0.009 | 0 | 0.85 | 0.95 | 0.86 | 0.87 | 0.86 | 0.77 |
| 11 | 4399281 | rs7934676 | 0.77 | 0.252 | -1.198 | 0.454 | -0.023 | -0.03 | -0.03 | 0 | 0.29 | 0.15 | 0.33 | 0.36 | 0.35 | 0.32 |
| 11 | 4400801 | rs11603547 | 0.374 | 0.401 | -1.077 | 0.347 | -0.06 | 0.048 | 0.048 | 0 | 0.27 | 0.03 | 0.23 | 0.17 | 0.33 | 0.14 |
| 11 | 4404192 | rs2052731 | -1.047 | 0.11 | 0.115 | 0.887 | 0.007 | 0.065 | 0.065 | 0 | 0.17 | 0.35 | 0.14 | 0.32 | 0.15 | 0.27 |
| 11 | 4406479 | rs4144331 | -1.077 | 0.063 | -0.147 | 0.737 | -0.001 | 0.016 | 0.016 | 0.439 | 0.11 | 0.2 | 0.07 | 0.21 | 0.1 | 0.2 |
| 11 | 4407725 | rs2269328 | -0.528 | -0.565 | -0.293 | 0.199 | 0.001 | 0 | 0 | 0 | 0.25 | 0.23 | 0.23 | 0.14 | 0.2 | 0.27 |
| 11 | 4409386 | rs2554933 | 0.022 | -0.189 | -0.823 | 0.118 | -0.008 | 0.016 | 0.016 | 0 | 0.53 | 0.68 | 0.43 | 0.44 | 0.48 | 0.59 |
| 11 | 4410752 | rs2855142 | 0.046 | -0.274 | -0.45 | -0.304 | -0.001 | 0.001 | 0.001 | 0 | 0.34 | 0.37 | 0.27 | 0.44 | 0.28 | 0.48 |
| 11 | 4411241 | rs926101 | 0.324 | -0.288 | -0.165 | -0.308 | 0 | 0.011 | 0.011 | 0 | 0.16 | 0.19 | 0.13 | 0.28 | 0.12 | 0.31 |
| 11 | 4413960 | rs928915 | 0.697 | -0.279 | -0.334 | -0.358 | -0.018 | 0.004 | 0.004 | 0 | 0.4 | 0.27 | 0.49 | 0.58 | 0.48 | 0.32 |
| 11 | 4415064 | rs12365876 | 0.87 | -0.3 | -0.27 | -0.442 | -0.009 | -0.104 | -0.104 | 0.074 | 0.23 | 0.31 | 0.16 | 0 | 0.21 | 0.11 |
| 11 | 4415079 | rs11032080 | 0.854 | -0.299 | -0.27 | -0.441 | -0.009 | -0.104 | -0.104 | 0 | 0.23 | 0.31 | 0.16 | 0 | 0.21 | 0.11 |
| 11 | 4430936 | rs1376679 | -0.132 | -0.025 | -0.523 | -0.005 | -0.001 | 0 | 0 | 0 | 0.4 | 0.58 | 0.51 | 0.69 | 0.44 | 0.44 |
| 11 | 4431327 | rs11032188 | 1.309 | 0.006 | -0.602 | 0.046 | -0.048 | -0.077 | -0.077 | 0.009 | 0.22 | 0.08 | 0.35 | 0.41 | 0.32 | 0.12 |
| 11 | 4431543 | rs1451656 | -0.788 | -0.304 | -0.61 | -0.395 | -0.003 | -0.01 | -0.01 | 0 | 0.83 | 0.74 | 0.78 | 0.87 | 0.81 | 0.79 |
| 11 | 4433414 | rs4910978 | 0.034 | -0.274 | -1.055 | -0.553 | 0.022 | 0.129 | 0.129 | 0 | 0.42 | 0.15 | 0.28 | 0.19 | 0.38 | 0.33 |
| 11 | 4436470 | rs1549489 | 0.514 | -0.06 | -0.438 | -0.285 | 0.011 | 0.075 | 0.075 | 0 | 0.57 | 0.24 | 0.64 | 0.75 | 0.59 | 0.64 |
| 11 | 4440617 | rs1376685 | 0.477 | -0.669 | 0.528 | -0.487 | -0.012 | -0.024 | -0.024 | 0.38 | 0.6 | 0.47 | 0.62 | 0.73 | 0.65 | 0.49 |
| 11 | 4440692 | rs1823633 | 0.199 | -0.675 | 0.566 | -0.454 | -0.013 | -0.012 | -0.012 | 0 | 0.65 | 0.56 | 0.77 | 0.73 | 0.71 | 0.55 |
| 11 | 4440838 | rs1823634 | 0.329 | -0.667 | 0.557 | -0.503 | -0.02 | -0.019 | -0.019 | 0.235 | 0.62 | 0.44 | 0.75 | 0.71 | 0.71 | 0.49 |
| 11 | 4443472 | rs1030392 | 0.551 | -0.559 | 0.349 | -0.438 | 0.029 | 0.058 | 0.058 | 0 | 0.57 | 0.25 | 0.59 | 0.73 | 0.66 | 0.43 |
| 11 | 4448649 | rs10836055 | -0.474 | -0.08 | -0.073 | 0.198 | -0.004 | -0.014 | -0.014 | 0.773 | 0.07 | 0.05 | 0.07 | 0.09 | 0.08 | 0.14 |
| 11 | 4454297 | rs4910980 | 0.367 | -0.177 | -0.274 | -0.015 | 0.049 | 0.075 | 0.075 | 0 | 0.5 | 0.4 | 0.66 | 0.73 | 0.65 | 0.42 |
| 11 | 4454913 | rs6578494 | 0.01 | -0.21 | -0.338 | 0.057 | 0.019 | 0.027 | 0.027 | 0 | 0.6 | 0.71 | 0.64 | 0.76 | 0.7 | 0.42 |
| 11 | 4455955 | rs10500601 | -0.608 | 1.132 | -0.549 | 0.869 | -0.057 | -0.074 | -0.074 | 0 | 0.21 | 0.05 | 0.23 | 0.36 | 0.34 | 0.21 |
| 11 | 4455985 | rs966496 | -1.505 | 0.896 | 0.292 | 0.529 | 0.021 | 0.009 | 0.009 | 0 | 0.32 | 0.51 | 0.36 | 0.37 | 0.4 | 0.31 |
| 11 | 4456562 | rs10500602 | -0.448 | 0.842 | 0.02 | 0.461 | 0.003 | -0.028 | -0.028 | 0.027 | 0.07 | 0.15 | 0.11 | 0.02 | 0.07 | 0.1 |
| 11 | 4458967 | rs4910981 | -1.811 | 0.81 | -0.411 | 0.805 | -0.038 | -0.033 | -0.033 | 0 | 0.42 | 0.75 | 0.48 | 0.59 | 0.47 | 0.57 |
| 11 | 4460348 | rs1863514 | -0.428 | 1.065 | -0.437 | 1.202 | -0.02 | -0.039 | -0.039 | 0 | 0.2 | 0.09 | 0.23 | 0.36 | 0.3 | 0.21 |
| 11 | 4465378 | rs331539 | 0.41 | 0.794 | 0.226 | 1.02 | 0.009 | 0.061 | 0.061 | 0 | 0.9 | 0.81 | 0.89 | 0.79 | 0.92 | 0.76 |
| 11 | 4466869 | rs12286769 | 0.191 | 0.862 | 0.074 | 0.791 | 0 | 0 | 0 | 0 | 0.55 | 0.55 | 0.5 | 0.42 | 0.47 | 0.43 |
| 11 | 4467638 | rs191353 | -0.927 | 0.549 | -0.213 | 0.527 | 0.017 | -0.038 | -0.038 | 0 | 0.58 | 0.84 | 0.69 | 0.63 | 0.65 | 0.65 |
| 11 | 4471473 | rs7934336 | -1.052 | 0.505 | 0.926 | 0.433 | 0.045 | 0.077 | 0.077 | 0 | 0.07 | 0.37 | 0.07 | 0.03 | 0.03 | 0.03 |
| 11 | 4472186 | rs11032298 | -0.312 | 0.699 | 0.619 | 0.414 | -0.008 | -0.012 | -0.012 | 0 | 0.06 | 0.04 | 0.07 | 0.17 | 0.09 | 0.12 |
| 11 | 4472880 | rs11032304 | -1.776 | 0.663 | 0.968 | 0.428 | 0.028 | 0.078 | 0.078 | 0 | 0.12 | 0.36 | 0.14 | 0.2 | 0.12 | 0.13 |
| 11 | 4473634 | rs1365317 | -1.688 | 0.573 | 1.176 | 0.435 | 0.048 | 0.096 | 0.096 | 0 | 0.1 | 0.34 | 0.11 | 0.17 | 0.09 | 0.17 |
| 11 | 4473720 | rs167948 | -1.153 | 0.999 | 0.641 | 0.382 | -0.04 | -0.043 | -0.043 | 0 | 0.37 | 0.76 | 0.48 | 0.58 | 0.45 | 0.43 |
| 11 | 4476699 | rs331535 | -1.715 | 0.856 | 0.486 | 0.161 | 0.052 | -0.006 | -0.006 | 0 | 0.52 | 0.75 | 0.58 | 0.6 | 0.62 | 0.61 |
| 11 | 4483678 | rs11032345 | 0.583 | 0.817 | 1.086 | 0.147 | 0.032 | 0.036 | 0.036 | 0 | 0.2 | 0.24 | 0.31 | 0.38 | 0.31 | 0.24 |
| 11 | 4485007 | rs11032347 | -0.846 | 0.812 | 0.986 | 0.122 | 0.002 | 0.005 | 0.005 | 0 | 0.27 | 0.26 | 0.2 | 0.22 | 0.26 | 0.34 |
| 11 | 4488230 | rs11032361 | -0.319 | 0.996 | 1.122 | 0.15 | 0.075 | 0.064 | 0.064 | 0 | 0.24 | 0.44 | 0.36 | 0.38 | 0.33 | 0.25 |
| 11 | 4491786 | rs331501 | 0.545 | 0.508 | -0.303 | 0.002 | -0.031 | 0.059 | 0.059 | 0 | 0.33 | 0.04 | 0.26 | 0.21 | 0.33 | 0.21 |
| 11 | 4492466 | rs11032378 | 0.029 | 1.017 | 0.689 | 0.319 | 0.013 | 0.01 | 0.01 | 0.146 | 0.21 | 0.22 | 0.33 | 0.38 | 0.33 | 0.24 |
| 11 | 4495028 | rs331503 | 0.192 | 1.028 | 0.187 | 0.334 | -0.052 | -0.063 | -0.063 | 0 | 0.19 | 0.12 | 0.3 | 0.42 | 0.31 | 0.23 |
| 11 | 4496846 | rs2278170 | 0.159 | 1.044 | 0.3 | 0.272 | -0.032 | -0.037 | -0.037 | 0.074 | 0.2 | 0.16 | 0.29 | 0.37 | 0.31 | 0.24 |
| 11 | 4496961 | rs2278172 | 0.009 | 1.025 | 0.533 | 0.272 | 0.019 | 0.022 | 0.022 | 0 | 0.21 | 0.24 | 0.31 | 0.37 | 0.32 | 0.24 |
| 11 | 4500031 | rs12796582 | -0.479 | 0.663 | 0.656 | -0.383 | 0.004 | -0.037 | -0.037 | 0 | 0.22 | 0.3 | 0.17 | 0.07 | 0.24 | 0.22 |
| 11 | 4500876 | rs7110328 | -2.313 | 0.362 | 0.973 | -0.093 | 0.011 | 0.04 | 0.04 | 1.076 | 0.06 | 0.62 | 0.1 | 0.05 | 0.03 | 0.03 |
| 11 | 4508405 | rs331508 | -0.726 | -0.43 | 0.705 | 0.011 | -0.006 | -0.001 | -0.001 | 0 | 0.35 | 0.55 | 0.22 | 0.32 | 0.3 | 0.44 |
| 11 | 4510285 | rs96489 | -0.734 | -0.437 | 0.729 | 0.017 | -0.006 | -0.001 | -0.001 | 0 | 0.35 | 0.55 | 0.22 | 0.32 | 0.3 | 0.44 |
| 11 | 4513664 | rs167947 | -0.729 | -0.448 | 1.012 | 0.011 | -0.023 | 0.004 | 0.004 | 0.072 | 0.34 | 0.44 | 0.21 | 0.32 | 0.3 | 0.44 |
| 11 | 4516365 | rs10160717 | -0.937 | -0.448 | 1.236 | 0.072 | 0.013 | 0.071 | 0.071 | 0 | 0.06 | 0.19 | 0.05 | 0.09 | 0.05 | 0.05 |
| 11 | 4516576 | rs10160399 | -1.315 | -0.481 | 1.277 | 0.056 | 0.009 | 0.068 | 0.068 | 0 | 0.07 | 0.23 | 0.07 | 0.09 | 0.06 | 0.05 |
| 11 | 4524438 | rs2412382 | -1.901 | -0.502 | 1.333 | 0.045 | 0.028 | 0.108 | 0.108 | 0.216 | 0.09 | 0.44 | 0.09 | 0.09 | 0.06 | 0.05 |
| 11 | 4527196 | rs2709193 | -0.086 | -0.449 | 0.717 | -0.02 | 0.012 | 0.004 | 0.004 | 0 | 0.27 | 0.1 | 0.15 | 0.23 | 0.25 | 0.39 |
| 11 | 4533118 | rs2566238 | -0.07 | -0.446 | 0.718 | -0.018 | 0.011 | 0.003 | 0.003 | 0 | 0.27 | 0.1 | 0.15 | 0.23 | 0.25 | 0.39 |
| 11 | 4534420 | rs2709152 | -0.21 | -0.45 | 0.73 | -0.02 | 0.015 | 0.007 | 0.007 | 0 | 0.28 | 0.12 | 0.15 | 0.23 | 0.25 | 0.39 |
| 11 | 4538034 | rs2709159 | -0.18 | -0.429 | 0.822 | -0.005 | 0.014 | 0.01 | 0.01 | 0 | 0.28 | 0.15 | 0.15 | 0.23 | 0.25 | 0.39 |
| 11 | 4539336 | rs2566208 | -0.189 | -0.509 | 0.747 | -0.018 | 0.003 | 0.006 | 0.006 | 0 | 0.28 | 0.1 | 0.16 | 0.23 | 0.27 | 0.37 |
| 11 | 4542813 | rs7131138 | 0.616 | -0.591 | 0.879 | -0.402 | 0.006 | -0.054 | -0.054 | 0.018 | 0.1 | 0.15 | 0.09 | 0 | 0.09 | 0.03 |
| 11 | 4545673 | rs2709135 | -1.021 | -0.781 | 1.095 | -0.407 | 0.009 | 0.017 | 0.017 | 0 | 0.45 | 0.54 | 0.28 | 0.27 | 0.38 | 0.43 |
| 11 | 4546132 | rs2657158 | -0.37 | -0.792 | 1.091 | -0.411 | -0.003 | 0.06 | 0.06 | 0 | 0.37 | 0.13 | 0.23 | 0.23 | 0.36 | 0.41 |
| 11 | 4546868 | rs2709136 | -1.206 | -0.799 | 1.049 | -0.431 | 0.031 | 0.035 | 0.035 | 0 | 0.47 | 0.61 | 0.3 | 0.32 | 0.39 | 0.44 |
| 11 | 4548949 | rs11823077 | -0.896 | -0.552 | 1.008 | -0.271 | 0.008 | 0.038 | 0.038 | 0 | 0.07 | 0.28 | 0.04 | 0.04 | 0.03 | 0.02 |

| | | | | | | | | | | | | | | | | |
|---|---|---|---|---|---|---|---|---|---|---|---|---|---|---|---|---|
| 11 | 4550729 | rs12282850 | -0.856 | -0.55 | 0.973 | -0.281 | 0.004 | 0.029 | 0.029 | 0 | 0.07 | 0.27 | 0.04 | 0.04 | 0.03 | 0.02 |
| 11 | 4552102 | rs2641426 | -0.809 | -0.702 | 0.796 | -0.325 | -0.01 | -0.013 | -0.013 | 0.09 | 0.43 | 0.47 | 0.26 | 0.3 | 0.36 | 0.43 |
| 11 | 4560369 | rs10836241 | -1.412 | -0.702 | 0.401 | -0.738 | -0.012 | 0.02 | 0.02 | 0 | 0.57 | 0.63 | 0.35 | 0.31 | 0.48 | 0.51 |
| 11 | 4563218 | rs12416860 | -0.069 | -0.685 | 0.524 | -0.769 | -0.016 | -0.025 | -0.025 | 0 | 0.15 | 0.26 | 0.09 | 0.07 | 0.11 | 0.11 |
| 11 | 4563728 | rs17220520 | 1.228 | -0.653 | 0.352 | -0.628 | -0.014 | 0.058 | 0.058 | 0 | 0.1 | 0 | 0.05 | 0 | 0.08 | 0.07 |
| 11 | 4565527 | rs12280424 | 0.244 | -0.703 | 0.402 | -0.766 | 0 | -0.011 | -0.011 | 0 | 0.13 | 0.15 | 0.1 | 0.04 | 0.12 | 0.05 |
| 11 | 4565825 | rs17220853 | 1.414 | -0.716 | 0.454 | -0.783 | -0.015 | 0.058 | 0.058 | 0 | 0.1 | 0 | 0.05 | 0 | 0.08 | 0.07 |
| 11 | 4566377 | rs2657168 | -0.468 | -0.77 | 0.537 | -0.756 | 0.023 | 0.037 | 0.037 | 0 | 0.44 | 0.29 | 0.23 | 0.26 | 0.36 | 0.48 |
| 11 | 4566711 | rs2709182 | -1.244 | -0.795 | 0.154 | -0.748 | -0.002 | 0.004 | 0.004 | 0 | 0.56 | 0.52 | 0.34 | 0.26 | 0.49 | 0.52 |
| 11 | 4568861 | rs12289941 | 0.519 | -0.722 | 0.045 | -0.702 | -0.001 | 0.007 | 0.007 | 0 | 0.12 | 0.1 | 0.1 | 0.04 | 0.11 | 0.05 |
| 11 | 4571782 | rs4910628 | 0.931 | -0.831 | 0.066 | -0.874 | 0.002 | -0.008 | -0.008 | 0 | 0.43 | 0.49 | 0.65 | 0.74 | 0.51 | 0.5 |
| 11 | 4573236 | rs2657147 | -0.541 | -0.836 | 0.001 | -0.853 | -0.006 | -0.012 | -0.012 | 0.422 | 0.46 | 0.42 | 0.26 | 0.27 | 0.38 | 0.48 |
| 11 | 4576666 | rs1394720 | -0.679 | -0.833 | -0.148 | -0.775 | -0.015 | -0.014 | -0.014 | 0 | 0.39 | 0.47 | 0.21 | 0.26 | 0.32 | 0.43 |
| 11 | 4578238 | rs2709175 | -0.911 | -0.67 | -0.153 | -0.58 | 0.006 | 0.013 | 0.013 | 0 | 0.48 | 0.49 | 0.29 | 0.27 | 0.41 | 0.47 |
| 11 | 4578709 | rs1505216 | 0.285 | -0.454 | -0.499 | -0.025 | 0.01 | -0.008 | -0.008 | 0 | 0.21 | 0.11 | 0.1 | 0.23 | 0.16 | 0.34 |
| 11 | 4587253 | rs11032827 | 0.051 | 0.416 | -0.616 | 0.437 | -0.029 | -0.034 | -0.034 | 0.515 | 0.29 | 0.23 | 0.48 | 0.61 | 0.37 | 0.35 |
| 11 | 4587940 | rs7129059 | -0.814 | -0.368 | 0.436 | 0.78 | 0.013 | 0.256 | 0.256 | 0.839 | 0.1 | 0.46 | 0.25 | 0.24 | 0.06 | 0.1 |
| 11 | 4589066 | rs2641399 | -1.011 | -1.136 | 0.278 | 0.607 | -0.021 | 0.041 | 0.041 | 0 | 0.28 | 0.42 | 0.33 | 0.52 | 0.19 | 0.41 |
| 11 | 4590639 | rs4910630 | 0.344 | -0.484 | -0.418 | 0.074 | 0.026 | 0.005 | 0.005 | 0 | 0.72 | 0.49 | 0.49 | 0.71 | 0.63 | 0.75 |
| 11 | 4591739 | rs10768096 | 0.046 | -0.598 | 0.168 | 0.029 | 0.012 | 0.006 | 0.006 | 0 | 0.76 | 0.73 | 0.52 | 0.72 | 0.63 | 0.75 |
| 11 | 4592277 | rs1505209 | 0.481 | -0.576 | -0.011 | 0.069 | -0.021 | -0.028 | -0.028 | 0 | 0.06 | 0.03 | 0.24 | 0.24 | 0.09 | 0.1 |
| 11 | 4592340 | rs1505208 | 1.584 | -0.986 | -0.513 | -0.391 | -0.14 | -0.077 | -0.077 | 0 | 0.43 | 0.05 | 0.34 | 0.42 | 0.45 | 0.4 |
| 11 | 4593052 | rs10836309 | 0.256 | -1.69 | -0.534 | -0.693 | -0.019 | 0.003 | 0.003 | 0 | 0.28 | 0.43 | 0.18 | 0.29 | 0.19 | 0.41 |
| 11 | 4594471 | rs12793041 | 1.39 | -1.54 | -0.569 | -0.654 | 0.004 | 0.016 | 0.016 | 0 | 0.07 | 0.04 | 0.06 | 0 | 0.07 | 0.02 |
| 11 | 4594545 | rs12795289 | 1.39 | -1.54 | -0.569 | -0.654 | 0.004 | 0.016 | 0.016 | 0 | 0.07 | 0.04 | 0.06 | 0 | 0.07 | 0.02 |
| 11 | 4594885 | rs2657148 | 0.277 | -1.618 | -0.408 | -0.487 | -0.011 | -0.036 | -0.036 | 0 | 0.31 | 0.09 | 0.29 | 0.34 | 0.28 | 0.45 |
| 11 | 4596449 | rs4459298 | -0.05 | -1.368 | 0.15 | -0.469 | -0.029 | 0.079 | 0.079 | 0 | 0.85 | 0.62 | 0.82 | 0.81 | 0.9 | 0.93 |
| 11 | 4596465 | rs4573672 | 0.702 | -1.454 | 0.09 | -0.533 | -0.048 | 0.001 | 0.001 | 0 | 0.78 | 0.55 | 0.76 | 0.81 | 0.83 | 0.92 |
| 11 | 4596897 | rs1505202 | 1.831 | -1.091 | -0.014 | -0.266 | -0.035 | -0.049 | -0.049 | 0 | 0.3 | 0.12 | 0.23 | 0.38 | 0.31 | 0.31 |
| 11 | 4597980 | rs2709155 | -0.962 | -0.972 | 0.23 | -0.405 | 0 | 0.018 | 0.018 | 0 | 0.43 | 0.35 | 0.41 | 0.37 | 0.41 | 0.5 |
| 11 | 4599221 | rs11032896 | 1.245 | -0.592 | 0.608 | -0.251 | 0 | -0.004 | -0.004 | 0 | 0.05 | 0.04 | 0.05 | 0 | 0.05 | 0.01 |
| 11 | 4601269 | rs1122778 | 0.135 | -0.634 | 0.958 | -0.291 | 0.022 | -0.071 | -0.071 | 0 | 0.89 | 0.67 | 0.87 | 1 | 0.9 | 0.97 |
| 11 | 4601302 | rs1000071 | 0.43 | -0.648 | 1.025 | -0.298 | 0.024 | -0.082 | -0.082 | 1.021 | 0.88 | 0.54 | 0.85 | 1 | 0.9 | 0.97 |
| 11 | 4601873 | rs2163284 | 0.43 | -0.648 | 1.038 | -0.298 | 0.027 | -0.082 | -0.082 | 3.27 | 0.88 | 0.55 | 0.85 | 1 | 0.9 | 0.97 |
| 11 | 4605490 | rs1346833 | 0.128 | -0.657 | 1.053 | -0.288 | 0.04 | -0.021 | -0.021 | 0.366 | 0.91 | 0.58 | 0.91 | 1 | 0.95 | 0.98 |
| 11 | 4606952 | rs7124305 | 1.057 | -0.633 | 1.154 | -0.277 | 0.004 | -0.031 | -0.031 | 0 | 0.06 | 0.1 | 0.05 | 0 | 0.04 | 0.01 |
| 11 | 4613203 | rs12146686 | 0.223 | -0.844 | 1.013 | -0.09 | -0.008 | -0.031 | -0.031 | 0.689 | 0.19 | 0.1 | 0.14 | 0.31 | 0.2 | 0.35 |
| 11 | 4613366 | rs905872 | 0.274 | -0.913 | 1.61 | 0.017 | 0.053 | -0.012 | -0.012 | 6.201 | 0.92 | 0.56 | 0.91 | 1 | 0.95 | 0.99 |
| 11 | 4620328 | rs12796784 | 0.866 | -0.866 | 1.106 | 0.042 | 0.003 | -0.03 | -0.03 | 0 | 0.06 | 0.1 | 0.05 | 0 | 0.04 | 0.01 |
| 11 | 4621323 | rs3750992 | 0.664 | -1.094 | 0.68 | 0.129 | 0.003 | 0.005 | 0.005 | 0 | 0.31 | 0.3 | 0.28 | 0.4 | 0.34 | 0.33 |
| 11 | 4623788 | rs931811 | 0.307 | -1.055 | 0.161 | 0.159 | -0.022 | 0.012 | 0.012 | 0 | 0.56 | 0.23 | 0.59 | 0.57 | 0.56 | 0.63 |
| 11 | 4624432 | rs17314159 | 0.708 | -0.919 | 0.337 | 0.449 | -0.024 | -0.028 | -0.028 | 0 | 0.28 | 0.15 | 0.26 | 0.4 | 0.34 | 0.33 |
| 11 | 4631352 | rs7950390 | -0.689 | -0.889 | 0.035 | -0.059 | 0.043 | -0.024 | -0.024 | 0.157 | 0.09 | 0.47 | 0.1 | 0 | 0.06 | 0.03 |
| 11 | 4639292 | rs11600417 | -0.421 | -0.774 | -0.221 | 0.22 | -0.004 | -0.02 | -0.02 | 0 | 0.64 | 0.38 | 0.67 | 0.54 | 0.62 | 0.66 |
| 11 | 4647770 | rs3861787 | -0.278 | -0.219 | 0.021 | -0.006 | 0.108 | 0.034 | 0.034 | 1.778 | 0.93 | 0.56 | 0.91 | 1 | 0.94 | 0.97 |
| 11 | 4658068 | rs10836397 | 0.1 | -0.376 | -1.401 | -0.036 | -0.06 | 0.036 | 0.036 | 0 | 0.41 | 0.15 | 0.37 | 0.32 | 0.51 | 0.46 |
| 11 | 4659085 | rs16923484 | 0.375 | -0.942 | -1.44 | -0.053 | 0.027 | 0.01 | 0.01 | 0 | 0.17 | 0.11 | 0.23 | 0.17 | 0.07 | 0.18 |
| 11 | 4659187 | rs4625457 | -0.866 | -0.95 | -1.632 | -0.193 | 0.015 | -0.034 | -0.034 | 0 | 0.67 | 0.27 | 0.67 | 0.48 | 0.68 | 0.74 |
| 11 | 4659419 | rs1908177 | 0.554 | -0.622 | -2.122 | -0.634 | -0.12 | 0.149 | 0.149 | 0.011 | 0.25 | 0.02 | 0.27 | 0 | 0.37 | 0.17 |
| 11 | 4659858 | rs11033126 | 0.333 | -0.617 | -2.017 | -0.559 | 0.006 | -0.015 | -0.015 | 0 | 0.27 | 0.22 | 0.18 | 0.31 | 0.22 | 0.4 |
| 11 | 4661285 | rs905871 | 1.292 | -0.361 | -1.883 | -0.587 | -0.089 | 0.103 | 0.103 | 0 | 0.18 | 0.02 | 0.2 | 0 | 0.3 | 0.13 |
| 11 | 4663543 | rs10768139 | -0.035 | -0.569 | -1.453 | -0.582 | -0.002 | 0.087 | 0.087 | 0 | 0.58 | 0.28 | 0.5 | 0.83 | 0.61 | 0.69 |
| 11 | 4664657 | rs7925271 | 0.882 | -0.044 | -0.582 | 0.048 | -0.002 | 0.004 | 0.004 | 0 | 0.27 | 0.28 | 0.35 | 0.62 | 0.28 | 0.28 |
| 11 | 4666074 | rs10768149 | 0.436 | 0.118 | -0.805 | -0.211 | -0.001 | 0.113 | 0.113 | 0 | 0.5 | 0.16 | 0.35 | 0.32 | 0.51 | 0.54 |
| 11 | 4666216 | rs10768150 | 0.307 | 0.103 | -0.391 | -0.218 | 0.005 | 0.08 | 0.08 | 0 | 0.51 | 0.23 | 0.36 | 0.32 | 0.51 | 0.54 |
| 11 | 4666810 | rs16924294 | -0.091 | 0.064 | -0.252 | 0.396 | 0 | 0.002 | 0.002 | 0 | 0.17 | 0.19 | 0.3 | 0.27 | 0.2 | 0.08 |
| 11 | 4667214 | rs7102992 | 0.57 | -0.255 | -0.442 | 0.331 | 0.003 | -0.017 | -0.017 | 0 | 0.3 | 0.27 | 0.36 | 0.6 | 0.26 | 0.38 |
| 11 | 4667594 | rs1394713 | -1.1 | -0.26 | 0.176 | 0.089 | 0.004 | -0.116 | -0.116 | 0 | 0.74 | 0.69 | 0.8 | 1 | 0.74 | 0.83 |
| 11 | 4667693 | rs7937180 | -0.632 | -0.153 | 0.162 | 0.14 | 0.001 | 0.017 | 0.017 | 0 | 0.48 | 0.45 | 0.45 | 0.37 | 0.49 | 0.55 |
| 11 | 4669916 | rs1857783 | -0.102 | -0.204 | 0.478 | 0.055 | -0.003 | 0.029 | 0.029 | 0 | 0.54 | 0.66 | 0.55 | 0.95 | 0.5 | 0.64 |
| 11 | 4670713 | rs7124317 | 0.149 | -0.186 | 0.282 | 0.064 | 0.001 | 0.034 | 0.034 | 0 | 0.52 | 0.39 | 0.53 | 0.94 | 0.5 | 0.64 |
| 11 | 4673106 | rs12280089 | -0.106 | -0.149 | -0.005 | 0.082 | -0.024 | 0.132 | 0.132 | 0 | 0.43 | 0.25 | 0.42 | 0.06 | 0.5 | 0.36 |
| 11 | 4673647 | rs4910524 | -1.102 | -0.179 | -0.098 | 0.191 | -0.048 | 0.036 | 0.036 | 0 | 0.74 | 0.63 | 0.67 | 0.38 | 0.85 | 0.73 |
| 11 | 4674575 | rs3817098 | -0.023 | -0.055 | -0.775 | -0.122 | -0.033 | 0.164 | 0.164 | 0 | 0.43 | 0.2 | 0.42 | 0.06 | 0.5 | 0.36 |
| 11 | 4677509 | rs12574901 | 1.104 | -0.267 | -1.09 | 0.472 | 0.01 | -0.061 | -0.061 | 0 | 0.22 | 0.16 | 0.29 | 0.44 | 0.22 | 0.26 |
| 11 | 4679286 | rs12281472 | 0.612 | -0.337 | -1.134 | -0.267 | -0.004 | 0.002 | 0.002 | 0 | 0.33 | 0.28 | 0.34 | 0.03 | 0.33 | 0.29 |
| 11 | 4679919 | rs10500610 | 0.866 | 0.104 | -1.005 | -0.009 | -0.042 | 0.063 | 0.063 | 0 | 0.11 | 0.02 | 0.09 | 0 | 0.15 | 0.07 |
| 11 | 4680467 | rs10500611 | 0.866 | 0.104 | -1.005 | -0.009 | -0.042 | 0.063 | 0.063 | 0 | 0.11 | 0.02 | 0.09 | 0 | 0.15 | 0.07 |
| 11 | 4682264 | rs10768156 | -0.193 | 0.075 | -0.891 | 0.33 | -0.014 | -0.04 | -0.04 | 0 | 0.62 | 0.25 | 0.53 | 0.45 | 0.6 | 0.59 |

| | | | | | | | | | | | | | | | | |
|---|---|---|---|---|---|---|---|---|---|---|---|---|---|---|---|---|
| 11 | 4682376 | rs10768157 | -0.209 | 0.089 | -0.919 | 0.337 | -0.012 | -0.035 | -0.035 | 0 | 0.62 | 0.27 | 0.53 | 0.45 | 0.6 | 0.59 |
| 11 | 4687238 | rs7124760 | -0.385 | 0.304 | -1.006 | -0.42 | -0.031 | 0.031 | 0.031 | 0 | 0.39 | 0.26 | 0.3 | 0.32 | 0.47 | 0.47 |
| 11 | 4693134 | rs11033275 | -0.466 | 0.074 | -1.397 | -0.79 | 0.06 | 0.052 | 0.052 | 0.177 | 0.56 | 0.27 | 0.46 | 0.32 | 0.69 | 0.59 |
| 11 | 4694308 | rs11033277 | 0.827 | 0.378 | -1.633 | -0.58 | -0.088 | -0.069 | -0.069 | 0 | 0.24 | 0.04 | 0.2 | 0.31 | 0.32 | 0.4 |
| 11 | 4694428 | rs10836452 | 0.698 | 0.035 | -1.541 | -0.397 | 0.019 | -0.066 | -0.066 | 4.968 | 0.24 | 0.06 | 0.31 | 0.65 | 0.18 | 0.25 |
| 11 | 4696875 | rs10836454 | -0.361 | 0.258 | -0.845 | -0.834 | -0.002 | -0.011 | -0.011 | 0 | 0.59 | 0.38 | 0.59 | 0.94 | 0.6 | 0.7 |
| 11 | 4700862 | rs1124113 | -0.414 | 0.094 | -0.669 | 0.089 | -0.005 | 0.02 | 0.02 | 0 | 0.08 | 0.13 | 0.15 | 0.22 | 0.05 | 0.05 |
| 11 | 4702108 | rs11033308 | 0.231 | 0.046 | -0.445 | 0.034 | -0.005 | -0.011 | -0.011 | 0.118 | 0.06 | 0.1 | 0.03 | 0.03 | 0.02 | 0.05 |
| 11 | 4703165 | rs1123991 | 1.253 | -0.042 | 0.187 | -0.079 | 0.012 | 0.006 | 0.006 | 0 | 0.81 | 0.53 | 0.77 | 0.82 | 0.83 | 0.89 |
| 11 | 4706179 | rs12223580 | 0.242 | -0.016 | -0.191 | 0.028 | -0.005 | -0.011 | -0.011 | 0 | 0.06 | 0.11 | 0.03 | 0.03 | 0.02 | 0.05 |
| 11 | 4707571 | rs10836470 | 0.288 | -0.813 | -1.018 | -0.154 | 0.037 | -0.068 | -0.068 | 0 | 0.25 | 0.12 | 0.17 | 0.58 | 0.16 | 0.37 |
| 11 | 4707755 | rs10742359 | 0.658 | -0.391 | -0.796 | -1.055 | 0.007 | 0.036 | 0.036 | 0 | 0.63 | 0.33 | 0.61 | 0.77 | 0.68 | 0.79 |
| 11 | 4709585 | rs1072894 | 0.543 | -0.434 | -0.767 | -0.657 | -0.009 | 0.003 | 0.003 | 0 | 0.59 | 0.46 | 0.68 | 0.42 | 0.69 | 0.69 |
| 11 | 4712353 | rs11033352 | -0.624 | -0.293 | -0.819 | -0.835 | 0.002 | -0.045 | -0.045 | 1.122 | 0.26 | 0.29 | 0.24 | 0.03 | 0.27 | 0.25 |
| 11 | 4713651 | rs4910526 | 1.008 | -0.536 | -0.878 | -0.584 | -0.029 | -0.088 | -0.088 | 1.241 | 0.18 | 0.03 | 0.14 | 0.36 | 0.14 | 0.24 |
| 11 | 4713811 | rs1505210 | 0.219 | -0.582 | -0.546 | -0.625 | -0.026 | -0.007 | -0.007 | 0 | 0.77 | 0.45 | 0.82 | 0.76 | 0.83 | 0.86 |
| 11 | 4714050 | rs7480497 | 0.381 | -0.514 | -0.415 | -0.627 | -0.003 | -0.002 | -0.002 | 0 | 0.71 | 0.28 | 0.75 | 0.76 | 0.78 | 0.85 |
| 11 | 4716066 | rs7108524 | -0.577 | -0.235 | -0.445 | -0.669 | -0.012 | -0.001 | -0.001 | 0 | 0.36 | 0.3 | 0.35 | 0.35 | 0.41 | 0.45 |
| 11 | 4716197 | rs10836480 | 1.334 | -0.238 | -0.716 | -0.644 | -0.03 | -0.074 | -0.074 | 2.285 | 0.17 | 0.03 | 0.14 | 0.31 | 0.17 | 0.22 |
| 11 | 4722142 | rs1032449 | 1.054 | -0.215 | -0.646 | -0.534 | -0.001 | -0.001 | -0.001 | 0.747 | 0.42 | 0.36 | 0.48 | 0.39 | 0.44 | 0.48 |
| 11 | 4724486 | rs7478759 | -0.524 | 0.014 | -0.432 | -0.358 | 0.001 | 0.216 | 0.216 | 0.972 | 0.06 | 0.37 | 0.08 | 0.26 | 0.02 | 0.01 |
| 11 | 4726878 | rs12271916 | -0.088 | 0.047 | -0.507 | -0.4 | -0.023 | 0.158 | 0.158 | 0.314 | 0.13 | 0.47 | 0.12 | 0.29 | 0.06 | 0.08 |
| 11 | 4729164 | rs1389463 | -0.853 | 0.224 | -0.755 | -0.353 | 0.025 | 0.008 | 0.008 | 0 | 0.52 | 0.71 | 0.4 | 0.6 | 0.42 | 0.39 |
| 11 | 4730320 | rs10836510 | 0.27 | 0.14 | -1.299 | -0.163 | 0.013 | 0.001 | 0.001 | 0 | 0.51 | 0.76 | 0.58 | 0.43 | 0.48 | 0.57 |
| 11 | 4731468 | rs4436535 | 0.391 | 0.219 | -1.41 | 0.159 | 0.02 | 0.01 | 0.01 | 0 | 0.69 | 0.72 | 0.77 | 0.71 | 0.76 | 0.85 |
| 11 | 4731583 | rs4471413 | -0.023 | 0.552 | -1.894 | 0.422 | 0.033 | -0.012 | -0.012 | 0 | 0.77 | 0.93 | 0.83 | 0.74 | 0.8 | 0.92 |
| 11 | 4732086 | rs4284391 | -0.023 | 0.467 | -1.894 | 0.422 | 0.044 | -0.012 | -0.012 | 0 | 0.77 | 0.93 | 0.82 | 0.72 | 0.81 | 0.92 |
| 11 | 4740665 | rs4498989 | 0.144 | 1.394 | -1.995 | 0.519 | -0.071 | -0.029 | -0.029 | 0.22 | 0.12 | 0.02 | 0.29 | 0.12 | 0.24 | 0.22 |
| 11 | 4741954 | rs7933187 | 0.599 | 0.843 | -1.783 | 0.463 | 0.012 | 0.01 | 0.01 | 0 | 0.41 | 0.57 | 0.24 | 0.36 | 0.25 | 0.35 |
| 11 | 4742926 | rs4331087 | 0.08 | 0.795 | -1.903 | 0.323 | -0.014 | -0.011 | -0.011 | 0 | 0.36 | 0.33 | 0.6 | 0.4 | 0.54 | 0.57 |
| 11 | 4745154 | rs10836532 | 0.51 | 0.758 | -1.898 | -0.453 | -0.001 | 0.01 | 0.01 | 0 | 0.42 | 0.54 | 0.27 | 0.2 | 0.35 | 0.34 |
| 11 | 4747352 | rs7102381 | -0.254 | 0.546 | -2.268 | 0.158 | -0.085 | -0.03 | -0.03 | 0.227 | 0.59 | 0.9 | 0.62 | 0.46 | 0.56 | 0.56 |
| 11 | 4748566 | rs4329682 | -0.331 | 0.532 | -2.148 | 0.021 | -0.06 | 0.016 | 0.016 | 0.137 | 0.6 | 0.84 | 0.6 | 0.39 | 0.58 | 0.59 |
| 11 | 4750773 | rs10836540 | 0.446 | -0.268 | -1.696 | -0.72 | 0.01 | 0.007 | 0.007 | 0 | 0.31 | 0.3 | 0.26 | 0.11 | 0.23 | 0.28 |
| 11 | 4752849 | rs7104808 | -0.919 | -0.444 | -0.721 | -0.648 | -0.018 | 0.048 | 0.048 | 0.116 | 0.58 | 0.75 | 0.59 | 0.2 | 0.57 | 0.57 |
| 11 | 4755505 | rs4375437 | -0.066 | -0.833 | -0.548 | -0.677 | -0.001 | 0 | 0 | 0.013 | 0.36 | 0.35 | 0.28 | 0.31 | 0.26 | 0.34 |
| 11 | 4757938 | rs10742372 | -0.765 | -0.906 | -0.07 | -0.881 | 0.004 | 0.004 | 0.004 | 0.261 | 0.47 | 0.57 | 0.39 | 0.31 | 0.4 | 0.35 |
| 11 | 4758152 | rs11033618 | 0.934 | -1.371 | -0.549 | -1.281 | 0.026 | 0.099 | 0.099 | 0 | 0.17 | 0.02 | 0.06 | 0 | 0.13 | 0.08 |
| 11 | 4759541 | rs4132539 | -0.36 | -1.436 | 0.132 | -1.267 | -0.027 | -0.065 | -0.065 | 0.07 | 0.17 | 0.32 | 0.13 | 0.07 | 0.15 | 0.05 |
| 11 | 4764043 | rs11033664 | -0.251 | -1.438 | 0.202 | -1.235 | -0.01 | 0.032 | 0.032 | 0 | 0.08 | 0.31 | 0.03 | 0.07 | 0.02 | 0.04 |
| 11 | 4772797 | rs7952595 | -0.738 | -2.213 | -0.199 | -1.679 | -0.032 | -0.089 | -0.089 | 0 | 0.4 | 0.47 | 0.21 | 0.07 | 0.29 | 0.16 |
| 11 | 4774410 | rs10836574 | -0.851 | -1.999 | -1.043 | -1.28 | -0.069 | -0.137 | -0.137 | 0 | 0.79 | 0.99 | 0.71 | 0.57 | 0.78 | 0.73 |
| 11 | 4777353 | rs10836579 | 1.335 | -2.226 | -0.421 | -1.768 | 0.021 | 0.028 | 0.028 | 0 | 0.17 | 0.09 | 0.07 | 0 | 0.13 | 0.07 |
| 11 | 4784679 | rs7947959 | -0.293 | -2.219 | -0.253 | -1.797 | -0.023 | -0.074 | -0.074 | 0.018 | 0.4 | 0.48 | 0.22 | 0.07 | 0.29 | 0.16 |
| 11 | 4790268 | rs1030726 | -0.037 | -2.497 | -0.327 | -2.134 | -0.045 | -0.147 | -0.147 | 0 | 0.29 | 0.46 | 0.2 | 0 | 0.27 | 0.07 |
| 11 | 4790308 | rs1030725 | -0.048 | -2.499 | -0.328 | -2.135 | -0.045 | -0.147 | -0.147 | 0.191 | 0.29 | 0.46 | 0.2 | 0 | 0.27 | 0.07 |
| 11 | 4790471 | rs1030723 | 0.936 | -2.653 | -0.401 | -2.225 | -0.017 | -0.111 | -0.111 | 0.32 | 0.15 | 0.44 | 0.11 | 0 | 0.13 | 0.05 |
| 11 | 4790575 | rs12788102 | 1.323 | -2.653 | -0.405 | -2.225 | 0.031 | 0.081 | 0.081 | 0.101 | 0.13 | 0.02 | 0.1 | 0 | 0.14 | 0.01 |
| 11 | 4790671 | rs11033795 | -0.246 | -2.639 | -0.396 | -2.218 | -0.045 | -0.147 | -0.147 | 0 | 0.29 | 0.46 | 0.2 | 0 | 0.27 | 0.07 |
| 11 | 4790857 | rs11033797 | -0.246 | -2.639 | -0.396 | -2.218 | -0.045 | -0.147 | -0.147 | 0 | 0.29 | 0.46 | 0.2 | 0 | 0.27 | 0.07 |
| 11 | 4790948 | rs11033800 | -0.246 | -2.639 | -0.396 | -2.218 | -0.045 | -0.147 | -0.147 | 0 | 0.29 | 0.46 | 0.2 | 0 | 0.27 | 0.07 |
| 11 | 4791111 | rs17324812 | 1.369 | -1.975 | -1.555 | -2.686 | -0.091 | 0.088 | 0.088 | 0 | 0.2 | 0.01 | 0.14 | 0.06 | 0.25 | 0.26 |
| 11 | 4793649 | rs11033816 | 1.076 | -2.065 | -0.467 | -2.32 | -0.016 | -0.096 | -0.096 | 0 | 0.15 | 0.29 | 0.1 | 0 | 0.13 | 0.05 |
| 11 | 4807054 | rs12801757 | 1.407 | -2.044 | -0.689 | -2.294 | 0.031 | 0.081 | 0.081 | 0.101 | 0.13 | 0.02 | 0.1 | 0 | 0.14 | 0.01 |
| 11 | 4811083 | rs12224171 | -0.804 | -1.996 | -0.548 | -1.455 | -0.016 | 0.025 | 0.025 | 0 | 0.82 | 0.71 | 0.78 | 0.77 | 0.85 | 0.89 |
| 11 | 4814850 | rs12805337 | -0.281 | -2.018 | -0.552 | -2.5 | -0.07 | -0.14 | -0.14 | 0 | 0.28 | 0.39 | 0.15 | 0.01 | 0.19 | 0.13 |
| 11 | 4815348 | rs10768242 | -0.517 | -1.897 | -0.714 | -2.923 | -0.006 | -0.071 | -0.071 | 0 | 0.37 | 0.55 | 0.19 | 0.01 | 0.25 | 0.2 |
| 11 | 4819009 | rs4910681 | -0.871 | -1.866 | -0.692 | -2.91 | 0.07 | 0.115 | 0.115 | 0 | 0.49 | 0.62 | 0.29 | 0.01 | 0.39 | 0.22 |
| 11 | 4821120 | rs7939275 | 1.382 | -1.872 | -0.842 | -2.914 | 0 | -0.008 | -0.008 | 0 | 0.2 | 0.15 | 0.12 | 0 | 0.2 | 0.08 |
| 11 | 4823689 | rs4382897 | -0.655 | -1.554 | -1.181 | -2.711 | 0.035 | 0.013 | 0.013 | 0 | 0.49 | 0.56 | 0.28 | 0.01 | 0.39 | 0.22 |
| 11 | 4824878 | rs2053116 | -0.678 | -1.566 | -1.186 | -2.717 | 0.036 | 0.015 | 0.015 | 0 | 0.49 | 0.56 | 0.28 | 0.01 | 0.39 | 0.22 |
| 11 | 4825225 | rs7941731 | -0.662 | -1.556 | -1.182 | -2.712 | 0.035 | 0.014 | 0.014 | 0 | 0.49 | 0.56 | 0.28 | 0.01 | 0.39 | 0.22 |
| 11 | 4839164 | rs12276131 | 0.377 | -0.592 | -1.904 | -2.219 | 0.005 | 0.176 | 0.176 | 0 | 0.51 | 0.21 | 0.36 | 0.12 | 0.51 | 0.59 |
| 11 | 4839847 | rs12276504 | 0.351 | -0.715 | -2.081 | -2.282 | -0.067 | 0.286 | 0.286 | 0 | 0.49 | 0.07 | 0.35 | 0.11 | 0.51 | 0.59 |
| 11 | 4840438 | rs17328039 | 0.836 | -0.492 | -2.526 | -2.733 | -0.029 | 0.11 | 0.11 | 0 | 0.28 | 0.06 | 0.21 | 0.04 | 0.27 | 0.32 |
| 11 | 4842023 | rs11034067 | 1.88 | -0.397 | -2.274 | -2.54 | 0.011 | 0.056 | 0.056 | 0.137 | 0.13 | 0.04 | 0.1 | 0 | 0.16 | 0.02 |
| 11 | 4848704 | rs10836729 | -0.587 | -0.648 | -1.662 | -2.776 | 0.012 | -0.019 | -0.019 | 0.004 | 0.73 | 0.64 | 0.52 | 0.12 | 0.69 | 0.69 |
| 11 | 4852009 | rs12420732 | 1.355 | -0.007 | -2.145 | -2.014 | -0.064 | -0.072 | -0.072 | 0 | 0.12 | 0.02 | 0.17 | 0.28 | 0.2 | 0.15 |
| 11 | 4852462 | rs7951380 | 1.753 | -0.384 | -1.673 | -2.646 | -0.015 | -0.064 | -0.064 | 0.028 | 0.11 | 0.24 | 0.05 | 0 | 0.05 | 0.09 |

| | | | | | | | | | | | | | | | | |
|---|---|---|---|---|---|---|---|---|---|---|---|---|---|---|---|---|
| 11 | 4852715 | rs17328663 | 0.524 | -0.453 | -1.785 | -2.648 | -0.078 | 0.263 | 0.263 | 0 | 0.5 | 0.07 | 0.34 | 0.13 | 0.5 | 0.59 |
| 11 | 4855124 | rs16906747 | 1.218 | -0.455 | -1.136 | -2.269 | -0.017 | -0.069 | -0.069 | 0 | 0.11 | 0.26 | 0.06 | 0 | 0.05 | 0.09 |
| 11 | 4862853 | rs11034170 | 0.367 | -0.43 | -1.263 | -2.371 | -0.004 | -0.055 | -0.055 | 0 | 0.48 | 0.36 | 0.26 | 0.1 | 0.39 | 0.6 |
| 11 | 4867824 | rs12795917 | 0.963 | -0.137 | -1.607 | -2.27 | 0.006 | 0.106 | 0.106 | 0.246 | 0.24 | 0.07 | 0.13 | 0.03 | 0.2 | 0.31 |
| 11 | 4869649 | rs12361955 | -0.121 | -0.034 | -1.464 | -2.264 | 0.017 | 0.005 | 0.005 | 0.867 | 0.47 | 0.34 | 0.26 | 0.07 | 0.4 | 0.4 |
| 11 | 4870298 | rs11601065 | 0.523 | 0.139 | -2.05 | -2.131 | -0.049 | 0.197 | 0.197 | 0 | 0.31 | 0.02 | 0.14 | 0.07 | 0.26 | 0.4 |
| 11 | 4893032 | rs12273277 | 0.62 | -0.113 | -1.735 | -1.721 | -0.13 | 0.22 | 0.22 | 0 | 0.35 | 0.03 | 0.23 | 0.08 | 0.39 | 0.43 |
| 11 | 4896254 | rs1865286 | 0.2 | -0.11 | -1.553 | -1.69 | -0.057 | 0.28 | 0.28 | 0.463 | 0.47 | 0.16 | 0.34 | 0.08 | 0.54 | 0.45 |
| 11 | 4901399 | rs11034467 | 0.064 | -0.124 | -1.485 | -1.713 | -0.038 | 0.262 | 0.262 | 0.306 | 0.48 | 0.24 | 0.35 | 0.08 | 0.54 | 0.45 |
| 11 | 4905202 | rs11034493 | 0.212 | -0.065 | -1.47 | -1.683 | -0.051 | 0.268 | 0.268 | 0.572 | 0.47 | 0.18 | 0.34 | 0.08 | 0.54 | 0.44 |
| 11 | 4910224 | rs2570591 | -0.541 | -0.06 | -1.457 | -1.679 | -0.038 | 0.265 | 0.265 | 0.318 | 0.53 | 0.81 | 0.67 | 0.92 | 0.48 | 0.56 |
| 11 | 4912192 | rs10836912 | 0.235 | -0.073 | -1.484 | -1.728 | -0.058 | 0.256 | 0.256 | 0 | 0.45 | 0.15 | 0.32 | 0.06 | 0.52 | 0.43 |
| 11 | 4913057 | rs10836914 | 0.368 | 0.052 | -1.019 | -0.898 | -0.012 | 0.11 | 0.11 | 0 | 0.11 | 0.21 | 0.23 | 0.59 | 0.08 | 0.07 |
| 11 | 4913314 | rs12577475 | 0.6 | 0.091 | -1.035 | -0.862 | -0.008 | 0.104 | 0.104 | 0 | 0.11 | 0.14 | 0.23 | 0.59 | 0.08 | 0.07 |
| 11 | 4914371 | rs2570586 | 0.742 | 0.191 | -0.535 | -0.913 | 0.039 | 0.085 | 0.085 | 0 | 0.13 | 0.25 | 0.3 | 0.32 | 0.17 | 0.15 |
| 11 | 4915072 | rs1816448 | 0.508 | 0.063 | -0.719 | -1.03 | -0.046 | 0.213 | 0.213 | 0 | 0.43 | 0.16 | 0.3 | 0.06 | 0.5 | 0.43 |
| 11 | 4919979 | rs11034573 | 0.26 | 0.114 | -0.839 | -0.804 | -0.013 | 0.093 | 0.093 | 0 | 0.12 | 0.16 | 0.16 | 0.59 | 0.08 | 0.07 |
| 11 | 4922016 | rs4910693 | 1.298 | 0.133 | -1.113 | -1.231 | -0.133 | 0.157 | 0.157 | 0 | 0.28 | 0.03 | 0.2 | 0.06 | 0.34 | 0.41 |
| 11 | 4925933 | rs10500632 | 1.587 | 0.007 | -0.768 | -1.247 | -0.027 | 0.028 | 0.028 | 0 | 0.08 | 0.03 | 0.07 | 0 | 0.12 | 0.08 |
| 11 | 4926740 | rs10768355 | -0.547 | -0.063 | -0.701 | -1.033 | 0.003 | 0.079 | 0.079 | 0 | 0.21 | 0.26 | 0.24 | 0.55 | 0.21 | 0.16 |
| 11 | 4928841 | rs7108225 | 1.624 | -0.013 | -0.807 | -1.035 | -0.027 | 0.028 | 0.028 | 0 | 0.08 | 0.03 | 0.07 | 0 | 0.12 | 0.08 |
| 11 | 4930763 | rs11034603 | -0.222 | -0.039 | -0.818 | -1.007 | -0.003 | -0.01 | -0.01 | 0 | 0.2 | 0.16 | 0.23 | 0.54 | 0.2 | 0.15 |
| 11 | 4931083 | rs16907254 | 1.466 | 0.145 | -1.435 | -1.025 | -0.034 | -0.043 | -0.043 | 0.138 | 0.09 | 0.02 | 0.15 | 0.36 | 0.11 | 0.09 |
| 11 | 4932775 | rs1368832 | 1.277 | -0.361 | -1.279 | -2.067 | 0.014 | 0.014 | 0.014 | 0 | 0.23 | 0.22 | 0.09 | 0.03 | 0.12 | 0.26 |
| 11 | 4934503 | rs11034615 | 1.348 | -0.031 | -1.574 | -2.054 | -0.13 | 0.156 | 0.156 | 0 | 0.28 | 0.02 | 0.2 | 0.06 | 0.34 | 0.41 |
| 11 | 4935470 | rs7112101 | -1.34 | -0.703 | -0.688 | -1.113 | 0.028 | 0.052 | 0.052 | 0 | 0.55 | 0.63 | 0.32 | 0.27 | 0.46 | 0.43 |
| 11 | 4935610 | rs12802922 | 1.167 | -0.373 | -1.414 | -1.067 | 0.016 | 0.079 | 0.079 | 1.057 | 0.13 | 0.02 | 0.09 | 0 | 0.14 | 0.02 |
| 11 | 4936608 | rs12419598 | 1.053 | -0.279 | -1.391 | -1.041 | -0.043 | 0.044 | 0.044 | 0 | 0.08 | 0.01 | 0.07 | 0 | 0.12 | 0.08 |
| 11 | 4936613 | rs16907312 | 0.195 | -1.198 | -0.812 | -0.637 | -0.061 | -0.054 | -0.054 | 0 | 0.24 | 0.34 | 0.1 | 0.05 | 0.12 | 0.28 |
| 11 | 4937040 | rs2595978 | -0.563 | -1.002 | -1.872 | -0.539 | 0.037 | 0.035 | 0.035 | 0 | 0.77 | 0.84 | 0.91 | 0.95 | 0.88 | 0.74 |
| 11 | 4941573 | rs17337649 | 0.79 | -0.952 | -1.979 | -0.523 | 0.014 | 0.082 | 0.082 | 2.166 | 0.13 | 0.01 | 0.1 | 0 | 0.15 | 0.02 |
| 11 | 4941904 | rs1433897 | 1.273 | -0.749 | -2.001 | -0.523 | 0.058 | 0.069 | 0.069 | 0 | 0.52 | 0.7 | 0.64 | 0.72 | 0.57 | 0.6 |
| 11 | 4942386 | rs11034658 | -1.181 | -0.129 | -2.31 | -0.244 | -0.029 | -0.001 | -0.001 | 0 | 0.24 | 0.07 | 0.26 | 0.23 | 0.31 | 0.14 |
| 11 | 4944986 | rs12796015 | 0.61 | -0.119 | -2.421 | -0.294 | 0.014 | 0.082 | 0.082 | 2.166 | 0.13 | 0.01 | 0.09 | 0 | 0.14 | 0.02 |
| 11 | 4945229 | rs10836954 | 0.325 | 0.345 | -2.096 | 0.081 | -0.001 | -0.026 | -0.026 | 0 | 0.06 | 0.01 | 0.02 | 0.22 | 0.07 | 0.04 |
| 11 | 4946056 | rs16907439 | -0.161 | 0.644 | -1.912 | 0.035 | 0.005 | -0.006 | -0.006 | 0 | 0.05 | 0.06 | 0.15 | 0.01 | 0.11 | 0.08 |
| 11 | 4946345 | rs11034685 | -0.495 | -0.421 | -2.201 | 0.079 | 0.117 | 0.058 | 0.058 | 0 | 0.4 | 0.12 | 0.2 | 0.27 | 0.32 | 0.33 |
| 11 | 4947313 | rs1469750 | 0.51 | 0.778 | -2.007 | 0.854 | -0.038 | -0.06 | -0.06 | 0.032 | 0.75 | 0.88 | 0.59 | 0.7 | 0.66 | 0.83 |
| 11 | 4947437 | rs11034689 | -0.569 | 0.758 | -2.003 | 0.807 | -0.027 | -0.049 | -0.049 | 0 | 0.17 | 0.11 | 0.37 | 0.33 | 0.23 | 0.15 |
| 11 | 4948560 | rs10500624 | 0.249 | 0.345 | -1.344 | 0.419 | -0.005 | 0.011 | 0.011 | 0.063 | 0.11 | 0.09 | 0.07 | 0.09 | 0.16 | 0.18 |
| 11 | 4951599 | rs840708 | -0.067 | 0.032 | -1.118 | -0.259 | -0.001 | -0.014 | -0.014 | 0 | 0.47 | 0.43 | 0.67 | 0.76 | 0.53 | 0.32 |
| 11 | 4951958 | rs840709 | -1.047 | -0.18 | -0.984 | -0.395 | 0.003 | 0.044 | 0.044 | 0 | 0.66 | 0.75 | 0.77 | 0.86 | 0.72 | 0.53 |
| 11 | 4953022 | rs16907494 | 1.704 | -0.266 | -1.18 | -0.667 | 0.002 | -0.014 | -0.014 | 0.557 | 0.18 | 0.16 | 0.09 | 0.01 | 0.12 | 0.08 |
| 11 | 4953462 | rs840716 | 0.703 | -0.326 | -0.779 | -0.268 | 0.035 | 0.199 | 0.199 | 0 | 0.07 | 0.16 | 0.02 | 0.25 | 0.06 | 0.03 |
| 11 | 4960032 | rs12288920 | 2.001 | -0.406 | -1.049 | -0.565 | -0.003 | 0.082 | 0.082 | 0 | 0.17 | 0.06 | 0.08 | 0.01 | 0.12 | 0.08 |
| 11 | 4961547 | rs17339295 | 0.953 | -0.348 | -0.924 | -0.53 | -0.001 | 0.001 | 0.001 | 0 | 0.13 | 0.14 | 0.09 | 0.08 | 0.17 | 0.18 |
| 11 | 4964103 | rs2595996 | -0.553 | -0.427 | -0.913 | -0.572 | -0.001 | -0.008 | -0.008 | 0 | 0.82 | 0.79 | 0.89 | 0.91 | 0.81 | 0.82 |
| 11 | 4964193 | rs2595995 | -1.152 | -0.392 | -0.907 | -0.552 | 0.001 | -0.015 | -0.015 | 0 | 0.87 | 0.8 | 0.91 | 0.91 | 0.84 | 0.82 |
| 11 | 4964302 | rs2570583 | -1.152 | -0.392 | -0.907 | -0.552 | 0.001 | -0.015 | -0.015 | 0 | 0.87 | 0.8 | 0.91 | 0.91 | 0.84 | 0.82 |
| 11 | 4964374 | rs2595994 | -1.152 | -0.392 | -0.907 | -0.552 | 0.001 | -0.015 | -0.015 | 0 | 0.87 | 0.8 | 0.91 | 0.91 | 0.84 | 0.82 |
| 11 | 4964627 | rs2595993 | 0.417 | -0.455 | -0.93 | -0.375 | 0.001 | -0.003 | -0.003 | 0 | 0.44 | 0.41 | 0.53 | 0.55 | 0.48 | 0.3 |
| 11 | 4964649 | rs2595992 | -0.922 | -0.161 | -0.782 | 0.124 | 0.001 | -0.015 | -0.015 | 0 | 0.87 | 0.8 | 0.91 | 0.91 | 0.84 | 0.82 |
| 11 | 4967061 | rs11034822 | 0.676 | -0.161 | -0.976 | 0.142 | -0.052 | -0.003 | -0.003 | 0.28 | 0.15 | 0.01 | 0.12 | 0.13 | 0.16 | 0.39 |
| 11 | 4979689 | rs11602673 | 0.946 | 0.07 | -0.816 | 0.136 | -0.005 | 0.012 | 0.012 | 0.09 | 0.11 | 0.08 | 0.07 | 0.09 | 0.16 | 0.18 |
| 11 | 4980153 | rs7105981 | -0.31 | -0.043 | -0.73 | 0.17 | 0.001 | -0.006 | -0.006 | 0 | 0.23 | 0.21 | 0.3 | 0.34 | 0.23 | 0.44 |
| 11 | 4980792 | rs3850509 | -0.56 | -0.273 | -0.68 | 0.235 | 0.001 | -0.001 | -0.001 | 0 | 0.36 | 0.34 | 0.38 | 0.43 | 0.39 | 0.62 |
| 11 | 4982916 | rs11034913 | -0.568 | -0.032 | -1.042 | 0.392 | -0.065 | 0.002 | 0.002 | 0.282 | 0.36 | 0.09 | 0.41 | 0.31 | 0.37 | 0.26 |
| 11 | 4983161 | rs11034915 | 0.939 | -0.262 | -0.609 | 0.492 | 0.019 | 0.051 | 0.051 | 0 | 0.53 | 0.27 | 0.4 | 0.38 | 0.59 | 0.67 |
| 11 | 4988619 | rs10734458 | 0.939 | -0.262 | -0.609 | 0.492 | 0.019 | 0.051 | 0.051 | 0 | 0.53 | 0.27 | 0.4 | 0.38 | 0.59 | 0.67 |
| 11 | 4991138 | rs11034951 | 0.859 | -0.264 | -0.475 | 0.552 | 0.004 | 0.01 | 0.01 | 0 | 0.53 | 0.43 | 0.41 | 0.4 | 0.59 | 0.67 |
| 11 | 4992282 | rs1378738 | 0.162 | -0.275 | -0.37 | 0.582 | -0.026 | 0.027 | 0.027 | 0.187 | 0.63 | 0.83 | 0.58 | 0.7 | 0.63 | 0.74 |
| 11 | 4993057 | rs1598507 | 0.782 | -0.152 | -0.453 | 0.66 | 0.004 | 0.01 | 0.01 | 0 | 0.53 | 0.43 | 0.41 | 0.4 | 0.59 | 0.67 |
| 11 | 4995686 | rs7102290 | -0.466 | 0.632 | -0.737 | 1.005 | -0.026 | 0.034 | 0.034 | 0.026 | 0.22 | 0.66 | 0.26 | 0.3 | 0.14 | 0.13 |
| 11 | 4995989 | rs2124447 | -0.466 | 0.632 | -0.737 | 1.005 | -0.026 | 0.034 | 0.034 | 0.026 | 0.22 | 0.66 | 0.25 | 0.31 | 0.14 | 0.13 |
| 11 | 5000043 | rs1455954 | 0.636 | 0.823 | -1.041 | 0.852 | 0 | -0.003 | -0.003 | 0 | 0.61 | 0.35 | 0.43 | 0.43 | 0.64 | 0.71 |
| 11 | 5005327 | rs12800414 | -0.174 | 1.373 | -0.734 | 1.39 | -0.001 | 0 | 0 | 0 | 0.16 | 0.1 | 0.12 | 0.09 | 0.23 | 0.23 |
| 11 | 5005685 | rs11034990 | -0.32 | 0.979 | -0.795 | 2.05 | -0.045 | -0.006 | -0.006 | 0 | 0.08 | 0.01 | 0.28 | 0.08 | 0.14 | 0.09 |
| 11 | 5010905 | rs2499953 | -1.071 | 0.053 | -0.089 | 1.907 | -0.028 | 0.209 | 0.209 | 0.02 | 0.09 | 0.28 | 0.16 | 0.28 | 0.03 | 0.05 |
| 11 | 5012894 | rs2499958 | -0.542 | 0.588 | -1.069 | 1.209 | -0.039 | 0.045 | 0.045 | 0 | 0.42 | 0.23 | 0.51 | 0.7 | 0.56 | 0.45 |

| | | | | | | | | | | | | | | | | |
|---|---|---|---|---|---|---|---|---|---|---|---|---|---|---|---|---|
| 11 | 5015571 | rs2500035 | 0.326 | 0.575 | -0.85 | 1.919 | -0.004 | -0.022 | -0.022 | 0 | 0.79 | 0.81 | 0.54 | 0.57 | 0.74 | 0.85 |
| 11 | 5016269 | rs11035046 | 0.384 | 0.609 | -1.014 | 0.752 | 0.02 | -0.102 | -0.102 | 0 | 0.34 | 0.38 | 0.18 | 0.07 | 0.35 | 0.29 |
| 11 | 5020189 | rs11035064 | 0.459 | 0.647 | -0.626 | 0.642 | 0.011 | -0.07 | -0.07 | 0.332 | 0.31 | 0.3 | 0.16 | 0.07 | 0.31 | 0.27 |
| 11 | 5020509 | rs11035066 | 0.459 | 0.647 | -0.626 | 0.642 | 0.011 | -0.07 | -0.07 | 0.332 | 0.31 | 0.3 | 0.16 | 0.07 | 0.31 | 0.27 |
| 11 | 5020799 | rs10768448 | 0.43 | 0.625 | -0.6 | 0.631 | 0.012 | -0.078 | -0.078 | 0 | 0.31 | 0.35 | 0.16 | 0.07 | 0.31 | 0.27 |
| 11 | 5022036 | rs2445293 | 0.339 | 0.829 | -0.769 | 0.791 | 0.02 | 0.114 | 0.114 | 0 | 0.06 | 0.14 | 0.11 | 0.27 | 0.1 | 0.06 |
| 11 | 5026200 | rs10837119 | 0.366 | 0.562 | -1.08 | 0.847 | 0.04 | 0.025 | 0.025 | 0 | 0.44 | 0.29 | 0.4 | 0.4 | 0.36 | 0.55 |
| 11 | 5026767 | rs2499955 | 0.347 | 0.662 | -0.915 | 1.071 | 0.025 | 0.207 | 0.207 | 0.123 | 0.89 | 0.74 | 0.86 | 0.54 | 0.85 | 0.89 |
| 11 | 5029457 | rs11035102 | -0.161 | 0.657 | -0.882 | 0.932 | -0.004 | -0.002 | -0.002 | 0 | 0.08 | 0.14 | 0.09 | 0.14 | 0.1 | 0.06 |
| 11 | 5037065 | rs17342537 | 0.784 | 0.663 | -1.035 | 0.852 | -0.024 | -0.004 | -0.004 | 1.356 | 0.13 | 0.03 | 0.28 | 0.09 | 0.14 | 0.07 |
| 11 | 5042941 | rs1378749 | -1.096 | 0.105 | -0.983 | 0.503 | -0.004 | 0.003 | 0.003 | 0 | 0.61 | 0.52 | 0.58 | 0.4 | 0.63 | 0.61 |
| 11 | 5045694 | rs1598512 | -0.401 | -0.086 | -0.828 | 0.424 | 0.004 | 0.001 | 0.001 | 0 | 0.53 | 0.34 | 0.53 | 0.38 | 0.58 | 0.41 |
| 11 | 5048058 | rs2445248 | 0.09 | -0.173 | -0.798 | 0.292 | 0 | -0.007 | -0.007 | 0 | 0.46 | 0.42 | 0.45 | 0.23 | 0.47 | 0.34 |
| 11 | 5053712 | rs2442421 | 0.069 | -0.211 | -0.812 | 0.273 | -0.002 | 0.006 | 0.006 | 0 | 0.46 | 0.43 | 0.45 | 0.23 | 0.47 | 0.35 |
| 11 | 5054643 | rs16908782 | 0.382 | -0.279 | -0.857 | 0.127 | -0.009 | -0.016 | -0.016 | 0 | 0.11 | 0.15 | 0.05 | 0.02 | 0.05 | 0.2 |
| 11 | 5055057 | rs2445263 | -0.636 | -0.259 | -0.875 | 0.246 | -0.004 | 0.016 | 0.016 | 0 | 0.57 | 0.54 | 0.49 | 0.26 | 0.51 | 0.55 |
| 11 | 5055875 | rs10768481 | 1.952 | 0.007 | -1.111 | -0.03 | -0.09 | -0.023 | -0.023 | 0 | 0.19 | 0.01 | 0.1 | 0.11 | 0.24 | 0.21 |
| 11 | 5057550 | rs16908907 | 0.751 | 0.035 | -0.957 | 0.004 | 0.007 | 0.006 | 0.006 | 0.296 | 0.11 | 0.11 | 0.04 | 0.02 | 0.05 | 0.2 |
| 11 | 5059064 | rs2445271 | 0.341 | 0.096 | -0.848 | 0.064 | 0.006 | -0.038 | -0.038 | 0 | 0.34 | 0.33 | 0.16 | 0.15 | 0.33 | 0.29 |
| 11 | 5066087 | rs1378736 | -1.154 | 0.064 | -0.941 | 0.017 | -0.006 | 0.017 | 0.017 | 0 | 0.57 | 0.59 | 0.5 | 0.26 | 0.51 | 0.55 |
| 11 | 5068137 | rs2500017 | -1.154 | 0.064 | -0.941 | 0.017 | -0.006 | 0.017 | 0.017 | 0 | 0.57 | 0.59 | 0.5 | 0.26 | 0.51 | 0.55 |
| 11 | 5068431 | rs17350764 | -0.261 | -0.015 | -0.924 | -0.086 | 0.007 | 0.014 | 0.014 | 0.523 | 0.23 | 0.25 | 0.34 | 0.11 | 0.18 | 0.26 |
| 11 | 5068661 | rs2500019 | -1.134 | 0.104 | -0.93 | 0.004 | -0.006 | 0.017 | 0.017 | 0 | 0.57 | 0.59 | 0.5 | 0.26 | 0.51 | 0.55 |
| 11 | 5069413 | rs2500027 | -1.134 | 0.104 | -0.93 | 0.004 | -0.006 | 0.017 | 0.017 | 0 | 0.57 | 0.59 | 0.5 | 0.26 | 0.51 | 0.55 |
| 11 | 5071626 | rs1463376 | 1.728 | 0.146 | -0.623 | -0.206 | 0.022 | -0.001 | -0.001 | 0 | 0.21 | 0.21 | 0.11 | 0.15 | 0.24 | 0.22 |
| 11 | 5072498 | rs2500036 | -1.287 | 0.184 | -0.634 | -0.161 | -0.006 | 0.016 | 0.016 | 0 | 0.57 | 0.59 | 0.5 | 0.26 | 0.51 | 0.55 |
| 11 | 5073101 | rs12574869 | -0.175 | 0.131 | -0.64 | -0.231 | 0.007 | 0.014 | 0.014 | 0.523 | 0.23 | 0.25 | 0.34 | 0.11 | 0.18 | 0.26 |
| 11 | 5074011 | rs1378747 | 1.599 | 0.127 | -0.642 | -0.231 | 0.017 | -0.004 | -0.004 | 4.415 | 0.21 | 0.21 | 0.11 | 0.15 | 0.24 | 0.22 |
| 11 | 5077653 | rs2500048 | 0.221 | 0.134 | -0.658 | -0.231 | 0.006 | -0.049 | -0.049 | 0.257 | 0.34 | 0.36 | 0.17 | 0.15 | 0.33 | 0.29 |
| 11 | 5078999 | rs9804487 | -1.28 | 0.169 | -0.708 | -0.193 | -0.007 | 0.019 | 0.019 | 0 | 0.57 | 0.6 | 0.5 | 0.26 | 0.51 | 0.55 |
| 11 | 5079353 | rs11035386 | 1.95 | 0.224 | -0.86 | 0.081 | 0.003 | -0.019 | -0.019 | 0 | 0.32 | 0.18 | 0.35 | 0.38 | 0.32 | 0.35 |
| 11 | 5081530 | rs7109504 | 1.512 | 0.287 | -0.834 | 0.02 | 0.007 | 0.008 | 0.008 | 0.095 | 0.12 | 0.06 | 0.05 | 0 | 0.09 | 0.07 |
| 11 | 5081942 | rs7116353 | -0.733 | 0.262 | -0.866 | 0.03 | -0.01 | 0.015 | 0.015 | 0 | 0.32 | 0.27 | 0.44 | 0.22 | 0.34 | 0.19 |
| 11 | 5082425 | rs1551487 | 1.063 | -0.193 | -0.681 | -0.278 | 0.002 | -0.019 | -0.019 | 0.092 | 0.13 | 0.12 | 0.06 | 0 | 0.09 | 0.07 |
| 11 | 5084742 | rs17351915 | -0.212 | -0.171 | -0.71 | -0.265 | 0.003 | 0.012 | 0.012 | 0 | 0.25 | 0.27 | 0.35 | 0.09 | 0.22 | 0.13 |
| 11 | 5085254 | rs16909520 | -0.896 | -0.211 | -0.679 | -0.292 | -0.004 | -0.022 | -0.022 | 0 | 0.36 | 0.46 | 0.4 | 0.11 | 0.27 | 0.34 |
| 11 | 5088772 | rs1378742 | 1.386 | -0.218 | -0.654 | -0.387 | 0.013 | -0.007 | -0.007 | 0 | 0.22 | 0.21 | 0.11 | 0.15 | 0.24 | 0.22 |
| 11 | 5089017 | rs2445311 | -0.709 | -0.235 | -0.502 | -0.313 | -0.017 | -0.061 | -0.061 | 0 | 0.25 | 0.33 | 0.11 | 0.02 | 0.13 | 0.28 |
| 11 | 5089436 | rs10768533 | 1.117 | -0.072 | -0.711 | -0.295 | -0.014 | -0.018 | -0.018 | 0 | 0.44 | 0.31 | 0.48 | 0.55 | 0.49 | 0.43 |
| 11 | 5090742 | rs1378743 | 0.115 | 0.185 | -0.755 | -0.635 | 0.004 | -0.04 | -0.04 | 0 | 0.28 | 0.35 | 0.13 | 0.14 | 0.21 | 0.22 |
| 11 | 5091332 | rs936143 | 2.435 | 0.197 | -0.677 | -0.634 | 0.044 | 0.04 | 0.04 | 0 | 0.13 | 0.16 | 0.05 | 0.13 | 0.13 | 0.15 |
| 11 | 5091633 | rs17352781 | 1.331 | 0.201 | -0.919 | -0.619 | -0.056 | 0.003 | 0.003 | 0 | 0.15 | 0.03 | 0.3 | 0.09 | 0.2 | 0.07 |
| 11 | 5091882 | rs4144717 | -0.494 | 0.106 | -0.704 | -0.464 | 0.004 | -0.014 | -0.014 | 0 | 0.43 | 0.42 | 0.43 | 0.22 | 0.41 | 0.3 |
| 11 | 5095383 | rs10837269 | -0.947 | 0.405 | -0.708 | -0.314 | -0.001 | 0.021 | 0.021 | 0 | 0.49 | 0.53 | 0.47 | 0.25 | 0.44 | 0.43 |
| 11 | 5096201 | rs2500056 | 0.162 | 1.02 | -0.563 | -0.266 | -0.006 | -0.005 | -0.005 | 0.583 | 0.21 | 0.26 | 0.14 | 0.2 | 0.23 | 0.2 |
| 11 | 5096331 | rs4144715 | 0.108 | 1.013 | -0.478 | -0.206 | 0 | 0 | 0 | 1.661 | 0.6 | 0.44 | 0.6 | 0.46 | 0.55 | 0.57 |
| 11 | 5096651 | rs10768545 | 0.682 | 1.01 | -0.728 | -0.2 | -0.024 | -0.049 | -0.049 | 0 | 0.15 | 0.05 | 0.23 | 0.33 | 0.17 | 0.18 |
| 11 | 5098553 | rs2445265 | -1.689 | 1.192 | -0.94 | -0.639 | -0.036 | -0.076 | -0.076 | 2.323 | 0.78 | 0.92 | 0.74 | 0.58 | 0.75 | 0.75 |
| 11 | 5098714 | rs10768550 | -0.227 | 1.08 | -0.536 | -0.686 | 0.05 | 0.132 | 0.132 | 0 | 0.22 | 0.37 | 0.31 | 0.39 | 0.26 | 0.22 |
| 11 | 5102476 | rs2500058 | -1.431 | 1.105 | -0.582 | -0.666 | -0.03 | -0.066 | -0.066 | 1.862 | 0.78 | 0.91 | 0.74 | 0.58 | 0.75 | 0.75 |
| 11 | 5105194 | rs10837299 | -1.126 | 0.995 | -0.494 | -0.772 | 0.007 | 0.028 | 0.028 | 0 | 0.32 | 0.48 | 0.36 | 0.48 | 0.35 | 0.27 |
| 11 | 5106155 | rs2500061 | 1.227 | 0.986 | -0.497 | -0.789 | 0.02 | 0.09 | 0.09 | 0 | 0.46 | 0.35 | 0.38 | 0.1 | 0.4 | 0.48 |
| 11 | 5109946 | rs10500616 | -0.493 | 1.017 | -0.343 | -0.73 | 0.017 | 0.049 | 0.049 | 0 | 0.25 | 0.28 | 0.31 | 0.39 | 0.28 | 0.24 |
| 11 | 5110068 | rs2499936 | -1.366 | 0.996 | -0.508 | -0.688 | -0.014 | -0.041 | -0.041 | 0.413 | 0.78 | 0.83 | 0.74 | 0.59 | 0.75 | 0.75 |
| 11 | 5115856 | rs1466266 | 0.515 | 0.606 | -0.492 | -0.666 | -0.035 | -0.112 | -0.112 | 0 | 0.18 | 0.03 | 0.23 | 0.38 | 0.17 | 0.18 |
| 11 | 5116109 | rs12280713 | -0.447 | 0.627 | -0.239 | -0.781 | 0.083 | 0.037 | 0.037 | 0 | 0.13 | 0.38 | 0.11 | 0.1 | 0.18 | 0.08 |
| 11 | 5120790 | rs10837328 | -0.852 | 0.525 | -0.723 | -0.767 | 0.017 | 0.067 | 0.067 | 0 | 0.31 | 0.41 | 0.34 | 0.48 | 0.35 | 0.26 |
| 11 | 5124833 | rs2445286 | 0.949 | 0.655 | -1.175 | -0.914 | 0.039 | 0.129 | 0.129 | 0 | 0.45 | 0.3 | 0.37 | 0.09 | 0.38 | 0.48 |
| 11 | 5127510 | rs2499945 | -1.003 | 0.803 | -1.156 | -1.047 | -0.04 | -0.056 | -0.056 | 0 | 0.75 | 0.9 | 0.72 | 0.57 | 0.71 | 0.73 |
| 11 | 5128534 | rs10837349 | 1.733 | 0.55 | -1.405 | -1.231 | -0.013 | 0.016 | 0.016 | 0 | 0.1 | 0 | 0.05 | 0.03 | 0.12 | 0.16 |
| 11 | 5132119 | rs2445296 | -1.195 | 0.901 | -0.43 | -0.659 | -0.041 | -0.063 | -0.063 | 0 | 0.78 | 0.92 | 0.73 | 0.57 | 0.73 | 0.75 |
| 11 | 5137846 | rs10837368 | 1.134 | 0.876 | -0.26 | -0.654 | 0.013 | 0.01 | 0.01 | 0 | 0.45 | 0.43 | 0.36 | 0.09 | 0.35 | 0.48 |
| 11 | 5138733 | rs4294557 | -0.081 | 0.617 | -0.134 | -0.327 | 0.001 | -0.062 | -0.062 | 0 | 0.21 | 0.11 | 0.24 | 0.38 | 0.17 | 0.19 |
| 11 | 5140977 | rs4512811 | -1.133 | 0.659 | -0.395 | -0.312 | 0.024 | 0.072 | 0.072 | 0 | 0.31 | 0.46 | 0.35 | 0.48 | 0.36 | 0.25 |
| 11 | 5143309 | rs11035765 | 1.083 | -0.128 | -1.079 | -0.26 | -0.051 | -0.069 | -0.069 | 0.175 | 0.11 | 0.01 | 0.22 | 0.38 | 0.15 | 0.15 |
| 11 | 5145686 | rs11035774 | 0.474 | -0.014 | -0.978 | -0.147 | -0.044 | -0.129 | -0.129 | 1.066 | 0.19 | 0.01 | 0.23 | 0.38 | 0.17 | 0.19 |
| 11 | 5146869 | rs12577357 | 0.824 | -0.569 | -0.77 | -0.533 | 0.046 | 0.053 | 0.053 | 0 | 0.35 | 0.29 | 0.29 | 0.09 | 0.25 | 0.46 |
| 11 | 5149332 | rs2499948 | 0.216 | 0.5 | -0.531 | 0.034 | -0.01 | -0.036 | -0.036 | 0.053 | 0.19 | 0.1 | 0.23 | 0.39 | 0.19 | 0.21 |
| 11 | 5153261 | rs2472530 | 0.292 | 0.335 | -0.387 | -0.125 | -0.069 | -0.128 | -0.128 | 0 | 0.19 | 0.01 | 0.23 | 0.39 | 0.2 | 0.22 |

| | | | | | | | | | | | | | | | | |
|---|---|---|---|---|---|---|---|---|---|---|---|---|---|---|---|---|
| 11 | 5157122 | rs11826290 | 1.43 | -0.303 | -0.529 | -0.583 | 0.056 | 0.089 | 0.089 | 0 | 0.29 | 0.14 | 0.26 | 0.03 | 0.24 | 0.34 |
| 11 | 5160096 | rs2472528 | 0.028 | -0.53 | -0.189 | -0.523 | 0.034 | -0.015 | -0.015 | 0 | 0.54 | 0.64 | 0.5 | 0.48 | 0.4 | 0.67 |
| 11 | 5167125 | rs2472523 | 0.673 | -0.197 | -0.493 | -0.413 | -0.147 | -0.105 | -0.105 | 1.833 | 0.3 | 0.06 | 0.29 | 0.43 | 0.37 | 0.26 |
| 11 | 5167247 | rs2472522 | 0.679 | -0.253 | -0.329 | -0.199 | -0.059 | -0.128 | -0.128 | 0 | 0.21 | 0.01 | 0.23 | 0.4 | 0.2 | 0.23 |
| 11 | 5172398 | rs7482505 | -0.025 | -0.681 | -0.032 | -0.221 | 0 | -0.004 | -0.004 | 0 | 0.56 | 0.52 | 0.56 | 0.51 | 0.51 | 0.58 |
| 11 | 5175477 | rs7940626 | 0.204 | -0.69 | -0.457 | -0.257 | 0.056 | 0.042 | 0.042 | 0 | 0.5 | 0.23 | 0.5 | 0.43 | 0.45 | 0.55 |
| 11 | 5180187 | rs10837461 | 0.423 | -0.402 | -0.513 | -0.052 | -0.054 | -0.126 | -0.126 | 0.787 | 0.22 | 0.01 | 0.23 | 0.4 | 0.21 | 0.27 |
| 11 | 5188844 | rs10734485 | 0.607 | -0.154 | 0.057 | 0.098 | -0.011 | 0.08 | 0.08 | 0 | 0.88 | 0.71 | 0.93 | 0.83 | 0.91 | 0.81 |
| 11 | 5190474 | rs4910717 | 1.379 | -0.26 | -0.068 | -0.055 | 0.041 | 0.08 | 0.08 | 0 | 0.27 | 0.14 | 0.27 | 0.03 | 0.24 | 0.27 |
| 11 | 5192095 | rs7481262 | -0.328 | -0.285 | 0.246 | -0.053 | -0.018 | -0.019 | -0.019 | 0.001 | 0.6 | 0.43 | 0.65 | 0.8 | 0.67 | 0.53 |
| 11 | 5192729 | rs4910539 | 0.203 | -0.241 | 0.13 | 0.017 | 0.009 | 0.141 | 0.141 | 0 | 0.9 | 0.68 | 0.94 | 0.83 | 0.92 | 0.81 |
| 11 | 5194582 | rs7480988 | 1.403 | -0.202 | -0.338 | -0.029 | 0.035 | 0.063 | 0.063 | 0 | 0.27 | 0.14 | 0.27 | 0.03 | 0.24 | 0.27 |
| 11 | 5198595 | rs6578559 | 0.995 | -0.151 | -0.48 | 0.041 | -0.067 | -0.113 | -0.113 | 0.98 | 0.2 | 0.01 | 0.22 | 0.35 | 0.2 | 0.27 |
| 11 | 5199208 | rs7927539 | -0.453 | -0.697 | 0.403 | -0.33 | -0.006 | -0.013 | -0.013 | 0 | 0.67 | 0.65 | 0.7 | 0.92 | 0.75 | 0.58 |
| 11 | 5200346 | rs7948698 | -0.416 | -0.656 | 0.245 | -0.309 | -0.022 | -0.037 | -0.037 | 0 | 0.67 | 0.61 | 0.7 | 0.9 | 0.75 | 0.58 |
| 11 | 5206294 | rs7934373 | -0.32 | -0.252 | -0.336 | -0.033 | 0.007 | 0.006 | 0.006 | 0 | 0.6 | 0.6 | 0.64 | 0.82 | 0.65 | 0.64 |
| 11 | 5209592 | rs11602279 | 0.355 | -0.142 | -0.707 | 0.358 | -0.03 | 0.061 | 0.061 | 0.105 | 0.86 | 0.44 | 0.91 | 0.83 | 0.92 | 0.82 |
| 11 | 5210250 | rs7117194 | 0.189 | 0.562 | -0.788 | -0.301 | 0.008 | 0.122 | 0.122 | 0 | 0.4 | 0.26 | 0.31 | 0.08 | 0.37 | 0.33 |
| 11 | 5223920 | rs11036227 | -0.388 | 1.637 | -0.389 | 1.402 | -0.095 | -0.07 | -0.07 | 0 | 0.16 | 0.02 | 0.16 | 0.26 | 0.29 | 0.15 |
| 11 | 5228708 | rs10837582 | -0.967 | 1.403 | 0.027 | 1.262 | -0.022 | -0.018 | -0.018 | 0 | 0.21 | 0.22 | 0.22 | 0.34 | 0.33 | 0.18 |
| 11 | 5230302 | rs7935470 | -1.649 | 1.095 | -0.134 | 0.506 | 0.001 | 0.004 | 0.004 | 0 | 0.43 | 0.57 | 0.52 | 0.52 | 0.56 | 0.45 |
| 11 | 5231897 | rs10837593 | 0.698 | 1.091 | 0.293 | 0.982 | 0.007 | 0.118 | 0.118 | 0 | 0.86 | 0.82 | 0.69 | 0.51 | 0.81 | 0.57 |
| 11 | 5233697 | rs6578584 | -1.502 | 0.911 | -0.012 | 0.84 | -0.016 | -0.021 | -0.021 | 0 | 0.25 | 0.44 | 0.3 | 0.18 | 0.24 | 0.31 |
| 11 | 5246512 | rs7110263 | 0.916 | 1.063 | -0.42 | 1.023 | -0.001 | 0.018 | 0.018 | 0 | 0.87 | 0.89 | 0.72 | 0.51 | 0.82 | 0.56 |
| 11 | 5247141 | rs1609812 | 0.754 | 1.022 | -0.41 | 1.034 | -0.001 | 0.026 | 0.026 | 0 | 0.87 | 0.88 | 0.73 | 0.51 | 0.83 | 0.57 |
| 11 | 5247726 | rs7946748 | -0.343 | 1.002 | -0.442 | 0.147 | -0.001 | 0.009 | 0.009 | 0 | 0.09 | 0.07 | 0.08 | 0.02 | 0.13 | 0.2 |
| 11 | 5249004 | rs11036364 | -0.352 | 0.752 | -0.602 | 0.145 | -0.028 | -0.018 | -0.018 | 0.072 | 0.43 | 0.29 | 0.49 | 0.44 | 0.47 | 0.39 |
| 11 | 5258162 | rs4320977 | 0.661 | 1.008 | 1.045 | -0.596 | -0.035 | 0.066 | 0.066 | 0 | 0.61 | 0.21 | 0.62 | 0.77 | 0.5 | 0.57 |
| 11 | 5258592 | rs4402323 | 0.4 | 1.073 | 1.246 | -0.585 | 0.027 | -0.033 | -0.033 | 0.053 | 0.64 | 0.52 | 0.65 | 0.77 | 0.5 | 0.57 |
| 11 | 5258989 | rs4910736 | 0.422 | 1.051 | 1.239 | -0.593 | 0.026 | -0.035 | -0.035 | 0 | 0.64 | 0.52 | 0.65 | 0.77 | 0.5 | 0.55 |
| 11 | 5268797 | rs10488676 | 0.884 | 0.926 | 1.065 | -0.676 | -0.024 | 0.087 | 0.087 | 0 | 0.58 | 0.17 | 0.59 | 0.76 | 0.49 | 0.55 |
| 11 | 5271671 | rs2855039 | -0.038 | 1.52 | 1.172 | -0.797 | -0.052 | 0.029 | 0.029 | 0.632 | 0.23 | 0.15 | 0.19 | 0.13 | 0.31 | 0.28 |
| 11 | 5273922 | rs10768707 | 0.791 | 1.597 | 1.219 | -0.799 | -0.03 | 0.012 | 0.012 | 0 | 0.86 | 0.92 | 0.84 | 0.9 | 0.79 | 0.83 |
| 11 | 5275343 | rs11036476 | 1.458 | 1.566 | 1.266 | -0.801 | -0.008 | 0.08 | 0.08 | 0 | 0.58 | 0.19 | 0.6 | 0.76 | 0.48 | 0.55 |
| 11 | 5279153 | rs5010979 | 1.017 | 1.581 | 1.423 | -0.772 | 0 | 0 | 0 | 0.122 | 0.85 | 0.85 | 0.83 | 0.89 | 0.79 | 0.83 |
| 11 | 5287156 | rs10768733 | 0.844 | 1.585 | 1.283 | -0.77 | -0.03 | 0.011 | 0.011 | 0 | 0.86 | 0.92 | 0.84 | 0.89 | 0.79 | 0.82 |
| 11 | 5287290 | rs4910740 | 1.204 | 1.361 | 1.444 | -0.77 | -0.001 | 0.027 | 0.027 | 0 | 0.59 | 0.34 | 0.6 | 0.76 | 0.47 | 0.56 |
| 11 | 5291532 | rs3759071 | 1.255 | 1.423 | 1.677 | -0.747 | -0.001 | 0.001 | 0.001 | 0 | 0.6 | 0.33 | 0.6 | 0.77 | 0.47 | 0.59 |
| 11 | 5298030 | rs10488675 | 0.888 | 0.743 | 0.725 | -1.404 | -0.054 | 0.105 | 0.105 | 0 | 0.24 | 0.01 | 0.19 | 0.06 | 0.28 | 0.23 |
| 11 | 5306509 | rs4910742 | -0.671 | 1.815 | 0.754 | -0.581 | -0.002 | -0.019 | -0.019 | 0 | 0.95 | 1 | 0.98 | 0.92 | 0.95 | 0.89 |
| 11 | 5309695 | rs16912979 | -0.5 | 1.892 | 0.935 | -0.507 | 0.054 | -0.022 | -0.022 | 0.034 | 0.25 | 0.52 | 0.25 | 0.21 | 0.31 | 0.25 |
| 11 | 5310150 | rs4910743 | 1.123 | 1.664 | 0.658 | -0.475 | -0.033 | 0.063 | 0.063 | 0 | 0.61 | 0.11 | 0.62 | 0.71 | 0.47 | 0.57 |
| 11 | 5312401 | rs4601817 | 0.993 | 1.759 | 0.822 | -0.363 | -0.003 | 0.025 | 0.025 | 0 | 0.6 | 0.3 | 0.48 | 0.71 | 0.45 | 0.56 |
| 11 | 5317553 | rs10837767 | 0.955 | 1.944 | 0.53 | -0.378 | 0.001 | -0.005 | -0.005 | 0 | 0.61 | 0.48 | 0.58 | 0.74 | 0.5 | 0.59 |
| 11 | 5321686 | rs6578599 | 0.193 | 2.172 | -0.728 | -0.205 | -0.03 | -0.001 | -0.001 | 0 | 0.88 | 0.95 | 0.89 | 0.9 | 0.76 | 0.8 |
| 11 | 5322737 | rs10837771 | 1.141 | 1.934 | -1.124 | -0.534 | -0.009 | 0.13 | 0.13 | 0 | 0.57 | 0.12 | 0.46 | 0.75 | 0.45 | 0.55 |
| 11 | 5323071 | rs7118113 | 1.054 | 1.826 | -0.291 | -0.221 | 0.048 | -0.03 | -0.03 | 0 | 0.8 | 0.42 | 0.74 | 0.88 | 0.69 | 0.78 |
| 11 | 5328361 | rs3898917 | 0.507 | 0.647 | -1.046 | 0.063 | 0.034 | 0.003 | 0.003 | 0 | 0.66 | 0.24 | 0.64 | 0.34 | 0.77 | 0.59 |
| 11 | 5328707 | rs10082566 | -0.238 | 0.361 | -0.623 | 0.738 | -0.012 | 0.012 | 0.012 | 0 | 0.14 | 0.21 | 0.15 | 0.22 | 0.1 | 0.15 |
| 11 | 5331522 | rs11036742 | 1.268 | 0.893 | -1.689 | -0.014 | 0.007 | 0.15 | 0.15 | 0 | 0.43 | 0.08 | 0.37 | 0.22 | 0.4 | 0.39 |
| 11 | 5332443 | rs7926089 | -0.644 | 0.823 | -0.786 | 0.091 | 0.038 | 0.289 | 0.289 | 0 | 0.89 | 0.79 | 0.85 | 0.55 | 0.91 | 0.77 |
| 11 | 5333124 | rs7930548 | 0.323 | 1.484 | -1.052 | -0.13 | 0.015 | 0.046 | 0.046 | 0.241 | 0.75 | 0.77 | 0.72 | 0.55 | 0.7 | 0.67 |
| 11 | 5335943 | rs11036766 | 1.168 | 1.517 | -0.901 | -0.197 | 0.004 | -0.002 | -0.002 | 0 | 0.59 | 0.52 | 0.55 | 0.43 | 0.5 | 0.53 |
| 11 | 5336475 | rs7942925 | 0.402 | 1.389 | -0.733 | -0.24 | -0.025 | 0.016 | 0.016 | 0 | 0.16 | 0.34 | 0.16 | 0.21 | 0.1 | 0.15 |
| 11 | 5336998 | rs7935554 | -0.284 | 1.477 | -0.641 | -0.146 | -0.006 | -0.029 | -0.029 | 0.359 | 0.3 | 0.47 | 0.31 | 0.21 | 0.3 | 0.24 |
| 11 | 5338211 | rs7118086 | -0.279 | 1.473 | -0.688 | -0.145 | -0.004 | -0.014 | -0.014 | 8.242 | 0.3 | 0.42 | 0.3 | 0.22 | 0.3 | 0.25 |
| 11 | 5338759 | rs2736567 | 1.799 | 1.48 | -0.744 | -0.161 | -0.011 | 0.141 | 0.141 | 0 | 0.42 | 0.07 | 0.37 | 0.21 | 0.4 | 0.38 |
| 11 | 5341509 | rs4910748 | 0.471 | 1.878 | -0.512 | 0.054 | 0.071 | 0.028 | 0.028 | 0 | 0.09 | 0.16 | 0.14 | 0.12 | 0.18 | 0.11 |
| 11 | 5343211 | rs11036801 | 1.552 | 1.909 | -0.521 | 0.077 | 0.023 | 0.218 | 0.218 | 0.018 | 0.51 | 0.2 | 0.33 | 0.2 | 0.46 | 0.43 |
| 11 | 5343533 | rs4910749 | 0.882 | 1.91 | -0.624 | 0.112 | 0.004 | 0.016 | 0.016 | 0 | 0.59 | 0.36 | 0.46 | 0.32 | 0.62 | 0.53 |
| 11 | 5343747 | rs3886223 | -0.538 | 1.896 | -0.56 | 0.096 | -0.007 | -0.018 | -0.018 | 0 | 0.3 | 0.44 | 0.3 | 0.22 | 0.29 | 0.24 |
| 11 | 5344159 | rs4433594 | -0.733 | 1.918 | -0.592 | 0.109 | 0.04 | 0.302 | 0.302 | 0 | 0.89 | 0.78 | 0.77 | 0.54 | 0.91 | 0.78 |
| 11 | 5344499 | rs7934005 | -0.733 | 1.918 | -0.592 | 0.109 | 0.04 | 0.302 | 0.302 | 0 | 0.89 | 0.78 | 0.77 | 0.54 | 0.91 | 0.78 |
| 11 | 5344561 | rs7933257 | -0.733 | 1.918 | -0.592 | 0.109 | 0.04 | 0.302 | 0.302 | 0 | 0.89 | 0.78 | 0.77 | 0.54 | 0.91 | 0.77 |
| 11 | 5344592 | rs11036814 | 0.867 | 1.928 | -0.597 | 0.105 | -0.052 | -0.04 | -0.04 | 2.029 | 0.08 | 0.01 | 0.12 | 0.12 | 0.17 | 0.11 |
| 11 | 5344681 | rs11036815 | -0.465 | 1.552 | -0.429 | 0.168 | 0.001 | 0.004 | 0.004 | 0 | 0.28 | 0.29 | 0.3 | 0.22 | 0.29 | 0.24 |
| 11 | 5344847 | rs10768793 | -0.64 | 1.531 | -0.59 | 0.208 | 0.04 | 0.302 | 0.302 | 0 | 0.89 | 0.78 | 0.77 | 0.54 | 0.91 | 0.78 |
| 11 | 5345128 | rs10837814 | 1.021 | 1.53 | -0.586 | 0.205 | 0.005 | 0.026 | 0.026 | 0 | 0.58 | 0.34 | 0.46 | 0.32 | 0.61 | 0.53 |
| 11 | 5345170 | rs7952293 | -0.273 | 1.404 | -0.609 | 0.544 | 0.04 | 0.302 | 0.302 | 0 | 0.89 | 0.79 | 0.77 | 0.54 | 0.91 | 0.78 |

| | | | | | | | | | | | | | | | | |
|---|---|---|---|---|---|---|---|---|---|---|---|---|---|---|---|---|
| 11 | 5345486 | rs4910750 | -0.251 | 1.402 | -0.61 | 0.544 | 0.038 | 0.299 | 0.299 | 0 | 0.89 | 0.78 | 0.77 | 0.54 | 0.91 | 0.78 |
| 11 | 5352500 | rs11036868 | -0.868 | 0.999 | -1.805 | -0.399 | -0.006 | 0.175 | 0.175 | 0.311 | 0.56 | 0.86 | 0.72 | 0.78 | 0.53 | 0.66 |
| 11 | 5352806 | rs11036870 | -0.412 | 0.828 | -1.587 | -0.211 | -0.011 | 0.075 | 0.075 | 0 | 0.46 | 0.32 | 0.45 | 0.2 | 0.45 | 0.42 |
| 11 | 5357048 | rs11036885 | -0.127 | 0.456 | -1.577 | 0.256 | 0.002 | -0.038 | -0.038 | 0.059 | 0.08 | 0.03 | 0.1 | 0.11 | 0.06 | 0.13 |
| 11 | 5357881 | rs4269953 | -0.69 | 0.467 | -1.479 | -0.2 | -0.001 | -0.009 | -0.009 | 0 | 0.18 | 0.72 | 0.22 | 0.07 | 0.15 | 0.15 |
| 11 | 5360615 | rs10837853 | -1.235 | 0.428 | -2.029 | -0.632 | -0.047 | 0.078 | 0.078 | 0 | 0.38 | 0.83 | 0.46 | 0.21 | 0.46 | 0.4 |
| 11 | 5364276 | rs4910551 | -0.772 | 0.401 | -0.602 | -0.802 | -0.018 | 0.046 | 0.046 | 0 | 0.33 | 0.26 | 0.4 | 0.18 | 0.44 | 0.39 |
| 11 | 5364450 | rs11036912 | -0.799 | 0.403 | -0.692 | -0.781 | -0.018 | 0.047 | 0.047 | 0 | 0.34 | 0.26 | 0.4 | 0.18 | 0.44 | 0.39 |
| 11 | 5364742 | rs11036913 | 1.149 | -0.412 | -1.372 | -0.456 | -0.09 | 0.03 | 0.03 | 0 | 0.4 | 0.08 | 0.44 | 0.24 | 0.43 | 0.41 |
| 11 | 5365428 | rs4326830 | -0.072 | -0.443 | -1.295 | -0.488 | 0.012 | -0.07 | -0.07 | 0.875 | 0.28 | 0.42 | 0.31 | 0.09 | 0.36 | 0.27 |
| 11 | 5366319 | rs5017236 | 1.105 | -0.435 | -1.739 | -0.436 | -0.115 | 0.034 | 0.034 | 0 | 0.41 | 0.08 | 0.44 | 0.25 | 0.45 | 0.4 |
| 11 | 5366991 | rs11036924 | -0.058 | 0.071 | -1.295 | -0.433 | -0.03 | 0.073 | 0.073 | 0 | 0.26 | 0.16 | 0.3 | 0.09 | 0.37 | 0.27 |
| 11 | 5369851 | rs7123112 | 0.158 | 0.341 | -1.149 | -0.302 | -0.004 | 0.009 | 0.009 | 0 | 0.19 | 0.19 | 0.24 | 0.11 | 0.28 | 0.22 |
| 11 | 5372147 | rs4439540 | -0.048 | 0.236 | -0.818 | -0.505 | 0.001 | -0.046 | -0.046 | 0 | 0.21 | 0.33 | 0.23 | 0.09 | 0.26 | 0.23 |
| 11 | 5372189 | rs4301813 | 0.777 | -0.668 | -1.358 | -0.496 | 0.088 | -0.061 | -0.061 | 0.834 | 0.34 | 0.13 | 0.16 | 0.34 | 0.18 | 0.34 |
| 11 | 5372751 | rs4910755 | 0.461 | -0.565 | -1.292 | -0.531 | 0.001 | -0.047 | -0.047 | 0 | 0.21 | 0.33 | 0.23 | 0.09 | 0.26 | 0.23 |
| 11 | 5372856 | rs4910756 | 1.508 | -0.586 | -1.284 | -0.555 | 0.012 | -0.052 | -0.052 | 0.032 | 0.13 | 0.23 | 0.16 | 0.02 | 0.21 | 0.12 |
| 11 | 5373006 | rs7483122 | 0.622 | -0.527 | -1.265 | -0.488 | 0.001 | -0.046 | -0.046 | 0 | 0.21 | 0.33 | 0.23 | 0.09 | 0.26 | 0.23 |
| 11 | 5373129 | rs5006887 | 0.622 | -0.527 | -1.265 | -0.488 | 0.001 | -0.046 | -0.046 | 0 | 0.21 | 0.33 | 0.23 | 0.09 | 0.26 | 0.23 |
| 11 | 5373242 | rs5006885 | 0.539 | -0.517 | -1.252 | -0.493 | 0.002 | -0.049 | -0.049 | 0 | 0.21 | 0.34 | 0.24 | 0.09 | 0.27 | 0.23 |
| 11 | 5373251 | rs5006884 | 0.625 | -0.518 | -1.252 | -0.493 | 0.001 | -0.046 | -0.046 | 0 | 0.21 | 0.33 | 0.23 | 0.09 | 0.26 | 0.23 |
| 11 | 5373334 | rs5006882 | 0.625 | -0.518 | -1.252 | -0.493 | 0.001 | -0.046 | -0.046 | 0 | 0.21 | 0.33 | 0.23 | 0.09 | 0.26 | 0.23 |
| 11 | 5373562 | rs5024042 | 0.638 | -0.513 | -1.25 | -0.49 | 0.001 | -0.046 | -0.046 | 0.033 | 0.2 | 0.33 | 0.23 | 0.09 | 0.26 | 0.22 |
| 11 | 5373646 | rs5024041 | -1.419 | -0.468 | -1.257 | -0.332 | 0.05 | 0.018 | 0.018 | 0 | 0.68 | 0.58 | 0.57 | 0.55 | 0.61 | 0.72 |
| 11 | 5374355 | rs4494328 | 0.796 | -0.527 | -1.353 | -0.321 | 0.093 | -0.079 | -0.079 | 0.062 | 0.35 | 0.13 | 0.16 | 0.39 | 0.18 | 0.35 |
| 11 | 5374857 | rs4492850 | 1.217 | -0.212 | -1.202 | -0.384 | -0.028 | 0.026 | 0.026 | 0.361 | 0.13 | 0.04 | 0.16 | 0.08 | 0.16 | 0.15 |
| 11 | 5376856 | rs2647601 | -0.35 | -0.336 | -1.145 | -0.377 | -0.001 | -0.005 | -0.005 | 0.378 | 0.33 | 0.36 | 0.38 | 0.16 | 0.4 | 0.37 |
| 11 | 5382834 | rs17270517 | -0.466 | -0.347 | -1.108 | -0.382 | 0.003 | -0.042 | -0.042 | 0.279 | 0.33 | 0.43 | 0.39 | 0.16 | 0.4 | 0.36 |
| 11 | 5384070 | rs17270818 | -0.419 | -0.339 | -1.146 | -0.378 | 0 | -0.013 | -0.013 | 0.209 | 0.33 | 0.37 | 0.39 | 0.16 | 0.4 | 0.36 |
| 11 | 5385932 | rs2647608 | 0.679 | -0.363 | -1.097 | -0.411 | -0.002 | 0.021 | 0.021 | 0 | 0.2 | 0.21 | 0.22 | 0.08 | 0.27 | 0.22 |
| 11 | 5386668 | rs10500635 | 0.512 | -0.365 | -1.096 | -0.412 | 0 | -0.052 | -0.052 | 0 | 0.21 | 0.34 | 0.24 | 0.08 | 0.27 | 0.22 |
| 11 | 5388488 | rs11601662 | 1.437 | -0.364 | -1.18 | -0.358 | -0.025 | 0.022 | 0.022 | 0.429 | 0.12 | 0.03 | 0.15 | 0.08 | 0.13 | 0.14 |
| 11 | 5394480 | rs2736549 | 0.692 | -0.648 | -1.254 | -0.132 | 0.059 | -0.039 | -0.039 | 0 | 0.35 | 0.19 | 0.18 | 0.36 | 0.19 | 0.35 |
| 11 | 5402554 | rs2723373 | -1.371 | -0.506 | -1.104 | -0.118 | 0.053 | 0.017 | 0.017 | 0 | 0.69 | 0.61 | 0.56 | 0.55 | 0.59 | 0.71 |
| 11 | 5404060 | rs2647549 | -1.202 | -0.54 | -1.067 | -0.134 | -0.006 | -0.002 | -0.002 | 0 | 0.67 | 0.39 | 0.54 | 0.55 | 0.59 | 0.71 |
| 11 | 5404735 | rs11037129 | 1.219 | -0.481 | -1.184 | -0.181 | -0.024 | 0.036 | 0.036 | 0.544 | 0.14 | 0.06 | 0.17 | 0.08 | 0.15 | 0.15 |
| 11 | 5410103 | rs2736533 | 1.079 | -1.197 | -1.359 | -0.174 | 0.003 | -0.019 | -0.019 | 0 | 0.43 | 0.37 | 0.41 | 0.22 | 0.45 | 0.28 |
| 11 | 5410934 | rs1498467 | -0.694 | -1.227 | -1.151 | -0.15 | 0.006 | 0.051 | 0.051 | 1.075 | 0.81 | 0.68 | 0.73 | 0.55 | 0.82 | 0.72 |
| 11 | 5411239 | rs1498469 | -0.733 | -1.225 | -1.178 | -0.148 | 0.002 | 0.024 | 0.024 | 0.951 | 0.81 | 0.74 | 0.73 | 0.55 | 0.82 | 0.73 |
| 11 | 5411398 | rs2736531 | 0.039 | -1.237 | -1.178 | -0.057 | 0.012 | -0.003 | -0.003 | 0.197 | 0.38 | 0.34 | 0.32 | 0.31 | 0.38 | 0.44 |
| 11 | 5411580 | rs10768907 | -0.627 | -1.038 | -1.181 | -0.016 | 0.002 | 0.024 | 0.024 | 0.951 | 0.81 | 0.74 | 0.73 | 0.55 | 0.82 | 0.73 |
| 11 | 5412591 | rs11037194 | -0.809 | -1.039 | -1.175 | -0.015 | -0.006 | -0.022 | -0.022 | 0 | 0.82 | 0.83 | 0.75 | 0.54 | 0.82 | 0.75 |
| 11 | 5413551 | rs2340324 | -0.627 | -1.036 | -1.181 | -0.015 | 0.002 | 0.024 | 0.024 | 0.951 | 0.81 | 0.74 | 0.73 | 0.55 | 0.82 | 0.73 |
| 11 | 5417034 | rs951747 | -0.704 | -1.299 | -1.115 | -0.059 | 0.002 | -0.002 | -0.002 | 0 | 0.57 | 0.67 | 0.43 | 0.47 | 0.49 | 0.56 |
| 11 | 5419559 | rs4432053 | 1.137 | -1.299 | -1.182 | -0.054 | -0.108 | 0.165 | 0.165 | 0 | 0.37 | 0.11 | 0.46 | 0.15 | 0.49 | 0.32 |
| 11 | 5420909 | rs6578634 | -0.997 | -1.3 | -1.167 | -0.042 | 0.001 | 0.146 | 0.146 | 0 | 0.95 | 0.87 | 0.91 | 0.62 | 0.98 | 0.9 |
| 11 | 5422198 | rs10838005 | 1.1 | -1.287 | -1.152 | -0.058 | -0.108 | 0.165 | 0.165 | 0 | 0.37 | 0.11 | 0.46 | 0.15 | 0.49 | 0.32 |
| 11 | 5424701 | rs16930511 | 2.018 | -1.72 | -1.359 | -0.074 | 0.025 | -0.041 | -0.041 | 0.002 | 0.08 | 0.01 | 0.04 | 0.18 | 0.05 | 0.12 |
| 11 | 5430603 | rs975115 | -1.607 | -1.755 | -1.201 | -0.293 | -0.048 | -0.001 | -0.001 | 0 | 0.8 | 0.61 | 0.87 | 0.85 | 0.89 | 0.88 |
| 11 | 5431986 | rs1566273 | 1.128 | -1.825 | -1.326 | -0.384 | -0.038 | 0.141 | 0.141 | 0 | 0.4 | 0.23 | 0.48 | 0.15 | 0.51 | 0.34 |
| 11 | 5432294 | rs10838051 | 1.263 | -1.759 | -1.41 | -0.336 | -0.107 | 0.167 | 0.167 | 0 | 0.38 | 0.11 | 0.46 | 0.15 | 0.49 | 0.33 |
| 11 | 5440287 | rs2647579 | 0.904 | -1.856 | -1.445 | -0.336 | 0.035 | -0.053 | -0.053 | 0 | 0.2 | 0.07 | 0.1 | 0.24 | 0.12 | 0.2 |
| 11 | 5442985 | rs10768949 | 1.225 | -1.576 | -1.318 | -0.451 | -0.088 | 0.155 | 0.155 | 0 | 0.37 | 0.14 | 0.47 | 0.15 | 0.49 | 0.34 |
| 11 | 5443190 | rs10768950 | 1.225 | -1.576 | -1.318 | -0.451 | -0.088 | 0.155 | 0.155 | 0 | 0.37 | 0.14 | 0.47 | 0.15 | 0.49 | 0.34 |
| 11 | 5443700 | rs2736588 | -0.483 | -1.963 | -1.238 | -0.347 | 0.016 | 0.04 | 0.04 | 0 | 0.41 | 0.28 | 0.4 | 0.7 | 0.4 | 0.53 |
| 11 | 5443739 | rs2736587 | -0.483 | -1.963 | -1.238 | -0.347 | 0.016 | 0.04 | 0.04 | 0 | 0.41 | 0.28 | 0.4 | 0.7 | 0.4 | 0.53 |
| 11 | 5443887 | rs10838093 | -0.483 | -1.963 | -1.238 | -0.347 | 0.016 | 0.04 | 0.04 | 0 | 0.41 | 0.28 | 0.4 | 0.7 | 0.4 | 0.53 |
| 11 | 5443963 | rs10838095 | -0.483 | -1.963 | -1.238 | -0.347 | 0.016 | 0.04 | 0.04 | 0 | 0.41 | 0.28 | 0.4 | 0.7 | 0.4 | 0.53 |
| 11 | 5444061 | rs2736586 | -0.483 | -1.963 | -1.238 | -0.347 | 0.016 | 0.04 | 0.04 | 0 | 0.41 | 0.28 | 0.4 | 0.7 | 0.4 | 0.53 |
| 11 | 5444136 | rs2647574 | -0.483 | -1.963 | -1.238 | -0.347 | 0.016 | 0.04 | 0.04 | 0 | 0.41 | 0.28 | 0.4 | 0.7 | 0.4 | 0.53 |
| 11 | 5444353 | rs2647573 | -0.483 | -1.963 | -1.238 | -0.347 | 0.016 | 0.04 | 0.04 | 0 | 0.41 | 0.28 | 0.4 | 0.7 | 0.4 | 0.53 |
| 11 | 5446474 | rs1472453 | 0.668 | -1.632 | -1.181 | -0.495 | 0.001 | -0.025 | -0.025 | 0.093 | 0.38 | 0.46 | 0.49 | 0.15 | 0.53 | 0.36 |
| 11 | 5448543 | rs11037417 | 0.128 | -1.551 | -1.332 | -0.171 | 0.009 | 0.035 | 0.035 | 0 | 0.18 | 0.26 | 0.25 | 0.44 | 0.25 | 0.24 |
| 11 | 5451875 | rs1498474 | 2.06 | -1.439 | -1.743 | -0.468 | -0.161 | 0.047 | 0.047 | 1.01 | 0.22 | 0.04 | 0.4 | 0.13 | 0.43 | 0.2 |
| 11 | 5455929 | rs1391619 | -0.364 | -2.283 | -0.554 | -0.267 | -0.035 | 0.018 | 0.018 | 0.159 | 0.75 | 0.76 | 0.87 | 0.73 | 0.87 | 0.81 |
| 11 | 5457598 | rs986772 | -1.299 | -2.43 | -0.439 | -0.312 | -0.023 | -0.016 | -0.016 | 1.06 | 0.91 | 0.86 | 0.92 | 0.91 | 0.95 | 0.9 |
| 11 | 5457631 | rs10838102 | 0.983 | -2.523 | -0.228 | -0.348 | 0.067 | 0.121 | 0.121 | 0 | 0.5 | 0.26 | 0.53 | 0.26 | 0.57 | 0.47 |
| 11 | 5461263 | rs10838107 | 0.964 | -2.648 | -0.2 | -0.342 | 0.098 | 0.133 | 0.133 | 0 | 0.5 | 0.21 | 0.53 | 0.25 | 0.59 | 0.47 |
| 11 | 5462085 | rs11037444 | 0.575 | -2.64 | -0.311 | -0.233 | -0.005 | -0.028 | -0.028 | 0 | 0.16 | 0.12 | 0.23 | 0.37 | 0.23 | 0.22 |

| | | | | | | | | | | | | | | | | |
|---|---|---|---|---|---|---|---|---|---|---|---|---|---|---|---|---|
| 11 | 5462374 | rs16930982 | 1.068 | -2.712 | -0.371 | -0.275 | 0.013 | -0.012 | -0.012 | 0 | 0.06 | 0.01 | 0.1 | 0.36 | 0.03 | 0.17 |
| 11 | 5462702 | rs16930998 | 1.068 | -2.712 | -0.371 | -0.275 | 0.013 | -0.012 | -0.012 | 0.129 | 0.06 | 0.01 | 0.1 | 0.36 | 0.03 | 0.17 |
| 11 | 5462905 | rs1498487 | -0.964 | -2.738 | -0.254 | -0.281 | 0.01 | -0.011 | -0.011 | 0 | 0.77 | 0.88 | 0.89 | 0.72 | 0.87 | 0.81 |
| 11 | 5463061 | rs4910793 | 1.751 | -2.574 | -0.421 | -0.228 | 0.007 | 0.004 | 0.004 | 0 | 0.08 | 0.05 | 0.07 | 0.09 | 0.05 | 0.1 |
| 11 | 5463425 | rs16931041 | 0.326 | -2.432 | -0.464 | -0.097 | -0.011 | -0.052 | -0.052 | 0 | 0.16 | 0.1 | 0.23 | 0.39 | 0.23 | 0.22 |
| 11 | 5469576 | rs10500641 | -0.491 | -2.472 | -0.371 | -0.149 | -0.034 | 0.02 | 0.02 | 0.131 | 0.75 | 0.77 | 0.87 | 0.72 | 0.87 | 0.81 |
| 11 | 5469959 | rs16931173 | 1.317 | -2.195 | -0.871 | -0.474 | -0.012 | 0.001 | 0.001 | 0 | 0.15 | 0.18 | 0.08 | 0.12 | 0.06 | 0.15 |
| 11 | 5470716 | rs956471 | 0.802 | -1.909 | -1.234 | -0.816 | -0.031 | 0.149 | 0.149 | 0 | 0.37 | 0.15 | 0.47 | 0.13 | 0.54 | 0.34 |
| 11 | 5470990 | rs4094102 | 0.714 | -1.926 | -1.308 | -0.888 | -0.037 | 0.071 | 0.071 | 0.052 | 0.14 | 0.03 | 0.18 | 0.02 | 0.23 | 0.07 |
| 11 | 5471746 | rs7948471 | 0.646 | -1.633 | -0.742 | -0.733 | 0.016 | -0.011 | -0.011 | 3.547 | 0.33 | 0.45 | 0.46 | 0.25 | 0.5 | 0.35 |
| 11 | 5474093 | rs2030090 | 0.624 | -1.741 | -0.332 | -0.771 | -0.006 | 0.003 | 0.003 | 0.842 | 0.33 | 0.71 | 0.47 | 0.26 | 0.5 | 0.35 |
| 11 | 5474838 | rs2030094 | 0.834 | -1.695 | -0.591 | -0.748 | 0.025 | -0.007 | -0.007 | 0.026 | 0.32 | 0.45 | 0.45 | 0.26 | 0.5 | 0.35 |
| 11 | 5475118 | rs12577167 | -0.343 | -1.965 | -1.319 | -0.697 | 0.078 | -0.069 | -0.069 | 0 | 0.24 | 0.07 | 0.16 | 0.36 | 0.13 | 0.28 |
| 11 | 5475597 | rs11037503 | 2.139 | -1.069 | -1.297 | -0.747 | -0.127 | 0.098 | 0.098 | 0.081 | 0.2 | 0.05 | 0.3 | 0.04 | 0.37 | 0.18 |
| 11 | 5477390 | rs2736578 | -0.395 | -1.423 | -1.224 | -0.809 | 0.069 | 0.003 | 0.003 | 0 | 0.39 | 0.22 | 0.36 | 0.43 | 0.31 | 0.37 |
| 11 | 5479891 | rs1603776 | 2.405 | -0.973 | -1.258 | -0.865 | -0.113 | 0.089 | 0.089 | 0.141 | 0.18 | 0.04 | 0.3 | 0.04 | 0.35 | 0.18 |
| 11 | 5480444 | rs11037542 | 1.358 | -1.11 | -1.308 | -0.907 | -0.003 | 0.049 | 0.049 | 0 | 0.13 | 0.05 | 0.05 | 0 | 0.11 | 0.13 |
| 11 | 5483235 | rs1498482 | 2.135 | -0.911 | -1.245 | -0.828 | -0.126 | 0.099 | 0.099 | 0.477 | 0.2 | 0.04 | 0.3 | 0.04 | 0.37 | 0.2 |
| 11 | 5484136 | rs1498481 | -0.035 | -1.192 | -0.212 | -0.626 | 0.014 | 0.002 | 0.002 | 0.017 | 0.58 | 0.39 | 0.67 | 0.47 | 0.68 | 0.55 |
| 11 | 5487596 | rs2647593 | -0.171 | -1.173 | -0.043 | -0.603 | -0.003 | 0 | 0 | 0 | 0.61 | 0.67 | 0.71 | 0.47 | 0.68 | 0.6 |
| 11 | 5488533 | rs12576842 | 0.339 | -1.441 | -0.353 | -0.46 | 0.034 | -0.071 | -0.071 | 3.373 | 0.22 | 0.06 | 0.16 | 0.36 | 0.12 | 0.29 |
| 11 | 5488840 | rs454465 | -0.554 | -1.439 | 0.204 | -0.379 | -0.031 | 0.015 | 0.015 | 0.078 | 0.68 | 0.49 | 0.77 | 0.69 | 0.79 | 0.75 |
| 11 | 5490077 | rs2736569 | 1.981 | -1.397 | 0.163 | -0.511 | 0.003 | 0.017 | 0.017 | 0 | 0.16 | 0.12 | 0.18 | 0.08 | 0.18 | 0.07 |
| 11 | 5490345 | rs449937 | -0.759 | -1.454 | 0.342 | -0.518 | -0.022 | 0.013 | 0.013 | 0 | 0.69 | 0.68 | 0.79 | 0.69 | 0.79 | 0.75 |
| 11 | 5491183 | rs7106927 | 0.123 | -1.362 | 0.319 | -0.459 | 0.055 | 0.013 | 0.013 | 0.121 | 0.52 | 0.32 | 0.59 | 0.6 | 0.61 | 0.68 |
| 11 | 5491568 | rs7106553 | 0.078 | -1.348 | 0.252 | -0.459 | 0.037 | 0.011 | 0.011 | 0.087 | 0.52 | 0.38 | 0.59 | 0.61 | 0.62 | 0.68 |
| 11 | 5493370 | rs425610 | -1.672 | -1.294 | 0.2 | -0.432 | 0.004 | 0.004 | 0.004 | 0 | 0.81 | 0.91 | 0.87 | 0.84 | 0.85 | 0.86 |
| 11 | 5493557 | rs416586 | 0.296 | -1.333 | 0.291 | -0.705 | -0.009 | -0.067 | -0.067 | 0 | 0.31 | 0.44 | 0.27 | 0.08 | 0.27 | 0.21 |
| 11 | 5496318 | rs438996 | -0.234 | -1.456 | 0.289 | -0.543 | 0.004 | 0.002 | 0.002 | 0 | 0.55 | 0.47 | 0.6 | 0.62 | 0.62 | 0.7 |
| 11 | 5496811 | rs2647543 | 0.568 | -1.184 | 0.373 | -0.571 | -0.022 | -0.007 | -0.007 | 0.285 | 0.39 | 0.3 | 0.51 | 0.61 | 0.5 | 0.51 |
| 11 | 5505149 | rs12283559 | 0.31 | -1.29 | 0.587 | -0.527 | 0.021 | 0.023 | 0.023 | 0 | 0.41 | 0.3 | 0.35 | 0.32 | 0.35 | 0.28 |
| 11 | 5506470 | rs2244431 | 1.29 | -1.055 | 0.003 | -0.328 | -0.008 | -0.011 | -0.011 | 0 | 0.11 | 0.1 | 0.1 | 0.22 | 0.13 | 0.18 |
| 11 | 5509689 | rs12577059 | 1.326 | -1.161 | 0.063 | -0.416 | 0.027 | 0.003 | 0.003 | 0 | 0.26 | 0.12 | 0.16 | 0.24 | 0.2 | 0.26 |
| 11 | 5510341 | rs404280 | -0.177 | -1.002 | -0.077 | -0.333 | 0.042 | 0.022 | 0.022 | 0 | 0.53 | 0.32 | 0.58 | 0.61 | 0.59 | 0.6 |
| 11 | 5510688 | rs7101919 | 2.631 | -0.521 | -0.554 | -0.48 | -0.052 | 0.065 | 0.065 | 0 | 0.19 | 0.06 | 0.28 | 0.04 | 0.31 | 0.16 |
| 11 | 5512590 | rs438994 | 1.106 | -0.462 | -0.636 | -0.209 | -0.033 | -0.039 | -0.039 | 0 | 0.11 | 0.06 | 0.12 | 0.19 | 0.17 | 0.19 |
| 11 | 5514892 | rs2879747 | 1.455 | -0.518 | -0.595 | -0.439 | -0.002 | 0.03 | 0.03 | 0 | 0.14 | 0.08 | 0.06 | 0 | 0.11 | 0.1 |
| 11 | 5517485 | rs366533 | -0.188 | -0.576 | -0.403 | -0.255 | 0.037 | 0.047 | 0.047 | 0 | 0.56 | 0.32 | 0.61 | 0.69 | 0.62 | 0.63 |
| 11 | 5518911 | rs397076 | 0.658 | -0.726 | -0.984 | -0.072 | -0.005 | -0.004 | -0.004 | 0 | 0.13 | 0.11 | 0.12 | 0.22 | 0.17 | 0.2 |
| 11 | 5518978 | rs435810 | -0.524 | -1.038 | -1.035 | -0.286 | -0.021 | -0.009 | -0.009 | 0 | 0.7 | 0.37 | 0.79 | 0.77 | 0.82 | 0.7 |
| 11 | 5525654 | rs10500645 | 0.75 | -0.433 | -2.009 | 0.354 | 0.01 | -0.103 | -0.103 | 0 | 0.21 | 0.04 | 0.27 | 0.39 | 0.18 | 0.23 |
| 11 | 5528633 | rs1060574 | 0.349 | 0.077 | -2.553 | -1.492 | -0.009 | 0.079 | 0.079 | 0 | 0.32 | 0.09 | 0.22 | 0.19 | 0.3 | 0.27 |
| 11 | 5528918 | rs2227271 | 0.668 | 0.724 | -2.194 | -1.409 | -0.078 | 0.096 | 0.096 | 0 | 0.24 | 0.06 | 0.33 | 0.05 | 0.37 | 0.2 |
| 11 | 5529139 | rs2234456 | 0.64 | 0.738 | -2.168 | -1.382 | -0.086 | 0.11 | 0.11 | 0 | 0.26 | 0.06 | 0.33 | 0.05 | 0.38 | 0.21 |
| 11 | 5529152 | rs2234455 | -0.183 | -0.099 | -1.415 | -0.517 | -0.013 | 0.077 | 0.077 | 0 | 0.21 | 0.01 | 0.24 | 0.09 | 0.21 | 0.1 |
| 11 | 5533402 | rs317780 | -0.467 | 0.383 | -0.84 | -0.627 | -0.016 | 0.11 | 0.11 | 0.059 | 0.43 | 0.28 | 0.36 | 0.1 | 0.51 | 0.23 |
| 11 | 5533606 | rs3763880 | 0.588 | -0.303 | -0.916 | -0.215 | 0.01 | -0.005 | -0.005 | 0 | 0.26 | 0.09 | 0.25 | 0.29 | 0.21 | 0.4 |
| 11 | 5534114 | rs10734521 | -0.274 | -0.454 | -0.069 | 0.551 | 0.004 | 0.03 | 0.03 | 0.298 | 0.53 | 0.39 | 0.6 | 0.66 | 0.47 | 0.66 |
| 11 | 5535007 | rs4910812 | -0.445 | -0.522 | 0.103 | 0.561 | 0 | 0.008 | 0.008 | 0.488 | 0.55 | 0.43 | 0.6 | 0.67 | 0.48 | 0.67 |
| 11 | 5535529 | rs10838245 | 0.584 | -0.596 | 0.241 | 0.783 | 0.002 | -0.007 | -0.007 | 0 | 0.27 | 0.29 | 0.33 | 0.37 | 0.26 | 0.26 |
| 11 | 5536415 | rs10769023 | 0.669 | -0.644 | 0.287 | 0.755 | 0.002 | -0.012 | -0.012 | 0 | 0.27 | 0.25 | 0.32 | 0.37 | 0.26 | 0.27 |
| 11 | 5536852 | rs2017434 | 0.407 | -0.645 | 0.288 | 0.754 | -0.002 | 0.004 | 0.004 | 2.906 | 0.28 | 0.35 | 0.35 | 0.38 | 0.26 | 0.27 |
| 11 | 5537045 | rs872751 | 0.407 | -0.645 | 0.288 | 0.754 | -0.002 | 0.004 | 0.004 | 2.906 | 0.28 | 0.35 | 0.35 | 0.38 | 0.26 | 0.27 |
| 11 | 5537161 | rs2047456 | 0.407 | -0.645 | 0.342 | 0.754 | 0 | -0.001 | -0.001 | 0 | 0.28 | 0.33 | 0.35 | 0.38 | 0.26 | 0.27 |
| 11 | 5539139 | rs979752 | 0.469 | -0.312 | 0.484 | 0.186 | -0.004 | -0.002 | -0.002 | 0.042 | 0.26 | 0.28 | 0.21 | 0.26 | 0.31 | 0.24 |
| 11 | 5539670 | rs7131665 | -0.419 | -0.336 | 0.689 | 0.224 | 0.001 | 0.004 | 0.004 | 0 | 0.53 | 0.61 | 0.55 | 0.61 | 0.57 | 0.5 |
| 11 | 5540466 | rs12418972 | 0.669 | -0.35 | 0.729 | 0.19 | -0.013 | 0.012 | 0.012 | 0.131 | 0.24 | 0.21 | 0.19 | 0.03 | 0.3 | 0.14 |
| 11 | 5541179 | rs1391615 | -0.479 | -0.347 | 0.901 | 0.194 | 0.006 | 0.025 | 0.025 | 0.019 | 0.54 | 0.68 | 0.56 | 0.62 | 0.57 | 0.51 |
| 11 | 5543705 | rs7945342 | 1.027 | -0.285 | 0.834 | 0.152 | 0 | -0.028 | -0.028 | 0 | 0.25 | 0.16 | 0.32 | 0.36 | 0.26 | 0.26 |
| 11 | 5546722 | rs10160557 | -0.033 | -0.525 | 0.763 | -0.287 | 0.032 | 0.145 | 0.145 | 0 | 0.5 | 0.36 | 0.43 | 0.08 | 0.57 | 0.29 |
| 11 | 5547530 | rs7932885 | 1.105 | -0.267 | 0.577 | -0.381 | -0.008 | 0.008 | 0.008 | 0 | 0.17 | 0.16 | 0.29 | 0.07 | 0.23 | 0.09 |
| 11 | 5547957 | rs4910815 | 0.135 | -0.631 | 0.603 | 0.048 | 0 | 0 | 0 | 0 | 0.35 | 0.37 | 0.31 | 0.38 | 0.33 | 0.33 |
| 11 | 5550630 | rs317773 | -0.282 | -0.576 | 0.544 | 0.101 | -0.002 | -0.002 | -0.002 | 0.433 | 0.42 | 0.66 | 0.37 | 0.63 | 0.37 | 0.47 |
| 11 | 5554825 | rs12270763 | 0.678 | -0.512 | 0.478 | 0.062 | -0.002 | 0 | 0 | 0.743 | 0.31 | 0.3 | 0.21 | 0.27 | 0.32 | 0.26 |
| 11 | 5555043 | rs10742709 | -0.67 | -0.457 | 0.553 | -0.137 | -0.003 | 0.001 | 0.001 | 0 | 0.57 | 0.62 | 0.64 | 0.45 | 0.6 | 0.48 |
| 11 | 5555551 | rs10838277 | -0.155 | -0.49 | 0.47 | -0.211 | 0.014 | 0.029 | 0.029 | 0 | 0.51 | 0.47 | 0.52 | 0.09 | 0.58 | 0.3 |
| 11 | 5565906 | rs7934354 | 0.816 | -0.493 | 0.662 | -0.177 | -0.012 | 0.29 | 0.29 | 1.939 | 0.06 | 0.16 | 0.11 | 0.37 | 0.02 | 0.17 |
| 11 | 5566030 | rs1995158 | -1.594 | -0.365 | 0.091 | -1.32 | -0.017 | 0.057 | 0.057 | 0 | 0.83 | 0.94 | 0.84 | 0.92 | 0.8 | 0.92 |
| 11 | 5566051 | rs1995157 | -0.052 | -0.372 | 0.105 | -1.327 | 0.018 | 0.047 | 0.047 | 0 | 0.51 | 0.45 | 0.51 | 0.09 | 0.58 | 0.3 |

| | | | | | | | | | | | | | | | | |
|---|---|---|---|---|---|---|---|---|---|---|---|---|---|---|---|---|
| 11 | 5566112 | rs1995156 | -0.013 | -0.375 | 0.103 | -1.328 | 0.017 | 0.038 | 0.038 | 0 | 0.51 | 0.46 | 0.51 | 0.09 | 0.58 | 0.3 |
| 11 | 5566365 | rs1566275 | -1.593 | -0.368 | 0.101 | -1.32 | -0.015 | 0.056 | 0.056 | 0 | 0.83 | 0.93 | 0.84 | 0.92 | 0.8 | 0.92 |
| 11 | 5566489 | rs10769054 | -0.088 | -0.334 | -0.019 | -1.303 | 0.016 | 0.035 | 0.035 | 0 | 0.51 | 0.47 | 0.51 | 0.09 | 0.58 | 0.3 |
| 11 | 5566789 | rs12798586 | 0.705 | -0.223 | -0.18 | -1.186 | -0.027 | 0.029 | 0.029 | 0 | 0.2 | 0.15 | 0.34 | 0.08 | 0.28 | 0.1 |
| 11 | 5569452 | rs431702 | -0.899 | -0.361 | -0.182 | -1.279 | -0.012 | 0.122 | 0.122 | 0 | 0.61 | 0.78 | 0.72 | 0.92 | 0.55 | 0.79 |
| 11 | 5572816 | rs401426 | -0.37 | -0.376 | 0.021 | -1.006 | 0.027 | 0.048 | 0.048 | 0 | 0.39 | 0.64 | 0.43 | 0.84 | 0.27 | 0.68 |
| 11 | 5575253 | rs431409 | 0.346 | -0.321 | 0.061 | -0.975 | -0.022 | -0.048 | -0.048 | 0.763 | 0.31 | 0.48 | 0.33 | 0.84 | 0.22 | 0.62 |
| 11 | 5577421 | rs12289789 | 1.379 | -0.059 | -0.745 | -0.961 | -0.004 | 0.052 | 0.052 | 0 | 0.23 | 0.17 | 0.15 | 0.02 | 0.22 | 0.14 |
| 11 | 5578343 | rs1541941 | -0.01 | -0.147 | -0.568 | -0.849 | -0.02 | -0.053 | -0.053 | 0 | 0.37 | 0.46 | 0.35 | 0.84 | 0.27 | 0.68 |
| 11 | 5581076 | rs12270701 | 1.091 | 0.17 | -0.791 | -0.824 | -0.005 | 0.028 | 0.028 | 0 | 0.23 | 0.18 | 0.15 | 0.02 | 0.22 | 0.14 |
| 11 | 5583533 | rs11038110 | 0.599 | 0.296 | -0.892 | -0.727 | 0.013 | -0.027 | -0.027 | 0 | 0.19 | 0.19 | 0.22 | 0.08 | 0.23 | 0.09 |
| 11 | 5584496 | rs7936389 | 0.691 | 0.145 | -1.57 | -0.263 | 0.003 | -0.014 | -0.014 | 0 | 0.21 | 0.07 | 0.16 | 0.23 | 0.17 | 0.36 |
| 11 | 5585744 | rs10838341 | 0.576 | 0.469 | -1.528 | -0.659 | -0.018 | 0.055 | 0.055 | 0 | 0.23 | 0.13 | 0.21 | 0.02 | 0.28 | 0.15 |
| 11 | 5586252 | rs10742719 | -0.146 | 0.422 | -1.832 | -1.169 | -0.038 | 0.155 | 0.155 | 0 | 0.59 | 0.81 | 0.67 | 0.98 | 0.48 | 0.7 |
| 11 | 5587111 | rs4604918 | -0.087 | 0.365 | -1.817 | -1.197 | -0.034 | 0.161 | 0.161 | 0 | 0.58 | 0.8 | 0.67 | 0.98 | 0.48 | 0.7 |
| 11 | 5588174 | rs9804698 | 0.361 | 0.69 | -2.024 | -1.415 | -0.061 | 0.116 | 0.116 | 0 | 0.31 | 0.12 | 0.27 | 0.02 | 0.42 | 0.16 |
| 11 | 5589862 | rs2341430 | -1.002 | 0.576 | -2.039 | -1.423 | 0.01 | 0.073 | 0.073 | 0 | 0.86 | 0.97 | 0.92 | 1 | 0.85 | 0.85 |
| 11 | 5590525 | rs4382942 | 0.266 | 0.077 | -2.418 | -0.824 | -0.001 | -0.003 | -0.003 | 1.41 | 0.36 | 0.63 | 0.55 | 0.62 | 0.33 | 0.46 |
| 11 | 5591634 | rs11038137 | 1.476 | 0.098 | -2.564 | -1.116 | -0.001 | -0.012 | -0.012 | 0 | 0.13 | 0.09 | 0.07 | 0.01 | 0.1 | 0.02 |
| 11 | 5592226 | rs11038141 | 0.771 | 0.198 | -2.443 | -0.685 | 0.007 | 0.002 | 0.002 | 0 | 0.2 | 0.11 | 0.1 | 0.22 | 0.15 | 0.16 |
| 11 | 5594093 | rs2341431 | 0.943 | 0.19 | -2.478 | -0.217 | -0.029 | -0.049 | -0.049 | 0 | 0.18 | 0.06 | 0.26 | 0.33 | 0.22 | 0.09 |
| 11 | 5597130 | rs10838362 | 1.449 | 0.596 | -2.426 | -1.012 | -0.022 | 0.049 | 0.049 | 0 | 0.25 | 0.14 | 0.21 | 0.01 | 0.28 | 0.15 |
| 11 | 5597633 | rs10500652 | 0.768 | 0.547 | -2.456 | -1.019 | 0.011 | 0.005 | 0.005 | 0 | 0.21 | 0.13 | 0.11 | 0.21 | 0.15 | 0.16 |
| 11 | 5597789 | rs7120684 | 0.308 | 0.507 | -2.123 | -1.055 | 0.001 | -0.011 | -0.011 | 0 | 0.36 | 0.58 | 0.56 | 0.72 | 0.33 | 0.53 |
| 11 | 5597948 | rs11038171 | -0.59 | 1 | -2.026 | -1.176 | 0.004 | 0.004 | 0.004 | 0 | 0.81 | 0.86 | 0.88 | 0.99 | 0.76 | 0.84 |
| 11 | 5602615 | rs2341434 | 0.067 | 0.874 | -2.145 | -1.247 | -0.015 | 0.096 | 0.096 | 0 | 0.56 | 0.74 | 0.67 | 0.98 | 0.48 | 0.69 |
| 11 | 5602679 | rs892336 | 0.067 | 0.874 | -2.145 | -1.247 | -0.015 | 0.096 | 0.096 | 0 | 0.56 | 0.74 | 0.67 | 0.98 | 0.48 | 0.69 |
| 11 | 5602790 | rs2163946 | 0.067 | 0.874 | -2.145 | -1.247 | -0.015 | 0.096 | 0.096 | 0 | 0.56 | 0.74 | 0.67 | 0.98 | 0.48 | 0.69 |
| 11 | 5602968 | rs10769086 | -0.116 | 0.665 | -2.063 | -0.399 | -0.015 | 0.102 | 0.102 | 0 | 0.82 | 0.97 | 0.88 | 0.99 | 0.76 | 0.84 |
| 11 | 5614275 | rs7117257 | -0.65 | 0.735 | -1.849 | 0.067 | -0.003 | 0.04 | 0.04 | 0 | 0.27 | 0.15 | 0.25 | 0.23 | 0.23 | 0.21 |
| 11 | 5617896 | rs10838406 | -0.417 | 0.968 | -1.019 | 0.814 | 0.009 | 0.052 | 0.052 | 0 | 0.08 | 0.18 | 0.05 | 0.18 | 0.07 | 0.14 |
| 11 | 5617987 | rs12272467 | 0.396 | 1.422 | -0.654 | 0.854 | 0.008 | 0.005 | 0.005 | 1.605 | 0.62 | 0.42 | 0.52 | 0.49 | 0.51 | 0.35 |
| 11 | 5618417 | rs7927012 | 0.416 | 1.41 | -0.477 | 0.846 | 0 | -0.001 | -0.001 | 0.013 | 0.65 | 0.35 | 0.57 | 0.49 | 0.6 | 0.36 |
| 11 | 5618502 | rs7936854 | 0.416 | 1.407 | -0.477 | 0.846 | 0 | -0.001 | -0.001 | 0 | 0.65 | 0.35 | 0.57 | 0.49 | 0.61 | 0.36 |
| 11 | 5622301 | rs6578665 | 0.22 | 1.116 | -0.351 | 0.886 | 0.006 | 0.067 | 0.067 | 0 | 0.08 | 0.2 | 0.05 | 0.18 | 0.06 | 0.12 |
| 11 | 5625603 | rs7108470 | -0.589 | 0.501 | -0.891 | 0.09 | 0.003 | 0.005 | 0.005 | 0 | 0.53 | 0.51 | 0.54 | 0.68 | 0.38 | 0.59 |
| 11 | 5625713 | rs10769118 | 1.146 | -0.21 | -1.22 | -0.397 | 0.036 | 0.033 | 0.033 | 0 | 0.21 | 0.11 | 0.08 | 0.02 | 0.13 | 0.11 |
| 11 | 5625847 | rs3751006 | -1.31 | 0.192 | -0.85 | -0.614 | 0.015 | -0.003 | -0.003 | 0 | 0.7 | 0.72 | 0.66 | 0.72 | 0.59 | 0.8 |
| 11 | 5626241 | rs1074353 | -1.093 | 0.072 | -0.842 | -0.644 | 0.005 | -0.006 | -0.006 | 0 | 0.65 | 0.68 | 0.62 | 0.72 | 0.57 | 0.8 |
| 11 | 5626899 | rs3763881 | -0.616 | -0.186 | -1.023 | -0.249 | -0.001 | 0.011 | 0.011 | 0 | 0.38 | 0.54 | 0.36 | 0.47 | 0.34 | 0.57 |
| 11 | 5628072 | rs7104061 | 0.493 | -0.274 | -0.881 | -0.282 | -0.002 | 0 | 0 | 0 | 0.27 | 0.26 | 0.26 | 0.25 | 0.2 | 0.18 |
| 11 | 5630635 | rs4910828 | -0.286 | 0.258 | -0.78 | -0.175 | 0.009 | 0.015 | 0.015 | 0 | 0.79 | 0.68 | 0.72 | 0.74 | 0.78 | 0.71 |
| 11 | 5630865 | rs11038336 | 0.469 | 0.111 | -1.048 | -0.556 | -0.012 | 0.014 | 0.014 | 0 | 0.07 | 0.02 | 0.07 | 0.05 | 0.06 | 0.15 |
| 11 | 5631479 | rs3740999 | 0.629 | 0.184 | -1.216 | -1.056 | -0.028 | 0.014 | 0.014 | 0 | 0.32 | 0.3 | 0.3 | 0.04 | 0.49 | 0.19 |
| 11 | 5633747 | rs7104648 | 0.803 | -0.869 | -1.191 | -0.542 | 0.005 | -0.01 | -0.01 | 0 | 0.17 | 0.13 | 0.15 | 0.29 | 0.15 | 0.17 |
| 11 | 5636039 | rs4910573 | -0.595 | -1.434 | -1.116 | -1.046 | 0.011 | 0.006 | 0.006 | 0 | 0.77 | 0.8 | 0.87 | 0.94 | 0.88 | 0.75 |
| 11 | 5639873 | rs10400271 | -0.233 | -1.792 | -0.862 | -0.836 | 0.009 | 0.023 | 0.023 | 0 | 0.4 | 0.36 | 0.49 | 0.75 | 0.37 | 0.54 |
| 11 | 5640497 | rs7109361 | -1.17 | -1.542 | -1.07 | -1.033 | -0.024 | 0.146 | 0.146 | 0 | 0.58 | 0.79 | 0.71 | 0.93 | 0.56 | 0.66 |
| 11 | 5642001 | rs7939159 | 0.187 | -1.355 | -0.486 | -0.571 | 0.011 | 0.021 | 0.021 | 0 | 0.23 | 0.3 | 0.22 | 0.29 | 0.3 | 0.19 |
| 11 | 5647033 | rs12274893 | 0.032 | -1.787 | -0.27 | -0.006 | 0.003 | -0.017 | -0.017 | 0 | 0.23 | 0.21 | 0.33 | 0.47 | 0.21 | 0.36 |
| 11 | 5652341 | rs12226152 | -0.48 | -1.814 | -0.224 | -0.404 | -0.023 | -0.045 | -0.045 | 0 | 0.86 | 0.85 | 0.95 | 1 | 0.95 | 0.86 |
| 11 | 5655233 | rs3824951 | 0.873 | -1.904 | -0.155 | -0.413 | 0.014 | 0.016 | 0.016 | 0 | 0.51 | 0.38 | 0.48 | 0.32 | 0.66 | 0.4 |
| 11 | 5659680 | rs11038476 | -0.388 | -2.209 | 0.053 | -0.437 | -0.007 | 0.002 | 0.002 | 0 | 0.33 | 0.43 | 0.48 | 0.65 | 0.23 | 0.44 |
| 11 | 5660023 | rs10838475 | -0.285 | -2.208 | 0.145 | -0.436 | 0.021 | -0.016 | -0.016 | 0.54 | 0.32 | 0.25 | 0.46 | 0.65 | 0.23 | 0.44 |
| 11 | 5661046 | rs7944669 | 0.861 | -2.272 | 0.18 | -0.479 | 0.037 | 0.017 | 0.017 | 0 | 0.53 | 0.35 | 0.48 | 0.35 | 0.72 | 0.43 |
| 11 | 5662319 | rs7108834 | -0.618 | -2.008 | 0.501 | -0.339 | -0.032 | -0.057 | -0.057 | 1.385 | 0.87 | 0.85 | 0.96 | 1 | 0.95 | 0.9 |
| 11 | 5663575 | rs12792498 | 0.793 | -1.992 | 0.379 | -0.451 | 0.005 | -0.035 | -0.035 | 0.02 | 0.37 | 0.38 | 0.37 | 0.1 | 0.52 | 0.28 |
| 11 | 5664684 | rs16933844 | 0.926 | -1.862 | 0.417 | -0.103 | 0.006 | 0.012 | 0.012 | 0 | 0.05 | 0.13 | 0.06 | 0.04 | 0.05 | 0.06 |
| 11 | 5664974 | rs7109861 | -0.172 | -1.703 | -0.193 | 0.036 | -0.06 | -0.094 | -0.094 | 0 | 0.86 | 0.67 | 0.94 | 1 | 0.95 | 0.9 |
| 11 | 5665225 | rs7106798 | -0.178 | -1.703 | -0.193 | 0.036 | -0.059 | -0.094 | -0.094 | 0 | 0.86 | 0.67 | 0.94 | 1 | 0.95 | 0.9 |
| 11 | 5667538 | rs16933888 | 0.4 | -1.313 | -0.305 | 0.674 | -0.004 | 0.067 | 0.067 | 0.009 | 0.11 | 0.16 | 0.22 | 0.39 | 0.08 | 0.12 |
| 11 | 5669109 | rs11038527 | 0.568 | -1.254 | -0.512 | 0.282 | 0.013 | -0.019 | -0.019 | 0.44 | 0.07 | 0.03 | 0.11 | 0.17 | 0.03 | 0.04 |
| 11 | 5669228 | rs4910832 | 0 | -1.131 | -0.395 | 0.186 | -0.014 | -0.003 | -0.003 | 0 | 0.82 | 0.74 | 0.84 | 0.84 | 0.88 | 0.88 |
| 11 | 5671623 | rs11038546 | -0.162 | -0.906 | -0.325 | 0.194 | 0.05 | 0.03 | 0.03 | 0 | 0.51 | 0.73 | 0.65 | 0.56 | 0.63 | 0.38 |
| 11 | 5672354 | rs7950311 | 0.422 | -0.833 | -0.417 | 0.248 | 0.01 | -0.055 | -0.055 | 0 | 0.4 | 0.52 | 0.42 | 0.13 | 0.54 | 0.27 |
| 11 | 5675972 | rs16933930 | 0.348 | -0.433 | -0.623 | 0.927 | 0 | 0.028 | 0.028 | 0 | 0.1 | 0.08 | 0.21 | 0.43 | 0.08 | 0.1 |
| 11 | 5683567 | rs7122620 | 0.125 | -0.586 | -0.401 | -0.203 | -0.002 | -0.038 | -0.038 | 0 | 0.47 | 0.5 | 0.44 | 0.11 | 0.6 | 0.3 |
| 11 | 5688940 | rs11038628 | -0.569 | -1.06 | 0.244 | 0.855 | -0.039 | 0.232 | 0.232 | 0.109 | 0.14 | 0.31 | 0.23 | 0.43 | 0.06 | 0.17 |
| 11 | 5689975 | rs12808061 | -0.228 | -0.438 | 0.437 | 0.492 | 0.057 | -0.057 | -0.057 | 0 | 0.1 | 0.35 | 0.08 | 0 | 0.12 | 0.08 |

| | | | | | | | | | | | | | | | | |
|---|---|---|---|---|---|---|---|---|---|---|---|---|---|---|---|---|
| 11 | 5690691 | rs12271333 | 0.278 | -0.782 | 0.094 | 0.376 | -0.006 | 0.006 | 0.006 | 0 | 0.17 | 0.23 | 0.2 | 0.19 | 0.14 | 0.13 |
| 11 | 5691214 | rs2015090 | -0.247 | -0.817 | 0.149 | 0.354 | -0.04 | 0.247 | 0.247 | 0.003 | 0.15 | 0.35 | 0.23 | 0.44 | 0.06 | 0.18 |
| 11 | 5691980 | rs904375 | -1.266 | -0.774 | -0.031 | 0.206 | -0.012 | 0.033 | 0.033 | 0.001 | 0.27 | 0.63 | 0.33 | 0.43 | 0.2 | 0.27 |
| 11 | 5693181 | rs2342390 | -0.535 | -0.819 | -0.116 | 0.216 | -0.007 | -0.003 | -0.003 | 0 | 0.81 | 0.71 | 0.79 | 0.83 | 0.85 | 0.86 |
| 11 | 5695743 | rs10838517 | -0.615 | -0.809 | -0.127 | 0.24 | -0.008 | 0 | 0 | 0 | 0.82 | 0.73 | 0.8 | 0.83 | 0.86 | 0.86 |
| 11 | 5696104 | rs10742766 | 1.352 | -0.822 | -0.118 | 0.215 | 0.036 | 0.036 | 0.036 | 0.211 | 0.55 | 0.12 | 0.47 | 0.4 | 0.66 | 0.6 |
| 11 | 5700340 | rs3740995 | -0.464 | -0.761 | 0.031 | 0.357 | -0.009 | -0.002 | -0.002 | 0.309 | 0.81 | 0.73 | 0.8 | 0.83 | 0.86 | 0.86 |
| 11 | 5700649 | rs10769175 | 0.41 | -0.608 | -0.256 | 0.241 | -0.01 | 0.002 | 0.002 | 0.318 | 0.67 | 0.39 | 0.57 | 0.4 | 0.79 | 0.68 |
| 11 | 5701001 | rs10838525 | 1.061 | -0.146 | -0.932 | -0.266 | -0.041 | 0.071 | 0.071 | 0 | 0.26 | 0.1 | 0.23 | 0.08 | 0.38 | 0.16 |
| 11 | 5701074 | rs11601507 | 0.776 | -0.24 | -1.068 | 0.09 | -0.004 | -0.03 | -0.03 | 0.066 | 0.08 | 0.01 | 0.06 | 0.09 | 0.07 | 0.23 |
| 11 | 5701883 | rs17305868 | 0.064 | -0.407 | -0.44 | 0.266 | -0.002 | 0.004 | 0.004 | 0 | 0.43 | 0.41 | 0.41 | 0.36 | 0.56 | 0.37 |
| 11 | 5702462 | rs4992800 | -0.596 | -0.5 | 0.057 | 0.692 | 0.006 | 0.163 | 0.163 | 0.095 | 0.93 | 0.75 | 0.87 | 0.78 | 0.97 | 0.94 |
| 11 | 5706311 | rs3802980 | 0.18 | -0.591 | -0.235 | 0.396 | -0.004 | -0.005 | -0.005 | 0.409 | 0.52 | 0.59 | 0.52 | 0.47 | 0.53 | 0.49 |
| 11 | 5710183 | rs885002 | -1.215 | -0.434 | 0.011 | 0.637 | 0.001 | 0.097 | 0.097 | 0 | 0.95 | 0.85 | 0.88 | 0.78 | 0.96 | 0.94 |
| 11 | 5712260 | rs1995162 | -0.184 | -0.493 | 0.016 | 0.639 | -0.002 | 0.035 | 0.035 | 0 | 0.09 | 0.12 | 0.14 | 0.27 | 0.06 | 0.18 |
| 11 | 5713658 | rs7112085 | 0.488 | -0.511 | -0.182 | 0.196 | 0.005 | 0.008 | 0.008 | 0 | 0.33 | 0.24 | 0.31 | 0.32 | 0.31 | 0.43 |
| 11 | 5716100 | rs16934565 | 1.368 | -0.202 | -0.022 | -0.222 | 0.016 | -0.044 | -0.044 | 0 | 0.09 | 0.1 | 0.08 | 0 | 0.12 | 0.01 |
| 11 | 5717063 | rs4910837 | 0.286 | -0.272 | -0.023 | -0.282 | -0.005 | 0.067 | 0.067 | 0.412 | 0.48 | 0.37 | 0.41 | 0.18 | 0.52 | 0.51 |
| 11 | 5717762 | rs10838543 | 0.253 | -0.237 | -0.028 | -0.288 | -0.004 | 0.069 | 0.069 | 0.32 | 0.48 | 0.37 | 0.42 | 0.18 | 0.52 | 0.51 |
| 11 | 5718517 | rs7935564 | -0.064 | -0.277 | -0.185 | -0.095 | -0.006 | -0.039 | -0.039 | 0 | 0.57 | 0.47 | 0.49 | 0.18 | 0.64 | 0.51 |
| 11 | 5719251 | rs10769180 | -0.841 | 0.214 | -0.601 | 0.16 | -0.008 | -0.002 | -0.002 | 0 | 0.74 | 0.77 | 0.71 | 0.75 | 0.65 | 0.83 |
| 11 | 5719667 | rs2291842 | -0.9 | -0.739 | -1.284 | 0.212 | 0.01 | 0.002 | 0.002 | 0 | 0.35 | 0.34 | 0.31 | 0.3 | 0.21 | 0.33 |
| 11 | 5719835 | rs2291844 | -0.228 | -0.604 | -2.489 | -0.317 | -0.047 | 0.031 | 0.031 | 0 | 0.39 | 0.08 | 0.47 | 0.64 | 0.38 | 0.37 |
| 11 | 5719848 | rs7950912 | -0.362 | -0.883 | -2.134 | -1.045 | -0.135 | 0.172 | 0.172 | 0.362 | 0.68 | 0.96 | 0.7 | 0.89 | 0.57 | 0.75 |
| 11 | 5722444 | rs4525262 | -0.036 | -0.714 | -0.925 | -0.359 | 0 | 0 | 0 | 0 | 0.54 | 0.6 | 0.61 | 0.45 | 0.59 | 0.53 |
| 11 | 5722948 | rs4290258 | -0.814 | -0.766 | -0.828 | -0.228 | 0.012 | -0.029 | -0.029 | 0 | 0.73 | 0.84 | 0.79 | 0.59 | 0.75 | 0.69 |
| 11 | 5729580 | rs2291845 | 1.659 | -0.554 | -1.073 | -0.368 | -0.124 | 0.1 | 0.1 | 0 | 0.19 | 0.02 | 0.15 | 0.07 | 0.31 | 0.19 |
| 11 | 5731749 | rs12419535 | 0.632 | -0.836 | -0.212 | 0.354 | 0.017 | -0.031 | -0.031 | 0 | 0.06 | 0.17 | 0.05 | 0 | 0.04 | 0.06 |
| 11 | 5740634 | rs12285602 | 0.916 | -1.013 | 0.28 | 0.453 | 0.003 | 0.06 | 0.06 | 0 | 0.1 | 0.27 | 0.23 | 0.2 | 0.08 | 0.08 |
| 11 | 5741845 | rs12287535 | -0.77 | -1.068 | 0.136 | 0.438 | -0.005 | -0.012 | -0.012 | 0.079 | 0.28 | 0.45 | 0.26 | 0.23 | 0.23 | 0.21 |
| 11 | 5744372 | rs4243963 | -0.337 | -1.1 | -0.119 | 0.444 | 0.008 | -0.016 | -0.016 | 0 | 0.68 | 0.81 | 0.78 | 0.58 | 0.72 | 0.66 |
| 11 | 5745637 | rs2047458 | 0.805 | -0.981 | -0.479 | 0.483 | 0 | -0.001 | -0.001 | 0.016 | 0.07 | 0.1 | 0.19 | 0.12 | 0.05 | 0.1 |
| 11 | 5747303 | rs11604371 | 0.295 | -0.795 | -0.226 | 0.244 | -0.006 | 0.006 | 0.006 | 0 | 0.6 | 0.62 | 0.7 | 0.49 | 0.65 | 0.61 |
| 11 | 5748222 | rs7933860 | 1.107 | -0.815 | -0.212 | 0.219 | 0.003 | -0.015 | -0.015 | 0 | 0.41 | 0.45 | 0.38 | 0.24 | 0.49 | 0.42 |
| 11 | 5748354 | rs7944807 | 0.269 | -0.798 | -0.203 | 0.236 | -0.006 | 0.006 | 0.006 | 0 | 0.6 | 0.61 | 0.7 | 0.49 | 0.65 | 0.61 |
| 11 | 5749836 | rs7930210 | 0.638 | -0.43 | -0.857 | 0.305 | 0.002 | -0.017 | -0.017 | 0 | 0.19 | 0.16 | 0.32 | 0.26 | 0.17 | 0.2 |
| 11 | 5755605 | rs1498552 | -0.773 | -0.731 | -1.256 | 0.482 | -0.012 | -0.091 | -0.091 | 0 | 0.86 | 0.99 | 0.85 | 0.71 | 0.86 | 0.84 |
| 11 | 5755682 | rs11038888 | 0.85 | -0.148 | -0.713 | -0.198 | -0.001 | 0 | 0 | 0 | 0.08 | 0.09 | 0.07 | 0.09 | 0.07 | 0.04 |
| 11 | 5758062 | rs7397032 | -0.27 | 0.075 | -0.672 | -0.255 | -0.019 | 0.021 | 0.021 | 0 | 0.79 | 0.66 | 0.66 | 0.75 | 0.83 | 0.8 |
| 11 | 5770840 | rs2343231 | -0.377 | 0.158 | -0.458 | 0.301 | -0.002 | -0.004 | -0.004 | 0.461 | 0.45 | 0.51 | 0.38 | 0.68 | 0.38 | 0.46 |
| 11 | 5776595 | rs12363178 | -1.211 | -0.487 | 0.155 | 0.69 | 0.024 | -0.011 | -0.011 | 0 | 0.23 | 0.15 | 0.15 | 0.23 | 0.16 | 0.22 |
| 11 | 5777096 | rs12363545 | -1.211 | -0.487 | 0.155 | 0.69 | 0.024 | -0.011 | -0.011 | 0.476 | 0.23 | 0.15 | 0.15 | 0.23 | 0.16 | 0.22 |
| 11 | 5777210 | rs7395852 | 1.016 | -0.487 | 0.121 | 0.682 | -0.021 | 0.021 | 0.021 | 0 | 0.75 | 0.63 | 0.72 | 0.67 | 0.81 | 0.75 |
| 11 | 5809477 | rs10838637 | 0.002 | -0.984 | 0.007 | 0.319 | -0.019 | -0.03 | -0.03 | 1.268 | 0.37 | 0.46 | 0.27 | 0.2 | 0.33 | 0.29 |
| 11 | 5809548 | rs7948009 | -0.031 | -1.141 | 0.485 | 0.279 | 0.032 | 0.04 | 0.04 | 0 | 0.6 | 0.25 | 0.59 | 0.71 | 0.64 | 0.69 |
| 11 | 5809673 | rs10769224 | 0.055 | -1.132 | 0.48 | 0.28 | -0.019 | -0.03 | -0.03 | 0 | 0.37 | 0.46 | 0.27 | 0.2 | 0.33 | 0.29 |
| 11 | 5809746 | rs10742787 | 0.055 | -1.132 | 0.474 | 0.28 | -0.019 | -0.03 | -0.03 | 0 | 0.37 | 0.46 | 0.27 | 0.2 | 0.33 | 0.29 |
| 11 | 5809811 | rs12365487 | -0.41 | -1.331 | 0.359 | 0.221 | 0.035 | -0.07 | -0.07 | 0 | 0.19 | 0.03 | 0.13 | 0.23 | 0.11 | 0.19 |
| 11 | 5809990 | rs11039038 | 0.458 | -1.396 | 0.329 | 0.261 | 0.009 | -0.052 | -0.052 | 0 | 0.09 | 0.01 | 0.07 | 0.22 | 0.07 | 0.12 |
| 11 | 5810080 | rs10742788 | 0.118 | -1.314 | 0.393 | 0.222 | -0.019 | -0.03 | -0.03 | 0.536 | 0.37 | 0.46 | 0.27 | 0.2 | 0.33 | 0.29 |
| 11 | 5821436 | rs7940624 | 0.118 | -1.316 | 0.501 | 0.218 | -0.023 | -0.037 | -0.037 | 0 | 0.37 | 0.47 | 0.27 | 0.2 | 0.33 | 0.29 |
| 11 | 5821573 | rs10047437 | -0.1 | -1.328 | 0.41 | 0.219 | 0.035 | 0.044 | 0.044 | 0 | 0.6 | 0.24 | 0.59 | 0.71 | 0.64 | 0.69 |
| 11 | 5827003 | rs11039100 | -0.555 | -1.393 | 0.318 | 0.252 | 0.033 | -0.067 | -0.067 | 0 | 0.17 | 0.05 | 0.12 | 0.23 | 0.11 | 0.18 |
| 11 | 5827452 | rs11607346 | 0.298 | -1.46 | 0.524 | 0.179 | -0.033 | -0.033 | -0.033 | 0 | 0.34 | 0.54 | 0.26 | 0.2 | 0.29 | 0.25 |
| 11 | 5841856 | rs4758435 | 0.925 | -1.19 | 0.435 | 0.108 | -0.012 | 0.018 | 0.018 | 0 | 0.76 | 0.72 | 0.71 | 0.67 | 0.82 | 0.77 |
| 11 | 5842356 | rs8181512 | 0.587 | -1.119 | 0.593 | 0.185 | 0.008 | -0.003 | -0.003 | 0 | 0.44 | 0.34 | 0.48 | 0.47 | 0.54 | 0.49 |
| 11 | 5844353 | rs10838688 | 0.311 | -1.209 | 0.584 | 0.498 | -0.007 | 0.193 | 0.193 | 0 | 0.89 | 0.68 | 0.78 | 0.69 | 0.9 | 0.89 |
| 11 | 5845004 | rs4758113 | 0.254 | -1.225 | -0.023 | 0.423 | 0.014 | -0.017 | -0.017 | 0.923 | 0.08 | 0.08 | 0.09 | 0.21 | 0.07 | 0.08 |
| 11 | 5850737 | rs7394956 | -1.569 | -1.244 | 0.185 | 0.34 | -0.019 | 0.02 | 0.02 | 0 | 0.29 | 0.4 | 0.23 | 0.43 | 0.18 | 0.29 |
| 11 | 5850768 | rs4910851 | -0.346 | -1.125 | 0.26 | 0.446 | -0.016 | 0.039 | 0.039 | 0 | 0.11 | 0.24 | 0.1 | 0.2 | 0.07 | 0.1 |
| 11 | 5850793 | rs12364796 | 0.255 | -0.998 | 0.096 | 0.42 | 0.005 | -0.049 | -0.049 | 0.184 | 0.09 | 0.01 | 0.07 | 0.22 | 0.08 | 0.15 |
| 11 | 5851596 | rs1377522 | -1.272 | -0.478 | 0.143 | 0.244 | 0.032 | 0.024 | 0.024 | 0 | 0.7 | 0.64 | 0.65 | 0.63 | 0.62 | 0.69 |
| 11 | 5862532 | rs10742809 | 0.346 | -0.994 | -0.198 | 0.033 | 0.018 | -0.037 | -0.037 | 0 | 0.36 | 0.22 | 0.3 | 0.46 | 0.34 | 0.34 |
| 11 | 5862780 | rs4362172 | 0.056 | -0.865 | -0.104 | 0.113 | 0.005 | -0.002 | -0.002 | 0 | 0.42 | 0.38 | 0.32 | 0.46 | 0.37 | 0.37 |
| 11 | 5863113 | rs10742810 | 0.056 | -0.865 | -0.104 | 0.113 | 0.005 | -0.002 | -0.002 | 0 | 0.42 | 0.38 | 0.32 | 0.46 | 0.37 | 0.37 |
| 11 | 5866626 | rs11039272 | 0.872 | -0.567 | -0.254 | 0.145 | -0.029 | -0.021 | -0.021 | 0 | 0.31 | 0.26 | 0.4 | 0.38 | 0.43 | 0.32 |
| 11 | 5867283 | rs7938036 | 0.844 | -0.577 | -0.186 | 0.159 | -0.019 | -0.014 | -0.014 | 0 | 0.32 | 0.29 | 0.41 | 0.38 | 0.43 | 0.32 |
| 11 | 5899862 | rs10838779 | 0.909 | -0.427 | -0.218 | 0.223 | -0.01 | -0.007 | -0.007 | 0 | 0.3 | 0.32 | 0.4 | 0.37 | 0.41 | 0.32 |
| 11 | 5904497 | rs10500657 | 0.187 | -0.834 | -0.104 | 0.009 | 0.01 | 0.022 | 0.022 | 0 | 0.74 | 0.86 | 0.76 | 0.83 | 0.8 | 0.7 |

| | | | | | | | | | | | | | | | | |
|---|---|---|---|---|---|---|---|---|---|---|---|---|---|---|---|---|
| 11 | 5906048 | rs4758168 | 0.187 | -0.836 | 0.141 | 0.008 | -0.003 | -0.008 | -0.008 | 0 | 0.74 | 0.81 | 0.75 | 0.83 | 0.8 | 0.69 |
| 11 | 5906073 | rs4757986 | 0.187 | -0.836 | 0.141 | 0.008 | -0.003 | -0.008 | -0.008 | 0 | 0.74 | 0.81 | 0.75 | 0.83 | 0.8 | 0.69 |
| 11 | 5906192 | rs1453435 | 0.129 | -0.839 | 0.085 | 0.007 | 0.012 | 0.028 | 0.028 | 0 | 0.74 | 0.88 | 0.76 | 0.83 | 0.8 | 0.69 |
| 11 | 5906291 | rs11823842 | 1.152 | -0.822 | 0.105 | 0.047 | -0.009 | -0.006 | -0.006 | 0 | 0.3 | 0.32 | 0.4 | 0.36 | 0.41 | 0.32 |
| 11 | 5908939 | rs1453433 | 0.383 | -1.093 | -0.081 | 0.204 | 0.013 | -0.038 | -0.038 | 0 | 0.11 | 0.05 | 0.09 | 0.23 | 0.08 | 0.09 |
| 11 | 5915809 | rs1453438 | -0.478 | -1.238 | 0.148 | 0.058 | 0.042 | 0.002 | 0.002 | 0 | 0.51 | 0.64 | 0.4 | 0.48 | 0.42 | 0.43 |
| 11 | 5920721 | rs10838795 | 1.164 | -1.339 | 0.39 | 0.038 | 0.007 | 0.018 | 0.018 | 0 | 0.28 | 0.19 | 0.21 | 0.24 | 0.29 | 0.23 |
| 11 | 5922222 | rs10160336 | 0.31 | -1.481 | 0.276 | -0.042 | 0.003 | 0.088 | 0.088 | 0 | 0.46 | 0.3 | 0.53 | 0.6 | 0.53 | 0.55 |
| 11 | 5922707 | rs7106300 | -0.308 | -1.452 | 0.372 | -0.027 | -0.011 | -0.02 | -0.02 | 0 | 0.91 | 0.89 | 0.95 | 0.99 | 0.95 | 0.92 |
| 11 | 5923082 | rs10500656 | 0.232 | -1.427 | 0.415 | -0.011 | 0 | -0.005 | -0.005 | 4.069 | 0.15 | 0.12 | 0.17 | 0.15 | 0.13 | 0.16 |
| 11 | 5925432 | rs11039518 | 0.253 | -1.319 | 0.354 | 0.049 | 0.01 | -0.02 | -0.02 | 0 | 0.12 | 0.11 | 0.1 | 0.25 | 0.09 | 0.09 |
| 11 | 5933112 | rs1453428 | 0.186 | -1.26 | 0.352 | 0.055 | -0.001 | -0.004 | -0.004 | 0 | 0.15 | 0.12 | 0.17 | 0.15 | 0.13 | 0.16 |
| 11 | 5934737 | rs4237755 | 0.018 | -1.291 | 0.4 | 0.067 | -0.02 | -0.032 | -0.032 | 0 | 0.89 | 0.81 | 0.93 | 0.99 | 0.95 | 0.92 |
| 11 | 5954809 | rs10769353 | 0.136 | -1.249 | 0.344 | 0.119 | 0.009 | -0.007 | -0.007 | 0 | 0.4 | 0.37 | 0.32 | 0.49 | 0.38 | 0.32 |
| 11 | 5959757 | rs3858526 | 0.88 | -1.193 | 0.367 | 0.08 | -0.01 | -0.016 | -0.016 | 0 | 0.3 | 0.4 | 0.25 | 0.24 | 0.29 | 0.23 |
| 11 | 5960036 | rs7946713 | 0.98 | -1.103 | 0.333 | 0.14 | -0.048 | -0.025 | -0.025 | 0 | 0.34 | 0.22 | 0.43 | 0.35 | 0.45 | 0.42 |
| 11 | 5961535 | rs11039767 | 0.906 | -0.848 | 0.341 | 0.197 | -0.017 | -0.02 | -0.02 | 0 | 0.35 | 0.27 | 0.42 | 0.41 | 0.41 | 0.42 |
| 11 | 5966415 | rs1901843 | 1.28 | -0.102 | 0.159 | 0.372 | 0 | 0.004 | 0.004 | 0 | 0.27 | 0.28 | 0.35 | 0.35 | 0.28 | 0.38 |
| 11 | 5967747 | rs1453431 | 0.314 | -0.114 | -0.116 | 0.32 | 0.005 | -0.022 | -0.022 | 0 | 0.13 | 0.14 | 0.09 | 0.08 | 0.14 | 0.16 |
| 11 | 5968728 | rs1840178 | -0.524 | -0.217 | -0.254 | 0.377 | 0.007 | 0.019 | 0.019 | 0 | 0.5 | 0.49 | 0.64 | 0.65 | 0.54 | 0.53 |
| 11 | 5969185 | rs10769378 | -0.607 | -0.061 | -0.719 | 0.479 | -0.056 | -0.063 | -0.063 | 0.108 | 0.21 | 0.06 | 0.28 | 0.3 | 0.26 | 0.14 |
| 11 | 5977678 | rs11039888 | -0.783 | -0.242 | -0.727 | 0.738 | -0.004 | -0.007 | -0.007 | 0 | 0.25 | 0.2 | 0.38 | 0.45 | 0.25 | 0.27 |
| 11 | 5979497 | rs7113402 | -0.914 | -0.307 | -0.592 | 0.708 | 0.006 | 0.03 | 0.03 | 0 | 0.26 | 0.29 | 0.39 | 0.45 | 0.26 | 0.27 |
| 11 | 5989293 | rs7113548 | 1.438 | -0.779 | -0.769 | 0.387 | 0.002 | 0.016 | 0.016 | 0.698 | 0.26 | 0.2 | 0.22 | 0.21 | 0.29 | 0.22 |
| 11 | 5992215 | rs4758290 | -0.417 | -0.672 | -0.764 | 0.414 | -0.001 | 0.001 | 0.001 | 0 | 0.53 | 0.5 | 0.61 | 0.67 | 0.54 | 0.5 |
| 11 | 5995159 | rs4758294 | -0.91 | -0.769 | -1.054 | 0.359 | -0.042 | -0.005 | -0.005 | 0.223 | 0.61 | 0.76 | 0.65 | 0.68 | 0.58 | 0.55 |
| 11 | 6003512 | rs10839170 | 0.695 | -0.173 | -0.666 | 0.292 | -0.002 | 0.031 | 0.031 | 0.011 | 0.29 | 0.18 | 0.25 | 0.21 | 0.32 | 0.21 |
| 11 | 6004097 | rs12575289 | 0.23 | -0.004 | -0.437 | 0.723 | -0.067 | -0.081 | -0.081 | 3.318 | 0.12 | 0.01 | 0.3 | 0.34 | 0.19 | 0.11 |
| 11 | 6005759 | rs1989213 | 1.162 | -0.514 | 0.096 | 0.022 | -0.027 | -0.054 | -0.054 | 0 | 0.75 | 0.42 | 0.82 | 0.87 | 0.81 | 0.76 |
| 11 | 6007272 | rs4237768 | -0.008 | -0.804 | 0.091 | 0.052 | -0.001 | -0.003 | -0.003 | 0 | 0.85 | 0.78 | 0.87 | 0.87 | 0.85 | 0.8 |
| 11 | 6007613 | rs4354673 | 0.53 | -0.556 | -0.338 | 0.182 | -0.042 | -0.056 | -0.056 | 0 | 0.75 | 0.42 | 0.82 | 0.87 | 0.82 | 0.76 |
| 11 | 6011761 | rs10769550 | 0.161 | -0.154 | -0.806 | 0.388 | 0.033 | 0.055 | 0.055 | 0 | 0.48 | 0.27 | 0.56 | 0.64 | 0.53 | 0.54 |
| 11 | 6013529 | rs4388872 | -0.597 | 0.067 | -0.803 | 0.278 | -0.01 | -0.035 | -0.035 | 0.053 | 0.32 | 0.26 | 0.49 | 0.54 | 0.4 | 0.21 |
| 11 | 6015194 | rs4758342 | 0.48 | -0.085 | -0.168 | 0.44 | -0.074 | -0.081 | -0.081 | 0 | 0.15 | 0.01 | 0.35 | 0.25 | 0.27 | 0.14 |
| 11 | 6016511 | rs7939215 | -0.693 | -0.134 | 0.082 | 0.456 | -0.005 | 0.01 | 0.01 | 0 | 0.83 | 0.79 | 0.86 | 0.72 | 0.88 | 0.92 |
| 11 | 6019989 | rs7947235 | 1.293 | -0.241 | -0.167 | -0.334 | -0.008 | -0.003 | -0.003 | 0.94 | 0.36 | 0.29 | 0.26 | 0.38 | 0.33 | 0.51 |
| 11 | 6020806 | rs997433 | 0.949 | -0.125 | -0.86 | -0.283 | -0.001 | 0.002 | 0.002 | 0 | 0.4 | 0.59 | 0.3 | 0.39 | 0.34 | 0.53 |
| 11 | 6021988 | rs10839219 | -0.151 | -0.016 | -1.231 | -0.744 | -0.02 | 0.09 | 0.09 | 0.074 | 0.26 | 0.06 | 0.19 | 0.08 | 0.27 | 0.25 |
| 11 | 6023581 | rs11040248 | -0.159 | -0.014 | -1.231 | -0.742 | -0.02 | 0.09 | 0.09 | 0.105 | 0.26 | 0.06 | 0.19 | 0.08 | 0.27 | 0.26 |
| 11 | 6023640 | rs11040249 | -0.155 | -0.004 | -1.226 | -0.737 | -0.02 | 0.09 | 0.09 | 0.027 | 0.26 | 0.06 | 0.19 | 0.08 | 0.27 | 0.27 |
| 11 | 6026661 | rs4573661 | -0.643 | -0.271 | -0.369 | -0.283 | -0.007 | 0.01 | 0.01 | 0 | 0.35 | 0.51 | 0.45 | 0.51 | 0.35 | 0.21 |
| 11 | 6028091 | rs6578717 | 0.2 | 0.172 | -0.159 | -1.385 | 0.016 | -0.077 | -0.077 | 0 | 0.15 | 0.32 | 0.19 | 0.01 | 0.18 | 0.14 |
| 11 | 6029540 | rs11040271 | 1.286 | 0.192 | -1.005 | -1.818 | -0.043 | 0.006 | 0.006 | 0 | 0.3 | 0.07 | 0.32 | 0.76 | 0.24 | 0.46 |
| 11 | 6030442 | rs4494290 | -0.185 | 0.134 | -0.535 | -1.988 | -0.013 | -0.052 | -0.052 | 0 | 0.67 | 0.63 | 0.77 | 0.94 | 0.67 | 0.76 |
| 11 | 6032001 | rs4317963 | -0.334 | 0.095 | -0.431 | -2.007 | -0.002 | -0.018 | -0.018 | 1.213 | 0.7 | 0.72 | 0.78 | 0.94 | 0.67 | 0.78 |
| 11 | 6038674 | rs7928574 | -0.528 | 0.166 | -0.471 | -2.102 | 0.005 | -0.064 | -0.064 | 0.007 | 0.34 | 0.59 | 0.45 | 0.18 | 0.39 | 0.26 |
| 11 | 6041247 | rs4351799 | 0.467 | 0.319 | -0.343 | -1.729 | 0.012 | -0.021 | -0.021 | 0 | 0.17 | 0.41 | 0.26 | 0.17 | 0.2 | 0.1 |
| 11 | 6041346 | rs4237774 | 0.112 | 0.242 | -0.801 | -1.726 | 0.001 | 0.001 | 0.001 | 0 | 0.7 | 0.78 | 0.79 | 0.94 | 0.67 | 0.78 |
| 11 | 6041406 | rs11040306 | 0.138 | 0.481 | -1.554 | -1.696 | -0.047 | 0.129 | 0.129 | 0 | 0.2 | 0.01 | 0.14 | 0.01 | 0.24 | 0.14 |
| 11 | 6044201 | rs4758368 | -0.192 | 0.587 | -1.339 | -1.695 | -0.054 | 0.114 | 0.114 | 0 | 0.29 | 0.12 | 0.36 | 0.05 | 0.38 | 0.29 |
| 11 | 6045931 | rs4758369 | -0.448 | 0.452 | -0.641 | -1.818 | -0.005 | -0.03 | -0.03 | 0 | 0.33 | 0.31 | 0.38 | 0.05 | 0.4 | 0.31 |
| 11 | 6048056 | rs11040335 | -0.402 | 0.508 | -0.646 | -1.816 | -0.01 | 0.003 | 0.003 | 0 | 0.32 | 0.26 | 0.37 | 0.05 | 0.4 | 0.31 |
| 11 | 6051317 | rs4758371 | 0.902 | 0.24 | -0.356 | -2.522 | 0.005 | 0.063 | 0.063 | 0 | 0.38 | 0.24 | 0.47 | 0.92 | 0.31 | 0.46 |
| 11 | 6051811 | rs7946925 | -0.01 | 0.215 | -0.098 | -1.998 | -0.002 | -0.035 | -0.035 | 0 | 0.22 | 0.23 | 0.13 | 0.03 | 0.24 | 0.17 |
| 11 | 6063985 | rs4420239 | 0.166 | 0.11 | -0.204 | -1.316 | 0.002 | 0.013 | 0.013 | 0 | 0.49 | 0.54 | 0.5 | 0.69 | 0.43 | 0.53 |
| 11 | 6070462 | rs5015541 | 0.215 | 0.276 | -0.336 | -1.28 | -0.001 | -0.004 | -0.004 | 0 | 0.48 | 0.49 | 0.49 | 0.69 | 0.42 | 0.53 |
| 11 | 6072356 | rs4625434 | 0.469 | 0.172 | -0.336 | -1.357 | -0.007 | 0.037 | 0.037 | 0.012 | 0.43 | 0.15 | 0.46 | 0.69 | 0.4 | 0.5 |
| 11 | 6075446 | rs10839311 | 0.044 | 0.396 | -0.723 | -1.92 | 0 | -0.028 | -0.028 | 0 | 0.52 | 0.53 | 0.65 | 0.95 | 0.45 | 0.59 |
| 11 | 6083952 | rs4564313 | 0.628 | 0.351 | -1.639 | -1.476 | -0.058 | -0.079 | -0.079 | 0 | 0.32 | 0.06 | 0.47 | 0.75 | 0.29 | 0.46 |
| 11 | 6084034 | rs12417160 | -0.833 | 0.623 | -1.123 | -1.996 | -0.008 | -0.03 | -0.03 | 0.496 | 0.26 | 0.6 | 0.26 | 0.14 | 0.23 | 0.21 |
| 11 | 6088712 | rs7930953 | -0.519 | 0.92 | -1.137 | -2.327 | 0 | -0.012 | -0.012 | 0 | 0.75 | 0.78 | 0.81 | 0.95 | 0.65 | 0.89 |
| 11 | 6089740 | rs6578729 | 0.459 | 0.795 | -1.223 | -2.469 | -0.011 | 0.008 | 0.008 | 0 | 0.12 | 0.09 | 0.09 | 0 | 0.14 | 0.08 |
| 11 | 6090372 | rs4353251 | 0.19 | 0.749 | -1.111 | -2.505 | 0.001 | 0.014 | 0.014 | 0 | 0.65 | 0.32 | 0.68 | 0.83 | 0.6 | 0.78 |
| 11 | 6107855 | rs11040663 | 1.095 | 0.244 | -1.217 | -2.814 | -0.085 | 0.244 | 0.244 | 0 | 0.48 | 0.1 | 0.61 | 0.9 | 0.49 | 0.67 |
| 11 | 6111054 | rs4328190 | -0.168 | 0.203 | -0.99 | -2.838 | -0.031 | -0.004 | -0.004 | 0 | 0.12 | 0.37 | 0.09 | 0.05 | 0.05 | 0.14 |
| 11 | 6113332 | rs7106139 | 1.104 | 0.239 | -1.2 | -2.813 | -0.085 | 0.243 | 0.243 | 0 | 0.48 | 0.1 | 0.61 | 0.9 | 0.49 | 0.67 |
| 11 | 6120908 | rs1903778 | -0.035 | 0.412 | -1.046 | -2.744 | -0.003 | -0.015 | -0.015 | 0 | 0.65 | 0.68 | 0.72 | 0.95 | 0.54 | 0.81 |
| 11 | 6129837 | rs1462983 | 1.524 | 0.371 | -1.47 | -2.772 | -0.056 | 0.175 | 0.175 | 0 | 0.45 | 0.1 | 0.6 | 0.88 | 0.45 | 0.64 |
| 11 | 6134006 | rs1381911 | 0.585 | 0.224 | -1.478 | -2.981 | 0.003 | -0.001 | -0.001 | 1.285 | 0.52 | 0.54 | 0.66 | 0.94 | 0.46 | 0.76 |

| | | | | | | | | | | | | | | | | |
|---|---|---|---|---|---|---|---|---|---|---|---|---|---|---|---|---|
| 11 | 6134904 | rs4758085 | 0.28 | 0.231 | -1.488 | -2.989 | -0.009 | 0.043 | 0.043 | 0 | 0.55 | 0.65 | 0.69 | 0.95 | 0.49 | 0.8 |
| 11 | 6136252 | rs1381912 | -0.26 | 0.162 | -1.423 | -2.991 | -0.027 | -0.018 | -0.018 | 1.106 | 0.13 | 0.4 | 0.1 | 0.05 | 0.05 | 0.12 |
| 11 | 6144998 | rs4758391 | 1.52 | 0.198 | -1.547 | -2.98 | -0.045 | 0.114 | 0.114 | 0 | 0.39 | 0.1 | 0.56 | 0.88 | 0.4 | 0.63 |
| 11 | 6146580 | rs7480133 | 0.394 | 0.362 | -1.461 | -2.828 | -0.01 | 0.037 | 0.037 | 0 | 0.55 | 0.63 | 0.69 | 0.95 | 0.49 | 0.8 |
| 11 | 6150738 | rs325637 | 0.203 | 0.5 | -1.506 | -2.766 | -0.021 | 0.071 | 0.071 | 0 | 0.58 | 0.7 | 0.7 | 0.95 | 0.5 | 0.8 |
| 11 | 6161724 | rs325661 | 0.849 | 0.82 | -1.538 | -2.74 | 0 | -0.01 | -0.01 | 0 | 0.22 | 0.21 | 0.11 | 0.04 | 0.21 | 0.1 |
| 11 | 6164700 | rs325664 | 0.03 | 0.388 | -1.196 | -2.399 | -0.031 | -0.012 | -0.012 | 0 | 0.87 | 0.6 | 0.9 | 0.95 | 0.95 | 0.91 |
| 11 | 6173346 | rs11826453 | 0.158 | 1.244 | -1.832 | -2.065 | -0.021 | 0.052 | 0.052 | 0 | 0.33 | 0.24 | 0.23 | 0.05 | 0.4 | 0.18 |
| 11 | 6176111 | rs325697 | -0.092 | 0.845 | -1.609 | -1.798 | -0.026 | 0.07 | 0.07 | 0.003 | 0.41 | 0.3 | 0.29 | 0.05 | 0.5 | 0.2 |
| 11 | 6179462 | rs325687 | -0.214 | 0.634 | -1.64 | -1.557 | 0.015 | 0.14 | 0.14 | 0.041 | 0.46 | 0.29 | 0.31 | 0.06 | 0.56 | 0.26 |
| 11 | 6189084 | rs325628 | -0.05 | 0.422 | -1.55 | -1.305 | 0.015 | 0.138 | 0.138 | 0.722 | 0.51 | 0.28 | 0.61 | 0.81 | 0.57 | 0.72 |
| 11 | 6189514 | rs12226204 | 0.055 | -0.01 | -0.976 | -1.402 | -0.022 | 0.05 | 0.05 | 0.365 | 0.05 | 0.14 | 0.03 | 0.17 | 0.02 | 0.06 |
| 11 | 6192443 | rs4505047 | 0.251 | -0.001 | -1.11 | -1.608 | -0.027 | 0.128 | 0.128 | 0.012 | 0.45 | 0.23 | 0.59 | 0.77 | 0.52 | 0.65 |
| 11 | 6193508 | rs10769670 | 0.225 | 0.041 | -1.003 | -1.608 | -0.024 | 0.197 | 0.197 | 0.88 | 0.47 | 0.16 | 0.58 | 0.78 | 0.53 | 0.66 |
| 11 | 6198688 | rs11601498 | 0.933 | -0.025 | -1.457 | -1.939 | -0.045 | 0.184 | 0.184 | 2.475 | 0.27 | 0.03 | 0.15 | 0.01 | 0.25 | 0.08 |
| 11 | 6203685 | rs10431016 | 0.081 | -0.023 | -1.046 | -1.94 | 0.002 | 0.009 | 0.009 | 0.977 | 0.34 | 0.27 | 0.2 | 0.2 | 0.31 | 0.16 |
| 11 | 6204316 | rs325648 | 1.288 | -0.148 | -1.158 | -1.023 | -0.016 | 0.047 | 0.047 | 0.071 | 0.73 | 0.41 | 0.63 | 0.62 | 0.77 | 0.5 |
| 11 | 6213501 | rs325611 | 0.496 | 0.211 | -1.013 | -0.847 | 0.023 | -0.092 | -0.092 | 0 | 0.82 | 0.46 | 0.79 | 0.97 | 0.83 | 0.82 |
| 11 | 6214230 | rs10839530 | 1.313 | 0.158 | -1.16 | -0.967 | 0.002 | 0.069 | 0.069 | 1.223 | 0.25 | 0.05 | 0.13 | 0.18 | 0.21 | 0.14 |
| 11 | 6217122 | rs869900 | 1.022 | 0.141 | -1.17 | -0.966 | -0.035 | 0.007 | 0.007 | 0.557 | 0.43 | 0.15 | 0.44 | 0.6 | 0.47 | 0.42 |
| 11 | 6219515 | rs1381914 | 1.347 | 0.111 | -1.183 | -0.988 | 0.01 | 0.056 | 0.056 | 1.19 | 0.25 | 0.07 | 0.13 | 0.16 | 0.21 | 0.14 |
| 11 | 6221427 | rs12417049 | 1.684 | -0.039 | -1.472 | -0.312 | 0.004 | -0.07 | -0.07 | 0 | 0.11 | 0 | 0.15 | 0.21 | 0.08 | 0.17 |
| 11 | 6222847 | rs16910016 | -0.418 | 0.253 | -0.758 | -0.545 | 0.014 | -0.036 | -0.036 | 0.104 | 0.08 | 0.25 | 0.06 | 0.01 | 0.06 | 0.02 |
| 11 | 6223837 | rs188980 | -0.522 | 0.251 | -0.752 | -0.546 | 0.023 | -0.032 | -0.032 | 3.988 | 0.08 | 0.28 | 0.06 | 0.01 | 0.06 | 0.02 |
| 11 | 6226115 | rs11040803 | 1.566 | 0.252 | -0.78 | -0.603 | -0.002 | 0.001 | 0.001 | 0 | 0.73 | 0.21 | 0.71 | 0.95 | 0.74 | 0.8 |
| 11 | 6227083 | rs10742968 | 2.127 | 0.306 | -0.802 | -0.537 | 0.003 | -0.01 | -0.01 | 0 | 0.68 | 0.33 | 0.67 | 0.8 | 0.68 | 0.52 |
| 11 | 6231731 | rs10769671 | 1.554 | 0.526 | -0.527 | -0.391 | 0.029 | 0.016 | 0.016 | 0.651 | 0.77 | 0.36 | 0.72 | 0.81 | 0.75 | 0.56 |
| 11 | 6234688 | rs4758399 | 1.54 | 0.584 | -0.47 | -0.852 | 0.004 | -0.028 | -0.028 | 0 | 0.79 | 0.23 | 0.78 | 0.94 | 0.77 | 0.84 |
| 11 | 6234753 | rs11040807 | 0.761 | 0.625 | -0.557 | -0.878 | -0.015 | 0.185 | 0.185 | 0.562 | 0.52 | 0.16 | 0.59 | 0.78 | 0.55 | 0.69 |
| 11 | 6239344 | rs3750944 | -0.466 | 0.883 | -0.061 | -0.836 | -0.002 | -0.004 | -0.004 | 0 | 0.45 | 0.62 | 0.43 | 0.5 | 0.54 | 0.5 |
| 11 | 6241350 | rs7114859 | 1.568 | 0.74 | 0.386 | -1.173 | 0.035 | -0.082 | -0.082 | 0 | 0.78 | 0.42 | 0.78 | 0.94 | 0.76 | 0.83 |
| 11 | 6241827 | rs1531480 | 1.18 | 0.758 | 0.462 | -1.17 | 0.04 | -0.055 | -0.055 | 0.204 | 0.83 | 0.38 | 0.81 | 0.96 | 0.82 | 0.85 |
| 11 | 6242337 | rs17238860 | -0.223 | 0.752 | 0.449 | -1.236 | 0.002 | 0.004 | 0.004 | 2.559 | 0.06 | 0.04 | 0.04 | 0.01 | 0.05 | 0.02 |
| 11 | 6243804 | rs3750945 | 1.204 | 0.77 | 0.464 | -1.224 | 0.036 | -0.061 | -0.061 | 0 | 0.83 | 0.36 | 0.81 | 0.96 | 0.82 | 0.85 |
| 11 | 6243982 | rs325606 | 0.903 | 0.771 | 0.475 | -1.224 | 0.078 | -0.049 | -0.049 | 0 | 0.85 | 0.54 | 0.83 | 0.96 | 0.82 | 0.85 |
| 11 | 6254397 | rs1564131 | 1.411 | 0.803 | 0.413 | -1.239 | 0.005 | 0.216 | 0.216 | 0.037 | 0.51 | 0.16 | 0.58 | 0.8 | 0.56 | 0.69 |
| 11 | 6259501 | rs325702 | 0.033 | 0.633 | 0.562 | -1.287 | 0.1 | 0.017 | 0.017 | 0 | 0.92 | 0.59 | 0.87 | 0.97 | 0.89 | 0.86 |
| 11 | 6259835 | rs1553746 | 0.899 | 0.506 | 0.181 | -1.221 | 0.008 | -0.077 | -0.077 | 0 | 0.78 | 0.34 | 0.79 | 0.95 | 0.8 | 0.83 |
| 11 | 6262715 | rs7129498 | -0.25 | 0.521 | 0.149 | -1.202 | 0.066 | -0.047 | -0.047 | 0 | 0.15 | 0.45 | 0.16 | 0.04 | 0.16 | 0.15 |
| 11 | 6270384 | rs2344608 | 1.075 | 0.504 | -0.153 | -1.184 | -0.004 | 0.193 | 0.193 | 0.059 | 0.52 | 0.16 | 0.6 | 0.83 | 0.56 | 0.68 |
| 11 | 6272677 | rs11040813 | 0.234 | 0.341 | -0.63 | -1.118 | 0.016 | 0.084 | 0.084 | 0 | 0.26 | 0.07 | 0.18 | 0.13 | 0.21 | 0.15 |
| 11 | 6273744 | rs10500661 | 0.304 | 0.291 | -0.649 | -1.082 | 0.009 | 0.012 | 0.012 | 0.019 | 0.26 | 0.17 | 0.15 | 0.13 | 0.19 | 0.13 |
| 11 | 6280248 | rs906895 | 1.056 | 0.324 | -0.854 | -1.001 | 0.001 | 0.189 | 0.189 | 0.004 | 0.51 | 0.19 | 0.61 | 0.79 | 0.57 | 0.67 |
| 11 | 6281400 | rs11040819 | 0.376 | 0.579 | -0.667 | -1.231 | 0.032 | -0.085 | -0.085 | 0 | 0.81 | 0.48 | 0.83 | 0.97 | 0.82 | 0.84 |
| 11 | 6282808 | rs2929183 | 0.304 | 0.674 | -0.779 | -1.183 | 0.052 | -0.096 | -0.096 | 0 | 0.79 | 0.49 | 0.8 | 0.94 | 0.76 | 0.82 |
| 11 | 6283452 | rs1396860 | 0.23 | 0.55 | -0.8 | -1.177 | 0.024 | 0.063 | 0.063 | 0 | 0.26 | 0.1 | 0.17 | 0.12 | 0.18 | 0.15 |
| 11 | 6288282 | rs2947029 | 0.477 | 0.611 | -0.847 | -0.613 | 0.008 | 0.068 | 0.068 | 0.27 | 0.51 | 0.38 | 0.65 | 0.78 | 0.57 | 0.65 |
| 11 | 6290756 | rs3901667 | -1.043 | 0.649 | -0.687 | -0.202 | 0.001 | 0.003 | 0.003 | 0.031 | 0.07 | 0.07 | 0.06 | 0.02 | 0.08 | 0.03 |
| 11 | 6293182 | rs1042047 | -0.941 | 0.782 | -0.58 | -0.132 | -0.002 | 0.001 | 0.001 | 0.032 | 0.44 | 0.59 | 0.54 | 0.46 | 0.38 | 0.4 |
| 11 | 6293210 | rs1042048 | -1.188 | 1.137 | -0.057 | -1.147 | 0.001 | -0.075 | -0.075 | 0 | 0.31 | 0.4 | 0.3 | 0.02 | 0.29 | 0.2 |
| 11 | 6294394 | rs6578750 | -1.309 | 0.784 | 0.152 | -0.902 | -0.018 | -0.041 | -0.041 | 0 | 0.21 | 0.59 | 0.17 | 0.18 | 0.19 | 0.07 |
| 11 | 6295483 | rs2880829 | -1.456 | 0.785 | 0.008 | -0.668 | -0.032 | 0.124 | 0.124 | 0.877 | 0.19 | 0.38 | 0.33 | 0.42 | 0.16 | 0.19 |
| 11 | 6298803 | rs10839538 | 1.874 | 1.078 | -0.249 | -0.765 | 0.005 | -0.008 | -0.008 | 0 | 0.54 | 0.18 | 0.36 | 0.56 | 0.54 | 0.64 |
| 11 | 6300016 | rs10769677 | 1.51 | 0.694 | -0.176 | -0.64 | 0.008 | -0.005 | -0.005 | 0 | 0.63 | 0.34 | 0.51 | 0.57 | 0.68 | 0.76 |
| 11 | 6301177 | rs7951875 | 1.026 | 0.718 | -0.295 | -0.791 | -0.047 | 0.031 | 0.031 | 0.865 | 0.77 | 0.62 | 0.66 | 0.75 | 0.84 | 0.81 |
| 11 | 6303396 | rs12296096 | -0.505 | 1.623 | -1.272 | -1.295 | -0.012 | 0.033 | 0.033 | 0 | 0.19 | 0.12 | 0.2 | 0 | 0.2 | 0.15 |
| 11 | 6304710 | rs11040840 | -0.168 | 1.928 | -1.529 | -1.17 | -0.049 | 0.055 | 0.055 | 0 | 0.13 | 0.05 | 0.08 | 0 | 0.18 | 0.06 |
| 11 | 6311226 | rs7104040 | 0.809 | 1.458 | -1.714 | -0.819 | -0.002 | -0.008 | -0.008 | 0 | 0.38 | 0.07 | 0.3 | 0.68 | 0.39 | 0.39 |
| 11 | 6311485 | rs6578752 | 0.292 | 1.57 | -1.357 | -0.952 | 0.005 | 0.025 | 0.025 | 0 | 0.53 | 0.4 | 0.4 | 0.69 | 0.5 | 0.46 |
| 11 | 6312769 | rs10742969 | 0.552 | 1.621 | -1.235 | -0.584 | 0.029 | 0.028 | 0.028 | 0.009 | 0.71 | 0.45 | 0.49 | 0.69 | 0.73 | 0.65 |
| 11 | 6313741 | rs10769684 | 0.374 | 1.687 | -1.938 | 0.102 | -0.031 | -0.004 | -0.004 | 0 | 0.61 | 0.27 | 0.45 | 0.68 | 0.63 | 0.54 |
| 11 | 6318959 | rs11040851 | -0.104 | 1.73 | -2.033 | 0.663 | -0.019 | -0.045 | -0.045 | 0 | 0.32 | 0.16 | 0.23 | 0.4 | 0.31 | 0.32 |
| 11 | 6319626 | rs11600546 | -0.727 | 2.115 | -1.919 | 0.736 | -0.046 | -0.039 | -0.039 | 0.215 | 0.09 | 0.02 | 0.17 | 0.13 | 0.15 | 0.08 |
| 11 | 6320703 | rs10839547 | -0.11 | 2.113 | -1.472 | 0.429 | 0.035 | -0.027 | -0.027 | 0.056 | 0.11 | 0.15 | 0.12 | 0.06 | 0.16 | 0.04 |
| 11 | 6322284 | rs10839548 | 0.07 | 1.498 | -2.631 | 0.539 | -0.094 | -0.081 | -0.081 | 0 | 0.48 | 0.05 | 0.45 | 0.47 | 0.52 | 0.42 |
| 11 | 6324061 | rs7930238 | -0.516 | 1.207 | -1.9 | 0.614 | -0.002 | 0.002 | 0.002 | 0 | 0.08 | 0.13 | 0.16 | 0.12 | 0.07 | 0.05 |
| 11 | 6324634 | rs4758405 | -0.175 | 1.078 | -1.785 | 0.594 | 0.011 | 0.001 | 0.001 | 0 | 0.52 | 0.63 | 0.48 | 0.43 | 0.45 | 0.34 |
| 11 | 6324678 | rs2344350 | 0.285 | 1.235 | -0.769 | -0.245 | 0.04 | -0.11 | -0.11 | 1.496 | 0.84 | 0.59 | 0.89 | 0.94 | 0.85 | 0.88 |
| 11 | 6324964 | rs4604857 | -0.2 | 1.021 | -0.464 | -0.075 | 0.003 | -0.009 | -0.009 | 0 | 0.4 | 0.45 | 0.37 | 0.35 | 0.37 | 0.36 |

| | | | | | | | | | | | | | | | | |
|---|---|---|---|---|---|---|---|---|---|---|---|---|---|---|---|---|
| 11 | 6326376 | rs4758406 | -0.716 | 0.788 | -0.307 | -0.595 | 0.003 | 0.023 | 0.023 | 0.767 | 0.77 | 0.83 | 0.69 | 0.85 | 0.82 | 0.84 |
| 11 | 6327406 | rs4758407 | 0.552 | 0.625 | -0.053 | -0.584 | 0.003 | 0.034 | 0.034 | 1.761 | 0.55 | 0.24 | 0.47 | 0.59 | 0.58 | 0.62 |
| 11 | 6334734 | rs10769687 | 0.913 | 0.741 | 0.758 | -0.548 | 0.003 | 0.006 | 0.006 | 0 | 0.5 | 0.42 | 0.44 | 0.59 | 0.5 | 0.6 |
| 11 | 6335562 | rs2634184 | 0.052 | 0.747 | 0.182 | -0.548 | -0.005 | -0.01 | -0.01 | 0 | 0.77 | 0.72 | 0.82 | 0.88 | 0.76 | 0.76 |
| 11 | 6336114 | rs11040858 | 0.32 | 0.56 | -0.558 | -0.338 | -0.026 | -0.037 | -0.037 | 3.103 | 0.27 | 0.05 | 0.37 | 0.29 | 0.25 | 0.17 |
| 11 | 6338507 | rs4237775 | 0.795 | 0.332 | -0.097 | -0.227 | -0.011 | 0.01 | 0.01 | 0 | 0.47 | 0.32 | 0.42 | 0.59 | 0.47 | 0.56 |
| 11 | 6340706 | rs1051992 | 0.971 | -0.15 | 0.289 | -0.211 | 0.023 | 0.039 | 0.039 | 0 | 0.5 | 0.39 | 0.47 | 0.62 | 0.58 | 0.61 |
| 11 | 6342329 | rs1488864 | 0.534 | -0.073 | 1.283 | 0.131 | -0.074 | -0.027 | -0.027 | 0 | 0.84 | 0.5 | 0.91 | 0.91 | 0.92 | 0.79 |
| 11 | 6347072 | rs10769688 | 1.146 | -0.042 | 0.95 | 0.133 | -0.039 | 0.008 | 0.008 | 0 | 0.8 | 0.42 | 0.84 | 0.75 | 0.87 | 0.74 |
| 11 | 6348639 | rs4758408 | 0.578 | -0.023 | 0.916 | 0.15 | -0.056 | -0.03 | -0.03 | 0 | 0.84 | 0.4 | 0.9 | 0.91 | 0.92 | 0.79 |
| 11 | 6349463 | rs2682121 | -1.151 | 0.042 | 1.033 | 0.148 | -0.006 | 0.027 | 0.027 | 0 | 0.13 | 0.18 | 0.11 | 0.26 | 0.12 | 0.21 |
| 11 | 6350791 | rs10839553 | -0.115 | 0.054 | 0.702 | 0.152 | -0.008 | -0.005 | -0.005 | 0.25 | 0.05 | 0.05 | 0.07 | 0.16 | 0.08 | 0.06 |
| 11 | 6351109 | rs2682119 | 0.054 | 0.074 | 0.737 | 0.149 | -0.025 | 0.035 | 0.035 | 0 | 0.08 | 0.2 | 0.04 | 0.09 | 0.04 | 0.15 |
| 11 | 6355016 | rs2171605 | 0.743 | 0.172 | -0.435 | 0.14 | -0.014 | 0.003 | 0.003 | 0 | 0.79 | 0.34 | 0.83 | 0.75 | 0.84 | 0.72 |
| 11 | 6362948 | rs4396271 | -0.377 | 0.869 | -0.657 | -0.1 | -0.009 | 0.005 | 0.005 | 0.285 | 0.25 | 0.12 | 0.41 | 0.25 | 0.25 | 0.16 |
| 11 | 6363763 | rs2634206 | 0.807 | 0.715 | -0.496 | -0.345 | 0.001 | 0.001 | 0.001 | 0 | 0.87 | 0.84 | 0.9 | 0.88 | 0.88 | 0.91 |
| 11 | 6365157 | rs2682111 | -0.189 | 1.255 | -0.581 | -0.857 | -0.012 | 0.018 | 0.018 | 0 | 0.72 | 0.78 | 0.64 | 0.84 | 0.65 | 0.63 |
| 11 | 6365704 | rs4758411 | 0.829 | 1.336 | -0.11 | -0.855 | 0.011 | 0.009 | 0.009 | 0 | 0.45 | 0.43 | 0.42 | 0.63 | 0.35 | 0.48 |
| 11 | 6370376 | rs7128818 | 1.04 | 1.624 | 0.149 | -0.663 | -0.001 | 0 | 0 | 0 | 0.56 | 0.47 | 0.57 | 0.57 | 0.47 | 0.58 |
| 11 | 6371908 | rs2682108 | 1.219 | 1.49 | 0.727 | -0.358 | 0.005 | -0.026 | -0.026 | 0 | 0.71 | 0.49 | 0.73 | 0.76 | 0.68 | 0.62 |
| 11 | 6373975 | rs2682107 | 0.516 | 1.409 | 0.523 | -0.004 | 0.007 | 0.01 | 0.01 | 0 | 0.8 | 0.71 | 0.79 | 0.76 | 0.76 | 0.73 |
| 11 | 6394176 | rs11040876 | -0.677 | 1.202 | 0.482 | -0.047 | 0.001 | 0 | 0 | 1.028 | 0.06 | 0.1 | 0.07 | 0.07 | 0.08 | 0.03 |
| 11 | 6397336 | rs2101469 | -1.331 | 1.646 | 0.583 | -0.065 | 0.005 | 0.001 | 0.001 | 0 | 0.11 | 0.08 | 0.16 | 0.17 | 0.21 | 0.04 |
| 11 | 6400407 | rs7932237 | -0.854 | 1.639 | 0.857 | -0.302 | -0.002 | -0.035 | -0.035 | 0.62 | 0.12 | 0.16 | 0.14 | 0 | 0.13 | 0.16 |
| 11 | 6405441 | rs2723646 | -1.643 | 1.718 | 1.573 | -0.257 | 0.082 | 0.075 | 0.075 | 0.002 | 0.16 | 0.33 | 0.22 | 0.24 | 0.25 | 0.08 |
| 11 | 6406055 | rs4758415 | -0.667 | 1.618 | 1.451 | -0.29 | -0.001 | -0.031 | -0.031 | 0.134 | 0.11 | 0.15 | 0.14 | 0 | 0.12 | 0.13 |
| 11 | 6409790 | rs11040883 | -0.758 | 1.615 | 1.483 | -0.292 | -0.001 | -0.04 | -0.04 | 1.379 | 0.11 | 0.16 | 0.15 | 0 | 0.13 | 0.13 |
| 11 | 6409818 | rs2723659 | -2.179 | 1.557 | 1.946 | -0.294 | 0.04 | -0.004 | -0.004 | 0.005 | 0.26 | 0.49 | 0.36 | 0.25 | 0.37 | 0.21 |
| 11 | 6410397 | rs1542705 | -2.434 | 1.289 | 1.741 | -0.434 | -0.002 | 0.004 | 0.004 | 0 | 0.32 | 0.64 | 0.39 | 0.25 | 0.4 | 0.27 |
| 11 | 6415392 | rs11601088 | -1.151 | 1.654 | 1.41 | -0.49 | 0.009 | 0.04 | 0.04 | 0 | 0.12 | 0.06 | 0.09 | 0 | 0.12 | 0.16 |
| 11 | 6415463 | rs1050239 | -1.677 | 1.52 | 1.527 | -0.133 | 0.003 | 0.002 | 0.002 | 0.087 | 0.14 | 0.11 | 0.21 | 0.15 | 0.21 | 0.1 |
| 11 | 6415882 | rs8164 | -1.348 | 1.441 | 1.579 | -0.163 | -0.01 | -0.061 | -0.061 | 0.358 | 0.12 | 0.29 | 0.11 | 0 | 0.12 | 0.17 |
| 11 | 6416959 | rs1800606 | 0.794 | 1.528 | 0.949 | -0.151 | 0.005 | 0.014 | 0.014 | 0 | 0.88 | 0.92 | 0.84 | 0.99 | 0.88 | 0.74 |
| 11 | 6428106 | rs1079199 | -1.549 | 1.938 | 1.118 | 0.013 | 0.058 | -0.005 | -0.005 | 0 | 0.22 | 0.48 | 0.29 | 0.14 | 0.3 | 0.17 |
| 11 | 6434317 | rs10839563 | -1.516 | 1.882 | 0.904 | 0.145 | 0.066 | 0.023 | 0.023 | 0.804 | 0.17 | 0.49 | 0.25 | 0.16 | 0.19 | 0.14 |
| 11 | 6436028 | rs1552513 | -0.919 | 1.754 | 0.716 | -0.654 | -0.001 | -0.007 | -0.007 | 0 | 0.37 | 0.35 | 0.46 | 0.19 | 0.43 | 0.46 |
| 11 | 6436704 | rs2682095 | 0.102 | 1.351 | 0.462 | -0.726 | -0.007 | 0.005 | 0.005 | 0 | 0.23 | 0.2 | 0.31 | 0.2 | 0.3 | 0.2 |
| 11 | 6441051 | rs2723664 | -0.968 | 1.038 | 0.528 | -1.013 | 0.005 | 0.04 | 0.04 | 0.334 | 0.49 | 0.37 | 0.59 | 0.32 | 0.52 | 0.52 |
| 11 | 6459938 | rs4758417 | 0.121 | 1.481 | 0.712 | -0.725 | -0.028 | -0.036 | -0.036 | 0 | 0.09 | 0 | 0.12 | 0.12 | 0.08 | 0.14 |
| 11 | 6460026 | rs10500671 | -0.244 | 1.361 | 1.175 | -0.73 | 0.092 | -0.059 | -0.059 | 1.934 | 0.15 | 0.36 | 0.2 | 0.06 | 0.22 | 0.23 |
| 11 | 6466076 | rs7113742 | -0.807 | 1.186 | 1.048 | -0.67 | 0.038 | -0.033 | -0.033 | 0 | 0.18 | 0.41 | 0.18 | 0.11 | 0.18 | 0.32 |
| 11 | 6468326 | rs1906656 | -1.115 | 1.149 | -0.153 | -0.833 | -0.001 | -0.001 | -0.001 | 0 | 0.6 | 0.73 | 0.65 | 0.43 | 0.57 | 0.67 |
| 11 | 6470057 | rs2723636 | -0.454 | 1.588 | -0.472 | -1.346 | 0.009 | -0.078 | -0.078 | 0.175 | 0.17 | 0.3 | 0.18 | 0.02 | 0.17 | 0.21 |
| 11 | 6471722 | rs17821052 | -0.766 | 1.321 | -1.055 | -0.573 | -0.016 | 0.052 | 0.052 | 0 | 0.23 | 0.11 | 0.25 | 0.11 | 0.23 | 0.26 |
| 11 | 6477831 | rs10606 | -0.898 | 1.338 | -0.527 | -0.683 | 0.013 | -0.074 | -0.074 | 1.613 | 0.18 | 0.3 | 0.18 | 0.02 | 0.18 | 0.22 |
| 11 | 6478619 | rs16913748 | -0.891 | 1.337 | -0.528 | -0.684 | 0.013 | -0.074 | -0.074 | 0 | 0.18 | 0.3 | 0.18 | 0.02 | 0.18 | 0.22 |
| 11 | 6494786 | rs11605777 | -1.263 | 1.837 | -0.443 | -0.803 | 0.019 | -0.035 | -0.035 | 0 | 0.38 | 0.46 | 0.48 | 0.17 | 0.51 | 0.3 |
| 11 | 6495600 | rs11605881 | -0.219 | 1.821 | -0.351 | -0.831 | 0.004 | -0.063 | -0.063 | 0.68 | 0.17 | 0.27 | 0.17 | 0.03 | 0.17 | 0.2 |
| 11 | 6496333 | rs11603308 | -0.305 | 1.804 | -0.239 | -0.84 | 0.009 | -0.089 | -0.089 | 0 | 0.18 | 0.35 | 0.17 | 0.03 | 0.17 | 0.2 |
| 11 | 6499215 | rs2303493 | -1.293 | 1.733 | -0.453 | -1.016 | -0.067 | 0.096 | 0.096 | 0 | 0.37 | 0.19 | 0.47 | 0.13 | 0.52 | 0.24 |
| 11 | 6503575 | rs1048121 | -0.386 | 1.573 | 0.481 | -1.205 | -0.011 | 0.024 | 0.024 | 0.528 | 0.17 | 0.13 | 0.16 | 0.03 | 0.18 | 0.21 |
| 11 | 6509393 | rs7924933 | -0.746 | 1.376 | 0.949 | -1.299 | 0.013 | -0.007 | -0.007 | 0 | 0.06 | 0.24 | 0.02 | 0 | 0.02 | 0.01 |
| 11 | 6510244 | rs2555162 | -1.83 | 0.841 | 0.269 | -1.803 | -0.001 | 0.025 | 0.025 | 0 | 0.62 | 0.86 | 0.66 | 0.27 | 0.68 | 0.56 |
| 11 | 6518785 | rs2555159 | -2.249 | -0.128 | -0.516 | -1.034 | 0.049 | -0.138 | -0.138 | 0 | 0.79 | 0.99 | 0.81 | 0.55 | 0.89 | 0.76 |
| 11 | 6518932 | rs17243759 | 0.676 | -0.279 | 0.127 | -1.459 | -0.015 | 0.014 | 0.014 | 0 | 0.24 | 0.14 | 0.32 | 0.13 | 0.35 | 0.22 |
| 11 | 6520503 | rs10769694 | -1.137 | -0.374 | 0.479 | -1.503 | 0.003 | 0.151 | 0.151 | 0 | 0.49 | 0.7 | 0.52 | 0.16 | 0.56 | 0.44 |
| 11 | 6523657 | rs2723620 | -0.577 | -0.357 | 0.501 | -1.493 | 0.006 | -0.033 | -0.033 | 0 | 0.43 | 0.46 | 0.5 | 0.16 | 0.54 | 0.43 |
| 11 | 6530106 | rs11605607 | 1.214 | -0.349 | 0.425 | -1.494 | -0.031 | 0.061 | 0.061 | 0.221 | 0.15 | 0.08 | 0.16 | 0.02 | 0.18 | 0.15 |
| 11 | 6530278 | rs10839568 | 0.696 | -0.338 | 0.459 | -1.478 | -0.015 | 0.014 | 0.014 | 0 | 0.24 | 0.14 | 0.32 | 0.13 | 0.35 | 0.22 |
| 11 | 6531702 | rs2555151 | -0.169 | -0.354 | 0.504 | -1.322 | -0.001 | 0.001 | 0.001 | 0 | 0.31 | 0.6 | 0.4 | 0.17 | 0.38 | 0.23 |
| 11 | 6531995 | rs4758422 | -0.153 | -0.334 | 0.633 | -1.315 | 0.019 | -0.037 | -0.037 | 0.202 | 0.29 | 0.42 | 0.39 | 0.17 | 0.37 | 0.23 |
| 11 | 6532519 | rs4758423 | 0.662 | -0.323 | 0.472 | -1.303 | -0.015 | 0.013 | 0.013 | 0 | 0.24 | 0.14 | 0.32 | 0.13 | 0.35 | 0.22 |
| 11 | 6533090 | rs2555149 | -1.138 | -0.378 | 0.385 | -1.294 | -0.005 | 0.153 | 0.153 | 0 | 0.49 | 0.78 | 0.55 | 0.23 | 0.53 | 0.45 |
| 11 | 6533231 | rs2555148 | -0.951 | -0.434 | 0.163 | -1.653 | 0.009 | 0.032 | 0.032 | 0 | 0.34 | 0.72 | 0.44 | 0.17 | 0.29 | 0.43 |
| 11 | 6541580 | rs2555143 | -1.56 | -0.382 | -0.034 | -1.559 | -0.028 | 0.03 | 0.03 | 0 | 0.6 | 0.86 | 0.6 | 0.28 | 0.64 | 0.52 |
| 11 | 6543931 | rs7931672 | 1.24 | -0.27 | 0.103 | -1.465 | -0.008 | -0.015 | -0.015 | 0 | 0.06 | 0.08 | 0.06 | 0.01 | 0.02 | 0.13 |
| 11 | 6549411 | rs2880963 | 0.798 | -0.091 | 0.169 | -1.492 | -0.001 | -0.024 | -0.024 | 0.04 | 0.19 | 0.18 | 0.17 | 0.03 | 0.18 | 0.22 |
| 11 | 6551470 | rs10839574 | 0.596 | -0.133 | 0.19 | -1.504 | -0.001 | -0.005 | -0.005 | 0 | 0.54 | 0.51 | 0.49 | 0.84 | 0.45 | 0.56 |
| 11 | 6553861 | rs11040912 | -0.99 | 0.79 | 0.399 | -0.472 | 0.098 | 0.161 | 0.161 | 0 | 0.1 | 0.34 | 0.28 | 0.24 | 0.13 | 0.21 |

| | | | | | | | | | | | | | | | | |
|---|---|---|---|---|---|---|---|---|---|---|---|---|---|---|---|---|
| 11 | 6554069 | rs4237778 | 0.975 | -0.001 | 0.074 | -1.52 | -0.002 | 0.051 | 0.051 | 0 | 0.57 | 0.32 | 0.6 | 0.76 | 0.52 | 0.68 |
| 11 | 6555365 | rs7103810 | -0.482 | -0.219 | 0.212 | -1.456 | -0.012 | -0.016 | -0.016 | 0.563 | 0.84 | 0.78 | 0.91 | 0.92 | 0.89 | 0.88 |
| 11 | 6555853 | rs10734609 | 0.628 | -0.288 | -0.427 | -1.521 | 0.001 | 0.001 | 0.001 | 0 | 0.72 | 0.36 | 0.73 | 0.82 | 0.81 | 0.72 |
| 11 | 6555914 | rs10742974 | 1.379 | -0.415 | -0.422 | -1.606 | -0.005 | 0.043 | 0.043 | 0 | 0.41 | 0.17 | 0.42 | 0.75 | 0.36 | 0.39 |
| 11 | 6556400 | rs11040913 | -0.043 | -0.157 | -0.118 | -1.37 | -0.008 | 0.01 | 0.01 | 0 | 0.26 | 0.16 | 0.19 | 0.08 | 0.36 | 0.22 |
| 11 | 6556895 | rs2344929 | 0.787 | -0.099 | -0.393 | -1.33 | -0.002 | 0.082 | 0.082 | 0 | 0.56 | 0.32 | 0.7 | 0.8 | 0.53 | 0.62 |
| 11 | 6560141 | rs11040914 | -0.545 | -0.095 | -0.301 | -1.163 | -0.025 | -0.016 | -0.016 | 0 | 0.85 | 0.66 | 0.92 | 0.92 | 0.91 | 0.89 |
| 11 | 6563565 | rs11604481 | -0.086 | 0.583 | -0.878 | -1.194 | 0.002 | 0.032 | 0.032 | 0 | 0.23 | 0.18 | 0.24 | 0.02 | 0.24 | 0.22 |
| 11 | 6563826 | rs11040915 | -0.531 | 0.932 | -0.84 | -0.814 | -0.005 | 0.006 | 0.006 | 0 | 0.27 | 0.17 | 0.3 | 0.14 | 0.36 | 0.31 |
| 11 | 6564384 | rs7103173 | -0.131 | 1.014 | -0.973 | -0.848 | -0.038 | 0.037 | 0.037 | 0 | 0.21 | 0.08 | 0.25 | 0.09 | 0.31 | 0.23 |
| 11 | 6564674 | rs2344931 | -0.052 | 0.951 | -0.909 | -0.538 | -0.012 | -0.005 | -0.005 | 0 | 0.69 | 0.76 | 0.65 | 0.64 | 0.57 | 0.68 |
| 11 | 6569896 | rs11606889 | -0.096 | 0.435 | -1.122 | -1.043 | -0.011 | 0.081 | 0.081 | 0 | 0.2 | 0.08 | 0.17 | 0.02 | 0.18 | 0.2 |
| 11 | 6577577 | rs2345345 | 0.45 | 0.327 | -0.752 | -0.754 | -0.002 | 0.021 | 0.021 | 0 | 0.59 | 0.66 | 0.64 | 0.78 | 0.57 | 0.64 |
| 11 | 6585007 | rs11040923 | 1.165 | -0.085 | -0.998 | -0.767 | -0.076 | 0.036 | 0.036 | 1.782 | 0.49 | 0.07 | 0.47 | 0.72 | 0.47 | 0.57 |
| 11 | 6585502 | rs7107072 | 1.079 | -0.338 | -0.592 | -0.981 | -0.009 | -0.034 | -0.034 | 0 | 0.83 | 0.24 | 0.83 | 0.98 | 0.93 | 0.87 |
| 11 | 6585670 | rs16915423 | 0.179 | -0.354 | 0.137 | -0.975 | 0.057 | 0.027 | 0.027 | 0 | 0.06 | 0.3 | 0.09 | 0.01 | 0.02 | 0.12 |
| 11 | 6585838 | rs11825255 | -1.199 | -0.338 | -0.019 | -0.978 | -0.026 | -0.071 | -0.071 | 4.967 | 0.12 | 0.55 | 0.13 | 0.01 | 0.04 | 0.12 |
| 11 | 6587228 | rs7947368 | -1.507 | -0.322 | -0.53 | -0.97 | 0.003 | -0.013 | -0.013 | 0.101 | 0.15 | 0.77 | 0.16 | 0.01 | 0.06 | 0.13 |
| 11 | 6588228 | rs10769699 | -0.156 | -0.165 | -0.687 | -0.858 | -0.099 | -0.063 | -0.063 | 0 | 0.28 | 0.07 | 0.31 | 0.22 | 0.41 | 0.22 |
| 11 | 6598183 | rs4758112 | 0.013 | -0.268 | -0.491 | -0.673 | 0 | -0.002 | -0.002 | 0 | 0.91 | 0.91 | 0.94 | 0.91 | 0.92 | 0.92 |
| 11 | 6609791 | rs10839586 | 1.448 | -0.548 | 0.043 | -0.801 | -0.048 | -0.034 | -0.034 | 2.305 | 0.81 | 0.45 | 0.81 | 0.9 | 0.89 | 0.89 |
| 11 | 6609990 | rs4423167 | 0.239 | -0.481 | -0.175 | -0.762 | -0.015 | -0.024 | -0.024 | 0.189 | 0.08 | 0.02 | 0.05 | 0.08 | 0.08 | 0.05 |
| 11 | 6618548 | rs11040930 | 1.645 | -0.579 | -0.427 | -1.036 | -0.02 | 0.068 | 0.068 | 1.207 | 0.48 | 0.13 | 0.42 | 0.73 | 0.48 | 0.58 |
| 11 | 6621516 | rs11040933 | -0.66 | -0.325 | -0.509 | -0.773 | -0.003 | 0.004 | 0.004 | 0.004 | 0.29 | 0.22 | 0.33 | 0.16 | 0.4 | 0.15 |
| 11 | 6622714 | rs2288283 | 1.189 | -0.543 | -0.04 | -0.589 | -0.045 | -0.049 | -0.049 | 0.064 | 0.78 | 0.39 | 0.79 | 0.91 | 0.86 | 0.87 |
| 11 | 6623918 | rs11606458 | -0.846 | -0.105 | -0.776 | -0.55 | -0.053 | 0.025 | 0.025 | 0 | 0.16 | 0.01 | 0.15 | 0.07 | 0.25 | 0.12 |
| 11 | 6624181 | rs4758438 | 0.567 | -0.762 | -0.932 | -0.044 | 0.022 | -0.014 | -0.014 | 0.295 | 0.25 | 0.16 | 0.17 | 0.33 | 0.14 | 0.14 |
| 11 | 6626675 | rs4758441 | -0.117 | 0.078 | -0.181 | -0.067 | -0.006 | 0.006 | 0.006 | 0.007 | 0.33 | 0.5 | 0.29 | 0.36 | 0.25 | 0.28 |
| 11 | 6633608 | rs2555173 | 0.154 | -0.265 | 0.035 | -0.024 | -0.046 | 0.015 | 0.015 | 0.267 | 0.29 | 0.45 | 0.21 | 0.33 | 0.16 | 0.24 |
| 11 | 6634149 | rs7488 | 0.402 | -0.141 | 0.172 | -0.238 | 0.003 | -0.011 | -0.011 | 0 | 0.82 | 0.8 | 0.78 | 0.91 | 0.75 | 0.8 |
| 11 | 6637504 | rs2072651 | 0.752 | -0.219 | 0.284 | -0.11 | 0.004 | -0.006 | -0.006 | 0 | 0.25 | 0.21 | 0.2 | 0.34 | 0.16 | 0.19 |
| 11 | 6640753 | rs1800732 | -0.661 | -0.352 | 0.306 | -0.236 | -0.01 | -0.018 | -0.018 | 0.046 | 0.06 | 0.26 | 0.03 | 0 | 0.01 | 0.05 |
| 11 | 6648424 | rs4758443 | 0.05 | -0.393 | -0.225 | -0.185 | -0.011 | -0.053 | -0.053 | 3.619 | 0.37 | 0.12 | 0.34 | 0.42 | 0.37 | 0.3 |
| 11 | 6652618 | rs2659871 | -0.595 | -0.271 | -0.228 | -0.137 | 0.003 | 0 | 0 | 0 | 0.46 | 0.59 | 0.42 | 0.43 | 0.38 | 0.38 |
| 11 | 6666054 | rs2555172 | -0.827 | -0.415 | -1.162 | -0.115 | 0.007 | -0.014 | -0.014 | 0 | 0.68 | 0.6 | 0.68 | 0.78 | 0.63 | 0.6 |
| 11 | 6666267 | rs7111291 | 0.008 | 0.013 | -1.902 | 0.07 | -0.132 | -0.114 | -0.114 | 0.064 | 0.4 | 0.04 | 0.41 | 0.51 | 0.47 | 0.34 |
| 11 | 6668328 | rs2063082 | -0.652 | 0.633 | -1.199 | 0.221 | -0.016 | 0.007 | 0.007 | 0 | 0.16 | 0.11 | 0.19 | 0.16 | 0.28 | 0.24 |
| 11 | 6670921 | rs4758114 | 0.933 | 0.173 | -1.694 | 0.154 | -0.093 | -0.097 | -0.097 | 0 | 0.35 | 0.06 | 0.26 | 0.39 | 0.36 | 0.3 |
| 11 | 6672457 | rs1876301 | -0.53 | 0.068 | -1.3 | 0.066 | -0.001 | 0.039 | 0.039 | 0.677 | 0.62 | 0.36 | 0.69 | 0.82 | 0.64 | 0.52 |
| 11 | 6673190 | rs12419946 | 0.812 | 0.464 | -1.323 | 0.245 | -0.092 | -0.081 | -0.081 | 0.855 | 0.37 | 0.06 | 0.26 | 0.38 | 0.36 | 0.3 |
| 11 | 6679132 | rs12294943 | -0.445 | 0.171 | -0.372 | 0.492 | 0.017 | 0.007 | 0.007 | 0 | 0.11 | 0.19 | 0.13 | 0.11 | 0.16 | 0.07 |
| 11 | 6683542 | rs937856 | 0.702 | -0.516 | -0.426 | 0.306 | 0.05 | -0.003 | -0.003 | 0 | 0.69 | 0.23 | 0.58 | 0.68 | 0.79 | 0.61 |
| 11 | 6684511 | rs10769704 | 0.531 | -0.536 | 0.475 | 0.341 | -0.051 | 0.033 | 0.033 | 0 | 0.75 | 0.53 | 0.65 | 0.68 | 0.83 | 0.62 |
| 11 | 6689064 | rs4758117 | 0.649 | -0.532 | 0.526 | 0.345 | -0.004 | 0 | 0 | 0 | 0.73 | 0.34 | 0.64 | 0.68 | 0.83 | 0.62 |
| 11 | 6695844 | rs7950601 | 0.005 | -0.414 | 0.587 | 0.186 | -0.018 | -0.088 | -0.088 | 0.842 | 0.85 | 0.63 | 0.92 | 0.99 | 0.91 | 0.77 |
| 11 | 6700602 | rs1111585 | -0.067 | -0.28 | 0.654 | 0.596 | -0.018 | 0.056 | 0.056 | 0 | 0.11 | 0.15 | 0.31 | 0.3 | 0.06 | 0.14 |
| 11 | 6700887 | rs7128249 | -1.163 | -0.357 | 0.991 | 0.36 | 0.031 | -0.028 | -0.028 | 0 | 0.09 | 0.46 | 0.12 | 0 | 0.08 | 0.02 |
| 11 | 6702409 | rs3741277 | -1.63 | -0.36 | 0.589 | 0.391 | -0.022 | 0.02 | 0.02 | 0 | 0.2 | 0.69 | 0.42 | 0.3 | 0.14 | 0.17 |
| 11 | 6703105 | rs3741276 | 0.137 | -0.356 | 0.566 | 0.395 | 0.018 | -0.037 | -0.037 | 0 | 0.1 | 0.03 | 0.31 | 0.31 | 0.06 | 0.14 |
| 11 | 6703892 | rs1567135 | 0.473 | -0.094 | -0.245 | 0.134 | -0.108 | -0.012 | -0.012 | 0.074 | 0.36 | 0.1 | 0.31 | 0.31 | 0.52 | 0.35 |
| 11 | 6705253 | rs7126152 | -1.172 | -0.12 | 0.527 | 0.391 | -0.015 | -0.058 | -0.058 | 0 | 0.1 | 0.67 | 0.12 | 0 | 0.08 | 0.02 |
| 11 | 6705464 | rs12296090 | -1.363 | -0.156 | 0.56 | 0.373 | -0.017 | -0.063 | -0.063 | 3.521 | 0.11 | 0.65 | 0.12 | 0 | 0.08 | 0.02 |
| 11 | 6707260 | rs4758124 | 0.317 | 0.223 | 0.442 | 0.4 | -0.011 | -0.046 | -0.046 | 3.091 | 0.11 | 0.03 | 0.31 | 0.31 | 0.1 | 0.14 |
| 11 | 6707887 | rs12573864 | -1.183 | 0.282 | 0.379 | 0.448 | -0.007 | 0.011 | 0.011 | 0.018 | 0.21 | 0.69 | 0.43 | 0.31 | 0.18 | 0.16 |
| 11 | 6710947 | rs4408289 | -0.577 | 0.703 | 0.482 | 0.178 | 0.015 | -0.011 | -0.011 | 1.702 | 0.75 | 0.61 | 0.77 | 0.84 | 0.72 | 0.83 |
| 11 | 6717557 | rs12794064 | 0.533 | 0.702 | 0.139 | 0.177 | -0.025 | -0.003 | -0.003 | 1.306 | 0.22 | 0.12 | 0.21 | 0.16 | 0.28 | 0.16 |
| 11 | 6717608 | rs1472836 | 0.278 | 0.677 | 0.354 | 0.316 | 0.004 | 0.045 | 0.045 | 0 | 0.72 | 0.56 | 0.52 | 0.5 | 0.72 | 0.52 |
| 11 | 6717724 | rs11040966 | 0.278 | 0.677 | 0.354 | 0.324 | 0.004 | 0.043 | 0.043 | 0 | 0.72 | 0.56 | 0.52 | 0.5 | 0.72 | 0.52 |
| 11 | 6717814 | rs10839592 | 0.664 | 0.717 | 0.236 | 0.418 | -0.049 | -0.083 | -0.083 | 0 | 0.12 | 0.01 | 0.36 | 0.41 | 0.16 | 0.21 |
| 11 | 6719027 | rs1466974 | 1.124 | 0.664 | 0.514 | 0.379 | 0.008 | 0.064 | 0.064 | 0 | 0.47 | 0.13 | 0.29 | 0.34 | 0.44 | 0.36 |
| 11 | 6719118 | rs10769708 | 0.48 | 0.687 | 0.53 | 0.392 | 0.005 | 0.046 | 0.046 | 0 | 0.72 | 0.56 | 0.51 | 0.5 | 0.71 | 0.52 |
| 11 | 6720186 | rs10839595 | -0.204 | 0.707 | 0.213 | 0.454 | -0.025 | -0.019 | -0.019 | 0.838 | 0.36 | 0.27 | 0.58 | 0.56 | 0.45 | 0.37 |
| 11 | 6727468 | rs1395558 | 0.37 | 0.563 | -0.285 | 0.063 | 0.007 | -0.01 | -0.01 | 0 | 0.61 | 0.29 | 0.47 | 0.5 | 0.67 | 0.48 |
| 11 | 6729454 | rs10839597 | 0.663 | 0.512 | -0.471 | 0.027 | 0.009 | 0.018 | 0.018 | 0 | 0.39 | 0.17 | 0.27 | 0.34 | 0.4 | 0.32 |
| 11 | 6729493 | rs10839598 | -0.946 | 0.73 | -0.106 | 0.069 | 0 | 0.002 | 0.002 | 0 | 0.42 | 0.51 | 0.54 | 0.5 | 0.41 | 0.55 |
| 11 | 6734053 | rs11040978 | -0.427 | 0.637 | -0.609 | 0.194 | -0.022 | 0.006 | 0.006 | 0 | 0.42 | 0.2 | 0.62 | 0.41 | 0.42 | 0.34 |
| 11 | 6735019 | rs17176958 | 0.779 | 0.75 | -0.835 | 0.043 | -0.012 | 0.081 | 0.081 | 0 | 0.19 | 0.07 | 0.16 | 0.06 | 0.19 | 0.11 |
| 11 | 6735869 | rs7937911 | -0.097 | 0.821 | -0.869 | 0.21 | -0.019 | -0.01 | -0.01 | 0 | 0.27 | 0.15 | 0.52 | 0.29 | 0.31 | 0.24 |
| 11 | 6737138 | rs3741274 | 0.56 | 0.824 | -0.864 | 0.216 | 0.006 | 0.038 | 0.038 | 0 | 0.07 | 0.08 | 0.36 | 0.23 | 0.12 | 0.12 |

| | | | | | | | | | | | | | | | | |
|---|---|---|---|---|---|---|---|---|---|---|---|---|---|---|---|---|
| 11 | 6737893 | rs11040980 | 0.56 | 0.824 | -0.864 | 0.216 | 0.006 | 0.038 | 0.038 | 0 | 0.07 | 0.08 | 0.36 | 0.23 | 0.12 | 0.12 |
| 11 | 6737979 | rs11040982 | 1.285 | 0.49 | -1.318 | 0.502 | 0.016 | -0.078 | -0.078 | 0 | 0.13 | 0.01 | 0.07 | 0.25 | 0.11 | 0.11 |
| 11 | 6738291 | rs11040983 | 0.617 | 1.114 | -0.994 | 0.537 | 0.006 | 0.038 | 0.038 | 0 | 0.07 | 0.08 | 0.36 | 0.23 | 0.12 | 0.12 |
| 11 | 6738411 | rs17262467 | 0.939 | 1.114 | -0.998 | 0.535 | -0.01 | 0.078 | 0.078 | 0 | 0.19 | 0.06 | 0.16 | 0.06 | 0.19 | 0.11 |
| 11 | 6738952 | rs10769716 | -0.193 | 1.114 | -1.013 | 0.532 | -0.009 | -0.022 | -0.022 | 0 | 0.23 | 0.13 | 0.46 | 0.36 | 0.23 | 0.22 |
| 11 | 6739407 | rs12284429 | 0.68 | 1.173 | -0.282 | 0.449 | -0.001 | -0.001 | -0.001 | 0 | 0.62 | 0.41 | 0.46 | 0.48 | 0.66 | 0.42 |
| 11 | 6740308 | rs12223353 | 1.003 | 0.991 | -0.388 | 0.55 | 0.004 | -0.04 | -0.04 | 0 | 0.42 | 0.21 | 0.29 | 0.42 | 0.47 | 0.3 |
| 11 | 6743665 | rs16917433 | 0.93 | 1.099 | -0.607 | 0.489 | -0.001 | 0.01 | 0.01 | 0 | 0.16 | 0.05 | 0.09 | 0.12 | 0.11 | 0.11 |
| 11 | 6743890 | rs7942744 | 0.239 | 1.064 | -0.123 | 0.493 | 0.012 | -0.008 | -0.008 | 0 | 0.61 | 0.48 | 0.4 | 0.62 | 0.58 | 0.45 |
| 11 | 6744362 | rs7942521 | -0.211 | 1.039 | -0.208 | 0.559 | -0.023 | 0.005 | 0.005 | 0 | 0.42 | 0.2 | 0.62 | 0.41 | 0.42 | 0.34 |
| 11 | 6749978 | rs10769719 | -0.398 | 0.892 | 0.401 | 0.598 | 0.023 | -0.016 | -0.016 | 0 | 0.88 | 0.7 | 0.93 | 0.91 | 0.89 | 0.68 |
| 11 | 6756331 | rs949901 | -0.459 | 0.86 | 0.408 | 0.598 | 0.022 | -0.014 | -0.014 | 0 | 0.88 | 0.73 | 0.94 | 0.91 | 0.89 | 0.68 |
| 11 | 6756574 | rs4475903 | 0.319 | 0.617 | 0.902 | 0.574 | -0.031 | -0.004 | -0.004 | 0 | 0.86 | 0.39 | 0.92 | 0.85 | 0.94 | 0.66 |
| 11 | 6758224 | rs7935848 | -0.38 | 0.677 | 0.272 | 0.494 | 0.009 | 0.041 | 0.041 | 0 | 0.56 | 0.27 | 0.69 | 0.67 | 0.54 | 0.45 |
| 11 | 6761413 | rs1506972 | 1.25 | 0.654 | 0.693 | 0.495 | -0.004 | -0.013 | -0.013 | 0 | 0.3 | 0.35 | 0.22 | 0.24 | 0.35 | 0.24 |
| 11 | 6766102 | rs7931228 | -0.01 | 1.026 | 0.534 | 0.865 | 0.008 | 0.024 | 0.024 | 0 | 0.48 | 0.17 | 0.33 | 0.45 | 0.41 | 0.33 |
| 11 | 6767609 | rs11040999 | 1.465 | 1.037 | 0.875 | 0.883 | 0 | 0.008 | 0.008 | 0 | 0.29 | 0.2 | 0.21 | 0.24 | 0.35 | 0.24 |
| 11 | 6768561 | rs1354803 | 1.498 | 1.001 | 0.536 | 0.819 | -0.044 | -0.028 | -0.028 | 0.61 | 0.13 | 0 | 0.09 | 0.13 | 0.11 | 0.11 |
| 11 | 6773439 | rs7928685 | 0.845 | 1.023 | 0.682 | 0.847 | -0.02 | -0.011 | -0.011 | 0 | 0.07 | 0.05 | 0.36 | 0.22 | 0.13 | 0.12 |
| 11 | 6777637 | rs10500662 | 1.675 | 0.981 | 0.891 | 0.84 | 0 | 0 | 0 | 0.279 | 0.29 | 0.26 | 0.22 | 0.22 | 0.34 | 0.24 |
| 11 | 6779071 | rs1506981 | -0.356 | 0.976 | 0.813 | 0.83 | 0.015 | 0.019 | 0.019 | 0 | 0.49 | 0.31 | 0.35 | 0.47 | 0.41 | 0.33 |
| 11 | 6779721 | rs12286483 | 1.848 | 0.982 | 0.616 | 0.821 | -0.007 | 0.025 | 0.025 | 0 | 0.28 | 0.14 | 0.21 | 0.15 | 0.34 | 0.19 |
| 11 | 6783013 | rs12795894 | 1.823 | 0.98 | 0.615 | 0.787 | -0.007 | 0.03 | 0.03 | 0 | 0.28 | 0.14 | 0.21 | 0.16 | 0.35 | 0.19 |
| 11 | 6783401 | rs4758132 | -0.725 | 0.996 | 0.766 | 0.791 | 0.001 | -0.017 | -0.017 | 0.007 | 0.87 | 0.71 | 0.93 | 0.91 | 0.9 | 0.69 |
| 11 | 6788642 | rs10839615 | 0.829 | 1.065 | 0.779 | 0.894 | -0.019 | -0.006 | -0.006 | 0 | 0.07 | 0.05 | 0.36 | 0.24 | 0.14 | 0.12 |
| 11 | 6790106 | rs7102536 | -1.674 | 0.835 | 0.653 | 0.583 | -0.013 | 0.051 | 0.051 | 0 | 0.82 | 0.9 | 0.84 | 0.93 | 0.81 | 0.88 |
| 11 | 6791151 | rs670151 | -1.352 | 0.898 | 0.819 | 0.681 | -0.006 | -0.009 | -0.009 | 0 | 0.63 | 0.67 | 0.42 | 0.53 | 0.51 | 0.64 |
| 11 | 6792058 | rs11041009 | 0.157 | 0.841 | 0.458 | 0.672 | -0.001 | 0.003 | 0.003 | 0 | 0.27 | 0.27 | 0.16 | 0.21 | 0.21 | 0.42 |
| 11 | 6793428 | rs593313 | 0.011 | 0.455 | 0.411 | 0.633 | 0.017 | 0.017 | 0.017 | 0 | 0.35 | 0.22 | 0.25 | 0.32 | 0.29 | 0.22 |
| 11 | 6794304 | rs555693 | 0.51 | 0.742 | 0.598 | 0.691 | -0.003 | -0.004 | -0.004 | 0 | 0.63 | 0.37 | 0.46 | 0.48 | 0.65 | 0.42 |
| 11 | 6804067 | rs1463956 | 1.637 | 0.551 | 0.375 | 0.653 | 0.01 | 0.02 | 0.02 | 0 | 0.3 | 0.16 | 0.21 | 0.18 | 0.33 | 0.18 |
| 11 | 6805524 | rs7126011 | 1.632 | 0.561 | 0.38 | 0.658 | 0.01 | 0.02 | 0.02 | 0 | 0.3 | 0.16 | 0.21 | 0.18 | 0.33 | 0.18 |
| 11 | 6805900 | rs11041021 | 0.518 | 0.682 | 0.349 | 0.699 | -0.002 | 0.002 | 0.002 | 0 | 0.21 | 0.17 | 0.12 | 0.19 | 0.17 | 0.3 |
| 11 | 6807243 | rs497681 | 0.355 | 0.811 | 0.495 | 0.851 | 0.027 | 0.011 | 0.011 | 0 | 0.67 | 0.53 | 0.49 | 0.63 | 0.63 | 0.52 |
| 11 | 6808368 | rs2741752 | 1.883 | 0.8 | 0.334 | 0.84 | -0.001 | 0.029 | 0.029 | 0 | 0.3 | 0.11 | 0.21 | 0.18 | 0.35 | 0.18 |
| 11 | 6809901 | rs11041026 | 0.89 | 0.735 | 0.361 | 0.898 | 0.002 | -0.006 | -0.006 | 0.013 | 0.15 | 0.12 | 0.17 | 0.27 | 0.14 | 0.09 |
| 11 | 6810464 | rs2595498 | 0.927 | 0.726 | 0.35 | 0.885 | 0.021 | -0.029 | -0.029 | 0 | 0.43 | 0.17 | 0.37 | 0.45 | 0.46 | 0.27 |
| 11 | 6810524 | rs11041027 | 1.039 | 0.731 | 0.145 | 0.906 | 0.001 | -0.046 | -0.046 | 0.117 | 0.14 | 0.06 | 0.16 | 0.27 | 0.14 | 0.09 |
| 11 | 6816778 | rs866347 | 0.108 | 0.699 | 0.442 | 0.516 | 0.079 | 0.04 | 0.04 | 0 | 0.84 | 0.48 | 0.59 | 0.82 | 0.8 | 0.81 |
| 11 | 6816875 | rs7122644 | -0.06 | 0.658 | 0.376 | 0.487 | 0.004 | -0.049 | -0.049 | 3.181 | 0.33 | 0.19 | 0.27 | 0.46 | 0.29 | 0.38 |
| 11 | 6817395 | rs604109 | -1.082 | 0.702 | 0.39 | 0.487 | -0.028 | -0.021 | -0.021 | 0 | 0.89 | 0.9 | 0.62 | 0.82 | 0.8 | 0.82 |
| 11 | 6817741 | rs2741740 | 1.876 | 0.629 | 0.374 | 0.445 | 0.002 | 0.028 | 0.028 | 0 | 0.3 | 0.15 | 0.22 | 0.18 | 0.35 | 0.18 |
| 11 | 6818254 | rs2659885 | 1.961 | 0.636 | 0.268 | 0.448 | 0.003 | 0.024 | 0.024 | 0 | 0.29 | 0.1 | 0.21 | 0.18 | 0.33 | 0.18 |
| 11 | 6819423 | rs502464 | 0.877 | 0.645 | 0.309 | 0.465 | -0.005 | -0.009 | -0.009 | 0 | 0.22 | 0.25 | 0.12 | 0.18 | 0.17 | 0.26 |
| 11 | 6819568 | rs2741845 | 1.942 | 0.647 | 0.265 | 0.462 | -0.026 | 0.015 | 0.015 | 0 | 0.29 | 0.06 | 0.21 | 0.18 | 0.35 | 0.18 |
| 11 | 6846072 | rs2595424 | 0.077 | 0.647 | 0.491 | 0.519 | -0.005 | -0.001 | -0.001 | 0 | 0.66 | 0.68 | 0.51 | 0.64 | 0.64 | 0.56 |
| 11 | 6848913 | rs7129692 | 0.332 | 0.608 | 0.29 | 0.521 | -0.02 | -0.032 | -0.032 | 0 | 0.25 | 0.52 | 0.13 | 0.18 | 0.17 | 0.25 |
| 11 | 6851301 | rs2741816 | 0.204 | 0.655 | 0.192 | 0.542 | -0.011 | 0.002 | 0.002 | 0 | 0.64 | 0.35 | 0.49 | 0.64 | 0.63 | 0.59 |
| 11 | 6851427 | rs11041043 | -0.279 | 0.605 | 0.016 | 0.556 | 0.007 | -0.035 | -0.035 | 0 | 0.35 | 0.25 | 0.28 | 0.46 | 0.31 | 0.41 |
| 11 | 6854547 | rs752708 | 0.19 | 0.549 | -0.025 | 0.638 | -0.001 | -0.002 | -0.002 | 0 | 0.26 | 0.31 | 0.25 | 0.39 | 0.24 | 0.35 |
| 11 | 6856817 | rs2741806 | 1.29 | 0.464 | 0.365 | 0.535 | -0.025 | -0.035 | -0.035 | 0 | 0.34 | 0.44 | 0.23 | 0.18 | 0.33 | 0.19 |
| 11 | 6865648 | rs2741803 | -0.766 | 0.649 | 0.359 | 0.644 | -0.012 | -0.01 | -0.01 | 0 | 0.89 | 0.9 | 0.62 | 0.82 | 0.81 | 0.81 |
| 11 | 6868131 | rs7117015 | -0.235 | 0.54 | 0.206 | 0.628 | -0.007 | 0.009 | 0.009 | 0 | 0.36 | 0.56 | 0.3 | 0.46 | 0.31 | 0.37 |
| 11 | 6879173 | rs2595471 | 0.78 | 0.488 | -0.015 | 0.519 | 0.011 | 0.032 | 0.032 | 0 | 0.52 | 0.32 | 0.31 | 0.36 | 0.5 | 0.44 |
| 11 | 6883912 | rs2741775 | 0.78 | 0.488 | -0.015 | 0.57 | 0.011 | 0.019 | 0.019 | 0 | 0.52 | 0.32 | 0.31 | 0.4 | 0.5 | 0.44 |
| 11 | 6885407 | rs2741767 | 0.753 | 0.993 | 0.352 | 0.634 | 0.021 | 0.039 | 0.039 | 0 | 0.5 | 0.41 | 0.4 | 0.27 | 0.44 | 0.28 |
| 11 | 6890136 | rs4300376 | -0.49 | 0.447 | 0.347 | 0.546 | -0.037 | -0.001 | -0.001 | 0 | 0.33 | 0.2 | 0.37 | 0.31 | 0.36 | 0.43 |
| 11 | 6891704 | rs10839632 | 0.723 | 0.665 | 0.327 | 0.508 | -0.022 | -0.03 | -0.03 | 0.888 | 0.25 | 0.21 | 0.29 | 0.41 | 0.33 | 0.36 |
| 11 | 6892866 | rs2741762 | 0.683 | 0.627 | 0.395 | 0.459 | -0.01 | 0.017 | 0.017 | 0 | 0.62 | 0.29 | 0.6 | 0.69 | 0.64 | 0.57 |
| 11 | 6895415 | rs1449138 | 0.492 | 0.651 | 0.38 | 0.468 | 0.004 | 0.019 | 0.019 | 0 | 0.49 | 0.23 | 0.58 | 0.43 | 0.5 | 0.4 |
| 11 | 6897251 | rs2741757 | -1.109 | 0.641 | 0.633 | 0.462 | 0.004 | 0.007 | 0.007 | 0 | 0.8 | 0.83 | 0.9 | 0.83 | 0.83 | 0.75 |
| 11 | 6897670 | rs10769728 | 0.687 | 0.676 | 0.659 | 0.482 | -0.021 | -0.028 | -0.028 | 0 | 0.25 | 0.2 | 0.27 | 0.4 | 0.31 | 0.35 |
| 11 | 6898540 | rs7938371 | -0.894 | 0.441 | 0.781 | 0.536 | 0.07 | 0.065 | 0.065 | 2.285 | 0.05 | 0.39 | 0.03 | 0 | 0 | 0 |
| 11 | 6898663 | rs10839635 | 0.761 | 0.456 | 0.651 | 0.544 | -0.024 | -0.029 | -0.029 | 2.285 | 0.25 | 0.2 | 0.29 | 0.4 | 0.33 | 0.36 |
| 11 | 6898758 | rs7102808 | 0.701 | 0.368 | 0.353 | 0.532 | -0.047 | -0.027 | -0.027 | 0 | 0.11 | 0.03 | 0.27 | 0.14 | 0.19 | 0.18 |
| 11 | 6901845 | rs2595445 | 0.209 | 0.418 | 0.64 | 0.59 | 0 | 0.001 | 0.001 | 0 | 0.52 | 0.45 | 0.6 | 0.52 | 0.5 | 0.44 |
| 11 | 6905024 | rs2595448 | 1.22 | 0.39 | 0.232 | 0.589 | 0.05 | 0.028 | 0.028 | 0 | 0.38 | 0.19 | 0.31 | 0.29 | 0.31 | 0.22 |
| 11 | 6911127 | rs2741829 | 0.413 | 0.426 | 0.112 | 0.613 | 0.002 | -0.007 | -0.007 | 0.62 | 0.6 | 0.4 | 0.42 | 0.37 | 0.48 | 0.42 |
| 11 | 6913127 | rs2741804 | 0.463 | 0.477 | 0.006 | 0.603 | -0.023 | 0.003 | 0.003 | 0.036 | 0.49 | 0.12 | 0.57 | 0.43 | 0.5 | 0.41 |

| | | | | | | | | | | | | | | | |
|---|---|---|---|---|---|---|---|---|---|---|---|---|---|---|---|
| 11 | 6913290 | rs1965209 | 0.574 | 0.764 | -0.141 | 0.722 | -0.05 | -0.024 | -0.024 | 0 | 0.12 | 0.03 | 0.26 | 0.14 | 0.19 | 0.19 |
| 11 | 6915109 | rs1449123 | 0.721 | 0.861 | -0.29 | 0.541 | 0.004 | 0.013 | 0.013 | 0 | 0.2 | 0.13 | 0.09 | 0.08 | 0.16 | 0.2 |
| 11 | 6915315 | rs1965211 | 0.066 | 1.452 | -0.037 | 1.314 | 0.038 | 0.056 | 0.056 | 0 | 0.84 | 0.66 | 0.71 | 0.76 | 0.81 | 0.77 |
| 11 | 6915546 | rs10769730 | -0.537 | 1.056 | -0.555 | 1.231 | 0 | 0.001 | 0.001 | 0.059 | 0.25 | 0.27 | 0.29 | 0.4 | 0.33 | 0.36 |
| 11 | 6919452 | rs10769731 | -0.377 | -0.246 | -0.472 | 0.973 | -0.034 | -0.002 | -0.002 | 0.807 | 0.32 | 0.56 | 0.32 | 0.33 | 0.18 | 0.35 |
| 11 | 6925675 | rs1470627 | -0.454 | 0.346 | -0.899 | 1.052 | 0.002 | 0 | 0 | 0.135 | 0.81 | 0.79 | 0.77 | 0.78 | 0.78 | 0.68 |
| 11 | 6933160 | rs12789816 | 0.34 | 0.503 | -0.922 | 1.028 | -0.04 | -0.05 | -0.05 | 0.561 | 0.17 | 0.04 | 0.17 | 0.21 | 0.16 | 0.32 |
| 11 | 6933289 | rs10769736 | 0.818 | 0.288 | -0.948 | 1.518 | 0.021 | 0.075 | 0.075 | 0 | 0.47 | 0.15 | 0.44 | 0.34 | 0.59 | 0.34 |
| 11 | 6935158 | rs4758146 | 0.082 | 0.001 | -0.526 | 1.408 | -0.011 | 0.005 | 0.005 | 0.168 | 0.32 | 0.64 | 0.33 | 0.45 | 0.19 | 0.34 |
| 11 | 6941739 | rs12288515 | -0.354 | 0.835 | -0.852 | 0.825 | 0.013 | -0.003 | -0.003 | 0 | 0.45 | 0.6 | 0.44 | 0.45 | 0.39 | 0.38 |
| 11 | 6942476 | rs10839658 | 0.152 | 0.829 | -0.831 | 0.887 | 0.057 | 0.063 | 0.063 | 0 | 0.52 | 0.27 | 0.49 | 0.34 | 0.65 | 0.34 |
| 11 | 6942628 | rs11605995 | 0.613 | 0.831 | -0.842 | 0.887 | -0.012 | 0.006 | 0.006 | 0.337 | 0.31 | 0.62 | 0.33 | 0.45 | 0.19 | 0.35 |
| 11 | 6942695 | rs12806437 | 0.733 | 0.964 | -0.942 | 0.828 | -0.043 | -0.05 | -0.05 | 0 | 0.17 | 0.04 | 0.17 | 0.21 | 0.16 | 0.32 |
| 11 | 6951734 | rs11820166 | 1.236 | 0.925 | -0.75 | 0.994 | -0.001 | -0.016 | -0.016 | 0 | 0.06 | 0.02 | 0.06 | 0.17 | 0.07 | 0.1 |
| 11 | 6953456 | rs2053835 | -0.197 | 1.103 | -0.308 | 0.921 | 0.04 | -0.016 | -0.016 | 0 | 0.05 | 0.19 | 0.07 | 0 | 0.06 | 0 |
| 11 | 6953858 | rs11041108 | 0.955 | 1.124 | -0.464 | 0.951 | -0.086 | -0.086 | -0.086 | 0 | 0.32 | 0.04 | 0.3 | 0.37 | 0.37 | 0.3 |
| 11 | 6958594 | rs1602569 | 0.453 | 1.225 | -0.231 | 0.91 | -0.012 | 0.004 | 0.004 | 0.546 | 0.42 | 0.51 | 0.44 | 0.45 | 0.33 | 0.52 |
| 11 | 6962957 | rs2857891 | 0.565 | 1.179 | -0.272 | 0.822 | -0.009 | -0.005 | -0.005 | 0.52 | 0.14 | 0.09 | 0.13 | 0 | 0.22 | 0.03 |
| 11 | 6963477 | rs1491826 | 0.639 | 1.189 | -0.226 | 0.865 | 0.045 | -0.035 | -0.035 | 0 | 0.75 | 0.57 | 0.75 | 0.82 | 0.68 | 0.82 |
| 11 | 6963730 | rs10839664 | 0.979 | 1.302 | -0.724 | 1.106 | -0.096 | -0.087 | -0.087 | 0 | 0.32 | 0.04 | 0.35 | 0.36 | 0.38 | 0.31 |
| 11 | 6964018 | rs1491827 | -0.173 | 1.417 | -0.343 | 1.201 | 0.001 | 0.001 | 0.001 | 17.394 | 0.92 | 0.92 | 0.94 | 0.81 | 0.95 | 0.85 |
| 11 | 6965652 | rs1907612 | 0.145 | 1.523 | -0.321 | 1.196 | 0.003 | 0.001 | 0.001 | 0 | 0.15 | 0.14 | 0.13 | 0.15 | 0.23 | 0.04 |
| 11 | 6966499 | rs1388534 | 0.038 | 1.539 | -0.047 | 1.253 | 0.009 | 0.093 | 0.093 | 2.672 | 0.92 | 0.72 | 0.93 | 0.82 | 0.95 | 0.88 |
| 11 | 6966708 | rs957748 | 0.612 | 1.555 | -0.241 | 1.2 | 0.005 | 0.014 | 0.014 | 8.283 | 0.77 | 0.73 | 0.81 | 0.7 | 0.72 | 0.83 |
| 11 | 6977175 | rs2239730 | -0.368 | 1.376 | 0.128 | 0.906 | 0.018 | 0.002 | 0.002 | 2.252 | 0.18 | 0.35 | 0.13 | 0.14 | 0.23 | 0.05 |
| 11 | 6980511 | rs2857901 | -0.333 | 1.389 | 0.115 | 0.87 | 0.019 | 0.003 | 0.003 | 2.176 | 0.18 | 0.34 | 0.13 | 0.14 | 0.23 | 0.05 |
| 11 | 6984001 | rs2857887 | 0.016 | 1.32 | -0.585 | 0.876 | -0.018 | -0.001 | -0.001 | 0 | 0.18 | 0.1 | 0.16 | 0.14 | 0.28 | 0.05 |
| 11 | 6985309 | rs1388525 | 0.922 | 1.566 | -0.937 | 0.779 | -0.059 | -0.079 | -0.079 | 0 | 0.32 | 0.04 | 0.33 | 0.35 | 0.32 | 0.33 |
| 11 | 6986426 | rs7117376 | -1.411 | 1.988 | 0.044 | 0.588 | 0.006 | 0.003 | 0.003 | 7.362 | 0.26 | 0.28 | 0.25 | 0.3 | 0.38 | 0.18 |
| 11 | 6993077 | rs980159 | 0.042 | 1.963 | 0.041 | 0.59 | 0.018 | 0.04 | 0.04 | 3.594 | 0.92 | 0.83 | 0.91 | 0.83 | 0.9 | 0.87 |
| 11 | 6993922 | rs2638110 | -0.662 | 1.692 | -0.409 | 0.337 | 0.006 | -0.002 | -0.002 | 0 | 0.19 | 0.21 | 0.17 | 0.14 | 0.28 | 0.05 |
| 11 | 6998597 | rs969652 | -0.705 | 1.917 | -0.609 | 0.281 | -0.015 | 0.004 | 0.004 | 0.236 | 0.21 | 0.16 | 0.18 | 0.16 | 0.33 | 0.08 |
| 11 | 7001786 | rs2857917 | -1.206 | 2.091 | -0.416 | -0.484 | -0.001 | -0.001 | -0.001 | 0 | 0.34 | 0.58 | 0.33 | 0.25 | 0.41 | 0.21 |
| 11 | 7008962 | rs11041124 | -0.557 | 1.696 | -0.473 | -0.331 | -0.009 | 0.042 | 0.042 | 0 | 0.21 | 0.12 | 0.2 | 0.08 | 0.23 | 0.11 |
| 11 | 7012482 | rs2515949 | -0.622 | 1.106 | -0.551 | -0.517 | -0.001 | -0.004 | -0.004 | 0.174 | 0.58 | 0.42 | 0.63 | 0.67 | 0.6 | 0.55 |
| 11 | 7014066 | rs11041127 | -0.518 | 0.845 | -0.375 | -0.311 | -0.001 | -0.019 | -0.019 | 0 | 0.16 | 0.14 | 0.16 | 0.08 | 0.15 | 0.1 |
| 11 | 7020454 | rs2857930 | -1.031 | 1.305 | -0.434 | -0.216 | -0.012 | 0.008 | 0.008 | 0 | 0.22 | 0.15 | 0.29 | 0.15 | 0.32 | 0.2 |
| 11 | 7020629 | rs1388526 | -0.013 | 0.864 | 0.633 | -0.477 | 0.016 | -0.006 | -0.006 | 3.124 | 0.94 | 0.73 | 0.97 | 1 | 0.98 | 0.98 |
| 11 | 7020687 | rs1352860 | 0.811 | 0.47 | 0.479 | -0.669 | -0.037 | -0.058 | -0.058 | 0 | 0.9 | 0.73 | 0.96 | 1 | 0.97 | 0.96 |
| 11 | 7022160 | rs2857919 | 0.866 | 0.475 | 0.467 | -0.667 | -0.036 | -0.057 | -0.057 | 0.015 | 0.9 | 0.73 | 0.96 | 1 | 0.97 | 0.96 |
| 11 | 7022360 | rs2239734 | -0.325 | 0.625 | 0.433 | -0.693 | -0.004 | -0.006 | -0.006 | 0 | 0.07 | 0.04 | 0.15 | 0.13 | 0.11 | 0.13 |
| 11 | 7022717 | rs1156526 | 2.011 | 0.063 | 0.663 | -0.462 | 0 | 0.002 | 0.002 | 0 | 0.61 | 0.42 | 0.64 | 0.71 | 0.62 | 0.65 |
| 11 | 7023688 | rs7932579 | 1.798 | 0.052 | 0.66 | -0.467 | 0 | -0.019 | -0.019 | 0.643 | 0.62 | 0.57 | 0.65 | 0.71 | 0.63 | 0.65 |
| 11 | 7037829 | rs6578820 | -1.556 | 0.082 | 0.699 | -0.46 | 0.002 | 0 | 0 | 0.608 | 0.3 | 0.35 | 0.33 | 0.29 | 0.35 | 0.31 |
| 11 | 7041763 | rs12418753 | 0.453 | 0.075 | 0.656 | -0.436 | 0.012 | -0.037 | -0.037 | 0.006 | 0.06 | 0 | 0.02 | 0.12 | 0.03 | 0.11 |
| 11 | 7049140 | rs10839692 | 1.724 | 0.045 | 0.763 | -0.47 | 0.001 | -0.004 | -0.004 | 0.052 | 0.64 | 0.42 | 0.65 | 0.71 | 0.62 | 0.67 |
| 11 | 7052875 | rs902736 | 1.154 | 0.002 | 1.381 | -0.518 | 0.021 | 0.001 | 0.001 | 0 | 0.7 | 0.55 | 0.66 | 0.71 | 0.65 | 0.7 |
| 11 | 7054279 | rs10839696 | 1.319 | -0.056 | 1.337 | -0.522 | 0.018 | -0.04 | -0.04 | 0.273 | 0.71 | 0.45 | 0.66 | 0.85 | 0.65 | 0.78 |
| 11 | 7054683 | rs10839697 | 1.697 | -0.056 | 1.324 | -0.523 | 0.001 | -0.004 | -0.004 | 0 | 0.64 | 0.43 | 0.65 | 0.71 | 0.62 | 0.67 |
| 11 | 7058885 | rs11041149 | -0.508 | 0.038 | 1.272 | -0.508 | 0.003 | 0.005 | 0.005 | 0 | 0.08 | 0.1 | 0.16 | 0.13 | 0.11 | 0.13 |
| 11 | 7061782 | rs969233 | -0.495 | 0.012 | 1.314 | -0.529 | 0.028 | -0.037 | -0.037 | 0 | 0.86 | 0.81 | 0.86 | 0.98 | 0.79 | 0.93 |
| 11 | 7065289 | rs6578821 | 1.026 | 0.447 | 1.235 | -0.655 | 0.036 | -0.021 | -0.021 | 0.121 | 0.75 | 0.65 | 0.67 | 0.85 | 0.64 | 0.79 |
| 11 | 7079038 | rs10839708 | 2.054 | 0.244 | 1.395 | -0.658 | 0 | 0.017 | 0.017 | 0 | 0.59 | 0.35 | 0.61 | 0.71 | 0.6 | 0.64 |
| 11 | 7091357 | rs10769762 | 1.099 | 0.251 | 1.306 | -0.714 | 0.041 | -0.095 | -0.095 | 0.907 | 0.73 | 0.53 | 0.67 | 0.92 | 0.65 | 0.87 |
| 11 | 7091569 | rs17280682 | 0.391 | 0.26 | 1.317 | -0.721 | -0.021 | 0.022 | 0.022 | 0 | 0.14 | 0.07 | 0.13 | 0.02 | 0.21 | 0.07 |
| 11 | 7092893 | rs11041157 | 1.508 | 0.249 | 1.323 | -0.557 | 0.008 | -0.042 | -0.042 | 0 | 0.67 | 0.43 | 0.64 | 0.9 | 0.63 | 0.84 |
| 11 | 7093083 | rs902738 | -0.794 | 0.257 | 1.317 | -0.558 | -0.007 | -0.024 | -0.024 | 0.095 | 0.05 | 0.11 | 0.04 | 0 | 0.03 | 0.03 |
| 11 | 7095322 | rs10769764 | 1.487 | 0.148 | 1.319 | -0.513 | 0.009 | -0.024 | -0.024 | 2.835 | 0.66 | 0.53 | 0.66 | 0.75 | 0.63 | 0.76 |
| 11 | 7113134 | rs6578828 | 1.015 | 0.314 | 1.363 | -0.34 | 0.011 | -0.026 | -0.026 | 2.273 | 0.86 | 0.63 | 0.81 | 0.91 | 0.87 | 0.93 |
| 11 | 7118082 | rs4277091 | 1.841 | 0.205 | 1.307 | -0.332 | -0.057 | -0.004 | -0.004 | 0.225 | 0.75 | 0.49 | 0.78 | 0.77 | 0.84 | 0.8 |
| 11 | 7119010 | rs4758162 | 1.467 | 0.295 | 1.351 | -0.419 | -0.021 | -0.08 | -0.08 | 0 | 0.82 | 0.49 | 0.79 | 0.93 | 0.87 | 0.92 |
| 11 | 7126726 | rs12283053 | 0.055 | 0.198 | 1.204 | -0.359 | -0.014 | -0.029 | -0.029 | 0.03 | 0.06 | 0.21 | 0.03 | 0 | 0.02 | 0.02 |
| 11 | 7132245 | rs9804679 | -1.976 | -0.204 | 0.933 | -0.618 | -0.009 | -0.018 | -0.018 | 0 | 0.39 | 0.43 | 0.34 | 0.24 | 0.35 | 0.26 |
| 11 | 7133804 | rs7102031 | -1.378 | -0.207 | 0.949 | -0.6 | -0.009 | -0.019 | -0.019 | 0 | 0.32 | 0.38 | 0.21 | 0.17 | 0.28 | 0.22 |
| 11 | 7137616 | rs1507625 | -0.818 | -0.21 | 0.868 | -0.584 | 0.002 | 0.014 | 0.014 | 0 | 0.25 | 0.17 | 0.18 | 0.17 | 0.26 | 0.2 |
| 11 | 7142133 | rs11041200 | -0.273 | -0.159 | 0.754 | -0.534 | 0 | 0 | 0 | 0 | 0.07 | 0.06 | 0.13 | 0.07 | 0.07 | 0.04 |
| 11 | 7162780 | rs11041213 | -1.147 | -0.174 | 0.782 | -0.547 | -0.02 | -0.031 | -0.031 | 0.037 | 0.14 | 0.27 | 0.16 | 0.07 | 0.09 | 0.06 |
| 11 | 7172160 | rs3105716 | 1.392 | -0.107 | 0.572 | -0.522 | 0.001 | 0.001 | 0.001 | 0 | 0.61 | 0.4 | 0.64 | 0.76 | 0.65 | 0.75 |
| 11 | 7177851 | rs10839732 | 1.001 | -0.243 | -0.459 | -0.406 | -0.085 | -0.009 | -0.009 | 0.291 | 0.27 | 0.02 | 0.22 | 0.26 | 0.3 | 0.27 |

| | | | | | | | | | | | | | | | | |
|---|---|---|---|---|---|---|---|---|---|---|---|---|---|---|---|---|
| 11 | 7180491 | rs10839734 | 0.275 | -0.2 | 0.355 | -0.102 | -0.004 | 0.038 | 0.038 | 0.577 | 0.48 | 0.29 | 0.51 | 0.5 | 0.49 | 0.5 |
| 11 | 7181575 | rs2346323 | -0.759 | -0.182 | 0.331 | 0.376 | -0.022 | 0.009 | 0.009 | 0 | 0.66 | 0.75 | 0.61 | 0.76 | 0.57 | 0.69 |
| 11 | 7181636 | rs10500681 | 0.755 | -0.369 | 0.083 | 0.402 | 0.007 | -0.045 | -0.045 | 0.385 | 0.08 | 0 | 0.03 | 0.27 | 0.05 | 0.13 |
| 11 | 7183800 | rs11041221 | 0.418 | -0.163 | 0.338 | -0.013 | 0.001 | 0.01 | 0.01 | 0 | 0.07 | 0.1 | 0.13 | 0.15 | 0.07 | 0.04 |
| 11 | 7184989 | rs10839736 | -0.426 | -0.178 | 0.475 | -0.044 | -0.011 | 0.005 | 0.005 | 1.987 | 0.57 | 0.67 | 0.57 | 0.49 | 0.51 | 0.56 |
| 11 | 7188457 | rs11041225 | 0.239 | 0.277 | -0.61 | 0.285 | 0.002 | 0.009 | 0.009 | 0 | 0.07 | 0.1 | 0.13 | 0.15 | 0.07 | 0.04 |
| 11 | 7189003 | rs10500683 | 0.202 | 0.385 | -0.503 | 0.264 | -0.011 | 0.035 | 0.035 | 0.748 | 0.48 | 0.28 | 0.52 | 0.5 | 0.49 | 0.53 |
| 11 | 7191722 | rs12270243 | 0.221 | 0.321 | -0.676 | 0.246 | 0.018 | 0.007 | 0.007 | 0.505 | 0.44 | 0.3 | 0.48 | 0.51 | 0.44 | 0.49 |
| 11 | 7195494 | rs7937849 | -0.831 | 0.382 | -0.404 | 0.04 | -0.009 | -0.029 | -0.029 | 0.063 | 0.17 | 0.39 | 0.12 | 0.13 | 0.16 | 0.16 |
| 11 | 7195508 | rs7951678 | -1.412 | 0.796 | -0.799 | 0.04 | 0.017 | -0.003 | -0.003 | 0 | 0.4 | 0.54 | 0.31 | 0.35 | 0.47 | 0.32 |
| 11 | 7196338 | rs12361949 | 0.34 | 1.587 | -1.553 | 0.265 | -0.05 | 0.059 | 0.059 | 0 | 0.13 | 0.07 | 0.15 | 0 | 0.21 | 0.03 |
| 11 | 7198054 | rs7131435 | -1.257 | 0.698 | -1.299 | 0.631 | -0.023 | -0.075 | -0.075 | 0 | 0.23 | 0.11 | 0.18 | 0.32 | 0.23 | 0.23 |
| 11 | 7207824 | rs2135871 | 0.121 | 0.247 | -0.638 | -0.224 | -0.009 | -0.046 | -0.046 | 0 | 0.78 | 0.69 | 0.73 | 0.88 | 0.79 | 0.78 |
| 11 | 7210964 | rs1507623 | 0.397 | 0.077 | -1.318 | 0.024 | -0.04 | -0.068 | -0.068 | 0 | 0.19 | 0.03 | 0.14 | 0.22 | 0.24 | 0.22 |
| 11 | 7212657 | rs1395908 | -0.906 | 0.273 | -1.327 | -0.116 | -0.013 | 0.013 | 0.013 | 0 | 0.8 | 0.86 | 0.72 | 0.96 | 0.74 | 0.75 |
| 11 | 7213503 | rs3905311 | 0.771 | 0.041 | -0.558 | -0.184 | -0.003 | 0.003 | 0.003 | 0 | 0.54 | 0.47 | 0.45 | 0.63 | 0.47 | 0.5 |
| 11 | 7216241 | rs1507628 | 1.192 | -0.317 | -0.816 | -0.217 | -0.019 | 0.091 | 0.091 | 0 | 0.43 | 0.11 | 0.33 | 0.3 | 0.39 | 0.34 |
| 11 | 7217025 | rs1911627 | -1.141 | -0.338 | -0.645 | -0.253 | -0.031 | 0.048 | 0.048 | 0 | 0.81 | 0.9 | 0.66 | 0.96 | 0.73 | 0.76 |
| 11 | 7232673 | rs7106726 | 0.646 | -0.491 | -0.602 | -0.099 | 0.008 | -0.013 | -0.013 | 0 | 0.49 | 0.22 | 0.38 | 0.51 | 0.45 | 0.44 |
| 11 | 7233442 | rs1857994 | -0.12 | -0.338 | -0.587 | -0.344 | 0.007 | -0.053 | -0.053 | 0 | 0.3 | 0.36 | 0.44 | 0.14 | 0.32 | 0.26 |
| 11 | 7236253 | rs16923546 | 0.939 | -0.419 | -0.562 | -0.112 | -0.041 | -0.074 | -0.074 | 0 | 0.17 | 0.03 | 0.13 | 0.27 | 0.22 | 0.21 |
| 11 | 7244215 | rs3812741 | -0.525 | 0.109 | -0.403 | -0.587 | -0.002 | 0.002 | 0.002 | 0 | 0.35 | 0.6 | 0.49 | 0.14 | 0.38 | 0.29 |
| 11 | 7244599 | rs1487841 | 0.883 | -0.364 | -1.057 | -0.507 | -0.034 | -0.128 | -0.128 | 0 | 0.38 | 0.04 | 0.25 | 0.52 | 0.37 | 0.47 |
| 11 | 7257474 | rs766498 | -1.713 | -0.333 | 0.674 | -0.893 | 0.018 | 0.095 | 0.095 | 2.609 | 0.08 | 0.48 | 0.12 | 0.12 | 0.03 | 0.05 |
| 11 | 7257798 | rs17283958 | 0.043 | -0.181 | 0.32 | -1.135 | -0.088 | 0.128 | 0.128 | 0 | 0.25 | 0.02 | 0.35 | 0.03 | 0.31 | 0.32 |
| 11 | 7260691 | rs6578839 | -1.89 | -0.425 | 1.442 | -0.261 | -0.011 | -0.021 | -0.021 | 0.083 | 0.09 | 0.44 | 0.04 | 0 | 0.01 | 0.04 |
| 11 | 7260983 | rs1487845 | -0.784 | -0.193 | 0.515 | -0.256 | -0.004 | 0.003 | 0.003 | 0 | 0.11 | 0.08 | 0.17 | 0.11 | 0.13 | 0.05 |
| 11 | 7261376 | rs11041270 | -1.876 | -0.109 | 0.756 | -0.261 | 0.011 | -0.025 | -0.025 | 0 | 0.17 | 0.35 | 0.2 | 0.13 | 0.18 | 0.08 |
| 11 | 7262248 | rs11041273 | -0.687 | -0.229 | 0.633 | -0.233 | 0.001 | 0.003 | 0.003 | 0.01 | 0.11 | 0.07 | 0.16 | 0.11 | 0.13 | 0.05 |
| 11 | 7262943 | rs1552463 | -1.377 | -0.089 | 0.696 | -0.163 | 0.079 | 0.029 | 0.029 | 0 | 0.05 | 0.25 | 0.03 | 0.01 | 0.05 | 0.03 |
| 11 | 7266724 | rs11041279 | -1.234 | -0.364 | 0.067 | -0.086 | -0.023 | -0.028 | -0.028 | 0 | 0.19 | 0.29 | 0.19 | 0.11 | 0.15 | 0.09 |
| 11 | 7272026 | rs2055708 | 0.735 | -0.47 | 0.279 | 0.354 | -0.037 | 0.029 | 0.029 | 1.362 | 0.77 | 0.56 | 0.82 | 0.7 | 0.85 | 0.75 |
| 11 | 7272363 | rs2135872 | -1.211 | -1.153 | 1.117 | 0.123 | -0.026 | -0.053 | -0.053 | 0 | 0.19 | 0.21 | 0.13 | 0.06 | 0.11 | 0.29 |
| 11 | 7272397 | rs2055709 | 1.642 | -0.624 | 0.43 | -0.126 | 0.009 | -0.004 | -0.004 | 0 | 0.53 | 0.41 | 0.56 | 0.48 | 0.6 | 0.49 |
| 11 | 7278278 | rs1487863 | 0.786 | -0.246 | 1.24 | -0.187 | 0.015 | 0.069 | 0.069 | 0 | 0.72 | 0.46 | 0.68 | 0.49 | 0.69 | 0.71 |
| 11 | 7278401 | rs1487864 | 0.108 | -0.034 | 1.443 | -0.193 | 0.021 | 0.044 | 0.044 | 0 | 0.85 | 0.72 | 0.76 | 0.74 | 0.83 | 0.78 |
| 11 | 7283249 | rs10839752 | -2.095 | 0.353 | 1.382 | -0.322 | 0.002 | 0.002 | 0.002 | 0 | 0.43 | 0.51 | 0.46 | 0.5 | 0.44 | 0.51 |
| 11 | 7284298 | rs2346804 | 0.944 | 0.34 | 1.343 | -0.332 | 0.007 | -0.024 | -0.024 | 0.033 | 0.69 | 0.57 | 0.61 | 0.75 | 0.71 | 0.56 |
| 11 | 7284357 | rs10743013 | 2.005 | 0.331 | 1.368 | -0.359 | -0.014 | -0.017 | -0.017 | 0 | 0.56 | 0.34 | 0.53 | 0.49 | 0.56 | 0.48 |
| 11 | 7286422 | rs10839753 | 1.098 | 0.451 | 1.495 | -0.206 | 0.046 | 0.128 | 0.128 | 0 | 0.76 | 0.51 | 0.68 | 0.5 | 0.71 | 0.73 |
| 11 | 7302406 | rs4757989 | 1.202 | 0.559 | 1.201 | -0.029 | 0.017 | 0.112 | 0.112 | 0.731 | 0.85 | 0.66 | 0.87 | 0.57 | 0.82 | 0.9 |
| 11 | 7306095 | rs16924357 | -0.929 | 0.555 | 1.266 | -0.014 | 0.002 | 0.007 | 0.007 | 0.07 | 0.05 | 0.14 | 0.05 | 0.09 | 0.05 | 0.02 |
| 11 | 7309061 | rs6578842 | 0.263 | 0.56 | 1.28 | -0.002 | 0.012 | 0.086 | 0.086 | 0.092 | 0.9 | 0.8 | 0.92 | 0.66 | 0.87 | 0.93 |
| 11 | 7309243 | rs11041297 | -0.755 | 0.594 | 0.851 | -0.407 | -0.069 | 0.096 | 0.096 | 0 | 0.28 | 0.06 | 0.34 | 0.1 | 0.33 | 0.43 |
| 11 | 7310138 | rs11041299 | 0.899 | 0.596 | 0.845 | -0.388 | -0.036 | -0.031 | -0.031 | 0 | 0.07 | 0 | 0.05 | 0.25 | 0.12 | 0.06 |
| 11 | 7310767 | rs4758170 | 1.795 | 0.545 | 1.004 | -0.418 | -0.002 | -0.002 | -0.002 | 0.206 | 0.56 | 0.52 | 0.53 | 0.49 | 0.49 | 0.48 |
| 11 | 7313866 | rs1487875 | 1.155 | 0.544 | 0.979 | -0.418 | -0.036 | -0.028 | -0.028 | 0.115 | 0.62 | 0.75 | 0.59 | 0.57 | 0.54 | 0.5 |
| 11 | 7320166 | rs1319309 | 1.617 | 0.535 | 0.937 | -0.429 | -0.011 | -0.012 | -0.012 | 0 | 0.57 | 0.58 | 0.54 | 0.48 | 0.49 | 0.48 |
| 11 | 7320535 | rs12799959 | -0.841 | 0.558 | 0.745 | -0.478 | -0.035 | 0.078 | 0.078 | 0 | 0.28 | 0.1 | 0.34 | 0.1 | 0.33 | 0.43 |
| 11 | 7321481 | rs755831 | -0.136 | 0.432 | 1.188 | -0.427 | 0 | -0.037 | -0.037 | 0 | 0.15 | 0.17 | 0.13 | 0.01 | 0.13 | 0.23 |
| 11 | 7325275 | rs11041303 | 0.306 | 0.505 | 1.151 | -0.184 | 0.001 | 0.046 | 0.046 | 0 | 0.08 | 0.06 | 0.05 | 0.34 | 0.12 | 0.07 |
| 11 | 7325445 | rs7948390 | -1.127 | 0.443 | 1.316 | -0.177 | 0.006 | 0.067 | 0.067 | 0 | 0.05 | 0.24 | 0.04 | 0.1 | 0.03 | 0.02 |
| 11 | 7326195 | rs2200650 | 1.82 | 0.44 | 0.667 | -0.98 | -0.023 | -0.091 | -0.091 | 0.232 | 0.39 | 0.1 | 0.42 | 0.47 | 0.37 | 0.41 |
| 11 | 7329763 | rs2220371 | 1.334 | 0.917 | 1.136 | -1.023 | 0 | -0.02 | -0.02 | 0 | 0.64 | 0.27 | 0.7 | 0.55 | 0.65 | 0.62 |
| 11 | 7329893 | rs7946495 | 0.732 | 0.995 | 1.091 | -1.052 | 0 | 0 | 0 | 2.599 | 0.72 | 0.67 | 0.77 | 0.74 | 0.71 | 0.65 |
| 11 | 7330704 | rs11041306 | -0.644 | 1.086 | 1.242 | -0.76 | 0.022 | 0.095 | 0.095 | 0.198 | 0.11 | 0.21 | 0.1 | 0.3 | 0.15 | 0.06 |
| 11 | 7335823 | rs7939239 | -0.326 | 1.377 | 0.942 | -1.225 | -0.023 | 0.047 | 0.047 | 0 | 0.18 | 0.08 | 0.2 | 0 | 0.23 | 0.2 |
| 11 | 7336536 | rs931378 | 0.19 | 1.393 | 1.277 | -1.231 | 0.013 | -0.069 | -0.069 | 0.017 | 0.81 | 0.72 | 0.8 | 1 | 0.77 | 0.8 |
| 11 | 7336916 | rs17215501 | 0.007 | 1.267 | 1.143 | -1.505 | -0.003 | 0.003 | 0.003 | 0 | 0.27 | 0.23 | 0.3 | 0.09 | 0.33 | 0.15 |
| 11 | 7341459 | rs1487856 | -1.268 | 1.21 | 1.244 | -1.431 | 0.002 | -0.05 | -0.05 | 0 | 0.27 | 0.64 | 0.27 | 0.07 | 0.29 | 0.23 |
| 11 | 7343638 | rs1017940 | -0.089 | 1.237 | 1.42 | -1.466 | 0.012 | -0.051 | -0.051 | 0 | 0.7 | 0.53 | 0.65 | 0.91 | 0.64 | 0.84 |
| 11 | 7349792 | rs10734617 | -0.304 | 1.23 | 1.479 | -1.463 | -0.001 | -0.002 | -0.002 | 0 | 0.73 | 0.68 | 0.69 | 0.91 | 0.68 | 0.85 |
| 11 | 7350851 | rs1487853 | 1.278 | 1.192 | 1.574 | -1.505 | 0.02 | 0.064 | 0.064 | 0 | 0.51 | 0.34 | 0.45 | 0.9 | 0.41 | 0.65 |
| 11 | 7351331 | rs10839766 | 0.31 | 1.287 | 1.238 | -1.213 | 0.018 | 0.08 | 0.08 | 0 | 0.09 | 0.13 | 0.05 | 0.27 | 0.13 | 0.16 |
| 11 | 7353097 | rs901755 | -0.16 | 1.212 | 1.033 | -1.283 | -0.017 | 0.029 | 0.029 | 0 | 0.19 | 0.12 | 0.2 | 0 | 0.23 | 0.2 |
| 11 | 7354137 | rs12792362 | 0.347 | 1.194 | 1.065 | -1.319 | -0.045 | 0.063 | 0.063 | 0.033 | 0.25 | 0.08 | 0.29 | 0.09 | 0.32 | 0.15 |
| 11 | 7354174 | rs973108 | -1.087 | 1.187 | 1.192 | -1.314 | 0.002 | 0.03 | 0.03 | 0 | 0.45 | 0.34 | 0.51 | 0.1 | 0.55 | 0.35 |
| 11 | 7355217 | rs1843984 | -0.435 | 1.054 | 0.647 | -1.21 | 0.009 | -0.046 | -0.046 | 0 | 0.33 | 0.5 | 0.37 | 0.17 | 0.38 | 0.18 |
| 11 | 7361787 | rs10430870 | 1.791 | 0.78 | -0.246 | -1.086 | 0.021 | -0.144 | -0.144 | 0 | 0.37 | 0.05 | 0.37 | 0.57 | 0.25 | 0.47 |

| | | | | | | | | | | | | | | | | |
|---|---|---|---|---|---|---|---|---|---|---|---|---|---|---|---|---|
| 11 | 7366506 | rs4758175 | -0.47 | 1.196 | 0.069 | -1.449 | -0.029 | 0.047 | 0.047 | 0 | 0.22 | 0.13 | 0.21 | 0.06 | 0.27 | 0.23 |
| 11 | 7367095 | rs10769780 | -0.47 | 1.196 | 0.069 | -1.449 | -0.029 | 0.047 | 0.047 | 0 | 0.22 | 0.13 | 0.21 | 0.07 | 0.27 | 0.23 |
| 11 | 7368862 | rs6578849 | 1.18 | 1.1 | 0.593 | -1.33 | -0.011 | 0.017 | 0.017 | 0 | 0.65 | 0.29 | 0.7 | 0.72 | 0.62 | 0.74 |
| 11 | 7369875 | rs4525211 | -1.336 | 0.977 | 0.816 | -1.405 | 0.002 | -0.009 | -0.009 | 0 | 0.51 | 0.48 | 0.56 | 0.18 | 0.66 | 0.49 |
| 11 | 7371849 | rs1487880 | 0.032 | 1.405 | 0.456 | -1.351 | -0.016 | 0.018 | 0.018 | 0 | 0.26 | 0.21 | 0.33 | 0.1 | 0.38 | 0.23 |
| 11 | 7372799 | rs6578851 | -0.849 | 0.827 | 0.556 | -1.143 | 0.025 | 0.038 | 0.038 | 0 | 0.92 | 0.81 | 0.95 | 0.83 | 0.89 | 0.97 |
| 11 | 7374594 | rs7933960 | -1.045 | 0.451 | 0.301 | -1.245 | 0.021 | 0.025 | 0.025 | 0.038 | 0.49 | 0.44 | 0.55 | 0.17 | 0.66 | 0.48 |
| 11 | 7375914 | rs10500696 | 0.563 | 1.014 | -0.135 | -1.155 | -0.012 | 0.023 | 0.023 | 0 | 0.09 | 0.03 | 0.07 | 0 | 0.1 | 0 |
| 11 | 7378221 | rs12800586 | -0.055 | 1.237 | -0.19 | -1.246 | -0.006 | 0.048 | 0.048 | 1.25 | 0.18 | 0.08 | 0.16 | 0.04 | 0.16 | 0.25 |
| 11 | 7384411 | rs977749 | 0.501 | 1.63 | -0.118 | -1.102 | -0.027 | 0.003 | 0.003 | 0.035 | 0.18 | 0.12 | 0.3 | 0.17 | 0.35 | 0.25 |
| 11 | 7385448 | rs6578855 | -0.685 | 1.442 | 0.733 | -1.28 | 0.043 | -0.048 | -0.048 | 0 | 0.3 | 0.46 | 0.39 | 0.17 | 0.46 | 0.26 |
| 11 | 7387401 | rs7937079 | 1.488 | 1.271 | 0.548 | -1.072 | 0.069 | 0.003 | 0.003 | 0 | 0.43 | 0.21 | 0.39 | 0.56 | 0.3 | 0.46 |
| 11 | 7388171 | rs7107520 | 0.31 | 1.359 | 0.446 | -1.039 | 0.076 | -0.013 | -0.013 | 0 | 0.19 | 0.4 | 0.3 | 0.17 | 0.35 | 0.25 |
| 11 | 7397799 | rs2346824 | 1.552 | 0.869 | -0.277 | -1.036 | 0.033 | -0.084 | -0.084 | 0 | 0.41 | 0.04 | 0.37 | 0.56 | 0.29 | 0.46 |
| 11 | 7398961 | rs7934287 | -0.963 | 0.877 | 0.492 | -1.204 | -0.003 | -0.122 | -0.122 | 0.056 | 0.31 | 0.44 | 0.28 | 0.04 | 0.27 | 0.25 |
| 11 | 7400623 | rs7933282 | 0.797 | 0.679 | 0.252 | -1.009 | 0.001 | 0.087 | 0.087 | 1.876 | 0.53 | 0.27 | 0.53 | 0.81 | 0.44 | 0.54 |
| 11 | 7400684 | rs10839774 | -0.063 | 0.936 | 0.177 | -1.189 | -0.016 | 0.031 | 0.031 | 2.428 | 0.27 | 0.23 | 0.2 | 0.05 | 0.38 | 0.23 |
| 11 | 7400790 | rs1023996 | 1.067 | 0.817 | -0.099 | -1.044 | -0.014 | 0.199 | 0.199 | 2.854 | 0.51 | 0.13 | 0.52 | 0.81 | 0.44 | 0.54 |
| 11 | 7402370 | rs16925527 | 0.174 | 0.559 | 0.278 | -0.614 | 0 | 0.07 | 0.07 | 0 | 0.13 | 0.28 | 0.06 | 0.27 | 0.1 | 0.1 |
| 11 | 7402760 | rs12361175 | 0.368 | 0.69 | -0.026 | -0.952 | -0.064 | 0.1 | 0.1 | 1.081 | 0.26 | 0.1 | 0.25 | 0.05 | 0.39 | 0.21 |
| 11 | 7403546 | rs11041339 | -0.298 | 0.64 | -0.118 | -1.043 | -0.001 | 0 | 0 | 0 | 0.22 | 0.17 | 0.13 | 0.26 | 0.2 | 0.12 |
| 11 | 7404901 | rs16925591 | 1.632 | 0.608 | -0.148 | -0.983 | 0.029 | -0.02 | -0.02 | 0 | 0.41 | 0.11 | 0.42 | 0.56 | 0.33 | 0.47 |
| 11 | 7411652 | rs1320041 | 0.005 | 0.282 | 0.514 | -0.59 | 0.006 | 0.041 | 0.041 | 0.01 | 0.91 | 0.75 | 0.78 | 0.86 | 0.93 | 0.77 |
| 11 | 7415972 | rs2346828 | 1.02 | 0.283 | 0.708 | -0.639 | -0.003 | 0.003 | 0.003 | 0 | 0.68 | 0.38 | 0.64 | 0.61 | 0.72 | 0.68 |
| 11 | 7419377 | rs11041350 | 0.16 | 0.285 | 0.797 | -0.562 | 0.012 | 0.037 | 0.037 | 0 | 0.13 | 0.34 | 0.07 | 0.17 | 0.1 | 0.1 |
| 11 | 7419615 | rs9804600 | 0.465 | 0.374 | 0.152 | -0.599 | -0.001 | 0.001 | 0.001 | 0.448 | 0.81 | 0.71 | 0.7 | 0.76 | 0.82 | 0.76 |
| 11 | 7421269 | rs11041352 | 0.101 | 0.14 | 0.056 | -0.481 | -0.018 | -0.014 | -0.014 | 0.901 | 0.17 | 0.3 | 0.07 | 0.16 | 0.11 | 0.14 |
| 11 | 7421720 | rs4264136 | -0.723 | 0.727 | -0.705 | -0.432 | -0.027 | 0.036 | 0.036 | 0 | 0.33 | 0.79 | 0.25 | 0.27 | 0.39 | 0.22 |
| 11 | 7426594 | rs4397827 | -0.733 | 0.897 | -1.114 | -0.682 | -0.028 | -0.06 | -0.06 | 0.198 | 0.84 | 0.94 | 0.8 | 0.76 | 0.82 | 0.71 |
| 11 | 7431727 | rs11522663 | 0.859 | 1.534 | -0.487 | -0.676 | 0.01 | -0.017 | -0.017 | 0 | 0.14 | 0.18 | 0.14 | 0.04 | 0.3 | 0.06 |
| 11 | 7432446 | rs2346833 | 0.28 | 1.001 | -1.273 | 0.048 | -0.005 | 0.008 | 0.008 | 0 | 0.17 | 0.13 | 0.39 | 0.46 | 0.23 | 0.22 |
| 11 | 7433790 | rs7124004 | -0.423 | 0.8 | -0.491 | -0.421 | 0.001 | -0.002 | -0.002 | 0 | 0.34 | 0.64 | 0.26 | 0.31 | 0.45 | 0.18 |
| 11 | 7434147 | rs7127462 | -0.636 | 0.679 | -1.307 | -0.881 | -0.025 | 0.092 | 0.092 | 0.021 | 0.37 | 0.8 | 0.26 | 0.21 | 0.42 | 0.17 |
| 11 | 7435295 | rs7938208 | -0.406 | 1.057 | -1.13 | -0.692 | -0.028 | 0.054 | 0.054 | 0.333 | 0.33 | 0.78 | 0.26 | 0.2 | 0.42 | 0.17 |
| 11 | 7435415 | rs7116485 | -0.481 | 0.686 | -0.641 | -0.191 | 0 | -0.023 | -0.023 | 0.001 | 0.1 | 0.1 | 0.08 | 0 | 0.11 | 0.01 |
| 11 | 7437599 | rs7946653 | -0.166 | 0.586 | -0.663 | -0.398 | -0.009 | 0.022 | 0.022 | 0 | 0.42 | 0.35 | 0.33 | 0.11 | 0.45 | 0.32 |
| 11 | 7437611 | rs7946655 | -0.744 | 0.698 | -0.059 | 0.571 | 0.041 | 0.028 | 0.028 | 0 | 0.49 | 0.72 | 0.57 | 0.45 | 0.57 | 0.48 |
| 11 | 7439093 | rs4757993 | 0.523 | 0.7 | -0.773 | 0.622 | -0.026 | 0.084 | 0.084 | 0 | 0.57 | 0.26 | 0.43 | 0.19 | 0.6 | 0.42 |
| 11 | 7440390 | rs11041365 | 1.22 | 0.48 | -0.783 | 0.613 | -0.027 | -0.005 | -0.005 | 0 | 0.11 | 0.02 | 0.11 | 0.11 | 0.13 | 0.11 |
| 11 | 7441272 | rs11041371 | 0.495 | 0.527 | -0.684 | 0.614 | -0.017 | 0.065 | 0.065 | 0 | 0.58 | 0.26 | 0.61 | 0.21 | 0.62 | 0.44 |
| 11 | 7441554 | rs12786812 | 0.527 | 0.542 | -0.596 | 0.878 | 0.002 | 0.016 | 0.016 | 0 | 0.14 | 0.14 | 0.13 | 0.29 | 0.15 | 0.12 |
| 11 | 7441952 | rs11041373 | 0.687 | 0.524 | -0.604 | 1.048 | 0.003 | 0.031 | 0.031 | 0 | 0.13 | 0.13 | 0.13 | 0.32 | 0.16 | 0.13 |
| 11 | 7442670 | rs12362353 | 0.266 | 0.454 | -0.629 | 0.908 | 0.003 | 0.038 | 0.038 | 0 | 0.16 | 0.25 | 0.14 | 0.3 | 0.16 | 0.12 |
| 11 | 7443183 | rs11601238 | 1.131 | 0.229 | -0.9 | 0.593 | -0.076 | 0.17 | 0.17 | 0 | 0.28 | 0.04 | 0.33 | 0.01 | 0.32 | 0.21 |
| 11 | 7444027 | rs1320042 | -0.536 | 0.114 | -0.635 | 0.957 | -0.005 | 0.072 | 0.072 | 0.34 | 0.75 | 0.44 | 0.73 | 0.54 | 0.8 | 0.61 |
| 11 | 7445905 | rs7118182 | -1.157 | 0.225 | -1.433 | 0.607 | -0.02 | -0.018 | -0.018 | 0 | 0.72 | 0.95 | 0.76 | 0.79 | 0.71 | 0.74 |
| 11 | 7446969 | rs10839781 | -0.3 | 0.147 | -1.064 | 1.01 | 0.046 | -0.073 | -0.073 | 0 | 0.3 | 0.82 | 0.3 | 0.52 | 0.18 | 0.32 |
| 11 | 7447857 | rs4757994 | -0.759 | 0.309 | -0.761 | 1.006 | 0.015 | -0.005 | -0.005 | 1.379 | 0.39 | 0.66 | 0.38 | 0.64 | 0.28 | 0.46 |
| 11 | 7448700 | rs7949007 | -0.591 | 0.152 | -0.67 | 0.971 | 0.012 | -0.024 | -0.024 | 0.6 | 0.32 | 0.71 | 0.3 | 0.64 | 0.26 | 0.39 |
| 11 | 7456307 | rs4417248 | 0.226 | 0.253 | -0.143 | 0.808 | 0.006 | -0.004 | -0.004 | 0 | 0.42 | 0.35 | 0.3 | 0.48 | 0.37 | 0.42 |
| 11 | 7456529 | rs4313569 | -0.015 | 0.256 | -0.313 | 0.866 | 0.012 | 0.001 | 0.001 | 0 | 0.48 | 0.63 | 0.36 | 0.48 | 0.47 | 0.43 |
| 11 | 7457741 | rs12575833 | -0.662 | 0.274 | -0.401 | 1.13 | 0.006 | 0.063 | 0.063 | 0 | 0.06 | 0.11 | 0.13 | 0.25 | 0.09 | 0.12 |
| 11 | 7458223 | rs4319495 | -0.296 | 0.259 | -0.674 | 0.563 | -0.019 | 0.076 | 0.076 | 0 | 0.54 | 0.75 | 0.49 | 0.73 | 0.56 | 0.55 |
| 11 | 7460958 | rs4072103 | 0.436 | 0.876 | -0.436 | 0.302 | -0.016 | 0.025 | 0.025 | 0 | 0.17 | 0.11 | 0.27 | 0.02 | 0.18 | 0.19 |
| 11 | 7461253 | rs4072144 | -0.353 | 0.775 | 0.103 | 0.95 | 0.007 | 0.012 | 0.012 | 1.836 | 0.11 | 0.14 | 0.19 | 0.23 | 0.19 | 0.06 |
| 11 | 7463443 | rs12271195 | -0.319 | 0.756 | 0.551 | 0.672 | 0.036 | -0.029 | -0.029 | 0 | 0.05 | 0.1 | 0.04 | 0 | 0.1 | 0.02 |
| 11 | 7464958 | rs10839783 | -0.905 | 0.499 | 1.466 | 0.832 | 0.029 | 0.082 | 0.082 | 0.311 | 0.07 | 0.33 | 0.1 | 0.12 | 0.06 | 0.05 |
| 11 | 7469119 | rs7130698 | -1.341 | -0.103 | 1.242 | 0.566 | 0.016 | 0.033 | 0.033 | 0.355 | 0.34 | 0.75 | 0.26 | 0.24 | 0.29 | 0.23 |
| 11 | 7469359 | rs7111363 | -0.674 | -0.103 | 1.867 | 0.537 | 0.049 | -0.041 | -0.041 | 0.891 | 0.08 | 0.27 | 0.07 | 0 | 0.1 | 0.03 |
| 11 | 7471878 | rs7942332 | -0.915 | -0.117 | 1.942 | 0.534 | 0.044 | -0.036 | -0.036 | 0 | 0.09 | 0.26 | 0.09 | 0.01 | 0.12 | 0.03 |
| 11 | 7472813 | rs7121843 | -1.578 | -0.167 | 1.835 | 0.664 | 0.022 | 0.044 | 0.044 | 0.115 | 0.34 | 0.83 | 0.27 | 0.28 | 0.3 | 0.23 |
| 11 | 7473121 | rs7106254 | -0.962 | -0.179 | 2.341 | 0.532 | -0.011 | -0.082 | -0.082 | 0.478 | 0.28 | 0.48 | 0.2 | 0.14 | 0.24 | 0.17 |
| 11 | 7473153 | rs10839786 | -1.248 | -0.188 | 2.446 | 0.532 | 0.01 | 0.008 | 0.008 | 0.662 | 0.3 | 0.69 | 0.22 | 0.24 | 0.19 | 0.2 |
| 11 | 7473381 | rs11041387 | -0.587 | 0.034 | 2.263 | 0.606 | 0.033 | 0.097 | 0.097 | 0.028 | 0.06 | 0.32 | 0.07 | 0.11 | 0.05 | 0.04 |
| 11 | 7473625 | rs4589277 | -0.977 | 0.133 | 1.783 | 0.55 | 0.046 | -0.036 | -0.036 | 5.406 | 0.09 | 0.27 | 0.09 | 0.01 | 0.12 | 0.03 |
| 11 | 7475440 | rs4758185 | 1.1 | -0.274 | 0.935 | 1.046 | 0.03 | 0.015 | 0.015 | 0 | 0.53 | 0.3 | 0.55 | 0.43 | 0.58 | 0.51 |
| 11 | 7475989 | rs11041390 | -0.945 | -0.303 | 0.972 | 0.73 | -0.019 | -0.048 | -0.048 | 0 | 0.35 | 0.52 | 0.2 | 0.15 | 0.25 | 0.29 |
| 11 | 7476061 | rs11041391 | -0.945 | -0.303 | 0.948 | 0.73 | -0.019 | -0.048 | -0.048 | 0 | 0.35 | 0.52 | 0.2 | 0.15 | 0.25 | 0.29 |
| 11 | 7476534 | rs11041393 | -0.905 | -0.297 | 0.931 | 0.321 | -0.021 | -0.09 | -0.09 | 0 | 0.34 | 0.49 | 0.2 | 0.06 | 0.25 | 0.26 |

| | | | | | | | | | | | | | | | | |
|---|---|---|---|---|---|---|---|---|---|---|---|---|---|---|---|---|
| 11 | 7482226 | rs7107529 | -0.675 | -0.09 | 0.276 | 0.491 | -0.018 | -0.051 | -0.051 | 0 | 0.29 | 0.57 | 0.17 | 0.13 | 0.26 | 0.23 |
| 11 | 7485711 | rs4628658 | 0.45 | 0.615 | -0.268 | 0.919 | -0.004 | -0.002 | -0.002 | 0.275 | 0.62 | 0.64 | 0.49 | 0.62 | 0.55 | 0.52 |
| 11 | 7488341 | rs4344466 | -0.064 | 0.63 | -0.607 | 0.986 | -0.014 | -0.009 | -0.009 | 0 | 0.85 | 0.94 | 0.85 | 0.79 | 0.84 | 0.86 |
| 11 | 7488872 | rs10732509 | 0.543 | 0.555 | -0.509 | 1.008 | 0.003 | 0.01 | 0.01 | 0 | 0.51 | 0.43 | 0.62 | 0.31 | 0.52 | 0.55 |
| 11 | 7488927 | rs10732510 | 0.524 | 0.54 | -0.515 | 1.023 | 0.003 | 0.009 | 0.009 | 0 | 0.51 | 0.44 | 0.63 | 0.31 | 0.52 | 0.55 |
| 11 | 7489440 | rs4408288 | -0.389 | 0.85 | -0.37 | 0.923 | -0.006 | 0.001 | 0.001 | 0 | 0.82 | 0.85 | 0.82 | 0.79 | 0.79 | 0.84 |
| 11 | 7489819 | rs4351800 | 0.105 | 0.675 | 0.337 | 0.725 | -0.005 | 0.01 | 0.01 | 0.394 | 0.55 | 0.62 | 0.65 | 0.31 | 0.57 | 0.57 |
| 11 | 7490548 | rs4073619 | 0.721 | 0.168 | -0.514 | 0.848 | -0.05 | 0.065 | 0.065 | 0.164 | 0.35 | 0.12 | 0.47 | 0.16 | 0.44 | 0.39 |
| 11 | 7491479 | rs11607516 | 0.612 | 0.289 | -0.045 | 0.889 | -0.011 | -0.001 | -0.001 | 0 | 0.18 | 0.22 | 0.15 | 0.15 | 0.11 | 0.19 |
| 11 | 7498835 | rs10769793 | -1.031 | 0.222 | -0.691 | 0.776 | -0.022 | -0.041 | -0.041 | 0.141 | 0.79 | 0.93 | 0.8 | 0.62 | 0.79 | 0.78 |
| 11 | 7499483 | rs4338502 | -1.181 | 0.224 | -0.146 | 1.158 | -0.019 | 0.071 | 0.071 | 0 | 0.51 | 0.85 | 0.47 | 0.61 | 0.43 | 0.53 |
| 11 | 7500103 | rs11821306 | 0.022 | 0.884 | 0.233 | 0.531 | 0 | -0.005 | -0.005 | 0 | 0.21 | 0.19 | 0.15 | 0.12 | 0.2 | 0.16 |
| 11 | 7501472 | rs12792158 | 0.104 | 0.925 | 0.006 | 0.551 | -0.014 | 0.019 | 0.019 | 0 | 0.2 | 0.13 | 0.14 | 0.12 | 0.2 | 0.16 |
| 11 | 7505333 | rs10839791 | 0.223 | 0.378 | 0.192 | 0.609 | -0.006 | 0.018 | 0.018 | 0 | 0.17 | 0.22 | 0.16 | 0.19 | 0.15 | 0.19 |
| 11 | 7505961 | rs10839793 | -0.589 | 0.445 | 0.323 | 0.274 | -0.013 | -0.015 | -0.015 | 0.114 | 0.34 | 0.37 | 0.24 | 0.16 | 0.21 | 0.3 |
| 11 | 7508289 | rs7949140 | -0.227 | 0.549 | 0.252 | -0.063 | -0.007 | -0.048 | -0.048 | 0 | 0.31 | 0.41 | 0.26 | 0.13 | 0.25 | 0.28 |
| 11 | 7508840 | rs11041416 | -0.878 | 1.621 | -0.099 | 0.178 | -0.006 | -0.004 | -0.004 | 0 | 0.12 | 0.13 | 0.23 | 0.16 | 0.24 | 0.17 |
| 11 | 7509682 | rs12803770 | -0.81 | 1.687 | -0.05 | 0.206 | 0.008 | 0.001 | 0.001 | 0 | 0.1 | 0.13 | 0.22 | 0.16 | 0.19 | 0.17 |
| 11 | 7512590 | rs6578868 | -0.357 | 1.497 | 0.007 | -0.354 | 0.003 | 0.018 | 0.018 | 0 | 0.44 | 0.58 | 0.33 | 0.16 | 0.42 | 0.36 |
| 11 | 7520533 | rs3923015 | -0.722 | 1.21 | 0.145 | -0.26 | 0.008 | -0.001 | -0.001 | 0 | 0.09 | 0.2 | 0.17 | 0.08 | 0.1 | 0.15 |
| 11 | 7523550 | rs10769795 | -0.951 | 1.14 | 0.203 | -0.504 | 0.001 | 0.003 | 0.003 | 0.019 | 0.55 | 0.63 | 0.52 | 0.34 | 0.61 | 0.52 |
| 11 | 7532175 | rs3884596 | -0.386 | 1.3 | 0.053 | -0.665 | -0.001 | -0.001 | -0.001 | 0.016 | 0.48 | 0.46 | 0.36 | 0.26 | 0.51 | 0.35 |
| 11 | 7533228 | rs4370912 | -0.325 | 1.282 | 0.059 | -0.628 | 0 | -0.009 | -0.009 | 0.242 | 0.47 | 0.51 | 0.36 | 0.24 | 0.51 | 0.34 |
| 11 | 7539678 | rs7102169 | -0.157 | 1.255 | 0.098 | -0.733 | 0 | -0.027 | -0.027 | 0 | 0.44 | 0.48 | 0.34 | 0.2 | 0.45 | 0.33 |
| 11 | 7542298 | rs4078258 | -0.955 | 1.307 | -0.256 | -0.736 | 0.018 | 0.072 | 0.072 | 0 | 0.58 | 0.9 | 0.58 | 0.27 | 0.61 | 0.52 |
| 11 | 7546402 | rs10839797 | -0.878 | 1.348 | -0.101 | -0.802 | 0.004 | 0.034 | 0.034 | 0 | 0.56 | 0.7 | 0.54 | 0.3 | 0.57 | 0.51 |
| 11 | 7555066 | rs4322372 | -0.174 | 1.094 | -0.256 | -0.756 | -0.009 | 0.026 | 0.026 | 0 | 0.42 | 0.67 | 0.35 | 0.21 | 0.46 | 0.32 |
| 11 | 7556057 | rs4757999 | -0.107 | 1.109 | -0.167 | -0.771 | 0.011 | -0.028 | -0.028 | 0.065 | 0.39 | 0.5 | 0.34 | 0.2 | 0.44 | 0.31 |
| 11 | 7556577 | rs12791447 | 0.408 | 0.71 | -0.683 | -0.217 | -0.006 | -0.021 | -0.021 | 0.677 | 0.08 | 0.01 | 0.2 | 0.12 | 0.06 | 0.16 |
| 11 | 7559930 | rs4758001 | -1.136 | 0.485 | -0.966 | -0.516 | 0.012 | -0.059 | -0.059 | 0 | 0.72 | 0.92 | 0.85 | 0.38 | 0.72 | 0.64 |
| 11 | 7561198 | rs6578869 | 0.258 | 0.387 | -0.34 | -0.665 | -0.014 | -0.045 | -0.045 | 0 | 0.24 | 0.43 | 0.16 | 0.11 | 0.24 | 0.22 |
| 11 | 7563487 | rs12788646 | -0.966 | 0.336 | -0.465 | 0.279 | -0.024 | 0.08 | 0.08 | 0.036 | 0.22 | 0.42 | 0.39 | 0.38 | 0.17 | 0.24 |
| 11 | 7563642 | rs7481343 | -0.752 | 0.268 | -1.221 | 0.209 | -0.018 | -0.126 | -0.126 | 0.569 | 0.74 | 0.99 | 0.7 | 0.41 | 0.76 | 0.65 |
| 11 | 7566887 | rs4076808 | -0.423 | 0.667 | -1.248 | -0.112 | -0.042 | 0.115 | 0.115 | 0 | 0.5 | 0.87 | 0.28 | 0.29 | 0.48 | 0.4 |
| 11 | 7567875 | rs10839803 | -0.348 | 0.731 | -0.784 | -0.212 | 0.021 | -0.066 | -0.066 | 1.944 | 0.23 | 0.52 | 0.14 | 0.12 | 0.21 | 0.2 |
| 11 | 7568001 | rs7105146 | 1.088 | 0.753 | -0.702 | -0.152 | -0.004 | -0.014 | -0.014 | 1.999 | 0.26 | 0.32 | 0.13 | 0.17 | 0.24 | 0.21 |
| 11 | 7570607 | rs7122137 | 0.047 | 0.818 | -0.717 | -0.201 | -0.011 | -0.019 | -0.019 | 0.738 | 0.38 | 0.44 | 0.17 | 0.25 | 0.33 | 0.37 |
| 11 | 7571426 | rs11041441 | 1.216 | 0.919 | -0.759 | -0.129 | -0.009 | -0.006 | -0.006 | 0 | 0.12 | 0 | 0.03 | 0.08 | 0.08 | 0.16 |
| 11 | 7576318 | rs7479626 | 0.533 | 0.757 | -0.43 | -0.141 | -0.003 | -0.034 | -0.034 | 3.969 | 0.29 | 0.5 | 0.15 | 0.16 | 0.26 | 0.19 |
| 11 | 7576350 | rs7481625 | 1.03 | 0.766 | -0.615 | -0.121 | -0.002 | -0.02 | -0.02 | 0 | 0.24 | 0.38 | 0.12 | 0.16 | 0.22 | 0.18 |
| 11 | 7578417 | rs7112087 | -0.392 | 1.018 | -1.347 | 0.171 | -0.028 | 0.023 | 0.023 | 0 | 0.48 | 0.81 | 0.28 | 0.38 | 0.46 | 0.38 |
| 11 | 7581032 | rs7478756 | -0.368 | 1.025 | -1.343 | 0.175 | -0.03 | 0.021 | 0.021 | 0 | 0.48 | 0.81 | 0.28 | 0.38 | 0.46 | 0.38 |
| 11 | 7583923 | rs7129193 | 0.444 | 1.342 | -0.964 | 0.383 | 0.004 | -0.003 | -0.003 | 0 | 0.15 | 0.16 | 0.09 | 0.13 | 0.19 | 0.17 |
| 11 | 7590356 | rs11041459 | 0.444 | 1.342 | -0.988 | 0.378 | 0.002 | -0.001 | -0.001 | 0.058 | 0.15 | 0.16 | 0.09 | 0.14 | 0.19 | 0.17 |
| 11 | 7593036 | rs6578875 | 0.411 | 1.016 | -0.846 | 0.292 | -0.003 | -0.003 | -0.003 | 0 | 0.33 | 0.58 | 0.17 | 0.25 | 0.26 | 0.22 |
| 11 | 7597280 | rs7131430 | -0.275 | 0.932 | -1.29 | 0.524 | -0.045 | -0.054 | -0.054 | 3.163 | 0.28 | 0.07 | 0.54 | 0.31 | 0.29 | 0.33 |
| 11 | 7599391 | rs6578882 | 0.03 | 0.852 | 0.067 | 0.448 | -0.001 | 0.001 | 0.001 | 0 | 0.42 | 0.36 | 0.28 | 0.29 | 0.4 | 0.36 |
| 11 | 7599454 | rs6578883 | -0.08 | 0.852 | 0.2 | 0.448 | 0 | -0.016 | -0.016 | 0.077 | 0.44 | 0.46 | 0.29 | 0.29 | 0.4 | 0.37 |
| 11 | 7603479 | rs12098959 | -0.671 | 0.636 | 0.26 | 0.621 | -0.007 | 0.035 | 0.035 | 0 | 0.16 | 0.25 | 0.24 | 0.27 | 0.13 | 0.22 |
| 11 | 7604023 | rs6578885 | -1.033 | 0.683 | 0.326 | 0.582 | -0.03 | -0.055 | -0.055 | 0 | 0.61 | 0.84 | 0.54 | 0.56 | 0.55 | 0.58 |
| 11 | 7605737 | rs11041463 | 0.901 | 0.774 | 0.369 | 0.486 | 0 | -0.002 | -0.002 | 0 | 0.32 | 0.31 | 0.24 | 0.25 | 0.3 | 0.21 |
| 11 | 7606125 | rs4758201 | 0.057 | 0.737 | -0.252 | 0.281 | -0.007 | -0.055 | -0.055 | 0 | 0.12 | 0.01 | 0.19 | 0.15 | 0.08 | 0.14 |
| 11 | 7607536 | rs7933428 | 0.434 | 0.866 | -0.06 | 0.342 | -0.001 | -0.004 | -0.004 | 0 | 0.11 | 0.11 | 0.06 | 0.05 | 0.12 | 0.13 |
| 11 | 7613349 | rs4472926 | -1.219 | 0.889 | -0.388 | 0.235 | -0.002 | 0.004 | 0.004 | 0.769 | 0.33 | 0.3 | 0.33 | 0.31 | 0.31 | 0.39 |
| 11 | 7615485 | rs11041469 | 0.635 | 0.752 | -0.221 | 0.573 | 0 | -0.036 | -0.036 | 0 | 0.34 | 0.21 | 0.31 | 0.46 | 0.38 | 0.37 |
| 11 | 7616191 | rs9787734 | -1.58 | 0.456 | 0.286 | 0.865 | 0.002 | -0.002 | -0.002 | 0 | 0.22 | 0.2 | 0.27 | 0.25 | 0.2 | 0.25 |
| 11 | 7617899 | rs4758003 | -1.455 | 0.369 | 0.317 | 0.899 | 0.005 | -0.003 | -0.003 | 0 | 0.22 | 0.2 | 0.27 | 0.25 | 0.17 | 0.24 |
| 11 | 7619262 | rs4758205 | 1.212 | 0.782 | 1.005 | 1.17 | 0.032 | 0.099 | 0.099 | 0 | 0.8 | 0.46 | 0.75 | 0.64 | 0.71 | 0.81 |
| 11 | 7619518 | rs4758206 | 0.924 | 0.132 | 1.47 | 0.185 | 0.005 | -0.041 | -0.041 | 10.068 | 0.88 | 0.64 | 0.92 | 0.96 | 0.91 | 0.94 |
| 11 | 7621867 | rs11041474 | -1.27 | 0.143 | 1.563 | 0.289 | 0.021 | 0.153 | 0.153 | 0 | 0.05 | 0.21 | 0.15 | 0.15 | 0.04 | 0.04 |
| 11 | 7626251 | rs12803437 | -0.846 | 0.365 | 1.509 | 0.239 | 0.037 | -0.052 | -0.052 | 0 | 0.1 | 0.28 | 0.08 | 0.01 | 0.11 | 0.04 |
| 11 | 7627786 | rs3892335 | -0.876 | -0.24 | 0.891 | -0.454 | 0.068 | 0.135 | 0.135 | 0 | 0.5 | 0.71 | 0.37 | 0.17 | 0.32 | 0.37 |
| 11 | 7628780 | rs12099341 | -1.192 | 0.597 | 1.146 | 0.08 | 0.059 | -0.052 | -0.052 | 0 | 0.1 | 0.42 | 0.09 | 0.01 | 0.11 | 0.04 |
| 11 | 7635723 | rs7129654 | -0.854 | 0.747 | 0.307 | 0.153 | 0.028 | -0.046 | -0.046 | 1.179 | 0.09 | 0.22 | 0.08 | 0.01 | 0.11 | 0.04 |
| 11 | 7641875 | rs7118058 | 0.634 | -0.644 | 0.794 | 0.704 | -0.062 | 0.058 | 0.058 | 0 | 0.8 | 0.61 | 0.81 | 0.71 | 0.91 | 0.75 |
| 11 | 7642031 | rs10769813 | -1.112 | -0.476 | 0.183 | -0.046 | 0.003 | 0.05 | 0.05 | 0 | 0.54 | 0.66 | 0.5 | 0.88 | 0.46 | 0.49 |
| 11 | 7646734 | rs10839825 | 0.78 | -0.273 | -0.027 | -0.82 | -0.033 | 0.06 | 0.06 | 0 | 0.35 | 0.22 | 0.45 | 0.1 | 0.5 | 0.42 |
| 11 | 7651309 | rs12576185 | 0.108 | -0.625 | -0.314 | -0.728 | -0.002 | -0.005 | -0.005 | 0 | 0.17 | 0.14 | 0.09 | 0.5 | 0.21 | 0.15 |
| 11 | 7653090 | rs12708352 | 0.208 | -0.878 | -0.307 | -0.91 | -0.048 | 0.105 | 0.105 | 0 | 0.27 | 0.1 | 0.26 | 0.05 | 0.26 | 0.11 |

| | | | | | | | | | | | | | | | | |
|---|---|---|---|---|---|---|---|---|---|---|---|---|---|---|---|---|
| 11 | 7653567 | rs7113054 | 0.013 | -1.049 | -0.143 | -0.736 | -0.001 | -0.001 | -0.001 | 0.109 | 0.18 | 0.16 | 0.1 | 0.59 | 0.21 | 0.16 |
| 11 | 7653867 | rs11041489 | -0.475 | -1.462 | -0.213 | -1.24 | -0.052 | -0.08 | -0.08 | 0.022 | 0.17 | 0.35 | 0.12 | 0.04 | 0.05 | 0.17 |
| 11 | 7657259 | rs4372441 | 1.005 | -1.528 | -0.171 | -0.635 | 0.003 | -0.007 | -0.007 | 10.722 | 0.67 | 0.36 | 0.78 | 0.46 | 0.75 | 0.74 |
| 11 | 7659293 | rs7119574 | 0.153 | -1.481 | -0.116 | -0.655 | 0.006 | -0.019 | -0.019 | 0 | 0.07 | 0.28 | 0.06 | 0.01 | 0.04 | 0.07 |
| 11 | 7663726 | rs4402283 | 0.276 | -1.618 | -0.032 | -0.758 | -0.033 | -0.123 | -0.123 | 0 | 0.8 | 0.49 | 0.88 | 0.97 | 0.89 | 0.87 |
| 11 | 7663779 | rs3861066 | -0.16 | -1.639 | -0.02 | -0.558 | -0.009 | 0.094 | 0.094 | 0.095 | 0.17 | 0.28 | 0.13 | 0.46 | 0.14 | 0.14 |
| 11 | 7663887 | rs4525226 | -0.662 | -1.799 | -0.251 | -0.705 | 0.001 | -0.004 | -0.004 | 0.226 | 0.51 | 0.55 | 0.48 | 0.85 | 0.43 | 0.48 |
| 11 | 7664432 | rs4130499 | 0.575 | -1.748 | -0.168 | -0.383 | 0.008 | -0.008 | -0.008 | 0.551 | 0.12 | 0.37 | 0.08 | 0.1 | 0.07 | 0.09 |
| 11 | 7667790 | rs7117146 | -0.67 | -1.697 | -0.259 | -0.428 | 0.002 | -0.033 | -0.033 | 0 | 0.66 | 0.81 | 0.65 | 0.64 | 0.7 | 0.65 |
| 11 | 7668524 | rs10839830 | 2.173 | -1.688 | -0.339 | -0.465 | -0.146 | 0.064 | 0.064 | 0.51 | 0.28 | 0.02 | 0.41 | 0.09 | 0.45 | 0.38 |
| 11 | 7668861 | rs6578890 | 0.508 | -1.719 | -0.244 | -0.475 | 0.001 | 0.009 | 0.009 | 0 | 0.47 | 0.43 | 0.52 | 0.12 | 0.57 | 0.51 |
| 11 | 7670160 | rs4421737 | -0.048 | -1.705 | -0.162 | -0.408 | -0.007 | 0.069 | 0.069 | 0.071 | 0.17 | 0.22 | 0.12 | 0.52 | 0.14 | 0.15 |
| 11 | 7670353 | rs4758009 | 0.661 | -1.698 | -0.154 | -0.433 | -0.009 | -0.001 | -0.001 | 1.189 | 0.45 | 0.23 | 0.54 | 0.61 | 0.59 | 0.53 |
| 11 | 7670752 | rs11607941 | -0.779 | -1.754 | -0.195 | -0.429 | -0.024 | -0.032 | -0.032 | 0 | 0.25 | 0.43 | 0.15 | 0.14 | 0.2 | 0.21 |
| 11 | 7673015 | rs5864 | -0.435 | -1.62 | -0.431 | -0.36 | 0.002 | -0.018 | -0.018 | 0 | 0.63 | 0.75 | 0.61 | 0.6 | 0.64 | 0.66 |
| 11 | 7676385 | rs10839833 | 0.186 | -1.554 | -0.581 | -0.413 | 0.033 | 0.076 | 0.076 | 0 | 0.5 | 0.31 | 0.52 | 0.24 | 0.62 | 0.56 |
| 11 | 7685465 | rs12793971 | -0.734 | -1.373 | -0.709 | -0.534 | 0 | -0.001 | -0.001 | 0.003 | 0.25 | 0.27 | 0.16 | 0.12 | 0.25 | 0.17 |
| 11 | 7686395 | rs1128643 | 2.054 | -1.382 | -1.017 | -0.548 | -0.098 | 0.023 | 0.023 | 0.313 | 0.27 | 0.04 | 0.37 | 0.12 | 0.38 | 0.39 |
| 11 | 7686596 | rs13172 | -1.231 | -1.288 | -0.722 | -0.336 | 0 | -0.001 | -0.001 | 0 | 0.73 | 0.84 | 0.67 | 0.77 | 0.76 | 0.75 |
| 11 | 7686674 | rs13854 | -1.255 | -1.283 | -0.727 | -0.335 | -0.001 | 0.001 | 0.001 | 0.111 | 0.73 | 0.85 | 0.67 | 0.77 | 0.76 | 0.75 |
| 11 | 7687715 | rs12801394 | -1.29 | -1.298 | -0.735 | -0.285 | -0.002 | 0.004 | 0.004 | 0.279 | 0.74 | 0.85 | 0.68 | 0.82 | 0.76 | 0.77 |
| 11 | 7692194 | rs7931064 | -0.309 | -1.373 | -0.507 | -0.301 | -0.01 | 0 | 0 | 0 | 0.19 | 0.47 | 0.13 | 0.13 | 0.15 | 0.17 |
| 11 | 7694762 | rs945888 | -0.864 | -1.591 | -0.79 | -0.31 | -0.001 | 0.024 | 0.024 | 0.537 | 0.68 | 0.79 | 0.64 | 0.81 | 0.7 | 0.68 |
| 11 | 7696338 | rs12292013 | -1.014 | -1.502 | -0.565 | -0.178 | -0.025 | -0.006 | -0.006 | 0 | 0.23 | 0.63 | 0.32 | 0.19 | 0.18 | 0.16 |
| 11 | 7700373 | rs11041529 | 0.518 | -1.234 | -0.798 | 0.094 | -0.029 | -0.027 | -0.027 | 0 | 0.1 | 0.04 | 0.06 | 0.45 | 0.14 | 0.07 |
| 11 | 7702063 | rs4758212 | -0.545 | -1.976 | -0.558 | -0.204 | -0.012 | -0.027 | -0.027 | 0 | 0.18 | 0.25 | 0.26 | 0.05 | 0.11 | 0.12 |
| 11 | 7707101 | rs12271537 | -0.698 | -2.042 | -0.592 | -0.169 | -0.033 | -0.007 | -0.007 | 0.051 | 0.22 | 0.56 | 0.19 | 0.19 | 0.17 | 0.13 |
| 11 | 7707958 | rs7113286 | -1.79 | -2.113 | -0.647 | -0.217 | 0.079 | 0.079 | 0.079 | 0.191 | 0.41 | 0.81 | 0.44 | 0.24 | 0.28 | 0.24 |
| 11 | 7708281 | rs7932813 | -0.938 | -1.885 | -0.444 | 0.132 | -0.033 | -0.002 | -0.002 | 0 | 0.22 | 0.56 | 0.19 | 0.19 | 0.17 | 0.13 |
| 11 | 7708857 | rs12792768 | -0.24 | -1.738 | -0.731 | 0.217 | -0.003 | 0.001 | 0.001 | 0 | 0.15 | 0.17 | 0.15 | 0.17 | 0.14 | 0.07 |
| 11 | 7709450 | rs11041534 | -1.177 | -1.348 | -0.909 | 0.383 | -0.019 | -0.068 | -0.068 | 0.06 | 0.16 | 0.07 | 0.14 | 0.44 | 0.15 | 0.13 |
| 11 | 7709769 | rs10734622 | -0.23 | -0.909 | -0.521 | -0.298 | -0.008 | -0.013 | -0.013 | 0.012 | 0.58 | 0.48 | 0.72 | 0.74 | 0.71 | 0.57 |
| 11 | 7711924 | rs10839843 | 0.866 | -1.245 | -0.456 | -0.125 | 0.044 | 0.017 | 0.017 | 0.006 | 0.44 | 0.17 | 0.57 | 0.59 | 0.62 | 0.5 |
| 11 | 7711956 | rs11820292 | -0.25 | -1.271 | -0.433 | -0.135 | -0.001 | 0.01 | 0.01 | 0 | 0.12 | 0.22 | 0.13 | 0.15 | 0.12 | 0.05 |
| 11 | 7712471 | rs4509745 | 0.779 | -1.151 | -0.465 | -0.069 | 0.044 | 0.024 | 0.024 | 0 | 0.45 | 0.23 | 0.58 | 0.59 | 0.62 | 0.49 |
| 11 | 7712661 | rs4520590 | -0.975 | -0.511 | -0.352 | 0.176 | -0.013 | -0.009 | -0.009 | 0.027 | 0.19 | 0.28 | 0.14 | 0.16 | 0.14 | 0.1 |
| 11 | 7714238 | rs10839846 | -0.763 | -0.456 | -0.364 | 0.01 | -0.051 | -0.105 | -0.105 | 1.088 | 0.18 | 0.48 | 0.15 | 0.03 | 0.1 | 0.12 |
| 11 | 7714414 | rs10839847 | -0.416 | -0.437 | -1.138 | 0.241 | 0.004 | -0.001 | -0.001 | 0.206 | 0.68 | 0.71 | 0.71 | 0.63 | 0.73 | 0.66 |
| 11 | 7714879 | rs4488184 | 0.255 | 0.931 | -2.017 | 0.53 | -0.107 | -0.063 | -0.063 | 0.34 | 0.33 | 0.08 | 0.36 | 0.29 | 0.39 | 0.49 |
| 11 | 7715079 | rs7117567 | 0.119 | 1.265 | -2.036 | 0.563 | -0.087 | -0.065 | -0.065 | 0 | 0.32 | 0.1 | 0.3 | 0.28 | 0.39 | 0.48 |
| 11 | 7715174 | rs11041537 | -0.469 | 1.237 | -1.431 | 0.931 | 0.007 | 0.037 | 0.037 | 0 | 0.23 | 0.25 | 0.17 | 0.49 | 0.26 | 0.28 |
| 11 | 7715253 | rs12419873 | -0.247 | 1.083 | -1.547 | 0.484 | -0.041 | -0.005 | -0.005 | 0 | 0.34 | 0.14 | 0.25 | 0.3 | 0.36 | 0.51 |
| 11 | 7721743 | rs12279973 | 0.464 | -0.161 | 0.788 | 0.561 | 0.021 | -0.035 | -0.035 | 0.067 | 0.65 | 0.26 | 0.65 | 0.46 | 0.69 | 0.69 |
| 11 | 7721870 | rs10839849 | 0.131 | -0.311 | 1.354 | 0.425 | -0.04 | -0.07 | -0.07 | 1.142 | 0.88 | 0.57 | 0.91 | 1 | 0.95 | 0.91 |
| 11 | 7725092 | rs3924747 | 0.112 | -0.209 | 0.814 | -0.1 | -0.108 | 0.115 | 0.115 | 1.366 | 0.24 | 0.03 | 0.13 | 0.05 | 0.31 | 0.08 |
| 11 | 7725173 | rs3924748 | -0.506 | -0.202 | 1.018 | -0.089 | -0.052 | 0.162 | 0.162 | 0.824 | 0.33 | 0.1 | 0.17 | 0.05 | 0.36 | 0.17 |
| 11 | 7727607 | rs10839852 | 0.207 | -0.167 | 1.112 | 0.026 | -0.001 | 0.007 | 0.007 | 0 | 0.22 | 0.23 | 0.25 | 0.53 | 0.25 | 0.2 |
| 11 | 7727706 | rs12364824 | 0.356 | -0.207 | 1.169 | 0.416 | -0.009 | -0.02 | -0.02 | 0.51 | 0.2 | 0.13 | 0.22 | 0.26 | 0.23 | 0.11 |
| 11 | 7727886 | rs7927138 | 1.611 | -0.306 | 1.052 | 0.318 | -0.011 | -0.027 | -0.027 | 0 | 0.39 | 0.21 | 0.51 | 0.42 | 0.37 | 0.61 |
| 11 | 7746352 | rs1459826 | 1.368 | -0.395 | 2.028 | 0.894 | 0.032 | -0.006 | -0.006 | 0 | 0.59 | 0.25 | 0.62 | 0.42 | 0.65 | 0.59 |
| 11 | 7747954 | rs12289723 | 1.36 | -0.349 | 2.021 | 0.931 | 0.036 | -0.008 | -0.008 | 0 | 0.59 | 0.22 | 0.62 | 0.42 | 0.65 | 0.59 |
| 11 | 7748532 | rs1384507 | 0.223 | -0.19 | 2.01 | 1.025 | -0.002 | 0.039 | 0.039 | 0 | 0.76 | 0.66 | 0.73 | 0.48 | 0.75 | 0.79 |
| 11 | 7750622 | rs4237761 | -0.266 | -0.177 | 1.801 | 1.087 | 0 | 0.001 | 0.001 | 0 | 0.22 | 0.26 | 0.25 | 0.26 | 0.23 | 0.11 |
| 11 | 7765884 | rs11821046 | -1.091 | -0.225 | 2.076 | 1.045 | 0.046 | 0.069 | 0.069 | 0 | 0.07 | 0.37 | 0.06 | 0.07 | 0.05 | 0.1 |
| 11 | 7775321 | rs7129903 | -1.405 | -0.34 | 1.938 | 1.045 | -0.056 | 0.004 | 0.004 | 0 | 0.18 | 0.48 | 0.1 | 0.19 | 0.1 | 0.26 |
| 11 | 7775909 | rs1384505 | 0.511 | 0.132 | 1.952 | 1.112 | -0.003 | 0.014 | 0.014 | 0.236 | 0.74 | 0.67 | 0.73 | 0.59 | 0.76 | 0.82 |
| 11 | 7780767 | rs4758233 | 0.321 | 0.131 | 1.984 | 1.111 | -0.001 | 0.013 | 0.013 | 0 | 0.76 | 0.7 | 0.73 | 0.59 | 0.76 | 0.82 |
| 11 | 7782422 | rs7939444 | 0.515 | 0.281 | 2.148 | 0.767 | -0.019 | 0.058 | 0.058 | 0.338 | 0.76 | 0.54 | 0.76 | 0.62 | 0.78 | 0.83 |
| 11 | 7784484 | rs7948490 | -0.897 | 0.291 | 2.295 | 0.732 | -0.047 | -0.016 | -0.016 | 0.337 | 0.14 | 0.39 | 0.08 | 0.11 | 0.06 | 0.08 |
| 11 | 7795895 | rs7110160 | 1.576 | 0.354 | 1.967 | 0.715 | 0.051 | -0.057 | -0.057 | 0.045 | 0.61 | 0.15 | 0.67 | 0.48 | 0.7 | 0.63 |
| 11 | 7799392 | rs7107954 | -0.533 | 0.568 | 2.279 | 0.679 | 0.004 | -0.014 | -0.014 | 2.694 | 0.22 | 0.29 | 0.28 | 0.12 | 0.27 | 0.11 |
| 11 | 7801547 | rs2063424 | -1.805 | 0.531 | 2.388 | 0.684 | 0.008 | 0.033 | 0.033 | 0 | 0.38 | 0.73 | 0.39 | 0.24 | 0.37 | 0.27 |
| 11 | 7801926 | rs12288289 | -0.594 | 0.598 | 2.561 | 0.783 | 0.02 | 0.063 | 0.063 | 0 | 0.05 | 0.23 | 0.05 | 0.11 | 0.03 | 0.09 |
| 11 | 7838441 | rs11500784 | -0.766 | 0.6 | 2.533 | 0.784 | 0.023 | 0.069 | 0.069 | 0 | 0.05 | 0.25 | 0.05 | 0.11 | 0.03 | 0.09 |
| 11 | 7848385 | rs735976 | 1.571 | 0.571 | 2.213 | 0.793 | 0.002 | -0.097 | -0.097 | 0 | 0.6 | 0.06 | 0.59 | 0.48 | 0.63 | 0.62 |
| 11 | 7849162 | rs2361056 | 1.571 | 0.571 | 2.213 | 0.793 | 0.002 | -0.097 | -0.097 | 1.188 | 0.6 | 0.06 | 0.59 | 0.48 | 0.63 | 0.62 |
| 11 | 7850345 | rs1482795 | 1.211 | 0.542 | 2.492 | 0.781 | -0.044 | 0.069 | 0.069 | 0 | 0.82 | 0.35 | 0.87 | 0.59 | 0.9 | 0.73 |
| 11 | 7850753 | rs1482796 | 0.999 | 0.597 | 2.544 | 0.824 | 0 | -0.011 | -0.011 | 0 | 0.7 | 0.22 | 0.63 | 0.49 | 0.67 | 0.68 |
| 11 | 7854578 | rs1828459 | -0.474 | 0.649 | 2.146 | 0.746 | 0.004 | -0.011 | -0.011 | 0.004 | 0.21 | 0.26 | 0.28 | 0.12 | 0.27 | 0.11 |

| | | | | | | | | | | | | | | | | |
|---|---|---|---|---|---|---|---|---|---|---|---|---|---|---|---|---|
| 11 | 7859075 | rs1037815 | 1.805 | 1.002 | 1.084 | 0.514 | -0.001 | -0.097 | -0.097 | 0.745 | 0.44 | 0.03 | 0.5 | 0.48 | 0.39 | 0.55 |
| 11 | 7870760 | rs17221305 | -0.032 | 0.904 | 1.27 | 0.473 | -0.017 | 0.016 | 0.016 | 0.209 | 0.19 | 0.14 | 0.26 | 0.11 | 0.27 | 0.11 |
| 11 | 7873366 | rs10839886 | 1.766 | 0.956 | 1.016 | 0.418 | 0.011 | -0.088 | -0.088 | 0 | 0.44 | 0.04 | 0.5 | 0.5 | 0.38 | 0.56 |
| 11 | 7878493 | rs962638 | 1.087 | 1.073 | 1.608 | 0.509 | -0.001 | -0.001 | -0.001 | 0 | 0.66 | 0.2 | 0.72 | 0.61 | 0.66 | 0.6 |
| 11 | 7879176 | rs7112002 | 0.295 | 1.148 | 2.009 | 0.508 | 0.027 | -0.03 | -0.03 | 0 | 0.77 | 0.68 | 0.85 | 0.88 | 0.7 | 0.91 |
| 11 | 7879804 | rs720892 | 0.258 | 0.883 | 1.956 | 0.527 | 0.01 | -0.005 | -0.005 | 6.818 | 0.15 | 0.21 | 0.23 | 0.12 | 0.2 | 0.2 |
| 11 | 7882544 | rs16933056 | 0.027 | 0.889 | 2.104 | 0.519 | 0.025 | -0.006 | -0.006 | 0 | 0.17 | 0.32 | 0.23 | 0.13 | 0.2 | 0.2 |
| 11 | 7890697 | rs7950307 | 0.57 | 0.959 | 2.232 | 0.478 | 0.064 | -0.072 | -0.072 | 0 | 0.75 | 0.52 | 0.83 | 0.88 | 0.69 | 0.91 |
| 11 | 7891693 | rs931238 | 0.037 | 0.899 | 1.803 | 0.449 | 0.002 | -0.013 | -0.013 | 0 | 0.1 | 0.07 | 0.03 | 0 | 0.06 | 0.07 |
| 11 | 7891959 | rs10839898 | 2.102 | 0.802 | 1.663 | 0.535 | 0.026 | -0.056 | -0.056 | 2.122 | 0.46 | 0.09 | 0.52 | 0.48 | 0.42 | 0.56 |
| 11 | 7892911 | rs4758247 | -0.061 | 0.922 | 1.946 | 0.729 | 0.024 | 0.074 | 0.074 | 0 | 0.82 | 0.6 | 0.74 | 0.72 | 0.79 | 0.76 |
| 11 | 7895876 | rs10500702 | -0.475 | 0.802 | 1.435 | 0.744 | -0.015 | 0.013 | 0.013 | 0 | 0.12 | 0.13 | 0.06 | 0.26 | 0.06 | 0.15 |
| 11 | 7896345 | rs10769843 | 2.15 | 0.772 | 1.214 | 0.643 | -0.007 | -0.076 | -0.076 | 0 | 0.45 | 0.03 | 0.51 | 0.45 | 0.41 | 0.51 |
| 11 | 7897196 | rs6578908 | 0.081 | 1.227 | 1.814 | 0.792 | 0.029 | 0.095 | 0.095 | 0 | 0.82 | 0.49 | 0.73 | 0.71 | 0.8 | 0.75 |
| 11 | 7900472 | rs11604937 | -1.306 | 1.226 | 1.731 | 0.527 | 0.01 | -0.059 | -0.059 | 1.667 | 0.25 | 0.29 | 0.17 | 0.02 | 0.31 | 0.25 |
| 11 | 7900940 | rs10743039 | 0.486 | 1.287 | 1.763 | 0.6 | 0.002 | 0.006 | 0.006 | 0 | 0.75 | 0.74 | 0.74 | 0.86 | 0.77 | 0.83 |
| 11 | 7901368 | rs2361065 | -0.473 | 1.293 | 1.744 | 0.602 | 0.003 | 0 | 0 | 0 | 0.83 | 0.8 | 0.76 | 0.86 | 0.8 | 0.9 |
| 11 | 7901644 | rs16933365 | 0.001 | 1.374 | 1.732 | 0.786 | 0.007 | 0.165 | 0.165 | 0 | 0.05 | 0.18 | 0.06 | 0.26 | 0.04 | 0.09 |
| 11 | 7902813 | rs16933394 | -0.837 | 1.372 | 1.555 | 0.751 | 0.094 | -0.016 | -0.016 | 1.905 | 0.19 | 0.37 | 0.16 | 0.17 | 0.28 | 0.2 |
| 11 | 7904380 | rs7928132 | 1.346 | 1.353 | 0.841 | 0.425 | -0.002 | 0.003 | 0.003 | 0 | 0.58 | 0.59 | 0.62 | 0.85 | 0.5 | 0.77 |
| 11 | 7904993 | rs4758026 | 0.302 | 1.287 | 0.202 | 0.434 | -0.006 | -0.023 | -0.023 | 0 | 0.73 | 0.6 | 0.72 | 0.86 | 0.76 | 0.72 |
| 11 | 7906288 | rs10839906 | -0.314 | 1.473 | -0.316 | 0.438 | -0.011 | 0 | 0 | 0 | 0.15 | 0.1 | 0.12 | 0.01 | 0.27 | 0.06 |
| 11 | 7907474 | rs4758249 | 0.975 | 0.625 | -0.594 | 0.798 | -0.018 | -0.045 | -0.045 | 0 | 0.51 | 0.09 | 0.53 | 0.46 | 0.49 | 0.53 |
| 11 | 7909574 | rs11607118 | -0.88 | 1.754 | 0.112 | 0.395 | 0.074 | -0.047 | -0.047 | 0 | 0.3 | 0.5 | 0.38 | 0.14 | 0.43 | 0.26 |
| 11 | 7910740 | rs11041644 | -1.366 | 2.287 | 0.289 | 0.805 | 0.092 | 0.045 | 0.045 | 0 | 0.06 | 0.18 | 0.21 | 0.14 | 0.17 | 0.15 |
| 11 | 7914606 | rs12788171 | -1.416 | 1.387 | 0.562 | 0.25 | -0.009 | -0.017 | -0.017 | 0.631 | 0.25 | 0.34 | 0.25 | 0.15 | 0.24 | 0.16 |
| 11 | 7922651 | rs10769847 | 0.708 | 1.2 | -0.262 | 0.195 | 0.012 | 0.147 | 0.147 | 0 | 0.51 | 0.18 | 0.46 | 0.76 | 0.55 | 0.51 |
| 11 | 7926432 | rs10769848 | 0.259 | 1.476 | 0.354 | 0.19 | 0.047 | 0.039 | 0.039 | 0.473 | 0.9 | 0.81 | 0.6 | 0.84 | 0.8 | 0.81 |
| 11 | 7930640 | rs10839916 | 0.734 | 1.065 | 0.462 | 0.393 | -0.017 | -0.007 | -0.007 | 0.765 | 0.57 | 0.61 | 0.49 | 0.53 | 0.55 | 0.54 |
| 11 | 7931710 | rs2360839 | 0.656 | 1.033 | 0.408 | 0.477 | 0 | 0.004 | 0.004 | 0.483 | 0.86 | 0.85 | 0.9 | 0.74 | 0.89 | 0.88 |
| 11 | 7933613 | rs12291194 | 0.391 | 1.205 | 0.667 | 0.389 | -0.033 | -0.027 | -0.027 | 0.011 | 0.07 | 0.02 | 0.33 | 0.22 | 0.17 | 0.18 |
| 11 | 7945818 | rs7109112 | -0.509 | 1.061 | 1.185 | 0.099 | -0.005 | -0.013 | -0.013 | 2.128 | 0.16 | 0.18 | 0.08 | 0.11 | 0.12 | 0.14 |
| 11 | 7949350 | rs4758258 | 0.548 | 1.484 | 1.055 | -0.055 | 0.004 | -0.003 | -0.003 | 0.005 | 0.8 | 0.75 | 0.85 | 0.76 | 0.8 | 0.81 |
| 11 | 7950669 | rs1397596 | 0.75 | 1.202 | 0.694 | -0.261 | -0.027 | 0.159 | 0.159 | 0 | 0.24 | 0.02 | 0.11 | 0 | 0.21 | 0.17 |
| 11 | 7957953 | rs10839947 | 1.422 | 1.147 | 1.087 | 0.163 | 0.013 | -0.026 | -0.026 | 0.409 | 0.45 | 0.29 | 0.37 | 0.5 | 0.41 | 0.44 |
| 11 | 7961130 | rs4758259 | -0.109 | 1.079 | 1.179 | 0.191 | -0.001 | -0.001 | -0.001 | 1.545 | 0.33 | 0.23 | 0.46 | 0.26 | 0.35 | 0.37 |
| 11 | 7961146 | rs9787855 | 1.206 | 0.831 | 1.04 | 0.151 | 0 | -0.011 | -0.011 | 0 | 0.44 | 0.39 | 0.39 | 0.5 | 0.43 | 0.37 |
| 11 | 7962235 | rs7942833 | 1.132 | 0.758 | 1.167 | 0.488 | 0.022 | 0.056 | 0.056 | 0 | 0.7 | 0.52 | 0.6 | 0.47 | 0.64 | 0.63 |
| 11 | 7963498 | rs1355384 | 0.03 | 0.582 | 1.233 | 0.169 | -0.021 | -0.025 | -0.025 | 0 | 0.32 | 0.44 | 0.33 | 0.2 | 0.27 | 0.27 |
| 11 | 7963691 | rs11041681 | 0.145 | 0.504 | 1.192 | 0.088 | -0.006 | -0.023 | -0.023 | 0 | 0.07 | 0.11 | 0.03 | 0.01 | 0.03 | 0.09 |
| 11 | 7964654 | rs9665778 | -0.037 | 0.618 | 1.06 | 0.162 | 0.01 | 0.019 | 0.019 | 0 | 0.94 | 0.91 | 0.94 | 0.84 | 0.9 | 0.88 |
| 11 | 7964667 | rs11041683 | -0.796 | 0.626 | 1.164 | 0.158 | 0.003 | -0.003 | -0.003 | 0 | 0.45 | 0.37 | 0.4 | 0.41 | 0.41 | 0.47 |
| 11 | 7968359 | rs878860 | 1.578 | 0.653 | 1.119 | 0.145 | -0.029 | 0.006 | 0.006 | 0 | 0.41 | 0.16 | 0.47 | 0.42 | 0.44 | 0.32 |
| 11 | 7968467 | rs2360838 | 1.254 | 0.65 | 1.238 | 0.144 | 0.013 | 0.049 | 0.049 | 0 | 0.72 | 0.45 | 0.8 | 0.63 | 0.72 | 0.59 |
| 11 | 7968562 | rs907203 | -1.633 | 0.66 | 1.147 | 0.157 | -0.001 | -0.002 | -0.002 | 0 | 0.54 | 0.7 | 0.48 | 0.49 | 0.46 | 0.62 |
| 11 | 7973233 | rs2136907 | -0.057 | 0.683 | 1.08 | 0.169 | -0.002 | 0.043 | 0.043 | 0 | 0.16 | 0.23 | 0.11 | 0.28 | 0.15 | 0.26 |
| 11 | 7974549 | rs11041688 | -0.226 | 0.71 | 1.06 | 0.167 | 0.049 | 0.132 | 0.132 | 0.35 | 0.9 | 0.79 | 0.73 | 0.72 | 0.81 | 0.84 |
| 11 | 7977065 | rs4304763 | 0.479 | 0.488 | 0.866 | 0.024 | -0.007 | -0.009 | -0.009 | 0 | 0.77 | 0.79 | 0.84 | 0.7 | 0.74 | 0.66 |
| 11 | 7977152 | rs4421735 | 1.666 | 0.515 | 0.876 | 0.08 | -0.039 | -0.004 | -0.004 | 0 | 0.41 | 0.14 | 0.46 | 0.41 | 0.44 | 0.33 |
| 11 | 7982094 | rs12363522 | 1.913 | 0.418 | 0.301 | -0.282 | -0.015 | 0.035 | 0.035 | 0 | 0.07 | 0.01 | 0.08 | 0 | 0.06 | 0.01 |
| 11 | 7983199 | rs9919582 | -0.631 | 0.417 | 0.674 | 0.06 | 0.012 | 0.059 | 0.059 | 0 | 0.39 | 0.28 | 0.21 | 0.22 | 0.32 | 0.35 |
| 11 | 7985439 | rs12289513 | 1.099 | 0.449 | 0.122 | 0.048 | 0.014 | 0.15 | 0.15 | 0 | 0.06 | 0.15 | 0.2 | 0.2 | 0.09 | 0.11 |
| 11 | 7987347 | rs1858279 | -0.65 | 0.467 | 0.113 | -0.213 | 0.019 | 0.036 | 0.036 | 0 | 0.4 | 0.25 | 0.21 | 0.29 | 0.32 | 0.41 |
| 11 | 7989181 | rs6578916 | 1.062 | 1.105 | -0.242 | 0.187 | -0.008 | 0.038 | 0.038 | 0 | 0.52 | 0.3 | 0.69 | 0.61 | 0.54 | 0.53 |
| 11 | 7997397 | rs12420107 | 0.392 | 1.164 | -0.423 | -0.295 | -0.002 | 0.014 | 0.014 | 0.497 | 0.18 | 0.11 | 0.11 | 0 | 0.17 | 0.06 |
| 11 | 7999184 | rs2884017 | 0.821 | 1.118 | -0.618 | -0.025 | -0.042 | -0.027 | -0.027 | 0 | 0.6 | 0.16 | 0.56 | 0.65 | 0.59 | 0.69 |
| 11 | 8004045 | rs10734627 | 0.432 | 1.114 | 0.265 | -0.682 | 0 | -0.038 | -0.038 | 0 | 0.93 | 0.75 | 0.95 | 1 | 0.95 | 0.99 |
| 11 | 8005470 | rs11041699 | -0.85 | 1.465 | 0.824 | -0.789 | -0.046 | -0.045 | -0.045 | 0 | 0.2 | 0.27 | 0.11 | 0.11 | 0.13 | 0.22 |
| 11 | 8006713 | rs4758265 | -0.111 | 1.602 | 0.399 | -1.113 | -0.01 | 0.007 | 0.007 | 0.218 | 0.74 | 0.76 | 0.79 | 0.9 | 0.65 | 0.87 |
| 11 | 8007515 | rs10769861 | -0.166 | 1.628 | 0.416 | -1.1 | -0.027 | 0.02 | 0.02 | 0.163 | 0.76 | 0.82 | 0.8 | 0.9 | 0.65 | 0.93 |
| 11 | 8010130 | rs12275353 | -0.529 | 1.319 | 1.113 | -1.318 | -0.001 | 0.02 | 0.02 | 0 | 0.1 | 0.22 | 0.11 | 0.17 | 0.1 | 0.18 |
| 11 | 8012077 | rs7951923 | 0.184 | 1.556 | 1.022 | -1.342 | 0.012 | -0.083 | -0.083 | 0 | 0.21 | 0.26 | 0.14 | 0 | 0.22 | 0.04 |
| 11 | 8013450 | rs7941782 | -0.694 | 1.344 | 1.237 | -1.453 | -0.05 | -0.05 | -0.05 | 0 | 0.25 | 0.33 | 0.13 | 0.13 | 0.14 | 0.25 |
| 11 | 8015146 | rs10839960 | -1.235 | 1.601 | 1.228 | -1.447 | 0.02 | 0.033 | 0.033 | 1.162 | 0.46 | 0.38 | 0.23 | 0.13 | 0.34 | 0.31 |
| 11 | 8015651 | rs4559682 | -0.808 | 1.914 | 1.344 | -1.43 | -0.008 | -0.058 | -0.058 | 0.009 | 0.3 | 0.57 | 0.19 | 0.15 | 0.28 | 0.3 |
| 11 | 8019790 | rs6578918 | -0.791 | 2.238 | 0.77 | -0.946 | 0.012 | 0.003 | 0.003 | 1.103 | 0.48 | 0.56 | 0.26 | 0.13 | 0.37 | 0.32 |
| 11 | 8022014 | rs4758038 | -0.894 | 2.314 | -0.038 | -0.924 | -0.009 | 0.164 | 0.164 | 0.621 | 0.51 | 0.73 | 0.31 | 0.15 | 0.48 | 0.32 |
| 11 | 8024468 | rs11041706 | 0.608 | 1.358 | -0.561 | -0.721 | -0.008 | 0.269 | 0.269 | 0.415 | 0.51 | 0.13 | 0.66 | 0.81 | 0.48 | 0.66 |
| 11 | 8037138 | rs2280730 | -0.587 | 2.067 | 0.884 | 0.324 | 0.001 | 0.001 | 0.001 | 0 | 0.36 | 0.39 | 0.53 | 0.52 | 0.37 | 0.36 |

| | | | | | | | | | | | | | | | | |
|---|---|---|---|---|---|---|---|---|---|---|---|---|---|---|---|---|
| 11 | 8037597 | rs2280728 | -1.096 | 2.657 | 1.089 | 0.34 | 0.014 | 0.037 | 0.037 | 0 | 0.43 | 0.63 | 0.6 | 0.72 | 0.55 | 0.49 |
| 11 | 8048210 | rs1609254 | -0.472 | 3.017 | 0.922 | 0.611 | 0.005 | 0.01 | 0.01 | 0 | 0.26 | 0.25 | 0.49 | 0.52 | 0.33 | 0.41 |
| 11 | 8053159 | rs4758272 | -0.545 | 2.878 | 1.107 | 0.821 | 0.001 | 0.038 | 0.038 | 0 | 0.27 | 0.31 | 0.48 | 0.5 | 0.29 | 0.4 |
| 11 | 8055832 | rs7952059 | -0.672 | 3.076 | 1.257 | 0.667 | 0.105 | 0.071 | 0.071 | 1.739 | 0.14 | 0.3 | 0.12 | 0.2 | 0.22 | 0.08 |
| 11 | 8059308 | rs12799847 | 0.577 | 2.292 | 0.286 | -0.399 | -0.049 | 0.099 | 0.099 | 0 | 0.22 | 0.01 | 0.19 | 0.11 | 0.19 | 0.24 |
| 11 | 8061926 | rs1406094 | 0.325 | 1.952 | 0.826 | 0.266 | 0 | 0.009 | 0.009 | 0 | 0.18 | 0.2 | 0.18 | 0.52 | 0.15 | 0.29 |
| 11 | 8062963 | rs1528130 | 0.97 | 2.149 | 0.645 | 0.214 | -0.074 | -0.053 | -0.053 | 0 | 0.64 | 0.12 | 0.68 | 0.46 | 0.62 | 0.65 |
| 11 | 8063419 | rs1918849 | -0.08 | 2.256 | 1.128 | 0.148 | 0.092 | -0.06 | -0.06 | 0 | 0.13 | 0.32 | 0.1 | 0.04 | 0.17 | 0.06 |
| 11 | 8067451 | rs2141322 | 0.425 | 2.242 | 0.985 | 0.141 | 0.106 | -0.097 | -0.097 | 0 | 0.83 | 0.52 | 0.88 | 0.96 | 0.76 | 0.93 |
| 11 | 8073236 | rs7949880 | 0.083 | 1.973 | 0.225 | 0.208 | 0.006 | -0.024 | -0.024 | 0 | 0.74 | 0.81 | 0.53 | 0.47 | 0.77 | 0.63 |
| 11 | 8076174 | rs10839972 | -0.347 | 1.788 | -0.154 | 0.865 | 0.025 | 0.1 | 0.1 | 0.118 | 0.79 | 0.47 | 0.78 | 0.72 | 0.78 | 0.86 |
| 11 | 8078704 | rs11041723 | 0.685 | 1.466 | -1.343 | 0.235 | -0.055 | 0.019 | 0.019 | 0.01 | 0.58 | 0.17 | 0.39 | 0.43 | 0.51 | 0.56 |
| 11 | 8088757 | rs7932687 | -0.699 | 0.93 | -1.566 | 0.047 | -0.062 | -0.065 | -0.065 | 0 | 0.45 | 0.87 | 0.33 | 0.46 | 0.5 | 0.5 |
| 11 | 8089121 | rs7113729 | -0.843 | 0.727 | -1.278 | -0.052 | 0.009 | -0.017 | -0.017 | 0 | 0.48 | 0.8 | 0.37 | 0.46 | 0.55 | 0.51 |
| 11 | 8089999 | rs7114039 | -0.725 | 1.248 | -0.86 | -0.173 | 0.001 | 0 | 0 | 0 | 0.39 | 0.56 | 0.31 | 0.42 | 0.43 | 0.48 |
| 11 | 8094682 | rs1406095 | -0.684 | 1.14 | -0.654 | 0.07 | 0.015 | 0.015 | 0.015 | 0.027 | 0.39 | 0.5 | 0.53 | 0.44 | 0.48 | 0.54 |
| 11 | 8097434 | rs3794018 | -1.086 | 0.841 | -1.606 | 0.144 | -0.122 | -0.164 | -0.164 | 0.902 | 0.58 | 0.97 | 0.65 | 0.52 | 0.58 | 0.63 |
| 11 | 8099189 | rs7926527 | 0.347 | 0.277 | -0.472 | 0.519 | -0.002 | 0 | 0 | 1.955 | 0.64 | 0.66 | 0.43 | 0.55 | 0.68 | 0.55 |
| 11 | 8099913 | rs4758287 | 0.305 | 0.332 | 0.045 | 0.748 | -0.013 | 0.026 | 0.026 | 0.794 | 0.71 | 0.59 | 0.54 | 0.59 | 0.75 | 0.69 |
| 11 | 8101173 | rs4758040 | 1.236 | 0.925 | 0.014 | 0.681 | -0.014 | -0.041 | -0.041 | 0.013 | 0.55 | 0.15 | 0.38 | 0.52 | 0.59 | 0.57 |
| 11 | 8103104 | rs11041733 | 0.951 | 1.044 | 0.18 | 0.706 | -0.01 | -0.044 | -0.044 | 0.132 | 0.61 | 0.25 | 0.41 | 0.56 | 0.64 | 0.58 |
| 11 | 8110479 | rs1881237 | -1.496 | 1.058 | 1.202 | 0.393 | 0.127 | 0.078 | 0.078 | 1.173 | 0.05 | 0.45 | 0.09 | 0 | 0.03 | 0 |
| 11 | 8111121 | rs1317500 | -0.704 | 1.039 | 0.549 | 0.332 | 0.001 | 0.033 | 0.033 | 0 | 0.09 | 0.04 | 0.04 | 0 | 0.05 | 0.06 |
| 11 | 8117518 | rs11041740 | -0.748 | 1.154 | 1.527 | 0.381 | 0.051 | -0.09 | -0.09 | 0.357 | 0.18 | 0.42 | 0.15 | 0.06 | 0.2 | 0.22 |
| 11 | 8118103 | rs2242501 | 1.206 | 1.115 | 1.526 | 1.153 | 0.001 | 0.008 | 0.008 | 0 | 0.75 | 0.32 | 0.57 | 0.66 | 0.77 | 0.71 |
| 11 | 8118472 | rs2242503 | 1.292 | 1.066 | 1.37 | 0.955 | -0.005 | 0.007 | 0.007 | 0.289 | 0.69 | 0.35 | 0.75 | 0.84 | 0.68 | 0.61 |
| 11 | 8120601 | rs16935778 | -1.779 | 1.214 | 1.201 | 0.957 | 0.042 | 0.008 | 0.008 | 0 | 0.08 | 0.18 | 0.11 | 0.08 | 0.12 | 0.16 |
| 11 | 8120902 | rs7481121 | 1.347 | 1.157 | 1.332 | 1.047 | -0.005 | 0.039 | 0.039 | 0 | 0.89 | 0.74 | 0.88 | 0.86 | 0.91 | 0.8 |
| 11 | 8121218 | rs1108277 | 1.925 | 0.961 | 0.736 | 0.971 | -0.059 | -0.063 | -0.063 | 0 | 0.6 | 0.11 | 0.47 | 0.54 | 0.63 | 0.53 |
| 11 | 8126235 | rs3750956 | -2.731 | 0.999 | 1.897 | 0.725 | -0.009 | -0.009 | -0.009 | 0 | 0.13 | 0.39 | 0.21 | 0.1 | 0.1 | 0.22 |
| 11 | 8126923 | rs11041742 | -0.567 | 0.933 | 1.969 | 1.441 | 0.014 | 0.06 | 0.06 | 0 | 0.2 | 0.28 | 0.34 | 0.42 | 0.21 | 0.27 |
| 11 | 8128158 | rs4340037 | -0.597 | 0.891 | 1.969 | 1.456 | 0.012 | 0.058 | 0.058 | 0 | 0.2 | 0.29 | 0.35 | 0.44 | 0.21 | 0.27 |
| 11 | 8129204 | rs1406093 | -1.99 | 0.89 | 2.092 | 1.423 | 0.004 | 0.018 | 0.018 | 0 | 0.09 | 0.26 | 0.18 | 0.1 | 0.08 | 0.18 |
| 11 | 8130176 | rs2141321 | -1.911 | 0.871 | 2.059 | 1.445 | 0.001 | 0.052 | 0.052 | 0.029 | 0.31 | 0.55 | 0.53 | 0.53 | 0.31 | 0.49 |
| 11 | 8149753 | rs1528133 | -1.655 | 0.826 | 2.004 | 1.369 | 0.012 | 0.01 | 0.01 | 0 | 0.05 | 0.15 | 0.09 | 0.06 | 0.07 | 0.12 |
| 11 | 8152080 | rs4758295 | -3.184 | 0.805 | 2.098 | 1.356 | -0.017 | -0.017 | -0.017 | 0 | 0.14 | 0.39 | 0.2 | 0.1 | 0.11 | 0.25 |
| 11 | 8154451 | rs11041759 | -3.022 | 0.795 | 2.246 | 1.394 | -0.008 | 0.004 | 0.004 | 0.131 | 0.12 | 0.4 | 0.19 | 0.1 | 0.09 | 0.21 |
| 11 | 8159892 | rs10839976 | -0.63 | 0.893 | 1.925 | 1.483 | 0.015 | 0.061 | 0.061 | 0 | 0.2 | 0.28 | 0.34 | 0.42 | 0.22 | 0.27 |
| 11 | 8172150 | rs7122904 | 1.359 | 0.918 | 0.95 | 1.039 | 0.003 | 0.013 | 0.013 | 0.331 | 0.39 | 0.32 | 0.29 | 0.27 | 0.41 | 0.21 |
| 11 | 8190057 | rs7945726 | -1.389 | 1.189 | 1.322 | 1.253 | -0.011 | -0.005 | -0.005 | 0.237 | 0.05 | 0.08 | 0.02 | 0.04 | 0.02 | 0.1 |
| 11 | 8201186 | rs7481683 | -1.508 | 1.879 | 1.921 | 1.348 | 0.06 | 0.053 | 0.053 | 0.016 | 0.26 | 0.46 | 0.39 | 0.42 | 0.32 | 0.42 |
| 11 | 8208348 | rs11041788 | -1.181 | 1.963 | 1.921 | 1.394 | 0.037 | 0.043 | 0.043 | 0 | 0.22 | 0.34 | 0.35 | 0.39 | 0.26 | 0.33 |
| 11 | 8208730 | rs4758309 | 1.515 | 1.803 | 1.726 | 1.421 | 0.002 | 0.007 | 0.007 | 0.034 | 0.69 | 0.37 | 0.6 | 0.55 | 0.7 | 0.57 |
| 11 | 8212242 | rs11041791 | -1.928 | 1.786 | 1.668 | 1.444 | 0.018 | 0.047 | 0.047 | 0 | 0.29 | 0.46 | 0.38 | 0.46 | 0.29 | 0.42 |
| 11 | 8217326 | rs3911309 | 1.998 | 2.068 | 0.912 | 1.222 | -0.042 | -0.032 | -0.032 | 0.719 | 0.62 | 0.23 | 0.53 | 0.5 | 0.58 | 0.5 |
| 11 | 8222286 | rs4636658 | 2.111 | 2.106 | 0.572 | 1.252 | -0.081 | -0.074 | -0.074 | 0 | 0.61 | 0.12 | 0.53 | 0.49 | 0.58 | 0.5 |
| 11 | 8225688 | rs12360753 | 1.738 | 1.469 | 0.866 | 1.395 | 0.01 | -0.045 | -0.045 | 2.401 | 0.7 | 0.16 | 0.62 | 0.59 | 0.76 | 0.72 |
| 11 | 8227964 | rs1881229 | 0.845 | 1.706 | 0.911 | 1.368 | -0.006 | 0.05 | 0.05 | 0.062 | 0.81 | 0.65 | 0.74 | 0.7 | 0.85 | 0.8 |
| 11 | 8228920 | rs4627068 | -1.724 | 1.846 | 0.326 | 1.266 | -0.005 | -0.014 | -0.014 | 0 | 0.23 | 0.76 | 0.36 | 0.34 | 0.22 | 0.19 |
| 11 | 8233876 | rs7951027 | -2.212 | 2.472 | 0.933 | 0.788 | 0.068 | -0.063 | -0.063 | 0.893 | 0.19 | 0.57 | 0.21 | 0.09 | 0.29 | 0.13 |
| 11 | 8238639 | rs4758051 | 1.937 | 1.936 | 0.023 | 1.087 | -0.064 | -0.094 | -0.094 | 2.531 | 0.58 | 0.07 | 0.51 | 0.42 | 0.57 | 0.46 |
| 11 | 8239818 | rs7122225 | -1.855 | 2.17 | 0.852 | 1.05 | 0.087 | -0.015 | -0.015 | 0.168 | 0.11 | 0.3 | 0.12 | 0.05 | 0.14 | 0.08 |
| 11 | 8242629 | rs7109806 | -1.767 | 2.105 | 1.162 | 1.059 | 0.061 | 0.021 | 0.021 | 0.483 | 0.1 | 0.41 | 0.1 | 0.06 | 0.08 | 0.07 |
| 11 | 8243026 | rs10840002 | 2.053 | 2.2 | 0.845 | 1.089 | 0.028 | -0.083 | -0.083 | 0.043 | 0.61 | 0.15 | 0.55 | 0.44 | 0.64 | 0.5 |
| 11 | 8243170 | rs6578942 | -1.631 | 2.162 | 1.117 | 0.624 | 0.046 | 0.017 | 0.017 | 0 | 0.08 | 0.3 | 0.09 | 0.06 | 0.07 | 0.07 |
| 11 | 8243798 | rs11041816 | 1.916 | 1.729 | -0.181 | 0.347 | -0.057 | 0.254 | 0.254 | 0 | 0.48 | 0.06 | 0.67 | 0.15 | 0.47 | 0.39 |
| 11 | 8246181 | rs1042359 | -1.937 | 1.781 | 1.132 | 0.421 | 0.029 | -0.047 | -0.047 | 0.196 | 0.16 | 0.38 | 0.12 | 0.06 | 0.14 | 0.12 |
| 11 | 8247131 | rs7933956 | -1.549 | 1.443 | 1.088 | 0.502 | 0.047 | 0.002 | 0.002 | 2.521 | 0.08 | 0.34 | 0.1 | 0.05 | 0.07 | 0.05 |
| 11 | 8250811 | rs4758317 | 1.398 | 1.68 | 0.778 | 0.924 | 0.012 | 0.01 | 0.01 | 0 | 0.69 | 0.6 | 0.52 | 0.42 | 0.56 | 0.73 |
| 11 | 8251438 | rs11041820 | -0.164 | 1.831 | 0.041 | 0.941 | -0.067 | -0.002 | -0.002 | 0 | 0.14 | 0.04 | 0.09 | 0.18 | 0.26 | 0.09 |
| 11 | 8252853 | rs110419 | 1.702 | 1.104 | 0.328 | 0.615 | -0.05 | -0.05 | -0.05 | 0 | 0.61 | 0.15 | 0.47 | 0.42 | 0.52 | 0.69 |
| 11 | 8261298 | rs413708 | 1.708 | 0.768 | 0.896 | -0.235 | -0.044 | -0.07 | -0.07 | 0 | 0.78 | 0.66 | 0.89 | 0.94 | 0.83 | 0.66 |
| 11 | 8262401 | rs3849994 | -1.197 | 0.883 | 1.209 | -0.393 | 0.041 | -0.07 | -0.07 | 0 | 0.11 | 0.44 | 0.1 | 0 | 0.06 | 0.03 |
| 11 | 8264183 | rs406281 | 2.008 | 1.236 | 0.922 | -0.637 | 0.001 | -0.022 | -0.022 | 0 | 0.68 | 0.32 | 0.75 | 0.96 | 0.7 | 0.82 |
| 11 | 8264350 | rs16937230 | -0.986 | 1.314 | 0.908 | -0.561 | -0.001 | -0.002 | -0.002 | 0 | 0.05 | 0.08 | 0.19 | 0.06 | 0.03 | 0.2 |
| 11 | 8264525 | rs484161 | 2.782 | 1.705 | 0.623 | -0.56 | 0.064 | 0.187 | 0.187 | 0.267 | 0.49 | 0.12 | 0.46 | 0.8 | 0.41 | 0.53 |
| 11 | 8264900 | rs437976 | 2.677 | 1.548 | 0.716 | -0.584 | 0.01 | 0.088 | 0.088 | 0 | 0.62 | 0.22 | 0.56 | 0.9 | 0.66 | 0.62 |
| 11 | 8265652 | rs12277079 | -2.268 | 1.306 | 1.464 | -0.314 | 0.028 | -0.054 | -0.054 | 0 | 0.15 | 0.46 | 0.28 | 0.06 | 0.09 | 0.22 |
| 11 | 8266488 | rs3794014 | 3.108 | 1.156 | 0.936 | 0.13 | -0.043 | -0.103 | -0.103 | 0.082 | 0.48 | 0.04 | 0.37 | 0.53 | 0.42 | 0.43 |

| | | | | | | | | | | | | | | | | |
|---|---|---|---|---|---|---|---|---|---|---|---|---|---|---|---|---|
| 11 | 8266559 | rs2061036 | -0.745 | 1.437 | 1.515 | 0.245 | 0.037 | -0.015 | -0.015 | 0.448 | 0.08 | 0.24 | 0.08 | 0.02 | 0.06 | 0.02 |
| 11 | 8266629 | rs2061037 | -2.403 | 1.432 | 1.337 | 0.233 | -0.089 | 0.072 | 0.072 | 0.097 | 0.33 | 0.8 | 0.41 | 0.18 | 0.42 | 0.35 |
| 11 | 8269405 | rs417210 | -1.677 | 1.355 | 1.409 | 0.234 | 0.036 | 0.043 | 0.043 | 0 | 0.24 | 0.35 | 0.28 | 0.37 | 0.33 | 0.24 |
| 11 | 8270244 | rs3794012 | 3.11 | 1.314 | 0.993 | 0.212 | -0.037 | -0.08 | -0.08 | 0.733 | 0.49 | 0.05 | 0.38 | 0.53 | 0.44 | 0.45 |
| 11 | 8271723 | rs2311011 | 2.473 | 1.289 | 1.197 | 0.163 | 0.004 | 0.018 | 0.018 | 0 | 0.72 | 0.29 | 0.62 | 0.91 | 0.72 | 0.66 |
| 11 | 8272605 | rs379951 | 1.462 | 1.337 | 0.886 | 0.216 | 0.064 | -0.073 | -0.073 | 0.002 | 0.69 | 0.54 | 0.77 | 0.87 | 0.58 | 0.81 |
| 11 | 8273973 | rs7949134 | 1.04 | 1.358 | 0.865 | 0.451 | 0.005 | 0.144 | 0.144 | 0 | 0.91 | 0.72 | 0.81 | 0.71 | 0.93 | 0.88 |
| 11 | 8274457 | rs11606296 | -0.715 | 1.2 | 0.549 | 0.343 | 0.047 | -0.008 | -0.008 | 0 | 0.08 | 0.26 | 0.08 | 0.02 | 0.06 | 0.03 |
| 11 | 8275835 | rs4237770 | 2.289 | 1.047 | 0.175 | 0.305 | -0.134 | -0.123 | -0.123 | 0 | 0.56 | 0.05 | 0.43 | 0.52 | 0.52 | 0.47 |
| 11 | 8276780 | rs7107078 | 2.139 | 0.829 | 0.368 | 0.425 | -0.033 | 0.043 | 0.043 | 0 | 0.77 | 0.34 | 0.6 | 0.66 | 0.8 | 0.6 |
| 11 | 8278197 | rs204938 | 1.077 | 1.017 | -0.41 | 0.197 | -0.055 | 0.083 | 0.083 | 0.012 | 0.63 | 0.24 | 0.56 | 0.8 | 0.59 | 0.57 |
| 11 | 8280938 | rs204937 | 1.102 | 1.194 | -0.561 | 0.243 | -0.066 | 0.176 | 0.176 | 0.569 | 0.58 | 0.13 | 0.69 | 0.85 | 0.56 | 0.75 |
| 11 | 8282786 | rs11041833 | 1.02 | 1.107 | -0.356 | 0.597 | -0.003 | -0.027 | -0.027 | 0.011 | 0.54 | 0.17 | 0.35 | 0.51 | 0.45 | 0.46 |
| 11 | 8291906 | rs7118748 | -1.771 | 1.242 | 0.494 | 1.189 | 0.001 | 0.081 | 0.081 | 0 | 0.21 | 0.58 | 0.45 | 0.41 | 0.2 | 0.35 |
| 11 | 8301191 | rs7104520 | -1.021 | 1.525 | 0.511 | 1.536 | 0.099 | 0.284 | 0.284 | 0 | 0.05 | 0.2 | 0.17 | 0.31 | 0.07 | 0.14 |
| 11 | 8304015 | rs4758322 | 0.016 | 1.056 | -0.66 | 1.019 | -0.012 | -0.06 | -0.06 | 0.194 | 0.33 | 0.14 | 0.34 | 0.39 | 0.33 | 0.33 |
| 11 | 8315496 | rs10160665 | -0.627 | 1.355 | -0.566 | 0.483 | -0.015 | -0.015 | -0.015 | 0 | 0.39 | 0.3 | 0.61 | 0.52 | 0.41 | 0.6 |
| 11 | 8320592 | rs1454441 | 0.233 | 1.288 | 0.042 | 0.635 | 0.006 | -0.007 | -0.007 | 0 | 0.23 | 0.19 | 0.3 | 0.16 | 0.25 | 0.3 |
| 11 | 8324546 | rs7109141 | 0.718 | 1.32 | 0.22 | 0.604 | 0.017 | -0.012 | -0.012 | 0 | 0.38 | 0.27 | 0.22 | 0.42 | 0.35 | 0.32 |
| 11 | 8326607 | rs10840011 | -0.536 | 1.274 | 0.358 | 0.574 | 0.014 | -0.005 | -0.005 | 0 | 0.55 | 0.32 | 0.73 | 0.55 | 0.59 | 0.64 |
| 11 | 8328097 | rs10840015 | -0.403 | 1.486 | 0.445 | 0.613 | 0.04 | -0.041 | -0.041 | 0 | 0.3 | 0.43 | 0.36 | 0.19 | 0.33 | 0.37 |
| 11 | 8329193 | rs4758330 | -0.12 | 1.356 | 0.254 | 0.654 | 0.018 | -0.029 | -0.029 | 6.685 | 0.27 | 0.33 | 0.33 | 0.17 | 0.28 | 0.34 |
| 11 | 8329230 | rs4758331 | -0.668 | 1.369 | 0.086 | 0.683 | 0.002 | -0.012 | -0.012 | 0 | 0.67 | 0.71 | 0.58 | 0.61 | 0.67 | 0.69 |
| 11 | 8329840 | rs4758332 | -0.146 | 1.356 | -0.028 | 0.836 | 0.015 | -0.005 | -0.005 | 0 | 0.28 | 0.3 | 0.33 | 0.22 | 0.3 | 0.35 |
| 11 | 8330352 | rs10840016 | 0.018 | 1.248 | 0.068 | 1.073 | -0.008 | 0.005 | 0.005 | 0 | 0.41 | 0.47 | 0.25 | 0.46 | 0.39 | 0.37 |
| 11 | 8331494 | rs897102 | 1.05 | 1.747 | 0.125 | 1.159 | 0.055 | 0.075 | 0.075 | 0 | 0.87 | 0.72 | 0.75 | 0.7 | 0.83 | 0.77 |
| 11 | 8332661 | rs874811 | 0.131 | 1.552 | -0.672 | 0.744 | -0.022 | 0.008 | 0.008 | 0 | 0.72 | 0.79 | 0.74 | 0.77 | 0.74 | 0.69 |
| 11 | 8332942 | rs897101 | -0.094 | 0.925 | -0.793 | 0.834 | 0 | -0.004 | -0.004 | 0 | 0.7 | 0.66 | 0.59 | 0.51 | 0.68 | 0.71 |
| 11 | 8333255 | rs1869551 | 0.14 | 0.44 | -1.089 | -0.231 | 0.001 | -0.003 | -0.003 | 0 | 0.72 | 0.71 | 0.69 | 0.88 | 0.68 | 0.77 |
| 11 | 8333826 | rs6578953 | -0.08 | 0.3 | -1.266 | 0.061 | 0.018 | -0.006 | -0.006 | 0 | 0.63 | 0.75 | 0.64 | 0.39 | 0.69 | 0.52 |
| 11 | 8334784 | rs11041869 | -0.123 | 0.416 | -1.277 | -0.461 | 0 | -0.013 | -0.013 | 0 | 0.14 | 0.1 | 0.08 | 0 | 0.14 | 0.09 |
| 11 | 8334841 | rs7102266 | -0.075 | 0.454 | -1.507 | -0.461 | -0.02 | 0.045 | 0.045 | 0 | 0.37 | 0.17 | 0.47 | 0.25 | 0.54 | 0.29 |
| 11 | 8337966 | rs2871610 | 0.108 | 0.41 | -1.506 | -0.111 | -0.019 | 0.01 | 0.01 | 0 | 0.47 | 0.27 | 0.33 | 0.45 | 0.47 | 0.65 |
| 11 | 8338357 | rs4758059 | 0.645 | 0.361 | -1.529 | 0.181 | 0.009 | 0.014 | 0.014 | 0 | 0.25 | 0.13 | 0.16 | 0.21 | 0.22 | 0.35 |
| 11 | 8339628 | rs12575529 | 0.434 | 0.168 | -1.225 | -0.258 | -0.012 | -0.036 | -0.036 | 0 | 0.3 | 0.2 | 0.53 | 0.53 | 0.35 | 0.21 |
| 11 | 8339894 | rs10840020 | -0.015 | 0.059 | -0.849 | -0.35 | -0.001 | -0.001 | -0.001 | 0 | 0.57 | 0.59 | 0.38 | 0.47 | 0.55 | 0.73 |
| 11 | 8340063 | rs10769895 | -0.369 | 0.278 | -1.771 | 0.058 | -0.014 | -0.005 | -0.005 | 1.069 | 0.46 | 0.26 | 0.56 | 0.49 | 0.52 | 0.51 |
| 11 | 8340724 | rs4758062 | 0.74 | 0.382 | -1.336 | 0.475 | 0.001 | 0.001 | 0.001 | 0 | 0.24 | 0.22 | 0.17 | 0.21 | 0.23 | 0.35 |
| 11 | 8341669 | rs4362135 | -0.301 | 0.685 | -1.104 | 0.799 | -0.013 | 0.006 | 0.006 | 0.656 | 0.73 | 0.4 | 0.73 | 0.72 | 0.79 | 0.85 |
| 11 | 8341686 | rs4412747 | -0.431 | 1.169 | -1.207 | 0.981 | -0.042 | -0.043 | -0.043 | 0.22 | 0.07 | 0.01 | 0.29 | 0.26 | 0.14 | 0.1 |
| 11 | 8343361 | rs1026345 | -0.49 | 0.34 | -0.707 | 0.078 | -0.018 | -0.034 | -0.034 | 0.579 | 0.34 | 0.55 | 0.25 | 0.24 | 0.28 | 0.37 |
| 11 | 8348340 | rs4274193 | -0.474 | 0.498 | -1.273 | 0.027 | -0.031 | 0.008 | 0.008 | 0 | 0.25 | 0.08 | 0.16 | 0.24 | 0.27 | 0.31 |
| 11 | 8348674 | rs10840024 | 0.278 | -0.073 | -0.773 | -0.127 | -0.017 | -0.006 | -0.006 | 0.284 | 0.19 | 0.35 | 0.18 | 0.17 | 0.13 | 0.18 |
| 11 | 8348725 | rs10840025 | 0.237 | -0.092 | -0.781 | -0.136 | -0.017 | -0.007 | -0.007 | 0 | 0.2 | 0.35 | 0.17 | 0.17 | 0.13 | 0.18 |
| 11 | 8349118 | rs12575952 | 0.577 | 0.103 | -0.932 | -0.243 | -0.001 | -0.01 | -0.01 | 0 | 0.21 | 0.07 | 0.14 | 0.24 | 0.2 | 0.26 |
| 11 | 8349506 | rs1026347 | 0.56 | -0.778 | -1.188 | -0.808 | 0.032 | 0.036 | 0.036 | 0 | 0.28 | 0.17 | 0.13 | 0.06 | 0.15 | 0.18 |
| 11 | 8350541 | rs756057 | 1.586 | -0.798 | -1.476 | -0.481 | -0.139 | -0.144 | -0.144 | 0 | 0.22 | 0.02 | 0.45 | 0.42 | 0.39 | 0.25 |
| 11 | 8351264 | rs11041884 | -0.749 | -1.12 | -0.983 | -0.534 | -0.059 | -0.087 | -0.087 | 0 | 0.69 | 0.55 | 0.84 | 0.94 | 0.85 | 0.82 |
| 11 | 8351657 | rs7124392 | -0.793 | -1.119 | -0.933 | -0.534 | -0.051 | -0.077 | -0.077 | 0 | 0.69 | 0.57 | 0.84 | 0.94 | 0.85 | 0.82 |
| 11 | 8352511 | rs10840026 | 0.644 | -0.91 | -0.96 | -0.456 | -0.029 | 0.042 | 0.042 | 0 | 0.22 | 0.39 | 0.16 | 0.28 | 0.19 | 0.24 |
| 11 | 8354463 | rs874360 | 0.901 | -0.798 | -1.184 | -0.376 | -0.155 | -0.126 | -0.126 | 0 | 0.26 | 0.04 | 0.46 | 0.43 | 0.47 | 0.3 |
| 11 | 8356996 | rs12279469 | 1.023 | -1.417 | -1.169 | -0.661 | 0.003 | 0.002 | 0.002 | 0 | 0.14 | 0.11 | 0.07 | 0.11 | 0.08 | 0.07 |
| 11 | 8357802 | rs10769901 | 1.348 | -1.389 | -0.62 | -0.643 | -0.046 | 0.001 | 0.001 | 0 | 0.17 | 0.32 | 0.09 | 0.17 | 0.08 | 0.16 |
| 11 | 8358049 | rs9804548 | 1.229 | -0.916 | -0.946 | -0.536 | -0.101 | -0.118 | -0.118 | 0 | 0.25 | 0.07 | 0.45 | 0.43 | 0.4 | 0.27 |
| 11 | 8359318 | rs10840027 | -0.145 | -1.556 | -0.832 | -0.702 | 0.028 | 0.026 | 0.026 | 0 | 0.69 | 0.77 | 0.81 | 0.83 | 0.76 | 0.77 |
| 11 | 8359495 | rs4758347 | -0.48 | -1.65 | -0.86 | -0.803 | -0.001 | -0.008 | -0.008 | 0.013 | 0.76 | 0.76 | 0.87 | 0.94 | 0.85 | 0.9 |
| 11 | 8359880 | rs4758348 | -0.427 | -1.462 | -0.665 | -0.684 | 0.026 | 0.03 | 0.03 | 0 | 0.72 | 0.79 | 0.88 | 0.94 | 0.83 | 0.83 |
| 11 | 8360804 | rs747190 | 0.675 | -1.225 | -0.644 | -0.546 | -0.043 | -0.037 | -0.037 | 0 | 0.31 | 0.24 | 0.49 | 0.44 | 0.48 | 0.32 |
| 11 | 8366256 | rs6578959 | 0.333 | -1.391 | -0.372 | -0.404 | 0.009 | 0.002 | 0.002 | 0.671 | 0.35 | 0.29 | 0.28 | 0.37 | 0.25 | 0.46 |
| 11 | 8368246 | rs7123904 | 0.407 | -1.273 | -0.222 | -0.339 | -0.009 | -0.001 | -0.001 | 0 | 0.33 | 0.4 | 0.3 | 0.37 | 0.25 | 0.44 |
| 11 | 8368490 | rs7113611 | 0.556 | -1.246 | -0.269 | -0.344 | 0.001 | -0.001 | -0.001 | 0.047 | 0.31 | 0.3 | 0.28 | 0.37 | 0.25 | 0.43 |
| 11 | 8369415 | rs897111 | 0.097 | -1.159 | -0.313 | -0.399 | -0.001 | -0.001 | -0.001 | 0.24 | 0.36 | 0.39 | 0.31 | 0.37 | 0.31 | 0.48 |
| 11 | 8370644 | rs4414207 | 0.875 | -1.022 | -0.524 | -0.5 | -0.001 | -0.008 | -0.008 | 0 | 0.05 | 0.06 | 0.03 | 0 | 0.06 | 0.05 |
| 11 | 8374548 | rs10840028 | 0.142 | -0.501 | -0.453 | -0.384 | 0.001 | -0.006 | -0.006 | 0 | 0.35 | 0.34 | 0.52 | 0.48 | 0.39 | 0.48 |
| 11 | 8378655 | rs4758350 | 0.739 | -0.303 | -0.737 | -0.309 | -0.1 | -0.06 | -0.06 | 0.398 | 0.29 | 0.11 | 0.54 | 0.56 | 0.51 | 0.33 |
| 11 | 8378752 | rs4430495 | -0.069 | -0.561 | -0.643 | -0.461 | 0 | 0 | 0 | 0.232 | 0.4 | 0.41 | 0.37 | 0.46 | 0.44 | 0.37 |
| 11 | 8380719 | rs4237772 | 0.01 | -0.496 | -0.396 | -0.571 | -0.002 | -0.002 | -0.002 | 0 | 0.6 | 0.44 | 0.57 | 0.64 | 0.6 | 0.6 |
| 11 | 8382106 | rs7931324 | -0.678 | -0.177 | 0.032 | -0.493 | -0.007 | -0.062 | -0.062 | 0.022 | 0.12 | 0.24 | 0.05 | 0 | 0.09 | 0.1 |
| 11 | 8384816 | rs2045140 | -0.832 | 0.207 | 0.087 | -0.387 | 0.006 | -0.091 | -0.091 | 0 | 0.3 | 0.42 | 0.23 | 0.13 | 0.26 | 0.32 |

| | | | | | | | | | | | | | | | | |
|---|---|---|---|---|---|---|---|---|---|---|---|---|---|---|---|---|
| 11 | 8388703 | rs7927814 | 0.65 | 0.514 | 0.327 | 0.349 | 0.09 | 0.075 | 0.075 | 0 | 0.93 | 0.79 | 0.85 | 0.88 | 0.82 | 0.89 |
| 11 | 8390147 | rs16937906 | 0.269 | 0.279 | -0.375 | 0.691 | 0.031 | 0.044 | 0.044 | 1.181 | 0.42 | 0.06 | 0.26 | 0.38 | 0.32 | 0.42 |
| 11 | 8391230 | rs897105 | -0.352 | 0.248 | -0.298 | 0.844 | 0.025 | -0.019 | -0.019 | 0 | 0.54 | 0.73 | 0.59 | 0.48 | 0.56 | 0.52 |
| 11 | 8392768 | rs4758063 | -0.136 | 0.148 | -0.12 | 0.925 | 0.007 | 0.057 | 0.057 | 0.453 | 0.51 | 0.66 | 0.59 | 0.6 | 0.53 | 0.51 |
| 11 | 8394169 | rs7106385 | 0.129 | 0.187 | -0.054 | 0.695 | 0 | -0.002 | -0.002 | 0 | 0.46 | 0.38 | 0.5 | 0.46 | 0.5 | 0.4 |
| 11 | 8398847 | rs1466234 | -0.097 | -0.295 | 0.108 | 0.276 | 0.023 | 0.043 | 0.043 | 0.185 | 0.49 | 0.37 | 0.65 | 0.75 | 0.61 | 0.47 |
| 11 | 8402917 | rs7108135 | -0.368 | -1.298 | 0.419 | -0.096 | -0.082 | -0.077 | -0.077 | 0.527 | 0.17 | 0.38 | 0.05 | 0.04 | 0.05 | 0.12 |
| 11 | 8403392 | rs7112091 | -0.787 | -1.856 | 0.036 | -0.43 | 0.036 | 0.045 | 0.045 | 0.071 | 0.49 | 0.59 | 0.37 | 0.32 | 0.36 | 0.57 |
| 11 | 8403699 | rs7112477 | -1.811 | -1.791 | -0.441 | -0.159 | -0.054 | -0.043 | -0.043 | 0 | 0.74 | 0.84 | 0.52 | 0.6 | 0.54 | 0.74 |
| 11 | 8404501 | rs16937956 | 0.039 | -1.771 | -0.553 | -0.127 | 0.041 | -0.037 | -0.037 | 0 | 0.45 | 0.23 | 0.42 | 0.53 | 0.39 | 0.51 |
| 11 | 8404535 | rs16937958 | 1.076 | -1.634 | -0.477 | -0.253 | 0.013 | 0.011 | 0.011 | 0 | 0.28 | 0.19 | 0.26 | 0.28 | 0.25 | 0.39 |
| 11 | 8416123 | rs7111218 | 0.278 | -1.435 | -0.197 | -0.31 | 0.001 | -0.078 | -0.078 | 0 | 0.17 | 0.37 | 0.13 | 0.04 | 0.15 | 0.17 |
| 11 | 8424590 | rs7929500 | -1.675 | -1.455 | -0.335 | -0.042 | -0.027 | -0.02 | -0.02 | 0.366 | 0.74 | 0.79 | 0.52 | 0.61 | 0.54 | 0.73 |
| 11 | 8430938 | rs4586148 | 1.079 | -1.694 | -0.506 | 0.016 | 0.017 | -0.082 | -0.082 | 0 | 0.13 | 0.01 | 0.1 | 0.25 | 0.08 | 0.06 |
| 11 | 8433864 | rs10500703 | 1.948 | -1.958 | -0.654 | -0.12 | -0.011 | 0.004 | 0.004 | 0.481 | 0.21 | 0.03 | 0.21 | 0.21 | 0.21 | 0.35 |
| 11 | 8434164 | rs7126521 | 1.508 | -2.044 | -0.465 | -0.205 | -0.003 | -0.042 | -0.042 | 0 | 0.12 | 0.22 | 0.05 | 0.04 | 0.1 | 0.11 |
| 11 | 8434693 | rs10840037 | 0.108 | -2.217 | -0.166 | -0.22 | -0.041 | 0.04 | 0.04 | 0 | 0.21 | 0.28 | 0.13 | 0.29 | 0.11 | 0.14 |
| 11 | 8435078 | rs3751096 | 1.236 | -2.122 | -0.152 | -0.175 | 0.006 | -0.001 | -0.001 | 0 | 0.27 | 0.11 | 0.26 | 0.29 | 0.25 | 0.38 |
| 11 | 8437827 | rs12808779 | 2.001 | -2.1 | -0.144 | -0.166 | 0.003 | 0.014 | 0.014 | 0.588 | 0.21 | 0.06 | 0.21 | 0.21 | 0.21 | 0.35 |
| 11 | 8442054 | rs7939568 | -0.597 | -2.035 | -0.048 | -0.08 | -0.051 | 0.004 | 0.004 | 0 | 0.3 | 0.33 | 0.14 | 0.29 | 0.14 | 0.18 |
| 11 | 8453594 | rs2121312 | -0.724 | -1.759 | 0.017 | -0.06 | -0.003 | 0.011 | 0.011 | 0 | 0.58 | 0.62 | 0.39 | 0.36 | 0.46 | 0.67 |
| 11 | 8461178 | rs7112153 | -1.035 | -1.617 | 0.437 | -0.013 | 0.014 | 0.022 | 0.022 | 0 | 0.65 | 0.64 | 0.47 | 0.54 | 0.5 | 0.68 |
| 11 | 8462719 | rs7127684 | -0.931 | -1.581 | 0.523 | -0.017 | 0.014 | 0.023 | 0.023 | 0 | 0.64 | 0.5 | 0.45 | 0.54 | 0.5 | 0.68 |
| 11 | 8483335 | rs1900273 | -1.221 | -1.317 | 0.352 | -0.008 | 0.008 | 0.013 | 0.013 | 0 | 0.68 | 0.71 | 0.48 | 0.54 | 0.5 | 0.69 |
| 11 | 8486310 | rs1446464 | -0.533 | -1.389 | 0.261 | -0.032 | 0.007 | -0.008 | -0.008 | 0 | 0.59 | 0.34 | 0.37 | 0.49 | 0.42 | 0.62 |
| 11 | 8525293 | rs7931834 | 1.255 | -1.422 | 0.145 | -0.098 | -0.005 | -0.009 | -0.009 | 0 | 0.24 | 0.35 | 0.25 | 0.26 | 0.23 | 0.36 |
| 11 | 8535922 | rs1992756 | -0.543 | -1.683 | 0.155 | -0.152 | 0.025 | 0.012 | 0.012 | 0.254 | 0.38 | 0.34 | 0.18 | 0.29 | 0.2 | 0.26 |
| 11 | 8551916 | rs7105027 | -0.552 | -1.672 | 0.146 | -0.134 | 0.028 | 0.013 | 0.013 | 0.047 | 0.38 | 0.34 | 0.18 | 0.29 | 0.21 | 0.26 |
| 11 | 8561002 | rs11041966 | -0.173 | -1.941 | -0.05 | -0.396 | 0.078 | -0.005 | -0.005 | 1.207 | 0.29 | 0.13 | 0.14 | 0.26 | 0.16 | 0.17 |
| 11 | 8564778 | rs1026021 | 1.627 | -1.899 | -0.104 | -0.356 | -0.043 | -0.003 | -0.003 | 0 | 0.21 | 0.01 | 0.2 | 0.2 | 0.22 | 0.35 |
| 11 | 8568207 | rs10840070 | -0.06 | -1.898 | -0.068 | -0.324 | -0.003 | -0.013 | -0.013 | 0 | 0.4 | 0.46 | 0.33 | 0.3 | 0.37 | 0.51 |
| 11 | 8588823 | rs9300091 | -1.471 | -1.448 | 0.104 | -0.056 | 0.029 | 0.04 | 0.04 | 0 | 0.69 | 0.65 | 0.47 | 0.53 | 0.5 | 0.69 |
| 11 | 8597866 | rs11041994 | -1.146 | -1.455 | 0.13 | -0.047 | 0.019 | 0.024 | 0.024 | 0 | 0.66 | 0.48 | 0.45 | 0.5 | 0.49 | 0.68 |
| 11 | 8599718 | rs12361165 | -0.028 | -1.455 | 0.12 | -0.096 | 0.015 | 0.005 | 0.005 | 0 | 0.08 | 0.29 | 0.09 | 0.05 | 0.08 | 0.07 |
| 11 | 8622855 | rs10840089 | 0.351 | -1.476 | -0.117 | 0.064 | -0.007 | 0.012 | 0.012 | 0.08 | 0.4 | 0.48 | 0.6 | 0.47 | 0.58 | 0.37 |
| 11 | 8623193 | rs10769924 | 1.143 | -1.383 | -0.069 | 0.179 | 0.024 | 0.027 | 0.027 | 0 | 0.32 | 0.34 | 0.53 | 0.41 | 0.5 | 0.31 |
| 11 | 8625817 | rs11042008 | 0.495 | -1.66 | -0.15 | 0.03 | -0.001 | 0.001 | 0.001 | 0 | 0.55 | 0.38 | 0.73 | 0.68 | 0.72 | 0.67 |
| 11 | 8639200 | rs4929923 | 0.078 | -1.387 | -0.263 | 0.681 | 0.001 | 0 | 0 | 0 | 0.55 | 0.55 | 0.59 | 0.4 | 0.62 | 0.52 |
| 11 | 8643989 | rs10840096 | -0.874 | -1.548 | -0.375 | 0.802 | -0.014 | 0.016 | 0.016 | 0 | 0.83 | 0.77 | 0.84 | 0.77 | 0.88 | 0.92 |
| 11 | 8646670 | rs7947230 | 1.284 | -1.608 | -0.189 | 0.768 | 0 | 0.001 | 0.001 | 0.109 | 0.22 | 0.26 | 0.21 | 0.36 | 0.19 | 0.34 |
| 11 | 8653359 | rs9633894 | -0.965 | -1.608 | -0.136 | 0.766 | -0.004 | 0.004 | 0.004 | 0 | 0.84 | 0.84 | 0.85 | 0.77 | 0.88 | 0.92 |
| 11 | 8657321 | rs10743084 | 1.49 | -1.618 | -0.191 | 0.779 | 0.007 | -0.027 | -0.027 | 0.038 | 0.21 | 0.15 | 0.19 | 0.36 | 0.19 | 0.35 |
| 11 | 8658485 | rs4929927 | 0.204 | -1.489 | -0.106 | 0.531 | -0.004 | -0.002 | -0.002 | 0 | 0.55 | 0.5 | 0.58 | 0.4 | 0.63 | 0.51 |
| 11 | 8662516 | rs11042023 | -0.065 | -1.436 | -0.165 | 0.475 | -0.004 | -0.002 | -0.002 | 2.49 | 0.58 | 0.52 | 0.58 | 0.4 | 0.63 | 0.51 |
| 11 | 8671369 | rs10840101 | 2.1 | -0.758 | -0.819 | 0.453 | -0.132 | -0.002 | -0.002 | 0.452 | 0.2 | 0.01 | 0.2 | 0.13 | 0.27 | 0.19 |
| 11 | 8678756 | rs885114 | 0.037 | -1.047 | -0.593 | 0.634 | 0 | -0.001 | -0.001 | 5.114 | 0.39 | 0.42 | 0.35 | 0.6 | 0.31 | 0.42 |
| 11 | 8694830 | rs10128597 | 0.199 | -0.961 | -1.156 | 0.677 | 0.04 | -0.018 | -0.018 | 0.257 | 0.37 | 0.16 | 0.32 | 0.6 | 0.32 | 0.43 |
| 11 | 8694999 | rs7121168 | 0.88 | -0.624 | -0.975 | 0.533 | 0 | -0.019 | -0.019 | 0 | 0.06 | 0.08 | 0.06 | 0 | 0.06 | 0.06 |
| 11 | 8699225 | rs2036423 | 0.085 | -1.067 | -1.742 | 0.479 | 0.038 | 0.029 | 0.029 | 0 | 0.5 | 0.3 | 0.36 | 0.61 | 0.42 | 0.5 |
| 11 | 8701836 | rs10840106 | 1.153 | -1.568 | -2.13 | 0.504 | -0.066 | 0.016 | 0.016 | 0 | 0.28 | 0.14 | 0.27 | 0.19 | 0.38 | 0.24 |
| 11 | 8703941 | rs10840108 | 0.666 | -2.072 | -2.224 | 0.66 | -0.003 | -0.001 | -0.001 | 0 | 0.37 | 0.39 | 0.29 | 0.33 | 0.24 | 0.37 |
| 11 | 8706628 | rs2073685 | 2.473 | -2.143 | -2.35 | 0.666 | 0.004 | -0.03 | -0.03 | 0 | 0.19 | 0.09 | 0.17 | 0.28 | 0.18 | 0.23 |
| 11 | 8708989 | rs7925000 | 0.64 | -2.17 | -2.367 | 0.639 | -0.003 | -0.001 | -0.001 | 0 | 0.36 | 0.39 | 0.28 | 0.34 | 0.24 | 0.36 |
| 11 | 8709551 | rs10840111 | 2.16 | -1.662 | -2.968 | 0.86 | 0.005 | -0.031 | -0.031 | 0.198 | 0.2 | 0.09 | 0.17 | 0.28 | 0.18 | 0.23 |
| 11 | 8710511 | rs10840112 | 1.499 | -1.439 | -2.814 | 0.751 | -0.183 | -0.032 | -0.032 | 4.154 | 0.28 | 0.03 | 0.26 | 0.18 | 0.38 | 0.24 |
| 11 | 8714325 | rs4929908 | 0.56 | -1.568 | -2.248 | 0.803 | -0.026 | -0.106 | -0.106 | 0.098 | 0.31 | 0.11 | 0.43 | 0.52 | 0.39 | 0.38 |
| 11 | 8735662 | rs931756 | -0.006 | -1.146 | -1.562 | 0.692 | -0.002 | -0.008 | -0.008 | 0.126 | 0.61 | 0.59 | 0.53 | 0.64 | 0.52 | 0.62 |
| 11 | 8736695 | rs10840114 | 0.632 | -0.812 | -2.144 | 0.838 | -0.125 | -0.13 | -0.13 | 0.509 | 0.32 | 0.03 | 0.31 | 0.37 | 0.45 | 0.28 |
| 11 | 8744219 | rs7105372 | -0.396 | -1.055 | -1.285 | 0.65 | 0.004 | -0.001 | -0.001 | 2.04 | 0.24 | 0.17 | 0.13 | 0.28 | 0.21 | 0.35 |
| 11 | 8753552 | rs905289 | -0.943 | -1.334 | -0.962 | 0.6 | 0 | 0 | 0 | 0 | 0.52 | 0.54 | 0.63 | 0.66 | 0.64 | 0.64 |
| 11 | 8753729 | rs905290 | 0.823 | -1.26 | -1.264 | 0.636 | -0.17 | -0.156 | -0.156 | 0.002 | 0.25 | 0.03 | 0.47 | 0.35 | 0.43 | 0.27 |
| 11 | 8758114 | rs10769947 | 0.685 | -1.314 | -1.087 | 0.717 | -0.098 | -0.067 | -0.067 | 2.011 | 0.26 | 0.14 | 0.47 | 0.3 | 0.43 | 0.27 |
| 11 | 8760156 | rs10840115 | 0.509 | -2.089 | -1.22 | 0.403 | 0.061 | -0.056 | -0.056 | 0 | 0.56 | 0.17 | 0.64 | 0.38 | 0.6 | 0.49 |
| 11 | 8762162 | rs12296005 | 1.628 | -1.517 | -0.835 | 0.706 | -0.018 | -0.053 | -0.053 | 0 | 0.08 | 0.01 | 0.05 | 0.22 | 0.1 | 0.21 |
| 11 | 8778352 | rs10769953 | -1.518 | -1.788 | -0.298 | 0.593 | -0.006 | -0.002 | -0.002 | 0 | 0.58 | 0.59 | 0.66 | 0.64 | 0.66 | 0.69 |
| 11 | 8786058 | rs10743087 | -0.72 | -1.868 | -0.104 | 0.511 | 0.074 | 0.045 | 0.045 | 0 | 0.51 | 0.34 | 0.61 | 0.64 | 0.63 | 0.64 |
| 11 | 8787620 | rs10743088 | -0.862 | -1.825 | 0.02 | 0.5 | 0.007 | 0.005 | 0.005 | 0.026 | 0.52 | 0.46 | 0.64 | 0.64 | 0.64 | 0.64 |
| 11 | 8791227 | rs2873297 | -0.708 | -1.86 | 0.065 | 0.492 | 0.084 | 0.051 | 0.051 | 0.007 | 0.51 | 0.32 | 0.62 | 0.64 | 0.64 | 0.64 |
| 11 | 8792622 | rs4929936 | 1.379 | -1.944 | 0.108 | 0.514 | 0.042 | 0.061 | 0.061 | 0 | 0.39 | 0.11 | 0.32 | 0.26 | 0.34 | 0.3 |

| | | | | | | | | | | | | | | | | |
|---|---|---|---|---|---|---|---|---|---|---|---|---|---|---|---|---|
| 11 | 8799749 | rs11042064 | 0.194 | -1.911 | 0.074 | 0.544 | 0.01 | -0.002 | -0.002 | 0 | 0.25 | 0.21 | 0.15 | 0.31 | 0.21 | 0.37 |
| 11 | 8807179 | rs2292046 | -0.752 | -1.946 | 0.115 | 0.546 | 0.054 | 0.036 | 0.036 | 0.034 | 0.51 | 0.37 | 0.62 | 0.64 | 0.63 | 0.64 |
| 11 | 8813956 | rs12421789 | -0.012 | -1.845 | 0.521 | 0.569 | 0.036 | 0.041 | 0.041 | 0.429 | 0.66 | 0.51 | 0.49 | 0.58 | 0.55 | 0.67 |
| 11 | 8815115 | rs10769958 | -0.056 | -1.845 | 0.521 | 0.569 | 0.039 | 0.044 | 0.044 | 0.438 | 0.66 | 0.52 | 0.49 | 0.58 | 0.55 | 0.67 |
| 11 | 8816495 | rs4929911 | 0.844 | -1.854 | 0.483 | 0.548 | -0.083 | -0.055 | -0.055 | 0 | 0.25 | 0.11 | 0.48 | 0.32 | 0.43 | 0.27 |
| 11 | 8817206 | rs6484484 | 0.143 | -1.787 | 0.571 | 0.583 | -0.035 | -0.016 | -0.016 | 0 | 0.32 | 0.25 | 0.5 | 0.32 | 0.45 | 0.32 |
| 11 | 8817580 | rs10500707 | 1.338 | -1.918 | 0.343 | 0.456 | 0.037 | 0.052 | 0.052 | 0 | 0.39 | 0.21 | 0.35 | 0.26 | 0.35 | 0.3 |
| 11 | 8819808 | rs3763917 | 0.07 | -1.845 | 0.315 | 0.48 | 0 | -0.001 | -0.001 | 0.238 | 0.26 | 0.26 | 0.14 | 0.32 | 0.2 | 0.36 |
| 11 | 8824388 | rs4929914 | -0.673 | -1.748 | 0.124 | 0.507 | -0.006 | -0.004 | -0.004 | 1.555 | 0.71 | 0.73 | 0.51 | 0.69 | 0.57 | 0.7 |
| 11 | 8824418 | rs4929915 | 1.873 | -1.907 | -0.006 | 0.331 | 0.02 | 0.086 | 0.086 | 0 | 0.32 | 0.15 | 0.24 | 0.07 | 0.29 | 0.2 |
| 11 | 8825966 | rs11042077 | -0.851 | -1.707 | -0.045 | 0.527 | -0.004 | -0.005 | -0.005 | 0 | 0.55 | 0.58 | 0.62 | 0.69 | 0.61 | 0.68 |
| 11 | 8826700 | rs11042078 | 0.348 | -1.173 | -0.047 | 0.658 | -0.06 | -0.028 | -0.028 | 1.174 | 0.27 | 0.13 | 0.48 | 0.3 | 0.41 | 0.28 |
| 11 | 8836030 | rs7101895 | 0.912 | -1.269 | 0.147 | 0.545 | -0.01 | 0.02 | 0.02 | 0.059 | 0.08 | 0.13 | 0.1 | 0.21 | 0.06 | 0.1 |
| 11 | 8836801 | rs10769962 | 0.12 | -1.376 | 0.549 | 0.474 | -0.042 | 0.089 | 0.089 | 0.324 | 0.87 | 0.59 | 0.86 | 0.79 | 0.93 | 0.89 |
| 11 | 8838660 | rs4929918 | 0.182 | -1.531 | 0.028 | 0.395 | -0.005 | -0.002 | -0.002 | 0.07 | 0.7 | 0.39 | 0.82 | 0.67 | 0.76 | 0.64 |
| 11 | 8844065 | rs7935617 | -0.229 | -1.481 | 0.144 | 0.378 | 0 | -0.001 | -0.001 | 0.2 | 0.28 | 0.31 | 0.15 | 0.34 | 0.26 | 0.37 |
| 11 | 8849495 | rs2568090 | 0.345 | -1.413 | -0.051 | 0.272 | -0.08 | -0.071 | -0.071 | 0.053 | 0.28 | 0.14 | 0.48 | 0.39 | 0.41 | 0.34 |
| 11 | 8852239 | rs2243566 | -0.263 | -1.435 | -0.079 | 0.183 | -0.039 | -0.005 | -0.005 | 0 | 0.36 | 0.27 | 0.58 | 0.58 | 0.47 | 0.42 |
| 11 | 8859384 | rs2568025 | 0.466 | -1.413 | -0.471 | 0.003 | 0.026 | -0.016 | -0.016 | 0 | 0.62 | 0.36 | 0.71 | 0.47 | 0.69 | 0.53 |
| 11 | 8862530 | rs7945705 | 0.448 | -1.123 | -0.926 | 0.071 | -0.187 | -0.131 | -0.131 | 1.036 | 0.26 | 0.02 | 0.46 | 0.33 | 0.41 | 0.3 |
| 11 | 8865363 | rs10840134 | 0.448 | -1.166 | -0.926 | 0.071 | -0.189 | -0.131 | -0.131 | 5.444 | 0.26 | 0.02 | 0.46 | 0.33 | 0.41 | 0.3 |
| 11 | 8874075 | rs4910068 | 1.717 | -0.978 | -0.492 | -0.14 | 0.026 | 0.137 | 0.137 | 0.021 | 0.33 | 0.09 | 0.23 | 0.05 | 0.27 | 0.19 |
| 11 | 8876852 | rs2742552 | -0.322 | -1.086 | -0.164 | -0.038 | 0.015 | 0.015 | 0.015 | 0 | 0.67 | 0.45 | 0.47 | 0.6 | 0.54 | 0.65 |
| 11 | 8879794 | rs10840135 | -0.488 | -1.096 | -0.175 | -0.032 | 0.023 | -0.023 | -0.023 | 0 | 0.33 | 0.24 | 0.23 | 0.55 | 0.27 | 0.47 |
| 11 | 8887097 | rs16906256 | -0.488 | -1.096 | -0.175 | -0.068 | 0.023 | -0.022 | -0.022 | 0 | 0.33 | 0.25 | 0.23 | 0.55 | 0.27 | 0.47 |
| 11 | 8893193 | rs2653565 | 1.395 | -1.192 | -0.296 | -0.096 | 0.034 | 0.074 | 0.074 | 0.076 | 0.42 | 0.11 | 0.27 | 0.27 | 0.35 | 0.44 |
| 11 | 8901824 | rs12223690 | 0.554 | -1.264 | -0.261 | -0.13 | -0.001 | 0.017 | 0.017 | 0.301 | 0.14 | 0.2 | 0.16 | 0.33 | 0.15 | 0.2 |
| 11 | 8905461 | rs1025438 | 1.738 | -0.994 | -0.206 | 0.024 | -0.011 | -0.045 | -0.045 | 0 | 0.08 | 0.01 | 0.04 | 0.22 | 0.08 | 0.25 |
| 11 | 8920704 | rs10840147 | 0.8 | -1.001 | 0.015 | -0.059 | -0.169 | -0.122 | -0.122 | 4.234 | 0.26 | 0.05 | 0.48 | 0.33 | 0.44 | 0.26 |
| 11 | 8923528 | rs10840148 | 0.807 | -0.969 | 0.022 | -0.035 | -0.163 | -0.12 | -0.12 | 0 | 0.25 | 0.05 | 0.47 | 0.33 | 0.42 | 0.26 |
| 11 | 8927518 | rs2568029 | 1.908 | -1.106 | -0.188 | -0.252 | 0.021 | 0.14 | 0.14 | 0.099 | 0.33 | 0.06 | 0.22 | 0.05 | 0.27 | 0.19 |
| 11 | 8933064 | rs1133833 | 2.064 | -1.152 | -0.29 | -0.344 | -0.023 | 0.232 | 0.232 | 0.245 | 0.32 | 0.01 | 0.21 | 0 | 0.27 | 0.17 |
| 11 | 8939887 | rs1560569 | 0.897 | -1.056 | -0.111 | -0.237 | -0.042 | -0.032 | -0.032 | 0 | 0.25 | 0.17 | 0.5 | 0.33 | 0.42 | 0.25 |
| 11 | 8940149 | rs2056903 | 1.242 | -1.133 | -0.096 | -0.284 | 0.022 | 0.14 | 0.14 | 0.182 | 0.4 | 0.11 | 0.25 | 0.22 | 0.35 | 0.46 |
| 11 | 8946950 | rs3751068 | -0.774 | -1.181 | 0.069 | -0.321 | 0.029 | 0.048 | 0.048 | 0.3 | 0.51 | 0.4 | 0.69 | 0.71 | 0.61 | 0.49 |
| 11 | 8959020 | rs3751063 | -1.087 | -1.178 | 0.086 | -0.325 | 0.055 | 0.086 | 0.086 | 0.326 | 0.54 | 0.29 | 0.69 | 0.71 | 0.64 | 0.51 |
| 11 | 8959545 | rs4909951 | -1.186 | -1.181 | 0.088 | -0.327 | 0.05 | 0.079 | 0.079 | 0.261 | 0.54 | 0.3 | 0.69 | 0.71 | 0.64 | 0.51 |
| 11 | 8962764 | rs2742502 | 2.161 | -1.198 | 0.026 | -0.362 | -0.029 | 0.236 | 0.236 | 0.263 | 0.32 | 0.01 | 0.21 | 0 | 0.28 | 0.17 |
| 11 | 8964319 | rs3763912 | -0.839 | -1.108 | 0.147 | -0.255 | 0.064 | 0.115 | 0.115 | 0.379 | 0.5 | 0.29 | 0.68 | 0.71 | 0.61 | 0.49 |
| 11 | 8964691 | rs2742497 | 1.338 | -1.115 | 0.142 | -0.277 | 0.024 | 0.145 | 0.145 | 0.206 | 0.4 | 0.12 | 0.25 | 0.22 | 0.35 | 0.46 |
| 11 | 8964938 | rs2455603 | -0.06 | -1.116 | 0.254 | -0.278 | 0.043 | 0.002 | 0.002 | 0 | 0.93 | 0.63 | 0.95 | 1 | 0.96 | 0.98 |
| 11 | 8975748 | rs2568085 | -0.379 | -0.983 | 0.129 | -0.219 | -0.04 | 0.062 | 0.062 | 0.063 | 0.28 | 0.44 | 0.24 | 0.45 | 0.21 | 0.27 |
| 11 | 8975776 | rs4929922 | 1.064 | -0.977 | 0.12 | -0.218 | -0.145 | -0.123 | -0.123 | 0 | 0.23 | 0.05 | 0.47 | 0.33 | 0.4 | 0.24 |
| 11 | 8981537 | rs2163625 | -0.82 | -1.107 | 0.206 | -0.213 | 0.026 | 0.049 | 0.049 | 0.307 | 0.51 | 0.41 | 0.7 | 0.75 | 0.61 | 0.49 |
| 11 | 8983284 | rs2116510 | -0.966 | -1.097 | 0.14 | -0.245 | -0.005 | -0.026 | -0.026 | 0.431 | 0.52 | 0.5 | 0.7 | 0.78 | 0.61 | 0.52 |
| 11 | 8986978 | rs2653569 | -0.923 | -0.976 | 0.247 | -0.083 | -0.035 | 0.043 | 0.043 | 0.065 | 0.32 | 0.5 | 0.25 | 0.44 | 0.25 | 0.29 |
| 11 | 9003306 | rs1123936 | -0.312 | -1.073 | 0.198 | -0.15 | 0.031 | 0.03 | 0.03 | 0 | 0.66 | 0.46 | 0.47 | 0.61 | 0.54 | 0.69 |
| 11 | 9007669 | rs3815991 | 0.061 | -1.132 | 0.263 | -0.164 | 0.014 | -0.036 | -0.036 | 0.006 | 0.24 | 0.13 | 0.2 | 0.38 | 0.21 | 0.23 |
| 11 | 9008674 | rs11042148 | 0.035 | -1.128 | 0.235 | -0.162 | 0.003 | -0.005 | -0.005 | 0.002 | 0.25 | 0.22 | 0.2 | 0.38 | 0.21 | 0.23 |
| 11 | 9010737 | rs11042149 | -0.284 | -1.028 | 0.126 | -0.28 | -0.009 | -0.008 | -0.008 | 0.609 | 0.65 | 0.36 | 0.47 | 0.61 | 0.54 | 0.69 |
| 11 | 9013639 | rs7930026 | 1.129 | -0.924 | -0.102 | -0.207 | -0.159 | -0.135 | -0.135 | 0.543 | 0.25 | 0.05 | 0.46 | 0.35 | 0.42 | 0.23 |
| 11 | 9024867 | rs6486061 | 0.572 | -0.985 | 0.071 | -0.357 | 0.016 | 0.022 | 0.022 | 0.057 | 0.48 | 0.47 | 0.31 | 0.28 | 0.37 | 0.55 |
| 11 | 9035027 | rs1984791 | 1.531 | -0.243 | -0.25 | -0.279 | -0.108 | -0.117 | -0.117 | 0.115 | 0.18 | 0.05 | 0.43 | 0.34 | 0.31 | 0.19 |
| 11 | 9038925 | rs7941341 | 1.004 | -1.91 | -0.586 | -0.784 | 0.041 | 0.145 | 0.145 | 0 | 0.35 | 0.16 | 0.24 | 0 | 0.28 | 0.2 |
| 11 | 9042331 | rs1136966 | -1.341 | -1.824 | -0.212 | -0.736 | 0.025 | 0.034 | 0.034 | 0.456 | 0.66 | 0.78 | 0.77 | 1 | 0.73 | 0.81 |
| 11 | 9042569 | rs1367 | 0.321 | -1.865 | -0.241 | -0.561 | -0.001 | 0 | 0 | 0 | 0.07 | 0.1 | 0.04 | 0.13 | 0.07 | 0.11 |
| 11 | 9044767 | rs932287 | -0.274 | -2.391 | -0.205 | -0.551 | 0.014 | 0.024 | 0.024 | 0.399 | 0.46 | 0.61 | 0.6 | 0.77 | 0.61 | 0.67 |
| 11 | 9048513 | rs2056902 | 0.741 | -2.403 | -0.403 | -0.366 | -0.001 | 0.001 | 0.001 | 0 | 0.1 | 0.11 | 0.06 | 0.21 | 0.1 | 0.26 |
| 11 | 9048668 | rs3829273 | 0.346 | -2.403 | -0.432 | -0.391 | -0.001 | -0.014 | -0.014 | 0 | 0.12 | 0.06 | 0.06 | 0.21 | 0.13 | 0.28 |
| 11 | 9051224 | rs1960180 | -0.882 | -2.608 | -0.473 | -0.578 | 0.037 | 0.067 | 0.067 | 1.513 | 0.57 | 0.69 | 0.72 | 0.79 | 0.64 | 0.56 |
| 11 | 9051406 | rs3751058 | 0.898 | -2.326 | -0.557 | -0.42 | 0 | -0.019 | -0.019 | 0 | 0.09 | 0.06 | 0.05 | 0.21 | 0.1 | 0.26 |
| 11 | 9054522 | rs7109896 | -0.103 | -2.733 | -0.558 | -0.476 | 0.027 | 0.022 | 0.022 | 0.026 | 0.54 | 0.59 | 0.41 | 0.31 | 0.38 | 0.38 |
| 11 | 9055970 | rs3794149 | -0.489 | -2.695 | -1.004 | -0.519 | -0.016 | -0.01 | -0.01 | 1.92 | 0.31 | 0.39 | 0.27 | 0.23 | 0.27 | 0.22 |
| 11 | 9059192 | rs1883100 | -1.713 | -2.354 | -0.561 | -0.121 | -0.014 | -0.037 | -0.037 | 3.763 | 0.64 | 0.63 | 0.76 | 0.9 | 0.74 | 0.77 |
| 11 | 9063570 | rs1883099 | 0.368 | -2.007 | -0.968 | 0.023 | -0.108 | -0.094 | -0.094 | 1.281 | 0.32 | 0.07 | 0.47 | 0.55 | 0.47 | 0.5 |
| 11 | 9064231 | rs3794147 | -0.139 | -1.893 | -0.537 | -0.282 | 0.002 | -0.002 | -0.002 | 0 | 0.09 | 0.15 | 0.08 | 0.12 | 0.12 | 0.11 |
| 11 | 9067400 | rs10840164 | -0.916 | -1.833 | -0.651 | -0.137 | -0.004 | -0.004 | -0.004 | 0 | 0.38 | 0.38 | 0.2 | 0.39 | 0.33 | 0.47 |
| 11 | 9068716 | rs6486112 | -0.204 | -1.93 | -0.63 | -0.127 | 0.001 | 0.001 | 0.001 | 0.389 | 0.49 | 0.54 | 0.69 | 0.61 | 0.62 | 0.44 |
| 11 | 9069046 | rs3751055 | -0.206 | -1.449 | -1.044 | 0.358 | 0.014 | -0.017 | -0.017 | 0 | 0.15 | 0.05 | 0.06 | 0.26 | 0.13 | 0.28 |

| | | | | | | | | | | | | | | | | |
|---|---|---|---|---|---|---|---|---|---|---|---|---|---|---|---|---|
| 11 | 9071106 | rs10840166 | 0.38 | -1.178 | -0.967 | 0.487 | -0.018 | -0.011 | -0.011 | 0 | 0.38 | 0.28 | 0.62 | 0.46 | 0.54 | 0.38 |
| 11 | 9071175 | rs10769988 | -0.996 | -1.712 | -1.08 | 0.31 | 0.001 | 0.005 | 0.005 | 2.009 | 0.73 | 0.71 | 0.77 | 0.89 | 0.75 | 0.89 |
| 11 | 9071236 | rs7106593 | 0.178 | -1.709 | -0.977 | 0.467 | 0.022 | 0.032 | 0.032 | 0.536 | 0.47 | 0.63 | 0.67 | 0.64 | 0.59 | 0.49 |
| 11 | 9072140 | rs3751052 | 0.062 | -1.751 | -1.008 | 0.485 | 0.045 | 0.045 | 0.045 | 0.161 | 0.5 | 0.63 | 0.7 | 0.63 | 0.61 | 0.53 |
| 11 | 9072944 | rs7126825 | -1.224 | -1.731 | -1.151 | 0.401 | -0.007 | -0.009 | -0.009 | 0 | 0.75 | 0.69 | 0.82 | 0.89 | 0.8 | 0.91 |
| 11 | 9074535 | rs3751051 | -0.014 | -1.326 | -1.824 | 0.432 | 0.001 | -0.001 | -0.001 | 0 | 0.24 | 0.03 | 0.11 | 0.26 | 0.19 | 0.38 |
| 11 | 9081926 | rs7112378 | -0.253 | -0.939 | -1.407 | 0.541 | 0.031 | -0.011 | -0.011 | 1.12 | 0.59 | 0.26 | 0.73 | 0.7 | 0.68 | 0.76 |
| 11 | 9086295 | rs6486125 | 0.388 | -1.255 | -0.632 | 0.68 | -0.024 | -0.013 | -0.013 | 0.02 | 0.62 | 0.56 | 0.78 | 0.63 | 0.7 | 0.5 |
| 11 | 9089120 | rs7107892 | 0.975 | -1.514 | -0.6 | 0.771 | -0.011 | 0.002 | 0.002 | 0 | 0.67 | 0.65 | 0.84 | 0.56 | 0.76 | 0.61 |
| 11 | 9090408 | rs11606516 | -0.686 | -0.952 | -0.949 | 0.586 | 0.002 | -0.004 | -0.004 | 0 | 0.1 | 0.06 | 0.03 | 0.14 | 0.07 | 0.1 |
| 11 | 9091244 | rs3751050 | -0.615 | -0.98 | -1.016 | 0.596 | 0.018 | 0.045 | 0.045 | 0 | 0.13 | 0.05 | 0.06 | 0.08 | 0.11 | 0.21 |
| 11 | 9093486 | rs7929797 | 0.492 | -0.732 | -0.998 | 0.631 | 0.002 | -0.013 | -0.013 | 0 | 0.86 | 0.89 | 0.82 | 0.78 | 0.87 | 0.76 |
| 11 | 9109287 | rs2647528 | -1.461 | -0.99 | -0.882 | 0.406 | 0.012 | 0.012 | 0.012 | 0 | 0.25 | 0.19 | 0.21 | 0.21 | 0.2 | 0.27 |
| 11 | 9114999 | rs4910440 | -0.074 | -0.488 | -1.457 | 0.36 | -0.024 | -0.018 | -0.018 | 0.148 | 0.54 | 0.81 | 0.59 | 0.45 | 0.47 | 0.58 |
| 11 | 9117898 | rs10840170 | 0.606 | -0.688 | -0.763 | -0.569 | 0.003 | -0.085 | -0.085 | 0 | 0.2 | 0.32 | 0.15 | 0.01 | 0.22 | 0.09 |
| 11 | 9127130 | rs1121629 | -0.094 | -1.199 | -0.546 | -0.736 | -0.012 | -0.014 | -0.014 | 0 | 0.16 | 0.24 | 0.07 | 0.07 | 0.11 | 0.15 |
| 11 | 9130147 | rs11042182 | 1.123 | -1.236 | -0.547 | -0.385 | -0.029 | -0.048 | -0.048 | 0 | 0.33 | 0.13 | 0.31 | 0.43 | 0.36 | 0.28 |
| 11 | 9133052 | rs3898554 | -0.286 | -1.495 | 0.221 | -0.504 | -0.008 | -0.06 | -0.06 | 0.435 | 0.37 | 0.45 | 0.32 | 0.11 | 0.36 | 0.28 |
| 11 | 9133878 | rs4910443 | 1.324 | -1.554 | 0.235 | -0.511 | -0.035 | 0.104 | 0.104 | 0.091 | 0.19 | 0.02 | 0.12 | 0.01 | 0.23 | 0.1 |
| 11 | 9142797 | rs7128469 | -1.075 | -1.925 | 0.274 | -0.889 | 0.043 | 0.163 | 0.163 | 0 | 0.5 | 0.73 | 0.4 | 0.14 | 0.45 | 0.38 |
| 11 | 9145807 | rs10769990 | -0.551 | -1.387 | 0.113 | -1.206 | 0 | 0.002 | 0.002 | 0 | 0.62 | 0.57 | 0.55 | 0.47 | 0.63 | 0.51 |
| 11 | 9159818 | rs1353729 | -0.013 | -1.485 | 0.348 | -1.392 | -0.049 | -0.073 | -0.073 | 0 | 0.2 | 0.43 | 0.21 | 0.1 | 0.13 | 0.18 |
| 11 | 9160070 | rs1353730 | -0.34 | -1.18 | 0.28 | -1.47 | -0.003 | -0.053 | -0.053 | 0 | 0.43 | 0.45 | 0.35 | 0.1 | 0.39 | 0.29 |
| 11 | 9162868 | rs10769993 | -0.136 | -1.05 | 0.28 | -1.328 | 0.004 | -0.001 | -0.001 | 0.06 | 0.38 | 0.62 | 0.35 | 0.1 | 0.36 | 0.25 |
| 11 | 9171775 | rs7123970 | -0.204 | -1.093 | 0.233 | -1.344 | 0.018 | 0.064 | 0.064 | 0 | 0.4 | 0.68 | 0.36 | 0.13 | 0.36 | 0.26 |
| 11 | 9185077 | rs9888257 | -0.636 | -1.077 | 0.376 | -1.287 | -0.019 | -0.042 | -0.042 | 3.266 | 0.24 | 0.3 | 0.22 | 0.11 | 0.19 | 0.24 |
| 11 | 9198010 | rs7925308 | -0.836 | -1.082 | 0.042 | -1.41 | 0.02 | 0.078 | 0.078 | 0 | 0.47 | 0.68 | 0.38 | 0.12 | 0.42 | 0.34 |
| 11 | 9206279 | rs12099067 | 0.517 | -1.075 | 0.254 | -1.436 | -0.01 | -0.009 | -0.009 | 1.304 | 0.16 | 0.2 | 0.17 | 0.11 | 0.12 | 0.15 |
| 11 | 9215691 | rs11042217 | 0.397 | -0.905 | 0.227 | -1.403 | 0 | -0.171 | -0.171 | 0.016 | 0.3 | 0.5 | 0.21 | 0.01 | 0.3 | 0.17 |
| 11 | 9232590 | rs12222684 | 1.603 | -0.807 | -0.22 | -1.161 | -0.003 | -0.038 | -0.038 | 0 | 0.24 | 0.11 | 0.22 | 0.39 | 0.25 | 0.2 |
| 11 | 9237540 | rs12360767 | 1.795 | -1.279 | -0.17 | -1.33 | -0.061 | 0.114 | 0.114 | 0.153 | 0.18 | 0.01 | 0.13 | 0.01 | 0.22 | 0.09 |
| 11 | 9258146 | rs11042231 | 1.487 | -1.217 | -0.049 | -1.334 | -0.022 | 0.07 | 0.07 | 0.057 | 0.2 | 0.09 | 0.13 | 0.01 | 0.22 | 0.09 |
| 11 | 9275898 | rs7116459 | -0.242 | -1.253 | 0.18 | -1.21 | 0.04 | -0.069 | -0.069 | 0 | 0.38 | 0.2 | 0.47 | 0.46 | 0.28 | 0.54 |
| 11 | 9276226 | rs7929621 | 0.403 | -1.347 | 0.423 | -1.26 | -0.003 | -0.14 | -0.14 | 0.47 | 0.3 | 0.56 | 0.21 | 0.01 | 0.3 | 0.17 |
| 11 | 9283449 | rs7936622 | 1.462 | -1.281 | 0.513 | -1.197 | 0 | -0.044 | -0.044 | 0.02 | 0.21 | 0.27 | 0.15 | 0.01 | 0.22 | 0.09 |
| 11 | 9304120 | rs4369406 | 1.152 | -1.017 | 0.424 | -1.064 | 0.026 | 0.037 | 0.037 | 0 | 0.35 | 0.44 | 0.36 | 0.51 | 0.45 | 0.28 |
| 11 | 9309405 | rs5014247 | 0.609 | -1.078 | 0.379 | -1.112 | 0.001 | 0 | 0 | 0 | 0.52 | 0.52 | 0.48 | 0.52 | 0.67 | 0.36 |
| 11 | 9310982 | rs11042267 | 1.218 | -1.057 | 0.087 | -1.1 | -0.023 | -0.029 | -0.029 | 0 | 0.34 | 0.23 | 0.34 | 0.51 | 0.45 | 0.28 |
| 11 | 9311128 | rs10770008 | 1.26 | -1.059 | 0.082 | -1.101 | -0.029 | -0.037 | -0.037 | 4.528 | 0.34 | 0.19 | 0.34 | 0.51 | 0.45 | 0.28 |
| 11 | 9311172 | rs7114748 | -0.15 | -1.19 | -0.061 | -1.124 | -0.009 | 0.008 | 0.008 | 0 | 0.66 | 0.64 | 0.64 | 0.64 | 0.76 | 0.55 |
| 11 | 9313814 | rs7103472 | 1.825 | -1.099 | -0.132 | -1.131 | -0.041 | 0.073 | 0.073 | 0 | 0.17 | 0.07 | 0.12 | 0.01 | 0.21 | 0.08 |
| 11 | 9323353 | rs4537761 | 0.579 | -1.102 | -0.051 | -1.037 | -0.031 | -0.022 | -0.022 | 0 | 0.16 | 0.33 | 0.17 | 0.11 | 0.11 | 0.19 |
| 11 | 9328743 | rs4282978 | 0.103 | -1.027 | -0.24 | -1.034 | 0.005 | 0.014 | 0.014 | 0 | 0.5 | 0.57 | 0.52 | 0.63 | 0.55 | 0.47 |
| 11 | 9352780 | rs7119300 | 0.805 | -0.274 | -0.075 | -0.574 | -0.005 | -0.019 | -0.019 | 0 | 0.69 | 0.33 | 0.72 | 0.85 | 0.74 | 0.69 |
| 11 | 9352951 | rs7131017 | 0.376 | 0.161 | -0.459 | -0.305 | -0.008 | -0.071 | -0.071 | 0 | 0.21 | 0.07 | 0.32 | 0.33 | 0.19 | 0.34 |
| 11 | 9363612 | rs7935301 | 0.982 | 0.417 | 0.258 | -0.613 | -0.005 | 0.021 | 0.021 | 1.494 | 0.79 | 0.42 | 0.7 | 0.77 | 0.81 | 0.73 |
| 11 | 9365378 | rs7483960 | 0.602 | 0.772 | 0.183 | -1.056 | 0.001 | -0.108 | -0.108 | 0 | 0.68 | 0.51 | 0.74 | 0.92 | 0.67 | 0.63 |
| 11 | 9366086 | rs7115641 | -0.042 | 0.927 | 0.611 | -0.245 | 0.033 | -0.039 | -0.039 | 0 | 0.91 | 0.68 | 0.93 | 1 | 0.92 | 0.92 |
| 11 | 9372626 | rs7950276 | -0.136 | 1.361 | 0.057 | -0.08 | 0.008 | -0.004 | -0.004 | 0 | 0.35 | 0.39 | 0.36 | 0.25 | 0.37 | 0.23 |
| 11 | 9376134 | rs10770022 | -0.42 | 1.003 | 0.25 | 0.152 | 0.017 | -0.02 | -0.02 | 0 | 0.61 | 0.71 | 0.58 | 0.42 | 0.62 | 0.46 |
| 11 | 9379005 | rs1578971 | 0.473 | 1.556 | 0.051 | -0.373 | -0.044 | 0.125 | 0.125 | 0 | 0.43 | 0.08 | 0.34 | 0.24 | 0.41 | 0.3 |
| 11 | 9389875 | rs4910048 | -0.043 | 1.259 | 1.067 | -0.303 | -0.001 | -0.01 | -0.01 | 1.318 | 0.53 | 0.56 | 0.43 | 0.29 | 0.53 | 0.4 |
| 11 | 9395649 | rs4625466 | -0.419 | 0.964 | 1.062 | -0.422 | -0.01 | -0.001 | -0.001 | 0.232 | 0.61 | 0.68 | 0.72 | 0.3 | 0.79 | 0.49 |
| 11 | 9406023 | rs6486307 | 0.019 | 1.305 | 1.818 | -0.627 | -0.005 | -0.005 | -0.005 | 0.189 | 0.4 | 0.41 | 0.34 | 0.7 | 0.33 | 0.54 |
| 11 | 9408426 | rs10840228 | -0.135 | 1.368 | 1.751 | -0.628 | 0.112 | 0.06 | 0.06 | 0.744 | 0.58 | 0.24 | 0.63 | 0.3 | 0.74 | 0.48 |
| 11 | 9416258 | rs11042329 | 0.4 | 1.236 | 2.021 | -0.654 | 0.078 | 0.016 | 0.016 | 0.189 | 0.35 | 0.2 | 0.29 | 0.7 | 0.19 | 0.48 |
| 11 | 9419206 | rs4910051 | 0.386 | 1.508 | 1.76 | -0.621 | 0.042 | 0.178 | 0.178 | 0 | 0.51 | 0.17 | 0.58 | 0.2 | 0.59 | 0.4 |
| 11 | 9429619 | rs7115477 | -0.038 | 1.498 | 1.825 | -0.619 | 0.031 | 0.068 | 0.068 | 0.07 | 0.56 | 0.29 | 0.61 | 0.19 | 0.65 | 0.46 |
| 11 | 9431886 | rs7480643 | -0.322 | 1.485 | 1.867 | -0.618 | -0.011 | -0.03 | -0.03 | 0 | 0.58 | 0.45 | 0.64 | 0.2 | 0.66 | 0.46 |
| 11 | 9451800 | rs7122252 | 0.202 | 1.589 | 1.79 | -0.613 | 0.009 | 0.045 | 0.045 | 0 | 0.5 | 0.38 | 0.49 | 0.3 | 0.51 | 0.4 |
| 11 | 9457344 | rs7104568 | 0.924 | 1.556 | 1.699 | -0.651 | -0.063 | 0.064 | 0.064 | 0 | 0.42 | 0.12 | 0.45 | 0.29 | 0.45 | 0.34 |
| 11 | 9473074 | rs6486361 | -0.161 | 1.663 | 1.897 | -0.597 | 0.037 | 0.035 | 0.035 | 0 | 0.55 | 0.65 | 0.57 | 0.3 | 0.67 | 0.43 |
| 11 | 9476564 | rs7944578 | -0.631 | 1.708 | 1.661 | -0.564 | 0.011 | 0.07 | 0.07 | 0 | 0.51 | 0.69 | 0.51 | 0.69 | 0.49 | 0.6 |
| 11 | 9483493 | rs10840243 | 0.395 | 1.605 | 1.569 | -0.564 | 0.006 | 0.078 | 0.078 | 0 | 0.48 | 0.21 | 0.48 | 0.32 | 0.52 | 0.4 |
| 11 | 9491474 | rs10840245 | 0.773 | 1.589 | 1.456 | -0.583 | -0.082 | 0.039 | 0.039 | 0 | 0.43 | 0.1 | 0.45 | 0.31 | 0.45 | 0.34 |
| 11 | 9502592 | rs2313090 | -1.671 | 1.692 | 1.671 | -0.373 | 0.139 | -0.019 | -0.019 | 0 | 0.07 | 0.31 | 0.07 | 0.01 | 0.14 | 0.09 |
| 11 | 9533701 | rs2290424 | 0.382 | 1.321 | 1.22 | -0.213 | -0.029 | 0.055 | 0.055 | 0 | 0.45 | 0.22 | 0.47 | 0.32 | 0.46 | 0.34 |
| 11 | 9539663 | rs10770037 | -0.041 | 1.224 | 0.912 | -0.224 | 0 | -0.002 | -0.002 | 0 | 0.5 | 0.49 | 0.51 | 0.29 | 0.51 | 0.36 |
| 11 | 9540323 | rs17258462 | 0.498 | 1.262 | 0.152 | -0.279 | -0.141 | 0.19 | 0.19 | 0 | 0.42 | 0.03 | 0.43 | 0.16 | 0.43 | 0.3 |

| | | | | | | | | | | | | | | | | |
|---|---|---|---|---|---|---|---|---|---|---|---|---|---|---|---|---|
| 11 | 9550673 | rs3763873 | -0.059 | 1.235 | 0.726 | -0.276 | 0.001 | 0.003 | 0.003 | 0 | 0.51 | 0.48 | 0.49 | 0.3 | 0.53 | 0.39 |
| 11 | 9553209 | rs10840253 | 0.267 | 1.265 | 0.662 | -0.162 | -0.007 | 0.023 | 0.023 | 0 | 0.46 | 0.4 | 0.49 | 0.35 | 0.46 | 0.35 |
| 11 | 9556117 | rs7102407 | -1.892 | 1.659 | 1.484 | -0.105 | 0.085 | -0.039 | -0.039 | 0 | 0.09 | 0.44 | 0.08 | 0 | 0.1 | 0.05 |
| 11 | 9568463 | rs10840257 | -2.461 | 1.977 | 1.356 | -0.256 | 0.114 | -0.103 | -0.103 | 0.233 | 0.15 | 0.47 | 0.16 | 0.01 | 0.23 | 0.18 |
| 11 | 9573518 | rs7951583 | -1.574 | 1.804 | 1.005 | -0.118 | 0.057 | -0.032 | -0.032 | 0.043 | 0.07 | 0.21 | 0.1 | 0.01 | 0.13 | 0.12 |
| 11 | 9587187 | rs10770039 | 0.346 | 1.157 | 0.261 | 0.237 | 0 | -0.001 | -0.001 | 2.782 | 0.66 | 0.59 | 0.69 | 0.43 | 0.7 | 0.59 |
| 11 | 9587478 | rs7937639 | 0.223 | 1.072 | 0.256 | 0.558 | 0.005 | -0.02 | -0.02 | 0 | 0.68 | 0.72 | 0.69 | 0.54 | 0.71 | 0.61 |
| 11 | 9588893 | rs10770040 | -0.43 | 0.955 | -0.123 | 1.219 | 0 | 0 | 0 | 0 | 0.82 | 0.81 | 0.88 | 0.72 | 0.9 | 0.7 |
| 11 | 9590024 | rs921678 | 0.992 | 1.418 | -0.367 | 0.964 | -0.057 | -0.068 | -0.068 | 0 | 0.63 | 0.78 | 0.71 | 0.51 | 0.56 | 0.48 |
| 11 | 9590389 | rs7939541 | 0.621 | 1.648 | -0.425 | 0.805 | -0.05 | -0.072 | -0.072 | 0 | 0.67 | 0.81 | 0.77 | 0.55 | 0.62 | 0.58 |
| 11 | 9593427 | rs3763872 | 1.244 | 1.678 | -0.213 | 1.007 | -0.02 | 0.017 | 0.017 | 0 | 0.58 | 0.6 | 0.65 | 0.67 | 0.51 | 0.55 |
| 11 | 9594827 | rs3763869 | -2.264 | 1.646 | -0.185 | 0.965 | 0.024 | 0.018 | 0.018 | 0 | 0.5 | 0.49 | 0.46 | 0.38 | 0.61 | 0.61 |
| 11 | 9599943 | rs3910384 | 1.729 | 1.628 | -0.296 | 0.942 | 0.011 | 0.014 | 0.014 | 0 | 0.52 | 0.5 | 0.55 | 0.61 | 0.43 | 0.41 |
| 11 | 9600158 | rs11042429 | -1.491 | 1.658 | -0.331 | 0.925 | -0.053 | 0.036 | 0.036 | 0 | 0.28 | 0.13 | 0.25 | 0.18 | 0.4 | 0.22 |
| 11 | 9603236 | rs4370932 | 1.867 | 1.501 | -0.206 | 0.972 | 0.013 | 0.004 | 0.004 | 0 | 0.49 | 0.47 | 0.51 | 0.42 | 0.39 | 0.39 |
| 11 | 9607032 | rs10770042 | -0.527 | 1.381 | -0.293 | 0.865 | -0.001 | -0.005 | -0.005 | 0 | 0.15 | 0.12 | 0.18 | 0.18 | 0.18 | 0.12 |
| 11 | 9610805 | rs1049403 | -0.432 | 1.395 | -0.364 | 0.872 | -0.004 | -0.013 | -0.013 | 0 | 0.14 | 0.09 | 0.17 | 0.18 | 0.18 | 0.12 |
| 11 | 9615459 | rs2872220 | 0.221 | 1.126 | -0.341 | 0.914 | 0.013 | -0.028 | -0.028 | 1.131 | 0.19 | 0.07 | 0.13 | 0.2 | 0.14 | 0.27 |
| 11 | 9622111 | rs7928738 | -0.7 | 1.474 | 0.053 | 0.823 | 0.042 | -0.062 | -0.062 | 0 | 0.14 | 0.27 | 0.11 | 0.01 | 0.25 | 0.11 |
| 11 | 9625196 | rs12420438 | -1.149 | 1.526 | -0.5 | 0.654 | 0.067 | 0.035 | 0.035 | 0.412 | 0.51 | 0.18 | 0.43 | 0.44 | 0.63 | 0.51 |
| 11 | 9627471 | rs10770043 | -0.871 | 1.447 | -0.492 | 0.611 | -0.014 | 0.014 | 0.014 | 0 | 0.45 | 0.36 | 0.4 | 0.44 | 0.47 | 0.48 |
| 11 | 9628802 | rs10840269 | -1.543 | 1.229 | -0.231 | 0.699 | 0.001 | 0 | 0 | 0 | 0.57 | 0.56 | 0.5 | 0.41 | 0.68 | 0.57 |
| 11 | 9645397 | rs7109545 | 0.831 | 1.213 | -0.385 | 0.731 | 0.062 | -0.01 | -0.01 | 0.625 | 0.46 | 0.73 | 0.53 | 0.52 | 0.34 | 0.48 |
| 11 | 9650585 | rs1865558 | 1 | 1.251 | -0.388 | 0.807 | -0.004 | 0 | 0 | 0.688 | 0.42 | 0.48 | 0.48 | 0.34 | 0.3 | 0.45 |
| 11 | 9659903 | rs11042453 | 1.207 | 1.218 | -0.263 | 1.196 | -0.026 | -0.015 | -0.015 | 0 | 0.4 | 0.39 | 0.45 | 0.26 | 0.28 | 0.39 |
| 11 | 9661603 | rs11042454 | 1.207 | 1.218 | -0.263 | 1.055 | -0.026 | 0.004 | 0.004 | 0.277 | 0.4 | 0.39 | 0.45 | 0.33 | 0.29 | 0.39 |
| 11 | 9663767 | rs11042458 | -0.449 | 1.416 | -0.431 | 1.118 | -0.076 | -0.01 | -0.01 | 0.102 | 0.42 | 0.12 | 0.38 | 0.38 | 0.46 | 0.47 |
| 11 | 9667828 | rs12275368 | -0.093 | 1.302 | -0.419 | 1.136 | -0.047 | -0.028 | -0.028 | 0 | 0.36 | 0.13 | 0.34 | 0.37 | 0.39 | 0.44 |
| 11 | 9679145 | rs12807017 | -0.42 | 1.485 | -0.127 | 0.889 | -0.044 | 0.044 | 0.044 | 0 | 0.14 | 0.07 | 0.12 | 0 | 0.26 | 0.1 |
| 11 | 9690013 | rs10840285 | 1.495 | 0.647 | -0.341 | 1.146 | -0.01 | -0.085 | -0.085 | 0.483 | 0.65 | 0.12 | 0.64 | 0.52 | 0.67 | 0.7 |
| 11 | 9691765 | rs12223505 | 1.448 | 0.676 | -0.163 | 1.174 | 0 | -0.052 | -0.052 | 0 | 0.66 | 0.2 | 0.64 | 0.52 | 0.67 | 0.69 |
| 11 | 9697344 | rs4548612 | 0.577 | 0.854 | 0.312 | 1.25 | 0.055 | 0.018 | 0.018 | 0 | 0.8 | 0.56 | 0.71 | 0.7 | 0.71 | 0.83 |
| 11 | 9710354 | rs1435463 | 1.187 | 0.86 | 0.596 | 1.25 | -0.003 | -0.012 | -0.012 | 0 | 0.69 | 0.28 | 0.65 | 0.54 | 0.68 | 0.69 |
| 11 | 9734945 | rs1390679 | -1.433 | 1.184 | 0.428 | 0.882 | 0.018 | -0.015 | -0.015 | 0 | 0.25 | 0.33 | 0.18 | 0.15 | 0.32 | 0.24 |
| 11 | 9737520 | rs379692 | -0.337 | 1.131 | 0.415 | 0.854 | -0.048 | 0.101 | 0.101 | 0 | 0.1 | 0.29 | 0.06 | 0.14 | 0.03 | 0.13 |
| 11 | 9737574 | rs397250 | -1.196 | 1.197 | 0.423 | 0.888 | 0.018 | -0.015 | -0.015 | 0 | 0.25 | 0.33 | 0.18 | 0.15 | 0.32 | 0.24 |
| 11 | 9746201 | rs1695613 | -1.316 | 1.223 | 0.426 | 0.907 | 0.02 | -0.013 | -0.013 | 0 | 0.25 | 0.33 | 0.18 | 0.17 | 0.32 | 0.24 |
| 11 | 9749645 | rs397686 | -1.316 | 1.223 | 0.426 | 0.907 | 0.02 | -0.013 | -0.013 | 0.004 | 0.25 | 0.33 | 0.18 | 0.17 | 0.32 | 0.24 |
| 11 | 9750658 | rs431480 | -1.391 | 1.266 | 0.192 | 0.3 | 0.001 | -0.026 | -0.026 | 0 | 0.45 | 0.47 | 0.4 | 0.24 | 0.47 | 0.37 |
| 11 | 9754221 | rs360157 | 0.413 | 1.11 | -0.249 | 0.135 | 0.005 | 0.004 | 0.004 | 0.097 | 0.59 | 0.46 | 0.51 | 0.37 | 0.59 | 0.5 |
| 11 | 9759713 | rs93138 | 0.398 | 1.079 | -0.253 | 0.131 | 0.005 | 0.004 | 0.004 | 0.186 | 0.59 | 0.47 | 0.51 | 0.37 | 0.59 | 0.5 |
| 11 | 9763094 | rs373894 | 1.425 | 0.664 | -0.065 | -0.109 | 0.021 | -0.013 | -0.013 | 0.701 | 0.38 | 0.31 | 0.39 | 0.38 | 0.3 | 0.38 |
| 11 | 9765348 | rs7939896 | -0.483 | 0.5 | 0.23 | 0.431 | 0.053 | 0.362 | 0.362 | 1.186 | 0.05 | 0.34 | 0.16 | 0.31 | 0.01 | 0.05 |
| 11 | 9772840 | rs1056593 | -1.109 | 0.526 | 0.257 | 0.378 | 0.018 | 0.158 | 0.158 | 0 | 0.1 | 0.31 | 0.19 | 0.31 | 0.07 | 0.07 |
| 11 | 9773568 | rs360137 | 0.577 | 0.535 | 0.076 | 0.356 | 0.002 | 0.002 | 0.002 | 0.059 | 0.59 | 0.44 | 0.52 | 0.39 | 0.6 | 0.5 |
| 11 | 9776567 | rs360140 | -0.342 | 0.554 | -0.618 | 0.225 | -0.027 | -0.003 | -0.003 | 0 | 0.69 | 0.82 | 0.71 | 0.68 | 0.66 | 0.57 |
| 11 | 9784297 | rs360120 | -0.047 | 0.036 | -0.773 | 0.083 | 0.02 | -0.019 | -0.019 | 0.023 | 0.81 | 0.87 | 0.9 | 0.68 | 0.9 | 0.72 |
| 11 | 9790817 | rs360147 | 1.297 | -0.342 | -0.381 | -0.236 | 0.016 | -0.005 | -0.005 | 0 | 0.38 | 0.62 | 0.39 | 0.33 | 0.28 | 0.38 |
| 11 | 9791579 | rs960909 | 2.011 | -0.343 | -0.186 | -0.236 | -0.006 | 0.009 | 0.009 | 0 | 0.31 | 0.29 | 0.35 | 0.33 | 0.27 | 0.34 |
| 11 | 9796699 | rs10770061 | 0.23 | 0.02 | -0.494 | -0.296 | -0.027 | -0.009 | -0.009 | 0.068 | 0.28 | 0.21 | 0.32 | 0.35 | 0.38 | 0.19 |
| 11 | 9800450 | rs1045634 | 0.304 | 0.006 | -0.518 | -0.298 | -0.035 | -0.014 | -0.014 | 0.017 | 0.27 | 0.19 | 0.32 | 0.35 | 0.38 | 0.19 |
| 11 | 9800601 | rs10118 | 0.732 | -0.028 | -0.484 | -0.327 | 0.001 | 0.004 | 0.004 | 0 | 0.6 | 0.58 | 0.67 | 0.67 | 0.64 | 0.52 |
| 11 | 9801451 | rs3829252 | 0.296 | 0.025 | -0.517 | -0.296 | -0.032 | -0.014 | -0.014 | 0.043 | 0.27 | 0.19 | 0.32 | 0.35 | 0.38 | 0.19 |
| 11 | 9802952 | rs3763862 | -0.338 | -0.472 | -0.409 | -0.119 | 0.041 | 0.052 | 0.052 | 0.767 | 0.44 | 0.27 | 0.52 | 0.35 | 0.63 | 0.35 |
| 11 | 9805523 | rs10770062 | 0.464 | -0.426 | -0.285 | -0.091 | -0.038 | 0.016 | 0.016 | 0 | 0.79 | 0.68 | 0.88 | 0.66 | 0.91 | 0.7 |
| 11 | 9809956 | rs360129 | -0.234 | -0.195 | -0.89 | 0.024 | 0.004 | -0.005 | -0.005 | 0.021 | 0.82 | 0.84 | 0.89 | 0.65 | 0.89 | 0.72 |
| 11 | 9827021 | rs421752 | 0.586 | 0.261 | -1.361 | -0.447 | -0.005 | 0.018 | 0.018 | 0 | 0.55 | 0.35 | 0.7 | 0.8 | 0.53 | 0.67 |
| 11 | 9830595 | rs11042500 | 0.047 | 0.405 | -1.708 | -0.908 | -0.013 | 0.119 | 0.119 | 0 | 0.42 | 0.18 | 0.31 | 0.01 | 0.54 | 0.28 |
| 11 | 9839856 | rs17268414 | -0.807 | 0.318 | -1.752 | -0.862 | 0.098 | 0.158 | 0.158 | 0.101 | 0.49 | 0.22 | 0.4 | 0.15 | 0.65 | 0.39 |
| 11 | 9841970 | rs12576763 | 0.851 | 0.373 | -1.758 | -0.872 | -0.029 | -0.034 | -0.034 | 0.043 | 0.07 | 0.01 | 0.09 | 0.13 | 0.1 | 0.1 |
| 11 | 9842436 | rs10770068 | -0.75 | 0.265 | -1.732 | -0.872 | 0.088 | 0.148 | 0.148 | 0.021 | 0.49 | 0.23 | 0.4 | 0.14 | 0.64 | 0.38 |
| 11 | 9842491 | rs10734650 | -0.546 | -0.063 | -1.746 | -0.498 | -0.013 | 0.042 | 0.042 | 0 | 0.82 | 0.77 | 0.68 | 0.32 | 0.88 | 0.69 |
| 11 | 9847690 | rs4910508 | 1.499 | 0.006 | -1.65 | -0.585 | -0.037 | -0.028 | -0.028 | 0 | 0.33 | 0.56 | 0.28 | 0.18 | 0.24 | 0.32 |
| 11 | 9856524 | rs4243951 | 0.652 | 0.121 | -1.475 | -0.562 | -0.002 | -0.006 | -0.006 | 0 | 0.72 | 0.69 | 0.57 | 0.16 | 0.75 | 0.55 |
| 11 | 9865856 | rs11603291 | 1.511 | 0.412 | -1.77 | -0.834 | -0.056 | 0.088 | 0.088 | 0.318 | 0.19 | 0.04 | 0.18 | 0 | 0.25 | 0.15 |
| 11 | 9867666 | rs745389 | -0.757 | 0.396 | -1.715 | -0.492 | -0.004 | 0.095 | 0.095 | 0 | 0.86 | 0.77 | 0.68 | 0.32 | 0.88 | 0.69 |
| 11 | 9869224 | rs1002718 | 1.169 | 0.541 | -1.765 | -0.837 | 0 | -0.019 | -0.019 | 0 | 0.48 | 0.51 | 0.46 | 0.16 | 0.5 | 0.4 |
| 11 | 9870622 | rs11042511 | 0.484 | 0.629 | -1.701 | -0.737 | -0.004 | -0.006 | -0.006 | 0 | 0.24 | 0.18 | 0.11 | 0 | 0.25 | 0.12 |
| 11 | 9875519 | rs360169 | -0.324 | 0.479 | -1.698 | -0.325 | 0.028 | 0.12 | 0.12 | 0 | 0.51 | 0.25 | 0.48 | 0.3 | 0.57 | 0.36 |

| | | | | | | | | | | | | | | | | |
|---|---|---|---|---|---|---|---|---|---|---|---|---|---|---|---|---|
| 11 | 9881475 | rs780382 | -0.327 | 0.439 | -1.8 | -0.3 | 0.001 | -0.004 | -0.004 | 2.33 | 0.65 | 0.7 | 0.73 | 0.6 | 0.62 | 0.61 |
| 11 | 9885849 | rs4910067 | 1.359 | 0.256 | -1.789 | -0.219 | 0.004 | 0.01 | 0.01 | 0 | 0.32 | 0.49 | 0.37 | 0.28 | 0.31 | 0.36 |
| 11 | 9890714 | rs12577195 | 1.409 | 0.46 | -1.907 | -0.014 | -0.008 | 0.005 | 0.005 | 0 | 0.06 | 0.04 | 0.1 | 0.16 | 0.11 | 0.14 |
| 11 | 9896939 | rs7124727 | -1.688 | 0.317 | -1.901 | -0.177 | 0.006 | -0.008 | -0.008 | 0 | 0.87 | 0.92 | 0.85 | 0.58 | 0.87 | 0.73 |
| 11 | 9900438 | rs1546541 | 1.795 | 0.331 | -1.833 | -0.825 | -0.02 | -0.059 | -0.059 | 0 | 0.27 | 0.49 | 0.28 | 0.12 | 0.2 | 0.22 |
| 11 | 9902592 | rs4910071 | 0.996 | 0.64 | -2.105 | -0.835 | -0.031 | 0.091 | 0.091 | 0 | 0.2 | 0.08 | 0.11 | 0 | 0.23 | 0.11 |
| 11 | 9908358 | rs10840315 | 1.564 | 0.682 | -2.099 | -0.737 | -0.095 | 0.109 | 0.109 | 0.435 | 0.18 | 0.02 | 0.18 | 0 | 0.29 | 0.17 |
| 11 | 9910728 | rs1996062 | -0.52 | 0.29 | -1.997 | -0.655 | 0.002 | 0.032 | 0.032 | 0 | 0.54 | 0.35 | 0.47 | 0.27 | 0.55 | 0.35 |
| 11 | 9912052 | rs10743120 | 0.381 | 0.131 | -2.036 | -0.425 | 0.026 | 0.039 | 0.039 | 0.036 | 0.52 | 0.63 | 0.54 | 0.24 | 0.61 | 0.48 |
| 11 | 9916603 | rs10840318 | -1.939 | 0.135 | -2.061 | -0.397 | 0.001 | 0.015 | 0.015 | 0 | 0.87 | 0.89 | 0.83 | 0.52 | 0.86 | 0.66 |
| 11 | 9920537 | rs7105337 | 1.15 | 0.218 | -2.096 | -0.398 | -0.007 | 0.07 | 0.07 | 0 | 0.46 | 0.67 | 0.48 | 0.17 | 0.51 | 0.39 |
| 11 | 9923714 | rs11608212 | 1.481 | -0.144 | -2.197 | -0.538 | -0.015 | 0.072 | 0.072 | 0 | 0.2 | 0.09 | 0.12 | 0 | 0.21 | 0.1 |
| 11 | 9924436 | rs11042540 | -0.813 | -0.132 | -2.086 | -0.417 | -0.006 | -0.055 | -0.055 | 0 | 0.67 | 0.79 | 0.72 | 0.51 | 0.65 | 0.56 |
| 11 | 9927904 | rs7931062 | 0.269 | -0.131 | -2.049 | -0.421 | 0.036 | 0.055 | 0.055 | 0.089 | 0.52 | 0.67 | 0.55 | 0.24 | 0.61 | 0.48 |
| 11 | 9929006 | rs16907248 | 1.565 | -0.296 | -1.897 | -0.353 | -0.006 | -0.003 | -0.003 | 0 | 0.15 | 0.11 | 0.17 | 0.27 | 0.04 | 0.08 |
| 11 | 9930529 | rs12796169 | 1.306 | 0.159 | -1.715 | -0.517 | -0.022 | -0.037 | -0.037 | 0 | 0.27 | 0.49 | 0.29 | 0.17 | 0.21 | 0.21 |
| 11 | 9938545 | rs7950655 | 0.223 | -0.148 | -2.023 | -0.672 | -0.009 | 0.058 | 0.058 | 0 | 0.45 | 0.66 | 0.48 | 0.18 | 0.52 | 0.39 |
| 11 | 9942432 | rs7945309 | 2.04 | -0.039 | -2.112 | -0.781 | -0.004 | -0.005 | -0.005 | 0 | 0.1 | 0.09 | 0.07 | 0 | 0.14 | 0.08 |
| 11 | 9948104 | rs4910083 | 0.255 | -0.08 | -2.1 | -0.809 | 0.003 | 0.033 | 0.033 | 0 | 0.48 | 0.61 | 0.38 | 0.18 | 0.53 | 0.4 |
| 11 | 9951257 | rs1512829 | -0.977 | -0.092 | -2.183 | -0.446 | -0.003 | -0.077 | -0.077 | 0 | 0.8 | 0.87 | 0.7 | 0.5 | 0.8 | 0.64 |
| 11 | 9951968 | rs10840327 | 1.349 | -0.599 | -1.912 | -0.882 | -0.067 | 0.084 | 0.084 | 0.107 | 0.17 | 0.04 | 0.17 | 0 | 0.28 | 0.17 |
| 11 | 9961079 | rs7939451 | -0.698 | -0.909 | -1.769 | -0.605 | -0.001 | -0.001 | -0.001 | 0 | 0.55 | 0.42 | 0.52 | 0.42 | 0.57 | 0.49 |
| 11 | 9973062 | rs10500715 | -0.27 | -0.548 | -1.494 | -0.718 | -0.017 | 0.015 | 0.015 | 0 | 0.33 | 0.25 | 0.42 | 0.19 | 0.44 | 0.35 |
| 11 | 9979761 | rs7952455 | 1.453 | -0.436 | -1.495 | -0.724 | -0.008 | 0.01 | 0.01 | 0 | 0.11 | 0.06 | 0.07 | 0 | 0.14 | 0.08 |
| 11 | 9981165 | rs4909917 | 0.622 | -0.406 | -1.471 | -0.675 | -0.008 | -0.001 | -0.001 | 0.557 | 0.21 | 0.17 | 0.35 | 0.19 | 0.3 | 0.3 |
| 11 | 9987228 | rs4576815 | 0.657 | -0.407 | -1.473 | -0.7 | -0.026 | -0.004 | -0.004 | 0 | 0.21 | 0.13 | 0.34 | 0.19 | 0.3 | 0.3 |
| 11 | 9995399 | rs11042575 | -0.704 | -0.615 | -1.498 | -0.871 | 0.01 | 0.097 | 0.097 | 0 | 0.5 | 0.26 | 0.48 | 0.3 | 0.55 | 0.44 |
| 11 | 10003829 | rs7106234 | 0.133 | -0.726 | -0.942 | -0.72 | -0.02 | -0.019 | -0.019 | 1.04 | 0.66 | 0.58 | 0.68 | 0.7 | 0.73 | 0.73 |
| 11 | 10024311 | rs10500716 | 0.233 | -0.747 | -1.645 | -0.578 | 0.006 | 0.013 | 0.013 | 0 | 0.16 | 0.17 | 0.27 | 0.27 | 0.2 | 0.15 |
| 11 | 10029016 | rs9888170 | 0.049 | -0.773 | -1.625 | -0.565 | 0.017 | 0.147 | 0.147 | 0.402 | 0.06 | 0.16 | 0.21 | 0.27 | 0.06 | 0.04 |
| 11 | 10047647 | rs17272751 | 0.435 | -0.851 | -1.907 | -0.773 | 0.036 | 0.142 | 0.142 | 0 | 0.23 | 0.02 | 0.06 | 0.02 | 0.14 | 0.15 |
| 11 | 10048319 | rs7925889 | -0.582 | -0.828 | -1.903 | -0.798 | 0.031 | 0.037 | 0.037 | 0 | 0.37 | 0.21 | 0.33 | 0.3 | 0.31 | 0.27 |
| 11 | 10058400 | rs16907410 | 0.035 | -0.56 | -1.384 | -0.186 | 0.001 | 0.003 | 0.003 | 0 | 0.15 | 0.12 | 0.26 | 0.28 | 0.2 | 0.14 |
| 11 | 10062082 | rs2896516 | -0.751 | -0.523 | -1.578 | -0.328 | -0.037 | 0.04 | 0.04 | 0.124 | 0.28 | 0.09 | 0.31 | 0.17 | 0.31 | 0.31 |
| 11 | 10093760 | rs7946995 | 0.444 | -0.534 | -0.971 | 0.039 | 0.008 | 0 | 0 | 0.503 | 0.22 | 0.24 | 0.36 | 0.19 | 0.3 | 0.3 |
| 11 | 10108009 | rs11042615 | 1.162 | -0.551 | -0.923 | 0.041 | -0.043 | 0.051 | 0.051 | 0.039 | 0.18 | 0.07 | 0.18 | 0.02 | 0.28 | 0.2 |
| 11 | 10108484 | rs11042617 | -0.628 | -0.598 | -0.95 | 0.109 | -0.004 | -0.004 | -0.004 | 0.084 | 0.36 | 0.32 | 0.61 | 0.47 | 0.48 | 0.4 |
| 11 | 10143942 | rs7129047 | 0.333 | -0.581 | -0.963 | -0.048 | 0.002 | 0.012 | 0.012 | 0 | 0.14 | 0.13 | 0.27 | 0.28 | 0.19 | 0.1 |
| 11 | 10172545 | rs16907505 | 0.51 | -0.949 | -1.198 | -0.279 | 0.055 | 0.103 | 0.103 | 0 | 0.18 | 0.02 | 0.04 | 0.02 | 0.08 | 0.13 |
| 11 | 10174447 | rs7129531 | -0.837 | -0.991 | -1.127 | -0.273 | 0.027 | 0.015 | 0.015 | 0 | 0.34 | 0.21 | 0.31 | 0.31 | 0.27 | 0.24 |
| 11 | 10186289 | rs17293593 | -0.354 | -0.95 | -1.149 | -0.333 | 0.05 | 0.093 | 0.093 | 0.135 | 0.3 | 0.12 | 0.11 | 0.02 | 0.21 | 0.2 |
| 11 | 10211233 | rs12288275 | 0.044 | -0.856 | -0.937 | -0.189 | 0.023 | 0.192 | 0.192 | 0.035 | 0.06 | 0.17 | 0.22 | 0.28 | 0.06 | 0.05 |
| 11 | 10217575 | rs2069214 | 0.103 | -0.855 | -0.943 | -0.189 | 0.023 | 0.191 | 0.191 | 0.171 | 0.06 | 0.17 | 0.22 | 0.28 | 0.06 | 0.05 |
| 11 | 10224357 | rs7929332 | 1.333 | -0.881 | -0.98 | -0.245 | -0.001 | -0.001 | -0.001 | 0 | 0.11 | 0.07 | 0.07 | 0 | 0.13 | 0.06 |
| 11 | 10226268 | rs2119991 | -0.19 | -0.891 | -0.967 | -0.216 | -0.002 | 0.042 | 0.042 | 0 | 0.17 | 0.24 | 0.28 | 0.29 | 0.18 | 0.11 |
| 11 | 10229851 | rs7938543 | 0.967 | -0.792 | -1.078 | -0.436 | -0.001 | 0.008 | 0.008 | 0 | 0.46 | 0.61 | 0.5 | 0.67 | 0.45 | 0.55 |
| 11 | 10234930 | rs12278680 | -0.032 | -0.906 | -1.074 | -0.427 | -0.024 | 0.027 | 0.027 | 0 | 0.17 | 0.23 | 0.25 | 0.28 | 0.11 | 0.1 |
| 11 | 10264122 | rs11042686 | 0.813 | -0.537 | -1.025 | -0.287 | -0.048 | -0.024 | -0.024 | 0 | 0.21 | 0.08 | 0.37 | 0.29 | 0.34 | 0.22 |
| 11 | 10267618 | rs11042689 | 0.807 | -0.532 | -1.023 | -0.296 | -0.048 | -0.026 | -0.026 | 0.185 | 0.21 | 0.08 | 0.37 | 0.29 | 0.34 | 0.22 |
| 11 | 10273128 | rs10840378 | -0.475 | -0.692 | -1.029 | -0.368 | -0.007 | -0.047 | -0.047 | 0 | 0.33 | 0.32 | 0.13 | 0.04 | 0.21 | 0.23 |
| 11 | 10275120 | rs1470260 | -0.174 | -0.657 | -1.006 | -0.351 | 0.031 | 0.035 | 0.035 | 0.025 | 0.31 | 0.22 | 0.12 | 0.04 | 0.21 | 0.23 |
| 11 | 10275343 | rs1442740 | -0.433 | -0.658 | -1.007 | -0.352 | -0.008 | -0.049 | -0.049 | 1.178 | 0.33 | 0.31 | 0.13 | 0.04 | 0.21 | 0.23 |
| 11 | 10294772 | rs7107290 | -1.235 | -0.621 | -1.005 | -0.345 | 0 | 0.007 | 0.007 | 0 | 0.54 | 0.39 | 0.5 | 0.33 | 0.55 | 0.45 |
| 11 | 10295144 | rs6484148 | -0.997 | -0.602 | -1.032 | -0.335 | 0.004 | 0.06 | 0.06 | 0 | 0.52 | 0.29 | 0.49 | 0.33 | 0.55 | 0.46 |
| 11 | 10296667 | rs2920151 | -0.164 | -0.653 | -1.027 | -0.446 | 0.042 | 0.07 | 0.07 | 0.034 | 0.3 | 0.18 | 0.11 | 0.04 | 0.21 | 0.23 |
| 11 | 10331311 | rs7944706 | 0.944 | -0.627 | -0.805 | -0.068 | -0.023 | -0.044 | -0.044 | 0 | 0.42 | 0.25 | 0.31 | 0.46 | 0.43 | 0.43 |
| 11 | 10333383 | rs11042727 | 0.822 | -0.154 | -1.445 | -0.414 | -0.024 | -0.029 | -0.029 | 0.126 | 0.05 | 0 | 0.03 | 0.14 | 0.07 | 0.1 |
| 11 | 10335700 | rs1450265 | 0.211 | -0.491 | -1.078 | -1.08 | -0.004 | -0.053 | -0.053 | 0 | 0.12 | 0.15 | 0.08 | 0 | 0.13 | 0.06 |
| 11 | 10337640 | rs7108385 | 0.21 | -0.494 | -1.084 | -1.082 | -0.004 | -0.05 | -0.05 | 0.663 | 0.12 | 0.15 | 0.08 | 0 | 0.13 | 0.06 |
| 11 | 10346994 | rs896043 | 0.749 | -0.398 | -1.248 | -1.565 | -0.001 | 0.009 | 0.009 | 0.039 | 0.11 | 0.05 | 0.07 | 0 | 0.13 | 0.06 |
| 11 | 10347130 | rs1450276 | -0.218 | -0.492 | -1.301 | -1.632 | 0.007 | 0.011 | 0.011 | 2.172 | 0.15 | 0.08 | 0.07 | 0 | 0.14 | 0.06 |
| 11 | 10353745 | rs11822548 | 0.324 | -0.135 | -1.415 | -1.421 | 0.042 | 0.048 | 0.048 | 0 | 0.19 | 0.08 | 0.04 | 0.02 | 0.09 | 0.14 |
| 11 | 10355181 | rs7941557 | 0.453 | -0.031 | -1.355 | -1.346 | 0.003 | -0.059 | -0.059 | 0 | 0.12 | 0.28 | 0.08 | 0.02 | 0.13 | 0.07 |
| 11 | 10368614 | rs2923098 | -0.897 | 0.534 | -1.196 | -1.125 | 0.001 | 0.003 | 0.003 | 0 | 0.49 | 0.39 | 0.3 | 0.04 | 0.48 | 0.41 |
| 11 | 10368976 | rs2044782 | 0.36 | 0.623 | -1.185 | -0.993 | -0.029 | 0.023 | 0.023 | 0.005 | 0.18 | 0.11 | 0.18 | 0 | 0.28 | 0.17 |
| 11 | 10370420 | rs1580004 | -0.207 | 0.543 | -0.885 | -0.217 | 0 | 0.003 | 0.003 | 0 | 0.53 | 0.45 | 0.44 | 0.27 | 0.51 | 0.51 |
| 11 | 10372959 | rs11042750 | -0.197 | 0.309 | -0.764 | -0.751 | -0.001 | 0.064 | 0.064 | 0 | 0.54 | 0.63 | 0.64 | 0.87 | 0.5 | 0.65 |
| 11 | 10375899 | rs2957705 | -0.757 | -0.256 | 0.208 | -0.275 | -0.045 | 0.027 | 0.027 | 0.189 | 0.23 | 0.37 | 0.32 | 0.69 | 0.15 | 0.35 |
| 11 | 10377258 | rs2957707 | 0.962 | -0.291 | 0.116 | -0.671 | -0.006 | -0.023 | -0.023 | 0.027 | 0.65 | 0.4 | 0.51 | 0.18 | 0.7 | 0.55 |

| | | | | | | | | | | | | | | | | |
|---|---|---|---|---|---|---|---|---|---|---|---|---|---|---|---|---|
| 11 | 10380828 | rs2218793 | -1.127 | -0.393 | 0.644 | -0.729 | -0.002 | -0.011 | -0.011 | 0 | 0.31 | 0.33 | 0.47 | 0.81 | 0.29 | 0.43 |
| 11 | 10387833 | rs10840395 | 0.648 | -0.599 | 0.809 | 0.487 | -0.04 | 0.029 | 0.029 | 0.331 | 0.75 | 0.4 | 0.84 | 0.7 | 0.86 | 0.66 |
| 11 | 10390581 | rs6484218 | -0.967 | -0.734 | 1.211 | 0.451 | -0.032 | 0.056 | 0.056 | 0.102 | 0.21 | 0.36 | 0.13 | 0.28 | 0.16 | 0.22 |
| 11 | 10391880 | rs10770107 | 0.776 | -0.768 | 1.262 | 0.442 | -0.043 | 0.058 | 0.058 | 0.55 | 0.77 | 0.43 | 0.85 | 0.72 | 0.84 | 0.77 |
| 11 | 10392437 | rs7128306 | 0.657 | -0.928 | 1.195 | 0.429 | -0.057 | 0.066 | 0.066 | 2.01 | 0.79 | 0.41 | 0.87 | 0.71 | 0.88 | 0.79 |
| 11 | 10393873 | rs11042759 | 0.399 | -0.947 | 0.724 | 0.374 | -0.047 | -0.101 | -0.101 | 0 | 0.14 | 0.01 | 0.35 | 0.39 | 0.19 | 0.14 |
| 11 | 10397340 | rs2923088 | 0.244 | -0.722 | 0.735 | -0.378 | 0.001 | -0.001 | -0.001 | 1.302 | 0.49 | 0.27 | 0.63 | 0.58 | 0.52 | 0.44 |
| 11 | 10397518 | rs2957661 | 1.091 | -0.842 | 0.898 | -0.493 | -0.012 | 0.009 | 0.009 | 5.303 | 0.28 | 0.26 | 0.26 | 0.13 | 0.34 | 0.33 |
| 11 | 10398518 | rs2923123 | -0.521 | -0.694 | 0.76 | -0.447 | 0.003 | 0.002 | 0.002 | 1.108 | 0.5 | 0.59 | 0.36 | 0.42 | 0.48 | 0.56 |
| 11 | 10410261 | rs16907765 | 0.305 | -0.739 | 0.443 | -0.438 | 0.035 | -0.06 | -0.06 | 2.902 | 0.15 | 0.05 | 0.05 | 0.29 | 0.08 | 0.2 |
| 11 | 10411717 | rs7934497 | -0.787 | -0.546 | 0.709 | -0.975 | 0.004 | -0.008 | -0.008 | 0 | 0.55 | 0.62 | 0.35 | 0.42 | 0.51 | 0.59 |
| 11 | 10414509 | rs10840400 | 0.084 | -0.448 | 0.694 | -0.852 | -0.004 | 0 | 0 | 0.357 | 0.51 | 0.26 | 0.65 | 0.56 | 0.58 | 0.45 |
| 11 | 10415804 | rs10743136 | 0.435 | 0.075 | 1.013 | -0.396 | 0.005 | 0.012 | 0.012 | 2.657 | 0.32 | 0.36 | 0.27 | 0.35 | 0.32 | 0.35 |
| 11 | 10417254 | rs4909926 | -0.52 | 0.111 | 1.155 | -0.501 | -0.003 | -0.038 | -0.038 | 0 | 0.47 | 0.46 | 0.68 | 0.82 | 0.51 | 0.51 |
| 11 | 10422641 | rs10770114 | 0.344 | 0.14 | 0.612 | -0.5 | 0.052 | 0.213 | 0.213 | 0 | 0.49 | 0.18 | 0.26 | 0.17 | 0.43 | 0.45 |
| 11 | 10424509 | rs12577015 | -0.526 | 0.433 | -0.061 | 0.233 | 0.001 | 0.036 | 0.036 | 2.705 | 0.17 | 0.16 | 0.42 | 0.48 | 0.19 | 0.18 |
| 11 | 10429860 | rs7103258 | -0.738 | 0.996 | -0.173 | -1.498 | 0.015 | -0.073 | -0.073 | 0 | 0.19 | 0.29 | 0.21 | 0.03 | 0.2 | 0.12 |
| 11 | 10429998 | rs12801986 | -0.175 | 1.154 | -0.298 | -1.488 | -0.028 | 0.026 | 0.026 | 0 | 0.06 | 0.03 | 0.04 | 0 | 0.1 | 0.02 |
| 11 | 10430899 | rs12226767 | -0.006 | 0.713 | -0.605 | -1.604 | 0.013 | -0.005 | -0.005 | 0 | 0.08 | 0.02 | 0.01 | 0.13 | 0.05 | 0.26 |
| 11 | 10431834 | rs7123414 | 0.698 | 1.119 | -0.34 | -2.105 | 0.008 | -0.046 | -0.046 | 0.556 | 0.1 | 0.16 | 0.08 | 0 | 0.12 | 0.07 |
| 11 | 10432370 | rs3741044 | -1.391 | 0.947 | -0.395 | -2.736 | 0.008 | 0.046 | 0.046 | 0.022 | 0.57 | 0.72 | 0.45 | 0.29 | 0.57 | 0.62 |
| 11 | 10433426 | rs10840406 | 0.296 | 0.627 | -0.449 | -2.721 | -0.033 | 0.02 | 0.02 | 0 | 0.12 | 0.05 | 0.05 | 0.13 | 0.14 | 0.28 |
| 11 | 10433795 | rs10840407 | -0.683 | 0.584 | -0.265 | -2.77 | 0 | -0.019 | -0.019 | 0 | 0.25 | 0.29 | 0.21 | 0.16 | 0.24 | 0.37 |
| 11 | 10433907 | rs10840408 | -0.683 | 0.584 | -0.265 | -2.77 | 0 | -0.019 | -0.019 | 0 | 0.25 | 0.29 | 0.21 | 0.16 | 0.24 | 0.37 |
| 11 | 10435751 | rs7109187 | -0.511 | 0.47 | 0.246 | -2.84 | 0.008 | -0.036 | -0.036 | 1.057 | 0.41 | 0.45 | 0.29 | 0.26 | 0.46 | 0.53 |
| 11 | 10437096 | rs10840410 | 1.738 | 0.281 | -0.404 | -3.281 | -0.015 | 0.074 | 0.074 | 0 | 0.22 | 0.08 | 0.18 | 0.04 | 0.25 | 0.16 |
| 11 | 10438541 | rs10840411 | 1.236 | 0.625 | -0.034 | -2.912 | 0 | 0.003 | 0.003 | 0 | 0.2 | 0.16 | 0.14 | 0.13 | 0.21 | 0.18 |
| 11 | 10440344 | rs12288723 | 0.221 | 0.531 | 0.028 | -2.935 | -0.011 | 0.014 | 0.014 | 0.459 | 0.29 | 0.17 | 0.18 | 0.26 | 0.33 | 0.43 |
| 11 | 10440668 | rs11042796 | 0.396 | 0.69 | 0.093 | -2.817 | 0.002 | -0.005 | -0.005 | 0 | 0.12 | 0.1 | 0.05 | 0.13 | 0.14 | 0.25 |
| 11 | 10452981 | rs1376000 | 0.299 | 0.422 | 0.456 | -2.811 | -0.001 | -0.026 | -0.026 | 2.362 | 0.3 | 0.4 | 0.29 | 0.16 | 0.28 | 0.26 |
| 11 | 10458596 | rs1450270 | -0.074 | 0.494 | 0.393 | -2.837 | 0.003 | -0.027 | -0.027 | 0 | 0.58 | 0.41 | 0.67 | 0.84 | 0.53 | 0.66 |
| 11 | 10464912 | rs10840415 | 0.99 | 0.455 | 0.364 | -2.681 | 0.003 | -0.022 | -0.022 | 0.119 | 0.13 | 0.22 | 0.17 | 0.06 | 0.14 | 0.13 |
| 11 | 10467383 | rs7939845 | -0.652 | 0.264 | 0.232 | -2.72 | 0.003 | 0.009 | 0.009 | 0 | 0.72 | 0.82 | 0.83 | 0.73 | 0.73 | 0.57 |
| 11 | 10470275 | rs4909928 | 0.255 | 0.417 | 0.618 | -2.924 | 0.048 | -0.07 | -0.07 | 0 | 0.17 | 0.4 | 0.43 | 0.06 | 0.22 | 0.2 |
| 11 | 10475967 | rs4909932 | -0.172 | 0.299 | 0.734 | -2.948 | -0.001 | -0.051 | -0.051 | 0 | 0.6 | 0.54 | 0.72 | 0.77 | 0.59 | 0.51 |
| 11 | 10476898 | rs899011 | -0.635 | 0.312 | 0.663 | -2.961 | 0.002 | 0.012 | 0.012 | 0 | 0.71 | 0.73 | 0.8 | 0.78 | 0.71 | 0.58 |
| 11 | 10477127 | rs899010 | 0.216 | 0.312 | 0.888 | -2.937 | -0.001 | -0.014 | -0.014 | 0 | 0.56 | 0.41 | 0.42 | 0.7 | 0.57 | 0.42 |
| 11 | 10478228 | rs2071020 | 0.584 | 0.32 | 0.748 | -2.86 | 0.007 | 0.065 | 0.065 | 0 | 0.47 | 0.33 | 0.35 | 0.69 | 0.45 | 0.35 |
| 11 | 10481216 | rs11820856 | 0.587 | 0.353 | 0.574 | -2.868 | -0.011 | 0.012 | 0.012 | 0.915 | 0.28 | 0.28 | 0.5 | 0.19 | 0.33 | 0.33 |
| 11 | 10486254 | rs10840419 | 0.618 | 0.135 | 0.092 | -3.09 | -0.005 | -0.009 | -0.009 | 2.524 | 0.23 | 0.17 | 0.2 | 0.67 | 0.25 | 0.27 |
| 11 | 10497467 | rs11042830 | 0.778 | 0.1 | -0.033 | -2.977 | -0.001 | -0.001 | -0.001 | 0 | 0.21 | 0.19 | 0.19 | 0.64 | 0.23 | 0.24 |
| 11 | 10499103 | rs4910143 | -0.064 | -0.06 | -0.164 | -3.289 | 0.007 | 0.026 | 0.026 | 0 | 0.55 | 0.41 | 0.7 | 0.74 | 0.58 | 0.45 |
| 11 | 10499334 | rs10770118 | -2.519 | -0.087 | -0.198 | -3.307 | -0.006 | 0.007 | 0.007 | 0.053 | 0.84 | 0.9 | 0.88 | 0.93 | 0.8 | 0.82 |
| 11 | 10499995 | rs3741043 | 1.503 | -0.166 | -0.213 | -3.324 | 0.024 | 0.007 | 0.007 | 0 | 0.1 | 0.06 | 0.05 | 0.16 | 0.05 | 0.26 |
| 11 | 10502422 | rs11042834 | 0.164 | 0.002 | -0.091 | -3.336 | -0.001 | -0.041 | -0.041 | 0.603 | 0.47 | 0.54 | 0.41 | 0.08 | 0.44 | 0.28 |
| 11 | 10503756 | rs16907852 | 1.154 | -0.051 | -0.355 | -3.198 | 0 | -0.087 | -0.087 | 0 | 0.13 | 0.33 | 0.09 | 0 | 0.1 | 0.06 |
| 11 | 10504008 | rs10840424 | -0.126 | 0.222 | -0.913 | -2.518 | 0.002 | 0.018 | 0.018 | 0 | 0.69 | 0.75 | 0.79 | 0.72 | 0.67 | 0.52 |
| 11 | 10507646 | rs12419888 | -0.251 | 0.216 | -0.877 | -2.417 | 0.019 | 0.154 | 0.154 | 0 | 0.48 | 0.22 | 0.32 | 0.25 | 0.45 | 0.53 |
| 11 | 10510272 | rs7125888 | 0.11 | 1.017 | -0.578 | -2.21 | -0.001 | -0.028 | -0.028 | 0 | 0.58 | 0.56 | 0.43 | 0.85 | 0.57 | 0.65 |
| 11 | 10511142 | rs12795565 | -1.602 | 0.7 | -0.267 | -1.738 | -0.011 | -0.081 | -0.081 | 1.715 | 0.12 | 0.38 | 0.1 | 0.01 | 0.1 | 0.08 |
| 11 | 10513317 | rs1466427 | 0.526 | 1.019 | -1.371 | -1.613 | -0.004 | -0.029 | -0.029 | 0.35 | 0.74 | 0.73 | 0.56 | 0.91 | 0.76 | 0.81 |
| 11 | 10514707 | rs1001118 | -0.928 | 1.185 | -1.135 | -1.037 | 0.004 | -0.037 | -0.037 | 0 | 0.12 | 0.15 | 0.07 | 0.01 | 0.1 | 0.09 |
| 11 | 10518860 | rs10840427 | -0.331 | 1.265 | -1.843 | -1.023 | -0.075 | 0.116 | 0.116 | 0 | 0.37 | 0.16 | 0.33 | 0.16 | 0.51 | 0.4 |
| 11 | 10518977 | rs11042854 | -0.038 | 0.971 | -1.955 | -1.426 | -0.008 | 0.069 | 0.069 | 0 | 0.58 | 0.73 | 0.68 | 0.78 | 0.5 | 0.6 |
| 11 | 10519316 | rs11042856 | -0.098 | 0.604 | -1.529 | -0.932 | -0.002 | -0.001 | -0.001 | 0.021 | 0.59 | 0.45 | 0.61 | 0.38 | 0.65 | 0.56 |
| 11 | 10521764 | rs3741041 | 0.075 | 0.776 | -2.201 | -1.025 | -0.094 | -0.075 | -0.075 | 0 | 0.33 | 0.05 | 0.4 | 0.69 | 0.41 | 0.38 |
| 11 | 10522120 | rs4910146 | -0.351 | 0.398 | -0.977 | -1.358 | 0 | -0.002 | -0.002 | 0 | 0.42 | 0.52 | 0.47 | 0.69 | 0.47 | 0.38 |
| 11 | 10523476 | rs7932321 | -1.05 | -0.535 | -0.578 | -1.889 | -0.015 | -0.004 | -0.004 | 0 | 0.82 | 0.76 | 0.94 | 0.86 | 0.9 | 0.9 |
| 11 | 10523519 | rs2071018 | 0.491 | -0.067 | 0.3 | -1.68 | -0.012 | -0.009 | -0.009 | 0 | 0.34 | 0.21 | 0.4 | 0.69 | 0.41 | 0.35 |
| 11 | 10533952 | rs12448 | 0.643 | -0.046 | 0.146 | -1.67 | -0.029 | 0.089 | 0.089 | 0 | 0.33 | 0.12 | 0.25 | 0.16 | 0.32 | 0.21 |
| 11 | 10534866 | rs10770123 | 0.974 | -0.262 | 0.351 | -1.912 | -0.052 | -0.003 | -0.003 | 0 | 0.37 | 0.22 | 0.6 | 0.7 | 0.51 | 0.46 |
| 11 | 10536116 | rs4146830 | -0.637 | -0.305 | 0.737 | -1.939 | -0.001 | -0.014 | -0.014 | 0 | 0.62 | 0.61 | 0.73 | 0.85 | 0.66 | 0.77 |
| 11 | 10538328 | rs11042868 | 0.971 | -0.302 | 0.666 | -1.918 | -0.037 | 0.124 | 0.124 | 0 | 0.32 | 0.09 | 0.24 | 0.12 | 0.32 | 0.21 |
| 11 | 10538816 | rs11602469 | 1.169 | -0.427 | 0.754 | -1.983 | -0.004 | 0.007 | 0.007 | 0 | 0.14 | 0.12 | 0.07 | 0.15 | 0.09 | 0.25 |
| 11 | 10540331 | rs1993818 | 1.041 | -0.381 | 0.754 | -1.985 | -0.063 | -0.007 | -0.007 | 0 | 0.37 | 0.2 | 0.59 | 0.69 | 0.51 | 0.46 |
| 11 | 10545404 | rs4243915 | -0.622 | -0.391 | 0.822 | -2.101 | -0.002 | -0.037 | -0.037 | 0 | 0.61 | 0.56 | 0.72 | 0.84 | 0.66 | 0.76 |
| 11 | 10553324 | rs10500727 | 0.88 | -0.399 | 0.761 | -2.07 | -0.03 | -0.066 | -0.066 | 0.007 | 0.11 | 0.26 | 0.06 | 0 | 0.06 | 0.06 |
| 11 | 10554630 | rs1350991 | -0.781 | -0.308 | 0.669 | -2.034 | -0.001 | 0.063 | 0.063 | 0 | 0.63 | 0.73 | 0.73 | 0.84 | 0.66 | 0.77 |
| 11 | 10558777 | rs7938782 | -0.361 | -0.356 | 0.772 | -1.916 | -0.044 | -0.021 | -0.021 | 0.107 | 0.25 | 0.35 | 0.13 | 0.14 | 0.15 | 0.32 |

| | | | | | | | | | | | | | | | | |
|---|---|---|---|---|---|---|---|---|---|---|---|---|---|---|---|---|
| 11 | 10567602 | rs10219205 | 0.274 | 0.292 | 0.978 | -2.001 | -0.012 | 0.021 | 0.021 | 1.311 | 0.41 | 0.33 | 0.62 | 0.71 | 0.53 | 0.47 |
| 11 | 10572232 | rs11042876 | 0.961 | 0.565 | 0.551 | -2.046 | -0.053 | -0.012 | -0.012 | 0.482 | 0.31 | 0.13 | 0.4 | 0.7 | 0.4 | 0.35 |
| 11 | 10577397 | rs728528 | 0.84 | 0.389 | 0.671 | -2.262 | -0.002 | 0.064 | 0.064 | 0.148 | 0.37 | 0.25 | 0.27 | 0.17 | 0.34 | 0.22 |
| 11 | 10579144 | rs7358373 | 0.854 | 0.83 | 0.593 | -2.195 | -0.038 | -0.006 | -0.006 | 0 | 0.32 | 0.15 | 0.4 | 0.71 | 0.4 | 0.36 |
| 11 | 10579636 | rs1062912 | 0.854 | 0.83 | 0.593 | -2.217 | -0.042 | -0.005 | -0.005 | 0 | 0.32 | 0.14 | 0.4 | 0.71 | 0.4 | 0.36 |
| 11 | 10594503 | rs11042882 | 0.575 | 0.4 | 0.355 | -2.159 | -0.032 | -0.001 | -0.001 | 0 | 0.32 | 0.2 | 0.41 | 0.71 | 0.4 | 0.35 |
| 11 | 10597081 | rs7115310 | 0.587 | 0.424 | 0.345 | -2.163 | -0.033 | 0 | 0 | 1.355 | 0.32 | 0.2 | 0.41 | 0.7 | 0.41 | 0.35 |
| 11 | 10597671 | rs11042883 | 0.737 | 0.439 | 0.032 | -2.16 | -0.137 | -0.062 | -0.062 | 0 | 0.31 | 0.03 | 0.39 | 0.71 | 0.41 | 0.35 |
| 11 | 10603648 | rs2288236 | 0.715 | 0.384 | 0.032 | -2.163 | -0.13 | -0.065 | -0.065 | 0.246 | 0.31 | 0.03 | 0.39 | 0.71 | 0.4 | 0.35 |
| 11 | 10614128 | rs11042889 | 1.071 | 0.368 | 0.248 | -2.187 | 0 | -0.012 | -0.012 | 0 | 0.07 | 0.06 | 0.18 | 0 | 0.07 | 0.06 |
| 11 | 10615473 | rs7126245 | -0.76 | 0.14 | 0.658 | -2.304 | 0.009 | 0.079 | 0.079 | 1.179 | 0.49 | 0.67 | 0.65 | 0.72 | 0.54 | 0.51 |
| 11 | 10619419 | rs11042892 | 1.239 | -0.001 | 0.64 | -2.438 | -0.016 | 0.094 | 0.094 | 2.808 | 0.37 | 0.18 | 0.26 | 0.17 | 0.36 | 0.23 |
| 11 | 10620629 | rs16927077 | -0.889 | 0.21 | 0.916 | -2.291 | 0.043 | -0.031 | -0.031 | 0 | 0.1 | 0.45 | 0.08 | 0.01 | 0.07 | 0.1 |
| 11 | 10623622 | rs749045 | -0.738 | 0.163 | 0.629 | -2.263 | 0.011 | 0.079 | 0.079 | 1.459 | 0.5 | 0.64 | 0.66 | 0.73 | 0.55 | 0.51 |
| 11 | 10624215 | rs10840444 | -0.395 | 0.109 | 0.697 | -2.277 | -0.001 | -0.004 | -0.004 | 0 | 0.47 | 0.4 | 0.64 | 0.73 | 0.55 | 0.48 |
| 11 | 10627711 | rs1017275 | -0.214 | -0.228 | 0.169 | -2.039 | 0.024 | 0.002 | 0.002 | 0 | 0.84 | 0.91 | 0.92 | 0.87 | 0.9 | 0.74 |
| 11 | 10632252 | rs964426 | -0.016 | 0.433 | 0.416 | -1.829 | 0.006 | -0.011 | -0.011 | 0 | 0.38 | 0.41 | 0.43 | 0.68 | 0.42 | 0.38 |
| 11 | 10632484 | rs964425 | 0.399 | 0.453 | 0.108 | -1.823 | -0.054 | -0.011 | -0.011 | 0 | 0.33 | 0.12 | 0.4 | 0.69 | 0.4 | 0.35 |
| 11 | 10638972 | rs4910163 | 0.86 | 0.583 | 0.142 | -1.751 | -0.021 | 0.036 | 0.036 | 0 | 0.07 | 0 | 0.17 | 0 | 0.07 | 0.07 |
| 11 | 10649161 | rs1425152 | 0.154 | 0.44 | -0.08 | -1.653 | -0.019 | 0.024 | 0.024 | 2.053 | 0.4 | 0.25 | 0.44 | 0.69 | 0.45 | 0.42 |
| 11 | 10650329 | rs2278125 | -0.372 | 0.449 | -0.256 | -1.626 | 0.002 | 0.006 | 0.006 | 3.21 | 0.51 | 0.53 | 0.64 | 0.72 | 0.54 | 0.51 |
| 11 | 10656687 | rs11042904 | 0.123 | 0.517 | -0.442 | -2.082 | 0.003 | -0.025 | -0.025 | 1.832 | 0.41 | 0.45 | 0.31 | 0.2 | 0.44 | 0.38 |
| 11 | 10658735 | rs1035691 | -0.366 | 0.509 | -0.467 | -2.375 | -0.006 | -0.087 | -0.087 | 0.218 | 0.62 | 0.57 | 0.62 | 0.97 | 0.66 | 0.81 |
| 11 | 10661951 | rs7944136 | 0.927 | 0.519 | -0.374 | -2.358 | 0 | -0.024 | -0.024 | 0 | 0.07 | 0.09 | 0.17 | 0 | 0.05 | 0.05 |
| 11 | 10666405 | rs2033049 | -0.045 | 0.504 | -0.467 | -2.542 | 0.001 | 0.022 | 0.022 | 0 | 0.49 | 0.44 | 0.48 | 0.2 | 0.49 | 0.46 |
| 11 | 10668522 | rs11042909 | 0.648 | 0.518 | -0.999 | -2.501 | -0.048 | -0.029 | -0.029 | 0.228 | 0.19 | 0.03 | 0.3 | 0.76 | 0.2 | 0.43 |
| 11 | 10670073 | rs4909943 | -0.277 | 0.508 | -0.918 | -2.533 | -0.017 | 0.025 | 0.025 | 1.029 | 0.27 | 0.15 | 0.48 | 0.76 | 0.26 | 0.49 |
| 11 | 10673739 | rs4909945 | -0.86 | 0.317 | -1.138 | -2.788 | -0.033 | 0.206 | 0.206 | 0.002 | 0.7 | 0.98 | 0.82 | 0.97 | 0.7 | 0.86 |
| 11 | 10682175 | rs10840460 | 0.688 | 0.482 | -0.87 | -2.494 | -0.04 | -0.004 | -0.004 | 0 | 0.17 | 0.04 | 0.34 | 0.79 | 0.2 | 0.43 |
| 11 | 10682248 | rs10770131 | -0.586 | 0.412 | -1.009 | -2.913 | -0.006 | 0.229 | 0.229 | 0.42 | 0.59 | 0.88 | 0.61 | 0.96 | 0.58 | 0.77 |
| 11 | 10688494 | rs10840463 | 0.704 | 0.447 | -0.556 | -2.826 | -0.027 | 0.004 | 0.004 | 0.028 | 0.17 | 0.09 | 0.34 | 0.79 | 0.2 | 0.43 |
| 11 | 10689262 | rs881870 | 0.588 | 0.191 | -0.264 | -3.221 | -0.003 | -0.059 | -0.059 | 0 | 0.31 | 0.36 | 0.21 | 0.16 | 0.32 | 0.31 |
| 11 | 10689820 | rs1390571 | -0.882 | 0.257 | -0.148 | -3.061 | 0.007 | 0.089 | 0.089 | 0 | 0.62 | 0.77 | 0.63 | 0.96 | 0.62 | 0.8 |
| 11 | 10691080 | rs903823 | -0.942 | 0.256 | -0.174 | -3.061 | -0.001 | 0.155 | 0.155 | 0.016 | 0.62 | 0.85 | 0.63 | 0.96 | 0.62 | 0.8 |
| 11 | 10691638 | rs1874444 | 0.969 | 0.362 | -0.323 | -3.014 | -0.026 | 0.003 | 0.003 | 0.044 | 0.17 | 0.09 | 0.35 | 0.79 | 0.2 | 0.43 |
| 11 | 10691721 | rs1874445 | -0.922 | 0.36 | -0.347 | -3.014 | 0.002 | -0.051 | -0.051 | 0 | 0.6 | 0.59 | 0.61 | 0.96 | 0.62 | 0.79 |
| 11 | 10693584 | rs7938027 | -0.245 | 0.223 | -0.133 | -2.612 | -0.001 | -0.048 | -0.048 | 0 | 0.42 | 0.43 | 0.26 | 0.17 | 0.41 | 0.37 |
| 11 | 10698073 | rs10770132 | -0.308 | 0.247 | -0.248 | -2.6 | -0.002 | -0.042 | -0.042 | 0 | 0.43 | 0.48 | 0.26 | 0.17 | 0.41 | 0.37 |
| 11 | 10699071 | rs903825 | -0.308 | 0.247 | -0.259 | -2.6 | -0.001 | -0.04 | -0.04 | 0.247 | 0.43 | 0.48 | 0.26 | 0.17 | 0.41 | 0.37 |
| 11 | 10702517 | rs12789511 | 0.116 | 0.49 | -0.069 | -2.316 | 0.005 | -0.026 | -0.026 | 0.06 | 0.26 | 0.28 | 0.17 | 0.16 | 0.31 | 0.32 |
| 11 | 10709168 | rs4910169 | 0.055 | 0.192 | -0.183 | -2.582 | 0.002 | 0.01 | 0.01 | 0.2 | 0.3 | 0.25 | 0.29 | 0.05 | 0.27 | 0.17 |
| 11 | 10711415 | rs4909947 | -0.277 | 0.188 | -0.25 | -2.922 | -0.001 | -0.038 | -0.038 | 0 | 0.53 | 0.53 | 0.43 | 0.12 | 0.5 | 0.27 |
| 11 | 10715124 | rs10770136 | -1.17 | 0.244 | -0.502 | -2.952 | -0.001 | -0.004 | -0.004 | 0 | 0.75 | 0.75 | 0.81 | 0.86 | 0.72 | 0.69 |
| 11 | 10715721 | rs10840473 | 1.457 | 0.229 | -0.444 | -2.961 | 0.007 | 0.04 | 0.04 | 0 | 0.22 | 0.13 | 0.12 | 0.06 | 0.22 | 0.1 |
| 11 | 10718606 | rs17420140 | 1.138 | 0.31 | -0.709 | -2.843 | -0.065 | -0.071 | -0.071 | 0 | 0.18 | 0.02 | 0.34 | 0.74 | 0.19 | 0.4 |
| 11 | 10719454 | rs10734656 | -1.1 | 0.2 | -0.519 | -2.777 | 0.004 | -0.02 | -0.02 | 2.731 | 0.75 | 0.73 | 0.81 | 0.84 | 0.72 | 0.69 |
| 11 | 10724434 | rs7129273 | 0.432 | 0.207 | -0.606 | -2.747 | -0.004 | -0.043 | -0.043 | 0 | 0.31 | 0.38 | 0.29 | 0.06 | 0.27 | 0.19 |
| 11 | 10732967 | rs7114199 | -0.84 | 0.592 | -0.926 | -2.384 | -0.003 | 0.001 | 0.001 | 0 | 0.76 | 0.78 | 0.81 | 0.84 | 0.72 | 0.68 |
| 11 | 10740549 | rs10840479 | -0.034 | 0.66 | -0.813 | -2.099 | -0.009 | 0.047 | 0.047 | 0 | 0.37 | 0.24 | 0.47 | 0.79 | 0.39 | 0.49 |
| 11 | 10745394 | rs11042937 | -0.657 | 0.642 | -1.313 | -1.657 | -0.003 | 0.136 | 0.136 | 0 | 0.56 | 0.78 | 0.71 | 0.8 | 0.52 | 0.58 |
| 11 | 10754074 | rs11042939 | -0.411 | 1.269 | -0.694 | 0.391 | 0.017 | 0.039 | 0.039 | 0 | 0.27 | 0.4 | 0.43 | 0.44 | 0.25 | 0.35 |
| 11 | 10757334 | rs4910184 | -0.063 | 1.22 | -1.18 | 0.692 | -0.042 | -0.036 | -0.036 | 0 | 0.61 | 0.78 | 0.55 | 0.4 | 0.45 | 0.47 |
| 11 | 10764618 | rs924111 | -0.029 | 1.215 | -0.705 | 0.853 | -0.001 | -0.002 | -0.002 | 0 | 0.4 | 0.31 | 0.4 | 0.43 | 0.41 | 0.47 |
| 11 | 10771479 | rs11042949 | -0.826 | 1.16 | 0.236 | 1.046 | 0.002 | 0.082 | 0.082 | 0 | 0.08 | 0.17 | 0.1 | 0.21 | 0.06 | 0.18 |
| 11 | 10772555 | rs2279696 | 0.325 | 1.128 | -0.081 | 1.184 | -0.015 | -0.029 | -0.029 | 0 | 0.42 | 0.16 | 0.41 | 0.4 | 0.43 | 0.51 |
| 11 | 10781155 | rs12418180 | -1.876 | 1.041 | 1.794 | 1.262 | 0.019 | 0.212 | 0.212 | 1.116 | 0.09 | 0.45 | 0.11 | 0.2 | 0.05 | 0.13 |
| 11 | 10786142 | rs3736326 | 0.875 | 1.718 | 1.466 | 0.759 | -0.003 | 0.004 | 0.004 | 0.028 | 0.65 | 0.32 | 0.76 | 0.78 | 0.6 | 0.74 |
| 11 | 10788376 | rs10219256 | -0.408 | 1.683 | 1.012 | 0.751 | -0.017 | 0.035 | 0.035 | 0 | 0.26 | 0.11 | 0.15 | 0.21 | 0.3 | 0.22 |
| 11 | 10791983 | rs2132517 | -2.128 | 1.442 | 1.092 | 0.951 | -0.001 | 0.041 | 0.041 | 0.745 | 0.13 | 0.71 | 0.25 | 0.23 | 0.12 | 0.16 |
| 11 | 10792326 | rs10500731 | -0.374 | 1.533 | 0.777 | 0.321 | 0.019 | -0.037 | -0.037 | 0 | 0.08 | 0.15 | 0.07 | 0 | 0.1 | 0.07 |
| 11 | 10795125 | rs17330399 | -0.615 | 1.844 | 0.541 | 0.497 | -0.074 | 0.009 | 0.009 | 0 | 0.23 | 0.01 | 0.14 | 0.18 | 0.27 | 0.15 |
| 11 | 10805459 | rs7103192 | 0.47 | 1.593 | 0.225 | 0.534 | -0.016 | 0.02 | 0.02 | 0 | 0.73 | 0.82 | 0.82 | 0.78 | 0.65 | 0.78 |
| 11 | 10813613 | rs7103582 | -1.321 | 1.471 | 0.698 | 0.887 | 0.013 | 0.028 | 0.028 | 0 | 0.24 | 0.31 | 0.4 | 0.61 | 0.36 | 0.33 |
| 11 | 10821055 | rs3736327 | 1.135 | 1.491 | 1.243 | 0.394 | 0.029 | 0.03 | 0.03 | 0.18 | 0.69 | 0.58 | 0.74 | 0.58 | 0.61 | 0.72 |
| 11 | 10823038 | rs7103202 | -0.232 | 1.484 | 1.253 | 0.373 | -0.001 | 0.003 | 0.003 | 0.563 | 0.07 | 0.04 | 0.02 | 0.01 | 0.06 | 0.03 |
| 11 | 10826256 | rs2242361 | -1.172 | 1.464 | 1.313 | 0.373 | 0.011 | 0.022 | 0.022 | 0 | 0.24 | 0.27 | 0.23 | 0.41 | 0.33 | 0.25 |
| 11 | 10846117 | rs7937695 | -1.129 | 1.409 | 1.473 | 0.382 | 0.02 | 0.042 | 0.042 | 0 | 0.24 | 0.29 | 0.27 | 0.42 | 0.34 | 0.24 |
| 11 | 10848430 | rs4910193 | -1.575 | 1.451 | 1.507 | 0.235 | 0.008 | 0.011 | 0.011 | 0 | 0.28 | 0.29 | 0.3 | 0.41 | 0.38 | 0.24 |
| 11 | 10849558 | rs4910198 | 1.882 | 1.524 | 1.533 | 0.252 | -0.045 | -0.018 | -0.018 | 0.046 | 0.63 | 0.17 | 0.62 | 0.59 | 0.55 | 0.73 |

| | | | | | | | | | | | | | | | | |
|---|---|---|---|---|---|---|---|---|---|---|---|---|---|---|---|---|
| 11 | 10849613 | rs2132522 | 1.225 | 1.695 | 1.797 | 0.268 | 0.081 | 0.029 | 0.029 | 0.118 | 0.94 | 0.59 | 0.93 | 1 | 0.95 | 0.99 |
| 11 | 10852822 | rs903836 | 1.334 | 1.695 | 1.796 | 0.268 | 0.072 | 0.021 | 0.021 | 0.273 | 0.93 | 0.58 | 0.93 | 1 | 0.95 | 0.99 |
| 11 | 10859590 | rs12418217 | -1.095 | 1.702 | 1.782 | 0.264 | -0.001 | -0.018 | -0.018 | 0.172 | 0.1 | 0.13 | 0.07 | 0.01 | 0.11 | 0.04 |
| 11 | 10860810 | rs11042979 | -1.739 | 1.701 | 1.774 | 0.264 | 0.024 | -0.101 | -0.101 | 0 | 0.14 | 0.51 | 0.11 | 0.01 | 0.11 | 0.04 |
| 11 | 10864595 | rs7950378 | -0.944 | 1.565 | 1.616 | 0.368 | 0.082 | 0.027 | 0.027 | 0.651 | 0.07 | 0.5 | 0.09 | 0 | 0.05 | 0.01 |
| 11 | 10865739 | rs771374 | -1.524 | 1.6 | 1.095 | 0.404 | -0.078 | -0.047 | -0.047 | 3.528 | 0.37 | 0.88 | 0.38 | 0.41 | 0.45 | 0.27 |
| 11 | 10877684 | rs11042983 | -1.024 | 2.358 | 0.607 | 0.868 | 0.033 | 0.034 | 0.034 | 0 | 0.26 | 0.33 | 0.29 | 0.44 | 0.41 | 0.28 |
| 11 | 10878223 | rs12292697 | -1.082 | 1.794 | 0.415 | 0.515 | -0.005 | 0.007 | 0.007 | 0 | 0.06 | 0.03 | 0.04 | 0.07 | 0.06 | 0.08 |
| 11 | 10887294 | rs7109947 | -1.68 | 1.626 | 0.95 | 0.181 | -0.002 | -0.08 | -0.08 | 0 | 0.16 | 0.31 | 0.1 | 0.03 | 0.14 | 0.16 |
| 11 | 10899436 | rs7125450 | -1.586 | 1.644 | 1.268 | 0.189 | 0.006 | -0.066 | -0.066 | 0.003 | 0.14 | 0.26 | 0.1 | 0.05 | 0.15 | 0.13 |
| 11 | 10900309 | rs11345 | 1.37 | 1.952 | 0.895 | 0.231 | 0.042 | -0.037 | -0.037 | 1.672 | 0.46 | 0.17 | 0.3 | 0.5 | 0.41 | 0.31 |
| 11 | 10901377 | rs16908491 | -0.391 | 2.047 | 0.849 | 0.255 | -0.002 | -0.03 | -0.03 | 0.003 | 0.08 | 0.14 | 0.06 | 0.03 | 0.07 | 0.08 |
| 11 | 10904067 | rs1554519 | 0.258 | 1.932 | 1.155 | 0.33 | -0.001 | -0.001 | -0.001 | 0 | 0.62 | 0.62 | 0.41 | 0.55 | 0.56 | 0.44 |
| 11 | 10909710 | rs2140214 | -0.026 | 1.858 | 1.287 | 0.602 | -0.032 | -0.02 | -0.02 | 2.11 | 0.7 | 0.8 | 0.74 | 0.65 | 0.64 | 0.55 |
| 11 | 10918161 | rs6484537 | 1.051 | 1.762 | 1.366 | 0.665 | -0.001 | 0.002 | 0.002 | 0 | 0.55 | 0.4 | 0.64 | 0.6 | 0.49 | 0.42 |
| 11 | 10918279 | rs2896567 | 0.96 | 1.758 | 1.393 | 0.64 | 0 | 0 | 0 | 0 | 0.57 | 0.5 | 0.65 | 0.6 | 0.5 | 0.43 |
| 11 | 10919211 | rs1533877 | 0.71 | 1.798 | 0.874 | 0.559 | 0.008 | -0.003 | -0.003 | 0.633 | 0.63 | 0.41 | 0.68 | 0.61 | 0.56 | 0.47 |
| 11 | 10922394 | rs6484539 | 0.835 | 1.503 | 0.874 | 0.788 | -0.003 | 0.011 | 0.011 | 0 | 0.55 | 0.36 | 0.64 | 0.6 | 0.49 | 0.42 |
| 11 | 10949045 | rs1386147 | 0.229 | 1.53 | -0.398 | 1.166 | -0.008 | -0.01 | -0.01 | 0 | 0.69 | 0.72 | 0.75 | 0.63 | 0.65 | 0.54 |
| 11 | 10955151 | rs636020 | -1.456 | 1.088 | -0.244 | 2.088 | -0.005 | 0.13 | 0.13 | 0 | 0.07 | 0.11 | 0.12 | 0.35 | 0.09 | 0.12 |
| 11 | 10958321 | rs6484550 | -0.455 | 0.993 | -0.255 | 1.795 | -0.002 | -0.001 | -0.001 | 5.094 | 0.44 | 0.56 | 0.47 | 0.54 | 0.44 | 0.44 |
| 11 | 10960273 | rs11043025 | 0.286 | 0.541 | -0.857 | 1.558 | 0.011 | -0.015 | -0.015 | 0 | 0.08 | 0.01 | 0.02 | 0.08 | 0.04 | 0.28 |
| 11 | 10960394 | rs652612 | -0.353 | 0.101 | -0.588 | 1.163 | -0.01 | 0.036 | 0.036 | 0 | 0.55 | 0.74 | 0.57 | 0.59 | 0.54 | 0.49 |
| 11 | 10962378 | rs557096 | 0.264 | 0.343 | -0.748 | 1.12 | -0.009 | 0.028 | 0.028 | 1.244 | 0.82 | 0.78 | 0.69 | 0.65 | 0.83 | 0.63 |
| 11 | 10963705 | rs523397 | 0.55 | 0.284 | -0.492 | 0.601 | -0.006 | -0.005 | -0.005 | 1.301 | 0.63 | 0.46 | 0.56 | 0.61 | 0.72 | 0.52 |
| 11 | 10963958 | rs2403433 | 0.279 | 0.308 | -0.792 | 0.554 | 0.005 | 0.002 | 0.002 | 2.605 | 0.11 | 0.14 | 0.09 | 0.12 | 0.15 | 0.11 |
| 11 | 10965371 | rs4910203 | 0.297 | 0.265 | -0.879 | 1.111 | -0.008 | 0.012 | 0.012 | 0.553 | 0.05 | 0.02 | 0.03 | 0.23 | 0.07 | 0.05 |
| 11 | 10965821 | rs7131542 | 1.158 | 0.263 | -0.804 | 1.064 | 0.003 | 0.009 | 0.009 | 0.068 | 0.07 | 0.08 | 0.06 | 0.12 | 0.09 | 0.07 |
| 11 | 10967638 | rs1601589 | 1.181 | 0.251 | -0.796 | 1.062 | -0.002 | -0.002 | -0.002 | 0 | 0.07 | 0.07 | 0.06 | 0.12 | 0.1 | 0.07 |
| 11 | 10970735 | rs1081557 | -0.508 | 0 | -1.182 | 0.56 | 0.012 | -0.012 | -0.012 | 0.016 | 0.42 | 0.71 | 0.31 | 0.51 | 0.43 | 0.34 |
| 11 | 10984880 | rs1601588 | 0.123 | -0.27 | -0.801 | 0.851 | 0 | 0.117 | 0.117 | 0.934 | 0.89 | 0.71 | 0.94 | 0.8 | 0.9 | 0.94 |
| 11 | 10992850 | rs4910210 | 1.182 | -0.155 | -0.766 | 0.883 | 0 | -0.003 | -0.003 | 0 | 0.17 | 0.17 | 0.33 | 0.32 | 0.14 | 0.4 |
| 11 | 10994559 | rs7112994 | 1.031 | -0.095 | -0.78 | 0.744 | 0.028 | 0.096 | 0.096 | 0.015 | 0.5 | 0.22 | 0.45 | 0.3 | 0.56 | 0.44 |
| 11 | 10997718 | rs7947494 | 0.15 | 0.035 | -0.942 | 0.662 | 0.025 | 0.007 | 0.007 | 0 | 0.54 | 0.31 | 0.68 | 0.5 | 0.54 | 0.69 |
| 11 | 10997949 | rs11603305 | 1.059 | 0.114 | -1.02 | 0.655 | 0 | 0.003 | 0.003 | 0 | 0.29 | 0.3 | 0.21 | 0.22 | 0.31 | 0.23 |
| 11 | 11004825 | rs7112391 | -1.033 | 0.216 | -0.84 | 0.719 | -0.032 | -0.034 | -0.034 | 0 | 0.71 | 0.86 | 0.7 | 0.72 | 0.68 | 0.74 |
| 11 | 11005966 | rs7123082 | -0.546 | 0.177 | -0.788 | 0.736 | 0.001 | 0.042 | 0.042 | 0.126 | 0.91 | 0.91 | 0.94 | 0.68 | 0.88 | 0.94 |
| 11 | 11007570 | rs17334036 | 1.757 | 0.226 | -0.917 | 0.622 | -0.069 | 0.034 | 0.034 | 0 | 0.22 | 0.06 | 0.25 | 0.12 | 0.25 | 0.21 |
| 11 | 11007971 | rs732392 | 0.406 | 0.195 | -0.669 | 0.634 | -0.001 | 0.002 | 0.002 | 0 | 0.33 | 0.29 | 0.44 | 0.37 | 0.29 | 0.47 |
| 11 | 11009775 | rs11043041 | 1.445 | 0.105 | -0.678 | 0.195 | -0.005 | 0.006 | 0.006 | 0.006 | 0.23 | 0.17 | 0.18 | 0.17 | 0.27 | 0.2 |
| 11 | 11011024 | rs11605707 | 1.469 | 0.14 | -0.143 | 0.123 | 0 | -0.028 | -0.028 | 0.005 | 0.06 | 0.17 | 0.03 | 0.01 | 0.04 | 0.03 |
| 11 | 11011562 | rs6484589 | 0.908 | 0.131 | -0.137 | 0.122 | -0.003 | 0.023 | 0.023 | 0 | 0.31 | 0.27 | 0.3 | 0.13 | 0.31 | 0.25 |
| 11 | 11014952 | rs7941808 | -0.208 | 0.096 | -0.005 | 0.278 | -0.001 | -0.004 | -0.004 | 0 | 0.6 | 0.49 | 0.65 | 0.66 | 0.6 | 0.69 |
| 11 | 11017279 | rs7117665 | -0.327 | 0.108 | -0.073 | 0.279 | 0 | -0.004 | -0.004 | 0 | 0.61 | 0.58 | 0.65 | 0.67 | 0.6 | 0.68 |
| 11 | 11020083 | rs12269906 | 1.682 | 0.197 | -0.216 | 0.174 | -0.002 | 0.004 | 0.004 | 0.75 | 0.16 | 0.12 | 0.1 | 0.03 | 0.14 | 0.07 |
| 11 | 11021312 | rs10743159 | 0.122 | 0.351 | 0.01 | 0.546 | -0.001 | 0.084 | 0.084 | 0.053 | 0.86 | 0.68 | 0.93 | 0.79 | 0.86 | 0.9 |
| 11 | 11027552 | rs2403375 | -0.106 | 0.398 | -0.052 | 0.608 | -0.001 | 0.006 | 0.006 | 0.025 | 0.59 | 0.57 | 0.62 | 0.55 | 0.55 | 0.67 |
| 11 | 11028685 | rs10743162 | 0.771 | 0.365 | -0.265 | 0.61 | -0.001 | 0.012 | 0.012 | 0 | 0.35 | 0.24 | 0.43 | 0.37 | 0.29 | 0.48 |
| 11 | 11029062 | rs7123407 | 0.641 | 0.665 | -0.298 | 0.245 | 0.003 | -0.006 | -0.006 | 0 | 0.38 | 0.34 | 0.47 | 0.49 | 0.34 | 0.49 |
| 11 | 11030667 | rs2171236 | -0.34 | 0.598 | -0.263 | 0.272 | 0.013 | 0.055 | 0.055 | 0 | 0.71 | 0.59 | 0.77 | 0.63 | 0.65 | 0.75 |
| 11 | 11032427 | rs11828788 | 1.155 | 0.725 | -0.14 | 0.355 | -0.003 | -0.03 | -0.03 | 0 | 0.06 | 0.19 | 0.03 | 0.01 | 0.04 | 0.03 |
| 11 | 11032988 | rs7121565 | 0.37 | 0.823 | -0.206 | 0.343 | -0.027 | 0.047 | 0.047 | 0 | 0.45 | 0.23 | 0.43 | 0.3 | 0.51 | 0.4 |
| 11 | 11033096 | rs7121789 | 2.035 | 0.793 | -0.469 | 0.333 | -0.083 | 0.019 | 0.019 | 0 | 0.22 | 0.02 | 0.25 | 0.12 | 0.25 | 0.21 |
| 11 | 11037124 | rs2129286 | 0.725 | 0.853 | -0.342 | 0.349 | 0.003 | -0.003 | -0.003 | 0 | 0.4 | 0.34 | 0.46 | 0.49 | 0.35 | 0.51 |
| 11 | 11037708 | rs4910217 | -0.008 | 0.862 | -0.307 | 0.357 | 0.009 | 0.033 | 0.033 | 0 | 0.48 | 0.57 | 0.53 | 0.68 | 0.45 | 0.56 |
| 11 | 11038332 | rs10770159 | 0.315 | 0.918 | -0.156 | 0.607 | 0.022 | 0.129 | 0.129 | 0.828 | 0.92 | 0.8 | 0.93 | 0.81 | 0.9 | 0.94 |
| 11 | 11040934 | rs1386155 | -0.176 | 1.014 | -0.496 | 0.639 | 0.012 | 0.007 | 0.007 | 0 | 0.67 | 0.53 | 0.67 | 0.68 | 0.64 | 0.71 |
| 11 | 11042958 | rs1486541 | 0.376 | 0.688 | -0.267 | 0.868 | 0.029 | 0.14 | 0.14 | 0.015 | 0.87 | 0.55 | 0.9 | 0.79 | 0.86 | 0.9 |
| 11 | 11046259 | rs10500737 | 1.107 | 1.044 | -1.216 | 0.441 | -0.056 | 0 | 0 | 0.47 | 0.18 | 0.02 | 0.21 | 0.12 | 0.18 | 0.2 |
| 11 | 11050931 | rs11601235 | 0.54 | 1.343 | -1.047 | 0.658 | -0.046 | -0.029 | -0.029 | 0 | 0.1 | 0 | 0.11 | 0.1 | 0.15 | 0.11 |
| 11 | 11051503 | rs1486536 | -0.918 | 0.754 | -0.739 | 0.953 | 0 | 0 | 0 | 2.227 | 0.77 | 0.75 | 0.77 | 0.73 | 0.83 | 0.81 |
| 11 | 11052290 | rs2403435 | -0.716 | 0.659 | -0.78 | 1.219 | 0.004 | -0.007 | -0.007 | 0 | 0.74 | 0.74 | 0.7 | 0.65 | 0.82 | 0.78 |
| 11 | 11053680 | rs2129283 | -0.107 | 0.66 | -0.124 | 1.066 | -0.012 | 0.035 | 0.035 | 0 | 0.67 | 0.54 | 0.66 | 0.59 | 0.72 | 0.73 |
| 11 | 11058190 | rs17335592 | 0.124 | 0.865 | -0.425 | 0.887 | 0.021 | 0.002 | 0.002 | 0 | 0.21 | 0.24 | 0.17 | 0.22 | 0.26 | 0.28 |
| 11 | 11060903 | rs1386158 | -0.073 | 0.447 | -0.44 | 0.761 | 0.005 | -0.001 | -0.001 | 0 | 0.56 | 0.49 | 0.61 | 0.61 | 0.59 | 0.7 |
| 11 | 11060923 | rs10770165 | 0.121 | 0.366 | 0.194 | 0.716 | -0.012 | 0.011 | 0.011 | 0 | 0.53 | 0.34 | 0.58 | 0.61 | 0.58 | 0.69 |
| 11 | 11065405 | rs7940981 | -1.658 | 0.267 | 0.022 | 0.802 | 0.003 | -0.005 | -0.005 | 0 | 0.8 | 0.81 | 0.82 | 0.75 | 0.88 | 0.83 |
| 11 | 11069843 | rs7946997 | -0.355 | 0.122 | 0.059 | 0.139 | 0.035 | -0.12 | -0.12 | 0.148 | 0.27 | 0.44 | 0.19 | 0.07 | 0.29 | 0.14 |
| 11 | 11076017 | rs11043067 | 0.943 | -0.131 | -0.585 | 0.219 | -0.105 | 0.097 | 0.097 | 0.313 | 0.21 | 0.02 | 0.15 | 0.04 | 0.26 | 0.09 |

| | | | | | | | | | | | | | | | | |
|---|---|---|---|---|---|---|---|---|---|---|---|---|---|---|---|---|
| 11 | 11076056 | rs1586381 | -1.039 | -0.757 | -0.326 | 0.708 | -0.062 | 0.046 | 0.046 | 0 | 0.7 | 0.52 | 0.72 | 0.64 | 0.84 | 0.76 |
| 11 | 11080548 | rs10840545 | 1.025 | -0.334 | -0.199 | 0.772 | 0 | 0.013 | 0.013 | 0.002 | 0.12 | 0.14 | 0.28 | 0.37 | 0.16 | 0.37 |
| 11 | 11082966 | rs12273047 | 0.979 | -0.524 | 0.204 | 0.562 | 0.015 | -0.045 | -0.045 | 0.419 | 0.19 | 0.19 | 0.16 | 0.04 | 0.23 | 0.07 |
| 11 | 11083800 | rs4909961 | 0.433 | -0.53 | 0.446 | 0.748 | 0.001 | 0 | 0 | 0.444 | 0.35 | 0.39 | 0.44 | 0.42 | 0.44 | 0.58 |
| 11 | 11084001 | rs274486 | -0.957 | -0.711 | 0.289 | 0.815 | -0.007 | -0.012 | -0.012 | 0.585 | 0.69 | 0.73 | 0.71 | 0.81 | 0.76 | 0.78 |
| 11 | 11084102 | rs10743172 | 0.549 | -0.735 | 0.156 | 0.82 | 0.021 | 0.02 | 0.02 | 0.176 | 0.36 | 0.54 | 0.45 | 0.43 | 0.45 | 0.58 |
| 11 | 11088504 | rs1394120 | 0.029 | -0.677 | 0.069 | 0.839 | -0.034 | -0.072 | -0.072 | 6.111 | 0.35 | 0.09 | 0.24 | 0.39 | 0.31 | 0.2 |
| 11 | 11088815 | rs1486531 | -0.819 | -0.58 | 0.172 | 0.823 | 0.024 | 0.02 | 0.02 | 0.143 | 0.65 | 0.46 | 0.55 | 0.58 | 0.55 | 0.43 |
| 11 | 11092386 | rs7938527 | 1.012 | -0.513 | 0.123 | 1.031 | -0.013 | 0.047 | 0.047 | 0 | 0.29 | 0.11 | 0.33 | 0.19 | 0.33 | 0.23 |
| 11 | 11092888 | rs1843676 | -0.17 | -0.498 | 0.039 | 1.008 | -0.015 | -0.024 | -0.024 | 0 | 0.38 | 0.12 | 0.26 | 0.38 | 0.34 | 0.23 |
| 11 | 11095452 | rs2896570 | -0.967 | -0.438 | 0.301 | 1.205 | 0.004 | 0 | 0 | 0 | 0.72 | 0.64 | 0.63 | 0.7 | 0.69 | 0.64 |
| 11 | 11098039 | rs2631235 | -0.421 | -0.316 | 0.032 | 1.308 | 0.012 | -0.023 | -0.023 | 0.245 | 0.4 | 0.29 | 0.26 | 0.52 | 0.34 | 0.44 |
| 11 | 11099460 | rs12289317 | 0.755 | -0.362 | -0.193 | 1.467 | -0.074 | -0.003 | -0.003 | 1.937 | 0.25 | 0.06 | 0.18 | 0.22 | 0.29 | 0.17 |
| 11 | 11102238 | rs274500 | -0.509 | -0.385 | 0.243 | 1.681 | -0.04 | 0.079 | 0.079 | 0 | 0.16 | 0.32 | 0.12 | 0.3 | 0.08 | 0.28 |
| 11 | 11102542 | rs274503 | 0.536 | -0.522 | -0.383 | 1.392 | 0.011 | -0.012 | -0.012 | 0.057 | 0.11 | 0.07 | 0.06 | 0.17 | 0.04 | 0.08 |
| 11 | 11105662 | rs274508 | -0.68 | -0.094 | -0.483 | 1.274 | 0 | 0 | 0 | 0 | 0.73 | 0.67 | 0.62 | 0.71 | 0.68 | 0.69 |
| 11 | 11108680 | rs274512 | -0.686 | -0.202 | -0.678 | 1.299 | 0.005 | 0 | 0 | 0 | 0.7 | 0.63 | 0.58 | 0.68 | 0.65 | 0.68 |
| 11 | 11109239 | rs1910505 | 1.113 | -0.211 | -1.227 | 1.201 | -0.038 | 0.025 | 0.025 | 0 | 0.25 | 0.06 | 0.31 | 0.2 | 0.31 | 0.22 |
| 11 | 11109296 | rs1910504 | 0.747 | -0.036 | -1.029 | 1.156 | 0 | 0.012 | 0.012 | 0 | 0.3 | 0.31 | 0.34 | 0.41 | 0.33 | 0.41 |
| 11 | 11111125 | rs924796 | 0.255 | 0.605 | -1.768 | 0.751 | -0.06 | 0.042 | 0.042 | 0.468 | 0.26 | 0.08 | 0.19 | 0.14 | 0.28 | 0.19 |
| 11 | 11114939 | rs11043078 | -0.009 | 0.795 | -1.572 | 0.754 | -0.014 | 0.01 | 0.01 | 0.111 | 0.29 | 0.23 | 0.42 | 0.26 | 0.34 | 0.31 |
| 11 | 11115392 | rs10219213 | -0.226 | 1.115 | -1.695 | 1.024 | -0.047 | -0.032 | -0.032 | 0.019 | 0.32 | 0.12 | 0.5 | 0.32 | 0.36 | 0.28 |
| 11 | 11115648 | rs7108095 | -0.458 | 0.366 | -1.101 | 0.559 | 0.013 | 0.019 | 0.019 | 0 | 0.54 | 0.43 | 0.39 | 0.32 | 0.59 | 0.46 |
| 11 | 11118343 | rs11043080 | -0.66 | 0.615 | -1.172 | 1.087 | -0.023 | -0.064 | -0.064 | 0 | 0.1 | 0.01 | 0.08 | 0.25 | 0.11 | 0.18 |
| 11 | 11119228 | rs4909963 | 0.303 | 0.569 | -0.537 | 0.429 | 0.007 | -0.036 | -0.036 | 2.362 | 0.9 | 0.8 | 0.94 | 0.97 | 0.9 | 0.96 |
| 11 | 11119860 | rs7936368 | -0.407 | 0.014 | -0.209 | 0.074 | -0.021 | -0.031 | -0.031 | 0 | 0.19 | 0.29 | 0.15 | 0.13 | 0.15 | 0.12 |
| 11 | 11120206 | rs17344770 | 0.565 | 0.33 | -1.217 | -0.481 | -0.07 | 0.088 | 0.088 | 0.241 | 0.3 | 0.05 | 0.33 | 0.12 | 0.34 | 0.2 |
| 11 | 11121557 | rs4482023 | -0.594 | -0.051 | -0.38 | -0.717 | -0.014 | -0.049 | -0.049 | 0.503 | 0.34 | 0.57 | 0.31 | 0.19 | 0.26 | 0.21 |
| 11 | 11123656 | rs7116607 | -0.496 | -0.093 | -0.302 | -0.743 | -0.004 | -0.021 | -0.021 | 0 | 0.32 | 0.37 | 0.3 | 0.19 | 0.25 | 0.2 |
| 11 | 11128017 | rs924795 | 0.78 | -0.157 | -0.344 | -0.081 | -0.001 | 0.008 | 0.008 | 0.003 | 0.41 | 0.34 | 0.42 | 0.32 | 0.41 | 0.25 |
| 11 | 11129275 | rs1503930 | -0.441 | -0.24 | -0.506 | 0.107 | -0.001 | -0.002 | -0.002 | 0 | 0.7 | 0.71 | 0.71 | 0.5 | 0.65 | 0.43 |
| 11 | 11129688 | rs11043086 | 0.004 | -0.397 | 0.05 | 0.063 | 0.008 | 0.005 | 0.005 | 0 | 0.19 | 0.13 | 0.24 | 0.19 | 0.16 | 0.16 |
| 11 | 11135795 | rs11043097 | -0.093 | -0.388 | -0.072 | 0.068 | 0.016 | 0.009 | 0.009 | 0 | 0.2 | 0.07 | 0.23 | 0.19 | 0.17 | 0.17 |
| 11 | 11143030 | rs6484689 | 0.222 | -0.448 | -0.04 | -0.008 | 0.009 | 0.013 | 0.013 | 0.004 | 0.17 | 0.13 | 0.24 | 0.12 | 0.15 | 0.11 |
| 11 | 11150830 | rs7103691 | 0.666 | -0.323 | 0.056 | 0.125 | -0.03 | 0.139 | 0.139 | 0 | 0.46 | 0.15 | 0.4 | 0.23 | 0.5 | 0.32 |
| 11 | 11151347 | rs7107340 | 0.712 | -0.354 | 0.044 | 0.098 | -0.049 | 0.139 | 0.139 | 0 | 0.44 | 0.11 | 0.4 | 0.23 | 0.48 | 0.31 |
| 11 | 11151466 | rs7122257 | 0.304 | -0.103 | -0.008 | 0.806 | 0.019 | -0.009 | -0.009 | 0 | 0.58 | 0.22 | 0.6 | 0.41 | 0.64 | 0.51 |
| 11 | 11152428 | rs1394118 | 0.284 | -0.219 | 0.65 | 0.987 | -0.012 | 0.007 | 0.007 | 0 | 0.78 | 0.57 | 0.76 | 0.75 | 0.83 | 0.77 |
| 11 | 11155517 | rs1503938 | -0.806 | -0.138 | 0.814 | 1.014 | 0.005 | 0.015 | 0.015 | 0 | 0.19 | 0.22 | 0.36 | 0.36 | 0.25 | 0.24 |
| 11 | 11155959 | rs11602104 | -0.154 | -0.161 | 0.755 | 0.979 | -0.001 | -0.001 | -0.001 | 0 | 0.14 | 0.11 | 0.19 | 0.18 | 0.13 | 0.08 |
| 11 | 11160916 | rs7127097 | 0.682 | -0.275 | 0.976 | 1.079 | -0.017 | 0.016 | 0.016 | 0 | 0.64 | 0.45 | 0.77 | 0.6 | 0.73 | 0.59 |
| 11 | 11161248 | rs7110986 | -2.109 | -0.233 | 1.3 | 1.101 | -0.007 | 0.091 | 0.091 | 0 | 0.53 | 0.71 | 0.58 | 0.76 | 0.5 | 0.62 |
| 11 | 11162347 | rs869872 | 1.752 | -0.314 | 0.987 | 1.037 | -0.107 | 0.149 | 0.149 | 0 | 0.45 | 0.06 | 0.4 | 0.24 | 0.5 | 0.31 |
| 11 | 11163649 | rs10743178 | -1.209 | -0.165 | 1.282 | 1.188 | 0.064 | 0.137 | 0.137 | 0 | 0.94 | 0.8 | 0.82 | 0.83 | 0.88 | 0.86 |
| 11 | 11163681 | rs10734661 | -1.253 | -0.164 | 1.282 | 1.189 | 0.066 | 0.14 | 0.14 | 0 | 0.94 | 0.8 | 0.82 | 0.83 | 0.88 | 0.86 |
| 11 | 11165364 | rs11825182 | -0.007 | -0.283 | 1.112 | 1.084 | 0 | -0.003 | -0.003 | 0.525 | 0.2 | 0.15 | 0.15 | 0.35 | 0.2 | 0.29 |
| 11 | 11166593 | rs7122220 | -1.678 | -0.387 | 1.42 | 0.841 | -0.001 | 0.046 | 0.046 | 0 | 0.29 | 0.62 | 0.43 | 0.46 | 0.29 | 0.38 |
| 11 | 11169808 | rs10840586 | -1.594 | -0.376 | 1.493 | 0.594 | 0.012 | -0.016 | -0.016 | 0 | 0.31 | 0.45 | 0.4 | 0.25 | 0.34 | 0.29 |
| 11 | 11172701 | rs1875775 | 0.245 | -0.419 | 1.659 | 0.483 | -0.022 | -0.073 | -0.073 | 1.041 | 0.84 | 0.44 | 0.8 | 0.96 | 0.91 | 0.93 |
| 11 | 11175628 | rs11043118 | 1.24 | -0.38 | 1.535 | 0.637 | 0 | 0.016 | 0.016 | 0 | 0.68 | 0.41 | 0.67 | 0.63 | 0.65 | 0.6 |
| 11 | 11178959 | rs11043121 | -0.768 | -0.21 | 1.243 | 0.657 | -0.003 | -0.004 | -0.004 | 0.687 | 0.16 | 0.17 | 0.23 | 0.18 | 0.27 | 0.23 |
| 11 | 11183179 | rs7129423 | -0.893 | -0.261 | 1.533 | 0.624 | 0.04 | 0.057 | 0.057 | 0 | 0.14 | 0.32 | 0.22 | 0.18 | 0.22 | 0.22 |
| 11 | 11184095 | rs11043124 | 0.005 | -0.239 | 1.474 | 0.612 | 0.061 | 0.009 | 0.009 | 0.172 | 0.06 | 0.26 | 0.16 | 0.02 | 0.06 | 0.04 |
| 11 | 11184373 | rs12417680 | 1.835 | -0.392 | 1.243 | 0.495 | -0.022 | 0.006 | 0.006 | 1.125 | 0.54 | 0.13 | 0.49 | 0.62 | 0.56 | 0.52 |
| 11 | 11184721 | rs7105012 | -0.759 | -0.323 | 1.133 | 0.5 | 0 | -0.029 | -0.029 | 0.099 | 0.21 | 0.26 | 0.25 | 0.15 | 0.2 | 0.21 |
| 11 | 11186717 | rs10770179 | -0.752 | -0.318 | 1.135 | 0.478 | 0 | -0.012 | -0.012 | 0 | 0.21 | 0.26 | 0.25 | 0.18 | 0.2 | 0.22 |
| 11 | 11189537 | rs885117 | -0.937 | -0.32 | 1.096 | 0.452 | -0.004 | -0.006 | -0.006 | 0 | 0.22 | 0.28 | 0.27 | 0.19 | 0.19 | 0.22 |
| 11 | 11189970 | rs2403461 | 0.324 | -0.299 | 0.897 | 0.451 | 0.041 | -0.012 | -0.012 | 0.459 | 0.05 | 0.18 | 0.16 | 0.02 | 0.05 | 0.04 |
| 11 | 11190381 | rs7929304 | 1.041 | -0.311 | 0.497 | 0.376 | 0.001 | -0.01 | -0.01 | 0 | 0.7 | 0.38 | 0.58 | 0.81 | 0.67 | 0.72 |
| 11 | 11191487 | rs12363497 | 1.525 | -0.25 | 0.444 | 0.391 | -0.005 | 0.03 | 0.03 | 0 | 0.54 | 0.31 | 0.47 | 0.64 | 0.53 | 0.54 |
| 11 | 11193070 | rs12364300 | -0.578 | 0.049 | 0.236 | 0.294 | -0.001 | -0.001 | -0.001 | 0 | 0.1 | 0.11 | 0.09 | 0.05 | 0.11 | 0.08 |
| 11 | 11194217 | rs2403464 | -0.594 | 0.128 | 0.218 | 0.346 | 0.01 | -0.019 | -0.019 | 0.17 | 0.22 | 0.37 | 0.41 | 0.15 | 0.26 | 0.2 |
| 11 | 11195213 | rs2722779 | 0.671 | 0.08 | -0.736 | 0.411 | -0.073 | -0.079 | -0.079 | 0 | 0.38 | 0.13 | 0.32 | 0.59 | 0.44 | 0.49 |
| 11 | 11198251 | rs2655103 | 0.072 | 0.094 | -0.278 | 0.431 | 0.05 | 0.078 | 0.078 | 0.034 | 0.92 | 0.69 | 0.91 | 0.88 | 0.93 | 0.92 |
| 11 | 11200858 | rs2655074 | 0.802 | 0.286 | -1.345 | 0.482 | -0.05 | -0.099 | -0.099 | 0 | 0.33 | 0.08 | 0.26 | 0.52 | 0.37 | 0.37 |
| 11 | 11201684 | rs7932734 | 0.16 | -0.102 | -1.272 | 0.242 | 0.037 | 0.031 | 0.031 | 0 | 0.29 | 0.15 | 0.16 | 0.17 | 0.2 | 0.28 |
| 11 | 11205233 | rs2722772 | 0.151 | -0.001 | -0.787 | 0.185 | -0.002 | 0.031 | 0.031 | 0 | 0.47 | 0.65 | 0.4 | 0.69 | 0.51 | 0.38 |
| 11 | 11206934 | rs1503932 | 0.952 | 0.298 | -1.112 | 0.218 | -0.059 | -0.087 | -0.087 | 0 | 0.29 | 0.08 | 0.23 | 0.5 | 0.39 | 0.27 |
| 11 | 11208173 | rs11043139 | 0.355 | 0.673 | -0.564 | 0.194 | 0.026 | 0.024 | 0.024 | 0 | 0.08 | 0.13 | 0.18 | 0.11 | 0.14 | 0.15 |

| | | | | | | | | | | | | | | | | |
|---|---|---|---|---|---|---|---|---|---|---|---|---|---|---|---|---|
| 11 | 11209551 | rs12271102 | 0.611 | 0.176 | -0.977 | -0.161 | 0.015 | 0.012 | 0.012 | 0 | 0.34 | 0.02 | 0.16 | 0.21 | 0.22 | 0.37 |
| 11 | 11211567 | rs7130366 | 0.078 | 0.245 | -0.478 | -0.147 | 0.007 | -0.023 | -0.023 | 0 | 0.42 | 0.19 | 0.39 | 0.37 | 0.36 | 0.53 |
| 11 | 11214588 | rs12280481 | 0.062 | 0.246 | -0.475 | -0.151 | 0.011 | -0.013 | -0.013 | 0 | 0.42 | 0.24 | 0.39 | 0.37 | 0.36 | 0.53 |
| 11 | 11215710 | rs11822107 | 0.148 | 0.154 | -0.151 | -0.259 | -0.012 | -0.064 | -0.064 | 1.202 | 0.13 | 0.2 | 0.1 | 0 | 0.12 | 0.1 |
| 11 | 11223236 | rs7114703 | -0.208 | 0.29 | -0.02 | 0.061 | 0.009 | -0.005 | -0.005 | 0 | 0.43 | 0.31 | 0.4 | 0.37 | 0.36 | 0.53 |
| 11 | 11226136 | rs17347113 | 0.71 | 0.623 | -0.318 | 0.123 | 0.025 | 0.01 | 0.01 | 0 | 0.07 | 0.1 | 0.2 | 0.11 | 0.16 | 0.15 |
| 11 | 11227291 | rs7924382 | 0.207 | 0.009 | -0.127 | -0.064 | 0.02 | 0.009 | 0.009 | 0 | 0.37 | 0.27 | 0.22 | 0.26 | 0.23 | 0.38 |
| 11 | 11232008 | rs11043167 | 0.168 | 0.032 | -0.487 | 0.106 | -0.02 | 0.035 | 0.035 | 0 | 0.42 | 0.27 | 0.34 | 0.28 | 0.48 | 0.41 |
| 11 | 11232260 | rs4910278 | 0.23 | 0.094 | -0.525 | 0.111 | -0.024 | 0.038 | 0.038 | 0 | 0.41 | 0.26 | 0.34 | 0.23 | 0.47 | 0.41 |
| 11 | 11234314 | rs7123620 | -0.073 | -0.236 | -0.768 | 0.211 | 0.016 | 0.016 | 0.016 | 0 | 0.6 | 0.27 | 0.43 | 0.29 | 0.63 | 0.51 |
| 11 | 11241056 | rs12287713 | -0.489 | 0.129 | -1.098 | -0.156 | -0.007 | 0.02 | 0.02 | 0 | 0.24 | 0.19 | 0.16 | 0.07 | 0.26 | 0.2 |
| 11 | 11245936 | rs11021708 | 1.11 | 0.466 | -1.587 | -0.185 | -0.08 | 0.01 | 0.01 | 0 | 0.19 | 0.01 | 0.29 | 0.07 | 0.23 | 0.07 |
| 11 | 11246376 | rs4341525 | -0.949 | 0.394 | -1.053 | 0.506 | 0.038 | 0.108 | 0.108 | 0 | 0.91 | 0.79 | 0.79 | 0.66 | 0.84 | 0.8 |
| 11 | 11247842 | rs17347854 | 1.039 | -0.031 | -1.465 | -0.086 | 0.003 | 0.031 | 0.031 | 0 | 0.18 | 0.07 | 0.12 | 0.06 | 0.16 | 0.13 |
| 11 | 11251630 | rs11021712 | 1.115 | 0.156 | -0.557 | 0.031 | 0.017 | -0.049 | -0.049 | 0 | 0.22 | 0.34 | 0.32 | 0.07 | 0.23 | 0.11 |
| 11 | 11252484 | rs11021715 | 0.079 | -0.789 | -0.739 | 0.276 | 0.049 | 0.006 | 0.006 | 0 | 0.43 | 0.24 | 0.26 | 0.39 | 0.34 | 0.43 |
| 11 | 11254348 | rs7940763 | 0.455 | -0.668 | -0.404 | -0.215 | -0.012 | -0.086 | -0.086 | 0.397 | 0.2 | 0.42 | 0.1 | 0.05 | 0.15 | 0.18 |
| 11 | 11255991 | rs4408307 | -0.385 | -0.776 | -0.763 | -0.054 | 0.02 | 0.005 | 0.005 | 0 | 0.54 | 0.74 | 0.57 | 0.49 | 0.55 | 0.47 |
| 11 | 11256491 | rs7112035 | 0.114 | -0.555 | -0.008 | -0.156 | -0.013 | -0.079 | -0.079 | 0 | 0.2 | 0.4 | 0.09 | 0.05 | 0.14 | 0.16 |
| 11 | 11258928 | rs10831552 | 0.114 | -0.693 | -0.46 | -0.145 | 0.011 | 0.041 | 0.041 | 0.058 | 0.21 | 0.12 | 0.11 | 0.05 | 0.18 | 0.2 |
| 11 | 11260163 | rs4910287 | -0.26 | -0.929 | -0.278 | 0.199 | -0.011 | 0.054 | 0.054 | 0 | 0.75 | 0.69 | 0.7 | 0.55 | 0.74 | 0.63 |
| 11 | 11261045 | rs11021718 | -0.878 | -1.134 | -0.21 | 0.152 | 0 | -0.021 | -0.021 | 0.002 | 0.34 | 0.52 | 0.35 | 0.27 | 0.37 | 0.38 |
| 11 | 11261567 | rs4076941 | -0.797 | -1.111 | -0.158 | 0.166 | 0.004 | -0.02 | -0.02 | 0.061 | 0.33 | 0.48 | 0.34 | 0.27 | 0.36 | 0.38 |
| 11 | 11269086 | rs4441010 | -0.359 | -1.349 | -0.233 | 0.102 | -0.024 | 0.029 | 0.029 | 0 | 0.71 | 0.69 | 0.69 | 0.4 | 0.77 | 0.53 |
| 11 | 11270037 | rs4501975 | 0.32 | -1.534 | -0.25 | -0.118 | -0.005 | -0.002 | -0.002 | 0 | 0.61 | 0.47 | 0.66 | 0.26 | 0.71 | 0.34 |
| 11 | 11270851 | rs10831556 | 0.252 | -1.532 | -0.312 | -0.111 | -0.022 | -0.014 | -0.014 | 0 | 0.62 | 0.53 | 0.66 | 0.26 | 0.71 | 0.34 |
| 11 | 11272817 | rs10500742 | 1.253 | -0.948 | -1.135 | -0.579 | -0.041 | 0.045 | 0.045 | 0 | 0.25 | 0.08 | 0.29 | 0.13 | 0.29 | 0.17 |
| 11 | 11273949 | rs4909973 | -0.012 | -0.807 | -1.036 | -0.667 | -0.006 | -0.006 | -0.006 | 0.683 | 0.55 | 0.53 | 0.61 | 0.37 | 0.63 | 0.41 |
| 11 | 11274698 | rs7935881 | -0.145 | -0.711 | -0.895 | -0.461 | 0.006 | -0.026 | -0.026 | 0 | 0.43 | 0.54 | 0.4 | 0.23 | 0.45 | 0.27 |
| 11 | 11280690 | rs4508201 | -0.137 | -0.48 | -1.011 | -0.311 | -0.004 | 0.043 | 0.043 | 0 | 0.58 | 0.72 | 0.5 | 0.22 | 0.56 | 0.31 |
| 11 | 11280805 | rs4265593 | 0.683 | -0.443 | -1.174 | -0.358 | -0.024 | 0.059 | 0.059 | 0 | 0.39 | 0.28 | 0.37 | 0.2 | 0.43 | 0.25 |
| 11 | 11280918 | rs4545549 | 0.626 | -0.388 | -1.103 | -0.318 | -0.009 | 0.027 | 0.027 | 0 | 0.41 | 0.36 | 0.37 | 0.21 | 0.44 | 0.25 |
| 11 | 11283088 | rs4243925 | 0.145 | -0.662 | -1.137 | -0.134 | 0.051 | 0.032 | 0.032 | 0 | 0.52 | 0.33 | 0.53 | 0.4 | 0.59 | 0.46 |
| 11 | 11285592 | rs12286046 | 1.426 | -0.296 | -1.223 | 0.056 | -0.01 | 0.018 | 0.018 | 0.002 | 0.13 | 0.01 | 0.09 | 0.05 | 0.1 | 0.05 |
| 11 | 11288590 | rs10831562 | 0.09 | -0.354 | -1.252 | 0.048 | -0.005 | 0.001 | 0.001 | 0 | 0.22 | 0.09 | 0.26 | 0.2 | 0.2 | 0.17 |
| 11 | 11288621 | rs10831563 | -0.125 | -0.539 | -1.449 | -0.223 | -0.003 | 0.057 | 0.057 | 0.043 | 0.34 | 0.23 | 0.18 | 0.2 | 0.3 | 0.3 |
| 11 | 11290071 | rs4910291 | -1.031 | -0.884 | -1.04 | 0.339 | -0.006 | 0.017 | 0.017 | 0.03 | 0.73 | 0.68 | 0.67 | 0.7 | 0.71 | 0.73 |
| 11 | 11290164 | rs10831565 | -0.591 | -0.669 | -1.413 | 0.188 | -0.013 | 0.029 | 0.029 | 0 | 0.44 | 0.33 | 0.52 | 0.37 | 0.44 | 0.42 |
| 11 | 11290399 | rs11602638 | 1.054 | -0.667 | -1.637 | 0.175 | -0.063 | -0.01 | -0.01 | 0 | 0.2 | 0.05 | 0.25 | 0.18 | 0.22 | 0.25 |
| 11 | 11292864 | rs12282401 | 0.472 | -0.67 | -1.491 | 0.24 | -0.014 | -0.028 | -0.028 | 0.865 | 0.23 | 0.07 | 0.23 | 0.24 | 0.22 | 0.16 |
| 11 | 11294109 | rs4910292 | 0.501 | -0.783 | -1.415 | 0.131 | -0.016 | 0.067 | 0.067 | 0.194 | 0.44 | 0.35 | 0.26 | 0.24 | 0.46 | 0.36 |
| 11 | 11294115 | rs4909976 | 0.518 | -0.782 | -1.412 | 0.131 | -0.018 | 0.071 | 0.071 | 0.241 | 0.44 | 0.33 | 0.26 | 0.24 | 0.46 | 0.36 |
| 11 | 11294339 | rs7123377 | -0.547 | -0.775 | -1.337 | 0.339 | -0.001 | 0.002 | 0.002 | 0.018 | 0.68 | 0.43 | 0.65 | 0.5 | 0.67 | 0.52 |
| 11 | 11304251 | rs10831571 | -0.41 | -0.693 | -1.321 | 0.249 | -0.007 | 0.017 | 0.017 | 0 | 0.43 | 0.38 | 0.51 | 0.37 | 0.4 | 0.41 |
| 11 | 11306531 | rs10831572 | 1.331 | -0.561 | -1.609 | 0.272 | -0.046 | 0.03 | 0.03 | 0 | 0.18 | 0.03 | 0.24 | 0.13 | 0.2 | 0.24 |
| 11 | 11306534 | rs10831573 | 0.04 | -0.612 | -1.634 | 0.267 | -0.055 | -0.035 | -0.035 | 0 | 0.28 | 0.14 | 0.43 | 0.35 | 0.32 | 0.35 |
| 11 | 11308084 | rs17348788 | 0.827 | -0.66 | -1.641 | 0.194 | -0.016 | 0.032 | 0.032 | 1.281 | 0.21 | 0.13 | 0.27 | 0.15 | 0.22 | 0.26 |
| 11 | 11310293 | rs10741540 | -0.366 | -1.012 | -2.003 | 0.557 | -0.019 | -0.086 | -0.086 | 1.155 | 0.4 | 0.12 | 0.65 | 0.47 | 0.57 | 0.49 |
| 11 | 11310310 | rs10765829 | -0.409 | -1.063 | -2.076 | 0.524 | -0.034 | -0.091 | -0.091 | 0 | 0.39 | 0.1 | 0.65 | 0.45 | 0.56 | 0.48 |
| 11 | 11310634 | rs4909977 | 2.287 | -1.105 | -2.113 | 0.407 | -0.041 | 0.067 | 0.067 | 0 | 0.22 | 0.06 | 0.11 | 0.07 | 0.24 | 0.09 |
| 11 | 11310926 | rs4910294 | -1.565 | -1.084 | -2.063 | 0.194 | 0.027 | -0.025 | -0.025 | 0 | 0.82 | 0.96 | 0.88 | 0.78 | 0.86 | 0.87 |
| 11 | 11311191 | rs12360863 | 1.628 | -0.714 | -1.867 | -0.059 | -0.061 | -0.034 | -0.034 | 0 | 0.1 | 0.02 | 0.2 | 0.11 | 0.22 | 0.08 |
| 11 | 11311488 | rs12417080 | 2.355 | -0.773 | -1.895 | -0.097 | 0.008 | 0.033 | 0.033 | 0.006 | 0.12 | 0.01 | 0.06 | 0.03 | 0.08 | 0.05 |
| 11 | 11311506 | rs16909341 | 1.854 | -0.322 | -1.618 | 0.195 | -0.042 | -0.051 | -0.051 | 0.096 | 0.08 | 0.01 | 0.18 | 0.21 | 0.12 | 0.11 |
| 11 | 11311743 | rs4910295 | 0.251 | -0.395 | -1.562 | 0.164 | -0.091 | -0.083 | -0.083 | 0 | 0.19 | 0.09 | 0.38 | 0.33 | 0.33 | 0.21 |
| 11 | 11314995 | rs12788849 | 1.334 | -0.808 | -1.279 | -0.163 | -0.026 | 0.046 | 0.046 | 0.239 | 0.24 | 0.07 | 0.14 | 0.13 | 0.26 | 0.11 |
| 11 | 11316049 | rs10831575 | -0.656 | -0.73 | -1.127 | -0.085 | 0.003 | 0.061 | 0.061 | 0 | 0.7 | 0.57 | 0.52 | 0.47 | 0.65 | 0.54 |
| 11 | 11316074 | rs11021762 | 0.31 | -0.817 | -1.443 | -0.206 | 0.021 | 0.043 | 0.043 | 0 | 0.43 | 0.33 | 0.23 | 0.17 | 0.35 | 0.25 |
| 11 | 11316413 | rs4910297 | -0.305 | -0.26 | -1.198 | 0.112 | 0.004 | 0.029 | 0.029 | 0 | 0.46 | 0.3 | 0.51 | 0.36 | 0.41 | 0.45 |
| 11 | 11317263 | rs4559684 | 0.87 | 0.309 | -0.859 | 0.337 | 0.017 | 0.062 | 0.062 | 0 | 0.46 | 0.33 | 0.49 | 0.25 | 0.47 | 0.22 |
| 11 | 11317353 | rs11021763 | 0.586 | 0.268 | -0.119 | 0.952 | 0.008 | 0.272 | 0.272 | 0 | 0.89 | 0.63 | 0.73 | 0.62 | 0.89 | 0.71 |
| 11 | 11317490 | rs4334005 | -0.607 | 0.726 | -0.325 | 0.572 | 0.023 | 0.012 | 0.012 | 0 | 0.33 | 0.54 | 0.37 | 0.39 | 0.36 | 0.39 |
| 11 | 11317948 | rs7114163 | -0.794 | 0.916 | -0.594 | 0.545 | -0.002 | -0.002 | -0.002 | 0 | 0.33 | 0.33 | 0.37 | 0.38 | 0.37 | 0.39 |
| 11 | 11318289 | rs11021765 | 2.475 | 0.629 | -1.082 | 0.313 | -0.034 | 0.027 | 0.027 | 0.005 | 0.33 | 0.04 | 0.26 | 0.18 | 0.32 | 0.17 |
| 11 | 11318463 | rs11601477 | 0.33 | 0.789 | -1.019 | 0.315 | -0.091 | -0.097 | -0.097 | 0.484 | 0.22 | 0.08 | 0.3 | 0.38 | 0.28 | 0.35 |
| 11 | 11318593 | rs4375435 | -0.493 | 1.132 | -0.897 | 0.331 | -0.006 | -0.008 | -0.008 | 0 | 0.31 | 0.29 | 0.34 | 0.38 | 0.29 | 0.38 |
| 11 | 11318964 | rs4075678 | 2.093 | 0.618 | -0.662 | 0.198 | 0.023 | 0.052 | 0.052 | 0 | 0.35 | 0.22 | 0.28 | 0.18 | 0.32 | 0.16 |
| 11 | 11319245 | rs4909979 | 0.174 | 0.727 | -0.654 | 0.276 | 0.001 | -0.007 | -0.007 | 0.207 | 0.13 | 0.16 | 0.24 | 0.04 | 0.17 | 0.07 |
| 11 | 11319396 | rs4909980 | 0.727 | 0.787 | -0.539 | 0.377 | -0.008 | 0.008 | 0.008 | 0.164 | 0.59 | 0.64 | 0.56 | 0.27 | 0.56 | 0.27 |

| | | | | | | | | | | | | | | | | |
|---|---|---|---|---|---|---|---|---|---|---|---|---|---|---|---|---|
| 11 | 11319947 | rs4384379 | -0.404 | 1.247 | -0.845 | 0.742 | -0.012 | -0.014 | -0.014 | 2.63 | 0.15 | 0.12 | 0.42 | 0.3 | 0.2 | 0.13 |
| 11 | 11321819 | rs4375432 | 0.194 | 0.561 | -0.632 | 0.477 | -0.007 | 0.01 | 0.01 | 0.217 | 0.63 | 0.51 | 0.78 | 0.51 | 0.68 | 0.37 |
| 11 | 11322309 | rs4910304 | 1.2 | 0.564 | -0.666 | 0.467 | 0.002 | 0.002 | 0.002 | 0.194 | 0.5 | 0.49 | 0.56 | 0.45 | 0.51 | 0.29 |
| 11 | 11325713 | rs4910306 | 1.07 | 0.805 | -0.479 | 0.706 | 0 | 0 | 0 | 0 | 0.46 | 0.51 | 0.56 | 0.47 | 0.49 | 0.29 |
| 11 | 11326544 | rs7126515 | 0.466 | 0.693 | -0.345 | 0.75 | -0.004 | 0.001 | 0.001 | 0 | 0.58 | 0.47 | 0.79 | 0.51 | 0.66 | 0.35 |
| 11 | 11330390 | rs4909984 | 0.754 | 0.956 | 0.352 | 1.015 | -0.011 | -0.019 | -0.019 | 0.054 | 0.75 | 0.6 | 0.88 | 0.82 | 0.77 | 0.85 |
| 11 | 11331723 | rs12275946 | 1.458 | 1.581 | 0.113 | 1.328 | -0.01 | -0.01 | -0.01 | 0 | 0.62 | 0.66 | 0.52 | 0.51 | 0.55 | 0.48 |
| 11 | 11333188 | rs4909985 | 0.889 | 1.768 | 0.082 | 1.365 | 0.011 | 0.033 | 0.033 | 0 | 0.7 | 0.62 | 0.59 | 0.52 | 0.67 | 0.55 |
| 11 | 11335515 | rs11021791 | -0.262 | 1.767 | -0.395 | 1.315 | -0.042 | 0.024 | 0.024 | 0.53 | 0.08 | 0 | 0.08 | 0.02 | 0.13 | 0.07 |
| 11 | 11336127 | rs11021795 | -1.084 | 1.753 | -0.49 | 1.823 | -0.052 | -0.076 | -0.076 | 0 | 0.12 | 0.04 | 0.44 | 0.47 | 0.18 | 0.17 |
| 11 | 11337115 | rs16909413 | -0.42 | 2.015 | -0.434 | 1.788 | -0.032 | -0.013 | -0.013 | 0.01 | 0.07 | 0.04 | 0.4 | 0.31 | 0.15 | 0.12 |
| 11 | 11337543 | rs12419084 | -1.314 | 1.839 | -0.282 | 1.185 | -0.005 | 0.025 | 0.025 | 0 | 0.27 | 0.21 | 0.16 | 0.03 | 0.27 | 0.24 |
| 11 | 11337655 | rs4910317 | 0.966 | 1.593 | 0.077 | 1.386 | 0.017 | 0.105 | 0.105 | 2.314 | 0.89 | 0.4 | 0.92 | 0.8 | 0.91 | 0.76 |
| 11 | 11337945 | rs6484804 | -2.134 | 1.617 | 0.726 | 1.256 | 0.013 | -0.003 | -0.003 | 0 | 0.33 | 0.6 | 0.57 | 0.3 | 0.32 | 0.31 |
| 11 | 11338178 | rs2403512 | -0.312 | 1.75 | 0.444 | 1.203 | -0.028 | -0.02 | -0.02 | 0 | 0.07 | 0.04 | 0.4 | 0.24 | 0.15 | 0.13 |
| 11 | 11338190 | rs17435455 | -0.063 | 1.673 | 0.693 | 1.051 | 0.006 | -0.032 | -0.032 | 0 | 0.09 | 0.11 | 0.05 | 0 | 0.11 | 0.08 |
| 11 | 11338267 | rs2403514 | 1.828 | 1.539 | 0.56 | 1.14 | -0.005 | -0.029 | -0.029 | 0.333 | 0.65 | 0.26 | 0.78 | 0.6 | 0.66 | 0.59 |
| 11 | 11340599 | rs4442542 | 0.152 | 1.52 | 0.596 | 1.123 | 0.062 | 0.165 | 0.165 | 0.144 | 0.94 | 0.8 | 0.96 | 0.8 | 0.91 | 0.78 |
| 11 | 11341186 | rs1471895 | -0.76 | 1.602 | 0.738 | 1.065 | -0.016 | -0.024 | -0.024 | 0 | 0.11 | 0.17 | 0.05 | 0.03 | 0.06 | 0.1 |
| 11 | 11341813 | rs10741544 | 1.692 | 1.536 | 0.587 | 1.139 | -0.004 | 0.064 | 0.064 | 0 | 0.75 | 0.61 | 0.83 | 0.57 | 0.77 | 0.62 |
| 11 | 11343477 | rs16909447 | -0.481 | 1.534 | -0.34 | 1.236 | -0.047 | -0.045 | -0.045 | 0.862 | 0.08 | 0.03 | 0.4 | 0.26 | 0.14 | 0.13 |
| 11 | 11345579 | rs4909987 | 1.856 | 1 | -0.302 | 0.676 | -0.001 | 0.006 | 0.006 | 0.46 | 0.53 | 0.26 | 0.39 | 0.51 | 0.53 | 0.42 |
| 11 | 11347323 | rs11021804 | 2.137 | 0.989 | -0.359 | 0.405 | 0.017 | 0.075 | 0.075 | 0 | 0.43 | 0.21 | 0.31 | 0.29 | 0.44 | 0.35 |
| 11 | 11348669 | rs17435705 | -0.014 | 1.093 | -0.332 | 0.568 | -0.031 | 0.067 | 0.067 | 0 | 0.11 | 0 | 0.04 | 0 | 0.1 | 0.09 |
| 11 | 11350331 | rs10831587 | 2.128 | 1.121 | -0.629 | 0.518 | -0.036 | 0.072 | 0.072 | 0.229 | 0.43 | 0.1 | 0.28 | 0.29 | 0.44 | 0.33 |
| 11 | 11353300 | rs10500745 | -1.594 | 1.37 | -0.042 | 0.884 | -0.013 | -0.023 | -0.023 | 0.397 | 0.28 | 0.23 | 0.52 | 0.48 | 0.35 | 0.47 |
| 11 | 11354346 | rs901553 | -1.953 | 1.149 | 0.07 | 0.898 | 0 | 0 | 0 | 2.571 | 0.37 | 0.34 | 0.45 | 0.58 | 0.36 | 0.49 |
| 11 | 11354393 | rs901554 | 1.081 | 1.124 | 0.253 | 1.062 | -0.011 | -0.011 | -0.011 | 0.502 | 0.87 | 0.78 | 0.92 | 0.9 | 0.88 | 0.92 |
| 11 | 11355344 | rs901555 | 1.06 | 1.125 | 0.254 | 1.061 | -0.011 | -0.009 | -0.009 | 0.86 | 0.87 | 0.78 | 0.92 | 0.9 | 0.88 | 0.92 |
| 11 | 11356753 | rs10430875 | -1.831 | 1.304 | 0.14 | 1.473 | -0.018 | 0.103 | 0.103 | 0.012 | 0.23 | 0.5 | 0.16 | 0.37 | 0.21 | 0.32 |
| 11 | 11359886 | rs4909989 | -0.869 | 1.764 | -0.632 | 1.85 | 0.01 | 0.026 | 0.026 | 0 | 0.1 | 0.11 | 0.43 | 0.26 | 0.15 | 0.17 |
| 11 | 11360588 | rs12225810 | -0.219 | 1.203 | -0.646 | 1.438 | -0.008 | 0.041 | 0.041 | 2.932 | 0.05 | 0.07 | 0.04 | 0.16 | 0.04 | 0.04 |
| 11 | 11360796 | rs11021810 | -1.05 | 1.489 | -0.917 | 0.906 | -0.016 | -0.036 | -0.036 | 0 | 0.17 | 0.09 | 0.08 | 0.24 | 0.2 | 0.3 |
| 11 | 11363248 | rs9888256 | 0.699 | 1.844 | -0.718 | -0.18 | -0.009 | -0.006 | -0.006 | 0 | 0.66 | 0.69 | 0.79 | 0.91 | 0.59 | 0.82 |
| 11 | 11363305 | rs9888138 | 1.085 | 2.294 | -0.365 | -0.496 | 0.07 | -0.095 | -0.095 | 0.749 | 0.7 | 0.51 | 0.73 | 0.89 | 0.56 | 0.8 |
| 11 | 11363558 | rs4909991 | -1.254 | 1.862 | -1.094 | -0.201 | -0.06 | -0.023 | -0.023 | 0 | 0.16 | 0.01 | 0.12 | 0.14 | 0.21 | 0.15 |
| 11 | 11364026 | rs7120774 | -1.157 | 1.721 | -0.229 | 0.449 | 0.017 | 0.079 | 0.079 | 0 | 0.22 | 0.44 | 0.34 | 0.34 | 0.25 | 0.2 |
| 11 | 11367614 | rs10831594 | 1.099 | 0.781 | 0.356 | -0.147 | -0.031 | -0.038 | -0.038 | 0 | 0.92 | 0.33 | 0.94 | 1 | 0.99 | 0.99 |
| 11 | 11368527 | rs4910325 | 0.631 | 0.087 | 0.522 | -0.409 | -0.026 | -0.022 | -0.022 | 0 | 0.83 | 0.74 | 0.91 | 0.91 | 0.94 | 0.87 |
| 11 | 11369194 | rs7109161 | -0.136 | 0.387 | -0.217 | -1.037 | -0.006 | 0.024 | 0.024 | 0 | 0.36 | 0.25 | 0.57 | 0.85 | 0.4 | 0.62 |
| 11 | 11373960 | rs2071461 | -0.128 | 0.36 | 0.038 | -0.191 | -0.012 | 0.004 | 0.004 | 0 | 0.68 | 0.41 | 0.63 | 0.66 | 0.73 | 0.77 |
| 11 | 11374269 | rs2071460 | 0.212 | 0.86 | -0.765 | -0.142 | -0.033 | -0.032 | -0.032 | 0 | 0.37 | 0.19 | 0.46 | 0.52 | 0.48 | 0.54 |
| 11 | 11378655 | rs6484839 | 0.065 | 0.916 | -0.865 | -1.221 | 0.001 | -0.091 | -0.091 | 0.123 | 0.36 | 0.54 | 0.27 | 0.02 | 0.41 | 0.13 |
| 11 | 11379260 | rs2035631 | -0.876 | 1.806 | -0.789 | 0.005 | 0.092 | 0.196 | 0.196 | 0 | 0.91 | 0.78 | 0.78 | 0.57 | 0.83 | 0.71 |
| 11 | 11386340 | rs12787100 | 0.724 | 1.293 | -1.291 | -0.707 | -0.043 | 0.169 | 0.169 | 0 | 0.29 | 0.1 | 0.21 | 0.03 | 0.31 | 0.17 |
| 11 | 11387713 | rs12794600 | 0.657 | 1.275 | -1.259 | -0.721 | -0.035 | 0.161 | 0.161 | 0 | 0.29 | 0.11 | 0.22 | 0.03 | 0.31 | 0.17 |
| 11 | 11387855 | rs17436486 | 0.623 | 1.259 | -1.059 | -0.728 | -0.013 | 0.103 | 0.103 | 0 | 0.29 | 0.16 | 0.22 | 0.03 | 0.31 | 0.17 |
| 11 | 11388125 | rs16909529 | 0.58 | 1.228 | -0.976 | -0.743 | -0.003 | 0.046 | 0.046 | 0 | 0.3 | 0.2 | 0.22 | 0.03 | 0.31 | 0.17 |
| 11 | 11390172 | rs1487131 | -0.241 | 0.95 | -1.502 | -0.619 | -0.001 | 0 | 0 | 0.311 | 0.67 | 0.66 | 0.62 | 0.44 | 0.65 | 0.45 |
| 11 | 11390585 | rs3993153 | -0.216 | -0.126 | -1.665 | -0.739 | -0.012 | -0.054 | -0.054 | 0 | 0.23 | 0.62 | 0.13 | 0.06 | 0.21 | 0.13 |
| 11 | 11391079 | rs12286476 | 0.236 | -0.177 | -1.454 | -0.532 | -0.022 | -0.051 | -0.051 | 0 | 0.17 | 0.59 | 0.1 | 0.07 | 0.16 | 0.08 |
| 11 | 11392321 | rs10765842 | 0.775 | -0.047 | -1.382 | -0.627 | -0.002 | -0.092 | -0.092 | 0 | 0.14 | 0.32 | 0.07 | 0 | 0.15 | 0.04 |
| 11 | 11392472 | rs12288572 | 0.531 | 0.103 | -2.25 | -0.134 | 0.014 | 0.061 | 0.061 | 0 | 0.18 | 0.01 | 0.12 | 0.07 | 0.12 | 0.17 |
| 11 | 11393268 | rs11021823 | 0.181 | 0.56 | -1.76 | 0.416 | -0.016 | -0.144 | -0.144 | 0.046 | 0.22 | 0.02 | 0.32 | 0.41 | 0.2 | 0.24 |
| 11 | 11395489 | rs10047474 | -0.028 | 0.617 | -1.546 | -0.314 | -0.005 | -0.004 | -0.004 | 0 | 0.62 | 0.63 | 0.39 | 0.49 | 0.52 | 0.6 |
| 11 | 11395762 | rs10831606 | 0.161 | 0.954 | -1.638 | -0.529 | 0.015 | 0.029 | 0.029 | 0 | 0.75 | 0.67 | 0.52 | 0.46 | 0.69 | 0.56 |
| 11 | 11397975 | rs12575708 | 0.503 | 1.218 | -1.618 | -0.901 | -0.011 | 0.009 | 0.009 | 0 | 0.1 | 0.09 | 0.11 | 0.04 | 0.18 | 0.06 |
| 11 | 11398285 | rs2129530 | 0.258 | 0.137 | -1.589 | -1.393 | 0.003 | 0.002 | 0.002 | 0 | 0.54 | 0.49 | 0.47 | 0.4 | 0.7 | 0.36 |
| 11 | 11398347 | rs2129531 | -0.467 | -0.08 | -1.747 | -1.396 | 0.005 | -0.013 | -0.013 | 0 | 0.69 | 0.69 | 0.55 | 0.51 | 0.76 | 0.47 |
| 11 | 11398725 | rs12290003 | 1.016 | 0.339 | -1.447 | -1.301 | 0.01 | 0 | 0 | 0 | 0.11 | 0.01 | 0.05 | 0.11 | 0.05 | 0.08 |
| 11 | 11399406 | rs7118622 | -0.555 | 0.541 | -0.925 | -1.629 | 0.073 | -0.12 | -0.12 | 0 | 0.2 | 0.47 | 0.18 | 0.04 | 0.29 | 0.11 |
| 11 | 11400518 | rs4910328 | 0.143 | -0.148 | -1.306 | -1.757 | -0.016 | 0.163 | 0.163 | 0.724 | 0.32 | 0.09 | 0.23 | 0.06 | 0.33 | 0.17 |
| 11 | 11401119 | rs10831607 | -0.91 | -0.304 | -1.515 | -1.651 | 0.001 | 0 | 0 | 0 | 0.66 | 0.63 | 0.46 | 0.36 | 0.63 | 0.44 |
| 11 | 11401510 | rs11021825 | 0.887 | -1.006 | -1.999 | -1.451 | 0.06 | -0.029 | -0.029 | 0 | 0.28 | 0.13 | 0.15 | 0.29 | 0.19 | 0.24 |
| 11 | 11403717 | rs10765844 | -0.295 | -0.193 | -1.47 | -0.88 | 0.007 | -0.003 | -0.003 | 0 | 0.62 | 0.44 | 0.39 | 0.36 | 0.52 | 0.41 |
| 11 | 11404814 | rs10831608 | -0.309 | 1.054 | -1.51 | -0.705 | -0.078 | 0.056 | 0.056 | 0 | 0.23 | 0.05 | 0.22 | 0.11 | 0.35 | 0.19 |
| 11 | 11407649 | rs1466610 | -0.026 | 1.103 | -0.547 | -0.539 | 0.034 | -0.087 | -0.087 | 0 | 0.83 | 0.61 | 0.89 | 0.96 | 0.83 | 0.82 |
| 11 | 11409067 | rs11021826 | -0.283 | 1.184 | -0.688 | 0.309 | -0.004 | 0.019 | 0.019 | 0 | 0.12 | 0.13 | 0.26 | 0.52 | 0.16 | 0.27 |
| 11 | 11409938 | rs4910330 | 1.114 | 0.776 | -1.003 | -0.293 | 0.071 | 0.011 | 0.011 | 0.467 | 0.39 | 0.19 | 0.32 | 0.37 | 0.22 | 0.3 |

| | | | | | | | | | | | | | | | | |
|---|---|---|---|---|---|---|---|---|---|---|---|---|---|---|---|---|
| 11 | 11410588 | rs7931994 | 0.95 | 0.438 | -1.205 | -0.762 | 0.018 | 0.016 | 0.016 | 1.127 | 0.45 | 0.36 | 0.25 | 0.28 | 0.24 | 0.36 |
| 11 | 11410725 | rs7945106 | 1.275 | 0.556 | -1.183 | -0.797 | -0.008 | -0.027 | -0.027 | 0.004 | 0.16 | 0.16 | 0.09 | 0.05 | 0.13 | 0.17 |
| 11 | 11412180 | rs4910331 | -0.515 | 0.757 | -0.868 | -0.783 | 0.006 | -0.028 | -0.028 | 0 | 0.69 | 0.67 | 0.7 | 0.9 | 0.5 | 0.73 |
| 11 | 11414338 | rs4910333 | 1.554 | 0.241 | -0.919 | -0.496 | -0.035 | -0.036 | -0.036 | 1.1 | 0.11 | 0.01 | 0.17 | 0.14 | 0.1 | 0.12 |
| 11 | 11416073 | rs12418191 | 0.862 | 0.074 | -0.879 | -0.154 | 0.056 | 0.012 | 0.012 | 0.037 | 0.37 | 0.2 | 0.25 | 0.33 | 0.24 | 0.28 |
| 11 | 11417009 | rs1872794 | 0.721 | -0.003 | -0.835 | -0.209 | 0.099 | -0.001 | -0.001 | 0 | 0.41 | 0.15 | 0.33 | 0.4 | 0.24 | 0.35 |
| 11 | 11417894 | rs4910336 | -0.118 | 0.225 | -0.658 | -0.245 | -0.002 | -0.016 | -0.016 | 0 | 0.42 | 0.39 | 0.39 | 0.15 | 0.64 | 0.28 |
| 11 | 11419394 | rs12574825 | 1.657 | 0.114 | -0.808 | -0.164 | 0.092 | 0.002 | 0.002 | 0.432 | 0.29 | 0.1 | 0.15 | 0.27 | 0.13 | 0.23 |
| 11 | 11420246 | rs17437465 | 0.84 | 0.56 | -0.276 | -0.171 | 0.005 | -0.025 | -0.025 | 0 | 0.08 | 0.12 | 0.11 | 0 | 0.12 | 0.02 |
| 11 | 11421090 | rs7929019 | 0.724 | 0.515 | -0.115 | -0.198 | 0.003 | 0.005 | 0.005 | 0 | 0.63 | 0.42 | 0.44 | 0.45 | 0.64 | 0.49 |
| 11 | 11422607 | rs10765846 | -0.942 | 0.506 | 0.005 | -0.101 | 0.001 | 0.162 | 0.162 | 0 | 0.85 | 0.69 | 0.75 | 0.55 | 0.88 | 0.62 |
| 11 | 11422750 | rs12421938 | 0.026 | 0.563 | -0.069 | -0.327 | 0.008 | -0.011 | -0.011 | 2.637 | 0.19 | 0.23 | 0.28 | 0.12 | 0.21 | 0.14 |
| 11 | 11424906 | rs723459 | 0.13 | 0.46 | -0.205 | -0.614 | 0 | 0.013 | 0.013 | 0 | 0.57 | 0.41 | 0.61 | 0.84 | 0.4 | 0.76 |
| 11 | 11425052 | rs901557 | 0.418 | 0.465 | -0.238 | -0.618 | -0.023 | 0.03 | 0.03 | 0 | 0.33 | 0.2 | 0.27 | 0.15 | 0.49 | 0.22 |
| 11 | 11426198 | rs4909996 | 0.602 | 0.255 | -0.074 | -0.228 | 0.018 | 0.129 | 0.129 | 0 | 0.77 | 0.48 | 0.63 | 0.54 | 0.77 | 0.56 |
| 11 | 11426939 | rs11021831 | 0.187 | 0.26 | 0.004 | -0.009 | 0.032 | 0.24 | 0.24 | 0 | 0.09 | 0.26 | 0.26 | 0.43 | 0.11 | 0.3 |
| 11 | 11427040 | rs7101677 | 0.061 | 0.117 | -0.111 | -0.487 | -0.027 | 0.123 | 0.123 | 0 | 0.43 | 0.22 | 0.32 | 0.15 | 0.56 | 0.33 |
| 11 | 11427577 | rs7935882 | -0.531 | 0.223 | 0.051 | -0.455 | 0.017 | -0.006 | -0.006 | 0 | 0.82 | 0.69 | 0.73 | 0.85 | 0.82 | 0.9 |
| 11 | 11427928 | rs7928557 | 0.071 | 0.259 | -0.004 | -0.441 | -0.03 | 0.097 | 0.097 | 0 | 0.44 | 0.27 | 0.29 | 0.17 | 0.53 | 0.35 |
| 11 | 11428085 | rs2171344 | -0.75 | 0.587 | -0.155 | -0.586 | 0.033 | 0.057 | 0.057 | 1.78 | 0.51 | 0.64 | 0.67 | 0.71 | 0.54 | 0.68 |
| 11 | 11429150 | rs11021836 | 0.323 | -0.049 | 0.163 | -0.174 | -0.026 | -0.008 | -0.008 | 0.161 | 0.38 | 0.45 | 0.43 | 0.69 | 0.27 | 0.54 |
| 11 | 11429355 | rs2403517 | -0.097 | -0.017 | 0.158 | -0.118 | 0.065 | 0.299 | 0.299 | 0 | 0.11 | 0.33 | 0.28 | 0.45 | 0.13 | 0.3 |
| 11 | 11429376 | rs7128890 | -0.022 | 0.01 | 0.031 | -0.201 | 0.018 | 0.037 | 0.037 | 0 | 0.53 | 0.36 | 0.47 | 0.31 | 0.61 | 0.43 |
| 11 | 11429532 | rs2009422 | 1.06 | -0.455 | 0.168 | 0.256 | -0.001 | 0.002 | 0.002 | 0 | 0.1 | 0.13 | 0.18 | 0.14 | 0.08 | 0.08 |
| 11 | 11432042 | rs7948809 | -0.218 | -0.528 | 0.337 | 0.236 | 0.001 | 0 | 0 | 0 | 0.43 | 0.42 | 0.41 | 0.4 | 0.33 | 0.4 |
| 11 | 11434915 | rs11601517 | 0.809 | -0.342 | 0.161 | 0.234 | 0.065 | -0.013 | -0.013 | 0 | 0.05 | 0.2 | 0.07 | 0 | 0.09 | 0.04 |
| 11 | 11438041 | rs1565029 | -0.373 | -0.563 | 0.145 | 0.226 | 0.11 | 0.041 | 0.041 | 0 | 0.46 | 0.79 | 0.47 | 0.65 | 0.28 | 0.56 |
| 11 | 11441775 | rs11021845 | 0.507 | -0.434 | 0.093 | 0.407 | -0.025 | 0.126 | 0.126 | 0 | 0.11 | 0.19 | 0.23 | 0.41 | 0.05 | 0.28 |
| 11 | 11444621 | rs12293713 | 1.001 | -0.145 | -0.454 | 0.139 | -0.039 | -0.038 | -0.038 | 0 | 0.08 | 0.01 | 0.05 | 0.13 | 0.13 | 0.1 |
| 11 | 11445303 | rs6484897 | -1.068 | -1.203 | -0.177 | 0.206 | 0.03 | 0.113 | 0.113 | 0 | 0.56 | 0.82 | 0.64 | 0.81 | 0.36 | 0.65 |
| 11 | 11445760 | rs12578026 | 0.585 | -1.284 | -0.132 | 0.248 | -0.035 | -0.01 | -0.01 | 0.042 | 0.3 | 0.33 | 0.17 | 0.27 | 0.15 | 0.25 |
| 11 | 11447174 | rs7111888 | -1.185 | -0.851 | -0.174 | 0.515 | -0.009 | 0.105 | 0.105 | 0 | 0.27 | 0.5 | 0.47 | 0.53 | 0.21 | 0.39 |
| 11 | 11448210 | rs2403520 | -0.071 | -1.049 | -0.754 | 0.955 | -0.003 | 0.027 | 0.027 | 2.266 | 0.12 | 0.15 | 0.19 | 0.34 | 0.06 | 0.28 |
| 11 | 11450673 | rs4910344 | -0.603 | -0.862 | -0.935 | 0.603 | 0.022 | -0.069 | -0.069 | 0 | 0.25 | 0.16 | 0.4 | 0.53 | 0.18 | 0.39 |
| 11 | 11452313 | rs1023982 | -0.107 | -0.517 | -0.675 | 0.328 | 0.015 | 0.009 | 0.009 | 0.039 | 0.28 | 0.21 | 0.16 | 0.24 | 0.16 | 0.24 |
| 11 | 11455876 | rs12363821 | -0.088 | -0.094 | -0.695 | 0.03 | 0.001 | -0.011 | -0.011 | 0 | 0.12 | 0.1 | 0.09 | 0.01 | 0.09 | 0.19 |
| 11 | 11456027 | rs7108734 | 0.876 | 0.03 | -0.433 | 0.23 | -0.004 | -0.008 | -0.008 | 0 | 0.75 | 0.6 | 0.5 | 0.79 | 0.77 | 0.7 |
| 11 | 11456749 | rs7946706 | -0.393 | -0.213 | -0.19 | 0.434 | -0.012 | 0.055 | 0.055 | 0 | 0.21 | 0.27 | 0.13 | 0.54 | 0.15 | 0.31 |
| 11 | 11459744 | rs11021853 | 0.377 | 0.781 | -0.319 | 0.192 | -0.017 | 0.02 | 0.02 | 0 | 0.12 | 0.07 | 0.11 | 0.01 | 0.21 | 0.18 |
| 11 | 11459990 | rs11021854 | 0.424 | 0.768 | -0.318 | 0.193 | -0.016 | 0.02 | 0.02 | 0 | 0.12 | 0.07 | 0.12 | 0.01 | 0.21 | 0.18 |
| 11 | 11461222 | rs4910001 | 0.668 | 0.579 | -0.248 | 0.102 | -0.001 | 0.018 | 0.018 | 0 | 0.15 | 0.08 | 0.25 | 0.1 | 0.13 | 0.08 |
| 11 | 11462298 | rs4910003 | -0.804 | 0.332 | -0.327 | 0.182 | -0.002 | -0.001 | -0.001 | 0 | 0.37 | 0.33 | 0.39 | 0.64 | 0.36 | 0.56 |
| 11 | 11466613 | rs12278731 | 0.721 | 0.371 | -0.82 | 0.186 | 0.001 | 0.009 | 0.009 | 0 | 0.58 | 0.18 | 0.52 | 0.38 | 0.58 | 0.39 |
| 11 | 11469752 | rs9988868 | 1.092 | 0.354 | -0.694 | 0.237 | -0.004 | 0.034 | 0.034 | 0.007 | 0.43 | 0.18 | 0.29 | 0.35 | 0.44 | 0.38 |
| 11 | 11472193 | rs11021857 | -0.057 | -0.221 | -0.395 | 0.89 | -0.004 | 0.054 | 0.054 | 0 | 0.12 | 0.16 | 0.04 | 0.44 | 0.07 | 0.26 |
| 11 | 11472617 | rs1487122 | -0.626 | -0.348 | -0.101 | 0.254 | -0.027 | -0.039 | -0.039 | 0 | 0.85 | 0.73 | 0.89 | 0.95 | 0.94 | 0.93 |
| 11 | 11473089 | rs11822766 | -0.346 | -0.053 | 0.188 | 0.009 | 0.024 | -0.084 | -0.084 | 0 | 0.21 | 0.39 | 0.18 | 0.09 | 0.24 | 0.19 |
| 11 | 11473108 | rs10831614 | 0.123 | -0.497 | 0.132 | 0.559 | -0.012 | 0.005 | 0.005 | 0 | 0.72 | 0.7 | 0.87 | 0.62 | 0.84 | 0.56 |
| 11 | 11473228 | rs12417203 | -0.169 | 0.039 | 0.083 | 0.07 | 0.005 | -0.049 | -0.049 | 0 | 0.22 | 0.24 | 0.13 | 0.07 | 0.24 | 0.17 |
| 11 | 11474462 | rs10831616 | 0.322 | -0.209 | -0.456 | 0.444 | 0 | -0.131 | -0.131 | 0.136 | 0.21 | 0.02 | 0.14 | 0.48 | 0.17 | 0.27 |
| 11 | 11475480 | rs1487121 | 1.32 | 0.508 | -0.032 | -0.358 | -0.003 | 0.025 | 0.025 | 0 | 0.6 | 0.35 | 0.6 | 0.83 | 0.62 | 0.59 |
| 11 | 11475827 | rs4910346 | -0.313 | 0.72 | -0.431 | -0.157 | 0.011 | -0.064 | -0.064 | 6.488 | 0.19 | 0.21 | 0.1 | 0.01 | 0.2 | 0.08 |
| 11 | 11478010 | rs4910006 | -0.775 | 0.506 | -0.654 | 0.349 | -0.003 | -0.001 | -0.001 | 0 | 0.07 | 0.13 | 0.04 | 0.09 | 0.07 | 0.16 |
| 11 | 11479866 | rs4910008 | -0.11 | 0.381 | -1.243 | 0.194 | 0.01 | -0.015 | -0.015 | 0 | 0.43 | 0.29 | 0.61 | 0.51 | 0.54 | 0.44 |
| 11 | 11482863 | rs2054840 | 0.347 | 0.344 | -1.416 | -0.331 | 0.016 | 0.068 | 0.068 | 0 | 0.27 | 0.12 | 0.18 | 0.07 | 0.23 | 0.16 |
| 11 | 11486540 | rs6484928 | 0.297 | 0.449 | -0.905 | 0.17 | -0.019 | 0.023 | 0.023 | 0 | 0.72 | 0.57 | 0.61 | 0.35 | 0.77 | 0.57 |
| 11 | 11487025 | rs12786876 | 0.322 | 1.063 | -1.433 | -0.32 | -0.045 | 0.048 | 0.048 | 0.65 | 0.15 | 0.07 | 0.26 | 0.01 | 0.27 | 0.11 |
| 11 | 11489347 | rs11021862 | 0.024 | 0.447 | -1.359 | 0.274 | 0.005 | -0.05 | -0.05 | 0.499 | 0.09 | 0 | 0.05 | 0.16 | 0.06 | 0.18 |
| 11 | 11490764 | rs11021865 | 0.342 | 0.525 | -1.164 | 0.41 | -0.001 | -0.003 | -0.003 | 0 | 0.11 | 0.07 | 0.1 | 0.18 | 0.11 | 0.12 |
| 11 | 11497253 | rs3741409 | 0.604 | 0.457 | -0.428 | -0.285 | 0.004 | -0.035 | -0.035 | 0 | 0.71 | 0.44 | 0.79 | 0.77 | 0.72 | 0.75 |
| 11 | 11498410 | rs4910353 | 1.061 | -0.306 | -0.724 | -0.703 | -0.011 | -0.011 | -0.011 | 0 | 0.15 | 0.2 | 0.07 | 0.05 | 0.1 | 0.12 |
| 11 | 11498983 | rs11606228 | 1.952 | 0.002 | -1.075 | -0.3 | -0.023 | -0.045 | -0.045 | 0 | 0.14 | 0.05 | 0.12 | 0.36 | 0.16 | 0.24 |
| 11 | 11499145 | rs6416093 | -1.001 | -0.16 | -0.595 | -0.467 | 0.084 | -0.043 | -0.043 | 0.603 | 0.89 | 0.61 | 0.9 | 0.98 | 0.85 | 0.95 |
| 11 | 11499853 | rs4910355 | 0.626 | -0.205 | -0.776 | -0.467 | 0.025 | 0.07 | 0.07 | 0.011 | 0.55 | 0.25 | 0.71 | 0.72 | 0.62 | 0.63 |
| 11 | 11500907 | rs11021873 | 1.44 | -0.701 | -1.032 | -0.767 | -0.046 | -0.037 | -0.037 | 0 | 0.25 | 0.02 | 0.19 | 0.22 | 0.26 | 0.28 |
| 11 | 11501118 | rs6484942 | -0.621 | -0.743 | -0.694 | -0.808 | -0.03 | -0.012 | -0.012 | 0 | 0.79 | 0.56 | 0.84 | 0.8 | 0.86 | 0.81 |
| 11 | 11501643 | rs11021874 | 0.055 | -0.555 | -0.788 | -0.715 | -0.004 | 0.002 | 0.002 | 0 | 0.42 | 0.35 | 0.59 | 0.57 | 0.52 | 0.45 |
| 11 | 11504349 | rs7118797 | 0.008 | -0.483 | -0.783 | -0.365 | 0.03 | 0.019 | 0.019 | 0 | 0.45 | 0.18 | 0.3 | 0.42 | 0.4 | 0.48 |
| 11 | 11506346 | rs1565028 | 0.858 | -0.481 | -0.829 | -0.259 | -0.022 | -0.048 | -0.048 | 0.031 | 0.32 | 0.2 | 0.21 | 0.42 | 0.36 | 0.37 |

| | | | | | | | | | | | | | | | | |
|---|---|---|---|---|---|---|---|---|---|---|---|---|---|---|---|---|
| 11 | 11507091 | rs7117475 | 0.246 | -0.642 | -0.832 | -0.305 | -0.007 | 0.008 | 0.008 | 0 | 0.19 | 0.22 | 0.13 | 0.3 | 0.14 | 0.22 |
| 11 | 11507157 | rs4910362 | -0.258 | -0.637 | -0.835 | -0.234 | -0.001 | 0.026 | 0.026 | 2.64 | 0.51 | 0.43 | 0.34 | 0.69 | 0.5 | 0.58 |
| 11 | 11509984 | rs7943717 | 0.798 | -0.351 | -0.596 | 0.049 | -0.01 | -0.019 | -0.019 | 0 | 0.32 | 0.25 | 0.22 | 0.42 | 0.36 | 0.37 |
| 11 | 11512224 | rs7116469 | -0.705 | -0.196 | -0.62 | -0.023 | -0.035 | -0.025 | -0.025 | 0 | 0.8 | 0.48 | 0.57 | 0.87 | 0.84 | 0.88 |
| 11 | 11512305 | rs6484951 | -0.789 | -0.139 | -0.511 | -0.033 | -0.002 | -0.022 | -0.022 | 0 | 0.81 | 0.59 | 0.57 | 0.87 | 0.81 | 0.87 |
| 11 | 11512738 | rs1386425 | -0.743 | -0.122 | -0.581 | -0.223 | 0 | 0 | 0 | 0.041 | 0.84 | 0.83 | 0.81 | 0.87 | 0.81 | 0.76 |
| 11 | 11512807 | rs1386424 | -0.52 | -0.12 | -0.681 | -0.218 | 0.004 | -0.003 | -0.003 | 0 | 0.33 | 0.35 | 0.59 | 0.28 | 0.38 | 0.37 |
| 11 | 11512926 | rs1386423 | -0.372 | -0.196 | -0.745 | -0.268 | 0 | 0.003 | 0.003 | 0 | 0.57 | 0.58 | 0.67 | 0.59 | 0.58 | 0.56 |
| 11 | 11515530 | rs4910364 | 0.257 | -0.116 | -0.645 | -0.738 | 0.005 | -0.001 | -0.001 | 0 | 0.27 | 0.3 | 0.53 | 0.26 | 0.32 | 0.33 |
| 11 | 11517147 | rs12417042 | 0.554 | -0.262 | -0.663 | -0.675 | 0.025 | 0.055 | 0.055 | 0 | 0.33 | 0.17 | 0.23 | 0.16 | 0.26 | 0.25 |
| 11 | 11519691 | rs10831623 | 0.681 | -0.095 | -0.651 | -0.603 | 0.004 | 0.03 | 0.03 | 0 | 0.32 | 0.21 | 0.19 | 0.18 | 0.29 | 0.22 |
| 11 | 11521053 | rs923259 | 0.692 | -0.07 | -0.639 | -0.59 | 0.004 | 0.029 | 0.029 | 0 | 0.32 | 0.21 | 0.19 | 0.18 | 0.28 | 0.22 |
| 11 | 11521285 | rs6484954 | -0.671 | -0.023 | -0.761 | -0.393 | -0.005 | -0.014 | -0.014 | 2.286 | 0.63 | 0.61 | 0.78 | 0.81 | 0.72 | 0.74 |
| 11 | 11525424 | rs10765861 | 0.237 | 0.087 | -0.869 | 0.061 | -0.016 | -0.015 | -0.015 | 0 | 0.34 | 0.29 | 0.41 | 0.59 | 0.42 | 0.4 |
| 11 | 11527167 | rs4910365 | 0.581 | -0.376 | -0.696 | 0.229 | -0.021 | 0.001 | 0.001 | 0.74 | 0.13 | 0.16 | 0.22 | 0.13 | 0.09 | 0.1 |
| 11 | 11529564 | rs11021881 | -0.093 | -0.713 | -0.736 | 0.177 | -0.029 | 0.006 | 0.006 | 0 | 0.24 | 0.26 | 0.3 | 0.28 | 0.13 | 0.25 |
| 11 | 11529828 | rs9943588 | -0.223 | 0.519 | -0.916 | -0.64 | -0.027 | 0.078 | 0.078 | 0 | 0.39 | 0.22 | 0.44 | 0.11 | 0.48 | 0.35 |
| 11 | 11530583 | rs7925530 | 1.153 | 0.303 | -1.44 | -0.58 | -0.009 | 0.066 | 0.066 | 0 | 0.17 | 0.02 | 0.05 | 0.03 | 0.13 | 0.21 |
| 11 | 11532294 | rs3934815 | -1.1 | 0.31 | -1.366 | -0.783 | 0.085 | 0.108 | 0.108 | 0 | 0.53 | 0.28 | 0.51 | 0.22 | 0.68 | 0.61 |
| 11 | 11535410 | rs7934617 | 0.943 | 0.434 | -1.249 | -0.94 | 0.007 | 0.025 | 0.025 | 0 | 0.21 | 0.18 | 0.09 | 0.04 | 0.15 | 0.25 |
| 11 | 11535629 | rs4550214 | 0.611 | 1.118 | -1.126 | -0.841 | -0.003 | 0.02 | 0.02 | 0 | 0.55 | 0.54 | 0.59 | 0.82 | 0.48 | 0.55 |
| 11 | 11535880 | rs7935113 | 1.005 | 0.952 | -1.14 | -0.827 | 0.004 | 0.006 | 0.006 | 0 | 0.21 | 0.22 | 0.09 | 0.04 | 0.15 | 0.25 |
| 11 | 11536487 | rs4077732 | -0.146 | 1.079 | -1.13 | -0.78 | -0.008 | 0.007 | 0.007 | 0 | 0.77 | 0.77 | 0.68 | 0.86 | 0.64 | 0.8 |
| 11 | 11537403 | rs4556537 | 1.157 | 1.154 | -1.208 | -0.639 | -0.058 | -0.036 | -0.036 | 0.036 | 0.11 | 0.01 | 0.12 | 0.11 | 0.17 | 0.18 |
| 11 | 11538434 | rs7946738 | 1.028 | 1.122 | -1.173 | -0.705 | 0.004 | 0.011 | 0.011 | 0 | 0.21 | 0.21 | 0.09 | 0.03 | 0.15 | 0.25 |
| 11 | 11540207 | rs7943075 | 0.234 | 1.114 | -0.746 | -0.746 | -0.001 | -0.005 | -0.005 | 1.632 | 0.59 | 0.53 | 0.72 | 0.73 | 0.6 | 0.48 |
| 11 | 11540560 | rs7122801 | 0.239 | 1.164 | -0.899 | -0.755 | -0.051 | 0.053 | 0.053 | 0 | 0.31 | 0.08 | 0.19 | 0.14 | 0.31 | 0.42 |
| 11 | 11540856 | rs7122263 | 0.076 | 1.028 | -0.96 | -0.821 | -0.046 | 0.063 | 0.063 | 0 | 0.32 | 0.08 | 0.21 | 0.14 | 0.33 | 0.42 |
| 11 | 11540861 | rs4520594 | -0.816 | 1.085 | -0.979 | -0.935 | 0.044 | 0.156 | 0.156 | 0 | 0.46 | 0.13 | 0.42 | 0.15 | 0.59 | 0.2 |
| 11 | 11542409 | rs4264144 | 0.485 | 1.218 | -0.934 | -0.879 | -0.03 | 0.017 | 0.017 | 0 | 0.14 | 0.08 | 0.26 | 0.01 | 0.27 | 0.11 |
| 11 | 11543021 | rs4128873 | 0.338 | 1.102 | -1.169 | -0.934 | -0.086 | 0.03 | 0.03 | 0 | 0.28 | 0.03 | 0.18 | 0.14 | 0.3 | 0.41 |
| 11 | 11543155 | rs4310606 | -1.238 | 1.006 | -0.916 | -0.869 | -0.014 | 0.028 | 0.028 | 0 | 0.91 | 0.72 | 0.93 | 0.88 | 0.91 | 0.91 |
| 11 | 11543971 | rs10831626 | 1.127 | 0.922 | -0.656 | -0.928 | 0.013 | 0.013 | 0.013 | 0 | 0.45 | 0.4 | 0.46 | 0.72 | 0.33 | 0.38 |
| 11 | 11546924 | rs4910370 | 0.362 | 0.983 | -0.765 | -0.971 | 0.053 | -0.071 | -0.071 | 0 | 0.75 | 0.53 | 0.65 | 0.87 | 0.63 | 0.79 |
| 11 | 11549650 | rs7107470 | 1.11 | 0.929 | -0.771 | -0.916 | 0.017 | 0.021 | 0.021 | 0 | 0.46 | 0.57 | 0.47 | 0.77 | 0.33 | 0.38 |
| 11 | 11552072 | rs7120582 | 0.309 | 1.201 | -0.829 | -0.818 | -0.003 | -0.017 | -0.017 | 0.008 | 0.15 | 0.13 | 0.27 | 0.01 | 0.28 | 0.12 |
| 11 | 11553643 | rs12361327 | -0.223 | 1.184 | -1.057 | -0.842 | -0.101 | 0.132 | 0.132 | 0.061 | 0.3 | 0.08 | 0.31 | 0.06 | 0.4 | 0.32 |
| 11 | 11555046 | rs4910379 | -0.345 | 0.97 | -0.698 | -0.685 | -0.005 | 0.022 | 0.022 | 0 | 0.67 | 0.69 | 0.8 | 0.81 | 0.7 | 0.56 |
| 11 | 11558051 | rs10831636 | 0.527 | 0.751 | -0.992 | -0.221 | 0.023 | 0.04 | 0.04 | 0.002 | 0.53 | 0.63 | 0.52 | 0.8 | 0.41 | 0.43 |
| 11 | 11562006 | rs4910382 | 0.988 | 0.477 | -1.915 | 0.146 | -0.038 | 0.005 | 0.005 | 0 | 0.38 | 0.46 | 0.4 | 0.47 | 0.25 | 0.25 |
| 11 | 11566037 | rs4497400 | 0.467 | 1.757 | -2.682 | 0.122 | -0.025 | 0.02 | 0.02 | 0 | 0.12 | 0.08 | 0.28 | 0.01 | 0.31 | 0.11 |
| 11 | 11568136 | rs4910014 | -0.736 | 1.689 | -2.827 | 0.45 | 0 | 0.007 | 0.007 | 0 | 0.83 | 0.76 | 0.86 | 0.65 | 0.86 | 0.77 |
| 11 | 11570895 | rs7943730 | 0.503 | 2.582 | -3.167 | 0.319 | -0.021 | 0.013 | 0.013 | 0 | 0.12 | 0.08 | 0.26 | 0.01 | 0.25 | 0.1 |
| 11 | 11574999 | rs11021924 | 0.552 | 2.7 | -3.352 | 0.272 | -0.072 | -0.037 | -0.037 | 0 | 0.13 | 0.02 | 0.13 | 0.11 | 0.24 | 0.2 |
| 11 | 11575457 | rs10765863 | 1.395 | 1.92 | -3.541 | 0.244 | -0.053 | -0.07 | -0.07 | 0 | 0.11 | 0.01 | 0.1 | 0.31 | 0.17 | 0.1 |
| 11 | 11576549 | rs11021927 | 0.067 | 1.495 | -3.204 | -0.401 | -0.102 | -0.013 | -0.013 | 0 | 0.3 | 0.02 | 0.19 | 0.17 | 0.35 | 0.41 |
| 11 | 11578262 | rs7126296 | -0.549 | 0.751 | -3.064 | -0.268 | -0.008 | 0.108 | 0.108 | 0 | 0.42 | 0.11 | 0.44 | 0.18 | 0.6 | 0.51 |
| 11 | 11589711 | rs4910384 | -0.412 | 0.3 | -2.728 | -0.002 | -0.05 | 0.108 | 0.108 | 0.061 | 0.43 | 0.19 | 0.41 | 0.24 | 0.48 | 0.57 |
| 11 | 11590291 | rs4910018 | 0.698 | 0.645 | -2.702 | -0.132 | -0.004 | -0.008 | -0.008 | 0 | 0.09 | 0.08 | 0.17 | 0 | 0.09 | 0.06 |
| 11 | 11590742 | rs9919639 | 0.757 | 0.334 | -2.702 | 0.209 | -0.093 | -0.001 | -0.001 | 0 | 0.22 | 0.01 | 0.16 | 0.13 | 0.29 | 0.37 |
| 11 | 11590956 | rs7124824 | 0.606 | -0.034 | -2.814 | 0.577 | 0.04 | -0.062 | -0.062 | 0 | 0.14 | 0.03 | 0.07 | 0.26 | 0.04 | 0.14 |
| 11 | 11590971 | rs9919560 | 0.295 | 0.78 | -2.374 | 0.178 | -0.101 | 0.012 | 0.012 | 0 | 0.23 | 0.01 | 0.18 | 0.13 | 0.3 | 0.41 |
| 11 | 11592433 | rs4472929 | 0.103 | 0.521 | -1.265 | 0.453 | -0.008 | -0.005 | -0.005 | 0.994 | 0.86 | 0.79 | 0.87 | 0.89 | 0.87 | 0.92 |
| 11 | 11592794 | rs9919579 | 0.745 | 0.544 | -0.947 | 0.531 | -0.001 | -0.008 | -0.008 | 2.021 | 0.63 | 0.37 | 0.66 | 0.4 | 0.59 | 0.71 |
| 11 | 11592911 | rs9919625 | 0.077 | 0.399 | -0.932 | 0.072 | -0.005 | -0.003 | -0.003 | 1.151 | 0.86 | 0.82 | 0.87 | 0.89 | 0.87 | 0.91 |
| 11 | 11593110 | rs9919632 | 0.065 | 0.403 | -0.931 | 0.074 | -0.005 | -0.003 | -0.003 | 0 | 0.86 | 0.82 | 0.87 | 0.89 | 0.87 | 0.91 |
| 11 | 11593982 | rs7118819 | -0.075 | 0.379 | -0.838 | -0.003 | -0.002 | -0.009 | -0.009 | 0 | 0.68 | 0.63 | 0.63 | 0.71 | 0.73 | 0.81 |
| 11 | 11596334 | rs7935223 | 0.651 | 0.045 | -1.488 | -0.398 | -0.023 | 0.084 | 0.084 | 0 | 0.27 | 0.01 | 0.12 | 0.11 | 0.19 | 0.13 |
| 11 | 11596478 | rs11021941 | 0.088 | -0.528 | -0.665 | -0.629 | -0.042 | -0.047 | -0.047 | 0 | 0.3 | 0.55 | 0.3 | 0.13 | 0.17 | 0.14 |
| 11 | 11598372 | rs7106005 | -0.388 | -0.59 | -0.072 | -0.68 | -0.003 | -0.004 | -0.004 | 0 | 0.68 | 0.39 | 0.67 | 0.87 | 0.79 | 0.85 |
| 11 | 11598835 | rs4910387 | 0.467 | 0.19 | -0.322 | -0.428 | 0 | 0 | 0 | 0.616 | 0.1 | 0.1 | 0.17 | 0.11 | 0.14 | 0.19 |
| 11 | 11599177 | rs10831644 | 0.158 | 0.223 | -0.203 | -0.374 | -0.01 | -0.005 | -0.005 | 0 | 0.31 | 0.33 | 0.39 | 0.63 | 0.46 | 0.52 |
| 11 | 11601314 | rs10831645 | 0.189 | 0.206 | -0.451 | -0.388 | -0.113 | -0.066 | -0.066 | 0.067 | 0.29 | 0.12 | 0.37 | 0.64 | 0.46 | 0.52 |
| 11 | 11601537 | rs7103589 | -0.616 | -0.005 | -0.113 | -0.486 | 0.009 | 0.03 | 0.03 | 0 | 0.54 | 0.63 | 0.66 | 0.74 | 0.61 | 0.61 |
| 11 | 11601761 | rs11607730 | 0.782 | -0.048 | -0.655 | -0.693 | -0.004 | -0.007 | -0.007 | 0 | 0.23 | 0.2 | 0.27 | 0.1 | 0.15 | 0.1 |
| 11 | 11603494 | rs4469886 | 0.397 | 0.466 | -1.069 | -0.046 | -0.118 | -0.039 | -0.039 | 0 | 0.25 | 0.06 | 0.31 | 0.29 | 0.37 | 0.39 |
| 11 | 11613554 | rs7102078 | 0.209 | -0.057 | -1.055 | -0.293 | -0.016 | -0.01 | -0.01 | 0.012 | 0.26 | 0.24 | 0.26 | 0.15 | 0.14 | 0.1 |
| 11 | 11615837 | rs10765871 | 0.267 | 0.636 | -1.712 | -0.154 | -0.082 | 0.108 | 0.108 | 0 | 0.47 | 0.12 | 0.39 | 0.71 | 0.5 | 0.59 |
| 11 | 11626445 | rs7928310 | -0.118 | 0.356 | -0.8 | 0.019 | -0.013 | -0.02 | -0.02 | 0.092 | 0.21 | 0.24 | 0.13 | 0.15 | 0.13 | 0.08 |

| | | | | | | | | | | | | | | | | |
|---|---|---|---|---|---|---|---|---|---|---|---|---|---|---|---|---|
| 11 | 11627021 | rs10765873 | -0.144 | 0.756 | -1.111 | 0.188 | -0.013 | -0.02 | -0.02 | 0.848 | 0.36 | 0.2 | 0.31 | 0.54 | 0.45 | 0.51 |
| 11 | 11631234 | rs10765874 | 0.209 | 0.397 | -1.056 | -0.089 | 0.002 | 0.007 | 0.007 | 0 | 0.87 | 0.92 | 0.77 | 0.87 | 0.88 | 0.9 |
| 11 | 11631429 | rs7104225 | -0.075 | 1 | -0.789 | 0.149 | -0.035 | -0.018 | -0.018 | 0 | 0.37 | 0.22 | 0.34 | 0.55 | 0.49 | 0.48 |
| 11 | 11644920 | rs4910023 | 0.325 | 0.34 | -0.347 | 0.085 | 0.017 | 0.042 | 0.042 | 0 | 0.42 | 0.26 | 0.29 | 0.33 | 0.36 | 0.39 |
| 11 | 11651418 | rs10765879 | -0.178 | 0.506 | 0 | 0.017 | 0 | -0.001 | -0.001 | 0 | 0.55 | 0.53 | 0.49 | 0.54 | 0.58 | 0.47 |
| 11 | 11651588 | rs10765880 | -0.358 | 0.059 | -0.256 | -0.241 | -0.005 | -0.012 | -0.012 | 0.012 | 0.71 | 0.69 | 0.53 | 0.6 | 0.66 | 0.6 |
| 11 | 11651785 | rs10831651 | -0.97 | 0.274 | -0.389 | -0.415 | 0 | -0.034 | -0.034 | 0 | 0.9 | 0.82 | 0.9 | 1 | 0.87 | 0.94 |
| 11 | 11652231 | rs11021958 | 1.723 | -0.069 | -0.651 | -0.488 | -0.042 | 0.053 | 0.053 | 0 | 0.35 | 0.08 | 0.28 | 0.25 | 0.35 | 0.31 |
| 11 | 11652453 | rs12790312 | 0.43 | -0.126 | -0.231 | -0.105 | 0.02 | 0.046 | 0.046 | 0.754 | 0.15 | 0 | 0.03 | 0.06 | 0.09 | 0.14 |
| 11 | 11654972 | rs7484121 | 0.14 | 0.159 | 0.246 | 0.152 | 0.06 | 0.039 | 0.039 | 0 | 0.12 | 0.29 | 0.18 | 0.18 | 0.2 | 0.07 |
| 11 | 11657895 | rs12283957 | -0.652 | -0.043 | -0.554 | 0.068 | -0.013 | -0.002 | -0.002 | 0.703 | 0.2 | 0.19 | 0.09 | 0.12 | 0.08 | 0.2 |
| 11 | 11658269 | rs10765882 | -1.656 | 0.169 | -0.642 | 0.968 | 0 | 0.007 | 0.007 | 0 | 0.47 | 0.43 | 0.53 | 0.6 | 0.5 | 0.65 |
| 11 | 11660815 | rs10831656 | -1.328 | 0.37 | -0.614 | 1.191 | -0.001 | 0 | 0 | 0 | 0.29 | 0.3 | 0.4 | 0.34 | 0.29 | 0.29 |
| 11 | 11665804 | rs11021969 | -0.655 | 0.51 | -0.943 | 1.282 | -0.009 | -0.006 | -0.006 | 0 | 0.11 | 0.1 | 0.34 | 0.3 | 0.21 | 0.23 |
| 11 | 11667155 | rs10741559 | -0.912 | 0.532 | -0.929 | 1.299 | -0.003 | -0.005 | -0.005 | 0 | 0.83 | 0.85 | 0.87 | 0.72 | 0.8 | 0.66 |
| 11 | 11668648 | rs12803005 | -0.127 | 0.203 | -1.038 | 0.758 | 0.05 | 0.068 | 0.068 | 0 | 0.17 | 0 | 0.05 | 0.04 | 0.08 | 0.09 |
| 11 | 11671956 | rs11021970 | 0.157 | 0.571 | -0.891 | 0.95 | 0.033 | 0.037 | 0.037 | 0.153 | 0.06 | 0.09 | 0.14 | 0.21 | 0.2 | 0.11 |
| 11 | 11673195 | rs11606440 | 0.109 | 0.302 | -0.979 | 0.782 | 0.037 | 0.046 | 0.046 | 0 | 0.14 | 0.07 | 0.05 | 0.04 | 0.08 | 0.09 |
| 11 | 11673754 | rs11606539 | 0.149 | 0.322 | -1.182 | 0.792 | 0.031 | 0.048 | 0.048 | 0 | 0.14 | 0 | 0.04 | 0.04 | 0.08 | 0.08 |
| 11 | 11680935 | rs7480726 | -1.356 | 1.318 | -0.481 | 1.349 | 0.011 | 0.018 | 0.018 | 0 | 0.14 | 0.2 | 0.36 | 0.3 | 0.23 | 0.24 |
| 11 | 11690685 | rs10765884 | -0.527 | 0.562 | -1.218 | 1.44 | -0.017 | -0.097 | -0.097 | 0 | 0.17 | 0.03 | 0.31 | 0.48 | 0.16 | 0.38 |
| 11 | 11691268 | rs4078206 | -0.691 | 0.705 | -1.263 | 1.42 | -0.019 | -0.091 | -0.091 | 0 | 0.19 | 0.03 | 0.34 | 0.56 | 0.19 | 0.41 |
| 11 | 11693599 | rs7101423 | -0.008 | 0.679 | -0.714 | 0.926 | 0.003 | 0.011 | 0.011 | 0 | 0.85 | 0.72 | 0.85 | 0.84 | 0.83 | 0.89 |
| 11 | 11693828 | rs7949993 | -0.357 | 0.689 | -0.708 | 0.874 | -0.001 | -0.019 | -0.019 | 0 | 0.87 | 0.87 | 0.89 | 1 | 0.84 | 0.95 |
| 11 | 11700694 | rs4910407 | -0.42 | 0.674 | -0.852 | -0.153 | 0.001 | 0.044 | 0.044 | 0 | 0.45 | 0.36 | 0.6 | 0.79 | 0.42 | 0.67 |
| 11 | 11700746 | rs11021985 | 0.742 | 0.855 | -1.58 | -0.478 | -0.096 | 0.095 | 0.095 | 0.144 | 0.16 | 0.02 | 0.18 | 0.01 | 0.29 | 0.08 |
| 11 | 11706291 | rs7945768 | 0.433 | 0.559 | -1.41 | -0.227 | -0.03 | -0.029 | -0.029 | 0 | 0.16 | 0.09 | 0.22 | 0.22 | 0.19 | 0.22 |
| 11 | 11711942 | rs6485128 | -0.191 | -0.152 | -0.991 | -0.298 | -0.01 | 0.008 | 0.008 | 2.064 | 0.43 | 0.34 | 0.67 | 0.6 | 0.49 | 0.56 |
| 11 | 11719619 | rs4910411 | 0.03 | -0.315 | -1.142 | -0.449 | 0.01 | 0.039 | 0.039 | 0 | 0.08 | 0.03 | 0.03 | 0 | 0.08 | 0.01 |
| 11 | 11720806 | rs4910412 | -0.325 | -0.311 | -1.194 | -0.505 | -0.001 | 0.052 | 0.052 | 0 | 0.1 | 0.03 | 0.04 | 0 | 0.11 | 0.04 |
| 11 | 11730134 | rs7948970 | -0.135 | -0.611 | -0.859 | -1.299 | -0.01 | -0.102 | -0.102 | 0 | 0.38 | 0.46 | 0.19 | 0.01 | 0.28 | 0.17 |
| 11 | 11731251 | rs4910027 | 0.244 | -0.894 | -1.616 | -1.024 | 0.006 | -0.056 | -0.056 | 0 | 0.23 | 0.04 | 0.44 | 0.73 | 0.23 | 0.48 |
| 11 | 11739732 | rs1869697 | 1.304 | -1.285 | -1.855 | -1.379 | 0.063 | 0.154 | 0.154 | 1.284 | 0.26 | 0.03 | 0.08 | 0.01 | 0.16 | 0.12 |
| 11 | 11740963 | rs1471013 | 0.489 | -1.453 | -2.126 | -0.616 | -0.002 | -0.085 | -0.085 | 0.654 | 0.21 | 0.04 | 0.22 | 0.39 | 0.25 | 0.31 |
| 11 | 11744511 | rs4910413 | -1.641 | -1.798 | -1.647 | -0.934 | 0 | 0 | 0 | 0.815 | 0.72 | 0.7 | 0.7 | 0.75 | 0.6 | 0.71 |
| 11 | 11746431 | rs16910005 | 0.889 | -1.818 | -1.269 | -0.861 | -0.003 | 0.035 | 0.035 | 0 | 0.1 | 0.1 | 0.28 | 0.34 | 0.07 | 0.22 |
| 11 | 11749653 | rs11606749 | 1.175 | -1.919 | -0.847 | -0.924 | -0.055 | -0.126 | -0.126 | 0 | 0.31 | 0.41 | 0.13 | 0.01 | 0.18 | 0.11 |
| 11 | 11750180 | rs767165 | -0.29 | -1.858 | -1.082 | -0.88 | 0.042 | 0.031 | 0.031 | 0 | 0.48 | 0.64 | 0.48 | 0.36 | 0.34 | 0.38 |
| 11 | 11759507 | rs11022012 | 0.192 | -1.838 | -1.31 | -1.277 | 0.02 | -0.048 | -0.048 | 0 | 0.26 | 0.03 | 0.44 | 0.73 | 0.24 | 0.48 |
| 11 | 11761476 | rs1455255 | -0.369 | -1.941 | -1.159 | -1.376 | 0.057 | 0.085 | 0.085 | 0.115 | 0.55 | 0.2 | 0.56 | 0.74 | 0.43 | 0.59 |
| 11 | 11761553 | rs11604515 | 1.212 | -1.861 | -1.075 | -1.347 | 0.035 | 0.028 | 0.028 | 0.662 | 0.29 | 0.17 | 0.12 | 0.01 | 0.18 | 0.11 |
| 11 | 11761782 | rs17446056 | 1.02 | -1.889 | -1.1 | -1.371 | 0.043 | 0.05 | 0.05 | 0.167 | 0.31 | 0.17 | 0.14 | 0.03 | 0.23 | 0.15 |
| 11 | 11762588 | rs882036 | 1.099 | -1.635 | -0.679 | -1.349 | 0.023 | -0.058 | -0.058 | 0 | 0.11 | 0.24 | 0.1 | 0 | 0.12 | 0.05 |
| 11 | 11763848 | rs897360 | -1.068 | -1.758 | -0.655 | -1.316 | -0.006 | -0.026 | -0.026 | 0.006 | 0.42 | 0.5 | 0.59 | 0.74 | 0.42 | 0.58 |
| 11 | 11765613 | rs11604688 | -0.808 | -1.132 | -0.507 | -0.94 | -0.004 | 0.001 | 0.001 | 0 | 0.67 | 0.66 | 0.69 | 0.76 | 0.59 | 0.66 |
| 11 | 11771889 | rs10500747 | -0.68 | -1.099 | -0.44 | -0.591 | 0.014 | -0.014 | -0.014 | 0.046 | 0.28 | 0.22 | 0.47 | 0.65 | 0.25 | 0.56 |
| 11 | 11774738 | rs1979235 | -0.936 | -0.817 | -0.452 | -0.608 | 0.02 | -0.016 | -0.016 | 0.021 | 0.7 | 0.65 | 0.69 | 0.83 | 0.59 | 0.72 |
| 11 | 11777749 | rs7939727 | 0.809 | -0.972 | -0.504 | -0.751 | -0.017 | 0.011 | 0.011 | 0 | 0.12 | 0.07 | 0.09 | 0 | 0.15 | 0.05 |
| 11 | 11779374 | rs874651 | 1.955 | -1.082 | -0.501 | -0.714 | 0.053 | 0.033 | 0.033 | 0.089 | 0.27 | 0.15 | 0.11 | 0.17 | 0.19 | 0.11 |
| 11 | 11782282 | rs9988795 | 1.256 | 0.485 | -0.136 | -0.349 | -0.06 | -0.005 | -0.005 | 0 | 0.2 | 0.06 | 0.18 | 0.16 | 0.24 | 0.24 |
| 11 | 11785883 | rs7951610 | 1.672 | 0.38 | -0.212 | 0.108 | -0.015 | -0.037 | -0.037 | 0.045 | 0.1 | 0.02 | 0.15 | 0.22 | 0.12 | 0.26 |
| 11 | 11787253 | rs11824377 | -0.701 | 0.579 | 0.062 | 0.052 | 0.023 | -0.017 | -0.017 | 0.097 | 0.7 | 0.65 | 0.7 | 0.83 | 0.58 | 0.73 |
| 11 | 11796897 | rs2054117 | -0.371 | 0.12 | -0.077 | 0.672 | -0.036 | -0.009 | -0.009 | 0 | 0.67 | 0.25 | 0.65 | 0.66 | 0.57 | 0.69 |
| 11 | 11798389 | rs10458871 | 0.008 | -0.006 | -0.237 | 0.716 | 0.007 | -0.114 | -0.114 | 0 | 0.21 | 0.04 | 0.42 | 0.49 | 0.19 | 0.51 |
| 11 | 11799517 | rs1455263 | 0.395 | -0.181 | 0.078 | 0.823 | 0.006 | 0.005 | 0.005 | 0 | 0.43 | 0.4 | 0.25 | 0.34 | 0.38 | 0.2 |
| 11 | 11799968 | rs1455264 | 0.288 | -0.497 | -0.201 | 0.81 | 0.009 | -0.025 | -0.025 | 0 | 0.07 | 0.02 | 0.27 | 0.28 | 0.06 | 0.24 |
| 11 | 11800559 | rs12416973 | 0.928 | -0.672 | -0.055 | 0.155 | 0.037 | 0.031 | 0.031 | 0.075 | 0.32 | 0.2 | 0.13 | 0.18 | 0.23 | 0.13 |
| 11 | 11800932 | rs7116774 | -0.342 | -0.332 | 0.386 | 0.211 | 0.037 | -0.023 | -0.023 | 0 | 0.08 | 0.36 | 0.06 | 0 | 0.05 | 0.03 |
| 11 | 11801733 | rs12574375 | 0.646 | -0.688 | 0.162 | 0.208 | 0.055 | 0.048 | 0.048 | 1.007 | 0.36 | 0.22 | 0.13 | 0.19 | 0.22 | 0.13 |
| 11 | 11802361 | rs1378357 | 0.646 | -0.645 | 0.205 | 0.227 | 0.052 | 0.048 | 0.048 | 0.622 | 0.36 | 0.22 | 0.14 | 0.19 | 0.21 | 0.14 |
| 11 | 11803296 | rs1466257 | 0.253 | 0.244 | 0.425 | 0.697 | 0.051 | 0.043 | 0.043 | 0 | 0.06 | 0.12 | 0.12 | 0.14 | 0.17 | 0.09 |
| 11 | 11805433 | rs10831672 | 0.132 | 0.16 | 0.356 | 0.983 | 0.063 | 0.022 | 0.022 | 0 | 0.45 | 0.27 | 0.23 | 0.44 | 0.31 | 0.32 |
| 11 | 11806317 | rs11022029 | -0.423 | -0.218 | 0.953 | 0.967 | 0.002 | 0.118 | 0.118 | 0 | 0.15 | 0.41 | 0.11 | 0.33 | 0.11 | 0.19 |
| 11 | 11808547 | rs16910122 | 1.668 | -0.233 | 0.745 | 0.901 | 0.057 | 0.032 | 0.032 | 0.954 | 0.28 | 0.01 | 0.1 | 0.17 | 0.2 | 0.13 |
| 11 | 11809438 | rs10500751 | -0.106 | -0.025 | 0.86 | 1.221 | 0.002 | 0.002 | 0.002 | 0 | 0.5 | 0.52 | 0.24 | 0.7 | 0.4 | 0.42 |
| 11 | 11811025 | rs12418830 | -0.143 | 0.188 | 0.551 | 1.124 | -0.008 | -0.022 | -0.022 | 0.008 | 0.21 | 0.16 | 0.42 | 0.3 | 0.21 | 0.33 |
| 11 | 11814526 | rs1455250 | -1.224 | 0.055 | 0.727 | 1.053 | 0.006 | 0.125 | 0.125 | 2.091 | 0.19 | 0.6 | 0.14 | 0.53 | 0.18 | 0.29 |
| 11 | 11814701 | rs16910152 | 0.957 | 0.01 | 0.663 | 1.007 | 0.095 | 0.096 | 0.096 | 1.056 | 0.43 | 0.13 | 0.48 | 0.26 | 0.32 | 0.34 |
| 11 | 11817406 | rs12416698 | 0.897 | 0.286 | 0.798 | 1.135 | 0.004 | 0.012 | 0.012 | 0.881 | 0.15 | 0.12 | 0.39 | 0.09 | 0.13 | 0.22 |

| | | | | | | | | | | | | | | | | |
|---|---|---|---|---|---|---|---|---|---|---|---|---|---|---|---|---|
| 11 | 11818120 | rs4376903 | -0.239 | 0.523 | 0.738 | 1.083 | 0.028 | 0.001 | 0.001 | 0 | 0.12 | 0.2 | 0.14 | 0.09 | 0.21 | 0.13 |
| 11 | 11820449 | rs10831676 | -0.442 | 0.363 | 0.711 | 1.041 | -0.017 | 0.015 | 0.015 | 0 | 0.64 | 0.72 | 0.62 | 0.78 | 0.49 | 0.61 |
| 11 | 11820816 | rs11022043 | -1.349 | 0.432 | 0.387 | 1.134 | 0.002 | 0.109 | 0.109 | 2.155 | 0.19 | 0.62 | 0.14 | 0.52 | 0.17 | 0.29 |
| 11 | 11822694 | rs4910420 | 1.006 | 0.498 | -0.091 | 1.157 | 0.002 | 0.01 | 0.01 | 0.525 | 0.14 | 0.08 | 0.38 | 0.09 | 0.12 | 0.19 |
| 11 | 11829799 | rs16910169 | 0.768 | 0.588 | -0.127 | 1.309 | -0.007 | 0.005 | 0.005 | 0 | 0.06 | 0.04 | 0.03 | 0.24 | 0.09 | 0.11 |
| 11 | 11830325 | rs734462 | -0.416 | 0.579 | -0.12 | 1.275 | -0.021 | -0.047 | -0.047 | 0 | 0.14 | 0.08 | 0.09 | 0.52 | 0.17 | 0.29 |
| 11 | 11832343 | rs1020679 | 0.67 | 0.652 | -0.109 | 1.235 | 0.054 | 0.027 | 0.027 | 0 | 0.06 | 0.11 | 0.11 | 0.09 | 0.18 | 0.12 |
| 11 | 11836332 | rs16910187 | 1.927 | 0.479 | -0.319 | 1.148 | 0.104 | 0.092 | 0.092 | 2.366 | 0.32 | 0.1 | 0.1 | 0.16 | 0.2 | 0.13 |
| 11 | 11840925 | rs12790557 | 0.789 | 0.615 | -0.271 | 1.284 | -0.005 | -0.017 | -0.017 | 1.466 | 0.08 | 0.07 | 0.06 | 0.28 | 0.09 | 0.17 |
| 11 | 11840965 | rs10831680 | -1.416 | 0.631 | -0.311 | 1.03 | -0.045 | 0.028 | 0.028 | 0 | 0.77 | 0.92 | 0.76 | 0.86 | 0.7 | 0.77 |
| 11 | 11844516 | rs1542379 | 0.509 | 0.593 | -0.292 | 1.072 | 0.039 | 0.015 | 0.015 | 0 | 0.06 | 0.09 | 0.11 | 0.09 | 0.18 | 0.1 |
| 11 | 11862213 | rs11022066 | 1.857 | 0.064 | -0.511 | 0.899 | 0.081 | 0.054 | 0.054 | 4.055 | 0.3 | 0.01 | 0.1 | 0.16 | 0.19 | 0.13 |
| 11 | 11863410 | rs12418935 | 0.385 | 0.439 | 0.01 | 1.152 | -0.006 | -0.005 | -0.005 | 2.425 | 0.15 | 0.19 | 0.39 | 0.09 | 0.1 | 0.21 |
| 11 | 11866815 | rs17450541 | 0.597 | 0.542 | -0.077 | 1.162 | -0.003 | 0.015 | 0.015 | 0.021 | 0.16 | 0.09 | 0.41 | 0.09 | 0.16 | 0.24 |
| 11 | 11868549 | rs7951773 | -0.214 | 0.81 | -0.033 | 1.418 | 0 | 0.022 | 0.022 | 0.603 | 0.13 | 0.13 | 0.14 | 0.51 | 0.27 | 0.29 |
| 11 | 11885841 | rs7128291 | -0.008 | 0.506 | -0.224 | 1.221 | -0.068 | 0.081 | 0.081 | 0.116 | 0.58 | 0.88 | 0.34 | 0.77 | 0.52 | 0.53 |
| 11 | 11893217 | rs12418890 | 0.815 | 0.54 | -0.206 | 1.231 | -0.006 | 0.017 | 0.017 | 0.032 | 0.15 | 0.08 | 0.4 | 0.09 | 0.16 | 0.23 |
| 11 | 11898743 | rs10831690 | -0.061 | 0.621 | -0.215 | 1.535 | -0.023 | -0.024 | -0.024 | 0.041 | 0.1 | 0.06 | 0.11 | 0.27 | 0.25 | 0.17 |
| 11 | 11913585 | rs11022079 | -1.441 | 0.572 | -0.116 | 1.218 | -0.069 | 0.013 | 0.013 | 0.026 | 0.78 | 0.96 | 0.76 | 0.86 | 0.7 | 0.77 |
| 11 | 11918698 | rs4360694 | 1.221 | 0.587 | -0.097 | 1.225 | -0.066 | 0.015 | 0.015 | 0.04 | 0.22 | 0.04 | 0.24 | 0.14 | 0.3 | 0.24 |
| 11 | 11943043 | rs7480906 | -0.456 | 0.306 | 0.219 | 1.255 | -0.005 | 0.009 | 0.009 | 0.127 | 0.46 | 0.38 | 0.71 | 0.32 | 0.48 | 0.57 |
| 11 | 11954786 | rs10741562 | 0.009 | 0.924 | 0.059 | 1.548 | -0.023 | -0.023 | -0.023 | 0 | 0.9 | 0.94 | 0.88 | 0.78 | 0.75 | 0.83 |
| 11 | 11971165 | rs10741564 | 1.128 | 0.79 | 0.672 | 1.606 | 0.125 | 0.186 | 0.186 | 2.301 | 0.8 | 0.51 | 0.81 | 0.49 | 0.69 | 0.64 |
| 11 | 11976628 | rs7933089 | 0.51 | 0.877 | 0.514 | 1.631 | -0.024 | -0.024 | -0.024 | 0 | 0.83 | 0.87 | 0.84 | 0.49 | 0.69 | 0.68 |
| 11 | 11979674 | rs1979687 | -0.249 | 0.75 | 0.448 | 1.412 | -0.017 | -0.01 | -0.01 | 0 | 0.36 | 0.27 | 0.31 | 0.44 | 0.4 | 0.39 |
| 11 | 11980397 | rs13903 | 0.031 | 1.076 | 0.4 | 1.31 | -0.024 | -0.023 | -0.023 | 0.168 | 0.9 | 0.94 | 0.89 | 0.79 | 0.75 | 0.84 |
| 11 | 11983885 | rs1472189 | 0.86 | 1.229 | 0.198 | 1.442 | -0.057 | 0.002 | 0.002 | 0.074 | 0.22 | 0.06 | 0.25 | 0.18 | 0.33 | 0.27 |
| 11 | 11986061 | rs3206824 | 0.395 | 1.184 | 0.692 | 1.54 | 0.063 | 0.035 | 0.035 | 0 | 0.9 | 0.82 | 0.88 | 0.79 | 0.76 | 0.84 |
| 11 | 11988442 | rs7480026 | -0.984 | 0.678 | 0.499 | 1.309 | -0.028 | 0.003 | 0.003 | 0.536 | 0.14 | 0.61 | 0.1 | 0.09 | 0.07 | 0.08 |
| 11 | 11994832 | rs1552796 | 0.452 | 0.904 | -0.711 | 1.416 | -0.141 | -0.024 | -0.024 | 0 | 0.34 | 0.04 | 0.62 | 0.3 | 0.44 | 0.45 |
| 11 | 11996757 | rs7396140 | 0.867 | 1.55 | 0.554 | 2.027 | 0.084 | 0.146 | 0.146 | 0 | 0.78 | 0.58 | 0.81 | 0.46 | 0.69 | 0.63 |
| 11 | 11997895 | rs11022099 | -0.924 | 1.385 | 0.417 | 1.175 | -0.003 | 0.027 | 0.027 | 0 | 0.19 | 0.41 | 0.12 | 0.24 | 0.14 | 0.2 |
| 11 | 12003930 | rs11022102 | 0.282 | -0.477 | 0.107 | 1.419 | 0.011 | -0.011 | -0.011 | 0 | 0.23 | 0.18 | 0.08 | 0.28 | 0.12 | 0.16 |
| 11 | 12003946 | rs11022103 | 0.63 | -0.525 | 0.015 | 1.583 | 0.048 | -0.01 | -0.01 | 0 | 0.21 | 0.01 | 0.05 | 0.13 | 0.1 | 0.09 |
| 11 | 12004699 | rs10765917 | 0.215 | 0.376 | -0.126 | 1.291 | -0.013 | -0.035 | -0.035 | 0.852 | 0.3 | 0.15 | 0.28 | 0.41 | 0.32 | 0.37 |
| 11 | 12004835 | rs12225759 | 0.355 | 0.062 | -0.335 | 0.995 | 0.065 | -0.011 | -0.011 | 0 | 0.44 | 0.3 | 0.26 | 0.51 | 0.28 | 0.41 |
| 11 | 12007778 | rs7113678 | 0 | 0.311 | -0.666 | 0.413 | 0.047 | 0.093 | 0.093 | 0 | 0.55 | 0.71 | 0.34 | 0.88 | 0.36 | 0.58 |
| 11 | 12008908 | rs12295349 | -0.658 | 1.461 | -0.374 | 0.796 | 0.032 | -0.008 | -0.008 | 0 | 0.23 | 0.39 | 0.19 | 0.16 | 0.29 | 0.16 |
| 11 | 12009569 | rs16910295 | 0.079 | 1.066 | -0.767 | 1.143 | -0.023 | -0.036 | -0.036 | 0 | 0.07 | 0.01 | 0.04 | 0.18 | 0.06 | 0.13 |
| 11 | 12010327 | rs1038909 | 0.309 | 1.302 | -0.516 | 0.613 | 0.007 | -0.013 | -0.013 | 0 | 0.7 | 0.65 | 0.68 | 0.82 | 0.65 | 0.61 |
| 11 | 12010496 | rs988666 | 0.359 | 1.231 | -0.603 | 0.56 | -0.012 | -0.006 | -0.006 | 0.379 | 0.36 | 0.33 | 0.48 | 0.57 | 0.43 | 0.36 |
| 11 | 12013617 | rs6485350 | -1.035 | 0.977 | -0.351 | 0.398 | -0.004 | 0.01 | 0.01 | 0 | 0.52 | 0.67 | 0.55 | 0.36 | 0.5 | 0.57 |
| 11 | 12013813 | rs7116879 | -0.033 | 0.847 | -0.149 | 0.403 | -0.011 | 0.036 | 0.036 | 0 | 0.16 | 0.28 | 0.3 | 0.18 | 0.11 | 0.32 |
| 11 | 12014668 | rs987663 | 0.216 | 0.767 | -0.543 | 0.242 | 0.028 | -0.061 | -0.061 | 1.316 | 0.22 | 0.3 | 0.19 | 0.05 | 0.32 | 0.18 |
| 11 | 12015730 | rs11022105 | -1.345 | 0.373 | -0.677 | 1.002 | -0.004 | 0.006 | 0.006 | 0 | 0.33 | 0.39 | 0.41 | 0.32 | 0.29 | 0.41 |
| 11 | 12016528 | rs952932 | -0.068 | 0.037 | -1.281 | 0.506 | 0.004 | -0.016 | -0.016 | 0.22 | 0.06 | 0.05 | 0.25 | 0 | 0.06 | 0.05 |
| 11 | 12016945 | rs12421658 | -0.007 | -0.156 | -1.422 | 0.746 | 0.02 | -0.03 | -0.03 | 0 | 0.14 | 0 | 0.05 | 0.13 | 0.08 | 0.07 |
| 11 | 12018220 | rs7107048 | 0.306 | 0.443 | -0.991 | 0.949 | -0.021 | -0.041 | -0.041 | 0 | 0.91 | 1 | 0.96 | 0.81 | 0.92 | 0.83 |
| 11 | 12020026 | rs3750938 | -1.461 | 0.317 | -0.128 | 0.888 | 0.009 | 0.084 | 0.084 | 0 | 0.24 | 0.49 | 0.38 | 0.24 | 0.2 | 0.4 |
| 11 | 12020115 | rs3750939 | -1.034 | 0.275 | 0.323 | 0.906 | 0.003 | 0.011 | 0.011 | 0 | 0.16 | 0.25 | 0.09 | 0.13 | 0.14 | 0.31 |
| 11 | 12020616 | rs7104941 | -1.709 | -0.055 | 0.657 | 0.908 | -0.011 | -0.025 | -0.025 | 0.093 | 0.19 | 0.36 | 0.33 | 0.09 | 0.15 | 0.28 |
| 11 | 12022898 | rs3750940 | -1.643 | -0.13 | 0.968 | 0.878 | 0 | 0 | 0 | 1.052 | 0.18 | 0.19 | 0.32 | 0.1 | 0.16 | 0.27 |
| 11 | 12027863 | rs4290212 | -0.032 | -0.705 | 1.585 | 0.312 | -0.026 | 0.025 | 0.025 | 4.239 | 0.18 | 0.39 | 0.09 | 0.13 | 0.13 | 0.1 |
| 11 | 12038680 | rs872574 | -0.96 | -0.794 | 1.588 | 0.312 | -0.001 | -0.002 | -0.002 | 0 | 0.16 | 0.23 | 0.32 | 0.1 | 0.15 | 0.26 |
| 11 | 12038874 | rs11022114 | 1.548 | -0.517 | 0.56 | 0.02 | -0.106 | -0.135 | -0.135 | 0.046 | 0.2 | 0.02 | 0.36 | 0.43 | 0.31 | 0.17 |
| 11 | 12039278 | rs2896596 | -0.409 | -0.523 | 0.67 | 0.001 | -0.001 | -0.002 | -0.002 | 0.108 | 0.16 | 0.22 | 0.33 | 0.1 | 0.15 | 0.26 |
| 11 | 12040595 | rs903012 | 0.744 | -0.54 | 0.66 | 0 | -0.001 | -0.001 | -0.001 | 0.128 | 0.17 | 0.16 | 0.07 | 0.14 | 0.13 | 0.08 |
| 11 | 12040870 | rs903008 | -0.14 | -1.405 | 0.613 | -0.496 | -0.022 | -0.011 | -0.011 | 0 | 0.3 | 0.32 | 0.12 | 0.16 | 0.17 | 0.22 |
| 11 | 12041496 | rs4500466 | -1.528 | -1.38 | 0.309 | -0.486 | 0.08 | 0.059 | 0.059 | 0.523 | 0.47 | 0.75 | 0.46 | 0.26 | 0.32 | 0.47 |
| 11 | 12043269 | rs12274783 | 0.513 | -1.011 | 0.245 | -0.333 | -0.016 | -0.001 | -0.001 | 0 | 0.19 | 0.3 | 0.08 | 0.13 | 0.14 | 0.08 |
| 11 | 12050652 | rs11022119 | 1.387 | -0.233 | 0.194 | -0.125 | -0.087 | -0.133 | -0.133 | 0.101 | 0.2 | 0.02 | 0.3 | 0.42 | 0.3 | 0.2 |
| 11 | 12051466 | rs923805 | -1.617 | -0.693 | 0.644 | -0.365 | 0 | -0.001 | -0.001 | 0.169 | 0.37 | 0.66 | 0.43 | 0.2 | 0.25 | 0.4 |
| 11 | 12053190 | rs16910389 | 0.014 | -0.408 | 0.21 | -0.281 | -0.004 | -0.016 | -0.016 | 0.07 | 0.11 | 0.09 | 0.32 | 0.17 | 0.11 | 0.22 |
| 11 | 12055477 | rs7395308 | -0.253 | -0.324 | 0.392 | -0.123 | 0.012 | 0.051 | 0.051 | 0 | 0.12 | 0.28 | 0.33 | 0.17 | 0.11 | 0.21 |
| 11 | 12064576 | rs11603598 | 0.469 | -0.459 | 0.345 | -0.332 | -0.008 | -0.032 | -0.032 | 0 | 0.08 | 0.09 | 0.03 | 0 | 0.04 | 0.06 |
| 11 | 12068184 | rs1994326 | -1.22 | -0.659 | 0.046 | -0.438 | -0.045 | -0.089 | -0.089 | 3.056 | 0.25 | 0.37 | 0.09 | 0.05 | 0.14 | 0.11 |
| 11 | 12068504 | rs12278824 | -0.159 | -0.614 | -0.026 | -0.384 | 0.001 | 0.036 | 0.036 | 0 | 0.32 | 0.47 | 0.37 | 0.47 | 0.35 | 0.31 |
| 11 | 12069248 | rs923171 | -0.787 | -0.37 | -0.384 | -0.568 | -0.028 | -0.08 | -0.08 | 0 | 0.21 | 0.25 | 0.06 | 0.01 | 0.12 | 0.1 |
| 11 | 12072099 | rs12417334 | -0.39 | 0.852 | -1.14 | -0.106 | 0.006 | -0.021 | -0.021 | 0.042 | 0.31 | 0.39 | 0.39 | 0.13 | 0.3 | 0.32 |

| | | | | | | | | | | | | | | | | |
|---|---|---|---|---|---|---|---|---|---|---|---|---|---|---|---|---|
| 11 | 12073518 | rs6485416 | -0.156 | 0.734 | -1.744 | 0.58 | 0.002 | -0.027 | -0.027 | 0 | 0.66 | 0.78 | 0.77 | 0.6 | 0.62 | 0.67 |
| 11 | 12075829 | rs7928952 | -0.928 | 0.924 | -1.591 | 1.032 | -0.001 | 0.044 | 0.044 | 0 | 0.22 | 0.29 | 0.25 | 0.36 | 0.21 | 0.38 |
| 11 | 12076162 | rs7946686 | -0.357 | 0.966 | -2.775 | 0.24 | -0.003 | 0.107 | 0.107 | 0.033 | 0.53 | 0.84 | 0.38 | 0.23 | 0.53 | 0.42 |
| 11 | 12076178 | rs6485420 | -0.523 | 0.954 | -2.591 | 0.787 | 0.034 | -0.075 | -0.075 | 0.782 | 0.73 | 0.88 | 0.52 | 0.37 | 0.8 | 0.58 |
| 11 | 12076251 | rs11022134 | -0.727 | 2.007 | -1.934 | 0.782 | -0.04 | -0.03 | -0.03 | 0 | 0.17 | 0.05 | 0.14 | 0.16 | 0.25 | 0.27 |
| 11 | 12078926 | rs12804449 | -0.937 | 1.377 | -1.713 | 0.428 | -0.008 | 0.004 | 0.004 | 0.012 | 0.15 | 0.12 | 0.13 | 0.11 | 0.2 | 0.29 |
| 11 | 12080552 | rs1872561 | 0.134 | 1.061 | -2.034 | 0.716 | -0.033 | -0.071 | -0.071 | 0.049 | 0.13 | 0.02 | 0.32 | 0.19 | 0.16 | 0.09 |
| 11 | 12082627 | rs11022141 | 0.013 | 0.699 | -2.066 | 0.18 | -0.002 | 0.013 | 0.013 | 0.398 | 0.3 | 0.21 | 0.23 | 0.17 | 0.28 | 0.22 |
| 11 | 12082861 | rs7478802 | -1.374 | 0.236 | -1.844 | -0.314 | -0.013 | 0.014 | 0.014 | 0.1 | 0.75 | 0.8 | 0.8 | 0.81 | 0.69 | 0.78 |
| 11 | 12083368 | rs1814174 | 0.389 | 0.098 | -1.19 | -0.201 | 0.004 | -0.02 | -0.02 | 0.214 | 0.2 | 0.18 | 0.41 | 0.58 | 0.22 | 0.35 |
| 11 | 12083579 | rs1159828 | 0.45 | -0.42 | -0.753 | -0.581 | -0.005 | 0.009 | 0.009 | 0 | 0.43 | 0.59 | 0.32 | 0.4 | 0.45 | 0.39 |
| 11 | 12083803 | rs11022142 | 1.11 | -0.411 | -0.838 | -0.359 | -0.03 | -0.12 | -0.12 | 0 | 0.17 | 0.05 | 0.39 | 0.58 | 0.22 | 0.35 |
| 11 | 12084910 | rs11603050 | -0.342 | -0.474 | -0.302 | -0.318 | -0.01 | 0.024 | 0.024 | 0 | 0.47 | 0.35 | 0.63 | 0.59 | 0.51 | 0.57 |
| 11 | 12093458 | rs2018347 | -0.352 | -0.657 | 0.242 | -0.518 | -0.02 | -0.107 | -0.107 | 4.146 | 0.18 | 0.38 | 0.09 | 0 | 0.13 | 0.08 |
| 11 | 12094658 | rs10831716 | -0.321 | -0.638 | -0.042 | -0.856 | 0.001 | 0.113 | 0.113 | 0.166 | 0.49 | 0.28 | 0.64 | 0.8 | 0.51 | 0.67 |
| 11 | 12096833 | rs11601664 | 0.245 | -0.684 | -0.396 | -0.339 | -0.034 | -0.09 | -0.09 | 0 | 0.25 | 0.09 | 0.37 | 0.41 | 0.3 | 0.17 |
| 11 | 12098277 | rs7948759 | 0.01 | -1.085 | 0.015 | -0.247 | -0.017 | -0.072 | -0.072 | 0.191 | 0.16 | 0.28 | 0.08 | 0 | 0.12 | 0.05 |
| 11 | 12098506 | rs12794303 | 0.242 | -1.158 | -0.3 | -0.267 | -0.084 | -0.067 | -0.067 | 0 | 0.4 | 0.11 | 0.54 | 0.48 | 0.44 | 0.32 |
| 11 | 12099176 | rs11605511 | -1.99 | -1.199 | -0.277 | -0.205 | -0.016 | 0.042 | 0.042 | 0 | 0.7 | 0.8 | 0.75 | 0.88 | 0.64 | 0.76 |
| 11 | 12101898 | rs17383023 | 1.55 | -1.205 | -0.403 | -0.162 | -0.058 | -0.086 | -0.086 | 0.16 | 0.17 | 0.02 | 0.33 | 0.28 | 0.22 | 0.13 |
| 11 | 12105342 | rs12577400 | 0.418 | -1.063 | -0.156 | -0.029 | -0.006 | 0.13 | 0.13 | 0 | 0.06 | 0.12 | 0.09 | 0.37 | 0.07 | 0.26 |
| 11 | 12108029 | rs7480668 | -0.895 | -0.939 | -0.438 | -0.646 | 0 | -0.004 | -0.004 | 0.778 | 0.58 | 0.45 | 0.63 | 0.43 | 0.56 | 0.37 |
| 11 | 12108793 | rs11022159 | -0.337 | -0.94 | -0.401 | -0.614 | -0.001 | 0.003 | 0.003 | 0 | 0.48 | 0.45 | 0.6 | 0.43 | 0.5 | 0.33 |
| 11 | 12110348 | rs7129030 | -1.624 | -0.85 | -0.273 | -0.425 | -0.026 | 0.055 | 0.055 | 0 | 0.7 | 0.81 | 0.75 | 0.88 | 0.62 | 0.77 |
| 11 | 12110677 | rs10831728 | 0.962 | -0.942 | -0.74 | -0.433 | -0.01 | 0.004 | 0.004 | 3.51 | 0.09 | 0.09 | 0.04 | 0.13 | 0.04 | 0.01 |
| 11 | 12113297 | rs11600108 | 0.481 | -0.278 | -0.708 | -0.464 | -0.003 | 0 | 0 | 0 | 0.39 | 0.45 | 0.45 | 0.4 | 0.31 | 0.16 |
| 11 | 12113908 | rs7950266 | -0.643 | 0.452 | -0.626 | 0.609 | -0.031 | 0.269 | 0.269 | 0 | 0.12 | 0.3 | 0.12 | 0.47 | 0.07 | 0.39 |
| 11 | 12114121 | rs4757190 | -0.506 | 1.353 | -1.426 | -0.425 | -0.064 | 0.086 | 0.086 | 0 | 0.18 | 0.06 | 0.18 | 0.02 | 0.23 | 0.22 |
| 11 | 12122094 | rs7111481 | 0.045 | 1.209 | -0.958 | 0.067 | -0.004 | 0.008 | 0.008 | 0 | 0.56 | 0.61 | 0.64 | 0.41 | 0.52 | 0.57 |
| 11 | 12122574 | rs7130896 | 0.015 | 1.459 | -0.294 | 0.402 | 0.002 | 0.049 | 0.049 | 0 | 0.7 | 0.61 | 0.65 | 0.51 | 0.75 | 0.71 |
| 11 | 12126352 | rs12803936 | 0.119 | 1.873 | -0.621 | 0.332 | -0.025 | -0.02 | -0.02 | 0.919 | 0.14 | 0.07 | 0.22 | 0.23 | 0.23 | 0.2 |
| 11 | 12127181 | rs2004915 | 0.004 | 1.514 | -0.057 | 0.805 | -0.003 | -0.007 | -0.007 | 0 | 0.27 | 0.25 | 0.31 | 0.34 | 0.25 | 0.35 |
| 11 | 12127479 | rs2015963 | 0.624 | 1.378 | 0.128 | 0.564 | -0.012 | 0.017 | 0.017 | 0 | 0.57 | 0.33 | 0.46 | 0.39 | 0.6 | 0.43 |
| 11 | 12129389 | rs3763820 | 0.023 | 1.359 | 0.378 | 0.642 | 0.035 | 0.103 | 0.103 | 0.215 | 0.85 | 0.62 | 0.77 | 0.73 | 0.85 | 0.82 |
| 11 | 12131800 | rs1485964 | 0.332 | 1.461 | -0.062 | 0.514 | -0.008 | 0.01 | 0.01 | 0 | 0.59 | 0.39 | 0.47 | 0.38 | 0.61 | 0.45 |
| 11 | 12132538 | rs2307072 | 0.36 | 1.304 | -0.177 | 0.318 | 0.007 | 0.018 | 0.018 | 0 | 0.78 | 0.44 | 0.73 | 0.72 | 0.78 | 0.74 |
| 11 | 12137982 | rs4756772 | 0.154 | 1.085 | -0.374 | 0.515 | -0.013 | -0.013 | -0.013 | 0.009 | 0.29 | 0.25 | 0.34 | 0.32 | 0.33 | 0.36 |
| 11 | 12138524 | rs10765923 | -0.112 | 0.562 | -0.352 | 0.101 | 0.029 | 0.043 | 0.043 | 0 | 0.48 | 0.34 | 0.41 | 0.39 | 0.45 | 0.35 |
| 11 | 12139815 | rs7932017 | -0.579 | -0.067 | -0.459 | -0.347 | -0.001 | -0.036 | -0.036 | 0 | 0.23 | 0.26 | 0.13 | 0.03 | 0.16 | 0.11 |
| 11 | 12147250 | rs10741566 | -1.418 | 0.39 | -0.083 | 0.838 | 0.016 | 0.133 | 0.133 | 0 | 0.08 | 0.26 | 0.14 | 0.23 | 0.09 | 0.31 |
| 11 | 12147395 | rs10765924 | -0.411 | -0.012 | -0.69 | -0.011 | 0.01 | 0.071 | 0.071 | 0.006 | 0.26 | 0.11 | 0.18 | 0.08 | 0.23 | 0.17 |
| 11 | 12147458 | rs4237703 | 0.08 | -0.019 | -0.658 | -0.015 | 0.013 | 0.065 | 0.065 | 0.002 | 0.74 | 0.88 | 0.83 | 0.91 | 0.77 | 0.83 |
| 11 | 12148752 | rs11022188 | -0.314 | -0.172 | -0.742 | 0.214 | -0.017 | -0.07 | -0.07 | 0 | 0.12 | 0.01 | 0.09 | 0.24 | 0.12 | 0.32 |
| 11 | 12149531 | rs2013486 | -0.35 | -0.439 | -0.941 | 0.041 | -0.016 | -0.059 | -0.059 | 0 | 0.2 | 0.04 | 0.28 | 0.28 | 0.22 | 0.38 |
| 11 | 12154464 | rs7108923 | -0.709 | -0.873 | 0.127 | 0.46 | -0.003 | 0.097 | 0.097 | 0 | 0.06 | 0.29 | 0.1 | 0.1 | 0.01 | 0.08 |
| 11 | 12155083 | rs977244 | -0.018 | -0.797 | -0.112 | 0.359 | -0.005 | -0.002 | -0.002 | 0 | 0.47 | 0.48 | 0.46 | 0.39 | 0.41 | 0.46 |
| 11 | 12155701 | rs10831740 | 0.515 | -0.838 | -0.235 | 0.454 | -0.001 | 0.039 | 0.039 | 0 | 0.39 | 0.18 | 0.35 | 0.28 | 0.4 | 0.37 |
| 11 | 12156414 | rs16910593 | 0.536 | -0.833 | 0.195 | 0.235 | 0.005 | -0.004 | -0.004 | 0 | 0.33 | 0.38 | 0.25 | 0.33 | 0.35 | 0.21 |
| 11 | 12156853 | rs1485957 | 0.534 | -0.838 | 0.193 | 0.228 | 0.004 | -0.004 | -0.004 | 0 | 0.33 | 0.38 | 0.25 | 0.33 | 0.35 | 0.21 |
| 11 | 12157657 | rs2171150 | -0.301 | -0.822 | 0.242 | 0.203 | 0 | 0 | 0 | 1.206 | 0.61 | 0.43 | 0.66 | 0.56 | 0.64 | 0.7 |
| 11 | 12159098 | rs9971381 | 0.314 | -0.829 | 0.084 | 0.056 | 0.009 | -0.027 | -0.027 | 0.521 | 0.4 | 0.55 | 0.29 | 0.31 | 0.39 | 0.23 |
| 11 | 12159661 | rs1994318 | 0.693 | -0.954 | -0.273 | 0.612 | 0.032 | -0.027 | -0.027 | 0 | 0.61 | 0.28 | 0.4 | 0.52 | 0.64 | 0.4 |
| 11 | 12160054 | rs901285 | -0.333 | -0.46 | -0.06 | 0.482 | -0.005 | -0.023 | -0.023 | 0 | 0.1 | 0.15 | 0.19 | 0.04 | 0.12 | 0.17 |
| 11 | 12161625 | rs11022193 | 0.414 | -0.31 | -0.299 | 0.799 | 0.001 | -0.001 | -0.001 | 0 | 0.11 | 0.08 | 0.12 | 0.23 | 0.09 | 0.07 |
| 11 | 12163479 | rs4757227 | -0.184 | -0.022 | -0.301 | 0.284 | -0.001 | -0.015 | -0.015 | 0 | 0.44 | 0.43 | 0.49 | 0.31 | 0.44 | 0.21 |
| 11 | 12166233 | rs988189 | 1.124 | 0.663 | -1.041 | 0.883 | -0.08 | -0.084 | -0.084 | 0 | 0.14 | 0.01 | 0.17 | 0.27 | 0.23 | 0.14 |
| 11 | 12169301 | rs4644628 | 0.047 | -0.268 | -0.327 | 0.581 | 0.002 | 0.02 | 0.02 | 0.614 | 0.45 | 0.31 | 0.51 | 0.61 | 0.54 | 0.68 |
| 11 | 12169820 | rs4756775 | -0.987 | -0.701 | 0.391 | 1.194 | -0.032 | 0.01 | 0.01 | 0.585 | 0.25 | 0.67 | 0.16 | 0.36 | 0.13 | 0.21 |
| 11 | 12170325 | rs10831742 | -1.115 | -0.525 | 0.687 | 1.335 | -0.046 | 0.039 | 0.039 | 0 | 0.29 | 0.52 | 0.16 | 0.63 | 0.16 | 0.28 |
| 11 | 12170617 | rs7102041 | 1.116 | 0.313 | 1.263 | 0.734 | 0.005 | -0.075 | -0.075 | 0 | 0.88 | 0.73 | 0.93 | 1 | 0.9 | 0.94 |
| 11 | 12172059 | rs9804570 | -1.918 | 0.949 | 1.618 | 1.042 | 0.035 | -0.024 | -0.024 | 0 | 0.17 | 0.29 | 0.5 | 0.1 | 0.28 | 0.15 |
| 11 | 12174566 | rs12270057 | -0.675 | 1.112 | 1.645 | 1.623 | -0.035 | -0.005 | -0.005 | 0.014 | 0.11 | 0.04 | 0.44 | 0.1 | 0.23 | 0.16 |
| 11 | 12176273 | rs1564946 | 1.939 | 0.828 | 1.898 | 1.629 | 0.018 | 0.152 | 0.152 | 0 | 0.93 | 0.79 | 0.96 | 0.75 | 0.93 | 0.94 |
| 11 | 12183036 | rs7131034 | 1.321 | 0.77 | 2.068 | 1.887 | 0.049 | 0.133 | 0.133 | 0.204 | 0.84 | 0.69 | 0.77 | 0.61 | 0.78 | 0.8 |
| 11 | 12185469 | rs901302 | 1.191 | 0.725 | 2.213 | 1.846 | 0.024 | 0.279 | 0.279 | 0 | 0.93 | 0.73 | 0.91 | 0.71 | 0.94 | 0.94 |
| 11 | 12185801 | rs2016398 | -0.477 | 0.911 | 2.238 | 2.014 | -0.061 | -0.031 | -0.031 | 0 | 0.1 | 0.01 | 0.2 | 0.1 | 0.2 | 0.13 |
| 11 | 12186296 | rs2010463 | -2.164 | 0.532 | 1.789 | 1.482 | -0.001 | -0.001 | -0.001 | 0 | 0.39 | 0.65 | 0.5 | 0.57 | 0.4 | 0.62 |
| 11 | 12189013 | rs10831744 | 1.463 | 0.561 | 2.242 | 1.558 | 0.019 | 0.093 | 0.093 | 0.214 | 0.81 | 0.37 | 0.72 | 0.62 | 0.79 | 0.8 |
| 11 | 12194228 | rs4471395 | -0.786 | 0.438 | 2.169 | 1.751 | -0.003 | 0.145 | 0.145 | 0 | 0.12 | 0.24 | 0.09 | 0.45 | 0.08 | 0.35 |

| | | | | | | | | | | | | | | | | |
|---|---|---|---|---|---|---|---|---|---|---|---|---|---|---|---|---|
| 11 | 12194961 | rs7950540 | 0.213 | 0.864 | 2.063 | 1.434 | 0.023 | -0.033 | -0.033 | 1.385 | 0.06 | 0.18 | 0.06 | 0 | 0.07 | 0.01 |
| 11 | 12197358 | rs7121956 | 1.93 | 0.774 | 2.195 | 1.403 | 0.011 | 0.013 | 0.013 | 0 | 0.65 | 0.47 | 0.49 | 0.59 | 0.6 | 0.68 |
| 11 | 12199308 | rs7117024 | 1.094 | 0.818 | 2.067 | 1.434 | -0.008 | -0.014 | -0.014 | 0 | 0.74 | 0.8 | 0.58 | 0.6 | 0.67 | 0.69 |
| 11 | 12199666 | rs12293251 | -0.507 | 0.887 | 2.161 | 1.536 | 0.011 | 0.198 | 0.198 | 0 | 0.12 | 0.31 | 0.09 | 0.44 | 0.08 | 0.35 |
| 11 | 12202276 | rs7130607 | 1.406 | 0.899 | 2.018 | 1.181 | 0.05 | 0.068 | 0.068 | 0 | 0.78 | 0.63 | 0.7 | 0.61 | 0.67 | 0.81 |
| 11 | 12204581 | rs7949360 | -1.627 | 0.842 | 1.964 | 1.038 | -0.012 | 0.016 | 0.016 | 0 | 0.24 | 0.18 | 0.42 | 0.11 | 0.32 | 0.26 |
| 11 | 12205793 | rs901291 | -2.373 | 0.782 | 1.913 | 0.959 | 0.021 | -0.054 | -0.054 | 1.596 | 0.34 | 0.5 | 0.49 | 0.16 | 0.4 | 0.27 |
| 11 | 12208393 | rs6485561 | -1.933 | 0.479 | 2.033 | 0.917 | -0.002 | -0.124 | -0.124 | 0 | 0.23 | 0.52 | 0.3 | 0.05 | 0.2 | 0.15 |
| 11 | 12216248 | rs17477837 | 0.085 | 0.842 | 1.328 | 1.152 | -0.004 | 0.031 | 0.031 | 0 | 0.08 | 0.03 | 0.19 | 0 | 0.08 | 0.1 |
| 11 | 12218497 | rs10831758 | -1.395 | 0.737 | 1.673 | 1.42 | -0.007 | 0.087 | 0.087 | 0 | 0.22 | 0.59 | 0.32 | 0.43 | 0.19 | 0.39 |
| 11 | 12219217 | rs1032151 | -1.971 | 0.552 | 1.513 | 1.32 | -0.001 | 0.015 | 0.015 | 0 | 0.3 | 0.69 | 0.46 | 0.53 | 0.29 | 0.42 |
| 11 | 12219904 | rs17477991 | -0.874 | 0.527 | 1.055 | 1.17 | -0.057 | 0.04 | 0.04 | 0 | 0.25 | 0.04 | 0.23 | 0.12 | 0.32 | 0.17 |
| 11 | 12221382 | rs10831759 | 0.297 | 0.547 | 1.303 | 1.185 | 0.01 | 0.012 | 0.012 | 0.665 | 0.08 | 0.18 | 0.16 | 0.09 | 0.1 | 0.03 |
| 11 | 12227844 | rs12283453 | -1.526 | 0.507 | 1.292 | 1.171 | -0.021 | 0.019 | 0.019 | 0 | 0.33 | 0.22 | 0.38 | 0.21 | 0.41 | 0.2 |
| 11 | 12229116 | rs871703 | -2.084 | 0.431 | 1.653 | 1.256 | -0.015 | 0.02 | 0.02 | 0 | 0.42 | 0.72 | 0.48 | 0.65 | 0.47 | 0.48 |
| 11 | 12229437 | rs16910765 | -0.626 | 0.421 | 1.622 | 1.309 | 0.042 | 0.401 | 0.401 | 0 | 0.08 | 0.41 | 0.09 | 0.44 | 0.06 | 0.29 |
| 11 | 12229655 | rs2279390 | -1.577 | 0.477 | 1.59 | 1.266 | 0.047 | 0.182 | 0.182 | 0 | 0.21 | 0.49 | 0.25 | 0.5 | 0.26 | 0.44 |
| 11 | 12230672 | rs12807026 | -0.673 | 0.436 | 1.172 | 0.99 | 0.002 | -0.002 | -0.002 | 2.21 | 0.14 | 0.11 | 0.15 | 0.06 | 0.2 | 0.16 |
| 11 | 12231277 | rs12787479 | -0.391 | 0.48 | 1.063 | 1.013 | -0.009 | 0.005 | 0.005 | 0 | 0.13 | 0.08 | 0.15 | 0.06 | 0.2 | 0.16 |
| 11 | 12231732 | rs12788362 | -1.025 | 0.415 | 0.93 | 1.044 | -0.05 | 0.039 | 0.039 | 0 | 0.25 | 0.05 | 0.22 | 0.12 | 0.32 | 0.17 |
| 11 | 12233728 | rs12576744 | -1.142 | 0.273 | 1.55 | 1.548 | -0.008 | 0.301 | 0.301 | 0 | 0.1 | 0.35 | 0.09 | 0.47 | 0.06 | 0.29 |
| 11 | 12237135 | rs2012580 | 0.618 | 0.622 | 1.121 | 1.74 | -0.004 | 0.095 | 0.095 | 0.023 | 0.87 | 0.79 | 0.81 | 0.56 | 0.88 | 0.9 |
| 11 | 12238025 | rs2306728 | 0.093 | 0.692 | 1.031 | 1.704 | 0.001 | 0.081 | 0.081 | 0.594 | 0.89 | 0.8 | 0.82 | 0.64 | 0.89 | 0.9 |
| 11 | 12238115 | rs2306729 | -0.062 | 0.73 | 0.835 | 1.505 | 0 | 0 | 0 | 0 | 0.09 | 0.1 | 0.16 | 0.09 | 0.1 | 0.04 |
| 11 | 12239114 | rs11827631 | -1.057 | 0.782 | 0.922 | 1.098 | -0.014 | 0.031 | 0.031 | 0 | 0.25 | 0.15 | 0.21 | 0.06 | 0.33 | 0.16 |
| 11 | 12239326 | rs11827638 | -1.06 | 0.805 | 0.93 | 1.108 | -0.015 | 0.031 | 0.031 | 0 | 0.25 | 0.15 | 0.21 | 0.06 | 0.33 | 0.16 |
| 11 | 12244468 | rs10831769 | -1.669 | 0.778 | 1.192 | 1.012 | 0.003 | -0.011 | -0.011 | 0.783 | 0.43 | 0.43 | 0.64 | 0.66 | 0.46 | 0.55 |
| 11 | 12248515 | rs2706643 | -1.052 | 1.083 | 1.738 | 1.406 | 0.084 | 0.042 | 0.042 | 0.057 | 0.13 | 0.32 | 0.16 | 0.16 | 0.17 | 0.12 |
| 11 | 12251732 | rs2641940 | -0.348 | 1.135 | 1.146 | 1.412 | 0.01 | 0.001 | 0.001 | 0.018 | 0.1 | 0.09 | 0.12 | 0.1 | 0.15 | 0.12 |
| 11 | 12253424 | rs1609930 | -1.095 | 0.959 | 1.482 | 1.569 | 0.12 | 0.272 | 0.272 | 0.087 | 0.12 | 0.42 | 0.15 | 0.4 | 0.17 | 0.13 |
| 11 | 12253548 | rs2131771 | -1.334 | 0.619 | 0.268 | 1.362 | -0.003 | -0.111 | -0.111 | 0.155 | 0.3 | 0.09 | 0.49 | 0.5 | 0.29 | 0.43 |
| 11 | 12255546 | rs2010576 | 1.619 | 0.588 | 0.408 | 1.261 | -0.006 | 0.003 | 0.003 | 0 | 0.6 | 0.71 | 0.38 | 0.41 | 0.57 | 0.46 |
| 11 | 12257458 | rs2246778 | -1.857 | 0.735 | 0.725 | 1.688 | -0.002 | -0.002 | -0.002 | 1.042 | 0.26 | 0.24 | 0.55 | 0.65 | 0.28 | 0.52 |
| 11 | 12257976 | rs3794083 | -0.602 | 0.756 | 0.298 | 1.731 | 0.003 | -0.051 | -0.051 | 0 | 0.15 | 0.08 | 0.42 | 0.53 | 0.14 | 0.41 |
| 11 | 12259447 | rs7939601 | 1.32 | 0.741 | 0.365 | 1.566 | 0.005 | 0.006 | 0.006 | 0 | 0.73 | 0.71 | 0.44 | 0.35 | 0.71 | 0.5 |
| 11 | 12260355 | rs11022262 | -1.177 | 0.75 | -0.468 | 0.884 | -0.063 | 0.036 | 0.036 | 0 | 0.19 | 0.02 | 0.2 | 0.09 | 0.25 | 0.14 |
| 11 | 12260712 | rs11604904 | -1.146 | 0.725 | -0.43 | 0.873 | -0.014 | 0.037 | 0.037 | 0 | 0.19 | 0.07 | 0.17 | 0.08 | 0.22 | 0.14 |
| 11 | 12262410 | rs2279613 | -1.805 | 0.709 | -0.217 | 1.502 | -0.016 | 0.002 | 0.002 | 0 | 0.33 | 0.2 | 0.44 | 0.28 | 0.39 | 0.42 |
| 11 | 12263047 | rs2279614 | -0.486 | 0.676 | -0.202 | 1.538 | 0.002 | 0.005 | 0.005 | 0 | 0.14 | 0.13 | 0.27 | 0.21 | 0.17 | 0.28 |
| 11 | 12264140 | rs880963 | 2.05 | 0.442 | -0.042 | 2.117 | -0.006 | 0 | 0 | 0 | 0.58 | 0.67 | 0.34 | 0.36 | 0.55 | 0.52 |
| 11 | 12265083 | rs12294434 | 0.249 | 0.442 | -0.221 | 2.064 | -0.003 | -0.003 | -0.003 | 0 | 0.07 | 0.02 | 0.09 | 0.06 | 0.08 | 0.06 |
| 11 | 12265542 | rs2270511 | -0.457 | 0.47 | -0.182 | 2.103 | 0.002 | 0.005 | 0.005 | 0 | 0.15 | 0.16 | 0.27 | 0.21 | 0.17 | 0.28 |
| 11 | 12266498 | rs12574429 | -0.687 | 0.521 | -0.186 | 1.715 | 0 | 0.019 | 0.019 | 0 | 0.17 | 0.1 | 0.17 | 0.05 | 0.19 | 0.11 |
| 11 | 12269885 | rs2706627 | 0.711 | 0.372 | -0.118 | 1.731 | 0.003 | 0.069 | 0.069 | 0 | 0.94 | 0.9 | 0.84 | 0.64 | 0.95 | 0.92 |
| 11 | 12271440 | rs1973386 | 1.828 | 0.072 | 0.172 | 1.792 | -0.001 | -0.021 | -0.021 | 0.021 | 0.54 | 0.56 | 0.37 | 0.13 | 0.55 | 0.4 |
| 11 | 12273566 | rs7946327 | 1.435 | 0.087 | 0.036 | 1.877 | 0.002 | 0.002 | 0.002 | 0 | 0.58 | 0.61 | 0.53 | 0.48 | 0.6 | 0.47 |
| 11 | 12274529 | rs2706660 | 0.936 | 0.137 | -0.165 | 2.091 | -0.004 | -0.022 | -0.022 | 0.609 | 0.78 | 0.85 | 0.69 | 0.58 | 0.76 | 0.8 |
| 11 | 12277705 | rs11022264 | -1.134 | -0.24 | -0.504 | 1.968 | 0.007 | -0.024 | -0.024 | 0 | 0.19 | 0.15 | 0.19 | 0.34 | 0.15 | 0.28 |
| 11 | 12277822 | rs17480838 | -0.155 | -0.185 | -0.675 | 2.009 | 0.007 | 0.005 | 0.005 | 0 | 0.1 | 0.06 | 0.07 | 0.07 | 0.09 | 0.15 |
| 11 | 12279260 | rs7116182 | -1.193 | -0.005 | -0.164 | 2.272 | 0.002 | 0.108 | 0.108 | 0 | 0.09 | 0.2 | 0.13 | 0.27 | 0.06 | 0.13 |
| 11 | 12279937 | rs2270513 | -1.578 | -0.212 | -0.325 | 2.112 | -0.002 | 0.02 | 0.02 | 0 | 0.32 | 0.33 | 0.44 | 0.5 | 0.33 | 0.44 |
| 11 | 12282908 | rs3794075 | -0.855 | -0.194 | -0.75 | 1.504 | 0.001 | 0.005 | 0.005 | 0 | 0.22 | 0.14 | 0.29 | 0.16 | 0.25 | 0.29 |
| 11 | 12283928 | rs10765933 | -0.732 | -0.277 | -0.728 | 1.565 | 0 | -0.002 | -0.002 | 0 | 0.21 | 0.16 | 0.29 | 0.23 | 0.23 | 0.32 |
| 11 | 12284498 | rs3812747 | -0.746 | -0.099 | -0.773 | 1.345 | -0.001 | 0.001 | 0.001 | 0 | 0.21 | 0.16 | 0.28 | 0.16 | 0.24 | 0.22 |
| 11 | 12284925 | rs2279616 | 0.089 | -0.555 | -0.978 | 1.581 | 0.001 | -0.001 | -0.001 | 0.013 | 0.82 | 0.83 | 0.82 | 0.75 | 0.85 | 0.77 |
| 11 | 12285090 | rs8808 | -0.564 | -1.107 | -1.344 | 2.031 | 0.03 | -0.091 | -0.091 | 0.649 | 0.15 | 0.04 | 0.1 | 0.48 | 0.11 | 0.17 |
| 11 | 12285952 | rs903208 | 0.513 | -0.766 | -1.914 | 1.104 | 0.007 | 0.024 | 0.024 | 0 | 0.49 | 0.1 | 0.34 | 0.61 | 0.57 | 0.49 |
| 11 | 12289058 | rs2706652 | -0.492 | -0.498 | -0.384 | 0.873 | 0.002 | -0.001 | -0.001 | 0.542 | 0.35 | 0.37 | 0.54 | 0.64 | 0.35 | 0.4 |
| 11 | 12289881 | rs11022267 | -0.767 | -0.716 | -0.417 | 0.311 | -0.001 | -0.007 | -0.007 | 0.005 | 0.3 | 0.3 | 0.38 | 0.14 | 0.26 | 0.27 |
| 11 | 12290812 | rs7121541 | 1.273 | -1.098 | -0.697 | -0.446 | -0.055 | 0.11 | 0.11 | 0 | 0.39 | 0.17 | 0.25 | 0.15 | 0.49 | 0.41 |
| 11 | 12291805 | rs963012 | 0.235 | -1.385 | -0.552 | -0.382 | -0.012 | 0.034 | 0.034 | 0 | 0.63 | 0.71 | 0.41 | 0.26 | 0.64 | 0.55 |
| 11 | 12293261 | rs7123104 | 0.402 | -1.333 | -0.482 | -0.602 | 0.011 | 0.029 | 0.029 | 0 | 0.62 | 0.65 | 0.43 | 0.26 | 0.65 | 0.55 |
| 11 | 12294363 | rs16910954 | -0.277 | -1.192 | 0.186 | -0.308 | -0.005 | 0.039 | 0.039 | 0.036 | 0.07 | 0.2 | 0.07 | 0.09 | 0.05 | 0.07 |
| 11 | 12294609 | rs11607178 | -0.87 | -1.189 | 0.266 | -0.368 | 0.002 | -0.027 | -0.027 | 1.884 | 0.17 | 0.27 | 0.14 | 0.09 | 0.15 | 0.11 |
| 11 | 12296409 | rs2706646 | -1.134 | -1.347 | 0.186 | -0.041 | -0.006 | -0.007 | -0.007 | 0 | 0.36 | 0.48 | 0.45 | 0.25 | 0.29 | 0.36 |
| 11 | 12296781 | rs7106205 | -1.842 | -1.782 | 0.394 | -0.602 | 0.007 | 0.01 | 0.01 | 0 | 0.49 | 0.5 | 0.65 | 0.82 | 0.4 | 0.54 |
| 11 | 12296810 | rs10765937 | -1.552 | -1.955 | 0.725 | -0.135 | 0.005 | 0.006 | 0.006 | 0 | 0.4 | 0.39 | 0.56 | 0.78 | 0.32 | 0.49 |
| 11 | 12298921 | rs1125742 | -0.418 | -2.089 | 0.508 | -0.012 | -0.012 | -0.018 | -0.018 | 0 | 0.26 | 0.13 | 0.47 | 0.7 | 0.28 | 0.45 |
| 11 | 12298990 | rs1125744 | -1.705 | -2.282 | 0.97 | -0.108 | 0.015 | 0.029 | 0.029 | 0 | 0.4 | 0.68 | 0.58 | 0.77 | 0.34 | 0.51 |

| | | | | | | | | | | | | | | | | |
|---|---|---|---|---|---|---|---|---|---|---|---|---|---|---|---|---|
| 11 | 12300654 | rs4757330 | -0.339 | -2.135 | 0.627 | -0.036 | -0.013 | -0.021 | -0.021 | 0 | 0.26 | 0.13 | 0.48 | 0.7 | 0.28 | 0.45 |
| 11 | 12301823 | rs7947131 | -0.292 | -2.192 | 0.657 | 0.385 | -0.011 | -0.048 | -0.048 | 0 | 0.24 | 0.16 | 0.47 | 0.57 | 0.27 | 0.4 |
| 11 | 12302760 | rs7940285 | -0.635 | -2.173 | 0.619 | 0.322 | -0.009 | -0.015 | -0.015 | 1.962 | 0.27 | 0.17 | 0.49 | 0.7 | 0.3 | 0.48 |
| 11 | 12303354 | rs4757332 | -1.725 | -2.118 | 0.829 | 0.239 | 0.006 | 0.008 | 0.008 | 0 | 0.39 | 0.42 | 0.55 | 0.78 | 0.33 | 0.52 |
| 11 | 12305051 | rs10741577 | -1.586 | -2.088 | 0.707 | 0.224 | 0.002 | 0.002 | 0.002 | 0 | 0.4 | 0.62 | 0.57 | 0.76 | 0.33 | 0.52 |
| 11 | 12306437 | rs4756813 | 1.486 | -2.41 | -0.01 | -0.019 | 0.065 | 0.075 | 0.075 | 0 | 0.52 | 0.22 | 0.39 | 0.3 | 0.64 | 0.44 |
| 11 | 12308500 | rs903209 | 0.346 | -2.461 | 0.332 | -0.026 | -0.04 | -0.062 | -0.062 | 0 | 0.81 | 0.69 | 0.92 | 0.93 | 0.91 | 0.86 |
| 11 | 12309002 | rs3736304 | 0.157 | -2.511 | 0.111 | 0.144 | -0.011 | 0.075 | 0.075 | 0 | 0.3 | 0.48 | 0.45 | 0.52 | 0.29 | 0.39 |
| 11 | 12309866 | rs7122652 | 0.196 | -2.469 | 0.146 | 0.064 | -0.048 | -0.072 | -0.072 | 0 | 0.8 | 0.64 | 0.91 | 0.93 | 0.91 | 0.85 |
| 11 | 12311730 | rs11606810 | -0.362 | -2.337 | 0.171 | 0.136 | -0.024 | -0.051 | -0.051 | 0.22 | 0.09 | 0.22 | 0.03 | 0.01 | 0.02 | 0.03 |
| 11 | 12312280 | rs1994902 | -1.289 | -2.373 | 0.171 | 0.176 | -0.027 | -0.016 | -0.016 | 0 | 0.14 | 0.25 | 0.14 | 0.09 | 0.05 | 0.07 |
| 11 | 12312347 | rs1389548 | 0.069 | -2.428 | 0.28 | 0.251 | 0 | 0.047 | 0.047 | 0 | 0.29 | 0.38 | 0.44 | 0.5 | 0.3 | 0.39 |
| 11 | 12312550 | rs1389549 | -1.277 | -2.322 | -0.048 | 0.331 | 0.027 | 0.019 | 0.019 | 0 | 0.45 | 0.71 | 0.6 | 0.6 | 0.37 | 0.48 |
| 11 | 12314551 | rs7127178 | -0.826 | -2.262 | -0.03 | 0.359 | 0.001 | -0.002 | -0.002 | 0 | 0.4 | 0.65 | 0.48 | 0.51 | 0.34 | 0.44 |
| 11 | 12315186 | rs10741578 | -0.915 | -2.208 | -0.155 | 0.387 | 0.01 | 0.006 | 0.006 | 0 | 0.43 | 0.64 | 0.59 | 0.6 | 0.37 | 0.47 |
| 11 | 12315891 | rs1493953 | -1.124 | -2.069 | -0.451 | -0.036 | 0 | 0 | 0 | 0 | 0.89 | 0.91 | 0.95 | 0.94 | 0.95 | 0.88 |
| 11 | 12317933 | rs11022280 | 1.421 | -1.96 | -0.594 | 0.037 | -0.014 | 0.139 | 0.139 | 0 | 0.43 | 0.13 | 0.34 | 0.18 | 0.55 | 0.35 |
| 11 | 12327308 | rs11022285 | -0.934 | -1.878 | -0.128 | 0.522 | -0.015 | 0.007 | 0.007 | 0 | 0.36 | 0.57 | 0.32 | 0.63 | 0.28 | 0.31 |
| 11 | 12336668 | rs7937533 | 0.481 | -1.901 | 0.133 | 0.676 | -0.003 | 0.101 | 0.101 | 0 | 0.2 | 0.35 | 0.18 | 0.42 | 0.21 | 0.2 |
| 11 | 12337522 | rs6485652 | 0.885 | -2.254 | 0.051 | 0.489 | 0.046 | -0.022 | -0.022 | 0 | 0.61 | 0.3 | 0.75 | 0.53 | 0.72 | 0.65 |
| 11 | 12341061 | rs7130018 | -0.497 | -2.195 | 0.2 | 0.518 | -0.016 | 0.03 | 0.03 | 0 | 0.29 | 0.53 | 0.19 | 0.42 | 0.23 | 0.19 |
| 11 | 12342923 | rs10831787 | 1.064 | -2.065 | 0.177 | 0.795 | 0 | 0.002 | 0.002 | 0 | 0.1 | 0.12 | 0.26 | 0.13 | 0.09 | 0.23 |
| 11 | 12343617 | rs7934505 | 1.006 | -2.337 | 0.281 | 0.714 | -0.002 | 0 | 0 | 0 | 0.61 | 0.45 | 0.63 | 0.31 | 0.69 | 0.58 |
| 11 | 12344362 | rs1493960 | 0.548 | -2.273 | 0.218 | 0.679 | -0.026 | 0.007 | 0.007 | 0 | 0.8 | 0.75 | 0.79 | 0.73 | 0.88 | 0.74 |
| 11 | 12345459 | rs903213 | -1.216 | -2.043 | 0.171 | 0.359 | -0.008 | -0.007 | -0.007 | 0.48 | 0.9 | 0.9 | 0.95 | 0.94 | 0.95 | 0.87 |
| 11 | 12347397 | rs7931843 | 0.989 | -1.963 | 0.146 | 0.438 | 0.028 | 0.028 | 0.028 | 0 | 0.55 | 0.41 | 0.4 | 0.33 | 0.64 | 0.43 |
| 11 | 12353977 | rs10831789 | 0.803 | -1.869 | 0.048 | 0.449 | -0.003 | 0.027 | 0.027 | 0 | 0.1 | 0.16 | 0.26 | 0.15 | 0.08 | 0.21 |
| 11 | 12362700 | rs4501962 | 0.055 | -1.984 | 0.098 | 0.288 | -0.027 | 0.019 | 0.019 | 0 | 0.63 | 0.56 | 0.66 | 0.44 | 0.71 | 0.63 |
| 11 | 12364308 | rs11022316 | 0.878 | -1.805 | -0.293 | 0.359 | 0.001 | 0.024 | 0.024 | 0 | 0.1 | 0.16 | 0.26 | 0.15 | 0.09 | 0.21 |
| 11 | 12365147 | rs2055102 | 0.954 | -1.75 | -0.68 | 0.295 | 0.071 | 0.125 | 0.125 | 0 | 0.49 | 0.21 | 0.37 | 0.29 | 0.61 | 0.39 |
| 11 | 12367459 | rs10831796 | 0.569 | -1.506 | -0.449 | 0.264 | -0.003 | 0.005 | 0.005 | 0 | 0.13 | 0.13 | 0.26 | 0.15 | 0.09 | 0.23 |
| 11 | 12367700 | rs10831797 | 0.827 | -0.733 | -0.956 | -0.091 | -0.1 | 0.017 | 0.017 | 0 | 0.37 | 0.12 | 0.28 | 0.32 | 0.48 | 0.35 |
| 11 | 12368388 | rs7113079 | -0.086 | -0.152 | -0.33 | 0.272 | -0.003 | 0.009 | 0.009 | 0 | 0.06 | 0.15 | 0.13 | 0.07 | 0.03 | 0.03 |
| 11 | 12368559 | rs16911142 | 0.127 | -0.115 | -0.497 | 0.263 | -0.005 | 0.009 | 0.009 | 0.018 | 0.05 | 0.01 | 0.12 | 0.05 | 0.04 | 0.04 |
| 11 | 12368645 | rs4757381 | 1.242 | -0.309 | -0.626 | 0.281 | -0.038 | -0.002 | -0.002 | 0 | 0.41 | 0.18 | 0.28 | 0.38 | 0.47 | 0.34 |
| 11 | 12369397 | rs4625438 | -0.071 | -0.414 | 0.357 | 0.543 | 0.038 | 0.146 | 0.146 | 0 | 0.14 | 0.3 | 0.17 | 0.4 | 0.19 | 0.16 |
| 11 | 12370967 | rs4237713 | -0.26 | -0.588 | 0.186 | 0.088 | 0.035 | 0.059 | 0.059 | 0 | 0.68 | 0.81 | 0.68 | 0.89 | 0.81 | 0.59 |
| 11 | 12371595 | rs7951657 | -0.582 | -0.399 | 0.445 | 0.176 | 0.065 | 0.277 | 0.277 | 0 | 0.17 | 0.48 | 0.21 | 0.5 | 0.21 | 0.21 |
| 11 | 12373680 | rs16911154 | 0.064 | -0.666 | 0.438 | 0.159 | -0.013 | -0.006 | -0.006 | 2.671 | 0.05 | 0.16 | 0.09 | 0.03 | 0.01 | 0.02 |
| 11 | 12373851 | rs10831800 | 0.774 | -0.592 | 0.272 | 0.225 | -0.003 | -0.004 | -0.004 | 0.801 | 0.52 | 0.51 | 0.49 | 0.38 | 0.6 | 0.4 |
| 11 | 12375443 | rs11022320 | -0.017 | -0.804 | 0.015 | -0.714 | 0.023 | 0.027 | 0.027 | 0 | 0.67 | 0.8 | 0.76 | 0.76 | 0.75 | 0.58 |
| 11 | 12375529 | rs7101744 | 0.861 | -0.808 | -0.134 | -1.095 | 0.104 | 0.173 | 0.173 | 0.001 | 0.54 | 0.16 | 0.63 | 0.77 | 0.7 | 0.48 |
| 11 | 12377438 | rs10741582 | 0.651 | -0.669 | 0.734 | -0.766 | 0.02 | 0.019 | 0.019 | 6.598 | 0.74 | 0.23 | 0.88 | 0.93 | 0.88 | 0.8 |
| 11 | 12377766 | rs7129480 | -0.29 | -0.595 | 0.983 | -0.729 | -0.023 | -0.029 | -0.029 | 0 | 0.82 | 0.76 | 0.93 | 0.93 | 0.9 | 0.84 |
| 11 | 12384466 | rs6485683 | -0.288 | -0.496 | -0.274 | -0.726 | -0.015 | -0.023 | -0.023 | 0.817 | 0.82 | 0.79 | 0.94 | 0.93 | 0.9 | 0.84 |
| 11 | 12393940 | rs12365064 | -0.007 | 0.044 | -0.909 | -0.593 | -0.05 | -0.014 | -0.014 | 0 | 0.09 | 0.01 | 0.1 | 0.05 | 0.14 | 0.07 |
| 11 | 12397634 | rs10734193 | 0.556 | -0.47 | -0.577 | -0.641 | 0.064 | 0.09 | 0.09 | 6.221 | 0.73 | 0.17 | 0.86 | 0.91 | 0.84 | 0.79 |
| 11 | 12398611 | rs4757405 | 0.387 | -0.463 | -0.427 | -0.657 | -0.001 | -0.013 | -0.013 | 0 | 0.76 | 0.28 | 0.88 | 0.92 | 0.86 | 0.82 |
| 11 | 12403836 | rs10219167 | 0.487 | -0.251 | -0.691 | -0.609 | 0.081 | 0.142 | 0.142 | 4.999 | 0.69 | 0.13 | 0.77 | 0.91 | 0.8 | 0.77 |
| 11 | 12415077 | rs4757420 | -1.889 | 1.071 | 0.629 | 0.184 | 0.042 | -0.038 | -0.038 | 0.26 | 0.15 | 0.41 | 0.1 | 0.1 | 0.17 | 0.13 |
| 11 | 12419452 | rs960806 | -1.786 | 0.869 | 0.805 | 0.226 | 0.04 | -0.031 | -0.031 | 0 | 0.09 | 0.52 | 0.06 | 0 | 0.07 | 0.03 |
| 11 | 12422990 | rs12275332 | -1.198 | 1.007 | 0.697 | 0.294 | 0.094 | 0.02 | 0.02 | 0.253 | 0.07 | 0.45 | 0.06 | 0 | 0.07 | 0.03 |
| 11 | 12423832 | rs11022340 | -0.862 | 1.394 | -0.596 | 1.086 | -0.027 | -0.065 | -0.065 | 0.086 | 0.16 | 0.06 | 0.18 | 0.53 | 0.22 | 0.19 |
| 11 | 12426303 | rs12292762 | -1.092 | 1.352 | -0.452 | 0.209 | -0.002 | -0.05 | -0.05 | 0.562 | 0.25 | 0.62 | 0.14 | 0.09 | 0.21 | 0.19 |
| 11 | 12429376 | rs10831809 | -0.837 | 1.923 | -1.236 | 0.78 | -0.077 | -0.079 | -0.079 | 0 | 0.26 | 0.09 | 0.32 | 0.46 | 0.38 | 0.15 |
| 11 | 12429493 | rs10831810 | -0.713 | 2.182 | -1.211 | 1.061 | -0.059 | -0.083 | -0.083 | 3.459 | 0.17 | 0.07 | 0.28 | 0.49 | 0.29 | 0.13 |
| 11 | 12431361 | rs4485089 | -1.198 | 1.894 | -0.738 | 0.913 | 0.001 | 0.002 | 0.002 | 0 | 0.26 | 0.71 | 0.22 | 0.36 | 0.24 | 0.34 |
| 11 | 12434458 | rs11022344 | -1.602 | 1.662 | -0.692 | 0.67 | -0.008 | 0.017 | 0.017 | 0 | 0.46 | 0.69 | 0.38 | 0.46 | 0.47 | 0.43 |
| 11 | 12434945 | rs11022346 | -1.278 | 2.023 | -0.18 | 0.676 | 0.006 | 0.01 | 0.01 | 0 | 0.37 | 0.49 | 0.38 | 0.49 | 0.43 | 0.27 |
| 11 | 12434982 | rs11022347 | -1.192 | 1.835 | -0.055 | 0.693 | 0 | 0.011 | 0.011 | 0 | 0.37 | 0.47 | 0.37 | 0.49 | 0.41 | 0.26 |
| 11 | 12435456 | rs11022348 | -1.46 | 1.8 | 0.103 | 0.679 | 0.011 | 0.067 | 0.067 | 0 | 0.2 | 0.49 | 0.16 | 0.34 | 0.16 | 0.27 |
| 11 | 12435603 | rs4375421 | -0.449 | 1.781 | 0.094 | 0.665 | -0.007 | 0.024 | 0.024 | 0.019 | 0.25 | 0.2 | 0.26 | 0.16 | 0.33 | 0.12 |
| 11 | 12437526 | rs4450154 | -1.603 | 1.589 | 0.18 | 0.77 | 0.012 | 0.064 | 0.064 | 0 | 0.21 | 0.49 | 0.16 | 0.35 | 0.16 | 0.26 |
| 11 | 12440284 | rs1002316 | -1.427 | 1.546 | -0.225 | 0.684 | -0.008 | 0.025 | 0.025 | 0 | 0.43 | 0.63 | 0.36 | 0.23 | 0.49 | 0.34 |
| 11 | 12443223 | rs4418789 | 1.27 | 1.449 | -0.297 | 0.927 | 0.001 | -0.017 | -0.017 | 0.174 | 0.56 | 0.24 | 0.55 | 0.46 | 0.56 | 0.73 |
| 11 | 12445357 | rs10765947 | 1.171 | 1.472 | -0.11 | 0.934 | -0.019 | 0.062 | 0.062 | 0.472 | 0.86 | 0.34 | 0.89 | 0.74 | 0.92 | 0.9 |
| 11 | 12446924 | rs10831815 | 2.106 | 1.51 | -0.098 | 0.927 | -0.01 | -0.011 | -0.011 | 0 | 0.49 | 0.11 | 0.5 | 0.49 | 0.48 | 0.55 |
| 11 | 12447602 | rs7946268 | -1.835 | 1.493 | 0.34 | 0.931 | 0 | -0.001 | -0.001 | 0 | 0.43 | 0.6 | 0.43 | 0.49 | 0.45 | 0.27 |
| 11 | 12450237 | rs952251 | -1.498 | 1.646 | 0.341 | 0.916 | -0.004 | 0.005 | 0.005 | 0.147 | 0.36 | 0.61 | 0.41 | 0.48 | 0.36 | 0.24 |

| | | | | | | | | | | | | | | | | |
|---|---|---|---|---|---|---|---|---|---|---|---|---|---|---|---|---|
| 11 | 12451704 | rs1344613 | -0.304 | 1.676 | 0.236 | 0.783 | -0.037 | 0.051 | 0.051 | 0 | 0.2 | 0.13 | 0.16 | 0.02 | 0.27 | 0.22 |
| 11 | 12451820 | rs6485735 | -2.385 | 1.677 | 0.247 | 0.815 | -0.018 | 0.055 | 0.055 | 0.232 | 0.29 | 0.56 | 0.31 | 0.53 | 0.23 | 0.21 |
| 11 | 12452744 | rs885467 | -1.991 | 1.386 | 0.366 | 0.934 | -0.042 | 0.118 | 0.118 | 1.738 | 0.23 | 0.56 | 0.29 | 0.53 | 0.16 | 0.2 |
| 11 | 12453041 | rs2081612 | -0.916 | 1.408 | 0.44 | 0.877 | 0.016 | 0.186 | 0.186 | 0.24 | 0.11 | 0.51 | 0.1 | 0.25 | 0.07 | 0.08 |
| 11 | 12454046 | rs1344615 | 0.139 | 1.478 | 0.338 | 0.69 | 0.027 | -0.045 | -0.045 | 0.556 | 0.91 | 0.78 | 0.94 | 0.98 | 0.93 | 0.83 |
| 11 | 12455682 | rs884272 | -0.674 | 1.401 | 0.274 | 0.749 | 0.002 | 0.009 | 0.009 | 0 | 0.26 | 0.13 | 0.32 | 0.24 | 0.29 | 0.16 |
| 11 | 12463318 | rs10741590 | 0.901 | 1.319 | 0.736 | 0.791 | 0.008 | -0.049 | -0.049 | 0 | 0.9 | 0.43 | 0.93 | 1 | 0.95 | 0.95 |
| 11 | 12470201 | rs7930590 | -1.651 | 1.461 | 0.472 | 1.006 | 0.052 | 0.057 | 0.057 | 0 | 0.17 | 0.5 | 0.16 | 0.25 | 0.15 | 0.28 |
| 11 | 12474098 | rs4756884 | 0.193 | 1.247 | -1.039 | 1.031 | -0.07 | -0.115 | -0.115 | 0.049 | 0.26 | 0.1 | 0.52 | 0.51 | 0.33 | 0.56 |
| 11 | 12476231 | rs1365403 | 0.416 | 0.656 | -1.391 | 0.57 | 0.001 | 0.001 | 0.001 | 0 | 0.12 | 0.12 | 0.07 | 0.04 | 0.1 | 0.08 |
| 11 | 12478598 | rs10741591 | -0.144 | 0.635 | -1.383 | 0.657 | -0.05 | -0.063 | -0.063 | 0.084 | 0.36 | 0.19 | 0.57 | 0.52 | 0.41 | 0.63 |
| 11 | 12479067 | rs11022360 | 0.869 | 0.171 | -1.413 | 0.659 | 0.012 | 0.019 | 0.019 | 0.055 | 0.38 | 0.19 | 0.23 | 0.27 | 0.37 | 0.14 |
| 11 | 12482568 | rs1426576 | 0.032 | 0.069 | -1.729 | 0.807 | -0.001 | -0.072 | -0.072 | 0 | 0.38 | 0.16 | 0.32 | 0.48 | 0.34 | 0.51 |
| 11 | 12483197 | rs10831827 | -0.748 | 0.252 | -1.513 | 0.45 | 0.034 | 0.054 | 0.054 | 0 | 0.86 | 0.68 | 0.63 | 0.73 | 0.85 | 0.79 |
| 11 | 12483337 | rs12226436 | -0.035 | 0.328 | -1.621 | 0.55 | 0.003 | -0.024 | -0.024 | 0 | 0.39 | 0.16 | 0.33 | 0.56 | 0.35 | 0.54 |
| 11 | 12484144 | rs11022361 | 0.44 | 0.516 | -1.483 | 0.265 | -0.071 | 0.13 | 0.13 | 0.281 | 0.43 | 0.15 | 0.27 | 0.18 | 0.49 | 0.25 |
| 11 | 12490545 | rs1991320 | -0.927 | 0.364 | -1.22 | 0.361 | 0.01 | -0.013 | -0.013 | 2.287 | 0.86 | 0.68 | 0.89 | 0.87 | 0.88 | 0.77 |
| 11 | 12492141 | rs10831829 | 0.581 | 0.64 | -2.286 | 0.438 | -0.098 | -0.003 | -0.003 | 0 | 0.33 | 0.02 | 0.21 | 0.18 | 0.38 | 0.15 |
| 11 | 12495699 | rs2288292 | 0.487 | 0.131 | -2.068 | 0.324 | 0.017 | 0.045 | 0.045 | 0.92 | 0.24 | 0.03 | 0.15 | 0.13 | 0.18 | 0.2 |
| 11 | 12497173 | rs2403627 | -0.215 | 0.081 | -1.731 | 0.643 | 0.038 | 0.004 | 0.004 | 0 | 0.42 | 0.3 | 0.28 | 0.39 | 0.33 | 0.4 |
| 11 | 12500768 | rs4536192 | -0.717 | 0.674 | -1.752 | 0.657 | 0.007 | 0.069 | 0.069 | 0 | 0.53 | 0.36 | 0.7 | 0.79 | 0.58 | 0.61 |
| 11 | 12500833 | rs12223442 | 1.228 | 0.933 | -1.949 | 0.553 | -0.057 | 0.085 | 0.085 | 0 | 0.24 | 0.07 | 0.18 | 0.08 | 0.3 | 0.18 |
| 11 | 12517482 | rs10765955 | 0.466 | 1.343 | -0.835 | 1.494 | -0.05 | -0.045 | -0.045 | 0 | 0.1 | 0.03 | 0.41 | 0.38 | 0.24 | 0.21 |
| 11 | 12525925 | rs11547363 | 0.692 | 0.71 | -0.764 | 0.828 | 0 | 0.005 | 0.005 | 0.491 | 0.13 | 0.17 | 0.08 | 0.15 | 0.11 | 0.17 |
| 11 | 12528489 | rs4277094 | -0.315 | 1.091 | -2.08 | 1.132 | -0.108 | -0.158 | -0.158 | 0 | 0.34 | 0.04 | 0.56 | 0.54 | 0.41 | 0.4 |
| 11 | 12530688 | rs11022378 | -0.339 | 0.934 | -1.974 | 1.064 | -0.111 | -0.092 | -0.092 | 0 | 0.37 | 0.1 | 0.59 | 0.59 | 0.45 | 0.44 |
| 11 | 12533508 | rs7941378 | -1.309 | 0.51 | -1.046 | 1.371 | -0.009 | 0.035 | 0.035 | 3.404 | 0.1 | 0.15 | 0.27 | 0.17 | 0.09 | 0.22 |
| 11 | 12534707 | rs2243707 | -0.797 | 0.162 | -1.651 | 0.682 | 0.02 | 0.083 | 0.083 | 0 | 0.63 | 0.79 | 0.79 | 0.8 | 0.64 | 0.73 |
| 11 | 12535663 | rs11022381 | 0.253 | 0.228 | -1.66 | 0.686 | -0.01 | -0.031 | -0.031 | 0 | 0.05 | 0.01 | 0.24 | 0.09 | 0.08 | 0.16 |
| 11 | 12539759 | rs871939 | -0.334 | -0.123 | -0.591 | 0.557 | 0.005 | 0.005 | 0.005 | 0.292 | 0.58 | 0.45 | 0.76 | 0.75 | 0.66 | 0.56 |
| 11 | 12542882 | rs933930 | -0.293 | 0.139 | -0.577 | 1.017 | 0 | 0.02 | 0.02 | 0.969 | 0.45 | 0.62 | 0.64 | 0.66 | 0.53 | 0.61 |
| 11 | 12543952 | rs12363780 | -0.501 | 0.478 | -0.755 | 1.224 | -0.016 | 0.034 | 0.034 | 1.445 | 0.28 | 0.14 | 0.26 | 0.16 | 0.29 | 0.28 |
| 11 | 12546292 | rs10831841 | -0.687 | 0.438 | -0.502 | 1.408 | -0.004 | -0.017 | -0.017 | 0.011 | 0.31 | 0.24 | 0.34 | 0.34 | 0.29 | 0.45 |
| 11 | 12547401 | rs12366151 | -0.055 | 0.201 | -0.016 | 0.879 | 0.013 | -0.05 | -0.05 | 0 | 0.23 | 0.29 | 0.17 | 0.08 | 0.27 | 0.27 |
| 11 | 12548017 | rs11022386 | -0.373 | 0.201 | -0.322 | 1.48 | -0.045 | -0.049 | -0.049 | 0 | 0.19 | 0.12 | 0.3 | 0.38 | 0.34 | 0.26 |
| 11 | 12551844 | rs7644 | -0.472 | 0.027 | -0.24 | 1.794 | 0.016 | -0.004 | -0.004 | 1.547 | 0.2 | 0.23 | 0.3 | 0.22 | 0.34 | 0.26 |
| 11 | 12554678 | rs10765959 | 0.662 | -0.851 | -0.124 | 1.235 | 0.009 | 0 | 0 | 0 | 0.56 | 0.65 | 0.52 | 0.55 | 0.37 | 0.47 |
| 11 | 12555184 | rs11022399 | 1.23 | -1.212 | -0.972 | 0.89 | 0.084 | -0.007 | -0.007 | 0 | 0.26 | 0.04 | 0.21 | 0.16 | 0.15 | 0.15 |
| 11 | 12557016 | rs3812760 | -0.458 | 0.258 | -0.279 | 1.346 | 0.019 | 0.004 | 0.004 | 0 | 0.86 | 0.86 | 0.83 | 0.76 | 0.68 | 0.82 |
| 11 | 12559202 | rs4757597 | 0.366 | 0.206 | -0.12 | 1.45 | 0.064 | 0.096 | 0.096 | 0 | 0.79 | 0.64 | 0.63 | 0.53 | 0.66 | 0.72 |
| 11 | 12559691 | rs4756926 | 0.289 | 0.499 | -0.43 | 1.045 | 0.003 | -0.021 | -0.021 | 0.062 | 0.64 | 0.75 | 0.65 | 0.54 | 0.39 | 0.52 |
| 11 | 12561933 | rs12421465 | 0.327 | 0.684 | -1.146 | -0.439 | 0.001 | -0.021 | -0.021 | 1.232 | 0.24 | 0.21 | 0.16 | 0.08 | 0.28 | 0.3 |
| 11 | 12562720 | rs1365406 | -0.044 | -0.856 | -1.731 | -0.123 | 0.014 | -0.02 | -0.02 | 0.701 | 0.42 | 0.64 | 0.37 | 0.52 | 0.6 | 0.52 |
| 11 | 12563953 | rs10741594 | 0.468 | -0.526 | -0.894 | -0.852 | -0.013 | -0.067 | -0.067 | 0.15 | 0.71 | 0.67 | 0.8 | 0.97 | 0.81 | 0.82 |
| 11 | 12564852 | rs10831847 | 0.605 | -0.294 | -0.821 | -0.655 | -0.013 | -0.027 | -0.027 | 1.439 | 0.76 | 0.68 | 0.83 | 0.85 | 0.82 | 0.75 |
| 11 | 12565175 | rs4113950 | -0.702 | -0.08 | -1.414 | -1.379 | 0.01 | 0.136 | 0.136 | 0 | 0.35 | 0.12 | 0.39 | 0.13 | 0.31 | 0.33 |
| 11 | 12566917 | rs10831850 | 0.697 | 0.598 | -2.082 | -1.969 | -0.144 | 0.116 | 0.116 | 0 | 0.23 | 0.03 | 0.2 | 0.04 | 0.34 | 0.19 |
| 11 | 12569384 | rs4353252 | -0.284 | -0.289 | -1.744 | -1.683 | 0.016 | 0.037 | 0.037 | 0 | 0.52 | 0.66 | 0.54 | 0.59 | 0.53 | 0.58 |
| 11 | 12569519 | rs2114391 | -2.081 | -0.677 | -1.79 | -1.808 | -0.001 | 0.018 | 0.018 | 0 | 0.88 | 0.93 | 0.92 | 1 | 0.88 | 0.97 |
| 11 | 12571019 | rs7948477 | -0.134 | -0.55 | -1.256 | -1.564 | -0.013 | 0.013 | 0.013 | 0 | 0.29 | 0.63 | 0.35 | 0.54 | 0.2 | 0.4 |
| 11 | 12575039 | rs1863584 | 0.143 | -0.324 | -0.296 | -1.52 | 0.01 | 0.026 | 0.026 | 0 | 0.47 | 0.38 | 0.46 | 0.4 | 0.46 | 0.41 |
| 11 | 12577353 | rs4505050 | 0.176 | -0.158 | -0.253 | -1.614 | 0.005 | -0.011 | -0.011 | 0 | 0.24 | 0.21 | 0.3 | 0.55 | 0.18 | 0.41 |
| 11 | 12578118 | rs5023635 | -0.255 | -0.172 | -0.297 | -1.811 | 0.017 | 0.11 | 0.11 | 0 | 0.48 | 0.24 | 0.5 | 0.65 | 0.51 | 0.6 |
| 11 | 12578182 | rs6485827 | -0.985 | -0.16 | -0.158 | -1.809 | -0.024 | 0.024 | 0.024 | 0 | 0.73 | 0.78 | 0.77 | 0.95 | 0.64 | 0.82 |
| 11 | 12580178 | rs1365404 | 0.162 | -0.172 | -0.155 | -1.777 | 0.002 | -0.004 | -0.004 | 0.321 | 0.25 | 0.25 | 0.3 | 0.6 | 0.18 | 0.43 |
| 11 | 12581045 | rs7120273 | -0.972 | -0.154 | -0.12 | -1.785 | -0.011 | -0.005 | -0.005 | 0 | 0.72 | 0.77 | 0.77 | 0.95 | 0.64 | 0.82 |
| 11 | 12589751 | rs7950684 | -0.523 | -0.185 | -0.14 | -1.75 | 0.003 | -0.005 | -0.005 | 0 | 0.76 | 0.8 | 0.7 | 0.4 | 0.82 | 0.57 |
| 11 | 12589844 | rs7935247 | 1.121 | -0.241 | -0.373 | -1.953 | -0.007 | 0 | 0 | 0 | 0.26 | 0.51 | 0.24 | 0.33 | 0.26 | 0.24 |
| 11 | 12596885 | rs4757634 | 0.243 | -0.367 | -0.607 | -2.022 | -0.063 | 0.329 | 0.329 | 0 | 0.45 | 0.06 | 0.44 | 0.07 | 0.55 | 0.32 |
| 11 | 12599138 | rs4756936 | 1.028 | -0.474 | -0.409 | -1.957 | -0.092 | 0.111 | 0.111 | 0 | 0.26 | 0.06 | 0.23 | 0.04 | 0.38 | 0.18 |
| 11 | 12611849 | rs10765970 | 0.852 | -0.157 | 0.063 | -1.483 | -0.002 | 0.066 | 0.066 | 0 | 0.15 | 0.26 | 0.24 | 0.6 | 0.12 | 0.36 |
| 11 | 12616210 | rs11022428 | 0.994 | -0.223 | -0.337 | -1.651 | 0.019 | 0.084 | 0.084 | 0 | 0.16 | 0.03 | 0.19 | 0.01 | 0.14 | 0.14 |
| 11 | 12619919 | rs7124295 | 0.863 | 0.103 | 0.046 | -1.475 | -0.031 | 0.038 | 0.038 | 0 | 0.34 | 0.22 | 0.29 | 0.05 | 0.4 | 0.24 |
| 11 | 12623871 | rs10831870 | 0.496 | 0.351 | 0.424 | -0.89 | 0.044 | 0.119 | 0.119 | 0.059 | 0.16 | 0.34 | 0.23 | 0.57 | 0.18 | 0.36 |
| 11 | 12624390 | rs12360881 | -0.07 | 0.311 | 0.442 | -0.711 | -0.053 | -0.063 | -0.063 | 0.117 | 0.28 | 0.36 | 0.3 | 0.13 | 0.21 | 0.18 |
| 11 | 12624678 | rs12277767 | 0.488 | 0.527 | -0.365 | -0.558 | 0.014 | 0.064 | 0.064 | 0 | 0.16 | 0.24 | 0.23 | 0.5 | 0.18 | 0.36 |
| 11 | 12626621 | rs7121444 | -0.437 | 0.626 | -0.603 | -0.789 | -0.006 | -0.014 | -0.014 | 0 | 0.42 | 0.45 | 0.52 | 0.62 | 0.41 | 0.53 |
| 11 | 12636791 | rs9919652 | -0.127 | 0.472 | -0.666 | -0.652 | 0.005 | 0.009 | 0.009 | 0 | 0.49 | 0.48 | 0.5 | 0.6 | 0.52 | 0.6 |
| 11 | 12638232 | rs4237723 | -0.022 | 1.017 | -0.45 | -1.099 | 0.011 | 0.022 | 0.022 | 0 | 0.53 | 0.34 | 0.46 | 0.09 | 0.42 | 0.36 |

| | | | | | | | | | | | | | | | | |
|---|---|---|---|---|---|---|---|---|---|---|---|---|---|---|---|---|
| 11 | 12641391 | rs7128566 | 0.5 | 1.628 | 0.134 | 0.489 | 0.067 | 0.154 | 0.154 | 0 | 0.92 | 0.82 | 0.83 | 0.64 | 0.81 | 0.83 |
| 11 | 12643136 | rs4757702 | 0.14 | 1.214 | -0.116 | 0.023 | 0.039 | 0.01 | 0.01 | 0.256 | 0.55 | 0.35 | 0.55 | 0.57 | 0.35 | 0.68 |
| 11 | 12650458 | rs7124836 | 0.478 | 1.655 | -0.064 | 0.536 | 0.091 | 0.189 | 0.189 | 0 | 0.92 | 0.75 | 0.82 | 0.65 | 0.81 | 0.83 |
| 11 | 12652968 | rs4288736 | -0.692 | 1.164 | -0.766 | 0.149 | -0.102 | 0.138 | 0.138 | 0 | 0.63 | 0.87 | 0.72 | 0.82 | 0.52 | 0.76 |
| 11 | 12655710 | rs11022458 | -0.102 | 0.496 | -0.673 | 0.888 | 0.002 | 0.016 | 0.016 | 0 | 0.13 | 0.1 | 0.17 | 0.34 | 0.1 | 0.11 |
| 11 | 12660201 | rs6485968 | -0.341 | 0.722 | -0.34 | 0.785 | 0.063 | 0.032 | 0.032 | 0.885 | 0.5 | 0.87 | 0.55 | 0.43 | 0.4 | 0.62 |
| 11 | 12662120 | rs10831886 | 0.124 | 0.455 | 0.19 | 0.293 | 0.037 | 0.045 | 0.045 | 0.122 | 0.39 | 0.3 | 0.44 | 0.24 | 0.26 | 0.53 |
| 11 | 12679245 | rs7931934 | -0.226 | 0.463 | 0.407 | 0.376 | 0.005 | 0.048 | 0.048 | 0 | 0.44 | 0.33 | 0.41 | 0.31 | 0.44 | 0.6 |
| 11 | 12680733 | rs10831894 | -0.31 | 0.465 | 0.606 | 0.377 | 0.003 | 0.017 | 0.017 | 0 | 0.45 | 0.41 | 0.42 | 0.31 | 0.44 | 0.6 |
| 11 | 12680889 | rs4757752 | -0.678 | 1.328 | -0.202 | 0.611 | -0.069 | -0.052 | -0.052 | 0 | 0.89 | 0.99 | 0.89 | 0.78 | 0.79 | 0.82 |
| 11 | 12691007 | rs4522149 | 0.043 | 0.773 | -0.106 | 0.731 | 0.023 | 0.016 | 0.016 | 0 | 0.07 | 0 | 0.06 | 0.46 | 0.04 | 0.12 |
| 11 | 12695171 | rs6485989 | -0.392 | 0.912 | 0.405 | 0.392 | 0.022 | 0.103 | 0.103 | 0.133 | 0.48 | 0.75 | 0.45 | 0.31 | 0.44 | 0.6 |
| 11 | 12723693 | rs6486032 | -0.535 | 0.799 | 0.836 | 0.482 | 0.02 | 0.041 | 0.041 | 0.141 | 0.49 | 0.63 | 0.39 | 0.3 | 0.38 | 0.65 |
| 11 | 12759834 | rs4553350 | -0.213 | 0.643 | 1.535 | 1.028 | -0.007 | -0.043 | -0.043 | 0.317 | 0.45 | 0.14 | 0.37 | 0.48 | 0.41 | 0.66 |
| 11 | 12772213 | rs4444062 | -1.014 | 1.193 | 1.478 | 1.073 | -0.058 | -0.045 | -0.045 | 0 | 0.91 | 0.99 | 0.9 | 0.78 | 0.79 | 0.84 |
| 11 | 12776467 | rs10831905 | -0.264 | 1.06 | 1.649 | 0.995 | 0.021 | -0.008 | -0.008 | 1.034 | 0.46 | 0.21 | 0.39 | 0.49 | 0.41 | 0.66 |
| 11 | 12778769 | rs7930077 | -0.607 | 1.064 | 1.753 | 0.94 | 0.016 | 0.038 | 0.038 | 0.044 | 0.49 | 0.62 | 0.39 | 0.29 | 0.39 | 0.66 |
| 11 | 12783911 | rs4415740 | -0.757 | 1.064 | 1.651 | 0.939 | 0.036 | 0.094 | 0.094 | 0.196 | 0.5 | 0.71 | 0.4 | 0.29 | 0.39 | 0.66 |
| 11 | 12810971 | rs1866714 | 1.317 | 0.982 | 1.54 | 0.692 | -0.03 | 0.183 | 0.183 | 0.022 | 0.36 | 0.07 | 0.42 | 0 | 0.36 | 0.13 |
| 11 | 12812419 | rs10831908 | -0.17 | 0.842 | 1.676 | 0.912 | 0 | 0.122 | 0.122 | 0 | 0.45 | 0.15 | 0.35 | 0.27 | 0.39 | 0.66 |
| 11 | 12822588 | rs10500759 | -0.555 | 0.965 | 1.797 | 0.39 | 0.012 | 0.042 | 0.042 | 0 | 0.49 | 0.26 | 0.37 | 0.4 | 0.43 | 0.67 |
| 11 | 12824581 | rs6486059 | 0.351 | 0.926 | 1.8 | 0.26 | 0.015 | 0.095 | 0.095 | 0 | 0.5 | 0.33 | 0.63 | 0.8 | 0.6 | 0.34 |
| 11 | 12835952 | rs11022512 | -0.874 | 0.498 | 0.988 | -0.371 | -0.007 | 0.19 | 0.19 | 0 | 0.48 | 0.19 | 0.36 | 0.21 | 0.41 | 0.67 |
| 11 | 12839527 | rs10466408 | 0.807 | -0.096 | 0.686 | -0.552 | -0.007 | 0.017 | 0.017 | 0 | 0.17 | 0.01 | 0.11 | 0.1 | 0.1 | 0.26 |
| 11 | 12841992 | rs10466378 | -1.111 | 0.165 | 0.755 | -0.518 | -0.027 | 0.191 | 0.191 | 0 | 0.45 | 0.16 | 0.35 | 0.21 | 0.4 | 0.63 |
| 11 | 12842990 | rs4757060 | 1.309 | -0.156 | 1.003 | -0.57 | 0.015 | 0.033 | 0.033 | 0.123 | 0.49 | 0.34 | 0.57 | 0.56 | 0.57 | 0.35 |
| 11 | 12843878 | rs10765992 | 0.5 | -0.139 | 0.984 | -0.65 | 0.005 | 0.015 | 0.015 | 0 | 0.66 | 0.71 | 0.69 | 0.68 | 0.68 | 0.56 |
| 11 | 12845835 | rs17409101 | -0.505 | -0.128 | 0.821 | -1.607 | -0.008 | 0.129 | 0.129 | 0 | 0.31 | 0.15 | 0.24 | 0.12 | 0.3 | 0.44 |
| 11 | 12865441 | rs10047490 | 0.758 | 0.004 | 0.669 | -1.378 | 0.003 | 0.023 | 0.023 | 0 | 0.15 | 0.05 | 0.11 | 0.1 | 0.1 | 0.2 |
| 11 | 12867171 | rs1866705 | 0.195 | 0.007 | 0.828 | -1.407 | -0.002 | -0.011 | -0.011 | 0 | 0.68 | 0.63 | 0.77 | 0.79 | 0.72 | 0.64 |
| 11 | 12868119 | rs1866707 | 0.555 | 0.012 | 0.632 | -1.405 | -0.003 | -0.013 | -0.013 | 0.319 | 0.65 | 0.36 | 0.74 | 0.79 | 0.72 | 0.58 |
| 11 | 12876666 | rs4757956 | 0.592 | 0.086 | -0.143 | -1.356 | 0.007 | 0.015 | 0.015 | 0 | 0.62 | 0.62 | 0.75 | 0.79 | 0.69 | 0.52 |
| 11 | 12886540 | rs10500764 | 1.577 | 0.568 | -1.152 | -1.46 | -0.024 | 0.219 | 0.219 | 0.011 | 0.34 | 0.04 | 0.4 | 0 | 0.32 | 0.15 |
| 11 | 12893947 | rs11022522 | 0.143 | 1.436 | -0.353 | -0.28 | -0.066 | -0.063 | -0.063 | 0 | 0.1 | 0.02 | 0.18 | 0.73 | 0.27 | 0.18 |
| 11 | 12894315 | rs7115063 | -1.819 | 1.199 | -0.059 | -0.517 | -0.001 | -0.038 | -0.038 | 0 | 0.43 | 0.59 | 0.4 | 0.96 | 0.53 | 0.62 |
| 11 | 12894758 | rs12289262 | -1.288 | 0.772 | -0.57 | -0.305 | -0.002 | 0.02 | 0.02 | 0 | 0.29 | 0.16 | 0.18 | 0.23 | 0.25 | 0.43 |
| 11 | 12895233 | rs12277651 | -0.263 | 0.568 | -0.726 | -0.379 | 0.005 | 0.01 | 0.01 | 0 | 0.17 | 0.12 | 0.09 | 0.1 | 0.09 | 0.3 |
| 11 | 12902966 | rs10766002 | -1.196 | 0.642 | -0.883 | -0.759 | 0.009 | 0.124 | 0.124 | 0 | 0.48 | 0.68 | 0.41 | 0.94 | 0.55 | 0.72 |
| 11 | 12903565 | rs2304734 | -0.081 | 1.352 | -0.935 | 0.732 | -0.047 | -0.019 | -0.019 | 0 | 0.08 | 0.01 | 0.12 | 0.47 | 0.16 | 0.08 |
| 11 | 12909146 | rs4757064 | -0.254 | 0.413 | -0.933 | -0.509 | 0.005 | 0.044 | 0.044 | 0 | 0.26 | 0.19 | 0.17 | 0.05 | 0.26 | 0.15 |
| 11 | 12911236 | rs10831916 | -0.547 | 0.09 | -0.405 | -0.532 | 0.003 | 0.044 | 0.044 | 1.673 | 0.19 | 0.39 | 0.18 | 0.69 | 0.18 | 0.27 |
| 11 | 12911769 | rs16911712 | -1.053 | 0.27 | -0.117 | -0.524 | 0.014 | 0.072 | 0.072 | 0.008 | 0.06 | 0.19 | 0.06 | 0.22 | 0.07 | 0.28 |
| 11 | 12912067 | rs4757963 | -0.788 | 0.227 | -0.233 | -0.496 | -0.001 | -0.047 | -0.047 | 0.007 | 0.45 | 0.47 | 0.38 | 0.89 | 0.48 | 0.49 |
| 11 | 12912104 | rs3937603 | -0.664 | 0.024 | -0.18 | -0.465 | -0.003 | -0.031 | -0.031 | 0 | 0.43 | 0.47 | 0.34 | 0.75 | 0.44 | 0.41 |
| 11 | 12915874 | rs10766003 | 1.092 | -0.049 | 0.671 | -0.075 | -0.01 | 0.065 | 0.065 | 0.4 | 0.86 | 0.49 | 0.88 | 0.8 | 0.86 | 0.77 |
| 11 | 12916329 | rs16911722 | -1.088 | -0.096 | 1.042 | 0.335 | -0.008 | 0.088 | 0.088 | 0 | 0.06 | 0.15 | 0.03 | 0.15 | 0.06 | 0.24 |
| 11 | 12919675 | rs16911732 | -1.548 | -0.066 | 1.374 | 0.943 | -0.013 | 0.039 | 0.039 | 0 | 0.11 | 0.24 | 0.07 | 0.15 | 0.11 | 0.25 |
| 11 | 12920271 | rs12364619 | 1.207 | 0.04 | 0.587 | -0.081 | -0.117 | 0.238 | 0.238 | 0.134 | 0.36 | 0.05 | 0.53 | 0 | 0.42 | 0.24 |
| 11 | 12923434 | rs2289436 | -1.998 | 0.471 | 1.702 | 0.721 | 0.006 | 0.223 | 0.223 | 0.079 | 0.15 | 0.53 | 0.18 | 0.58 | 0.18 | 0.15 |
| 11 | 12928226 | rs4319500 | 0.97 | 0.651 | 1.606 | 0.595 | 0.012 | 0.019 | 0.019 | 0 | 0.72 | 0.38 | 0.72 | 0.36 | 0.66 | 0.68 |
| 11 | 12933933 | rs17411627 | -0.218 | 0.876 | 1.064 | 0.008 | -0.041 | 0.065 | 0.065 | 0.103 | 0.18 | 0.08 | 0.14 | 0.06 | 0.22 | 0.13 |
| 11 | 12935451 | rs1866709 | 0.29 | 0.841 | 1.196 | 0.04 | 0.007 | -0.008 | -0.008 | 0 | 0.79 | 0.67 | 0.8 | 0.81 | 0.75 | 0.79 |
| 11 | 12940196 | rs747113 | 1.224 | 0.511 | 1.057 | -0.296 | -0.002 | -0.003 | -0.003 | 0 | 0.61 | 0.37 | 0.58 | 0.58 | 0.57 | 0.68 |
| 11 | 12941423 | rs12291766 | -0.611 | 0.399 | 0.982 | -0.338 | 0.006 | 0.004 | 0.004 | 0 | 0.11 | 0.08 | 0.04 | 0 | 0.07 | 0.08 |
| 11 | 12945058 | rs12792409 | -1.985 | 0.49 | 1.395 | -0.336 | 0.008 | 0.124 | 0.124 | 0 | 0.21 | 0.4 | 0.17 | 0.38 | 0.23 | 0.17 |
| 11 | 12946203 | rs745361 | -1.081 | 0.469 | 1.501 | -0.348 | -0.019 | -0.071 | -0.071 | 0 | 0.13 | 0.27 | 0.06 | 0 | 0.07 | 0.07 |
| 11 | 12948464 | rs2581566 | 0.74 | 0.562 | 1.49 | -0.394 | 0.013 | 0.026 | 0.026 | 0.221 | 0.72 | 0.65 | 0.63 | 0.59 | 0.63 | 0.78 |
| 11 | 12950553 | rs16911799 | 0.814 | 0.583 | 1.52 | -0.383 | 0.02 | 0.041 | 0.041 | 0.167 | 0.72 | 0.61 | 0.63 | 0.59 | 0.63 | 0.78 |
| 11 | 12957618 | rs4369386 | -0.36 | 0.669 | 1.52 | -0.169 | 0.022 | 0.152 | 0.152 | 0 | 0.08 | 0.14 | 0.11 | 0.39 | 0.17 | 0.11 |
| 11 | 12958051 | rs12807171 | -1.08 | 0.444 | 1.285 | -0.549 | 0.004 | 0.077 | 0.077 | 1.122 | 0.36 | 0.21 | 0.33 | 0.05 | 0.35 | 0.19 |
| 11 | 12960082 | rs16911828 | -0.26 | 0.796 | 1.488 | 0.182 | 0.003 | -0.002 | -0.002 | 0 | 0.1 | 0.09 | 0.05 | 0 | 0.07 | 0.06 |
| 11 | 12966547 | rs16911855 | -0.74 | 1.621 | 1.79 | 0.747 | 0.049 | 0.245 | 0.245 | 0.011 | 0.1 | 0.28 | 0.12 | 0.37 | 0.17 | 0.11 |
| 11 | 12980357 | rs10766011 | -1.583 | 1.515 | 1.71 | 0.704 | 0.045 | 0.049 | 0.049 | 0 | 0.29 | 0.53 | 0.41 | 0.42 | 0.41 | 0.25 |
| 11 | 12987192 | rs4450153 | -0.974 | 0.657 | 1.593 | -0.195 | 0.022 | -0.103 | -0.103 | 0.367 | 0.2 | 0.41 | 0.23 | 0.04 | 0.2 | 0.14 |
| 11 | 12987342 | rs4436527 | -0.67 | 0.661 | 1.371 | -0.193 | -0.007 | 0.007 | 0.007 | 0 | 0.18 | 0.14 | 0.2 | 0.04 | 0.2 | 0.14 |
| 11 | 12993149 | rs10831930 | -0.55 | 0.5 | 0.342 | 0.933 | -0.003 | 0.066 | 0.066 | 0.191 | 0.23 | 0.52 | 0.26 | 0.33 | 0.22 | 0.13 |
| 11 | 12994112 | rs12295316 | -0.828 | -0.068 | -0.262 | -0.13 | 0.01 | 0.004 | 0.004 | 0 | 0.21 | 0.2 | 0.18 | 0.01 | 0.18 | 0.12 |
| 11 | 13002186 | rs7929650 | -0.798 | -0.467 | -0.622 | 0.57 | -0.009 | 0.021 | 0.021 | 0 | 0.14 | 0.21 | 0.1 | 0.23 | 0.11 | 0.08 |
| 11 | 13005461 | rs7942018 | 0.236 | 0.154 | -0.975 | 0.123 | -0.021 | -0.018 | -0.018 | 0 | 0.65 | 0.75 | 0.64 | 0.42 | 0.59 | 0.56 |

| | | | | | | | | | | | | | | | | |
|---|---|---|---|---|---|---|---|---|---|---|---|---|---|---|---|---|
| 11 | 13006033 | rs10831931 | 0.515 | 0.085 | -0.49 | 0.105 | 0.005 | 0.001 | 0.001 | 0.177 | 0.63 | 0.57 | 0.6 | 0.41 | 0.58 | 0.56 |
| 11 | 13006721 | rs3935745 | 0.253 | -0.18 | -0.517 | -0.766 | -0.006 | -0.002 | -0.002 | 0 | 0.35 | 0.35 | 0.2 | 0.29 | 0.28 | 0.28 |
| 11 | 13007397 | rs3935878 | 0.173 | -0.185 | -0.5 | -0.779 | -0.027 | -0.01 | -0.01 | 0 | 0.36 | 0.42 | 0.2 | 0.29 | 0.28 | 0.28 |
| 11 | 13008591 | rs7114499 | 1.429 | -0.149 | -0.539 | -0.995 | -0.01 | -0.065 | -0.065 | 0 | 0.21 | 0.22 | 0.12 | 0 | 0.18 | 0.06 |
| 11 | 13009700 | rs7934110 | 1.108 | -0.239 | -0.684 | -1.079 | 0.007 | -0.103 | -0.103 | 1.046 | 0.31 | 0.44 | 0.43 | 0.13 | 0.31 | 0.3 |
| 11 | 13011522 | rs11022589 | 0.703 | -0.326 | -0.759 | -1.072 | -0.022 | -0.087 | -0.087 | 0 | 0.26 | 0.38 | 0.18 | 0.07 | 0.23 | 0.19 |
| 11 | 13011847 | rs4757071 | -0.662 | 0.079 | -0.755 | -0.077 | 0.001 | 0.001 | 0.001 | 1.08 | 0.52 | 0.52 | 0.41 | 0.62 | 0.43 | 0.41 |
| 11 | 13015732 | rs4485088 | -0.302 | 0.177 | -0.636 | -0.683 | 0.005 | 0.006 | 0.006 | 0 | 0.42 | 0.41 | 0.35 | 0.36 | 0.38 | 0.35 |
| 11 | 13019218 | rs10500769 | 0.881 | 0.337 | -0.683 | -0.734 | 0.009 | -0.073 | -0.073 | 0.153 | 0.33 | 0.42 | 0.44 | 0.13 | 0.33 | 0.28 |
| 11 | 13020422 | rs10766022 | -0.713 | -0.035 | -0.611 | -0.597 | -0.011 | -0.026 | -0.026 | 0 | 0.75 | 0.83 | 0.8 | 0.5 | 0.71 | 0.63 |
| 11 | 13021135 | rs10831938 | -0.726 | -0.035 | -0.61 | -0.597 | -0.018 | -0.04 | -0.04 | 0 | 0.75 | 0.86 | 0.8 | 0.5 | 0.71 | 0.63 |
| 11 | 13025736 | rs7951763 | -0.753 | -0.033 | -0.672 | -0.597 | -0.037 | -0.072 | -0.072 | 0.868 | 0.76 | 0.9 | 0.8 | 0.5 | 0.71 | 0.63 |
| 11 | 13026298 | rs4411272 | 0.877 | 0.249 | -0.856 | -0.779 | -0.011 | -0.043 | -0.043 | 0 | 0.29 | 0.4 | 0.26 | 0.21 | 0.24 | 0.21 |
| 11 | 13033155 | rs1141090 | 0.044 | 0.252 | -1.158 | -0.782 | 0.005 | 0.04 | 0.04 | 0 | 0.42 | 0.3 | 0.47 | 0.22 | 0.41 | 0.3 |
| 11 | 13049812 | rs4237696 | -0.259 | 0.154 | -1.451 | -0.873 | -0.022 | 0.112 | 0.112 | 0 | 0.54 | 0.22 | 0.52 | 0.28 | 0.51 | 0.48 |
| 11 | 13053972 | rs17506253 | -0.038 | 0.163 | -1.222 | -1.35 | 0.006 | 0.112 | 0.112 | 0.353 | 0.55 | 0.76 | 0.44 | 0.28 | 0.47 | 0.4 |
| 11 | 13057028 | rs12273457 | -0.409 | 0.154 | -1.5 | -1.231 | -0.017 | 0.046 | 0.046 | 0 | 0.44 | 0.66 | 0.62 | 0.37 | 0.46 | 0.41 |
| 11 | 13061030 | rs4757096 | 0.408 | 0.546 | -1.629 | -0.858 | -0.048 | 0.014 | 0.014 | 0 | 0.19 | 0.06 | 0.31 | 0.15 | 0.22 | 0.24 |
| 11 | 13063882 | rs2727378 | -0.529 | 0.74 | -1.437 | -0.655 | -0.036 | -0.053 | -0.053 | 0 | 0.76 | 0.88 | 0.79 | 0.44 | 0.69 | 0.65 |
| 11 | 13071286 | rs11022618 | 0.131 | 0.213 | -0.393 | -1.179 | 0.002 | 0.021 | 0.021 | 0 | 0.4 | 0.32 | 0.43 | 0.22 | 0.37 | 0.3 |
| 11 | 13074670 | rs2583122 | -0.362 | 0.078 | -0.059 | -1.172 | -0.009 | -0.02 | -0.02 | 0 | 0.71 | 0.77 | 0.74 | 0.43 | 0.64 | 0.53 |
| 11 | 13075300 | rs11022621 | 1.09 | -0.103 | -0.237 | -1.665 | -0.017 | -0.004 | -0.004 | 0 | 0.24 | 0.25 | 0.16 | 0.21 | 0.18 | 0.12 |
| 11 | 13083053 | rs2727409 | -1.762 | 0.147 | -0.559 | -1.667 | -0.019 | 0.046 | 0.046 | 0 | 0.86 | 0.94 | 0.88 | 0.96 | 0.82 | 0.92 |
| 11 | 13085602 | rs12294180 | 1.067 | -0.181 | -0.493 | -1.771 | 0.041 | 0.026 | 0.026 | 0.023 | 0.32 | 0.2 | 0.16 | 0.26 | 0.24 | 0.15 |
| 11 | 13086274 | rs11022626 | 0.561 | 0.009 | -0.302 | -1.628 | 0.001 | 0.033 | 0.033 | 0 | 0.17 | 0.2 | 0.15 | 0.59 | 0.18 | 0.41 |
| 11 | 13086849 | rs12799420 | 2.147 | -0.081 | -0.26 | -2.001 | 0.014 | 0.003 | 0.003 | 0 | 0.18 | 0.13 | 0.07 | 0 | 0.13 | 0.06 |
| 11 | 13087360 | rs12281796 | 0.749 | 0.331 | 0.56 | -1.373 | 0.023 | -0.058 | -0.058 | 0 | 0.17 | 0.25 | 0.28 | 0.01 | 0.19 | 0.18 |
| 11 | 13087407 | rs16911996 | -0.214 | 0.311 | 0.367 | -1.122 | 0.038 | -0.001 | -0.001 | 0.474 | 0.07 | 0.16 | 0.06 | 0.08 | 0.1 | 0.15 |
| 11 | 13088435 | rs17416058 | 0.918 | 0.203 | 0.083 | -1.137 | 0.003 | 0.01 | 0.01 | 0 | 0.18 | 0.17 | 0.22 | 0.08 | 0.16 | 0.18 |
| 11 | 13090871 | rs10831950 | 1.08 | 0.095 | -0.045 | -0.81 | 0.009 | 0.017 | 0.017 | 0 | 0.12 | 0.12 | 0.09 | 0.49 | 0.06 | 0.31 |
| 11 | 13090942 | rs2727405 | -0.286 | 0.228 | 0.445 | -1.042 | -0.013 | 0.03 | 0.03 | 0 | 0.32 | 0.45 | 0.34 | 0.52 | 0.28 | 0.43 |
| 11 | 13096116 | rs11022633 | -0.792 | 0.345 | 0.435 | -1.354 | 0.004 | 0.059 | 0.059 | 0 | 0.49 | 0.34 | 0.54 | 0.08 | 0.47 | 0.39 |
| 11 | 13096415 | rs1441329 | -2.207 | 0.276 | 0.158 | -1.379 | -0.005 | 0.154 | 0.154 | 0 | 0.69 | 0.88 | 0.8 | 0.97 | 0.68 | 0.86 |
| 11 | 13097871 | rs17416453 | 0.684 | 0.364 | 0.085 | -1.453 | -0.003 | 0.047 | 0.047 | 0 | 0.16 | 0.06 | 0.08 | 0.03 | 0.14 | 0.13 |
| 11 | 13098051 | rs10831953 | -0.239 | 0.343 | 0.126 | -1.473 | -0.056 | 0.117 | 0.117 | 0 | 0.32 | 0.12 | 0.34 | 0.07 | 0.33 | 0.3 |
| 11 | 13099336 | rs10831954 | 1.134 | 0.282 | 0.136 | -1.271 | -0.042 | 0.051 | 0.051 | 0 | 0.15 | 0.06 | 0.27 | 0.03 | 0.19 | 0.17 |
| 11 | 13099808 | rs2727400 | -1.655 | 0.013 | 0.47 | -0.991 | 0.008 | 0.005 | 0.005 | 0 | 0.67 | 0.76 | 0.72 | 0.69 | 0.65 | 0.82 |
| 11 | 13102476 | rs2727398 | -0.981 | 0.097 | 0.43 | -1.251 | 0.021 | 0.059 | 0.059 | 0 | 0.55 | 0.66 | 0.64 | 0.17 | 0.58 | 0.49 |
| 11 | 13106910 | rs977169 | -0.879 | 0.254 | 0.525 | -1.147 | 0 | -0.019 | -0.019 | 0.57 | 0.52 | 0.51 | 0.57 | 0.17 | 0.5 | 0.48 |
| 11 | 13119308 | rs10766036 | -0.364 | 0.106 | 0.077 | -1.009 | -0.022 | -0.036 | -0.036 | 0 | 0.37 | 0.48 | 0.38 | 0.81 | 0.27 | 0.5 |
| 11 | 13122553 | rs7126761 | 1.263 | 0.372 | -0.107 | -0.542 | 0.016 | -0.051 | -0.051 | 0.107 | 0.1 | 0.03 | 0.05 | 0.49 | 0.06 | 0.3 |
| 11 | 13127473 | rs7103853 | -1.077 | 0.553 | -0.062 | -0.827 | 0.011 | 0.061 | 0.061 | 0 | 0.88 | 0.81 | 0.85 | 0.38 | 0.83 | 0.59 |
| 11 | 13127893 | rs6486088 | -0.132 | 0.447 | -0.104 | -1.108 | -0.013 | -0.011 | -0.011 | 0.436 | 0.37 | 0.47 | 0.37 | 0.32 | 0.31 | 0.24 |
| 11 | 13129269 | rs1441327 | 1.025 | 0.347 | -0.165 | -1.179 | -0.053 | 0.3 | 0.3 | 0.175 | 0.46 | 0.16 | 0.47 | 0.06 | 0.51 | 0.35 |
| 11 | 13129539 | rs1441323 | -0.694 | 0.357 | -0.099 | -1.139 | 0.004 | 0.063 | 0.063 | 0 | 0.83 | 0.74 | 0.84 | 0.38 | 0.82 | 0.59 |
| 11 | 13129912 | rs729620 | 0.902 | 0.716 | -0.872 | -1.951 | -0.049 | 0.057 | 0.057 | 0.293 | 0.15 | 0.08 | 0.26 | 0.02 | 0.18 | 0.2 |
| 11 | 13130170 | rs2896628 | 0.827 | 0.637 | -0.777 | -2 | -0.002 | -0.029 | -0.029 | 0 | 0.21 | 0.16 | 0.21 | 0.04 | 0.17 | 0.14 |
| 11 | 13131002 | rs4757108 | -0.352 | 0.579 | -1.05 | -1.773 | -0.008 | -0.077 | -0.077 | 0.106 | 0.67 | 0.48 | 0.69 | 0.1 | 0.68 | 0.48 |
| 11 | 13132504 | rs7928742 | -0.427 | 0.805 | -0.616 | -1.28 | 0.004 | -0.064 | -0.064 | 0 | 0.37 | 0.39 | 0.48 | 0.07 | 0.36 | 0.34 |
| 11 | 13135055 | rs10766043 | 0.218 | 0.629 | -0.821 | -0.992 | 0.025 | 0.131 | 0.131 | 0.143 | 0.49 | 0.32 | 0.53 | 0.16 | 0.53 | 0.42 |
| 11 | 13137171 | rs4756748 | 0.172 | 0.76 | -0.183 | -0.705 | -0.004 | -0.023 | -0.023 | 0 | 0.7 | 0.49 | 0.67 | 0.74 | 0.74 | 0.76 |
| 11 | 13137352 | rs7124054 | -0.297 | 0.335 | -0.021 | -1.048 | 0.005 | -0.04 | -0.04 | 0 | 0.07 | 0.2 | 0.05 | 0.01 | 0.06 | 0.09 |
| 11 | 13138160 | rs4756751 | -0.049 | 0.375 | 0.22 | -0.881 | 0 | -0.072 | -0.072 | 0.037 | 0.71 | 0.52 | 0.71 | 0.83 | 0.74 | 0.76 |
| 11 | 13140242 | rs1348150 | -0.152 | 0.143 | 0.582 | 0.079 | 0.009 | -0.018 | -0.018 | 0.56 | 0.71 | 0.75 | 0.75 | 0.53 | 0.78 | 0.63 |
| 11 | 13142080 | rs17418644 | 0.744 | 0.403 | 0.717 | 0.098 | -0.007 | 0.052 | 0.052 | 3.157 | 0.2 | 0.1 | 0.2 | 0.1 | 0.18 | 0.09 |
| 11 | 13144514 | rs11824425 | -0.322 | 0.37 | 1.047 | 0.067 | 0.048 | 0.004 | 0.004 | 1.505 | 0.09 | 0.27 | 0.1 | 0.07 | 0.1 | 0.16 |
| 11 | 13144810 | rs7938273 | -0.128 | 0.371 | 1.059 | 0.067 | 0.02 | -0.008 | -0.008 | 1.188 | 0.08 | 0.17 | 0.08 | 0.07 | 0.09 | 0.16 |
| 11 | 13145416 | rs2119561 | 0.047 | 0.369 | 1.051 | 0.071 | -0.004 | 0.01 | 0.01 | 0.747 | 0.7 | 0.67 | 0.73 | 0.53 | 0.76 | 0.62 |
| 11 | 13146288 | rs11022673 | 1.238 | 0.357 | 1.042 | 0.078 | 0.016 | -0.004 | -0.004 | 0 | 0.09 | 0.04 | 0.05 | 0.37 | 0.06 | 0.29 |
| 11 | 13147538 | rs7342278 | -0.215 | 0.398 | 1.043 | 0.06 | 0.028 | -0.005 | -0.005 | 5.206 | 0.08 | 0.2 | 0.09 | 0.07 | 0.09 | 0.16 |
| 11 | 13159838 | rs10831981 | 0.726 | 0.344 | 0.814 | -0.063 | 0.007 | -0.004 | -0.004 | 0 | 0.61 | 0.35 | 0.6 | 0.38 | 0.66 | 0.46 |
| 11 | 13163618 | rs10766048 | 1.528 | 0.308 | 0.469 | -0.298 | -0.04 | -0.009 | -0.009 | 0 | 0.45 | 0.15 | 0.5 | 0.39 | 0.53 | 0.36 |
| 11 | 13163667 | rs10766049 | 1.55 | 0.307 | 0.41 | -0.298 | -0.058 | -0.023 | -0.023 | 0 | 0.45 | 0.13 | 0.5 | 0.39 | 0.53 | 0.36 |
| 11 | 13164269 | rs11022681 | -1.526 | 0.295 | 0.563 | -0.21 | 0.007 | 0.007 | 0.007 | 0.042 | 0.5 | 0.39 | 0.42 | 0.56 | 0.47 | 0.63 |
| 11 | 13164586 | rs4114383 | 0.93 | 0.867 | -0.056 | -0.328 | -0.048 | -0.041 | -0.041 | 0 | 0.15 | 0.04 | 0.25 | 0.23 | 0.19 | 0.19 |
| 11 | 13164896 | rs11022685 | 0.338 | 0.245 | 0.025 | -0.268 | 0.002 | -0.059 | -0.059 | 3.453 | 0.17 | 0.06 | 0.11 | 0.45 | 0.15 | 0.45 |
| 11 | 13166047 | rs7926443 | 0.063 | 0.054 | 0.276 | -0.491 | -0.011 | -0.066 | -0.066 | 0.445 | 0.16 | 0.24 | 0.1 | 0.01 | 0.13 | 0.1 |
| 11 | 13171946 | rs11022693 | 1.545 | 0.31 | -0.571 | -0.312 | -0.016 | 0.009 | 0.009 | 0.202 | 0.29 | 0.01 | 0.21 | 0.14 | 0.26 | 0.17 |
| 11 | 13181590 | rs10500774 | -1.62 | 0.283 | 0.111 | -0.046 | -0.009 | -0.014 | -0.014 | 0 | 0.2 | 0.74 | 0.15 | 0.1 | 0.11 | 0.13 |

| | | | | | | | | | | | | | | | | |
|---|---|---|---|---|---|---|---|---|---|---|---|---|---|---|---|---|
| 11 | 13181618 | rs3849161 | -1.947 | 0.43 | -0.613 | -0.278 | -0.051 | 0.122 | 0.122 | 0.743 | 0.57 | 0.85 | 0.6 | 0.76 | 0.53 | 0.69 |
| 11 | 13185226 | rs16924750 | -0.884 | 0.463 | -0.4 | -0.249 | -0.015 | -0.029 | -0.029 | 1.162 | 0.16 | 0.69 | 0.14 | 0.09 | 0.11 | 0.11 |
| 11 | 13187977 | rs16912297 | -0.553 | 0.444 | 0.221 | -0.257 | 0.011 | -0.003 | -0.003 | 0 | 0.13 | 0.44 | 0.11 | 0.09 | 0.1 | 0.11 |
| 11 | 13188142 | rs16912299 | -0.43 | 0.42 | 0.367 | -0.279 | 0.042 | 0.043 | 0.043 | 0 | 0.09 | 0.36 | 0.08 | 0.09 | 0.08 | 0.1 |
| 11 | 13189586 | rs6486097 | -1.781 | 0.401 | 0.428 | -0.296 | -0.022 | -0.04 | -0.04 | 0 | 0.17 | 0.61 | 0.12 | 0.09 | 0.1 | 0.13 |
| 11 | 13191293 | rs1481871 | 0.161 | 0.492 | 0.327 | -0.25 | -0.017 | 0.019 | 0.019 | 0 | 0.61 | 0.8 | 0.52 | 0.33 | 0.57 | 0.44 |
| 11 | 13191527 | rs7935099 | -0.101 | 0.523 | 0.184 | 0.104 | -0.018 | -0.023 | -0.023 | 0 | 0.1 | 0.04 | 0.29 | 0.16 | 0.12 | 0.2 |
| 11 | 13192706 | rs4756756 | 1.674 | 0.6 | 0.162 | 0.108 | -0.028 | 0.139 | 0.139 | 0.235 | 0.47 | 0.2 | 0.43 | 0.25 | 0.49 | 0.31 |
| 11 | 13194858 | rs7933806 | -0.892 | 0.148 | 0.023 | 0.34 | 0.012 | 0.013 | 0.013 | 0.757 | 0.11 | 0.58 | 0.1 | 0.09 | 0.08 | 0.1 |
| 11 | 13197316 | rs10831990 | -0.328 | 0.15 | -0.478 | 0.467 | -0.059 | -0.184 | -0.184 | 0.233 | 0.28 | 0.02 | 0.18 | 0.5 | 0.3 | 0.37 |
| 11 | 13206160 | rs748923 | 0.228 | 0.054 | 0.528 | 0.971 | 0.059 | 0.333 | 0.333 | 0.036 | 0.09 | 0.3 | 0.12 | 0.39 | 0.12 | 0.23 |
| 11 | 13209260 | rs4757122 | 1.032 | -0.407 | 0.02 | 0.532 | 0.046 | 0.052 | 0.052 | 0 | 0.53 | 0.33 | 0.55 | 0.36 | 0.61 | 0.43 |
| 11 | 13211208 | rs16912392 | -0.659 | -0.681 | 0.047 | 0.501 | -0.001 | 0.033 | 0.033 | 0.289 | 0.23 | 0.15 | 0.22 | 0.06 | 0.2 | 0.23 |
| 11 | 13218438 | rs1080775 | 0.395 | -0.863 | 0.874 | 1.077 | 0.02 | -0.026 | -0.026 | 2.087 | 0.75 | 0.84 | 0.77 | 0.41 | 0.79 | 0.66 |
| 11 | 13227981 | rs10831997 | 0.597 | -0.848 | 1.092 | 1.095 | -0.055 | 0.056 | 0.056 | 0.239 | 0.73 | 0.53 | 0.76 | 0.41 | 0.79 | 0.67 |
| 11 | 13228643 | rs10734210 | -0.343 | 0.165 | 1.179 | 1.695 | 0.029 | 0.25 | 0.25 | 0 | 0.9 | 0.78 | 0.85 | 0.57 | 0.86 | 0.79 |
| 11 | 13232738 | rs2170436 | 1.035 | -0.242 | 1.17 | 0.708 | 0.005 | 0.01 | 0.01 | 0.633 | 0.67 | 0.29 | 0.62 | 0.81 | 0.7 | 0.75 |
| 11 | 13237709 | rs11022710 | 0.812 | -0.216 | 1.384 | 0.697 | 0.003 | 0.004 | 0.004 | 1.671 | 0.73 | 0.27 | 0.67 | 0.81 | 0.75 | 0.77 |
| 11 | 13239330 | rs11022713 | 0.3 | 0.217 | 1.07 | 1.056 | 0.018 | 0.095 | 0.095 | 1.265 | 0.13 | 0.18 | 0.32 | 0.46 | 0.2 | 0.28 |
| 11 | 13239500 | rs7112005 | 0.542 | -0.472 | 0.964 | 0.742 | -0.006 | -0.012 | -0.012 | 0 | 0.77 | 0.66 | 0.71 | 0.81 | 0.76 | 0.77 |
| 11 | 13250399 | rs10832000 | -0.484 | -0.396 | 0.788 | 0.769 | -0.019 | 0 | 0 | 0 | 0.19 | 0.07 | 0.27 | 0.16 | 0.24 | 0.2 |
| 11 | 13252041 | rs12795870 | 0.032 | -0.489 | 0.649 | 0.657 | 0.004 | 0.003 | 0.003 | 0 | 0.08 | 0.06 | 0.06 | 0 | 0.05 | 0.04 |
| 11 | 13254504 | rs7104311 | -0.098 | -0.399 | 0.853 | 0.979 | 0.016 | 0.228 | 0.228 | 0 | 0.82 | 0.54 | 0.67 | 0.49 | 0.82 | 0.69 |
| 11 | 13255971 | rs7112233 | 0.048 | -0.361 | -0.031 | 0.998 | -0.001 | 0.011 | 0.011 | 0 | 0.16 | 0.16 | 0.31 | 0.51 | 0.18 | 0.31 |
| 11 | 13260882 | rs12795287 | 1.365 | -0.289 | -0.354 | 0.669 | 0.003 | 0.113 | 0.113 | 0 | 0.53 | 0.13 | 0.34 | 0.32 | 0.55 | 0.46 |
| 11 | 13261390 | rs11607529 | 1.419 | -0.283 | -0.504 | 0.672 | -0.011 | 0.113 | 0.113 | 0 | 0.53 | 0.08 | 0.34 | 0.32 | 0.55 | 0.46 |
| 11 | 13267554 | rs1384014 | 0.498 | -1.093 | 0.325 | 1.233 | -0.009 | 0.036 | 0.036 | 0 | 0.74 | 0.72 | 0.64 | 0.56 | 0.78 | 0.64 |
| 11 | 13269536 | rs1481881 | 1.514 | -2.019 | 0.144 | 0.626 | 0.038 | 0.115 | 0.115 | 0 | 0.48 | 0.25 | 0.45 | 0.28 | 0.59 | 0.45 |
| 11 | 13269548 | rs1481880 | 0.355 | -2.121 | 0.655 | 0.745 | -0.002 | 0.014 | 0.014 | 0 | 0.72 | 0.74 | 0.63 | 0.51 | 0.74 | 0.65 |
| 11 | 13269792 | rs10832006 | -0.086 | -2.212 | 0.634 | 0.683 | -0.004 | -0.029 | -0.029 | 0 | 0.21 | 0.18 | 0.35 | 0.49 | 0.23 | 0.3 |
| 11 | 13272029 | rs11022724 | 0.13 | -2.244 | 0.902 | 0.633 | 0.023 | -0.019 | -0.019 | 0.423 | 0.2 | 0.14 | 0.15 | 0.23 | 0.15 | 0.2 |
| 11 | 13276329 | rs10832008 | 0.043 | -2.241 | 0.989 | 0.632 | 0.022 | -0.013 | -0.013 | 0.23 | 0.21 | 0.16 | 0.15 | 0.23 | 0.15 | 0.2 |
| 11 | 13278027 | rs10766066 | 0.573 | -2.265 | 1.209 | 0.621 | -0.005 | 0.034 | 0.034 | 0 | 0.71 | 0.69 | 0.62 | 0.51 | 0.74 | 0.64 |
| 11 | 13280556 | rs4756763 | 1.518 | -2.139 | 0.955 | 0.556 | 0.033 | 0.105 | 0.105 | 0 | 0.48 | 0.28 | 0.45 | 0.28 | 0.59 | 0.44 |
| 11 | 13288698 | rs7942486 | 1.54 | -2.271 | 1.231 | 0.746 | 0.006 | 0.024 | 0.024 | 0.019 | 0.45 | 0.34 | 0.42 | 0.32 | 0.57 | 0.4 |
| 11 | 13293905 | rs900145 | 0.905 | -2.002 | 1.677 | 1.111 | -0.016 | 0.023 | 0.023 | 0 | 0.65 | 0.44 | 0.56 | 0.56 | 0.72 | 0.63 |
| 11 | 13297789 | rs7126225 | -0.402 | -1.843 | 1.087 | 1.146 | -0.007 | -0.018 | -0.018 | 0.268 | 0.94 | 0.92 | 0.98 | 1 | 0.97 | 0.95 |
| 11 | 13312254 | rs7123257 | 0.348 | -1.865 | 0.841 | 1.139 | -0.024 | -0.045 | -0.045 | 0 | 0.92 | 0.75 | 0.97 | 1 | 0.97 | 0.95 |
| 11 | 13312969 | rs7107287 | -0.317 | -1.977 | 0.832 | 1.165 | 0.001 | -0.015 | -0.015 | 0.12 | 0.25 | 0.24 | 0.4 | 0.44 | 0.24 | 0.32 |
| 11 | 13313096 | rs10741615 | -0.346 | -1.962 | 0.745 | 1.172 | -0.021 | -0.037 | -0.037 | 0 | 0.93 | 0.89 | 0.98 | 1 | 0.97 | 0.95 |
| 11 | 13314307 | rs12290622 | 0.699 | -1.887 | 0.517 | 1.215 | -0.025 | 0.042 | 0.042 | 0 | 0.66 | 0.51 | 0.57 | 0.56 | 0.72 | 0.64 |
| 11 | 13318139 | rs7950226 | -0.295 | -1.128 | -0.69 | 1.458 | -0.062 | -0.04 | -0.04 | 0 | 0.43 | 0.14 | 0.56 | 0.62 | 0.46 | 0.45 |
| 11 | 13319894 | rs7949336 | -0.933 | -1.659 | -0.079 | 1.866 | -0.015 | -0.109 | -0.109 | 1.059 | 0.24 | 0.11 | 0.39 | 0.48 | 0.23 | 0.3 |
| 11 | 13321343 | rs10832020 | -0.165 | -1.642 | -0.097 | 1.908 | -0.014 | -0.104 | -0.104 | 0.574 | 0.18 | 0.03 | 0.18 | 0.28 | 0.16 | 0.25 |
| 11 | 13329139 | rs4757143 | -0.273 | -1.67 | 0.114 | 1.413 | 0.011 | -0.005 | -0.005 | 0.01 | 0.18 | 0.12 | 0.15 | 0.18 | 0.14 | 0.24 |
| 11 | 13331226 | rs4757144 | 1.627 | -1.886 | -0.04 | 1.433 | -0.006 | -0.007 | -0.007 | 0 | 0.49 | 0.05 | 0.44 | 0.41 | 0.59 | 0.44 |
| 11 | 13331324 | rs4757145 | -1.473 | -1.562 | 0.618 | 1.673 | 0.007 | 0.002 | 0.002 | 0 | 0.42 | 0.47 | 0.5 | 0.58 | 0.38 | 0.54 |
| 11 | 13333851 | rs10741616 | -1.455 | -1.572 | 0.622 | 1.673 | 0.009 | 0.002 | 0.002 | 0 | 0.42 | 0.41 | 0.5 | 0.58 | 0.38 | 0.54 |
| 11 | 13340867 | rs7126796 | 0.151 | -1.493 | 0.91 | 1.67 | -0.017 | 0 | 0 | 0 | 0.81 | 0.68 | 0.82 | 0.82 | 0.85 | 0.77 |
| 11 | 13345115 | rs11022765 | -0.507 | -1.315 | 0.837 | 1.924 | -0.006 | -0.037 | -0.037 | 0 | 0.23 | 0.17 | 0.32 | 0.38 | 0.23 | 0.31 |
| 11 | 13349781 | rs10766077 | -1.583 | -1.322 | 1.033 | 1.607 | 0.01 | 0.005 | 0.005 | 0 | 0.4 | 0.33 | 0.45 | 0.63 | 0.35 | 0.53 |
| 11 | 13350131 | rs1982350 | 1.179 | -1.322 | 0.876 | 1.607 | -0.009 | -0.006 | -0.006 | 0 | 0.6 | 0.49 | 0.54 | 0.37 | 0.65 | 0.47 |
| 11 | 13350747 | rs7924734 | -0.777 | -1.018 | 0.495 | 1.75 | -0.007 | -0.045 | -0.045 | 0 | 0.23 | 0.17 | 0.35 | 0.43 | 0.24 | 0.31 |
| 11 | 13355770 | rs6486121 | 1.205 | -0.694 | 0.757 | 1.229 | 0.018 | 0.044 | 0.044 | 0 | 0.59 | 0.4 | 0.49 | 0.16 | 0.66 | 0.39 |
| 11 | 13357183 | rs10832027 | 0.585 | -0.736 | 1.068 | 1.434 | 0.008 | -0.009 | -0.009 | 0 | 0.67 | 0.78 | 0.57 | 0.35 | 0.7 | 0.54 |
| 11 | 13362291 | rs16912751 | -0.636 | -0.824 | 1.04 | 1.481 | 0.011 | 0.15 | 0.15 | 0 | 0.08 | 0.25 | 0.08 | 0.19 | 0.05 | 0.14 |
| 11 | 13362815 | rs7937060 | 1.334 | -0.659 | 0.375 | 1.456 | 0.013 | 0.052 | 0.052 | 0 | 0.58 | 0.39 | 0.38 | 0.16 | 0.64 | 0.4 |
| 11 | 13364200 | rs1562438 | -0.436 | -0.2 | -0.555 | 1.615 | -0.051 | -0.128 | -0.128 | 0 | 0.23 | 0.05 | 0.49 | 0.45 | 0.25 | 0.39 |
| 11 | 13364413 | rs1562437 | -0.335 | -0.161 | -0.537 | 1.438 | -0.011 | 0.01 | 0.01 | 0 | 0.9 | 0.86 | 0.96 | 0.82 | 0.94 | 0.95 |
| 11 | 13364712 | rs1026071 | -0.83 | 0.491 | -0.719 | 1.375 | -0.013 | -0.059 | -0.059 | 0.193 | 0.23 | 0.12 | 0.49 | 0.48 | 0.26 | 0.38 |
| 11 | 13373764 | rs11022775 | -1.454 | 0.185 | -0.141 | 0.694 | -0.015 | -0.023 | -0.023 | 0 | 0.13 | 0.35 | 0.13 | 0.09 | 0.08 | 0.21 |
| 11 | 13379973 | rs2290037 | -0.926 | 0.411 | -0.539 | 1.046 | -0.008 | -0.032 | -0.032 | 1.759 | 0.05 | 0 | 0.06 | 0.15 | 0.06 | 0.09 |
| 11 | 13383682 | rs1868049 | -0.49 | 0.212 | -1.053 | 1.45 | 0.002 | -0.071 | -0.071 | 0 | 0.15 | 0.04 | 0.12 | 0.54 | 0.14 | 0.18 |
| 11 | 13385316 | rs3789327 | -0.021 | 0.42 | -1.076 | 0.335 | -0.004 | 0.021 | 0.021 | 0.031 | 0.56 | 0.66 | 0.62 | 0.32 | 0.47 | 0.63 |
| 11 | 13390860 | rs11022778 | 0.479 | 0.132 | -1.099 | -0.07 | 0.05 | 0.141 | 0.141 | 0 | 0.43 | 0.18 | 0.31 | 0.16 | 0.34 | 0.43 |
| 11 | 13392213 | rs4757151 | 0.013 | 0.278 | -1.695 | 0.06 | -0.022 | 0.04 | 0.04 | 0 | 0.5 | 0.17 | 0.52 | 0.39 | 0.51 | 0.54 |
| 11 | 13396166 | rs11600996 | 1.029 | -0.172 | -2.175 | -0.523 | -0.044 | 0.133 | 0.133 | 1.626 | 0.51 | 0.09 | 0.34 | 0.26 | 0.56 | 0.56 |
| 11 | 13396909 | rs11022780 | 0.942 | -0.239 | -1.175 | -0.559 | 0.009 | 0.085 | 0.085 | 1.201 | 0.52 | 0.25 | 0.35 | 0.26 | 0.56 | 0.66 |
| 11 | 13399791 | rs10832030 | -0.45 | -0.426 | -0.383 | -0.518 | -0.045 | 0.117 | 0.117 | 0 | 0.77 | 0.65 | 0.76 | 0.5 | 0.85 | 0.83 |

| | | | | | | | | | | | | | | | | |
|---|---|---|---|---|---|---|---|---|---|---|---|---|---|---|---|---|
| 11 | 13402409 | rs11022781 | 0.878 | -0.397 | -0.196 | -0.524 | 0 | 0.003 | 0.003 | 0.892 | 0.54 | 0.45 | 0.37 | 0.26 | 0.57 | 0.66 |
| 11 | 13403043 | rs969485 | 0.285 | -0.241 | -0.239 | -0.587 | -0.031 | 0.031 | 0.031 | 0 | 0.64 | 0.51 | 0.67 | 0.43 | 0.72 | 0.77 |
| 11 | 13405899 | rs11022783 | 1.026 | -0.396 | -0.383 | 0.383 | -0.005 | -0.082 | -0.082 | 0 | 0.15 | 0.01 | 0.08 | 0.25 | 0.14 | 0.11 |
| 11 | 13410371 | rs17452383 | 0.904 | -0.413 | -0.364 | 0.186 | -0.006 | 0.028 | 0.028 | 0 | 0.12 | 0.08 | 0.08 | 0.06 | 0.12 | 0.06 |
| 11 | 13411344 | rs10832032 | -1.183 | -0.421 | -0.17 | 0.177 | -0.003 | 0.021 | 0.021 | 0.599 | 0.41 | 0.4 | 0.6 | 0.74 | 0.41 | 0.3 |
| 11 | 13422274 | rs2896633 | 0.518 | -0.293 | -0.103 | 0.263 | -0.012 | -0.008 | -0.008 | 0 | 0.65 | 0.52 | 0.45 | 0.32 | 0.69 | 0.72 |
| 11 | 13425833 | rs4757153 | 0.059 | -0.298 | -0.114 | 0.306 | -0.037 | -0.037 | -0.037 | 0 | 0.11 | 0.05 | 0.3 | 0.16 | 0.15 | 0.11 |
| 11 | 13428961 | rs7951183 | -0.447 | -0.299 | -0.103 | 0.306 | -0.043 | 0.109 | 0.109 | 0 | 0.77 | 0.65 | 0.76 | 0.49 | 0.84 | 0.83 |
| 11 | 13441741 | rs2403664 | 1.319 | -0.299 | -0.132 | 0.287 | 0.016 | 0.099 | 0.099 | 0.885 | 0.51 | 0.27 | 0.36 | 0.25 | 0.58 | 0.66 |
| 11 | 13446295 | rs11022817 | 1.429 | -0.311 | -0.149 | 0.281 | 0.014 | 0.128 | 0.128 | 1.22 | 0.51 | 0.19 | 0.35 | 0.25 | 0.58 | 0.66 |
| 11 | 13452904 | rs11022824 | -0.061 | -0.201 | -0.248 | 0.289 | -0.019 | -0.019 | -0.019 | 0.842 | 0.11 | 0.08 | 0.3 | 0.17 | 0.15 | 0.11 |
| 11 | 13481096 | rs4356206 | 0.759 | -0.211 | -0.245 | 0.264 | -0.006 | 0.032 | 0.032 | 0 | 0.13 | 0.07 | 0.08 | 0.07 | 0.12 | 0.07 |
| 11 | 13483590 | rs2028976 | -0.17 | -0.229 | -0.193 | 0.341 | -0.008 | -0.009 | -0.009 | 0 | 0.11 | 0.1 | 0.3 | 0.17 | 0.15 | 0.1 |
| 11 | 13496731 | rs10766083 | 0.792 | -0.34 | -0.083 | 0.239 | 0.015 | 0.043 | 0.043 | 2.623 | 0.59 | 0.19 | 0.39 | 0.31 | 0.66 | 0.6 |
| 11 | 13498641 | rs10766084 | 0.914 | -0.336 | -0.106 | 0.214 | 0.027 | 0.035 | 0.035 | 3.223 | 0.6 | 0.19 | 0.41 | 0.31 | 0.72 | 0.63 |
| 11 | 13514263 | rs6254 | 1.673 | 0.122 | 0.029 | -0.066 | -0.069 | 0.054 | 0.054 | 0 | 0.25 | 0.07 | 0.22 | 0.12 | 0.36 | 0.19 |
| 11 | 13516194 | rs192802 | -0.69 | 0.101 | 0.333 | -0.069 | -0.003 | -0.007 | -0.007 | 0 | 0.37 | 0.43 | 0.57 | 0.69 | 0.34 | 0.28 |
| 11 | 13533893 | rs307222 | 0.449 | 0.057 | 0.447 | 0.003 | -0.026 | 0.07 | 0.07 | 0 | 0.22 | 0.29 | 0.24 | 0.52 | 0.15 | 0.17 |
| 11 | 13569182 | rs7107207 | 1.667 | 0.016 | 0.535 | -0.038 | -0.013 | 0.017 | 0.017 | 0 | 0.25 | 0.22 | 0.22 | 0.06 | 0.36 | 0.16 |
| 11 | 13571211 | rs4757167 | -0.659 | 0.042 | 0.336 | -0.033 | -0.004 | -0.007 | -0.007 | 0 | 0.37 | 0.42 | 0.57 | 0.69 | 0.34 | 0.28 |
| 11 | 13573825 | rs10500784 | 1.728 | 0.056 | 0.069 | -0.04 | -0.069 | 0.11 | 0.11 | 0 | 0.24 | 0.07 | 0.2 | 0.06 | 0.36 | 0.16 |
| 11 | 13574634 | rs10832063 | 1.647 | 0.089 | 0.084 | -0.034 | 0.028 | 0.046 | 0.046 | 1.341 | 0.24 | 0.06 | 0.11 | 0.13 | 0.17 | 0.33 |
| 11 | 13583784 | rs10500786 | 0.671 | 0.16 | 0.065 | -0.021 | -0.082 | 0.096 | 0.096 | 0 | 0.36 | 0.15 | 0.29 | 0.17 | 0.48 | 0.26 |
| 11 | 13584585 | rs11022886 | -0.236 | 0.214 | 0.511 | 0.362 | -0.046 | -0.036 | -0.036 | 0 | 0.14 | 0.06 | 0.33 | 0.17 | 0.19 | 0.11 |
| 11 | 13617608 | rs2036417 | 1.027 | 0.191 | 0.622 | 0.573 | 0.002 | -0.006 | -0.006 | 0 | 0.61 | 0.33 | 0.51 | 0.57 | 0.64 | 0.66 |
| 11 | 13618804 | rs7125774 | 0.657 | 0.174 | 0.721 | 0.563 | -0.001 | 0.003 | 0.003 | 1.968 | 0.64 | 0.59 | 0.53 | 0.57 | 0.64 | 0.77 |
| 11 | 13624089 | rs7948130 | -0.332 | 0.222 | 0.073 | 0.629 | -0.039 | 0 | 0 | 0 | 0.25 | 0.53 | 0.14 | 0.26 | 0.16 | 0.26 |
| 11 | 13625439 | rs1502243 | -0.319 | 0.659 | -0.55 | 0.646 | -0.054 | -0.038 | -0.038 | 0 | 0.14 | 0.06 | 0.33 | 0.18 | 0.2 | 0.08 |
| 11 | 13625619 | rs1502242 | 0.024 | 0.157 | -0.405 | 0.713 | -0.007 | -0.004 | -0.004 | 0.89 | 0.1 | 0.06 | 0.19 | 0.36 | 0.1 | 0.12 |
| 11 | 13628747 | rs10832085 | -1.486 | -0.001 | -0.841 | 0.084 | 0.02 | 0.078 | 0.078 | 0 | 0.51 | 0.66 | 0.65 | 0.86 | 0.43 | 0.51 |
| 11 | 13628890 | rs17455429 | -0.256 | 0.164 | -1.408 | 0.57 | -0.014 | -0.026 | -0.026 | 1.813 | 0.12 | 0.06 | 0.26 | 0.2 | 0.15 | 0.09 |
| 11 | 13629018 | rs10832087 | -0.636 | 0.137 | -1.222 | 0.567 | 0.018 | -0.025 | -0.025 | 0.259 | 0.36 | 0.24 | 0.38 | 0.53 | 0.32 | 0.31 |
| 11 | 13631160 | rs7120428 | -0.309 | 0.803 | -1.056 | 0.204 | -0.005 | -0.007 | -0.007 | 0 | 0.1 | 0.08 | 0.08 | 0 | 0.15 | 0.17 |
| 11 | 13634872 | rs7939526 | 0.463 | 1.051 | -0.794 | -0.547 | 0.003 | -0.045 | -0.045 | 0 | 0.14 | 0.19 | 0.13 | 0.03 | 0.16 | 0.26 |
| 11 | 13635897 | rs10832089 | -1.403 | 1.623 | -0.399 | -0.513 | 0.004 | -0.013 | -0.013 | 0 | 0.41 | 0.56 | 0.45 | 0.26 | 0.5 | 0.57 |
| 11 | 13640479 | rs1580873 | 0.963 | 1.448 | -0.333 | -0.538 | 0 | -0.029 | -0.029 | 0 | 0.15 | 0.16 | 0.12 | 0.03 | 0.15 | 0.26 |
| 11 | 13654078 | rs905278 | 1.374 | 1.046 | -1.218 | -0.252 | -0.021 | -0.076 | -0.076 | 0 | 0.16 | 0.01 | 0.27 | 0.21 | 0.17 | 0.08 |
| 11 | 13656086 | rs7927727 | 0.127 | 1.555 | -0.92 | -0.484 | -0.008 | 0.003 | 0.003 | 0 | 0.24 | 0.19 | 0.27 | 0.19 | 0.34 | 0.27 |
| 11 | 13658238 | rs10832097 | 1.693 | 1.009 | -0.564 | -0.182 | 0.016 | 0.022 | 0.022 | 1.183 | 0.37 | 0.17 | 0.19 | 0.27 | 0.28 | 0.31 |
| 11 | 13659797 | rs12420514 | 1.626 | 1.175 | -0.507 | -0.026 | -0.005 | -0.007 | -0.007 | 0.088 | 0.1 | 0.08 | 0.08 | 0 | 0.15 | 0.17 |
| 11 | 13660543 | rs12787381 | 1.456 | 1.168 | -0.482 | 0.032 | -0.004 | -0.004 | -0.004 | 0 | 0.14 | 0.1 | 0.19 | 0.19 | 0.18 | 0.11 |
| 11 | 13661321 | rs12223081 | 1.688 | 0.888 | -0.825 | -0.111 | -0.01 | -0.025 | -0.025 | 0.005 | 0.07 | 0 | 0.05 | 0.26 | 0.06 | 0.17 |
| 11 | 13680829 | rs35510318 | 1.515 | 0.94 | -0.432 | -0.225 | 0.002 | -0.029 | -0.029 | 0.132 | 0.11 | 0.13 | 0.09 | 0 | 0.15 | 0.17 |
| 11 | 13689583 | rs2173994 | -0.321 | 0.404 | -0.549 | -0.367 | 0 | 0.004 | 0.004 | 0 | 0.46 | 0.56 | 0.48 | 0.27 | 0.54 | 0.58 |
| 11 | 13698833 | rs10500796 | 1.507 | 0.407 | -0.523 | -0.383 | 0.01 | 0.005 | 0.005 | 0 | 0.15 | 0.16 | 0.2 | 0.2 | 0.18 | 0.11 |
| 11 | 13700271 | rs7129719 | -0.362 | 0.374 | -0.536 | -0.389 | 0 | 0.005 | 0.005 | 0 | 0.46 | 0.55 | 0.48 | 0.27 | 0.54 | 0.58 |
| 11 | 13724572 | rs10500797 | 1.351 | 0.277 | -0.686 | -0.384 | 0.004 | 0.002 | 0.002 | 1.935 | 0.33 | 0.17 | 0.18 | 0.29 | 0.27 | 0.3 |
| 11 | 13731446 | rs7944263 | 0.481 | 0.588 | -0.631 | -0.732 | -0.002 | -0.013 | -0.013 | 0 | 0.29 | 0.27 | 0.22 | 0.04 | 0.32 | 0.44 |
| 11 | 13731642 | rs17535089 | 1.38 | 0.605 | -0.635 | -0.715 | 0 | -0.001 | -0.001 | 0.402 | 0.14 | 0.12 | 0.19 | 0.19 | 0.18 | 0.11 |
| 11 | 13739556 | rs7926942 | 0.288 | 0.413 | -0.691 | -0.914 | 0 | -0.048 | -0.048 | 0 | 0.3 | 0.32 | 0.23 | 0.04 | 0.32 | 0.44 |
| 11 | 13758789 | rs1866948 | 1.276 | 0.344 | -1.024 | -0.428 | -0.043 | 0.021 | 0.021 | 0.583 | 0.1 | 0.01 | 0.32 | 0.03 | 0.12 | 0.09 |
| 11 | 13761491 | rs17535767 | 0.235 | 0.407 | -1.232 | -0.442 | -0.007 | 0.002 | 0.002 | 0 | 0.22 | 0.1 | 0.14 | 0.17 | 0.21 | 0.27 |
| 11 | 13762105 | rs310082 | -0.414 | -0.037 | -0.602 | -0.146 | 0 | 0 | 0 | 0.397 | 0.47 | 0.53 | 0.6 | 0.47 | 0.46 | 0.39 |
| 11 | 13764295 | rs723789 | 0.442 | -0.222 | -0.361 | -0.068 | 0.005 | -0.059 | -0.059 | 0 | 0.33 | 0.13 | 0.23 | 0.45 | 0.31 | 0.3 |
| 11 | 13767027 | rs1730833 | 0.213 | -0.586 | -0.201 | -0.078 | 0.008 | -0.013 | -0.013 | 0.348 | 0.35 | 0.33 | 0.24 | 0.45 | 0.31 | 0.31 |
| 11 | 13770831 | rs10766109 | 0.24 | -0.469 | -0.269 | -0.092 | -0.002 | 0.019 | 0.019 | 0 | 0.25 | 0.26 | 0.21 | 0.44 | 0.26 | 0.29 |
| 11 | 13782568 | rs7926320 | 0.218 | -0.374 | -0.451 | -0.235 | -0.002 | 0.027 | 0.027 | 0 | 0.6 | 0.36 | 0.41 | 0.28 | 0.59 | 0.53 |
| 11 | 13786793 | rs894203 | 0.723 | -0.553 | -0.942 | -0.242 | -0.016 | 0.143 | 0.143 | 0 | 0.49 | 0.21 | 0.3 | 0.25 | 0.5 | 0.43 |
| 11 | 13795424 | rs11604748 | 1.485 | -0.526 | -0.637 | -0.3 | 0.017 | -0.026 | -0.026 | 0 | 0.11 | 0.16 | 0.17 | 0.03 | 0.11 | 0.11 |
| 11 | 13798118 | rs16913144 | 1.864 | -0.601 | -1.089 | -0.563 | -0.038 | -0.05 | -0.05 | 0.031 | 0.13 | 0.01 | 0.27 | 0.15 | 0.16 | 0.08 |
| 11 | 13800622 | rs7949225 | -0.433 | -0.803 | -0.674 | 0.135 | 0 | 0.005 | 0.005 | 0 | 0.35 | 0.37 | 0.37 | 0.48 | 0.37 | 0.38 |
| 11 | 13802396 | rs12282103 | 2.695 | -0.741 | -1.436 | 0.094 | -0.027 | 0.058 | 0.058 | 0 | 0.1 | 0 | 0.05 | 0 | 0.11 | 0.06 |
| 11 | 13804757 | rs310045 | 0.891 | -0.722 | -1.244 | 0.23 | 0.016 | 0.135 | 0.135 | 0 | 0.51 | 0.25 | 0.49 | 0.23 | 0.56 | 0.44 |
| 11 | 13805483 | rs7108248 | 0.373 | -0.525 | -1.323 | 0.431 | 0.001 | -0.061 | -0.061 | 0.255 | 0.23 | 0.13 | 0.2 | 0.46 | 0.25 | 0.27 |
| 11 | 13810627 | rs310057 | 0.208 | -0.631 | -1.016 | -0.092 | -0.03 | 0.04 | 0.04 | 0 | 0.82 | 0.58 | 0.85 | 0.79 | 0.9 | 0.78 |
| 11 | 13812002 | rs7937076 | 0.74 | -0.227 | -1.571 | -0.358 | -0.028 | 0.121 | 0.121 | 0.209 | 0.45 | 0.08 | 0.47 | 0.28 | 0.48 | 0.42 |
| 11 | 13813234 | rs2119673 | 0.168 | -0.453 | -1.077 | -0.028 | 0 | 0 | 0 | 0 | 0.55 | 0.54 | 0.53 | 0.46 | 0.6 | 0.61 |
| 11 | 13814730 | rs7942935 | -0.104 | -0.065 | -0.916 | 0.219 | 0.001 | -0.006 | -0.006 | 0 | 0.61 | 0.66 | 0.57 | 0.46 | 0.64 | 0.63 |
| 11 | 13821182 | rs10832115 | 0.365 | 0.008 | -0.943 | 0.188 | 0 | 0.005 | 0.005 | 0 | 0.51 | 0.53 | 0.4 | 0.41 | 0.52 | 0.56 |

| | | | | | | | | | | | | | | | |
|---|---|---|---|---|---|---|---|---|---|---|---|---|---|---|---|
| 11 | 13824801 | rs10832116 | 1.021 | -0.139 | -1.029 | 0.133 | -0.024 | 0.036 | 0.036 | 0 | 0.38 | 0.11 | 0.27 | 0.31 | 0.43 | 0.41 |
| 11 | 13850832 | rs12798238 | 1.606 | 0.003 | -1.06 | 0.125 | -0.037 | -0.016 | -0.016 | 0.798 | 0.07 | 0.01 | 0.14 | 0.06 | 0.09 | 0.06 |
| 11 | 13854726 | rs12146602 | 0.477 | -0.028 | -1.015 | -0.004 | -0.02 | 0.023 | 0.023 | 0.397 | 0.11 | 0.26 | 0.12 | 0.1 | 0.07 | 0.15 |
| 11 | 13859998 | rs1507535 | -0.354 | -0.082 | -0.841 | -0.068 | -0.001 | 0 | 0 | 0 | 0.47 | 0.43 | 0.58 | 0.59 | 0.45 | 0.43 |
| 11 | 13863404 | rs1507534 | -1.447 | -0.113 | -0.941 | -0.052 | -0.003 | 0.041 | 0.041 | 0 | 0.61 | 0.83 | 0.73 | 0.69 | 0.57 | 0.59 |
| 11 | 13863474 | rs1507533 | -0.633 | -0.077 | -0.905 | -0.141 | 0 | 0.005 | 0.005 | 0 | 0.49 | 0.56 | 0.62 | 0.59 | 0.48 | 0.43 |
| 11 | 13864260 | rs11022984 | 0.413 | -0.107 | -1.66 | 0.212 | -0.072 | -0.067 | -0.067 | 0 | 0.32 | 0.03 | 0.38 | 0.34 | 0.32 | 0.29 |
| 11 | 13867368 | rs17537538 | 1.494 | -0.203 | -1.734 | -0.622 | -0.036 | 0.086 | 0.086 | 0 | 0.29 | 0.09 | 0.23 | 0.14 | 0.39 | 0.41 |
| 11 | 13867727 | rs10500799 | 0.012 | -0.003 | -1.75 | -0.544 | -0.097 | -0.155 | -0.155 | 0 | 0.35 | 0.07 | 0.42 | 0.45 | 0.38 | 0.38 |
| 11 | 13871193 | rs7121237 | -1.179 | 0.012 | -1.556 | -0.384 | 0.006 | 0.044 | 0.044 | 0 | 0.6 | 0.78 | 0.72 | 0.69 | 0.57 | 0.57 |
| 11 | 13875937 | rs4086666 | 1.253 | 0.871 | -1.52 | -0.003 | -0.025 | 0.059 | 0.059 | 0 | 0.11 | 0.03 | 0.06 | 0 | 0.12 | 0.09 |
| 11 | 13876119 | rs4756771 | 0.1 | 0.928 | -1.036 | 0.636 | -0.034 | -0.132 | -0.132 | 0 | 0.17 | 0.04 | 0.33 | 0.45 | 0.21 | 0.3 |
| 11 | 13881941 | rs5011495 | 0.005 | 0.735 | -1.013 | 0.661 | -0.058 | -0.139 | -0.139 | 0 | 0.18 | 0.04 | 0.37 | 0.45 | 0.23 | 0.3 |
| 11 | 13883385 | rs6486150 | -0.699 | 0.389 | -0.102 | -0.148 | 0.071 | 0.025 | 0.025 | 0 | 0.05 | 0.29 | 0.04 | 0 | 0.03 | 0.06 |
| 11 | 13893667 | rs11022995 | -0.043 | 0.485 | -1.144 | -0.505 | -0.001 | 0.027 | 0.027 | 0 | 0.6 | 0.7 | 0.71 | 0.87 | 0.55 | 0.51 |
| 11 | 13894144 | rs7130288 | -0.176 | 0.463 | -2.128 | 0.274 | -0.017 | -0.101 | -0.101 | 0 | 0.57 | 0.1 | 0.47 | 0.5 | 0.55 | 0.71 |
| 11 | 13894472 | rs990081 | -0.13 | 0.34 | -1.553 | 0.896 | 0.065 | -0.001 | -0.001 | 0 | 0.5 | 0.3 | 0.47 | 0.47 | 0.44 | 0.42 |
| 11 | 13895577 | rs10766117 | -0.033 | 0.508 | -0.565 | 1.235 | 0.003 | 0 | 0 | 0.933 | 0.52 | 0.44 | 0.38 | 0.25 | 0.43 | 0.4 |
| 11 | 13897617 | rs7116819 | 0.539 | -0.226 | -0.578 | 0.867 | -0.03 | -0.023 | -0.023 | 0 | 0.15 | 0.18 | 0.06 | 0.06 | 0.08 | 0.06 |
| 11 | 13900090 | rs11022997 | 0.624 | 0.858 | -0.878 | 1.021 | -0.028 | -0.021 | -0.021 | 0 | 0.07 | 0.04 | 0.09 | 0.25 | 0.16 | 0.12 |
| 11 | 13901466 | rs1503314 | 0.041 | 1.151 | -0.678 | 0.94 | -0.009 | -0.011 | -0.011 | 0.22 | 0.77 | 0.78 | 0.74 | 0.43 | 0.67 | 0.58 |
| 11 | 13903788 | rs17538757 | 1.66 | 1.404 | -0.342 | 0.664 | 0.016 | -0.035 | -0.035 | 0 | 0.12 | 0.2 | 0.12 | 0.01 | 0.13 | 0.04 |
| 11 | 13911295 | rs4757205 | 1.041 | 1.186 | -0.569 | 0.55 | -0.005 | 0.089 | 0.089 | 0 | 0.46 | 0.17 | 0.54 | 0.35 | 0.46 | 0.46 |
| 11 | 13912746 | rs1393878 | 0.704 | 1.336 | -0.39 | 0.82 | -0.03 | -0.003 | -0.003 | 0.046 | 0.07 | 0.04 | 0.17 | 0.32 | 0.15 | 0.16 |
| 11 | 13913003 | rs3898059 | 0.36 | 0.831 | -0.172 | 0.232 | -0.007 | 0.063 | 0.063 | 0.594 | 0.19 | 0.07 | 0.14 | 0.03 | 0.17 | 0.08 |
| 11 | 13917687 | rs7105593 | 1.04 | 1.046 | 0.068 | 0.325 | 0.028 | -0.05 | -0.05 | 0 | 0.13 | 0.25 | 0.12 | 0.01 | 0.13 | 0.04 |
| 11 | 13925137 | rs12416997 | 0.149 | 0.778 | -0.174 | 0.201 | -0.022 | -0.03 | -0.03 | 0.048 | 0.13 | 0.15 | 0.04 | 0.03 | 0.06 | 0.08 |
| 11 | 13925480 | rs12271326 | -0.564 | 0.669 | 0.309 | 0.113 | -0.038 | -0.084 | -0.084 | 0 | 0.16 | 0.47 | 0.07 | 0.03 | 0.08 | 0.08 |
| 11 | 13925737 | rs10832136 | 0.927 | 1.298 | 0.114 | 0.216 | -0.055 | -0.048 | -0.048 | 0.044 | 0.09 | 0.01 | 0.25 | 0.18 | 0.16 | 0.11 |
| 11 | 13929322 | rs7121835 | -0.248 | 1.218 | 0.195 | 0.198 | -0.031 | -0.071 | -0.071 | 0 | 0.16 | 0.39 | 0.06 | 0.03 | 0.08 | 0.08 |
| 11 | 13934928 | rs4471396 | 0.515 | 1.114 | 0.256 | 0.322 | 0.035 | 0.008 | 0.008 | 0 | 0.63 | 0.24 | 0.78 | 0.67 | 0.75 | 0.83 |
| 11 | 13940895 | rs7925707 | -0.227 | 1.101 | 0.289 | 0.312 | -0.033 | -0.08 | -0.08 | 0.221 | 0.16 | 0.44 | 0.06 | 0.03 | 0.08 | 0.08 |
| 11 | 13942849 | rs12361502 | -0.403 | 1.148 | 0.136 | 0.339 | -0.003 | 0.023 | 0.023 | 0.148 | 0.76 | 0.39 | 0.82 | 0.71 | 0.81 | 0.92 |
| 11 | 13946865 | rs1027837 | 0.884 | 1.125 | -0.029 | 0.296 | -0.004 | -0.004 | -0.004 | 0 | 0.55 | 0.44 | 0.54 | 0.49 | 0.59 | 0.73 |
| 11 | 13947545 | rs4757208 | -0.097 | 1.431 | -0.053 | 0.512 | -0.038 | -0.056 | -0.056 | 0.557 | 0.72 | 0.86 | 0.61 | 0.53 | 0.67 | 0.81 |
| 11 | 13949272 | rs1354900 | 0.856 | 1.325 | -0.368 | -0.049 | -0.001 | -0.027 | -0.027 | 0 | 0.17 | 0.17 | 0.17 | 0.02 | 0.23 | 0.15 |
| 11 | 13950967 | rs7932860 | 1.144 | 1.065 | -0.466 | -0.108 | 0.001 | -0.002 | -0.002 | 0.085 | 0.4 | 0.43 | 0.4 | 0.47 | 0.38 | 0.6 |
| 11 | 13952585 | rs16913462 | -0.443 | 0.488 | -0.418 | -0.156 | 0 | -0.065 | -0.065 | 0 | 0.3 | 0.35 | 0.21 | 0.06 | 0.29 | 0.21 |
| 11 | 13953404 | rs16913463 | 1.49 | 0.444 | -0.936 | 0.086 | -0.008 | -0.039 | -0.039 | 0 | 0.11 | 0 | 0.18 | 0.17 | 0.13 | 0.18 |
| 11 | 13954368 | rs4757211 | 1.773 | 0.659 | -0.843 | 0.076 | -0.038 | 0.033 | 0.033 | 0 | 0.28 | 0.15 | 0.16 | 0.21 | 0.33 | 0.36 |
| 11 | 13961461 | rs2403712 | 0.45 | 0.808 | -0.516 | 0.11 | -0.056 | -0.064 | -0.064 | 0 | 0.15 | 0.4 | 0.05 | 0.04 | 0.06 | 0.06 |
| 11 | 13961949 | rs11023028 | 1.501 | 0.823 | -0.575 | 0.2 | -0.022 | -0.048 | -0.048 | 0 | 0.08 | 0 | 0.22 | 0.18 | 0.07 | 0.1 |
| 11 | 13962364 | rs7120733 | 1.094 | 0.835 | -0.578 | 0.206 | -0.02 | -0.04 | -0.04 | 0 | 0.4 | 0.23 | 0.41 | 0.47 | 0.46 | 0.6 |
| 11 | 13962704 | rs6486160 | 0.386 | 0.545 | -0.399 | -0.051 | -0.056 | -0.064 | -0.064 | 2.406 | 0.15 | 0.4 | 0.05 | 0.04 | 0.06 | 0.06 |
| 11 | 13963368 | rs10832148 | -0.156 | 0.761 | -0.437 | 0.204 | -0.003 | 0.003 | 0.003 | 0 | 0.63 | 0.66 | 0.68 | 0.68 | 0.59 | 0.77 |
| 11 | 13965040 | rs1919299 | 0.34 | 0.815 | -0.489 | 0.208 | 0.002 | 0.015 | 0.015 | 0.013 | 0.53 | 0.44 | 0.64 | 0.64 | 0.56 | 0.74 |
| 11 | 13966344 | rs2049723 | 0.368 | 1.276 | -0.445 | 0.562 | -0.009 | -0.052 | -0.052 | 0 | 0.21 | 0.08 | 0.17 | 0.32 | 0.19 | 0.1 |
| 11 | 13967222 | rs1528663 | -1.041 | 1.517 | -0.217 | 0.387 | 0.014 | -0.084 | -0.084 | 0 | 0.84 | 0.69 | 0.82 | 0.98 | 0.79 | 0.88 |
| 11 | 13975534 | rs7107108 | -0.901 | 1.522 | -0.337 | 0.389 | 0.014 | -0.101 | -0.101 | 0 | 0.83 | 0.62 | 0.82 | 0.98 | 0.79 | 0.88 |
| 11 | 13978737 | rs732989 | 1.258 | 1.624 | -0.777 | 0.578 | -0.007 | -0.031 | -0.031 | 0 | 0.13 | 0.08 | 0.12 | 0.29 | 0.12 | 0.05 |
| 11 | 13990280 | rs1528658 | 0.723 | 1.972 | -1.021 | 0.488 | 0.039 | 0.026 | 0.026 | 0 | 0.13 | 0.22 | 0.25 | 0.2 | 0.21 | 0.17 |
| 11 | 13993498 | rs1881516 | 0.963 | 0.986 | -1.662 | 0.045 | 0.034 | 0.051 | 0.051 | 0 | 0.37 | 0.22 | 0.17 | 0.17 | 0.24 | 0.39 |
| 11 | 13996155 | rs1819084 | -0.843 | 1.447 | -1.479 | 0.422 | -0.02 | -0.08 | -0.08 | 0 | 0.23 | 0.09 | 0.28 | 0.47 | 0.26 | 0.24 |
| 11 | 13996226 | rs1830390 | -0.527 | 1.342 | -1.101 | -0.346 | -0.015 | -0.027 | -0.027 | 0 | 0.11 | 0.01 | 0.16 | 0.17 | 0.14 | 0.18 |
| 11 | 13996686 | rs1852755 | 0.474 | 1.745 | -0.331 | -0.662 | 0.072 | -0.137 | -0.137 | 0 | 0.81 | 0.52 | 0.8 | 0.99 | 0.71 | 0.86 |
| 11 | 13997219 | rs17463035 | -1.493 | 1.546 | -0.969 | -0.299 | -0.017 | 0.015 | 0.015 | 0 | 0.09 | 0.04 | 0.07 | 0.07 | 0.11 | 0.18 |
| 11 | 13997808 | rs12575169 | -0.843 | 1.971 | -1.225 | -0.51 | -0.064 | 0.023 | 0.023 | 0 | 0.12 | 0.01 | 0.12 | 0.05 | 0.22 | 0.17 |
| 11 | 14000241 | rs2697844 | 0.161 | 1.789 | -1.584 | -0.662 | -0.031 | -0.016 | -0.016 | 0.014 | 0.32 | 0.18 | 0.55 | 0.7 | 0.37 | 0.38 |
| 11 | 14002819 | rs2697846 | 0.062 | 0.959 | -1.122 | -1.464 | -0.003 | -0.001 | -0.001 | 0.034 | 0.64 | 0.8 | 0.76 | 0.7 | 0.6 | 0.48 |
| 11 | 14006744 | rs2618511 | 0.024 | 0.934 | -1.093 | -1.444 | -0.001 | -0.004 | -0.004 | 0 | 0.65 | 0.81 | 0.77 | 0.7 | 0.61 | 0.48 |
| 11 | 14021639 | rs2618516 | 0.063 | 0.836 | -1.047 | -1.468 | 0.003 | 0 | 0 | 0 | 0.64 | 0.74 | 0.76 | 0.7 | 0.62 | 0.48 |
| 11 | 14022014 | rs2697852 | -0.168 | 0.663 | -1.178 | -1.491 | -0.001 | 0 | 0 | 0 | 0.78 | 0.8 | 0.85 | 0.82 | 0.71 | 0.67 |
| 11 | 14029854 | rs1919309 | 0.708 | 0.199 | -0.35 | -1.976 | -0.006 | 0.03 | 0.03 | 0 | 0.45 | 0.32 | 0.63 | 0.7 | 0.5 | 0.43 |
| 11 | 14031713 | rs10500789 | 1.739 | 0.199 | -0.381 | -1.928 | 0.003 | 0.004 | 0.004 | 0 | 0.15 | 0.11 | 0.11 | 0 | 0.12 | 0.05 |
| 11 | 14033524 | rs1528667 | 1.252 | 0.383 | -0.517 | -1.862 | -0.067 | -0.032 | -0.032 | 0 | 0.31 | 0.12 | 0.55 | 0.7 | 0.39 | 0.38 |
| 11 | 14047447 | rs2012072 | -0.386 | 0.241 | -0.554 | -1.665 | -0.023 | -0.073 | -0.073 | 0 | 0.34 | 0.06 | 0.22 | 0.33 | 0.37 | 0.51 |
| 11 | 14048480 | rs10832164 | 0.756 | 0.094 | -0.279 | -1.784 | -0.006 | 0.018 | 0.018 | 0 | 0.46 | 0.28 | 0.62 | 0.67 | 0.51 | 0.42 |
| 11 | 14058186 | rs10766134 | -0.38 | 0.169 | -0.355 | -1.722 | -0.006 | -0.067 | -0.067 | 0 | 0.35 | 0.09 | 0.23 | 0.33 | 0.37 | 0.52 |
| 11 | 14058278 | rs12283632 | 0.053 | 0.272 | -0.286 | -1.621 | 0 | -0.001 | -0.001 | 0 | 0.22 | 0.2 | 0.16 | 0.18 | 0.3 | 0.33 |

| | | | | | | | | | | | | | | | | |
|---|---|---|---|---|---|---|---|---|---|---|---|---|---|---|---|---|
| 11 | 14066486 | rs10832169 | -0.664 | 0.246 | -0.453 | -1.663 | 0.007 | 0.042 | 0.042 | 0 | 0.51 | 0.2 | 0.33 | 0.33 | 0.5 | 0.58 |
| 11 | 14068251 | rs4756773 | 1.165 | 0.329 | -0.614 | -1.665 | -0.09 | -0.072 | -0.072 | 0 | 0.32 | 0.1 | 0.54 | 0.67 | 0.39 | 0.38 |
| 11 | 14072663 | rs2403708 | 0.119 | 0.42 | -0.5 | -1.43 | -0.032 | -0.023 | -0.023 | 0 | 0.2 | 0.05 | 0.15 | 0.2 | 0.3 | 0.33 |
| 11 | 14080925 | rs2697823 | 0.355 | 0.094 | -0.354 | -1.508 | 0.027 | 0.04 | 0.04 | 0 | 0.3 | 0.19 | 0.18 | 0.13 | 0.2 | 0.25 |
| 11 | 14089431 | rs2697825 | 0.272 | 0.093 | -0.148 | -1.523 | 0.014 | 0.021 | 0.021 | 0 | 0.31 | 0.25 | 0.19 | 0.13 | 0.2 | 0.25 |
| 11 | 14089499 | rs1919297 | -0.12 | 0.115 | -0.603 | -1.858 | -0.037 | 0.004 | 0.004 | 0 | 0.25 | 0.06 | 0.19 | 0.2 | 0.33 | 0.42 |
| 11 | 14089741 | rs1528646 | -0.107 | 0.118 | -0.602 | -1.857 | -0.038 | 0.004 | 0.004 | 0 | 0.25 | 0.06 | 0.19 | 0.2 | 0.33 | 0.42 |
| 11 | 14090461 | rs2697827 | -0.934 | 0.141 | 0.008 | -1.835 | -0.027 | 0.012 | 0.012 | 0 | 0.57 | 0.71 | 0.41 | 0.33 | 0.52 | 0.66 |
| 11 | 14093889 | rs12365614 | 1.238 | 0.23 | -0.056 | -1.953 | -0.048 | -0.046 | -0.046 | 0 | 0.28 | 0.13 | 0.5 | 0.67 | 0.36 | 0.29 |
| 11 | 14097139 | rs2178242 | 1.292 | 1.208 | -0.181 | -1.376 | -0.005 | -0.072 | -0.072 | 3.461 | 0.25 | 0.12 | 0.41 | 0.55 | 0.26 | 0.24 |
| 11 | 14097161 | rs11023075 | 0.296 | 0.708 | -0.23 | -1.423 | -0.051 | -0.066 | -0.066 | 0.015 | 0.22 | 0.06 | 0.17 | 0.31 | 0.31 | 0.41 |
| 11 | 14102118 | rs2697833 | -1.19 | 0.855 | 0.002 | -1.461 | -0.002 | -0.151 | -0.151 | 1.962 | 0.81 | 0.49 | 0.82 | 1 | 0.84 | 0.92 |
| 11 | 14102228 | rs7102409 | 1.421 | 0.662 | -0.13 | -1.602 | 0.04 | -0.022 | -0.022 | 0.437 | 0.17 | 0.04 | 0.1 | 0.15 | 0.08 | 0.2 |
| 11 | 14103114 | rs11023081 | 1.459 | 0.701 | -0.188 | -1.61 | -0.027 | -0.06 | -0.06 | 0.016 | 0.26 | 0.16 | 0.44 | 0.55 | 0.32 | 0.26 |
| 11 | 14103801 | rs1881512 | 1.536 | 0.716 | -0.232 | -1.602 | -0.035 | -0.073 | -0.073 | 2.665 | 0.25 | 0.12 | 0.44 | 0.55 | 0.32 | 0.25 |
| 11 | 14105497 | rs6486166 | -0.466 | 0.679 | -0.207 | -1.602 | -0.033 | -0.044 | -0.044 | 0 | 0.64 | 0.48 | 0.72 | 0.85 | 0.76 | 0.73 |
| 11 | 14113759 | rs7116296 | -0.686 | 0.775 | -0.509 | -1.561 | -0.007 | -0.007 | -0.007 | 0.146 | 0.55 | 0.37 | 0.38 | 0.45 | 0.52 | 0.67 |
| 11 | 14114044 | rs7941345 | -1.279 | 1.319 | -0.13 | -1.532 | 0 | -0.028 | -0.028 | 0 | 0.83 | 0.83 | 0.86 | 1 | 0.84 | 0.92 |
| 11 | 14114318 | rs12795982 | -0.177 | 1.234 | -0.199 | -1.025 | -0.039 | -0.053 | -0.053 | 0 | 0.21 | 0.06 | 0.17 | 0.31 | 0.31 | 0.41 |
| 11 | 14114334 | rs6486167 | 0.655 | 1.137 | 0.03 | -1.338 | -0.004 | -0.091 | -0.091 | 0 | 0.18 | 0.29 | 0.11 | 0 | 0.13 | 0.06 |
| 11 | 14124311 | rs1609341 | 0.546 | 1.475 | 0.002 | -0.994 | 0.034 | -0.053 | -0.053 | 0 | 0.77 | 0.56 | 0.77 | 0.85 | 0.66 | 0.62 |
| 11 | 14127493 | rs1528654 | 0.232 | 1.399 | -0.522 | -0.651 | 0.002 | -0.004 | -0.004 | 2.974 | 0.38 | 0.41 | 0.51 | 0.25 | 0.41 | 0.35 |
| 11 | 14128875 | rs10734220 | 0.129 | 1.797 | -0.517 | -0.434 | 0.004 | -0.001 | -0.001 | 0 | 0.86 | 0.85 | 0.83 | 0.88 | 0.71 | 0.79 |
| 11 | 14133293 | rs1919302 | 0.417 | 1.756 | -0.501 | -0.406 | -0.002 | 0 | 0 | 0 | 0.76 | 0.76 | 0.78 | 0.83 | 0.7 | 0.75 |
| 11 | 14134514 | rs4757226 | 0.429 | 1.32 | -0.678 | -0.126 | 0.008 | -0.021 | -0.021 | 0 | 0.35 | 0.23 | 0.32 | 0.56 | 0.36 | 0.33 |
| 11 | 14156176 | rs10832187 | 0.338 | 1.261 | -0.461 | 0.028 | 0.002 | -0.014 | -0.014 | 0.371 | 0.41 | 0.26 | 0.38 | 0.52 | 0.44 | 0.39 |
| 11 | 14159137 | rs7941383 | 0.216 | 1.221 | -0.36 | 0.088 | 0.005 | -0.011 | -0.011 | 0 | 0.42 | 0.32 | 0.38 | 0.54 | 0.44 | 0.39 |
| 11 | 14169984 | rs10741636 | 0.243 | 1.194 | -0.141 | -0.018 | 0.001 | -0.001 | -0.001 | 0 | 0.42 | 0.36 | 0.39 | 0.55 | 0.45 | 0.39 |
| 11 | 14185693 | rs10832223 | 0.724 | 0.985 | -0.295 | -0.331 | -0.001 | -0.004 | -0.004 | 0 | 0.25 | 0.23 | 0.26 | 0.12 | 0.16 | 0.18 |
| 11 | 14186021 | rs11023121 | 0.724 | 0.985 | -0.295 | -0.331 | -0.001 | -0.004 | -0.004 | 0 | 0.25 | 0.23 | 0.26 | 0.12 | 0.16 | 0.18 |
| 11 | 14198018 | rs11602192 | -0.761 | 1.299 | -0.457 | -0.215 | -0.025 | 0.003 | 0.003 | 0 | 0.86 | 0.91 | 0.86 | 0.87 | 0.82 | 0.82 |
| 11 | 14206679 | rs6486178 | -0.674 | 1.284 | -0.22 | -0.225 | 0.005 | -0.002 | -0.002 | 1.802 | 0.85 | 0.83 | 0.86 | 0.87 | 0.82 | 0.82 |
| 11 | 14213229 | rs7480396 | 0.546 | 1.112 | -0.242 | -0.102 | -0.001 | -0.004 | -0.004 | 0.177 | 0.42 | 0.33 | 0.37 | 0.56 | 0.46 | 0.39 |
| 11 | 14224883 | rs4756776 | -0.145 | 0.657 | -0.583 | -0.262 | 0.005 | 0.044 | 0.044 | 0 | 0.4 | 0.23 | 0.39 | 0.24 | 0.38 | 0.45 |
| 11 | 14224997 | rs4757240 | -0.553 | 0.827 | 0.044 | 0.046 | 0.001 | 0.001 | 0.001 | 0 | 0.48 | 0.54 | 0.54 | 0.42 | 0.45 | 0.52 |
| 11 | 14225465 | rs10766169 | 0.843 | 0.603 | -0.132 | -0.09 | 0.002 | 0.012 | 0.012 | 0 | 0.27 | 0.19 | 0.28 | 0.13 | 0.22 | 0.21 |
| 11 | 14233522 | rs6486182 | -0.842 | 0.489 | -0.143 | -0.041 | -0.003 | 0 | 0 | 0.01 | 0.55 | 0.6 | 0.6 | 0.44 | 0.49 | 0.59 |
| 11 | 14239961 | rs10766175 | 0.059 | 0.439 | -0.048 | 0.001 | 0.008 | 0.015 | 0.015 | 0 | 0.4 | 0.23 | 0.47 | 0.35 | 0.35 | 0.35 |
| 11 | 14241685 | rs4128525 | 0.024 | 0.426 | -0.058 | -0.005 | 0.001 | 0.009 | 0.009 | 0 | 0.39 | 0.17 | 0.46 | 0.35 | 0.35 | 0.35 |
| 11 | 14244240 | rs10832235 | 1.085 | 0.403 | -0.033 | 0.022 | 0 | 0.002 | 0.002 | 0 | 0.12 | 0.04 | 0.1 | 0.1 | 0.14 | 0.23 |
| 11 | 14244966 | rs10766176 | 0.688 | 0.298 | 0.064 | -0.026 | 0.001 | 0.001 | 0.001 | 0 | 0.32 | 0.26 | 0.33 | 0.16 | 0.25 | 0.27 |
| 11 | 14245688 | rs11023150 | 0.756 | 0.251 | -0.11 | -0.183 | -0.014 | -0.023 | -0.023 | 0 | 0.32 | 0.28 | 0.33 | 0.51 | 0.39 | 0.33 |
| 11 | 14246296 | rs1025412 | 0.077 | 0.39 | -0.56 | -0.225 | 0.003 | -0.006 | -0.006 | 0 | 0.46 | 0.5 | 0.45 | 0.62 | 0.52 | 0.57 |
| 11 | 14253101 | rs6486183 | -0.643 | 0.406 | -1.125 | -0.152 | -0.005 | 0.015 | 0.015 | 0.013 | 0.81 | 0.86 | 0.86 | 0.89 | 0.77 | 0.84 |
| 11 | 14253351 | rs7103919 | 0.861 | 0.392 | -1.137 | -0.148 | 0.017 | 0.011 | 0.011 | 0 | 0.33 | 0.19 | 0.4 | 0.26 | 0.24 | 0.27 |
| 11 | 14253824 | rs1123875 | -0.359 | 0.484 | -1.327 | -0.108 | -0.005 | 0.014 | 0.014 | 0.014 | 0.81 | 0.87 | 0.87 | 0.89 | 0.78 | 0.84 |
| 11 | 14254570 | rs4756781 | -0.057 | 0.325 | -1.402 | -0.192 | -0.003 | 0.024 | 0.024 | 0.006 | 0.79 | 0.86 | 0.87 | 0.89 | 0.76 | 0.84 |
| 11 | 14254606 | rs4756782 | 0.581 | 0.417 | -1.361 | 0.242 | 0.001 | 0.003 | 0.003 | 0 | 0.12 | 0.02 | 0.11 | 0.1 | 0.16 | 0.22 |
| 11 | 14255818 | rs10766177 | -0.268 | 0.819 | -1.708 | 0.474 | -0.04 | 0.016 | 0.016 | 0 | 0.41 | 0.2 | 0.48 | 0.33 | 0.51 | 0.42 |
| 11 | 14256184 | rs12291225 | -0.635 | 0.831 | -1.872 | 1.28 | -0.022 | -0.115 | -0.115 | 0 | 0.22 | 0.04 | 0.16 | 0.45 | 0.28 | 0.31 |
| 11 | 14256444 | rs1864658 | -0.772 | 1.006 | -1.773 | 1.238 | -0.034 | -0.099 | -0.099 | 3.903 | 0.24 | 0.07 | 0.28 | 0.49 | 0.32 | 0.34 |
| 11 | 14262375 | rs7116756 | -0.191 | 0.928 | -1.385 | 0.948 | 0 | 0 | 0 | 0 | 0.73 | 0.65 | 0.77 | 0.71 | 0.73 | 0.71 |
| 11 | 14264916 | rs2303973 | 1.146 | 0.922 | -1.315 | 1.045 | 0.005 | 0.033 | 0.033 | 2.155 | 0.51 | 0.56 | 0.56 | 0.24 | 0.41 | 0.41 |
| 11 | 14264979 | rs2303974 | -0.321 | 0.924 | -1.392 | 1.244 | -0.021 | -0.077 | -0.077 | 0 | 0.23 | 0.09 | 0.25 | 0.5 | 0.32 | 0.34 |
| 11 | 14273677 | rs731274 | -0.261 | 0.863 | -0.81 | 0.951 | -0.009 | 0.017 | 0.017 | 0 | 0.14 | 0.24 | 0.1 | 0.19 | 0.13 | 0.1 |
| 11 | 14275605 | rs2278347 | 0.818 | 0.479 | -1.384 | 0.976 | -0.012 | -0.024 | -0.024 | 0 | 0.16 | 0.01 | 0.18 | 0.12 | 0.18 | 0.18 |
| 11 | 14276931 | rs11023163 | -1.455 | 0.639 | -1.046 | 0.781 | 0.02 | -0.032 | -0.032 | 0.247 | 0.83 | 0.69 | 0.85 | 0.92 | 0.79 | 0.84 |
| 11 | 14285900 | rs4756786 | -0.669 | 0.268 | -0.869 | 1.238 | -0.004 | -0.044 | -0.044 | 0.147 | 0.3 | 0.08 | 0.27 | 0.37 | 0.34 | 0.34 |
| 11 | 14292377 | rs4756787 | -1.702 | 0.341 | -0.78 | 1.168 | 0.035 | 0.03 | 0.03 | 0 | 0.88 | 0.64 | 0.9 | 0.87 | 0.88 | 0.88 |
| 11 | 14297872 | rs4757245 | -0.464 | 0.307 | -1.024 | 1.169 | -0.002 | -0.042 | -0.042 | 0 | 0.3 | 0.09 | 0.28 | 0.37 | 0.34 | 0.35 |
| 11 | 14302533 | rs11023175 | 0.865 | 0.398 | -0.909 | 1.262 | -0.004 | -0.055 | -0.055 | 0 | 0.09 | 0.02 | 0.04 | 0.19 | 0.07 | 0.09 |
| 11 | 14325401 | rs2970335 | 1.647 | 0.431 | -0.84 | 1.18 | -0.003 | 0.009 | 0.009 | 0 | 0.41 | 0.37 | 0.51 | 0.32 | 0.37 | 0.36 |
| 11 | 14340504 | rs10832242 | -0.525 | 0.342 | -0.87 | 1.185 | -0.008 | -0.029 | -0.029 | 0 | 0.33 | 0.09 | 0.31 | 0.37 | 0.38 | 0.42 |
| 11 | 14342761 | rs11023186 | 0.272 | 0.388 | -0.75 | 1.224 | -0.006 | 0.011 | 0.011 | 0 | 0.14 | 0.2 | 0.09 | 0.19 | 0.13 | 0.1 |
| 11 | 14360435 | rs2970332 | 0.121 | 0.398 | -0.723 | 1.444 | -0.005 | 0.098 | 0.098 | 0.344 | 0.74 | 0.46 | 0.76 | 0.6 | 0.75 | 0.77 |
| 11 | 14362060 | rs4756788 | 0.718 | 0.46 | -0.941 | 1.084 | -0.048 | -0.077 | -0.077 | 0 | 0.13 | 0.01 | 0.09 | 0.26 | 0.16 | 0.16 |
| 11 | 14396485 | rs7937890 | -1.871 | 0.6 | -0.656 | 0.825 | -0.006 | 0.014 | 0.014 | 0 | 0.6 | 0.7 | 0.51 | 0.68 | 0.62 | 0.62 |
| 11 | 14409815 | rs11023203 | -0.426 | 0.454 | -0.883 | 0.722 | 0.009 | -0.035 | -0.035 | 1.035 | 0.36 | 0.1 | 0.29 | 0.4 | 0.37 | 0.39 |
| 11 | 14410383 | rs7125877 | 1.716 | 0.37 | -0.56 | 0.631 | 0.009 | -0.043 | -0.043 | 0 | 0.11 | 0.27 | 0.08 | 0.04 | 0.1 | 0.08 |

| | | | | | | | | | | | | | | | | |
|---|---|---|---|---|---|---|---|---|---|---|---|---|---|---|---|---|
| 11 | 14421101 | rs9667144 | -0.32 | 0.441 | -0.69 | 0.7 | 0.021 | -0.039 | -0.039 | 0 | 0.36 | 0.1 | 0.29 | 0.4 | 0.35 | 0.4 |
| 11 | 14424235 | rs12421378 | -0.125 | 0.331 | -0.54 | 0.669 | -0.002 | 0.013 | 0.013 | 0 | 0.15 | 0.22 | 0.16 | 0.17 | 0.15 | 0.11 |
| 11 | 14436733 | rs4382869 | -0.598 | 0.134 | -0.251 | 0.521 | -0.008 | 0.003 | 0.003 | 0.278 | 0.26 | 0.61 | 0.25 | 0.29 | 0.26 | 0.21 |
| 11 | 14447443 | rs12419533 | 0.892 | 0.102 | -0.344 | 0.506 | -0.005 | -0.055 | -0.055 | 0 | 0.1 | 0.02 | 0.1 | 0.18 | 0.08 | 0.09 |
| 11 | 14450531 | rs12295888 | -0.29 | 0.693 | -0.499 | 0.464 | -0.003 | -0.03 | -0.03 | 0.256 | 0.37 | 0.1 | 0.47 | 0.4 | 0.4 | 0.44 |
| 11 | 14478224 | rs10832275 | -0.218 | 0.661 | -0.504 | 0.449 | -0.009 | -0.038 | -0.038 | 0.174 | 0.36 | 0.09 | 0.48 | 0.4 | 0.4 | 0.44 |
| 11 | 14484590 | rs2597203 | 1.713 | 0.632 | -0.207 | 0.442 | 0.02 | -0.005 | -0.005 | 6.866 | 0.11 | 0.31 | 0.09 | 0.12 | 0.12 | 0.1 |
| 11 | 14504463 | rs7121538 | 0.304 | 0.501 | -0.259 | 0.444 | -0.009 | 0.024 | 0.024 | 0 | 0.15 | 0.24 | 0.15 | 0.18 | 0.14 | 0.1 |
| 11 | 14505962 | rs2597193 | -0.28 | 0.528 | -0.67 | 0.406 | -0.007 | 0.01 | 0.01 | 0.182 | 0.26 | 0.55 | 0.24 | 0.3 | 0.25 | 0.21 |
| 11 | 14508698 | rs11023238 | 0.543 | 0.424 | -0.863 | 0.377 | -0.046 | 0.082 | 0.082 | 1.436 | 0.25 | 0.01 | 0.41 | 0.09 | 0.29 | 0.3 |
| 11 | 14510736 | rs2575859 | -2.207 | 0.404 | -0.796 | 0.247 | -0.012 | 0.011 | 0.011 | 0 | 0.63 | 0.74 | 0.73 | 0.7 | 0.65 | 0.64 |
| 11 | 14512060 | rs2597189 | -2.515 | 0.095 | -0.818 | -0.043 | -0.042 | -0.022 | -0.022 | 0.202 | 0.71 | 0.88 | 0.76 | 0.7 | 0.69 | 0.66 |
| 11 | 14516906 | rs2597188 | -1.953 | 0.241 | -0.771 | 0.05 | -0.012 | 0.011 | 0.011 | 0 | 0.63 | 0.75 | 0.73 | 0.7 | 0.65 | 0.64 |
| 11 | 14536956 | rs11023246 | -0.377 | 0.04 | -0.843 | 0.14 | -0.004 | -0.036 | -0.036 | 0.144 | 0.37 | 0.09 | 0.48 | 0.4 | 0.4 | 0.44 |
| 11 | 14573552 | rs16930278 | 1.324 | 0.046 | -0.634 | -0.175 | -0.004 | -0.002 | -0.002 | 0 | 0.07 | 0.04 | 0.04 | 0 | 0.06 | 0.02 |
| 11 | 14579376 | rs12286408 | -0.274 | -0.089 | -0.734 | 0.121 | 0.003 | -0.012 | -0.012 | 0 | 0.35 | 0.09 | 0.47 | 0.38 | 0.38 | 0.44 |
| 11 | 14581762 | rs2597212 | -1.848 | -0.233 | -0.628 | 0.086 | -0.007 | 0.002 | 0.002 | 0 | 0.62 | 0.68 | 0.67 | 0.68 | 0.56 | 0.62 |
| 11 | 14583526 | rs2575843 | 1.628 | -0.263 | -0.53 | 0.067 | -0.001 | -0.028 | -0.028 | 0 | 0.11 | 0.22 | 0.07 | 0.05 | 0.09 | 0.07 |
| 11 | 14603548 | rs2575850 | -0.981 | -0.278 | -0.542 | 0.105 | 0.011 | -0.006 | -0.006 | 0 | 0.47 | 0.42 | 0.55 | 0.54 | 0.48 | 0.52 |
| 11 | 14632570 | rs1403247 | -1.935 | -0.27 | -0.564 | 0.118 | -0.007 | 0.002 | 0.002 | 0.744 | 0.62 | 0.69 | 0.67 | 0.68 | 0.57 | 0.62 |
| 11 | 14636316 | rs10741651 | -0.981 | -0.233 | -0.743 | 0.14 | 0.01 | -0.006 | -0.006 | 0.316 | 0.47 | 0.44 | 0.56 | 0.54 | 0.48 | 0.52 |
| 11 | 14637391 | rs10500802 | 1.23 | -0.191 | -0.779 | 0.257 | -0.008 | -0.056 | -0.056 | 0 | 0.09 | 0.01 | 0.05 | 0.29 | 0.09 | 0.14 |
| 11 | 14644426 | rs7931291 | -2.127 | -0.196 | -0.769 | 0.207 | -0.034 | 0.001 | 0.001 | 0.912 | 0.64 | 0.78 | 0.69 | 0.68 | 0.58 | 0.62 |
| 11 | 14648826 | rs10500803 | 1.131 | 0.123 | -0.951 | 0.379 | -0.005 | -0.055 | -0.055 | 0 | 0.09 | 0 | 0.05 | 0.29 | 0.08 | 0.14 |
| 11 | 14657389 | rs10832288 | 1.149 | 0.346 | -0.741 | 0.298 | -0.001 | 0.011 | 0.011 | 0.086 | 0.12 | 0.3 | 0.08 | 0.16 | 0.1 | 0.07 |
| 11 | 14659511 | rs10458874 | -0.954 | 0.332 | -0.762 | 0.262 | 0.009 | -0.006 | -0.006 | 0 | 0.47 | 0.44 | 0.56 | 0.54 | 0.48 | 0.52 |
| 11 | 14660826 | rs10766188 | -0.157 | 0.386 | -0.883 | 0.292 | 0 | -0.017 | -0.017 | 0 | 0.34 | 0.09 | 0.47 | 0.38 | 0.38 | 0.44 |
| 11 | 14662047 | rs7109770 | 0.911 | 0.662 | -0.584 | 0.572 | -0.001 | 0.008 | 0.008 | 0.113 | 0.12 | 0.24 | 0.07 | 0.16 | 0.1 | 0.07 |
| 11 | 14669802 | rs4757261 | -0.179 | 0.215 | -0.997 | 0.413 | -0.021 | 0.016 | 0.016 | 0 | 0.41 | 0.1 | 0.5 | 0.38 | 0.44 | 0.46 |
| 11 | 14678245 | rs11023293 | 1.219 | 0.104 | -0.768 | 0.468 | -0.007 | 0.013 | 0.013 | 0.096 | 0.11 | 0.24 | 0.07 | 0.16 | 0.09 | 0.07 |
| 11 | 14679017 | rs1403248 | -1.836 | -0.141 | -0.428 | 0.447 | 0 | 0 | 0 | 0.382 | 0.68 | 0.71 | 0.7 | 0.68 | 0.62 | 0.65 |
| 11 | 14679247 | rs1879888 | -0.795 | -0.148 | -0.353 | 0.441 | -0.002 | -0.003 | -0.003 | 0 | 0.53 | 0.46 | 0.59 | 0.54 | 0.53 | 0.55 |
| 11 | 14680514 | rs11023294 | 1.348 | -0.173 | -0.353 | 0.425 | -0.007 | 0.013 | 0.013 | 0.096 | 0.11 | 0.24 | 0.07 | 0.16 | 0.09 | 0.07 |
| 11 | 14685078 | rs11529568 | 1.348 | -0.173 | -0.353 | 0.425 | -0.007 | 0.013 | 0.013 | 0.096 | 0.11 | 0.24 | 0.07 | 0.16 | 0.09 | 0.07 |
| 11 | 14690192 | rs12283049 | 1.411 | -0.336 | -0.632 | 0.183 | -0.083 | 0.068 | 0.068 | 0.434 | 0.16 | 0.01 | 0.38 | 0.06 | 0.25 | 0.19 |
| 11 | 14690962 | rs7119073 | -1.877 | -0.157 | -0.36 | 0.434 | 0 | 0 | 0 | 0.768 | 0.68 | 0.71 | 0.7 | 0.68 | 0.62 | 0.65 |
| 11 | 14696325 | rs1533852 | 1.258 | -0.157 | -0.357 | 0.44 | -0.009 | 0.018 | 0.018 | 0.079 | 0.12 | 0.32 | 0.08 | 0.16 | 0.09 | 0.07 |
| 11 | 14698246 | rs11023304 | 1.12 | -0.157 | -0.357 | 0.44 | -0.011 | 0.019 | 0.019 | 0.13 | 0.12 | 0.36 | 0.09 | 0.16 | 0.09 | 0.08 |
| 11 | 14698458 | rs12417788 | 0.618 | -0.16 | -0.359 | 0.452 | -0.036 | -0.007 | -0.007 | 0 | 0.15 | 0.25 | 0.11 | 0.13 | 0.09 | 0.1 |
| 11 | 14702589 | rs16930437 | 3.121 | -0.197 | -0.391 | 0.417 | -0.001 | 0.027 | 0.027 | 0 | 0.05 | 0 | 0.02 | 0 | 0.04 | 0.02 |
| 11 | 14709324 | rs7938266 | 0.062 | -0.192 | -0.389 | 0.409 | -0.022 | 0.015 | 0.015 | 0 | 0.4 | 0.1 | 0.5 | 0.38 | 0.44 | 0.46 |
| 11 | 14710823 | rs11023307 | 0.935 | -0.186 | -0.377 | 0.416 | -0.007 | 0.013 | 0.013 | 0.096 | 0.11 | 0.24 | 0.07 | 0.16 | 0.09 | 0.07 |
| 11 | 14726242 | rs7942142 | 0.718 | -0.183 | -0.378 | 0.416 | -0.013 | 0.02 | 0.02 | 0.213 | 0.12 | 0.36 | 0.09 | 0.16 | 0.09 | 0.08 |
| 11 | 14736259 | rs7109368 | 0.847 | -0.181 | -0.378 | 0.418 | -0.009 | 0.018 | 0.018 | 0.079 | 0.12 | 0.32 | 0.08 | 0.16 | 0.09 | 0.08 |
| 11 | 14745554 | rs7947041 | -0.664 | -0.199 | -0.281 | 0.703 | -0.055 | 0.107 | 0.107 | 0.01 | 0.83 | 0.66 | 0.87 | 0.7 | 0.91 | 0.84 |
| 11 | 14747427 | rs10832294 | 0.083 | -0.089 | -0.286 | 0.228 | -0.026 | 0.016 | 0.016 | 0 | 0.4 | 0.1 | 0.5 | 0.38 | 0.44 | 0.46 |
| 11 | 14748052 | rs10741654 | -1.922 | -0.09 | -0.279 | 0.231 | 0 | 0 | 0 | 0 | 0.68 | 0.71 | 0.7 | 0.67 | 0.62 | 0.65 |
| 11 | 14750914 | rs7117383 | 0.67 | -0.094 | -0.271 | 0.234 | -0.011 | 0.017 | 0.017 | 0.1 | 0.12 | 0.36 | 0.09 | 0.16 | 0.09 | 0.08 |
| 11 | 14756272 | rs11826363 | 0.701 | -0.108 | -0.274 | 0.236 | -0.011 | 0.017 | 0.017 | 0 | 0.12 | 0.32 | 0.08 | 0.16 | 0.09 | 0.08 |
| 11 | 14757095 | rs16930498 | 0.633 | -0.09 | -0.32 | 0.215 | -0.036 | 0 | 0 | 0.472 | 0.15 | 0.25 | 0.12 | 0.13 | 0.09 | 0.1 |
| 11 | 14757412 | rs16930500 | 0.633 | -0.09 | -0.32 | 0.215 | -0.036 | 0 | 0 | 0.192 | 0.15 | 0.25 | 0.12 | 0.14 | 0.09 | 0.1 |
| 11 | 14769392 | rs10832299 | 0.039 | 0.077 | -0.427 | 0.215 | -0.023 | 0.018 | 0.018 | 0 | 0.4 | 0.1 | 0.5 | 0.38 | 0.44 | 0.46 |
| 11 | 14774591 | rs1007392 | 0.07 | 0.077 | -0.428 | 0.215 | -0.026 | 0.013 | 0.013 | 0 | 0.4 | 0.1 | 0.5 | 0.38 | 0.44 | 0.46 |
| 11 | 14778269 | rs16930516 | 0.638 | 0.061 | -0.4 | 0.215 | -0.036 | -0.002 | -0.002 | 0.7 | 0.15 | 0.25 | 0.12 | 0.13 | 0.09 | 0.1 |
| 11 | 14781621 | rs1451678 | -0.865 | 0.056 | -0.387 | 0.217 | -0.003 | -0.003 | -0.003 | 0 | 0.53 | 0.46 | 0.59 | 0.54 | 0.53 | 0.55 |
| 11 | 14798806 | rs6486202 | -2.12 | 0.003 | -0.351 | 0.179 | -0.013 | -0.006 | -0.006 | 0 | 0.7 | 0.79 | 0.71 | 0.68 | 0.62 | 0.65 |
| 11 | 14806266 | rs1542291 | -2.12 | 0.003 | -0.351 | 0.179 | -0.013 | -0.006 | -0.006 | 0 | 0.7 | 0.79 | 0.71 | 0.68 | 0.62 | 0.65 |
| 11 | 14810762 | rs4757268 | -1.911 | 0.042 | -0.387 | 0.198 | 0.001 | 0 | 0 | 0 | 0.68 | 0.71 | 0.7 | 0.68 | 0.62 | 0.65 |
| 11 | 14813608 | rs11602976 | 0.984 | 0.04 | -0.383 | 0.198 | -0.008 | -0.036 | -0.036 | 0 | 0.11 | 0.25 | 0.07 | 0.05 | 0.09 | 0.07 |
| 11 | 14815233 | rs10832302 | 0.963 | 0.042 | -0.382 | 0.173 | -0.008 | 0.014 | 0.014 | 0.067 | 0.11 | 0.25 | 0.07 | 0.16 | 0.09 | 0.08 |
| 11 | 14825245 | rs10741656 | -1.909 | 0.044 | -0.384 | 0.173 | 0.001 | 0 | 0 | 0.248 | 0.68 | 0.71 | 0.7 | 0.68 | 0.62 | 0.65 |
| 11 | 14825704 | rs3740823 | -1.909 | 0.044 | -0.384 | 0.173 | 0.001 | 0 | 0 | 0.003 | 0.68 | 0.71 | 0.7 | 0.68 | 0.62 | 0.65 |
| 11 | 14826809 | rs12419657 | 0.622 | 0.03 | -0.397 | 0.178 | -0.036 | 0.001 | 0.001 | 0 | 0.15 | 0.25 | 0.12 | 0.14 | 0.09 | 0.1 |
| 11 | 14834904 | rs7944633 | 0.898 | 0.152 | -0.532 | 0.199 | -0.008 | 0.012 | 0.012 | 0.067 | 0.11 | 0.24 | 0.07 | 0.16 | 0.09 | 0.08 |
| 11 | 14836192 | rs4757269 | -0.061 | 0.211 | -0.671 | 0.201 | -0.025 | 0.015 | 0.015 | 0 | 0.4 | 0.1 | 0.51 | 0.38 | 0.45 | 0.46 |
| 11 | 14852490 | rs1037378 | -1.048 | 0.405 | -0.562 | 0.193 | 0.003 | -0.005 | -0.005 | 0 | 0.54 | 0.44 | 0.61 | 0.54 | 0.57 | 0.55 |
| 11 | 14855438 | rs11023346 | 0.769 | 0.443 | -0.622 | 0.166 | 0.001 | 0.006 | 0.006 | 0 | 0.12 | 0.21 | 0.08 | 0.17 | 0.12 | 0.1 |
| 11 | 14861517 | rs11023350 | 0 | 0.528 | -0.649 | 0.132 | -0.024 | 0.019 | 0.019 | 0 | 0.41 | 0.11 | 0.52 | 0.37 | 0.45 | 0.45 |
| 11 | 14863083 | rs11821380 | 0.662 | 0.527 | -0.632 | 0.155 | 0.002 | 0.01 | 0.01 | 0 | 0.12 | 0.28 | 0.09 | 0.17 | 0.12 | 0.09 |

| | | | | | | | | | | | | | | | | |
|---|---|---|---|---|---|---|---|---|---|---|---|---|---|---|---|---|
| 11 | 14865666 | rs2167453 | 0.532 | 0.527 | -0.631 | 0.154 | 0.002 | 0.011 | 0.011 | 0.001 | 0.13 | 0.32 | 0.09 | 0.18 | 0.12 | 0.1 |
| 11 | 14881256 | rs6486205 | -1.945 | 0.424 | -0.539 | 0.103 | 0.005 | 0.002 | 0.002 | 0.817 | 0.69 | 0.67 | 0.72 | 0.68 | 0.66 | 0.66 |
| 11 | 14881771 | rs7116978 | -1.935 | 0.422 | -0.54 | 0.103 | 0.005 | 0.002 | 0.002 | 0.846 | 0.69 | 0.67 | 0.72 | 0.68 | 0.66 | 0.66 |
| 11 | 14887830 | rs10832312 | 0.658 | 0.456 | -0.519 | 0.166 | 0.003 | 0.015 | 0.015 | 0 | 0.12 | 0.3 | 0.09 | 0.18 | 0.12 | 0.09 |
| 11 | 14894220 | rs1868997 | -1.926 | 0.51 | -0.679 | 0.095 | 0.005 | 0.002 | 0.002 | 0.713 | 0.69 | 0.67 | 0.72 | 0.68 | 0.66 | 0.66 |
| 11 | 14903636 | rs11023374 | 0.831 | 0.431 | -0.516 | 0.371 | -0.088 | 0.112 | 0.112 | 0.06 | 0.22 | 0.01 | 0.43 | 0.06 | 0.29 | 0.21 |
| 11 | 14909461 | rs7935792 | 0.44 | 0.438 | 0.012 | 0.86 | 0.02 | 0.055 | 0.055 | 0 | 0.07 | 0.2 | 0.06 | 0.17 | 0.07 | 0.08 |
| 11 | 14910234 | rs1993116 | -1.664 | 0.395 | 0.06 | 0.857 | -0.013 | -0.002 | -0.002 | 0 | 0.67 | 0.78 | 0.72 | 0.68 | 0.61 | 0.65 |
| 11 | 14910273 | rs10500804 | 0.292 | 0.409 | 0.065 | 0.885 | -0.022 | 0.023 | 0.023 | 0.022 | 0.42 | 0.1 | 0.52 | 0.37 | 0.45 | 0.45 |
| 11 | 14912417 | rs7129781 | 0.5 | 0.436 | 0.081 | 0.889 | 0.019 | 0.054 | 0.054 | 0 | 0.07 | 0.2 | 0.06 | 0.17 | 0.07 | 0.08 |
| 11 | 14913575 | rs12794714 | 0.259 | 0.464 | -0.07 | 0.89 | -0.022 | 0.024 | 0.024 | 0.032 | 0.42 | 0.1 | 0.51 | 0.37 | 0.45 | 0.44 |
| 11 | 14915310 | rs2060793 | -1.409 | 0.568 | -0.046 | 0.801 | 0 | 0 | 0 | 0.816 | 0.65 | 0.66 | 0.7 | 0.68 | 0.62 | 0.64 |
| 11 | 14918216 | rs1562902 | -1.263 | 0.245 | -0.191 | 0.965 | 0 | -0.003 | -0.003 | 0.576 | 0.56 | 0.52 | 0.67 | 0.62 | 0.56 | 0.57 |
| 11 | 14918308 | rs16930625 | 0.003 | 0.308 | -0.484 | 1.053 | -0.014 | 0.004 | 0.004 | 0 | 0.12 | 0.19 | 0.29 | 0.13 | 0.08 | 0.09 |
| 11 | 14921880 | rs10766197 | -0.305 | 0.357 | -0.494 | 1.079 | -0.043 | 0.022 | 0.022 | 0.038 | 0.41 | 0.1 | 0.36 | 0.36 | 0.47 | 0.45 |
| 11 | 14936943 | rs12416759 | -0.423 | 0.261 | -0.654 | 0.411 | 0.004 | -0.003 | -0.003 | 0.245 | 0.1 | 0.04 | 0.27 | 0.11 | 0.08 | 0.07 |
| 11 | 14950820 | rs7111531 | -0.956 | 0.464 | -0.574 | -0.142 | -0.022 | 0.045 | 0.045 | 0 | 0.33 | 0.2 | 0.57 | 0.14 | 0.43 | 0.31 |
| 11 | 14952338 | rs10741661 | -1.041 | 0.306 | 0.086 | -0.173 | 0.005 | -0.024 | -0.024 | 0 | 0.35 | 0.38 | 0.6 | 0.15 | 0.44 | 0.33 |
| 11 | 14963244 | rs11023389 | -0.875 | 0.402 | 0.127 | -0.127 | -0.028 | 0.048 | 0.048 | 0 | 0.32 | 0.19 | 0.57 | 0.14 | 0.43 | 0.31 |
| 11 | 14967610 | rs1874432 | 0.999 | 0.196 | 0.499 | 0.343 | 0.011 | -0.005 | -0.005 | 0 | 0.63 | 0.44 | 0.4 | 0.66 | 0.56 | 0.62 |
| 11 | 14971607 | rs11023392 | -0.005 | 0.328 | 0.294 | 0.214 | -0.02 | 0.082 | 0.082 | 0 | 0.14 | 0.01 | 0.11 | 0 | 0.14 | 0.05 |
| 11 | 14988998 | rs5242 | -1.207 | 0.706 | 0.775 | -0.029 | 0.061 | -0.016 | -0.016 | 0 | 0.19 | 0.38 | 0.47 | 0.16 | 0.3 | 0.29 |
| 11 | 14998061 | rs10832338 | -0.472 | 0.717 | 0.655 | -0.028 | -0.031 | 0.003 | 0.003 | 0 | 0.15 | 0.06 | 0.42 | 0.14 | 0.28 | 0.25 |
| 11 | 15000384 | rs7948017 | -1.497 | 0.806 | 0.908 | -0.061 | 0.069 | -0.032 | -0.032 | 0 | 0.21 | 0.44 | 0.49 | 0.15 | 0.34 | 0.29 |
| 11 | 15014179 | rs12293178 | -0.757 | 0.878 | 0.866 | -0.052 | 0.021 | -0.006 | -0.006 | 0.071 | 0.16 | 0.21 | 0.43 | 0.14 | 0.25 | 0.26 |
| 11 | 15029576 | rs2029877 | -0.71 | 0.89 | 0.847 | -0.032 | -0.005 | 0.001 | 0.001 | 0.66 | 0.15 | 0.12 | 0.42 | 0.14 | 0.24 | 0.26 |
| 11 | 15094057 | rs10832355 | -0.654 | 0.905 | 0.79 | -0.053 | -0.086 | -0.023 | -0.023 | 0.087 | 0.15 | 0.02 | 0.42 | 0.14 | 0.25 | 0.25 |
| 11 | 15097002 | rs11603873 | -0.679 | 0.613 | 0.827 | -0.052 | 0.001 | 0 | 0 | 2.366 | 0.07 | 0.05 | 0.11 | 0.07 | 0.09 | 0.09 |
| 11 | 15099143 | rs2282654 | 0.457 | 0.645 | 0.839 | -0.07 | 0.039 | -0.007 | -0.007 | 0 | 0.83 | 0.7 | 0.56 | 0.85 | 0.74 | 0.72 |
| 11 | 15099695 | rs12274674 | -0.516 | 0.664 | 0.56 | -0.062 | -0.029 | 0.005 | 0.005 | 1.018 | 0.16 | 0.08 | 0.42 | 0.14 | 0.26 | 0.25 |
| 11 | 15111580 | rs2226917 | 1.458 | 0.63 | 0.596 | -0.072 | 0.036 | -0.026 | -0.026 | 0 | 0.73 | 0.45 | 0.44 | 0.78 | 0.65 | 0.62 |
| 11 | 15113058 | rs7127660 | 1.458 | 0.63 | 0.575 | -0.072 | 0.035 | -0.025 | -0.025 | 1.416 | 0.73 | 0.44 | 0.44 | 0.78 | 0.65 | 0.62 |
| 11 | 15120775 | rs11023431 | -0.855 | 0.668 | 0.276 | -0.037 | -0.034 | 0.003 | 0.003 | 3.079 | 0.17 | 0.07 | 0.44 | 0.17 | 0.28 | 0.3 |
| 11 | 15121130 | rs756152 | 1.401 | 0.471 | 0.62 | -0.176 | 0.022 | -0.029 | -0.029 | 0 | 0.7 | 0.43 | 0.42 | 0.78 | 0.62 | 0.62 |
| 11 | 15126447 | rs10766209 | -0.601 | 0.626 | 0.297 | -0.116 | -0.03 | 0.007 | 0.007 | 2.341 | 0.16 | 0.07 | 0.43 | 0.13 | 0.25 | 0.29 |
| 11 | 15126792 | rs4757283 | 0.712 | 0.434 | 0.761 | -0.108 | 0.024 | -0.013 | -0.013 | 0 | 0.81 | 0.65 | 0.55 | 0.85 | 0.78 | 0.68 |
| 11 | 15130597 | rs10766210 | -0.027 | -0.205 | 0.221 | 0.026 | -0.001 | 0 | 0 | 0 | 0.61 | 0.63 | 0.61 | 0.38 | 0.58 | 0.56 |
| 11 | 15139288 | rs872172 | 0.182 | -0.43 | 0.178 | -0.087 | 0.014 | 0.033 | 0.033 | 0 | 0.55 | 0.29 | 0.58 | 0.35 | 0.53 | 0.45 |
| 11 | 15149784 | rs4492803 | -0.22 | -0.415 | -0.013 | -0.031 | -0.001 | 0 | 0 | 0.001 | 0.6 | 0.62 | 0.62 | 0.38 | 0.55 | 0.53 |
| 11 | 15164117 | rs4757292 | 0.724 | 0.055 | -2.586 | -0.301 | -0.151 | 0.063 | 0.063 | 0.003 | 0.32 | 0.03 | 0.5 | 0.16 | 0.36 | 0.24 |
| 11 | 15164461 | rs4757293 | 0.482 | 0.088 | -1.884 | 0.037 | 0.03 | 0.068 | 0.068 | 0 | 0.51 | 0.23 | 0.57 | 0.34 | 0.52 | 0.44 |
| 11 | 15166201 | rs12226609 | -0.526 | 0.118 | -1.828 | 0.476 | -0.005 | -0.111 | -0.111 | 0 | 0.2 | 0.01 | 0.1 | 0.36 | 0.18 | 0.25 |
| 11 | 15168771 | rs1792538 | -0.271 | 0.798 | -1.647 | -0.855 | -0.016 | 0.057 | 0.057 | 0 | 0.31 | 0.22 | 0.31 | 0.13 | 0.35 | 0.36 |
| 11 | 15168799 | rs1792537 | -0.211 | 0.552 | -1.213 | -1.573 | -0.003 | -0.017 | -0.017 | 0 | 0.71 | 0.67 | 0.68 | 0.23 | 0.71 | 0.67 |
| 11 | 15170235 | rs12275748 | 0.393 | 0.776 | -1.135 | -1.267 | 0.007 | 0.006 | 0.006 | 0 | 0.06 | 0.08 | 0.07 | 0.12 | 0.12 | 0.04 |
| 11 | 15173964 | rs7104355 | 0.306 | 0.483 | -0.368 | -1.325 | 0.014 | -0.04 | -0.04 | 0 | 0.2 | 0.08 | 0.23 | 0.65 | 0.16 | 0.26 |
| 11 | 15175985 | rs10832368 | 0.331 | 0.22 | -0.185 | -1.952 | -0.005 | 0.023 | 0.023 | 0 | 0.37 | 0.35 | 0.34 | 0.2 | 0.4 | 0.33 |
| 11 | 15177259 | rs1714348 | 0.617 | 0.171 | -0.226 | -2.117 | -0.034 | 0.088 | 0.088 | 0 | 0.37 | 0.23 | 0.41 | 0.15 | 0.44 | 0.43 |
| 11 | 15177950 | rs1714347 | -0.203 | -0.022 | -0.048 | -2.019 | -0.02 | 0.016 | 0.016 | 0 | 0.76 | 0.68 | 0.75 | 0.36 | 0.82 | 0.73 |
| 11 | 15180923 | rs1714345 | -0.169 | 0.292 | -0.203 | -2.122 | 0.038 | 0.066 | 0.066 | 0 | 0.49 | 0.71 | 0.61 | 0.31 | 0.61 | 0.58 |
| 11 | 15181696 | rs1792571 | 0.323 | 0.237 | -0.309 | -2.054 | -0.011 | -0.022 | -0.022 | 0.308 | 0.67 | 0.45 | 0.66 | 0.24 | 0.72 | 0.71 |
| 11 | 15182729 | rs1109328 | -0.489 | 1.055 | -0.737 | -2.671 | -0.023 | 0.055 | 0.055 | 0 | 0.69 | 0.83 | 0.64 | 0.85 | 0.62 | 0.66 |
| 11 | 15185729 | rs1612783 | -0.22 | 0.478 | -1.02 | -3.18 | 0.042 | 0.121 | 0.121 | 0 | 0.53 | 0.27 | 0.62 | 0.85 | 0.38 | 0.64 |
| 11 | 15187704 | rs1540160 | -0.251 | 0.475 | -0.951 | -3.192 | 0.023 | 0.089 | 0.089 | 0.034 | 0.54 | 0.35 | 0.64 | 0.86 | 0.4 | 0.65 |
| 11 | 15189510 | rs1792532 | 1.226 | 0.24 | -1.264 | -3.261 | 0.007 | 0.017 | 0.017 | 0 | 0.16 | 0.09 | 0.23 | 0.13 | 0.13 | 0.22 |
| 11 | 15191412 | rs4597047 | 0.615 | 0.08 | -1.623 | -3.445 | 0.048 | 0.015 | 0.015 | 0 | 0.29 | 0.15 | 0.32 | 0.69 | 0.17 | 0.34 |
| 11 | 15192049 | rs1627020 | -0.577 | 0.407 | -1.357 | -3.599 | 0.042 | -0.08 | -0.08 | 0.872 | 0.86 | 0.46 | 0.87 | 0.98 | 0.86 | 0.91 |
| 11 | 15195816 | rs1714379 | 0.269 | 0.101 | -1.367 | -3.562 | 0.014 | 0.063 | 0.063 | 0 | 0.27 | 0.1 | 0.28 | 0.16 | 0.21 | 0.36 |
| 11 | 15198772 | rs1624731 | -0.789 | 0.368 | -1.275 | -3.591 | 0.004 | -0.046 | -0.046 | 0 | 0.61 | 0.45 | 0.71 | 0.93 | 0.52 | 0.7 |
| 11 | 15201184 | rs2679632 | 0.032 | 0.066 | -1.39 | -3.233 | -0.007 | 0.05 | 0.05 | 0 | 0.24 | 0.11 | 0.28 | 0.17 | 0.23 | 0.31 |
| 11 | 15208263 | rs1714373 | 0.124 | 0.019 | -1.411 | -3.297 | 0.003 | 0.051 | 0.051 | 0 | 0.24 | 0.1 | 0.23 | 0.16 | 0.21 | 0.3 |
| 11 | 15209391 | rs1792574 | 0.046 | 0.125 | -1.332 | -3.226 | -0.006 | 0.054 | 0.054 | 0 | 0.25 | 0.11 | 0.28 | 0.17 | 0.23 | 0.31 |
| 11 | 15216227 | rs11023467 | 0.869 | 0.257 | -0.582 | -3.521 | 0.001 | -0.073 | -0.073 | 0 | 0.33 | 0.43 | 0.25 | 0.1 | 0.31 | 0.26 |
| 11 | 15216692 | rs1108010 | 1.412 | 0.165 | -0.335 | -3.546 | -0.007 | 0.043 | 0.043 | 0 | 0.31 | 0.19 | 0.24 | 0.1 | 0.31 | 0.26 |
| 11 | 15216886 | rs1540149 | -1.236 | 0.315 | -0.319 | -3.665 | -0.004 | -0.005 | -0.005 | 0 | 0.87 | 0.92 | 0.88 | 0.98 | 0.76 | 0.9 |
| 11 | 15220073 | rs1540151 | -0.782 | -0.141 | -0.074 | -3.495 | 0.007 | 0.055 | 0.055 | 0 | 0.54 | 0.37 | 0.62 | 0.87 | 0.44 | 0.65 |
| 11 | 15221092 | rs1540154 | -0.317 | 0.025 | -0.53 | -3.381 | 0.012 | 0.033 | 0.033 | 0 | 0.31 | 0.22 | 0.31 | 0.23 | 0.25 | 0.42 |
| 11 | 15223735 | rs4756800 | 0.199 | -0.151 | -0.463 | -2.398 | 0.003 | 0.038 | 0.038 | 0 | 0.31 | 0.24 | 0.37 | 0.74 | 0.28 | 0.37 |
| 11 | 15225295 | rs4757298 | 0.422 | -0.354 | -0.406 | -2.396 | 0.018 | 0.024 | 0.024 | 0.207 | 0.29 | 0.18 | 0.32 | 0.73 | 0.23 | 0.35 |

| | | | | | | | | | | | | | | | | |
|---|---|---|---|---|---|---|---|---|---|---|---|---|---|---|---|---|
| 11 | 15227989 | rs1945628 | -0.216 | -0.858 | 0.032 | -2.218 | -0.04 | -0.017 | -0.017 | 0 | 0.31 | 0.58 | 0.28 | 0.71 | 0.22 | 0.32 |
| 11 | 15228054 | rs4343008 | 1.096 | -0.674 | -0.55 | -2.687 | -0.061 | 0.138 | 0.138 | 0 | 0.35 | 0.08 | 0.37 | 0.11 | 0.36 | 0.35 |
| 11 | 15228450 | rs1792551 | -1.462 | -0.348 | -0.28 | -2.883 | -0.03 | 0.095 | 0.095 | 0 | 0.72 | 0.89 | 0.73 | 0.96 | 0.68 | 0.74 |
| 11 | 15228940 | rs939052 | 0.741 | -0.216 | -0.346 | -2.89 | -0.042 | 0.032 | 0.032 | 0.001 | 0.22 | 0.13 | 0.3 | 0.16 | 0.37 | 0.22 |
| 11 | 15232698 | rs1945630 | 0.873 | -0.612 | -0.136 | -2.851 | 0.005 | 0.006 | 0.006 | 0.007 | 0.32 | 0.28 | 0.32 | 0.73 | 0.24 | 0.36 |
| 11 | 15237493 | rs4757301 | 0.542 | -0.437 | -0.08 | -2.79 | 0.033 | 0.051 | 0.051 | 0.27 | 0.34 | 0.2 | 0.32 | 0.73 | 0.26 | 0.43 |
| 11 | 15239354 | rs939048 | 0.641 | -0.119 | -0.114 | -2.672 | -0.045 | 0.027 | 0.027 | 0.017 | 0.22 | 0.13 | 0.3 | 0.18 | 0.38 | 0.23 |
| 11 | 15243593 | rs16931247 | -0.705 | -0.244 | 0.125 | -2.661 | 0.005 | -0.013 | -0.013 | 0 | 0.09 | 0.36 | 0.08 | 0.06 | 0.03 | 0.05 |
| 11 | 15245118 | rs1714336 | -0.339 | -0.16 | 0.066 | -2.72 | 0.061 | -0.063 | -0.063 | 3.596 | 0.3 | 0.47 | 0.38 | 0.24 | 0.41 | 0.28 |
| 11 | 15247879 | rs1714338 | -0.528 | -0.281 | 0 | -2.608 | -0.011 | -0.112 | -0.112 | 0 | 0.63 | 0.56 | 0.69 | 0.97 | 0.67 | 0.71 |
| 11 | 15250018 | rs1792565 | -0.157 | -0.236 | -0.567 | -2.456 | 0.025 | 0.078 | 0.078 | 0.157 | 0.56 | 0.27 | 0.62 | 0.91 | 0.63 | 0.66 |
| 11 | 15258633 | rs16931277 | 0.564 | -0.316 | -0.151 | -1.855 | -0.05 | -0.052 | -0.052 | 0.031 | 0.09 | 0.2 | 0.02 | 0 | 0.02 | 0.05 |
| 11 | 15260124 | rs2156827 | 0.485 | 0.037 | -0.287 | -2.111 | -0.012 | 0.013 | 0.013 | 0 | 0.63 | 0.39 | 0.64 | 0.75 | 0.53 | 0.68 |
| 11 | 15261668 | rs2271831 | 0.778 | -0.088 | -0.428 | -2.305 | -0.038 | 0.136 | 0.136 | 0 | 0.57 | 0.25 | 0.55 | 0.77 | 0.46 | 0.65 |
| 11 | 15261749 | rs2271829 | -0.468 | 0.282 | -0.151 | -2.268 | 0.032 | -0.033 | -0.033 | 0 | 0.75 | 0.73 | 0.72 | 0.87 | 0.61 | 0.85 |
| 11 | 15267597 | rs1054952 | 0.334 | 0.188 | -0.311 | -2.482 | 0.005 | -0.031 | -0.031 | 0 | 0.25 | 0.3 | 0.26 | 0.1 | 0.29 | 0.27 |
| 11 | 15267727 | rs1945624 | -0.78 | 0.172 | -0.392 | -2.444 | 0.009 | -0.012 | -0.012 | 1.098 | 0.75 | 0.78 | 0.73 | 0.87 | 0.61 | 0.85 |
| 11 | 15274077 | rs11821916 | 1.398 | 0.328 | -0.285 | -2.226 | -0.004 | 0.017 | 0.017 | 0.093 | 0.17 | 0.1 | 0.11 | 0.01 | 0.16 | 0.12 |
| 11 | 15274990 | rs939047 | 0.229 | 0.671 | -0.382 | -2.294 | -0.025 | 0.084 | 0.084 | 0.045 | 0.57 | 0.34 | 0.59 | 0.77 | 0.54 | 0.59 |
| 11 | 15275242 | rs939046 | 1.187 | 0.354 | -1.007 | -2.398 | 0.031 | 0.084 | 0.084 | 0.508 | 0.4 | 0.1 | 0.33 | 0.7 | 0.34 | 0.45 |
| 11 | 15279421 | rs1945607 | 0.545 | 0.671 | -0.939 | -2.311 | 0.018 | 0.187 | 0.187 | 0 | 0.51 | 0.19 | 0.5 | 0.75 | 0.43 | 0.59 |
| 11 | 15287899 | rs12273915 | 1.387 | 0.446 | -0.944 | -2.249 | -0.052 | 0.016 | 0.016 | 0 | 0.1 | 0.02 | 0.16 | 0.06 | 0.13 | 0.1 |
| 11 | 15289832 | rs10766226 | -1.407 | -0.081 | -0.568 | -2.114 | 0.063 | 0.013 | 0.013 | 0 | 0.82 | 0.67 | 0.75 | 0.77 | 0.77 | 0.83 |
| 11 | 15290883 | rs12279544 | 0.484 | -0.108 | -0.581 | -2.048 | -0.019 | -0.097 | -0.097 | 0 | 0.23 | 0.32 | 0.13 | 0.02 | 0.19 | 0.18 |
| 11 | 15298929 | rs10741668 | -0.178 | -0.194 | -0.378 | -1.368 | 0.017 | -0.007 | -0.007 | 0 | 0.76 | 0.73 | 0.65 | 0.74 | 0.67 | 0.77 |
| 11 | 15301137 | rs12289726 | 0.204 | -1.078 | -0.263 | -1.128 | -0.025 | -0.086 | -0.086 | 2.548 | 0.14 | 0.3 | 0.15 | 0 | 0.12 | 0.09 |
| 11 | 15301762 | rs11826924 | 0.527 | -1.179 | -1.058 | -0.701 | 0.094 | -0.014 | -0.014 | 2.978 | 0.31 | 0.09 | 0.24 | 0.62 | 0.14 | 0.37 |
| 11 | 15310981 | rs10766227 | -0.329 | -0.933 | -0.808 | -0.719 | 0.063 | 0.047 | 0.047 | 0.693 | 0.47 | 0.67 | 0.41 | 0.62 | 0.27 | 0.46 |
| 11 | 15319228 | rs1343841 | -0.289 | -0.095 | -0.819 | -0.312 | 0.013 | 0.005 | 0.005 | 0 | 0.63 | 0.5 | 0.56 | 0.59 | 0.61 | 0.67 |
| 11 | 15321099 | rs11023514 | 0.474 | -0.633 | -0.726 | 0.068 | 0.008 | -0.003 | -0.003 | 0 | 0.16 | 0.11 | 0.1 | 0.51 | 0.09 | 0.27 |
| 11 | 15322956 | rs1320211 | 0.165 | 0.461 | -0.508 | -0.028 | 0.006 | 0.003 | 0.003 | 0.348 | 0.59 | 0.49 | 0.56 | 0.56 | 0.53 | 0.55 |
| 11 | 15323955 | rs4145786 | 0.85 | 1.298 | -0.509 | 0.02 | -0.053 | -0.045 | -0.045 | 0 | 0.15 | 0.07 | 0.14 | 0.2 | 0.21 | 0.18 |
| 11 | 15327994 | rs7942153 | -0.507 | 1.522 | -0.132 | 0.137 | 0.084 | 0.119 | 0.119 | 0.061 | 0.91 | 0.83 | 0.67 | 0.74 | 0.78 | 0.87 |
| 11 | 15329304 | rs10832400 | 0.639 | 1.005 | -0.049 | -0.287 | 0.045 | 0.078 | 0.078 | 0 | 0.44 | 0.3 | 0.34 | 0.16 | 0.35 | 0.4 |
| 11 | 15332750 | rs11023519 | -0.731 | 0.849 | 0.057 | -0.289 | -0.003 | -0.008 | -0.008 | 0 | 0.74 | 0.77 | 0.59 | 0.4 | 0.68 | 0.7 |
| 11 | 15341413 | rs4548599 | -1.111 | 0.938 | 0.18 | -0.373 | -0.008 | -0.037 | -0.037 | 0 | 0.55 | 0.54 | 0.64 | 0.84 | 0.65 | 0.59 |
| 11 | 15345573 | rs4334003 | 0.462 | 1.002 | 0.151 | -0.315 | -0.003 | -0.009 | -0.009 | 0 | 0.26 | 0.22 | 0.41 | 0.59 | 0.32 | 0.3 |
| 11 | 15350605 | rs7117265 | 0.351 | 0.714 | -0.639 | -0.361 | -0.012 | -0.005 | -0.005 | 0.308 | 0.23 | 0.19 | 0.17 | 0.2 | 0.23 | 0.26 |
| 11 | 15361578 | rs11023534 | -0.906 | 0.677 | -0.65 | -0.431 | 0.059 | 0.02 | 0.02 | 0.115 | 0.29 | 0.48 | 0.46 | 0.31 | 0.41 | 0.26 |
| 11 | 15366827 | rs3936095 | -1.539 | 0.617 | -0.657 | -0.44 | 0.05 | 0.014 | 0.014 | 0.87 | 0.52 | 0.65 | 0.62 | 0.51 | 0.64 | 0.51 |
| 11 | 15369216 | rs7930357 | 0.799 | 0.634 | -0.69 | -0.468 | 0.002 | -0.024 | -0.024 | 0 | 0.06 | 0.1 | 0.05 | 0.01 | 0.07 | 0.05 |
| 11 | 15372794 | rs4237705 | -1.942 | 0.575 | -0.724 | -0.463 | 0.013 | 0.023 | 0.023 | 0 | 0.7 | 0.78 | 0.72 | 0.85 | 0.73 | 0.69 |
| 11 | 15375848 | rs11023542 | 1.806 | 0.608 | -0.725 | -0.381 | 0.053 | -0.078 | -0.078 | 0 | 0.17 | 0.04 | 0.09 | 0.34 | 0.1 | 0.17 |
| 11 | 15382631 | rs3925090 | -0.196 | 0.774 | -0.606 | -0.438 | 0.083 | 0.068 | 0.068 | 0.861 | 0.22 | 0.37 | 0.41 | 0.3 | 0.34 | 0.21 |
| 11 | 15391027 | rs12794836 | -1.905 | 0.727 | -0.703 | -0.487 | 0.016 | 0.03 | 0.03 | 0 | 0.7 | 0.8 | 0.72 | 0.85 | 0.73 | 0.69 |
| 11 | 15398601 | rs7932410 | 0.395 | 1.08 | -0.713 | -0.248 | -0.033 | -0.11 | -0.11 | 0 | 0.33 | 0.12 | 0.23 | 0.54 | 0.3 | 0.38 |
| 11 | 15401144 | rs4500467 | -1.882 | 1.016 | -0.587 | -0.285 | 0.029 | 0.089 | 0.089 | 0 | 0.68 | 0.86 | 0.73 | 0.95 | 0.71 | 0.71 |
| 11 | 15419972 | rs12290103 | -0.876 | 1.278 | -0.319 | 0.027 | 0.052 | 0.056 | 0.056 | 0.66 | 0.27 | 0.54 | 0.46 | 0.42 | 0.39 | 0.29 |
| 11 | 15422400 | rs4450156 | -0.218 | 0.907 | -0.263 | 0.023 | 0.056 | 0.047 | 0.047 | 0.748 | 0.21 | 0.27 | 0.39 | 0.31 | 0.33 | 0.2 |
| 11 | 15423130 | rs11023563 | -0.453 | 0.905 | -0.292 | 0.039 | 0.068 | 0.06 | 0.06 | 0.502 | 0.22 | 0.34 | 0.4 | 0.31 | 0.34 | 0.2 |
| 11 | 15438256 | rs6486245 | -0.607 | 0.988 | -0.376 | -0.048 | -0.004 | 0.015 | 0.015 | 0 | 0.42 | 0.59 | 0.54 | 0.75 | 0.45 | 0.45 |
| 11 | 15441149 | rs4360688 | -1.094 | 0.933 | -0.903 | -0.072 | 0.007 | 0.12 | 0.12 | 0 | 0.51 | 0.78 | 0.56 | 0.75 | 0.46 | 0.45 |
| 11 | 15444484 | rs4237706 | -1.947 | 1.211 | -0.752 | -0.044 | 0.069 | 0.081 | 0.081 | 0 | 0.68 | 0.86 | 0.78 | 0.96 | 0.8 | 0.73 |
| 11 | 15444505 | rs4237707 | -1.285 | 2.057 | -0.117 | 0.757 | -0.002 | 0.049 | 0.049 | 0 | 0.44 | 0.69 | 0.6 | 0.75 | 0.54 | 0.47 |
| 11 | 15445575 | rs4291657 | -0.042 | 1.617 | -0.2 | 0.68 | 0.213 | 0.319 | 0.319 | 0.125 | 0.09 | 0.32 | 0.34 | 0.27 | 0.22 | 0.12 |
| 11 | 15446814 | rs4418790 | -1.708 | 1.663 | -0.207 | 0.66 | 0.002 | 0.044 | 0.044 | 0 | 0.49 | 0.66 | 0.59 | 0.65 | 0.53 | 0.4 |
| 11 | 15448583 | rs7937161 | -0.398 | 1.562 | 0.106 | 1.319 | 0.205 | 0.317 | 0.317 | 0.12 | 0.09 | 0.31 | 0.34 | 0.26 | 0.21 | 0.12 |
| 11 | 15448704 | rs11602776 | 0.381 | 1.277 | -0.395 | 1.191 | 0.017 | -0.028 | -0.028 | 0.355 | 0.18 | 0.14 | 0.12 | 0.35 | 0.14 | 0.19 |
| 11 | 15452247 | rs4237709 | -0.368 | 1.53 | -0.32 | 0.894 | 0.004 | 0.009 | 0.009 | 0 | 0.33 | 0.53 | 0.47 | 0.61 | 0.37 | 0.31 |
| 11 | 15455777 | rs4756808 | -0.331 | 1.713 | -0.583 | 0.811 | -0.038 | -0.042 | -0.042 | 0 | 0.13 | 0.06 | 0.18 | 0.19 | 0.22 | 0.14 |
| 11 | 15456468 | rs11023571 | 0.316 | 1.789 | -0.22 | 0.482 | 0.006 | 0.013 | 0.013 | 0.196 | 0.42 | 0.36 | 0.43 | 0.23 | 0.36 | 0.43 |
| 11 | 15457467 | rs7114467 | 0.553 | 1.722 | 0.047 | 0.425 | -0.027 | -0.051 | -0.051 | 0.504 | 0.38 | 0.19 | 0.35 | 0.57 | 0.4 | 0.4 |
| 11 | 15460101 | rs4522153 | -0.203 | 1.134 | 0.379 | -0.026 | 0.015 | 0.037 | 0.037 | 0.008 | 0.55 | 0.46 | 0.55 | 0.76 | 0.62 | 0.55 |
| 11 | 15461116 | rs4396274 | -0.94 | 2.15 | 0.514 | 0.64 | 0.02 | 0.015 | 0.015 | 0.193 | 0.08 | 0.1 | 0.17 | 0.13 | 0.11 | 0.11 |
| 11 | 15465860 | rs10734234 | 1.287 | 2.023 | 0.727 | 0.591 | 0.068 | 0.03 | 0.03 | 0 | 0.89 | 0.71 | 0.86 | 0.82 | 0.89 | 0.85 |
| 11 | 15471048 | rs7949682 | 0.978 | 2.917 | 1 | 0.633 | 0.062 | 0.033 | 0.033 | 0 | 0.85 | 0.71 | 0.76 | 0.76 | 0.79 | 0.84 |
| 11 | 15487237 | rs7936919 | -1.203 | 3.478 | 0.971 | 0.846 | 0 | -0.001 | -0.001 | 0 | 0.17 | 0.2 | 0.2 | 0.22 | 0.28 | 0.24 |
| 11 | 15491837 | rs7947152 | 0.851 | 3.567 | 0.986 | 0.844 | 0.048 | 0.024 | 0.024 | 0 | 0.85 | 0.73 | 0.76 | 0.77 | 0.78 | 0.84 |
| 11 | 15497017 | rs10832444 | 2.146 | 3.371 | 1.221 | 0.72 | 0.012 | 0.009 | 0.009 | 0 | 0.65 | 0.38 | 0.54 | 0.55 | 0.5 | 0.6 |

| | | | | | | | | | | | | | | | | |
|---|---|---|---|---|---|---|---|---|---|---|---|---|---|---|---|---|
| 11 | 15499846 | rs6416166 | 1.162 | 3.622 | 1.143 | 0.799 | 0.041 | 0.023 | 0.023 | 0 | 0.81 | 0.73 | 0.66 | 0.66 | 0.66 | 0.76 |
| 11 | 15503208 | rs7938802 | -1.001 | 3.592 | 1.169 | 0.794 | 0 | -0.001 | -0.001 | 0.258 | 0.12 | 0.1 | 0.1 | 0.12 | 0.15 | 0.16 |
| 11 | 15514889 | rs10766261 | 1.3 | 3.615 | 1.174 | 0.804 | 0.095 | 0.056 | 0.056 | 0 | 0.8 | 0.62 | 0.65 | 0.67 | 0.66 | 0.76 |
| 11 | 15540871 | rs7113468 | 0.923 | 3.379 | 1.254 | 0.883 | 0.064 | 0.048 | 0.048 | 0 | 0.83 | 0.7 | 0.68 | 0.67 | 0.71 | 0.78 |
| 11 | 15554358 | rs11023623 | -0.94 | 3.345 | 1.239 | 0.875 | -0.004 | -0.005 | -0.005 | 0 | 0.11 | 0.07 | 0.09 | 0.12 | 0.16 | 0.16 |
| 11 | 15555070 | rs2351096 | -0.178 | 3.55 | 1.277 | 0.89 | 0.097 | 0.15 | 0.15 | 0 | 0.95 | 0.82 | 0.83 | 0.8 | 0.92 | 0.9 |
| 11 | 15558038 | rs10832471 | 2.111 | 3.531 | 1.308 | 0.868 | 0.065 | 0.047 | 0.047 | 0 | 0.7 | 0.45 | 0.57 | 0.55 | 0.55 | 0.62 |
| 11 | 15560186 | rs2351097 | -0.992 | 3.497 | 1.264 | 0.846 | -0.002 | 0.017 | 0.017 | 0 | 0.12 | 0.09 | 0.09 | 0.06 | 0.15 | 0.16 |
| 11 | 15575772 | rs7936873 | 1.215 | 3.406 | 1.338 | 0.801 | 0.066 | 0.073 | 0.073 | 0 | 0.82 | 0.67 | 0.64 | 0.61 | 0.71 | 0.76 |
| 11 | 15578457 | rs2351044 | 0.999 | 2.387 | 0.937 | -0.134 | -0.043 | -0.057 | -0.057 | 0 | 0.29 | 0.12 | 0.42 | 0.54 | 0.37 | 0.39 |
| 11 | 15580347 | rs2351046 | 2.35 | 3.135 | 1.609 | 0.648 | 0.014 | 0.009 | 0.009 | 0.028 | 0.67 | 0.38 | 0.54 | 0.55 | 0.56 | 0.6 |
| 11 | 15584211 | rs2351050 | 2.274 | 3.162 | 1.609 | 0.585 | 0.054 | 0.022 | 0.022 | 0 | 0.68 | 0.46 | 0.62 | 0.58 | 0.56 | 0.6 |
| 11 | 15591042 | rs17438762 | -0.407 | 3.066 | 1.403 | 0.536 | -0.033 | -0.002 | -0.002 | 0 | 0.07 | 0 | 0.07 | 0.06 | 0.13 | 0.11 |
| 11 | 15601040 | rs2351058 | 1.101 | 2.932 | 1.646 | 0.695 | 0.071 | 0.056 | 0.056 | 0 | 0.83 | 0.72 | 0.71 | 0.64 | 0.71 | 0.78 |
| 11 | 15609588 | rs7117096 | 1.799 | 2.954 | 1.659 | 0.681 | 0.113 | 0.08 | 0.08 | 0 | 0.77 | 0.58 | 0.64 | 0.58 | 0.6 | 0.69 |
| 11 | 15610823 | rs4756814 | 2.627 | 2.896 | 1.707 | 0.669 | 0.023 | 0.015 | 0.015 | 0.16 | 0.72 | 0.36 | 0.65 | 0.58 | 0.62 | 0.66 |
| 11 | 15610834 | rs7111186 | -1.361 | 2.675 | 1.684 | 0.672 | -0.01 | -0.02 | -0.02 | 0.533 | 0.05 | 0.09 | 0.02 | 0 | 0.02 | 0.05 |
| 11 | 15612068 | rs11023648 | -0.612 | 2.754 | 1.642 | 0.706 | -0.03 | -0.025 | -0.025 | 0 | 0.11 | 0.08 | 0.15 | 0.18 | 0.21 | 0.15 |
| 11 | 15615679 | rs729418 | -0.866 | 2.76 | 1.666 | 0.709 | -0.013 | -0.011 | -0.011 | 0 | 0.12 | 0.12 | 0.16 | 0.18 | 0.21 | 0.15 |
| 11 | 15616694 | rs11023650 | 2.13 | 2.69 | 1.605 | 0.67 | 0.029 | 0.016 | 0.016 | 0.139 | 0.7 | 0.39 | 0.63 | 0.58 | 0.59 | 0.64 |
| 11 | 15619533 | rs10832482 | 1.426 | 2.833 | 1.383 | 0.74 | 0.078 | 0.068 | 0.068 | 0 | 0.81 | 0.63 | 0.72 | 0.63 | 0.71 | 0.78 |
| 11 | 15620907 | rs2351059 | -1.293 | 1.95 | 1.059 | 0.511 | -0.001 | -0.008 | -0.008 | 0 | 0.09 | 0.13 | 0.07 | 0.07 | 0.13 | 0.14 |
| 11 | 15626189 | rs10160745 | 0.926 | 1.917 | 1.332 | 0.646 | 0.07 | 0.097 | 0.097 | 0 | 0.91 | 0.68 | 0.85 | 0.8 | 0.92 | 0.97 |
| 11 | 15631476 | rs7927948 | -1.168 | 1.444 | 0.553 | 1.154 | -0.01 | 0.113 | 0.113 | 0.325 | 0.09 | 0.17 | 0.27 | 0.38 | 0.04 | 0.24 |
| 11 | 15634227 | rs11601272 | 1.19 | 1.291 | -0.097 | 0.79 | -0.003 | -0.017 | -0.017 | 0 | 0.4 | 0.13 | 0.34 | 0.37 | 0.45 | 0.44 |
| 11 | 15640818 | rs1124115 | 0.253 | 0.931 | 0.239 | 0.746 | 0.01 | -0.032 | -0.032 | 0 | 0.63 | 0.51 | 0.65 | 0.78 | 0.6 | 0.8 |
| 11 | 15649614 | rs7107683 | -1.425 | 1.053 | 0.527 | 0.914 | 0.007 | 0.187 | 0.187 | 0.289 | 0.11 | 0.31 | 0.29 | 0.38 | 0.06 | 0.24 |
| 11 | 15651472 | rs931282 | -0.196 | 1.059 | 0.421 | 0.93 | -0.002 | 0.104 | 0.104 | 0 | 0.07 | 0.12 | 0.25 | 0.38 | 0.04 | 0.23 |
| 11 | 15664512 | rs11023674 | 1.021 | 0.943 | 0.521 | 0.898 | -0.001 | 0.025 | 0.025 | 0 | 0.51 | 0.34 | 0.38 | 0.38 | 0.54 | 0.56 |
| 11 | 15686485 | rs4757348 | -0.799 | 1.051 | 0.527 | 0.814 | 0 | 0.002 | 0.002 | 0 | 0.3 | 0.33 | 0.37 | 0.39 | 0.23 | 0.31 |
| 11 | 15692228 | rs7128738 | -0.219 | 0.953 | 0.384 | 0.844 | 0.011 | 0.006 | 0.006 | 0 | 0.8 | 0.69 | 0.75 | 0.77 | 0.75 | 0.83 |
| 11 | 15693576 | rs1484425 | -0.145 | 0.942 | 0.275 | 0.849 | -0.001 | -0.04 | -0.04 | 0.037 | 0.25 | 0.14 | 0.34 | 0.38 | 0.2 | 0.3 |
| 11 | 15694529 | rs12363664 | -0.105 | 0.942 | 0.269 | 0.849 | -0.004 | -0.053 | -0.053 | 0.771 | 0.25 | 0.11 | 0.34 | 0.38 | 0.2 | 0.3 |
| 11 | 15704877 | rs10766276 | -0.022 | 0.933 | 0.261 | 0.848 | -0.017 | -0.081 | -0.081 | 0.023 | 0.24 | 0.06 | 0.33 | 0.38 | 0.2 | 0.3 |
| 11 | 15709984 | rs11023685 | -0.697 | 0.955 | 0.291 | 0.826 | 0 | -0.012 | -0.012 | 2 | 0.29 | 0.26 | 0.36 | 0.38 | 0.23 | 0.32 |
| 11 | 15713477 | rs747784 | -0.014 | 0.953 | 0.196 | 0.812 | -0.017 | -0.083 | -0.083 | 0 | 0.24 | 0.06 | 0.33 | 0.39 | 0.2 | 0.3 |
| 11 | 15713556 | rs747783 | -0.419 | 0.965 | 0.196 | 0.923 | -0.01 | -0.039 | -0.039 | 3.239 | 0.28 | 0.13 | 0.32 | 0.34 | 0.23 | 0.32 |
| 11 | 15714623 | rs4757353 | -0.122 | 1.019 | 0.245 | 0.808 | 0.014 | 0.005 | 0.005 | 0 | 0.79 | 0.64 | 0.74 | 0.77 | 0.75 | 0.83 |
| 11 | 15724868 | rs12365254 | 0.609 | 0.902 | 0.169 | 0.606 | -0.002 | 0.017 | 0.017 | 0 | 0.51 | 0.41 | 0.38 | 0.38 | 0.53 | 0.52 |
| 11 | 15730356 | rs1994214 | 0.559 | 1.004 | -0.013 | 0.635 | -0.002 | 0.033 | 0.033 | 0 | 0.5 | 0.32 | 0.38 | 0.38 | 0.53 | 0.47 |
| 11 | 15732077 | rs11023691 | 0.358 | 1.038 | 0.015 | 0.809 | 0 | 0.009 | 0.009 | 0.011 | 0.54 | 0.38 | 0.41 | 0.37 | 0.57 | 0.5 |
| 11 | 15735653 | rs3886607 | -0.742 | 1.092 | -0.182 | 0.943 | 0 | -0.002 | -0.002 | 0 | 0.47 | 0.52 | 0.6 | 0.62 | 0.44 | 0.51 |
| 11 | 15745640 | rs1484419 | 1.132 | 1.192 | 0.426 | 1.866 | 0.056 | 0.19 | 0.19 | 0.12 | 0.86 | 0.43 | 0.81 | 0.67 | 0.82 | 0.83 |
| 11 | 15752803 | rs1484444 | 1.281 | 0.448 | -0.791 | 1.597 | 0 | -0.001 | -0.001 | 0 | 0.6 | 0.4 | 0.71 | 0.56 | 0.65 | 0.66 |
| 11 | 15754676 | rs12361668 | -0.409 | 0.375 | -0.825 | 1.938 | 0.041 | -0.004 | -0.004 | 0 | 0.26 | 0.01 | 0.1 | 0.18 | 0.18 | 0.17 |
| 11 | 15755356 | rs1484446 | 1.316 | 0.522 | -0.948 | 1.749 | -0.047 | 0.039 | 0.039 | 0 | 0.42 | 0.21 | 0.37 | 0.33 | 0.41 | 0.44 |
| 11 | 15758914 | rs11023696 | -0.482 | 0.84 | 0.172 | 2.067 | 0.163 | 0.287 | 0.287 | 0.339 | 0.08 | 0.47 | 0.1 | 0.22 | 0.12 | 0.12 |
| 11 | 15770297 | rs1484439 | -1.487 | 0.788 | 0.173 | 1.759 | 0 | 0.005 | 0.005 | 1.247 | 0.44 | 0.61 | 0.54 | 0.59 | 0.6 | 0.51 |
| 11 | 15770954 | rs961776 | -1.278 | 0.96 | 0.751 | 2.013 | 0.042 | 0.102 | 0.102 | 0.509 | 0.23 | 0.42 | 0.23 | 0.39 | 0.34 | 0.2 |
| 11 | 15772315 | rs7128920 | 0.275 | 1.015 | 0.645 | 2.008 | 0.006 | -0.014 | -0.014 | 0 | 0.12 | 0.05 | 0.06 | 0.12 | 0.09 | 0.18 |
| 11 | 15776274 | rs1601251 | -0.821 | 0.912 | 0.978 | 2.069 | -0.044 | -0.071 | -0.071 | 0 | 0.19 | 0.09 | 0.19 | 0.4 | 0.34 | 0.19 |
| 11 | 15776786 | rs1882133 | -2.107 | 0.922 | 1.117 | 2.067 | 0.042 | 0.039 | 0.039 | 3.439 | 0.32 | 0.51 | 0.48 | 0.47 | 0.51 | 0.32 |
| 11 | 15779599 | rs1385164 | -0.437 | 0.97 | 1.384 | 1.97 | 0.055 | -0.009 | -0.009 | 0 | 0.1 | 0.18 | 0.26 | 0.08 | 0.17 | 0.14 |
| 11 | 15781027 | rs7129677 | -2.454 | 0.72 | 1.65 | 1.869 | -0.017 | 0.017 | 0.017 | 1.326 | 0.44 | 0.76 | 0.54 | 0.6 | 0.57 | 0.49 |
| 11 | 15799460 | rs1484433 | -1.044 | 1.007 | 1.801 | 1.927 | 0.061 | 0.035 | 0.035 | 1.686 | 0.13 | 0.23 | 0.27 | 0.19 | 0.2 | 0.28 |
| 11 | 15801938 | rs1545896 | 1.216 | 0.816 | 1.755 | 2.184 | 0.027 | 0.06 | 0.06 | 0.244 | 0.76 | 0.66 | 0.61 | 0.44 | 0.67 | 0.75 |
| 11 | 15803262 | rs11023717 | 0.947 | 0.839 | 1.768 | 2.196 | 0.01 | 0.023 | 0.023 | 0.136 | 0.78 | 0.76 | 0.62 | 0.44 | 0.67 | 0.75 |
| 11 | 15805225 | rs4462300 | -0.544 | 0.947 | 1.381 | 2.199 | -0.048 | -0.069 | -0.069 | 0 | 0.16 | 0.05 | 0.19 | 0.39 | 0.3 | 0.15 |
| 11 | 15815555 | rs4757365 | -0.902 | 0.901 | 1.537 | 2.388 | -0.052 | -0.065 | -0.065 | 0 | 0.19 | 0.05 | 0.17 | 0.37 | 0.31 | 0.18 |
| 11 | 15816667 | rs7103463 | -0.161 | 0.47 | 2.178 | 1.794 | -0.043 | -0.062 | -0.062 | 0 | 0.11 | 0.43 | 0.08 | 0 | 0.04 | 0.03 |
| 11 | 15821026 | rs923031 | -1.535 | 0.693 | 1.728 | 1.884 | -0.071 | -0.068 | -0.068 | 0 | 0.3 | 0.13 | 0.48 | 0.56 | 0.52 | 0.49 |
| 11 | 15829043 | rs11023727 | -1.526 | 0.674 | 1.715 | 1.875 | -0.071 | -0.064 | -0.064 | 0 | 0.3 | 0.15 | 0.48 | 0.56 | 0.51 | 0.49 |
| 11 | 15830495 | rs1837607 | -1.792 | 0.082 | 2.239 | 1.509 | 0.013 | 0.004 | 0.004 | 0 | 0.27 | 0.44 | 0.52 | 0.26 | 0.28 | 0.42 |
| 11 | 15835975 | rs3959930 | -1.667 | 0.024 | 2.527 | 1.478 | -0.002 | -0.001 | -0.001 | 0 | 0.27 | 0.24 | 0.51 | 0.26 | 0.28 | 0.42 |
| 11 | 15838137 | rs1837606 | -2.06 | 0.056 | 1.939 | 1.494 | 0 | 0.001 | 0.001 | 0.168 | 0.3 | 0.67 | 0.55 | 0.26 | 0.28 | 0.44 |
| 11 | 15844653 | rs2615043 | 1.215 | -0.019 | 1.633 | 0.819 | -0.002 | -0.008 | -0.008 | 0 | 0.69 | 0.57 | 0.45 | 0.25 | 0.71 | 0.45 |
| 11 | 15848847 | rs1440708 | 1.223 | -0.054 | 2.1 | 0.802 | -0.001 | -0.007 | -0.007 | 0 | 0.69 | 0.6 | 0.46 | 0.25 | 0.71 | 0.45 |
| 11 | 15855895 | rs10832522 | -1.703 | -0.089 | 2.291 | 0.734 | -0.004 | 0.009 | 0.009 | 0.188 | 0.29 | 0.22 | 0.53 | 0.75 | 0.29 | 0.55 |
| 11 | 15858203 | rs7112764 | -0.755 | -0.29 | 2.492 | 0.613 | -0.017 | -0.034 | -0.034 | 0 | 0.06 | 0.16 | 0.04 | 0 | 0.03 | 0.01 |

| | | | | | | | | | | | | | | | | |
|---|---|---|---|---|---|---|---|---|---|---|---|---|---|---|---|---|
| 11 | 15863328 | rs1465893 | -1.55 | -0.281 | 2.103 | 0.602 | 0 | -0.002 | -0.002 | 0.785 | 0.27 | 0.32 | 0.53 | 0.76 | 0.27 | 0.51 |
| 11 | 15866420 | rs7109948 | -0.625 | -0.271 | 2.128 | 0.473 | -0.011 | -0.029 | -0.029 | 0 | 0.06 | 0.18 | 0.03 | 0 | 0.03 | 0.01 |
| 11 | 15872496 | rs11023743 | -1.398 | -0.393 | 2.462 | 0.837 | -0.007 | -0.028 | -0.028 | 0.974 | 0.07 | 0.31 | 0.05 | 0 | 0.03 | 0.01 |
| 11 | 15873904 | rs7124200 | -0.776 | -0.39 | 2.371 | 0.838 | -0.012 | -0.03 | -0.03 | 0 | 0.06 | 0.19 | 0.04 | 0 | 0.03 | 0.01 |
| 11 | 15877983 | rs11023744 | 0.35 | 0.161 | 1.072 | -0.427 | -0.123 | 0.139 | 0.139 | 0 | 0.22 | 0.03 | 0.18 | 0.01 | 0.34 | 0.21 |
| 11 | 15879763 | rs10832525 | -0.62 | -0.07 | 2.09 | 0.254 | 0.025 | 0.055 | 0.055 | 0.075 | 0.17 | 0.22 | 0.46 | 0.59 | 0.27 | 0.35 |
| 11 | 15880212 | rs11828021 | 0.158 | -0.248 | 1.844 | -0.047 | -0.016 | -0.02 | -0.02 | 0 | 0.05 | 0.06 | 0.02 | 0 | 0.02 | 0.01 |
| 11 | 15880494 | rs12293229 | -0.557 | -0.126 | 2.112 | -0.035 | 0.02 | -0.006 | -0.006 | 0 | 0.06 | 0.28 | 0.05 | 0 | 0.02 | 0.01 |
| 11 | 15886799 | rs1866821 | 1.413 | -0.322 | 2.092 | -0.134 | -0.005 | -0.032 | -0.032 | 0 | 0.67 | 0.6 | 0.46 | 0.23 | 0.69 | 0.45 |
| 11 | 15890865 | rs2582321 | 0.188 | -0.126 | 1.883 | 0.008 | -0.006 | -0.001 | -0.001 | 0.321 | 0.9 | 0.92 | 0.99 | 0.86 | 0.96 | 0.9 |
| 11 | 15892588 | rs12364139 | -0.827 | -0.03 | 1.644 | 0.057 | 0.005 | 0.011 | 0.011 | 0 | 0.2 | 0.22 | 0.46 | 0.61 | 0.27 | 0.34 |
| 11 | 15893286 | rs1440707 | -0.792 | -0.029 | 1.644 | 0.058 | 0.005 | 0.012 | 0.012 | 0 | 0.2 | 0.23 | 0.46 | 0.61 | 0.27 | 0.33 |
| 11 | 15914404 | rs1440703 | 0.294 | -0.112 | 0.794 | -0.209 | -0.028 | -0.028 | -0.028 | 0 | 0.07 | 0.03 | 0.08 | 0.22 | 0.15 | 0.12 |
| 11 | 15915779 | rs2008923 | 1.279 | -0.601 | 1.171 | 0.017 | 0.006 | -0.007 | -0.007 | 0 | 0.7 | 0.64 | 0.53 | 0.27 | 0.68 | 0.57 |
| 11 | 15921283 | rs11023768 | -1.4 | -0.505 | 0.773 | 0.065 | 0.002 | 0.005 | 0.005 | 0.605 | 0.27 | 0.3 | 0.51 | 0.63 | 0.29 | 0.43 |
| 11 | 15923562 | rs722317 | 1.364 | -0.738 | 0.007 | -0.745 | -0.088 | 0.247 | 0.247 | 0 | 0.4 | 0.09 | 0.27 | 0.01 | 0.45 | 0.27 |
| 11 | 15930472 | rs10832529 | -1.405 | -1.103 | 1.26 | 0.645 | -0.03 | 0.21 | 0.21 | 0 | 0.16 | 0.42 | 0.16 | 0.57 | 0.08 | 0.38 |
| 11 | 15931747 | rs11023775 | 1.695 | -1.049 | 0.864 | 0.454 | -0.135 | 0.238 | 0.238 | 0 | 0.45 | 0.05 | 0.36 | 0.09 | 0.52 | 0.37 |
| 11 | 15932777 | rs2615040 | 1.42 | -1.004 | 1.382 | 0.474 | -0.006 | 0.134 | 0.134 | 0 | 0.49 | 0.33 | 0.4 | 0.09 | 0.52 | 0.38 |
| 11 | 15937067 | rs11023779 | 0.414 | -0.972 | 1.435 | 0.837 | -0.006 | 0.048 | 0.048 | 0 | 0.07 | 0.04 | 0.07 | 0.49 | 0.06 | 0.2 |
| 11 | 15941099 | rs10458877 | -1.623 | -0.988 | 1.816 | 0.831 | -0.028 | 0.18 | 0.18 | 0 | 0.16 | 0.4 | 0.16 | 0.58 | 0.08 | 0.38 |
| 11 | 15942032 | rs16932318 | 0.146 | -0.959 | 1.605 | 0.834 | 0.002 | 0.177 | 0.177 | 0 | 0.07 | 0.13 | 0.08 | 0.49 | 0.06 | 0.2 |
| 11 | 15946803 | rs2582295 | 0.363 | -1.001 | 1.369 | 0.665 | -0.008 | 0.016 | 0.016 | 0 | 0.63 | 0.79 | 0.55 | 0.66 | 0.6 | 0.74 |
| 11 | 15955113 | rs12361765 | 1.494 | -0.96 | 1.062 | 0.681 | -0.066 | 0.27 | 0.27 | 0 | 0.48 | 0.14 | 0.39 | 0.09 | 0.52 | 0.37 |
| 11 | 15955304 | rs2615018 | 0.305 | -0.763 | 1.119 | 0.743 | 0.006 | -0.009 | -0.009 | 0 | 0.65 | 0.66 | 0.57 | 0.68 | 0.6 | 0.77 |
| 11 | 15962351 | rs1866822 | 1.474 | -1.535 | 1.274 | 1.265 | -0.03 | 0.001 | 0.001 | 0 | 0.72 | 0.36 | 0.79 | 0.26 | 0.9 | 0.63 |
| 11 | 15972672 | rs16932353 | -1.297 | -0.561 | 1.128 | 1.323 | -0.034 | 0.033 | 0.033 | 0 | 0.09 | 0.22 | 0.08 | 0.09 | 0.01 | 0.11 |
| 11 | 15993057 | rs4539287 | -0.583 | 0.374 | 0.182 | 1.858 | -0.163 | -0.043 | -0.043 | 2.291 | 0.3 | 0.03 | 0.43 | 0.25 | 0.41 | 0.29 |
| 11 | 16003297 | rs4237711 | -1.031 | -0.341 | 0.602 | 1.34 | -0.005 | 0.006 | 0.006 | 0 | 0.43 | 0.32 | 0.51 | 0.35 | 0.52 | 0.46 |
| 11 | 16004562 | rs10832539 | -0.323 | -0.027 | 1.388 | 1.542 | 0.068 | 0.012 | 0.012 | 0.414 | 0.29 | 0.43 | 0.46 | 0.26 | 0.4 | 0.36 |
| 11 | 16007047 | rs11023813 | -0.689 | 0.004 | 1.272 | 1.224 | -0.001 | 0 | 0 | 0 | 0.36 | 0.33 | 0.53 | 0.27 | 0.53 | 0.38 |
| 11 | 16015766 | rs10766293 | 0.221 | 0.489 | 1.406 | 1.509 | -0.051 | -0.022 | -0.022 | 0 | 0.23 | 0.13 | 0.41 | 0.24 | 0.38 | 0.28 |
| 11 | 16022866 | rs11023821 | -0.277 | 0.472 | 1.442 | 1.528 | 0.021 | 0.018 | 0.018 | 0 | 0.28 | 0.35 | 0.57 | 0.61 | 0.4 | 0.33 |
| 11 | 16026842 | rs7942880 | -0.872 | 0.315 | 1.309 | 1.199 | 0.001 | -0.002 | -0.002 | 0 | 0.4 | 0.44 | 0.69 | 0.61 | 0.59 | 0.39 |
| 11 | 16027418 | rs4437995 | 0.397 | 0.529 | 1.313 | 1.531 | -0.125 | -0.07 | -0.07 | 0 | 0.23 | 0.03 | 0.4 | 0.24 | 0.38 | 0.28 |
| 11 | 16041305 | rs7941496 | 0.241 | 0.316 | 1.55 | 1.303 | 0.007 | -0.009 | -0.009 | 0 | 0.67 | 0.63 | 0.34 | 0.25 | 0.49 | 0.64 |
| 11 | 16072004 | rs12577495 | -1.429 | 0.493 | 1.52 | 1.018 | 0.044 | 0.43 | 0.43 | 0 | 0.05 | 0.27 | 0.18 | 0.38 | 0.02 | 0.05 |
| 11 | 16105390 | rs10766295 | 0.477 | 0.629 | 1.397 | 1.233 | -0.122 | -0.064 | -0.064 | 0 | 0.22 | 0.03 | 0.39 | 0.24 | 0.39 | 0.28 |
| 11 | 16118679 | rs2014828 | -0.576 | 0.3 | 1.53 | 1.3 | 0.065 | 0.031 | 0.031 | 0 | 0.31 | 0.46 | 0.65 | 0.61 | 0.51 | 0.36 |
| 11 | 16133413 | rs4617548 | -0.573 | 0.303 | 1.532 | 1.301 | 0.065 | 0.031 | 0.031 | 0 | 0.31 | 0.46 | 0.65 | 0.61 | 0.51 | 0.36 |
| 11 | 16200444 | rs1455104 | -0.029 | 0.48 | 1.039 | 1.234 | 0.024 | -0.001 | -0.001 | 0 | 0.72 | 0.65 | 0.52 | 0.75 | 0.51 | 0.7 |
| 11 | 16221145 | rs6486278 | -0.31 | 0.486 | 0.862 | 1.237 | -0.026 | -0.003 | -0.003 | 0 | 0.73 | 0.77 | 0.53 | 0.75 | 0.51 | 0.69 |
| 11 | 16221683 | rs1455118 | 0.065 | 0.57 | 0.604 | 0.73 | 0.04 | 0.02 | 0.02 | 0 | 0.68 | 0.59 | 0.35 | 0.38 | 0.49 | 0.64 |
| 11 | 16225741 | rs7125634 | 0.171 | 0.572 | 0.43 | 0.718 | 0.046 | 0.02 | 0.02 | 0.074 | 0.67 | 0.49 | 0.35 | 0.38 | 0.49 | 0.64 |
| 11 | 16238087 | rs10832558 | 0.163 | 1.041 | -0.067 | 0.673 | -0.062 | -0.065 | -0.065 | 0 | 0.21 | 0.02 | 0.34 | 0.23 | 0.26 | 0.25 |
| 11 | 16240252 | rs10766305 | -0.539 | 0.414 | -0.369 | 0 | -0.001 | -0.001 | -0.001 | 0.107 | 0.52 | 0.45 | 0.56 | 0.48 | 0.5 | 0.64 |
| 11 | 16242054 | rs10832565 | -0.53 | 0.412 | -0.371 | -0.001 | -0.001 | -0.001 | -0.001 | 0 | 0.52 | 0.45 | 0.56 | 0.48 | 0.5 | 0.64 |
| 11 | 16242575 | rs10832567 | -0.53 | 0.41 | -0.37 | -0.001 | -0.001 | -0.001 | -0.001 | 0.027 | 0.52 | 0.45 | 0.56 | 0.48 | 0.5 | 0.64 |
| 11 | 16248020 | rs2351958 | -0.325 | 0.4 | -0.259 | -0.277 | 0.001 | 0 | 0 | 1.072 | 0.5 | 0.45 | 0.54 | 0.5 | 0.42 | 0.64 |
| 11 | 16248204 | rs2351959 | -0.293 | -0.109 | -0.207 | -0.536 | 0.022 | -0.042 | -0.042 | 1 | 0.92 | 0.8 | 0.93 | 1 | 0.9 | 0.97 |
| 11 | 16259405 | rs1949481 | 0.083 | 0.379 | -0.502 | -0.6 | -0.01 | 0.005 | 0.005 | 0.084 | 0.45 | 0.37 | 0.42 | 0.47 | 0.54 | 0.33 |
| 11 | 16263349 | rs12291520 | 1.061 | 0.153 | -0.709 | -0.694 | 0.002 | 0.001 | 0.001 | 0 | 0.12 | 0.12 | 0.16 | 0.32 | 0.09 | 0.12 |
| 11 | 16264711 | rs12418887 | 1.069 | 0.226 | -0.345 | -0.872 | 0.009 | 0.022 | 0.022 | 0.3 | 0.23 | 0.15 | 0.1 | 0 | 0.18 | 0.2 |
| 11 | 16266740 | rs1595373 | -1.328 | 0.26 | -0.293 | -0.693 | -0.001 | 0.005 | 0.005 | 0 | 0.79 | 0.75 | 0.64 | 0.76 | 0.81 | 0.75 |
| 11 | 16284626 | rs2351962 | -1.339 | 0.259 | -0.381 | -0.694 | -0.001 | 0 | 0 | 0 | 0.79 | 0.77 | 0.64 | 0.76 | 0.81 | 0.75 |
| 11 | 16285686 | rs10832577 | 0.765 | 0.237 | -0.398 | -0.765 | -0.016 | -0.025 | -0.025 | 0 | 0.37 | 0.27 | 0.36 | 0.47 | 0.44 | 0.3 |
| 11 | 16307700 | rs1156725 | -1.107 | 0.36 | -0.46 | -0.553 | 0.001 | -0.005 | -0.005 | 0 | 0.77 | 0.76 | 0.64 | 0.76 | 0.8 | 0.75 |
| 11 | 16308668 | rs12800659 | 0.999 | 0.359 | -0.531 | -0.553 | -0.031 | -0.081 | -0.081 | 0.902 | 0.22 | 0.03 | 0.34 | 0.24 | 0.2 | 0.25 |
| 11 | 16327983 | rs12420808 | 1 | 0.29 | -0.506 | -0.782 | 0.012 | 0.024 | 0.024 | 0.191 | 0.23 | 0.14 | 0.1 | 0 | 0.18 | 0.2 |
| 11 | 16329009 | rs297367 | 0.214 | 0.44 | -0.371 | -0.674 | 0 | 0.002 | 0.002 | 0 | 0.57 | 0.5 | 0.71 | 0.72 | 0.64 | 0.55 |
| 11 | 16351041 | rs4757392 | 0.842 | 0.535 | -0.499 | -0.606 | -0.019 | -0.027 | -0.027 | 0.089 | 0.34 | 0.28 | 0.35 | 0.47 | 0.44 | 0.3 |
| 11 | 16355566 | rs297347 | -0.787 | 0.616 | -0.207 | -0.55 | 0.001 | 0.089 | 0.089 | 0 | 0.89 | 0.79 | 0.89 | 0.72 | 0.92 | 0.78 |
| 11 | 16379226 | rs7127006 | -0.483 | 0.641 | -0.471 | -1.064 | -0.002 | -0.089 | -0.089 | 0 | 0.31 | 0.39 | 0.18 | 0 | 0.29 | 0.23 |
| 11 | 16381813 | rs7929530 | 0.016 | 0.522 | -0.721 | -0.545 | 0.001 | -0.022 | -0.022 | 0 | 0.19 | 0.17 | 0.1 | 0 | 0.16 | 0.18 |
| 11 | 16383430 | rs631163 | -0.284 | 0.805 | -0.526 | 0.148 | -0.001 | 0.016 | 0.016 | 0 | 0.35 | 0.58 | 0.39 | 0.51 | 0.35 | 0.36 |
| 11 | 16383722 | rs1866109 | 0.168 | 1.094 | -0.552 | -0.25 | 0.009 | -0.039 | -0.039 | 0 | 0.62 | 0.52 | 0.71 | 0.81 | 0.57 | 0.66 |
| 11 | 16394068 | rs1370465 | -0.154 | 1.347 | -1.516 | -0.552 | -0.104 | 0.098 | 0.098 | 0.281 | 0.31 | 0.02 | 0.24 | 0.1 | 0.39 | 0.22 |
| 11 | 16415904 | rs1018143 | -0.898 | 1.175 | -0.933 | -0.119 | -0.001 | -0.004 | -0.004 | 0.419 | 0.59 | 0.6 | 0.65 | 0.49 | 0.6 | 0.52 |
| 11 | 16418711 | rs1393926 | 0.623 | 0.952 | -0.448 | -0.059 | -0.001 | -0.02 | -0.02 | 0 | 0.19 | 0.16 | 0.34 | 0.33 | 0.16 | 0.18 |

| | | | | | | | | | | | | | | | | |
|---|---|---|---|---|---|---|---|---|---|---|---|---|---|---|---|---|
| 11 | 16427972 | rs2174163 | 0.73 | 0.971 | -0.193 | -0.074 | -0.003 | -0.012 | -0.012 | 0 | 0.41 | 0.33 | 0.35 | 0.51 | 0.4 | 0.48 |
| 11 | 16437678 | rs7121140 | 0.961 | 1.907 | -0.175 | -0.328 | 0.022 | -0.03 | -0.03 | 0.093 | 0.36 | 0.28 | 0.29 | 0.51 | 0.27 | 0.46 |
| 11 | 16438411 | rs7932807 | 0.513 | 1.877 | 0.576 | -0.351 | -0.008 | 0.056 | 0.056 | 0.01 | 0.35 | 0.23 | 0.28 | 0.14 | 0.41 | 0.3 |
| 11 | 16443567 | rs2174164 | 0.731 | 1.788 | 0.561 | -0.373 | -0.017 | -0.06 | -0.06 | 0 | 0.19 | 0.08 | 0.34 | 0.33 | 0.19 | 0.19 |
| 11 | 16445649 | rs17470624 | 0.581 | 1.758 | 0.582 | -0.347 | 0.018 | -0.016 | -0.016 | 0 | 0.05 | 0.04 | 0.06 | 0 | 0.14 | 0.02 |
| 11 | 16455794 | rs16933090 | -0.595 | 1.755 | 0.979 | -0.349 | 0.039 | -0.021 | -0.021 | 0.046 | 0.08 | 0.16 | 0.07 | 0.03 | 0.14 | 0.06 |
| 11 | 16469759 | rs7929629 | 0.972 | 1.461 | 1.141 | -0.345 | 0.028 | -0.062 | -0.062 | 0 | 0.35 | 0.18 | 0.28 | 0.51 | 0.27 | 0.46 |
| 11 | 16474547 | rs10832608 | 0.523 | 1.47 | 1.248 | -0.347 | -0.01 | 0.007 | 0.007 | 0 | 0.39 | 0.46 | 0.31 | 0.53 | 0.27 | 0.5 |
| 11 | 16480518 | rs10832610 | 0.387 | 1.125 | 1.241 | -0.333 | -0.001 | -0.007 | -0.007 | 0 | 0.21 | 0.2 | 0.35 | 0.32 | 0.19 | 0.19 |
| 11 | 16485301 | rs1354329 | 0.81 | 1.526 | 0.927 | -0.602 | -0.001 | -0.043 | -0.043 | 0 | 0.28 | 0.3 | 0.23 | 0.03 | 0.3 | 0.19 |
| 11 | 16487065 | rs7118874 | 1.13 | 1.472 | 1.037 | -0.424 | 0.06 | -0.047 | -0.047 | 0 | 0.36 | 0.21 | 0.22 | 0.52 | 0.18 | 0.44 |
| 11 | 16494570 | rs7945007 | 0.293 | 1.7 | 1.146 | -0.404 | 0.016 | 0.002 | 0.002 | 0 | 0.1 | 0.16 | 0.13 | 0.1 | 0.18 | 0.13 |
| 11 | 16499623 | rs7937011 | -0.106 | 1.69 | 1.133 | -0.38 | -0.001 | -0.001 | -0.001 | 0 | 0.51 | 0.51 | 0.43 | 0.67 | 0.51 | 0.62 |
| 11 | 16504966 | rs3924790 | -0.133 | 1.687 | 1.237 | -0.381 | 0 | -0.001 | -0.001 | 2.161 | 0.52 | 0.54 | 0.44 | 0.67 | 0.51 | 0.62 |
| 11 | 16506090 | rs10832616 | 0.88 | 1.684 | 1.055 | -0.342 | -0.018 | -0.067 | -0.067 | 2.572 | 0.19 | 0.05 | 0.33 | 0.3 | 0.19 | 0.19 |
| 11 | 16510682 | rs7109165 | 0.393 | 1.662 | 1.227 | -0.34 | 0.02 | -0.001 | -0.001 | 0 | 0.1 | 0.19 | 0.14 | 0.09 | 0.18 | 0.13 |
| 11 | 16533793 | rs7924783 | -0.444 | 1.664 | 1.246 | -0.336 | 0 | 0.002 | 0.002 | 0 | 0.3 | 0.31 | 0.48 | 0.45 | 0.36 | 0.34 |
| 11 | 16534415 | rs12802656 | -0.115 | 1.665 | 1.249 | -0.433 | 0.009 | -0.045 | -0.045 | 0 | 0.39 | 0.5 | 0.37 | 0.19 | 0.48 | 0.34 |
| 11 | 16541049 | rs12799418 | 0.266 | 1.743 | 1.208 | -0.375 | -0.024 | 0.061 | 0.061 | 0.019 | 0.36 | 0.19 | 0.34 | 0.13 | 0.48 | 0.32 |
| 11 | 16582920 | rs4603269 | 0.889 | 1.681 | 1.145 | -0.323 | 0.036 | -0.018 | -0.018 | 0.885 | 0.38 | 0.32 | 0.22 | 0.49 | 0.18 | 0.47 |
| 11 | 16591594 | rs4593976 | 1.164 | 1.712 | 0.844 | -0.331 | -0.031 | 0.103 | 0.103 | 0.09 | 0.27 | 0.07 | 0.21 | 0.03 | 0.3 | 0.19 |
| 11 | 16592120 | rs11023990 | 0.947 | 1.51 | 0.824 | -0.262 | -0.049 | -0.111 | -0.111 | 0.098 | 0.18 | 0.01 | 0.33 | 0.32 | 0.18 | 0.17 |
| 11 | 16608600 | rs12576360 | 0.71 | 1.459 | 0.974 | -0.277 | -0.014 | 0.004 | 0.004 | 0.32 | 0.39 | 0.44 | 0.25 | 0.55 | 0.18 | 0.49 |
| 11 | 16614806 | rs4757416 | 1.07 | 1.492 | 0.982 | -0.288 | -0.034 | -0.088 | -0.088 | 0.014 | 0.17 | 0.03 | 0.32 | 0.32 | 0.19 | 0.17 |
| 11 | 16615280 | rs4757417 | -0.021 | 1.473 | 0.995 | -0.307 | -0.001 | -0.021 | -0.021 | 0 | 0.57 | 0.47 | 0.57 | 0.87 | 0.37 | 0.66 |
| 11 | 16634707 | rs9988855 | 0.76 | 1.489 | 0.996 | -0.273 | 0.009 | -0.007 | -0.007 | 0 | 0.39 | 0.39 | 0.24 | 0.55 | 0.18 | 0.48 |
| 11 | 16669707 | rs7481887 | 0.589 | 1.642 | 0.979 | -0.345 | -0.042 | 0.06 | 0.06 | 0.009 | 0.31 | 0.17 | 0.27 | 0.03 | 0.43 | 0.21 |
| 11 | 16673384 | rs3921675 | 0.157 | 1.66 | 1.007 | -0.3 | 0.024 | -0.005 | -0.005 | 0 | 0.11 | 0.24 | 0.15 | 0.09 | 0.2 | 0.13 |
| 11 | 16686989 | rs10766338 | 1.105 | 1.666 | 0.915 | -0.245 | -0.03 | -0.075 | -0.075 | 0.005 | 0.17 | 0.04 | 0.32 | 0.32 | 0.19 | 0.18 |
| 11 | 16691743 | rs11024012 | 0.817 | 1.596 | 0.948 | -0.294 | 0.009 | -0.006 | -0.006 | 0 | 0.39 | 0.39 | 0.24 | 0.49 | 0.19 | 0.47 |
| 11 | 16692990 | rs7128484 | -0.334 | 1.612 | 0.964 | -0.313 | 0.02 | 0.026 | 0.026 | 0 | 0.88 | 0.75 | 0.85 | 0.85 | 0.8 | 0.85 |
| 11 | 16699145 | rs1501452 | 0.762 | 1.749 | 0.888 | -0.339 | -0.073 | 0.123 | 0.123 | 0.123 | 0.29 | 0.07 | 0.25 | 0.03 | 0.4 | 0.21 |
| 11 | 16730733 | rs1566579 | -0.397 | 1.599 | 0.904 | -0.336 | 0.021 | 0.018 | 0.018 | 0.188 | 0.5 | 0.39 | 0.61 | 0.36 | 0.62 | 0.39 |
| 11 | 16735491 | rs11024021 | 0.512 | 1.632 | 0.591 | -0.336 | -0.011 | -0.011 | -0.011 | 0 | 0.32 | 0.23 | 0.28 | 0.03 | 0.43 | 0.21 |
| 11 | 16745643 | rs1392998 | 0.669 | 1.652 | 0.495 | -0.326 | -0.04 | 0.051 | 0.051 | 0 | 0.31 | 0.14 | 0.27 | 0.03 | 0.43 | 0.21 |
| 11 | 16748143 | rs2352663 | 1.106 | 1.631 | 0.144 | -0.29 | -0.029 | -0.082 | -0.082 | 0.209 | 0.17 | 0.04 | 0.32 | 0.33 | 0.18 | 0.18 |
| 11 | 16758246 | rs3809095 | 1.327 | 1.658 | 0.126 | -0.309 | -0.083 | 0.128 | 0.128 | 0.196 | 0.24 | 0.01 | 0.2 | 0.03 | 0.28 | 0.2 |
| 11 | 16759078 | rs11024029 | 0.994 | 0.951 | 0.336 | -0.075 | -0.029 | -0.083 | -0.083 | 0 | 0.17 | 0.04 | 0.32 | 0.33 | 0.18 | 0.17 |
| 11 | 16760244 | rs1846936 | -0.365 | 0.441 | 0.788 | 0.206 | 0.007 | -0.003 | -0.003 | 0 | 0.89 | 0.87 | 0.86 | 0.9 | 0.81 | 0.87 |
| 11 | 16783824 | rs7127877 | 0.048 | 0.065 | 0.724 | 0.09 | -0.009 | 0.001 | 0.001 | 0.164 | 0.39 | 0.6 | 0.26 | 0.49 | 0.19 | 0.46 |
| 11 | 16787777 | rs10832678 | 0.474 | -0.394 | -0.09 | 0.371 | 0.031 | -0.039 | -0.039 | 0 | 0.15 | 0.1 | 0.08 | 0.37 | 0.04 | 0.22 |
| 11 | 16787922 | rs7128359 | 0.311 | -0.296 | 0.26 | 0.101 | -0.024 | -0.025 | -0.025 | 0 | 0.26 | 0.59 | 0.19 | 0.19 | 0.15 | 0.31 |
| 11 | 16789091 | rs4757431 | 0.34 | 0.281 | 0.146 | 0.494 | 0.038 | 0.003 | 0.003 | 0 | 0.86 | 0.64 | 0.83 | 0.88 | 0.8 | 0.85 |
| 11 | 16791151 | rs10766348 | -0.052 | -0.277 | -0.631 | 0.781 | 0.031 | -0.038 | -0.038 | 0.225 | 0.15 | 0.09 | 0.08 | 0.37 | 0.04 | 0.23 |
| 11 | 16796707 | rs10832680 | 0.697 | 0.172 | -1.444 | 0.894 | -0.106 | -0.082 | -0.082 | 0 | 0.27 | 0.02 | 0.3 | 0.38 | 0.34 | 0.26 |
| 11 | 16798672 | rs217766 | -0.237 | 0.666 | -0.496 | 0.858 | 0.094 | 0.024 | 0.024 | 0 | 0.1 | 0.26 | 0.15 | 0.07 | 0.22 | 0.06 |
| 11 | 16799044 | rs217765 | -0.038 | 0.546 | -0.104 | 1.103 | 0.001 | 0.001 | 0.001 | 0 | 0.38 | 0.39 | 0.46 | 0.49 | 0.56 | 0.45 |
| 11 | 16800105 | rs217762 | -0.787 | 0.211 | -0.791 | 0.795 | -0.012 | -0.042 | -0.042 | 0.043 | 0.68 | 0.53 | 0.74 | 0.76 | 0.75 | 0.68 |
| 11 | 16806087 | rs4756864 | -0.556 | 0.099 | -1.321 | 0.664 | 0.018 | -0.008 | -0.008 | 0 | 0.38 | 0.33 | 0.38 | 0.35 | 0.3 | 0.29 |
| 11 | 16806160 | rs217756 | -1.442 | -0.007 | -1.708 | 0.537 | -0.02 | 0.046 | 0.046 | 0 | 0.83 | 0.92 | 0.88 | 0.92 | 0.84 | 0.9 |
| 11 | 16817072 | rs11024041 | 0.99 | -0.197 | -1.791 | 0.52 | -0.003 | -0.056 | -0.056 | 0 | 0.18 | 0.08 | 0.26 | 0.28 | 0.17 | 0.17 |
| 11 | 16818754 | rs217738 | 0.094 | -0.042 | -1.788 | 0.286 | -0.007 | 0.038 | 0.038 | 0.077 | 0.14 | 0.05 | 0.07 | 0 | 0.14 | 0.07 |
| 11 | 16821166 | rs216486 | 0.988 | 0.044 | -1.581 | 0.8 | -0.018 | 0.019 | 0.019 | 0 | 0.29 | 0.18 | 0.22 | 0.3 | 0.33 | 0.37 |
| 11 | 16823736 | rs216489 | -0.002 | 0.308 | -1.333 | 0.692 | 0.018 | -0.005 | -0.005 | 0 | 0.65 | 0.57 | 0.61 | 0.65 | 0.64 | 0.66 |
| 11 | 16833461 | rs10832683 | 0.922 | 0.619 | -2.035 | 0.806 | -0.046 | -0.078 | -0.078 | 0 | 0.26 | 0.08 | 0.27 | 0.36 | 0.32 | 0.21 |
| 11 | 16834031 | rs216493 | -0.822 | 0.322 | -1.757 | 1.032 | 0.008 | 0.001 | 0.001 | 0 | 0.8 | 0.8 | 0.82 | 0.82 | 0.83 | 0.87 |
| 11 | 16834510 | rs216495 | -0.321 | 0.65 | -1.86 | 0.996 | -0.029 | 0.01 | 0.01 | 0 | 0.7 | 0.76 | 0.69 | 0.75 | 0.62 | 0.66 |
| 11 | 16835984 | rs10766349 | -0.114 | 0.132 | -1.422 | 1.611 | 0.002 | 0.015 | 0.015 | 0 | 0.08 | 0.1 | 0.23 | 0.2 | 0.11 | 0.09 |
| 11 | 16836571 | rs216496 | 0.245 | -0.066 | -1.675 | 1.609 | -0.005 | 0.008 | 0.008 | 0 | 0.12 | 0.07 | 0.07 | 0.24 | 0.05 | 0.31 |
| 11 | 16837904 | rs10832685 | -0.277 | 0.714 | -1.764 | 1.329 | -0.005 | -0.012 | -0.012 | 0 | 0.08 | 0.05 | 0.23 | 0.21 | 0.11 | 0.1 |
| 11 | 16842787 | rs1027617 | -0.05 | 1.009 | -1.882 | 0.812 | -0.058 | 0.013 | 0.013 | 0 | 0.43 | 0.21 | 0.41 | 0.36 | 0.48 | 0.41 |
| 11 | 16844409 | rs217755 | -0.936 | 1.293 | -1.315 | 0.837 | 0.003 | 0.007 | 0.007 | 0 | 0.22 | 0.24 | 0.26 | 0.34 | 0.21 | 0.36 |
| 11 | 16844924 | rs4756865 | 0.407 | 0.687 | -1.699 | 0.319 | 0.019 | -0.054 | -0.054 | 0.69 | 0.3 | 0.17 | 0.42 | 0.48 | 0.34 | 0.24 |
| 11 | 16847317 | rs440212 | -0.004 | 0.901 | -0.157 | 0.07 | 0.069 | 0.064 | 0.064 | 0 | 0.11 | 0.31 | 0.2 | 0.16 | 0.14 | 0.13 |
| 11 | 16848077 | rs390974 | 0.009 | 0.899 | -0.158 | 0.05 | 0.071 | 0.062 | 0.062 | 0 | 0.11 | 0.31 | 0.2 | 0.16 | 0.14 | 0.13 |
| 11 | 16851951 | rs402982 | -0.006 | 0.901 | -0.157 | 0.051 | 0.069 | 0.06 | 0.06 | 0 | 0.11 | 0.31 | 0.2 | 0.16 | 0.14 | 0.13 |
| 11 | 16852672 | rs740624 | 0.23 | 1.019 | -0.646 | -0.618 | -0.021 | 0.077 | 0.077 | 0 | 0.46 | 0.22 | 0.53 | 0.7 | 0.49 | 0.49 |
| 11 | 16853714 | rs407932 | 0.709 | 1.015 | -0.433 | -0.618 | -0.011 | -0.01 | -0.01 | 2.098 | 0.09 | 0.07 | 0.19 | 0.16 | 0.13 | 0.11 |
| 11 | 16858229 | rs383623 | -0.62 | 1.012 | -0.414 | -0.637 | -0.005 | -0.033 | -0.033 | 0 | 0.57 | 0.54 | 0.73 | 0.86 | 0.63 | 0.62 |

| | | | | | | | | | | | | | | | | |
|---|---|---|---|---|---|---|---|---|---|---|---|---|---|---|---|---|
| 11 | 16863087 | rs452745 | 0.508 | 1.025 | -0.453 | -0.614 | 0.069 | 0.06 | 0.06 | 0 | 0.11 | 0.31 | 0.2 | 0.16 | 0.14 | 0.13 |
| 11 | 16863130 | rs369819 | 1.535 | 0.53 | -0.704 | -0.836 | 0.014 | -0.029 | -0.029 | 0.198 | 0.05 | 0 | 0.03 | 0.08 | 0.01 | 0.11 |
| 11 | 16864694 | rs11024054 | 0.843 | 0.333 | -0.897 | -1.172 | -0.046 | 0.06 | 0.06 | 0 | 0.18 | 0.06 | 0.13 | 0.05 | 0.25 | 0.12 |
| 11 | 16865033 | rs376456 | -0.164 | 0.41 | -0.78 | -1.034 | 0.011 | 0.034 | 0.034 | 0 | 0.58 | 0.66 | 0.75 | 0.86 | 0.63 | 0.62 |
| 11 | 16872440 | rs366590 | -0.799 | 0.684 | -1.172 | -0.811 | -0.05 | -0.022 | -0.022 | 0 | 0.24 | 0.11 | 0.27 | 0.29 | 0.32 | 0.21 |
| 11 | 16876747 | rs366052 | -0.004 | 0.199 | -1.033 | -1.4 | 0.022 | 0.085 | 0.085 | 0 | 0.57 | 0.31 | 0.59 | 0.81 | 0.65 | 0.58 |
| 11 | 16877497 | rs11024058 | 1.07 | 0.043 | -1.151 | -1.547 | -0.058 | 0.058 | 0.058 | 0 | 0.17 | 0.04 | 0.13 | 0.06 | 0.23 | 0.11 |
| 11 | 16878455 | rs392981 | -1.255 | -0.05 | -0.611 | -1.553 | -0.053 | -0.045 | -0.045 | 0 | 0.8 | 0.67 | 0.76 | 0.9 | 0.9 | 0.72 |
| 11 | 16878698 | rs393633 | 1.106 | 0.125 | -0.798 | -1.545 | -0.03 | -0.026 | -0.026 | 0 | 0.09 | 0.03 | 0.18 | 0.16 | 0.13 | 0.1 |
| 11 | 16883854 | rs376152 | -0.594 | 0.019 | -0.734 | -1.552 | 0.004 | 0.005 | 0.005 | 0 | 0.74 | 0.31 | 0.72 | 0.87 | 0.88 | 0.69 |
| 11 | 16889745 | rs415300 | -0.623 | 0.038 | -0.791 | -1.611 | -0.014 | 0.024 | 0.024 | 0 | 0.35 | 0.26 | 0.34 | 0.25 | 0.39 | 0.25 |
| 11 | 16889852 | rs11024063 | 0.868 | 0.044 | -0.92 | -1.603 | -0.041 | 0.053 | 0.053 | 0 | 0.44 | 0.24 | 0.39 | 0.66 | 0.51 | 0.47 |
| 11 | 16894877 | rs370161 | -1.153 | 0.177 | -1.409 | -1.689 | 0.006 | 0.031 | 0.031 | 0 | 0.44 | 0.37 | 0.44 | 0.25 | 0.55 | 0.27 |
| 11 | 16896925 | rs17772101 | 1.158 | 0.209 | -1.447 | -1.734 | -0.048 | 0.054 | 0.054 | 0 | 0.17 | 0.05 | 0.13 | 0.06 | 0.23 | 0.11 |
| 11 | 16897496 | rs11024066 | 1.824 | 0.024 | -1.533 | -1.753 | -0.057 | -0.091 | -0.091 | 0 | 0.32 | 0.07 | 0.3 | 0.64 | 0.34 | 0.37 |
| 11 | 16902268 | rs381815 | -0.542 | -0.005 | -1.418 | -1.704 | -0.014 | 0.011 | 0.011 | 0 | 0.23 | 0.19 | 0.29 | 0.16 | 0.3 | 0.13 |
| 11 | 16912116 | rs393625 | -1.155 | -0.077 | -1.42 | -1.68 | -0.075 | -0.046 | -0.046 | 0 | 0.77 | 0.5 | 0.74 | 0.88 | 0.89 | 0.64 |
| 11 | 16917219 | rs11024074 | -1.047 | 0.195 | -1.43 | -1.571 | 0.008 | -0.013 | -0.013 | 1.169 | 0.27 | 0.33 | 0.31 | 0.19 | 0.31 | 0.16 |
| 11 | 16920570 | rs10832696 | 1.446 | 0.421 | -1.656 | -1.402 | -0.018 | 0.018 | 0.018 | 0 | 0.05 | 0.02 | 0.06 | 0 | 0.13 | 0.02 |
| 11 | 16924324 | rs7109209 | 0.874 | -0.25 | -1.677 | -1.345 | -0.021 | -0.06 | -0.06 | 0 | 0.37 | 0.07 | 0.3 | 0.64 | 0.35 | 0.38 |
| 11 | 16935949 | rs16933680 | 0.192 | 0.705 | -1.12 | -1.578 | -0.059 | -0.034 | -0.034 | 0 | 0.13 | 0.02 | 0.21 | 0.15 | 0.17 | 0.08 |
| 11 | 16937968 | rs4757444 | 0.724 | 0.324 | -0.666 | -1.714 | 0.001 | -0.004 | -0.004 | 0 | 0.54 | 0.5 | 0.54 | 0.77 | 0.42 | 0.74 |
| 11 | 16952482 | rs16933684 | -0.13 | 0.216 | -0.5 | -1.165 | 0.003 | 0.002 | 0.002 | 0 | 0.13 | 0.17 | 0.2 | 0.15 | 0.14 | 0.08 |
| 11 | 16952894 | rs7117795 | -0.075 | 0.279 | -0.741 | -1.694 | -0.019 | 0.025 | 0.025 | 0 | 0.69 | 0.73 | 0.77 | 0.91 | 0.61 | 0.85 |
| 11 | 16955369 | rs4757447 | -0.417 | 0.188 | -0.113 | -1.738 | 0.032 | -0.098 | -0.098 | 0 | 0.32 | 0.5 | 0.25 | 0.09 | 0.39 | 0.17 |
| 11 | 16959611 | rs177563 | -0.453 | 0.199 | -0.196 | -1.699 | -0.008 | -0.008 | -0.008 | 0.348 | 0.88 | 0.92 | 0.8 | 0.85 | 0.86 | 0.93 |
| 11 | 16967712 | rs10832704 | 0.186 | 0.267 | -0.488 | -1.716 | -0.091 | 0.096 | 0.096 | 0 | 0.23 | 0.08 | 0.18 | 0.05 | 0.36 | 0.12 |
| 11 | 16973883 | rs4756878 | 0.783 | -0.003 | -0.322 | -1.608 | 0.004 | 0.052 | 0.052 | 0 | 0.55 | 0.35 | 0.54 | 0.76 | 0.47 | 0.75 |
| 11 | 16990290 | rs6486330 | 0.153 | -0.196 | -0.832 | -1.686 | -0.001 | -0.02 | -0.02 | 0 | 0.46 | 0.39 | 0.46 | 0.85 | 0.49 | 0.54 |
| 11 | 16990389 | rs6486331 | -0.57 | 0.359 | -1.206 | -1.207 | 0 | 0.015 | 0.015 | 0 | 0.53 | 0.73 | 0.54 | 0.41 | 0.42 | 0.62 |
| 11 | 16992940 | rs1544506 | -0.083 | -0.412 | -1.557 | -1.668 | 0.047 | 0.061 | 0.061 | 0 | 0.57 | 0.71 | 0.59 | 0.27 | 0.36 | 0.51 |
| 11 | 16994581 | rs7130301 | -0.48 | -0.244 | -1.69 | -1.605 | 0 | -0.003 | -0.003 | 0 | 0.8 | 0.78 | 0.83 | 0.87 | 0.66 | 0.88 |
| 11 | 16994792 | rs10832706 | 0.668 | -0.043 | -2.226 | -1.584 | -0.084 | 0.054 | 0.054 | 0 | 0.16 | 0.05 | 0.16 | 0.1 | 0.33 | 0.09 |
| 11 | 16997462 | rs3927931 | 0.338 | -0.519 | -1.709 | -1.105 | -0.03 | -0.034 | -0.034 | 0.662 | 0.29 | 0.33 | 0.29 | 0.12 | 0.13 | 0.36 |
| 11 | 16999029 | rs11024097 | 0.407 | 0.316 | -1.718 | -0.733 | -0.105 | -0.09 | -0.09 | 0 | 0.23 | 0.05 | 0.24 | 0.62 | 0.31 | 0.37 |
| 11 | 16999882 | rs713503 | -0.012 | 0.567 | -1.032 | -1.278 | -0.005 | -0.016 | -0.016 | 0 | 0.58 | 0.42 | 0.61 | 0.86 | 0.47 | 0.76 |
| 11 | 17001584 | rs4757467 | -0.042 | 0.838 | -0.915 | -1.136 | -0.003 | -0.016 | -0.016 | 0 | 0.68 | 0.7 | 0.7 | 0.96 | 0.6 | 0.74 |
| 11 | 17004150 | rs4757470 | 0.721 | 0.537 | -0.84 | -0.476 | -0.007 | -0.001 | -0.001 | 0.021 | 0.4 | 0.45 | 0.29 | 0.45 | 0.31 | 0.31 |
| 11 | 17005654 | rs7103326 | 0.848 | 0.311 | -0.89 | -0.017 | 0.008 | -0.009 | -0.009 | 0 | 0.07 | 0.06 | 0.12 | 0.14 | 0.01 | 0.1 |
| 11 | 17008605 | rs11024102 | 0.794 | 0.524 | -0.95 | -0.13 | -0.104 | -0.125 | -0.125 | 0 | 0.16 | 0.02 | 0.27 | 0.37 | 0.26 | 0.33 |
| 11 | 17008644 | rs10832709 | 0.447 | 0.517 | -0.933 | -0.128 | 0 | 0 | 0 | 0 | 0.47 | 0.5 | 0.41 | 0.58 | 0.33 | 0.41 |
| 11 | 17022030 | rs7940499 | -0.259 | 0.435 | -0.819 | -0.47 | 0.003 | -0.109 | -0.109 | 0.946 | 0.64 | 0.51 | 0.73 | 0.96 | 0.58 | 0.74 |
| 11 | 17025740 | rs4757484 | -0.272 | 0.465 | -0.82 | -0.471 | -0.002 | -0.11 | -0.11 | 0.284 | 0.65 | 0.52 | 0.73 | 0.96 | 0.61 | 0.75 |
| 11 | 17027329 | rs10741719 | -0.305 | 0.474 | -0.807 | -0.48 | -0.001 | -0.126 | -0.126 | 0.008 | 0.65 | 0.54 | 0.73 | 0.97 | 0.61 | 0.75 |
| 11 | 17037793 | rs7130826 | -0.45 | 0.618 | -0.796 | 0.068 | -0.016 | -0.031 | -0.031 | 0.499 | 0.24 | 0.21 | 0.39 | 0.53 | 0.25 | 0.41 |
| 11 | 17085192 | rs7949405 | -0.514 | 0.561 | -0.21 | 0.375 | 0.015 | -0.014 | -0.014 | 0 | 0.24 | 0.35 | 0.4 | 0.8 | 0.24 | 0.42 |
| 11 | 17089005 | rs11024145 | 0.092 | -0.37 | 0.318 | 0.958 | -0.02 | 0.082 | 0.082 | 0 | 0.08 | 0.13 | 0.17 | 0.42 | 0.01 | 0.16 |
| 11 | 17096211 | rs1541533 | 1.248 | -0.887 | -0.267 | 0.329 | -0.003 | 0 | 0 | 0.166 | 0.1 | 0 | 0.14 | 0.07 | 0.07 | 0.11 |
| 11 | 17099613 | rs16933823 | 0.183 | -1.479 | -0.421 | -0.92 | 0.003 | -0.033 | -0.033 | 0 | 0.27 | 0.22 | 0.18 | 0.02 | 0.19 | 0.17 |
| 11 | 17111272 | rs10832733 | -1.079 | -1.315 | -0.472 | -1.001 | -0.004 | 0.033 | 0.033 | 0 | 0.57 | 0.62 | 0.62 | 0.81 | 0.5 | 0.6 |
| 11 | 17118687 | rs11604561 | 1.203 | -1.778 | -0.549 | -1.277 | 0.009 | 0.011 | 0.011 | 0 | 0.15 | 0.1 | 0.11 | 0.01 | 0.12 | 0.06 |
| 11 | 17157366 | rs11603765 | -0.748 | -1.869 | -0.712 | -1.326 | -0.003 | -0.057 | -0.057 | 0 | 0.33 | 0.27 | 0.3 | 0.02 | 0.21 | 0.13 |
| 11 | 17191019 | rs214936 | 1.004 | -1.734 | -1.72 | -1.338 | -0.004 | 0.035 | 0.035 | 0.013 | 0.43 | 0.37 | 0.38 | 0.18 | 0.5 | 0.38 |
| 11 | 17194584 | rs214933 | 0.691 | -1.654 | -1.69 | -1.32 | -0.028 | 0.005 | 0.005 | 0.064 | 0.23 | 0.19 | 0.31 | 0.8 | 0.29 | 0.48 |
| 11 | 17207433 | rs542274 | -0.699 | -1.696 | -1.791 | -1.301 | -0.005 | 0.022 | 0.022 | 0.529 | 0.42 | 0.59 | 0.52 | 0.81 | 0.38 | 0.55 |
| 11 | 17247708 | rs7949699 | -1.283 | -1.547 | -1.752 | -1.151 | -0.004 | 0.034 | 0.034 | 0 | 0.58 | 0.63 | 0.61 | 0.81 | 0.5 | 0.63 |
| 11 | 17250019 | rs10832747 | -1.268 | -1.551 | -1.755 | -1.153 | -0.003 | 0.035 | 0.035 | 0 | 0.58 | 0.63 | 0.61 | 0.81 | 0.5 | 0.62 |
| 11 | 17260116 | rs12577525 | 1.072 | -1.349 | -1.771 | -0.91 | -0.128 | -0.14 | -0.14 | 0 | 0.2 | 0.02 | 0.25 | 0.38 | 0.28 | 0.34 |
| 11 | 17267029 | rs1488933 | -0.454 | -1.388 | -1.687 | -1.005 | -0.003 | 0.002 | 0.002 | 0.181 | 0.37 | 0.57 | 0.4 | 0.8 | 0.36 | 0.54 |
| 11 | 17291499 | rs214097 | -1.184 | -1.493 | -2.02 | -1.14 | -0.001 | 0.003 | 0.003 | 0 | 0.63 | 0.64 | 0.66 | 0.82 | 0.57 | 0.64 |
| 11 | 17300844 | rs7127347 | -0.243 | -1.038 | -1.448 | -0.594 | -0.021 | 0.159 | 0.159 | 0 | 0.17 | 0.36 | 0.2 | 0.43 | 0.11 | 0.21 |
| 11 | 17313521 | rs214105 | 1.081 | -0.805 | -1.774 | -0.707 | -0.116 | -0.129 | -0.129 | 0 | 0.19 | 0.02 | 0.18 | 0.36 | 0.28 | 0.31 |
| 11 | 17316029 | rs1330 | 0.182 | -0.995 | -1.772 | -0.524 | -0.055 | -0.051 | -0.051 | 0 | 0.33 | 0.14 | 0.35 | 0.37 | 0.33 | 0.37 |
| 11 | 17320797 | rs10741725 | -0.824 | -0.988 | -1.622 | -0.56 | -0.001 | 0.067 | 0.067 | 0.806 | 0.51 | 0.65 | 0.56 | 0.81 | 0.42 | 0.6 |
| 11 | 17337230 | rs10466382 | 1.514 | -0.734 | -2.078 | -0.801 | -0.021 | -0.015 | -0.015 | 2.155 | 0.14 | 0.1 | 0.15 | 0.51 | 0.3 | 0.2 |
| 11 | 17339127 | rs2634462 | -1.042 | -1.148 | -1.979 | -1.089 | -0.017 | -0.041 | -0.041 | 0.031 | 0.67 | 0.65 | 0.69 | 0.88 | 0.71 | 0.65 |
| 11 | 17369517 | rs7112244 | 0.75 | -1.031 | -1.954 | -0.947 | 0.064 | 0.06 | 0.06 | 0.062 | 0.05 | 0.41 | 0.02 | 0 | 0 | 0.01 |
| 11 | 17370701 | rs2354867 | 0.319 | -1.062 | -2.298 | -0.985 | -0.047 | -0.087 | -0.087 | 0 | 0.24 | 0.13 | 0.26 | 0.53 | 0.37 | 0.25 |
| 11 | 17390855 | rs2018218 | -1.039 | -1.194 | -2.465 | -1.195 | 0.001 | -0.006 | -0.006 | 0 | 0.45 | 0.65 | 0.46 | 0.54 | 0.42 | 0.31 |

| | | | | | | | | | | | | | | | | |
|---|---|---|---|---|---|---|---|---|---|---|---|---|---|---|---|---|
| 11 | 17393644 | rs12791318 | 1.449 | -1.436 | -2.665 | -1.156 | -0.026 | -0.03 | -0.03 | 1.443 | 0.28 | 0.31 | 0.23 | 0.13 | 0.22 | 0.26 |
| 11 | 17394597 | rs4756887 | -0.907 | -1.387 | -2.615 | -1.121 | 0.001 | -0.006 | -0.006 | 0 | 0.46 | 0.66 | 0.46 | 0.54 | 0.42 | 0.31 |
| 11 | 17395760 | rs11024272 | -0.888 | -1.383 | -2.615 | -1.119 | 0.001 | -0.006 | -0.006 | 0.619 | 0.45 | 0.65 | 0.46 | 0.54 | 0.42 | 0.31 |
| 11 | 17396517 | rs10832781 | 0.678 | -1.281 | -2.645 | -1.064 | -0.035 | -0.063 | -0.063 | 1.603 | 0.24 | 0.16 | 0.26 | 0.53 | 0.37 | 0.24 |
| 11 | 17405333 | rs10832785 | -0.398 | -1.252 | -2.646 | -1.326 | 0.002 | 0.014 | 0.014 | 2.084 | 0.36 | 0.56 | 0.38 | 0.57 | 0.36 | 0.33 |
| 11 | 17405347 | rs1002227 | 1.807 | -1.244 | -2.689 | -1.309 | -0.042 | -0.06 | -0.06 | 4.145 | 0.16 | 0.08 | 0.18 | 0.56 | 0.31 | 0.26 |
| 11 | 17406051 | rs4757512 | 0.571 | -1.435 | -2.707 | -1.409 | -0.013 | -0.051 | -0.051 | 0 | 0.38 | 0.39 | 0.32 | 0.08 | 0.27 | 0.27 |
| 11 | 17408025 | rs2285676 | -0.374 | -0.99 | -2.712 | -1.377 | 0.002 | 0.009 | 0.009 | 2.628 | 0.37 | 0.59 | 0.39 | 0.58 | 0.38 | 0.34 |
| 11 | 17408630 | rs5215 | -1.52 | -0.949 | -2.723 | -1.353 | -0.122 | -0.116 | -0.116 | 0 | 0.74 | 0.95 | 0.7 | 0.66 | 0.65 | 0.6 |
| 11 | 17409572 | rs5219 | -1.532 | -0.987 | -2.609 | -1.23 | -0.138 | -0.131 | -0.131 | 0 | 0.74 | 0.98 | 0.71 | 0.66 | 0.65 | 0.6 |
| 11 | 17411163 | rs886288 | -0.484 | -1.079 | -2.62 | -1.284 | 0.002 | 0.008 | 0.008 | 2.478 | 0.38 | 0.58 | 0.44 | 0.58 | 0.39 | 0.34 |
| 11 | 17411821 | rs2074314 | -1.478 | -0.892 | -2.615 | -1.185 | -0.138 | -0.128 | -0.128 | 0 | 0.74 | 0.98 | 0.71 | 0.66 | 0.65 | 0.6 |
| 11 | 17412960 | rs7124355 | -1.739 | -0.816 | -2.625 | -1.212 | -0.103 | -0.118 | -0.118 | 0.108 | 0.77 | 0.97 | 0.73 | 0.66 | 0.69 | 0.61 |
| 11 | 17414570 | rs8192690 | 1.217 | -1.088 | -2.476 | -1.119 | 0.006 | 0.04 | 0.04 | 0 | 0.09 | 0.03 | 0.03 | 0 | 0.05 | 0.04 |
| 11 | 17414715 | rs1109591 | -0.554 | -1.136 | -2.473 | -1.141 | 0.011 | 0.01 | 0.01 | 0.02 | 0.53 | 0.46 | 0.5 | 0.13 | 0.34 | 0.37 |
| 11 | 17417337 | rs8192691 | 1.095 | -1.074 | -2.526 | -1.248 | -0.02 | 0.065 | 0.065 | 0 | 0.12 | 0.02 | 0.16 | 0.01 | 0.06 | 0.05 |
| 11 | 17418023 | rs757111 | 1.525 | -1.07 | -2.526 | -1.246 | -0.026 | 0.057 | 0.057 | 0 | 0.11 | 0.01 | 0.15 | 0.01 | 0.06 | 0.05 |
| 11 | 17418477 | rs757110 | -1.686 | -1.002 | -2.57 | -1.187 | -0.134 | -0.135 | -0.135 | 0 | 0.73 | 0.98 | 0.69 | 0.64 | 0.65 | 0.58 |
| 11 | 17419279 | rs1799859 | -0.097 | -1.107 | -2.482 | -1.404 | 0.089 | 0.206 | 0.206 | 0 | 0.45 | 0.77 | 0.42 | 0.09 | 0.27 | 0.29 |
| 11 | 17421366 | rs16934019 | 1.325 | -1.106 | -2.378 | -1.357 | 0.075 | 0.071 | 0.071 | 0 | 0.05 | 0.38 | 0.02 | 0 | 0 | 0.01 |
| 11 | 17421714 | rs4148640 | -0.321 | -1.037 | -2.598 | -1.346 | 0.11 | 0.255 | 0.255 | 0 | 0.48 | 0.82 | 0.48 | 0.12 | 0.28 | 0.35 |
| 11 | 17421790 | rs2074312 | 1.955 | -1.057 | -2.623 | -1.3 | -0.039 | -0.056 | -0.056 | 0.241 | 0.15 | 0.06 | 0.18 | 0.57 | 0.31 | 0.2 |
| 11 | 17421860 | rs2074311 | -0.939 | -0.996 | -2.574 | -1.259 | -0.082 | -0.034 | -0.034 | 0 | 0.64 | 0.88 | 0.66 | 0.68 | 0.6 | 0.55 |
| 11 | 17421886 | rs2074310 | -1.347 | -0.421 | -2.282 | -0.832 | -0.111 | -0.087 | -0.087 | 0 | 0.73 | 0.96 | 0.69 | 0.7 | 0.65 | 0.62 |
| 11 | 17422397 | rs12422139 | 0.241 | -0.147 | -1.146 | -0.163 | -0.028 | 0.066 | 0.066 | 0 | 0.12 | 0.01 | 0.16 | 0.01 | 0.06 | 0.07 |
| 11 | 17424395 | rs4757513 | -0.689 | -0.08 | -0.383 | 0.548 | -0.008 | 0.016 | 0.016 | 0.092 | 0.09 | 0.43 | 0.08 | 0.03 | 0.01 | 0.05 |
| 11 | 17424711 | rs2067043 | -0.009 | 0.656 | -1.423 | 0.887 | -0.031 | -0.045 | -0.045 | 2.271 | 0.16 | 0.09 | 0.18 | 0.56 | 0.31 | 0.23 |
| 11 | 17427363 | rs4148636 | -1.211 | -0.866 | -1.237 | -0.248 | 0.028 | 0.026 | 0.026 | 4.949 | 0.23 | 0.79 | 0.15 | 0.05 | 0.11 | 0.13 |
| 11 | 17427557 | rs739688 | 1.074 | -0.877 | -2.174 | -0.368 | 0.018 | 0.082 | 0.082 | 0 | 0.49 | 0.11 | 0.59 | 0.66 | 0.6 | 0.39 |
| 11 | 17427862 | rs2077655 | -0.579 | -0.906 | -2.116 | -0.376 | 0.028 | 0.027 | 0.027 | 4.959 | 0.23 | 0.79 | 0.15 | 0.05 | 0.11 | 0.13 |
| 11 | 17429358 | rs1319447 | 0.634 | -1.038 | -1.205 | -0.406 | 0.008 | 0.042 | 0.042 | 0 | 0.58 | 0.34 | 0.64 | 0.72 | 0.61 | 0.45 |
| 11 | 17429872 | rs4148634 | 0.227 | -1.083 | -0.276 | -0.449 | 0.021 | -0.04 | -0.04 | 0.383 | 0.08 | 0.27 | 0.06 | 0.02 | 0.1 | 0.06 |
| 11 | 17429922 | rs2106865 | 0.563 | -1.111 | -0.222 | -0.42 | 0 | 0.001 | 0.001 | 0 | 0.61 | 0.63 | 0.66 | 0.72 | 0.61 | 0.46 |
| 11 | 17430933 | rs2237992 | -0.119 | -1.106 | -0.36 | -0.411 | -0.054 | -0.005 | -0.005 | 0.224 | 0.7 | 0.97 | 0.72 | 0.74 | 0.71 | 0.52 |
| 11 | 17431210 | rs7947462 | -0.119 | -1.106 | -0.36 | -0.411 | -0.054 | -0.005 | -0.005 | 0 | 0.7 | 0.96 | 0.72 | 0.74 | 0.71 | 0.52 |
| 11 | 17433088 | rs7926934 | 0.082 | -1.012 | -0.118 | -0.36 | 0.013 | -0.047 | -0.047 | 0 | 0.09 | 0.29 | 0.06 | 0.02 | 0.1 | 0.06 |
| 11 | 17433862 | rs10766398 | 1.272 | -1.171 | -0.03 | -0.351 | 0.049 | 0.085 | 0.085 | 0 | 0.49 | 0.2 | 0.59 | 0.64 | 0.59 | 0.38 |
| 11 | 17434284 | rs1805036 | 0.155 | -1.219 | -0.003 | -0.345 | 0.012 | -0.047 | -0.047 | 0.956 | 0.09 | 0.29 | 0.06 | 0.02 | 0.1 | 0.06 |
| 11 | 17435214 | rs2074309 | 1.255 | -1.217 | -0.113 | -0.339 | 0.05 | 0.085 | 0.085 | 0 | 0.49 | 0.2 | 0.59 | 0.64 | 0.59 | 0.38 |
| 11 | 17436936 | rs4148626 | -0.763 | -1.305 | -0.307 | -0.43 | 0.003 | 0.008 | 0.008 | 0 | 0.82 | 0.98 | 0.89 | 0.92 | 0.84 | 0.85 |
| 11 | 17436951 | rs12293228 | -0.621 | -1.344 | 0.087 | -0.457 | 0.009 | -0.071 | -0.071 | 13.358 | 0.12 | 0.52 | 0.07 | 0.02 | 0.1 | 0.07 |
| 11 | 17437785 | rs4148625 | 1.723 | -1.289 | -0.002 | -0.44 | 0.045 | 0.087 | 0.087 | 0 | 0.49 | 0.17 | 0.59 | 0.64 | 0.59 | 0.38 |
| 11 | 17437826 | rs2074316 | -0.279 | -1.213 | 0.203 | -0.372 | -0.001 | 0.001 | 0.001 | 0 | 0.19 | 0.15 | 0.23 | 0.27 | 0.16 | 0.41 |
| 11 | 17438890 | rs2074315 | -1.268 | -1.206 | 0.19 | -0.33 | -0.008 | -0.01 | -0.01 | 0 | 0.28 | 0.53 | 0.29 | 0.29 | 0.25 | 0.47 |
| 11 | 17440229 | rs2237991 | -1.134 | -1.094 | -0.33 | -0.363 | 0.008 | 0.01 | 0.01 | 0.056 | 0.3 | 0.73 | 0.3 | 0.3 | 0.26 | 0.47 |
| 11 | 17440473 | rs916829 | -0.771 | -1.089 | -0.489 | -0.371 | 0.003 | 0.011 | 0.011 | 0 | 0.82 | 0.98 | 0.89 | 0.92 | 0.84 | 0.85 |
| 11 | 17441228 | rs2299640 | -0.164 | -1.069 | -0.305 | -0.456 | 0.007 | -0.077 | -0.077 | 0 | 0.14 | 0.3 | 0.1 | 0.02 | 0.16 | 0.07 |
| 11 | 17441385 | rs2299639 | -1.038 | -1.149 | -0.361 | -0.483 | 0.006 | 0.013 | 0.013 | 0 | 0.3 | 0.7 | 0.29 | 0.23 | 0.22 | 0.38 |
| 11 | 17441828 | rs4757517 | -1.239 | -1.135 | -0.32 | -0.423 | 0.007 | 0.04 | 0.04 | 0.152 | 0.36 | 0.7 | 0.32 | 0.23 | 0.31 | 0.41 |
| 11 | 17442460 | rs929235 | 0.327 | -1.068 | -0.318 | -0.381 | 0.004 | -0.07 | -0.07 | 3.69 | 0.14 | 0.24 | 0.1 | 0.03 | 0.16 | 0.08 |
| 11 | 17444024 | rs4148624 | 0.144 | -1.093 | -0.324 | -0.323 | 0.008 | -0.081 | -0.081 | 0 | 0.15 | 0.36 | 0.11 | 0.04 | 0.17 | 0.08 |
| 11 | 17444134 | rs2237988 | -0.115 | -0.806 | -1.274 | -0.536 | -0.017 | -0.045 | -0.045 | 0 | 0.76 | 0.97 | 0.82 | 0.8 | 0.75 | 0.72 |
| 11 | 17444590 | rs11024284 | 0.555 | -0.721 | -1.132 | -0.496 | 0.003 | -0.055 | -0.055 | 0 | 0.14 | 0.23 | 0.1 | 0.04 | 0.16 | 0.08 |
| 11 | 17445026 | rs916827 | 1.291 | -0.457 | -1.238 | -0.323 | -0.059 | -0.047 | -0.047 | 0 | 0.4 | 0.14 | 0.54 | 0.56 | 0.42 | 0.3 |
| 11 | 17448704 | rs1799854 | 1.442 | -0.595 | -1.314 | -0.388 | -0.066 | -0.061 | -0.061 | 3.203 | 0.36 | 0.14 | 0.54 | 0.55 | 0.42 | 0.32 |
| 11 | 17449002 | rs4148622 | 0.059 | -0.656 | -1.09 | -0.562 | 0.018 | 0.03 | 0.03 | 0 | 0.68 | 0.78 | 0.83 | 0.89 | 0.73 | 0.79 |
| 11 | 17451427 | rs4148621 | 0.235 | -0.183 | -1.099 | -0.153 | -0.019 | -0.013 | -0.013 | 0.252 | 0.1 | 0.02 | 0.12 | 0.08 | 0.14 | 0.12 |
| 11 | 17452580 | rs2074308 | 0.43 | -0.208 | -1.171 | 0.02 | -0.028 | -0.04 | -0.04 | 0.061 | 0.11 | 0.03 | 0.11 | 0.2 | 0.13 | 0.31 |
| 11 | 17453443 | rs4148618 | 0.115 | -0.133 | -1.031 | -0.651 | -0.01 | 0.017 | 0.017 | 0.077 | 0.17 | 0.09 | 0.11 | 0.06 | 0.22 | 0.08 |
| 11 | 17455371 | rs4148617 | -1.183 | -0.767 | -0.231 | -0.988 | -0.006 | -0.068 | -0.068 | 0 | 0.31 | 0.61 | 0.19 | 0.07 | 0.26 | 0.17 |
| 11 | 17458136 | rs7108578 | 0.215 | -0.471 | -0.276 | -0.782 | -0.01 | 0.019 | 0.019 | 0.065 | 0.17 | 0.1 | 0.15 | 0.07 | 0.22 | 0.08 |
| 11 | 17458299 | rs7119439 | 0.233 | -0.47 | -0.276 | -0.781 | -0.01 | 0.019 | 0.019 | 0.077 | 0.17 | 0.1 | 0.15 | 0.07 | 0.22 | 0.08 |
| 11 | 17462220 | rs886293 | -0.725 | -0.447 | -0.713 | -0.38 | -0.007 | -0.048 | -0.048 | 0.092 | 0.76 | 0.88 | 0.71 | 0.38 | 0.73 | 0.58 |
| 11 | 17462304 | rs2283261 | 0.619 | -0.776 | -0.471 | -0.774 | 0.016 | 0.08 | 0.08 | 0 | 0.48 | 0.71 | 0.4 | 0.27 | 0.39 | 0.32 |
| 11 | 17462727 | rs757109 | 0.626 | -0.783 | -0.474 | -0.778 | 0.015 | 0.079 | 0.079 | 0 | 0.48 | 0.71 | 0.4 | 0.27 | 0.39 | 0.32 |
| 11 | 17466268 | rs7117162 | 0.889 | -0.802 | -0.322 | -0.785 | 0.012 | 0.032 | 0.032 | 0 | 0.45 | 0.63 | 0.38 | 0.26 | 0.36 | 0.24 |
| 11 | 17466523 | rs2040653 | 0.984 | -0.871 | 0.079 | -0.818 | 0.003 | 0.009 | 0.009 | 0 | 0.44 | 0.38 | 0.36 | 0.26 | 0.36 | 0.24 |
| 11 | 17468317 | rs7950189 | -0.382 | -0.832 | -0.276 | -0.821 | -0.051 | 0.028 | 0.028 | 0 | 0.59 | 0.84 | 0.57 | 0.3 | 0.51 | 0.5 |
| 11 | 17473146 | rs722341 | 1.473 | -0.724 | -0.376 | -0.868 | -0.021 | 0.012 | 0.012 | 0.121 | 0.1 | 0.01 | 0.15 | 0.03 | 0.12 | 0.11 |

| | | | | | | | | | | | | | | | | |
|---|---|---|---|---|---|---|---|---|---|---|---|---|---|---|---|---|
| 11 | 17473437 | rs2283258 | 2.393 | -1.124 | -0.726 | -1.028 | -0.114 | -0.118 | -0.118 | 4.349 | 0.24 | 0.03 | 0.3 | 0.33 | 0.31 | 0.17 |
| 11 | 17475283 | rs2073586 | 1.321 | -1.281 | -0.343 | -1.082 | -0.053 | -0.02 | -0.02 | 1.152 | 0.36 | 0.16 | 0.33 | 0.35 | 0.38 | 0.29 |
| 11 | 17475443 | rs2073585 | 0.359 | -1.368 | 0.08 | -1.125 | -0.001 | -0.002 | -0.002 | 0 | 0.46 | 0.54 | 0.38 | 0.35 | 0.39 | 0.31 |
| 11 | 17476516 | rs2237976 | 0.039 | -1.438 | 0.156 | -1.161 | 0.006 | 0.018 | 0.018 | 0 | 0.47 | 0.63 | 0.4 | 0.34 | 0.39 | 0.3 |
| 11 | 17477830 | rs2237975 | 0.77 | -1.438 | 0.09 | -1.125 | 0.008 | 0.029 | 0.029 | 0 | 0.41 | 0.25 | 0.35 | 0.33 | 0.35 | 0.21 |
| 11 | 17478157 | rs2237974 | 1.422 | -1.428 | 0.097 | -1.119 | -0.014 | 0 | 0 | 0 | 0.34 | 0.22 | 0.34 | 0.33 | 0.35 | 0.2 |
| 11 | 17480835 | rs7105391 | 0.425 | -1.464 | 0.336 | -1.109 | -0.004 | -0.006 | -0.006 | 0 | 0.44 | 0.49 | 0.37 | 0.37 | 0.35 | 0.21 |
| 11 | 17481948 | rs886291 | 0.44 | -1.463 | 0.333 | -1.108 | -0.004 | -0.005 | -0.005 | 0 | 0.44 | 0.49 | 0.37 | 0.37 | 0.35 | 0.21 |
| 11 | 17482329 | rs886289 | 2.364 | -1.539 | 0.043 | -1.158 | -0.11 | -0.123 | -0.123 | 0 | 0.24 | 0.02 | 0.3 | 0.37 | 0.31 | 0.17 |
| 11 | 17484731 | rs4148609 | 1.002 | -1.291 | 0.341 | -0.914 | 0.007 | 0.014 | 0.014 | 0 | 0.42 | 0.3 | 0.35 | 0.37 | 0.35 | 0.21 |
| 11 | 17486737 | rs3819521 | 1.639 | -1.248 | 0.292 | -0.893 | -0.011 | -0.006 | -0.006 | 0 | 0.35 | 0.27 | 0.33 | 0.37 | 0.35 | 0.21 |
| 11 | 17487278 | rs3819517 | 0.611 | -1.246 | 0.469 | -0.929 | 0.002 | 0.011 | 0.011 | 0 | 0.46 | 0.62 | 0.39 | 0.38 | 0.39 | 0.3 |
| 11 | 17487445 | rs11024295 | 0.64 | -1.246 | 0.449 | -0.929 | 0.001 | 0.005 | 0.005 | 0 | 0.46 | 0.59 | 0.39 | 0.38 | 0.39 | 0.3 |
| 11 | 17489106 | rs11826760 | 1.713 | -1.249 | 0.414 | -0.942 | -0.009 | -0.004 | -0.004 | 0 | 0.35 | 0.3 | 0.33 | 0.37 | 0.35 | 0.21 |
| 11 | 17489267 | rs4148608 | 1.713 | -1.249 | 0.414 | -0.942 | -0.009 | -0.004 | -0.004 | 0 | 0.35 | 0.3 | 0.33 | 0.37 | 0.35 | 0.21 |
| 11 | 17489279 | rs4148607 | 1.713 | -1.249 | 0.414 | -0.907 | -0.009 | -0.005 | -0.005 | 0.03 | 0.35 | 0.3 | 0.33 | 0.37 | 0.35 | 0.21 |
| 11 | 17489363 | rs4148603 | 0.335 | -1.235 | 0.274 | -0.899 | 0.008 | 0.032 | 0.032 | 0 | 0.47 | 0.72 | 0.39 | 0.38 | 0.39 | 0.3 |
| 11 | 17489445 | rs2283257 | 0.83 | -1.09 | 0.035 | -1.048 | -0.019 | 0.054 | 0.054 | 0 | 0.18 | 0.06 | 0.15 | 0.06 | 0.2 | 0.07 |
| 11 | 17491831 | rs11024298 | 0.807 | -1.126 | 0.216 | -1.014 | -0.046 | -0.049 | -0.049 | 0 | 0.09 | 0.29 | 0.04 | 0 | 0.01 | 0.01 |
| 11 | 17492664 | rs2237967 | 0.109 | -1.076 | -0.254 | -0.998 | -0.025 | -0.018 | -0.018 | 0 | 0.59 | 0.8 | 0.56 | 0.41 | 0.51 | 0.53 |
| 11 | 17495051 | rs2283255 | -0.328 | -0.918 | -0.202 | -0.662 | 0.007 | -0.036 | -0.036 | 0.173 | 0.62 | 0.71 | 0.57 | 0.46 | 0.62 | 0.37 |
| 11 | 17495180 | rs2283254 | 0.635 | -0.83 | 0.056 | -0.284 | 0.003 | 0.004 | 0.004 | 0 | 0.35 | 0.41 | 0.35 | 0.35 | 0.34 | 0.21 |
| 11 | 17495430 | rs2283253 | 0.153 | -0.596 | -0.423 | -0.267 | 0.004 | -0.005 | -0.005 | 1.102 | 0.43 | 0.61 | 0.39 | 0.41 | 0.39 | 0.28 |
| 11 | 17499547 | rs3758953 | -1.065 | -0.84 | -0.098 | -0.334 | 0.002 | -0.001 | -0.001 | 1.864 | 0.57 | 0.59 | 0.59 | 0.41 | 0.5 | 0.62 |
| 11 | 17499885 | rs2188966 | -0.333 | -0.993 | -0.417 | -0.332 | -0.024 | 0.005 | 0.005 | 1.357 | 0.38 | 0.22 | 0.5 | 0.32 | 0.4 | 0.44 |
| 11 | 17501063 | rs3758949 | -0.331 | -0.965 | -0.403 | -0.301 | -0.02 | 0.005 | 0.005 | 1.078 | 0.39 | 0.23 | 0.51 | 0.33 | 0.4 | 0.44 |
| 11 | 17501210 | rs3758947 | 1.152 | -1.499 | -0.587 | -0.356 | -0.051 | -0.091 | -0.091 | 0.161 | 0.17 | 0.01 | 0.34 | 0.26 | 0.18 | 0.27 |
| 11 | 17506874 | rs1076204 | -0.67 | -1.4 | -0.216 | -0.517 | 0.001 | 0.001 | 0.001 | 0.56 | 0.45 | 0.42 | 0.5 | 0.41 | 0.45 | 0.52 |
| 11 | 17508213 | rs12278956 | -0.284 | -1.67 | -0.107 | -0.521 | -0.038 | 0.011 | 0.011 | 0 | 0.27 | 0.34 | 0.22 | 0.3 | 0.14 | 0.19 |
| 11 | 17510231 | rs11606229 | -0.512 | -1.699 | -0.066 | -0.706 | -0.028 | -0.115 | -0.115 | 0 | 0.34 | 0.41 | 0.15 | 0.04 | 0.24 | 0.21 |
| 11 | 17510419 | rs11603988 | 1.441 | -1.702 | -0.576 | -0.637 | -0.04 | -0.1 | -0.1 | 0 | 0.18 | 0.02 | 0.35 | 0.3 | 0.18 | 0.27 |
| 11 | 17510565 | rs4757527 | -0.524 | -1.721 | -0.385 | -0.682 | -0.028 | -0.114 | -0.114 | 0 | 0.34 | 0.41 | 0.15 | 0.04 | 0.24 | 0.21 |
| 11 | 17511427 | rs7129639 | 0.883 | -1.222 | -0.254 | -0.411 | 0.012 | 0.007 | 0.007 | 0 | 0.45 | 0.31 | 0.63 | 0.58 | 0.6 | 0.55 |
| 11 | 17513147 | rs17703233 | 1.117 | -1.059 | -0.314 | -0.357 | -0.011 | 0.025 | 0.025 | 0.097 | 0.1 | 0.06 | 0.05 | 0.02 | 0.09 | 0.04 |
| 11 | 17513943 | rs12576915 | 1 | -1.351 | 0.037 | -0.043 | 0.044 | -0.053 | -0.053 | 0.798 | 0.19 | 0.09 | 0.17 | 0.3 | 0.09 | 0.11 |
| 11 | 17515143 | rs4757536 | -0.552 | -1.284 | 0.117 | -0.069 | 0.007 | -0.009 | -0.009 | 0.041 | 0.75 | 0.71 | 0.8 | 0.62 | 0.88 | 0.83 |
| 11 | 17515668 | rs1055581 | 0.677 | -1.094 | -0.047 | -0.03 | 0.048 | -0.02 | -0.02 | 0.278 | 0.33 | 0.43 | 0.32 | 0.3 | 0.46 | 0.37 |
| 11 | 17515693 | rs1055577 | 0.708 | -1.087 | -0.191 | -0.009 | 0.033 | -0.017 | -0.017 | 0 | 0.32 | 0.39 | 0.32 | 0.29 | 0.46 | 0.36 |
| 11 | 17515833 | rs1055574 | 0.708 | -1.087 | -0.191 | -0.072 | 0.033 | -0.013 | -0.013 | 0 | 0.32 | 0.39 | 0.32 | 0.29 | 0.46 | 0.37 |
| 11 | 17516038 | rs2072227 | 0.686 | -1.089 | -0.192 | -0.074 | 0.032 | -0.014 | -0.014 | 0 | 0.33 | 0.39 | 0.32 | 0.29 | 0.46 | 0.36 |
| 11 | 17517879 | rs2072228 | -0.058 | -0.946 | -0.685 | 0.072 | -0.01 | 0.032 | 0.032 | 0.425 | 0.45 | 0.29 | 0.49 | 0.32 | 0.46 | 0.54 |
| 11 | 17518909 | rs2072231 | -0.111 | -1.544 | -0.423 | 0.199 | -0.017 | 0.008 | 0.008 | 0.176 | 0.22 | 0.2 | 0.19 | 0.31 | 0.1 | 0.11 |
| 11 | 17520339 | rs2270024 | 0.018 | -1.011 | -0.849 | 0.226 | 0.038 | 0.014 | 0.014 | 0 | 0.55 | 0.39 | 0.68 | 0.63 | 0.66 | 0.61 |
| 11 | 17521825 | rs11024311 | -1.015 | -1.119 | -0.637 | -0.252 | -0.004 | -0.029 | -0.029 | 0 | 0.13 | 0.11 | 0.06 | 0 | 0.08 | 0.11 |
| 11 | 17522086 | rs878208 | -1.957 | -1.615 | 0.31 | -0.751 | -0.057 | -0.153 | -0.153 | 0 | 0.24 | 0.52 | 0.12 | 0.05 | 0.13 | 0.28 |
| 11 | 17522810 | rs2072234 | -0.781 | -1.859 | -0.293 | -1.168 | -0.017 | 0.064 | 0.064 | 0.08 | 0.59 | 0.81 | 0.39 | 0.34 | 0.45 | 0.49 |
| 11 | 17523289 | rs12270644 | -0.686 | -1.866 | 0.129 | -1.186 | -0.026 | -0.06 | -0.06 | 0 | 0.16 | 0.21 | 0.11 | 0.01 | 0.08 | 0.03 |
| 11 | 17523474 | rs17703528 | 0.337 | -1.753 | 0.151 | -1.123 | -0.004 | 0.009 | 0.009 | 0 | 0.07 | 0.06 | 0.04 | 0.04 | 0.05 | 0.23 |
| 11 | 17530463 | rs1006099 | -1.703 | -2.032 | 0.365 | -0.418 | -0.028 | 0.032 | 0.032 | 0 | 0.28 | 0.49 | 0.26 | 0.29 | 0.2 | 0.38 |
| 11 | 17530484 | rs7104083 | -0.012 | -2.265 | 0.484 | -0.335 | -0.012 | 0.024 | 0.024 | 0.427 | 0.39 | 0.27 | 0.6 | 0.61 | 0.52 | 0.51 |
| 11 | 17532597 | rs1076311 | -0.018 | -2.266 | 0.491 | -0.412 | -0.023 | 0.04 | 0.04 | 0.317 | 0.39 | 0.2 | 0.6 | 0.62 | 0.53 | 0.52 |
| 11 | 17534413 | rs7107406 | 0.438 | -2.23 | 0.481 | -0.414 | -0.001 | 0.007 | 0.007 | 0 | 0.07 | 0.08 | 0.04 | 0.04 | 0.05 | 0.26 |
| 11 | 17535273 | rs16934329 | 0.735 | -2.259 | 0.578 | -0.427 | -0.019 | -0.026 | -0.026 | 0 | 0.08 | 0.11 | 0.08 | 0.03 | 0.05 | 0.02 |
| 11 | 17537033 | rs7119876 | -1.522 | -2.333 | 0.699 | -0.471 | -0.009 | -0.029 | -0.029 | 0 | 0.23 | 0.3 | 0.19 | 0.09 | 0.13 | 0.31 |
| 11 | 17537263 | rs963401 | -1.864 | -2.347 | 0.837 | -0.452 | 0.008 | 0.027 | 0.027 | 0 | 0.64 | 0.75 | 0.81 | 0.72 | 0.66 | 0.83 |
| 11 | 17538048 | rs2237960 | 0.322 | -2.528 | 0.826 | -0.357 | -0.004 | 0.073 | 0.073 | 0 | 0.1 | 0.14 | 0.13 | 0.23 | 0.1 | 0.11 |
| 11 | 17538539 | rs7128086 | 0.016 | -2.552 | 0.804 | -0.307 | 0.006 | 0.174 | 0.174 | 0 | 0.11 | 0.3 | 0.14 | 0.25 | 0.1 | 0.11 |
| 11 | 17539212 | rs10766408 | -0.731 | -2.569 | 0.766 | -0.484 | -0.002 | -0.01 | -0.01 | 0 | 0.48 | 0.53 | 0.66 | 0.68 | 0.58 | 0.78 |
| 11 | 17539562 | rs12793661 | -0.733 | -2.319 | 0.458 | -0.428 | 0.012 | 0.035 | 0.035 | 0 | 0.13 | 0.1 | 0.1 | 0.05 | 0.08 | 0.29 |
| 11 | 17540079 | rs11603262 | -0.263 | -2.19 | 0.514 | -0.582 | 0.003 | 0.003 | 0.003 | 0 | 0.4 | 0.44 | 0.57 | 0.46 | 0.5 | 0.43 |
| 11 | 17541798 | rs7129173 | -1.434 | -2.502 | 0.685 | -0.987 | -0.079 | -0.101 | -0.101 | 0 | 0.19 | 0.49 | 0.17 | 0.06 | 0.08 | 0.2 |
| 11 | 17542439 | rs2240487 | -0.694 | -2.623 | 0.257 | -1.05 | 0.025 | 0.098 | 0.098 | 0 | 0.51 | 0.3 | 0.65 | 0.67 | 0.53 | 0.53 |
| 11 | 17542649 | rs2041032 | -1.392 | -2.474 | 0.357 | -1.1 | -0.003 | -0.022 | -0.022 | 0 | 0.57 | 0.46 | 0.61 | 0.7 | 0.54 | 0.77 |
| 11 | 17546378 | rs4756898 | -0.951 | -2.076 | -0.076 | -1.14 | -0.164 | -0.163 | -0.163 | 0.051 | 0.76 | 0.98 | 0.66 | 0.6 | 0.65 | 0.62 |
| 11 | 17549409 | rs2355021 | -0.105 | -2.031 | 0.303 | -1.119 | 0.016 | -0.02 | -0.02 | 0.227 | 0.11 | 0.24 | 0.11 | 0.06 | 0.11 | 0.28 |
| 11 | 17550138 | rs2108333 | -0.712 | -2.116 | -0.02 | -1.17 | -0.112 | -0.104 | -0.104 | 0.507 | 0.7 | 0.9 | 0.64 | 0.61 | 0.63 | 0.6 |
| 11 | 17553105 | rs2041027 | 0.73 | -2.214 | 0.104 | -1.311 | 0.017 | -0.005 | -0.005 | 0 | 0.47 | 0.38 | 0.45 | 0.52 | 0.47 | 0.29 |
| 11 | 17557017 | rs2237959 | 0.811 | -2.179 | 0.042 | -1.393 | 0.006 | -0.002 | -0.002 | 0 | 0.46 | 0.42 | 0.45 | 0.54 | 0.47 | 0.29 |
| 11 | 17560127 | rs972676 | 0.213 | -1.681 | 0.342 | -1.229 | -0.005 | -0.006 | -0.006 | 0 | 0.45 | 0.68 | 0.49 | 0.58 | 0.48 | 0.54 |

| | | | | | | | | | | | | | | | | |
|---|---|---|---|---|---|---|---|---|---|---|---|---|---|---|---|---|
| 11 | 17562113 | rs10741729 | 1.038 | -1.533 | 0.229 | -0.934 | 0.006 | 0.03 | 0.03 | 0 | 0.33 | 0.44 | 0.4 | 0.56 | 0.39 | 0.27 |
| 11 | 17572284 | rs7102164 | -0.643 | -1.598 | -0.396 | -1.578 | -0.017 | -0.107 | -0.107 | 0.782 | 0.46 | 0.47 | 0.21 | 0.04 | 0.26 | 0.38 |
| 11 | 17572402 | rs7119071 | -0.253 | -0.966 | -0.348 | -1.064 | 0 | 0 | 0 | 0 | 0.53 | 0.49 | 0.51 | 0.6 | 0.56 | 0.58 |
| 11 | 17573329 | rs10766410 | 0.684 | -0.719 | -0.743 | -0.662 | 0.005 | 0.008 | 0.008 | 0 | 0.34 | 0.4 | 0.41 | 0.56 | 0.44 | 0.27 |
| 11 | 17574588 | rs4757543 | 0.59 | 0.544 | -0.951 | 0.026 | -0.075 | -0.091 | -0.091 | 0 | 0.77 | 0.9 | 0.58 | 0.47 | 0.65 | 0.62 |
| 11 | 17577223 | rs4757544 | -0.856 | -0.683 | -0.265 | -1.001 | -0.015 | -0.038 | -0.038 | 0 | 0.23 | 0.22 | 0.12 | 0.05 | 0.2 | 0.21 |
| 11 | 17584135 | rs11024322 | -1.206 | -0.397 | 0.118 | -0.976 | -0.008 | -0.055 | -0.055 | 0 | 0.19 | 0.23 | 0.11 | 0 | 0.14 | 0.08 |
| 11 | 17586381 | rs757985 | -1.999 | -0.696 | 0.93 | -0.8 | -0.018 | -0.025 | -0.025 | 4.141 | 0.17 | 0.67 | 0.21 | 0.09 | 0.15 | 0.17 |
| 11 | 17586522 | rs4757547 | -1.771 | -0.688 | 1.762 | -0.77 | 0.003 | 0.007 | 0.007 | 0.166 | 0.13 | 0.32 | 0.13 | 0.08 | 0.13 | 0.07 |
| 11 | 17590046 | rs11024323 | -1.929 | -0.024 | 1.677 | -0.864 | -0.003 | 0.006 | 0.006 | 0 | 0.32 | 0.26 | 0.23 | 0.31 | 0.37 | 0.3 |
| 11 | 17593115 | rs2240491 | -2.362 | -0.03 | 2.509 | -0.918 | -0.004 | 0.023 | 0.023 | 0.269 | 0.38 | 0.67 | 0.33 | 0.32 | 0.42 | 0.37 |
| 11 | 17593709 | rs7106548 | -2.414 | -0.023 | 2.529 | -0.9 | -0.002 | 0.025 | 0.025 | 0.027 | 0.39 | 0.67 | 0.32 | 0.32 | 0.42 | 0.37 |
| 11 | 17597822 | rs7104990 | -2.292 | 0.167 | 2.691 | -0.854 | 0.009 | -0.041 | -0.041 | 1.469 | 0.37 | 0.49 | 0.31 | 0.33 | 0.42 | 0.37 |
| 11 | 17599093 | rs10832804 | -2.306 | 0.232 | 2.714 | -0.871 | 0.007 | -0.032 | -0.032 | 0 | 0.36 | 0.48 | 0.3 | 0.27 | 0.42 | 0.4 |
| 11 | 17601219 | rs11024327 | -1.77 | 0.081 | 2.526 | -1.001 | 0 | -0.018 | -0.018 | 0 | 0.16 | 0.12 | 0.09 | 0 | 0.16 | 0.09 |
| 11 | 17607572 | rs7924620 | -1.903 | 0.741 | 3.105 | -0.761 | 0.078 | -0.011 | -0.011 | 0 | 0.15 | 0.42 | 0.15 | 0.1 | 0.17 | 0.1 |
| 11 | 17608675 | rs7106021 | -2.794 | 0.693 | 3.252 | -0.796 | 0.022 | -0.044 | -0.044 | 1.863 | 0.19 | 0.57 | 0.21 | 0.1 | 0.19 | 0.18 |
| 11 | 17609294 | rs11024329 | -2.027 | 0.694 | 3.302 | -0.795 | 0.079 | -0.01 | -0.01 | 0 | 0.15 | 0.42 | 0.15 | 0.1 | 0.17 | 0.1 |
| 11 | 17611668 | rs7947947 | -1.173 | 0.844 | 3.244 | -0.774 | -0.007 | -0.01 | -0.01 | 1.477 | 0.13 | 0.08 | 0.09 | 0.22 | 0.16 | 0.23 |
| 11 | 17612486 | rs10832816 | -2.905 | 0.845 | 3.27 | -0.778 | 0.024 | -0.003 | -0.003 | 1.242 | 0.28 | 0.53 | 0.24 | 0.33 | 0.34 | 0.33 |
| 11 | 17612776 | rs720459 | -3.158 | 0.843 | 3.201 | -0.779 | -0.004 | 0.003 | 0.003 | 0 | 0.32 | 0.68 | 0.31 | 0.33 | 0.36 | 0.41 |
| 11 | 17615203 | rs11024333 | -1.368 | 0.648 | 2.929 | -0.479 | -0.009 | -0.013 | -0.013 | 0 | 0.13 | 0.08 | 0.09 | 0.22 | 0.17 | 0.23 |
| 11 | 17618298 | rs7126777 | -1.868 | 0.649 | 3.172 | -0.48 | -0.001 | -0.001 | -0.001 | 0 | 0.16 | 0.15 | 0.14 | 0.22 | 0.19 | 0.31 |
| 11 | 17625937 | rs7107432 | -2.275 | 0.732 | 3.048 | -0.429 | 0.017 | 0.068 | 0.068 | 0.012 | 0.18 | 0.33 | 0.17 | 0.34 | 0.19 | 0.32 |
| 11 | 17631091 | rs740318 | -2.004 | 0.551 | 2.439 | -1.04 | -0.001 | -0.006 | -0.006 | 0.451 | 0.18 | 0.36 | 0.1 | 0.21 | 0.17 | 0.35 |
| 11 | 17656686 | rs11024350 | -1.192 | 0.75 | 1.176 | -0.761 | -0.004 | -0.011 | -0.011 | 0 | 0.12 | 0.06 | 0.08 | 0.24 | 0.14 | 0.22 |
| 11 | 17667439 | rs11024357 | -1.837 | 1.25 | 1.065 | -1.327 | -0.022 | 0.03 | 0.03 | 0 | 0.2 | 0.14 | 0.13 | 0.07 | 0.26 | 0.28 |
| 11 | 17667561 | rs11024358 | -1.813 | 1.257 | 1.068 | -1.324 | -0.022 | 0.028 | 0.028 | 0 | 0.19 | 0.14 | 0.13 | 0.07 | 0.26 | 0.28 |
| 11 | 17670293 | rs7107666 | -2.884 | 1.063 | 0.941 | -1.472 | -0.055 | 0.149 | 0.149 | 0.192 | 0.39 | 0.86 | 0.41 | 0.16 | 0.42 | 0.55 |
| 11 | 17672112 | rs869108 | 1.645 | 0.864 | 1.611 | -1.649 | 0.007 | -0.085 | -0.085 | 0 | 0.75 | 0.64 | 0.86 | 0.96 | 0.74 | 0.73 |
| 11 | 17676066 | rs9633896 | 2.54 | 0.798 | 1.613 | -1.524 | -0.016 | 0.144 | 0.144 | 0.227 | 0.65 | 0.13 | 0.64 | 0.88 | 0.67 | 0.54 |
| 11 | 17676406 | rs12575678 | -1.87 | 0.789 | 2.227 | -1.545 | -0.022 | -0.022 | -0.022 | 0 | 0.14 | 0.34 | 0.15 | 0.09 | 0.1 | 0.15 |
| 11 | 17692011 | rs7483847 | 1.786 | 0.801 | 2.2 | -1.526 | 0.049 | -0.133 | -0.133 | 1.757 | 0.81 | 0.58 | 0.8 | 1 | 0.78 | 0.71 |
| 11 | 17702336 | rs10766416 | 1.719 | 0.957 | 2.079 | -1.448 | 0.08 | -0.032 | -0.032 | 0 | 0.86 | 0.57 | 0.71 | 0.91 | 0.8 | 0.74 |
| 11 | 17702641 | rs3858493 | 1.755 | 0.956 | 2.073 | -1.448 | 0.071 | -0.034 | -0.034 | 0.086 | 0.85 | 0.6 | 0.72 | 0.91 | 0.79 | 0.74 |
| 11 | 17705368 | rs3858497 | -0.781 | 0.864 | 1.485 | -1.519 | -0.017 | 0.062 | 0.062 | 0 | 0.11 | 0.01 | 0.05 | 0 | 0.1 | 0.16 |
| 11 | 17708585 | rs10766419 | 2.785 | 0.861 | 1.509 | -1.566 | -0.021 | 0.155 | 0.155 | 0.117 | 0.65 | 0.14 | 0.61 | 0.88 | 0.65 | 0.54 |
| 11 | 17722192 | rs4757570 | 1.898 | 0.654 | 1.705 | -1.749 | 0.042 | -0.126 | -0.126 | 0.037 | 0.82 | 0.52 | 0.75 | 0.95 | 0.76 | 0.73 |
| 11 | 17723296 | rs3858510 | -2.03 | 0.608 | 1.81 | -1.751 | -0.025 | -0.104 | -0.104 | 0 | 0.18 | 0.35 | 0.08 | 0.01 | 0.11 | 0.2 |
| 11 | 17726374 | rs4757574 | 1.849 | 0.673 | 1.422 | -1.788 | 0.036 | -0.124 | -0.124 | 0.035 | 0.81 | 0.49 | 0.75 | 0.96 | 0.75 | 0.73 |
| 11 | 17738564 | rs2249104 | -1.844 | 0.539 | 1.026 | -1.739 | 0.012 | 0.003 | 0.003 | 0 | 0.18 | 0.16 | 0.06 | 0.01 | 0.09 | 0.28 |
| 11 | 17739729 | rs7938556 | -0.801 | 0.563 | 1.21 | -1.503 | -0.024 | -0.029 | -0.029 | 0 | 0.06 | 0.13 | 0.02 | 0 | 0.01 | 0.1 |
| 11 | 17740778 | rs3911833 | -1.812 | 0.705 | 0.941 | -1.452 | 0.039 | -0.084 | -0.084 | 0 | 0.14 | 0.32 | 0.22 | 0.03 | 0.2 | 0.23 |
| 11 | 17756213 | rs28724581 | 1.658 | 0.552 | -0.478 | -0.935 | 0.034 | 0.044 | 0.044 | 0 | 0.54 | 0.08 | 0.6 | 0.61 | 0.66 | 0.37 |
| 11 | 17758587 | rs7926805 | -1.718 | 0.324 | 0.274 | -1.353 | -0.006 | -0.029 | -0.029 | 0.134 | 0.35 | 0.58 | 0.2 | 0.16 | 0.24 | 0.45 |
| 11 | 17760904 | rs4757587 | 1.037 | 0.442 | 0.38 | -1.025 | -0.013 | -0.026 | -0.026 | 0 | 0.83 | 0.4 | 0.84 | 0.83 | 0.84 | 0.72 |
| 11 | 17769755 | rs1012105 | 0.951 | 0.627 | -0.516 | -1.725 | 0.015 | 0.17 | 0.17 | 0.089 | 0.55 | 0.22 | 0.76 | 0.88 | 0.64 | 0.67 |
| 11 | 17771168 | rs7124411 | -0.322 | 0.742 | 0.396 | -1.931 | 0.07 | -0.071 | -0.071 | 0 | 0.13 | 0.3 | 0.11 | 0.02 | 0.18 | 0.1 |
| 11 | 17771487 | rs10734255 | -0.122 | 0.67 | 0.516 | -2.16 | -0.014 | -0.077 | -0.077 | 0 | 0.81 | 0.75 | 0.93 | 1 | 0.89 | 0.82 |
| 11 | 17774473 | rs757511 | 2.309 | 1.249 | -0.05 | -1.856 | -0.094 | 0.074 | 0.074 | 0 | 0.42 | 0.06 | 0.6 | 0.69 | 0.47 | 0.34 |
| 11 | 17775029 | rs10832857 | -1.502 | 1.232 | 0.63 | -1.873 | 0.021 | -0.066 | -0.066 | 0 | 0.26 | 0.37 | 0.16 | 0.1 | 0.29 | 0.19 |
| 11 | 17775799 | rs2299636 | -0.698 | 1.111 | 0.137 | -1.73 | 0.003 | 0.005 | 0.005 | 0 | 0.14 | 0.14 | 0.05 | 0.1 | 0.1 | 0.09 |
| 11 | 17790227 | rs917032 | 0.692 | 0.921 | 0.252 | -1.812 | 0.001 | -0.15 | -0.15 | 0.906 | 0.66 | 0.51 | 0.83 | 0.98 | 0.71 | 0.72 |
| 11 | 17791822 | rs7946133 | -0.952 | 0.717 | -0.195 | -1.054 | 0.003 | 0.006 | 0.006 | 0 | 0.13 | 0.12 | 0.06 | 0.1 | 0.1 | 0.09 |
| 11 | 17803711 | rs1236205 | -1.438 | 0.589 | -0.006 | -1.508 | -0.007 | -0.083 | -0.083 | 0 | 0.28 | 0.4 | 0.13 | 0 | 0.21 | 0.21 |
| 11 | 17804998 | rs999420 | 1.728 | 1.394 | -0.89 | -1.515 | -0.033 | 0.148 | 0.148 | 0.075 | 0.49 | 0.08 | 0.52 | 0.7 | 0.41 | 0.34 |
| 11 | 17809343 | rs1236207 | -0.234 | 1.101 | -0.568 | -1.618 | 0.002 | -0.025 | -0.025 | 0.012 | 0.19 | 0.22 | 0.06 | 0 | 0.12 | 0.18 |
| 11 | 17809639 | rs10788 | -0.214 | 1.095 | -0.569 | -1.621 | 0.002 | -0.022 | -0.022 | 0 | 0.19 | 0.22 | 0.07 | 0 | 0.12 | 0.18 |
| 11 | 17809724 | rs1528 | -0.214 | 1.095 | -0.569 | -1.621 | 0.002 | -0.022 | -0.022 | 0 | 0.19 | 0.22 | 0.07 | 0 | 0.12 | 0.18 |
| 11 | 17814359 | rs11024400 | -0.363 | 1.214 | -0.35 | -1.638 | -0.006 | -0.061 | -0.061 | 0.564 | 0.2 | 0.31 | 0.07 | 0 | 0.13 | 0.19 |
| 11 | 17816412 | rs4756919 | -0.559 | 1.555 | 0.041 | -1.645 | -0.002 | -0.002 | -0.002 | 0.02 | 0.89 | 0.94 | 0.87 | 0.86 | 0.83 | 0.83 |
| 11 | 17820177 | rs2283250 | 2.168 | 1.669 | -0.152 | -1.726 | -0.019 | 0.157 | 0.157 | 0.195 | 0.52 | 0.08 | 0.61 | 0.72 | 0.51 | 0.38 |
| 11 | 17820848 | rs11024405 | -1.283 | 1.664 | -0.06 | -1.748 | 0.006 | -0.028 | -0.028 | 0.155 | 0.28 | 0.33 | 0.12 | 0 | 0.2 | 0.22 |
| 11 | 17821182 | rs3802970 | 0.922 | 1.677 | 0.007 | -1.734 | 0.032 | 0.01 | 0.01 | 0.457 | 0.78 | 0.48 | 0.72 | 0.72 | 0.67 | 0.56 |
| 11 | 17823296 | rs11024408 | -0.227 | 1.557 | -0.193 | -1.701 | 0.008 | 0.01 | 0.01 | 0 | 0.18 | 0.14 | 0.06 | 0 | 0.12 | 0.18 |
| 11 | 17824372 | rs2283249 | -0.994 | 1.514 | 0.068 | -1.677 | -0.002 | 0.009 | 0.009 | 0 | 0.26 | 0.26 | 0.19 | 0.13 | 0.28 | 0.44 |
| 11 | 17825099 | rs2299633 | -1.696 | 1.47 | 0.023 | -1.653 | 0 | -0.012 | -0.012 | 0.916 | 0.35 | 0.42 | 0.25 | 0.13 | 0.36 | 0.47 |
| 11 | 17832433 | rs2189593 | -0.547 | 1.283 | -0.412 | -1.714 | -0.005 | 0.099 | 0.099 | 0 | 0.22 | 0.08 | 0.1 | 0.05 | 0.21 | 0.31 |
| 11 | 17833064 | rs2237949 | -1.42 | 1.05 | 0.282 | -1.384 | 0.076 | 0.295 | 0.295 | 4.036 | 0.05 | 0.45 | 0.11 | 0.16 | 0 | 0.1 |

| | | | | | | | | | | | | | | | | |
|---|---|---|---|---|---|---|---|---|---|---|---|---|---|---|---|---|
| 11 | 17835042 | rs1548528 | 1.614 | 1.367 | 0.151 | -1.532 | -0.001 | -0.015 | -0.015 | 0 | 0.55 | 0.5 | 0.68 | 0.87 | 0.48 | 0.42 |
| 11 | 17835483 | rs4756921 | -0.144 | 1.389 | 0.168 | -1.548 | 0.002 | -0.031 | -0.031 | 0 | 0.88 | 0.92 | 0.91 | 1 | 0.82 | 0.92 |
| 11 | 17842951 | rs11821847 | -1.062 | 1.294 | 0.147 | -1.419 | -0.002 | 0.053 | 0.053 | 0 | 0.33 | 0.26 | 0.23 | 0.15 | 0.35 | 0.51 |
| 11 | 17844037 | rs4757591 | -1.115 | 1.29 | 0.178 | -1.414 | 0.001 | 0.023 | 0.023 | 0 | 0.33 | 0.32 | 0.23 | 0.15 | 0.35 | 0.5 |
| 11 | 17847381 | rs2107427 | -1.037 | 1.305 | 0.086 | -1.406 | 0 | 0.027 | 0.027 | 0 | 0.33 | 0.34 | 0.23 | 0.14 | 0.35 | 0.5 |
| 11 | 17851342 | rs6486399 | 1.339 | 1.048 | -0.107 | -1.374 | 0 | -0.026 | -0.026 | 0 | 0.53 | 0.46 | 0.66 | 0.85 | 0.45 | 0.41 |
| 11 | 17859904 | rs10832861 | 0.902 | 0.42 | -1.54 | -0.538 | -0.114 | -0.076 | -0.076 | 0 | 0.41 | 0.06 | 0.58 | 0.47 | 0.46 | 0.45 |
| 11 | 17863925 | rs16934735 | -0.494 | -0.004 | -1.999 | -0.507 | 0.045 | 0.066 | 0.066 | 0 | 0.29 | 0.06 | 0.16 | 0.14 | 0.2 | 0.2 |
| 11 | 17867235 | rs1029499 | -0.373 | -0.042 | -1.776 | 0.178 | 0.035 | -0.062 | -0.062 | 0 | 0.27 | 0.2 | 0.18 | 0.39 | 0.21 | 0.24 |
| 11 | 17868673 | rs1015445 | 1.443 | -0.265 | -1.923 | 0.302 | -0.078 | -0.021 | -0.021 | 0.013 | 0.38 | 0.11 | 0.45 | 0.42 | 0.42 | 0.38 |
| 11 | 17872749 | rs2237939 | -0.481 | -0.219 | -1.13 | 0.162 | 0.017 | -0.028 | -0.028 | 0 | 0.29 | 0.24 | 0.18 | 0.4 | 0.23 | 0.25 |
| 11 | 17877603 | rs12419230 | 0.034 | -0.01 | -0.991 | 0.407 | -0.008 | -0.045 | -0.045 | 0 | 0.65 | 0.15 | 0.7 | 0.59 | 0.62 | 0.59 |
| 11 | 17878912 | rs4756925 | -0.371 | -0.026 | -0.398 | 0.357 | 0.005 | -0.05 | -0.05 | 0 | 0.69 | 0.56 | 0.77 | 0.85 | 0.65 | 0.64 |
| 11 | 17883386 | rs1015896 | 0.318 | 0.032 | -0.716 | 0.355 | -0.03 | -0.04 | -0.04 | 0.888 | 0.62 | 0.14 | 0.67 | 0.6 | 0.57 | 0.58 |
| 11 | 17893079 | rs549686 | -0.416 | 0.21 | -0.634 | 0.236 | 0.001 | -0.022 | -0.022 | 0 | 0.62 | 0.64 | 0.64 | 0.85 | 0.56 | 0.62 |
| 11 | 17894411 | rs512819 | 0.995 | 0.128 | -0.575 | 0.223 | -0.008 | -0.021 | -0.021 | 0 | 0.35 | 0.22 | 0.44 | 0.47 | 0.33 | 0.38 |
| 11 | 17894731 | rs537393 | 0.684 | 0.131 | -0.534 | 0.214 | 0 | -0.002 | -0.002 | 0 | 0.38 | 0.38 | 0.5 | 0.74 | 0.37 | 0.42 |
| 11 | 17897324 | rs2237930 | -0.247 | 0.326 | -0.46 | 0.272 | 0 | -0.032 | -0.032 | 0 | 0.6 | 0.58 | 0.64 | 0.85 | 0.56 | 0.62 |
| 11 | 17904501 | rs2107426 | -0.082 | 0.179 | -0.734 | 0.109 | -0.021 | 0.136 | 0.136 | 0 | 0.23 | 0.02 | 0.13 | 0.02 | 0.19 | 0.2 |
| 11 | 17904634 | rs121704 | -1.349 | 0.385 | -0.859 | 0.163 | -0.022 | -0.075 | -0.075 | 0.208 | 0.66 | 0.86 | 0.57 | 0.49 | 0.65 | 0.69 |
| 11 | 17908706 | rs501996 | 0.374 | 0.441 | -0.402 | -0.042 | -0.002 | -0.018 | -0.018 | 2.287 | 0.37 | 0.37 | 0.37 | 0.26 | 0.39 | 0.45 |
| 11 | 17910664 | rs7119188 | -0.438 | 0.447 | -0.231 | -0.042 | -0.008 | 0.037 | 0.037 | 0 | 0.25 | 0.16 | 0.17 | 0.03 | 0.25 | 0.22 |
| 11 | 17923676 | rs550460 | -0.869 | 0.508 | -0.148 | 0.214 | 0.008 | -0.019 | -0.019 | 0 | 0.61 | 0.42 | 0.51 | 0.27 | 0.58 | 0.67 |
| 11 | 17925059 | rs542350 | -0.34 | 0.511 | -0.084 | 0.636 | -0.02 | -0.032 | -0.032 | 1.197 | 0.43 | 0.81 | 0.46 | 0.48 | 0.47 | 0.51 |
| 11 | 17930293 | rs529191 | -0.105 | 0.576 | -0.185 | 0.061 | -0.002 | 0.005 | 0.005 | 0 | 0.41 | 0.53 | 0.44 | 0.27 | 0.47 | 0.49 |
| 11 | 17934383 | rs2237924 | -0.717 | -0.643 | -0.617 | 0.268 | 0.009 | -0.066 | -0.066 | 0 | 0.19 | 0.02 | 0.13 | 0.25 | 0.17 | 0.23 |
| 11 | 17935204 | rs4757603 | -0.19 | -0.53 | -0.274 | 0.081 | 0 | 0.071 | 0.071 | 0 | 0.54 | 0.25 | 0.56 | 0.76 | 0.55 | 0.49 |
| 11 | 17942831 | rs739916 | 1.207 | -0.592 | -0.248 | 0.041 | -0.016 | -0.025 | -0.025 | 0 | 0.35 | 0.19 | 0.43 | 0.51 | 0.39 | 0.26 |
| 11 | 17943805 | rs4141243 | 1.168 | -0.601 | -0.189 | 0.036 | -0.053 | -0.077 | -0.077 | 0 | 0.35 | 0.12 | 0.42 | 0.51 | 0.39 | 0.26 |
| 11 | 17943876 | rs2283240 | -0.384 | -0.574 | 0.103 | 0.003 | 0.001 | 0.028 | 0.028 | 0 | 0.55 | 0.31 | 0.56 | 0.74 | 0.56 | 0.49 |
| 11 | 17945020 | rs10832866 | 0.225 | -0.608 | 0.435 | -0.208 | 0.03 | 0.084 | 0.084 | 0.059 | 0.15 | 0.39 | 0.24 | 0.23 | 0.17 | 0.38 |
| 11 | 17960851 | rs758437 | -0.185 | -0.656 | 0.139 | -0.178 | 0.015 | 0.018 | 0.018 | 0 | 0.18 | 0.05 | 0.13 | 0.08 | 0.16 | 0.17 |
| 11 | 17961609 | rs12277387 | 0.062 | -0.68 | 0.421 | -0.21 | 0 | -0.002 | -0.002 | 0 | 0.55 | 0.4 | 0.55 | 0.59 | 0.56 | 0.43 |
| 11 | 17981047 | rs211146 | -1.741 | -0.63 | 0.399 | -0.211 | -0.08 | -0.099 | -0.099 | 0 | 0.65 | 0.92 | 0.58 | 0.49 | 0.6 | 0.74 |
| 11 | 17988860 | rs211095 | -1.803 | -0.632 | 0.531 | -0.227 | -0.017 | -0.024 | -0.024 | 0 | 0.64 | 0.81 | 0.58 | 0.49 | 0.61 | 0.74 |
| 11 | 18001888 | rs211114 | -0.508 | -0.598 | 0.625 | -0.253 | 0.001 | 0.013 | 0.013 | 0 | 0.45 | 0.69 | 0.45 | 0.4 | 0.44 | 0.57 |
| 11 | 18008376 | rs623143 | 0.136 | -0.598 | 0.607 | -0.255 | 0.024 | 0.071 | 0.071 | 0.003 | 0.15 | 0.38 | 0.24 | 0.22 | 0.17 | 0.38 |
| 11 | 18016873 | rs211130 | -0.449 | -0.57 | 0.383 | -0.252 | 0.003 | 0.029 | 0.029 | 0 | 0.45 | 0.75 | 0.45 | 0.4 | 0.45 | 0.57 |
| 11 | 18035436 | rs6486402 | 0.562 | -0.386 | 0.557 | -0.138 | -0.002 | 0 | 0 | 0 | 0.12 | 0.14 | 0.27 | 0.23 | 0.17 | 0.37 |
| 11 | 18038118 | rs12292915 | -0.037 | -0.532 | 0.449 | -0.26 | -0.01 | -0.004 | -0.004 | 0.392 | 0.6 | 0.22 | 0.51 | 0.58 | 0.57 | 0.43 |
| 11 | 18045987 | rs7933505 | 1.701 | -0.676 | 0.172 | -0.267 | -0.13 | -0.132 | -0.132 | 0.762 | 0.34 | 0.04 | 0.37 | 0.48 | 0.39 | 0.27 |
| 11 | 18046493 | rs211102 | -0.635 | -0.693 | 0.601 | -0.255 | 0.024 | 0.107 | 0.107 | 0 | 0.86 | 0.68 | 0.72 | 0.76 | 0.83 | 0.63 |
| 11 | 18055920 | rs17794760 | -0.49 | -0.721 | 0.151 | -0.301 | 0.04 | 0.061 | 0.061 | 0 | 0.25 | 0.06 | 0.15 | 0.1 | 0.18 | 0.17 |
| 11 | 18056225 | rs172423 | 1.29 | -0.633 | 0.229 | -0.271 | -0.041 | 0.04 | 0.04 | 0.634 | 0.23 | 0.03 | 0.16 | 0.18 | 0.26 | 0.18 |
| 11 | 18058341 | rs211107 | 1.582 | -0.664 | 0.321 | -0.482 | -0.037 | -0.047 | -0.047 | 0.017 | 0.34 | 0.21 | 0.38 | 0.52 | 0.4 | 0.29 |
| 11 | 18059292 | rs169806 | 1.671 | -0.653 | -0.013 | -0.477 | -0.134 | -0.153 | -0.153 | 0 | 0.34 | 0.04 | 0.37 | 0.52 | 0.4 | 0.29 |
| 11 | 18059929 | rs10832876 | 1.113 | -0.51 | 0.223 | -0.521 | -0.036 | 0.072 | 0.072 | 0.246 | 0.23 | 0.04 | 0.16 | 0.13 | 0.26 | 0.17 |
| 11 | 18068163 | rs7130929 | -0.676 | -0.456 | 0.583 | -0.502 | 0.018 | 0.043 | 0.043 | 0 | 0.49 | 0.44 | 0.34 | 0.21 | 0.43 | 0.37 |
| 11 | 18072273 | rs1079785 | -0.727 | -0.486 | 0.864 | -0.517 | 0.038 | 0.103 | 0.103 | 0 | 0.51 | 0.7 | 0.35 | 0.21 | 0.43 | 0.37 |
| 11 | 18073039 | rs7950705 | -0.692 | -0.486 | 0.753 | -0.517 | 0.016 | 0.036 | 0.036 | 0 | 0.49 | 0.45 | 0.34 | 0.21 | 0.43 | 0.37 |
| 11 | 18073142 | rs517005 | 1.389 | -0.396 | 0.209 | -0.732 | -0.131 | -0.148 | -0.148 | 0 | 0.35 | 0.04 | 0.29 | 0.51 | 0.39 | 0.24 |
| 11 | 18088946 | rs7943884 | 0.521 | -0.553 | 0.941 | -1.745 | -0.011 | -0.097 | -0.097 | 0 | 0.27 | 0.45 | 0.19 | 0.12 | 0.26 | 0.21 |
| 11 | 18105302 | rs3741199 | -1.269 | -1.141 | 1.036 | -1.731 | 0.011 | 0.002 | 0.002 | 0 | 0.49 | 0.55 | 0.28 | 0.15 | 0.41 | 0.31 |
| 11 | 18105871 | rs6486406 | -0.804 | -1.221 | 0.665 | -1.794 | 0.011 | 0.003 | 0.003 | 0 | 0.43 | 0.64 | 0.25 | 0.15 | 0.35 | 0.29 |
| 11 | 18117516 | rs7937884 | -0.461 | -0.932 | 0.598 | -1.475 | -0.018 | -0.041 | -0.041 | 0 | 0.16 | 0.27 | 0.07 | 0.01 | 0.09 | 0.08 |
| 11 | 18135642 | rs2468782 | 0.884 | -0.704 | 0.386 | -1.362 | -0.021 | -0.094 | -0.094 | 0 | 0.78 | 0.62 | 0.88 | 0.98 | 0.82 | 0.91 |
| 11 | 18138153 | rs2468849 | 0.345 | -0.63 | 0.51 | -1.125 | -0.031 | -0.079 | -0.079 | 0 | 0.85 | 0.63 | 0.94 | 0.99 | 0.9 | 0.94 |
| 11 | 18154120 | rs2468842 | 0.174 | -0.31 | 0.168 | -0.854 | 0.015 | 0.06 | 0.06 | 0 | 0.25 | 0.09 | 0.14 | 0.13 | 0.22 | 0.21 |
| 11 | 18159254 | rs4274188 | -0.093 | -0.336 | 0.672 | -0.866 | -0.034 | -0.097 | -0.097 | 0.879 | 0.72 | 0.54 | 0.84 | 0.87 | 0.78 | 0.79 |
| 11 | 18164687 | rs2445162 | -0.61 | -0.217 | -0.154 | -0.962 | 0.003 | -0.005 | -0.005 | 0 | 0.37 | 0.65 | 0.26 | 0.15 | 0.31 | 0.3 |
| 11 | 18170688 | rs2468826 | 0.164 | -0.144 | -0.436 | -1.762 | 0.003 | -0.008 | -0.008 | 1.656 | 0.62 | 0.68 | 0.76 | 0.95 | 0.6 | 0.61 |
| 11 | 18177494 | rs17567204 | 0.074 | -0.097 | -0.499 | -1.866 | -0.052 | 0.178 | 0.178 | 0 | 0.29 | 0.05 | 0.18 | 0.04 | 0.32 | 0.19 |
| 11 | 18178448 | rs11024502 | 0.772 | -0.041 | -1.07 | -1.504 | -0.015 | -0.093 | -0.093 | 0.488 | 0.28 | 0.09 | 0.36 | 0.51 | 0.27 | 0.23 |
| 11 | 18192072 | rs2246304 | 0.844 | -0.787 | -0.628 | -1.886 | -0.007 | -0.014 | -0.014 | 0 | 0.26 | 0.26 | 0.34 | 0.44 | 0.31 | 0.34 |
| 11 | 18192239 | rs2860278 | 1.191 | -0.688 | -0.59 | -1.835 | -0.005 | -0.005 | -0.005 | 0 | 0.24 | 0.25 | 0.34 | 0.44 | 0.31 | 0.34 |
| 11 | 18193107 | rs7107957 | 1.037 | -0.742 | -0.661 | -1.873 | -0.049 | 0.177 | 0.177 | 0 | 0.29 | 0.04 | 0.18 | 0.04 | 0.31 | 0.19 |
| 11 | 18195348 | rs11024532 | 1.166 | -0.917 | -0.595 | -2.031 | -0.029 | 0.137 | 0.137 | 0 | 0.24 | 0.05 | 0.14 | 0.03 | 0.26 | 0.18 |
| 11 | 18200030 | rs2445192 | -0.297 | -1.013 | -0.478 | -2.049 | -0.039 | -0.081 | -0.081 | 4.701 | 0.85 | 0.7 | 0.91 | 0.98 | 0.89 | 0.76 |
| 11 | 18211343 | rs11024542 | 1.159 | -0.958 | -0.581 | -2.046 | -0.006 | 0.104 | 0.104 | 0 | 0.24 | 0.1 | 0.15 | 0.03 | 0.27 | 0.18 |

| | | | | | | | | | | | | | | | | |
|---|---|---|---|---|---|---|---|---|---|---|---|---|---|---|---|---|
| 11 | 18212384 | rs4417246 | 0.682 | -0.874 | -0.591 | -2.013 | -0.019 | -0.067 | -0.067 | 0 | 0.35 | 0.11 | 0.4 | 0.51 | 0.33 | 0.24 |
| 11 | 18213135 | rs2198706 | 1.077 | -0.823 | -0.811 | -1.928 | -0.056 | -0.096 | -0.096 | 0 | 0.24 | 0.14 | 0.34 | 0.44 | 0.3 | 0.34 |
| 11 | 18214205 | rs12365565 | 0.387 | -0.924 | -0.242 | -2.245 | 0.007 | 0.014 | 0.014 | 0 | 0.26 | 0.25 | 0.16 | 0.04 | 0.27 | 0.18 |
| 11 | 18218678 | rs16935005 | -0.51 | -1.24 | -0.019 | -1.468 | -0.005 | -0.021 | -0.021 | 0 | 0.13 | 0.16 | 0.06 | 0.01 | 0.08 | 0.13 |
| 11 | 18220910 | rs2468812 | 0.338 | -0.887 | -0.282 | -0.742 | 0.002 | -0.08 | -0.08 | 1.012 | 0.28 | 0.12 | 0.43 | 0.57 | 0.26 | 0.35 |
| 11 | 18225742 | rs2468802 | 0.293 | -0.807 | -0.24 | -0.993 | 0.008 | -0.012 | -0.012 | 0 | 0.68 | 0.23 | 0.73 | 0.62 | 0.75 | 0.68 |
| 11 | 18228871 | rs11024552 | 0.487 | -0.691 | -0.542 | -0.889 | -0.045 | -0.092 | -0.092 | 0 | 0.2 | 0.04 | 0.2 | 0.27 | 0.19 | 0.13 |
| 11 | 18231289 | rs4757627 | 0.51 | -0.621 | -0.023 | -1.256 | -0.047 | 0.163 | 0.163 | 0.21 | 0.41 | 0.18 | 0.3 | 0.05 | 0.49 | 0.33 |
| 11 | 18235412 | rs12364058 | 0.597 | -0.53 | 0.024 | -1.151 | -0.024 | 0.068 | 0.068 | 0 | 0.4 | 0.22 | 0.31 | 0.05 | 0.49 | 0.33 |
| 11 | 18246943 | rs7935726 | 0.374 | -0.676 | 0.044 | -1.066 | -0.011 | 0.017 | 0.017 | 0 | 0.43 | 0.38 | 0.32 | 0.05 | 0.52 | 0.34 |
| 11 | 18253176 | rs2460827 | 0.241 | -0.749 | 0.058 | -0.867 | -0.031 | 0.037 | 0.037 | 1.903 | 0.91 | 0.7 | 0.96 | 0.89 | 0.97 | 0.96 |
| 11 | 18258282 | rs1378609 | 0.265 | -0.737 | 0.063 | -0.862 | -0.033 | 0.034 | 0.034 | 1.443 | 0.91 | 0.7 | 0.96 | 0.9 | 0.97 | 0.96 |
| 11 | 18258928 | rs2168363 | 0.213 | -0.24 | -0.776 | -0.824 | -0.012 | 0.081 | 0.081 | 0.212 | 0.43 | 0.32 | 0.34 | 0.05 | 0.54 | 0.35 |
| 11 | 18264571 | rs1520887 | -0.193 | -0.227 | -0.681 | 0.167 | -0.014 | 0.025 | 0.025 | 2.784 | 0.3 | 0.51 | 0.31 | 0.35 | 0.24 | 0.14 |
| 11 | 18264756 | rs1520885 | 0.24 | -0.399 | -1.201 | 0.806 | -0.003 | -0.11 | -0.11 | 0.013 | 0.21 | 0.07 | 0.22 | 0.49 | 0.18 | 0.29 |
| 11 | 18267027 | rs2468844 | 0.562 | -0.979 | -0.199 | -0.12 | -0.07 | -0.073 | -0.073 | 0.432 | 0.76 | 0.61 | 0.78 | 0.93 | 0.89 | 0.7 |
| 11 | 18270669 | rs3825023 | -1.413 | -0.74 | 0.003 | 0.113 | -0.041 | -0.011 | -0.011 | 0 | 0.14 | 0.49 | 0.18 | 0.14 | 0.08 | 0.2 |
| 11 | 18271929 | rs7933280 | -0.908 | -0.859 | 0.066 | 0.018 | -0.067 | -0.051 | -0.051 | 0.384 | 0.12 | 0.49 | 0.05 | 0.06 | 0.03 | 0.15 |
| 11 | 18275661 | rs10832911 | 0.05 | -0.334 | -0.67 | 0.205 | 0.006 | -0.002 | -0.002 | 0 | 0.61 | 0.42 | 0.73 | 0.46 | 0.67 | 0.6 |
| 11 | 18278098 | rs2045272 | -0.425 | 0.561 | -0.452 | 0.041 | 0.028 | -0.007 | -0.007 | 0 | 0.29 | 0.33 | 0.45 | 0.22 | 0.46 | 0.29 |
| 11 | 18278912 | rs7950019 | -1.025 | 0.072 | -0.539 | 0.113 | 0.078 | 0.075 | 0.075 | 0.005 | 0.48 | 0.63 | 0.63 | 0.65 | 0.69 | 0.64 |
| 11 | 18280635 | rs1993373 | -1.032 | 0.069 | -0.526 | 0.111 | 0.079 | 0.075 | 0.075 | 0.008 | 0.49 | 0.63 | 0.63 | 0.65 | 0.69 | 0.64 |
| 11 | 18282051 | rs11603089 | 0.792 | 1.405 | -0.687 | 0.118 | -0.041 | -0.041 | -0.041 | 2.163 | 0.1 | 0.03 | 0.25 | 0.19 | 0.19 | 0.12 |
| 11 | 18295810 | rs11024600 | -0.56 | -0.231 | -0.665 | -0.081 | 0.004 | -0.019 | -0.019 | 0.643 | 0.35 | 0.71 | 0.33 | 0.41 | 0.34 | 0.41 |
| 11 | 18300954 | rs1046615 | -0.523 | -0.518 | -0.379 | -0.06 | -0.007 | 0.028 | 0.028 | 0.06 | 0.33 | 0.42 | 0.3 | 0.38 | 0.32 | 0.32 |
| 11 | 18303404 | rs2271997 | 0.589 | -0.602 | -0.505 | -0.183 | -0.025 | -0.007 | -0.007 | 1.092 | 0.29 | 0.21 | 0.43 | 0.33 | 0.51 | 0.48 |
| 11 | 18305333 | rs2049129 | 0.034 | -0.45 | -0.261 | -0.765 | -0.005 | -0.112 | -0.112 | 0.083 | 0.21 | 0.27 | 0.11 | 0 | 0.17 | 0.05 |
| 11 | 18306399 | rs11024603 | 0.113 | -0.406 | -0.284 | -0.665 | 0.003 | 0.044 | 0.044 | 1.384 | 0.2 | 0.24 | 0.14 | 0.34 | 0.2 | 0.19 |
| 11 | 18319180 | rs7128017 | 0.892 | -0.186 | 0.079 | -0.042 | -0.009 | -0.003 | -0.003 | 0.444 | 0.13 | 0.19 | 0.17 | 0.12 | 0.13 | 0.15 |
| 11 | 18325146 | rs2403254 | 0.447 | -0.213 | -0.014 | -0.083 | -0.101 | -0.1 | -0.1 | 0.239 | 0.31 | 0.09 | 0.51 | 0.59 | 0.54 | 0.5 |
| 11 | 18342081 | rs10741741 | -0.92 | -0.337 | -0.005 | -0.198 | 0.06 | 0.095 | 0.095 | 0 | 0.48 | 0.31 | 0.71 | 0.96 | 0.69 | 0.68 |
| 11 | 18343694 | rs4150530 | 1.098 | -0.357 | -0.022 | -0.197 | -0.008 | -0.001 | -0.001 | 0 | 0.12 | 0.2 | 0.16 | 0.13 | 0.13 | 0.13 |
| 11 | 18344064 | rs3802967 | 0.591 | -0.33 | -0.026 | -0.15 | -0.113 | -0.114 | -0.114 | 0 | 0.3 | 0.07 | 0.51 | 0.59 | 0.54 | 0.4 |
| 11 | 18357180 | rs4150579 | -0.824 | -0.405 | 0.115 | -1.175 | 0.048 | 0.046 | 0.046 | 0 | 0.5 | 0.38 | 0.72 | 0.96 | 0.71 | 0.69 |
| 11 | 18363569 | rs4150606 | -0.022 | 0.553 | -0.647 | -0.45 | -0.066 | 0.122 | 0.122 | 0 | 0.35 | 0.12 | 0.55 | 0.84 | 0.57 | 0.54 |
| 11 | 18400973 | rs7946331 | 0.398 | 0.449 | -1.152 | 0.407 | 0.026 | 0.03 | 0.03 | 0 | 0.47 | 0.26 | 0.58 | 0.57 | 0.59 | 0.63 |
| 11 | 18415687 | rs2403278 | 0.405 | 0.481 | -1.155 | 0.438 | 0.023 | 0.024 | 0.024 | 0 | 0.47 | 0.23 | 0.57 | 0.58 | 0.58 | 0.63 |
| 11 | 18417197 | rs7932887 | 0.511 | 0.122 | -0.22 | 0.096 | -0.009 | -0.075 | -0.075 | 0.138 | 0.12 | 0.31 | 0.06 | 0 | 0.09 | 0.03 |
| 11 | 18419534 | rs1830426 | 1.36 | -0.23 | -0.006 | -0.14 | 0.003 | 0.023 | 0.023 | 0 | 0.73 | 0.42 | 0.73 | 0.65 | 0.73 | 0.74 |
| 11 | 18421683 | rs11601413 | 0.507 | -0.181 | 0.194 | -0.089 | 0.002 | -0.006 | -0.006 | 2.034 | 0.09 | 0.1 | 0.14 | 0.06 | 0.12 | 0.11 |
| 11 | 18422029 | rs12274651 | 0.801 | -0.185 | 0.206 | -0.105 | -0.008 | -0.073 | -0.073 | 0 | 0.12 | 0.31 | 0.06 | 0 | 0.09 | 0.03 |
| 11 | 18429467 | rs3758682 | 1.167 | -0.203 | 0.146 | -0.065 | 0 | 0.015 | 0.015 | 0 | 0.73 | 0.4 | 0.73 | 0.65 | 0.73 | 0.74 |
| 11 | 18435700 | rs10832934 | 1.3 | 0.603 | -0.308 | 0.153 | -0.052 | -0.028 | -0.028 | 0 | 0.11 | 0.01 | 0.18 | 0.15 | 0.17 | 0.13 |
| 11 | 18439212 | rs2037868 | -0.253 | 0.313 | 0.143 | 0.32 | 0.05 | 0.174 | 0.174 | 0 | 0.93 | 0.83 | 0.92 | 0.68 | 0.91 | 0.88 |
| 11 | 18451096 | rs3740713 | 1.544 | 0.119 | -0.043 | 0.159 | 0.015 | 0.06 | 0.06 | 1.965 | 0.18 | 0.01 | 0.14 | 0.06 | 0.12 | 0.08 |
| 11 | 18470487 | rs16935432 | 1.208 | 0.152 | 0.019 | 0.178 | 0.012 | 0.011 | 0.011 | 0.9 | 0.21 | 0.16 | 0.15 | 0.06 | 0.13 | 0.08 |
| 11 | 18472133 | rs1881717 | -0.724 | 0.978 | 0.494 | 0.918 | 0.048 | 0.173 | 0.173 | 0 | 0.07 | 0.17 | 0.08 | 0.33 | 0.09 | 0.12 |
| 11 | 18475100 | rs2721123 | -1.955 | 0.895 | 0.628 | 0.838 | 0.013 | 0.045 | 0.045 | 0 | 0.25 | 0.58 | 0.26 | 0.39 | 0.27 | 0.26 |
| 11 | 18476729 | rs2698559 | -0.922 | 1.181 | 0.215 | 1.235 | 0.051 | 0.172 | 0.172 | 0 | 0.07 | 0.17 | 0.08 | 0.3 | 0.09 | 0.12 |
| 11 | 18503159 | rs2279900 | -0.709 | 1.138 | -0.417 | 1.681 | -0.02 | -0.066 | -0.066 | 1.747 | 0.16 | 0.07 | 0.24 | 0.44 | 0.2 | 0.4 |
| 11 | 18503737 | rs3781640 | -0.617 | 1.122 | -0.431 | 1.673 | -0.027 | -0.066 | -0.066 | 0.005 | 0.14 | 0.01 | 0.18 | 0.24 | 0.14 | 0.29 |
| 11 | 18509034 | rs11024677 | -0.745 | 1.118 | -0.433 | 1.658 | -0.022 | -0.064 | -0.064 | 0 | 0.15 | 0.02 | 0.18 | 0.24 | 0.14 | 0.29 |
| 11 | 18509993 | rs16935476 | -0.89 | 1.249 | -0.313 | 1.495 | 0.011 | 0.018 | 0.018 | 0 | 0.25 | 0.33 | 0.33 | 0.29 | 0.27 | 0.31 |
| 11 | 18528815 | rs2658547 | -0.431 | 1.273 | -0.22 | 1.466 | 0 | -0.002 | -0.002 | 0.023 | 0.9 | 0.86 | 0.89 | 0.93 | 0.89 | 0.92 |
| 11 | 18529108 | rs10500836 | -1.64 | 1.2 | -0.18 | 1.497 | 0.002 | 0 | 0 | 0.316 | 0.31 | 0.32 | 0.47 | 0.66 | 0.36 | 0.63 |
| 11 | 18530047 | rs10832939 | -0.351 | 1.203 | -0.18 | 1.513 | 0.008 | 0.043 | 0.043 | 0 | 0.15 | 0.24 | 0.23 | 0.22 | 0.16 | 0.22 |
| 11 | 18536215 | rs12574333 | -0.538 | 1.223 | -0.242 | 1.527 | -0.02 | -0.067 | -0.067 | 0.254 | 0.16 | 0.07 | 0.24 | 0.44 | 0.2 | 0.4 |
| 11 | 18552899 | rs12222154 | -0.158 | 1.243 | -0.281 | 1.464 | -0.032 | -0.07 | -0.07 | 0 | 0.14 | 0.01 | 0.18 | 0.24 | 0.14 | 0.29 |
| 11 | 18562982 | rs2271079 | 0.168 | 1.284 | -0.334 | 1.449 | -0.004 | 0.002 | 0.002 | 0 | 0.1 | 0.09 | 0.1 | 0.07 | 0.11 | 0.08 |
| 11 | 18575406 | rs4756949 | -1.865 | 0.821 | 0.033 | 1.62 | 0.008 | -0.005 | -0.005 | 0 | 0.32 | 0.39 | 0.48 | 0.67 | 0.36 | 0.63 |
| 11 | 18582814 | rs12418726 | -0.715 | 0.861 | 0.037 | 1.649 | -0.002 | 0.007 | 0.007 | 0 | 0.16 | 0.12 | 0.24 | 0.44 | 0.2 | 0.41 |
| 11 | 18586122 | rs10741747 | -1.812 | 0.803 | -0.21 | 1.453 | 0.008 | -0.005 | -0.005 | 6.116 | 0.32 | 0.4 | 0.48 | 0.66 | 0.36 | 0.63 |
| 11 | 18597312 | rs11024719 | 0.171 | 0.625 | -0.618 | 1.272 | -0.018 | -0.03 | -0.03 | 0 | 0.06 | 0.01 | 0.13 | 0.17 | 0.06 | 0.23 |
| 11 | 18600322 | rs10500835 | -0.868 | 0.636 | -0.604 | 1.309 | -0.022 | -0.072 | -0.072 | 0.828 | 0.15 | 0.01 | 0.18 | 0.26 | 0.14 | 0.29 |
| 11 | 18620047 | rs4436523 | -0.845 | 0.657 | -0.478 | 1.108 | 0.002 | 0.03 | 0.03 | 0 | 0.18 | 0.28 | 0.23 | 0.27 | 0.16 | 0.27 |
| 11 | 18626834 | rs17494675 | 0.437 | 0.543 | -0.79 | 1.056 | 0.007 | -0.027 | -0.027 | 0 | 0.1 | 0.01 | 0.04 | 0.08 | 0.06 | 0.07 |
| 11 | 18628418 | rs17579600 | 0.494 | 0.544 | -0.789 | 1.057 | 0.006 | -0.027 | -0.027 | 0 | 0.1 | 0.01 | 0.04 | 0.08 | 0.06 | 0.07 |
| 11 | 18628730 | rs10832948 | 0.739 | 0.621 | -1.577 | 0.681 | -0.06 | -0.014 | -0.014 | 0 | 0.42 | 0.04 | 0.44 | 0.35 | 0.37 | 0.35 |
| 11 | 18629304 | rs7934205 | 0.285 | 0.527 | -1.231 | 0.583 | 0.002 | 0.007 | 0.007 | 0 | 0.52 | 0.44 | 0.49 | 0.37 | 0.44 | 0.37 |

| | | | | | | | | | | | | | | | | |
|---|---|---|---|---|---|---|---|---|---|---|---|---|---|---|---|---|
| 11 | 18631527 | rs11024731 | 0.821 | 0.75 | -1.581 | 0.677 | -0.033 | 0.076 | 0.076 | 0 | 0.27 | 0.1 | 0.22 | 0.14 | 0.33 | 0.19 |
| 11 | 18633971 | rs11605097 | 0.529 | 0.757 | -2.708 | 0.657 | -0.007 | 0.058 | 0.058 | 0 | 0.48 | 0.08 | 0.45 | 0.35 | 0.42 | 0.36 |
| 11 | 18636481 | rs16935599 | 1.206 | 0.861 | -2.507 | 0.474 | -0.003 | -0.013 | -0.013 | 0.114 | 0.05 | 0.03 | 0.02 | 0 | 0.05 | 0.01 |
| 11 | 18638853 | rs10500833 | 0.918 | 1.093 | -2.672 | 0.673 | 0.005 | 0.023 | 0.023 | 0 | 0.06 | 0.06 | 0.16 | 0.19 | 0.07 | 0.23 |
| 11 | 18641561 | rs10832951 | 1.586 | 1.029 | -2.885 | 0.622 | -0.052 | -0.005 | -0.005 | 0 | 0.27 | 0.03 | 0.23 | 0.22 | 0.23 | 0.12 |
| 11 | 18643758 | rs10766487 | -0.531 | 1.147 | -2.068 | 0.933 | 0.013 | 0.03 | 0.03 | 0 | 0.1 | 0.16 | 0.18 | 0.19 | 0.11 | 0.25 |
| 11 | 18671991 | rs1550868 | 0.37 | 1.089 | -2.545 | 0.313 | -0.092 | 0.049 | 0.049 | 0.008 | 0.23 | 0.02 | 0.21 | 0.15 | 0.31 | 0.18 |
| 11 | 18687025 | rs12787397 | 0.326 | 1.045 | -2.544 | 0.314 | -0.088 | 0.052 | 0.052 | 0.02 | 0.23 | 0.02 | 0.21 | 0.15 | 0.32 | 0.18 |
| 11 | 18689653 | rs12295403 | 0.001 | 0.978 | -2.472 | 0.523 | -0.091 | 0.013 | 0.013 | 0 | 0.33 | 0.05 | 0.25 | 0.25 | 0.38 | 0.24 |
| 11 | 18690219 | rs4757695 | 0.895 | 0.94 | -2.499 | 0.614 | 0.011 | 0.094 | 0.094 | 0 | 0.48 | 0.18 | 0.46 | 0.31 | 0.44 | 0.41 |
| 11 | 18690389 | rs12789252 | -0.202 | 0.514 | -2.211 | -0.203 | -0.091 | 0.018 | 0.018 | 0 | 0.33 | 0.05 | 0.25 | 0.25 | 0.38 | 0.25 |
| 11 | 18696407 | rs2403303 | 0.162 | 1.023 | -1.893 | 0.102 | 0.005 | 0.058 | 0.058 | 0 | 0.57 | 0.29 | 0.67 | 0.74 | 0.56 | 0.75 |
| 11 | 18702509 | rs4757697 | -0.661 | 1.489 | -1.816 | 0.486 | -0.065 | -0.033 | -0.033 | 0 | 0.19 | 0.05 | 0.25 | 0.19 | 0.24 | 0.29 |
| 11 | 18719433 | rs11024765 | -0.514 | 1.583 | -1.275 | 0.074 | 0.006 | -0.012 | -0.012 | 0 | 0.29 | 0.25 | 0.39 | 0.18 | 0.38 | 0.17 |
| 11 | 18719929 | rs12282271 | -0.692 | 1.244 | -0.679 | 0.369 | 0.003 | -0.004 | -0.004 | 0.343 | 0.36 | 0.53 | 0.49 | 0.29 | 0.43 | 0.29 |
| 11 | 18722791 | rs733460 | 0.601 | 0.217 | -1.901 | -0.198 | 0.047 | 0.103 | 0.103 | 0.075 | 0.33 | 0.04 | 0.19 | 0.11 | 0.19 | 0.27 |
| 11 | 18724558 | rs14792 | 0.395 | 0.799 | -1.71 | 0.654 | -0.002 | 0.006 | 0.006 | 0 | 0.11 | 0.08 | 0.14 | 0.33 | 0.16 | 0.24 |
| 11 | 18728650 | rs2289965 | 0.557 | 1.185 | -2.14 | 0.175 | -0.134 | 0.009 | 0.009 | 1.87 | 0.21 | 0.02 | 0.34 | 0.13 | 0.4 | 0.16 |
| 11 | 18730895 | rs11024768 | -0.394 | 0.147 | -1.113 | 0.129 | 0.008 | 0.009 | 0.009 | 0 | 0.47 | 0.45 | 0.65 | 0.76 | 0.72 | 0.57 |
| 11 | 18732846 | rs4537730 | -0.585 | 0.024 | -0.761 | -0.003 | 0.057 | 0.072 | 0.072 | 0 | 0.52 | 0.7 | 0.68 | 0.81 | 0.74 | 0.57 |
| 11 | 18735396 | rs10741752 | -1.222 | -0.039 | -0.791 | -0.087 | 0.087 | 0.097 | 0.097 | 0 | 0.59 | 0.85 | 0.71 | 0.81 | 0.78 | 0.58 |
| 11 | 18735947 | rs7125943 | -1.11 | -0.103 | -0.819 | -0.119 | 0.082 | 0.088 | 0.088 | 0 | 0.6 | 0.85 | 0.71 | 0.81 | 0.78 | 0.58 |
| 11 | 18737095 | rs4424652 | -1.22 | -0.071 | -0.63 | -0.104 | 0.079 | 0.084 | 0.084 | 0 | 0.59 | 0.83 | 0.7 | 0.81 | 0.77 | 0.58 |
| 11 | 18738281 | rs10766494 | -1.236 | -0.092 | -0.642 | -0.114 | 0.079 | 0.084 | 0.084 | 0 | 0.59 | 0.83 | 0.7 | 0.81 | 0.77 | 0.58 |
| 11 | 18738784 | rs10832973 | 0.177 | 0.259 | -0.553 | 0.011 | -0.011 | -0.011 | -0.011 | 4.09 | 0.36 | 0.34 | 0.5 | 0.44 | 0.56 | 0.4 |
| 11 | 18739358 | rs4561190 | -0.114 | -0.078 | -0.586 | -0.06 | -0.013 | -0.006 | -0.006 | 0 | 0.9 | 0.8 | 0.89 | 0.94 | 0.94 | 0.88 |
| 11 | 18742043 | rs12421865 | -0.266 | 0.652 | -0.832 | 0.045 | -0.024 | -0.018 | -0.018 | 0 | 0.13 | 0.08 | 0.31 | 0.19 | 0.21 | 0.14 |
| 11 | 18742570 | rs2278734 | 0.461 | 0.452 | -1.093 | -0.234 | -0.001 | 0.023 | 0.023 | 0 | 0.41 | 0.21 | 0.49 | 0.32 | 0.36 | 0.47 |
| 11 | 18744695 | rs1160219 | 0.784 | 1.16 | -0.599 | 0.173 | 0.027 | 0.037 | 0.037 | 0.387 | 0.43 | 0.31 | 0.23 | 0.13 | 0.22 | 0.41 |
| 11 | 18745177 | rs1979050 | -0.217 | 1.821 | -0.119 | 0.826 | 0.003 | 0.001 | 0.001 | 0.169 | 0.31 | 0.33 | 0.3 | 0.33 | 0.42 | 0.32 |
| 11 | 18747959 | rs4237729 | 0.642 | 2.091 | 0.287 | 0.89 | 0.043 | -0.008 | -0.008 | 0 | 0.83 | 0.73 | 0.82 | 0.82 | 0.71 | 0.76 |
| 11 | 18748046 | rs748944 | 1.206 | 1.856 | 0.527 | 1.186 | 0.013 | 0.044 | 0.044 | 0.132 | 0.46 | 0.3 | 0.34 | 0.17 | 0.36 | 0.36 |
| 11 | 18748819 | rs873670 | -1.286 | 1.807 | 0.734 | 1.43 | 0.031 | 0.031 | 0.031 | 0 | 0.52 | 0.71 | 0.66 | 0.68 | 0.63 | 0.51 |
| 11 | 18751041 | rs1550870 | -0.179 | 1.969 | 0.621 | 1.744 | -0.007 | 0 | 0 | 0 | 0.31 | 0.32 | 0.31 | 0.26 | 0.44 | 0.25 |
| 11 | 18754679 | rs6483524 | 1.443 | 1.83 | 0.453 | 1.489 | 0.03 | 0.1 | 0.1 | 0 | 0.43 | 0.16 | 0.32 | 0.15 | 0.35 | 0.36 |
| 11 | 18757355 | rs4756961 | 0.488 | 1.762 | 0.48 | 1.44 | 0.023 | 0.018 | 0.018 | 0.066 | 0.62 | 0.52 | 0.67 | 0.56 | 0.52 | 0.61 |
| 11 | 18762983 | rs4756962 | 1.292 | 1.876 | -0.335 | 1.564 | 0.011 | -0.021 | -0.021 | 0.901 | 0.46 | 0.03 | 0.56 | 0.37 | 0.41 | 0.52 |
| 11 | 18764113 | rs2278732 | 1.543 | 1.865 | -0.328 | 1.65 | -0.016 | -0.001 | -0.001 | 0 | 0.41 | 0.02 | 0.49 | 0.32 | 0.38 | 0.41 |
| 11 | 18766647 | rs10766500 | 0.265 | 1.952 | 0.113 | 1.504 | 0.132 | 0.009 | 0.009 | 2.428 | 0.83 | 0.52 | 0.74 | 0.81 | 0.7 | 0.76 |
| 11 | 18768068 | rs1867876 | -0.926 | 1.884 | 0.097 | 1.584 | -0.01 | 0.009 | 0.009 | 2.082 | 0.42 | 0.5 | 0.24 | 0.46 | 0.29 | 0.35 |
| 11 | 18770822 | rs921082 | -0.585 | 1.689 | 0.123 | 1.608 | 0.001 | 0.014 | 0.014 | 2.295 | 0.37 | 0.49 | 0.25 | 0.42 | 0.29 | 0.25 |
| 11 | 18771614 | rs930617 | 0.21 | 1.784 | 0.214 | 1.638 | -0.005 | -0.015 | -0.015 | 0 | 0.18 | 0.28 | 0.09 | 0.15 | 0.13 | 0.08 |
| 11 | 18775682 | rs4757710 | 0.354 | 2.013 | 0.121 | 1.529 | 0.119 | -0.001 | -0.001 | 0 | 0.82 | 0.53 | 0.77 | 0.81 | 0.72 | 0.79 |
| 11 | 18777171 | rs755796 | 0.342 | 1.976 | 0.124 | 1.523 | 0.116 | 0.007 | 0.007 | 3.158 | 0.82 | 0.53 | 0.77 | 0.81 | 0.72 | 0.79 |
| 11 | 18780344 | rs10832979 | -0.409 | 2.05 | 0.172 | 1.507 | 0.154 | -0.037 | -0.037 | 0 | 0.88 | 0.59 | 0.85 | 0.96 | 0.78 | 0.92 |
| 11 | 18781866 | rs1106866 | 0.056 | 2.083 | 0.017 | 1.614 | -0.016 | 0.001 | 0.001 | 2.073 | 0.23 | 0.34 | 0.11 | 0.22 | 0.17 | 0.2 |
| 11 | 18782131 | rs1106865 | 0.863 | 2.253 | 0.159 | 1.666 | -0.002 | -0.031 | -0.031 | 0 | 0.4 | 0.03 | 0.5 | 0.34 | 0.39 | 0.36 |
| 11 | 18782408 | rs7941067 | -0.228 | 2.15 | 0.455 | 1.455 | -0.003 | 0.005 | 0.005 | 2.32 | 0.43 | 0.5 | 0.27 | 0.47 | 0.32 | 0.43 |
| 11 | 18782916 | rs1867875 | -0.118 | 2.206 | -0.004 | 1.35 | 0.118 | -0.04 | -0.04 | 0 | 0.11 | 0.33 | 0.14 | 0.03 | 0.22 | 0.09 |
| 11 | 18782967 | rs7106657 | 0.192 | 1.15 | -0.888 | 1.736 | 0.012 | -0.075 | -0.075 | 0 | 0.46 | 0.08 | 0.58 | 0.51 | 0.46 | 0.49 |
| 11 | 18787044 | rs4272766 | 0.968 | 1.326 | 0.726 | 2.039 | 0.129 | 0.463 | 0.463 | 0.091 | 0.94 | 0.6 | 0.82 | 0.75 | 0.93 | 0.97 |
| 11 | 18795058 | rs4757717 | -0.116 | 1.145 | -0.323 | -0.156 | 0.013 | 0.104 | 0.104 | 0.042 | 0.42 | 0.22 | 0.55 | 0.84 | 0.39 | 0.59 |
| 11 | 18796480 | rs4073915 | 0.277 | 1.246 | -0.777 | -0.932 | 0.016 | 0.121 | 0.121 | 0 | 0.31 | 0.13 | 0.23 | 0.03 | 0.29 | 0.13 |
| 11 | 18802503 | rs11024792 | -0.303 | 1.427 | 0.32 | -0.915 | 0.056 | -0.077 | -0.077 | 0 | 0.23 | 0.36 | 0.19 | 0.11 | 0.33 | 0.25 |
| 11 | 18804524 | rs7131023 | -0.866 | 1.194 | 0.876 | -1.034 | 0.025 | -0.046 | -0.046 | 0 | 0.26 | 0.37 | 0.17 | 0.17 | 0.31 | 0.32 |
| 11 | 18804934 | rs4757718 | 0.724 | 0.837 | 0.781 | -1.206 | -0.017 | -0.042 | -0.042 | 0.528 | 0.42 | 0.46 | 0.54 | 0.72 | 0.33 | 0.52 |
| 11 | 18808355 | rs7117716 | 0.507 | 0.542 | 1.011 | -1.42 | 0.022 | 0.035 | 0.035 | 0 | 0.48 | 0.46 | 0.45 | 0.7 | 0.33 | 0.55 |
| 11 | 18808557 | rs10832986 | -0.007 | 0.867 | 0.734 | -1.67 | -0.079 | 0.055 | 0.055 | 0 | 0.15 | 0.01 | 0.11 | 0.07 | 0.19 | 0.22 |
| 11 | 18809135 | rs7950455 | 0.489 | -0.044 | 0.511 | -1.079 | -0.044 | 0.03 | 0.03 | 0.112 | 0.13 | 0.25 | 0.11 | 0.16 | 0.04 | 0.04 |
| 11 | 18809542 | rs12577551 | 1.198 | 0.031 | -0.089 | -1.495 | -0.017 | 0.086 | 0.086 | 0.953 | 0.5 | 0.18 | 0.54 | 0.62 | 0.52 | 0.67 |
| 11 | 18815295 | rs7932938 | -0.52 | 0.159 | 0.522 | -1.476 | 0.011 | -0.016 | -0.016 | 0 | 0.35 | 0.35 | 0.45 | 0.27 | 0.52 | 0.42 |
| 11 | 18816245 | rs7129128 | -0.178 | 0.21 | 0.706 | -1.463 | 0.057 | -0.012 | -0.012 | 0.01 | 0.19 | 0.28 | 0.26 | 0.19 | 0.31 | 0.27 |
| 11 | 18822037 | rs4627050 | 0.625 | -0.012 | 0.796 | -1.38 | 0.001 | -0.001 | -0.001 | 0 | 0.57 | 0.55 | 0.67 | 0.73 | 0.61 | 0.79 |
| 11 | 18824315 | rs11024797 | 1.067 | 0.153 | 0.602 | -1.322 | -0.011 | 0.021 | 0.021 | 0 | 0.16 | 0.1 | 0.25 | 0.1 | 0.22 | 0.15 |
| 11 | 18843054 | rs4756970 | -0.601 | 0.835 | 0.724 | -1.324 | 0.04 | -0.008 | -0.008 | 0 | 0.16 | 0.29 | 0.21 | 0.17 | 0.22 | 0.25 |
| 11 | 18843233 | rs4756971 | 0.476 | 0.41 | -0.167 | -0.717 | 0.019 | 0.004 | 0.004 | 0 | 0.49 | 0.35 | 0.53 | 0.52 | 0.55 | 0.66 |
| 11 | 18857747 | rs10832998 | 0.956 | 0.191 | -0.048 | -0.79 | -0.009 | 0.031 | 0.031 | 0 | 0.52 | 0.28 | 0.45 | 0.57 | 0.51 | 0.55 |
| 11 | 18858316 | rs7948997 | -0.274 | 0.228 | 0.105 | -0.807 | -0.009 | -0.007 | -0.007 | 0 | 0.7 | 0.61 | 0.67 | 0.73 | 0.75 | 0.8 |
| 11 | 18858540 | rs3935980 | -0.078 | 0.243 | 0.107 | -0.801 | 0.049 | -0.003 | -0.003 | 0 | 0.16 | 0.31 | 0.2 | 0.16 | 0.21 | 0.25 |

| | | | | | | | | | | | | | | | | |
|---|---|---|---|---|---|---|---|---|---|---|---|---|---|---|---|---|
| 11 | 18860716 | rs11500050 | 0.833 | 0.042 | -0.014 | -0.733 | -0.009 | 0.031 | 0.031 | 0 | 0.53 | 0.28 | 0.45 | 0.57 | 0.51 | 0.55 |
| 11 | 18866536 | rs4643051 | -1.244 | 0.088 | 0.117 | -0.699 | 0.012 | 0.032 | 0.032 | 0 | 0.51 | 0.67 | 0.58 | 0.63 | 0.52 | 0.57 |
| 11 | 18866999 | rs6483541 | -1.524 | 0.043 | 0.171 | -0.684 | -0.002 | -0.06 | -0.06 | 0.261 | 0.88 | 0.8 | 0.92 | 1 | 0.86 | 0.98 |
| 11 | 18869581 | rs5019888 | 0.122 | 0.245 | 0.072 | -0.708 | -0.027 | 0.002 | 0.002 | 0 | 0.28 | 0.13 | 0.34 | 0.27 | 0.31 | 0.23 |
| 11 | 18870823 | rs4756973 | -0.566 | 0.15 | 0.28 | -0.324 | -0.006 | -0.045 | -0.045 | 0.318 | 0.24 | 0.42 | 0.25 | 0.16 | 0.26 | 0.24 |
| 11 | 18871028 | rs4757729 | -0.863 | 0.131 | 0.27 | -0.202 | -0.006 | 0.02 | 0.02 | 0 | 0.31 | 0.44 | 0.29 | 0.38 | 0.34 | 0.35 |
| 11 | 18871881 | rs11024838 | -0.03 | 0.194 | -0.003 | -0.126 | 0 | 0 | 0 | 0.027 | 0.2 | 0.22 | 0.14 | 0.21 | 0.21 | 0.14 |
| 11 | 18882221 | rs6483543 | 1.912 | 0.265 | -0.065 | 0.02 | -0.001 | 0.014 | 0.014 | 0 | 0.08 | 0.07 | 0.05 | 0.21 | 0.07 | 0.11 |
| 11 | 18885423 | rs7936877 | 0.714 | 0.262 | -0.022 | -0.022 | 0 | 0.007 | 0.007 | 0 | 0.14 | 0.15 | 0.1 | 0.2 | 0.13 | 0.09 |
| 11 | 18887345 | rs7944558 | 1.983 | 0.258 | -0.061 | -0.012 | -0.001 | 0.025 | 0.025 | 0 | 0.08 | 0.09 | 0.05 | 0.21 | 0.07 | 0.11 |
| 11 | 18887813 | rs1483096 | 0.775 | 0.228 | 0.148 | -0.066 | -0.001 | 0.03 | 0.03 | 0 | 0.66 | 0.51 | 0.69 | 0.44 | 0.68 | 0.59 |
| 11 | 18890206 | rs11024850 | 1.94 | 0.312 | -0.424 | -0.19 | -0.046 | 0.026 | 0.026 | 0 | 0.15 | 0.03 | 0.27 | 0.09 | 0.2 | 0.14 |
| 11 | 18896034 | rs2086698 | 1.637 | 0.108 | -0.339 | -0.221 | 0.006 | 0.008 | 0.008 | 0.297 | 0.51 | 0.25 | 0.43 | 0.54 | 0.54 | 0.55 |
| 11 | 18901853 | rs4757732 | 0.19 | 0.082 | -0.212 | -0.217 | -0.019 | 0.001 | 0.001 | 0.19 | 0.84 | 0.61 | 0.82 | 0.84 | 0.89 | 0.79 |
| 11 | 18903330 | rs12363574 | 2.639 | 0.143 | -1.051 | -0.256 | -0.055 | -0.057 | -0.057 | 0 | 0.33 | 0.07 | 0.31 | 0.34 | 0.33 | 0.42 |
| 11 | 18903604 | rs7946938 | 1.288 | 0.159 | -0.749 | 0.002 | 0.002 | -0.002 | -0.002 | 0.736 | 0.67 | 0.29 | 0.7 | 0.64 | 0.75 | 0.7 |
| 11 | 18906829 | rs11024866 | 1.397 | 0.346 | -0.746 | 0.092 | -0.003 | -0.018 | -0.018 | 0 | 0.13 | 0.08 | 0.08 | 0.2 | 0.11 | 0.09 |
| 11 | 18911346 | rs1483098 | 1.623 | 0.095 | -0.786 | 0.16 | -0.008 | 0.029 | 0.029 | 0.197 | 0.45 | 0.26 | 0.41 | 0.33 | 0.5 | 0.46 |
| 11 | 18929815 | rs958061 | -0.188 | 0.067 | -1.104 | 0.177 | -0.061 | -0.093 | -0.093 | 0 | 0.36 | 0.1 | 0.38 | 0.53 | 0.38 | 0.36 |
| 11 | 18933049 | rs1817503 | 1.553 | 0.05 | -1.074 | 0.168 | -0.01 | 0.028 | 0.028 | 0.124 | 0.44 | 0.2 | 0.38 | 0.33 | 0.47 | 0.45 |
| 11 | 18935038 | rs1026898 | 1.362 | 0.279 | -1.142 | 0.535 | 0 | 0.006 | 0.006 | 0 | 0.08 | 0.07 | 0.05 | 0.24 | 0.08 | 0.13 |
| 11 | 18936554 | rs11600150 | 0.097 | 0.352 | -1.174 | 0.506 | -0.106 | -0.062 | -0.062 | 0 | 0.28 | 0.03 | 0.34 | 0.29 | 0.32 | 0.23 |
| 11 | 18939666 | rs10833018 | -0.44 | 0.285 | -1.129 | 0.46 | -0.071 | -0.093 | -0.093 | 0 | 0.36 | 0.1 | 0.38 | 0.53 | 0.38 | 0.36 |
| 11 | 18949220 | rs7939600 | -0.533 | -0.131 | -0.219 | 0.619 | -0.027 | 0.017 | 0.017 | 0 | 0.15 | 0.25 | 0.07 | 0.21 | 0.08 | 0.16 |
| 11 | 18950092 | rs11024884 | 0.78 | 0.385 | -0.447 | 0.864 | 0 | 0.006 | 0.006 | 0.042 | 0.08 | 0.08 | 0.05 | 0.21 | 0.07 | 0.14 |
| 11 | 18956690 | rs2004384 | 0.4 | 0.565 | -0.872 | 0.418 | 0.007 | 0.028 | 0.028 | 0.023 | 0.43 | 0.25 | 0.35 | 0.3 | 0.4 | 0.41 |
| 11 | 18973083 | rs11024906 | 0.817 | 0.616 | -1.452 | 0.46 | -0.049 | -0.051 | -0.051 | 0 | 0.39 | 0.03 | 0.24 | 0.33 | 0.36 | 0.41 |
| 11 | 18974860 | rs1842921 | 0.546 | 0.59 | -1.206 | 0.849 | 0.006 | -0.059 | -0.059 | 0 | 0.43 | 0.16 | 0.27 | 0.5 | 0.41 | 0.5 |
| 11 | 18979904 | rs2007048 | 0.583 | 0.271 | -0.055 | 0.701 | 0.007 | -0.054 | -0.054 | 0 | 0.43 | 0.17 | 0.27 | 0.5 | 0.41 | 0.51 |
| 11 | 18982634 | rs4757741 | -0.08 | 0.734 | -0.11 | 0.819 | -0.061 | 0.009 | 0.009 | 0.413 | 0.36 | 0.12 | 0.56 | 0.31 | 0.44 | 0.42 |
| 11 | 18986882 | rs4757742 | -0.463 | 0.715 | 0.225 | 0.832 | -0.019 | -0.014 | -0.014 | 0.353 | 0.41 | 0.31 | 0.6 | 0.48 | 0.5 | 0.5 |
| 11 | 18988572 | rs1842920 | -0.454 | 0.716 | 0.225 | 0.832 | -0.019 | -0.014 | -0.014 | 0 | 0.41 | 0.31 | 0.6 | 0.48 | 0.5 | 0.5 |
| 11 | 18989826 | rs1842917 | 0.66 | 0.993 | -0.206 | 0.805 | -0.016 | 0.025 | 0.025 | 0.087 | 0.24 | 0.1 | 0.33 | 0.19 | 0.25 | 0.33 |
| 11 | 18993465 | rs7946550 | 1.622 | 0.84 | -0.102 | 0.92 | -0.044 | -0.042 | -0.042 | 0 | 0.39 | 0.03 | 0.24 | 0.33 | 0.35 | 0.42 |
| 11 | 19009495 | rs6483561 | -0.68 | 0.862 | 0.465 | 0.931 | -0.009 | -0.005 | -0.005 | 0 | 0.42 | 0.38 | 0.61 | 0.48 | 0.5 | 0.5 |
| 11 | 19021421 | rs2170655 | -0.455 | 0.882 | 0.307 | 0.941 | -0.027 | -0.02 | -0.02 | 0 | 0.4 | 0.29 | 0.6 | 0.47 | 0.49 | 0.48 |
| 11 | 19044777 | rs7118407 | 1.245 | 0.732 | 0.019 | 1.044 | -0.002 | 0.008 | 0.008 | 0.67 | 0.39 | 0.11 | 0.24 | 0.33 | 0.36 | 0.43 |
| 11 | 19051872 | rs7111562 | -0.645 | 1.174 | 0.961 | 1.356 | 0.006 | 0.006 | 0.006 | 0 | 0.12 | 0.2 | 0.12 | 0.2 | 0.14 | 0.09 |
| 11 | 19055921 | rs11024956 | -1.201 | 1.254 | 1.146 | 1.341 | 0.038 | 0.068 | 0.068 | 0 | 0.14 | 0.17 | 0.15 | 0.34 | 0.2 | 0.16 |
| 11 | 19057767 | rs11024958 | -1.521 | 0.886 | 0.93 | 1.014 | -0.001 | -0.002 | -0.002 | 0 | 0.39 | 0.65 | 0.4 | 0.43 | 0.4 | 0.26 |
| 11 | 19059114 | rs6483573 | -1.595 | 0.936 | 1.114 | 1.126 | 0.096 | 0.188 | 0.188 | 1.611 | 0.06 | 0.34 | 0.05 | 0.14 | 0.06 | 0.07 |
| 11 | 19072151 | rs7109068 | -1.503 | 0.94 | 1.037 | 1.092 | 0.024 | 0.076 | 0.076 | 0 | 0.19 | 0.41 | 0.17 | 0.31 | 0.19 | 0.16 |
| 11 | 19076150 | rs7126062 | -0.524 | 0.884 | 1.37 | 0.945 | 0 | 0.013 | 0.013 | 0.156 | 0.24 | 0.42 | 0.27 | 0.25 | 0.26 | 0.18 |
| 11 | 19076431 | rs7126313 | -0.406 | 0.874 | 1.441 | 0.969 | 0.001 | 0.011 | 0.011 | 0.015 | 0.24 | 0.29 | 0.27 | 0.26 | 0.26 | 0.18 |
| 11 | 19077765 | rs10833049 | -0.64 | 0.857 | 1.426 | 0.961 | -0.007 | 0.016 | 0.016 | 0.903 | 0.26 | 0.53 | 0.28 | 0.27 | 0.26 | 0.18 |
| 11 | 19079292 | rs7116312 | -0.57 | 0.755 | 1.295 | 0.918 | 0.001 | 0.007 | 0.007 | 0.144 | 0.26 | 0.54 | 0.29 | 0.25 | 0.27 | 0.18 |
| 11 | 19079584 | rs7102329 | -1.238 | 0.68 | 1.792 | 1.141 | 0.016 | 0.015 | 0.015 | 0 | 0.16 | 0.43 | 0.14 | 0.17 | 0.13 | 0.08 |
| 11 | 19080944 | rs10833052 | 0.48 | 0.005 | 1.097 | 1.363 | 0 | 0 | 0 | 0 | 0.65 | 0.46 | 0.66 | 0.65 | 0.62 | 0.52 |
| 11 | 19083477 | rs1503502 | -1.391 | -0.016 | 0.826 | 1.405 | 0.032 | -0.013 | -0.013 | 0 | 0.42 | 0.78 | 0.37 | 0.46 | 0.41 | 0.46 |
| 11 | 19088750 | rs10833053 | 1.249 | -0.007 | 0.621 | 1.399 | 0.024 | -0.045 | -0.045 | 0 | 0.56 | 0.12 | 0.62 | 0.53 | 0.59 | 0.54 |
| 11 | 19089384 | rs11024974 | -0.647 | 0.131 | 1.278 | 2.007 | -0.01 | -0.012 | -0.012 | 0.138 | 0.19 | 0.26 | 0.15 | 0.14 | 0.19 | 0.25 |
| 11 | 19094723 | rs16936091 | -0.891 | 0.227 | 1.016 | 2.37 | 0.001 | -0.018 | -0.018 | 0 | 0.08 | 0.04 | 0.08 | 0.19 | 0.05 | 0.15 |
| 11 | 19096159 | rs905552 | 2.463 | 0.071 | 0.802 | 2.294 | 0.036 | -0.167 | -0.167 | 0 | 0.58 | 0.03 | 0.63 | 0.49 | 0.68 | 0.55 |
| 11 | 19097685 | rs2062819 | 0.955 | -0.061 | 1.215 | 2.274 | -0.014 | -0.01 | -0.01 | 0 | 0.79 | 0.31 | 0.82 | 0.86 | 0.81 | 0.77 |
| 11 | 19097912 | rs16936098 | -2.031 | -0.023 | 1.417 | 2.245 | -0.026 | 0.05 | 0.05 | 0.033 | 0.2 | 0.31 | 0.19 | 0.37 | 0.13 | 0.22 |
| 11 | 19100736 | rs10766527 | 1.824 | -0.582 | 1.795 | 2.134 | -0.052 | 0.099 | 0.099 | 0.369 | 0.78 | 0.49 | 0.8 | 0.63 | 0.87 | 0.78 |
| 11 | 19101491 | rs868337 | -0.739 | -0.542 | 1.742 | 2.156 | 0.001 | 0.036 | 0.036 | 0 | 0.08 | 0.06 | 0.13 | 0.37 | 0.05 | 0.15 |
| 11 | 19105332 | rs10833056 | -0.441 | -0.513 | 1.893 | 2.145 | 0.001 | 0.027 | 0.027 | 0 | 0.14 | 0.25 | 0.14 | 0.21 | 0.18 | 0.19 |
| 11 | 19106045 | rs11024979 | -1.994 | -0.4 | 1.658 | 2.222 | -0.009 | 0.001 | 0.001 | 0.098 | 0.19 | 0.2 | 0.13 | 0.2 | 0.13 | 0.21 |
| 11 | 19106267 | rs11024980 | -2.188 | -0.398 | 1.651 | 2.227 | -0.035 | 0.081 | 0.081 | 0.151 | 0.2 | 0.32 | 0.18 | 0.38 | 0.13 | 0.21 |
| 11 | 19108683 | rs16936115 | -1.131 | -0.017 | 1.626 | 2.151 | -0.02 | 0.024 | 0.024 | 0.588 | 0.11 | 0.2 | 0.1 | 0.17 | 0.07 | 0.07 |
| 11 | 19111756 | rs7926950 | 0.946 | 0.005 | 1.655 | 2.093 | -0.016 | 0.001 | 0.001 | 0 | 0.81 | 0.52 | 0.83 | 0.8 | 0.83 | 0.77 |
| 11 | 19115179 | rs11024995 | -1.244 | 0.164 | 1.497 | 2.042 | -0.006 | -0.054 | -0.054 | 0 | 0.12 | 0.18 | 0.07 | 0 | 0.13 | 0.07 |
| 11 | 19117778 | rs11025000 | -0.366 | 0.171 | 1.398 | 2.2 | -0.007 | -0.016 | -0.016 | 7.369 | 0.06 | 0.03 | 0.08 | 0.22 | 0.05 | 0.15 |
| 11 | 19121231 | rs11025007 | -0.759 | 0.162 | 1.405 | 2.165 | -0.002 | 0.024 | 0.024 | 0.346 | 0.11 | 0.03 | 0.06 | 0 | 0.13 | 0.07 |
| 11 | 19122590 | rs12223284 | -0.62 | 0.166 | 1.427 | 2.262 | 0 | 0.073 | 0.073 | 0 | 0.07 | 0.06 | 0.13 | 0.39 | 0.05 | 0.15 |
| 11 | 19126858 | rs10833061 | -2.127 | 0.158 | 1.494 | 2.255 | -0.006 | 0.045 | 0.045 | 0.051 | 0.19 | 0.26 | 0.2 | 0.39 | 0.18 | 0.2 |
| 11 | 19128052 | rs10833062 | -2.043 | 0.158 | 1.486 | 2.26 | -0.003 | 0.004 | 0.004 | 0.033 | 0.19 | 0.23 | 0.16 | 0.22 | 0.18 | 0.19 |
| 11 | 19130858 | rs11025013 | -2.24 | 0.153 | 1.531 | 2.24 | -0.009 | 0.06 | 0.06 | 0.027 | 0.2 | 0.28 | 0.21 | 0.39 | 0.18 | 0.19 |

| | | | | | | | | | | | | | | | | |
|---|---|---|---|---|---|---|---|---|---|---|---|---|---|---|---|---|
| 11 | 19133909 | rs11608243 | -1.718 | 0.289 | 1.54 | 2.07 | -0.005 | -0.002 | -0.002 | 0 | 0.17 | 0.14 | 0.17 | 0.17 | 0.2 | 0.2 |
| 11 | 19135804 | rs4267049 | 0.674 | 0.166 | 1.817 | 1.999 | 0.006 | -0.02 | -0.02 | 0 | 0.94 | 0.79 | 0.95 | 0.99 | 0.97 | 0.95 |
| 11 | 19136906 | rs11025018 | -1.937 | 0.158 | 1.557 | 2.075 | 0.004 | 0.046 | 0.046 | 0.089 | 0.18 | 0.26 | 0.22 | 0.33 | 0.2 | 0.2 |
| 11 | 19137947 | rs1546329 | 2.569 | 0.038 | 1.137 | 2.138 | -0.022 | -0.098 | -0.098 | 0.405 | 0.6 | 0.03 | 0.63 | 0.39 | 0.65 | 0.54 |
| 11 | 19138127 | rs11025021 | -0.415 | -0.083 | 1.496 | 2.048 | -0.012 | 0.064 | 0.064 | 0 | 0.13 | 0.26 | 0.09 | 0.26 | 0.12 | 0.22 |
| 11 | 19140385 | rs11025026 | 1.446 | 0.015 | 0.695 | 1.76 | -0.009 | -0.009 | -0.009 | 0.959 | 0.72 | 0.46 | 0.78 | 0.73 | 0.7 | 0.65 |
| 11 | 19145974 | rs1503506 | 1.45 | 0.076 | 0.658 | 1.813 | -0.001 | -0.001 | -0.001 | 0.204 | 0.72 | 0.36 | 0.78 | 0.73 | 0.7 | 0.65 |
| 11 | 19167850 | rs2271001 | 1.591 | 0.07 | 0.656 | 1.81 | 0.005 | 0.005 | 0.005 | 0.367 | 0.71 | 0.33 | 0.78 | 0.73 | 0.7 | 0.65 |
| 11 | 19182348 | rs3802798 | 1.544 | 0.098 | 0.526 | 1.856 | 0.009 | -0.022 | -0.022 | 0 | 0.69 | 0.32 | 0.68 | 0.52 | 0.68 | 0.55 |
| 11 | 19184044 | rs1393954 | 0.539 | 0.223 | 0.401 | 1.82 | -0.021 | 0.083 | 0.083 | 0 | 0.82 | 0.55 | 0.86 | 0.73 | 0.85 | 0.74 |
| 11 | 19188122 | rs2242458 | 1.507 | 0.398 | -0.767 | 1.403 | -0.038 | -0.163 | -0.163 | 0.412 | 0.6 | 0.03 | 0.62 | 0.48 | 0.61 | 0.58 |
| 11 | 19189752 | rs11823088 | -0.143 | 0.658 | -0.193 | 1.44 | 0.012 | 0.037 | 0.037 | 0 | 0.05 | 0.08 | 0.1 | 0.15 | 0.09 | 0.05 |
| 11 | 19198018 | rs1503501 | 1.041 | 0.644 | -0.552 | 1.064 | 0.011 | -0.037 | -0.037 | 3.77 | 0.6 | 0.27 | 0.58 | 0.49 | 0.6 | 0.59 |
| 11 | 19203165 | rs1991249 | 0.698 | 0.766 | -0.014 | 1.332 | 0.021 | 0.085 | 0.085 | 0 | 0.78 | 0.62 | 0.74 | 0.51 | 0.75 | 0.72 |
| 11 | 19205132 | rs9888224 | -1.132 | -0.419 | 0.102 | 0.358 | -0.04 | -0.063 | -0.063 | 1.645 | 0.21 | 0.32 | 0.16 | 0.06 | 0.13 | 0.13 |
| 11 | 19205798 | rs7130460 | -1.549 | -0.448 | 0.688 | 0.155 | -0.057 | -0.111 | -0.111 | 0 | 0.24 | 0.52 | 0.17 | 0.06 | 0.13 | 0.14 |
| 11 | 19205980 | rs7102888 | -1.674 | -0.523 | 0.768 | -0.268 | -0.033 | -0.11 | -0.11 | 0.007 | 0.3 | 0.5 | 0.2 | 0.1 | 0.22 | 0.22 |
| 11 | 19207405 | rs10833069 | 1.523 | -0.198 | -0.828 | 0.195 | -0.165 | -0.161 | -0.161 | 0 | 0.4 | 0.03 | 0.4 | 0.51 | 0.48 | 0.4 |
| 11 | 19208319 | rs3781799 | -0.172 | -0.052 | -0.539 | 0.501 | 0.007 | 0.012 | 0.012 | 0 | 0.1 | 0.15 | 0.07 | 0.1 | 0.12 | 0.07 |
| 11 | 19208573 | rs1077687 | 1.59 | -0.396 | -1.28 | 0.292 | -0.14 | -0.131 | -0.131 | 0 | 0.37 | 0.04 | 0.47 | 0.47 | 0.45 | 0.39 |
| 11 | 19209050 | rs10833071 | 0.385 | -0.177 | -0.703 | 0.565 | -0.019 | -0.031 | -0.031 | 0 | 0.15 | 0.05 | 0.27 | 0.22 | 0.16 | 0.31 |
| 11 | 19223143 | rs10741762 | 0.561 | -0.474 | 0.252 | 0.053 | -0.052 | -0.088 | -0.088 | 0 | 0.74 | 0.53 | 0.85 | 0.91 | 0.8 | 0.89 |
| 11 | 19224718 | rs10741763 | 1.824 | -0.448 | 0.084 | 0.385 | -0.048 | 0.001 | 0.001 | 0.096 | 0.43 | 0.14 | 0.47 | 0.44 | 0.47 | 0.44 |
| 11 | 19227355 | rs10766536 | -0.4 | -0.345 | 0.394 | 0.509 | 0.043 | 0.059 | 0.059 | 1.557 | 0.17 | 0.31 | 0.1 | 0.25 | 0.2 | 0.16 |
| 11 | 19230165 | rs11025056 | 0.14 | -0.474 | 0.758 | 0.7 | -0.006 | -0.003 | -0.003 | 1.321 | 0.11 | 0.19 | 0.05 | 0.09 | 0.07 | 0.08 |
| 11 | 19234797 | rs11025059 | -1.422 | -0.499 | 1.016 | 0.69 | 0.009 | 0.051 | 0.051 | 0.486 | 0.22 | 0.41 | 0.17 | 0.25 | 0.22 | 0.1 |
| 11 | 19235154 | rs12146588 | 0.102 | -0.468 | 0.912 | 0.708 | 0.002 | 0.03 | 0.03 | 2.601 | 0.08 | 0.08 | 0.07 | 0.24 | 0.13 | 0.08 |
| 11 | 19238380 | rs2553974 | 1.727 | -0.595 | 0.797 | 0.49 | -0.004 | 0 | 0 | 0 | 0.62 | 0.42 | 0.78 | 0.66 | 0.68 | 0.81 |
| 11 | 19240538 | rs998263 | -1.803 | -0.606 | 0.817 | 0.494 | -0.003 | 0.009 | 0.009 | 0.578 | 0.27 | 0.54 | 0.2 | 0.25 | 0.28 | 0.12 |
| 11 | 19244377 | rs924693 | -1.082 | -0.522 | 0.473 | 0.489 | -0.017 | -0.04 | -0.04 | 0 | 0.31 | 0.28 | 0.4 | 0.47 | 0.35 | 0.38 |
| 11 | 19245604 | rs2292061 | 2.165 | -0.566 | 0.504 | 0.424 | -0.021 | 0.015 | 0.015 | 0 | 0.44 | 0.25 | 0.48 | 0.44 | 0.51 | 0.52 |
| 11 | 19245854 | rs793278 | 0.048 | -0.684 | 0.757 | 0.163 | -0.006 | 0.004 | 0.004 | 0 | 0.79 | 0.79 | 0.68 | 0.78 | 0.83 | 0.71 |
| 11 | 19246590 | rs793276 | -0.529 | -0.628 | 0.71 | 0.198 | -0.008 | -0.029 | -0.029 | 0 | 0.83 | 0.93 | 0.7 | 0.78 | 0.83 | 0.71 |
| 11 | 19247767 | rs7936229 | -0.751 | -0.578 | 0.52 | 0.205 | -0.019 | -0.028 | -0.028 | 0 | 0.18 | 0.25 | 0.09 | 0.09 | 0.13 | 0.09 |
| 11 | 19250164 | rs10500844 | 1.121 | -0.407 | 0.325 | 0.361 | 0.01 | 0.008 | 0.008 | 0.017 | 0.56 | 0.44 | 0.58 | 0.68 | 0.69 | 0.6 |
| 11 | 19251704 | rs12273302 | -0.498 | -0.026 | 0.345 | 0.591 | -0.016 | -0.01 | -0.01 | 0 | 0.12 | 0.19 | 0.04 | 0.09 | 0.04 | 0.08 |
| 11 | 19251968 | rs1863382 | -0.875 | 0.739 | 0.245 | 0.991 | 0.021 | 0.063 | 0.063 | 0.583 | 0.07 | 0.12 | 0.04 | 0.17 | 0.08 | 0.13 |
| 11 | 19252427 | rs793252 | -0.75 | 1.017 | 0.004 | 0.611 | 0.011 | 0.015 | 0.015 | 0 | 0.33 | 0.42 | 0.45 | 0.46 | 0.39 | 0.43 |
| 11 | 19264497 | rs793283 | -0.369 | 1.152 | 0.173 | 0.1 | 0.018 | -0.046 | -0.046 | 0 | 0.17 | 0.3 | 0.13 | 0.1 | 0.24 | 0.09 |
| 11 | 19265226 | rs7129281 | 0.727 | 0.845 | -0.222 | -0.021 | 0.018 | -0.058 | -0.058 | 0 | 0.34 | 0.22 | 0.31 | 0.52 | 0.24 | 0.46 |
| 11 | 19268555 | rs1698897 | 0.016 | 0.815 | 0.046 | -0.078 | 0.019 | -0.059 | -0.059 | 0 | 0.19 | 0.38 | 0.13 | 0.11 | 0.24 | 0.09 |
| 11 | 19270031 | rs1587704 | 0.752 | 0.727 | -0.398 | -0.009 | 0.023 | -0.06 | -0.06 | 0 | 0.35 | 0.21 | 0.31 | 0.52 | 0.25 | 0.48 |
| 11 | 19278691 | rs2553970 | -0.034 | 0.708 | 0.595 | -0.463 | 0.028 | -0.057 | -0.057 | 0.102 | 0.82 | 0.51 | 0.86 | 0.89 | 0.76 | 0.91 |
| 11 | 19297340 | rs1502987 | -0.388 | 0.687 | 0.445 | -0.448 | 0.023 | -0.056 | -0.056 | 0.25 | 0.18 | 0.39 | 0.14 | 0.11 | 0.24 | 0.09 |
| 11 | 19306336 | rs11025088 | -0.223 | 0.649 | -0.024 | -0.615 | 0 | 0.003 | 0.003 | 0 | 0.38 | 0.32 | 0.41 | 0.15 | 0.4 | 0.22 |
| 11 | 19308744 | rs7111578 | 0.01 | 0.649 | -0.618 | -0.571 | -0.004 | -0.035 | -0.035 | 1.354 | 0.61 | 0.43 | 0.56 | 0.85 | 0.61 | 0.78 |
| 11 | 19311030 | rs11025093 | -0.386 | 0.656 | -1.112 | -0.068 | -0.046 | -0.015 | -0.015 | 0.495 | 0.14 | 0.01 | 0.18 | 0.16 | 0.17 | 0.17 |
| 11 | 19313016 | rs1648222 | -0.302 | -0.01 | -0.846 | -0.357 | 0.009 | 0.02 | 0.02 | 0 | 0.49 | 0.59 | 0.47 | 0.49 | 0.59 | 0.44 |
| 11 | 19314614 | rs11025095 | -0.6 | -0.66 | -1.348 | -0.847 | 0.017 | 0.021 | 0.021 | 0.123 | 0.58 | 0.96 | 0.56 | 0.48 | 0.69 | 0.54 |
| 11 | 19323503 | rs793247 | 0.176 | -1.022 | -0.964 | -0.734 | 0.031 | 0.01 | 0.01 | 0.025 | 0.63 | 0.9 | 0.6 | 0.48 | 0.7 | 0.66 |
| 11 | 19331010 | rs4757787 | -1.047 | -1.148 | -0.164 | -0.81 | 0.033 | 0.029 | 0.029 | 0 | 0.45 | 0.71 | 0.46 | 0.51 | 0.32 | 0.39 |
| 11 | 19335282 | rs17500922 | 0.15 | -1.003 | -0.565 | -0.734 | 0 | 0.019 | 0.019 | 0 | 0.09 | 0.04 | 0.05 | 0 | 0.09 | 0.05 |
| 11 | 19338029 | rs10734274 | 0.056 | -1.015 | -0.281 | -0.74 | -0.01 | -0.057 | -0.057 | 0.178 | 0.89 | 0.79 | 0.94 | 1 | 0.91 | 0.95 |
| 11 | 19344878 | rs1393696 | 0.614 | -0.964 | -0.524 | -0.752 | 0.013 | 0.014 | 0.014 | 0 | 0.57 | 0.58 | 0.57 | 0.49 | 0.67 | 0.62 |
| 11 | 19345331 | rs7940014 | 0.17 | -0.958 | -0.557 | -0.749 | -0.004 | -0.063 | -0.063 | 1.39 | 0.29 | 0.04 | 0.35 | 0.51 | 0.22 | 0.3 |
| 11 | 19349935 | rs7116214 | -0.001 | -0.936 | -0.518 | -0.744 | 0.002 | -0.016 | -0.016 | 0 | 0.87 | 0.84 | 0.94 | 1 | 0.89 | 0.92 |
| 11 | 19350262 | rs11025101 | 0.114 | -0.945 | -0.477 | -0.583 | 0.016 | -0.013 | -0.013 | 0.383 | 0.29 | 0.18 | 0.36 | 0.51 | 0.22 | 0.31 |
| 11 | 19352288 | rs10766542 | 0.787 | -0.969 | -0.395 | -0.524 | 0 | -0.001 | -0.001 | 0.574 | 0.56 | 0.48 | 0.55 | 0.43 | 0.67 | 0.61 |
| 11 | 19352616 | rs16936547 | 0.049 | -0.949 | -1.05 | -0.472 | 0.006 | 0.02 | 0.02 | 0 | 0.11 | 0.05 | 0.06 | 0.06 | 0.11 | 0.08 |
| 11 | 19355372 | rs4756995 | 0.035 | -0.791 | -1.07 | -0.314 | 0.005 | -0.057 | -0.057 | 0 | 0.29 | 0.05 | 0.35 | 0.51 | 0.21 | 0.3 |
| 11 | 19371907 | rs4393295 | -0.059 | -0.352 | -0.983 | 0.046 | 0.003 | -0.018 | -0.018 | 0 | 0.67 | 0.72 | 0.62 | 0.53 | 0.71 | 0.67 |
| 11 | 19372943 | rs10833098 | 0.053 | -0.34 | -1.031 | 0.045 | -0.033 | -0.082 | -0.082 | 0 | 0.3 | 0.08 | 0.36 | 0.47 | 0.29 | 0.33 |
| 11 | 19374222 | rs1354268 | -0.096 | -0.54 | -1.032 | 0.168 | -0.054 | -0.093 | -0.093 | 0.012 | 0.3 | 0.04 | 0.37 | 0.51 | 0.29 | 0.33 |
| 11 | 19374244 | rs1354269 | -0.923 | -0.603 | -0.795 | 0.088 | 0.004 | 0.011 | 0.011 | 0 | 0.43 | 0.42 | 0.44 | 0.55 | 0.38 | 0.4 |
| 11 | 19379879 | rs2702705 | 0.173 | -0.086 | -0.828 | 0.482 | -0.013 | -0.034 | -0.034 | 0 | 0.21 | 0.15 | 0.35 | 0.44 | 0.27 | 0.14 |
| 11 | 19383628 | rs4551739 | 0.607 | 0.168 | -1.611 | 0.239 | -0.029 | 0.017 | 0.017 | 0 | 0.13 | 0.01 | 0.21 | 0.13 | 0.15 | 0.15 |
| 11 | 19391182 | rs17593306 | -0.408 | 0.248 | -0.924 | 0.757 | 0.001 | 0.001 | 0.001 | 0 | 0.11 | 0.09 | 0.05 | 0.04 | 0.1 | 0.07 |
| 11 | 19399302 | rs2729843 | -0.712 | 0.337 | -0.917 | 0.877 | -0.008 | -0.022 | -0.022 | 2.081 | 0.31 | 0.23 | 0.4 | 0.49 | 0.37 | 0.21 |
| 11 | 19401373 | rs2702703 | -0.715 | -0.062 | -0.611 | 0.752 | 0.016 | -0.012 | -0.012 | 0 | 0.42 | 0.66 | 0.24 | 0.52 | 0.32 | 0.46 |

| | | | | | | | | | | | | | | | | |
|---|---|---|---|---|---|---|---|---|---|---|---|---|---|---|---|---|
| 11 | 19412661 | rs4073508 | 0.445 | -0.335 | -0.044 | 0.67 | 0.034 | -0.03 | -0.03 | 0 | 0.58 | 0.26 | 0.79 | 0.52 | 0.71 | 0.6 |
| 11 | 19416483 | rs10833101 | -0.467 | -0.612 | -0.867 | 0.279 | 0.045 | 0.058 | 0.058 | 0 | 0.15 | 0 | 0.04 | 0.09 | 0.05 | 0.16 |
| 11 | 19420097 | rs4353249 | -0.318 | 0.024 | -0.282 | 0.547 | -0.003 | 0.004 | 0.004 | 0.062 | 0.39 | 0.39 | 0.42 | 0.52 | 0.34 | 0.29 |
| 11 | 19422628 | rs10833106 | 0.576 | 0.147 | -1.059 | -0.138 | -0.004 | 0.103 | 0.103 | 0 | 0.27 | 0.06 | 0.15 | 0.16 | 0.25 | 0.4 |
| 11 | 19423117 | rs2729910 | -0.771 | 0.471 | -0.822 | 0.408 | 0.001 | 0.06 | 0.06 | 0.045 | 0.73 | 0.48 | 0.76 | 0.57 | 0.77 | 0.76 |
| 11 | 19423358 | rs2702671 | 0.147 | 0.501 | -0.725 | -0.002 | 0.025 | 0.075 | 0.075 | 0 | 0.4 | 0.26 | 0.34 | 0.25 | 0.36 | 0.46 |
| 11 | 19425092 | rs2702676 | -0.205 | 0.647 | -0.227 | 0.87 | 0.007 | 0.212 | 0.212 | 0 | 0.89 | 0.68 | 0.82 | 0.5 | 0.89 | 0.78 |
| 11 | 19427092 | rs12364788 | -0.157 | 0.966 | -0.303 | -0.033 | 0.032 | -0.043 | -0.043 | 1.565 | 0.21 | 0.31 | 0.34 | 0.1 | 0.32 | 0.16 |
| 11 | 19430197 | rs2702630 | -0.417 | 0.542 | -0.498 | 0.142 | 0 | -0.008 | -0.008 | 0.909 | 0.64 | 0.62 | 0.72 | 0.76 | 0.63 | 0.66 |
| 11 | 19440132 | rs2702736 | -0.857 | 0.198 | 0.083 | 0.248 | 0.001 | 0.001 | 0.001 | 0 | 0.64 | 0.69 | 0.77 | 0.7 | 0.7 | 0.71 |
| 11 | 19440246 | rs11025130 | 0.414 | -0.252 | 0.399 | 0.849 | -0.023 | 0.076 | 0.076 | 1.665 | 0.11 | 0.21 | 0.17 | 0.27 | 0.07 | 0.13 |
| 11 | 19440493 | rs2702738 | -1.28 | -0.322 | 0.532 | 0.739 | -0.006 | 0.006 | 0.006 | 0 | 0.66 | 0.84 | 0.79 | 0.7 | 0.69 | 0.71 |
| 11 | 19441522 | rs4515943 | -1.101 | -0.316 | 0.542 | 0.756 | -0.001 | -0.003 | -0.003 | 0 | 0.64 | 0.66 | 0.77 | 0.7 | 0.68 | 0.71 |
| 11 | 19442926 | rs4462314 | -1.191 | -0.341 | 0.719 | 0.746 | -0.001 | -0.003 | -0.003 | 0 | 0.65 | 0.62 | 0.78 | 0.7 | 0.69 | 0.71 |
| 11 | 19445475 | rs10766556 | 0.379 | -0.033 | 0.396 | 0.615 | 0.008 | 0.037 | 0.037 | 0 | 0.38 | 0.14 | 0.36 | 0.32 | 0.36 | 0.48 |
| 11 | 19446705 | rs11025133 | 1.526 | -0.254 | 0.053 | 0.261 | 0.023 | -0.004 | -0.004 | 0 | 0.08 | 0.01 | 0.11 | 0.09 | 0.05 | 0.2 |
| 11 | 19447873 | rs2702639 | -0.561 | -0.032 | 0.302 | 0.323 | 0.037 | 0.004 | 0.004 | 0 | 0.51 | 0.69 | 0.54 | 0.55 | 0.43 | 0.6 |
| 11 | 19450893 | rs4550206 | 1.815 | 0.238 | -0.199 | 0.21 | -0.024 | -0.01 | -0.01 | 0.904 | 0.31 | 0.08 | 0.16 | 0.28 | 0.25 | 0.22 |
| 11 | 19451020 | rs11025135 | 0.292 | 0.613 | 0.018 | 0.065 | -0.075 | -0.022 | -0.022 | 0 | 0.13 | 0.02 | 0.21 | 0.11 | 0.25 | 0.09 |
| 11 | 19460084 | rs4757809 | 1.788 | 0.507 | -0.214 | -0.046 | -0.058 | 0.08 | 0.08 | 0 | 0.17 | 0.01 | 0.1 | 0.06 | 0.2 | 0.16 |
| 11 | 19466002 | rs10766560 | 1.384 | 0.454 | 0.308 | 0.083 | 0.018 | -0.021 | -0.021 | 0 | 0.38 | 0.21 | 0.3 | 0.54 | 0.27 | 0.42 |
| 11 | 19466720 | rs2632010 | 0.473 | 0.588 | 0.206 | 0.08 | -0.018 | 0.001 | 0.001 | 0 | 0.8 | 0.82 | 0.69 | 0.83 | 0.64 | 0.79 |
| 11 | 19466820 | rs11025138 | -0.394 | 0.617 | 0.254 | 0.098 | 0 | 0.001 | 0.001 | 0 | 0.17 | 0.15 | 0.25 | 0.11 | 0.29 | 0.11 |
| 11 | 19468769 | rs4756999 | -0.856 | 0.263 | 0.119 | 0.21 | 0.025 | 0.023 | 0.023 | 0 | 0.53 | 0.37 | 0.6 | 0.41 | 0.65 | 0.48 |
| 11 | 19483904 | rs2200568 | 0.623 | 0.073 | 1.018 | 0.488 | -0.022 | 0.012 | 0.012 | 0 | 0.65 | 0.44 | 0.53 | 0.6 | 0.68 | 0.66 |
| 11 | 19486268 | rs12271178 | -0.245 | 0.07 | 1.07 | 0.374 | 0.005 | 0.002 | 0.002 | 0 | 0.24 | 0.2 | 0.29 | 0.24 | 0.22 | 0.22 |
| 11 | 19489018 | rs10766561 | 1.39 | 0.13 | 0.818 | 0.442 | -0.05 | -0.009 | -0.009 | 1.052 | 0.34 | 0.18 | 0.24 | 0.33 | 0.4 | 0.4 |
| 11 | 19492634 | rs2729878 | 1.145 | 0.064 | 1.136 | 0.37 | 0.001 | -0.017 | -0.017 | 3.026 | 0.84 | 0.58 | 0.79 | 0.9 | 0.83 | 0.89 |
| 11 | 19493284 | rs11820210 | 0.448 | -0.007 | 1.081 | 0.381 | 0.006 | -0.001 | -0.001 | 0 | 0.32 | 0.24 | 0.29 | 0.34 | 0.27 | 0.31 |
| 11 | 19493779 | rs1565041 | 0.144 | 0.182 | 1.241 | 0.363 | -0.001 | 0.014 | 0.014 | 0 | 0.9 | 0.73 | 0.84 | 0.9 | 0.9 | 0.94 |
| 11 | 19494407 | rs2729881 | -0.868 | 0.084 | 1.425 | 0.461 | 0.001 | 0 | 0 | 0.539 | 0.33 | 0.3 | 0.44 | 0.33 | 0.34 | 0.28 |
| 11 | 19495611 | rs2702725 | -0.257 | 0.134 | 1.388 | 0.441 | -0.02 | -0.003 | -0.003 | 0 | 0.77 | 0.7 | 0.82 | 0.77 | 0.82 | 0.79 |
| 11 | 19498686 | rs2729884 | 0.407 | 0.133 | 1.523 | 0.461 | 0.003 | 0 | 0 | 1.094 | 0.47 | 0.55 | 0.6 | 0.44 | 0.54 | 0.47 |
| 11 | 19505545 | rs11025151 | 0.559 | 0.045 | 1.412 | 0.467 | 0.011 | -0.007 | -0.007 | 0.005 | 0.31 | 0.17 | 0.22 | 0.32 | 0.27 | 0.31 |
| 11 | 19510887 | rs2632020 | 0.554 | -0.609 | 1.271 | 0.639 | 0.017 | -0.009 | -0.009 | 0 | 0.45 | 0.31 | 0.58 | 0.44 | 0.54 | 0.47 |
| 11 | 19511587 | rs11604718 | -0.614 | -0.604 | 1.199 | 0.629 | 0.031 | 0.001 | 0.001 | 0 | 0.52 | 0.3 | 0.39 | 0.56 | 0.46 | 0.53 |
| 11 | 19512670 | rs12418988 | 0.659 | -0.609 | 1.079 | 0.634 | 0.023 | -0.028 | -0.028 | 0 | 0.18 | 0.06 | 0.15 | 0.23 | 0.16 | 0.18 |
| 11 | 19518595 | rs11025158 | -0.092 | -0.521 | 0.926 | 0.686 | -0.003 | -0.001 | -0.001 | 0 | 0.1 | 0.13 | 0.22 | 0.11 | 0.15 | 0.07 |
| 11 | 19522864 | rs2702656 | -0.572 | -0.754 | 1.03 | 0.699 | -0.039 | 0.003 | 0.003 | 0 | 0.79 | 0.59 | 0.82 | 0.78 | 0.84 | 0.82 |
| 11 | 19523388 | rs2403441 | 1.467 | -0.763 | 0.964 | 0.696 | -0.066 | -0.026 | -0.026 | 0 | 0.34 | 0.12 | 0.39 | 0.34 | 0.39 | 0.41 |
| 11 | 19524048 | rs10766565 | 1.3 | -0.728 | 0.942 | 0.708 | -0.035 | -0.009 | -0.009 | 0 | 0.35 | 0.19 | 0.4 | 0.34 | 0.39 | 0.41 |
| 11 | 19525056 | rs4757815 | 0.301 | -0.73 | 0.942 | 0.686 | 0.009 | 0.002 | 0.002 | 0.107 | 0.34 | 0.21 | 0.19 | 0.33 | 0.31 | 0.36 |
| 11 | 19528822 | rs4423163 | 1.114 | -0.62 | 0.967 | 0.743 | -0.045 | -0.015 | -0.015 | 0 | 0.34 | 0.16 | 0.4 | 0.34 | 0.39 | 0.41 |
| 11 | 19532506 | rs1055543 | -0.184 | -0.59 | 0.821 | 0.732 | 0 | 0.002 | 0.002 | 0.214 | 0.44 | 0.36 | 0.42 | 0.44 | 0.46 | 0.41 |
| 11 | 19533093 | rs1565040 | -0.176 | -0.372 | 0.403 | 0.998 | -0.045 | -0.002 | -0.002 | 0 | 0.79 | 0.56 | 0.82 | 0.78 | 0.84 | 0.82 |
| 11 | 19538163 | rs4757818 | 0.963 | 0.106 | -0.243 | 0.891 | -0.113 | -0.041 | -0.041 | 0 | 0.33 | 0.07 | 0.41 | 0.28 | 0.37 | 0.4 |
| 11 | 19538185 | rs4757819 | 0.921 | 0.094 | -0.197 | 0.877 | -0.096 | -0.033 | -0.033 | 0 | 0.33 | 0.1 | 0.41 | 0.28 | 0.37 | 0.4 |
| 11 | 19540109 | rs2702735 | -1.384 | -0.675 | 1.326 | 1.066 | -0.026 | -0.009 | -0.009 | 0 | 0.23 | 0.51 | 0.25 | 0.2 | 0.2 | 0.13 |
| 11 | 19541784 | rs2632057 | -1.547 | -0.665 | 1.259 | 1.214 | -0.015 | -0.008 | -0.008 | 0 | 0.4 | 0.52 | 0.5 | 0.38 | 0.4 | 0.38 |
| 11 | 19546520 | rs2702646 | -1.652 | -0.648 | 1.453 | 1.198 | -0.009 | -0.006 | -0.006 | 0 | 0.41 | 0.59 | 0.5 | 0.38 | 0.39 | 0.38 |
| 11 | 19547027 | rs11605946 | -0.55 | 0.176 | 1.2 | 1.362 | -0.013 | -0.017 | -0.017 | 0 | 0.19 | 0.03 | 0.26 | 0.19 | 0.2 | 0.24 |
| 11 | 19553185 | rs12803842 | -2.382 | 0.378 | 2.239 | 1.539 | 0.001 | 0.001 | 0.001 | 2.835 | 0.21 | 0.74 | 0.2 | 0.16 | 0.18 | 0.22 |
| 11 | 19554417 | rs7117077 | -2.373 | 0.239 | 2.238 | 1.539 | 0.001 | 0.001 | 0.001 | 2.783 | 0.21 | 0.74 | 0.19 | 0.16 | 0.18 | 0.22 |
| 11 | 19559749 | rs10500854 | -1.989 | 0.173 | 2.653 | 1.531 | 0.006 | 0.022 | 0.022 | 0 | 0.1 | 0.19 | 0.13 | 0.15 | 0.1 | 0.02 |
| 11 | 19561445 | rs16936939 | -1.731 | 0.175 | 2.707 | 1.418 | -0.035 | -0.084 | -0.084 | 0 | 0.13 | 0.59 | 0.07 | 0 | 0.08 | 0.2 |
| 11 | 19564549 | rs11025177 | -1.598 | 0.32 | 2.082 | 1.691 | 0.027 | 0.093 | 0.093 | 0 | 0.09 | 0.22 | 0.13 | 0.19 | 0.1 | 0.02 |
| 11 | 19566243 | rs7950505 | -1.649 | 0.354 | 1.269 | 1.74 | -0.008 | 0.008 | 0.008 | 0.817 | 0.2 | 0.66 | 0.2 | 0.19 | 0.18 | 0.17 |
| 11 | 19569563 | rs874426 | 1.512 | 0.542 | 1.322 | 2.272 | -0.008 | 0.051 | 0.051 | 1.345 | 0.77 | 0.38 | 0.74 | 0.43 | 0.83 | 0.73 |
| 11 | 19574197 | rs10741778 | -2.184 | 0.84 | 1.405 | 2.259 | 0.002 | 0.021 | 0.021 | 0.061 | 0.25 | 0.67 | 0.25 | 0.5 | 0.27 | 0.45 |
| 11 | 19578952 | rs10766569 | -2.282 | 0.895 | 1.799 | 2.25 | 0.039 | 0.137 | 0.137 | 0.047 | 0.24 | 0.45 | 0.25 | 0.5 | 0.28 | 0.45 |
| 11 | 19579055 | rs10766570 | -1.241 | 1.115 | 1.572 | 2.415 | -0.073 | -0.072 | -0.072 | 0 | 0.11 | 0.01 | 0.17 | 0.44 | 0.22 | 0.25 |
| 11 | 19590448 | rs4338501 | -1.739 | 0.975 | 0.18 | 2.23 | 0 | 0 | 0 | 0 | 0.22 | 0.19 | 0.23 | 0.35 | 0.28 | 0.33 |
| 11 | 19609944 | rs10766573 | 1.382 | 0.508 | -0.485 | 1.501 | 0.001 | 0.017 | 0.017 | 7.113 | 0.8 | 0.07 | 0.76 | 0.85 | 0.85 | 0.76 |
| 11 | 19614747 | rs7931824 | -1.639 | 0.448 | 1.513 | 1.455 | 0.023 | -0.023 | -0.023 | 0 | 0.13 | 0.34 | 0.14 | 0.09 | 0.13 | 0.15 |
| 11 | 19619540 | rs7930497 | -1.885 | 0.353 | 2.208 | 1.456 | 0.069 | 0.12 | 0.12 | 2.751 | 0.05 | 0.39 | 0.06 | 0.03 | 0.01 | 0.08 |
| 11 | 19619944 | rs1471903 | -0.238 | 1.266 | 0.921 | 1.863 | -0.132 | -0.138 | -0.138 | 0 | 0.21 | 0.02 | 0.39 | 0.4 | 0.39 | 0.33 |
| 11 | 19620227 | rs11025189 | 1.856 | 0.695 | 0.977 | 1.699 | 0.087 | 0.005 | 0.005 | 9.47 | 0.81 | 0.07 | 0.89 | 0.86 | 0.97 | 0.84 |
| 11 | 19623277 | rs11025190 | -1.517 | 0.691 | 1.781 | 1.699 | -0.054 | 0.048 | 0.048 | 2.366 | 0.13 | 0.54 | 0.06 | 0.12 | 0.03 | 0.12 |
| 11 | 19625940 | rs10741784 | 1.915 | 0.666 | 1.566 | 1.736 | 0.091 | -0.029 | -0.029 | 0 | 0.81 | 0.07 | 0.84 | 0.81 | 0.95 | 0.79 |

| | | | | | | | | | | | | | | | | |
|---|---|---|---|---|---|---|---|---|---|---|---|---|---|---|---|---|
| 11 | 19627527 | rs10833134 | 1.842 | 0.668 | 1.586 | 1.713 | 0.065 | -0.015 | -0.015 | 0 | 0.81 | 0.11 | 0.84 | 0.8 | 0.95 | 0.79 |
| 11 | 19635453 | rs17599506 | 1.675 | 0.259 | 0.669 | 1.228 | -0.055 | -0.007 | -0.007 | 1.315 | 0.41 | 0.02 | 0.3 | 0.27 | 0.36 | 0.36 |
| 11 | 19637385 | rs11025194 | -0.291 | 0.261 | 0.567 | 1.147 | -0.191 | -0.17 | -0.17 | 0.274 | 0.34 | 0.02 | 0.47 | 0.59 | 0.5 | 0.41 |
| 11 | 19648600 | rs4757829 | 1.865 | 0.081 | 1.847 | 1.281 | 0.013 | 0.186 | 0.186 | 6.593 | 0.93 | 0.41 | 0.88 | 0.87 | 0.99 | 0.87 |
| 11 | 19650087 | rs2017056 | 1.649 | 0.08 | 1.894 | 1.131 | 0.041 | 0.148 | 0.148 | 7.34 | 0.94 | 0.45 | 0.94 | 0.94 | 0.99 | 0.91 |
| 11 | 19674471 | rs1987420 | 1.466 | 0.292 | 2.243 | 0.972 | 0.121 | 0.136 | 0.136 | 2.903 | 0.95 | 0.53 | 0.93 | 0.94 | 0.94 | 0.87 |
| 11 | 19697364 | rs2216997 | 1.54 | -0.023 | 1.919 | 0.786 | -0.05 | -0.047 | -0.047 | 0 | 0.8 | 0.34 | 0.89 | 0.89 | 0.91 | 0.77 |
| 11 | 19700177 | rs2012651 | 1.941 | -0.066 | 1.916 | 0.837 | 0.012 | 0.012 | 0.012 | 5.197 | 0.77 | 0.19 | 0.88 | 0.86 | 0.9 | 0.73 |
| 11 | 19700324 | rs4267050 | 1.583 | -0.042 | 1.795 | 0.847 | -0.066 | -0.026 | -0.026 | 0 | 0.8 | 0.4 | 0.89 | 0.82 | 0.9 | 0.73 |
| 11 | 19702039 | rs12288601 | -1.099 | -0.024 | 1.752 | 0.841 | -0.016 | -0.034 | -0.034 | 0.581 | 0.14 | 0.2 | 0.05 | 0 | 0.03 | 0.08 |
| 11 | 19702427 | rs7931452 | -1.184 | -0.021 | 1.672 | 0.851 | -0.066 | -0.061 | -0.061 | 0 | 0.16 | 0.36 | 0.06 | 0.07 | 0.04 | 0.11 |
| 11 | 19702865 | rs10741791 | -0.086 | -0.016 | 0.294 | 0.651 | -0.098 | -0.041 | -0.041 | 0.447 | 0.4 | 0.05 | 0.58 | 0.64 | 0.55 | 0.43 |
| 11 | 19707839 | rs7130790 | 1.278 | -0.181 | 1.264 | 0.596 | 0.114 | 0.145 | 0.145 | 1.387 | 0.95 | 0.65 | 0.94 | 0.9 | 0.92 | 0.89 |
| 11 | 19709674 | rs7938698 | -1.67 | -0.161 | 1.224 | 0.594 | -0.058 | -0.044 | -0.044 | 1.534 | 0.2 | 0.51 | 0.15 | 0.15 | 0.1 | 0.18 |
| 11 | 19711993 | rs1364795 | 1.788 | -0.162 | 1.063 | 0.599 | 0.035 | 0.016 | 0.016 | 4.838 | 0.74 | 0.13 | 0.73 | 0.78 | 0.82 | 0.71 |
| 11 | 19712486 | rs10741792 | 1.295 | 0.584 | 1.151 | 0.761 | 0.109 | 0.099 | 0.099 | 1.202 | 0.95 | 0.67 | 0.89 | 0.94 | 0.93 | 0.89 |
| 11 | 19730635 | rs890136 | 1.069 | 0.244 | 0.438 | 0.877 | -0.002 | 0.004 | 0.004 | 0 | 0.8 | 0.76 | 0.84 | 0.75 | 0.82 | 0.7 |
| 11 | 19735987 | rs2278132 | 0.032 | 0.424 | -0.197 | 0.222 | -0.045 | 0.035 | 0.035 | 1.097 | 0.57 | 0.88 | 0.33 | 0.33 | 0.49 | 0.51 |
| 11 | 19736996 | rs1319701 | 0.083 | 0.26 | 0.041 | 0.133 | -0.005 | 0.035 | 0.035 | 1.413 | 0.57 | 0.72 | 0.34 | 0.36 | 0.51 | 0.5 |
| 11 | 19743250 | rs7118901 | -0.121 | 0.129 | -0.228 | 0.247 | -0.067 | -0.045 | -0.045 | 0.205 | 0.61 | 0.88 | 0.41 | 0.44 | 0.51 | 0.59 |
| 11 | 19748584 | rs11025224 | -1.987 | -0.257 | 0.103 | 0.096 | -0.023 | 0.094 | 0.094 | 4.177 | 0.14 | 0.63 | 0.12 | 0.35 | 0.09 | 0.27 |
| 11 | 19748992 | rs1364792 | -1.896 | -0.254 | 0.25 | -0.05 | -0.02 | 0.077 | 0.077 | 3.805 | 0.13 | 0.6 | 0.12 | 0.26 | 0.08 | 0.25 |
| 11 | 19756763 | rs1035699 | -0.147 | -0.216 | 1.51 | -0.048 | -0.029 | 0.028 | 0.028 | 1.683 | 0.6 | 0.76 | 0.39 | 0.35 | 0.51 | 0.56 |
| 11 | 19756789 | rs1425227 | 1.028 | 0.339 | 1.356 | -0.65 | -0.007 | -0.067 | -0.067 | 0 | 0.42 | 0.51 | 0.22 | 0.07 | 0.31 | 0.27 |
| 11 | 19757220 | rs1820351 | -0.763 | 0.611 | 0.934 | -0.375 | 0.018 | -0.055 | -0.055 | 0 | 0.11 | 0.2 | 0.09 | 0.02 | 0.13 | 0.05 |
| 11 | 19765013 | rs1346028 | 0.3 | 0.322 | 1.052 | -0.124 | 0.002 | 0.003 | 0.003 | 0.495 | 0.51 | 0.46 | 0.3 | 0.35 | 0.38 | 0.53 |
| 11 | 19765839 | rs11025234 | -0.793 | 0.342 | 1.216 | -0.131 | 0.002 | -0.005 | -0.005 | 0.034 | 0.1 | 0.12 | 0.1 | 0.08 | 0.13 | 0.18 |
| 11 | 19770390 | rs10741799 | -0.246 | 0.33 | 1.373 | -0.168 | -0.01 | 0.006 | 0.006 | 0.116 | 0.59 | 0.59 | 0.42 | 0.37 | 0.48 | 0.67 |
| 11 | 19770798 | rs10833162 | -0.532 | 0.308 | 1.38 | -0.186 | 0.005 | -0.032 | -0.032 | 0 | 0.08 | 0.13 | 0.12 | 0.02 | 0.09 | 0.14 |
| 11 | 19770940 | rs10766589 | 0.212 | 0.29 | 1.506 | -0.188 | -0.002 | -0.049 | -0.049 | 1.755 | 0.53 | 0.43 | 0.33 | 0.1 | 0.4 | 0.45 |
| 11 | 19770987 | rs10766590 | 0.212 | 0.29 | 1.506 | -0.188 | -0.002 | -0.049 | -0.049 | 1.385 | 0.53 | 0.43 | 0.29 | 0.1 | 0.4 | 0.45 |
| 11 | 19771054 | rs4757842 | 0.085 | 0.327 | 1.04 | -0.149 | -0.051 | -0.028 | -0.028 | 1.601 | 0.39 | 0.24 | 0.54 | 0.57 | 0.48 | 0.26 |
| 11 | 19774304 | rs11025239 | -0.235 | 0.253 | 0.753 | -0.239 | 0.034 | 0.157 | 0.157 | 0 | 0.05 | 0.15 | 0.08 | 0.27 | 0.08 | 0.23 |
| 11 | 19776660 | rs10741800 | -0.84 | -0.096 | 0.415 | -1.195 | 0.029 | 0.05 | 0.05 | 0.951 | 0.54 | 0.65 | 0.77 | 0.92 | 0.69 | 0.67 |
| 11 | 19782191 | rs7118473 | -0.289 | -0.584 | -0.221 | -1.559 | 0.039 | 0.067 | 0.067 | 1.584 | 0.48 | 0.66 | 0.41 | 0.17 | 0.35 | 0.38 |
| 11 | 19786640 | rs11025246 | -0.759 | -0.808 | -0.393 | -1.814 | 0.033 | 0.078 | 0.078 | 0 | 0.55 | 0.69 | 0.47 | 0.14 | 0.4 | 0.39 |
| 11 | 19787053 | rs11819786 | -0.572 | -0.548 | -0.228 | -1.689 | 0.053 | 0.098 | 0.098 | 1.153 | 0.51 | 0.68 | 0.45 | 0.14 | 0.35 | 0.38 |
| 11 | 19789118 | rs4757843 | 0.731 | -0.656 | -0.394 | -1.803 | 0.059 | 0.083 | 0.083 | 3.197 | 0.3 | 0.1 | 0.19 | 0.14 | 0.17 | 0.16 |
| 11 | 19790118 | rs2004082 | -0.499 | -0.558 | -0.151 | -1.615 | 0.012 | 0.016 | 0.016 | 2.108 | 0.47 | 0.59 | 0.38 | 0.14 | 0.32 | 0.28 |
| 11 | 19790494 | rs2004083 | -0.103 | -0.34 | -0.579 | -1.415 | 0.02 | 0.059 | 0.059 | 3.501 | 0.41 | 0.3 | 0.35 | 0.15 | 0.3 | 0.27 |
| 11 | 19801267 | rs2896583 | -0.408 | 0.005 | -0.664 | -1.413 | 0.002 | -0.034 | -0.034 | 0.478 | 0.59 | 0.58 | 0.67 | 0.83 | 0.56 | 0.7 |
| 11 | 19801679 | rs10833168 | -0.829 | -0.196 | -0.451 | -0.063 | -0.005 | -0.05 | -0.05 | 0 | 0.14 | 0.07 | 0.35 | 0.59 | 0.19 | 0.35 |
| 11 | 19808063 | rs10766594 | -0.754 | -0.183 | -0.204 | -0.447 | 0 | 0.004 | 0.004 | 0.246 | 0.15 | 0.16 | 0.33 | 0.71 | 0.17 | 0.38 |
| 11 | 19811187 | rs7931362 | -0.457 | -0.133 | -0.221 | -0.858 | -0.004 | -0.058 | -0.058 | 0 | 0.11 | 0.2 | 0.07 | 0.01 | 0.11 | 0.05 |
| 11 | 19817809 | rs918988 | -0.758 | -0.047 | -0.418 | -0.746 | -0.007 | -0.014 | -0.014 | 0 | 0.41 | 0.51 | 0.6 | 0.73 | 0.41 | 0.57 |
| 11 | 19838663 | rs1425235 | 0.492 | -0.139 | 0.08 | -0.867 | -0.023 | -0.123 | -0.123 | 0.146 | 0.82 | 0.65 | 0.9 | 1 | 0.88 | 0.85 |
| 11 | 19841363 | rs7944986 | 0.674 | -0.442 | -0.347 | -0.846 | 0.009 | 0.01 | 0.01 | 0.81 | 0.71 | 0.27 | 0.79 | 0.88 | 0.84 | 0.78 |
| 11 | 19842316 | rs2896584 | -0.204 | -0.262 | 0.042 | -0.762 | 0.004 | -0.022 | -0.022 | 0.806 | 0.84 | 0.69 | 0.9 | 0.89 | 0.89 | 0.87 |
| 11 | 19843574 | rs11025273 | 0.76 | 0.329 | -1.142 | -0.417 | -0.046 | -0.017 | -0.017 | 0 | 0.43 | 0.19 | 0.59 | 0.55 | 0.5 | 0.44 |
| 11 | 19843968 | rs4757851 | 0.444 | 0.351 | -1.051 | -0.387 | -0.009 | 0.031 | 0.031 | 1.029 | 0.5 | 0.23 | 0.59 | 0.55 | 0.55 | 0.53 |
| 11 | 19848022 | rs6483616 | 0.302 | 0.4 | -0.356 | -0.229 | -0.008 | -0.006 | -0.006 | 0 | 0.63 | 0.55 | 0.72 | 0.65 | 0.67 | 0.59 |
| 11 | 19848918 | rs11025276 | -0.005 | 0.268 | -0.401 | -0.304 | -0.025 | -0.051 | -0.051 | 1.888 | 0.27 | 0.06 | 0.19 | 0.34 | 0.28 | 0.31 |
| 11 | 19853393 | rs11025279 | -0.223 | 0.355 | -0.517 | -0.159 | 0.004 | 0.003 | 0.003 | 0.022 | 0.42 | 0.64 | 0.39 | 0.36 | 0.34 | 0.19 |
| 11 | 19854088 | rs6483617 | -0.101 | 0.363 | -0.54 | -0.158 | 0.006 | 0.003 | 0.003 | 0 | 0.4 | 0.67 | 0.38 | 0.35 | 0.34 | 0.19 |
| 11 | 19855883 | rs1348349 | -0.723 | 0.945 | -1.144 | -0.781 | -0.004 | -0.007 | -0.007 | 0 | 0.64 | 0.81 | 0.64 | 0.38 | 0.62 | 0.33 |
| 11 | 19855954 | rs11025281 | 0.404 | 0.667 | -0.858 | -0.734 | -0.001 | 0 | 0 | 0 | 0.15 | 0.18 | 0.22 | 0.23 | 0.11 | 0.07 |
| 11 | 19860949 | rs7925366 | 0.818 | 1.172 | -1.024 | -0.869 | -0.001 | -0.046 | -0.046 | 0 | 0.16 | 0.2 | 0.13 | 0.01 | 0.16 | 0.09 |
| 11 | 19861308 | rs17531257 | -0.441 | 1.685 | -1.257 | 0.051 | -0.039 | -0.048 | -0.048 | 0 | 0.08 | 0.01 | 0.14 | 0.19 | 0.15 | 0.14 |
| 11 | 19861995 | rs1443118 | -0.132 | 1.421 | -1.104 | 0.005 | -0.007 | 0.001 | 0.001 | 1.666 | 0.58 | 0.69 | 0.57 | 0.44 | 0.52 | 0.28 |
| 11 | 19870164 | rs730348 | -0.029 | 1.272 | -1.508 | 0.222 | -0.045 | -0.056 | -0.056 | 0 | 0.66 | 0.85 | 0.57 | 0.51 | 0.56 | 0.66 |
| 11 | 19870947 | rs873764 | 0.11 | 0.908 | -1.17 | 0.053 | -0.001 | 0.005 | 0.005 | 0.083 | 0.54 | 0.62 | 0.5 | 0.41 | 0.43 | 0.6 |
| 11 | 19871270 | rs12287043 | 1.826 | 1.234 | -1.35 | 0.182 | -0.026 | -0.043 | -0.043 | 0 | 0.12 | 0.02 | 0.23 | 0.28 | 0.15 | 0.07 |
| 11 | 19871312 | rs2165802 | 1.152 | 0.868 | -1.271 | -0.31 | -0.001 | -0.007 | -0.007 | 0 | 0.15 | 0.17 | 0.15 | 0 | 0.08 | 0.07 |
| 11 | 19872846 | rs2249178 | 1.407 | 0.624 | -1.505 | -0.533 | 0.022 | -0.107 | -0.107 | 0.32 | 0.25 | 0.07 | 0.19 | 0.36 | 0.21 | 0.43 |
| 11 | 19873489 | rs2707110 | -0.211 | 0.609 | -1.075 | -0.468 | -0.002 | -0.002 | -0.002 | 0 | 0.48 | 0.57 | 0.35 | 0.34 | 0.3 | 0.53 |
| 11 | 19874227 | rs2625316 | -0.767 | 0.786 | -1.156 | -0.488 | -0.016 | -0.016 | -0.016 | 2.944 | 0.62 | 0.75 | 0.53 | 0.46 | 0.51 | 0.61 |
| 11 | 19875168 | rs716133 | 0.632 | 0.435 | -1.021 | -0.408 | -0.029 | -0.037 | -0.037 | 0 | 0.22 | 0.6 | 0.15 | 0.12 | 0.17 | 0.1 |
| 11 | 19885687 | rs2707094 | -0.674 | 0.417 | -0.997 | -0.421 | -0.007 | -0.06 | -0.06 | 0 | 0.7 | 0.57 | 0.75 | 0.87 | 0.76 | 0.86 |
| 11 | 19886909 | rs11025301 | 2.003 | 0.48 | -0.991 | -0.352 | -0.001 | 0 | 0 | 0 | 0.14 | 0.16 | 0.11 | 0.11 | 0.16 | 0.04 |

| | | | | | | | | | | | | | | | | |
|---|---|---|---|---|---|---|---|---|---|---|---|---|---|---|---|---|
| 11 | 19889078 | rs2585788 | 1.453 | 0.631 | -1.65 | -0.276 | -0.004 | -0.046 | -0.046 | 0.38 | 0.23 | 0.11 | 0.22 | 0.36 | 0.24 | 0.46 |
| 11 | 19891290 | rs894548 | 1.399 | 0.84 | -1.843 | -0.304 | -0.06 | -0.055 | -0.055 | 0 | 0.12 | 0.01 | 0.14 | 0.17 | 0.23 | 0.14 |
| 11 | 19891456 | rs2585789 | 1.541 | 0.271 | -1.442 | -0.09 | -0.031 | -0.097 | -0.097 | 0 | 0.21 | 0.03 | 0.18 | 0.32 | 0.23 | 0.46 |
| 11 | 19893621 | rs7934251 | -0.09 | 0.345 | -1.344 | -0.116 | 0.007 | 0.005 | 0.005 | 0.69 | 0.58 | 0.62 | 0.62 | 0.61 | 0.67 | 0.44 |
| 11 | 19894277 | rs2585752 | -1.233 | 0.496 | -1.597 | -0.101 | -0.007 | -0.061 | -0.061 | 0.018 | 0.84 | 0.78 | 0.86 | 1 | 0.92 | 0.94 |
| 11 | 19898874 | rs2218329 | 0.419 | 0.756 | -1.999 | 0.367 | -0.075 | -0.056 | -0.056 | 0 | 0.37 | 0.18 | 0.48 | 0.49 | 0.49 | 0.35 |
| 11 | 19903757 | rs2625312 | 0.112 | 0.639 | -1.519 | 1.002 | -0.016 | -0.109 | -0.109 | 0 | 0.24 | 0.04 | 0.18 | 0.34 | 0.24 | 0.46 |
| 11 | 19904490 | rs12575413 | -0.56 | 0.605 | -0.631 | 0.909 | -0.002 | 0.013 | 0.013 | 0 | 0.22 | 0.25 | 0.32 | 0.34 | 0.18 | 0.13 |
| 11 | 19906795 | rs2028603 | 0.815 | 0.721 | -0.305 | 0.732 | 0.003 | -0.026 | -0.026 | 0 | 0.65 | 0.46 | 0.55 | 0.26 | 0.66 | 0.45 |
| 11 | 19907563 | rs2625303 | 0.055 | 0.737 | -0.653 | 1.408 | -0.031 | -0.095 | -0.095 | 0 | 0.18 | 0.06 | 0.17 | 0.42 | 0.21 | 0.43 |
| 11 | 19908690 | rs2707096 | 0.648 | 0.678 | -0.214 | 1.346 | 0.021 | 0.06 | 0.06 | 0 | 0.87 | 0.79 | 0.77 | 0.71 | 0.9 | 0.91 |
| 11 | 19911733 | rs2707098 | 0.17 | 0.49 | -0.636 | 1.492 | -0.006 | -0.08 | -0.08 | 0 | 0.16 | 0.06 | 0.16 | 0.39 | 0.18 | 0.39 |
| 11 | 19913145 | rs2625291 | 1.097 | -0.194 | 0.151 | 1.099 | -0.011 | 0.033 | 0.033 | 0.113 | 0.82 | 0.38 | 0.73 | 0.71 | 0.89 | 0.85 |
| 11 | 19916098 | rs7935182 | -0.099 | -0.705 | -0.088 | 0.075 | -0.003 | 0.007 | 0.007 | 0 | 0.35 | 0.32 | 0.25 | 0.15 | 0.36 | 0.25 |
| 11 | 19922410 | rs4757857 | 0.942 | -1.28 | -0.787 | 0.233 | -0.147 | 0.021 | 0.021 | 0 | 0.38 | 0.05 | 0.5 | 0.31 | 0.53 | 0.37 |
| 11 | 19923122 | rs2707107 | -0.543 | -1.313 | -0.35 | 0.468 | -0.076 | 0.165 | 0.165 | 0.498 | 0.22 | 0.49 | 0.18 | 0.45 | 0.11 | 0.35 |
| 11 | 19923487 | rs2707106 | -0.795 | -1.424 | -0.514 | 0.19 | -0.02 | 0.001 | 0.001 | 0 | 0.37 | 0.47 | 0.34 | 0.55 | 0.26 | 0.44 |
| 11 | 19924341 | rs967543 | -0.356 | -1.463 | -0.33 | 0.196 | -0.056 | 0.124 | 0.124 | 0.528 | 0.22 | 0.41 | 0.17 | 0.44 | 0.11 | 0.34 |
| 11 | 19926420 | rs2028606 | -0.782 | -1.595 | -0.637 | -0.137 | 0.003 | 0.002 | 0.002 | 0 | 0.42 | 0.39 | 0.38 | 0.58 | 0.29 | 0.44 |
| 11 | 19927215 | rs2028607 | -1.121 | -1.319 | -0.309 | 0.351 | -0.019 | 0.018 | 0.018 | 0 | 0.29 | 0.34 | 0.22 | 0.5 | 0.15 | 0.32 |
| 11 | 19928344 | rs2028608 | -0.922 | -1.324 | -0.407 | 0.237 | 0.004 | 0.041 | 0.041 | 0 | 0.58 | 0.32 | 0.67 | 0.76 | 0.64 | 0.69 |
| 11 | 19928395 | rs2028609 | 0.536 | -1.386 | -0.268 | 0.202 | -0.05 | 0.007 | 0.007 | 0 | 0.34 | 0.18 | 0.48 | 0.35 | 0.51 | 0.37 |
| 11 | 19931051 | rs7131523 | -1.061 | -1.202 | -0.478 | 0.398 | 0.04 | -0.05 | -0.05 | 0 | 0.27 | 0.16 | 0.2 | 0.4 | 0.14 | 0.3 |
| 11 | 19933123 | rs10766603 | 1.024 | -0.884 | -0.526 | 0.461 | -0.158 | -0.035 | -0.035 | 1.023 | 0.28 | 0.02 | 0.39 | 0.27 | 0.47 | 0.33 |
| 11 | 19935832 | rs2707100 | -1.045 | -1.395 | 0.066 | 0.494 | -0.086 | 0.148 | 0.148 | 0.036 | 0.21 | 0.42 | 0.15 | 0.43 | 0.08 | 0.41 |
| 11 | 19936835 | rs10833190 | -1.218 | -1.371 | -0.619 | 0.533 | -0.003 | 0.001 | 0.001 | 0 | 0.26 | 0.2 | 0.18 | 0.42 | 0.14 | 0.3 |
| 11 | 19937855 | rs10833192 | 0.438 | -0.615 | -1.102 | 0.367 | -0.133 | -0.064 | -0.064 | 0 | 0.29 | 0.05 | 0.39 | 0.33 | 0.47 | 0.4 |
| 11 | 19946239 | rs920719 | -0.852 | -0.571 | -0.791 | 0.486 | 0.006 | 0.056 | 0.056 | 0 | 0.51 | 0.38 | 0.47 | 0.7 | 0.58 | 0.66 |
| 11 | 19947749 | rs7118085 | 0.021 | -0.741 | -0.907 | 0.507 | -0.021 | 0.057 | 0.057 | 0 | 0.15 | 0.19 | 0.17 | 0.35 | 0.1 | 0.28 |
| 11 | 19948073 | rs2707081 | 0.977 | -0.801 | -1.065 | 0.433 | -0.021 | 0.108 | 0.108 | 0 | 0.43 | 0.17 | 0.45 | 0.27 | 0.4 | 0.25 |
| 11 | 19952664 | rs4757862 | -1.107 | -0.238 | -1.527 | 0.825 | 0.014 | 0.053 | 0.053 | 0 | 0.52 | 0.37 | 0.53 | 0.65 | 0.59 | 0.66 |
| 11 | 19953492 | rs10833197 | -0.071 | -0.209 | -1.577 | 1.072 | -0.158 | -0.036 | -0.036 | 0 | 0.37 | 0.04 | 0.34 | 0.33 | 0.51 | 0.38 |
| 11 | 19955500 | rs1372989 | 0.276 | -0.496 | -0.998 | 1.507 | -0.043 | 0.14 | 0.14 | 0 | 0.84 | 0.65 | 0.79 | 0.64 | 0.91 | 0.73 |
| 11 | 19956427 | rs894557 | -0.604 | -0.376 | -1.408 | 1.342 | -0.028 | 0.093 | 0.093 | 0 | 0.16 | 0.27 | 0.19 | 0.42 | 0.09 | 0.37 |
| 11 | 19960147 | rs12807015 | 0.21 | 0.56 | -1.897 | 1.147 | -0.14 | -0.077 | -0.077 | 3.789 | 0.26 | 0.03 | 0.32 | 0.29 | 0.42 | 0.29 |
| 11 | 19960636 | rs10766606 | -0.068 | 0.41 | -1.801 | 0.929 | -0.13 | -0.134 | -0.134 | 0 | 0.33 | 0.06 | 0.47 | 0.45 | 0.48 | 0.32 |
| 11 | 19962033 | rs2585769 | -0.242 | 0.539 | -1.525 | 0.972 | -0.023 | -0.027 | -0.027 | 0 | 0.4 | 0.29 | 0.51 | 0.48 | 0.49 | 0.5 |
| 11 | 19963642 | rs11025333 | -0.951 | 0.116 | -1.77 | 1.21 | 0.013 | -0.072 | -0.072 | 0.306 | 0.26 | 0.07 | 0.25 | 0.53 | 0.2 | 0.31 |
| 11 | 19970507 | rs12284679 | -0.672 | 0.637 | -1.805 | 1.02 | -0.031 | 0.002 | 0.002 | 0 | 0.06 | 0 | 0.09 | 0.02 | 0.08 | 0.21 |
| 11 | 19971439 | rs2018644 | 0.482 | 0.536 | -1.043 | 0.944 | -0.001 | -0.002 | -0.002 | 0 | 0.93 | 0.94 | 0.9 | 0.98 | 0.92 | 0.85 |
| 11 | 19972603 | rs11025336 | -0.721 | 0.32 | -1.498 | 1.892 | -0.032 | -0.113 | -0.113 | 0.219 | 0.16 | 0.01 | 0.15 | 0.37 | 0.16 | 0.22 |
| 11 | 19973768 | rs7131051 | 0.761 | 0.278 | -2.116 | 1.472 | -0.065 | -0.01 | -0.01 | 0.169 | 0.35 | 0.13 | 0.42 | 0.3 | 0.37 | 0.37 |
| 11 | 19975876 | rs2165800 | -0.008 | -0.361 | -1.342 | 1.284 | 0.055 | 0.067 | 0.067 | 0.419 | 0.55 | 0.23 | 0.61 | 0.69 | 0.64 | 0.7 |
| 11 | 19975906 | rs2165801 | 0.063 | -0.497 | -1.103 | 1.139 | 0.005 | -0.014 | -0.014 | 0 | 0.92 | 0.88 | 0.9 | 0.98 | 0.92 | 0.85 |
| 11 | 19977594 | rs730000 | 0.351 | -0.515 | -0.682 | 1.286 | -0.026 | -0.006 | -0.006 | 0 | 0.37 | 0.58 | 0.3 | 0.34 | 0.28 | 0.15 |
| 11 | 19977718 | rs729999 | -0.054 | -0.388 | -0.829 | 1.36 | 0.039 | 0.025 | 0.025 | 0.38 | 0.55 | 0.27 | 0.6 | 0.65 | 0.64 | 0.68 |
| 11 | 19983180 | rs10833206 | 1.119 | -0.041 | -0.776 | 1.358 | -0.027 | 0.12 | 0.12 | 1.056 | 0.46 | 0.13 | 0.46 | 0.3 | 0.54 | 0.4 |
| 11 | 19984811 | rs10766609 | 0.798 | -0.057 | 0.033 | 1.399 | -0.002 | -0.012 | -0.012 | 0 | 0.53 | 0.49 | 0.5 | 0.33 | 0.55 | 0.46 |
| 11 | 19985501 | rs7950088 | 0.769 | -0.108 | -0.135 | 1.365 | -0.01 | 0.074 | 0.074 | 1.325 | 0.56 | 0.77 | 0.52 | 0.32 | 0.56 | 0.46 |
| 11 | 19987362 | rs7106847 | 0.662 | -0.055 | -0.471 | 1.937 | -0.059 | -0.126 | -0.126 | 0.058 | 0.82 | 0.98 | 0.78 | 0.63 | 0.79 | 0.65 |
| 11 | 19988412 | rs2403547 | -0.77 | 0.039 | 0.323 | 1.839 | -0.008 | 0.007 | 0.007 | 0 | 0.33 | 0.44 | 0.32 | 0.35 | 0.3 | 0.3 |
| 11 | 19988487 | rs4447157 | 0.014 | 0.094 | 0.202 | 2.172 | -0.026 | -0.018 | -0.018 | 0.189 | 0.93 | 0.95 | 0.87 | 0.75 | 0.88 | 0.84 |
| 11 | 19988569 | rs4429037 | 0.743 | -0.097 | 0.224 | 1.566 | 0.001 | -0.052 | -0.052 | 0 | 0.73 | 0.89 | 0.66 | 0.43 | 0.71 | 0.59 |
| 11 | 19990087 | rs4757872 | 1.433 | -0.069 | 0.544 | 0.605 | 0.003 | 0.013 | 0.013 | 0 | 0.68 | 0.46 | 0.72 | 0.41 | 0.71 | 0.65 |
| 11 | 19990226 | rs10766610 | 2.432 | -0.115 | 0.39 | 0.56 | -0.09 | 0.262 | 0.262 | 0.145 | 0.47 | 0.08 | 0.46 | 0.16 | 0.52 | 0.39 |
| 11 | 19990758 | rs2625300 | 0.499 | -0.047 | 0.415 | 0.659 | -0.055 | -0.099 | -0.099 | 0.239 | 0.86 | 0.95 | 0.82 | 0.51 | 0.83 | 0.72 |
| 11 | 19992797 | rs2251973 | 0.935 | -0.17 | 0.652 | 0.579 | 0 | -0.014 | -0.014 | 0.163 | 0.11 | 0.19 | 0.07 | 0.05 | 0.08 | 0.04 |
| 11 | 19993776 | rs7933147 | 1.945 | -0.215 | 0.803 | 0.338 | 0.003 | 0.001 | 0.001 | 0 | 0.52 | 0.44 | 0.53 | 0.18 | 0.57 | 0.45 |
| 11 | 19995837 | rs11604874 | 0.38 | 0.181 | 0.818 | 0.674 | 0.002 | -0.003 | -0.003 | 0 | 0.16 | 0.14 | 0.19 | 0.23 | 0.15 | 0.19 |
| 11 | 19998919 | rs2625295 | 0.544 | 0.652 | 1.542 | 0.912 | 0.018 | -0.057 | -0.057 | 0 | 0.83 | 0.63 | 0.85 | 0.92 | 0.85 | 0.89 |
| 11 | 20003227 | rs7931593 | -1.065 | 0.655 | 1.208 | 0.915 | 0.012 | -0.051 | -0.051 | 0 | 0.19 | 0.32 | 0.16 | 0.08 | 0.19 | 0.14 |
| 11 | 20004148 | rs7129191 | -1.112 | 0.651 | 1.207 | 0.913 | 0.011 | -0.052 | -0.052 | 0 | 0.19 | 0.32 | 0.16 | 0.08 | 0.19 | 0.16 |
| 11 | 20005452 | rs2568118 | -1.275 | 0.553 | 0.65 | 0.94 | -0.005 | -0.037 | -0.037 | 0 | 0.25 | 0.19 | 0.2 | 0.53 | 0.21 | 0.28 |
| 11 | 20007542 | rs10833213 | 1.753 | 0.55 | 0.691 | 0.932 | -0.002 | 0 | 0 | 0 | 0.63 | 0.37 | 0.65 | 0.4 | 0.66 | 0.56 |
| 11 | 20008532 | rs7130917 | -0.594 | 0.469 | 0.297 | 0.936 | -0.011 | -0.056 | -0.056 | 0 | 0.18 | 0.33 | 0.09 | 0.09 | 0.12 | 0.07 |
| 11 | 20009445 | rs17614100 | 0.51 | 0.496 | 0.304 | 0.946 | -0.032 | 0.027 | 0.027 | 0 | 0.06 | 0.01 | 0.09 | 0.03 | 0.1 | 0.12 |
| 11 | 20009559 | rs12099330 | -0.565 | 0.495 | 0.304 | 0.946 | -0.01 | -0.055 | -0.055 | 0 | 0.18 | 0.33 | 0.09 | 0.09 | 0.12 | 0.07 |
| 11 | 20009681 | rs12099071 | 1.307 | 0.548 | 0.333 | 0.999 | 0.025 | 0.125 | 0.125 | 0 | 0.77 | 0.5 | 0.72 | 0.49 | 0.77 | 0.62 |
| 11 | 20010351 | rs2625322 | -1.287 | 0.646 | 0.052 | 1.052 | -0.021 | -0.053 | -0.053 | 0 | 0.2 | 0.15 | 0.24 | 0.51 | 0.22 | 0.37 |

| | | | | | | | | | | | | | | | | |
|---|---|---|---|---|---|---|---|---|---|---|---|---|---|---|---|---|
| 11 | 20011154 | rs2584847 | -0.679 | 0.424 | 0.149 | 1.16 | -0.004 | -0.065 | -0.065 | 1.363 | 0.16 | 0.1 | 0.15 | 0.49 | 0.12 | 0.26 |
| 11 | 20011537 | rs2568135 | -0.539 | 0.577 | 0.21 | 1.244 | -0.001 | -0.016 | -0.016 | 0 | 0.16 | 0.15 | 0.15 | 0.49 | 0.12 | 0.26 |
| 11 | 20016559 | rs7111951 | 0.42 | 0.654 | 0.117 | 1.248 | -0.002 | 0.021 | 0.021 | 0 | 0.81 | 0.76 | 0.84 | 0.51 | 0.87 | 0.74 |
| 11 | 20016618 | rs7112354 | -0.393 | 0.706 | -0.393 | 0.943 | 0.008 | -0.033 | -0.033 | 0.496 | 0.14 | 0.26 | 0.09 | 0.08 | 0.12 | 0.07 |
| 11 | 20019137 | rs4237746 | 0.653 | 1.155 | -1.524 | 1.159 | -0.056 | 0.011 | 0.011 | 0 | 0.56 | 0.17 | 0.49 | 0.42 | 0.59 | 0.52 |
| 11 | 20019657 | rs10500866 | -0.581 | 1.525 | -0.839 | 1.78 | -0.015 | -0.017 | -0.017 | 0 | 0.12 | 0.09 | 0.19 | 0.32 | 0.16 | 0.14 |
| 11 | 20020097 | rs4757879 | -0.291 | 1.041 | -0.455 | 1.108 | -0.004 | -0.002 | -0.002 | 0.434 | 0.42 | 0.6 | 0.63 | 0.52 | 0.46 | 0.57 |
| 11 | 20020564 | rs11601815 | -1.49 | 0.423 | 0.649 | 0.671 | -0.027 | -0.068 | -0.068 | 0.47 | 0.17 | 0.47 | 0.1 | 0.04 | 0.09 | 0.19 |
| 11 | 20020884 | rs10741808 | 0.965 | 0.521 | 0.666 | 0.796 | -0.011 | -0.016 | -0.016 | 0 | 0.77 | 0.64 | 0.85 | 0.78 | 0.82 | 0.93 |
| 11 | 20023619 | rs2584840 | 1.165 | 0.847 | 0.564 | 0.991 | 0.007 | 0.012 | 0.012 | 0 | 0.89 | 0.86 | 0.71 | 0.85 | 0.86 | 0.81 |
| 11 | 20024139 | rs16937344 | -0.855 | 0.662 | 0.434 | 0.878 | 0 | -0.001 | -0.001 | 0 | 0.07 | 0.08 | 0.02 | 0.09 | 0.06 | 0.13 |
| 11 | 20030430 | rs11606785 | -0.523 | 0.718 | 0.306 | 0.989 | -0.021 | -0.033 | -0.033 | 0 | 0.06 | 0.01 | 0.03 | 0.11 | 0.07 | 0.14 |
| 11 | 20031390 | rs12286318 | -1.449 | 0.46 | 1.072 | 1.352 | -0.025 | 0.031 | 0.031 | 2.937 | 0.21 | 0.4 | 0.14 | 0.29 | 0.15 | 0.06 |
| 11 | 20031865 | rs11605275 | -0.773 | 0.566 | 0.621 | 1.437 | 0 | -0.004 | -0.004 | 0 | 0.08 | 0.08 | 0.03 | 0.09 | 0.06 | 0.13 |
| 11 | 20037120 | rs11025356 | -0.576 | 0.562 | 0.593 | 1.426 | -0.019 | -0.065 | -0.065 | 0 | 0.13 | 0.02 | 0.1 | 0.29 | 0.12 | 0.06 |
| 11 | 20043439 | rs2584834 | -1.63 | 0.884 | 0.625 | 1.228 | -0.003 | -0.003 | -0.003 | 0 | 0.12 | 0.1 | 0.09 | 0.15 | 0.14 | 0.17 |
| 11 | 20044696 | rs2449501 | -1.667 | 0.7 | 1.08 | 0.923 | -0.004 | -0.021 | -0.021 | 0 | 0.17 | 0.31 | 0.21 | 0.1 | 0.15 | 0.18 |
| 11 | 20046407 | rs2568131 | -1.609 | 0.521 | 1.279 | 0.855 | -0.027 | -0.015 | -0.015 | 0 | 0.16 | 0.44 | 0.08 | 0.1 | 0.09 | 0.13 |
| 11 | 20046848 | rs12577511 | -0.712 | 0.797 | 1.037 | 1.123 | -0.002 | -0.007 | -0.007 | 0 | 0.13 | 0.11 | 0.1 | 0.19 | 0.14 | 0.06 |
| 11 | 20048273 | rs12807657 | -1.838 | 0.915 | 1.213 | 1.104 | 0.012 | -0.002 | -0.002 | 0 | 0.06 | 0.12 | 0.05 | 0.04 | 0.07 | 0.05 |
| 11 | 20048928 | rs7101488 | -1.451 | 0.747 | 1.515 | 1.305 | 0.006 | 0.055 | 0.055 | 0 | 0.08 | 0.14 | 0.08 | 0.2 | 0.09 | 0.05 |
| 11 | 20049859 | rs4757880 | 1.336 | -0.195 | 1.185 | 0.581 | 0.032 | 0.018 | 0.018 | 0.5 | 0.54 | 0.25 | 0.7 | 0.55 | 0.67 | 0.67 |
| 11 | 20050052 | rs920671 | 1.924 | -0.239 | 1.994 | 0.586 | 0.034 | 0.091 | 0.091 | 0.469 | 0.84 | 0.6 | 0.87 | 0.77 | 0.84 | 0.89 |
| 11 | 20050205 | rs2006636 | 1.473 | -0.232 | 2.32 | 0.619 | 0.051 | 0.103 | 0.103 | 0 | 0.87 | 0.63 | 0.88 | 0.8 | 0.87 | 0.93 |
| 11 | 20054591 | rs11025359 | -1.597 | -0.234 | 2.136 | 0.595 | 0.026 | 0.054 | 0.054 | 0 | 0.12 | 0.26 | 0.11 | 0.19 | 0.12 | 0.07 |
| 11 | 20055361 | rs12807369 | -0.531 | -0.283 | 1.849 | 0.486 | 0.025 | 0.064 | 0.064 | 2.78 | 0.3 | 0.13 | 0.13 | 0.22 | 0.22 | 0.27 |
| 11 | 20055663 | rs12276043 | 1.639 | -0.308 | 1.956 | 0.487 | 0.028 | 0.029 | 0.029 | 0.216 | 0.52 | 0.27 | 0.69 | 0.54 | 0.62 | 0.63 |
| 11 | 20056273 | rs7131550 | -1.935 | -0.418 | 2.04 | 0.435 | 0.017 | 0.016 | 0.016 | 0.263 | 0.48 | 0.62 | 0.29 | 0.46 | 0.39 | 0.38 |
| 11 | 20057158 | rs4757882 | -0.247 | -0.454 | 1.781 | 0.368 | 0.02 | 0.057 | 0.057 | 2.482 | 0.28 | 0.12 | 0.12 | 0.17 | 0.19 | 0.24 |
| 11 | 20058093 | rs10741810 | 0.399 | -0.561 | 1.729 | 0.262 | -0.005 | -0.007 | -0.007 | 0.868 | 0.68 | 0.4 | 0.83 | 0.74 | 0.8 | 0.77 |
| 11 | 20060147 | rs7126277 | -0.618 | 0.833 | 1.073 | -0.256 | -0.001 | -0.011 | -0.011 | 0 | 0.39 | 0.34 | 0.22 | 0.2 | 0.43 | 0.27 |
| 11 | 20062704 | rs12795296 | 1.343 | 0.777 | 0.842 | -0.18 | 0.026 | 0.009 | 0.009 | 0.025 | 0.45 | 0.31 | 0.62 | 0.53 | 0.39 | 0.58 |
| 11 | 20063760 | rs10833229 | 0.353 | 0.826 | 0.896 | -0.14 | 0 | -0.007 | -0.007 | 0.111 | 0.59 | 0.45 | 0.72 | 0.73 | 0.56 | 0.66 |
| 11 | 20065673 | rs3802799 | 1.398 | 0.449 | -0.289 | -0.423 | -0.04 | -0.14 | -0.14 | 0.018 | 0.17 | 0.02 | 0.27 | 0.45 | 0.22 | 0.47 |
| 11 | 20067064 | rs7125647 | -0.562 | 0.629 | -0.263 | -0.874 | 0.006 | -0.058 | -0.058 | 0.258 | 0.61 | 0.6 | 0.66 | 0.83 | 0.54 | 0.7 |
| 11 | 20068489 | rs4757026 | 0.086 | 0.582 | -0.226 | -0.809 | 0.003 | -0.061 | -0.061 | 0 | 0.45 | 0.45 | 0.32 | 0.19 | 0.54 | 0.33 |
| 11 | 20078496 | rs11025365 | 0.02 | 0.409 | -0.403 | 0.181 | -0.022 | 0.043 | 0.043 | 0 | 0.23 | 0.33 | 0.51 | 0.55 | 0.2 | 0.49 |
| 11 | 20079333 | rs1372796 | 0.074 | 0.646 | -0.95 | -0.28 | -0.002 | -0.037 | -0.037 | 0.004 | 0.12 | 0.13 | 0.07 | 0 | 0.08 | 0.06 |
| 11 | 20086764 | rs1978947 | -0.181 | 1.093 | -1.141 | 0.361 | -0.016 | -0.065 | -0.065 | 0 | 0.26 | 0.13 | 0.4 | 0.51 | 0.33 | 0.57 |
| 11 | 20087212 | rs12785600 | 0.458 | 0.965 | -0.962 | -0.334 | -0.016 | 0.083 | 0.083 | 0.502 | 0.55 | 0.32 | 0.42 | 0.21 | 0.49 | 0.37 |
| 11 | 20089944 | rs1442710 | -0.668 | 0.515 | -0.366 | -0.34 | -0.018 | 0.069 | 0.069 | 0.151 | 0.23 | 0.35 | 0.29 | 0.55 | 0.19 | 0.17 |
| 11 | 20090650 | rs11025367 | -0.083 | 0.81 | 0.12 | -0.653 | 0.018 | -0.009 | -0.009 | 0.049 | 0.12 | 0.3 | 0.21 | 0.08 | 0.14 | 0.2 |
| 11 | 20090841 | rs11824188 | -1.132 | 0.88 | -0.422 | -0.959 | -0.003 | -0.007 | -0.007 | 0 | 0.38 | 0.54 | 0.53 | 0.65 | 0.39 | 0.37 |
| 11 | 20091148 | rs7951443 | -0.056 | 0.324 | -0.503 | -1.36 | -0.037 | 0.06 | 0.06 | 0 | 0.22 | 0.41 | 0.17 | 0.35 | 0.14 | 0.27 |
| 11 | 20091724 | rs6483639 | 0.219 | 0.56 | -0.453 | -1.545 | -0.015 | -0.014 | -0.014 | 0 | 0.3 | 0.44 | 0.28 | 0.28 | 0.26 | 0.46 |
| 11 | 20091919 | rs6483640 | -0.904 | 0.667 | -0.602 | -1.659 | -0.001 | 0.035 | 0.035 | 0 | 0.52 | 0.65 | 0.68 | 0.93 | 0.49 | 0.68 |
| 11 | 20098298 | rs11025378 | 0.098 | 0.467 | -0.395 | -1.377 | -0.022 | 0.002 | 0.002 | 0 | 0.3 | 0.53 | 0.4 | 0.33 | 0.25 | 0.47 |
| 11 | 20098670 | rs10734289 | -0.57 | 1.318 | -0.078 | -0.977 | 0.002 | 0.011 | 0.011 | 0 | 0.25 | 0.27 | 0.38 | 0.65 | 0.29 | 0.19 |
| 11 | 20099399 | rs6483641 | -0.531 | 0.56 | -0.355 | -1.301 | 0.004 | 0.007 | 0.007 | 0 | 0.85 | 0.87 | 0.88 | 0.96 | 0.87 | 0.92 |
| 11 | 20099541 | rs6483642 | -0.485 | 0.587 | -0.327 | -1.276 | 0.002 | 0.002 | 0.002 | 0 | 0.86 | 0.88 | 0.89 | 0.96 | 0.89 | 0.92 |
| 11 | 20100381 | rs4757031 | -1.021 | 0.582 | -0.332 | -1.318 | 0.001 | 0.012 | 0.012 | 0 | 0.87 | 0.91 | 0.89 | 0.96 | 0.89 | 0.95 |
| 11 | 20100472 | rs10741815 | -0.701 | 0.583 | -0.333 | -1.318 | 0.001 | 0.001 | 0.001 | 0 | 0.86 | 0.89 | 0.89 | 0.96 | 0.88 | 0.95 |
| 11 | 20101744 | rs1867115 | -0.721 | 0.583 | -0.419 | -1.304 | 0.001 | 0.043 | 0.043 | 0 | 0.86 | 0.96 | 0.89 | 0.96 | 0.88 | 0.95 |
| 11 | 20102251 | rs7103409 | 0.401 | 0.516 | -0.232 | -1.316 | 0.004 | 0.021 | 0.021 | 0 | 0.59 | 0.37 | 0.46 | 0.34 | 0.6 | 0.72 |
| 11 | 20104669 | rs2028570 | 0.719 | 0.878 | -0.185 | -1.097 | 0 | 0.065 | 0.065 | 0.026 | 0.56 | 0.23 | 0.43 | 0.31 | 0.58 | 0.72 |
| 11 | 20107179 | rs7124576 | -0.764 | 1.484 | -0.315 | -0.934 | -0.005 | -0.059 | -0.059 | 0.037 | 0.81 | 0.73 | 0.85 | 0.96 | 0.83 | 0.92 |
| 11 | 20107602 | rs11025380 | 0.282 | 1.761 | -0.446 | -1.054 | -0.049 | 0.051 | 0.051 | 0 | 0.16 | 0.05 | 0.1 | 0.04 | 0.21 | 0.14 |
| 11 | 20108492 | rs11025381 | 1.235 | 1.708 | 0.172 | -0.328 | -0.004 | 0.017 | 0.017 | 0 | 0.56 | 0.47 | 0.41 | 0.32 | 0.51 | 0.73 |
| 11 | 20108966 | rs2005163 | -0.001 | 1.756 | -0.023 | -0.207 | 0.011 | 0.067 | 0.067 | 0 | 0.18 | 0.24 | 0.37 | 0.59 | 0.22 | 0.14 |
| 11 | 20112373 | rs2289560 | -0.017 | 1.278 | 0.195 | -0.142 | 0.018 | 0.017 | 0.017 | 0.161 | 0.23 | 0.25 | 0.43 | 0.63 | 0.31 | 0.17 |
| 11 | 20112417 | rs2243624 | -0.103 | 1.334 | -0.63 | -0.114 | 0.066 | 0.064 | 0.064 | 0.355 | 0.24 | 0.46 | 0.45 | 0.63 | 0.31 | 0.17 |
| 11 | 20112492 | rs1837971 | -0.139 | 0.878 | -0.465 | 0.279 | -0.002 | 0.004 | 0.004 | 0 | 0.57 | 0.48 | 0.42 | 0.33 | 0.51 | 0.74 |
| 11 | 20119298 | rs2289564 | -0.726 | 0.705 | -0.36 | 1.021 | -0.004 | -0.004 | -0.004 | 0 | 0.16 | 0.12 | 0.1 | 0.21 | 0.21 | 0.41 |
| 11 | 20119312 | rs2289563 | -0.849 | 0.822 | -0.376 | 1.131 | -0.008 | -0.01 | -0.01 | 0 | 0.16 | 0.12 | 0.1 | 0.29 | 0.22 | 0.41 |
| 11 | 20119506 | rs2289561 | -0.845 | 0.711 | -0.369 | 1.139 | -0.004 | -0.01 | -0.01 | 0 | 0.16 | 0.12 | 0.11 | 0.29 | 0.23 | 0.41 |
| 11 | 20120853 | rs10741818 | -0.692 | 0.59 | -0.359 | 1.153 | -0.004 | -0.01 | -0.01 | 0 | 0.15 | 0.11 | 0.12 | 0.29 | 0.24 | 0.41 |
| 11 | 20124914 | rs2243411 | 0.975 | 0.092 | 0.806 | 1.213 | -0.014 | 0.026 | 0.026 | 0 | 0.94 | 0.78 | 0.97 | 0.96 | 0.99 | 0.95 |
| 11 | 20132086 | rs7102814 | 1.185 | -0.289 | 1.127 | 2.069 | -0.015 | 0.052 | 0.052 | 5.004 | 0.81 | 0.31 | 0.69 | 0.59 | 0.88 | 0.84 |
| 11 | 20133413 | rs4757893 | 1.178 | -0.366 | 2.008 | 2.173 | -0.017 | 0.053 | 0.053 | 3.978 | 0.81 | 0.32 | 0.71 | 0.57 | 0.88 | 0.84 |

| | | | | | | | | | | | | | | | | |
|---|---|---|---|---|---|---|---|---|---|---|---|---|---|---|---|---|
| 11 | 20134433 | rs1867125 | 1.239 | -0.428 | 2.678 | 2.258 | 0.045 | 0.471 | 0.471 | 0 | 0.91 | 0.59 | 0.74 | 0.57 | 0.92 | 0.86 |
| 11 | 20134869 | rs10766627 | 2.159 | -0.831 | 2.194 | 2.215 | 0.016 | -0.005 | -0.005 | 0.02 | 0.73 | 0.26 | 0.66 | 0.37 | 0.84 | 0.8 |
| 11 | 20135750 | rs10833247 | -1.952 | -0.72 | 2.532 | 2.132 | 0.047 | 0 | 0 | 0.814 | 0.07 | 0.3 | 0.05 | 0 | 0.05 | 0.02 |
| 11 | 20138670 | rs2249878 | -2.36 | -0.706 | 2.346 | 2.28 | 0.011 | 0.328 | 0.328 | 0 | 0.08 | 0.27 | 0.25 | 0.37 | 0.08 | 0.06 |
| 11 | 20139937 | rs3802803 | -1.799 | -0.525 | 2.165 | 2.364 | 0.02 | 0.232 | 0.232 | 0 | 0.06 | 0.14 | 0.23 | 0.37 | 0.1 | 0.08 |
| 11 | 20140471 | rs894445 | 1.689 | -0.823 | 2.39 | 2.363 | -0.045 | 0.033 | 0.033 | 0.992 | 0.84 | 0.44 | 0.84 | 0.75 | 0.91 | 0.89 |
| 11 | 20141839 | rs2246192 | -2.093 | -0.279 | 1.496 | 2.022 | -0.016 | -0.018 | -0.018 | 0 | 0.26 | 0.61 | 0.38 | 0.77 | 0.21 | 0.21 |
| 11 | 20148569 | rs831457 | 0.438 | -0.1 | 0.86 | 1.67 | -0.01 | 0 | 0 | 0 | 0.85 | 0.81 | 0.89 | 0.88 | 0.89 | 0.71 |
| 11 | 20150777 | rs4757895 | 0.965 | -1.131 | 0.698 | 1.916 | 0.043 | -0.029 | -0.029 | 0 | 0.61 | 0.33 | 0.71 | 0.54 | 0.76 | 0.5 |
| 11 | 20154539 | rs1348303 | 1.098 | -0.353 | -0.314 | 0.935 | -0.044 | 0.067 | 0.067 | 0 | 0.23 | 0.12 | 0.18 | 0.09 | 0.31 | 0.22 |
| 11 | 20156166 | rs831467 | -0.91 | -0.19 | 0.048 | 1.123 | 0.002 | 0 | 0 | 0 | 0.68 | 0.67 | 0.6 | 0.61 | 0.65 | 0.75 |
| 11 | 20157726 | rs7927739 | 0.763 | -0.322 | -0.525 | 1.035 | -0.007 | -0.003 | -0.003 | 0 | 0.17 | 0.1 | 0.28 | 0.2 | 0.2 | 0.16 |
| 11 | 20158569 | rs1520912 | 0.348 | -0.571 | -0.1 | 1.172 | 0.024 | -0.049 | -0.049 | 0 | 0.28 | 0.25 | 0.28 | 0.47 | 0.19 | 0.21 |
| 11 | 20159583 | rs1520905 | 0.238 | -0.439 | 0.188 | 1.166 | -0.02 | 0.038 | 0.038 | 0.014 | 0.19 | 0.43 | 0.21 | 0.19 | 0.14 | 0.34 |
| 11 | 20159603 | rs1568444 | 0.339 | -0.41 | -0.022 | 1.114 | 0.026 | -0.037 | -0.037 | 0.284 | 0.28 | 0.25 | 0.21 | 0.41 | 0.18 | 0.2 |
| 11 | 20160965 | rs961746 | -0.466 | -0.394 | -0.016 | 1.097 | 0.005 | 0.007 | 0.007 | 0 | 0.33 | 0.51 | 0.48 | 0.38 | 0.37 | 0.5 |
| 11 | 20161913 | rs831470 | -0.493 | -0.579 | -0.332 | 0.879 | 0 | -0.001 | -0.001 | 0 | 0.33 | 0.66 | 0.41 | 0.39 | 0.31 | 0.49 |
| 11 | 20161931 | rs831471 | -0.686 | -0.287 | -0.451 | 0.3 | -0.004 | 0.108 | 0.108 | 0.181 | 0.57 | 0.77 | 0.68 | 0.8 | 0.46 | 0.67 |
| 11 | 20162806 | rs831473 | -0.172 | -0.261 | -0.15 | 0.657 | -0.01 | 0.013 | 0.013 | 0 | 0.15 | 0.29 | 0.18 | 0.13 | 0.12 | 0.39 |
| 11 | 20163318 | rs12280260 | 0.602 | 0.527 | -1.579 | 0.052 | -0.06 | 0.014 | 0.014 | 0.138 | 0.27 | 0.03 | 0.27 | 0.17 | 0.28 | 0.17 |
| 11 | 20164332 | rs11025411 | -0.06 | 0.342 | -1.333 | 0.38 | -0.001 | -0.01 | -0.01 | 0 | 0.19 | 0.15 | 0.15 | 0.3 | 0.19 | 0.4 |
| 11 | 20165500 | rs1442717 | -1.298 | 0.036 | -1.102 | 0.094 | -0.026 | 0.035 | 0.035 | 0.19 | 0.7 | 0.62 | 0.63 | 0.53 | 0.76 | 0.72 |
| 11 | 20166668 | rs11025413 | 1.398 | 0.331 | -1.306 | 0.396 | -0.01 | -0.02 | -0.02 | 0 | 0.12 | 0.08 | 0.11 | 0.2 | 0.15 | 0.38 |
| 11 | 20168139 | rs1520906 | 0.742 | 0.78 | -1.245 | -0.122 | -0.11 | 0.03 | 0.03 | 0.121 | 0.32 | 0.04 | 0.32 | 0.17 | 0.37 | 0.21 |
| 11 | 20174299 | rs831460 | -0.535 | 0.742 | -0.683 | -0.133 | 0.004 | -0.024 | -0.024 | 0 | 0.17 | 0.28 | 0.11 | 0.12 | 0.17 | 0.1 |
| 11 | 20175202 | rs831461 | -0.507 | 0.713 | -0.663 | -0.23 | 0.013 | 0.034 | 0.034 | 0.199 | 0.5 | 0.55 | 0.46 | 0.31 | 0.54 | 0.32 |
| 11 | 20176986 | rs10833256 | 0.592 | 0.835 | -1.27 | -0.268 | -0.144 | 0.017 | 0.017 | 0.006 | 0.34 | 0.03 | 0.33 | 0.2 | 0.39 | 0.23 |
| 11 | 20178717 | rs831464 | -0.259 | 0.291 | -0.321 | -0.241 | -0.031 | -0.064 | -0.064 | 0 | 0.14 | 0.24 | 0.08 | 0.02 | 0.08 | 0.02 |
| 11 | 20179231 | rs2139751 | 0.031 | 0.968 | -0.899 | 0.092 | -0.005 | 0.099 | 0.099 | 0 | 0.49 | 0.26 | 0.46 | 0.68 | 0.48 | 0.62 |
| 11 | 20184958 | rs10833259 | 0.387 | 0.577 | -1.353 | 0.355 | 0.008 | 0.04 | 0.04 | 0.152 | 0.36 | 0.12 | 0.23 | 0.27 | 0.32 | 0.52 |
| 11 | 20188883 | rs1520903 | 0.962 | 1.071 | -1.014 | 0.68 | -0.074 | -0.108 | -0.108 | 0.007 | 0.21 | 0.05 | 0.36 | 0.35 | 0.31 | 0.17 |
| 11 | 20191933 | rs1402376 | -0.074 | 0.274 | -0.559 | -0.181 | -0.004 | -0.015 | -0.015 | 0 | 0.6 | 0.58 | 0.7 | 0.78 | 0.67 | 0.72 |
| 11 | 20195292 | rs1520910 | -0.197 | 0.143 | -0.191 | -0.126 | -0.009 | -0.02 | -0.02 | 0 | 0.74 | 0.53 | 0.78 | 0.77 | 0.76 | 0.81 |
| 11 | 20196522 | rs11821968 | 0.541 | -0.062 | -0.188 | 0.005 | -0.009 | -0.008 | -0.008 | 0.714 | 0.18 | 0.22 | 0.15 | 0.13 | 0.13 | 0.11 |
| 11 | 20199347 | rs10766633 | -1.848 | 0.24 | -0.404 | -0.15 | -0.015 | 0.011 | 0.011 | 0 | 0.94 | 0.98 | 0.94 | 1 | 0.89 | 0.97 |
| 11 | 20199610 | rs7941540 | -1.267 | 0.268 | -0.45 | -0.228 | -0.05 | 0.014 | 0.014 | 0 | 0.71 | 0.86 | 0.62 | 0.76 | 0.61 | 0.79 |
| 11 | 20201557 | rs7949454 | 0.701 | -0.037 | -0.323 | -0.519 | -0.023 | 0.016 | 0.016 | 2.348 | 0.11 | 0.49 | 0.07 | 0.09 | 0.06 | 0.08 |
| 11 | 20202134 | rs4757040 | 0.801 | -0.017 | -0.498 | -0.487 | -0.001 | -0.012 | -0.012 | 0 | 0.4 | 0.21 | 0.34 | 0.4 | 0.38 | 0.52 |
| 11 | 20205484 | rs1004786 | 0.827 | 0.216 | -0.501 | -0.478 | -0.001 | -0.001 | -0.001 | 0 | 0.17 | 0.18 | 0.14 | 0.13 | 0.12 | 0.12 |
| 11 | 20209972 | rs7113122 | 0.094 | 0.292 | -0.57 | -0.513 | -0.001 | 0.012 | 0.012 | 0 | 0.23 | 0.2 | 0.2 | 0.14 | 0.23 | 0.15 |
| 11 | 20213955 | rs11025430 | -0.558 | 0.573 | -0.39 | -0.205 | 0.03 | 0 | 0 | 0.12 | 0.65 | 0.46 | 0.62 | 0.64 | 0.58 | 0.7 |
| 11 | 20219834 | rs11025436 | 1.648 | 0.467 | -0.75 | -0.343 | 0.003 | -0.024 | -0.024 | 0 | 0.21 | 0.14 | 0.28 | 0.41 | 0.16 | 0.47 |
| 11 | 20222703 | rs11025438 | 0.351 | 0.686 | -0.65 | -0.462 | 0.009 | 0.028 | 0.028 | 0 | 0.52 | 0.33 | 0.52 | 0.63 | 0.43 | 0.67 |
| 11 | 20224852 | rs11529659 | 0.927 | 1.049 | -1.289 | 0.021 | -0.059 | -0.067 | -0.067 | 0.095 | 0.25 | 0.08 | 0.24 | 0.34 | 0.33 | 0.19 |
| 11 | 20229993 | rs9943572 | -0.711 | 1.144 | -0.711 | 0.006 | 0.031 | 0 | 0 | 0 | 0.13 | 0.22 | 0.17 | 0.14 | 0.19 | 0.08 |
| 11 | 20231630 | rs7128588 | 1.554 | 0.637 | -1.256 | -0.108 | 0.004 | -0.016 | -0.016 | 0 | 0.21 | 0.16 | 0.23 | 0.35 | 0.17 | 0.47 |
| 11 | 20232335 | rs4757045 | -0.15 | 0.338 | -0.823 | -0.243 | 0.013 | -0.016 | -0.016 | 0 | 0.6 | 0.33 | 0.44 | 0.52 | 0.66 | 0.4 |
| 11 | 20235409 | rs11025443 | 1.413 | 0.373 | -1.399 | -0.328 | -0.04 | -0.048 | -0.048 | 0 | 0.26 | 0.12 | 0.27 | 0.37 | 0.37 | 0.2 |
| 11 | 20236780 | rs11025444 | 0.104 | 0.455 | -1.045 | -0.606 | 0.018 | 0.04 | 0.04 | 0 | 0.31 | 0.12 | 0.17 | 0.21 | 0.25 | 0.2 |
| 11 | 20237064 | rs10766635 | 0.45 | 0.417 | -0.977 | -0.673 | -0.001 | -0.005 | -0.005 | 0 | 0.71 | 0.32 | 0.58 | 0.81 | 0.69 | 0.81 |
| 11 | 20238149 | rs7127406 | 1.479 | 0.54 | -0.962 | -0.573 | -0.033 | 0.032 | 0.032 | 0 | 0.41 | 0.19 | 0.41 | 0.6 | 0.44 | 0.62 |
| 11 | 20243053 | rs16905844 | -0.481 | 0.74 | 0.212 | 0.042 | 0.088 | -0.004 | -0.004 | 0.464 | 0.09 | 0.26 | 0.12 | 0.07 | 0.13 | 0.05 |
| 11 | 20245723 | rs10500872 | -0.261 | 0.339 | -0.857 | 0.027 | 0.022 | 0.001 | 0.001 | 0.245 | 0.21 | 0.11 | 0.12 | 0.2 | 0.17 | 0.16 |
| 11 | 20245859 | rs10766637 | 1.234 | 0.309 | -0.845 | -0.016 | 0.001 | 0.084 | 0.084 | 0.834 | 0.53 | 0.15 | 0.46 | 0.61 | 0.53 | 0.67 |
| 11 | 20246269 | rs11025448 | -0.401 | -0.084 | -0.269 | 0.358 | 0.011 | 0.007 | 0.007 | 0 | 0.16 | 0.11 | 0.09 | 0.12 | 0.12 | 0.1 |
| 11 | 20252106 | rs1589379 | 0.954 | 0.346 | 0.346 | 0.525 | 0.007 | 0.005 | 0.005 | 0.038 | 0.73 | 0.56 | 0.63 | 0.29 | 0.71 | 0.76 |
| 11 | 20253599 | rs1353649 | 0.656 | 0.202 | 0.612 | 0.685 | -0.016 | 0.046 | 0.046 | 0 | 0.72 | 0.55 | 0.66 | 0.47 | 0.73 | 0.88 |
| 11 | 20261282 | rs1553075 | -0.183 | 0.197 | 0.692 | 1.055 | -0.032 | 0.07 | 0.07 | 0 | 0.8 | 0.62 | 0.75 | 0.66 | 0.82 | 0.94 |
| 11 | 20262014 | rs1607840 | 0.525 | 0.494 | 0.421 | 0.638 | -0.021 | -0.014 | -0.014 | 0 | 0.41 | 0.32 | 0.27 | 0.41 | 0.43 | 0.38 |
| 11 | 20264424 | rs4757913 | 1.064 | 0.439 | 0.372 | 0.559 | 0.005 | -0.037 | -0.037 | 0.842 | 0.29 | 0.29 | 0.37 | 0.13 | 0.29 | 0.48 |
| 11 | 20266222 | rs10458884 | -0.006 | 0.495 | 0.346 | 0.572 | 0.026 | 0.015 | 0.015 | 0 | 0.5 | 0.31 | 0.37 | 0.52 | 0.53 | 0.47 |
| 11 | 20269390 | rs1520896 | -0.447 | 0.334 | 0.608 | 0.808 | -0.035 | 0.072 | 0.072 | 0 | 0.8 | 0.58 | 0.75 | 0.66 | 0.82 | 0.94 |
| 11 | 20269732 | rs10766639 | 1.222 | 0.403 | 0.445 | 0.604 | -0.049 | 0.106 | 0.106 | 0.318 | 0.3 | 0.08 | 0.16 | 0.12 | 0.31 | 0.26 |
| 11 | 20272436 | rs10741823 | 1.287 | 0.444 | 0.457 | 0.624 | -0.054 | 0.104 | 0.104 | 0.435 | 0.3 | 0.08 | 0.16 | 0.12 | 0.32 | 0.26 |
| 11 | 20275946 | rs1520895 | 0.146 | 0.429 | 0.719 | 0.71 | 0.003 | -0.017 | -0.017 | 1.091 | 0.45 | 0.45 | 0.54 | 0.3 | 0.47 | 0.64 |
| 11 | 20278865 | rs10833273 | 1.549 | 0.418 | 0.589 | 0.309 | -0.05 | 0.092 | 0.092 | 0.014 | 0.28 | 0.06 | 0.15 | 0.11 | 0.31 | 0.26 |
| 11 | 20282368 | rs871401 | 0.722 | 0.53 | 0.703 | 0.425 | 0.009 | -0.053 | -0.053 | 2.397 | 0.36 | 0.38 | 0.43 | 0.19 | 0.36 | 0.57 |
| 11 | 20284420 | rs4757915 | 0.226 | 0.57 | 0.602 | 0.413 | 0.022 | 0.133 | 0.133 | 1.237 | 0.47 | 0.27 | 0.33 | 0.13 | 0.46 | 0.28 |
| 11 | 20286574 | rs7950594 | -0.463 | 0.393 | 0.638 | 0.458 | 0.003 | 0.002 | 0.002 | 0.157 | 0.49 | 0.51 | 0.59 | 0.53 | 0.5 | 0.67 |

| | | | | | | | | | | | | | | | | |
|---|---|---|---|---|---|---|---|---|---|---|---|---|---|---|---|---|
| 11 | 20289037 | rs4757917 | 0.623 | 0.44 | 0.286 | 0.32 | -0.001 | 0.004 | 0.004 | 0.184 | 0.33 | 0.29 | 0.41 | 0.19 | 0.33 | 0.58 |
| 11 | 20290931 | rs2403575 | 0.385 | 0.43 | 0.343 | 0.379 | -0.004 | -0.046 | -0.046 | 0.87 | 0.36 | 0.43 | 0.43 | 0.19 | 0.33 | 0.58 |
| 11 | 20292093 | rs10766642 | 0.205 | 0.744 | -0.287 | 0.588 | -0.002 | 0.001 | 0.001 | 0.067 | 0.35 | 0.32 | 0.19 | 0.12 | 0.35 | 0.26 |
| 11 | 20297638 | rs12786945 | -0.889 | 0.673 | -0.517 | 2.007 | -0.002 | -0.013 | -0.013 | 0 | 0.1 | 0.09 | 0.13 | 0.34 | 0.1 | 0.1 |
| 11 | 20299723 | rs1874692 | -0.421 | 0.529 | -0.257 | 2.002 | -0.013 | -0.056 | -0.056 | 0.241 | 0.35 | 0.15 | 0.35 | 0.45 | 0.31 | 0.59 |
| 11 | 20305097 | rs11529173 | -0.427 | 0.508 | -0.565 | 2.008 | -0.012 | -0.024 | -0.024 | 0.002 | 0.28 | 0.14 | 0.38 | 0.43 | 0.32 | 0.17 |
| 11 | 20305479 | rs10766646 | 0.842 | 0.283 | 0.238 | 1.898 | -0.002 | 0.005 | 0.005 | 0 | 0.39 | 0.53 | 0.32 | 0.4 | 0.39 | 0.32 |
| 11 | 20310244 | rs7112050 | -1.092 | 0.262 | 0.505 | 1.927 | -0.002 | 0.011 | 0.011 | 0 | 0.59 | 0.51 | 0.55 | 0.51 | 0.59 | 0.63 |
| 11 | 20310498 | rs10458885 | -0.23 | 0.343 | 0.675 | 1.991 | -0.003 | -0.002 | -0.002 | 0.081 | 0.27 | 0.24 | 0.26 | 0.34 | 0.31 | 0.12 |
| 11 | 20313272 | rs12787276 | -0.01 | 0.502 | 0.617 | 2.044 | -0.025 | 0.075 | 0.075 | 0 | 0.24 | 0.06 | 0.21 | 0.14 | 0.26 | 0.1 |
| 11 | 20316000 | rs7929873 | -0.176 | 0.545 | 0.797 | 2.093 | -0.006 | 0.002 | 0.002 | 0 | 0.63 | 0.41 | 0.61 | 0.34 | 0.69 | 0.4 |
| 11 | 20328070 | rs11822785 | 0.242 | 0.489 | 0.9 | 1.974 | -0.001 | 0.036 | 0.036 | 0 | 0.06 | 0.11 | 0.17 | 0.1 | 0.06 | 0.08 |
| 11 | 20328482 | rs7106374 | 0.143 | 0.468 | 1.017 | 2.204 | -0.004 | -0.004 | -0.004 | 0 | 0.36 | 0.37 | 0.39 | 0.44 | 0.37 | 0.57 |
| 11 | 20328774 | rs11606925 | 0.325 | 0.404 | 1.031 | 2.19 | -0.003 | 0 | 0 | 0 | 0.34 | 0.36 | 0.29 | 0.14 | 0.37 | 0.5 |
| 11 | 20328840 | rs904182 | -0.057 | 0.326 | 0.509 | 1.842 | -0.032 | 0.087 | 0.087 | 0 | 0.26 | 0.07 | 0.22 | 0.13 | 0.3 | 0.08 |
| 11 | 20329227 | rs883051 | -1.522 | 0.225 | 0.813 | 1.887 | 0.001 | 0.026 | 0.026 | 0.061 | 0.69 | 0.79 | 0.81 | 0.85 | 0.69 | 0.74 |
| 11 | 20331241 | rs4757925 | -0.768 | 0.374 | 1.153 | 2.208 | 0.017 | 0.095 | 0.095 | 0 | 0.08 | 0.28 | 0.09 | 0.09 | 0.05 | 0.07 |
| 11 | 20332435 | rs2132964 | 0.773 | 0.292 | 1.046 | 2.194 | 0.003 | -0.007 | -0.007 | 0 | 0.64 | 0.43 | 0.48 | 0.45 | 0.68 | 0.78 |
| 11 | 20335391 | rs10430897 | -0.064 | 0.495 | 0.867 | 1.797 | -0.001 | 0 | 0 | 3.417 | 0.36 | 0.4 | 0.38 | 0.41 | 0.44 | 0.55 |
| 11 | 20336470 | rs11025485 | 0.234 | 0.39 | 1.274 | 1.87 | -0.002 | 0.051 | 0.051 | 0 | 0.06 | 0.16 | 0.16 | 0.08 | 0.04 | 0.07 |
| 11 | 20337228 | rs10766651 | 0.748 | 0.27 | 1.025 | 1.814 | -0.001 | 0.11 | 0.11 | 0.396 | 0.45 | 0.26 | 0.43 | 0.76 | 0.45 | 0.39 |
| 11 | 20337341 | rs10430900 | -0.388 | 0.18 | 1.098 | 1.9 | -0.001 | 0.063 | 0.063 | 0 | 0.1 | 0.18 | 0.13 | 0.3 | 0.09 | 0.08 |
| 11 | 20341067 | rs7938228 | 0.352 | 0.136 | 1.114 | 1.624 | 0.001 | -0.016 | -0.016 | 0 | 0.18 | 0.13 | 0.14 | 0.02 | 0.18 | 0.06 |
| 11 | 20342934 | rs1874690 | 1.436 | 0.138 | 1.223 | 1.765 | 0.024 | -0.048 | -0.048 | 0 | 0.37 | 0.26 | 0.31 | 0.48 | 0.33 | 0.31 |
| 11 | 20343746 | rs10500876 | 2.017 | 0.141 | 1.151 | 1.831 | -0.036 | 0.047 | 0.047 | 0.452 | 0.33 | 0.05 | 0.19 | 0.14 | 0.32 | 0.25 |
| 11 | 20348349 | rs4757930 | 0.147 | 0.058 | 1.286 | 1.754 | 0.002 | -0.024 | -0.024 | 0 | 0.31 | 0.32 | 0.35 | 0.11 | 0.37 | 0.5 |
| 11 | 20348932 | rs4757931 | -0.884 | 0.057 | 1.333 | 1.826 | -0.003 | 0.038 | 0.038 | 0 | 0.15 | 0.19 | 0.26 | 0.36 | 0.13 | 0.14 |
| 11 | 20349670 | rs16906047 | 0.069 | 0.049 | 1.258 | 1.852 | -0.003 | -0.002 | -0.002 | 1.774 | 0.09 | 0.07 | 0.12 | 0.28 | 0.09 | 0.08 |
| 11 | 20351011 | rs4757053 | 1.554 | 0.043 | 1.251 | 1.825 | 0.043 | 0.063 | 0.063 | 0 | 0.59 | 0.18 | 0.52 | 0.25 | 0.67 | 0.74 |
| 11 | 20355297 | rs10766653 | 1.66 | -0.307 | 1.101 | 1.619 | 0.027 | 0.09 | 0.09 | 1.921 | 0.39 | 0.19 | 0.2 | 0.14 | 0.34 | 0.26 |
| 11 | 20356444 | rs7128099 | 1.538 | 0.303 | 1.23 | 1.71 | -0.011 | 0.057 | 0.057 | 0 | 0.78 | 0.49 | 0.67 | 0.28 | 0.83 | 0.77 |
| 11 | 20357928 | rs7928187 | 0.402 | 0.43 | 0.672 | 1.73 | -0.039 | 0.07 | 0.07 | 0.002 | 0.27 | 0.13 | 0.24 | 0.12 | 0.33 | 0.47 |
| 11 | 20358702 | rs7929060 | -0.141 | 0.414 | 0.571 | 1.868 | -0.021 | 0.026 | 0.026 | 0 | 0.78 | 0.66 | 0.73 | 0.65 | 0.82 | 0.89 |
| 11 | 20359771 | rs952619 | 0.93 | 0.428 | 0.488 | 1.872 | -0.012 | 0.016 | 0.016 | 0 | 0.67 | 0.47 | 0.6 | 0.35 | 0.72 | 0.8 |
| 11 | 20362474 | rs10766656 | 1.471 | 0.667 | 0.232 | 1.62 | -0.028 | -0.096 | -0.096 | 0.229 | 0.41 | 0.12 | 0.3 | 0.43 | 0.41 | 0.33 |
| 11 | 20365604 | rs10833300 | -0.443 | 0.654 | 0.3 | 1.483 | 0.008 | -0.02 | -0.02 | 0 | 0.35 | 0.57 | 0.44 | 0.21 | 0.39 | 0.53 |
| 11 | 20372321 | rs10833308 | 1.294 | 0.594 | -0.126 | 1.47 | -0.034 | -0.083 | -0.083 | 2.969 | 0.42 | 0.13 | 0.31 | 0.48 | 0.41 | 0.33 |
| 11 | 20372789 | rs7121349 | 0.851 | 0.613 | 0.498 | 1.875 | -0.011 | 0.024 | 0.024 | 0 | 0.71 | 0.57 | 0.58 | 0.67 | 0.73 | 0.8 |
| 11 | 20378100 | rs2403583 | -1.343 | 0.787 | 0.367 | 2.216 | 0.004 | 0.038 | 0.038 | 0 | 0.19 | 0.25 | 0.39 | 0.41 | 0.19 | 0.14 |
| 11 | 20378124 | rs11025523 | -0.075 | 0.846 | -0.239 | 1.471 | -0.001 | 0.021 | 0.021 | 0 | 0.18 | 0.06 | 0.1 | 0.14 | 0.21 | 0.35 |
| 11 | 20386301 | rs10833312 | 1.007 | 0.858 | 0.393 | 1.68 | -0.002 | 0 | 0 | 0 | 0.59 | 0.37 | 0.51 | 0.52 | 0.55 | 0.45 |
| 11 | 20397200 | rs10500879 | -0.99 | 0.321 | 1.066 | 1.29 | 0.005 | -0.043 | -0.043 | 0 | 0.1 | 0.31 | 0.18 | 0.02 | 0.08 | 0.02 |
| 11 | 20401007 | rs10766662 | 1.463 | 0.327 | 1.931 | 2.452 | 0.012 | 0.163 | 0.163 | 0.669 | 0.84 | 0.57 | 0.54 | 0.41 | 0.86 | 0.87 |
| 11 | 20401835 | rs7934001 | 1.153 | 0.321 | 1.87 | 2.484 | 0.015 | 0.256 | 0.256 | 1.751 | 0.92 | 0.75 | 0.8 | 0.43 | 0.93 | 0.89 |
| 11 | 20405135 | rs16906210 | -2.044 | 0.259 | 2.029 | 2.717 | -0.014 | 0.253 | 0.253 | 2.521 | 0.06 | 0.27 | 0.08 | 0.3 | 0.01 | 0.04 |
| 11 | 20408926 | rs11025542 | -1.824 | 0.272 | 2.033 | 2.745 | 0.011 | 0.132 | 0.132 | 0.082 | 0.08 | 0.27 | 0.14 | 0.23 | 0.05 | 0.07 |
| 11 | 20419028 | rs7934372 | 2.305 | -0.017 | 1.702 | 2.366 | -0.046 | 0.094 | 0.094 | 0.222 | 0.74 | 0.5 | 0.52 | 0.41 | 0.82 | 0.81 |
| 11 | 20433483 | rs7479049 | 2.12 | -0.023 | 1.706 | 2.137 | -0.041 | 0.279 | 0.279 | 2.025 | 0.84 | 0.5 | 0.57 | 0.46 | 0.93 | 0.88 |
| 11 | 20435108 | rs10734291 | 2.097 | -0.006 | 1.705 | 2.137 | -0.026 | 0.281 | 0.281 | 2.577 | 0.84 | 0.5 | 0.57 | 0.46 | 0.92 | 0.88 |
| 11 | 20443875 | rs11025559 | 0.144 | 0.035 | 1.702 | 1.811 | 0.001 | -0.008 | -0.008 | 0.308 | 0.18 | 0.29 | 0.11 | 0.19 | 0.18 | 0.32 |
| 11 | 20449607 | rs7480750 | 2.66 | 0.287 | 1.219 | 1.516 | -0.128 | 0.106 | 0.106 | 2.473 | 0.41 | 0.03 | 0.28 | 0.15 | 0.46 | 0.31 |
| 11 | 20458498 | rs6483683 | 1.291 | 0.202 | 1.439 | 1.596 | 0.014 | 0.428 | 0.428 | 2.443 | 0.9 | 0.55 | 0.71 | 0.47 | 0.94 | 0.89 |
| 11 | 20475560 | rs10833328 | 1.557 | 0.212 | 1.28 | 1.553 | -0.023 | 0.144 | 0.144 | 0 | 0.79 | 0.56 | 0.66 | 0.42 | 0.83 | 0.83 |
| 11 | 20483748 | rs3758803 | 2.053 | 0.261 | 1.161 | 1.464 | 0.041 | 0.129 | 0.129 | 0.301 | 0.5 | 0.19 | 0.39 | 0.22 | 0.57 | 0.42 |
| 11 | 20488453 | rs1036923 | 1.633 | 0.202 | 1.223 | 1.567 | -0.036 | 0.291 | 0.291 | 1.803 | 0.85 | 0.53 | 0.58 | 0.45 | 0.93 | 0.88 |
| 11 | 20495405 | rs2120309 | 1.763 | 0.254 | 1.213 | 1.543 | -0.007 | 0.18 | 0.18 | 0.503 | 0.8 | 0.52 | 0.54 | 0.45 | 0.82 | 0.86 |
| 11 | 20498814 | rs7946612 | 1.251 | 0.258 | 1.322 | 1.559 | 0.011 | 0.253 | 0.253 | 0.826 | 0.89 | 0.68 | 0.71 | 0.47 | 0.89 | 0.89 |
| 11 | 20499412 | rs7115830 | 2.342 | 0.269 | 1.254 | 1.439 | 0 | 0.079 | 0.079 | 0.099 | 0.47 | 0.26 | 0.37 | 0.22 | 0.51 | 0.4 |
| 11 | 20509740 | rs12420525 | -0.024 | 0.254 | 1.189 | 1.571 | 0 | 0.034 | 0.034 | 0 | 0.09 | 0.02 | 0.16 | 0.02 | 0.07 | 0.07 |
| 11 | 20513260 | rs11025583 | -0.463 | 0.131 | 1.3 | 1.568 | -0.006 | -0.021 | -0.021 | 0 | 0.25 | 0.31 | 0.14 | 0.19 | 0.25 | 0.42 |
| 11 | 20520461 | rs2403600 | 1.657 | 0.121 | 1.285 | 1.847 | -0.011 | 0.399 | 0.399 | 1.508 | 0.89 | 0.58 | 0.73 | 0.48 | 0.94 | 0.89 |
| 11 | 20522290 | rs7395007 | 1.194 | 0.172 | 1.3 | 1.872 | -0.003 | 0.358 | 0.358 | 1.305 | 0.91 | 0.67 | 0.73 | 0.48 | 0.94 | 0.89 |
| 11 | 20522915 | rs10741842 | 1.428 | 0.201 | 1.184 | 1.866 | -0.034 | 0.23 | 0.23 | 0.868 | 0.82 | 0.54 | 0.57 | 0.47 | 0.88 | 0.87 |
| 11 | 20534286 | rs11025592 | -1.198 | 0.454 | 1.173 | 1.859 | -0.013 | 0.02 | 0.02 | 0 | 0.32 | 0.48 | 0.26 | 0.41 | 0.29 | 0.47 |
| 11 | 20537099 | rs2120306 | 1.16 | 0.335 | 1.154 | 1.801 | 0.018 | 0.122 | 0.122 | 0 | 0.83 | 0.66 | 0.62 | 0.37 | 0.85 | 0.8 |
| 11 | 20543270 | rs7122152 | -1.067 | 0.492 | 1.065 | 1.983 | 0.015 | 0.038 | 0.038 | 0.004 | 0.11 | 0.18 | 0.23 | 0.26 | 0.12 | 0.16 |
| 11 | 20545376 | rs7131043 | 0.851 | -0.734 | 1.255 | 1.409 | -0.04 | 0.034 | 0.034 | 0 | 0.7 | 0.41 | 0.7 | 0.45 | 0.81 | 0.62 |
| 11 | 20545926 | rs12787810 | 0.994 | -0.75 | 1.174 | 1.414 | -0.061 | 0.058 | 0.058 | 0 | 0.7 | 0.46 | 0.7 | 0.45 | 0.81 | 0.61 |
| 11 | 20546869 | rs11025602 | -0.612 | -0.54 | 1.084 | 1.368 | -0.001 | 0.003 | 0.003 | 0 | 0.18 | 0.14 | 0.12 | 0.15 | 0.19 | 0.35 |

| | | | | | | | | | | | | | | | | |
|---|---|---|---|---|---|---|---|---|---|---|---|---|---|---|---|---|
| 11 | 20553689 | rs7112553 | 0.905 | -0.461 | 1.463 | 1.379 | -0.005 | 0.09 | 0.09 | 0.019 | 0.78 | 0.66 | 0.73 | 0.49 | 0.79 | 0.62 |
| 11 | 20554790 | rs11025608 | 2.231 | -0.427 | 0.825 | 0.763 | -0.063 | 0.091 | 0.091 | 0 | 0.44 | 0.13 | 0.39 | 0.22 | 0.56 | 0.36 |
| 11 | 20557198 | rs16906402 | -1.381 | 0.16 | 1.293 | 1.275 | 0.007 | -0.023 | -0.023 | 1.03 | 0.23 | 0.26 | 0.15 | 0.15 | 0.26 | 0.44 |
| 11 | 20557535 | rs7129522 | -1.294 | 0.16 | 1.293 | 1.274 | 0.009 | -0.023 | -0.023 | 0 | 0.23 | 0.26 | 0.15 | 0.15 | 0.25 | 0.44 |
| 11 | 20561991 | rs2403602 | -0.99 | 0.126 | 1.432 | 1.185 | -0.02 | -0.075 | -0.075 | 0.768 | 0.21 | 0.47 | 0.12 | 0.12 | 0.15 | 0.34 |
| 11 | 20563041 | rs7924410 | 1.443 | 0.108 | 1.396 | 1.205 | 0.001 | 0.064 | 0.064 | 0.14 | 0.71 | 0.45 | 0.67 | 0.46 | 0.73 | 0.52 |
| 11 | 20566376 | rs1867306 | -1.423 | 0.172 | 1.173 | 0.897 | 0 | 0 | 0 | 0 | 0.27 | 0.28 | 0.31 | 0.32 | 0.26 | 0.42 |
| 11 | 20566453 | rs1894160 | -1.359 | 0.17 | 1.117 | 0.889 | 0 | 0 | 0 | 0.121 | 0.26 | 0.27 | 0.3 | 0.32 | 0.26 | 0.38 |
| 11 | 20573959 | rs2165735 | -1.593 | 0.231 | 0.938 | 0.811 | 0.001 | 0 | 0 | 0 | 0.3 | 0.39 | 0.34 | 0.33 | 0.27 | 0.42 |
| 11 | 20574079 | rs1465968 | 1.432 | 1.147 | 0.644 | 1.033 | 0.01 | 0.004 | 0.004 | 0.223 | 0.66 | 0.53 | 0.47 | 0.33 | 0.65 | 0.53 |
| 11 | 20579466 | rs4331088 | -1.083 | 1.222 | 0.713 | 1.148 | 0.056 | 0.295 | 0.295 | 0.217 | 0.05 | 0.24 | 0.2 | 0.33 | 0.08 | 0.06 |
| 11 | 20588922 | rs4923013 | -0.15 | 1.328 | 0.113 | 0.914 | -0.016 | 0.017 | 0.017 | 0 | 0.17 | 0.09 | 0.11 | 0.13 | 0.17 | 0.29 |
| 11 | 20589324 | rs7115019 | -1.849 | 0.947 | 0.604 | 0.938 | 0.011 | 0.136 | 0.136 | 0.881 | 0.23 | 0.52 | 0.46 | 0.54 | 0.24 | 0.26 |
| 11 | 20594783 | rs7929521 | -1.921 | 0.773 | 0.264 | 0.185 | -0.002 | 0.006 | 0.006 | 0.692 | 0.38 | 0.68 | 0.4 | 0.35 | 0.37 | 0.3 |
| 11 | 20594890 | rs12418677 | -1.039 | 0.851 | 0.631 | 0.432 | 0.079 | 0.338 | 0.338 | 0 | 0.07 | 0.32 | 0.21 | 0.33 | 0.1 | 0.06 |
| 11 | 20602331 | rs11025637 | -0.528 | 0.788 | 0.36 | 0.162 | -0.001 | -0.032 | -0.032 | 0.009 | 0.15 | 0.21 | 0.09 | 0.02 | 0.13 | 0.16 |
| 11 | 20602578 | rs7126169 | 1.752 | 0.883 | -0.606 | -0.218 | -0.017 | -0.112 | -0.112 | 0.022 | 0.35 | 0.01 | 0.28 | 0.35 | 0.31 | 0.35 |
| 11 | 20605665 | rs10458887 | 1.704 | 0.894 | -0.25 | -0.257 | 0.02 | -0.006 | -0.006 | 0.001 | 0.36 | 0.25 | 0.3 | 0.35 | 0.31 | 0.38 |
| 11 | 20608950 | rs2012543 | -0.473 | 1.254 | -0.146 | -0.092 | 0.003 | 0.002 | 0.002 | 0 | 0.7 | 0.71 | 0.51 | 0.35 | 0.61 | 0.6 |
| 11 | 20609379 | rs10833362 | 0.318 | 1.059 | -0.675 | -0.227 | -0.083 | -0.002 | -0.002 | 0 | 0.3 | 0.09 | 0.34 | 0.29 | 0.4 | 0.36 |
| 11 | 20611698 | rs16906495 | 1.34 | 1.101 | -0.625 | -0.437 | -0.001 | -0.017 | -0.017 | 0 | 0.1 | 0.12 | 0.06 | 0 | 0.11 | 0.09 |
| 11 | 20612269 | rs7131598 | -1.74 | 0.835 | -0.566 | -0.556 | -0.02 | -0.004 | -0.004 | 0 | 0.58 | 0.54 | 0.68 | 0.66 | 0.73 | 0.64 |
| 11 | 20618589 | rs894747 | 0.682 | 0.982 | -0.435 | -0.623 | 0.009 | 0.014 | 0.014 | 0.795 | 0.69 | 0.42 | 0.64 | 0.62 | 0.65 | 0.72 |
| 11 | 20622975 | rs1443547 | 0.93 | 0.88 | -0.441 | -0.552 | 0.004 | 0.002 | 0.002 | 0 | 0.48 | 0.45 | 0.38 | 0.37 | 0.38 | 0.43 |
| 11 | 20623023 | rs2241941 | -0.249 | 1.398 | -0.742 | -0.641 | 0.027 | 0.026 | 0.026 | 0 | 0.23 | 0.33 | 0.32 | 0.36 | 0.35 | 0.27 |
| 11 | 20623042 | rs1443548 | -1.288 | 1.199 | -0.968 | -0.174 | -0.003 | -0.003 | -0.003 | 0.164 | 0.78 | 0.87 | 0.75 | 0.75 | 0.79 | 0.77 |
| 11 | 20630163 | rs1373165 | 0.481 | 0.872 | -0.608 | -0.081 | -0.005 | -0.003 | -0.003 | 0.022 | 0.85 | 0.85 | 0.9 | 0.98 | 0.82 | 0.87 |
| 11 | 20638059 | rs7942430 | 0.241 | 0.2 | -0.721 | -0.127 | 0.002 | 0.002 | 0.002 | 0.823 | 0.57 | 0.39 | 0.51 | 0.32 | 0.67 | 0.7 |
| 11 | 20638648 | rs11025652 | 1.577 | -0.007 | -0.758 | -0.17 | 0.009 | 0.039 | 0.039 | 0 | 0.21 | 0.11 | 0.14 | 0.15 | 0.19 | 0.18 |
| 11 | 20639864 | rs7119457 | 1.009 | 0.035 | -0.956 | -0.447 | -0.01 | 0.06 | 0.06 | 0 | 0.63 | 0.78 | 0.69 | 0.8 | 0.63 | 0.74 |
| 11 | 20641676 | rs10766703 | -0.291 | -0.033 | -0.955 | -0.433 | 0.01 | 0.011 | 0.011 | 1.077 | 0.64 | 0.69 | 0.72 | 0.67 | 0.7 | 0.76 |
| 11 | 20643434 | rs2001982 | 0.355 | 0.123 | -0.727 | -0.31 | -0.001 | -0.002 | -0.002 | 2.814 | 0.52 | 0.5 | 0.66 | 0.64 | 0.6 | 0.71 |
| 11 | 20643784 | rs7942096 | 1.454 | 0.117 | 0.145 | 0.017 | 0 | -0.001 | -0.001 | 0 | 0.41 | 0.36 | 0.42 | 0.33 | 0.45 | 0.56 |
| 11 | 20646348 | rs12803296 | 0.798 | -0.267 | -0.223 | -0.033 | 0.004 | 0.095 | 0.095 | 0 | 0.29 | 0.11 | 0.18 | 0.13 | 0.26 | 0.2 |
| 11 | 20647669 | rs11025661 | 0.058 | -0.306 | -0.066 | 0.069 | -0.012 | 0.025 | 0.025 | 0.359 | 0.16 | 0.2 | 0.14 | 0.23 | 0.1 | 0.06 |
| 11 | 20648020 | rs11025664 | 0.004 | -0.289 | -0.236 | 0.114 | 0.004 | 0.002 | 0.002 | 0.022 | 0.16 | 0.13 | 0.25 | 0.59 | 0.12 | 0.1 |
| 11 | 20648380 | rs1805091 | 0.806 | -0.288 | -0.27 | -0.061 | 0.013 | 0.107 | 0.107 | 0 | 0.28 | 0.12 | 0.18 | 0.09 | 0.25 | 0.2 |
| 11 | 20649404 | rs3819252 | -0.285 | -0.092 | -0.139 | 0.056 | 0.034 | 0.049 | 0.049 | 2.15 | 0.43 | 0.23 | 0.31 | 0.32 | 0.35 | 0.26 |
| 11 | 20652620 | rs2276431 | -0.258 | -0.091 | -0.188 | 0.057 | 0.035 | 0.054 | 0.054 | 2.212 | 0.43 | 0.19 | 0.31 | 0.31 | 0.35 | 0.26 |
| 11 | 20653783 | rs7110148 | 0.42 | 0.432 | -0.185 | -0.107 | -0.062 | 0.072 | 0.072 | 0 | 0.12 | 0.01 | 0.13 | 0 | 0.19 | 0.15 |
| 11 | 20655239 | rs4923601 | -1.626 | 0.312 | 0.321 | -0.047 | -0.001 | -0.002 | -0.002 | 0.836 | 0.69 | 0.77 | 0.65 | 0.69 | 0.66 | 0.58 |
| 11 | 20657494 | rs11025670 | 0.249 | 0.102 | 0.733 | 0.055 | -0.003 | -0.033 | -0.033 | 0 | 0.36 | 0.36 | 0.21 | 0.09 | 0.3 | 0.24 |
| 11 | 20658679 | rs2000959 | 1.506 | 0.193 | 0.04 | 0.146 | -0.024 | -0.025 | -0.025 | 0 | 0.32 | 0.14 | 0.43 | 0.35 | 0.36 | 0.51 |
| 11 | 20664403 | rs4923627 | -0.202 | 1.3 | -0.181 | -1.057 | -0.028 | 0.017 | 0.017 | 0.655 | 0.23 | 0.2 | 0.21 | 0 | 0.3 | 0.2 |
| 11 | 20669971 | rs1617769 | -0.941 | 0.437 | -0.09 | -0.76 | -0.007 | -0.039 | -0.039 | 0 | 0.84 | 0.81 | 0.92 | 1 | 0.9 | 0.85 |
| 11 | 20673867 | rs2276433 | 0.791 | 0.291 | -0.19 | -0.827 | -0.031 | 0.165 | 0.165 | 0 | 0.46 | 0.15 | 0.56 | 0.82 | 0.44 | 0.54 |
| 11 | 20675645 | rs17298481 | 0.856 | 0.345 | -0.293 | -0.978 | -0.001 | -0.01 | -0.01 | 0 | 0.06 | 0.05 | 0.05 | 0 | 0.06 | 0.01 |
| 11 | 20676068 | rs1792974 | -0.653 | 0.521 | -0.099 | -0.626 | -0.008 | -0.044 | -0.044 | 0.376 | 0.84 | 0.79 | 0.92 | 1 | 0.9 | 0.85 |
| 11 | 20676253 | rs2276432 | 0.415 | 1.632 | 0.625 | 0.276 | -0.012 | -0.047 | -0.047 | 2.452 | 0.17 | 0.24 | 0.12 | 0.05 | 0.16 | 0.2 |
| 11 | 20678604 | rs17232722 | 0.228 | 1.492 | -0.037 | 0.279 | -0.1 | 0.106 | 0.106 | 0 | 0.17 | 0.02 | 0.22 | 0.01 | 0.29 | 0.09 |
| 11 | 20680283 | rs7127159 | 0.102 | 1.538 | 0.112 | 0.511 | -0.003 | 0.027 | 0.027 | 0 | 0.08 | 0.12 | 0.11 | 0.16 | 0.07 | 0.08 |
| 11 | 20680404 | rs1792961 | -0.137 | 1.754 | 0.511 | -0.021 | 0 | -0.036 | -0.036 | 0 | 0.8 | 0.79 | 0.76 | 0.99 | 0.69 | 0.92 |
| 11 | 20681439 | rs10466424 | 0.97 | 1.621 | -0.014 | 0.062 | -0.02 | 0.084 | 0.084 | 0.663 | 0.59 | 0.22 | 0.59 | 0.8 | 0.47 | 0.71 |
| 11 | 20681572 | rs12789398 | -0.734 | 0.903 | 0.273 | 0.622 | 0.023 | -0.07 | -0.07 | 0.016 | 0.18 | 0.37 | 0.14 | 0.06 | 0.18 | 0.19 |
| 11 | 20692208 | rs2282661 | -0.054 | 0.807 | -1.472 | 1.038 | -0.06 | 0.007 | 0.007 | 1.307 | 0.43 | 0.08 | 0.45 | 0.35 | 0.48 | 0.32 |
| 11 | 20699857 | rs1793002 | -0.138 | 0.496 | -0.953 | 1.536 | 0.009 | -0.023 | -0.023 | 0 | 0.83 | 0.88 | 0.84 | 0.54 | 0.86 | 0.76 |
| 11 | 20700336 | rs1615231 | -0.138 | 0.516 | -0.953 | 1.536 | 0.011 | -0.023 | -0.023 | 0 | 0.83 | 0.88 | 0.84 | 0.54 | 0.87 | 0.78 |
| 11 | 20702034 | rs1519717 | -0.885 | 1.133 | -0.6 | 1.42 | 0.025 | -0.005 | -0.005 | 0 | 0.07 | 0.19 | 0.1 | 0.08 | 0.13 | 0.11 |
| 11 | 20706934 | rs1793000 | -0.246 | 1.074 | -0.657 | 1.421 | 0.01 | -0.028 | -0.028 | 4.264 | 0.83 | 0.88 | 0.84 | 0.53 | 0.86 | 0.72 |
| 11 | 20707280 | rs7481267 | 1.322 | 1.26 | -0.64 | 0.656 | -0.029 | -0.026 | -0.026 | 0 | 0.54 | 0.17 | 0.54 | 0.44 | 0.51 | 0.51 |
| 11 | 20707292 | rs870195 | -0.281 | 0.836 | 0.275 | 0.522 | 0.008 | -0.112 | -0.112 | 0 | 0.2 | 0.31 | 0.12 | 0 | 0.2 | 0.1 |
| 11 | 20718728 | rs1792984 | -0.298 | 0.85 | 0.33 | 0.529 | 0.008 | -0.096 | -0.096 | 0 | 0.19 | 0.28 | 0.11 | 0 | 0.2 | 0.1 |
| 11 | 20730761 | rs1715282 | -0.075 | 0.88 | 0.335 | 1.168 | 0.006 | -0.024 | -0.024 | 0 | 0.81 | 0.87 | 0.81 | 0.53 | 0.82 | 0.68 |
| 11 | 20736800 | rs7111828 | -1.468 | 1.097 | 0.445 | 0.867 | 0.008 | -0.009 | -0.009 | 0.409 | 0.09 | 0.14 | 0.07 | 0.05 | 0.14 | 0.11 |
| 11 | 20737138 | rs1611930 | -1.158 | 0.786 | 1.188 | 1.051 | -0.024 | 0.078 | 0.078 | 0 | 0.25 | 0.5 | 0.31 | 0.45 | 0.22 | 0.33 |
| 11 | 20743433 | rs1614419 | 1.41 | 1.056 | 1.411 | 0.808 | 0.068 | -0.008 | -0.008 | 0 | 0.9 | 0.71 | 0.94 | 0.94 | 0.86 | 0.89 |
| 11 | 20749657 | rs9630161 | 0.02 | 0.997 | 1.285 | 1.211 | 0 | -0.063 | -0.063 | 0 | 0.17 | 0.06 | 0.23 | 0.31 | 0.18 | 0.31 |
| 11 | 20752657 | rs7104232 | -1.149 | 1.008 | 1.298 | 1.189 | 0.007 | -0.001 | -0.001 | 0 | 0.07 | 0.11 | 0.05 | 0.06 | 0.14 | 0.11 |
| 11 | 20754440 | rs11025724 | -0.957 | 1.01 | 1.289 | 1.19 | 0.006 | -0.001 | -0.001 | 0 | 0.07 | 0.07 | 0.05 | 0.06 | 0.14 | 0.11 |

| | | | | | | | | | | | | | | | | |
|---|---|---|---|---|---|---|---|---|---|---|---|---|---|---|---|---|
| 11 | 20754814 | rs10766719 | 1.613 | 0.977 | 1.664 | 1.202 | -0.017 | -0.026 | -0.026 | 1.427 | 0.66 | 0.15 | 0.63 | 0.61 | 0.64 | 0.56 |
| 11 | 20755417 | rs7116986 | -1.222 | 1.02 | 1.678 | 1.169 | 0.016 | -0.003 | -0.003 | 0 | 0.08 | 0.09 | 0.05 | 0.06 | 0.14 | 0.11 |
| 11 | 20762771 | rs2403620 | 1.119 | 0.869 | 2.083 | 1.133 | 0.022 | 0.03 | 0.03 | 0 | 0.72 | 0.59 | 0.69 | 0.63 | 0.66 | 0.58 |
| 11 | 20763999 | rs10766721 | 1.2 | 0.859 | 2.13 | 1.128 | 0.028 | 0.037 | 0.037 | 0 | 0.72 | 0.54 | 0.68 | 0.63 | 0.66 | 0.57 |
| 11 | 20766299 | rs7111284 | -0.579 | 0.837 | 2.316 | 1.24 | -0.016 | 0.015 | 0.015 | 0 | 0.19 | 0.28 | 0.26 | 0.25 | 0.16 | 0.3 |
| 11 | 20766506 | rs1401792 | 1.88 | 0.803 | 2.334 | 1.221 | -0.018 | -0.018 | -0.018 | 0 | 0.66 | 0.13 | 0.63 | 0.64 | 0.64 | 0.55 |
| 11 | 20775758 | rs1554368 | 1.844 | 0.828 | 2.492 | 1.058 | 0.117 | 0.033 | 0.033 | 0 | 0.89 | 0.65 | 0.83 | 0.88 | 0.82 | 0.87 |
| 11 | 20776391 | rs1554367 | 2.409 | 0.827 | 2.432 | 1.091 | 0.093 | 0.014 | 0.014 | 1.48 | 0.87 | 0.44 | 0.82 | 0.88 | 0.82 | 0.85 |
| 11 | 20778649 | rs1356737 | 1.165 | 0.918 | 2.577 | 1.117 | 0.07 | 0.031 | 0.031 | 0 | 0.93 | 0.87 | 0.86 | 0.91 | 0.85 | 0.88 |
| 11 | 20781368 | rs908944 | 1.378 | 0.864 | 2.672 | 1.256 | 0.008 | 0.006 | 0.006 | 0 | 0.7 | 0.36 | 0.58 | 0.69 | 0.64 | 0.56 |
| 11 | 20782829 | rs437797 | 1.102 | 0.909 | 2.647 | 1.285 | 0.081 | -0.001 | -0.001 | 0 | 0.93 | 0.86 | 0.86 | 0.96 | 0.84 | 0.89 |
| 11 | 20784543 | rs4922623 | 1.405 | 0.775 | 2.607 | 1.215 | 0.018 | 0.019 | 0.019 | 0 | 0.71 | 0.4 | 0.6 | 0.69 | 0.66 | 0.55 |
| 11 | 20785239 | rs11025746 | -0.387 | 0.781 | 2.534 | 1.297 | 0.002 | -0.006 | -0.006 | 0 | 0.18 | 0.19 | 0.22 | 0.26 | 0.19 | 0.32 |
| 11 | 20787791 | rs11025748 | 1.352 | 0.791 | 2.279 | 1.285 | -0.001 | -0.001 | -0.001 | 0 | 0.69 | 0.31 | 0.57 | 0.67 | 0.64 | 0.56 |
| 11 | 20789657 | rs327025 | 1.536 | 0.914 | 2.409 | 1.053 | 0.071 | 0.006 | 0.006 | 0 | 0.93 | 0.85 | 0.92 | 0.95 | 0.84 | 0.89 |
| 11 | 20792419 | rs327028 | 0.829 | 0.575 | 2 | 1.267 | 0.031 | 0.06 | 0.06 | 0 | 0.94 | 0.84 | 0.94 | 0.9 | 0.91 | 0.94 |
| 11 | 20793645 | rs1949523 | 2.551 | 0.578 | 1.626 | 1.161 | -0.012 | -0.002 | -0.002 | 0 | 0.77 | 0.23 | 0.82 | 0.81 | 0.73 | 0.77 |
| 11 | 20804971 | rs2280360 | -2.708 | 0.583 | 1.863 | 1.137 | 0.08 | 0.027 | 0.027 | 0 | 0.2 | 0.45 | 0.15 | 0.16 | 0.26 | 0.26 |
| 11 | 20805335 | rs2280363 | -2.708 | 0.583 | 1.863 | 1.137 | 0.08 | 0.027 | 0.027 | 0 | 0.2 | 0.45 | 0.15 | 0.16 | 0.26 | 0.26 |
| 11 | 20809264 | rs1519736 | 1.676 | -0.04 | 0.782 | 0.511 | 0.013 | 0.091 | 0.091 | 0.045 | 0.65 | 0.15 | 0.74 | 0.87 | 0.68 | 0.66 |
| 11 | 20822918 | rs4923055 | 1.498 | 0.249 | 1.067 | 0.695 | -0.001 | 0.006 | 0.006 | 0 | 0.71 | 0.31 | 0.78 | 0.87 | 0.71 | 0.69 |
| 11 | 20839874 | rs7130897 | 1.583 | -0.585 | 1.636 | 0.427 | -0.029 | -0.085 | -0.085 | 0 | 0.78 | 0.41 | 0.85 | 0.96 | 0.85 | 0.78 |
| 11 | 20840492 | rs1401794 | 2.126 | -0.627 | 1.438 | 1.226 | -0.003 | 0.002 | 0.002 | 0.048 | 0.67 | 0.4 | 0.55 | 0.68 | 0.72 | 0.6 |
| 11 | 20840726 | rs10500883 | 2.299 | -0.719 | 1.471 | 1.178 | 0.018 | -0.008 | -0.008 | 0 | 0.66 | 0.28 | 0.54 | 0.68 | 0.72 | 0.61 |
| 11 | 20842974 | rs1018335 | 2.237 | -0.8 | 2.499 | 1.445 | -0.013 | 0.059 | 0.059 | 0 | 0.72 | 0.52 | 0.49 | 0.63 | 0.73 | 0.68 |
| 11 | 20843445 | rs7925869 | 2.15 | -0.8 | 2.488 | 1.427 | -0.01 | 0.048 | 0.048 | 0 | 0.73 | 0.58 | 0.5 | 0.64 | 0.75 | 0.68 |
| 11 | 20843934 | rs435001 | 1.207 | -0.809 | 2.524 | 1.394 | -0.01 | -0.078 | -0.078 | 0 | 0.84 | 0.66 | 0.9 | 0.97 | 0.87 | 0.89 |
| 11 | 20846161 | rs2403635 | 3.078 | -0.954 | 2.118 | 1.298 | 0.022 | -0.015 | -0.015 | 0 | 0.61 | 0.32 | 0.49 | 0.63 | 0.67 | 0.58 |
| 11 | 20851337 | rs1914984 | -0.806 | -0.732 | 1.87 | 1.431 | 0.001 | -0.004 | -0.004 | 0.754 | 0.06 | 0.06 | 0.04 | 0.04 | 0.08 | 0.09 |
| 11 | 20853197 | rs2680989 | -1.601 | -0.687 | 1.865 | 1.561 | -0.002 | -0.013 | -0.013 | 0 | 0.12 | 0.08 | 0.33 | 0.33 | 0.13 | 0.21 |
| 11 | 20854167 | rs7933216 | -1.995 | -0.807 | 2.544 | 1.609 | -0.01 | 0.217 | 0.217 | 0 | 0.11 | 0.33 | 0.26 | 0.3 | 0.08 | 0.18 |
| 11 | 20854395 | rs11025776 | -3.623 | -0.942 | 2.884 | 1.52 | -0.009 | 0.008 | 0.008 | 0 | 0.33 | 0.58 | 0.42 | 0.37 | 0.3 | 0.37 |
| 11 | 20855440 | rs12270521 | -2.451 | -0.836 | 3.015 | 1.581 | -0.028 | 0.12 | 0.12 | 0 | 0.13 | 0.27 | 0.25 | 0.31 | 0.08 | 0.17 |
| 11 | 20856545 | rs10833397 | -1.45 | -0.831 | 3.037 | 1.589 | -0.008 | 0.154 | 0.154 | 0 | 0.1 | 0.22 | 0.25 | 0.3 | 0.07 | 0.15 |
| 11 | 20860026 | rs10833398 | -2.558 | -0.832 | 3.154 | 1.56 | -0.002 | -0.039 | -0.039 | 0 | 0.14 | 0.26 | 0.12 | 0.03 | 0.14 | 0.18 |
| 11 | 20863059 | rs4923099 | -3.444 | -0.83 | 3.141 | 1.527 | -0.014 | 0.013 | 0.013 | 0.186 | 0.27 | 0.59 | 0.37 | 0.33 | 0.21 | 0.33 |
| 11 | 20869554 | rs2278318 | -1.962 | -0.922 | 3.054 | 1.718 | -0.022 | 0.14 | 0.14 | 0 | 0.12 | 0.27 | 0.25 | 0.32 | 0.07 | 0.16 |
| 11 | 20870736 | rs7927476 | 2.198 | -0.968 | 3.316 | 1.695 | -0.043 | 0.154 | 0.154 | 0.463 | 0.83 | 0.48 | 0.72 | 0.68 | 0.89 | 0.83 |
| 11 | 20871243 | rs1346690 | 2.867 | -1.024 | 3.191 | 1.658 | -0.026 | 0.057 | 0.057 | 0.027 | 0.76 | 0.42 | 0.66 | 0.65 | 0.8 | 0.75 |
| 11 | 20874438 | rs7121400 | -2.429 | -0.843 | 2.654 | 1.726 | -0.027 | 0.076 | 0.076 | 0.024 | 0.16 | 0.29 | 0.27 | 0.32 | 0.11 | 0.17 |
| 11 | 20878176 | rs1017324 | 2.144 | -0.817 | 2.201 | 1.285 | -0.002 | -0.003 | -0.003 | 0 | 0.69 | 0.3 | 0.7 | 0.75 | 0.74 | 0.72 |
| 11 | 20887699 | rs6483735 | 2.073 | -0.841 | 2.629 | 1.254 | -0.018 | -0.022 | -0.022 | 1.252 | 0.73 | 0.42 | 0.74 | 0.77 | 0.78 | 0.74 |
| 11 | 20888625 | rs919476 | -1.429 | -0.86 | 2.839 | 1.332 | 0.002 | 0.11 | 0.11 | 0 | 0.1 | 0.28 | 0.15 | 0.19 | 0.07 | 0.17 |
| 11 | 20892763 | rs7109004 | 2.249 | -0.391 | 1.862 | 1.967 | 0.004 | -0.04 | -0.04 | 0 | 0.68 | 0.18 | 0.7 | 0.63 | 0.71 | 0.72 |
| 11 | 20894098 | rs10766737 | 2.188 | -0.638 | 1.897 | 2.004 | 0.018 | -0.034 | -0.034 | 0 | 0.68 | 0.2 | 0.67 | 0.62 | 0.74 | 0.73 |
| 11 | 20898390 | rs10766739 | 2.313 | -0.683 | 1.552 | 2.053 | 0.016 | -0.082 | -0.082 | 0 | 0.67 | 0.14 | 0.57 | 0.58 | 0.73 | 0.73 |
| 11 | 20899163 | rs10741857 | 1.062 | -0.61 | 1.734 | 1.728 | 0.001 | -0.008 | -0.008 | 0 | 0.86 | 0.84 | 0.87 | 0.92 | 0.83 | 0.91 |
| 11 | 20903707 | rs4922666 | 1.745 | -0.559 | 1.912 | 1.574 | -0.011 | -0.063 | -0.063 | 0 | 0.78 | 0.65 | 0.85 | 0.93 | 0.84 | 0.89 |
| 11 | 20905942 | rs11025798 | 1.685 | -0.672 | 0.586 | 1.699 | -0.136 | 0.062 | 0.062 | 0.248 | 0.41 | 0.03 | 0.31 | 0.28 | 0.43 | 0.27 |
| 11 | 20908913 | rs7128766 | -0.983 | -1.423 | 1.967 | 2.027 | 0.016 | 0.161 | 0.161 | 0 | 0.1 | 0.3 | 0.25 | 0.19 | 0.07 | 0.19 |
| 11 | 20909301 | rs10766740 | 2.356 | -1.391 | 1.592 | 1.95 | 0.01 | -0.02 | -0.02 | 0 | 0.69 | 0.28 | 0.59 | 0.58 | 0.75 | 0.66 |
| 11 | 20910614 | rs1549717 | 1.156 | -1.3 | 0.985 | 1.737 | 0.003 | 0.043 | 0.043 | 0 | 0.83 | 0.89 | 0.88 | 0.94 | 0.86 | 0.88 |
| 11 | 20925039 | rs10766743 | 2.292 | -1.361 | 0.675 | 2.102 | 0.025 | -0.169 | -0.169 | 0 | 0.66 | 0.05 | 0.58 | 0.51 | 0.75 | 0.66 |
| 11 | 20926256 | rs1429793 | -1.513 | -1.338 | 1.283 | 2.076 | -0.006 | 0.106 | 0.106 | 0.043 | 0.18 | 0.43 | 0.32 | 0.34 | 0.16 | 0.29 |
| 11 | 20930384 | rs1594254 | 2.095 | -1.324 | 1.441 | 1.992 | -0.019 | 0.009 | 0.009 | 0.299 | 0.73 | 0.4 | 0.62 | 0.67 | 0.79 | 0.68 |
| 11 | 20930997 | rs7116897 | 0.677 | -1.306 | 1.157 | 1.913 | -0.006 | -0.03 | -0.03 | 0 | 0.92 | 0.85 | 0.94 | 1 | 0.94 | 0.98 |
| 11 | 20931291 | rs10833411 | 1.721 | -1.231 | 0.701 | 2.002 | 0.035 | -0.017 | -0.017 | 0 | 0.7 | 0.08 | 0.59 | 0.68 | 0.79 | 0.68 |
| 11 | 20933965 | rs1465604 | 1.748 | -1.322 | 0.819 | 1.908 | -0.024 | -0.015 | -0.015 | 0 | 0.72 | 0.37 | 0.61 | 0.74 | 0.8 | 0.68 |
| 11 | 20934448 | rs2082080 | 2.03 | -1.197 | 0.261 | 1.808 | 0.034 | -0.127 | -0.127 | 0.729 | 0.66 | 0.08 | 0.57 | 0.51 | 0.74 | 0.66 |
| 11 | 20939655 | rs3740874 | -1.388 | -1.222 | 0.389 | 1.293 | -0.014 | -0.036 | -0.036 | 0 | 0.07 | 0 | 0.02 | 0.11 | 0.06 | 0.03 |
| 11 | 20942507 | rs1429781 | 1.677 | -1.361 | 0.594 | 1.159 | -0.012 | -0.017 | -0.017 | 1.536 | 0.74 | 0.29 | 0.84 | 0.82 | 0.82 | 0.87 |
| 11 | 20949763 | rs4363596 | 0.816 | -2.183 | 0.799 | 0.509 | -0.017 | -0.081 | -0.081 | 0 | 0.7 | 0.64 | 0.85 | 1 | 0.84 | 0.85 |
| 11 | 20958343 | rs4923177 | 1.198 | -2.555 | 1.296 | 0.469 | -0.056 | -0.093 | -0.093 | 0.177 | 0.69 | 0.47 | 0.84 | 0.91 | 0.81 | 0.83 |
| 11 | 20959429 | rs1880087 | 1.937 | -2.674 | 1.472 | 0.947 | 0.053 | -0.009 | -0.009 | 0 | 0.6 | 0.25 | 0.7 | 0.56 | 0.74 | 0.71 |
| 11 | 20960904 | rs4923183 | 1.954 | -2.754 | 1.5 | 0.938 | 0.062 | -0.01 | -0.01 | 0 | 0.6 | 0.24 | 0.7 | 0.56 | 0.75 | 0.71 |
| 11 | 20962109 | rs7943038 | 1.039 | -2.709 | 1.353 | 0.917 | -0.031 | -0.1 | -0.1 | 0 | 0.71 | 0.63 | 0.84 | 1 | 0.84 | 0.84 |
| 11 | 20963548 | rs2896621 | -1.081 | -2.674 | 1.3 | 1.093 | -0.012 | -0.035 | -0.035 | 0 | 0.1 | 0.03 | 0.05 | 0.29 | 0.1 | 0.14 |
| 11 | 20963571 | rs2403645 | 0.125 | -2.716 | 1.434 | 1.061 | -0.004 | -0.003 | -0.003 | 0.41 | 0.06 | 0.03 | 0.04 | 0.2 | 0.06 | 0.11 |
| 11 | 20966234 | rs7930660 | 0.905 | -2.736 | 1.564 | 0.811 | -0.101 | -0.173 | -0.173 | 0.546 | 0.74 | 0.48 | 0.88 | 1 | 0.9 | 0.87 |

| | | | | | | | | | | | | | | | |
|---|---|---|---|---|---|---|---|---|---|---|---|---|---|---|---|
| 11 | 20973270 | rs6416031 | 0.069 | -2.417 | 0.916 | 0.928 | 0.023 | 0.011 | 0.011 | 2.151 | 0.85 | 0.9 | 0.92 | 1 | 0.93 | 0.9 |
| 11 | 20978633 | rs16907140 | -0.983 | -1.728 | 0.383 | 1.972 | 0 | 0.054 | 0.054 | 0 | 0.1 | 0.08 | 0.22 | 0.46 | 0.08 | 0.14 |
| 11 | 20980037 | rs11025836 | -1.109 | -2.511 | -0.055 | 1.226 | 0.036 | -0.009 | -0.009 | 0.055 | 0.3 | 0.19 | 0.14 | 0.35 | 0.2 | 0.24 |
| 11 | 20980514 | rs7108916 | -2.013 | -2.651 | 0.141 | 1.173 | 0.036 | 0.002 | 0.002 | 0.238 | 0.44 | 0.34 | 0.4 | 0.62 | 0.3 | 0.32 |
| 11 | 20981894 | rs2280584 | -2.07 | -2.692 | 0.594 | 1.161 | -0.014 | -0.002 | -0.002 | 0 | 0.44 | 0.48 | 0.35 | 0.61 | 0.31 | 0.33 |
| 11 | 20983410 | rs7105756 | -1.805 | -2.743 | 0.796 | 1.152 | 0.005 | -0.004 | -0.004 | 0 | 0.4 | 0.41 | 0.3 | 0.51 | 0.24 | 0.27 |
| 11 | 20984077 | rs7109380 | -1.864 | -2.749 | 0.599 | 1.126 | -0.028 | 0.002 | 0.002 | 0.659 | 0.4 | 0.52 | 0.33 | 0.6 | 0.28 | 0.31 |
| 11 | 20989375 | rs11025839 | -0.788 | -2.68 | 0.48 | 1.147 | -0.003 | 0.008 | 0.008 | 0.013 | 0.14 | 0.18 | 0.2 | 0.26 | 0.11 | 0.09 |
| 11 | 20992263 | rs2003175 | -0.368 | -2.704 | 0.539 | 1.178 | -0.002 | 0.11 | 0.11 | 0 | 0.1 | 0.26 | 0.18 | 0.26 | 0.08 | 0.07 |
| 11 | 20995172 | rs10833419 | -0.814 | -2.762 | 0.319 | 1.065 | -0.009 | -0.001 | -0.001 | 0 | 0.26 | 0.26 | 0.1 | 0.26 | 0.13 | 0.19 |
| 11 | 20999558 | rs1400373 | -0.801 | -2.699 | 0.313 | 1.052 | -0.002 | 0 | 0 | 0 | 0.26 | 0.25 | 0.1 | 0.26 | 0.14 | 0.19 |
| 11 | 21002718 | rs2040316 | -1.601 | -2.249 | -0.249 | 0.643 | -0.043 | 0.058 | 0.058 | 0 | 0.61 | 0.8 | 0.48 | 0.77 | 0.55 | 0.59 |
| 11 | 21003275 | rs7482901 | 0.098 | -2.67 | -0.286 | 0.926 | -0.003 | 0.044 | 0.044 | 0 | 0.08 | 0.11 | 0.17 | 0.28 | 0.08 | 0.07 |
| 11 | 21005890 | rs10833425 | -1.324 | -2.759 | -0.329 | 0.826 | -0.022 | 0.224 | 0.224 | 0.229 | 0.19 | 0.51 | 0.22 | 0.52 | 0.16 | 0.16 |
| 11 | 21008120 | rs10833426 | -0.753 | -2.736 | -0.501 | 0.694 | 0.065 | 0.075 | 0.075 | 12.35 | 0.23 | 0.03 | 0.11 | 0.1 | 0.13 | 0.16 |
| 11 | 21010668 | rs7939928 | -0.161 | -2.557 | -0.36 | 0.765 | 0 | 0.004 | 0.004 | 0 | 0.07 | 0.14 | 0.06 | 0.09 | 0.07 | 0.07 |
| 11 | 21011821 | rs7122648 | -2.235 | -2.758 | -0.681 | 0.653 | 0 | 0.001 | 0.001 | 0.446 | 0.51 | 0.54 | 0.35 | 0.62 | 0.32 | 0.34 |
| 11 | 21012786 | rs10833428 | -1.325 | -2.766 | -0.674 | 0.7 | -0.009 | -0.002 | -0.002 | 1.569 | 0.38 | 0.41 | 0.16 | 0.35 | 0.22 | 0.26 |
| 11 | 21014765 | rs1914243 | -0.76 | -1.827 | -0.393 | 0.937 | 0.014 | -0.022 | -0.022 | 0 | 0.25 | 0.15 | 0.25 | 0.36 | 0.19 | 0.16 |
| 11 | 21016033 | rs10766761 | -1.268 | -1.781 | -0.494 | 0.717 | -0.021 | -0.003 | -0.003 | 0 | 0.43 | 0.51 | 0.32 | 0.61 | 0.3 | 0.27 |
| 11 | 21016295 | rs1400369 | -1.094 | -1.261 | -0.533 | 0.581 | -0.019 | 0.011 | 0.011 | 0 | 0.34 | 0.37 | 0.27 | 0.43 | 0.23 | 0.23 |
| 11 | 21017397 | rs2896623 | -0.658 | -1.178 | -1.237 | -0.257 | 0.015 | 0.086 | 0.086 | 0 | 0.22 | 0.05 | 0.09 | 0.02 | 0.16 | 0.3 |
| 11 | 21018940 | rs4923240 | -1.769 | -1.394 | -1.366 | 0.027 | 0.02 | 0.004 | 0.004 | 0 | 0.54 | 0.77 | 0.44 | 0.59 | 0.41 | 0.53 |
| 11 | 21020242 | rs1400367 | -1.016 | -1.961 | -0.524 | -0.386 | 0.008 | -0.011 | -0.011 | 3.335 | 0.32 | 0.27 | 0.15 | 0.02 | 0.23 | 0.31 |
| 11 | 21020560 | rs1400366 | -0.968 | -2.086 | -0.333 | -0.45 | -0.03 | -0.091 | -0.091 | 0.136 | 0.28 | 0.62 | 0.13 | 0.02 | 0.16 | 0.29 |
| 11 | 21021337 | rs4644637 | -2.134 | -2.291 | -0.197 | -0.634 | -0.013 | -0.068 | -0.068 | 0.262 | 0.29 | 0.66 | 0.17 | 0.02 | 0.19 | 0.22 |
| 11 | 21023900 | rs17232918 | -0.269 | -2.277 | -0.232 | -0.621 | 0.01 | 0.053 | 0.053 | 0 | 0.13 | 0.02 | 0.07 | 0 | 0.1 | 0.05 |
| 11 | 21024896 | rs7949652 | -1.26 | -2.284 | -0.214 | -0.618 | 0.008 | 0.027 | 0.027 | 0.408 | 0.22 | 0.13 | 0.12 | 0.03 | 0.17 | 0.22 |
| 11 | 21030704 | rs1516773 | -1.111 | -2.262 | -0.177 | -0.618 | 0.023 | -0.001 | -0.001 | 0.52 | 0.26 | 0.18 | 0.11 | 0 | 0.15 | 0.17 |
| 11 | 21032150 | rs17232925 | -0.198 | -2.224 | -0.192 | -0.593 | 0.011 | 0.043 | 0.043 | 0 | 0.13 | 0.03 | 0.07 | 0 | 0.1 | 0.04 |
| 11 | 21032271 | rs7941875 | -1.366 | -1.997 | -0.008 | -0.435 | -0.021 | -0.053 | -0.053 | 0.662 | 0.18 | 0.72 | 0.12 | 0.03 | 0.1 | 0.17 |
| 11 | 21034351 | rs11025874 | 1.329 | -1.101 | -0.457 | -0.265 | -0.077 | -0.089 | -0.089 | 0 | 0.14 | 0.01 | 0.14 | 0.29 | 0.23 | 0.17 |
| 11 | 21041759 | rs11025878 | -1.062 | -1.817 | -0.41 | -0.632 | -0.002 | -0.056 | -0.056 | 0.363 | 0.41 | 0.6 | 0.2 | 0.03 | 0.25 | 0.33 |
| 11 | 21044896 | rs7122403 | -1.487 | -0.902 | 0.38 | 0.578 | -0.017 | -0.02 | -0.02 | 0 | 0.08 | 0.59 | 0.05 | 0 | 0.01 | 0.01 |
| 11 | 21047605 | rs1914249 | 0.742 | -0.405 | -0.383 | 1.379 | -0.001 | 0.033 | 0.033 | 0 | 0.89 | 0.87 | 0.77 | 0.69 | 0.92 | 0.84 |
| 11 | 21049960 | rs16907292 | -0.42 | -0.573 | -0.709 | 1.818 | 0 | -0.018 | -0.018 | 0 | 0.07 | 0 | 0.1 | 0.35 | 0.04 | 0.1 |
| 11 | 21052712 | rs2063916 | -0.608 | -0.507 | -0.916 | 1.573 | -0.012 | 0.019 | 0.019 | 0 | 0.15 | 0.04 | 0.2 | 0.1 | 0.18 | 0.24 |
| 11 | 21053214 | rs4922728 | -1.509 | -0.654 | -0.208 | 1.371 | -0.007 | 0.013 | 0.013 | 0 | 0.26 | 0.36 | 0.21 | 0.39 | 0.22 | 0.33 |
| 11 | 21061196 | rs7937542 | 0.688 | -0.478 | -0.609 | 0.947 | 0.013 | 0.078 | 0.078 | 0.104 | 0.49 | 0.21 | 0.54 | 0.3 | 0.51 | 0.45 |
| 11 | 21062689 | rs11025886 | -0.572 | -0.684 | -0.916 | 1.264 | 0.005 | -0.062 | -0.062 | 0 | 0.1 | 0 | 0.08 | 0.18 | 0.05 | 0.11 |
| 11 | 21063785 | rs16907306 | -0.229 | -0.985 | -1.08 | 1.021 | 0.006 | 0.039 | 0.039 | 0 | 0.19 | 0.03 | 0.21 | 0.17 | 0.18 | 0.29 |
| 11 | 21064035 | rs12803840 | 1.66 | -1.211 | -1.292 | 0.865 | -0.028 | 0.042 | 0.042 | 0 | 0.36 | 0.17 | 0.48 | 0.23 | 0.4 | 0.32 |
| 11 | 21064322 | rs12577018 | -1.055 | -0.974 | -0.12 | 1.497 | -0.004 | 0.008 | 0.008 | 0 | 0.12 | 0.27 | 0.11 | 0.13 | 0.12 | 0.16 |
| 11 | 21064356 | rs16907316 | 0.007 | -0.812 | -0.082 | 1.355 | 0.033 | 0.019 | 0.019 | 0 | 0.16 | 0.04 | 0.16 | 0.17 | 0.09 | 0.26 |
| 11 | 21065744 | rs1516761 | -0.736 | -0.2 | 0.96 | 2.134 | 0.065 | 0.182 | 0.182 | 0 | 0.18 | 0.52 | 0.28 | 0.51 | 0.27 | 0.17 |
| 11 | 21069877 | rs940199 | -1.04 | -0.283 | 0.898 | 2.169 | 0.04 | 0.1 | 0.1 | 0 | 0.2 | 0.53 | 0.27 | 0.61 | 0.27 | 0.22 |
| 11 | 21075331 | rs10833447 | 0.972 | -0.166 | 0.121 | 2.226 | -0.05 | -0.045 | -0.045 | 0 | 0.09 | 0.01 | 0.08 | 0.35 | 0.16 | 0.1 |
| 11 | 21076108 | rs10833448 | -1.016 | -0.171 | 0.593 | 1.755 | 0.021 | 0.089 | 0.089 | 0 | 0.21 | 0.59 | 0.28 | 0.61 | 0.26 | 0.22 |
| 11 | 21076967 | rs7951965 | -0.55 | -0.045 | 0.589 | 1.643 | 0.035 | 0.246 | 0.246 | 0.034 | 0.18 | 0.51 | 0.12 | 0.41 | 0.17 | 0.14 |
| 11 | 21079942 | rs7120935 | -0.915 | -0.003 | -0.717 | 0.378 | -0.023 | 0.123 | 0.123 | 0 | 0.58 | 0.85 | 0.63 | 0.82 | 0.56 | 0.69 |
| 11 | 21081277 | rs1516757 | -1.31 | 0.152 | -1.194 | -0.182 | -0.067 | 0.089 | 0.089 | 0 | 0.83 | 0.95 | 0.82 | 1 | 0.73 | 0.89 |
| 11 | 21081891 | rs9666195 | -0.09 | 0.629 | -0.921 | 0.056 | 0.003 | -0.011 | -0.011 | 0 | 0.39 | 0.68 | 0.3 | 0.53 | 0.35 | 0.4 |
| 11 | 21082675 | rs1914247 | -0.002 | 0.494 | -0.403 | 0.25 | 0.001 | 0.023 | 0.023 | 0 | 0.24 | 0.27 | 0.33 | 0.38 | 0.22 | 0.42 |
| 11 | 21082995 | rs1516755 | 0.04 | 0.475 | -0.376 | 0.255 | 0 | 0.017 | 0.017 | 0 | 0.25 | 0.26 | 0.33 | 0.38 | 0.22 | 0.42 |
| 11 | 21090426 | rs7948905 | -0.011 | 0.476 | -0.287 | 0.399 | 0.001 | -0.002 | -0.002 | 0 | 0.57 | 0.57 | 0.48 | 0.63 | 0.49 | 0.49 |
| 11 | 21094330 | rs1516747 | 0.325 | 0.218 | -0.442 | 0.215 | 0.051 | 0.024 | 0.024 | 0 | 0.31 | 0.47 | 0.36 | 0.44 | 0.42 | 0.27 |
| 11 | 21094956 | rs7924855 | -0.323 | 0.163 | -0.59 | 0.193 | -0.002 | 0.001 | 0.001 | 0 | 0.6 | 0.56 | 0.59 | 0.62 | 0.61 | 0.54 |
| 11 | 21099887 | rs10833455 | 0.217 | 0.387 | -0.493 | 0.226 | 0.062 | 0.027 | 0.027 | 0 | 0.31 | 0.53 | 0.35 | 0.44 | 0.41 | 0.26 |
| 11 | 21104020 | rs1894054 | -0.657 | 0.305 | -0.748 | 0.229 | 0.008 | 0.021 | 0.021 | 0 | 0.62 | 0.77 | 0.47 | 0.7 | 0.43 | 0.57 |
| 11 | 21107429 | rs7110500 | -0.132 | -0.144 | -0.582 | 0.119 | -0.024 | 0.008 | 0.008 | 0.265 | 0.07 | 0.13 | 0.02 | 0.09 | 0.01 | 0.12 |
| 11 | 21109584 | rs12418446 | 1.439 | -0.353 | -0.832 | 0.074 | 0.039 | -0.004 | -0.004 | 0 | 0.24 | 0.03 | 0.2 | 0.17 | 0.17 | 0.19 |
| 11 | 21114308 | rs2138599 | 0.429 | -0.42 | -0.155 | 0.072 | 0.003 | 0.002 | 0.002 | 0 | 0.33 | 0.34 | 0.23 | 0.25 | 0.17 | 0.36 |
| 11 | 21121959 | rs11025915 | -0.252 | -0.246 | -0.582 | 0.115 | 0.038 | -0.031 | -0.031 | 0.253 | 0.08 | 0 | 0.01 | 0.09 | 0.02 | 0.14 |
| 11 | 21129269 | rs2040315 | 0 | -0.213 | 0.048 | 0.217 | -0.029 | -0.006 | -0.006 | 0 | 0.39 | 0.48 | 0.29 | 0.31 | 0.24 | 0.45 |
| 11 | 21136563 | rs7928566 | -0.462 | 0.339 | -0.015 | 0.376 | -0.004 | -0.005 | -0.005 | 0 | 0.74 | 0.84 | 0.62 | 0.67 | 0.71 | 0.7 |
| 11 | 21141191 | rs11025927 | 0.311 | 0.184 | 0.007 | -0.004 | -0.003 | 0.002 | 0.002 | 0 | 0.14 | 0.12 | 0.09 | 0.06 | 0.15 | 0.11 |
| 11 | 21148724 | rs12282210 | 1.478 | -0.298 | 0.347 | 0.122 | -0.002 | 0 | 0 | 0.121 | 0.3 | 0.3 | 0.27 | 0.19 | 0.19 | 0.24 |
| 11 | 21157762 | rs1607098 | -0.353 | -0.255 | 0.442 | 0.163 | 0 | -0.012 | -0.012 | 0 | 0.69 | 0.78 | 0.61 | 0.55 | 0.71 | 0.58 |
| 11 | 21158204 | rs10766768 | -0.314 | -0.306 | 0.753 | 0.127 | 0.004 | -0.02 | -0.02 | 0.556 | 0.43 | 0.53 | 0.35 | 0.35 | 0.52 | 0.37 |

| | | | | | | | | | | | | | | | | |
|---|---|---|---|---|---|---|---|---|---|---|---|---|---|---|---|---|
| 11 | 21158287 | rs10766769 | -0.314 | -0.306 | 0.753 | 0.127 | 0.004 | -0.02 | -0.02 | 1.499 | 0.43 | 0.53 | 0.35 | 0.36 | 0.52 | 0.37 |
| 11 | 21159558 | rs1260149 | 1.46 | -0.305 | 0.805 | 0.133 | -0.015 | -0.006 | -0.006 | 0 | 0.3 | 0.31 | 0.27 | 0.2 | 0.19 | 0.24 |
| 11 | 21159590 | rs10833465 | -0.005 | -0.084 | 0.821 | 0.066 | -0.003 | -0.002 | -0.002 | 0 | 0.48 | 0.52 | 0.37 | 0.36 | 0.35 | 0.47 |
| 11 | 21164694 | rs1984369 | 1.157 | -0.056 | 0.788 | 0.053 | -0.007 | 0.002 | 0.002 | 0 | 0.25 | 0.25 | 0.24 | 0.3 | 0.36 | 0.24 |
| 11 | 21165137 | rs11025944 | 0.707 | -0.046 | 0.704 | 0.053 | -0.001 | -0.001 | -0.001 | 0 | 0.14 | 0.13 | 0.09 | 0.06 | 0.16 | 0.11 |
| 11 | 21167078 | rs1453991 | 1.1 | -0.455 | 0.627 | 0.125 | 0.015 | -0.004 | -0.004 | 0.396 | 0.34 | 0.28 | 0.27 | 0.3 | 0.2 | 0.35 |
| 11 | 21167190 | rs1614424 | 1.03 | -0.44 | 0.582 | 0.133 | 0.003 | -0.002 | -0.002 | 0.221 | 0.34 | 0.31 | 0.27 | 0.3 | 0.2 | 0.35 |
| 11 | 21168295 | rs1791810 | -0.288 | 0.069 | 0.399 | 0.481 | -0.002 | -0.003 | -0.003 | 0 | 0.74 | 0.8 | 0.61 | 0.66 | 0.71 | 0.7 |
| 11 | 21171018 | rs12788723 | -0.173 | 0.436 | 0.568 | 0.065 | 0.006 | -0.015 | -0.015 | 0 | 0.14 | 0.14 | 0.17 | 0.06 | 0.17 | 0.11 |
| 11 | 21172069 | rs1628441 | -1.018 | -0.49 | 0.498 | -0.107 | -0.001 | -0.006 | -0.006 | 0 | 0.44 | 0.48 | 0.51 | 0.58 | 0.38 | 0.54 |
| 11 | 21173654 | rs7113280 | -0.45 | -0.595 | 0.489 | -0.135 | 0.008 | -0.031 | -0.031 | 0 | 0.35 | 0.19 | 0.45 | 0.54 | 0.3 | 0.49 |
| 11 | 21174849 | rs1631571 | -0.795 | -0.674 | 0.296 | -0.37 | -0.014 | 0.003 | 0.003 | 0 | 0.74 | 0.83 | 0.8 | 0.69 | 0.67 | 0.72 |
| 11 | 21175537 | rs10766770 | 1.035 | -0.683 | 0.546 | -0.398 | 0.03 | 0.066 | 0.066 | 0.256 | 0.37 | 0.26 | 0.31 | 0.12 | 0.34 | 0.25 |
| 11 | 21177159 | rs1453997 | 0.89 | -0.647 | 0.723 | -0.093 | 0.026 | 0.032 | 0.032 | 0.099 | 0.39 | 0.34 | 0.31 | 0.18 | 0.33 | 0.34 |
| 11 | 21182718 | rs11025951 | 0.039 | -0.521 | 0.822 | 0.06 | 0.001 | -0.007 | -0.007 | 0 | 0.09 | 0.12 | 0.05 | 0.04 | 0.08 | 0.05 |
| 11 | 21183993 | rs1453998 | -0.267 | -0.441 | 1.023 | 0.157 | 0.031 | 0.006 | 0.006 | 0 | 0.75 | 0.65 | 0.79 | 0.67 | 0.65 | 0.75 |
| 11 | 21184296 | rs10766771 | -1.786 | -0.32 | 0.911 | 0.154 | 0.003 | 0.001 | 0.001 | 0 | 0.85 | 0.83 | 0.85 | 0.7 | 0.76 | 0.79 |
| 11 | 21184613 | rs1454000 | -1.752 | -0.298 | 0.723 | 0.166 | -0.011 | -0.009 | -0.009 | 0 | 0.86 | 0.88 | 0.86 | 0.7 | 0.76 | 0.8 |
| 11 | 21186719 | rs4923385 | 0.421 | -0.429 | 1.007 | 0.512 | 0 | 0.006 | 0.006 | 0 | 0.27 | 0.25 | 0.44 | 0.48 | 0.26 | 0.34 |
| 11 | 21186995 | rs7119634 | -0.241 | -0.417 | 0.927 | 0.507 | 0 | 0.007 | 0.007 | 0 | 0.54 | 0.57 | 0.39 | 0.23 | 0.45 | 0.41 |
| 11 | 21189157 | rs10500895 | 0.614 | -0.396 | 0.661 | 0.523 | -0.013 | -0.064 | -0.064 | 2.13 | 0.25 | 0.11 | 0.42 | 0.48 | 0.25 | 0.34 |
| 11 | 21191016 | rs10500896 | 0.816 | -0.207 | 0.473 | 0.59 | 0.017 | 0.043 | 0.043 | 0 | 0.4 | 0.34 | 0.32 | 0.12 | 0.35 | 0.25 |
| 11 | 21192129 | rs7940204 | 0.734 | -0.182 | 0.176 | 0.602 | 0.02 | 0.054 | 0.054 | 0 | 0.4 | 0.33 | 0.32 | 0.12 | 0.35 | 0.25 |
| 11 | 21192532 | rs10833472 | 0.02 | -0.196 | -0.038 | 0.617 | 0.001 | -0.003 | -0.003 | 0.279 | 0.49 | 0.51 | 0.39 | 0.15 | 0.45 | 0.3 |
| 11 | 21195352 | rs10500898 | 0.51 | 0.379 | -0.112 | 0.815 | 0.005 | 0.008 | 0.008 | 0 | 0.42 | 0.4 | 0.33 | 0.2 | 0.35 | 0.35 |
| 11 | 21199804 | rs4923403 | -0.379 | 1.003 | 0.019 | 1.144 | -0.02 | -0.002 | -0.002 | 0 | 0.78 | 0.87 | 0.82 | 0.7 | 0.7 | 0.74 |
| 11 | 21205090 | rs7130084 | -1.428 | 0.684 | 1.026 | 2.004 | 0.049 | 0.108 | 0.108 | 0 | 0.16 | 0.51 | 0.16 | 0.28 | 0.16 | 0.15 |
| 11 | 21207667 | rs12575552 | -1.289 | 0.446 | 0.637 | 2.573 | 0.001 | 0.089 | 0.089 | 0 | 0.08 | 0.08 | 0.21 | 0.49 | 0.13 | 0.13 |
| 11 | 21210363 | rs7115088 | -1.789 | 0.372 | 0.895 | 2.556 | 0.001 | 0.143 | 0.143 | 0 | 0.1 | 0.17 | 0.22 | 0.51 | 0.14 | 0.15 |
| 11 | 21213962 | rs17233103 | -0.832 | 0.404 | 0.838 | 2.362 | -0.022 | -0.015 | -0.015 | 1.033 | 0.05 | 0.01 | 0.06 | 0.05 | 0.08 | 0.08 |
| 11 | 21214340 | rs11823347 | -0.482 | 0.244 | 0.844 | 2.601 | 0.006 | -0.017 | -0.017 | 0 | 0.11 | 0.05 | 0.07 | 0.12 | 0.06 | 0.07 |
| 11 | 21215845 | rs1791847 | 1.423 | 0.132 | 1.625 | 2.558 | -0.004 | 0.017 | 0.017 | 0.017 | 0.82 | 0.68 | 0.84 | 0.83 | 0.86 | 0.87 |
| 11 | 21216852 | rs1670644 | 2.04 | -0.091 | 1.119 | 2.863 | 0.023 | -0.055 | -0.055 | 1.144 | 0.72 | 0.17 | 0.7 | 0.46 | 0.8 | 0.71 |
| 11 | 21227564 | rs16907672 | -0.874 | -0.093 | 0.313 | 2.787 | 0.01 | -0.009 | -0.009 | 1.035 | 0.11 | 0.07 | 0.07 | 0.12 | 0.06 | 0.06 |
| 11 | 21234023 | rs11025965 | -1.368 | 0.125 | -0.258 | 2.623 | 0.012 | -0.025 | -0.025 | 0 | 0.16 | 0.05 | 0.11 | 0.17 | 0.09 | 0.14 |
| 11 | 21234245 | rs1945327 | -1.021 | 0.373 | 0.177 | 3.265 | 0.002 | 0.124 | 0.124 | 0 | 0.13 | 0.16 | 0.26 | 0.49 | 0.11 | 0.15 |
| 11 | 21235289 | rs12272954 | -0.224 | 0.45 | -0.729 | 2.499 | -0.042 | -0.064 | -0.064 | 2.968 | 0.22 | 0.06 | 0.24 | 0.24 | 0.24 | 0.25 |
| 11 | 21240934 | rs1791869 | -0.4 | 0.458 | -0.624 | 2.467 | -0.014 | -0.012 | -0.012 | 0 | 0.23 | 0.14 | 0.25 | 0.19 | 0.24 | 0.19 |
| 11 | 21244703 | rs1453988 | -0.433 | 0.441 | -0.745 | 2.317 | -0.03 | -0.031 | -0.031 | 0 | 0.27 | 0.09 | 0.26 | 0.22 | 0.27 | 0.21 |
| 11 | 21251200 | rs1670638 | -0.455 | 0.568 | -0.582 | 1.663 | -0.006 | -0.009 | -0.009 | 0 | 0.27 | 0.18 | 0.27 | 0.23 | 0.27 | 0.21 |
| 11 | 21258878 | rs1454013 | 0.069 | 0.507 | -0.125 | 1.178 | 0 | -0.001 | -0.001 | 0 | 0.43 | 0.55 | 0.4 | 0.59 | 0.39 | 0.49 |
| 11 | 21259545 | rs2085028 | 0.353 | 0.314 | 0.245 | 1.283 | -0.005 | -0.003 | -0.003 | 0 | 0.34 | 0.32 | 0.3 | 0.37 | 0.36 | 0.39 |
| 11 | 21260658 | rs7109435 | -0.253 | 0.709 | -0.314 | 0.448 | -0.005 | -0.009 | -0.009 | 0.247 | 0.21 | 0.18 | 0.2 | 0.03 | 0.27 | 0.16 |
| 11 | 21262032 | rs11025990 | -0.443 | 0.567 | 0.015 | 0.889 | 0 | 0.001 | 0.001 | 0.02 | 0.61 | 0.36 | 0.64 | 0.57 | 0.61 | 0.67 |
| 11 | 21262344 | rs7117679 | -0.054 | 0.154 | -0.179 | 0.782 | -0.009 | 0.008 | 0.008 | 0 | 0.3 | 0.3 | 0.24 | 0.41 | 0.22 | 0.38 |
| 11 | 21262460 | rs12805365 | -0.613 | 1.579 | -0.159 | 0.607 | 0.031 | 0.017 | 0.017 | 0 | 0.16 | 0.25 | 0.18 | 0.22 | 0.23 | 0.3 |
| 11 | 21262591 | rs1454001 | 0.347 | 0.451 | -1.03 | -0.042 | 0.074 | -0.029 | -0.029 | 0 | 0.44 | 0.26 | 0.41 | 0.52 | 0.31 | 0.55 |
| 11 | 21263171 | rs10766778 | 0.185 | 0.883 | -0.76 | -0.826 | -0.009 | -0.089 | -0.089 | 0 | 0.73 | 0.41 | 0.84 | 0.99 | 0.78 | 0.89 |
| 11 | 21263842 | rs1945331 | -0.251 | 1.495 | -1.413 | -0.622 | -0.058 | 0.014 | 0.014 | 0.055 | 0.11 | 0.01 | 0.08 | 0.05 | 0.16 | 0.05 |
| 11 | 21264300 | rs4923452 | -0.549 | 1.262 | -0.973 | -0.978 | -0.002 | 0.008 | 0.008 | 0.787 | 0.3 | 0.24 | 0.17 | 0.01 | 0.29 | 0.16 |
| 11 | 21267570 | rs16907774 | -0.363 | 1.259 | -0.981 | 0.009 | 0.008 | 0.018 | 0.018 | 0 | 0.13 | 0.14 | 0.21 | 0.26 | 0.18 | 0.16 |
| 11 | 21268909 | rs1791862 | -0.506 | 1.104 | -1.351 | -0.303 | -0.09 | 0.093 | 0.093 | 0 | 0.29 | 0.11 | 0.24 | 0.1 | 0.45 | 0.13 |
| 11 | 21274168 | rs1629381 | -0.111 | 0.801 | -0.97 | 0.145 | -0.001 | 0.003 | 0.003 | 0 | 0.88 | 0.89 | 0.9 | 0.8 | 0.93 | 0.91 |
| 11 | 21274683 | rs10833482 | 0.18 | -0.47 | -0.663 | 0.499 | -0.046 | 0.104 | 0.104 | 0 | 0.2 | 0.4 | 0.22 | 0.35 | 0.11 | 0.39 |
| 11 | 21276725 | rs1454004 | -0.326 | -0.448 | -0.619 | 0.471 | -0.006 | 0.011 | 0.011 | 0 | 0.88 | 0.86 | 0.9 | 0.8 | 0.93 | 0.92 |
| 11 | 21277189 | rs10766779 | 0.354 | -0.74 | -0.502 | 0.239 | 0 | 0.002 | 0.002 | 0 | 0.54 | 0.55 | 0.57 | 0.59 | 0.47 | 0.76 |
| 11 | 21280519 | rs1628540 | 0.267 | -0.895 | -0.529 | 0.122 | -0.037 | 0.046 | 0.046 | 0 | 0.33 | 0.19 | 0.28 | 0.16 | 0.47 | 0.16 |
| 11 | 21281105 | rs723468 | 0.703 | -0.83 | -0.85 | 0.128 | -0.018 | 0.002 | 0.002 | 0 | 0.06 | 0 | 0.07 | 0.04 | 0.06 | 0.03 |
| 11 | 21282111 | rs10466386 | 1.63 | -1.07 | -1.015 | 0.083 | -0.074 | 0.045 | 0.045 | 0 | 0.31 | 0.02 | 0.33 | 0.18 | 0.32 | 0.28 |
| 11 | 21284695 | rs1670629 | -0.121 | -0.922 | -0.709 | 0.269 | -0.006 | 0.013 | 0.013 | 0 | 0.84 | 0.81 | 0.84 | 0.77 | 0.9 | 0.89 |
| 11 | 21285045 | rs1791827 | -1.111 | -0.976 | -0.692 | 0.266 | -0.008 | 0.044 | 0.044 | 0 | 0.89 | 0.84 | 0.9 | 0.8 | 0.93 | 0.91 |
| 11 | 21302821 | rs1791872 | -0.16 | -1.551 | -0.895 | 0.086 | 0.002 | 0.009 | 0.009 | 0 | 0.4 | 0.52 | 0.51 | 0.56 | 0.58 | 0.61 |
| 11 | 21306113 | rs4151054 | 0.163 | -1.232 | -0.833 | 0.305 | -0.097 | 0.101 | 0.101 | 0 | 0.36 | 0.13 | 0.3 | 0.21 | 0.49 | 0.19 |
| 11 | 21308077 | rs10833492 | 0.69 | -1.317 | -0.754 | 0.335 | 0.007 | 0 | 0 | 0 | 0.48 | 0.47 | 0.54 | 0.53 | 0.44 | 0.69 |
| 11 | 21308199 | rs7127622 | 0.294 | -1.256 | -1.075 | 0.363 | -0.115 | 0.095 | 0.095 | 0 | 0.35 | 0.1 | 0.29 | 0.2 | 0.48 | 0.2 |
| 11 | 21310513 | rs7113875 | -1.302 | -1.183 | -0.867 | 0.495 | -0.017 | 0.075 | 0.075 | 0 | 0.9 | 0.8 | 0.91 | 0.8 | 0.96 | 0.92 |
| 11 | 21316281 | rs6483761 | -1.371 | -1.112 | -1.004 | 0.5 | -0.016 | 0.056 | 0.056 | 0 | 0.91 | 0.84 | 0.92 | 0.79 | 0.96 | 0.93 |
| 11 | 21321010 | rs11820583 | 0.045 | -1.366 | -1.338 | 0.378 | -0.035 | 0.057 | 0.057 | 0 | 0.2 | 0.31 | 0.22 | 0.32 | 0.12 | 0.41 |
| 11 | 21321305 | rs10833498 | 1.604 | -1.313 | -1.47 | 0.295 | -0.069 | 0.034 | 0.034 | 0 | 0.3 | 0.03 | 0.31 | 0.19 | 0.31 | 0.26 |

| | | | | | | | | | | | | | | | | |
|---|---|---|---|---|---|---|---|---|---|---|---|---|---|---|---|---|
| 11 | 21325338 | rs936008 | 0.377 | -1.101 | -1.217 | 0.338 | -0.111 | 0.081 | 0.081 | 0 | 0.33 | 0.09 | 0.29 | 0.2 | 0.48 | 0.18 |
| 11 | 21327123 | rs10833500 | 0.465 | -1.131 | -1.105 | 0.343 | -0.027 | 0.047 | 0.047 | 0 | 0.17 | 0.26 | 0.2 | 0.31 | 0.1 | 0.39 |
| 11 | 21327159 | rs1531005 | -0.097 | -0.642 | -0.855 | 0.536 | 0.011 | 0.004 | 0.004 | 0 | 0.51 | 0.41 | 0.49 | 0.47 | 0.57 | 0.58 |
| 11 | 21339877 | rs1453987 | 0.255 | -0.46 | -1.09 | 0.746 | -0.009 | 0.025 | 0.025 | 0 | 0.46 | 0.35 | 0.47 | 0.35 | 0.52 | 0.44 |
| 11 | 21343939 | rs4923528 | -0.315 | -0.427 | -1.554 | 1.102 | -0.006 | 0.034 | 0.034 | 0.03 | 0.87 | 0.86 | 0.9 | 0.67 | 0.92 | 0.81 |
| 11 | 21345606 | rs11603932 | 0.642 | 0.009 | -1.996 | 0.507 | -0.057 | 0.036 | 0.036 | 0.001 | 0.26 | 0.03 | 0.31 | 0.17 | 0.27 | 0.21 |
| 11 | 21346365 | rs4399327 | -0.198 | 0.559 | -1.703 | 0.691 | -0.006 | -0.002 | -0.002 | 0 | 0.58 | 0.43 | 0.59 | 0.55 | 0.52 | 0.55 |
| 11 | 21350620 | rs10833507 | 0.268 | 0.798 | -2.109 | 0.443 | -0.017 | 0.033 | 0.033 | 0 | 0.25 | 0.06 | 0.28 | 0.15 | 0.28 | 0.25 |
| 11 | 21351945 | rs1945425 | 0.91 | 0.696 | -2.388 | 0.551 | -0.019 | -0.054 | -0.054 | 1.878 | 0.17 | 0.01 | 0.09 | 0.15 | 0.17 | 0.08 |
| 11 | 21355150 | rs10833509 | -0.41 | 1.079 | -1.251 | 0.485 | 0.028 | 0.013 | 0.013 | 0 | 0.2 | 0.28 | 0.39 | 0.29 | 0.24 | 0.24 |
| 11 | 21355784 | rs12421240 | 0.207 | 0.345 | -1.242 | 0.719 | -0.017 | -0.037 | -0.037 | 0 | 0.12 | 0.06 | 0.24 | 0.25 | 0.12 | 0.13 |
| 11 | 21357846 | rs10437585 | -0.474 | 0.253 | -0.99 | 0.759 | 0.005 | 0.007 | 0.007 | 0.129 | 0.47 | 0.37 | 0.67 | 0.42 | 0.51 | 0.56 |
| 11 | 21358546 | rs4597066 | -0.433 | 0.305 | -0.978 | 0.375 | 0.003 | 0.003 | 0.003 | 0.466 | 0.47 | 0.38 | 0.67 | 0.44 | 0.5 | 0.56 |
| 11 | 21360104 | rs1557461 | -0.79 | -0.17 | -1.023 | -0.227 | -0.007 | -0.01 | -0.01 | 0 | 0.77 | 0.81 | 0.75 | 0.72 | 0.81 | 0.74 |
| 11 | 21362764 | rs11026033 | 0.721 | -0.221 | -1.381 | -0.461 | -0.038 | 0.086 | 0.086 | 0 | 0.16 | 0.02 | 0.13 | 0.05 | 0.19 | 0.12 |
| 11 | 21363327 | rs2040322 | 0.365 | -0.708 | -1.36 | -0.568 | -0.004 | -0.015 | -0.015 | 0 | 0.4 | 0.66 | 0.53 | 0.52 | 0.44 | 0.45 |
| 11 | 21365903 | rs12222217 | 1.05 | -0.378 | -1.76 | -0.413 | -0.04 | -0.05 | -0.05 | 1.27 | 0.23 | 0.08 | 0.24 | 0.27 | 0.2 | 0.25 |
| 11 | 21366526 | rs10833513 | 0.951 | -0.41 | -1.606 | -0.361 | -0.039 | -0.049 | -0.049 | 0 | 0.24 | 0.09 | 0.25 | 0.28 | 0.2 | 0.26 |
| 11 | 21367530 | rs11026038 | -0.409 | -0.246 | -1.708 | -0.231 | 0.025 | 0.035 | 0.035 | 0 | 0.39 | 0.24 | 0.31 | 0.28 | 0.38 | 0.34 |
| 11 | 21368523 | rs16907971 | 0.752 | 0.293 | -1.829 | 0.469 | 0.007 | -0.056 | -0.056 | 0.064 | 0.16 | 0 | 0.07 | 0.15 | 0.15 | 0.11 |
| 11 | 21368871 | rs12296077 | -0.1 | 0.624 | -1.217 | 0.477 | 0.013 | 0.015 | 0.015 | 0 | 0.29 | 0.35 | 0.4 | 0.44 | 0.4 | 0.33 |
| 11 | 21372044 | rs950253 | 0.072 | 0.869 | -1.73 | 0.501 | -0.013 | -0.01 | -0.01 | 0 | 0.27 | 0.24 | 0.4 | 0.41 | 0.39 | 0.34 |
| 11 | 21376439 | rs10500904 | -0.236 | 0.574 | -1.805 | 0.679 | -0.015 | -0.01 | -0.01 | 0.128 | 0.06 | 0.13 | 0.1 | 0.05 | 0.03 | 0.14 |
| 11 | 21377151 | rs7126959 | -0.007 | 0.917 | -2.464 | 0.789 | -0.003 | -0.011 | -0.011 | 0 | 0.28 | 0.22 | 0.3 | 0.28 | 0.27 | 0.3 |
| 11 | 21378056 | rs12786220 | -0.83 | 1.186 | -2.495 | 0.651 | -0.083 | -0.056 | -0.056 | 0 | 0.23 | 0.01 | 0.16 | 0.21 | 0.28 | 0.22 |
| 11 | 21383866 | rs8181537 | -0.812 | 1.126 | -2.792 | 0.527 | -0.101 | -0.031 | -0.031 | 0 | 0.32 | 0.02 | 0.18 | 0.27 | 0.39 | 0.38 |
| 11 | 21384100 | rs7926858 | -0.759 | 1.345 | -3.149 | 0.388 | -0.017 | 0.072 | 0.072 | 0 | 0.48 | 0.23 | 0.53 | 0.37 | 0.57 | 0.48 |
| 11 | 21384701 | rs7104612 | -0.413 | 0.798 | -2.996 | 0.254 | 0 | -0.008 | -0.008 | 0 | 0.76 | 0.82 | 0.59 | 0.68 | 0.75 | 0.84 |
| 11 | 21386502 | rs1945405 | -0.207 | 0.092 | -3.58 | -0.386 | 0.02 | 0.02 | 0.02 | 0 | 0.56 | 0.77 | 0.43 | 0.6 | 0.62 | 0.69 |
| 11 | 21387464 | rs1945403 | -0.382 | 0.006 | -3.543 | -0.51 | 0.033 | 0.012 | 0.012 | 0 | 0.58 | 0.79 | 0.47 | 0.61 | 0.66 | 0.71 |
| 11 | 21388542 | rs4923569 | -0.364 | -0.2 | -2.812 | -0.369 | 0.003 | 0 | 0 | 0 | 0.48 | 0.55 | 0.38 | 0.53 | 0.6 | 0.58 |
| 11 | 21389217 | rs4488195 | -1.312 | -0.644 | -3.052 | -0.619 | -0.005 | -0.012 | -0.012 | 0.77 | 0.83 | 0.86 | 0.86 | 0.94 | 0.92 | 0.89 |
| 11 | 21390901 | rs1945454 | 1.462 | -0.685 | -3.134 | -0.618 | -0.02 | -0.043 | -0.043 | 0.094 | 0.27 | 0.09 | 0.4 | 0.35 | 0.28 | 0.16 |
| 11 | 21391498 | rs7949183 | 0.347 | -0.423 | -2.089 | 0.021 | -0.002 | 0.088 | 0.088 | 0 | 0.15 | 0.26 | 0.13 | 0.32 | 0.1 | 0.21 |
| 11 | 21391795 | rs7934374 | -1.35 | 0.272 | -2.396 | -0.043 | -0.07 | 0.004 | 0.004 | 1.581 | 0.81 | 0.95 | 0.84 | 0.85 | 0.7 | 0.74 |
| 11 | 21392487 | rs4151056 | 1.322 | 0.3 | -2.059 | 0.209 | 0.01 | 0.121 | 0.121 | 0 | 0.09 | 0.22 | 0.07 | 0.27 | 0.07 | 0.11 |
| 11 | 21393859 | rs7120820 | 0.746 | 0.273 | -1.965 | 0.161 | 0.021 | 0.002 | 0.002 | 0 | 0.45 | 0.31 | 0.6 | 0.41 | 0.36 | 0.26 |
| 11 | 21394295 | rs1945453 | 0.171 | 0.256 | -0.712 | 0.364 | -0.009 | 0 | 0 | 0 | 0.15 | 0.13 | 0.19 | 0.11 | 0.08 | 0.11 |
| 11 | 21395259 | rs12798947 | -0.446 | 0.556 | -0.456 | 0.368 | -0.002 | 0.002 | 0.002 | 0 | 0.19 | 0.17 | 0.14 | 0.16 | 0.22 | 0.25 |
| 11 | 21396218 | rs16908013 | -0.554 | 0.383 | -0.234 | 0.627 | -0.005 | -0.004 | -0.004 | 0 | 0.23 | 0.23 | 0.21 | 0.24 | 0.21 | 0.29 |
| 11 | 21396360 | rs10833520 | -0.106 | 1.011 | -0.036 | 0.834 | 0.004 | 0 | 0 | 0 | 0.11 | 0.15 | 0.11 | 0.14 | 0.21 | 0.22 |
| 11 | 21400116 | rs7105978 | -0.98 | 0.34 | 0.596 | 1.186 | 0.041 | 0.121 | 0.121 | 0 | 0.15 | 0.42 | 0.2 | 0.25 | 0.13 | 0.19 |
| 11 | 21401298 | rs4244544 | -1.401 | 0.214 | 0.258 | 1.077 | -0.006 | 0.025 | 0.025 | 0 | 0.68 | 0.87 | 0.65 | 0.75 | 0.65 | 0.66 |
| 11 | 21401516 | rs11026059 | -1.152 | 0.518 | 0.542 | 1.381 | 0.037 | 0.085 | 0.085 | 0 | 0.23 | 0.48 | 0.25 | 0.39 | 0.26 | 0.41 |
| 11 | 21408571 | rs10766793 | 0.026 | 0.357 | 0.404 | 1.183 | -0.003 | 0.002 | 0.002 | 0 | 0.47 | 0.37 | 0.45 | 0.47 | 0.53 | 0.67 |
| 11 | 21411124 | rs10833521 | 0.903 | 0.406 | 0.206 | 1.152 | -0.005 | -0.028 | -0.028 | 0.465 | 0.34 | 0.16 | 0.35 | 0.38 | 0.36 | 0.39 |
| 11 | 21411321 | rs10833522 | 1.38 | 0.388 | 0.165 | 1.172 | -0.017 | -0.014 | -0.014 | 0 | 0.3 | 0.11 | 0.34 | 0.31 | 0.34 | 0.31 |
| 11 | 21411891 | rs12146653 | 1.211 | 0.004 | -0.048 | 0.896 | -0.047 | -0.068 | -0.068 | 0 | 0.14 | 0.01 | 0.09 | 0.2 | 0.14 | 0.11 |
| 11 | 21412957 | rs10160607 | -0.069 | 0.313 | 0.238 | 0.887 | -0.005 | 0.001 | 0.001 | 0 | 0.47 | 0.36 | 0.45 | 0.48 | 0.53 | 0.67 |
| 11 | 21413780 | rs7109477 | 0.352 | 0.616 | 0.127 | 1.155 | -0.047 | -0.046 | -0.046 | 2.107 | 0.55 | 0.18 | 0.61 | 0.45 | 0.56 | 0.48 |
| 11 | 21415127 | rs10500907 | 1.295 | 1.055 | -0.05 | 0.709 | -0.011 | -0.015 | -0.015 | 0 | 0.3 | 0.22 | 0.36 | 0.36 | 0.35 | 0.36 |
| 11 | 21416589 | rs10734296 | -0.246 | 1.279 | -0.43 | 0.614 | 0.061 | 0.045 | 0.045 | 0 | 0.51 | 0.73 | 0.61 | 0.63 | 0.61 | 0.7 |
| 11 | 21420224 | rs7101928 | 0.473 | 1.462 | -0.511 | 0.276 | 0.002 | -0.002 | -0.002 | 0.991 | 0.61 | 0.42 | 0.67 | 0.61 | 0.61 | 0.61 |
| 11 | 21421753 | rs12796542 | -1.612 | 1.596 | -0.638 | 0.378 | -0.005 | 0.003 | 0.003 | 0 | 0.13 | 0.12 | 0.1 | 0.12 | 0.18 | 0.2 |
| 11 | 21424273 | rs12791761 | -1.193 | 1.034 | -0.579 | 0.351 | -0.002 | -0.006 | -0.006 | 0.009 | 0.06 | 0.06 | 0.03 | 0 | 0.06 | 0.03 |
| 11 | 21425763 | rs10833525 | -1.137 | 0.483 | 0.16 | 0.407 | -0.027 | 0.012 | 0.012 | 0 | 0.16 | 0.42 | 0.11 | 0.16 | 0.09 | 0.29 |
| 11 | 21425793 | rs2226912 | -1.441 | 1.425 | 0.396 | 0.072 | 0.029 | -0.061 | -0.061 | 0 | 0.22 | 0.42 | 0.15 | 0.11 | 0.21 | 0.13 |
| 11 | 21429135 | rs1945436 | 0.184 | 1.257 | -0.71 | -0.397 | -0.029 | 0.102 | 0.102 | 0 | 0.43 | 0.19 | 0.33 | 0.21 | 0.46 | 0.42 |
| 11 | 21429653 | rs10833526 | -0.209 | 1.178 | -0.596 | 0.249 | -0.016 | 0.002 | 0.002 | 0 | 0.29 | 0.21 | 0.37 | 0.31 | 0.35 | 0.32 |
| 11 | 21429888 | rs4531470 | 0.236 | 1.449 | -0.843 | 0.205 | -0.061 | -0.069 | -0.069 | 0.266 | 0.19 | 0.03 | 0.28 | 0.27 | 0.22 | 0.24 |
| 11 | 21434022 | rs2105791 | 0.476 | 1.387 | -0.585 | 0.13 | -0.001 | 0 | 0 | 0 | 0.72 | 0.71 | 0.62 | 0.69 | 0.66 | 0.67 |
| 11 | 21434464 | rs11026074 | 0.131 | 1.663 | -0.806 | 0.121 | -0.021 | 0.02 | 0.02 | 0.01 | 0.05 | 0.02 | 0.09 | 0 | 0.12 | 0.03 |
| 11 | 21434708 | rs883128 | -0.689 | 1.56 | -0.878 | 0.283 | -0.038 | -0.031 | -0.031 | 0 | 0.07 | 0 | 0.04 | 0.11 | 0.12 | 0.17 |
| 11 | 21436232 | rs11026076 | 0.468 | 1.067 | -0.747 | -0.154 | -0.021 | 0.052 | 0.052 | 2.014 | 0.41 | 0.25 | 0.34 | 0.27 | 0.48 | 0.44 |
| 11 | 21439476 | rs10833529 | -0.608 | 1.342 | -0.604 | 0.353 | 0 | 0 | 0 | 0.322 | 0.37 | 0.67 | 0.5 | 0.45 | 0.48 | 0.5 |
| 11 | 21443083 | rs1945428 | -0.271 | 0.821 | -0.612 | 0.638 | 0.041 | 0.019 | 0.019 | 0.031 | 0.49 | 0.62 | 0.39 | 0.47 | 0.39 | 0.5 |
| 11 | 21446808 | rs1945424 | 0.126 | 0.689 | -0.415 | 0.604 | 0.017 | 0.108 | 0.108 | 1.289 | 0.43 | 0.22 | 0.24 | 0.24 | 0.39 | 0.47 |
| 11 | 21450509 | rs1001199 | 0.373 | -0.139 | -0.153 | 0.766 | -0.002 | 0.001 | 0.001 | 0 | 0.34 | 0.34 | 0.27 | 0.41 | 0.26 | 0.39 |
| 11 | 21451466 | rs1557464 | 0.119 | -0.033 | -0.043 | 0.817 | -0.007 | -0.001 | -0.001 | 0 | 0.38 | 0.37 | 0.28 | 0.41 | 0.27 | 0.4 |

| | | | | | | | | | | | | | | | | |
|---|---|---|---|---|---|---|---|---|---|---|---|---|---|---|---|---|
| 11 | 21464433 | rs17305470 | 0.263 | -0.199 | -0.594 | 0.753 | -0.088 | 0.093 | 0.093 | 0.149 | 0.28 | 0.01 | 0.12 | 0.1 | 0.3 | 0.29 |
| 11 | 21468342 | rs10833534 | -0.925 | 0.109 | -0.174 | 1.386 | -0.018 | 0.119 | 0.119 | 0.42 | 0.55 | 0.88 | 0.56 | 0.79 | 0.45 | 0.58 |
| 11 | 21469913 | rs1945410 | 0.436 | 0.345 | -0.102 | 1.361 | -0.041 | -0.051 | -0.051 | 0 | 0.45 | 0.2 | 0.53 | 0.49 | 0.46 | 0.44 |
| 11 | 21471818 | rs1945408 | 0.231 | 0.577 | 0.398 | 1.547 | 0.073 | 0.137 | 0.137 | 2.363 | 0.88 | 0.48 | 0.77 | 0.77 | 0.84 | 0.91 |
| 11 | 21473003 | rs1955066 | -0.22 | 0.466 | 0.44 | 1.545 | 0.045 | 0.301 | 0.301 | 0.408 | 0.06 | 0.44 | 0.12 | 0.2 | 0.01 | 0.03 |
| 11 | 21478022 | rs7950933 | 1.244 | 0.526 | 0.266 | 1.595 | 0.019 | 0.017 | 0.017 | 0.183 | 0.78 | 0.42 | 0.64 | 0.72 | 0.69 | 0.77 |
| 11 | 21478922 | rs10833538 | 0.375 | 0.458 | 0.26 | 1.59 | 0.005 | 0.064 | 0.064 | 0.805 | 0.41 | 0.18 | 0.24 | 0.28 | 0.38 | 0.45 |
| 11 | 21479238 | rs10766816 | -0.346 | 0.489 | 0.287 | 1.55 | 0.049 | -0.022 | -0.022 | 0.031 | 0.92 | 0.8 | 0.91 | 0.98 | 0.85 | 0.93 |
| 11 | 21480151 | rs7115839 | 0.329 | 0.411 | 0.262 | 1.57 | 0.007 | 0.108 | 0.108 | 0.626 | 0.41 | 0.19 | 0.24 | 0.24 | 0.38 | 0.45 |
| 11 | 21480256 | rs7116093 | 0.783 | 0.459 | 0.363 | 1.622 | 0.046 | 0.108 | 0.108 | 0.989 | 0.85 | 0.49 | 0.72 | 0.72 | 0.81 | 0.84 |
| 11 | 21480796 | rs1945415 | -1.681 | 0.482 | 0.149 | 1.616 | 0.032 | -0.005 | -0.005 | 0.665 | 0.95 | 0.92 | 0.92 | 0.98 | 0.85 | 0.96 |
| 11 | 21481485 | rs11026097 | 0.355 | 0.472 | 0.104 | 1.604 | 0.004 | 0.108 | 0.108 | 0.87 | 0.41 | 0.19 | 0.24 | 0.24 | 0.4 | 0.46 |
| 11 | 21484115 | rs10833542 | 0.813 | 0.811 | -0.484 | 1.27 | -0.087 | 0.089 | 0.089 | 0 | 0.39 | 0.06 | 0.35 | 0.21 | 0.39 | 0.36 |
| 11 | 21490773 | rs4923611 | -1.297 | 1.077 | -0.218 | 1.335 | 0.067 | 0.007 | 0.007 | 2.125 | 0.94 | 0.88 | 0.89 | 0.97 | 0.85 | 0.93 |
| 11 | 21496533 | rs7924766 | 0.76 | 1.017 | -0.254 | 1.286 | 0.026 | -0.043 | -0.043 | 0.273 | 0.78 | 0.42 | 0.71 | 0.91 | 0.68 | 0.8 |
| 11 | 21496683 | rs4922828 | 0.749 | 1.023 | -0.251 | 1.29 | 0.026 | -0.043 | -0.043 | 3.386 | 0.78 | 0.42 | 0.71 | 0.91 | 0.68 | 0.8 |
| 11 | 21497779 | rs10833547 | -0.141 | 0.784 | -0.742 | 1.808 | 0.025 | -0.007 | -0.007 | 0 | 0.51 | 0.64 | 0.44 | 0.53 | 0.44 | 0.57 |
| 11 | 21516600 | rs10766824 | 0.061 | -0.235 | -0.268 | 2.194 | -0.036 | 0.049 | 0.049 | 0 | 0.85 | 0.8 | 0.79 | 0.65 | 0.95 | 0.72 |
| 11 | 21517163 | rs12576288 | -0.036 | 0.383 | 0.06 | 2.378 | 0.028 | 0.065 | 0.065 | 0 | 0.11 | 0.22 | 0.21 | 0.26 | 0.2 | 0.08 |
| 11 | 21522529 | rs10219188 | 0.708 | 0.149 | -0.623 | 1.731 | -0.07 | 0.07 | 0.07 | 0 | 0.38 | 0.19 | 0.3 | 0.23 | 0.48 | 0.45 |
| 11 | 21523125 | rs11026113 | -1.056 | 0.219 | -0.513 | 2.01 | -0.001 | 0.007 | 0.007 | 0 | 0.24 | 0.31 | 0.4 | 0.6 | 0.26 | 0.36 |
| 11 | 21531851 | rs11601433 | 0.428 | 0.306 | -0.459 | 1.388 | 0.013 | -0.01 | -0.01 | 0 | 0.11 | 0.16 | 0.18 | 0.06 | 0.2 | 0.07 |
| 11 | 21532049 | rs11026117 | -1.367 | 0.045 | -0.316 | 1.477 | -0.003 | 0.006 | 0.006 | 0 | 0.27 | 0.38 | 0.41 | 0.63 | 0.26 | 0.37 |
| 11 | 21532167 | rs11606073 | 0.395 | 0.405 | -0.49 | 1.086 | 0.007 | -0.005 | -0.005 | 0 | 0.1 | 0.13 | 0.17 | 0.06 | 0.21 | 0.07 |
| 11 | 21532867 | rs4074586 | 0.015 | 0.24 | -0.627 | 0.863 | 0.037 | 0.082 | 0.082 | 0 | 0.45 | 0.33 | 0.45 | 0.23 | 0.62 | 0.48 |
| 11 | 21533812 | rs11026120 | 0.107 | 0.527 | -0.62 | 0.985 | -0.068 | 0.065 | 0.065 | 0.014 | 0.41 | 0.11 | 0.31 | 0.32 | 0.44 | 0.54 |
| 11 | 21534059 | rs4550216 | -0.113 | 0.367 | -0.073 | 0.967 | -0.003 | 0.036 | 0.036 | 0 | 0.55 | 0.4 | 0.59 | 0.65 | 0.56 | 0.45 |
| 11 | 21536070 | rs10833552 | 0.764 | 0.214 | -0.422 | 1.316 | 0.015 | -0.019 | -0.019 | 0.305 | 0.36 | 0.2 | 0.29 | 0.37 | 0.32 | 0.21 |
| 11 | 21536391 | rs6483775 | -0.387 | 0.315 | -0.306 | 1.323 | 0.01 | -0.01 | -0.01 | 0 | 0.88 | 0.85 | 0.83 | 0.93 | 0.82 | 0.93 |
| 11 | 21536418 | rs7130553 | -0.217 | 0.201 | 0.226 | 1.345 | 0 | 0 | 0 | 0.509 | 0.51 | 0.48 | 0.47 | 0.53 | 0.5 | 0.72 |
| 11 | 21536710 | rs7952026 | 0.588 | 0.556 | -0.221 | 1.436 | -0.121 | -0.002 | -0.002 | 1.112 | 0.34 | 0.08 | 0.41 | 0.31 | 0.44 | 0.42 |
| 11 | 21537020 | rs11026122 | -0.695 | 1.254 | 0.21 | 1.846 | 0.052 | 0.117 | 0.117 | 0 | 0.19 | 0.48 | 0.34 | 0.3 | 0.24 | 0.23 |
| 11 | 21537527 | rs7104588 | 0.761 | 0.814 | -0.213 | 1.23 | 0.004 | 0 | 0 | 0 | 0.38 | 0.24 | 0.21 | 0.32 | 0.34 | 0.29 |
| 11 | 21539055 | rs6483778 | 0.249 | 0.566 | -0.297 | 1.374 | 0.009 | -0.006 | -0.006 | 0 | 0.13 | 0.14 | 0.22 | 0.14 | 0.11 | 0.2 |
| 11 | 21539845 | rs4923633 | 0.813 | 0.838 | -0.508 | 1.39 | -0.103 | -0.001 | -0.001 | 0.112 | 0.29 | 0.05 | 0.37 | 0.27 | 0.37 | 0.27 |
| 11 | 21542570 | rs10766829 | -0.299 | 0.327 | -0.028 | 1.326 | 0.004 | 0.006 | 0.006 | 0 | 0.45 | 0.32 | 0.56 | 0.45 | 0.56 | 0.49 |
| 11 | 21546136 | rs4614448 | -0.364 | 0.19 | 0.513 | 1.301 | -0.012 | 0.166 | 0.166 | 0.449 | 0.88 | 0.52 | 0.93 | 0.77 | 0.95 | 0.77 |
| 11 | 21560094 | rs7927068 | -0.328 | 0.106 | 0.328 | 1.202 | 0.003 | 0.03 | 0.03 | 0.011 | 0.47 | 0.33 | 0.43 | 0.38 | 0.55 | 0.5 |
| 11 | 21562763 | rs4923643 | -0.269 | 0.313 | 0.754 | 1.358 | -0.027 | 0.047 | 0.047 | 0.759 | 0.16 | 0.37 | 0.24 | 0.21 | 0.12 | 0.2 |
| 11 | 21574121 | rs7933730 | 1 | 0.008 | 0.371 | 1.071 | -0.016 | -0.005 | -0.005 | 0.286 | 0.37 | 0.14 | 0.36 | 0.32 | 0.35 | 0.24 |
| 11 | 21578367 | rs4319515 | 0.551 | -0.035 | 0.377 | 1.017 | -0.075 | 0.061 | 0.061 | 0 | 0.35 | 0.11 | 0.27 | 0.26 | 0.46 | 0.34 |
| 11 | 21579442 | rs4262719 | 0.718 | -0.218 | 0.646 | 1.122 | -0.007 | 0.008 | 0.008 | 0.68 | 0.74 | 0.29 | 0.64 | 0.58 | 0.81 | 0.58 |
| 11 | 21581005 | rs4343020 | 0.287 | -0.205 | 1.014 | 1.156 | -0.021 | 0.001 | 0.001 | 0 | 0.15 | 0.22 | 0.18 | 0.14 | 0.11 | 0.19 |
| 11 | 21581665 | rs7119475 | -0.425 | -0.111 | 0.96 | 1.195 | -0.004 | 0.126 | 0.126 | 0 | 0.89 | 0.6 | 0.88 | 0.81 | 0.94 | 0.79 |
| 11 | 21582217 | rs11026133 | -0.634 | -0.065 | 0.581 | 1.217 | 0.001 | 0.119 | 0.119 | 0 | 0.9 | 0.69 | 0.83 | 0.72 | 0.92 | 0.78 |
| 11 | 21586339 | rs10766833 | -0.085 | -0.037 | 0.296 | 1.16 | -0.025 | 0.05 | 0.05 | 0 | 0.79 | 0.7 | 0.71 | 0.65 | 0.83 | 0.59 |
| 11 | 21587215 | rs11604321 | 0.866 | 0.038 | -0.601 | 1.06 | -0.007 | -0.009 | -0.009 | 0 | 0.37 | 0.16 | 0.35 | 0.31 | 0.35 | 0.21 |
| 11 | 21587453 | rs12576212 | 1.204 | 0.315 | -1.131 | 0.609 | -0.053 | -0.021 | -0.021 | 0 | 0.18 | 0.03 | 0.09 | 0.2 | 0.22 | 0.2 |
| 11 | 21588442 | rs4922560 | 0.628 | -0.278 | -0.559 | 0.672 | -0.008 | -0.034 | -0.034 | 0 | 0.37 | 0.16 | 0.39 | 0.39 | 0.33 | 0.23 |
| 11 | 21600567 | rs4922849 | -0.579 | -0.662 | -0.156 | 0.669 | -0.027 | 0.001 | 0.001 | 3.76 | 0.2 | 0.69 | 0.25 | 0.23 | 0.1 | 0.26 |
| 11 | 21605753 | rs1945569 | -0.581 | -0.41 | -0.682 | 0.651 | -0.057 | -0.074 | -0.074 | 0 | 0.58 | 0.94 | 0.62 | 0.55 | 0.43 | 0.45 |
| 11 | 21609366 | rs11602745 | 0.412 | -0.387 | -0.241 | 0.496 | 0.012 | -0.012 | -0.012 | 0 | 0.3 | 0.2 | 0.29 | 0.31 | 0.22 | 0.2 |
| 11 | 21610969 | rs1459894 | 0.29 | -0.793 | 0.038 | 0.3 | -0.021 | -0.002 | -0.002 | 0 | 0.17 | 0.22 | 0.19 | 0.16 | 0.12 | 0.26 |
| 11 | 21611029 | rs898364 | -0.37 | -0.837 | 0.141 | 0.189 | -0.008 | -0.016 | -0.016 | 0 | 0.22 | 0.31 | 0.27 | 0.16 | 0.25 | 0.28 |
| 11 | 21611168 | rs744046 | -1.292 | -0.823 | -0.463 | 0.386 | -0.092 | -0.008 | -0.008 | 0 | 0.57 | 0.94 | 0.66 | 0.59 | 0.49 | 0.56 |
| 11 | 21616465 | rs2199234 | -0.802 | -0.768 | -0.003 | 0.334 | 0.019 | 0.009 | 0.009 | 0 | 0.48 | 0.69 | 0.61 | 0.54 | 0.44 | 0.42 |
| 11 | 21617059 | rs11608012 | -0.802 | -0.759 | -0.003 | 0.334 | 0.023 | 0.009 | 0.009 | 0 | 0.48 | 0.69 | 0.61 | 0.54 | 0.43 | 0.42 |
| 11 | 21617550 | rs1459892 | -0.653 | -0.744 | 0.066 | 0.342 | 0.019 | 0.003 | 0.003 | 0 | 0.47 | 0.64 | 0.6 | 0.54 | 0.43 | 0.42 |
| 11 | 21624588 | rs1599203 | -0.68 | -0.867 | 0.108 | 0.378 | 0.01 | 0 | 0 | 0 | 0.47 | 0.65 | 0.6 | 0.54 | 0.44 | 0.42 |
| 11 | 21625552 | rs1459885 | 0.937 | -0.912 | -0.041 | 0.371 | -0.103 | -0.077 | -0.077 | 0 | 0.43 | 0.06 | 0.33 | 0.43 | 0.52 | 0.49 |
| 11 | 21625898 | rs1945549 | 0.348 | -0.536 | 0.578 | 0.619 | -0.021 | 0.017 | 0.017 | 1.273 | 0.08 | 0.19 | 0.14 | 0.15 | 0.07 | 0.1 |
| 11 | 21632174 | rs4565893 | -1.037 | -0.432 | 0.288 | 0.603 | 0.02 | 0.006 | 0.006 | 0 | 0.48 | 0.7 | 0.62 | 0.54 | 0.44 | 0.43 |
| 11 | 21638463 | rs2032373 | -0.892 | 0.155 | 0.193 | 0.523 | 0.006 | 0.003 | 0.003 | 0 | 0.54 | 0.56 | 0.62 | 0.45 | 0.59 | 0.44 |
| 11 | 21641576 | rs4457740 | -0.55 | 0.094 | -0.035 | 0.705 | -0.001 | 0.006 | 0.006 | 0 | 0.45 | 0.63 | 0.45 | 0.46 | 0.45 | 0.43 |
| 11 | 21641954 | rs3958160 | 0.11 | 0.153 | 0.072 | 0.29 | -0.013 | 0.031 | 0.031 | 0 | 0.34 | 0.24 | 0.22 | 0.2 | 0.38 | 0.24 |
| 11 | 21649811 | rs7931276 | 0.257 | 0.355 | 0.496 | 0.514 | 0.006 | 0.009 | 0.009 | 0.023 | 0.11 | 0.13 | 0.24 | 0.12 | 0.13 | 0.09 |
| 11 | 21652727 | rs990339 | -1.707 | 0.323 | 0.632 | 0.426 | 0.008 | -0.019 | -0.019 | 0 | 0.27 | 0.53 | 0.49 | 0.25 | 0.33 | 0.21 |
| 11 | 21653988 | rs7129096 | -1.165 | -0.127 | 0.455 | 0.471 | 0.005 | -0.031 | -0.031 | 0.172 | 0.16 | 0.37 | 0.25 | 0.13 | 0.19 | 0.12 |
| 11 | 21654066 | rs7129113 | -0.688 | -0.363 | 0.394 | 0.646 | -0.024 | 0.002 | 0.002 | 1.847 | 0.1 | 0.24 | 0.16 | 0.13 | 0.07 | 0.1 |

| | | | | | | | | | | | | | | | | |
|---|---|---|---|---|---|---|---|---|---|---|---|---|---|---|---|---|
| 11 | 21658844 | rs4492823 | 0.112 | -0.413 | -0.106 | 0.559 | 0.008 | 0.044 | 0.044 | 1.036 | 0.48 | 0.57 | 0.31 | 0.33 | 0.51 | 0.47 |
| 11 | 21660464 | rs10766838 | -0.485 | -0.565 | -0.354 | 0.859 | 0.006 | -0.016 | -0.016 | 0 | 0.64 | 0.69 | 0.46 | 0.58 | 0.64 | 0.71 |
| 11 | 21663111 | rs1380509 | 0.147 | -0.444 | 0.19 | 0.889 | 0 | 0 | 0 | 0.107 | 0.55 | 0.55 | 0.44 | 0.45 | 0.54 | 0.57 |
| 11 | 21666116 | rs11026144 | 0.703 | -0.378 | 0.065 | 0.912 | -0.003 | 0 | 0 | 0 | 0.13 | 0.05 | 0.09 | 0.12 | 0.14 | 0.12 |
| 11 | 21667094 | rs1380510 | 0.923 | -0.286 | 0.178 | 0.708 | 0.011 | -0.005 | -0.005 | 0 | 0.16 | 0.24 | 0.1 | 0.15 | 0.19 | 0.28 |
| 11 | 21667334 | rs764949 | -0.977 | -0.422 | 0.583 | 0.902 | -0.008 | -0.013 | -0.013 | 0 | 0.73 | 0.77 | 0.61 | 0.67 | 0.68 | 0.8 |
| 11 | 21668395 | rs1380511 | -0.049 | -0.377 | 0.632 | 0.942 | -0.001 | -0.001 | -0.001 | 0 | 0.6 | 0.61 | 0.52 | 0.56 | 0.55 | 0.69 |
| 11 | 21669202 | rs1945546 | -0.639 | -0.573 | 0.774 | 0.938 | -0.015 | 0.001 | 0.001 | 0 | 0.2 | 0.25 | 0.25 | 0.22 | 0.18 | 0.18 |
| 11 | 21669219 | rs1945547 | -0.763 | -0.446 | 0.634 | 0.823 | 0.007 | 0.036 | 0.036 | 0 | 0.71 | 0.65 | 0.55 | 0.53 | 0.68 | 0.7 |
| 11 | 21669293 | rs10833583 | 0.656 | -0.505 | 0.551 | 0.8 | -0.017 | -0.011 | -0.011 | 0 | 0.13 | 0.02 | 0.08 | 0.12 | 0.13 | 0.12 |
| 11 | 21674597 | rs1459868 | -0.933 | -0.286 | 0.886 | 0.875 | -0.012 | -0.014 | -0.014 | 0 | 0.73 | 0.75 | 0.61 | 0.67 | 0.67 | 0.83 |
| 11 | 21680488 | rs2061500 | -0.65 | -0.226 | 0.593 | 0.593 | 0.014 | 0.06 | 0.06 | 0.02 | 0.71 | 0.58 | 0.54 | 0.52 | 0.67 | 0.7 |
| 11 | 21686312 | rs1459865 | 0.393 | -0.447 | 0.312 | 0.326 | 0.007 | 0.056 | 0.056 | 2.829 | 0.5 | 0.35 | 0.3 | 0.33 | 0.48 | 0.54 |
| 11 | 21706014 | rs11026160 | -1.09 | -0.908 | 0.513 | 0.527 | -0.115 | -0.009 | -0.009 | 0 | 0.66 | 0.94 | 0.59 | 0.72 | 0.57 | 0.65 |
| 11 | 21712019 | rs4278510 | -1.111 | -0.92 | 0.508 | 0.522 | -0.115 | -0.01 | -0.01 | 0 | 0.66 | 0.94 | 0.59 | 0.72 | 0.57 | 0.65 |
| 11 | 21720698 | rs7933853 | -0.731 | -1.054 | 1.283 | 0.718 | 0.008 | 0.001 | 0.001 | 0.784 | 0.6 | 0.49 | 0.37 | 0.42 | 0.56 | 0.52 |
| 11 | 21724734 | rs4492824 | -0.807 | -1.043 | 1.702 | 0.731 | 0.016 | 0.005 | 0.005 | 1.047 | 0.61 | 0.57 | 0.37 | 0.41 | 0.56 | 0.52 |
| 11 | 21727820 | rs4244479 | -0.984 | -1.041 | 1.719 | 0.731 | -0.026 | -0.013 | -0.013 | 0 | 0.63 | 0.78 | 0.39 | 0.41 | 0.56 | 0.52 |
| 11 | 21730707 | rs2726125 | -0.567 | -1.032 | 1.341 | 0.736 | -0.062 | -0.022 | -0.022 | 0.021 | 0.58 | 0.22 | 0.34 | 0.41 | 0.56 | 0.52 |
| 11 | 21738115 | rs3886894 | 0.537 | -1.017 | 1.394 | 0.69 | -0.012 | -0.006 | -0.006 | 0 | 0.38 | 0.29 | 0.61 | 0.58 | 0.44 | 0.48 |
| 11 | 21738350 | rs7104861 | 0.196 | -1.028 | 1.339 | 0.684 | -0.023 | -0.005 | -0.005 | 0.827 | 0.42 | 0.69 | 0.65 | 0.59 | 0.44 | 0.48 |
| 11 | 21738451 | rs1945359 | 0.854 | -0.944 | 1.143 | 0.745 | -0.084 | 0.011 | 0.011 | 1.534 | 0.34 | 0.09 | 0.41 | 0.28 | 0.43 | 0.35 |
| 11 | 21752014 | rs7119311 | 0.427 | -0.694 | 0.929 | -0.24 | -0.009 | -0.003 | -0.003 | 0.21 | 0.41 | 0.59 | 0.64 | 0.59 | 0.44 | 0.48 |
| 11 | 21753111 | rs1459873 | 0.481 | -0.683 | 0.698 | -0.234 | -0.001 | -0.001 | -0.001 | 0 | 0.41 | 0.54 | 0.63 | 0.59 | 0.44 | 0.48 |
| 11 | 21758947 | rs1599196 | 0.682 | -0.575 | -0.338 | -0.181 | -0.018 | -0.01 | -0.01 | 0.063 | 0.37 | 0.26 | 0.61 | 0.59 | 0.44 | 0.48 |
| 11 | 21759043 | rs985660 | 0.223 | -0.677 | -0.997 | -0.223 | 0.048 | 0.017 | 0.017 | 0 | 0.25 | 0.1 | 0.12 | 0.15 | 0.18 | 0.15 |
| 11 | 21762062 | rs2611744 | 0.55 | 0.227 | -0.466 | -0.115 | -0.009 | 0.018 | 0.018 | 2.182 | 0.05 | 0.03 | 0.05 | 0 | 0.08 | 0.08 |
| 11 | 21763627 | rs7114439 | -0.148 | 0.304 | -0.116 | 0.218 | -0.015 | -0.008 | -0.008 | 0.291 | 0.38 | 0.32 | 0.61 | 0.59 | 0.44 | 0.48 |
| 11 | 21772385 | rs2000949 | -0.145 | 0.313 | -0.621 | 0.205 | 0.001 | 0.003 | 0.003 | 1.802 | 0.54 | 0.45 | 0.43 | 0.45 | 0.5 | 0.46 |
| 11 | 21775739 | rs2840375 | -0.371 | -0.276 | -0.417 | -0.455 | -0.014 | -0.047 | -0.047 | 0 | 0.85 | 0.81 | 0.94 | 1 | 0.91 | 0.98 |
| 11 | 21788258 | rs7936787 | 0.772 | -0.941 | -0.076 | -0.571 | -0.007 | -0.006 | -0.006 | 0 | 0.72 | 0.36 | 0.88 | 0.9 | 0.85 | 0.87 |
| 11 | 21788732 | rs1459891 | 0.766 | -0.948 | -0.502 | -1.146 | -0.001 | 0.019 | 0.019 | 0 | 0.12 | 0.05 | 0.11 | 0 | 0.08 | 0.07 |
| 11 | 21795746 | rs10766875 | 0.926 | -0.905 | -0.189 | -0.175 | 0.019 | -0.005 | -0.005 | 0 | 0.7 | 0.29 | 0.77 | 0.68 | 0.85 | 0.76 |
| 11 | 21796279 | rs12289507 | 1.062 | -0.08 | -0.686 | -0.474 | -0.002 | 0.013 | 0.013 | 0.024 | 0.54 | 0.23 | 0.6 | 0.58 | 0.5 | 0.64 |
| 11 | 21802981 | rs7934550 | 0.334 | -0.334 | -0.098 | -0.349 | -0.038 | -0.035 | -0.035 | 0 | 0.73 | 0.45 | 0.89 | 0.9 | 0.85 | 0.87 |
| 11 | 21823202 | rs1945339 | 0.313 | -0.313 | -0.086 | -0.388 | -0.036 | -0.033 | -0.033 | 1.021 | 0.73 | 0.45 | 0.89 | 0.9 | 0.85 | 0.87 |
| 11 | 21829711 | rs12419270 | -0.049 | -0.38 | 0.022 | -0.399 | -0.005 | 0.002 | 0.002 | 0 | 0.1 | 0.11 | 0.03 | 0.1 | 0.06 | 0.11 |
| 11 | 21830364 | rs10833635 | 0.465 | -0.377 | 0.019 | -0.406 | 0.007 | 0.008 | 0.008 | 0.875 | 0.73 | 0.31 | 0.88 | 0.9 | 0.85 | 0.87 |
| 11 | 21832897 | rs10833637 | -0.246 | -0.16 | -0.354 | -0.434 | -0.002 | 0.012 | 0.012 | 0.146 | 0.1 | 0.12 | 0.28 | 0.23 | 0.07 | 0.14 |
| 11 | 21841206 | rs11026263 | -1.103 | -0.204 | -0.402 | -0.449 | 0.008 | -0.015 | -0.015 | 0.236 | 0.19 | 0.16 | 0.3 | 0.33 | 0.13 | 0.25 |
| 11 | 21846314 | rs1945360 | 0.877 | -0.425 | 0.273 | -0.63 | -0.002 | 0 | 0 | 1.216 | 0.65 | 0.41 | 0.62 | 0.67 | 0.78 | 0.74 |
| 11 | 21846894 | rs10741912 | -1.237 | -0.426 | 0.499 | -0.657 | -0.01 | -0.038 | -0.038 | 0 | 0.85 | 0.83 | 0.94 | 1 | 0.91 | 0.98 |
| 11 | 21848074 | rs16908606 | 0.316 | -0.439 | 0.403 | -0.657 | 0 | 0.021 | 0.021 | 0 | 0.09 | 0.05 | 0.16 | 0.01 | 0.07 | 0.02 |
| 11 | 21853264 | rs1032230 | 0.834 | -0.443 | 0.395 | -0.648 | -0.006 | -0.014 | -0.014 | 0 | 0.66 | 0.41 | 0.73 | 0.89 | 0.78 | 0.85 |
| 11 | 21864596 | rs2029299 | -1.342 | -0.423 | 0.971 | -0.646 | -0.035 | -0.078 | -0.078 | 0.008 | 0.85 | 0.75 | 0.94 | 1 | 0.91 | 0.98 |
| 11 | 21866461 | rs12420752 | -0.156 | -0.4 | 0.822 | -0.603 | -0.006 | 0.036 | 0.036 | 0 | 0.11 | 0.12 | 0.14 | 0.32 | 0.06 | 0.22 |
| 11 | 21869165 | rs11026296 | -0.65 | -0.134 | 0.688 | -0.756 | -0.014 | -0.017 | -0.017 | 0 | 0.27 | 0.26 | 0.32 | 0.41 | 0.41 | 0.35 |
| 11 | 21874253 | rs7122488 | 0.628 | -0.063 | 0.621 | -0.715 | -0.019 | -0.001 | -0.001 | 0 | 0.65 | 0.44 | 0.62 | 0.67 | 0.78 | 0.74 |
| 11 | 21882858 | rs4617585 | 1.041 | -0.118 | 0.322 | -0.652 | 0.002 | 0.01 | 0.01 | 0 | 0.49 | 0.39 | 0.45 | 0.58 | 0.43 | 0.57 |
| 11 | 21891843 | rs2403686 | -1.632 | 0.475 | 0.108 | -0.195 | -0.006 | 0.006 | 0.006 | 0 | 0.41 | 0.69 | 0.48 | 0.38 | 0.46 | 0.42 |
| 11 | 21894814 | rs4922945 | 0.321 | 0.856 | -0.35 | -0.039 | -0.07 | 0.026 | 0.026 | 0 | 0.16 | 0.03 | 0.14 | 0.09 | 0.3 | 0.12 |
| 11 | 21896880 | rs11026318 | -0.915 | 0.229 | 0.421 | 0.361 | -0.016 | 0.056 | 0.056 | 0.038 | 0.14 | 0.33 | 0.28 | 0.19 | 0.09 | 0.14 |
| 11 | 21905241 | rs2129728 | -0.642 | 0.588 | -0.225 | 0.368 | 0.003 | 0.012 | 0.012 | 0 | 0.08 | 0.12 | 0.04 | 0.1 | 0.08 | 0.16 |
| 11 | 21909377 | rs1487824 | -0.263 | 0.211 | -1.053 | 0.116 | 0.011 | 0.018 | 0.018 | 0 | 0.28 | 0.1 | 0.21 | 0.26 | 0.27 | 0.19 |
| 11 | 21915389 | rs10766893 | 0.083 | 0.738 | -0.917 | 0.166 | 0.046 | 0.045 | 0.045 | 0 | 0.26 | 0.34 | 0.43 | 0.49 | 0.41 | 0.39 |
| 11 | 21915497 | rs10766894 | 0.083 | 0.738 | -0.917 | 0.166 | 0.046 | 0.045 | 0.045 | 0 | 0.26 | 0.34 | 0.43 | 0.49 | 0.41 | 0.39 |
| 11 | 21919310 | rs12272451 | 0.947 | 0.118 | -1.344 | -0.194 | 0.02 | 0.122 | 0.122 | 0.141 | 0.26 | 0.05 | 0.11 | 0.02 | 0.2 | 0.15 |
| 11 | 21929071 | rs1487819 | -0.863 | -0.326 | -0.638 | 0.19 | -0.021 | 0.067 | 0.067 | 0 | 0.84 | 0.66 | 0.75 | 0.7 | 0.87 | 0.75 |
| 11 | 21929882 | rs2403695 | 0.633 | -0.277 | -0.368 | 0.149 | 0.01 | 0.009 | 0.009 | 0 | 0.25 | 0.26 | 0.42 | 0.47 | 0.42 | 0.38 |
| 11 | 21934390 | rs4922599 | -0.389 | -0.446 | -0.258 | 0.043 | 0.01 | -0.007 | -0.007 | 0 | 0.56 | 0.57 | 0.54 | 0.46 | 0.6 | 0.56 |
| 11 | 21934916 | rs12277777 | 1.348 | -0.482 | -0.273 | 0.01 | 0.043 | 0.116 | 0.116 | 0.105 | 0.25 | 0.05 | 0.09 | 0.02 | 0.17 | 0.14 |
| 11 | 21936062 | rs426984 | 1.325 | -0.499 | -0.272 | 0.003 | 0.005 | 0.016 | 0.016 | 0 | 0.28 | 0.08 | 0.21 | 0.24 | 0.27 | 0.19 |
| 11 | 21936644 | rs453700 | -1.151 | -0.626 | 0.056 | 0.056 | -0.002 | -0.003 | -0.003 | 0 | 0.86 | 0.83 | 0.91 | 0.9 | 0.88 | 0.83 |
| 11 | 21936812 | rs1487815 | 0.632 | -0.601 | -0.123 | 0.007 | 0.009 | -0.004 | -0.004 | 0.086 | 0.3 | 0.23 | 0.12 | 0.05 | 0.19 | 0.18 |
| 11 | 21945004 | rs2009912 | 0.041 | -0.543 | -0.114 | 0.059 | 0.012 | 0.011 | 0.011 | 0 | 0.54 | 0.42 | 0.63 | 0.72 | 0.68 | 0.56 |
| 11 | 21952288 | rs324172 | -0.377 | -0.535 | -0.154 | 0.043 | 0.013 | 0.01 | 0.01 | 0 | 0.58 | 0.6 | 0.73 | 0.89 | 0.69 | 0.65 |
| 11 | 21968847 | rs324258 | 1.215 | -0.487 | -0.25 | 0.088 | 0.006 | 0.017 | 0.017 | 0 | 0.28 | 0.09 | 0.21 | 0.24 | 0.27 | 0.19 |
| 11 | 21983094 | rs170489 | -1.06 | -0.459 | -0.252 | -0.328 | -0.008 | -0.044 | -0.044 | 0 | 0.84 | 0.68 | 0.89 | 0.96 | 0.85 | 0.83 |
| 11 | 21983447 | rs2198999 | -0.442 | -0.476 | -0.263 | 0.464 | 0.013 | -0.004 | -0.004 | 0 | 0.58 | 0.61 | 0.67 | 0.51 | 0.61 | 0.64 |

| | | | | | | | | | | | | | | | | |
|---|---|---|---|---|---|---|---|---|---|---|---|---|---|---|---|---|
| 11 | 21983941 | rs1386735 | 0.683 | -0.506 | -0.211 | 0.474 | 0.068 | -0.015 | -0.015 | 0 | 0.26 | 0.36 | 0.41 | 0.28 | 0.42 | 0.38 |
| 11 | 21984078 | rs324217 | 1.387 | -0.51 | -0.216 | 0.474 | 0.01 | -0.123 | -0.123 | 0.569 | 0.27 | 0.07 | 0.22 | 0.45 | 0.24 | 0.19 |
| 11 | 21984498 | rs10833683 | 0.357 | -0.511 | -0.205 | 0.484 | 0.079 | 0.039 | 0.039 | 0 | 0.28 | 0.41 | 0.55 | 0.46 | 0.42 | 0.46 |
| 11 | 21984582 | rs4922606 | -0.229 | -0.521 | -0.213 | 0.449 | 0.009 | 0.015 | 0.015 | 0 | 0.54 | 0.55 | 0.51 | 0.29 | 0.59 | 0.51 |
| 11 | 21984785 | rs324220 | 0.262 | -0.51 | -0.241 | 0.46 | 0.012 | -0.001 | -0.001 | 0 | 0.57 | 0.29 | 0.33 | 0.5 | 0.43 | 0.37 |
| 11 | 21987037 | rs4550219 | 0.332 | -0.479 | -0.281 | 0.478 | 0.022 | 0.011 | 0.011 | 0 | 0.56 | 0.24 | 0.39 | 0.52 | 0.43 | 0.38 |
| 11 | 21987561 | rs324228 | 0.956 | -0.211 | -0.366 | 0.565 | 0.016 | -0.103 | -0.103 | 1.498 | 0.28 | 0.08 | 0.23 | 0.48 | 0.25 | 0.23 |
| 11 | 21993126 | rs7110604 | 0.274 | 0.097 | -0.172 | 0.142 | 0.091 | -0.024 | -0.024 | 0.151 | 0.27 | 0.41 | 0.41 | 0.29 | 0.42 | 0.38 |
| 11 | 21997038 | rs1825836 | 0.176 | 0.154 | -0.162 | -0.002 | 0.017 | -0.009 | -0.009 | 0.982 | 0.29 | 0.3 | 0.4 | 0.27 | 0.45 | 0.42 |
| 11 | 22001998 | rs324204 | -0.349 | -0.309 | -0.308 | -0.391 | -0.018 | -0.064 | -0.064 | 0.076 | 0.64 | 0.55 | 0.81 | 0.95 | 0.75 | 0.73 |
| 11 | 22007739 | rs11026358 | 0.707 | -0.167 | -0.72 | 0.203 | 0.001 | -0.003 | -0.003 | 0 | 0.05 | 0.02 | 0.02 | 0.11 | 0.03 | 0.01 |
| 11 | 22010721 | rs324159 | 0.368 | 0.177 | -1.02 | -0.482 | 0.005 | 0.151 | 0.151 | 12.943 | 0.4 | 0.16 | 0.21 | 0.08 | 0.36 | 0.23 |
| 11 | 22016670 | rs324193 | 0.571 | 0.307 | -0.987 | -0.657 | 0.01 | 0.102 | 0.102 | 12.596 | 0.4 | 0.26 | 0.27 | 0.11 | 0.36 | 0.22 |
| 11 | 22024184 | rs324178 | -0.368 | -0.442 | -0.74 | -0.288 | 0.034 | -0.011 | -0.011 | 0 | 0.67 | 0.48 | 0.55 | 0.66 | 0.59 | 0.56 |
| 11 | 22030566 | rs10734310 | -1.071 | -0.001 | -0.28 | -0.423 | -0.002 | 0.001 | 0.001 | 1.506 | 0.78 | 0.77 | 0.68 | 0.78 | 0.7 | 0.71 |
| 11 | 22037445 | rs11026380 | 0.251 | -0.066 | -0.513 | 0.075 | 0.049 | -0.104 | -0.104 | 3.906 | 0.25 | 0.1 | 0.2 | 0.41 | 0.22 | 0.26 |
| 11 | 22059879 | rs1157385 | -0.034 | -0.035 | -0.179 | 0.07 | 0.016 | 0.002 | 0.002 | 0 | 0.1 | 0.14 | 0.11 | 0.11 | 0.11 | 0.15 |
| 11 | 22062310 | rs1350395 | -0.425 | 0.014 | -0.308 | 0.061 | 0.048 | 0.064 | 0.064 | 2.979 | 0.73 | 0.49 | 0.51 | 0.51 | 0.66 | 0.55 |
| 11 | 22065266 | rs11026390 | 0.227 | -0.089 | -0.304 | 0.16 | 0.049 | -0.116 | -0.116 | 0.06 | 0.24 | 0.1 | 0.2 | 0.41 | 0.22 | 0.26 |
| 11 | 22067536 | rs10500921 | 1.329 | -0.125 | -0.704 | -0.16 | -0.021 | 0.023 | 0.023 | 0.024 | 0.12 | 0 | 0.06 | 0.07 | 0.12 | 0.14 |
| 11 | 22083652 | rs11026398 | -0.134 | 0.299 | -0.098 | 0.364 | 0.035 | -0.078 | -0.078 | 0.175 | 0.27 | 0.1 | 0.27 | 0.52 | 0.25 | 0.33 |
| 11 | 22105030 | rs11026407 | 0.262 | 0.753 | 0.409 | 0.558 | 0.043 | -0.013 | -0.013 | 0.057 | 0.68 | 0.46 | 0.55 | 0.67 | 0.59 | 0.55 |
| 11 | 22106815 | rs10833700 | 0.262 | 0.753 | 0.409 | 0.558 | 0.043 | -0.013 | -0.013 | 0 | 0.68 | 0.46 | 0.55 | 0.67 | 0.59 | 0.55 |
| 11 | 22116783 | rs7934260 | -1.276 | 0.407 | 1.221 | 1.363 | 0.119 | 0.215 | 0.215 | 0.005 | 0.1 | 0.49 | 0.21 | 0.21 | 0.09 | 0.16 |
| 11 | 22120723 | rs4442550 | -0.659 | 0.292 | 0.904 | 1.326 | 0.026 | -0.071 | -0.071 | 0.473 | 0.24 | 0.1 | 0.21 | 0.36 | 0.21 | 0.24 |
| 11 | 22123113 | rs7358426 | -1.53 | 0.356 | 0.813 | 1.343 | 0.011 | -0.008 | -0.008 | 0.948 | 0.35 | 0.76 | 0.42 | 0.57 | 0.3 | 0.4 |
| 11 | 22141171 | rs10833705 | -1.519 | 0.83 | 0.86 | 1.116 | 0.075 | 0.16 | 0.16 | 1.862 | 0.12 | 0.53 | 0.21 | 0.2 | 0.09 | 0.17 |
| 11 | 22178677 | rs4244488 | 0.865 | 0.507 | 0.174 | 0.354 | 0 | -0.015 | -0.015 | 0.9 | 0.58 | 0.44 | 0.41 | 0.21 | 0.6 | 0.47 |
| 11 | 22207300 | rs10833715 | -1.31 | 0.293 | 0.409 | 1.407 | -0.003 | 0.02 | 0.02 | 1.443 | 0.23 | 0.7 | 0.37 | 0.44 | 0.15 | 0.24 |
| 11 | 22207495 | rs11026452 | -0.422 | 0.3 | 0.372 | 1.452 | -0.024 | -0.033 | -0.033 | 0 | 0.16 | 0.06 | 0.32 | 0.21 | 0.15 | 0.16 |
| 11 | 22221653 | rs11026460 | -0.173 | 0.242 | 0.706 | 1.214 | -0.004 | 0.003 | 0.003 | 2.907 | 0.13 | 0.08 | 0.24 | 0.13 | 0.1 | 0.13 |
| 11 | 22227418 | rs6416032 | 0.082 | 0.317 | 0.88 | 1.277 | -0.008 | 0.067 | 0.067 | 1.476 | 0.82 | 0.64 | 0.7 | 0.58 | 0.8 | 0.82 |
| 11 | 22232365 | rs7102678 | -0.259 | 0.287 | 0.587 | 1.302 | -0.005 | -0.09 | -0.09 | 0.685 | 0.83 | 0.95 | 0.73 | 0.58 | 0.8 | 0.82 |
| 11 | 22242729 | rs4312063 | -0.312 | 0.289 | 0.588 | 1.295 | -0.007 | -0.087 | -0.087 | 0.104 | 0.83 | 0.95 | 0.73 | 0.59 | 0.79 | 0.82 |
| 11 | 22249612 | rs10766928 | 0.686 | 0.231 | 0.849 | 1.26 | -0.031 | 0.052 | 0.052 | 1.029 | 0.78 | 0.44 | 0.67 | 0.36 | 0.79 | 0.74 |
| 11 | 22257284 | rs4271385 | -0.688 | 0.255 | 0.734 | 1.271 | 0.053 | 0.419 | 0.419 | 2.216 | 0.06 | 0.54 | 0.05 | 0.23 | 0 | 0.08 |
| 11 | 22266469 | rs7941608 | -0.688 | 0.255 | 0.724 | 1.28 | 0.052 | 0.408 | 0.408 | 2.216 | 0.06 | 0.55 | 0.05 | 0.22 | 0.01 | 0.08 |
| 11 | 22277852 | rs11026477 | 0.128 | 0.461 | -0.192 | 1.043 | -0.062 | -0.026 | -0.026 | 0 | 0.15 | 0.01 | 0.24 | 0.15 | 0.14 | 0.15 |
| 11 | 22293946 | rs7126673 | -1.326 | 0.271 | -0.208 | 1.169 | -0.009 | 0.021 | 0.021 | 5.428 | 0.17 | 0.75 | 0.12 | 0.23 | 0.1 | 0.19 |
| 11 | 22295434 | rs7395682 | 0.115 | 0.735 | -0.511 | 1.256 | -0.006 | -0.085 | -0.085 | 0.215 | 0.83 | 0.95 | 0.72 | 0.58 | 0.79 | 0.82 |
| 11 | 22316942 | rs7128422 | 0.698 | 0.804 | -0.619 | 0.885 | -0.025 | -0.061 | -0.061 | 0.413 | 0.74 | 0.86 | 0.64 | 0.54 | 0.67 | 0.69 |
| 11 | 22326069 | rs4570579 | 0.004 | 0.636 | -0.489 | 0.798 | -0.002 | 0.076 | 0.076 | 0 | 0.89 | 0.87 | 0.73 | 0.52 | 0.85 | 0.86 |
| 11 | 22327542 | rs11026499 | 0.3 | 0.104 | -0.461 | 0.355 | 0 | 0 | 0 | 0.024 | 0.11 | 0.1 | 0.21 | 0.12 | 0.04 | 0.17 |
| 11 | 22334571 | rs11026505 | -0.564 | 0.248 | 0.205 | 0.234 | 0.044 | 0.049 | 0.049 | 0 | 0.14 | 0.33 | 0.18 | 0.23 | 0.18 | 0.25 |
| 11 | 22344665 | rs11026510 | 0.16 | 0.488 | 0.223 | 0.006 | 0.004 | 0.009 | 0.009 | 0 | 0.15 | 0.11 | 0.24 | 0.11 | 0.1 | 0.2 |
| 11 | 22345050 | rs7103652 | 0.247 | 0.535 | 0.244 | -0.012 | -0.002 | -0.015 | -0.015 | 0 | 0.11 | 0.15 | 0.15 | 0 | 0.17 | 0.12 |
| 11 | 22348844 | rs10833743 | -1.744 | 0.283 | 0.682 | -0.063 | 0.002 | -0.001 | -0.001 | 0 | 0.33 | 0.59 | 0.43 | 0.32 | 0.29 | 0.48 |
| 11 | 22351468 | rs11026518 | 0.135 | 0.341 | 0.512 | -0.257 | 0.004 | 0.007 | 0.007 | 0 | 0.15 | 0.1 | 0.24 | 0.12 | 0.11 | 0.2 |
| 11 | 22352006 | rs4362145 | 1.323 | 0.581 | -0.152 | -1.094 | 0.001 | 0.106 | 0.106 | 0 | 0.3 | 0.12 | 0.15 | 0 | 0.27 | 0.12 |
| 11 | 22358720 | rs1155821 | -0.152 | 1.149 | 0.053 | -0.202 | 0.007 | -0.029 | -0.029 | 0.75 | 0.15 | 0.21 | 0.15 | 0.01 | 0.25 | 0.16 |
| 11 | 22359279 | rs2593696 | -0.126 | 1.142 | 0.064 | -0.193 | 0.007 | -0.029 | -0.029 | 0 | 0.15 | 0.21 | 0.15 | 0.01 | 0.24 | 0.16 |
| 11 | 22362296 | rs1447514 | 0.212 | 1.163 | -0.031 | -0.175 | -0.006 | 0.007 | 0.007 | 0 | 0.14 | 0.09 | 0.24 | 0.11 | 0.11 | 0.16 |
| 11 | 22364744 | rs2246710 | -0.359 | 1.239 | 0.2 | -0.2 | 0.015 | -0.041 | -0.041 | 0.587 | 0.15 | 0.24 | 0.16 | 0.01 | 0.24 | 0.16 |
| 11 | 22368301 | rs2593693 | -0.547 | 1.242 | -0.147 | 0.116 | 0.068 | 0.074 | 0.074 | 0.419 | 0.19 | 0.5 | 0.18 | 0.31 | 0.25 | 0.29 |
| 11 | 22371256 | rs2665691 | 0.249 | 0.664 | -0.414 | 0.204 | 0 | 0 | 0 | 3.44 | 0.67 | 0.66 | 0.48 | 0.38 | 0.74 | 0.61 |
| 11 | 22376572 | rs10500923 | -0.145 | 0.692 | -0.313 | 0.402 | -0.014 | 0.011 | 0.011 | 0 | 0.1 | 0.06 | 0.05 | 0.05 | 0.11 | 0.13 |
| 11 | 22378057 | rs11026529 | -0.318 | 0.614 | -0.601 | 1.156 | -0.012 | -0.069 | -0.069 | 0 | 0.11 | 0.01 | 0.14 | 0.48 | 0.13 | 0.15 |
| 11 | 22379184 | rs11026531 | -1.951 | 0.493 | -0.332 | 1.175 | -0.003 | 0.027 | 0.027 | 0 | 0.23 | 0.28 | 0.2 | 0.52 | 0.25 | 0.28 |
| 11 | 22383413 | rs11026533 | -0.469 | 0.745 | -0.633 | 1.257 | -0.028 | -0.091 | -0.091 | 0 | 0.12 | 0.01 | 0.15 | 0.48 | 0.16 | 0.16 |
| 11 | 22384166 | rs2290046 | -0.404 | 0.716 | -0.175 | 0.536 | 0.007 | -0.018 | -0.018 | 0 | 0.11 | 0.19 | 0.05 | 0.05 | 0.11 | 0.13 |
| 11 | 22387745 | rs10833748 | 0.256 | 0.512 | -0.869 | 0.74 | -0.012 | -0.056 | -0.056 | 0 | 0.37 | 0.22 | 0.44 | 0.49 | 0.41 | 0.38 |
| 11 | 22388769 | rs7107169 | -0.041 | -0.016 | 0.589 | 0.401 | 0.002 | -0.002 | -0.002 | 0 | 0.1 | 0.32 | 0.06 | 0.05 | 0.06 | 0.15 |
| 11 | 22389031 | rs2593688 | -1.006 | 0.075 | 0.781 | 0.556 | 0.009 | -0.009 | -0.009 | 0 | 0.6 | 0.74 | 0.69 | 0.55 | 0.59 | 0.59 |
| 11 | 22391419 | rs2593685 | -1.119 | -0.043 | 0.851 | 0.498 | -0.021 | -0.064 | -0.064 | 5.149 | 0.62 | 0.92 | 0.7 | 0.55 | 0.59 | 0.59 |
| 11 | 22395570 | rs12280547 | -0.56 | -0.092 | 0.549 | 0.31 | 0.002 | -0.001 | -0.001 | 0 | 0.34 | 0.34 | 0.48 | 0.28 | 0.42 | 0.37 |
| 11 | 22400124 | rs1979072 | 0.218 | -0.192 | 0.662 | 0.311 | -0.015 | 0.015 | 0.015 | 0.385 | 0.24 | 0.41 | 0.2 | 0.27 | 0.17 | 0.22 |
| 11 | 22400269 | rs1979073 | 0.233 | -0.191 | 0.653 | 0.311 | -0.014 | 0.015 | 0.015 | 0.562 | 0.24 | 0.41 | 0.2 | 0.27 | 0.17 | 0.22 |
| 11 | 22400813 | rs11026539 | 0.82 | -0.22 | 0.506 | 0.402 | -0.041 | 0.01 | 0.01 | 1.096 | 0.2 | 0.01 | 0.16 | 0.12 | 0.17 | 0.19 |
| 11 | 22403421 | rs11026541 | 0.773 | -0.219 | 0.607 | 0.287 | -0.005 | -0.042 | -0.042 | 0 | 0.2 | 0.09 | 0.17 | 0.27 | 0.17 | 0.22 |

| | | | | | | | | | | | | | | | | |
|---|---|---|---|---|---|---|---|---|---|---|---|---|---|---|---|---|
| 11 | 22418380 | rs956349 | -0.623 | -0.201 | 0.848 | 0.24 | 0.011 | -0.006 | -0.006 | 1.358 | 0.35 | 0.36 | 0.49 | 0.29 | 0.45 | 0.38 |
| 11 | 22418570 | rs726859 | -1.474 | -0.207 | 0.881 | 0.241 | 0.027 | -0.001 | -0.001 | 0.367 | 0.57 | 0.72 | 0.68 | 0.55 | 0.61 | 0.6 |
| 11 | 22420124 | rs7952608 | 0.664 | -0.157 | 1.467 | 0.162 | -0.01 | -0.012 | -0.012 | 0.399 | 0.09 | 0.17 | 0.05 | 0.05 | 0.06 | 0.15 |
| 11 | 22421282 | rs2665716 | -0.755 | -0.105 | 1.232 | 0.193 | -0.011 | 0.008 | 0.008 | 0 | 0.36 | 0.64 | 0.51 | 0.29 | 0.45 | 0.37 |
| 11 | 22430874 | rs2593665 | -0.755 | -0.102 | 1.224 | 0.195 | -0.006 | 0.003 | 0.003 | 0 | 0.36 | 0.64 | 0.51 | 0.28 | 0.45 | 0.37 |
| 11 | 22434494 | rs2665672 | -0.734 | -0.101 | 1.231 | 0.195 | 0.003 | -0.003 | -0.003 | 0 | 0.36 | 0.58 | 0.51 | 0.29 | 0.45 | 0.37 |
| 11 | 22435111 | rs3106185 | -1.676 | 0.052 | 1.202 | 0.224 | -0.011 | -0.074 | -0.074 | 0.276 | 0.62 | 0.92 | 0.71 | 0.53 | 0.62 | 0.6 |
| 11 | 22436189 | rs932085 | -1.346 | -0.653 | 1.421 | 0.241 | -0.003 | -0.012 | -0.012 | 0 | 0.52 | 0.73 | 0.61 | 0.5 | 0.45 | 0.59 |
| 11 | 22436299 | rs12793010 | -0.155 | -0.822 | 1.511 | 0.175 | 0.017 | 0 | 0 | 0.401 | 0.32 | 0.08 | 0.21 | 0.28 | 0.23 | 0.38 |
| 11 | 22437287 | rs2671117 | 0.099 | -0.794 | 1.838 | 0.186 | 0.048 | 0.06 | 0.06 | 0 | 0.18 | 0.48 | 0.37 | 0.22 | 0.22 | 0.21 |
| 11 | 22442119 | rs2671126 | 0.063 | -0.797 | 1.816 | 0.185 | 0.038 | 0.048 | 0.048 | 0 | 0.18 | 0.51 | 0.38 | 0.21 | 0.22 | 0.21 |
| 11 | 22444436 | rs340960 | -1.467 | -0.824 | 1.86 | 0.189 | 0.002 | -0.002 | -0.002 | 0.066 | 0.52 | 0.6 | 0.6 | 0.5 | 0.45 | 0.59 |
| 11 | 22445120 | rs340959 | 0.165 | -0.813 | 1.849 | 0.19 | 0.053 | 0.066 | 0.066 | 0 | 0.17 | 0.45 | 0.37 | 0.22 | 0.22 | 0.21 |
| 11 | 22446863 | rs340957 | -1.307 | -0.817 | 1.568 | 0.191 | 0.006 | -0.004 | -0.004 | 0 | 0.51 | 0.66 | 0.6 | 0.5 | 0.45 | 0.59 |
| 11 | 22453193 | rs1868070 | -0.014 | -0.861 | 1.433 | 0.154 | 0.027 | 0.035 | 0.035 | 0 | 0.18 | 0.57 | 0.38 | 0.22 | 0.22 | 0.21 |
| 11 | 22455894 | rs2665680 | -0.184 | -0.868 | 1.046 | 0.347 | -0.006 | -0.084 | -0.084 | 0 | 0.68 | 0.95 | 0.79 | 0.63 | 0.76 | 0.62 |
| 11 | 22457449 | rs2457664 | -0.062 | -0.783 | 1.2 | -0.14 | 0.02 | 0.025 | 0.025 | 0 | 0.19 | 0.59 | 0.37 | 0.21 | 0.22 | 0.21 |
| 11 | 22465753 | rs2671116 | -1.419 | -0.778 | 1.193 | -0.071 | 0.011 | -0.007 | -0.007 | 0.076 | 0.49 | 0.69 | 0.59 | 0.49 | 0.43 | 0.58 |
| 11 | 22466364 | rs2593676 | 0.098 | -0.782 | 1.222 | -0.066 | 0.041 | 0.057 | 0.057 | 0 | 0.17 | 0.55 | 0.37 | 0.22 | 0.2 | 0.21 |
| 11 | 22467554 | rs2665675 | -1.385 | -0.754 | 0.829 | -0.061 | 0.006 | -0.012 | -0.012 | 0 | 0.51 | 0.74 | 0.6 | 0.49 | 0.44 | 0.58 |
| 11 | 22475813 | rs2671122 | -0.815 | -0.409 | 0.253 | 0.108 | 0 | -0.006 | -0.006 | 0.644 | 0.23 | 0.87 | 0.41 | 0.22 | 0.21 | 0.21 |
| 11 | 22477620 | rs720092 | -0.365 | -0.477 | 0.21 | -0.053 | 0.01 | -0.009 | -0.009 | 0 | 0.27 | 0.04 | 0.18 | 0.22 | 0.17 | 0.25 |
| 11 | 22486266 | rs1996163 | -0.543 | -0.203 | 0.604 | -0.032 | 0.001 | -0.009 | -0.009 | 0 | 0.27 | 0.32 | 0.23 | 0.2 | 0.22 | 0.25 |
| 11 | 22489981 | rs340938 | 1.263 | -0.489 | 0.214 | -0.167 | 0.016 | 0.075 | 0.075 | 0 | 0.47 | 0.18 | 0.65 | 0.71 | 0.62 | 0.58 |
| 11 | 22490228 | rs11026577 | -0.224 | -0.655 | 0.105 | -0.278 | -0.072 | 0.065 | 0.065 | 0 | 0.32 | 0.03 | 0.24 | 0.16 | 0.3 | 0.23 |
| 11 | 22490475 | rs6483853 | -1.066 | -0.733 | 0.385 | -0.039 | -0.02 | -0.008 | -0.008 | 0.223 | 0.2 | 0.74 | 0.1 | 0.12 | 0.08 | 0.19 |
| 11 | 22495330 | rs11026578 | -0.197 | -0.702 | 0.52 | -0.024 | 0.005 | 0.004 | 0.004 | 1.306 | 0.14 | 0.17 | 0.05 | 0.07 | 0.07 | 0.16 |
| 11 | 22499539 | rs1514081 | 1.793 | -0.701 | 0.315 | 0.071 | -0.127 | -0.118 | -0.118 | 0.077 | 0.38 | 0.06 | 0.4 | 0.5 | 0.48 | 0.4 |
| 11 | 22501906 | rs1514077 | -1.755 | -1.183 | 1.147 | -0.15 | 0.016 | 0.019 | 0.019 | 0 | 0.47 | 0.63 | 0.29 | 0.31 | 0.34 | 0.42 |
| 11 | 22516207 | rs11026587 | 0.334 | -1.168 | 0.562 | -0.112 | 0.007 | -0.006 | -0.006 | 0 | 0.11 | 0.07 | 0.04 | 0.13 | 0.07 | 0.19 |
| 11 | 22519082 | rs1028067 | -0.52 | -1.108 | -0.009 | -0.097 | -0.055 | 0.023 | 0.023 | 0.01 | 0.29 | 0.03 | 0.2 | 0.19 | 0.26 | 0.22 |
| 11 | 22525081 | rs925633 | -0.055 | -0.948 | -0.042 | -0.103 | -0.024 | -0.02 | -0.02 | 0.788 | 0.86 | 0.71 | 0.94 | 0.93 | 0.92 | 0.84 |
| 11 | 22525721 | rs974837 | 1.319 | -0.926 | -0.739 | -0.092 | 0.049 | 0.066 | 0.066 | 0 | 0.52 | 0.22 | 0.7 | 0.69 | 0.67 | 0.58 |
| 11 | 22533052 | rs11026604 | -0.359 | -0.48 | -0.641 | 0.201 | -0.014 | -0.011 | -0.011 | 0 | 0.14 | 0.24 | 0.06 | 0.07 | 0.07 | 0.16 |
| 11 | 22533437 | rs10833764 | 1.197 | -0.525 | -0.725 | 0.153 | 0.017 | 0.034 | 0.034 | 0 | 0.57 | 0.2 | 0.71 | 0.69 | 0.67 | 0.6 |
| 11 | 22535344 | rs16909585 | -0.973 | -0.284 | -0.393 | 0.223 | 0.002 | 0.002 | 0.002 | 1.313 | 0.14 | 0.18 | 0.05 | 0.07 | 0.07 | 0.16 |
| 11 | 22536864 | rs10833766 | 0.603 | 0.729 | -0.491 | 0.301 | -0.001 | -0.001 | -0.001 | 0.473 | 0.81 | 0.84 | 0.85 | 0.84 | 0.8 | 0.91 |
| 11 | 22537165 | rs12273019 | -1.054 | -0.085 | -0.314 | 0.359 | -0.007 | -0.01 | -0.01 | 0 | 0.32 | 0.64 | 0.46 | 0.19 | 0.23 | 0.25 |
| 11 | 22537674 | rs7120717 | -0.953 | -0.545 | -0.243 | 1.129 | 0.002 | -0.016 | -0.016 | 0 | 0.28 | 0.22 | 0.28 | 0.54 | 0.29 | 0.24 |
| 11 | 22540139 | rs16927135 | -0.781 | -0.517 | 1.076 | 1.045 | -0.002 | -0.011 | -0.011 | 0 | 0.13 | 0.36 | 0.14 | 0.11 | 0.1 | 0.04 |
| 11 | 22551767 | rs1514084 | 0.357 | -0.86 | 1.049 | 1.116 | 0.009 | -0.024 | -0.024 | 0 | 0.68 | 0.72 | 0.72 | 0.51 | 0.71 | 0.77 |
| 11 | 22559731 | rs10833776 | 1.557 | -0.96 | 0.798 | 1.221 | 0.001 | 0.038 | 0.038 | 0.442 | 0.52 | 0.05 | 0.54 | 0.34 | 0.61 | 0.64 |
| 11 | 22560585 | rs4923008 | 0.863 | -0.989 | 1.76 | 1.108 | 0 | -0.026 | -0.026 | 0 | 0.86 | 0.42 | 0.84 | 0.93 | 0.9 | 0.96 |
| 11 | 22567197 | rs1399103 | -1.135 | -0.96 | 1.366 | 1.126 | -0.004 | 0.035 | 0.035 | 0.453 | 0.14 | 0.42 | 0.15 | 0.17 | 0.1 | 0.14 |
| 11 | 22580526 | rs16909659 | -0.096 | -1.086 | 1.076 | 1.593 | 0.006 | -0.045 | -0.045 | 0 | 0.2 | 0.05 | 0.1 | 0.2 | 0.2 | 0.09 |
| 11 | 22587033 | rs6483870 | -0.957 | -1.755 | 1.678 | 1.096 | -0.013 | -0.066 | -0.066 | 0 | 0.19 | 0.21 | 0.11 | 0.01 | 0.13 | 0.13 |
| 11 | 22591011 | rs10833781 | -1.745 | -1.651 | 1.073 | 1.288 | 0.033 | 0.026 | 0.026 | 0 | 0.46 | 0.61 | 0.37 | 0.3 | 0.37 | 0.25 |
| 11 | 22592845 | rs7950479 | -2.136 | -1.565 | 0.696 | 1.609 | 0.068 | 0.096 | 0.096 | 0.014 | 0.52 | 0.85 | 0.47 | 0.69 | 0.4 | 0.39 |
| 11 | 22598840 | rs10833782 | -1.731 | -1.573 | 0.707 | 1.512 | 0.063 | 0.05 | 0.05 | 0 | 0.47 | 0.66 | 0.38 | 0.3 | 0.38 | 0.25 |
| 11 | 22599604 | rs11026668 | 0.331 | -1.581 | 0.693 | 1.52 | 0.006 | -0.05 | -0.05 | 0 | 0.2 | 0.05 | 0.1 | 0.2 | 0.2 | 0.09 |
| 11 | 22602387 | rs11026669 | -1.117 | -1.567 | 0.785 | 1.426 | -0.033 | -0.056 | -0.056 | 0.962 | 0.24 | 0.52 | 0.26 | 0.1 | 0.16 | 0.14 |
| 11 | 22608265 | rs16909724 | -0.349 | -1.508 | 0.764 | 1.602 | 0.023 | -0.059 | -0.059 | 0 | 0.25 | 0.18 | 0.17 | 0.5 | 0.22 | 0.22 |
| 11 | 22613389 | rs1355889 | 1.903 | -1.543 | 0.577 | 1.569 | 0.021 | 0.062 | 0.062 | 0 | 0.47 | 0.05 | 0.52 | 0.3 | 0.59 | 0.6 |
| 11 | 22613488 | rs1355888 | 1.894 | -1.543 | 0.6 | 1.569 | 0.055 | 0.091 | 0.091 | 0 | 0.47 | 0.08 | 0.52 | 0.3 | 0.59 | 0.6 |
| 11 | 22618626 | rs11607117 | 0.298 | -1.491 | 0.503 | 1.599 | -0.012 | -0.069 | -0.069 | 0 | 0.2 | 0.03 | 0.09 | 0.2 | 0.2 | 0.09 |
| 11 | 22641217 | rs448101 | 1.767 | -1.422 | 0.514 | 1.561 | 0.069 | 0.103 | 0.103 | 0 | 0.48 | 0.11 | 0.52 | 0.3 | 0.59 | 0.61 |
| 11 | 22646025 | rs4447177 | 0.765 | -1.617 | 0.618 | 1.536 | -0.026 | -0.037 | -0.037 | 0 | 0.75 | 0.34 | 0.76 | 0.88 | 0.85 | 0.82 |
| 11 | 22647366 | rs3740615 | 0.276 | -1.603 | 0.201 | 1.652 | -0.002 | -0.042 | -0.042 | 0 | 0.19 | 0.04 | 0.09 | 0.2 | 0.2 | 0.05 |
| 11 | 22648806 | rs7112345 | -1.171 | -1.458 | 0.325 | 0.878 | -0.066 | -0.141 | -0.141 | 0.413 | 0.24 | 0.46 | 0.22 | 0.04 | 0.15 | 0.15 |
| 11 | 22648843 | rs4442551 | 0.154 | -1.385 | -0.978 | 0.919 | 0.011 | -0.056 | -0.056 | 0 | 0.2 | 0.07 | 0.14 | 0.26 | 0.2 | 0.05 |
| 11 | 22657331 | rs337484 | 1.181 | -0.861 | -0.244 | 0.451 | 0.033 | 0.037 | 0.037 | 0 | 0.74 | 0.22 | 0.75 | 0.88 | 0.85 | 0.82 |
| 11 | 22689884 | rs11026723 | 0.629 | 0.385 | -0.93 | 0.783 | -0.044 | 0.033 | 0.033 | 0.203 | 0.59 | 0.13 | 0.54 | 0.63 | 0.59 | 0.45 |
| 11 | 22690204 | rs680436 | -2.151 | -0.023 | 0.668 | 0.491 | -0.01 | -0.02 | -0.02 | 0 | 0.13 | 0.37 | 0.07 | 0.07 | 0.07 | 0.17 |
| 11 | 22696029 | rs1479896 | -1.549 | 0.049 | 0.877 | 0.569 | -0.001 | -0.016 | -0.016 | 0 | 0.17 | 0.25 | 0.12 | 0.14 | 0.17 | 0.27 |
| 11 | 22696451 | rs922571 | 2.331 | 0.234 | 0.531 | 0.428 | 0.012 | 0.055 | 0.055 | 0.471 | 0.64 | 0.17 | 0.61 | 0.75 | 0.67 | 0.55 |
| 11 | 22696692 | rs380241 | -1.009 | 0.28 | 0.866 | 0.417 | 0.009 | -0.025 | -0.025 | 0 | 0.09 | 0.17 | 0.14 | 0.04 | 0.1 | 0.14 |
| 11 | 22697623 | rs7117792 | 1.984 | 0.316 | 0.595 | 0.45 | 0.011 | 0.05 | 0.05 | 0.461 | 0.65 | 0.19 | 0.62 | 0.75 | 0.67 | 0.55 |
| 11 | 22700825 | rs443661 | -2.323 | 0.324 | 0.85 | 0.431 | -0.003 | -0.022 | -0.022 | 1.844 | 0.33 | 0.61 | 0.3 | 0.18 | 0.3 | 0.43 |
| 11 | 22711144 | rs91294 | -1.54 | 0.332 | -0.132 | 0.205 | -0.001 | -0.035 | -0.035 | 0 | 0.17 | 0.18 | 0.16 | 0.01 | 0.14 | 0.23 |

| | | | | | | | | | | | | | | | | |
|---|---|---|---|---|---|---|---|---|---|---|---|---|---|---|---|---|
| 11 | 22717120 | rs1628924 | -0.865 | 0.03 | -0.216 | 1.081 | -0.002 | 0.004 | 0.004 | 0 | 0.24 | 0.25 | 0.11 | 0.33 | 0.17 | 0.2 |
| 11 | 22721656 | rs7952704 | 1.201 | -0.498 | 0.246 | 1.494 | 0.054 | -0.004 | -0.004 | 0 | 0.66 | 0.19 | 0.82 | 0.39 | 0.78 | 0.69 |
| 11 | 22729464 | rs337454 | -0.618 | -0.534 | 0.958 | 1.759 | -0.012 | 0.023 | 0.023 | 0 | 0.25 | 0.28 | 0.12 | 0.36 | 0.17 | 0.2 |
| 11 | 22730694 | rs440991 | -0.575 | -0.559 | 1.036 | 1.838 | 0.003 | -0.005 | -0.005 | 0 | 0.24 | 0.2 | 0.1 | 0.35 | 0.17 | 0.19 |
| 11 | 22747745 | rs337448 | 1.048 | -0.637 | 2.269 | 1.752 | 0.057 | 0.053 | 0.053 | 2.757 | 0.94 | 0.57 | 0.96 | 1 | 0.99 | 0.98 |
| 11 | 22752092 | rs10833804 | -1.299 | 0.288 | 1.95 | 1.453 | 0.001 | 0 | 0 | 0 | 0.32 | 0.4 | 0.29 | 0.6 | 0.34 | 0.32 |
| 11 | 22753580 | rs11026758 | -0.669 | 0.31 | 1.945 | 1.489 | -0.021 | -0.02 | -0.02 | 2.606 | 0.08 | 0.06 | 0.17 | 0.18 | 0.17 | 0.13 |
| 11 | 22754297 | rs11026760 | 1.679 | 0.291 | 1.941 | 1.411 | -0.002 | 0.016 | 0.016 | 0.67 | 0.61 | 0.13 | 0.67 | 0.4 | 0.65 | 0.65 |
| 11 | 22765079 | rs7929747 | 1.373 | 0.326 | 2.372 | 1.44 | 0.003 | 0.007 | 0.007 | 0.229 | 0.63 | 0.27 | 0.69 | 0.41 | 0.66 | 0.65 |
| 11 | 22779484 | rs10766966 | -1.201 | 0.356 | 2.193 | 1.452 | 0 | 0 | 0 | 1.391 | 0.31 | 0.31 | 0.28 | 0.59 | 0.34 | 0.34 |
| 11 | 22786413 | rs10734321 | 1.988 | 0.42 | 2.53 | 1.558 | 0.059 | 0.108 | 0.108 | 1.137 | 0.85 | 0.5 | 0.79 | 0.77 | 0.82 | 0.83 |
| 11 | 22789765 | rs12226338 | -1.536 | 0.448 | 2.45 | 1.583 | 0.01 | 0.019 | 0.019 | 0 | 0.11 | 0.16 | 0.17 | 0.23 | 0.18 | 0.15 |
| 11 | 22790693 | rs2896652 | -1.777 | 0.038 | 2.33 | 1.338 | -0.003 | -0.008 | -0.008 | 0 | 0.39 | 0.58 | 0.42 | 0.57 | 0.39 | 0.36 |
| 11 | 22792091 | rs17306223 | -0.441 | 0.073 | 2.22 | 1.366 | -0.029 | -0.025 | -0.025 | 0 | 0.08 | 0.03 | 0.16 | 0.15 | 0.17 | 0.15 |
| 11 | 22793221 | rs11026784 | 1.455 | 0.336 | 2.547 | 1.34 | 0.003 | -0.001 | -0.001 | 0 | 0.59 | 0.33 | 0.49 | 0.5 | 0.6 | 0.62 |
| 11 | 22793809 | rs1499242 | -0.905 | 0.326 | 2.812 | 1.337 | -0.021 | 0.021 | 0.021 | 0 | 0.3 | 0.38 | 0.24 | 0.35 | 0.22 | 0.21 |
| 11 | 22793906 | rs4307722 | 1.706 | 0.317 | 2.723 | 1.33 | 0.002 | -0.009 | -0.009 | 0.009 | 0.57 | 0.16 | 0.48 | 0.5 | 0.6 | 0.62 |
| 11 | 22798431 | rs1499244 | 0.808 | 1.812 | 2.365 | 0.963 | 0 | -0.006 | -0.006 | 0 | 0.62 | 0.37 | 0.58 | 0.58 | 0.62 | 0.72 |
| 11 | 22803712 | rs7129006 | -1.398 | 1.733 | 2.164 | 0.92 | 0 | 0.002 | 0.002 | 0 | 0.43 | 0.46 | 0.55 | 0.57 | 0.55 | 0.39 |
| 11 | 22812535 | rs2041466 | -0.617 | 1.303 | 0.875 | 0.395 | 0.006 | -0.008 | -0.008 | 0 | 0.37 | 0.57 | 0.45 | 0.32 | 0.45 | 0.28 |
| 11 | 22813463 | rs2109681 | -0.442 | 1.335 | 0.709 | 0.352 | -0.013 | 0.019 | 0.019 | 0.831 | 0.34 | 0.3 | 0.35 | 0.21 | 0.43 | 0.27 |
| 11 | 22825559 | rs1544570 | -0.411 | 1.147 | -0.292 | 0.364 | -0.124 | 0.028 | 0.028 | 0 | 0.31 | 0.05 | 0.4 | 0.16 | 0.42 | 0.36 |
| 11 | 22828470 | rs7129430 | 1.123 | 1.154 | -0.411 | 0.051 | -0.047 | 0.119 | 0.119 | 0 | 0.54 | 0.16 | 0.43 | 0.77 | 0.44 | 0.64 |
| 11 | 22828912 | rs16910194 | 0.297 | 1.06 | 0.492 | -0.014 | 0.017 | -0.068 | -0.068 | 0 | 0.13 | 0.33 | 0.13 | 0.04 | 0.16 | 0.05 |
| 11 | 22832056 | rs10833823 | 1.675 | 0.86 | -0.494 | 0.402 | 0.02 | -0.12 | -0.12 | 0 | 0.41 | 0.04 | 0.35 | 0.49 | 0.3 | 0.36 |
| 11 | 22843220 | rs1019216 | 0.748 | 0.95 | 0.068 | 0.187 | 0 | -0.014 | -0.014 | 0 | 0.12 | 0.16 | 0.11 | 0.04 | 0.17 | 0.05 |
| 11 | 22854616 | rs7112397 | -0.717 | 0.964 | -0.176 | 0.229 | 0.002 | -0.018 | -0.018 | 1.048 | 0.43 | 0.51 | 0.54 | 0.21 | 0.56 | 0.34 |
| 11 | 22860940 | rs7940960 | -0.109 | 0.8 | -0.283 | 0.666 | 0.013 | -0.002 | -0.002 | 0 | 0.65 | 0.45 | 0.54 | 0.68 | 0.55 | 0.63 |
| 11 | 22881002 | rs3213706 | 0.467 | 0.705 | -0.313 | 0.801 | 0.037 | 0.056 | 0.056 | 0 | 0.5 | 0.32 | 0.42 | 0.63 | 0.36 | 0.6 |
| 11 | 22881341 | rs2240942 | -0.917 | 1.265 | 0.386 | 1.243 | 0.008 | 0.027 | 0.027 | 0 | 0.15 | 0.22 | 0.09 | 0.28 | 0.14 | 0.3 |
| 11 | 22881542 | rs1860678 | -0.962 | 1.25 | 0.418 | 1.256 | 0.008 | 0.035 | 0.035 | 0 | 0.16 | 0.26 | 0.1 | 0.28 | 0.14 | 0.3 |
| 11 | 22881687 | rs2240941 | -0.348 | 0.905 | 0.184 | 1.172 | -0.03 | -0.044 | -0.044 | 0 | 0.1 | 0.05 | 0.07 | 0.21 | 0.13 | 0.28 |
| 11 | 22887435 | rs7946387 | -1.688 | 0.452 | 0.493 | 1.028 | 0.002 | -0.002 | -0.002 | 0 | 0.59 | 0.67 | 0.62 | 0.54 | 0.74 | 0.61 |
| 11 | 22903273 | rs12575930 | 0.293 | 0.867 | 0.428 | 0.944 | -0.043 | -0.019 | -0.019 | 0 | 0.3 | 0.11 | 0.37 | 0.3 | 0.38 | 0.28 |
| 11 | 22904119 | rs2429773 | -0.567 | 0.951 | 0.824 | 0.943 | 0.031 | 0.045 | 0.045 | 0.395 | 0.14 | 0.41 | 0.12 | 0.19 | 0.16 | 0.29 |
| 11 | 22913265 | rs11600550 | 0.353 | 0.854 | 0.906 | 0.949 | -0.031 | -0.012 | -0.012 | 0 | 0.3 | 0.15 | 0.37 | 0.3 | 0.38 | 0.27 |
| 11 | 22916992 | rs887567 | -1.599 | 0.544 | 0.956 | 0.982 | -0.01 | 0.001 | 0.001 | 0 | 0.58 | 0.61 | 0.62 | 0.54 | 0.74 | 0.61 |
| 11 | 22937497 | rs2470332 | -0.426 | 0.687 | 1.131 | 1.065 | -0.005 | -0.003 | -0.003 | 0 | 0.12 | 0.17 | 0.1 | 0.18 | 0.18 | 0.27 |
| 11 | 22944542 | rs2159669 | -0.428 | 0.715 | 1.197 | 1.153 | 0.016 | -0.005 | -0.005 | 0 | 0.35 | 0.43 | 0.44 | 0.26 | 0.44 | 0.28 |
| 11 | 22946149 | rs10766983 | 1.799 | 0.263 | 0.937 | 0.991 | 0.093 | -0.054 | -0.054 | 0 | 0.35 | 0.1 | 0.3 | 0.44 | 0.19 | 0.39 |
| 11 | 22947196 | rs2403778 | 0.136 | 0.312 | 0.929 | 0.881 | -0.035 | 0.006 | 0.006 | 0 | 0.33 | 0.26 | 0.42 | 0.28 | 0.44 | 0.28 |
| 11 | 22958481 | rs4565895 | 0.844 | 0.478 | 0.821 | 0.605 | -0.02 | 0.068 | 0.068 | 0.022 | 0.16 | 0.04 | 0.12 | 0.04 | 0.18 | 0.04 |
| 11 | 22974588 | rs2433178 | -0.119 | 0.588 | 1.391 | 0.762 | 0.078 | 0.035 | 0.035 | 0 | 0.12 | 0.31 | 0.12 | 0.17 | 0.19 | 0.24 |
| 11 | 23000413 | rs7934814 | 0.755 | 0.445 | 1.426 | 0.784 | 0.045 | 0.009 | 0.009 | 0.421 | 0.54 | 0.27 | 0.42 | 0.46 | 0.38 | 0.43 |
| 11 | 23000971 | rs2403793 | -0.276 | 0.394 | 1.509 | 0.69 | 0.103 | 0.06 | 0.06 | 0 | 0.13 | 0.39 | 0.14 | 0.18 | 0.19 | 0.24 |
| 11 | 23020640 | rs3909703 | 1.577 | -0.063 | 1.204 | 0.708 | 0.073 | -0.015 | -0.015 | 0 | 0.36 | 0.14 | 0.3 | 0.42 | 0.21 | 0.4 |
| 11 | 23020845 | rs11026887 | 0.348 | -0.046 | 1.09 | 0.707 | 0.033 | 0.007 | 0.007 | 0.061 | 0.54 | 0.3 | 0.43 | 0.47 | 0.38 | 0.44 |
| 11 | 23027542 | rs12270677 | -1.422 | 0.46 | 0.993 | 0.532 | 0.041 | 0.142 | 0.142 | 0.929 | 0.07 | 0.27 | 0.07 | 0.17 | 0.05 | 0.11 |
| 11 | 23027613 | rs2433474 | 0.214 | 0.58 | 0.272 | 0.183 | -0.002 | -0.006 | -0.006 | 0 | 0.18 | 0.12 | 0.14 | 0.04 | 0.22 | 0.05 |
| 11 | 23034703 | rs2433479 | -1.081 | -0.035 | 0.432 | 0.514 | 0.038 | 0.007 | 0.007 | 0 | 0.59 | 0.73 | 0.65 | 0.37 | 0.74 | 0.51 |
| 11 | 23036819 | rs10430910 | 1.13 | -0.036 | 0.321 | 0.69 | 0.072 | -0.025 | -0.025 | 0 | 0.36 | 0.14 | 0.3 | 0.42 | 0.2 | 0.41 |
| 11 | 23037293 | rs10833903 | 0.677 | 0.721 | 0.376 | 0.154 | -0.028 | 0.006 | 0.006 | 0 | 0.2 | 0.14 | 0.31 | 0.01 | 0.41 | 0.15 |
| 11 | 23044153 | rs3909702 | 1.307 | 0.42 | -0.146 | 0.339 | 0.063 | -0.002 | -0.002 | 0.519 | 0.35 | 0.14 | 0.29 | 0.36 | 0.21 | 0.39 |
| 11 | 23061291 | rs10833917 | 0.631 | 0.965 | -0.045 | 0.421 | -0.129 | 0.116 | 0.116 | 0.228 | 0.19 | 0.04 | 0.37 | 0.01 | 0.41 | 0.16 |
| 11 | 23061510 | rs7939038 | -0.285 | 0.96 | 0.291 | 0.484 | 0.068 | -0.013 | -0.013 | 0 | 0.73 | 0.61 | 0.55 | 0.78 | 0.54 | 0.75 |
| 11 | 23070714 | rs6483890 | -0.596 | 0.857 | 0.083 | 0.53 | 0.011 | -0.002 | -0.002 | 0 | 0.76 | 0.78 | 0.66 | 0.79 | 0.58 | 0.7 |
| 11 | 23083059 | rs957830 | -0.419 | 0.56 | -0.28 | 0.535 | -0.003 | 0 | 0 | 1.156 | 0.43 | 0.51 | 0.34 | 0.42 | 0.38 | 0.39 |
| 11 | 23094511 | rs2433518 | -0.714 | 0.794 | -1.332 | 0.575 | -0.04 | 0.001 | 0.001 | 0.159 | 0.77 | 0.85 | 0.66 | 0.78 | 0.58 | 0.78 |
| 11 | 23095871 | rs2195030 | 1.307 | 0.542 | -1.467 | 0.588 | 0.065 | -0.023 | -0.023 | 3.443 | 0.32 | 0.11 | 0.28 | 0.36 | 0.2 | 0.33 |
| 11 | 23096397 | rs2468574 | -0.877 | 0.614 | -1.441 | 0.587 | -0.026 | 0.001 | 0.001 | 0.041 | 0.78 | 0.85 | 0.66 | 0.78 | 0.58 | 0.78 |
| 11 | 23099792 | rs2433578 | -0.429 | 0.578 | -1.497 | 0.606 | 0.021 | -0.001 | -0.001 | 0 | 0.45 | 0.74 | 0.38 | 0.42 | 0.39 | 0.44 |
| 11 | 23102000 | rs11026966 | 0.193 | 1.132 | -1.725 | 0.719 | -0.064 | 0.001 | 0.001 | 0.022 | 0.23 | 0.11 | 0.33 | 0.22 | 0.42 | 0.22 |
| 11 | 23108987 | rs2468585 | -0.673 | 0.341 | -1.293 | 0.965 | 0.003 | -0.019 | -0.019 | 1.552 | 0.27 | 0.13 | 0.18 | 0.29 | 0.24 | 0.13 |
| 11 | 23118929 | rs2433567 | 0.641 | 0.164 | -1.012 | 1.036 | 0.002 | -0.007 | -0.007 | 0 | 0.85 | 0.84 | 0.92 | 0.73 | 0.9 | 0.68 |
| 11 | 23122704 | rs10833940 | -0.163 | 0.365 | -0.852 | 0.651 | 0.004 | 0.015 | 0.015 | 1.052 | 0.35 | 0.49 | 0.38 | 0.49 | 0.36 | 0.21 |
| 11 | 23125286 | rs7112320 | -1.245 | 0.264 | -0.369 | 0.758 | 0 | 0.011 | 0.011 | 0 | 0.49 | 0.64 | 0.47 | 0.59 | 0.46 | 0.51 |
| 11 | 23125291 | rs10833941 | -0.095 | 0.262 | -0.37 | 0.747 | 0 | 0.012 | 0.012 | 0.019 | 0.14 | 0.05 | 0.07 | 0.11 | 0.1 | 0.29 |
| 11 | 23125374 | rs10833942 | -0.095 | 0.262 | -0.37 | 0.747 | 0 | 0.012 | 0.012 | 0.019 | 0.14 | 0.05 | 0.07 | 0.11 | 0.1 | 0.29 |
| 11 | 23132261 | rs7119653 | -0.203 | 0.241 | -0.365 | 0.906 | 0.007 | -0.013 | -0.013 | 0.631 | 0.36 | 0.59 | 0.4 | 0.65 | 0.36 | 0.31 |

| | | | | | | | | | | | | | | | | |
|---|---|---|---|---|---|---|---|---|---|---|---|---|---|---|---|---|
| 11 | 23136879 | rs2468662 | 1.165 | -0.08 | -0.995 | -0.065 | -0.052 | 0.099 | 0.099 | 0 | 0.23 | 0.06 | 0.3 | 0.01 | 0.35 | 0.18 |
| 11 | 23140031 | rs2433455 | -0.509 | -0.058 | -0.974 | -0.03 | 0.001 | -0.001 | -0.001 | 1.032 | 0.86 | 0.85 | 0.92 | 0.82 | 0.9 | 0.78 |
| 11 | 23140654 | rs3110275 | -0.346 | -0.065 | -0.953 | -0.034 | -0.001 | 0 | 0 | 0.182 | 0.86 | 0.82 | 0.91 | 0.82 | 0.9 | 0.7 |
| 11 | 23145520 | rs971535 | 0.414 | -0.085 | -0.998 | -0.072 | 0.002 | 0.064 | 0.064 | 0 | 0.5 | 0.24 | 0.52 | 0.25 | 0.54 | 0.47 |
| 11 | 23155225 | rs11027001 | 0.158 | -0.183 | -1.037 | -0.121 | 0.002 | 0.002 | 0.002 | 0.43 | 0.38 | 0.61 | 0.4 | 0.57 | 0.37 | 0.31 |
| 11 | 23155251 | rs12421611 | 1.228 | -0.171 | -0.85 | -0.354 | 0.019 | 0.003 | 0.003 | 0.013 | 0.26 | 0.1 | 0.2 | 0.25 | 0.19 | 0.28 |
| 11 | 23160403 | rs7933654 | -0.396 | -0.031 | -0.826 | -0.168 | 0.001 | -0.002 | -0.002 | 0.29 | 0.87 | 0.85 | 0.92 | 0.82 | 0.9 | 0.76 |
| 11 | 23161082 | rs7936974 | 0.457 | -0.117 | -0.915 | -0.426 | 0 | 0.006 | 0.006 | 0 | 0.51 | 0.38 | 0.54 | 0.33 | 0.55 | 0.52 |
| 11 | 23161451 | rs10767012 | 0.056 | -0.074 | -0.959 | -0.405 | 0.014 | -0.001 | -0.001 | 1.945 | 0.37 | 0.58 | 0.4 | 0.63 | 0.4 | 0.27 |
| 11 | 23161499 | rs7115663 | -0.694 | -0.126 | -0.834 | -0.257 | 0 | 0 | 0 | 0 | 0.5 | 0.6 | 0.56 | 0.67 | 0.49 | 0.47 |
| 11 | 23163357 | rs4497407 | -0.326 | -0.538 | -0.617 | -1.011 | 0.002 | 0.004 | 0.004 | 0 | 0.49 | 0.56 | 0.37 | 0.2 | 0.56 | 0.51 |
| 11 | 23164031 | rs4578382 | 1.082 | -0.636 | -0.469 | -1.007 | -0.003 | -0.016 | -0.016 | 0.316 | 0.08 | 0.06 | 0.03 | 0 | 0.04 | 0.01 |
| 11 | 23168804 | rs11027007 | 0.2 | -0.519 | -0.597 | -0.623 | -0.014 | -0.038 | -0.038 | 0 | 0.18 | 0.12 | 0.2 | 0.35 | 0.18 | 0.1 |
| 11 | 23169829 | rs16910726 | 1.51 | -0.52 | -0.589 | -0.616 | 0.02 | -0.021 | -0.021 | 0.548 | 0.1 | 0.05 | 0.13 | 0.34 | 0.08 | 0.07 |
| 11 | 23170844 | rs4472938 | 1.004 | -0.546 | -0.567 | -1.077 | 0.017 | 0.037 | 0.037 | 0 | 0.3 | 0.24 | 0.2 | 0.03 | 0.31 | 0.32 |
| 11 | 23171592 | rs4077895 | 0.249 | -0.579 | -0.565 | -1.095 | 0.027 | 0.019 | 0.019 | 0 | 0.39 | 0.31 | 0.23 | 0.03 | 0.34 | 0.32 |
| 11 | 23175316 | rs10160246 | -0.483 | -0.422 | -0.537 | -0.817 | 0.002 | 0.043 | 0.043 | 0 | 0.49 | 0.42 | 0.32 | 0.37 | 0.52 | 0.55 |
| 11 | 23183529 | rs11027021 | -0.716 | -0.365 | -0.168 | -0.986 | 0.046 | -0.079 | -0.079 | 0.968 | 0.22 | 0.42 | 0.25 | 0.1 | 0.19 | 0.18 |
| 11 | 23190252 | rs16910749 | 1.401 | -0.341 | -0.306 | -1.024 | -0.01 | -0.031 | -0.031 | 1.007 | 0.08 | 0.09 | 0.03 | 0 | 0.03 | 0.01 |
| 11 | 23192677 | rs1871953 | 0.319 | -0.331 | -0.33 | -1.028 | -0.006 | -0.014 | -0.014 | 0 | 0.16 | 0.14 | 0.09 | 0.01 | 0.14 | 0.03 |
| 11 | 23193095 | rs2449446 | 1.111 | -0.369 | -0.366 | -1.048 | 0.018 | 0.035 | 0.035 | 0 | 0.3 | 0.25 | 0.2 | 0.03 | 0.31 | 0.31 |
| 11 | 23196241 | rs11828289 | 0.497 | -0.174 | -0.141 | -0.869 | 0.075 | -0.06 | -0.06 | 0.268 | 0.11 | 0.23 | 0.2 | 0.01 | 0.16 | 0.06 |
| 11 | 23202785 | rs16910800 | -0.405 | -0.264 | -0.161 | -0.919 | 0.036 | -0.108 | -0.108 | 0.655 | 0.2 | 0.29 | 0.23 | 0.01 | 0.19 | 0.06 |
| 11 | 23214366 | rs17234274 | 0.481 | -0.412 | -0.709 | -0.939 | -0.017 | 0.006 | 0.006 | 0 | 0.39 | 0.35 | 0.45 | 0.31 | 0.44 | 0.38 |
| 11 | 23214880 | rs10833965 | -0.186 | -0.425 | -0.705 | -1.165 | 0.004 | -0.04 | -0.04 | 1.389 | 0.54 | 0.46 | 0.61 | 0.98 | 0.6 | 0.69 |
| 11 | 23216479 | rs2468642 | 0.426 | -0.056 | -0.923 | -0.941 | -0.003 | -0.037 | -0.037 | 0 | 0.35 | 0.27 | 0.35 | 0.02 | 0.37 | 0.31 |
| 11 | 23217814 | rs1384464 | 1.063 | -0.021 | -0.931 | -0.945 | -0.001 | -0.053 | -0.053 | 0.002 | 0.27 | 0.2 | 0.28 | 0.01 | 0.26 | 0.28 |
| 11 | 23219311 | rs12099049 | 1.258 | 0.15 | -1.98 | -0.774 | -0.017 | 0.015 | 0.015 | 0 | 0.08 | 0.06 | 0.06 | 0.01 | 0.11 | 0.03 |
| 11 | 23220093 | rs11027052 | -0.172 | 0.179 | -1.984 | -0.491 | 0.002 | -0.069 | -0.069 | 1.158 | 0.16 | 0.33 | 0.22 | 0.04 | 0.18 | 0.12 |
| 11 | 23223423 | rs1004730 | -0.035 | 0.265 | -1.942 | -0.434 | 0.006 | -0.068 | -0.068 | 0.042 | 0.16 | 0.33 | 0.22 | 0.04 | 0.17 | 0.11 |
| 11 | 23224901 | rs2170597 | 0.831 | 0.018 | -1.839 | -1.013 | 0.019 | -0.015 | -0.015 | 3.865 | 0.06 | 0.22 | 0.06 | 0.04 | 0.09 | 0.07 |
| 11 | 23226964 | rs1384487 | -1.347 | -0.018 | -1.973 | -0.939 | 0.02 | 0.121 | 0.121 | 0 | 0.77 | 0.96 | 0.85 | 0.99 | 0.78 | 0.75 |
| 11 | 23232884 | rs11602836 | 0.67 | 0.532 | -1.413 | -0.128 | -0.099 | -0.148 | -0.148 | 0 | 0.24 | 0.02 | 0.23 | 0.35 | 0.27 | 0.38 |
| 11 | 23233501 | rs12280639 | -0.509 | 0.196 | -0.265 | -0.463 | 0.009 | -0.063 | -0.063 | 0 | 0.15 | 0.31 | 0.22 | 0.04 | 0.17 | 0.11 |
| 11 | 23243241 | rs1564628 | 0.128 | -0.005 | -0.596 | -0.652 | 0.005 | 0.015 | 0.015 | 0 | 0.6 | 0.64 | 0.61 | 0.86 | 0.61 | 0.62 |
| 11 | 23243454 | rs1384479 | 0.129 | -0.031 | -0.421 | -0.636 | 0.001 | -0.007 | -0.007 | 0 | 0.6 | 0.63 | 0.62 | 0.86 | 0.61 | 0.62 |
| 11 | 23244871 | rs10833989 | 0.054 | 0.001 | -0.385 | -0.488 | 0 | -0.002 | -0.002 | 0 | 0.1 | 0.07 | 0.22 | 0.11 | 0.08 | 0.13 |
| 11 | 23245030 | rs16910901 | 0.885 | -0.829 | -0.065 | 0.33 | -0.033 | 0.082 | 0.082 | 0 | 0.1 | 0.18 | 0.14 | 0.35 | 0.07 | 0.06 |
| 11 | 23245183 | rs12362091 | 0.816 | -1.314 | 0.117 | 0.02 | 0 | -0.019 | -0.019 | 4.061 | 0.26 | 0.23 | 0.2 | 0 | 0.29 | 0.18 |
| 11 | 23246812 | rs1482728 | 0.351 | -0.758 | 0.484 | 0.533 | 0.013 | 0.077 | 0.077 | 0 | 0.81 | 0.73 | 0.66 | 0.55 | 0.78 | 0.6 |
| 11 | 23247163 | rs922858 | 1.122 | -0.794 | 0.416 | 0.49 | 0 | -0.026 | -0.026 | 0 | 0.26 | 0.24 | 0.2 | 0 | 0.29 | 0.18 |
| 11 | 23247620 | rs11027076 | -0.165 | -0.848 | 0.415 | 0.587 | -0.018 | 0.042 | 0.042 | 0 | 0.2 | 0.26 | 0.36 | 0.46 | 0.16 | 0.19 |
| 11 | 23251330 | rs1384483 | 1.577 | -0.49 | -0.377 | 0.029 | -0.054 | -0.007 | -0.007 | 5.392 | 0.26 | 0.02 | 0.12 | 0.15 | 0.24 | 0.29 |
| 11 | 23255522 | rs1072786 | 0.056 | -0.626 | 0.129 | -0.113 | 0.014 | 0.025 | 0.025 | 0 | 0.47 | 0.41 | 0.46 | 0.78 | 0.4 | 0.53 |
| 11 | 23256291 | rs11027081 | -0.096 | -0.65 | 0.181 | -0.138 | 0.005 | 0 | 0 | 0.088 | 0.51 | 0.49 | 0.53 | 0.17 | 0.59 | 0.42 |
| 11 | 23259140 | rs11027083 | -0.736 | -0.438 | 0.712 | 0.222 | 0.009 | 0.009 | 0.009 | 0 | 0.31 | 0.42 | 0.42 | 0.4 | 0.33 | 0.16 |
| 11 | 23259274 | rs12283577 | 0.917 | -0.989 | 0.795 | 0.489 | -0.03 | -0.015 | -0.015 | 0.135 | 0.72 | 0.56 | 0.59 | 0.17 | 0.78 | 0.58 |
| 11 | 23259338 | rs10833992 | 0.219 | -0.86 | 0.714 | 0.443 | -0.02 | 0.108 | 0.108 | 0 | 0.16 | 0.25 | 0.21 | 0.47 | 0.13 | 0.15 |
| 11 | 23259685 | rs10833993 | 1.086 | -0.855 | 0.714 | 0.255 | -0.031 | 0.065 | 0.065 | 0 | 0.11 | 0.17 | 0.15 | 0.37 | 0.08 | 0.05 |
| 11 | 23259996 | rs721320 | -0.718 | -0.936 | 0.884 | 0.511 | -0.039 | 0.145 | 0.145 | 0 | 0.24 | 0.4 | 0.51 | 0.52 | 0.22 | 0.32 |
| 11 | 23260397 | rs10833994 | 1.153 | -0.97 | 0.773 | 0.541 | 0.036 | -0.043 | -0.043 | 0.729 | 0.11 | 0.05 | 0.13 | 0.36 | 0.08 | 0.05 |
| 11 | 23262395 | rs17306503 | 0.005 | -0.738 | 0.788 | 0.553 | 0.012 | 0 | 0 | 0 | 0.18 | 0.17 | 0.3 | 0.16 | 0.22 | 0.31 |
| 11 | 23263048 | rs17306510 | 0.62 | -0.75 | 0.506 | 0.517 | -0.005 | -0.011 | -0.011 | 0 | 0.15 | 0.11 | 0.21 | 0 | 0.16 | 0.09 |
| 11 | 23268171 | rs989169 | -0.426 | -0.768 | 0.45 | 0.548 | 0.007 | -0.026 | -0.026 | 0 | 0.26 | 0.13 | 0.33 | 0.36 | 0.24 | 0.15 |
| 11 | 23269184 | rs10834000 | -0.417 | -0.762 | 0.452 | 0.55 | 0.006 | -0.026 | -0.026 | 0 | 0.25 | 0.13 | 0.33 | 0.36 | 0.24 | 0.15 |
| 11 | 23276124 | rs12361664 | 1.143 | -0.965 | 0.753 | 0.566 | -0.011 | -0.014 | -0.014 | 0.558 | 0.62 | 0.5 | 0.48 | 0.21 | 0.73 | 0.7 |
| 11 | 23286024 | rs7936937 | 1.165 | -0.965 | 0.87 | 0.566 | -0.02 | -0.024 | -0.024 | 0 | 0.62 | 0.4 | 0.47 | 0.21 | 0.72 | 0.7 |
| 11 | 23288275 | rs12417735 | 0.787 | -0.932 | 0.838 | 0.574 | -0.018 | 0.018 | 0.018 | 0 | 0.14 | 0.08 | 0.21 | 0 | 0.16 | 0.09 |
| 11 | 23289329 | rs7121330 | -1.317 | -0.944 | 0.887 | 0.621 | -0.004 | -0.003 | -0.003 | 1.372 | 0.34 | 0.41 | 0.52 | 0.73 | 0.27 | 0.25 |
| 11 | 23292409 | rs2062142 | -1.427 | -0.851 | 0.848 | 0.605 | 0.002 | -0.009 | -0.009 | 0 | 0.37 | 0.4 | 0.61 | 0.88 | 0.34 | 0.47 |
| 11 | 23292949 | rs10834017 | -1.437 | -0.777 | 0.805 | 0.623 | -0.015 | -0.087 | -0.087 | 0 | 0.37 | 0.5 | 0.62 | 0.88 | 0.34 | 0.47 |
| 11 | 23294641 | rs10500951 | 0.611 | -0.776 | 0.766 | 0.636 | -0.001 | 0.005 | 0.005 | 0 | 0.09 | 0.12 | 0.11 | 0.16 | 0.07 | 0.22 |
| 11 | 23297414 | rs1994118 | -1.24 | -0.746 | 0.776 | 0.468 | 0.006 | -0.013 | -0.013 | 0 | 0.34 | 0.35 | 0.54 | 0.52 | 0.31 | 0.37 |
| 11 | 23304889 | rs12575780 | 0.168 | -0.338 | 0.601 | 0.781 | 0.024 | 0.096 | 0.096 | 0 | 0.16 | 0.23 | 0.29 | 0.37 | 0.2 | 0.2 |
| 11 | 23330293 | rs7931071 | -0.218 | -0.473 | 0.088 | 0.134 | -0.003 | 0.004 | 0.004 | 0 | 0.51 | 0.57 | 0.45 | 0.39 | 0.5 | 0.45 |
| 11 | 23337714 | rs7931150 | -0.324 | -0.549 | 0.025 | 0.09 | -0.006 | 0 | 0 | 0 | 0.52 | 0.6 | 0.46 | 0.39 | 0.51 | 0.45 |
| 11 | 23356139 | rs11027153 | 0.957 | -0.644 | 0.31 | 0.136 | 0 | 0.001 | 0.001 | 0 | 0.29 | 0.22 | 0.16 | 0.08 | 0.3 | 0.26 |
| 11 | 23362010 | rs11027159 | 0.591 | -0.704 | 0.526 | 0.264 | -0.004 | 0.033 | 0.033 | 0 | 0.33 | 0.4 | 0.23 | 0.41 | 0.33 | 0.34 |
| 11 | 23363861 | rs4307718 | -1.084 | -0.89 | 0.474 | 0.192 | -0.003 | 0.002 | 0.002 | 0 | 0.94 | 0.91 | 0.99 | 0.95 | 0.99 | 0.94 |

| | | | | | | | | | | | | | | | | |
|---|---|---|---|---|---|---|---|---|---|---|---|---|---|---|---|---|
| 11 | 23363924 | rs11027160 | 0.659 | -0.843 | 0.597 | 0.203 | -0.003 | 0.022 | 0.022 | 0 | 0.33 | 0.37 | 0.23 | 0.41 | 0.33 | 0.34 |
| 11 | 23372511 | rs4617580 | -0.467 | -1.34 | 0.405 | 0.3 | 0.013 | -0.019 | -0.019 | 3.029 | 0.82 | 0.83 | 0.85 | 0.58 | 0.92 | 0.89 |
| 11 | 23374739 | rs11027171 | 1.228 | -1.391 | 0.633 | -0.296 | -0.002 | -0.02 | -0.02 | 0 | 0.29 | 0.28 | 0.16 | 0.08 | 0.3 | 0.26 |
| 11 | 23376126 | rs6483921 | 0.407 | -1.313 | 0.568 | -0.49 | -0.002 | -0.015 | -0.015 | 0.26 | 0.44 | 0.37 | 0.37 | 0.08 | 0.46 | 0.36 |
| 11 | 23376448 | rs11027173 | 2.103 | -0.981 | 0.389 | -0.801 | -0.019 | 0.06 | 0.06 | 0 | 0.2 | 0.08 | 0.14 | 0 | 0.21 | 0.15 |
| 11 | 23376777 | rs12288166 | 0.533 | -0.904 | 0.599 | -0.698 | -0.01 | -0.003 | -0.003 | 0 | 0.14 | 0.13 | 0.17 | 0.17 | 0.21 | 0.29 |
| 11 | 23386182 | rs7113386 | 0.645 | -0.948 | 0.644 | -0.786 | -0.002 | 0.03 | 0.03 | 0 | 0.45 | 0.5 | 0.38 | 0.08 | 0.45 | 0.36 |
| 11 | 23386882 | rs7114349 | -1.355 | -0.967 | 0.498 | -0.682 | -0.003 | -0.001 | -0.001 | 0.049 | 0.94 | 0.91 | 0.99 | 0.93 | 1 | 0.94 |
| 11 | 23388974 | rs12417697 | 1.067 | -0.873 | 0.521 | -0.812 | -0.004 | -0.007 | -0.007 | 0 | 0.16 | 0.15 | 0.22 | 0 | 0.16 | 0.1 |
| 11 | 23389010 | rs7119607 | 1.495 | -0.908 | 0.586 | -0.781 | -0.002 | -0.029 | -0.029 | 0 | 0.29 | 0.29 | 0.16 | 0.08 | 0.3 | 0.26 |
| 11 | 23399673 | rs7103438 | 1.322 | -1.002 | 0.72 | -0.83 | -0.002 | -0.027 | -0.027 | 0 | 0.3 | 0.34 | 0.15 | 0.08 | 0.3 | 0.27 |
| 11 | 23410903 | rs1585579 | 0.088 | -0.986 | 0.833 | -0.811 | 0.003 | -0.025 | -0.025 | 0 | 0.45 | 0.53 | 0.31 | 0.24 | 0.46 | 0.54 |
| 11 | 23412515 | rs4300392 | -0.473 | -0.961 | 0.871 | -0.8 | -0.001 | -0.004 | -0.004 | 0 | 0.6 | 0.62 | 0.52 | 0.25 | 0.61 | 0.64 |
| 11 | 23415864 | rs10834062 | 0.581 | -0.982 | 0.923 | -0.809 | 0.001 | 0.008 | 0.008 | 0 | 0.35 | 0.27 | 0.46 | 0.75 | 0.38 | 0.36 |
| 11 | 23416345 | rs1381720 | 0.538 | -0.964 | 0.904 | -0.851 | -0.01 | 0 | 0 | 0 | 0.14 | 0.13 | 0.15 | 0.16 | 0.16 | 0.27 |
| 11 | 23416801 | rs1381719 | -0.714 | -1.002 | 0.923 | -0.914 | -0.002 | -0.055 | -0.055 | 0.726 | 0.8 | 0.79 | 0.77 | 1 | 0.84 | 0.9 |
| 11 | 23420155 | rs6483934 | -1.568 | -0.927 | 0.806 | -1.03 | -0.006 | -0.056 | -0.056 | 0 | 0.69 | 0.55 | 0.84 | 0.92 | 0.69 | 0.73 |
| 11 | 23420932 | rs7929370 | 1.808 | -1.064 | 0.577 | -1.023 | -0.042 | 0.075 | 0.075 | 0 | 0.2 | 0.06 | 0.15 | 0 | 0.27 | 0.16 |
| 11 | 23422446 | rs10834065 | 0.81 | -0.92 | 0.681 | -0.956 | 0.015 | -0.018 | -0.018 | 0.633 | 0.16 | 0.11 | 0.3 | 0.76 | 0.12 | 0.2 |
| 11 | 23423825 | rs11027200 | 1.799 | -1.137 | 0.231 | -1.103 | -0.038 | 0.077 | 0.077 | 0 | 0.2 | 0.05 | 0.16 | 0 | 0.26 | 0.15 |
| 11 | 23425775 | rs11825990 | 0.37 | -0.829 | 0.661 | -1.076 | 0.004 | -0.018 | -0.018 | 0 | 0.42 | 0.39 | 0.31 | 0.23 | 0.49 | 0.54 |
| 11 | 23425890 | rs10767046 | 0.933 | -0.712 | 0.426 | -0.491 | 0.016 | -0.022 | -0.022 | 3.778 | 0.14 | 0.1 | 0.25 | 0.4 | 0.11 | 0.12 |
| 11 | 23429071 | rs12284538 | 0.2 | -0.154 | 0.476 | -0.767 | -0.014 | 0 | 0 | 0 | 0.12 | 0.1 | 0.14 | 0.12 | 0.21 | 0.28 |
| 11 | 23437553 | rs1471365 | -0.699 | -0.5 | 0.41 | -1.04 | -0.005 | -0.057 | -0.057 | 0 | 0.56 | 0.48 | 0.65 | 0.92 | 0.64 | 0.68 |
| 11 | 23439096 | rs17234449 | 0.414 | -0.442 | 0.378 | -1.062 | -0.013 | 0 | 0 | 0 | 0.11 | 0.1 | 0.14 | 0.12 | 0.21 | 0.28 |
| 11 | 23443409 | rs10734333 | -1.05 | -0.577 | 0.526 | -0.885 | -0.006 | -0.055 | -0.055 | 0.094 | 0.82 | 0.71 | 0.79 | 1 | 0.88 | 0.92 |
| 11 | 23448534 | rs2199471 | -0.413 | -0.57 | 0.333 | -0.747 | -0.002 | 0 | 0 | 0 | 0.54 | 0.43 | 0.6 | 0.53 | 0.63 | 0.6 |
| 11 | 23457793 | rs4548619 | -0.647 | -0.597 | 0.484 | -0.671 | -0.022 | -0.139 | -0.139 | 0 | 0.81 | 0.48 | 0.77 | 1 | 0.88 | 0.92 |
| 11 | 23458658 | rs1580284 | 0.567 | -0.331 | 0.045 | -0.566 | 0.013 | -0.052 | -0.052 | 0.026 | 0.21 | 0.1 | 0.22 | 0.4 | 0.24 | 0.13 |
| 11 | 23460111 | rs10834086 | 0.698 | 0.272 | 0.052 | -0.54 | -0.013 | -0.048 | -0.048 | 3.125 | 0.2 | 0.11 | 0.24 | 0.4 | 0.26 | 0.15 |
| 11 | 23460491 | rs7107238 | -0.468 | 0.08 | 0.589 | -0.762 | -0.028 | -0.118 | -0.118 | 0 | 0.78 | 0.6 | 0.76 | 1 | 0.86 | 0.89 |
| 11 | 23461597 | rs980452 | -0.939 | 0.079 | 0.626 | -0.761 | -0.015 | -0.089 | -0.089 | 0.343 | 0.81 | 0.66 | 0.77 | 1 | 0.87 | 0.92 |
| 11 | 23466073 | rs11027216 | 1 | 0.145 | 0.501 | -0.735 | -0.012 | -0.044 | -0.044 | 0 | 0.19 | 0.09 | 0.23 | 0.39 | 0.24 | 0.14 |
| 11 | 23470627 | rs11027220 | 1.776 | 0.186 | 0.55 | -0.955 | 0 | -0.038 | -0.038 | 0 | 0.2 | 0.19 | 0.22 | 0 | 0.19 | 0.1 |
| 11 | 23470846 | rs10767062 | -1.633 | 0.16 | 0.584 | -0.974 | 0 | -0.026 | -0.026 | 1.499 | 0.82 | 0.82 | 0.79 | 1 | 0.87 | 0.92 |
| 11 | 23471025 | rs10834089 | 0.557 | 0.367 | 0.478 | -0.903 | -0.014 | -0.014 | -0.014 | 0 | 0.46 | 0.2 | 0.36 | 0.48 | 0.49 | 0.44 |
| 11 | 23475803 | rs11027223 | 2.041 | 0.363 | 0.333 | -0.911 | -0.017 | 0.053 | 0.053 | 0 | 0.24 | 0.06 | 0.12 | 0.08 | 0.23 | 0.29 |
| 11 | 23477604 | rs7935052 | 0.687 | -0.207 | 0.726 | -0.687 | -0.011 | -0.01 | -0.01 | 0 | 0.47 | 0.23 | 0.36 | 0.48 | 0.49 | 0.44 |
| 11 | 23478455 | rs10500956 | 0.7 | -0.128 | 0.547 | -0.586 | 0.02 | 0.003 | 0.003 | 0 | 0.09 | 0.13 | 0.13 | 0.13 | 0.17 | 0.27 |
| 11 | 23482052 | rs7110960 | -0.001 | -0.435 | 0.463 | -0.5 | 0.016 | 0.06 | 0.06 | 0 | 0.68 | 0.49 | 0.59 | 0.48 | 0.68 | 0.54 |
| 11 | 23482944 | rs12146457 | 1.531 | -0.342 | -0.175 | -0.547 | -0.016 | 0.127 | 0.127 | 0 | 0.19 | 0.03 | 0.21 | 0 | 0.18 | 0.1 |
| 11 | 23489969 | rs10834096 | 0.532 | -0.266 | -0.033 | -0.39 | -0.039 | -0.035 | -0.035 | 0 | 0.45 | 0.15 | 0.35 | 0.47 | 0.49 | 0.44 |
| 11 | 23494273 | rs4923097 | -0.429 | -0.236 | 0.327 | -0.404 | -0.003 | -0.002 | -0.002 | 0.197 | 0.72 | 0.7 | 0.65 | 0.87 | 0.69 | 0.65 |
| 11 | 23494877 | rs7110439 | 0.364 | -0.136 | 0.207 | -0.529 | 0 | -0.001 | -0.001 | 0 | 0.45 | 0.44 | 0.38 | 0.48 | 0.36 | 0.45 |
| 11 | 23504057 | rs11027245 | 1.574 | -0.035 | 0.243 | -0.545 | -0.009 | -0.018 | -0.018 | 0.81 | 0.2 | 0.21 | 0.1 | 0.08 | 0.16 | 0.23 |
| 11 | 23513736 | rs1462546 | 0.016 | 0.126 | -0.061 | -0.145 | 0 | 0 | 0 | 0 | 0.67 | 0.64 | 0.57 | 0.35 | 0.66 | 0.56 |
| 11 | 23515684 | rs7117598 | -0.5 | 0.129 | -0.2 | -0.297 | -0.005 | 0.006 | 0.006 | 0 | 0.7 | 0.71 | 0.63 | 0.76 | 0.68 | 0.64 |
| 11 | 23517131 | rs11027262 | 0.908 | -0.888 | -0.229 | -0.067 | 0.017 | 0.037 | 0.037 | 0 | 0.2 | 0.11 | 0.21 | 0 | 0.17 | 0.14 |
| 11 | 23517187 | rs11027263 | 1.276 | -0.859 | -0.397 | -0.051 | 0.003 | 0.036 | 0.036 | 0 | 0.2 | 0.08 | 0.1 | 0.09 | 0.2 | 0.23 |
| 11 | 23517596 | rs1879033 | -0.092 | -0.965 | 0.588 | 0.854 | -0.021 | 0.064 | 0.064 | 0 | 0.24 | 0.34 | 0.28 | 0.42 | 0.21 | 0.21 |
| 11 | 23518405 | rs1350948 | -0.793 | -0.813 | 0.107 | 0.744 | 0.019 | 0.041 | 0.041 | 0 | 0.8 | 0.86 | 0.78 | 1 | 0.84 | 0.87 |
| 11 | 23520512 | rs4923101 | -0.149 | -1.034 | 0.058 | 0.958 | 0.025 | -0.03 | -0.03 | 5.452 | 0.75 | 0.81 | 0.67 | 0.35 | 0.81 | 0.62 |
| 11 | 23521491 | rs7120514 | 0.468 | -1.207 | -0.014 | 0.767 | 0.013 | 0.027 | 0.027 | 0.555 | 0.15 | 0.06 | 0.17 | 0 | 0.09 | 0.07 |
| 11 | 23522689 | rs9299969 | -0.129 | -0.828 | 0.226 | 1.567 | 0.027 | 0.005 | 0.005 | 0 | 0.77 | 0.82 | 0.72 | 0.75 | 0.83 | 0.71 |
| 11 | 23523142 | rs7949875 | -0.014 | -0.82 | 0.285 | 0.962 | 0.014 | -0.014 | -0.014 | 0 | 0.75 | 0.79 | 0.66 | 0.35 | 0.81 | 0.63 |
| 11 | 23523475 | rs7949958 | 0.122 | -0.8 | 0.236 | 0.972 | -0.025 | 0.016 | 0.016 | 0 | 0.73 | 0.63 | 0.65 | 0.35 | 0.81 | 0.63 |
| 11 | 23531129 | rs11826112 | -1.167 | -0.352 | 0.568 | 1.635 | 0.003 | 0.309 | 0.309 | 0.473 | 0.07 | 0.21 | 0.15 | 0.59 | 0.04 | 0.18 |
| 11 | 23532943 | rs7926648 | -0.33 | -0.588 | 0.211 | 1.32 | 0.009 | 0.007 | 0.007 | 0.03 | 0.38 | 0.43 | 0.42 | 0.27 | 0.27 | 0.32 |
| 11 | 23533754 | rs1850824 | 0.917 | -0.812 | 0.494 | 1.489 | -0.015 | 0.312 | 0.312 | 3.027 | 0.89 | 0.61 | 0.85 | 0.61 | 0.94 | 0.82 |
| 11 | 23533902 | rs1850822 | -0.534 | -0.492 | -0.013 | 0.998 | -0.016 | -0.003 | -0.003 | 0.019 | 0.43 | 0.57 | 0.52 | 0.63 | 0.29 | 0.46 |
| 11 | 23536654 | rs11027268 | 0.221 | -0.021 | -0.595 | 0.92 | -0.009 | -0.004 | -0.004 | 0.007 | 0.17 | 0.07 | 0.16 | 0.15 | 0.22 | 0.33 |
| 11 | 23536736 | rs1462545 | -0.626 | -0.129 | -0.469 | 0.532 | 0.001 | -0.007 | -0.007 | 0.881 | 0.56 | 0.54 | 0.56 | 0.67 | 0.56 | 0.64 |
| 11 | 23540006 | rs11027271 | -0.108 | -0.276 | -0.068 | 0.713 | 0.034 | 0.079 | 0.079 | 0 | 0.45 | 0.29 | 0.38 | 0.21 | 0.36 | 0.26 |
| 11 | 23540492 | rs10834123 | -1.01 | -0.274 | 0.019 | 0.88 | -0.033 | -0.057 | -0.057 | 0.012 | 0.69 | 0.79 | 0.61 | 0.43 | 0.64 | 0.7 |
| 11 | 23540793 | rs12224604 | -1.031 | -0.273 | 0.029 | 0.783 | -0.026 | -0.009 | -0.009 | 0 | 0.7 | 0.79 | 0.61 | 0.67 | 0.65 | 0.77 |
| 11 | 23543826 | rs7931521 | -1.147 | -0.32 | -0.099 | 0.908 | -0.078 | -0.036 | -0.036 | 0 | 0.71 | 0.92 | 0.63 | 0.7 | 0.65 | 0.78 |
| 11 | 23545552 | rs10767079 | -1.107 | -0.322 | 0.074 | 0.912 | -0.048 | -0.02 | -0.02 | 0 | 0.71 | 0.89 | 0.63 | 0.69 | 0.65 | 0.78 |
| 11 | 23546058 | rs1462564 | 0.313 | -0.41 | 0.041 | 1.116 | -0.036 | -0.048 | -0.048 | 0 | 0.26 | 0.1 | 0.21 | 0.31 | 0.31 | 0.5 |
| 11 | 23547384 | rs16911459 | 1.323 | -0.729 | 0.345 | 1.124 | -0.018 | 0.071 | 0.071 | 0.094 | 0.09 | 0.22 | 0.04 | 0.22 | 0.05 | 0.08 |

| | | | | | | | | | | | | | | | | |
|---|---|---|---|---|---|---|---|---|---|---|---|---|---|---|---|---|
| 11 | 23548673 | rs11027285 | -1.499 | -0.889 | 0.363 | 0.961 | -0.012 | -0.007 | -0.007 | 0 | 0.73 | 0.82 | 0.64 | 0.68 | 0.67 | 0.8 |
| 11 | 23549065 | rs10500962 | 1.213 | -0.841 | 0.144 | 1.107 | -0.056 | 0.011 | 0.011 | 0.358 | 0.18 | 0.06 | 0.19 | 0.15 | 0.28 | 0.4 |
| 11 | 23552720 | rs7110620 | -0.266 | -1.339 | 0.252 | 0.926 | 0.005 | -0.006 | -0.006 | 0 | 0.31 | 0.28 | 0.21 | 0.43 | 0.19 | 0.21 |
| 11 | 23559112 | rs1599613 | 0.813 | -1.333 | 0.223 | 0.942 | -0.003 | -0.002 | -0.002 | 0 | 0.19 | 0.24 | 0.2 | 0.18 | 0.18 | 0.16 |
| 11 | 23564329 | rs7122954 | -0.503 | -1.327 | 0.089 | 0.805 | 0.002 | 0.001 | 0.001 | 0.046 | 0.41 | 0.37 | 0.39 | 0.31 | 0.32 | 0.3 |
| 11 | 23564493 | rs7122789 | -0.386 | -1.313 | 0.105 | 0.817 | 0.018 | 0.047 | 0.047 | 0 | 0.4 | 0.26 | 0.36 | 0.19 | 0.31 | 0.26 |
| 11 | 23564843 | rs11027305 | -1.277 | -1.296 | -0.084 | 0.535 | 0.01 | 0.006 | 0.006 | 0.094 | 0.5 | 0.55 | 0.43 | 0.54 | 0.37 | 0.38 |
| 11 | 23571242 | rs9943612 | 0.742 | -1.326 | -0.08 | 0.375 | -0.002 | 0.059 | 0.059 | 1.91 | 0.19 | 0.31 | 0.22 | 0.31 | 0.19 | 0.2 |
| 11 | 23572959 | rs11819873 | -0.248 | -1.329 | -0.227 | -0.068 | 0.018 | -0.005 | -0.005 | 0 | 0.29 | 0.25 | 0.21 | 0.01 | 0.17 | 0.11 |
| 11 | 23574446 | rs10741985 | -0.552 | -0.801 | -0.261 | 0.604 | 0.008 | 0.002 | 0.002 | 0.069 | 0.44 | 0.38 | 0.46 | 0.56 | 0.34 | 0.38 |
| 11 | 23575384 | rs1519423 | 0.38 | -0.299 | -0.285 | 0.667 | 0.032 | 0.033 | 0.033 | 0.394 | 0.3 | 0.34 | 0.39 | 0.5 | 0.43 | 0.34 |
| 11 | 23577442 | rs11027313 | -0.476 | -1.26 | -0.152 | -0.409 | -0.018 | -0.034 | -0.034 | 0 | 0.45 | 0.49 | 0.35 | 0.2 | 0.3 | 0.47 |
| 11 | 23581968 | rs11027318 | -0.054 | -1.192 | -0.248 | -0.404 | 0.017 | 0.023 | 0.023 | 0 | 0.41 | 0.36 | 0.33 | 0.18 | 0.3 | 0.43 |
| 11 | 23582397 | rs7110068 | -0.08 | -1.273 | -0.275 | -0.425 | 0.02 | 0.03 | 0.03 | 0 | 0.41 | 0.35 | 0.34 | 0.17 | 0.3 | 0.43 |
| 11 | 23583863 | rs1401615 | -1.134 | -0.925 | -0.653 | -0.929 | 0.007 | -0.001 | -0.001 | 2.675 | 0.78 | 0.76 | 0.73 | 0.73 | 0.72 | 0.77 |
| 11 | 23584367 | rs7925363 | -0.812 | -1.613 | -0.546 | -1.173 | 0.05 | 0.031 | 0.031 | 0.386 | 0.51 | 0.74 | 0.46 | 0.54 | 0.34 | 0.35 |
| 11 | 23585386 | rs1467238 | -0.429 | -1.459 | -0.017 | -1.483 | -0.067 | -0.134 | -0.134 | 0.095 | 0.19 | 0.49 | 0.16 | 0.01 | 0.05 | 0.05 |
| 11 | 23585898 | rs7110525 | 1.788 | -0.752 | -0.449 | -0.979 | -0.01 | -0.022 | -0.022 | 0 | 0.06 | 0.11 | 0.02 | 0 | 0.01 | 0.01 |
| 11 | 23590764 | rs4606474 | -0.095 | -0.784 | -0.705 | -1.102 | -0.012 | 0.082 | 0.082 | 0 | 0.39 | 0.23 | 0.31 | 0.24 | 0.45 | 0.39 |
| 11 | 23590788 | rs1005743 | -0.089 | -0.777 | -0.702 | -1.099 | -0.013 | 0.081 | 0.081 | 0 | 0.38 | 0.22 | 0.31 | 0.24 | 0.45 | 0.39 |
| 11 | 23591733 | rs7932853 | -0.139 | -0.805 | -0.714 | -1.113 | -0.011 | 0.085 | 0.085 | 0 | 0.39 | 0.22 | 0.31 | 0.24 | 0.45 | 0.39 |
| 11 | 23592020 | rs11027331 | -0.714 | -0.937 | -0.561 | -0.921 | -0.006 | 0.032 | 0.032 | 4.685 | 0.54 | 0.36 | 0.45 | 0.64 | 0.57 | 0.58 |
| 11 | 23596570 | rs10500964 | 1.141 | -0.928 | -0.544 | -0.864 | 0.006 | -0.058 | -0.058 | 0.029 | 0.15 | 0.06 | 0.13 | 0.38 | 0.12 | 0.16 |
| 11 | 23596802 | rs11027351 | -0.634 | -0.947 | -0.563 | -0.94 | -0.007 | 0.03 | 0.03 | 6.055 | 0.54 | 0.36 | 0.45 | 0.63 | 0.55 | 0.58 |
| 11 | 23601081 | rs7107114 | 0.939 | -0.868 | -0.493 | -0.781 | 0 | 0.019 | 0.019 | 10.098 | 0.16 | 0.14 | 0.14 | 0.39 | 0.12 | 0.19 |
| 11 | 23618613 | rs7128154 | 1.603 | -0.801 | -0.577 | -0.906 | 0.002 | -0.007 | -0.007 | 0 | 0.09 | 0.08 | 0.02 | 0 | 0.04 | 0.02 |
| 11 | 23618856 | rs16911592 | 1.241 | -0.827 | -0.719 | -0.848 | -0.017 | -0.113 | -0.113 | 0 | 0.15 | 0.01 | 0.13 | 0.38 | 0.12 | 0.16 |
| 11 | 23622247 | rs11823466 | -1.216 | -0.845 | -0.375 | -1.113 | 0.032 | -0.007 | -0.007 | 0 | 0.68 | 0.58 | 0.52 | 0.67 | 0.64 | 0.58 |
| 11 | 23628652 | rs7939820 | -1.316 | -0.801 | -0.537 | -1.092 | 0.039 | -0.003 | -0.003 | 0 | 0.7 | 0.52 | 0.51 | 0.67 | 0.65 | 0.6 |
| 11 | 23628945 | rs7950168 | 0.11 | -0.643 | -0.768 | -0.986 | -0.015 | -0.022 | -0.022 | 0 | 0.18 | 0.24 | 0.07 | 0.05 | 0.1 | 0.12 |
| 11 | 23639325 | rs11027397 | 0.54 | -0.434 | -1.06 | -0.913 | 0.014 | 0.033 | 0.033 | 0 | 0.16 | 0.08 | 0.06 | 0.05 | 0.12 | 0.08 |
| 11 | 23645618 | rs11500117 | 1.217 | -0.335 | -0.787 | -0.996 | -0.008 | -0.059 | -0.059 | 0 | 0.1 | 0.24 | 0.03 | 0 | 0.04 | 0.02 |
| 11 | 23660635 | rs7941380 | -0.262 | 0.104 | -1.678 | -0.895 | 0.024 | -0.006 | -0.006 | 0 | 0.46 | 0.64 | 0.22 | 0.46 | 0.29 | 0.26 |
| 11 | 23667516 | rs1519414 | 0.019 | 0.619 | -2.016 | -1.33 | -0.033 | 0.03 | 0.03 | 0.932 | 0.25 | 0.06 | 0.3 | 0.23 | 0.34 | 0.4 |
| 11 | 23671985 | rs11027415 | -0.932 | 0.653 | -1.305 | -1.072 | 0.013 | 0.042 | 0.042 | 0.122 | 0.16 | 0.07 | 0.07 | 0.04 | 0.13 | 0.08 |
| 11 | 23673754 | rs10500966 | -0.657 | 1.752 | -1.115 | -1.266 | -0.024 | 0.005 | 0.005 | 0.433 | 0.2 | 0.13 | 0.13 | 0.01 | 0.28 | 0.16 |
| 11 | 23676139 | rs10741988 | 0.041 | 1.715 | -1.224 | -1.424 | -0.012 | 0.048 | 0.048 | 0.663 | 0.82 | 0.87 | 0.88 | 0.99 | 0.81 | 0.94 |
| 11 | 23678817 | rs11027419 | -0.927 | 1.827 | -1.13 | -1.61 | -0.006 | -0.003 | -0.003 | 0 | 0.07 | 0.05 | 0.06 | 0.01 | 0.12 | 0.08 |
| 11 | 23679692 | rs10767101 | 1.462 | 1.248 | -0.623 | -2.211 | -0.019 | 0.052 | 0.052 | 1.893 | 0.62 | 0.25 | 0.75 | 0.96 | 0.62 | 0.72 |
| 11 | 23682502 | rs10500967 | 0.548 | 0.732 | 0.254 | -2.149 | -0.001 | -0.041 | -0.041 | 0 | 0.74 | 0.73 | 0.85 | 0.99 | 0.75 | 0.89 |
| 11 | 23685952 | rs11027423 | 0.5 | 0.304 | -0.689 | -2.011 | -0.107 | -0.095 | -0.095 | 0.016 | 0.28 | 0.04 | 0.52 | 0.68 | 0.37 | 0.43 |
| 11 | 23690735 | rs4923118 | 0.212 | -0.286 | 0.192 | -2.169 | -0.019 | 0.051 | 0.051 | 1.289 | 0.78 | 0.85 | 0.88 | 0.99 | 0.77 | 0.9 |
| 11 | 23694031 | rs12281955 | -1.694 | -0.378 | 0.444 | -2.182 | 0.003 | -0.009 | -0.009 | 0 | 0.33 | 0.7 | 0.19 | 0.05 | 0.27 | 0.21 |
| 11 | 23699123 | rs7127739 | 1.407 | -0.577 | 0.547 | -2.28 | 0.026 | 0.129 | 0.129 | 5.767 | 0.63 | 0.18 | 0.79 | 0.95 | 0.73 | 0.79 |
| 11 | 23706463 | rs10741989 | 0.342 | -0.565 | 0.571 | -2.275 | -0.004 | 0.023 | 0.023 | 2.026 | 0.75 | 0.81 | 0.87 | 1 | 0.77 | 0.9 |
| 11 | 23706483 | rs11027433 | 1.393 | -0.566 | 0.601 | -2.275 | 0.024 | 0.105 | 0.105 | 5.422 | 0.64 | 0.2 | 0.8 | 0.95 | 0.73 | 0.78 |
| 11 | 23715283 | rs10741991 | 1.303 | -0.554 | 0.652 | -2.269 | 0.016 | 0.05 | 0.05 | 4.881 | 0.64 | 0.24 | 0.81 | 0.95 | 0.73 | 0.79 |
| 11 | 23718743 | rs12272427 | -1.547 | -0.269 | 0.832 | -2.07 | -0.02 | -0.024 | -0.024 | 0 | 0.11 | 0.56 | 0.06 | 0.05 | 0.04 | 0.11 |
| 11 | 23723766 | rs4922665 | 0.658 | -0.273 | 1.147 | -2.094 | -0.001 | -0.144 | -0.144 | 0.393 | 0.73 | 0.49 | 0.84 | 0.98 | 0.77 | 0.85 |
| 11 | 23726195 | rs10834179 | 1.068 | -0.323 | 0.96 | -2.107 | 0.009 | 0.06 | 0.06 | 3.834 | 0.67 | 0.19 | 0.8 | 0.98 | 0.76 | 0.81 |
| 11 | 23727350 | rs11027449 | -0.527 | -0.317 | 0.949 | -2.14 | -0.03 | 0.075 | 0.075 | 2.726 | 0.23 | 0.16 | 0.12 | 0 | 0.23 | 0.1 |
| 11 | 23741440 | rs10834184 | 0.991 | -0.275 | 0.527 | -2.115 | 0.008 | 0.031 | 0.031 | 4.571 | 0.68 | 0.2 | 0.81 | 0.99 | 0.76 | 0.87 |
| 11 | 23760361 | rs9666772 | -0.539 | -0.612 | 0.322 | -1.943 | 0.012 | 0.049 | 0.049 | 1.318 | 0.23 | 0.15 | 0.11 | 0 | 0.17 | 0.06 |
| 11 | 23762416 | rs7947895 | -1.137 | -0.665 | 0.685 | -1.97 | -0.037 | -0.169 | -0.169 | 0 | 0.28 | 0.52 | 0.15 | 0 | 0.18 | 0.07 |
| 11 | 23765762 | rs11600566 | -0.446 | -0.379 | -0.334 | -1.82 | 0 | 0.102 | 0.102 | 1.532 | 0.22 | 0.09 | 0.1 | 0 | 0.17 | 0.06 |
| 11 | 23773031 | rs12275084 | -0.48 | -0.38 | -0.206 | -1.821 | 0.009 | 0.036 | 0.036 | 1.515 | 0.22 | 0.13 | 0.11 | 0 | 0.18 | 0.07 |
| 11 | 23776103 | rs11820608 | -1.045 | -0.564 | 0.4 | -2.073 | -0.001 | -0.045 | -0.045 | 0.03 | 0.1 | 0.53 | 0.07 | 0.01 | 0.06 | 0.05 |
| 11 | 23777459 | rs4317983 | -1.301 | -0.616 | 0.338 | -2.151 | 0.012 | -0.017 | -0.017 | 1.165 | 0.33 | 0.69 | 0.17 | 0.01 | 0.22 | 0.16 |
| 11 | 23779099 | rs11827258 | -1.196 | -0.677 | 0.582 | -1.96 | -0.001 | -0.044 | -0.044 | 0 | 0.1 | 0.54 | 0.07 | 0.01 | 0.05 | 0.05 |
| 11 | 23780120 | rs4589296 | 0.288 | -0.648 | -0.568 | -2.401 | -0.046 | 0.147 | 0.147 | 0.032 | 0.45 | 0.2 | 0.66 | 0.78 | 0.5 | 0.43 |
| 11 | 23783338 | rs4923127 | -0.573 | -0.822 | 0.106 | -2.334 | -0.012 | 0.042 | 0.042 | 0 | 0.62 | 0.73 | 0.74 | 0.79 | 0.6 | 0.65 |
| 11 | 23784536 | rs4601775 | -0.718 | -0.884 | 0.076 | -2.385 | -0.014 | 0.031 | 0.031 | 0 | 0.64 | 0.73 | 0.74 | 0.8 | 0.6 | 0.65 |
| 11 | 23801412 | rs11027491 | 0.069 | -0.8 | 0.695 | -2.135 | 0.007 | -0.029 | -0.029 | 0.062 | 0.06 | 0.14 | 0.07 | 0 | 0.06 | 0.02 |
| 11 | 23803362 | rs4418809 | -1.963 | -0.937 | 1.215 | -2.198 | 0.025 | 0.023 | 0.023 | 4.397 | 0.05 | 0.36 | 0.03 | 0 | 0 | 0 |
| 11 | 23804832 | rs7118415 | -1.881 | -1.578 | 0.93 | -1.417 | -0.047 | 0.054 | 0.054 | 0.077 | 0.24 | 0.67 | 0.41 | 0.58 | 0.14 | 0.24 |
| 11 | 23815456 | rs6483989 | 1.452 | -1.471 | 0.818 | -1.35 | 0.047 | 0.044 | 0.044 | 3.936 | 0.95 | 0.61 | 0.97 | 1 | 1 | 0.99 |
| 11 | 23816444 | rs4586161 | 0.537 | -1.667 | -0.153 | -1.653 | -0.077 | 0.129 | 0.129 | 0.29 | 0.29 | 0.06 | 0.2 | 0.11 | 0.35 | 0.38 |
| 11 | 23820633 | rs7111329 | -0.588 | -1.552 | 0.503 | -0.667 | -0.012 | 0.077 | 0.077 | 0 | 0.9 | 0.89 | 0.77 | 0.53 | 0.95 | 0.85 |
| 11 | 23833286 | rs7113510 | -0.048 | -1.56 | 0.651 | -0.661 | -0.03 | 0.109 | 0.109 | 0 | 0.88 | 0.81 | 0.69 | 0.44 | 0.93 | 0.81 |

| | | | | | | | | | | | | | | | | |
|---|---|---|---|---|---|---|---|---|---|---|---|---|---|---|---|---|
| 11 | 23841971 | rs4362144 | -0.153 | -1.51 | 0.814 | -0.643 | -0.037 | 0.218 | 0.218 | 0 | 0.12 | 0.26 | 0.26 | 0.49 | 0.08 | 0.14 |
| 11 | 23844975 | rs11027540 | 0.515 | -1.486 | 0.69 | -0.626 | -0.004 | 0.072 | 0.072 | 0 | 0.09 | 0.09 | 0.23 | 0.46 | 0.08 | 0.14 |
| 11 | 23847957 | rs11027542 | 1.226 | -1.454 | 0.25 | -1.072 | -0.111 | -0.104 | -0.104 | 2.796 | 0.3 | 0.04 | 0.33 | 0.32 | 0.4 | 0.36 |
| 11 | 23857767 | rs11027549 | -1.286 | -1.709 | 1.803 | -0.205 | -0.008 | 0.019 | 0.019 | 0.522 | 0.06 | 0.25 | 0.1 | 0.07 | 0.01 | 0.05 |
| 11 | 23859991 | rs11027552 | -0.751 | -1.722 | 1.789 | -0.213 | -0.009 | 0.093 | 0.093 | 0 | 0.13 | 0.17 | 0.3 | 0.47 | 0.14 | 0.16 |
| 11 | 23861558 | rs11027556 | -0.834 | -1.771 | 1.786 | -0.316 | -0.004 | -0.024 | -0.024 | 0 | 0.09 | 0.1 | 0.07 | 0 | 0.07 | 0.03 |
| 11 | 23861993 | rs2178219 | -2.484 | -1.795 | 1.903 | -0.314 | -0.03 | 0.033 | 0.033 | 0.017 | 0.27 | 0.66 | 0.43 | 0.58 | 0.17 | 0.23 |
| 11 | 23864400 | rs4267072 | -2.053 | -1.788 | 1.938 | -0.312 | -0.047 | 0.077 | 0.077 | 0 | 0.25 | 0.61 | 0.42 | 0.54 | 0.17 | 0.22 |
| 11 | 23867685 | rs10834213 | -2.589 | -1.438 | 2.224 | -1.062 | -0.056 | -0.051 | -0.051 | 0 | 0.15 | 0.57 | 0.18 | 0.1 | 0.06 | 0.09 |
| 11 | 23875164 | rs11027570 | -1.397 | -1.395 | 2.207 | -1.072 | -0.017 | -0.028 | -0.028 | 0 | 0.1 | 0.17 | 0.14 | 0.06 | 0.06 | 0.08 |
| 11 | 23883355 | rs7123551 | -2.234 | -1.42 | 2.413 | -1.084 | -0.037 | -0.062 | -0.062 | 0 | 0.13 | 0.47 | 0.16 | 0.06 | 0.06 | 0.09 |
| 11 | 23888384 | rs13377220 | -1.518 | -1.431 | 2.396 | -1.091 | -0.007 | -0.037 | -0.037 | 0 | 0.09 | 0.38 | 0.08 | 0 | 0.04 | 0.02 |
| 11 | 23890569 | rs10834219 | -2.614 | -1.453 | 2.082 | -1.095 | -0.059 | -0.059 | -0.059 | 0.002 | 0.16 | 0.63 | 0.19 | 0.1 | 0.06 | 0.09 |
| 11 | 23890660 | rs11027579 | -0.793 | -1.386 | 1.768 | -1.083 | -0.014 | -0.037 | -0.037 | 0 | 0.07 | 0.2 | 0.06 | 0 | 0.04 | 0.02 |
| 11 | 23901749 | rs10400331 | -1.782 | -1.417 | 1.135 | -1.079 | -0.077 | -0.072 | -0.072 | 0 | 0.17 | 0.55 | 0.18 | 0.09 | 0.06 | 0.09 |
| 11 | 23915409 | rs4316508 | 0.013 | -0.545 | 0.099 | -1.401 | -0.06 | 0.12 | 0.12 | 1.232 | 0.22 | 0.04 | 0.15 | 0.04 | 0.27 | 0.25 |
| 11 | 23922318 | rs10500970 | -0.163 | -0.664 | 1.317 | -1.436 | 0.005 | -0.018 | -0.018 | 0 | 0.09 | 0.19 | 0.06 | 0.05 | 0.09 | 0.08 |
| 11 | 23922815 | rs12291881 | -1.626 | -0.573 | 1.516 | -1.682 | 0.014 | -0.094 | -0.094 | 0.426 | 0.19 | 0.51 | 0.19 | 0.05 | 0.19 | 0.14 |
| 11 | 23926630 | rs10834232 | -1.296 | -0.727 | 1.671 | -1.73 | -0.006 | 0 | 0 | 2.726 | 0.42 | 0.62 | 0.34 | 0.09 | 0.46 | 0.39 |
| 11 | 23938197 | rs2141396 | 0.293 | -0.891 | 1.931 | -1.458 | 0.001 | -0.088 | -0.088 | 0.561 | 0.71 | 0.56 | 0.77 | 0.87 | 0.69 | 0.6 |
| 11 | 23940407 | rs7941280 | 0.636 | -0.897 | 1.984 | -1.355 | -0.022 | 0.042 | 0.042 | 0.689 | 0.94 | 0.79 | 0.98 | 0.92 | 0.99 | 0.91 |
| 11 | 23945841 | rs3992998 | 0.873 | -0.889 | 1.699 | -1.382 | -0.016 | -0.013 | -0.013 | 0 | 0.84 | 0.71 | 0.9 | 0.88 | 0.9 | 0.8 |
| 11 | 23948464 | rs4627077 | 0.147 | -1.041 | 0.958 | -1.527 | -0.043 | 0.122 | 0.122 | 2.076 | 0.25 | 0.06 | 0.2 | 0.06 | 0.3 | 0.32 |
| 11 | 23948850 | rs10742015 | 0.612 | -0.963 | 1.225 | -1.429 | -0.001 | -0.043 | -0.043 | 0 | 0.61 | 0.41 | 0.68 | 0.82 | 0.6 | 0.52 |
| 11 | 23950320 | rs1919221 | 1.02 | -0.836 | 0.673 | -1.347 | 0.02 | 0.135 | 0.135 | 2.627 | 0.56 | 0.24 | 0.67 | 0.82 | 0.59 | 0.49 |
| 11 | 23955079 | rs7130113 | 1.366 | -0.798 | 1.208 | -0.673 | -0.033 | 0.01 | 0.01 | 0.898 | 0.84 | 0.34 | 0.87 | 0.82 | 0.92 | 0.84 |
| 11 | 23973802 | rs4923145 | -1.357 | -0.83 | 1.904 | -0.642 | -0.01 | 0.052 | 0.052 | 0.054 | 0.12 | 0.37 | 0.09 | 0.18 | 0.08 | 0.17 |
| 11 | 23991218 | rs6484002 | -1.971 | -0.822 | 1.537 | 0.278 | -0.03 | 0.074 | 0.074 | 1.14 | 0.23 | 0.52 | 0.19 | 0.64 | 0.2 | 0.31 |
| 11 | 23994642 | rs6484003 | -1.701 | -0.827 | 1.548 | 0.296 | -0.021 | 0.065 | 0.065 | 0.705 | 0.15 | 0.5 | 0.11 | 0.19 | 0.08 | 0.18 |
| 11 | 23995576 | rs2176788 | -1.4 | -0.803 | 1.33 | 0.308 | -0.016 | 0.057 | 0.057 | 0.396 | 0.14 | 0.42 | 0.11 | 0.19 | 0.08 | 0.18 |
| 11 | 23997478 | rs16911971 | 0.606 | -0.556 | 0.821 | 0.417 | -0.003 | 0.027 | 0.027 | 0 | 0.08 | 0.03 | 0.08 | 0.46 | 0.12 | 0.13 |
| 11 | 23999115 | rs1915056 | -0.024 | -0.765 | 0.438 | -0.247 | -0.007 | 0.111 | 0.111 | 0.002 | 0.29 | 0.11 | 0.2 | 0.05 | 0.3 | 0.26 |
| 11 | 24015283 | rs7931562 | -1.105 | -0.582 | 0.917 | 0.693 | -0.002 | -0.01 | -0.01 | 1.093 | 0.14 | 0.16 | 0.13 | 0.56 | 0.2 | 0.2 |
| 11 | 24016876 | rs1949541 | -1.708 | -0.633 | 1.153 | 0.55 | -0.018 | 0.069 | 0.069 | 0.151 | 0.22 | 0.46 | 0.19 | 0.62 | 0.22 | 0.33 |
| 11 | 24033297 | rs7110905 | -1.877 | -0.411 | 1.223 | 0.412 | 0.048 | 0.059 | 0.059 | 0 | 0.09 | 0.39 | 0.09 | 0.11 | 0.08 | 0.12 |
| 11 | 24038287 | rs10500973 | -0.479 | -0.431 | 0.483 | 0.51 | -0.001 | 0.009 | 0.009 | 0.3 | 0.3 | 0.65 | 0.25 | 0.5 | 0.33 | 0.3 |
| 11 | 24048346 | rs2403884 | -0.294 | 0.835 | -0.024 | -0.097 | 0 | 0 | 0 | 2.026 | 0.38 | 0.62 | 0.51 | 0.45 | 0.38 | 0.5 |
| 11 | 24060691 | rs7949813 | 0.502 | 0.824 | 0.093 | -0.123 | -0.004 | -0.018 | -0.018 | 0 | 0.28 | 0.2 | 0.46 | 0.4 | 0.33 | 0.41 |
| 11 | 24061085 | rs4412756 | 0.226 | 0.849 | -0.172 | -0.107 | -0.009 | -0.032 | -0.032 | 0 | 0.29 | 0.15 | 0.45 | 0.41 | 0.33 | 0.43 |
| 11 | 24073794 | rs1996652 | -0.343 | 0.835 | -0.172 | -0.141 | 0.004 | 0 | 0 | 0 | 0.4 | 0.54 | 0.54 | 0.45 | 0.45 | 0.53 |
| 11 | 24074207 | rs11027689 | 0.419 | 0.838 | -0.331 | -0.142 | -0.025 | -0.031 | -0.031 | 0 | 0.32 | 0.13 | 0.49 | 0.41 | 0.38 | 0.45 |
| 11 | 24081147 | rs12361319 | -0.338 | 0.593 | -0.472 | -0.18 | -0.002 | 0.001 | 0.001 | 0 | 0.43 | 0.58 | 0.55 | 0.45 | 0.46 | 0.56 |
| 11 | 24084200 | rs2946782 | 0.581 | 0.222 | -1.004 | -0.449 | -0.056 | 0.144 | 0.144 | 0 | 0.25 | 0.03 | 0.18 | 0.01 | 0.25 | 0.17 |
| 11 | 24085740 | rs2139518 | -0.741 | 0.457 | -0.501 | 0.268 | 0.001 | 0.012 | 0.012 | 0 | 0.48 | 0.65 | 0.57 | 0.46 | 0.49 | 0.57 |
| 11 | 24087672 | rs12275191 | -0.589 | -0.104 | -1.664 | 0.968 | 0.018 | -0.077 | -0.077 | 0 | 0.35 | 0.16 | 0.3 | 0.42 | 0.37 | 0.42 |
| 11 | 24087714 | rs12294573 | -0.34 | -0.783 | -1.596 | 1.048 | 0.035 | -0.051 | -0.051 | 0 | 0.31 | 0.15 | 0.28 | 0.37 | 0.32 | 0.39 |
| 11 | 24087801 | rs12275238 | -0.448 | -0.978 | -1.763 | 0.766 | 0.059 | -0.075 | -0.075 | 0.369 | 0.34 | 0.1 | 0.27 | 0.38 | 0.31 | 0.36 |
| 11 | 24088689 | rs2957787 | 0.452 | -0.662 | -1.136 | 0.496 | 0.001 | -0.015 | -0.015 | 0.16 | 0.64 | 0.69 | 0.45 | 0.53 | 0.65 | 0.48 |
| 11 | 24088858 | rs2403885 | -0.327 | -0.689 | -1.111 | 0.601 | -0.014 | -0.045 | -0.045 | 0 | 0.22 | 0.11 | 0.48 | 0.42 | 0.25 | 0.26 |
| 11 | 24089953 | rs10767156 | 0.911 | -1.01 | -0.823 | 0.462 | 0 | -0.001 | -0.001 | 0 | 0.6 | 0.41 | 0.7 | 0.4 | 0.65 | 0.62 |
| 11 | 24091682 | rs10834273 | 0.143 | -0.534 | -1.166 | 0.738 | -0.055 | -0.13 | -0.13 | 3.519 | 0.18 | 0.01 | 0.16 | 0.4 | 0.19 | 0.19 |
| 11 | 24099773 | rs1915063 | 0.838 | -0.537 | -0.442 | 0.381 | -0.001 | 0.009 | 0.009 | 0 | 0.55 | 0.34 | 0.36 | 0.36 | 0.55 | 0.47 |
| 11 | 24100977 | rs1879228 | 0.2 | -0.314 | -0.635 | 0.314 | -0.001 | -0.001 | -0.001 | 0.006 | 0.67 | 0.65 | 0.75 | 0.32 | 0.69 | 0.69 |
| 11 | 24108934 | rs2139515 | -0.279 | -0.227 | -0.705 | 1.016 | -0.001 | 0.026 | 0.026 | 0 | 0.93 | 0.9 | 0.94 | 0.75 | 0.91 | 0.87 |
| 11 | 24110315 | rs12791495 | -0.053 | -0.305 | -1.005 | 0.406 | -0.032 | 0.048 | 0.048 | 0 | 0.46 | 0.23 | 0.35 | 0.29 | 0.55 | 0.57 |
| 11 | 24116195 | rs16912086 | 0.511 | 0.12 | -1.595 | 0.599 | -0.093 | -0.019 | -0.019 | 0.266 | 0.33 | 0.08 | 0.25 | 0.27 | 0.41 | 0.38 |
| 11 | 24130487 | rs1915061 | -0.011 | -0.253 | -0.888 | 0.475 | 0.003 | 0 | 0 | 0 | 0.4 | 0.39 | 0.27 | 0.45 | 0.32 | 0.22 |
| 11 | 24131110 | rs2176789 | -0.522 | -0.339 | -0.777 | 0.475 | 0.003 | 0.01 | 0.01 | 0 | 0.07 | 0.04 | 0.07 | 0.02 | 0.1 | 0.14 |
| 11 | 24132249 | rs16912094 | -0.959 | -0.61 | -0.798 | 0.439 | 0.014 | 0.01 | 0.01 | 0 | 0.12 | 0.06 | 0.06 | 0.05 | 0.1 | 0.19 |
| 11 | 24132555 | rs7121343 | 1.101 | -1.388 | -1.116 | 0.586 | 0.1 | 0.049 | 0.049 | 0 | 0.53 | 0.19 | 0.6 | 0.32 | 0.65 | 0.4 |
| 11 | 24144286 | rs2957762 | -0.163 | -1.566 | -0.122 | 0.833 | 0.009 | -0.036 | -0.036 | 0.052 | 0.21 | 0.12 | 0.19 | 0.4 | 0.2 | 0.13 |
| 11 | 24145350 | rs7930560 | -0.622 | -0.877 | 0.646 | 0.796 | 0.057 | 0.061 | 0.061 | 0 | 0.14 | 0.37 | 0.4 | 0.23 | 0.15 | 0.25 |
| 11 | 24156271 | rs2403895 | -0.175 | -1.654 | 0.648 | 0.673 | 0.008 | -0.031 | -0.031 | 1.66 | 0.21 | 0.12 | 0.19 | 0.4 | 0.2 | 0.13 |
| 11 | 24157765 | rs2403896 | 0.854 | -1.718 | 0.825 | 0.67 | -0.012 | 0.006 | 0.006 | 0 | 0.64 | 0.65 | 0.73 | 0.55 | 0.7 | 0.62 |
| 11 | 24162541 | rs2403900 | 0.95 | -1.848 | 0.741 | 0.58 | 0.004 | -0.003 | -0.003 | 0 | 0.63 | 0.7 | 0.73 | 0.54 | 0.69 | 0.59 |
| 11 | 24164611 | rs2947738 | -0.207 | -1.901 | 1.176 | 0.684 | -0.031 | 0.052 | 0.052 | 0 | 0.17 | 0.25 | 0.13 | 0.42 | 0.12 | 0.1 |
| 11 | 24168134 | rs7938053 | 1.346 | -2.042 | 1.502 | 0.623 | 0.019 | -0.008 | -0.008 | 0 | 0.63 | 0.42 | 0.71 | 0.53 | 0.7 | 0.64 |
| 11 | 24172032 | rs2947745 | 0.637 | -1.792 | 1.293 | 0.532 | -0.012 | -0.061 | -0.061 | 0 | 0.8 | 0.67 | 0.83 | 0.95 | 0.82 | 0.74 |
| 11 | 24175046 | rs2957776 | 0.728 | -1.791 | 1.349 | 0.553 | -0.021 | -0.067 | -0.067 | 0 | 0.79 | 0.57 | 0.82 | 0.94 | 0.82 | 0.74 |

| | | | | | | | | | | | | | | | | |
|---|---|---|---|---|---|---|---|---|---|---|---|---|---|---|---|---|
| 11 | 24175352 | rs7107397 | -0.177 | -1.8 | 1.106 | 1.019 | -0.001 | 0.039 | 0.039 | 0 | 0.09 | 0.13 | 0.13 | 0.32 | 0.07 | 0.19 |
| 11 | 24177400 | rs1320356 | -0.967 | -1.815 | 1.394 | 0.712 | -0.005 | -0.034 | -0.034 | 1.528 | 0.19 | 0.24 | 0.16 | 0.03 | 0.17 | 0.13 |
| 11 | 24177461 | rs1320357 | -0.344 | -2.163 | 1.464 | 0.764 | -0.021 | 0.03 | 0.03 | 0 | 0.19 | 0.27 | 0.13 | 0.43 | 0.13 | 0.15 |
| 11 | 24178134 | rs2403903 | 0.77 | -1.829 | 1.627 | 0.491 | -0.022 | -0.103 | -0.103 | 0 | 0.79 | 0.56 | 0.82 | 0.96 | 0.83 | 0.7 |
| 11 | 24192136 | rs4614453 | -0.278 | -1.82 | 0.436 | 0.334 | -0.005 | 0.017 | 0.017 | 0 | 0.88 | 0.93 | 0.92 | 0.94 | 0.89 | 0.88 |
| 11 | 24192528 | rs4923161 | -0.179 | -2.646 | 0.566 | 0.335 | -0.057 | 0.067 | 0.067 | 0 | 0.21 | 0.41 | 0.15 | 0.39 | 0.1 | 0.24 |
| 11 | 24192686 | rs4923162 | 0.133 | -2.782 | 0.268 | 0.476 | 0.008 | -0.057 | -0.057 | 0 | 0.79 | 0.83 | 0.82 | 0.53 | 0.83 | 0.69 |
| 11 | 24193340 | rs12575283 | 1.639 | -2.634 | 0.114 | -0.312 | -0.001 | 0.077 | 0.077 | 0 | 0.4 | 0.22 | 0.53 | 0.11 | 0.58 | 0.32 |
| 11 | 24196119 | rs11027743 | -0.631 | -2.051 | 0.207 | 0.218 | 0.005 | 0.004 | 0.004 | 0 | 0.18 | 0.12 | 0.14 | 0.03 | 0.14 | 0.12 |
| 11 | 24208560 | rs7129631 | -0.706 | -2.146 | 0.538 | 0.55 | -0.03 | -0.008 | -0.008 | 0 | 0.16 | 0.21 | 0.15 | 0.09 | 0.1 | 0.21 |
| 11 | 24210175 | rs7937322 | -0.003 | -1.798 | 0.556 | 1.435 | -0.001 | -0.051 | -0.051 | 0.155 | 0.2 | 0.14 | 0.22 | 0.43 | 0.15 | 0.23 |
| 11 | 24211321 | rs11827778 | -1.662 | -1.99 | 1.238 | 1.204 | -0.047 | 0.039 | 0.039 | 0.583 | 0.25 | 0.54 | 0.22 | 0.3 | 0.16 | 0.31 |
| 11 | 24231257 | rs11819948 | -0.654 | -2.011 | 1.358 | 1.208 | -0.012 | -0.002 | -0.002 | 0.012 | 0.18 | 0.19 | 0.15 | 0.13 | 0.1 | 0.22 |
| 11 | 24236096 | rs7106204 | -1.009 | -2.014 | 1.488 | 1.21 | -0.059 | -0.011 | -0.011 | 0 | 0.2 | 0.39 | 0.17 | 0.14 | 0.12 | 0.24 |
| 11 | 24246904 | rs7951530 | -2.157 | -2.504 | 1.536 | 1.021 | 0.044 | -0.02 | -0.02 | 0 | 0.3 | 0.72 | 0.23 | 0.34 | 0.16 | 0.3 |
| 11 | 24248274 | rs7119181 | -0.954 | -2.27 | 1.589 | 1.04 | 0.019 | 0.149 | 0.149 | 0 | 0.07 | 0.23 | 0.06 | 0.21 | 0.06 | 0.09 |
| 11 | 24248805 | rs7941806 | -0.747 | -2.332 | 1.411 | 1.013 | -0.005 | 0.043 | 0.043 | 0 | 0.05 | 0.22 | 0.09 | 0.05 | 0.01 | 0.03 |
| 11 | 24257425 | rs11027786 | -1.305 | -2.412 | 1.562 | 1.05 | -0.03 | 0.157 | 0.157 | 1.277 | 0.1 | 0.48 | 0.07 | 0.19 | 0.02 | 0.06 |
| 11 | 24259879 | rs7107321 | -1.805 | -2.571 | 1.499 | 0.991 | -0.039 | 0.006 | 0.006 | 1.015 | 0.26 | 0.57 | 0.15 | 0.28 | 0.13 | 0.27 |
| 11 | 24267222 | rs11821633 | -2.268 | -2.634 | 1.356 | 0.981 | -0.055 | 0.033 | 0.033 | 1.571 | 0.18 | 0.64 | 0.16 | 0.27 | 0.06 | 0.12 |
| 11 | 24269719 | rs1020277 | 2.251 | -2.577 | 0.357 | 0.405 | -0.127 | 0.094 | 0.094 | 0.063 | 0.33 | 0.06 | 0.37 | 0.15 | 0.48 | 0.32 |
| 11 | 24280281 | rs7104694 | -1.415 | -2.638 | 0.441 | 0.489 | -0.031 | 0.006 | 0.006 | 0.287 | 0.28 | 0.62 | 0.23 | 0.33 | 0.15 | 0.3 |
| 11 | 24290777 | rs12365463 | 2.304 | -2.545 | 0.016 | 0.417 | -0.2 | 0.058 | 0.058 | 0.145 | 0.33 | 0.02 | 0.36 | 0.15 | 0.49 | 0.32 |
| 11 | 24305935 | rs11827033 | 0.334 | -2.446 | 0.422 | 0.571 | -0.011 | -0.009 | -0.009 | 0.077 | 0.15 | 0.14 | 0.09 | 0.09 | 0.11 | 0.2 |
| 11 | 24308924 | rs16925305 | 1.264 | -2.447 | 0.133 | 0.568 | 0.017 | 0.008 | 0.008 | 0 | 0.11 | 0.05 | 0.07 | 0.07 | 0.09 | 0.14 |
| 11 | 24316872 | rs10834309 | 0.522 | -2.501 | -0.008 | 0.711 | -0.003 | -0.054 | -0.054 | 0.619 | 0.23 | 0.14 | 0.23 | 0.41 | 0.18 | 0.26 |
| 11 | 24321827 | rs12801367 | -0.841 | -2.358 | 0.311 | 0.648 | -0.064 | 0.065 | 0.065 | 0.083 | 0.23 | 0.41 | 0.21 | 0.33 | 0.14 | 0.29 |
| 11 | 24333824 | rs2012346 | 1.016 | -1.518 | 0.051 | 0.474 | -0.001 | -0.001 | -0.001 | 0 | 0.45 | 0.56 | 0.36 | 0.39 | 0.48 | 0.4 |
| 11 | 24335156 | rs10834311 | 0.458 | -1.463 | -0.244 | 0.658 | -0.037 | -0.069 | -0.069 | 0 | 0.22 | 0.09 | 0.38 | 0.41 | 0.24 | 0.28 |
| 11 | 24345162 | rs7104550 | -0.45 | -2.18 | 0.445 | 0.36 | -0.08 | 0.021 | 0.021 | 0.072 | 0.23 | 0.45 | 0.14 | 0.26 | 0.11 | 0.22 |
| 11 | 24346093 | rs7946392 | 0.075 | -2.132 | 0.114 | 0.301 | -0.021 | -0.039 | -0.039 | 1.786 | 0.63 | 0.79 | 0.61 | 0.46 | 0.63 | 0.6 |
| 11 | 24356464 | rs7950920 | -0.064 | -2.107 | 0.436 | 0.326 | -0.039 | 0.025 | 0.025 | 0.05 | 0.2 | 0.36 | 0.13 | 0.27 | 0.12 | 0.23 |
| 11 | 24359362 | rs1822339 | 1.955 | -2.308 | 0.102 | 0.084 | -0.186 | 0.084 | 0.084 | 0.372 | 0.34 | 0.02 | 0.44 | 0.15 | 0.52 | 0.36 |
| 11 | 24359477 | rs1822338 | 0.758 | -2.218 | 0.212 | 0.156 | -0.002 | -0.011 | -0.011 | 0 | 0.52 | 0.46 | 0.6 | 0.22 | 0.62 | 0.45 |
| 11 | 24362286 | rs11027822 | 1.367 | -2.068 | -0.174 | 0.253 | -0.04 | 0.065 | 0.065 | 0.226 | 0.4 | 0.23 | 0.53 | 0.2 | 0.53 | 0.39 |
| 11 | 24364459 | rs12796137 | -0.174 | -2.268 | -0.042 | 0.201 | -0.072 | -0.004 | -0.004 | 0 | 0.27 | 0.47 | 0.14 | 0.27 | 0.14 | 0.24 |
| 11 | 24365908 | rs7119079 | 0.652 | -1.823 | -0.327 | 0.384 | -0.015 | -0.032 | -0.032 | 0.089 | 0.65 | 0.7 | 0.6 | 0.47 | 0.65 | 0.61 |
| 11 | 24372661 | rs12362261 | 0.775 | -1.546 | -0.092 | 0.376 | 0.003 | 0.013 | 0.013 | 0 | 0.67 | 0.61 | 0.61 | 0.47 | 0.68 | 0.63 |
| 11 | 24373079 | rs1372956 | 1.792 | -1.871 | -0.818 | -0.08 | -0.162 | 0.116 | 0.116 | 1.252 | 0.38 | 0.03 | 0.4 | 0.15 | 0.55 | 0.37 |
| 11 | 24377105 | rs11027828 | 0.828 | -2.154 | 0.16 | -0.016 | 0.001 | -0.02 | -0.02 | 0 | 0.63 | 0.47 | 0.57 | 0.27 | 0.68 | 0.58 |
| 11 | 24378843 | rs10834318 | -0.635 | -2.388 | 0.208 | -0.111 | -0.049 | -0.044 | -0.044 | 0 | 0.36 | 0.48 | 0.21 | 0.16 | 0.19 | 0.26 |
| 11 | 24383954 | rs1596279 | 0.391 | -2.447 | 0.243 | 0.145 | -0.001 | -0.002 | -0.002 | 0 | 0.64 | 0.64 | 0.61 | 0.51 | 0.69 | 0.59 |
| 11 | 24399097 | rs12577327 | 1.139 | -2.487 | 0.507 | 0.087 | 0.035 | 0.043 | 0.043 | 0 | 0.2 | 0.08 | 0.08 | 0.07 | 0.13 | 0.14 |
| 11 | 24399805 | rs7940888 | 0.664 | -2.491 | 0.689 | 0.122 | -0.013 | -0.003 | -0.003 | 0 | 0.23 | 0.24 | 0.18 | 0.15 | 0.14 | 0.17 |
| 11 | 24403620 | rs11027857 | 0.953 | -2.418 | 0.842 | 0.163 | 0.003 | -0.007 | -0.007 | 0 | 0.44 | 0.53 | 0.45 | 0.35 | 0.56 | 0.43 |
| 11 | 24405921 | rs11027859 | 2.15 | -2.36 | 0.292 | 0.222 | -0.2 | 0.071 | 0.071 | 0.227 | 0.32 | 0.02 | 0.37 | 0.15 | 0.53 | 0.34 |
| 11 | 24406140 | rs11027860 | 1.493 | -2.322 | 0.412 | 0.09 | -0.075 | -0.006 | -0.006 | 0.023 | 0.37 | 0.17 | 0.42 | 0.35 | 0.55 | 0.42 |
| 11 | 24407108 | rs7122189 | 0.127 | -1.681 | 1.064 | 0.789 | -0.017 | -0.035 | -0.035 | 0.17 | 0.15 | 0.08 | 0.21 | 0.38 | 0.14 | 0.27 |
| 11 | 24412621 | rs4923170 | -1.62 | -2.127 | 1.461 | 0.364 | 0.011 | 0.004 | 0.004 | 0.228 | 0.47 | 0.61 | 0.44 | 0.58 | 0.37 | 0.52 |
| 11 | 24419481 | rs9299979 | 0.417 | -2.136 | 1.916 | 0.326 | -0.023 | -0.018 | -0.018 | 0 | 0.8 | 0.56 | 0.78 | 0.88 | 0.87 | 0.78 |
| 11 | 24429419 | rs4633474 | 0.035 | -2.147 | 2.043 | 0.303 | -0.012 | -0.014 | -0.014 | 0 | 0.1 | 0.24 | 0.1 | 0.05 | 0.06 | 0.05 |
| 11 | 24433663 | rs7105586 | 1.315 | -1.95 | 2.072 | -0.017 | -0.001 | -0.039 | -0.039 | 0 | 0.07 | 0.14 | 0.03 | 0 | 0.05 | 0.03 |
| 11 | 24435073 | rs7113296 | 0.45 | -1.999 | 2.007 | 0.025 | -0.001 | 0 | 0 | 8.298 | 0.2 | 0.18 | 0.16 | 0.44 | 0.18 | 0.26 |
| 11 | 24435202 | rs4307720 | 0.155 | -1.983 | 2.058 | 0.019 | -0.003 | 0.027 | 0.027 | 6.42 | 0.72 | 0.57 | 0.79 | 0.57 | 0.76 | 0.71 |
| 11 | 24440965 | rs7924982 | 1.511 | -1.944 | 1.761 | -0.065 | -0.057 | -0.063 | -0.063 | 0 | 0.43 | 0.12 | 0.5 | 0.44 | 0.57 | 0.41 |
| 11 | 24446873 | rs16912551 | -1.167 | -1.828 | 1.958 | -0.056 | -0.015 | -0.025 | -0.025 | 1.203 | 0.17 | 0.36 | 0.19 | 0.08 | 0.11 | 0.13 |
| 11 | 24452761 | rs12795773 | -0.848 | -1.768 | 1.939 | 0.057 | -0.002 | 0.004 | 0.004 | 4.987 | 0.43 | 0.39 | 0.37 | 0.54 | 0.37 | 0.55 |
| 11 | 24457272 | rs11027908 | 0.299 | -1.757 | 1.918 | 0.06 | -0.004 | -0.024 | -0.024 | 0.265 | 0.3 | 0.21 | 0.25 | 0.5 | 0.27 | 0.44 |
| 11 | 24457755 | rs7102470 | -0.278 | -1.704 | 2.002 | 0.021 | -0.009 | -0.028 | -0.028 | 0 | 0.13 | 0.21 | 0.12 | 0.05 | 0.1 | 0.12 |
| 11 | 24469451 | rs6484051 | -0.324 | -1.746 | 1.663 | 0.004 | 0.003 | -0.001 | -0.001 | 2.087 | 0.41 | 0.35 | 0.32 | 0.5 | 0.34 | 0.51 |
| 11 | 24475606 | rs10767190 | 1.931 | -1.656 | 1.318 | -0.073 | -0.114 | -0.115 | -0.115 | 0 | 0.36 | 0.05 | 0.43 | 0.42 | 0.51 | 0.38 |
| 11 | 24477706 | rs12276835 | -1.411 | -1.625 | 1.666 | 0.02 | 0.022 | 0.009 | 0.009 | 2.071 | 0.5 | 0.6 | 0.44 | 0.55 | 0.38 | 0.56 |
| 11 | 24480040 | rs10834345 | 0.372 | -1.655 | 1.636 | -0.001 | -0.006 | -0.031 | -0.031 | 0 | 0.31 | 0.19 | 0.27 | 0.49 | 0.28 | 0.43 |
| 11 | 24483140 | rs10834346 | 1.514 | -1.427 | 1.232 | 0.119 | -0.091 | -0.078 | -0.078 | 0 | 0.4 | 0.11 | 0.46 | 0.42 | 0.53 | 0.39 |
| 11 | 24488141 | rs10742039 | 0.122 | -0.998 | 1.593 | -0.088 | -0.006 | -0.035 | -0.035 | 0 | 0.89 | 0.71 | 0.89 | 0.98 | 0.91 | 0.95 |
| 11 | 24493876 | rs10500979 | 0.663 | -0.977 | 1.328 | -0.075 | -0.001 | -0.002 | -0.002 | 0 | 0.05 | 0.04 | 0.09 | 0.06 | 0.04 | 0.05 |
| 11 | 24499370 | rs7938928 | 1.164 | -0.989 | 1.325 | -0.076 | -0.027 | -0.053 | -0.053 | 0 | 0.81 | 0.5 | 0.79 | 0.92 | 0.87 | 0.9 |
| 11 | 24504903 | rs10219185 | 1.001 | -0.843 | 1.221 | -0.037 | -0.021 | -0.048 | -0.048 | 0 | 0.82 | 0.53 | 0.79 | 0.93 | 0.87 | 0.9 |
| 11 | 24511240 | rs4923190 | 1.273 | -0.94 | 1.153 | -0.118 | -0.031 | -0.076 | -0.076 | 0 | 0.75 | 0.54 | 0.77 | 0.91 | 0.82 | 0.87 |

| | | | | | | | | | | | | | | | | |
|---|---|---|---|---|---|---|---|---|---|---|---|---|---|---|---|---|
| 11 | 24513821 | rs12223724 | 0.994 | -1.164 | 0.396 | -0.073 | 0.006 | 0.075 | 0.075 | 1.788 | 0.49 | 0.09 | 0.52 | 0.25 | 0.6 | 0.43 |
| 11 | 24515523 | rs7925550 | -0.829 | -1.404 | 1.11 | -0.227 | -0.036 | -0.063 | -0.063 | 0 | 0.15 | 0.38 | 0.15 | 0.03 | 0.09 | 0.07 |
| 11 | 24517243 | rs11027951 | 0.819 | -1.383 | 1.348 | 0.183 | -0.003 | -0.006 | -0.006 | 0 | 0.25 | 0.23 | 0.26 | 0.5 | 0.26 | 0.42 |
| 11 | 24518441 | rs1509616 | -0.642 | -1.381 | 1.146 | 0.18 | 0.016 | 0.061 | 0.061 | 0.19 | 0.72 | 0.52 | 0.72 | 0.5 | 0.74 | 0.57 |
| 11 | 24520669 | rs9667545 | -1.376 | -1.219 | 1.16 | 0.303 | -0.054 | 0.007 | 0.007 | 0.231 | 0.24 | 0.46 | 0.2 | 0.25 | 0.15 | 0.16 |
| 11 | 24521666 | rs7943454 | -1.731 | -0.999 | 0.634 | 0.239 | -0.068 | 0.031 | 0.031 | 0.483 | 0.63 | 0.88 | 0.58 | 0.78 | 0.51 | 0.67 |
| 11 | 24527958 | rs7927293 | -0.075 | -0.793 | 0.892 | 0.345 | 0.005 | 0.014 | 0.014 | 1.066 | 0.35 | 0.42 | 0.29 | 0.47 | 0.31 | 0.46 |
| 11 | 24533104 | rs1995837 | -0.66 | -1.508 | 1.161 | 0.211 | 0.012 | 0.048 | 0.048 | 0 | 0.1 | 0.19 | 0.15 | 0.21 | 0.08 | 0.12 |
| 11 | 24537624 | rs11027967 | -0.845 | -1.707 | 1.227 | -0.082 | -0.07 | -0.046 | -0.046 | 10.426 | 0.16 | 0.38 | 0.08 | 0.07 | 0.07 | 0.06 |
| 11 | 24538279 | rs11027970 | -1.471 | -1.729 | 1.316 | -0.138 | -0.014 | -0.045 | -0.045 | 0 | 0.16 | 0.4 | 0.15 | 0.09 | 0.09 | 0.09 |
| 11 | 24539314 | rs1318297 | 0.258 | -1.655 | 0.836 | -0.086 | -0.028 | -0.008 | -0.008 | 0 | 0.1 | 0.28 | 0.06 | 0.06 | 0.05 | 0.04 |
| 11 | 24542153 | rs883002 | -0.444 | -1.501 | 0.619 | -0.03 | -0.029 | -0.034 | -0.034 | 0 | 0.06 | 0.18 | 0.02 | 0 | 0.01 | 0.02 |
| 11 | 24543242 | rs11824978 | 0.554 | -1.125 | -0.311 | -0.08 | -0.086 | -0.095 | -0.095 | 0 | 0.36 | 0.13 | 0.45 | 0.52 | 0.45 | 0.6 |
| 11 | 24545353 | rs7102196 | -1.275 | -1.158 | 0.58 | -0.273 | -0.045 | -0.055 | -0.055 | 0.143 | 0.25 | 0.51 | 0.19 | 0.16 | 0.15 | 0.14 |
| 11 | 24549045 | rs10500982 | -1.185 | -1.064 | 0.731 | -0.242 | -0.038 | -0.058 | -0.058 | 1.299 | 0.12 | 0.36 | 0.17 | 0.02 | 0.08 | 0.08 |
| 11 | 24551110 | rs4244514 | -1.961 | -1.175 | 0.943 | -0.227 | -0.053 | -0.053 | -0.053 | 0.118 | 0.18 | 0.54 | 0.28 | 0.07 | 0.13 | 0.24 |
| 11 | 24552429 | rs11027980 | -1.652 | -1.216 | 0.979 | -0.283 | -0.063 | -0.088 | -0.088 | 0 | 0.15 | 0.53 | 0.19 | 0.02 | 0.08 | 0.08 |
| 11 | 24553333 | rs1396850 | -0.205 | -1.333 | 0.101 | -0.776 | 0.003 | -0.006 | -0.006 | 4.207 | 0.45 | 0.56 | 0.39 | 0.15 | 0.37 | 0.24 |
| 11 | 24553440 | rs10767205 | -0.591 | -0.88 | 0.63 | -0.536 | -0.01 | -0.03 | -0.03 | 0.269 | 0.09 | 0.27 | 0.16 | 0.01 | 0.08 | 0.08 |
| 11 | 24554271 | rs10834369 | -0.048 | -0.605 | 0.859 | -0.211 | 0.017 | 0.025 | 0.025 | 0 | 0.11 | 0.18 | 0.21 | 0.14 | 0.12 | 0.19 |
| 11 | 24557125 | rs10834370 | -0.13 | -0.87 | 0.738 | -0.758 | 0.003 | -0.005 | -0.005 | 0 | 0.47 | 0.54 | 0.41 | 0.16 | 0.39 | 0.25 |
| 11 | 24558167 | rs1588346 | -0.275 | -0.933 | 0.706 | -0.868 | -0.001 | -0.047 | -0.047 | 0 | 0.13 | 0.24 | 0.19 | 0.02 | 0.14 | 0.1 |
| 11 | 24564217 | rs4491224 | -0.232 | -1.065 | 0.723 | -0.518 | 0.006 | 0.009 | 0.009 | 0 | 0.54 | 0.41 | 0.59 | 0.77 | 0.61 | 0.74 |
| 11 | 24570856 | rs2403959 | -0.146 | -1.154 | 0.688 | -0.393 | 0.006 | 0.009 | 0.009 | 0 | 0.46 | 0.57 | 0.42 | 0.21 | 0.39 | 0.3 |
| 11 | 24573538 | rs12221725 | 0.264 | -1.149 | 0.671 | -0.362 | 0.018 | -0.003 | -0.003 | 0 | 0.13 | 0.18 | 0.21 | 0.09 | 0.12 | 0.2 |
| 11 | 24575390 | rs16912792 | 0.058 | -1.223 | 0.655 | -0.371 | 0.029 | 0.103 | 0.103 | 0 | 0.44 | 0.29 | 0.38 | 0.16 | 0.39 | 0.24 |
| 11 | 24577183 | rs1509605 | 0.869 | -1.275 | 0.458 | -0.381 | -0.11 | -0.098 | -0.098 | 0.015 | 0.39 | 0.12 | 0.37 | 0.59 | 0.48 | 0.47 |
| 11 | 24577558 | rs10834385 | -0.21 | -1.263 | 0.52 | -0.406 | 0.006 | 0.008 | 0.008 | 0 | 0.53 | 0.43 | 0.58 | 0.72 | 0.6 | 0.67 |
| 11 | 24579598 | rs12797112 | 1.399 | -1.281 | 0.328 | -0.425 | -0.023 | 0.07 | 0.07 | 0 | 0.3 | 0.03 | 0.2 | 0.13 | 0.26 | 0.16 |
| 11 | 24586870 | rs12289378 | 1.407 | -1.301 | 0.115 | -0.6 | -0.084 | -0.093 | -0.093 | 0 | 0.29 | 0.11 | 0.34 | 0.62 | 0.38 | 0.41 |
| 11 | 24593626 | rs1509589 | -0.344 | -1.306 | 0.258 | -0.541 | -0.002 | -0.007 | -0.007 | 0.012 | 0.54 | 0.44 | 0.59 | 0.79 | 0.63 | 0.71 |
| 11 | 24599638 | rs7115380 | -0.281 | -1.16 | 0.337 | -0.27 | -0.002 | -0.003 | -0.003 | 0.575 | 0.55 | 0.42 | 0.61 | 0.73 | 0.63 | 0.72 |
| 11 | 24606921 | rs10742044 | -0.692 | -1.049 | 0.445 | -0.232 | -0.01 | 0.011 | 0.011 | 0 | 0.56 | 0.63 | 0.59 | 0.32 | 0.49 | 0.48 |
| 11 | 24607717 | rs9633851 | -0.633 | -1.066 | 0.432 | -0.247 | -0.012 | 0.016 | 0.016 | 0 | 0.55 | 0.64 | 0.58 | 0.35 | 0.49 | 0.42 |
| 11 | 24610019 | rs1858156 | 0.526 | -1.054 | 0.206 | -0.29 | -0.038 | -0.024 | -0.024 | 0.331 | 0.42 | 0.22 | 0.4 | 0.6 | 0.51 | 0.52 |
| 11 | 24610850 | rs11028040 | 1.078 | -0.84 | -0.019 | -0.251 | -0.002 | 0.02 | 0.02 | 0 | 0.29 | 0.04 | 0.17 | 0.19 | 0.23 | 0.16 |
| 11 | 24613989 | rs10834401 | -0.538 | -0.324 | 0.914 | -0.241 | 0.021 | 0.009 | 0.009 | 0 | 0.13 | 0.17 | 0.21 | 0.11 | 0.12 | 0.2 |
| 11 | 24618295 | rs16912906 | -0.92 | -0.491 | 0.579 | -0.387 | 0 | -0.016 | -0.016 | 0.219 | 0.2 | 0.24 | 0.2 | 0.03 | 0.21 | 0.13 |
| 11 | 24630127 | rs3992982 | -1.162 | -0.427 | -0.071 | -0.761 | -0.001 | -0.021 | -0.021 | 2.328 | 0.28 | 0.3 | 0.35 | 0.05 | 0.26 | 0.19 |
| 11 | 24635152 | rs982036 | -0.746 | -0.343 | -0.354 | -0.954 | -0.001 | -0.002 | -0.002 | 0.405 | 0.43 | 0.43 | 0.25 | 0.28 | 0.33 | 0.35 |
| 11 | 24635763 | rs7110784 | 1.095 | -0.382 | -0.268 | -0.903 | -0.045 | -0.022 | -0.022 | 0 | 0.46 | 0.15 | 0.54 | 0.56 | 0.57 | 0.43 |
| 11 | 24636501 | rs10834405 | 0.866 | -0.625 | -0.42 | -1.038 | -0.058 | -0.069 | -0.069 | 2.698 | 0.21 | 0.01 | 0.13 | 0.26 | 0.17 | 0.2 |
| 11 | 24639566 | rs2136481 | 0.903 | -0.594 | -0.425 | -1.04 | -0.059 | -0.072 | -0.072 | 0 | 0.2 | 0.01 | 0.13 | 0.26 | 0.17 | 0.2 |
| 11 | 24640213 | rs1543012 | 1.341 | -0.665 | -0.346 | -1.059 | -0.049 | -0.019 | -0.019 | 2.904 | 0.46 | 0.15 | 0.55 | 0.57 | 0.57 | 0.43 |
| 11 | 24641934 | rs12418943 | 0.881 | -0.613 | -0.358 | -1.031 | -0.062 | -0.077 | -0.077 | 1.092 | 0.2 | 0.01 | 0.13 | 0.26 | 0.18 | 0.2 |
| 11 | 24642587 | rs906882 | -0.715 | -0.656 | -0.292 | -1.047 | -0.001 | -0.003 | -0.003 | 0 | 0.43 | 0.43 | 0.24 | 0.28 | 0.33 | 0.35 |
| 11 | 24648325 | rs4923200 | -0.032 | -0.631 | 0.237 | -1.057 | -0.005 | -0.007 | -0.007 | 0 | 0.27 | 0.17 | 0.27 | 0.33 | 0.26 | 0.31 |
| 11 | 24649296 | rs12285329 | -0.478 | -0.693 | 0.163 | -1.143 | -0.035 | -0.108 | -0.108 | 0 | 0.27 | 0.42 | 0.13 | 0.02 | 0.18 | 0.16 |
| 11 | 24649362 | rs7124697 | -1.413 | -0.699 | 0.484 | -1.147 | 0.009 | 0.024 | 0.024 | 0 | 0.49 | 0.59 | 0.39 | 0.35 | 0.41 | 0.44 |
| 11 | 24654654 | rs16912971 | -0.245 | -0.72 | 0.303 | -1.098 | -0.002 | -0.032 | -0.032 | 0 | 0.25 | 0.29 | 0.12 | 0.01 | 0.18 | 0.16 |
| 11 | 24657338 | rs4923202 | 0.935 | -0.698 | 0.298 | -0.955 | -0.018 | -0.066 | -0.066 | 0 | 0.17 | 0.06 | 0.12 | 0.32 | 0.15 | 0.25 |
| 11 | 24664261 | rs1995841 | -0.929 | -0.714 | 0.651 | -0.902 | 0.011 | 0.008 | 0.008 | 0 | 0.44 | 0.4 | 0.25 | 0.35 | 0.34 | 0.43 |
| 11 | 24665985 | rs3903036 | -0.709 | -0.792 | 0.625 | -0.92 | 0.001 | -0.006 | -0.006 | 1.484 | 0.29 | 0.32 | 0.16 | 0.09 | 0.19 | 0.27 |
| 11 | 24666607 | rs870383 | 0.785 | -0.268 | 0.32 | -1.283 | -0.017 | 0.061 | 0.061 | 0 | 0.19 | 0.1 | 0.32 | 0.04 | 0.22 | 0.12 |
| 11 | 24668637 | rs12576803 | 0.803 | -0.138 | 0.663 | -1.17 | 0 | 0.014 | 0.014 | 0 | 0.2 | 0.24 | 0.13 | 0.35 | 0.17 | 0.26 |
| 11 | 24669006 | rs7119980 | -0.447 | -0.031 | 0.418 | -1.671 | -0.004 | -0.116 | -0.116 | 0.289 | 0.66 | 0.63 | 0.55 | 0.96 | 0.67 | 0.83 |
| 11 | 24678322 | rs12573950 | 0.699 | 0.12 | 0.272 | -1.312 | 0.001 | -0.005 | -0.005 | 0 | 0.21 | 0.19 | 0.16 | 0.34 | 0.18 | 0.25 |
| 11 | 24681532 | rs3923615 | -0.62 | 0.038 | 0.203 | -1.552 | 0.011 | 0.006 | 0.006 | 0 | 0.46 | 0.64 | 0.52 | 0.43 | 0.41 | 0.45 |
| 11 | 24681659 | rs4561214 | 0.792 | 0.006 | 0.225 | -1.536 | 0 | 0.002 | 0.002 | 0 | 0.22 | 0.2 | 0.16 | 0.36 | 0.18 | 0.26 |
| 11 | 24683117 | rs4465366 | -1.093 | 0.032 | 0.163 | -1.634 | -0.009 | -0.01 | -0.01 | 0.079 | 0.59 | 0.69 | 0.63 | 0.46 | 0.53 | 0.5 |
| 11 | 24683854 | rs4531473 | 1.198 | -0.273 | -0.064 | -1.774 | -0.001 | -0.001 | -0.001 | 0 | 0.33 | 0.35 | 0.36 | 0.54 | 0.41 | 0.45 |
| 11 | 24686598 | rs4592420 | -0.159 | -0.303 | 0.063 | -1.769 | -0.019 | 0.01 | 0.01 | 0 | 0.67 | 0.55 | 0.79 | 0.65 | 0.75 | 0.67 |
| 11 | 24690002 | rs10834419 | 1.035 | -0.293 | -0.102 | -1.806 | -0.008 | 0.042 | 0.042 | 0 | 0.21 | 0.15 | 0.33 | 0.09 | 0.23 | 0.17 |
| 11 | 24690099 | rs10834420 | 0.777 | -0.298 | -0.095 | -1.816 | -0.013 | 0.01 | 0.01 | 0 | 0.45 | 0.35 | 0.45 | 0.58 | 0.52 | 0.5 |
| 11 | 24690524 | rs10767233 | 0.344 | -0.1 | -0.586 | -1.491 | -0.016 | 0.137 | 0.137 | 0.264 | 0.34 | 0.15 | 0.42 | 0.09 | 0.34 | 0.22 |
| 11 | 24691773 | rs7944845 | 0.262 | 0.055 | -0.229 | -0.858 | 0 | 0.001 | 0.001 | 0 | 0.22 | 0.21 | 0.16 | 0.35 | 0.18 | 0.26 |
| 11 | 24699102 | rs12577048 | 0.772 | 0.1 | -0.317 | -0.888 | 0 | -0.005 | -0.005 | 0.587 | 0.18 | 0.13 | 0.12 | 0.26 | 0.16 | 0.18 |
| 11 | 24700545 | rs7118149 | 0.628 | -0.102 | -0.145 | -1.476 | -0.019 | 0.007 | 0.007 | 0 | 0.44 | 0.31 | 0.49 | 0.56 | 0.52 | 0.5 |
| 11 | 24700655 | rs10834426 | 0.733 | 0.045 | -0.302 | -1.283 | 0 | -0.004 | -0.004 | 0 | 0.21 | 0.17 | 0.15 | 0.35 | 0.18 | 0.25 |

| | | | | | | | | | | | | | | | | |
|---|---|---|---|---|---|---|---|---|---|---|---|---|---|---|---|---|
| 11 | 24715305 | rs4500487 | 0.158 | -0.021 | -0.117 | -1.298 | -0.002 | 0.007 | 0.007 | 0 | 0.25 | 0.25 | 0.17 | 0.35 | 0.2 | 0.3 |
| 11 | 24718860 | rs7949406 | -0.465 | -0.038 | -0.432 | -1.59 | -0.017 | 0.034 | 0.034 | 2.422 | 0.54 | 0.29 | 0.53 | 0.38 | 0.5 | 0.44 |
| 11 | 24719920 | rs7395433 | -0.846 | -0.035 | -0.265 | -1.688 | 0.003 | -0.007 | -0.007 | 0 | 0.92 | 0.81 | 0.96 | 0.95 | 0.93 | 0.94 |
| 11 | 24721154 | rs7484277 | 0.255 | 0.01 | -0.74 | -1.706 | -0.001 | 0.138 | 0.138 | 1.14 | 0.34 | 0.15 | 0.37 | 0.03 | 0.32 | 0.17 |
| 11 | 24722672 | rs12798775 | 0.338 | 0.12 | -0.625 | -0.967 | -0.001 | -0.026 | -0.026 | 0 | 0.25 | 0.16 | 0.2 | 0.38 | 0.24 | 0.29 |
| 11 | 24722990 | rs7109593 | 0.313 | 0.087 | -0.619 | -1.006 | 0 | -0.018 | -0.018 | 1.157 | 0.25 | 0.16 | 0.2 | 0.36 | 0.23 | 0.29 |
| 11 | 24724857 | rs4604884 | 0.347 | 0.131 | -0.702 | -1.174 | 0 | 0.001 | 0.001 | 0 | 0.38 | 0.38 | 0.5 | 0.57 | 0.44 | 0.46 |
| 11 | 24724888 | rs4359207 | 0.358 | 0.245 | -0.52 | -1.212 | 0.022 | 0.061 | 0.061 | 1.417 | 0.3 | 0.45 | 0.44 | 0.57 | 0.37 | 0.42 |
| 11 | 24726825 | rs12099324 | 0.283 | 0.06 | -0.568 | -1.435 | -0.005 | 0.006 | 0.006 | 0 | 0.13 | 0.1 | 0.06 | 0 | 0.14 | 0.14 |
| 11 | 24726922 | rs12790301 | -0.16 | -0.152 | -0.077 | -1.062 | -0.003 | -0.002 | -0.002 | 0 | 0.25 | 0.24 | 0.15 | 0.21 | 0.2 | 0.24 |
| 11 | 24733266 | rs2716456 | -0.278 | 0.381 | -0.093 | -0.964 | 0.001 | -0.006 | -0.006 | 0 | 0.34 | 0.32 | 0.5 | 0.76 | 0.45 | 0.45 |
| 11 | 24733553 | rs2631441 | -1.011 | -0.027 | 0.534 | -0.287 | -0.001 | 0.103 | 0.103 | 2.487 | 0.1 | 0.41 | 0.19 | 0.14 | 0.04 | 0.16 |
| 11 | 24734804 | rs2029192 | -1.184 | -0.212 | 0.395 | -0.378 | -0.05 | 0.039 | 0.039 | 2.674 | 0.16 | 0.52 | 0.32 | 0.16 | 0.07 | 0.27 |
| 11 | 24736493 | rs10834444 | 0.752 | 0.026 | -0.281 | -0.831 | -0.027 | 0.041 | 0.041 | 0.423 | 0.32 | 0.2 | 0.26 | 0.1 | 0.38 | 0.23 |
| 11 | 24739804 | rs10834449 | 0.078 | -0.341 | -0.426 | -0.425 | 0.029 | -0.002 | -0.002 | 0 | 0.56 | 0.32 | 0.51 | 0.53 | 0.66 | 0.45 |
| 11 | 24740434 | rs2716458 | -0.495 | -0.101 | -0.806 | -0.188 | -0.037 | 0.048 | 0.048 | 2.163 | 0.15 | 0.46 | 0.31 | 0.16 | 0.07 | 0.26 |
| 11 | 24741384 | rs2631438 | 0.275 | 0.196 | -1.292 | -0.043 | -0.009 | 0.056 | 0.056 | 0 | 0.11 | 0.27 | 0.29 | 0.16 | 0.06 | 0.27 |
| 11 | 24744592 | rs1994373 | 0.326 | 0.478 | -2.118 | 0.266 | -0.066 | -0.175 | -0.175 | 0 | 0.24 | 0.03 | 0.24 | 0.42 | 0.3 | 0.2 |
| 11 | 24749376 | rs2631433 | -0.629 | 0.294 | -1.677 | 0.012 | -0.038 | -0.055 | -0.055 | 0 | 0.74 | 0.97 | 0.85 | 0.7 | 0.74 | 0.72 |
| 11 | 24750211 | rs10834460 | -0.18 | 0.481 | -1.06 | -0.44 | 0.014 | 0.044 | 0.044 | 0 | 0.49 | 0.62 | 0.52 | 0.24 | 0.59 | 0.36 |
| 11 | 24750454 | rs2716460 | 0.076 | 0.499 | -0.817 | 0.214 | -0.006 | 0.01 | 0.01 | 0.094 | 0.63 | 0.44 | 0.61 | 0.54 | 0.67 | 0.5 |
| 11 | 24750980 | rs2716461 | -0.389 | 0.825 | -1.062 | -0.179 | 0.007 | 0.005 | 0.005 | 0 | 0.55 | 0.62 | 0.65 | 0.45 | 0.58 | 0.37 |
| 11 | 24751744 | rs4923205 | 0.218 | 0.777 | -1.62 | -0.17 | 0.009 | -0.062 | -0.062 | 3.024 | 0.4 | 0.12 | 0.31 | 0.53 | 0.36 | 0.61 |
| 11 | 24751859 | rs4923206 | 0.126 | 1.583 | -1.468 | -0.115 | 0.025 | -0.024 | -0.024 | 0.017 | 0.06 | 0.1 | 0.06 | 0.01 | 0.13 | 0.04 |
| 11 | 24751895 | rs10500989 | 0 | 1.265 | -2.005 | -0.097 | -0.107 | -0.166 | -0.166 | 1.781 | 0.23 | 0.04 | 0.39 | 0.52 | 0.3 | 0.19 |
| 11 | 24753430 | rs16913118 | 0.04 | 0.482 | -1.414 | -0.785 | -0.015 | -0.124 | -0.124 | 0.165 | 0.25 | 0.39 | 0.2 | 0 | 0.18 | 0.14 |
| 11 | 24755890 | rs2716449 | -0.327 | 0.971 | -2.425 | -0.022 | -0.007 | -0.038 | -0.038 | 2.028 | 0.81 | 0.85 | 0.9 | 0.61 | 0.78 | 0.67 |
| 11 | 24756924 | rs2631431 | 0.007 | 0.881 | -2.685 | -1.298 | 0.012 | 0.13 | 0.13 | 0.078 | 0.43 | 0.76 | 0.37 | 0.03 | 0.41 | 0.29 |
| 11 | 24757325 | rs4923207 | 0.375 | 1.4 | -2.661 | -0.547 | -0.086 | 0.033 | 0.033 | 0 | 0.15 | 0.01 | 0.26 | 0.12 | 0.22 | 0.14 |
| 11 | 24758432 | rs17235001 | -0.138 | 0.644 | -2.384 | -0.42 | -0.036 | 0.08 | 0.08 | 0 | 0.13 | 0.01 | 0.09 | 0 | 0.14 | 0.13 |
| 11 | 24758856 | rs1906099 | 0.1 | 0.263 | -1.636 | 0.555 | -0.031 | 0.146 | 0.146 | 0.166 | 0.79 | 0.53 | 0.73 | 0.56 | 0.87 | 0.76 |
| 11 | 24759043 | rs1564994 | -0.127 | 0.496 | -2.224 | 0.465 | 0.023 | -0.049 | -0.049 | 0 | 0.29 | 0.2 | 0.35 | 0.55 | 0.24 | 0.31 |
| 11 | 24759664 | rs11028130 | -0.621 | 1.227 | -1.812 | 0.762 | -0.047 | -0.075 | -0.075 | 4.491 | 0.18 | 0.07 | 0.17 | 0.28 | 0.23 | 0.22 |
| 11 | 24770644 | rs12791552 | -1.051 | 1.109 | -1.389 | 0.287 | -0.034 | 0.038 | 0.038 | 0 | 0.08 | 0.01 | 0.06 | 0.01 | 0.14 | 0.05 |
| 11 | 24774037 | rs10834475 | -1.406 | 0.311 | -0.599 | 0.807 | -0.004 | 0.086 | 0.086 | 0.01 | 0.22 | 0.33 | 0.22 | 0.45 | 0.22 | 0.35 |
| 11 | 24774506 | rs10834476 | -1.267 | 0.237 | -0.36 | 0.81 | 0.002 | 0.061 | 0.061 | 0.068 | 0.17 | 0.23 | 0.18 | 0.42 | 0.2 | 0.24 |
| 11 | 24774864 | rs2716441 | 0.974 | -0.399 | -0.388 | -0.148 | -0.003 | -0.007 | -0.007 | 0.219 | 0.42 | 0.38 | 0.46 | 0.44 | 0.56 | 0.47 |
| 11 | 24777925 | rs12803470 | -0.519 | -0.425 | -0.274 | -0.185 | 0.015 | -0.034 | -0.034 | 0 | 0.07 | 0.11 | 0.06 | 0.01 | 0.11 | 0.02 |
| 11 | 24778111 | rs2716548 | -0.556 | -0.632 | -0.127 | -0.321 | 0.028 | 0.069 | 0.069 | 0.42 | 0.65 | 0.82 | 0.68 | 0.89 | 0.76 | 0.81 |
| 11 | 24779345 | rs2631413 | -0.593 | -0.668 | -0.125 | -0.339 | 0.03 | 0.069 | 0.069 | 0.097 | 0.64 | 0.78 | 0.68 | 0.89 | 0.76 | 0.81 |
| 11 | 24780891 | rs10450584 | -2.268 | -0.737 | 0.131 | -0.246 | -0.014 | 0.091 | 0.091 | 0.438 | 0.24 | 0.39 | 0.23 | 0.43 | 0.22 | 0.33 |
| 11 | 24782186 | rs12284066 | -2.061 | -0.732 | 0.186 | -0.426 | 0.007 | -0.093 | -0.093 | 0 | 0.16 | 0.38 | 0.12 | 0.02 | 0.15 | 0.13 |
| 11 | 24786653 | rs12281860 | -0.889 | -0.936 | 0.076 | -0.382 | -0.003 | -0.027 | -0.027 | 0 | 0.08 | 0.07 | 0.06 | 0.01 | 0.06 | 0.02 |
| 11 | 24793160 | rs2931360 | 1.06 | -1.079 | 0.146 | -0.417 | 0.002 | -0.001 | -0.001 | 0 | 0.45 | 0.43 | 0.48 | 0.46 | 0.6 | 0.58 |
| 11 | 24794324 | rs1007719 | 0.396 | -1.069 | 0.4 | -0.227 | 0.055 | -0.016 | -0.016 | 0 | 0.55 | 0.74 | 0.57 | 0.48 | 0.68 | 0.62 |
| 11 | 24795275 | rs2716539 | 0.943 | -1.066 | 0.404 | -0.212 | 0.001 | 0 | 0 | 0 | 0.51 | 0.48 | 0.53 | 0.48 | 0.65 | 0.6 |
| 11 | 24800404 | rs1486699 | -0.335 | -1.052 | 0.49 | 0.027 | -0.002 | 0.06 | 0.06 | 0 | 0.1 | 0.11 | 0.12 | 0.39 | 0.09 | 0.17 |
| 11 | 24801648 | rs1486698 | 0.118 | -0.992 | 0.462 | -0.023 | -0.009 | -0.034 | -0.034 | 0 | 0.62 | 0.59 | 0.67 | 0.87 | 0.76 | 0.79 |
| 11 | 24803895 | rs10834484 | -0.491 | -0.967 | 0.463 | 0.085 | 0 | 0.037 | 0.037 | 0 | 0.11 | 0.11 | 0.13 | 0.39 | 0.1 | 0.17 |
| 11 | 24808750 | rs2716471 | 0.095 | -0.805 | 0.606 | 0.108 | -0.012 | -0.029 | -0.029 | 0 | 0.65 | 0.63 | 0.7 | 0.87 | 0.79 | 0.8 |
| 11 | 24816638 | rs1486681 | -0.079 | -0.736 | 0.65 | 0.195 | 0.018 | 0.028 | 0.028 | 1.946 | 0.69 | 0.71 | 0.77 | 0.88 | 0.78 | 0.85 |
| 11 | 24819089 | rs12799525 | -0.043 | -0.686 | 0.381 | 0.19 | 0.025 | 0.141 | 0.141 | 1.98 | 0.28 | 0.08 | 0.2 | 0.06 | 0.2 | 0.13 |
| 11 | 24819568 | rs729092 | -0.773 | -0.64 | 0.459 | 0.449 | -0.03 | 0.031 | 0.031 | 0.545 | 0.37 | 0.51 | 0.33 | 0.5 | 0.28 | 0.28 |
| 11 | 24822669 | rs11028183 | -1.118 | 0.208 | 0.336 | 1.266 | 0.013 | 0.285 | 0.285 | 0.189 | 0.05 | 0.17 | 0.09 | 0.43 | 0.06 | 0.1 |
| 11 | 24831746 | rs2716473 | 0.643 | 0.539 | 0.117 | 1.025 | 0.001 | -0.029 | -0.029 | 0.629 | 0.66 | 0.28 | 0.76 | 0.5 | 0.69 | 0.71 |
| 11 | 24832009 | rs2716474 | 0.667 | 1.056 | 0.438 | 1.11 | 0.018 | 0.026 | 0.026 | 0 | 0.67 | 0.47 | 0.74 | 0.65 | 0.65 | 0.7 |
| 11 | 24832707 | rs2716475 | 0.39 | 1.126 | 0.429 | 1.076 | 0.018 | 0.025 | 0.025 | 0 | 0.7 | 0.54 | 0.76 | 0.66 | 0.67 | 0.71 |
| 11 | 24835665 | rs12801073 | 0.072 | 1.291 | 0.476 | 1.108 | 0 | 0 | 0 | 0 | 0.17 | 0.16 | 0.12 | 0.21 | 0.22 | 0.15 |
| 11 | 24837488 | rs11028189 | -0.501 | 1.165 | -0.04 | -0.137 | -0.022 | 0.039 | 0.039 | 0.008 | 0.18 | 0.1 | 0.16 | 0 | 0.2 | 0.07 |
| 11 | 24837801 | rs10834489 | -0.686 | 1.313 | -0.218 | 0.133 | -0.079 | -0.021 | -0.021 | 2.901 | 0.33 | 0.13 | 0.37 | 0.63 | 0.42 | 0.39 |
| 11 | 24838571 | rs2716479 | 0.948 | 0.843 | 0.17 | 0.288 | 0 | 0 | 0 | 0 | 0.53 | 0.45 | 0.48 | 0.35 | 0.44 | 0.52 |
| 11 | 24839058 | rs11028190 | 0.027 | 0.655 | -0.508 | 0.496 | -0.137 | -0.151 | -0.151 | 0 | 0.24 | 0.02 | 0.3 | 0.5 | 0.33 | 0.3 |
| 11 | 24840073 | rs1486688 | 0 | -0.244 | 0.135 | 0.429 | 0.007 | 0.007 | 0.007 | 0.404 | 0.53 | 0.53 | 0.47 | 0.35 | 0.4 | 0.55 |
| 11 | 24840482 | rs2631511 | 0.219 | -0.31 | -0.113 | 0.783 | 0.015 | -0.001 | -0.001 | 0.583 | 0.74 | 0.8 | 0.74 | 0.71 | 0.8 | 0.66 |
| 11 | 24848824 | rs2631506 | 1.167 | 1.03 | 0.808 | 0.852 | 0.107 | 0.112 | 0.112 | 0.017 | 0.86 | 0.55 | 0.76 | 0.77 | 0.83 | 0.74 |
| 11 | 24850630 | rs17307187 | -0.883 | 1.093 | 1.054 | 0.89 | 0.012 | 0.007 | 0.007 | 0 | 0.08 | 0.11 | 0.17 | 0.13 | 0.12 | 0.23 |
| 11 | 24858636 | rs1486708 | 1.796 | 1.07 | 0.787 | 0.416 | -0.037 | 0.067 | 0.067 | 3.18 | 0.62 | 0.18 | 0.6 | 0.77 | 0.65 | 0.6 |
| 11 | 24859125 | rs11028209 | -0.712 | 1.196 | 0.858 | 0.5 | 0.008 | 0.009 | 0.009 | 0 | 0.1 | 0.12 | 0.2 | 0.23 | 0.17 | 0.26 |
| 11 | 24861214 | rs2716508 | -0.715 | 1.177 | 1.71 | 0.408 | 0.014 | -0.023 | -0.023 | 0 | 0.14 | 0.23 | 0.15 | 0.06 | 0.13 | 0.12 |

| | | | | | | | | | | | | | | | | |
|---|---|---|---|---|---|---|---|---|---|---|---|---|---|---|---|---|
| 11 | 24862072 | rs2716509 | 0.926 | 1.14 | 2.078 | 0.387 | 0.011 | 0.004 | 0.004 | 0 | 0.95 | 0.76 | 0.97 | 1 | 0.99 | 0.98 |
| 11 | 24863089 | rs10834496 | 2.065 | 0.867 | 1.858 | 0.206 | -0.002 | 0 | 0 | 0.42 | 0.67 | 0.36 | 0.61 | 0.71 | 0.68 | 0.57 |
| 11 | 24868957 | rs2631468 | -0.83 | 1.063 | 1.576 | 0.36 | 0.012 | -0.023 | -0.023 | 0 | 0.14 | 0.22 | 0.15 | 0.06 | 0.13 | 0.15 |
| 11 | 24869249 | rs2716522 | -0.738 | 1.08 | 1.617 | 0.363 | 0.015 | -0.026 | -0.026 | 0 | 0.14 | 0.23 | 0.16 | 0.06 | 0.13 | 0.12 |
| 11 | 24876622 | rs2631445 | -0.77 | 1.112 | 1.596 | 0.358 | 0.016 | -0.026 | -0.026 | 0 | 0.14 | 0.22 | 0.15 | 0.06 | 0.13 | 0.12 |
| 11 | 24879336 | rs11028231 | -1.002 | 1.107 | 1.65 | 0.374 | 0.009 | -0.029 | -0.029 | 0 | 0.15 | 0.23 | 0.16 | 0.06 | 0.13 | 0.12 |
| 11 | 24879829 | rs1872720 | 2.19 | 1.079 | 1.835 | 0.411 | -0.016 | 0.001 | 0.001 | 3.855 | 0.64 | 0.25 | 0.6 | 0.64 | 0.63 | 0.6 |
| 11 | 24884719 | rs1960472 | 0.561 | 1.083 | 1.855 | 0.412 | 0.021 | 0.015 | 0.015 | 2.22 | 0.85 | 0.7 | 0.82 | 0.84 | 0.86 | 0.88 |
| 11 | 24885339 | rs2403986 | 2.072 | 1.065 | 1.538 | 0.342 | -0.016 | 0.001 | 0.001 | 3.855 | 0.64 | 0.25 | 0.6 | 0.64 | 0.63 | 0.59 |
| 11 | 24891999 | rs16913433 | -0.966 | 0.639 | 1.234 | 0.049 | -0.006 | -0.025 | -0.025 | 0 | 0.1 | 0.25 | 0.03 | 0.04 | 0.07 | 0.03 |
| 11 | 24892439 | rs10834519 | -0.915 | 0.742 | 0.678 | 0.228 | 0.007 | -0.012 | -0.012 | 0.956 | 0.13 | 0.19 | 0.16 | 0.07 | 0.14 | 0.12 |
| 11 | 24902164 | rs1486683 | -0.321 | 0.209 | 0.845 | -0.078 | 0.008 | 0.017 | 0.017 | 0 | 0.46 | 0.35 | 0.49 | 0.68 | 0.54 | 0.53 |
| 11 | 24905957 | rs1386231 | -0.301 | 0.191 | 0.837 | -0.086 | 0.007 | 0.016 | 0.016 | 0 | 0.46 | 0.36 | 0.49 | 0.68 | 0.54 | 0.53 |
| 11 | 24906515 | rs749150 | 0.355 | -0.091 | 1.059 | -0.282 | 0.034 | 0.067 | 0.067 | 0 | 0.48 | 0.34 | 0.45 | 0.24 | 0.44 | 0.41 |
| 11 | 24908541 | rs10834527 | -0.05 | -0.079 | 1.057 | -0.286 | 0.007 | 0.017 | 0.017 | 0 | 0.46 | 0.35 | 0.49 | 0.68 | 0.54 | 0.53 |
| 11 | 24911153 | rs5000667 | -0.459 | -0.177 | 1.276 | -0.263 | 0.017 | 0.026 | 0.026 | 0 | 0.49 | 0.38 | 0.51 | 0.69 | 0.55 | 0.56 |
| 11 | 24911312 | rs1386234 | 0.486 | -0.285 | 1.505 | -0.34 | 0.032 | 0.082 | 0.082 | 0 | 0.46 | 0.32 | 0.44 | 0.23 | 0.44 | 0.42 |
| 11 | 24934662 | rs7478820 | -0.891 | -0.843 | 1.228 | -0.976 | 0.003 | 0.057 | 0.057 | 0.14 | 0.57 | 0.66 | 0.56 | 0.82 | 0.54 | 0.56 |
| 11 | 24955222 | rs7481329 | -0.281 | -0.544 | 1.24 | -0.949 | 0.005 | -0.03 | -0.03 | 1.203 | 0.42 | 0.47 | 0.5 | 0.75 | 0.48 | 0.54 |
| 11 | 24960510 | rs7481112 | -0.426 | -0.642 | 1.293 | -1.035 | 0.001 | -0.033 | -0.033 | 0 | 0.43 | 0.52 | 0.49 | 0.82 | 0.48 | 0.54 |
| 11 | 24969228 | rs7481402 | -0.198 | -0.729 | 1.292 | -0.904 | 0.015 | -0.038 | -0.038 | 0 | 0.39 | 0.46 | 0.45 | 0.73 | 0.47 | 0.48 |
| 11 | 24972495 | rs7483842 | 0.26 | -0.62 | 1.205 | -0.924 | -0.001 | -0.026 | -0.026 | 0.365 | 0.56 | 0.49 | 0.5 | 0.19 | 0.51 | 0.45 |
| 11 | 24986010 | rs6484088 | 0.262 | -0.667 | 1.072 | -0.984 | -0.134 | 0.025 | 0.025 | 0 | 0.32 | 0.05 | 0.42 | 0.68 | 0.48 | 0.46 |
| 11 | 25029641 | rs1446190 | -0.315 | -0.634 | 0.715 | -1.24 | -0.084 | 0.211 | 0.211 | 0 | 0.38 | 0.07 | 0.51 | 0.87 | 0.51 | 0.56 |
| 11 | 25039983 | rs12274531 | -0.301 | -0.654 | 0.902 | -1.253 | -0.044 | 0.123 | 0.123 | 0 | 0.64 | 0.81 | 0.49 | 0.14 | 0.5 | 0.54 |
| 11 | 25041972 | rs1562264 | -0.264 | -0.671 | 0.894 | -1.108 | -0.054 | 0.098 | 0.098 | 0 | 0.65 | 0.82 | 0.49 | 0.24 | 0.51 | 0.54 |
| 11 | 25042503 | rs7112294 | -0.32 | -0.67 | 0.894 | -1.108 | -0.051 | 0.1 | 0.1 | 0 | 0.65 | 0.81 | 0.49 | 0.24 | 0.51 | 0.54 |
| 11 | 25047871 | rs4503492 | -0.266 | -0.672 | 0.914 | -1.109 | -0.042 | 0.092 | 0.092 | 0 | 0.64 | 0.79 | 0.49 | 0.24 | 0.51 | 0.54 |
| 11 | 25054314 | rs11823079 | -0.275 | -0.675 | 0.927 | -1.11 | -0.039 | 0.087 | 0.087 | 0 | 0.64 | 0.78 | 0.49 | 0.24 | 0.51 | 0.54 |
| 11 | 25064297 | rs1596854 | 0.425 | -0.646 | 1.079 | -1.115 | 0.001 | -0.02 | -0.02 | 0 | 0.55 | 0.43 | 0.44 | 0.09 | 0.48 | 0.44 |
| 11 | 25066819 | rs4923233 | 0.5 | -0.678 | 1.07 | -1.138 | 0.002 | -0.004 | -0.004 | 0 | 0.54 | 0.46 | 0.44 | 0.19 | 0.49 | 0.45 |
| 11 | 25077951 | rs12279520 | 0.495 | -0.593 | 1.232 | -1.225 | 0 | -0.042 | -0.042 | 0.036 | 0.55 | 0.46 | 0.44 | 0.08 | 0.5 | 0.44 |
| 11 | 25078894 | rs1867754 | 0.057 | -0.632 | 0.812 | -1.238 | -0.151 | 0.123 | 0.123 | 0 | 0.33 | 0.05 | 0.48 | 0.77 | 0.47 | 0.48 |
| 11 | 25088584 | rs1446160 | 1.111 | -0.635 | 1.046 | -1.228 | -0.03 | -0.005 | -0.005 | 0 | 0.9 | 0.67 | 0.93 | 0.9 | 0.97 | 0.92 |
| 11 | 25088712 | rs1838773 | 1.188 | -0.799 | 0.933 | -1.373 | 0.004 | 0.141 | 0.141 | 0.441 | 0.47 | 0.26 | 0.41 | 0.08 | 0.48 | 0.43 |
| 11 | 25096309 | rs7108394 | 0.508 | -0.399 | 0.238 | -1.164 | -0.004 | 0.04 | 0.04 | 0 | 0.57 | 0.62 | 0.46 | 0.13 | 0.5 | 0.45 |
| 11 | 25098691 | rs2166413 | -0.181 | 0.469 | -0.058 | -1.135 | -0.082 | 0.116 | 0.116 | 0 | 0.32 | 0.09 | 0.43 | 0.78 | 0.4 | 0.44 |
| 11 | 25099453 | rs11028406 | -0.342 | 0.496 | 0.009 | -1.101 | -0.095 | 0.115 | 0.115 | 0 | 0.34 | 0.12 | 0.47 | 0.78 | 0.46 | 0.48 |
| 11 | 25100071 | rs12420723 | 0.544 | 0.842 | -0.078 | -0.168 | -0.034 | -0.037 | -0.037 | 0.109 | 0.1 | 0.02 | 0.22 | 0.36 | 0.17 | 0.12 |
| 11 | 25100385 | rs11028409 | 0.188 | 0.379 | 0.282 | -1.01 | 0.001 | -0.021 | -0.021 | 0 | 0.05 | 0.17 | 0.03 | 0.01 | 0.02 | 0.07 |
| 11 | 25102331 | rs17050 | -0.458 | 0.396 | 0.147 | -1.238 | -0.035 | 0.057 | 0.057 | 0 | 0.36 | 0.24 | 0.48 | 0.87 | 0.46 | 0.5 |
| 11 | 25102415 | rs1374133 | -0.504 | 0.401 | 0.004 | -1.229 | -0.055 | 0.09 | 0.09 | 0 | 0.35 | 0.19 | 0.47 | 0.87 | 0.46 | 0.5 |
| 11 | 25105591 | rs1010404 | 0.42 | 0.614 | -0.178 | -1.101 | -0.001 | -0.037 | -0.037 | 0 | 0.58 | 0.55 | 0.47 | 0.12 | 0.51 | 0.43 |
| 11 | 25113781 | rs11602337 | 0.696 | 0.314 | -1.571 | -1.576 | -0.018 | 0.168 | 0.168 | 0 | 0.3 | 0.05 | 0.21 | 0.05 | 0.25 | 0.3 |
| 11 | 25114454 | rs4923243 | 0.75 | -0.072 | -2.035 | -1.967 | 0.011 | 0.03 | 0.03 | 0 | 0.11 | 0.05 | 0.05 | 0 | 0.08 | 0.09 |
| 11 | 25115305 | rs11028418 | 1.327 | 0.004 | -2.326 | -2.345 | -0.015 | 0.048 | 0.048 | 0 | 0.19 | 0.1 | 0.17 | 0.05 | 0.22 | 0.15 |
| 11 | 25117751 | rs12805702 | 1.178 | 0.334 | -1.969 | -2.459 | -0.006 | 0.089 | 0.089 | 0 | 0.33 | 0.17 | 0.26 | 0.06 | 0.31 | 0.31 |
| 11 | 25120420 | rs11028423 | 1.392 | 0.329 | -1.992 | -2.442 | -0.015 | 0.049 | 0.049 | 0 | 0.19 | 0.1 | 0.17 | 0.05 | 0.22 | 0.15 |
| 11 | 25120833 | rs1446184 | 1.392 | 0.329 | -2.008 | -2.443 | -0.016 | 0.052 | 0.052 | 0 | 0.19 | 0.1 | 0.17 | 0.05 | 0.22 | 0.15 |
| 11 | 25132070 | rs7928682 | 0.477 | 0.347 | -1.954 | -2.432 | 0.006 | 0.142 | 0.142 | 0.301 | 0.53 | 0.27 | 0.43 | 0.11 | 0.53 | 0.46 |
| 11 | 25134179 | rs11028434 | 1.165 | 0.413 | -2.247 | -1.878 | -0.022 | -0.081 | -0.081 | 0 | 0.17 | 0.03 | 0.12 | 0.34 | 0.17 | 0.28 |
| 11 | 25134742 | rs12787764 | 1.172 | 0.32 | -2.052 | -3.304 | -0.004 | 0.084 | 0.084 | 0 | 0.33 | 0.17 | 0.26 | 0.09 | 0.31 | 0.31 |
| 11 | 25140302 | rs12805962 | 1.373 | 0.348 | -2.079 | -3.288 | -0.044 | 0.143 | 0.143 | 0 | 0.32 | 0.07 | 0.25 | 0.09 | 0.31 | 0.32 |
| 11 | 25144319 | rs11824385 | 1.119 | 0.371 | -2.023 | -3.315 | -0.001 | -0.02 | -0.02 | 0 | 0.21 | 0.2 | 0.18 | 0.02 | 0.22 | 0.14 |
| 11 | 25150091 | rs11600287 | 0.987 | 0.184 | -1.728 | -3.509 | -0.008 | 0.275 | 0.275 | 0.206 | 0.51 | 0.16 | 0.42 | 0.1 | 0.53 | 0.5 |
| 11 | 25154378 | rs7125050 | 0.499 | 0.142 | -1.217 | -3.524 | 0 | -0.023 | -0.023 | 0 | 0.54 | 0.48 | 0.45 | 0.11 | 0.53 | 0.5 |
| 11 | 25157728 | rs7943819 | 0.874 | 0.283 | -2.342 | -3.433 | -0.006 | 0.256 | 0.256 | 0.181 | 0.52 | 0.17 | 0.45 | 0.12 | 0.53 | 0.51 |
| 11 | 25162545 | rs17307542 | -0.024 | 0.326 | -1.496 | -2.802 | -0.006 | 0.059 | 0.059 | 0 | 0.1 | 0 | 0.05 | 0 | 0.08 | 0.09 |
| 11 | 25171374 | rs12790361 | 0.013 | 0.247 | -1.494 | -2.789 | -0.013 | 0.066 | 0.066 | 0.198 | 0.11 | 0.01 | 0.07 | 0 | 0.09 | 0.07 |
| 11 | 25172009 | rs11028464 | 0.078 | 0.309 | -1.369 | -3.246 | -0.007 | 0.023 | 0.023 | 0.738 | 0.23 | 0.15 | 0.17 | 0.08 | 0.25 | 0.35 |
| 11 | 25172401 | rs11028466 | -0.093 | 0.256 | -1.202 | -3.268 | -0.001 | -0.002 | -0.002 | 0 | 0.24 | 0.23 | 0.18 | 0.08 | 0.25 | 0.36 |
| 11 | 25175032 | rs12416792 | -0.1 | 0.315 | -1.327 | -3.354 | -0.041 | 0.058 | 0.058 | 0 | 0.3 | 0.16 | 0.23 | 0.09 | 0.4 | 0.42 |
| 11 | 25180965 | rs1584186 | -1.319 | -0.136 | -1.257 | -3.85 | 0 | -0.026 | -0.026 | 0 | 0.79 | 0.79 | 0.82 | 0.99 | 0.82 | 0.91 |
| 11 | 25181297 | rs1584184 | 0.481 | -0.463 | -1.44 | -4.104 | -0.047 | -0.151 | -0.151 | 0 | 0.24 | 0.52 | 0.14 | 0 | 0.12 | 0.08 |
| 11 | 25181838 | rs7126778 | -1.108 | -0.276 | -1.312 | -3.781 | -0.006 | -0.087 | -0.087 | 0 | 0.78 | 0.67 | 0.81 | 0.99 | 0.81 | 0.91 |
| 11 | 25186505 | rs10834614 | 0.452 | 0.232 | -1.78 | -3.082 | -0.007 | 0.061 | 0.061 | 0.547 | 0.24 | 0.05 | 0.44 | 0.9 | 0.28 | 0.42 |
| 11 | 25196853 | rs10742072 | -0.219 | 0.34 | -1.531 | -3.239 | 0.063 | 0.218 | 0.218 | 1.752 | 0.54 | 0.18 | 0.68 | 0.98 | 0.69 | 0.83 |
| 11 | 25196998 | rs12281577 | 0.288 | 0.844 | -1.763 | -3.434 | -0.018 | 0.033 | 0.033 | 0.107 | 0.18 | 0.09 | 0.11 | 0.06 | 0.18 | 0.3 |
| 11 | 25197109 | rs12282883 | -0.469 | 0.524 | -0.692 | -3.728 | -0.006 | -0.065 | -0.065 | 0.606 | 0.58 | 0.44 | 0.4 | 0.07 | 0.59 | 0.49 |

| | | | | | | | | | | | | | | | | |
|---|---|---|---|---|---|---|---|---|---|---|---|---|---|---|---|---|
| 11 | 25197463 | rs10458889 | -0.123 | -0.04 | -0.519 | -4.053 | 0 | -0.027 | -0.027 | 0.386 | 0.59 | 0.64 | 0.66 | 0.92 | 0.49 | 0.57 |
| 11 | 25202338 | rs11028494 | 1.174 | -0.246 | -0.409 | -3.993 | -0.054 | -0.056 | -0.056 | 0.817 | 0.12 | 0.22 | 0.03 | 0 | 0.01 | 0.01 |
| 11 | 25208539 | rs2194907 | 1.35 | -0.251 | -0.4 | -3.996 | -0.037 | -0.038 | -0.038 | 0 | 0.11 | 0.17 | 0.02 | 0 | 0.01 | 0.01 |
| 11 | 25209982 | rs2217032 | -1.137 | -0.019 | -0.278 | -4.187 | -0.001 | 0.029 | 0.029 | 0.13 | 0.44 | 0.38 | 0.27 | 0.06 | 0.45 | 0.39 |
| 11 | 25219752 | rs12278154 | 0.145 | 0.004 | -0.516 | -4.126 | -0.065 | 0.047 | 0.047 | 1.039 | 0.18 | 0.04 | 0.14 | 0.06 | 0.31 | 0.33 |
| 11 | 25220054 | rs10767342 | 1.505 | -0.185 | -0.605 | -4.251 | -0.018 | 0.134 | 0.134 | 0.018 | 0.31 | 0.06 | 0.52 | 0.92 | 0.36 | 0.49 |
| 11 | 25235091 | rs4335492 | -1.24 | 0.118 | -0.206 | -3.842 | -0.007 | 0.035 | 0.035 | 0.594 | 0.43 | 0.34 | 0.31 | 0.11 | 0.43 | 0.42 |
| 11 | 25236950 | rs17241923 | 0.643 | 0.118 | -0.291 | -3.86 | -0.002 | -0.003 | -0.003 | 1.289 | 0.13 | 0.12 | 0.08 | 0 | 0.12 | 0.09 |
| 11 | 25237942 | rs11028518 | -0.659 | 0.13 | -0.265 | -3.861 | -0.043 | 0.064 | 0.064 | 0 | 0.29 | 0.13 | 0.23 | 0.11 | 0.32 | 0.33 |
| 11 | 25242350 | rs7109318 | -0.403 | -0.584 | -0.04 | -3.437 | -0.004 | -0.044 | -0.044 | 0.116 | 0.22 | 0.23 | 0.11 | 0 | 0.13 | 0.09 |
| 11 | 25244001 | rs17307708 | -0.049 | -0.233 | -0.166 | -3.576 | 0.01 | 0.056 | 0.056 | 0 | 0.37 | 0.23 | 0.24 | 0.01 | 0.31 | 0.15 |
| 11 | 25256554 | rs2162164 | -1.116 | -0.044 | -0.187 | -3.582 | -0.001 | -0.067 | -0.067 | 0.193 | 0.57 | 0.53 | 0.39 | 0.1 | 0.54 | 0.38 |
| 11 | 25258687 | rs11028528 | 0.09 | 0.145 | -1.069 | -3.596 | -0.096 | 0.137 | 0.137 | 1.226 | 0.32 | 0.1 | 0.26 | 0.1 | 0.4 | 0.29 |
| 11 | 25262125 | rs7927431 | 0.477 | 0.228 | -0.671 | -3.246 | -0.015 | 0.151 | 0.151 | 0.084 | 0.41 | 0.19 | 0.59 | 0.93 | 0.46 | 0.63 |
| 11 | 25270335 | rs2042680 | -0.402 | 0 | -0.619 | -3.084 | 0.006 | -0.029 | -0.029 | 0 | 0.49 | 0.54 | 0.33 | 0.01 | 0.41 | 0.21 |
| 11 | 25274897 | rs12289156 | -0.835 | -0.855 | -0.942 | -2.747 | 0.016 | 0.011 | 0.011 | 0.564 | 0.24 | 0.19 | 0.09 | 0.07 | 0.09 | 0.23 |
| 11 | 25275069 | rs11028539 | -0.149 | 0.164 | -1.611 | -3.652 | -0.047 | 0.274 | 0.274 | 0.157 | 0.44 | 0.15 | 0.36 | 0.07 | 0.52 | 0.42 |
| 11 | 25275875 | rs10437619 | -0.501 | 0.232 | -1.415 | -3.589 | -0.002 | 0.018 | 0.018 | 0.835 | 0.41 | 0.37 | 0.27 | 0.08 | 0.39 | 0.27 |
| 11 | 25277449 | rs717607 | -0.215 | 0.695 | -1.075 | -3.676 | -0.039 | 0.051 | 0.051 | 0 | 0.23 | 0.13 | 0.21 | 0.06 | 0.34 | 0.19 |
| 11 | 25284208 | rs10767347 | -0.22 | 0.751 | -0.585 | -3.658 | -0.002 | -0.003 | -0.003 | 0 | 0.81 | 0.79 | 0.8 | 0.94 | 0.74 | 0.8 |
| 11 | 25287389 | rs7942193 | -0.04 | 0.682 | 0.037 | -3.687 | 0.052 | -0.054 | -0.054 | 0 | 0.8 | 0.67 | 0.77 | 0.94 | 0.67 | 0.78 |
| 11 | 25291015 | rs11028546 | -1.692 | 0.329 | 0.641 | -3.052 | 0.003 | 0.062 | 0.062 | 0.021 | 0.08 | 0.38 | 0.04 | 0.06 | 0.02 | 0.07 |
| 11 | 25315971 | rs2114076 | 0.187 | 0.329 | 0.312 | -3.229 | -0.014 | 0.015 | 0.015 | 0 | 0.2 | 0.17 | 0.23 | 0.06 | 0.26 | 0.23 |
| 11 | 25316492 | rs2059541 | 0.072 | 0.275 | 0.81 | -3.244 | 0.081 | -0.09 | -0.09 | 0.831 | 0.77 | 0.53 | 0.72 | 0.94 | 0.67 | 0.74 |
| 11 | 25323589 | rs7102124 | -0.079 | 0.264 | 0.785 | -3.249 | -0.013 | 0.011 | 0.011 | 0 | 0.22 | 0.19 | 0.26 | 0.06 | 0.33 | 0.26 |
| 11 | 25332979 | rs1425822 | -0.163 | 0.26 | 0.791 | -3.249 | 0.033 | -0.063 | -0.063 | 1.503 | 0.78 | 0.61 | 0.75 | 0.94 | 0.74 | 0.77 |
| 11 | 25337780 | rs11028585 | -0.88 | 0.189 | 0.874 | -3.263 | -0.019 | 0.02 | 0.02 | 0 | 0.05 | 0.18 | 0.02 | 0.05 | 0.01 | 0.07 |
| 11 | 25338988 | rs12363488 | 0.984 | 0.18 | 0.885 | -3.275 | 0.002 | -0.01 | -0.01 | 0.203 | 0.7 | 0.31 | 0.69 | 0.89 | 0.65 | 0.68 |
| 11 | 25340131 | rs7129251 | -1.134 | 0.021 | 0.83 | -3.177 | -0.018 | 0.026 | 0.026 | 0 | 0.06 | 0.2 | 0.02 | 0.05 | 0.01 | 0.07 |
| 11 | 25349335 | rs12223666 | 0.159 | 0.268 | 0.674 | -3.205 | -0.016 | 0.022 | 0.022 | 0 | 0.19 | 0.17 | 0.22 | 0.06 | 0.21 | 0.23 |
| 11 | 25362364 | rs17307813 | 0.196 | 0.365 | 0.492 | -3.017 | -0.04 | 0.043 | 0.043 | 0 | 0.19 | 0.09 | 0.22 | 0.06 | 0.23 | 0.24 |
| 11 | 25372142 | rs10834656 | 0.639 | 0.324 | 0.711 | -3.007 | 0.022 | -0.015 | -0.015 | 0 | 0.72 | 0.42 | 0.7 | 0.86 | 0.68 | 0.67 |
| 11 | 25375512 | rs2404060 | 0.848 | 0.248 | 0.512 | -2.972 | 0.008 | 0.031 | 0.031 | 0.492 | 0.92 | 0.73 | 0.95 | 0.94 | 0.91 | 0.92 |
| 11 | 25377046 | rs11028633 | 0.177 | 0.236 | 0.561 | -3.003 | -0.004 | 0.003 | 0.003 | 0 | 0.2 | 0.17 | 0.23 | 0.08 | 0.23 | 0.25 |
| 11 | 25377220 | rs11028634 | -0.253 | 0.052 | 0.165 | -3.269 | -0.047 | 0.045 | 0.045 | 0 | 0.24 | 0.09 | 0.25 | 0.08 | 0.28 | 0.28 |
| 11 | 25378706 | rs4459251 | 0.32 | 0.073 | 0.215 | -3.124 | 0.023 | -0.015 | -0.015 | 0 | 0.72 | 0.56 | 0.71 | 0.86 | 0.68 | 0.67 |
| 11 | 25385677 | rs12417317 | 1.237 | -0.041 | -1.085 | -3.587 | -0.021 | 0.067 | 0.067 | 0 | 0.38 | 0.06 | 0.51 | 0.86 | 0.41 | 0.48 |
| 11 | 25388752 | rs11028640 | 0.833 | -0.325 | -0.613 | -3.211 | -0.05 | 0.109 | 0.109 | 0 | 0.57 | 0.11 | 0.61 | 0.86 | 0.58 | 0.6 |
| 11 | 25391785 | rs7935930 | -0.602 | -0.381 | 0.824 | -2.852 | 0.042 | -0.067 | -0.067 | 0 | 0.11 | 0.38 | 0.1 | 0 | 0.1 | 0.07 |
| 11 | 25394425 | rs2404071 | 0.031 | -1.005 | 0.337 | -2.903 | 0.02 | 0.019 | 0.019 | 0 | 0.89 | 0.9 | 0.95 | 0.97 | 0.96 | 0.85 |
| 11 | 25396639 | rs10834664 | 0.277 | -0.877 | 0.554 | -2.276 | 0.014 | -0.004 | -0.004 | 0 | 0.17 | 0.21 | 0.14 | 0.09 | 0.22 | 0.19 |
| 11 | 25397438 | rs10767368 | 0.964 | -0.854 | 0.766 | -2.251 | -0.001 | -0.017 | -0.017 | 0 | 0.7 | 0.61 | 0.8 | 0.88 | 0.73 | 0.65 |
| 11 | 25399796 | rs10742088 | 0.938 | -0.962 | 1.172 | -2.274 | -0.007 | -0.036 | -0.036 | 0 | 0.68 | 0.44 | 0.79 | 0.88 | 0.72 | 0.65 |
| 11 | 25400627 | rs10834666 | 1.662 | -1.322 | 0.783 | -2.545 | 0.018 | 0.2 | 0.2 | 0 | 0.53 | 0.11 | 0.69 | 0.88 | 0.6 | 0.57 |
| 11 | 25407911 | rs7111550 | 0.659 | -1.08 | 0.839 | -2.298 | -0.003 | 0.027 | 0.027 | 0 | 0.71 | 0.78 | 0.82 | 0.88 | 0.73 | 0.65 |
| 11 | 25417222 | rs7127730 | 0.629 | -1.067 | 0.76 | -2.315 | -0.002 | -0.022 | -0.022 | 0 | 0.7 | 0.61 | 0.81 | 0.88 | 0.73 | 0.65 |
| 11 | 25431743 | rs12807415 | 0.117 | -1.124 | 0.68 | -2.266 | 0.008 | 0.008 | 0.008 | 0 | 0.11 | 0.1 | 0.05 | 0.03 | 0.05 | 0.16 |
| 11 | 25433466 | rs10767383 | 0.854 | -1.105 | 0.957 | -2.349 | -0.004 | -0.044 | -0.044 | 0 | 0.69 | 0.46 | 0.79 | 0.88 | 0.72 | 0.66 |
| 11 | 25437677 | rs7107962 | 0.684 | -1.071 | 0.824 | -2.349 | -0.048 | -0.007 | -0.007 | 0.304 | 0.15 | 0.06 | 0.11 | 0.09 | 0.21 | 0.16 |
| 11 | 25438153 | rs4444035 | 1.704 | -1.286 | 0.732 | -2.581 | 0.002 | 0.201 | 0.201 | 0 | 0.53 | 0.09 | 0.69 | 0.88 | 0.6 | 0.58 |
| 11 | 25453443 | rs12287650 | -0.152 | -1.29 | 0.884 | -2.576 | -0.032 | -0.117 | -0.117 | 0 | 0.18 | 0.56 | 0.11 | 0 | 0.13 | 0.07 |
| 11 | 25453482 | rs5017351 | -0.592 | -1.281 | 0.783 | -2.575 | 0.009 | -0.004 | -0.004 | 0.831 | 0.82 | 0.78 | 0.86 | 0.91 | 0.77 | 0.81 |
| 11 | 25455664 | rs2618799 | -1.412 | -1.364 | 0.772 | -2.588 | -0.008 | -0.04 | -0.04 | 0 | 0.3 | 0.67 | 0.17 | 0.03 | 0.19 | 0.26 |
| 11 | 25456008 | rs7935271 | 1.675 | -1.309 | 0.52 | -2.569 | 0.012 | 0.197 | 0.197 | 0 | 0.53 | 0.11 | 0.69 | 0.88 | 0.59 | 0.58 |
| 11 | 25457869 | rs2953117 | 0.254 | -1.113 | 0.875 | -2.448 | -0.014 | -0.095 | -0.095 | 0 | 0.15 | 0.33 | 0.09 | 0 | 0.12 | 0.07 |
| 11 | 25467085 | rs11028689 | 0.727 | -0.995 | 0.449 | -2.224 | -0.054 | -0.029 | -0.029 | 9.736 | 0.14 | 0.04 | 0.11 | 0.1 | 0.21 | 0.17 |
| 11 | 25498699 | rs11028708 | 1.254 | -1.278 | 0.281 | -2.429 | 0.012 | 0.17 | 0.17 | 0 | 0.54 | 0.13 | 0.7 | 0.87 | 0.6 | 0.68 |
| 11 | 25501832 | rs10501008 | 0.663 | -1.352 | 0.76 | -2.43 | 0.017 | 0.002 | 0.002 | 0.45 | 0.16 | 0.22 | 0.11 | 0.1 | 0.19 | 0.13 |
| 11 | 25526050 | rs10767395 | 1.456 | -1.315 | 0.325 | -2.479 | 0.006 | 0.194 | 0.194 | 0 | 0.52 | 0.15 | 0.7 | 0.87 | 0.58 | 0.59 |
| 11 | 25528202 | rs10767396 | -0.68 | -1.363 | 0.48 | -2.51 | 0.013 | -0.048 | -0.048 | 3.556 | 0.33 | 0.47 | 0.22 | 0.1 | 0.35 | 0.27 |
| 11 | 25529663 | rs2028614 | 0.273 | -1.278 | 0.562 | -2.429 | -0.047 | -0.057 | -0.057 | 0 | 0.86 | 0.72 | 0.93 | 0.97 | 0.93 | 0.87 |
| 11 | 25529987 | rs9633819 | 1.864 | -1.682 | -0.199 | -2.817 | -0.002 | 0.051 | 0.051 | 0 | 0.18 | 0.01 | 0.14 | 0.06 | 0.16 | 0.14 |
| 11 | 25545164 | rs1036886 | 1.729 | -1.196 | 0.1 | -1.738 | -0.113 | -0.081 | -0.081 | 0.002 | 0.37 | 0.05 | 0.33 | 0.63 | 0.46 | 0.38 |
| 11 | 25552964 | rs10501013 | 0.699 | -0.936 | 0.522 | -1.755 | -0.059 | -0.034 | -0.034 | 0.333 | 0.14 | 0.01 | 0.1 | 0.1 | 0.19 | 0.15 |
| 11 | 25555059 | rs17242293 | -0.858 | -0.987 | 0.551 | -1.781 | -0.004 | 0.014 | 0.014 | 0 | 0.25 | 0.12 | 0.15 | 0.13 | 0.24 | 0.28 |
| 11 | 25558538 | rs10834705 | 1.426 | -0.995 | 0.434 | -1.871 | 0.026 | 0.093 | 0.093 | 0.006 | 0.65 | 0.14 | 0.77 | 0.89 | 0.72 | 0.69 |
| 11 | 25562027 | rs11028760 | 0.495 | -0.765 | 0.46 | -1.758 | -0.04 | -0.01 | -0.01 | 0 | 0.15 | 0.05 | 0.1 | 0.09 | 0.19 | 0.15 |
| 11 | 25563417 | rs2165818 | -0.379 | -0.788 | 0.652 | -1.771 | 0.01 | -0.014 | -0.014 | 0 | 0.2 | 0.32 | 0.13 | 0.09 | 0.2 | 0.18 |
| 11 | 25581074 | rs7120268 | -1.87 | -0.844 | 0.806 | -1.788 | -0.008 | -0.022 | -0.022 | 0.789 | 0.08 | 0.59 | 0.06 | 0 | 0.02 | 0.03 |

| | | | | | | | | | | | | | | | | |
|---|---|---|---|---|---|---|---|---|---|---|---|---|---|---|---|---|
| 11 | 25582331 | rs10767405 | 1.504 | -0.945 | 0.679 | -1.897 | 0.059 | 0.15 | 0.15 | 0 | 0.66 | 0.11 | 0.81 | 0.97 | 0.76 | 0.73 |
| 11 | 25585180 | rs1561937 | 1.47 | -0.954 | 0.624 | -1.896 | 0.042 | 0.043 | 0.043 | 1.544 | 0.79 | 0.13 | 0.87 | 0.97 | 0.91 | 0.84 |
| 11 | 25599986 | rs11028773 | 0.801 | -0.838 | 0.464 | -1.866 | -0.061 | -0.028 | -0.028 | 0 | 0.14 | 0.01 | 0.1 | 0.09 | 0.18 | 0.13 |
| 11 | 25605844 | rs2896754 | -2.138 | -0.867 | 0.577 | -1.869 | -0.034 | -0.051 | -0.051 | 0 | 0.1 | 0.66 | 0.06 | 0 | 0.02 | 0.03 |
| 11 | 25610986 | rs2120148 | -0.114 | -0.904 | 0.581 | -1.889 | -0.009 | -0.014 | -0.014 | 0 | 0.11 | 0.11 | 0.05 | 0.03 | 0.07 | 0.13 |
| 11 | 25616540 | rs16914165 | -1.976 | -0.839 | 0.619 | -1.848 | -0.022 | -0.041 | -0.041 | 0 | 0.09 | 0.64 | 0.06 | 0 | 0.02 | 0.03 |
| 11 | 25630365 | rs1443160 | -0.018 | -0.709 | 0.475 | -1.814 | -0.005 | -0.123 | -0.123 | 3.112 | 0.82 | 0.46 | 0.89 | 1 | 0.85 | 0.89 |
| 11 | 25636886 | rs882257 | -0.018 | -0.731 | 0.475 | -1.814 | -0.008 | -0.123 | -0.123 | 0.221 | 0.82 | 0.46 | 0.89 | 1 | 0.85 | 0.89 |
| 11 | 25637259 | rs882258 | 1.916 | -0.755 | -0.181 | -1.811 | 0.003 | 0.198 | 0.198 | 0 | 0.51 | 0.1 | 0.55 | 0.84 | 0.58 | 0.6 |
| 11 | 25637280 | rs894578 | -0.402 | -0.851 | -0.059 | -1.76 | 0.004 | -0.101 | -0.101 | 0 | 0.84 | 0.67 | 0.9 | 1 | 0.85 | 0.89 |
| 11 | 25642425 | rs7102465 | -0.405 | -0.59 | -0.698 | -1.531 | -0.016 | -0.023 | -0.023 | 0 | 0.12 | 0.15 | 0.06 | 0.03 | 0.07 | 0.13 |
| 11 | 25651226 | rs10834729 | -1.503 | -0.259 | -1.006 | -1.305 | 0.004 | 0.013 | 0.013 | 0 | 0.45 | 0.62 | 0.4 | 0.18 | 0.39 | 0.39 |
| 11 | 25651414 | rs11606269 | -0.238 | 0.25 | -1.662 | -0.358 | -0.01 | -0.011 | -0.011 | 1.45 | 0.16 | 0.1 | 0.26 | 0.13 | 0.18 | 0.14 |
| 11 | 25662734 | rs4356180 | -1.023 | -0.18 | -1.6 | -0.513 | 0.01 | 0.01 | 0.01 | 0 | 0.12 | 0.13 | 0.04 | 0.05 | 0.06 | 0.15 |
| 11 | 25664199 | rs10742108 | -0.153 | 0.219 | -2.186 | -0.917 | 0.067 | 0.244 | 0.244 | 0 | 0.51 | 0.16 | 0.51 | 0.88 | 0.59 | 0.65 |
| 11 | 25679555 | rs7126145 | -0.521 | 1.325 | -1.844 | -0.127 | -0.031 | 0.023 | 0.023 | 0.279 | 0.15 | 0.07 | 0.13 | 0.1 | 0.22 | 0.29 |
| 11 | 25680364 | rs12577727 | -0.704 | 1.285 | -1.88 | 1.052 | -0.022 | -0.037 | -0.037 | 0 | 0.11 | 0.06 | 0.15 | 0.52 | 0.11 | 0.15 |
| 11 | 25682233 | rs1493661 | 0.008 | 0.785 | -0.882 | 0.557 | 0.03 | 0.02 | 0.02 | 2.651 | 0.93 | 0.61 | 0.95 | 1 | 0.98 | 0.94 |
| 11 | 25684596 | rs11028833 | -0.456 | 1.033 | -1.567 | 1.285 | -0.023 | -0.028 | -0.028 | 0 | 0.1 | 0.05 | 0.14 | 0.42 | 0.1 | 0.14 |
| 11 | 25687417 | rs1389412 | 0.664 | 0.772 | -2.437 | 0.297 | -0.101 | 0.104 | 0.104 | 0.472 | 0.46 | 0.08 | 0.6 | 0.7 | 0.49 | 0.51 |
| 11 | 25688809 | rs11607275 | 0.099 | 0.676 | -2.367 | -0.122 | -0.036 | 0.154 | 0.154 | 0 | 0.27 | 0.06 | 0.2 | 0.01 | 0.28 | 0.1 |
| 11 | 25696461 | rs1602916 | 1.103 | 0.39 | -2.414 | 0.608 | -0.027 | 0.02 | 0.02 | 0.294 | 0.46 | 0.14 | 0.52 | 0.44 | 0.56 | 0.41 |
| 11 | 25700470 | rs1908161 | 0.422 | 0.34 | -1 | 0.957 | 0.04 | 0.504 | 0.504 | 0.698 | 0.93 | 0.66 | 0.9 | 0.63 | 0.95 | 0.95 |
| 11 | 25700712 | rs17308279 | 0.564 | 0.196 | -1.237 | 0.614 | 0 | 0.139 | 0.139 | 0.218 | 0.21 | 0.01 | 0.14 | 0.01 | 0.16 | 0.04 |
| 11 | 25700989 | rs1493664 | 1.614 | 0.454 | -1.256 | 0.904 | -0.018 | -0.025 | -0.025 | 0 | 0.48 | 0.14 | 0.54 | 0.56 | 0.56 | 0.46 |
| 11 | 25701351 | rs1159087 | 0.325 | 0.406 | -1.127 | 0.843 | 0.021 | 0.488 | 0.488 | 2.839 | 0.93 | 0.67 | 0.92 | 0.64 | 0.97 | 0.96 |
| 11 | 25706158 | rs4923292 | 0.081 | 0.412 | -0.337 | -0.018 | 0.009 | -0.004 | -0.004 | 2.866 | 0.13 | 0.16 | 0.17 | 0.13 | 0.18 | 0.31 |
| 11 | 25710774 | rs1493670 | 1.124 | 0.338 | -0.217 | 0.156 | -0.023 | 0.062 | 0.062 | 0.318 | 0.82 | 0.38 | 0.86 | 0.76 | 0.88 | 0.82 |
| 11 | 25712300 | rs1493669 | 1.004 | 0.313 | -0.254 | 0.095 | 0.043 | 0.024 | 0.024 | 1.853 | 0.6 | 0.29 | 0.72 | 0.75 | 0.74 | 0.76 |
| 11 | 25717654 | rs4923297 | 0.234 | 0.336 | 0.202 | 0.122 | 0.013 | 0.169 | 0.169 | 0.126 | 0.94 | 0.79 | 0.93 | 0.82 | 0.95 | 0.96 |
| 11 | 25718943 | rs1602919 | 0.35 | 0.324 | 0.175 | 0.092 | 0 | 0 | 0 | 0 | 0.12 | 0.11 | 0.16 | 0.13 | 0.16 | 0.29 |
| 11 | 25720706 | rs7113321 | -0.591 | 0.233 | 0.171 | 0.071 | -0.045 | -0.112 | -0.112 | 0 | 0.33 | 0.47 | 0.2 | 0.07 | 0.2 | 0.19 |
| 11 | 25724484 | rs2172582 | 0.061 | 0.259 | 0.127 | 0.1 | 0.014 | 0.168 | 0.168 | 0 | 0.95 | 0.81 | 0.93 | 0.82 | 0.95 | 0.96 |
| 11 | 25730008 | rs1353128 | 0.73 | 0.245 | 0.247 | 0.091 | -0.004 | -0.001 | -0.001 | 0 | 0.62 | 0.43 | 0.75 | 0.74 | 0.75 | 0.77 |
| 11 | 25753192 | rs12420001 | 1.571 | 0.352 | 0.084 | 0.064 | 0.001 | 0.036 | 0.036 | 0 | 0.48 | 0.17 | 0.53 | 0.62 | 0.58 | 0.47 |
| 11 | 25753430 | rs1038934 | 0.559 | 0.475 | 0.143 | 0.181 | 0.018 | 0.039 | 0.039 | 0 | 0.81 | 0.71 | 0.74 | 0.68 | 0.78 | 0.66 |
| 11 | 25755528 | rs10501021 | -0.573 | 0.159 | 0.102 | -0.097 | -0.041 | -0.094 | -0.094 | 0 | 0.29 | 0.37 | 0.18 | 0.06 | 0.17 | 0.19 |
| 11 | 25755821 | rs10501022 | -0.235 | 0.104 | 0.256 | 0.016 | -0.056 | -0.099 | -0.099 | 0.504 | 0.24 | 0.4 | 0.15 | 0.06 | 0.13 | 0.15 |
| 11 | 25756735 | rs11823995 | -0.369 | 0.138 | 0.311 | 0.022 | -0.01 | -0.031 | -0.031 | 0 | 0.26 | 0.3 | 0.15 | 0.07 | 0.16 | 0.15 |
| 11 | 25764292 | rs11028895 | 0.414 | 0.277 | 0.253 | 0.17 | 0.015 | 0.051 | 0.051 | 0 | 0.83 | 0.71 | 0.76 | 0.69 | 0.8 | 0.66 |
| 11 | 25773322 | rs2349308 | 0.414 | 0.274 | 0.252 | 0.168 | 0.015 | 0.051 | 0.051 | 0 | 0.83 | 0.7 | 0.76 | 0.69 | 0.8 | 0.66 |
| 11 | 25777213 | rs12274588 | -0.537 | 0.137 | 0.292 | 0.074 | -0.058 | -0.117 | -0.117 | 0 | 0.28 | 0.48 | 0.17 | 0.07 | 0.16 | 0.15 |
| 11 | 25779039 | rs1493663 | 0.459 | 0.189 | 0.333 | 0.188 | 0.015 | 0.052 | 0.052 | 0 | 0.83 | 0.7 | 0.76 | 0.69 | 0.8 | 0.68 |
| 11 | 25786560 | rs1389420 | 0.44 | 0.19 | 0.333 | 0.188 | 0.016 | 0.052 | 0.052 | 0 | 0.83 | 0.7 | 0.76 | 0.69 | 0.8 | 0.68 |
| 11 | 25787631 | rs6484176 | -0.525 | 0.466 | -0.011 | -0.302 | -0.017 | -0.07 | -0.07 | 0 | 0.36 | 0.52 | 0.22 | 0.08 | 0.26 | 0.17 |
| 11 | 25791846 | rs1389423 | 1.467 | 0.446 | -0.041 | -0.246 | -0.042 | 0.021 | 0.021 | 3.716 | 0.46 | 0.17 | 0.52 | 0.62 | 0.51 | 0.51 |
| 11 | 25791864 | rs2307140 | 0.513 | 0.469 | 0.021 | -0.238 | -0.003 | 0.001 | 0.001 | 0 | 0.11 | 0.1 | 0.17 | 0.12 | 0.16 | 0.28 |
| 11 | 25792678 | rs1389424 | 1.42 | 0.424 | 0.033 | -0.203 | -0.034 | 0.024 | 0.024 | 3.705 | 0.46 | 0.18 | 0.52 | 0.62 | 0.51 | 0.51 |
| 11 | 25794462 | rs1156762 | -1.274 | 0.392 | 0.176 | -0.257 | -0.003 | 0.043 | 0.043 | 0.66 | 0.48 | 0.34 | 0.45 | 0.31 | 0.48 | 0.43 |
| 11 | 25798364 | rs10767453 | -1.316 | 0.428 | 0.128 | -0.243 | -0.003 | 0.051 | 0.051 | 0.897 | 0.48 | 0.34 | 0.43 | 0.31 | 0.48 | 0.42 |
| 11 | 25801048 | rs10834801 | -1.253 | 0.436 | 0.199 | -0.23 | 0 | -0.001 | -0.001 | 0 | 0.48 | 0.44 | 0.44 | 0.31 | 0.48 | 0.43 |
| 11 | 25802244 | rs4922741 | 0.078 | 0.659 | 0.426 | -0.264 | -0.017 | 0.022 | 0.022 | 0 | 0.11 | 0.06 | 0.15 | 0.11 | 0.15 | 0.28 |
| 11 | 25808617 | rs10458890 | 0.08 | 0.843 | -0.148 | -0.274 | -0.017 | 0.024 | 0.024 | 0 | 0.11 | 0.05 | 0.15 | 0.08 | 0.15 | 0.28 |
| 11 | 25813830 | rs10834804 | 0.08 | 0.843 | -0.148 | -0.273 | -0.017 | 0.022 | 0.022 | 0 | 0.11 | 0.06 | 0.15 | 0.11 | 0.16 | 0.28 |
| 11 | 25817694 | rs4923307 | -1.575 | 0.519 | 0.329 | -0.1 | -0.006 | 0.09 | 0.09 | 0.034 | 0.17 | 0.42 | 0.29 | 0.29 | 0.16 | 0.3 |
| 11 | 25818836 | rs1532286 | 1.349 | 0.549 | -0.45 | -0.415 | -0.003 | 0.061 | 0.061 | 0.215 | 0.54 | 0.22 | 0.5 | 0.68 | 0.57 | 0.56 |
| 11 | 25819061 | rs11028937 | -0.305 | 0.183 | -0.523 | -0.586 | -0.003 | 0.006 | 0.006 | 0 | 0.13 | 0.19 | 0.1 | 0.19 | 0.11 | 0.24 |
| 11 | 25821550 | rs7481377 | 0.153 | 0.184 | -0.689 | -0.876 | -0.019 | 0.022 | 0.022 | 0 | 0.08 | 0.03 | 0.12 | 0.02 | 0.09 | 0.09 |
| 11 | 25823936 | rs11823887 | 0.192 | 0.478 | -0.532 | -1.538 | 0 | -0.047 | -0.047 | 0 | 0.22 | 0.2 | 0.14 | 0.02 | 0.25 | 0.17 |
| 11 | 25826554 | rs327483 | 0.552 | 0.5 | -0.777 | -1.533 | -0.008 | 0.004 | 0.004 | 0 | 0.25 | 0.21 | 0.29 | 0.25 | 0.24 | 0.25 |
| 11 | 25835890 | rs10767461 | 0.766 | 0.402 | -0.28 | -1.656 | 0.002 | 0.087 | 0.087 | 0 | 0.51 | 0.35 | 0.53 | 0.73 | 0.51 | 0.56 |
| 11 | 25837926 | rs1372274 | 0.766 | 0.402 | -0.28 | -1.656 | 0.002 | 0.087 | 0.087 | 0 | 0.51 | 0.35 | 0.53 | 0.73 | 0.51 | 0.57 |
| 11 | 25839087 | rs7116993 | 0.811 | 0.365 | -0.314 | -1.692 | 0.005 | 0.086 | 0.086 | 0 | 0.49 | 0.35 | 0.52 | 0.73 | 0.51 | 0.56 |
| 11 | 25841229 | rs7111524 | 1.255 | 0.428 | -0.425 | -1.696 | -0.049 | 0.043 | 0.043 | 0 | 0.2 | 0.05 | 0.27 | 0.13 | 0.21 | 0.16 |
| 11 | 25843698 | rs11028958 | 0.16 | 0.576 | -0.212 | -1.642 | 0.149 | 0.026 | 0.026 | 0.173 | 0.93 | 0.59 | 0.92 | 1 | 0.89 | 0.96 |
| 11 | 25845567 | rs1372270 | 1.375 | -0.058 | -1.064 | -1.88 | -0.028 | 0.099 | 0.099 | 0 | 0.2 | 0.03 | 0.19 | 0.02 | 0.2 | 0.1 |
| 11 | 25846889 | rs12418699 | 1.81 | -0.023 | -1.02 | -1.197 | -0.071 | -0.169 | -0.169 | 0 | 0.23 | 0.03 | 0.21 | 0.5 | 0.27 | 0.41 |
| 11 | 25851394 | rs1596071 | 0.203 | -0.418 | -0.595 | -1.322 | -0.007 | -0.016 | -0.016 | 0 | 0.29 | 0.37 | 0.24 | 0.11 | 0.25 | 0.23 |
| 11 | 25860349 | rs7131574 | 1.161 | -0.251 | -0.629 | -1.379 | -0.046 | 0.058 | 0.058 | 0 | 0.25 | 0.1 | 0.3 | 0.14 | 0.31 | 0.2 |

| | | | | | | | | | | | | | | | | |
|---|---|---|---|---|---|---|---|---|---|---|---|---|---|---|---|---|
| 11 | 25860834 | rs1372264 | -1.289 | -0.333 | -0.622 | -1.46 | -0.025 | 0.049 | 0.049 | 0 | 0.8 | 0.54 | 0.78 | 0.74 | 0.85 | 0.82 |
| 11 | 25864177 | rs1441504 | -1.105 | -0.105 | -0.45 | -1.476 | -0.001 | -0.011 | -0.011 | 1.257 | 0.56 | 0.52 | 0.55 | 0.25 | 0.57 | 0.43 |
| 11 | 25874463 | rs767689 | -0.025 | -0.178 | -0.42 | -1.421 | -0.011 | -0.025 | -0.025 | 0 | 0.31 | 0.41 | 0.26 | 0.11 | 0.25 | 0.23 |
| 11 | 25878681 | rs2033979 | -0.206 | -0.185 | -0.338 | -1.615 | 0.028 | 0.138 | 0.138 | 0 | 0.48 | 0.29 | 0.54 | 0.17 | 0.55 | 0.34 |
| 11 | 25878875 | rs2033977 | -0.789 | -0.146 | -0.238 | -1.565 | 0.005 | 0.011 | 0.011 | 0 | 0.55 | 0.45 | 0.57 | 0.21 | 0.6 | 0.43 |
| 11 | 25880093 | rs2859991 | 0.486 | 0.141 | -0.134 | -1.236 | 0.008 | 0.013 | 0.013 | 0 | 0.28 | 0.33 | 0.43 | 0.42 | 0.34 | 0.29 |
| 11 | 25886485 | rs11603501 | 0.606 | -0.037 | -0.718 | -1.395 | -0.075 | -0.058 | -0.058 | 0 | 0.42 | 0.12 | 0.42 | 0.52 | 0.47 | 0.5 |
| 11 | 25889500 | rs10767474 | 0.645 | 0.053 | -0.292 | -1.511 | 0.006 | -0.04 | -0.04 | 0 | 0.27 | 0.32 | 0.32 | 0.15 | 0.32 | 0.23 |
| 11 | 25895955 | rs11028988 | 1.232 | 0.073 | -0.346 | -1.509 | -0.053 | 0.056 | 0.056 | 0 | 0.22 | 0.09 | 0.3 | 0.15 | 0.3 | 0.22 |
| 11 | 25901964 | rs7108065 | 0.915 | -0.166 | -0.324 | -1.477 | 0.033 | -0.06 | -0.06 | 0.638 | 0.23 | 0.08 | 0.19 | 0.3 | 0.17 | 0.21 |
| 11 | 25902183 | rs7108465 | 1.521 | -0.163 | -0.344 | -1.572 | -0.006 | 0.126 | 0.126 | 0 | 0.19 | 0.01 | 0.09 | 0.02 | 0.15 | 0.15 |
| 11 | 25902929 | rs1441516 | 0.121 | 0.008 | -0.185 | -1.6 | 0.064 | 0.035 | 0.035 | 0 | 0.92 | 0.64 | 0.87 | 0.96 | 0.92 | 0.93 |
| 11 | 25905509 | rs7930215 | 1.373 | -0.157 | -0.272 | -1.544 | -0.003 | 0.11 | 0.11 | 0 | 0.2 | 0.02 | 0.1 | 0.02 | 0.17 | 0.15 |
| 11 | 25906066 | rs10834836 | -0.056 | -0.238 | -0.308 | -1.63 | 0.009 | 0.044 | 0.044 | 0 | 0.5 | 0.39 | 0.55 | 0.18 | 0.55 | 0.37 |
| 11 | 25916225 | rs12223678 | -0.072 | -0.304 | -0.35 | -1.52 | 0.032 | 0.051 | 0.051 | 0 | 0.47 | 0.26 | 0.43 | 0.34 | 0.41 | 0.35 |
| 11 | 25917182 | rs4550176 | -0.068 | -0.316 | -0.441 | -1.527 | 0.025 | 0.053 | 0.053 | 0 | 0.47 | 0.15 | 0.42 | 0.34 | 0.41 | 0.35 |
| 11 | 25924376 | rs4923316 | 1.221 | -0.304 | -0.466 | -1.508 | -0.009 | -0.062 | -0.062 | 0 | 0.25 | 0.11 | 0.31 | 0.33 | 0.24 | 0.2 |
| 11 | 25929959 | rs1441519 | 0.078 | -0.33 | -0.264 | -1.498 | 0.024 | 0.021 | 0.021 | 0 | 0.48 | 0.32 | 0.53 | 0.63 | 0.58 | 0.6 |
| 11 | 25931262 | rs16914877 | 1.234 | -0.699 | -0.417 | -1.465 | -0.04 | -0.121 | -0.121 | 0 | 0.23 | 0.05 | 0.21 | 0.48 | 0.27 | 0.39 |
| 11 | 25944931 | rs10834851 | 1.455 | -0.461 | -0.389 | -1.501 | -0.02 | 0.049 | 0.049 | 0 | 0.22 | 0.11 | 0.28 | 0.13 | 0.21 | 0.21 |
| 11 | 25946859 | rs2033976 | -0.122 | -0.495 | -0.362 | -1.464 | 0.001 | 0.041 | 0.041 | 0 | 0.48 | 0.32 | 0.46 | 0.33 | 0.5 | 0.36 |
| 11 | 25951917 | rs12803523 | 0.706 | -0.477 | -0.302 | -1.398 | 0.018 | 0.025 | 0.025 | 0 | 0.22 | 0.12 | 0.11 | 0.02 | 0.16 | 0.14 |
| 11 | 25958308 | rs10400315 | 0.916 | -0.282 | -0.105 | -1.201 | -0.001 | 0 | 0 | 0 | 0.24 | 0.19 | 0.22 | 0.5 | 0.26 | 0.39 |
| 11 | 25959832 | rs4922749 | -0.722 | -0.286 | -0.114 | -1.206 | -0.005 | 0.016 | 0.016 | 0 | 0.7 | 0.4 | 0.73 | 0.46 | 0.71 | 0.56 |
| 11 | 25964920 | rs12282385 | 0.617 | -0.051 | -0.074 | -1.303 | 0.004 | -0.025 | -0.025 | 0.123 | 0.27 | 0.29 | 0.28 | 0.07 | 0.33 | 0.18 |
| 11 | 25966124 | rs7125344 | -0.899 | -0.206 | -0.055 | -1.225 | -0.002 | 0.011 | 0.011 | 0 | 0.7 | 0.69 | 0.76 | 0.5 | 0.72 | 0.6 |
| 11 | 25967152 | rs1441494 | -1.466 | -0.202 | -0.075 | -1.207 | -0.001 | 0.01 | 0.01 | 0 | 0.76 | 0.75 | 0.78 | 0.51 | 0.75 | 0.62 |
| 11 | 25973247 | rs12275759 | 0.719 | -0.188 | -0.127 | -1.278 | -0.004 | -0.03 | -0.03 | 0 | 0.21 | 0.22 | 0.13 | 0.06 | 0.17 | 0.19 |
| 11 | 25973647 | rs11029042 | 0.247 | -0.403 | -0.141 | -1.304 | -0.021 | 0.127 | 0.127 | 0 | 0.39 | 0.17 | 0.33 | 0.06 | 0.44 | 0.24 |
| 11 | 25975003 | rs7111070 | 0.82 | -0.394 | 0.068 | -1.131 | -0.004 | -0.011 | -0.011 | 0 | 0.26 | 0.24 | 0.39 | 0.42 | 0.27 | 0.28 |
| 11 | 25978136 | rs7931950 | 1.132 | -0.55 | 0.059 | -1.108 | -0.012 | 0.02 | 0.02 | 0 | 0.22 | 0.17 | 0.24 | 0.04 | 0.3 | 0.13 |
| 11 | 25982048 | rs328466 | 0.573 | -0.461 | 0.279 | -0.949 | -0.009 | 0.029 | 0.029 | 0 | 0.29 | 0.61 | 0.27 | 0.52 | 0.24 | 0.45 |
| 11 | 25983215 | rs328467 | 1.036 | -0.453 | 0.157 | -0.942 | -0.007 | 0.084 | 0.084 | 0 | 0.26 | 0.41 | 0.23 | 0.49 | 0.23 | 0.4 |
| 11 | 25983309 | rs328468 | 1.114 | -0.441 | -0.253 | -0.934 | -0.001 | 0.013 | 0.013 | 0 | 0.25 | 0.27 | 0.22 | 0.49 | 0.23 | 0.4 |
| 11 | 25989302 | rs10767478 | 0.223 | -0.401 | -0.427 | -1.002 | 0.003 | 0.207 | 0.207 | 0 | 0.46 | 0.11 | 0.51 | 0.19 | 0.58 | 0.36 |
| 11 | 26009964 | rs12277478 | -0.057 | -0.56 | -0.202 | -0.895 | 0 | 0.032 | 0.032 | 0 | 0.08 | 0.34 | 0.06 | 0.04 | 0.03 | 0.06 |
| 11 | 26013925 | rs7928222 | 1.051 | -0.559 | -0.312 | -0.968 | -0.001 | 0.013 | 0.013 | 0 | 0.25 | 0.27 | 0.22 | 0.49 | 0.23 | 0.41 |
| 11 | 26014763 | rs724027 | 1.214 | -0.252 | -1.242 | -1.244 | -0.08 | 0.104 | 0.104 | 0 | 0.21 | 0.02 | 0.21 | 0.04 | 0.29 | 0.13 |
| 11 | 26015219 | rs11029066 | -0.038 | -0.851 | -0.518 | -0.527 | -0.001 | 0 | 0 | 0 | 0.34 | 0.63 | 0.28 | 0.53 | 0.27 | 0.48 |
| 11 | 26018501 | rs12360762 | 0.048 | 0.227 | -1.11 | -0.815 | 0.021 | 0.191 | 0.191 | 0 | 0.46 | 0.17 | 0.52 | 0.18 | 0.57 | 0.38 |
| 11 | 26027163 | rs10767484 | -0.589 | 0.584 | -0.137 | 0.36 | -0.007 | 0.093 | 0.093 | 0 | 0.1 | 0.15 | 0.19 | 0.29 | 0.07 | 0.26 |
| 11 | 26027235 | rs1542715 | -0.606 | 0.773 | -0.693 | 0.412 | -0.061 | -0.149 | -0.149 | 0 | 0.21 | 0.03 | 0.26 | 0.4 | 0.24 | 0.24 |
| 11 | 26027540 | rs1542714 | -1.283 | 0.581 | -0.579 | 0.228 | -0.006 | -0.101 | -0.101 | 0.07 | 0.3 | 0.09 | 0.34 | 0.47 | 0.3 | 0.35 |
| 11 | 26035814 | rs10834882 | -0.276 | 0.679 | -0.244 | 0.338 | -0.008 | 0.134 | 0.134 | 0 | 0.1 | 0.18 | 0.2 | 0.29 | 0.07 | 0.27 |
| 11 | 26038634 | rs1532107 | 1.933 | 0.616 | 0.178 | 0.096 | 0.013 | 0.055 | 0.055 | 0 | 0.54 | 0.39 | 0.41 | 0.2 | 0.56 | 0.36 |
| 11 | 26038760 | rs1387487 | 0.228 | 0.458 | -0.279 | 0.15 | 0.017 | 0.02 | 0.02 | 0.353 | 0.1 | 0.06 | 0.08 | 0.04 | 0.08 | 0.12 |
| 11 | 26041706 | rs10501033 | 2.085 | 0.805 | -0.222 | 0.128 | -0.001 | 0.016 | 0.016 | 0 | 0.52 | 0.45 | 0.4 | 0.2 | 0.51 | 0.33 |
| 11 | 26041936 | rs7126809 | -0.873 | 1.264 | -0.251 | 0.483 | -0.12 | -0.161 | -0.161 | 0.788 | 0.23 | 0.02 | 0.28 | 0.43 | 0.31 | 0.26 |
| 11 | 26044992 | rs12363688 | 0.878 | 1.267 | -0.118 | -0.7 | 0.024 | -0.039 | -0.039 | 0 | 0.65 | 0.53 | 0.49 | 0.25 | 0.6 | 0.48 |
| 11 | 26045487 | rs10834893 | -0.652 | 0.399 | -0.765 | -0.299 | 0.006 | -0.01 | -0.01 | 1.028 | 0.23 | 0.11 | 0.4 | 0.7 | 0.19 | 0.47 |
| 11 | 26045601 | rs7932614 | -0.657 | 0.715 | -0.523 | -0.499 | -0.001 | -0.023 | -0.023 | 0 | 0.43 | 0.44 | 0.61 | 0.81 | 0.4 | 0.63 |
| 11 | 26061440 | rs2130234 | 0.789 | 0.587 | -0.279 | -0.405 | -0.003 | -0.058 | -0.058 | 0 | 0.81 | 0.74 | 0.88 | 0.96 | 0.84 | 0.88 |
| 11 | 26070407 | rs2349798 | 0.768 | 0.507 | 0.403 | -0.452 | -0.003 | -0.087 | -0.087 | 0.082 | 0.83 | 0.61 | 0.89 | 0.95 | 0.86 | 0.9 |
| 11 | 26079373 | rs7927115 | 0.764 | 0.505 | 0.402 | -0.453 | -0.002 | -0.087 | -0.087 | 0.784 | 0.83 | 0.6 | 0.89 | 0.95 | 0.86 | 0.9 |
| 11 | 26087226 | rs7930886 | 0.448 | 0.511 | 0.595 | -0.492 | 0 | -0.011 | -0.011 | 0.022 | 0.94 | 0.78 | 0.98 | 0.99 | 0.99 | 0.93 |
| 11 | 26091038 | rs11029127 | -0.889 | 0.466 | 0.476 | -0.498 | 0.006 | -0.041 | -0.041 | 0 | 0.12 | 0.18 | 0.09 | 0.03 | 0.13 | 0.03 |
| 11 | 26093556 | rs17242991 | -0.957 | 0.403 | 0.455 | -0.53 | 0.004 | -0.049 | -0.049 | 0 | 0.12 | 0.14 | 0.08 | 0 | 0.12 | 0.03 |
| 11 | 26095784 | rs11029134 | 0.915 | 0.385 | -0.843 | -0.435 | -0.091 | -0.055 | -0.055 | 0 | 0.37 | 0.02 | 0.46 | 0.32 | 0.42 | 0.38 |
| 11 | 26117392 | rs7117360 | 0.553 | 0.372 | -0.081 | 0.057 | 0.007 | 0.002 | 0.002 | 0 | 0.3 | 0.34 | 0.3 | 0.34 | 0.35 | 0.4 |
| 11 | 26143738 | rs3108814 | -0.888 | 0.24 | -0.122 | 0.182 | 0.009 | 0.083 | 0.083 | 0.058 | 0.51 | 0.8 | 0.42 | 0.36 | 0.51 | 0.52 |
| 11 | 26160068 | rs4391777 | -1.516 | 0.261 | 0.352 | 0.414 | -0.019 | 0.013 | 0.013 | 0 | 0.1 | 0.34 | 0.05 | 0.09 | 0.04 | 0.14 |
| 11 | 26172770 | rs4072100 | 0.351 | 0.71 | -0.348 | 0.492 | -0.08 | -0.015 | -0.015 | 0 | 0.18 | 0.04 | 0.19 | 0.18 | 0.28 | 0.24 |
| 11 | 26178913 | rs3108813 | -0.29 | 0.502 | -0.392 | 1.191 | -0.018 | -0.048 | -0.048 | 0 | 0.92 | 1 | 0.9 | 0.72 | 0.95 | 0.9 |
| 11 | 26188592 | rs12786982 | -0.922 | 0.583 | -0.119 | 0.329 | -0.01 | 0.003 | 0.003 | 0 | 0.31 | 0.26 | 0.26 | 0.31 | 0.39 | 0.33 |
| 11 | 26221061 | rs11601570 | -0.797 | 0.626 | -0.097 | 0.316 | -0.019 | 0.019 | 0.019 | 0 | 0.3 | 0.24 | 0.25 | 0.19 | 0.37 | 0.29 |
| 11 | 26244793 | rs11029398 | -0.903 | 0.613 | -0.098 | 0.341 | -0.007 | 0.002 | 0.002 | 0.033 | 0.31 | 0.28 | 0.26 | 0.31 | 0.39 | 0.34 |
| 11 | 26249239 | rs11605752 | -0.774 | 0.625 | -0.379 | 0.236 | -0.061 | 0.053 | 0.053 | 0 | 0.3 | 0.15 | 0.25 | 0.18 | 0.37 | 0.29 |
| 11 | 26258632 | rs17243196 | 0.074 | 0.875 | -0.421 | 0.283 | -0.099 | -0.016 | -0.016 | 2.993 | 0.21 | 0.06 | 0.21 | 0.18 | 0.32 | 0.27 |
| 11 | 26272253 | rs2045702 | -0.855 | 0.832 | -0.084 | 0.124 | -0.012 | 0.012 | 0.012 | 0 | 0.29 | 0.27 | 0.25 | 0.15 | 0.35 | 0.29 |

| | | | | | | | | | | | | | | | | |
|---|---|---|---|---|---|---|---|---|---|---|---|---|---|---|---|---|
| 11 | 26276143 | rs12224061 | -0.868 | 0.761 | -0.044 | 0.146 | 0.002 | -0.001 | -0.001 | 0 | 0.31 | 0.36 | 0.26 | 0.27 | 0.37 | 0.39 |
| 11 | 26287657 | rs7105867 | -1.799 | 0.538 | 0.739 | 0.133 | 0.011 | -0.027 | -0.027 | 0 | 0.18 | 0.48 | 0.16 | 0.13 | 0.2 | 0.1 |
| 11 | 26300330 | rs10834940 | -1.001 | 0.588 | 0.721 | 0.13 | 0.037 | 0.019 | 0.019 | 0 | 0.11 | 0.3 | 0.11 | 0.12 | 0.16 | 0.06 |
| 11 | 26303474 | rs7930530 | -1.673 | 0.549 | 0.753 | 0.105 | 0.051 | 0.006 | 0.006 | 0 | 0.14 | 0.47 | 0.14 | 0.12 | 0.17 | 0.08 |
| 11 | 26303605 | rs7945497 | -1.308 | 0.335 | 0.872 | 0.282 | 0.02 | 0.095 | 0.095 | 0 | 0.07 | 0.27 | 0.05 | 0.12 | 0.06 | 0.06 |
| 11 | 26344166 | rs2350208 | -0.845 | 0.26 | 0.129 | -0.084 | 0 | -0.004 | -0.004 | 0.005 | 0.43 | 0.53 | 0.34 | 0.27 | 0.47 | 0.36 |
| 11 | 26353121 | rs7117246 | 0.949 | 0.061 | -0.713 | 0.429 | -0.039 | -0.072 | -0.072 | 0 | 0.43 | 0.06 | 0.56 | 0.45 | 0.45 | 0.5 |
| 11 | 26353951 | rs17243259 | 0.191 | 0.303 | -0.536 | 0.452 | -0.01 | 0.019 | 0.019 | 0.349 | 0.31 | 0.21 | 0.21 | 0.16 | 0.31 | 0.3 |
| 11 | 26355874 | rs11029525 | 1.075 | -0.137 | -1.129 | 0.156 | -0.05 | -0.096 | -0.096 | 0 | 0.31 | 0.11 | 0.52 | 0.41 | 0.37 | 0.43 |
| 11 | 26356083 | rs1381186 | 1.304 | 0.026 | -1.254 | -0.082 | -0.025 | -0.094 | -0.094 | 0 | 0.27 | 0.11 | 0.48 | 0.41 | 0.3 | 0.4 |
| 11 | 26357170 | rs7103837 | 0.511 | 0.012 | -0.919 | -0.291 | 0.013 | -0.01 | -0.01 | 0 | 0.37 | 0.43 | 0.29 | 0.33 | 0.39 | 0.32 |
| 11 | 26357364 | rs4922755 | 0.103 | -0.02 | -0.966 | -0.271 | -0.026 | 0.034 | 0.034 | 0 | 0.09 | 0.05 | 0.06 | 0.04 | 0.12 | 0.06 |
| 11 | 26359576 | rs2045705 | -0.688 | -0.105 | -0.847 | -0.332 | 0 | 0.01 | 0.01 | 0.13 | 0.25 | 0.22 | 0.18 | 0.15 | 0.25 | 0.34 |
| 11 | 26360665 | rs12793737 | 0.657 | 0.186 | -0.945 | -0.054 | 0.007 | 0 | 0 | 0 | 0.31 | 0.38 | 0.23 | 0.32 | 0.3 | 0.2 |
| 11 | 26361216 | rs1015267 | -0.335 | 0.133 | -1.129 | -0.148 | -0.016 | -0.033 | -0.033 | 0 | 0.37 | 0.43 | 0.25 | 0.26 | 0.32 | 0.36 |
| 11 | 26361574 | rs10834964 | 0.264 | 0.22 | -1.263 | -0.128 | 0 | -0.001 | -0.001 | 0 | 0.35 | 0.37 | 0.27 | 0.35 | 0.37 | 0.22 |
| 11 | 26366279 | rs4923350 | 0.667 | -0.053 | -0.741 | -0.143 | -0.006 | -0.024 | -0.024 | 0 | 0.58 | 0.49 | 0.7 | 0.74 | 0.61 | 0.6 |
| 11 | 26376106 | rs11605942 | -0.835 | -0.267 | -0.748 | 0.175 | -0.005 | -0.002 | -0.002 | 0 | 0.2 | 0.14 | 0.1 | 0.25 | 0.13 | 0.25 |
| 11 | 26380169 | rs10834970 | 0.371 | -0.255 | -0.924 | 0.044 | 0.022 | 0.05 | 0.05 | 0 | 0.52 | 0.33 | 0.56 | 0.63 | 0.57 | 0.47 |
| 11 | 26380805 | rs10834971 | -0.675 | -0.27 | -0.833 | 0.752 | 0.007 | -0.016 | -0.016 | 0 | 0.14 | 0.12 | 0.06 | 0.21 | 0.08 | 0.25 |
| 11 | 26380992 | rs2349814 | -0.275 | -0.266 | -0.376 | 0.706 | -0.007 | 0.027 | 0.027 | 0 | 0.15 | 0.19 | 0.12 | 0.3 | 0.13 | 0.15 |
| 11 | 26381742 | rs4923351 | 0.663 | -0.097 | -0.706 | 0.209 | 0.018 | 0.03 | 0.03 | 0 | 0.49 | 0.17 | 0.59 | 0.55 | 0.6 | 0.4 |
| 11 | 26382402 | rs7950410 | 0.266 | 0.587 | 0.376 | -0.068 | 0.016 | -0.044 | -0.044 | 0.584 | 0.87 | 0.76 | 0.8 | 0.98 | 0.83 | 0.94 |
| 11 | 26390566 | rs716701 | 1.661 | 0.532 | 0.366 | -0.143 | -0.008 | 0.011 | 0.011 | 0.685 | 0.62 | 0.29 | 0.61 | 0.67 | 0.6 | 0.6 |
| 11 | 26395548 | rs922246 | -0.412 | 0.503 | 0.549 | -0.168 | -0.002 | 0 | 0 | 0 | 0.17 | 0.12 | 0.1 | 0.18 | 0.19 | 0.28 |
| 11 | 26398012 | rs12806876 | 0.382 | 0.524 | 0.622 | -0.213 | -0.003 | 0.002 | 0.002 | 0 | 0.09 | 0.09 | 0.19 | 0.02 | 0.14 | 0.06 |
| 11 | 26407220 | rs12283791 | -1.574 | 0.108 | 1.139 | -0.092 | -0.006 | 0.024 | 0.024 | 0 | 0.25 | 0.43 | 0.17 | 0.3 | 0.23 | 0.34 |
| 11 | 26410587 | rs7480892 | -1.586 | 0.108 | 1.132 | -0.092 | -0.006 | 0.024 | 0.024 | 0 | 0.25 | 0.43 | 0.17 | 0.3 | 0.23 | 0.34 |
| 11 | 26427359 | rs7102859 | 1.457 | 0.072 | 0.808 | -0.149 | 0.001 | 0.003 | 0.003 | 0 | 0.71 | 0.27 | 0.81 | 0.72 | 0.73 | 0.72 |
| 11 | 26438486 | rs7113405 | -1.847 | 0.035 | 1.079 | -0.164 | -0.001 | -0.001 | -0.001 | 0 | 0.28 | 0.64 | 0.18 | 0.28 | 0.25 | 0.37 |
| 11 | 26442126 | rs2126183 | -1.477 | 0.005 | 1.53 | -0.164 | 0.008 | -0.025 | -0.025 | 0 | 0.24 | 0.37 | 0.15 | 0.18 | 0.24 | 0.34 |
| 11 | 26442211 | rs2169339 | -0.547 | -0.002 | 1.426 | -0.168 | 0.001 | -0.001 | -0.001 | 0 | 0.17 | 0.16 | 0.1 | 0.18 | 0.19 | 0.31 |
| 11 | 26446191 | rs10834976 | 2.177 | 0.007 | 1.569 | -0.028 | -0.019 | 0.021 | 0.021 | 1.713 | 0.87 | 0.41 | 0.91 | 0.89 | 0.93 | 0.94 |
| 11 | 26448386 | rs4394800 | -2.346 | 0.027 | 1.266 | -0.011 | 0 | 0.056 | 0.056 | 1.07 | 0.11 | 0.48 | 0.08 | 0.11 | 0.06 | 0.06 |
| 11 | 26489594 | rs1493739 | 1.347 | 0.101 | 1.12 | -0.183 | 0.01 | -0.003 | -0.003 | 0 | 0.68 | 0.53 | 0.68 | 0.68 | 0.62 | 0.53 |
| 11 | 26492563 | rs11029574 | -1.242 | 0.15 | 1.22 | -0.092 | -0.023 | 0.013 | 0.013 | 0.496 | 0.26 | 0.15 | 0.26 | 0.2 | 0.32 | 0.42 |
| 11 | 26499535 | rs11029577 | -1.36 | 0.129 | 1.385 | -0.105 | -0.004 | 0.004 | 0.004 | 0.435 | 0.26 | 0.23 | 0.26 | 0.2 | 0.33 | 0.42 |
| 11 | 26499618 | rs1355130 | 0.142 | 0.155 | 1.226 | -0.132 | -0.031 | 0.014 | 0.014 | 0 | 0.09 | 0.01 | 0.17 | 0.04 | 0.12 | 0.07 |
| 11 | 26500028 | rs4505017 | -1.413 | 0.072 | 1.408 | -0.169 | 0.003 | 0.067 | 0.067 | 0 | 0.07 | 0.2 | 0.05 | 0.13 | 0.06 | 0.05 |
| 11 | 26500273 | rs1509170 | 2.169 | 0.121 | 1.496 | -0.161 | -0.002 | 0.006 | 0.006 | 0.201 | 0.62 | 0.29 | 0.63 | 0.66 | 0.6 | 0.52 |
| 11 | 26514931 | rs1995800 | 1.031 | 0.345 | 1.304 | -0.09 | 0.014 | 0.047 | 0.047 | 0.009 | 0.78 | 0.63 | 0.84 | 0.69 | 0.76 | 0.6 |
| 11 | 26516176 | rs11029582 | -0.629 | 0.371 | 1.265 | -0.089 | -0.007 | 0 | 0 | 0.007 | 0.16 | 0.11 | 0.11 | 0.16 | 0.21 | 0.3 |
| 11 | 26516202 | rs11029583 | -0.629 | 0.371 | 1.265 | -0.089 | -0.007 | 0 | 0 | 0.007 | 0.16 | 0.11 | 0.11 | 0.16 | 0.21 | 0.3 |
| 11 | 26521821 | rs7929544 | 1.418 | 0.436 | 0.963 | -0.06 | 0.003 | -0.003 | -0.003 | 0 | 0.66 | 0.44 | 0.68 | 0.68 | 0.64 | 0.56 |
| 11 | 26527171 | rs10767545 | 1.208 | 0.523 | 0.869 | -0.131 | 0.011 | -0.041 | -0.041 | 0.227 | 0.68 | 0.48 | 0.69 | 0.79 | 0.64 | 0.59 |
| 11 | 26530798 | rs17243350 | 0.452 | 0.491 | 0.542 | -0.197 | -0.036 | 0.062 | 0.062 | 0 | 0.11 | 0 | 0.16 | 0.01 | 0.14 | 0.04 |
| 11 | 26538774 | rs7126899 | 1.576 | 0.52 | 0.712 | -0.211 | 0.002 | -0.012 | -0.012 | 0 | 0.67 | 0.38 | 0.67 | 0.79 | 0.63 | 0.61 |
| 11 | 26543031 | rs10834998 | -0.059 | 0.366 | 0.244 | -0.114 | -0.016 | -0.108 | -0.108 | 0 | 0.14 | 0.01 | 0.25 | 0.53 | 0.12 | 0.11 |
| 11 | 26544374 | rs372878 | 0.848 | 0.607 | 0.799 | -0.429 | 0.037 | 0.002 | 0.002 | 0 | 0.82 | 0.57 | 0.85 | 0.8 | 0.79 | 0.65 |
| 11 | 26554127 | rs4923362 | 1.083 | 1.065 | 0.197 | -0.815 | 0.007 | -0.036 | -0.036 | 0 | 0.65 | 0.47 | 0.74 | 0.8 | 0.6 | 0.62 |
| 11 | 26555141 | rs293939 | 0.657 | 0.593 | 0.615 | -0.835 | 0.007 | -0.048 | -0.048 | 0 | 0.65 | 0.55 | 0.75 | 0.83 | 0.62 | 0.68 |
| 11 | 26558873 | rs375562 | -0.373 | 0.52 | 0.667 | -0.871 | 0.002 | -0.043 | -0.043 | 0 | 0.1 | 0.19 | 0.07 | 0.03 | 0.08 | 0.12 |
| 11 | 26567487 | rs2703403 | 0.123 | 0.437 | 0.592 | -0.923 | 0.011 | -0.045 | -0.045 | 2.032 | 0.69 | 0.55 | 0.7 | 0.81 | 0.66 | 0.69 |
| 11 | 26568966 | rs2663168 | 0.123 | 0.437 | 0.592 | -0.923 | 0.011 | -0.045 | -0.045 | 0 | 0.69 | 0.55 | 0.71 | 0.81 | 0.67 | 0.69 |
| 11 | 26575491 | rs293969 | 0.578 | 0.341 | 0.42 | -0.812 | 0 | -0.014 | -0.014 | 0 | 0.78 | 0.75 | 0.82 | 0.84 | 0.79 | 0.8 |
| 11 | 26576191 | rs12418975 | 1.29 | 0.273 | 0.344 | -0.929 | -0.011 | 0.083 | 0.083 | 0 | 0.48 | 0.27 | 0.55 | 0.65 | 0.51 | 0.48 |
| 11 | 26576725 | rs10501053 | 0.246 | 0.255 | 0.484 | -0.855 | -0.001 | -0.011 | -0.011 | 0 | 0.27 | 0.29 | 0.26 | 0.17 | 0.26 | 0.28 |
| 11 | 26576968 | rs17243440 | 0.595 | 0.346 | 0.387 | -0.789 | -0.003 | 0.01 | 0.01 | 0 | 0.23 | 0.15 | 0.17 | 0.15 | 0.25 | 0.27 |
| 11 | 26579493 | rs293966 | 0.568 | 0.543 | 0.149 | -0.752 | 0.04 | -0.011 | -0.011 | 0.382 | 0.92 | 0.72 | 0.93 | 0.97 | 0.93 | 0.87 |
| 11 | 26580120 | rs17309195 | -0.136 | 0.605 | -0.462 | -0.596 | -0.029 | -0.01 | -0.01 | 0 | 0.14 | 0.02 | 0.11 | 0.15 | 0.13 | 0.08 |
| 11 | 26584445 | rs381348 | -0.324 | 0.258 | 0.009 | 0.55 | -0.006 | 0.021 | 0.021 | 0 | 0.68 | 0.67 | 0.8 | 0.48 | 0.73 | 0.66 |
| 11 | 26584625 | rs444178 | 0.29 | 0.234 | 0.058 | 0.472 | -0.005 | 0.01 | 0.01 | 0 | 0.62 | 0.57 | 0.75 | 0.44 | 0.66 | 0.56 |
| 11 | 26585014 | rs502093 | 0.27 | 0.242 | 0.002 | 0.476 | -0.004 | 0.007 | 0.007 | 0 | 0.62 | 0.62 | 0.75 | 0.44 | 0.66 | 0.56 |
| 11 | 26587133 | rs293980 | -0.359 | 0.294 | 0.028 | 0.553 | -0.001 | 0.002 | 0.002 | 0 | 0.68 | 0.71 | 0.8 | 0.48 | 0.73 | 0.66 |
| 11 | 26590126 | rs293974 | 0.053 | 0.112 | -0.123 | 1.039 | 0.008 | -0.001 | -0.001 | 0 | 0.32 | 0.19 | 0.19 | 0.3 | 0.28 | 0.4 |
| 11 | 26592361 | rs10430847 | 0.633 | 0.26 | -0.489 | 1.597 | -0.021 | -0.036 | -0.036 | 0 | 0.08 | 0.01 | 0.05 | 0.35 | 0.09 | 0.29 |
| 11 | 26592685 | rs293971 | -0.19 | 0.124 | 0.141 | 1.385 | 0.002 | -0.011 | -0.011 | 0 | 0.95 | 0.8 | 0.87 | 0.98 | 0.98 | 0.98 |
| 11 | 26596402 | rs2131691 | 0.226 | 0.279 | -0.359 | 0.761 | 0.001 | 0.003 | 0.003 | 0 | 0.53 | 0.52 | 0.38 | 0.54 | 0.53 | 0.63 |
| 11 | 26599911 | rs7119386 | 0.835 | 0.252 | -0.611 | 0.49 | 0 | 0.004 | 0.004 | 0 | 0.36 | 0.34 | 0.26 | 0.39 | 0.36 | 0.48 |

| | | | | | | | | | | | | | | | | |
|---|---|---|---|---|---|---|---|---|---|---|---|---|---|---|---|---|
| 11 | 26600371 | rs293983 | 0.374 | 0.782 | -1.406 | 0.279 | -0.017 | 0.021 | 0.021 | 0 | 0.25 | 0.13 | 0.37 | 0.17 | 0.31 | 0.24 |
| 11 | 26603926 | rs12793968 | 0.856 | 0.737 | -1.072 | 0.409 | -0.044 | -0.004 | -0.004 | 0 | 0.15 | 0.01 | 0.11 | 0.13 | 0.16 | 0.14 |
| 11 | 26605331 | rs12295638 | 0.064 | 0.682 | -0.745 | 0.8 | -0.005 | -0.003 | -0.003 | 2.206 | 0.11 | 0.1 | 0.09 | 0.34 | 0.09 | 0.08 |
| 11 | 26606690 | rs211066 | -0.937 | 0.272 | -0.312 | 0.322 | -0.023 | -0.025 | -0.025 | 0 | 0.26 | 0.4 | 0.34 | 0.13 | 0.18 | 0.21 |
| 11 | 26608497 | rs7945560 | 0.984 | 0.928 | -0.353 | 0.14 | 0 | 0.005 | 0.005 | 0 | 0.13 | 0.12 | 0.15 | 0.33 | 0.24 | 0.11 |
| 11 | 26611665 | rs11606418 | 1.412 | 0.937 | -0.367 | 0.13 | -0.015 | -0.019 | -0.019 | 0 | 0.11 | 0.08 | 0.13 | 0.32 | 0.21 | 0.09 |
| 11 | 26611796 | rs10767555 | 0.795 | 0.871 | -0.32 | -0.477 | 0.012 | 0.007 | 0.007 | 0 | 0.49 | 0.44 | 0.3 | 0.39 | 0.41 | 0.52 |
| 11 | 26619658 | rs10835008 | 0.073 | 0.801 | -0.293 | -0.543 | -0.016 | -0.049 | -0.049 | 0.895 | 0.15 | 0.22 | 0.09 | 0.04 | 0.08 | 0.07 |
| 11 | 26625685 | rs11029633 | -0.262 | 0.982 | -0.382 | -0.637 | 0 | 0.026 | 0.026 | 0 | 0.32 | 0.23 | 0.2 | 0.18 | 0.27 | 0.26 |
| 11 | 26635432 | rs293934 | -1.636 | 0.984 | 0.023 | -0.617 | 0.002 | -0.022 | -0.022 | 0 | 0.9 | 0.84 | 0.84 | 0.97 | 0.92 | 0.93 |
| 11 | 26636926 | rs10835019 | 0.285 | 0.942 | 0.137 | -0.608 | 0 | -0.017 | -0.017 | 1.931 | 0.77 | 0.53 | 0.53 | 0.86 | 0.8 | 0.8 |
| 11 | 26642580 | rs294006 | 1.244 | 1.008 | 0.103 | -0.532 | 0.007 | 0.033 | 0.033 | 0 | 0.1 | 0.18 | 0.27 | 0.11 | 0.08 | 0.12 |
| 11 | 26642796 | rs11029648 | 1.895 | 1.041 | -0.273 | -0.488 | 0.002 | -0.07 | -0.07 | 0 | 0.34 | 0.08 | 0.19 | 0.36 | 0.33 | 0.45 |
| 11 | 26642975 | rs7930308 | 1.26 | 1.09 | -0.326 | -0.494 | -0.012 | 0.051 | 0.051 | 0.055 | 0.45 | 0.29 | 0.33 | 0.68 | 0.53 | 0.54 |
| 11 | 26643186 | rs7107869 | 1.649 | 0.931 | -0.273 | -0.389 | 0.018 | -0.022 | -0.022 | 1.524 | 0.34 | 0.22 | 0.21 | 0.36 | 0.33 | 0.45 |
| 11 | 26647318 | rs10767563 | -1.13 | 0.423 | 0.01 | -0.096 | 0.003 | 0.029 | 0.029 | 0 | 0.94 | 0.93 | 0.74 | 0.89 | 0.94 | 0.89 |
| 11 | 26648185 | rs1389455 | 0.034 | 0.402 | 0.061 | -0.174 | 0.011 | 0.021 | 0.021 | 0 | 0.83 | 0.63 | 0.57 | 0.86 | 0.86 | 0.82 |
| 11 | 26655658 | rs7943086 | 1.78 | 0.566 | -0.375 | -0.376 | -0.003 | 0.002 | 0.002 | 0.032 | 0.41 | 0.25 | 0.27 | 0.36 | 0.42 | 0.49 |
| 11 | 26656563 | rs7946841 | 1.78 | 0.566 | -0.375 | -0.376 | -0.003 | 0.002 | 0.002 | 0.032 | 0.41 | 0.23 | 0.25 | 0.36 | 0.42 | 0.47 |
| 11 | 26659406 | rs6484224 | 1.769 | 0.559 | -0.375 | -0.376 | -0.002 | 0.003 | 0.003 | 0.034 | 0.41 | 0.23 | 0.25 | 0.36 | 0.42 | 0.47 |
| 11 | 26661690 | rs7110072 | -0.796 | 0.555 | -0.112 | -0.326 | 0.054 | 0.143 | 0.143 | 0 | 0.93 | 0.72 | 0.73 | 0.9 | 0.94 | 0.89 |
| 11 | 26665695 | rs17243517 | 1.249 | 0.537 | -0.442 | -0.35 | -0.023 | -0.03 | -0.03 | 0.627 | 0.1 | 0.01 | 0.08 | 0.14 | 0.09 | 0.17 |
| 11 | 26666902 | rs4923372 | -1.054 | 0.48 | -0.293 | -0.442 | 0 | -0.001 | -0.001 | 5.04 | 0.37 | 0.39 | 0.39 | 0.51 | 0.41 | 0.34 |
| 11 | 26667604 | rs2298 | 1.188 | 0.45 | -0.249 | -0.443 | 0.002 | 0.004 | 0.004 | 0.3 | 0.6 | 0.44 | 0.42 | 0.68 | 0.69 | 0.58 |
| 11 | 26669844 | rs7928737 | 1.178 | 0.45 | -0.261 | -0.443 | 0.002 | 0.004 | 0.004 | 0.271 | 0.6 | 0.44 | 0.42 | 0.68 | 0.69 | 0.58 |
| 11 | 26674033 | rs10734381 | -0.035 | 0.423 | -0.233 | -0.467 | 0.03 | 0.049 | 0.049 | 0 | 0.86 | 0.64 | 0.7 | 0.9 | 0.89 | 0.86 |
| 11 | 26674408 | rs7947969 | 1.079 | 0.441 | -0.335 | -0.503 | -0.01 | -0.038 | -0.038 | 0.448 | 0.64 | 0.31 | 0.52 | 0.53 | 0.64 | 0.71 |
| 11 | 26674513 | rs7934757 | 1.412 | 0.422 | -0.351 | -0.522 | -0.004 | -0.008 | -0.008 | 1.562 | 0.6 | 0.5 | 0.43 | 0.68 | 0.69 | 0.58 |
| 11 | 26674768 | rs6484237 | 2.007 | 0.423 | -0.442 | -0.522 | -0.02 | 0.002 | 0.002 | 0 | 0.43 | 0.19 | 0.28 | 0.36 | 0.47 | 0.48 |
| 11 | 26677947 | rs10835051 | 0.471 | 0.45 | -0.401 | -0.499 | -0.01 | -0.027 | -0.027 | 0.395 | 0.7 | 0.33 | 0.55 | 0.57 | 0.67 | 0.76 |
| 11 | 26682387 | rs3802753 | -0.009 | 0.461 | -0.484 | -0.506 | -0.008 | 0.019 | 0.019 | 0 | 0.2 | 0.12 | 0.3 | 0.17 | 0.19 | 0.24 |
| 11 | 26682434 | rs3802754 | 0.896 | 0.489 | -0.48 | -0.492 | 0.032 | 0.067 | 0.067 | 0 | 0.17 | 0.31 | 0.15 | 0.33 | 0.22 | 0.11 |
| 11 | 26684142 | rs17309300 | 1.44 | 0.493 | -0.627 | -0.503 | -0.007 | 0.015 | 0.015 | 0 | 0.09 | 0.04 | 0.21 | 0.03 | 0.09 | 0.07 |
| 11 | 26684617 | rs3550 | 2.207 | 0.405 | -0.803 | -0.515 | -0.092 | -0.076 | -0.076 | 0 | 0.41 | 0.03 | 0.24 | 0.36 | 0.43 | 0.46 |
| 11 | 26686075 | rs10501062 | 0.88 | 0.443 | -0.633 | -0.435 | 0.015 | 0.036 | 0.036 | 0 | 0.17 | 0.26 | 0.14 | 0.33 | 0.22 | 0.11 |
| 11 | 26686565 | rs10501063 | 1.193 | 0.392 | -0.756 | -0.441 | -0.025 | -0.029 | -0.029 | 0.027 | 0.1 | 0.01 | 0.08 | 0.14 | 0.09 | 0.17 |
| 11 | 26691664 | rs10835054 | 3.031 | 0.121 | -1.196 | -0.718 | 0.023 | 0.088 | 0.088 | 0.033 | 0.22 | 0.02 | 0.13 | 0.04 | 0.17 | 0.11 |
| 11 | 26691876 | rs7932573 | -0.588 | 0.512 | -0.003 | 0.049 | -0.001 | 0 | 0 | 0 | 0.27 | 0.28 | 0.22 | 0.47 | 0.31 | 0.28 |
| 11 | 26692435 | rs11029666 | 2.248 | 0.524 | 0.086 | 0.037 | 0.001 | 0.015 | 0.015 | 0 | 0.43 | 0.28 | 0.29 | 0.35 | 0.47 | 0.48 |
| 11 | 26692481 | rs7128239 | -0.624 | 0.567 | 0.011 | 0.142 | 0 | 0.001 | 0.001 | 0 | 0.26 | 0.28 | 0.22 | 0.47 | 0.31 | 0.28 |
| 11 | 26692733 | rs7114378 | -0.652 | 0.567 | 0.011 | 0.142 | -0.001 | 0 | 0 | 0 | 0.27 | 0.28 | 0.22 | 0.47 | 0.31 | 0.28 |
| 11 | 26692742 | rs7128823 | -0.613 | 0.574 | 0.014 | 0.146 | 0 | 0.001 | 0.001 | 0 | 0.26 | 0.28 | 0.22 | 0.47 | 0.32 | 0.28 |
| 11 | 26693686 | rs4312025 | -0.567 | 0.463 | 0.086 | 0.116 | -0.007 | -0.005 | -0.005 | 0.209 | 0.18 | 0.23 | 0.12 | 0.18 | 0.12 | 0.21 |
| 11 | 26694812 | rs4278475 | -0.564 | 0.548 | 0.37 | 0.188 | 0.048 | 0.131 | 0.131 | 0 | 0.93 | 0.75 | 0.81 | 0.89 | 0.95 | 0.88 |
| 11 | 26694842 | rs4551716 | 1.702 | 0.529 | 0.27 | 0.075 | 0.005 | 0.023 | 0.023 | 0 | 0.5 | 0.44 | 0.31 | 0.35 | 0.49 | 0.48 |
| 11 | 26695195 | rs12286201 | -0.491 | 0.578 | 0.278 | 0.153 | 0.002 | -0.053 | -0.053 | 0 | 0.09 | 0.28 | 0.06 | 0 | 0.06 | 0.02 |
| 11 | 26696343 | rs4244525 | 1.545 | 0.845 | 0.394 | 0.389 | -0.007 | -0.001 | -0.001 | 0.02 | 0.58 | 0.23 | 0.5 | 0.38 | 0.57 | 0.54 |
| 11 | 26697017 | rs10835056 | 1.102 | 0.938 | 0.41 | 0.499 | -0.004 | 0.014 | 0.014 | 0.228 | 0.72 | 0.51 | 0.62 | 0.72 | 0.76 | 0.64 |
| 11 | 26697300 | rs10835057 | 1.519 | 0.811 | 0.386 | 0.406 | 0.001 | 0.001 | 0.001 | 0 | 0.57 | 0.45 | 0.49 | 0.43 | 0.54 | 0.53 |
| 11 | 26698005 | rs12420230 | -0.537 | 0.577 | 0.326 | 0.422 | -0.021 | -0.052 | -0.052 | 0 | 0.13 | 0.02 | 0.23 | 0.13 | 0.09 | 0.14 |
| 11 | 26701094 | rs11029672 | 1.924 | 0.596 | 0.158 | 0.45 | 0.017 | -0.005 | -0.005 | 0 | 0.5 | 0.22 | 0.42 | 0.43 | 0.47 | 0.56 |
| 11 | 26701172 | rs10767567 | 0.712 | 0.6 | 0.232 | 0.652 | -0.01 | 0.025 | 0.025 | 0 | 0.85 | 0.75 | 0.76 | 0.87 | 0.91 | 0.87 |
| 11 | 26702222 | rs10767568 | 0.54 | 0.864 | 0.074 | 0.661 | 0 | 0 | 0 | 0 | 0.69 | 0.67 | 0.78 | 0.66 | 0.7 | 0.83 |
| 11 | 26708727 | rs10767570 | 1.195 | 0.734 | -0.151 | 0.527 | 0.012 | 0.018 | 0.018 | 0 | 0.55 | 0.32 | 0.37 | 0.35 | 0.63 | 0.53 |
| 11 | 26715618 | rs12270727 | -1.238 | 0.404 | 0.318 | 0.464 | 0 | 0.058 | 0.058 | 0 | 0.08 | 0.32 | 0.19 | 0.12 | 0.04 | 0.11 |
| 11 | 26716217 | rs10767571 | 1.723 | 0.71 | -0.157 | 0.459 | -0.104 | -0.048 | -0.048 | 1.178 | 0.48 | 0.05 | 0.32 | 0.39 | 0.54 | 0.49 |
| 11 | 26717345 | rs11822508 | -0.072 | 0.732 | 0.045 | 0.401 | 0.046 | -0.052 | -0.052 | 1.36 | 0.09 | 0.25 | 0.09 | 0 | 0.12 | 0.04 |
| 11 | 26720410 | rs4300357 | 1.228 | 0.527 | -0.141 | 0.423 | 0.014 | 0.02 | 0.02 | 0.15 | 0.77 | 0.21 | 0.72 | 0.82 | 0.84 | 0.82 |
| 11 | 26720959 | rs11029680 | -0.875 | 0.214 | 0.363 | 0.825 | 0 | 0.002 | 0.002 | 0 | 0.31 | 0.34 | 0.41 | 0.43 | 0.3 | 0.33 |
| 11 | 26722399 | rs10835061 | 1.884 | 0.118 | 0.252 | 1.215 | 0 | -0.008 | -0.008 | 0 | 0.56 | 0.05 | 0.39 | 0.3 | 0.67 | 0.55 |
| 11 | 26722523 | rs4477399 | -0.735 | 0.204 | 0.347 | 0.912 | -0.018 | -0.054 | -0.054 | 0.623 | 0.27 | 0.15 | 0.46 | 0.41 | 0.27 | 0.3 |
| 11 | 26730322 | rs4923381 | -0.038 | 0.222 | 0.246 | 0.981 | -0.073 | -0.047 | -0.047 | 0.132 | 0.18 | 0.02 | 0.31 | 0.26 | 0.2 | 0.19 |
| 11 | 26732923 | rs4322349 | -0.862 | 0.196 | 0.399 | 0.802 | -0.013 | -0.048 | -0.048 | 0.723 | 0.27 | 0.15 | 0.46 | 0.41 | 0.27 | 0.29 |
| 11 | 26736428 | rs10767574 | 0.536 | 0.142 | 0.767 | 0.782 | -0.009 | 0.061 | 0.061 | 0.023 | 0.91 | 0.4 | 0.9 | 0.87 | 0.96 | 0.9 |
| 11 | 26742602 | rs11029687 | 1.723 | 0.203 | -0.634 | 0.799 | -0.002 | -0.073 | -0.073 | 0 | 0.56 | 0.06 | 0.39 | 0.36 | 0.67 | 0.56 |
| 11 | 26742676 | rs12807847 | 0.869 | 0.033 | -0.396 | 0.88 | -0.006 | -0.001 | -0.001 | 0 | 0.84 | 0.21 | 0.85 | 0.78 | 0.94 | 0.88 |
| 11 | 26747108 | rs4328172 | 1.48 | 0.597 | -0.865 | 0.679 | -0.092 | -0.045 | -0.045 | 0 | 0.48 | 0.05 | 0.32 | 0.36 | 0.53 | 0.52 |
| 11 | 26749737 | rs7946266 | -1.528 | 0.651 | -0.025 | 0.855 | -0.001 | 0.003 | 0.003 | 0 | 0.38 | 0.46 | 0.52 | 0.43 | 0.34 | 0.44 |
| 11 | 26750200 | rs7936032 | -0.539 | 0.301 | 0.186 | 1.116 | 0.015 | -0.022 | -0.022 | 0 | 0.16 | 0.12 | 0.15 | 0.22 | 0.08 | 0.15 |

| | | | | | | | | | | | | | | | | |
|---|---|---|---|---|---|---|---|---|---|---|---|---|---|---|---|---|
| 11 | 26751042 | rs12274162 | -0.845 | 0.815 | 0.353 | 0.709 | 0.039 | 0.021 | 0.021 | 0 | 0.06 | 0.26 | 0.06 | 0.06 | 0.07 | 0.11 |
| 11 | 26756407 | rs10835063 | 0.177 | 0.531 | -1.31 | 0.929 | -0.034 | -0.147 | -0.147 | 2.031 | 0.21 | 0.01 | 0.2 | 0.41 | 0.22 | 0.3 |
| 11 | 26758401 | rs4539270 | 0.493 | 0.745 | -1.052 | 0.401 | 0 | -0.001 | -0.001 | 0 | 0.71 | 0.32 | 0.67 | 0.83 | 0.76 | 0.82 |
| 11 | 26761225 | rs4420216 | -0.91 | 0.826 | -1.024 | 0.502 | -0.017 | -0.035 | -0.035 | 0.016 | 0.22 | 0.16 | 0.46 | 0.56 | 0.27 | 0.29 |
| 11 | 26762851 | rs12098963 | 0.288 | 0.558 | -0.878 | 0.814 | 0.001 | 0.051 | 0.051 | 0 | 0.07 | 0.05 | 0.17 | 0.4 | 0.06 | 0.21 |
| 11 | 26763016 | rs10501065 | 0.418 | 0.412 | -1.213 | 0.186 | 0.025 | -0.014 | -0.014 | 3.579 | 0.58 | 0.22 | 0.69 | 0.57 | 0.67 | 0.52 |
| 11 | 26766933 | rs16916200 | 0.214 | 0.845 | -1.19 | 0.106 | 0.006 | 0.009 | 0.009 | 0 | 0.08 | 0.09 | 0.23 | 0.17 | 0.09 | 0.06 |
| 11 | 26767433 | rs7926655 | 0.173 | 0.723 | -1.153 | -0.026 | 0.001 | 0 | 0 | 0 | 0.12 | 0.1 | 0.04 | 0.05 | 0.08 | 0.14 |
| 11 | 26767952 | rs10501066 | 0.377 | 0.75 | -1.21 | 0.017 | 0 | 0.001 | 0.001 | 0 | 0.08 | 0.07 | 0.23 | 0.17 | 0.09 | 0.06 |
| 11 | 26770926 | rs10128713 | -0.548 | 0.739 | -0.998 | -0.026 | 0 | -0.003 | -0.003 | 0 | 0.37 | 0.63 | 0.48 | 0.55 | 0.32 | 0.39 |
| 11 | 26770983 | rs10128596 | 0.642 | 0.692 | -1.616 | 0.036 | 0.011 | -0.064 | -0.064 | 0 | 0.24 | 0.12 | 0.21 | 0.35 | 0.23 | 0.3 |
| 11 | 26771794 | rs10501069 | 0.609 | 0.677 | -1.665 | -0.017 | -0.024 | 0.031 | 0.031 | 0 | 0.5 | 0.13 | 0.46 | 0.4 | 0.58 | 0.47 |
| 11 | 26774813 | rs4922771 | 0.281 | 0.58 | -1.878 | 0.149 | -0.049 | -0.036 | -0.036 | 0.256 | 0.37 | 0.05 | 0.32 | 0.33 | 0.35 | 0.33 |
| 11 | 26782020 | rs10835070 | -0.643 | 0.357 | -0.867 | 0.648 | -0.003 | 0.02 | 0.02 | 0 | 0.15 | 0.19 | 0.16 | 0.3 | 0.13 | 0.3 |
| 11 | 26786121 | rs7945103 | 0.625 | 0.462 | -1.057 | 0.438 | 0.025 | 0.023 | 0.023 | 0.116 | 0.34 | 0.15 | 0.18 | 0.28 | 0.29 | 0.25 |
| 11 | 26792264 | rs1017602 | 0.145 | 0.638 | -0.035 | 0.351 | 0.114 | 0.092 | 0.092 | 0 | 0.94 | 0.72 | 0.77 | 0.89 | 0.91 | 0.92 |
| 11 | 26792747 | rs7935084 | -0.133 | 0.667 | -0.058 | 0.365 | 0.125 | 0.103 | 0.103 | 0 | 0.94 | 0.73 | 0.78 | 0.89 | 0.91 | 0.92 |
| 11 | 26793052 | rs1530778 | 0.787 | 0.594 | -0.424 | 0.234 | -0.148 | -0.022 | -0.022 | 0 | 0.38 | 0.03 | 0.38 | 0.28 | 0.44 | 0.34 |
| 11 | 26803448 | rs10767578 | 0.572 | 0.386 | 0.383 | 0.059 | 0.131 | 0.108 | 0.108 | 0.008 | 0.93 | 0.65 | 0.89 | 0.88 | 0.9 | 0.93 |
| 11 | 26815537 | rs10742160 | 0.75 | 0.374 | 0.462 | 0.053 | 0.131 | 0.107 | 0.107 | 0.009 | 0.93 | 0.64 | 0.89 | 0.88 | 0.9 | 0.93 |
| 11 | 26819928 | rs10835072 | 0.389 | 0.448 | 0.551 | 0.091 | 0.004 | 0.017 | 0.017 | 0 | 0.47 | 0.22 | 0.43 | 0.4 | 0.52 | 0.47 |
| 11 | 26822461 | rs12417428 | 0.218 | 0.301 | 0.37 | 0.2 | -0.013 | -0.069 | -0.069 | 0.831 | 0.1 | 0.01 | 0.29 | 0.27 | 0.1 | 0.18 |
| 11 | 26824992 | rs9633856 | 0.882 | 0.473 | 0.413 | -0.317 | -0.159 | -0.105 | -0.105 | 0 | 0.39 | 0.01 | 0.39 | 0.35 | 0.47 | 0.35 |
| 11 | 26835113 | rs10835074 | 1.08 | 0.157 | 0.673 | -0.443 | 0.001 | -0.041 | -0.041 | 0.312 | 0.68 | 0.25 | 0.5 | 0.44 | 0.68 | 0.65 |
| 11 | 26843292 | rs12363428 | 0.918 | 0.545 | 1.026 | -0.226 | 0.058 | 0.127 | 0.127 | 0.201 | 0.33 | 0.11 | 0.14 | 0.04 | 0.24 | 0.23 |
| 11 | 26850692 | rs2060651 | -0.71 | 0.685 | 1.505 | -0.148 | -0.003 | -0.003 | -0.003 | 0.22 | 0.57 | 0.64 | 0.57 | 0.57 | 0.46 | 0.6 |
| 11 | 26856571 | rs7122599 | 0.826 | 0.87 | 1.443 | -0.054 | -0.073 | -0.039 | -0.039 | 0 | 0.38 | 0.12 | 0.34 | 0.34 | 0.44 | 0.35 |
| 11 | 26857146 | rs4307673 | 0.327 | 0.964 | 1.747 | 0.043 | 0.019 | -0.006 | -0.006 | 0 | 0.94 | 0.85 | 0.93 | 0.95 | 0.95 | 0.89 |
| 11 | 26863615 | rs1545433 | 0.597 | 0.964 | 1.657 | 0.039 | 0.007 | 0.017 | 0.017 | 0 | 0.94 | 0.87 | 0.93 | 0.91 | 0.95 | 0.89 |
| 11 | 26874580 | rs7124231 | -0.1 | 0.937 | 1.393 | -0.118 | 0.03 | 0.101 | 0.101 | 0 | 0.47 | 0.57 | 0.34 | 0.21 | 0.41 | 0.41 |
| 11 | 26876833 | rs7102662 | 0.514 | 0.918 | 1.543 | -0.127 | 0.031 | 0.017 | 0.017 | 0.051 | 0.39 | 0.28 | 0.18 | 0.03 | 0.27 | 0.27 |
| 11 | 26880991 | rs1375979 | -0.031 | 1.035 | 1.178 | 0.059 | 0.001 | 0.05 | 0.05 | 0 | 0.82 | 0.66 | 0.66 | 0.57 | 0.83 | 0.73 |
| 11 | 26884756 | rs1375978 | 0.205 | 1.039 | 1.201 | 0.048 | 0.004 | 0.007 | 0.007 | 0 | 0.46 | 0.45 | 0.32 | 0.21 | 0.41 | 0.41 |
| 11 | 26889460 | rs12799405 | 0.94 | 1.087 | 0.921 | 0.081 | -0.102 | -0.076 | -0.076 | 0 | 0.34 | 0.07 | 0.32 | 0.36 | 0.42 | 0.34 |
| 11 | 26903063 | rs1375977 | -0.469 | 0.715 | 1.572 | -0.694 | 0.025 | 0.081 | 0.081 | 0 | 0.47 | 0.55 | 0.34 | 0.22 | 0.41 | 0.41 |
| 11 | 26936695 | rs12282988 | -3.016 | 1.464 | 2.83 | 0.53 | 0.033 | 0.023 | 0.023 | 0.108 | 0.14 | 0.58 | 0.19 | 0.18 | 0.15 | 0.17 |
| 11 | 26938505 | rs10835084 | -1.971 | 1.521 | 2.508 | 0.558 | 0.052 | 0.043 | 0.043 | 0 | 0.09 | 0.17 | 0.15 | 0.18 | 0.14 | 0.12 |
| 11 | 26958573 | rs12574167 | -2.093 | 1.247 | 2.46 | 0.593 | 0.092 | 0.101 | 0.101 | 0 | 0.1 | 0.29 | 0.13 | 0.18 | 0.12 | 0.11 |
| 11 | 26962816 | rs12279221 | -2.728 | 0.926 | 2.325 | 0.618 | 0.016 | 0.033 | 0.033 | 0 | 0.14 | 0.6 | 0.2 | 0.22 | 0.14 | 0.18 |
| 11 | 26970603 | rs959044 | -1.357 | 1.014 | 2.589 | 0.664 | 0.072 | 0.084 | 0.084 | 0 | 0.05 | 0.26 | 0.07 | 0.09 | 0.04 | 0.09 |
| 11 | 26986106 | rs12286947 | -1.39 | 0.956 | 2.293 | 0.65 | 0.057 | 0.067 | 0.067 | 0 | 0.06 | 0.27 | 0.09 | 0.12 | 0.08 | 0.06 |
| 11 | 27007293 | rs12362707 | -0.588 | 0.798 | 2.173 | 0.392 | -0.006 | -0.041 | -0.041 | 0 | 0.08 | 0.11 | 0.04 | 0 | 0.05 | 0.02 |
| 11 | 27014032 | rs7106676 | -1.968 | 1.016 | 2.188 | 0.795 | 0.068 | 0.109 | 0.109 | 0 | 0.08 | 0.43 | 0.11 | 0.11 | 0.06 | 0.14 |
| 11 | 27015219 | rs16916399 | -0.879 | 1.045 | 1.361 | 1.181 | -0.025 | -0.061 | -0.061 | 0 | 0.1 | 0.01 | 0.16 | 0.29 | 0.09 | 0.15 |
| 11 | 27018019 | rs2350934 | -2.269 | 0.77 | 1.314 | 0.445 | -0.005 | 0.008 | 0.008 | 0 | 0.16 | 0.65 | 0.2 | 0.2 | 0.13 | 0.21 |
| 11 | 27018626 | rs11029769 | -1.394 | 0.848 | 0.958 | 0.613 | 0.108 | 0.24 | 0.24 | 0 | 0.06 | 0.44 | 0.09 | 0.17 | 0.03 | 0.1 |
| 11 | 27019873 | rs11029771 | -0.797 | 1.095 | -0.465 | 0.189 | -0.006 | 0.022 | 0.022 | 0.029 | 0.32 | 0.72 | 0.33 | 0.21 | 0.34 | 0.38 |
| 11 | 27020461 | rs10835093 | -0.187 | 1.242 | -0.533 | 0.807 | -0.006 | -0.008 | -0.008 | 0 | 0.06 | 0.08 | 0.24 | 0.17 | 0.1 | 0.06 |
| 11 | 27020684 | rs4922781 | 0.236 | 0.608 | -0.628 | -0.541 | 0.003 | -0.017 | -0.017 | 0 | 0.31 | 0.27 | 0.13 | 0.03 | 0.26 | 0.22 |
| 11 | 27024192 | rs1020372 | -0.516 | 1.125 | -0.96 | 0.177 | -0.023 | -0.071 | -0.071 | 0 | 0.16 | 0.1 | 0.34 | 0.41 | 0.17 | 0.2 |
| 11 | 27025520 | rs10501071 | 0.845 | 0.888 | -0.343 | -0.101 | -0.016 | -0.117 | -0.117 | 0 | 0.25 | 0.37 | 0.12 | 0.01 | 0.23 | 0.16 |
| 11 | 27025877 | rs1822891 | -0.732 | 0.856 | -0.28 | -0.099 | -0.012 | -0.058 | -0.058 | 0 | 0.18 | 0.15 | 0.36 | 0.42 | 0.18 | 0.23 |
| 11 | 27027201 | rs10767590 | -0.318 | 1.04 | -0.444 | -0.905 | 0.017 | 0.028 | 0.028 | 0 | 0.5 | 0.4 | 0.73 | 0.92 | 0.54 | 0.5 |
| 11 | 27028245 | rs6484257 | 0.967 | 1.067 | -0.156 | -0.835 | -0.008 | -0.017 | -0.017 | 0 | 0.32 | 0.25 | 0.37 | 0.5 | 0.36 | 0.27 |
| 11 | 27029560 | rs11029778 | 0.462 | 0.717 | -0.436 | -0.836 | 0.002 | 0.004 | 0.004 | 0 | 0.24 | 0.18 | 0.15 | 0.07 | 0.22 | 0.34 |
| 11 | 27030676 | rs16916435 | 1.064 | 0.833 | -0.448 | -0.585 | -0.018 | -0.043 | -0.043 | 0 | 0.08 | 0.01 | 0.22 | 0.17 | 0.08 | 0.06 |
| 11 | 27034437 | rs1445325 | -0.749 | 1.113 | -0.57 | -0.716 | -0.011 | -0.013 | -0.013 | 0 | 0.64 | 0.31 | 0.59 | 0.51 | 0.54 | 0.69 |
| 11 | 27034956 | rs10835101 | 0.646 | 0.89 | -0.854 | -0.961 | -0.002 | -0.044 | -0.044 | 1.586 | 0.2 | 0.19 | 0.14 | 0.04 | 0.18 | 0.14 |
| 11 | 27036066 | rs7109303 | 0.007 | 0.935 | -0.935 | -0.972 | -0.017 | -0.102 | -0.102 | 0.665 | 0.44 | 0.03 | 0.44 | 0.46 | 0.37 | 0.54 |
| 11 | 27039674 | rs7116404 | 0.612 | 0.81 | -0.737 | -1.026 | 0 | -0.016 | -0.016 | 0 | 0.2 | 0.15 | 0.14 | 0.04 | 0.18 | 0.12 |
| 11 | 27051962 | rs12226573 | 0.578 | 0.808 | -1.03 | -1.13 | -0.022 | 0.1 | 0.1 | 0 | 0.33 | 0.02 | 0.18 | 0.19 | 0.27 | 0.4 |
| 11 | 27052661 | rs980804 | 0.514 | 0.879 | -0.606 | -0.988 | -0.003 | -0.005 | -0.005 | 0 | 0.38 | 0.66 | 0.46 | 0.51 | 0.44 | 0.31 |
| 11 | 27060940 | rs7939644 | -0.172 | 0.328 | -0.946 | -1.298 | -0.014 | -0.046 | -0.046 | 1.75 | 0.67 | 0.82 | 0.65 | 0.56 | 0.7 | 0.5 |
| 11 | 27061217 | rs11029795 | -0.294 | 0.295 | -0.673 | -1.473 | -0.002 | -0.033 | -0.033 | 0.15 | 0.39 | 0.57 | 0.37 | 0.12 | 0.33 | 0.28 |
| 11 | 27062338 | rs10767592 | 0.134 | 0.316 | -0.726 | -1.406 | 0.008 | 0.006 | 0.006 | 0 | 0.49 | 0.66 | 0.48 | 0.52 | 0.51 | 0.42 |
| 11 | 27062975 | rs11029796 | 0.644 | 0.24 | -0.182 | -1.304 | 0.03 | 0.019 | 0.019 | 0 | 0.07 | 0.28 | 0.17 | 0.07 | 0.06 | 0.07 |
| 11 | 27063867 | rs12385796 | 0.445 | 0.258 | -0.465 | -1.294 | -0.032 | -0.081 | -0.081 | 0.883 | 0.19 | 0.43 | 0.22 | 0.08 | 0.14 | 0.19 |
| 11 | 27075551 | rs1031176 | -1.083 | 0.21 | -0.483 | -0.984 | -0.025 | -0.048 | -0.048 | 0 | 0.89 | 0.97 | 0.77 | 0.75 | 0.88 | 0.85 |
| 11 | 27076977 | rs2305095 | 0.895 | 0.21 | -0.501 | -0.973 | -0.043 | -0.07 | -0.07 | 1.247 | 0.11 | 0.01 | 0.23 | 0.26 | 0.12 | 0.15 |

| | | | | | | | | | | | | | | | | |
|---|---|---|---|---|---|---|---|---|---|---|---|---|---|---|---|---|
| 11 | 27078246 | rs11029812 | 0.394 | 0.095 | -0.458 | -1.041 | -0.031 | 0.016 | 0.016 | 0 | 0.27 | 0.05 | 0.27 | 0.26 | 0.23 | 0.38 |
| 11 | 27079310 | rs7119888 | -0.636 | 0.079 | 0.31 | -0.856 | 0 | 0.012 | 0.012 | 0 | 0.67 | 0.36 | 0.69 | 0.56 | 0.63 | 0.72 |
| 11 | 27085406 | rs2351069 | 0.665 | 0.143 | -0.253 | -0.821 | -0.009 | -0.035 | -0.035 | 1.069 | 0.3 | 0.24 | 0.27 | 0.44 | 0.37 | 0.26 |
| 11 | 27092316 | rs10835111 | 0.599 | -0.277 | -0.311 | -0.726 | -0.011 | 0.061 | 0.061 | 0 | 0.27 | 0.58 | 0.24 | 0.44 | 0.27 | 0.26 |
| 11 | 27096926 | rs7113071 | -0.759 | -0.76 | -0.719 | -1.433 | 0.053 | 0.089 | 0.089 | 0 | 0.78 | 0.93 | 0.93 | 0.93 | 0.9 | 0.66 |
| 11 | 27100432 | rs11029827 | 0.65 | -0.735 | -0.489 | -1.752 | 0.009 | -0.128 | -0.128 | 0 | 0.18 | 0.4 | 0.12 | 0 | 0.18 | 0.08 |
| 11 | 27104501 | rs1020269 | 0.289 | -0.91 | -0.48 | -1.839 | 0.054 | 0.103 | 0.103 | 0 | 0.57 | 0.26 | 0.8 | 0.92 | 0.71 | 0.59 |
| 11 | 27109732 | rs2015372 | -0.114 | -0.982 | -0.52 | -1.863 | 0.003 | -0.017 | -0.017 | 0 | 0.31 | 0.68 | 0.17 | 0.05 | 0.22 | 0.19 |
| 11 | 27114001 | rs987687 | 1.141 | -0.92 | -0.355 | -1.834 | -0.003 | -0.007 | -0.007 | 0 | 0.16 | 0.11 | 0.09 | 0 | 0.18 | 0.07 |
| 11 | 27123970 | rs4278476 | 1.084 | -0.852 | -0.992 | -1.803 | -0.003 | -0.013 | -0.013 | 0.374 | 0.16 | 0.11 | 0.09 | 0 | 0.19 | 0.07 |
| 11 | 27124364 | rs4539271 | -1.792 | -0.834 | -1.262 | -1.779 | 0.012 | 0.069 | 0.069 | 0 | 0.88 | 0.99 | 0.97 | 0.98 | 0.94 | 0.76 |
| 11 | 27134451 | rs7951152 | -1.781 | -0.853 | -1.295 | -1.789 | 0.012 | 0.069 | 0.069 | 0 | 0.88 | 0.99 | 0.97 | 0.98 | 0.94 | 0.79 |
| 11 | 27141676 | rs1899506 | 0.814 | -0.701 | -1.445 | -1.445 | 0.003 | 0.041 | 0.041 | 0 | 0.46 | 0.23 | 0.67 | 0.66 | 0.58 | 0.42 |
| 11 | 27145005 | rs3781677 | 1.097 | -0.479 | -1.554 | -1.255 | -0.047 | -0.07 | -0.07 | 0 | 0.11 | 0 | 0.13 | 0.26 | 0.13 | 0.15 |
| 11 | 27145929 | rs1550617 | -0.514 | -0.557 | -1.355 | -1.363 | 0.001 | 0.003 | 0.003 | 0 | 0.39 | 0.3 | 0.27 | 0.31 | 0.36 | 0.33 |
| 11 | 27147270 | rs1044273 | 1.055 | -0.52 | -1.343 | -1.301 | -0.003 | -0.017 | -0.017 | 0 | 0.16 | 0.11 | 0.09 | 0 | 0.19 | 0.07 |
| 11 | 27154853 | rs1584010 | 1.419 | -0.601 | -1.313 | -0.914 | -0.061 | -0.176 | -0.176 | 0 | 0.24 | 0.01 | 0.32 | 0.5 | 0.29 | 0.27 |
| 11 | 27154995 | rs1348331 | 1.291 | -0.48 | -1.137 | -1.023 | -0.011 | -0.04 | -0.04 | 0.192 | 0.13 | 0.14 | 0.03 | 0.03 | 0.06 | 0.23 |
| 11 | 27162750 | rs10835119 | 1.173 | -0.425 | -1.269 | -0.877 | -0.051 | 0.07 | 0.07 | 0 | 0.28 | 0.11 | 0.35 | 0.14 | 0.35 | 0.19 |
| 11 | 27167641 | rs1972922 | 0.561 | -0.615 | -1.027 | -0.926 | 0.022 | 0.019 | 0.019 | 0 | 0.57 | 0.3 | 0.74 | 0.66 | 0.67 | 0.47 |
| 11 | 27175455 | rs1442920 | 0.362 | -0.407 | -0.75 | -0.278 | -0.025 | 0.042 | 0.042 | 0.003 | 0.75 | 0.55 | 0.84 | 0.7 | 0.83 | 0.75 |
| 11 | 27177204 | rs16916620 | 0.487 | -0.703 | -0.448 | -0.696 | -0.05 | -0.041 | -0.041 | 0.96 | 0.13 | 0.42 | 0.06 | 0.05 | 0.04 | 0.11 |
| 11 | 27189492 | rs7931240 | 0.024 | 0.145 | -1.152 | -0.506 | -0.031 | 0.12 | 0.12 | 0.053 | 0.28 | 0.06 | 0.31 | 0.11 | 0.3 | 0.17 |
| 11 | 27191577 | rs7925848 | 0.083 | 0.565 | -0.562 | 0.117 | 0.004 | -0.01 | -0.01 | 0 | 0.46 | 0.31 | 0.47 | 0.49 | 0.43 | 0.42 |
| 11 | 27192715 | rs12277945 | -0.955 | 0.685 | 0.283 | -0.188 | 0.064 | -0.057 | -0.057 | 0.401 | 0.11 | 0.39 | 0.09 | 0 | 0.14 | 0.04 |
| 11 | 27194627 | rs920698 | -0.1 | 0.125 | 0.551 | -0.231 | -0.011 | -0.028 | -0.028 | 0 | 0.89 | 0.77 | 0.95 | 0.93 | 0.91 | 0.91 |
| 11 | 27194855 | rs725014 | 0.841 | -0.42 | 0.909 | 0.045 | -0.038 | 0.048 | 0.048 | 0.732 | 0.82 | 0.38 | 0.89 | 0.73 | 0.88 | 0.78 |
| 11 | 27198361 | rs7938462 | 1.039 | -1.053 | -0.07 | -0.408 | 0.067 | 0.009 | 0.009 | 0.192 | 0.53 | 0.15 | 0.66 | 0.61 | 0.62 | 0.6 |
| 11 | 27200995 | rs11606133 | 0.735 | -1.251 | -0.036 | -0.526 | -0.021 | -0.011 | -0.011 | 0.004 | 0.13 | 0.05 | 0.16 | 0.12 | 0.14 | 0.1 |
| 11 | 27201105 | rs7115994 | 0.191 | -1.356 | 0.505 | -0.573 | -0.017 | -0.069 | -0.069 | 0 | 0.17 | 0.24 | 0.05 | 0 | 0.14 | 0.1 |
| 11 | 27209910 | rs7952355 | 0.569 | -1.386 | 0.651 | -0.57 | -0.002 | -0.013 | -0.013 | 0 | 0.67 | 0.4 | 0.77 | 0.88 | 0.72 | 0.8 |
| 11 | 27212251 | rs4923427 | 1.504 | -1.698 | 0.202 | -0.912 | 0.043 | 0.008 | 0.008 | 0.181 | 0.47 | 0.15 | 0.64 | 0.61 | 0.57 | 0.57 |
| 11 | 27212541 | rs7395215 | 0.247 | -1.634 | 0.373 | -0.888 | -0.007 | -0.022 | -0.022 | 0.535 | 0.68 | 0.43 | 0.77 | 0.88 | 0.73 | 0.81 |
| 11 | 27215155 | rs7944449 | -0.175 | -1.555 | 0.317 | -0.962 | 0 | -0.001 | -0.001 | 4.535 | 0.18 | 0.17 | 0.12 | 0.27 | 0.15 | 0.21 |
| 11 | 27218677 | rs1442933 | -0.601 | -1.571 | 0.36 | -0.904 | 0.008 | 0.001 | 0.001 | 0 | 0.31 | 0.22 | 0.29 | 0.29 | 0.29 | 0.31 |
| 11 | 27219795 | rs7939668 | 1.054 | -1.516 | 0.234 | -0.86 | 0.078 | 0.122 | 0.122 | 0.007 | 0.5 | 0.2 | 0.65 | 0.71 | 0.58 | 0.6 |
| 11 | 27220728 | rs11029863 | 0.596 | -0.677 | -0.144 | -0.797 | -0.074 | -0.12 | -0.12 | 0.109 | 0.14 | 0.01 | 0.37 | 0.47 | 0.19 | 0.18 |
| 11 | 27225418 | rs1964434 | -0.28 | -0.989 | 0.415 | -0.786 | -0.014 | 0.018 | 0.018 | 0 | 0.19 | 0.24 | 0.13 | 0.26 | 0.17 | 0.23 |
| 11 | 27226591 | rs10835128 | -0.233 | -1.117 | -0.138 | -1.436 | -0.004 | -0.038 | -0.038 | 0 | 0.27 | 0.17 | 0.42 | 0.56 | 0.23 | 0.23 |
| 11 | 27229587 | rs7930038 | 1.101 | -1.49 | -0.206 | -1.741 | -0.06 | -0.05 | -0.05 | 0.005 | 0.38 | 0.15 | 0.42 | 0.44 | 0.49 | 0.62 |
| 11 | 27230938 | rs1442931 | 0.006 | -1.704 | 0.404 | -1.908 | -0.069 | -0.178 | -0.178 | 0.688 | 0.68 | 0.46 | 0.87 | 1 | 0.75 | 0.85 |
| 11 | 27232408 | rs10767616 | 0.014 | -1.703 | 0.423 | -1.907 | -0.068 | -0.175 | -0.175 | 0 | 0.68 | 0.46 | 0.87 | 1 | 0.75 | 0.85 |
| 11 | 27232497 | rs10767617 | -1.13 | -1.905 | 0.826 | -2.009 | -0.044 | -0.09 | -0.09 | 0 | 0.86 | 0.64 | 0.95 | 1 | 0.91 | 0.92 |
| 11 | 27235715 | rs4597027 | 0.332 | -1.771 | 0.336 | -2.046 | 0.003 | -0.027 | -0.027 | 0 | 0.18 | 0.19 | 0.08 | 0 | 0.16 | 0.08 |
| 11 | 27236316 | rs4923434 | 0.332 | -1.771 | 0.336 | -2.046 | 0.003 | -0.027 | -0.027 | 0 | 0.18 | 0.19 | 0.08 | 0 | 0.16 | 0.08 |
| 11 | 27240841 | rs1442926 | -1.163 | -1.689 | 0.454 | -1.902 | -0.035 | -0.073 | -0.073 | 0 | 0.87 | 0.72 | 0.95 | 1 | 0.91 | 0.92 |
| 11 | 27249097 | rs11029878 | 0.761 | -1.839 | 0.198 | -1.742 | -0.041 | -0.122 | -0.122 | 0 | 0.14 | 0.01 | 0.36 | 0.48 | 0.15 | 0.18 |
| 11 | 27249688 | rs6484280 | -1.048 | -1.704 | 0.535 | -1.648 | -0.039 | -0.078 | -0.078 | 1.139 | 0.86 | 0.71 | 0.95 | 1 | 0.91 | 0.93 |
| 11 | 27255158 | rs7395420 | -1.143 | -1.693 | 0.337 | -1.643 | -0.031 | -0.066 | -0.066 | 0 | 0.87 | 0.73 | 0.95 | 1 | 0.91 | 0.93 |
| 11 | 27258899 | rs10835138 | 1.115 | -1.419 | 0.233 | -1.763 | -0.006 | -0.023 | -0.023 | 0 | 0.4 | 0.19 | 0.46 | 0.45 | 0.54 | 0.61 |
| 11 | 27260264 | rs10742171 | -0.697 | -1.596 | 0.245 | -1.884 | 0.017 | 0.011 | 0.011 | 0 | 0.82 | 0.86 | 0.91 | 1 | 0.82 | 0.93 |
| 11 | 27260806 | rs10835141 | -0.45 | -1.598 | 0.397 | -1.857 | -0.021 | 0.003 | 0.003 | 0 | 0.41 | 0.5 | 0.45 | 0.55 | 0.29 | 0.31 |
| 11 | 27260838 | rs4639892 | 0.131 | -1.48 | 0.064 | -1.811 | 0 | -0.002 | -0.002 | 0 | 0.27 | 0.22 | 0.4 | 0.55 | 0.2 | 0.24 |
| 11 | 27261193 | rs3892729 | 1.223 | -1.122 | -0.307 | -1.663 | -0.092 | -0.065 | -0.065 | 0 | 0.37 | 0.11 | 0.42 | 0.45 | 0.48 | 0.6 |
| 11 | 27262287 | rs5004411 | -0.17 | -1.206 | 0.08 | -1.413 | -0.013 | 0.08 | 0.08 | 0.075 | 0.29 | 0.47 | 0.43 | 0.46 | 0.21 | 0.44 |
| 11 | 27269546 | rs11029892 | 0.857 | -0.319 | -0.619 | -1.634 | -0.054 | 0.072 | 0.072 | 0 | 0.2 | 0.09 | 0.23 | 0.04 | 0.3 | 0.15 |
| 11 | 27271285 | rs7928078 | -0.117 | -0.442 | -0.503 | -1.789 | -0.02 | 0.019 | 0.019 | 0 | 0.29 | 0.23 | 0.28 | 0.04 | 0.39 | 0.19 |
| 11 | 27276450 | rs7948420 | -1.254 | -0.671 | -0.568 | -1.757 | 0.011 | -0.029 | -0.029 | 0 | 0.62 | 0.78 | 0.72 | 0.5 | 0.6 | 0.63 |
| 11 | 27295678 | rs10835149 | 1.773 | -0.719 | -0.822 | -1.696 | -0.079 | 0.078 | 0.078 | 0 | 0.26 | 0.02 | 0.18 | 0.15 | 0.31 | 0.2 |
| 11 | 27303937 | rs7126943 | -1.363 | -0.386 | -0.083 | -1.535 | 0.011 | -0.039 | -0.039 | 0 | 0.61 | 0.7 | 0.7 | 0.48 | 0.6 | 0.62 |
| 11 | 27311814 | rs7479309 | -0.669 | -0.401 | 0.15 | -1.601 | 0.004 | 0 | 0 | 0 | 0.49 | 0.4 | 0.66 | 0.47 | 0.5 | 0.59 |
| 11 | 27316394 | rs7483162 | -1.462 | -0.397 | 0.119 | -1.595 | 0.001 | -0.002 | -0.002 | 0.565 | 0.59 | 0.56 | 0.72 | 0.47 | 0.59 | 0.62 |
| 11 | 27318099 | rs11029926 | 0.169 | -0.41 | 0.257 | -1.659 | -0.003 | 0.01 | 0.01 | 0.011 | 0.3 | 0.26 | 0.44 | 0.43 | 0.21 | 0.44 |
| 11 | 27319188 | rs11029928 | 1.094 | -0.324 | -0.19 | -1.665 | -0.025 | 0.033 | 0.033 | 0 | 0.2 | 0.13 | 0.22 | 0.04 | 0.28 | 0.15 |
| 11 | 27335009 | rs1478682 | 0.164 | -0.41 | -0.361 | -1.426 | -0.034 | 0.051 | 0.051 | 0 | 0.28 | 0.22 | 0.28 | 0.04 | 0.37 | 0.18 |
| 11 | 27336872 | rs9299997 | -1.251 | -0.273 | -0.771 | -1.3 | 0.001 | -0.003 | -0.003 | 0.102 | 0.6 | 0.58 | 0.71 | 0.47 | 0.59 | 0.62 |
| 11 | 27337843 | rs2045935 | -0.105 | -0.265 | -1.061 | -0.433 | -0.013 | -0.11 | -0.11 | 0 | 0.27 | 0.07 | 0.41 | 0.43 | 0.21 | 0.44 |
| 11 | 27350042 | rs10835162 | -0.152 | -0.366 | -0.537 | -0.409 | 0.005 | -0.044 | -0.044 | 0.484 | 0.27 | 0.17 | 0.42 | 0.44 | 0.21 | 0.43 |
| 11 | 27355031 | rs3929244 | -0.152 | -0.525 | -0.399 | -0.344 | 0 | -0.004 | -0.004 | 0 | 0.3 | 0.33 | 0.46 | 0.5 | 0.26 | 0.41 |

| | | | | | | | | | | | | | | | | |
|---|---|---|---|---|---|---|---|---|---|---|---|---|---|---|---|---|
| 11 | 27356095 | rs7937220 | 0.839 | -0.597 | -0.399 | -0.341 | -0.02 | 0.002 | 0.002 | 0.206 | 0.1 | 0.22 | 0.07 | 0.09 | 0.04 | 0.04 |
| 11 | 27362917 | rs12282639 | 1.211 | -1.018 | -0.328 | -0.499 | -0.028 | -0.029 | -0.029 | 0 | 0.06 | 0.11 | 0.01 | 0 | 0 | 0 |
| 11 | 27372239 | rs10835169 | 1.212 | -0.434 | -0.979 | -1.032 | -0.089 | 0.074 | 0.074 | 0 | 0.22 | 0.06 | 0.26 | 0.07 | 0.34 | 0.14 |
| 11 | 27373436 | rs7935021 | -0.269 | -0.311 | -0.227 | -0.753 | 0.012 | 0.04 | 0.04 | 0.529 | 0.67 | 0.46 | 0.49 | 0.5 | 0.69 | 0.58 |
| 11 | 27381899 | rs16916901 | 1.193 | -0.578 | -0.197 | -0.55 | -0.019 | 0.003 | 0.003 | 0.13 | 0.1 | 0.22 | 0.07 | 0.09 | 0.04 | 0.04 |
| 11 | 27385939 | rs1038101 | 1.949 | -0.648 | -0.575 | -0.786 | -0.037 | 0.032 | 0.032 | 0 | 0.09 | 0 | 0.08 | 0.03 | 0.12 | 0.15 |
| 11 | 27386508 | rs7937671 | -0.267 | -1.003 | -0.181 | -0.872 | -0.008 | -0.002 | -0.002 | 0 | 0.57 | 0.58 | 0.78 | 0.58 | 0.7 | 0.57 |
| 11 | 27388504 | rs1046568 | -0.242 | -1.012 | -0.166 | -0.851 | -0.006 | -0.001 | -0.001 | 0 | 0.57 | 0.58 | 0.78 | 0.58 | 0.7 | 0.57 |
| 11 | 27388910 | rs2447995 | 0.022 | -0.671 | -0.414 | -0.99 | -0.004 | 0.005 | 0.005 | 0 | 0.29 | 0.27 | 0.32 | 0.16 | 0.36 | 0.17 |
| 11 | 27393813 | rs2448001 | -0.176 | -0.704 | -0.326 | -0.969 | 0.003 | 0.001 | 0.001 | 0.076 | 0.46 | 0.49 | 0.27 | 0.45 | 0.37 | 0.44 |
| 11 | 27398845 | rs10835171 | 0.761 | -0.564 | -0.25 | -0.976 | 0 | 0.02 | 0.02 | 2.593 | 0.35 | 0.39 | 0.49 | 0.5 | 0.31 | 0.42 |
| 11 | 27400717 | rs2472625 | -0.744 | -0.56 | -0.278 | -0.995 | 0.011 | 0.023 | 0.023 | 2.972 | 0.62 | 0.54 | 0.47 | 0.5 | 0.63 | 0.57 |
| 11 | 27401190 | rs10835173 | 1.683 | -0.582 | -0.385 | -0.956 | -0.022 | -0.055 | -0.055 | 4.99 | 0.26 | 0.14 | 0.44 | 0.41 | 0.28 | 0.4 |
| 11 | 27406844 | rs6484295 | 0.296 | -0.892 | -0.39 | -1.081 | 0.011 | -0.003 | -0.003 | 0 | 0.38 | 0.37 | 0.22 | 0.45 | 0.31 | 0.43 |
| 11 | 27408417 | rs7935151 | 0.836 | -0.607 | 0.163 | -1.185 | -0.008 | 0.028 | 0.028 | 0 | 0.62 | 0.36 | 0.66 | 0.85 | 0.59 | 0.84 |
| 11 | 27415403 | rs11029992 | 0.474 | -0.571 | 0.286 | -1.129 | 0.031 | -0.036 | -0.036 | 0.43 | 0.2 | 0.25 | 0.24 | 0.05 | 0.32 | 0.13 |
| 11 | 27417226 | rs11029994 | -1.366 | -0.729 | -0.058 | -1.2 | -0.024 | 0.093 | 0.093 | 0.034 | 0.39 | 0.74 | 0.35 | 0.15 | 0.41 | 0.15 |
| 11 | 27417696 | rs11029995 | -0.613 | -0.702 | -0.052 | -1.164 | 0.014 | -0.02 | -0.02 | 0 | 0.29 | 0.36 | 0.29 | 0.15 | 0.38 | 0.14 |
| 11 | 27426391 | rs7936621 | -1.92 | -0.967 | -1.181 | -0.871 | -0.002 | 0.008 | 0.008 | 0.046 | 0.6 | 0.84 | 0.5 | 0.34 | 0.62 | 0.43 |
| 11 | 27433832 | rs4457709 | -1.574 | -1.381 | -1.047 | -0.454 | 0.048 | 0.07 | 0.07 | 3.121 | 0.42 | 0.82 | 0.27 | 0.29 | 0.31 | 0.29 |
| 11 | 27444517 | rs4128868 | 0.038 | -1.246 | -0.948 | -0.341 | 0 | -0.012 | -0.012 | 0 | 0.21 | 0.16 | 0.15 | 0.28 | 0.21 | 0.26 |
| 11 | 27451759 | rs12271278 | -1.051 | -1.226 | -0.601 | -0.546 | -0.016 | -0.075 | -0.075 | 0 | 0.22 | 0.66 | 0.14 | 0.01 | 0.12 | 0.03 |
| 11 | 27456059 | rs4514364 | 0.853 | -0.893 | -0.982 | -0.065 | -0.002 | 0.004 | 0.004 | 0 | 0.69 | 0.7 | 0.69 | 0.74 | 0.68 | 0.87 |
| 11 | 27464328 | rs4357647 | 0.501 | -1.003 | -1.042 | -0.034 | -0.029 | 0.004 | 0.004 | 0.062 | 0.8 | 0.68 | 0.87 | 0.78 | 0.9 | 0.94 |
| 11 | 27465591 | rs4074516 | 2.41 | -1.405 | -1.682 | -0.001 | -0.085 | -0.111 | -0.111 | 0.332 | 0.36 | 0.06 | 0.46 | 0.41 | 0.41 | 0.56 |
| 11 | 27471596 | rs4923445 | 1.27 | -1.286 | -1.558 | 0.184 | -0.015 | -0.034 | -0.034 | 0 | 0.16 | 0.07 | 0.1 | 0.26 | 0.17 | 0.26 |
| 11 | 27472777 | rs7945211 | 1.54 | -1.452 | -1.555 | 0.225 | -0.098 | -0.065 | -0.065 | 0 | 0.48 | 0.07 | 0.55 | 0.41 | 0.51 | 0.59 |
| 11 | 27473981 | rs4542364 | 2.125 | -1.548 | -1.614 | -0.068 | -0.081 | -0.056 | -0.056 | 1.496 | 0.37 | 0.06 | 0.47 | 0.6 | 0.4 | 0.56 |
| 11 | 27480827 | rs11030014 | 0.142 | -1.332 | -0.525 | 0.415 | 0.018 | -0.052 | -0.052 | 0 | 0.22 | 0.43 | 0.29 | 0.15 | 0.21 | 0.27 |
| 11 | 27482352 | rs4617543 | 0.06 | -1.467 | -0.65 | 0.803 | -0.044 | 0.143 | 0.143 | 0 | 0.87 | 0.67 | 0.78 | 0.65 | 0.94 | 0.9 |
| 11 | 27487992 | rs11030016 | 1.356 | -1.43 | -0.828 | 0.701 | 0.048 | -0.107 | -0.107 | 0.022 | 0.57 | 0.07 | 0.47 | 0.51 | 0.72 | 0.58 |
| 11 | 27495259 | rs4923447 | 0.483 | -1.322 | -0.544 | 0.709 | -0.05 | 0.001 | 0.001 | 0 | 0.84 | 0.66 | 0.81 | 0.87 | 0.94 | 0.89 |
| 11 | 27496161 | rs10767643 | -0.667 | -1.292 | -0.551 | 0.729 | -0.027 | 0.035 | 0.035 | 0 | 0.9 | 0.74 | 0.82 | 0.87 | 0.95 | 0.91 |
| 11 | 27505954 | rs12786130 | 1.318 | -1.231 | -0.604 | 0.7 | -0.003 | -0.013 | -0.013 | 0 | 0.73 | 0.65 | 0.64 | 0.83 | 0.78 | 0.83 |
| 11 | 27507902 | rs4922787 | 1.18 | -1.202 | -0.602 | 0.728 | 0 | 0.001 | 0.001 | 0 | 0.74 | 0.73 | 0.65 | 0.82 | 0.78 | 0.83 |
| 11 | 27508681 | rs11030024 | -0.294 | -1.182 | -0.652 | 0.76 | -0.005 | 0.019 | 0.019 | 5.565 | 0.26 | 0.59 | 0.33 | 0.37 | 0.22 | 0.32 |
| 11 | 27510830 | rs11030026 | -0.101 | -0.912 | -0.894 | 0.721 | -0.046 | 0.033 | 0.033 | 0.004 | 0.1 | 0.01 | 0.15 | 0.04 | 0.14 | 0.05 |
| 11 | 27515109 | rs7104230 | 1.974 | -1.071 | -1.024 | 0.861 | -0.101 | -0.062 | -0.062 | 1.747 | 0.46 | 0.07 | 0.3 | 0.44 | 0.55 | 0.54 |
| 11 | 27517670 | rs6484306 | 1.477 | -1.121 | -0.362 | 0.978 | -0.024 | 0.043 | 0.043 | 0 | 0.69 | 0.47 | 0.61 | 0.59 | 0.78 | 0.81 |
| 11 | 27518123 | rs16917051 | -1.29 | -1.144 | -0.273 | 1.037 | -0.037 | 0.051 | 0.051 | 0 | 0.11 | 0.19 | 0.05 | 0.25 | 0.02 | 0.07 |
| 11 | 27518920 | rs7934816 | -0.971 | -1.062 | -0.235 | 1.08 | -0.034 | 0.066 | 0.066 | 0 | 0.1 | 0.19 | 0.05 | 0.25 | 0.02 | 0.07 |
| 11 | 27541995 | rs10835189 | 2.174 | -1.082 | -0.362 | 1.065 | -0.078 | -0.061 | -0.061 | 0.553 | 0.46 | 0.07 | 0.29 | 0.44 | 0.52 | 0.54 |
| 11 | 27547379 | rs7127239 | 0.659 | -0.984 | -0.198 | 1.074 | 0.022 | 0.031 | 0.031 | 0.155 | 0.79 | 0.58 | 0.61 | 0.78 | 0.8 | 0.88 |
| 11 | 27571603 | rs1304100 | 0.032 | -0.987 | -0.398 | 1.065 | 0.021 | -0.062 | -0.062 | 0 | 0.25 | 0.42 | 0.31 | 0.15 | 0.25 | 0.27 |
| 11 | 27582949 | rs11030048 | 2.181 | -0.974 | -0.48 | 0.851 | -0.069 | -0.056 | -0.056 | 0.087 | 0.46 | 0.07 | 0.3 | 0.44 | 0.52 | 0.54 |
| 11 | 27583129 | rs7481311 | 0.648 | -0.854 | -0.354 | 0.903 | 0.008 | 0.01 | 0.01 | 0.167 | 0.8 | 0.67 | 0.63 | 0.78 | 0.79 | 0.88 |
| 11 | 27618265 | rs10835201 | -1.433 | -0.665 | -0.38 | 1.019 | 0.015 | 0.052 | 0.052 | 0 | 0.23 | 0.48 | 0.38 | 0.31 | 0.22 | 0.14 |
| 11 | 27623611 | rs7949590 | 1.7 | -0.761 | -0.404 | 1.012 | -0.052 | -0.029 | -0.029 | 0 | 0.53 | 0.1 | 0.31 | 0.44 | 0.53 | 0.59 |
| 11 | 27628412 | rs10734394 | -0.103 | -0.641 | -0.17 | 0.703 | 0.02 | -0.009 | -0.009 | 0 | 0.75 | 0.58 | 0.69 | 0.75 | 0.75 | 0.72 |
| 11 | 27633409 | rs7939810 | 1.073 | -0.767 | -0.101 | 0.068 | 0 | -0.001 | -0.001 | 0 | 0.59 | 0.44 | 0.4 | 0.28 | 0.64 | 0.63 |
| 11 | 27635319 | rs1387144 | 1.069 | -0.673 | -0.09 | 0.14 | 0.002 | 0.002 | 0.002 | 0.496 | 0.61 | 0.42 | 0.64 | 0.41 | 0.6 | 0.37 |
| 11 | 27636885 | rs1488830 | -0.597 | -0.908 | 0.133 | 0.28 | 0.01 | 0.085 | 0.085 | 0 | 0.79 | 0.72 | 0.82 | 0.51 | 0.78 | 0.74 |
| 11 | 27637014 | rs1488831 | 0.094 | -0.869 | 0.074 | 0.128 | -0.008 | -0.014 | -0.014 | 1.307 | 0.09 | 0.16 | 0.04 | 0.07 | 0.06 | 0.3 |
| 11 | 27646745 | rs6416056 | 0.694 | -0.907 | -0.136 | 0.089 | 0.003 | -0.012 | -0.012 | 0 | 0.66 | 0.66 | 0.78 | 0.48 | 0.71 | 0.39 |
| 11 | 27647285 | rs4074134 | 0.274 | -0.816 | -0.353 | 0.056 | -0.008 | -0.034 | -0.034 | 1.952 | 0.21 | 0.16 | 0.18 | 0.44 | 0.22 | 0.27 |
| 11 | 27662826 | rs7130131 | 0.041 | -0.802 | -0.321 | -0.26 | -0.011 | -0.025 | -0.025 | 0 | 0.09 | 0.19 | 0.04 | 0.05 | 0.06 | 0.29 |
| 11 | 27665543 | rs11030096 | 1.832 | -0.888 | -0.55 | -0.454 | -0.018 | 0.012 | 0.012 | 0 | 0.43 | 0.11 | 0.3 | 0.3 | 0.47 | 0.3 |
| 11 | 27665788 | rs7126752 | 0.556 | -0.917 | -0.343 | -0.583 | -0.042 | -0.033 | -0.033 | 0.042 | 0.09 | 0.2 | 0.03 | 0.03 | 0.01 | 0 |
| 11 | 27667202 | rs925946 | 0.733 | -0.898 | -0.266 | -0.611 | 0.008 | -0.044 | -0.044 | 1.646 | 0.76 | 0.71 | 0.75 | 0.94 | 0.71 | 0.63 |
| 11 | 27672252 | rs10767658 | 0.733 | -0.898 | -0.266 | -0.611 | 0.008 | -0.044 | -0.044 | 1.27 | 0.76 | 0.71 | 0.75 | 0.94 | 0.71 | 0.63 |
| 11 | 27675712 | rs1519480 | 1.432 | -0.876 | -0.341 | -0.544 | -0.016 | 0.071 | 0.071 | 0.003 | 0.65 | 0.16 | 0.47 | 0.75 | 0.7 | 0.61 |
| 11 | 27677041 | rs7124442 | 1.244 | -0.857 | -0.33 | -0.579 | 0.006 | -0.129 | -0.129 | 0 | 0.67 | 0.49 | 0.73 | 0.91 | 0.71 | 0.6 |
| 11 | 27679916 | rs6265 | 1.356 | -0.751 | -0.463 | -0.471 | -0.056 | -0.12 | -0.12 | 0 | 0.15 | 0.01 | 0.15 | 0.49 | 0.2 | 0.2 |
| 11 | 27684517 | rs11030104 | 0.907 | -0.79 | -0.509 | -0.423 | -0.068 | -0.153 | -0.153 | 0 | 0.19 | 0.02 | 0.17 | 0.49 | 0.22 | 0.26 |
| 11 | 27695464 | rs11030108 | 0.7 | -0.716 | -0.831 | -0.413 | 0.014 | -0.054 | -0.054 | 0 | 0.77 | 0.73 | 0.76 | 0.94 | 0.71 | 0.62 |
| 11 | 27695910 | rs10835210 | 2.229 | -1.091 | -1.121 | -0.672 | -0.081 | -0.013 | -0.013 | 0 | 0.34 | 0.06 | 0.26 | 0.26 | 0.44 | 0.27 |
| 11 | 27700125 | rs7103411 | -0.557 | -0.202 | -0.85 | -0.658 | -0.026 | -0.122 | -0.122 | 0.647 | 0.77 | 0.93 | 0.83 | 0.51 | 0.77 | 0.7 |
| 11 | 27701365 | rs10835211 | -0.265 | 0.033 | -0.748 | -0.634 | -0.033 | 0.029 | 0.029 | 0 | 0.14 | 0.07 | 0.2 | 0.01 | 0.23 | 0.09 |
| 11 | 27728102 | rs11030119 | -1.45 | 0.01 | -0.687 | -0.607 | 0.005 | -0.044 | -0.044 | 0 | 0.25 | 0.32 | 0.23 | 0.06 | 0.28 | 0.38 |

| | | | | | | | | | | | | | | | | |
|---|---|---|---|---|---|---|---|---|---|---|---|---|---|---|---|---|
| 11 | 27731983 | rs7934165 | 1.56 | 0.041 | -0.85 | -0.541 | -0.002 | 0.001 | 0.001 | 0 | 0.44 | 0.44 | 0.57 | 0.41 | 0.47 | 0.32 |
| 11 | 27734420 | rs962369 | -1.435 | 0.93 | -1.539 | -0.463 | -0.023 | 0.029 | 0.029 | 0 | 0.22 | 0.16 | 0.22 | 0.06 | 0.28 | 0.38 |
| 11 | 27744859 | rs12273363 | -0.306 | 0.868 | -1.598 | -0.46 | -0.015 | 0.012 | 0.012 | 0 | 0.11 | 0.07 | 0.18 | 0.01 | 0.18 | 0.08 |
| 11 | 27745764 | rs908867 | -0.415 | 0.385 | -1.394 | -0.144 | -0.001 | -0.002 | -0.002 | 0 | 0.11 | 0.1 | 0.05 | 0.05 | 0.1 | 0.3 |
| 11 | 27749725 | rs1491850 | -0.726 | 0.62 | -1.464 | 0.167 | -0.031 | -0.031 | -0.031 | 0 | 0.37 | 0.24 | 0.37 | 0.48 | 0.43 | 0.36 |
| 11 | 27750586 | rs2049048 | -1.445 | 0.856 | -1.123 | -0.284 | 0.043 | -0.002 | -0.002 | 0.081 | 0.08 | 0.23 | 0.14 | 0.04 | 0.14 | 0.18 |
| 11 | 27752763 | rs1491851 | -0.349 | 0.651 | -2.398 | -0.672 | -0.107 | 0.078 | 0.078 | 0 | 0.38 | 0.09 | 0.37 | 0.28 | 0.54 | 0.4 |
| 11 | 27755298 | rs7482257 | -1.162 | 0.302 | -2.393 | -1.005 | 0.01 | 0.054 | 0.054 | 0 | 0.55 | 0.37 | 0.43 | 0.28 | 0.63 | 0.57 |
| 11 | 27757622 | rs1157659 | -0.064 | 0.573 | -2.576 | -0.767 | -0.088 | 0.06 | 0.06 | 0 | 0.36 | 0.09 | 0.33 | 0.27 | 0.48 | 0.35 |
| 11 | 27773337 | rs2131060 | -0.388 | 0.54 | -2.042 | -0.558 | 0.012 | 0.045 | 0.045 | 0 | 0.13 | 0.02 | 0.04 | 0 | 0.09 | 0.14 |
| 11 | 27776622 | rs2172229 | -0.724 | 0.617 | -2.096 | -0.541 | 0.008 | 0.085 | 0.085 | 0 | 0.49 | 0.35 | 0.47 | 0.26 | 0.56 | 0.52 |
| 11 | 27778592 | rs11030129 | 0.028 | 0.772 | -2.696 | -0.284 | 0.005 | -0.012 | -0.012 | 3.068 | 0.44 | 0.3 | 0.41 | 0.51 | 0.42 | 0.38 |
| 11 | 27781979 | rs728633 | 0.343 | 1.199 | -2.936 | -0.94 | -0.073 | 0.103 | 0.103 | 0 | 0.18 | 0.02 | 0.1 | 0.01 | 0.21 | 0.26 |
| 11 | 27786474 | rs11030134 | -0.085 | 0.845 | -2.488 | -0.252 | -0.019 | 0.064 | 0.064 | 0 | 0.37 | 0.11 | 0.35 | 0.26 | 0.38 | 0.31 |
| 11 | 27787783 | rs10501091 | -0.147 | 0.867 | -2.479 | -0.236 | -0.006 | 0.062 | 0.062 | 0 | 0.38 | 0.17 | 0.35 | 0.26 | 0.38 | 0.31 |
| 11 | 27791859 | rs10835221 | -0.262 | 0.838 | -2.176 | 0.505 | -0.004 | -0.023 | -0.023 | 0 | 0.42 | 0.73 | 0.47 | 0.51 | 0.4 | 0.38 |
| 11 | 27792023 | rs7933739 | -0.274 | 0.869 | -2.035 | 0.686 | 0.009 | 0.005 | 0.005 | 0 | 0.27 | 0.34 | 0.34 | 0.31 | 0.31 | 0.55 |
| 11 | 27794045 | rs12419289 | -0.654 | 0.029 | -2.031 | 1.207 | 0.033 | 0.012 | 0.012 | 0 | 0.4 | 0.73 | 0.35 | 0.39 | 0.33 | 0.45 |
| 11 | 27799994 | rs10835222 | -0.625 | 0.028 | -1.833 | 1.193 | 0.006 | 0.002 | 0.002 | 0 | 0.38 | 0.59 | 0.34 | 0.39 | 0.34 | 0.44 |
| 11 | 27808814 | rs1519481 | -0.508 | -0.95 | -1.284 | 2.029 | -0.05 | 0.094 | 0.094 | 0 | 0.23 | 0.52 | 0.19 | 0.43 | 0.12 | 0.38 |
| 11 | 27818706 | rs1491864 | -0.857 | -0.98 | -1.312 | 2.032 | -0.013 | 0.012 | 0.012 | 0 | 0.25 | 0.68 | 0.2 | 0.43 | 0.12 | 0.39 |
| 11 | 27828980 | rs10835227 | -0.074 | -0.62 | -0.075 | 1.929 | 0.01 | 0.041 | 0.041 | 0 | 0.85 | 0.67 | 0.8 | 0.73 | 0.87 | 0.73 |
| 11 | 27847646 | rs722361 | -0.308 | -0.884 | 0.362 | 2.101 | -0.047 | 0.143 | 0.143 | 0 | 0.21 | 0.41 | 0.18 | 0.49 | 0.12 | 0.37 |
| 11 | 27850631 | rs1826333 | 1.207 | -0.963 | 0.368 | 2.208 | -0.002 | -0.003 | -0.003 | 0 | 0.64 | 0.36 | 0.62 | 0.18 | 0.75 | 0.36 |
| 11 | 27866783 | rs1352864 | -0.081 | -0.594 | 0.223 | 2.321 | 0.002 | 0.034 | 0.034 | 0 | 0.86 | 0.78 | 0.81 | 0.67 | 0.87 | 0.72 |
| 11 | 27871543 | rs12797855 | 0.967 | -0.703 | 0.255 | 2.615 | 0.002 | 0.048 | 0.048 | 0 | 0.06 | 0.07 | 0.06 | 0.25 | 0.07 | 0.04 |
| 11 | 27874482 | rs6484335 | -0.113 | -0.742 | 0.155 | 2.133 | 0 | 0.01 | 0.01 | 0 | 0.86 | 0.81 | 0.81 | 0.67 | 0.88 | 0.73 |
| 11 | 27885449 | rs2638116 | -0.962 | -0.682 | 0.085 | 2.164 | -0.009 | -0.014 | -0.014 | 0.452 | 0.9 | 0.94 | 0.82 | 0.67 | 0.89 | 0.74 |
| 11 | 27890362 | rs2638122 | -0.145 | -0.725 | 0.14 | 2.107 | 0 | -0.001 | -0.001 | 0 | 0.86 | 0.83 | 0.81 | 0.68 | 0.87 | 0.71 |
| 11 | 27893387 | rs1491845 | -0.062 | -0.78 | 0.217 | 2.079 | -0.002 | 0.014 | 0.014 | 0 | 0.85 | 0.77 | 0.8 | 0.68 | 0.87 | 0.7 |
| 11 | 27898208 | rs1038840 | 1.546 | -1.095 | 0.339 | 1.965 | -0.012 | -0.016 | -0.016 | 0 | 0.63 | 0.42 | 0.62 | 0.19 | 0.74 | 0.34 |
| 11 | 27906952 | rs4923503 | 0.109 | -0.842 | 0.024 | 1.822 | -0.002 | 0.014 | 0.014 | 0 | 0.85 | 0.75 | 0.81 | 0.7 | 0.88 | 0.7 |
| 11 | 27911646 | rs2726836 | -0.249 | -0.801 | -0.065 | 1.842 | 0 | -0.003 | -0.003 | 0 | 0.87 | 0.85 | 0.82 | 0.7 | 0.88 | 0.71 |
| 11 | 27928591 | rs7937195 | -0.001 | -1.119 | 0.203 | 2.012 | -0.043 | 0.068 | 0.068 | 0.338 | 0.15 | 0.2 | 0.12 | 0.37 | 0.06 | 0.16 |
| 11 | 27930079 | rs4923506 | 0.357 | -1.108 | 0.278 | 1.951 | 0 | 0 | 0 | 0.083 | 0.8 | 0.76 | 0.82 | 0.71 | 0.82 | 0.67 |
| 11 | 27932280 | rs4923507 | 1.327 | -1.378 | 0.394 | 1.953 | -0.054 | 0.058 | 0.058 | 0 | 0.7 | 0.51 | 0.72 | 0.35 | 0.82 | 0.56 |
| 11 | 27946346 | rs10835237 | -0.227 | -1.341 | 0.372 | 1.942 | -0.067 | 0.119 | 0.119 | 0 | 0.16 | 0.33 | 0.12 | 0.36 | 0.06 | 0.16 |
| 11 | 27958820 | rs2726832 | 0.131 | -1.23 | 0.163 | 1.877 | -0.001 | -0.009 | -0.009 | 0 | 0.12 | 0.1 | 0.15 | 0.28 | 0.12 | 0.26 |
| 11 | 27970856 | rs1404950 | 1.263 | -1.433 | 0.585 | 1.986 | -0.04 | 0.035 | 0.035 | 0.07 | 0.71 | 0.47 | 0.72 | 0.34 | 0.82 | 0.59 |
| 11 | 27972606 | rs2726829 | 0.062 | -1.417 | 0.577 | 1.972 | -0.001 | 0.001 | 0.001 | 0 | 0.13 | 0.13 | 0.15 | 0.28 | 0.12 | 0.26 |
| 11 | 27978450 | rs1464896 | 1.347 | -1.873 | -0.46 | 0.948 | 0.025 | 0.112 | 0.112 | 0 | 0.3 | 0.08 | 0.16 | 0.11 | 0.28 | 0.16 |
| 11 | 27984145 | rs12799489 | 1.024 | -1.486 | 0.079 | 1.456 | -0.038 | 0.033 | 0.033 | 0.106 | 0.71 | 0.46 | 0.72 | 0.34 | 0.82 | 0.59 |
| 11 | 27990119 | rs10501093 | 0.631 | -0.843 | -0.179 | 1.475 | -0.036 | -0.042 | -0.042 | 0 | 0.1 | 0 | 0.07 | 0.17 | 0.11 | 0.24 |
| 11 | 28005579 | rs10742181 | -0.45 | -0.638 | -0.013 | 1.485 | 0.002 | 0.013 | 0.013 | 0 | 0.87 | 0.84 | 0.81 | 0.72 | 0.86 | 0.68 |
| 11 | 28008972 | rs4244533 | 1.021 | -0.999 | 0.358 | 1.691 | -0.029 | 0.076 | 0.076 | 0.023 | 0.74 | 0.51 | 0.77 | 0.35 | 0.81 | 0.55 |
| 11 | 28013323 | rs12417122 | -0.406 | -1.049 | 0.378 | 1.69 | -0.045 | 0.175 | 0.175 | 0 | 0.12 | 0.3 | 0.05 | 0.37 | 0.05 | 0.12 |
| 11 | 28017166 | rs11030182 | 0.401 | -0.948 | 0.206 | 1.747 | -0.001 | 0.004 | 0.004 | 0 | 0.13 | 0.11 | 0.18 | 0.28 | 0.14 | 0.33 |
| 11 | 28022326 | rs10835255 | 0.969 | -1.017 | 0.18 | 1.642 | -0.033 | 0.085 | 0.085 | 0.016 | 0.74 | 0.53 | 0.77 | 0.35 | 0.81 | 0.55 |
| 11 | 28031454 | rs12285612 | 0.424 | -0.94 | -0.013 | 1.558 | -0.005 | -0.013 | -0.013 | 0.289 | 0.13 | 0.09 | 0.18 | 0.28 | 0.14 | 0.33 |
| 11 | 28037253 | rs10835259 | 1.265 | -1.728 | -0.257 | 0.957 | 0.055 | 0.122 | 0.122 | 0.777 | 0.51 | 0.22 | 0.61 | 0.24 | 0.56 | 0.44 |
| 11 | 28046681 | rs10835263 | 0.477 | -1.82 | 0.067 | 1.06 | -0.037 | 0.112 | 0.112 | 0 | 0.11 | 0.16 | 0.04 | 0.4 | 0.05 | 0.13 |
| 11 | 28057957 | rs10458896 | -0.661 | -1.865 | 0.121 | 1.028 | 0.011 | -0.012 | -0.012 | 0.292 | 0.36 | 0.64 | 0.2 | 0.48 | 0.3 | 0.23 |
| 11 | 28070798 | rs11030196 | 2.814 | -1.048 | -0.201 | 0.856 | -0.036 | -0.02 | -0.02 | 0 | 0.06 | 0.01 | 0.1 | 0.07 | 0.11 | 0.04 |
| 11 | 28104605 | rs7937778 | 0.456 | -1.492 | 0.03 | 0.704 | -0.022 | -0.078 | -0.078 | 0 | 0.24 | 0.47 | 0.17 | 0.08 | 0.25 | 0.1 |
| 11 | 28110151 | rs12272419 | 0.546 | -1.467 | 0.017 | 0.728 | -0.037 | 0.114 | 0.114 | 0 | 0.11 | 0.16 | 0.04 | 0.4 | 0.05 | 0.13 |
| 11 | 28129982 | rs3812738 | -0.561 | -1.471 | -0.001 | 0.7 | 0.008 | -0.01 | -0.01 | 0 | 0.35 | 0.64 | 0.2 | 0.47 | 0.3 | 0.23 |
| 11 | 28133420 | rs17310069 | 1.77 | -1.453 | -0.071 | 0.784 | -0.051 | -0.031 | -0.031 | 0 | 0.1 | 0 | 0.1 | 0.17 | 0.13 | 0.3 |
| 11 | 28160335 | rs12273897 | 0.558 | -1.438 | 0.002 | 0.61 | -0.037 | 0.112 | 0.112 | 0 | 0.11 | 0.16 | 0.04 | 0.4 | 0.05 | 0.13 |
| 11 | 28174813 | rs960357 | 1.53 | -1.793 | -0.092 | 0.43 | 0.006 | 0.282 | 0.282 | 0.227 | 0.46 | 0.06 | 0.54 | 0.09 | 0.55 | 0.33 |
| 11 | 28175245 | rs7931847 | 0.719 | -1.463 | 0.28 | 0.283 | -0.006 | -0.028 | -0.028 | 0.139 | 0.23 | 0.29 | 0.16 | 0.07 | 0.25 | 0.1 |
| 11 | 28177723 | rs12796293 | 0.396 | -1.439 | -0.049 | 0.295 | -0.02 | -0.075 | -0.075 | 0.058 | 0.24 | 0.45 | 0.17 | 0.07 | 0.25 | 0.1 |
| 11 | 28182581 | rs11030231 | 0.951 | -1.422 | -0.059 | 0.317 | -0.005 | -0.012 | -0.012 | 0 | 0.13 | 0.1 | 0.18 | 0.28 | 0.14 | 0.32 |
| 11 | 28189639 | rs1483858 | -0.574 | -1.414 | -0.064 | 0.324 | 0.005 | -0.007 | -0.007 | 10.722 | 0.35 | 0.62 | 0.2 | 0.47 | 0.3 | 0.23 |
| 11 | 28237516 | rs7127104 | 1.596 | -1.246 | 0.275 | 0.494 | 0.057 | 0.129 | 0.129 | 0.767 | 0.51 | 0.2 | 0.61 | 0.24 | 0.56 | 0.44 |
| 11 | 28248121 | rs17244387 | 1.517 | -1.239 | 0.245 | 0.501 | -0.031 | 0.01 | 0.01 | 0 | 0.05 | 0.01 | 0.09 | 0.02 | 0.17 | 0.01 |
| 11 | 28255053 | rs7949057 | -0.228 | -1.225 | 0.355 | 0.56 | -0.02 | -0.076 | -0.076 | 0 | 0.24 | 0.45 | 0.17 | 0.07 | 0.25 | 0.11 |
| 11 | 28292078 | rs1032091 | -1.783 | -1.081 | -0.011 | 0.604 | 0.055 | 0.118 | 0.118 | 0.767 | 0.49 | 0.76 | 0.39 | 0.76 | 0.44 | 0.56 |
| 11 | 28360360 | rs11030281 | -0.51 | -0.752 | 0.477 | 1.055 | -0.047 | 0.136 | 0.136 | 0 | 0.12 | 0.22 | 0.04 | 0.4 | 0.05 | 0.13 |
| 11 | 28381599 | rs11030292 | -0.473 | -0.323 | 0.629 | 0.942 | -0.013 | 0.123 | 0.123 | 0 | 0.12 | 0.17 | 0.05 | 0.39 | 0.07 | 0.2 |

| | | | | | | | | | | | | | | | | |
|---|---|---|---|---|---|---|---|---|---|---|---|---|---|---|---|---|
| 11 | 28416646 | rs4923535 | 1.464 | -0.231 | 0.337 | 0.451 | -0.007 | 0.155 | 0.155 | 1.984 | 0.48 | 0.19 | 0.59 | 0.24 | 0.51 | 0.35 |
| 11 | 28421841 | rs7931626 | -1.488 | -0.278 | 0.976 | 0.415 | -0.003 | -0.015 | -0.015 | 1.638 | 0.46 | 0.46 | 0.38 | 0.76 | 0.46 | 0.61 |
| 11 | 28446722 | rs7932458 | 1.669 | -0.292 | 0.897 | 0.41 | -0.009 | 0.15 | 0.15 | 2.075 | 0.48 | 0.19 | 0.59 | 0.24 | 0.51 | 0.35 |
| 11 | 28466083 | rs7941325 | -0.905 | -0.378 | 0.994 | 0.391 | -0.004 | -0.052 | -0.052 | 0 | 0.26 | 0.39 | 0.17 | 0.07 | 0.3 | 0.18 |
| 11 | 28469325 | rs7110074 | 0.619 | -0.371 | 0.943 | 0.359 | -0.007 | 0.001 | 0.001 | 0 | 0.11 | 0.1 | 0.12 | 0.08 | 0.22 | 0.11 |
| 11 | 28507389 | rs4923542 | -1.739 | -0.316 | 0.892 | 0.561 | 0.003 | -0.006 | -0.006 | 0 | 0.39 | 0.58 | 0.22 | 0.47 | 0.36 | 0.38 |
| 11 | 28510100 | rs2582905 | -2.539 | -0.381 | 0.936 | 0.521 | -0.02 | 0.133 | 0.133 | 1.802 | 0.54 | 0.84 | 0.42 | 0.76 | 0.49 | 0.66 |
| 11 | 28511643 | rs4075635 | -0.13 | -0.376 | 0.988 | 0.526 | -0.001 | -0.013 | -0.013 | 0 | 0.21 | 0.24 | 0.15 | 0.08 | 0.27 | 0.15 |
| 11 | 28512458 | rs871004 | -1.062 | -0.064 | 0.695 | 0.731 | 0.003 | -0.006 | -0.006 | 0 | 0.39 | 0.58 | 0.22 | 0.47 | 0.36 | 0.38 |
| 11 | 28518811 | rs10835348 | 1.013 | 0.113 | 0.503 | 0.808 | -0.03 | -0.065 | -0.065 | 0 | 0.12 | 0.01 | 0.18 | 0.29 | 0.13 | 0.28 |
| 11 | 28532658 | rs2585807 | -0.793 | 0.043 | 0.542 | 0.79 | -0.032 | -0.059 | -0.059 | 0 | 0.86 | 0.97 | 0.82 | 0.7 | 0.85 | 0.68 |
| 11 | 28537265 | rs10501099 | 1.142 | 0.115 | 0.627 | 0.757 | -0.007 | 0.081 | 0.081 | 1.421 | 0.47 | 0.32 | 0.6 | 0.21 | 0.5 | 0.32 |
| 11 | 28548229 | rs990845 | 1.168 | 0.161 | 0.265 | -0.437 | -0.012 | 0.102 | 0.102 | 1.583 | 0.47 | 0.29 | 0.6 | 0.22 | 0.5 | 0.32 |
| 11 | 28549266 | rs4598653 | 1.305 | 0.163 | 0.229 | -0.436 | -0.057 | 0.155 | 0.155 | 1.909 | 0.45 | 0.12 | 0.59 | 0.22 | 0.5 | 0.32 |
| 11 | 28575986 | rs1596297 | 1.18 | 0.188 | 0.217 | -0.424 | -0.013 | 0.103 | 0.103 | 1.59 | 0.46 | 0.28 | 0.6 | 0.22 | 0.5 | 0.32 |
| 11 | 28578605 | rs11030378 | -0.67 | 0.01 | -0.092 | -0.582 | -0.015 | -0.118 | -0.118 | 0 | 0.34 | 0.48 | 0.21 | 0.08 | 0.32 | 0.26 |
| 11 | 28580338 | rs12419325 | -0.029 | -0.072 | -0.297 | 0.001 | -0.01 | -0.033 | -0.033 | 0 | 0.17 | 0.09 | 0.19 | 0.68 | 0.18 | 0.33 |
| 11 | 28581430 | rs1596300 | 0.452 | -0.507 | -0.089 | -0.76 | -0.016 | -0.094 | -0.094 | 0.004 | 0.61 | 0.45 | 0.79 | 0.93 | 0.7 | 0.69 |
| 11 | 28583814 | rs12294302 | 0.108 | -0.516 | -0.088 | -0.507 | 0.001 | 0.007 | 0.007 | 0 | 0.21 | 0.16 | 0.09 | 0.09 | 0.15 | 0.25 |
| 11 | 28602221 | rs2585817 | -0.958 | -0.707 | 0.422 | -0.393 | -0.003 | -0.011 | -0.011 | 0 | 0.53 | 0.52 | 0.36 | 0.77 | 0.38 | 0.64 |
| 11 | 28622910 | rs3885071 | 0.281 | -0.817 | 0.235 | -0.499 | 0.006 | -0.002 | -0.002 | 0 | 0.11 | 0.08 | 0.03 | 0 | 0.03 | 0.04 |
| 11 | 28623873 | rs7931440 | 0.87 | -0.847 | 0.373 | -0.527 | -0.007 | 0.092 | 0.092 | 0.089 | 0.43 | 0.15 | 0.62 | 0.22 | 0.61 | 0.36 |
| 11 | 28627504 | rs10835363 | -0.934 | -0.891 | 0.791 | -0.601 | 0.005 | 0.014 | 0.014 | 0 | 0.52 | 0.41 | 0.35 | 0.79 | 0.4 | 0.67 |
| 11 | 28648737 | rs11030393 | -1.141 | -0.926 | 0.764 | -0.598 | 0.01 | 0.054 | 0.054 | 0 | 0.57 | 0.66 | 0.38 | 0.81 | 0.37 | 0.73 |
| 11 | 28658030 | rs12577364 | -0.996 | -0.804 | 0.731 | -0.584 | -0.002 | -0.023 | -0.023 | 0 | 0.52 | 0.51 | 0.35 | 0.81 | 0.37 | 0.7 |
| 11 | 28659827 | rs3935075 | 0.512 | -0.902 | 0.805 | -0.485 | -0.016 | -0.007 | -0.007 | 0 | 0.08 | 0.08 | 0.11 | 0.07 | 0.03 | 0.05 |
| 11 | 28660205 | rs4337023 | -1.033 | -0.895 | 0.773 | -0.487 | 0 | -0.016 | -0.016 | 0 | 0.52 | 0.5 | 0.36 | 0.81 | 0.37 | 0.7 |
| 11 | 28676838 | rs4923554 | -0.449 | -0.752 | 0.602 | -0.384 | 0.014 | 0.057 | 0.057 | 0.137 | 0.44 | 0.36 | 0.25 | 0.72 | 0.34 | 0.65 |
| 11 | 28683234 | rs1344299 | 0.248 | -0.337 | 0.625 | -0.213 | 0.016 | 0.014 | 0.014 | 0 | 0.13 | 0.21 | 0.13 | 0.15 | 0.2 | 0.1 |
| 11 | 28687018 | rs11030406 | -0.506 | -0.529 | 0.455 | 0.734 | 0.001 | 0.215 | 0.215 | 0 | 0.08 | 0.21 | 0.05 | 0.44 | 0.06 | 0.12 |
| 11 | 28697941 | rs2355997 | 0.282 | -0.597 | 0.358 | 0.782 | 0.065 | 0.625 | 0.625 | 0.392 | 0.93 | 0.55 | 0.92 | 0.55 | 0.97 | 0.93 |
| 11 | 28697951 | rs2355996 | 0.146 | -0.834 | 0.224 | 0.423 | -0.018 | -0.003 | -0.003 | 0 | 0.08 | 0.09 | 0.11 | 0.12 | 0.03 | 0.05 |
| 11 | 28697984 | rs7940384 | 0.748 | -0.837 | -0.23 | 0.038 | 0.018 | 0.141 | 0.141 | 0 | 0.47 | 0.22 | 0.58 | 0.23 | 0.6 | 0.33 |
| 11 | 28700423 | rs10767741 | -0.061 | -0.966 | -0.27 | -0.025 | 0 | -0.001 | -0.001 | 0 | 0.58 | 0.44 | 0.71 | 0.35 | 0.63 | 0.39 |
| 11 | 28700544 | rs11030410 | 0.5 | -0.445 | -0.451 | 0.123 | 0.017 | 0.148 | 0.148 | 0 | 0.47 | 0.21 | 0.58 | 0.23 | 0.59 | 0.33 |
| 11 | 28710946 | rs11030411 | -0.811 | -0.275 | 0.339 | 1.158 | 0.074 | 0.664 | 0.664 | 0.17 | 0.07 | 0.45 | 0.08 | 0.43 | 0.03 | 0.07 |
| 11 | 28711791 | rs578440 | 1.076 | -0.463 | 0.692 | 1.129 | 0.055 | 0.534 | 0.534 | 0 | 0.92 | 0.52 | 0.93 | 0.51 | 0.96 | 0.92 |
| 11 | 28714139 | rs939505 | 0.456 | -0.669 | 0.125 | 0.068 | 0.026 | 0.031 | 0.031 | 0 | 0.66 | 0.25 | 0.66 | 0.3 | 0.76 | 0.74 |
| 11 | 28721187 | rs481554 | 0.882 | -0.335 | -0.505 | -0.523 | -0.016 | 0.158 | 0.158 | 0 | 0.37 | 0.14 | 0.28 | 0.09 | 0.3 | 0.51 |
| 11 | 28723710 | rs12360565 | 0.734 | -0.291 | -0.482 | -0.496 | -0.083 | 0.06 | 0.06 | 0 | 0.27 | 0.08 | 0.38 | 0.2 | 0.45 | 0.24 |
| 11 | 28726564 | rs528016 | -0.484 | -0.336 | -0.208 | -0.472 | 0.092 | 0.048 | 0.048 | 0.073 | 0.95 | 0.62 | 0.97 | 1 | 0.98 | 0.98 |
| 11 | 28732417 | rs520869 | 0.625 | -0.375 | -0.269 | -0.532 | 0.003 | 0.147 | 0.147 | 0 | 0.38 | 0.17 | 0.29 | 0.09 | 0.31 | 0.51 |
| 11 | 28732442 | rs1879445 | -0.552 | -0.356 | -0.426 | -0.525 | -0.002 | -0.042 | -0.042 | 1.059 | 0.56 | 0.45 | 0.68 | 0.91 | 0.66 | 0.46 |
| 11 | 28739440 | rs580377 | 0.779 | -0.27 | -0.713 | -0.418 | -0.02 | 0.166 | 0.166 | 0 | 0.37 | 0.13 | 0.28 | 0.09 | 0.31 | 0.51 |
| 11 | 28741136 | rs10835395 | 0.141 | 0.308 | -0.87 | -0.046 | -0.075 | 0.062 | 0.062 | 0 | 0.27 | 0.1 | 0.38 | 0.2 | 0.44 | 0.23 |
| 11 | 28743235 | rs560596 | 0.337 | -0.027 | -0.645 | 0.327 | -0.004 | 0.012 | 0.012 | 0.434 | 0.55 | 0.36 | 0.38 | 0.38 | 0.48 | 0.7 |
| 11 | 28744244 | rs530349 | 0.145 | -0.071 | -0.291 | 0.152 | 0 | -0.001 | -0.001 | 0 | 0.6 | 0.6 | 0.43 | 0.37 | 0.53 | 0.67 |
| 11 | 28746318 | rs1878533 | 0.596 | -0.128 | -0.407 | 0.689 | -0.092 | -0.091 | -0.091 | 0.643 | 0.26 | 0.06 | 0.41 | 0.38 | 0.4 | 0.27 |
| 11 | 28749414 | rs1914475 | 0.602 | -0.175 | -0.024 | 0.708 | -0.092 | -0.068 | -0.068 | 0 | 0.26 | 0.06 | 0.41 | 0.34 | 0.4 | 0.26 |
| 11 | 28751437 | rs10835397 | 0.56 | -0.218 | 0.01 | 0.701 | -0.016 | -0.001 | -0.001 | 0 | 0.58 | 0.28 | 0.41 | 0.41 | 0.53 | 0.69 |
| 11 | 28753841 | rs502249 | 0.568 | -0.204 | 0.025 | 0.701 | -0.027 | -0.004 | -0.004 | 0 | 0.58 | 0.2 | 0.4 | 0.41 | 0.53 | 0.67 |
| 11 | 28763321 | rs7948120 | 1.216 | -0.608 | -0.581 | 0.262 | 0.025 | 0.085 | 0.085 | 0 | 0.32 | 0.13 | 0.22 | 0.14 | 0.24 | 0.45 |
| 11 | 28764514 | rs548797 | 0.36 | -0.31 | -0.279 | 0.384 | -0.039 | -0.029 | -0.029 | 0 | 0.6 | 0.17 | 0.4 | 0.42 | 0.57 | 0.69 |
| 11 | 28786347 | rs1517573 | 0.314 | -0.213 | 0.017 | 0.448 | 0.032 | 0.017 | 0.017 | 0 | 0.28 | 0.4 | 0.43 | 0.34 | 0.36 | 0.27 |
| 11 | 28808408 | rs2090492 | 0.134 | -0.235 | -0.199 | 0.448 | -0.038 | -0.044 | -0.044 | 0 | 0.62 | 0.17 | 0.43 | 0.41 | 0.6 | 0.7 |
| 11 | 28826913 | rs11030448 | -0.098 | 0.092 | -0.065 | 0.378 | 0.014 | 0.068 | 0.068 | 0 | 0.12 | 0.23 | 0.22 | 0.25 | 0.15 | 0.04 |
| 11 | 28828762 | rs1949419 | 1.128 | -0.015 | 0.03 | 0.318 | -0.005 | 0.002 | 0.002 | 0.458 | 0.79 | 0.25 | 0.71 | 0.75 | 0.82 | 0.95 |
| 11 | 28838085 | rs11030451 | 1.205 | 0.14 | -0.233 | 0.245 | -0.062 | 0.056 | 0.056 | 0 | 0.19 | 0.03 | 0.14 | 0.08 | 0.27 | 0.2 |
| 11 | 28841147 | rs12293036 | 0.209 | 0.061 | 0.689 | 0.847 | 0.024 | 0.049 | 0.049 | 0.047 | 0.23 | 0.39 | 0.34 | 0.35 | 0.25 | 0.27 |
| 11 | 28847155 | rs2945090 | -0.142 | 0.116 | 0.363 | 0.942 | 0.037 | 0.115 | 0.115 | 0 | 0.89 | 0.7 | 0.78 | 0.75 | 0.86 | 0.97 |
| 11 | 28850329 | rs7108408 | 0.655 | 0.173 | -0.244 | 0.888 | -0.031 | -0.044 | -0.044 | 0.074 | 0.21 | 0.09 | 0.31 | 0.35 | 0.26 | 0.28 |
| 11 | 28854844 | rs10835415 | 0.696 | 0.02 | -0.363 | 0.756 | -0.033 | -0.01 | -0.01 | 0 | 0.6 | 0.13 | 0.41 | 0.39 | 0.59 | 0.73 |
| 11 | 28858714 | rs973556 | 0.66 | 0.021 | -0.248 | 0.743 | -0.022 | -0.001 | -0.001 | 0 | 0.6 | 0.2 | 0.41 | 0.39 | 0.59 | 0.73 |
| 11 | 28858778 | rs1400752 | 0.66 | 0.021 | -0.248 | 0.743 | -0.022 | -0.001 | -0.001 | 2.267 | 0.6 | 0.2 | 0.41 | 0.39 | 0.59 | 0.73 |
| 11 | 28859158 | rs1400751 | -0.916 | 0.253 | 0.348 | 0.465 | 0.029 | -0.116 | -0.116 | 0.78 | 0.15 | 0.5 | 0.15 | 0 | 0.15 | 0.03 |
| 11 | 28867779 | rs1517569 | -0.776 | 0.184 | 0.325 | 0.607 | 0.01 | 0.002 | 0.002 | 0 | 0.36 | 0.55 | 0.46 | 0.35 | 0.4 | 0.27 |
| 11 | 28871669 | rs11030473 | 0.651 | 0.316 | -0.053 | 0.693 | -0.035 | -0.067 | -0.067 | 1.855 | 0.21 | 0.08 | 0.31 | 0.35 | 0.23 | 0.23 |
| 11 | 28876111 | rs16918370 | 0.59 | 0.281 | 0.556 | 0.676 | -0.005 | -0.01 | -0.01 | 0.144 | 0.21 | 0.24 | 0.32 | 0.35 | 0.23 | 0.23 |
| 11 | 28876280 | rs12795950 | -0.041 | 0.334 | 0.539 | 0.606 | 0.003 | -0.024 | -0.024 | 0.051 | 0.09 | 0.13 | 0.11 | 0 | 0.16 | 0.02 |

| | | | | | | | | | | | | | | | | |
|---|---|---|---|---|---|---|---|---|---|---|---|---|---|---|---|---|
| 11 | 28882115 | rs4282967 | -0.768 | 0.238 | 0.663 | 0.614 | -0.008 | -0.002 | -0.002 | 0.025 | 0.35 | 0.34 | 0.45 | 0.35 | 0.4 | 0.25 |
| 11 | 28884379 | rs12802872 | -0.78 | 0.253 | 0.459 | 0.621 | -0.034 | -0.016 | -0.016 | 0 | 0.34 | 0.22 | 0.44 | 0.35 | 0.4 | 0.25 |
| 11 | 28887766 | rs11604366 | -0.788 | 0.251 | 0.474 | 0.668 | -0.033 | -0.009 | -0.009 | 0 | 0.34 | 0.22 | 0.43 | 0.33 | 0.41 | 0.26 |
| 11 | 28890210 | rs12364789 | -0.232 | 0.083 | 0.519 | 0.833 | 0.005 | 0.05 | 0.05 | 0.443 | 0.73 | 0.53 | 0.53 | 0.41 | 0.73 | 0.74 |
| 11 | 28891471 | rs7938748 | -0.243 | 0.084 | 0.519 | 0.834 | 0.005 | 0.051 | 0.051 | 0.409 | 0.74 | 0.53 | 0.53 | 0.41 | 0.73 | 0.74 |
| 11 | 28898961 | rs11030496 | 1.114 | -0.397 | -0.307 | 0.276 | -0.014 | 0.03 | 0.03 | 0 | 0.18 | 0.02 | 0.12 | 0.08 | 0.12 | 0.24 |
| 11 | 28927943 | rs4133156 | 0.122 | -0.129 | -0.175 | 0.308 | 0.017 | 0.008 | 0.008 | 0.044 | 0.63 | 0.41 | 0.42 | 0.41 | 0.56 | 0.72 |
| 11 | 28933682 | rs10501106 | 1.371 | -0.297 | -0.38 | 0.277 | -0.004 | -0.026 | -0.026 | 0.342 | 0.05 | 0 | 0.02 | 0.22 | 0.03 | 0.04 |
| 11 | 28937491 | rs991489 | 0.005 | -0.074 | -0.29 | 0.284 | 0.006 | 0.01 | 0.01 | 0.036 | 0.64 | 0.42 | 0.45 | 0.41 | 0.62 | 0.72 |
| 11 | 28943475 | rs7130212 | 0.128 | -0.081 | -0.413 | 0.484 | -0.081 | -0.061 | -0.061 | 0.094 | 0.31 | 0.12 | 0.41 | 0.34 | 0.41 | 0.26 |
| 11 | 28944386 | rs2959112 | -0.047 | -0.076 | -0.234 | 0.51 | -0.016 | -0.013 | -0.013 | 0 | 0.32 | 0.3 | 0.4 | 0.34 | 0.37 | 0.26 |
| 11 | 28945896 | rs2959111 | 0.152 | -0.079 | -0.263 | 0.609 | -0.019 | -0.011 | -0.011 | 0 | 0.31 | 0.26 | 0.38 | 0.32 | 0.34 | 0.26 |
| 11 | 28948098 | rs1433692 | -0.178 | -0.09 | -0.132 | 0.467 | -0.005 | -0.003 | -0.003 | 0.024 | 0.67 | 0.6 | 0.57 | 0.66 | 0.59 | 0.74 |
| 11 | 28948495 | rs4923583 | 0.58 | -0.5 | -0.169 | 0.174 | 0.011 | 0.014 | 0.014 | 0 | 0.26 | 0.16 | 0.18 | 0.14 | 0.17 | 0.4 |
| 11 | 28955149 | rs2959108 | -1.267 | -0.195 | -0.419 | 0.318 | -0.012 | -0.032 | -0.032 | 0.13 | 0.77 | 0.8 | 0.69 | 0.66 | 0.75 | 0.76 |
| 11 | 28955342 | rs2959107 | -0.527 | -0.201 | -0.419 | 0.318 | -0.019 | -0.023 | -0.023 | 0 | 0.69 | 0.74 | 0.62 | 0.66 | 0.66 | 0.75 |
| 11 | 28956624 | rs11030513 | -0.049 | -0.207 | -0.431 | 0.309 | 0.007 | 0.009 | 0.009 | 0.022 | 0.64 | 0.44 | 0.46 | 0.41 | 0.63 | 0.72 |
| 11 | 28962881 | rs934463 | 1.089 | -0.188 | -0.634 | 0.187 | -0.064 | 0.038 | 0.038 | 0.022 | 0.19 | 0.02 | 0.16 | 0.07 | 0.2 | 0.24 |
| 11 | 28977527 | rs1120073 | 0.838 | -0.071 | -0.155 | 0.476 | 0.001 | 0.107 | 0.107 | 0 | 0.85 | 0.45 | 0.76 | 0.75 | 0.88 | 0.88 |
| 11 | 28979318 | rs2356274 | -1.213 | 0.062 | -0.399 | 0.466 | -0.028 | -0.055 | -0.055 | 0 | 0.78 | 0.9 | 0.68 | 0.66 | 0.74 | 0.76 |
| 11 | 28988703 | rs986253 | -1.159 | -0.021 | -0.386 | 0.416 | -0.02 | -0.047 | -0.047 | 0.437 | 0.77 | 0.87 | 0.69 | 0.66 | 0.75 | 0.76 |
| 11 | 29001308 | rs10835451 | 0.32 | -0.061 | 0.029 | 0.519 | 0.009 | 0.041 | 0.041 | 0.399 | 0.27 | 0.39 | 0.47 | 0.58 | 0.28 | 0.26 |
| 11 | 29002237 | rs715190 | 0.998 | -0.052 | -0.098 | 0.548 | -0.022 | -0.052 | -0.052 | 0 | 0.23 | 0.11 | 0.31 | 0.34 | 0.25 | 0.24 |
| 11 | 29016502 | rs7126178 | 0.304 | -0.08 | -0.017 | 0.517 | 0.015 | 0.006 | 0.006 | 0.106 | 0.63 | 0.42 | 0.41 | 0.41 | 0.56 | 0.63 |
| 11 | 29020150 | rs11030540 | 0.807 | 0.006 | -0.042 | 0.554 | -0.007 | -0.024 | -0.024 | 0 | 0.23 | 0.16 | 0.32 | 0.35 | 0.25 | 0.24 |
| 11 | 29026450 | rs12416708 | -0.655 | 0.195 | -0.018 | 0.538 | 0.031 | 0.084 | 0.084 | 1.436 | 0.73 | 0.46 | 0.5 | 0.44 | 0.71 | 0.77 |
| 11 | 29035497 | rs11030546 | -0.121 | 0.17 | -0.067 | 0.508 | -0.021 | -0.03 | -0.03 | 0 | 0.32 | 0.19 | 0.51 | 0.58 | 0.34 | 0.33 |
| 11 | 29037522 | rs768082 | 1.153 | -0.1 | 0.098 | 0.227 | 0 | 0.01 | 0.01 | 0 | 0.32 | 0.25 | 0.23 | 0.28 | 0.35 | 0.23 |
| 11 | 29044894 | rs11030551 | 1.092 | 0.102 | 0.097 | 0.587 | -0.003 | -0.003 | -0.003 | 0.282 | 0.22 | 0.19 | 0.32 | 0.34 | 0.24 | 0.22 |
| 11 | 29045588 | rs11030553 | 0.245 | 0.104 | 0.108 | 0.579 | 0.028 | 0.015 | 0.015 | 0.231 | 0.65 | 0.44 | 0.42 | 0.42 | 0.56 | 0.67 |
| 11 | 29046543 | rs17662858 | 1.097 | 0.096 | 0.107 | 0.581 | -0.003 | -0.003 | -0.003 | 0.046 | 0.22 | 0.19 | 0.32 | 0.34 | 0.24 | 0.22 |
| 11 | 29049401 | rs1368735 | 0.26 | 0.095 | 0.107 | 0.579 | 0.017 | 0.018 | 0.018 | 0.279 | 0.65 | 0.44 | 0.45 | 0.42 | 0.63 | 0.67 |
| 11 | 29049887 | rs7938205 | 1.085 | 0.095 | 0.082 | 0.582 | -0.003 | -0.003 | -0.003 | 0 | 0.22 | 0.19 | 0.32 | 0.34 | 0.24 | 0.22 |
| 11 | 29054545 | rs1897697 | 1.028 | 0.038 | -0.026 | 0.56 | 0.026 | 0.069 | 0.069 | 0 | 0.84 | 0.48 | 0.72 | 0.77 | 0.8 | 0.88 |
| 11 | 29059115 | rs7105259 | 0.222 | 0.042 | -0.034 | 0.563 | 0.027 | 0.016 | 0.016 | 0.295 | 0.65 | 0.43 | 0.42 | 0.42 | 0.56 | 0.67 |
| 11 | 29063396 | rs7123034 | 0.367 | 0.051 | -0.029 | 0.566 | 0.021 | -0.006 | -0.006 | 0.093 | 0.63 | 0.42 | 0.41 | 0.39 | 0.56 | 0.66 |
| 11 | 29083746 | rs10742213 | 0.71 | -0.164 | -0.353 | 0.301 | -0.045 | -0.083 | -0.083 | 4.508 | 0.27 | 0.06 | 0.33 | 0.34 | 0.27 | 0.23 |
| 11 | 29086045 | rs7114955 | -0.562 | -0.074 | -0.045 | 0.089 | -0.005 | -0.081 | -0.081 | 0 | 0.13 | 0.34 | 0.09 | 0 | 0.1 | 0.07 |
| 11 | 29106959 | rs10767785 | -0.53 | -0.138 | -0.52 | 0.168 | -0.008 | -0.001 | -0.001 | 0.092 | 0.38 | 0.32 | 0.52 | 0.57 | 0.37 | 0.34 |
| 11 | 29107555 | rs10767786 | -0.53 | -0.138 | -0.52 | 0.157 | -0.008 | -0.004 | -0.004 | 0.712 | 0.38 | 0.32 | 0.52 | 0.56 | 0.37 | 0.34 |
| 11 | 29115116 | rs2055015 | 0.051 | 0.323 | -0.659 | 0.469 | -0.038 | -0.073 | -0.073 | 0 | 0.18 | 0.08 | 0.25 | 0.28 | 0.19 | 0.17 |
| 11 | 29135383 | rs2933170 | -0.267 | 0.474 | -0.567 | 0.502 | -0.001 | 0.001 | 0.001 | 0 | 0.25 | 0.33 | 0.42 | 0.23 | 0.26 | 0.3 |
| 11 | 29137386 | rs16918650 | 1.709 | 0.396 | -0.675 | 0.57 | -0.04 | -0.052 | -0.052 | 0.034 | 0.1 | 0.05 | 0.08 | 0.2 | 0.13 | 0.11 |
| 11 | 29138255 | rs2933171 | -0.642 | 0.346 | -0.583 | 0.294 | 0.004 | -0.002 | -0.002 | 0 | 0.83 | 0.84 | 0.8 | 0.85 | 0.77 | 0.68 |
| 11 | 29141778 | rs2933168 | -0.584 | 0.339 | -0.393 | 0.349 | 0.006 | 0.032 | 0.032 | 0 | 0.36 | 0.56 | 0.48 | 0.43 | 0.33 | 0.4 |
| 11 | 29147594 | rs2958638 | -0.011 | 0.247 | -0.301 | 0.274 | -0.017 | 0.01 | 0.01 | 0 | 0.25 | 0.51 | 0.39 | 0.23 | 0.2 | 0.29 |
| 11 | 29162136 | rs1602565 | 2.123 | 0.224 | -0.364 | 0.261 | -0.004 | -0.023 | -0.023 | 0 | 0.13 | 0.02 | 0.19 | 0.13 | 0.11 | 0.15 |
| 11 | 29167659 | rs4923593 | 1.974 | -0.22 | -0.372 | -0.032 | -0.003 | 0.042 | 0.042 | 0 | 0.41 | 0.2 | 0.31 | 0.42 | 0.44 | 0.25 |
| 11 | 29172397 | rs2035841 | 2.118 | -0.136 | -0.264 | 0.162 | -0.04 | -0.052 | -0.052 | 0 | 0.1 | 0.04 | 0.08 | 0.2 | 0.13 | 0.11 |
| 11 | 29175395 | rs10767787 | -0.417 | -0.12 | -0.157 | -0.016 | 0.004 | 0.03 | 0.03 | 0 | 0.36 | 0.56 | 0.48 | 0.44 | 0.33 | 0.4 |
| 11 | 29182459 | rs617346 | 1.774 | -0.056 | -0.137 | -0.063 | -0.008 | -0.002 | -0.002 | 0 | 0.16 | 0.3 | 0.1 | 0.14 | 0.14 | 0.16 |
| 11 | 29191130 | rs8181513 | 1.226 | 0.161 | -0.431 | -0.031 | -0.031 | -0.024 | -0.024 | 0.096 | 0.07 | 0.01 | 0.26 | 0.07 | 0.12 | 0.16 |
| 11 | 29195137 | rs1491818 | 1.459 | 0.187 | -0.453 | -0.276 | 0.02 | 0.001 | 0.001 | 0 | 0.34 | 0.22 | 0.22 | 0.46 | 0.3 | 0.22 |
| 11 | 29195732 | rs2055014 | -0.304 | 0.15 | -0.101 | -0.165 | -0.004 | 0.017 | 0.017 | 0 | 0.34 | 0.62 | 0.33 | 0.45 | 0.28 | 0.38 |
| 11 | 29196242 | rs11030600 | 0.961 | 0.088 | 0.276 | -0.259 | 0.028 | 0.016 | 0.016 | 0 | 0.06 | 0.26 | 0.03 | 0.01 | 0.04 | 0.05 |
| 11 | 29197549 | rs2467592 | 0.429 | 0.09 | -0.051 | -0.225 | -0.035 | -0.022 | -0.022 | 0.663 | 0.22 | 0.57 | 0.13 | 0.16 | 0.15 | 0.23 |
| 11 | 29202440 | rs7938219 | 2.813 | 0.281 | -0.232 | -0.125 | -0.001 | -0.019 | -0.019 | 0 | 0.12 | 0.02 | 0.08 | 0.14 | 0.11 | 0.12 |
| 11 | 29204531 | rs1994721 | 0.635 | 0.196 | 0.021 | -0.16 | -0.025 | -0.008 | -0.008 | 0 | 0.2 | 0.49 | 0.12 | 0.15 | 0.15 | 0.23 |
| 11 | 29205697 | rs2467589 | 2.753 | 0.25 | -0.164 | -0.132 | -0.002 | -0.019 | -0.019 | 0 | 0.12 | 0.03 | 0.08 | 0.14 | 0.11 | 0.12 |
| 11 | 29211299 | rs7939402 | 0.536 | 0.463 | 0.146 | -0.182 | 0.037 | 0.018 | 0.018 | 0 | 0.07 | 0.34 | 0.04 | 0.01 | 0.04 | 0.05 |
| 11 | 29213968 | rs2933175 | -0.449 | 0.675 | -0.41 | -0.169 | -0.005 | 0.002 | 0.002 | 0 | 0.77 | 0.81 | 0.6 | 0.88 | 0.63 | 0.64 |
| 11 | 29218767 | rs2467590 | 0.852 | -0.03 | -0.808 | -0.033 | 0.006 | -0.067 | -0.067 | 0 | 0.43 | 0.09 | 0.28 | 0.58 | 0.39 | 0.31 |
| 11 | 29221294 | rs591032 | -0.97 | -0.057 | -0.164 | -0.103 | -0.002 | -0.001 | -0.001 | 0 | 0.54 | 0.49 | 0.67 | 0.45 | 0.6 | 0.61 |
| 11 | 29224006 | rs2467594 | -1.397 | -0.316 | -0.42 | -0.296 | -0.012 | -0.076 | -0.076 | 0 | 0.63 | 0.89 | 0.72 | 0.46 | 0.64 | 0.71 |
| 11 | 29228396 | rs2467599 | 1.053 | -0.391 | -0.641 | -0.261 | -0.045 | -0.061 | -0.061 | 0 | 0.3 | 0.02 | 0.2 | 0.4 | 0.28 | 0.19 |
| 11 | 29231552 | rs11030613 | -0.109 | 0.132 | -0.055 | -0.237 | -0.06 | 0.058 | 0.058 | 1.101 | 0.29 | 0.11 | 0.39 | 0.13 | 0.41 | 0.35 |
| 11 | 29239598 | rs1491797 | -0.923 | -0.065 | 0.381 | 0.141 | 0.022 | -0.001 | -0.001 | 0 | 0.56 | 0.34 | 0.55 | 0.36 | 0.64 | 0.57 |
| 11 | 29249635 | rs10767795 | -0.931 | 0.062 | 0.157 | 0.309 | 0.011 | 0.007 | 0.007 | 0 | 0.52 | 0.85 | 0.66 | 0.45 | 0.55 | 0.65 |
| 11 | 29250158 | rs1502447 | -0.905 | 0.059 | 0.316 | 0.318 | 0.016 | 0.015 | 0.015 | 0 | 0.52 | 0.76 | 0.66 | 0.45 | 0.54 | 0.65 |

| | | | | | | | | | | | | | | | | |
|---|---|---|---|---|---|---|---|---|---|---|---|---|---|---|---|---|
| 11 | 29258947 | rs10835498 | -0.223 | 0.069 | 0.192 | 0.155 | -0.014 | 0.065 | 0.065 | 0.13 | 0.4 | 0.27 | 0.44 | 0.14 | 0.46 | 0.37 |
| 11 | 29272080 | rs10742217 | -0.133 | 0.282 | 0.252 | 0.427 | 0.002 | -0.016 | -0.016 | 0 | 0.36 | 0.41 | 0.44 | 0.13 | 0.41 | 0.34 |
| 11 | 29275928 | rs7943620 | 1.086 | 0.332 | 0.398 | 0.572 | 0.016 | 0.075 | 0.075 | 0.155 | 0.14 | 0.29 | 0.1 | 0.21 | 0.13 | 0.14 |
| 11 | 29276505 | rs16918880 | 1.535 | 0.411 | -0.283 | 0.61 | -0.013 | -0.028 | -0.028 | 0.21 | 0.12 | 0.06 | 0.08 | 0.21 | 0.13 | 0.14 |
| 11 | 29289848 | rs4459336 | -0.971 | 0.366 | -0.841 | 0.162 | 0.003 | 0.013 | 0.013 | 0 | 0.54 | 0.84 | 0.67 | 0.44 | 0.54 | 0.74 |
| 11 | 29290569 | rs7929751 | -0.766 | 0.325 | -0.808 | 0.191 | 0.016 | 0.043 | 0.043 | 0 | 0.5 | 0.84 | 0.67 | 0.45 | 0.52 | 0.7 |
| 11 | 29293934 | rs17609863 | -0.105 | 0.39 | -0.269 | 0.291 | -0.002 | -0.006 | -0.006 | 0 | 0.08 | 0.2 | 0.05 | 0.06 | 0.06 | 0.25 |
| 11 | 29299709 | rs4447203 | 0.693 | 0.526 | -0.812 | 0.253 | -0.021 | -0.017 | -0.017 | 0.604 | 0.23 | 0.13 | 0.2 | 0.41 | 0.27 | 0.09 |
| 11 | 29303887 | rs17513662 | 0.935 | 0.356 | -0.695 | 0.217 | 0 | 0 | 0 | 0.381 | 0.06 | 0.05 | 0.03 | 0.05 | 0.06 | 0.1 |
| 11 | 29304247 | rs17513684 | 1.828 | 0.043 | -1.052 | 0.096 | -0.023 | -0.006 | -0.006 | 0.005 | 0.21 | 0.01 | 0.14 | 0.16 | 0.18 | 0.21 |
| 11 | 29312419 | rs10835507 | 0.282 | -0.095 | -0.099 | 0.311 | -0.003 | 0.004 | 0.004 | 0.802 | 0.3 | 0.23 | 0.39 | 0.17 | 0.33 | 0.28 |
| 11 | 29312842 | rs12417717 | -0.271 | -0.078 | -0.159 | 0.369 | -0.003 | -0.007 | -0.007 | 0 | 0.42 | 0.36 | 0.6 | 0.38 | 0.45 | 0.39 |
| 11 | 29316153 | rs12792909 | 0.862 | -0.065 | -0.137 | 0.411 | -0.001 | 0 | 0 | 0 | 0.07 | 0.07 | 0.03 | 0.07 | 0.06 | 0.1 |
| 11 | 29332489 | rs7481118 | 0.045 | -0.106 | 0.031 | 0.39 | -0.003 | 0.002 | 0.002 | 1.599 | 0.89 | 0.62 | 0.94 | 0.92 | 0.94 | 0.73 |
| 11 | 29346876 | rs10835511 | -0.852 | 0.028 | -0.648 | 0.456 | 0 | 0.004 | 0.004 | 0.051 | 0.92 | 0.85 | 0.96 | 0.93 | 0.94 | 0.9 |
| 11 | 29350878 | rs17297770 | 0.417 | 0.24 | -1.317 | 0.347 | 0.004 | 0.03 | 0.03 | 0.015 | 0.51 | 0.21 | 0.38 | 0.53 | 0.57 | 0.35 |
| 11 | 29353459 | rs12226886 | 1.39 | 0.403 | -1.123 | 0.678 | -0.002 | -0.042 | -0.042 | 0 | 0.12 | 0.07 | 0.21 | 0.27 | 0.12 | 0.11 |
| 11 | 29355932 | rs11030665 | 0.301 | 0.516 | -0.956 | 0.006 | -0.008 | 0.009 | 0.009 | 0 | 0.47 | 0.31 | 0.35 | 0.54 | 0.5 | 0.35 |
| 11 | 29360384 | rs11030667 | -0.333 | 0.751 | -0.663 | -0.314 | 0 | 0.003 | 0.003 | 0 | 0.53 | 0.45 | 0.6 | 0.66 | 0.45 | 0.74 |
| 11 | 29362171 | rs12808270 | -0.302 | 0.933 | -1.087 | -0.244 | -0.004 | 0.013 | 0.013 | 0 | 0.21 | 0.17 | 0.25 | 0.13 | 0.24 | 0.32 |
| 11 | 29363716 | rs1341296 | -0.212 | 0.828 | -0.566 | -0.258 | 0 | 0.008 | 0.008 | 0 | 0.54 | 0.54 | 0.61 | 0.66 | 0.45 | 0.74 |
| 11 | 29370702 | rs2793476 | 0.332 | 0.807 | -1.18 | -0.397 | -0.023 | 0.019 | 0.019 | 0 | 0.35 | 0.1 | 0.22 | 0.35 | 0.36 | 0.23 |
| 11 | 29376704 | rs2711816 | 0.306 | 1.266 | -1.356 | -0.105 | 0.034 | 0.086 | 0.086 | 0 | 0.5 | 0.81 | 0.38 | 0.59 | 0.38 | 0.55 |
| 11 | 29380194 | rs11030671 | 1.282 | 1.238 | -1.38 | -0.132 | -0.008 | 0.017 | 0.017 | 0 | 0.09 | 0.03 | 0.13 | 0.06 | 0.09 | 0.19 |
| 11 | 29385080 | rs2711823 | 0.412 | 0.946 | -0.778 | -0.124 | 0.014 | -0.005 | -0.005 | 0 | 0.47 | 0.64 | 0.28 | 0.48 | 0.39 | 0.53 |
| 11 | 29385292 | rs2793472 | 1.241 | 0.776 | -1.152 | 0.191 | -0.036 | -0.046 | -0.046 | 0 | 0.22 | 0.1 | 0.17 | 0.26 | 0.26 | 0.17 |
| 11 | 29402234 | rs11030679 | 1.242 | 0.808 | -1.184 | 0.138 | -0.037 | 0.04 | 0.04 | 0 | 0.14 | 0.04 | 0.34 | 0.05 | 0.19 | 0.24 |
| 11 | 29403471 | rs1397870 | 1.277 | 0.736 | -1.419 | 0.03 | 0.004 | -0.014 | -0.014 | 0 | 0.23 | 0.13 | 0.13 | 0.34 | 0.22 | 0.11 |
| 11 | 29404798 | rs792514 | 1.81 | 0.967 | -0.984 | 0.463 | -0.061 | -0.06 | -0.06 | 0 | 0.17 | 0.01 | 0.11 | 0.19 | 0.19 | 0.15 |
| 11 | 29412441 | rs10767806 | -1.121 | 0.829 | 0.184 | 0.477 | 0.12 | 0.228 | 0.228 | 1.708 | 0.94 | 0.64 | 0.84 | 0.89 | 0.94 | 0.97 |
| 11 | 29413728 | rs792520 | 0.622 | 0.805 | -0.474 | 0.373 | -0.001 | -0.001 | -0.001 | 2.14 | 0.75 | 0.27 | 0.46 | 0.77 | 0.71 | 0.61 |
| 11 | 29414130 | rs979756 | 1.241 | 0.969 | -0.429 | 0.311 | -0.024 | 0.034 | 0.034 | 0 | 0.14 | 0.05 | 0.34 | 0.05 | 0.19 | 0.23 |
| 11 | 29425920 | rs2356579 | 0.648 | 0.816 | -0.828 | 0.328 | -0.006 | -0.023 | -0.023 | 0 | 0.55 | 0.25 | 0.3 | 0.58 | 0.45 | 0.45 |
| 11 | 29428392 | rs973851 | 0.622 | 0.848 | -0.813 | 0.344 | -0.008 | -0.024 | -0.024 | 0.056 | 0.56 | 0.25 | 0.3 | 0.58 | 0.45 | 0.45 |
| 11 | 29449099 | rs2779154 | -0.258 | 0.311 | -0.392 | 0.775 | 0.001 | 0.003 | 0.003 | 0 | 0.7 | 0.3 | 0.65 | 0.63 | 0.68 | 0.68 |
| 11 | 29455138 | rs329545 | 0.485 | -0.453 | -0.727 | 0.746 | -0.006 | 0.006 | 0.006 | 0 | 0.41 | 0.46 | 0.37 | 0.43 | 0.36 | 0.4 |
| 11 | 29460313 | rs2667433 | 0.976 | -0.028 | -1.363 | 0.697 | -0.035 | -0.071 | -0.071 | 0 | 0.11 | 0.01 | 0.12 | 0.33 | 0.17 | 0.08 |
| 11 | 29465015 | rs634740 | -0.611 | -0.298 | -1.148 | 0.473 | 0.022 | 0.025 | 0.025 | 0 | 0.59 | 0.78 | 0.7 | 0.67 | 0.67 | 0.78 |
| 11 | 29470621 | rs2225571 | 0.413 | -0.388 | -1.333 | 0.477 | -0.053 | -0.048 | -0.048 | 0 | 0.22 | 0.07 | 0.17 | 0.24 | 0.29 | 0.18 |
| 11 | 29472815 | rs907333 | -1.737 | -0.43 | -1.357 | 0.481 | -0.002 | 0 | 0 | 0.627 | 0.79 | 0.82 | 0.82 | 0.84 | 0.88 | 0.93 |
| 11 | 29475113 | rs10767809 | 1.072 | -0.401 | -1.935 | 0.311 | -0.118 | -0.087 | -0.087 | 0 | 0.17 | 0.01 | 0.3 | 0.25 | 0.28 | 0.28 |
| 11 | 29475619 | rs2667431 | 2.765 | -0.902 | -2.139 | 0.04 | 0.004 | -0.065 | -0.065 | 0 | 0.17 | 0.03 | 0.1 | 0.32 | 0.16 | 0.21 |
| 11 | 29476307 | rs541109 | 0.15 | -1.127 | -1.469 | -0.101 | 0.001 | 0.004 | 0.004 | 0 | 0.34 | 0.34 | 0.21 | 0.24 | 0.31 | 0.28 |
| 11 | 29477079 | rs12224306 | 1.157 | -0.998 | -1.712 | -0.116 | -0.041 | -0.027 | -0.027 | 1.96 | 0.18 | 0.1 | 0.3 | 0.25 | 0.28 | 0.28 |
| 11 | 29477653 | rs10488789 | 1.824 | -1.382 | -1.422 | -0.329 | -0.012 | -0.028 | -0.028 | 0.619 | 0.06 | 0.14 | 0.22 | 0 | 0.05 | 0.04 |
| 11 | 29478337 | rs10488790 | -1.53 | -1.327 | -1.745 | -0.361 | -0.002 | 0 | 0 | 0 | 0.79 | 0.82 | 0.89 | 0.84 | 0.88 | 0.93 |
| 11 | 29479973 | rs2667428 | 0.716 | -0.476 | -1.596 | -0.084 | -0.031 | -0.069 | -0.069 | 0 | 0.19 | 0.02 | 0.14 | 0.32 | 0.24 | 0.22 |
| 11 | 29485514 | rs12292242 | 0.719 | -0.848 | -1.415 | -0.018 | 0.001 | -0.011 | -0.011 | 0.002 | 0.12 | 0.13 | 0.08 | 0.01 | 0.1 | 0.1 |
| 11 | 29488271 | rs584041 | 0.654 | -0.432 | -1.333 | 0.466 | 0.012 | -0.051 | -0.051 | 0 | 0.18 | 0.06 | 0.1 | 0.34 | 0.16 | 0.22 |
| 11 | 29490531 | rs546771 | 0.482 | -0.41 | -1.283 | 0.368 | 0.006 | -0.01 | -0.01 | 0.054 | 0.19 | 0.17 | 0.11 | 0.33 | 0.16 | 0.22 |
| 11 | 29492526 | rs534880 | -0.358 | 0.076 | -1.311 | 0.23 | -0.028 | -0.063 | -0.063 | 0.095 | 0.78 | 0.93 | 0.86 | 0.7 | 0.76 | 0.81 |
| 11 | 29493216 | rs7116943 | -0.162 | -0.171 | -0.757 | -0.445 | 0.001 | 0.001 | 0.001 | 0 | 0.16 | 0.19 | 0.11 | 0.12 | 0.12 | 0.16 |
| 11 | 29500160 | rs1511518 | -0.601 | -0.11 | -0.615 | -0.656 | -0.001 | -0.006 | -0.006 | 0 | 0.37 | 0.62 | 0.56 | 0.15 | 0.41 | 0.48 |
| 11 | 29507866 | rs688610 | 0.588 | -0.273 | -0.81 | -0.651 | -0.025 | -0.043 | -0.043 | 0 | 0.13 | 0.33 | 0.08 | 0.05 | 0.06 | 0.1 |
| 11 | 29512184 | rs481303 | -0.134 | 0.223 | -1.444 | -0.374 | 0.024 | 0.01 | 0.01 | 0.19 | 0.46 | 0.65 | 0.35 | 0.57 | 0.34 | 0.47 |
| 11 | 29514307 | rs7126351 | 0.826 | 0.302 | -1.433 | -0.4 | -0.018 | 0.018 | 0.018 | 0 | 0.14 | 0.08 | 0.23 | 0.06 | 0.22 | 0.18 |
| 11 | 29517230 | rs645286 | 0.106 | -0.733 | -0.762 | 0.232 | -0.038 | 0.054 | 0.054 | 2.212 | 0.32 | 0.47 | 0.24 | 0.47 | 0.22 | 0.22 |
| 11 | 29521581 | rs2779149 | 0.023 | -0.63 | -1.003 | 0.284 | -0.034 | 0.039 | 0.039 | 1.294 | 0.33 | 0.51 | 0.25 | 0.47 | 0.22 | 0.34 |
| 11 | 29523877 | rs2779148 | -0.51 | -0.101 | -1.19 | 0.342 | 0.034 | 0.014 | 0.014 | 1.071 | 0.53 | 0.86 | 0.57 | 0.57 | 0.38 | 0.52 |
| 11 | 29527661 | rs1009070 | -0.484 | 0.284 | -0.718 | 0.362 | -0.01 | -0.028 | -0.028 | 0 | 0.07 | 0.22 | 0.22 | 0 | 0.05 | 0.04 |
| 11 | 29530247 | rs1231637 | -0.281 | 0.539 | -1.714 | 0.55 | -0.019 | 0.008 | 0.008 | 0.151 | 0.56 | 0.87 | 0.58 | 0.59 | 0.42 | 0.55 |
| 11 | 29532586 | rs11030732 | 0.31 | 0.798 | -2.479 | 0.33 | -0.029 | 0.093 | 0.093 | 0.493 | 0.61 | 0.09 | 0.72 | 0.75 | 0.65 | 0.59 |
| 11 | 29536437 | rs1123563 | 0.332 | 0.775 | -2.49 | 0.451 | -0.032 | 0.076 | 0.076 | 0.533 | 0.61 | 0.08 | 0.72 | 0.72 | 0.65 | 0.57 |
| 11 | 29541532 | rs10835534 | -0.479 | 0.736 | -2.263 | 0.298 | -0.006 | 0.003 | 0.003 | 0 | 0.12 | 0.09 | 0.12 | 0.13 | 0.15 | 0.11 |
| 11 | 29545690 | rs2137063 | -0.919 | 1.205 | -1.656 | 1.081 | -0.056 | -0.089 | -0.089 | 1.057 | 0.87 | 0.99 | 0.85 | 0.73 | 0.82 | 0.89 |
| 11 | 29552091 | rs10742224 | -1.109 | 1.192 | -0.433 | 0.771 | 0.038 | -0.029 | -0.029 | 1.452 | 0.18 | 0.43 | 0.18 | 0.13 | 0.21 | 0.16 |
| 11 | 29559404 | rs1511512 | -1.26 | 1.129 | -0.201 | 0.725 | 0.041 | -0.041 | -0.041 | 0 | 0.19 | 0.52 | 0.19 | 0.13 | 0.21 | 0.16 |
| 11 | 29568315 | rs7929790 | -1.115 | 1.129 | -0.206 | 0.725 | 0.037 | -0.033 | -0.033 | 0.046 | 0.18 | 0.43 | 0.17 | 0.13 | 0.21 | 0.16 |
| 11 | 29575377 | rs1511516 | -1.189 | 0.957 | -0.417 | 0.626 | 0.02 | 0.035 | 0.035 | 0.828 | 0.38 | 0.95 | 0.24 | 0.29 | 0.27 | 0.43 |

| | | | | | | | | | | | | | | | | |
|---|---|---|---|---|---|---|---|---|---|---|---|---|---|---|---|---|
| 11 | 29591246 | rs326751 | -0.348 | 0.898 | -0.267 | 0.595 | 0.05 | 0.061 | 0.061 | 0.851 | 0.26 | 0.84 | 0.13 | 0.14 | 0.15 | 0.31 |
| 11 | 29598857 | rs326753 | -0.729 | 0.904 | -0.148 | 0.647 | -0.007 | -0.01 | -0.01 | 0.249 | 0.06 | 0.67 | 0.04 | 0 | 0.01 | 0.04 |
| 11 | 29606266 | rs326765 | -1.323 | 0.917 | -0.116 | 0.674 | -0.008 | -0.011 | -0.011 | 0 | 0.17 | 0.76 | 0.15 | 0.13 | 0.12 | 0.14 |
| 11 | 29609480 | rs326769 | -0.701 | 0.893 | 0.045 | 0.652 | 0.002 | -0.001 | -0.001 | 0 | 0.06 | 0.64 | 0.04 | 0 | 0.01 | 0.04 |
| 11 | 29610778 | rs1947305 | -0.934 | 0.974 | -0.019 | 0.655 | -0.014 | 0.019 | 0.019 | 0 | 0.11 | 0.09 | 0.15 | 0.05 | 0.18 | 0.13 |
| 11 | 29623372 | rs326706 | 0.231 | 1.24 | 0.526 | 0.667 | -0.002 | -0.001 | -0.001 | 0 | 0.21 | 0.23 | 0.11 | 0.23 | 0.19 | 0.2 |
| 11 | 29637764 | rs1361643 | -1.305 | 1.137 | 0.505 | 0.528 | 0.013 | -0.083 | -0.083 | 0 | 0.15 | 0.34 | 0.16 | 0.02 | 0.18 | 0.11 |
| 11 | 29638345 | rs326788 | 0.083 | 1.095 | 0.516 | 0.981 | -0.003 | -0.002 | -0.002 | 0 | 0.22 | 0.24 | 0.11 | 0.22 | 0.18 | 0.28 |
| 11 | 29646947 | rs10835549 | 0.101 | 1.102 | 0.706 | 0.957 | 0 | -0.01 | -0.01 | 0.319 | 0.07 | 0.07 | 0.1 | 0.01 | 0.08 | 0.09 |
| 11 | 29649814 | rs7944673 | -1.844 | 1.084 | 0.796 | 0.959 | 0.014 | -0.052 | -0.052 | 0.01 | 0.18 | 0.68 | 0.19 | 0.08 | 0.21 | 0.15 |
| 11 | 29651299 | rs704617 | -1.458 | 1.349 | 0.615 | 0.946 | -0.08 | -0.024 | -0.024 | 0 | 0.4 | 0.95 | 0.31 | 0.39 | 0.42 | 0.44 |
| 11 | 29659444 | rs2447045 | -0.019 | 1.446 | 0.942 | 0.89 | -0.001 | -0.014 | -0.014 | 1.922 | 0.23 | 0.31 | 0.2 | 0.22 | 0.22 | 0.28 |
| 11 | 29661845 | rs4614422 | 0.966 | 1.377 | -0.004 | 1.052 | -0.043 | -0.08 | -0.08 | 0 | 0.14 | 0.01 | 0.21 | 0.2 | 0.17 | 0.12 |
| 11 | 29663086 | rs704616 | -1.972 | 1.132 | -0.614 | 0.641 | -0.032 | 0.134 | 0.134 | 1.001 | 0.68 | 0.97 | 0.7 | 0.91 | 0.7 | 0.82 |
| 11 | 29675343 | rs697311 | -0.808 | 0.941 | -0.257 | 1.023 | -0.032 | -0.031 | -0.031 | 0.07 | 0.37 | 0.95 | 0.34 | 0.38 | 0.33 | 0.33 |
| 11 | 29680576 | rs324299 | -1.139 | 1.009 | 0.504 | 1.097 | -0.008 | -0.005 | -0.005 | 0.176 | 0.39 | 0.64 | 0.39 | 0.44 | 0.45 | 0.44 |
| 11 | 29693578 | rs324293 | -1.055 | 0.986 | 0.57 | 1.054 | -0.001 | -0.017 | -0.017 | 0.326 | 0.38 | 0.76 | 0.35 | 0.44 | 0.35 | 0.34 |
| 11 | 29699270 | rs10835555 | -0.983 | 1.08 | 0.747 | 1.068 | 0.006 | -0.008 | -0.008 | 0 | 0.13 | 0.14 | 0.17 | 0.07 | 0.21 | 0.12 |
| 11 | 29710982 | rs326740 | -0.369 | 1.066 | 1.078 | 1.041 | 0.001 | 0.001 | 0.001 | 0 | 0.3 | 0.29 | 0.45 | 0.31 | 0.32 | 0.29 |
| 11 | 29757305 | rs10160238 | -0.319 | 1.105 | 0.367 | 0.859 | -0.02 | 0.02 | 0.02 | 0 | 0.1 | 0.05 | 0.15 | 0.02 | 0.18 | 0.1 |
| 11 | 29772443 | rs1154975 | -1.817 | 0.913 | 0.365 | 1.023 | 0.005 | 0.063 | 0.063 | 0 | 0.54 | 0.75 | 0.69 | 0.7 | 0.55 | 0.67 |
| 11 | 29774810 | rs1002728 | -0.751 | 0.902 | 0.864 | 1.255 | 0.017 | -0.005 | -0.005 | 0 | 0.36 | 0.5 | 0.5 | 0.32 | 0.41 | 0.4 |
| 11 | 29776185 | rs722530 | 0.571 | 0.76 | 0.974 | 1.213 | 0 | 0.006 | 0.006 | 0 | 0.21 | 0.23 | 0.19 | 0.3 | 0.21 | 0.25 |
| 11 | 29782559 | rs10835564 | -0.025 | 0.704 | 1.07 | 1.192 | -0.005 | 0.039 | 0.039 | 3.641 | 0.16 | 0.21 | 0.18 | 0.39 | 0.13 | 0.27 |
| 11 | 29795497 | rs1322507 | -0.117 | 0.724 | 0.752 | 1.21 | -0.002 | -0.013 | -0.013 | 0 | 0.17 | 0.11 | 0.21 | 0.34 | 0.18 | 0.25 |
| 11 | 29795917 | rs10835566 | 0.132 | 0.79 | 0.6 | 0.996 | -0.002 | 0.008 | 0.008 | 0 | 0.08 | 0.05 | 0.11 | 0.02 | 0.09 | 0.08 |
| 11 | 29795932 | rs10835567 | 0.099 | 0.788 | 0.624 | 0.994 | 0 | -0.006 | -0.006 | 0.025 | 0.08 | 0.07 | 0.11 | 0.02 | 0.09 | 0.08 |
| 11 | 29800788 | rs4278493 | 0.391 | 0.747 | 0.489 | 1.218 | -0.047 | -0.003 | -0.003 | 0.363 | 0.1 | 0.01 | 0.11 | 0.1 | 0.14 | 0.17 |
| 11 | 29806844 | rs1590287 | -1.867 | 0.26 | 0.634 | 0.97 | -0.001 | 0.003 | 0.003 | 1.992 | 0.37 | 0.57 | 0.37 | 0.47 | 0.34 | 0.41 |
| 11 | 29814400 | rs417424 | 2.183 | 0.159 | -0.094 | 0.979 | -0.062 | 0.06 | 0.06 | 0.989 | 0.42 | 0.08 | 0.43 | 0.26 | 0.47 | 0.34 |
| 11 | 29815530 | rs10835581 | -0.463 | 0.261 | 0.053 | 0.992 | -0.005 | -0.012 | -0.012 | 0 | 0.21 | 0.14 | 0.23 | 0.38 | 0.24 | 0.28 |
| 11 | 29821636 | rs322622 | -1.083 | 0.356 | -0.465 | 0.81 | -0.002 | -0.021 | -0.021 | 1.605 | 0.81 | 0.87 | 0.81 | 0.74 | 0.82 | 0.75 |
| 11 | 29822057 | rs11030828 | -0.62 | 0.336 | -0.453 | 0.417 | -0.004 | -0.041 | -0.041 | 1.861 | 0.13 | 0.2 | 0.13 | 0.04 | 0.12 | 0.11 |
| 11 | 29822605 | rs441703 | 0.622 | 0.448 | -0.776 | 0.738 | -0.023 | 0.09 | 0.09 | 4.834 | 0.54 | 0.15 | 0.54 | 0.29 | 0.57 | 0.42 |
| 11 | 29823335 | rs322623 | 0.36 | 0.077 | -0.415 | 0.911 | -0.001 | -0.001 | -0.001 | 0 | 0.61 | 0.6 | 0.63 | 0.31 | 0.61 | 0.55 |
| 11 | 29826143 | rs1336820 | 0.429 | -0.826 | -0.955 | 1.091 | 0.022 | -0.096 | -0.096 | 0.169 | 0.14 | 0.01 | 0.21 | 0.31 | 0.08 | 0.2 |
| 11 | 29827214 | rs9300005 | 0.401 | -1.361 | -0.393 | 0.516 | 0.007 | 0.01 | 0.01 | 0 | 0.38 | 0.28 | 0.26 | 0.26 | 0.35 | 0.42 |
| 11 | 29845532 | rs535806 | -1.318 | -1.34 | -0.696 | 0.54 | -0.053 | 0.063 | 0.063 | 1.324 | 0.64 | 0.9 | 0.54 | 0.79 | 0.47 | 0.69 |
| 11 | 29861453 | rs12291179 | 1.312 | -1.556 | -0.428 | 0.376 | -0.003 | -0.03 | -0.03 | 0 | 0.14 | 0.15 | 0.07 | 0.01 | 0.14 | 0.19 |
| 11 | 29863909 | rs1415118 | -1.468 | -1.304 | -0.541 | 0.31 | -0.089 | 0.058 | 0.058 | 0.697 | 0.67 | 0.89 | 0.57 | 0.81 | 0.52 | 0.7 |
| 11 | 29872433 | rs11030846 | 1.296 | -1.592 | -0.397 | 0.593 | 0.023 | -0.079 | -0.079 | 0 | 0.14 | 0.06 | 0.21 | 0.35 | 0.1 | 0.22 |
| 11 | 29887903 | rs3120601 | -1.455 | -1.639 | -0.056 | 0.568 | -0.037 | 0.01 | 0.01 | 0.632 | 0.74 | 0.65 | 0.88 | 0.72 | 0.83 | 0.79 |
| 11 | 29888148 | rs12421390 | 1.089 | -1.635 | -0.04 | 0.502 | 0.001 | -0.016 | -0.016 | 0 | 0.16 | 0.15 | 0.16 | 0.01 | 0.19 | 0.23 |
| 11 | 29895210 | rs10501118 | 0.897 | -1.672 | -0.071 | 0.528 | 0.036 | -0.086 | -0.086 | 0.005 | 0.17 | 0.1 | 0.23 | 0.51 | 0.11 | 0.25 |
| 11 | 29897097 | rs10501120 | 0.9 | -1.663 | -0.072 | 0.486 | 0.034 | -0.103 | -0.103 | 0 | 0.17 | 0.04 | 0.22 | 0.36 | 0.1 | 0.23 |
| 11 | 29897759 | rs1746942 | 1.033 | -1.574 | -0.016 | 0.333 | -0.038 | 0.096 | 0.096 | 1.062 | 0.37 | 0.12 | 0.47 | 0.19 | 0.53 | 0.31 |
| 11 | 29902645 | rs1764528 | 0.211 | -1.54 | 0.009 | 0.713 | 0.097 | 0.114 | 0.114 | 1.801 | 0.54 | 0.28 | 0.63 | 0.21 | 0.72 | 0.54 |
| 11 | 29941778 | rs554651 | 1.177 | -0.688 | -0.63 | 0.837 | 0.001 | -0.001 | -0.001 | 0 | 0.21 | 0.14 | 0.1 | 0.24 | 0.16 | 0.17 |
| 11 | 29948481 | rs12576044 | 0.361 | 0.448 | -1.012 | 0.939 | -0.073 | 0.054 | 0.054 | 0 | 0.31 | 0.1 | 0.37 | 0.2 | 0.36 | 0.3 |
| 11 | 29964840 | rs1425170 | 0.193 | 0.761 | -1.605 | 0.992 | -0.057 | -0.115 | -0.115 | 0 | 0.24 | 0.03 | 0.31 | 0.32 | 0.25 | 0.26 |
| 11 | 29969227 | rs568970 | 0.036 | 0.131 | -0.758 | 0.455 | 0.002 | 0.005 | 0.005 | 0.021 | 0.4 | 0.31 | 0.25 | 0.26 | 0.39 | 0.38 |
| 11 | 29990325 | rs500745 | 0.356 | 0.312 | -0.585 | 0.999 | 0.005 | 0.058 | 0.058 | 1.405 | 0.74 | 0.58 | 0.47 | 0.56 | 0.71 | 0.8 |
| 11 | 29994465 | rs2993051 | 0.554 | 0.005 | -0.996 | 0.805 | -0.004 | -0.035 | -0.035 | 5.501 | 0.74 | 0.78 | 0.49 | 0.4 | 0.69 | 0.71 |
| 11 | 30004238 | rs10835606 | -0.222 | 0.148 | -1.026 | 1.235 | -0.01 | -0.022 | -0.022 | 0 | 0.06 | 0 | 0.14 | 0.27 | 0.06 | 0.1 |
| 11 | 30012458 | rs533448 | 0.845 | 0.642 | -1.815 | -0.307 | -0.108 | 0.096 | 0.096 | 0 | 0.22 | 0.02 | 0.21 | 0.05 | 0.3 | 0.25 |
| 11 | 30023175 | rs569724 | 0.828 | 0.204 | -2.136 | -0.597 | -0.056 | -0.057 | -0.057 | 0 | 0.19 | 0.06 | 0.16 | 0.24 | 0.25 | 0.15 |
| 11 | 30027074 | rs524373 | 0.848 | 0.197 | -2.144 | -0.6 | -0.075 | -0.074 | -0.074 | 0 | 0.19 | 0.02 | 0.15 | 0.24 | 0.25 | 0.15 |
| 11 | 30033191 | rs3802914 | 0.022 | -0.336 | -1.442 | -1.04 | -0.021 | -0.086 | -0.086 | 0 | 0.2 | 0.32 | 0.18 | 0.04 | 0.17 | 0.11 |
| 11 | 30035236 | rs1323860 | -0.683 | -0.527 | -1.416 | -1.182 | 0 | -0.012 | -0.012 | 0 | 0.42 | 0.46 | 0.37 | 0.27 | 0.44 | 0.28 |
| 11 | 30036177 | rs10835608 | 0.716 | -0.792 | -1.243 | -1.262 | -0.004 | 0.004 | 0.004 | 0 | 0.29 | 0.27 | 0.4 | 0.63 | 0.24 | 0.36 |
| 11 | 30047512 | rs10767831 | -0.76 | -0.759 | -1.273 | -1.321 | 0.047 | 0.053 | 0.053 | 0 | 0.52 | 0.67 | 0.62 | 0.69 | 0.43 | 0.49 |
| 11 | 30049635 | rs11030921 | 0.948 | -0.464 | -1.429 | -1.05 | -0.019 | 0.071 | 0.071 | 0 | 0.12 | 0.01 | 0.13 | 0 | 0.13 | 0.07 |
| 11 | 30054459 | rs7944308 | -0.416 | -0.305 | -1.055 | -0.977 | 0.005 | -0.024 | -0.024 | 0 | 0.61 | 0.51 | 0.65 | 0.75 | 0.57 | 0.73 |
| 11 | 30056918 | rs1425172 | -0.294 | -0.455 | -0.725 | -0.176 | -0.006 | 0.005 | 0.005 | 0 | 0.29 | 0.27 | 0.4 | 0.65 | 0.23 | 0.36 |
| 11 | 30065448 | rs542926 | -0.197 | 0.225 | -0.646 | -1.089 | 0 | 0.072 | 0.072 | 0 | 0.43 | 0.19 | 0.56 | 0.72 | 0.4 | 0.4 |
| 11 | 30092081 | rs1436094 | 0.759 | 0.647 | -0.196 | -1.714 | -0.028 | 0.038 | 0.038 | 0 | 0.34 | 0.04 | 0.21 | 0.23 | 0.34 | 0.32 |
| 11 | 30096930 | rs665153 | -0.073 | 0.681 | 0.261 | -1.705 | 0.089 | -0.127 | -0.127 | 0.09 | 0.83 | 0.5 | 0.81 | 0.98 | 0.76 | 0.75 |
| 11 | 30100728 | rs12277573 | 0.285 | 0.822 | 0.377 | -1.75 | -0.014 | 0.011 | 0.011 | 0.29 | 0.07 | 0.04 | 0.06 | 0.02 | 0.14 | 0.19 |
| 11 | 30111109 | rs1436093 | 1.143 | 0.768 | 0.39 | -1.777 | 0.012 | -0.08 | -0.08 | 0 | 0.78 | 0.26 | 0.77 | 0.95 | 0.74 | 0.72 |

| | | | | | | | | | | | | | | | | |
|---|---|---|---|---|---|---|---|---|---|---|---|---|---|---|---|---|
| 11 | 30134926 | rs1222164 | -0.699 | 0.865 | 0.228 | -1.824 | 0.084 | -0.034 | -0.034 | 0 | 0.94 | 0.79 | 0.93 | 0.99 | 0.86 | 0.94 |
| 11 | 30135595 | rs1369810 | 0.938 | 0.671 | -0.071 | -1.865 | -0.024 | 0.039 | 0.039 | 0.744 | 0.35 | 0.04 | 0.21 | 0.23 | 0.34 | 0.32 |
| 11 | 30137328 | rs1436096 | -0.307 | 0.182 | 0.118 | -1.732 | -0.001 | -0.008 | -0.008 | 1.926 | 0.5 | 0.47 | 0.6 | 0.76 | 0.42 | 0.57 |
| 11 | 30144366 | rs997083 | 0.028 | 0.14 | 0.242 | -1.724 | -0.002 | -0.02 | -0.02 | 0 | 0.46 | 0.45 | 0.58 | 0.76 | 0.41 | 0.44 |
| 11 | 30144581 | rs16920207 | -0.282 | 0.349 | 0.564 | -1.223 | 0.048 | 0.045 | 0.045 | 0 | 0.05 | 0.29 | 0.05 | 0.08 | 0.04 | 0.04 |
| 11 | 30147285 | rs4923621 | 0.496 | 0.98 | 0.022 | -1.551 | -0.035 | 0.021 | 0.021 | 2.817 | 0.35 | 0.18 | 0.41 | 0.25 | 0.42 | 0.28 |
| 11 | 30164732 | rs10835621 | 0.313 | 0.741 | 0.681 | -1.681 | 0.008 | -0.015 | -0.015 | 7.638 | 0.36 | 0.41 | 0.25 | 0.24 | 0.37 | 0.38 |
| 11 | 30165896 | rs930321 | 0.406 | 0.716 | 0.892 | -1.695 | -0.018 | 0.017 | 0.017 | 5.717 | 0.34 | 0.2 | 0.24 | 0.24 | 0.37 | 0.37 |
| 11 | 30166240 | rs1898174 | 0.3 | 0.716 | 0.876 | -1.695 | 0.008 | -0.015 | -0.015 | 0 | 0.36 | 0.41 | 0.25 | 0.24 | 0.37 | 0.38 |
| 11 | 30167146 | rs16925531 | 0.631 | 0.404 | 0.611 | -1.752 | -0.004 | 0.048 | 0.048 | 0.147 | 0.19 | 0.23 | 0.41 | 0.62 | 0.19 | 0.23 |
| 11 | 30169600 | rs601937 | 0.503 | 0.098 | 0.707 | -1.884 | -0.009 | -0.04 | -0.04 | 0 | 0.11 | 0.17 | 0.07 | 0 | 0.09 | 0.05 |
| 11 | 30169856 | rs663173 | 0.32 | 0.133 | 0.637 | -1.845 | 0.003 | -0.015 | -0.015 | 0 | 0.38 | 0.56 | 0.27 | 0.24 | 0.37 | 0.38 |
| 11 | 30170373 | rs3908935 | 0.513 | 0.19 | 0.54 | -1.817 | 0.008 | -0.015 | -0.015 | 0.34 | 0.36 | 0.41 | 0.25 | 0.24 | 0.37 | 0.38 |
| 11 | 30177010 | rs3858479 | 0.834 | 0.171 | 0.362 | -1.812 | -0.021 | 0.009 | 0.009 | 4.294 | 0.32 | 0.19 | 0.24 | 0.24 | 0.37 | 0.25 |
| 11 | 30206532 | rs11030997 | 0.966 | 0.626 | -1.17 | -1.612 | -0.083 | -0.004 | -0.004 | 0 | 0.3 | 0.08 | 0.23 | 0.24 | 0.37 | 0.25 |
| 11 | 30206846 | rs2682344 | -0.362 | 0.166 | -0.491 | -1.194 | -0.005 | -0.032 | -0.032 | 0 | 0.12 | 0.16 | 0.07 | 0 | 0.09 | 0.06 |
| 11 | 30214713 | rs961605 | -0.613 | 0.119 | -0.989 | -0.989 | 0.086 | 0.054 | 0.054 | 0 | 0.48 | 0.73 | 0.38 | 0.63 | 0.4 | 0.56 |
| 11 | 30226528 | rs11031006 | 1.224 | -0.276 | -1.212 | -1.214 | 0.007 | 0.034 | 0.034 | 0 | 0.12 | 0.05 | 0.1 | 0.03 | 0.14 | 0.1 |
| 11 | 30234394 | rs601681 | -0.79 | 0.664 | -0.725 | -1.453 | 0.02 | 0.016 | 0.016 | 0 | 0.49 | 0.55 | 0.39 | 0.64 | 0.41 | 0.56 |
| 11 | 30243247 | rs543401 | -0.733 | 0.684 | -0.761 | -1.417 | 0.022 | 0.025 | 0.025 | 0 | 0.5 | 0.56 | 0.52 | 0.67 | 0.45 | 0.58 |
| 11 | 30243868 | rs1782507 | -1.158 | 0.689 | -1.077 | -1.782 | -0.03 | 0.091 | 0.091 | 0.246 | 0.69 | 0.82 | 0.65 | 0.9 | 0.67 | 0.7 |
| 11 | 30251625 | rs639403 | -0.358 | 0.668 | -1.067 | -1.773 | 0.022 | 0.025 | 0.025 | 0 | 0.5 | 0.56 | 0.52 | 0.67 | 0.45 | 0.58 |
| 11 | 30255185 | rs6169 | -0.538 | 0.651 | -1.039 | -1.783 | 0.063 | 0.082 | 0.082 | 0 | 0.52 | 0.76 | 0.53 | 0.67 | 0.45 | 0.59 |
| 11 | 30255982 | rs676349 | -0.333 | 0.639 | -1.053 | -1.78 | 0.022 | 0.026 | 0.026 | 0 | 0.5 | 0.56 | 0.52 | 0.67 | 0.45 | 0.58 |
| 11 | 30258989 | rs615577 | -0.355 | 0.639 | -1.054 | -1.779 | 0.021 | 0.025 | 0.025 | 0 | 0.5 | 0.56 | 0.52 | 0.67 | 0.45 | 0.58 |
| 11 | 30273750 | rs10734405 | -1.852 | 0.691 | -1.053 | -1.734 | -0.05 | -0.042 | -0.042 | 0 | 0.83 | 0.95 | 0.89 | 0.77 | 0.78 | 0.89 |
| 11 | 30310992 | rs1933343 | -1.809 | 0.673 | -1.002 | -1.739 | -0.043 | -0.037 | -0.037 | 0 | 0.83 | 0.93 | 0.88 | 0.77 | 0.78 | 0.89 |
| 11 | 30334223 | rs1856142 | 0.182 | 0.54 | -0.836 | -1.83 | -0.041 | 0.008 | 0.008 | 0 | 0.36 | 0.46 | 0.29 | 0.61 | 0.28 | 0.46 |
| 11 | 30342542 | rs4083926 | 0.878 | 0.135 | -0.39 | -2.065 | -0.011 | 0.022 | 0.022 | 0.261 | 0.23 | 0.22 | 0.22 | 0.6 | 0.2 | 0.31 |
| 11 | 30343925 | rs1222220 | -1.169 | 0.568 | -0.562 | -1.814 | -0.035 | 0.076 | 0.076 | 0.275 | 0.69 | 0.81 | 0.54 | 0.87 | 0.67 | 0.7 |
| 11 | 30353725 | rs1765139 | 1.319 | 0.608 | -0.403 | -1.691 | -0.014 | -0.013 | -0.013 | 0 | 0.18 | 0.12 | 0.12 | 0.23 | 0.22 | 0.1 |
| 11 | 30361377 | rs1222208 | 1.515 | 0.636 | -0.572 | -1.678 | -0.052 | -0.046 | -0.046 | 0 | 0.17 | 0.05 | 0.11 | 0.23 | 0.22 | 0.11 |
| 11 | 30378559 | rs1765142 | -0.882 | 0.522 | -0.696 | -1.456 | -0.041 | 0.064 | 0.064 | 0.017 | 0.72 | 0.82 | 0.64 | 0.9 | 0.66 | 0.7 |
| 11 | 30389710 | rs528292 | -1.425 | 0.214 | -0.766 | -1.106 | -0.048 | -0.049 | -0.049 | 0 | 0.78 | 0.95 | 0.87 | 0.76 | 0.75 | 0.88 |
| 11 | 30393360 | rs640245 | 0.248 | 0.173 | -0.784 | -0.914 | -0.038 | 0.094 | 0.094 | 0.053 | 0.4 | 0.25 | 0.46 | 0.18 | 0.45 | 0.39 |
| 11 | 30394734 | rs2183848 | -0.2 | -0.023 | -0.565 | -0.736 | 0.028 | -0.02 | -0.02 | 0 | 0.38 | 0.65 | 0.3 | 0.54 | 0.3 | 0.49 |
| 11 | 30397010 | rs522089 | -0.459 | 0.084 | -0.643 | -0.644 | -0.014 | 0.019 | 0.019 | 0 | 0.69 | 0.61 | 0.73 | 0.43 | 0.71 | 0.6 |
| 11 | 30405254 | rs1765132 | 1.123 | 0.065 | -1.151 | -0.842 | 0.023 | -0.007 | -0.007 | 0.23 | 0.16 | 0.08 | 0.2 | 0.17 | 0.11 | 0.13 |
| 11 | 30412174 | rs592328 | 0.564 | 0.404 | -0.934 | -1.036 | 0.021 | -0.09 | -0.09 | 0 | 0.3 | 0.4 | 0.3 | 0.07 | 0.35 | 0.33 |
| 11 | 30422068 | rs2065418 | 0.327 | 0.339 | -1.089 | -0.982 | 0.004 | -0.006 | -0.006 | 0.271 | 0.38 | 0.13 | 0.34 | 0.34 | 0.35 | 0.25 |
| 11 | 30432176 | rs553951 | -0.587 | -0.008 | -0.909 | -0.851 | -0.022 | 0.026 | 0.026 | 0 | 0.68 | 0.55 | 0.64 | 0.41 | 0.71 | 0.58 |
| 11 | 30435051 | rs567056 | 0.48 | 0.157 | -1.029 | -0.988 | 0.005 | -0.047 | -0.047 | 0.095 | 0.29 | 0.32 | 0.3 | 0.06 | 0.36 | 0.32 |
| 11 | 30435585 | rs2282300 | 0.899 | 0.119 | -1.495 | -0.546 | -0.057 | -0.011 | -0.011 | 0 | 0.22 | 0.05 | 0.14 | 0.17 | 0.24 | 0.13 |
| 11 | 30436400 | rs10767843 | 0.523 | -0.239 | -0.963 | -0.314 | 0.01 | -0.001 | -0.001 | 0.746 | 0.17 | 0.14 | 0.2 | 0.17 | 0.11 | 0.12 |
| 11 | 30439707 | rs16920647 | 0.682 | -0.236 | -1.079 | -0.138 | 0.038 | -0.021 | -0.021 | 0.032 | 0.16 | 0.05 | 0.08 | 0.18 | 0.11 | 0.13 |
| 11 | 30440708 | rs474376 | 0.469 | -0.862 | -0.782 | -0.591 | -0.002 | 0 | 0 | 0 | 0.65 | 0.31 | 0.7 | 0.79 | 0.79 | 0.67 |
| 11 | 30441946 | rs597053 | 1.306 | -0.201 | -1.495 | -0.823 | -0.116 | 0.081 | 0.081 | 0 | 0.2 | 0.02 | 0.17 | 0.06 | 0.29 | 0.24 |
| 11 | 30443248 | rs11605215 | 1.289 | -0.122 | -1.67 | 0.148 | -0.052 | -0.088 | -0.088 | 0 | 0.16 | 0.02 | 0.27 | 0.31 | 0.22 | 0.16 |
| 11 | 30447998 | rs704660 | -0.279 | -0.592 | -0.835 | 0.076 | 0.008 | 0.038 | 0.038 | 0 | 0.45 | 0.67 | 0.29 | 0.24 | 0.4 | 0.38 |
| 11 | 30449743 | rs10835665 | 0.657 | -0.702 | -0.343 | 0.096 | -0.076 | -0.004 | -0.004 | 1.396 | 0.21 | 0.44 | 0.11 | 0.18 | 0.11 | 0.14 |
| 11 | 30452181 | rs514644 | -0.351 | -0.8 | -0.379 | 0.422 | -0.037 | 0.022 | 0.022 | 0 | 0.32 | 0.66 | 0.29 | 0.41 | 0.18 | 0.42 |
| 11 | 30455242 | rs808182 | 0.495 | -0.778 | -0.391 | 0.447 | -0.007 | 0.044 | 0.044 | 0 | 0.11 | 0.22 | 0.18 | 0.23 | 0.07 | 0.28 |
| 11 | 30456390 | rs11031084 | 1.143 | -0.683 | -0.555 | 0.359 | -0.109 | 0.09 | 0.09 | 0 | 0.23 | 0.04 | 0.23 | 0.06 | 0.32 | 0.26 |
| 11 | 30456545 | rs980561 | 1.143 | -0.683 | -0.555 | 0.359 | -0.109 | 0.09 | 0.09 | 0 | 0.23 | 0.04 | 0.23 | 0.06 | 0.32 | 0.26 |
| 11 | 30456817 | rs980562 | 1.366 | -0.626 | -0.925 | 0.379 | -0.097 | 0.094 | 0.094 | 0 | 0.21 | 0.03 | 0.16 | 0.06 | 0.29 | 0.24 |
| 11 | 30457877 | rs808183 | 0.54 | -0.523 | -0.839 | 0.495 | 0 | 0.002 | 0.002 | 0 | 0.1 | 0.12 | 0.06 | 0.2 | 0.06 | 0.28 |
| 11 | 30459058 | rs504181 | -0.988 | -0.495 | -0.945 | 0.488 | -0.037 | -0.034 | -0.034 | 0.183 | 0.57 | 0.87 | 0.54 | 0.48 | 0.5 | 0.68 |
| 11 | 30469383 | rs567688 | 0.567 | -0.46 | -0.797 | 0.557 | -0.003 | 0.013 | 0.013 | 0 | 0.1 | 0.13 | 0.07 | 0.2 | 0.06 | 0.28 |
| 11 | 30471861 | rs557957 | -1.033 | -0.531 | -0.848 | 0.506 | -0.043 | -0.037 | -0.037 | 0 | 0.57 | 0.88 | 0.54 | 0.48 | 0.5 | 0.68 |
| 11 | 30475477 | rs529444 | 0.469 | -0.646 | -0.659 | 0.543 | -0.007 | 0.06 | 0.06 | 0 | 0.11 | 0.22 | 0.18 | 0.23 | 0.07 | 0.28 |
| 11 | 30479911 | rs10437628 | 0.384 | -0.765 | -0.573 | 0.407 | -0.084 | -0.014 | -0.014 | 3.195 | 0.23 | 0.56 | 0.12 | 0.18 | 0.11 | 0.14 |
| 11 | 30489027 | rs1026477 | 0.547 | -0.614 | -0.805 | 0.402 | -0.029 | 0 | 0 | 0 | 0.45 | 0.17 | 0.65 | 0.58 | 0.53 | 0.35 |
| 11 | 30489044 | rs7115139 | 0.177 | -0.872 | -0.423 | 0.472 | -0.065 | -0.012 | -0.012 | 0.213 | 0.23 | 0.6 | 0.12 | 0.18 | 0.12 | 0.15 |
| 11 | 30489776 | rs11031104 | 0.231 | -0.838 | -0.248 | 0.488 | -0.078 | -0.01 | -0.01 | 0 | 0.23 | 0.45 | 0.12 | 0.18 | 0.12 | 0.14 |
| 11 | 30504621 | rs2242144 | 0.932 | -0.276 | -0.658 | 0.779 | 0.004 | 0.01 | 0.01 | 0 | 0.15 | 0.12 | 0.28 | 0.3 | 0.21 | 0.16 |
| 11 | 30507095 | rs16920793 | -0.24 | -0.559 | 0.02 | 0.611 | 0.008 | 0.033 | 0.033 | 0.042 | 0.22 | 0.53 | 0.38 | 0.3 | 0.24 | 0.21 |
| 11 | 30508108 | rs545610 | -1.307 | -0.395 | -0.475 | -0.035 | -0.093 | 0.083 | 0.083 | 0.304 | 0.74 | 0.95 | 0.7 | 0.93 | 0.66 | 0.76 |
| 11 | 30511811 | rs1336516 | 0.831 | -0.581 | -0.558 | 0.123 | 0.034 | -0.017 | -0.017 | 0.567 | 0.41 | 0.18 | 0.24 | 0.42 | 0.35 | 0.27 |
| 11 | 30517996 | rs1458727 | 0.122 | -0.476 | -0.698 | 0.2 | 0.006 | 0.144 | 0.144 | 0 | 0.22 | 0.43 | 0.39 | 0.5 | 0.24 | 0.31 |

| | | | | | | | | | | | | | | | | |
|---|---|---|---|---|---|---|---|---|---|---|---|---|---|---|---|---|
| 11 | 30520163 | rs11031118 | 1.456 | -0.245 | -1.051 | 0.064 | -0.122 | 0.106 | 0.106 | 0 | 0.22 | 0.02 | 0.17 | 0.04 | 0.32 | 0.25 |
| 11 | 30528944 | rs1026479 | 1.203 | 0.405 | -0.892 | 0.399 | 0.006 | 0.03 | 0.03 | 0 | 0.16 | 0.13 | 0.3 | 0.46 | 0.26 | 0.29 |
| 11 | 30531483 | rs7944883 | 1.231 | -0.122 | -0.794 | 0.164 | 0.005 | 0.006 | 0.006 | 0 | 0.3 | 0.29 | 0.19 | 0.23 | 0.24 | 0.15 |
| 11 | 30532992 | rs6484487 | 0.081 | -0.119 | -0.796 | 0.15 | -0.002 | 0.009 | 0.009 | 0 | 0.47 | 0.42 | 0.56 | 0.7 | 0.52 | 0.46 |
| 11 | 30537157 | rs12419132 | 1.99 | -0.192 | -0.927 | 0.135 | 0.03 | -0.032 | -0.032 | 0.954 | 0.16 | 0.05 | 0.11 | 0.21 | 0.12 | 0.15 |
| 11 | 30541384 | rs531353 | -1.145 | 0.163 | -0.814 | 0.084 | -0.104 | 0.07 | 0.07 | 2.738 | 0.76 | 0.98 | 0.74 | 0.91 | 0.71 | 0.76 |
| 11 | 30541876 | rs7113764 | 1.2 | 0.262 | -0.641 | 0.171 | 0.006 | 0.031 | 0.031 | 0 | 0.16 | 0.13 | 0.3 | 0.46 | 0.25 | 0.29 |
| 11 | 30547438 | rs11031122 | 1.515 | 0.212 | -0.717 | -0.254 | -0.092 | 0.084 | 0.084 | 0.221 | 0.2 | 0.01 | 0.12 | 0.05 | 0.25 | 0.23 |
| 11 | 30552900 | rs6484488 | 1.089 | 0.102 | -0.767 | 0.054 | 0.003 | 0.013 | 0.013 | 0 | 0.17 | 0.12 | 0.26 | 0.42 | 0.25 | 0.3 |
| 11 | 30553329 | rs11031125 | 1.478 | 0.033 | -0.748 | 0.158 | -0.047 | 0.076 | 0.076 | 0.38 | 0.2 | 0.05 | 0.13 | 0.05 | 0.25 | 0.23 |
| 11 | 30554404 | rs559337 | -0.023 | -0.057 | -0.475 | 0.09 | 0.021 | 0.038 | 0.038 | 0.051 | 0.28 | 0.51 | 0.33 | 0.42 | 0.32 | 0.45 |
| 11 | 30555332 | rs811210 | -0.789 | -0.057 | -0.974 | 0.149 | 0.047 | 0.024 | 0.024 | 0 | 0.52 | 0.74 | 0.53 | 0.48 | 0.59 | 0.7 |
| 11 | 30564774 | rs537839 | -0.771 | -0.044 | -1.067 | 0.162 | 0.036 | 0.019 | 0.019 | 0 | 0.51 | 0.67 | 0.53 | 0.48 | 0.59 | 0.7 |
| 11 | 30567671 | rs10835687 | 0.985 | 0.175 | -1.517 | -0.166 | -0.087 | 0.083 | 0.083 | 0.323 | 0.2 | 0.01 | 0.12 | 0.05 | 0.25 | 0.23 |
| 11 | 30571347 | rs10767859 | 0.084 | -0.131 | -1.775 | -0.336 | -0.048 | -0.031 | -0.031 | 0 | 0.31 | 0.13 | 0.37 | 0.64 | 0.41 | 0.44 |
| 11 | 30574638 | rs16920925 | -0.135 | -0.606 | -1.19 | -0.498 | -0.007 | 0.016 | 0.016 | 0 | 0.13 | 0.4 | 0.11 | 0.13 | 0.1 | 0.09 |
| 11 | 30579485 | rs521069 | -0.103 | -0.537 | -1.756 | -0.702 | 0.018 | 0.005 | 0.005 | 0 | 0.35 | 0.36 | 0.57 | 0.66 | 0.45 | 0.37 |
| 11 | 30584613 | rs1374616 | 1.141 | -0.683 | -2.482 | -0.475 | -0.007 | 0.006 | 0.006 | 3.006 | 0.19 | 0.01 | 0.1 | 0.12 | 0.14 | 0.24 |
| 11 | 30592926 | rs533824 | -0.978 | -0.354 | -1.889 | -0.226 | 0.008 | 0.042 | 0.042 | 0.178 | 0.88 | 0.99 | 0.96 | 0.97 | 0.92 | 0.94 |
| 11 | 30599745 | rs3824982 | 0.243 | -0.103 | -1.931 | -0.054 | -0.035 | -0.045 | -0.045 | 0 | 0.19 | 0.05 | 0.14 | 0.19 | 0.23 | 0.29 |
| 11 | 30600383 | rs3781852 | -0.424 | -0.408 | -1.067 | -0.352 | 0.002 | -0.001 | -0.001 | 0 | 0.24 | 0.56 | 0.18 | 0.19 | 0.26 | 0.29 |
| 11 | 30602252 | rs4922556 | 0.96 | -0.624 | -1.506 | -0.475 | 0.001 | 0.011 | 0.011 | 2.799 | 0.21 | 0.12 | 0.15 | 0.16 | 0.2 | 0.14 |
| 11 | 30602310 | rs2045528 | 0.205 | -0.65 | -1.21 | -0.401 | 0.017 | 0.012 | 0.012 | 0.381 | 0.4 | 0.26 | 0.26 | 0.32 | 0.38 | 0.52 |
| 11 | 30605923 | rs809462 | 0.065 | -0.097 | -0.947 | 0.046 | 0.026 | 0.05 | 0.05 | 0 | 0.26 | 0.32 | 0.45 | 0.54 | 0.36 | 0.34 |
| 11 | 30613628 | rs12281369 | 0.504 | -0.095 | -0.981 | 0.103 | -0.002 | 0.001 | 0.001 | 0.162 | 0.22 | 0.2 | 0.16 | 0.18 | 0.23 | 0.29 |
| 11 | 30613825 | rs7119684 | 0.441 | -0.097 | -0.982 | 0.102 | -0.002 | 0.001 | 0.001 | 0.148 | 0.23 | 0.24 | 0.16 | 0.18 | 0.23 | 0.29 |
| 11 | 30618816 | rs1374612 | -0.013 | 0.06 | -0.617 | 0.412 | 0.044 | 0.076 | 0.076 | 0 | 0.26 | 0.47 | 0.44 | 0.47 | 0.32 | 0.34 |
| 11 | 30625358 | rs2593549 | -0.459 | -0.193 | -1.093 | 0.187 | 0.001 | 0.03 | 0.03 | 0 | 0.5 | 0.69 | 0.67 | 0.65 | 0.46 | 0.51 |
| 11 | 30626660 | rs294377 | 0.051 | -1.15 | -1.461 | 0.002 | 0.03 | 0.037 | 0.037 | 0 | 0.7 | 0.96 | 0.77 | 0.79 | 0.77 | 0.77 |
| 11 | 30635384 | rs2871802 | 1.387 | -0.52 | -1.355 | 0.088 | -0.016 | 0.006 | 0.006 | 0.526 | 0.21 | 0.13 | 0.2 | 0.15 | 0.29 | 0.19 |
| 11 | 30640248 | rs4922559 | 0.873 | -0.698 | -1.552 | 0.026 | 0.004 | -0.005 | -0.005 | 0.339 | 0.2 | 0.02 | 0.17 | 0.18 | 0.15 | 0.11 |
| 11 | 30647282 | rs294348 | -0.498 | -0.786 | -1.638 | -0.109 | -0.019 | 0.046 | 0.046 | 2.995 | 0.89 | 0.99 | 0.93 | 0.95 | 0.88 | 0.75 |
| 11 | 30647845 | rs294349 | 0.782 | -1.161 | -1.384 | 0.456 | -0.066 | -0.102 | -0.102 | 1.986 | 0.38 | 0.1 | 0.49 | 0.47 | 0.42 | 0.43 |
| 11 | 30653141 | rs7931108 | 0.337 | -1.164 | -0.493 | 0.476 | 0.015 | -0.016 | -0.016 | 0 | 0.28 | 0.52 | 0.24 | 0.21 | 0.31 | 0.2 |
| 11 | 30658439 | rs427996 | 0.063 | -0.918 | 0.025 | 0.801 | -0.047 | -0.081 | -0.081 | 0 | 0.38 | 0.12 | 0.49 | 0.47 | 0.42 | 0.43 |
| 11 | 30658947 | rs16921067 | -0.112 | -0.933 | 0.293 | 0.789 | 0.05 | 0.016 | 0.016 | 0.107 | 0.06 | 0.24 | 0.03 | 0 | 0.02 | 0 |
| 11 | 30661857 | rs294354 | -0.196 | -0.603 | -0.151 | 0.612 | -0.001 | 0.001 | 0.001 | 0 | 0.41 | 0.48 | 0.52 | 0.57 | 0.42 | 0.43 |
| 11 | 30687010 | rs1447267 | 0.6 | -0.724 | 0.642 | 1.012 | -0.014 | 0.168 | 0.168 | 0 | 0.82 | 0.6 | 0.78 | 0.51 | 0.85 | 0.71 |
| 11 | 30691539 | rs1447265 | -1.289 | -0.825 | 0.822 | 0.188 | -0.002 | -0.012 | -0.012 | 0 | 0.16 | 0.19 | 0.07 | 0.09 | 0.1 | 0.34 |
| 11 | 30707698 | rs294363 | 1.787 | -1.151 | 1.171 | 0.142 | 0.013 | 0.004 | 0.004 | 0 | 0.64 | 0.27 | 0.64 | 0.63 | 0.81 | 0.54 |
| 11 | 30709418 | rs294364 | 1.872 | -1.145 | 1.207 | 0.145 | 0.042 | 0.011 | 0.011 | 0 | 0.63 | 0.19 | 0.64 | 0.63 | 0.81 | 0.54 |
| 11 | 30724208 | rs169803 | 0.564 | -1.023 | 1.191 | 0.154 | -0.043 | 0.044 | 0.044 | 0.02 | 0.8 | 0.39 | 0.7 | 0.72 | 0.91 | 0.88 |
| 11 | 30729398 | rs435026 | -0.186 | -0.983 | 1.366 | 0.174 | -0.004 | 0.011 | 0.011 | 0 | 0.15 | 0.17 | 0.27 | 0.28 | 0.08 | 0.11 |
| 11 | 30736810 | rs292434 | -0.407 | -0.922 | 1.043 | 0.125 | -0.001 | -0.013 | -0.013 | 0 | 0.11 | 0.08 | 0.06 | 0.02 | 0.09 | 0.22 |
| 11 | 30741502 | rs4922844 | 0.161 | -1.041 | 1.064 | 0.35 | -0.025 | 0.07 | 0.07 | 0 | 0.83 | 0.64 | 0.72 | 0.72 | 0.92 | 0.89 |
| 11 | 30747467 | rs1374610 | 1.048 | -1.085 | 0.976 | -0.099 | -0.024 | -0.006 | -0.006 | 0 | 0.77 | 0.3 | 0.68 | 0.74 | 0.9 | 0.81 |
| 11 | 30749936 | rs795235 | -0.306 | -0.535 | 0.669 | 0.139 | -0.021 | 0.011 | 0.011 | 0 | 0.86 | 0.78 | 0.81 | 0.8 | 0.93 | 0.91 |
| 11 | 30750391 | rs618585 | -0.738 | -0.358 | -0.04 | -0.123 | -0.003 | 0.023 | 0.023 | 0 | 0.13 | 0.09 | 0.15 | 0.03 | 0.11 | 0.2 |
| 11 | 30751955 | rs685270 | -0.175 | -0.266 | -0.267 | -0.034 | -0.003 | -0.011 | -0.011 | 0 | 0.4 | 0.32 | 0.33 | 0.49 | 0.4 | 0.57 |
| 11 | 30760335 | rs3925584 | 0.963 | -0.31 | -0.496 | -0.063 | -0.141 | -0.091 | -0.091 | 1.586 | 0.37 | 0.06 | 0.42 | 0.33 | 0.49 | 0.26 |
| 11 | 30761310 | rs689434 | 0.014 | -0.96 | 0.553 | -0.351 | -0.052 | -0.037 | -0.037 | 0 | 0.79 | 0.64 | 0.76 | 0.82 | 0.89 | 0.83 |
| 11 | 30773173 | rs911269 | 0.973 | -0.959 | 0.197 | -0.355 | 0.004 | 0.004 | 0.004 | 0 | 0.68 | 0.41 | 0.62 | 0.73 | 0.81 | 0.66 |
| 11 | 30773321 | rs795232 | -1.026 | 0.081 | -0.112 | -0.012 | -0.007 | -0.023 | -0.023 | 0 | 0.11 | 0.18 | 0.14 | 0.09 | 0.08 | 0.17 |
| 11 | 30780863 | rs682398 | -0.039 | 0.308 | -0.427 | 0.091 | 0.016 | 0.045 | 0.045 | 0 | 0.49 | 0.27 | 0.36 | 0.6 | 0.43 | 0.51 |
| 11 | 30780931 | rs638196 | 0.433 | -0.044 | 0.658 | -0.209 | -0.021 | -0.087 | -0.087 | 0 | 0.85 | 0.5 | 0.9 | 0.98 | 0.89 | 0.87 |
| 11 | 30796603 | rs1232052 | -0.853 | -0.881 | 1.324 | -0.265 | -0.063 | -0.061 | -0.061 | 0 | 0.24 | 0.5 | 0.24 | 0.13 | 0.09 | 0.11 |
| 11 | 30810059 | rs607165 | -0.703 | -0.437 | 1.527 | -0.059 | 0.002 | 0.008 | 0.008 | 0 | 0.42 | 0.4 | 0.5 | 0.64 | 0.39 | 0.46 |
| 11 | 30824829 | rs4520568 | -0.615 | -0.473 | 1.543 | -0.087 | -0.006 | 0.021 | 0.021 | 0 | 0.42 | 0.24 | 0.55 | 0.66 | 0.39 | 0.48 |
| 11 | 30838689 | rs978751 | 0.602 | -0.529 | 2.451 | -0.107 | -0.002 | -0.003 | -0.003 | 0 | 0.56 | 0.45 | 0.42 | 0.33 | 0.61 | 0.52 |
| 11 | 30849863 | rs650602 | 0.63 | -0.596 | 2.663 | -0.14 | 0.009 | 0.017 | 0.017 | 0 | 0.54 | 0.34 | 0.41 | 0.33 | 0.61 | 0.52 |
| 11 | 30852782 | rs1005019 | 0.842 | -0.623 | 2.516 | -0.108 | 0.032 | 0.087 | 0.087 | 0 | 0.52 | 0.14 | 0.4 | 0.3 | 0.61 | 0.51 |
| 11 | 30854905 | rs1015541 | -0.214 | -0.704 | 2.719 | -0.072 | 0.029 | 0.099 | 0.099 | 1.281 | 0.74 | 0.42 | 0.63 | 0.45 | 0.7 | 0.64 |
| 11 | 30855237 | rs546181 | -0.637 | -0.716 | 2.8 | -0.077 | 0.004 | 0.016 | 0.016 | 0 | 0.77 | 0.72 | 0.65 | 0.45 | 0.7 | 0.64 |
| 11 | 30871743 | rs273553 | -0.315 | -0.74 | 2.939 | -0.066 | 0.018 | 0.055 | 0.055 | 0.564 | 0.75 | 0.59 | 0.65 | 0.45 | 0.7 | 0.64 |
| 11 | 30873739 | rs11031225 | 0.927 | -0.756 | 2.991 | -0.076 | 0.031 | 0.071 | 0.071 | 0 | 0.52 | 0.23 | 0.41 | 0.3 | 0.61 | 0.51 |
| 11 | 30875674 | rs273595 | 0.258 | -0.754 | 2.689 | -0.075 | 0.043 | 0.027 | 0.027 | 0 | 0.19 | 0.04 | 0.21 | 0.15 | 0.09 | 0.13 |
| 11 | 30887928 | rs273584 | 0.086 | -0.78 | 2.723 | -0.09 | 0.024 | 0.018 | 0.018 | 0 | 0.21 | 0.11 | 0.22 | 0.15 | 0.09 | 0.13 |
| 11 | 30918425 | rs273565 | -0.628 | -0.694 | 2.771 | -0.024 | -0.01 | -0.016 | -0.016 | 0 | 0.78 | 0.78 | 0.66 | 0.45 | 0.7 | 0.63 |
| 11 | 30930374 | rs175399 | -0.31 | -0.754 | 2.896 | -0.053 | 0.024 | 0.073 | 0.073 | 0.498 | 0.75 | 0.54 | 0.64 | 0.45 | 0.7 | 0.63 |

| | | | | | | | | | | | | | | | | |
|---|---|---|---|---|---|---|---|---|---|---|---|---|---|---|---|---|
| 11 | 30935629 | rs273611 | 0.321 | -0.835 | 2.669 | -0.107 | 0.046 | 0.029 | 0.029 | 0.013 | 0.19 | 0.04 | 0.21 | 0.16 | 0.09 | 0.11 |
| 11 | 30941272 | rs273608 | -0.417 | -0.797 | 2.713 | -0.049 | -0.004 | -0.004 | -0.004 | 0 | 0.76 | 0.72 | 0.65 | 0.45 | 0.69 | 0.61 |
| 11 | 30980983 | rs163867 | 0.516 | -0.456 | 2.522 | -0.103 | 0.008 | 0.022 | 0.022 | 0 | 0.57 | 0.66 | 0.44 | 0.29 | 0.61 | 0.49 |
| 11 | 30992875 | rs11031254 | 0.76 | -0.154 | 1.377 | -1.04 | -0.033 | 0.075 | 0.075 | 0 | 0.27 | 0.1 | 0.19 | 0.02 | 0.33 | 0.24 |
| 11 | 31000774 | rs7933758 | 0.77 | -0.175 | 1.296 | -1.05 | -0.042 | 0.094 | 0.094 | 0 | 0.27 | 0.09 | 0.19 | 0.02 | 0.33 | 0.24 |
| 11 | 31010962 | rs290108 | -0.033 | -0.211 | 1.149 | -0.544 | 0.006 | 0.028 | 0.028 | 0 | 0.57 | 0.72 | 0.44 | 0.29 | 0.6 | 0.48 |
| 11 | 31027219 | rs459581 | 0.235 | -0.45 | 0.402 | -0.458 | 0.024 | 0.018 | 0.018 | 0 | 0.2 | 0.12 | 0.21 | 0.17 | 0.1 | 0.15 |
| 11 | 31041739 | rs290741 | -0.206 | -1.004 | -0.106 | 0.023 | 0.043 | -0.038 | -0.038 | 1.044 | 0.17 | 0.06 | 0.18 | 0.29 | 0.08 | 0.15 |
| 11 | 31044673 | rs288460 | -0.905 | -0.999 | -0.266 | -0.407 | -0.033 | 0.011 | 0.011 | 0 | 0.58 | 0.79 | 0.51 | 0.41 | 0.49 | 0.43 |
| 11 | 31048097 | rs288462 | -0.781 | -1.047 | 0.565 | -0.5 | -0.002 | 0.002 | 0.002 | 0 | 0.57 | 0.61 | 0.49 | 0.34 | 0.49 | 0.43 |
| 11 | 31056014 | rs288470 | 0.501 | -1.057 | 0.541 | -0.506 | 0.043 | -0.038 | -0.038 | 0.021 | 0.17 | 0.06 | 0.18 | 0.29 | 0.08 | 0.16 |
| 11 | 31063901 | rs288463 | -0.886 | -1.054 | 0.621 | -0.504 | -0.014 | 0.021 | 0.021 | 0 | 0.57 | 0.7 | 0.5 | 0.34 | 0.49 | 0.43 |
| 11 | 31064570 | rs11824421 | 0.404 | -1.18 | 1.24 | -0.566 | -0.007 | 0.01 | 0.01 | 0 | 0.37 | 0.34 | 0.3 | 0.05 | 0.41 | 0.27 |
| 11 | 31066467 | rs601305 | 0.046 | -1.168 | 1.074 | -0.56 | -0.003 | -0.002 | -0.002 | 2.72 | 0.18 | 0.18 | 0.18 | 0.29 | 0.08 | 0.16 |
| 11 | 31113273 | rs587492 | -0.964 | -1.142 | 0.822 | -0.506 | -0.027 | 0.033 | 0.033 | 0 | 0.58 | 0.77 | 0.51 | 0.34 | 0.49 | 0.43 |
| 11 | 31167203 | rs208104 | -0.366 | -1.142 | 1.143 | -0.572 | 0.01 | -0.085 | -0.085 | 0 | 0.61 | 0.51 | 0.56 | 0.93 | 0.58 | 0.72 |
| 11 | 31195157 | rs208085 | -0.43 | -1.163 | 1.182 | -0.582 | 0 | -0.03 | -0.03 | 0 | 0.6 | 0.41 | 0.55 | 0.93 | 0.58 | 0.72 |
| 11 | 31210771 | rs16921914 | 1.482 | -1.149 | 0.699 | 0.199 | -0.08 | -0.118 | -0.118 | 0 | 0.16 | 0.01 | 0.26 | 0.38 | 0.26 | 0.28 |
| 11 | 31225707 | rs471211 | -0.667 | -1.047 | 0.837 | -0.023 | 0.011 | -0.086 | -0.086 | 0 | 0.61 | 0.51 | 0.56 | 0.93 | 0.58 | 0.72 |
| 11 | 31229804 | rs286651 | -0.999 | -1.04 | 0.768 | -0.026 | -0.056 | 0.122 | 0.122 | 0 | 0.65 | 0.87 | 0.7 | 0.95 | 0.58 | 0.72 |
| 11 | 31230540 | rs12364868 | 0.711 | -1.351 | 0.657 | -0.116 | -0.002 | 0.001 | 0.001 | 0 | 0.27 | 0.34 | 0.25 | 0.31 | 0.24 | 0.31 |
| 11 | 31231534 | rs2774406 | -1.124 | -1.339 | 0.588 | -0.142 | -0.054 | 0.121 | 0.121 | 0 | 0.65 | 0.86 | 0.7 | 0.95 | 0.58 | 0.72 |
| 11 | 31263112 | rs2616812 | -1.078 | -1.261 | 0.59 | -0.111 | -0.048 | 0.14 | 0.14 | 0.017 | 0.63 | 0.86 | 0.69 | 0.95 | 0.57 | 0.69 |
| 11 | 31271585 | rs1330714 | 0.675 | -1.225 | 0.583 | -0.086 | -0.004 | 0.001 | 0.001 | 0 | 0.27 | 0.34 | 0.25 | 0.31 | 0.24 | 0.31 |
| 11 | 31283377 | rs7116699 | -0.838 | -1.239 | 0.616 | -0.088 | 0.005 | -0.076 | -0.076 | 0 | 0.59 | 0.51 | 0.56 | 0.93 | 0.57 | 0.69 |
| 11 | 31286472 | rs11824587 | 0.746 | -1.143 | 0.557 | -0.058 | 0 | -0.001 | -0.001 | 0 | 0.26 | 0.23 | 0.24 | 0.31 | 0.24 | 0.31 |
| 11 | 31309818 | rs11031342 | 0.272 | -1.02 | 0.205 | 0.149 | -0.039 | -0.08 | -0.08 | 1.986 | 0.3 | 0.07 | 0.3 | 0.62 | 0.33 | 0.38 |
| 11 | 31324298 | rs2616792 | -0.627 | -1.012 | 0.204 | 0.164 | -0.031 | -0.079 | -0.079 | 0 | 0.7 | 0.94 | 0.7 | 0.38 | 0.68 | 0.63 |
| 11 | 31327316 | rs2616815 | -0.627 | -1.012 | 0.204 | 0.164 | -0.031 | -0.079 | -0.079 | 1.247 | 0.7 | 0.94 | 0.7 | 0.38 | 0.67 | 0.63 |
| 11 | 31385819 | rs7108340 | 0.498 | -0.801 | 0.36 | -0.049 | 0.002 | -0.004 | -0.004 | 0 | 0.3 | 0.34 | 0.27 | 0.31 | 0.29 | 0.33 |
| 11 | 31390733 | rs11031367 | 0.387 | -0.803 | 0.424 | -0.052 | -0.002 | 0.001 | 0.001 | 0 | 0.3 | 0.26 | 0.27 | 0.31 | 0.29 | 0.33 |
| 11 | 31404484 | rs578666 | -0.779 | -0.876 | 0.115 | -0.174 | -0.024 | -0.075 | -0.075 | 0 | 0.7 | 0.94 | 0.7 | 0.38 | 0.67 | 0.62 |
| 11 | 31418602 | rs7129979 | 1.444 | -0.476 | -1.007 | -0.643 | -0.076 | -0.033 | -0.033 | 2.282 | 0.2 | 0.01 | 0.14 | 0.24 | 0.21 | 0.17 |
| 11 | 31424603 | rs6484503 | 0.417 | -0.32 | -0.274 | -0.465 | 0.003 | -0.083 | -0.083 | 0 | 0.38 | 0.49 | 0.42 | 0.07 | 0.38 | 0.29 |
| 11 | 31424823 | rs6484504 | -2.29 | -0.4 | -0.457 | -0.248 | -0.069 | -0.112 | -0.112 | 0.649 | 0.84 | 0.98 | 0.74 | 0.62 | 0.74 | 0.72 |
| 11 | 31425196 | rs12272996 | 0.282 | -0.491 | -0.384 | -0.269 | 0 | -0.028 | -0.028 | 0.769 | 0.4 | 0.6 | 0.43 | 0.07 | 0.38 | 0.29 |
| 11 | 31428184 | rs11031375 | 1.305 | -0.759 | -0.396 | 0.014 | 0.035 | -0.044 | -0.044 | 0 | 0.15 | 0.05 | 0.04 | 0.24 | 0.07 | 0.09 |
| 11 | 31466339 | rs1223119 | -0.048 | -0.438 | -0.238 | -0.356 | 0.003 | 0.004 | 0.004 | 0 | 0.34 | 0.69 | 0.38 | 0.31 | 0.25 | 0.37 |
| 11 | 31478058 | rs1223067 | -0.082 | -0.463 | -0.212 | -0.369 | 0.022 | 0.028 | 0.028 | 0 | 0.36 | 0.8 | 0.4 | 0.31 | 0.25 | 0.37 |
| 11 | 31479370 | rs1237428 | 0.001 | -0.485 | 0.014 | -0.379 | 0.014 | 0.018 | 0.018 | 0 | 0.35 | 0.75 | 0.4 | 0.31 | 0.25 | 0.37 |
| 11 | 31486211 | rs11031384 | 1.711 | -0.37 | -0.191 | -0.481 | -0.085 | 0.084 | 0.084 | 0.01 | 0.26 | 0.03 | 0.23 | 0.06 | 0.32 | 0.19 |
| 11 | 31488890 | rs1223070 | 0.115 | -0.369 | -0.143 | -0.434 | 0.018 | 0.023 | 0.023 | 0 | 0.36 | 0.76 | 0.39 | 0.31 | 0.25 | 0.37 |
| 11 | 31548522 | rs2973129 | 0.054 | -0.395 | -0.008 | -0.438 | 0.024 | 0.027 | 0.027 | 0 | 0.36 | 0.78 | 0.4 | 0.32 | 0.25 | 0.37 |
| 11 | 31573838 | rs2093613 | 0.515 | -0.391 | 0.125 | -0.469 | -0.003 | -0.004 | -0.004 | 0 | 0.32 | 0.36 | 0.26 | 0.32 | 0.24 | 0.37 |
| 11 | 31589199 | rs2996464 | 1.629 | -0.314 | -0.238 | -0.475 | -0.025 | 0.051 | 0.051 | 0 | 0.27 | 0.14 | 0.23 | 0.04 | 0.32 | 0.19 |
| 11 | 31622534 | rs1232186 | 0.091 | -0.282 | -0.447 | -0.353 | -0.022 | -0.023 | -0.023 | 2.956 | 0.37 | 0.14 | 0.37 | 0.62 | 0.43 | 0.44 |
| 11 | 31636481 | rs986527 | 1.165 | -0.229 | -0.461 | -0.138 | -0.115 | -0.153 | -0.153 | 0 | 0.23 | 0.02 | 0.34 | 0.44 | 0.38 | 0.35 |
| 11 | 31649475 | rs4922570 | -0.243 | -0.358 | 0.267 | 0.039 | 0.008 | 0.006 | 0.006 | 0 | 0.65 | 0.3 | 0.61 | 0.66 | 0.75 | 0.63 |
| 11 | 31664340 | rs2146568 | 1.473 | -0.481 | 0.116 | -0.122 | -0.017 | -0.049 | -0.049 | 0.064 | 0.13 | 0 | 0.08 | 0.23 | 0.13 | 0.17 |
| 11 | 31742816 | rs7950909 | 0.273 | -0.311 | 0.481 | -0.043 | -0.009 | -0.014 | -0.014 | 0 | 0.33 | 0.52 | 0.27 | 0.32 | 0.26 | 0.37 |
| 11 | 31768884 | rs7104512 | 0.262 | -0.295 | 0.405 | -0.019 | -0.008 | -0.009 | -0.009 | 0 | 0.33 | 0.56 | 0.29 | 0.33 | 0.26 | 0.38 |
| 11 | 31781829 | rs1014122 | 0.391 | -0.185 | 0.255 | 0.023 | -0.008 | -0.012 | -0.012 | 0 | 0.31 | 0.5 | 0.28 | 0.32 | 0.26 | 0.39 |
| 11 | 31793051 | rs10835818 | 0.277 | -0.147 | 0.107 | 0.1 | -0.006 | -0.006 | -0.006 | 0 | 0.32 | 0.37 | 0.28 | 0.33 | 0.28 | 0.39 |
| 11 | 31796560 | rs2863231 | -0.223 | 0.42 | -0.42 | 0.543 | -0.054 | -0.115 | -0.115 | 0 | 0.76 | 0.93 | 0.6 | 0.45 | 0.69 | 0.69 |
| 11 | 31800874 | rs1541315 | -0.035 | 0.434 | -0.486 | 0.359 | -0.053 | -0.13 | -0.13 | 0 | 0.75 | 0.94 | 0.59 | 0.46 | 0.68 | 0.69 |
| 11 | 31802357 | rs2177482 | 0.56 | -0.232 | -0.469 | -0.086 | -0.029 | 0.098 | 0.098 | 0.22 | 0.33 | 0.12 | 0.27 | 0.04 | 0.37 | 0.2 |
| 11 | 31807524 | rs3026401 | 0.394 | -0.395 | 0.302 | 0.486 | -0.02 | 0.041 | 0.041 | 0 | 0.71 | 0.58 | 0.62 | 0.5 | 0.79 | 0.74 |
| 11 | 31809070 | rs662702 | 0.065 | -0.468 | 0.249 | 0.602 | -0.038 | 0.001 | 0.001 | 0 | 0.18 | 0.33 | 0.16 | 0.18 | 0.06 | 0.09 |
| 11 | 31812582 | rs2071754 | 0.487 | -0.439 | 0.162 | 0.56 | -0.043 | 0.093 | 0.093 | 0 | 0.71 | 0.47 | 0.61 | 0.49 | 0.81 | 0.75 |
| 11 | 31813509 | rs3026390 | 0.276 | 0.221 | 0.125 | 0.623 | -0.001 | 0.005 | 0.005 | 0.061 | 0.46 | 0.32 | 0.39 | 0.46 | 0.53 | 0.61 |
| 11 | 31830657 | rs3026354 | -0.769 | 0.376 | 0.834 | 1.015 | 0.007 | 0.08 | 0.08 | 0 | 0.12 | 0.23 | 0.24 | 0.3 | 0.14 | 0.18 |
| 11 | 31835418 | rs7942007 | -0.795 | 0.378 | 1.036 | 1.001 | 0.002 | 0.037 | 0.037 | 0 | 0.12 | 0.19 | 0.24 | 0.3 | 0.16 | 0.19 |
| 11 | 31838128 | rs4440995 | -1.042 | 0.373 | 0.682 | 0.983 | 0.009 | 0.102 | 0.102 | 0 | 0.14 | 0.32 | 0.24 | 0.3 | 0.16 | 0.18 |
| 11 | 31871538 | rs1524143 | -0.815 | 0.45 | 0.234 | 0.522 | -0.001 | 0.001 | 0.001 | 0 | 0.35 | 0.37 | 0.38 | 0.21 | 0.4 | 0.32 |
| 11 | 31872707 | rs7935771 | -0.715 | 0.402 | 0.233 | 0.497 | -0.001 | 0.001 | 0.001 | 0 | 0.35 | 0.37 | 0.39 | 0.21 | 0.4 | 0.38 |
| 11 | 31878000 | rs761453 | -1.174 | 0.385 | -0.356 | 0.711 | -0.021 | -0.094 | -0.094 | 0.099 | 0.8 | 0.95 | 0.73 | 0.58 | 0.79 | 0.68 |
| 11 | 31885579 | rs686699 | 0.964 | -0.069 | -0.001 | 0.598 | 0.001 | -0.069 | -0.069 | 0.526 | 0.4 | 0.46 | 0.28 | 0.11 | 0.38 | 0.27 |
| 11 | 31891303 | rs677874 | -1.036 | 0.108 | -0.286 | 0.646 | -0.006 | -0.084 | -0.084 | 1.53 | 0.78 | 0.93 | 0.73 | 0.58 | 0.78 | 0.61 |
| 11 | 31892255 | rs11031497 | -0.724 | 0.234 | 0.26 | 0.493 | -0.027 | 0.025 | 0.025 | 0 | 0.34 | 0.2 | 0.37 | 0.21 | 0.41 | 0.38 |

| | | | | | | | | | | | | | | | | |
|---|---|---|---|---|---|---|---|---|---|---|---|---|---|---|---|---|
| 11 | 31900314 | rs11031509 | -0.889 | 0.161 | 0.171 | 0.432 | 0.004 | 0.037 | 0.037 | 0 | 0.32 | 0.16 | 0.31 | 0.15 | 0.31 | 0.36 |
| 11 | 31902533 | rs7106566 | -0.553 | 0.221 | 0.161 | 0.449 | 0.022 | -0.024 | -0.024 | 0 | 0.26 | 0.12 | 0.2 | 0.26 | 0.2 | 0.33 |
| 11 | 31904417 | rs10835824 | 0.092 | 0.747 | 0.476 | 0.706 | 0.02 | 0.066 | 0.066 | 0 | 0.11 | 0.18 | 0.26 | 0.37 | 0.17 | 0.05 |
| 11 | 31905534 | rs652722 | -1.357 | 0.47 | 1.176 | 0.542 | -0.012 | 0.023 | 0.023 | 0 | 0.31 | 0.49 | 0.35 | 0.4 | 0.27 | 0.21 |
| 11 | 31910743 | rs12221893 | 0.139 | 0.966 | 1.507 | 0.79 | 0.008 | 0.038 | 0.038 | 0 | 0.1 | 0.11 | 0.25 | 0.37 | 0.17 | 0.05 |
| 11 | 31913642 | rs604900 | -2.644 | 0.731 | 1.631 | 0.702 | -0.02 | 0.058 | 0.058 | 0 | 0.46 | 0.79 | 0.56 | 0.67 | 0.49 | 0.42 |
| 11 | 31917599 | rs673946 | -1.923 | 0.824 | 1.626 | 0.687 | 0.024 | 0.015 | 0.015 | 0 | 0.33 | 0.5 | 0.43 | 0.4 | 0.39 | 0.31 |
| 11 | 31921805 | rs11031526 | 0.578 | 0.798 | 1.152 | 0.662 | 0.019 | 0.051 | 0.051 | 0 | 0.14 | 0.03 | 0.07 | 0.03 | 0.09 | 0.24 |
| 11 | 31922680 | rs682572 | -2.485 | 0.839 | 1.123 | 0.725 | -0.018 | 0.069 | 0.069 | 0.011 | 0.48 | 0.77 | 0.57 | 0.68 | 0.52 | 0.51 |
| 11 | 31922718 | rs12419837 | -0.748 | 0.995 | 1.2 | 1.473 | 0 | 0.009 | 0.009 | 0 | 0.13 | 0.13 | 0.13 | 0.28 | 0.1 | 0.19 |
| 11 | 31936515 | rs17720050 | -1.787 | 1.057 | 0.918 | 0.813 | -0.02 | -0.038 | -0.038 | 0 | 0.23 | 0.11 | 0.38 | 0.65 | 0.3 | 0.24 |
| 11 | 31941035 | rs2014429 | 1.376 | 1.334 | 0.996 | 0.204 | 0.046 | 0.106 | 0.106 | 0.006 | 0.77 | 0.55 | 0.6 | 0.38 | 0.7 | 0.74 |
| 11 | 31945989 | rs10835826 | -0.998 | 1.263 | 0.318 | 0.36 | -0.008 | -0.026 | -0.026 | 0 | 0.16 | 0.09 | 0.27 | 0.62 | 0.19 | 0.26 |
| 11 | 31976902 | rs224604 | 0.541 | 1.247 | 0.946 | -0.224 | 0.031 | -0.002 | -0.002 | 0 | 0.64 | 0.47 | 0.45 | 0.33 | 0.58 | 0.7 |
| 11 | 31979424 | rs224597 | 0.294 | 0.995 | 1.007 | -0.234 | 0.012 | 0.008 | 0.008 | 0 | 0.67 | 0.56 | 0.52 | 0.33 | 0.7 | 0.76 |
| 11 | 31983665 | rs797370 | 0.223 | 1.025 | 0.884 | -0.226 | 0.005 | 0.004 | 0.004 | 0 | 0.68 | 0.66 | 0.52 | 0.34 | 0.7 | 0.76 |
| 11 | 31985157 | rs224644 | 0.111 | 1.047 | 0.74 | -0.305 | 0.003 | 0.003 | 0.003 | 0 | 0.7 | 0.72 | 0.53 | 0.34 | 0.7 | 0.76 |
| 11 | 31993727 | rs12796318 | -0.335 | 0.764 | -0.201 | -0.502 | -0.002 | 0.05 | 0.05 | 0 | 0.16 | 0.01 | 0.1 | 0.04 | 0.13 | 0.28 |
| 11 | 32016633 | rs224674 | -0.995 | 1.05 | 0.757 | -0.35 | 0.07 | -0.009 | -0.009 | 0 | 0.07 | 0.35 | 0.09 | 0 | 0.11 | 0.01 |
| 11 | 32019404 | rs224670 | -0.089 | 1.205 | -0.12 | -0.097 | -0.087 | -0.095 | -0.095 | 0 | 0.76 | 0.99 | 0.63 | 0.7 | 0.77 | 0.83 |
| 11 | 32020344 | rs1003715 | 1.161 | 1.067 | 0.251 | -0.463 | -0.007 | 0.018 | 0.018 | 0 | 0.57 | 0.34 | 0.33 | 0.3 | 0.54 | 0.55 |
| 11 | 32027444 | rs224663 | -1.626 | 0.758 | 0.862 | -0.923 | -0.017 | -0.078 | -0.078 | 0 | 0.17 | 0.58 | 0.17 | 0.04 | 0.16 | 0.26 |
| 11 | 32033826 | rs10734409 | -0.262 | 1.429 | -0.017 | -1.027 | -0.009 | -0.016 | -0.016 | 0 | 0.7 | 0.71 | 0.72 | 0.95 | 0.56 | 0.69 |
| 11 | 32036899 | rs1033506 | 0.315 | 1.314 | -0.133 | -0.737 | 0.012 | 0.013 | 0.013 | 0 | 0.5 | 0.5 | 0.38 | 0.6 | 0.41 | 0.35 |
| 11 | 32039609 | rs10767910 | 0.39 | 0.989 | -0.815 | -0.416 | -0.016 | -0.002 | -0.002 | 0 | 0.41 | 0.17 | 0.35 | 0.59 | 0.43 | 0.29 |
| 11 | 32042563 | rs1995622 | -0.816 | 0.844 | -0.699 | -0.48 | 0.014 | -0.11 | -0.11 | 0.19 | 0.6 | 0.47 | 0.68 | 0.93 | 0.53 | 0.6 |
| 11 | 32044946 | rs1506525 | -0.602 | 1.01 | -0.69 | -0.209 | -0.003 | 0.06 | 0.06 | 0.033 | 0.2 | 0.3 | 0.4 | 0.35 | 0.18 | 0.34 |
| 11 | 32053419 | rs11031557 | 0.116 | 0.849 | -1.432 | 0.039 | 0.003 | -0.007 | -0.007 | 0 | 0.15 | 0.12 | 0.32 | 0.26 | 0.12 | 0.13 |
| 11 | 32058968 | rs10767913 | -0.361 | 0.985 | -1.502 | 0.395 | -0.004 | 0.003 | 0.003 | 0 | 0.42 | 0.2 | 0.36 | 0.58 | 0.42 | 0.29 |
| 11 | 32061633 | rs11031560 | -0.267 | 0.936 | -1.778 | 0.713 | -0.025 | -0.094 | -0.094 | 0 | 0.3 | 0.12 | 0.2 | 0.56 | 0.32 | 0.2 |
| 11 | 32062536 | rs2294908 | 0.226 | 0.624 | -1.021 | 0.986 | -0.02 | 0.039 | 0.039 | 0 | 0.69 | 0.41 | 0.69 | 0.6 | 0.77 | 0.61 |
| 11 | 32063229 | rs224612 | 0.824 | 2.216 | -0.866 | 0.82 | 0.049 | 0.002 | 0.002 | 0 | 0.9 | 0.88 | 0.85 | 0.89 | 0.73 | 0.92 |
| 11 | 32064649 | rs224618 | 1.149 | 1.962 | -0.796 | 0.753 | 0.013 | 0 | 0 | 0 | 0.89 | 0.84 | 0.9 | 0.88 | 0.84 | 0.92 |
| 11 | 32065256 | rs224619 | 0.546 | 1.802 | -1.58 | 0.741 | -0.031 | -0.012 | -0.012 | 0 | 0.6 | 0.77 | 0.6 | 0.58 | 0.49 | 0.6 |
| 11 | 32068176 | rs2658577 | 0.09 | 1.789 | -1.621 | 0.776 | -0.026 | -0.02 | -0.02 | 0 | 0.91 | 0.97 | 0.91 | 0.85 | 0.84 | 0.94 |
| 11 | 32068785 | rs7938472 | -0.025 | 1.805 | -1.495 | 0.905 | -0.04 | -0.017 | -0.017 | 0 | 0.29 | 0.19 | 0.31 | 0.32 | 0.36 | 0.34 |
| 11 | 32069802 | rs2745886 | 0.353 | 1.764 | -1.415 | 0.886 | -0.01 | -0.008 | -0.008 | 0.235 | 0.9 | 0.93 | 0.91 | 0.85 | 0.84 | 0.93 |
| 11 | 32069871 | rs2063016 | -0.009 | 1.749 | -1.433 | 0.88 | -0.04 | -0.015 | -0.015 | 0.396 | 0.29 | 0.19 | 0.31 | 0.32 | 0.36 | 0.34 |
| 11 | 32070352 | rs4922888 | -0.091 | 0.665 | -1.787 | -0.179 | 0.032 | 0.022 | 0.022 | 0 | 0.6 | 0.73 | 0.56 | 0.31 | 0.65 | 0.53 |
| 11 | 32071695 | rs2745882 | 0.914 | -0.002 | -1.937 | -0.137 | 0.013 | 0.016 | 0.016 | 0 | 0.11 | 0.07 | 0.06 | 0.05 | 0.04 | 0.12 |
| 11 | 32073206 | rs10835832 | 0.811 | 0.062 | -1.908 | 0.009 | 0.035 | -0.106 | -0.106 | 0 | 0.19 | 0.05 | 0.28 | 0.48 | 0.15 | 0.27 |
| 11 | 32073229 | rs7949970 | 0.699 | 0.133 | -0.858 | -0.09 | 0.014 | 0.003 | 0.003 | 0 | 0.31 | 0.36 | 0.32 | 0.32 | 0.36 | 0.34 |
| 11 | 32075089 | rs7929409 | 0.138 | -0.272 | -0.846 | 0.173 | 0.006 | -0.009 | -0.009 | 0 | 0.71 | 0.73 | 0.61 | 0.44 | 0.75 | 0.58 |
| 11 | 32077032 | rs224638 | -0.582 | -0.083 | -0.588 | 0.033 | -0.017 | 0.024 | 0.024 | 0.086 | 0.06 | 0.19 | 0.03 | 0.06 | 0.01 | 0.14 |
| 11 | 32077503 | rs224637 | -0.161 | 0.461 | -0.839 | -0.498 | -0.002 | -0.002 | -0.002 | 0 | 0.44 | 0.6 | 0.34 | 0.18 | 0.48 | 0.41 |
| 11 | 32079221 | rs12420599 | 1.163 | 0.629 | -1.327 | 0.426 | -0.08 | -0.104 | -0.104 | 0 | 0.16 | 0.02 | 0.27 | 0.41 | 0.21 | 0.28 |
| 11 | 32080615 | rs224634 | -0.086 | -0.599 | -1.397 | -0.247 | 0.047 | -0.04 | -0.04 | 5.399 | 0.36 | 0.25 | 0.33 | 0.49 | 0.3 | 0.43 |
| 11 | 32085297 | rs224633 | -0.674 | -0.996 | -0.59 | -0.534 | 0.005 | -0.003 | -0.003 | 0.003 | 0.39 | 0.63 | 0.33 | 0.5 | 0.3 | 0.44 |
| 11 | 32085420 | rs224632 | 0.106 | -0.883 | -1.249 | -1.112 | -0.027 | 0.158 | 0.158 | 0.883 | 0.62 | 0.16 | 0.71 | 0.99 | 0.64 | 0.78 |
| 11 | 32086617 | rs224631 | 0.725 | -2.285 | -1.013 | -0.305 | 0.02 | -0.014 | -0.014 | 0 | 0.19 | 0.1 | 0.09 | 0.23 | 0.14 | 0.21 |
| 11 | 32087108 | rs224630 | -1.499 | -2.342 | -1.041 | -0.43 | 0.089 | -0.036 | -0.036 | 1.47 | 0.54 | 0.85 | 0.37 | 0.49 | 0.35 | 0.44 |
| 11 | 32092617 | rs11031570 | 1.063 | -1.55 | -0.504 | 0.332 | 0.013 | -0.065 | -0.065 | 1.469 | 0.13 | 0.04 | 0.2 | 0.27 | 0.13 | 0.21 |
| 11 | 32101281 | rs477838 | -0.658 | -1.495 | -0.069 | 0.322 | -0.009 | 0.009 | 0.009 | 0.002 | 0.35 | 0.42 | 0.34 | 0.51 | 0.31 | 0.44 |
| 11 | 32107384 | rs610376 | 0.719 | -1.448 | -0.3 | 0.488 | -0.001 | -0.044 | -0.044 | 0.529 | 0.15 | 0.07 | 0.22 | 0.26 | 0.16 | 0.22 |
| 11 | 32109174 | rs581573 | 0.218 | -1.593 | 0.056 | 0.418 | -0.005 | 0.01 | 0.01 | 0 | 0.18 | 0.22 | 0.23 | 0.26 | 0.16 | 0.23 |
| 11 | 32112143 | rs7109100 | 0.148 | -1.854 | 0.465 | 0.419 | -0.042 | 0.072 | 0.072 | 0 | 0.21 | 0.44 | 0.15 | 0.26 | 0.17 | 0.25 |
| 11 | 32119174 | rs223070 | 0.746 | -1.795 | 0.218 | 0.497 | 0.002 | -0.037 | -0.037 | 0 | 0.16 | 0.08 | 0.18 | 0.28 | 0.16 | 0.2 |
| 11 | 32122003 | rs2281411 | -0.205 | -1.868 | 0.202 | -0.027 | 0 | 0.014 | 0.014 | 0 | 0.53 | 0.56 | 0.48 | 0.68 | 0.52 | 0.6 |
| 11 | 32125855 | rs223058 | -1.256 | -1.744 | 0.151 | -0.124 | -0.002 | -0.009 | -0.009 | 0 | 0.7 | 0.76 | 0.66 | 0.96 | 0.68 | 0.8 |
| 11 | 32133448 | rs7123826 | 1.377 | -1.804 | -0.012 | 0.008 | -0.073 | -0.065 | -0.065 | 0.718 | 0.28 | 0.08 | 0.33 | 0.39 | 0.34 | 0.35 |
| 11 | 32134761 | rs689176 | -0.811 | -1.859 | 0.133 | -0.135 | -0.001 | -0.079 | -0.079 | 0 | 0.66 | 0.57 | 0.69 | 0.96 | 0.66 | 0.79 |
| 11 | 32136436 | rs4922579 | 0.421 | -2.134 | 0.359 | -0.017 | 0.001 | 0 | 0 | 0 | 0.43 | 0.43 | 0.43 | 0.57 | 0.48 | 0.52 |
| 11 | 32144681 | rs110688 | -1.311 | -2.043 | 0.446 | -0.182 | -0.006 | 0.006 | 0.006 | 0 | 0.72 | 0.81 | 0.71 | 0.95 | 0.68 | 0.8 |
| 11 | 32149398 | rs1605271 | 0.101 | -2.006 | 0.576 | -0.019 | 0.002 | 0.014 | 0.014 | 0 | 0.48 | 0.45 | 0.52 | 0.67 | 0.51 | 0.53 |
| 11 | 32158713 | rs10742265 | -0.826 | -1.991 | 0.478 | -0.166 | -0.001 | -0.011 | -0.011 | 0 | 0.64 | 0.68 | 0.62 | 0.86 | 0.65 | 0.71 |
| 11 | 32163222 | rs1010038 | -0.626 | -1.813 | 0.329 | 0.199 | -0.013 | 0.032 | 0.032 | 0 | 0.32 | 0.54 | 0.29 | 0.46 | 0.31 | 0.38 |
| 11 | 32164835 | rs2236000 | -0.595 | -1.809 | 0.362 | 0.201 | -0.009 | 0.022 | 0.022 | 0 | 0.32 | 0.41 | 0.29 | 0.46 | 0.31 | 0.38 |
| 11 | 32168691 | rs7950374 | -0.444 | -1.701 | -0.401 | -0.034 | 0.006 | 0.03 | 0.03 | 0 | 0.53 | 0.46 | 0.53 | 0.85 | 0.56 | 0.71 |
| 11 | 32173989 | rs11031614 | 1.122 | -1.404 | -0.31 | 0.286 | -0.032 | -0.073 | -0.073 | 1.72 | 0.13 | 0.03 | 0.17 | 0.25 | 0.17 | 0.19 |

| | | | | | | | | | | | | | | | | |
|---|---|---|---|---|---|---|---|---|---|---|---|---|---|---|---|---|
| 11 | 32179877 | rs1033584 | 0.157 | -1.392 | -0.285 | 0.292 | -0.002 | 0.001 | 0.001 | 0 | 0.17 | 0.22 | 0.18 | 0.26 | 0.18 | 0.2 |
| 11 | 32196933 | rs11031630 | 1.19 | -1.424 | -0.789 | 0.274 | -0.005 | -0.03 | -0.03 | 0 | 0.12 | 0.04 | 0.09 | 0.21 | 0.13 | 0.19 |
| 11 | 32197320 | rs17668735 | 1.016 | -1.523 | -0.775 | 0.065 | 0.011 | 0.008 | 0.008 | 0.861 | 0.12 | 0.04 | 0.12 | 0.06 | 0.1 | 0.08 |
| 11 | 32199488 | rs10742267 | -0.059 | -1.525 | -0.726 | 0.258 | 0.008 | -0.091 | -0.091 | 0.554 | 0.27 | 0.14 | 0.27 | 0.47 | 0.29 | 0.4 |
| 11 | 32202065 | rs6484552 | -0.782 | -1.674 | -0.933 | 0.216 | 0.034 | -0.022 | -0.022 | 1.659 | 0.39 | 0.76 | 0.34 | 0.47 | 0.33 | 0.43 |
| 11 | 32203846 | rs6484553 | -1.356 | -1.098 | -1.583 | 0.447 | -0.085 | -0.142 | -0.142 | 2.981 | 0.76 | 0.99 | 0.78 | 0.58 | 0.76 | 0.69 |
| 11 | 32208950 | rs10835855 | 0.583 | -1.741 | -0.774 | 1.186 | 0.01 | -0.013 | -0.013 | 0 | 0.14 | 0.08 | 0.09 | 0.39 | 0.08 | 0.23 |
| 11 | 32211032 | rs10835856 | 0.131 | -1.573 | -0.959 | 0.658 | -0.014 | -0.15 | -0.15 | 0 | 0.3 | 0.08 | 0.22 | 0.47 | 0.29 | 0.47 |
| 11 | 32215128 | rs10767917 | -1.283 | -1.958 | -0.452 | 0.33 | 0.008 | -0.003 | -0.003 | 0 | 0.55 | 0.62 | 0.42 | 0.51 | 0.45 | 0.56 |
| 11 | 32219777 | rs10767919 | -1 | -1.995 | -0.096 | 0.322 | -0.001 | -0.001 | -0.001 | 0 | 0.51 | 0.53 | 0.42 | 0.51 | 0.44 | 0.56 |
| 11 | 32219851 | rs10835861 | 0.456 | -1.807 | -0.301 | 0.401 | 0.003 | -0.12 | -0.12 | 0 | 0.3 | 0.08 | 0.22 | 0.43 | 0.28 | 0.47 |
| 11 | 32222212 | rs4922905 | -0.84 | -1.223 | -0.013 | 0.477 | 0.009 | 0.013 | 0.013 | 0 | 0.56 | 0.63 | 0.43 | 0.64 | 0.45 | 0.57 |
| 11 | 32224611 | rs4922906 | -0.836 | -0.875 | -1.621 | 0.054 | 0.003 | -0.041 | -0.041 | 0 | 0.57 | 0.86 | 0.44 | 0.53 | 0.47 | 0.57 |
| 11 | 32224732 | rs10767921 | -0.24 | -0.599 | -1.35 | -0.604 | 0.025 | -0.063 | -0.063 | 0.317 | 0.3 | 0.13 | 0.22 | 0.42 | 0.26 | 0.44 |
| 11 | 32226445 | rs2418579 | 0.039 | 0.118 | -0.015 | -1.579 | 0.008 | -0.094 | -0.094 | 0.452 | 0.82 | 0.46 | 0.85 | 0.95 | 0.81 | 0.91 |
| 11 | 32231059 | rs7948785 | -0.574 | 1.046 | -0.455 | -1.942 | -0.006 | 0.005 | 0.005 | 0 | 0.16 | 0.14 | 0.16 | 0.04 | 0.2 | 0.12 |
| 11 | 32232204 | rs10767923 | 0.449 | 0.894 | 0.373 | -1.957 | 0.026 | -0.057 | -0.057 | 0 | 0.69 | 0.61 | 0.68 | 0.91 | 0.57 | 0.78 |
| 11 | 32242706 | rs1467459 | 0.617 | 0.33 | -0.06 | -2.356 | 0.028 | 0.085 | 0.085 | 0 | 0.44 | 0.28 | 0.5 | 0.79 | 0.34 | 0.5 |
| 11 | 32247542 | rs10835872 | -0.198 | 0.524 | 0.386 | -2.455 | -0.019 | -0.067 | -0.067 | 1.047 | 0.21 | 0.41 | 0.17 | 0.11 | 0.19 | 0.24 |
| 11 | 32249249 | rs10767924 | -1.155 | 0.523 | -0.611 | -2.866 | -0.048 | 0.103 | 0.103 | 0 | 0.75 | 0.94 | 0.84 | 0.94 | 0.69 | 0.87 |
| 11 | 32251787 | rs11031650 | 1.251 | 0.317 | -0.826 | -3.009 | -0.059 | 0.029 | 0.029 | 0.284 | 0.33 | 0.04 | 0.47 | 0.76 | 0.33 | 0.46 |
| 11 | 32253286 | rs1998003 | -1.494 | 0.407 | -0.828 | -3.011 | -0.061 | 0.09 | 0.09 | 0 | 0.76 | 0.95 | 0.84 | 0.9 | 0.68 | 0.88 |
| 11 | 32254978 | rs7110087 | -0.671 | 0.331 | -0.601 | -3.128 | -0.038 | -0.091 | -0.091 | 0 | 0.28 | 0.49 | 0.19 | 0.08 | 0.13 | 0.21 |
| 11 | 32257346 | rs11031651 | 1.241 | 0.393 | -1.565 | -3.056 | -0.028 | -0.017 | -0.017 | 0 | 0.09 | 0.19 | 0.04 | 0.04 | 0.03 | 0.04 |
| 11 | 32263545 | rs11031657 | 0.898 | 0.81 | -2.431 | -2.908 | 0.002 | -0.011 | -0.011 | 0 | 0.11 | 0.16 | 0.13 | 0.06 | 0.16 | 0.17 |
| 11 | 32263818 | rs11031658 | 0.338 | 0.817 | -2.371 | -2.854 | -0.034 | 0.145 | 0.145 | 2.368 | 0.44 | 0.21 | 0.6 | 0.82 | 0.5 | 0.64 |
| 11 | 32266453 | rs1033628 | -0.564 | 1.552 | -1.476 | -2.071 | -0.061 | 0.091 | 0.091 | 0 | 0.76 | 0.95 | 0.84 | 0.9 | 0.69 | 0.88 |
| 11 | 32272327 | rs11031665 | 0.227 | 1.712 | -0.491 | -1.242 | 0.004 | -0.016 | -0.016 | 0 | 0.89 | 0.83 | 0.88 | 0.94 | 0.84 | 0.8 |
| 11 | 32273170 | rs12417249 | -0.792 | 1.551 | -0.897 | -1.174 | -0.031 | 0.053 | 0.053 | 0 | 0.2 | 0.07 | 0.2 | 0.09 | 0.22 | 0.21 |
| 11 | 32276662 | rs16923002 | -1.079 | 1.303 | 0.006 | -0.958 | 0.048 | -0.09 | -0.09 | 0.584 | 0.15 | 0.37 | 0.18 | 0.03 | 0.19 | 0.18 |
| 11 | 32279713 | rs2418589 | -0.672 | 0.96 | -0.143 | -0.631 | -0.006 | 0.02 | 0.02 | 0 | 0.17 | 0.27 | 0.33 | 0.7 | 0.16 | 0.38 |
| 11 | 32284173 | rs6484559 | -0.432 | 1.293 | -0.416 | -1.691 | 0.037 | -0.074 | -0.074 | 0 | 0.69 | 0.61 | 0.64 | 0.84 | 0.63 | 0.67 |
| 11 | 32288129 | rs6484560 | 0.04 | 0.676 | -0.86 | -2.243 | -0.004 | -0.015 | -0.015 | 0 | 0.52 | 0.53 | 0.51 | 0.72 | 0.42 | 0.52 |
| 11 | 32289894 | rs6484562 | 0.779 | 0.561 | -1.328 | -2.372 | 0.045 | 0.109 | 0.109 | 0 | 0.35 | 0.17 | 0.18 | 0.02 | 0.28 | 0.14 |
| 11 | 32293377 | rs7105928 | -0.211 | 1.072 | -1.197 | -2.107 | -0.066 | 0.095 | 0.095 | 0 | 0.71 | 0.9 | 0.66 | 0.89 | 0.67 | 0.73 |
| 11 | 32295312 | rs7130589 | 0.532 | 0.861 | -1.03 | -2.432 | 0.009 | 0.047 | 0.047 | 0 | 0.42 | 0.29 | 0.33 | 0.18 | 0.35 | 0.42 |
| 11 | 32296109 | rs11031688 | 0.091 | 0.806 | -1.145 | -2.798 | 0.019 | 0.038 | 0.038 | 0 | 0.2 | 0.15 | 0.06 | 0.05 | 0.14 | 0.17 |
| 11 | 32302441 | rs764670 | -0.898 | 1.26 | -1.117 | -2.516 | -0.089 | -0.004 | -0.004 | 0 | 0.83 | 0.99 | 0.83 | 0.89 | 0.71 | 0.92 |
| 11 | 32302799 | rs764669 | -0.869 | 0.787 | -0.342 | -2.516 | -0.006 | -0.019 | -0.019 | 0.007 | 0.4 | 0.56 | 0.49 | 0.72 | 0.36 | 0.5 |
| 11 | 32308979 | rs11823078 | 0.071 | 0.787 | 0.043 | -2.562 | 0.007 | -0.006 | -0.006 | 0 | 0.21 | 0.21 | 0.07 | 0.05 | 0.15 | 0.17 |
| 11 | 32309418 | rs12574511 | 0.641 | 0.754 | 0.252 | -2.556 | -0.004 | -0.008 | -0.008 | 0 | 0.18 | 0.13 | 0.4 | 0.67 | 0.21 | 0.33 |
| 11 | 32309976 | rs1474844 | -0.76 | 0.851 | 0.287 | -2.514 | 0.002 | 0.007 | 0.007 | 0 | 0.41 | 0.65 | 0.51 | 0.72 | 0.37 | 0.53 |
| 11 | 32320110 | rs763583 | -1.609 | 0.796 | 0.512 | -2.624 | 0 | -0.002 | -0.002 | 0.498 | 0.61 | 0.65 | 0.67 | 0.81 | 0.69 | 0.66 |
| 11 | 32327017 | rs12224688 | 0.705 | 0.609 | 0.421 | -2.084 | -0.058 | -0.006 | -0.006 | 0 | 0.16 | 0.01 | 0.12 | 0.1 | 0.22 | 0.07 |
| 11 | 32331591 | rs10835884 | -0.209 | 0.208 | 0.837 | -2.095 | -0.018 | -0.103 | -0.103 | 0.583 | 0.23 | 0.37 | 0.11 | 0.04 | 0.17 | 0.22 |
| 11 | 32332499 | rs11031708 | 0.062 | 0.17 | 1.078 | -2.202 | 0 | 0.001 | 0.001 | 0.805 | 0.22 | 0.23 | 0.41 | 0.68 | 0.23 | 0.37 |
| 11 | 32332995 | rs742996 | -0.132 | 0.08 | 0.867 | -2.198 | -0.017 | -0.103 | -0.103 | 0.232 | 0.23 | 0.37 | 0.11 | 0.04 | 0.17 | 0.22 |
| 11 | 32334100 | rs12361455 | 0.009 | 0.101 | 0.812 | -2.196 | -0.004 | -0.11 | -0.11 | 0 | 0.22 | 0.4 | 0.12 | 0.04 | 0.19 | 0.22 |
| 11 | 32338224 | rs910514 | -1.44 | 0.007 | 0.37 | -2.184 | 0.025 | 0.083 | 0.083 | 0.073 | 0.47 | 0.76 | 0.55 | 0.72 | 0.42 | 0.6 |
| 11 | 32340688 | rs7934939 | 0.359 | -0.006 | 0.062 | -2.093 | -0.033 | -0.06 | -0.06 | 0 | 0.19 | 0.06 | 0.4 | 0.68 | 0.22 | 0.37 |
| 11 | 32342517 | rs7121219 | 0.539 | -0.037 | 0.199 | -2.183 | 0.004 | 0.018 | 0.018 | 0 | 0.16 | 0.13 | 0.07 | 0.03 | 0.13 | 0.21 |
| 11 | 32346834 | rs11031716 | -0.079 | -0.037 | 0.467 | -2.182 | -0.025 | -0.107 | -0.107 | 0.114 | 0.21 | 0.32 | 0.08 | 0.03 | 0.14 | 0.22 |
| 11 | 32358979 | rs2144327 | 0.377 | -0.034 | 0.11 | -1.712 | -0.006 | -0.025 | -0.025 | 0.838 | 0.2 | 0.11 | 0.37 | 0.66 | 0.22 | 0.37 |
| 11 | 32361685 | rs10767930 | -1.148 | -0.169 | 0.535 | -1.813 | -0.002 | -0.025 | -0.025 | 0 | 0.28 | 0.69 | 0.16 | 0.07 | 0.2 | 0.24 |
| 11 | 32365430 | rs11031731 | -0.22 | -0.19 | 0.445 | -1.78 | -0.017 | -0.061 | -0.061 | 0 | 0.21 | 0.32 | 0.09 | 0.07 | 0.15 | 0.22 |
| 11 | 32367735 | rs11031734 | 0.747 | 0.121 | -0.422 | -1.853 | -0.071 | 0.006 | 0.006 | 2.579 | 0.16 | 0.01 | 0.11 | 0.11 | 0.22 | 0.07 |
| 11 | 32368744 | rs2207548 | -1.013 | 0.171 | -0.031 | -1.692 | 0.001 | -0.004 | -0.004 | 0.022 | 0.42 | 0.57 | 0.22 | 0.16 | 0.39 | 0.3 |
| 11 | 32374199 | rs4922591 | -1.908 | 0.094 | -0.49 | -1.652 | -0.049 | 0.071 | 0.071 | 0 | 0.64 | 0.87 | 0.55 | 0.75 | 0.59 | 0.71 |
| 11 | 32375669 | rs1884232 | -1.6 | -0.345 | -0.293 | -1.553 | -0.006 | 0.153 | 0.153 | 0 | 0.49 | 0.83 | 0.45 | 0.72 | 0.49 | 0.55 |
| 11 | 32378306 | rs6484569 | -0.753 | -0.552 | 0.309 | -1.792 | 0.002 | -0.005 | -0.005 | 0 | 0.32 | 0.62 | 0.37 | 0.68 | 0.36 | 0.44 |
| 11 | 32380916 | rs2418899 | -0.177 | -0.583 | 0.595 | -1.854 | 0 | 0.012 | 0.012 | 0 | 0.14 | 0.14 | 0.27 | 0.63 | 0.14 | 0.33 |
| 11 | 32381969 | rs7109881 | -0.417 | -0.592 | 0.576 | -1.897 | -0.004 | -0.027 | -0.027 | 0 | 0.17 | 0.19 | 0.08 | 0.03 | 0.14 | 0.12 |
| 11 | 32382167 | rs12285677 | 0.299 | -0.406 | 1.044 | -1.699 | 0.014 | -0.043 | -0.043 | 0.257 | 0.16 | 0.27 | 0.1 | 0.06 | 0.22 | 0.11 |
| 11 | 32386419 | rs11031746 | -1.069 | -1.634 | 1.205 | -1.436 | -0.007 | 0.018 | 0.018 | 0 | 0.2 | 0.25 | 0.27 | 0.64 | 0.15 | 0.33 |
| 11 | 32386914 | rs3858439 | -0.943 | -1.64 | 1.248 | -1.438 | -0.002 | 0.005 | 0.005 | 0 | 0.19 | 0.21 | 0.27 | 0.64 | 0.15 | 0.32 |
| 11 | 32387919 | rs7105538 | -1.565 | -1.738 | 1.423 | -1.521 | -0.006 | 0.001 | 0.001 | 0 | 0.3 | 0.39 | 0.3 | 0.64 | 0.25 | 0.41 |
| 11 | 32399200 | rs10835893 | -2.074 | -1.89 | 1.751 | -1.78 | 0.004 | 0.004 | 0.004 | 0 | 0.42 | 0.6 | 0.41 | 0.72 | 0.33 | 0.54 |
| 11 | 32400053 | rs12277905 | 1.296 | -1.913 | 1.869 | -1.788 | -0.021 | -0.013 | -0.013 | 0.008 | 0.08 | 0.17 | 0.02 | 0.04 | 0.03 | 0.1 |
| 11 | 32401036 | rs7946388 | -1.666 | -1.895 | 1.969 | -1.779 | -0.014 | -0.011 | -0.011 | 0.67 | 0.32 | 0.39 | 0.37 | 0.68 | 0.27 | 0.44 |

| | | | | | | | | | | | | | | | | |
|---|---|---|---|---|---|---|---|---|---|---|---|---|---|---|---|---|
| 11 | 32401414 | rs10835894 | -1.602 | -1.897 | 2.061 | -1.78 | -0.001 | -0.001 | -0.001 | 0 | 0.32 | 0.32 | 0.36 | 0.68 | 0.27 | 0.44 |
| 11 | 32405355 | rs12574749 | -1.848 | -1.896 | 2.029 | -1.779 | -0.016 | -0.019 | -0.019 | 0.351 | 0.33 | 0.43 | 0.39 | 0.68 | 0.3 | 0.45 |
| 11 | 32408693 | rs5030335 | -0.55 | -1.938 | 1.954 | -1.774 | -0.014 | -0.06 | -0.06 | 0 | 0.17 | 0.26 | 0.07 | 0.02 | 0.12 | 0.12 |
| 11 | 32410002 | rs5030320 | -2.649 | -2.038 | 1.956 | -1.852 | 0.071 | 0.111 | 0.111 | 0.044 | 0.45 | 0.79 | 0.43 | 0.72 | 0.33 | 0.54 |
| 11 | 32410380 | rs5030316 | -0.567 | -1.978 | 2.006 | -1.812 | -0.025 | -0.079 | -0.079 | 0 | 0.16 | 0.29 | 0.06 | 0.02 | 0.09 | 0.11 |
| 11 | 32410516 | rs5030315 | -0.703 | -1.979 | 2.02 | -1.812 | -0.029 | -0.091 | -0.091 | 0 | 0.16 | 0.33 | 0.07 | 0.02 | 0.09 | 0.11 |
| 11 | 32412040 | rs5030302 | -0.841 | -1.977 | 2.027 | -1.812 | -0.018 | -0.098 | -0.098 | 0.2 | 0.17 | 0.35 | 0.09 | 0.02 | 0.12 | 0.12 |
| 11 | 32417432 | rs5030280 | -0.248 | -1.961 | 1.918 | -1.773 | -0.001 | -0.03 | -0.03 | 0.968 | 0.14 | 0.07 | 0.3 | 0.66 | 0.14 | 0.33 |
| 11 | 32417447 | rs5030279 | 1.284 | -1.983 | 1.914 | -1.806 | -0.022 | -0.013 | -0.013 | 0 | 0.08 | 0.16 | 0.02 | 0.04 | 0.03 | 0.09 |
| 11 | 32417945 | rs16754 | -0.744 | -1.989 | 1.847 | -1.808 | 0.017 | -0.007 | -0.007 | 0 | 0.22 | 0.12 | 0.32 | 0.7 | 0.17 | 0.41 |
| 11 | 32424604 | rs6484577 | -2.124 | -1.953 | 1.919 | -1.783 | 0.027 | 0.039 | 0.039 | 0 | 0.41 | 0.67 | 0.41 | 0.72 | 0.32 | 0.54 |
| 11 | 32426704 | rs10767935 | -1.488 | -1.949 | 1.841 | -1.772 | -0.01 | -0.003 | -0.003 | 0 | 0.29 | 0.34 | 0.35 | 0.68 | 0.23 | 0.44 |
| 11 | 32435529 | rs2900740 | 1.283 | -1.908 | 1.737 | -1.746 | -0.021 | -0.015 | -0.015 | 0.025 | 0.08 | 0.16 | 0.02 | 0.04 | 0.03 | 0.09 |
| 11 | 32438477 | rs5030207 | -1.905 | -1.947 | 1.512 | -1.756 | -0.009 | -0.011 | -0.011 | 0.043 | 0.39 | 0.55 | 0.37 | 0.72 | 0.26 | 0.54 |
| 11 | 32438684 | rs5030203 | -0.319 | -1.961 | 1.124 | -1.67 | -0.002 | -0.029 | -0.029 | 0 | 0.16 | 0.07 | 0.3 | 0.65 | 0.18 | 0.34 |
| 11 | 32438753 | rs3818055 | -2.201 | -1.849 | 1.03 | -1.672 | 0.071 | 0.108 | 0.108 | 0.32 | 0.43 | 0.82 | 0.43 | 0.72 | 0.32 | 0.55 |
| 11 | 32444020 | rs2418910 | -1.765 | -1.955 | 0.87 | -1.916 | -0.015 | -0.027 | -0.027 | 0 | 0.41 | 0.52 | 0.44 | 0.74 | 0.3 | 0.57 |
| 11 | 32446985 | rs12293750 | 1.391 | -1.935 | 0.905 | -1.908 | -0.026 | -0.016 | -0.016 | 0.045 | 0.08 | 0.12 | 0.02 | 0.03 | 0.02 | 0.1 |
| 11 | 32449799 | rs5030166 | -1.35 | -2.134 | 0.972 | -2.134 | -0.047 | -0.164 | -0.164 | 0.023 | 0.27 | 0.49 | 0.14 | 0.03 | 0.18 | 0.19 |
| 11 | 32453536 | rs3858449 | -0.478 | -2.161 | 0.461 | -2.102 | -0.006 | 0.011 | 0.011 | 0.067 | 0.19 | 0.22 | 0.33 | 0.69 | 0.17 | 0.38 |
| 11 | 32453622 | rs7936152 | -1.433 | -2.204 | 0.48 | -2.128 | 0.006 | -0.021 | -0.021 | 0 | 0.35 | 0.66 | 0.16 | 0.07 | 0.18 | 0.28 |
| 11 | 32458219 | rs2301250 | -2.516 | -2.078 | 0.459 | -2.071 | 0.058 | 0.102 | 0.102 | 0 | 0.54 | 0.75 | 0.48 | 0.75 | 0.35 | 0.66 |
| 11 | 32460464 | rs6508 | -0.774 | -2.237 | 0.701 | -2.088 | -0.031 | -0.088 | -0.088 | 0.189 | 0.13 | 0.44 | 0.07 | 0.01 | 0.07 | 0.05 |
| 11 | 32461090 | rs7946086 | -1.611 | -2.213 | 0.64 | -2.009 | -0.005 | -0.021 | -0.021 | 0.046 | 0.42 | 0.41 | 0.42 | 0.86 | 0.27 | 0.61 |
| 11 | 32464168 | rs4980794 | -2.037 | -2.103 | 0.066 | -2.14 | 0.048 | 0.06 | 0.06 | 0 | 0.43 | 0.72 | 0.44 | 0.72 | 0.27 | 0.57 |
| 11 | 32466649 | rs3858451 | -0.287 | -2.142 | -0.04 | -2.087 | 0.01 | -0.076 | -0.076 | 0.517 | 0.19 | 0.06 | 0.31 | 0.65 | 0.16 | 0.41 |
| 11 | 32473626 | rs3858454 | -3.091 | -2.128 | -0.039 | -2.191 | -0.007 | 0.098 | 0.098 | 0 | 0.59 | 0.81 | 0.52 | 0.76 | 0.39 | 0.7 |
| 11 | 32481177 | rs7927401 | -1.58 | -1.872 | 0.208 | -2.069 | 0.027 | 0.025 | 0.025 | 0 | 0.36 | 0.71 | 0.18 | 0.09 | 0.22 | 0.26 |
| 11 | 32494962 | rs3858461 | -3.252 | -1.071 | 0.241 | -1.788 | -0.07 | 0.119 | 0.119 | 0.378 | 0.55 | 0.9 | 0.54 | 0.73 | 0.45 | 0.55 |
| 11 | 32495550 | rs10835921 | -0.217 | -1.073 | 0.226 | -1.773 | -0.011 | -0.081 | -0.081 | 0 | 0.17 | 0.05 | 0.31 | 0.65 | 0.15 | 0.3 |
| 11 | 32498355 | rs2067666 | -0.914 | -0.91 | 0.388 | -1.818 | 0.002 | 0.088 | 0.088 | 0 | 0.21 | 0.05 | 0.11 | 0.05 | 0.15 | 0.2 |
| 11 | 32499100 | rs12276814 | -1.286 | -0.908 | 0.464 | -1.787 | -0.022 | -0.065 | -0.065 | 0 | 0.14 | 0.71 | 0.09 | 0.02 | 0.09 | 0.05 |
| 11 | 32500324 | rs17748949 | 0.227 | -0.828 | 0.14 | -1.704 | 0.013 | 0.072 | 0.072 | 0 | 0.14 | 0.03 | 0.05 | 0.04 | 0.07 | 0.14 |
| 11 | 32509232 | rs12272563 | -2.177 | -0.616 | 0.647 | -1.608 | -0.001 | -0.075 | -0.075 | 1.355 | 0.23 | 0.69 | 0.18 | 0.03 | 0.23 | 0.12 |
| 11 | 32515997 | rs10835931 | -0.557 | -0.706 | 0.698 | -1.274 | -0.005 | 0.035 | 0.035 | 0 | 0.17 | 0.24 | 0.32 | 0.67 | 0.16 | 0.26 |
| 11 | 32517406 | rs11031821 | -1.301 | -0.299 | 0.754 | -1.332 | 0.018 | -0.047 | -0.047 | 1.194 | 0.17 | 0.3 | 0.14 | 0.03 | 0.23 | 0.11 |
| 11 | 32517428 | rs11031822 | -0.48 | -0.278 | 0.272 | -1.33 | 0.006 | 0.077 | 0.077 | 0.385 | 0.16 | 0.28 | 0.33 | 0.67 | 0.18 | 0.27 |
| 11 | 32518070 | rs7113149 | -0.068 | -0.274 | -0.608 | -1.393 | 0.02 | 0.021 | 0.021 | 0 | 0.19 | 0.11 | 0.07 | 0.04 | 0.1 | 0.14 |
| 11 | 32521562 | rs7103780 | -1.818 | 0.067 | -0.773 | -1.144 | -0.095 | 0.1 | 0.1 | 0 | 0.55 | 0.88 | 0.55 | 0.74 | 0.5 | 0.52 |
| 11 | 32521972 | rs11031829 | -1.145 | 1.215 | 0.118 | -0.364 | -0.041 | 0.009 | 0.009 | 0 | 0.07 | 0 | 0.07 | 0.02 | 0.14 | 0.07 |
| 11 | 32524630 | rs10835933 | -1.111 | 1.226 | 0.123 | -0.359 | -0.041 | 0.009 | 0.009 | 0 | 0.07 | 0 | 0.07 | 0.02 | 0.14 | 0.07 |
| 11 | 32530384 | rs4963212 | -1.367 | 0.962 | 0.693 | -0.053 | 0.014 | 0.039 | 0.039 | 0.04 | 0.2 | 0.23 | 0.35 | 0.66 | 0.22 | 0.34 |
| 11 | 32549413 | rs17749506 | -0.721 | 1.23 | 0.53 | -0.23 | -0.032 | 0.027 | 0.027 | 0 | 0.05 | 0 | 0.07 | 0 | 0.12 | 0.02 |
| 11 | 32549624 | rs12285345 | -1.434 | 1.395 | 1.055 | -0.519 | 0.081 | -0.102 | -0.102 | 0 | 0.16 | 0.38 | 0.15 | 0.01 | 0.25 | 0.04 |
| 11 | 32553135 | rs10835941 | -0.811 | 1.261 | 0.502 | -0.102 | -0.011 | -0.052 | -0.052 | 0.696 | 0.18 | 0.1 | 0.15 | 0.64 | 0.18 | 0.32 |
| 11 | 32554805 | rs12226638 | -1.653 | 1.231 | 1.163 | -0.184 | 0.023 | 0 | 0 | 0.471 | 0.36 | 0.58 | 0.34 | 0.66 | 0.46 | 0.37 |
| 11 | 32571594 | rs10767949 | -1.591 | 1.186 | 0.678 | -0.166 | -0.032 | -0.002 | -0.002 | 0 | 0.35 | 0.71 | 0.35 | 0.66 | 0.42 | 0.36 |
| 11 | 32576698 | rs1872706 | -1.081 | 1.086 | 0.673 | -0.053 | 0.003 | 0.004 | 0.004 | 0.206 | 0.26 | 0.27 | 0.23 | 0.65 | 0.31 | 0.34 |
| 11 | 32583695 | rs7112453 | -1.059 | 0.728 | 0.058 | -0.049 | -0.012 | 0.007 | 0.007 | 0 | 0.33 | 0.43 | 0.26 | 0.63 | 0.3 | 0.34 |
| 11 | 32586375 | rs2467584 | 0.916 | 0.507 | -1.151 | -0.092 | -0.022 | 0.078 | 0.078 | 0 | 0.52 | 0.19 | 0.62 | 0.35 | 0.54 | 0.56 |
| 11 | 32593954 | rs169323 | -0.553 | 0.445 | -1.368 | -0.736 | 0.055 | 0.129 | 0.129 | 0 | 0.49 | 0.74 | 0.35 | 0.23 | 0.41 | 0.3 |
| 11 | 32596111 | rs12788831 | 0.014 | 0.057 | -1.242 | -0.878 | 0.006 | -0.002 | -0.002 | 0 | 0.13 | 0.12 | 0.1 | 0.16 | 0.08 | 0.1 |
| 11 | 32600674 | rs11031883 | -0.62 | 0.159 | -0.868 | -1.077 | -0.012 | -0.005 | -0.005 | 0 | 0.17 | 0.21 | 0.12 | 0.16 | 0.12 | 0.15 |
| 11 | 32600937 | rs201839 | -1.223 | 0.122 | -1.002 | -1.264 | 0.019 | 0.144 | 0.144 | 0 | 0.53 | 0.79 | 0.37 | 0.23 | 0.45 | 0.34 |
| 11 | 32603285 | rs201887 | -1.305 | 0.338 | -1.752 | -1.543 | -0.127 | -0.019 | -0.019 | 0.007 | 0.65 | 0.95 | 0.56 | 0.28 | 0.6 | 0.63 |
| 11 | 32604045 | rs201884 | 1.091 | -0.207 | -0.722 | -2.045 | -0.001 | -0.012 | -0.012 | 0 | 0.25 | 0.23 | 0.21 | 0.07 | 0.28 | 0.16 |
| 11 | 32608473 | rs201881 | -0.475 | -0.363 | -0.186 | -2.047 | 0.002 | -0.002 | -0.002 | 0 | 0.44 | 0.53 | 0.3 | 0.17 | 0.4 | 0.3 |
| 11 | 32615001 | rs552512 | -1.034 | 0.071 | -0.138 | -2.435 | 0.004 | 0.023 | 0.023 | 0 | 0.59 | 0.67 | 0.59 | 0.23 | 0.67 | 0.65 |
| 11 | 32622694 | rs11031889 | 1.661 | 0.139 | -0.16 | -2.441 | -0.03 | 0.03 | 0.03 | 0 | 0.06 | 0.01 | 0.11 | 0.01 | 0.16 | 0.08 |
| 11 | 32636914 | rs7116407 | 1.454 | 0.051 | -0.452 | -2.201 | 0 | 0.001 | 0.001 | 0 | 0.07 | 0.07 | 0.03 | 0.13 | 0.04 | 0.03 |
| 11 | 32637182 | rs201904 | -1.473 | -0.024 | -0.231 | -2.26 | 0.001 | 0.025 | 0.025 | 0 | 0.69 | 0.8 | 0.63 | 0.23 | 0.76 | 0.71 |
| 11 | 32639192 | rs201905 | -0.136 | -0.026 | -0.139 | -2.292 | 0.004 | 0.003 | 0.003 | 0 | 0.48 | 0.49 | 0.32 | 0.17 | 0.45 | 0.33 |
| 11 | 32658926 | rs185689 | 1.845 | -0.028 | -0.155 | -2.3 | -0.002 | 0 | 0 | 0 | 0.24 | 0.2 | 0.21 | 0.07 | 0.28 | 0.16 |
| 11 | 32662981 | rs7115208 | 0.223 | 0.007 | -0.184 | -2.244 | 0.016 | -0.033 | -0.033 | 0 | 0.2 | 0.28 | 0.31 | 0.06 | 0.31 | 0.38 |
| 11 | 32677070 | rs12801320 | 1.546 | -0.004 | -0.195 | -2.23 | 0.002 | -0.012 | -0.012 | 0 | 0.13 | 0.16 | 0.19 | 0.05 | 0.15 | 0.3 |
| 11 | 32680927 | rs7130754 | 0.416 | 0.038 | -0.201 | -2.237 | -0.005 | 0 | 0 | 0 | 0.19 | 0.17 | 0.3 | 0.06 | 0.32 | 0.38 |
| 11 | 32681698 | rs2748353 | -0.054 | -0.047 | -0.16 | -2.275 | -0.002 | -0.026 | -0.026 | 0 | 0.47 | 0.41 | 0.32 | 0.17 | 0.45 | 0.33 |
| 11 | 32691569 | rs11031930 | 2.19 | -0.042 | -0.22 | -2.255 | -0.015 | -0.031 | -0.031 | 0 | 0.05 | 0 | 0.02 | 0.13 | 0.04 | 0.03 |
| 11 | 32699987 | rs10742285 | -1.989 | -0.059 | -0.195 | -2.252 | -0.001 | -0.006 | -0.006 | 0 | 0.76 | 0.79 | 0.8 | 0.93 | 0.72 | 0.84 |

| | | | | | | | | | | | | | | | | |
|---|---|---|---|---|---|---|---|---|---|---|---|---|---|---|---|---|
| 11 | 32720150 | rs11031940 | -0.33 | -0.059 | -0.149 | -2.256 | 0.003 | 0.002 | 0.002 | 0 | 0.53 | 0.6 | 0.68 | 0.83 | 0.55 | 0.67 |
| 11 | 32731192 | rs2087085 | 1.602 | -0.004 | -0.284 | -2.251 | 0.002 | -0.01 | -0.01 | 0.527 | 0.13 | 0.16 | 0.19 | 0.05 | 0.15 | 0.3 |
| 11 | 32746276 | rs11031951 | 1.985 | 0.023 | -0.348 | -2.235 | -0.03 | 0.03 | 0.03 | 0 | 0.06 | 0.01 | 0.11 | 0.01 | 0.16 | 0.07 |
| 11 | 32775602 | rs11031964 | 2.268 | -0.047 | -0.345 | -2.222 | -0.047 | -0.105 | -0.105 | 0.228 | 0.23 | 0.03 | 0.34 | 0.64 | 0.2 | 0.26 |
| 11 | 32781250 | rs926735 | 1.525 | -0.017 | -0.316 | -2.299 | -0.063 | -0.007 | -0.007 | 0 | 0.29 | 0.04 | 0.36 | 0.77 | 0.24 | 0.29 |
| 11 | 32782993 | rs7107642 | 1.202 | 0.041 | -0.189 | -2.183 | 0.009 | 0.007 | 0.007 | 0 | 0.13 | 0.09 | 0.07 | 0.1 | 0.07 | 0.11 |
| 11 | 32795973 | rs7933005 | 0.249 | 0.128 | -0.146 | -2.19 | 0.009 | -0.021 | -0.021 | 0 | 0.2 | 0.24 | 0.3 | 0.06 | 0.31 | 0.38 |
| 11 | 32801143 | rs4756244 | -0.181 | 0.076 | -0.107 | -2.281 | -0.002 | -0.022 | -0.022 | 0.354 | 0.52 | 0.56 | 0.68 | 0.83 | 0.55 | 0.66 |
| 11 | 32805169 | rs198264 | -2.634 | -0.059 | -0.475 | -2.682 | -0.06 | 0.056 | 0.056 | 0 | 0.9 | 0.99 | 0.88 | 1 | 0.79 | 0.98 |
| 11 | 32811380 | rs2419766 | 0 | 0.181 | -0.234 | -2.44 | 0.011 | 0.035 | 0.035 | 0 | 0.52 | 0.48 | 0.67 | 0.83 | 0.55 | 0.66 |
| 11 | 32812529 | rs1509891 | -0.077 | 0.163 | -0.283 | -2.454 | -0.003 | -0.029 | -0.029 | 0 | 0.52 | 0.53 | 0.68 | 0.83 | 0.55 | 0.66 |
| 11 | 32814473 | rs10767966 | -1.668 | 0.322 | -0.225 | -2.454 | 0.001 | -0.03 | -0.03 | 0 | 0.74 | 0.73 | 0.79 | 0.93 | 0.71 | 0.83 |
| 11 | 32815580 | rs960389 | -0.103 | 0.396 | -0.267 | -2.383 | -0.002 | -0.025 | -0.025 | 0 | 0.52 | 0.55 | 0.69 | 0.83 | 0.56 | 0.66 |
| 11 | 32822434 | rs7102857 | -0.146 | 0.405 | -0.255 | -2.378 | 0 | -0.016 | -0.016 | 0 | 0.53 | 0.58 | 0.68 | 0.83 | 0.57 | 0.66 |
| 11 | 32830362 | rs12789110 | 1.471 | 0.423 | -0.064 | -2.128 | 0 | -0.021 | -0.021 | 0 | 0.09 | 0.07 | 0.03 | 0 | 0.08 | 0.08 |
| 11 | 32830564 | rs12787998 | 0.433 | 0.404 | -0.019 | -2.156 | 0 | -0.04 | -0.04 | 0 | 0.13 | 0.13 | 0.05 | 0 | 0.12 | 0.12 |
| 11 | 32834416 | rs197697 | -0.125 | 0.339 | -0.075 | -1.832 | -0.005 | -0.001 | -0.001 | 0 | 0.55 | 0.59 | 0.66 | 0.47 | 0.51 | 0.68 |
| 11 | 32837363 | rs7936055 | 1.847 | 0.474 | -0.091 | -1.819 | 0.009 | -0.069 | -0.069 | 0 | 0.25 | 0.15 | 0.16 | 0.53 | 0.19 | 0.12 |
| 11 | 32841636 | rs11827537 | 1.835 | 0.508 | -0.015 | -1.818 | -0.001 | -0.011 | -0.011 | 0 | 0.1 | 0.16 | 0.18 | 0.05 | 0.13 | 0.25 |
| 11 | 32843875 | rs197706 | -1.841 | 0.519 | -0.004 | -1.817 | 0.027 | -0.026 | -0.026 | 0 | 0.94 | 0.92 | 0.88 | 0.99 | 0.83 | 0.93 |
| 11 | 32851008 | rs12363973 | 1.819 | 0.45 | -0.027 | -1.79 | -0.001 | -0.011 | -0.011 | 0 | 0.1 | 0.16 | 0.18 | 0.05 | 0.12 | 0.25 |
| 11 | 32855043 | rs12800054 | 0.854 | 0.458 | -0.057 | -1.81 | -0.002 | -0.008 | -0.008 | 0 | 0.12 | 0.07 | 0.05 | 0 | 0.12 | 0.12 |
| 11 | 32860138 | rs11605633 | 1.46 | 0.361 | 0.243 | -1.905 | -0.022 | -0.006 | -0.006 | 0 | 0.31 | 0.13 | 0.4 | 0.31 | 0.32 | 0.32 |
| 11 | 32862393 | rs7944652 | 0.436 | 0.46 | 0.257 | -1.857 | 0 | 0 | 0 | 0 | 0.45 | 0.42 | 0.49 | 0.41 | 0.4 | 0.43 |
| 11 | 32868623 | rs11032017 | -1.905 | 0.507 | 0.241 | -1.857 | 0.017 | -0.02 | -0.02 | 0 | 0.94 | 0.95 | 0.89 | 0.99 | 0.83 | 0.93 |
| 11 | 32874926 | rs11032025 | 0.369 | 0.46 | 0.281 | -1.819 | 0.005 | 0.007 | 0.007 | 0 | 0.44 | 0.35 | 0.48 | 0.41 | 0.4 | 0.43 |
| 11 | 32875597 | rs7933966 | -0.325 | 0.485 | 0.126 | -1.753 | -0.023 | -0.009 | -0.009 | 0.864 | 0.57 | 0.73 | 0.67 | 0.49 | 0.52 | 0.68 |
| 11 | 32882358 | rs7118737 | 0.834 | 0.557 | 0.088 | -1.71 | -0.002 | -0.008 | -0.008 | 0 | 0.12 | 0.07 | 0.05 | 0 | 0.12 | 0.12 |
| 11 | 32886090 | rs12364134 | 1.464 | 0.559 | 0.111 | -1.657 | 0.001 | -0.007 | -0.007 | 0.51 | 0.1 | 0.16 | 0.17 | 0.07 | 0.13 | 0.26 |
| 11 | 32895664 | rs10767971 | -0.85 | 0.631 | -0.176 | -1.583 | -0.019 | -0.033 | -0.033 | 0.291 | 0.65 | 0.76 | 0.7 | 0.49 | 0.61 | 0.77 |
| 11 | 32903263 | rs4755454 | -0.047 | 0.422 | -0.182 | -1.556 | 0.003 | 0.008 | 0.008 | 0 | 0.49 | 0.61 | 0.63 | 0.49 | 0.49 | 0.67 |
| 11 | 32911737 | rs652963 | 0.139 | 0.284 | 0.247 | -1.549 | -0.007 | -0.086 | -0.086 | 0 | 0.85 | 0.71 | 0.87 | 1 | 0.83 | 0.92 |
| 11 | 32916005 | rs2786800 | 0.784 | 0.206 | -0.022 | -1.606 | -0.038 | -0.146 | -0.146 | 0 | 0.78 | 0.49 | 0.84 | 1 | 0.81 | 0.89 |
| 11 | 32917505 | rs2746037 | 0.759 | 0.209 | -0.02 | -1.605 | -0.037 | -0.145 | -0.145 | 0 | 0.78 | 0.49 | 0.84 | 1 | 0.81 | 0.89 |
| 11 | 32937406 | rs2024636 | 1.529 | 0.613 | -0.324 | -1.417 | -0.039 | -0.115 | -0.115 | 0 | 0.22 | 0.04 | 0.16 | 0.64 | 0.19 | 0.14 |
| 11 | 32938165 | rs4756359 | 1.024 | 0.324 | -0.055 | -1.441 | -0.028 | 0.07 | 0.07 | 0 | 0.41 | 0.21 | 0.44 | 0.24 | 0.41 | 0.46 |
| 11 | 32955122 | rs2297781 | 0.578 | 0.182 | 0.26 | -1.051 | -0.003 | 0 | 0 | 0 | 0.07 | 0.08 | 0.08 | 0.06 | 0.06 | 0.12 |
| 11 | 32956620 | rs7481878 | 1.518 | 0.39 | 0.182 | -0.952 | -0.033 | -0.161 | -0.161 | 0.241 | 0.79 | 0.55 | 0.85 | 1 | 0.81 | 0.88 |
| 11 | 33005960 | rs7944318 | 1.532 | 0.43 | -0.242 | -0.914 | -0.006 | -0.158 | -0.158 | 0 | 0.78 | 0.57 | 0.81 | 1 | 0.76 | 0.88 |
| 11 | 33036294 | rs11826269 | -0.495 | 1.003 | -1.304 | 0.896 | -0.011 | -0.039 | -0.039 | 0 | 0.15 | 0.1 | 0.1 | 0.37 | 0.15 | 0.3 |
| 11 | 33039093 | rs4755351 | -0.557 | 0.995 | -1.287 | 0.889 | -0.011 | -0.031 | -0.031 | 0 | 0.15 | 0.11 | 0.1 | 0.34 | 0.15 | 0.3 |
| 11 | 33041338 | rs7105777 | 1.047 | 0.848 | -1.23 | 0.482 | -0.008 | -0.008 | -0.008 | 2.403 | 0.64 | 0.62 | 0.61 | 0.54 | 0.64 | 0.34 |
| 11 | 33046610 | rs17336103 | 1.287 | 0.672 | -1.906 | 0.406 | -0.045 | -0.005 | -0.005 | 0 | 0.18 | 0.01 | 0.13 | 0.12 | 0.22 | 0.1 |
| 11 | 33053107 | rs966191 | 1.315 | 0.333 | -1.15 | 0.208 | 0.002 | 0.011 | 0.011 | 1.091 | 0.4 | 0.44 | 0.46 | 0.42 | 0.41 | 0.25 |
| 11 | 33054562 | rs2273536 | 1.296 | 0.292 | -1.152 | 0.208 | 0 | 0.01 | 0.01 | 1.288 | 0.4 | 0.44 | 0.45 | 0.42 | 0.4 | 0.25 |
| 11 | 33065394 | rs1064005 | 1.214 | 0.301 | -1.168 | 0.212 | 0 | 0.01 | 0.01 | 1.191 | 0.4 | 0.46 | 0.45 | 0.42 | 0.4 | 0.25 |
| 11 | 33065547 | rs7926030 | 1.278 | 0.245 | -0.843 | 0.185 | 0 | 0.003 | 0.003 | 1.675 | 0.39 | 0.4 | 0.44 | 0.42 | 0.4 | 0.25 |
| 11 | 33066571 | rs1033717 | -0.092 | 0.328 | -0.959 | 0.198 | 0.008 | -0.038 | -0.038 | 0 | 0.35 | 0.22 | 0.23 | 0.46 | 0.34 | 0.39 |
| 11 | 33075970 | rs6484637 | 1.224 | 0.37 | -0.981 | 0.199 | -0.002 | 0.009 | 0.009 | 1.167 | 0.4 | 0.48 | 0.45 | 0.42 | 0.39 | 0.25 |
| 11 | 33076191 | rs2273544 | 0.255 | 0.33 | -0.99 | 0.148 | 0.003 | -0.033 | -0.033 | 0.324 | 0.73 | 0.65 | 0.67 | 0.88 | 0.74 | 0.65 |
| 11 | 33079580 | rs2273549 | 1.134 | 0.519 | -0.753 | 0.448 | 0.016 | 0.025 | 0.025 | 0 | 0.19 | 0.07 | 0.13 | 0.12 | 0.19 | 0.1 |
| 11 | 33091567 | rs2024967 | -0.301 | 0.587 | -0.701 | 0.324 | 0.024 | -0.004 | -0.004 | 0 | 0.88 | 0.7 | 0.76 | 0.89 | 0.88 | 0.75 |
| 11 | 33097267 | rs7127111 | -0.233 | 0.366 | -1.454 | 0.329 | 0.006 | -0.063 | -0.063 | 0.358 | 0.36 | 0.14 | 0.22 | 0.46 | 0.34 | 0.4 |
| 11 | 33097360 | rs7127229 | -0.273 | 0.622 | -1.333 | 0.468 | -0.019 | -0.05 | -0.05 | 0 | 0.36 | 0.17 | 0.25 | 0.46 | 0.39 | 0.4 |
| 11 | 33102251 | rs10836020 | -0.225 | 0.487 | -1.311 | 0.527 | -0.001 | -0.057 | -0.057 | 0 | 0.34 | 0.17 | 0.22 | 0.47 | 0.34 | 0.41 |
| 11 | 33103277 | rs12365818 | 0.929 | 0.22 | -1.394 | 0.37 | 0.005 | 0.013 | 0.013 | 0 | 0.19 | 0.03 | 0.12 | 0.12 | 0.19 | 0.09 |
| 11 | 33106616 | rs3758741 | 1.065 | 0.312 | -1.51 | 0.64 | -0.05 | 0.022 | 0.022 | 1.783 | 0.38 | 0.1 | 0.25 | 0.21 | 0.4 | 0.23 |
| 11 | 33112903 | rs6484644 | -0.024 | 0.243 | -1.454 | 0.837 | 0.058 | 0.051 | 0.051 | 0.339 | 0.56 | 0.2 | 0.62 | 0.61 | 0.67 | 0.59 |
| 11 | 33118274 | rs7103953 | 1.25 | 0.266 | -1.203 | 0.848 | -0.05 | 0.025 | 0.025 | 1.783 | 0.38 | 0.1 | 0.27 | 0.22 | 0.4 | 0.23 |
| 11 | 33118888 | rs10836021 | -0.158 | 0.236 | -1.214 | 0.836 | 0.053 | 0.043 | 0.043 | 0.245 | 0.58 | 0.19 | 0.62 | 0.61 | 0.68 | 0.61 |
| 11 | 33125046 | rs10836023 | 0.476 | 0.247 | -1.224 | 0.825 | -0.037 | -0.053 | -0.053 | 0 | 0.15 | 0.04 | 0.32 | 0.39 | 0.22 | 0.31 |
| 11 | 33128256 | rs6484648 | -0.167 | 0.248 | -1.223 | 0.849 | 0.053 | 0.038 | 0.038 | 0.245 | 0.58 | 0.19 | 0.62 | 0.6 | 0.68 | 0.61 |
| 11 | 33132864 | rs3117655 | 0.89 | 0.247 | -1.224 | 0.792 | -0.041 | 0.053 | 0.053 | 2.107 | 0.41 | 0.15 | 0.29 | 0.22 | 0.45 | 0.29 |
| 11 | 33143126 | rs429439 | -0.836 | 0.061 | -0.769 | 0.7 | 0.023 | -0.008 | -0.008 | 0.182 | 0.66 | 0.81 | 0.68 | 0.61 | 0.72 | 0.62 |
| 11 | 33162009 | rs1028564 | -0.053 | 0.122 | -0.701 | 0.805 | 0.053 | 0.043 | 0.043 | 0.245 | 0.58 | 0.19 | 0.62 | 0.61 | 0.68 | 0.61 |
| 11 | 33182278 | rs11032185 | 0.581 | 0.123 | -0.423 | 0.795 | 0.11 | 0.055 | 0.055 | 0 | 0.06 | 0.42 | 0.05 | 0 | 0.04 | 0.01 |
| 11 | 33183017 | rs3736213 | -0.055 | 0.108 | -0.473 | 0.825 | 0.052 | 0.04 | 0.04 | 0.255 | 0.58 | 0.19 | 0.62 | 0.61 | 0.68 | 0.61 |
| 11 | 33183073 | rs3808985 | 1.667 | -0.027 | -0.954 | 0.514 | 0 | -0.024 | -0.024 | 0 | 0.17 | 0.01 | 0.07 | 0.11 | 0.11 | 0.13 |
| 11 | 33190513 | rs2901036 | 0.89 | 0.118 | -0.958 | 0.522 | -0.044 | -0.059 | -0.059 | 1.147 | 0.15 | 0.04 | 0.33 | 0.39 | 0.23 | 0.31 |

| | | | | | | | | | | | | | | | | |
|---|---|---|---|---|---|---|---|---|---|---|---|---|---|---|---|---|
| 11 | 33202539 | rs7118193 | 2.055 | 0.109 | -0.95 | 0.511 | -0.048 | 0.019 | 0.019 | 0.293 | 0.2 | 0.06 | 0.18 | 0.1 | 0.28 | 0.1 |
| 11 | 33211190 | rs11032198 | 0.859 | 0.164 | -0.938 | 0.549 | -0.041 | -0.055 | -0.055 | 0.048 | 0.15 | 0.04 | 0.33 | 0.39 | 0.24 | 0.31 |
| 11 | 33222902 | rs7131287 | -0.838 | 0.176 | -0.889 | 0.452 | 0.015 | 0.009 | 0.009 | 0 | 0.62 | 0.61 | 0.67 | 0.62 | 0.7 | 0.62 |
| 11 | 33233486 | rs10768001 | -1.088 | 0.175 | -0.905 | 0.453 | 0.022 | 0.01 | 0.01 | 0 | 0.64 | 0.7 | 0.68 | 0.62 | 0.72 | 0.62 |
| 11 | 33270782 | rs267442 | -1.216 | 0.379 | -0.737 | 0.537 | 0.012 | 0 | 0 | 0 | 0.66 | 0.71 | 0.68 | 0.62 | 0.72 | 0.63 |
| 11 | 33281328 | rs267449 | -1.94 | 0.345 | -0.788 | 0.612 | 0.008 | -0.046 | -0.046 | 0 | 0.75 | 0.99 | 0.86 | 0.72 | 0.79 | 0.88 |
| 11 | 33288446 | rs590224 | -1.653 | 0.521 | -0.383 | 0.697 | 0.005 | 0 | 0 | 0 | 0.64 | 0.61 | 0.68 | 0.61 | 0.72 | 0.63 |
| 11 | 33291462 | rs10836046 | -1.581 | 0.522 | -0.373 | 0.67 | -0.01 | -0.002 | -0.002 | 0 | 0.63 | 0.54 | 0.67 | 0.61 | 0.72 | 0.63 |
| 11 | 33302060 | rs1005934 | 0.279 | 0.532 | -0.395 | 0.675 | -0.04 | -0.055 | -0.055 | 4.655 | 0.15 | 0.04 | 0.33 | 0.42 | 0.23 | 0.33 |
| 11 | 33302474 | rs16924091 | 0.634 | 0.538 | -0.377 | 0.67 | 0.094 | 0.04 | 0.04 | 0.002 | 0.05 | 0.31 | 0.04 | 0 | 0.04 | 0.01 |
| 11 | 33310265 | rs4388862 | 0.781 | 0.58 | -0.945 | 0.799 | -0.022 | -0.06 | -0.06 | 1.518 | 0.12 | 0.02 | 0.27 | 0.39 | 0.13 | 0.29 |
| 11 | 33312458 | rs11607288 | 0.133 | 0.033 | -0.928 | 0.855 | -0.035 | 0.06 | 0.06 | 1.688 | 0.37 | 0.1 | 0.26 | 0.18 | 0.4 | 0.22 |
| 11 | 33314798 | rs10734415 | -1.256 | 0.038 | -0.925 | 0.864 | -0.01 | -0.002 | -0.002 | 1.793 | 0.63 | 0.55 | 0.67 | 0.61 | 0.72 | 0.63 |
| 11 | 33317202 | rs4755562 | -0.11 | -0.044 | -0.902 | 0.974 | -0.039 | 0.056 | 0.056 | 2.057 | 0.4 | 0.14 | 0.29 | 0.22 | 0.44 | 0.29 |
| 11 | 33345888 | rs2068825 | 1.019 | -0.124 | -0.937 | 0.765 | -0.006 | -0.009 | -0.009 | 0 | 0.16 | 0.01 | 0.07 | 0.1 | 0.11 | 0.13 |
| 11 | 33370884 | rs4756641 | 0.517 | 0.354 | -0.762 | 1.275 | -0.038 | 0.081 | 0.081 | 2.057 | 0.4 | 0.14 | 0.29 | 0.19 | 0.44 | 0.28 |
| 11 | 33376095 | rs4755571 | 0.612 | 0.231 | -0.881 | 1.357 | -0.033 | 0.061 | 0.061 | 1.709 | 0.38 | 0.1 | 0.26 | 0.18 | 0.4 | 0.22 |
| 11 | 33378209 | rs10600 | 0.132 | -0.636 | -0.381 | 1.4 | -0.006 | -0.002 | -0.002 | 0 | 0.62 | 0.54 | 0.66 | 0.61 | 0.7 | 0.62 |
| 11 | 33387008 | rs10836057 | -1.173 | -0.187 | -0.377 | 1.872 | 0.005 | -0.018 | -0.018 | 0 | 0.11 | 0.07 | 0.14 | 0.27 | 0.09 | 0.15 |
| 11 | 33387545 | rs11032239 | -1.173 | -0.187 | -0.377 | 1.881 | 0.005 | -0.017 | -0.017 | 0 | 0.11 | 0.08 | 0.14 | 0.28 | 0.09 | 0.16 |
| 11 | 33390547 | rs11032242 | -1.682 | -0.446 | 0.067 | 1.762 | -0.039 | 0.097 | 0.097 | 0 | 0.19 | 0.42 | 0.17 | 0.33 | 0.11 | 0.25 |
| 11 | 33394460 | rs2615924 | 0.537 | -0.172 | 0.056 | 1.581 | -0.017 | 0.084 | 0.084 | 0 | 0.74 | 0.61 | 0.66 | 0.47 | 0.8 | 0.61 |
| 11 | 33399991 | rs2761207 | 0.217 | 0.488 | -0.033 | 1.602 | 0.018 | 0.07 | 0.07 | 0 | 0.78 | 0.75 | 0.67 | 0.44 | 0.75 | 0.59 |
| 11 | 33401635 | rs7933529 | -0.625 | 0.449 | -0.273 | 1.565 | -0.004 | -0.012 | -0.012 | 0 | 0.09 | 0.09 | 0.13 | 0.12 | 0.09 | 0.14 |
| 11 | 33402402 | rs2753413 | 0.666 | 0.337 | -0.158 | 1.461 | 0.015 | 0.022 | 0.022 | 0 | 0.7 | 0.66 | 0.54 | 0.37 | 0.68 | 0.46 |
| 11 | 33406776 | rs3898926 | 0.517 | -0.043 | -0.962 | 1.042 | -0.1 | -0.035 | -0.035 | 0 | 0.42 | 0.09 | 0.38 | 0.36 | 0.54 | 0.37 |
| 11 | 33412704 | rs16924257 | 0.002 | 0.054 | -0.891 | 1.076 | -0.022 | -0.028 | -0.028 | 0 | 0.07 | 0 | 0.12 | 0.08 | 0.07 | 0.1 |
| 11 | 33416070 | rs7929907 | 0.169 | 0.155 | -1.341 | 0.868 | 0.018 | -0.027 | -0.027 | 1.107 | 0.5 | 0.13 | 0.5 | 0.45 | 0.61 | 0.5 |
| 11 | 33428255 | rs2615919 | -0.35 | 0.663 | -0.725 | 0.361 | 0.018 | -0.02 | -0.02 | 0 | 0.32 | 0.64 | 0.49 | 0.23 | 0.46 | 0.38 |
| 11 | 33433623 | rs2615945 | -0.645 | 1.198 | -0.574 | -0.088 | 0.014 | -0.011 | -0.011 | 0.225 | 0.24 | 0.24 | 0.33 | 0.11 | 0.41 | 0.28 |
| 11 | 33435125 | rs12282223 | -0.079 | 0.558 | -0.534 | 0.371 | -0.011 | 0.029 | 0.029 | 0 | 0.31 | 0.4 | 0.29 | 0.52 | 0.26 | 0.4 |
| 11 | 33436631 | rs10836061 | -0.027 | 0.7 | -0.333 | 0.378 | 0 | -0.002 | -0.002 | 0 | 0.3 | 0.28 | 0.27 | 0.53 | 0.27 | 0.4 |
| 11 | 33439135 | rs2615943 | -0.868 | 0.375 | -0.482 | 0.335 | -0.004 | -0.002 | -0.002 | 0 | 0.56 | 0.57 | 0.62 | 0.64 | 0.71 | 0.7 |
| 11 | 33443587 | rs4756683 | -1.037 | -0.09 | -0.606 | 0.264 | 0.035 | 0.018 | 0.018 | 0 | 0.62 | 0.77 | 0.65 | 0.66 | 0.74 | 0.69 |
| 11 | 33443726 | rs4756684 | -1.245 | -0.497 | -0.913 | 0.284 | 0.02 | 0.008 | 0.008 | 0.108 | 0.68 | 0.8 | 0.68 | 0.7 | 0.77 | 0.75 |
| 11 | 33449347 | rs2761201 | -0.188 | -1.372 | -0.736 | -0.156 | 0.001 | -0.001 | -0.001 | 0 | 0.41 | 0.58 | 0.36 | 0.46 | 0.33 | 0.4 |
| 11 | 33459596 | rs2615906 | 1.246 | -1.578 | -0.339 | 0.138 | 0.006 | 0.074 | 0.074 | 0 | 0.19 | 0.32 | 0.24 | 0.3 | 0.18 | 0.17 |
| 11 | 33466642 | rs2023571 | -1.89 | -1.198 | -0.817 | -0.171 | -0.021 | -0.034 | -0.034 | 0 | 0.64 | 0.77 | 0.57 | 0.56 | 0.61 | 0.7 |
| 11 | 33471804 | rs7131170 | 0.707 | -1.259 | -0.783 | -0.232 | -0.044 | 0.062 | 0.062 | 0 | 0.22 | 0.07 | 0.2 | 0.1 | 0.28 | 0.33 |
| 11 | 33472945 | rs1998305 | -0.244 | -1.283 | -0.677 | -0.204 | 0.006 | -0.002 | -0.002 | 3.68 | 0.41 | 0.41 | 0.44 | 0.39 | 0.47 | 0.5 |
| 11 | 33474665 | rs16924343 | 0.18 | -1.132 | -0.577 | -0.218 | -0.033 | -0.023 | -0.023 | 0 | 0.23 | 0.31 | 0.12 | 0.15 | 0.13 | 0.2 |
| 11 | 33476703 | rs7943651 | -0.433 | -0.639 | -0.766 | -0.253 | -0.102 | -0.038 | -0.038 | 0 | 0.35 | 0.08 | 0.41 | 0.38 | 0.45 | 0.49 |
| 11 | 33477930 | rs911585 | 0.973 | -0.24 | -0.003 | 0.358 | -0.004 | -0.009 | -0.009 | 0 | 0.15 | 0.15 | 0.24 | 0.3 | 0.19 | 0.16 |
| 11 | 33498468 | rs10836066 | 1.338 | -0.33 | -0.032 | -0.129 | 0.002 | 0.074 | 0.074 | 0.29 | 0.35 | 0.18 | 0.23 | 0.02 | 0.33 | 0.26 |
| 11 | 33518390 | rs2756277 | -0.374 | 0.319 | 0.748 | 0.929 | -0.048 | -0.009 | -0.009 | 0.015 | 0.75 | 0.87 | 0.62 | 0.69 | 0.65 | 0.68 |
| 11 | 33520628 | rs2615939 | -0.332 | 0.313 | 0.753 | 0.945 | -0.076 | -0.034 | -0.034 | 0 | 0.75 | 0.9 | 0.63 | 0.68 | 0.65 | 0.68 |
| 11 | 33522613 | rs2615938 | -0.329 | 0.314 | 0.799 | 0.961 | -0.04 | -0.02 | -0.02 | 0.532 | 0.74 | 0.86 | 0.62 | 0.66 | 0.65 | 0.67 |
| 11 | 33525405 | rs2756276 | 0.491 | 0.251 | 1.288 | 1.09 | 0.002 | 0.011 | 0.011 | 0.008 | 0.55 | 0.43 | 0.34 | 0.33 | 0.44 | 0.44 |
| 11 | 33530851 | rs10836069 | 1.165 | 0.244 | 1.577 | 1.238 | 0.003 | 0.01 | 0.01 | 0 | 0.17 | 0.14 | 0.25 | 0.25 | 0.21 | 0.08 |
| 11 | 33532393 | rs2615935 | 0.44 | 0.236 | 1.62 | 1.192 | 0 | -0.003 | -0.003 | 0 | 0.56 | 0.47 | 0.35 | 0.45 | 0.43 | 0.57 |
| 11 | 33541163 | rs12273774 | 0.929 | 0.24 | 1.785 | 0.974 | 0.001 | 0.014 | 0.014 | 0 | 0.17 | 0.14 | 0.24 | 0.32 | 0.21 | 0.2 |
| 11 | 33559186 | rs2615929 | 1.156 | 0.234 | 1.849 | 0.816 | 0.014 | -0.011 | -0.011 | 0 | 0.43 | 0.39 | 0.26 | 0.02 | 0.35 | 0.27 |
| 11 | 33563598 | rs2076624 | 0.339 | 0.148 | 1.662 | 0.956 | -0.005 | 0.076 | 0.076 | 0 | 0.09 | 0.13 | 0.08 | 0.31 | 0.06 | 0.13 |
| 11 | 33564123 | rs2076623 | -0.471 | 0.181 | 1.575 | 0.806 | -0.067 | -0.029 | -0.029 | 0.283 | 0.73 | 0.89 | 0.61 | 0.66 | 0.62 | 0.62 |
| 11 | 33578510 | rs1883916 | -0.51 | 0.212 | 1.937 | 0.846 | 0.004 | 0.013 | 0.013 | 0 | 0.28 | 0.47 | 0.35 | 0.64 | 0.26 | 0.29 |
| 11 | 33581262 | rs2281381 | 0.453 | -0.076 | 1.955 | 0.768 | 0.008 | 0.013 | 0.013 | 0 | 0.52 | 0.45 | 0.32 | 0.33 | 0.39 | 0.39 |
| 11 | 33581306 | rs2281380 | 0.563 | -0.081 | 1.946 | 0.765 | 0.007 | 0.011 | 0.011 | 0 | 0.52 | 0.47 | 0.32 | 0.33 | 0.39 | 0.39 |
| 11 | 33585978 | rs11032297 | 0.072 | -0.098 | 1.781 | 0.906 | -0.009 | 0.069 | 0.069 | 0 | 0.11 | 0.17 | 0.08 | 0.32 | 0.06 | 0.16 |
| 11 | 33590268 | rs10836075 | 1.062 | -0.101 | 2.002 | 0.412 | 0.047 | 0.113 | 0.113 | 0.948 | 0.41 | 0.26 | 0.27 | 0.02 | 0.32 | 0.26 |
| 11 | 33599446 | rs7128963 | -0.407 | 0.285 | 1.423 | 1.146 | -0.002 | -0.005 | -0.005 | 0 | 0.23 | 0.78 | 0.29 | 0.36 | 0.23 | 0.15 |
| 11 | 33604885 | rs2273117 | 0.906 | 0.341 | 1.336 | 1.17 | -0.036 | -0.05 | -0.05 | 0 | 0.16 | 0.08 | 0.24 | 0.36 | 0.22 | 0.13 |
| 11 | 33616112 | rs7939586 | 0.926 | 0.341 | 1.359 | 1.17 | -0.03 | -0.039 | -0.039 | 0 | 0.16 | 0.1 | 0.25 | 0.36 | 0.22 | 0.12 |
| 11 | 33617771 | rs761435 | -0.565 | 0.373 | 1.144 | 1.121 | -0.06 | -0.014 | -0.014 | 2.807 | 0.74 | 0.95 | 0.64 | 0.72 | 0.61 | 0.6 |
| 11 | 33628117 | rs2076625 | -0.472 | 0.227 | 1.639 | 1.028 | -0.001 | -0.001 | -0.001 | 0 | 0.25 | 0.74 | 0.29 | 0.36 | 0.23 | 0.16 |
| 11 | 33630792 | rs4755750 | -0.279 | 0.422 | 0.869 | 1.121 | -0.035 | 0.015 | 0.015 | 0.296 | 0.72 | 0.92 | 0.64 | 0.75 | 0.61 | 0.6 |
| 11 | 33631423 | rs2076622 | 0.423 | 0.284 | 0.765 | 1.046 | -0.003 | -0.003 | -0.003 | 0.001 | 0.52 | 0.56 | 0.38 | 0.38 | 0.39 | 0.49 |
| 11 | 33638388 | rs10768011 | 0.496 | 0.285 | 0.601 | 0.97 | -0.004 | 0.003 | 0.003 | 0.959 | 0.58 | 0.38 | 0.51 | 0.38 | 0.51 | 0.39 |
| 11 | 33638992 | rs10836082 | 1.273 | 0.04 | 0.456 | 0.974 | 0.025 | 0.079 | 0.079 | 0.482 | 0.37 | 0.15 | 0.31 | 0.11 | 0.31 | 0.29 |
| 11 | 33640104 | rs768474 | -0.542 | 0.046 | 0.283 | 1.156 | -0.086 | -0.034 | -0.034 | 0 | 0.74 | 0.97 | 0.64 | 0.75 | 0.63 | 0.63 |

| | | | | | | | | | | | | | | | | |
|---|---|---|---|---|---|---|---|---|---|---|---|---|---|---|---|---|
| 11 | 33643373 | rs4755765 | 0.448 | -0.021 | 0.931 | 1.426 | 0 | 0.007 | 0.007 | 0.34 | 0.63 | 0.53 | 0.59 | 0.58 | 0.58 | 0.48 |
| 11 | 33644514 | rs4755771 | 0.481 | 0.088 | 0.922 | 1.421 | -0.003 | -0.009 | -0.009 | 0 | 0.62 | 0.64 | 0.59 | 0.53 | 0.56 | 0.44 |
| 11 | 33645458 | rs7114092 | 0.034 | 0.155 | 0.507 | 1.399 | -0.081 | -0.114 | -0.114 | 0 | 0.67 | 0.92 | 0.62 | 0.54 | 0.58 | 0.48 |
| 11 | 33653780 | rs4333997 | 0.28 | 0.146 | 0.912 | 1.444 | 0.001 | 0.003 | 0.003 | 0.125 | 0.64 | 0.58 | 0.59 | 0.55 | 0.57 | 0.48 |
| 11 | 33654867 | rs11605684 | 1.317 | -0.071 | 0.665 | 1.161 | 0.02 | 0.065 | 0.065 | 0.253 | 0.37 | 0.16 | 0.32 | 0.11 | 0.33 | 0.29 |
| 11 | 33665166 | rs1073655 | -0.553 | -0.014 | 0.467 | 1.255 | -0.039 | -0.001 | -0.001 | 0 | 0.72 | 0.94 | 0.61 | 0.72 | 0.59 | 0.57 |
| 11 | 33666319 | rs7949170 | -0.53 | -0.031 | 0.455 | 1.26 | -0.038 | -0.005 | -0.005 | 0 | 0.72 | 0.94 | 0.61 | 0.72 | 0.6 | 0.57 |
| 11 | 33670718 | rs4755795 | -0.309 | 0.047 | 0.953 | 1.3 | 0.002 | 0.004 | 0.004 | 0 | 0.31 | 0.37 | 0.29 | 0.61 | 0.28 | 0.3 |
| 11 | 33671867 | rs7948932 | -0.498 | -0.613 | 1.759 | 1.366 | 0.012 | 0.017 | 0.017 | 0 | 0.54 | 0.63 | 0.57 | 0.54 | 0.38 | 0.64 |
| 11 | 33676609 | rs10768015 | -0.146 | -0.653 | 1.655 | 1.188 | 0.039 | 0.019 | 0.019 | 0 | 0.49 | 0.34 | 0.4 | 0.5 | 0.37 | 0.49 |
| 11 | 33680249 | rs11032332 | 0.907 | -0.966 | 1.69 | 1.232 | 0.025 | -0.02 | -0.02 | 0 | 0.35 | 0.21 | 0.28 | 0.33 | 0.27 | 0.37 |
| 11 | 33680613 | rs2295064 | 0.839 | -0.963 | 1.592 | 1.233 | 0.021 | -0.014 | -0.014 | 0 | 0.36 | 0.24 | 0.28 | 0.34 | 0.27 | 0.37 |
| 11 | 33681621 | rs763851 | -0.087 | -0.914 | 1.594 | 1.25 | 0.011 | 0.007 | 0.007 | 0 | 0.47 | 0.41 | 0.38 | 0.51 | 0.34 | 0.47 |
| 11 | 33686605 | rs11032335 | 0.775 | -0.809 | 1.129 | 1.216 | 0.017 | -0.023 | -0.023 | 0.008 | 0.35 | 0.2 | 0.29 | 0.33 | 0.27 | 0.37 |
| 11 | 33692315 | rs3741021 | 0.839 | -0.573 | 1.278 | 1.163 | 0.005 | -0.003 | -0.003 | 0 | 0.35 | 0.3 | 0.28 | 0.34 | 0.27 | 0.35 |
| 11 | 33692422 | rs3741020 | 0.648 | -0.573 | 1.276 | 1.164 | 0.003 | -0.002 | -0.002 | 0 | 0.37 | 0.32 | 0.3 | 0.34 | 0.31 | 0.35 |
| 11 | 33696361 | rs10742304 | -0.833 | -0.438 | 0.83 | 1.235 | -0.036 | 0.003 | 0.003 | 0 | 0.58 | 0.88 | 0.46 | 0.56 | 0.45 | 0.63 |
| 11 | 33705172 | rs4142964 | -1.093 | -0.609 | 1.133 | 0.928 | -0.031 | 0.062 | 0.062 | 0 | 0.14 | 0.37 | 0.11 | 0.22 | 0.07 | 0.26 |
| 11 | 33706353 | rs1009448 | -1.025 | -0.593 | 1.197 | 0.935 | -0.027 | 0.037 | 0.037 | 0 | 0.13 | 0.36 | 0.12 | 0.17 | 0.07 | 0.1 |
| 11 | 33706877 | rs7127596 | -1.627 | -0.244 | 0.897 | 1.055 | 0.013 | 0.059 | 0.059 | 0 | 0.14 | 0.41 | 0.13 | 0.22 | 0.13 | 0.24 |
| 11 | 33707552 | rs10768019 | -1.274 | -0.158 | 0.738 | 1.094 | 0.015 | 0.053 | 0.053 | 0 | 0.13 | 0.32 | 0.13 | 0.22 | 0.14 | 0.25 |
| 11 | 33708184 | rs11032342 | -0.083 | -0.163 | 0.511 | 1.097 | -0.004 | 0.037 | 0.037 | 0 | 0.08 | 0.19 | 0.09 | 0.17 | 0.05 | 0.08 |
| 11 | 33709273 | rs831620 | -0.536 | -0.214 | 0.404 | 1.308 | 0.003 | 0.003 | 0.003 | 0 | 0.48 | 0.55 | 0.36 | 0.6 | 0.41 | 0.59 |
| 11 | 33710945 | rs11032343 | -1.272 | -0.02 | -0.259 | 1.3 | -0.005 | 0.017 | 0.017 | 0 | 0.15 | 0.27 | 0.13 | 0.22 | 0.14 | 0.25 |
| 11 | 33714641 | rs2746600 | -0.127 | 0.05 | -0.669 | 1.034 | 0.026 | 0.014 | 0.014 | 0 | 0.46 | 0.4 | 0.35 | 0.57 | 0.34 | 0.47 |
| 11 | 33718980 | rs831608 | 0.022 | 0.861 | -1.327 | 1.422 | -0.078 | -0.108 | -0.108 | 3.178 | 0.8 | 0.97 | 0.67 | 0.57 | 0.77 | 0.68 |
| 11 | 33719850 | rs831612 | -0.272 | 0.563 | -1.633 | 0.76 | -0.003 | -0.038 | -0.038 | 0 | 0.32 | 0.26 | 0.25 | 0.45 | 0.29 | 0.39 |
| 11 | 33724780 | rs704697 | -0.056 | -0.073 | -1.369 | 0.004 | 0.001 | 0.005 | 0.005 | 0 | 0.52 | 0.44 | 0.4 | 0.54 | 0.58 | 0.58 |
| 11 | 33724992 | rs1047581 | 1.061 | -0.218 | -0.838 | -0.096 | 0.014 | 0.071 | 0.071 | 0 | 0.33 | 0.21 | 0.2 | 0.1 | 0.33 | 0.2 |
| 11 | 33727515 | rs12272660 | 0.791 | -0.487 | -0.474 | -0.24 | -0.018 | -0.023 | -0.023 | 0 | 0.07 | 0.13 | 0.02 | 0 | 0.01 | 0.03 |
| 11 | 33727705 | rs11585 | 0.519 | -0.348 | -0.406 | 0.5 | 0.005 | 0.12 | 0.12 | 0 | 0.09 | 0.19 | 0.05 | 0.19 | 0.07 | 0.08 |
| 11 | 33728614 | rs831618 | -0.036 | -0.319 | -0.389 | 0.406 | 0.001 | 0.005 | 0.005 | 0 | 0.52 | 0.44 | 0.41 | 0.53 | 0.58 | 0.58 |
| 11 | 33730060 | rs11032350 | 1.504 | -0.376 | -0.531 | 0.233 | -0.005 | 0.137 | 0.137 | 0 | 0.31 | 0.07 | 0.19 | 0.09 | 0.33 | 0.15 |
| 11 | 33731406 | rs842 | 1.395 | -0.366 | -0.525 | 0.239 | 0.01 | 0.112 | 0.112 | 0 | 0.32 | 0.13 | 0.19 | 0.1 | 0.33 | 0.2 |
| 11 | 33740780 | rs831633 | -0.707 | -0.26 | 0.199 | 0.52 | -0.005 | -0.011 | -0.011 | 0 | 0.61 | 0.74 | 0.46 | 0.5 | 0.62 | 0.61 |
| 11 | 33743019 | rs831630 | -1.155 | -0.031 | 0.376 | 0.499 | 0.022 | 0.089 | 0.089 | 0.338 | 0.28 | 0.54 | 0.25 | 0.4 | 0.3 | 0.46 |
| 11 | 33743534 | rs704700 | 0.848 | 0.07 | 0.331 | 0.644 | -0.012 | -0.011 | -0.011 | 0 | 0.11 | 0.11 | 0.18 | 0.24 | 0.18 | 0.29 |
| 11 | 33747333 | rs861256 | -0.691 | -0.105 | 0.891 | 0.677 | -0.004 | -0.016 | -0.016 | 0 | 0.61 | 0.68 | 0.46 | 0.48 | 0.63 | 0.59 |
| 11 | 33752403 | rs1738548 | 0.982 | -0.006 | 0.91 | 0.696 | 0.049 | 0.036 | 0.036 | 0 | 0.38 | 0.19 | 0.31 | 0.28 | 0.34 | 0.3 |
| 11 | 33754477 | rs10768024 | -0.534 | 0.307 | 1.161 | 0.946 | -0.014 | 0.019 | 0.019 | 0 | 0.15 | 0.26 | 0.07 | 0.16 | 0.09 | 0.11 |
| 11 | 33757770 | rs2273121 | 0.787 | 0.226 | 0.988 | 0.939 | -0.055 | 0.024 | 0.024 | 0 | 0.09 | 0.02 | 0.1 | 0.02 | 0.23 | 0.15 |
| 11 | 33758638 | rs831603 | 0.073 | 0.186 | 0.986 | 0.827 | 0.002 | 0.056 | 0.056 | 0 | 0.55 | 0.28 | 0.45 | 0.32 | 0.63 | 0.52 |
| 11 | 33763193 | rs831592 | -1.05 | 0.231 | 1.059 | 0.872 | -0.001 | 0 | 0 | 0.439 | 0.69 | 0.69 | 0.51 | 0.48 | 0.69 | 0.62 |
| 11 | 33766630 | rs831596 | 0.838 | 0.244 | 1.064 | 0.867 | 0.048 | 0.07 | 0.07 | 0 | 0.45 | 0.28 | 0.37 | 0.29 | 0.44 | 0.37 |
| 11 | 33773352 | rs7103573 | -1.259 | 0.192 | 1.081 | 0.843 | 0.037 | -0.004 | -0.004 | 0 | 0.23 | 0.41 | 0.14 | 0.19 | 0.25 | 0.25 |
| 11 | 33775424 | rs1522085 | -0.953 | 0.261 | 1.039 | 0.973 | -0.001 | -0.001 | -0.001 | 0 | 0.69 | 0.69 | 0.51 | 0.48 | 0.69 | 0.62 |
| 11 | 33778729 | rs12224154 | 0.739 | 0.397 | 0.906 | 1.1 | -0.118 | -0.121 | -0.121 | 0.881 | 0.3 | 0.02 | 0.46 | 0.44 | 0.3 | 0.37 |
| 11 | 33779673 | rs1112841 | 0.709 | 0.404 | 0.931 | 1.042 | -0.019 | -0.051 | -0.051 | 0 | 0.31 | 0.17 | 0.49 | 0.52 | 0.31 | 0.37 |
| 11 | 33796413 | rs2272067 | -0.475 | 0.692 | 1.29 | 1.325 | -0.005 | 0.031 | 0.031 | 0 | 0.12 | 0.21 | 0.07 | 0.28 | 0.08 | 0.11 |
| 11 | 33797408 | rs3817442 | -0.05 | 0.73 | 1.244 | 1.261 | -0.012 | -0.038 | -0.038 | 0 | 0.42 | 0.19 | 0.55 | 0.53 | 0.5 | 0.56 |
| 11 | 33799472 | rs2177121 | -0.626 | 0.728 | 1.287 | 1.353 | -0.004 | 0.184 | 0.184 | 0.125 | 0.13 | 0.32 | 0.21 | 0.37 | 0.08 | 0.12 |
| 11 | 33802300 | rs11032383 | 0.04 | 0.8 | 1.177 | 1.383 | -0.075 | -0.033 | -0.033 | 0 | 0.42 | 0.07 | 0.41 | 0.44 | 0.5 | 0.56 |
| 11 | 33804010 | rs11032385 | 0.722 | 0.877 | 1.151 | 1.367 | -0.051 | 0.031 | 0.031 | 0 | 0.09 | 0.02 | 0.09 | 0.02 | 0.2 | 0.16 |
| 11 | 33804557 | rs884710 | 0.747 | 0.875 | 1.151 | 1.374 | -0.073 | -0.075 | -0.075 | 0 | 0.32 | 0.03 | 0.32 | 0.41 | 0.3 | 0.39 |
| 11 | 33815975 | rs11601836 | 0.37 | 0.901 | 1.481 | 1.53 | 0.062 | 0.029 | 0.029 | 0 | 0.1 | 0.18 | 0.12 | 0.11 | 0.2 | 0.16 |
| 11 | 33826317 | rs17038 | 1.145 | 0.759 | 1.429 | 1.278 | 0.004 | 0.011 | 0.011 | 1.133 | 0.75 | 0.28 | 0.68 | 0.6 | 0.72 | 0.7 |
| 11 | 33828847 | rs939804 | 0.97 | 0.517 | 1.06 | 1.149 | 0.042 | 0.091 | 0.091 | 0 | 0.4 | 0.09 | 0.31 | 0.18 | 0.37 | 0.31 |
| 11 | 33830115 | rs7950864 | -0.681 | 0.78 | 1.545 | 1.499 | -0.005 | 0.13 | 0.13 | 0.051 | 0.12 | 0.22 | 0.2 | 0.37 | 0.08 | 0.11 |
| 11 | 33838377 | rs12223394 | -1.265 | 0.811 | 1.088 | 1.473 | -0.017 | 0.089 | 0.089 | 0 | 0.14 | 0.22 | 0.29 | 0.6 | 0.08 | 0.2 |
| 11 | 33840893 | rs11032401 | -0.133 | 0.915 | 1.105 | 1.559 | -0.001 | -0.02 | -0.02 | 0 | 0.58 | 0.46 | 0.45 | 0.23 | 0.62 | 0.57 |
| 11 | 33841739 | rs7946109 | -1.264 | 0.975 | 0.663 | 1.458 | 0.054 | 0.115 | 0.115 | 0.423 | 0.24 | 0.55 | 0.31 | 0.45 | 0.27 | 0.25 |
| 11 | 33852673 | rs4756053 | 0.233 | 0.524 | 0.45 | 1.492 | 0.104 | 0.093 | 0.093 | 0 | 0.51 | 0.3 | 0.33 | 0.25 | 0.39 | 0.39 |
| 11 | 33859314 | rs3924417 | -0.761 | 0.495 | 0.223 | 1.536 | -0.004 | -0.019 | -0.019 | 0 | 0.44 | 0.56 | 0.63 | 0.83 | 0.35 | 0.58 |
| 11 | 33864261 | rs7938708 | -1.419 | 0.56 | 0.082 | 2.282 | 0.003 | 0.007 | 0.007 | 0 | 0.11 | 0.13 | 0.11 | 0.28 | 0.04 | 0.18 |
| 11 | 33865882 | rs4130603 | -0.167 | 0.721 | 0.219 | 1.484 | -0.019 | 0.002 | 0.002 | 0.192 | 0.07 | 0.08 | 0.01 | 0.07 | 0.01 | 0.02 |
| 11 | 33879547 | rs7121669 | -1.119 | 2.129 | 0.479 | 1.658 | 0.142 | 0.089 | 0.089 | 0 | 0.06 | 0.16 | 0.12 | 0.21 | 0.19 | 0.09 |
| 11 | 33879892 | rs7106112 | -0.811 | 2.067 | -0.009 | 1.795 | -0.014 | -0.015 | -0.015 | 0 | 0.34 | 0.33 | 0.25 | 0.5 | 0.4 | 0.36 |
| 11 | 33880429 | rs7447 | 0.004 | 2.044 | -0.059 | 1.938 | -0.014 | 0.044 | 0.044 | 0.48 | 0.25 | 0.08 | 0.12 | 0.13 | 0.19 | 0.17 |
| 11 | 33881016 | rs3740617 | 0.76 | 1.904 | 0.785 | 1.864 | 0.027 | 0.029 | 0.029 | 0 | 0.64 | 0.47 | 0.73 | 0.49 | 0.6 | 0.61 |

| | | | | | | | | | | | | | | | | |
|---|---|---|---|---|---|---|---|---|---|---|---|---|---|---|---|---|
| 11 | 33885319 | rs3781575 | -1.768 | 2.013 | 0.913 | 1.839 | 0.17 | 0.235 | 0.235 | 0 | 0.11 | 0.43 | 0.13 | 0.3 | 0.18 | 0.21 |
| 11 | 33886080 | rs2273797 | -0.247 | 2.047 | 1.404 | 1.855 | 0.011 | -0.005 | -0.005 | 0 | 0.29 | 0.27 | 0.12 | 0.28 | 0.18 | 0.28 |
| 11 | 33886294 | rs2038602 | -1.035 | 1.816 | 1.378 | 1.721 | 0.006 | 0.001 | 0.001 | 0 | 0.39 | 0.58 | 0.28 | 0.41 | 0.42 | 0.38 |
| 11 | 33892656 | rs11606830 | -0.139 | 1.04 | 1.03 | 1.332 | 0.003 | 0.008 | 0.008 | 0 | 0.28 | 0.21 | 0.13 | 0.2 | 0.22 | 0.29 |
| 11 | 33899956 | rs3758638 | -0.494 | 0.494 | 1.217 | 1.46 | -0.026 | 0.005 | 0.005 | 0.012 | 0.38 | 0.5 | 0.25 | 0.43 | 0.24 | 0.35 |
| 11 | 33902612 | rs3740619 | -0.406 | 0.339 | 0.818 | 1.391 | 0.018 | 0.008 | 0.008 | 0 | 0.37 | 0.34 | 0.21 | 0.36 | 0.22 | 0.34 |
| 11 | 33903814 | rs1885523 | 0.268 | 0.537 | -0.109 | 1.15 | 0.023 | 0.015 | 0.015 | 0.942 | 0.3 | 0.34 | 0.49 | 0.41 | 0.47 | 0.51 |
| 11 | 33903882 | rs4307693 | 0.062 | 0.271 | -0.578 | 0.891 | 0.02 | 0.016 | 0.016 | 0 | 0.33 | 0.2 | 0.19 | 0.24 | 0.21 | 0.33 |
| 11 | 33906826 | rs3781578 | -0.207 | 1.009 | -0.606 | 1.172 | -0.001 | 0.002 | 0.002 | 0.215 | 0.19 | 0.16 | 0.26 | 0.34 | 0.25 | 0.34 |
| 11 | 33907955 | rs4756077 | -0.047 | 0.841 | -1.121 | 0.506 | -0.027 | -0.033 | -0.033 | 0 | 0.78 | 0.88 | 0.65 | 0.66 | 0.73 | 0.75 |
| 11 | 33908547 | rs911817 | 0.346 | 0.327 | -0.816 | 0.284 | -0.005 | -0.004 | -0.004 | 0 | 0.77 | 0.77 | 0.64 | 0.66 | 0.76 | 0.74 |
| 11 | 33908609 | rs883495 | 0.569 | 0.089 | 0.06 | 0.161 | 0.007 | 0 | 0 | 0 | 0.74 | 0.53 | 0.63 | 0.66 | 0.75 | 0.74 |
| 11 | 33912790 | rs11032440 | 0.439 | -0.303 | -0.465 | -0.38 | -0.013 | 0.001 | 0.001 | 0 | 0.39 | 0.27 | 0.39 | 0.56 | 0.48 | 0.51 |
| 11 | 33913568 | rs2273799 | 0.415 | -0.288 | -0.473 | -0.422 | -0.014 | 0.012 | 0.012 | 0 | 0.39 | 0.26 | 0.39 | 0.58 | 0.48 | 0.51 |
| 11 | 33914428 | rs3758640 | 0.176 | -0.397 | -0.331 | -0.774 | -0.002 | 0.002 | 0.002 | 0 | 0.64 | 0.62 | 0.57 | 0.67 | 0.64 | 0.64 |
| 11 | 33915891 | rs3758642 | 1.089 | -1.093 | -0.458 | -0.568 | -0.002 | 0.072 | 0.072 | 0 | 0.5 | 0.26 | 0.36 | 0.6 | 0.57 | 0.48 |
| 11 | 33917594 | rs750780 | 0.262 | -0.933 | -0.15 | -0.35 | -0.02 | -0.005 | -0.005 | 0 | 0.73 | 0.46 | 0.61 | 0.7 | 0.78 | 0.72 |
| 11 | 33924787 | rs941940 | 0.449 | -0.644 | -0.417 | -0.251 | -0.003 | -0.001 | -0.001 | 0 | 0.64 | 0.67 | 0.5 | 0.6 | 0.65 | 0.56 |
| 11 | 33925975 | rs7941248 | -0.195 | -1.358 | -0.548 | 0.109 | 0.041 | -0.015 | -0.015 | 0 | 0.61 | 0.79 | 0.59 | 0.5 | 0.7 | 0.65 |
| 11 | 33929348 | rs7946026 | 0.738 | -1.321 | -1.268 | 0.21 | 0.001 | -0.149 | -0.149 | 0 | 0.46 | 0.03 | 0.51 | 0.42 | 0.65 | 0.57 |
| 11 | 33931867 | rs7119405 | -0.954 | -0.962 | 0.144 | -0.112 | 0.002 | -0.069 | -0.069 | 0 | 0.11 | 0.38 | 0.06 | 0 | 0.05 | 0.06 |
| 11 | 33943644 | rs2181165 | 0.602 | -0.861 | -0.729 | 0.3 | -0.001 | -0.015 | -0.015 | 0.225 | 0.43 | 0.18 | 0.46 | 0.53 | 0.56 | 0.52 |
| 11 | 33945335 | rs7109636 | -0.547 | -1.237 | -0.322 | -0.426 | -0.009 | -0.061 | -0.061 | 1.129 | 0.17 | 0.23 | 0.11 | 0 | 0.13 | 0.14 |
| 11 | 33949342 | rs7103973 | -0.037 | -1.023 | -0.281 | 0.278 | 0.015 | -0.008 | -0.008 | 0 | 0.64 | 0.7 | 0.49 | 0.37 | 0.69 | 0.64 |
| 11 | 33954193 | rs1176363 | -0.769 | -0.844 | -0.128 | 0.277 | -0.001 | -0.042 | -0.042 | 0.179 | 0.1 | 0.53 | 0.07 | 0 | 0.06 | 0.05 |
| 11 | 33956132 | rs1176359 | 0.192 | -0.492 | -0.67 | 0.726 | 0.01 | -0.01 | -0.01 | 2.218 | 0.67 | 0.71 | 0.52 | 0.36 | 0.7 | 0.65 |
| 11 | 33958297 | rs10836131 | -0.941 | -0.092 | -0.501 | 0.78 | 0 | 0.011 | 0.011 | 0 | 0.28 | 0.36 | 0.48 | 0.64 | 0.3 | 0.34 |
| 11 | 33959436 | rs7945752 | 0.139 | -0.038 | -0.75 | 0.223 | 0.002 | 0.002 | 0.002 | 0.048 | 0.62 | 0.44 | 0.51 | 0.3 | 0.69 | 0.7 |
| 11 | 33961756 | rs7119163 | 0.556 | 0.157 | -1.114 | 0.05 | 0.015 | 0.081 | 0.081 | 0.027 | 0.49 | 0.3 | 0.41 | 0.27 | 0.52 | 0.61 |
| 11 | 33963848 | rs10742308 | 0.092 | 0.36 | -0.949 | 0.112 | -0.014 | 0.03 | 0.03 | 0 | 0.85 | 0.92 | 0.89 | 0.9 | 0.8 | 0.9 |
| 11 | 33971407 | rs10768037 | 0.477 | 0.35 | -1.133 | -0.431 | 0.014 | 0.11 | 0.11 | 0 | 0.52 | 0.68 | 0.6 | 0.75 | 0.46 | 0.53 |
| 11 | 33982201 | rs11032467 | 0.724 | 0.18 | -1.646 | 0.421 | 0.012 | -0.087 | -0.087 | 0.973 | 0.12 | 0 | 0.2 | 0.31 | 0.08 | 0.12 |
| 11 | 33982853 | rs2176611 | -0.115 | 0.447 | -0.734 | -0.229 | -0.007 | 0.007 | 0.007 | 0 | 0.63 | 0.73 | 0.67 | 0.75 | 0.55 | 0.58 |
| 11 | 33985242 | rs2421466 | 0.986 | 0.545 | -1.277 | -0.097 | -0.007 | 0.049 | 0.049 | 0.018 | 0.09 | 0.01 | 0.05 | 0 | 0.08 | 0.04 |
| 11 | 33992938 | rs2421467 | 0.176 | 0.562 | -1.112 | -0.134 | -0.03 | 0.136 | 0.136 | 0 | 0.56 | 0.84 | 0.64 | 0.75 | 0.48 | 0.54 |
| 11 | 33995732 | rs7941690 | 0.329 | 0.665 | -0.139 | -0.248 | 0.005 | -0.067 | -0.067 | 0 | 0.84 | 0.59 | 0.91 | 0.96 | 0.85 | 0.86 |
| 11 | 33995797 | rs10836136 | 0.106 | 0.976 | -0.373 | -0.687 | 0.014 | -0.079 | -0.079 | 1.152 | 0.6 | 0.49 | 0.66 | 0.89 | 0.47 | 0.62 |
| 11 | 33999742 | rs7123907 | 0.002 | 0.929 | -0.758 | -0.34 | -0.004 | 0.008 | 0.008 | 0.019 | 0.58 | 0.37 | 0.63 | 0.82 | 0.46 | 0.6 |
| 11 | 34000018 | rs10836139 | -0.365 | 1.114 | -0.704 | 0.792 | -0.006 | -0.012 | -0.012 | 0 | 0.2 | 0.13 | 0.29 | 0.26 | 0.2 | 0.17 |
| 11 | 34011084 | rs11032476 | 0.359 | 0.759 | -0.771 | 1.112 | 0.023 | -0.05 | -0.05 | 0 | 0.08 | 0 | 0.19 | 0.23 | 0.06 | 0.15 |
| 11 | 34013886 | rs4756093 | -0.423 | 1.75 | -0.014 | 1.091 | 0.043 | 0.058 | 0.058 | 0 | 0.25 | 0.34 | 0.27 | 0.37 | 0.35 | 0.29 |
| 11 | 34019537 | rs12285326 | -0.157 | 1.896 | 0.061 | 0.795 | -0.037 | -0.061 | -0.061 | 3.483 | 0.22 | 0.1 | 0.21 | 0.39 | 0.3 | 0.27 |
| 11 | 34024580 | rs11601630 | 0.2 | 1.991 | 0.264 | 0.361 | 0.015 | -0.058 | -0.058 | 0 | 0.12 | 0.18 | 0.1 | 0 | 0.13 | 0.06 |
| 11 | 34028130 | rs10501131 | 0.646 | 1.961 | 0.306 | 0.362 | 0.002 | -0.034 | -0.034 | 0.524 | 0.1 | 0.1 | 0.05 | 0 | 0.08 | 0.06 |
| 11 | 34032330 | rs7121633 | 0.837 | 1.81 | 1.28 | 0.368 | -0.007 | -0.004 | -0.004 | 0 | 0.63 | 0.4 | 0.67 | 0.59 | 0.57 | 0.65 |
| 11 | 34033151 | rs10768046 | 0.567 | 1.914 | 1.413 | 0.107 | 0 | -0.001 | -0.001 | 1.386 | 0.67 | 0.42 | 0.76 | 0.69 | 0.69 | 0.68 |
| 11 | 34039096 | rs11032492 | 0.158 | 2.013 | 1.385 | 0.126 | -0.006 | -0.015 | -0.015 | 0.057 | 0.21 | 0.16 | 0.17 | 0.31 | 0.24 | 0.26 |
| 11 | 34042300 | rs10836149 | 0.762 | 2.046 | 1.728 | 0.091 | 0.01 | -0.002 | -0.002 | 2.529 | 0.66 | 0.4 | 0.72 | 0.69 | 0.58 | 0.65 |
| 11 | 34042996 | rs7950733 | 0.066 | 2.051 | 1.573 | 0.077 | -0.006 | -0.014 | -0.014 | 0.037 | 0.22 | 0.17 | 0.17 | 0.33 | 0.23 | 0.26 |
| 11 | 34053873 | rs7120520 | 0.76 | 1.997 | 1.759 | 0.074 | 0.014 | -0.001 | -0.001 | 0 | 0.66 | 0.42 | 0.71 | 0.67 | 0.58 | 0.64 |
| 11 | 34056207 | rs12279034 | -0.283 | 2.03 | 1.189 | 0.191 | -0.007 | -0.006 | -0.006 | 0 | 0.23 | 0.22 | 0.21 | 0.33 | 0.34 | 0.3 |
| 11 | 34067121 | rs4497383 | -0.583 | 1.731 | 0.944 | -0.178 | -0.012 | -0.019 | -0.019 | 0 | 0.29 | 0.18 | 0.23 | 0.33 | 0.33 | 0.34 |
| 11 | 34072011 | rs12418412 | 0.086 | 1.76 | 1.27 | -0.904 | 0.048 | -0.065 | -0.065 | 0.36 | 0.12 | 0.21 | 0.09 | 0 | 0.18 | 0.09 |
| 11 | 34091679 | rs16925114 | 0.425 | 1.59 | 1.218 | -0.949 | -0.009 | -0.063 | -0.063 | 0.618 | 0.12 | 0.18 | 0.05 | 0 | 0.08 | 0.07 |
| 11 | 34097209 | rs10047428 | -0.259 | 1.572 | 1.688 | -0.964 | 0.008 | -0.092 | -0.092 | 0.081 | 0.87 | 0.56 | 0.93 | 1 | 0.92 | 0.93 |
| 11 | 34100008 | rs17699517 | 0.563 | 1.59 | 1.432 | -0.955 | -0.003 | -0.03 | -0.03 | 1.144 | 0.11 | 0.11 | 0.05 | 0 | 0.08 | 0.07 |
| 11 | 34108305 | rs10836154 | 0.222 | 1.604 | 1.387 | -1.005 | -0.006 | 0.041 | 0.041 | 0 | 0.08 | 0.01 | 0.05 | 0 | 0.09 | 0.06 |
| 11 | 34123814 | rs16925167 | 0.344 | 1.702 | 1.609 | -0.884 | -0.007 | -0.063 | -0.063 | 0 | 0.12 | 0.18 | 0.05 | 0 | 0.08 | 0.07 |
| 11 | 34126170 | rs16925170 | 0.344 | 1.681 | 1.609 | -0.884 | -0.009 | -0.063 | -0.063 | 2.987 | 0.12 | 0.18 | 0.05 | 0 | 0.08 | 0.07 |
| 11 | 34136028 | rs2957481 | -0.451 | 1.682 | 1.6 | -0.883 | -0.008 | -0.071 | -0.071 | 0.144 | 0.88 | 0.74 | 0.94 | 1 | 0.92 | 0.93 |
| 11 | 34141528 | rs2957509 | -0.428 | 1.663 | 1.638 | -0.883 | 0 | -0.079 | -0.079 | 0.373 | 0.88 | 0.65 | 0.94 | 1 | 0.92 | 0.93 |
| 11 | 34189564 | rs4756111 | -0.166 | 0.87 | 1.318 | -0.942 | -0.026 | -0.094 | -0.094 | 0 | 0.86 | 0.71 | 0.84 | 1 | 0.92 | 0.9 |
| 11 | 34190423 | rs1569774 | -0.042 | 0.454 | 0.503 | -1.348 | 0.01 | 0.107 | 0.107 | 0.305 | 0.46 | 0.22 | 0.49 | 0.71 | 0.42 | 0.51 |
| 11 | 34190762 | rs2982608 | -0.859 | 0.399 | 0.725 | -2.072 | -0.018 | 0.028 | 0.028 | 0.164 | 0.61 | 0.35 | 0.71 | 0.88 | 0.51 | 0.68 |
| 11 | 34192092 | rs2957489 | 0.212 | 0.293 | 0.556 | -2.1 | -0.004 | -0.043 | -0.043 | 0 | 0.18 | 0.16 | 0.16 | 0 | 0.1 | 0.14 |
| 11 | 34196441 | rs7121208 | -0.222 | 0.086 | 0.676 | -2.176 | -0.002 | -0.05 | -0.05 | 0 | 0.51 | 0.52 | 0.61 | 0.89 | 0.45 | 0.61 |
| 11 | 34196484 | rs2467383 | 0.018 | 0.059 | 0.52 | -2.182 | 0.004 | 0.113 | 0.113 | 0.565 | 0.47 | 0.3 | 0.39 | 0.08 | 0.54 | 0.39 |
| 11 | 34201657 | rs1321017 | 0.012 | 0.098 | 0.393 | -2.171 | 0.004 | 0.042 | 0.042 | 0.747 | 0.47 | 0.36 | 0.39 | 0.11 | 0.54 | 0.38 |
| 11 | 34202953 | rs2467377 | -0.263 | 0.032 | 0.247 | -2.191 | -0.044 | -0.119 | -0.119 | 0 | 0.69 | 0.58 | 0.75 | 0.97 | 0.83 | 0.78 |

| | | | | | | | | | | | | | | | | |
|---|---|---|---|---|---|---|---|---|---|---|---|---|---|---|---|---|
| 11 | 34203862 | rs2143868 | -0.154 | 0.234 | -0.335 | -2.038 | -0.001 | 0.034 | 0.034 | 0.147 | 0.6 | 0.66 | 0.45 | 0.13 | 0.57 | 0.42 |
| 11 | 34204396 | rs4756113 | -0.249 | 0.216 | -1.03 | -1.759 | 0 | 0.081 | 0.081 | 0.019 | 0.38 | 0.24 | 0.48 | 0.81 | 0.39 | 0.53 |
| 11 | 34212883 | rs9666645 | 0.627 | -0.063 | -0.525 | -1.546 | 0.019 | 0.03 | 0.03 | 0.758 | 0.62 | 0.3 | 0.71 | 0.97 | 0.72 | 0.71 |
| 11 | 34215606 | rs2473904 | -0.364 | -0.066 | -0.512 | -1.134 | -0.016 | -0.069 | -0.069 | 0 | 0.26 | 0.3 | 0.24 | 0.03 | 0.18 | 0.22 |
| 11 | 34218556 | rs2957497 | 0.105 | 0.573 | -0.549 | 0.17 | 0 | 0.001 | 0.001 | 0.062 | 0.63 | 0.42 | 0.5 | 0.5 | 0.61 | 0.67 |
| 11 | 34220504 | rs2467372 | 0.413 | 0.564 | -0.454 | 0.524 | 0.018 | 0.031 | 0.031 | 0 | 0.8 | 0.71 | 0.82 | 0.74 | 0.73 | 0.64 |
| 11 | 34227327 | rs10768051 | -0.398 | -0.076 | -1.444 | 0.601 | 0.016 | -0.075 | -0.075 | 0 | 0.32 | 0.07 | 0.53 | 0.57 | 0.27 | 0.34 |
| 11 | 34228306 | rs2746633 | -0.947 | -0.265 | -1.372 | 1.239 | 0.01 | -0.05 | -0.05 | 0 | 0.25 | 0.15 | 0.21 | 0.39 | 0.22 | 0.31 |
| 11 | 34234145 | rs1883959 | -0.765 | 0.098 | -0.399 | 1.321 | 0.018 | 0.053 | 0.053 | 0 | 0.12 | 0.23 | 0.25 | 0.25 | 0.16 | 0.16 |
| 11 | 34236890 | rs7120814 | -1.184 | 0.021 | 0.488 | 1.266 | 0.05 | 0.123 | 0.123 | 0 | 0.14 | 0.49 | 0.26 | 0.25 | 0.16 | 0.16 |
| 11 | 34237292 | rs2746628 | 0.977 | -0.296 | 0.279 | 0.45 | 0.006 | 0.029 | 0.029 | 0 | 0.61 | 0.31 | 0.6 | 0.75 | 0.65 | 0.63 |
| 11 | 34240676 | rs911498 | 0.032 | -0.568 | 0.088 | 0.647 | -0.01 | -0.013 | -0.013 | 0.161 | 0.68 | 0.62 | 0.82 | 0.86 | 0.8 | 0.72 |
| 11 | 34248586 | rs6484701 | 0.765 | -0.353 | -0.015 | 0.674 | -0.008 | -0.001 | -0.001 | 0 | 0.55 | 0.51 | 0.57 | 0.62 | 0.64 | 0.57 |
| 11 | 34251855 | rs2957523 | 0.828 | 0.314 | -0.017 | 0.85 | 0.025 | 0.118 | 0.118 | 0.014 | 0.76 | 0.6 | 0.63 | 0.59 | 0.72 | 0.65 |
| 11 | 34251991 | rs1033752 | 0.689 | 0.321 | -0.053 | 0.854 | 0.009 | 0.047 | 0.047 | 0.016 | 0.78 | 0.72 | 0.64 | 0.59 | 0.72 | 0.65 |
| 11 | 34253421 | rs4756118 | 0.315 | 0.296 | -0.846 | 0.126 | 0 | -0.091 | -0.091 | 0.077 | 0.31 | 0.1 | 0.34 | 0.46 | 0.23 | 0.32 |
| 11 | 34254039 | rs17255930 | 1.183 | 0.443 | -0.634 | 0.295 | -0.038 | -0.014 | -0.014 | 0 | 0.06 | 0 | 0.07 | 0.06 | 0.14 | 0.09 |
| 11 | 34254625 | rs1983756 | -1.147 | 0.025 | -0.375 | 0.529 | -0.024 | 0.048 | 0.048 | 0 | 0.93 | 0.85 | 0.95 | 0.89 | 0.99 | 0.93 |
| 11 | 34256153 | rs6484702 | 1.615 | 0.317 | -0.927 | 0.054 | -0.007 | -0.013 | -0.013 | 0 | 0.23 | 0.2 | 0.19 | 0.08 | 0.35 | 0.19 |
| 11 | 34256377 | rs4756119 | 0.022 | 0.65 | -1.24 | 0.997 | -0.005 | -0.088 | -0.088 | 0.698 | 0.2 | 0.1 | 0.19 | 0.42 | 0.15 | 0.22 |
| 11 | 34257064 | rs10768053 | -0.699 | 0.708 | -0.611 | 1.07 | -0.032 | -0.031 | -0.031 | 0.329 | 0.22 | 0.11 | 0.44 | 0.3 | 0.33 | 0.33 |
| 11 | 34257916 | rs10836163 | -0.784 | 0.685 | -0.574 | 1.139 | -0.014 | -0.02 | -0.02 | 0 | 0.23 | 0.15 | 0.47 | 0.34 | 0.31 | 0.34 |
| 11 | 34259820 | rs9787727 | -0.684 | 0.951 | -0.646 | 1.056 | -0.076 | -0.024 | -0.024 | 0 | 0.14 | 0.01 | 0.3 | 0.16 | 0.22 | 0.14 |
| 11 | 34265140 | rs2755176 | 0.804 | 0.426 | -0.158 | 1.094 | -0.007 | -0.016 | -0.016 | 0 | 0.63 | 0.72 | 0.46 | 0.55 | 0.59 | 0.57 |
| 11 | 34266683 | rs7105914 | 1.068 | 0.349 | -0.253 | 1.057 | 0 | 0.001 | 0.001 | 0 | 0.53 | 0.36 | 0.39 | 0.55 | 0.55 | 0.54 |
| 11 | 34271143 | rs10836168 | -0.76 | 0.231 | -0.363 | 1.053 | -0.016 | 0.003 | 0.003 | 0 | 0.15 | 0.04 | 0.14 | 0.14 | 0.17 | 0.17 |
| 11 | 34271712 | rs7941113 | -0.923 | -0.014 | 0.187 | 1.034 | -0.015 | -0.003 | -0.003 | 0.132 | 0.13 | 0.3 | 0.08 | 0.07 | 0.11 | 0.13 |
| 11 | 34275289 | rs10734423 | -1.985 | -0.031 | 0.859 | 1.096 | -0.01 | -0.007 | -0.007 | 0 | 0.3 | 0.56 | 0.23 | 0.22 | 0.28 | 0.32 |
| 11 | 34278873 | rs10836174 | -2.167 | 0.004 | 1.142 | 0.967 | -0.013 | -0.025 | -0.025 | 0 | 0.29 | 0.44 | 0.22 | 0.2 | 0.27 | 0.32 |
| 11 | 34280911 | rs12289607 | -1.08 | -0.335 | 0.972 | 0.794 | -0.053 | -0.021 | -0.021 | 0 | 0.12 | 0.33 | 0.03 | 0.05 | 0.03 | 0.04 |
| 11 | 34282229 | rs10836178 | -1.927 | -0.287 | 0.66 | 0.843 | -0.018 | -0.019 | -0.019 | 0 | 0.28 | 0.59 | 0.19 | 0.19 | 0.2 | 0.23 |
| 11 | 34282979 | rs16925360 | 0.178 | -0.21 | 0.531 | 0.769 | -0.001 | 0.017 | 0.017 | 0.402 | 0.08 | 0.03 | 0.05 | 0.02 | 0.07 | 0.08 |
| 11 | 34283861 | rs2611145 | 2.022 | 0.005 | 0.414 | 1.359 | 0.014 | 0.034 | 0.034 | 0.301 | 0.53 | 0.33 | 0.46 | 0.39 | 0.53 | 0.62 |
| 11 | 34290063 | rs2755167 | -0.85 | -0.035 | 0.848 | 1.153 | -0.017 | -0.039 | -0.039 | 0 | 0.1 | 0.49 | 0.05 | 0 | 0.02 | 0 |
| 11 | 34297505 | rs12361605 | -0.214 | 0.582 | 0.923 | 1.374 | -0.002 | 0 | 0 | 0 | 0.09 | 0.1 | 0.23 | 0.02 | 0.12 | 0.09 |
| 11 | 34297823 | rs7110436 | 1.911 | 0.538 | 0.628 | 1.711 | -0.017 | -0.003 | -0.003 | 0 | 0.59 | 0.26 | 0.29 | 0.48 | 0.51 | 0.62 |
| 11 | 34302118 | rs1967745 | -1.581 | 0.873 | 0.926 | 1.443 | 0.054 | -0.044 | -0.044 | 0.006 | 0.18 | 0.49 | 0.32 | 0.07 | 0.21 | 0.16 |
| 11 | 34309519 | rs1998276 | 0.395 | 0.831 | 0.3 | 1.544 | 0.008 | 0.036 | 0.036 | 0.531 | 0.79 | 0.73 | 0.61 | 0.65 | 0.72 | 0.87 |
| 11 | 34311031 | rs2420520 | 0.602 | 0.626 | -0.232 | 1.442 | 0.018 | 0.022 | 0.022 | 2.578 | 0.69 | 0.44 | 0.51 | 0.67 | 0.61 | 0.77 |
| 11 | 34312019 | rs5008984 | -1.094 | 0.103 | -0.718 | 1.481 | 0.008 | -0.014 | -0.014 | 0 | 0.1 | 0.04 | 0.05 | 0.16 | 0.09 | 0.29 |
| 11 | 34312648 | rs2611127 | -1.14 | -0.571 | -0.471 | 2.157 | -0.033 | 0.111 | 0.111 | 0 | 0.15 | 0.22 | 0.21 | 0.48 | 0.08 | 0.26 |
| 11 | 34313693 | rs2755149 | -0.534 | -0.44 | -1.085 | 0.798 | -0.004 | 0.006 | 0.006 | 0 | 0.58 | 0.61 | 0.65 | 0.64 | 0.57 | 0.54 |
| 11 | 34314249 | rs911500 | -1.009 | -0.351 | -1.668 | 0.551 | -0.118 | 0.115 | 0.115 | 0 | 0.71 | 0.98 | 0.83 | 0.9 | 0.69 | 0.87 |
| 11 | 34314373 | rs4254051 | 0.803 | -0.208 | -2.056 | -0.04 | -0.004 | -0.095 | -0.095 | 2.456 | 0.31 | 0.47 | 0.32 | 0.04 | 0.28 | 0.22 |
| 11 | 34318636 | rs2755147 | -0.24 | -0.091 | -2.903 | -0.217 | -0.022 | 0.04 | 0.04 | 0.407 | 0.54 | 0.77 | 0.5 | 0.63 | 0.47 | 0.45 |
| 11 | 34318749 | rs7939713 | 0.756 | 0.259 | -2.535 | 0.164 | -0.003 | 0.067 | 0.067 | 0 | 0.11 | 0.19 | 0.17 | 0.25 | 0.12 | 0.33 |
| 11 | 34320849 | rs11032604 | 0.648 | 0.407 | -2.453 | 0.669 | -0.041 | -0.088 | -0.088 | 0 | 0.17 | 0.04 | 0.29 | 0.46 | 0.24 | 0.2 |
| 11 | 34321193 | rs2755145 | -0.729 | 0.501 | -2.145 | -0.089 | -0.077 | 0.086 | 0.086 | 0 | 0.73 | 0.97 | 0.85 | 0.87 | 0.72 | 0.88 |
| 11 | 34321466 | rs2180949 | -0.709 | 0.194 | -1.188 | 0.578 | 0.001 | -0.003 | -0.003 | 0.527 | 0.25 | 0.2 | 0.35 | 0.24 | 0.25 | 0.41 |
| 11 | 34324297 | rs4755367 | -0.508 | 0.376 | -0.813 | 0.1 | -0.004 | 0.014 | 0.014 | 0 | 0.59 | 0.67 | 0.59 | 0.7 | 0.5 | 0.73 |
| 11 | 34326471 | rs1267476 | -0.41 | 0.323 | -0.52 | 0.088 | -0.034 | 0.082 | 0.082 | 0 | 0.6 | 0.78 | 0.59 | 0.79 | 0.5 | 0.75 |
| 11 | 34327576 | rs2611093 | 0.256 | -0.165 | -0.243 | 0.529 | -0.025 | 0.027 | 0.027 | 0 | 0.13 | 0.26 | 0.05 | 0.18 | 0.07 | 0.13 |
| 11 | 34328563 | rs2955957 | 0.238 | -0.169 | -0.244 | 0.361 | -0.025 | 0.067 | 0.067 | 0 | 0.13 | 0.26 | 0.05 | 0.27 | 0.07 | 0.14 |
| 11 | 34333973 | rs2077441 | 0.27 | 0.177 | -1.623 | 0.399 | -0.003 | 0.039 | 0.039 | 0 | 0.18 | 0.19 | 0.22 | 0.43 | 0.17 | 0.4 |
| 11 | 34337564 | rs2611140 | -0.296 | 0.941 | -2.195 | -0.551 | -0.102 | 0.112 | 0.112 | 0 | 0.62 | 0.88 | 0.61 | 0.79 | 0.54 | 0.77 |
| 11 | 34338766 | rs1267471 | 0.168 | 0.885 | -1.892 | -0.695 | -0.006 | 0.049 | 0.049 | 3.184 | 0.81 | 0.93 | 0.91 | 0.97 | 0.83 | 0.92 |
| 11 | 34343483 | rs1267468 | -0.173 | 1.058 | -2.354 | -1.024 | -0.048 | 0.128 | 0.128 | 0.775 | 0.23 | 0.04 | 0.22 | 0.05 | 0.23 | 0.25 |
| 11 | 34352436 | rs2755148 | 0.245 | 0.885 | -3.435 | -1.773 | -0.055 | 0.208 | 0.208 | 0.472 | 0.52 | 0.14 | 0.44 | 0.16 | 0.52 | 0.55 |
| 11 | 34352852 | rs11032618 | 0.596 | 1.072 | -3.458 | -0.568 | -0.002 | 0.018 | 0.018 | 0.332 | 0.07 | 0.08 | 0.08 | 0.36 | 0.08 | 0.03 |
| 11 | 34353226 | rs4756135 | 0.993 | 1.045 | -3.722 | -0.949 | -0.095 | 0.166 | 0.166 | 0 | 0.27 | 0.05 | 0.3 | 0.04 | 0.34 | 0.27 |
| 11 | 34357036 | rs1886657 | 1.504 | 1.087 | -2.677 | -0.458 | 0.032 | 0.05 | 0.05 | 0 | 0.08 | 0.22 | 0.07 | 0.13 | 0.07 | 0.08 |
| 11 | 34357080 | rs10836186 | 0.819 | 1.033 | -2.561 | -0.284 | -0.007 | 0.062 | 0.062 | 0.064 | 0.07 | 0.12 | 0.06 | 0.37 | 0.05 | 0.02 |
| 11 | 34359571 | rs7104735 | -0.204 | 0.718 | -1.942 | -0.048 | 0.011 | -0.014 | -0.014 | 0 | 0.28 | 0.39 | 0.28 | 0.19 | 0.25 | 0.19 |
| 11 | 34362999 | rs2611143 | 1.612 | 0.626 | -2.08 | -0.019 | 0.016 | 0.046 | 0.046 | 0 | 0.26 | 0.07 | 0.16 | 0.12 | 0.21 | 0.28 |
| 11 | 34364133 | rs926623 | -0.62 | 0.805 | -1.446 | 0.247 | 0.005 | 0.028 | 0.028 | 0.023 | 0.33 | 0.49 | 0.41 | 0.35 | 0.34 | 0.36 |
| 11 | 34369054 | rs742635 | 1.238 | 0.88 | -1.13 | 0.665 | -0.014 | -0.021 | -0.021 | 0 | 0.07 | 0.04 | 0.05 | 0.17 | 0.09 | 0.08 |
| 11 | 34370668 | rs2078165 | -1.483 | 0.857 | -1.427 | 0.3 | -0.074 | 0.223 | 0.223 | 0 | 0.46 | 0.9 | 0.55 | 0.86 | 0.45 | 0.4 |
| 11 | 34370840 | rs2078164 | -1.48 | 0.861 | -1.426 | 0.566 | -0.075 | 0.162 | 0.162 | 0 | 0.46 | 0.9 | 0.55 | 0.8 | 0.45 | 0.38 |
| 11 | 34373443 | rs1883736 | -0.126 | 0.664 | -0.855 | 0.793 | 0.031 | 0.256 | 0.256 | 0.397 | 0.09 | 0.4 | 0.09 | 0.37 | 0.07 | 0.02 |

| | | | | | | | | | | | | | | | |
|---|---|---|---|---|---|---|---|---|---|---|---|---|---|---|---|
| 11 | 34376317 | rs742634 | 0.27 | 0.665 | -0.478 | 0.87 | 0.063 | 0.363 | 0.363 | 0.822 | 0.07 | 0.32 | 0.09 | 0.38 | 0.06 | 0.02 |
| 11 | 34376882 | rs2745915 | 1.108 | 0.439 | -0.968 | 0.514 | 0.028 | 0.064 | 0.064 | 0 | 0.28 | 0.07 | 0.15 | 0.14 | 0.23 | 0.32 |
| 11 | 34379220 | rs3740795 | 0.504 | 0.602 | -0.642 | 0.427 | 0.057 | 0.345 | 0.345 | 0.654 | 0.06 | 0.29 | 0.09 | 0.37 | 0.06 | 0.02 |
| 11 | 34401083 | rs2746167 | 1.316 | 0.345 | -0.545 | 0.195 | 0.001 | 0.001 | 0.001 | 0 | 0.26 | 0.23 | 0.16 | 0.16 | 0.21 | 0.28 |
| 11 | 34404491 | rs10219333 | -0.002 | 0.935 | -0.067 | 0.189 | 0.093 | 0.168 | 0.168 | 0 | 0.1 | 0.26 | 0.12 | 0.33 | 0.18 | 0.04 |
| 11 | 34405407 | rs2746152 | 1.359 | 0.361 | -0.067 | 0.222 | 0.022 | 0.027 | 0.027 | 0 | 0.25 | 0.14 | 0.15 | 0.16 | 0.21 | 0.28 |
| 11 | 34409822 | rs997704 | 0.519 | 0.33 | -0.34 | 0.036 | 0.022 | 0.013 | 0.013 | 0 | 0.46 | 0.37 | 0.43 | 0.37 | 0.38 | 0.49 |
| 11 | 34410017 | rs11032644 | 0.217 | 0.195 | -0.614 | 0.042 | -0.008 | -0.007 | -0.007 | 2.642 | 0.15 | 0.21 | 0.13 | 0.07 | 0.09 | 0.09 |
| 11 | 34416247 | rs2746157 | -0.365 | 0.326 | -0.676 | 0.036 | 0.006 | -0.012 | -0.012 | 0.041 | 0.22 | 0.21 | 0.28 | 0.27 | 0.16 | 0.2 |
| 11 | 34417143 | rs2103711 | 0.523 | 0.613 | -0.218 | -0.059 | -0.001 | 0 | 0 | 0.686 | 0.46 | 0.52 | 0.45 | 0.46 | 0.38 | 0.47 |
| 11 | 34420930 | rs7928060 | -0.393 | 0.675 | -0.392 | 0.005 | 0.015 | -0.001 | -0.001 | 0 | 0.76 | 0.92 | 0.69 | 0.79 | 0.79 | 0.81 |
| 11 | 34423616 | rs10836214 | -0.514 | 0.71 | -0.188 | 0.146 | 0.015 | 0.212 | 0.212 | 1.013 | 0.13 | 0.51 | 0.09 | 0.31 | 0.08 | 0.04 |
| 11 | 34427724 | rs12574586 | 1.584 | 0.661 | -0.441 | 0.046 | 0.035 | 0.015 | 0.015 | 0 | 0.31 | 0.19 | 0.24 | 0.24 | 0.26 | 0.37 |
| 11 | 34428362 | rs5026710 | 0.883 | 0.786 | -0.389 | -0.068 | 0.02 | 0.06 | 0.06 | 0 | 0.56 | 0.32 | 0.44 | 0.28 | 0.63 | 0.65 |
| 11 | 34428509 | rs12289239 | 0.799 | 0.783 | -0.2 | 0.02 | 0.029 | 0.291 | 0.291 | 0 | 0.07 | 0.28 | 0.05 | 0.31 | 0.04 | 0.02 |
| 11 | 34431488 | rs12804689 | 0.949 | 0.808 | -0.266 | -0.074 | -0.04 | 0.039 | 0.039 | 0 | 0.25 | 0.18 | 0.2 | 0.04 | 0.37 | 0.28 |
| 11 | 34434323 | rs7946153 | 0.928 | 0.74 | -0.304 | -0.052 | -0.011 | 0.005 | 0.005 | 0 | 0.37 | 0.42 | 0.29 | 0.54 | 0.3 | 0.39 |
| 11 | 34434761 | rs12294378 | 0.928 | 0.74 | -0.304 | -0.052 | -0.011 | 0.005 | 0.005 | 0 | 0.37 | 0.42 | 0.29 | 0.54 | 0.3 | 0.39 |
| 11 | 34436686 | rs873466 | 1.041 | 0.756 | -0.152 | -0.075 | 0.008 | -0.042 | -0.042 | 0 | 0.25 | 0.27 | 0.2 | 0.04 | 0.37 | 0.28 |
| 11 | 34437027 | rs910091 | 0.29 | 0.756 | -0.153 | -0.073 | 0.003 | -0.005 | -0.005 | 1.388 | 0.63 | 0.68 | 0.49 | 0.59 | 0.67 | 0.67 |
| 11 | 34438558 | rs10488735 | 0.698 | 0.638 | -0.142 | 0.067 | 0.012 | 0.026 | 0.026 | 0 | 0.13 | 0.23 | 0.21 | 0.21 | 0.12 | 0.14 |
| 11 | 34440025 | rs208668 | 0.127 | 0.63 | -0.131 | -0.012 | 0.021 | 0.059 | 0.059 | 0 | 0.5 | 0.65 | 0.5 | 0.75 | 0.42 | 0.53 |
| 11 | 34440923 | rs7131473 | 0.449 | 0.912 | 0.112 | 0.394 | 0.027 | 0.287 | 0.287 | 0 | 0.07 | 0.28 | 0.05 | 0.31 | 0.04 | 0.02 |
| 11 | 34445032 | rs1408039 | 0.82 | 0.878 | 0.062 | 0.317 | -0.034 | 0.08 | 0.08 | 1.158 | 0.14 | 0.26 | 0.11 | 0.31 | 0.09 | 0.03 |
| 11 | 34447586 | rs11032695 | 1.125 | 1.268 | -0.239 | -0.025 | -0.058 | 0.065 | 0.065 | 0 | 0.22 | 0.12 | 0.18 | 0.04 | 0.35 | 0.28 |
| 11 | 34448269 | rs2223676 | 0.938 | 1.267 | -0.131 | 0.032 | -0.038 | 0.089 | 0.089 | 1.17 | 0.15 | 0.3 | 0.11 | 0.3 | 0.09 | 0.03 |
| 11 | 34449260 | rs16925514 | 1.299 | 1.291 | -0.031 | -0.053 | 0.001 | -0.029 | -0.029 | 0 | 0.06 | 0.13 | 0.04 | 0 | 0.04 | 0.02 |
| 11 | 34450461 | rs12295136 | 1.001 | 1.278 | 0.105 | -0.059 | 0.03 | -0.021 | -0.021 | 0 | 0.06 | 0.25 | 0.04 | 0 | 0.04 | 0.02 |
| 11 | 34453965 | rs208678 | 1.695 | 1.332 | 0.079 | 0.056 | 0.005 | 0.031 | 0.031 | 0 | 0.07 | 0.08 | 0.18 | 0.22 | 0.07 | 0.13 |
| 11 | 34454061 | rs208679 | 1.652 | 1.33 | 0.094 | 0.055 | 0.009 | 0.046 | 0.046 | 0 | 0.07 | 0.11 | 0.18 | 0.22 | 0.07 | 0.13 |
| 11 | 34454147 | rs7118388 | -0.511 | 1.286 | 0.075 | 0.023 | -0.013 | 0.018 | 0.018 | 0.008 | 0.44 | 0.66 | 0.34 | 0.35 | 0.49 | 0.33 |
| 11 | 34455309 | rs7944397 | 0.121 | 1.27 | 0.068 | 0.034 | -0.049 | 0.08 | 0.08 | 1.01 | 0.21 | 0.49 | 0.16 | 0.3 | 0.14 | 0.05 |
| 11 | 34455941 | rs208682 | -0.397 | 1.176 | -0.215 | -0.074 | -0.01 | -0.028 | -0.028 | 0 | 0.81 | 0.97 | 0.77 | 0.8 | 0.84 | 0.81 |
| 11 | 34456108 | rs12807961 | 1.32 | 1.259 | -0.206 | -0.151 | -0.061 | 0.068 | 0.068 | 0 | 0.22 | 0.12 | 0.18 | 0.04 | 0.35 | 0.28 |
| 11 | 34456149 | rs12808450 | 1.32 | 1.259 | -0.206 | -0.151 | -0.061 | 0.068 | 0.068 | 0 | 0.22 | 0.12 | 0.18 | 0.04 | 0.35 | 0.28 |
| 11 | 34456188 | rs554518 | -1.186 | 1.174 | -0.252 | -0.175 | -0.052 | 0.035 | 0.035 | 0 | 0.9 | 0.99 | 0.79 | 1 | 0.85 | 0.95 |
| 11 | 34456208 | rs1107573 | 1.442 | 1.42 | -0.047 | -0.089 | 0.001 | -0.027 | -0.027 | 0 | 0.06 | 0.12 | 0.03 | 0 | 0.04 | 0.02 |
| 11 | 34457654 | rs10836234 | 1.225 | 1.438 | -0.09 | -0.084 | -0.061 | 0.069 | 0.069 | 0 | 0.22 | 0.12 | 0.18 | 0.04 | 0.35 | 0.28 |
| 11 | 34459717 | rs769214 | 0.547 | 1.673 | -0.057 | -0.026 | -0.008 | -0.017 | -0.017 | 0 | 0.58 | 0.56 | 0.46 | 0.27 | 0.67 | 0.64 |
| 11 | 34460231 | rs1001179 | 1.928 | 2.103 | -0.611 | -0.069 | -0.069 | 0.089 | 0.089 | 0.012 | 0.17 | 0.02 | 0.12 | 0.03 | 0.23 | 0.25 |
| 11 | 34460704 | rs10836235 | 1.3 | 1.223 | -0.366 | 0.141 | 0.027 | -0.043 | -0.043 | 0 | 0.14 | 0.09 | 0.11 | 0.3 | 0.09 | 0.03 |
| 11 | 34460711 | rs12270780 | 1.642 | 1.304 | -0.218 | 0.112 | -0.024 | -0.02 | -0.02 | 0 | 0.31 | 0.38 | 0.26 | 0.23 | 0.27 | 0.34 |
| 11 | 34460769 | rs11604331 | 1.604 | 1.347 | -0.254 | 0.107 | -0.091 | 0.097 | 0.097 | 0 | 0.21 | 0.06 | 0.18 | 0.03 | 0.35 | 0.29 |
| 11 | 34461568 | rs11032699 | 0.715 | 1.248 | -0.183 | 0.123 | 0.008 | 0.016 | 0.016 | 0 | 0.58 | 0.6 | 0.48 | 0.26 | 0.66 | 0.65 |
| 11 | 34461665 | rs12272630 | 0.652 | 1.248 | -0.126 | 0.114 | 0.075 | 0.009 | 0.009 | 0 | 0.07 | 0.34 | 0.06 | 0 | 0.04 | 0.02 |
| 11 | 34461703 | rs11032700 | 0.431 | 1.241 | -0.13 | 0.115 | 0.023 | 0.062 | 0.062 | 0 | 0.59 | 0.77 | 0.48 | 0.27 | 0.66 | 0.65 |
| 11 | 34463760 | rs524154 | -0.861 | 0.86 | -0.061 | 0.184 | 0.008 | 0.011 | 0.011 | 0 | 0.41 | 0.4 | 0.47 | 0.74 | 0.29 | 0.34 |
| 11 | 34463872 | rs17881482 | 1.714 | 0.872 | -0.074 | 0.161 | 0.028 | 0.019 | 0.019 | 0 | 0.29 | 0.16 | 0.24 | 0.23 | 0.26 | 0.34 |
| 11 | 34464748 | rs494024 | -1.798 | 0.872 | -0.075 | 0.158 | -0.133 | 0.126 | 0.126 | 0 | 0.78 | 0.97 | 0.77 | 0.97 | 0.6 | 0.7 |
| 11 | 34467146 | rs2179625 | 1.716 | 0.895 | -0.045 | 0.155 | 0.025 | 0.015 | 0.015 | 0 | 0.29 | 0.1 | 0.23 | 0.23 | 0.26 | 0.34 |
| 11 | 34467272 | rs484214 | -0.741 | 0.812 | -0.033 | 0.159 | 0.034 | 0.05 | 0.05 | 0 | 0.4 | 0.28 | 0.47 | 0.74 | 0.29 | 0.34 |
| 11 | 34467490 | rs482322 | -0.751 | 0.812 | -0.033 | 0.159 | 0.035 | 0.051 | 0.051 | 0 | 0.4 | 0.28 | 0.47 | 0.74 | 0.29 | 0.34 |
| 11 | 34467538 | rs524167 | -1.748 | 0.812 | -0.033 | 0.159 | -0.106 | 0.109 | 0.109 | 0 | 0.77 | 0.94 | 0.77 | 0.97 | 0.6 | 0.7 |
| 11 | 34467646 | rs480575 | -0.818 | 0.812 | -0.034 | 0.16 | 0.035 | 0.053 | 0.053 | 0 | 0.39 | 0.22 | 0.47 | 0.73 | 0.29 | 0.34 |
| 11 | 34467848 | rs2300182 | 0.786 | 0.785 | -0.072 | 0.196 | 0.004 | -0.037 | -0.037 | 0 | 0.17 | 0.03 | 0.19 | 0.2 | 0.12 | 0.18 |
| 11 | 34468123 | rs11032702 | 0.705 | 0.793 | -0.064 | 0.2 | 0.078 | 0.011 | 0.011 | 0 | 0.07 | 0.36 | 0.06 | 0 | 0.04 | 0.02 |
| 11 | 34468639 | rs560807 | -1.821 | 0.796 | -0.133 | 0.201 | -0.133 | 0.126 | 0.126 | 0 | 0.78 | 0.97 | 0.77 | 0.97 | 0.6 | 0.7 |
| 11 | 34468936 | rs2284369 | -0.302 | 0.777 | -0.105 | 0.258 | 0.056 | -0.102 | -0.102 | 0 | 0.32 | 0.14 | 0.29 | 0.5 | 0.22 | 0.21 |
| 11 | 34469295 | rs2284368 | -0.302 | 0.777 | -0.105 | 0.258 | 0.056 | -0.102 | -0.102 | 0.868 | 0.32 | 0.14 | 0.29 | 0.5 | 0.22 | 0.21 |
| 11 | 34469775 | rs1408036 | 1.134 | 0.777 | -0.105 | 0.258 | 0.034 | -0.049 | -0.049 | 0.858 | 0.15 | 0.1 | 0.1 | 0.3 | 0.09 | 0.03 |
| 11 | 34470679 | rs769218 | -0.268 | 0.775 | -0.105 | 0.257 | 0.056 | -0.102 | -0.102 | 0.011 | 0.32 | 0.14 | 0.29 | 0.5 | 0.22 | 0.21 |
| 11 | 34471066 | rs17884086 | 1.338 | 0.8 | -0.074 | 0.252 | 0.005 | -0.027 | -0.027 | 0 | 0.06 | 0.13 | 0.04 | 0 | 0.04 | 0.02 |
| 11 | 34472774 | rs12269988 | 1.338 | 0.449 | -0.042 | 0.338 | 0.004 | -0.026 | -0.026 | 0 | 0.06 | 0.13 | 0.04 | 0 | 0.04 | 0.02 |
| 11 | 34473422 | rs2076556 | 0.753 | 0.483 | -0.765 | 0.377 | 0.004 | -0.036 | -0.036 | 0 | 0.17 | 0.03 | 0.19 | 0.2 | 0.12 | 0.18 |
| 11 | 34476539 | rs2300181 | 1.666 | 0.109 | -0.815 | 0.368 | 0.028 | 0.019 | 0.019 | 0 | 0.29 | 0.16 | 0.24 | 0.23 | 0.26 | 0.34 |
| 11 | 34477125 | rs7933285 | 1.653 | 0.109 | -0.815 | 0.368 | 0.029 | 0.019 | 0.019 | 0 | 0.29 | 0.16 | 0.24 | 0.23 | 0.26 | 0.34 |
| 11 | 34478073 | rs2073058 | 1.663 | 0.117 | -0.811 | 0.372 | 0.028 | 0.019 | 0.019 | 0 | 0.29 | 0.16 | 0.24 | 0.23 | 0.26 | 0.34 |
| 11 | 34481095 | rs10836244 | -0.281 | 0.092 | -0.799 | 0.416 | 0.055 | -0.093 | -0.093 | 0.111 | 0.32 | 0.15 | 0.29 | 0.5 | 0.22 | 0.21 |
| 11 | 34482908 | rs769217 | -0.281 | 0.092 | -0.799 | 0.416 | 0.055 | -0.093 | -0.093 | 0 | 0.32 | 0.14 | 0.29 | 0.5 | 0.21 | 0.21 |

| | | | | | | | | | | | | | | | | |
|---|---|---|---|---|---|---|---|---|---|---|---|---|---|---|---|---|
| 11 | 34483147 | rs7113917 | -0.281 | 0.092 | -0.799 | 0.416 | 0.055 | -0.093 | -0.093 | 0 | 0.32 | 0.14 | 0.29 | 0.5 | 0.22 | 0.21 |
| 11 | 34484542 | rs2284367 | -0.281 | 0.092 | -0.808 | 0.416 | 0.055 | -0.093 | -0.093 | 0.928 | 0.32 | 0.14 | 0.29 | 0.5 | 0.22 | 0.21 |
| 11 | 34484973 | rs2284365 | -0.551 | 0.089 | -0.786 | 0.281 | 0.011 | -0.013 | -0.013 | 0 | 0.33 | 0.32 | 0.3 | 0.51 | 0.22 | 0.21 |
| 11 | 34486604 | rs4755374 | 0.713 | -0.088 | -0.808 | 0.299 | 0.005 | -0.036 | -0.036 | 0 | 0.17 | 0.04 | 0.18 | 0.2 | 0.12 | 0.18 |
| 11 | 34487729 | rs511895 | -1.715 | -0.032 | -0.829 | 0.243 | -0.107 | 0.11 | 0.11 | 0 | 0.77 | 0.94 | 0.77 | 0.97 | 0.6 | 0.7 |
| 11 | 34488830 | rs6484722 | -0.276 | -0.058 | -0.825 | 0.254 | 0.055 | -0.093 | -0.093 | 1.373 | 0.32 | 0.14 | 0.29 | 0.5 | 0.22 | 0.21 |
| 11 | 34489252 | rs10488736 | 0.906 | -0.056 | -0.797 | 0.251 | -0.013 | -0.019 | -0.019 | 0 | 0.37 | 0.53 | 0.29 | 0.23 | 0.31 | 0.36 |
| 11 | 34490537 | rs2420388 | -0.291 | -0.043 | -0.96 | 0.277 | 0.052 | -0.079 | -0.079 | 0.885 | 0.32 | 0.17 | 0.29 | 0.5 | 0.22 | 0.21 |
| 11 | 34491680 | rs7947841 | 1.162 | -0.039 | -0.969 | 0.244 | 0.026 | -0.037 | -0.037 | 0.429 | 0.15 | 0.1 | 0.1 | 0.3 | 0.09 | 0.03 |
| 11 | 34491941 | rs499406 | -1.773 | 0.005 | -0.992 | 0.213 | -0.133 | 0.127 | 0.127 | 0 | 0.78 | 0.97 | 0.77 | 0.97 | 0.6 | 0.7 |
| 11 | 34492309 | rs16925614 | 0.732 | -0.039 | -1.023 | 0.249 | 0.005 | -0.04 | -0.04 | 0 | 0.17 | 0.04 | 0.18 | 0.2 | 0.12 | 0.18 |
| 11 | 34492443 | rs17881488 | 1.109 | -0.044 | -1.014 | 0.258 | 0.025 | -0.036 | -0.036 | 0 | 0.15 | 0.12 | 0.11 | 0.3 | 0.09 | 0.03 |
| 11 | 34493638 | rs7104301 | 1.692 | -0.042 | -1.014 | 0.213 | 0.029 | 0.022 | 0.022 | 0 | 0.29 | 0.16 | 0.23 | 0.23 | 0.26 | 0.34 |
| 11 | 34493801 | rs475043 | -1.651 | 0.344 | -0.898 | 0.382 | -0.136 | 0.129 | 0.129 | 0 | 0.78 | 0.97 | 0.77 | 0.97 | 0.6 | 0.7 |
| 11 | 34495358 | rs4756151 | 0.863 | 0.161 | -0.796 | 0.462 | -0.003 | 0.011 | 0.011 | 0 | 0.15 | 0.19 | 0.19 | 0.21 | 0.12 | 0.15 |
| 11 | 34500079 | rs1535721 | 0.35 | 0.083 | -0.824 | 0.473 | 0.006 | -0.014 | -0.014 | 0 | 0.69 | 0.3 | 0.65 | 0.45 | 0.78 | 0.82 |
| 11 | 34500687 | rs1323690 | 0.556 | 0.059 | -0.891 | 0.456 | 0.06 | -0.007 | -0.007 | 0.008 | 0.45 | 0.19 | 0.41 | 0.42 | 0.37 | 0.5 |
| 11 | 34502042 | rs7949972 | 0.46 | 0.069 | -0.876 | 0.462 | 0.023 | 0.005 | 0.005 | 0 | 0.47 | 0.38 | 0.42 | 0.43 | 0.37 | 0.5 |
| 11 | 34502345 | rs2231828 | 1.27 | 0.29 | -0.913 | 0.346 | -0.145 | 0.138 | 0.138 | 0 | 0.23 | 0.03 | 0.23 | 0.03 | 0.41 | 0.32 |
| 11 | 34504659 | rs3781708 | -1.547 | 0.31 | -0.831 | 0.418 | 0.008 | 0.035 | 0.035 | 0 | 0.92 | 0.9 | 0.81 | 0.77 | 0.92 | 0.87 |
| 11 | 34507407 | rs3758732 | 0.848 | 0.29 | -1.397 | 0.51 | 0.002 | -0.02 | -0.02 | 0 | 0.14 | 0.1 | 0.19 | 0.24 | 0.12 | 0.15 |
| 11 | 34508792 | rs11032722 | 1.169 | 0.329 | -1.457 | 0.225 | -0.14 | 0.134 | 0.134 | 0 | 0.23 | 0.03 | 0.23 | 0.03 | 0.4 | 0.31 |
| 11 | 34513571 | rs11032727 | -0.962 | 0.332 | -1.463 | 0.369 | 0.006 | -0.002 | -0.002 | 0 | 0.54 | 0.47 | 0.54 | 0.51 | 0.62 | 0.48 |
| 11 | 34513932 | rs1000768 | -0.186 | 0.5 | -1.474 | 0.224 | -0.005 | 0.012 | 0.012 | 0.007 | 0.39 | 0.35 | 0.43 | 0.24 | 0.53 | 0.46 |
| 11 | 34514941 | rs737254 | 0.409 | 1.213 | -1.061 | -0.067 | -0.007 | -0.015 | -0.015 | 0.535 | 0.25 | 0.22 | 0.24 | 0.04 | 0.41 | 0.31 |
| 11 | 34521649 | rs16930094 | -0.101 | 0.288 | -1.257 | 0.693 | 0.007 | -0.01 | -0.01 | 0 | 0.15 | 0.13 | 0.11 | 0.27 | 0.1 | 0.03 |
| 11 | 34528766 | rs766826 | -0.127 | 0.482 | -1.382 | 0.582 | 0.045 | -0.032 | -0.032 | 0.001 | 0.41 | 0.25 | 0.4 | 0.47 | 0.36 | 0.49 |
| 11 | 34532993 | rs11606405 | -0.315 | 1.683 | -1.087 | 0.271 | -0.016 | 0.011 | 0.011 | 0 | 0.08 | 0.06 | 0.1 | 0 | 0.19 | 0.03 |
| 11 | 34534035 | rs836145 | -0.6 | 1.537 | -1.263 | 0.257 | 0 | -0.002 | -0.002 | 0 | 0.34 | 0.7 | 0.43 | 0.65 | 0.33 | 0.29 |
| 11 | 34535131 | rs10836251 | -0.095 | 1.308 | -0.687 | -0.103 | 0.003 | -0.069 | -0.069 | 0 | 0.21 | 0.27 | 0.19 | 0 | 0.24 | 0.04 |
| 11 | 34537316 | rs836141 | -1.52 | 1.334 | -1.507 | -0.623 | 0.008 | 0.078 | 0.078 | 0.291 | 0.48 | 0.8 | 0.39 | 0.68 | 0.43 | 0.39 |
| 11 | 34538237 | rs11032736 | -1.12 | 1.635 | -1.393 | -0.446 | -0.102 | 0.104 | 0.104 | 0 | 0.27 | 0.11 | 0.3 | 0.06 | 0.43 | 0.19 |
| 11 | 34538539 | rs17271097 | 0.157 | 1.62 | -1.222 | -0.458 | -0.006 | 0.026 | 0.026 | 0 | 0.14 | 0.08 | 0.07 | 0.04 | 0.15 | 0.14 |
| 11 | 34540469 | rs3847600 | -1.04 | 1.453 | -1.084 | -0.422 | -0.068 | 0.066 | 0.066 | 0 | 0.28 | 0.17 | 0.3 | 0.06 | 0.43 | 0.19 |
| 11 | 34543942 | rs11032742 | -1.151 | 1.425 | -0.391 | 0.409 | -0.008 | 0.003 | 0.003 | 0 | 0.31 | 0.3 | 0.32 | 0.25 | 0.44 | 0.3 |
| 11 | 34545317 | rs3901901 | -1.288 | 1.411 | -0.357 | 0.402 | -0.017 | 0.007 | 0.007 | 0 | 0.32 | 0.28 | 0.31 | 0.25 | 0.43 | 0.29 |
| 11 | 34548906 | rs836135 | -0.278 | 1.504 | 0.154 | 0.867 | 0.025 | 0.226 | 0.226 | 0.029 | 0.1 | 0.34 | 0.14 | 0.41 | 0.12 | 0.06 |
| 11 | 34555191 | rs836132 | -0.18 | 1.501 | 0.169 | 0.86 | 0.002 | 0.08 | 0.08 | 0 | 0.1 | 0.14 | 0.13 | 0.42 | 0.12 | 0.08 |
| 11 | 34555676 | rs12788347 | -0.047 | 1.416 | 0.294 | 0.651 | 0.001 | -0.01 | -0.01 | 4.074 | 0.15 | 0.15 | 0.06 | 0.03 | 0.13 | 0.14 |
| 11 | 34560674 | rs3886899 | 0.756 | 0.883 | -0.236 | 0.663 | -0.018 | -0.073 | -0.073 | 0 | 0.11 | 0 | 0.22 | 0.31 | 0.12 | 0.24 |
| 11 | 34562176 | rs836125 | -0.251 | 1.143 | 0.273 | 0.565 | 0.013 | 0.169 | 0.169 | 0.01 | 0.1 | 0.26 | 0.14 | 0.42 | 0.12 | 0.08 |
| 11 | 34562247 | rs836124 | -1.47 | 1.24 | -0.097 | 0.525 | 0.013 | 0.004 | 0.004 | 0.609 | 0.38 | 0.56 | 0.43 | 0.48 | 0.52 | 0.28 |
| 11 | 34566510 | rs10836255 | 0.264 | 1.716 | -0.677 | 1.243 | -0.059 | -0.059 | -0.059 | 0 | 0.11 | 0.02 | 0.14 | 0.38 | 0.24 | 0.11 |
| 11 | 34568598 | rs836116 | -1.616 | 0.24 | -0.637 | 0.612 | 0.013 | -0.001 | -0.001 | 2.077 | 0.59 | 0.77 | 0.5 | 0.64 | 0.62 | 0.37 |
| 11 | 34570208 | rs836109 | -0.486 | -0.494 | 0.324 | 0.858 | -0.046 | 0.135 | 0.135 | 0.288 | 0.21 | 0.42 | 0.16 | 0.61 | 0.14 | 0.1 |
| 11 | 34571836 | rs3847603 | -0.31 | -0.522 | 0.59 | 0.838 | -0.011 | 0.044 | 0.044 | 0.434 | 0.19 | 0.18 | 0.13 | 0.59 | 0.15 | 0.1 |
| 11 | 34573817 | rs836104 | -0.537 | -0.512 | 0.351 | 0.806 | -0.046 | 0.115 | 0.115 | 0.184 | 0.21 | 0.42 | 0.17 | 0.64 | 0.15 | 0.1 |
| 11 | 34578340 | rs836098 | -1.111 | -0.299 | 0.351 | 0.555 | -0.036 | 0.042 | 0.042 | 0 | 0.35 | 0.43 | 0.21 | 0.58 | 0.27 | 0.25 |
| 11 | 34583335 | rs10836257 | -0.712 | 0.237 | 0.235 | 0.06 | 0.025 | 0.025 | 0.025 | 1.57 | 0.27 | 0.34 | 0.45 | 0.51 | 0.42 | 0.26 |
| 11 | 34584382 | rs16925781 | -0.865 | -0.098 | 0.897 | -0.819 | 0.004 | -0.015 | -0.015 | 0 | 0.17 | 0.13 | 0.06 | 0.01 | 0.1 | 0.11 |
| 11 | 34584938 | rs7942802 | 1.314 | -0.267 | 0.741 | -0.953 | -0.033 | -0.097 | -0.097 | 0.049 | 0.75 | 0.34 | 0.87 | 0.99 | 0.87 | 0.87 |
| 11 | 34585751 | rs1453391 | 0.972 | 0.482 | 1.11 | -0.603 | 0.045 | -0.126 | -0.126 | 1.736 | 0.77 | 0.6 | 0.81 | 0.99 | 0.69 | 0.88 |
| 11 | 34586444 | rs7947487 | 1.15 | 0.445 | 0.954 | -0.619 | 0.015 | -0.083 | -0.083 | 0 | 0.73 | 0.66 | 0.79 | 0.96 | 0.66 | 0.88 |
| 11 | 34588886 | rs1037457 | 0.747 | 0.805 | 1.137 | -0.207 | 0.021 | -0.042 | -0.042 | 0 | 0.8 | 0.75 | 0.83 | 0.95 | 0.71 | 0.9 |
| 11 | 34590268 | rs11602149 | -1.532 | 0.685 | 1.318 | -0.342 | 0.007 | -0.043 | -0.043 | 0 | 0.25 | 0.27 | 0.18 | 0.05 | 0.31 | 0.1 |
| 11 | 34595608 | rs1530966 | -1.518 | 0.705 | 1.753 | -0.353 | 0.039 | -0.111 | -0.111 | 2.192 | 0.26 | 0.45 | 0.19 | 0.05 | 0.31 | 0.1 |
| 11 | 34597605 | rs12797608 | -1.326 | 0.695 | 1.759 | -0.357 | -0.052 | 0.084 | 0.084 | 0 | 0.24 | 0.08 | 0.16 | 0.05 | 0.3 | 0.1 |
| 11 | 34598118 | rs713510 | -1.539 | 0.701 | 1.784 | -0.354 | 0.001 | -0.027 | -0.027 | 0 | 0.25 | 0.27 | 0.18 | 0.05 | 0.31 | 0.1 |
| 11 | 34605435 | rs5028798 | 0.087 | 0.449 | 1.436 | 0.095 | -0.024 | 0.12 | 0.12 | 0 | 0.77 | 0.55 | 0.65 | 0.52 | 0.8 | 0.67 |
| 11 | 34609541 | rs10768081 | 1.907 | 0.439 | 1.228 | 0.135 | -0.117 | -0.097 | -0.097 | 0 | 0.5 | 0.03 | 0.33 | 0.42 | 0.48 | 0.58 |
| 11 | 34614555 | rs2084969 | 0.049 | 0.419 | 1.165 | 0.144 | -0.024 | 0.133 | 0.133 | 0 | 0.77 | 0.53 | 0.66 | 0.51 | 0.79 | 0.67 |
| 11 | 34618756 | rs2045040 | 1.513 | 0.569 | 0.345 | 0.235 | -0.112 | -0.097 | -0.097 | 0 | 0.5 | 0.03 | 0.33 | 0.42 | 0.47 | 0.58 |
| 11 | 34621004 | rs1024276 | 1.463 | 0.608 | 0.374 | 0.245 | -0.032 | -0.031 | -0.031 | 0 | 0.51 | 0.12 | 0.33 | 0.42 | 0.47 | 0.58 |
| 11 | 34622955 | rs286953 | -0.93 | -0.359 | 0.828 | 0.845 | -0.016 | 0.068 | 0.068 | 0 | 0.31 | 0.4 | 0.49 | 0.52 | 0.29 | 0.36 |
| 11 | 34637669 | rs10836265 | 1.333 | -0.382 | -0.18 | 0.763 | 0.013 | -0.092 | -0.092 | 0 | 0.51 | 0.04 | 0.41 | 0.51 | 0.62 | 0.53 |
| 11 | 34640304 | rs11032786 | -0.189 | -0.144 | 0.943 | 1.044 | 0.003 | 0.141 | 0.141 | 0 | 0.06 | 0.08 | 0.09 | 0.49 | 0.08 | 0.27 |
| 11 | 34642668 | rs286925 | 1.455 | -0.364 | 1.344 | 0.995 | -0.013 | 0.029 | 0.029 | 0 | 0.75 | 0.65 | 0.72 | 0.38 | 0.83 | 0.51 |
| 11 | 34642823 | rs717582 | 1.218 | -0.412 | 1.463 | 1.011 | 0.004 | -0.002 | -0.002 | 1.787 | 0.58 | 0.38 | 0.55 | 0.54 | 0.64 | 0.58 |
| 11 | 34643721 | rs286921 | 0.792 | -0.433 | 1.468 | 1.005 | -0.002 | 0.03 | 0.03 | 0 | 0.84 | 0.81 | 0.87 | 0.41 | 0.86 | 0.57 |

| | | | | | | | | | | | | | | | | |
|---|---|---|---|---|---|---|---|---|---|---|---|---|---|---|---|---|
| 11 | 34644483 | rs443816 | -0.984 | -0.446 | 1.577 | 1.002 | -0.006 | 0.011 | 0.011 | 0 | 0.33 | 0.46 | 0.4 | 0.37 | 0.31 | 0.27 |
| 11 | 34645203 | rs16925915 | -0.256 | -0.519 | 1.473 | 1.027 | 0.001 | 0.123 | 0.123 | 0 | 0.07 | 0.08 | 0.09 | 0.5 | 0.09 | 0.28 |
| 11 | 34648615 | rs10488740 | 1.809 | -0.506 | 1.456 | 0.536 | 0.008 | 0.036 | 0.036 | 0 | 0.51 | 0.35 | 0.47 | 0.04 | 0.54 | 0.29 |
| 11 | 34650764 | rs10768085 | 1.315 | -0.666 | 1.572 | 0.872 | 0 | -0.001 | -0.001 | 0 | 0.6 | 0.58 | 0.63 | 0.79 | 0.63 | 0.69 |
| 11 | 34653771 | rs17358295 | -0.303 | -0.555 | 0.663 | 0.863 | -0.005 | 0.019 | 0.019 | 0 | 0.12 | 0.07 | 0.2 | 0.03 | 0.11 | 0.07 |
| 11 | 34655363 | rs286913 | -0.111 | -0.456 | 0.664 | 1 | 0.003 | 0.001 | 0.001 | 0 | 0.91 | 0.92 | 0.96 | 0.91 | 0.95 | 0.84 |
| 11 | 34655936 | rs968251 | -0.182 | -0.427 | 0.575 | 1.097 | -0.001 | 0.132 | 0.132 | 0 | 0.06 | 0.06 | 0.07 | 0.63 | 0.06 | 0.31 |
| 11 | 34656664 | rs7945550 | 1.045 | -0.42 | 0.495 | 0.996 | -0.011 | -0.008 | -0.008 | 0 | 0.59 | 0.52 | 0.57 | 0.67 | 0.64 | 0.61 |
| 11 | 34657144 | rs286912 | -0.894 | 0.082 | 0.05 | 1.117 | 0 | 0.001 | 0.001 | 0 | 0.31 | 0.3 | 0.38 | 0.24 | 0.31 | 0.23 |
| 11 | 34660618 | rs12366193 | -1.335 | -0.045 | 0.31 | 1.811 | -0.028 | 0.088 | 0.088 | 0 | 0.23 | 0.4 | 0.17 | 0.49 | 0.15 | 0.49 |
| 11 | 34662120 | rs3794155 | -1.275 | 0.047 | 0.457 | 1.713 | -0.01 | -0.006 | -0.006 | 1.284 | 0.14 | 0.27 | 0.1 | 0.09 | 0.08 | 0.32 |
| 11 | 34662961 | rs286905 | 0.943 | 0.773 | 0.473 | 2.334 | 0.041 | 0.132 | 0.132 | 0.306 | 0.87 | 0.72 | 0.79 | 0.62 | 0.85 | 0.85 |
| 11 | 34665925 | rs3751074 | -0.978 | 0.604 | 0.11 | 2.183 | -0.002 | 0.044 | 0.044 | 0.097 | 0.12 | 0.18 | 0.2 | 0.29 | 0.1 | 0.14 |
| 11 | 34666063 | rs3751075 | -1.064 | 0.678 | -0.124 | 2.201 | -0.006 | -0.011 | -0.011 | 0 | 0.13 | 0.08 | 0.2 | 0.27 | 0.14 | 0.14 |
| 11 | 34666850 | rs3751076 | -0.968 | 0.976 | -0.791 | 1.898 | -0.058 | -0.049 | -0.049 | 0.284 | 0.21 | 0.04 | 0.19 | 0.22 | 0.23 | 0.41 |
| 11 | 34672161 | rs286899 | -0.664 | 0.809 | -0.21 | 1.512 | 0.012 | -0.003 | -0.003 | 0 | 0.08 | 0.17 | 0.06 | 0.09 | 0.11 | 0.1 |
| 11 | 34672889 | rs11032801 | -1.055 | 0.933 | -0.395 | 2.477 | 0.002 | 0.123 | 0.123 | 0 | 0.06 | 0.06 | 0.19 | 0.42 | 0.07 | 0.18 |
| 11 | 34681761 | rs7940320 | -0.191 | 0.419 | -1.204 | 1.699 | 0.002 | 0.047 | 0.047 | 1.436 | 0.24 | 0.04 | 0.33 | 0.13 | 0.19 | 0.13 |
| 11 | 34682767 | rs286887 | -0.202 | 0.456 | -1.266 | 1.967 | -0.007 | -0.011 | -0.011 | 0 | 0.09 | 0.06 | 0.08 | 0.33 | 0.12 | 0.29 |
| 11 | 34686005 | rs704730 | -0.024 | 0.448 | -1.272 | 1.947 | -0.004 | 0.07 | 0.07 | 2.142 | 0.23 | 0.03 | 0.32 | 0.09 | 0.19 | 0.12 |
| 11 | 34689296 | rs286883 | -0.26 | 0.515 | -1.187 | 2.024 | -0.008 | -0.016 | -0.016 | 0 | 0.09 | 0.06 | 0.08 | 0.33 | 0.12 | 0.28 |
| 11 | 34699269 | rs7112608 | 1.269 | 0.167 | -0.255 | 1.318 | 0.004 | 0.02 | 0.02 | 0 | 0.53 | 0.33 | 0.32 | 0.13 | 0.53 | 0.38 |
| 11 | 34702158 | rs286873 | 0.105 | 0.169 | -0.139 | 1.682 | -0.003 | 0.008 | 0.008 | 0 | 0.09 | 0.09 | 0.07 | 0.31 | 0.11 | 0.29 |
| 11 | 34702640 | rs704738 | 0.028 | 0.128 | 0.342 | 1.685 | 0.015 | 0.042 | 0.042 | 1.797 | 0.25 | 0.13 | 0.35 | 0.13 | 0.2 | 0.13 |
| 11 | 34725028 | rs1901830 | 0.467 | -0.023 | 1.682 | 1.634 | -0.001 | 0.002 | 0.002 | 0.155 | 0.64 | 0.63 | 0.57 | 0.53 | 0.7 | 0.58 |
| 11 | 34745964 | rs2027495 | -0.432 | -0.041 | 1.839 | 1.637 | -0.034 | -0.022 | -0.022 | 0.701 | 0.08 | 0.04 | 0.23 | 0.4 | 0.16 | 0.2 |
| 11 | 34746360 | rs1361646 | 0.361 | -0.067 | 1.873 | 1.648 | 0.01 | -0.043 | -0.043 | 0 | 0.66 | 0.77 | 0.57 | 0.52 | 0.7 | 0.58 |
| 11 | 34748502 | rs11032829 | 0.172 | -0.03 | 1.828 | 1.681 | -0.002 | 0.02 | 0.02 | 0 | 0.09 | 0.1 | 0.08 | 0.35 | 0.11 | 0.29 |
| 11 | 34752562 | rs7121272 | 1.116 | -0.245 | 2.008 | 1.369 | 0.001 | 0.006 | 0.006 | 0 | 0.55 | 0.58 | 0.33 | 0.17 | 0.53 | 0.4 |
| 11 | 34753375 | rs947501 | 0.318 | -0.242 | 1.927 | 1.395 | 0.01 | -0.018 | -0.018 | 0 | 0.65 | 0.74 | 0.57 | 0.56 | 0.69 | 0.6 |
| 11 | 34753747 | rs11032833 | -0.773 | -0.192 | 1.679 | 1.387 | 0.011 | 0.102 | 0.102 | 0 | 0.1 | 0.14 | 0.25 | 0.39 | 0.16 | 0.21 |
| 11 | 34768948 | rs10742323 | -0.048 | -0.345 | 1.882 | 1.052 | -0.023 | -0.076 | -0.076 | 0 | 0.72 | 0.63 | 0.68 | 0.95 | 0.8 | 0.89 |
| 11 | 34770494 | rs12277584 | -1.224 | -0.201 | 1.998 | 1.288 | 0.029 | 0.205 | 0.205 | 0 | 0.17 | 0.47 | 0.18 | 0.43 | 0.2 | 0.33 |
| 11 | 34776331 | rs7924717 | -0.438 | -0.203 | 2.036 | 1.294 | 0.019 | 0.232 | 0.232 | 0 | 0.11 | 0.34 | 0.13 | 0.43 | 0.11 | 0.31 |
| 11 | 34776532 | rs7939918 | -0.166 | -0.122 | 1.334 | 0.216 | 0.054 | 0.036 | 0.036 | 1.853 | 0.33 | 0.49 | 0.32 | 0.43 | 0.46 | 0.39 |
| 11 | 34778351 | rs7110293 | 1.017 | -0.504 | 1.421 | -0.011 | -0.026 | -0.049 | -0.049 | 0 | 0.86 | 0.43 | 0.71 | 0.95 | 0.87 | 0.91 |
| 11 | 34796804 | rs2136509 | -0.425 | -0.306 | 1.429 | -0.032 | -0.024 | -0.021 | -0.021 | 0 | 0.38 | 0.71 | 0.35 | 0.44 | 0.48 | 0.4 |
| 11 | 34805210 | rs1396885 | 0.39 | -0.296 | 1.244 | -0.006 | 0.007 | 0.007 | 0.007 | 0.659 | 0.31 | 0.36 | 0.28 | 0.44 | 0.39 | 0.38 |
| 11 | 34806466 | rs11032873 | 1.041 | -0.422 | 0.825 | -0.06 | -0.052 | -0.132 | -0.132 | 0.059 | 0.23 | 0.01 | 0.28 | 0.39 | 0.21 | 0.22 |
| 11 | 34811180 | rs10501135 | 0.439 | -0.308 | 1.097 | 0.07 | -0.039 | -0.033 | -0.033 | 0 | 0.3 | 0.22 | 0.27 | 0.4 | 0.39 | 0.36 |
| 11 | 34816058 | rs7130756 | 0.647 | -0.364 | 1.319 | 0.088 | -0.007 | 0.072 | 0.072 | 0.622 | 0.93 | 0.63 | 0.78 | 0.89 | 0.96 | 0.92 |
| 11 | 34816531 | rs1858167 | 0.437 | -0.453 | 1.071 | 0.011 | -0.109 | -0.115 | -0.115 | 1.616 | 0.55 | 0.05 | 0.44 | 0.49 | 0.48 | 0.52 |
| 11 | 34821948 | rs525202 | -0.207 | -0.422 | 1.898 | -0.456 | -0.015 | -0.022 | -0.022 | 0 | 0.6 | 0.46 | 0.61 | 0.77 | 0.68 | 0.6 |
| 11 | 34829410 | rs474636 | 0.57 | -0.354 | 2.041 | -0.383 | -0.001 | -0.018 | -0.018 | 0 | 0.9 | 0.7 | 0.73 | 0.94 | 0.87 | 0.91 |
| 11 | 34851961 | rs473885 | -0.371 | -0.408 | 1.96 | -0.458 | 0.021 | 0.031 | 0.031 | 0 | 0.62 | 0.65 | 0.62 | 0.82 | 0.69 | 0.62 |
| 11 | 34853382 | rs535719 | 0.226 | -0.398 | 1.933 | -0.457 | 0.064 | 0.12 | 0.12 | 0 | 0.54 | 0.25 | 0.54 | 0.76 | 0.6 | 0.58 |
| 11 | 34854662 | rs502857 | -0.098 | -0.391 | 1.888 | -0.469 | 0.015 | 0.019 | 0.019 | 0 | 0.6 | 0.37 | 0.61 | 0.78 | 0.69 | 0.61 |
| 11 | 34857707 | rs731726 | 1.449 | -0.861 | 1.166 | -0.942 | 0.021 | 0.049 | 0.049 | 0 | 0.13 | 0.01 | 0.07 | 0.05 | 0.08 | 0.15 |
| 11 | 34857834 | rs731727 | -0.407 | -0.804 | 1.385 | -0.932 | 0.023 | 0.042 | 0.042 | 0 | 0.34 | 0.24 | 0.18 | 0.05 | 0.27 | 0.31 |
| 11 | 34858084 | rs967751 | 0.31 | -1.042 | 1.672 | -0.524 | 0 | 0.081 | 0.081 | 0 | 0.89 | 0.69 | 0.73 | 0.82 | 0.88 | 0.89 |
| 11 | 34859944 | rs12292993 | -0.104 | -1.049 | 1.561 | -0.806 | -0.006 | -0.019 | -0.019 | 0 | 0.08 | 0.32 | 0.05 | 0.02 | 0.03 | 0.08 |
| 11 | 34862880 | rs1553759 | -0.26 | -1.138 | 1.614 | -0.775 | -0.031 | -0.026 | -0.026 | 2.991 | 0.76 | 0.64 | 0.66 | 0.82 | 0.81 | 0.72 |
| 11 | 34867695 | rs2163374 | 0.326 | -0.976 | 1.259 | -1.077 | 0.015 | 0.051 | 0.051 | 1.093 | 0.49 | 0.39 | 0.45 | 0.82 | 0.43 | 0.65 |
| 11 | 34867968 | rs1549791 | -0.306 | -0.971 | 1.226 | -1.076 | -0.002 | 0.075 | 0.075 | 0.122 | 0.56 | 0.71 | 0.5 | 0.84 | 0.46 | 0.72 |
| 11 | 34871556 | rs2915180 | -0.818 | -0.975 | 1.23 | -1.072 | -0.019 | -0.017 | -0.017 | 0.57 | 0.79 | 0.73 | 0.68 | 0.84 | 0.82 | 0.84 |
| 11 | 34877761 | rs2941061 | -0.843 | -1.006 | 1.189 | -1.066 | -0.006 | -0.005 | -0.005 | 0 | 0.79 | 0.8 | 0.68 | 0.84 | 0.82 | 0.84 |
| 11 | 34879009 | rs7120509 | 0.255 | -0.857 | 1.069 | -0.912 | -0.005 | -0.021 | -0.021 | 0.14 | 0.08 | 0.3 | 0.05 | 0.02 | 0.03 | 0.08 |
| 11 | 34885510 | rs2915236 | 0.494 | -0.795 | 0.58 | -0.868 | 0.035 | 0.135 | 0.135 | 0.559 | 0.46 | 0.27 | 0.42 | 0.79 | 0.37 | 0.6 |
| 11 | 34886417 | rs2941054 | 1.016 | -0.724 | 0.548 | -0.929 | 0.024 | 0.089 | 0.089 | 1.721 | 0.42 | 0.26 | 0.39 | 0.81 | 0.34 | 0.52 |
| 11 | 34889989 | rs3962032 | 0.402 | -0.771 | 0.573 | -0.925 | -0.075 | 0.095 | 0.095 | 1.629 | 0.32 | 0.14 | 0.25 | 0.09 | 0.47 | 0.21 |
| 11 | 34892293 | rs2915233 | -0.217 | -0.923 | 0.821 | -0.693 | -0.052 | -0.039 | -0.039 | 0.506 | 0.75 | 0.45 | 0.65 | 0.82 | 0.8 | 0.73 |
| 11 | 34895100 | rs2033359 | 0.032 | -0.613 | 0.597 | -0.558 | 0 | 0.004 | 0.004 | 0 | 0.1 | 0.17 | 0.08 | 0.09 | 0.12 | 0.11 |
| 11 | 34902521 | rs2956086 | -0.01 | -0.82 | 0.793 | -0.689 | -0.03 | -0.029 | -0.029 | 0 | 0.73 | 0.64 | 0.64 | 0.81 | 0.77 | 0.65 |
| 11 | 34903671 | rs6484746 | 0.173 | -0.852 | 1.097 | -0.7 | 0.041 | 0.065 | 0.065 | 0 | 0.07 | 0.28 | 0.07 | 0.07 | 0.09 | 0.03 |
| 11 | 34910077 | rs1571134 | 0.764 | -0.859 | 1.013 | -0.707 | 0.008 | 0.006 | 0.006 | 0.018 | 0.66 | 0.35 | 0.57 | 0.74 | 0.68 | 0.62 |
| 11 | 34910397 | rs1977420 | 0.931 | -0.629 | 0.826 | -0.929 | 0.022 | 0.057 | 0.057 | 1.581 | 0.44 | 0.31 | 0.4 | 0.74 | 0.36 | 0.52 |
| 11 | 34914137 | rs2915223 | 0.663 | -0.631 | 0.848 | -0.901 | 0.008 | 0.006 | 0.006 | 0.018 | 0.66 | 0.36 | 0.57 | 0.74 | 0.68 | 0.62 |
| 11 | 34918375 | rs2986429 | 0.163 | -0.636 | 0.936 | -0.899 | -0.036 | -0.017 | -0.017 | 0 | 0.71 | 0.51 | 0.6 | 0.75 | 0.71 | 0.7 |
| 11 | 34922603 | rs2915216 | 1.092 | -0.645 | 0.881 | -0.914 | 0.021 | 0.072 | 0.072 | 1.181 | 0.42 | 0.24 | 0.4 | 0.74 | 0.35 | 0.52 |

| | | | | | | | | | | | | | | | | |
|---|---|---|---|---|---|---|---|---|---|---|---|---|---|---|---|---|
| 11 | 34934652 | rs2956116 | -0.791 | -0.663 | 0.997 | -0.911 | 0.006 | 0.005 | 0.005 | 0 | 0.78 | 0.84 | 0.68 | 0.82 | 0.8 | 0.73 |
| 11 | 34935661 | rs984690 | -0.775 | -0.663 | 1.019 | -0.911 | -0.005 | -0.004 | -0.004 | 0 | 0.78 | 0.79 | 0.67 | 0.82 | 0.8 | 0.73 |
| 11 | 34937138 | rs3794170 | -0.479 | -0.672 | 1.084 | -0.925 | 0.014 | 0.023 | 0.023 | 0 | 0.13 | 0.44 | 0.1 | 0.09 | 0.12 | 0.11 |
| 11 | 34937330 | rs3763933 | -0.479 | -0.672 | 1.084 | -0.925 | 0.014 | 0.023 | 0.023 | 0 | 0.13 | 0.44 | 0.1 | 0.09 | 0.12 | 0.11 |
| 11 | 34937813 | rs2956114 | 1.146 | -0.677 | 1.058 | -0.953 | 0.02 | 0.071 | 0.071 | 1.116 | 0.42 | 0.22 | 0.4 | 0.74 | 0.35 | 0.52 |
| 11 | 34938269 | rs1049306 | -0.593 | -0.76 | 0.993 | -0.783 | 0.013 | 0.022 | 0.022 | 0 | 0.13 | 0.44 | 0.1 | 0.09 | 0.12 | 0.11 |
| 11 | 34938310 | rs1049307 | -0.568 | -0.76 | 0.993 | -0.783 | 0.014 | 0.023 | 0.023 | 0 | 0.13 | 0.44 | 0.11 | 0.09 | 0.12 | 0.11 |
| 11 | 34941494 | rs724714 | -0.696 | -0.765 | 0.965 | -0.713 | -0.005 | -0.003 | -0.003 | 0 | 0.78 | 0.79 | 0.67 | 0.82 | 0.8 | 0.73 |
| 11 | 34944453 | rs2956103 | 0.822 | -0.762 | 0.879 | -0.712 | 0.018 | 0.011 | 0.011 | 0.042 | 0.65 | 0.31 | 0.57 | 0.73 | 0.68 | 0.6 |
| 11 | 34945449 | rs16926479 | 0.866 | -0.569 | 0.818 | -0.585 | -0.007 | -0.018 | -0.018 | 0 | 0.06 | 0.15 | 0.03 | 0.02 | 0.03 | 0.08 |
| 11 | 34949502 | rs12283472 | 0.733 | -0.499 | 0.72 | -0.534 | 0.006 | 0.015 | 0.015 | 0 | 0.05 | 0.08 | 0.05 | 0.07 | 0.09 | 0.03 |
| 11 | 34968327 | rs496620 | 0.26 | -0.619 | 0.425 | -0.604 | 0.002 | 0 | 0 | 0 | 0.68 | 0.71 | 0.6 | 0.73 | 0.7 | 0.66 |
| 11 | 34968523 | rs614668 | -0.251 | -0.62 | 0.652 | -0.581 | -0.008 | -0.002 | -0.002 | 0 | 0.74 | 0.72 | 0.64 | 0.76 | 0.74 | 0.68 |
| 11 | 34969112 | rs11539202 | 1.668 | -0.419 | 0.481 | -0.752 | -0.132 | 0.14 | 0.14 | 0 | 0.21 | 0.03 | 0.15 | 0 | 0.32 | 0.09 |
| 11 | 34979548 | rs7119807 | -0.271 | -1.161 | 1.545 | 0.068 | 0.001 | -0.016 | -0.016 | 0 | 0.08 | 0.32 | 0.06 | 0.02 | 0.03 | 0.08 |
| 11 | 34987357 | rs511776 | -0.257 | -1.157 | 1.374 | -0.005 | 0.006 | 0.004 | 0.004 | 0 | 0.78 | 0.84 | 0.69 | 0.81 | 0.81 | 0.71 |
| 11 | 34991727 | rs497582 | 1.191 | -1.136 | 1.299 | 0.005 | 0.025 | 0.02 | 0.02 | 0.063 | 0.64 | 0.32 | 0.56 | 0.74 | 0.71 | 0.61 |
| 11 | 35024965 | rs1326947 | -0.02 | -0.906 | 0.329 | 0.131 | -0.007 | -0.005 | -0.005 | 0 | 0.77 | 0.77 | 0.69 | 0.81 | 0.81 | 0.71 |
| 11 | 35033098 | rs11032973 | -0.829 | -0.454 | 0.188 | 0.294 | -0.004 | 0 | 0 | 0 | 0.1 | 0.22 | 0.09 | 0.06 | 0.09 | 0.09 |
| 11 | 35036029 | rs1035732 | -1.092 | -0.652 | 0.084 | 0.052 | -0.017 | -0.062 | -0.062 | 0.042 | 0.16 | 0.29 | 0.11 | 0.03 | 0.13 | 0.26 |
| 11 | 35043180 | rs2421763 | -1.036 | -0.65 | -0.278 | 0.918 | -0.031 | -0.081 | -0.081 | 0 | 0.14 | 0.01 | 0.12 | 0.23 | 0.17 | 0.33 |
| 11 | 35043231 | rs497476 | 0.082 | -1.768 | -0.704 | 0.504 | 0.014 | -0.014 | -0.014 | 0.006 | 0.35 | 0.62 | 0.41 | 0.5 | 0.2 | 0.34 |
| 11 | 35048359 | rs2785162 | -0.289 | -1.761 | -0.686 | 0.464 | 0.051 | -0.028 | -0.028 | 1.619 | 0.38 | 0.7 | 0.41 | 0.52 | 0.21 | 0.43 |
| 11 | 35052489 | rs2785160 | 0.06 | -1.741 | -0.514 | 0.417 | -0.004 | 0.001 | 0.001 | 0 | 0.34 | 0.59 | 0.41 | 0.5 | 0.21 | 0.34 |
| 11 | 35054185 | rs2732521 | -0.305 | -1.727 | -0.577 | 0.421 | 0.038 | -0.023 | -0.023 | 0 | 0.38 | 0.67 | 0.42 | 0.52 | 0.23 | 0.44 |
| 11 | 35055259 | rs7113469 | -0.234 | -1.746 | -0.653 | 0.406 | 0 | -0.02 | -0.02 | 0 | 0.14 | 0.12 | 0.12 | 0.23 | 0.16 | 0.33 |
| 11 | 35057285 | rs2785158 | -1.441 | -1.729 | -0.644 | 0.367 | 0.043 | 0.182 | 0.182 | 0 | 0.53 | 0.9 | 0.54 | 0.75 | 0.39 | 0.76 |
| 11 | 35060090 | rs11032980 | 1.695 | -2.082 | -1.182 | -0.368 | 0.022 | 0.081 | 0.081 | 4.285 | 0.16 | 0.02 | 0.06 | 0 | 0.09 | 0.08 |
| 11 | 35069058 | rs2785153 | 0.35 | -1.995 | -0.757 | -0.112 | 0.024 | -0.021 | -0.021 | 0 | 0.36 | 0.67 | 0.42 | 0.5 | 0.21 | 0.36 |
| 11 | 35074000 | rs737503 | 1.093 | -2.575 | 0.09 | -0.083 | 0.008 | 0.008 | 0.008 | 0 | 0.12 | 0 | 0.25 | 0.1 | 0.07 | 0.12 |
| 11 | 35074665 | rs693163 | -1.933 | -1.831 | -0.133 | -0.012 | -0.075 | -0.014 | -0.014 | 0.002 | 0.7 | 0.91 | 0.71 | 0.69 | 0.56 | 0.81 |
| 11 | 35076452 | rs11032981 | 2.132 | -2.073 | -0.164 | -0.38 | 0.03 | 0.063 | 0.063 | 0.953 | 0.14 | 0.03 | 0.05 | 0 | 0.07 | 0.02 |
| 11 | 35077756 | rs2162159 | -0.256 | -1.82 | 0.187 | 0.266 | 0.021 | -0.009 | -0.009 | 0.246 | 0.28 | 0.2 | 0.34 | 0.33 | 0.18 | 0.45 |
| 11 | 35079922 | rs2553774 | -0.584 | -1.777 | 0.041 | 0.217 | -0.005 | -0.001 | -0.001 | 0 | 0.32 | 0.65 | 0.37 | 0.32 | 0.19 | 0.45 |
| 11 | 35084592 | rs2732552 | -1.086 | -1.661 | 0.01 | 0.26 | -0.005 | 0.026 | 0.026 | 0 | 0.62 | 0.31 | 0.65 | 0.75 | 0.55 | 0.72 |
| 11 | 35088399 | rs2732549 | -1.979 | -1.715 | -0.051 | 0.235 | -0.134 | -0.02 | -0.02 | 0 | 0.71 | 0.97 | 0.71 | 0.78 | 0.56 | 0.81 |
| 11 | 35088683 | rs2732547 | -1.163 | -1.611 | 0.071 | 0.287 | -0.001 | 0.004 | 0.004 | 0 | 0.62 | 0.36 | 0.65 | 0.75 | 0.55 | 0.72 |
| 11 | 35090546 | rs11032988 | -0.277 | -1.733 | -0.166 | 0.328 | 0.049 | -0.006 | -0.006 | 1.39 | 0.3 | 0.12 | 0.32 | 0.3 | 0.19 | 0.47 |
| 11 | 35095324 | rs2553828 | 0.598 | -1.613 | -0.078 | 0.404 | -0.042 | -0.055 | -0.055 | 0 | 0.32 | 0.18 | 0.33 | 0.45 | 0.35 | 0.25 |
| 11 | 35099386 | rs2553825 | 0.52 | -1.653 | 0.102 | 0.525 | 0.002 | -0.009 | -0.009 | 0 | 0.12 | 0.1 | 0.08 | 0.2 | 0.11 | 0.29 |
| 11 | 35099975 | rs1425801 | -0.561 | -1.558 | 0.267 | 0.28 | 0.003 | 0.026 | 0.026 | 0 | 0.5 | 0.41 | 0.45 | 0.64 | 0.47 | 0.64 |
| 11 | 35100877 | rs378517 | -0.665 | -1.367 | 0.345 | 0.329 | 0 | -0.003 | -0.003 | 0 | 0.56 | 0.58 | 0.64 | 0.53 | 0.44 | 0.55 |
| 11 | 35105495 | rs353605 | -1.082 | -1.412 | -0.156 | 0.168 | -0.001 | 0 | 0 | 0.484 | 0.43 | 0.56 | 0.39 | 0.37 | 0.31 | 0.7 |
| 11 | 35106925 | rs12790065 | 0.81 | -1.059 | -1.811 | 0.338 | -0.013 | 0.057 | 0.057 | 0 | 0.2 | 0.01 | 0.27 | 0.09 | 0.15 | 0.23 |
| 11 | 35109675 | rs1365057 | 0.656 | -0.487 | -1.241 | 0.813 | 0.001 | -0.01 | -0.01 | 0.537 | 0.26 | 0.24 | 0.31 | 0.4 | 0.24 | 0.16 |
| 11 | 35110118 | rs7131078 | -0.217 | 0.165 | -1.371 | 1.07 | -0.001 | 0.015 | 0.015 | 0 | 0.12 | 0.12 | 0.06 | 0.25 | 0.13 | 0.32 |
| 11 | 35120741 | rs353590 | -0.877 | 0.357 | -2.117 | 0.616 | -0.177 | -0.027 | -0.027 | 0 | 0.71 | 0.98 | 0.7 | 0.76 | 0.55 | 0.86 |
| 11 | 35121226 | rs353589 | -0.114 | -0.711 | -1.897 | 0.083 | -0.004 | -0.015 | -0.015 | 0 | 0.33 | 0.59 | 0.33 | 0.12 | 0.18 | 0.37 |
| 11 | 35122661 | rs353587 | -0.706 | -1.173 | -1.836 | -0.046 | 0.04 | 0.105 | 0.105 | 0.151 | 0.48 | 0.83 | 0.42 | 0.67 | 0.39 | 0.63 |
| 11 | 35123051 | rs1559759 | 0.276 | -0.703 | -2.018 | 0.65 | -0.011 | -0.026 | -0.026 | 0 | 0.14 | 0.1 | 0.09 | 0.32 | 0.17 | 0.34 |
| 11 | 35123195 | rs1559758 | 0.095 | -0.698 | -1.854 | 0.242 | -0.008 | 0.003 | 0.003 | 0 | 0.36 | 0.26 | 0.36 | 0.63 | 0.37 | 0.49 |
| 11 | 35124040 | rs4756190 | 0.876 | -0.216 | -1.876 | 0.153 | -0.083 | 0.023 | 0.023 | 0 | 0.29 | 0.1 | 0.29 | 0.24 | 0.41 | 0.2 |
| 11 | 35124560 | rs353582 | -0.152 | -1.154 | -1.107 | -0.398 | 0.041 | 0.048 | 0.048 | 0 | 0.45 | 0.7 | 0.47 | 0.71 | 0.29 | 0.6 |
| 11 | 35129094 | rs7106681 | 0.149 | -1.184 | -0.627 | 0.177 | -0.003 | 0.013 | 0.013 | 0 | 0.08 | 0.11 | 0.05 | 0.18 | 0.06 | 0.2 |
| 11 | 35131751 | rs2732578 | 0.874 | -1.54 | -0.526 | 0.104 | 0.007 | -0.031 | -0.031 | 1.314 | 0.27 | 0.2 | 0.38 | 0.4 | 0.23 | 0.21 |
| 11 | 35132267 | rs404977 | -0.399 | -1.636 | 0.393 | -0.028 | 0.012 | 0.021 | 0.021 | 0 | 0.41 | 0.66 | 0.47 | 0.68 | 0.31 | 0.46 |
| 11 | 35139072 | rs353565 | 0.304 | -1.609 | 0.908 | 0.044 | -0.002 | 0.006 | 0.006 | 0 | 0.31 | 0.62 | 0.42 | 0.46 | 0.25 | 0.26 |
| 11 | 35150272 | rs353573 | 0.333 | -1.715 | 1.241 | -0.005 | -0.008 | 0.021 | 0.021 | 0 | 0.32 | 0.41 | 0.41 | 0.45 | 0.26 | 0.27 |
| 11 | 35151341 | rs353570 | -0.284 | -1.767 | 1.277 | -0.107 | 0.042 | -0.033 | -0.033 | 0 | 0.4 | 0.73 | 0.44 | 0.47 | 0.28 | 0.4 |
| 11 | 35156500 | rs353560 | -1.365 | -1.633 | 0.84 | -0.095 | -0.005 | 0.024 | 0.024 | 0.457 | 0.11 | 0.5 | 0.05 | 0.1 | 0.03 | 0.16 |
| 11 | 35158401 | rs1425802 | 0.261 | -1.584 | 0.102 | 0.007 | 0.03 | -0.054 | -0.054 | 0.044 | 0.34 | 0.16 | 0.42 | 0.4 | 0.26 | 0.36 |
| 11 | 35161076 | rs3751031 | 0.069 | -1.177 | 0.155 | -0.103 | -0.014 | -0.052 | -0.052 | 0.046 | 0.09 | 0.01 | 0.05 | 0.19 | 0.07 | 0.2 |
| 11 | 35161670 | rs353558 | -1.826 | -0.759 | -0.154 | -0.628 | 0.007 | 0.002 | 0.002 | 0 | 0.78 | 0.8 | 0.77 | 0.74 | 0.66 | 0.86 |
| 11 | 35163114 | rs11033008 | 0.013 | -0.719 | -0.559 | -0.894 | -0.05 | 0.114 | 0.114 | 0.018 | 0.19 | 0.03 | 0.18 | 0.01 | 0.2 | 0.19 |
| 11 | 35167359 | rs11033010 | -1.036 | -0.697 | -0.089 | -1.087 | 0.001 | 0.022 | 0.022 | 0.63 | 0.34 | 0.29 | 0.32 | 0.19 | 0.31 | 0.37 |
| 11 | 35172109 | rs3794125 | 0.979 | -1.062 | 0.138 | -1.311 | -0.01 | 0 | 0 | 0 | 0.13 | 0.07 | 0.14 | 0.09 | 0.04 | 0.06 |
| 11 | 35174136 | rs353620 | -1.402 | -1.061 | 0.412 | -1.421 | -0.003 | -0.014 | -0.014 | 0 | 0.4 | 0.54 | 0.39 | 0.19 | 0.34 | 0.29 |
| 11 | 35174595 | rs353619 | -0.454 | -1.002 | 0.554 | -1.395 | 0.005 | -0.034 | -0.034 | 0 | 0.23 | 0.38 | 0.22 | 0.1 | 0.21 | 0.21 |
| 11 | 35177479 | rs353615 | -1.252 | -0.912 | 0.54 | -1.259 | 0.003 | 0.003 | 0.003 | 0 | 0.37 | 0.62 | 0.35 | 0.22 | 0.28 | 0.29 |

| | | | | | | | | | | | | | | | | |
|---|---|---|---|---|---|---|---|---|---|---|---|---|---|---|---|---|
| 11 | 35177771 | rs7952493 | 0.035 | -0.948 | 0.538 | -1.516 | -0.038 | -0.018 | -0.018 | 0 | 0.19 | 0.23 | 0.16 | 0.09 | 0.06 | 0.09 |
| 11 | 35178638 | rs12280381 | 0.138 | -0.907 | 0.443 | -1.566 | -0.01 | -0.002 | -0.002 | 0.009 | 0.18 | 0.18 | 0.15 | 0.09 | 0.06 | 0.09 |
| 11 | 35179651 | rs353612 | -0.264 | -0.206 | 0.852 | -1.353 | 0.012 | -0.039 | -0.039 | 0.947 | 0.19 | 0.32 | 0.17 | 0.04 | 0.22 | 0.19 |
| 11 | 35183838 | rs193276 | -0.788 | -0.147 | 0.88 | -1.286 | 0.016 | -0.004 | -0.004 | 0 | 0.18 | 0.33 | 0.3 | 0.13 | 0.23 | 0.21 |
| 11 | 35184364 | rs353639 | -1.202 | -0.235 | 1.161 | -1.299 | 0.021 | -0.013 | -0.013 | 0 | 0.2 | 0.58 | 0.31 | 0.13 | 0.23 | 0.21 |
| 11 | 35184527 | rs353638 | -1.005 | -0.263 | 0.958 | -1.29 | 0.036 | -0.007 | -0.007 | 0 | 0.19 | 0.45 | 0.3 | 0.13 | 0.22 | 0.2 |
| 11 | 35185056 | rs353634 | -2.405 | -0.107 | 0.105 | -1.332 | -0.089 | -0.05 | -0.05 | 0 | 0.83 | 0.99 | 0.86 | 0.79 | 0.71 | 0.87 |
| 11 | 35185294 | rs102518 | -1.07 | -0.497 | 0.922 | -1.666 | 0.014 | -0.002 | -0.002 | 0 | 0.18 | 0.33 | 0.3 | 0.13 | 0.22 | 0.2 |
| 11 | 35188191 | rs353630 | -1.352 | -0.502 | 0.808 | -1.733 | 0.011 | -0.107 | -0.107 | 0 | 0.26 | 0.51 | 0.32 | 0.05 | 0.25 | 0.21 |
| 11 | 35189148 | rs3794113 | 0.509 | 0.172 | 0.552 | -1.331 | -0.011 | -0.011 | -0.011 | 0 | 0.07 | 0.03 | 0.08 | 0.08 | 0.06 | 0.22 |
| 11 | 35189245 | rs353626 | -1.623 | 0.122 | 0.795 | -1.388 | -0.003 | -0.006 | -0.006 | 0 | 0.24 | 0.4 | 0.36 | 0.21 | 0.21 | 0.39 |
| 11 | 35192114 | rs353646 | 0.381 | 0.286 | 0.245 | -1.317 | 0.001 | -0.009 | -0.009 | 0 | 0.1 | 0.18 | 0.25 | 0.04 | 0.12 | 0.13 |
| 11 | 35192279 | rs3794109 | 0.115 | 0.43 | -0.324 | -1.293 | 0.024 | 0.002 | 0.002 | 0 | 0.1 | 0.18 | 0.12 | 0.09 | 0.15 | 0.24 |
| 11 | 35198108 | rs4756196 | -1.436 | 0.231 | -0.711 | -1.39 | -0.037 | 0.029 | 0.029 | 0 | 0.58 | 0.76 | 0.63 | 0.29 | 0.47 | 0.52 |
| 11 | 35201842 | rs1071695 | -0.592 | -0.43 | -1.301 | -1.135 | 0.049 | 0.077 | 0.077 | 0 | 0.31 | 0.15 | 0.22 | 0.07 | 0.17 | 0.1 |
| 11 | 35210798 | rs996076 | 0.175 | 0.635 | -0.999 | -0.959 | 0 | 0.008 | 0.008 | 0 | 0.52 | 0.43 | 0.44 | 0.86 | 0.55 | 0.71 |
| 11 | 35213126 | rs7934770 | -0.348 | 0.665 | -1.308 | -1.287 | -0.017 | 0.111 | 0.111 | 0.004 | 0.64 | 0.93 | 0.53 | 0.88 | 0.71 | 0.78 |
| 11 | 35213378 | rs10488810 | -0.048 | 0.875 | -1.299 | -1.018 | -0.005 | -0.003 | -0.003 | 2.195 | 0.08 | 0.06 | 0.09 | 0.08 | 0.09 | 0.14 |
| 11 | 35215726 | rs10128562 | 0.489 | 0.954 | -1.051 | -1.014 | -0.002 | 0.006 | 0.006 | 0 | 0.44 | 0.37 | 0.36 | 0.77 | 0.51 | 0.59 |
| 11 | 35223945 | rs713330 | -0.002 | 0.523 | -0.804 | -1.353 | -0.04 | 0.162 | 0.162 | 0.001 | 0.68 | 0.92 | 0.66 | 0.93 | 0.66 | 0.86 |
| 11 | 35226155 | rs9666607 | -0.026 | 0.363 | -0.8 | -1.348 | -0.019 | 0.158 | 0.158 | 0 | 0.68 | 0.92 | 0.69 | 0.93 | 0.68 | 0.86 |
| 11 | 35230605 | rs2421826 | 0.37 | -0.376 | -0.18 | -1.249 | 0.002 | -0.002 | -0.002 | 0 | 0.62 | 0.7 | 0.76 | 0.85 | 0.61 | 0.8 |
| 11 | 35238459 | rs1547059 | -0.001 | -0.513 | -0.127 | -0.721 | 0.005 | -0.024 | -0.024 | 3.876 | 0.67 | 0.73 | 0.81 | 0.99 | 0.68 | 0.85 |
| 11 | 35241229 | rs2295756 | 0.877 | -0.435 | -0.54 | -1.297 | -0.017 | 0.017 | 0.017 | 0 | 0.37 | 0.21 | 0.35 | 0.58 | 0.34 | 0.52 |
| 11 | 35241989 | rs7945310 | -0.238 | -0.326 | -0.601 | -0.511 | 0.002 | 0.001 | 0.001 | 0 | 0.61 | 0.66 | 0.64 | 0.57 | 0.64 | 0.79 |
| 11 | 35244058 | rs7116432 | 0.617 | -0.063 | -0.901 | -0.28 | -0.012 | 0.036 | 0.036 | 0 | 0.32 | 0.23 | 0.5 | 0.81 | 0.33 | 0.56 |
| 11 | 35244574 | rs11607862 | 0.842 | -0.034 | -1.423 | -0.114 | -0.071 | -0.144 | -0.144 | 0.774 | 0.29 | 0.03 | 0.32 | 0.36 | 0.3 | 0.49 |
| 11 | 35244643 | rs11033031 | 0.545 | 0.198 | -0.871 | 0.146 | -0.029 | 0.031 | 0.031 | 0 | 0.11 | 0.04 | 0.14 | 0.05 | 0.14 | 0.07 |
| 11 | 35245907 | rs10128586 | -0.639 | 0.089 | -0.551 | 0.146 | 0.004 | -0.018 | -0.018 | 0.894 | 0.18 | 0.29 | 0.16 | 0.13 | 0.17 | 0.1 |
| 11 | 35249670 | rs12419062 | -0.559 | -0.247 | -1.064 | 0.103 | 0.032 | 0.025 | 0.025 | 0.101 | 0.42 | 0.71 | 0.49 | 0.41 | 0.36 | 0.23 |
| 11 | 35251318 | rs8193 | 0.862 | 0 | -1.539 | 0.11 | -0.043 | -0.12 | -0.12 | 0 | 0.4 | 0.07 | 0.36 | 0.42 | 0.35 | 0.65 |
| 11 | 35253133 | rs10836347 | 0.98 | -0.09 | -1.188 | -0.012 | -0.028 | 0.018 | 0.018 | 0 | 0.11 | 0.04 | 0.14 | 0.07 | 0.14 | 0.07 |
| 11 | 35253272 | rs13347 | 0.032 | -0.428 | -1.237 | -0.064 | 0.01 | 0.003 | 0.003 | 0 | 0.31 | 0.7 | 0.35 | 0.34 | 0.22 | 0.16 |
| 11 | 35256693 | rs7115638 | 0.762 | -0.864 | -0.914 | 0.047 | -0.008 | 0.007 | 0.007 | 0 | 0.46 | 0.36 | 0.4 | 0.37 | 0.45 | 0.53 |
| 11 | 35257316 | rs7119757 | -0.697 | -0.87 | -0.987 | 0.042 | -0.06 | 0.061 | 0.061 | 0.263 | 0.81 | 0.95 | 0.84 | 0.91 | 0.71 | 0.78 |
| 11 | 35264055 | rs2421823 | 0.34 | -1.04 | -0.007 | 0.1 | -0.006 | 0 | 0 | 0 | 0.59 | 0.49 | 0.58 | 0.53 | 0.61 | 0.61 |
| 11 | 35265444 | rs10836355 | 0.897 | -1.081 | 0.058 | -0.09 | 0.014 | 0.047 | 0.047 | 0 | 0.51 | 0.33 | 0.52 | 0.61 | 0.52 | 0.6 |
| 11 | 35266944 | rs736374 | 2.017 | -1.523 | -0.348 | -0.397 | -0.085 | -0.133 | -0.133 | 0 | 0.3 | 0.04 | 0.34 | 0.48 | 0.35 | 0.5 |
| 11 | 35267855 | rs10768118 | 1.234 | -1.366 | -0.166 | -0.244 | -0.006 | -0.024 | -0.024 | 0 | 0.4 | 0.24 | 0.37 | 0.48 | 0.38 | 0.52 |
| 11 | 35270601 | rs12287327 | 1.387 | -1.515 | 0.209 | -0.22 | 0.021 | -0.013 | -0.013 | 0 | 0.18 | 0.14 | 0.22 | 0.23 | 0.1 | 0.13 |
| 11 | 35271419 | rs10836356 | 0.603 | -1.327 | 0.169 | -0.173 | -0.009 | 0.002 | 0.002 | 0 | 0.48 | 0.34 | 0.43 | 0.48 | 0.47 | 0.54 |
| 11 | 35273268 | rs3088168 | 1.123 | -1.308 | 0.05 | -0.159 | -0.05 | -0.079 | -0.079 | 0 | 0.39 | 0.12 | 0.36 | 0.48 | 0.38 | 0.52 |
| 11 | 35274829 | rs1043101 | 1.104 | -1.313 | 0.11 | -0.161 | -0.039 | -0.066 | -0.066 | 1.321 | 0.39 | 0.13 | 0.36 | 0.48 | 0.38 | 0.52 |
| 11 | 35276059 | rs12360706 | 1.711 | -1.353 | 0.183 | -0.206 | -0.007 | -0.002 | -0.002 | 0.131 | 0.11 | 0.07 | 0.15 | 0.13 | 0.14 | 0.07 |
| 11 | 35277270 | rs3812778 | 1.807 | -1.36 | 0.19 | -0.207 | -0.026 | -0.016 | -0.016 | 0 | 0.11 | 0.04 | 0.15 | 0.13 | 0.14 | 0.07 |
| 11 | 35278959 | rs7936950 | -1.781 | -1.53 | 0.391 | -0.339 | -0.08 | 0.054 | 0.054 | 0.973 | 0.88 | 0.99 | 0.87 | 0.97 | 0.73 | 0.92 |
| 11 | 35279301 | rs12294045 | 0.631 | -1.645 | 0.601 | -0.34 | 0.036 | -0.003 | -0.003 | 0 | 0.25 | 0.16 | 0.28 | 0.23 | 0.19 | 0.16 |
| 11 | 35280852 | rs10768122 | 1.272 | -1.554 | 0.451 | -0.285 | -0.039 | -0.066 | -0.066 | 0 | 0.39 | 0.13 | 0.36 | 0.48 | 0.38 | 0.52 |
| 11 | 35282096 | rs4508184 | 1.712 | -1.511 | 0.419 | -0.338 | -0.007 | -0.004 | -0.004 | 0.277 | 0.11 | 0.07 | 0.15 | 0.13 | 0.14 | 0.07 |
| 11 | 35282334 | rs1570216 | 1.712 | -1.511 | 0.419 | -0.338 | -0.007 | -0.004 | -0.004 | 0 | 0.11 | 0.07 | 0.15 | 0.13 | 0.14 | 0.07 |
| 11 | 35292312 | rs4756204 | 1.145 | -1.609 | 0.448 | 0.059 | -0.037 | -0.061 | -0.061 | 0 | 0.39 | 0.14 | 0.36 | 0.43 | 0.38 | 0.52 |
| 11 | 35300905 | rs3794097 | 1.001 | -1.625 | 0.41 | 0.114 | -0.046 | -0.034 | -0.034 | 0.458 | 0.17 | 0.06 | 0.23 | 0.18 | 0.23 | 0.09 |
| 11 | 35306345 | rs10836364 | 1.257 | -1.651 | 0.407 | 0.242 | -0.048 | -0.061 | -0.061 | 0 | 0.39 | 0.1 | 0.36 | 0.42 | 0.38 | 0.52 |
| 11 | 35307148 | rs3763888 | 1.098 | -1.646 | 0.483 | 0.244 | -0.001 | -0.004 | -0.004 | 0 | 0.41 | 0.35 | 0.37 | 0.42 | 0.38 | 0.52 |
| 11 | 35308783 | rs3818275 | 0.172 | -1.648 | 0.542 | 0.254 | -0.001 | -0.003 | -0.003 | 0 | 0.59 | 0.45 | 0.63 | 0.71 | 0.64 | 0.64 |
| 11 | 35314173 | rs2281634 | 1.133 | -1.763 | 0.732 | 0.289 | -0.004 | -0.002 | -0.002 | 0.625 | 0.21 | 0.17 | 0.19 | 0.22 | 0.06 | 0.14 |
| 11 | 35317212 | rs6484776 | -1.056 | -1.646 | 0.779 | 0.191 | -0.093 | 0.057 | 0.057 | 2.296 | 0.82 | 0.98 | 0.85 | 0.92 | 0.71 | 0.78 |
| 11 | 35317353 | rs3794091 | 0.912 | -1.567 | 0.849 | 0.248 | -0.002 | 0.009 | 0.009 | 0 | 0.42 | 0.5 | 0.39 | 0.52 | 0.38 | 0.55 |
| 11 | 35321585 | rs3847607 | 0.204 | -1.59 | 0.212 | 0.241 | 0.025 | 0.046 | 0.046 | 0 | 0.58 | 0.25 | 0.63 | 0.71 | 0.64 | 0.65 |
| 11 | 35324660 | rs3794089 | 1.148 | -1.575 | 0.156 | 0.255 | -0.032 | -0.082 | -0.082 | 5.083 | 0.41 | 0.11 | 0.38 | 0.52 | 0.38 | 0.56 |
| 11 | 35327748 | rs752949 | 0.772 | -1.493 | 0.221 | 0.251 | -0.014 | -0.005 | -0.005 | 0 | 0.17 | 0.14 | 0.25 | 0.19 | 0.26 | 0.09 |
| 11 | 35329615 | rs3794087 | 0.588 | -1.402 | 0.005 | 0.208 | -0.05 | -0.015 | -0.015 | 0 | 0.2 | 0.07 | 0.24 | 0.19 | 0.26 | 0.09 |
| 11 | 35331642 | rs3903687 | 0.494 | -1.395 | -0.072 | 0.104 | 0.029 | 0.059 | 0.059 | 0 | 0.58 | 0.17 | 0.61 | 0.71 | 0.62 | 0.65 |
| 11 | 35334668 | rs17379710 | 1.205 | -1.245 | 0.378 | 0.065 | 0.013 | 0.037 | 0.037 | 0 | 0.53 | 0.3 | 0.54 | 0.64 | 0.44 | 0.66 |
| 11 | 35338743 | rs2273688 | -0.885 | -1.024 | 0.318 | 0.067 | -0.031 | 0.023 | 0.023 | 0.38 | 0.86 | 0.93 | 0.85 | 0.92 | 0.73 | 0.8 |
| 11 | 35347929 | rs3847615 | -0.924 | -1.029 | 0.5 | 0.064 | 0.002 | -0.003 | -0.003 | 0 | 0.85 | 0.85 | 0.84 | 0.93 | 0.73 | 0.8 |
| 11 | 35348532 | rs7102331 | 0.044 | -1.037 | 0.579 | 0.084 | -0.004 | 0.002 | 0.002 | 0.028 | 0.77 | 0.76 | 0.81 | 0.83 | 0.7 | 0.76 |
| 11 | 35353111 | rs4756211 | 0.619 | -0.964 | 0.345 | 0.125 | -0.019 | 0 | 0 | 0 | 0.19 | 0.12 | 0.24 | 0.19 | 0.25 | 0.08 |
| 11 | 35353944 | rs11033073 | 1.521 | -1.175 | 0.221 | 0.121 | -0.037 | -0.057 | -0.057 | 0.785 | 0.34 | 0.13 | 0.36 | 0.44 | 0.37 | 0.52 |

| | | | | | | | | | | | | | | | | |
|---|---|---|---|---|---|---|---|---|---|---|---|---|---|---|---|---|
| 11 | 35360016 | rs3847618 | 1.174 | -1.291 | 0.294 | -0.021 | -0.001 | 0 | 0 | 0 | 0.19 | 0.16 | 0.18 | 0.13 | 0.07 | 0.16 |
| 11 | 35362489 | rs10768129 | -1.09 | -1.174 | 0.217 | 0.125 | -0.037 | 0.01 | 0.01 | 0.574 | 0.86 | 0.94 | 0.85 | 0.92 | 0.73 | 0.8 |
| 11 | 35363078 | rs7942698 | 1.423 | -1.293 | 0.512 | 0.093 | -0.003 | -0.01 | -0.01 | 0.399 | 0.37 | 0.33 | 0.4 | 0.51 | 0.38 | 0.52 |
| 11 | 35364435 | rs3847621 | 0.386 | -1.347 | 0.537 | 0.084 | 0.003 | 0.003 | 0.003 | 0 | 0.56 | 0.45 | 0.62 | 0.7 | 0.62 | 0.6 |
| 11 | 35365938 | rs4756214 | 1.625 | -1.289 | 0.534 | 0.119 | -0.009 | -0.021 | -0.021 | 0.634 | 0.35 | 0.24 | 0.4 | 0.51 | 0.38 | 0.52 |
| 11 | 35375280 | rs10768131 | 0.243 | -1.26 | 0.365 | 0.098 | -0.082 | 0.029 | 0.029 | 0.052 | 0.64 | 0.86 | 0.64 | 0.73 | 0.49 | 0.72 |
| 11 | 35375388 | rs16927393 | 1.16 | -1.352 | 0.58 | 0.167 | -0.006 | 0.033 | 0.033 | 0 | 0.07 | 0.19 | 0.04 | 0.09 | 0.04 | 0.04 |
| 11 | 35375958 | rs4756219 | 0.476 | -1.225 | 0.435 | 0.235 | -0.024 | 0.006 | 0.006 | 0 | 0.22 | 0.08 | 0.21 | 0.19 | 0.24 | 0.08 |
| 11 | 35380789 | rs1320443 | 0.252 | -1.483 | 0.561 | 0.318 | -0.031 | 0.002 | 0.002 | 0 | 0.27 | 0.34 | 0.22 | 0.21 | 0.11 | 0.21 |
| 11 | 35382124 | rs4756222 | 0.504 | -0.425 | 0.496 | -0.053 | -0.004 | 0.006 | 0.006 | 0.175 | 0.59 | 0.63 | 0.59 | 0.73 | 0.44 | 0.72 |
| 11 | 35384432 | rs1923298 | -0.723 | 0.122 | 0.547 | -0.213 | 0.002 | 0.028 | 0.028 | 0 | 0.18 | 0.39 | 0.16 | 0.17 | 0.17 | 0.25 |
| 11 | 35387697 | rs11033097 | -0.854 | 0.383 | 0.485 | -0.367 | -0.006 | -0.015 | -0.015 | 0 | 0.3 | 0.41 | 0.22 | 0.19 | 0.27 | 0.28 |
| 11 | 35387874 | rs11033098 | -0.872 | 0.362 | 0.724 | -0.421 | -0.01 | -0.029 | -0.029 | 0 | 0.31 | 0.44 | 0.23 | 0.19 | 0.27 | 0.28 |
| 11 | 35389487 | rs4287314 | -0.249 | 1.32 | 1.022 | -0.853 | 0.029 | -0.061 | -0.061 | 0 | 0.83 | 0.67 | 0.83 | 0.95 | 0.76 | 0.92 |
| 11 | 35392601 | rs1885345 | -0.663 | 1.277 | 0.759 | -0.857 | -0.036 | 0.041 | 0.041 | 0 | 0.86 | 0.95 | 0.85 | 0.95 | 0.76 | 0.92 |
| 11 | 35394729 | rs9704794 | -0.558 | 1.171 | 0.731 | -0.817 | 0.001 | 0.003 | 0.003 | 0 | 0.29 | 0.29 | 0.22 | 0.21 | 0.27 | 0.27 |
| 11 | 35394834 | rs3847623 | 1.332 | 1.129 | 0.781 | -0.814 | -0.016 | 0.091 | 0.091 | 0 | 0.53 | 0.24 | 0.6 | 0.74 | 0.49 | 0.65 |
| 11 | 35399110 | rs911561 | 1.184 | 1.188 | 0.676 | -0.785 | 0.003 | -0.015 | -0.015 | 0 | 0.55 | 0.51 | 0.62 | 0.73 | 0.49 | 0.65 |
| 11 | 35399975 | rs10836386 | -0.96 | 1.284 | -0.002 | -0.702 | 0.005 | 0.018 | 0.018 | 0 | 0.46 | 0.39 | 0.44 | 0.33 | 0.55 | 0.36 |
| 11 | 35402408 | rs10836387 | -0.839 | 1.412 | 0.16 | -0.586 | -0.001 | 0.001 | 0.001 | 0 | 0.41 | 0.44 | 0.34 | 0.33 | 0.42 | 0.39 |
| 11 | 35403124 | rs4593985 | 1.185 | 0.872 | -0.301 | -0.324 | 0.054 | -0.005 | -0.005 | 0 | 0.41 | 0.18 | 0.39 | 0.6 | 0.3 | 0.44 |
| 11 | 35403189 | rs7119850 | -0.22 | 0.837 | 0.025 | -0.14 | -0.004 | -0.045 | -0.045 | 0.008 | 0.16 | 0.21 | 0.11 | 0.03 | 0.14 | 0.12 |
| 11 | 35408126 | rs11827417 | 0.063 | 0.894 | -0.213 | 0.086 | -0.002 | -0.004 | -0.004 | 0 | 0.11 | 0.07 | 0.16 | 0.12 | 0.14 | 0.16 |
| 11 | 35409341 | rs7108848 | -0.435 | 1.06 | -0.014 | -0.185 | 0 | -0.057 | -0.057 | 0 | 0.18 | 0.27 | 0.14 | 0.03 | 0.17 | 0.12 |
| 11 | 35413047 | rs10768132 | 1.268 | 0.901 | -0.372 | 0.624 | -0.017 | 0.006 | 0.006 | 0 | 0.45 | 0.2 | 0.42 | 0.43 | 0.41 | 0.42 |
| 11 | 35414366 | rs1923304 | -0.303 | 0.702 | -0.153 | 0.757 | -0.034 | -0.035 | -0.035 | 0 | 0.12 | 0.01 | 0.16 | 0.11 | 0.16 | 0.11 |
| 11 | 35420858 | rs16927464 | -1.547 | 0.416 | 0.29 | 0.933 | -0.012 | 0.022 | 0.022 | 0 | 0.22 | 0.66 | 0.25 | 0.34 | 0.2 | 0.23 |
| 11 | 35428375 | rs2145906 | 0.478 | 0.803 | 0.872 | 0.577 | 0.014 | -0.007 | -0.007 | 0 | 0.65 | 0.43 | 0.64 | 0.64 | 0.61 | 0.61 |
| 11 | 35430474 | rs4144755 | -0.246 | 0.814 | 0.761 | 0.578 | -0.034 | 0.017 | 0.017 | 0 | 0.73 | 0.83 | 0.69 | 0.77 | 0.63 | 0.69 |
| 11 | 35432417 | rs1885342 | -0.246 | 0.814 | 0.761 | 0.578 | -0.034 | 0.017 | 0.017 | 0 | 0.73 | 0.83 | 0.69 | 0.77 | 0.63 | 0.69 |
| 11 | 35435433 | rs11033115 | -0.334 | 0.511 | 0.919 | 0.64 | 0.013 | 0.102 | 0.102 | 0.158 | 0.06 | 0.28 | 0.03 | 0.13 | 0.02 | 0.08 |
| 11 | 35438179 | rs1923292 | 0.988 | 0.725 | 0.644 | 0.651 | 0.015 | -0.004 | -0.004 | 3.531 | 0.51 | 0.13 | 0.54 | 0.56 | 0.41 | 0.46 |
| 11 | 35439103 | rs1923293 | -1.113 | 0.793 | 1.03 | 0.659 | 0.054 | 0.116 | 0.116 | 0.286 | 0.07 | 0.36 | 0.04 | 0.12 | 0.04 | 0.1 |
| 11 | 35439816 | rs1923294 | 0.496 | 0.713 | 0.203 | 0.754 | 0.003 | 0 | 0 | 0.131 | 0.54 | 0.54 | 0.56 | 0.51 | 0.4 | 0.43 |
| 11 | 35442608 | rs1923295 | 0.444 | 0.893 | -0.083 | 0.769 | 0.002 | 0.005 | 0.005 | 0 | 0.57 | 0.66 | 0.58 | 0.56 | 0.49 | 0.44 |
| 11 | 35443444 | rs16927467 | -0.827 | 0.456 | 0.013 | 0.65 | -0.008 | -0.002 | -0.002 | 1.742 | 0.13 | 0.12 | 0.05 | 0.12 | 0.07 | 0.11 |
| 11 | 35445574 | rs4755410 | 0.648 | 0.524 | -0.231 | 0.612 | -0.017 | 0.023 | 0.023 | 0 | 0.61 | 0.31 | 0.51 | 0.72 | 0.55 | 0.58 |
| 11 | 35451976 | rs1173923 | 0.64 | 0.563 | -0.01 | 0.626 | -0.009 | 0.015 | 0.015 | 0 | 0.61 | 0.32 | 0.51 | 0.73 | 0.56 | 0.59 |
| 11 | 35453719 | rs7767 | 0.661 | 0.612 | 0.002 | 0.637 | -0.018 | 0.028 | 0.028 | 0 | 0.61 | 0.3 | 0.51 | 0.72 | 0.56 | 0.59 |
| 11 | 35458946 | rs599338 | -0.871 | 0.563 | 0.393 | 0.624 | -0.004 | 0.004 | 0.004 | 0 | 0.37 | 0.63 | 0.49 | 0.25 | 0.44 | 0.29 |
| 11 | 35460843 | rs644755 | -1.799 | -0.269 | -0.499 | 0.865 | -0.02 | 0.003 | 0.003 | 0.046 | 0.68 | 0.87 | 0.73 | 0.68 | 0.66 | 0.7 |
| 11 | 35472451 | rs11033125 | 0.513 | 0.23 | -0.04 | 0.647 | 0.005 | -0.015 | -0.015 | 0 | 0.2 | 0.22 | 0.34 | 0.1 | 0.28 | 0.19 |
| 11 | 35479735 | rs1516988 | -1.596 | -0.036 | -0.229 | 0.691 | -0.008 | 0.02 | 0.02 | 0.184 | 0.53 | 0.72 | 0.55 | 0.58 | 0.5 | 0.44 |
| 11 | 35481588 | rs7943168 | 1.149 | -0.064 | -0.213 | 0.619 | -0.022 | 0.022 | 0.022 | 0 | 0.08 | 0.05 | 0.05 | 0 | 0.13 | 0.03 |
| 11 | 35486237 | rs7112485 | 0.103 | 0.034 | 0.247 | 0.847 | 0.036 | -0.021 | -0.021 | 0 | 0.23 | 0.47 | 0.38 | 0.21 | 0.29 | 0.24 |
| 11 | 35487642 | rs10501140 | -2.108 | 0.069 | 0.181 | 0.808 | 0.001 | 0.001 | 0.001 | 0 | 0.67 | 0.75 | 0.73 | 0.62 | 0.67 | 0.61 |
| 11 | 35487715 | rs10836401 | -0.552 | 0.223 | 0.206 | 0.835 | 0.015 | -0.027 | -0.027 | 0.019 | 0.29 | 0.53 | 0.43 | 0.25 | 0.31 | 0.26 |
| 11 | 35487857 | rs11602786 | 0.861 | 0.472 | 0.147 | 0.929 | -0.048 | 0.006 | 0.006 | 0 | 0.13 | 0.01 | 0.17 | 0.05 | 0.16 | 0.16 |
| 11 | 35489891 | rs6484787 | -0.821 | 0.436 | 0.69 | 1.32 | -0.003 | -0.007 | -0.007 | 0 | 0.57 | 0.49 | 0.64 | 0.57 | 0.62 | 0.56 |
| 11 | 35491891 | rs7933964 | 0.837 | 0.727 | 1.265 | 1.819 | -0.008 | -0.005 | -0.005 | 0.195 | 0.05 | 0.09 | 0.05 | 0.04 | 0.02 | 0.02 |
| 11 | 35492351 | rs16927500 | 0.119 | 0.732 | 1.217 | 1.821 | 0.003 | 0.009 | 0.009 | 2.459 | 0.07 | 0.26 | 0.06 | 0.04 | 0.02 | 0.02 |
| 11 | 35495674 | rs12799002 | -1.552 | 0.789 | 0.628 | 1.851 | 0.02 | 0.055 | 0.055 | 0 | 0.26 | 0.58 | 0.2 | 0.34 | 0.29 | 0.3 |
| 11 | 35495951 | rs16927508 | -1.037 | 1.169 | 0.728 | 1.824 | 0.074 | 0.115 | 0.115 | 0 | 0.19 | 0.47 | 0.17 | 0.29 | 0.26 | 0.29 |
| 11 | 35497059 | rs7106288 | -0.588 | 1.192 | 0.541 | 1.452 | 0.079 | -0.072 | -0.072 | 0 | 0.17 | 0.36 | 0.16 | 0.07 | 0.25 | 0.19 |
| 11 | 35500592 | rs4456236 | -0.833 | 1.226 | 0.184 | 1.489 | 0.074 | 0.178 | 0.178 | 0 | 0.19 | 0.45 | 0.2 | 0.38 | 0.26 | 0.27 |
| 11 | 35501843 | rs592268 | -0.163 | 1.042 | -0.454 | 1.014 | 0 | 0.001 | 0.001 | 0 | 0.58 | 0.45 | 0.45 | 0.5 | 0.57 | 0.53 |
| 11 | 35504853 | rs4756234 | 0.061 | 0.822 | -1.34 | 0.38 | -0.1 | 0.088 | 0.088 | 0 | 0.29 | 0.06 | 0.29 | 0.15 | 0.42 | 0.32 |
| 11 | 35505197 | rs686971 | 0.177 | 0.242 | -1.232 | 0.542 | 0 | -0.009 | -0.009 | 0.051 | 0.75 | 0.76 | 0.72 | 0.58 | 0.8 | 0.68 |
| 11 | 35507700 | rs621363 | -0.409 | 0.204 | -1.226 | 0.678 | -0.039 | -0.058 | -0.058 | 0 | 0.82 | 0.97 | 0.84 | 0.76 | 0.85 | 0.72 |
| 11 | 35508918 | rs649640 | 0.417 | -0.325 | -1.15 | 0.49 | -0.042 | 0.182 | 0.182 | 0.511 | 0.19 | 0.49 | 0.22 | 0.37 | 0.12 | 0.18 |
| 11 | 35510907 | rs10768140 | 0.6 | -0.38 | -1.694 | 0.367 | 0.029 | 0.006 | 0.006 | 0.738 | 0.61 | 0.11 | 0.59 | 0.4 | 0.71 | 0.53 |
| 11 | 35512220 | rs4142611 | 1.3 | -0.121 | -1.738 | 0.104 | -0.051 | 0.153 | 0.153 | 0 | 0.31 | 0.05 | 0.22 | 0.07 | 0.29 | 0.2 |
| 11 | 35518995 | rs694961 | 0.548 | 0.464 | -1.186 | 0.415 | 0.024 | -0.007 | -0.007 | 5.75 | 0.47 | 0.69 | 0.44 | 0.44 | 0.39 | 0.35 |
| 11 | 35519169 | rs581513 | 0.294 | 0.718 | -0.435 | 1.059 | 0.008 | 0.036 | 0.036 | 0 | 0.82 | 0.77 | 0.74 | 0.6 | 0.82 | 0.71 |
| 11 | 35522638 | rs7946272 | 0.636 | 1.076 | -0.561 | 0.653 | 0.001 | 0 | 0 | 0 | 0.15 | 0.16 | 0.27 | 0.27 | 0.23 | 0.19 |
| 11 | 35526523 | rs11033151 | -1.089 | 0.484 | -0.316 | 0.707 | 0 | -0.012 | -0.012 | 0 | 0.41 | 0.56 | 0.52 | 0.71 | 0.41 | 0.51 |
| 11 | 35528204 | rs581660 | -0.455 | 0.337 | -0.23 | 0.833 | 0.004 | 0.003 | 0.003 | 0 | 0.27 | 0.4 | 0.33 | 0.34 | 0.31 | 0.3 |
| 11 | 35532452 | rs769149 | 0.338 | 0.023 | -0.258 | 0.358 | 0.002 | 0.065 | 0.065 | 0 | 0.17 | 0.3 | 0.13 | 0.21 | 0.11 | 0.22 |
| 11 | 35532656 | rs645079 | 0.607 | 0.021 | -0.353 | 0.399 | -0.014 | -0.013 | -0.013 | 3.611 | 0.11 | 0.19 | 0.05 | 0.06 | 0.08 | 0.11 |

| | | | | | | | | | | | | | | | | |
|---|---|---|---|---|---|---|---|---|---|---|---|---|---|---|---|---|
| 11 | 35533895 | rs11602690 | 0.657 | 0.02 | -0.455 | 0.375 | 0.007 | -0.002 | -0.002 | 0 | 0.1 | 0.01 | 0.04 | 0.07 | 0.08 | 0.12 |
| 11 | 35535317 | rs11033157 | -0.364 | 0.084 | -0.329 | 0.418 | 0.002 | 0.004 | 0.004 | 0 | 0.29 | 0.33 | 0.46 | 0.64 | 0.33 | 0.38 |
| 11 | 35536881 | rs10734439 | -0.48 | 0.083 | -0.312 | 0.384 | 0.012 | 0.03 | 0.03 | 0 | 0.3 | 0.42 | 0.47 | 0.64 | 0.33 | 0.39 |
| 11 | 35538592 | rs589475 | -0.613 | 0.062 | -0.359 | 0.307 | -0.003 | -0.008 | -0.008 | 0 | 0.34 | 0.67 | 0.42 | 0.53 | 0.34 | 0.42 |
| 11 | 35539269 | rs650950 | -2.263 | 0.05 | -0.575 | 0.124 | -0.025 | 0.203 | 0.203 | 0 | 0.57 | 0.82 | 0.65 | 0.97 | 0.53 | 0.74 |
| 11 | 35539344 | rs656009 | -1.418 | -0.079 | -0.41 | 0.312 | 0.025 | 0.044 | 0.044 | 0 | 0.42 | 0.64 | 0.38 | 0.7 | 0.3 | 0.56 |
| 11 | 35540398 | rs1760315 | -1.353 | -0.034 | -0.09 | 0.341 | -0.002 | -0.006 | -0.006 | 0 | 0.41 | 0.54 | 0.37 | 0.7 | 0.3 | 0.56 |
| 11 | 35540677 | rs16926171 | 0.465 | -0.1 | -0.015 | 0.424 | -0.004 | 0.044 | 0.044 | 0 | 0.14 | 0.17 | 0.19 | 0.37 | 0.1 | 0.2 |
| 11 | 35541465 | rs695121 | -1.399 | -0.085 | 0.001 | 0.386 | 0.004 | 0.006 | 0.006 | 0 | 0.4 | 0.58 | 0.37 | 0.7 | 0.29 | 0.56 |
| 11 | 35542658 | rs637585 | -1.005 | 0.363 | 0.191 | 0.565 | -0.023 | 0.045 | 0.045 | 0 | 0.27 | 0.4 | 0.18 | 0.33 | 0.19 | 0.37 |
| 11 | 35544493 | rs610966 | -1.969 | 0.925 | -0.395 | 0.005 | -0.025 | 0.014 | 0.014 | 0 | 0.61 | 0.54 | 0.64 | 0.51 | 0.68 | 0.68 |
| 11 | 35544737 | rs16927618 | -0.21 | 1.264 | -0.625 | 0.05 | -0.004 | -0.005 | -0.005 | 0 | 0.2 | 0.19 | 0.21 | 0.03 | 0.22 | 0.19 |
| 11 | 35544832 | rs12800242 | 0.324 | 1.237 | -0.682 | 0.035 | 0.009 | 0.019 | 0.019 | 0 | 0.12 | 0.07 | 0.06 | 0.06 | 0.1 | 0.1 |
| 11 | 35545094 | rs12803861 | -1.219 | 1.257 | -0.688 | 0.051 | 0 | -0.011 | -0.011 | 0.26 | 0.33 | 0.33 | 0.27 | 0.09 | 0.32 | 0.29 |
| 11 | 35546361 | rs7942841 | 0.044 | 2.045 | -0.863 | 0.645 | -0.055 | -0.019 | -0.019 | 0.126 | 0.16 | 0.01 | 0.26 | 0.19 | 0.2 | 0.2 |
| 11 | 35546732 | rs17795186 | -1.247 | 2.066 | 0.338 | 1.018 | -0.01 | 0.013 | 0.013 | 0 | 0.2 | 0.13 | 0.2 | 0.09 | 0.23 | 0.19 |
| 11 | 35551866 | rs506859 | -1.65 | 1.639 | 1.152 | 0.997 | 0.01 | -0.097 | -0.097 | 0.154 | 0.14 | 0.39 | 0.07 | 0 | 0.13 | 0.11 |
| 11 | 35553793 | rs1144589 | 1.305 | 2.081 | 2.107 | 2.286 | -0.008 | 0.006 | 0.006 | 0.195 | 0.76 | 0.8 | 0.67 | 0.87 | 0.7 | 0.79 |
| 11 | 35558748 | rs661993 | 1.121 | 2.164 | 2.152 | 2.543 | 0.018 | -0.001 | -0.001 | 1.205 | 0.87 | 0.84 | 0.73 | 0.91 | 0.79 | 0.89 |
| 11 | 35558904 | rs586910 | -1.312 | 1.972 | 1.938 | 2.354 | -0.026 | 0.069 | 0.069 | 0 | 0.14 | 0.21 | 0.17 | 0.32 | 0.11 | 0.17 |
| 11 | 35560205 | rs645267 | -1.168 | 2.192 | 1.915 | 1.998 | 0.043 | 0.057 | 0.057 | 0 | 0.13 | 0.35 | 0.14 | 0.18 | 0.13 | 0.12 |
| 11 | 35560298 | rs644814 | -1.816 | 2.176 | 1.903 | 1.99 | -0.013 | -0.016 | -0.016 | 0.552 | 0.25 | 0.36 | 0.19 | 0.21 | 0.23 | 0.23 |
| 11 | 35561585 | rs697317 | -2.479 | 2.25 | 1.64 | 2.383 | -0.023 | -0.006 | -0.006 | 0 | 0.41 | 0.68 | 0.5 | 0.42 | 0.47 | 0.38 |
| 11 | 35563367 | rs623230 | -1.868 | 2.129 | 1.524 | 2.364 | -0.007 | -0.026 | -0.026 | 0 | 0.27 | 0.43 | 0.22 | 0.22 | 0.25 | 0.27 |
| 11 | 35566223 | rs687460 | -1.142 | 2.303 | 1.18 | 2.824 | 0.043 | 0.083 | 0.083 | 0 | 0.08 | 0.13 | 0.22 | 0.25 | 0.16 | 0.11 |
| 11 | 35566613 | rs685734 | -1.35 | 2.189 | 1.194 | 2.77 | 0.024 | 0.029 | 0.029 | 0.043 | 0.11 | 0.13 | 0.23 | 0.22 | 0.18 | 0.12 |
| 11 | 35568275 | rs17294960 | -1.049 | 2.306 | 1.234 | 2.678 | 0.026 | 0.006 | 0.006 | 0 | 0.09 | 0.12 | 0.16 | 0.07 | 0.15 | 0.12 |
| 11 | 35568911 | rs11603781 | -0.324 | 2.749 | 0.955 | 2.891 | -0.03 | -0.029 | -0.029 | 0 | 0.05 | 0.01 | 0.11 | 0.11 | 0.13 | 0.07 |
| 11 | 35570378 | rs12577501 | -1.807 | 2.513 | 0.681 | 3.06 | -0.006 | -0.08 | -0.08 | 0 | 0.25 | 0.02 | 0.13 | 0.31 | 0.2 | 0.19 |
| 11 | 35571914 | rs10128679 | -1.205 | 2.646 | 1.244 | 3.234 | 0.065 | 0.137 | 0.137 | 0 | 0.07 | 0.21 | 0.19 | 0.29 | 0.16 | 0.08 |
| 11 | 35573109 | rs12290869 | -1.121 | 2.202 | 1.25 | 3.243 | 0.053 | 0.139 | 0.139 | 0 | 0.07 | 0.21 | 0.17 | 0.29 | 0.14 | 0.07 |
| 11 | 35574050 | rs12365267 | -0.873 | 2.214 | 1.911 | 3.15 | 0.07 | 0.021 | 0.021 | 0.227 | 0.1 | 0.24 | 0.17 | 0.07 | 0.16 | 0.13 |
| 11 | 35575775 | rs12361794 | -0.596 | 2.104 | 1.681 | 3.168 | 0.003 | 0.014 | 0.014 | 0 | 0.12 | 0.01 | 0.07 | 0.08 | 0.1 | 0.1 |
| 11 | 35575974 | rs10836416 | -2.968 | 2.101 | 1.988 | 3.151 | -0.005 | 0.015 | 0.015 | 0.006 | 0.44 | 0.61 | 0.49 | 0.67 | 0.49 | 0.39 |
| 11 | 35576921 | rs11033178 | -1.998 | 2.073 | 1.742 | 3.206 | -0.07 | -0.118 | -0.118 | 0 | 0.3 | 0.04 | 0.24 | 0.42 | 0.33 | 0.25 |
| 11 | 35579889 | rs3858472 | -2.863 | 2.131 | 1.869 | 3.038 | -0.001 | -0.001 | -0.001 | 0 | 0.43 | 0.57 | 0.48 | 0.67 | 0.51 | 0.4 |
| 11 | 35580703 | rs10836419 | -1.554 | 2.17 | 1.186 | 3.198 | 0.033 | 0.111 | 0.111 | 0.017 | 0.11 | 0.19 | 0.19 | 0.41 | 0.19 | 0.13 |
| 11 | 35582259 | rs10836424 | -1.222 | 2.147 | 1.025 | 3.144 | -0.007 | -0.039 | -0.039 | 0.128 | 0.2 | 0.01 | 0.11 | 0.18 | 0.16 | 0.11 |
| 11 | 35586059 | rs3911014 | -2.725 | 2.141 | 1.155 | 3.13 | -0.02 | 0.032 | 0.032 | 0.207 | 0.42 | 0.26 | 0.46 | 0.67 | 0.51 | 0.4 |
| 11 | 35588932 | rs3858475 | -2.083 | 1.811 | 0.923 | 2.731 | -0.021 | -0.011 | -0.011 | 0 | 0.31 | 0.14 | 0.27 | 0.26 | 0.31 | 0.27 |
| 11 | 35593491 | rs3980449 | -1.724 | 1.895 | 0.891 | 2.621 | -0.041 | -0.044 | -0.044 | 0 | 0.15 | 0.06 | 0.19 | 0.26 | 0.2 | 0.24 |
| 11 | 35596889 | rs11033190 | -0.955 | 1.855 | 1.315 | 2.631 | -0.058 | -0.05 | -0.05 | 0 | 0.1 | 0.01 | 0.18 | 0.12 | 0.16 | 0.16 |
| 11 | 35597154 | rs659401 | -2.05 | 1.526 | 1.842 | 2.57 | -0.049 | 0.051 | 0.051 | 0 | 0.26 | 0.51 | 0.17 | 0.46 | 0.16 | 0.16 |
| 11 | 35602337 | rs10768154 | -1.722 | 1.541 | 1.637 | 2.479 | 0.022 | -0.017 | -0.017 | 0 | 0.22 | 0.05 | 0.09 | 0.29 | 0.16 | 0.16 |
| 11 | 35608275 | rs678679 | -1.733 | 1.551 | 1.937 | 2.21 | -0.049 | 0.07 | 0.07 | 0 | 0.18 | 0.38 | 0.14 | 0.37 | 0.12 | 0.16 |
| 11 | 35611601 | rs1447007 | -2.065 | 1.905 | 1.479 | 2.362 | -0.138 | -0.073 | -0.073 | 0 | 0.23 | 0.02 | 0.32 | 0.2 | 0.35 | 0.23 |
| 11 | 35623147 | rs506418 | -2.839 | 1.76 | 1.85 | 2.304 | 0.006 | 0.009 | 0.009 | 0 | 0.4 | 0.48 | 0.45 | 0.56 | 0.45 | 0.42 |
| 11 | 35623483 | rs12806627 | -0.424 | 1.831 | 1.804 | 2.337 | -0.033 | -0.024 | -0.024 | 0 | 0.07 | 0.02 | 0.12 | 0.08 | 0.16 | 0.08 |
| 11 | 35624868 | rs645052 | -2.656 | 1.813 | 1.887 | 2.379 | 0.003 | 0.004 | 0.004 | 0 | 0.37 | 0.41 | 0.44 | 0.44 | 0.44 | 0.36 |
| 11 | 35625278 | rs697320 | -2.483 | 1.535 | 1.912 | 1.995 | -0.013 | 0.034 | 0.034 | 0 | 0.33 | 0.49 | 0.33 | 0.48 | 0.29 | 0.34 |
| 11 | 35629494 | rs579687 | -1.191 | 1.578 | 1.761 | 1.759 | -0.05 | -0.02 | -0.02 | 0 | 0.15 | 0.03 | 0.19 | 0.12 | 0.19 | 0.15 |
| 11 | 35631174 | rs697321 | -2.133 | 1.573 | 1.973 | 1.849 | -0.013 | 0.03 | 0.03 | 0 | 0.32 | 0.51 | 0.32 | 0.45 | 0.28 | 0.31 |
| 11 | 35635460 | rs697325 | -2.152 | 1.528 | 1.533 | 1.833 | -0.008 | 0.01 | 0.01 | 0 | 0.33 | 0.65 | 0.31 | 0.45 | 0.27 | 0.3 |
| 11 | 35635800 | rs697316 | -2.019 | 1.771 | 0.999 | 1.656 | 0.018 | -0.014 | -0.014 | 0 | 0.35 | 0.74 | 0.3 | 0.46 | 0.29 | 0.24 |
| 11 | 35638772 | rs697315 | -1.481 | 2.173 | 1.147 | 1.507 | 0.033 | 0.002 | 0.002 | 0.084 | 0.26 | 0.63 | 0.32 | 0.29 | 0.36 | 0.2 |
| 11 | 35644794 | rs521081 | -0.831 | 1.951 | 1.073 | 1.24 | 0.048 | -0.024 | -0.024 | 0 | 0.08 | 0.23 | 0.1 | 0.02 | 0.11 | 0.06 |
| 11 | 35650073 | rs10836429 | -1.5 | 1.842 | 0.619 | 1.645 | -0.051 | -0.038 | -0.038 | 0 | 0.38 | 0.17 | 0.57 | 0.53 | 0.47 | 0.37 |
| 11 | 35657763 | rs16927798 | -0.995 | 1.939 | 1.136 | 1.399 | 0.019 | -0.015 | -0.015 | 0.021 | 0.14 | 0.27 | 0.12 | 0.06 | 0.17 | 0.07 |
| 11 | 35666886 | rs520850 | -1.318 | 1.409 | 1.543 | 1.573 | 0.011 | 0.111 | 0.111 | 0.19 | 0.06 | 0.34 | 0.12 | 0.13 | 0.01 | 0.02 |
| 11 | 35679996 | rs10466459 | -0.918 | 1.376 | 1.611 | 1.55 | -0.009 | -0.013 | -0.013 | 0 | 0.05 | 0.07 | 0.01 | 0 | 0.01 | 0 |
| 11 | 35683884 | rs4755422 | -1.622 | 1.348 | 2.112 | 1.543 | -0.028 | 0.036 | 0.036 | 0 | 0.06 | 0.18 | 0.1 | 0.13 | 0.01 | 0.02 |
| 11 | 35689490 | rs628673 | -1.212 | 1.348 | 2.171 | 1.543 | -0.018 | 0.056 | 0.056 | 0.01 | 0.05 | 0.19 | 0.1 | 0.13 | 0.01 | 0.02 |
| 11 | 35753511 | rs11033260 | -1.38 | 2.33 | 2.528 | 2.579 | 0.07 | 0.097 | 0.097 | 0 | 0.19 | 0.32 | 0.21 | 0.41 | 0.31 | 0.36 |
| 11 | 35754464 | rs7129170 | -0.173 | 2.509 | 2.336 | 2.584 | -0.056 | -0.015 | -0.015 | 0.023 | 0.12 | 0 | 0.11 | 0.09 | 0.16 | 0.08 |
| 11 | 35759888 | rs1323856 | -0.779 | 2.721 | 2.469 | 2.914 | 0.083 | 0.054 | 0.054 | 0 | 0.07 | 0.22 | 0.08 | 0.07 | 0.11 | 0.06 |
| 11 | 35775813 | rs2985152 | 1.279 | 2.088 | 1.865 | 2.641 | -0.003 | -0.01 | -0.01 | 0 | 0.71 | 0.79 | 0.56 | 0.71 | 0.7 | 0.79 |
| 11 | 35802040 | rs2147094 | 0.121 | 2.073 | 1.885 | 2.631 | -0.001 | 0 | 0 | 0 | 0.85 | 0.82 | 0.66 | 0.78 | 0.87 | 0.87 |
| 11 | 35823818 | rs671783 | 1.985 | 2.195 | 1.893 | 2.74 | 0.006 | -0.005 | -0.005 | 1.42 | 0.6 | 0.34 | 0.41 | 0.56 | 0.65 | 0.71 |
| 11 | 35841160 | rs17735796 | -0.583 | 2.382 | 1.463 | 3.01 | -0.055 | -0.063 | -0.063 | 1.644 | 0.12 | 0.01 | 0.1 | 0.28 | 0.18 | 0.17 |

| | | | | | | | | | | | | | | | | |
|---|---|---|---|---|---|---|---|---|---|---|---|---|---|---|---|---|
| 11 | 35841576 | rs10501143 | -1.351 | 2.363 | 1.54 | 2.493 | 0 | 0.001 | 0.001 | 0 | 0.18 | 0.15 | 0.19 | 0.21 | 0.18 | 0.15 |
| 11 | 35842766 | rs12226613 | -1.331 | 2.35 | 1.535 | 2.304 | 0 | 0.022 | 0.022 | 0 | 0.18 | 0.15 | 0.1 | 0.09 | 0.18 | 0.14 |
| 11 | 35844930 | rs11033288 | -1.871 | 2.49 | 1.571 | 2.455 | 0.082 | 0.129 | 0.129 | 0 | 0.21 | 0.36 | 0.3 | 0.57 | 0.31 | 0.39 |
| 11 | 35849841 | rs11033289 | -1.689 | 2.202 | 0.865 | 1.972 | -0.008 | 0.051 | 0.051 | 0 | 0.19 | 0.1 | 0.12 | 0.08 | 0.22 | 0.18 |
| 11 | 35853091 | rs11033291 | -0.622 | 2.206 | 1.298 | 2.111 | 0.023 | 0.058 | 0.058 | 0 | 0.15 | 0.35 | 0.17 | 0.2 | 0.15 | 0.16 |
| 11 | 35854776 | rs2957939 | -1.333 | 2.21 | 1.39 | 2.4 | 0.005 | 0.105 | 0.105 | 0.073 | 0.11 | 0.21 | 0.05 | 0.31 | 0.1 | 0.1 |
| 11 | 35858686 | rs7937174 | -1.64 | 2.388 | 1.19 | 2.186 | 0.055 | 0.036 | 0.036 | 0 | 0.25 | 0.41 | 0.32 | 0.34 | 0.36 | 0.31 |
| 11 | 35859844 | rs623374 | 2.624 | 2.439 | 0.731 | 1.824 | -0.004 | 0.146 | 0.146 | 0.083 | 0.52 | 0.05 | 0.52 | 0.28 | 0.41 | 0.47 |
| 11 | 35861029 | rs1408154 | -0.748 | 2.634 | 0.845 | 1.874 | 0.038 | 0.009 | 0.009 | 0 | 0.13 | 0.18 | 0.16 | 0.16 | 0.21 | 0.18 |
| 11 | 35861618 | rs2956340 | 1.174 | 2.354 | 1.072 | 1.924 | 0.004 | -0.042 | -0.042 | 0 | 0.83 | 0.71 | 0.87 | 0.92 | 0.8 | 0.83 |
| 11 | 35862496 | rs666110 | 1.599 | 2.358 | 0.112 | 1.974 | 0 | -0.008 | -0.008 | 1.418 | 0.65 | 0.25 | 0.68 | 0.42 | 0.62 | 0.65 |
| 11 | 35865302 | rs11033300 | -1.296 | 2.326 | -0.825 | 1.813 | -0.02 | 0.031 | 0.031 | 0.013 | 0.15 | 0.05 | 0.1 | 0.08 | 0.21 | 0.17 |
| 11 | 35866085 | rs2956343 | -1.132 | 2.236 | -0.825 | 1.652 | -0.005 | 0.039 | 0.039 | 0 | 0.26 | 0.4 | 0.17 | 0.38 | 0.24 | 0.17 |
| 11 | 35868300 | rs676184 | -0.283 | 1.535 | -1.876 | 1.171 | 0.023 | -0.036 | -0.036 | 0 | 0.13 | 0 | 0.16 | 0.14 | 0.07 | 0.27 |
| 11 | 35869257 | rs6484799 | -1.141 | 2.283 | -1.48 | 1.513 | -0.002 | -0.002 | -0.002 | 0.019 | 0.27 | 0.32 | 0.22 | 0.54 | 0.29 | 0.3 |
| 11 | 35869732 | rs679270 | 1.792 | 1.91 | -1.523 | 1.317 | 0.027 | 0.178 | 0.178 | 0.111 | 0.48 | 0.03 | 0.5 | 0.23 | 0.38 | 0.43 |
| 11 | 35871007 | rs4756258 | -0.842 | 1.747 | -0.801 | 2.09 | -0.005 | 0.04 | 0.04 | 0.118 | 0.2 | 0.29 | 0.14 | 0.38 | 0.17 | 0.11 |
| 11 | 35871679 | rs10836458 | -1.314 | 2.022 | -1.015 | 1.832 | -0.025 | -0.024 | -0.024 | 0 | 0.1 | 0.1 | 0.15 | 0.23 | 0.23 | 0.24 |
| 11 | 35873934 | rs7106732 | -1.035 | 1.033 | -1.024 | 1.571 | 0.006 | 0.003 | 0.003 | 1.351 | 0.37 | 0.54 | 0.49 | 0.63 | 0.47 | 0.44 |
| 11 | 35874466 | rs11827345 | -0.763 | -0.365 | -1.539 | 2.217 | 0.036 | -0.077 | -0.077 | 1.24 | 0.24 | 0.12 | 0.18 | 0.43 | 0.16 | 0.29 |
| 11 | 35875537 | rs635828 | 1.481 | -0.197 | -1.445 | 2.119 | 0.022 | 0.046 | 0.046 | 0.01 | 0.54 | 0.16 | 0.62 | 0.34 | 0.59 | 0.4 |
| 11 | 35875576 | rs12226262 | -0.628 | -0.381 | -1.544 | 2.332 | -0.003 | -0.017 | -0.017 | 0 | 0.23 | 0.02 | 0.18 | 0.22 | 0.18 | 0.32 |
| 11 | 35879571 | rs671068 | 0.77 | -0.574 | -1.549 | 2.414 | -0.01 | -0.03 | -0.03 | 0 | 0.74 | 0.94 | 0.82 | 0.78 | 0.76 | 0.8 |
| 11 | 35879671 | rs671503 | 1.986 | -0.509 | -1.504 | 2.454 | 0.037 | 0.03 | 0.03 | 0.041 | 0.54 | 0.16 | 0.62 | 0.35 | 0.6 | 0.39 |
| 11 | 35879951 | rs672597 | 0.233 | -0.371 | -1.624 | 2.034 | -0.025 | -0.044 | -0.044 | 0.363 | 0.8 | 0.99 | 0.84 | 0.81 | 0.81 | 0.92 |
| 11 | 35886919 | rs684909 | 1.316 | -0.264 | -0.851 | 1.842 | 0.013 | 0.009 | 0.009 | 0 | 0.57 | 0.42 | 0.67 | 0.6 | 0.62 | 0.54 |
| 11 | 35893012 | rs1473844 | 1.178 | -0.274 | -0.873 | 2.065 | 0.047 | -0.033 | -0.033 | 0.013 | 0.61 | 0.24 | 0.7 | 0.53 | 0.71 | 0.5 |
| 11 | 35910376 | rs671658 | 0.853 | 0.115 | -0.377 | 2.024 | 0.014 | -0.011 | -0.011 | 1.052 | 0.73 | 0.21 | 0.75 | 0.75 | 0.79 | 0.84 |
| 11 | 35912362 | rs593542 | 0.307 | 0.049 | 0.73 | 2.04 | -0.006 | -0.024 | -0.024 | 0 | 0.8 | 0.5 | 0.79 | 0.9 | 0.83 | 0.91 |
| 11 | 35913571 | rs16928055 | -0.372 | -0.066 | 0.106 | 2.301 | -0.037 | -0.077 | -0.077 | 0.189 | 0.13 | 0.01 | 0.12 | 0.27 | 0.13 | 0.24 |
| 11 | 35917089 | rs7119112 | 1.125 | -0.533 | 0.249 | 2.111 | 0.037 | -0.053 | -0.053 | 0 | 0.68 | 0.14 | 0.66 | 0.63 | 0.77 | 0.79 |
| 11 | 35932164 | rs2048568 | 1.085 | -0.405 | 0.103 | 2.287 | 0.027 | -0.075 | -0.075 | 0.171 | 0.69 | 0.15 | 0.62 | 0.56 | 0.75 | 0.74 |
| 11 | 35941082 | rs12419834 | -1.791 | 0.13 | 1.226 | 2.653 | 0.034 | 0.175 | 0.175 | 0 | 0.17 | 0.5 | 0.27 | 0.38 | 0.17 | 0.14 |
| 11 | 35945048 | rs7932394 | -2.066 | 0.72 | 1.209 | 2.807 | 0.002 | 0.003 | 0.003 | 0 | 0.3 | 0.31 | 0.3 | 0.51 | 0.36 | 0.37 |
| 11 | 35947565 | rs2063581 | -2.039 | 0.709 | 1.27 | 2.815 | -0.012 | -0.014 | -0.014 | 0 | 0.29 | 0.29 | 0.3 | 0.51 | 0.35 | 0.37 |
| 11 | 35957322 | rs11033335 | -1.241 | 0.738 | 1.016 | 3.059 | -0.034 | -0.034 | -0.034 | 0 | 0.16 | 0.13 | 0.24 | 0.39 | 0.29 | 0.31 |
| 11 | 35968743 | rs10219322 | -1.779 | 0.759 | 0.873 | 2.932 | 0.026 | 0.063 | 0.063 | 0.802 | 0.13 | 0.22 | 0.22 | 0.44 | 0.23 | 0.33 |
| 11 | 35971750 | rs263096 | -1.849 | -0.448 | 0.846 | 2.619 | -0.015 | -0.001 | -0.001 | 0 | 0.21 | 0.73 | 0.18 | 0.21 | 0.11 | 0.18 |
| 11 | 35977163 | rs16928142 | -1.346 | -1.013 | 0.383 | 2.916 | -0.017 | -0.116 | -0.116 | 1.149 | 0.16 | 0 | 0.19 | 0.4 | 0.16 | 0.26 |
| 11 | 35980394 | rs263088 | -0.905 | -1.277 | 1.415 | 2.55 | -0.036 | 0.048 | 0.048 | 0.11 | 0.1 | 0.29 | 0.04 | 0.16 | 0.03 | 0.18 |
| 11 | 35983221 | rs11033346 | -0.942 | -1.936 | 1.049 | 2.208 | 0.01 | -0.011 | -0.011 | 0 | 0.37 | 0.24 | 0.45 | 0.6 | 0.35 | 0.47 |
| 11 | 35985584 | rs2984699 | -2.118 | -2.149 | 1.38 | 2.057 | 0.046 | 0.13 | 0.13 | 0 | 0.52 | 0.84 | 0.52 | 0.77 | 0.4 | 0.66 |
| 11 | 35986927 | rs2984701 | -1.841 | -2.159 | 1.468 | 2.067 | 0.083 | 0.091 | 0.091 | 0 | 0.5 | 0.81 | 0.51 | 0.67 | 0.38 | 0.57 |
| 11 | 35990164 | rs10768172 | -2.073 | -2.374 | 1.2 | 1.998 | 0.037 | 0.087 | 0.087 | 0.059 | 0.47 | 0.9 | 0.51 | 0.71 | 0.39 | 0.66 |
| 11 | 35991050 | rs2901651 | -1.14 | -2.382 | 1.643 | 2.002 | -0.001 | -0.001 | -0.001 | 0 | 0.34 | 0.36 | 0.46 | 0.64 | 0.37 | 0.56 |
| 11 | 35991881 | rs7119978 | -2.146 | -2.098 | 1.172 | 2.035 | 0.037 | 0.091 | 0.091 | 0.01 | 0.52 | 0.9 | 0.51 | 0.71 | 0.4 | 0.66 |
| 11 | 36001396 | rs4755432 | -1.214 | -1.984 | 1.145 | 2.049 | 0.001 | 0.034 | 0.034 | 0 | 0.39 | 0.24 | 0.46 | 0.7 | 0.39 | 0.56 |
| 11 | 36013229 | rs2146359 | -0.712 | -2.235 | 0.548 | 1.897 | 0.004 | -0.069 | -0.069 | 0 | 0.36 | 0.13 | 0.45 | 0.53 | 0.35 | 0.5 |
| 11 | 36013671 | rs982859 | -0.848 | -2.398 | 0.676 | 1.829 | 0.017 | -0.034 | -0.034 | 0.138 | 0.39 | 0.28 | 0.43 | 0.49 | 0.36 | 0.58 |
| 11 | 36020797 | rs12274406 | -2.003 | -2.521 | 0.392 | 1.229 | 0.074 | -0.047 | -0.047 | 0.014 | 0.53 | 0.92 | 0.39 | 0.45 | 0.37 | 0.67 |
| 11 | 36025118 | rs262418 | 2.568 | -1.983 | 0.928 | 1.548 | -0.013 | 0.058 | 0.058 | 1.947 | 0.85 | 0.32 | 0.8 | 0.73 | 0.92 | 0.81 |
| 11 | 36028475 | rs1322737 | -0.495 | -1.938 | 0.953 | 1.309 | 0.017 | -0.01 | -0.01 | 0.562 | 0.36 | 0.25 | 0.3 | 0.37 | 0.33 | 0.55 |
| 11 | 36029375 | rs262407 | 1.127 | -1.919 | 0.814 | 1.297 | 0.071 | 0.025 | 0.025 | 0.207 | 0.94 | 0.68 | 0.85 | 0.98 | 0.96 | 0.96 |
| 11 | 36029527 | rs11033374 | -1.218 | -1.74 | 0.571 | 1.404 | -0.007 | 0.002 | 0.002 | 0.012 | 0.44 | 0.53 | 0.45 | 0.52 | 0.37 | 0.66 |
| 11 | 36033069 | rs7947746 | -0.992 | -1.652 | 0.323 | 1.198 | -0.008 | 0.001 | 0.001 | 0 | 0.4 | 0.43 | 0.44 | 0.41 | 0.36 | 0.59 |
| 11 | 36041260 | rs262426 | 1.015 | -1.655 | -0.177 | 0.969 | 0.023 | 0.009 | 0.009 | 1.138 | 0.45 | 0.19 | 0.61 | 0.37 | 0.55 | 0.37 |
| 11 | 36041753 | rs1016215 | -0.528 | -1.878 | 0.535 | 0.982 | 0.013 | 0.003 | 0.003 | 0 | 0.37 | 0.24 | 0.3 | 0.4 | 0.37 | 0.39 |
| 11 | 36044106 | rs7927259 | -1 | -1.886 | 0.658 | 0.977 | -0.02 | 0.011 | 0.011 | 0 | 0.12 | 0.21 | 0.07 | 0.11 | 0.07 | 0.16 |
| 11 | 36048840 | rs995328 | -0.869 | -1.884 | 0.611 | 0.975 | -0.006 | 0.002 | 0.002 | 0 | 0.11 | 0.14 | 0.07 | 0.11 | 0.07 | 0.16 |
| 11 | 36052212 | rs262440 | 1.116 | -1.898 | 0.644 | 0.919 | 0.049 | 0.004 | 0.004 | 0.173 | 0.5 | 0.34 | 0.62 | 0.47 | 0.57 | 0.43 |
| 11 | 36054985 | rs374544 | 0.826 | -2.018 | 0.623 | 0.56 | 0.057 | -0.015 | -0.015 | 0 | 0.52 | 0.29 | 0.64 | 0.45 | 0.58 | 0.38 |
| 11 | 36056922 | rs262453 | 0.289 | -2.106 | 1.05 | 0.392 | -0.016 | -0.07 | -0.07 | 0.028 | 0.8 | 0.5 | 0.86 | 0.92 | 0.81 | 0.85 |
| 11 | 36059983 | rs262450 | -0.033 | -2.026 | 0.911 | 0.486 | 0.003 | 0.001 | 0.001 | 0 | 0.81 | 0.71 | 0.83 | 0.83 | 0.77 | 0.76 |
| 11 | 36062969 | rs882366 | -0.583 | -2.091 | 1.162 | 0.635 | -0.037 | -0.003 | -0.003 | 0 | 0.27 | 0.38 | 0.11 | 0.29 | 0.18 | 0.29 |
| 11 | 36066656 | rs11033386 | -0.224 | -1.417 | 0.684 | 0.803 | -0.005 | -0.031 | -0.031 | 0.88 | 0.1 | 0.07 | 0.15 | 0.22 | 0.08 | 0.22 |
| 11 | 36068766 | rs2173554 | 0.448 | -0.595 | 0.55 | 0.513 | -0.009 | -0.007 | -0.007 | 1.059 | 0.1 | 0.07 | 0.34 | 0.13 | 0.13 | 0.12 |
| 11 | 36069793 | rs1499514 | 0.348 | -0.664 | 0.614 | 0.6 | -0.003 | -0.001 | -0.001 | 0.724 | 0.17 | 0.22 | 0.06 | 0.24 | 0.15 | 0.24 |
| 11 | 36071595 | rs7122453 | 0.036 | -0.67 | 0.556 | 0.706 | 0 | 0.015 | 0.015 | 0 | 0.19 | 0.22 | 0.23 | 0.47 | 0.16 | 0.34 |
| 11 | 36073654 | rs1499512 | 1.563 | -0.83 | 0.437 | 0.644 | 0.01 | -0.032 | -0.032 | 0 | 0.57 | 0.26 | 0.51 | 0.41 | 0.65 | 0.47 |

| | | | | | | | | | | | | | | | | |
|---|---|---|---|---|---|---|---|---|---|---|---|---|---|---|---|---|
| 11 | 36077902 | rs13377557 | 0.475 | -0.613 | 0.536 | 0.747 | 0.008 | -0.019 | -0.019 | 0.061 | 0.09 | 0.04 | 0.06 | 0.12 | 0.06 | 0.14 |
| 11 | 36078171 | rs4756265 | 1.772 | -0.439 | 0.41 | 0.519 | -0.029 | -0.023 | -0.023 | 0 | 0.39 | 0.16 | 0.37 | 0.36 | 0.43 | 0.33 |
| 11 | 36078941 | rs11033397 | 0.617 | -0.395 | -0.128 | 0.55 | 0.004 | 0.011 | 0.011 | 0 | 0.21 | 0.18 | 0.15 | 0.17 | 0.23 | 0.16 |
| 11 | 36079538 | rs904559 | -1.323 | -0.355 | -0.028 | 0.415 | -0.001 | 0.007 | 0.007 | 0 | 0.49 | 0.59 | 0.55 | 0.52 | 0.5 | 0.53 |
| 11 | 36081144 | rs4265577 | 0.356 | -0.028 | -0.808 | 0.5 | 0.015 | 0.033 | 0.033 | 0 | 0.29 | 0.18 | 0.2 | 0.18 | 0.29 | 0.3 |
| 11 | 36087853 | rs1499511 | -1.356 | 0.212 | -1.317 | 0.388 | -0.045 | -0.04 | -0.04 | 0 | 0.62 | 0.87 | 0.64 | 0.66 | 0.57 | 0.67 |
| 11 | 36088509 | rs1499509 | -0.597 | 0.633 | -0.601 | 0.302 | -0.003 | 0.001 | 0.001 | 0 | 0.17 | 0.21 | 0.06 | 0.31 | 0.14 | 0.22 |
| 11 | 36090420 | rs17799503 | -0.702 | 0.904 | -0.857 | 0.122 | -0.047 | 0.049 | 0.049 | 3.37 | 0.1 | 0.01 | 0.18 | 0 | 0.12 | 0.06 |
| 11 | 36090919 | rs7952591 | -1.078 | 0.7 | -0.573 | 0.064 | 0.005 | -0.058 | -0.058 | 0.581 | 0.15 | 0.22 | 0.24 | 0 | 0.17 | 0.13 |
| 11 | 36091888 | rs2173553 | 0.376 | 0.836 | -1.262 | 0.779 | -0.034 | -0.074 | -0.074 | 0.628 | 0.79 | 0.94 | 0.79 | 0.6 | 0.74 | 0.76 |
| 11 | 36092555 | rs4545528 | -0.898 | 0.41 | -0.378 | 1.079 | -0.005 | 0.076 | 0.076 | 0 | 0.18 | 0.29 | 0.18 | 0.44 | 0.18 | 0.3 |
| 11 | 36093636 | rs7126914 | -0.292 | 0.439 | -0.425 | 1.141 | -0.015 | -0.046 | -0.046 | 0 | 0.14 | 0.05 | 0.17 | 0.37 | 0.17 | 0.2 |
| 11 | 36093840 | rs1392067 | -0.267 | 0.439 | -0.425 | 1.134 | -0.015 | -0.045 | -0.045 | 0 | 0.14 | 0.05 | 0.17 | 0.36 | 0.17 | 0.2 |
| 11 | 36093992 | rs1392066 | -0.725 | -0.218 | -0.772 | -0.122 | 0.011 | 0.001 | 0.001 | 0 | 0.75 | 0.76 | 0.8 | 0.97 | 0.86 | 0.78 |
| 11 | 36096165 | rs904551 | 0.442 | -0.491 | -1.61 | -0.206 | 0.037 | -0.073 | -0.073 | 0 | 0.29 | 0.07 | 0.23 | 0.34 | 0.24 | 0.17 |
| 11 | 36096312 | rs10430962 | 0.841 | -0.318 | -1.465 | -0.108 | -0.011 | -0.044 | -0.044 | 0 | 0.14 | 0.06 | 0.16 | 0.36 | 0.17 | 0.21 |
| 11 | 36102894 | rs7928546 | 1.292 | -0.792 | -0.728 | -0.545 | -0.008 | -0.04 | -0.04 | 0 | 0.2 | 0.2 | 0.07 | 0.01 | 0.1 | 0.19 |
| 11 | 36103228 | rs1138807 | 0.008 | -1.157 | -0.363 | -0.755 | 0.006 | 0 | 0 | 0 | 0.4 | 0.41 | 0.4 | 0.35 | 0.32 | 0.22 |
| 11 | 36103381 | rs3740962 | -0.221 | -0.922 | -0.232 | -0.571 | 0.003 | -0.014 | -0.014 | 0 | 0.39 | 0.17 | 0.52 | 0.63 | 0.61 | 0.6 |
| 11 | 36104536 | rs10836488 | 0.514 | -0.238 | 0.147 | -0.264 | 0.011 | 0.047 | 0.047 | 0 | 0.15 | 0.17 | 0.17 | 0.37 | 0.19 | 0.18 |
| 11 | 36108497 | rs2030834 | -0.083 | -0.555 | 0.012 | -0.444 | -0.003 | 0.001 | 0.001 | 0 | 0.7 | 0.73 | 0.76 | 0.62 | 0.69 | 0.72 |
| 11 | 36111616 | rs1917899 | 0.116 | -0.238 | -0.603 | -0.702 | 0.027 | -0.064 | -0.064 | 0 | 0.29 | 0.48 | 0.26 | 0.08 | 0.34 | 0.16 |
| 11 | 36111766 | rs1917900 | -0.187 | -0.586 | -1.196 | -0.298 | 0.011 | -0.017 | -0.017 | 0.38 | 0.73 | 0.88 | 0.79 | 0.63 | 0.73 | 0.74 |
| 11 | 36113666 | rs4756270 | -0.355 | -0.17 | -1.332 | 0 | 0.018 | 0.011 | 0.011 | 0 | 0.46 | 0.72 | 0.56 | 0.41 | 0.51 | 0.48 |
| 11 | 36114157 | rs16928390 | -1.069 | -0.325 | -1.547 | 0.225 | 0.024 | 0.039 | 0.039 | 0 | 0.28 | 0.15 | 0.32 | 0.21 | 0.23 | 0.29 |
| 11 | 36115705 | rs11033417 | -0.59 | 0.242 | -1.251 | 0.569 | -0.003 | 0.033 | 0.033 | 0 | 0.24 | 0.08 | 0.31 | 0.16 | 0.24 | 0.29 |
| 11 | 36117340 | rs10836489 | -0.982 | 0.245 | -0.369 | 1.209 | -0.009 | 0.056 | 0.056 | 0 | 0.15 | 0.32 | 0.1 | 0.32 | 0.13 | 0.22 |
| 11 | 36121110 | rs4756275 | -0.565 | 0.28 | -0.438 | 1.056 | -0.005 | 0.014 | 0.014 | 1.188 | 0.32 | 0.45 | 0.4 | 0.47 | 0.28 | 0.3 |
| 11 | 36124199 | rs1961758 | 0.043 | 0.627 | -0.77 | 0.516 | -0.011 | 0.018 | 0.018 | 0 | 0.07 | 0.03 | 0.06 | 0.01 | 0.09 | 0.04 |
| 11 | 36125711 | rs11033427 | 0.983 | 0.899 | -1.684 | 0.396 | -0.071 | -0.087 | -0.087 | 0.635 | 0.21 | 0.01 | 0.15 | 0.21 | 0.23 | 0.28 |
| 11 | 36125835 | rs10768177 | -0.541 | 0.286 | -0.799 | -0.071 | -0.026 | -0.073 | -0.073 | 1.468 | 0.88 | 0.66 | 0.93 | 1 | 0.95 | 0.94 |
| 11 | 36126923 | rs7928494 | 0.578 | -0.156 | -0.36 | 0.141 | -0.005 | 0.001 | 0.001 | 0 | 0.4 | 0.29 | 0.43 | 0.73 | 0.43 | 0.49 |
| 11 | 36132273 | rs7104875 | -0.041 | -0.094 | -0.201 | 0.071 | -0.001 | 0.003 | 0.003 | 3.784 | 0.26 | 0.25 | 0.33 | 0.17 | 0.25 | 0.23 |
| 11 | 36134670 | rs4756281 | 0.998 | 0.48 | -0.256 | 0.732 | -0.056 | -0.072 | -0.072 | 0.005 | 0.21 | 0.06 | 0.19 | 0.27 | 0.26 | 0.25 |
| 11 | 36136580 | rs4755435 | 0.169 | 0.5 | -0.049 | 0.706 | 0.016 | 0.076 | 0.076 | 0 | 0.54 | 0.28 | 0.57 | 0.31 | 0.61 | 0.53 |
| 11 | 36137453 | rs12276510 | 0.102 | 0.179 | -0.35 | 0.64 | -0.018 | 0.126 | 0.126 | 0 | 0.1 | 0.19 | 0.08 | 0.33 | 0.07 | 0.1 |
| 11 | 36139934 | rs10501144 | -0.275 | 0.46 | -0.286 | 0.547 | 0.027 | 0.012 | 0.012 | 0 | 0.58 | 0.27 | 0.72 | 0.63 | 0.7 | 0.62 |
| 11 | 36141825 | rs7107244 | -0.139 | 0.622 | -0.458 | 0.345 | -0.033 | -0.02 | -0.02 | 0 | 0.41 | 0.24 | 0.52 | 0.43 | 0.48 | 0.45 |
| 11 | 36142727 | rs10768183 | -0.179 | 0.628 | -0.505 | 0.342 | -0.01 | -0.004 | -0.004 | 0 | 0.41 | 0.32 | 0.53 | 0.44 | 0.48 | 0.45 |
| 11 | 36145076 | rs10501145 | -0.079 | 0.765 | -0.125 | 0.129 | 0 | -0.005 | -0.005 | 0 | 0.46 | 0.45 | 0.54 | 0.67 | 0.48 | 0.5 |
| 11 | 36147644 | rs7117000 | 0.444 | 1.025 | 0.355 | 0.243 | 0.01 | 0.053 | 0.053 | 0 | 0.14 | 0.22 | 0.19 | 0.24 | 0.18 | 0.18 |
| 11 | 36150591 | rs11606633 | 0.831 | 1.17 | -0.035 | -0.141 | -0.029 | 0.105 | 0.105 | 0 | 0.24 | 0.05 | 0.17 | 0.03 | 0.22 | 0.18 |
| 11 | 36153721 | rs4450151 | -1.578 | 1.497 | 0.503 | -0.331 | 0.023 | -0.054 | -0.054 | 0 | 0.86 | 0.78 | 0.69 | 1 | 0.8 | 0.84 |
| 11 | 36154375 | rs11033448 | 1.764 | 1.393 | 0.166 | 0.036 | 0.001 | -0.089 | -0.089 | 1.293 | 0.26 | 0.07 | 0.17 | 0.36 | 0.23 | 0.23 |
| 11 | 36155578 | rs2422178 | -0.292 | 1.536 | 0.457 | 0.028 | -0.006 | -0.006 | -0.006 | 0 | 0.44 | 0.48 | 0.31 | 0.39 | 0.34 | 0.39 |
| 11 | 36155717 | rs12417545 | 1.805 | 1.51 | 0.385 | 0 | -0.051 | -0.095 | -0.095 | 2.091 | 0.25 | 0.07 | 0.19 | 0.37 | 0.28 | 0.24 |
| 11 | 36155730 | rs16928442 | 1.189 | 1.398 | 0.685 | -0.102 | 0.005 | -0.047 | -0.047 | 0 | 0.09 | 0.2 | 0.07 | 0 | 0.08 | 0.02 |
| 11 | 36155789 | rs11033450 | -0.184 | 1.378 | 0.564 | 0.08 | 0.001 | 0 | 0 | 0 | 0.42 | 0.43 | 0.3 | 0.43 | 0.34 | 0.39 |
| 11 | 36156051 | rs11033452 | -0.56 | 1.404 | 0.563 | 0.125 | -0.027 | 0.069 | 0.069 | 0.34 | 0.21 | 0.44 | 0.16 | 0.35 | 0.17 | 0.21 |
| 11 | 36156124 | rs11033453 | -1.132 | 1.494 | 0.368 | 0.203 | -0.034 | 0.029 | 0.029 | 0 | 0.59 | 0.74 | 0.5 | 0.65 | 0.51 | 0.58 |
| 11 | 36158084 | rs11828868 | 1.057 | 1.279 | 0.433 | 0.354 | 0.001 | 0.013 | 0.013 | 0 | 0.14 | 0.11 | 0.2 | 0.29 | 0.21 | 0.18 |
| 11 | 36158694 | rs867718 | -1.196 | 1.373 | 0.543 | 0.146 | 0.022 | 0.07 | 0.07 | 1.235 | 0.51 | 0.75 | 0.66 | 0.65 | 0.58 | 0.58 |
| 11 | 36159947 | rs9645614 | -1.204 | 1.302 | 0.637 | 0.159 | 0 | -0.028 | -0.028 | 2.997 | 0.74 | 0.75 | 0.82 | 0.94 | 0.79 | 0.81 |
| 11 | 36161425 | rs6484818 | -0.451 | 1.277 | 0.856 | 0.392 | -0.002 | 0.051 | 0.051 | 0 | 0.26 | 0.43 | 0.31 | 0.59 | 0.29 | 0.37 |
| 11 | 36161604 | rs6484820 | -1.195 | 1.487 | 0.707 | 0.332 | -0.001 | 0.003 | 0.003 | 0 | 0.38 | 0.71 | 0.39 | 0.63 | 0.42 | 0.41 |
| 11 | 36162728 | rs10836505 | -0.651 | 1.488 | 1.165 | 0.718 | -0.001 | 0.011 | 0.011 | 0 | 0.26 | 0.32 | 0.3 | 0.59 | 0.29 | 0.37 |
| 11 | 36162848 | rs11033457 | 0.674 | 1.435 | 0.742 | 0.901 | -0.001 | -0.049 | -0.049 | 0 | 0.08 | 0 | 0.08 | 0.28 | 0.08 | 0.16 |
| 11 | 36169869 | rs10768195 | -0.038 | 1.414 | 0.834 | 0.589 | 0.038 | 0.008 | 0.008 | 0 | 0.49 | 0.71 | 0.37 | 0.62 | 0.43 | 0.5 |
| 11 | 36173252 | rs955249 | -0.272 | 1.325 | 0.822 | 0.371 | -0.002 | -0.016 | -0.016 | 0 | 0.61 | 0.61 | 0.55 | 0.93 | 0.65 | 0.68 |
| 11 | 36175645 | rs2422179 | 0.168 | 1.413 | 0.356 | 0.526 | 0.012 | 0.037 | 0.037 | 0 | 0.53 | 0.39 | 0.46 | 0.64 | 0.6 | 0.53 |
| 11 | 36176258 | rs7111826 | -0.122 | 1.432 | 0.089 | 0.481 | 0.016 | 0.018 | 0.018 | 1.779 | 0.23 | 0.41 | 0.39 | 0.29 | 0.25 | 0.29 |
| 11 | 36177369 | rs10836509 | 0.983 | 1.607 | -0.259 | 0.567 | -0.02 | -0.044 | -0.044 | 0 | 0.15 | 0.07 | 0.21 | 0.29 | 0.22 | 0.2 |
| 11 | 36177551 | rs904556 | -0.225 | 1.573 | -0.249 | 0.512 | 0.03 | 0.157 | 0.157 | 0 | 0.59 | 0.21 | 0.54 | 0.92 | 0.71 | 0.69 |
| 11 | 36177734 | rs2422180 | 1.222 | 1.566 | -0.256 | 0.521 | -0.038 | -0.004 | -0.004 | 0 | 0.34 | 0.14 | 0.23 | 0.35 | 0.38 | 0.32 |
| 11 | 36178090 | rs12576603 | 0.571 | 1.513 | -0.12 | 0.453 | -0.022 | -0.057 | -0.057 | 2.515 | 0.09 | 0 | 0.1 | 0.28 | 0.1 | 0.15 |
| 11 | 36178765 | rs4756288 | 1.065 | 1.365 | -0.037 | 0.452 | 0.031 | -0.091 | -0.091 | 0 | 0.15 | 0.4 | 0.29 | 0.01 | 0.15 | 0.13 |
| 11 | 36178902 | rs7924470 | 0.44 | 1.352 | -1.211 | 0.519 | -0.02 | 0.075 | 0.075 | 0.034 | 0.49 | 0.21 | 0.43 | 0.64 | 0.55 | 0.52 |
| 11 | 36179252 | rs2133522 | -0.803 | 0.473 | -0.969 | 0.209 | 0.014 | 0.033 | 0.033 | 0 | 0.77 | 0.89 | 0.86 | 0.93 | 0.86 | 0.82 |
| 11 | 36180755 | rs11033470 | -0.024 | 0.871 | -0.952 | 0.761 | -0.073 | -0.025 | -0.025 | 0 | 0.42 | 0.15 | 0.33 | 0.65 | 0.5 | 0.47 |

| | | | | | | | | | | | | | | | | |
|---|---|---|---|---|---|---|---|---|---|---|---|---|---|---|---|---|
| 11 | 36182280 | rs7951966 | 0.859 | 0.418 | 0.197 | 1.12 | -0.008 | 0.031 | 0.031 | 0.835 | 0.75 | 0.41 | 0.63 | 0.64 | 0.77 | 0.76 |
| 11 | 36183297 | rs12269916 | -0.997 | 0.414 | 0.949 | 1.178 | -0.013 | 0.162 | 0.162 | 0 | 0.12 | 0.33 | 0.17 | 0.32 | 0.08 | 0.12 |
| 11 | 36187878 | rs4755439 | 0.384 | 0.364 | 1.037 | 1.141 | 0.016 | 0.135 | 0.135 | 0.555 | 0.81 | 0.53 | 0.67 | 0.64 | 0.8 | 0.81 |
| 11 | 36199843 | rs7121939 | 1.115 | 0.296 | 1.16 | 1.256 | 0.002 | 0.245 | 0.245 | 0.025 | 0.88 | 0.48 | 0.8 | 0.67 | 0.91 | 0.87 |
| 11 | 36201710 | rs16928480 | -0.342 | 0.599 | 1.051 | 0.813 | 0.026 | 0.018 | 0.018 | 0 | 0.06 | 0.14 | 0.06 | 0.03 | 0.06 | 0.09 |
| 11 | 36202307 | rs4755440 | 0.42 | 0.469 | 1.449 | 0.787 | 0.001 | 0.003 | 0.003 | 0.097 | 0.79 | 0.65 | 0.64 | 0.76 | 0.76 | 0.82 |
| 11 | 36203519 | rs10501146 | 0.691 | 0.519 | 1.069 | 0.554 | -0.001 | 0.001 | 0.001 | 0 | 0.3 | 0.29 | 0.24 | 0.34 | 0.26 | 0.25 |
| 11 | 36212439 | rs11607726 | 0.644 | 0.487 | 0.951 | 0.542 | -0.015 | -0.015 | -0.015 | 0 | 0.48 | 0.19 | 0.38 | 0.42 | 0.49 | 0.53 |
| 11 | 36215572 | rs1526054 | 0.1 | 0.368 | 1.027 | 0.351 | 0.001 | 0.004 | 0.004 | 0 | 0.36 | 0.41 | 0.39 | 0.52 | 0.36 | 0.35 |
| 11 | 36215938 | rs12574525 | -0.195 | 0.409 | 1.006 | 0.37 | 0 | 0.014 | 0.014 | 0 | 0.07 | 0.08 | 0.15 | 0.21 | 0.11 | 0.12 |
| 11 | 36221890 | rs11601619 | -0.36 | 0.373 | 0.969 | 0.372 | -0.001 | -0.007 | -0.007 | 0 | 0.07 | 0.05 | 0.15 | 0.22 | 0.1 | 0.06 |
| 11 | 36222477 | rs11033492 | 0.523 | 0.332 | 0.984 | 0.342 | 0 | -0.005 | -0.005 | 0.044 | 0.3 | 0.35 | 0.25 | 0.3 | 0.26 | 0.29 |
| 11 | 36226559 | rs871429 | 0.036 | 0.356 | 0.906 | 0.355 | 0.001 | 0.005 | 0.005 | 0 | 0.36 | 0.41 | 0.4 | 0.52 | 0.37 | 0.35 |
| 11 | 36233751 | rs1374376 | -0.19 | 0.432 | 0.654 | 0.374 | 0.005 | 0.002 | 0.002 | 0.606 | 0.2 | 0.32 | 0.37 | 0.23 | 0.23 | 0.18 |
| 11 | 36235024 | rs12420852 | 0.419 | 0.519 | 0.084 | 0.491 | -0.021 | -0.027 | -0.027 | 0 | 0.05 | 0 | 0.15 | 0.22 | 0.1 | 0.06 |
| 11 | 36235363 | rs10836520 | 1.351 | 0.427 | -0.051 | 0.298 | -0.028 | -0.105 | -0.105 | 0 | 0.36 | 0.11 | 0.28 | 0.42 | 0.39 | 0.37 |
| 11 | 36237675 | rs11605941 | -0.248 | 0.559 | 0.478 | 0.399 | -0.021 | -0.026 | -0.026 | 0 | 0.05 | 0 | 0.15 | 0.19 | 0.1 | 0.1 |
| 11 | 36244190 | rs12275029 | -0.821 | 0.491 | 0.566 | 0.245 | -0.015 | 0.049 | 0.049 | 5.341 | 0.23 | 0.15 | 0.29 | 0.05 | 0.25 | 0.26 |
| 11 | 36251046 | rs4755444 | 1.469 | 0.112 | 0.818 | 0.488 | 0 | 0 | 0 | 0.431 | 0.7 | 0.34 | 0.55 | 0.73 | 0.69 | 0.67 |
| 11 | 36255087 | rs16928593 | -0.502 | 0.098 | 0.474 | 0.463 | 0.001 | 0.002 | 0.002 | 6.549 | 0.2 | 0.21 | 0.24 | 0.05 | 0.17 | 0.24 |
| 11 | 36258285 | rs10501147 | 1.034 | 0.124 | 0.327 | 0.538 | -0.021 | 0.005 | 0.005 | 4.699 | 0.11 | 0.03 | 0.17 | 0.11 | 0.14 | 0.13 |
| 11 | 36269071 | rs17814201 | 0.969 | 0.15 | 0.289 | 0.571 | -0.015 | 0.007 | 0.007 | 4.358 | 0.11 | 0.04 | 0.17 | 0.11 | 0.14 | 0.13 |
| 11 | 36270057 | rs10400279 | -0.416 | 0.272 | 0.431 | 0.579 | -0.005 | -0.006 | -0.006 | 0 | 0.09 | 0.15 | 0.06 | 0.05 | 0.06 | 0.12 |
| 11 | 36272348 | rs7945114 | 1.106 | 0.053 | 0.883 | 0.495 | -0.011 | -0.01 | -0.01 | 0 | 0.69 | 0.44 | 0.58 | 0.72 | 0.71 | 0.67 |
| 11 | 36273754 | rs7114323 | 0.146 | -0.082 | 0.825 | 0.275 | -0.001 | -0.002 | -0.002 | 0 | 0.78 | 0.71 | 0.77 | 0.83 | 0.81 | 0.75 |
| 11 | 36275074 | rs10768202 | 0.575 | 0.013 | 0.548 | 0.319 | -0.025 | -0.049 | -0.049 | 0 | 0.73 | 0.5 | 0.74 | 0.83 | 0.79 | 0.72 |
| 11 | 36285390 | rs12417278 | -0.153 | 0.414 | 0.153 | 0.535 | 0.004 | 0.001 | 0.001 | 0 | 0.29 | 0.27 | 0.16 | 0.35 | 0.28 | 0.25 |
| 11 | 36286635 | rs746890 | -0.682 | 0.526 | -0.091 | 0.246 | -0.013 | -0.008 | -0.008 | 0.094 | 0.28 | 0.31 | 0.13 | 0.26 | 0.21 | 0.19 |
| 11 | 36288752 | rs7109505 | 1.752 | -0.376 | -0.433 | 0.598 | -0.027 | 0 | 0 | 1.294 | 0.35 | 0.16 | 0.47 | 0.31 | 0.45 | 0.34 |
| 11 | 36290278 | rs2289989 | -0.182 | -0.473 | 0.023 | 0.838 | -0.003 | 0.015 | 0.015 | 0 | 0.35 | 0.34 | 0.35 | 0.26 | 0.33 | 0.46 |
| 11 | 36292065 | rs4756305 | -0.305 | -0.469 | 0.167 | 0.83 | -0.001 | 0.002 | 0.002 | 0 | 0.36 | 0.38 | 0.36 | 0.27 | 0.33 | 0.46 |
| 11 | 36293048 | rs7107433 | -2.116 | -0.469 | 0.217 | 0.832 | -0.017 | 0.004 | 0.004 | 1.129 | 0.64 | 0.78 | 0.52 | 0.69 | 0.55 | 0.66 |
| 11 | 36294052 | rs3812766 | 0.866 | -0.462 | 0.255 | 0.851 | -0.013 | 0.014 | 0.014 | 0.08 | 0.19 | 0.25 | 0.09 | 0.25 | 0.14 | 0.07 |
| 11 | 36301180 | rs7128260 | 0.378 | -0.622 | 0.099 | 0.731 | -0.037 | 0.04 | 0.04 | 0.034 | 0.22 | 0.05 | 0.13 | 0.16 | 0.19 | 0.32 |
| 11 | 36310677 | rs12418067 | 0.944 | -0.535 | 0.454 | 0.836 | -0.015 | 0.017 | 0.017 | 0.043 | 0.19 | 0.25 | 0.08 | 0.25 | 0.14 | 0.06 |
| 11 | 36311014 | rs7939948 | -1.436 | -0.549 | 0.514 | 0.827 | -0.003 | 0 | 0 | 0 | 0.55 | 0.56 | 0.43 | 0.51 | 0.47 | 0.53 |
| 11 | 36311570 | rs2289977 | 1.311 | -0.485 | 0.82 | 0.785 | 0.006 | -0.008 | -0.008 | 0.138 | 0.16 | 0.1 | 0.05 | 0.25 | 0.11 | 0.06 |
| 11 | 36313749 | rs12282145 | 0.412 | -0.519 | 0.583 | 0.746 | 0.001 | 0.002 | 0.002 | 0.166 | 0.09 | 0.07 | 0.04 | 0.03 | 0.07 | 0.12 |
| 11 | 36315560 | rs12292771 | 0.562 | -0.478 | 0.673 | 0.923 | -0.025 | 0.126 | 0.126 | 0.066 | 0.19 | 0.37 | 0.12 | 0.39 | 0.12 | 0.07 |
| 11 | 36316168 | rs6484834 | 0.668 | -0.773 | 0.423 | 0.556 | 0 | 0 | 0 | 0 | 0.54 | 0.54 | 0.57 | 0.53 | 0.56 | 0.4 |
| 11 | 36318335 | rs11033537 | 1.28 | -0.866 | 0.252 | 0.772 | -0.014 | 0.041 | 0.041 | 0 | 0.15 | 0.22 | 0.13 | 0.24 | 0.1 | 0.06 |
| 11 | 36320195 | rs10836529 | -0.7 | -1.069 | 0.254 | 0.757 | -0.004 | 0.012 | 0.012 | 0 | 0.38 | 0.49 | 0.43 | 0.52 | 0.35 | 0.28 |
| 11 | 36320992 | rs1562327 | 1.814 | -1.139 | 0.158 | 0.508 | -0.051 | 0.023 | 0.023 | 0.09 | 0.35 | 0.15 | 0.41 | 0.28 | 0.46 | 0.34 |
| 11 | 36322149 | rs11033541 | 1.171 | -1.223 | 0.262 | 0.6 | -0.015 | 0.027 | 0.027 | 0 | 0.16 | 0.23 | 0.13 | 0.24 | 0.11 | 0.05 |
| 11 | 36324593 | rs12292704 | -0.401 | -1.147 | -0.111 | -0.053 | -0.002 | 0.001 | 0.001 | 0 | 0.39 | 0.53 | 0.45 | 0.53 | 0.35 | 0.3 |
| 11 | 36324962 | rs11033542 | 1.008 | -0.965 | -0.335 | -0.7 | -0.042 | 0.042 | 0.042 | 2.047 | 0.3 | 0.13 | 0.34 | 0.18 | 0.39 | 0.26 |
| 11 | 36325764 | rs12294763 | 0.713 | -0.051 | 0.133 | -0.31 | -0.029 | -0.009 | -0.009 | 0 | 0.19 | 0.14 | 0.28 | 0.2 | 0.31 | 0.24 |
| 11 | 36329290 | rs10836533 | 0.722 | -0.327 | 0.464 | -0.447 | 0 | 0.004 | 0.004 | 0.338 | 0.53 | 0.42 | 0.41 | 0.68 | 0.38 | 0.46 |
| 11 | 36329456 | rs11033546 | -0.792 | -0.34 | 0.764 | -0.763 | -0.014 | 0.017 | 0.017 | 0 | 0.11 | 0.31 | 0.07 | 0.12 | 0.09 | 0.12 |
| 11 | 36334103 | rs9971501 | -1.255 | -0.158 | 0.554 | -0.785 | 0 | -0.009 | -0.009 | 0 | 0.89 | 0.89 | 0.79 | 0.99 | 0.86 | 0.83 |
| 11 | 36336263 | rs2218565 | 1.339 | -0.23 | 0.758 | -0.871 | 0.007 | 0.031 | 0.031 | 0.112 | 0.45 | 0.35 | 0.37 | 0.77 | 0.33 | 0.41 |
| 11 | 36349396 | rs11033553 | -0.004 | -0.215 | 0.978 | -0.929 | -0.006 | -0.042 | -0.042 | 3.382 | 0.14 | 0.28 | 0.09 | 0.05 | 0.11 | 0.11 |
| 11 | 36353471 | rs12419146 | -1.112 | -0.285 | 1.268 | -1.066 | 0.014 | -0.025 | -0.025 | 0 | 0.07 | 0.28 | 0.04 | 0 | 0.04 | 0.03 |
| 11 | 36357037 | rs1900337 | -0.263 | -0.339 | 1.248 | -1.134 | -0.016 | 0.022 | 0.022 | 0.417 | 0.68 | 0.74 | 0.53 | 0.89 | 0.52 | 0.59 |
| 11 | 36358876 | rs4755449 | 0.824 | -0.371 | 1.295 | -1.095 | -0.019 | 0.066 | 0.066 | 1.137 | 0.56 | 0.31 | 0.44 | 0.8 | 0.44 | 0.51 |
| 11 | 36361918 | rs7101640 | -0.972 | -0.479 | 1.428 | -1.033 | -0.039 | -0.023 | -0.023 | 0 | 0.22 | 0.53 | 0.14 | 0.14 | 0.15 | 0.15 |
| 11 | 36363575 | rs4755450 | 1.275 | -0.428 | 0.257 | -1.025 | 0.01 | 0.058 | 0.058 | 0.153 | 0.44 | 0.25 | 0.37 | 0.75 | 0.33 | 0.41 |
| 11 | 36367117 | rs11033564 | 0.374 | -0.247 | -0.256 | -0.943 | 0.001 | 0.008 | 0.008 | 0 | 0.12 | 0.1 | 0.07 | 0.05 | 0.11 | 0.11 |
| 11 | 36375707 | rs7105283 | 0.546 | -0.272 | -0.141 | -0.99 | -0.016 | -0.041 | -0.041 | 0 | 0.75 | 0.5 | 0.66 | 0.88 | 0.75 | 0.67 |
| 11 | 36376603 | rs10501151 | 0.742 | 0.281 | -0.994 | -0.955 | -0.074 | 0.026 | 0.026 | 0.281 | 0.19 | 0.04 | 0.24 | 0.09 | 0.33 | 0.17 |
| 11 | 36379532 | rs7129794 | 0.249 | 0.197 | -0.791 | -0.78 | 0.001 | 0.001 | 0.001 | 0 | 0.13 | 0.14 | 0.07 | 0.09 | 0.12 | 0.11 |
| 11 | 36381265 | rs567260 | -0.491 | 0.495 | -1.098 | -0.816 | -0.011 | 0.017 | 0.017 | 0 | 0.84 | 0.88 | 0.75 | 0.96 | 0.8 | 0.72 |
| 11 | 36384112 | rs2162195 | -0.455 | 0.15 | -1.027 | -0.754 | 0.006 | 0.025 | 0.025 | 0 | 0.23 | 0.2 | 0.28 | 0.05 | 0.21 | 0.32 |
| 11 | 36385236 | rs17615296 | 0.858 | 0.358 | -1.196 | -0.421 | 0.001 | -0.027 | -0.027 | 0 | 0.38 | 0.15 | 0.34 | 0.71 | 0.31 | 0.37 |
| 11 | 36386755 | rs10501153 | 0.142 | 0.926 | -1.21 | -0.485 | -0.086 | 0.053 | 0.053 | 0 | 0.21 | 0.03 | 0.23 | 0.08 | 0.33 | 0.2 |
| 11 | 36388162 | rs7929961 | 0.252 | 0.942 | -1.145 | -0.44 | -0.076 | 0.052 | 0.052 | 0 | 0.21 | 0.03 | 0.22 | 0.08 | 0.3 | 0.18 |
| 11 | 36391882 | rs525634 | -0.094 | 0.249 | -0.333 | 0.213 | 0.013 | -0.006 | -0.006 | 0 | 0.54 | 0.26 | 0.61 | 0.46 | 0.62 | 0.66 |
| 11 | 36395345 | rs2554012 | -0.22 | -0.239 | -0.225 | 0.558 | -0.024 | 0.027 | 0.027 | 0.048 | 0.63 | 0.49 | 0.65 | 0.46 | 0.71 | 0.72 |
| 11 | 36399331 | rs3812774 | -1.007 | -0.506 | -0.742 | 0.284 | 0.031 | 0.02 | 0.02 | 0 | 0.13 | 0.03 | 0.06 | 0.1 | 0.07 | 0.08 |

| | | | | | | | | | | | | | | | | |
|---|---|---|---|---|---|---|---|---|---|---|---|---|---|---|---|---|
| 11 | 36399514 | rs1425936 | -0.195 | -1.083 | -1.708 | -0.149 | -0.004 | 0.04 | 0.04 | 0 | 0.24 | 0.02 | 0.16 | 0.13 | 0.18 | 0.26 |
| 11 | 36401004 | rs1425939 | 0.103 | -1.346 | -1.762 | 0.556 | 0.002 | -0.004 | -0.004 | 0 | 0.12 | 0.14 | 0.18 | 0.29 | 0.08 | 0.22 |
| 11 | 36402006 | rs11608110 | 0.096 | -1.425 | -1.888 | 0.569 | 0.014 | -0.005 | -0.005 | 0 | 0.08 | 0.02 | 0.03 | 0.09 | 0.04 | 0.06 |
| 11 | 36404730 | rs330255 | -0.416 | -2.123 | -1.463 | 0.246 | 0.027 | 0.05 | 0.05 | 0 | 0.73 | 0.84 | 0.81 | 0.9 | 0.81 | 0.7 |
| 11 | 36407102 | rs404079 | 1.105 | -2.527 | -0.613 | 0.994 | -0.006 | 0 | 0 | 2.357 | 0.48 | 0.49 | 0.59 | 0.44 | 0.67 | 0.41 |
| 11 | 36407403 | rs10836545 | 0.297 | -2.548 | -0.617 | 1.061 | 0.007 | -0.011 | -0.011 | 0 | 0.14 | 0.12 | 0.17 | 0.29 | 0.08 | 0.22 |
| 11 | 36408045 | rs446134 | 0.25 | -2.489 | -0.545 | 0.937 | -0.004 | -0.008 | -0.008 | 0 | 0.62 | 0.64 | 0.77 | 0.81 | 0.76 | 0.63 |
| 11 | 36409580 | rs7931347 | 1.11 | -2.584 | -0.08 | 1.067 | 0.035 | 0.01 | 0.01 | 0 | 0.48 | 0.35 | 0.57 | 0.42 | 0.67 | 0.41 |
| 11 | 36409826 | rs606442 | 0.09 | -2.561 | -0.109 | 1.081 | -0.017 | 0.063 | 0.063 | 0 | 0.15 | 0.24 | 0.19 | 0.39 | 0.09 | 0.22 |
| 11 | 36411034 | rs11601958 | 0.814 | -2.367 | 0.323 | 0.975 | 0.076 | 0.001 | 0.001 | 0 | 0.46 | 0.22 | 0.56 | 0.43 | 0.67 | 0.41 |
| 11 | 36412655 | rs4077044 | 0.187 | -1.235 | 0.088 | 0.889 | 0.002 | 0.003 | 0.003 | 0 | 0.51 | 0.45 | 0.5 | 0.38 | 0.6 | 0.37 |
| 11 | 36419929 | rs12785381 | -1.214 | -0.554 | 0.915 | 0.763 | 0.061 | 0.021 | 0.021 | 2.423 | 0.07 | 0.31 | 0.06 | 0.05 | 0.09 | 0.11 |
| 11 | 36421759 | rs10836549 | -1.119 | -0.616 | 1.345 | 0.708 | -0.007 | -0.016 | -0.016 | 0 | 0.09 | 0.24 | 0.07 | 0.09 | 0.06 | 0.09 |
| 11 | 36422172 | rs330260 | 2.077 | -0.699 | 1.191 | 0.772 | -0.009 | -0.008 | -0.008 | 0 | 0.8 | 0.38 | 0.8 | 0.8 | 0.82 | 0.77 |
| 11 | 36422438 | rs7933726 | -0.997 | -0.688 | 1.217 | 0.731 | 0 | 0 | 0 | 0 | 0.09 | 0.09 | 0.12 | 0.16 | 0.09 | 0.12 |
| 11 | 36422792 | rs330261 | 1.31 | -0.71 | 1.184 | 0.663 | 0.06 | -0.004 | -0.004 | 0 | 0.9 | 0.58 | 0.93 | 0.97 | 0.91 | 0.89 |
| 11 | 36427893 | rs10836550 | -2.026 | -0.486 | 1.069 | 0.658 | 0.054 | -0.016 | -0.016 | 0 | 0.13 | 0.53 | 0.1 | 0.06 | 0.14 | 0.13 |
| 11 | 36429041 | rs7103677 | -0.385 | -0.47 | -0.306 | 0.409 | -0.01 | 0.023 | 0.023 | 0 | 0.18 | 0 | 0.12 | 0.1 | 0.19 | 0.1 |
| 11 | 36437709 | rs765539 | -1.509 | -0.449 | 1.149 | 0.589 | 0.026 | -0.017 | -0.017 | 0 | 0.13 | 0.35 | 0.05 | 0.11 | 0.09 | 0.18 |
| 11 | 36438075 | rs1365120 | 2.102 | -0.512 | 1.581 | 1.047 | 0.02 | 0.056 | 0.056 | 0.039 | 0.88 | 0.34 | 0.86 | 0.87 | 0.9 | 0.8 |
| 11 | 36444395 | rs7951020 | -0.496 | -0.085 | 0.534 | 1.063 | -0.003 | -0.018 | -0.018 | 0 | 0.28 | 0.26 | 0.36 | 0.57 | 0.32 | 0.36 |
| 11 | 36445197 | rs10501155 | -1.026 | -0.211 | 0.854 | 0.938 | 0.006 | 0.01 | 0.01 | 0 | 0.35 | 0.44 | 0.39 | 0.61 | 0.39 | 0.42 |
| 11 | 36446216 | rs7107720 | 1.17 | 0.105 | 0.969 | 0.747 | 0 | -0.006 | -0.006 | 0 | 0.55 | 0.41 | 0.49 | 0.26 | 0.56 | 0.39 |
| 11 | 36446712 | rs331446 | -0.777 | 0.31 | 0.77 | 1.518 | 0.015 | 0.146 | 0.146 | 0 | 0.17 | 0.36 | 0.36 | 0.52 | 0.22 | 0.31 |
| 11 | 36448379 | rs744895 | -0.394 | 0.319 | 0.74 | 1.509 | 0.01 | 0.072 | 0.072 | 0 | 0.15 | 0.32 | 0.27 | 0.4 | 0.2 | 0.23 |
| 11 | 36448523 | rs744894 | 0.143 | 0.385 | 0.584 | 1.375 | 0.017 | -0.011 | -0.011 | 0 | 0.05 | 0.08 | 0.03 | 0.04 | 0.06 | 0.05 |
| 11 | 36451458 | rs12418202 | 0.145 | 0.312 | 0.38 | 1.345 | -0.028 | -0.042 | -0.042 | 0 | 0.18 | 0.01 | 0.1 | 0.18 | 0.13 | 0.23 |
| 11 | 36452850 | rs751926 | 1.832 | 0.257 | 0.897 | 1.257 | -0.005 | 0.032 | 0.032 | 0 | 0.55 | 0.31 | 0.48 | 0.26 | 0.58 | 0.4 |
| 11 | 36454231 | rs331485 | -0.674 | 0.179 | 0.737 | 0.848 | -0.006 | 0.005 | 0.005 | 0 | 0.87 | 0.74 | 0.84 | 0.91 | 0.88 | 0.84 |
| 11 | 36457253 | rs10836561 | -0.756 | 0.387 | 0.15 | 1.006 | 0.014 | 0.033 | 0.033 | 0 | 0.19 | 0.29 | 0.28 | 0.44 | 0.26 | 0.28 |
| 11 | 36458617 | rs7929580 | 0.167 | 0.019 | 0.191 | 0.855 | -0.021 | -0.011 | -0.011 | 0 | 0.8 | 0.36 | 0.78 | 0.87 | 0.86 | 0.82 |
| 11 | 36458777 | rs577054 | -0.699 | 0.212 | 0.145 | 1.097 | 0.009 | 0.05 | 0.05 | 0 | 0.16 | 0.24 | 0.27 | 0.47 | 0.23 | 0.25 |
| 11 | 36459130 | rs4756316 | 1.028 | 0.113 | 0.474 | 0.746 | -0.001 | -0.001 | -0.001 | 0.004 | 0.63 | 0.4 | 0.68 | 0.77 | 0.7 | 0.6 |
| 11 | 36459378 | rs331482 | -1.128 | -0.072 | 0.428 | 1.209 | 0.006 | -0.01 | -0.01 | 0 | 0.32 | 0.27 | 0.21 | 0.43 | 0.26 | 0.47 |
| 11 | 36460081 | rs7930554 | -0.366 | 0.081 | 0.364 | 1.204 | 0.009 | 0.096 | 0.096 | 2.119 | 0.16 | 0.28 | 0.15 | 0.41 | 0.2 | 0.27 |
| 11 | 36460958 | rs331478 | -0.435 | 0.023 | 0.066 | 0.956 | 0.016 | 0.05 | 0.05 | 0 | 0.8 | 0.89 | 0.85 | 0.94 | 0.85 | 0.76 |
| 11 | 36465453 | rs10742373 | 1.262 | 0.234 | 0.396 | 1.165 | 0.001 | 0.01 | 0.01 | 0 | 0.59 | 0.48 | 0.66 | 0.47 | 0.58 | 0.43 |
| 11 | 36465719 | rs10742374 | 0.747 | 0.115 | 0.475 | 1.251 | -0.003 | 0 | 0 | 0 | 0.64 | 0.63 | 0.75 | 0.57 | 0.66 | 0.49 |
| 11 | 36466141 | rs4756321 | 0.975 | 0.14 | 0.376 | 1.045 | -0.011 | 0.022 | 0.022 | 0 | 0.62 | 0.49 | 0.68 | 0.47 | 0.65 | 0.45 |
| 11 | 36469758 | rs2458927 | 0.733 | -0.131 | 0.292 | 0.675 | -0.008 | -0.05 | -0.05 | 0 | 0.7 | 0.52 | 0.8 | 0.84 | 0.72 | 0.66 |
| 11 | 36470815 | rs10501156 | 0.655 | -0.106 | -0.088 | 0.633 | 0.004 | 0.059 | 0.059 | 0 | 0.14 | 0.02 | 0.09 | 0.02 | 0.12 | 0.17 |
| 11 | 36472800 | rs3740958 | 0.039 | -0.15 | 0.123 | 1.038 | 0 | 0.03 | 0.03 | 0.558 | 0.13 | 0.17 | 0.11 | 0.38 | 0.13 | 0.22 |
| 11 | 36473784 | rs11600757 | 0.827 | -0.073 | 0.073 | 0.908 | -0.011 | 0.005 | 0.005 | 0 | 0.11 | 0.06 | 0.08 | 0.07 | 0.16 | 0.12 |
| 11 | 36475317 | rs890259 | -1.483 | -0.854 | -0.086 | 0.506 | 0.01 | -0.002 | -0.002 | 0 | 0.49 | 0.6 | 0.32 | 0.54 | 0.41 | 0.53 |
| 11 | 36478114 | rs10836563 | 0.058 | -0.128 | 0.121 | -0.155 | 0.018 | 0.005 | 0.005 | 0 | 0.21 | 0.32 | 0.29 | 0.27 | 0.25 | 0.15 |
| 11 | 36479789 | rs11033637 | 0.983 | -0.335 | -0.262 | -0.257 | -0.018 | 0.055 | 0.055 | 0.057 | 0.57 | 0.22 | 0.55 | 0.66 | 0.55 | 0.69 |
| 11 | 36480683 | rs9630172 | 1.033 | -0.834 | -0.029 | -0.742 | -0.018 | -0.032 | -0.032 | 0 | 0.12 | 0.19 | 0.07 | 0 | 0.07 | 0.02 |
| 11 | 36480986 | rs6484858 | 0.946 | -0.294 | -0.16 | -0.537 | 0.011 | 0.1 | 0.1 | 0 | 0.58 | 0.23 | 0.74 | 0.78 | 0.62 | 0.68 |
| 11 | 36486064 | rs331436 | 0.783 | -0.267 | 0.465 | -0.469 | 0.001 | -0.02 | -0.02 | 0 | 0.59 | 0.47 | 0.62 | 0.8 | 0.59 | 0.7 |
| 11 | 36487597 | rs11033641 | 0.557 | -0.207 | 0.484 | -0.578 | -0.007 | -0.032 | -0.032 | 0.906 | 0.18 | 0.21 | 0.07 | 0.04 | 0.13 | 0.15 |
| 11 | 36488796 | rs331432 | -0.505 | -0.214 | 0.372 | -0.566 | -0.007 | 0.004 | 0.004 | 0 | 0.78 | 0.83 | 0.7 | 0.84 | 0.73 | 0.85 |
| 11 | 36491389 | rs981982 | 0.43 | -0.379 | 0.307 | -0.32 | -0.002 | -0.014 | -0.014 | 1.951 | 0.17 | 0.17 | 0.06 | 0.04 | 0.13 | 0.15 |
| 11 | 36510071 | rs3740961 | 0.805 | -0.117 | -0.571 | 0.307 | -0.02 | -0.111 | -0.111 | 0.449 | 0.15 | 0.03 | 0.29 | 0.53 | 0.12 | 0.29 |
| 11 | 36513786 | rs5030472 | 0.888 | 0.054 | -0.189 | -0.367 | -0.006 | 0.018 | 0.018 | 0 | 0.1 | 0.05 | 0.05 | 0.06 | 0.11 | 0.13 |
| 11 | 36533753 | rs5030411 | 0.825 | -0.05 | 0.033 | -0.888 | -0.036 | 0.085 | 0.085 | 0.104 | 0.54 | 0.13 | 0.58 | 0.71 | 0.6 | 0.64 |
| 11 | 36533885 | rs5030409 | 1.048 | -0.042 | 0.087 | -1.273 | 0.016 | 0.036 | 0.036 | 0 | 0.17 | 0.09 | 0.06 | 0.06 | 0.11 | 0.14 |
| 11 | 36540549 | rs331455 | 0.21 | 0.107 | 0.328 | -1.363 | -0.01 | -0.038 | -0.038 | 0 | 0.69 | 0.65 | 0.82 | 0.9 | 0.75 | 0.71 |
| 11 | 36559278 | rs12419174 | 0.205 | 0.406 | 0.316 | -0.939 | -0.001 | -0.004 | -0.004 | 0 | 0.11 | 0.1 | 0.09 | 0.01 | 0.12 | 0.07 |
| 11 | 36579290 | rs11033693 | -1.389 | 0.445 | 0.386 | -0.79 | -0.016 | 0.005 | 0.005 | 0.946 | 0.26 | 0.81 | 0.33 | 0.2 | 0.25 | 0.21 |
| 11 | 36582111 | rs12225867 | 0.51 | 0.414 | 1.095 | -0.786 | 0.035 | 0.024 | 0.024 | 0 | 0.12 | 0.2 | 0.2 | 0.15 | 0.15 | 0.08 |
| 11 | 36587017 | rs3758873 | -1.527 | 0.374 | 1.012 | -0.815 | -0.013 | 0.006 | 0.006 | 0.535 | 0.27 | 0.81 | 0.33 | 0.2 | 0.25 | 0.2 |
| 11 | 36589622 | rs872053 | 0.762 | 0.445 | 1.09 | -0.878 | -0.001 | -0.008 | -0.008 | 0 | 0.08 | 0.08 | 0.06 | 0.01 | 0.08 | 0.06 |
| 11 | 36595600 | rs3740955 | -1.117 | 0.099 | 0.686 | -1.04 | 0.006 | 0.109 | 0.109 | 0 | 0.43 | 0.77 | 0.61 | 0.76 | 0.36 | 0.52 |
| 11 | 36597313 | rs2227973 | 1.2 | -0.022 | 0.732 | -0.946 | 0.021 | -0.055 | -0.055 | 0 | 0.18 | 0.08 | 0.28 | 0.57 | 0.12 | 0.3 |
| 11 | 36605134 | rs7480202 | 0.433 | -0.026 | 0.968 | -0.974 | -0.005 | -0.01 | -0.01 | 0 | 0.12 | 0.16 | 0.1 | 0.04 | 0.11 | 0.13 |
| 11 | 36612090 | rs1818545 | -0.609 | -0.09 | 1.421 | -0.966 | -0.012 | 0.001 | 0.001 | 0 | 0.33 | 0.63 | 0.41 | 0.6 | 0.23 | 0.44 |
| 11 | 36617989 | rs867801 | 0.018 | 0.014 | 1.393 | -1.056 | -0.019 | -0.039 | -0.039 | 0.616 | 0.14 | 0.33 | 0.11 | 0.04 | 0.11 | 0.1 |
| 11 | 36618883 | rs867803 | 0.769 | 0.016 | 1.388 | -1.058 | -0.007 | -0.028 | -0.028 | 0 | 0.11 | 0.19 | 0.07 | 0.01 | 0.09 | 0.08 |
| 11 | 36625604 | rs10501158 | 0.796 | 0.016 | 1.382 | -1.06 | -0.005 | -0.027 | -0.027 | 1.5 | 0.11 | 0.16 | 0.07 | 0.01 | 0.09 | 0.06 |

| | | | | | | | | | | | | | | | | |
|---|---|---|---|---|---|---|---|---|---|---|---|---|---|---|---|---|
| 11 | 36633302 | rs10836576 | -0.522 | 0.11 | 1.457 | -1.073 | 0.054 | 0.005 | 0.005 | 0 | 0.22 | 0.47 | 0.31 | 0.18 | 0.25 | 0.16 |
| 11 | 36655089 | rs3758874 | 1.41 | -0.077 | 1.204 | -1.109 | 0.01 | -0.03 | -0.03 | 0 | 0.16 | 0.1 | 0.28 | 0.58 | 0.12 | 0.31 |
| 11 | 36657272 | rs7128419 | -0.044 | 0.055 | 1.128 | -1.252 | -0.019 | -0.04 | -0.04 | 1.27 | 0.14 | 0.34 | 0.11 | 0.04 | 0.11 | 0.1 |
| 11 | 36687767 | rs10836586 | 0.002 | 0.309 | 0.567 | -1.284 | 0.002 | -0.038 | -0.038 | 0 | 0.09 | 0.17 | 0.09 | 0.02 | 0.08 | 0.02 |
| 11 | 36707480 | rs1391544 | 0.02 | 0.313 | 0.521 | -1.282 | 0.001 | -0.038 | -0.038 | 2.576 | 0.09 | 0.17 | 0.09 | 0.02 | 0.08 | 0.02 |
| 11 | 36717383 | rs7123291 | 1.033 | 0.361 | 0.309 | -1.408 | -0.022 | 0.197 | 0.197 | 1.582 | 0.51 | 0.16 | 0.43 | 0.22 | 0.55 | 0.47 |
| 11 | 36719758 | rs332438 | -0.977 | 0.217 | 1.058 | -1.346 | -0.018 | -0.114 | -0.114 | 0 | 0.21 | 0.51 | 0.18 | 0.03 | 0.17 | 0.1 |
| 11 | 36721017 | rs332435 | -0.102 | 0.212 | 1.164 | -1.348 | 0.002 | -0.035 | -0.035 | 1.077 | 0.09 | 0.22 | 0.1 | 0.02 | 0.08 | 0.02 |
| 11 | 36721248 | rs332433 | -1.097 | 0.2 | 1.165 | -1.36 | -0.019 | -0.111 | -0.111 | 0.854 | 0.21 | 0.54 | 0.19 | 0.03 | 0.17 | 0.1 |
| 11 | 36726818 | rs10836589 | -0.072 | 0.221 | 1.075 | -1.342 | 0.012 | -0.034 | -0.034 | 0 | 0.09 | 0.23 | 0.1 | 0.02 | 0.08 | 0.02 |
| 11 | 36728261 | rs12294719 | -1.163 | 0.208 | 1.099 | -1.338 | -0.02 | -0.097 | -0.097 | 0 | 0.22 | 0.52 | 0.2 | 0.05 | 0.18 | 0.14 |
| 11 | 36729545 | rs11033751 | -0.692 | 0.248 | 0.987 | -1.374 | -0.006 | -0.08 | -0.08 | 0 | 0.19 | 0.34 | 0.18 | 0.02 | 0.17 | 0.08 |
| 11 | 36730884 | rs1533588 | -0.985 | 0.279 | 1.015 | -1.384 | -0.008 | -0.103 | -0.103 | 0 | 0.21 | 0.41 | 0.18 | 0.03 | 0.18 | 0.1 |
| 11 | 36737903 | rs16929335 | 0.364 | 0.371 | 0.579 | -1.494 | -0.007 | -0.056 | -0.056 | 0 | 0.24 | 0.33 | 0.18 | 0.11 | 0.21 | 0.2 |
| 11 | 36738257 | rs11033758 | 1.097 | 0.4 | 0.085 | -1.585 | -0.089 | 0.141 | 0.141 | 0.299 | 0.38 | 0.03 | 0.35 | 0.16 | 0.42 | 0.32 |
| 11 | 36738897 | rs7102442 | 1.359 | 0.524 | 0.735 | -1.093 | -0.013 | 0.099 | 0.099 | 0 | 0.46 | 0.28 | 0.48 | 0.16 | 0.54 | 0.38 |
| 11 | 36741120 | rs7948751 | -1.129 | 0.546 | 0.628 | -0.72 | -0.011 | 0.052 | 0.052 | 0 | 0.81 | 0.74 | 0.73 | 0.45 | 0.87 | 0.64 |
| 11 | 36741502 | rs7949234 | -1.335 | 0.564 | 0.45 | -0.7 | 0.011 | -0.033 | -0.033 | 0.137 | 0.82 | 0.85 | 0.75 | 0.46 | 0.87 | 0.65 |
| 11 | 36741709 | rs10836590 | 1.441 | 0.669 | 0.374 | -0.822 | -0.056 | 0.204 | 0.204 | 0 | 0.45 | 0.16 | 0.48 | 0.16 | 0.54 | 0.37 |
| 11 | 36745881 | rs10768222 | 0.062 | 0.555 | 0.734 | -0.709 | -0.037 | 0.024 | 0.024 | 0 | 0.68 | 0.49 | 0.67 | 0.36 | 0.8 | 0.53 |
| 11 | 36750204 | rs4756339 | 0.636 | 0.766 | 0.892 | -0.606 | 0.069 | 0.056 | 0.056 | 0 | 0.11 | 0.24 | 0.11 | 0.16 | 0.17 | 0.09 |
| 11 | 36756359 | rs1589177 | 0.137 | 0.752 | 1.051 | -0.614 | 0.132 | 0.099 | 0.099 | 0 | 0.13 | 0.44 | 0.12 | 0.16 | 0.18 | 0.08 |
| 11 | 36759552 | rs1518746 | 1.286 | 0.74 | 0.597 | -0.747 | -0.08 | 0.223 | 0.223 | 0.016 | 0.44 | 0.12 | 0.47 | 0.16 | 0.53 | 0.36 |
| 11 | 36759882 | rs11033782 | 1.056 | 0.762 | 0.77 | -0.728 | 0.033 | 0.016 | 0.016 | 0 | 0.1 | 0.13 | 0.09 | 0.14 | 0.16 | 0.07 |
| 11 | 36760325 | rs1022367 | 0.775 | 0.686 | 0.895 | -0.755 | 0.019 | 0.017 | 0.017 | 0 | 0.55 | 0.39 | 0.58 | 0.31 | 0.7 | 0.45 |
| 11 | 36761612 | rs4480515 | -0.353 | 0.615 | 0.798 | -0.937 | -0.002 | -0.002 | -0.002 | 0 | 0.72 | 0.71 | 0.65 | 0.3 | 0.71 | 0.57 |
| 11 | 36763813 | rs964524 | 1.357 | 0.388 | 0.783 | -0.933 | -0.038 | 0.166 | 0.166 | 0.012 | 0.44 | 0.19 | 0.47 | 0.14 | 0.53 | 0.36 |
| 11 | 36764142 | rs964526 | 1.374 | 0.384 | 0.678 | -0.937 | -0.087 | 0.233 | 0.233 | 0.022 | 0.44 | 0.12 | 0.47 | 0.14 | 0.53 | 0.36 |
| 11 | 36764288 | rs964527 | -0.247 | 0.484 | 0.892 | -0.867 | 0.013 | 0.012 | 0.012 | 0 | 0.7 | 0.5 | 0.64 | 0.3 | 0.71 | 0.57 |
| 11 | 36766061 | rs11033786 | -0.804 | 0.03 | 0.787 | -0.847 | -0.003 | -0.003 | -0.003 | 0 | 0.26 | 0.25 | 0.15 | 0.14 | 0.18 | 0.19 |
| 11 | 36766225 | rs4756345 | -1.282 | 0.269 | 0.451 | -1.035 | -0.004 | -0.013 | -0.013 | 0 | 0.34 | 0.33 | 0.25 | 0.15 | 0.31 | 0.23 |
| 11 | 36768765 | rs7930272 | -0.024 | 0.373 | 0.591 | -0.655 | 0.031 | -0.011 | -0.011 | 1.313 | 0.67 | 0.52 | 0.57 | 0.26 | 0.62 | 0.5 |
| 11 | 36770392 | rs11033790 | 1.004 | 0.477 | 0.06 | -0.558 | -0.001 | 0.061 | 0.061 | 0 | 0.42 | 0.31 | 0.39 | 0.13 | 0.46 | 0.32 |
| 11 | 36776618 | rs7478973 | -0.14 | 0.636 | -0.101 | -0.05 | 0.001 | -0.041 | -0.041 | 0 | 0.71 | 0.83 | 0.63 | 0.33 | 0.72 | 0.5 |
| 11 | 36780585 | rs11033794 | 0.511 | 0.602 | -0.187 | -0.325 | -0.001 | 0.071 | 0.071 | 0 | 0.58 | 0.81 | 0.53 | 0.26 | 0.59 | 0.42 |
| 11 | 36781155 | rs11603790 | -0.567 | 0.434 | -0.268 | -0.256 | -0.023 | -0.034 | -0.034 | 0 | 0.17 | 0.05 | 0.15 | 0.14 | 0.18 | 0.13 |
| 11 | 36785787 | rs7940927 | -0.537 | 0.589 | -0.056 | -0.242 | -0.003 | -0.075 | -0.075 | 0 | 0.75 | 0.89 | 0.66 | 0.4 | 0.75 | 0.55 |
| 11 | 36805997 | rs925922 | 0.425 | 0.713 | 0.418 | -0.066 | 0.049 | -0.005 | -0.005 | 0 | 0.11 | 0.2 | 0.2 | 0.09 | 0.16 | 0.08 |
| 11 | 36810334 | rs7928723 | -0.025 | -0.177 | 0.928 | -0.194 | -0.001 | 0.003 | 0.003 | 0 | 0.53 | 0.56 | 0.38 | 0.29 | 0.52 | 0.44 |
| 11 | 36812881 | rs4755461 | 0.772 | -0.241 | 1.102 | -0.224 | 0.003 | 0.012 | 0.012 | 0 | 0.42 | 0.37 | 0.25 | 0.14 | 0.41 | 0.31 |
| 11 | 36825618 | rs7117886 | -0.453 | -0.26 | 1.001 | -0.239 | -0.035 | 0.01 | 0.01 | 0 | 0.58 | 0.76 | 0.46 | 0.3 | 0.53 | 0.46 |
| 11 | 36827869 | rs1518754 | -0.171 | -0.185 | 1.036 | -0.189 | 0 | 0.018 | 0.018 | 0 | 0.11 | 0.19 | 0.13 | 0.16 | 0.11 | 0.13 |
| 11 | 36834398 | rs7119238 | 0.69 | -0.173 | 0.982 | -0.184 | 0.003 | 0.028 | 0.028 | 0 | 0.05 | 0.22 | 0.08 | 0.02 | 0.01 | 0.01 |
| 11 | 36859326 | rs11033833 | -0.278 | 0.255 | 0.374 | -0.126 | 0 | 0.009 | 0.009 | 0 | 0.12 | 0.16 | 0.13 | 0.16 | 0.11 | 0.13 |
| 11 | 36863551 | rs10836622 | -0.477 | 0.337 | 0.468 | 0.133 | 0.001 | -0.008 | -0.008 | 0 | 0.47 | 0.47 | 0.62 | 0.71 | 0.48 | 0.56 |
| 11 | 36873565 | rs7104698 | 0.422 | 0.267 | 0.478 | 1.014 | 0.007 | 0.08 | 0.08 | 0.057 | 0.84 | 0.35 | 0.73 | 0.69 | 0.87 | 0.79 |
| 11 | 36874579 | rs986996 | 0.244 | 0.13 | -0.625 | 0.445 | -0.119 | 0.134 | 0.134 | 0 | 0.33 | 0.1 | 0.45 | 0.13 | 0.52 | 0.27 |
| 11 | 36877740 | rs12222306 | 0.378 | -0.263 | -0.475 | 0.48 | 0.07 | -0.011 | -0.011 | 0.054 | 0.47 | 0.11 | 0.27 | 0.56 | 0.33 | 0.53 |
| 11 | 36884595 | rs7935463 | 0.447 | -0.408 | -0.711 | 0.483 | 0.052 | -0.048 | -0.048 | 0 | 0.46 | 0.07 | 0.26 | 0.55 | 0.33 | 0.52 |
| 11 | 36889740 | rs7934277 | 0.822 | -0.659 | -0.863 | 0.685 | 0.044 | 0.164 | 0.164 | 0.587 | 0.37 | 0.08 | 0.12 | 0.13 | 0.26 | 0.35 |
| 11 | 36892152 | rs10836628 | -0.173 | -0.244 | -0.419 | 0.996 | -0.006 | 0.111 | 0.111 | 0.064 | 0.09 | 0.12 | 0.14 | 0.42 | 0.07 | 0.17 |
| 11 | 36892288 | rs12272065 | -0.173 | -0.244 | -0.419 | 0.975 | -0.006 | 0.112 | 0.112 | 0.064 | 0.09 | 0.12 | 0.14 | 0.43 | 0.07 | 0.17 |
| 11 | 36896281 | rs4517489 | 0.013 | -0.332 | -0.274 | 0.446 | 0.007 | 0.008 | 0.008 | 0.015 | 0.53 | 0.48 | 0.45 | 0.71 | 0.34 | 0.64 |
| 11 | 36896591 | rs4434979 | -0.483 | 0.034 | -0.233 | 0.585 | -0.001 | 0 | 0 | 0 | 0.44 | 0.42 | 0.59 | 0.66 | 0.59 | 0.55 |
| 11 | 36897721 | rs10836630 | -1.317 | 0.277 | -0.011 | 0.878 | -0.009 | 0.012 | 0.012 | 0 | 0.26 | 0.38 | 0.38 | 0.75 | 0.24 | 0.46 |
| 11 | 36897737 | rs7924804 | 0.787 | 0.102 | 0.003 | 1.49 | -0.012 | 0.036 | 0.036 | 0 | 0.87 | 0.83 | 0.94 | 0.65 | 0.92 | 0.67 |
| 11 | 36898350 | rs4304747 | -0.226 | 0.215 | -1.105 | 0.394 | -0.016 | 0.186 | 0.186 | 0.13 | 0.53 | 0.85 | 0.59 | 0.77 | 0.47 | 0.7 |
| 11 | 36898373 | rs7928357 | -1.386 | 0.609 | -0.032 | 0.281 | 0.1 | 0.003 | 0.003 | 0.151 | 0.06 | 0.33 | 0.06 | 0 | 0.06 | 0.01 |
| 11 | 36898750 | rs7120395 | 0.966 | 0.399 | -1.066 | 0.964 | -0.088 | -0.16 | -0.16 | 0 | 0.63 | 0.09 | 0.66 | 0.57 | 0.61 | 0.46 |
| 11 | 36906202 | rs4307696 | 0.119 | 0.402 | 0.001 | 1.534 | 0.019 | 0.107 | 0.107 | 0.693 | 0.93 | 0.73 | 0.97 | 0.85 | 0.96 | 0.67 |
| 11 | 36923515 | rs2127729 | 1.165 | 0.985 | -0.232 | 1.099 | -0.05 | -0.034 | -0.034 | 0 | 0.34 | 0.05 | 0.4 | 0.32 | 0.36 | 0.24 |
| 11 | 36938773 | rs1383349 | 1.204 | 0.907 | -0.283 | 1.151 | -0.027 | -0.121 | -0.121 | 0 | 0.68 | 0.12 | 0.74 | 0.57 | 0.73 | 0.49 |
| 11 | 36951734 | rs1480547 | -0.384 | 0.93 | -0.183 | 1.258 | 0.009 | 0.042 | 0.042 | 0 | 0.19 | 0.2 | 0.19 | 0.29 | 0.21 | 0.15 |
| 11 | 36952592 | rs10836640 | 0.696 | 1.048 | -0.249 | 1.229 | -0.06 | 0.122 | 0.122 | 0 | 0.37 | 0.06 | 0.34 | 0.24 | 0.37 | 0.28 |
| 11 | 36960292 | rs1825015 | 0.071 | 1.069 | 0.071 | 1.311 | -0.001 | -0.007 | -0.007 | 0 | 0.16 | 0.11 | 0.18 | 0.28 | 0.18 | 0.14 |
| 11 | 36962971 | rs1383348 | -0.205 | 0.708 | -0.28 | 1.043 | 0.009 | -0.014 | -0.014 | 0 | 0.46 | 0.21 | 0.49 | 0.52 | 0.54 | 0.35 |
| 11 | 36967260 | rs1480550 | 0.628 | 0.705 | -0.038 | 1.1 | 0 | 0.084 | 0.084 | 4.936 | 0.9 | 0.31 | 0.92 | 0.84 | 0.95 | 0.68 |
| 11 | 36969003 | rs764848 | 0.932 | 0.838 | -0.355 | 0.918 | -0.086 | 0.071 | 0.071 | 0 | 0.28 | 0.02 | 0.3 | 0.22 | 0.31 | 0.2 |
| 11 | 36969254 | rs764849 | 0.624 | 0.797 | -0.351 | 0.9 | 0.011 | 0.035 | 0.035 | 0 | 0.1 | 0.04 | 0.05 | 0.01 | 0.06 | 0.07 |

| | | | | | | | | | | | | | | | | |
|---|---|---|---|---|---|---|---|---|---|---|---|---|---|---|---|---|
| 11 | 36970447 | rs10836642 | 0.832 | 0.781 | -0.382 | 0.891 | -0.086 | 0.073 | 0.073 | 0 | 0.29 | 0.02 | 0.3 | 0.22 | 0.33 | 0.2 |
| 11 | 36975569 | rs10836643 | 1.651 | 0.656 | 0.117 | 1.028 | 0.009 | 0.004 | 0.004 | 0.777 | 0.36 | 0.32 | 0.41 | 0.33 | 0.34 | 0.26 |
| 11 | 36982289 | rs4756360 | 0.105 | 0.832 | 0.22 | 1.37 | -0.005 | 0.074 | 0.074 | 0 | 0.14 | 0.18 | 0.15 | 0.26 | 0.14 | 0.12 |
| 11 | 36983770 | rs1600591 | 0.523 | 1.165 | -0.093 | 1.197 | 0.125 | 0.194 | 0.194 | 1.315 | 0.91 | 0.57 | 0.93 | 0.82 | 0.91 | 0.67 |
| 11 | 36983798 | rs1600590 | -0.167 | 1.227 | -0.413 | 1.18 | 0.126 | 0.115 | 0.115 | 0.183 | 0.93 | 0.69 | 0.94 | 0.88 | 0.92 | 0.85 |
| 11 | 36985037 | rs10836645 | 0.52 | 1.202 | -0.617 | 1.059 | -0.115 | 0.088 | 0.088 | 0.004 | 0.31 | 0.02 | 0.32 | 0.23 | 0.37 | 0.22 |
| 11 | 36987205 | rs4756362 | 0.3 | 1.029 | 0.002 | 1.268 | 0.094 | 0.163 | 0.163 | 0.598 | 0.92 | 0.68 | 0.94 | 0.81 | 0.92 | 0.68 |
| 11 | 36988302 | rs10836648 | -0.552 | 0.952 | -1.009 | 0.801 | -0.013 | 0.08 | 0.08 | 0 | 0.44 | 0.29 | 0.49 | 0.3 | 0.55 | 0.31 |
| 11 | 36989149 | rs11033890 | 1.484 | 1.245 | -1.323 | 0.997 | -0.065 | -0.136 | -0.136 | 0 | 0.62 | 0.08 | 0.68 | 0.54 | 0.67 | 0.49 |
| 11 | 36994253 | rs433374 | 0.205 | 1.448 | -0.629 | 0.28 | 0.002 | -0.02 | -0.02 | 0 | 0.83 | 0.81 | 0.87 | 0.91 | 0.82 | 0.87 |
| 11 | 36995113 | rs413460 | 0.979 | 1.528 | -0.997 | 0.296 | 0.06 | 0.079 | 0.079 | 0.125 | 0.46 | 0.29 | 0.34 | 0.3 | 0.36 | 0.45 |
| 11 | 36999042 | rs1849342 | 1.9 | 1.459 | -1.094 | 0.265 | 0.046 | 0.038 | 0.038 | 0.307 | 0.36 | 0.05 | 0.27 | 0.25 | 0.26 | 0.27 |
| 11 | 36999385 | rs1871684 | 0.717 | 1.516 | -1.153 | 0.396 | -0.068 | -0.109 | -0.109 | 0.307 | 0.25 | 0.1 | 0.47 | 0.49 | 0.37 | 0.2 |
| 11 | 37002821 | rs10768240 | 0.545 | 1.529 | -1.158 | 0.39 | -0.086 | -0.112 | -0.112 | 0 | 0.27 | 0.1 | 0.5 | 0.5 | 0.42 | 0.23 |
| 11 | 37008440 | rs2199895 | 1.995 | 1.447 | -1.152 | 0.388 | 0.019 | 0.012 | 0.012 | 0.317 | 0.35 | 0.02 | 0.27 | 0.25 | 0.26 | 0.27 |
| 11 | 37009621 | rs1119048 | 1.951 | 1.496 | -1.14 | 0.387 | 0.035 | 0.036 | 0.036 | 0.268 | 0.36 | 0.05 | 0.27 | 0.26 | 0.27 | 0.27 |
| 11 | 37014014 | rs4756365 | 0.768 | 1.26 | -1.117 | 0.406 | -0.067 | -0.106 | -0.106 | 0.747 | 0.25 | 0.1 | 0.47 | 0.48 | 0.37 | 0.2 |
| 11 | 37015305 | rs12419135 | 0.336 | 0.637 | -0.768 | 0.174 | 0.003 | 0.039 | 0.039 | 0 | 0.09 | 0.02 | 0.04 | 0.01 | 0.06 | 0.06 |
| 11 | 37020611 | rs333822 | 0.982 | 0.794 | -0.526 | 0.391 | -0.001 | 0.001 | 0.001 | 0.171 | 0.71 | 0.39 | 0.81 | 0.8 | 0.73 | 0.65 |
| 11 | 37043351 | rs333805 | 0.423 | 0.754 | -1.051 | 0.759 | -0.057 | -0.178 | -0.178 | 1.221 | 0.23 | 0.03 | 0.39 | 0.51 | 0.26 | 0.31 |
| 11 | 37044130 | rs333808 | 0.973 | 0.728 | -1.184 | 0.376 | -0.005 | 0.042 | 0.042 | 0 | 0.62 | 0.29 | 0.71 | 0.77 | 0.62 | 0.57 |
| 11 | 37044901 | rs12788604 | 1.724 | 0.77 | -1.251 | 0.322 | -0.062 | 0.063 | 0.063 | 0.013 | 0.38 | 0.06 | 0.33 | 0.26 | 0.37 | 0.27 |
| 11 | 37045724 | rs333810 | 1.003 | 0.782 | -1.228 | 0.359 | -0.032 | -0.173 | -0.173 | 2.63 | 0.21 | 0.03 | 0.36 | 0.52 | 0.21 | 0.28 |
| 11 | 37046609 | rs7129384 | -0.292 | 0.58 | -0.709 | 0.282 | 0.002 | -0.016 | -0.016 | 0.989 | 0.22 | 0.24 | 0.17 | 0.13 | 0.24 | 0.16 |
| 11 | 37051399 | rs7128101 | 1.46 | 0.627 | -0.795 | 0.261 | -0.046 | 0.09 | 0.09 | 0 | 0.4 | 0.13 | 0.35 | 0.25 | 0.41 | 0.29 |
| 11 | 37060502 | rs7129592 | 1.444 | 0.511 | -0.615 | 0.246 | -0.021 | 0.005 | 0.005 | 0.399 | 0.61 | 0.11 | 0.72 | 0.7 | 0.63 | 0.49 |
| 11 | 37060662 | rs16929844 | 0.11 | 0.429 | -0.36 | 0.113 | 0.017 | -0.048 | -0.048 | 0.268 | 0.09 | 0.29 | 0.06 | 0 | 0.07 | 0.06 |
| 11 | 37061671 | rs4442530 | 0.763 | 0.417 | -0.663 | 0.269 | 0.007 | 0.028 | 0.028 | 0.017 | 0.64 | 0.28 | 0.75 | 0.76 | 0.67 | 0.6 |
| 11 | 37063291 | rs2089917 | 0.805 | 0.433 | -0.77 | 0.276 | 0.007 | 0.035 | 0.035 | 0.042 | 0.63 | 0.22 | 0.75 | 0.77 | 0.67 | 0.61 |
| 11 | 37080974 | rs333786 | 1.217 | 0.26 | -0.664 | 0.249 | 0 | 0.049 | 0.049 | 0.087 | 0.41 | 0.26 | 0.34 | 0.22 | 0.38 | 0.25 |
| 11 | 37083616 | rs12788923 | -0.787 | 0.358 | -0.523 | 0.148 | 0.015 | -0.047 | -0.047 | 0.248 | 0.32 | 0.45 | 0.25 | 0.2 | 0.36 | 0.35 |
| 11 | 37090499 | rs10768243 | 0.757 | 0.327 | -0.856 | 0.345 | -0.04 | -0.175 | -0.175 | 0 | 0.22 | 0.02 | 0.39 | 0.47 | 0.23 | 0.24 |
| 11 | 37100764 | rs12289536 | 1.464 | 0.233 | -0.68 | -0.063 | -0.01 | 0.038 | 0.038 | 0 | 0.38 | 0.23 | 0.34 | 0.24 | 0.38 | 0.33 |
| 11 | 37102463 | rs10768245 | -0.273 | 0.293 | -1.102 | -0.055 | 0.012 | -0.001 | -0.001 | 0 | 0.69 | 0.78 | 0.58 | 0.37 | 0.7 | 0.54 |
| 11 | 37106860 | rs11033931 | 1.521 | 0.267 | -0.971 | -0.23 | 0.005 | -0.051 | -0.051 | 0 | 0.38 | 0.42 | 0.35 | 0.2 | 0.38 | 0.25 |
| 11 | 37108723 | rs1512371 | -0.36 | 0.316 | -1.011 | -0.172 | 0.006 | -0.002 | -0.002 | 0.305 | 0.7 | 0.78 | 0.57 | 0.36 | 0.69 | 0.56 |
| 11 | 37109186 | rs1512372 | 0.105 | 0.625 | -1.586 | -0.139 | -0.064 | 0.096 | 0.096 | 1.183 | 0.17 | 0.02 | 0.13 | 0.02 | 0.2 | 0.12 |
| 11 | 37111867 | rs1160627 | -0.334 | 0.657 | -1.381 | 0.069 | -0.001 | -0.027 | -0.027 | 0 | 0.31 | 0.36 | 0.23 | 0.12 | 0.32 | 0.21 |
| 11 | 37113368 | rs1858417 | -0.305 | 0.712 | -1.356 | 0.088 | 0 | -0.018 | -0.018 | 0 | 0.31 | 0.36 | 0.23 | 0.16 | 0.31 | 0.31 |
| 11 | 37115656 | rs1512369 | -0.531 | 0.675 | -1.165 | 0.882 | -0.07 | -0.146 | -0.146 | 0.172 | 0.26 | 0.02 | 0.43 | 0.55 | 0.31 | 0.33 |
| 11 | 37115843 | rs1512368 | -0.535 | 0.592 | -1.662 | 0.812 | -0.039 | -0.13 | -0.13 | 0 | 0.26 | 0.03 | 0.41 | 0.56 | 0.28 | 0.34 |
| 11 | 37116815 | rs746517 | -0.107 | 0.8 | -1.011 | 0.38 | 0.005 | -0.003 | -0.003 | 0 | 0.51 | 0.42 | 0.45 | 0.53 | 0.46 | 0.57 |
| 11 | 37119431 | rs4756369 | 0.661 | 0.991 | -0.093 | -0.068 | 0.049 | -0.042 | -0.042 | 0 | 0.87 | 0.53 | 0.88 | 0.95 | 0.89 | 0.91 |
| 11 | 37119712 | rs17651845 | 0.546 | 1.491 | -0.982 | -0.557 | -0.057 | 0.145 | 0.145 | 0 | 0.33 | 0.07 | 0.27 | 0.1 | 0.37 | 0.2 |
| 11 | 37126189 | rs428215 | -0.269 | 1.924 | -0.346 | 0.101 | 0.061 | 0.145 | 0.145 | 0 | 0.07 | 0.11 | 0.09 | 0.28 | 0.12 | 0.13 |
| 11 | 37131842 | rs10501169 | 0.252 | 1.345 | -0.472 | -0.375 | -0.001 | -0.022 | -0.022 | 0 | 0.16 | 0.16 | 0.07 | 0.01 | 0.15 | 0.17 |
| 11 | 37133420 | rs7946463 | 0.659 | 1.871 | -0.021 | 0.001 | 0.09 | 0.141 | 0.141 | 0.065 | 0.81 | 0.54 | 0.8 | 0.67 | 0.78 | 0.79 |
| 11 | 37134171 | rs10768246 | 0.584 | 1.641 | -0.639 | -0.637 | -0.014 | 0.051 | 0.051 | 0 | 0.33 | 0.21 | 0.28 | 0.1 | 0.36 | 0.2 |
| 11 | 37140316 | rs12578014 | -0.553 | 1.887 | -0.775 | -0.16 | -0.021 | -0.01 | -0.01 | 0.081 | 0.15 | 0.06 | 0.12 | 0.14 | 0.18 | 0.16 |
| 11 | 37140672 | rs334980 | -0.296 | 1.319 | -0.261 | -0.115 | 0 | 0 | 0 | 0 | 0.45 | 0.44 | 0.54 | 0.75 | 0.48 | 0.51 |
| 11 | 37140732 | rs334979 | -0.754 | 1.402 | -0.197 | -0.183 | -0.002 | 0.002 | 0.002 | 0 | 0.52 | 0.57 | 0.63 | 0.79 | 0.56 | 0.56 |
| 11 | 37141853 | rs11033958 | 0.056 | 0.989 | -0.254 | 0.033 | -0.004 | 0.003 | 0.003 | 0 | 0.09 | 0.09 | 0.08 | 0.04 | 0.16 | 0.13 |
| 11 | 37142093 | rs334978 | -0.165 | 0.232 | -0.163 | 0.679 | 0.002 | -0.004 | -0.004 | 0.123 | 0.34 | 0.3 | 0.42 | 0.6 | 0.29 | 0.3 |
| 11 | 37145360 | rs1983257 | -1.233 | 0.824 | 0.339 | 1.159 | 0.036 | 0.11 | 0.11 | 0 | 0.15 | 0.34 | 0.27 | 0.27 | 0.17 | 0.2 |
| 11 | 37145651 | rs2704917 | -0.278 | 0.794 | -0.827 | 1.268 | -0.046 | -0.138 | -0.138 | 0.31 | 0.25 | 0.04 | 0.15 | 0.53 | 0.27 | 0.3 |
| 11 | 37150239 | rs2614171 | -1.024 | 0.181 | -0.764 | 0.555 | -0.058 | 0.144 | 0.144 | 1.915 | 0.41 | 0.17 | 0.52 | 0.9 | 0.51 | 0.56 |
| 11 | 37152778 | rs2614170 | -0.67 | 0.222 | -0.823 | 1.068 | -0.048 | -0.008 | -0.008 | 2.021 | 0.33 | 0.13 | 0.41 | 0.7 | 0.43 | 0.5 |
| 11 | 37154319 | rs2662201 | -0.865 | 0.094 | -0.814 | 1.117 | -0.061 | 0.061 | 0.061 | 3.074 | 0.35 | 0.16 | 0.5 | 0.81 | 0.48 | 0.54 |
| 11 | 37154519 | rs11033967 | 1.606 | -0.727 | -1.209 | 0.65 | -0.107 | 0.144 | 0.144 | 0.582 | 0.45 | 0.06 | 0.42 | 0.16 | 0.45 | 0.32 |
| 11 | 37159857 | rs11822786 | -0.466 | -1.084 | -0.219 | 0.688 | -0.069 | -0.088 | -0.088 | 0 | 0.15 | 0.5 | 0.12 | 0.04 | 0.06 | 0.16 |
| 11 | 37172103 | rs12292339 | -1.015 | -0.949 | -0.058 | 0.822 | -0.078 | -0.031 | -0.031 | 0 | 0.17 | 0.52 | 0.13 | 0.14 | 0.07 | 0.19 |
| 11 | 37182945 | rs1563951 | -0.177 | -0.864 | 0.222 | 0.726 | -0.04 | -0.049 | -0.049 | 0 | 0.12 | 0.46 | 0.1 | 0.04 | 0.05 | 0.16 |
| 11 | 37187241 | rs716365 | 0.309 | -0.6 | 0.102 | 1.143 | -0.026 | -0.034 | -0.034 | 0 | 0.23 | 0.16 | 0.38 | 0.46 | 0.4 | 0.24 |
| 11 | 37193333 | rs4756372 | -0.778 | -0.611 | -0.538 | 0.968 | -0.06 | -0.014 | -0.014 | 1.026 | 0.2 | 0.39 | 0.14 | 0.23 | 0.09 | 0.4 |
| 11 | 37200795 | rs2662199 | -0.013 | -0.441 | -0.623 | 0.966 | 0.016 | 0.017 | 0.017 | 1.306 | 0.67 | 0.43 | 0.57 | 0.47 | 0.54 | 0.71 |
| 11 | 37205935 | rs10768256 | -0.913 | -0.497 | -0.232 | 0.88 | -0.055 | 0.01 | 0.01 | 0 | 0.19 | 0.37 | 0.14 | 0.25 | 0.09 | 0.4 |
| 11 | 37217499 | rs7130755 | -0.927 | -0.287 | -0.435 | 0.645 | -0.001 | 0.094 | 0.094 | 1.576 | 0.49 | 0.69 | 0.57 | 0.79 | 0.52 | 0.66 |
| 11 | 37222358 | rs11034021 | -0.859 | -0.76 | -0.096 | 1.143 | -0.044 | 0.065 | 0.065 | 1.128 | 0.11 | 0.6 | 0.28 | 0.17 | 0.05 | 0.1 |
| 11 | 37232526 | rs2100819 | -0.906 | -0.818 | -0.113 | 0.659 | 0.016 | 0.037 | 0.037 | 2.043 | 0.55 | 0.62 | 0.54 | 0.77 | 0.45 | 0.68 |

| | | | | | | | | | | | | | | | | |
|---|---|---|---|---|---|---|---|---|---|---|---|---|---|---|---|---|
| 11 | 37235843 | rs2100818 | -0.418 | -0.99 | 0.04 | 1.247 | 0.002 | 0 | 0 | 0.654 | 0.46 | 0.53 | 0.47 | 0.6 | 0.41 | 0.47 |
| 11 | 37249967 | rs4756376 | 0.362 | -0.98 | -0.342 | 1.588 | -0.009 | -0.114 | -0.114 | 0 | 0.34 | 0.05 | 0.26 | 0.49 | 0.31 | 0.37 |
| 11 | 37252050 | rs9633876 | -0.56 | -1.086 | 0.436 | 1.602 | -0.024 | 0.03 | 0.03 | 0.235 | 0.1 | 0.34 | 0.22 | 0.09 | 0.05 | 0.09 |
| 11 | 37256343 | rs11034037 | -0.836 | -1.268 | 0.1 | 1.173 | 0 | -0.001 | -0.001 | 0 | 0.6 | 0.47 | 0.61 | 0.64 | 0.49 | 0.78 |
| 11 | 37283532 | rs6484915 | -0.436 | -1.352 | 0.161 | 1.019 | 0.029 | 0.009 | 0.009 | 2.21 | 0.54 | 0.33 | 0.54 | 0.61 | 0.45 | 0.63 |
| 11 | 37285494 | rs11034056 | -0.761 | -1.349 | -0.047 | 1.047 | -0.002 | 0.003 | 0.003 | 0.287 | 0.61 | 0.49 | 0.55 | 0.64 | 0.48 | 0.63 |
| 11 | 37287828 | rs2177216 | -0.782 | -1.347 | -0.046 | 1.048 | -0.002 | -0.004 | -0.004 | 0 | 0.61 | 0.49 | 0.57 | 0.67 | 0.48 | 0.66 |
| 11 | 37288175 | rs1343828 | 0.032 | -1.427 | -0.112 | 1.024 | 0.011 | 0.02 | 0.02 | 0 | 0.16 | 0.01 | 0.07 | 0.07 | 0.12 | 0.17 |
| 11 | 37291017 | rs1949703 | -1.308 | -1.413 | -0.05 | 1.024 | 0.007 | 0 | 0 | 1.669 | 0.66 | 0.62 | 0.64 | 0.71 | 0.52 | 0.81 |
| 11 | 37302238 | rs10836719 | -0.619 | -1.401 | -0.165 | 1.075 | -0.02 | 0.041 | 0.041 | 0.585 | 0.1 | 0.4 | 0.22 | 0.09 | 0.05 | 0.09 |
| 11 | 37306300 | rs7927846 | 0.989 | -1.404 | -0.244 | 1.06 | -0.001 | -0.102 | -0.102 | 0 | 0.34 | 0.06 | 0.26 | 0.48 | 0.31 | 0.37 |
| 11 | 37313489 | rs7934426 | -0.249 | -1.147 | -0.092 | 1.047 | 0.009 | -0.069 | -0.069 | 1.528 | 0.53 | 0.09 | 0.38 | 0.58 | 0.44 | 0.65 |
| 11 | 37317728 | rs7129803 | -1.167 | -1.063 | 0.263 | 1.075 | -0.039 | 0.072 | 0.072 | 0.382 | 0.11 | 0.43 | 0.24 | 0.13 | 0.05 | 0.12 |
| 11 | 37325465 | rs10836727 | -0.995 | -1.019 | 0.252 | 1.051 | -0.012 | 0.051 | 0.051 | 0.979 | 0.11 | 0.47 | 0.24 | 0.1 | 0.07 | 0.09 |
| 11 | 37327721 | rs1916060 | -0.01 | -1.174 | -0.528 | 1.015 | 0.044 | -0.086 | -0.086 | 0 | 0.49 | 0.07 | 0.32 | 0.52 | 0.42 | 0.5 |
| 11 | 37339216 | rs1522677 | -0.462 | -0.773 | -0.531 | 0.889 | -0.076 | -0.048 | -0.048 | 0 | 0.61 | 0.09 | 0.41 | 0.6 | 0.51 | 0.71 |
| 11 | 37344446 | rs4756394 | 0.714 | -0.984 | -0.669 | 0.89 | 0.006 | -0.101 | -0.101 | 1.779 | 0.41 | 0.07 | 0.34 | 0.5 | 0.38 | 0.53 |
| 11 | 37346956 | rs10768265 | -1.122 | -0.59 | 0.319 | 1.035 | -0.05 | -0.003 | -0.003 | 0 | 0.69 | 0.84 | 0.66 | 0.69 | 0.54 | 0.78 |
| 11 | 37350828 | rs1522679 | 0.242 | -0.786 | 0.647 | 0.996 | 0.013 | 0.003 | 0.003 | 0 | 0.55 | 0.31 | 0.4 | 0.61 | 0.47 | 0.56 |
| 11 | 37354863 | rs2177212 | 0.279 | -0.559 | 0.792 | 0.993 | -0.029 | 0 | 0 | 0 | 0.55 | 0.14 | 0.42 | 0.62 | 0.51 | 0.61 |
| 11 | 37361820 | rs10836728 | -0.388 | -0.459 | 0.711 | 0.908 | -0.008 | 0.002 | 0.002 | 0 | 0.63 | 0.39 | 0.45 | 0.68 | 0.53 | 0.7 |
| 11 | 37362281 | rs1403264 | -0.922 | -0.45 | 0.255 | 0.876 | -0.148 | -0.009 | -0.009 | 0.912 | 0.69 | 0.93 | 0.68 | 0.75 | 0.55 | 0.78 |
| 11 | 37378324 | rs11034107 | -0.436 | -0.531 | 0.397 | 0.959 | 0.033 | -0.03 | -0.03 | 1.098 | 0.06 | 0.01 | 0.01 | 0.08 | 0.02 | 0.13 |
| 11 | 37396461 | rs1822933 | 0.447 | -0.678 | 0.286 | 0.805 | -0.022 | -0.009 | -0.009 | 0 | 0.53 | 0.22 | 0.44 | 0.59 | 0.52 | 0.56 |
| 11 | 37400375 | rs12224887 | -0.192 | -0.763 | 0.312 | 0.764 | 0.004 | -0.002 | -0.002 | 0 | 0.1 | 0.1 | 0.02 | 0.09 | 0.03 | 0.15 |
| 11 | 37406069 | rs10836736 | 0.741 | -0.754 | 0.311 | 0.745 | -0.089 | -0.056 | -0.056 | 0.352 | 0.51 | 0.05 | 0.44 | 0.6 | 0.52 | 0.57 |
| 11 | 37426627 | rs987827 | 1.181 | -0.99 | 0.054 | 0.477 | -0.061 | -0.141 | -0.141 | 0 | 0.4 | 0.04 | 0.52 | 0.58 | 0.4 | 0.44 |
| 11 | 37433074 | rs7942072 | -0.657 | -0.953 | 0.455 | 0.558 | -0.085 | 0.025 | 0.025 | 0.033 | 0.66 | 0.85 | 0.72 | 0.76 | 0.57 | 0.79 |
| 11 | 37439348 | rs1822934 | 0.69 | -0.97 | 0.775 | 0.548 | -0.041 | -0.006 | -0.006 | 0 | 0.49 | 0.11 | 0.61 | 0.61 | 0.52 | 0.6 |
| 11 | 37449297 | rs7113598 | -0.786 | -0.871 | 0.781 | 0.415 | -0.144 | -0.006 | -0.006 | 0.037 | 0.68 | 0.93 | 0.72 | 0.73 | 0.57 | 0.78 |
| 11 | 37461620 | rs961772 | 0.68 | -0.947 | 0.905 | 0.42 | -0.045 | 0.03 | 0.03 | 0.28 | 0.49 | 0.11 | 0.63 | 0.64 | 0.52 | 0.62 |
| 11 | 37461807 | rs12226252 | -0.946 | -0.963 | 1.28 | 0.263 | -0.065 | -0.006 | -0.006 | 0 | 0.14 | 0.32 | 0.04 | 0.09 | 0.03 | 0.15 |
| 11 | 37476792 | rs2053669 | 0.722 | -0.977 | 1.216 | 0.254 | -0.074 | 0.047 | 0.047 | 0.247 | 0.49 | 0.08 | 0.63 | 0.65 | 0.52 | 0.62 |
| 11 | 37477609 | rs12418089 | 0.693 | -0.981 | 1.47 | 0.253 | -0.001 | 0.048 | 0.048 | 0.053 | 0.51 | 0.31 | 0.66 | 0.65 | 0.54 | 0.63 |
| 11 | 37481054 | rs11034154 | -0.686 | -0.999 | 1.507 | 0.27 | -0.03 | -0.003 | -0.003 | 0 | 0.13 | 0.19 | 0.03 | 0.09 | 0.03 | 0.15 |
| 11 | 37484940 | rs2166283 | 0.548 | -0.894 | 1.567 | 0.271 | -0.001 | 0.004 | 0.004 | 0 | 0.52 | 0.41 | 0.67 | 0.69 | 0.54 | 0.64 |
| 11 | 37489822 | rs11606085 | 0.643 | -0.865 | 0.618 | 0.391 | -0.004 | 0.057 | 0.057 | 0.209 | 0.49 | 0.22 | 0.65 | 0.61 | 0.52 | 0.62 |
| 11 | 37501843 | rs10836745 | -0.557 | -0.436 | -0.222 | 0.479 | -0.147 | -0.004 | -0.004 | 0.008 | 0.68 | 0.91 | 0.67 | 0.73 | 0.58 | 0.79 |
| 11 | 37509065 | rs11034171 | 0.112 | 0.214 | -0.42 | 0.273 | -0.103 | -0.031 | -0.031 | 0 | 0.2 | 0.02 | 0.21 | 0.21 | 0.31 | 0.18 |
| 11 | 37516205 | rs10742390 | -2.053 | -0.096 | -0.316 | 0.017 | 0.014 | 0.059 | 0.059 | 4.01 | 0.5 | 0.69 | 0.4 | 0.38 | 0.52 | 0.5 |
| 11 | 37516664 | rs11034174 | 0 | 0.012 | 0.098 | 0.024 | -0.014 | 0.009 | 0.009 | 0.621 | 0.22 | 0.17 | 0.22 | 0.21 | 0.32 | 0.18 |
| 11 | 37518784 | rs12287198 | -1.705 | -0.101 | 0.144 | -0.045 | -0.008 | -0.036 | -0.036 | 0 | 0.26 | 0.48 | 0.18 | 0.16 | 0.2 | 0.33 |
| 11 | 37519542 | rs10836747 | -0.304 | -0.01 | -0.084 | -0.02 | -0.001 | -0.002 | -0.002 | 0 | 0.1 | 0.06 | 0.09 | 0.11 | 0.11 | 0.17 |
| 11 | 37521486 | rs4630265 | -1.427 | -0.102 | 0.014 | 0.011 | -0.002 | 0.022 | 0.022 | 1.064 | 0.41 | 0.68 | 0.32 | 0.32 | 0.41 | 0.35 |
| 11 | 37524085 | rs10501178 | -0.442 | -0.599 | -0.091 | -0.125 | -0.002 | 0.019 | 0.019 | 0 | 0.12 | 0.07 | 0.05 | 0.06 | 0.08 | 0.12 |
| 11 | 37526394 | rs1545342 | -2.238 | -0.655 | -0.16 | -0.132 | 0.007 | 0.065 | 0.065 | 0.19 | 0.51 | 0.78 | 0.39 | 0.37 | 0.44 | 0.44 |
| 11 | 37528194 | rs1545344 | 0.047 | -0.723 | -0.09 | -0.273 | -0.009 | -0.001 | -0.001 | 0 | 0.15 | 0.14 | 0.14 | 0.14 | 0.2 | 0.2 |
| 11 | 37532517 | rs4356200 | -2.23 | -0.825 | 0.298 | -0.358 | -0.028 | -0.018 | -0.018 | 0.082 | 0.38 | 0.53 | 0.26 | 0.27 | 0.31 | 0.31 |
| 11 | 37547750 | rs7948504 | 0.453 | -0.804 | 0.284 | -0.354 | -0.006 | 0.002 | 0.002 | 0 | 0.07 | 0.05 | 0.08 | 0.06 | 0.09 | 0.12 |
| 11 | 37555803 | rs7108561 | -1.215 | -0.884 | 0.388 | -0.357 | -0.111 | -0.034 | -0.034 | 0 | 0.24 | 0.39 | 0.12 | 0.14 | 0.1 | 0.11 |
| 11 | 37556022 | rs12291143 | -1.904 | -0.859 | 0.28 | -0.339 | -0.063 | -0.041 | -0.041 | 0.022 | 0.29 | 0.5 | 0.17 | 0.17 | 0.15 | 0.14 |
| 11 | 37556897 | rs1461917 | 0.418 | -0.839 | 0.22 | -0.336 | -0.025 | -0.002 | -0.002 | 0 | 0.14 | 0.06 | 0.11 | 0.13 | 0.18 | 0.19 |
| 11 | 37560570 | rs10768268 | -3.342 | -0.845 | 0.222 | -0.334 | 0.007 | 0.05 | 0.05 | 0.168 | 0.51 | 0.77 | 0.39 | 0.39 | 0.44 | 0.47 |
| 11 | 37561201 | rs10836750 | -0.346 | -1.007 | 0.561 | -0.352 | 0.004 | 0.003 | 0.003 | 0.528 | 0.17 | 0.12 | 0.04 | 0.13 | 0.04 | 0.09 |
| 11 | 37564368 | rs12271313 | 0.482 | -0.839 | 0.359 | -0.279 | 0.017 | 0.003 | 0.003 | 0 | 0.07 | 0.14 | 0.09 | 0.09 | 0.09 | 0.13 |
| 11 | 37564954 | rs12271598 | 0.482 | -0.839 | 0.364 | -0.279 | 0.018 | 0.003 | 0.003 | 0 | 0.07 | 0.14 | 0.09 | 0.09 | 0.09 | 0.13 |
| 11 | 37565039 | rs12278705 | 0.358 | -0.84 | 0.36 | -0.279 | 0.016 | 0.009 | 0.009 | 0 | 0.07 | 0.14 | 0.09 | 0.11 | 0.09 | 0.16 |
| 11 | 37569819 | rs10836754 | -1.78 | -0.873 | 0.344 | -0.312 | -0.036 | -0.029 | -0.029 | 2.399 | 0.29 | 0.61 | 0.18 | 0.15 | 0.15 | 0.12 |
| 11 | 37573427 | rs7931655 | -2.808 | -0.705 | 0.241 | -0.228 | 0.022 | 0.061 | 0.061 | 0.061 | 0.5 | 0.74 | 0.37 | 0.36 | 0.42 | 0.46 |
| 11 | 37577216 | rs3980290 | -2.357 | -1.318 | 0.364 | -0.187 | 0.026 | 0.034 | 0.034 | 0.594 | 0.43 | 0.68 | 0.29 | 0.28 | 0.33 | 0.31 |
| 11 | 37579996 | rs7342222 | -2.448 | -1.477 | 1.141 | 0.141 | -0.002 | 0 | 0 | 0.477 | 0.35 | 0.59 | 0.24 | 0.24 | 0.28 | 0.31 |
| 11 | 37590232 | rs1461898 | -2.09 | -1.429 | 1.154 | 0.177 | -0.035 | -0.016 | -0.016 | 0 | 0.31 | 0.48 | 0.18 | 0.23 | 0.21 | 0.29 |
| 11 | 37591199 | rs7932763 | -0.882 | -1.335 | 1.168 | -0.071 | -0.02 | -0.05 | -0.05 | 0 | 0.2 | 0.41 | 0.17 | 0.1 | 0.19 | 0.21 |
| 11 | 37630303 | rs10836807 | -1.841 | -1.363 | 1.158 | -0.105 | -0.028 | -0.021 | -0.021 | 0.184 | 0.28 | 0.52 | 0.18 | 0.21 | 0.21 | 0.27 |
| 11 | 37631526 | rs4528301 | -0.751 | -1.37 | 1.301 | -0.116 | -0.016 | -0.046 | -0.046 | 0 | 0.19 | 0.33 | 0.15 | 0.08 | 0.17 | 0.18 |
| 11 | 37646906 | rs1381422 | -1.826 | -1.335 | 1.094 | -0.082 | -0.043 | 0.005 | 0.005 | 0 | 0.15 | 0.42 | 0.12 | 0.14 | 0.11 | 0.1 |
| 11 | 37651249 | rs1599505 | -1.371 | -1.364 | 1.036 | -0.098 | -0.038 | 0.023 | 0.023 | 0 | 0.13 | 0.46 | 0.06 | 0.14 | 0.05 | 0.1 |
| 11 | 37654158 | rs7938719 | -2.127 | -1.458 | 0.949 | -0.153 | -0.002 | -0.001 | -0.001 | 1.345 | 0.31 | 0.62 | 0.18 | 0.23 | 0.21 | 0.3 |
| 11 | 37667462 | rs767289 | -3.044 | -1.356 | 0.852 | -0.088 | 0.046 | 0.028 | 0.028 | 0.311 | 0.43 | 0.77 | 0.32 | 0.38 | 0.37 | 0.44 |

| | | | | | | | | | | | | | | | | |
|---|---|---|---|---|---|---|---|---|---|---|---|---|---|---|---|---|
| 11 | 37669581 | rs12362125 | -2.598 | -1.326 | 0.875 | -0.044 | 0.026 | 0.023 | 0.023 | 1.022 | 0.36 | 0.69 | 0.25 | 0.25 | 0.28 | 0.32 |
| 11 | 37670386 | rs11034323 | -0.641 | -0.926 | 0.983 | 0.137 | 0.004 | 0.007 | 0.007 | 1.736 | 0.19 | 0.16 | 0.13 | 0.1 | 0.17 | 0.21 |
| 11 | 37675392 | rs1073480 | 0.311 | -1.268 | 0.512 | -0.241 | -0.004 | -0.031 | -0.031 | 0.269 | 0.75 | 0.69 | 0.83 | 0.87 | 0.77 | 0.73 |
| 11 | 37683652 | rs17665586 | 0.091 | -1.085 | 0.09 | -0.017 | 0.014 | -0.036 | -0.036 | 0 | 0.07 | 0 | 0.01 | 0.11 | 0.02 | 0.08 |
| 11 | 37688427 | rs7939975 | 0.661 | -0.692 | 0.463 | -0.018 | -0.002 | 0.009 | 0.009 | 0.223 | 0.62 | 0.68 | 0.64 | 0.69 | 0.55 | 0.57 |
| 11 | 37688849 | rs10501184 | 1.737 | -0.665 | -0.128 | -0.366 | -0.096 | -0.081 | -0.081 | 0 | 0.42 | 0.08 | 0.54 | 0.49 | 0.43 | 0.39 |
| 11 | 37698742 | rs12281581 | -0.443 | -1.132 | 0.157 | -0.365 | -0.011 | -0.031 | -0.031 | 0 | 0.13 | 0.19 | 0.04 | 0.02 | 0.06 | 0.02 |
| 11 | 37700125 | rs1461903 | 1.58 | -0.669 | -0.218 | -0.216 | -0.087 | -0.079 | -0.079 | 0 | 0.41 | 0.08 | 0.53 | 0.49 | 0.42 | 0.38 |
| 11 | 37706893 | rs10400338 | -0.326 | -0.436 | 0.622 | -0.798 | 0.014 | -0.037 | -0.037 | 0 | 0.07 | 0.15 | 0.07 | 0 | 0.09 | 0.02 |
| 11 | 37757910 | rs7938114 | -0.085 | -0.446 | 0.58 | -0.841 | -0.006 | -0.051 | -0.051 | 0 | 0.1 | 0.15 | 0.08 | 0 | 0.1 | 0.02 |
| 11 | 37758393 | rs1515038 | -0.07 | -0.775 | -0.015 | -0.553 | 0.01 | -0.117 | -0.117 | 0.459 | 0.2 | 0.01 | 0.22 | 0.58 | 0.17 | 0.4 |
| 11 | 37768477 | rs11034401 | 2.04 | 0.327 | 0.824 | -1.37 | -0.065 | 0.21 | 0.21 | 0 | 0.61 | 0.11 | 0.67 | 0.88 | 0.56 | 0.73 |
| 11 | 37793480 | rs292801 | -0.417 | 0.391 | 0.556 | -1.369 | -0.011 | 0.006 | 0.006 | 0 | 0.11 | 0.06 | 0.12 | 0.1 | 0.14 | 0.08 |
| 11 | 37808290 | rs7105398 | -0.424 | 0.316 | 0.502 | -1.318 | 0.064 | -0.038 | -0.038 | 0 | 0.12 | 0.29 | 0.11 | 0.05 | 0.2 | 0.14 |
| 11 | 37825890 | rs292752 | -0.06 | 0.049 | -0.653 | -1.411 | 0.016 | 0.008 | 0.008 | 0 | 0.88 | 0.92 | 0.93 | 1 | 0.91 | 0.97 |
| 11 | 37832410 | rs292744 | -0.007 | 0.053 | -0.638 | -1.409 | -0.004 | -0.027 | -0.027 | 0 | 0.87 | 0.87 | 0.92 | 1 | 0.91 | 0.97 |
| 11 | 37834436 | rs11034430 | 0.74 | 0.322 | -1.023 | -1.154 | -0.054 | 0.204 | 0.204 | 0 | 0.61 | 0.13 | 0.67 | 0.88 | 0.57 | 0.73 |
| 11 | 37848312 | rs292754 | 1.452 | 0.059 | 0.688 | -0.375 | -0.012 | -0.004 | -0.004 | 0 | 0.86 | 0.26 | 0.8 | 0.82 | 0.88 | 0.75 |
| 11 | 37873350 | rs292774 | 1.599 | 0.017 | 1.5 | -0.395 | -0.01 | -0.005 | -0.005 | 0.051 | 0.86 | 0.24 | 0.79 | 0.82 | 0.88 | 0.76 |
| 11 | 37883072 | rs10836878 | -0.881 | -0.477 | 1.357 | -0.576 | 0.013 | 0.003 | 0.003 | 0 | 0.11 | 0.07 | 0.11 | 0.14 | 0.08 | 0.18 |
| 11 | 37891119 | rs2705413 | 1.367 | -0.509 | 1.895 | -0.591 | -0.042 | -0.006 | -0.006 | 0 | 0.85 | 0.46 | 0.8 | 0.82 | 0.89 | 0.79 |
| 11 | 37908227 | rs2705423 | 1.795 | -0.536 | 1.754 | -0.604 | -0.023 | -0.01 | -0.01 | 0 | 0.82 | 0.26 | 0.78 | 0.8 | 0.89 | 0.79 |
| 11 | 37909208 | rs12222979 | -0.567 | -0.5 | 1.677 | -0.587 | 0.011 | -0.001 | -0.001 | 0 | 0.09 | 0.07 | 0.1 | 0.15 | 0.07 | 0.15 |
| 11 | 37919311 | rs10836887 | -2.079 | -0.434 | 1.809 | -0.601 | -0.039 | -0.016 | -0.016 | 0 | 0.16 | 0.61 | 0.22 | 0.19 | 0.12 | 0.21 |
| 11 | 37929587 | rs11034498 | -0.55 | -0.373 | 1.493 | -0.556 | 0.007 | -0.002 | -0.002 | 3.724 | 0.09 | 0.07 | 0.11 | 0.15 | 0.08 | 0.15 |
| 11 | 37938425 | rs11034506 | -2.137 | -0.563 | 2.045 | -0.652 | 0.013 | 0.004 | 0.004 | 0.546 | 0.07 | 0.3 | 0.03 | 0 | 0.02 | 0.05 |
| 11 | 37941821 | rs16930850 | -0.923 | -0.569 | 2.466 | -0.676 | 0.004 | 0.038 | 0.038 | 0.255 | 0.11 | 0.36 | 0.14 | 0.19 | 0.09 | 0.18 |
| 11 | 37944909 | rs1478762 | 0.845 | -0.545 | 0.985 | -0.962 | -0.182 | -0.043 | -0.043 | 0.875 | 0.36 | 0.02 | 0.39 | 0.36 | 0.4 | 0.41 |
| 11 | 37945831 | rs16930853 | -0.362 | -0.206 | 1.679 | -1.222 | -0.003 | 0.003 | 0.003 | 0.214 | 0.1 | 0.15 | 0.12 | 0.19 | 0.09 | 0.18 |
| 11 | 37950922 | rs2618291 | 1.584 | -0.222 | 1.574 | -1.23 | -0.027 | -0.014 | -0.014 | 0 | 0.82 | 0.29 | 0.83 | 0.81 | 0.89 | 0.77 |
| 11 | 37954517 | rs4503507 | -0.888 | 0.692 | 1.738 | -1.528 | 0.03 | -0.073 | -0.073 | 0.048 | 0.17 | 0.29 | 0.16 | 0.03 | 0.19 | 0.07 |
| 11 | 37955876 | rs2618294 | 1.438 | 0.62 | 1.734 | -1.499 | 0 | -0.04 | -0.04 | 0 | 0.77 | 0.28 | 0.79 | 0.94 | 0.79 | 0.87 |
| 11 | 37968043 | rs10836904 | -1.748 | 0.621 | 1.715 | -1.54 | 0.011 | -0.125 | -0.125 | 0 | 0.22 | 0.55 | 0.19 | 0.03 | 0.2 | 0.12 |
| 11 | 37968683 | rs7927134 | -1.033 | 0.72 | 1.721 | -1.491 | 0.048 | -0.099 | -0.099 | 0 | 0.18 | 0.42 | 0.18 | 0.03 | 0.19 | 0.07 |
| 11 | 37975643 | rs12286680 | -1.076 | 0.702 | 1.71 | -1.491 | 0.049 | -0.106 | -0.106 | 0 | 0.18 | 0.5 | 0.18 | 0.03 | 0.19 | 0.07 |
| 11 | 37977241 | rs10836905 | -0.764 | 0.712 | 1.432 | -1.486 | 0.013 | -0.047 | -0.047 | 0.078 | 0.16 | 0.19 | 0.16 | 0.03 | 0.19 | 0.07 |
| 11 | 37984194 | rs4408282 | 1.331 | 0.869 | 1.514 | -1.516 | 0.01 | -0.062 | -0.062 | 0 | 0.77 | 0.31 | 0.77 | 0.94 | 0.76 | 0.89 |
| 11 | 37994667 | rs1488524 | -0.593 | 0.921 | 1.04 | -1.455 | -0.027 | 0.037 | 0.037 | 0 | 0.15 | 0.09 | 0.14 | 0.03 | 0.19 | 0.06 |
| 11 | 38002226 | rs10836913 | 1.42 | 0.427 | 0.371 | -2.002 | 0.043 | 0.207 | 0.207 | 0 | 0.45 | 0.1 | 0.52 | 0.8 | 0.39 | 0.58 |
| 11 | 38002718 | rs2129930 | -0.483 | 1.503 | 0.847 | -1.607 | 0.025 | -0.05 | -0.05 | 0.788 | 0.15 | 0.34 | 0.16 | 0.05 | 0.18 | 0.06 |
| 11 | 38015477 | rs11034550 | 0.278 | 1.463 | 0.184 | -1.744 | -0.085 | 0.073 | 0.073 | 0 | 0.17 | 0.04 | 0.18 | 0 | 0.32 | 0.15 |
| 11 | 38017896 | rs984783 | -0.743 | 0.816 | 0.599 | -1.143 | 0.022 | -0.045 | -0.045 | 0 | 0.15 | 0.31 | 0.16 | 0.05 | 0.18 | 0.06 |
| 11 | 38040340 | rs11034562 | -0.724 | 0.501 | 0.468 | -1.189 | 0.012 | 0.052 | 0.052 | 0 | 0.23 | 0.1 | 0.14 | 0.12 | 0.2 | 0.23 |
| 11 | 38048445 | rs10836928 | 0.397 | 1.84 | 0.343 | -1.35 | -0.066 | 0.06 | 0.06 | 0 | 0.1 | 0.02 | 0.14 | 0.03 | 0.26 | 0.1 |
| 11 | 38051454 | rs1679407 | -1.45 | 1.527 | 0.714 | -1.422 | 0.061 | 0.05 | 0.05 | 0.009 | 0.09 | 0.48 | 0.11 | 0.06 | 0.08 | 0.08 |
| 11 | 38070462 | rs12296091 | -1.119 | 1.439 | 1.122 | -1.403 | 0.009 | 0.017 | 0.017 | 0 | 0.08 | 0.22 | 0.09 | 0.06 | 0.07 | 0.08 |
| 11 | 38072445 | rs821027 | -1.214 | 1.405 | 1.346 | -1.419 | 0.038 | 0.04 | 0.04 | 0 | 0.06 | 0.38 | 0.06 | 0.04 | 0.04 | 0.05 |
| 11 | 38073809 | rs11034572 | -1.791 | 1.409 | 1.334 | -1.418 | 0.052 | 0.065 | 0.065 | 0 | 0.09 | 0.51 | 0.12 | 0.06 | 0.07 | 0.08 |
| 11 | 38097003 | rs11034580 | -0.158 | 1.079 | 1.093 | -1.585 | -0.059 | 0.046 | 0.046 | 0 | 0.14 | 0.04 | 0.15 | 0.03 | 0.29 | 0.12 |
| 11 | 38125567 | rs821011 | -2.021 | 0.448 | 1.61 | -1.4 | 0.007 | 0.017 | 0.017 | 0.098 | 0.1 | 0.64 | 0.13 | 0.06 | 0.07 | 0.08 |
| 11 | 38128635 | rs11034587 | 0.705 | 0.229 | 1.033 | -1.099 | -0.013 | -0.027 | -0.027 | 2.801 | 0.05 | 0 | 0.02 | 0.08 | 0.06 | 0.15 |
| 11 | 38173409 | rs2127541 | 1.545 | -0.842 | 0.546 | -1.752 | -0.001 | 0.171 | 0.171 | 0.517 | 0.43 | 0.1 | 0.55 | 0.81 | 0.38 | 0.45 |
| 11 | 38182666 | rs820903 | -0.212 | -1.041 | 0.513 | -1.764 | 0.058 | 0.052 | 0.052 | 0 | 0.78 | 0.93 | 0.87 | 0.88 | 0.87 | 0.75 |
| 11 | 38200445 | rs7947901 | 1.441 | -0.792 | 1.155 | -1.543 | 0.045 | 0.057 | 0.057 | 0.448 | 0.92 | 0.45 | 0.89 | 0.95 | 0.94 | 0.92 |
| 11 | 38205546 | rs10742414 | 0.765 | -0.953 | 1.311 | -1.64 | -0.003 | -0.012 | -0.012 | 0 | 0.92 | 0.55 | 0.96 | 1 | 0.97 | 0.93 |
| 11 | 38211902 | rs6484977 | 1.222 | -0.955 | 1.303 | -1.643 | -0.036 | -0.035 | -0.035 | 0 | 0.89 | 0.38 | 0.9 | 0.97 | 0.94 | 0.91 |
| 11 | 38231760 | rs11034619 | 1.722 | -1.105 | 0.633 | -1.039 | -0.041 | 0.017 | 0.017 | 1.43 | 0.41 | 0.05 | 0.42 | 0.69 | 0.36 | 0.36 |
| 11 | 38232995 | rs1564054 | 1.02 | -1.165 | 1.066 | -0.981 | -0.013 | -0.012 | -0.012 | 0 | 0.89 | 0.56 | 0.92 | 0.97 | 0.94 | 0.91 |
| 11 | 38264006 | rs7128697 | 0.22 | -0.808 | 0.219 | -0.286 | -0.138 | 0.112 | 0.112 | 0.295 | 0.18 | 0.02 | 0.17 | 0.03 | 0.39 | 0.19 |
| 11 | 38285501 | rs10836945 | 2.005 | -1.239 | 1.089 | -0.024 | -0.02 | 0.142 | 0.142 | 0.849 | 0.91 | 0.31 | 0.9 | 0.79 | 0.95 | 0.77 |
| 11 | 38291744 | rs2085970 | 1.064 | -1.602 | 0.744 | -0.197 | 0.008 | -0.041 | -0.041 | 1.592 | 0.44 | 0.05 | 0.56 | 0.62 | 0.37 | 0.33 |
| 11 | 38306943 | rs7945710 | -1.644 | -1.505 | 0.991 | 0.167 | 0.082 | 0.134 | 0.134 | 0.913 | 0.05 | 0.47 | 0.04 | 0.04 | 0.01 | 0.07 |
| 11 | 38314253 | rs6484988 | 1.918 | -1.506 | 0.777 | 0.175 | -0.017 | 0.157 | 0.157 | 0.99 | 0.91 | 0.32 | 0.9 | 0.8 | 0.95 | 0.77 |
| 11 | 38340284 | rs1583443 | 0.962 | -1.966 | 0.328 | -0.183 | 0.059 | -0.035 | -0.035 | 1.552 | 0.37 | 0.11 | 0.53 | 0.62 | 0.31 | 0.33 |
| 11 | 38345032 | rs1897931 | 2.173 | -1.484 | 1.237 | 0.596 | -0.017 | 0.157 | 0.157 | 0.99 | 0.91 | 0.32 | 0.9 | 0.8 | 0.95 | 0.77 |
| 11 | 38349608 | rs1435154 | 0.685 | -1.485 | 1.186 | 0.613 | -0.038 | 0.282 | 0.282 | 2.355 | 0.89 | 0.53 | 0.83 | 0.74 | 0.96 | 0.82 |
| 11 | 38353300 | rs1435155 | 1.245 | -1.512 | 1.171 | 0.9 | -0.067 | 0.261 | 0.261 | 2.854 | 0.86 | 0.46 | 0.82 | 0.67 | 0.94 | 0.72 |
| 11 | 38377892 | rs1435143 | 0.832 | -1.502 | 0.784 | 0.888 | -0.048 | 0.222 | 0.222 | 1.493 | 0.88 | 0.47 | 0.81 | 0.74 | 0.95 | 0.81 |
| 11 | 38384838 | rs958846 | 1.869 | -1.813 | 0.59 | 0.738 | -0.067 | 0.049 | 0.049 | 1.903 | 0.78 | 0.33 | 0.77 | 0.67 | 0.92 | 0.66 |

| | | | | | | | | | | | | | | | | |
|---|---|---|---|---|---|---|---|---|---|---|---|---|---|---|---|---|
| 11 | 38396884 | rs9633825 | 0.691 | -1.364 | 0.565 | 0.861 | -0.04 | 0.244 | 0.244 | 0.923 | 0.88 | 0.57 | 0.81 | 0.74 | 0.95 | 0.8 |
| 11 | 38409443 | rs1369322 | 0.847 | -1.263 | -0.119 | 0.342 | 0.009 | 0.005 | 0.005 | 1.025 | 0.54 | 0.35 | 0.55 | 0.64 | 0.48 | 0.47 |
| 11 | 38423206 | rs2901881 | 0.284 | -1.549 | -0.728 | 0.035 | -0.092 | 0.099 | 0.099 | 0 | 0.26 | 0.09 | 0.23 | 0.03 | 0.46 | 0.19 |
| 11 | 38427410 | rs10836964 | 1.692 | -2.133 | -0.998 | 0.239 | 0 | -0.011 | -0.011 | 0 | 0.42 | 0.06 | 0.46 | 0.42 | 0.42 | 0.26 |
| 11 | 38428141 | rs7929709 | 0.604 | -2.203 | -0.874 | 0.291 | -0.033 | 0.045 | 0.045 | 0.301 | 0.77 | 0.31 | 0.72 | 0.52 | 0.9 | 0.6 |
| 11 | 38429425 | rs10768377 | 1.727 | -2.171 | -0.921 | 0.267 | 0 | -0.011 | -0.011 | 0 | 0.42 | 0.06 | 0.46 | 0.42 | 0.41 | 0.26 |
| 11 | 38445715 | rs17621211 | 0.353 | -2.195 | -0.813 | 0.225 | -0.045 | 0.045 | 0.045 | 0 | 0.27 | 0.15 | 0.25 | 0.03 | 0.49 | 0.2 |
| 11 | 38453246 | rs7116317 | 1.23 | -2.24 | -0.816 | 0.259 | 0.087 | 0.051 | 0.051 | 1.447 | 0.48 | 0.11 | 0.47 | 0.42 | 0.4 | 0.32 |
| 11 | 38458567 | rs7107634 | 1.282 | -2.213 | -0.848 | 0.258 | 0.06 | 0.023 | 0.023 | 1.57 | 0.48 | 0.06 | 0.47 | 0.42 | 0.4 | 0.32 |
| 11 | 38460043 | rs1550061 | 1.697 | -2.255 | -0.85 | 0.257 | 0.014 | -0.026 | -0.026 | 3.202 | 0.41 | 0.06 | 0.45 | 0.42 | 0.38 | 0.26 |
| 11 | 38464918 | rs11605064 | 0.477 | -2.376 | -0.879 | 0.177 | -0.137 | 0.179 | 0.179 | 0 | 0.27 | 0.03 | 0.24 | 0.03 | 0.49 | 0.2 |
| 11 | 38468689 | rs1435158 | 0.692 | -2.512 | -0.766 | 0.31 | -0.054 | 0.061 | 0.061 | 0.361 | 0.77 | 0.35 | 0.73 | 0.51 | 0.89 | 0.6 |
| 11 | 38473382 | rs10501199 | 1.73 | -2.449 | -0.824 | 0.305 | 0.048 | 0.016 | 0.016 | 2.096 | 0.41 | 0.14 | 0.45 | 0.42 | 0.36 | 0.25 |
| 11 | 38476586 | rs11034744 | 1.784 | -2.459 | -0.862 | 0.299 | 0.03 | -0.01 | -0.01 | 0 | 0.41 | 0.08 | 0.45 | 0.42 | 0.36 | 0.25 |
| 11 | 38477541 | rs920041 | 1.784 | -2.459 | -0.862 | 0.299 | 0.03 | -0.01 | -0.01 | 0.205 | 0.41 | 0.08 | 0.45 | 0.42 | 0.36 | 0.25 |
| 11 | 38477555 | rs10836987 | 0.376 | -2.298 | -0.777 | 0.135 | -0.126 | 0.183 | 0.183 | 0 | 0.27 | 0.03 | 0.24 | 0.03 | 0.5 | 0.2 |
| 11 | 38486300 | rs11034752 | -0.496 | -2.312 | -0.608 | 0.144 | -0.002 | 0.012 | 0.012 | 0 | 0.36 | 0.33 | 0.28 | 0.09 | 0.54 | 0.34 |
| 11 | 38497601 | rs1030774 | 1.229 | -2.631 | -0.774 | 0.672 | 0.054 | -0.025 | -0.025 | 0 | 0.71 | 0.19 | 0.72 | 0.64 | 0.87 | 0.65 |
| 11 | 38509330 | rs10836993 | -1.234 | -2.478 | -0.736 | 0.654 | 0.04 | 0.03 | 0.03 | 0 | 0.48 | 0.63 | 0.48 | 0.39 | 0.59 | 0.5 |
| 11 | 38517941 | rs2861190 | 1.534 | -2.476 | -0.916 | 0.688 | 0.036 | -0.073 | -0.073 | 0 | 0.43 | 0.08 | 0.48 | 0.55 | 0.36 | 0.36 |
| 11 | 38527379 | rs9633880 | 1.612 | -2.517 | -0.911 | 0.68 | 0.033 | -0.073 | -0.073 | 0 | 0.41 | 0.09 | 0.48 | 0.54 | 0.34 | 0.37 |
| 11 | 38527606 | rs10742430 | 0.77 | -2.571 | -0.783 | 0.675 | -0.035 | 0.013 | 0.013 | 2.124 | 0.79 | 0.32 | 0.78 | 0.64 | 0.91 | 0.69 |
| 11 | 38530527 | rs2861192 | 0.66 | -2.208 | -0.549 | 0.903 | -0.063 | 0.084 | 0.084 | 3.934 | 0.85 | 0.35 | 0.8 | 0.64 | 0.93 | 0.72 |
| 11 | 38547214 | rs7933866 | -0.81 | -2.209 | -0.529 | 0.902 | -0.075 | 0.147 | 0.147 | 2.345 | 0.14 | 0.58 | 0.19 | 0.36 | 0.06 | 0.27 |
| 11 | 38560367 | rs10837001 | 1.02 | -2.202 | -0.656 | 0.826 | 0.046 | -0.029 | -0.029 | 0.026 | 0.54 | 0.15 | 0.45 | 0.54 | 0.41 | 0.46 |
| 11 | 38566424 | rs1026833 | -0.577 | -2.137 | -0.641 | 0.843 | -0.012 | 0.165 | 0.165 | 0 | 0.37 | 0.14 | 0.37 | 0.09 | 0.58 | 0.33 |
| 11 | 38571985 | rs931164 | -0.72 | -2.055 | -0.555 | 0.908 | -0.064 | 0.167 | 0.167 | 3.111 | 0.14 | 0.57 | 0.19 | 0.37 | 0.07 | 0.25 |
| 11 | 38574857 | rs7950759 | 0.479 | -1.985 | -0.957 | 0.912 | -0.047 | 0.064 | 0.064 | 4.595 | 0.85 | 0.32 | 0.8 | 0.65 | 0.93 | 0.75 |
| 11 | 38577786 | rs7934284 | -0.561 | -2.041 | -1.051 | 0.856 | -0.02 | 0.172 | 0.172 | 0 | 0.37 | 0.12 | 0.37 | 0.09 | 0.58 | 0.32 |
| 11 | 38586444 | rs10837011 | 1.339 | -2.003 | -1.063 | 0.876 | 0.09 | -0.01 | -0.01 | 0.873 | 0.49 | 0.09 | 0.43 | 0.55 | 0.35 | 0.43 |
| 11 | 38592842 | rs11034859 | 1.524 | -2.019 | -1.076 | 1.078 | 0.069 | 0.019 | 0.019 | 1.451 | 0.47 | 0.08 | 0.38 | 0.41 | 0.34 | 0.35 |
| 11 | 38597421 | rs1481292 | 0.513 | -1.89 | -0.839 | 1.031 | -0.056 | 0.085 | 0.085 | 4.94 | 0.85 | 0.34 | 0.8 | 0.64 | 0.93 | 0.74 |
| 11 | 38604075 | rs1481294 | -0.081 | -1.87 | -1.068 | 1.097 | -0.049 | 0.043 | 0.043 | 0 | 0.31 | 0.15 | 0.44 | 0.18 | 0.53 | 0.33 |
| 11 | 38605473 | rs10742431 | 0.878 | -1.649 | -0.401 | 1.325 | -0.04 | 0.021 | 0.021 | 5.453 | 0.86 | 0.27 | 0.89 | 0.73 | 0.94 | 0.81 |
| 11 | 38611186 | rs7108795 | -0.546 | -1.273 | -0.508 | 1.414 | 0.004 | -0.001 | -0.001 | 0 | 0.07 | 0.04 | 0.07 | 0.14 | 0.07 | 0.13 |
| 11 | 38620095 | rs1552115 | 0.456 | -1.046 | 0.157 | 1.506 | 0.017 | 0.092 | 0.092 | 0.943 | 0.88 | 0.39 | 0.89 | 0.77 | 0.9 | 0.96 |
| 11 | 38629231 | rs1038258 | 1.049 | -1.235 | -0.474 | 1.344 | 0.022 | -0.009 | -0.009 | 0 | 0.57 | 0.09 | 0.62 | 0.44 | 0.44 | 0.6 |
| 11 | 38632770 | rs12788032 | 0.129 | -1.26 | -0.474 | 1.102 | -0.112 | 0.112 | 0.112 | 0 | 0.25 | 0.06 | 0.2 | 0.03 | 0.44 | 0.15 |
| 11 | 38664589 | rs1585961 | 0.616 | -1.057 | 0.14 | 1.679 | -0.003 | 0.137 | 0.137 | 1.215 | 0.86 | 0.37 | 0.87 | 0.66 | 0.89 | 0.86 |
| 11 | 38670343 | rs1383741 | 0.554 | -1.048 | 0.882 | 1.684 | 0.019 | 0.221 | 0.221 | 0.893 | 0.87 | 0.51 | 0.88 | 0.66 | 0.9 | 0.86 |
| 11 | 38676579 | rs2291204 | -0.597 | -1.058 | 1.373 | 1.671 | 0.045 | 0.253 | 0.253 | 0.873 | 0.12 | 0.45 | 0.1 | 0.29 | 0.1 | 0.06 |
| 11 | 38681306 | rs1354429 | 0.701 | -1.075 | 1.314 | 1.662 | -0.003 | 0.116 | 0.116 | 1.887 | 0.86 | 0.36 | 0.89 | 0.71 | 0.9 | 0.94 |
| 11 | 38683984 | rs1904967 | -1.734 | -1.285 | 1.175 | 1.648 | -0.022 | 0.084 | 0.084 | 0.385 | 0.2 | 0.53 | 0.17 | 0.33 | 0.18 | 0.28 |
| 11 | 38687306 | rs12271338 | -1.663 | -1.299 | 1.173 | 1.613 | -0.021 | 0.034 | 0.034 | 0.298 | 0.2 | 0.42 | 0.17 | 0.27 | 0.18 | 0.26 |
| 11 | 38707653 | rs3817089 | 0.463 | -0.949 | 0.872 | 1.575 | 0.061 | 0.037 | 0.037 | 0.126 | 0.77 | 0.65 | 0.78 | 0.67 | 0.57 | 0.64 |
| 11 | 38788615 | rs11034952 | 0.428 | -0.357 | 0.875 | 1.637 | 0.07 | 0.021 | 0.021 | 0 | 0.79 | 0.66 | 0.78 | 0.73 | 0.6 | 0.66 |
| 11 | 38817699 | rs10768414 | 0.804 | -0.339 | 1.235 | 1.656 | 0.066 | 0.08 | 0.08 | 0 | 0.74 | 0.59 | 0.76 | 0.54 | 0.6 | 0.58 |
| 11 | 38831942 | rs2068461 | -1.015 | -0.342 | 1.182 | 1.67 | 0 | -0.002 | -0.002 | 0 | 0.24 | 0.23 | 0.21 | 0.35 | 0.38 | 0.31 |
| 11 | 38841059 | rs7928323 | -1.015 | -0.342 | 1.179 | 1.67 | -0.005 | -0.005 | -0.005 | 0 | 0.24 | 0.22 | 0.21 | 0.35 | 0.38 | 0.31 |
| 11 | 38845174 | rs10837055 | 0.936 | -0.397 | 1.268 | 1.68 | 0.076 | 0.093 | 0.093 | 0 | 0.73 | 0.55 | 0.76 | 0.54 | 0.6 | 0.57 |
| 11 | 38860022 | rs11034972 | -0.721 | -0.348 | 0.904 | 1.615 | -0.134 | -0.009 | -0.009 | 0.062 | 0.22 | 0.04 | 0.18 | 0.17 | 0.38 | 0.26 |
| 11 | 38881682 | rs2555524 | -1.501 | -0.42 | 0.931 | 1.388 | 0.033 | 0.048 | 0.048 | 0 | 0.28 | 0.38 | 0.22 | 0.44 | 0.36 | 0.38 |
| 11 | 38903213 | rs1706963 | -0.503 | -0.425 | 0.934 | 1.414 | 0.001 | 0.296 | 0.296 | 0 | 0.06 | 0.28 | 0.04 | 0.27 | 0.01 | 0.18 |
| 11 | 38908347 | rs12785451 | 0.514 | -0.314 | 0.558 | 1.205 | -0.096 | 0.071 | 0.071 | 0 | 0.13 | 0.02 | 0.13 | 0.05 | 0.33 | 0.07 |
| 11 | 38908483 | rs11034982 | 0.785 | -0.401 | 0.579 | 1.265 | 0.01 | -0.035 | -0.035 | 0 | 0.06 | 0 | 0.05 | 0.11 | 0.03 | 0.11 |
| 11 | 38914113 | rs1155099 | -1.12 | -0.619 | 0.64 | 0.764 | -0.091 | 0.084 | 0.084 | 0 | 0.24 | 0.1 | 0.27 | 0.11 | 0.46 | 0.25 |
| 11 | 38915575 | rs7122369 | 0.137 | -0.695 | 1.413 | 0.87 | 0.079 | -0.015 | -0.015 | 0.09 | 0.81 | 0.36 | 0.8 | 0.84 | 0.67 | 0.8 |
| 11 | 38916274 | rs11605619 | -0.787 | -1.415 | 0.624 | 0.749 | 0.011 | -0.017 | -0.017 | 0 | 0.31 | 0.18 | 0.23 | 0.32 | 0.29 | 0.32 |
| 11 | 38922321 | rs1706979 | 0.168 | -0.854 | 1.567 | 0.901 | 0.082 | -0.003 | -0.003 | 0.053 | 0.15 | 0.26 | 0.17 | 0.16 | 0.34 | 0.2 |
| 11 | 38922899 | rs10837066 | -0.088 | -0.853 | 1.729 | 0.879 | 0.171 | -0.004 | -0.004 | 0 | 0.17 | 0.44 | 0.19 | 0.16 | 0.34 | 0.2 |
| 11 | 38950513 | rs2912573 | 0.05 | -0.81 | 1.658 | 0.691 | 0.103 | -0.019 | -0.019 | 0 | 0.81 | 0.39 | 0.8 | 0.83 | 0.66 | 0.79 |
| 11 | 38979245 | rs7116065 | 0.823 | -0.863 | 1.46 | 0.724 | -0.019 | 0.001 | 0.001 | 0 | 0.74 | 0.21 | 0.76 | 0.77 | 0.63 | 0.7 |
| 11 | 38998451 | rs7936255 | 0.011 | -0.616 | 1.18 | 0.258 | 0.139 | -0.127 | -0.127 | 0.02 | 0.82 | 0.43 | 0.83 | 0.94 | 0.66 | 0.93 |
| 11 | 39008571 | rs10837089 | 0.847 | 0.095 | 0.929 | 0.425 | -0.016 | 0.018 | 0.018 | 0.055 | 0.72 | 0.23 | 0.71 | 0.82 | 0.6 | 0.75 |
| 11 | 39011026 | rs11605135 | 2.182 | -0.441 | 0.375 | 0.467 | -0.074 | -0.043 | -0.043 | 0 | 0.37 | 0.02 | 0.49 | 0.37 | 0.37 | 0.37 |
| 11 | 39011336 | rs12274346 | -1.071 | -0.417 | 0.586 | 0.277 | -0.005 | -0.011 | -0.011 | 0 | 0.31 | 0.7 | 0.31 | 0.12 | 0.41 | 0.24 |
| 11 | 39033555 | rs1461630 | 0.508 | -0.296 | 0.478 | 0.255 | -0.062 | 0.064 | 0.064 | 0 | 0.18 | 0.07 | 0.14 | 0.05 | 0.29 | 0.07 |
| 11 | 39043546 | rs6485050 | -0.929 | -0.349 | 0.65 | 0.266 | 0.034 | 0.003 | 0.003 | 2.096 | 0.11 | 0.45 | 0.15 | 0.07 | 0.11 | 0.16 |
| 11 | 39050892 | rs1599459 | 2.186 | -0.503 | 0.279 | 0.297 | -0.082 | -0.051 | -0.051 | 0.811 | 0.36 | 0.02 | 0.45 | 0.38 | 0.39 | 0.38 |

| | | | | | | | | | | | | | | | | |
|---|---|---|---|---|---|---|---|---|---|---|---|---|---|---|---|---|
| 11 | 39065712 | rs11035061 | 1.965 | -0.371 | 0.601 | 0.123 | -0.017 | 0.058 | 0.058 | 0.038 | 0.6 | 0.26 | 0.63 | 0.76 | 0.56 | 0.52 |
| 11 | 39110776 | rs10768455 | 1.038 | 0.063 | 0.628 | 0.347 | 0.052 | 0.089 | 0.089 | 0.329 | 0.94 | 0.61 | 0.94 | 0.98 | 1 | 0.9 |
| 11 | 39116538 | rs16932859 | 0.356 | 0.119 | 0.335 | 0.367 | -0.005 | -0.007 | -0.007 | 0 | 0.06 | 0.04 | 0.12 | 0.13 | 0.11 | 0.12 |
| 11 | 39124659 | rs7121190 | 0.217 | 0.127 | 0.323 | 0.323 | -0.074 | 0.08 | 0.08 | 0 | 0.18 | 0.08 | 0.14 | 0.05 | 0.29 | 0.11 |
| 11 | 39136760 | rs1079149 | 0.272 | 0.107 | 0.417 | 0.493 | -0.005 | -0.006 | -0.006 | 0.142 | 0.06 | 0.04 | 0.14 | 0.14 | 0.11 | 0.12 |
| 11 | 39139342 | rs12420328 | 0.272 | 0.107 | 0.417 | 0.493 | -0.005 | -0.006 | -0.006 | 0 | 0.06 | 0.04 | 0.14 | 0.14 | 0.11 | 0.12 |
| 11 | 39163635 | rs7120739 | 1.167 | 0.061 | 0.613 | 0.475 | 0.008 | 0.062 | 0.062 | 0.183 | 0.89 | 0.66 | 0.83 | 0.84 | 0.89 | 0.73 |
| 11 | 39179160 | rs16932986 | 0.279 | 0.074 | 0.523 | 0.479 | -0.005 | -0.007 | -0.007 | 0 | 0.06 | 0.04 | 0.14 | 0.14 | 0.11 | 0.12 |
| 11 | 39184056 | rs10742464 | 0.956 | 0.12 | 0.595 | 0.395 | 0.025 | -0.027 | -0.027 | 0 | 0.7 | 0.39 | 0.68 | 0.74 | 0.6 | 0.62 |
| 11 | 39199019 | rs12417280 | -1.196 | 0.052 | 0.676 | 0.634 | 0.008 | 0.06 | 0.06 | 0.053 | 0.1 | 0.29 | 0.16 | 0.2 | 0.11 | 0.22 |
| 11 | 39212217 | rs12420552 | -2.058 | 0.036 | 0.816 | 0.654 | 0.004 | 0.082 | 0.082 | 0.583 | 0.13 | 0.45 | 0.18 | 0.2 | 0.11 | 0.26 |
| 11 | 39229850 | rs1488942 | -1.679 | 0.064 | 0.789 | 0.691 | 0.012 | 0.09 | 0.09 | 0.448 | 0.12 | 0.4 | 0.17 | 0.2 | 0.11 | 0.21 |
| 11 | 39234089 | rs1906679 | -0.493 | 0.112 | 0.757 | 0.671 | 0.009 | 0.03 | 0.03 | 0.182 | 0.07 | 0.16 | 0.15 | 0.13 | 0.11 | 0.11 |
| 11 | 39261260 | rs11035169 | -0.543 | 0.152 | 0.626 | 0.622 | 0.012 | 0.034 | 0.034 | 0.113 | 0.08 | 0.17 | 0.15 | 0.13 | 0.11 | 0.11 |
| 11 | 39263969 | rs12285179 | -0.543 | 0.152 | 0.626 | 0.622 | 0.012 | 0.034 | 0.034 | 0.113 | 0.08 | 0.17 | 0.15 | 0.13 | 0.11 | 0.11 |
| 11 | 39266095 | rs7924427 | 0.17 | -0.127 | 0.113 | 0.291 | -0.018 | 0.017 | 0.017 | 0 | 0.15 | 0.01 | 0.1 | 0.08 | 0.13 | 0.21 |
| 11 | 39280571 | rs2030088 | 1.784 | 0.477 | 0.281 | 0.194 | 0.002 | 0.003 | 0.003 | 0.752 | 0.63 | 0.34 | 0.64 | 0.67 | 0.69 | 0.44 |
| 11 | 39280831 | rs11035198 | 1.705 | 0.084 | 0.002 | -0.158 | -0.011 | -0.044 | -0.044 | 0 | 0.35 | 0.15 | 0.54 | 0.45 | 0.41 | 0.39 |
| 11 | 39282575 | rs10837134 | -0.005 | 0.474 | 0.303 | -0.059 | 0.001 | 0.001 | 0.001 | 0.521 | 0.36 | 0.34 | 0.26 | 0.37 | 0.39 | 0.15 |
| 11 | 39283877 | rs4755505 | 0.636 | 0.596 | 0.447 | -0.638 | 0.039 | -0.123 | -0.123 | 0 | 0.73 | 0.54 | 0.81 | 0.88 | 0.67 | 0.74 |
| 11 | 39286042 | rs4604853 | -0.797 | 0.586 | 0.469 | -0.637 | -0.003 | -0.006 | -0.006 | 0 | 0.82 | 0.86 | 0.9 | 0.93 | 0.78 | 0.83 |
| 11 | 39289766 | rs7118523 | 0.717 | 0.603 | 0.432 | -0.635 | 0.033 | -0.113 | -0.113 | 3.61 | 0.72 | 0.48 | 0.81 | 0.88 | 0.67 | 0.74 |
| 11 | 39299257 | rs11035204 | 0.364 | 0.624 | 0.338 | -0.612 | -0.012 | 0.015 | 0.015 | 0 | 0.07 | 0.05 | 0.06 | 0.05 | 0.11 | 0.08 |
| 11 | 39299569 | rs1532513 | 0.57 | 0.626 | 0.322 | -0.608 | 0.04 | -0.118 | -0.118 | 0 | 0.73 | 0.54 | 0.81 | 0.88 | 0.67 | 0.74 |
| 11 | 39309289 | rs17725930 | 0.771 | 0.189 | 0.062 | -0.552 | -0.046 | 0.064 | 0.064 | 0 | 0.21 | 0.06 | 0.21 | 0.07 | 0.27 | 0.12 |
| 11 | 39310021 | rs7124898 | -0.789 | 0.167 | 0.292 | -0.534 | 0.006 | -0.03 | -0.03 | 0.042 | 0.32 | 0.38 | 0.31 | 0.14 | 0.37 | 0.34 |
| 11 | 39310422 | rs17726019 | -0.694 | 0.166 | 0.341 | -0.526 | 0.01 | -0.024 | -0.024 | 0 | 0.11 | 0.27 | 0.09 | 0.07 | 0.11 | 0.17 |
| 11 | 39314653 | rs1982988 | 0.692 | 0.237 | 0.298 | -0.505 | -0.024 | 0.045 | 0.045 | 0 | 0.21 | 0.11 | 0.21 | 0.08 | 0.27 | 0.17 |
| 11 | 39319703 | rs10444236 | 0.03 | -0.174 | 1.33 | 0.307 | -0.004 | 0.039 | 0.039 | 0.006 | 0.22 | 0.3 | 0.21 | 0.26 | 0.16 | 0.18 |
| 11 | 39322214 | rs7944994 | -0.807 | -0.176 | 1.42 | -0.072 | 0 | -0.001 | -0.001 | 0 | 0.35 | 0.35 | 0.34 | 0.42 | 0.26 | 0.35 |
| 11 | 39325576 | rs10734472 | -1.137 | -0.121 | 1.597 | 0.014 | 0.003 | -0.008 | -0.008 | 0.296 | 0.43 | 0.38 | 0.41 | 0.47 | 0.36 | 0.45 |
| 11 | 39327563 | rs12807582 | 0.983 | -0.145 | 1.511 | 0.014 | 0.009 | -0.008 | -0.008 | 0 | 0.55 | 0.37 | 0.57 | 0.53 | 0.63 | 0.55 |
| 11 | 39335783 | rs10768471 | 1.139 | -0.072 | 2.053 | -0.298 | 0.078 | 0.105 | 0.105 | 0.725 | 0.95 | 0.44 | 0.95 | 0.99 | 1 | 0.88 |
| 11 | 39353212 | rs10837146 | 1.149 | -0.062 | 1.856 | -0.373 | 0.011 | 0.008 | 0.008 | 0.869 | 0.73 | 0.22 | 0.85 | 0.92 | 0.82 | 0.66 |
| 11 | 39390707 | rs7105155 | 1.135 | 0.083 | 1.078 | 0.3 | -0.004 | -0.012 | -0.012 | 0 | 0.93 | 0.61 | 0.96 | 1 | 0.97 | 1 |
| 11 | 39393388 | rs7117622 | 1.049 | 0.16 | 0.649 | -0.072 | -0.011 | -0.049 | -0.049 | 0 | 0.8 | 0.65 | 0.88 | 0.93 | 0.8 | 0.84 |
| 11 | 39394070 | rs10768476 | 1.038 | 0.195 | 0.711 | -0.083 | -0.006 | -0.033 | -0.033 | 1.728 | 0.8 | 0.71 | 0.88 | 0.93 | 0.8 | 0.82 |
| 11 | 39400626 | rs994587 | 1.932 | 0.355 | 0.212 | 0.464 | -0.003 | -0.091 | -0.091 | 0.056 | 0.61 | 0.1 | 0.43 | 0.57 | 0.6 | 0.49 |
| 11 | 39424358 | rs4494281 | -0.153 | 0.178 | 0.417 | 0.296 | -0.002 | -0.004 | -0.004 | 0 | 0.08 | 0.08 | 0.02 | 0.05 | 0.03 | 0.04 |
| 11 | 39426409 | rs12279967 | -1.266 | 0.155 | 0.935 | 0.267 | -0.03 | -0.029 | -0.029 | 0 | 0.12 | 0.26 | 0.04 | 0.08 | 0.05 | 0.18 |
| 11 | 39434429 | rs12294840 | -0.23 | 0.209 | 0.826 | 0.286 | -0.018 | -0.021 | -0.021 | 0.001 | 0.08 | 0.14 | 0.03 | 0.05 | 0.04 | 0.04 |
| 11 | 39436698 | rs11035257 | 0.286 | 0.214 | 0.63 | 0.641 | -0.057 | -0.063 | -0.063 | 0.096 | 0.1 | 0 | 0.31 | 0.25 | 0.18 | 0.15 |
| 11 | 39439328 | rs11035260 | -0.049 | 0.012 | 0.657 | 0.573 | -0.003 | -0.004 | -0.004 | 0 | 0.08 | 0.06 | 0.02 | 0.05 | 0.03 | 0.04 |
| 11 | 39442174 | rs6485094 | 1.65 | 0.746 | 1.141 | 0.769 | -0.001 | -0.001 | -0.001 | 0 | 0.66 | 0.3 | 0.42 | 0.58 | 0.57 | 0.63 |
| 11 | 39445016 | rs4237674 | 1.597 | 0.848 | 1.142 | 0.771 | -0.001 | -0.001 | -0.001 | 2.356 | 0.66 | 0.3 | 0.42 | 0.58 | 0.56 | 0.63 |
| 11 | 39456990 | rs11035274 | -0.204 | 1.245 | 1.276 | 1.114 | -0.015 | -0.009 | -0.009 | 0 | 0.11 | 0.07 | 0.33 | 0.28 | 0.22 | 0.22 |
| 11 | 39474379 | rs10742494 | -2.349 | 0.729 | 1.665 | 0.917 | -0.01 | 0.02 | 0.02 | 2.375 | 0.19 | 0.7 | 0.33 | 0.18 | 0.19 | 0.19 |
| 11 | 39484014 | rs11035287 | -2.339 | 0.697 | 1.765 | 0.894 | -0.007 | 0.028 | 0.028 | 1.228 | 0.18 | 0.65 | 0.32 | 0.18 | 0.2 | 0.19 |
| 11 | 39522637 | rs4628651 | -2.25 | 0.588 | 1.813 | 1.088 | -0.012 | 0.047 | 0.047 | 0.325 | 0.2 | 0.67 | 0.34 | 0.23 | 0.2 | 0.23 |
| 11 | 39535809 | rs7946013 | 1.659 | 0.838 | 1.815 | 1.111 | 0.031 | 0.061 | 0.061 | 0 | 0.81 | 0.36 | 0.63 | 0.81 | 0.71 | 0.78 |
| 11 | 39539470 | rs3884617 | -1.078 | 0.877 | 1.773 | 0.871 | 0.012 | -0.024 | -0.024 | 0 | 0.12 | 0.1 | 0.12 | 0.02 | 0.26 | 0.07 |
| 11 | 39567316 | rs4237676 | 0.873 | 0.383 | 1.075 | 0.572 | -0.001 | 0.008 | 0.008 | 0 | 0.62 | 0.31 | 0.55 | 0.76 | 0.56 | 0.5 |
| 11 | 39604010 | rs4592392 | -1.996 | 0.25 | 1.501 | 0.973 | 0.027 | 0.083 | 0.083 | 0.156 | 0.1 | 0.49 | 0.23 | 0.08 | 0.07 | 0.19 |
| 11 | 39605868 | rs4456239 | -1.884 | 0.089 | 2.129 | 1.49 | 0.011 | 0.318 | 0.318 | 0.224 | 0.07 | 0.41 | 0.2 | 0.19 | 0.03 | 0.2 |
| 11 | 39606277 | rs4315042 | 0.8 | -0.514 | 1.785 | 0.65 | -0.021 | -0.002 | -0.002 | 0 | 0.61 | 0.43 | 0.56 | 0.7 | 0.69 | 0.69 |
| 11 | 39611165 | rs10501215 | -0.436 | -0.653 | 1.731 | 0.578 | 0.022 | 0.077 | 0.077 | 1.31 | 0.74 | 0.87 | 0.78 | 0.92 | 0.77 | 0.95 |
| 11 | 39611400 | rs10837180 | 1.018 | -0.657 | 2.126 | 0.668 | 0.008 | 0 | 0 | 0.172 | 0.6 | 0.33 | 0.55 | 0.7 | 0.7 | 0.69 |
| 11 | 39620131 | rs7932035 | 0.008 | -0.762 | 2.436 | 0.712 | 0.014 | 0.037 | 0.037 | 0.463 | 0.74 | 0.82 | 0.81 | 0.89 | 0.8 | 0.92 |
| 11 | 39623896 | rs7112347 | 0.602 | -0.775 | 2.466 | 0.712 | -0.003 | 0 | 0 | 0 | 0.68 | 0.32 | 0.62 | 0.72 | 0.79 | 0.72 |
| 11 | 39632367 | rs16933573 | -2.215 | -0.775 | 2.411 | 0.706 | 0.003 | 0.231 | 0.231 | 0 | 0.1 | 0.55 | 0.21 | 0.2 | 0.04 | 0.25 |
| 11 | 39632506 | rs12416853 | 0.315 | -0.784 | 2.363 | 0.678 | 0.023 | 0.065 | 0.065 | 0.707 | 0.21 | 0.06 | 0.16 | 0.09 | 0.17 | 0.03 |
| 11 | 39633353 | rs1477869 | -0.501 | -0.777 | 2.339 | 0.681 | 0.017 | 0.031 | 0.031 | 0.739 | 0.78 | 0.84 | 0.83 | 0.91 | 0.83 | 0.97 |
| 11 | 39635775 | rs10837188 | 0.522 | -0.778 | 2.351 | 0.688 | 0.002 | -0.002 | -0.002 | 0 | 0.69 | 0.31 | 0.62 | 0.72 | 0.79 | 0.85 |
| 11 | 39647400 | rs17648172 | 0.346 | 0.127 | 1.356 | 0.61 | -0.124 | 0.004 | 0.004 | 0 | 0.2 | 0.01 | 0.32 | 0.15 | 0.36 | 0.16 |
| 11 | 39652635 | rs10837198 | -0.035 | -0.545 | 2.167 | 0.996 | 0.022 | 0.047 | 0.047 | 1.037 | 0.22 | 0.13 | 0.17 | 0.09 | 0.17 | 0.03 |
| 11 | 39662187 | rs7930822 | -0.084 | -0.533 | 2.165 | 0.987 | 0.005 | 0.011 | 0.011 | 0.08 | 0.77 | 0.8 | 0.82 | 0.91 | 0.81 | 0.83 |
| 11 | 39666972 | rs1477870 | 0.515 | -0.599 | 2.024 | 1.15 | -0.034 | 0.018 | 0.018 | 0.016 | 0.69 | 0.43 | 0.62 | 0.67 | 0.79 | 0.73 |
| 11 | 39667955 | rs1354926 | 0.575 | -0.558 | 1.754 | 1.142 | -0.024 | 0.011 | 0.011 | 0.032 | 0.68 | 0.38 | 0.62 | 0.67 | 0.79 | 0.73 |
| 11 | 39671205 | rs10837204 | 0.878 | -0.868 | 1.085 | 1.322 | 0.049 | -0.011 | -0.011 | 0 | 0.66 | 0.18 | 0.67 | 0.68 | 0.79 | 0.67 |

| | | | | | | | | | | | | | | | | |
|---|---|---|---|---|---|---|---|---|---|---|---|---|---|---|---|---|
| 11 | 39672274 | rs1354925 | 0.842 | -0.871 | 1.24 | 1.285 | 0.007 | -0.002 | -0.002 | 0 | 0.66 | 0.27 | 0.67 | 0.72 | 0.8 | 0.67 |
| 11 | 39673927 | rs11035388 | -0.2 | -0.639 | 1.002 | 0.536 | 0.009 | 0.004 | 0.004 | 0.149 | 0.2 | 0.13 | 0.11 | 0.01 | 0.14 | 0.04 |
| 11 | 39688047 | rs17727783 | -0.453 | -0.573 | 1.631 | 0.469 | -0.008 | -0.078 | -0.078 | 0 | 0.22 | 0.29 | 0.11 | 0.01 | 0.15 | 0.04 |
| 11 | 39698786 | rs11035405 | -0.972 | -1.001 | 0.704 | 0.952 | 0.034 | -0.01 | -0.01 | 0 | 0.38 | 0.19 | 0.37 | 0.43 | 0.28 | 0.31 |
| 11 | 39700963 | rs1702619 | -1.62 | -0.608 | 1.872 | 0.848 | 0.025 | 0.086 | 0.086 | 1.141 | 0.13 | 0.48 | 0.08 | 0.17 | 0.12 | 0.24 |
| 11 | 39709068 | rs769864 | -1.574 | -0.628 | 2.07 | 0.839 | 0.023 | 0.077 | 0.077 | 0.592 | 0.13 | 0.53 | 0.08 | 0.17 | 0.12 | 0.24 |
| 11 | 39717326 | rs11035412 | -1.156 | -0.636 | 2.398 | 0.835 | 0.053 | 0.112 | 0.112 | 0.001 | 0.1 | 0.32 | 0.07 | 0.17 | 0.12 | 0.08 |
| 11 | 39726391 | rs17728870 | 0.693 | -0.355 | 2.021 | 0.58 | -0.03 | 0.028 | 0.028 | 0 | 0.06 | 0 | 0.05 | 0 | 0.08 | 0.02 |
| 11 | 39728091 | rs1993836 | 1.464 | -1.009 | 2.258 | 0.653 | 0.025 | 0.089 | 0.089 | 0.597 | 0.6 | 0.12 | 0.69 | 0.73 | 0.67 | 0.64 |
| 11 | 39729692 | rs769825 | -0.021 | -1.016 | 2.386 | 0.621 | -0.017 | -0.102 | -0.102 | 0 | 0.77 | 0.66 | 0.87 | 0.99 | 0.85 | 0.95 |
| 11 | 39740536 | rs16933754 | 0.041 | -1.005 | 2.507 | 0.637 | 0.006 | 0.007 | 0.007 | 0 | 0.05 | 0.11 | 0.11 | 0.1 | 0.06 | 0.1 |
| 11 | 39740619 | rs1676681 | -2.176 | -1.012 | 2.57 | 0.643 | 0.025 | 0.08 | 0.08 | 0.076 | 0.17 | 0.48 | 0.18 | 0.26 | 0.18 | 0.31 |
| 11 | 39740695 | rs1676680 | -1.215 | -1.013 | 2.591 | 0.642 | 0.035 | 0.086 | 0.086 | 0.041 | 0.11 | 0.34 | 0.07 | 0.17 | 0.12 | 0.08 |
| 11 | 39741702 | rs10837218 | 1.516 | -1.017 | 2.487 | 0.637 | 0.023 | 0.091 | 0.091 | 0.669 | 0.6 | 0.11 | 0.69 | 0.73 | 0.67 | 0.64 |
| 11 | 39743574 | rs7939395 | 1.071 | -1.015 | 2.499 | 0.636 | 0.011 | 0.028 | 0.028 | 1.957 | 0.66 | 0.24 | 0.8 | 0.83 | 0.73 | 0.87 |
| 11 | 39746905 | rs11035428 | 1.519 | -1.101 | 2.282 | 0.733 | 0 | 0.067 | 0.067 | 1.141 | 0.6 | 0.06 | 0.69 | 0.72 | 0.69 | 0.65 |
| 11 | 39749305 | rs769866 | 0.67 | -0.87 | 2.36 | 0.602 | -0.011 | -0.076 | -0.076 | 0 | 0.69 | 0.41 | 0.74 | 0.89 | 0.75 | 0.71 |
| 11 | 39753414 | rs1676670 | 1.133 | -0.827 | 2.112 | 0.613 | 0.022 | 0.049 | 0.049 | 1.895 | 0.65 | 0.22 | 0.81 | 0.83 | 0.75 | 0.74 |
| 11 | 39772957 | rs1395919 | 1.879 | -0.848 | 1.76 | 1.098 | -0.021 | -0.056 | -0.056 | 0 | 0.6 | 0.07 | 0.69 | 0.63 | 0.66 | 0.62 |
| 11 | 39774191 | rs11602882 | -0.02 | -1.026 | 1.903 | 0.981 | 0.014 | 0.074 | 0.074 | 0.765 | 0.19 | 0.06 | 0.09 | 0.01 | 0.13 | 0.05 |
| 11 | 39774565 | rs10837243 | -0.807 | -0.986 | 2.062 | 1.066 | 0.062 | 0.205 | 0.205 | 0 | 0.08 | 0.23 | 0.06 | 0.26 | 0.11 | 0.09 |
| 11 | 39797971 | rs10837259 | 0.465 | -1.032 | 2.017 | 0.736 | -0.039 | -0.134 | -0.134 | 0 | 0.75 | 0.39 | 0.87 | 0.99 | 0.87 | 0.92 |
| 11 | 39800072 | rs991293 | 0.465 | -1.031 | 2.017 | 0.736 | -0.039 | -0.134 | -0.134 | 0.802 | 0.75 | 0.39 | 0.87 | 0.99 | 0.87 | 0.92 |
| 11 | 39810064 | rs7939125 | 1.306 | -0.937 | 1.383 | 1.07 | 0.007 | 0.021 | 0.021 | 0.021 | 0.64 | 0.22 | 0.71 | 0.72 | 0.68 | 0.66 |
| 11 | 39818824 | rs12223410 | -1.547 | -0.974 | 2.698 | 1.201 | 0.023 | 0.123 | 0.123 | 0.421 | 0.13 | 0.22 | 0.18 | 0.34 | 0.19 | 0.17 |
| 11 | 39839671 | rs7935404 | -1.837 | -1.084 | 2.875 | 1.137 | 0.007 | 0.058 | 0.058 | 0.393 | 0.14 | 0.19 | 0.18 | 0.33 | 0.19 | 0.18 |
| 11 | 39840315 | rs7102927 | -0.231 | -1.308 | 2.73 | 0.939 | 0.004 | -0.022 | -0.022 | 0 | 0.8 | 0.82 | 0.91 | 0.99 | 0.87 | 0.93 |
| 11 | 39851288 | rs930996 | 1.166 | -1.269 | 2.914 | 0.971 | -0.011 | -0.008 | -0.008 | 0 | 0.65 | 0.56 | 0.72 | 0.69 | 0.69 | 0.75 |
| 11 | 39865056 | rs4492793 | 1.449 | -1.251 | 2.766 | 1.001 | 0.015 | 0.01 | 0.01 | 0.108 | 0.63 | 0.32 | 0.7 | 0.7 | 0.69 | 0.66 |
| 11 | 39876405 | rs12224631 | -0.423 | -1.227 | 2.663 | 0.891 | 0.006 | 0.035 | 0.035 | 0 | 0.05 | 0.06 | 0.05 | 0.15 | 0.08 | 0.09 |
| 11 | 39877553 | rs10768552 | 1.149 | -1.211 | 2.692 | 0.891 | -0.015 | -0.026 | -0.026 | 0 | 0.66 | 0.54 | 0.81 | 0.77 | 0.7 | 0.76 |
| 11 | 39893958 | rs12364614 | -0.657 | -1.444 | 1.629 | 0.677 | 0.014 | 0.111 | 0.111 | 0.029 | 0.25 | 0.08 | 0.11 | 0.04 | 0.19 | 0.09 |
| 11 | 39894373 | rs11035554 | 1.454 | -1.671 | 1.566 | 0.95 | 0.04 | -0.005 | -0.005 | 0 | 0.64 | 0.09 | 0.81 | 0.74 | 0.77 | 0.67 |
| 11 | 39894911 | rs4756552 | 1.568 | -1.663 | 2.134 | 0.935 | 0.064 | 0.211 | 0.211 | 2.95 | 0.94 | 0.65 | 0.96 | 0.85 | 0.96 | 0.89 |
| 11 | 39902443 | rs10768556 | 1.975 | -1.664 | 2.273 | 0.961 | 0.06 | 0.219 | 0.219 | 3.961 | 0.93 | 0.56 | 0.95 | 0.85 | 0.96 | 0.89 |
| 11 | 39915234 | rs2199409 | -1.092 | -1.729 | 2.173 | 0.868 | -0.027 | -0.043 | -0.043 | 0.017 | 0.3 | 0.57 | 0.15 | 0.16 | 0.19 | 0.14 |
| 11 | 39916430 | rs1461788 | -1.212 | -1.777 | 2.178 | 0.834 | -0.027 | -0.051 | -0.051 | 0.09 | 0.31 | 0.57 | 0.15 | 0.11 | 0.19 | 0.19 |
| 11 | 39924543 | rs12360555 | 1.437 | -1.769 | 1.965 | 0.883 | 0.046 | 0 | 0 | 1.846 | 0.64 | 0.09 | 0.81 | 0.7 | 0.78 | 0.75 |
| 11 | 39928302 | rs12362799 | -0.548 | -1.759 | 2.296 | 0.767 | 0.018 | 0.099 | 0.099 | 0.006 | 0.25 | 0.08 | 0.1 | 0.04 | 0.18 | 0.09 |
| 11 | 39928935 | rs11035594 | -1.134 | -1.73 | 2.393 | 0.774 | -0.007 | -0.02 | -0.02 | 0.114 | 0.29 | 0.25 | 0.13 | 0.11 | 0.19 | 0.15 |
| 11 | 39933968 | rs10837301 | 1.497 | -1.326 | 2.593 | 1.079 | 0.104 | 0.238 | 0.238 | 3.553 | 0.95 | 0.61 | 0.95 | 0.86 | 0.97 | 0.95 |
| 11 | 39951362 | rs6485159 | -0.915 | -1.841 | 1.663 | 0.608 | -0.019 | -0.026 | -0.026 | 0 | 0.32 | 0.29 | 0.13 | 0.17 | 0.2 | 0.12 |
| 11 | 39962707 | rs7927729 | -0.977 | -1.777 | 1.859 | 0.571 | -0.037 | -0.055 | -0.055 | 0 | 0.32 | 0.39 | 0.13 | 0.17 | 0.21 | 0.16 |
| 11 | 39966521 | rs7943388 | -0.667 | -1.774 | 1.845 | 0.567 | -0.007 | -0.056 | -0.056 | 0 | 0.28 | 0.22 | 0.1 | 0 | 0.14 | 0.11 |
| 11 | 39981948 | rs10837311 | -1.007 | -1.759 | 1.496 | 0.607 | -0.03 | -0.041 | -0.041 | 0 | 0.32 | 0.36 | 0.13 | 0.15 | 0.2 | 0.11 |
| 11 | 39997451 | rs7483173 | -1.653 | -2.302 | 1.245 | 1.307 | -0.007 | 0 | 0 | 0 | 0.47 | 0.48 | 0.38 | 0.53 | 0.27 | 0.37 |
| 11 | 39998287 | rs12286970 | -0.919 | -2.23 | 1.127 | 1.406 | 0.013 | 0.041 | 0.041 | 0 | 0.06 | 0.11 | 0.05 | 0.18 | 0.09 | 0.04 |
| 11 | 40000800 | rs10837317 | -1.695 | -2.442 | 1.135 | 1.636 | -0.004 | 0 | 0 | 0 | 0.46 | 0.45 | 0.38 | 0.52 | 0.27 | 0.37 |
| 11 | 40004506 | rs7111832 | 0.388 | -2.527 | 1.118 | 1.628 | 0.013 | -0.006 | -0.006 | 1.008 | 0.11 | 0.08 | 0.23 | 0.35 | 0.03 | 0.2 |
| 11 | 40013964 | rs7952564 | 0.241 | -2.529 | 1.184 | 1.627 | -0.014 | 0.034 | 0.034 | 0 | 0.11 | 0.13 | 0.23 | 0.34 | 0.04 | 0.2 |
| 11 | 40015352 | rs9888247 | 1.775 | -2.422 | 1.567 | 1.473 | 0.064 | 0.031 | 0.031 | 0 | 0.49 | 0.36 | 0.52 | 0.41 | 0.7 | 0.53 |
| 11 | 40016567 | rs7483295 | -0.45 | -2.459 | 1.58 | 1.381 | 0.056 | 0.054 | 0.054 | 0 | 0.29 | 0.12 | 0.1 | 0 | 0.14 | 0.11 |
| 11 | 40029625 | rs968845 | 1.53 | -1.966 | 1.436 | 1.683 | -0.026 | 0.103 | 0.103 | 0 | 0.77 | 0.47 | 0.61 | 0.4 | 0.83 | 0.6 |
| 11 | 40032861 | rs16934228 | -0.879 | -1.228 | 1.04 | 1.575 | -0.027 | -0.026 | -0.026 | 0 | 0.05 | 0.01 | 0.06 | 0.13 | 0.13 | 0.06 |
| 11 | 40041290 | rs6485172 | 1.458 | -1.164 | 1.592 | 2.075 | 0.119 | 0.296 | 0.296 | 0 | 0.9 | 0.61 | 0.78 | 0.61 | 0.83 | 0.84 |
| 11 | 40047754 | rs4755538 | 1.571 | -1.884 | 1.103 | 1.622 | 0.041 | 0.005 | 0.005 | 0 | 0.51 | 0.29 | 0.54 | 0.46 | 0.64 | 0.53 |
| 11 | 40056052 | rs2861776 | -0.581 | -1.926 | 0.567 | 1.491 | 0.052 | 0.047 | 0.047 | 0 | 0.32 | 0.21 | 0.18 | 0.15 | 0.16 | 0.25 |
| 11 | 40067242 | rs11035659 | -0.482 | -1.91 | 0.289 | 1.499 | 0.08 | 0.069 | 0.069 | 0 | 0.31 | 0.1 | 0.17 | 0.15 | 0.16 | 0.23 |
| 11 | 40071723 | rs1031778 | -0.463 | -1.9 | 0.3 | 1.476 | 0.072 | 0.068 | 0.068 | 0 | 0.31 | 0.13 | 0.18 | 0.15 | 0.16 | 0.21 |
| 11 | 40073820 | rs10768576 | 1.382 | -1.347 | 0.752 | 1.805 | 0.069 | 0.192 | 0.192 | 0 | 0.9 | 0.73 | 0.75 | 0.59 | 0.83 | 0.84 |
| 11 | 40076546 | rs1462183 | -1.076 | -1.396 | -0.209 | 1.139 | 0.008 | 0.023 | 0.023 | 0 | 0.56 | 0.47 | 0.51 | 0.76 | 0.4 | 0.6 |
| 11 | 40079008 | rs7950925 | -0.778 | -1.44 | -0.183 | 1.641 | 0.061 | 0.056 | 0.056 | 0 | 0.5 | 0.38 | 0.39 | 0.36 | 0.35 | 0.47 |
| 11 | 40080055 | rs11035665 | -0.22 | -1.497 | -0.236 | 1.652 | 0.058 | 0.076 | 0.076 | 0 | 0.43 | 0.34 | 0.26 | 0.16 | 0.25 | 0.42 |
| 11 | 40082660 | rs16934294 | -0.803 | -1.359 | -0.134 | 1.75 | -0.006 | 0.04 | 0.04 | 0 | 0.12 | 0.33 | 0.15 | 0.21 | 0.1 | 0.12 |
| 11 | 40082790 | rs10837327 | -0.71 | -1.042 | -0.322 | 1.69 | 0 | 0.075 | 0.075 | 0 | 0.07 | 0.1 | 0.12 | 0.41 | 0.05 | 0.13 |
| 11 | 40089635 | rs4756571 | -1.19 | 0.252 | -0.474 | 0.725 | 0.04 | -0.041 | -0.041 | 0 | 0.73 | 0.65 | 0.67 | 0.86 | 0.66 | 0.68 |
| 11 | 40092369 | rs10501217 | 0.195 | 0.355 | -1.194 | 0.746 | -0.073 | -0.001 | -0.001 | 0 | 0.16 | 0.01 | 0.16 | 0.1 | 0.26 | 0.09 |
| 11 | 40093494 | rs7929966 | 0.235 | 0.346 | -0.625 | 1.308 | -0.001 | -0.001 | -0.001 | 0 | 0.32 | 0.35 | 0.36 | 0.43 | 0.35 | 0.5 |
| 11 | 40094796 | rs11035673 | -1.019 | -0.322 | -0.216 | 1.276 | 0.016 | 0.092 | 0.092 | 0 | 0.08 | 0.27 | 0.04 | 0.17 | 0.07 | 0.08 |

| | | | | | | | | | | | | | | | | |
|---|---|---|---|---|---|---|---|---|---|---|---|---|---|---|---|---|
| 11 | 40096025 | rs1381571 | 0.605 | 0.167 | -0.558 | 0.931 | -0.053 | 0.04 | 0.04 | 0.031 | 0.28 | 0.17 | 0.33 | 0.15 | 0.43 | 0.32 |
| 11 | 40105120 | rs12279098 | -0.942 | -0.704 | 0.088 | 1.842 | -0.025 | 0.182 | 0.182 | 0 | 0.06 | 0.16 | 0.07 | 0.3 | 0.01 | 0.07 |
| 11 | 40105881 | rs16934335 | -1.881 | -0.83 | 0.465 | 1.704 | -0.027 | 0.181 | 0.181 | 0 | 0.1 | 0.31 | 0.1 | 0.3 | 0.03 | 0.1 |
| 11 | 40111384 | rs4756573 | 0.294 | -1.366 | -0.002 | 1.697 | -0.022 | -0.014 | -0.014 | 0 | 0.57 | 0.5 | 0.56 | 0.69 | 0.68 | 0.62 |
| 11 | 40117479 | rs10837342 | 0.757 | -1.358 | -0.015 | 1.731 | 0.112 | 0.059 | 0.059 | 0 | 0.5 | 0.25 | 0.44 | 0.37 | 0.67 | 0.54 |
| 11 | 40118333 | rs3109378 | -0.836 | -0.822 | 0.631 | 1.781 | -0.005 | 0.267 | 0.267 | 0.903 | 0.06 | 0.29 | 0.08 | 0.3 | 0 | 0.07 |
| 11 | 40118598 | rs3109384 | 0.491 | -0.774 | 0.124 | 1.681 | -0.029 | -0.018 | -0.018 | 0 | 0.71 | 0.56 | 0.59 | 0.76 | 0.77 | 0.74 |
| 11 | 40133483 | rs10837344 | -0.907 | -0.294 | -0.446 | 1.808 | 0.012 | -0.096 | -0.096 | 0 | 0.25 | 0.08 | 0.27 | 0.36 | 0.17 | 0.17 |
| 11 | 40134083 | rs10501219 | -0.692 | -0.366 | -0.483 | 1.891 | 0.06 | -0.07 | -0.07 | 0 | 0.18 | 0.08 | 0.2 | 0.34 | 0.05 | 0.14 |
| 11 | 40135242 | rs11035693 | -0.944 | -0.343 | -0.152 | 1.975 | 0.01 | -0.015 | -0.015 | 0 | 0.2 | 0.21 | 0.21 | 0.39 | 0.06 | 0.17 |
| 11 | 40137543 | rs2953310 | -1.899 | 0.02 | -0.058 | 1.697 | 0.016 | 0.031 | 0.031 | 0 | 0.44 | 0.7 | 0.51 | 0.61 | 0.37 | 0.24 |
| 11 | 40142377 | rs7943560 | 1.235 | 0.711 | -0.21 | 1.427 | 0.003 | 0.013 | 0.013 | 0 | 0.6 | 0.33 | 0.52 | 0.33 | 0.68 | 0.6 |
| 11 | 40142665 | rs7930525 | 1.359 | 0.287 | 0 | 1.528 | -0.001 | -0.001 | -0.001 | 0 | 0.62 | 0.38 | 0.56 | 0.33 | 0.71 | 0.6 |
| 11 | 40147696 | rs16934385 | -0.079 | 0.821 | 0.109 | 1.738 | 0.002 | 0.03 | 0.03 | 0 | 0.06 | 0.06 | 0.1 | 0.18 | 0.07 | 0.02 |
| 11 | 40149684 | rs1381567 | -1.708 | 1.434 | 0.251 | 1.586 | 0.011 | 0.017 | 0.017 | 0 | 0.18 | 0.21 | 0.18 | 0.28 | 0.23 | 0.06 |
| 11 | 40150122 | rs10837346 | -0.146 | 1.369 | 0.418 | 1.848 | -0.005 | -0.007 | -0.007 | 0.224 | 0.22 | 0.21 | 0.19 | 0.24 | 0.2 | 0.11 |
| 11 | 40150291 | rs12577341 | -1.018 | 1.353 | 0.693 | 1.751 | 0.019 | 0.044 | 0.044 | 0 | 0.31 | 0.5 | 0.38 | 0.52 | 0.28 | 0.14 |
| 11 | 40156059 | rs7936167 | -1.732 | 1.084 | 0.607 | 1.726 | 0.023 | 0.155 | 0.155 | 2.001 | 0.15 | 0.42 | 0.22 | 0.29 | 0.14 | 0.05 |
| 11 | 40158154 | rs7929592 | -1.19 | 0.807 | 0.564 | 1.205 | 0.02 | 0.07 | 0.07 | 1.832 | 0.09 | 0.36 | 0.12 | 0.11 | 0.06 | 0.03 |
| 11 | 40161972 | rs7952540 | 0.462 | 0.772 | -0.518 | 0.665 | -0.001 | -0.002 | -0.002 | 0 | 0.3 | 0.2 | 0.25 | 0.31 | 0.34 | 0.34 |
| 11 | 40174630 | rs10160235 | -0.123 | 0.616 | -0.649 | 0.318 | 0.073 | 0.022 | 0.022 | 0 | 0.49 | 0.3 | 0.52 | 0.44 | 0.68 | 0.58 |
| 11 | 40175794 | rs1841704 | 0.767 | 0.486 | -0.306 | 0.213 | -0.003 | -0.026 | -0.026 | 0 | 0.05 | 0.09 | 0.03 | 0 | 0.03 | 0.01 |
| 11 | 40180572 | rs11035719 | 0.19 | 0.989 | -0.39 | 0.12 | 0.019 | 0.003 | 0.003 | 0.764 | 0.05 | 0.08 | 0.14 | 0.08 | 0.09 | 0.19 |
| 11 | 40181413 | rs1551833 | -0.774 | 0.209 | -0.307 | -0.312 | -0.041 | 0.018 | 0.018 | 0 | 0.64 | 0.53 | 0.59 | 0.45 | 0.74 | 0.62 |
| 11 | 40185254 | rs3109382 | -1.643 | 0.749 | -0.482 | -0.407 | -0.016 | -0.015 | -0.015 | 0 | 0.7 | 0.8 | 0.78 | 0.67 | 0.63 | 0.65 |
| 11 | 40187882 | rs10768582 | 1.381 | 0.776 | -0.412 | -0.454 | -0.003 | -0.003 | -0.003 | 0 | 0.3 | 0.24 | 0.21 | 0.3 | 0.34 | 0.34 |
| 11 | 40189231 | rs11035722 | 0.6 | 0.755 | -0.432 | -0.444 | -0.004 | -0.001 | -0.001 | 0.069 | 0.38 | 0.34 | 0.37 | 0.4 | 0.45 | 0.55 |
| 11 | 40191609 | rs4623893 | -1.012 | 0.15 | -0.004 | -0.412 | -0.017 | -0.016 | -0.016 | 0 | 0.83 | 0.69 | 0.85 | 0.85 | 0.87 | 0.73 |
| 11 | 40192526 | rs4429048 | -0.615 | 0.231 | -0.419 | -0.526 | -0.004 | -0.012 | -0.012 | 0 | 0.32 | 0.31 | 0.25 | 0.1 | 0.37 | 0.26 |
| 11 | 40192854 | rs4237677 | 0.579 | 0.214 | -0.647 | -0.158 | 0.012 | -0.033 | -0.033 | 0 | 0.22 | 0.13 | 0.29 | 0.43 | 0.17 | 0.15 |
| 11 | 40195513 | rs4074148 | 0.59 | 0.41 | -0.671 | -0.233 | -0.001 | 0 | 0 | 0 | 0.05 | 0.03 | 0.13 | 0.06 | 0.07 | 0.15 |
| 11 | 40195564 | rs4076390 | 1.29 | 0.432 | -1.063 | -0.268 | -0.068 | -0.07 | -0.07 | 0 | 0.29 | 0.04 | 0.2 | 0.34 | 0.33 | 0.39 |
| 11 | 40196093 | rs7926722 | -1.03 | 0.689 | -0.214 | -0.457 | 0.023 | -0.039 | -0.039 | 0.009 | 0.37 | 0.5 | 0.49 | 0.27 | 0.45 | 0.26 |
| 11 | 40205141 | rs4319506 | 0.333 | 0.647 | -0.086 | -0.792 | 0.001 | -0.054 | -0.054 | 0 | 0.63 | 0.62 | 0.7 | 0.89 | 0.65 | 0.72 |
| 11 | 40205400 | rs16934423 | -0.071 | 0.41 | 0.071 | -0.785 | -0.035 | -0.05 | -0.05 | 0 | 0.09 | 0.22 | 0.03 | 0 | 0.03 | 0.04 |
| 11 | 40205503 | rs6485187 | -0.31 | 0.09 | -0.199 | -0.344 | -0.032 | 0.04 | 0.04 | 0.015 | 0.33 | 0.63 | 0.28 | 0.4 | 0.24 | 0.22 |
| 11 | 40207084 | rs17477151 | 0.88 | 0.469 | -0.46 | -0.15 | -0.032 | -0.003 | -0.003 | 0 | 0.09 | 0.01 | 0.12 | 0.07 | 0.1 | 0.09 |
| 11 | 40207923 | rs4075903 | 0.595 | 0.856 | -0.09 | -0.046 | -0.002 | -0.002 | -0.002 | 0.008 | 0.41 | 0.22 | 0.45 | 0.47 | 0.43 | 0.55 |
| 11 | 40209856 | rs7930773 | -0.503 | 0.989 | -0.055 | 0.013 | -0.006 | -0.037 | -0.037 | 0 | 0.59 | 0.71 | 0.66 | 0.57 | 0.6 | 0.83 |
| 11 | 40211252 | rs7924805 | -0.397 | 1.011 | -0.035 | -0.011 | -0.035 | -0.024 | -0.024 | 0 | 0.58 | 0.71 | 0.63 | 0.57 | 0.54 | 0.82 |
| 11 | 40212151 | rs10734479 | -0.907 | 0.793 | 0.21 | -0.106 | -0.019 | 0.007 | 0.007 | 0 | 0.59 | 0.75 | 0.53 | 0.52 | 0.48 | 0.5 |
| 11 | 40212372 | rs7942248 | -0.676 | 1.404 | 0.091 | 0.047 | -0.013 | -0.015 | -0.015 | 0 | 0.66 | 0.67 | 0.63 | 0.55 | 0.57 | 0.82 |
| 11 | 40216951 | rs10768584 | -0.394 | 1.362 | 0.149 | 0.007 | -0.006 | 0.068 | 0.068 | 0.052 | 0.48 | 0.37 | 0.4 | 0.58 | 0.48 | 0.33 |
| 11 | 40217821 | rs10768586 | -0.42 | 1.324 | 0.133 | -0.008 | -0.004 | 0.071 | 0.071 | 0.043 | 0.49 | 0.36 | 0.4 | 0.59 | 0.47 | 0.32 |
| 11 | 40218057 | rs10837363 | 0.679 | 1.434 | 0.057 | 0.034 | 0.012 | -0.029 | -0.029 | 0 | 0.33 | 0.24 | 0.3 | 0.5 | 0.3 | 0.18 |
| 11 | 40219825 | rs4073087 | -0.33 | 1.53 | 0.089 | 0.053 | -0.006 | 0.054 | 0.054 | 0.019 | 0.47 | 0.34 | 0.42 | 0.54 | 0.5 | 0.23 |
| 11 | 40221414 | rs7106026 | 0.518 | 1.585 | -0.134 | -0.101 | -0.011 | 0.012 | 0.012 | 0.013 | 0.14 | 0.11 | 0.1 | 0.09 | 0.17 | 0.05 |
| 11 | 40221507 | rs7106166 | -0.115 | 0.983 | 0.601 | -0.048 | -0.008 | 0.058 | 0.058 | 0.721 | 0.47 | 0.32 | 0.53 | 0.4 | 0.52 | 0.66 |
| 11 | 40222380 | rs12363681 | -0.027 | 0.739 | 0.611 | -0.043 | 0.009 | 0.053 | 0.053 | 0.337 | 0.46 | 0.31 | 0.49 | 0.37 | 0.45 | 0.64 |
| 11 | 40224135 | rs4509742 | -0.53 | 0.364 | 0.853 | -0.211 | -0.023 | -0.07 | -0.07 | 0.002 | 0.6 | 0.6 | 0.66 | 0.76 | 0.65 | 0.53 |
| 11 | 40226381 | rs4369393 | 0.154 | 0.518 | 0.531 | -0.025 | 0.029 | 0.104 | 0.104 | 0 | 0.52 | 0.24 | 0.58 | 0.63 | 0.53 | 0.5 |
| 11 | 40227204 | rs11035737 | 0.511 | 0.453 | 0.36 | -0.036 | 0.03 | 0.05 | 0.05 | 0 | 0.35 | 0.09 | 0.23 | 0.33 | 0.33 | 0.44 |
| 11 | 40229937 | rs10742530 | -0.015 | 0.591 | 0.533 | 0.046 | 0.004 | 0.005 | 0.005 | 0 | 0.55 | 0.48 | 0.62 | 0.62 | 0.58 | 0.42 |
| 11 | 40238179 | rs976933 | 1.661 | 0.772 | 1.461 | 0.174 | 0.071 | 0.028 | 0.028 | 0 | 0.9 | 0.54 | 0.85 | 0.95 | 0.91 | 0.87 |
| 11 | 40246485 | rs16934480 | -0.416 | 0.818 | 1.058 | 0.2 | -0.019 | -0.023 | -0.023 | 0.913 | 0.16 | 0.06 | 0.29 | 0.18 | 0.17 | 0.25 |
| 11 | 40247102 | rs7933522 | 2.01 | 0.706 | 1.3 | 0.111 | -0.015 | -0.014 | -0.014 | 0 | 0.88 | 0.29 | 0.87 | 0.95 | 0.94 | 0.87 |
| 11 | 40260495 | rs1452473 | 0.496 | 0.738 | 1.701 | 0.127 | 0.14 | 0.141 | 0.141 | 0 | 0.95 | 0.67 | 0.89 | 0.95 | 0.94 | 0.98 |
| 11 | 40269665 | rs7930449 | -0.311 | 0.741 | 1.489 | 0.152 | -0.013 | -0.01 | -0.01 | 2.402 | 0.15 | 0.07 | 0.3 | 0.17 | 0.17 | 0.23 |
| 11 | 40274933 | rs1452465 | 1.752 | 0.677 | 1.521 | 0.119 | 0.015 | 0.018 | 0.018 | 0.409 | 0.72 | 0.22 | 0.57 | 0.78 | 0.77 | 0.63 |
| 11 | 40277888 | rs999004 | 1.692 | 0.572 | 1.78 | -0.064 | -0.019 | -0.045 | -0.045 | 0.482 | 0.8 | 0.4 | 0.8 | 0.9 | 0.83 | 0.8 |
| 11 | 40280436 | rs1020579 | 2.136 | 0.583 | 1.718 | -0.06 | 0 | 0.045 | 0.045 | 0 | 0.63 | 0.22 | 0.47 | 0.73 | 0.64 | 0.52 |
| 11 | 40285275 | rs7944315 | 1.269 | 0.559 | 1.976 | -0.042 | 0.012 | -0.043 | -0.043 | 0 | 0.82 | 0.64 | 0.78 | 0.91 | 0.81 | 0.8 |
| 11 | 40287160 | rs16934661 | -0.584 | 0.811 | 1.669 | 0.007 | 0.005 | 0.003 | 0.003 | 0.362 | 0.17 | 0.2 | 0.32 | 0.2 | 0.23 | 0.25 |
| 11 | 40301193 | rs1901583 | 1.689 | 0.869 | 1.088 | -0.191 | 0 | 0.021 | 0.021 | 0.02 | 0.71 | 0.27 | 0.55 | 0.79 | 0.67 | 0.61 |
| 11 | 40306157 | rs10501227 | -0.547 | 0.962 | 0.702 | -0.026 | 0.024 | 0.012 | 0.012 | 0 | 0.05 | 0.14 | 0.09 | 0.02 | 0.06 | 0.03 |
| 11 | 40310789 | rs998447 | -2.183 | 1.044 | 0.693 | -0.032 | 0.011 | -0.014 | -0.014 | 0 | 0.35 | 0.4 | 0.53 | 0.28 | 0.44 | 0.43 |
| 11 | 40316673 | rs992797 | -2.19 | 0.889 | 0.724 | 0.108 | 0.009 | -0.014 | -0.014 | 0 | 0.38 | 0.5 | 0.54 | 0.33 | 0.45 | 0.45 |
| 11 | 40318617 | rs11035770 | -0.456 | 1.014 | 0.956 | 0.092 | -0.026 | -0.004 | -0.004 | 0 | 0.17 | 0.03 | 0.3 | 0.14 | 0.19 | 0.27 |
| 11 | 40321916 | rs2167662 | 0.859 | 0.727 | 1.185 | 0.148 | 0.058 | 0.051 | 0.051 | 1.921 | 0.95 | 0.72 | 0.91 | 0.95 | 0.95 | 0.92 |

| | | | | | | | | | | | | | | | | |
|---|---|---|---|---|---|---|---|---|---|---|---|---|---|---|---|---|
| 11 | 40329182 | rs896618 | 1.479 | 0.654 | 1.138 | 0.138 | -0.001 | -0.014 | -0.014 | 1.99 | 0.69 | 0.41 | 0.46 | 0.78 | 0.68 | 0.61 |
| 11 | 40348304 | rs4755547 | 0.787 | 0.75 | 0.293 | 0.283 | 0.055 | 0.057 | 0.057 | 0 | 0.93 | 0.72 | 0.88 | 0.94 | 0.93 | 0.97 |
| 11 | 40357253 | rs10501230 | -0.839 | 0.657 | -0.363 | 0.255 | -0.022 | 0.008 | 0.008 | 0.992 | 0.2 | 0.06 | 0.36 | 0.14 | 0.24 | 0.28 |
| 11 | 40357510 | rs1158362 | -1.293 | 0.535 | -0.362 | 0.219 | -0.003 | -0.008 | -0.008 | 0.005 | 0.29 | 0.33 | 0.46 | 0.19 | 0.28 | 0.32 |
| 11 | 40362590 | rs12785682 | -0.195 | 0.154 | -1.337 | 0.327 | 0.007 | -0.032 | -0.032 | 0.155 | 0.25 | 0.07 | 0.14 | 0.26 | 0.18 | 0.21 |
| 11 | 40370150 | rs7932379 | 0.738 | 0.146 | -2.263 | 0.015 | -0.034 | 0.023 | 0.023 | 1.081 | 0.25 | 0.05 | 0.29 | 0.15 | 0.33 | 0.28 |
| 11 | 40370881 | rs11035794 | 0.576 | 0.157 | -2.195 | 0.025 | 0.016 | -0.028 | -0.028 | 0 | 0.25 | 0.07 | 0.14 | 0.26 | 0.18 | 0.2 |
| 11 | 40373075 | rs1303850 | -0.525 | 0.226 | -1.752 | -0.443 | 0 | 0.052 | 0.052 | 0.713 | 0.63 | 0.78 | 0.66 | 0.84 | 0.58 | 0.58 |
| 11 | 40374155 | rs11601600 | -0.159 | 0.336 | -0.833 | -0.724 | -0.004 | 0.034 | 0.034 | 0.009 | 0.56 | 0.37 | 0.56 | 0.84 | 0.5 | 0.54 |
| 11 | 40376628 | rs1303932 | 1.064 | 0.264 | -0.531 | -0.575 | -0.02 | -0.058 | -0.058 | 0 | 0.3 | 0.19 | 0.41 | 0.54 | 0.32 | 0.33 |
| 11 | 40382104 | rs4397839 | -0.142 | 0.094 | 0.14 | -0.586 | 0 | -0.024 | -0.024 | 0 | 0.55 | 0.45 | 0.56 | 0.8 | 0.5 | 0.53 |
| 11 | 40388022 | rs7103883 | 0.607 | -0.121 | -0.008 | -0.879 | 0.01 | -0.008 | -0.008 | 0.246 | 0.43 | 0.33 | 0.38 | 0.44 | 0.42 | 0.47 |
| 11 | 40388879 | rs11035807 | 0.806 | -0.159 | -0.099 | -0.891 | -0.041 | -0.085 | -0.085 | 0 | 0.41 | 0.06 | 0.37 | 0.42 | 0.42 | 0.47 |
| 11 | 40389399 | rs12576114 | 0.074 | -0.18 | -0.081 | -0.871 | -0.031 | -0.061 | -0.061 | 0 | 0.39 | 0.18 | 0.47 | 0.53 | 0.41 | 0.34 |
| 11 | 40405192 | rs9651608 | -0.078 | -0.171 | 0.197 | -0.864 | 0.048 | -0.043 | -0.043 | 0 | 0.08 | 0.29 | 0.1 | 0 | 0.09 | 0.04 |
| 11 | 40405631 | rs10768599 | -0.596 | -0.113 | -0.173 | -0.853 | 0.038 | -0.113 | -0.113 | 1.28 | 0.82 | 0.56 | 0.85 | 0.99 | 0.81 | 0.81 |
| 11 | 40409913 | rs11605725 | 1.646 | -0.185 | -0.52 | -0.835 | -0.018 | 0.052 | 0.052 | 0 | 0.1 | 0.01 | 0.04 | 0.01 | 0.08 | 0.15 |
| 11 | 40423092 | rs920443 | 0.092 | -0.217 | -0.635 | -0.858 | -0.033 | -0.048 | -0.048 | 0 | 0.41 | 0.19 | 0.47 | 0.54 | 0.42 | 0.35 |
| 11 | 40426092 | rs16934802 | 0.222 | -0.117 | -0.605 | -0.558 | -0.063 | -0.066 | -0.066 | 0 | 0.37 | 0.13 | 0.42 | 0.47 | 0.4 | 0.34 |
| 11 | 40426864 | rs7946870 | 0.442 | -0.271 | -0.394 | -0.237 | -0.036 | -0.097 | -0.097 | 0 | 0.39 | 0.07 | 0.37 | 0.42 | 0.41 | 0.46 |
| 11 | 40438810 | rs1440105 | 0.456 | -0.266 | -0.407 | -0.235 | -0.021 | -0.078 | -0.078 | 0.018 | 0.39 | 0.11 | 0.37 | 0.42 | 0.41 | 0.45 |
| 11 | 40440844 | rs6485196 | 0.447 | -0.265 | -0.407 | -0.234 | -0.02 | -0.078 | -0.078 | 1.58 | 0.39 | 0.12 | 0.37 | 0.42 | 0.41 | 0.45 |
| 11 | 40448662 | rs7935540 | 0.073 | -0.383 | -0.095 | -0.253 | 0.004 | -0.001 | -0.001 | 0 | 0.45 | 0.54 | 0.44 | 0.51 | 0.42 | 0.46 |
| 11 | 40456042 | rs7945771 | 0.074 | -0.387 | -0.072 | -0.257 | 0 | -0.001 | -0.001 | 0 | 0.44 | 0.43 | 0.43 | 0.48 | 0.42 | 0.46 |
| 11 | 40456268 | rs1992315 | 0.19 | -0.382 | -0.167 | -0.255 | 0.028 | -0.038 | -0.038 | 0.139 | 0.43 | 0.23 | 0.41 | 0.48 | 0.42 | 0.46 |
| 11 | 40462553 | rs11035846 | 0.443 | -0.287 | -0.183 | -0.236 | -0.025 | -0.088 | -0.088 | 0 | 0.39 | 0.06 | 0.34 | 0.41 | 0.38 | 0.45 |
| 11 | 40468065 | rs7944171 | 0.165 | -0.26 | -0.119 | -0.239 | 0.023 | -0.021 | -0.021 | 0 | 0.43 | 0.31 | 0.39 | 0.48 | 0.39 | 0.45 |
| 11 | 40476768 | rs1440123 | 0.352 | -0.326 | -0.167 | -0.271 | 0.009 | -0.049 | -0.049 | 0.227 | 0.4 | 0.23 | 0.38 | 0.48 | 0.38 | 0.43 |
| 11 | 40491232 | rs2083420 | 0.575 | -0.318 | -0.475 | -0.257 | 0.001 | 0.01 | 0.01 | 0 | 0.34 | 0.34 | 0.45 | 0.56 | 0.36 | 0.37 |
| 11 | 40498570 | rs2119179 | 0.04 | -0.463 | -0.923 | -0.349 | 0.014 | 0 | 0 | 0 | 0.17 | 0.11 | 0.14 | 0.16 | 0.12 | 0.06 |
| 11 | 40506912 | rs7117433 | 0.995 | -0.432 | -0.876 | -0.342 | -0.004 | -0.019 | -0.019 | 0 | 0.34 | 0.22 | 0.44 | 0.56 | 0.36 | 0.37 |
| 11 | 40508635 | rs17484798 | 0.984 | -0.603 | -0.957 | -0.359 | -0.027 | -0.031 | -0.031 | 0.281 | 0.06 | 0.01 | 0.05 | 0.14 | 0.1 | 0.06 |
| 11 | 40510064 | rs17484924 | 0.363 | -0.675 | -0.94 | -0.41 | 0.019 | 0.006 | 0.006 | 0 | 0.16 | 0.05 | 0.04 | 0.12 | 0.1 | 0.07 |
| 11 | 40511611 | rs1039038 | -0.369 | -0.589 | -0.866 | -0.493 | 0.001 | -0.017 | -0.017 | 0.302 | 0.54 | 0.45 | 0.61 | 0.72 | 0.49 | 0.49 |
| 11 | 40524159 | rs1440107 | -0.442 | -0.555 | -0.903 | -0.866 | -0.003 | -0.001 | -0.001 | 0 | 0.62 | 0.38 | 0.72 | 0.81 | 0.53 | 0.53 |
| 11 | 40530245 | rs17406736 | 0.952 | -0.728 | -1.121 | -0.832 | 0.012 | -0.011 | -0.011 | 0 | 0.16 | 0.04 | 0.06 | 0.13 | 0.11 | 0.07 |
| 11 | 40532449 | rs17406814 | 2.002 | -0.268 | -1.352 | -1.062 | -0.038 | -0.019 | -0.019 | 0 | 0.12 | 0.01 | 0.1 | 0.14 | 0.21 | 0.13 |
| 11 | 40532950 | rs12574103 | 1.024 | -0.567 | -1.183 | -0.9 | 0.012 | -0.011 | -0.011 | 0.172 | 0.16 | 0.04 | 0.06 | 0.13 | 0.11 | 0.07 |
| 11 | 40538734 | rs17407151 | 0.506 | -0.539 | -1.056 | -0.827 | -0.006 | 0.016 | 0.016 | 0 | 0.42 | 0.23 | 0.54 | 0.63 | 0.4 | 0.42 |
| 11 | 40545629 | rs7129451 | 0.675 | -0.523 | -1.134 | -0.819 | -0.026 | 0.005 | 0.005 | 0 | 0.4 | 0.16 | 0.53 | 0.63 | 0.4 | 0.42 |
| 11 | 40547616 | rs11035895 | 2.158 | -0.493 | -1.419 | -0.821 | -0.022 | 0.034 | 0.034 | 0.709 | 0.18 | 0.01 | 0.2 | 0.13 | 0.18 | 0.18 |
| 11 | 40556645 | rs16934914 | 0.115 | -0.553 | -1.104 | -0.85 | -0.033 | -0.002 | -0.002 | 0 | 0.19 | 0.36 | 0.17 | 0.15 | 0.11 | 0.07 |
| 11 | 40557004 | rs1440112 | 0.881 | -0.373 | -1.22 | -1.061 | -0.048 | 0.011 | 0.011 | 0.05 | 0.21 | 0.1 | 0.21 | 0.18 | 0.37 | 0.32 |
| 11 | 40569126 | rs10837435 | 0.508 | -0.234 | -1.215 | -0.796 | -0.126 | -0.058 | -0.058 | 0 | 0.38 | 0.06 | 0.53 | 0.63 | 0.41 | 0.42 |
| 11 | 40569225 | rs7481977 | 0.426 | -0.314 | -1.109 | -0.782 | -0.044 | -0.002 | -0.002 | 0.232 | 0.4 | 0.15 | 0.54 | 0.63 | 0.4 | 0.42 |
| 11 | 40578791 | rs4755553 | 0.111 | -0.31 | -0.861 | -0.785 | 0 | -0.012 | -0.012 | 0.792 | 0.43 | 0.45 | 0.64 | 0.65 | 0.39 | 0.41 |
| 11 | 40579533 | rs7945461 | -0.813 | -0.302 | -0.866 | -0.817 | 0.014 | -0.041 | -0.041 | 0.681 | 0.62 | 0.51 | 0.73 | 0.79 | 0.52 | 0.52 |
| 11 | 40580159 | rs11035918 | 0.866 | -0.013 | -1.046 | -0.646 | -0.044 | -0.1 | -0.1 | 0 | 0.2 | 0.06 | 0.33 | 0.5 | 0.22 | 0.23 |
| 11 | 40586402 | rs10837436 | 0.277 | -0.072 | -1.042 | -0.67 | -0.016 | -0.057 | -0.057 | 0 | 0.24 | 0.13 | 0.37 | 0.53 | 0.25 | 0.26 |
| 11 | 40587040 | rs10837438 | 0.842 | 0.008 | -1.016 | -0.624 | -0.024 | -0.06 | -0.06 | 0 | 0.2 | 0.11 | 0.34 | 0.5 | 0.23 | 0.23 |
| 11 | 40591800 | rs7944117 | -0.238 | -0.445 | -0.745 | -0.799 | 0.016 | 0.023 | 0.023 | 0 | 0.41 | 0.35 | 0.3 | 0.24 | 0.31 | 0.28 |
| 11 | 40597707 | rs6485204 | 1.532 | -0.398 | -0.919 | -0.787 | -0.004 | 0.049 | 0.049 | 0.867 | 0.19 | 0.04 | 0.19 | 0.11 | 0.17 | 0.18 |
| 11 | 40600161 | rs1390177 | 0.958 | -0.586 | -0.897 | -0.48 | -0.016 | -0.016 | -0.016 | 0.088 | 0.07 | 0 | 0.11 | 0.06 | 0.06 | 0.05 |
| 11 | 40602448 | rs7124249 | -0.129 | -0.695 | -0.723 | -0.549 | 0.05 | 0.06 | 0.06 | 0.007 | 0.38 | 0.21 | 0.27 | 0.26 | 0.27 | 0.29 |
| 11 | 40606832 | rs4755554 | -0.224 | -0.662 | -0.619 | -0.534 | -0.032 | -0.04 | -0.04 | 0 | 0.41 | 0.45 | 0.29 | 0.26 | 0.27 | 0.29 |
| 11 | 40615445 | rs4756605 | -0.16 | -0.58 | -0.986 | -0.501 | 0.065 | 0.057 | 0.057 | 0.016 | 0.39 | 0.13 | 0.29 | 0.32 | 0.28 | 0.29 |
| 11 | 40616921 | rs7927392 | 1.479 | -0.493 | -0.148 | -0.566 | 0.048 | 0.18 | 0.18 | 0 | 0.16 | 0.3 | 0.25 | 0.48 | 0.2 | 0.21 |
| 11 | 40617461 | rs12278621 | 0.721 | -0.314 | -0.264 | -0.842 | -0.009 | -0.004 | -0.004 | 0 | 0.28 | 0.37 | 0.26 | 0.23 | 0.25 | 0.4 |
| 11 | 40618475 | rs1994976 | 0.61 | -0.279 | -0.915 | -1.054 | -0.017 | 0.127 | 0.127 | 0.201 | 0.33 | 0.05 | 0.35 | 0.15 | 0.33 | 0.27 |
| 11 | 40619212 | rs7121049 | -0.178 | -0.276 | -0.486 | -0.964 | -0.001 | -0.001 | -0.001 | 0 | 0.62 | 0.63 | 0.62 | 0.41 | 0.58 | 0.68 |
| 11 | 40619828 | rs1027388 | -0.35 | -0.31 | -0.31 | -1.095 | -0.002 | 0.117 | 0.117 | 0 | 0.52 | 0.25 | 0.44 | 0.24 | 0.52 | 0.34 |
| 11 | 40626074 | rs17488852 | 1.119 | -0.582 | -0.534 | -1.347 | -0.027 | 0.068 | 0.068 | 0 | 0.19 | 0.01 | 0.17 | 0.06 | 0.18 | 0.17 |
| 11 | 40639344 | rs16934985 | 0.882 | -0.291 | -0.022 | -0.953 | -0.011 | 0.004 | 0.004 | 0 | 0.28 | 0.36 | 0.28 | 0.27 | 0.25 | 0.41 |
| 11 | 40640360 | rs2861934 | 0.417 | -0.281 | -0.19 | -0.947 | 0.004 | -0.035 | -0.035 | 0 | 0.42 | 0.53 | 0.51 | 0.73 | 0.45 | 0.62 |
| 11 | 40641079 | rs11035962 | 0.976 | -0.011 | -0.754 | -0.809 | -0.035 | 0.015 | 0.015 | 0 | 0.14 | 0.04 | 0.1 | 0.11 | 0.18 | 0.08 |
| 11 | 40642327 | rs2132215 | 2.088 | 0.117 | -0.691 | -0.627 | -0.035 | -0.05 | -0.05 | 0 | 0.14 | 0.07 | 0.22 | 0.46 | 0.2 | 0.21 |
| 11 | 40648282 | rs11035972 | 0.941 | -0.302 | -0.63 | -0.716 | -0.003 | 0.007 | 0.007 | 0 | 0.09 | 0.09 | 0.12 | 0.25 | 0.15 | 0.26 |
| 11 | 40658277 | rs10742544 | -0.906 | -0.163 | -0.752 | -0.991 | 0.006 | 0.127 | 0.127 | 1.999 | 0.67 | 0.89 | 0.68 | 0.85 | 0.7 | 0.76 |
| 11 | 40661364 | rs2172860 | 0.085 | -0.165 | -0.916 | -0.846 | -0.005 | -0.013 | -0.013 | 0 | 0.51 | 0.15 | 0.58 | 0.6 | 0.54 | 0.48 |

| | | | | | | | | | | | | | | | | |
|---|---|---|---|---|---|---|---|---|---|---|---|---|---|---|---|---|
| 11 | 40667312 | rs11035989 | 0.934 | 0.098 | -0.503 | -0.339 | -0.03 | -0.056 | -0.056 | 0.096 | 0.19 | 0.13 | 0.27 | 0.45 | 0.24 | 0.24 |
| 11 | 40668583 | rs4756614 | 1.357 | 0.074 | -0.582 | -0.42 | -0.027 | 0.033 | 0.033 | 0.283 | 0.22 | 0.01 | 0.26 | 0.14 | 0.18 | 0.23 |
| 11 | 40669349 | rs16935026 | -0.352 | 0.006 | -0.277 | -0.481 | 0.004 | -0.032 | -0.032 | 0.634 | 0.21 | 0.26 | 0.13 | 0.11 | 0.21 | 0.11 |
| 11 | 40673063 | rs7394597 | 0.258 | 0.03 | -0.325 | -0.469 | -0.055 | -0.055 | -0.055 | 0 | 0.41 | 0.17 | 0.52 | 0.59 | 0.42 | 0.47 |
| 11 | 40677496 | rs6485219 | -1.037 | 0.136 | -0.218 | -0.407 | 0.009 | 0.004 | 0.004 | 0 | 0.6 | 0.38 | 0.65 | 0.7 | 0.62 | 0.57 |
| 11 | 40678140 | rs1495317 | 0.118 | 0.201 | -0.389 | -0.4 | -0.028 | -0.025 | -0.025 | 2.235 | 0.43 | 0.25 | 0.55 | 0.59 | 0.43 | 0.49 |
| 11 | 40679479 | rs11603172 | -0.223 | 0.816 | 0.052 | -0.17 | 0.007 | -0.013 | -0.013 | 0.249 | 0.17 | 0.16 | 0.11 | 0.11 | 0.2 | 0.08 |
| 11 | 40692912 | rs7949003 | 0.778 | 0.83 | -0.384 | 0.179 | -0.059 | -0.091 | -0.091 | 0 | 0.18 | 0.07 | 0.29 | 0.47 | 0.25 | 0.22 |
| 11 | 40695391 | rs10742547 | -0.692 | 0.529 | -0.23 | 0.059 | 0.019 | 0.009 | 0.009 | 0.38 | 0.37 | 0.51 | 0.41 | 0.61 | 0.42 | 0.32 |
| 11 | 40698650 | rs10742548 | -0.344 | 0.873 | -0.894 | 0.21 | 0.006 | 0.004 | 0.004 | 0 | 0.32 | 0.38 | 0.4 | 0.6 | 0.42 | 0.3 |
| 11 | 40707037 | rs4755560 | -0.429 | 0.674 | -1.256 | 0.336 | 0.001 | -0.001 | -0.001 | 0 | 0.68 | 0.69 | 0.71 | 0.71 | 0.63 | 0.6 |
| 11 | 40708010 | rs7939606 | 0.69 | 0.704 | -1.358 | 0.418 | -0.103 | -0.09 | -0.09 | 0 | 0.49 | 0.05 | 0.58 | 0.56 | 0.47 | 0.52 |
| 11 | 40709637 | rs11036017 | -0.236 | 0.718 | -1.296 | 0.421 | -0.002 | -0.018 | -0.018 | 1.796 | 0.17 | 0.61 | 0.12 | 0.14 | 0.14 | 0.08 |
| 11 | 40711439 | rs11036021 | -0.008 | 0.937 | -2.193 | 0.616 | -0.104 | -0.091 | -0.091 | 0 | 0.49 | 0.05 | 0.58 | 0.57 | 0.47 | 0.52 |
| 11 | 40736710 | rs11036035 | -0.772 | 1.256 | -0.743 | 0.559 | 0.037 | 0.008 | 0.008 | 0 | 0.12 | 0.3 | 0.12 | 0.09 | 0.17 | 0.16 |
| 11 | 40738299 | rs12366050 | -0.821 | 1.002 | -0.97 | 0.474 | -0.002 | 0.001 | 0.001 | 0 | 0.53 | 0.57 | 0.51 | 0.54 | 0.53 | 0.49 |
| 11 | 40742299 | rs10837484 | -0.767 | 0.766 | -1.042 | -0.352 | -0.008 | 0.046 | 0.046 | 0 | 0.42 | 0.57 | 0.32 | 0.1 | 0.45 | 0.3 |
| 11 | 40744768 | rs4132508 | -0.479 | 0.758 | -0.628 | -0.362 | 0.007 | -0.068 | -0.068 | 0.288 | 0.38 | 0.39 | 0.28 | 0.1 | 0.42 | 0.28 |
| 11 | 40756460 | rs11036050 | 0.863 | 0.365 | -1.17 | -0.841 | -0.007 | 0.026 | 0.026 | 0.328 | 0.38 | 0.22 | 0.26 | 0.21 | 0.46 | 0.29 |
| 11 | 40764768 | rs10837494 | 0.483 | 0.35 | -1.194 | -0.868 | 0 | 0.064 | 0.064 | 0.125 | 0.46 | 0.25 | 0.3 | 0.21 | 0.5 | 0.33 |
| 11 | 40764896 | rs11036058 | 0.772 | 0.407 | -1.113 | -0.86 | 0.02 | -0.013 | -0.013 | 0 | 0.14 | 0.2 | 0.16 | 0.07 | 0.2 | 0.17 |
| 11 | 40767022 | rs10742554 | -1.32 | 0.254 | -1.094 | -0.745 | -0.001 | 0.001 | 0.001 | 1.92 | 0.79 | 0.74 | 0.72 | 0.7 | 0.83 | 0.73 |
| 11 | 40769111 | rs11036061 | 0.053 | 0.127 | -1.139 | -0.826 | -0.013 | -0.016 | -0.016 | 0.003 | 0.6 | 0.45 | 0.45 | 0.3 | 0.69 | 0.49 |
| 11 | 40772512 | rs4550210 | -1.416 | 0.17 | -1.192 | -0.816 | -0.001 | 0 | 0 | 1.892 | 0.8 | 0.74 | 0.72 | 0.71 | 0.82 | 0.73 |
| 11 | 40772616 | rs4643070 | -1.591 | 0.099 | -0.976 | -0.851 | 0.001 | -0.043 | -0.043 | 2.283 | 0.81 | 0.89 | 0.74 | 0.69 | 0.82 | 0.72 |
| 11 | 40781345 | rs11036066 | 0.4 | 0.126 | -0.855 | -0.811 | -0.003 | 0.015 | 0.015 | 0 | 0.19 | 0.25 | 0.27 | 0.41 | 0.13 | 0.23 |
| 11 | 40783291 | rs7106315 | 1.024 | 0.061 | -0.89 | -0.88 | -0.018 | 0.008 | 0.008 | 0 | 0.13 | 0.07 | 0.14 | 0.08 | 0.2 | 0.17 |
| 11 | 40790356 | rs11036068 | 0.514 | -0.184 | -0.807 | -1.051 | -0.003 | 0.003 | 0.003 | 0 | 0.22 | 0.2 | 0.18 | 0.09 | 0.27 | 0.18 |
| 11 | 40790624 | rs4382893 | 0.106 | -0.308 | -0.393 | -0.452 | 0.019 | 0.014 | 0.014 | 0.551 | 0.67 | 0.73 | 0.49 | 0.3 | 0.72 | 0.52 |
| 11 | 40791501 | rs4132214 | 0.645 | -0.365 | -0.444 | -0.268 | 0.027 | -0.088 | -0.088 | 0 | 0.15 | 0.06 | 0.22 | 0.41 | 0.09 | 0.19 |
| 11 | 40801197 | rs10837502 | 0.434 | 0.033 | -0.02 | -0.819 | -0.008 | -0.019 | -0.019 | 0 | 0.54 | 0.44 | 0.41 | 0.24 | 0.66 | 0.45 |
| 11 | 40808616 | rs12277565 | -1.416 | 0.081 | 1.418 | -0.34 | 0.029 | 0.098 | 0.098 | 0 | 0.11 | 0.35 | 0.13 | 0.17 | 0.1 | 0.15 |
| 11 | 40809093 | rs4296032 | -2.467 | -0.016 | 1.653 | -0.496 | -0.003 | 0.017 | 0.017 | 0.018 | 0.18 | 0.55 | 0.19 | 0.16 | 0.15 | 0.22 |
| 11 | 40816057 | rs4340035 | -1.36 | -0.005 | 1.383 | -0.495 | -0.004 | -0.043 | -0.043 | 0 | 0.08 | 0.18 | 0.06 | 0 | 0.06 | 0.08 |
| 11 | 40817882 | rs12277750 | -1.158 | 0.077 | 1.139 | -0.419 | 0.007 | 0.041 | 0.041 | 0 | 0.09 | 0.18 | 0.11 | 0.17 | 0.1 | 0.15 |
| 11 | 40828597 | rs7480771 | 1.549 | 0.057 | 1.108 | -0.728 | -0.005 | -0.122 | -0.122 | 0 | 0.84 | 0.5 | 0.86 | 0.99 | 0.87 | 0.78 |
| 11 | 40830685 | rs4491209 | 1.998 | -0.171 | -0.01 | -0.819 | 0.039 | 0.071 | 0.071 | 0.173 | 0.67 | 0.12 | 0.66 | 0.81 | 0.74 | 0.61 |
| 11 | 40831698 | rs7479005 | 1.993 | 0.218 | -0.032 | -0.625 | 0.003 | 0.124 | 0.124 | 0.102 | 0.6 | 0.13 | 0.66 | 0.87 | 0.68 | 0.6 |
| 11 | 40840931 | rs7943949 | 1.329 | 0.199 | 1.382 | -0.396 | 0.067 | -0.052 | -0.052 | 0.515 | 0.9 | 0.59 | 0.92 | 0.99 | 0.9 | 0.86 |
| 11 | 40854095 | rs11036094 | -1.289 | 0.059 | 1.345 | -0.43 | 0.007 | -0.006 | -0.006 | 0.155 | 0.14 | 0.09 | 0.1 | 0.13 | 0.11 | 0.15 |
| 11 | 40859924 | rs7128152 | -1.669 | 0.247 | 1.543 | -0.488 | -0.024 | -0.018 | -0.018 | 0 | 0.23 | 0.36 | 0.19 | 0.13 | 0.13 | 0.22 |
| 11 | 40864899 | rs11036097 | 1.63 | -0.042 | 0.82 | -0.461 | 0.013 | 0.032 | 0.032 | 0 | 0.69 | 0.22 | 0.73 | 0.86 | 0.78 | 0.65 |
| 11 | 40875131 | rs10430982 | 1.775 | -0.03 | 0.825 | -0.473 | 0.019 | 0.044 | 0.044 | 0 | 0.67 | 0.22 | 0.73 | 0.86 | 0.76 | 0.64 |
| 11 | 40884510 | rs12288111 | -1.792 | -0.658 | 1.31 | -0.32 | -0.069 | -0.036 | -0.036 | 0.215 | 0.25 | 0.52 | 0.19 | 0.15 | 0.15 | 0.26 |
| 11 | 40884545 | rs12281223 | -2.449 | -0.465 | 1.067 | 0.046 | 0.033 | 0.016 | 0.016 | 0.576 | 0.37 | 0.81 | 0.37 | 0.32 | 0.31 | 0.42 |
| 11 | 40886521 | rs4755563 | 1.312 | -0.567 | 1.295 | 0.123 | -0.01 | -0.003 | -0.003 | 0 | 0.75 | 0.28 | 0.8 | 0.83 | 0.88 | 0.7 |
| 11 | 40888726 | rs7948597 | -1.517 | -0.458 | 1.214 | 0.064 | 0.006 | -0.032 | -0.032 | 0 | 0.08 | 0.13 | 0.09 | 0 | 0.14 | 0.09 |
| 11 | 40891793 | rs10430983 | 2.382 | -0.59 | 1.072 | 0.135 | 0.041 | -0.015 | -0.015 | 0 | 0.62 | 0.09 | 0.64 | 0.66 | 0.7 | 0.52 |
| 11 | 40904848 | rs11036136 | -0.701 | -0.806 | 1.436 | 0.132 | 0.01 | -0.02 | -0.02 | 0 | 0.1 | 0.04 | 0.14 | 0.15 | 0.06 | 0.12 |
| 11 | 40915304 | rs11036142 | -1.204 | -0.921 | 2.275 | -0.025 | -0.013 | -0.041 | -0.041 | 0.507 | 0.1 | 0.18 | 0.04 | 0 | 0.07 | 0.05 |
| 11 | 40915339 | rs10768645 | 1.268 | -0.861 | 2.443 | 0.105 | -0.029 | 0.017 | 0.017 | 0 | 0.87 | 0.71 | 0.84 | 0.88 | 0.94 | 0.82 |
| 11 | 40918036 | rs4482016 | 1.198 | -0.854 | 2.446 | 0.109 | -0.026 | 0.02 | 0.02 | 0 | 0.87 | 0.71 | 0.85 | 0.88 | 0.94 | 0.82 |
| 11 | 40921383 | rs11036146 | -0.738 | -0.829 | 2.321 | 0.121 | 0.007 | -0.011 | -0.011 | 0 | 0.1 | 0.04 | 0.14 | 0.12 | 0.06 | 0.12 |
| 11 | 40927453 | rs979221 | 0.79 | -1.024 | 2.67 | 0.001 | -0.002 | -0.04 | -0.04 | 0 | 0.91 | 0.7 | 0.97 | 1 | 0.97 | 0.84 |
| 11 | 40939009 | rs1038892 | -1.327 | -1.002 | 2.383 | 0.047 | -0.027 | 0.027 | 0.027 | 0 | 0.11 | 0.25 | 0.14 | 0.12 | 0.05 | 0.17 |
| 11 | 40946450 | rs10501242 | -0.549 | -0.961 | 2.194 | 0.067 | 0.013 | -0.022 | -0.022 | 0.533 | 0.09 | 0.03 | 0.13 | 0.12 | 0.05 | 0.1 |
| 11 | 40961066 | rs10768653 | 0.177 | -1.131 | 1.208 | 0.108 | 0.021 | -0.002 | -0.002 | 0 | 0.44 | 0.32 | 0.44 | 0.43 | 0.34 | 0.26 |
| 11 | 40966294 | rs11604079 | 0.953 | -0.988 | 1.026 | 0.077 | 0.014 | 0.008 | 0.008 | 0 | 0.34 | 0.18 | 0.29 | 0.32 | 0.28 | 0.16 |
| 11 | 40981020 | rs355230 | 0.296 | -0.865 | 1.189 | 0.244 | -0.01 | 0.023 | 0.023 | 0 | 0.84 | 0.81 | 0.74 | 0.71 | 0.89 | 0.83 |
| 11 | 40981334 | rs355231 | 0.747 | -0.914 | 1.207 | 0.19 | -0.009 | 0.014 | 0.014 | 0 | 0.82 | 0.77 | 0.72 | 0.71 | 0.86 | 0.83 |
| 11 | 40985242 | rs11036160 | 1.169 | -0.998 | 1.323 | 0.07 | 0.005 | 0.095 | 0.095 | 0.15 | 0.33 | 0.14 | 0.2 | 0.15 | 0.28 | 0.15 |
| 11 | 40989912 | rs10837535 | 1.169 | -1.013 | 1.435 | 0.063 | 0.006 | 0.093 | 0.093 | 0.15 | 0.33 | 0.13 | 0.2 | 0.15 | 0.28 | 0.15 |
| 11 | 40993375 | rs1585552 | 1.069 | -1.018 | 1.433 | 0.479 | -0.067 | 0.03 | 0.03 | 0 | 0.36 | 0.15 | 0.48 | 0.31 | 0.53 | 0.34 |
| 11 | 41001237 | rs10768656 | 1.441 | -1.026 | 1.401 | 0.1 | -0.141 | -0.116 | -0.116 | 2.735 | 0.32 | 0.05 | 0.41 | 0.38 | 0.42 | 0.41 |
| 11 | 41004199 | rs10837539 | 1.266 | -0.965 | 1.462 | -0.13 | 0.008 | 0.096 | 0.096 | 0.184 | 0.33 | 0.13 | 0.21 | 0.15 | 0.29 | 0.15 |
| 11 | 41005467 | rs355214 | 0.008 | -0.962 | 1.675 | -0.13 | 0.024 | 0.042 | 0.042 | 0.261 | 0.47 | 0.62 | 0.27 | 0.15 | 0.35 | 0.36 |
| 11 | 41015571 | rs355247 | 0.072 | -1.108 | 1.593 | -0.395 | -0.002 | 0 | 0 | 0 | 0.71 | 0.7 | 0.66 | 0.7 | 0.78 | 0.75 |
| 11 | 41019181 | rs355249 | -1.393 | -1.146 | 1.9 | -0.432 | 0.021 | -0.091 | -0.091 | 0 | 0.14 | 0.33 | 0.1 | 0 | 0.18 | 0.13 |
| 11 | 41020215 | rs7941735 | 1.688 | -1.164 | 1.821 | -0.438 | -0.113 | -0.097 | -0.097 | 0.177 | 0.35 | 0.07 | 0.41 | 0.56 | 0.45 | 0.51 |

| | | | | | | | | | | | | | | | | |
|---|---|---|---|---|---|---|---|---|---|---|---|---|---|---|---|---|
| 11 | 41021781 | rs355256 | -0.497 | -1.088 | 1.846 | -0.492 | -0.005 | -0.068 | -0.068 | 0 | 0.33 | 0.5 | 0.23 | 0.14 | 0.33 | 0.25 |
| 11 | 41021965 | rs10837541 | 1.279 | -1.096 | 1.777 | -0.473 | -0.004 | -0.009 | -0.009 | 0 | 0.19 | 0.21 | 0.13 | 0.14 | 0.15 | 0.12 |
| 11 | 41024138 | rs16935255 | -0.154 | -0.859 | 1.696 | -0.514 | 0.002 | -0.024 | -0.024 | 0 | 0.1 | 0.11 | 0.09 | 0 | 0.17 | 0.13 |
| 11 | 41024966 | rs355240 | -0.816 | -0.898 | 1.724 | -0.524 | -0.001 | -0.03 | -0.03 | 0 | 0.37 | 0.57 | 0.25 | 0.14 | 0.36 | 0.3 |
| 11 | 41025766 | rs355241 | 0.653 | -0.853 | 1.486 | -0.497 | -0.002 | -0.001 | -0.001 | 0 | 0.47 | 0.48 | 0.51 | 0.56 | 0.6 | 0.63 |
| 11 | 41029732 | rs10837544 | 1.457 | -0.815 | 1.222 | -0.486 | -0.067 | -0.051 | -0.051 | 0.129 | 0.37 | 0.16 | 0.42 | 0.56 | 0.44 | 0.5 |
| 11 | 41035085 | rs16935277 | -0.348 | -0.747 | 1.325 | -0.478 | 0.011 | -0.036 | -0.036 | 0 | 0.1 | 0.11 | 0.09 | 0 | 0.17 | 0.13 |
| 11 | 41039084 | rs7125467 | 0.287 | -0.833 | 1.462 | -0.494 | 0.013 | 0.005 | 0.005 | 0 | 0.53 | 0.44 | 0.52 | 0.55 | 0.62 | 0.63 |
| 11 | 41044219 | rs4539314 | 0.748 | -1.127 | 1.549 | -0.496 | -0.008 | -0.004 | -0.004 | 0 | 0.62 | 0.44 | 0.62 | 0.68 | 0.71 | 0.63 |
| 11 | 41049352 | rs7937902 | 0.366 | -1.128 | 1.554 | -0.494 | -0.024 | -0.004 | -0.004 | 0 | 0.66 | 0.51 | 0.63 | 0.68 | 0.74 | 0.64 |
| 11 | 41052020 | rs1350914 | 1.298 | -1.07 | 1.382 | -0.444 | 0.003 | 0.008 | 0.008 | 0 | 0.19 | 0.15 | 0.13 | 0.13 | 0.15 | 0.11 |
| 11 | 41055270 | rs1381583 | 1.143 | -0.964 | 1.533 | -0.39 | 0.012 | 0.022 | 0.022 | 0 | 0.57 | 0.35 | 0.55 | 0.68 | 0.59 | 0.61 |
| 11 | 41057577 | rs4631866 | 1.137 | -1.171 | 1.512 | -0.494 | -0.003 | -0.004 | -0.004 | 0 | 0.4 | 0.41 | 0.5 | 0.54 | 0.56 | 0.57 |
| 11 | 41059281 | rs10837555 | 1.24 | -0.826 | 1.711 | -0.454 | 0.011 | 0.022 | 0.022 | 0 | 0.57 | 0.38 | 0.55 | 0.68 | 0.59 | 0.62 |
| 11 | 41059708 | rs1462225 | 0.763 | -0.827 | 1.728 | -0.441 | -0.024 | -0.013 | -0.013 | 0 | 0.63 | 0.55 | 0.63 | 0.68 | 0.71 | 0.68 |
| 11 | 41059740 | rs1462224 | 0.455 | -0.754 | 1.357 | -0.194 | -0.01 | 0 | 0 | 0 | 0.67 | 0.57 | 0.59 | 0.68 | 0.67 | 0.73 |
| 11 | 41064232 | rs355220 | 0.253 | -0.573 | 0.978 | -0.163 | -0.01 | -0.026 | -0.026 | 0 | 0.05 | 0.11 | 0.02 | 0 | 0.03 | 0.03 |
| 11 | 41066277 | rs1462220 | 0.916 | -0.715 | 0.701 | -0.024 | -0.03 | 0.017 | 0.017 | 0 | 0.46 | 0.2 | 0.44 | 0.37 | 0.49 | 0.52 |
| 11 | 41066640 | rs6485244 | -0.556 | -0.692 | 1.241 | 0.001 | 0.044 | 0.074 | 0.074 | 0 | 0.15 | 0.33 | 0.12 | 0.33 | 0.21 | 0.29 |
| 11 | 41069095 | rs1381577 | 1.434 | -0.918 | 1.174 | -0.121 | -0.032 | -0.016 | -0.016 | 0 | 0.42 | 0.29 | 0.43 | 0.57 | 0.47 | 0.52 |
| 11 | 41070348 | rs7951025 | 0.636 | -0.941 | 1.181 | -0.125 | 0.043 | 0.031 | 0.031 | 0 | 0.53 | 0.37 | 0.52 | 0.69 | 0.66 | 0.75 |
| 11 | 41074215 | rs9943482 | 0.782 | -0.692 | 1.124 | -0.14 | 0.05 | 0.051 | 0.051 | 0 | 0.51 | 0.28 | 0.51 | 0.36 | 0.6 | 0.54 |
| 11 | 41074546 | rs16935312 | -0.753 | -0.687 | 1.119 | -0.154 | -0.003 | -0.011 | -0.011 | 0 | 0.12 | 0.13 | 0.1 | 0 | 0.16 | 0.07 |
| 11 | 41074667 | rs1350917 | 1.637 | -0.682 | 1.096 | -0.127 | -0.071 | -0.016 | -0.016 | 0 | 0.39 | 0.15 | 0.41 | 0.36 | 0.44 | 0.47 |
| 11 | 41074787 | rs355223 | -0.026 | -0.742 | 1.448 | -0.191 | 0.038 | -0.003 | -0.003 | 0 | 0.35 | 0.19 | 0.42 | 0.31 | 0.29 | 0.18 |
| 11 | 41075561 | rs4488182 | -0.542 | -0.87 | 1.198 | -0.29 | 0.006 | 0.033 | 0.033 | 0 | 0.25 | 0.13 | 0.15 | 0.21 | 0.24 | 0.12 |
| 11 | 41076836 | rs355260 | -0.534 | -0.862 | 1.51 | -0.277 | -0.018 | 0.008 | 0.008 | 0 | 0.31 | 0.37 | 0.4 | 0.31 | 0.27 | 0.25 |
| 11 | 41081964 | rs10837562 | 1.154 | -0.815 | 1.464 | -0.27 | 0.018 | 0.002 | 0.002 | 0.099 | 0.49 | 0.37 | 0.46 | 0.48 | 0.52 | 0.65 |
| 11 | 41083620 | rs10501248 | -0.583 | -0.806 | 1.558 | -0.257 | -0.021 | 0.015 | 0.015 | 0 | 0.38 | 0.42 | 0.44 | 0.52 | 0.3 | 0.24 |
| 11 | 41086726 | rs7115895 | -0.853 | -0.813 | 1.466 | -0.26 | -0.006 | 0.008 | 0.008 | 0 | 0.43 | 0.43 | 0.5 | 0.52 | 0.43 | 0.3 |
| 11 | 41089465 | rs2902125 | -0.468 | -0.808 | 1.472 | -0.248 | 0.003 | 0.026 | 0.026 | 0 | 0.29 | 0.43 | 0.44 | 0.31 | 0.34 | 0.25 |
| 11 | 41096261 | rs1462216 | 1.814 | -0.847 | 1.23 | -0.306 | -0.09 | -0.041 | -0.041 | 0 | 0.39 | 0.11 | 0.4 | 0.38 | 0.44 | 0.46 |
| 11 | 41097249 | rs6485247 | -0.128 | -0.811 | 1.438 | -0.267 | -0.016 | 0.014 | 0.014 | 0 | 0.73 | 0.38 | 0.6 | 0.7 | 0.77 | 0.8 |
| 11 | 41100409 | rs10837565 | 0.654 | -0.795 | 1.39 | -0.253 | 0.003 | 0.001 | 0.001 | 0 | 0.09 | 0.13 | 0.04 | 0.11 | 0.08 | 0.18 |
| 11 | 41103092 | rs12806432 | -0.174 | -0.911 | 1.725 | -0.14 | -0.012 | 0.027 | 0.027 | 0 | 0.24 | 0.32 | 0.38 | 0.3 | 0.23 | 0.2 |
| 11 | 41104950 | rs11036223 | 0.919 | -0.894 | 0.339 | -0.115 | 0.018 | -0.013 | -0.013 | 0 | 0.14 | 0.06 | 0.06 | 0.21 | 0.09 | 0.05 |
| 11 | 41105242 | rs1350915 | 0.911 | -0.769 | 0.112 | -0.166 | 0.028 | -0.023 | -0.023 | 0 | 0.54 | 0.27 | 0.48 | 0.49 | 0.56 | 0.69 |
| 11 | 41119492 | rs10501249 | -0.325 | -0.33 | 0.517 | -0.307 | 0.005 | 0.021 | 0.021 | 0 | 0.07 | 0.03 | 0.02 | 0 | 0.04 | 0.05 |
| 11 | 41133162 | rs7106877 | -0.788 | -0.136 | 0.801 | -0.108 | 0.028 | -0.038 | -0.038 | 0 | 0.07 | 0.13 | 0.07 | 0 | 0.12 | 0.08 |
| 11 | 41133555 | rs10742576 | 0.565 | -0.205 | 0.304 | -0.064 | -0.013 | -0.039 | -0.039 | 0 | 0.76 | 0.43 | 0.84 | 0.82 | 0.79 | 0.87 |
| 11 | 41139222 | rs10501250 | -0.703 | 0.01 | -0.128 | -0.05 | -0.003 | -0.015 | -0.015 | 1.517 | 0.05 | 0.04 | 0.02 | 0 | 0.04 | 0.02 |
| 11 | 41142855 | rs12293450 | -0.287 | 0.384 | -0.728 | 0.199 | -0.029 | -0.076 | -0.076 | 0.173 | 0.18 | 0.06 | 0.38 | 0.35 | 0.2 | 0.31 |
| 11 | 41145927 | rs2902117 | -0.308 | 0.545 | -0.696 | -0.11 | -0.011 | 0.022 | 0.022 | 0 | 0.2 | 0.14 | 0.16 | 0.08 | 0.25 | 0.11 |
| 11 | 41148131 | rs9787820 | 1.836 | 0.693 | -0.798 | -0.145 | 0.005 | -0.022 | -0.022 | 0.086 | 0.49 | 0.21 | 0.4 | 0.47 | 0.47 | 0.44 |
| 11 | 41148433 | rs11036247 | -0.549 | 0.499 | -0.32 | 0.29 | 0.005 | 0.027 | 0.027 | 0 | 0.2 | 0.24 | 0.4 | 0.46 | 0.23 | 0.39 |
| 11 | 41154796 | rs10837579 | -0.842 | 0.055 | -0.644 | 0.26 | -0.026 | -0.085 | -0.085 | 0 | 0.31 | 0.1 | 0.45 | 0.4 | 0.32 | 0.31 |
| 11 | 41159081 | rs12285634 | 0.739 | 0.166 | 0.86 | 0.344 | 0.037 | 0.115 | 0.115 | 1.139 | 0.79 | 0.55 | 0.55 | 0.63 | 0.78 | 0.81 |
| 11 | 41162903 | rs717112 | -0.158 | 0.175 | 0.631 | 0.401 | -0.001 | 0.009 | 0.009 | 2.257 | 0.15 | 0.16 | 0.42 | 0.37 | 0.19 | 0.17 |
| 11 | 41164213 | rs10837583 | -0.113 | -0.786 | 0.329 | -0.157 | 0.006 | 0.007 | 0.007 | 0 | 0.16 | 0.13 | 0.08 | 0.06 | 0.17 | 0.08 |
| 11 | 41173522 | rs7933937 | -0.036 | -0.67 | 0.616 | -0.092 | 0.039 | 0.012 | 0.012 | 0 | 0.95 | 0.77 | 0.98 | 1 | 0.97 | 0.98 |
| 11 | 41180155 | rs10768669 | 0.061 | -0.588 | 0.44 | 0.069 | -0.001 | 0.013 | 0.013 | 2.793 | 0.14 | 0.15 | 0.41 | 0.36 | 0.17 | 0.21 |
| 11 | 41187089 | rs4755570 | -0.07 | -1.108 | 0.496 | -0.102 | -0.015 | -0.01 | -0.01 | 0 | 0.14 | 0.2 | 0.07 | 0.13 | 0.1 | 0.14 |
| 11 | 41195670 | rs1386404 | 0.359 | -1.018 | 0.233 | -0.05 | -0.012 | -0.031 | -0.031 | 0 | 0.14 | 0.1 | 0.4 | 0.36 | 0.17 | 0.21 |
| 11 | 41200764 | rs2035628 | 1.807 | -1.227 | -0.455 | -0.74 | -0.121 | -0.101 | -0.101 | 0.017 | 0.34 | 0.06 | 0.32 | 0.45 | 0.41 | 0.49 |
| 11 | 41209500 | rs12786091 | 1.09 | -1.403 | 0.013 | -0.643 | 0.031 | -0.025 | -0.025 | 0 | 0.61 | 0.21 | 0.47 | 0.43 | 0.68 | 0.58 |
| 11 | 41212227 | rs11036281 | 1.191 | -1.462 | 0.1 | -0.738 | 0.001 | -0.013 | -0.013 | 0 | 0.11 | 0.08 | 0.06 | 0 | 0.13 | 0.02 |
| 11 | 41212670 | rs7943825 | -0.733 | -1.582 | 0.112 | -0.699 | -0.039 | -0.003 | -0.003 | 0.124 | 0.21 | 0.37 | 0.11 | 0.19 | 0.15 | 0.17 |
| 11 | 41221274 | rs990200 | 0.901 | -1.53 | -0.291 | -0.581 | 0.036 | 0.043 | 0.043 | 0.012 | 0.65 | 0.24 | 0.81 | 0.75 | 0.71 | 0.79 |
| 11 | 41225858 | rs2902115 | 1.488 | -1.573 | -0.488 | -0.361 | 0.023 | 0.039 | 0.039 | 0.054 | 0.52 | 0.09 | 0.39 | 0.34 | 0.53 | 0.64 |
| 11 | 41226338 | rs11036286 | 0.471 | -1.445 | -0.095 | -0.312 | 0 | 0.027 | 0.027 | 2.39 | 0.14 | 0.15 | 0.42 | 0.4 | 0.18 | 0.16 |
| 11 | 41226389 | rs16935437 | 0.619 | -1.444 | -0.216 | -0.307 | -0.011 | -0.037 | -0.037 | 0.08 | 0.14 | 0.07 | 0.41 | 0.4 | 0.18 | 0.15 |
| 11 | 41228774 | rs7122101 | 0.374 | -1.476 | -0.08 | -0.411 | -0.005 | 0.004 | 0.004 | 0 | 0.12 | 0.14 | 0.07 | 0.21 | 0.11 | 0.09 |
| 11 | 41230701 | rs11036290 | 1.316 | -1.442 | -0.482 | -0.419 | -0.002 | -0.019 | -0.019 | 0 | 0.08 | 0.02 | 0.03 | 0.18 | 0.1 | 0.07 |
| 11 | 41232860 | rs1021270 | 1.83 | -1.491 | -0.87 | -0.701 | -0.127 | -0.085 | -0.085 | 0 | 0.35 | 0.04 | 0.31 | 0.38 | 0.41 | 0.45 |
| 11 | 41235144 | rs4325297 | -1.498 | -1.498 | -0.672 | -0.458 | 0.042 | 0.071 | 0.071 | 0.475 | 0.47 | 0.87 | 0.59 | 0.65 | 0.44 | 0.36 |
| 11 | 41257362 | rs1352230 | 0.037 | -0.586 | -0.308 | 0.267 | -0.001 | 0.028 | 0.028 | 2.53 | 0.14 | 0.14 | 0.42 | 0.43 | 0.18 | 0.16 |
| 11 | 41259180 | rs729210 | -0.324 | -0.352 | -0.673 | 0.331 | -0.051 | -0.093 | -0.093 | 0 | 0.15 | 0.05 | 0.42 | 0.44 | 0.21 | 0.2 |
| 11 | 41261033 | rs12290967 | 1.263 | -1.015 | -0.75 | -0.278 | -0.045 | 0 | 0 | 0 | 0.36 | 0.18 | 0.34 | 0.36 | 0.45 | 0.4 |
| 11 | 41265670 | rs10501254 | 0.739 | -1.057 | -0.048 | -0.261 | 0.006 | 0.076 | 0.076 | 0.035 | 0.12 | 0.15 | 0.41 | 0.42 | 0.16 | 0.13 |

| | | | | | | | | | | | | | | | | |
|---|---|---|---|---|---|---|---|---|---|---|---|---|---|---|---|---|
| 11 | 41271950 | rs7105234 | 1.036 | -1.128 | -0.146 | -0.328 | -0.049 | -0.006 | -0.006 | 0 | 0.41 | 0.17 | 0.36 | 0.39 | 0.5 | 0.48 |
| 11 | 41277361 | rs7948297 | 0.335 | -1.632 | -0.362 | -0.734 | 0.054 | 0.117 | 0.117 | 0.012 | 0.25 | 0.09 | 0.1 | 0 | 0.14 | 0.22 |
| 11 | 41284958 | rs16935503 | 0.197 | -1.629 | -0.004 | -0.703 | -0.014 | -0.032 | -0.032 | 0 | 0.22 | 0.46 | 0.13 | 0.2 | 0.2 | 0.17 |
| 11 | 41286448 | rs10501259 | 0.322 | -1.627 | -0.164 | -0.709 | 0.052 | 0.137 | 0.137 | 0 | 0.25 | 0.08 | 0.09 | 0 | 0.13 | 0.22 |
| 11 | 41294454 | rs10837607 | 0.894 | -1.517 | -0.024 | -0.562 | 0.008 | 0.09 | 0.09 | 0.015 | 0.12 | 0.16 | 0.42 | 0.42 | 0.16 | 0.13 |
| 11 | 41299498 | rs10837613 | 0.144 | -1.584 | 0.196 | -0.679 | -0.013 | -0.032 | -0.032 | 0 | 0.22 | 0.43 | 0.12 | 0.2 | 0.2 | 0.16 |
| 11 | 41301051 | rs4625446 | -1.978 | -1.603 | 0.162 | -0.68 | -0.048 | -0.009 | -0.009 | 0 | 0.63 | 0.81 | 0.65 | 0.64 | 0.55 | 0.6 |
| 11 | 41309275 | rs2922037 | -0.582 | -1.72 | 0.535 | -0.825 | -0.026 | -0.016 | -0.016 | 0 | 0.81 | 0.61 | 0.89 | 0.81 | 0.84 | 0.84 |
| 11 | 41312360 | rs12285478 | 0.345 | -1.822 | 0.439 | -0.912 | 0.055 | 0.106 | 0.106 | 0 | 0.25 | 0.09 | 0.1 | 0 | 0.13 | 0.22 |
| 11 | 41316326 | rs2922054 | -0.401 | -1.489 | 0.744 | -0.627 | -0.03 | -0.017 | -0.017 | 0 | 0.8 | 0.53 | 0.89 | 0.81 | 0.84 | 0.84 |
| 11 | 41334238 | rs2922036 | 0.057 | -1.513 | 0.671 | -0.641 | -0.01 | -0.011 | -0.011 | 0.007 | 0.77 | 0.61 | 0.87 | 0.78 | 0.78 | 0.75 |
| 11 | 41338695 | rs12363107 | 0.835 | -1.366 | 0.301 | -0.23 | -0.006 | 0.019 | 0.019 | 0.045 | 0.12 | 0.1 | 0.3 | 0.41 | 0.16 | 0.14 |
| 11 | 41345282 | rs9971500 | 0.016 | -1.872 | -0.034 | -0.754 | 0.041 | 0.061 | 0.061 | 0 | 0.22 | 0.1 | 0.08 | 0 | 0.13 | 0.23 |
| 11 | 41348409 | rs11036358 | 0.809 | -1.945 | 0.429 | -0.878 | 0.003 | -0.016 | -0.016 | 0 | 0.16 | 0.16 | 0.09 | 0.01 | 0.16 | 0.12 |
| 11 | 41349594 | rs2958847 | 1.598 | -2.046 | 0.261 | -0.937 | -0.069 | 0.005 | 0.005 | 0 | 0.4 | 0.11 | 0.46 | 0.37 | 0.47 | 0.34 |
| 11 | 41360862 | rs12270847 | 1.077 | -2.2 | 0.424 | -1.069 | 0.014 | 0.015 | 0.015 | 0.09 | 0.14 | 0.08 | 0.07 | 0 | 0.1 | 0.19 |
| 11 | 41375122 | rs1588038 | 0.287 | -1.881 | 0.718 | -0.771 | -0.019 | -0.04 | -0.04 | 0.106 | 0.62 | 0.53 | 0.79 | 0.78 | 0.66 | 0.57 |
| 11 | 41379763 | rs11036368 | 0.342 | -1.901 | 0.586 | -0.746 | -0.014 | -0.059 | -0.059 | 0.02 | 0.19 | 0.1 | 0.28 | 0.4 | 0.19 | 0.22 |
| 11 | 41384850 | rs1037161 | 1.558 | -1.939 | 0.638 | -0.729 | -0.088 | -0.023 | -0.023 | 0 | 0.4 | 0.1 | 0.48 | 0.39 | 0.48 | 0.34 |
| 11 | 41386393 | rs2922047 | 1.37 | -1.855 | 0.568 | -0.687 | -0.027 | 0.006 | 0.006 | 0 | 0.41 | 0.22 | 0.5 | 0.39 | 0.48 | 0.34 |
| 11 | 41389802 | rs11036372 | 0.259 | -1.827 | 0.425 | -0.682 | -0.011 | -0.049 | -0.049 | 0 | 0.19 | 0.12 | 0.28 | 0.38 | 0.18 | 0.21 |
| 11 | 41389942 | rs2922048 | 0.625 | -1.784 | 0.677 | -0.582 | -0.01 | -0.023 | -0.023 | 0 | 0.61 | 0.43 | 0.78 | 0.78 | 0.66 | 0.56 |
| 11 | 41396875 | rs2922049 | -0.038 | -1.675 | 0.302 | -0.474 | -0.022 | -0.037 | -0.037 | 0 | 0.68 | 0.59 | 0.82 | 0.81 | 0.73 | 0.65 |
| 11 | 41401550 | rs2939753 | 0.87 | -0.207 | -0.008 | -0.238 | 0.023 | 0.033 | 0.033 | 0 | 0.27 | 0.31 | 0.37 | 0.39 | 0.33 | 0.34 |
| 11 | 41407333 | rs1506716 | -0.491 | -0.037 | -0.217 | -0.652 | 0.009 | 0.018 | 0.018 | 0 | 0.42 | 0.41 | 0.31 | 0.22 | 0.41 | 0.47 |
| 11 | 41411565 | rs10837639 | -0.466 | -0.03 | -0.275 | -0.649 | 0.016 | 0.04 | 0.04 | 0 | 0.42 | 0.34 | 0.31 | 0.22 | 0.41 | 0.47 |
| 11 | 41413943 | rs10837640 | -0.583 | -0.33 | -0.34 | -0.529 | -0.011 | -0.01 | -0.01 | 0.413 | 0.6 | 0.66 | 0.57 | 0.54 | 0.5 | 0.4 |
| 11 | 41414793 | rs2939763 | -1.614 | -0.457 | -0.202 | -1.08 | 0.002 | -0.021 | -0.021 | 0 | 0.94 | 0.9 | 0.96 | 1 | 0.93 | 0.89 |
| 11 | 41417982 | rs2939766 | 1.288 | -0.574 | -0.519 | -0.928 | -0.043 | -0.027 | -0.027 | 0 | 0.31 | 0.14 | 0.29 | 0.58 | 0.41 | 0.54 |
| 11 | 41420388 | rs2922021 | 0.104 | -0.952 | -0.513 | -1.059 | 0.011 | 0.017 | 0.017 | 0 | 0.8 | 0.85 | 0.87 | 0.85 | 0.83 | 0.81 |
| 11 | 41421949 | rs2956774 | -0.356 | -0.961 | -0.581 | -1.069 | 0.005 | 0.005 | 0.005 | 0 | 0.84 | 0.92 | 0.89 | 0.86 | 0.85 | 0.84 |
| 11 | 41424075 | rs986189 | -2.042 | -0.889 | -0.529 | -1.047 | -0.001 | 0.001 | 0.001 | 2.774 | 0.94 | 0.93 | 0.96 | 1 | 0.93 | 0.89 |
| 11 | 41429355 | rs7130734 | 0.116 | -1.145 | 0.108 | -1.289 | 0 | 0.003 | 0.003 | 0 | 0.39 | 0.58 | 0.47 | 0.29 | 0.34 | 0.24 |
| 11 | 41430746 | rs11036390 | 0.419 | -1.051 | 0.119 | -1.267 | -0.002 | -0.023 | -0.023 | 0 | 0.37 | 0.51 | 0.45 | 0.28 | 0.33 | 0.22 |
| 11 | 41438442 | rs1868029 | 0.662 | -0.935 | 0.157 | -1.181 | -0.016 | 0.023 | 0.023 | 0.028 | 0.44 | 0.3 | 0.42 | 0.56 | 0.49 | 0.58 |
| 11 | 41441656 | rs16935698 | 1.104 | -0.856 | 0.191 | -1.103 | 0 | -0.028 | -0.028 | 0 | 0.08 | 0.03 | 0.06 | 0.15 | 0.08 | 0.05 |
| 11 | 41443472 | rs2939755 | -0.542 | -0.99 | 0.134 | -1.178 | -0.029 | 0.083 | 0.083 | 0 | 0.61 | 0.8 | 0.54 | 0.71 | 0.59 | 0.66 |
| 11 | 41444545 | rs996082 | 0.246 | -0.962 | 0.216 | -1.135 | -0.001 | 0.008 | 0.008 | 0 | 0.48 | 0.47 | 0.43 | 0.56 | 0.48 | 0.6 |
| 11 | 41451266 | rs1506712 | -0.345 | -0.777 | 0.095 | -1.362 | 0.002 | -0.003 | -0.003 | 0 | 0.49 | 0.5 | 0.46 | 0.21 | 0.39 | 0.28 |
| 11 | 41453508 | rs11036399 | -1.111 | -0.651 | 0.097 | -1.432 | -0.002 | 0.019 | 0.019 | 0.688 | 0.6 | 0.61 | 0.54 | 0.22 | 0.58 | 0.4 |
| 11 | 41454639 | rs10768685 | -1.841 | -0.629 | -0.034 | -1.322 | -0.04 | -0.01 | -0.01 | 0.747 | 0.69 | 0.86 | 0.61 | 0.35 | 0.64 | 0.46 |
| 11 | 41464670 | rs1001952 | 0.609 | -0.753 | -0.241 | -1.471 | -0.077 | 0.096 | 0.096 | 0 | 0.37 | 0.03 | 0.3 | 0.2 | 0.3 | 0.25 |
| 11 | 41468936 | rs7105487 | -1.227 | -0.621 | 0.127 | -1.357 | 0.007 | -0.006 | -0.006 | 0 | 0.62 | 0.55 | 0.52 | 0.35 | 0.48 | 0.37 |
| 11 | 41469843 | rs7106617 | -1.578 | -0.672 | 0.058 | -1.29 | 0 | -0.001 | -0.001 | 0 | 0.67 | 0.62 | 0.58 | 0.36 | 0.64 | 0.44 |
| 11 | 41470267 | rs7124437 | -1.908 | -0.599 | -0.343 | -1.288 | -0.067 | -0.039 | -0.039 | 0.163 | 0.7 | 0.93 | 0.61 | 0.36 | 0.65 | 0.45 |
| 11 | 41504551 | rs4411275 | -1.135 | -0.954 | 0.549 | -1.433 | -0.034 | -0.086 | -0.086 | 0.053 | 0.87 | 0.68 | 0.93 | 1 | 0.94 | 0.98 |
| 11 | 41504686 | rs7120811 | 0.156 | -0.872 | 0.684 | -1.489 | 0.04 | -0.028 | -0.028 | 0.156 | 0.05 | 0.17 | 0.07 | 0 | 0.16 | 0.06 |
| 11 | 41506412 | rs1506714 | 1.072 | -0.934 | 0.466 | -1.661 | 0 | -0.009 | -0.009 | 1.781 | 0.38 | 0.35 | 0.47 | 0.77 | 0.42 | 0.61 |
| 11 | 41510850 | rs7109873 | 0.425 | -0.927 | 0.354 | -1.657 | -0.01 | 0.052 | 0.052 | 0.097 | 0.46 | 0.64 | 0.55 | 0.77 | 0.58 | 0.68 |
| 11 | 41518672 | rs1911165 | 0.086 | -0.879 | 0.398 | -1.612 | 0.06 | 0.045 | 0.045 | 0 | 0.06 | 0.21 | 0.08 | 0.11 | 0.16 | 0.08 |
| 11 | 41519485 | rs12361464 | -0.593 | -0.918 | 0.417 | -1.546 | -0.005 | 0.094 | 0.094 | 0.627 | 0.52 | 0.29 | 0.57 | 0.24 | 0.41 | 0.31 |
| 11 | 41521691 | rs7118044 | 0.721 | -0.901 | -0.095 | -1.616 | -0.076 | 0.021 | 0.021 | 0 | 0.34 | 0.03 | 0.39 | 0.24 | 0.27 | 0.25 |
| 11 | 41522530 | rs10837649 | 1.83 | -0.709 | 0.144 | -1.526 | -0.014 | -0.003 | -0.003 | 0 | 0.31 | 0.24 | 0.3 | 0.65 | 0.37 | 0.56 |
| 11 | 41532026 | rs11823636 | -0.994 | -0.624 | 0.388 | -1.334 | 0.002 | -0.11 | -0.11 | 0.001 | 0.15 | 0.41 | 0.13 | 0 | 0.21 | 0.11 |
| 11 | 41532609 | rs10501267 | 1.167 | -0.599 | 0.055 | -1.605 | -0.026 | 0.026 | 0.026 | 0 | 0.12 | 0.01 | 0.09 | 0.04 | 0.1 | 0.1 |
| 11 | 41533025 | rs7131668 | -0.248 | -0.611 | 0.518 | -1.276 | -0.011 | 0.049 | 0.049 | 0.095 | 0.46 | 0.69 | 0.43 | 0.65 | 0.58 | 0.66 |
| 11 | 41535969 | rs11036434 | 1.199 | -0.616 | 0.466 | -1.272 | -0.006 | -0.002 | -0.002 | 0.11 | 0.31 | 0.28 | 0.3 | 0.65 | 0.37 | 0.51 |
| 11 | 41541964 | rs11036436 | -0.063 | -0.367 | 0.706 | -1.112 | -0.003 | 0.013 | 0.013 | 0.158 | 0.46 | 0.6 | 0.42 | 0.65 | 0.58 | 0.66 |
| 11 | 41543073 | rs10768696 | 0.007 | -0.301 | 0.72 | -1.115 | -0.007 | 0.021 | 0.021 | 0.171 | 0.46 | 0.4 | 0.4 | 0.65 | 0.57 | 0.66 |
| 11 | 41554681 | rs10837668 | -0.091 | -0.371 | 0.28 | -1.098 | -0.009 | 0.048 | 0.048 | 0.184 | 0.46 | 0.68 | 0.43 | 0.65 | 0.58 | 0.66 |
| 11 | 41560989 | rs1457329 | -0.206 | -0.231 | 0.404 | -1.135 | 0.035 | -0.027 | -0.027 | 0 | 0.05 | 0.1 | 0.06 | 0 | 0.16 | 0.11 |
| 11 | 41563764 | rs7938549 | -0.663 | -0.977 | 0.25 | -0.765 | 0.006 | -0.019 | -0.019 | 0 | 0.6 | 0.74 | 0.49 | 0.6 | 0.65 | 0.7 |
| 11 | 41566028 | rs10768703 | -0.372 | -1.217 | 0.83 | -1.268 | 0.005 | 0.02 | 0.02 | 0 | 0.1 | 0.28 | 0.15 | 0.12 | 0.06 | 0.1 |
| 11 | 41573627 | rs1902637 | 0.381 | -1.459 | 0.519 | -1.154 | 0.036 | 0.013 | 0.013 | 0 | 0.4 | 0.21 | 0.42 | 0.34 | 0.3 | 0.27 |
| 11 | 41576981 | rs1457319 | -0.07 | -1.521 | 0.523 | -1.179 | -0.001 | -0.001 | -0.001 | 0 | 0.44 | 0.51 | 0.55 | 0.45 | 0.33 | 0.36 |
| 11 | 41577807 | rs7123780 | -0.007 | -1.53 | 0.299 | -1.183 | 0.038 | -0.008 | -0.008 | 0 | 0.45 | 0.69 | 0.57 | 0.45 | 0.33 | 0.36 |
| 11 | 41580712 | rs11036457 | -0.337 | -1.76 | 0.725 | -0.963 | -0.036 | -0.016 | -0.016 | 0 | 0.14 | 0.24 | 0.07 | 0.1 | 0.06 | 0.08 |
| 11 | 41582587 | rs12788207 | -0.027 | -1.562 | 0.768 | -1.128 | 0.006 | -0.005 | -0.005 | 0 | 0.41 | 0.44 | 0.32 | 0.63 | 0.45 | 0.52 |
| 11 | 41583033 | rs10837679 | -0.795 | -1.804 | 0.8 | -1.307 | -0.015 | -0.065 | -0.065 | 0 | 0.88 | 0.76 | 0.83 | 1 | 0.93 | 0.91 |

| | | | | | | | | | | | | | | | | |
|---|---|---|---|---|---|---|---|---|---|---|---|---|---|---|---|---|
| 11 | 41586140 | rs4756656 | 0.479 | -1.803 | 0.674 | -1.306 | -0.004 | 0.034 | 0.034 | 0 | 0.47 | 0.32 | 0.5 | 0.37 | 0.48 | 0.38 |
| 11 | 41587763 | rs11036463 | 1.02 | -1.4 | 0.807 | -1.077 | -0.006 | 0.016 | 0.016 | 0 | 0.43 | 0.22 | 0.46 | 0.37 | 0.4 | 0.24 |
| 11 | 41588707 | rs11036464 | -0.221 | -1.319 | 0.822 | -1.008 | 0 | -0.001 | -0.001 | 0 | 0.42 | 0.53 | 0.33 | 0.63 | 0.45 | 0.52 |
| 11 | 41594447 | rs1350320 | 1 | -1.171 | 0.429 | -0.947 | 0.004 | -0.004 | -0.004 | 0 | 0.71 | 0.33 | 0.7 | 0.9 | 0.79 | 0.69 |
| 11 | 41595069 | rs1457326 | 0.577 | -1.17 | 0.618 | -0.946 | -0.016 | -0.039 | -0.039 | 0 | 0.74 | 0.68 | 0.73 | 0.9 | 0.79 | 0.69 |
| 11 | 41598896 | rs11036468 | 1.056 | -1.122 | -0.278 | -0.893 | -0.03 | -0.042 | -0.042 | 0 | 0.27 | 0.21 | 0.25 | 0.53 | 0.39 | 0.44 |
| 11 | 41599967 | rs10501269 | 0.202 | -0.814 | -0.165 | -0.799 | 0.002 | 0.007 | 0.007 | 0 | 0.45 | 0.42 | 0.47 | 0.36 | 0.4 | 0.24 |
| 11 | 41600172 | rs4755576 | 0.837 | -1.029 | -0.013 | -0.595 | 0.001 | -0.005 | -0.005 | 0 | 0.94 | 0.94 | 0.86 | 0.99 | 0.96 | 0.88 |
| 11 | 41606671 | rs1463083 | 1.358 | -1.397 | 0.05 | -0.756 | 0.016 | 0.028 | 0.028 | 0 | 0.66 | 0.32 | 0.63 | 0.86 | 0.72 | 0.75 |
| 11 | 41611349 | rs1463068 | 0.081 | -1.47 | 0.186 | -0.813 | -0.003 | -0.021 | -0.021 | 0 | 0.95 | 0.9 | 0.87 | 0.99 | 0.95 | 0.93 |
| 11 | 41612018 | rs878084 | 1.138 | -1.571 | 0.126 | -0.877 | 0.007 | -0.003 | -0.003 | 0 | 0.89 | 0.9 | 0.82 | 0.99 | 0.91 | 0.92 |
| 11 | 41614306 | rs7930633 | 2.387 | -1.991 | -0.172 | -1.086 | 0.075 | 0.179 | 0.179 | 0.084 | 0.55 | 0.04 | 0.56 | 0.82 | 0.68 | 0.65 |
| 11 | 41616494 | rs10768709 | 1.539 | -1.449 | 0.539 | -0.837 | 0.015 | 0.008 | 0.008 | 0 | 0.77 | 0.84 | 0.71 | 0.98 | 0.82 | 0.86 |
| 11 | 41629903 | rs11036477 | 2.284 | -0.914 | 1.285 | -0.219 | -0.008 | -0.001 | -0.001 | 0 | 0.8 | 0.31 | 0.71 | 0.81 | 0.81 | 0.76 |
| 11 | 41631862 | rs1463075 | 0.453 | -0.898 | 1.669 | -0.219 | 0.035 | -0.009 | -0.009 | 0 | 0.94 | 0.78 | 0.85 | 0.99 | 0.95 | 0.9 |
| 11 | 41632353 | rs1463076 | 2.394 | -0.902 | 1.66 | -0.204 | -0.008 | 0 | 0 | 0 | 0.8 | 0.32 | 0.71 | 0.81 | 0.81 | 0.76 |
| 11 | 41633538 | rs931029 | -0.971 | -0.911 | 2.111 | -0.2 | 0.019 | 0.046 | 0.046 | 0.009 | 0.07 | 0.28 | 0.06 | 0.08 | 0.07 | 0.08 |
| 11 | 41639081 | rs1463079 | 2.171 | -0.914 | 2.071 | -0.197 | -0.011 | 0.036 | 0.036 | 0 | 0.84 | 0.46 | 0.78 | 0.83 | 0.86 | 0.81 |
| 11 | 41653695 | rs7109792 | 2.372 | -0.845 | 1.764 | -0.146 | -0.007 | 0.002 | 0.002 | 0 | 0.8 | 0.34 | 0.71 | 0.8 | 0.81 | 0.76 |
| 11 | 41662510 | rs11820725 | -1.02 | -0.927 | 1.546 | -0.124 | -0.003 | 0.015 | 0.015 | 0 | 0.08 | 0.22 | 0.05 | 0.08 | 0.07 | 0.09 |
| 11 | 41683052 | rs899035 | 1.885 | -0.933 | 1.579 | -0.122 | 0.004 | -0.025 | -0.025 | 0 | 0.84 | 0.52 | 0.77 | 0.89 | 0.84 | 0.78 |
| 11 | 41684258 | rs7113382 | 1.783 | -0.938 | 1.385 | -0.124 | 0.016 | -0.02 | -0.02 | 0 | 0.84 | 0.57 | 0.78 | 0.89 | 0.84 | 0.78 |
| 11 | 41685519 | rs10742596 | -0.016 | -0.913 | 1.336 | -0.105 | 0.002 | -0.005 | -0.005 | 0 | 0.93 | 0.95 | 0.96 | 0.92 | 0.93 | 0.91 |
| 11 | 41710829 | rs2199556 | 2.147 | -1.188 | 1.119 | -0.243 | 0.051 | 0.048 | 0.048 | 0.268 | 0.64 | 0.11 | 0.64 | 0.77 | 0.72 | 0.66 |
| 11 | 41711851 | rs1381939 | 0.558 | -0.691 | 1.558 | -0.051 | 0.017 | 0.04 | 0.04 | 0 | 0.92 | 0.74 | 0.95 | 0.92 | 0.93 | 0.91 |
| 11 | 41712470 | rs16935853 | -1.02 | -0.587 | 0.581 | -0.022 | 0.015 | -0.022 | -0.022 | 0.923 | 0.05 | 0.15 | 0.11 | 0.01 | 0.05 | 0.09 |
| 11 | 41723364 | rs7482552 | -0.832 | -0.22 | 0.301 | -0.043 | 0.004 | -0.014 | -0.014 | 0 | 0.05 | 0.09 | 0.06 | 0.01 | 0.1 | 0.09 |
| 11 | 41727989 | rs287761 | 0.923 | -0.667 | 0.386 | 0.447 | -0.007 | 0.013 | 0.013 | 0 | 0.69 | 0.59 | 0.65 | 0.57 | 0.72 | 0.75 |
| 11 | 41731763 | rs287756 | 1.86 | -0.641 | 0.295 | 0.404 | -0.002 | 0 | 0 | 0.56 | 0.56 | 0.49 | 0.54 | 0.46 | 0.59 | 0.57 |
| 11 | 41731790 | rs10837702 | 1.969 | -0.759 | 0.472 | 0.346 | 0.019 | -0.054 | -0.054 | 0 | 0.54 | 0.12 | 0.52 | 0.46 | 0.59 | 0.57 |
| 11 | 41737203 | rs287746 | -1.624 | -0.756 | 0.581 | 0.394 | 0.008 | 0.016 | 0.016 | 0.161 | 0.15 | 0.42 | 0.13 | 0.11 | 0.13 | 0.18 |
| 11 | 41741527 | rs287733 | -1.081 | -0.721 | 0.537 | 0.398 | 0.059 | 0.093 | 0.093 | 0 | 0.09 | 0.43 | 0.08 | 0.09 | 0.09 | 0.09 |
| 11 | 41742472 | rs479434 | 1.405 | -0.49 | 0.515 | 0.416 | 0.005 | -0.006 | -0.006 | 0.026 | 0.63 | 0.59 | 0.6 | 0.54 | 0.66 | 0.66 |
| 11 | 41750510 | rs1960350 | -1.223 | -0.528 | 0.425 | 0.467 | 0.024 | -0.033 | -0.033 | 0 | 0.3 | 0.22 | 0.32 | 0.45 | 0.23 | 0.25 |
| 11 | 41754073 | rs11825544 | -2.354 | -1.146 | 1.187 | 0.052 | -0.053 | -0.084 | -0.084 | 0 | 0.13 | 0.59 | 0.07 | 0 | 0.04 | 0.06 |
| 11 | 41757604 | rs12289055 | -1.722 | -1.147 | 1.885 | 0.052 | -0.01 | -0.041 | -0.041 | 0 | 0.09 | 0.4 | 0.06 | 0 | 0.04 | 0.03 |
| 11 | 41769608 | rs11036536 | -0.647 | -1.899 | 0.997 | 0.024 | 0.036 | -0.014 | -0.014 | 0 | 0.37 | 0.15 | 0.47 | 0.64 | 0.29 | 0.39 |
| 11 | 41781425 | rs11605567 | 1.68 | -1.668 | 0.851 | 0.095 | -0.007 | 0.136 | 0.136 | 0 | 0.45 | 0.14 | 0.39 | 0.22 | 0.59 | 0.44 |
| 11 | 41786402 | rs10837719 | 1.465 | -1.833 | 0.729 | 0.184 | 0.115 | 0.067 | 0.067 | 0 | 0.5 | 0.15 | 0.45 | 0.36 | 0.66 | 0.55 |
| 11 | 41799496 | rs723562 | -1.026 | -1.564 | 1.166 | 0.352 | -0.006 | -0.032 | -0.032 | 0.027 | 0.08 | 0.24 | 0.04 | 0 | 0.03 | 0.03 |
| 11 | 41806690 | rs10768739 | -2.001 | -1.527 | 0.423 | 0.287 | 0.02 | 0.05 | 0.05 | 0 | 0.53 | 0.84 | 0.59 | 0.63 | 0.4 | 0.5 |
| 11 | 41807281 | rs7128034 | -1.904 | -1.472 | 0.739 | 0.033 | -0.02 | -0.079 | -0.079 | 0 | 0.16 | 0.61 | 0.09 | 0.01 | 0.09 | 0.1 |
| 11 | 41819536 | rs1385379 | -1.216 | -1.552 | 1.136 | 0.33 | 0.027 | 0.005 | 0.005 | 0 | 0.42 | 0.34 | 0.53 | 0.63 | 0.34 | 0.42 |
| 11 | 41828613 | rs16936003 | 0.123 | -1.588 | 1.097 | 0.295 | -0.014 | -0.033 | -0.033 | 0 | 0.07 | 0.1 | 0.03 | 0 | 0.04 | 0.01 |
| 11 | 41830523 | rs10837748 | -1.116 | -1.489 | 1.093 | 0.33 | 0.025 | 0.002 | 0.002 | 0 | 0.41 | 0.29 | 0.53 | 0.63 | 0.34 | 0.42 |
| 11 | 41839645 | rs2606421 | -2.233 | -1.582 | 0.992 | 0.273 | -0.01 | 0.136 | 0.136 | 0 | 0.55 | 0.82 | 0.62 | 0.77 | 0.43 | 0.59 |
| 11 | 41842341 | rs12797436 | -1.046 | -1.247 | 1.585 | 0.145 | 0.002 | -0.023 | -0.023 | 0.241 | 0.09 | 0.14 | 0.05 | 0.01 | 0.08 | 0.08 |
| 11 | 41845688 | rs12794557 | -0.995 | -1.431 | 1.507 | 0.18 | 0.015 | 0.046 | 0.046 | 0 | 0.31 | 0.19 | 0.23 | 0.12 | 0.26 | 0.3 |
| 11 | 41859366 | rs9783353 | -2.273 | -1.438 | 1.542 | 0.227 | 0.037 | 0.095 | 0.095 | 0 | 0.46 | 0.72 | 0.32 | 0.27 | 0.36 | 0.47 |
| 11 | 41872033 | rs11600495 | -0.848 | -1.4 | 1.692 | 0.249 | -0.003 | 0.004 | 0.004 | 3.395 | 0.09 | 0.1 | 0.05 | 0.01 | 0.08 | 0.08 |
| 11 | 41879455 | rs7103550 | -1.453 | -1.348 | 1.72 | 0.324 | -0.022 | -0.027 | -0.027 | 0 | 0.36 | 0.43 | 0.26 | 0.25 | 0.28 | 0.4 |
| 11 | 41883445 | rs7120856 | -1.16 | -1.413 | 1.292 | 0.18 | 0.007 | 0.006 | 0.006 | 0 | 0.32 | 0.24 | 0.24 | 0.12 | 0.26 | 0.3 |
| 11 | 41893605 | rs11036615 | -0.676 | -1.334 | 1.123 | 0.448 | 0 | 0.071 | 0.071 | 0 | 0.24 | 0.1 | 0.2 | 0.12 | 0.22 | 0.29 |
| 11 | 41894463 | rs291712 | 0.349 | -1.334 | 1.209 | 0.448 | 0.004 | 0.046 | 0.046 | 0 | 0.76 | 0.86 | 0.79 | 0.88 | 0.78 | 0.71 |
| 11 | 41897826 | rs4756680 | -2.057 | -1.498 | 1.333 | 0.903 | 0.016 | 0.001 | 0.001 | 0 | 0.37 | 0.77 | 0.29 | 0.35 | 0.31 | 0.46 |
| 11 | 41898104 | rs4756681 | -0.431 | -1.504 | 1.419 | 0.873 | -0.005 | 0.054 | 0.054 | 0 | 0.2 | 0.09 | 0.19 | 0.12 | 0.2 | 0.25 |
| 11 | 41898126 | rs12793795 | -0.2 | -1.51 | 1.54 | 0.873 | -0.001 | -0.002 | -0.002 | 0 | 0.06 | 0.05 | 0.04 | 0.01 | 0.06 | 0.04 |
| 11 | 41899112 | rs7122740 | -2.09 | -1.515 | 2.123 | 0.877 | 0 | 0 | 0 | 0 | 0.36 | 0.63 | 0.29 | 0.35 | 0.31 | 0.46 |
| 11 | 41906381 | rs12788548 | -1.582 | -1.519 | 2.347 | 0.876 | 0.005 | 0.06 | 0.06 | 2.207 | 0.09 | 0.13 | 0.05 | 0.23 | 0.09 | 0.16 |
| 11 | 41906525 | rs10768764 | 0.926 | -1.522 | 2.416 | 0.871 | -0.016 | -0.073 | -0.073 | 0.013 | 0.73 | 0.43 | 0.76 | 0.88 | 0.77 | 0.7 |
| 11 | 41909745 | rs2036830 | 1.821 | -1.519 | 2.386 | 0.878 | 0.009 | 0.003 | 0.003 | 0 | 0.64 | 0.3 | 0.71 | 0.65 | 0.69 | 0.54 |
| 11 | 41915366 | rs9300039 | -1.728 | -1.335 | 2.429 | 0.94 | 0.004 | 0.054 | 0.054 | 1.251 | 0.1 | 0.13 | 0.05 | 0.23 | 0.09 | 0.15 |
| 11 | 41916166 | rs11036622 | -1.787 | -1.341 | 2.432 | 0.957 | 0.003 | 0.052 | 0.052 | 1.08 | 0.1 | 0.13 | 0.05 | 0.23 | 0.09 | 0.15 |
| 11 | 41925201 | rs7949300 | -2.978 | -1.395 | 2.797 | 0.91 | 0.003 | 0.099 | 0.099 | 1.127 | 0.16 | 0.47 | 0.1 | 0.23 | 0.12 | 0.19 |
| 11 | 41926369 | rs2019626 | -3.415 | -1.395 | 2.637 | 0.9 | -0.01 | 0.021 | 0.021 | 1.633 | 0.17 | 0.66 | 0.11 | 0.23 | 0.12 | 0.21 |
| 11 | 41932593 | rs7105457 | -2.772 | -1.376 | 2.525 | 0.91 | -0.002 | 0.038 | 0.038 | 2.193 | 0.15 | 0.23 | 0.07 | 0.23 | 0.12 | 0.18 |
| 11 | 41947768 | rs2176002 | 1.154 | -1.379 | 2.481 | 0.687 | 0.001 | -0.034 | -0.034 | 4.926 | 0.72 | 0.4 | 0.75 | 0.8 | 0.73 | 0.67 |
| 11 | 41955190 | rs10768772 | -0.571 | -1.321 | 2.403 | 0.732 | 0.005 | 0.003 | 0.003 | 0 | 0.89 | 0.92 | 0.91 | 0.87 | 0.92 | 0.79 |
| 11 | 41955855 | rs1913599 | 1.241 | -1.446 | 2.726 | 0.767 | -0.011 | 0.002 | 0.002 | 0 | 0.75 | 0.45 | 0.8 | 0.73 | 0.79 | 0.63 |

| | | | | | | | | | | | | | | | | |
|---|---|---|---|---|---|---|---|---|---|---|---|---|---|---|---|---|
| 11 | 41956646 | rs4489731 | 1.641 | -1.445 | 2.565 | 0.768 | 0.013 | 0.005 | 0.005 | 0 | 0.72 | 0.27 | 0.78 | 0.73 | 0.79 | 0.6 |
| 11 | 41958077 | rs4559679 | 1.622 | -1.428 | 2.571 | 0.776 | 0.012 | 0.005 | 0.005 | 0 | 0.72 | 0.26 | 0.78 | 0.73 | 0.79 | 0.61 |
| 11 | 41958375 | rs7110692 | 1.023 | -1.421 | 2.588 | 0.772 | -0.005 | -0.018 | -0.018 | 0 | 0.77 | 0.36 | 0.79 | 0.84 | 0.81 | 0.71 |
| 11 | 41965205 | rs7939620 | 0.335 | -1.41 | 2.647 | 0.774 | 0.007 | -0.05 | -0.05 | 0.289 | 0.82 | 0.45 | 0.82 | 0.9 | 0.84 | 0.74 |
| 11 | 41965909 | rs11036636 | 2.195 | -1.419 | 2.564 | 0.769 | 0.006 | 0.026 | 0.026 | 0 | 0.66 | 0.21 | 0.73 | 0.72 | 0.71 | 0.56 |
| 11 | 41967671 | rs7112663 | 1.119 | -1.392 | 2.568 | 0.801 | -0.005 | -0.016 | -0.016 | 0.187 | 0.77 | 0.34 | 0.79 | 0.84 | 0.81 | 0.71 |
| 11 | 41968107 | rs6485298 | 0.234 | -1.371 | 2.699 | 0.796 | 0.026 | -0.052 | -0.052 | 0.72 | 0.83 | 0.59 | 0.82 | 0.9 | 0.84 | 0.77 |
| 11 | 41972550 | rs2713820 | 1.078 | -1.366 | 2.234 | 1.445 | -0.005 | 0.031 | 0.031 | 0 | 0.76 | 0.33 | 0.61 | 0.51 | 0.8 | 0.7 |
| 11 | 41978556 | rs2672546 | 0.168 | -1.375 | 2.134 | 1.259 | 0 | 0.076 | 0.076 | 0 | 0.82 | 0.78 | 0.65 | 0.57 | 0.84 | 0.76 |
| 11 | 42008948 | rs11036649 | -1.774 | -1.398 | 2.209 | 1.181 | 0.004 | 0.032 | 0.032 | 1.197 | 0.11 | 0.25 | 0.07 | 0.17 | 0.09 | 0.18 |
| 11 | 42012945 | rs7127839 | 0.88 | -1.499 | 2.416 | 1.171 | -0.011 | 0.13 | 0.13 | 0 | 0.77 | 0.54 | 0.61 | 0.58 | 0.81 | 0.71 |
| 11 | 42014693 | rs11036652 | -1.899 | -1.5 | 2.39 | 1.167 | 0.005 | 0.041 | 0.041 | 1.105 | 0.11 | 0.3 | 0.07 | 0.17 | 0.09 | 0.18 |
| 11 | 42016120 | rs971744 | 1.81 | -1.496 | 1.997 | 1.094 | 0.012 | -0.019 | -0.019 | 0 | 0.66 | 0.26 | 0.54 | 0.41 | 0.71 | 0.53 |
| 11 | 42025245 | rs1514815 | 1.491 | -1.306 | 1.758 | 1.03 | 0.026 | -0.052 | -0.052 | 2.774 | 0.68 | 0.23 | 0.6 | 0.51 | 0.76 | 0.66 |
| 11 | 42026659 | rs2862451 | 1.982 | -1.352 | 1.766 | 0.996 | 0.012 | -0.039 | -0.039 | 0 | 0.63 | 0.21 | 0.54 | 0.42 | 0.69 | 0.55 |
| 11 | 42032191 | rs7103501 | 0.85 | -1.56 | 1.918 | 1.095 | -0.013 | 0.068 | 0.068 | 0 | 0.76 | 0.69 | 0.63 | 0.48 | 0.81 | 0.67 |
| 11 | 42047336 | rs9787747 | 1.491 | -2.059 | 1.184 | 0.746 | 0.019 | -0.024 | -0.024 | 0 | 0.62 | 0.31 | 0.56 | 0.45 | 0.7 | 0.58 |
| 11 | 42064674 | rs2465363 | 0.307 | -2.062 | 1.422 | 0.505 | -0.001 | 0.016 | 0.016 | 0 | 0.77 | 0.78 | 0.65 | 0.64 | 0.8 | 0.77 |
| 11 | 42065028 | rs11036670 | -0.649 | -1.93 | 1.492 | 0.38 | 0.017 | 0.021 | 0.021 | 0 | 0.06 | 0.17 | 0.1 | 0.07 | 0.05 | 0.05 |
| 11 | 42069186 | rs16936245 | -2.009 | -2.013 | 1.704 | 0.322 | 0.011 | 0 | 0 | 0.179 | 0.1 | 0.31 | 0.12 | 0.07 | 0.08 | 0.11 |
| 11 | 42070045 | rs2465362 | 1.896 | -2.041 | 1.624 | 0.176 | 0.008 | -0.043 | -0.043 | 0.414 | 0.59 | 0.27 | 0.48 | 0.53 | 0.65 | 0.54 |
| 11 | 42083016 | rs11036673 | -0.868 | -1.994 | 1.637 | 0.174 | 0 | 0.012 | 0.012 | 0 | 0.1 | 0.05 | 0.06 | 0.05 | 0.09 | 0.14 |
| 11 | 42091353 | rs4553355 | -1.072 | -2.004 | 1.673 | 0.17 | -0.004 | -0.017 | -0.017 | 0 | 0.1 | 0.15 | 0.1 | 0.04 | 0.07 | 0.14 |
| 11 | 42093134 | rs2931814 | 2.381 | -2.054 | 1.685 | 0.169 | -0.006 | -0.073 | -0.073 | 1.35 | 0.55 | 0.13 | 0.46 | 0.53 | 0.62 | 0.52 |
| 11 | 42101548 | rs2200724 | 1.042 | -2.035 | 1.728 | 0.178 | -0.001 | 0.008 | 0.008 | 0 | 0.68 | 0.42 | 0.58 | 0.57 | 0.72 | 0.68 |
| 11 | 42109065 | rs10837773 | 2.946 | -2.827 | 1.072 | -0.182 | -0.18 | -0.017 | -0.017 | 0.367 | 0.36 | 0.02 | 0.32 | 0.28 | 0.52 | 0.4 |
| 11 | 42110038 | rs11605864 | -0.84 | -2.15 | 1.616 | -0.105 | 0.036 | -0.022 | -0.022 | 0.202 | 0.07 | 0.18 | 0.06 | 0 | 0.08 | 0.05 |
| 11 | 42115286 | rs1488610 | -0.071 | -2.195 | 1.524 | -0.099 | -0.017 | -0.071 | -0.071 | 2.919 | 0.2 | 0.08 | 0.33 | 0.39 | 0.2 | 0.22 |
| 11 | 42130431 | rs12365941 | 1.881 | -2.237 | 1.533 | -0.177 | -0.01 | -0.078 | -0.078 | 0 | 0.56 | 0.15 | 0.46 | 0.52 | 0.63 | 0.51 |
| 11 | 42151229 | rs7930659 | 1.049 | -2.949 | 2.093 | -0.256 | -0.045 | -0.043 | -0.043 | 0 | 0.91 | 0.71 | 0.97 | 0.98 | 0.99 | 0.89 |
| 11 | 42154191 | rs11036712 | -0.677 | -3.006 | 2.039 | -0.173 | -0.002 | 0.02 | 0.02 | 0 | 0.21 | 0.26 | 0.29 | 0.41 | 0.17 | 0.19 |
| 11 | 42165001 | rs966536 | 2.108 | -3.086 | 1.913 | -0.272 | 0.037 | -0.044 | -0.044 | 0 | 0.59 | 0.2 | 0.6 | 0.55 | 0.7 | 0.6 |
| 11 | 42167038 | rs1601905 | 1.425 | -2.942 | 1.772 | -0.13 | -0.03 | 0.031 | 0.031 | 0 | 0.7 | 0.48 | 0.68 | 0.57 | 0.83 | 0.67 |
| 11 | 42185849 | rs4375424 | 1.624 | -2.924 | 1.618 | -0.203 | 0.019 | -0.018 | -0.018 | 0 | 0.67 | 0.33 | 0.64 | 0.55 | 0.78 | 0.66 |
| 11 | 42191577 | rs11036743 | -0.942 | -2.881 | 1.389 | -0.192 | -0.013 | 0.021 | 0.021 | 0 | 0.23 | 0.27 | 0.26 | 0.43 | 0.11 | 0.28 |
| 11 | 42196881 | rs4755594 | -1.683 | -2.869 | 1.355 | -0.188 | -0.038 | 0.043 | 0.043 | 0 | 0.31 | 0.5 | 0.32 | 0.45 | 0.2 | 0.33 |
| 11 | 42206273 | rs16936409 | 0.545 | -2.796 | 1.131 | -0.163 | -0.026 | -0.023 | -0.023 | 0 | 0.05 | 0.07 | 0.01 | 0.02 | 0 | 0.11 |
| 11 | 42211231 | rs16936417 | -1.308 | -2.746 | 0.502 | -0.098 | -0.021 | 0.044 | 0.044 | 0 | 0.26 | 0.34 | 0.3 | 0.44 | 0.17 | 0.29 |
| 11 | 42217564 | rs1011340 | -0.081 | -2.504 | -0.159 | 0.014 | -0.001 | -0.106 | -0.106 | 0.238 | 0.17 | 0.03 | 0.25 | 0.42 | 0.11 | 0.16 |
| 11 | 42234380 | rs11036765 | -0.403 | -2.386 | -1.644 | 0.084 | 0.006 | -0.052 | -0.052 | 0 | 0.2 | 0.12 | 0.28 | 0.42 | 0.17 | 0.18 |
| 11 | 42239022 | rs9300041 | -1.303 | -1.832 | -1.9 | -0.201 | -0.071 | -0.03 | -0.03 | 0.639 | 0.57 | 0.97 | 0.56 | 0.67 | 0.39 | 0.48 |
| 11 | 42246718 | rs7480010 | 1.246 | -1.976 | -1.419 | -0.927 | 0.101 | 0.188 | 0.188 | 0.688 | 0.51 | 0.04 | 0.71 | 0.8 | 0.72 | 0.58 |
| 11 | 42247461 | rs12364934 | -1.121 | -2.612 | -0.404 | -1.076 | -0.051 | -0.012 | -0.012 | 0 | 0.22 | 0.37 | 0.15 | 0.17 | 0.1 | 0.22 |
| 11 | 42250690 | rs16936434 | -0.572 | -2.678 | -0.39 | -1.047 | 0.023 | -0.016 | -0.016 | 0.273 | 0.12 | 0.1 | 0.06 | 0.17 | 0.06 | 0.12 |
| 11 | 42257092 | rs16936445 | -1.106 | -2.682 | -0.403 | -1.048 | 0.032 | -0.036 | -0.036 | 0 | 0.1 | 0.03 | 0.05 | 0.17 | 0.05 | 0.13 |
| 11 | 42266666 | rs12285569 | -0.772 | -2.552 | -0.116 | -1.092 | -0.011 | -0.088 | -0.088 | 0 | 0.17 | 0.64 | 0.12 | 0 | 0.14 | 0.11 |
| 11 | 42278390 | rs12791857 | -1.205 | -2.647 | -0.217 | -1.135 | 0.033 | 0.007 | 0.007 | 0 | 0.22 | 0.16 | 0.16 | 0.17 | 0.15 | 0.23 |
| 11 | 42279633 | rs7358398 | 0.866 | -2.637 | -0.187 | -1.118 | -0.02 | -0.002 | -0.002 | 0.009 | 0.77 | 0.7 | 0.84 | 0.79 | 0.89 | 0.73 |
| 11 | 42285225 | rs10837802 | 2.129 | -2.579 | -0.31 | -1.143 | 0.111 | 0.146 | 0.146 | 0.926 | 0.54 | 0.04 | 0.71 | 0.79 | 0.73 | 0.58 |
| 11 | 42293243 | rs4619139 | -0.812 | -2.344 | 1.053 | -0.466 | 0.059 | 0.032 | 0.032 | 0.07 | 0.06 | 0.36 | 0.03 | 0 | 0.02 | 0.02 |
| 11 | 42297268 | rs10742620 | 1.859 | -2.348 | 0.209 | -0.439 | 0.119 | 0.05 | 0.05 | 0.341 | 0.66 | 0.15 | 0.8 | 0.78 | 0.86 | 0.7 |
| 11 | 42305375 | rs12364729 | -1.221 | -1.648 | 0.328 | -0.151 | 0.027 | -0.003 | -0.003 | 0 | 0.18 | 0.16 | 0.14 | 0.17 | 0.09 | 0.18 |
| 11 | 42307166 | rs11036800 | -1.427 | -0.595 | 0.909 | 0.07 | 0.03 | -0.003 | -0.003 | 0 | 0.08 | 0.49 | 0.05 | 0 | 0.03 | 0.03 |
| 11 | 42310568 | rs7480538 | -1.37 | 0.048 | 0.422 | 0.429 | 0.023 | 0.036 | 0.036 | 0 | 0.09 | 0.15 | 0.12 | 0.13 | 0.14 | 0.17 |
| 11 | 42337746 | rs4373908 | 0.226 | -0.094 | 0.599 | -0.609 | -0.001 | -0.012 | -0.012 | 0 | 0.54 | 0.47 | 0.49 | 0.77 | 0.54 | 0.38 |
| 11 | 42348239 | rs10768794 | -0.377 | -0.331 | 1.639 | 0.551 | -0.042 | 0.167 | 0.167 | 0 | 0.38 | 0.12 | 0.26 | 0.15 | 0.41 | 0.53 |
| 11 | 42350265 | rs6485312 | 0.133 | -0.317 | 2.163 | 0.594 | 0.061 | 0.03 | 0.03 | 0.774 | 0.94 | 0.66 | 0.9 | 1 | 0.96 | 0.95 |
| 11 | 42351047 | rs10768795 | -0.689 | -0.65 | 1.797 | 0.431 | -0.005 | 0.127 | 0.127 | 0 | 0.44 | 0.16 | 0.42 | 0.23 | 0.44 | 0.57 |
| 11 | 42385511 | rs11036859 | 0.769 | -0.728 | 2.11 | 1.158 | 0.009 | 0.044 | 0.044 | 0 | 0.5 | 0.47 | 0.48 | 0.23 | 0.51 | 0.35 |
| 11 | 42405924 | rs10837837 | -0.155 | -0.946 | 1.7 | 0.917 | 0.001 | -0.022 | -0.022 | 0 | 0.26 | 0.14 | 0.3 | 0.65 | 0.28 | 0.23 |
| 11 | 42414918 | rs1841773 | -0.909 | 0.631 | 2.084 | 0.661 | 0.018 | -0.004 | -0.004 | 0 | 0.29 | 0.37 | 0.42 | 0.71 | 0.41 | 0.29 |
| 11 | 42416190 | rs7105232 | -0.2 | 0.614 | 2.087 | 0.665 | -0.009 | -0.025 | -0.025 | 2.505 | 0.23 | 0.13 | 0.31 | 0.66 | 0.28 | 0.23 |
| 11 | 42418255 | rs1508528 | -0.057 | 0.616 | 2.164 | 0.643 | -0.006 | -0.021 | -0.021 | 0 | 0.23 | 0.14 | 0.29 | 0.66 | 0.27 | 0.23 |
| 11 | 42420411 | rs11036878 | 0.477 | 0.477 | 2.263 | 0.481 | -0.01 | -0.015 | -0.015 | 0.632 | 0.84 | 0.52 | 0.88 | 0.85 | 0.88 | 0.62 |
| 11 | 42424649 | rs1157545 | 1.019 | 0.486 | 2.266 | 0.464 | 0 | -0.001 | -0.001 | 0 | 0.61 | 0.39 | 0.58 | 0.19 | 0.61 | 0.39 |
| 11 | 42427853 | rs1903727 | 1.705 | 0.492 | 1.157 | -0.518 | 0.026 | 0.147 | 0.147 | 0.065 | 0.32 | 0.13 | 0.22 | 0.05 | 0.27 | 0.16 |
| 11 | 42429389 | rs11036882 | -0.056 | 0.436 | 1.284 | -0.451 | 0 | -0.006 | -0.006 | 0 | 0.14 | 0.33 | 0.11 | 0.14 | 0.12 | 0.37 |
| 11 | 42430569 | rs2219699 | -0.78 | 0.726 | 1.019 | -0.765 | 0.018 | -0.051 | -0.051 | 0 | 0.33 | 0.52 | 0.53 | 0.77 | 0.35 | 0.31 |
| 11 | 42431121 | rs12362863 | 0.16 | 0.7 | 0.736 | -0.775 | -0.06 | 0.109 | 0.109 | 0.038 | 0.31 | 0.09 | 0.23 | 0.17 | 0.35 | 0.53 |

| | | | | | | | | | | | | | | | | |
|---|---|---|---|---|---|---|---|---|---|---|---|---|---|---|---|---|
| 11 | 42432492 | rs1508521 | 1.672 | 0.801 | 0.821 | -0.675 | 0.005 | 0.006 | 0.006 | 0 | 0.34 | 0.37 | 0.32 | 0.55 | 0.43 | 0.21 |
| 11 | 42434026 | rs7106357 | -0.528 | 0.716 | 0.728 | -0.661 | -0.012 | 0.001 | 0.001 | 0 | 0.3 | 0.39 | 0.44 | 0.27 | 0.21 | 0.24 |
| 11 | 42435777 | rs7949101 | -1.73 | 0.235 | 0.05 | -0.912 | 0.004 | 0.076 | 0.076 | 0.933 | 0.74 | 0.86 | 0.79 | 0.94 | 0.74 | 0.84 |
| 11 | 42437741 | rs1355052 | -1.253 | 0.324 | -0.062 | -1.119 | 0 | 0.069 | 0.069 | 0.213 | 0.76 | 0.61 | 0.81 | 0.45 | 0.74 | 0.84 |
| 11 | 42443899 | rs4756730 | 0.323 | 0.26 | -0.845 | -1.529 | -0.08 | 0.046 | 0.046 | 0 | 0.26 | 0.05 | 0.27 | 0.1 | 0.28 | 0.11 |
| 11 | 42444628 | rs10837849 | 1.403 | 0.18 | -1.28 | -1.567 | -0.085 | 0.106 | 0.106 | 0 | 0.32 | 0.03 | 0.22 | 0.17 | 0.34 | 0.46 |
| 11 | 42448612 | rs12789680 | 0.238 | -0.003 | -0.599 | -1.38 | -0.006 | -0.038 | -0.038 | 0.023 | 0.14 | 0.15 | 0.12 | 0.01 | 0.1 | 0.16 |
| 11 | 42450746 | rs3861059 | -1.016 | 0.166 | -0.583 | -1.074 | 0.001 | 0.023 | 0.023 | 0.012 | 0.45 | 0.58 | 0.59 | 0.28 | 0.4 | 0.38 |
| 11 | 42458508 | rs7103942 | -0.885 | 0.173 | -0.603 | -1.042 | -0.003 | -0.01 | -0.01 | 0 | 0.43 | 0.48 | 0.43 | 0.29 | 0.36 | 0.36 |
| 11 | 42458745 | rs7104098 | -0.892 | 0.164 | -0.898 | -0.827 | 0.004 | -0.002 | -0.002 | 0 | 0.62 | 0.57 | 0.66 | 0.32 | 0.53 | 0.52 |
| 11 | 42458937 | rs3849982 | -0.658 | 0.133 | -1.426 | -0.836 | -0.024 | 0.113 | 0.113 | 0 | 0.4 | 0.21 | 0.29 | 0.14 | 0.41 | 0.24 |
| 11 | 42459273 | rs7115681 | 0.23 | -0.19 | -1.112 | -0.493 | 0.003 | -0.002 | -0.002 | 0 | 0.36 | 0.34 | 0.38 | 0.63 | 0.36 | 0.3 |
| 11 | 42461789 | rs16936634 | 0.813 | -0.48 | -1.194 | -0.871 | 0.016 | 0.016 | 0.016 | 0 | 0.16 | 0.09 | 0.12 | 0.09 | 0.13 | 0.15 |
| 11 | 42464898 | rs7933132 | -1.393 | -0.515 | -0.775 | -1.048 | 0.039 | -0.091 | -0.091 | 2.34 | 0.87 | 0.62 | 0.89 | 1 | 0.88 | 0.99 |
| 11 | 42466106 | rs11036893 | 0.613 | -0.508 | -1.113 | -1.051 | -0.035 | -0.087 | -0.087 | 1.455 | 0.3 | 0.12 | 0.3 | 0.52 | 0.32 | 0.22 |
| 11 | 42469313 | rs1381782 | 2.288 | -0.364 | -1.118 | -1.272 | -0.044 | 0.052 | 0.052 | 0 | 0.27 | 0.12 | 0.35 | 0.13 | 0.31 | 0.43 |
| 11 | 42485667 | rs2169545 | 2.276 | -0.555 | -1.046 | -1.2 | -0.042 | 0.05 | 0.05 | 0 | 0.27 | 0.13 | 0.35 | 0.13 | 0.31 | 0.43 |
| 11 | 42488495 | rs10734503 | 0.071 | -0.392 | -0.779 | -0.957 | -0.017 | -0.085 | -0.085 | 0.323 | 0.7 | 0.52 | 0.78 | 0.9 | 0.75 | 0.83 |
| 11 | 42498183 | rs6485317 | -1.303 | -0.382 | -1.024 | -0.954 | 0.043 | -0.104 | -0.104 | 0 | 0.86 | 0.57 | 0.89 | 1 | 0.88 | 0.99 |
| 11 | 42499299 | rs7130621 | 0.676 | -0.4 | -1.342 | -0.933 | -0.011 | 0.078 | 0.078 | 1.945 | 0.54 | 0.25 | 0.54 | 0.26 | 0.54 | 0.64 |
| 11 | 42500196 | rs10768817 | 1.412 | -0.23 | -1.287 | -0.744 | -0.017 | 0.066 | 0.066 | 1.19 | 0.37 | 0.2 | 0.43 | 0.19 | 0.41 | 0.49 |
| 11 | 42504862 | rs2199482 | 0.212 | -0.481 | -0.548 | -0.34 | -0.008 | -0.031 | -0.031 | 0 | 0.69 | 0.45 | 0.75 | 0.83 | 0.74 | 0.8 |
| 11 | 42509547 | rs2063218 | -0.145 | -0.396 | -0.186 | 0.196 | -0.008 | -0.009 | -0.009 | 0 | 0.81 | 0.72 | 0.84 | 0.88 | 0.82 | 0.83 |
| 11 | 42513173 | rs12279506 | -0.083 | -0.395 | -0.195 | 0.584 | -0.007 | 0.006 | 0.006 | 0 | 0.81 | 0.72 | 0.82 | 0.73 | 0.81 | 0.81 |
| 11 | 42514486 | rs1026644 | -0.166 | -0.403 | -0.141 | 0.564 | 0.001 | 0.001 | 0.001 | 0 | 0.81 | 0.76 | 0.85 | 0.88 | 0.81 | 0.83 |
| 11 | 42519684 | rs1870911 | -0.281 | -0.411 | -0.134 | 0.56 | 0.006 | 0.009 | 0.009 | 0 | 0.82 | 0.81 | 0.85 | 0.88 | 0.81 | 0.83 |
| 11 | 42520092 | rs4755609 | -0.01 | -0.45 | -0.237 | 0.509 | -0.005 | -0.054 | -0.054 | 0 | 0.23 | 0.13 | 0.28 | 0.46 | 0.24 | 0.21 |
| 11 | 42520431 | rs4294547 | -0.541 | -0.511 | 0.215 | 0.377 | -0.005 | 0.01 | 0.01 | 0 | 0.27 | 0.56 | 0.33 | 0.74 | 0.26 | 0.33 |
| 11 | 42523366 | rs2862573 | -0.074 | -0.491 | 0.06 | 0.397 | 0 | -0.001 | -0.001 | 1.146 | 0.91 | 0.88 | 0.92 | 0.92 | 0.91 | 0.88 |
| 11 | 42543277 | rs2862571 | 1.517 | -0.662 | 0.241 | 0.289 | 0.014 | 0.027 | 0.027 | 0 | 0.55 | 0.4 | 0.54 | 0.27 | 0.58 | 0.59 |
| 11 | 42552055 | rs10837869 | 0.79 | -0.386 | 0.38 | 0.485 | -0.004 | 0.005 | 0.005 | 0 | 0.67 | 0.46 | 0.61 | 0.34 | 0.68 | 0.64 |
| 11 | 42566600 | rs10768832 | -0.484 | -0.377 | 0.374 | 0.397 | -0.056 | -0.134 | -0.134 | 0 | 0.8 | 0.66 | 0.92 | 0.99 | 0.9 | 0.86 |
| 11 | 42574260 | rs7947115 | 1.321 | -0.254 | 0.409 | 0.532 | -0.001 | -0.001 | -0.001 | 0 | 0.59 | 0.45 | 0.56 | 0.4 | 0.57 | 0.6 |
| 11 | 42579592 | rs4445620 | 1.612 | -0.287 | 0.362 | 0.502 | 0.013 | 0.068 | 0.068 | 0 | 0.57 | 0.29 | 0.54 | 0.18 | 0.56 | 0.49 |
| 11 | 42587300 | rs10837880 | -0.096 | -0.243 | 0.525 | 0.639 | -0.001 | -0.045 | -0.045 | 0 | 0.25 | 0.15 | 0.31 | 0.45 | 0.26 | 0.22 |
| 11 | 42595662 | rs12362425 | 2.101 | -0.157 | 0.031 | 0.24 | -0.113 | 0.125 | 0.125 | 0.22 | 0.42 | 0.03 | 0.42 | 0.15 | 0.42 | 0.45 |
| 11 | 42599548 | rs4331079 | -1.037 | -0.043 | 0.42 | 0.604 | -0.004 | -0.048 | -0.048 | 0.023 | 0.33 | 0.19 | 0.36 | 0.51 | 0.37 | 0.28 |
| 11 | 42618719 | rs4755162 | 1.605 | -0.098 | 0.511 | 0.601 | 0.016 | 0.077 | 0.077 | 0 | 0.56 | 0.22 | 0.53 | 0.24 | 0.58 | 0.57 |
| 11 | 42623993 | rs10742634 | -1.554 | -0.093 | 0.639 | 0.637 | 0.001 | 0 | 0 | 0 | 0.41 | 0.43 | 0.43 | 0.55 | 0.42 | 0.39 |
| 11 | 42641136 | rs4755163 | 1.578 | -0.04 | 0.371 | 0.606 | 0.015 | 0.116 | 0.116 | 0.037 | 0.56 | 0.21 | 0.54 | 0.15 | 0.57 | 0.49 |
| 11 | 42642595 | rs4306288 | 0.601 | -0.039 | 0.515 | 0.65 | 0.011 | 0.081 | 0.081 | 0.031 | 0.84 | 0.46 | 0.86 | 0.75 | 0.83 | 0.75 |
| 11 | 42645071 | rs7127452 | 1.174 | -0.045 | 0.43 | 0.411 | 0.011 | 0.002 | 0.002 | 0.014 | 0.62 | 0.34 | 0.7 | 0.37 | 0.68 | 0.57 |
| 11 | 42647155 | rs4584565 | 1.179 | -0.043 | 0.431 | 0.412 | 0.011 | 0.003 | 0.003 | 0.018 | 0.62 | 0.34 | 0.7 | 0.37 | 0.68 | 0.57 |
| 11 | 42648398 | rs7932533 | -0.196 | -0.047 | 0.597 | 0.424 | 0.036 | 0.082 | 0.082 | 0 | 0.92 | 0.69 | 0.95 | 0.91 | 0.95 | 0.88 |
| 11 | 42649538 | rs11604506 | 0.526 | -0.098 | 0.495 | 0.387 | 0.002 | -0.002 | -0.002 | 0 | 0.08 | 0.13 | 0.07 | 0.07 | 0.12 | 0.07 |
| 11 | 42650289 | rs4755620 | -0.813 | -0.394 | 0.699 | 0.654 | 0.005 | -0.016 | -0.016 | 0.243 | 0.3 | 0.27 | 0.24 | 0.54 | 0.27 | 0.32 |
| 11 | 42654042 | rs4567442 | 0.417 | -0.426 | 0.735 | 0.65 | 0.041 | 0.091 | 0.091 | 0.161 | 0.91 | 0.61 | 0.94 | 0.91 | 0.95 | 0.88 |
| 11 | 42655301 | rs12422064 | 0.965 | -0.354 | 0.557 | 0.628 | 0.012 | 0.003 | 0.003 | 0.013 | 0.62 | 0.34 | 0.7 | 0.37 | 0.68 | 0.57 |
| 11 | 42663357 | rs7114882 | -1.104 | 0.395 | 0.717 | 0.818 | -0.002 | 0.006 | 0.006 | 0 | 0.16 | 0.23 | 0.26 | 0.69 | 0.25 | 0.35 |
| 11 | 42663401 | rs7129198 | -0.484 | 0.382 | 0.695 | 0.894 | 0.001 | 0.01 | 0.01 | 0 | 0.06 | 0.12 | 0.07 | 0.16 | 0.08 | 0.14 |
| 11 | 42673024 | rs7118795 | 1.601 | 0.188 | 0.273 | 0.5 | -0.002 | 0.114 | 0.114 | 0.871 | 0.51 | 0.12 | 0.52 | 0.24 | 0.57 | 0.49 |
| 11 | 42680319 | rs4755624 | 1.479 | 0.272 | 0.237 | 0.517 | 0.002 | 0.087 | 0.087 | 0.382 | 0.53 | 0.21 | 0.52 | 0.24 | 0.57 | 0.49 |
| 11 | 42691021 | rs4326780 | 0.203 | 0.268 | -0.034 | 0.969 | 0.043 | 0.173 | 0.173 | 0 | 0.86 | 0.65 | 0.77 | 0.52 | 0.8 | 0.74 |
| 11 | 42696239 | rs7111587 | -0.106 | 0.148 | -0.159 | 0.619 | -0.049 | 0.052 | 0.052 | 0 | 0.13 | 0.01 | 0.1 | 0.02 | 0.14 | 0.14 |
| 11 | 42705646 | rs4565889 | 1.556 | 0.181 | -0.125 | 0.776 | 0.002 | 0.001 | 0.001 | 0 | 0.58 | 0.32 | 0.6 | 0.32 | 0.61 | 0.47 |
| 11 | 42715836 | rs10837908 | 0.5 | 0.441 | 0.196 | 1.304 | 0.027 | 0.206 | 0.206 | 0 | 0.84 | 0.6 | 0.76 | 0.47 | 0.81 | 0.72 |
| 11 | 42722587 | rs11037018 | -0.195 | 0.521 | -0.365 | 1.317 | -0.018 | -0.023 | -0.023 | 0 | 0.11 | 0.1 | 0.19 | 0.45 | 0.18 | 0.21 |
| 11 | 42726949 | rs4256962 | -0.551 | 0.511 | -0.153 | 0.869 | 0.032 | -0.033 | -0.033 | 0 | 0.82 | 0.64 | 0.88 | 0.91 | 0.87 | 0.84 |
| 11 | 42730635 | rs10837917 | -0.573 | 0.517 | -0.15 | 0.872 | 0.034 | -0.031 | -0.031 | 0 | 0.82 | 0.64 | 0.88 | 0.91 | 0.87 | 0.84 |
| 11 | 42736694 | rs3924155 | 0.112 | 0.56 | -0.218 | 0.902 | -0.001 | 0.008 | 0.008 | 0 | 0.77 | 0.4 | 0.84 | 0.74 | 0.85 | 0.8 |
| 11 | 42737996 | rs12278579 | -0.32 | 0.17 | -0.33 | 0.427 | -0.053 | 0.025 | 0.025 | 0 | 0.14 | 0.39 | 0.09 | 0.18 | 0.06 | 0.07 |
| 11 | 42745078 | rs11037036 | -1.081 | 0.273 | -0.751 | 0.672 | -0.002 | 0.016 | 0.016 | 0 | 0.23 | 0.34 | 0.27 | 0.6 | 0.23 | 0.2 |
| 11 | 42745494 | rs7947641 | 0.675 | -0.03 | -1.197 | 0.135 | -0.022 | 0.087 | 0.087 | 0 | 0.14 | 0.02 | 0.1 | 0 | 0.11 | 0.06 |
| 11 | 42747982 | rs4755646 | -0.87 | 0.212 | -1.029 | 0.29 | 0.015 | -0.107 | -0.107 | 0 | 0.8 | 0.51 | 0.82 | 0.99 | 0.84 | 0.83 |
| 11 | 42754350 | rs7127925 | 1.139 | 0.284 | -1.182 | 0.356 | -0.057 | 0.071 | 0.071 | 0 | 0.54 | 0.06 | 0.51 | 0.24 | 0.58 | 0.51 |
| 11 | 42756151 | rs7105806 | -0.214 | 0.284 | -1.032 | 0.465 | 0.018 | 0.149 | 0.149 | 0 | 0.13 | 0.33 | 0.23 | 0.51 | 0.19 | 0.16 |
| 11 | 42765063 | rs11822413 | -0.803 | 0.041 | -1.05 | 0.445 | -0.014 | 0.021 | 0.021 | 0 | 0.23 | 0.34 | 0.23 | 0.59 | 0.15 | 0.2 |
| 11 | 42767612 | rs10837919 | 0.294 | 0.385 | -0.876 | 0.573 | 0.022 | -0.028 | -0.028 | 0 | 0.6 | 0.33 | 0.6 | 0.45 | 0.68 | 0.55 |
| 11 | 42767819 | rs10837921 | 0.642 | 0.406 | -0.94 | 0.498 | 0.03 | 0.063 | 0.063 | 0.006 | 0.55 | 0.11 | 0.56 | 0.25 | 0.66 | 0.49 |

| | | | | | | | | | | | | | | | | |
|---|---|---|---|---|---|---|---|---|---|---|---|---|---|---|---|---|
| 11 | 42768045 | rs10837923 | 0.604 | 0.427 | -0.907 | 0.508 | 0.041 | 0.064 | 0.064 | 0 | 0.55 | 0.23 | 0.56 | 0.25 | 0.66 | 0.49 |
| 11 | 42771556 | rs7125369 | -0.648 | 0.855 | -0.619 | 0.343 | 0.004 | 0.013 | 0.013 | 0 | 0.07 | 0.17 | 0.06 | 0.1 | 0.04 | 0.07 |
| 11 | 42786155 | rs12286647 | 0.372 | 1.089 | -0.804 | 0.065 | 0.008 | -0.006 | -0.006 | 0.359 | 0.12 | 0.17 | 0.12 | 0.08 | 0.19 | 0.12 |
| 11 | 42795097 | rs7928165 | -1.152 | 1.061 | -0.913 | -0.126 | 0 | -0.011 | -0.011 | 0 | 0.36 | 0.4 | 0.33 | 0.24 | 0.39 | 0.4 |
| 11 | 42811366 | rs4408298 | -1.365 | 0.985 | -0.958 | -0.093 | 0.004 | -0.013 | -0.013 | 0 | 0.39 | 0.51 | 0.37 | 0.33 | 0.43 | 0.41 |
| 11 | 42837715 | rs4391804 | 1.05 | 0.744 | -0.996 | -0.013 | -0.012 | 0.031 | 0.031 | 0 | 0.61 | 0.3 | 0.55 | 0.69 | 0.58 | 0.58 |
| 11 | 42844441 | rs4472923 | -0.957 | 0.66 | -0.843 | 0.164 | 0.006 | -0.043 | -0.043 | 0.402 | 0.32 | 0.41 | 0.25 | 0.23 | 0.35 | 0.24 |
| 11 | 42847003 | rs7118856 | -0.399 | 0.697 | -0.796 | 0.222 | 0.021 | 0.003 | 0.003 | 0 | 0.87 | 0.71 | 0.78 | 0.89 | 0.87 | 0.83 |
| 11 | 42848592 | rs11603601 | 1.035 | 0.761 | -2.035 | 0.233 | -0.079 | -0.103 | -0.103 | 0.009 | 0.27 | 0.01 | 0.28 | 0.27 | 0.31 | 0.34 |
| 11 | 42848816 | rs10768893 | 0.657 | 0.586 | -1.018 | 0.074 | -0.006 | 0.005 | 0.005 | 0 | 0.74 | 0.39 | 0.79 | 0.67 | 0.73 | 0.69 |
| 11 | 42849616 | rs7105434 | 0.158 | 0.138 | -1.017 | 0.326 | -0.037 | -0.028 | -0.028 | 0.175 | 0.74 | 0.51 | 0.84 | 0.8 | 0.81 | 0.78 |
| 11 | 42856020 | rs7118693 | -0.296 | 0.968 | -1.954 | 0.651 | -0.08 | -0.067 | -0.067 | 1.576 | 0.28 | 0.04 | 0.34 | 0.32 | 0.35 | 0.23 |
| 11 | 42856868 | rs6485361 | -0.193 | 0.61 | -1.819 | 0.419 | -0.01 | -0.015 | -0.015 | 0 | 0.54 | 0.25 | 0.58 | 0.51 | 0.53 | 0.46 |
| 11 | 42859152 | rs10837943 | -0.037 | -0.269 | -1.003 | -0.146 | -0.061 | -0.107 | -0.107 | 0 | 0.82 | 0.43 | 0.86 | 0.97 | 0.92 | 0.92 |
| 11 | 42864284 | rs7938181 | -0.571 | -0.449 | -1.156 | -0.066 | 0.005 | -0.012 | -0.012 | 4.683 | 0.78 | 0.83 | 0.76 | 0.71 | 0.77 | 0.82 |
| 11 | 42864437 | rs7951701 | 0.242 | -0.347 | -0.456 | -0.328 | -0.019 | -0.012 | -0.012 | 1.085 | 0.64 | 0.47 | 0.62 | 0.69 | 0.66 | 0.76 |
| 11 | 42867577 | rs4132365 | 0.279 | -0.389 | -0.414 | -0.348 | -0.012 | -0.008 | -0.008 | 0 | 0.64 | 0.42 | 0.62 | 0.69 | 0.66 | 0.76 |
| 11 | 42868302 | rs7117480 | 1.01 | -0.257 | -0.931 | -1.075 | -0.026 | 0.049 | 0.049 | 0 | 0.16 | 0.06 | 0.1 | 0 | 0.17 | 0.06 |
| 11 | 42874407 | rs4469879 | 1.647 | -0.283 | -1.204 | -0.784 | -0.017 | -0.022 | -0.022 | 0.089 | 0.25 | 0.16 | 0.26 | 0.38 | 0.32 | 0.43 |
| 11 | 42876624 | rs7129336 | 0.432 | -0.536 | -0.806 | -1.033 | -0.001 | 0.03 | 0.03 | 0 | 0.45 | 0.28 | 0.43 | 0.65 | 0.43 | 0.69 |
| 11 | 42878498 | rs7930355 | -0.207 | -0.986 | -0.776 | -1.083 | -0.001 | -0.026 | -0.026 | 0.014 | 0.24 | 0.24 | 0.15 | 0.06 | 0.22 | 0.09 |
| 11 | 42879073 | rs4132802 | 0.771 | -0.936 | -0.892 | -0.994 | -0.026 | 0.014 | 0.014 | 0 | 0.41 | 0.22 | 0.39 | 0.63 | 0.47 | 0.67 |
| 11 | 42879811 | rs12283338 | -0.656 | -0.819 | -0.821 | -0.853 | 0.032 | 0.053 | 0.053 | 0 | 0.51 | 0.73 | 0.52 | 0.35 | 0.45 | 0.28 |
| 11 | 42885511 | rs7937072 | 0.992 | -0.441 | -0.789 | -0.629 | 0.001 | -0.001 | -0.001 | 0.168 | 0.12 | 0.11 | 0.05 | 0.06 | 0.07 | 0.05 |
| 11 | 42889289 | rs11037114 | 0.235 | -0.567 | -0.518 | -0.765 | -0.003 | 0 | 0 | 0.799 | 0.45 | 0.43 | 0.39 | 0.45 | 0.53 | 0.6 |
| 11 | 42913353 | rs1532096 | 0.096 | -0.876 | -0.733 | -0.995 | 0.037 | 0.065 | 0.065 | 0 | 0.23 | 0.11 | 0.12 | 0.01 | 0.13 | 0.07 |
| 11 | 42916859 | rs4572134 | -1.322 | -1.472 | -0.593 | -1.106 | 0.016 | -0.004 | -0.004 | 0.076 | 0.46 | 0.4 | 0.36 | 0.47 | 0.43 | 0.38 |
| 11 | 42924732 | rs964188 | 1.193 | -0.824 | -0.569 | -0.728 | 0.001 | 0.001 | 0.001 | 0 | 0.29 | 0.24 | 0.31 | 0.36 | 0.36 | 0.32 |
| 11 | 42925407 | rs6485383 | -0.925 | -0.945 | -0.366 | -0.815 | -0.003 | -0.001 | -0.001 | 0.382 | 0.45 | 0.59 | 0.36 | 0.46 | 0.41 | 0.39 |
| 11 | 42929685 | rs989357 | -1.578 | -0.958 | -0.406 | -0.822 | 0 | 0.016 | 0.016 | 0.075 | 0.75 | 0.84 | 0.67 | 0.82 | 0.78 | 0.73 |
| 11 | 42929738 | rs11037133 | 0.352 | -0.921 | -0.415 | -0.889 | 0.029 | 0.064 | 0.064 | 0 | 0.27 | 0.17 | 0.17 | 0.01 | 0.2 | 0.08 |
| 11 | 42933187 | rs7124810 | -0.07 | -0.915 | -0.407 | -0.874 | 0.029 | 0.007 | 0.007 | 0 | 0.32 | 0.21 | 0.23 | 0.28 | 0.25 | 0.24 |
| 11 | 42935188 | rs16937119 | -0.829 | -0.953 | -0.378 | -0.868 | -0.018 | 0.008 | 0.008 | 0 | 0.4 | 0.56 | 0.31 | 0.46 | 0.33 | 0.38 |
| 11 | 42941263 | rs1967081 | -0.48 | -0.984 | -0.365 | -0.948 | -0.001 | -0.004 | -0.004 | 0 | 0.6 | 0.42 | 0.54 | 0.63 | 0.6 | 0.58 |
| 11 | 42941438 | rs1489196 | 0.383 | -0.968 | -0.35 | -0.934 | 0.015 | 0.017 | 0.017 | 0.124 | 0.15 | 0.41 | 0.13 | 0.18 | 0.18 | 0.15 |
| 11 | 42942016 | rs11037140 | 1.285 | -0.956 | -0.407 | -0.926 | 0.036 | 0.055 | 0.055 | 0 | 0.2 | 0.1 | 0.11 | 0.02 | 0.1 | 0.08 |
| 11 | 42942712 | rs1352524 | 0.701 | -0.811 | -0.278 | -0.767 | 0.015 | 0.027 | 0.027 | 0 | 0.13 | 0.36 | 0.13 | 0.18 | 0.17 | 0.15 |
| 11 | 42948480 | rs10742656 | -1.742 | -0.643 | -0.468 | -0.665 | 0 | 0.015 | 0.015 | 0.075 | 0.75 | 0.83 | 0.67 | 0.82 | 0.78 | 0.72 |
| 11 | 42952281 | rs10768901 | 1.318 | -0.843 | -1.093 | -0.665 | 0 | 0 | 0 | 0 | 0.26 | 0.2 | 0.3 | 0.27 | 0.34 | 0.3 |
| 11 | 42954601 | rs10742658 | 0.651 | -0.933 | -1.099 | -0.614 | 0.007 | 0.001 | 0.001 | 0 | 0.33 | 0.32 | 0.36 | 0.35 | 0.45 | 0.35 |
| 11 | 42955957 | rs12577738 | 1.175 | -0.66 | -1.365 | -0.164 | -0.001 | -0.001 | -0.001 | 0 | 0.11 | 0.11 | 0.1 | 0.19 | 0.17 | 0.15 |
| 11 | 42957368 | rs4584564 | -1.406 | -0.55 | -1.47 | -0.101 | -0.019 | -0.016 | -0.016 | 0 | 0.77 | 0.99 | 0.71 | 0.82 | 0.83 | 0.77 |
| 11 | 42958117 | rs16937161 | 1.172 | -0.181 | -0.812 | 0.294 | -0.001 | 0.042 | 0.042 | 0 | 0.05 | 0.03 | 0.07 | 0.28 | 0.04 | 0.16 |
| 11 | 42959909 | rs1489205 | -0.605 | 0.332 | -0.627 | 0.522 | -0.002 | -0.003 | -0.003 | 0.031 | 0.31 | 0.32 | 0.26 | 0.45 | 0.38 | 0.31 |
| 11 | 42968875 | rs2046769 | 0.939 | -0.118 | -0.958 | 0.073 | -0.041 | -0.01 | -0.01 | 0 | 0.25 | 0.13 | 0.35 | 0.28 | 0.35 | 0.29 |
| 11 | 42969116 | rs11037169 | 1.927 | -0.644 | -1.009 | -0.268 | 0.044 | 0.069 | 0.069 | 0.015 | 0.21 | 0.11 | 0.09 | 0 | 0.12 | 0.05 |
| 11 | 42969584 | rs4237639 | -1.214 | -0.617 | -0.538 | -0.248 | 0.045 | 0.07 | 0.07 | 0 | 0.59 | 0.76 | 0.61 | 0.82 | 0.73 | 0.7 |
| 11 | 42971040 | rs11037175 | 0.753 | -0.617 | -0.333 | 0 | -0.003 | 0.006 | 0.006 | 0 | 0.27 | 0.24 | 0.21 | 0.25 | 0.32 | 0.25 |
| 11 | 42973253 | rs10837979 | 1.921 | -0.633 | -0.5 | -0.133 | -0.032 | -0.02 | -0.02 | 0 | 0.18 | 0.09 | 0.3 | 0.26 | 0.31 | 0.24 |
| 11 | 42977439 | rs7939657 | 0.079 | -0.655 | -0.406 | -0.121 | -0.002 | 0.015 | 0.015 | 0.271 | 0.46 | 0.39 | 0.45 | 0.28 | 0.47 | 0.34 |
| 11 | 42979811 | rs12363299 | 1.77 | -0.707 | -0.313 | -0.17 | -0.025 | -0.075 | -0.075 | 0 | 0.22 | 0.25 | 0.1 | 0.01 | 0.11 | 0.05 |
| 11 | 42983369 | rs10128702 | 1.732 | -0.509 | -0.413 | -0.063 | 0 | -0.018 | -0.018 | 0 | 0.09 | 0.07 | 0.03 | 0 | 0.08 | 0.07 |
| 11 | 42985037 | rs12292153 | -0.129 | -0.532 | -0.227 | 0.195 | 0.002 | 0.011 | 0.011 | 0.247 | 0.33 | 0.37 | 0.25 | 0.55 | 0.37 | 0.41 |
| 11 | 42987088 | rs4256967 | -0.139 | -0.537 | -0.229 | 0.193 | 0.002 | 0.01 | 0.01 | 0.26 | 0.33 | 0.37 | 0.25 | 0.55 | 0.37 | 0.41 |
| 11 | 42988930 | rs4267059 | -1.328 | -0.735 | -0.354 | 0.239 | 0.048 | 0.079 | 0.079 | 0 | 0.62 | 0.85 | 0.62 | 0.82 | 0.74 | 0.73 |
| 11 | 42989183 | rs4272772 | -1.35 | -0.709 | -0.461 | 0.246 | 0.052 | 0.097 | 0.097 | 0 | 0.62 | 0.89 | 0.62 | 0.82 | 0.74 | 0.73 |
| 11 | 42992520 | rs975104 | 1.303 | -0.125 | -0.955 | 0.518 | -0.023 | -0.014 | -0.014 | 0.261 | 0.2 | 0.15 | 0.3 | 0.25 | 0.32 | 0.25 |
| 11 | 42994299 | rs6485389 | -0.682 | -0.351 | -1.244 | 0.515 | -0.029 | -0.14 | -0.14 | 0 | 0.68 | 0.92 | 0.65 | 0.53 | 0.72 | 0.53 |
| 11 | 43001206 | rs7131154 | -1.544 | -0.271 | -0.918 | 0.22 | 0.031 | 0.104 | 0.104 | 0 | 0.64 | 0.92 | 0.62 | 0.83 | 0.74 | 0.75 |
| 11 | 43005220 | rs1489200 | -1.467 | -0.329 | -0.571 | 0.191 | 0.035 | 0.083 | 0.083 | 0 | 0.63 | 0.82 | 0.62 | 0.83 | 0.74 | 0.74 |
| 11 | 43008961 | rs1565251 | -0.65 | -0.199 | -0.119 | 0.399 | -0.002 | -0.004 | -0.004 | 0.336 | 0.41 | 0.57 | 0.34 | 0.61 | 0.42 | 0.49 |
| 11 | 43013511 | rs11037204 | 1.491 | -0.282 | 0.115 | 0.763 | 0.004 | 0.04 | 0.04 | 0 | 0.12 | 0.15 | 0.15 | 0.32 | 0.16 | 0.16 |
| 11 | 43017772 | rs7952412 | -2.142 | -0.28 | 0.008 | 0.518 | 0.03 | 0.099 | 0.099 | 0 | 0.64 | 0.91 | 0.62 | 0.83 | 0.74 | 0.75 |
| 11 | 43028129 | rs4307709 | -0.918 | -0.271 | 0.005 | 0.507 | -0.002 | -0.005 | -0.005 | 0 | 0.41 | 0.57 | 0.34 | 0.63 | 0.42 | 0.49 |
| 11 | 43029294 | rs7929118 | -1.87 | -0.374 | -0.231 | 0.264 | 0.033 | 0.102 | 0.102 | 0 | 0.64 | 0.87 | 0.62 | 0.85 | 0.74 | 0.74 |
| 11 | 43040945 | rs7120544 | -1.221 | -0.482 | -0.326 | 0.191 | 0.059 | 0.126 | 0.126 | 0 | 0.57 | 0.78 | 0.61 | 0.83 | 0.71 | 0.69 |
| 11 | 43045671 | rs262314 | 0.048 | -0.387 | 0.146 | 0.537 | -0.015 | -0.034 | -0.034 | 0 | 0.14 | 0.29 | 0.08 | 0.03 | 0.12 | 0.08 |
| 11 | 43049826 | rs11037217 | 1.038 | -0.348 | 0.237 | 0.856 | 0.004 | 0.023 | 0.023 | 0 | 0.12 | 0.16 | 0.15 | 0.27 | 0.16 | 0.15 |
| 11 | 43050524 | rs262328 | 0.021 | -0.373 | 0.19 | 0.839 | -0.019 | -0.045 | -0.045 | 0 | 0.14 | 0.33 | 0.08 | 0.03 | 0.12 | 0.07 |

| | | | | | | | | | | | | | | | | |
|---|---|---|---|---|---|---|---|---|---|---|---|---|---|---|---|---|
| 11 | 43055629 | rs371614 | 0.824 | -0.244 | 0.276 | 0.982 | 0.019 | 0.071 | 0.071 | 0 | 0.12 | 0.26 | 0.16 | 0.27 | 0.16 | 0.15 |
| 11 | 43061065 | rs2162167 | 1.129 | -0.185 | -0.271 | 1.193 | -0.039 | -0.06 | -0.06 | 0 | 0.1 | 0.02 | 0.09 | 0.18 | 0.13 | 0.19 |
| 11 | 43063682 | rs262322 | 0.649 | -0.654 | -0.317 | 0.756 | 0.012 | 0.011 | 0.011 | 0.577 | 0.12 | 0.11 | 0.03 | 0.03 | 0.05 | 0.06 |
| 11 | 43077155 | rs1425852 | -0.34 | -0.769 | -0.284 | 0.582 | 0.001 | -0.003 | -0.003 | 0 | 0.66 | 0.59 | 0.66 | 0.74 | 0.58 | 0.71 |
| 11 | 43079138 | rs2217038 | 2.28 | -0.52 | -0.633 | 0.288 | -0.018 | 0.013 | 0.013 | 0 | 0.13 | 0.09 | 0.16 | 0.04 | 0.25 | 0.14 |
| 11 | 43086094 | rs10501293 | 2.168 | -1.026 | -0.672 | 0.204 | -0.005 | 0.094 | 0.094 | 0 | 0.31 | 0.08 | 0.37 | 0.14 | 0.25 | 0.26 |
| 11 | 43087966 | rs1365083 | -2.165 | -1.015 | -0.511 | 0.209 | 0.001 | -0.026 | -0.026 | 0 | 0.94 | 0.9 | 0.94 | 1 | 0.91 | 0.99 |
| 11 | 43095376 | rs1159528 | 0.137 | -1.083 | -0.422 | 0.203 | -0.075 | -0.112 | -0.112 | 0.509 | 0.76 | 0.53 | 0.89 | 0.96 | 0.86 | 0.89 |
| 11 | 43099328 | rs11037234 | 1.51 | -0.978 | -1.023 | 0.558 | -0.044 | -0.035 | -0.035 | 0.013 | 0.21 | 0.12 | 0.27 | 0.32 | 0.32 | 0.27 |
| 11 | 43099947 | rs1895824 | 0.611 | -1.139 | -0.937 | 0.596 | -0.012 | -0.003 | -0.003 | 0.554 | 0.67 | 0.42 | 0.75 | 0.67 | 0.71 | 0.74 |
| 11 | 43101260 | rs7935833 | -0.243 | -1.013 | -0.828 | 0.504 | 0.001 | 0.029 | 0.029 | 1.258 | 0.78 | 0.51 | 0.78 | 0.71 | 0.76 | 0.8 |
| 11 | 43103439 | rs12575607 | 1.521 | -0.969 | -1.39 | 0.561 | -0.033 | -0.053 | -0.053 | 0 | 0.1 | 0.02 | 0.13 | 0.29 | 0.16 | 0.16 |
| 11 | 43105620 | rs9645617 | 2.11 | -0.933 | -1.352 | 0.562 | -0.003 | 0.045 | 0.045 | 0 | 0.31 | 0.1 | 0.37 | 0.22 | 0.26 | 0.27 |
| 11 | 43116022 | rs1425851 | 0.452 | -0.741 | -0.861 | 0.773 | -0.001 | -0.003 | -0.003 | 0 | 0.69 | 0.33 | 0.69 | 0.63 | 0.65 | 0.71 |
| 11 | 43124987 | rs4755186 | -0.072 | -0.705 | -0.784 | 0.715 | -0.009 | 0.003 | 0.003 | 2.33 | 0.76 | 0.4 | 0.77 | 0.71 | 0.76 | 0.81 |
| 11 | 43132078 | rs16937261 | 0.897 | -0.747 | -1.094 | 0.627 | 0.004 | -0.01 | -0.01 | 0 | 0.1 | 0.1 | 0.04 | 0.21 | 0.07 | 0.17 |
| 11 | 43135973 | rs11606803 | 1.06 | -0.777 | -1.151 | 0.611 | 0.007 | -0.043 | -0.043 | 0 | 0.59 | 0.17 | 0.65 | 0.56 | 0.59 | 0.57 |
| 11 | 43137342 | rs10838016 | 0.446 | -0.591 | -0.905 | 0.216 | 0.002 | 0.017 | 0.017 | 1.838 | 0.68 | 0.28 | 0.69 | 0.79 | 0.66 | 0.73 |
| 11 | 43138659 | rs7113868 | 0.399 | -0.222 | -0.771 | -0.124 | -0.016 | 0.03 | 0.03 | 0 | 0.16 | 0.07 | 0.12 | 0 | 0.18 | 0.12 |
| 11 | 43141144 | rs7941240 | -0.656 | -0.772 | -0.817 | -0.324 | -0.009 | -0.01 | -0.01 | 0.837 | 0.6 | 0.49 | 0.75 | 0.73 | 0.67 | 0.8 |
| 11 | 43141324 | rs2162171 | 1.995 | -0.839 | -0.998 | -0.238 | 0.001 | -0.002 | -0.002 | 0 | 0.15 | 0.11 | 0.31 | 0.18 | 0.12 | 0.31 |
| 11 | 43141663 | rs12785997 | 2.471 | -0.684 | -1.099 | -0.002 | -0.036 | -0.041 | -0.041 | 0 | 0.14 | 0.05 | 0.2 | 0.32 | 0.26 | 0.2 |
| 11 | 43142892 | rs2862667 | -1.63 | -0.985 | -0.699 | -0.273 | 0.001 | -0.001 | -0.001 | 0 | 0.69 | 0.69 | 0.78 | 0.74 | 0.69 | 0.84 |
| 11 | 43144664 | rs11037255 | -1.081 | -0.977 | -0.719 | -0.263 | -0.004 | -0.005 | -0.005 | 1.111 | 0.64 | 0.57 | 0.76 | 0.74 | 0.67 | 0.83 |
| 11 | 43157244 | rs11607331 | 1.349 | -0.887 | -0.949 | 0.173 | -0.06 | -0.079 | -0.079 | 0 | 0.19 | 0.05 | 0.26 | 0.43 | 0.32 | 0.28 |
| 11 | 43157346 | rs10838024 | 0.619 | -1.659 | -0.63 | 0.114 | -0.008 | -0.003 | -0.003 | 0 | 0.22 | 0.23 | 0.34 | 0.18 | 0.14 | 0.33 |
| 11 | 43161000 | rs11037262 | 0.283 | -1.277 | -0.853 | 0.175 | -0.026 | -0.006 | -0.006 | 0.008 | 0.12 | 0.23 | 0.05 | 0.12 | 0.05 | 0.11 |
| 11 | 43167522 | rs11601828 | 0.754 | -1.009 | -1.075 | 0.295 | 0.006 | -0.006 | -0.006 | 0 | 0.12 | 0.03 | 0.08 | 0.1 | 0.09 | 0.1 |
| 11 | 43168059 | rs11037268 | 0.889 | -1.01 | -0.855 | 0.222 | -0.022 | -0.039 | -0.039 | 0.406 | 0.09 | 0.13 | 0.02 | 0.01 | 0.03 | 0.05 |
| 11 | 43190388 | rs10768931 | -0.939 | -1.272 | -1.194 | 0.205 | 0.025 | 0 | 0 | 0.377 | 0.71 | 0.8 | 0.84 | 0.76 | 0.82 | 0.86 |
| 11 | 43193487 | rs11037309 | 1.757 | -1.457 | -1.582 | 0.184 | 0.006 | -0.022 | -0.022 | 1.66 | 0.1 | 0 | 0.26 | 0.1 | 0.08 | 0.16 |
| 11 | 43194814 | rs11037310 | 1.985 | -1.31 | -1.401 | 0.37 | 0 | 0.012 | 0.012 | 0 | 0.08 | 0.11 | 0.09 | 0.17 | 0.05 | 0.14 |
| 11 | 43196352 | rs1544872 | 1.32 | -1.197 | -1.448 | 0.358 | 0 | -0.013 | -0.013 | 0 | 0.12 | 0.09 | 0.11 | 0.21 | 0.12 | 0.17 |
| 11 | 43196391 | rs1496238 | 2.021 | -1.179 | -1.422 | 0.331 | 0 | 0.013 | 0.013 | 0 | 0.08 | 0.12 | 0.09 | 0.17 | 0.05 | 0.14 |
| 11 | 43196401 | rs2201615 | 2.209 | -1.164 | -1.446 | 0.292 | -0.001 | -0.001 | -0.001 | 0.291 | 0.05 | 0.05 | 0.04 | 0.12 | 0.05 | 0.1 |
| 11 | 43196656 | rs2220956 | 2.209 | -1.164 | -1.446 | 0.292 | -0.001 | -0.001 | -0.001 | 0 | 0.05 | 0.05 | 0.04 | 0.12 | 0.05 | 0.1 |
| 11 | 43197227 | rs1548003 | -2.335 | -1.591 | -1.654 | -0.035 | 0.02 | 0.077 | 0.077 | 0 | 0.78 | 0.92 | 0.9 | 1 | 0.85 | 0.97 |
| 11 | 43199535 | rs11037317 | 2.335 | -1.261 | -1.538 | 0.197 | -0.001 | -0.001 | -0.001 | 0 | 0.05 | 0.05 | 0.04 | 0.12 | 0.05 | 0.1 |
| 11 | 43199669 | rs7942141 | 1.95 | -1.38 | -1.625 | 0.042 | 0.018 | 0.025 | 0.025 | 0.082 | 0.08 | 0.05 | 0.02 | 0 | 0.03 | 0.05 |
| 11 | 43199938 | rs890249 | -0.429 | -1.368 | -1.605 | 0.151 | 0.008 | -0.018 | -0.018 | 0.244 | 0.28 | 0.27 | 0.21 | 0.39 | 0.2 | 0.35 |
| 11 | 43201243 | rs746474 | -1.383 | -1.188 | -1.717 | 0.337 | 0.029 | 0 | 0 | 0 | 0.71 | 0.81 | 0.84 | 0.76 | 0.81 | 0.86 |
| 11 | 43201350 | rs12808373 | 2.085 | 0.38 | -1.833 | 0.406 | -0.058 | -0.042 | -0.042 | 0 | 0.15 | 0.04 | 0.2 | 0.26 | 0.32 | 0.23 |
| 11 | 43206167 | rs11821780 | 0.617 | 0.152 | -1.216 | 0.176 | 0.009 | -0.045 | -0.045 | 0 | 0.16 | 0.19 | 0.15 | 0 | 0.22 | 0.11 |
| 11 | 43209934 | rs1425858 | 1.583 | -0.098 | -1.319 | 0.854 | -0.006 | -0.019 | -0.019 | 0 | 0.07 | 0.02 | 0.07 | 0.1 | 0.07 | 0.07 |
| 11 | 43213599 | rs11037328 | 1.363 | 0 | -1.108 | 0.768 | -0.001 | -0.006 | -0.006 | 0 | 0.05 | 0.04 | 0.04 | 0.12 | 0.05 | 0.1 |
| 11 | 43216769 | rs11037341 | 0.437 | 0.001 | -1.106 | 0.885 | 0.001 | 0 | 0 | 0.026 | 0.12 | 0.13 | 0.11 | 0.28 | 0.08 | 0.22 |
| 11 | 43218424 | rs11037342 | 1.339 | 0.514 | -1.508 | 0.805 | -0.059 | -0.026 | -0.026 | 2.956 | 0.17 | 0.05 | 0.2 | 0.22 | 0.32 | 0.28 |
| 11 | 43218491 | rs17591338 | 1.821 | -0.187 | -2.061 | -0.023 | 0.022 | 0.092 | 0.092 | 0 | 0.18 | 0.03 | 0.06 | 0 | 0.1 | 0.05 |
| 11 | 43221969 | rs12226499 | 2.018 | -0.153 | -1.985 | 0.233 | 0.003 | -0.006 | -0.006 | 0 | 0.06 | 0.05 | 0.1 | 0.17 | 0.03 | 0.09 |
| 11 | 43235460 | rs17591394 | 1.026 | 0.044 | -1.79 | -0.018 | 0.03 | -0.021 | -0.021 | 0.131 | 0.14 | 0.18 | 0.15 | 0.04 | 0.26 | 0.08 |
| 11 | 43235761 | rs7115932 | 1.057 | -0.644 | -1.833 | 0.091 | 0.036 | 0.006 | 0.006 | 0.013 | 0.31 | 0.14 | 0.18 | 0.24 | 0.17 | 0.23 |
| 11 | 43236061 | rs12365397 | 2.483 | -0.562 | -1.845 | 0.174 | -0.061 | -0.027 | -0.027 | 0 | 0.17 | 0.05 | 0.2 | 0.22 | 0.32 | 0.28 |
| 11 | 43238542 | rs7934456 | 0.036 | -0.68 | -1.514 | 0.217 | -0.002 | 0.002 | 0.002 | 0 | 0.42 | 0.39 | 0.33 | 0.29 | 0.43 | 0.29 |
| 11 | 43239836 | rs1353464 | 2.434 | -0.89 | -1.443 | 0.227 | 0.001 | -0.004 | -0.004 | 0 | 0.07 | 0.05 | 0.04 | 0.1 | 0.07 | 0.07 |
| 11 | 43239926 | rs11037361 | 2.421 | -0.947 | -1.364 | 0.222 | -0.001 | -0.001 | -0.001 | 0 | 0.06 | 0.06 | 0.04 | 0.11 | 0.05 | 0.1 |
| 11 | 43242587 | rs1353465 | -0.675 | -1.014 | -1.149 | 0.217 | 0 | -0.001 | -0.001 | 5.222 | 0.56 | 0.57 | 0.41 | 0.5 | 0.54 | 0.46 |
| 11 | 43242743 | rs1353466 | 1.329 | -1.069 | -1.158 | 0.186 | -0.002 | -0.019 | -0.019 | 0.092 | 0.09 | 0.05 | 0.11 | 0.24 | 0.07 | 0.16 |
| 11 | 43244906 | rs11037366 | -0.658 | -0.994 | -1.085 | 0.219 | 0.001 | -0.001 | -0.001 | 0 | 0.56 | 0.57 | 0.41 | 0.5 | 0.54 | 0.46 |
| 11 | 43256235 | rs7113638 | -0.225 | -0.904 | -1.01 | 0.207 | 0 | -0.001 | -0.001 | 0 | 0.48 | 0.52 | 0.37 | 0.41 | 0.47 | 0.39 |
| 11 | 43260324 | rs6485409 | -1.194 | -1.278 | -0.91 | 0.262 | -0.025 | 0.018 | 0.018 | 0 | 0.86 | 0.76 | 0.87 | 0.83 | 0.95 | 0.89 |
| 11 | 43262685 | rs7938844 | 0.781 | -1.013 | -1.286 | 0.239 | 0.029 | -0.003 | -0.003 | 0.016 | 0.28 | 0.19 | 0.18 | 0.24 | 0.17 | 0.21 |
| 11 | 43264321 | rs735984 | 0.998 | -1.014 | -1.264 | 0.261 | 0.027 | -0.006 | -0.006 | 0.523 | 0.27 | 0.19 | 0.18 | 0.24 | 0.16 | 0.2 |
| 11 | 43266834 | rs1353461 | 0.169 | -0.869 | -1.203 | 0.162 | -0.001 | -0.001 | -0.001 | 0 | 0.45 | 0.6 | 0.63 | 0.6 | 0.56 | 0.63 |
| 11 | 43267879 | rs12223779 | 1.603 | -0.616 | -0.888 | 0.608 | -0.002 | 0.01 | 0.01 | 0 | 0.08 | 0.05 | 0.1 | 0.18 | 0.03 | 0.12 |
| 11 | 43268827 | rs10838078 | 0.4 | -0.794 | -0.94 | 0.483 | -0.022 | 0.003 | 0.003 | 0 | 0.14 | 0.24 | 0.07 | 0.12 | 0.06 | 0.12 |
| 11 | 43278127 | rs12223066 | 0.387 | -0.773 | -1.123 | 0.439 | -0.018 | -0.024 | -0.024 | 0 | 0.23 | 0.11 | 0.22 | 0.26 | 0.27 | 0.22 |
| 11 | 43281629 | rs12270977 | 1.381 | -0.647 | -0.819 | 0.69 | 0.016 | 0.032 | 0.032 | 0 | 0.07 | 0.03 | 0.01 | 0 | 0.02 | 0.02 |
| 11 | 43283377 | rs11037390 | 0.842 | -0.531 | -0.643 | 0.846 | 0.01 | 0.074 | 0.074 | 0 | 0.07 | 0.23 | 0.05 | 0.12 | 0.04 | 0.11 |
| 11 | 43285357 | rs11037391 | 0.233 | -0.612 | -1.026 | 0.723 | -0.002 | 0.001 | 0.001 | 0 | 0.1 | 0.09 | 0.11 | 0.17 | 0.05 | 0.1 |

| | | | | | | | | | | | | | | | | |
|---|---|---|---|---|---|---|---|---|---|---|---|---|---|---|---|---|
| 11 | 43285881 | rs1496231 | -0.39 | -0.676 | -1.162 | 1.071 | 0.002 | -0.003 | -0.003 | 1.091 | 0.13 | 0.14 | 0.07 | 0.22 | 0.1 | 0.11 |
| 11 | 43290063 | rs2172998 | 0.891 | -0.508 | -1.782 | 0.558 | -0.043 | 0.033 | 0.033 | 0 | 0.31 | 0.14 | 0.25 | 0.28 | 0.37 | 0.34 |
| 11 | 43290350 | rs10838086 | -1.197 | -0.797 | -1.306 | 0.813 | 0.004 | -0.004 | -0.004 | 0.133 | 0.3 | 0.65 | 0.27 | 0.57 | 0.19 | 0.36 |
| 11 | 43300073 | rs1496236 | -0.886 | -0.607 | -0.946 | 0.926 | -0.005 | 0.05 | 0.05 | 0 | 0.85 | 0.82 | 0.82 | 0.65 | 0.91 | 0.75 |
| 11 | 43303925 | rs770770 | 0.308 | -0.814 | -1.371 | -0.093 | 0.023 | 0.043 | 0.043 | 0 | 0.51 | 0.61 | 0.48 | 0.41 | 0.65 | 0.47 |
| 11 | 43307196 | rs770765 | 0.49 | -0.692 | -1.338 | -0.756 | 0.017 | -0.076 | -0.076 | 0 | 0.25 | 0.49 | 0.18 | 0.12 | 0.27 | 0.13 |
| 11 | 43307811 | rs7107014 | -0.854 | -0.288 | -1.783 | -0.866 | -0.015 | 0.1 | 0.1 | 0 | 0.56 | 0.76 | 0.56 | 0.19 | 0.53 | 0.38 |
| 11 | 43310658 | rs1552942 | 0.807 | 0.06 | -1.727 | -0.647 | -0.023 | 0.051 | 0.051 | 0 | 0.22 | 0.14 | 0.16 | 0.12 | 0.27 | 0.12 |
| 11 | 43319979 | rs4338513 | 0.51 | 0.153 | -1.619 | -0.648 | -0.01 | 0.008 | 0.008 | 0 | 0.14 | 0.14 | 0.33 | 0.06 | 0.2 | 0.23 |
| 11 | 43334249 | rs11823539 | -0.128 | -0.318 | -1.659 | -0.933 | -0.013 | -0.077 | -0.077 | 0 | 0.17 | 0.56 | 0.31 | 0.03 | 0.12 | 0.14 |
| 11 | 43338968 | rs11037414 | 1.753 | -0.232 | -2.386 | -1.172 | -0.044 | -0.004 | -0.004 | 0 | 0.2 | 0.05 | 0.2 | 0.19 | 0.28 | 0.31 |
| 11 | 43344751 | rs11607472 | 3.09 | -0.256 | -2.614 | -1.513 | -0.03 | 0.08 | 0.08 | 0 | 0.14 | 0.01 | 0.08 | 0 | 0.11 | 0.03 |
| 11 | 43379506 | rs11037420 | 0.582 | -0.118 | -2.761 | -1.459 | -0.015 | 0.065 | 0.065 | 0 | 0.38 | 0.1 | 0.38 | 0.75 | 0.39 | 0.57 |
| 11 | 43381417 | rs11037421 | 2.408 | 0.177 | -2.768 | -1.554 | -0.026 | 0.04 | 0.04 | 0.065 | 0.07 | 0.01 | 0.09 | 0 | 0.12 | 0.06 |
| 11 | 43384448 | rs11605997 | -0.424 | -0.197 | -2.505 | -1.358 | 0.008 | 0.078 | 0.078 | 0 | 0.54 | 0.7 | 0.58 | 0.18 | 0.56 | 0.31 |
| 11 | 43384933 | rs903550 | -0.545 | -0.199 | -2.501 | -1.359 | -0.001 | 0.115 | 0.115 | 0 | 0.55 | 0.78 | 0.58 | 0.18 | 0.56 | 0.31 |
| 11 | 43388161 | rs4132524 | 2.582 | -0.277 | -2.624 | -1.488 | -0.015 | 0.109 | 0.109 | 0 | 0.18 | 0.03 | 0.09 | 0 | 0.12 | 0.04 |
| 11 | 43393699 | rs2132190 | -2.281 | -0.234 | -2.602 | -1.446 | 0.002 | -0.001 | -0.001 | 3.634 | 0.93 | 0.87 | 0.96 | 0.93 | 0.95 | 0.88 |
| 11 | 43408419 | rs4755711 | 1.392 | -0.296 | -2.637 | -1.013 | -0.066 | -0.137 | -0.137 | 0 | 0.32 | 0.06 | 0.27 | 0.47 | 0.36 | 0.47 |
| 11 | 43415570 | rs11827196 | 1.084 | -0.081 | -2.208 | -0.911 | -0.007 | -0.022 | -0.022 | 0.01 | 0.09 | 0.13 | 0.02 | 0 | 0.03 | 0.03 |
| 11 | 43432046 | rs7113471 | -0.247 | -0.055 | -2.233 | -0.926 | 0.006 | -0.042 | -0.042 | 23.486 | 0.33 | 0.58 | 0.45 | 0.14 | 0.38 | 0.21 |
| 11 | 43450090 | rs17594022 | 1.652 | -0.009 | -2.114 | -0.887 | 0.006 | -0.002 | -0.002 | 0 | 0.07 | 0.09 | 0.01 | 0 | 0.03 | 0.03 |
| 11 | 43480174 | rs17509794 | 2.155 | 0.472 | -2.359 | -0.565 | -0.062 | 0.084 | 0.084 | 0 | 0.14 | 0.01 | 0.12 | 0 | 0.23 | 0.05 |
| 11 | 43496797 | rs12418410 | 1.702 | 0.238 | -1.897 | 0.092 | -0.004 | 0.081 | 0.081 | 0 | 0.05 | 0.03 | 0.11 | 0.28 | 0.03 | 0.1 |
| 11 | 43499928 | rs11037447 | 1.702 | 0.238 | -1.897 | 0.09 | -0.004 | 0.079 | 0.079 | 0 | 0.05 | 0.03 | 0.11 | 0.28 | 0.03 | 0.1 |
| 11 | 43509169 | rs17510173 | 2.643 | 0.454 | -1.976 | -0.236 | -0.009 | 0.001 | 0.001 | 0 | 0.05 | 0.02 | 0.26 | 0.03 | 0.1 | 0.11 |
| 11 | 43511407 | rs900465 | 0.87 | 0.021 | -1.987 | -0.617 | -0.019 | -0.103 | -0.103 | 0 | 0.25 | 0.51 | 0.16 | 0.04 | 0.18 | 0.1 |
| 11 | 43514530 | rs11037452 | 1.314 | 0.309 | -2.378 | -0.283 | -0.064 | -0.021 | -0.021 | 0 | 0.26 | 0.06 | 0.25 | 0.29 | 0.34 | 0.39 |
| 11 | 43521662 | rs11037453 | 0.506 | 0.538 | -1.717 | -0.475 | -0.002 | -0.005 | -0.005 | 0 | 0.07 | 0.09 | 0.04 | 0.04 | 0.06 | 0.1 |
| 11 | 43521746 | rs1471672 | -0.705 | 0.477 | -1.855 | -0.625 | -0.006 | 0.03 | 0.03 | 0 | 0.38 | 0.68 | 0.49 | 0.18 | 0.43 | 0.3 |
| 11 | 43532023 | rs4755193 | 0.167 | -0.015 | -1.62 | -0.208 | -0.02 | 0.081 | 0.081 | 0 | 0.79 | 0.64 | 0.81 | 0.62 | 0.85 | 0.71 |
| 11 | 43534247 | rs7929418 | 0.524 | 0.28 | -2.418 | -0.342 | -0.016 | -0.11 | -0.11 | 0 | 0.57 | 0.11 | 0.62 | 0.51 | 0.6 | 0.62 |
| 11 | 43536288 | rs10768957 | 0.357 | 0.728 | -2.26 | -0.149 | -0.115 | -0.129 | -0.129 | 0 | 0.39 | 0.08 | 0.53 | 0.53 | 0.48 | 0.58 |
| 11 | 43536586 | rs12277553 | 0.071 | 0.637 | -2.002 | -0.168 | 0.021 | 0 | 0 | 0 | 0.13 | 0.48 | 0.1 | 0.09 | 0.11 | 0.18 |
| 11 | 43539013 | rs7941216 | 0.32 | 0.694 | -2.361 | -0.071 | -0.124 | -0.139 | -0.139 | 0.648 | 0.38 | 0.07 | 0.53 | 0.51 | 0.48 | 0.57 |
| 11 | 43543339 | rs878511 | -0.38 | 1.054 | -1.718 | 0.041 | 0.088 | -0.011 | -0.011 | 0 | 0.14 | 0.47 | 0.14 | 0.09 | 0.21 | 0.2 |
| 11 | 43544831 | rs10838120 | 0.95 | 0.663 | -2.45 | 0.097 | -0.024 | -0.065 | -0.065 | 0 | 0.18 | 0.12 | 0.24 | 0.41 | 0.23 | 0.19 |
| 11 | 43545235 | rs11607570 | 1.041 | 0.373 | -2.685 | -0.37 | 0.006 | 0.162 | 0.162 | 0.043 | 0.29 | 0.07 | 0.15 | 0.04 | 0.22 | 0.12 |
| 11 | 43545756 | rs16937556 | 1.022 | 0.356 | -2.683 | -0.368 | 0.003 | 0.163 | 0.163 | 0.057 | 0.29 | 0.07 | 0.15 | 0.04 | 0.22 | 0.12 |
| 11 | 43546698 | rs7119999 | 0.131 | 0.819 | -2.032 | 0.421 | -0.062 | -0.081 | -0.081 | 0.058 | 0.32 | 0.09 | 0.28 | 0.42 | 0.39 | 0.42 |
| 11 | 43550070 | rs10838121 | 0.638 | 0.755 | -1.734 | -0.011 | -0.049 | 0.005 | 0.005 | 0.025 | 0.2 | 0.04 | 0.2 | 0.16 | 0.27 | 0.19 |
| 11 | 43551518 | rs16937562 | -0.383 | 0.462 | -0.771 | 0.26 | 0.049 | 0.045 | 0.045 | 0.181 | 0.08 | 0.45 | 0.06 | 0.06 | 0.05 | 0.13 |
| 11 | 43554976 | rs900467 | -0.436 | 0.545 | -1.701 | 0.176 | -0.002 | 0.025 | 0.025 | 0 | 0.15 | 0.06 | 0.06 | 0.07 | 0.13 | 0.2 |
| 11 | 43559991 | rs10400240 | 0.088 | 0.856 | -1.914 | 0.356 | -0.035 | -0.001 | -0.001 | 0 | 0.38 | 0.23 | 0.37 | 0.41 | 0.53 | 0.42 |
| 11 | 43569964 | rs3740981 | 0.167 | 0.927 | -2.193 | 0.198 | -0.097 | 0.074 | 0.074 | 0.122 | 0.33 | 0.02 | 0.23 | 0.11 | 0.34 | 0.09 |
| 11 | 43572788 | rs4755197 | -0.234 | 0.908 | -2.299 | 0.853 | -0.107 | -0.117 | -0.117 | 0.047 | 0.39 | 0.03 | 0.32 | 0.5 | 0.39 | 0.27 |
| 11 | 43575766 | rs11037472 | 0.161 | 0.707 | -1.2 | 1.424 | -0.005 | 0.058 | 0.058 | 0 | 0.05 | 0.01 | 0.09 | 0.41 | 0.05 | 0.17 |
| 11 | 43590892 | rs11037478 | -0.466 | 0.606 | -0.485 | 0.523 | 0 | -0.01 | -0.01 | 0 | 0.11 | 0.22 | 0.06 | 0.11 | 0.08 | 0.14 |
| 11 | 43593325 | rs1483130 | 0.031 | 0.736 | -0.603 | 0.269 | 0.004 | 0.011 | 0.011 | 0 | 0.4 | 0.34 | 0.53 | 0.33 | 0.4 | 0.56 |
| 11 | 43595031 | rs4755200 | -0.023 | 0.934 | -0.365 | 1.034 | 0.015 | 0.193 | 0.193 | 0 | 0.07 | 0.13 | 0.11 | 0.47 | 0.09 | 0.13 |
| 11 | 43596350 | rs11606426 | -0.563 | 0.773 | -0.473 | -0.024 | 0.003 | -0.001 | -0.001 | 0.206 | 0.09 | 0.11 | 0.07 | 0.13 | 0.1 | 0.13 |
| 11 | 43604093 | rs2679051 | -0.336 | 0.279 | -0.315 | -0.802 | 0.01 | -0.032 | -0.032 | 0 | 0.37 | 0.5 | 0.61 | 0.73 | 0.42 | 0.48 |
| 11 | 43605304 | rs2679053 | -0.516 | -0.096 | -0.73 | -1.421 | 0 | -0.006 | -0.006 | 0 | 0.61 | 0.6 | 0.7 | 0.73 | 0.59 | 0.55 |
| 11 | 43617546 | rs11037491 | 0.818 | -0.066 | -0.818 | -1.655 | 0.008 | 0.127 | 0.127 | 0.689 | 0.24 | 0.07 | 0.09 | 0 | 0.17 | 0.08 |
| 11 | 43619938 | rs4755202 | 0.121 | -0.379 | -0.764 | -1.818 | -0.039 | 0.106 | 0.106 | 0 | 0.39 | 0.17 | 0.6 | 0.76 | 0.42 | 0.47 |
| 11 | 43628749 | rs4755720 | -0.63 | -0.356 | -0.579 | -1.799 | 0.004 | -0.046 | -0.046 | 0.052 | 0.65 | 0.55 | 0.71 | 0.79 | 0.6 | 0.56 |
| 11 | 43637563 | rs2902373 | 0.732 | -0.336 | -0.707 | -1.853 | 0.018 | 0.068 | 0.068 | 0.719 | 0.27 | 0.16 | 0.13 | 0 | 0.22 | 0.11 |
| 11 | 43643624 | rs10838139 | 0.328 | -0.364 | -0.714 | -1.783 | -0.046 | 0.085 | 0.085 | 0 | 0.37 | 0.17 | 0.58 | 0.74 | 0.41 | 0.43 |
| 11 | 43653833 | rs2862996 | -0.996 | -0.579 | -0.446 | -1.827 | 0.031 | -0.039 | -0.039 | 0 | 0.76 | 0.6 | 0.75 | 0.8 | 0.66 | 0.69 |
| 11 | 43664670 | rs7940743 | -0.731 | -0.609 | -0.252 | -1.75 | -0.048 | -0.046 | -0.046 | 0 | 0.89 | 0.71 | 0.96 | 0.97 | 0.98 | 0.88 |
| 11 | 43671734 | rs11037537 | 1.136 | -0.335 | -0.69 | -1.653 | -0.087 | 0.005 | 0.005 | 0.276 | 0.19 | 0.01 | 0.38 | 0.14 | 0.25 | 0.25 |
| 11 | 43686618 | rs6485438 | 0.627 | -0.399 | -0.216 | -1.487 | 0.026 | -0.024 | -0.024 | 0 | 0.24 | 0.34 | 0.25 | 0.22 | 0.34 | 0.3 |
| 11 | 43686685 | rs6485440 | -0.367 | -0.658 | -0.173 | -1.552 | -0.053 | -0.029 | -0.029 | 0 | 0.88 | 0.63 | 0.95 | 0.94 | 0.98 | 0.88 |
| 11 | 43688411 | rs11037545 | 0.924 | -0.749 | -0.361 | -1.763 | 0.012 | 0.129 | 0.129 | 0.786 | 0.26 | 0.08 | 0.12 | 0 | 0.22 | 0.11 |
| 11 | 43689610 | rs10838146 | -0.108 | -0.701 | -0.353 | -1.741 | -0.019 | -0.103 | -0.103 | 0.053 | 0.62 | 0.55 | 0.82 | 0.96 | 0.75 | 0.74 |
| 11 | 43694016 | rs10501303 | 0.564 | -0.678 | -0.368 | -1.361 | -0.013 | 0 | 0 | 0 | 0.12 | 0.1 | 0.1 | 0.13 | 0.04 | 0.13 |
| 11 | 43695374 | rs7944241 | 0.551 | -0.461 | -0.415 | -1.745 | -0.043 | 0.087 | 0.087 | 0 | 0.37 | 0.16 | 0.57 | 0.75 | 0.41 | 0.43 |
| 11 | 43696917 | rs7116641 | 0.843 | -0.435 | -0.231 | -1.766 | 0.024 | -0.017 | -0.017 | 0 | 0.23 | 0.31 | 0.25 | 0.22 | 0.35 | 0.3 |
| 11 | 43698499 | rs7928182 | 0.609 | -0.485 | -0.311 | -1.768 | -0.048 | 0.074 | 0.074 | 0 | 0.37 | 0.13 | 0.57 | 0.72 | 0.4 | 0.42 |

| | | | | | | | | | | | | | | | | |
|---|---|---|---|---|---|---|---|---|---|---|---|---|---|---|---|---|
| 11 | 43698956 | rs11037556 | -0.034 | -0.565 | -0.268 | -1.76 | -0.025 | -0.117 | -0.117 | 0.006 | 0.6 | 0.52 | 0.83 | 0.96 | 0.76 | 0.74 |
| 11 | 43713989 | rs7106967 | 0.017 | -0.673 | -0.368 | -1.842 | -0.001 | -0.005 | -0.005 | 0 | 0.45 | 0.46 | 0.36 | 0.23 | 0.54 | 0.44 |
| 11 | 43717275 | rs11037567 | 1.181 | -0.713 | -0.536 | -1.948 | 0.011 | 0.041 | 0.041 | 1.514 | 0.2 | 0.13 | 0.09 | 0 | 0.17 | 0.11 |
| 11 | 43721500 | rs4359199 | 0.181 | -0.604 | -0.224 | -1.563 | -0.001 | 0.017 | 0.017 | 0 | 0.46 | 0.58 | 0.37 | 0.26 | 0.55 | 0.44 |
| 11 | 43727577 | rs7932905 | 0.313 | -0.604 | -0.228 | -1.623 | 0 | -0.025 | -0.025 | 0 | 0.46 | 0.51 | 0.36 | 0.23 | 0.54 | 0.45 |
| 11 | 43740422 | rs6485450 | -0.372 | -0.894 | -0.284 | -1.556 | -0.043 | -0.066 | -0.066 | 0 | 0.81 | 0.72 | 0.93 | 0.97 | 0.94 | 0.86 |
| 11 | 43748673 | rs6485453 | 0.318 | -0.211 | -0.304 | -1.425 | -0.006 | 0.053 | 0.053 | 0 | 0.53 | 0.58 | 0.37 | 0.26 | 0.53 | 0.54 |
| 11 | 43749199 | rs7928422 | -0.413 | -0.113 | -0.332 | -1.327 | -0.029 | 0.055 | 0.055 | 0 | 0.37 | 0.24 | 0.36 | 0.73 | 0.4 | 0.38 |
| 11 | 43774274 | rs10838160 | -0.29 | 0.033 | -0.256 | -1.14 | -0.02 | 0.015 | 0.015 | 0 | 0.35 | 0.24 | 0.35 | 0.67 | 0.39 | 0.38 |
| 11 | 43788330 | rs11037611 | -0.197 | -0.013 | -0.239 | -1.069 | -0.019 | 0.029 | 0.029 | 0 | 0.35 | 0.25 | 0.35 | 0.72 | 0.4 | 0.38 |
| 11 | 43791865 | rs4755737 | -0.27 | 0.008 | -0.044 | -1.147 | -0.04 | 0.024 | 0.024 | 0.005 | 0.36 | 0.16 | 0.36 | 0.67 | 0.43 | 0.41 |
| 11 | 43803113 | rs4755740 | -0.134 | -0.015 | -0.078 | -1.155 | -0.1 | -0.012 | -0.012 | 2.128 | 0.33 | 0.05 | 0.33 | 0.67 | 0.39 | 0.38 |
| 11 | 43803254 | rs4755741 | -0.414 | -0.121 | 0.04 | -1.515 | 0.068 | 0.002 | 0.002 | 0.601 | 0.78 | 0.59 | 0.53 | 0.74 | 0.68 | 0.66 |
| 11 | 43820719 | rs10838173 | -0.117 | -0.031 | -0.036 | -1.459 | -0.031 | 0.02 | 0.02 | 0 | 0.34 | 0.2 | 0.35 | 0.67 | 0.39 | 0.38 |
| 11 | 43832505 | rs10400343 | -0.243 | -0.027 | 0.01 | -1.442 | 0.07 | 0.029 | 0.029 | 0.265 | 0.77 | 0.54 | 0.53 | 0.7 | 0.67 | 0.66 |
| 11 | 43851308 | rs11037662 | 1.39 | 0.326 | -0.306 | -1.441 | -0.044 | -0.089 | -0.089 | 0 | 0.12 | 0.01 | 0.18 | 0.53 | 0.14 | 0.17 |
| 11 | 43852491 | rs6485471 | 0.036 | 0.262 | -0.049 | -1.806 | -0.031 | 0.021 | 0.021 | 0 | 0.34 | 0.2 | 0.35 | 0.68 | 0.39 | 0.38 |
| 11 | 43864278 | rs2176598 | 0.057 | 1.584 | 0.551 | -0.806 | 0.081 | -0.025 | -0.025 | 0 | 0.82 | 0.62 | 0.58 | 0.85 | 0.74 | 0.79 |
| 11 | 43873539 | rs1518820 | -0.24 | 1.538 | 0.666 | -0.804 | 0.052 | 0.09 | 0.09 | 0 | 0.05 | 0.25 | 0.05 | 0.11 | 0.06 | 0.21 |
| 11 | 43876698 | rs11555762 | -1.003 | 1.536 | 0.503 | -0.814 | 0.067 | -0.003 | -0.003 | 0.239 | 0.22 | 0.42 | 0.46 | 0.26 | 0.32 | 0.34 |
| 11 | 43879540 | rs11037685 | -0.78 | 1.571 | -0.298 | -0.951 | -0.015 | 0.01 | 0.01 | 0.571 | 0.21 | 0.17 | 0.46 | 0.19 | 0.36 | 0.16 |
| 11 | 43884515 | rs1914771 | 0.022 | 0.364 | 0.219 | -0.998 | -0.049 | -0.081 | -0.081 | 0 | 0.8 | 0.62 | 0.82 | 0.94 | 0.9 | 0.94 |
| 11 | 43885537 | rs12804822 | -0.072 | 0.422 | -0.022 | -0.912 | -0.012 | -0.016 | -0.016 | 1.763 | 0.21 | 0.1 | 0.42 | 0.24 | 0.23 | 0.44 |
| 11 | 43886365 | rs2863000 | 0.473 | 0.443 | 0.015 | -0.869 | -0.053 | -0.105 | -0.105 | 0 | 0.75 | 0.55 | 0.61 | 0.92 | 0.87 | 0.88 |
| 11 | 43890633 | rs12364347 | -0.286 | 0.242 | 0.017 | -1.048 | 0.006 | -0.011 | -0.011 | 0.507 | 0.24 | 0.1 | 0.27 | 0.26 | 0.22 | 0.43 |
| 11 | 43890799 | rs11037690 | 2.149 | 0.347 | -0.234 | -1.117 | -0.059 | -0.109 | -0.109 | 0 | 0.32 | 0.04 | 0.21 | 0.52 | 0.34 | 0.2 |
| 11 | 43891694 | rs2862999 | 0.221 | 0.47 | -0.062 | -1.349 | -0.003 | -0.019 | -0.019 | 0.764 | 0.76 | 0.31 | 0.77 | 0.93 | 0.84 | 0.91 |
| 11 | 43893658 | rs10838191 | -0.343 | 0.472 | -0.084 | -1.334 | -0.05 | 0.034 | 0.034 | 0.067 | 0.36 | 0.19 | 0.35 | 0.31 | 0.46 | 0.51 |
| 11 | 43899601 | rs2434467 | 0.473 | 0.469 | 0 | -1.333 | -0.008 | -0.023 | -0.023 | 0 | 0.6 | 0.48 | 0.44 | 0.74 | 0.66 | 0.48 |
| 11 | 43903600 | rs11037703 | 1.833 | 0.468 | -0.104 | -1.345 | -0.04 | -0.053 | -0.053 | 0.556 | 0.35 | 0.06 | 0.21 | 0.6 | 0.35 | 0.34 |
| 11 | 43903842 | rs12806569 | 1.086 | 0.442 | -0.017 | -1.283 | -0.016 | -0.022 | -0.022 | 0 | 0.17 | 0.07 | 0.21 | 0.21 | 0.2 | 0.37 |
| 11 | 43905096 | rs2271820 | 0.009 | 0.449 | -0.013 | -1.278 | -0.006 | -0.001 | -0.001 | 0.898 | 0.23 | 0.14 | 0.42 | 0.24 | 0.24 | 0.47 |
| 11 | 43908230 | rs7101434 | -0.318 | 0.687 | 0.228 | -1.09 | 0.064 | -0.018 | -0.018 | 0.004 | 0.07 | 0.31 | 0.05 | 0 | 0.07 | 0.03 |
| 11 | 43912925 | rs1518818 | 0.21 | 0.704 | 0.124 | -1.013 | -0.006 | -0.006 | -0.006 | 0.178 | 0.21 | 0.14 | 0.42 | 0.24 | 0.23 | 0.43 |
| 11 | 43913386 | rs7116503 | -0.685 | 0.678 | 0.156 | -1.033 | -0.026 | -0.099 | -0.099 | 0 | 0.83 | 0.61 | 0.86 | 0.98 | 0.9 | 0.94 |
| 11 | 43914538 | rs1554316 | 0.871 | 0.703 | 0.101 | -1.018 | -0.015 | -0.023 | -0.023 | 0 | 0.17 | 0.06 | 0.21 | 0.21 | 0.2 | 0.37 |
| 11 | 43919778 | rs2434474 | 0.673 | 0.718 | 0.155 | -1.014 | -0.007 | -0.012 | -0.012 | 0 | 0.6 | 0.47 | 0.39 | 0.69 | 0.67 | 0.51 |
| 11 | 43920006 | rs7107050 | 0.194 | 0.746 | 0.046 | -0.926 | -0.006 | -0.006 | -0.006 | 0.039 | 0.21 | 0.14 | 0.42 | 0.24 | 0.23 | 0.43 |
| 11 | 43934717 | rs10734520 | 0.667 | 0.807 | 0.103 | -1.058 | 0.001 | 0.005 | 0.005 | 0 | 0.6 | 0.38 | 0.66 | 0.84 | 0.64 | 0.8 |
| 11 | 43936945 | rs10768994 | 1.097 | 0.815 | 0.339 | -0.992 | 0 | 0.003 | 0.003 | 0 | 0.44 | 0.41 | 0.46 | 0.62 | 0.43 | 0.42 |
| 11 | 43940644 | rs1048928 | 0.843 | 0.712 | 0.168 | -1.414 | 0.003 | 0.076 | 0.076 | 0 | 0.6 | 0.3 | 0.65 | 0.84 | 0.62 | 0.79 |
| 11 | 43942186 | rs2292891 | -0.396 | 0.711 | 0.276 | -1.436 | 0.008 | -0.025 | -0.025 | 0.384 | 0.23 | 0.31 | 0.21 | 0.14 | 0.27 | 0.16 |
| 11 | 43942994 | rs883856 | -0.539 | 0.698 | 0.216 | -1.432 | 0.016 | -0.036 | -0.036 | 0.015 | 0.08 | 0.2 | 0.23 | 0.02 | 0.08 | 0.08 |
| 11 | 43946967 | rs1973717 | 0.661 | 0.794 | 0.165 | -1.387 | 0 | -0.066 | -0.066 | 0.059 | 0.63 | 0.44 | 0.72 | 0.88 | 0.64 | 0.8 |
| 11 | 43947990 | rs12797684 | 0.678 | 0.763 | 0.163 | -1.389 | 0 | -0.066 | -0.066 | 0 | 0.63 | 0.46 | 0.72 | 0.88 | 0.63 | 0.8 |
| 11 | 43949008 | rs2434484 | -1.021 | 1.258 | -0.187 | -0.827 | 0.001 | -0.006 | -0.006 | 0 | 0.27 | 0.29 | 0.26 | 0.16 | 0.33 | 0.21 |
| 11 | 43949511 | rs7103602 | -0.007 | 1.254 | -0.58 | -0.527 | 0.04 | -0.044 | -0.044 | 0 | 0.78 | 0.62 | 0.61 | 0.84 | 0.73 | 0.81 |
| 11 | 43952762 | rs11037740 | -0.262 | 0.777 | -1.471 | -0.089 | -0.021 | -0.008 | -0.008 | 0.016 | 0.55 | 0.31 | 0.32 | 0.46 | 0.53 | 0.35 |
| 11 | 43953774 | rs7120772 | -0.937 | 0.801 | -0.955 | -0.498 | -0.002 | -0.014 | -0.014 | 0.365 | 0.18 | 0.17 | 0.13 | 0 | 0.19 | 0.04 |
| 11 | 43955081 | rs7128816 | -1.017 | 0.507 | -0.94 | -0.512 | 0.001 | -0.024 | -0.024 | 0 | 0.19 | 0.19 | 0.13 | 0 | 0.18 | 0.03 |
| 11 | 43956936 | rs6485477 | -0.373 | 0.379 | -0.761 | -0.214 | 0.004 | -0.003 | -0.003 | 0 | 0.42 | 0.58 | 0.64 | 0.62 | 0.48 | 0.62 |
| 11 | 43957859 | rs11037747 | 0.296 | -0.08 | -0.657 | -0.807 | -0.001 | 0 | 0 | 0.048 | 0.33 | 0.31 | 0.23 | 0.38 | 0.33 | 0.28 |
| 11 | 43960386 | rs6485479 | -0.987 | 0.312 | -0.625 | -0.439 | 0.003 | 0.003 | 0.003 | 0 | 0.75 | 0.75 | 0.79 | 0.69 | 0.67 | 0.71 |
| 11 | 43960532 | rs6485480 | 0.588 | -0.028 | -0.442 | -0.797 | 0.009 | -0.008 | -0.008 | 0 | 0.26 | 0.2 | 0.13 | 0 | 0.2 | 0.1 |
| 11 | 43970363 | rs10768996 | 0.192 | -0.129 | -0.306 | -0.42 | 0.001 | 0.003 | 0.003 | 0 | 0.52 | 0.58 | 0.48 | 0.68 | 0.58 | 0.5 |
| 11 | 43972609 | rs7126420 | 0.225 | -0.431 | -0.822 | -0.391 | 0.062 | 0.079 | 0.079 | 0 | 0.34 | 0.13 | 0.34 | 0.16 | 0.24 | 0.38 |
| 11 | 43975742 | rs1401317 | 0.668 | -0.51 | -0.586 | -0.645 | 0.046 | 0.05 | 0.05 | 0 | 0.29 | 0.16 | 0.13 | 0 | 0.18 | 0.14 |
| 11 | 43980440 | rs630211 | -0.66 | -0.159 | -0.641 | -0.165 | 0.01 | 0.007 | 0.007 | 0.147 | 0.5 | 0.42 | 0.7 | 0.54 | 0.53 | 0.66 |
| 11 | 43985156 | rs2863028 | -0.408 | 0.134 | -0.635 | -0.006 | 0.004 | 0.007 | 0.007 | 0.079 | 0.46 | 0.38 | 0.45 | 0.43 | 0.53 | 0.48 |
| 11 | 43989324 | rs12417085 | 0.69 | 0.264 | -0.936 | -0.006 | 0.007 | 0.023 | 0.023 | 0 | 0.09 | 0.04 | 0.14 | 0.02 | 0.06 | 0.11 |
| 11 | 43991355 | rs2434487 | 0.394 | 0.407 | -0.828 | 0.142 | -0.007 | 0.009 | 0.009 | 0 | 0.93 | 0.94 | 0.73 | 0.75 | 0.89 | 0.79 |
| 11 | 43993024 | rs7932326 | -0.567 | 0.337 | -1.025 | -0.432 | 0.017 | 0.091 | 0.091 | 0 | 0.56 | 0.72 | 0.76 | 0.82 | 0.58 | 0.7 |
| 11 | 43993199 | rs16937773 | 0.563 | 1.147 | -0.867 | 0.031 | -0.013 | -0.011 | -0.011 | 0.393 | 0.2 | 0.12 | 0.18 | 0.23 | 0.22 | 0.19 |
| 11 | 44003258 | rs2863030 | 0.021 | 0.395 | -0.842 | -0.04 | 0.001 | -0.011 | -0.011 | 0 | 0.11 | 0.09 | 0.06 | 0 | 0.11 | 0.1 |
| 11 | 44004135 | rs933302 | 0.483 | 0.714 | -1.222 | 0.281 | -0.036 | -0.086 | -0.086 | 0 | 0.14 | 0.01 | 0.21 | 0.28 | 0.12 | 0.22 |
| 11 | 44006007 | rs1557709 | 1.102 | 0.735 | -0.78 | 0.05 | -0.009 | 0.018 | 0.018 | 0 | 0.14 | 0.1 | 0.12 | 0.02 | 0.16 | 0.09 |
| 11 | 44006539 | rs2902389 | 0.873 | 0.551 | -0.217 | 0.404 | 0 | -0.001 | -0.001 | 0.541 | 0.47 | 0.47 | 0.45 | 0.47 | 0.48 | 0.27 |
| 11 | 44009414 | rs11037774 | -0.169 | 0.734 | -0.428 | 0.609 | -0.012 | -0.074 | -0.074 | 0 | 0.68 | 0.85 | 0.6 | 0.54 | 0.66 | 0.46 |
| 11 | 44014531 | rs7103415 | 0.558 | 0.822 | -0.116 | 0.51 | -0.016 | -0.024 | -0.024 | 1.565 | 0.57 | 0.79 | 0.53 | 0.48 | 0.53 | 0.37 |

| | | | | | | | | | | | | | | | | |
|---|---|---|---|---|---|---|---|---|---|---|---|---|---|---|---|---|
| 11 | 44015075 | rs11037782 | -0.655 | 0.784 | -0.232 | 0.605 | -0.015 | -0.051 | -0.051 | 0 | 0.27 | 0.13 | 0.29 | 0.35 | 0.3 | 0.37 |
| 11 | 44015866 | rs4595529 | -0.489 | 0.779 | -0.295 | 0.601 | -0.08 | -0.112 | -0.112 | 0.117 | 0.25 | 0.02 | 0.28 | 0.33 | 0.29 | 0.35 |
| 11 | 44016605 | rs11037788 | 1.126 | 0.842 | 0.466 | 0.741 | 0.003 | 0.069 | 0.069 | 0 | 0.52 | 0.37 | 0.42 | 0.09 | 0.51 | 0.24 |
| 11 | 44017314 | rs756956 | 1.201 | 0.827 | 0.536 | 0.67 | 0.002 | 0.114 | 0.114 | 0 | 0.5 | 0.32 | 0.41 | 0.06 | 0.49 | 0.23 |
| 11 | 44018822 | rs4755224 | -0.886 | 0.701 | 0.622 | 0.762 | 0.02 | 0.296 | 0.296 | 1.807 | 0.1 | 0.52 | 0.14 | 0.39 | 0.06 | 0.19 |
| 11 | 44019643 | rs728648 | -0.043 | 0.793 | 0.477 | 0.806 | -0.011 | 0.054 | 0.054 | 0.019 | 0.74 | 0.38 | 0.69 | 0.46 | 0.78 | 0.55 |
| 11 | 44019715 | rs10838212 | -0.003 | 0.812 | 0.785 | 0.873 | 0.027 | 0.209 | 0.209 | 0.095 | 0.08 | 0.27 | 0.1 | 0.31 | 0.06 | 0.13 |
| 11 | 44023161 | rs11037794 | 1.102 | 0.838 | 0.654 | 0.736 | 0.008 | 0.065 | 0.065 | 0 | 0.51 | 0.35 | 0.43 | 0.11 | 0.52 | 0.25 |
| 11 | 44025878 | rs2074048 | 0.656 | 0.898 | 0.774 | 0.91 | -0.002 | -0.005 | -0.005 | 0 | 0.16 | 0.15 | 0.25 | 0.35 | 0.18 | 0.32 |
| 11 | 44026030 | rs2074049 | 0.511 | 0.913 | 0.853 | 0.899 | -0.005 | 0.006 | 0.006 | 0 | 0.6 | 0.47 | 0.53 | 0.44 | 0.59 | 0.39 |
| 11 | 44028547 | rs7932879 | 1.825 | 1.135 | 0.162 | 0.37 | -0.093 | 0.126 | 0.126 | 0 | 0.28 | 0.02 | 0.29 | 0.05 | 0.3 | 0.12 |
| 11 | 44028907 | rs7943127 | -1.182 | 1.645 | 0.988 | 1.088 | 0.004 | -0.005 | -0.005 | 0 | 0.3 | 0.37 | 0.38 | 0.73 | 0.32 | 0.49 |
| 11 | 44029310 | rs11037800 | -0.286 | 1.639 | 0.994 | 0.825 | 0 | 0.01 | 0.01 | 0 | 0.22 | 0.23 | 0.28 | 0.4 | 0.26 | 0.35 |
| 11 | 44032746 | rs7942915 | 0.603 | 1.579 | 0.795 | 0.463 | 0.001 | 0.002 | 0.002 | 0 | 0.53 | 0.56 | 0.44 | 0.15 | 0.55 | 0.28 |
| 11 | 44032955 | rs7931200 | -0.776 | 1.376 | 0.878 | 0.762 | -0.008 | -0.001 | -0.001 | 0 | 0.34 | 0.48 | 0.39 | 0.68 | 0.29 | 0.49 |
| 11 | 44033602 | rs7109590 | 0.379 | 1.25 | 0.623 | 0.96 | -0.003 | -0.051 | -0.051 | 0.059 | 0.24 | 0.09 | 0.17 | 0.29 | 0.2 | 0.28 |
| 11 | 44036397 | rs17795583 | 1.363 | 1.421 | 0.546 | 0.795 | -0.016 | 0.009 | 0.009 | 0 | 0.06 | 0 | 0.04 | 0.02 | 0.08 | 0.03 |
| 11 | 44036866 | rs11037803 | 0.529 | 1.379 | 0.792 | 0.849 | 0.002 | -0.014 | -0.014 | 0 | 0.24 | 0.19 | 0.18 | 0.31 | 0.19 | 0.27 |
| 11 | 44036919 | rs929213 | -0.381 | 1.381 | 0.941 | 0.855 | -0.003 | -0.003 | -0.003 | 0 | 0.78 | 0.76 | 0.76 | 0.78 | 0.77 | 0.71 |
| 11 | 44038444 | rs11037804 | 1.344 | 1.405 | 1.077 | 0.855 | -0.015 | 0.041 | 0.041 | 0 | 0.37 | 0.3 | 0.37 | 0.12 | 0.42 | 0.18 |
| 11 | 44038877 | rs10838216 | 0.832 | 1.402 | 1.15 | 0.888 | 0.002 | 0.044 | 0.044 | 0 | 0.15 | 0.23 | 0.2 | 0.29 | 0.15 | 0.21 |
| 11 | 44038970 | rs10769002 | -0.178 | 1.398 | 1.191 | 0.893 | -0.009 | 0.041 | 0.041 | 0 | 0.69 | 0.6 | 0.6 | 0.49 | 0.7 | 0.5 |
| 11 | 44039024 | rs885628 | -1.47 | 1.399 | 0.999 | 0.897 | 0.007 | -0.001 | -0.001 | 0 | 0.86 | 0.91 | 0.8 | 0.85 | 0.85 | 0.75 |
| 11 | 44041903 | rs10769004 | 0.388 | 1.4 | 1.005 | 0.891 | -0.003 | -0.001 | -0.001 | 0 | 0.54 | 0.52 | 0.58 | 0.45 | 0.57 | 0.41 |
| 11 | 44046351 | rs7127815 | 0.381 | 1.411 | 1.011 | 0.897 | -0.004 | -0.001 | -0.001 | 0.142 | 0.54 | 0.52 | 0.58 | 0.45 | 0.57 | 0.41 |
| 11 | 44047632 | rs7938319 | 0.738 | 1.407 | 0.879 | 0.873 | 0.002 | -0.024 | -0.024 | 2.389 | 0.24 | 0.16 | 0.17 | 0.31 | 0.19 | 0.27 |
| 11 | 44048650 | rs1073369 | -1.301 | 1.464 | 0.726 | 0.595 | 0.001 | 0 | 0 | 4.027 | 0.85 | 0.88 | 0.8 | 0.86 | 0.85 | 0.75 |
| 11 | 44049526 | rs7101977 | 0.904 | 1.549 | 0.629 | 0.375 | 0.004 | -0.03 | -0.03 | 4.472 | 0.39 | 0.44 | 0.38 | 0.16 | 0.42 | 0.2 |
| 11 | 44052466 | rs9888287 | 1.157 | 1.199 | 0.254 | 0.389 | 0.002 | 0.046 | 0.046 | 0 | 0.37 | 0.27 | 0.34 | 0.15 | 0.34 | 0.17 |
| 11 | 44052598 | rs2074050 | 0.763 | 1.364 | -0.297 | 0.631 | -0.017 | -0.011 | -0.011 | 0.414 | 0.16 | 0.08 | 0.21 | 0.2 | 0.22 | 0.21 |
| 11 | 44053497 | rs6485492 | -0.853 | 0.586 | -0.048 | 1.19 | 0.001 | -0.023 | -0.023 | 0.591 | 0.86 | 0.91 | 0.79 | 0.77 | 0.83 | 0.72 |
| 11 | 44055493 | rs10838223 | 0.122 | 0.482 | 0.025 | 1.119 | -0.004 | 0.013 | 0.013 | 0 | 0.26 | 0.28 | 0.18 | 0.41 | 0.2 | 0.31 |
| 11 | 44055694 | rs10838226 | 0.147 | 0.833 | -0.118 | 1.267 | 0.002 | -0.029 | -0.029 | 0 | 0.83 | 0.91 | 0.81 | 0.75 | 0.81 | 0.71 |
| 11 | 44059419 | rs7122031 | 0.893 | 0.77 | -0.253 | 1.15 | -0.003 | 0.016 | 0.016 | 0 | 0.77 | 0.63 | 0.7 | 0.65 | 0.73 | 0.62 |
| 11 | 44066354 | rs3094395 | -1.06 | 0.883 | 0.032 | 0.904 | 0.02 | 0.044 | 0.044 | 0 | 0.09 | 0.27 | 0.09 | 0.14 | 0.11 | 0.08 |
| 11 | 44077227 | rs12284962 | -0.815 | 0.969 | -0.512 | 0.639 | 0 | -0.001 | -0.001 | 0 | 0.1 | 0.1 | 0.1 | 0.06 | 0.12 | 0.03 |
| 11 | 44080210 | rs2074051 | -0.511 | 0.233 | -1.28 | -0.23 | 0.03 | 0.076 | 0.076 | 0 | 0.71 | 0.9 | 0.59 | 0.84 | 0.78 | 0.77 |
| 11 | 44081027 | rs4755775 | 0.645 | 0.29 | -1.832 | -0.322 | -0.017 | -0.006 | -0.006 | 0 | 0.13 | 0.05 | 0.08 | 0.1 | 0.14 | 0.11 |
| 11 | 44081716 | rs178501 | 0.705 | -0.256 | -0.279 | -0.545 | -0.046 | -0.014 | -0.014 | 0 | 0.35 | 0.2 | 0.34 | 0.62 | 0.45 | 0.5 |
| 11 | 44082931 | rs178503 | 0.225 | -0.244 | -0.218 | -0.527 | -0.001 | 0 | 0 | 0 | 0.42 | 0.44 | 0.39 | 0.63 | 0.51 | 0.52 |
| 11 | 44087311 | rs178512 | 0.765 | 0.647 | -0.598 | 0.365 | 0.056 | 0.046 | 0.046 | 0 | 0.15 | 0.24 | 0.24 | 0.36 | 0.27 | 0.42 |
| 11 | 44089161 | rs178514 | -1.325 | -0.362 | -1.599 | -0.327 | 0.013 | 0.056 | 0.056 | 0 | 0.73 | 0.99 | 0.6 | 0.87 | 0.82 | 0.77 |
| 11 | 44091399 | rs7951555 | 0.81 | -0.121 | -1.328 | -0.346 | 0.012 | -0.046 | -0.046 | 0 | 0.2 | 0.1 | 0.08 | 0.3 | 0.17 | 0.16 |
| 11 | 44093297 | rs178519 | -1.077 | -0.305 | -1.351 | -0.795 | 0.026 | 0.137 | 0.137 | 0 | 0.68 | 0.91 | 0.86 | 0.92 | 0.76 | 0.74 |
| 11 | 44094416 | rs7935243 | -0.283 | -0.18 | -0.7 | -0.92 | 0 | -0.006 | -0.006 | 0 | 0.34 | 0.36 | 0.24 | 0.18 | 0.37 | 0.22 |
| 11 | 44094621 | rs3094390 | 1.065 | 0.108 | -0.167 | -0.418 | 0.012 | 0.144 | 0.144 | 0 | 0.11 | 0.22 | 0.09 | 0.31 | 0.11 | 0.2 |
| 11 | 44098000 | rs178524 | 0.51 | -0.291 | -0.383 | -0.851 | -0.024 | -0.042 | -0.042 | 0.253 | 0.22 | 0.09 | 0.51 | 0.41 | 0.28 | 0.38 |
| 11 | 44098751 | rs2074041 | 1.247 | -0.35 | -0.348 | -1.127 | 0.007 | 0.014 | 0.014 | 0 | 0.16 | 0.11 | 0.1 | 0.12 | 0.12 | 0.11 |
| 11 | 44098985 | rs2074043 | -0.578 | -0.425 | 0.219 | -1.234 | 0 | 0.001 | 0.001 | 0 | 0.44 | 0.56 | 0.31 | 0.47 | 0.46 | 0.38 |
| 11 | 44099588 | rs3134907 | 0.369 | -0.422 | 0.304 | -1.222 | 0.009 | 0.025 | 0.025 | 0 | 0.27 | 0.33 | 0.21 | 0.36 | 0.34 | 0.28 |
| 11 | 44099858 | rs178525 | -1.497 | -0.389 | 0.299 | -1.184 | 0.029 | 0.064 | 0.064 | 0 | 0.67 | 0.83 | 0.84 | 0.92 | 0.75 | 0.79 |
| 11 | 44101807 | rs113080 | -1.433 | -0.359 | 0.131 | -1.135 | 0 | -0.011 | -0.011 | 0 | 0.67 | 0.65 | 0.83 | 0.87 | 0.74 | 0.78 |
| 11 | 44104981 | rs3107275 | -0.438 | -0.358 | 0.145 | -1.025 | -0.002 | -0.004 | -0.004 | 0.028 | 0.43 | 0.41 | 0.3 | 0.47 | 0.46 | 0.35 |
| 11 | 44110690 | rs11037858 | 0.764 | -0.267 | -0.694 | -1.169 | -0.047 | 0.059 | 0.059 | 0 | 0.19 | 0.04 | 0.12 | 0.05 | 0.25 | 0.06 |
| 11 | 44115742 | rs3814767 | 0.405 | -0.494 | -0.503 | -1.197 | -0.006 | -0.009 | -0.009 | 6.079 | 0.23 | 0.17 | 0.5 | 0.4 | 0.28 | 0.4 |
| 11 | 44117122 | rs3923808 | -0.526 | -0.707 | -0.4 | -0.741 | 0.052 | 0.151 | 0.151 | 0 | 0.47 | 0.19 | 0.68 | 0.74 | 0.61 | 0.54 |
| 11 | 44126330 | rs7938320 | 0.234 | -0.836 | -0.066 | -0.697 | -0.032 | -0.044 | -0.044 | 0 | 0.19 | 0.35 | 0.11 | 0.14 | 0.14 | 0.19 |
| 11 | 44131354 | rs10838235 | -0.572 | -0.807 | -0.048 | -0.694 | 0.004 | 0.013 | 0.013 | 0 | 0.47 | 0.63 | 0.72 | 0.77 | 0.55 | 0.59 |
| 11 | 44134141 | rs7111879 | -0.454 | -1.343 | -0.216 | -0.677 | 0.001 | -0.007 | -0.007 | 0 | 0.46 | 0.47 | 0.71 | 0.75 | 0.57 | 0.55 |
| 11 | 44134916 | rs11037869 | 1.13 | -1.018 | -0.661 | -0.681 | -0.048 | -0.008 | -0.008 | 0 | 0.09 | 0.01 | 0.08 | 0.05 | 0.17 | 0.03 |
| 11 | 44142934 | rs7935138 | 0.819 | -0.995 | -0.583 | -0.961 | -0.03 | -0.038 | -0.038 | 0 | 0.19 | 0.09 | 0.48 | 0.41 | 0.3 | 0.41 |
| 11 | 44182017 | rs3935072 | -0.319 | -1.685 | -0.167 | -1.409 | -0.004 | 0.006 | 0.006 | 0 | 0.37 | 0.39 | 0.31 | 0.45 | 0.38 | 0.23 |
| 11 | 44198964 | rs903511 | -1.294 | -1.755 | -0.047 | -1.52 | -0.01 | -0.049 | -0.049 | 0.061 | 0.86 | 0.53 | 0.91 | 0.95 | 0.91 | 0.84 |
| 11 | 44207456 | rs2067787 | 0.272 | -1.91 | -0.134 | -1.673 | -0.014 | 0.035 | 0.035 | 0 | 0.26 | 0.18 | 0.17 | 0.05 | 0.33 | 0.07 |
| 11 | 44219286 | rs4755233 | 0.052 | -1.904 | -0.161 | -1.625 | 0.014 | 0.024 | 0.024 | 0.03 | 0.64 | 0.48 | 0.45 | 0.53 | 0.62 | 0.44 |
| 11 | 44238178 | rs2863054 | 0.423 | -1.775 | 0.268 | -1.532 | 0.002 | 0 | 0 | 0 | 0.53 | 0.46 | 0.38 | 0.39 | 0.55 | 0.35 |
| 11 | 44248728 | rs2034716 | -2.241 | -1.877 | 0.336 | -1.624 | -0.002 | -0.02 | -0.02 | 0 | 0.86 | 0.86 | 0.95 | 0.98 | 0.89 | 0.87 |
| 11 | 44249325 | rs869915 | 1.624 | -1.908 | 0.339 | -1.61 | -0.03 | -0.043 | -0.043 | 0 | 0.19 | 0.06 | 0.46 | 0.42 | 0.27 | 0.4 |
| 11 | 44250895 | rs16937862 | 1.683 | -1.918 | 0.41 | -1.612 | -0.007 | -0.004 | -0.004 | 0 | 0.12 | 0.15 | 0.07 | 0.11 | 0.06 | 0.11 |

| | | | | | | | | | | | | | | | | |
|---|---|---|---|---|---|---|---|---|---|---|---|---|---|---|---|---|
| 11 | 44252958 | rs11037908 | 2.661 | -2.155 | 0.21 | -1.832 | 0.012 | 0.081 | 0.081 | 0 | 0.2 | 0.03 | 0.07 | 0.07 | 0.14 | 0.18 |
| 11 | 44253403 | rs1113132 | 1.787 | -1.918 | 0.393 | -1.637 | -0.029 | -0.041 | -0.041 | 0 | 0.18 | 0.06 | 0.46 | 0.43 | 0.27 | 0.39 |
| 11 | 44255614 | rs11037909 | 1.678 | -1.888 | 0.364 | -1.622 | -0.003 | 0 | 0 | 0 | 0.19 | 0.11 | 0.46 | 0.43 | 0.27 | 0.4 |
| 11 | 44260804 | rs11037911 | 1.359 | -1.78 | 0.201 | -1.758 | -0.002 | -0.001 | -0.001 | 0 | 0.18 | 0.19 | 0.13 | 0.04 | 0.25 | 0.04 |
| 11 | 44272522 | rs7122149 | 1.163 | -1.821 | 0.149 | -1.569 | -0.002 | 0.075 | 0.075 | 0 | 0.34 | 0.14 | 0.22 | 0.18 | 0.3 | 0.27 |
| 11 | 44273424 | rs7126447 | 2.494 | -1.981 | 0.556 | -1.602 | -0.011 | -0.04 | -0.04 | 0.171 | 0.1 | 0.15 | 0.05 | 0 | 0.08 | 0.02 |
| 11 | 44273442 | rs7126360 | -1.583 | -1.71 | 0.361 | -1.404 | -0.013 | -0.047 | -0.047 | 0 | 0.85 | 0.73 | 0.92 | 0.95 | 0.89 | 0.85 |
| 11 | 44283722 | rs897005 | 0.064 | -1.541 | -0.105 | -1.281 | -0.008 | -0.006 | -0.006 | 0 | 0.71 | 0.71 | 0.81 | 0.75 | 0.65 | 0.77 |
| 11 | 44284705 | rs7942612 | 0.947 | -1.305 | -0.412 | -0.934 | 0.009 | -0.01 | -0.01 | 0 | 0.13 | 0.09 | 0.07 | 0.29 | 0.07 | 0.17 |
| 11 | 44285887 | rs4755797 | -0.556 | -1.327 | -0.101 | -1.31 | 0.027 | -0.074 | -0.074 | 0 | 0.74 | 0.52 | 0.81 | 0.93 | 0.69 | 0.94 |
| 11 | 44286566 | rs3802805 | 0.843 | -1.688 | -0.628 | -1.351 | -0.03 | -0.061 | -0.061 | 0 | 0.27 | 0.12 | 0.52 | 0.53 | 0.33 | 0.41 |
| 11 | 44287048 | rs11037921 | 0.037 | -1.331 | -0.147 | -1.365 | 0.005 | 0.014 | 0.014 | 0 | 0.28 | 0.32 | 0.21 | 0.35 | 0.32 | 0.23 |
| 11 | 44287292 | rs7925657 | -0.35 | -1.555 | -0.458 | -1.544 | -0.002 | 0 | 0 | 0 | 0.42 | 0.39 | 0.26 | 0.38 | 0.43 | 0.36 |
| 11 | 44287681 | rs4755800 | 0.092 | -1.218 | -0.365 | -0.792 | 0 | 0.002 | 0.002 | 0 | 0.62 | 0.55 | 0.75 | 0.63 | 0.62 | 0.76 |
| 11 | 44289071 | rs12419361 | 1.31 | -0.548 | -0.898 | -1.568 | 0.022 | 0.021 | 0.021 | 0.029 | 0.13 | 0.05 | 0.09 | 0.14 | 0.12 | 0.14 |
| 11 | 44290066 | rs12291270 | 0.264 | 0.045 | -0.564 | -1.415 | -0.026 | 0.01 | 0.01 | 0 | 0.14 | 0.11 | 0.11 | 0.07 | 0.24 | 0.06 |
| 11 | 44290126 | rs7108942 | -0.179 | -0.255 | -0.406 | -1.514 | -0.05 | -0.108 | -0.108 | 0 | 0.64 | 0.55 | 0.78 | 0.88 | 0.76 | 0.8 |
| 11 | 44291484 | rs7949067 | 0.147 | -0.027 | -0.569 | -1.083 | 0 | -0.017 | -0.017 | 0.917 | 0.51 | 0.5 | 0.67 | 0.81 | 0.52 | 0.74 |
| 11 | 44292747 | rs12288455 | -0.852 | -0.552 | -0.194 | -1.016 | -0.028 | -0.124 | -0.124 | 0 | 0.22 | 0.34 | 0.13 | 0.01 | 0.16 | 0.17 |
| 11 | 44293434 | rs10838248 | -0.5 | -1.391 | -0.206 | -0.462 | 0.001 | 0 | 0 | 0.064 | 0.31 | 0.27 | 0.22 | 0.28 | 0.21 | 0.21 |
| 11 | 44293529 | rs10838249 | -0.57 | -1.481 | -0.089 | -0.5 | -0.011 | -0.002 | -0.002 | 1.031 | 0.31 | 0.3 | 0.23 | 0.28 | 0.24 | 0.2 |
| 11 | 44296002 | rs7927029 | -1.063 | -0.695 | 0.134 | -0.892 | 0 | -0.004 | -0.004 | 4.265 | 0.56 | 0.51 | 0.65 | 0.36 | 0.56 | 0.31 |
| 11 | 44296591 | rs10769027 | -0.316 | -0.771 | 0.303 | -1.019 | -0.011 | -0.027 | -0.027 | 0.667 | 0.95 | 0.86 | 0.98 | 1 | 0.95 | 0.98 |
| 11 | 44296828 | rs11037927 | -0.8 | -0.673 | 0.184 | -1.189 | 0.004 | 0.035 | 0.035 | 0.041 | 0.5 | 0.32 | 0.6 | 0.36 | 0.52 | 0.26 |
| 11 | 44298621 | rs10769029 | 0.947 | -1.107 | 0.061 | -0.685 | -0.03 | -0.014 | -0.014 | 0 | 0.8 | 0.34 | 0.89 | 0.83 | 0.89 | 0.73 |
| 11 | 44302311 | rs2002991 | 0.141 | -0.474 | 0.041 | -0.18 | 0.031 | 0.018 | 0.018 | 0 | 0.75 | 0.62 | 0.6 | 0.69 | 0.69 | 0.74 |
| 11 | 44304276 | rs12575691 | 1.772 | -0.459 | -0.535 | -0.215 | -0.145 | -0.166 | -0.166 | 1.732 | 0.33 | 0.04 | 0.28 | 0.44 | 0.38 | 0.5 |
| 11 | 44305553 | rs9971516 | 0.497 | -1.17 | -0.483 | -0.334 | 0.02 | 0.063 | 0.063 | 0 | 0.18 | 0.07 | 0.1 | 0 | 0.14 | 0.03 |
| 11 | 44306381 | rs7939723 | -0.199 | -0.815 | -0.255 | -0.016 | -0.055 | -0.016 | -0.016 | 0 | 0.34 | 0.42 | 0.27 | 0.31 | 0.24 | 0.19 |
| 11 | 44306882 | rs4755240 | 0.337 | -0.026 | -0.024 | -0.263 | 0.027 | 0.03 | 0.03 | 0.538 | 0.79 | 0.56 | 0.64 | 0.75 | 0.74 | 0.77 |
| 11 | 44311968 | rs10838251 | 0.759 | 0.114 | -0.48 | -0.076 | 0.001 | 0.001 | 0.001 | 0 | 0.73 | 0.6 | 0.59 | 0.73 | 0.68 | 0.9 |
| 11 | 44315390 | rs1828656 | -0.48 | -0.049 | -0.463 | 0.369 | -0.007 | -0.006 | -0.006 | 0 | 0.43 | 0.44 | 0.53 | 0.33 | 0.45 | 0.11 |
| 11 | 44316919 | rs3853292 | 0.361 | 0.051 | -0.634 | 0.443 | -0.004 | 0.005 | 0.005 | 0 | 0.9 | 0.89 | 0.93 | 0.82 | 0.94 | 0.93 |
| 11 | 44317477 | rs10838253 | -0.466 | 0.04 | -0.568 | 0.431 | -0.002 | -0.002 | -0.002 | 0 | 0.43 | 0.53 | 0.53 | 0.33 | 0.44 | 0.11 |
| 11 | 44323193 | rs4755809 | 1.002 | 0.08 | -0.611 | 0.315 | 0.013 | -0.016 | -0.016 | 0 | 0.47 | 0.29 | 0.39 | 0.51 | 0.49 | 0.82 |
| 11 | 44325700 | rs2054029 | -0.584 | 0.262 | -0.605 | 0.285 | -0.003 | -0.003 | -0.003 | 1.927 | 0.42 | 0.39 | 0.52 | 0.31 | 0.45 | 0.11 |
| 11 | 44338255 | rs4568996 | 0.606 | 0.796 | -1.42 | -0.028 | -0.003 | 0.031 | 0.031 | 0.006 | 0.49 | 0.34 | 0.4 | 0.56 | 0.48 | 0.8 |
| 11 | 44340085 | rs12796759 | -0.006 | 1.081 | -2.328 | 0.021 | -0.091 | -0.032 | -0.032 | 0 | 0.16 | 0.01 | 0.34 | 0.13 | 0.28 | 0.05 |
| 11 | 44340618 | rs1454075 | -0.546 | 0.958 | -1.925 | 0.126 | -0.001 | 0.034 | 0.034 | 0 | 0.51 | 0.73 | 0.6 | 0.47 | 0.54 | 0.29 |
| 11 | 44343675 | rs11037953 | 0.608 | 1.083 | -2.507 | 0.225 | -0.095 | -0.118 | -0.118 | 0 | 0.39 | 0.03 | 0.33 | 0.53 | 0.37 | 0.65 |
| 11 | 44343856 | rs1869479 | 0.446 | 1.274 | -2.485 | -0.044 | -0.135 | 0.008 | 0.008 | 0.968 | 0.44 | 0.05 | 0.37 | 0.69 | 0.48 | 0.65 |
| 11 | 44344213 | rs4755811 | -0.833 | 1.367 | -2.011 | -0.591 | -0.042 | 0.086 | 0.086 | 0 | 0.29 | 0.14 | 0.2 | 0.03 | 0.35 | 0.14 |
| 11 | 44344634 | rs1377779 | 0.566 | 0.97 | -0.696 | 0.239 | 0.013 | 0.016 | 0.016 | 0.651 | 0.77 | 0.52 | 0.77 | 0.67 | 0.77 | 0.78 |
| 11 | 44348056 | rs2220885 | 0.664 | 0.551 | -1.161 | 0.359 | -0.027 | -0.02 | -0.02 | 0 | 0.53 | 0.23 | 0.4 | 0.49 | 0.52 | 0.56 |
| 11 | 44352394 | rs903514 | -0.504 | 0.236 | -0.916 | 0.177 | -0.05 | -0.01 | -0.01 | 0.009 | 0.35 | 0.18 | 0.52 | 0.29 | 0.42 | 0.3 |
| 11 | 44354596 | rs11037957 | -0.936 | 0.345 | -0.184 | 0.11 | 0 | -0.002 | -0.002 | 0 | 0.46 | 0.51 | 0.67 | 0.35 | 0.52 | 0.41 |
| 11 | 44357711 | rs10838264 | -0.001 | 0.476 | -0.738 | 0.209 | -0.035 | -0.01 | -0.01 | 0 | 0.3 | 0.2 | 0.51 | 0.28 | 0.41 | 0.29 |
| 11 | 44360545 | rs753688 | 0.141 | 0.493 | -0.58 | 0.587 | 0.019 | 0.035 | 0.035 | 0 | 0.48 | 0.21 | 0.29 | 0.65 | 0.43 | 0.55 |
| 11 | 44368892 | rs10838267 | -0.002 | 0.495 | 0.321 | 0.987 | -0.072 | 0.028 | 0.028 | 0.106 | 0.46 | 0.08 | 0.37 | 0.29 | 0.52 | 0.38 |
| 11 | 44369477 | rs3853291 | -0.131 | 0.541 | 0.671 | 1.014 | -0.001 | 0.032 | 0.032 | 0.018 | 0.48 | 0.31 | 0.38 | 0.29 | 0.52 | 0.38 |
| 11 | 44370034 | rs2863089 | -0.339 | 0.614 | 0.275 | 1.045 | -0.002 | 0.028 | 0.028 | 0.213 | 0.53 | 0.7 | 0.41 | 0.36 | 0.54 | 0.44 |
| 11 | 44375944 | rs11601884 | -0.688 | 0.886 | 0.087 | 2.121 | -0.013 | 0.044 | 0.044 | 0 | 0.06 | 0.01 | 0.09 | 0.45 | 0.06 | 0.19 |
| 11 | 44378010 | rs10838269 | -1.572 | 1.295 | 0.102 | 1.339 | -0.017 | 0.008 | 0.008 | 0 | 0.22 | 0.15 | 0.21 | 0.86 | 0.3 | 0.42 |
| 11 | 44384928 | rs10838271 | -1.468 | 1.156 | 0.128 | 1.343 | -0.005 | 0.005 | 0.005 | 0 | 0.21 | 0.14 | 0.2 | 0.86 | 0.26 | 0.41 |
| 11 | 44388505 | rs10769041 | -0.98 | 1.143 | 0.003 | 1.63 | -0.032 | -0.086 | -0.086 | 0 | 0.16 | 0.05 | 0.16 | 0.66 | 0.23 | 0.32 |
| 11 | 44389241 | rs11037969 | -1.015 | 1.191 | 0.208 | 0.35 | 0 | -0.001 | -0.001 | 0 | 0.1 | 0.13 | 0.3 | 0.09 | 0.09 | 0.11 |
| 11 | 44396537 | rs1158267 | 1.081 | 0.909 | 1.204 | 0.029 | 0.004 | -0.045 | -0.045 | 0 | 0.83 | 0.54 | 0.64 | 0.91 | 0.88 | 0.84 |
| 11 | 44398031 | rs922017 | -1.289 | 1.367 | 0.785 | 0.243 | -0.002 | 0.002 | 0.002 | 0 | 0.23 | 0.22 | 0.26 | 0.86 | 0.34 | 0.41 |
| 11 | 44402503 | rs4755821 | 1.125 | 1.06 | 1.175 | 0.276 | 0.037 | -0.041 | -0.041 | 0 | 0.92 | 0.74 | 0.94 | 1 | 0.94 | 0.99 |
| 11 | 44407358 | rs2123824 | -1.111 | 0.965 | 1.121 | 0.585 | -0.003 | 0.045 | 0.045 | 0 | 0.07 | 0.18 | 0.27 | 0.16 | 0.06 | 0.09 |
| 11 | 44421567 | rs10742707 | 1.721 | 1.016 | 1.858 | 0.247 | 0.088 | -0.044 | -0.044 | 0 | 0.9 | 0.55 | 0.89 | 1 | 0.9 | 0.99 |
| 11 | 44426290 | rs1037735 | 1.11 | 1.029 | 1.914 | 0.254 | 0.012 | -0.031 | -0.031 | 0 | 0.93 | 0.9 | 0.92 | 1 | 0.9 | 0.99 |
| 11 | 44430715 | rs897012 | 1.668 | 0.971 | 1.651 | 0.105 | 0.026 | -0.077 | -0.077 | 0 | 0.77 | 0.52 | 0.73 | 0.92 | 0.72 | 0.83 |
| 11 | 44432418 | rs6485523 | 0.9 | 1.072 | 1.059 | -0.003 | -0.001 | -0.017 | -0.017 | 0 | 0.88 | 0.9 | 0.92 | 1 | 0.83 | 0.96 |
| 11 | 44439439 | rs1012329 | -1.622 | 1.029 | 1.092 | -0.01 | -0.001 | -0.021 | -0.021 | 0 | 0.13 | 0.12 | 0.12 | 0 | 0.23 | 0.04 |
| 11 | 44440047 | rs11037982 | -1.735 | 1.006 | 1.146 | -0.021 | 0.005 | -0.034 | -0.034 | 0 | 0.14 | 0.16 | 0.12 | 0 | 0.23 | 0.04 |
| 11 | 44440970 | rs7936636 | -2.425 | 0.865 | 1.156 | 0.056 | -0.002 | 0.002 | 0.002 | 0 | 0.22 | 0.21 | 0.39 | 0.19 | 0.29 | 0.14 |
| 11 | 44445437 | rs11037988 | -1.678 | 0.917 | 1.105 | 0.078 | 0.001 | -0.025 | -0.025 | 0 | 0.14 | 0.18 | 0.12 | 0 | 0.22 | 0.03 |
| 11 | 44449519 | rs4755247 | 1.027 | 0.953 | 1.247 | 0.082 | 0.008 | -0.05 | -0.05 | 0.755 | 0.88 | 0.75 | 0.91 | 1 | 0.83 | 0.97 |

| | | | | | | | | | | | | | | | | |
|---|---|---|---|---|---|---|---|---|---|---|---|---|---|---|---|---|
| 11 | 44450995 | rs898019 | -2.127 | 0.827 | 1.142 | 0.219 | 0.003 | 0.008 | 0.008 | 1.073 | 0.34 | 0.68 | 0.34 | 0.66 | 0.34 | 0.41 |
| 11 | 44451792 | rs1902961 | -0.968 | 0.627 | 1.056 | -0.702 | -0.02 | -0.068 | -0.068 | 0 | 0.37 | 0.43 | 0.59 | 0.87 | 0.33 | 0.5 |
| 11 | 44452378 | rs10769048 | 1.643 | 1.09 | 1.102 | -0.761 | -0.013 | 0.234 | 0.234 | 0 | 0.49 | 0.2 | 0.27 | 0.11 | 0.45 | 0.46 |
| 11 | 44454921 | rs4755249 | -0.848 | 1.144 | 1.091 | -0.737 | 0.028 | -0.043 | -0.043 | 0 | 0.14 | 0.24 | 0.13 | 0.02 | 0.21 | 0.04 |
| 11 | 44456087 | rs1379877 | -0.571 | 1.075 | 1.46 | -0.674 | -0.006 | -0.006 | -0.006 | 1.654 | 0.32 | 0.32 | 0.34 | 0.7 | 0.29 | 0.41 |
| 11 | 44458019 | rs7926989 | -1.945 | 1.065 | 1.392 | -0.736 | 0.004 | 0.092 | 0.092 | 0 | 0.51 | 0.62 | 0.72 | 0.9 | 0.56 | 0.56 |
| 11 | 44459083 | rs11038009 | -1.201 | 1.117 | 1.283 | -0.274 | 0.013 | -0.008 | -0.008 | 0 | 0.16 | 0.23 | 0.34 | 0.19 | 0.22 | 0.11 |
| 11 | 44469452 | rs813632 | -1.248 | 0.992 | 1.219 | -0.215 | 0.003 | 0.001 | 0.001 | 0 | 0.16 | 0.22 | 0.34 | 0.22 | 0.21 | 0.13 |
| 11 | 44472378 | rs782411 | -1.225 | 1.104 | 1.215 | -0.186 | 0.004 | 0.001 | 0.001 | 0 | 0.16 | 0.21 | 0.34 | 0.22 | 0.21 | 0.13 |
| 11 | 44476521 | rs724411 | -0.114 | 0.555 | 0.545 | -1.296 | -0.005 | -0.01 | -0.01 | 1.454 | 0.45 | 0.43 | 0.43 | 0.69 | 0.39 | 0.53 |
| 11 | 44479082 | rs7947798 | 0.399 | 0.167 | 0.74 | -1.318 | 0.01 | 0.014 | 0.014 | 0 | 0.25 | 0.19 | 0.24 | 0.11 | 0.15 | 0.26 |
| 11 | 44480971 | rs1379881 | 0.798 | 1.427 | 0.071 | -1.539 | -0.023 | 0.032 | 0.032 | 0 | 0.06 | 0.01 | 0.11 | 0 | 0.22 | 0.05 |
| 11 | 44484876 | rs1379879 | -0.637 | 1.406 | 0.219 | -1.55 | 0.071 | -0.028 | -0.028 | 0.208 | 0.93 | 0.83 | 0.88 | 1 | 0.78 | 0.95 |
| 11 | 44492699 | rs11038025 | 1.201 | 1.191 | 0.098 | -1.789 | 0.075 | 0.095 | 0.095 | 0 | 0.3 | 0.12 | 0.12 | 0.1 | 0.17 | 0.27 |
| 11 | 44492899 | rs357917 | -0.677 | 1.35 | 0.208 | -1.804 | -0.005 | 0.02 | 0.02 | 0 | 0.62 | 0.62 | 0.76 | 0.9 | 0.61 | 0.69 |
| 11 | 44495910 | rs984144 | 0.61 | 1.536 | 0.152 | -1.761 | 0.003 | -0.006 | -0.006 | 0 | 0.08 | 0.08 | 0.06 | 0.02 | 0.12 | 0.03 |
| 11 | 44496100 | rs357924 | -0.552 | 1.442 | 0.143 | -1.723 | 0.089 | -0.035 | -0.035 | 0 | 0.92 | 0.8 | 0.87 | 1 | 0.77 | 0.95 |
| 11 | 44501126 | rs357938 | -1.31 | 0.955 | 0.455 | -1.823 | 0.007 | 0.002 | 0.002 | 0 | 0.69 | 0.72 | 0.86 | 0.91 | 0.83 | 0.75 |
| 11 | 44502787 | rs357936 | -0.286 | 1.624 | 0.431 | -1.307 | 0 | 0.001 | 0.001 | 0 | 0.13 | 0.14 | 0.31 | 0.21 | 0.17 | 0.14 |
| 11 | 44506198 | rs6485532 | 0.786 | 0.836 | -0.017 | -1.757 | 0.003 | -0.006 | -0.006 | 0 | 0.62 | 0.37 | 0.46 | 0.72 | 0.49 | 0.67 |
| 11 | 44506360 | rs7937346 | 0.777 | 0.556 | -0.205 | -1.642 | -0.018 | -0.018 | -0.018 | 0 | 0.16 | 0.23 | 0.1 | 0.09 | 0.12 | 0.14 |
| 11 | 44509386 | rs1240751 | -0.965 | 0.734 | -0.361 | -0.98 | 0.006 | -0.007 | -0.007 | 0 | 0.41 | 0.48 | 0.55 | 0.39 | 0.59 | 0.46 |
| 11 | 44513660 | rs6485535 | -0.371 | 0.648 | -0.642 | -0.333 | 0.003 | 0.011 | 0.011 | 0 | 0.53 | 0.6 | 0.44 | 0.34 | 0.59 | 0.47 |
| 11 | 44513752 | rs11607361 | 0.755 | 0.622 | -0.618 | -0.405 | -0.012 | 0.025 | 0.025 | 0 | 0.22 | 0.14 | 0.15 | 0.09 | 0.27 | 0.15 |
| 11 | 44515975 | rs1529530 | -1.262 | 0.588 | -0.805 | -0.221 | -0.001 | 0 | 0 | 0 | 0.63 | 0.63 | 0.73 | 0.52 | 0.66 | 0.64 |
| 11 | 44516235 | rs10838297 | 0.217 | 0.443 | -0.778 | -0.065 | -0.001 | 0.002 | 0.002 | 0 | 0.32 | 0.34 | 0.19 | 0.31 | 0.33 | 0.37 |
| 11 | 44518879 | rs4755840 | -1 | 0.011 | -0.843 | 0.202 | 0 | -0.019 | -0.019 | 0 | 0.62 | 0.68 | 0.74 | 0.52 | 0.66 | 0.62 |
| 11 | 44521492 | rs4755258 | -0.629 | -0.215 | -1.043 | 0.726 | 0.007 | 0.011 | 0.011 | 0 | 0.53 | 0.4 | 0.61 | 0.57 | 0.58 | 0.61 |
| 11 | 44522373 | rs1560295 | -0.205 | -0.407 | -0.601 | 0.927 | 0.001 | -0.002 | -0.002 | 0 | 0.56 | 0.57 | 0.6 | 0.48 | 0.42 | 0.63 |
| 11 | 44522529 | rs1560293 | 0.418 | -0.42 | -0.894 | 0.863 | 0.028 | 0.076 | 0.076 | 0 | 0.45 | 0.21 | 0.31 | 0.21 | 0.37 | 0.37 |
| 11 | 44526825 | rs7111599 | 0.106 | -0.032 | -0.485 | 0.621 | 0.013 | 0.024 | 0.024 | 0 | 0.53 | 0.41 | 0.61 | 0.6 | 0.45 | 0.62 |
| 11 | 44527505 | rs7950827 | 0.063 | 0.145 | -0.186 | 0.041 | -0.007 | -0.012 | -0.012 | 0 | 0.72 | 0.47 | 0.83 | 0.75 | 0.77 | 0.73 |
| 11 | 44528786 | rs12295488 | 0.652 | 0.609 | -0.087 | -0.095 | -0.016 | -0.049 | -0.049 | 4.435 | 0.09 | 0.18 | 0.03 | 0 | 0.03 | 0.01 |
| 11 | 44528947 | rs357922 | -0.949 | 0.573 | -0.686 | -0.051 | 0.057 | 0.092 | 0.092 | 0 | 0.5 | 0.66 | 0.43 | 0.26 | 0.61 | 0.43 |
| 11 | 44535670 | rs782404 | -0.273 | 1.638 | -1.1 | 0.562 | 0.016 | -0.002 | -0.002 | 0.031 | 0.25 | 0.31 | 0.23 | 0.25 | 0.35 | 0.3 |
| 11 | 44541295 | rs11038044 | -0.497 | 1.288 | -1.33 | 0.58 | -0.058 | 0.05 | 0.05 | 0 | 0.13 | 0.02 | 0.14 | 0.02 | 0.19 | 0.03 |
| 11 | 44550047 | rs2863104 | -0.822 | 1.552 | -0.194 | 1.798 | 0.063 | 0.182 | 0.182 | 1.67 | 0.14 | 0.35 | 0.22 | 0.54 | 0.22 | 0.21 |
| 11 | 44553121 | rs10838303 | -1.114 | 1.52 | 0.003 | 1.427 | 0.055 | 0.126 | 0.126 | 1.12 | 0.19 | 0.34 | 0.28 | 0.55 | 0.26 | 0.21 |
| 11 | 44554064 | rs7116194 | 0.288 | 1.396 | -0.739 | 0.332 | -0.034 | 0.027 | 0.027 | 0.463 | 0.29 | 0.13 | 0.37 | 0.22 | 0.35 | 0.33 |
| 11 | 44558374 | rs4755844 | -0.901 | 0.955 | -0.367 | 0.2 | -0.003 | -0.005 | -0.005 | 0 | 0.48 | 0.51 | 0.65 | 0.76 | 0.62 | 0.57 |
| 11 | 44559513 | rs7115558 | 0.556 | 0.981 | -0.367 | 0.184 | 0.008 | 0.01 | 0.01 | 0.194 | 0.52 | 0.53 | 0.35 | 0.24 | 0.38 | 0.43 |
| 11 | 44559898 | rs16937996 | 0.796 | -0.328 | -0.8 | -0.601 | 0.029 | 0.006 | 0.006 | 0 | 0.27 | 0.19 | 0.15 | 0.15 | 0.17 | 0.2 |
| 11 | 44560201 | rs12295769 | 0.119 | -0.078 | -0.761 | -0.874 | -0.005 | -0.058 | -0.058 | 0.026 | 0.26 | 0.32 | 0.23 | 0.08 | 0.23 | 0.26 |
| 11 | 44562224 | rs7127975 | -0.204 | -0.19 | -0.557 | -0.861 | -0.011 | -0.02 | -0.02 | 0 | 0.89 | 0.78 | 0.91 | 0.94 | 0.93 | 0.9 |
| 11 | 44573235 | rs10838304 | -0.698 | -0.131 | -0.88 | -1.165 | 0.001 | 0.037 | 0.037 | 0 | 0.77 | 0.71 | 0.64 | 0.65 | 0.75 | 0.66 |
| 11 | 44576263 | rs1991550 | 0.516 | -0.281 | -0.601 | -1.56 | 0.01 | 0.005 | 0.005 | 0 | 0.5 | 0.24 | 0.47 | 0.57 | 0.54 | 0.44 |
| 11 | 44580741 | rs9943566 | -0.536 | -0.239 | -0.639 | -2.485 | -0.015 | 0.049 | 0.049 | 0 | 0.57 | 0.68 | 0.45 | 0.2 | 0.51 | 0.41 |
| 11 | 44581593 | rs12282890 | 0.329 | -0.259 | -0.565 | -2.584 | -0.005 | -0.047 | -0.047 | 0.05 | 0.46 | 0.41 | 0.32 | 0.14 | 0.4 | 0.38 |
| 11 | 44585933 | rs934178 | -0.527 | -0.213 | -0.683 | -2.524 | -0.018 | 0.061 | 0.061 | 0 | 0.56 | 0.67 | 0.45 | 0.19 | 0.51 | 0.4 |
| 11 | 44588754 | rs12362531 | -0.41 | -0.215 | -0.663 | -2.525 | -0.002 | -0.003 | -0.003 | 0 | 0.55 | 0.54 | 0.44 | 0.19 | 0.51 | 0.41 |
| 11 | 44589841 | rs10838307 | -0.801 | -0.113 | -0.9 | -2.488 | -0.017 | 0.05 | 0.05 | 0 | 0.62 | 0.75 | 0.52 | 0.2 | 0.55 | 0.43 |
| 11 | 44595224 | rs7126437 | 0.82 | -0.178 | -0.831 | -2.545 | -0.024 | -0.091 | -0.091 | 0.039 | 0.3 | 0.47 | 0.29 | 0.11 | 0.27 | 0.2 |
| 11 | 44596817 | rs17541041 | 1.179 | 0.335 | -0.65 | -2.193 | 0 | -0.018 | -0.018 | 0 | 0.17 | 0.14 | 0.06 | 0.03 | 0.16 | 0.17 |
| 11 | 44598235 | rs7123132 | -0.377 | 0.38 | -0.85 | -1.844 | -0.007 | 0.042 | 0.042 | 0 | 0.59 | 0.73 | 0.52 | 0.2 | 0.54 | 0.4 |
| 11 | 44598323 | rs7109092 | -0.377 | 0.38 | -0.85 | -1.844 | -0.007 | 0.042 | 0.042 | 0 | 0.59 | 0.73 | 0.52 | 0.2 | 0.54 | 0.4 |
| 11 | 44603328 | rs730129 | 0.366 | 0.629 | -0.702 | -1.589 | 0.01 | -0.039 | -0.039 | 0.211 | 0.71 | 0.59 | 0.74 | 0.85 | 0.67 | 0.77 |
| 11 | 44604181 | rs2863107 | -0.291 | 0.846 | -1.015 | -1.211 | 0 | 0.01 | 0.01 | 8.365 | 0.87 | 0.87 | 0.85 | 0.94 | 0.89 | 0.96 |
| 11 | 44607332 | rs1981992 | 0.085 | 1.223 | -1.464 | -1.57 | -0.046 | 0.085 | 0.085 | 0.432 | 0.34 | 0.12 | 0.48 | 0.81 | 0.4 | 0.51 |
| 11 | 44613671 | rs12364180 | 0.454 | 0.907 | -1.547 | -2.035 | 0.01 | 0.09 | 0.09 | 0.121 | 0.29 | 0.13 | 0.16 | 0.05 | 0.23 | 0.24 |
| 11 | 44614583 | rs7932712 | 0.145 | 1.081 | -1.163 | -1.286 | -0.002 | 0.003 | 0.003 | 0 | 0.61 | 0.63 | 0.73 | 0.63 | 0.63 | 0.6 |
| 11 | 44615178 | rs17613700 | 1.721 | 1.048 | -1.412 | -1.351 | 0.015 | 0.039 | 0.039 | 0.006 | 0.19 | 0.09 | 0.06 | 0.03 | 0.13 | 0.23 |
| 11 | 44615430 | rs6485550 | 0.798 | 0.93 | -1.04 | -1.425 | -0.011 | 0.024 | 0.024 | 0 | 0.32 | 0.27 | 0.32 | 0.09 | 0.39 | 0.24 |
| 11 | 44615699 | rs7129118 | 0.064 | 1.243 | -0.799 | -0.88 | 0.131 | 0.196 | 0.196 | 0.824 | 0.1 | 0.31 | 0.1 | 0.29 | 0.17 | 0.14 |
| 11 | 44616829 | rs891307 | -1.091 | 0.621 | -1.618 | -0.799 | 0.021 | 0.022 | 0.022 | 12.984 | 0.89 | 0.96 | 0.88 | 0.94 | 0.93 | 0.97 |
| 11 | 44617142 | rs10838312 | 0.34 | 0.759 | -1.655 | -0.674 | -0.025 | -0.14 | -0.14 | 1.581 | 0.29 | 0.06 | 0.39 | 0.53 | 0.28 | 0.4 |
| 11 | 44623897 | rs10838313 | 0.736 | 0.901 | -1.456 | -1.011 | -0.079 | 0.036 | 0.036 | 0 | 0.15 | 0.01 | 0.21 | 0.06 | 0.23 | 0.13 |
| 11 | 44627712 | rs3781751 | 0.221 | 1.473 | -0.832 | -0.407 | 0.023 | 0.045 | 0.045 | 4.307 | 0.16 | 0.17 | 0.17 | 0.3 | 0.25 | 0.06 |
| 11 | 44630049 | rs3781753 | -0.734 | 0.779 | -1.265 | -0.859 | -0.008 | 0.166 | 0.166 | 0 | 0.56 | 0.81 | 0.65 | 0.86 | 0.55 | 0.55 |
| 11 | 44631655 | rs11601257 | -0.337 | 0.601 | -1.311 | -0.718 | -0.004 | -0.01 | -0.01 | 0 | 0.37 | 0.65 | 0.31 | 0.39 | 0.41 | 0.14 |

| | | | | | | | | | | | | | | | | |
|---|---|---|---|---|---|---|---|---|---|---|---|---|---|---|---|---|
| 11 | 44632954 | rs919445 | 1.032 | -0.118 | -1.686 | -0.25 | -0.024 | -0.117 | -0.117 | 0.554 | 0.18 | 0.05 | 0.32 | 0.48 | 0.15 | 0.41 |
| 11 | 44633068 | rs3781759 | 0.538 | -0.408 | -1.301 | -0.293 | -0.016 | -0.027 | -0.027 | 0.426 | 0.2 | 0.28 | 0.13 | 0.08 | 0.16 | 0.08 |
| 11 | 44636547 | rs7935166 | 0.787 | 0.031 | -1.96 | 0.332 | -0.033 | -0.133 | -0.133 | 0.088 | 0.18 | 0.01 | 0.32 | 0.47 | 0.13 | 0.4 |
| 11 | 44636839 | rs883752 | 0.628 | 0.418 | -1.704 | 0.149 | -0.061 | -0.064 | -0.064 | 0 | 0.25 | 0.1 | 0.17 | 0.27 | 0.34 | 0.28 |
| 11 | 44640016 | rs2303861 | 0.1 | -0.069 | -0.698 | -0.087 | 0.005 | -0.019 | -0.019 | 0 | 0.76 | 0.75 | 0.71 | 0.87 | 0.72 | 0.8 |
| 11 | 44640100 | rs12421108 | -0.628 | -0.37 | -0.91 | -0.307 | 0.017 | 0.011 | 0.011 | 7.689 | 0.14 | 0.09 | 0.08 | 0.01 | 0.08 | 0.03 |
| 11 | 44640268 | rs1139971 | 0.293 | -0.087 | -0.448 | 0.47 | -0.003 | 0.004 | 0.004 | 5.697 | 0.63 | 0.39 | 0.63 | 0.5 | 0.66 | 0.84 |
| 11 | 44644926 | rs11038083 | 0.271 | 0.13 | -0.714 | -0.039 | 0.003 | 0.008 | 0.008 | 1.632 | 0.15 | 0.1 | 0.15 | 0.09 | 0.11 | 0.08 |
| 11 | 44649122 | rs7122659 | 0.323 | 0.214 | -1.407 | -0.222 | -0.014 | 0.014 | 0.014 | 0.558 | 0.18 | 0.05 | 0.09 | 0.13 | 0.15 | 0.27 |
| 11 | 44649133 | rs4755852 | -0.746 | -0.033 | -1.389 | -0.449 | 0.007 | 0.026 | 0.026 | 0.578 | 0.83 | 0.9 | 0.85 | 0.91 | 0.88 | 0.92 |
| 11 | 44649231 | rs4755854 | 0.11 | 0.681 | -1.371 | 0.083 | -0.016 | -0.026 | -0.026 | 0.914 | 0.7 | 0.73 | 0.78 | 0.47 | 0.63 | 0.61 |
| 11 | 44650927 | rs6485554 | -0.057 | 0.538 | -0.939 | -0.287 | 0.01 | 0.075 | 0.075 | 0 | 0.57 | 0.7 | 0.78 | 0.84 | 0.64 | 0.7 |
| 11 | 44652154 | rs10769062 | 0.309 | 0.58 | -0.565 | -0.253 | -0.002 | -0.002 | -0.002 | 0 | 0.57 | 0.4 | 0.7 | 0.49 | 0.55 | 0.57 |
| 11 | 44653664 | rs714679 | 0.513 | -0.511 | -0.348 | -0.025 | 0.003 | 0.002 | 0.002 | 0 | 0.51 | 0.53 | 0.69 | 0.49 | 0.58 | 0.6 |
| 11 | 44654385 | rs7113954 | -0.116 | -0.351 | -0.206 | 0.153 | -0.005 | 0.018 | 0.018 | 0 | 0.2 | 0.23 | 0.1 | 0.36 | 0.14 | 0.13 |
| 11 | 44657500 | rs12420889 | -0.712 | -0.18 | -0.122 | 0.215 | 0.002 | 0.272 | 0.272 | 0.25 | 0.05 | 0.19 | 0.05 | 0.35 | 0.02 | 0.1 |
| 11 | 44664135 | rs10742713 | 0.785 | -0.314 | -0.263 | -0.053 | -0.015 | -0.013 | -0.013 | 0 | 0.39 | 0.27 | 0.36 | 0.45 | 0.54 | 0.52 |
| 11 | 44664975 | rs4604873 | -0.087 | -0.244 | -0.52 | -0.233 | -0.001 | -0.011 | -0.011 | 0 | 0.62 | 0.61 | 0.59 | 0.84 | 0.67 | 0.69 |
| 11 | 44671894 | rs4399316 | -0.437 | -0.192 | -0.642 | -0.204 | 0.022 | 0.07 | 0.07 | 0 | 0.66 | 0.8 | 0.63 | 0.84 | 0.7 | 0.7 |
| 11 | 44674234 | rs10769065 | -0.457 | -0.131 | -0.844 | -0.174 | 0.017 | 0.067 | 0.067 | 0 | 0.66 | 0.81 | 0.62 | 0.83 | 0.69 | 0.7 |
| 11 | 44675888 | rs11821326 | -0.821 | -0.016 | -0.392 | -0.053 | 0.005 | -0.022 | -0.022 | 0 | 0.05 | 0.22 | 0.04 | 0.01 | 0.03 | 0.01 |
| 11 | 44690081 | rs10732499 | -0.387 | 0.204 | -0.744 | 0.059 | -0.003 | -0.005 | -0.005 | 0.081 | 0.57 | 0.66 | 0.52 | 0.56 | 0.55 | 0.53 |
| 11 | 44691091 | rs7932189 | -0.527 | -0.118 | -0.617 | 0.112 | 0.003 | -0.006 | -0.006 | 0 | 0.59 | 0.65 | 0.55 | 0.57 | 0.61 | 0.55 |
| 11 | 44691479 | rs7935663 | -0.653 | -0.99 | -0.301 | -0.171 | -0.009 | -0.008 | -0.008 | 0 | 0.2 | 0.28 | 0.2 | 0.15 | 0.13 | 0.1 |
| 11 | 44698662 | rs7120821 | 0.745 | -1.255 | -0.194 | -0.485 | -0.003 | -0.01 | -0.01 | 0 | 0.75 | 0.77 | 0.83 | 0.9 | 0.8 | 0.94 |
| 11 | 44702928 | rs12294859 | 0.432 | -1.779 | 0.486 | -0.713 | -0.012 | 0.004 | 0.004 | 0 | 0.92 | 0.89 | 0.91 | 0.9 | 0.97 | 0.99 |
| 11 | 44704008 | rs10838330 | -0.533 | -2.654 | 0.966 | -0.546 | -0.026 | 0.069 | 0.069 | 0 | 0.15 | 0.22 | 0.35 | 0.61 | 0.07 | 0.31 |
| 11 | 44704275 | rs10742717 | 1.178 | -2.579 | 1.191 | -0.952 | -0.011 | -0.064 | -0.064 | 0 | 0.66 | 0.54 | 0.76 | 0.89 | 0.7 | 0.91 |
| 11 | 44705461 | rs4589285 | -2.037 | -2.865 | 1.286 | -1.045 | 0.032 | 0.072 | 0.072 | 0 | 0.47 | 0.67 | 0.53 | 0.75 | 0.37 | 0.44 |
| 11 | 44706160 | rs11038106 | -1.881 | -2.826 | 1.301 | -1.073 | 0.058 | 0.102 | 0.102 | 0 | 0.45 | 0.76 | 0.52 | 0.74 | 0.33 | 0.49 |
| 11 | 44710825 | rs10838333 | 1.926 | -3.069 | 1.326 | -1.237 | 0.08 | 0.192 | 0.192 | 0 | 0.49 | 0.26 | 0.4 | 0.16 | 0.66 | 0.5 |
| 11 | 44710934 | rs4755868 | -0.864 | -3.198 | 1.64 | -1.217 | -0.026 | 0.005 | 0.005 | 0 | 0.1 | 0.21 | 0.04 | 0.11 | 0.04 | 0.06 |
| 11 | 44711162 | rs4755869 | -0.064 | -3.29 | 1.722 | -1.268 | -0.018 | 0.037 | 0.037 | 0 | 0.87 | 0.86 | 0.65 | 0.37 | 0.94 | 0.67 |
| 11 | 44713928 | rs11605105 | 1.282 | -3.214 | 1.459 | -1.509 | -0.099 | -0.087 | -0.087 | 9.251 | 0.71 | 0.52 | 0.53 | 0.16 | 0.89 | 0.6 |
| 11 | 44716705 | rs10838336 | 2.387 | -3.027 | 0.82 | -1.487 | -0.002 | 0.26 | 0.26 | 0.123 | 0.45 | 0.1 | 0.37 | 0.15 | 0.62 | 0.48 |
| 11 | 44719357 | rs7933067 | -0.419 | -2.296 | 1.129 | -1.04 | -0.01 | -0.006 | -0.006 | 0.035 | 0.07 | 0.09 | 0.02 | 0.06 | 0.02 | 0.04 |
| 11 | 44719970 | rs4643068 | -0.805 | -1.957 | 1.398 | -0.899 | 0.02 | -0.094 | -0.094 | 0 | 0.18 | 0.27 | 0.12 | 0 | 0.21 | 0.1 |
| 11 | 44725188 | rs7479467 | 0.888 | -2.035 | 0.947 | -0.323 | -0.025 | 0.003 | 0.003 | 0.389 | 0.93 | 0.86 | 0.93 | 0.91 | 0.99 | 0.98 |
| 11 | 44729180 | rs10769071 | 1.48 | -2.934 | 1.446 | -0.972 | -0.076 | -0.097 | -0.097 | 0.03 | 0.71 | 0.5 | 0.84 | 0.9 | 0.89 | 0.9 |
| 11 | 44729754 | rs4237647 | 2.227 | -3.191 | 1.02 | -1.467 | 0.066 | 0.178 | 0.178 | 0.329 | 0.46 | 0.14 | 0.69 | 0.79 | 0.67 | 0.7 |
| 11 | 44732574 | rs4237648 | 1.016 | -3.295 | 1.407 | -1.555 | -0.009 | -0.072 | -0.072 | 0.325 | 0.75 | 0.45 | 0.81 | 0.87 | 0.79 | 0.85 |
| 11 | 44733505 | rs7120723 | 1.041 | -3.122 | 1.443 | -1.249 | 0.011 | 0.019 | 0.019 | 0 | 0.66 | 0.29 | 0.73 | 0.76 | 0.75 | 0.82 |
| 11 | 44734819 | rs4755874 | 0.153 | -3.3 | 1.529 | -1.376 | -0.008 | -0.001 | -0.001 | 0 | 0.92 | 0.86 | 0.92 | 0.9 | 0.96 | 0.97 |
| 11 | 44735776 | rs7951107 | 1.571 | -3.339 | 1.517 | -1.365 | -0.011 | -0.005 | -0.005 | 0.099 | 0.8 | 0.76 | 0.59 | 0.21 | 0.87 | 0.71 |
| 11 | 44737646 | rs6485566 | -1.058 | -3.427 | 1.858 | -1.459 | -0.034 | -0.084 | -0.084 | 0 | 0.35 | 0.46 | 0.19 | 0.11 | 0.23 | 0.24 |
| 11 | 44741200 | rs4322381 | 2.299 | -3.349 | 1.804 | -1.421 | 0.027 | 0.197 | 0.197 | 0.278 | 0.44 | 0.17 | 0.39 | 0.1 | 0.64 | 0.47 |
| 11 | 44749884 | rs4755880 | -0.83 | -2.87 | 1.805 | -1.09 | 0 | 0.047 | 0.047 | 0 | 0.09 | 0.1 | 0.32 | 0.69 | 0.08 | 0.26 |
| 11 | 44758398 | rs4755281 | 0.788 | -2.87 | 2 | -1.078 | -0.039 | -0.035 | -0.035 | 2.076 | 0.82 | 0.68 | 0.89 | 0.9 | 0.94 | 0.91 |
| 11 | 44764946 | rs4755283 | -1.515 | -2.751 | 1.775 | -1.029 | -0.038 | 0.004 | 0.004 | 0.066 | 0.25 | 0.37 | 0.38 | 0.71 | 0.14 | 0.34 |
| 11 | 44776732 | rs10769076 | -0.803 | -2.716 | 1.166 | -1.156 | -0.04 | 0.006 | 0.006 | 0 | 0.2 | 0.36 | 0.41 | 0.77 | 0.1 | 0.34 |
| 11 | 44780925 | rs1665150 | -0.534 | -2.66 | -0.097 | -1.226 | 0.018 | 0.046 | 0.046 | 0 | 0.27 | 0.16 | 0.14 | 0.07 | 0.22 | 0.16 |
| 11 | 44781631 | rs4394833 | -1.156 | -1.768 | -0.355 | -0.869 | 0.024 | 0.196 | 0.196 | 0.184 | 0.56 | 0.83 | 0.61 | 0.88 | 0.36 | 0.53 |
| 11 | 44787201 | rs704664 | 1.023 | -1.009 | 0.804 | 0.11 | 0.07 | 0.181 | 0.181 | 2.211 | 0.91 | 0.49 | 0.92 | 0.78 | 0.95 | 0.95 |
| 11 | 44788115 | rs10838353 | 0.855 | -1.167 | 0.045 | 0.636 | -0.002 | 0.007 | 0.007 | 0 | 0.73 | 0.36 | 0.54 | 0.34 | 0.76 | 0.85 |
| 11 | 44788542 | rs4755890 | 0.099 | -1.162 | -0.055 | 0.663 | -0.042 | 0.03 | 0.03 | 0 | 0.17 | 0.01 | 0.11 | 0.04 | 0.18 | 0.05 |
| 11 | 44796843 | rs751705 | 1.33 | -1.5 | -0.603 | -0.271 | -0.006 | 0.13 | 0.13 | 0 | 0.48 | 0.29 | 0.4 | 0.06 | 0.54 | 0.44 |
| 11 | 44797486 | rs704662 | 0.361 | -1.513 | -0.363 | -0.199 | -0.002 | 0.039 | 0.039 | 0 | 0.18 | 0.07 | 0.13 | 0.04 | 0.18 | 0.05 |
| 11 | 44800106 | rs12787702 | 1.395 | -1.2 | 0.461 | -0.203 | -0.025 | 0.196 | 0.196 | 0 | 0.46 | 0.23 | 0.38 | 0.05 | 0.52 | 0.44 |
| 11 | 44807313 | rs1670308 | -0.92 | -1.108 | 0.859 | 0.225 | 0.023 | 0.259 | 0.259 | 0 | 0.09 | 0.23 | 0.32 | 0.53 | 0.05 | 0.1 |
| 11 | 44811462 | rs11038144 | -0.152 | -1.077 | 0.421 | 0.028 | -0.013 | -0.001 | -0.001 | 0.043 | 0.24 | 0.09 | 0.13 | 0.22 | 0.23 | 0.37 |
| 11 | 44819999 | rs835855 | 0.924 | -1.463 | 0.601 | -0.458 | 0.002 | 0.004 | 0.004 | 0 | 0.61 | 0.4 | 0.51 | 0.12 | 0.69 | 0.52 |
| 11 | 44827034 | rs7120459 | 1.318 | -1.385 | 0.333 | -0.625 | -0.006 | 0.003 | 0.003 | 0 | 0.45 | 0.42 | 0.37 | 0.08 | 0.53 | 0.39 |
| 11 | 44830294 | rs17778931 | 0.284 | -1.5 | 0.04 | -0.672 | -0.037 | 0.106 | 0.106 | 0.035 | 0.21 | 0.01 | 0.07 | 0.06 | 0.19 | 0.34 |
| 11 | 44830973 | rs1670317 | -1.635 | -1.705 | 0.83 | -0.19 | 0.009 | 0.009 | 0.009 | 0 | 0.45 | 0.35 | 0.44 | 0.67 | 0.31 | 0.52 |
| 11 | 44834194 | rs2166452 | 2.092 | -0.886 | 0.287 | -1.137 | -0.075 | 0.159 | 0.159 | 0.008 | 0.32 | 0.08 | 0.3 | 0.06 | 0.41 | 0.39 |
| 11 | 44835771 | rs1001292 | -0.182 | -0.831 | 0.654 | -0.775 | 0.006 | 0.026 | 0.026 | 0.007 | 0.27 | 0.13 | 0.37 | 0.84 | 0.23 | 0.48 |
| 11 | 44844974 | rs12790857 | 1.566 | -0.283 | 0.01 | -0.68 | -0.081 | 0.019 | 0.019 | 0 | 0.16 | 0.01 | 0.19 | 0.05 | 0.24 | 0.06 |
| 11 | 44846431 | rs835826 | -1.792 | -0.557 | 0.429 | -0.994 | -0.004 | -0.117 | -0.117 | 1.855 | 0.59 | 0.5 | 0.65 | 0.94 | 0.57 | 0.6 |
| 11 | 44848821 | rs10838364 | -0.752 | -0.671 | 0.292 | -0.812 | 0.013 | 0.024 | 0.024 | 0 | 0.35 | 0.23 | 0.15 | 0.22 | 0.25 | 0.44 |

| | | | | | | | | | | | | | | | | |
|---|---|---|---|---|---|---|---|---|---|---|---|---|---|---|---|---|
| 11 | 44849275 | rs16938094 | -0.237 | -0.527 | 0.283 | -0.717 | 0.002 | 0.003 | 0.003 | 0 | 0.19 | 0.14 | 0.08 | 0.14 | 0.12 | 0.18 |
| 11 | 44849820 | rs835833 | 0.875 | -0.393 | 0.372 | -0.834 | -0.004 | 0.014 | 0.014 | 0 | 0.58 | 0.38 | 0.42 | 0.24 | 0.6 | 0.71 |
| 11 | 44850616 | rs11038169 | -0.172 | -0.558 | -0.099 | -0.683 | 0.003 | 0.005 | 0.005 | 0 | 0.19 | 0.12 | 0.08 | 0.14 | 0.12 | 0.19 |
| 11 | 44851822 | rs11601331 | 2.317 | -0.399 | -0.463 | -0.872 | -0.015 | 0.029 | 0.029 | 0 | 0.06 | 0 | 0.09 | 0 | 0.07 | 0.02 |
| 11 | 44855165 | rs1470849 | 0.217 | -0.34 | -0.162 | -0.887 | 0 | 0 | 0 | 0 | 0.17 | 0.14 | 0.07 | 0.14 | 0.12 | 0.19 |
| 11 | 44856227 | rs7124825 | -0.622 | -0.457 | -0.238 | -1.111 | 0.007 | -0.034 | -0.034 | 2.794 | 0.4 | 0.49 | 0.56 | 0.79 | 0.39 | 0.3 |
| 11 | 44861654 | rs835782 | -1.294 | -0.596 | -0.31 | -1.302 | -0.002 | -0.004 | -0.004 | 0 | 0.83 | 0.84 | 0.93 | 0.93 | 0.88 | 0.77 |
| 11 | 44862229 | rs987632 | 0.212 | -0.625 | -0.153 | -1.305 | -0.001 | -0.002 | -0.002 | 0 | 0.2 | 0.31 | 0.09 | 0.14 | 0.13 | 0.2 |
| 11 | 44863818 | rs835784 | 0.787 | -0.448 | -0.293 | -1.093 | -0.002 | -0.009 | -0.009 | 0 | 0.47 | 0.5 | 0.69 | 0.75 | 0.5 | 0.54 |
| 11 | 44866986 | rs7110434 | 1.465 | 0.081 | -0.734 | -0.685 | -0.085 | 0.043 | 0.043 | 0 | 0.17 | 0.01 | 0.16 | 0.04 | 0.25 | 0.05 |
| 11 | 44871181 | rs2218993 | -0.462 | -0.197 | -0.268 | -0.379 | -0.014 | 0.052 | 0.052 | 0 | 0.24 | 0.47 | 0.42 | 0.69 | 0.13 | 0.22 |
| 11 | 44872253 | rs835799 | 0.421 | 0.07 | -0.715 | -0.915 | 0.02 | -0.052 | -0.052 | 1.469 | 0.26 | 0.35 | 0.28 | 0.12 | 0.36 | 0.39 |
| 11 | 44872419 | rs835800 | -1.221 | -0.352 | -0.76 | -0.882 | -0.033 | -0.002 | -0.002 | 0 | 0.63 | 0.48 | 0.52 | 0.31 | 0.75 | 0.59 |
| 11 | 44872877 | rs4755903 | 1.192 | 0.173 | -1.107 | -1.283 | 0.03 | -0.054 | -0.054 | 0 | 0.2 | 0.31 | 0.24 | 0.08 | 0.3 | 0.26 |
| 11 | 44873532 | rs861354 | -0.113 | 0.313 | -1.211 | -1.192 | 0 | 0.006 | 0.006 | 0 | 0.42 | 0.36 | 0.43 | 0.17 | 0.62 | 0.35 |
| 11 | 44874510 | rs17784515 | 1.595 | 0.643 | -1.698 | -1.014 | -0.099 | 0.053 | 0.053 | 0 | 0.18 | 0.01 | 0.17 | 0.04 | 0.3 | 0.05 |
| 11 | 44874638 | rs835808 | -0.694 | 0.089 | -1.219 | -0.765 | 0.032 | 0.051 | 0.051 | 0 | 0.63 | 0.74 | 0.84 | 0.89 | 0.75 | 0.55 |
| 11 | 44880092 | rs7935135 | 0.61 | 0.789 | -1.307 | -0.043 | -0.016 | -0.003 | -0.003 | 0 | 0.16 | 0.08 | 0.09 | 0.15 | 0.15 | 0.17 |
| 11 | 44880141 | rs11038178 | 0.09 | 0.855 | -0.909 | -0.052 | -0.001 | -0.021 | -0.021 | 2.335 | 0.21 | 0.19 | 0.17 | 0.06 | 0.24 | 0.23 |
| 11 | 44881997 | rs12802362 | 0.181 | 1.22 | -0.929 | 0.766 | 0.001 | -0.001 | -0.001 | 0 | 0.18 | 0.21 | 0.18 | 0.1 | 0.26 | 0.06 |
| 11 | 44886199 | rs835750 | -0.206 | -0.19 | -1.088 | 1.716 | 0.01 | -0.001 | -0.001 | 0 | 0.45 | 0.42 | 0.34 | 0.42 | 0.3 | 0.46 |
| 11 | 44886554 | rs1466199 | -0.15 | -0.093 | -1.554 | 1.216 | -0.008 | 0.002 | 0.002 | 0 | 0.43 | 0.27 | 0.28 | 0.5 | 0.45 | 0.47 |
| 11 | 44888411 | rs835744 | 0.349 | 0.189 | -2.315 | 0.506 | -0.116 | -0.007 | -0.007 | 0.007 | 0.35 | 0.07 | 0.37 | 0.32 | 0.48 | 0.43 |
| 11 | 44890010 | rs11602990 | 1.403 | 0.177 | -2.337 | 0.212 | -0.084 | 0.077 | 0.077 | 0 | 0.14 | 0.02 | 0.12 | 0 | 0.26 | 0.05 |
| 11 | 44890761 | rs835743 | 0.887 | 0.467 | -2.547 | 0.075 | -0.031 | -0.003 | -0.003 | 0 | 0.24 | 0.06 | 0.12 | 0.19 | 0.21 | 0.24 |
| 11 | 44891374 | rs11038182 | 0.756 | 0.766 | -2.001 | 0.356 | -0.014 | -0.021 | -0.021 | 0.003 | 0.11 | 0.07 | 0.23 | 0.24 | 0.23 | 0.2 |
| 11 | 44892829 | rs11604664 | 0.234 | 0.636 | -1.825 | -0.26 | -0.043 | 0.052 | 0.052 | 0.006 | 0.18 | 0.1 | 0.14 | 0.03 | 0.29 | 0.09 |
| 11 | 44897700 | rs1993205 | -0.739 | -0.179 | -1.408 | -0.322 | -0.006 | -0.034 | -0.034 | 0 | 0.58 | 0.51 | 0.78 | 0.8 | 0.68 | 0.76 |
| 11 | 44900373 | rs4755910 | -0.178 | 0.85 | -1.738 | 0.344 | -0.063 | 0.008 | 0.008 | 1.122 | 0.26 | 0.17 | 0.42 | 0.22 | 0.52 | 0.26 |
| 11 | 44901903 | rs835839 | 0.711 | 0.715 | -1.609 | 0.429 | -0.014 | -0.011 | -0.011 | 0.837 | 0.14 | 0.06 | 0.09 | 0.2 | 0.16 | 0.14 |
| 11 | 44907443 | rs3740792 | 0.417 | 0.024 | -1.42 | 0.897 | -0.018 | -0.057 | -0.057 | 1.297 | 0.1 | 0.01 | 0.07 | 0.18 | 0.1 | 0.34 |
| 11 | 44909493 | rs835768 | 0.172 | 0.585 | -1.198 | 0.066 | -0.087 | 0.067 | 0.067 | 0.449 | 0.3 | 0.11 | 0.24 | 0.2 | 0.43 | 0.19 |
| 11 | 44913682 | rs1670314 | 0.428 | -0.377 | 0.035 | 0.769 | -0.034 | 0.048 | 0.048 | 0 | 0.72 | 0.47 | 0.62 | 0.46 | 0.84 | 0.56 |
| 11 | 44915802 | rs1665139 | 0.365 | -0.258 | -0.231 | 0.807 | -0.041 | 0.074 | 0.074 | 0 | 0.72 | 0.47 | 0.64 | 0.49 | 0.84 | 0.59 |
| 11 | 44918572 | rs835864 | -0.317 | 0.34 | -0.689 | 0.078 | -0.007 | 0.041 | 0.041 | 2.289 | 0.58 | 0.37 | 0.66 | 0.74 | 0.57 | 0.79 |
| 11 | 44920626 | rs10838382 | -0.46 | 0.026 | 0.071 | 0.311 | 0.026 | 0.139 | 0.139 | 0 | 0.05 | 0.23 | 0.11 | 0.15 | 0.02 | 0.08 |
| 11 | 44921283 | rs2863131 | 0.034 | 0.419 | -0.911 | -0.039 | -0.129 | -0.112 | -0.112 | 2.469 | 0.27 | 0.03 | 0.42 | 0.63 | 0.38 | 0.56 |
| 11 | 44923975 | rs10838384 | -0.735 | -0.265 | -0.518 | -0.696 | -0.014 | -0.022 | -0.022 | 0 | 0.29 | 0.33 | 0.29 | 0.15 | 0.21 | 0.23 |
| 11 | 44926820 | rs7118905 | -0.138 | 0.075 | -0.725 | -0.724 | 0 | -0.016 | -0.016 | 0.003 | 0.11 | 0.08 | 0.07 | 0 | 0.11 | 0.05 |
| 11 | 44930635 | rs4755296 | -0.723 | -0.131 | -0.072 | -0.898 | 0 | 0.022 | 0.022 | 0 | 0.49 | 0.61 | 0.7 | 0.83 | 0.49 | 0.77 |
| 11 | 44935693 | rs835984 | 0.655 | -0.445 | -0.358 | -1.432 | -0.019 | 0.129 | 0.129 | 0.342 | 0.39 | 0.2 | 0.25 | 0 | 0.42 | 0.14 |
| 11 | 44937943 | rs1552299 | -0.58 | -0.451 | 0.174 | -0.999 | -0.003 | 0.08 | 0.08 | 0 | 0.49 | 0.62 | 0.69 | 0.79 | 0.48 | 0.76 |
| 11 | 44940828 | rs2291334 | 0.588 | -0.057 | -1.21 | -0.805 | -0.095 | -0.128 | -0.128 | 0 | 0.22 | 0.03 | 0.42 | 0.56 | 0.33 | 0.52 |
| 11 | 44940873 | rs11038214 | -0.584 | -0.414 | -0.325 | -0.699 | 0 | -0.017 | -0.017 | 0 | 0.19 | 0.25 | 0.16 | 0.06 | 0.14 | 0.15 |
| 11 | 44949238 | rs835989 | 0.337 | -0.589 | -0.607 | -0.318 | 0.009 | -0.016 | -0.016 | 0 | 0.82 | 0.9 | 0.93 | 0.79 | 0.89 | 0.89 |
| 11 | 44952930 | rs3740789 | 0.388 | -0.558 | -0.372 | -0.469 | 0.006 | 0.026 | 0.026 | 0 | 0.08 | 0.03 | 0.1 | 0.02 | 0.06 | 0.08 |
| 11 | 44959620 | rs12419970 | 1.125 | -0.126 | -1.369 | -1.231 | -0.102 | 0.154 | 0.154 | 0 | 0.24 | 0.02 | 0.21 | 0 | 0.35 | 0.09 |
| 11 | 44961128 | rs10769103 | 0.646 | -0.605 | -1.634 | -1.126 | -0.02 | -0.023 | -0.023 | 0.673 | 0.34 | 0.18 | 0.44 | 0.64 | 0.41 | 0.67 |
| 11 | 44961840 | rs900851 | 0.315 | -1.073 | -0.955 | -1.285 | -0.032 | -0.035 | -0.035 | 0 | 0.7 | 0.57 | 0.86 | 0.81 | 0.82 | 0.88 |
| 11 | 44965372 | rs16933 | 0.953 | -0.132 | -1.05 | 0.039 | -0.014 | 0.006 | 0.006 | 0 | 0.3 | 0.21 | 0.2 | 0.28 | 0.37 | 0.12 |
| 11 | 44968756 | rs4755920 | -0.265 | 0.034 | -1.506 | 0.045 | 0.029 | -0.008 | -0.008 | 0 | 0.49 | 0.22 | 0.53 | 0.5 | 0.44 | 0.65 |
| 11 | 44973082 | rs1994209 | 0.305 | -0.211 | -1.709 | 0.129 | -0.06 | -0.048 | -0.048 | 0 | 0.27 | 0.11 | 0.54 | 0.62 | 0.41 | 0.49 |
| 11 | 44974800 | rs1872276 | 0.295 | -0.438 | -2.136 | 0.182 | -0.169 | -0.184 | -0.184 | 0.24 | 0.28 | 0.03 | 0.59 | 0.5 | 0.45 | 0.51 |
| 11 | 44976681 | rs11038225 | 0.295 | -0.65 | -1.627 | -0.056 | 0.057 | 0.063 | 0.063 | 1.177 | 0.22 | 0.06 | 0.09 | 0.1 | 0.1 | 0.25 |
| 11 | 44978176 | rs4755921 | 0.218 | 0.177 | -1.253 | 0.444 | 0.003 | 0.029 | 0.029 | 0 | 0.53 | 0.34 | 0.3 | 0.34 | 0.46 | 0.37 |
| 11 | 44982802 | rs4755922 | 0.507 | -0.201 | -1.846 | 0.731 | 0.008 | -0.008 | -0.008 | 0 | 0.63 | 0.09 | 0.67 | 0.7 | 0.71 | 0.66 |
| 11 | 44985620 | rs7106313 | -1.054 | -0.785 | -0.909 | 1 | -0.007 | -0.001 | -0.001 | 0.976 | 0.31 | 0.63 | 0.28 | 0.29 | 0.24 | 0.19 |
| 11 | 44993507 | rs10838394 | -1.538 | -1.381 | -1.182 | 1.597 | 0.078 | -0.045 | -0.045 | 0 | 0.42 | 0.84 | 0.36 | 0.54 | 0.31 | 0.4 |
| 11 | 44994060 | rs10838395 | -1.301 | -1.526 | -0.366 | 1.501 | 0.013 | -0.023 | -0.023 | 0.368 | 0.28 | 0.23 | 0.22 | 0.43 | 0.19 | 0.36 |
| 11 | 44995441 | rs3802785 | -1.073 | -1.548 | -0.333 | 1.324 | 0.024 | 0.023 | 0.023 | 0.95 | 0.25 | 0.14 | 0.2 | 0.22 | 0.18 | 0.21 |
| 11 | 44996996 | rs17181389 | -1.297 | -1.531 | 0.145 | 1.582 | 0.001 | -0.002 | -0.002 | 0 | 0.27 | 0.27 | 0.39 | 0.45 | 0.19 | 0.35 |
| 11 | 44999605 | rs2279359 | -1.248 | -1.549 | -0.014 | 1.491 | 0.023 | -0.031 | -0.031 | 0.067 | 0.27 | 0.17 | 0.39 | 0.42 | 0.19 | 0.27 |
| 11 | 45001728 | rs12291203 | -0.401 | -1.553 | 0.18 | 1.375 | -0.009 | -0.027 | -0.027 | 0 | 0.1 | 0.64 | 0.09 | 0.06 | 0.06 | 0.03 |
| 11 | 45009133 | rs10838397 | -1.684 | -1.531 | -0.067 | 1.438 | 0.057 | -0.043 | -0.043 | 0 | 0.38 | 0.81 | 0.49 | 0.5 | 0.26 | 0.39 |
| 11 | 45013151 | rs11606362 | -0.649 | -1.977 | 0.435 | 1.215 | -0.044 | -0.009 | -0.009 | 0 | 0.19 | 0.33 | 0.11 | 0.23 | 0.12 | 0.15 |
| 11 | 45014551 | rs10838398 | -0.968 | -1.647 | 0.596 | 1.051 | -0.029 | 0.028 | 0.028 | 0 | 0.21 | 0.36 | 0.34 | 0.24 | 0.14 | 0.25 |
| 11 | 45015345 | rs4755302 | -1.189 | -1.599 | 0.619 | 1.05 | -0.044 | 0.04 | 0.04 | 0.013 | 0.23 | 0.5 | 0.35 | 0.24 | 0.14 | 0.25 |
| 11 | 45016421 | rs2902421 | 0.343 | -1.345 | -0.611 | -0.102 | 0.054 | -0.036 | -0.036 | 2.599 | 0.43 | 0.13 | 0.55 | 0.55 | 0.65 | 0.56 |
| 11 | 45023786 | rs4755303 | 0.406 | -0.234 | -0.138 | -1.063 | -0.024 | 0.033 | 0.033 | 0.656 | 0.18 | 0.09 | 0.13 | 0.09 | 0.22 | 0.08 |

| | | | | | | | | | | | | | | | | |
|---|---|---|---|---|---|---|---|---|---|---|---|---|---|---|---|---|
| 11 | 45027780 | rs2863140 | 1.438 | 0.098 | -0.07 | -1.565 | -0.025 | 0.026 | 0.026 | 1.035 | 0.45 | 0.2 | 0.44 | 0.68 | 0.5 | 0.55 |
| 11 | 45030050 | rs2046657 | -0.771 | 0.053 | 0.791 | -1.616 | 0.004 | -0.022 | -0.022 | 0 | 0.8 | 0.7 | 0.85 | 0.91 | 0.78 | 0.92 |
| 11 | 45033168 | rs11038268 | -0.607 | -0.09 | 0.946 | -1.653 | -0.006 | 0.003 | 0.003 | 0.358 | 0.26 | 0.29 | 0.37 | 0.21 | 0.23 | 0.36 |
| 11 | 45044671 | rs1484398 | 1.549 | -0.012 | 0.693 | -1.707 | -0.007 | 0.025 | 0.025 | 0.902 | 0.46 | 0.27 | 0.44 | 0.68 | 0.49 | 0.55 |
| 11 | 45045189 | rs17093 | 0.323 | -0.001 | 0.57 | -1.751 | -0.004 | 0.021 | 0.021 | 0.362 | 0.21 | 0.2 | 0.14 | 0.08 | 0.22 | 0.08 |
| 11 | 45048588 | rs1817653 | 0.592 | -0.044 | 0.753 | -1.399 | -0.021 | -0.041 | -0.041 | 0 | 0.08 | 0.21 | 0.05 | 0.03 | 0.07 | 0.01 |
| 11 | 45049150 | rs901720 | -0.959 | 0.082 | 0.504 | -1.305 | -0.01 | 0.012 | 0.012 | 0 | 0.23 | 0.26 | 0.39 | 0.26 | 0.18 | 0.33 |
| 11 | 45051827 | rs754970 | 0.477 | 0.504 | 0.559 | -1.774 | -0.001 | -0.016 | -0.016 | 0 | 0.57 | 0.54 | 0.55 | 0.83 | 0.6 | 0.6 |
| 11 | 45052398 | rs10838405 | 1.291 | 0.742 | 0.767 | -1.848 | 0.001 | 0.042 | 0.042 | 0.043 | 0.43 | 0.25 | 0.44 | 0.73 | 0.42 | 0.56 |
| 11 | 45058125 | rs1872277 | 0.382 | 0.78 | 0.702 | -2.002 | -0.002 | -0.012 | -0.012 | 0.063 | 0.62 | 0.59 | 0.73 | 0.9 | 0.58 | 0.89 |
| 11 | 45058482 | rs12361918 | -0.231 | 0.742 | 0.743 | -2.058 | -0.004 | -0.02 | -0.02 | 0.146 | 0.15 | 0.24 | 0.1 | 0.1 | 0.15 | 0.06 |
| 11 | 45060507 | rs11038279 | -0.237 | 0.678 | 0.859 | -2.06 | -0.003 | 0.009 | 0.009 | 0.03 | 0.15 | 0.12 | 0.1 | 0.1 | 0.17 | 0.06 |
| 11 | 45062196 | rs900859 | 0.523 | 0.686 | 0.779 | -2.068 | -0.004 | 0.002 | 0.002 | 4.419 | 0.59 | 0.55 | 0.72 | 0.9 | 0.55 | 0.87 |
| 11 | 45062339 | rs7125249 | -0.06 | 0.656 | 0.856 | -1.981 | -0.004 | 0.011 | 0.011 | 1.328 | 0.16 | 0.19 | 0.28 | 0.17 | 0.12 | 0.33 |
| 11 | 45063947 | rs10838407 | 0.908 | 0.673 | 0.707 | -2.005 | -0.004 | -0.006 | -0.006 | 0 | 0.68 | 0.53 | 0.62 | 0.74 | 0.71 | 0.61 |
| 11 | 45064490 | rs1484404 | 1.111 | 0.651 | 0.661 | -1.869 | -0.001 | -0.012 | -0.012 | 0.534 | 0.45 | 0.46 | 0.49 | 0.73 | 0.46 | 0.55 |
| 11 | 45068136 | rs9645639 | 0.281 | 0.531 | 0.15 | -1.865 | -0.008 | 0.02 | 0.02 | 0 | 0.61 | 0.67 | 0.57 | 0.84 | 0.59 | 0.61 |
| 11 | 45071979 | rs10769108 | 1.151 | 0.698 | -0.226 | -2.466 | 0.026 | 0.243 | 0.243 | 2.736 | 0.51 | 0.23 | 0.65 | 0.89 | 0.44 | 0.8 |
| 11 | 45073858 | rs1351848 | 1.239 | 0.85 | -0.311 | -2.377 | 0.019 | 0.329 | 0.329 | 2.715 | 0.5 | 0.13 | 0.65 | 0.88 | 0.44 | 0.8 |
| 11 | 45075826 | rs11038286 | -1.18 | 0.818 | 0.594 | -1.815 | 0 | -0.034 | -0.034 | 0 | 0.17 | 0.17 | 0.11 | 0.02 | 0.2 | 0.04 |
| 11 | 45076544 | rs7483442 | -0.552 | 1.072 | 1.212 | -1.525 | 0.033 | -0.054 | -0.054 | 0 | 0.1 | 0.21 | 0.08 | 0.02 | 0.16 | 0.02 |
| 11 | 45077007 | rs12788080 | -2.02 | 1.139 | 1.73 | -1.612 | 0.058 | -0.079 | -0.079 | 0 | 0.18 | 0.4 | 0.14 | 0.08 | 0.23 | 0.11 |
| 11 | 45077911 | rs11602755 | -2.183 | 1.116 | 1.862 | -1.623 | 0.063 | -0.093 | -0.093 | 0 | 0.19 | 0.44 | 0.15 | 0.08 | 0.23 | 0.11 |
| 11 | 45081399 | rs11605047 | -1.787 | 1.057 | 1.632 | -1.774 | 0.055 | -0.097 | -0.097 | 1.365 | 0.14 | 0.44 | 0.09 | 0.02 | 0.17 | 0.03 |
| 11 | 45084531 | rs7951911 | -2.144 | 0.804 | 1.029 | -1.754 | -0.032 | -0.105 | -0.105 | 2.962 | 0.19 | 0.59 | 0.1 | 0.04 | 0.1 | 0.09 |
| 11 | 45085525 | rs2863147 | -2.3 | 0.808 | 0.178 | -1.997 | -0.053 | 0.166 | 0.166 | 1.594 | 0.35 | 0.86 | 0.25 | 0.04 | 0.41 | 0.17 |
| 11 | 45093614 | rs4755312 | -2.245 | -0.275 | 0.759 | -1.325 | -0.001 | -0.067 | -0.067 | 1.206 | 0.13 | 0.69 | 0.11 | 0.03 | 0.1 | 0.09 |
| 11 | 45098100 | rs10466475 | -0.397 | -0.175 | 0.453 | -1.273 | 0.021 | -0.029 | -0.029 | 0 | 0.05 | 0.12 | 0.05 | 0 | 0.07 | 0.08 |
| 11 | 45101538 | rs4755957 | -1.567 | -0.185 | -0.045 | -1.272 | -0.003 | -0.038 | -0.038 | 0 | 0.19 | 0.75 | 0.11 | 0.02 | 0.09 | 0.09 |
| 11 | 45106508 | rs10769111 | -0.576 | 0.158 | -0.306 | -0.556 | -0.008 | 0.015 | 0.015 | 1.606 | 0.35 | 0.47 | 0.49 | 0.62 | 0.31 | 0.42 |
| 11 | 45107700 | rs4755955 | -1.162 | 0.49 | -0.096 | -0.771 | 0.044 | -0.002 | -0.002 | 0 | 0.11 | 0.26 | 0.35 | 0.06 | 0.15 | 0.09 |
| 11 | 45112870 | rs12363087 | -0.371 | 0.734 | -0.688 | -0.899 | -0.039 | 0.001 | 0.001 | 0.077 | 0.09 | 0.01 | 0.09 | 0.06 | 0.14 | 0.09 |
| 11 | 45117171 | rs11038296 | -0.001 | -1.013 | -1.008 | -1.682 | 0.076 | 0.07 | 0.07 | 1.283 | 0.27 | 0.11 | 0.1 | 0.01 | 0.11 | 0.11 |
| 11 | 45117657 | rs2077427 | 0.276 | -0.564 | -0.797 | -1.5 | 0.027 | 0.015 | 0.015 | 0.218 | 0.19 | 0.1 | 0.09 | 0 | 0.11 | 0.08 |
| 11 | 45118495 | rs4994000 | 0.354 | -0.567 | -0.864 | -1.502 | 0.036 | 0.035 | 0.035 | 0.255 | 0.18 | 0.07 | 0.09 | 0 | 0.11 | 0.08 |
| 11 | 45119181 | rs747614 | -0.668 | 0.25 | -0.802 | -1.734 | -0.004 | -0.083 | -0.083 | 0 | 0.35 | 0.58 | 0.22 | 0.03 | 0.3 | 0.22 |
| 11 | 45128302 | rs3740787 | -0.791 | 0.482 | -1.498 | -1.806 | -0.044 | 0.25 | 0.25 | 4.604 | 0.39 | 0.93 | 0.52 | 0.06 | 0.37 | 0.31 |
| 11 | 45140524 | rs11038308 | 0.619 | -0.114 | -1.196 | -1.233 | 0.011 | 0.101 | 0.101 | 5.131 | 0.54 | 0.19 | 0.38 | 0.27 | 0.56 | 0.54 |
| 11 | 45140744 | rs7122974 | 0.982 | -0.374 | -0.337 | -1.49 | -0.016 | -0.057 | -0.057 | 0 | 0.19 | 0.24 | 0.1 | 0 | 0.12 | 0.08 |
| 11 | 45141700 | rs10838418 | 1.087 | -0.319 | -0.344 | -1.433 | -0.005 | 0.148 | 0.148 | 5.939 | 0.52 | 0.07 | 0.38 | 0.27 | 0.56 | 0.54 |
| 11 | 45142471 | rs7131444 | -0.289 | -0.307 | 0.538 | -1.388 | -0.015 | -0.099 | -0.099 | 0.228 | 0.32 | 0.48 | 0.43 | 0.05 | 0.29 | 0.22 |
| 11 | 45152825 | rs7929875 | -0.125 | -0.258 | 0.476 | -0.953 | -0.003 | 0.015 | 0.015 | 0.193 | 0.89 | 0.88 | 0.83 | 0.36 | 0.86 | 0.77 |
| 11 | 45161460 | rs1487718 | 1.172 | -0.243 | 0.364 | -0.305 | -0.01 | 0.001 | 0.001 | 5.823 | 0.53 | 0.07 | 0.4 | 0.44 | 0.56 | 0.55 |
| 11 | 45173290 | rs4755953 | -0.183 | -0.032 | 0.453 | 0.008 | -0.013 | 0.007 | 0.007 | 0.039 | 0.89 | 0.92 | 0.85 | 0.5 | 0.86 | 0.77 |
| 11 | 45174530 | rs1487721 | 1.044 | -0.147 | 0.49 | -0.061 | 0.008 | 0.065 | 0.065 | 5.266 | 0.53 | 0.18 | 0.4 | 0.35 | 0.56 | 0.54 |
| 11 | 45175323 | rs4755952 | -0.607 | 0.855 | -0.363 | 0.046 | -0.082 | 0.099 | 0.099 | 0.606 | 0.41 | 0.87 | 0.55 | 0.21 | 0.49 | 0.23 |
| 11 | 45186029 | rs6485597 | -0.382 | 0.473 | 0.438 | 0.253 | -0.022 | 0.206 | 0.206 | 0.194 | 0.92 | 0.82 | 0.9 | 0.48 | 0.97 | 0.84 |
| 11 | 45188860 | rs2863156 | 0.813 | 0.333 | 0.911 | -0.119 | -0.022 | -0.077 | -0.077 | 1.312 | 0.21 | 0.32 | 0.12 | 0 | 0.14 | 0.07 |
| 11 | 45192009 | rs12362992 | 0.874 | 0.316 | 0.282 | -0.647 | -0.04 | 0.041 | 0.041 | 3.639 | 0.23 | 0.14 | 0.45 | 0.08 | 0.37 | 0.32 |
| 11 | 45203848 | rs1078931 | 1.346 | 0.009 | 0.52 | -0.556 | -0.013 | -0.056 | -0.056 | 0 | 0.2 | 0.26 | 0.12 | 0 | 0.13 | 0.07 |
| 11 | 45204071 | rs1078930 | -0.877 | -0.045 | 0.617 | -0.428 | -0.01 | 0.115 | 0.115 | 0.048 | 0.92 | 0.87 | 0.91 | 0.48 | 0.97 | 0.84 |
| 11 | 45207851 | rs7480089 | 1.38 | -0.061 | 0.703 | -0.461 | -0.025 | -0.075 | -0.075 | 4.546 | 0.2 | 0.29 | 0.12 | 0 | 0.13 | 0.07 |
| 11 | 45208897 | rs3758728 | -0.75 | 0.1 | 0.208 | -0.336 | -0.01 | 0.11 | 0.11 | 0.018 | 0.92 | 0.87 | 0.91 | 0.47 | 0.97 | 0.84 |
| 11 | 45218043 | rs10769125 | 0.195 | 0.017 | 0.06 | -0.527 | 0.038 | 0.031 | 0.031 | 3.423 | 0.63 | 0.23 | 0.44 | 0.34 | 0.73 | 0.68 |
| 11 | 45227567 | rs17723470 | 1.124 | 0.208 | -0.14 | -0.753 | -0.038 | 0.069 | 0.069 | 0.887 | 0.23 | 0.09 | 0.17 | 0.04 | 0.28 | 0.28 |
| 11 | 45227830 | rs7940871 | 0.705 | 0.321 | 0.03 | -0.795 | -0.017 | 0.015 | 0.015 | 0.118 | 0.25 | 0.16 | 0.19 | 0.04 | 0.31 | 0.28 |
| 11 | 45228817 | rs2863164 | 0.157 | 0.381 | -0.103 | -0.604 | -0.033 | -0.035 | -0.035 | 0.425 | 0.07 | 0.02 | 0.33 | 0.12 | 0.14 | 0.09 |
| 11 | 45230061 | rs7128207 | 0.104 | 0.218 | 0.122 | -0.679 | 0.003 | 0.022 | 0.022 | 0.257 | 0.46 | 0.23 | 0.6 | 0.39 | 0.56 | 0.5 |
| 11 | 45234877 | rs11038356 | -0.795 | 0.143 | 0.322 | -0.691 | -0.02 | 0.135 | 0.135 | 0.596 | 0.91 | 0.81 | 0.9 | 0.46 | 0.97 | 0.84 |
| 11 | 45239959 | rs6416127 | -0.473 | 0.125 | 0.205 | -0.719 | -0.009 | 0.001 | 0.001 | 4.046 | 0.69 | 0.31 | 0.77 | 0.46 | 0.84 | 0.77 |
| 11 | 45241581 | rs3758726 | 0.556 | 0.28 | 0.084 | -0.536 | -0.036 | 0.077 | 0.077 | 0 | 0.38 | 0.15 | 0.26 | 0.27 | 0.43 | 0.4 |
| 11 | 45242383 | rs3781703 | 0.102 | 0.315 | -0.109 | -0.495 | -0.033 | -0.035 | -0.035 | 0.068 | 0.07 | 0.02 | 0.33 | 0.14 | 0.14 | 0.09 |
| 11 | 45246570 | rs3740784 | 0.102 | 0.315 | -0.109 | -0.502 | -0.033 | -0.035 | -0.035 | 0 | 0.07 | 0.02 | 0.32 | 0.14 | 0.14 | 0.09 |
| 11 | 45247314 | rs11038360 | 0.518 | 0.245 | 0.027 | -0.944 | -0.071 | 0.084 | 0.084 | 0 | 0.38 | 0.08 | 0.26 | 0.27 | 0.42 | 0.4 |
| 11 | 45250732 | rs2863171 | 0.593 | 0.313 | 0.561 | -0.704 | -0.035 | -0.132 | -0.132 | 0 | 0.25 | 0.4 | 0.14 | 0 | 0.16 | 0.11 |
| 11 | 45259944 | rs7127746 | 0.187 | 0.56 | 0.393 | -0.468 | -0.002 | 0.053 | 0.053 | 2.749 | 0.51 | 0.41 | 0.37 | 0.29 | 0.54 | 0.48 |
| 11 | 45262126 | rs8929 | 0.237 | 0.56 | 0.547 | -0.494 | 0.003 | 0.085 | 0.085 | 2.768 | 0.49 | 0.33 | 0.35 | 0.3 | 0.51 | 0.48 |
| 11 | 45262242 | rs2863173 | 0.672 | 0.487 | 0.948 | -0.286 | -0.002 | 0.122 | 0.122 | 0 | 0.81 | 0.56 | 0.57 | 0.4 | 0.85 | 0.85 |
| 11 | 45262256 | rs2902427 | 0.596 | 0.476 | 1.119 | -0.291 | -0.003 | 0.089 | 0.089 | 0 | 0.82 | 0.63 | 0.58 | 0.4 | 0.85 | 0.85 |

| | | | | | | | | | | | | | | | | |
|---|---|---|---|---|---|---|---|---|---|---|---|---|---|---|---|---|
| 11 | 45263258 | rs7110580 | 0.395 | 0.48 | 1.042 | -0.304 | -0.002 | 0.089 | 0.089 | 2.505 | 0.5 | 0.34 | 0.36 | 0.3 | 0.54 | 0.48 |
| 11 | 45263340 | rs3180446 | 0.924 | 0.51 | 1.032 | -0.286 | -0.01 | 0.047 | 0.047 | 3.571 | 0.31 | 0.18 | 0.21 | 0.1 | 0.31 | 0.37 |
| 11 | 45263694 | rs6416129 | 0.359 | 0.802 | 1.434 | 0.19 | 0 | 0.021 | 0.021 | 0 | 0.85 | 0.79 | 0.6 | 0.4 | 0.86 | 0.85 |
| 11 | 45264277 | rs2863174 | -0.055 | 0.81 | 1.401 | 0.206 | 0 | 0.004 | 0.004 | 0 | 0.87 | 0.86 | 0.6 | 0.4 | 0.88 | 0.86 |
| 11 | 45272453 | rs2863178 | 0.35 | 0.784 | 1.457 | 0.05 | -0.005 | 0.031 | 0.031 | 1.797 | 0.41 | 0.35 | 0.31 | 0.3 | 0.46 | 0.42 |
| 11 | 45273511 | rs7103871 | 0.588 | 0.73 | 1.438 | 0.011 | -0.005 | 0.045 | 0.045 | 0 | 0.82 | 0.71 | 0.7 | 0.39 | 0.85 | 0.83 |
| 11 | 45274894 | rs10838436 | 1.004 | 0.676 | 1.503 | -0.025 | -0.035 | 0.159 | 0.159 | 5.983 | 0.37 | 0.13 | 0.37 | 0.09 | 0.38 | 0.41 |
| 11 | 45276308 | rs3816205 | 0.804 | -0.012 | 1.915 | 0.367 | -0.005 | 0.082 | 0.082 | 0 | 0.82 | 0.66 | 0.7 | 0.39 | 0.85 | 0.83 |
| 11 | 45278317 | rs12362444 | 0.965 | 0.203 | 1.604 | 0.215 | -0.043 | 0.236 | 0.236 | 4.834 | 0.43 | 0.1 | 0.43 | 0.09 | 0.51 | 0.43 |
| 11 | 45280653 | rs751198 | 0.888 | 0.186 | 1.667 | 0.229 | 0.005 | 0.19 | 0.19 | 4.614 | 0.45 | 0.17 | 0.45 | 0.09 | 0.54 | 0.43 |
| 11 | 45284774 | rs10742747 | 1.172 | 0.036 | 1.951 | 0.444 | -0.014 | 0.107 | 0.107 | 0.179 | 0.8 | 0.48 | 0.69 | 0.38 | 0.85 | 0.83 |
| 11 | 45285803 | rs1000664 | 0.815 | 0.047 | 1.877 | 0.451 | -0.005 | 0.065 | 0.065 | 0.009 | 0.82 | 0.67 | 0.7 | 0.39 | 0.84 | 0.83 |
| 11 | 45290804 | rs4755936 | 0.527 | 0.103 | 1.472 | 0.973 | -0.037 | -0.027 | -0.027 | 0 | 0.06 | 0 | 0.19 | 0.22 | 0.12 | 0.08 |
| 11 | 45297599 | rs7124508 | 0.901 | 0.144 | 1.542 | 0.693 | -0.006 | 0.098 | 0.098 | 4.29 | 0.44 | 0.14 | 0.53 | 0.26 | 0.54 | 0.45 |
| 11 | 45303377 | rs2863182 | -0.304 | 0.088 | 1.74 | 0.669 | 0.039 | 0.046 | 0.046 | 2.309 | 0.43 | 0.68 | 0.26 | 0.3 | 0.33 | 0.43 |
| 11 | 45303711 | rs1077490 | 0.337 | 0.092 | 1.812 | 0.671 | -0.001 | -0.001 | -0.001 | 0 | 0.35 | 0.35 | 0.23 | 0.29 | 0.3 | 0.4 |
| 11 | 45305682 | rs12363398 | 0.526 | 0.073 | 1.843 | 0.662 | 0.012 | 0.012 | 0.012 | 1.15 | 0.33 | 0.27 | 0.23 | 0.29 | 0.3 | 0.4 |
| 11 | 45310410 | rs4755306 | -0.193 | 0.076 | 1.845 | 0.657 | 0.008 | 0.008 | 0.008 | 2.821 | 0.59 | 0.39 | 0.75 | 0.7 | 0.67 | 0.56 |
| 11 | 45316657 | rs11038387 | -0.708 | 0.089 | 1.635 | 0.648 | 0.011 | -0.014 | -0.014 | 0.906 | 0.06 | 0.25 | 0.02 | 0.01 | 0.02 | 0.03 |
| 11 | 45319167 | rs7118408 | -1.231 | 0.123 | 1.429 | 0.644 | 0.002 | 0.072 | 0.072 | 4.719 | 0.55 | 0.73 | 0.48 | 0.74 | 0.49 | 0.61 |
| 11 | 45320055 | rs11038388 | 0.331 | 0.248 | 1.345 | 0.796 | -0.007 | -0.011 | -0.011 | 0.292 | 0.35 | 0.38 | 0.26 | 0.27 | 0.33 | 0.47 |
| 11 | 45322191 | rs7131174 | 0.013 | 0.31 | 0.983 | 0.827 | 0.009 | 0.013 | 0.013 | 4.718 | 0.4 | 0.37 | 0.26 | 0.29 | 0.35 | 0.5 |
| 11 | 45330221 | rs11038391 | -0.244 | 0.377 | 0.497 | 0.957 | -0.002 | 0.019 | 0.019 | 0 | 0.12 | 0.11 | 0.2 | 0.45 | 0.13 | 0.11 |
| 11 | 45335053 | rs7926308 | -0.012 | -0.049 | 0.595 | 0.059 | -0.024 | -0.107 | -0.107 | 0 | 0.28 | 0.62 | 0.15 | 0.01 | 0.19 | 0.18 |
| 11 | 45337343 | rs10838444 | -0.117 | 0.042 | 0.475 | 0.456 | 0.013 | 0.016 | 0.016 | 1.247 | 0.37 | 0.32 | 0.21 | 0.28 | 0.32 | 0.45 |
| 11 | 45343012 | rs9667484 | -0.288 | -0.067 | 0.516 | 0.481 | 0.003 | -0.017 | -0.017 | 0 | 0.82 | 0.87 | 0.88 | 0.69 | 0.82 | 0.76 |
| 11 | 45343932 | rs7478974 | 0.029 | -0.135 | 0.64 | 0.42 | 0.001 | -0.003 | -0.003 | 0 | 0.79 | 0.84 | 0.87 | 0.69 | 0.81 | 0.74 |
| 11 | 45348366 | rs7483751 | 0.45 | -0.23 | 0.92 | 0.454 | 0.07 | 0.053 | 0.053 | 3.667 | 0.55 | 0.25 | 0.72 | 0.66 | 0.66 | 0.52 |
| 11 | 45351106 | rs4755316 | 0.051 | -0.001 | 1.27 | 0.771 | -0.013 | 0.096 | 0.096 | 0.311 | 0.8 | 0.67 | 0.86 | 0.6 | 0.81 | 0.75 |
| 11 | 45352879 | rs4755974 | -0.759 | 0.05 | 0.838 | 0.788 | 0 | -0.015 | -0.015 | 1.788 | 0.85 | 0.9 | 0.88 | 0.58 | 0.83 | 0.78 |
| 11 | 45358483 | rs7121660 | -1.093 | 0.987 | 0.686 | 0.713 | 0.051 | 0.052 | 0.052 | 0.25 | 0.24 | 0.44 | 0.3 | 0.32 | 0.29 | 0.2 |
| 11 | 45359591 | rs6485621 | 0.59 | 1.265 | 0.298 | 0.439 | 0.022 | 0.095 | 0.095 | 8.901 | 0.44 | 0.1 | 0.43 | 0.25 | 0.37 | 0.36 |
| 11 | 45370488 | rs11038408 | 0.563 | 1.129 | 0.764 | 0.3 | -0.016 | -0.154 | -0.154 | 0.084 | 0.27 | 0.43 | 0.15 | 0.01 | 0.22 | 0.24 |
| 11 | 45379892 | rs10838456 | 0.563 | 1.19 | 0.864 | 0.296 | -0.016 | -0.156 | -0.156 | 0 | 0.27 | 0.43 | 0.15 | 0.01 | 0.22 | 0.24 |
| 11 | 45381346 | rs4755319 | 0.7 | 1.225 | 0.638 | 0.313 | -0.007 | -0.099 | -0.099 | 0 | 0.26 | 0.34 | 0.15 | 0.01 | 0.22 | 0.24 |
| 11 | 45381670 | rs4755320 | 0.838 | 1.045 | 0.505 | 0.322 | -0.01 | -0.062 | -0.062 | 0.552 | 0.25 | 0.29 | 0.11 | 0 | 0.15 | 0.23 |
| 11 | 45389225 | rs11601710 | -0.501 | 1.699 | 0.4 | 0.972 | 0.05 | 0.075 | 0.075 | 0.036 | 0.05 | 0.12 | 0.24 | 0.27 | 0.18 | 0.1 |
| 11 | 45394284 | rs7950659 | -0.958 | 1.561 | 0.411 | 0.846 | 0.033 | 0.041 | 0.041 | 7.885 | 0.56 | 0.76 | 0.57 | 0.64 | 0.64 | 0.59 |
| 11 | 45397885 | rs6485622 | -0.8 | 1.541 | 0.69 | 0.919 | 0.004 | -0.008 | -0.008 | 0 | 0.53 | 0.62 | 0.37 | 0.49 | 0.55 | 0.53 |
| 11 | 45398217 | rs7930399 | 1.551 | 1.989 | 1.383 | 1.295 | 0.006 | 0.053 | 0.053 | 0 | 0.57 | 0.3 | 0.48 | 0.25 | 0.43 | 0.55 |
| 11 | 45401413 | rs896334 | 1.127 | 2.099 | 1.489 | 1.136 | 0.058 | 0.07 | 0.07 | 0.183 | 0.82 | 0.62 | 0.69 | 0.62 | 0.74 | 0.84 |
| 11 | 45403404 | rs6485623 | 0.625 | 2.068 | 1.449 | 1.14 | 0.063 | 0.044 | 0.044 | 0 | 0.84 | 0.64 | 0.7 | 0.74 | 0.76 | 0.86 |
| 11 | 45404318 | rs11038428 | -0.254 | 2.044 | 1.24 | 1.419 | 0.016 | 0.022 | 0.022 | 0 | 0.24 | 0.28 | 0.21 | 0.26 | 0.31 | 0.29 |
| 11 | 45406048 | rs7127704 | 0.49 | 2.148 | 1.231 | 1.514 | 0.022 | 0.029 | 0.029 | 0 | 0.86 | 0.7 | 0.74 | 0.75 | 0.83 | 0.87 |
| 11 | 45414183 | rs1545466 | 1.298 | 1.366 | 1.146 | 1.148 | 0 | 0 | 0 | 0 | 0.59 | 0.4 | 0.53 | 0.37 | 0.52 | 0.6 |
| 11 | 45415061 | rs896330 | -0.468 | 1.315 | 0.651 | 1.05 | 0 | -0.013 | -0.013 | 0 | 0.91 | 0.88 | 0.94 | 1 | 0.94 | 0.96 |
| 11 | 45417089 | rs2863706 | 0.17 | 1.199 | 0.658 | 1.344 | 0.003 | -0.003 | -0.003 | 0 | 0.88 | 0.85 | 0.82 | 0.86 | 0.93 | 0.92 |
| 11 | 45419113 | rs11038440 | 1.453 | 1.27 | 1.006 | 1.38 | 0.002 | -0.002 | -0.002 | 0.219 | 0.58 | 0.4 | 0.54 | 0.37 | 0.52 | 0.61 |
| 11 | 45420899 | rs2044873 | 0.814 | 1.241 | 0.987 | 1.343 | -0.012 | 0.018 | 0.018 | 1.237 | 0.85 | 0.76 | 0.81 | 0.75 | 0.92 | 0.91 |
| 11 | 45429381 | rs1037363 | 0.789 | 1.357 | 0.987 | 1.312 | 0.029 | 0.019 | 0.019 | 0.329 | 0.85 | 0.63 | 0.77 | 0.86 | 0.86 | 0.91 |
| 11 | 45438374 | rs10742752 | 0.537 | 1.4 | 0.658 | 1.404 | -0.025 | -0.015 | -0.015 | 0.382 | 0.66 | 0.8 | 0.78 | 0.63 | 0.62 | 0.67 |
| 11 | 45446724 | rs7943688 | -0.427 | 1.298 | 0.557 | 1.421 | -0.02 | -0.03 | -0.03 | 0.341 | 0.27 | 0.16 | 0.19 | 0.37 | 0.33 | 0.29 |
| 11 | 45458867 | rs12286627 | -0.092 | 0.74 | 0.674 | 1.061 | 0.002 | -0.001 | -0.001 | 0 | 0.11 | 0.14 | 0.2 | 0.25 | 0.08 | 0.08 |
| 11 | 45460738 | rs4755986 | 0.615 | 0.854 | 1.107 | 0.971 | 0.068 | -0.046 | -0.046 | 0.221 | 0.92 | 0.79 | 0.92 | 1 | 0.88 | 0.95 |
| 11 | 45463152 | rs921455 | 1.731 | 0.844 | 0.895 | 0.898 | -0.001 | -0.002 | -0.002 | 0 | 0.74 | 0.32 | 0.68 | 0.75 | 0.77 | 0.87 |
| 11 | 45472839 | rs6485626 | 0.626 | 0.663 | 1.171 | 1.182 | -0.004 | -0.025 | -0.025 | 0 | 0.95 | 0.86 | 0.98 | 1 | 0.97 | 0.97 |
| 11 | 45473135 | rs4755988 | 1.82 | 0.746 | 1.428 | 1.202 | -0.005 | 0.015 | 0.015 | 3.361 | 0.77 | 0.33 | 0.69 | 0.72 | 0.81 | 0.87 |
| 11 | 45474823 | rs1530868 | 1.664 | 0.751 | 1.682 | 1.195 | 0.104 | -0.017 | -0.017 | 0.82 | 0.92 | 0.65 | 0.93 | 1 | 0.91 | 0.96 |
| 11 | 45478082 | rs1825934 | 1.588 | 0.749 | 1.722 | 1.21 | 0.004 | 0.08 | 0.08 | 1.372 | 0.8 | 0.5 | 0.71 | 0.72 | 0.82 | 0.87 |
| 11 | 45481150 | rs7931858 | -0.637 | 0.737 | 1.698 | 1.209 | -0.001 | 0.009 | 0.009 | 0.061 | 0.13 | 0.18 | 0.22 | 0.28 | 0.09 | 0.1 |
| 11 | 45484440 | rs1565222 | 1.627 | 0.726 | 1.783 | 1.199 | -0.008 | 0.045 | 0.045 | 2.792 | 0.79 | 0.4 | 0.7 | 0.72 | 0.82 | 0.87 |
| 11 | 45493769 | rs10742759 | 0.804 | 0.657 | 1.747 | 1.306 | -0.024 | 0.017 | 0.017 | 0.029 | 0.84 | 0.68 | 0.89 | 0.81 | 0.9 | 0.9 |
| 11 | 45494063 | rs11038484 | 0.26 | 0.724 | 1.02 | 1.341 | -0.118 | -0.151 | -0.151 | 0 | 0.27 | 0.01 | 0.21 | 0.44 | 0.36 | 0.37 |
| 11 | 45494703 | rs7115683 | 1.817 | 0.465 | 1.747 | 1.218 | -0.01 | 0.035 | 0.035 | 1.248 | 0.78 | 0.42 | 0.69 | 0.75 | 0.82 | 0.87 |
| 11 | 45500427 | rs4755996 | 1.834 | 0.533 | 1.655 | 1.249 | -0.007 | 0.029 | 0.029 | 2.232 | 0.78 | 0.35 | 0.7 | 0.73 | 0.81 | 0.89 |
| 11 | 45526697 | rs7947921 | 0.616 | 0.659 | 1.398 | 0.759 | -0.002 | -0.001 | -0.001 | 0 | 0.95 | 0.9 | 0.97 | 0.95 | 0.96 | 0.96 |
| 11 | 45528116 | rs1488665 | 0.977 | 0.284 | 1.029 | 0.507 | -0.004 | -0.003 | -0.003 | 0 | 0.78 | 0.7 | 0.87 | 0.86 | 0.83 | 0.9 |
| 11 | 45535213 | rs875021 | -2.331 | 0.372 | 1.342 | 0.76 | 0.057 | 0.018 | 0.018 | 0 | 0.1 | 0.35 | 0.09 | 0.09 | 0.13 | 0.05 |
| 11 | 45539520 | rs6485628 | 1.763 | 0.334 | 1.122 | 0.729 | -0.009 | -0.01 | -0.01 | 0 | 0.79 | 0.53 | 0.85 | 0.83 | 0.81 | 0.89 |

| | | | | | | | | | | | | | | | | |
|---|---|---|---|---|---|---|---|---|---|---|---|---|---|---|---|---|
| 11 | 45540603 | rs1873059 | 0.341 | 0.308 | 0.548 | 0.712 | 0.009 | -0.002 | -0.002 | 0 | 0.52 | 0.48 | 0.54 | 0.5 | 0.56 | 0.54 |
| 11 | 45542342 | rs12417911 | 0.499 | 0.302 | 0.496 | 0.764 | 0.019 | -0.064 | -0.064 | 1.279 | 0.35 | 0.15 | 0.36 | 0.41 | 0.3 | 0.38 |
| 11 | 45545583 | rs4755327 | 0.384 | 0.316 | 0.448 | 0.51 | 0.027 | 0.007 | 0.007 | 1.337 | 0.36 | 0.16 | 0.35 | 0.29 | 0.3 | 0.35 |
| 11 | 45546703 | rs12417465 | 0.071 | 0.334 | 0.539 | 0.52 | 0.006 | 0.005 | 0.005 | 0 | 0.55 | 0.49 | 0.56 | 0.62 | 0.58 | 0.55 |
| 11 | 45548263 | rs11038514 | -0.414 | 0.592 | 0.956 | 0.347 | -0.002 | -0.001 | -0.001 | 0 | 0.11 | 0.16 | 0.08 | 0.08 | 0.08 | 0.06 |
| 11 | 45550010 | rs985651 | 1.618 | 1.134 | 0.793 | 0.271 | 0.04 | -0.026 | -0.026 | 0.009 | 0.78 | 0.35 | 0.8 | 0.84 | 0.73 | 0.88 |
| 11 | 45554951 | rs896344 | 1.696 | 1.138 | 0.629 | 0.276 | 0.019 | -0.019 | -0.019 | 0 | 0.77 | 0.28 | 0.79 | 0.83 | 0.73 | 0.85 |
| 11 | 45565539 | rs2046735 | 0.712 | 1.039 | 0.425 | 0.095 | 0.001 | -0.006 | -0.006 | 0 | 0.73 | 0.67 | 0.8 | 0.81 | 0.74 | 0.82 |
| 11 | 45573797 | rs1488674 | -0.218 | 0.933 | 1.312 | 0.007 | -0.03 | -0.013 | -0.013 | 0 | 0.8 | 0.71 | 0.9 | 0.88 | 0.9 | 0.96 |
| 11 | 45575697 | rs1565224 | 1.583 | 1.004 | 1.523 | -0.076 | -0.015 | 0.035 | 0.035 | 2.477 | 0.67 | 0.18 | 0.74 | 0.82 | 0.67 | 0.8 |
| 11 | 45575813 | rs11038540 | -2.576 | 1.046 | 1.548 | -0.071 | 0.197 | -0.017 | -0.017 | 1.985 | 0.12 | 0.51 | 0.15 | 0.07 | 0.22 | 0.15 |
| 11 | 45576034 | rs10160816 | -1.759 | 1.044 | 1.434 | -0.06 | 0.022 | -0.007 | -0.007 | 0.089 | 0.09 | 0.17 | 0.1 | 0.07 | 0.16 | 0.14 |
| 11 | 45576417 | rs4756005 | 0.593 | 0.779 | 1.352 | 0.026 | 0.007 | 0.008 | 0.008 | 5.049 | 0.75 | 0.25 | 0.83 | 0.86 | 0.82 | 0.94 |
| 11 | 45580099 | rs16938306 | -1.913 | 0.932 | 1.202 | -0.291 | 0.09 | -0.063 | -0.063 | 0.621 | 0.16 | 0.49 | 0.15 | 0.06 | 0.2 | 0.18 |
| 11 | 45585234 | rs6485633 | -1.196 | 0.554 | 0.271 | -0.447 | 0.016 | 0.026 | 0.026 | 0 | 0.75 | 0.85 | 0.71 | 0.9 | 0.78 | 0.71 |
| 11 | 45586169 | rs2129969 | 1.238 | 0.782 | 0.077 | -0.265 | -0.066 | 0.042 | 0.042 | 0.44 | 0.42 | 0.13 | 0.44 | 0.7 | 0.44 | 0.52 |
| 11 | 45590328 | rs17195752 | -0.748 | 0.927 | 0.153 | -0.146 | -0.04 | -0.03 | -0.03 | 0 | 0.08 | 0.01 | 0.12 | 0.12 | 0.16 | 0.13 |
| 11 | 45596965 | rs10838488 | 1.047 | 0.847 | 0.225 | -0.185 | -0.026 | 0.042 | 0.042 | 0.379 | 0.44 | 0.21 | 0.45 | 0.69 | 0.45 | 0.49 |
| 11 | 45600662 | rs275586 | -0.862 | 0.969 | 0.471 | -0.195 | 0.023 | 0.034 | 0.034 | 0 | 0.73 | 0.85 | 0.72 | 0.9 | 0.78 | 0.68 |
| 11 | 45606983 | rs1488682 | 0.276 | 1.053 | 0.687 | -0.083 | -0.012 | 0.02 | 0.02 | 0 | 0.62 | 0.36 | 0.55 | 0.78 | 0.55 | 0.55 |
| 11 | 45612867 | rs1107449 | 0.064 | 1.051 | 0.807 | -0.079 | -0.001 | -0.002 | -0.002 | 2.601 | 0.64 | 0.42 | 0.58 | 0.79 | 0.62 | 0.56 |
| 11 | 45618861 | rs7127267 | 0.154 | 1.026 | 1.004 | -0.056 | 0.005 | -0.01 | -0.01 | 0 | 0.63 | 0.47 | 0.57 | 0.78 | 0.55 | 0.54 |
| 11 | 45631953 | rs4755330 | 2.119 | 1.582 | 1.613 | 0.18 | 0.12 | 0.017 | 0.017 | 0.311 | 0.85 | 0.47 | 0.83 | 0.87 | 0.78 | 0.85 |
| 11 | 45635476 | rs12576823 | -1.629 | 1.551 | 1.443 | 0.208 | 0.055 | 0.025 | 0.025 | 0.345 | 0.1 | 0.21 | 0.13 | 0.12 | 0.15 | 0.14 |
| 11 | 45636937 | rs1562459 | -1.629 | 1.551 | 1.443 | 0.208 | 0.055 | 0.025 | 0.025 | 0.345 | 0.1 | 0.21 | 0.13 | 0.12 | 0.15 | 0.14 |
| 11 | 45638478 | rs12577749 | -1.601 | 0.975 | 1.503 | 0.269 | 0.04 | 0.05 | 0.05 | 0.399 | 0.08 | 0.2 | 0.1 | 0.12 | 0.1 | 0.12 |
| 11 | 45640918 | rs4756006 | -0.117 | 0.452 | 0.88 | -0.023 | 0.015 | 0.016 | 0.016 | 4.988 | 0.2 | 0.16 | 0.11 | 0.1 | 0.12 | 0.05 |
| 11 | 45660457 | rs4755331 | 2.125 | 1.138 | 1.617 | 1.211 | 0.153 | 0.348 | 0.348 | 0 | 0.89 | 0.52 | 0.79 | 0.67 | 0.82 | 0.89 |
| 11 | 45663568 | rs1348965 | -0.537 | 0.89 | 1.317 | 0.847 | 0.054 | 0.115 | 0.115 | 4.565 | 0.55 | 0.71 | 0.61 | 0.84 | 0.65 | 0.6 |
| 11 | 45666866 | rs10769158 | 0.688 | 0.844 | 1.374 | 0.83 | -0.049 | -0.041 | -0.041 | 0 | 0.41 | 0.18 | 0.4 | 0.51 | 0.47 | 0.49 |
| 11 | 45668114 | rs1447569 | 0.222 | 0.831 | 1.344 | 0.813 | 0.068 | 0.164 | 0.164 | 4.109 | 0.45 | 0.21 | 0.38 | 0.17 | 0.35 | 0.4 |
| 11 | 45670395 | rs3740702 | -0.636 | 0.906 | 1.35 | 0.889 | -0.03 | 0.032 | 0.032 | 0 | 0.05 | 0 | 0.15 | 0.33 | 0.1 | 0.1 |
| 11 | 45672261 | rs3802762 | 2.754 | 0.977 | 1.445 | 0.927 | 0.076 | 0.194 | 0.194 | 0.579 | 0.88 | 0.4 | 0.78 | 0.67 | 0.83 | 0.88 |
| 11 | 45672725 | rs4148903 | 2.516 | 0.976 | 1.577 | 0.93 | 0.134 | 0.288 | 0.288 | 0.795 | 0.89 | 0.47 | 0.78 | 0.68 | 0.83 | 0.89 |
| 11 | 45672747 | rs4148902 | 3.144 | 0.975 | 1.528 | 0.915 | 0.067 | 0.18 | 0.18 | 1.47 | 0.87 | 0.39 | 0.78 | 0.67 | 0.83 | 0.88 |
| 11 | 45672926 | rs4148901 | 0.72 | 0.923 | 1.479 | 0.883 | -0.042 | -0.04 | -0.04 | 0.415 | 0.41 | 0.19 | 0.4 | 0.49 | 0.47 | 0.46 |
| 11 | 45673014 | rs4148900 | 0.182 | 0.918 | 1.474 | 0.879 | 0.073 | 0.173 | 0.173 | 3.152 | 0.46 | 0.21 | 0.38 | 0.18 | 0.35 | 0.41 |
| 11 | 45674170 | rs2277272 | 3.369 | 0.957 | 1.489 | 0.879 | 0.052 | 0.149 | 0.149 | 1.155 | 0.86 | 0.39 | 0.77 | 0.67 | 0.82 | 0.87 |
| 11 | 45674599 | rs4148898 | 3.09 | 0.958 | 1.49 | 0.88 | 0.064 | 0.173 | 0.173 | 0.561 | 0.87 | 0.39 | 0.78 | 0.67 | 0.83 | 0.88 |
| 11 | 45676823 | rs1026078 | 0.687 | 0.932 | 1.467 | 0.898 | -0.039 | -0.023 | -0.023 | 0 | 0.42 | 0.2 | 0.4 | 0.48 | 0.47 | 0.49 |
| 11 | 45677029 | rs4148897 | 0.266 | 0.918 | 1.467 | 0.872 | 0.065 | 0.194 | 0.194 | 3.417 | 0.45 | 0.12 | 0.38 | 0.18 | 0.35 | 0.39 |
| 11 | 45678820 | rs2012019 | 0.46 | 0.883 | 1.715 | 0.921 | 0.006 | 0.004 | 0.004 | 3.031 | 0.43 | 0.46 | 0.4 | 0.48 | 0.47 | 0.52 |
| 11 | 45679197 | rs895729 | -0.27 | 0.771 | 1.682 | 0.814 | 0.015 | 0.041 | 0.041 | 0.372 | 0.53 | 0.58 | 0.57 | 0.82 | 0.58 | 0.62 |
| 11 | 45684117 | rs895733 | 0.685 | 0.97 | 1.619 | 0.832 | -0.029 | -0.038 | -0.038 | 0 | 0.42 | 0.19 | 0.38 | 0.48 | 0.45 | 0.5 |
| 11 | 45684254 | rs895734 | 0.099 | 0.832 | 1.635 | 0.845 | 0.026 | 0.184 | 0.184 | 0.256 | 0.48 | 0.19 | 0.55 | 0.82 | 0.57 | 0.6 |
| 11 | 45690568 | rs12420703 | 0.765 | 0.681 | 1.649 | 0.868 | -0.045 | -0.04 | -0.04 | 0 | 0.41 | 0.19 | 0.39 | 0.48 | 0.47 | 0.5 |
| 11 | 45690907 | rs10838503 | -1.599 | 0.723 | 1.644 | 0.901 | 0.033 | 0.217 | 0.217 | 1.142 | 0.11 | 0.33 | 0.19 | 0.33 | 0.11 | 0.1 |
| 11 | 45691118 | rs11038608 | 0.453 | 0.707 | 1.307 | 0.744 | 0.041 | 0.158 | 0.158 | 0.545 | 0.41 | 0.13 | 0.37 | 0.18 | 0.35 | 0.36 |
| 11 | 45696552 | rs11038611 | -0.806 | 0.73 | 0.871 | 1.103 | -0.038 | -0.055 | -0.055 | 0 | 0.1 | 0 | 0.16 | 0.34 | 0.12 | 0.11 |
| 11 | 45698971 | rs4755335 | -0.259 | 0.869 | 1.065 | 0.785 | 0.034 | 0.063 | 0.063 | 0 | 0.52 | 0.57 | 0.6 | 0.81 | 0.64 | 0.59 |
| 11 | 45699134 | rs4756016 | -0.456 | 0.864 | 0.754 | 0.791 | 0.021 | 0.084 | 0.084 | 0.045 | 0.54 | 0.68 | 0.59 | 0.79 | 0.58 | 0.58 |
| 11 | 45704923 | rs3926466 | -0.954 | 0.914 | 0.064 | 1.161 | -0.042 | -0.088 | -0.088 | 0 | 0.12 | 0 | 0.26 | 0.37 | 0.13 | 0.11 |
| 11 | 45706162 | rs7929114 | 0.259 | 0.911 | 0.535 | 0.723 | -0.009 | 0 | 0 | 0 | 0.4 | 0.32 | 0.32 | 0.44 | 0.47 | 0.49 |
| 11 | 45706453 | rs10769165 | 0.337 | 1.114 | 0.386 | 0.799 | 0.015 | -0.006 | -0.006 | 0 | 0.37 | 0.41 | 0.3 | 0.35 | 0.5 | 0.5 |
| 11 | 45707377 | rs10838505 | -1.631 | 1.365 | 0.601 | 1.408 | -0.034 | -0.048 | -0.048 | 0.063 | 0.14 | 0.07 | 0.23 | 0.2 | 0.18 | 0.13 |
| 11 | 45709976 | rs1447575 | -0.667 | 1.142 | 1.226 | 1.545 | -0.004 | 0.156 | 0.156 | 0 | 0.13 | 0.29 | 0.12 | 0.38 | 0.09 | 0.09 |
| 11 | 45710652 | rs1447576 | -0.645 | 1.144 | 1.227 | 1.547 | -0.003 | 0.158 | 0.158 | 0 | 0.12 | 0.29 | 0.13 | 0.38 | 0.09 | 0.09 |
| 11 | 45711030 | rs2666895 | -0.857 | 1.116 | 1.684 | 1.533 | 0.011 | 0.283 | 0.283 | 0.002 | 0.13 | 0.5 | 0.14 | 0.38 | 0.09 | 0.09 |
| 11 | 45717929 | rs10838506 | -1.908 | 1.282 | 1.489 | 1.238 | 0.013 | 0.011 | 0.011 | 0 | 0.18 | 0.23 | 0.34 | 0.22 | 0.23 | 0.34 |
| 11 | 45719649 | rs921167 | -2.056 | 1.262 | 1.771 | 1.21 | 0.062 | 0.048 | 0.048 | 0.048 | 0.19 | 0.45 | 0.35 | 0.23 | 0.22 | 0.35 |
| 11 | 45719881 | rs7130134 | 0.049 | 1.269 | 1.61 | 1.47 | -0.003 | 0.06 | 0.06 | 0 | 0.1 | 0.11 | 0.1 | 0.39 | 0.08 | 0.08 |
| 11 | 45730783 | rs756379 | -2.638 | 1.057 | 1.787 | 1.212 | -0.001 | -0.001 | -0.001 | 0 | 0.36 | 0.64 | 0.5 | 0.61 | 0.35 | 0.44 |
| 11 | 45737527 | rs2955677 | -2.121 | 1.094 | 2.086 | 1.19 | 0.08 | 0.038 | 0.038 | 0 | 0.18 | 0.42 | 0.37 | 0.21 | 0.23 | 0.35 |
| 11 | 45744189 | rs2593731 | -1.598 | 1.092 | 2.058 | 1.212 | -0.006 | 0.176 | 0.176 | 0.224 | 0.2 | 0.44 | 0.18 | 0.4 | 0.18 | 0.1 |
| 11 | 45757401 | rs2666900 | -2.58 | 1 | 1.892 | 1.21 | -0.002 | 0.002 | 0.002 | 0.096 | 0.36 | 0.45 | 0.47 | 0.61 | 0.32 | 0.44 |
| 11 | 45758944 | rs11038626 | -0.646 | 1.17 | 1.525 | 1.008 | -0.011 | -0.02 | -0.02 | 0 | 0.07 | 0.03 | 0.19 | 0.18 | 0.09 | 0.11 |
| 11 | 45762996 | rs2593753 | -2.733 | 1.175 | 1.648 | 0.886 | 0.015 | -0.026 | -0.026 | 0 | 0.27 | 0.49 | 0.36 | 0.21 | 0.26 | 0.37 |
| 11 | 45767146 | rs2666908 | -2.139 | 1.767 | 2.049 | 1.055 | 0.111 | 0.01 | 0.01 | 0 | 0.05 | 0.31 | 0.05 | 0 | 0.06 | 0.02 |
| 11 | 45774997 | rs12365302 | -1.881 | 1.81 | 2.036 | 1.185 | 0.046 | 0.014 | 0.014 | 0 | 0.12 | 0.24 | 0.11 | 0.14 | 0.16 | 0.15 |

| | | | | | | | | | | | | | | | | |
|---|---|---|---|---|---|---|---|---|---|---|---|---|---|---|---|---|
| 11 | 45776968 | rs1562455 | -1.773 | 1.822 | 1.847 | 1.163 | -0.016 | 0.134 | 0.134 | 2.895 | 0.23 | 0.51 | 0.36 | 0.49 | 0.17 | 0.33 |
| 11 | 45777525 | rs1530634 | -2.01 | 2.129 | 1.865 | 1.147 | 0.058 | 0.003 | 0.003 | 0 | 0.13 | 0.23 | 0.12 | 0.14 | 0.21 | 0.14 |
| 11 | 45781188 | rs7941885 | -1.689 | 1.412 | 1.871 | 1.201 | -0.02 | 0.012 | 0.012 | 0 | 0.12 | 0.25 | 0.08 | 0.14 | 0.06 | 0.13 |
| 11 | 45793023 | rs2666890 | -0.327 | 1.529 | 2.043 | 1.346 | -0.009 | 0.121 | 0.121 | 0 | 0.11 | 0.17 | 0.24 | 0.46 | 0.08 | 0.11 |
| 11 | 45794136 | rs11038647 | -2.234 | 1.606 | 2.261 | 1.273 | 0.026 | 0.017 | 0.017 | 2.031 | 0.14 | 0.52 | 0.13 | 0.15 | 0.13 | 0.15 |
| 11 | 45795670 | rs2467287 | -1.2 | 1.791 | 2.143 | 1.39 | -0.009 | 0.083 | 0.083 | 0 | 0.17 | 0.26 | 0.28 | 0.47 | 0.14 | 0.15 |
| 11 | 45811278 | rs4756024 | -1.701 | 1.821 | 2.018 | 1.412 | 0.016 | 0.153 | 0.153 | 0.611 | 0.21 | 0.45 | 0.35 | 0.48 | 0.21 | 0.15 |
| 11 | 45828332 | rs11823366 | -2.389 | 2.185 | 1.957 | 0.703 | 0.208 | 0.048 | 0.048 | 4.137 | 0.07 | 0.47 | 0.09 | 0.01 | 0.1 | 0.03 |
| 11 | 45832509 | rs7130656 | -2.047 | 2.437 | 1.948 | 0.665 | 0.159 | -0.012 | -0.012 | 0 | 0.05 | 0.23 | 0.07 | 0 | 0.14 | 0.02 |
| 11 | 45833960 | rs4756026 | -0.271 | 2.58 | 1.121 | 0.148 | -0.049 | 0.049 | 0.049 | 0 | 0.19 | 0.02 | 0.15 | 0.1 | 0.24 | 0.15 |
| 11 | 45834198 | rs7943306 | 0.695 | 1.838 | 1.294 | 0.408 | -0.042 | 0.106 | 0.106 | 1.043 | 0.8 | 0.54 | 0.83 | 0.41 | 0.88 | 0.7 |
| 11 | 45839889 | rs12805422 | 2.075 | 1.946 | 0.945 | 0.403 | -0.117 | 0.164 | 0.164 | 1.538 | 0.53 | 0.05 | 0.51 | 0.26 | 0.53 | 0.51 |
| 11 | 45843023 | rs10742769 | -0.513 | 1.887 | 0.9 | 0.455 | -0.009 | -0.069 | -0.069 | 0.017 | 0.89 | 0.99 | 0.9 | 0.46 | 0.94 | 0.88 |
| 11 | 45846619 | rs11038673 | 1.639 | 1.859 | 1.637 | 0.368 | -0.003 | -0.013 | -0.013 | 0 | 0.6 | 0.55 | 0.58 | 0.28 | 0.64 | 0.63 |
| 11 | 45852646 | rs2090602 | 1.089 | 1.843 | 1.629 | 0.373 | -0.024 | 0.068 | 0.068 | 0.795 | 0.8 | 0.59 | 0.71 | 0.39 | 0.87 | 0.81 |
| 11 | 45870177 | rs10838524 | 2.451 | 1.835 | 1.421 | 0.425 | -0.113 | 0.186 | 0.186 | 1.347 | 0.51 | 0.04 | 0.51 | 0.26 | 0.52 | 0.5 |
| 11 | 45873091 | rs11605924 | 2.51 | 1.854 | 1.426 | 0.421 | -0.062 | 0.193 | 0.193 | 0.682 | 0.51 | 0.04 | 0.49 | 0.26 | 0.49 | 0.49 |
| 11 | 45874264 | rs11038689 | -0.045 | 1.874 | 1.686 | 0.445 | -0.005 | 0.014 | 0.014 | 0.099 | 0.19 | 0.12 | 0.12 | 0.12 | 0.22 | 0.18 |
| 11 | 45877043 | rs4756035 | -0.104 | 1.844 | 1.81 | 0.485 | -0.012 | 0.096 | 0.096 | 0.085 | 0.88 | 0.82 | 0.88 | 0.48 | 0.93 | 0.88 |
| 11 | 45877529 | rs2292913 | -0.104 | 1.844 | 1.81 | 0.485 | -0.012 | 0.096 | 0.096 | 0.085 | 0.88 | 0.82 | 0.88 | 0.48 | 0.93 | 0.88 |
| 11 | 45891418 | rs7123390 | -0.141 | 1.841 | 1.731 | 0.459 | -0.011 | 0.018 | 0.018 | 0.055 | 0.2 | 0.13 | 0.14 | 0.13 | 0.25 | 0.18 |
| 11 | 45891508 | rs4755345 | -0.014 | 1.743 | 1.805 | 0.521 | -0.015 | 0.121 | 0.121 | 0.055 | 0.87 | 0.78 | 0.88 | 0.48 | 0.93 | 0.88 |
| 11 | 45903194 | rs10838527 | -0.984 | 1.792 | 1.967 | 0.472 | 0.124 | -0.01 | -0.01 | 1.482 | 0.06 | 0.26 | 0.06 | 0 | 0.12 | 0.01 |
| 11 | 45903613 | rs2292910 | 1.179 | 1.752 | 1.765 | 0.491 | 0.001 | 0.003 | 0.003 | 0 | 0.65 | 0.63 | 0.73 | 0.33 | 0.68 | 0.58 |
| 11 | 45905605 | rs3824872 | 1.984 | 1.458 | 1.947 | 0.986 | -0.001 | -0.009 | -0.009 | 0 | 0.73 | 0.23 | 0.64 | 0.42 | 0.75 | 0.79 |
| 11 | 45906830 | rs1554338 | -2.39 | 0.764 | 2.084 | 1.182 | 0.029 | 0.077 | 0.077 | 1.041 | 0.09 | 0.36 | 0.19 | 0.1 | 0.06 | 0.16 |
| 11 | 45913652 | rs1533614 | 0.579 | 0.608 | 1.625 | 1.368 | -0.015 | 0.18 | 0.18 | 0 | 0.88 | 0.75 | 0.88 | 0.53 | 0.94 | 0.89 |
| 11 | 45929961 | rs7925824 | 0.871 | 0.421 | 2.112 | 1.283 | -0.012 | 0.404 | 0.404 | 1.818 | 0.9 | 0.56 | 0.69 | 0.46 | 0.95 | 0.9 |
| 11 | 45949534 | rs939105 | 1.811 | 0.122 | 1.684 | 0.886 | -0.028 | 0.021 | 0.021 | 3.64 | 0.79 | 0.34 | 0.63 | 0.4 | 0.9 | 0.74 |
| 11 | 45958637 | rs7934681 | 0.899 | 0.346 | 1.886 | 0.666 | -0.001 | -0.071 | -0.071 | 0 | 0.88 | 0.73 | 0.93 | 1 | 0.9 | 0.89 |
| 11 | 45970084 | rs2959103 | 0.867 | 0.345 | 1.93 | 0.666 | -0.004 | -0.069 | -0.069 | 0 | 0.88 | 0.76 | 0.93 | 1 | 0.9 | 0.89 |
| 11 | 46010847 | rs12418498 | 0.268 | 0.435 | 1.185 | 0.255 | -0.022 | 0.038 | 0.038 | 0 | 0.09 | 0.01 | 0.06 | 0 | 0.15 | 0.03 |
| 11 | 46018139 | rs2959101 | -2.116 | 0.775 | 0.223 | -0.596 | -0.039 | 0.276 | 0.276 | 1.697 | 0.5 | 0.97 | 0.55 | 0.81 | 0.45 | 0.54 |
| 11 | 46033104 | rs17787346 | 0.093 | 0.814 | 0.434 | -0.463 | -0.027 | 0.015 | 0.015 | 0 | 0.11 | 0 | 0.05 | 0.04 | 0.08 | 0.09 |
| 11 | 46057427 | rs7103299 | -0.329 | 0.678 | 0.861 | -0.406 | -0.01 | -0.05 | -0.05 | 0 | 0.1 | 0.17 | 0.06 | 0 | 0.09 | 0.1 |
| 11 | 46072537 | rs11038752 | -0.027 | 1.026 | 0.806 | -0.563 | -0.006 | -0.001 | -0.001 | 0 | 0.12 | 0.08 | 0.08 | 0.15 | 0.18 | 0.16 |
| 11 | 46086727 | rs3929339 | 1.03 | 0.866 | 1.117 | -0.334 | -0.062 | 0.158 | 0.158 | 0.516 | 0.82 | 0.51 | 0.64 | 0.4 | 0.89 | 0.78 |
| 11 | 46090550 | rs10501316 | 0.052 | 0.888 | 0.719 | -0.325 | -0.005 | -0.001 | -0.001 | 0 | 0.12 | 0.08 | 0.08 | 0.15 | 0.18 | 0.16 |
| 11 | 46122023 | rs4756055 | 1.779 | 0.559 | 0.788 | -0.342 | -0.022 | 0.099 | 0.099 | 1.029 | 0.58 | 0.15 | 0.49 | 0.25 | 0.57 | 0.57 |
| 11 | 46134807 | rs10838566 | -0.107 | 0.757 | 0.794 | -0.066 | -0.022 | 0.311 | 0.311 | 0.03 | 0.06 | 0.22 | 0.29 | 0.55 | 0.02 | 0.06 |
| 11 | 46145881 | rs1125082 | -0.615 | 1.163 | 0.26 | -0.49 | -0.032 | 0.014 | 0.014 | 0 | 0.22 | 0.13 | 0.15 | 0.17 | 0.32 | 0.22 |
| 11 | 46151490 | rs10769190 | 0.693 | 0.813 | 0.535 | -0.699 | -0.008 | -0.07 | -0.07 | 0 | 0.78 | 0.68 | 0.87 | 0.97 | 0.79 | 0.81 |
| 11 | 46155938 | rs7120013 | 1.757 | 0.704 | 0.383 | -0.618 | 0.008 | 0.116 | 0.116 | 0.105 | 0.49 | 0.23 | 0.46 | 0.27 | 0.44 | 0.54 |
| 11 | 46157060 | rs10769191 | 1.011 | 0.645 | 0.967 | -0.556 | -0.035 | 0.059 | 0.059 | 1.127 | 0.72 | 0.55 | 0.64 | 0.43 | 0.77 | 0.76 |
| 11 | 46157546 | rs12800259 | -0.059 | 0.717 | 1.046 | -0.588 | -0.001 | 0 | 0 | 0 | 0.19 | 0.18 | 0.14 | 0.15 | 0.29 | 0.19 |
| 11 | 46160551 | rs7118159 | 1.745 | 0.737 | 0.969 | -0.562 | 0.018 | 0.103 | 0.103 | 0.063 | 0.49 | 0.31 | 0.47 | 0.27 | 0.45 | 0.54 |
| 11 | 46169659 | rs11038784 | -1.511 | 0.953 | 1.225 | -0.787 | -0.002 | 0.005 | 0.005 | 0 | 0.4 | 0.32 | 0.27 | 0.25 | 0.53 | 0.42 |
| 11 | 46170469 | rs11038785 | 0.254 | 0.906 | 1.312 | -0.808 | 0 | 0.001 | 0.001 | 0 | 0.17 | 0.12 | 0.13 | 0.17 | 0.3 | 0.19 |
| 11 | 46171404 | rs3935366 | 2.111 | 0.767 | 1.46 | -0.85 | 0.016 | 0.095 | 0.095 | 0 | 0.51 | 0.37 | 0.48 | 0.23 | 0.43 | 0.48 |
| 11 | 46179854 | rs2029483 | 0.789 | 1.108 | 2.154 | -1.437 | 0.039 | -0.004 | -0.004 | 0 | 0.94 | 0.72 | 0.96 | 1 | 0.96 | 0.92 |
| 11 | 46185344 | rs2130474 | 1.702 | 0.912 | 1.682 | -1.786 | -0.025 | 0.109 | 0.109 | 0.024 | 0.56 | 0.29 | 0.7 | 0.75 | 0.51 | 0.55 |
| 11 | 46188225 | rs12282151 | 1.696 | 0.84 | 1.502 | -1.76 | -0.026 | 0.106 | 0.106 | 0.031 | 0.56 | 0.29 | 0.7 | 0.75 | 0.51 | 0.55 |
| 11 | 46197588 | rs11038803 | -0.427 | 0.918 | 1.538 | -1.754 | -0.022 | 0.02 | 0.02 | 0 | 0.23 | 0.13 | 0.16 | 0.17 | 0.31 | 0.21 |
| 11 | 46198841 | rs11038804 | -0.909 | 1.01 | 1.839 | -1.841 | 0.006 | -0.084 | -0.084 | 0 | 0.16 | 0.33 | 0.11 | 0.02 | 0.15 | 0.06 |
| 11 | 46206877 | rs11038808 | -0.384 | 1.35 | 1.917 | -1.536 | -0.022 | 0.02 | 0.02 | 0 | 0.22 | 0.12 | 0.16 | 0.17 | 0.31 | 0.2 |
| 11 | 46210259 | rs7103380 | 0.855 | 1.239 | 1.759 | -1.694 | 0 | -0.004 | -0.004 | 0 | 0.68 | 0.7 | 0.77 | 0.77 | 0.57 | 0.58 |
| 11 | 46227161 | rs10734545 | 1.125 | 1.256 | 1.886 | -1.595 | 0.01 | -0.081 | -0.081 | 0 | 0.84 | 0.54 | 0.9 | 0.98 | 0.87 | 0.85 |
| 11 | 46231502 | rs7121257 | -0.635 | 1.186 | 1.908 | -1.622 | 0.005 | -0.039 | -0.039 | 0 | 0.11 | 0.26 | 0.06 | 0.02 | 0.09 | 0.07 |
| 11 | 46231998 | rs12283653 | -1.575 | 1.434 | 1.665 | -1.541 | -0.011 | 0.016 | 0.016 | 0 | 0.36 | 0.65 | 0.27 | 0.27 | 0.43 | 0.38 |
| 11 | 46235626 | rs2863720 | 1.697 | 1.414 | 1.374 | -1.511 | -0.014 | 0.064 | 0.064 | 0.06 | 0.57 | 0.35 | 0.7 | 0.73 | 0.48 | 0.51 |
| 11 | 46237731 | rs4755349 | -0.673 | 1.537 | 1.333 | -1.481 | -0.037 | 0.023 | 0.023 | 0 | 0.25 | 0.12 | 0.19 | 0.19 | 0.37 | 0.24 |
| 11 | 46245361 | rs7124891 | 1.228 | 1.26 | 1.147 | -1.166 | 0.018 | -0.074 | -0.074 | 0 | 0.85 | 0.52 | 0.9 | 0.98 | 0.87 | 0.86 |
| 11 | 46260223 | rs7113478 | -0.936 | 1.373 | 0.509 | -1.441 | 0.012 | -0.072 | -0.072 | 0 | 0.36 | 0.55 | 0.23 | 0.16 | 0.41 | 0.39 |
| 11 | 46260696 | rs7116965 | -0.161 | 0.989 | 0.539 | -1.406 | 0.005 | 0.012 | 0.012 | 0 | 0.15 | 0.11 | 0.07 | 0.05 | 0.13 | 0.15 |
| 11 | 46262022 | rs11038830 | -0.514 | 0.796 | 0.636 | -1.001 | 0 | 0.018 | 0.018 | 0.034 | 0.13 | 0.21 | 0.09 | 0.12 | 0.11 | 0.1 |
| 11 | 46264906 | rs7926726 | -0.618 | 1.572 | 0.811 | -1.196 | 0.008 | -0.054 | -0.054 | 0 | 0.22 | 0.3 | 0.17 | 0.05 | 0.26 | 0.16 |
| 11 | 46268489 | rs7128258 | -0.806 | 1.677 | 0.526 | -1.214 | -0.008 | -0.082 | -0.082 | 0 | 0.24 | 0.41 | 0.15 | 0.12 | 0.24 | 0.26 |
| 11 | 46272585 | rs10838587 | -0.633 | 1.747 | 0.299 | -1.035 | 0 | -0.009 | -0.009 | 0 | 0.23 | 0.27 | 0.16 | 0.18 | 0.27 | 0.22 |
| 11 | 46274143 | rs10838591 | -1.112 | 1.734 | -0.197 | -0.913 | -0.001 | -0.007 | -0.007 | 0 | 0.42 | 0.62 | 0.32 | 0.24 | 0.51 | 0.43 |

| | | | | | | | | | | | | | | | | |
|---|---|---|---|---|---|---|---|---|---|---|---|---|---|---|---|---|
| 11 | 46279815 | rs6485671 | -1.42 | 1.383 | -0.215 | -0.509 | -0.002 | 0.001 | 0.001 | 6.811 | 0.17 | 0.81 | 0.13 | 0.11 | 0.15 | 0.11 |
| 11 | 46281835 | rs7131201 | -1.758 | 1.481 | -0.647 | -0.498 | -0.015 | 0.003 | 0.003 | 3.449 | 0.26 | 0.88 | 0.21 | 0.17 | 0.25 | 0.24 |
| 11 | 46297631 | rs10437653 | -1.372 | 1.523 | -1.287 | -1.388 | -0.037 | 0.3 | 0.3 | 0.365 | 0.49 | 0.92 | 0.34 | 0.15 | 0.51 | 0.44 |
| 11 | 46304368 | rs11038851 | -0.659 | 1.712 | -0.249 | -1.339 | 0.04 | -0.07 | -0.07 | 0.058 | 0.17 | 0.28 | 0.11 | 0.05 | 0.19 | 0.25 |
| 11 | 46308549 | rs7127254 | -1.631 | 1.19 | -0.063 | -1.421 | 0 | -0.002 | -0.002 | 3.064 | 0.28 | 0.7 | 0.17 | 0.14 | 0.26 | 0.31 |
| 11 | 46309968 | rs6485672 | -1.733 | 1.113 | -0.781 | -1.412 | -0.077 | 0.081 | 0.081 | 0.299 | 0.5 | 0.96 | 0.33 | 0.25 | 0.51 | 0.51 |
| 11 | 46332893 | rs4625425 | -1.099 | 0.976 | 0.077 | -1.219 | 0.007 | -0.077 | -0.077 | 0 | 0.2 | 0.5 | 0.12 | 0.06 | 0.18 | 0.26 |
| 11 | 46342834 | rs7476 | -1.521 | 0.911 | -0.655 | -1.262 | -0.017 | 0 | 0 | 0 | 0.39 | 0.94 | 0.23 | 0.25 | 0.33 | 0.5 |
| 11 | 46348324 | rs11038863 | -0.998 | 0.92 | 0.292 | -1.273 | 0.019 | -0.065 | -0.065 | 0 | 0.18 | 0.34 | 0.11 | 0.06 | 0.18 | 0.25 |
| 11 | 46370767 | rs876701 | -1.276 | 0.636 | 1.137 | -1.302 | -0.009 | -0.034 | -0.034 | 0 | 0.19 | 0.33 | 0.1 | 0.05 | 0.17 | 0.15 |
| 11 | 46378708 | rs10838599 | 0.366 | 1.119 | 0.255 | -1.85 | -0.084 | 0.063 | 0.063 | 0 | 0.15 | 0.01 | 0.23 | 0.03 | 0.27 | 0.09 |
| 11 | 46393757 | rs3740976 | 1.039 | 0.625 | 0.344 | -1.654 | -0.017 | 0 | 0 | 0 | 0.14 | 0.04 | 0.09 | 0.08 | 0.14 | 0.2 |
| 11 | 46396479 | rs2046768 | -1.085 | 0.558 | 0.521 | -1.676 | -0.02 | -0.06 | -0.06 | 0 | 0.21 | 0.46 | 0.11 | 0.06 | 0.17 | 0.15 |
| 11 | 46454313 | rs16938506 | -1.45 | -0.027 | 0.71 | -0.91 | -0.032 | -0.082 | -0.082 | 0.056 | 0.23 | 0.6 | 0.12 | 0.05 | 0.17 | 0.15 |
| 11 | 46469900 | rs7951579 | -1.87 | -0.087 | 0.706 | -0.939 | 0.043 | 0.08 | 0.08 | 1.505 | 0.26 | 0.87 | 0.14 | 0.05 | 0.18 | 0.15 |
| 11 | 46520302 | rs17197116 | -0.023 | -0.074 | 0.822 | -0.938 | -0.001 | -0.001 | -0.001 | 0 | 0.09 | 0.1 | 0.06 | 0.05 | 0.08 | 0.07 |
| 11 | 46534546 | rs3802890 | -1.67 | -0.108 | 0.747 | -0.976 | 0.036 | 0.128 | 0.128 | 0.159 | 0.4 | 0.91 | 0.23 | 0.13 | 0.31 | 0.35 |
| 11 | 46551918 | rs10160701 | -1.434 | -0.147 | 0.874 | -0.977 | -0.032 | -0.083 | -0.083 | 3.458 | 0.23 | 0.6 | 0.12 | 0.05 | 0.17 | 0.15 |
| 11 | 46560680 | rs11819869 | -1.414 | 0.029 | 0.813 | -1.038 | -0.033 | -0.083 | -0.083 | 0 | 0.23 | 0.6 | 0.12 | 0.05 | 0.17 | 0.15 |
| 11 | 46596674 | rs7481312 | -0.807 | 0.04 | 0.732 | -1.047 | -0.013 | -0.053 | -0.053 | 0 | 0.2 | 0.41 | 0.1 | 0.05 | 0.17 | 0.14 |
| 11 | 46622392 | rs7125907 | -0.797 | -0.003 | 0.704 | -1.058 | -0.007 | -0.026 | -0.026 | 0 | 0.2 | 0.29 | 0.1 | 0.05 | 0.17 | 0.14 |
| 11 | 46631181 | rs35623865 | -1.192 | -0.001 | 0.689 | -1.057 | -0.022 | -0.062 | -0.062 | 0 | 0.22 | 0.43 | 0.11 | 0.05 | 0.17 | 0.15 |
| 11 | 46640503 | rs7484002 | -0.815 | -0.001 | 0.689 | -1.057 | -0.016 | -0.053 | -0.053 | 0.043 | 0.2 | 0.41 | 0.1 | 0.05 | 0.17 | 0.14 |
| 11 | 46644143 | rs12292425 | -0.843 | 0.001 | 0.679 | -1.057 | -0.017 | -0.058 | -0.058 | 0 | 0.2 | 0.42 | 0.1 | 0.05 | 0.17 | 0.14 |
| 11 | 46646030 | rs7122039 | -1.168 | 0.016 | 0.616 | -1.05 | -0.022 | -0.062 | -0.062 | 0.002 | 0.22 | 0.43 | 0.11 | 0.05 | 0.18 | 0.15 |
| 11 | 46676357 | rs10769204 | 1.236 | 0.106 | 0.233 | -1.01 | 0.038 | 0.125 | 0.125 | 0.145 | 0.6 | 0.09 | 0.77 | 0.87 | 0.69 | 0.66 |
| 11 | 46686260 | rs10838610 | 1.273 | 0.108 | 0.227 | -1.01 | 0.039 | 0.128 | 0.128 | 0.124 | 0.6 | 0.09 | 0.77 | 0.87 | 0.69 | 0.65 |
| 11 | 46691737 | rs4606447 | 0.034 | -0.691 | -0.851 | -0.225 | 0.078 | -0.075 | -0.075 | 0 | 0.61 | 0.12 | 0.69 | 0.38 | 0.73 | 0.71 |
| 11 | 46692870 | rs4752926 | 0.54 | -0.665 | -0.88 | -0.283 | 0.022 | 0.089 | 0.089 | 0.056 | 0.47 | 0.08 | 0.6 | 0.27 | 0.59 | 0.51 |
| 11 | 46701728 | rs10838612 | 2.053 | -0.928 | -0.529 | 0.063 | 0.018 | -0.138 | -0.138 | 0.44 | 0.16 | 0 | 0.19 | 0.6 | 0.1 | 0.17 |
| 11 | 46702691 | rs4319473 | 0.047 | -0.888 | -0.396 | -0.033 | 0.004 | 0.02 | 0.02 | 0 | 0.16 | 0.12 | 0.08 | 0.05 | 0.16 | 0.12 |
| 11 | 46723603 | rs10769205 | 0.289 | -0.907 | -0.618 | 0.022 | 0.079 | -0.01 | -0.01 | 0 | 0.6 | 0.12 | 0.59 | 0.31 | 0.71 | 0.7 |
| 11 | 46734384 | rs11038977 | 2.026 | -0.866 | -0.634 | 0.091 | 0.006 | -0.14 | -0.14 | 1.158 | 0.16 | 0.01 | 0.28 | 0.6 | 0.12 | 0.17 |
| 11 | 46742426 | rs3136435 | -0.038 | -0.775 | -0.299 | 0.031 | 0.024 | 0.11 | 0.11 | 0.225 | 0.1 | 0.4 | 0.07 | 0.1 | 0.08 | 0.05 |
| 11 | 46743135 | rs3136439 | 1.236 | -0.754 | -0.475 | 0.019 | -0.01 | 0.054 | 0.054 | 0 | 0.1 | 0.02 | 0.03 | 0 | 0.08 | 0.07 |
| 11 | 46745861 | rs3136456 | 0.81 | -0.739 | -0.416 | 0.033 | -0.001 | 0.007 | 0.007 | 0 | 0.07 | 0.09 | 0.05 | 0.1 | 0.08 | 0.05 |
| 11 | 46749648 | rs5898 | 1.244 | -0.789 | -0.466 | 0.052 | -0.01 | 0.052 | 0.052 | 0 | 0.09 | 0.02 | 0.03 | 0 | 0.08 | 0.07 |
| 11 | 46784697 | rs7928445 | 0.503 | -0.85 | -0.519 | 0.05 | -0.008 | -0.083 | -0.083 | 0 | 0.13 | 0.34 | 0.05 | 0 | 0.09 | 0.07 |
| 11 | 46787573 | rs10734548 | -0.441 | -0.916 | -0.825 | 0.187 | -0.009 | 0.006 | 0.006 | 0 | 0.68 | 0.4 | 0.65 | 0.39 | 0.79 | 0.76 |
| 11 | 46798631 | rs6485690 | 0.195 | -0.572 | -0.858 | 0.166 | 0.051 | 0.015 | 0.015 | 0 | 0.59 | 0.13 | 0.56 | 0.29 | 0.68 | 0.71 |
| 11 | 46810916 | rs11038993 | 0.808 | -0.441 | -1.05 | 0.104 | -0.079 | 0.137 | 0.137 | 0 | 0.44 | 0.07 | 0.46 | 0.18 | 0.51 | 0.37 |
| 11 | 46812227 | rs7940578 | -0.306 | -0.621 | -0.493 | 0.486 | -0.015 | 0.013 | 0.013 | 0 | 0.68 | 0.41 | 0.65 | 0.39 | 0.79 | 0.76 |
| 11 | 46823681 | rs12575721 | 0.614 | -0.434 | -0.649 | 0.516 | -0.014 | -0.016 | -0.016 | 0.043 | 0.07 | 0.03 | 0.07 | 0.1 | 0.11 | 0.05 |
| 11 | 46842045 | rs6485696 | 0.388 | -0.599 | -0.761 | 0.448 | 0.052 | 0.018 | 0.018 | 0 | 0.59 | 0.12 | 0.56 | 0.29 | 0.68 | 0.71 |
| 11 | 46849360 | rs1007738 | -0.272 | -0.653 | -0.478 | 0.47 | -0.01 | 0.007 | 0.007 | 0 | 0.68 | 0.42 | 0.65 | 0.39 | 0.79 | 0.76 |
| 11 | 46872873 | rs10769211 | -0.285 | -0.661 | -0.451 | 0.466 | -0.007 | 0.004 | 0.004 | 0 | 0.68 | 0.4 | 0.65 | 0.39 | 0.79 | 0.76 |
| 11 | 46875933 | rs6485700 | -0.58 | -0.759 | -0.227 | 0.407 | -0.009 | -0.086 | -0.086 | 3.021 | 0.87 | 0.62 | 0.95 | 1 | 0.91 | 0.93 |
| 11 | 46890165 | rs3816614 | -0.172 | -0.739 | -0.517 | 0.187 | -0.006 | 0.003 | 0.003 | 0 | 0.68 | 0.4 | 0.65 | 0.39 | 0.79 | 0.76 |
| 11 | 46897446 | rs2306033 | 2.023 | -0.744 | -0.832 | 0.201 | 0.005 | -0.128 | -0.128 | 0 | 0.15 | 0 | 0.28 | 0.61 | 0.12 | 0.17 |
| 11 | 46898771 | rs6485702 | 0.397 | -0.616 | -0.783 | 0.152 | 0.052 | 0.015 | 0.015 | 0 | 0.59 | 0.13 | 0.57 | 0.29 | 0.68 | 0.71 |
| 11 | 46916693 | rs3751097 | 0.352 | -0.358 | -0.611 | 0.181 | -0.002 | 0.001 | 0.001 | 0 | 0.08 | 0.08 | 0.07 | 0.05 | 0.11 | 0.05 |
| 11 | 46917983 | rs898604 | -0.948 | -0.823 | -0.307 | 0.242 | -0.035 | 0.059 | 0.059 | 0 | 0.78 | 0.63 | 0.69 | 0.38 | 0.86 | 0.81 |
| 11 | 46929777 | rs7102325 | 0.043 | -0.689 | -0.194 | -0.252 | -0.017 | -0.091 | -0.091 | 0 | 0.14 | 0.56 | 0.06 | 0 | 0.09 | 0.07 |
| 11 | 46945227 | rs10838635 | -0.001 | -0.701 | -0.607 | -0.094 | 0.075 | -0.039 | -0.039 | 0 | 0.65 | 0.15 | 0.64 | 0.33 | 0.77 | 0.74 |
| 11 | 46947366 | rs17787912 | 0.871 | -0.067 | -1.329 | -0.825 | -0.069 | 0.09 | 0.09 | 0 | 0.14 | 0.01 | 0.14 | 0 | 0.23 | 0.03 |
| 11 | 46957808 | rs7945558 | -0.317 | -0.252 | -1.207 | -0.748 | 0.079 | -0.038 | -0.038 | 0 | 0.65 | 0.15 | 0.64 | 0.33 | 0.77 | 0.74 |
| 11 | 46962474 | rs6485713 | 0.029 | -0.355 | -0.991 | -0.799 | 0.02 | 0.017 | 0.017 | 6.376 | 0.05 | 0.66 | 0.04 | 0 | 0 | 0 |
| 11 | 46963196 | rs2035067 | 0.507 | -0.45 | -1.009 | -0.846 | -0.025 | -0.027 | -0.027 | 0 | 0.93 | 0.3 | 0.95 | 1 | 0.99 | 1 |
| 11 | 46982481 | rs10769225 | -0.929 | -0.446 | -0.977 | -0.844 | 0.036 | -0.045 | -0.045 | 1.264 | 0.71 | 0.86 | 0.69 | 0.33 | 0.78 | 0.74 |
| 11 | 46984803 | rs10769226 | -0.997 | -0.453 | -0.942 | -0.847 | 0.036 | -0.086 | -0.086 | 0 | 0.72 | 0.95 | 0.69 | 0.33 | 0.78 | 0.74 |
| 11 | 46998512 | rs4548577 | -0.974 | -0.452 | -0.957 | -0.846 | 0.036 | -0.089 | -0.089 | 0 | 0.72 | 0.95 | 0.69 | 0.33 | 0.78 | 0.74 |
| 11 | 47008379 | rs7935346 | -0.312 | -0.453 | -0.995 | -0.847 | 0.048 | -0.018 | -0.018 | 0 | 0.66 | 0.24 | 0.64 | 0.33 | 0.78 | 0.74 |
| 11 | 47038220 | rs4494268 | -2.254 | -0.414 | -1.092 | -0.801 | 0.008 | -0.13 | -0.13 | 0 | 0.82 | 1 | 0.72 | 0.33 | 0.87 | 0.8 |
| 11 | 47044249 | rs7124648 | -0.777 | -0.533 | -1.113 | -0.881 | 0.053 | -0.044 | -0.044 | 2.912 | 0.69 | 0.86 | 0.69 | 0.33 | 0.78 | 0.74 |
| 11 | 47072805 | rs7103581 | -0.129 | -0.381 | -1.16 | -0.979 | 0.013 | 0.009 | 0.009 | 0 | 0.61 | 0.67 | 0.61 | 0.28 | 0.68 | 0.7 |
| 11 | 47081732 | rs11039074 | -0.177 | -0.383 | -1.162 | -0.98 | 0.027 | 0.018 | 0.018 | 0 | 0.61 | 0.74 | 0.61 | 0.28 | 0.68 | 0.7 |
| 11 | 47100796 | rs7946709 | 0.18 | -0.46 | -1.164 | -1.017 | 0.071 | 0.044 | 0.044 | 0 | 0.56 | 0.1 | 0.57 | 0.28 | 0.68 | 0.7 |
| 11 | 47110652 | rs7945315 | -0.186 | -0.472 | -1.125 | -0.987 | 0.01 | 0.006 | 0.006 | 0 | 0.61 | 0.66 | 0.61 | 0.28 | 0.68 | 0.7 |
| 11 | 47127153 | rs7117404 | 2.269 | -0.274 | -1.911 | -1.613 | -0.038 | 0.086 | 0.086 | 0.068 | 0.17 | 0.01 | 0.07 | 0.03 | 0.13 | 0.06 |

| | | | | | | | | | | | | | | | | |
|---|---|---|---|---|---|---|---|---|---|---|---|---|---|---|---|---|
| 11 | 47127549 | rs17790336 | 1.015 | -0.284 | -1.453 | -1.165 | -0.002 | 0.003 | 0.003 | 0 | 0.09 | 0.08 | 0.07 | 0.05 | 0.1 | 0.04 |
| 11 | 47127706 | rs17790342 | 1.015 | -0.284 | -1.452 | -1.165 | -0.003 | 0.003 | 0.003 | 0 | 0.09 | 0.07 | 0.07 | 0.05 | 0.1 | 0.04 |
| 11 | 47151897 | rs17790390 | 1.046 | -0.285 | -1.453 | -1.165 | -0.002 | 0.002 | 0.002 | 0 | 0.09 | 0.08 | 0.07 | 0.05 | 0.1 | 0.04 |
| 11 | 47152273 | rs10838660 | -0.018 | -0.301 | -1.405 | -1.182 | 0.074 | 0.045 | 0.045 | 0 | 0.56 | 0.11 | 0.57 | 0.28 | 0.67 | 0.7 |
| 11 | 47168254 | rs1352307 | 1.82 | -0.297 | -1.498 | -1.188 | -0.012 | 0.052 | 0.052 | 0 | 0.09 | 0 | 0.02 | 0 | 0.08 | 0.06 |
| 11 | 47175327 | rs752849 | 0.616 | 0.016 | -1.48 | -1.185 | 0.018 | -0.064 | -0.064 | 0 | 0.23 | 0.59 | 0.17 | 0.07 | 0.23 | 0.37 |
| 11 | 47176005 | rs747650 | 0.277 | -0.02 | -1.516 | -1.203 | 0.022 | 0.013 | 0.013 | 0 | 0.39 | 0.28 | 0.39 | 0.72 | 0.32 | 0.29 |
| 11 | 47179143 | rs2291444 | -1.175 | -0.071 | -1.538 | -1.224 | -0.061 | 0.076 | 0.076 | 0 | 0.61 | 0.89 | 0.56 | 0.79 | 0.56 | 0.66 |
| 11 | 47179829 | rs1060573 | -0.529 | -0.026 | -1.536 | -1.156 | 0.013 | 0.008 | 0.008 | 0 | 0.6 | 0.67 | 0.61 | 0.28 | 0.68 | 0.7 |
| 11 | 47182926 | rs3740694 | 1.55 | 0.355 | -1.718 | -1.184 | -0.041 | 0.015 | 0.015 | 0 | 0.16 | 0.01 | 0.08 | 0.07 | 0.16 | 0.37 |
| 11 | 47184117 | rs10838662 | 1.264 | 0.084 | -1.654 | -1.139 | 0.042 | -0.074 | -0.074 | 0.178 | 0.28 | 0.07 | 0.31 | 0.67 | 0.22 | 0.25 |
| 11 | 47185680 | rs11039114 | 1.213 | 0.167 | -1.67 | -1.144 | -0.083 | 0.029 | 0.029 | 0 | 0.19 | 0.02 | 0.13 | 0.07 | 0.23 | 0.37 |
| 11 | 47188411 | rs3740691 | -0.326 | 0.263 | -1.516 | -1.13 | 0 | -0.007 | -0.007 | 0 | 0.33 | 0.69 | 0.24 | 0.12 | 0.34 | 0.42 |
| 11 | 47188592 | rs3740690 | 0.971 | 0.257 | -1.729 | -1.136 | -0.102 | 0.059 | 0.059 | 0 | 0.37 | 0.05 | 0.44 | 0.21 | 0.44 | 0.33 |
| 11 | 47196982 | rs2279439 | 0.846 | -0.052 | -1.152 | -0.531 | 0.039 | -0.059 | -0.059 | 0.033 | 0.28 | 0.11 | 0.31 | 0.67 | 0.23 | 0.25 |
| 11 | 47201924 | rs11039119 | 0.012 | 0.019 | -1.12 | -0.55 | 0.016 | 0.013 | 0.013 | 0 | 0.38 | 0.23 | 0.38 | 0.72 | 0.34 | 0.3 |
| 11 | 47203068 | rs7940240 | -0.918 | -0.036 | -1.171 | -0.505 | 0.014 | -0.016 | -0.016 | 0 | 0.71 | 0.78 | 0.68 | 0.33 | 0.76 | 0.75 |
| 11 | 47203904 | rs10838664 | 0.768 | -0.025 | -1.231 | -0.499 | 0.029 | -0.044 | -0.044 | 0 | 0.28 | 0.13 | 0.31 | 0.67 | 0.24 | 0.25 |
| 11 | 47234718 | rs2029298 | -0.331 | 0.063 | -1.01 | -0.539 | 0.001 | 0.001 | 0.001 | 0 | 0.61 | 0.55 | 0.6 | 0.29 | 0.66 | 0.7 |
| 11 | 47246397 | rs7395581 | -0.417 | 0.168 | -0.758 | -0.475 | -0.011 | -0.012 | -0.012 | 0.711 | 0.66 | 0.48 | 0.62 | 0.22 | 0.7 | 0.55 |
| 11 | 47249294 | rs2957873 | -0.975 | 0.096 | -0.815 | -0.39 | -0.043 | 0.054 | 0.054 | 0 | 0.74 | 0.5 | 0.69 | 0.27 | 0.81 | 0.67 |
| 11 | 47260319 | rs901746 | -0.247 | 0.023 | -0.588 | -0.376 | -0.005 | -0.005 | -0.005 | 0.294 | 0.35 | 0.62 | 0.35 | 0.78 | 0.3 | 0.46 |
| 11 | 47261260 | rs11988 | 0.453 | 0.21 | -1.421 | -1.314 | -0.082 | 0.138 | 0.138 | 0 | 0.29 | 0.07 | 0.2 | 0.02 | 0.38 | 0.24 |
| 11 | 47266808 | rs2242261 | 1.484 | 0.187 | -1.307 | -1.196 | -0.06 | 0.117 | 0.117 | 0 | 0.23 | 0.46 | 0.26 | 0.68 | 0.15 | 0.2 |
| 11 | 47270255 | rs2167079 | 0.095 | 0.196 | -1.308 | -1.2 | -0.005 | -0.005 | -0.005 | 3.052 | 0.35 | 0.62 | 0.35 | 0.78 | 0.3 | 0.45 |
| 11 | 47275064 | rs10838681 | 0.547 | 0.215 | -1.348 | -1.177 | -0.024 | 0.011 | 0.011 | 0 | 0.31 | 0.49 | 0.33 | 0.73 | 0.26 | 0.32 |
| 11 | 47278938 | rs3758674 | 0.552 | 0.207 | -1.355 | -1.173 | -0.026 | -0.006 | -0.006 | 0.049 | 0.12 | 0.04 | 0.07 | 0.1 | 0.15 | 0.26 |
| 11 | 47280653 | rs12221497 | 0.552 | 0.207 | -1.355 | -1.173 | -0.026 | -0.006 | -0.006 | 0 | 0.12 | 0.04 | 0.07 | 0.1 | 0.15 | 0.26 |
| 11 | 47280762 | rs11039155 | 2.145 | -0.031 | -1.439 | -1.249 | -0.004 | 0.106 | 0.106 | 0 | 0.17 | 0 | 0.13 | 0.01 | 0.14 | 0.16 |
| 11 | 47282024 | rs2279238 | 1.409 | 0.089 | -1.263 | -1.142 | -0.052 | 0.11 | 0.11 | 0 | 0.22 | 0.42 | 0.26 | 0.67 | 0.15 | 0.2 |
| 11 | 47286290 | rs7120118 | 0.064 | 0.117 | -1.282 | -1.169 | -0.015 | -0.014 | -0.014 | 0 | 0.34 | 0.52 | 0.34 | 0.77 | 0.3 | 0.46 |
| 11 | 47290984 | rs1449627 | 0.065 | 0.097 | -1.329 | -0.914 | -0.013 | 0.005 | 0.005 | 0 | 0.35 | 0.52 | 0.34 | 0.73 | 0.31 | 0.46 |
| 11 | 47298360 | rs326214 | -0.145 | -0.122 | -1.318 | -0.961 | 0.012 | 0.001 | 0.001 | 0 | 0.62 | 0.31 | 0.64 | 0.27 | 0.69 | 0.54 |
| 11 | 47303275 | rs326217 | 1.188 | -0.108 | -1.43 | -1.042 | 0.002 | 0.001 | 0.001 | 0 | 0.33 | 0.25 | 0.44 | 0.25 | 0.3 | 0.3 |
| 11 | 47306585 | rs1051006 | 1.648 | -0.065 | -1.491 | -0.973 | -0.043 | 0.122 | 0.122 | 0 | 0.22 | 0.39 | 0.26 | 0.62 | 0.16 | 0.2 |
| 11 | 47306628 | rs3736101 | 1.306 | 0.022 | -1.748 | -0.985 | -0.026 | 0 | 0 | 0 | 0.09 | 0.03 | 0.07 | 0.05 | 0.12 | 0.12 |
| 11 | 47312689 | rs11039165 | 1.296 | 0.107 | -1.913 | -1.087 | -0.099 | 0.065 | 0.065 | 0 | 0.18 | 0.01 | 0.15 | 0.02 | 0.27 | 0.15 |
| 11 | 47312892 | rs10838687 | 0.962 | -0.067 | -1.746 | -0.926 | -0.045 | 0.088 | 0.088 | 0 | 0.27 | 0.49 | 0.27 | 0.67 | 0.2 | 0.33 |
| 11 | 47318157 | rs749067 | 0.331 | -0.058 | -1.816 | -0.936 | -0.142 | 0.144 | 0.144 | 0 | 0.29 | 0.02 | 0.2 | 0.02 | 0.39 | 0.25 |
| 11 | 47339180 | rs4752979 | 0.568 | 0.095 | -1.296 | -0.486 | -0.047 | 0.095 | 0.095 | 0 | 0.26 | 0.49 | 0.27 | 0.67 | 0.2 | 0.34 |
| 11 | 47346145 | rs2290148 | 0.946 | 0.063 | -1.267 | -0.439 | -0.061 | 0.162 | 0.162 | 0 | 0.23 | 0.47 | 0.26 | 0.62 | 0.16 | 0.2 |
| 11 | 47354787 | rs1052373 | -0.248 | 0.194 | -1.336 | -0.524 | -0.014 | 0.007 | 0.007 | 0 | 0.35 | 0.51 | 0.33 | 0.72 | 0.31 | 0.46 |
| 11 | 47359706 | rs11570094 | 1.359 | -0.136 | -1.81 | -0.429 | -0.02 | -0.047 | -0.047 | 0.008 | 0.31 | 0.04 | 0.42 | 0.27 | 0.29 | 0.32 |
| 11 | 47362339 | rs10769253 | 0.922 | -0.181 | -0.96 | 0.167 | -0.041 | 0.157 | 0.157 | 0 | 0.21 | 0.39 | 0.26 | 0.6 | 0.17 | 0.2 |
| 11 | 47366095 | rs11570067 | 0.193 | 0.333 | -1.519 | -0.114 | -0.017 | 0.009 | 0.009 | 0 | 0.11 | 0.09 | 0.09 | 0.05 | 0.14 | 0.05 |
| 11 | 47369443 | rs11570058 | 0.314 | 0.345 | -1.63 | -0.159 | -0.033 | 0.033 | 0.033 | 0 | 0.1 | 0.03 | 0.08 | 0.02 | 0.14 | 0.05 |
| 11 | 47370041 | rs3729989 | 0.127 | 0.319 | -1.601 | -0.091 | -0.021 | 0.021 | 0.021 | 0 | 0.11 | 0.04 | 0.08 | 0.03 | 0.14 | 0.05 |
| 11 | 47370607 | rs2697920 | 0.147 | 0.219 | -2.083 | 0.164 | 0.029 | -0.05 | -0.05 | 0 | 0.57 | 0.15 | 0.6 | 0.49 | 0.65 | 0.48 |
| 11 | 47380593 | rs3740689 | 0.527 | -0.347 | -2.154 | -0.13 | 0.04 | 0.06 | 0.06 | 0 | 0.49 | 0.08 | 0.51 | 0.48 | 0.57 | 0.64 |
| 11 | 47385923 | rs10838698 | -1.521 | 0.224 | -1.646 | -0.044 | 0.024 | 0.017 | 0.017 | 0 | 0.63 | 0.75 | 0.56 | 0.7 | 0.67 | 0.67 |
| 11 | 47394305 | rs896817 | -0.684 | 0.057 | -1.397 | -0.173 | -0.061 | 0.001 | 0.001 | 0 | 0.33 | 0.15 | 0.25 | 0.37 | 0.38 | 0.5 |
| 11 | 47410951 | rs4752993 | -1.793 | 0.062 | -1.327 | -0.122 | 0.017 | 0.014 | 0.014 | 0 | 0.63 | 0.86 | 0.58 | 0.72 | 0.67 | 0.67 |
| 11 | 47428565 | rs4752999 | 1.6 | -0.036 | -1.347 | -0.169 | 0.003 | 0.011 | 0.011 | 0 | 0.35 | 0.12 | 0.42 | 0.26 | 0.33 | 0.33 |
| 11 | 47431529 | rs753812 | 1.402 | -0.116 | -0.642 | -0.174 | -0.028 | 0.079 | 0.079 | 0 | 0.17 | 0.36 | 0.21 | 0.3 | 0.14 | 0.13 |
| 11 | 47432303 | rs755553 | -1.563 | -0.114 | -0.505 | -0.172 | -0.003 | -0.004 | -0.004 | 0 | 0.61 | 0.58 | 0.56 | 0.74 | 0.67 | 0.67 |
| 11 | 47434986 | rs2293576 | -0.024 | 0.004 | -0.696 | -0.211 | -0.135 | -0.101 | -0.101 | 0 | 0.26 | 0.01 | 0.21 | 0.4 | 0.31 | 0.48 |
| 11 | 47441513 | rs10838708 | 1.092 | 0.28 | -0.648 | -0.237 | -0.049 | 0.052 | 0.052 | 0 | 0.4 | 0.18 | 0.5 | 0.24 | 0.45 | 0.36 |
| 11 | 47449072 | rs12292911 | -0.576 | 0.051 | -0.317 | -0.149 | -0.052 | -0.005 | -0.005 | 0 | 0.33 | 0.14 | 0.26 | 0.4 | 0.37 | 0.51 |
| 11 | 47449544 | rs7105122 | 0.059 | -0.375 | -0.059 | 0.073 | -0.005 | 0.044 | 0.044 | 0 | 0.28 | 0.44 | 0.3 | 0.34 | 0.29 | 0.16 |
| 11 | 47454701 | rs10742805 | -0.143 | -0.36 | -0.337 | 0.091 | -0.002 | 0.005 | 0.005 | 0 | 0.71 | 0.35 | 0.69 | 0.66 | 0.71 | 0.84 |
| 11 | 47460306 | rs7126210 | -0.138 | -0.36 | -0.327 | 0.098 | -0.003 | 0.006 | 0.006 | 0 | 0.71 | 0.35 | 0.69 | 0.66 | 0.71 | 0.84 |
| 11 | 47468569 | rs3824867 | -0.091 | -0.4 | -0.359 | 0.078 | 0.009 | -0.009 | -0.009 | 0 | 0.69 | 0.26 | 0.68 | 0.66 | 0.7 | 0.84 |
| 11 | 47474146 | rs12361415 | -0.001 | -0.293 | -0.627 | 0.103 | -0.099 | -0.065 | -0.065 | 0 | 0.24 | 0.01 | 0.18 | 0.33 | 0.26 | 0.26 |
| 11 | 47487740 | rs1044269 | -0.91 | -0.42 | -0.358 | 0.124 | -0.041 | 0.042 | 0.042 | 0.006 | 0.79 | 0.32 | 0.78 | 0.71 | 0.85 | 0.87 |
| 11 | 47495202 | rs4752839 | 1.377 | -0.456 | -0.009 | 0.113 | -0.026 | 0.144 | 0.144 | 0 | 0.16 | 0.49 | 0.2 | 0.28 | 0.14 | 0.12 |
| 11 | 47500267 | rs2242081 | -0.767 | -0.342 | -0.145 | 0.106 | -0.066 | -0.024 | -0.024 | 0 | 0.36 | 0.11 | 0.32 | 0.44 | 0.43 | 0.53 |
| 11 | 47529037 | rs17726787 | 0.756 | -0.304 | -0.14 | 0.087 | -0.012 | 0.01 | 0.01 | 0 | 0.1 | 0.04 | 0.1 | 0.05 | 0.15 | 0.04 |
| 11 | 47529947 | rs7124681 | 1.138 | -0.346 | -0.124 | 0.079 | 0.005 | 0.059 | 0.059 | 0 | 0.43 | 0.21 | 0.45 | 0.27 | 0.42 | 0.35 |
| 11 | 47539697 | rs4752845 | 2.006 | -0.4 | -0.184 | -0.03 | -0.051 | -0.017 | -0.017 | 0.413 | 0.35 | 0.05 | 0.4 | 0.27 | 0.36 | 0.33 |

| | | | | | | | | | | | | | | | | |
|---|---|---|---|---|---|---|---|---|---|---|---|---|---|---|---|---|
| 11 | 47540007 | rs9666924 | -1.118 | -0.405 | -0.176 | -0.027 | -0.043 | 0.051 | 0.051 | 0 | 0.79 | 0.36 | 0.78 | 0.71 | 0.85 | 0.87 |
| 11 | 47554337 | rs10742814 | -0.726 | -0.407 | -0.18 | -0.028 | -0.066 | -0.019 | -0.019 | 0 | 0.36 | 0.11 | 0.32 | 0.43 | 0.43 | 0.53 |
| 11 | 47583121 | rs12798346 | 1.963 | -0.398 | -0.188 | -0.023 | -0.052 | -0.018 | -0.018 | 0.411 | 0.35 | 0.05 | 0.4 | 0.27 | 0.36 | 0.33 |
| 11 | 47600438 | rs2280231 | 0.204 | -0.38 | -0.178 | 0.001 | -0.097 | -0.057 | -0.057 | 0 | 0.24 | 0.01 | 0.17 | 0.32 | 0.25 | 0.25 |
| 11 | 47606984 | rs10742816 | -1.609 | -0.389 | 0.06 | 0.018 | -0.034 | 0.09 | 0.09 | 0 | 0.82 | 0.57 | 0.79 | 0.71 | 0.86 | 0.87 |
| 11 | 47648042 | rs4752856 | 2.031 | -0.428 | -0.026 | -0.097 | -0.056 | -0.022 | -0.022 | 0 | 0.34 | 0.05 | 0.4 | 0.27 | 0.35 | 0.32 |
| 11 | 47659135 | rs7118178 | 0.204 | -0.427 | -0.029 | -0.088 | -0.097 | -0.057 | -0.057 | 0.004 | 0.24 | 0.01 | 0.17 | 0.32 | 0.26 | 0.25 |
| 11 | 47663049 | rs10838738 | 1.927 | -0.562 | -0.005 | -0.1 | -0.051 | -0.017 | -0.017 | 0 | 0.35 | 0.05 | 0.4 | 0.28 | 0.35 | 0.33 |
| 11 | 47674101 | rs2869024 | 0.676 | -0.409 | 0.19 | -0.09 | -0.036 | 0.001 | 0.001 | 0 | 0.62 | 0.8 | 0.67 | 0.57 | 0.57 | 0.47 |
| 11 | 47679980 | rs10838740 | -0.855 | -0.384 | 0.225 | -0.094 | -0.042 | -0.004 | -0.004 | 0 | 0.37 | 0.18 | 0.33 | 0.43 | 0.43 | 0.53 |
| 11 | 47681083 | rs11608017 | 1.392 | -0.432 | 0.085 | -0.076 | -0.005 | 0.031 | 0.031 | 0 | 0.06 | 0 | 0.03 | 0 | 0.06 | 0.02 |
| 11 | 47681370 | rs1056387 | -1.021 | -0.4 | 0.111 | -0.042 | -0.047 | 0.001 | 0.001 | 0 | 0.38 | 0.15 | 0.32 | 0.43 | 0.44 | 0.53 |
| 11 | 47689343 | rs10742817 | -0.771 | -0.361 | 0.072 | -0.043 | -0.065 | -0.014 | -0.014 | 0 | 0.36 | 0.11 | 0.32 | 0.43 | 0.43 | 0.53 |
| 11 | 47694699 | rs4539273 | 0.791 | -0.365 | 0.111 | -0.045 | 0 | -0.001 | -0.001 | 0 | 0.61 | 0.64 | 0.67 | 0.57 | 0.57 | 0.47 |
| 11 | 47699398 | rs7930612 | 0.779 | -0.306 | 0.107 | -0.05 | -0.004 | 0.003 | 0.003 | 0 | 0.1 | 0.06 | 0.1 | 0.05 | 0.15 | 0.04 |
| 11 | 47701528 | rs12286721 | 0.688 | -0.269 | 0.194 | 0.025 | 0 | -0.001 | -0.001 | 0 | 0.61 | 0.64 | 0.67 | 0.57 | 0.57 | 0.47 |
| 11 | 47720228 | rs11039345 | -0.832 | -0.284 | 0.029 | 0.032 | -0.074 | -0.012 | -0.012 | 0 | 0.38 | 0.11 | 0.32 | 0.43 | 0.44 | 0.53 |
| 11 | 47729180 | rs10769288 | -0.535 | -0.171 | -0.037 | 0.019 | -0.094 | -0.023 | -0.023 | 0.004 | 0.33 | 0.06 | 0.25 | 0.39 | 0.37 | 0.43 |
| 11 | 47733494 | rs1017730 | 0.681 | -0.486 | 0.047 | 0.014 | -0.091 | -0.076 | -0.076 | 0.075 | 0.16 | 0.01 | 0.15 | 0.32 | 0.24 | 0.18 |
| 11 | 47746962 | rs11039358 | 0.532 | -0.611 | 0.569 | 0.166 | -0.007 | -0.042 | -0.042 | 0 | 0.08 | 0.13 | 0.04 | 0 | 0.07 | 0.02 |
| 11 | 47747809 | rs11039359 | 1.52 | -0.608 | 0.187 | 0.103 | 0.011 | 0.057 | 0.057 | 0.078 | 0.1 | 0 | 0.05 | 0 | 0.08 | 0.01 |
| 11 | 47807774 | rs2290850 | 1.483 | 0.032 | 0.749 | 0.767 | -0.052 | -0.015 | -0.015 | 0 | 0.35 | 0.04 | 0.41 | 0.27 | 0.36 | 0.32 |
| 11 | 47856114 | rs10838774 | -0.399 | 0.142 | -0.181 | 0.852 | -0.108 | -0.037 | -0.037 | 0 | 0.37 | 0.08 | 0.32 | 0.44 | 0.43 | 0.53 |
| 11 | 47868474 | rs7942031 | -0.165 | 0.069 | 0.271 | 0.926 | -0.045 | 0.024 | 0.024 | 0.005 | 0.26 | 0.54 | 0.26 | 0.29 | 0.21 | 0.15 |
| 11 | 47907641 | rs4752801 | 1.301 | 0.686 | -0.129 | 0.785 | -0.002 | 0.009 | 0.009 | 0 | 0.36 | 0.24 | 0.41 | 0.25 | 0.38 | 0.32 |
| 11 | 47929846 | rs6485795 | 0.117 | 0.669 | -0.388 | 0.487 | -0.1 | -0.057 | -0.057 | 0 | 0.26 | 0.02 | 0.22 | 0.36 | 0.28 | 0.49 |
| 11 | 47931557 | rs10838790 | 0.317 | -0.143 | 0.055 | 0.51 | 0.018 | -0.01 | -0.01 | 0 | 0.87 | 0.7 | 0.88 | 0.93 | 0.85 | 0.96 |
| 11 | 47940925 | rs747782 | -0.208 | -0.161 | 0.193 | 0.825 | -0.035 | 0.002 | 0.002 | 0 | 0.24 | 0.42 | 0.17 | 0.26 | 0.19 | 0.16 |
| 11 | 47953477 | rs1228020 | 0.058 | -0.066 | 0.184 | 0.833 | 0.017 | -0.001 | -0.001 | 0 | 0.89 | 0.73 | 0.88 | 0.92 | 0.86 | 0.94 |
| 11 | 47969152 | rs1681630 | 0.869 | -0.247 | 0.223 | 0.828 | 0.015 | 0.008 | 0.008 | 0 | 0.62 | 0.23 | 0.7 | 0.66 | 0.67 | 0.78 |
| 11 | 47976882 | rs7936879 | 0.25 | -0.82 | 0.134 | 0.535 | 0.005 | -0.055 | -0.055 | 0.593 | 0.2 | 0.01 | 0.1 | 0.32 | 0.12 | 0.22 |
| 11 | 48004369 | rs7946766 | 0.296 | -0.408 | 1.003 | 0.34 | -0.015 | -0.018 | -0.018 | 1.207 | 0.16 | 0.24 | 0.12 | 0.13 | 0.13 | 0.13 |
| 11 | 48011369 | rs7111419 | -0.587 | -0.166 | 0.942 | 0.445 | -0.007 | 0.02 | 0.02 | 0 | 0.07 | 0.14 | 0.05 | 0.08 | 0.05 | 0.04 |
| 11 | 48050995 | rs7130876 | -0.624 | -0.313 | 0.773 | 0.237 | -0.01 | -0.025 | -0.025 | 1.249 | 0.22 | 0.29 | 0.15 | 0.14 | 0.19 | 0.15 |
| 11 | 48051442 | rs6485802 | -1.664 | -0.301 | 0.338 | 0.22 | 0.032 | 0.062 | 0.062 | 1.113 | 0.36 | 0.94 | 0.27 | 0.24 | 0.29 | 0.18 |
| 11 | 48058313 | rs1113480 | -1.793 | 0.095 | 0.391 | 0.134 | 0 | 0.071 | 0.071 | 2.041 | 0.35 | 0.94 | 0.25 | 0.23 | 0.29 | 0.18 |
| 11 | 48066048 | rs12099166 | -0.143 | 0.135 | 0.46 | 0.115 | -0.021 | -0.018 | -0.018 | 0 | 0.06 | 0.01 | 0.04 | 0.08 | 0.07 | 0.04 |
| 11 | 48085189 | rs11601310 | -2.413 | 0.059 | 0.345 | 0.009 | -0.002 | 0.032 | 0.032 | 3.862 | 0.26 | 0.94 | 0.18 | 0.2 | 0.21 | 0.13 |
| 11 | 48085968 | rs7943570 | -1.187 | 0.059 | 1.072 | 0.018 | -0.019 | -0.019 | -0.019 | 0 | 0.13 | 0.61 | 0.09 | 0.12 | 0.08 | 0.06 |
| 11 | 48097480 | rs10734561 | 1.046 | 0.12 | 1.096 | 0.006 | -0.011 | -0.045 | -0.045 | 0 | 0.7 | 0.46 | 0.78 | 0.8 | 0.74 | 0.82 |
| 11 | 48102806 | rs6485806 | 1.022 | 0.129 | 0.865 | 0.003 | 0.007 | 0.038 | 0.038 | 0.463 | 0.71 | 0.19 | 0.77 | 0.83 | 0.75 | 0.82 |
| 11 | 48102867 | rs6485807 | 1.312 | 0.138 | 0.727 | 0.013 | 0.005 | 0.103 | 0.103 | 0.95 | 0.64 | 0.11 | 0.73 | 0.8 | 0.69 | 0.79 |
| 11 | 48113232 | rs1910364 | 0.548 | 0.196 | 1.12 | 0.228 | -0.002 | -0.034 | -0.034 | 0.152 | 0.76 | 0.6 | 0.83 | 0.8 | 0.79 | 0.84 |
| 11 | 48117873 | rs1503190 | 0.621 | 0.185 | 1.22 | 0.223 | -0.005 | -0.031 | -0.031 | 0 | 0.76 | 0.47 | 0.81 | 0.8 | 0.79 | 0.84 |
| 11 | 48120936 | rs4752894 | 1.53 | -0.291 | 0.701 | 0.109 | -0.053 | 0.006 | 0.006 | 0.089 | 0.44 | 0.06 | 0.64 | 0.48 | 0.59 | 0.57 |
| 11 | 48132197 | rs4356181 | -0.109 | -0.282 | 0.762 | 0.098 | 0.043 | 0.019 | 0.019 | 0 | 0.25 | 0.08 | 0.12 | 0.32 | 0.16 | 0.24 |
| 11 | 48133526 | rs1875696 | 0.66 | -0.262 | 0.827 | 0.1 | -0.002 | -0.032 | -0.032 | 0.236 | 0.72 | 0.42 | 0.78 | 0.8 | 0.75 | 0.81 |
| 11 | 48141843 | rs4752898 | 0.441 | -0.223 | 1.007 | 0.105 | 0.015 | -0.021 | -0.021 | 0 | 0.91 | 0.83 | 0.94 | 0.96 | 0.91 | 0.96 |
| 11 | 48145375 | rs1566734 | -0.278 | -0.269 | 0.837 | 0.108 | 0.046 | 0.025 | 0.025 | 0 | 0.26 | 0.11 | 0.12 | 0.32 | 0.16 | 0.25 |
| 11 | 48146622 | rs1503185 | -0.403 | -0.255 | 0.905 | 0.103 | -0.012 | -0.007 | -0.007 | 0 | 0.27 | 0.31 | 0.13 | 0.32 | 0.16 | 0.25 |
| 11 | 48148925 | rs905476 | -0.265 | -0.225 | 0.65 | 0.122 | 0.045 | 0.024 | 0.024 | 0 | 0.26 | 0.1 | 0.12 | 0.32 | 0.16 | 0.25 |
| 11 | 48162305 | rs7124275 | 1.032 | 0.071 | 0.843 | 0.32 | -0.003 | -0.002 | -0.002 | 0 | 0.55 | 0.46 | 0.68 | 0.45 | 0.67 | 0.4 |
| 11 | 48164721 | rs1566729 | -0.081 | 0.095 | 0.783 | 0.313 | -0.008 | -0.046 | -0.046 | 0 | 0.8 | 0.68 | 0.86 | 0.85 | 0.83 | 0.87 |
| 11 | 48166267 | rs4752904 | 1.47 | 0.137 | 0.376 | 0.565 | -0.06 | -0.084 | -0.084 | 0 | 0.47 | 0.06 | 0.59 | 0.38 | 0.58 | 0.39 |
| 11 | 48182237 | rs1039481 | 0.624 | 0.303 | 0.757 | 0.602 | 0.021 | -0.017 | -0.017 | 0 | 0.73 | 0.55 | 0.8 | 0.77 | 0.75 | 0.81 |
| 11 | 48190777 | rs12798703 | -0.69 | 0.394 | 0.643 | 0.644 | 0.019 | 0.013 | 0.013 | 0.016 | 0.08 | 0.13 | 0.05 | 0.06 | 0.08 | 0.04 |
| 11 | 48198286 | rs12575187 | -0.271 | 0.293 | 0.534 | 0.669 | 0.028 | 0.001 | 0.001 | 0 | 0.23 | 0.13 | 0.17 | 0.35 | 0.16 | 0.41 |
| 11 | 48200931 | rs10838818 | -0.248 | 0.294 | 0.523 | 0.654 | 0.03 | -0.019 | -0.019 | 0.73 | 0.23 | 0.09 | 0.17 | 0.39 | 0.16 | 0.41 |
| 11 | 48203984 | rs9633889 | 1.108 | 0.397 | 0.728 | 0.591 | 0.069 | 0.005 | 0.005 | 0.038 | 0.88 | 0.66 | 0.9 | 0.94 | 0.87 | 0.93 |
| 11 | 48215853 | rs924714 | -0.378 | 0.29 | 0.533 | 0.586 | 0.004 | -0.008 | -0.008 | 0.222 | 0.24 | 0.03 | 0.13 | 0.3 | 0.17 | 0.25 |
| 11 | 48225680 | rs7940877 | 1.156 | 0.317 | 0.701 | 0.577 | -0.008 | -0.005 | -0.005 | 0 | 0.6 | 0.49 | 0.71 | 0.55 | 0.7 | 0.47 |
| 11 | 48238549 | rs11606506 | -0.511 | 0.273 | 0.687 | 0.593 | 0.041 | -0.001 | -0.001 | 0 | 0.28 | 0.17 | 0.19 | 0.39 | 0.17 | 0.46 |
| 11 | 48238880 | rs10838833 | 0.553 | 0.228 | 0.844 | 0.573 | -0.01 | -0.002 | -0.002 | 0 | 0.15 | 0.29 | 0.12 | 0.16 | 0.12 | 0.12 |
| 11 | 48239071 | rs12292056 | -0.662 | 0.279 | 0.821 | 0.59 | 0.04 | 0.028 | 0.028 | 0.002 | 0.07 | 0.2 | 0.06 | 0.06 | 0.08 | 0.04 |
| 11 | 48241821 | rs1018136 | 0.677 | 0.226 | 0.814 | 0.576 | -0.013 | 0 | 0 | 0 | 0.64 | 0.55 | 0.74 | 0.55 | 0.75 | 0.49 |
| 11 | 48264183 | rs1393791 | 0.654 | 0.191 | 0.8 | 0.589 | -0.012 | 0.002 | 0.002 | 0 | 0.65 | 0.55 | 0.74 | 0.55 | 0.75 | 0.5 |
| 11 | 48264370 | rs1354293 | 0.795 | 0.182 | 0.899 | 0.585 | -0.022 | 0.002 | 0.002 | 0 | 0.63 | 0.48 | 0.74 | 0.55 | 0.75 | 0.5 |
| 11 | 48284477 | rs871249 | -0.951 | 0.236 | 0.9 | 0.56 | 0.005 | -0.002 | -0.002 | 0 | 0.35 | 0.32 | 0.24 | 0.45 | 0.25 | 0.5 |
| 11 | 48285906 | rs16905753 | 0.51 | 0.275 | 0.969 | 0.549 | -0.01 | -0.001 | -0.001 | 0 | 0.15 | 0.32 | 0.13 | 0.16 | 0.13 | 0.12 |

| | | | | | | | | | | | | | | | | |
|---|---|---|---|---|---|---|---|---|---|---|---|---|---|---|---|---|
| 11 | 48286256 | rs10838852 | 1.528 | 0.257 | 0.975 | 0.603 | -0.031 | -0.042 | -0.042 | 0 | 0.45 | 0.16 | 0.58 | 0.38 | 0.57 | 0.35 |
| 11 | 48287931 | rs12799818 | 0.077 | 0.256 | 1.015 | 0.608 | 0.009 | 0.004 | 0.004 | 0 | 0.06 | 0.06 | 0.05 | 0.06 | 0.08 | 0.04 |
| 11 | 48319974 | rs2128677 | -0.977 | 0.286 | 0.906 | 0.611 | -0.012 | 0.003 | 0.003 | 0 | 0.36 | 0.45 | 0.26 | 0.45 | 0.25 | 0.5 |
| 11 | 48326773 | rs1483123 | 1.51 | 0.311 | 0.853 | 0.62 | -0.031 | -0.042 | -0.042 | 1.223 | 0.45 | 0.16 | 0.58 | 0.38 | 0.57 | 0.35 |
| 11 | 48331739 | rs1905284 | -0.423 | 0.366 | 0.848 | 0.645 | 0.053 | 0.035 | 0.035 | 0 | 0.07 | 0.21 | 0.07 | 0.06 | 0.08 | 0.04 |
| 11 | 48333823 | rs10769344 | -1.101 | 0.745 | 0.693 | 0.574 | -0.013 | 0.002 | 0.002 | 0 | 0.36 | 0.45 | 0.26 | 0.45 | 0.26 | 0.5 |
| 11 | 48336287 | rs10769346 | -1.101 | 0.745 | 0.693 | 0.574 | -0.013 | 0.002 | 0.002 | 0 | 0.36 | 0.45 | 0.26 | 0.45 | 0.26 | 0.5 |
| 11 | 48337132 | rs11605627 | -0.322 | 0.724 | 0.672 | 0.56 | 0.041 | -0.018 | -0.018 | 0 | 0.27 | 0.09 | 0.18 | 0.39 | 0.17 | 0.46 |
| 11 | 48347140 | rs10838874 | -0.942 | 0.754 | 0.634 | 0.664 | -0.013 | 0.006 | 0.006 | 0 | 0.36 | 0.45 | 0.26 | 0.46 | 0.26 | 0.5 |
| 11 | 48375604 | rs717897 | 0.039 | 0.741 | 0.74 | 0.576 | 0.005 | -0.026 | -0.026 | 0 | 0.81 | 0.61 | 0.84 | 0.84 | 0.82 | 0.85 |
| 11 | 48376650 | rs1483118 | 0.125 | 0.725 | 0.892 | 0.568 | 0.005 | -0.035 | -0.035 | 0.092 | 0.8 | 0.48 | 0.83 | 0.84 | 0.82 | 0.85 |
| 11 | 48449889 | rs588700 | -0.097 | 0.713 | 0.937 | 0.589 | 0.006 | -0.02 | -0.02 | 0 | 0.81 | 0.67 | 0.85 | 0.84 | 0.82 | 0.85 |
| 11 | 48457439 | rs666165 | -0.064 | 0.718 | 0.909 | 0.573 | 0.024 | -0.028 | -0.028 | 0.188 | 0.81 | 0.52 | 0.83 | 0.84 | 0.82 | 0.86 |
| 11 | 48476690 | rs650021 | -0.528 | 0.737 | 0.827 | 0.672 | 0.012 | -0.004 | -0.004 | 3.073 | 0.28 | 0.26 | 0.24 | 0.39 | 0.18 | 0.45 |
| 11 | 48512810 | rs4882087 | -0.548 | 0.861 | 0.89 | 0.701 | 0.077 | 0.058 | 0.058 | 0.041 | 0.08 | 0.32 | 0.07 | 0.07 | 0.08 | 0.04 |
| 11 | 48516606 | rs7944287 | -0.741 | 0.731 | 0.792 | 0.726 | 0.042 | -0.012 | -0.012 | 0 | 0.82 | 1 | 0.92 | 0.77 | 0.91 | 0.8 |
| 11 | 48518893 | rs7395662 | 1.157 | 0.763 | 0.802 | 0.724 | 0.015 | 0.004 | 0.004 | 0 | 0.53 | 0.38 | 0.59 | 0.44 | 0.64 | 0.41 |
| 11 | 48558249 | rs4882110 | 1.682 | 0.777 | 0.77 | 0.702 | -0.101 | -0.079 | -0.079 | 0.889 | 0.44 | 0.05 | 0.53 | 0.37 | 0.56 | 0.36 |
| 11 | 48581765 | rs10838956 | 1.167 | 0.74 | 0.783 | 0.706 | 0.016 | 0.004 | 0.004 | 0 | 0.53 | 0.38 | 0.59 | 0.44 | 0.64 | 0.41 |
| 11 | 48605015 | rs7120801 | 0.393 | 0.708 | 0.78 | 0.716 | 0.012 | -0.004 | -0.004 | 1.221 | 0.72 | 0.74 | 0.76 | 0.61 | 0.82 | 0.55 |
| 11 | 48609936 | rs7944230 | 0.393 | 0.708 | 0.941 | 0.716 | -0.004 | 0 | 0 | 0 | 0.72 | 0.69 | 0.76 | 0.61 | 0.82 | 0.55 |
| 11 | 48646903 | rs10838987 | 0.154 | 0.67 | 0.671 | 0.71 | -0.009 | -0.017 | -0.017 | 2.638 | 0.07 | 0.03 | 0.15 | 0.16 | 0.08 | 0.26 |
| 11 | 48647107 | rs11039885 | 1.721 | 1.149 | 0.646 | 0.614 | -0.067 | -0.056 | -0.056 | 0.073 | 0.43 | 0.09 | 0.52 | 0.37 | 0.53 | 0.35 |
| 11 | 48650491 | rs12281162 | 0.544 | 0.982 | 0.748 | 0.605 | -0.009 | 0.004 | 0.004 | 0 | 0.15 | 0.33 | 0.13 | 0.16 | 0.12 | 0.11 |
| 11 | 48694871 | rs1588069 | -0.062 | 0.978 | 0.744 | 0.603 | 0.009 | -0.019 | -0.019 | 0 | 0.18 | 0.3 | 0.15 | 0.16 | 0.18 | 0.14 |
| 11 | 48889126 | rs11040106 | -1.416 | 0.886 | 0.752 | 0.587 | 0.005 | 0 | 0 | 0 | 0.45 | 0.42 | 0.39 | 0.56 | 0.36 | 0.6 |
| 11 | 48942781 | rs12293923 | 0.554 | 0.748 | 0.653 | 0.417 | -0.01 | -0.002 | -0.002 | 0 | 0.15 | 0.3 | 0.13 | 0.16 | 0.12 | 0.11 |
| 11 | 48987539 | rs1809986 | 1.103 | 0.619 | 0.333 | 0.586 | 0.029 | -0.004 | -0.004 | 0.197 | 0.54 | 0.33 | 0.6 | 0.43 | 0.64 | 0.4 |
| 11 | 49000558 | rs11040198 | -0.969 | 0.546 | 0.123 | 0.624 | 0.054 | -0.04 | -0.04 | 0.198 | 0.29 | 0.12 | 0.24 | 0.4 | 0.18 | 0.47 |
| 11 | 49009877 | rs2865908 | 0.637 | 0.322 | 0.041 | 0.51 | 0.012 | -0.015 | -0.015 | 0 | 0.77 | 0.81 | 0.78 | 0.6 | 0.83 | 0.59 |
| 11 | 49010832 | rs10431006 | 0.155 | 0.441 | 0.07 | 0.596 | -0.028 | -0.046 | -0.046 | 0 | 0.08 | 0 | 0.15 | 0.19 | 0.09 | 0.26 |
| 11 | 49030187 | rs10769558 | 0.827 | 0.382 | 0.052 | 0.541 | 0.038 | -0.013 | -0.013 | 0.641 | 0.7 | 0.81 | 0.76 | 0.6 | 0.82 | 0.53 |
| 11 | 49053601 | rs2696923 | 0.694 | 0.325 | -0.011 | 0.525 | 0.05 | -0.029 | -0.029 | 0 | 0.71 | 0.85 | 0.76 | 0.6 | 0.82 | 0.51 |
| 11 | 49079350 | rs1819409 | 0.609 | 0.759 | 0.4 | 0.667 | 0.005 | 0.009 | 0.009 | 0 | 0.15 | 0.47 | 0.14 | 0.16 | 0.14 | 0.1 |
| 11 | 49132307 | rs12797843 | 0.409 | 0.917 | 0.035 | 0.863 | -0.016 | -0.016 | -0.016 | 0.084 | 0.07 | 0.03 | 0.05 | 0.1 | 0.1 | 0.05 |
| 11 | 49146066 | rs10839224 | 0.378 | 0.807 | 0.297 | 0.925 | 0.03 | 0.072 | 0.072 | 0 | 0.07 | 0.2 | 0.15 | 0.16 | 0.09 | 0.27 |
| 11 | 49157399 | rs11040265 | -0.315 | 0.52 | -0.061 | 0.8 | 0.045 | 0.003 | 0.003 | 0 | 0.17 | 0.1 | 0.09 | 0.22 | 0.09 | 0.22 |
| 11 | 49162948 | rs10839229 | 0.115 | 0.598 | 0.125 | 0.903 | 0.043 | 0.093 | 0.093 | 0 | 0.08 | 0.3 | 0.16 | 0.16 | 0.09 | 0.27 |
| 11 | 49170774 | rs16906158 | 0.123 | 0.593 | 0.119 | 0.932 | 0.043 | 0.097 | 0.097 | 0 | 0.08 | 0.3 | 0.16 | 0.15 | 0.09 | 0.3 |
| 11 | 49176227 | rs34033751 | 0.094 | 0.498 | -0.105 | 1.22 | 0.046 | -0.043 | -0.043 | 0 | 0.15 | 0.04 | 0.16 | 0.28 | 0.08 | 0.2 |
| 11 | 49214048 | rs588458 | -1.339 | 0.682 | -0.143 | 0.876 | -0.061 | -0.041 | -0.041 | 0 | 0.37 | 0.81 | 0.39 | 0.42 | 0.41 | 0.48 |
| 11 | 49242990 | rs679470 | -0.554 | 0.658 | 0.055 | 0.96 | -0.003 | 0.018 | 0.018 | 0 | 0.24 | 0.6 | 0.29 | 0.32 | 0.23 | 0.4 |
| 11 | 49250912 | rs650826 | 0.578 | 0.687 | 0.152 | 0.901 | -0.003 | -0.006 | -0.006 | 1.551 | 0.15 | 0.31 | 0.13 | 0.15 | 0.14 | 0.11 |
| 11 | 49253662 | rs683680 | 0.54 | 0.688 | 0.132 | 0.901 | -0.003 | -0.006 | -0.006 | 0 | 0.15 | 0.35 | 0.13 | 0.15 | 0.14 | 0.11 |
| 11 | 49257082 | rs632220 | 0.54 | 0.688 | 0.132 | 0.901 | -0.003 | -0.006 | -0.006 | 0 | 0.15 | 0.35 | 0.13 | 0.15 | 0.14 | 0.11 |
| 11 | 49280857 | rs7124266 | -0.985 | 0.714 | 0.118 | 0.907 | 0.053 | -0.002 | -0.002 | 0 | 0.27 | 0.5 | 0.22 | 0.25 | 0.32 | 0.21 |
| 11 | 49296825 | rs7117025 | 0.43 | 0.63 | 0.043 | 0.879 | -0.001 | -0.005 | -0.005 | 4.638 | 0.16 | 0.41 | 0.14 | 0.15 | 0.13 | 0.12 |
| 11 | 49297915 | rs1164657 | -0.432 | 0.708 | -0.042 | 0.932 | 0.006 | 0.002 | 0.002 | 0 | 0.1 | 0.12 | 0.06 | 0.09 | 0.13 | 0.06 |
| 11 | 49304358 | rs1164681 | -0.45 | 0.43 | -0.202 | 1.303 | 0.043 | -0.046 | -0.046 | 0.009 | 0.85 | 0.94 | 0.91 | 0.71 | 0.92 | 0.79 |
| 11 | 49311134 | rs7939300 | 0.237 | 0.577 | -0.08 | 1.236 | 0.029 | 0.078 | 0.078 | 0 | 0.09 | 0.25 | 0.16 | 0.17 | 0.09 | 0.28 |
| 11 | 49357043 | rs3862342 | 0.352 | 0.598 | -0.135 | 1.259 | 0.037 | 0.018 | 0.018 | 0 | 0.74 | 0.48 | 0.78 | 0.72 | 0.72 | 0.8 |
| 11 | 49370004 | rs1917325 | 1.089 | 0.532 | -0.409 | 1.256 | -0.07 | -0.063 | -0.063 | 0 | 0.63 | 0.16 | 0.62 | 0.55 | 0.59 | 0.51 |
| 11 | 49373782 | rs12797853 | 0.674 | 0.605 | -0.482 | 1.227 | -0.006 | -0.006 | -0.006 | 0 | 0.07 | 0.05 | 0.05 | 0.1 | 0.1 | 0.05 |
| 11 | 49377760 | rs1917321 | 1.642 | 0.462 | -0.649 | 1.163 | -0.112 | 0.057 | 0.057 | 0.299 | 0.47 | 0.03 | 0.52 | 0.29 | 0.51 | 0.29 |
| 11 | 49401125 | rs7102702 | 0.544 | 0.513 | -0.393 | 0.934 | 0.033 | -0.002 | -0.002 | 0 | 0.7 | 0.59 | 0.79 | 0.72 | 0.68 | 0.78 |
| 11 | 49426062 | rs12292560 | 0.201 | 0.282 | -0.402 | 1.036 | 0.04 | -0.029 | -0.029 | 0.206 | 0.14 | 0.07 | 0.09 | 0.25 | 0.08 | 0.22 |
| 11 | 49447850 | rs7358352 | 0.152 | 0.886 | 0.071 | 0.922 | -0.006 | 0.011 | 0.011 | 0 | 0.16 | 0.32 | 0.13 | 0.19 | 0.13 | 0.12 |
| 11 | 49458055 | rs7113075 | -0.02 | 0.947 | 0.155 | 1.012 | -0.001 | 0.067 | 0.067 | 0 | 0.09 | 0.24 | 0.16 | 0.17 | 0.08 | 0.28 |
| 11 | 49464833 | rs7934591 | -0.019 | 0.947 | 0.154 | 1.012 | -0.001 | 0.067 | 0.067 | 0 | 0.09 | 0.24 | 0.16 | 0.16 | 0.08 | 0.28 |
| 11 | 49470419 | rs1846285 | -1.238 | 1.18 | 0.157 | 0.946 | 0.065 | 0.034 | 0.034 | 0.044 | 0.15 | 0.42 | 0.2 | 0.16 | 0.18 | 0.32 |
| 11 | 49473034 | rs1811672 | -0.256 | 1.076 | -0.083 | 1.012 | -0.006 | -0.002 | -0.002 | 0 | 0.88 | 0.92 | 0.94 | 0.91 | 0.87 | 0.92 |
| 11 | 49477266 | rs6485981 | -0.196 | 0.959 | 0.163 | 1.017 | 0.018 | 0.115 | 0.115 | 0 | 0.9 | 0.68 | 0.85 | 0.84 | 0.93 | 0.72 |
| 11 | 49488707 | rs12222221 | 0.035 | 0.964 | 0.146 | 1.022 | 0.007 | 0.089 | 0.089 | 0 | 0.1 | 0.3 | 0.16 | 0.16 | 0.08 | 0.28 |
| 11 | 49493736 | rs11040387 | 0.462 | 0.904 | 0.078 | 1.021 | -0.003 | -0.021 | -0.021 | 0 | 0.13 | 0.04 | 0.1 | 0.19 | 0.12 | 0.12 |
| 11 | 49498923 | rs10839287 | 0.294 | 1.038 | 0.12 | 1.018 | 0.073 | 0.067 | 0.067 | 0 | 0.87 | 0.61 | 0.81 | 0.84 | 0.86 | 0.68 |
| 11 | 49527345 | rs10839295 | 1.376 | 1.088 | -0.174 | 0.942 | -0.052 | -0.038 | -0.038 | 0 | 0.58 | 0.19 | 0.6 | 0.56 | 0.58 | 0.48 |
| 11 | 49537617 | rs10839296 | -1.079 | 1.138 | -0.059 | 0.917 | 0.006 | 0.001 | 0.001 | 0 | 0.29 | 0.4 | 0.21 | 0.28 | 0.28 | 0.2 |
| 11 | 49542457 | rs937430 | -1.177 | 1.137 | -0.052 | 0.928 | 0.005 | 0.002 | 0.002 | 0 | 0.3 | 0.43 | 0.21 | 0.28 | 0.28 | 0.2 |
| 11 | 49546908 | rs12098986 | 0.122 | 1.078 | -0.013 | 0.934 | -0.009 | 0.011 | 0.011 | 0 | 0.16 | 0.32 | 0.13 | 0.19 | 0.13 | 0.12 |

| | | | | | | | | | | | | | | | | |
|---|---|---|---|---|---|---|---|---|---|---|---|---|---|---|---|---|
| 11 | 49557880 | rs7111711 | 0.674 | 1.103 | -0.044 | 0.93 | 0.005 | 0.003 | 0.003 | 0 | 0.71 | 0.6 | 0.79 | 0.72 | 0.73 | 0.8 |
| 11 | 49572109 | rs1909220 | 0.731 | 1.097 | -0.059 | 0.913 | 0.005 | 0.001 | 0.001 | 0 | 0.71 | 0.57 | 0.79 | 0.72 | 0.73 | 0.8 |
| 11 | 49595912 | rs3862350 | 1.353 | 1.094 | -0.152 | 0.931 | -0.053 | -0.039 | -0.039 | 0 | 0.58 | 0.19 | 0.6 | 0.56 | 0.58 | 0.48 |
| 11 | 49596079 | rs11040432 | -0.103 | 1.259 | 0.017 | 0.959 | 0.011 | 0.096 | 0.096 | 0 | 0.1 | 0.32 | 0.16 | 0.16 | 0.08 | 0.28 |
| 11 | 49627833 | rs1909218 | -0.306 | 1.325 | -0.48 | 0.891 | -0.027 | -0.001 | -0.001 | 0 | 0.88 | 0.93 | 0.92 | 0.92 | 0.84 | 0.93 |
| 11 | 49637416 | rs7125677 | -0.298 | 1.314 | -0.485 | 0.891 | -0.027 | -0.003 | -0.003 | 0.016 | 0.88 | 0.93 | 0.92 | 0.92 | 0.84 | 0.93 |
| 11 | 55836620 | rs11227133 | 0.75 | 0.523 | -0.01 | -0.114 | 0.001 | -0.008 | -0.008 | 0 | 0.22 | 0.16 | 0.14 | 0.24 | 0.22 | 0.19 |
| 11 | 55839278 | rs11227145 | 2.16 | 0.508 | 0.026 | -0.122 | 0.002 | -0.012 | -0.012 | 0.01 | 0.12 | 0.05 | 0.1 | 0.14 | 0.12 | 0.12 |
| 11 | 55849535 | rs4939020 | -0.185 | 0.62 | 0.057 | -0.157 | -0.005 | 0.003 | 0.003 | 0 | 0.72 | 0.69 | 0.8 | 0.76 | 0.69 | 0.75 |
| 11 | 55853708 | rs1481999 | 0.597 | 0.615 | 0.054 | -0.165 | -0.005 | -0.004 | -0.004 | 0 | 0.45 | 0.46 | 0.52 | 0.62 | 0.37 | 0.57 |
| 11 | 55856293 | rs7935319 | -0.242 | 0.603 | 0.059 | -0.167 | -0.009 | 0.005 | 0.005 | 0 | 0.73 | 0.75 | 0.81 | 0.76 | 0.69 | 0.75 |
| 11 | 55857698 | rs10895996 | 1.257 | 0.5 | 0.03 | -0.081 | 0.002 | -0.003 | -0.003 | 0.144 | 0.15 | 0.12 | 0.18 | 0.24 | 0.11 | 0.36 |
| 11 | 55862565 | rs7122627 | 0.715 | 0.575 | -0.122 | -0.141 | 0.001 | -0.008 | -0.008 | 1.33 | 0.22 | 0.16 | 0.14 | 0.24 | 0.22 | 0.19 |
| 11 | 55864161 | rs2460192 | 1.886 | 0.647 | -0.219 | -0.256 | -0.005 | -0.056 | -0.056 | 0.039 | 0.17 | 0.22 | 0.09 | 0 | 0.15 | 0.02 |
| 11 | 55864963 | rs2449155 | -0.734 | 0.618 | -0.249 | -0.106 | 0.022 | -0.024 | -0.024 | 0 | 0.95 | 0.91 | 0.94 | 1 | 0.91 | 0.95 |
| 11 | 55884690 | rs1482009 | 2.006 | 0.545 | -0.309 | -0.075 | 0.004 | -0.01 | -0.01 | 0 | 0.12 | 0.05 | 0.1 | 0.14 | 0.12 | 0.12 |
| 11 | 55901283 | rs7949386 | -0.086 | 0.56 | -0.189 | -0.043 | -0.004 | 0.02 | 0.02 | 0 | 0.55 | 0.57 | 0.56 | 0.71 | 0.46 | 0.65 |
| 11 | 55904751 | rs1947923 | 0.412 | 0.68 | -0.21 | -0.011 | 0.023 | -0.024 | -0.024 | 0 | 0.05 | 0.09 | 0.05 | 0 | 0.09 | 0.05 |
| 11 | 55920262 | rs1871848 | 1.683 | 0.668 | -0.285 | 0.027 | -0.008 | -0.005 | -0.005 | 0 | 0.08 | 0.04 | 0.03 | 0.09 | 0.09 | 0.08 |
| 11 | 55929181 | rs11227400 | 1.679 | 0.625 | -0.525 | -0.094 | -0.029 | -0.117 | -0.117 | 0 | 0.25 | 0.03 | 0.3 | 0.38 | 0.23 | 0.2 |
| 11 | 55934440 | rs4300408 | -0.23 | 0.62 | -0.239 | -0.076 | 0 | -0.004 | -0.004 | 0 | 0.55 | 0.44 | 0.55 | 0.71 | 0.47 | 0.65 |
| 11 | 55937954 | rs4578407 | -1.005 | 0.631 | -0.242 | -0.087 | -0.003 | 0.006 | 0.006 | 0 | 0.73 | 0.76 | 0.72 | 0.84 | 0.68 | 0.83 |
| 11 | 55949458 | rs4616058 | -0.197 | 0.628 | -0.28 | 0.112 | 0 | -0.004 | -0.004 | 0 | 0.55 | 0.44 | 0.55 | 0.71 | 0.47 | 0.65 |
| 11 | 55951305 | rs4519111 | 1.552 | 0.632 | -0.429 | 0.183 | -0.026 | -0.114 | -0.114 | 0.342 | 0.26 | 0.03 | 0.3 | 0.38 | 0.23 | 0.2 |
| 11 | 55954348 | rs11227462 | 0.909 | 0.776 | -0.417 | 0.161 | -0.044 | -0.033 | -0.033 | 0 | 0.13 | 0.02 | 0.19 | 0.16 | 0.2 | 0.15 |
| 11 | 55956773 | rs17150147 | 1.474 | 0.701 | -0.244 | 0.15 | 0 | 0 | 0 | 0.472 | 0.1 | 0.14 | 0.04 | 0.09 | 0.09 | 0.08 |
| 11 | 55972861 | rs11227513 | 1.828 | 0.712 | -0.294 | 0.156 | -0.008 | -0.007 | -0.007 | 0 | 0.08 | 0.04 | 0.03 | 0.09 | 0.09 | 0.08 |
| 11 | 55977308 | rs7935265 | -0.79 | 0.706 | -0.243 | 0.083 | -0.003 | 0.006 | 0.006 | 0 | 0.73 | 0.76 | 0.72 | 0.83 | 0.68 | 0.83 |
| 11 | 55979266 | rs7130251 | 0.195 | 0.639 | -0.18 | 0.029 | -0.003 | 0.067 | 0.067 | 0 | 0.53 | 0.3 | 0.54 | 0.71 | 0.46 | 0.65 |
| 11 | 56000451 | rs11227598 | 2.265 | 0.731 | -0.135 | 0.06 | 0.005 | -0.002 | -0.002 | 0 | 0.13 | 0.06 | 0.1 | 0.13 | 0.12 | 0.12 |
| 11 | 56000471 | rs11227599 | 0.866 | 0.729 | -0.122 | 0.06 | -0.007 | 0.009 | 0.009 | 0 | 0.18 | 0.19 | 0.16 | 0.12 | 0.21 | 0.17 |
| 11 | 56000600 | rs3919907 | 1.756 | 0.749 | -0.186 | 0.083 | -0.022 | -0.116 | -0.116 | 0.52 | 0.26 | 0.03 | 0.3 | 0.38 | 0.23 | 0.2 |
| 11 | 56003099 | rs10896137 | 0.675 | 0.878 | 0.001 | -0.014 | 0.014 | -0.024 | -0.024 | 0 | 0.19 | 0.32 | 0.17 | 0.13 | 0.21 | 0.17 |
| 11 | 56003560 | rs2128137 | -2.452 | 0.88 | 0.044 | -0.037 | -0.01 | -0.051 | -0.051 | 0 | 0.86 | 0.79 | 0.92 | 1 | 0.88 | 0.99 |
| 11 | 56003589 | rs10896138 | 0.671 | 0.904 | -0.084 | -0.007 | 0.013 | -0.024 | -0.024 | 0 | 0.19 | 0.32 | 0.17 | 0.13 | 0.21 | 0.17 |
| 11 | 56025434 | rs11227667 | 0.913 | 0.943 | -0.117 | -0.365 | -0.046 | -0.033 | -0.033 | 0 | 0.13 | 0.02 | 0.19 | 0.16 | 0.2 | 0.17 |
| 11 | 56027115 | rs10160392 | 0.498 | 0.876 | -0.081 | -0.348 | 0.021 | -0.025 | -0.025 | 0 | 0.05 | 0.12 | 0.06 | 0 | 0.09 | 0.06 |
| 11 | 56036787 | rs11227719 | 0.591 | 0.78 | -0.187 | -0.328 | 0.027 | 0.011 | 0.011 | 0 | 0.44 | 0.19 | 0.51 | 0.62 | 0.37 | 0.58 |
| 11 | 56068101 | rs11227802 | 0.127 | 1.003 | -0.137 | -0.278 | 0 | -0.006 | -0.006 | 0 | 0.55 | 0.45 | 0.65 | 0.74 | 0.47 | 0.65 |
| 11 | 56071557 | rs2870434 | -0.356 | 1.036 | -0.152 | -0.302 | 0.024 | -0.045 | -0.045 | 0.188 | 0.69 | 0.61 | 0.76 | 0.86 | 0.58 | 0.77 |
| 11 | 56077823 | rs11227836 | 0.832 | 1.009 | -0.39 | -0.269 | -0.041 | -0.03 | -0.03 | 0.902 | 0.13 | 0.03 | 0.18 | 0.17 | 0.2 | 0.15 |
| 11 | 56078325 | rs10791907 | -0.447 | 0.999 | -0.272 | -0.127 | -0.005 | 0.022 | 0.022 | 0.189 | 0.61 | 0.7 | 0.63 | 0.71 | 0.56 | 0.71 |
| 11 | 56086147 | rs960193 | -0.926 | 1.036 | -0.297 | -0.149 | -0.004 | 0.008 | 0.008 | 0 | 0.73 | 0.76 | 0.82 | 0.87 | 0.68 | 0.83 |
| 11 | 56104457 | rs12222334 | 0.838 | 0.842 | -0.309 | -0.075 | 0.009 | -0.013 | -0.013 | 0 | 0.18 | 0.1 | 0.28 | 0.23 | 0.14 | 0.37 |
| 11 | 56107439 | rs1586006 | -0.412 | 0.943 | -0.239 | -0.359 | 0.029 | -0.069 | -0.069 | 0 | 0.67 | 0.48 | 0.74 | 0.86 | 0.58 | 0.77 |
| 11 | 56113516 | rs1905055 | -1.019 | 1.011 | -0.346 | -0.361 | -0.004 | 0.008 | 0.008 | 0 | 0.73 | 0.75 | 0.81 | 0.86 | 0.67 | 0.83 |
| 11 | 56113593 | rs10896272 | 0.324 | 0.987 | -0.49 | -0.326 | 0.033 | 0.038 | 0.038 | 0 | 0.45 | 0.21 | 0.6 | 0.64 | 0.37 | 0.58 |
| 11 | 56115212 | rs17614415 | 1.794 | 0.966 | -0.526 | -0.25 | -0.009 | -0.007 | -0.007 | 0.838 | 0.08 | 0.04 | 0.03 | 0.09 | 0.09 | 0.08 |
| 11 | 56122057 | rs2128133 | 1.263 | 0.896 | -0.574 | -0.265 | -0.024 | 0.019 | 0.019 | 0 | 0.16 | 0.01 | 0.16 | 0.09 | 0.13 | 0.11 |
| 11 | 56128081 | rs10896290 | 0.69 | 1.086 | -0.31 | -0.268 | -0.004 | 0.008 | 0.008 | 0 | 0.26 | 0.25 | 0.19 | 0.14 | 0.33 | 0.17 |
| 11 | 56128524 | rs7928704 | -2.022 | 1.015 | -0.418 | -0.247 | 0.004 | -0.003 | -0.003 | 0 | 0.88 | 0.94 | 0.9 | 0.88 | 0.88 | 0.89 |
| 11 | 56128772 | rs7938038 | 0.52 | 1.022 | -0.331 | -0.265 | 0.032 | 0.037 | 0.037 | 0 | 0.45 | 0.18 | 0.59 | 0.64 | 0.37 | 0.58 |
| 11 | 56137454 | rs6591243 | -1.963 | 0.991 | -0.324 | -0.183 | 0.004 | -0.003 | -0.003 | 0 | 0.88 | 0.94 | 0.9 | 0.88 | 0.88 | 0.89 |
| 11 | 56141251 | rs12804193 | 1.585 | 1.121 | -0.16 | -0.169 | -0.009 | -0.007 | -0.007 | 0 | 0.08 | 0.04 | 0.03 | 0.09 | 0.09 | 0.08 |
| 11 | 56143424 | rs12788990 | 1.209 | 1.117 | -0.016 | -0.171 | 0.022 | 0.02 | 0.02 | 0 | 0.09 | 0.17 | 0.04 | 0.09 | 0.09 | 0.08 |
| 11 | 56143977 | rs12272403 | -0.086 | 1.181 | -0.052 | -0.273 | 0.018 | -0.064 | -0.064 | 0 | 0.32 | 0.55 | 0.24 | 0.14 | 0.36 | 0.22 |
| 11 | 56149723 | rs949361 | 0.576 | 1.104 | -0.231 | -0.159 | -0.001 | 0.002 | 0.002 | 0.013 | 0.25 | 0.25 | 0.16 | 0.14 | 0.27 | 0.17 |
| 11 | 56162014 | rs17543899 | 0.873 | 1.195 | -0.153 | -0.07 | 0.034 | 0.007 | 0.007 | 0 | 0.1 | 0.19 | 0.07 | 0.09 | 0.16 | 0.08 |
| 11 | 56163562 | rs11228236 | -0.087 | 1.25 | -0.336 | -0.184 | 0.005 | -0.003 | -0.003 | 0 | 0.73 | 0.72 | 0.79 | 0.78 | 0.64 | 0.74 |
| 11 | 56175534 | rs17150526 | 1.549 | 0.975 | -0.485 | -0.152 | -0.033 | -0.148 | -0.148 | 0 | 0.26 | 0.02 | 0.31 | 0.41 | 0.23 | 0.21 |
| 11 | 56184888 | rs998544 | 0.512 | 1.058 | -0.255 | -0.24 | 0.031 | 0.037 | 0.037 | 0 | 0.45 | 0.18 | 0.59 | 0.64 | 0.37 | 0.58 |
| 11 | 56185159 | rs7930678 | 0.48 | 0.986 | -0.25 | -0.225 | -0.007 | -0.064 | -0.064 | 0.043 | 0.28 | 0.54 | 0.19 | 0.14 | 0.27 | 0.17 |
| 11 | 56185224 | rs12785840 | 1.102 | 1.147 | -0.223 | -0.108 | 0.028 | 0.011 | 0.011 | 0 | 0.1 | 0.19 | 0.07 | 0.09 | 0.14 | 0.08 |
| 11 | 56185345 | rs6591324 | -0.487 | 1.294 | -0.238 | -0.116 | 0.004 | -0.002 | -0.002 | 0 | 0.77 | 0.74 | 0.83 | 0.78 | 0.72 | 0.8 |
| 11 | 56202667 | rs1945242 | -0.257 | 1.087 | 0.092 | -0.042 | 0.122 | 0 | 0 | 0.094 | 0.06 | 0.26 | 0.06 | 0 | 0.09 | 0.05 |
| 11 | 56209204 | rs661090 | -1.042 | 0.914 | -0.068 | -0.065 | 0.004 | -0.018 | -0.018 | 1.653 | 0.81 | 0.94 | 0.78 | 0.77 | 0.84 | 0.63 |
| 11 | 56228448 | rs1945238 | 2.148 | 0.748 | 0.041 | -0.684 | -0.01 | -0.063 | -0.063 | 0 | 0.13 | 0.22 | 0.08 | 0 | 0.12 | 0.03 |
| 11 | 56229310 | rs7924520 | 0.044 | 0.712 | -0.215 | -0.743 | -0.002 | -0.028 | -0.028 | 0.122 | 0.33 | 0.6 | 0.26 | 0.13 | 0.34 | 0.2 |
| 11 | 56235061 | rs688740 | 0.667 | 1.055 | -0.499 | -0.757 | -0.048 | -0.002 | -0.002 | 0.063 | 0.14 | 0.03 | 0.12 | 0.13 | 0.22 | 0.16 |

| | | | | | | | | | | | | | | | | |
|---|---|---|---|---|---|---|---|---|---|---|---|---|---|---|---|---|
| 11 | 56239881 | rs1945206 | 1.985 | 1.03 | -0.483 | -1.024 | 0.004 | 0.001 | 0.001 | 0 | 0.13 | 0.08 | 0.1 | 0.13 | 0.12 | 0.11 |
| 11 | 56241490 | rs637202 | 0.58 | 1.102 | -0.453 | -0.985 | -0.048 | -0.001 | -0.001 | 0 | 0.14 | 0.03 | 0.12 | 0.13 | 0.22 | 0.16 |
| 11 | 56242354 | rs675120 | -0.332 | 1.08 | -0.452 | -0.982 | 0.003 | -0.002 | -0.002 | 0 | 0.2 | 0.17 | 0.18 | 0.13 | 0.32 | 0.21 |
| 11 | 56250625 | rs11228609 | 2.021 | 1.017 | -0.5 | -0.99 | -0.011 | -0.094 | -0.094 | 0.679 | 0.14 | 0.33 | 0.09 | 0 | 0.13 | 0.03 |
| 11 | 56256864 | rs618594 | 0.481 | 1.263 | -0.815 | -0.845 | -0.048 | -0.001 | -0.001 | 0 | 0.15 | 0.03 | 0.12 | 0.13 | 0.22 | 0.16 |
| 11 | 56257302 | rs616683 | -0.128 | 1.292 | -0.823 | -0.853 | -0.086 | 0.018 | 0.018 | 0 | 0.19 | 0.03 | 0.16 | 0.13 | 0.32 | 0.21 |
| 11 | 56269410 | rs12577186 | 1.596 | 0.93 | -0.777 | -0.755 | 0.005 | -0.012 | -0.012 | 0 | 0.07 | 0.03 | 0.02 | 0.1 | 0.06 | 0.08 |
| 11 | 56294275 | rs658845 | -1.941 | 0.713 | -1.196 | -0.911 | -0.039 | -0.143 | -0.143 | 0 | 0.79 | 0.97 | 0.6 | 0.48 | 0.78 | 0.57 |
| 11 | 56294431 | rs1945196 | 2.19 | 0.618 | -1.182 | -1.027 | 0.001 | 0 | 0 | 0.66 | 0.12 | 0.11 | 0.11 | 0.13 | 0.12 | 0.11 |
| 11 | 56294818 | rs637912 | 0.542 | 0.529 | -1.037 | -1.138 | 0.003 | -0.003 | -0.003 | 0.27 | 0.33 | 0.64 | 0.23 | 0.13 | 0.25 | 0.15 |
| 11 | 56299930 | rs588805 | 0.136 | 0.574 | -1.06 | -1.164 | 0.008 | 0.035 | 0.035 | 0 | 0.38 | 0.65 | 0.29 | 0.13 | 0.35 | 0.21 |
| 11 | 56300082 | rs11606988 | 1.251 | 0.59 | -1.121 | -1.305 | 0.001 | 0.004 | 0.004 | 0 | 0.19 | 0.15 | 0.19 | 0.1 | 0.16 | 0.12 |
| 11 | 56309583 | rs641778 | 0.781 | 0.699 | -0.997 | -1.176 | -0.042 | -0.014 | -0.014 | 0 | 0.14 | 0.03 | 0.1 | 0.16 | 0.2 | 0.16 |
| 11 | 56310222 | rs628524 | -0.837 | 0.697 | -0.995 | -1.185 | 0.003 | -0.003 | -0.003 | 0.002 | 0.67 | 0.36 | 0.77 | 0.88 | 0.75 | 0.85 |
| 11 | 56312630 | rs11228641 | 2.26 | 0.698 | -1.016 | -1.184 | 0.001 | 0 | 0 | 5.62 | 0.12 | 0.11 | 0.11 | 0.13 | 0.12 | 0.11 |
| 11 | 56326343 | rs10736677 | -0.54 | 0.75 | -1.112 | -1.135 | 0.007 | 0.034 | 0.034 | 0 | 0.62 | 0.35 | 0.72 | 0.87 | 0.65 | 0.79 |
| 11 | 56330546 | rs11228656 | 0.662 | 0.783 | -1.128 | -1.106 | -0.004 | 0.004 | 0.004 | 0.019 | 0.4 | 0.19 | 0.6 | 0.62 | 0.39 | 0.55 |
| 11 | 56342048 | rs10896486 | 0.655 | 0.858 | -1.03 | -1.044 | -0.042 | -0.017 | -0.017 | 0 | 0.14 | 0.03 | 0.1 | 0.16 | 0.2 | 0.16 |
| 11 | 56344260 | rs10896488 | 0.655 | 0.857 | -1.041 | -1.045 | -0.042 | -0.017 | -0.017 | 0 | 0.14 | 0.03 | 0.1 | 0.16 | 0.2 | 0.16 |
| 11 | 56366884 | rs12797276 | 1.151 | 0.731 | -1.27 | -1.379 | 0.006 | 0.007 | 0.007 | 0 | 0.21 | 0.16 | 0.14 | 0.12 | 0.15 | 0.15 |
| 11 | 56370106 | rs11228676 | -0.35 | 1.017 | -0.636 | -1.281 | 0.008 | 0.035 | 0.035 | 0 | 0.62 | 0.35 | 0.71 | 0.88 | 0.65 | 0.79 |
| 11 | 56370355 | rs1397049 | 1.773 | 0.87 | -0.34 | -1.313 | -0.018 | -0.078 | -0.078 | 1.085 | 0.18 | 0.25 | 0.1 | 0 | 0.14 | 0.04 |
| 11 | 56370425 | rs11228677 | 1.218 | 1.018 | -0.512 | -1.221 | 0.017 | -0.012 | -0.012 | 0 | 0.36 | 0.19 | 0.56 | 0.62 | 0.32 | 0.53 |
| 11 | 56387773 | rs12418301 | 1.677 | 0.69 | -0.213 | -1.101 | -0.007 | -0.002 | -0.002 | 0 | 0.13 | 0.21 | 0.11 | 0.12 | 0.12 | 0.11 |
| 11 | 56406968 | rs906956 | 0.533 | 0.759 | -0.286 | -1.157 | 0.011 | 0.086 | 0.086 | 0.023 | 0.47 | 0.28 | 0.62 | 0.72 | 0.45 | 0.63 |
| 11 | 56409603 | rs11606499 | 1.661 | 0.715 | -0.311 | -1.171 | -0.021 | -0.086 | -0.086 | 0 | 0.19 | 0.26 | 0.1 | 0 | 0.14 | 0.04 |
| 11 | 56413925 | rs11228693 | 1.908 | 0.767 | -0.324 | -1.132 | -0.004 | 0 | 0 | 0 | 0.12 | 0.2 | 0.11 | 0.13 | 0.12 | 0.11 |
| 11 | 56418826 | rs11228699 | 1.126 | 1.003 | -0.439 | -1.108 | -0.041 | 0.017 | 0.017 | 0 | 0.1 | 0.02 | 0.08 | 0.05 | 0.16 | 0.06 |
| 11 | 56419115 | rs11228700 | 1.049 | 0.877 | -0.441 | -1.133 | -0.003 | 0.001 | 0.001 | 0 | 0.4 | 0.24 | 0.6 | 0.62 | 0.39 | 0.56 |
| 11 | 56421197 | rs1397056 | -0.242 | 0.425 | -0.048 | -0.874 | 0.037 | 0.052 | 0.052 | 1.822 | 0.41 | 0.68 | 0.26 | 0.22 | 0.31 | 0.22 |
| 11 | 56431216 | rs11228710 | -0.014 | 0.371 | -0.312 | -0.866 | 0.053 | 0.075 | 0.075 | 2.403 | 0.59 | 0.27 | 0.74 | 0.79 | 0.69 | 0.78 |
| 11 | 56438675 | rs11228713 | 1.091 | 0.473 | -0.232 | -0.693 | -0.013 | 0.014 | 0.014 | 0.713 | 0.08 | 0.2 | 0.04 | 0.09 | 0.05 | 0.08 |
| 11 | 56440236 | rs11228719 | -0.032 | 0.609 | -0.526 | -0.751 | 0.041 | 0.068 | 0.068 | 1.11 | 0.57 | 0.31 | 0.73 | 0.78 | 0.66 | 0.75 |
| 11 | 56441825 | rs10896503 | 1.379 | 0.414 | -0.429 | -0.758 | -0.005 | -0.002 | -0.002 | 0.544 | 0.13 | 0.24 | 0.11 | 0.12 | 0.12 | 0.11 |
| 11 | 56444138 | rs10896504 | 0.324 | 0.67 | -1.185 | -0.563 | -0.016 | -0.028 | -0.028 | 1.34 | 0.08 | 0.02 | 0.03 | 0.16 | 0.08 | 0.15 |
| 11 | 56452103 | rs11228726 | 0.597 | 0.774 | -1.2 | -0.56 | -0.052 | 0.023 | 0.023 | 0.147 | 0.45 | 0.17 | 0.65 | 0.61 | 0.51 | 0.61 |
| 11 | 56466099 | rs1397048 | -0.261 | 0.714 | -1.085 | -0.971 | 0.026 | 0.108 | 0.108 | 0.026 | 0.46 | 0.72 | 0.31 | 0.23 | 0.41 | 0.26 |
| 11 | 56467085 | rs10896513 | -0.261 | 0.714 | -1.085 | -0.971 | 0.026 | 0.108 | 0.108 | 0.026 | 0.46 | 0.72 | 0.31 | 0.23 | 0.41 | 0.26 |
| 11 | 56473046 | rs619748 | -0.005 | 0.852 | -1.02 | -0.663 | -0.02 | 0.004 | 0.004 | 0 | 0.08 | 0.02 | 0.04 | 0.07 | 0.08 | 0.11 |
| 11 | 56481768 | rs12361463 | 1.42 | 0.959 | -1.505 | -0.847 | -0.04 | -0.103 | -0.103 | 0 | 0.22 | 0.05 | 0.49 | 0.61 | 0.23 | 0.5 |
| 11 | 56482335 | rs545457 | -0.209 | 0.768 | -1.235 | -1.348 | 0.001 | 0.024 | 0.024 | 0 | 0.63 | 0.68 | 0.77 | 0.82 | 0.64 | 0.77 |
| 11 | 56483227 | rs11228739 | 2.053 | 0.691 | -1.196 | -1.31 | -0.006 | 0.018 | 0.018 | 0 | 0.06 | 0.13 | 0.02 | 0.09 | 0.05 | 0.05 |
| 11 | 56489004 | rs511107 | 0.318 | 0.663 | -0.724 | -1.471 | 0.016 | 0.073 | 0.073 | 0 | 0.5 | 0.33 | 0.65 | 0.7 | 0.53 | 0.67 |
| 11 | 56505081 | rs600778 | 0.284 | 0.574 | -0.42 | -1.541 | 0.003 | 0.021 | 0.021 | 0 | 0.52 | 0.53 | 0.66 | 0.71 | 0.53 | 0.67 |
| 11 | 56507170 | rs12285909 | 1.496 | 0.629 | -0.462 | -1.633 | 0 | -0.054 | -0.054 | 0 | 0.23 | 0.2 | 0.14 | 0.01 | 0.24 | 0.07 |
| 11 | 56507230 | rs630816 | 0.097 | 0.557 | -0.457 | -1.61 | 0.003 | 0.036 | 0.036 | 0 | 0.53 | 0.58 | 0.67 | 0.71 | 0.54 | 0.67 |
| 11 | 56508541 | rs11228759 | 2.029 | 0.617 | -0.652 | -1.597 | -0.005 | -0.007 | -0.007 | 0 | 0.09 | 0.04 | 0.1 | 0.12 | 0.1 | 0.11 |
| 11 | 56508686 | rs11228760 | 2.029 | 0.617 | -0.652 | -1.597 | -0.005 | -0.007 | -0.007 | 0 | 0.09 | 0.04 | 0.1 | 0.12 | 0.1 | 0.11 |
| 11 | 56510623 | rs513873 | 0.67 | 0.64 | -0.657 | -1.604 | -0.003 | -0.003 | -0.003 | 0 | 0.28 | 0.22 | 0.16 | 0.09 | 0.3 | 0.16 |
| 11 | 56510694 | rs1397053 | 1.303 | 0.568 | -0.647 | -1.626 | -0.001 | -0.049 | -0.049 | 0.437 | 0.24 | 0.2 | 0.13 | 0.01 | 0.21 | 0.07 |
| 11 | 56511160 | rs11228763 | 1.658 | 0.865 | -0.611 | -1.272 | 0.001 | 0 | 0 | 0 | 0.1 | 0.08 | 0.1 | 0.12 | 0.1 | 0.11 |
| 11 | 56514523 | rs11228764 | 0.073 | 0.838 | -0.654 | -1.247 | 0.021 | 0.031 | 0.031 | 0 | 0.32 | 0.22 | 0.17 | 0.17 | 0.26 | 0.17 |
| 11 | 56517109 | rs12223748 | 1.577 | 0.93 | -0.693 | -1.158 | 0.001 | 0 | 0 | 0 | 0.09 | 0.07 | 0.1 | 0.11 | 0.09 | 0.1 |
| 11 | 56517141 | rs17638028 | 0.718 | 0.807 | -0.788 | -1.27 | 0.013 | 0.077 | 0.077 | 0.008 | 0.23 | 0.06 | 0.13 | 0.09 | 0.21 | 0.11 |
| 11 | 56518769 | rs17550900 | 0.696 | 0.805 | -0.788 | -1.271 | 0.014 | 0.076 | 0.076 | 0.01 | 0.23 | 0.06 | 0.13 | 0.09 | 0.21 | 0.11 |
| 11 | 56526100 | rs10501357 | 1.028 | 0.863 | -0.574 | -1.191 | -0.004 | -0.001 | -0.001 | 0 | 0.16 | 0.13 | 0.05 | 0.09 | 0.1 | 0.07 |
| 11 | 56542928 | rs1943639 | -0.199 | 0.87 | -0.492 | -0.892 | 0.011 | 0.026 | 0.026 | 0 | 0.5 | 0.38 | 0.34 | 0.3 | 0.48 | 0.23 |
| 11 | 56544017 | rs1015195 | 1.563 | 1.037 | -0.483 | -0.868 | 0.008 | -0.046 | -0.046 | 0 | 0.28 | 0.23 | 0.45 | 0.51 | 0.25 | 0.51 |
| 11 | 56544745 | rs518385 | -0.047 | 1.077 | -0.447 | -0.941 | -0.001 | 0 | 0 | 0 | 0.43 | 0.52 | 0.35 | 0.28 | 0.47 | 0.26 |
| 11 | 56549319 | rs522662 | -0.731 | 1.084 | -0.49 | -0.931 | 0 | 0 | 0 | 0.017 | 0.63 | 0.39 | 0.68 | 0.8 | 0.61 | 0.84 |
| 11 | 56562425 | rs556566 | -0.141 | 1.027 | -0.571 | -0.807 | 0.01 | -0.005 | -0.005 | 0 | 0.31 | 0.17 | 0.18 | 0.3 | 0.26 | 0.27 |
| 11 | 56563462 | rs510456 | -0.572 | 1.067 | -0.445 | -0.864 | 0.002 | 0.009 | 0.009 | 0 | 0.62 | 0.38 | 0.68 | 0.8 | 0.61 | 0.84 |
| 11 | 56566495 | rs558434 | -0.755 | 1.089 | -0.411 | -0.852 | 0 | 0.005 | 0.005 | 0 | 0.65 | 0.66 | 0.71 | 0.8 | 0.61 | 0.84 |
| 11 | 56576321 | rs575227 | -0.287 | 1.306 | -0.841 | -0.746 | -0.056 | 0.01 | 0.01 | 0 | 0.12 | 0.03 | 0.1 | 0.1 | 0.17 | 0.16 |
| 11 | 56590550 | rs513995 | 0.846 | 1.197 | -0.802 | -0.705 | -0.019 | -0.009 | -0.009 | 0.123 | 0.07 | 0.02 | 0.04 | 0.1 | 0.07 | 0.1 |
| 11 | 56605222 | rs4938842 | 1.361 | 1.159 | -0.834 | -0.812 | -0.005 | 0.023 | 0.023 | 0 | 0.16 | 0.07 | 0.11 | 0.09 | 0.18 | 0.1 |
| 11 | 56608758 | rs652689 | -2.02 | 1.154 | -0.842 | -0.894 | -0.001 | -0.003 | -0.003 | 0 | 0.82 | 0.8 | 0.82 | 0.88 | 0.8 | 0.94 |
| 11 | 56610412 | rs764855 | 0.181 | 1.247 | -0.709 | -0.935 | 0.008 | 0.023 | 0.023 | 0.059 | 0.54 | 0.61 | 0.61 | 0.7 | 0.44 | 0.68 |
| 11 | 56611798 | rs679124 | -1.107 | 1.395 | -1.082 | -0.859 | -0.004 | 0.002 | 0.002 | 0 | 0.77 | 0.75 | 0.76 | 0.88 | 0.7 | 0.88 |

| | | | | | | | | | | | | | | | | |
|---|---|---|---|---|---|---|---|---|---|---|---|---|---|---|---|---|
| 11 | 56617544 | rs560518 | 0.345 | 1.292 | -1.474 | -0.478 | -0.019 | 0.015 | 0.015 | 0 | 0.07 | 0.02 | 0.04 | 0.04 | 0.07 | 0.1 |
| 11 | 56621020 | rs624662 | 0.681 | 1.298 | -1.456 | -0.457 | -0.018 | 0.009 | 0.009 | 0 | 0.06 | 0.02 | 0.04 | 0.04 | 0.07 | 0.1 |
| 11 | 56621491 | rs1938739 | 1.086 | 0.922 | -1.393 | -0.129 | 0.005 | -0.05 | -0.05 | 0.71 | 0.27 | 0.18 | 0.42 | 0.48 | 0.24 | 0.52 |
| 11 | 56623156 | rs7126523 | 0.175 | 1.057 | -1.072 | -0.108 | -0.01 | 0.005 | 0.005 | 0 | 0.14 | 0.14 | 0.16 | 0.11 | 0.19 | 0.16 |
| 11 | 56623492 | rs7107985 | -0.253 | 0.477 | -0.935 | 0.113 | 0.004 | 0.002 | 0.002 | 0.076 | 0.16 | 0.11 | 0.07 | 0.09 | 0.11 | 0.07 |
| 11 | 56626151 | rs7942208 | 0.064 | 1.531 | -0.932 | 0.261 | -0.006 | -0.001 | -0.001 | 0 | 0.11 | 0.1 | 0.07 | 0.14 | 0.18 | 0.16 |
| 11 | 56626311 | rs7119763 | 0.449 | 1.373 | -0.876 | -0.273 | -0.004 | -0.014 | -0.014 | 0.163 | 0.54 | 0.45 | 0.67 | 0.71 | 0.45 | 0.65 |
| 11 | 56629376 | rs657554 | 1.115 | 1.86 | -0.827 | 0.208 | 0.046 | -0.013 | -0.013 | 0 | 0.53 | 0.27 | 0.57 | 0.54 | 0.41 | 0.52 |
| 11 | 56630864 | rs7118940 | 1.098 | 1.333 | -1.25 | 0.272 | 0 | 0.005 | 0.005 | 0 | 0.23 | 0.21 | 0.21 | 0.09 | 0.23 | 0.09 |
| 11 | 56633934 | rs551339 | 0.486 | 1.464 | -0.906 | 0.266 | 0.013 | -0.015 | -0.015 | 0.03 | 0.65 | 0.45 | 0.78 | 0.77 | 0.58 | 0.71 |
| 11 | 56634316 | rs11228824 | 0.679 | 1.517 | -0.654 | 0.495 | 0.005 | 0.014 | 0.014 | 0 | 0.12 | 0.16 | 0.24 | 0.23 | 0.17 | 0.19 |
| 11 | 56643586 | rs1938746 | 0.486 | 1.463 | -0.723 | 0.337 | -0.004 | 0.008 | 0.008 | 0 | 0.3 | 0.21 | 0.25 | 0.16 | 0.31 | 0.14 |
| 11 | 56645710 | rs1938747 | 0.814 | 2.164 | -0.242 | 0.631 | -0.013 | -0.015 | -0.015 | 0 | 0.08 | 0.05 | 0.16 | 0.19 | 0.12 | 0.15 |
| 11 | 56650949 | rs1783466 | 0.555 | 1.712 | 0.077 | 0.003 | 0.004 | -0.025 | -0.025 | 0 | 0.07 | 0.28 | 0.04 | 0 | 0.04 | 0.01 |
| 11 | 56652705 | rs4939097 | 0.12 | 1.973 | -0.109 | 0.195 | 0.002 | 0 | 0 | 0.444 | 0.14 | 0.12 | 0.08 | 0.16 | 0.2 | 0.18 |
| 11 | 56659738 | rs1792500 | -0.913 | 1.878 | 0.258 | 0.147 | 0.025 | -0.042 | -0.042 | 0 | 0.23 | 0.4 | 0.13 | 0.16 | 0.24 | 0.2 |
| 11 | 56664357 | rs17640885 | -0.434 | 2.106 | 0.246 | 0.191 | 0.019 | -0.012 | -0.012 | 4.955 | 0.18 | 0.18 | 0.12 | 0.16 | 0.29 | 0.2 |
| 11 | 56665272 | rs7943283 | -1.165 | 2.04 | 0.225 | -0.028 | 0.006 | -0.016 | -0.016 | 1.06 | 0.28 | 0.27 | 0.25 | 0.16 | 0.38 | 0.3 |
| 11 | 56670810 | rs1792509 | -1.974 | 2.369 | 0.116 | 0.798 | 0 | 0.001 | 0.001 | 0.44 | 0.39 | 0.33 | 0.38 | 0.65 | 0.56 | 0.48 |
| 11 | 56672481 | rs12271904 | 0.499 | 2.462 | 0.086 | 0.937 | 0.003 | 0.006 | 0.006 | 0 | 0.1 | 0.09 | 0.13 | 0 | 0.09 | 0.1 |
| 11 | 56675823 | rs17641328 | 0.217 | 2.467 | 0.455 | 0.955 | 0.002 | 0.004 | 0.004 | 0.022 | 0.08 | 0.14 | 0.05 | 0.08 | 0.07 | 0.05 |
| 11 | 56677622 | rs1792513 | -2.579 | 1.525 | 0.1 | 0.516 | 0.082 | 0.039 | 0.039 | 2.968 | 0.69 | 0.88 | 0.64 | 0.74 | 0.83 | 0.6 |
| 11 | 56681750 | rs12272961 | -0.701 | 1.699 | 0.103 | 0.437 | 0.007 | -0.017 | -0.017 | 0 | 0.27 | 0.27 | 0.25 | 0.16 | 0.38 | 0.3 |
| 11 | 56683750 | rs1793426 | -0.723 | 1.752 | 0.119 | 0.458 | 0.007 | -0.017 | -0.017 | 0 | 0.27 | 0.28 | 0.25 | 0.16 | 0.38 | 0.29 |
| 11 | 56690570 | rs17151364 | 0.611 | 1.491 | 0.139 | 0.546 | 0.003 | 0.006 | 0.006 | 0 | 0.1 | 0.1 | 0.13 | 0 | 0.08 | 0.1 |
| 11 | 56691213 | rs17151375 | 0.819 | 1.491 | 0.136 | 0.546 | 0.003 | 0.016 | 0.016 | 0 | 0.09 | 0.04 | 0.12 | 0 | 0.08 | 0.1 |
| 11 | 56694692 | rs1622278 | -1.021 | 2.119 | 0.103 | 0.387 | -0.011 | 0.01 | 0.01 | 0.85 | 0.31 | 0.27 | 0.3 | 0.16 | 0.45 | 0.35 |
| 11 | 56697294 | rs1793429 | 0.484 | 2.153 | 0.526 | 0.817 | 0 | 0.02 | 0.02 | 2.696 | 0.08 | 0.06 | 0.12 | 0 | 0.08 | 0.1 |
| 11 | 56718765 | rs1894122 | 0.266 | 2.203 | 0.602 | 1.054 | 0.001 | 0.072 | 0.072 | 0 | 0.06 | 0.04 | 0.11 | 0.39 | 0.12 | 0.15 |
| 11 | 56719413 | rs7104992 | -0.685 | 2.057 | 0.973 | 1.001 | 0.017 | 0.04 | 0.04 | 0 | 0.31 | 0.44 | 0.32 | 0.48 | 0.33 | 0.24 |
| 11 | 56727619 | rs1793455 | 0.438 | 2.31 | 1.246 | 1.299 | 0 | 0.019 | 0.019 | 2.633 | 0.08 | 0.06 | 0.12 | 0 | 0.08 | 0.1 |
| 11 | 56738678 | rs7117365 | -0.34 | 2.31 | 1.234 | 1.3 | -0.019 | -0.029 | -0.029 | 0.553 | 0.25 | 0.4 | 0.21 | 0.1 | 0.21 | 0.1 |
| 11 | 56742853 | rs1945996 | 0.428 | 2.383 | 1.174 | 1.344 | -0.016 | 0.039 | 0.039 | 2.98 | 0.08 | 0 | 0.12 | 0 | 0.08 | 0.1 |
| 11 | 56755518 | rs7130724 | 0.095 | 2.399 | 1.27 | 1.351 | -0.003 | -0.032 | -0.032 | 0.346 | 0.09 | 0.14 | 0.13 | 0 | 0.08 | 0.1 |
| 11 | 56756664 | rs2853083 | -2.959 | 2.482 | 1.038 | 1.273 | 0.009 | 0.093 | 0.093 | 9.272 | 0.59 | 0.95 | 0.56 | 0.71 | 0.62 | 0.58 |
| 11 | 56774260 | rs7942314 | 0.737 | 2.481 | 1.338 | 1.289 | -0.003 | 0 | 0 | 0.023 | 0.17 | 0.19 | 0.17 | 0.1 | 0.17 | 0.07 |
| 11 | 56797089 | rs602991 | -1.452 | 2.52 | 1.329 | 1.345 | -0.005 | 0.005 | 0.005 | 4.805 | 0.4 | 0.59 | 0.43 | 0.57 | 0.4 | 0.37 |
| 11 | 56800970 | rs17562197 | -1.277 | 2.572 | 1.36 | 1.356 | 0.001 | -0.001 | -0.001 | 0 | 0.21 | 0.23 | 0.13 | 0.22 | 0.25 | 0.25 |
| 11 | 56805122 | rs11228882 | -0.687 | 2.583 | 1.249 | 1.348 | -0.008 | 0.012 | 0.012 | 0 | 0.17 | 0.13 | 0.11 | 0.12 | 0.21 | 0.18 |
| 11 | 56832642 | rs550124 | 2.052 | 2.534 | 1.156 | 1.412 | 0.039 | 0.098 | 0.098 | 8.279 | 0.42 | 0.13 | 0.45 | 0.32 | 0.39 | 0.42 |
| 11 | 56840338 | rs655949 | 1.808 | 2.316 | 1.341 | 1.555 | 0.037 | 0.166 | 0.166 | 4.433 | 0.49 | 0.13 | 0.47 | 0.32 | 0.45 | 0.55 |
| 11 | 56852098 | rs553086 | -0.046 | 2.123 | 1.921 | 1.339 | -0.015 | -0.004 | -0.004 | 0 | 0.76 | 0.68 | 0.79 | 0.86 | 0.8 | 0.89 |
| 11 | 56869386 | rs527174 | 1.86 | 2.215 | 0.761 | 1.375 | 0.01 | 0.124 | 0.124 | 4.657 | 0.49 | 0.08 | 0.47 | 0.35 | 0.44 | 0.56 |
| 11 | 56881013 | rs11228916 | 0.095 | 2.161 | 0.927 | 1.386 | 0.003 | 0.003 | 0.003 | 0 | 0.21 | 0.11 | 0.19 | 0.11 | 0.2 | 0.12 |
| 11 | 56886867 | rs2000737 | -1.331 | 2.53 | 1.653 | 1.746 | 0.024 | -0.056 | -0.056 | 0.053 | 0.19 | 0.33 | 0.2 | 0.08 | 0.22 | 0.14 |
| 11 | 56890350 | rs97624 | 1.936 | 2.61 | 0.802 | 1.765 | -0.029 | -0.045 | -0.045 | 0 | 0.7 | 0.23 | 0.67 | 0.46 | 0.65 | 0.68 |
| 11 | 56893050 | rs7110024 | -0.254 | 2.657 | 0.73 | 1.219 | -0.001 | -0.001 | -0.001 | 0.791 | 0.11 | 0.12 | 0.09 | 0.07 | 0.15 | 0.08 |
| 11 | 56905911 | rs524288 | 1.793 | 2.384 | 0.277 | 1.341 | -0.029 | -0.034 | -0.034 | 0 | 0.7 | 0.23 | 0.67 | 0.52 | 0.65 | 0.69 |
| 11 | 56905977 | rs3017313 | -0.905 | 1.695 | 0.628 | 1.159 | 0.014 | 0.032 | 0.032 | 0.033 | 0.1 | 0.21 | 0.13 | 0.12 | 0.1 | 0.09 |
| 11 | 56910484 | rs648409 | 0.381 | 1.49 | 0.119 | 1.228 | 0.006 | -0.02 | -0.02 | 0 | 0.75 | 0.85 | 0.81 | 0.67 | 0.79 | 0.84 |
| 11 | 56914157 | rs6591407 | -0.425 | 1.97 | 0.072 | 0.942 | -0.017 | 0.013 | 0.013 | 0 | 0.13 | 0.1 | 0.13 | 0.11 | 0.19 | 0.1 |
| 11 | 56921414 | rs10792076 | -1.513 | 2.327 | 0.024 | 0.954 | -0.027 | -0.006 | -0.006 | 1.344 | 0.2 | 0.13 | 0.25 | 0.26 | 0.3 | 0.22 |
| 11 | 56923361 | rs11228925 | 0.46 | 2.362 | 0.057 | 1.034 | -0.011 | -0.006 | -0.006 | 0 | 0.07 | 0.03 | 0.08 | 0.1 | 0.11 | 0.08 |
| 11 | 56925021 | rs3017569 | 0.334 | 2.246 | 0.858 | 0.936 | 0.054 | 0.022 | 0.022 | 0 | 0.06 | 0.28 | 0.1 | 0.01 | 0.05 | 0.04 |
| 11 | 56930712 | rs612948 | 2.372 | 2.057 | 0.411 | 0.913 | -0.029 | 0.031 | 0.031 | 1.27 | 0.45 | 0.1 | 0.43 | 0.4 | 0.43 | 0.53 |
| 11 | 56933839 | rs1941036 | -0.571 | 2.036 | 0.51 | 0.639 | -0.018 | 0.003 | 0.003 | 0 | 0.14 | 0.1 | 0.17 | 0.16 | 0.2 | 0.19 |
| 11 | 56940379 | rs10896573 | -2.052 | 2.072 | 0.669 | 0.621 | -0.04 | 0.02 | 0.02 | 0 | 0.29 | 0.19 | 0.34 | 0.25 | 0.42 | 0.31 |
| 11 | 56947528 | rs2077190 | -0.228 | 1.987 | 0.803 | 0.715 | 0.009 | -0.025 | -0.025 | 0 | 0.78 | 0.87 | 0.81 | 0.61 | 0.83 | 0.83 |
| 11 | 56947790 | rs2077189 | -2.338 | 1.934 | 0.992 | 0.668 | -0.011 | 0.009 | 0.009 | 0 | 0.34 | 0.69 | 0.38 | 0.25 | 0.43 | 0.31 |
| 11 | 56949691 | rs12364102 | 0.347 | 2.022 | 0.962 | 0.656 | -0.03 | 0.002 | 0.002 | 0 | 0.09 | 0.01 | 0.09 | 0.07 | 0.11 | 0.07 |
| 11 | 56950264 | rs609763 | -0.266 | 2.003 | 0.966 | 0.671 | 0.009 | -0.023 | -0.023 | 0 | 0.78 | 0.87 | 0.81 | 0.68 | 0.83 | 0.84 |
| 11 | 56951765 | rs567766 | -0.279 | 2.006 | 0.968 | 0.672 | 0.009 | -0.022 | -0.022 | 0.155 | 0.78 | 0.87 | 0.81 | 0.68 | 0.83 | 0.84 |
| 11 | 56952427 | rs10896578 | 2.555 | 1.997 | 0.974 | 0.674 | 0.01 | 0.003 | 0.003 | 1.568 | 0.43 | 0.12 | 0.43 | 0.36 | 0.4 | 0.52 |
| 11 | 56954846 | rs716745 | -1.508 | 2.058 | 1.048 | 0.681 | -0.05 | 0.029 | 0.029 | 0 | 0.25 | 0.14 | 0.26 | 0.19 | 0.38 | 0.27 |
| 11 | 56955091 | rs948851 | -0.045 | 2.068 | 0.906 | 0.684 | 0.009 | -0.025 | -0.025 | 0 | 0.22 | 0.13 | 0.19 | 0.39 | 0.17 | 0.17 |
| 11 | 56955677 | rs11820215 | -0.06 | 2.068 | 0.906 | 0.684 | 0.009 | -0.025 | -0.025 | 0 | 0.22 | 0.13 | 0.19 | 0.39 | 0.17 | 0.16 |
| 11 | 56956516 | rs2903267 | 1.987 | 2.047 | 0.908 | 0.668 | -0.011 | 0.011 | 0.011 | 0 | 0.66 | 0.3 | 0.62 | 0.75 | 0.57 | 0.69 |
| 11 | 56957418 | rs1943489 | 1.954 | 2.048 | 0.909 | 0.668 | -0.011 | 0.01 | 0.01 | 0 | 0.66 | 0.3 | 0.62 | 0.75 | 0.57 | 0.69 |
| 11 | 56959336 | rs1893678 | 0.312 | 2.128 | 0.88 | 0.693 | -0.033 | -0.002 | -0.002 | 0.002 | 0.08 | 0.01 | 0.09 | 0.08 | 0.11 | 0.07 |

| | | | | | | | | | | | | | | | | |
|---|---|---|---|---|---|---|---|---|---|---|---|---|---|---|---|---|
| 11 | 56966801 | rs4939111 | 1.825 | 2.074 | 0.956 | 0.677 | 0.008 | -0.006 | -0.006 | 0 | 0.68 | 0.32 | 0.62 | 0.75 | 0.57 | 0.69 |
| 11 | 56966815 | rs10501366 | -0.607 | 2.125 | 0.597 | 0.747 | -0.042 | 0.014 | 0.014 | 0 | 0.14 | 0.08 | 0.17 | 0.11 | 0.27 | 0.2 |
| 11 | 56969455 | rs17145573 | 0.651 | 1.644 | 0.456 | 0.966 | -0.023 | -0.061 | -0.061 | 0 | 0.1 | 0 | 0.12 | 0.3 | 0.09 | 0.07 |
| 11 | 56988451 | rs4442567 | 0.581 | 1.526 | 0.826 | 0.854 | -0.052 | 0.08 | 0.08 | 0 | 0.75 | 0.56 | 0.78 | 0.68 | 0.83 | 0.83 |
| 11 | 56990597 | rs11604352 | -1.276 | 1.435 | 0.646 | 0.663 | -0.036 | 0.047 | 0.047 | 0 | 0.29 | 0.41 | 0.24 | 0.39 | 0.21 | 0.17 |
| 11 | 56997434 | rs4938854 | -1.324 | 1.625 | 0.766 | 0.707 | -0.018 | 0.015 | 0.015 | 0 | 0.3 | 0.38 | 0.27 | 0.3 | 0.26 | 0.2 |
| 11 | 56997832 | rs721607 | 2.107 | 1.547 | 0.236 | 0.632 | 0.047 | -0.099 | -0.099 | 0 | 0.4 | 0.09 | 0.37 | 0.51 | 0.32 | 0.51 |
| 11 | 57001826 | rs2282623 | 2.205 | 1.548 | 0.248 | 0.633 | 0.034 | -0.046 | -0.046 | 0 | 0.38 | 0.08 | 0.37 | 0.4 | 0.31 | 0.5 |
| 11 | 57001911 | rs2282624 | -2.119 | 1.622 | 0.596 | 0.599 | 0.006 | 0.001 | 0.001 | 0 | 0.55 | 0.53 | 0.52 | 0.58 | 0.64 | 0.47 |
| 11 | 57002111 | rs746886 | 2.302 | 1.62 | 0.141 | 0.534 | 0.03 | -0.049 | -0.049 | 0 | 0.37 | 0.08 | 0.37 | 0.4 | 0.31 | 0.5 |
| 11 | 57004344 | rs948847 | 1.448 | 1.74 | 1.042 | -0.013 | 0.007 | -0.005 | -0.005 | 1.751 | 0.62 | 0.38 | 0.64 | 0.71 | 0.56 | 0.71 |
| 11 | 57004577 | rs11544374 | -0.365 | 2.031 | 0.783 | -0.107 | -0.068 | -0.01 | -0.01 | 0 | 0.2 | 0.08 | 0.21 | 0.18 | 0.29 | 0.21 |
| 11 | 57005067 | rs9943582 | 1.185 | 2.019 | 1.007 | -0.126 | -0.022 | 0.01 | 0.01 | 0.47 | 0.67 | 0.7 | 0.68 | 0.78 | 0.58 | 0.73 |
| 11 | 57005587 | rs10501367 | 0.304 | 1.984 | 1.043 | -0.088 | 0.015 | 0.002 | 0.002 | 0 | 0.78 | 0.71 | 0.78 | 0.8 | 0.71 | 0.79 |
| 11 | 57005964 | rs7119375 | 0.078 | 1.996 | 0.864 | -0.083 | -0.05 | -0.017 | -0.017 | 0 | 0.79 | 0.86 | 0.78 | 0.8 | 0.71 | 0.79 |
| 11 | 57012275 | rs7928905 | 2.365 | 1.786 | 0.757 | -0.557 | -0.047 | 0.103 | 0.103 | 1.167 | 0.52 | 0.07 | 0.61 | 0.69 | 0.54 | 0.66 |
| 11 | 57013248 | rs599947 | -1.798 | 1.821 | 0.997 | -0.547 | 0.026 | -0.032 | -0.032 | 0.756 | 0.37 | 0.53 | 0.3 | 0.24 | 0.43 | 0.27 |
| 11 | 57014757 | rs2511030 | -1.38 | 1.629 | 1.019 | -0.483 | 0.034 | -0.023 | -0.023 | 0 | 0.28 | 0.51 | 0.23 | 0.23 | 0.31 | 0.23 |
| 11 | 57015932 | rs17151797 | -0.65 | 1.601 | 1.127 | -0.494 | -0.036 | 0.002 | 0.002 | 0 | 0.1 | 0.28 | 0.07 | 0.07 | 0.03 | 0.06 |
| 11 | 57021799 | rs4384399 | -1.475 | 1.725 | 1.238 | -0.468 | 0.004 | -0.018 | -0.018 | 0 | 0.29 | 0.41 | 0.22 | 0.22 | 0.29 | 0.25 |
| 11 | 57022972 | rs7130252 | -0.836 | 1.786 | 1.229 | -0.455 | 0.02 | -0.005 | -0.005 | 7.355 | 0.23 | 0.41 | 0.18 | 0.22 | 0.25 | 0.21 |
| 11 | 57037040 | rs1892963 | -0.52 | 1.793 | 1.161 | -0.424 | -0.035 | -0.004 | -0.004 | 0 | 0.22 | 0.15 | 0.17 | 0.22 | 0.27 | 0.21 |
| 11 | 57046836 | rs11228984 | -1.237 | 1.873 | 1.336 | -0.369 | 0.005 | -0.022 | -0.022 | 0 | 0.33 | 0.45 | 0.26 | 0.21 | 0.32 | 0.28 |
| 11 | 57051835 | rs4939131 | 2.093 | 1.879 | 1.174 | -0.394 | -0.069 | 0.089 | 0.089 | 3.58 | 0.52 | 0.07 | 0.63 | 0.7 | 0.56 | 0.66 |
| 11 | 57058050 | rs17573746 | 2.055 | 1.911 | 1.188 | -0.378 | -0.068 | 0.085 | 0.085 | 3.226 | 0.53 | 0.07 | 0.63 | 0.7 | 0.56 | 0.66 |
| 11 | 57060562 | rs12792781 | -1.206 | 1.91 | 1.31 | -0.371 | -0.014 | 0.036 | 0.036 | 0.286 | 0.33 | 0.26 | 0.25 | 0.21 | 0.33 | 0.28 |
| 11 | 57066448 | rs11228988 | -0.971 | 2.009 | 1.297 | -0.335 | 0 | -0.01 | -0.01 | 0 | 0.33 | 0.46 | 0.25 | 0.28 | 0.31 | 0.27 |
| 11 | 57073233 | rs11602501 | -2.077 | 2.008 | 1.266 | -0.343 | -0.03 | 0.08 | 0.08 | 2.172 | 0.46 | 0.84 | 0.36 | 0.3 | 0.44 | 0.33 |
| 11 | 57077464 | rs10896602 | -1.061 | 2.006 | 1.243 | -0.357 | 0.002 | -0.009 | -0.009 | 0 | 0.34 | 0.45 | 0.25 | 0.28 | 0.33 | 0.28 |
| 11 | 57082896 | rs10792086 | -1.339 | 1.554 | 1.326 | -0.356 | 0.009 | -0.056 | -0.056 | 0 | 0.14 | 0.27 | 0.13 | 0.02 | 0.14 | 0.06 |
| 11 | 57084029 | rs11228997 | -0.561 | 1.597 | 1.305 | -0.343 | 0.002 | -0.028 | -0.028 | 0 | 0.11 | 0.16 | 0.08 | 0.02 | 0.1 | 0.04 |
| 11 | 57085213 | rs10896604 | -0.338 | 1.607 | 1.159 | -0.338 | -0.007 | 0.017 | 0.017 | 0 | 0.1 | 0.1 | 0.07 | 0.02 | 0.11 | 0.04 |
| 11 | 57096655 | rs10792088 | -1.178 | 1.614 | 1.146 | -0.345 | -0.002 | -0.007 | -0.007 | 0.492 | 0.14 | 0.15 | 0.12 | 0.02 | 0.14 | 0.05 |
| 11 | 57097924 | rs10896605 | -1.178 | 1.614 | 1.146 | -0.345 | -0.001 | -0.007 | -0.007 | 0 | 0.14 | 0.15 | 0.12 | 0.02 | 0.14 | 0.05 |
| 11 | 57102096 | rs2230649 | -1.547 | 1.596 | 1.142 | -0.353 | 0.023 | -0.081 | -0.081 | 0.207 | 0.16 | 0.39 | 0.13 | 0.02 | 0.15 | 0.07 |
| 11 | 57103797 | rs4432037 | -0.604 | 2.004 | 1.06 | -0.452 | 0.007 | -0.031 | -0.031 | 0 | 0.3 | 0.35 | 0.25 | 0.17 | 0.33 | 0.26 |
| 11 | 57104492 | rs7110052 | -1.506 | 1.85 | 0.996 | -0.366 | 0.025 | -0.071 | -0.071 | 0 | 0.15 | 0.35 | 0.13 | 0.02 | 0.15 | 0.07 |
| 11 | 57108911 | rs10896608 | -0.415 | 1.752 | 0.347 | -0.311 | -0.012 | 0.036 | 0.036 | 0 | 0.1 | 0.06 | 0.07 | 0.02 | 0.1 | 0.04 |
| 11 | 57111051 | rs12419699 | -0.124 | 1.66 | 0.212 | -0.279 | -0.025 | -0.03 | -0.03 | 0 | 0.07 | 0.01 | 0.06 | 0.07 | 0.1 | 0.04 |
| 11 | 57111693 | rs10732882 | -2.063 | 1.794 | 0.329 | -0.364 | 0.023 | 0.067 | 0.067 | 0.065 | 0.53 | 0.74 | 0.44 | 0.31 | 0.59 | 0.36 |
| 11 | 57117385 | rs2276042 | 1.764 | 1.476 | 0.172 | -0.359 | 0.018 | 0.059 | 0.059 | 0.317 | 0.45 | 0.13 | 0.54 | 0.67 | 0.39 | 0.62 |
| 11 | 57120047 | rs3781902 | 0.899 | 1.7 | 0.206 | -0.383 | -0.019 | 0.057 | 0.057 | 0 | 0.58 | 0.15 | 0.65 | 0.69 | 0.55 | 0.7 |
| 11 | 57126708 | rs4938863 | 0.51 | 1.708 | 0.714 | -0.427 | 0.066 | -0.063 | -0.063 | 5.495 | 0.87 | 0.62 | 0.91 | 0.98 | 0.86 | 0.95 |
| 11 | 57133371 | rs894450 | 1.476 | 1.582 | 0.21 | -0.211 | 0.005 | 0.025 | 0.025 | 0.195 | 0.47 | 0.08 | 0.51 | 0.64 | 0.41 | 0.64 |
| 11 | 57137371 | rs2276039 | 1.597 | 1.632 | 0.251 | -0.137 | 0.006 | -0.001 | -0.001 | 0 | 0.45 | 0.08 | 0.48 | 0.63 | 0.38 | 0.62 |
| 11 | 57137424 | rs2276038 | 1.437 | 1.577 | 0.236 | -0.37 | -0.002 | 0.059 | 0.059 | 0.421 | 0.44 | 0.07 | 0.49 | 0.72 | 0.39 | 0.63 |
| 11 | 57145633 | rs7107352 | -0.604 | 1.933 | 1.135 | -0.214 | 0.035 | -0.026 | -0.026 | 0 | 0.07 | 0.13 | 0.06 | 0.01 | 0.11 | 0.08 |
| 11 | 57148731 | rs10400305 | 1.609 | 1.859 | 1.632 | -0.295 | 0.032 | -0.097 | -0.097 | 6.554 | 0.84 | 0.43 | 0.88 | 0.98 | 0.83 | 0.89 |
| 11 | 57156438 | rs3741089 | -1.47 | 2.206 | 1.541 | 0.054 | -0.02 | 0.006 | 0.006 | 1.308 | 0.32 | 0.25 | 0.26 | 0.23 | 0.46 | 0.26 |
| 11 | 57157405 | rs490358 | 0.487 | 2.274 | 1.535 | 0.016 | 0.034 | 0.006 | 0.006 | 0.02 | 0.79 | 0.62 | 0.83 | 0.77 | 0.74 | 0.86 |
| 11 | 57158427 | rs7106575 | -0.016 | 2.316 | 1.282 | -0.083 | 0.007 | -0.001 | -0.001 | 0.749 | 0.05 | 0.05 | 0.07 | 0.05 | 0.12 | 0.07 |
| 11 | 57160838 | rs634737 | 0.69 | 2.294 | 1.386 | -0.086 | 0.057 | -0.003 | -0.003 | 0 | 0.77 | 0.48 | 0.83 | 0.77 | 0.74 | 0.85 |
| 11 | 57162716 | rs555097 | 1.255 | 2.28 | 1.376 | -0.091 | 0.062 | -0.013 | -0.013 | 0 | 0.71 | 0.43 | 0.75 | 0.73 | 0.62 | 0.78 |
| 11 | 57166661 | rs3851114 | 1.099 | 2.012 | 0.849 | -0.191 | -0.016 | 0.016 | 0.016 | 0 | 0.64 | 0.3 | 0.71 | 0.73 | 0.59 | 0.77 |
| 11 | 57171123 | rs10792095 | -0.955 | 1.658 | 0.942 | 0.05 | 0.055 | 0.031 | 0.031 | 0 | 0.06 | 0.17 | 0.09 | 0.12 | 0.12 | 0.08 |
| 11 | 57174584 | rs1049273 | -0.567 | 1.711 | 0.513 | -0.247 | 0.008 | -0.001 | -0.001 | 0.823 | 0.05 | 0.05 | 0.08 | 0.05 | 0.12 | 0.07 |
| 11 | 57184964 | rs7948027 | -1.607 | 1.515 | 0.753 | -0.407 | 0.044 | -0.047 | -0.047 | 0 | 0.27 | 0.53 | 0.23 | 0.16 | 0.32 | 0.27 |
| 11 | 57190420 | rs7122734 | -1.339 | 1.351 | 1.139 | -0.444 | 0.016 | -0.058 | -0.058 | 0.616 | 0.23 | 0.51 | 0.16 | 0.1 | 0.22 | 0.21 |
| 11 | 57193653 | rs12422015 | -1.374 | 1.145 | 0.971 | -0.55 | -0.001 | 0.002 | 0.002 | 0 | 0.23 | 0.25 | 0.15 | 0.09 | 0.23 | 0.26 |
| 11 | 57202184 | rs2250887 | -1.236 | 1.237 | 1.033 | -0.55 | -0.003 | 0.004 | 0.004 | 0 | 0.22 | 0.22 | 0.16 | 0.1 | 0.24 | 0.27 |
| 11 | 57203009 | rs11229030 | 1.659 | 0.831 | 1.267 | -0.867 | -0.001 | 0 | 0 | 0 | 0.64 | 0.33 | 0.72 | 0.84 | 0.62 | 0.68 |
| 11 | 57207414 | rs2581925 | 1.402 | 1.298 | 0.914 | -0.722 | 0.035 | -0.065 | -0.065 | 0 | 0.7 | 0.47 | 0.76 | 0.85 | 0.64 | 0.69 |
| 11 | 57212580 | rs2439451 | 0.864 | 1.244 | 0.275 | -0.623 | 0.002 | -0.006 | -0.006 | 0.222 | 0.82 | 0.79 | 0.82 | 0.91 | 0.78 | 0.78 |
| 11 | 57248003 | rs2729353 | 0.847 | 1.25 | 0.307 | -0.82 | 0.001 | -0.012 | -0.012 | 0.096 | 0.8 | 0.74 | 0.75 | 0.89 | 0.78 | 0.78 |
| 11 | 57249121 | rs2955849 | 0.721 | 0.979 | 0.069 | -1.14 | -0.003 | 0.026 | 0.026 | 0 | 0.42 | 0.29 | 0.47 | 0.7 | 0.43 | 0.55 |
| 11 | 57252328 | rs2848634 | -0.007 | 0.905 | -0.152 | -1.127 | 0.002 | -0.013 | -0.013 | 0 | 0.8 | 0.73 | 0.75 | 0.89 | 0.77 | 0.78 |
| 11 | 57262993 | rs2729385 | 0.9 | 1.289 | -0.607 | -0.694 | -0.025 | -0.145 | -0.145 | 0.027 | 0.33 | 0.03 | 0.38 | 0.56 | 0.33 | 0.42 |
| 11 | 57270509 | rs2649667 | -0.138 | 1.224 | -0.259 | -0.916 | -0.007 | 0.014 | 0.014 | 0 | 0.81 | 0.81 | 0.75 | 0.9 | 0.77 | 0.78 |
| 11 | 57272612 | rs4939146 | -1.511 | 1.158 | 0.458 | -1.056 | 0.049 | 0.021 | 0.021 | 2.376 | 0.1 | 0.54 | 0.09 | 0.08 | 0.08 | 0.04 |

| | | | | | | | | | | | | | | | | |
|---|---|---|---|---|---|---|---|---|---|---|---|---|---|---|---|---|
| 11 | 57274044 | rs7933797 | -0.472 | 1.282 | 0.951 | -1.028 | 0.065 | 0.037 | 0.037 | 0 | 0.05 | 0.14 | 0.07 | 0.08 | 0.08 | 0.02 |
| 11 | 57290961 | rs4939148 | -0.238 | 0.384 | 0.757 | -1.537 | 0 | -0.014 | -0.014 | 0 | 0.49 | 0.53 | 0.25 | 0.15 | 0.35 | 0.23 |
| 11 | 57291801 | rs4938869 | -0.858 | 0.204 | 0.547 | -1.84 | 0.031 | -0.083 | -0.083 | 0 | 0.19 | 0.44 | 0.17 | 0.07 | 0.23 | 0.08 |
| 11 | 57295864 | rs10896623 | -0.541 | 0.194 | 0.63 | -1.845 | -0.002 | -0.017 | -0.017 | 0.268 | 0.48 | 0.53 | 0.25 | 0.15 | 0.35 | 0.24 |
| 11 | 57296659 | rs10792101 | -0.846 | 0.188 | 0.632 | -1.85 | 0.032 | -0.083 | -0.083 | 0 | 0.19 | 0.44 | 0.17 | 0.07 | 0.23 | 0.08 |
| 11 | 57297271 | rs17453436 | 1.06 | 0.191 | 0.457 | -1.784 | 0.094 | 0.097 | 0.097 | 0 | 0.3 | 0.09 | 0.08 | 0.07 | 0.12 | 0.15 |
| 11 | 57306011 | rs514385 | 1.006 | 0.569 | 1.039 | -1.759 | 0.006 | -0.011 | -0.011 | 1.275 | 0.35 | 0.36 | 0.5 | 0.76 | 0.44 | 0.53 |
| 11 | 57312754 | rs2649659 | 0.973 | 0.534 | 1.014 | -1.775 | 0.014 | -0.023 | -0.023 | 0 | 0.35 | 0.4 | 0.51 | 0.75 | 0.44 | 0.53 |
| 11 | 57319339 | rs5847 | 0.567 | 0.491 | 0.996 | -1.851 | 0.024 | -0.061 | -0.061 | 0.257 | 0.66 | 0.52 | 0.59 | 0.8 | 0.55 | 0.69 |
| 11 | 57321811 | rs2234409 | 0.925 | 0.549 | 1.034 | -1.828 | 0.015 | -0.023 | -0.023 | 6.258 | 0.35 | 0.38 | 0.5 | 0.76 | 0.44 | 0.53 |
| 11 | 57327446 | rs2848624 | 0.997 | 0.565 | 1.067 | -1.833 | 0.008 | -0.011 | -0.011 | 0.518 | 0.34 | 0.36 | 0.5 | 0.75 | 0.44 | 0.53 |
| 11 | 57329402 | rs4938871 | -0.804 | 0.938 | 1.46 | -1.532 | 0.041 | -0.066 | -0.066 | 0 | 0.17 | 0.34 | 0.16 | 0.08 | 0.23 | 0.08 |
| 11 | 57330273 | rs1115172 | -0.804 | 0.938 | 1.46 | -1.532 | 0.041 | -0.066 | -0.066 | 0.062 | 0.17 | 0.34 | 0.16 | 0.08 | 0.23 | 0.08 |
| 11 | 57339950 | rs7124330 | -0.59 | 0.966 | 1.383 | -1.518 | 0.012 | -0.029 | -0.029 | 0 | 0.16 | 0.21 | 0.15 | 0.08 | 0.23 | 0.08 |
| 11 | 57346874 | rs2511983 | -0.837 | 0.958 | 0.902 | -1.535 | -0.019 | 0.025 | 0.025 | 0 | 0.84 | 0.9 | 0.75 | 0.88 | 0.79 | 0.77 |
| 11 | 57352562 | rs2649662 | -0.561 | 0.903 | 0.89 | -1.56 | -0.005 | 0.021 | 0.021 | 0 | 0.81 | 0.84 | 0.74 | 0.88 | 0.79 | 0.77 |
| 11 | 57352655 | rs7112126 | -0.897 | 0.984 | 0.917 | -1.515 | 0.041 | -0.064 | -0.064 | 0 | 0.17 | 0.37 | 0.16 | 0.08 | 0.24 | 0.08 |
| 11 | 57360416 | rs2729376 | 0.476 | 0.946 | 0.298 | -1.774 | -0.001 | -0.022 | -0.022 | 0.088 | 0.62 | 0.61 | 0.6 | 0.85 | 0.51 | 0.6 |
| 11 | 57362803 | rs3758919 | 0.782 | -0.094 | -0.188 | -2.08 | 0.035 | 0.015 | 0.015 | 0.2 | 0.2 | 0.09 | 0.06 | 0.09 | 0.08 | 0.11 |
| 11 | 57366656 | rs1005511 | 0.813 | 0.517 | -0.04 | -2.154 | -0.01 | -0.031 | -0.031 | 0 | 0.4 | 0.51 | 0.51 | 0.77 | 0.34 | 0.46 |
| 11 | 57367222 | rs1005510 | 0.844 | 0.514 | -0.06 | -2.155 | -0.009 | -0.03 | -0.03 | 0 | 0.4 | 0.48 | 0.51 | 0.76 | 0.34 | 0.46 |
| 11 | 57369730 | rs11229063 | 0.357 | 0.8 | -0.431 | -2.24 | -0.002 | 0.047 | 0.047 | 0.495 | 0.26 | 0.09 | 0.16 | 0.1 | 0.26 | 0.16 |
| 11 | 57377992 | rs2508443 | 0.224 | 0.793 | -0.198 | -2.279 | -0.04 | 0.075 | 0.075 | 0 | 0.31 | 0.18 | 0.31 | 0.13 | 0.41 | 0.36 |
| 11 | 57381519 | rs1557522 | 0.274 | 0.771 | -0.381 | -2.273 | -0.082 | 0.124 | 0.124 | 0 | 0.31 | 0.11 | 0.3 | 0.14 | 0.41 | 0.36 |
| 11 | 57381989 | rs4926 | 0.172 | 0.594 | -0.133 | -2.037 | -0.004 | 0.046 | 0.046 | 0.563 | 0.26 | 0.09 | 0.17 | 0.09 | 0.27 | 0.16 |
| 11 | 57389737 | rs11607122 | -0.162 | 0.592 | 0.02 | -1.871 | -0.001 | -0.002 | -0.002 | 0 | 0.28 | 0.27 | 0.18 | 0.13 | 0.27 | 0.18 |
| 11 | 57390302 | rs10896632 | -0.168 | 0.559 | 0.021 | -1.915 | -0.001 | -0.006 | -0.006 | 0.169 | 0.28 | 0.27 | 0.18 | 0.09 | 0.27 | 0.18 |
| 11 | 57396825 | rs7102271 | -0.165 | 0.572 | 0.054 | -1.908 | -0.004 | -0.023 | -0.023 | 0 | 0.28 | 0.31 | 0.18 | 0.09 | 0.27 | 0.18 |
| 11 | 57408382 | rs731384 | 0.295 | 0.631 | -0.351 | -1.995 | -0.025 | 0.027 | 0.027 | 0 | 0.22 | 0.08 | 0.17 | 0.09 | 0.27 | 0.24 |
| 11 | 57412150 | rs1798177 | -0.39 | 0.5 | -0.903 | -2.576 | -0.05 | 0.178 | 0.178 | 0.058 | 0.64 | 0.88 | 0.69 | 0.89 | 0.58 | 0.68 |
| 11 | 57420780 | rs1783983 | -0.075 | 0.298 | -0.127 | -2.483 | -0.001 | -0.027 | -0.027 | 0 | 0.61 | 0.61 | 0.68 | 0.88 | 0.6 | 0.68 |
| 11 | 57420885 | rs1631684 | -0.338 | 0.295 | -0.134 | -2.492 | -0.03 | 0.155 | 0.155 | 0.056 | 0.63 | 0.85 | 0.69 | 0.88 | 0.6 | 0.68 |
| 11 | 57424000 | rs1613592 | -0.101 | 0.277 | -0.152 | -2.478 | -0.001 | -0.03 | -0.03 | 3.507 | 0.61 | 0.61 | 0.68 | 0.88 | 0.6 | 0.68 |
| 11 | 57424040 | rs10896633 | 0.479 | 0.193 | -0.581 | -2.596 | -0.03 | 0.052 | 0.052 | 0.066 | 0.26 | 0.08 | 0.17 | 0.09 | 0.3 | 0.28 |
| 11 | 57425182 | rs2278108 | 1.815 | 0.021 | -0.773 | -2.68 | -0.004 | 0.007 | 0.007 | 0.064 | 0.09 | 0.02 | 0.04 | 0.05 | 0.06 | 0.03 |
| 11 | 57425699 | rs11229087 | 0.952 | -0.29 | -0.465 | -2.84 | -0.014 | 0.025 | 0.025 | 0 | 0.3 | 0.15 | 0.49 | 0.78 | 0.3 | 0.38 |
| 11 | 57434122 | rs499188 | 0.119 | -0.056 | -0.107 | -2.739 | 0.005 | -0.027 | -0.027 | 0 | 0.29 | 0.34 | 0.18 | 0.1 | 0.31 | 0.27 |
| 11 | 57456244 | rs549022 | 1.63 | 0.384 | -0.094 | -2.87 | 0 | -0.001 | -0.001 | 0 | 0.24 | 0.24 | 0.47 | 0.8 | 0.25 | 0.35 |
| 11 | 57460224 | rs12801694 | 0.981 | 0.149 | -0.134 | -2.984 | 0.04 | 0.098 | 0.098 | 0 | 0.31 | 0.17 | 0.25 | 0.08 | 0.26 | 0.27 |
| 11 | 57463500 | rs1783978 | -0.23 | 0.192 | -0.084 | -2.936 | 0.005 | 0.013 | 0.013 | 0.057 | 0.53 | 0.58 | 0.66 | 0.9 | 0.56 | 0.63 |
| 11 | 57468129 | rs1783979 | -0.401 | 0.195 | -0.083 | -2.935 | -0.007 | 0.214 | 0.214 | 0.53 | 0.56 | 0.82 | 0.67 | 0.9 | 0.56 | 0.63 |
| 11 | 57470844 | rs544446 | 1.715 | 0.207 | -0.097 | -2.928 | 0 | -0.001 | -0.001 | 0 | 0.24 | 0.24 | 0.47 | 0.8 | 0.25 | 0.35 |
| 11 | 57477889 | rs1647405 | 0.416 | 0.243 | -0.014 | -2.928 | 0.005 | -0.026 | -0.026 | 0 | 0.3 | 0.34 | 0.18 | 0.1 | 0.31 | 0.27 |
| 11 | 57479848 | rs1424752 | 1.715 | 0.201 | -0.015 | -2.928 | 0 | -0.001 | -0.001 | 0.21 | 0.24 | 0.24 | 0.47 | 0.8 | 0.25 | 0.35 |
| 11 | 57488160 | rs480933 | 1.715 | 0.201 | -0.015 | -2.928 | 0 | -0.001 | -0.001 | 0 | 0.24 | 0.24 | 0.47 | 0.8 | 0.25 | 0.35 |
| 11 | 57496820 | rs11607056 | 0.94 | 0.169 | 0.027 | -2.942 | -0.034 | 0.049 | 0.049 | 0.417 | 0.26 | 0.09 | 0.17 | 0.1 | 0.31 | 0.25 |
| 11 | 57498232 | rs2847308 | -0.666 | 0.172 | 0.028 | -2.941 | 0.004 | -0.026 | -0.026 | 0.144 | 0.7 | 0.66 | 0.82 | 0.9 | 0.69 | 0.73 |
| 11 | 57501182 | rs2956981 | -0.126 | 0.172 | 0.03 | -2.94 | 0.006 | 0.014 | 0.014 | 0.174 | 0.47 | 0.42 | 0.34 | 0.11 | 0.44 | 0.38 |
| 11 | 57510294 | rs9420 | -0.733 | 0.135 | 0.039 | -2.483 | 0.005 | -0.028 | -0.028 | 0 | 0.71 | 0.66 | 0.82 | 0.9 | 0.7 | 0.73 |
| 11 | 57529429 | rs11570176 | 0.046 | 0.149 | -0.244 | -2.52 | -0.002 | 0.212 | 0.212 | 0.427 | 0.44 | 0.18 | 0.33 | 0.1 | 0.43 | 0.37 |
| 11 | 57550785 | rs12362406 | 1.058 | 0.045 | -0.335 | -2.594 | 0.028 | 0.174 | 0.174 | 0.13 | 0.3 | 0.04 | 0.22 | 0.08 | 0.25 | 0.27 |
| 11 | 57555770 | rs11570181 | 0.223 | 0.192 | 0.004 | -2.243 | -0.001 | 0.213 | 0.213 | 0.369 | 0.44 | 0.18 | 0.33 | 0.1 | 0.43 | 0.37 |
| 11 | 57563991 | rs10896644 | 0.23 | 0.199 | 0.007 | -2.24 | -0.002 | 0.212 | 0.212 | 0.427 | 0.44 | 0.18 | 0.33 | 0.1 | 0.43 | 0.37 |
| 11 | 57564246 | rs11570195 | 2.847 | -0.006 | -0.017 | -2.349 | -0.024 | -0.025 | -0.025 | 0 | 0.05 | 0.06 | 0.01 | 0 | 0 | 0 |
| 11 | 57570430 | rs653889 | 0.177 | 0.339 | 0.029 | -2.222 | -0.004 | 0.213 | 0.213 | 0.451 | 0.44 | 0.18 | 0.33 | 0.1 | 0.43 | 0.37 |
| 11 | 57585662 | rs708228 | 0.782 | 0.256 | -0.141 | -2.101 | -0.034 | 0.05 | 0.05 | 0.424 | 0.26 | 0.09 | 0.17 | 0.1 | 0.31 | 0.25 |
| 11 | 57593840 | rs606499 | -0.228 | 0.267 | -0.119 | -2.035 | 0.023 | 0.067 | 0.067 | 0.12 | 0.5 | 0.33 | 0.64 | 0.89 | 0.57 | 0.6 |
| 11 | 57600285 | rs12807250 | 0.606 | 0.177 | -0.41 | -1.65 | -0.034 | 0.045 | 0.045 | 0.412 | 0.26 | 0.09 | 0.17 | 0.1 | 0.31 | 0.25 |
| 11 | 57667222 | rs12790660 | 0.507 | 0.253 | -0.417 | -1.855 | -0.037 | 0.054 | 0.054 | 0.04 | 0.24 | 0.09 | 0.15 | 0.07 | 0.3 | 0.27 |
| 11 | 57684680 | rs591322 | -0.294 | -0.161 | -0.079 | -1.632 | -0.01 | 0.182 | 0.182 | 1.675 | 0.43 | 0.19 | 0.33 | 0.11 | 0.44 | 0.38 |
| 11 | 57696234 | rs609154 | -0.971 | -0.342 | -0.085 | -1.645 | 0 | 0 | 0 | 0.375 | 0.64 | 0.42 | 0.59 | 0.8 | 0.72 | 0.83 |
| 11 | 57703762 | rs7926999 | -0.073 | -0.318 | -0.129 | -1.496 | -0.028 | 0.151 | 0.151 | 2.709 | 0.44 | 0.21 | 0.35 | 0.2 | 0.47 | 0.4 |
| 11 | 57705324 | rs1397273 | 0.746 | -0.316 | -0.005 | -1.494 | -0.003 | -0.016 | -0.016 | 0 | 0.35 | 0.53 | 0.41 | 0.2 | 0.28 | 0.15 |
| 11 | 57709146 | rs10792118 | -0.345 | -0.325 | -0.007 | -1.432 | -0.001 | 0.108 | 0.108 | 4.498 | 0.57 | 0.72 | 0.66 | 0.87 | 0.56 | 0.6 |
| 11 | 57709850 | rs10792119 | 1.405 | -0.315 | -0.007 | -1.422 | -0.002 | -0.014 | -0.014 | 0 | 0.2 | 0.18 | 0.24 | 0.59 | 0.25 | 0.44 |
| 11 | 57713781 | rs4245215 | 0.129 | -0.27 | -0.162 | -1.48 | -0.016 | 0.169 | 0.169 | 4.453 | 0.42 | 0.21 | 0.34 | 0.12 | 0.44 | 0.4 |
| 11 | 57731052 | rs7102929 | 0.913 | -0.121 | 0.07 | -1.122 | 0.002 | -0.035 | -0.035 | 0 | 0.31 | 0.29 | 0.17 | 0.08 | 0.28 | 0.14 |
| 11 | 57734088 | rs12804093 | 0.656 | -0.12 | 0.054 | -1.147 | 0.003 | -0.017 | -0.017 | 0 | 0.33 | 0.32 | 0.19 | 0.16 | 0.31 | 0.15 |
| 11 | 57743931 | rs7121128 | 1.078 | -0.115 | -0.096 | -1.144 | -0.021 | 0.064 | 0.064 | 0.333 | 0.29 | 0.12 | 0.16 | 0.1 | 0.28 | 0.13 |

| | | | | | | | | | | | | | | | | |
|---|---|---|---|---|---|---|---|---|---|---|---|---|---|---|---|---|
| 11 | 57751001 | rs12418547 | 0.565 | -0.4 | 0.009 | -1.25 | -0.037 | 0.008 | 0.008 | 0.665 | 0.14 | 0.33 | 0.27 | 0.12 | 0.06 | 0.03 |
| 11 | 57752099 | rs1510418 | -0.609 | -0.352 | -0.001 | -1.302 | 0.002 | 0.014 | 0.014 | 1.893 | 0.5 | 0.57 | 0.59 | 0.31 | 0.48 | 0.42 |
| 11 | 57753790 | rs11229194 | -1.179 | -0.075 | 0.057 | -1.071 | 0.003 | 0.015 | 0.015 | 0 | 0.79 | 0.76 | 0.75 | 0.34 | 0.72 | 0.55 |
| 11 | 57761232 | rs4939173 | 1.3 | -0.074 | -0.035 | -1.052 | -0.002 | -0.009 | -0.009 | 0 | 0.2 | 0.17 | 0.24 | 0.58 | 0.25 | 0.44 |
| 11 | 57779208 | rs7926462 | -1.108 | -0.165 | -0.38 | -1.331 | 0.007 | 0.178 | 0.178 | 0 | 0.51 | 0.82 | 0.66 | 0.8 | 0.56 | 0.62 |
| 11 | 57782028 | rs10431127 | -0.197 | 0.008 | -0.202 | -1.224 | 0.001 | -0.001 | -0.001 | 0 | 0.35 | 0.34 | 0.37 | 0.67 | 0.5 | 0.58 |
| 11 | 57785231 | rs4277129 | 1.037 | 0.061 | -0.178 | -1.306 | -0.001 | 0.001 | 0.001 | 0.011 | 0.21 | 0.21 | 0.25 | 0.67 | 0.27 | 0.46 |
| 11 | 57786238 | rs7103284 | -0.077 | -0.276 | -0.159 | -1.56 | 0.001 | 0.01 | 0.01 | 0 | 0.43 | 0.41 | 0.45 | 0.75 | 0.46 | 0.71 |
| 11 | 57786845 | rs10896687 | -0.891 | -0.266 | -0.147 | -1.586 | -0.004 | -0.015 | -0.015 | 0 | 0.56 | 0.53 | 0.57 | 0.77 | 0.66 | 0.84 |
| 11 | 57788450 | rs7128781 | 1.244 | -0.107 | -0.052 | -1.474 | 0.002 | -0.016 | -0.016 | 0.034 | 0.13 | 0.13 | 0.12 | 0.02 | 0.2 | 0.13 |
| 11 | 57790824 | rs2441950 | -0.296 | -0.132 | -0.053 | -1.487 | 0.024 | -0.015 | -0.015 | 0.651 | 0.37 | 0.49 | 0.39 | 0.69 | 0.49 | 0.6 |
| 11 | 57790851 | rs2866831 | 1.187 | -0.083 | -0.095 | -1.45 | 0.002 | -0.016 | -0.016 | 0.969 | 0.13 | 0.13 | 0.12 | 0.02 | 0.2 | 0.13 |
| 11 | 57796009 | rs2441956 | -0.911 | -0.425 | -0.001 | -1.282 | 0.01 | 0.014 | 0.014 | 0 | 0.77 | 0.81 | 0.79 | 0.93 | 0.81 | 0.75 |
| 11 | 57798723 | rs921135 | 0.679 | -0.306 | -0.086 | -1.257 | 0.011 | 0.019 | 0.019 | 0 | 0.24 | 0.32 | 0.27 | 0.68 | 0.29 | 0.46 |
| 11 | 57810680 | rs7118536 | 1.44 | -0.317 | -0.043 | -1.179 | -0.011 | -0.026 | -0.026 | 0 | 0.2 | 0.16 | 0.23 | 0.59 | 0.26 | 0.44 |
| 11 | 57825450 | rs11229240 | 1.252 | -0.377 | -0.169 | -1.277 | 0.008 | 0.069 | 0.069 | 0 | 0.2 | 0.04 | 0.14 | 0.09 | 0.16 | 0.22 |
| 11 | 57831996 | rs2514202 | -0.828 | -0.288 | 0.008 | -1.102 | 0.011 | 0.018 | 0.018 | 0.003 | 0.76 | 0.68 | 0.73 | 0.32 | 0.71 | 0.54 |
| 11 | 57835508 | rs1868882 | 1.073 | -0.542 | 0.133 | -1.067 | -0.028 | 0.025 | 0.025 | 3.816 | 0.12 | 0.27 | 0.32 | 0.13 | 0.06 | 0.05 |
| 11 | 57836249 | rs2443447 | -0.093 | -0.618 | 0.136 | -1.235 | 0.001 | 0.005 | 0.005 | 0 | 0.56 | 0.57 | 0.65 | 0.26 | 0.55 | 0.33 |
| 11 | 57837848 | rs2441976 | 0.528 | -0.629 | 0.185 | -1.121 | -0.045 | 0.036 | 0.036 | 5.523 | 0.14 | 0.41 | 0.34 | 0.13 | 0.06 | 0.06 |
| 11 | 57847510 | rs12793317 | 1.159 | -0.535 | -0.619 | -1.124 | 0.016 | 0.068 | 0.068 | 0 | 0.19 | 0.04 | 0.14 | 0.1 | 0.16 | 0.22 |
| 11 | 57854731 | rs3974323 | 1.365 | -0.263 | -0.507 | -0.93 | -0.08 | -0.145 | -0.145 | 0 | 0.19 | 0.01 | 0.17 | 0.57 | 0.25 | 0.43 |
| 11 | 57864710 | rs2514209 | 0.461 | -0.508 | 0.083 | -0.821 | -0.044 | 0.021 | 0.021 | 1.851 | 0.14 | 0.29 | 0.32 | 0.13 | 0.06 | 0.05 |
| 11 | 57871866 | rs1031360 | -0.061 | -0.457 | -0.116 | -1.199 | 0.014 | 0.039 | 0.039 | 0 | 0.52 | 0.34 | 0.47 | 0.7 | 0.61 | 0.78 |
| 11 | 57872281 | rs1031232 | 1.007 | -0.483 | -0.101 | -1.277 | 0.002 | -0.02 | -0.02 | 0 | 0.16 | 0.14 | 0.17 | 0.03 | 0.2 | 0.13 |
| 11 | 57876221 | rs1447170 | 0.05 | -0.563 | 0.005 | -1.304 | -0.054 | 0.041 | 0.041 | 3.556 | 0.81 | 0.46 | 0.63 | 0.79 | 0.9 | 0.92 |
| 11 | 57876436 | rs7935428 | 1.392 | -0.497 | -0.218 | -1.305 | -0.05 | 0.081 | 0.081 | 0 | 0.28 | 0.08 | 0.15 | 0.09 | 0.3 | 0.14 |
| 11 | 57880392 | rs1451326 | 0.856 | -0.362 | -0.179 | -1.236 | -0.023 | -0.03 | -0.03 | 0 | 0.37 | 0.2 | 0.31 | 0.64 | 0.41 | 0.64 |
| 11 | 57884611 | rs1447168 | 0.791 | -0.352 | -0.177 | -1.217 | -0.023 | -0.029 | -0.029 | 0 | 0.38 | 0.2 | 0.32 | 0.64 | 0.42 | 0.64 |
| 11 | 57889566 | rs11601841 | 1.344 | -0.413 | -0.199 | -1.253 | -0.02 | 0.057 | 0.057 | 0 | 0.28 | 0.12 | 0.15 | 0.09 | 0.29 | 0.14 |
| 11 | 57893004 | rs1447167 | -0.572 | -0.46 | 0.163 | -1.267 | 0.015 | -0.029 | -0.029 | 0 | 0.81 | 0.74 | 0.81 | 0.89 | 0.77 | 0.86 |
| 11 | 57893374 | rs2441965 | 0.83 | -0.533 | 0.362 | -1.257 | -0.039 | 0.015 | 0.015 | 1.848 | 0.14 | 0.27 | 0.32 | 0.13 | 0.06 | 0.05 |
| 11 | 57893967 | rs3853676 | 0.983 | -0.456 | 0.334 | -1.28 | -0.03 | 0.036 | 0.036 | 0 | 0.31 | 0.16 | 0.18 | 0.19 | 0.32 | 0.17 |
| 11 | 57895573 | rs921445 | 1.354 | -0.463 | 0.305 | -1.285 | -0.04 | 0.061 | 0.061 | 0 | 0.28 | 0.09 | 0.16 | 0.12 | 0.29 | 0.15 |
| 11 | 57899255 | rs7120442 | 1.311 | -0.453 | 0.522 | -1.292 | -0.018 | 0.058 | 0.058 | 0 | 0.28 | 0.14 | 0.16 | 0.09 | 0.29 | 0.14 |
| 11 | 57900983 | rs1447166 | 0.052 | -0.508 | 0.489 | -1.325 | 0.012 | 0.062 | 0.062 | 0 | 0.51 | 0.43 | 0.63 | 0.8 | 0.48 | 0.68 |
| 11 | 57901176 | rs1868881 | 0.974 | -0.447 | 0.376 | -1.311 | -0.003 | 0.007 | 0.007 | 0 | 0.32 | 0.26 | 0.2 | 0.17 | 0.32 | 0.16 |
| 11 | 57902114 | rs921443 | -1.171 | -0.172 | 0.011 | -1.17 | 0.011 | -0.049 | -0.049 | 0 | 0.84 | 0.81 | 0.84 | 0.97 | 0.8 | 0.88 |
| 11 | 57909502 | rs10896705 | 0.688 | -0.302 | -0.317 | -1.161 | -0.068 | -0.068 | -0.068 | 0.129 | 0.38 | 0.1 | 0.31 | 0.64 | 0.42 | 0.64 |
| 11 | 57915680 | rs1447181 | -0.085 | -0.098 | 0.39 | -0.921 | 0.032 | -0.044 | -0.044 | 0.352 | 0.81 | 0.64 | 0.81 | 0.89 | 0.76 | 0.87 |
| 11 | 57918692 | rs2441987 | -0.502 | -0.319 | -0.085 | -1.019 | 0.009 | -0.002 | -0.002 | 0 | 0.3 | 0.31 | 0.52 | 0.27 | 0.38 | 0.22 |
| 11 | 57922678 | rs2443426 | -0.611 | -0.367 | -0.106 | -1.069 | -0.002 | 0.001 | 0.001 | 0 | 0.33 | 0.29 | 0.53 | 0.3 | 0.4 | 0.23 |
| 11 | 57926106 | rs7130768 | -0.507 | -0.609 | -0.287 | -1.304 | -0.005 | 0.016 | 0.016 | 0 | 0.45 | 0.4 | 0.54 | 0.66 | 0.52 | 0.7 |
| 11 | 57930210 | rs964720 | 2.108 | -0.415 | -0.499 | -1.263 | -0.053 | -0.073 | -0.073 | 1.175 | 0.11 | 0.02 | 0.13 | 0.53 | 0.16 | 0.42 |
| 11 | 57934197 | rs2443439 | -0.105 | -0.611 | -0.307 | -1.41 | 0.008 | -0.011 | -0.011 | 0 | 0.34 | 0.45 | 0.54 | 0.31 | 0.39 | 0.23 |
| 11 | 57934480 | rs2122801 | -1.429 | -0.675 | -0.296 | -1.466 | -0.002 | -0.022 | -0.022 | 0 | 0.62 | 0.57 | 0.8 | 0.88 | 0.67 | 0.79 |
| 11 | 57935523 | rs1349480 | -0.138 | -0.546 | -0.296 | -1.397 | 0.009 | -0.015 | -0.015 | 0 | 0.34 | 0.45 | 0.54 | 0.27 | 0.39 | 0.22 |
| 11 | 57948952 | rs2443443 | -1.571 | -0.764 | -0.02 | -1.503 | -0.002 | 0.071 | 0.071 | 2.012 | 0.63 | 0.81 | 0.81 | 0.87 | 0.66 | 0.78 |
| 11 | 57951001 | rs10896706 | 0.776 | -1.123 | -0.703 | -1.645 | -0.05 | -0.157 | -0.157 | 0 | 0.25 | 0.06 | 0.24 | 0.58 | 0.27 | 0.55 |
| 11 | 57951574 | rs1451319 | 0.849 | -1.22 | -0.73 | -1.72 | -0.02 | -0.006 | -0.006 | 0 | 0.24 | 0.23 | 0.26 | 0.22 | 0.15 | 0.09 |
| 11 | 57956832 | rs1376486 | 0.727 | -1.021 | -0.62 | -1.811 | 0.059 | -0.015 | -0.015 | 0 | 0.13 | 0.26 | 0.29 | 0.07 | 0.24 | 0.13 |
| 11 | 57958775 | rs7120468 | 0.427 | -0.964 | -1.326 | -1.779 | 0.005 | 0.014 | 0.014 | 0 | 0.34 | 0.22 | 0.38 | 0.75 | 0.31 | 0.51 |
| 11 | 57958972 | rs1451317 | 0.448 | -0.965 | -1.326 | -1.779 | 0.005 | 0.013 | 0.013 | 0 | 0.33 | 0.22 | 0.38 | 0.75 | 0.31 | 0.51 |
| 11 | 57965017 | rs10896710 | -0.566 | -1.176 | -1.451 | -1.884 | 0.041 | 0.087 | 0.087 | 0 | 0.51 | 0.28 | 0.5 | 0.79 | 0.43 | 0.65 |
| 11 | 57965070 | rs4092189 | -0.537 | -1.177 | -1.451 | -1.884 | 0.042 | 0.089 | 0.089 | 0 | 0.51 | 0.28 | 0.5 | 0.79 | 0.43 | 0.65 |
| 11 | 57967467 | rs1376483 | -0.453 | -0.868 | -1.29 | -1.734 | 0.03 | 0.072 | 0.072 | 0 | 0.47 | 0.28 | 0.5 | 0.79 | 0.42 | 0.65 |
| 11 | 57969448 | rs1451315 | -0.453 | -0.868 | -1.29 | -1.734 | 0.03 | 0.072 | 0.072 | 0 | 0.47 | 0.28 | 0.5 | 0.79 | 0.43 | 0.65 |
| 11 | 57972160 | rs1947298 | 0.407 | -0.87 | -1.296 | -1.736 | 0.004 | 0.014 | 0.014 | 0 | 0.34 | 0.22 | 0.39 | 0.75 | 0.31 | 0.51 |
| 11 | 57973199 | rs1823598 | 1.438 | -0.815 | -1.281 | -1.762 | -0.007 | 0.002 | 0.002 | 0 | 0.11 | 0.09 | 0.27 | 0.07 | 0.24 | 0.13 |
| 11 | 57976907 | rs11229282 | 1.978 | -0.836 | -1.273 | -1.783 | 0.018 | 0.024 | 0.024 | 0 | 0.13 | 0.06 | 0.11 | 0.04 | 0.11 | 0.14 |
| 11 | 57982229 | rs1966836 | -1.142 | -0.794 | -1.243 | -1.782 | 0.003 | 0.008 | 0.008 | 1.204 | 0.58 | 0.38 | 0.76 | 0.87 | 0.66 | 0.78 |
| 11 | 57986934 | rs4245217 | -0.646 | -1.222 | -1.228 | -1.473 | 0.031 | 0.083 | 0.083 | 0 | 0.47 | 0.28 | 0.54 | 0.8 | 0.42 | 0.65 |
| 11 | 57987034 | rs4939188 | 0.786 | -1.113 | -1.078 | -1.326 | 0.014 | -0.005 | -0.005 | 0.28 | 0.21 | 0.12 | 0.29 | 0.22 | 0.16 | 0.08 |
| 11 | 57988265 | rs11229294 | 0.494 | -0.596 | -1.258 | -1.398 | 0.022 | 0.049 | 0.049 | 0 | 0.2 | 0.04 | 0.11 | 0.1 | 0.2 | 0.14 |
| 11 | 57988674 | rs4939190 | -0.376 | -0.301 | -0.785 | -1.513 | 0.002 | -0.036 | -0.036 | 0 | 0.57 | 0.52 | 0.62 | 0.86 | 0.52 | 0.64 |
| 11 | 57988853 | rs10792147 | 0.44 | -0.161 | -0.837 | -1.431 | -0.001 | -0.007 | -0.007 | 0 | 0.37 | 0.41 | 0.53 | 0.79 | 0.37 | 0.51 |
| 11 | 57990372 | rs4938888 | -0.09 | -0.284 | -0.623 | -1.317 | -0.028 | -0.052 | -0.052 | 0.133 | 0.28 | 0.45 | 0.19 | 0.12 | 0.23 | 0.19 |
| 11 | 57990457 | rs10896717 | 0.718 | -0.3 | -0.446 | -1.331 | 0.005 | 0.004 | 0.004 | 0 | 0.21 | 0.21 | 0.08 | 0.08 | 0.15 | 0.16 |
| 11 | 57991093 | rs1451327 | -0.116 | -0.38 | -0.507 | -1.578 | -0.067 | 0.095 | 0.095 | 1.899 | 0.39 | 0.16 | 0.31 | 0.15 | 0.45 | 0.33 |

| | | | | | | | | | | | | | | | | |
|---|---|---|---|---|---|---|---|---|---|---|---|---|---|---|---|---|
| 11 | 57992068 | rs12276038 | -1.417 | -0.423 | -0.427 | -1.491 | -0.006 | 0 | 0 | 0 | 0.66 | 0.61 | 0.5 | 0.27 | 0.67 | 0.53 |
| 11 | 57994955 | rs7927345 | 0.647 | -0.189 | -0.361 | -1.325 | -0.011 | -0.03 | -0.03 | 0.252 | 0.22 | 0.33 | 0.12 | 0.12 | 0.18 | 0.17 |
| 11 | 57995330 | rs17555381 | 0.811 | 0.103 | -0.776 | -1.54 | -0.006 | 0.072 | 0.072 | 0.329 | 0.25 | 0.09 | 0.12 | 0.07 | 0.21 | 0.18 |
| 11 | 58004159 | rs7933284 | 0.433 | 0.11 | -0.324 | -1.29 | -0.009 | -0.012 | -0.012 | 0.046 | 0.22 | 0.3 | 0.21 | 0.16 | 0.18 | 0.17 |
| 11 | 58008980 | rs1938659 | -0.037 | 0.154 | -0.448 | -1.404 | -0.047 | 0.058 | 0.058 | 0.5 | 0.64 | 0.8 | 0.71 | 0.85 | 0.56 | 0.68 |
| 11 | 58022390 | rs7934043 | -0.067 | 0.149 | -0.482 | -1.416 | -0.084 | 0.073 | 0.073 | 0.139 | 0.66 | 0.86 | 0.72 | 0.85 | 0.56 | 0.69 |
| 11 | 58032553 | rs11229323 | 1.121 | 0.099 | -0.402 | -1.442 | 0.007 | 0.017 | 0.017 | 0 | 0.2 | 0.15 | 0.19 | 0.12 | 0.16 | 0.16 |
| 11 | 58034543 | rs10792156 | 1.139 | 0.07 | -0.191 | -1.439 | -0.007 | -0.005 | -0.005 | 3.174 | 0.43 | 0.48 | 0.49 | 0.7 | 0.38 | 0.52 |
| 11 | 58036382 | rs10896733 | 1.405 | 0.075 | -0.227 | -1.436 | 0.014 | 0.006 | 0.006 | 0 | 0.39 | 0.23 | 0.48 | 0.69 | 0.38 | 0.51 |
| 11 | 58036405 | rs10896734 | 1.076 | 0.095 | -0.292 | -1.387 | 0.008 | 0.016 | 0.016 | 0 | 0.2 | 0.15 | 0.19 | 0.12 | 0.16 | 0.16 |
| 11 | 58040621 | rs10501370 | 2.078 | 0.151 | -0.423 | -1.428 | 0.002 | 0.014 | 0.014 | 0 | 0.07 | 0.04 | 0.03 | 0 | 0.05 | 0.03 |
| 11 | 58051078 | rs7107808 | 0.368 | 0.295 | -0.32 | -1.265 | -0.004 | -0.023 | -0.023 | 0 | 0.25 | 0.38 | 0.23 | 0.16 | 0.22 | 0.18 |
| 11 | 58053501 | rs947785 | 1.436 | 0.295 | -0.472 | -1.274 | 0.01 | 0.001 | 0.001 | 0 | 0.39 | 0.27 | 0.48 | 0.69 | 0.38 | 0.52 |
| 11 | 58059421 | rs10896742 | -0.198 | 0.438 | -0.74 | -1.383 | -0.055 | 0.036 | 0.036 | 0.055 | 0.72 | 0.86 | 0.77 | 0.86 | 0.64 | 0.71 |
| 11 | 58063166 | rs4459322 | 0.247 | 0.74 | -0.392 | -0.951 | 0.001 | 0.018 | 0.018 | 0 | 0.23 | 0.17 | 0.21 | 0.12 | 0.21 | 0.18 |
| 11 | 58069704 | rs10896745 | 0.913 | 0.539 | -0.409 | -0.983 | 0.017 | 0.037 | 0.037 | 0 | 0.43 | 0.29 | 0.5 | 0.73 | 0.41 | 0.53 |
| 11 | 58081152 | rs4939204 | 0.936 | 0.539 | -0.406 | -0.986 | 0.018 | 0.041 | 0.041 | 0 | 0.43 | 0.26 | 0.5 | 0.73 | 0.41 | 0.52 |
| 11 | 58086755 | rs2000486 | -0.06 | 0.89 | -0.261 | -1.066 | 0.011 | -0.022 | -0.022 | 0 | 0.69 | 0.58 | 0.73 | 0.86 | 0.63 | 0.7 |
| 11 | 58090384 | rs3887001 | -0.084 | 0.875 | -0.182 | -1.078 | -0.019 | 0.02 | 0.02 | 0 | 0.7 | 0.75 | 0.73 | 0.86 | 0.63 | 0.7 |
| 11 | 58090614 | rs11229362 | 1.478 | 0.741 | -0.372 | -1.163 | -0.045 | -0.04 | -0.04 | 0 | 0.12 | 0.01 | 0.17 | 0.12 | 0.17 | 0.13 |
| 11 | 58095004 | rs4938893 | 0.354 | 0.937 | -0.036 | -0.937 | 0.022 | -0.04 | -0.04 | 0.432 | 0.67 | 0.44 | 0.71 | 0.85 | 0.61 | 0.7 |
| 11 | 58112951 | rs1938651 | -0.172 | 0.984 | -0.112 | -0.911 | -0.041 | 0.031 | 0.031 | 0.028 | 0.72 | 0.84 | 0.74 | 0.86 | 0.63 | 0.7 |
| 11 | 58113741 | rs10896758 | -1 | 1.024 | -0.161 | -0.942 | -0.042 | 0.006 | 0.006 | 0 | 0.9 | 0.95 | 0.83 | 0.94 | 0.77 | 0.87 |
| 11 | 58114984 | rs4261301 | 0.853 | 0.891 | -0.065 | -0.976 | 0.009 | 0.029 | 0.029 | 0 | 0.44 | 0.33 | 0.51 | 0.73 | 0.43 | 0.51 |
| 11 | 58116532 | rs1938637 | 0.242 | 0.946 | 0.033 | -0.77 | 0.009 | -0.012 | -0.012 | 0 | 0.71 | 0.65 | 0.73 | 0.86 | 0.63 | 0.7 |
| 11 | 58127643 | rs1938670 | -0.555 | 0.949 | 0.055 | -0.769 | -0.007 | 0.006 | 0.006 | 0 | 0.29 | 0.28 | 0.27 | 0.14 | 0.37 | 0.3 |
| 11 | 58129053 | rs1938672 | -0.306 | 0.95 | 0.03 | -0.766 | -0.043 | 0.032 | 0.032 | 0.006 | 0.28 | 0.16 | 0.26 | 0.14 | 0.37 | 0.3 |
| 11 | 58135650 | rs10896763 | -0.5 | 0.968 | -0.021 | -0.756 | -0.009 | 0.008 | 0.008 | 0 | 0.29 | 0.27 | 0.27 | 0.14 | 0.37 | 0.3 |
| 11 | 58136902 | rs10896764 | -0.548 | 0.969 | -0.04 | -0.755 | 0.009 | -0.012 | -0.012 | 0 | 0.29 | 0.36 | 0.27 | 0.14 | 0.37 | 0.29 |
| 11 | 58170291 | rs11229410 | -0.536 | 0.853 | -0.05 | -0.752 | 0.007 | -0.012 | -0.012 | 0.306 | 0.29 | 0.35 | 0.27 | 0.14 | 0.36 | 0.29 |
| 11 | 58170342 | rs11229411 | -0.536 | 0.853 | -0.05 | -0.752 | 0.007 | -0.012 | -0.012 | 0.142 | 0.29 | 0.35 | 0.27 | 0.14 | 0.36 | 0.29 |
| 11 | 58170469 | rs11229412 | -0.536 | 0.884 | -0.05 | -0.752 | 0.008 | -0.012 | -0.012 | 0 | 0.29 | 0.35 | 0.27 | 0.14 | 0.36 | 0.29 |
| 11 | 58170738 | rs11229413 | -0.536 | 0.813 | -0.05 | -0.752 | 0.007 | -0.012 | -0.012 | 0 | 0.29 | 0.35 | 0.27 | 0.14 | 0.36 | 0.29 |
| 11 | 58180063 | rs11229426 | -0.174 | 0.473 | -0.105 | -0.563 | -0.002 | -0.004 | -0.004 | 0.022 | 0.25 | 0.27 | 0.2 | 0.14 | 0.21 | 0.27 |
| 11 | 58190136 | rs4298923 | -1.081 | 0.486 | -0.192 | -0.652 | 0.002 | 0.064 | 0.064 | 0 | 0.52 | 0.74 | 0.45 | 0.27 | 0.44 | 0.46 |
| 11 | 58191203 | rs11229438 | 0.859 | 0.439 | -0.151 | -0.694 | -0.007 | 0.042 | 0.042 | 0 | 0.23 | 0.07 | 0.2 | 0.12 | 0.21 | 0.18 |
| 11 | 58192611 | rs11229439 | 1.249 | 0.452 | -0.161 | -0.704 | -0.007 | 0.02 | 0.02 | 0 | 0.21 | 0.04 | 0.19 | 0.12 | 0.19 | 0.18 |
| 11 | 58194192 | rs11229440 | 0.901 | 0.463 | -0.144 | -0.704 | -0.008 | 0.041 | 0.041 | 0 | 0.23 | 0.07 | 0.2 | 0.12 | 0.21 | 0.18 |
| 11 | 58194697 | rs4581466 | -0.934 | 0.462 | -0.042 | -0.704 | 0.009 | 0.047 | 0.047 | 0 | 0.51 | 0.61 | 0.44 | 0.27 | 0.44 | 0.46 |
| 11 | 58202974 | rs17474904 | 1.82 | 0.008 | -0.732 | -0.692 | 0.038 | -0.004 | -0.004 | 0.296 | 0.2 | 0.01 | 0.21 | 0.14 | 0.14 | 0.08 |
| 11 | 58205540 | rs10792168 | -1.244 | 0.043 | -0.699 | -0.673 | -0.165 | -0.165 | -0.165 | 0 | 0.74 | 0.98 | 0.68 | 0.41 | 0.58 | 0.54 |
| 11 | 58207203 | rs11229457 | 0.217 | 0.058 | -0.577 | -0.713 | -0.002 | -0.004 | -0.004 | 1.561 | 0.25 | 0.27 | 0.2 | 0.14 | 0.21 | 0.27 |
| 11 | 58208025 | rs11229459 | 0.217 | 0.058 | -0.577 | -0.713 | -0.002 | -0.004 | -0.004 | 0 | 0.25 | 0.27 | 0.2 | 0.14 | 0.21 | 0.27 |
| 11 | 58208343 | rs11229460 | 0.217 | 0.058 | -0.577 | -0.713 | -0.002 | -0.004 | -0.004 | 0 | 0.25 | 0.27 | 0.2 | 0.14 | 0.21 | 0.27 |
| 11 | 58209418 | rs7929868 | 0.217 | 0.058 | -0.577 | -0.713 | -0.002 | -0.004 | -0.004 | 2.188 | 0.25 | 0.27 | 0.2 | 0.14 | 0.21 | 0.27 |
| 11 | 58226251 | rs11229472 | 0.858 | 0.062 | -0.58 | -0.718 | -0.003 | -0.014 | -0.014 | 0 | 0.26 | 0.29 | 0.22 | 0.12 | 0.21 | 0.19 |
| 11 | 58230672 | rs11601742 | 1.225 | 0.037 | -0.595 | -0.705 | 0.003 | 0.024 | 0.024 | 0 | 0.17 | 0.07 | 0.09 | 0.08 | 0.14 | 0.17 |
| 11 | 58232813 | rs2515350 | -1.307 | -0.248 | -0.478 | -0.732 | -0.061 | -0.048 | -0.048 | 0 | 0.75 | 0.44 | 0.73 | 0.86 | 0.84 | 0.91 |
| 11 | 58266248 | rs11229499 | -0.457 | -0.17 | -0.46 | -0.506 | 0.007 | 0.019 | 0.019 | 0 | 0.48 | 0.41 | 0.41 | 0.3 | 0.42 | 0.46 |
| 11 | 58274155 | rs11229505 | 0.452 | -0.114 | -0.412 | -0.547 | -0.004 | 0.001 | 0.001 | 0.004 | 0.24 | 0.14 | 0.2 | 0.19 | 0.23 | 0.28 |
| 11 | 58274370 | rs2515366 | 0.347 | -0.086 | -0.432 | -0.487 | 0.004 | 0.085 | 0.085 | 0 | 0.48 | 0.27 | 0.56 | 0.72 | 0.56 | 0.55 |
| 11 | 58287319 | rs7952094 | -0.606 | -0.06 | -0.363 | -0.55 | 0.006 | 0.015 | 0.015 | 0 | 0.5 | 0.57 | 0.42 | 0.3 | 0.42 | 0.45 |
| 11 | 58292976 | rs10896788 | 0.416 | -0.049 | -0.349 | -0.555 | -0.003 | 0.001 | 0.001 | 0 | 0.24 | 0.15 | 0.2 | 0.18 | 0.23 | 0.27 |
| 11 | 58336851 | rs11229529 | 1.767 | 0.12 | -0.618 | -0.552 | -0.072 | 0.116 | 0.116 | 0 | 0.18 | 0.01 | 0.14 | 0 | 0.24 | 0.07 |
| 11 | 58339124 | rs10896794 | 0.435 | 0.191 | -0.533 | -0.405 | -0.004 | 0.001 | 0.001 | 0.004 | 0.24 | 0.14 | 0.2 | 0.18 | 0.23 | 0.27 |
| 11 | 58366561 | rs10896795 | 0.363 | 0.133 | -0.745 | -0.353 | -0.003 | 0.005 | 0.005 | 0.116 | 0.26 | 0.16 | 0.22 | 0.18 | 0.25 | 0.28 |
| 11 | 58371283 | rs7102510 | 0.375 | 0.139 | -0.743 | -0.35 | -0.003 | 0.004 | 0.004 | 0.15 | 0.26 | 0.16 | 0.22 | 0.18 | 0.25 | 0.28 |
| 11 | 58376735 | rs7945889 | 1.066 | 0.108 | -0.734 | -0.372 | 0 | -0.001 | -0.001 | 0 | 0.23 | 0.25 | 0.2 | 0.12 | 0.18 | 0.18 |
| 11 | 58378424 | rs8373 | 0.404 | 0.11 | -0.775 | -0.371 | -0.003 | 0.004 | 0.004 | 0.15 | 0.26 | 0.16 | 0.22 | 0.18 | 0.25 | 0.28 |
| 11 | 58381548 | rs12419118 | 1.578 | 0.08 | -0.784 | -0.337 | -0.023 | -0.008 | -0.008 | 0 | 0.22 | 0.26 | 0.23 | 0.13 | 0.12 | 0.07 |
| 11 | 58386177 | rs3168135 | 0.046 | 1.012 | -0.32 | -0.036 | -0.004 | 0.005 | 0.005 | 0.15 | 0.26 | 0.16 | 0.22 | 0.18 | 0.25 | 0.28 |
| 11 | 58386650 | rs11229545 | 0.046 | 1.012 | -0.32 | -0.036 | -0.004 | 0.005 | 0.005 | 0.15 | 0.26 | 0.16 | 0.22 | 0.18 | 0.25 | 0.28 |
| 11 | 58391501 | rs1800169 | 1.202 | 1.017 | -0.345 | -0.041 | 0.009 | 0.024 | 0.024 | 0 | 0.2 | 0.04 | 0.18 | 0.12 | 0.16 | 0.17 |
| 11 | 58392951 | rs2515362 | -1.042 | 0.863 | -0.3 | 0.067 | -0.003 | 0.046 | 0.046 | 0 | 0.52 | 0.69 | 0.45 | 0.31 | 0.45 | 0.47 |
| 11 | 58394154 | rs550942 | -1.381 | 1.023 | -1.233 | 0.721 | -0.023 | -0.006 | -0.006 | 0 | 0.95 | 0.99 | 0.93 | 0.71 | 0.83 | 0.92 |
| 11 | 58403833 | rs10501372 | 0.81 | -0.264 | -0.958 | -0.223 | 0 | 0 | 0 | 0 | 0.08 | 0.06 | 0.12 | 0.06 | 0.08 | 0.09 |
| 11 | 58423297 | rs11601296 | 0.492 | -0.222 | -0.876 | -0.272 | -0.005 | 0.005 | 0.005 | 0 | 0.18 | 0.1 | 0.1 | 0.12 | 0.18 | 0.18 |
| 11 | 58424478 | rs10896810 | -1.013 | -0.201 | -0.973 | -0.258 | 0.004 | 0.019 | 0.019 | 0 | 0.49 | 0.41 | 0.42 | 0.3 | 0.43 | 0.45 |
| 11 | 58427583 | rs10750901 | -0.541 | -0.71 | -0.944 | -0.057 | 0.009 | 0.034 | 0.034 | 0 | 0.29 | 0.15 | 0.3 | 0.18 | 0.26 | 0.27 |

| | | | | | | | | | | | | | | | | |
|---|---|---|---|---|---|---|---|---|---|---|---|---|---|---|---|---|
| 11 | 58427589 | rs10792180 | -0.537 | -0.689 | -1.029 | -0.047 | 0.008 | 0.024 | 0.024 | 0 | 0.29 | 0.2 | 0.31 | 0.18 | 0.26 | 0.27 |
| 11 | 58438655 | rs12798524 | -0.429 | -1.105 | -0.671 | 0.059 | 0.017 | 0.014 | 0.014 | 0.01 | 0.28 | 0.21 | 0.27 | 0.21 | 0.19 | 0.24 |
| 11 | 58439593 | rs4939214 | 0.584 | -1.172 | -0.723 | 0.044 | -0.001 | -0.001 | -0.001 | 0 | 0.18 | 0.15 | 0.23 | 0.12 | 0.12 | 0.1 |
| 11 | 58440720 | rs1014968 | 0.597 | -1.164 | -0.712 | 0.048 | -0.002 | -0.003 | -0.003 | 0 | 0.18 | 0.16 | 0.23 | 0.11 | 0.12 | 0.08 |
| 11 | 58441903 | rs12794523 | 0.222 | -0.864 | -0.856 | 0.206 | 0.014 | 0.003 | 0.003 | 0 | 0.1 | 0.03 | 0.04 | 0.09 | 0.06 | 0.14 |
| 11 | 58452608 | rs517679 | 0.616 | -0.21 | -0.854 | 0.514 | -0.079 | -0.022 | -0.022 | 0 | 0.38 | 0.1 | 0.4 | 0.31 | 0.46 | 0.34 |
| 11 | 58452810 | rs686883 | 0.154 | 0.838 | 0.506 | 1.186 | 0.015 | 0.043 | 0.043 | 1.342 | 0.8 | 0.71 | 0.8 | 0.56 | 0.72 | 0.62 |
| 11 | 58453529 | rs609036 | 0.912 | 0.852 | 0.285 | 1.189 | 0.012 | 0.061 | 0.061 | 1.094 | 0.72 | 0.46 | 0.7 | 0.52 | 0.67 | 0.57 |
| 11 | 58455331 | rs948559 | -0.947 | 0.529 | 0.071 | 0.761 | -0.009 | -0.021 | -0.021 | 0 | 0.07 | 0.16 | 0.1 | 0.03 | 0.07 | 0.07 |
| 11 | 58460726 | rs11229578 | -1.158 | 1.143 | 0.205 | 1.306 | 0.047 | 0.073 | 0.073 | 0 | 0.08 | 0.21 | 0.27 | 0.26 | 0.16 | 0.22 |
| 11 | 58468173 | rs619945 | -0.396 | 1.651 | -0.343 | 1.309 | -0.035 | 0 | 0 | 0.831 | 0.22 | 0.06 | 0.25 | 0.17 | 0.26 | 0.27 |
| 11 | 58473596 | rs614561 | 1.234 | 0.727 | 0.249 | 2.053 | 0.012 | 0.105 | 0.105 | 0 | 0.85 | 0.7 | 0.68 | 0.55 | 0.82 | 0.66 |
| 11 | 58478039 | rs675757 | 1.348 | 0.667 | 0.39 | 2.024 | 0.01 | 0.105 | 0.105 | 0 | 0.84 | 0.67 | 0.68 | 0.54 | 0.82 | 0.66 |
| 11 | 58526064 | rs1938717 | -0.895 | 0.741 | 0.468 | 2.121 | 0.032 | 0.053 | 0.053 | 0 | 0.09 | 0.19 | 0.27 | 0.26 | 0.16 | 0.22 |
| 11 | 58528909 | rs1938714 | 1.769 | 0.728 | 0.416 | 2.128 | 0.012 | 0.104 | 0.104 | 0.006 | 0.82 | 0.65 | 0.68 | 0.51 | 0.8 | 0.62 |
| 11 | 58529143 | rs12417075 | -1.153 | 0.729 | 0.42 | 2.125 | 0.035 | 0.057 | 0.057 | 0.119 | 0.1 | 0.22 | 0.27 | 0.26 | 0.16 | 0.22 |
| 11 | 58553771 | rs10896825 | -2.078 | 0.73 | 0.415 | 2.099 | 0.014 | 0.108 | 0.108 | 0 | 0.18 | 0.36 | 0.32 | 0.49 | 0.2 | 0.39 |
| 11 | 58572797 | rs1943254 | -1.988 | 0.731 | 0.358 | 2.1 | 0.01 | 0.08 | 0.08 | 0 | 0.17 | 0.29 | 0.32 | 0.49 | 0.2 | 0.39 |
| 11 | 58575572 | rs10896833 | 0.124 | 0.732 | 0.356 | 2.12 | 0.012 | -0.009 | -0.009 | 1.338 | 0.05 | 0.01 | 0.04 | 0.19 | 0.02 | 0.12 |
| 11 | 58600796 | rs10896848 | -2.101 | 0.705 | 0.349 | 2.064 | 0.009 | 0.076 | 0.076 | 0 | 0.17 | 0.3 | 0.32 | 0.48 | 0.2 | 0.39 |
| 11 | 58600885 | rs10896849 | -2.138 | 0.705 | 0.349 | 2.064 | 0.008 | 0.074 | 0.074 | 0 | 0.18 | 0.3 | 0.32 | 0.48 | 0.2 | 0.39 |
| 11 | 58602309 | rs11229651 | -1.988 | 0.706 | 0.349 | 2.064 | 0.009 | 0.071 | 0.071 | 0 | 0.17 | 0.29 | 0.32 | 0.48 | 0.2 | 0.39 |
| 11 | 58610656 | rs11229658 | -2.083 | 0.707 | 0.43 | 2.065 | 0.012 | 0.092 | 0.092 | 0 | 0.17 | 0.32 | 0.32 | 0.48 | 0.2 | 0.39 |
| 11 | 58625585 | rs663578 | 1.956 | 0.389 | 1.124 | 1.726 | -0.008 | 0.02 | 0.02 | 0.514 | 0.81 | 0.4 | 0.78 | 0.79 | 0.84 | 0.69 |
| 11 | 58658875 | rs7933339 | 1.11 | 0.237 | 0.898 | 1.867 | -0.018 | 0.125 | 0.125 | 0.744 | 0.89 | 0.61 | 0.91 | 0.83 | 0.96 | 0.84 |
| 11 | 58661146 | rs2226794 | 0.836 | 0.221 | 0.893 | 1.788 | -0.022 | 0.111 | 0.111 | 0.098 | 0.91 | 0.63 | 0.92 | 0.86 | 0.98 | 0.88 |
| 11 | 58663660 | rs1125442 | 0.32 | -0.169 | -0.239 | 1.173 | 0.021 | -0.011 | -0.011 | 0 | 0.33 | 0.12 | 0.28 | 0.33 | 0.33 | 0.35 |
| 11 | 58666446 | rs6591499 | 0.643 | -0.04 | 0.726 | 1.347 | -0.004 | 0.077 | 0.077 | 1.767 | 0.9 | 0.65 | 0.92 | 0.84 | 0.96 | 0.87 |
| 11 | 58668565 | rs2156349 | 1.05 | -0.001 | 1.274 | 1.748 | 0.033 | 0.187 | 0.187 | 0.29 | 0.86 | 0.53 | 0.8 | 0.69 | 0.87 | 0.84 |
| 11 | 58670553 | rs17578360 | -0.163 | -0.167 | 0.801 | 1.453 | -0.007 | 0.014 | 0.014 | 0 | 0.09 | 0.01 | 0.06 | 0.05 | 0.08 | 0.09 |
| 11 | 58675461 | rs7126489 | 0.741 | -0.056 | 1.379 | 1.596 | 0.006 | 0.09 | 0.09 | 0.488 | 0.85 | 0.36 | 0.79 | 0.69 | 0.87 | 0.84 |
| 11 | 58677472 | rs1893628 | -0.74 | -0.063 | 1.954 | 1.591 | 0.063 | 0.216 | 0.216 | 0 | 0.11 | 0.39 | 0.19 | 0.31 | 0.13 | 0.15 |
| 11 | 58678387 | rs10896877 | -0.61 | -0.063 | 2.018 | 1.59 | 0.02 | 0.102 | 0.102 | 0 | 0.11 | 0.22 | 0.18 | 0.31 | 0.13 | 0.15 |
| 11 | 58678460 | rs503591 | 0.574 | -0.408 | 1.261 | 1.443 | 0.048 | 0.086 | 0.086 | 0 | 0.4 | 0.15 | 0.3 | 0.26 | 0.32 | 0.34 |
| 11 | 58687627 | rs11229695 | -0.071 | -0.39 | 1.396 | 1.462 | 0.043 | 0.168 | 0.168 | 0 | 0.11 | 0.33 | 0.19 | 0.31 | 0.13 | 0.16 |
| 11 | 58690549 | rs532318 | -0.866 | -0.35 | 1.375 | 1.482 | -0.009 | 0 | 0 | 0 | 0.55 | 0.75 | 0.51 | 0.6 | 0.46 | 0.56 |
| 11 | 58696996 | rs17578583 | 0.767 | -0.34 | 0.971 | 1.339 | -0.027 | -0.042 | -0.042 | 0.682 | 0.07 | 0 | 0.12 | 0.14 | 0.11 | 0.19 |
| 11 | 58697805 | rs12270298 | 0.377 | -0.359 | 1.244 | 1.462 | -0.002 | 0.005 | 0.005 | 0 | 0.09 | 0.06 | 0.12 | 0.04 | 0.12 | 0.1 |
| 11 | 58701011 | rs480196 | -0.721 | -0.37 | 1.315 | 1.47 | -0.001 | -0.001 | -0.001 | 0.124 | 0.52 | 0.49 | 0.5 | 0.57 | 0.51 | 0.51 |
| 11 | 58709446 | rs578610 | 0.298 | -0.372 | 1.082 | 1.46 | 0.017 | 0.089 | 0.089 | 0 | 0.4 | 0.16 | 0.31 | 0.26 | 0.37 | 0.34 |
| 11 | 58710580 | rs7120597 | 0.707 | -0.368 | 1.22 | 1.475 | 0 | -0.001 | -0.001 | 0 | 0.45 | 0.47 | 0.49 | 0.41 | 0.48 | 0.43 |
| 11 | 58715285 | rs565355 | 0.335 | -0.463 | 0.96 | 1.472 | 0.007 | 0.084 | 0.084 | 0.118 | 0.39 | 0.12 | 0.3 | 0.26 | 0.37 | 0.34 |
| 11 | 58724149 | rs517039 | 0.324 | -0.442 | 0.798 | 1.488 | 0.007 | 0.081 | 0.081 | 0.01 | 0.39 | 0.15 | 0.31 | 0.26 | 0.37 | 0.33 |
| 11 | 58747617 | rs485025 | 0.256 | -0.544 | 0.786 | 1.421 | 0.026 | 0.09 | 0.09 | 0 | 0.41 | 0.18 | 0.29 | 0.26 | 0.37 | 0.35 |
| 11 | 58748120 | rs480352 | -1.248 | -0.436 | 0.689 | 1.222 | -0.001 | -0.002 | -0.002 | 0.37 | 0.59 | 0.46 | 0.54 | 0.64 | 0.59 | 0.63 |
| 11 | 58757947 | rs505054 | -1.243 | -0.373 | 0.696 | 1.117 | 0.002 | 0.002 | 0.002 | 0 | 0.62 | 0.5 | 0.56 | 0.64 | 0.61 | 0.71 |
| 11 | 58761391 | rs489810 | 0.149 | -0.315 | 0.752 | 1.671 | 0.023 | -0.021 | -0.021 | 0 | 0.43 | 0.27 | 0.43 | 0.51 | 0.4 | 0.56 |
| 11 | 58766266 | rs7946908 | -0.814 | -0.294 | 0.823 | 1.504 | 0.008 | 0.028 | 0.028 | 0 | 0.2 | 0.21 | 0.26 | 0.4 | 0.21 | 0.28 |
| 11 | 58773879 | rs11229725 | -0.281 | -0.262 | 0.843 | 1.516 | -0.003 | 0.004 | 0.004 | 0 | 0.17 | 0.1 | 0.2 | 0.09 | 0.2 | 0.14 |
| 11 | 58776177 | rs567224 | -1.086 | -0.172 | 0.792 | 1.594 | 0.003 | 0.006 | 0.006 | 0 | 0.62 | 0.47 | 0.56 | 0.63 | 0.6 | 0.71 |
| 11 | 58807241 | rs11602551 | 1.08 | -0.158 | 0.35 | 1.423 | -0.012 | -0.016 | -0.016 | 0 | 0.35 | 0.3 | 0.42 | 0.37 | 0.39 | 0.29 |
| 11 | 58820988 | rs3740948 | -0.392 | 0.026 | 0.236 | 1.396 | -0.013 | -0.006 | -0.006 | 1.067 | 0.15 | 0.07 | 0.2 | 0.16 | 0.19 | 0.14 |
| 11 | 58834388 | rs568468 | 0.203 | 0.167 | 0.213 | 1.431 | 0 | 0.001 | 0.001 | 0 | 0.5 | 0.52 | 0.54 | 0.59 | 0.45 | 0.42 |
| 11 | 58834900 | rs563191 | 0.485 | 0.205 | 0.14 | 1.434 | 0.007 | 0.013 | 0.013 | 0 | 0.47 | 0.42 | 0.47 | 0.27 | 0.43 | 0.33 |
| 11 | 58846985 | rs1943147 | 0.365 | 0.499 | -0.17 | 1.191 | 0.006 | -0.001 | -0.001 | 0 | 0.1 | 0.11 | 0.18 | 0.11 | 0.18 | 0.11 |
| 11 | 58863775 | rs7939969 | -0.551 | 0.226 | -0.301 | 1.43 | -0.015 | 0.021 | 0.021 | 0 | 0.64 | 0.9 | 0.78 | 0.63 | 0.64 | 0.59 |
| 11 | 58872960 | rs10896919 | -0.56 | 0.211 | -0.251 | 1.427 | -0.01 | 0.023 | 0.023 | 0 | 0.64 | 0.89 | 0.78 | 0.63 | 0.64 | 0.59 |
| 11 | 58877531 | rs10792219 | 0.69 | 0.62 | -0.261 | 1.348 | 0.005 | 0 | 0 | 0 | 0.1 | 0.08 | 0.18 | 0.11 | 0.2 | 0.12 |
| 11 | 58877909 | rs4409856 | 0.33 | 0.594 | 0.184 | 1.614 | 0.008 | 0.005 | 0.005 | 0 | 0.5 | 0.49 | 0.52 | 0.34 | 0.43 | 0.38 |
| 11 | 58883915 | rs10437720 | -0.453 | 0.709 | 0.21 | 1.724 | 0.13 | 0.143 | 0.143 | 0.082 | 0.13 | 0.4 | 0.25 | 0.27 | 0.2 | 0.21 |
| 11 | 58889191 | rs10896924 | 0.627 | 0.836 | 0.115 | 1.633 | 0.005 | 0 | 0 | 0 | 0.1 | 0.08 | 0.18 | 0.11 | 0.2 | 0.12 |
| 11 | 58910926 | rs11229840 | -0.351 | 0.816 | 0.262 | 1.608 | 0.124 | 0.166 | 0.166 | 0.1 | 0.13 | 0.39 | 0.25 | 0.29 | 0.2 | 0.22 |
| 11 | 58921832 | rs11229848 | -0.313 | 0.834 | 0.149 | 1.606 | 0.099 | 0.149 | 0.149 | 0.019 | 0.13 | 0.37 | 0.25 | 0.29 | 0.19 | 0.21 |
| 11 | 58923529 | rs587617 | -0.615 | 0.552 | -0.193 | 1.509 | -0.05 | 0.016 | 0.016 | 0 | 0.64 | 0.93 | 0.77 | 0.63 | 0.63 | 0.59 |
| 11 | 58929387 | rs517845 | 0.428 | 0.841 | -0.182 | 1.438 | 0.004 | 0.002 | 0.002 | 0 | 0.5 | 0.51 | 0.5 | 0.34 | 0.41 | 0.37 |
| 11 | 58930744 | rs637541 | -0.263 | 0.61 | -0.625 | 0.767 | 0.022 | 0.011 | 0.011 | 0 | 0.51 | 0.65 | 0.67 | 0.49 | 0.45 | 0.51 |
| 11 | 58938892 | rs11229864 | 0.675 | 0.486 | -0.958 | 0.958 | -0.012 | -0.074 | -0.074 | 0 | 0.17 | 0.01 | 0.08 | 0.29 | 0.16 | 0.12 |
| 11 | 58942848 | rs6591507 | -0.81 | 0.202 | -0.692 | 1.111 | -0.021 | -0.052 | -0.052 | 0 | 0.32 | 0.74 | 0.49 | 0.45 | 0.29 | 0.49 |
| 11 | 58951440 | rs656163 | 0.552 | -0.062 | -0.032 | 0.916 | -0.001 | -0.001 | -0.001 | 0.021 | 0.52 | 0.53 | 0.47 | 0.36 | 0.55 | 0.51 |

| | | | | | | | | | | | | | | | | |
|---|---|---|---|---|---|---|---|---|---|---|---|---|---|---|---|---|
| 11 | 58953775 | rs11229868 | 1.082 | -0.241 | -0.112 | 0.984 | 0.005 | -0.081 | -0.081 | 1.516 | 0.15 | 0 | 0.05 | 0.31 | 0.11 | 0.11 |
| 11 | 58966176 | rs10501375 | 0.822 | -0.261 | 0.109 | 1.047 | 0.026 | -0.039 | -0.039 | 0 | 0.16 | 0.07 | 0.05 | 0.32 | 0.12 | 0.11 |
| 11 | 58972669 | rs3740946 | 0.982 | -0.261 | 0.112 | 1.041 | 0.021 | -0.065 | -0.065 | 0 | 0.15 | 0.01 | 0.05 | 0.32 | 0.12 | 0.15 |
| 11 | 58973731 | rs1048444 | -0.045 | -0.242 | 0.172 | 1.025 | 0.031 | 0.023 | 0.023 | 0 | 0.67 | 0.51 | 0.49 | 0.66 | 0.63 | 0.62 |
| 11 | 58977768 | rs5029315 | 0.891 | -0.247 | 0.14 | 1.02 | 0.009 | 0.086 | 0.086 | 0 | 0.49 | 0.36 | 0.42 | 0.26 | 0.5 | 0.42 |
| 11 | 58978684 | rs7926933 | 1.135 | -0.16 | 0.041 | 1.082 | 0.019 | -0.062 | -0.062 | 0 | 0.15 | 0.01 | 0.04 | 0.3 | 0.12 | 0.11 |
| 11 | 58978940 | rs544864 | 0.541 | -0.02 | 0.297 | 0.935 | -0.028 | -0.035 | -0.035 | 0 | 0.22 | 0.12 | 0.4 | 0.29 | 0.29 | 0.33 |
| 11 | 58981586 | rs2000875 | 0.014 | -0.01 | 0.331 | 0.951 | 0.039 | 0.028 | 0.028 | 0 | 0.68 | 0.54 | 0.5 | 0.66 | 0.63 | 0.62 |
| 11 | 58984717 | rs1938593 | 0.255 | -0.011 | 0.321 | 0.944 | 0.012 | 0.017 | 0.017 | 0 | 0.66 | 0.47 | 0.48 | 0.57 | 0.61 | 0.57 |
| 11 | 59020419 | rs11229889 | 0.389 | -0.075 | 0.007 | 1.012 | 0.012 | -0.019 | -0.019 | 0 | 0.19 | 0.13 | 0.07 | 0.43 | 0.14 | 0.21 |
| 11 | 59022668 | rs4938914 | -0.518 | 0.012 | 0.223 | 0.975 | 0.053 | -0.027 | -0.027 | 0 | 0.88 | 0.56 | 0.9 | 0.96 | 0.92 | 0.95 |
| 11 | 59025022 | rs12280107 | 0.261 | 0.113 | -0.011 | 1.014 | 0.005 | 0.007 | 0.007 | 0 | 0.65 | 0.44 | 0.48 | 0.57 | 0.61 | 0.57 |
| 11 | 59026901 | rs11229895 | 0.706 | 0.17 | -0.049 | 0.99 | -0.001 | -0.001 | -0.001 | 0 | 0.22 | 0.17 | 0.4 | 0.28 | 0.29 | 0.32 |
| 11 | 59029287 | rs6591518 | -1.39 | 0.18 | -0.442 | 0.975 | 0.026 | -0.016 | -0.016 | 0.01 | 0.92 | 0.82 | 0.91 | 0.95 | 0.93 | 0.95 |
| 11 | 59029485 | rs11229898 | 0.129 | 0.213 | -0.461 | 1.014 | -0.002 | -0.002 | -0.002 | 0 | 0.65 | 0.41 | 0.46 | 0.55 | 0.61 | 0.53 |
| 11 | 59030490 | rs10896957 | 0.73 | 0.445 | -0.843 | 1.064 | 0.021 | -0.026 | -0.026 | 0 | 0.16 | 0.1 | 0.05 | 0.31 | 0.12 | 0.11 |
| 11 | 59031974 | rs17582175 | 1.299 | 0.975 | -1.195 | 0.83 | -0.061 | -0.002 | -0.002 | 0 | 0.22 | 0.05 | 0.27 | 0.18 | 0.29 | 0.12 |
| 11 | 59035047 | rs7119212 | -0.672 | 1.389 | -0.291 | 1.199 | 0.024 | -0.019 | -0.019 | 0.019 | 0.92 | 0.81 | 0.91 | 0.95 | 0.93 | 0.96 |
| 11 | 59036693 | rs10896958 | -0.12 | 1.075 | -0.809 | 1.355 | 0.048 | 0.028 | 0.028 | 0 | 0.68 | 0.55 | 0.5 | 0.67 | 0.63 | 0.63 |
| 11 | 59037867 | rs7949688 | -0.567 | 0.548 | -1.018 | 1.741 | 0.029 | -0.088 | -0.088 | 0.474 | 0.17 | 0.03 | 0.09 | 0.42 | 0.13 | 0.17 |
| 11 | 59041725 | rs7926943 | 0.916 | 1.573 | -0.274 | 1.177 | 0.046 | 0.035 | 0.035 | 0 | 0.76 | 0.54 | 0.76 | 0.72 | 0.74 | 0.74 |
| 11 | 59065162 | rs1945589 | 0.744 | 1.751 | -0.073 | 1.283 | 0.034 | 0.035 | 0.035 | 0 | 0.78 | 0.64 | 0.78 | 0.72 | 0.75 | 0.74 |
| 11 | 59100216 | rs10736696 | 1.133 | 1.97 | 0.361 | 1.204 | 0.06 | 0.037 | 0.037 | 0 | 0.76 | 0.57 | 0.75 | 0.72 | 0.72 | 0.73 |
| 11 | 59101682 | rs4938916 | 1.011 | 2.013 | 0.382 | 1.211 | 0.021 | 0 | 0 | 0 | 0.78 | 0.69 | 0.81 | 0.8 | 0.75 | 0.8 |
| 11 | 59123814 | rs4939283 | 1.411 | 1.975 | 0.361 | 1.075 | 0.056 | 0.016 | 0.016 | 0 | 0.75 | 0.57 | 0.75 | 0.74 | 0.72 | 0.73 |
| 11 | 59130899 | rs4939284 | 1.138 | 1.973 | 0.312 | 1.043 | 0.033 | -0.02 | -0.02 | 0 | 0.77 | 0.6 | 0.79 | 0.83 | 0.76 | 0.81 |
| 11 | 59134367 | rs881834 | 1.138 | 1.962 | 0.312 | 1.044 | 0.035 | -0.022 | -0.022 | 0 | 0.77 | 0.6 | 0.79 | 0.83 | 0.76 | 0.81 |
| 11 | 59138860 | rs1357374 | 1.44 | 1.963 | 0.313 | 1.045 | 0.048 | 0.009 | 0.009 | 0 | 0.75 | 0.6 | 0.74 | 0.74 | 0.72 | 0.71 |
| 11 | 59147075 | rs1996858 | 1.034 | 1.91 | 0.4 | 1.036 | 0.019 | -0.013 | -0.013 | 0 | 0.77 | 0.7 | 0.81 | 0.83 | 0.76 | 0.81 |
| 11 | 59160868 | rs7933143 | 1.395 | 1.917 | 0.478 | 1.065 | 0.051 | 0.012 | 0.012 | 0 | 0.74 | 0.51 | 0.74 | 0.74 | 0.73 | 0.73 |
| 11 | 59166402 | rs11607526 | -1.174 | 1.915 | 0.464 | 1.061 | -0.069 | 0.049 | 0.049 | 0 | 0.2 | 0.04 | 0.13 | 0.1 | 0.24 | 0.15 |
| 11 | 59169295 | rs11604216 | -1.478 | 1.908 | 0.484 | 1.062 | -0.034 | 0.005 | 0.005 | 0 | 0.22 | 0.13 | 0.18 | 0.19 | 0.27 | 0.22 |
| 11 | 59179022 | rs7128953 | 1.552 | 1.911 | 0.441 | 1.057 | 0.051 | 0.012 | 0.012 | 0 | 0.74 | 0.51 | 0.74 | 0.74 | 0.73 | 0.73 |
| 11 | 59184192 | rs1879929 | 1.443 | 1.912 | 0.42 | 1.08 | 0.034 | 0.01 | 0.01 | 0 | 0.75 | 0.59 | 0.74 | 0.74 | 0.73 | 0.73 |
| 11 | 59189912 | rs1453547 | -1.17 | 2.028 | 0.09 | 0.926 | -0.103 | 0.017 | 0.017 | 0 | 0.19 | 0.02 | 0.13 | 0.11 | 0.25 | 0.15 |
| 11 | 59190499 | rs11602535 | -1.646 | 2.184 | 0.209 | 0.958 | -0.017 | 0.019 | 0.019 | 0 | 0.23 | 0.21 | 0.16 | 0.12 | 0.27 | 0.17 |
| 11 | 59209058 | rs12794700 | -1.131 | 2.009 | 0.428 | 0.895 | -0.103 | 0.017 | 0.017 | 0 | 0.19 | 0.03 | 0.13 | 0.11 | 0.25 | 0.14 |
| 11 | 59210550 | rs7947825 | 2.161 | 1.925 | 0.802 | 0.875 | 0.061 | 0.16 | 0.16 | 1.421 | 0.91 | 0.48 | 0.72 | 0.87 | 0.95 | 0.89 |
| 11 | 59211188 | rs6591536 | 1.77 | 1.926 | 0.802 | 0.854 | 0.037 | -0.002 | -0.002 | 0 | 0.72 | 0.44 | 0.58 | 0.76 | 0.69 | 0.74 |
| 11 | 59211421 | rs17591107 | -1.019 | 1.93 | 0.653 | 0.829 | -0.103 | 0.016 | 0.016 | 0 | 0.19 | 0.02 | 0.13 | 0.11 | 0.25 | 0.14 |
| 11 | 59220879 | rs11229957 | -1.372 | 1.935 | 0.689 | 0.84 | -0.112 | -0.022 | -0.022 | 0 | 0.22 | 0.04 | 0.23 | 0.17 | 0.28 | 0.21 |
| 11 | 59223326 | rs1453546 | 0.166 | 2.749 | -0.029 | 0.378 | -0.092 | 0.027 | 0.027 | 0 | 0.15 | 0.01 | 0.3 | 0.13 | 0.29 | 0.19 |
| 11 | 59224608 | rs1453544 | -0.854 | 2.282 | 0.323 | 0.525 | -0.103 | 0.016 | 0.016 | 0 | 0.19 | 0.02 | 0.13 | 0.11 | 0.25 | 0.14 |
| 11 | 59225221 | rs1453541 | -1.723 | 1.973 | 0.167 | 0.291 | 0.002 | -0.001 | -0.001 | 0 | 0.26 | 0.32 | 0.27 | 0.18 | 0.3 | 0.23 |
| 11 | 59225472 | rs17500478 | -0.794 | 2.009 | 0.181 | 0.308 | -0.103 | 0.016 | 0.016 | 0 | 0.19 | 0.03 | 0.13 | 0.11 | 0.25 | 0.14 |
| 11 | 59230701 | rs2867748 | -1.637 | 2.006 | 0.186 | 0.155 | 0.004 | -0.005 | -0.005 | 0 | 0.26 | 0.32 | 0.27 | 0.15 | 0.29 | 0.23 |
| 11 | 59239839 | rs7111674 | -1.815 | 1.325 | 0.647 | 0.522 | 0.035 | 0.04 | 0.04 | 0.459 | 0.07 | 0.24 | 0.14 | 0.07 | 0.04 | 0.09 |
| 11 | 59240474 | rs7126271 | -2.012 | 1.329 | 0.647 | 0.523 | 0.051 | 0.056 | 0.056 | 0.962 | 0.07 | 0.31 | 0.14 | 0.07 | 0.04 | 0.09 |
| 11 | 59241429 | rs10750923 | 1.556 | 1.332 | 0.675 | 0.548 | 0.065 | 0.134 | 0.134 | 0.762 | 0.91 | 0.51 | 0.72 | 0.88 | 0.95 | 0.88 |
| 11 | 59254395 | rs1545527 | -0.536 | 0.618 | -0.795 | 0.062 | 0.017 | 0.026 | 0.026 | 0 | 0.4 | 0.34 | 0.23 | 0.27 | 0.3 | 0.41 |
| 11 | 59256054 | rs12791616 | 0.144 | 0.37 | -1.212 | -0.073 | -0.018 | -0.014 | -0.014 | 2.443 | 0.14 | 0.01 | 0.1 | 0.14 | 0.1 | 0.12 |
| 11 | 59257636 | rs7116107 | -0.847 | 0.393 | -0.43 | -0.064 | 0.009 | 0.015 | 0.015 | 0 | 0.41 | 0.43 | 0.23 | 0.28 | 0.31 | 0.4 |
| 11 | 59265036 | rs10792246 | 0.495 | 0.68 | -0.887 | -0.618 | -0.061 | -0.045 | -0.045 | 0.306 | 0.27 | 0.11 | 0.46 | 0.63 | 0.42 | 0.27 |
| 11 | 59268721 | rs11229974 | -0.235 | 0.439 | -0.758 | -0.497 | 0.011 | 0.045 | 0.045 | 0 | 0.38 | 0.15 | 0.21 | 0.28 | 0.31 | 0.4 |
| 11 | 59271637 | rs7120079 | -0.317 | 0.428 | -0.7 | -0.503 | 0.024 | 0.043 | 0.043 | 0 | 0.4 | 0.29 | 0.22 | 0.28 | 0.31 | 0.4 |
| 11 | 59281654 | rs11229983 | -0.236 | 0.477 | -0.782 | -0.502 | 0.023 | 0.045 | 0.045 | 0 | 0.38 | 0.14 | 0.21 | 0.28 | 0.3 | 0.39 |
| 11 | 59291045 | rs1453531 | -0.403 | 0.615 | -0.861 | -0.509 | 0.013 | 0.02 | 0.02 | 0 | 0.42 | 0.41 | 0.23 | 0.28 | 0.31 | 0.39 |
| 11 | 59303602 | rs11229994 | 1.056 | 1.139 | -1.132 | 0.065 | -0.034 | -0.035 | -0.035 | 0.168 | 0.1 | 0 | 0.06 | 0.15 | 0.12 | 0.03 |
| 11 | 59338663 | rs17153910 | -0.94 | 1.239 | -0.452 | -0.06 | 0.084 | 0.043 | 0.043 | 1.112 | 0.06 | 0.39 | 0.17 | 0.07 | 0.06 | 0.04 |
| 11 | 59342178 | rs8280 | -1.224 | 1.23 | -0.419 | -0.064 | 0.095 | 0.05 | 0.05 | 1.99 | 0.07 | 0.48 | 0.18 | 0.07 | 0.06 | 0.04 |
| 11 | 59346568 | rs536910 | -1.224 | 1.23 | -0.419 | -0.064 | 0.095 | 0.05 | 0.05 | 1.99 | 0.07 | 0.48 | 0.18 | 0.07 | 0.06 | 0.04 |
| 11 | 59359841 | rs2186411 | 0.112 | 1.718 | -0.659 | 0.436 | -0.032 | -0.069 | -0.069 | 0.018 | 0.23 | 0.11 | 0.42 | 0.48 | 0.34 | 0.28 |
| 11 | 59367448 | rs4938923 | -1.26 | 1.375 | -1.195 | -0.032 | -0.015 | 0.061 | 0.061 | 0 | 0.84 | 0.95 | 0.91 | 0.96 | 0.85 | 0.8 |
| 11 | 59377543 | rs11603417 | 0.882 | 1.275 | -1.467 | -0.161 | -0.051 | -0.037 | -0.037 | 0.443 | 0.15 | 0.01 | 0.11 | 0.18 | 0.13 | 0.11 |
| 11 | 59389113 | rs11230020 | -0.569 | 1.272 | -1.408 | -0.176 | 0.101 | 0.049 | 0.049 | 1.798 | 0.07 | 0.5 | 0.18 | 0.07 | 0.06 | 0.04 |
| 11 | 59407375 | rs12280606 | -0.511 | 1.256 | -1.415 | -0.181 | 0.09 | 0.04 | 0.04 | 1.395 | 0.07 | 0.48 | 0.18 | 0.07 | 0.06 | 0.04 |
| 11 | 59424804 | rs11606058 | 0.889 | 1.246 | -1.405 | -0.186 | -0.051 | -0.04 | -0.04 | 0 | 0.15 | 0.01 | 0.11 | 0.18 | 0.13 | 0.11 |
| 11 | 59431837 | rs1938690 | -0.628 | 1.419 | -1.176 | -0.293 | 0.09 | 0.046 | 0.046 | 1.395 | 0.07 | 0.48 | 0.18 | 0.07 | 0.06 | 0.03 |
| 11 | 59445500 | rs11607283 | 2.872 | 1.034 | -1.579 | -0.741 | -0.009 | 0.037 | 0.037 | 0.018 | 0.14 | 0.01 | 0.05 | 0.03 | 0.1 | 0.07 |

| | | | | | | | | | | | | | | | | |
|---|---|---|---|---|---|---|---|---|---|---|---|---|---|---|---|---|
| 11 | 59448068 | rs7116484 | -1.107 | 1.196 | -1.13 | -0.274 | -0.001 | -0.001 | -0.001 | 0.305 | 0.48 | 0.59 | 0.43 | 0.49 | 0.41 | 0.45 |
| 11 | 59479582 | rs2927185 | 1.762 | 1.48 | -1.117 | 0.151 | -0.014 | -0.025 | -0.025 | 1.823 | 0.12 | 0.05 | 0.09 | 0.26 | 0.16 | 0.05 |
| 11 | 59480969 | rs472177 | -1.087 | 1.488 | -0.518 | 0.535 | 0.087 | 0.043 | 0.043 | 1.258 | 0.07 | 0.48 | 0.18 | 0.07 | 0.06 | 0.05 |
| 11 | 59483693 | rs518427 | -1.853 | 1.54 | -0.534 | 0.622 | -0.001 | -0.003 | -0.003 | 0 | 0.48 | 0.58 | 0.43 | 0.42 | 0.41 | 0.46 |
| 11 | 59487281 | rs485803 | -1.161 | 1.957 | 0.202 | 0.574 | 0.037 | 0.017 | 0.017 | 0 | 0.05 | 0.28 | 0.17 | 0.06 | 0.05 | 0.04 |
| 11 | 59491673 | rs11230061 | -0.786 | 2.015 | 0.174 | 0.558 | -0.039 | 0.09 | 0.09 | 0 | 0.2 | 0.04 | 0.12 | 0.03 | 0.18 | 0.22 |
| 11 | 59495792 | rs498099 | -1.407 | 2.1 | 0.658 | 1.053 | 0.073 | 0.053 | 0.053 | 0 | 0.05 | 0.36 | 0.17 | 0.07 | 0.05 | 0.04 |
| 11 | 59525741 | rs529270 | -1.053 | 2.116 | 0.608 | 1.054 | 0.056 | 0.022 | 0.022 | 0 | 0.05 | 0.3 | 0.17 | 0.06 | 0.06 | 0.04 |
| 11 | 59536179 | rs516912 | -1.519 | 2.106 | 0.691 | 0.989 | 0.09 | 0.036 | 0.036 | 0.133 | 0.06 | 0.46 | 0.14 | 0.06 | 0.06 | 0.04 |
| 11 | 59549840 | rs7952024 | -0.985 | 2.166 | 0.228 | 1.018 | 0.03 | -0.004 | -0.004 | 0.009 | 0.05 | 0.24 | 0.12 | 0.06 | 0.06 | 0.04 |
| 11 | 59586423 | rs692985 | -0.452 | 1.931 | -0.389 | 1.301 | -0.029 | 0.02 | 0.02 | 0.021 | 0.13 | 0.04 | 0.11 | 0.06 | 0.13 | 0.2 |
| 11 | 59611362 | rs35792306 | -0.968 | 1.795 | -0.013 | 1.805 | -0.026 | -0.031 | -0.031 | 0 | 0.05 | 0 | 0.03 | 0.1 | 0.08 | 0.07 |
| 11 | 59623433 | rs1042613 | -0.837 | 1.478 | -0.009 | 1.235 | 0.056 | 0.001 | 0.001 | 0.171 | 0.2 | 0.55 | 0.16 | 0.16 | 0.23 | 0.27 |
| 11 | 59629305 | rs557564 | -0.94 | 1.477 | 0.387 | 1.001 | 0.033 | -0.008 | -0.008 | 0.411 | 0.21 | 0.34 | 0.16 | 0.16 | 0.26 | 0.25 |
| 11 | 59633493 | rs526934 | -0.335 | 1.466 | -0.434 | 0.796 | -0.022 | 0.007 | 0.007 | 0 | 0.75 | 0.89 | 0.77 | 0.83 | 0.72 | 0.77 |
| 11 | 59659006 | rs12277676 | 0.04 | 1.395 | -0.733 | 0.439 | -0.008 | 0.103 | 0.103 | 0 | 0.32 | 0.1 | 0.28 | 0.11 | 0.34 | 0.2 |
| 11 | 59697671 | rs514088 | 1.293 | 0.42 | -1.202 | 0.889 | -0.009 | -0.08 | -0.08 | 0 | 0.21 | 0.01 | 0.42 | 0.28 | 0.18 | 0.28 |
| 11 | 59708244 | rs7925914 | 0.073 | 0.448 | -0.654 | 0.88 | -0.007 | 0.001 | 0.001 | 0 | 0.54 | 0.33 | 0.64 | 0.43 | 0.54 | 0.5 |
| 11 | 59711390 | rs627461 | -0.213 | 0.317 | -0.294 | 1.063 | 0.007 | 0.023 | 0.023 | 0.856 | 0.69 | 0.49 | 0.73 | 0.63 | 0.67 | 0.64 |
| 11 | 59738667 | rs12419726 | 0.932 | 0.015 | -0.697 | 0.749 | -0.019 | 0.039 | 0.039 | 0 | 0.07 | 0 | 0.03 | 0 | 0.09 | 0.03 |
| 11 | 59742575 | rs17154321 | -0.148 | 0.16 | -0.443 | 1.185 | -0.003 | -0.035 | -0.035 | 0.671 | 0.34 | 0.24 | 0.56 | 0.41 | 0.32 | 0.4 |
| 11 | 59746376 | rs11230115 | 0.365 | -0.388 | -0.027 | 1.449 | -0.007 | 0.037 | 0.037 | 0 | 0.1 | 0.11 | 0.09 | 0.31 | 0.08 | 0.14 |
| 11 | 59748663 | rs7935186 | -0.492 | -0.724 | 0.389 | 1.343 | -0.029 | 0.061 | 0.061 | 0 | 0.7 | 0.5 | 0.48 | 0.62 | 0.76 | 0.68 |
| 11 | 59756311 | rs662065 | 0.22 | -0.694 | 0.516 | 1.323 | -0.022 | 0.044 | 0.044 | 0 | 0.31 | 0.55 | 0.52 | 0.4 | 0.24 | 0.34 |
| 11 | 59764628 | rs512888 | 0.419 | -0.618 | 0.891 | 1.653 | -0.004 | 0.013 | 0.013 | 0 | 0.11 | 0.09 | 0.08 | 0.33 | 0.09 | 0.15 |
| 11 | 59767000 | rs538112 | 0.553 | -0.627 | 0.883 | 1.649 | 0 | 0.004 | 0.004 | 0 | 0.1 | 0.06 | 0.08 | 0.32 | 0.08 | 0.13 |
| 11 | 59767621 | rs637131 | -0.375 | -0.656 | 1.097 | 1.635 | -0.021 | 0.194 | 0.194 | 0 | 0.14 | 0.39 | 0.16 | 0.43 | 0.09 | 0.23 |
| 11 | 59767788 | rs530771 | -0.725 | -0.804 | 0.821 | 1.509 | 0.043 | 0.038 | 0.038 | 0 | 0.43 | 0.72 | 0.61 | 0.72 | 0.33 | 0.5 |
| 11 | 59772730 | rs559335 | -0.585 | -0.511 | 0.434 | 1.385 | -0.017 | 0.072 | 0.072 | 0.464 | 0.22 | 0.32 | 0.21 | 0.49 | 0.18 | 0.37 |
| 11 | 59784606 | rs567708 | 0.289 | -0.312 | 0.206 | 1.21 | -0.001 | 0.002 | 0.002 | 1.709 | 0.12 | 0.14 | 0.12 | 0.17 | 0.11 | 0.26 |
| 11 | 59785647 | rs12421628 | 0.63 | -0.637 | 0.028 | 1.228 | 0.021 | -0.047 | -0.047 | 0 | 0.25 | 0.11 | 0.42 | 0.3 | 0.21 | 0.18 |
| 11 | 59796124 | rs559744 | -0.681 | -0.637 | 0.539 | 1.122 | -0.009 | 0.002 | 0.002 | 0 | 0.41 | 0.58 | 0.58 | 0.48 | 0.33 | 0.49 |
| 11 | 59799013 | rs1144745 | -0.708 | -0.631 | 0.524 | 1.133 | -0.013 | 0.005 | 0.005 | 0 | 0.4 | 0.5 | 0.56 | 0.47 | 0.32 | 0.49 |
| 11 | 59807380 | rs575699 | -0.38 | -0.163 | 0.482 | 1.095 | 0.01 | 0.057 | 0.057 | 1.974 | 0.15 | 0.38 | 0.14 | 0.18 | 0.11 | 0.27 |
| 11 | 59807806 | rs1144747 | 0.315 | -0.084 | 0.023 | 1.139 | -0.002 | -0.016 | -0.016 | 0 | 0.12 | 0.11 | 0.12 | 0.17 | 0.11 | 0.25 |
| 11 | 59808676 | rs1286386 | -0.416 | -0.093 | -0.415 | 0.628 | -0.001 | 0 | 0 | 0 | 0.57 | 0.59 | 0.6 | 0.52 | 0.46 | 0.52 |
| 11 | 59814287 | rs2276044 | -0.115 | 0.172 | -0.917 | 0.689 | 0.004 | -0.034 | -0.034 | 0 | 0.27 | 0.17 | 0.46 | 0.41 | 0.27 | 0.18 |
| 11 | 59815139 | rs10501381 | 0.107 | 0.203 | -0.966 | 0.682 | 0.033 | -0.059 | -0.059 | 0 | 0.25 | 0.1 | 0.44 | 0.41 | 0.19 | 0.16 |
| 11 | 59815517 | rs663925 | 0.752 | 0.546 | -1.328 | 0.29 | -0.048 | 0.017 | 0.017 | 0.588 | 0.34 | 0.19 | 0.21 | 0.35 | 0.36 | 0.52 |
| 11 | 59815640 | rs1151102 | 0.687 | 0.532 | -1.112 | 0.283 | -0.028 | 0.015 | 0.015 | 0.658 | 0.34 | 0.25 | 0.22 | 0.35 | 0.36 | 0.52 |
| 11 | 59823461 | rs12419784 | 0.832 | 0.556 | -1.289 | 0.438 | -0.035 | -0.106 | -0.106 | 0 | 0.21 | 0.05 | 0.44 | 0.41 | 0.27 | 0.17 |
| 11 | 59824526 | rs571760 | 0.102 | 0.235 | -0.95 | -0.143 | -0.004 | -0.025 | -0.025 | 0 | 0.57 | 0.5 | 0.65 | 0.77 | 0.55 | 0.67 |
| 11 | 59826857 | rs11824940 | 1.467 | 0.491 | -1.162 | 0.072 | -0.03 | -0.083 | -0.083 | 0.434 | 0.18 | 0.07 | 0.45 | 0.44 | 0.27 | 0.12 |
| 11 | 59827800 | rs11230135 | 1.169 | 0.827 | -1.414 | -0.105 | -0.046 | -0.013 | -0.013 | 0 | 0.11 | 0 | 0.06 | 0.08 | 0.15 | 0.12 |
| 11 | 59829741 | rs506846 | 0.242 | 0.59 | -0.94 | -0.01 | -0.002 | -0.003 | -0.003 | 0 | 0.61 | 0.43 | 0.68 | 0.71 | 0.61 | 0.67 |
| 11 | 59829804 | rs506053 | -0.201 | 0.54 | -0.693 | -0.055 | 0.002 | -0.017 | -0.017 | 0 | 0.63 | 0.66 | 0.69 | 0.81 | 0.58 | 0.67 |
| 11 | 59837097 | rs528823 | 1.297 | 0.493 | -0.838 | -0.063 | 0.003 | 0.03 | 0.03 | 0.135 | 0.35 | 0.29 | 0.18 | 0.23 | 0.27 | 0.51 |
| 11 | 59839787 | rs7106518 | 0.802 | 0.552 | -0.896 | 0.012 | -0.024 | -0.082 | -0.082 | 0 | 0.25 | 0.13 | 0.48 | 0.48 | 0.31 | 0.15 |
| 11 | 59844700 | rs498100 | 1.971 | 0.647 | -1.062 | -0.397 | -0.012 | 0.027 | 0.027 | 0 | 0.06 | 0.01 | 0.04 | 0 | 0.06 | 0.02 |
| 11 | 59845829 | rs543695 | 0.74 | 0.618 | -0.781 | -0.172 | 0.009 | 0.039 | 0.039 | 0.434 | 0.4 | 0.32 | 0.2 | 0.27 | 0.31 | 0.54 |
| 11 | 59849325 | rs1151107 | 0.184 | 0.691 | -0.736 | -0.256 | 0.005 | -0.045 | -0.045 | 0 | 0.62 | 0.52 | 0.67 | 0.81 | 0.57 | 0.67 |
| 11 | 59852078 | rs558788 | 0.74 | 0.586 | -0.92 | -0.194 | 0.002 | 0.066 | 0.066 | 0.209 | 0.35 | 0.23 | 0.18 | 0.22 | 0.27 | 0.52 |
| 11 | 59855385 | rs573790 | -0.223 | -0.002 | -0.349 | 0.535 | 0.007 | 0.043 | 0.043 | 0 | 0.71 | 0.5 | 0.4 | 0.46 | 0.64 | 0.76 |
| 11 | 59855483 | rs574700 | -0.717 | -0.227 | -0.164 | 0.551 | 0.013 | 0.106 | 0.106 | 2.635 | 0.07 | 0.26 | 0.02 | 0.18 | 0.04 | 0.04 |
| 11 | 59855711 | rs1441585 | -0.401 | -0.194 | -0.121 | 0.567 | 0.011 | 0.093 | 0.093 | 4.165 | 0.06 | 0.21 | 0.02 | 0.18 | 0.04 | 0.05 |
| 11 | 59856028 | rs1441586 | 0.222 | -0.286 | -0.235 | 0.377 | 0 | -0.008 | -0.008 | 0 | 0.55 | 0.52 | 0.32 | 0.38 | 0.46 | 0.57 |
| 11 | 59860178 | rs502581 | 0.411 | -0.297 | -0.267 | 0.394 | -0.005 | 0.053 | 0.053 | 0 | 0.53 | 0.3 | 0.31 | 0.38 | 0.46 | 0.57 |
| 11 | 59863104 | rs569108 | -0.357 | -0.291 | -0.233 | 0.492 | 0.01 | 0.091 | 0.091 | 4.293 | 0.07 | 0.21 | 0.02 | 0.18 | 0.04 | 0.05 |
| 11 | 59863253 | rs512555 | 0.061 | -0.262 | -0.234 | 0.506 | 0.005 | 0.061 | 0.061 | 5.157 | 0.06 | 0.13 | 0.02 | 0.18 | 0.04 | 0.05 |
| 11 | 59867581 | rs12576889 | 1.065 | -0.3 | -0.31 | 0.678 | 0.008 | -0.025 | -0.025 | 0.216 | 0.05 | 0.01 | 0.03 | 0.11 | 0.04 | 0.05 |
| 11 | 59903287 | rs525794 | 0.199 | -0.152 | 0.111 | 0.384 | 0 | -0.011 | -0.011 | 0.426 | 0.55 | 0.57 | 0.33 | 0.38 | 0.46 | 0.56 |
| 11 | 59914726 | rs583296 | 0.295 | -0.143 | -0.179 | 0.388 | -0.009 | 0.058 | 0.058 | 0 | 0.54 | 0.31 | 0.31 | 0.38 | 0.46 | 0.58 |
| 11 | 59918837 | rs684961 | 0.211 | -0.14 | -0.08 | 0.391 | -0.001 | 0.004 | 0.004 | 0 | 0.55 | 0.5 | 0.33 | 0.38 | 0.46 | 0.58 |
| 11 | 59924959 | rs920573 | 0.737 | -0.175 | -0.175 | 0.271 | -0.03 | 0.221 | 0.221 | 0 | 0.48 | 0.17 | 0.29 | 0.2 | 0.42 | 0.54 |
| 11 | 59930966 | rs664034 | -0.45 | -0.456 | 0.108 | 0.41 | 0.001 | 0.02 | 0.02 | 0 | 0.67 | 0.65 | 0.39 | 0.47 | 0.61 | 0.73 |
| 11 | 59936979 | rs667897 | 0.229 | -0.277 | 0.123 | 0.281 | 0.001 | 0.015 | 0.015 | 0 | 0.54 | 0.47 | 0.33 | 0.38 | 0.47 | 0.59 |
| 11 | 59939307 | rs610932 | -0.7 | -0.748 | 0.195 | 0.343 | -0.001 | 0.014 | 0.014 | 0 | 0.48 | 0.5 | 0.69 | 0.62 | 0.56 | 0.42 |
| 11 | 59945745 | rs12453 | 0.858 | -0.841 | 0.201 | 0.271 | -0.045 | 0.168 | 0.168 | 0 | 0.44 | 0.16 | 0.27 | 0.23 | 0.4 | 0.52 |
| 11 | 59946302 | rs598862 | 0.745 | -0.782 | 0.26 | 0.321 | -0.007 | 0.068 | 0.068 | 4.246 | 0.91 | 0.7 | 0.96 | 0.85 | 0.96 | 0.94 |

| | | | | | | | | | | | | | | | | |
|---|---|---|---|---|---|---|---|---|---|---|---|---|---|---|---|---|
| 11 | 59962166 | rs7926344 | 0.836 | -0.845 | 0.009 | 0.228 | -0.079 | 0.187 | 0.187 | 0.01 | 0.43 | 0.1 | 0.26 | 0.22 | 0.41 | 0.51 |
| 11 | 59975078 | rs7929589 | 0.755 | -0.745 | 0.076 | 0.516 | -0.036 | 0.09 | 0.09 | 0 | 0.44 | 0.16 | 0.41 | 0.33 | 0.4 | 0.54 |
| 11 | 59977646 | rs11230193 | -0.124 | -0.837 | 0.276 | 0.697 | -0.01 | 0.003 | 0.003 | 0 | 0.14 | 0.25 | 0.07 | 0.11 | 0.13 | 0.14 |
| 11 | 59978698 | rs4939319 | -0.124 | -0.837 | 0.276 | 0.697 | -0.01 | 0.003 | 0.003 | 0 | 0.14 | 0.25 | 0.07 | 0.11 | 0.13 | 0.14 |
| 11 | 59980598 | rs7929057 | 0.257 | -0.84 | 0.309 | 0.696 | -0.003 | 0.001 | 0.001 | 0 | 0.13 | 0.14 | 0.06 | 0.1 | 0.13 | 0.14 |
| 11 | 59988353 | rs11230197 | -0.111 | -0.835 | 0.27 | 0.698 | -0.01 | 0.002 | 0.002 | 0 | 0.14 | 0.25 | 0.07 | 0.1 | 0.13 | 0.15 |
| 11 | 59997666 | rs600550 | 0.458 | -0.866 | 0.265 | 0.726 | 0.004 | 0.031 | 0.031 | 0 | 0.32 | 0.41 | 0.46 | 0.41 | 0.39 | 0.26 |
| 11 | 60000326 | rs668134 | -0.15 | -0.823 | 0.021 | 0.704 | -0.001 | 0.008 | 0.008 | 0 | 0.9 | 0.83 | 0.95 | 0.85 | 0.93 | 0.93 |
| 11 | 60005268 | rs611267 | -0.716 | -0.853 | 0.008 | 0.613 | -0.001 | 0.002 | 0.002 | 0 | 0.47 | 0.78 | 0.55 | 0.52 | 0.53 | 0.4 |
| 11 | 60013117 | rs7127662 | -1.377 | -0.827 | 0.158 | 0.672 | -0.009 | 0.017 | 0.017 | 0 | 0.25 | 0.42 | 0.13 | 0.25 | 0.21 | 0.21 |
| 11 | 60021948 | rs1582763 | 1.069 | -0.855 | -0.231 | 0.459 | -0.099 | 0.167 | 0.167 | 0 | 0.39 | 0.03 | 0.37 | 0.15 | 0.36 | 0.48 |
| 11 | 60025565 | rs6591559 | -1.099 | -0.759 | -0.048 | 0.526 | -0.022 | 0.108 | 0.108 | 0 | 0.54 | 0.84 | 0.6 | 0.67 | 0.6 | 0.47 |
| 11 | 60028940 | rs1530914 | -1.1 | -0.759 | -0.049 | 0.529 | -0.024 | 0.117 | 0.117 | 0 | 0.54 | 0.85 | 0.6 | 0.71 | 0.6 | 0.47 |
| 11 | 60034429 | rs4938933 | -1.001 | -0.731 | 0.06 | 0.516 | -0.003 | 0.074 | 0.074 | 0 | 0.53 | 0.69 | 0.59 | 0.71 | 0.6 | 0.47 |
| 11 | 60035550 | rs12421663 | 1.52 | -0.693 | 0.18 | 0.771 | -0.021 | -0.009 | -0.009 | 0 | 0.19 | 0.12 | 0.4 | 0.26 | 0.32 | 0.14 |
| 11 | 60039058 | rs1117067 | 0.226 | -0.816 | 0.366 | 0.712 | -0.001 | 0 | 0 | 0 | 0.12 | 0.14 | 0.07 | 0.1 | 0.13 | 0.14 |
| 11 | 60042973 | rs11230228 | -1.082 | -1.085 | 0.356 | 0.604 | -0.017 | 0.016 | 0.016 | 0.022 | 0.36 | 0.59 | 0.2 | 0.44 | 0.29 | 0.36 |
| 11 | 60044880 | rs10897021 | -0.133 | -0.858 | 0.383 | 0.599 | 0.007 | -0.009 | -0.009 | 3.024 | 0.21 | 0.19 | 0.09 | 0.3 | 0.14 | 0.22 |
| 11 | 60047410 | rs1426248 | 0.924 | -0.948 | 0.117 | 0.197 | -0.001 | -0.016 | -0.016 | 0 | 0.27 | 0.27 | 0.15 | 0.16 | 0.21 | 0.32 |
| 11 | 60049098 | rs2017549 | 0.734 | -1.055 | 0.075 | 0.271 | -0.026 | -0.016 | -0.016 | 0 | 0.4 | 0.3 | 0.7 | 0.44 | 0.53 | 0.33 |
| 11 | 60050125 | rs10792260 | 0.06 | -1.01 | 0.117 | 0.256 | 0.041 | 0.012 | 0.012 | 0 | 0.51 | 0.37 | 0.77 | 0.55 | 0.65 | 0.45 |
| 11 | 60055651 | rs10897024 | 0.531 | -0.927 | -0.195 | 0.413 | -0.045 | -0.062 | -0.062 | 0 | 0.3 | 0.13 | 0.48 | 0.39 | 0.43 | 0.26 |
| 11 | 60057715 | rs4939331 | 0.701 | -1.035 | -0.203 | 0.268 | -0.005 | 0.066 | 0.066 | 0 | 0.3 | 0.13 | 0.15 | 0.16 | 0.25 | 0.33 |
| 11 | 60059810 | rs10750931 | -0.207 | -0.615 | 0.375 | 1.162 | 0.006 | 0.006 | 0.006 | 0 | 0.22 | 0.18 | 0.09 | 0.13 | 0.17 | 0.17 |
| 11 | 60063596 | rs11230235 | 0.051 | -0.587 | 0.34 | 1.179 | -0.048 | -0.059 | -0.059 | 0 | 0.3 | 0.13 | 0.48 | 0.39 | 0.43 | 0.26 |
| 11 | 60070176 | rs6591561 | -0.908 | -0.251 | 0.305 | 1.042 | 0.005 | -0.002 | -0.002 | 0 | 0.38 | 0.35 | 0.18 | 0.38 | 0.33 | 0.34 |
| 11 | 60076105 | rs10681 | -0.336 | -0.254 | 0.299 | 1.058 | 0.025 | 0.015 | 0.015 | 0 | 0.32 | 0.18 | 0.15 | 0.24 | 0.29 | 0.3 |
| 11 | 60076693 | rs11824734 | 1.194 | -0.28 | 0.022 | 0.989 | -0.011 | 0.077 | 0.077 | 0 | 0.41 | 0.27 | 0.38 | 0.17 | 0.36 | 0.47 |
| 11 | 60081274 | rs7108879 | -0.217 | -0.572 | -0.314 | 1.262 | 0.022 | 0.062 | 0.062 | 0 | 0.27 | 0.44 | 0.46 | 0.42 | 0.35 | 0.18 |
| 11 | 60084069 | rs10897029 | 0.004 | -0.51 | -0.806 | 1.249 | 0.046 | -0.033 | -0.033 | 0 | 0.22 | 0.1 | 0.1 | 0.27 | 0.19 | 0.26 |
| 11 | 60099894 | rs10897048 | -0.636 | 0.757 | -0.257 | 1.24 | 0.013 | 0.026 | 0.026 | 0 | 0.24 | 0.3 | 0.45 | 0.39 | 0.34 | 0.18 |
| 11 | 60101453 | rs10750935 | -0.649 | 0.839 | -0.211 | 1.28 | 0.017 | 0.037 | 0.037 | 0 | 0.25 | 0.35 | 0.46 | 0.39 | 0.34 | 0.17 |
| 11 | 60101593 | rs4506675 | -0.638 | 0.839 | -0.212 | 1.28 | 0.017 | 0.036 | 0.036 | 0 | 0.25 | 0.34 | 0.45 | 0.4 | 0.34 | 0.17 |
| 11 | 60102384 | rs2304935 | -0.11 | 0.929 | -0.5 | 1.313 | 0.007 | 0.005 | 0.005 | 0 | 0.2 | 0.23 | 0.42 | 0.25 | 0.31 | 0.13 |
| 11 | 60102507 | rs2304933 | -0.1 | 0.949 | -0.491 | 1.323 | 0.007 | 0.006 | 0.006 | 0 | 0.2 | 0.23 | 0.42 | 0.26 | 0.31 | 0.13 |
| 11 | 60102631 | rs10792268 | -0.086 | 0.96 | -0.486 | 1.328 | 0.008 | 0.006 | 0.006 | 0 | 0.2 | 0.23 | 0.42 | 0.26 | 0.31 | 0.13 |
| 11 | 60107105 | rs2165832 | -0.162 | 0.421 | -0.785 | 0.884 | 0.001 | 0.001 | 0.001 | 0 | 0.48 | 0.48 | 0.37 | 0.57 | 0.44 | 0.34 |
| 11 | 60107470 | rs2289612 | -0.383 | 0.455 | -1.091 | 0.869 | -0.031 | 0.013 | 0.013 | 0 | 0.39 | 0.26 | 0.28 | 0.39 | 0.49 | 0.35 |
| 11 | 60107660 | rs11230281 | -0.14 | -0.836 | -1.384 | 0.593 | 0.083 | -0.004 | -0.004 | 0 | 0.51 | 0.39 | 0.67 | 0.5 | 0.72 | 0.43 |
| 11 | 60108278 | rs10212 | -0.471 | -0.465 | -0.448 | 1.189 | 0.045 | 0.122 | 0.122 | 1.109 | 0.18 | 0.56 | 0.25 | 0.38 | 0.24 | 0.19 |
| 11 | 60108714 | rs2289610 | 0.997 | -0.478 | -0.012 | 1.364 | 0.048 | 0.036 | 0.036 | 0.116 | 0.11 | 0.17 | 0.15 | 0.24 | 0.24 | 0.1 |
| 11 | 60111152 | rs12787263 | 1.216 | -0.883 | -0.387 | 0.972 | -0.107 | -0.008 | -0.008 | 0 | 0.23 | 0.04 | 0.37 | 0.16 | 0.33 | 0.2 |
| 11 | 60112812 | rs11230284 | -0.313 | -0.78 | -0.16 | 1.003 | 0.082 | 0.102 | 0.102 | 0.168 | 0.19 | 0.51 | 0.18 | 0.29 | 0.26 | 0.18 |
| 11 | 60117641 | rs17533662 | 1.419 | -0.861 | -0.443 | 0.915 | -0.022 | -0.011 | -0.011 | 0 | 0.06 | 0 | 0.03 | 0.07 | 0.07 | 0.16 |
| 11 | 60122857 | rs4939346 | 0.45 | -1.678 | -0.016 | 0.675 | 0.059 | 0.076 | 0.076 | 0.481 | 0.47 | 0.31 | 0.32 | 0.25 | 0.33 | 0.34 |
| 11 | 60130796 | rs1443246 | -0.895 | -1.301 | 0.29 | 0.967 | 0.08 | 0.097 | 0.097 | 0.066 | 0.8 | 0.57 | 0.75 | 0.66 | 0.73 | 0.8 |
| 11 | 60132164 | rs7952244 | 0.31 | -1.328 | 0.359 | 0.952 | 0.038 | 0.04 | 0.04 | 0.818 | 0.73 | 0.43 | 0.71 | 0.6 | 0.66 | 0.61 |
| 11 | 60152584 | rs950802 | -0.006 | -1.681 | 0.22 | 0.873 | -0.002 | 0 | 0 | 1.663 | 0.52 | 0.46 | 0.36 | 0.44 | 0.34 | 0.39 |
| 11 | 60156484 | rs12793970 | 1.224 | -1.189 | 0.22 | 0.793 | -0.032 | -0.047 | -0.047 | 0 | 0.27 | 0.14 | 0.38 | 0.35 | 0.33 | 0.28 |
| 11 | 60157166 | rs2233253 | -0.282 | -1.174 | 0.292 | 0.799 | 0.06 | 0.06 | 0.06 | 0.16 | 0.73 | 0.56 | 0.71 | 0.6 | 0.65 | 0.61 |
| 11 | 60159708 | rs8181557 | 1.398 | -1.101 | 0.365 | 0.926 | -0.018 | -0.009 | -0.009 | 0 | 0.17 | 0.12 | 0.16 | 0.24 | 0.27 | 0.17 |
| 11 | 60160838 | rs4938939 | 0.507 | -1.156 | 0.329 | 0.904 | 0.043 | 0.051 | 0.051 | 0 | 0.45 | 0.33 | 0.32 | 0.25 | 0.33 | 0.3 |
| 11 | 60161651 | rs1892 | -1.234 | -0.818 | 0.481 | 0.96 | -0.002 | -0.002 | -0.002 | 0.18 | 0.82 | 0.84 | 0.84 | 0.72 | 0.73 | 0.8 |
| 11 | 60163635 | rs3816270 | -1.349 | -0.802 | 0.418 | 0.967 | -0.007 | -0.007 | -0.007 | 0 | 0.82 | 0.86 | 0.83 | 0.72 | 0.73 | 0.8 |
| 11 | 60166455 | rs10897056 | -1.347 | -0.856 | 0.457 | 0.967 | 0.064 | 0.046 | 0.046 | 0.578 | 0.82 | 0.67 | 0.76 | 0.71 | 0.73 | 0.84 |
| 11 | 60167744 | rs2304938 | 1.475 | -0.891 | 0.254 | 0.98 | -0.005 | -0.012 | -0.012 | 0 | 0.07 | 0.03 | 0.04 | 0.11 | 0.08 | 0.23 |
| 11 | 60168927 | rs2197237 | -1.006 | -0.9 | 0.287 | 0.706 | 0.083 | 0.027 | 0.027 | 0.401 | 0.8 | 0.51 | 0.83 | 0.76 | 0.73 | 0.81 |
| 11 | 60173107 | rs4939352 | -1.107 | -0.898 | 0.295 | 0.692 | 0.081 | 0.058 | 0.058 | 0.364 | 0.8 | 0.54 | 0.75 | 0.71 | 0.73 | 0.81 |
| 11 | 60173360 | rs4938941 | -0.117 | -0.907 | 0.185 | 0.692 | 0.056 | 0.025 | 0.025 | 0.686 | 0.73 | 0.48 | 0.71 | 0.64 | 0.66 | 0.61 |
| 11 | 60179489 | rs7950677 | 1.96 | -0.892 | 0.187 | 0.693 | 0 | 0.004 | 0.004 | 0 | 0.07 | 0.05 | 0.1 | 0.16 | 0.08 | 0.13 |
| 11 | 60180335 | rs7926219 | 0.579 | -0.937 | 0.222 | 0.834 | 0.004 | -0.005 | -0.005 | 1.173 | 0.62 | 0.52 | 0.65 | 0.34 | 0.63 | 0.39 |
| 11 | 60181627 | rs1443241 | -0.786 | -0.942 | 0.235 | 0.902 | 0.017 | 0.021 | 0.021 | 1.059 | 0.73 | 0.58 | 0.73 | 0.66 | 0.72 | 0.7 |
| 11 | 60184191 | rs3825020 | 0.611 | -0.944 | 0.169 | 0.884 | 0.004 | -0.005 | -0.005 | 0.566 | 0.62 | 0.52 | 0.65 | 0.34 | 0.63 | 0.39 |
| 11 | 60185775 | rs1838093 | 0.867 | -0.818 | 0.071 | 0.752 | 0.004 | 0.008 | 0.008 | 0 | 0.38 | 0.38 | 0.32 | 0.2 | 0.34 | 0.22 |
| 11 | 60191241 | rs2120181 | 1.162 | -0.528 | -0.14 | 0.969 | -0.017 | -0.007 | -0.007 | 0.211 | 0.25 | 0.14 | 0.18 | 0.29 | 0.26 | 0.26 |
| 11 | 60193409 | rs4938944 | 0.469 | -0.696 | -0.077 | 0.909 | 0.025 | 0.007 | 0.007 | 0.996 | 0.28 | 0.41 | 0.27 | 0.31 | 0.31 | 0.27 |
| 11 | 60199046 | rs10897061 | 1.138 | -0.637 | -0.126 | 1.002 | -0.023 | 0.002 | 0.002 | 0.843 | 0.28 | 0.16 | 0.19 | 0.27 | 0.31 | 0.29 |
| 11 | 60200446 | rs1530354 | -0.076 | -0.429 | -0.427 | 0.496 | 0.004 | 0.005 | 0.005 | 0.094 | 0.54 | 0.53 | 0.66 | 0.73 | 0.61 | 0.62 |
| 11 | 60201527 | rs17542525 | 1.634 | -0.072 | -0.948 | 0.681 | -0.027 | -0.015 | -0.015 | 0 | 0.05 | 0 | 0.03 | 0.07 | 0.07 | 0.17 |

| | | | | | | | | | | | | | | | | |
|---|---|---|---|---|---|---|---|---|---|---|---|---|---|---|---|---|
| 11 | 60202598 | rs11820229 | 1.799 | -0.197 | -0.796 | 0.513 | -0.011 | 0.001 | 0.001 | 0 | 0.22 | 0.15 | 0.4 | 0.2 | 0.29 | 0.19 |
| 11 | 60212297 | rs2120182 | 0.13 | -0.346 | -1.199 | 0.359 | 0.016 | 0.023 | 0.023 | 0 | 0.39 | 0.36 | 0.29 | 0.19 | 0.31 | 0.22 |
| 11 | 60215066 | rs12787520 | 1.017 | 0.191 | -1.496 | 0.517 | -0.06 | 0.012 | 0.012 | 0.001 | 0.25 | 0.07 | 0.41 | 0.19 | 0.32 | 0.19 |
| 11 | 60220947 | rs2000636 | -0.05 | -0.382 | -0.953 | 0.722 | -0.003 | 0 | 0 | 0.165 | 0.6 | 0.6 | 0.6 | 0.44 | 0.66 | 0.53 |
| 11 | 60222883 | rs11230333 | 1.467 | -0.322 | -0.84 | 1.432 | -0.004 | 0.081 | 0.081 | 0 | 0.05 | 0.09 | 0.05 | 0.22 | 0.04 | 0.09 |
| 11 | 60223018 | rs4939362 | -0.685 | -0.505 | -1.299 | 1.301 | -0.046 | -0.007 | -0.007 | 0 | 0.66 | 0.88 | 0.72 | 0.67 | 0.6 | 0.6 |
| 11 | 60232325 | rs4939364 | 0.082 | -1.527 | -0.989 | 1.071 | -0.007 | 0.018 | 0.018 | 0 | 0.68 | 0.59 | 0.62 | 0.65 | 0.68 | 0.56 |
| 11 | 60232766 | rs4633490 | -1.346 | -1.993 | -0.06 | 1.194 | -0.002 | 0.001 | 0.001 | 0 | 0.6 | 0.66 | 0.65 | 0.66 | 0.51 | 0.58 |
| 11 | 60236272 | rs1051756 | -1.224 | -1.997 | 0.036 | 1.192 | -0.014 | 0.01 | 0.01 | 0 | 0.6 | 0.74 | 0.67 | 0.66 | 0.51 | 0.58 |
| 11 | 60237234 | rs1051461 | 0.96 | -2.3 | -0.157 | 1.056 | -0.054 | -0.006 | -0.006 | 2.524 | 0.27 | 0.04 | 0.38 | 0.26 | 0.31 | 0.22 |
| 11 | 60237410 | rs3802957 | 1.398 | -2.297 | -0.044 | 1.115 | 0.006 | -0.004 | -0.004 | 0 | 0.09 | 0.04 | 0.03 | 0.19 | 0.04 | 0.08 |
| 11 | 60237991 | rs1051593 | -0.945 | -2.221 | 0.009 | 1.068 | 0.001 | 0.001 | 0.001 | 0 | 0.52 | 0.57 | 0.5 | 0.46 | 0.43 | 0.39 |
| 11 | 60238054 | rs1051594 | -0.945 | -2.249 | 0.009 | 1.068 | 0.002 | 0.001 | 0.001 | 0 | 0.52 | 0.57 | 0.5 | 0.46 | 0.43 | 0.39 |
| 11 | 60238138 | rs3802954 | 0.831 | -2.037 | -0.09 | 1.154 | -0.019 | -0.025 | -0.025 | 0.059 | 0.06 | 0.02 | 0.03 | 0.14 | 0.07 | 0.19 |
| 11 | 60239723 | rs7124085 | 0.315 | -2.02 | -0.075 | 1.168 | -0.001 | 0.007 | 0.007 | 0 | 0.07 | 0.09 | 0.14 | 0.21 | 0.07 | 0.19 |
| 11 | 60246207 | rs17543260 | 0.758 | -2 | -0.069 | 1.117 | 0.001 | 0.004 | 0.004 | 0 | 0.07 | 0.08 | 0.03 | 0.14 | 0.07 | 0.19 |
| 11 | 60264947 | rs2298552 | -1.438 | -2.124 | -0.017 | 0.929 | -0.01 | 0.006 | 0.006 | 0 | 0.6 | 0.66 | 0.66 | 0.66 | 0.51 | 0.58 |
| 11 | 60265002 | rs2298553 | -1.33 | -2.752 | 0.507 | 1.144 | -0.001 | 0 | 0 | 0.7 | 0.53 | 0.53 | 0.49 | 0.48 | 0.43 | 0.44 |
| 11 | 60268765 | rs3819092 | 1.186 | -2.564 | 0.21 | 0.97 | -0.011 | -0.03 | -0.03 | 0 | 0.05 | 0 | 0.09 | 0.14 | 0.07 | 0.14 |
| 11 | 60269240 | rs2155710 | 0.001 | -2.744 | 0.531 | 1.012 | -0.065 | -0.012 | -0.012 | 0 | 0.3 | 0.06 | 0.38 | 0.3 | 0.34 | 0.24 |
| 11 | 60274393 | rs2298558 | -1.188 | -2.57 | 0.996 | 1.142 | 0.004 | 0.008 | 0.008 | 0 | 0.48 | 0.58 | 0.6 | 0.35 | 0.42 | 0.37 |
| 11 | 60274640 | rs2298559 | -0.95 | -2.505 | 0.904 | 1.19 | 0.009 | 0.02 | 0.02 | 0 | 0.46 | 0.38 | 0.6 | 0.37 | 0.42 | 0.38 |
| 11 | 60276161 | rs4939387 | 0.228 | -2.064 | 1.177 | 1.662 | 0.01 | -0.016 | -0.016 | 0 | 0.13 | 0.08 | 0.11 | 0.13 | 0.09 | 0.19 |
| 11 | 60282019 | rs10750940 | 1.271 | -2.192 | 1.588 | 1.737 | -0.07 | 0.032 | 0.032 | 0 | 0.72 | 0.52 | 0.7 | 0.72 | 0.88 | 0.67 |
| 11 | 60283312 | rs12285606 | 0.873 | -2.095 | 1.546 | 1.747 | -0.01 | -0.015 | -0.015 | 0 | 0.07 | 0.04 | 0.01 | 0.01 | 0.03 | 0.04 |
| 11 | 60291413 | rs6591595 | 1.112 | -2.142 | 1.65 | 1.773 | -0.048 | 0.022 | 0.022 | 0 | 0.73 | 0.61 | 0.69 | 0.72 | 0.88 | 0.78 |
| 11 | 60296879 | rs11230364 | 0.634 | -2.558 | 1.682 | 1.205 | 0.011 | -0.023 | -0.023 | 2.185 | 0.12 | 0.06 | 0.1 | 0.13 | 0.08 | 0.18 |
| 11 | 60297968 | rs2186919 | 1.952 | -2.703 | 1.761 | 1.093 | -0.013 | 0.001 | 0.001 | 0 | 0.39 | 0.26 | 0.29 | 0.42 | 0.54 | 0.4 |
| 11 | 60298541 | rs4939396 | -1.179 | -2.635 | 1.707 | 1.132 | 0.006 | -0.007 | -0.007 | 0.408 | 0.24 | 0.19 | 0.3 | 0.28 | 0.12 | 0.22 |
| 11 | 60311515 | rs2226681 | -0.351 | -2.538 | 1.574 | 1.004 | 0 | 0.002 | 0.002 | 0 | 0.34 | 0.31 | 0.39 | 0.3 | 0.33 | 0.38 |
| 11 | 60335964 | rs2847212 | 0.944 | -3.099 | 1.922 | 1.424 | -0.049 | -0.019 | -0.019 | 0.039 | 0.77 | 0.54 | 0.84 | 0.87 | 0.87 | 0.77 |
| 11 | 60362261 | rs4939416 | -1.34 | -3.07 | 1.91 | 1.421 | -0.048 | -0.02 | -0.02 | 0 | 0.23 | 0.45 | 0.15 | 0.13 | 0.13 | 0.23 |
| 11 | 60363961 | rs11230420 | 0.393 | -2.776 | 1.75 | 1.593 | -0.002 | 0.004 | 0.004 | 0.125 | 0.08 | 0.04 | 0.17 | 0.15 | 0.03 | 0.04 |
| 11 | 60366513 | rs7927817 | -0.549 | -2.825 | 1.848 | 0.923 | -0.025 | -0.005 | -0.005 | 0 | 0.19 | 0.34 | 0.12 | 0.12 | 0.12 | 0.19 |
| 11 | 60379083 | rs17627710 | 0.035 | -2.879 | 1.408 | 0.88 | -0.027 | 0.037 | 0.037 | 0.149 | 0.26 | 0.01 | 0.29 | 0.19 | 0.22 | 0.2 |
| 11 | 60382046 | rs11230437 | 0.56 | -2.865 | 1.467 | 0.855 | -0.008 | -0.012 | -0.012 | 7.02 | 0.08 | 0.05 | 0.01 | 0.01 | 0.03 | 0.03 |
| 11 | 60387600 | rs1994457 | -2.084 | -2.836 | 1.565 | 0.953 | 0.028 | -0.009 | -0.009 | 0.093 | 0.56 | 0.71 | 0.62 | 0.5 | 0.37 | 0.57 |
| 11 | 60394379 | rs474347 | -1.208 | -3.088 | 1.722 | 1.054 | -0.049 | -0.002 | -0.002 | 3.776 | 0.27 | 0.43 | 0.29 | 0.2 | 0.15 | 0.23 |
| 11 | 60400154 | rs482386 | -2.082 | -2.96 | 1.104 | 0.956 | 0.042 | -0.005 | -0.005 | 0.389 | 0.55 | 0.7 | 0.61 | 0.5 | 0.34 | 0.56 |
| 11 | 60442427 | rs12576721 | 0.014 | -2.447 | 0.382 | 1.059 | -0.031 | 0.043 | 0.043 | 0.004 | 0.24 | 0.01 | 0.29 | 0.16 | 0.22 | 0.2 |
| 11 | 60451943 | rs647376 | -0.214 | -2.278 | 0.646 | 1.039 | 0.005 | -0.008 | -0.008 | 0.857 | 0.85 | 0.86 | 0.89 | 0.86 | 0.89 | 0.8 |
| 11 | 60473821 | rs1567083 | 1.137 | -2.228 | 0.744 | 1.023 | -0.053 | 0.004 | 0.004 | 0.242 | 0.71 | 0.41 | 0.68 | 0.77 | 0.85 | 0.74 |
| 11 | 60495124 | rs612342 | -0.449 | -1.506 | -0.281 | 1.03 | 0.01 | 0.009 | 0.009 | 0 | 0.25 | 0.14 | 0.31 | 0.24 | 0.22 | 0.24 |
| 11 | 60504436 | rs2077227 | -0.997 | -0.925 | 0.244 | 1.128 | 0.005 | -0.023 | -0.023 | 0.102 | 0.14 | 0.1 | 0.1 | 0.2 | 0.11 | 0.19 |
| 11 | 60517887 | rs7948556 | -1.297 | -1.184 | 0.292 | 0.937 | -0.029 | 0.017 | 0.017 | 0.25 | 0.31 | 0.43 | 0.31 | 0.34 | 0.2 | 0.27 |
| 11 | 60518182 | rs10897109 | 1.003 | -1.185 | 0.349 | 0.348 | -0.008 | -0.006 | -0.006 | 1.221 | 0.76 | 0.76 | 0.76 | 0.87 | 0.86 | 0.86 |
| 11 | 60519980 | rs7127081 | -1.211 | -0.674 | 0.845 | 0.627 | 0.019 | 0.102 | 0.102 | 0 | 0.1 | 0.45 | 0.1 | 0.14 | 0.11 | 0.06 |
| 11 | 60521225 | rs10736710 | 0.277 | -0.145 | -0.325 | 0.58 | 0 | 0.003 | 0.003 | 0 | 0.75 | 0.73 | 0.78 | 0.74 | 0.75 | 0.9 |
| 11 | 60523815 | rs12364352 | -0.961 | -0.5 | 0.167 | 0.51 | -0.015 | 0.065 | 0.065 | 0 | 0.11 | 0.4 | 0.11 | 0.14 | 0.12 | 0.07 |
| 11 | 60525786 | rs7930940 | 0.266 | 0.789 | -0.976 | 0.134 | -0.036 | 0.029 | 0.029 | 0 | 0.69 | 0.76 | 0.73 | 0.74 | 0.65 | 0.85 |
| 11 | 60528162 | rs11230466 | -0.209 | 0.342 | -1.498 | -0.238 | 0.011 | 0.107 | 0.107 | 0.441 | 0.43 | 0.21 | 0.28 | 0.21 | 0.36 | 0.23 |
| 11 | 60531346 | rs1032939 | 0.08 | 0.758 | -2.035 | 0.172 | -0.156 | -0.071 | -0.071 | 0.241 | 0.3 | 0.06 | 0.48 | 0.63 | 0.48 | 0.51 |
| 11 | 60536854 | rs7117409 | 0.002 | -0.076 | -1.681 | -0.455 | 0.014 | 0.024 | 0.024 | 1.472 | 0.31 | 0.19 | 0.15 | 0.09 | 0.21 | 0.2 |
| 11 | 60537069 | rs4939451 | -0.177 | 0.551 | -1.243 | -0.7 | 0.006 | -0.052 | -0.052 | 0.062 | 0.34 | 0.45 | 0.48 | 0.82 | 0.33 | 0.56 |
| 11 | 60537578 | rs4939452 | -0.247 | 0.219 | -1.508 | -0.394 | -0.002 | -0.003 | -0.003 | 0 | 0.47 | 0.32 | 0.39 | 0.47 | 0.53 | 0.52 |
| 11 | 60540224 | rs11230472 | 0.853 | -0.035 | -1.745 | -1.325 | 0.01 | 0.121 | 0.121 | 2.11 | 0.27 | 0.06 | 0.13 | 0.04 | 0.21 | 0.06 |
| 11 | 60542953 | rs7120524 | 0.033 | 0.38 | -1.817 | -2.055 | -0.049 | 0.075 | 0.075 | 0.221 | 0.32 | 0.24 | 0.25 | 0.11 | 0.42 | 0.32 |
| 11 | 60543909 | rs2276324 | -0.168 | 0.324 | -1.319 | -2.453 | 0.005 | -0.021 | -0.021 | 1.057 | 0.27 | 0.31 | 0.21 | 0.13 | 0.33 | 0.29 |
| 11 | 60544505 | rs7946001 | 0.855 | 0.77 | -0.572 | -2.222 | -0.002 | -0.05 | -0.05 | 0.01 | 0.37 | 0.38 | 0.25 | 0.13 | 0.35 | 0.19 |
| 11 | 60547604 | rs2089343 | 0.757 | 0.416 | -0.707 | -1.753 | -0.008 | -0.053 | -0.053 | 0.001 | 0.16 | 0.04 | 0.4 | 0.67 | 0.16 | 0.35 |
| 11 | 60559045 | rs630084 | 0.827 | 0.568 | -0.31 | -1.9 | -0.002 | 0 | 0 | 0 | 0.09 | 0.08 | 0.18 | 0.05 | 0.11 | 0.06 |
| 11 | 60559805 | rs931861 | 0.629 | 0.294 | 0.559 | -2.129 | 0.021 | -0.063 | -0.063 | 1.6 | 0.8 | 0.48 | 0.71 | 0.89 | 0.79 | 0.88 |
| 11 | 60561613 | rs751026 | -0.399 | 0.257 | 0.265 | -2.602 | 0.001 | 0.129 | 0.129 | 0 | 0.51 | 0.29 | 0.51 | 0.85 | 0.5 | 0.79 |
| 11 | 60562641 | rs4939456 | 0.147 | 0.192 | 0.445 | -2.631 | 0.015 | 0.124 | 0.124 | 0 | 0.42 | 0.19 | 0.26 | 0.23 | 0.38 | 0.36 |
| 11 | 60562975 | rs4939457 | 0.405 | 0.15 | 0.356 | -2.652 | -0.025 | 0.184 | 0.184 | 0 | 0.39 | 0.1 | 0.23 | 0.16 | 0.37 | 0.36 |
| 11 | 60563383 | rs10466665 | 1.067 | 0.256 | 0.876 | -2.536 | 0.038 | -0.002 | -0.002 | 0.298 | 0.11 | 0.25 | 0.21 | 0.11 | 0.12 | 0.06 |
| 11 | 60566619 | rs11230482 | -0.543 | 0.201 | 0.887 | -2.512 | 0.033 | 0.045 | 0.045 | 0.111 | 0.07 | 0.28 | 0.05 | 0.06 | 0.05 | 0.05 |
| 11 | 60566750 | rs1032936 | 2.014 | 0.191 | 0.589 | -2.562 | -0.03 | 0.107 | 0.107 | 1.272 | 0.28 | 0.14 | 0.2 | 0.08 | 0.29 | 0.08 |
| 11 | 60567230 | rs11230483 | -0.395 | 0.069 | 0.727 | -2.351 | 0.014 | 0.117 | 0.117 | 0.193 | 0.48 | 0.29 | 0.49 | 0.81 | 0.45 | 0.75 |

| | | | | | | | | | | | | | | | | |
|---|---|---|---|---|---|---|---|---|---|---|---|---|---|---|---|---|
| 11 | 60567589 | rs10792292 | -0.395 | 0.069 | 0.727 | -2.351 | 0.014 | 0.117 | 0.117 | 0.193 | 0.48 | 0.29 | 0.49 | 0.81 | 0.45 | 0.75 |
| 11 | 60568155 | rs592637 | 0.109 | 0.061 | 0.859 | -2.352 | 0.007 | 0.039 | 0.039 | 0.102 | 0.51 | 0.62 | 0.5 | 0.19 | 0.55 | 0.25 |
| 11 | 60568200 | rs10792293 | -0.4 | 0.067 | 0.758 | -2.533 | 0.012 | 0.143 | 0.143 | 0.188 | 0.48 | 0.26 | 0.49 | 0.85 | 0.45 | 0.75 |
| 11 | 60568701 | rs10792295 | -0.41 | 0.078 | 0.754 | -2.527 | 0.015 | 0.139 | 0.139 | 0.117 | 0.48 | 0.28 | 0.49 | 0.85 | 0.45 | 0.75 |
| 11 | 60568768 | rs10750945 | -0.402 | 0.082 | 0.664 | -2.525 | 0.014 | 0.157 | 0.157 | 0.139 | 0.48 | 0.25 | 0.49 | 0.85 | 0.45 | 0.75 |
| 11 | 60595005 | rs7930400 | -0.986 | -0.002 | -0.135 | -2.543 | 0.028 | 0.145 | 0.145 | 0 | 0.5 | 0.32 | 0.62 | 0.86 | 0.57 | 0.75 |
| 11 | 60595190 | rs10792298 | 0.401 | 0.226 | 0.204 | -2.638 | 0.011 | -0.059 | -0.059 | 0 | 0.36 | 0.46 | 0.42 | 0.81 | 0.4 | 0.64 |
| 11 | 60598483 | rs7942904 | 1.146 | -0.107 | 0.188 | -2.612 | 0.022 | 0.001 | 0.001 | 0 | 0.11 | 0.25 | 0.23 | 0.11 | 0.1 | 0.06 |
| 11 | 60599009 | rs7112307 | 1.103 | 0.256 | 0.034 | -2.625 | 0.027 | 0.073 | 0.073 | 0 | 0.26 | 0.11 | 0.12 | 0.12 | 0.21 | 0.21 |
| 11 | 60602839 | rs1944778 | 1.664 | 0.213 | -0.205 | -2.741 | -0.026 | 0.159 | 0.159 | 1.7 | 0.3 | 0.06 | 0.2 | 0.04 | 0.31 | 0.12 |
| 11 | 60603209 | rs484509 | -0.898 | 0.4 | 0.424 | -2.219 | 0.024 | 0.003 | 0.003 | 0 | 0.89 | 0.73 | 0.77 | 0.89 | 0.9 | 0.94 |
| 11 | 60603298 | rs1940590 | 0.993 | 0.4 | 0.406 | -2.219 | 0.015 | 0 | 0 | 0 | 0.1 | 0.2 | 0.23 | 0.11 | 0.1 | 0.07 |
| 11 | 60603647 | rs582246 | -0.659 | 0.411 | 0.233 | -2.214 | 0.031 | 0.005 | 0.005 | 0.104 | 0.88 | 0.67 | 0.76 | 0.89 | 0.9 | 0.94 |
| 11 | 60622522 | rs4939469 | -0.069 | -0.081 | -0.354 | -2.226 | -0.002 | -0.028 | -0.028 | 0.483 | 0.12 | 0.12 | 0.07 | 0.03 | 0.12 | 0.21 |
| 11 | 60624496 | rs575042 | -0.713 | -0.304 | -0.636 | -2.568 | 0.019 | 0.114 | 0.114 | 0 | 0.58 | 0.77 | 0.73 | 0.92 | 0.59 | 0.73 |
| 11 | 60637164 | rs489172 | -0.478 | 0.369 | -0.09 | -1.502 | 0.016 | 0.061 | 0.061 | 2.38 | 0.88 | 0.84 | 0.82 | 0.4 | 0.82 | 0.58 |
| 11 | 60638461 | rs10897122 | 0.031 | 0.371 | -0.25 | -1.588 | -0.079 | -0.098 | -0.098 | 0.369 | 0.15 | 0.03 | 0.31 | 0.63 | 0.22 | 0.45 |
| 11 | 60640227 | rs7946859 | 0.614 | -0.193 | -0.151 | -2.176 | 0.004 | 0.022 | 0.022 | 0 | 0.41 | 0.32 | 0.4 | 0.33 | 0.39 | 0.39 |
| 11 | 60640480 | rs2072098 | 0.653 | -0.238 | -0.063 | -2.163 | -0.033 | 0.014 | 0.014 | 0 | 0.4 | 0.11 | 0.39 | 0.32 | 0.39 | 0.38 |
| 11 | 60645214 | rs514524 | 1.515 | -0.381 | -0.192 | -2.355 | -0.031 | 0.182 | 0.182 | 2.808 | 0.29 | 0.03 | 0.19 | 0.04 | 0.3 | 0.05 |
| 11 | 60657518 | rs7126832 | 0.742 | -0.311 | -0.04 | -2.313 | -0.025 | 0.026 | 0.026 | 0 | 0.4 | 0.12 | 0.39 | 0.32 | 0.39 | 0.38 |
| 11 | 60663629 | rs4939474 | -0.386 | -0.339 | 0.617 | -2.335 | 0 | 0.004 | 0.004 | 0 | 0.52 | 0.6 | 0.46 | 0.33 | 0.48 | 0.45 |
| 11 | 60666116 | rs3763841 | 0.661 | -0.316 | 0.574 | -2.23 | -0.019 | 0.029 | 0.029 | 0 | 0.4 | 0.15 | 0.39 | 0.32 | 0.39 | 0.38 |
| 11 | 60666341 | rs3763840 | 0.661 | -0.316 | 0.574 | -2.23 | -0.019 | 0.029 | 0.029 | 0 | 0.4 | 0.14 | 0.39 | 0.32 | 0.39 | 0.38 |
| 11 | 60669197 | rs476182 | -1.686 | -0.303 | 0.522 | -2.227 | -0.008 | -0.022 | -0.022 | 0 | 0.94 | 0.81 | 0.96 | 1 | 0.97 | 0.99 |
| 11 | 60676289 | rs10897124 | 0.384 | -0.269 | 0.447 | -2.217 | -0.079 | -0.099 | -0.099 | 0 | 0.15 | 0.03 | 0.31 | 0.63 | 0.22 | 0.45 |
| 11 | 60689445 | rs555835 | -0.748 | -0.29 | 0.64 | -2.238 | -0.005 | -0.088 | -0.088 | 0 | 0.58 | 0.58 | 0.74 | 0.95 | 0.61 | 0.88 |
| 11 | 60697643 | rs2074417 | 0.361 | -0.488 | 0.616 | -1.947 | 0.003 | 0.026 | 0.026 | 0 | 0.41 | 0.25 | 0.4 | 0.32 | 0.39 | 0.43 |
| 11 | 60703403 | rs524523 | 1.167 | -0.494 | 0.707 | -2.035 | -0.001 | -0.015 | -0.015 | 0.585 | 0.06 | 0.23 | 0.05 | 0.01 | 0.03 | 0.01 |
| 11 | 60704213 | rs2469887 | -0.062 | -0.474 | 0.627 | -2.004 | -0.001 | -0.002 | -0.002 | 0 | 0.47 | 0.47 | 0.45 | 0.32 | 0.42 | 0.44 |
| 11 | 60711887 | rs3016367 | 0.198 | -0.358 | 0.204 | -1.87 | -0.01 | 0.022 | 0.022 | 0 | 0.58 | 0.67 | 0.5 | 0.32 | 0.49 | 0.48 |
| 11 | 60712672 | rs485639 | 0.417 | -0.288 | -0.223 | -1.822 | -0.01 | 0.06 | 0.06 | 0 | 0.53 | 0.3 | 0.46 | 0.31 | 0.51 | 0.49 |
| 11 | 60715712 | rs566353 | 0.66 | -0.304 | -0.826 | -1.606 | -0.044 | 0.089 | 0.089 | 0 | 0.47 | 0.09 | 0.43 | 0.31 | 0.43 | 0.41 |
| 11 | 60723898 | rs509581 | -0.237 | -0.377 | 0.344 | -1.408 | 0.011 | -0.008 | -0.008 | 0.051 | 0.08 | 0.4 | 0.06 | 0.01 | 0.03 | 0.01 |
| 11 | 60725609 | rs3016369 | 0.205 | -0.573 | -0.734 | -1.784 | 0.013 | 0.072 | 0.072 | 0 | 0.79 | 0.94 | 0.92 | 1 | 0.86 | 0.91 |
| 11 | 60728321 | rs473756 | 1.255 | -0.466 | -1.144 | -2.169 | -0.028 | 0.234 | 0.234 | 0 | 0.46 | 0.16 | 0.65 | 0.89 | 0.51 | 0.78 |
| 11 | 60728850 | rs12422200 | 0.245 | -0.543 | -0.216 | -2.199 | 0.009 | -0.125 | -0.125 | 0 | 0.22 | 0.4 | 0.17 | 0.01 | 0.26 | 0.07 |
| 11 | 60733626 | rs491600 | 0.947 | -0.764 | 0.001 | -2.071 | 0.027 | 0.045 | 0.045 | 0 | 0.65 | 0.31 | 0.78 | 0.9 | 0.76 | 0.86 |
| 11 | 60735780 | rs636833 | 0.51 | -0.82 | 0.01 | -2.023 | -0.042 | -0.083 | -0.083 | 0 | 0.69 | 0.62 | 0.84 | 0.9 | 0.81 | 0.88 |
| 11 | 60738832 | rs1859206 | -0.352 | -0.585 | -0.666 | -1.84 | -0.013 | -0.03 | -0.03 | 3.396 | 0.91 | 0.88 | 0.97 | 0.99 | 0.95 | 0.93 |
| 11 | 60745771 | rs11230542 | 0.231 | -0.422 | -0.874 | -1.796 | -0.021 | 0.072 | 0.072 | 0 | 0.12 | 0.02 | 0.05 | 0 | 0.1 | 0.01 |
| 11 | 60748939 | rs3019548 | 0.37 | -0.394 | -0.526 | -1.771 | 0.002 | 0 | 0 | 0 | 0.67 | 0.35 | 0.74 | 0.89 | 0.68 | 0.86 |
| 11 | 60750048 | rs2905506 | -0.796 | -0.146 | -0.513 | -1.661 | -0.006 | 0.013 | 0.013 | 0 | 0.82 | 0.86 | 0.87 | 0.98 | 0.75 | 0.9 |
| 11 | 60751569 | rs11230548 | 0.775 | 0.757 | -1.036 | -0.89 | 0.013 | 0.06 | 0.06 | 0.236 | 0.14 | 0.14 | 0.48 | 0.55 | 0.22 | 0.4 |
| 11 | 60751923 | rs3019544 | -1.116 | 0.166 | -0.765 | -1.192 | -0.007 | 0.038 | 0.038 | 0 | 0.83 | 0.9 | 0.89 | 0.98 | 0.79 | 0.91 |
| 11 | 60756588 | rs2905509 | 0.607 | 0.006 | -0.786 | -1.27 | 0.011 | 0.047 | 0.047 | 0 | 0.66 | 0.25 | 0.72 | 0.88 | 0.65 | 0.85 |
| 11 | 60757633 | rs7125778 | -0.52 | -0.152 | -0.14 | -0.996 | -0.031 | -0.009 | -0.009 | 0.192 | 0.15 | 0.48 | 0.14 | 0.1 | 0.1 | 0.05 |
| 11 | 60760530 | rs11230553 | -0.301 | -0.198 | -0.596 | -0.743 | -0.004 | 0.01 | 0.01 | 0.015 | 0.27 | 0.27 | 0.21 | 0.28 | 0.25 | 0.25 |
| 11 | 60760971 | rs929230 | 1.672 | -0.583 | -1.516 | -1.681 | -0.016 | 0.168 | 0.168 | 0 | 0.27 | 0.01 | 0.12 | 0.04 | 0.23 | 0.07 |
| 11 | 60763134 | rs11230555 | 0.634 | -0.662 | -1.473 | -1.646 | -0.001 | -0.044 | -0.044 | 0 | 0.22 | 0.26 | 0.18 | 0.09 | 0.23 | 0.06 |
| 11 | 60763832 | rs916811 | 1.503 | -0.421 | -1.954 | -1.607 | -0.105 | -0.131 | -0.131 | 0 | 0.16 | 0.03 | 0.48 | 0.58 | 0.27 | 0.46 |
| 11 | 60765553 | rs7929471 | -0.797 | -0.63 | -1.339 | -1.713 | -0.023 | 0.017 | 0.017 | 0 | 0.3 | 0.43 | 0.18 | 0.28 | 0.22 | 0.36 |
| 11 | 60768044 | rs2905517 | -0.511 | -0.241 | -1.136 | -1.618 | -0.002 | 0.021 | 0.021 | 0 | 0.44 | 0.34 | 0.69 | 0.88 | 0.52 | 0.71 |
| 11 | 60772090 | rs2237997 | -1.17 | -0.264 | -1.155 | -1.565 | 0.035 | 0.036 | 0.036 | 0 | 0.55 | 0.66 | 0.78 | 0.75 | 0.63 | 0.67 |
| 11 | 60776209 | rs11230563 | -0.697 | -0.183 | -1.035 | -1.677 | 0.004 | 0.015 | 0.015 | 0 | 0.39 | 0.61 | 0.3 | 0.17 | 0.36 | 0.21 |
| 11 | 60783108 | rs2074229 | -0.342 | -0.194 | -1.243 | -1.571 | -0.004 | 0.005 | 0.005 | 0 | 0.41 | 0.34 | 0.7 | 0.83 | 0.51 | 0.71 |
| 11 | 60790839 | rs3016173 | 0.696 | -0.255 | -1.182 | -1.535 | 0.006 | 0.03 | 0.03 | 0 | 0.55 | 0.32 | 0.7 | 0.78 | 0.55 | 0.61 |
| 11 | 60797703 | rs2905524 | -0.019 | -0.091 | -1.092 | -1.282 | 0.005 | 0.082 | 0.082 | 0 | 0.82 | 0.49 | 0.87 | 0.8 | 0.83 | 0.72 |
| 11 | 60801310 | rs999297 | 1.268 | 0.232 | -1.275 | -1.093 | -0.106 | -0.13 | -0.13 | 0.03 | 0.16 | 0.02 | 0.5 | 0.56 | 0.27 | 0.42 |
| 11 | 60803588 | rs3016171 | 0.828 | 0.194 | -1.225 | -1.111 | -0.043 | -0.058 | -0.058 | 0.004 | 0.18 | 0.14 | 0.51 | 0.56 | 0.27 | 0.42 |
| 11 | 60806747 | rs12293191 | 0.041 | -0.036 | -0.542 | -1.125 | -0.006 | -0.073 | -0.073 | 0 | 0.22 | 0.37 | 0.18 | 0.11 | 0.2 | 0.06 |
| 11 | 60810757 | rs175102 | 1.154 | 0.01 | -0.966 | -0.926 | -0.085 | -0.033 | -0.033 | 0.948 | 0.52 | 0.03 | 0.64 | 0.64 | 0.53 | 0.57 |
| 11 | 60824609 | rs175110 | -0.746 | 0.155 | -0.262 | -1.031 | -0.001 | -0.001 | -0.001 | 0.056 | 0.35 | 0.35 | 0.23 | 0.26 | 0.35 | 0.35 |
| 11 | 60825214 | rs175115 | 0.834 | 0.153 | -0.35 | -1.052 | -0.001 | 0.005 | 0.005 | 0 | 0.57 | 0.41 | 0.7 | 0.74 | 0.54 | 0.59 |
| 11 | 60829178 | rs175119 | -0.37 | 0.255 | -0.234 | -1.99 | -0.004 | -0.023 | -0.023 | 0.045 | 0.2 | 0.14 | 0.15 | 0 | 0.23 | 0.08 |
| 11 | 60830367 | rs3016175 | 0.754 | -0.038 | -0.256 | -2.016 | -0.022 | -0.025 | -0.025 | 0 | 0.83 | 0.71 | 0.9 | 0.88 | 0.85 | 0.83 |
| 11 | 60832282 | rs650258 | 0.951 | 0.125 | -0.267 | -2.108 | 0 | -0.029 | -0.029 | 0 | 0.67 | 0.65 | 0.76 | 0.88 | 0.63 | 0.76 |
| 11 | 60833624 | rs175126 | 0.513 | 0.197 | -0.681 | -2.154 | 0 | -0.003 | -0.003 | 0.635 | 0.48 | 0.57 | 0.63 | 0.74 | 0.45 | 0.68 |
| 11 | 60834043 | rs11601370 | -0.182 | 0.778 | -1.236 | -1.795 | -0.031 | 0.039 | 0.039 | 0 | 0.07 | 0 | 0.05 | 0 | 0.13 | 0.05 |

| | | | | | | | | | | | | | | | | |
|---|---|---|---|---|---|---|---|---|---|---|---|---|---|---|---|---|
| 11 | 60835322 | rs175129 | 0.7 | 0.526 | -1.091 | -1.748 | -0.05 | 0.036 | 0.036 | 0 | 0.39 | 0.16 | 0.6 | 0.66 | 0.43 | 0.65 |
| 11 | 60835782 | rs175130 | 0.505 | 0.815 | -0.298 | -1.947 | 0.024 | -0.07 | -0.07 | 0 | 0.75 | 0.56 | 0.78 | 0.86 | 0.74 | 0.76 |
| 11 | 60836117 | rs11230584 | -1.404 | 0.571 | -0.305 | -1.767 | -0.002 | -0.015 | -0.015 | 0 | 0.17 | 0.25 | 0.11 | 0.13 | 0.15 | 0.18 |
| 11 | 60836519 | rs175131 | -0.646 | 0.562 | -0.499 | -1.924 | 0.002 | 0.004 | 0.004 | 0.652 | 0.1 | 0.05 | 0.03 | 0.08 | 0.09 | 0.06 |
| 11 | 60839578 | rs2905501 | 0.692 | 0.973 | -0.144 | -2.174 | 0.01 | -0.032 | -0.032 | 0.089 | 0.92 | 0.84 | 0.94 | 0.99 | 0.92 | 0.86 |
| 11 | 60846825 | rs11230586 | -0.565 | 1.01 | -0.392 | -2.405 | -0.007 | 0.014 | 0.014 | 0 | 0.08 | 0.04 | 0.05 | 0 | 0.1 | 0.05 |
| 11 | 60848143 | rs3016176 | -1.677 | 0.577 | 0.407 | -2.463 | -0.001 | -0.015 | -0.015 | 0.112 | 0.14 | 0.53 | 0.1 | 0.08 | 0.1 | 0.19 |
| 11 | 60850652 | rs2905503 | -1.565 | 0.885 | 0.443 | -2.5 | 0.027 | -0.017 | -0.017 | 0 | 0.14 | 0.39 | 0.1 | 0.08 | 0.12 | 0.18 |
| 11 | 60858031 | rs509936 | 0.756 | 0.783 | -1.045 | -3.253 | -0.123 | 0.205 | 0.205 | 0 | 0.4 | 0.03 | 0.54 | 0.91 | 0.42 | 0.64 |
| 11 | 60859791 | rs175133 | 0.125 | 0.737 | -0.667 | -3.494 | 0.004 | 0.036 | 0.036 | 0 | 0.5 | 0.42 | 0.61 | 0.93 | 0.51 | 0.71 |
| 11 | 60859990 | rs3019565 | -1.786 | 0.45 | -0.339 | -3.576 | -0.004 | -0.029 | -0.029 | 0 | 0.26 | 0.74 | 0.17 | 0.08 | 0.21 | 0.27 |
| 11 | 60860889 | rs440879 | 0.782 | 0.794 | -0.158 | -3.631 | 0.002 | 0.102 | 0.102 | 0 | 0.41 | 0.33 | 0.57 | 0.92 | 0.4 | 0.66 |
| 11 | 60872073 | rs3862666 | -1.389 | 0.966 | 0.422 | -3.547 | 0.039 | -0.047 | -0.047 | 0.767 | 0.16 | 0.31 | 0.16 | 0.07 | 0.21 | 0.16 |
| 11 | 60875029 | rs3862667 | -1.798 | 0.882 | 0.476 | -3.553 | 0.016 | -0.052 | -0.052 | 0 | 0.18 | 0.55 | 0.18 | 0.08 | 0.21 | 0.16 |
| 11 | 60878662 | rs522073 | 0.769 | 0.695 | 0.107 | -3.782 | -0.017 | 0.138 | 0.138 | 0 | 0.42 | 0.27 | 0.57 | 0.92 | 0.44 | 0.67 |
| 11 | 60886913 | rs2241002 | -1.333 | 0.693 | 0.658 | -3.432 | 0.008 | -0.048 | -0.048 | 0 | 0.15 | 0.31 | 0.14 | 0.06 | 0.15 | 0.15 |
| 11 | 60892606 | rs637186 | -0.815 | 0.833 | 0.461 | -3.614 | -0.016 | 0.029 | 0.029 | 0 | 0.94 | 0.99 | 0.95 | 1 | 0.92 | 0.96 |
| 11 | 60901112 | rs666886 | 1.151 | 0.145 | -0.047 | -4.061 | 0.066 | 0.129 | 0.129 | 1.12 | 0.2 | 0 | 0.05 | 0 | 0.08 | 0.02 |
| 11 | 60903335 | rs10792303 | 0.455 | 1.415 | 0.578 | -4.002 | 0 | -0.08 | -0.08 | 0 | 0.52 | 0.56 | 0.67 | 0.99 | 0.55 | 0.79 |
| 11 | 60909581 | rs595158 | 0.413 | 1.634 | 0.598 | -3.982 | -0.001 | -0.092 | -0.092 | 0 | 0.53 | 0.55 | 0.67 | 0.99 | 0.54 | 0.79 |
| 11 | 60917528 | rs11230611 | -0.965 | 1.65 | 0.533 | -3.717 | 0.03 | -0.024 | -0.024 | 0 | 0.05 | 0.11 | 0.08 | 0 | 0.1 | 0.02 |
| 11 | 60941495 | rs17155752 | -0.86 | 1.567 | 0.538 | -3.709 | 0.022 | -0.028 | -0.028 | 0 | 0.05 | 0.12 | 0.06 | 0 | 0.07 | 0.08 |
| 11 | 60943778 | rs590494 | 0.408 | 1.796 | 0.677 | -3.904 | 0 | -0.053 | -0.053 | 0 | 0.51 | 0.46 | 0.67 | 0.99 | 0.56 | 0.73 |
| 11 | 60970554 | rs12799829 | 0.519 | 1.56 | 0.918 | -3.613 | -0.001 | 0.058 | 0.058 | 0 | 0.35 | 0.2 | 0.25 | 0 | 0.33 | 0.17 |
| 11 | 61017951 | rs4939511 | -2.879 | 2.135 | 2.996 | -2.173 | 0.035 | -0.016 | -0.016 | 0 | 0.13 | 0.3 | 0.13 | 0.09 | 0.15 | 0.07 |
| 11 | 61046610 | rs7111608 | -3.917 | 1.674 | 2.546 | -2.262 | -0.023 | -0.071 | -0.071 | 0 | 0.18 | 0.69 | 0.13 | 0.03 | 0.14 | 0.18 |
| 11 | 61077636 | rs11822158 | -3.464 | 1.727 | 2.383 | -2.229 | -0.042 | -0.016 | -0.016 | 0 | 0.09 | 0.59 | 0.06 | 0.03 | 0.01 | 0.15 |
| 11 | 61078988 | rs11230666 | -3.438 | 1.727 | 2.41 | -2.229 | -0.035 | -0.008 | -0.008 | 0 | 0.09 | 0.57 | 0.06 | 0.03 | 0.01 | 0.15 |
| 11 | 61085499 | rs4939513 | -4.088 | 1.733 | 2.406 | -2.234 | -0.022 | -0.07 | -0.07 | 0 | 0.18 | 0.7 | 0.13 | 0.03 | 0.14 | 0.18 |
| 11 | 61100075 | rs2512811 | 2.954 | 1.251 | 2.5 | -1.837 | -0.042 | -0.016 | -0.016 | 2.505 | 0.91 | 0.41 | 0.94 | 0.97 | 0.99 | 0.84 |
| 11 | 61106525 | rs2512809 | 3.666 | 1.263 | 2.495 | -1.845 | -0.055 | -0.045 | -0.045 | 0 | 0.89 | 0.35 | 0.92 | 0.97 | 0.98 | 0.83 |
| 11 | 61108974 | rs2260655 | 3.305 | 1.243 | 2.503 | -1.833 | -0.051 | -0.031 | -0.031 | 0.299 | 0.9 | 0.4 | 0.94 | 0.97 | 0.99 | 0.83 |
| 11 | 61114616 | rs3972757 | 2.931 | 1.258 | 2.467 | -1.842 | -0.031 | -0.008 | -0.008 | 0 | 0.91 | 0.44 | 0.94 | 0.97 | 0.99 | 0.84 |
| 11 | 61126858 | rs4939517 | 2.954 | 1.266 | 2.333 | -1.834 | -0.009 | -0.069 | -0.069 | 0.001 | 0.85 | 0.32 | 0.87 | 0.97 | 0.87 | 0.82 |
| 11 | 61134532 | rs1037448 | 2.356 | 1.132 | 2.279 | -1.867 | -0.023 | -0.096 | -0.096 | 0 | 0.81 | 0.32 | 0.87 | 0.99 | 0.87 | 0.82 |
| 11 | 61152630 | rs4453253 | 1.636 | 1.254 | 2.359 | -1.775 | 0.002 | -0.001 | -0.001 | 0.938 | 0.95 | 0.74 | 0.95 | 1 | 0.99 | 0.88 |
| 11 | 61165280 | rs3741265 | 0.817 | 1.339 | 2.408 | -1.738 | 0.006 | -0.052 | -0.052 | 0.444 | 0.87 | 0.74 | 0.88 | 0.98 | 0.87 | 0.85 |
| 11 | 61166306 | rs7607 | 2.028 | 1.352 | 2.428 | -1.752 | -0.012 | -0.067 | -0.067 | 0 | 0.81 | 0.3 | 0.84 | 0.97 | 0.85 | 0.81 |
| 11 | 61173336 | rs921635 | 0.808 | 1.35 | 2.438 | -1.741 | 0.006 | -0.052 | -0.052 | 0 | 0.87 | 0.75 | 0.88 | 0.98 | 0.87 | 0.85 |
| 11 | 61179285 | rs3019198 | 2.006 | 1.162 | 2.328 | -1.702 | -0.012 | -0.067 | -0.067 | 4.834 | 0.81 | 0.3 | 0.84 | 0.97 | 0.85 | 0.81 |
| 11 | 61179728 | rs2860519 | 2.022 | 1.162 | 2.328 | -1.702 | -0.012 | -0.067 | -0.067 | 0 | 0.81 | 0.3 | 0.84 | 0.97 | 0.85 | 0.81 |
| 11 | 61183388 | rs17704641 | -0.315 | 1.171 | 2.224 | -1.684 | -0.024 | 0.051 | 0.051 | 0 | 0.11 | 0 | 0.07 | 0.01 | 0.12 | 0.13 |
| 11 | 61196289 | rs2943807 | 2.285 | 0.79 | 2.768 | -1.141 | -0.011 | -0.067 | -0.067 | 0 | 0.81 | 0.3 | 0.84 | 0.97 | 0.85 | 0.81 |
| 11 | 61202912 | rs2943806 | 1.07 | 0.703 | 2.812 | -1.136 | 0.002 | -0.043 | -0.043 | 0 | 0.87 | 0.77 | 0.88 | 0.98 | 0.87 | 0.85 |
| 11 | 61213920 | rs17702 | 1.07 | 0.703 | 2.812 | -1.136 | 0.002 | -0.043 | -0.043 | 0 | 0.87 | 0.77 | 0.88 | 0.98 | 0.87 | 0.85 |
| 11 | 61213999 | rs6632 | -3.319 | 0.548 | 2.76 | -1.082 | 0.074 | 0.076 | 0.076 | 0.384 | 0.05 | 0.47 | 0.04 | 0.01 | 0.01 | 0.03 |
| 11 | 61249383 | rs3019200 | 1.63 | 0.698 | 2.474 | -1.113 | 0.041 | -0.104 | -0.104 | 0 | 0.83 | 0.5 | 0.85 | 0.97 | 0.83 | 0.84 |
| 11 | 61253200 | rs3019187 | 2.21 | 0.578 | 2.424 | -1.059 | -0.022 | -0.09 | -0.09 | 2.542 | 0.75 | 0.39 | 0.77 | 0.97 | 0.84 | 0.75 |
| 11 | 61258468 | rs2943813 | 2.333 | 0.722 | 2.406 | -1.002 | -0.004 | -0.05 | -0.05 | 0 | 0.74 | 0.28 | 0.75 | 0.97 | 0.8 | 0.75 |
| 11 | 61259751 | rs7947046 | -2.835 | 0.838 | 1.757 | -0.903 | 0.007 | -0.034 | -0.034 | 0.504 | 0.1 | 0.24 | 0.13 | 0.01 | 0.07 | 0.18 |
| 11 | 61260882 | rs2924446 | 0.743 | 0.967 | 1.797 | -0.854 | 0.002 | -0.063 | -0.063 | 0 | 0.87 | 0.82 | 0.91 | 1 | 0.87 | 0.94 |
| 11 | 61263640 | rs11230715 | -1.793 | 0.764 | 1.626 | -0.238 | 0.001 | 0.002 | 0.002 | 0 | 0.06 | 0.07 | 0.11 | 0.01 | 0.03 | 0.14 |
| 11 | 61266827 | rs11230719 | -1.798 | 0.8 | 1.629 | -0.111 | -0.001 | 0.001 | 0.001 | 0 | 0.06 | 0.07 | 0.11 | 0.05 | 0.04 | 0.14 |
| 11 | 61271195 | rs12805182 | -2.988 | 0.623 | 1.687 | -0.132 | 0.021 | -0.015 | -0.015 | 0.334 | 0.09 | 0.23 | 0.13 | 0.05 | 0.07 | 0.18 |
| 11 | 61272309 | rs11230723 | -1.712 | 0.653 | 0.879 | 0.983 | -0.006 | -0.05 | -0.05 | 0 | 0.1 | 0.02 | 0.26 | 0.43 | 0.06 | 0.27 |
| 11 | 61277885 | rs10897165 | -2.154 | 0.567 | 0.894 | 1.059 | -0.014 | -0.048 | -0.048 | 0 | 0.12 | 0.07 | 0.28 | 0.45 | 0.11 | 0.32 |
| 11 | 61278246 | rs751984 | -2.379 | 0.564 | 0.957 | 1.065 | -0.001 | 0.018 | 0.018 | 0 | 0.13 | 0.19 | 0.29 | 0.46 | 0.12 | 0.33 |
| 11 | 61282504 | rs3741260 | -1.84 | 0.653 | 0.858 | 0.768 | -0.004 | 0.002 | 0.002 | 0.348 | 0.12 | 0.1 | 0.34 | 0.44 | 0.16 | 0.33 |
| 11 | 61291077 | rs3862736 | -2.421 | -0.185 | 1.743 | 0.025 | 0.009 | -0.031 | -0.031 | 0 | 0.09 | 0.39 | 0.05 | 0 | 0.03 | 0.1 |
| 11 | 61291922 | rs10897167 | -2.546 | -0.196 | 1.739 | 0.02 | 0.005 | -0.037 | -0.037 | 0 | 0.1 | 0.45 | 0.05 | 0 | 0.03 | 0.1 |
| 11 | 61303610 | rs3018736 | -2.502 | -0.548 | 1.326 | 0.158 | -0.022 | -0.002 | -0.002 | 0 | 0.16 | 0.81 | 0.14 | 0.2 | 0.11 | 0.25 |
| 11 | 61342696 | rs887646 | -2.415 | -0.24 | 2.225 | -0.128 | 0.008 | -0.008 | -0.008 | 0 | 0.09 | 0.47 | 0.05 | 0.01 | 0.03 | 0.08 |
| 11 | 61371283 | rs174868 | 1.77 | 0.147 | 1.657 | -0.121 | 0.067 | 0.009 | 0.009 | 3.861 | 0.92 | 0.56 | 0.94 | 0.99 | 0.94 | 0.91 |
| 11 | 61376970 | rs198746 | -0.021 | -0.789 | 0.812 | -1.116 | -0.04 | -0.133 | -0.133 | 0 | 0.68 | 0.49 | 0.85 | 0.99 | 0.8 | 0.88 |
| 11 | 61379646 | rs2429862 | 0 | -0.654 | 0.876 | -1.405 | -0.021 | -0.108 | -0.108 | 0.007 | 0.66 | 0.64 | 0.81 | 0.95 | 0.74 | 0.86 |
| 11 | 61388090 | rs198750 | -2.41 | -0.51 | 1.525 | -1.033 | 0.006 | -0.02 | -0.02 | 0 | 0.13 | 0.54 | 0.13 | 0.05 | 0.12 | 0.11 |
| 11 | 61399965 | rs1791786 | 0.741 | -0.044 | 1.177 | -1.551 | 0.008 | -0.072 | -0.072 | 1.138 | 0.77 | 0.67 | 0.64 | 0.91 | 0.75 | 0.63 |
| 11 | 61400154 | rs11820769 | -1.571 | -0.164 | 1.345 | -1.469 | 0.001 | 0.01 | 0.01 | 0 | 0.06 | 0.21 | 0.04 | 0.03 | 0.05 | 0.03 |
| 11 | 61408449 | rs1812458 | 1.011 | 0.358 | 0.815 | -2.089 | 0.012 | 0.002 | 0.002 | 0 | 0.35 | 0.21 | 0.21 | 0.02 | 0.28 | 0.19 |

| | | | | | | | | | | | | | | | | |
|---|---|---|---|---|---|---|---|---|---|---|---|---|---|---|---|---|
| 11 | 61409247 | rs1692116 | -0.746 | 0.493 | 0.19 | -2.277 | -0.006 | -0.035 | -0.035 | 2.142 | 0.14 | 0.31 | 0.09 | 0.03 | 0.15 | 0.06 |
| 11 | 61413328 | rs2453712 | 0.273 | 0.45 | 0.241 | -2.269 | -0.002 | -0.059 | -0.059 | 1.475 | 0.1 | 0.22 | 0.05 | 0 | 0.1 | 0.02 |
| 11 | 61417472 | rs1692120 | 0.315 | 0.592 | 0.216 | -2.281 | 0.011 | 0.01 | 0.01 | 0 | 0.5 | 0.45 | 0.3 | 0.04 | 0.45 | 0.25 |
| 11 | 61442492 | rs7925523 | -0.111 | 1.361 | 0.108 | -0.447 | -0.002 | 0.014 | 0.014 | 0 | 0.13 | 0.1 | 0.08 | 0.25 | 0.16 | 0.11 |
| 11 | 61446425 | rs7931563 | -1.5 | 1.592 | 0.127 | -0.378 | -0.001 | -0.001 | -0.001 | 0.486 | 0.31 | 0.34 | 0.43 | 0.42 | 0.39 | 0.48 |
| 11 | 61446983 | rs872456 | 0.156 | 1.44 | 0.175 | -0.058 | 0.006 | 0.029 | 0.029 | 0 | 0.05 | 0.12 | 0.05 | 0.15 | 0.06 | 0.08 |
| 11 | 61450330 | rs11230781 | 0.87 | 1.211 | -0.236 | -0.433 | -0.048 | 0.215 | 0.215 | 1.837 | 0.51 | 0.12 | 0.51 | 0.21 | 0.52 | 0.49 |
| 11 | 61462424 | rs3931642 | -0.403 | 1.389 | -0.221 | -0.416 | -0.041 | -0.006 | -0.006 | 0 | 0.17 | 0.1 | 0.33 | 0.19 | 0.24 | 0.34 |
| 11 | 61463388 | rs198456 | -1.571 | 1.32 | 0.113 | -0.506 | 0.008 | -0.004 | -0.004 | 0 | 0.56 | 0.67 | 0.75 | 0.94 | 0.63 | 0.76 |
| 11 | 61464525 | rs17827816 | -1.32 | 1.315 | 0.083 | -0.42 | 0.01 | 0.021 | 0.021 | 0 | 0.31 | 0.41 | 0.43 | 0.44 | 0.39 | 0.48 |
| 11 | 61469473 | rs198450 | 0.26 | 1.14 | -0.045 | -0.369 | 0.001 | -0.074 | -0.074 | 0.05 | 0.63 | 0.14 | 0.58 | 0.47 | 0.67 | 0.58 |
| 11 | 61479322 | rs198437 | 0.986 | 1.174 | -0.074 | -0.425 | -0.068 | 0.218 | 0.218 | 2.704 | 0.51 | 0.07 | 0.5 | 0.21 | 0.52 | 0.49 |
| 11 | 61484981 | rs198432 | 0.588 | 1.21 | -0.034 | -0.25 | -0.002 | -0.013 | -0.013 | 0 | 0.23 | 0.23 | 0.32 | 0.47 | 0.25 | 0.27 |
| 11 | 61487690 | rs198430 | 0.621 | 1.437 | -0.05 | -0.269 | 0 | -0.012 | -0.012 | 0.478 | 0.23 | 0.23 | 0.32 | 0.46 | 0.24 | 0.28 |
| 11 | 61488968 | rs2903825 | -0.065 | 1.43 | -0.022 | -0.237 | 0 | 0.026 | 0.026 | 0 | 0.14 | 0.14 | 0.08 | 0.26 | 0.15 | 0.11 |
| 11 | 61490880 | rs9735635 | -0.065 | 1.43 | -0.022 | -0.237 | 0 | 0.026 | 0.026 | 0 | 0.14 | 0.13 | 0.08 | 0.26 | 0.15 | 0.11 |
| 11 | 61492039 | rs12285167 | -0.465 | 1.477 | -0.18 | -0.289 | -0.042 | -0.005 | -0.005 | 0 | 0.17 | 0.1 | 0.32 | 0.19 | 0.22 | 0.36 |
| 11 | 61494327 | rs4963243 | 0.517 | 1.45 | -0.186 | -0.225 | -0.042 | -0.075 | -0.075 | 0 | 0.11 | 0.02 | 0.06 | 0.26 | 0.15 | 0.12 |
| 11 | 61500487 | rs198448 | -0.485 | 1.478 | -0.144 | -0.219 | 0 | 0 | 0 | 0 | 0.37 | 0.37 | 0.42 | 0.71 | 0.39 | 0.39 |
| 11 | 61505168 | rs198444 | -0.682 | 1.413 | -0.186 | -0.191 | -0.003 | 0.018 | 0.018 | 0.03 | 0.4 | 0.36 | 0.5 | 0.8 | 0.43 | 0.41 |
| 11 | 61514085 | rs2852786 | 1.127 | 1.525 | -0.661 | -0.053 | -0.016 | -0.056 | -0.056 | 0 | 0.66 | 0.22 | 0.64 | 0.58 | 0.65 | 0.51 |
| 11 | 61516910 | rs3741255 | 0.016 | 1.7 | -0.892 | -0.015 | -0.059 | -0.016 | -0.016 | 1.433 | 0.11 | 0.06 | 0.21 | 0.14 | 0.16 | 0.31 |
| 11 | 61520668 | rs569258 | 1.345 | 1.426 | -0.786 | 0.214 | 0.023 | 0.062 | 0.062 | 0.389 | 0.39 | 0.2 | 0.22 | 0.24 | 0.34 | 0.31 |
| 11 | 61521621 | rs198464 | 0.718 | 1.36 | -0.593 | -0.058 | -0.001 | 0 | 0 | 0 | 0.54 | 0.39 | 0.3 | 0.38 | 0.49 | 0.33 |
| 11 | 61523735 | rs2238003 | -0.49 | 1.809 | -0.884 | -0.014 | -0.096 | -0.085 | -0.085 | 0.018 | 0.15 | 0.02 | 0.35 | 0.4 | 0.26 | 0.38 |
| 11 | 61524119 | rs198462 | 0.835 | 1.663 | -0.783 | -0.072 | 0.002 | 0 | 0 | 0.32 | 0.55 | 0.43 | 0.31 | 0.38 | 0.49 | 0.37 |
| 11 | 61525730 | rs198476 | 0.813 | 1.706 | -0.849 | -0.02 | 0.003 | -0.001 | -0.001 | 2.411 | 0.56 | 0.42 | 0.44 | 0.41 | 0.5 | 0.38 |
| 11 | 61526071 | rs198475 | 0.081 | 0.848 | -0.667 | -0.238 | -0.027 | -0.036 | -0.036 | 0 | 0.71 | 0.5 | 0.78 | 0.76 | 0.76 | 0.75 |
| 11 | 61534010 | rs2071213 | -1.298 | 1.324 | -0.691 | 0.258 | -0.001 | 0.001 | 0.001 | 0.485 | 0.33 | 0.48 | 0.37 | 0.58 | 0.31 | 0.65 |
| 11 | 61539691 | rs628993 | -0.403 | 1.259 | -1.2 | 0.669 | -0.028 | -0.089 | -0.089 | 0.467 | 0.12 | 0 | 0.05 | 0.37 | 0.09 | 0.43 |
| 11 | 61547237 | rs108499 | -0.779 | 2.48 | -1.921 | -0.329 | -0.12 | -0.11 | -0.11 | 0 | 0.21 | 0.02 | 0.58 | 0.56 | 0.33 | 0.12 |
| 11 | 61548559 | rs509360 | 0.575 | 1.743 | -2.726 | -1.414 | -0.008 | -0.008 | -0.008 | 0 | 0.6 | 0.08 | 0.79 | 0.57 | 0.71 | 0.24 |
| 11 | 61551356 | rs174535 | -1.29 | 1.97 | -2.012 | -1.115 | -0.049 | -0.045 | -0.045 | 0.014 | 0.26 | 0.15 | 0.6 | 0.57 | 0.35 | 0.16 |
| 11 | 61557803 | rs102275 | -2.013 | 1.526 | -1.335 | -1.358 | -0.001 | -0.001 | -0.001 | 0 | 0.34 | 0.7 | 0.65 | 0.57 | 0.36 | 0.16 |
| 11 | 61560081 | rs174538 | -0.348 | 2.075 | -1.534 | -1.168 | -0.108 | -0.109 | -0.109 | 0 | 0.21 | 0.02 | 0.57 | 0.56 | 0.31 | 0.12 |
| 11 | 61560261 | rs412334 | 1.747 | 2.074 | -1.773 | -1.421 | -0.022 | 0.02 | 0.02 | 0 | 0.06 | 0.01 | 0.08 | 0 | 0.16 | 0.01 |
| 11 | 61569830 | rs174546 | -1.108 | 1.809 | -1.352 | -1.396 | -0.116 | -0.11 | -0.11 | 3.221 | 0.24 | 0.02 | 0.59 | 0.57 | 0.35 | 0.14 |
| 11 | 61570783 | rs174547 | -1.108 | 1.809 | -1.352 | -1.396 | -0.116 | -0.11 | -0.11 | 0.044 | 0.24 | 0.02 | 0.59 | 0.57 | 0.35 | 0.14 |
| 11 | 61571478 | rs174550 | -1.108 | 1.809 | -1.352 | -1.396 | -0.116 | -0.11 | -0.11 | 0 | 0.24 | 0.02 | 0.59 | 0.57 | 0.35 | 0.14 |
| 11 | 61597212 | rs174570 | -0.594 | 1.865 | -1.305 | -1.352 | -0.039 | -0.027 | -0.027 | 3.304 | 0.08 | 0.01 | 0.49 | 0.56 | 0.16 | 0.07 |
| 11 | 61597972 | rs1535 | -1.57 | 1.749 | -1.109 | -1.424 | -0.043 | -0.041 | -0.041 | 2.322 | 0.26 | 0.11 | 0.59 | 0.57 | 0.35 | 0.14 |
| 11 | 61603237 | rs2727270 | -1.105 | 1.818 | -1.202 | -1.288 | -0.035 | -0.056 | -0.056 | 0 | 0.09 | 0.01 | 0.26 | 0.42 | 0.14 | 0.06 |
| 11 | 61603510 | rs174576 | -2.033 | 1.704 | -0.556 | -1.355 | -0.001 | -0.001 | -0.001 | 0.191 | 0.29 | 0.25 | 0.61 | 0.57 | 0.36 | 0.14 |
| 11 | 61604814 | rs174577 | -2.078 | 1.698 | -0.488 | -1.358 | 0.01 | 0.01 | 0.01 | 0.977 | 0.29 | 0.35 | 0.62 | 0.57 | 0.36 | 0.14 |
| 11 | 61609750 | rs174583 | -1.97 | 1.615 | -0.651 | -1.373 | -0.005 | -0.005 | -0.005 | 1.534 | 0.31 | 0.24 | 0.61 | 0.58 | 0.36 | 0.15 |
| 11 | 61614023 | rs17156442 | -1.406 | 1.586 | -0.295 | -1.345 | -0.007 | -0.051 | -0.051 | 0 | 0.09 | 0.27 | 0.04 | 0.01 | 0.05 | 0.04 |
| 11 | 61616012 | rs2851682 | -0.274 | 1.656 | -0.566 | -1.248 | -0.026 | -0.001 | -0.001 | 0 | 0.05 | 0 | 0.25 | 0.41 | 0.11 | 0.05 |
| 11 | 61618831 | rs174593 | -1.223 | 1.552 | -0.53 | -1.808 | -0.004 | 0.007 | 0.007 | 1.097 | 0.25 | 0.18 | 0.36 | 0.16 | 0.26 | 0.13 |
| 11 | 61624705 | rs498793 | -1.532 | 1.18 | -0.583 | -2.251 | 0.011 | 0.043 | 0.043 | 0.842 | 0.55 | 0.64 | 0.62 | 0.92 | 0.57 | 0.69 |
| 11 | 61627881 | rs174611 | -0.795 | 1.167 | -0.559 | -1.976 | -0.095 | 0.125 | 0.125 | 0.405 | 0.21 | 0.05 | 0.2 | 0.01 | 0.29 | 0.08 |
| 11 | 61629322 | rs174618 | -1.919 | 0.789 | 0.267 | -2.263 | 0 | -0.121 | -0.121 | 0 | 0.41 | 0.6 | 0.3 | 0.03 | 0.43 | 0.29 |
| 11 | 61638044 | rs7104849 | -0.037 | 0.666 | 0.212 | -2.049 | -0.01 | 0.011 | 0.011 | 0 | 0.94 | 0.92 | 0.9 | 0.99 | 0.88 | 0.97 |
| 11 | 61639573 | rs174448 | 1.314 | 0.559 | 0.435 | -1.54 | -0.001 | -0.001 | -0.001 | 0.933 | 0.65 | 0.58 | 0.38 | 0.7 | 0.62 | 0.65 |
| 11 | 61640379 | rs174449 | 1.579 | 0.523 | 0.489 | -1.562 | 0 | -0.002 | -0.002 | 1.248 | 0.63 | 0.34 | 0.37 | 0.68 | 0.62 | 0.58 |
| 11 | 61641542 | rs174450 | 1.985 | 0.566 | 0.304 | -1.423 | 0.008 | 0.075 | 0.075 | 0 | 0.52 | 0.21 | 0.31 | 0.61 | 0.53 | 0.56 |
| 11 | 61647288 | rs7942717 | 0.802 | 0.186 | 0.262 | -1.397 | -0.004 | -0.02 | -0.02 | 0.432 | 0.86 | 0.82 | 0.94 | 0.99 | 0.91 | 0.9 |
| 11 | 61655305 | rs1000778 | 1.379 | 0.211 | 0.186 | -1.332 | 0.024 | 0.022 | 0.022 | 0.232 | 0.65 | 0.22 | 0.44 | 0.69 | 0.73 | 0.61 |
| 11 | 61656117 | rs174455 | 1.696 | 0.256 | -0.309 | -1.327 | -0.028 | 0.009 | 0.009 | 0.004 | 0.59 | 0.03 | 0.33 | 0.59 | 0.61 | 0.57 |
| 11 | 61660704 | rs528285 | 0.634 | 0.65 | -0.036 | -1.059 | -0.005 | -0.017 | -0.017 | 0 | 0.69 | 0.72 | 0.47 | 0.44 | 0.65 | 0.69 |
| 11 | 61664992 | rs13966 | -1.431 | 0.945 | 0.283 | -0.539 | 0 | -0.039 | -0.039 | 0.091 | 0.46 | 0.48 | 0.64 | 0.86 | 0.52 | 0.72 |
| 11 | 61669946 | rs3815045 | -0.559 | 0.855 | -0.535 | -0.131 | 0.011 | -0.006 | -0.006 | 0.382 | 0.05 | 0 | 0.1 | 0.33 | 0.03 | 0.3 |
| 11 | 61671956 | rs174472 | -0.883 | 1.015 | -0.366 | -0.903 | -0.014 | 0.087 | 0.087 | 0 | 0.4 | 0.25 | 0.59 | 0.85 | 0.42 | 0.69 |
| 11 | 61687454 | rs11230827 | -1.24 | 0.458 | 0.285 | 0.108 | -0.023 | 0.046 | 0.046 | 0 | 0.08 | 0.23 | 0.11 | 0.16 | 0.03 | 0.4 |
| 11 | 61705521 | rs2736601 | -1.322 | 0.944 | -0.051 | 0.644 | 0.007 | 0.111 | 0.111 | 0 | 0.05 | 0.07 | 0.21 | 0.52 | 0.07 | 0.1 |
| 11 | 61710822 | rs2727269 | 0.4 | 0.429 | -0.26 | -0.4 | 0.006 | -0.064 | -0.064 | 0 | 0.42 | 0.38 | 0.33 | 0.16 | 0.5 | 0.12 |
| 11 | 61711475 | rs2521572 | 0.806 | 0.214 | 0.007 | 0.562 | -0.002 | 0.1 | 0.1 | 0 | 0.94 | 0.86 | 0.81 | 0.49 | 0.96 | 0.88 |
| 11 | 61722645 | rs1109748 | -1.379 | -0.116 | -0.004 | 0.455 | -0.003 | 0.007 | 0.007 | 0 | 0.14 | 0.08 | 0.28 | 0.65 | 0.07 | 0.57 |
| 11 | 61727791 | rs195157 | 1.159 | 0.188 | 0.503 | 0.068 | 0.055 | -0.014 | -0.014 | 0 | 0.89 | 0.49 | 0.78 | 0.97 | 0.9 | 0.97 |
| 11 | 61731233 | rs2668897 | -1.474 | 0.255 | 0.64 | 0.548 | 0.015 | 0.135 | 0.135 | 0 | 0.08 | 0.22 | 0.23 | 0.65 | 0.09 | 0.14 |

| | | | | | | | | | | | | | | | | |
|---|---|---|---|---|---|---|---|---|---|---|---|---|---|---|---|---|
| 11 | 61737244 | rs3758977 | -1.02 | -0.237 | 0.459 | -0.414 | -0.012 | -0.066 | -0.066 | 0.021 | 0.4 | 0.52 | 0.49 | 0.88 | 0.35 | 0.71 |
| 11 | 61744342 | rs195445 | -1.015 | -0.161 | 0.265 | -1.008 | 0.01 | -0.035 | -0.035 | 0 | 0.72 | 0.68 | 0.83 | 0.88 | 0.71 | 0.78 |
| 11 | 61745694 | rs17185574 | 0.64 | -0.318 | -0.081 | -0.942 | -0.051 | 0.159 | 0.159 | 0 | 0.23 | 0.01 | 0.13 | 0 | 0.23 | 0.18 |
| 11 | 61746269 | rs10897193 | -1.065 | -0.319 | 0.298 | -0.493 | 0.005 | 0.096 | 0.096 | 0.137 | 0.09 | 0.18 | 0.24 | 0.63 | 0.09 | 0.15 |
| 11 | 61759714 | rs7115826 | -1.283 | -0.325 | 0.72 | -0.467 | 0.112 | 0.215 | 0.215 | 1.444 | 0.06 | 0.54 | 0.11 | 0.13 | 0.06 | 0.02 |
| 11 | 61761552 | rs10792322 | 0.934 | -0.464 | 0.45 | -0.777 | -0.015 | -0.071 | -0.071 | 0.938 | 0.3 | 0.11 | 0.31 | 0.52 | 0.33 | 0.15 |
| 11 | 61766174 | rs12575056 | -0.852 | -0.471 | 0.638 | -0.713 | -0.014 | 0.109 | 0.109 | 0 | 0.13 | 0.3 | 0.24 | 0.35 | 0.11 | 0.43 |
| 11 | 61768706 | rs4963442 | 0.339 | -0.485 | 0.374 | -0.933 | 0.001 | 0.004 | 0.004 | 0 | 0.6 | 0.37 | 0.59 | 0.65 | 0.63 | 0.39 |
| 11 | 61770839 | rs11605994 | 1.039 | -0.365 | 0.104 | -1.102 | -0.033 | 0.083 | 0.083 | 0 | 0.21 | 0.06 | 0.12 | 0.04 | 0.23 | 0.14 |
| 11 | 61781117 | rs7935835 | 1.22 | -0.379 | 0.146 | -0.836 | -0.052 | 0.058 | 0.058 | 0 | 0.28 | 0.12 | 0.24 | 0.15 | 0.37 | 0.06 |
| 11 | 61783433 | rs17156751 | 0.862 | -0.489 | 0.126 | -0.668 | 0.018 | -0.003 | -0.003 | 0 | 0.06 | 0.03 | 0.14 | 0.26 | 0.04 | 0.39 |
| 11 | 61785538 | rs3855281 | -0.524 | -0.585 | 0.449 | -0.776 | 0.003 | 0.01 | 0.01 | 0 | 0.61 | 0.42 | 0.57 | 0.58 | 0.56 | 0.55 |
| 11 | 61789317 | rs10897207 | 1.241 | -0.538 | 0.062 | -0.783 | -0.063 | 0.062 | 0.062 | 0 | 0.29 | 0.1 | 0.25 | 0.15 | 0.39 | 0.06 |
| 11 | 61789941 | rs10792324 | 0.666 | -0.323 | 0.251 | -0.754 | -0.001 | -0.001 | -0.001 | 0 | 0.25 | 0.25 | 0.26 | 0.49 | 0.25 | 0.24 |
| 11 | 61790662 | rs10501388 | 0.676 | -0.294 | 0.093 | -0.75 | 0.003 | 0.008 | 0.008 | 0 | 0.38 | 0.47 | 0.41 | 0.42 | 0.44 | 0.45 |
| 11 | 61793535 | rs17706767 | 0.351 | -0.185 | -0.593 | -0.792 | -0.081 | 0.182 | 0.182 | 0 | 0.31 | 0.03 | 0.19 | 0.08 | 0.31 | 0.3 |
| 11 | 61795055 | rs10792327 | 0.839 | -0.163 | -0.511 | -0.695 | -0.028 | -0.045 | -0.045 | 0 | 0.37 | 0.13 | 0.38 | 0.43 | 0.43 | 0.46 |
| 11 | 61800812 | rs11230882 | -0.028 | -0.025 | -0.571 | -0.61 | -0.002 | 0.002 | 0.002 | 0.053 | 0.67 | 0.33 | 0.5 | 0.32 | 0.65 | 0.74 |
| 11 | 61801006 | rs12786494 | 0.038 | 0.149 | -1.227 | -0.529 | -0.079 | 0.165 | 0.165 | 0 | 0.32 | 0.04 | 0.19 | 0.12 | 0.31 | 0.31 |
| 11 | 61808092 | rs3019624 | 0.312 | 0.078 | -1.041 | -0.426 | -0.03 | 0.101 | 0.101 | 0.05 | 0.23 | 0.11 | 0.13 | 0.06 | 0.21 | 0.28 |
| 11 | 61808272 | rs3019626 | -0.693 | 0.23 | -1.029 | -0.136 | -0.038 | -0.127 | -0.127 | 0 | 0.77 | 0.94 | 0.87 | 0.56 | 0.77 | 0.78 |
| 11 | 61809207 | rs522090 | -0.646 | 0.251 | -0.18 | -0.272 | 0.054 | 0.215 | 0.215 | 0 | 0.09 | 0.33 | 0.21 | 0.27 | 0.11 | 0.04 |
| 11 | 61809694 | rs12277563 | -0.245 | -0.055 | -0.607 | -0.134 | -0.003 | -0.003 | -0.003 | 0 | 0.52 | 0.51 | 0.46 | 0.38 | 0.53 | 0.45 |
| 11 | 61815803 | rs4963452 | -0.685 | -0.294 | -1.087 | 0.014 | 0.021 | -0.052 | -0.052 | 0 | 0.21 | 0.15 | 0.47 | 0.43 | 0.16 | 0.35 |
| 11 | 61817200 | rs259885 | -0.879 | 0.176 | -0.997 | -0.476 | -0.012 | -0.02 | -0.02 | 0 | 0.66 | 0.71 | 0.77 | 0.55 | 0.56 | 0.77 |
| 11 | 61823386 | rs11230889 | -0.076 | 0.118 | -1.457 | -0.553 | 0.021 | 0.017 | 0.017 | 0 | 0.24 | 0.13 | 0.44 | 0.21 | 0.17 | 0.34 |
| 11 | 61827748 | rs10897213 | -1.101 | 0.292 | -0.859 | -0.839 | 0.01 | 0.022 | 0.022 | 1.279 | 0.63 | 0.5 | 0.75 | 0.57 | 0.55 | 0.63 |
| 11 | 61829542 | rs12793785 | 0.935 | 0.39 | -0.784 | -1.13 | 0.001 | -0.027 | -0.027 | 0.786 | 0.13 | 0.12 | 0.14 | 0 | 0.17 | 0.17 |
| 11 | 61829795 | rs449397 | -1.474 | 0.396 | -1.017 | -1.103 | 0.016 | 0.03 | 0.03 | 0.664 | 0.64 | 0.56 | 0.76 | 0.57 | 0.56 | 0.63 |
| 11 | 61830850 | rs11607063 | 0.429 | 0.362 | -1.379 | -1.694 | -0.024 | 0.069 | 0.069 | 1.826 | 0.22 | 0.01 | 0.12 | 0.13 | 0.19 | 0.1 |
| 11 | 61835423 | rs2926520 | 0.217 | 0.385 | -0.926 | -1.482 | 0.007 | -0.03 | -0.03 | 0 | 0.94 | 0.81 | 0.95 | 1 | 0.95 | 0.98 |
| 11 | 61840413 | rs12802587 | 0.295 | 0.631 | -1.354 | -1.278 | -0.038 | 0.042 | 0.042 | 0.195 | 0.13 | 0.05 | 0.14 | 0 | 0.17 | 0.17 |
| 11 | 61869513 | rs11603446 | -0.112 | -0.129 | -1.492 | -0.73 | 0.025 | 0.056 | 0.056 | 1.488 | 0.23 | 0.03 | 0.12 | 0.14 | 0.17 | 0.07 |
| 11 | 61875611 | rs259871 | 0.544 | 0.224 | -0.891 | -0.629 | 0.002 | -0.026 | -0.026 | 0 | 0.92 | 0.87 | 0.81 | 0.98 | 0.93 | 0.91 |
| 11 | 61878515 | rs7934431 | -0.402 | 0.56 | -1.365 | -1.538 | -0.023 | 0.128 | 0.128 | 0.05 | 0.4 | 0.23 | 0.4 | 0.13 | 0.39 | 0.24 |
| 11 | 61879689 | rs11230906 | 1.339 | 0.529 | -1.981 | -2.389 | -0.032 | 0.093 | 0.093 | 1.559 | 0.18 | 0.02 | 0.11 | 0 | 0.26 | 0.04 |
| 11 | 61884849 | rs11606620 | 0.151 | 0.564 | -1.949 | -2.344 | -0.025 | 0.137 | 0.137 | 0.05 | 0.4 | 0.23 | 0.4 | 0.11 | 0.39 | 0.24 |
| 11 | 61885073 | rs2264278 | -0.389 | 0.414 | -0.953 | -1.103 | -0.01 | 0.204 | 0.204 | 0.179 | 0.9 | 0.8 | 0.91 | 0.49 | 0.95 | 0.74 |
| 11 | 61890812 | rs17707648 | 0.672 | 0.475 | -1.061 | -1.133 | -0.001 | -0.001 | -0.001 | 0 | 0.05 | 0.04 | 0.18 | 0.01 | 0.05 | 0.06 |
| 11 | 61892667 | rs11230916 | 0.393 | 0.221 | -1.237 | -1.227 | -0.083 | -0.131 | -0.131 | 0 | 0.28 | 0.02 | 0.35 | 0.38 | 0.27 | 0.44 |
| 11 | 61893356 | rs1675129 | 0.36 | 0.205 | -0.316 | -1.295 | -0.002 | -0.08 | -0.08 | 0 | 0.58 | 0.53 | 0.38 | 0.1 | 0.67 | 0.25 |
| 11 | 61894482 | rs1628349 | 0.302 | 0.046 | 0.253 | -0.909 | -0.008 | 0.29 | 0.29 | 0.168 | 0.88 | 0.66 | 0.75 | 0.48 | 0.94 | 0.69 |
| 11 | 61906374 | rs1675126 | -0.247 | 0.204 | 0.126 | -0.904 | 0 | 0.088 | 0.088 | 0.504 | 0.92 | 0.91 | 0.9 | 0.49 | 0.91 | 0.72 |
| 11 | 61906864 | rs3781974 | -0.182 | 0.352 | 0.24 | -1.238 | 0.001 | -0.01 | -0.01 | 0 | 0.32 | 0.22 | 0.39 | 0.36 | 0.27 | 0.45 |
| 11 | 61907969 | rs12795611 | 0.066 | 0.358 | 0.292 | -1.178 | 0.002 | -0.014 | -0.014 | 0 | 0.28 | 0.19 | 0.36 | 0.35 | 0.22 | 0.43 |
| 11 | 61908672 | rs1675123 | 1.15 | 0.352 | 0.867 | -1.053 | 0.008 | 0.194 | 0.194 | 0.288 | 0.81 | 0.52 | 0.72 | 0.49 | 0.85 | 0.63 |
| 11 | 61916802 | rs1792886 | 0.294 | 0.163 | 0.841 | -0.945 | 0 | 0.027 | 0.027 | 0 | 0.86 | 0.86 | 0.89 | 0.49 | 0.9 | 0.7 |
| 11 | 61917002 | rs1792888 | 0.71 | 0.138 | 1.041 | -0.959 | -0.024 | 0.25 | 0.25 | 0.002 | 0.83 | 0.59 | 0.86 | 0.49 | 0.89 | 0.71 |
| 11 | 61940106 | rs1675090 | -0.135 | -0.04 | 0.172 | -1.45 | 0.021 | 0.021 | 0.021 | 0.09 | 0.5 | 0.66 | 0.55 | 0.43 | 0.5 | 0.56 |
| 11 | 61945388 | rs10897222 | 0.855 | -0.086 | 0.061 | -1.648 | 0.015 | 0.159 | 0.159 | 0.442 | 0.35 | 0.07 | 0.35 | 0.06 | 0.4 | 0.17 |
| 11 | 61947706 | rs12789264 | 0.969 | 0.138 | 0.115 | -1.134 | -0.021 | 0.036 | 0.036 | 0 | 0.26 | 0.15 | 0.16 | 0.18 | 0.25 | 0.15 |
| 11 | 61957621 | rs2232937 | 1.066 | 0.073 | 0.196 | -1.25 | 0.007 | 0.109 | 0.109 | 0.442 | 0.34 | 0.09 | 0.35 | 0.05 | 0.4 | 0.16 |
| 11 | 61958043 | rs12800538 | 0.911 | 0.034 | 0.19 | -1.124 | -0.017 | 0.033 | 0.033 | 0 | 0.26 | 0.16 | 0.16 | 0.18 | 0.25 | 0.17 |
| 11 | 61960809 | rs7951859 | -0.592 | 0.063 | 0.396 | -1.127 | -0.002 | -0.007 | -0.007 | 0.045 | 0.69 | 0.33 | 0.65 | 0.8 | 0.75 | 0.75 |
| 11 | 61961614 | rs12293403 | 0.532 | 0.029 | 0.191 | -0.79 | 0.002 | 0.013 | 0.013 | 0 | 0.11 | 0.09 | 0.07 | 0.4 | 0.09 | 0.32 |
| 11 | 61963632 | rs1792876 | -0.555 | 0.153 | 0.528 | -0.909 | 0.019 | 0.057 | 0.057 | 0 | 0.73 | 0.52 | 0.62 | 0.64 | 0.73 | 0.66 |
| 11 | 61969430 | rs1729374 | -0.561 | 0.19 | 0.445 | -0.871 | 0.007 | 0.025 | 0.025 | 0 | 0.73 | 0.61 | 0.63 | 0.64 | 0.73 | 0.66 |
| 11 | 61971537 | rs2463838 | -0.29 | 0.166 | 0.523 | -0.667 | -0.003 | 0 | 0 | 2.112 | 0.4 | 0.53 | 0.42 | 0.6 | 0.34 | 0.54 |
| 11 | 61988127 | rs1729388 | 0.269 | 0.31 | 0.133 | -1.278 | 0.002 | -0.001 | -0.001 | 0 | 0.44 | 0.34 | 0.43 | 0.13 | 0.47 | 0.24 |
| 11 | 61999746 | rs10736719 | 0.474 | 0.29 | -0.358 | -1.275 | -0.046 | 0.063 | 0.063 | 0 | 0.36 | 0.09 | 0.42 | 0.71 | 0.34 | 0.63 |
| 11 | 62003572 | rs1729404 | -0.137 | 0.164 | 0.485 | -1.401 | 0.02 | -0.015 | -0.015 | 0 | 0.78 | 0.54 | 0.67 | 0.81 | 0.76 | 0.79 |
| 11 | 62010863 | rs2232950 | 0.153 | 0.191 | 0.596 | -1.336 | 0.003 | 0.061 | 0.061 | 0 | 0.39 | 0.23 | 0.43 | 0.79 | 0.34 | 0.63 |
| 11 | 62017553 | rs1528726 | 0.375 | 0.187 | 0.203 | -1.172 | -0.019 | -0.019 | -0.019 | 0 | 0.13 | 0.11 | 0.27 | 0.2 | 0.18 | 0.2 |
| 11 | 62017763 | rs1528727 | -0.601 | 0.413 | 0.4 | -1.423 | 0.001 | -0.026 | -0.026 | 0.029 | 0.57 | 0.51 | 0.51 | 0.28 | 0.55 | 0.36 |
| 11 | 62018940 | rs2511345 | -0.002 | 0.444 | 0.403 | -1.282 | 0.006 | 0.016 | 0.016 | 0 | 0.15 | 0.24 | 0.27 | 0.27 | 0.18 | 0.2 |
| 11 | 62023301 | rs10501333 | 0.3 | 0.449 | 0.316 | -1.246 | -0.001 | 0.002 | 0.002 | 0 | 0.41 | 0.34 | 0.47 | 0.71 | 0.45 | 0.63 |
| 11 | 62028465 | rs11231002 | 1.417 | 0.789 | -0.981 | 0.001 | -0.063 | -0.008 | -0.008 | 0 | 0.38 | 0.03 | 0.29 | 0.32 | 0.36 | 0.48 |
| 11 | 62032425 | rs11602047 | -0.337 | 0.838 | -1.064 | -0.046 | -0.035 | 0.106 | 0.106 | 0 | 0.32 | 0.06 | 0.17 | 0.17 | 0.28 | 0.22 |
| 11 | 62032990 | rs17157079 | 0.931 | 0.826 | -1.183 | -0.076 | -0.081 | -0.158 | -0.158 | 0 | 0.22 | 0.04 | 0.3 | 0.4 | 0.31 | 0.29 |

| | | | | | | | | | | | | | | | | |
|---|---|---|---|---|---|---|---|---|---|---|---|---|---|---|---|---|
| 11 | 62035677 | rs12800927 | 0.372 | 0.778 | -0.999 | -0.235 | -0.038 | -0.003 | -0.003 | 0 | 0.15 | 0.01 | 0.1 | 0.14 | 0.16 | 0.13 |
| 11 | 62037583 | rs11603383 | -0.656 | 0.761 | -1.014 | -0.254 | -0.065 | 0.109 | 0.109 | 0 | 0.3 | 0.03 | 0.16 | 0.14 | 0.28 | 0.18 |
| 11 | 62040519 | rs17709552 | 1.324 | 0.77 | -0.696 | -0.331 | -0.029 | 0.085 | 0.085 | 0 | 0.15 | 0.01 | 0.06 | 0 | 0.12 | 0.05 |
| 11 | 62043929 | rs7925016 | -0.098 | 0.704 | -0.482 | -0.074 | 0 | 0 | 0 | 0 | 0.81 | 0.82 | 0.72 | 0.76 | 0.81 | 0.88 |
| 11 | 62053687 | rs10750972 | -0.112 | 0.6 | -0.216 | 0.005 | -0.006 | -0.011 | -0.011 | 1.2 | 0.78 | 0.38 | 0.72 | 0.86 | 0.84 | 0.93 |
| 11 | 62061349 | rs11231017 | 1.576 | 0.85 | -0.557 | 0.18 | -0.06 | -0.004 | -0.004 | 0 | 0.36 | 0.07 | 0.29 | 0.34 | 0.36 | 0.41 |
| 11 | 62062509 | rs11231018 | 1.57 | 0.864 | -0.514 | 0.192 | -0.054 | 0 | 0 | 0.024 | 0.36 | 0.08 | 0.29 | 0.34 | 0.36 | 0.41 |
| 11 | 62064715 | rs954237 | 0.644 | 0.684 | 0.184 | 0.392 | -0.001 | -0.009 | -0.009 | 0 | 0.63 | 0.56 | 0.78 | 0.8 | 0.67 | 0.8 |
| 11 | 62068526 | rs17157147 | 0.063 | 2.174 | 0.905 | 0.863 | -0.019 | -0.02 | -0.02 | 0 | 0.22 | 0.17 | 0.3 | 0.3 | 0.3 | 0.37 |
| 11 | 62068685 | rs1406381 | 0.873 | 2.196 | 0.953 | 0.915 | -0.011 | 0.077 | 0.077 | 0 | 0.83 | 0.39 | 0.72 | 0.7 | 0.82 | 0.86 |
| 11 | 62069365 | rs2178260 | 1.835 | 2.123 | 0.577 | 0.826 | -0.013 | 0.021 | 0.021 | 0.118 | 0.44 | 0.12 | 0.33 | 0.4 | 0.41 | 0.44 |
| 11 | 62080042 | rs11231027 | 1.812 | 2.104 | 0.547 | 0.826 | -0.017 | 0.017 | 0.017 | 0.08 | 0.44 | 0.1 | 0.33 | 0.4 | 0.4 | 0.44 |
| 11 | 62081181 | rs1528730 | -0.908 | 2.081 | 0.744 | 0.748 | -0.002 | -0.04 | -0.04 | 0.12 | 0.83 | 0.8 | 0.9 | 1 | 0.88 | 0.95 |
| 11 | 62083908 | rs2463833 | 0.641 | 2.172 | 0.861 | 0.836 | 0.001 | -0.003 | -0.003 | 0 | 0.59 | 0.45 | 0.6 | 0.69 | 0.59 | 0.57 |
| 11 | 62092267 | rs2513081 | -1.039 | 2.167 | 0.921 | 1.099 | -0.009 | 0.097 | 0.097 | 0 | 0.16 | 0.35 | 0.28 | 0.36 | 0.19 | 0.14 |
| 11 | 62096355 | rs7941992 | 2.015 | 2.172 | 0.914 | 1.097 | -0.025 | 0.013 | 0.013 | 0 | 0.44 | 0.09 | 0.33 | 0.4 | 0.41 | 0.44 |
| 11 | 62098735 | rs2513747 | -0.864 | 2.171 | 1.035 | 1.048 | -0.017 | -0.091 | -0.091 | 0 | 0.82 | 0.71 | 0.89 | 1 | 0.88 | 0.95 |
| 11 | 62103420 | rs2463822 | -1.158 | 2.166 | 1.06 | 1.045 | 0.006 | -0.002 | -0.002 | 0 | 0.84 | 0.87 | 0.91 | 1 | 0.88 | 0.95 |
| 11 | 62104937 | rs2246920 | -0.779 | 2.169 | 1.051 | 1.048 | -0.02 | -0.082 | -0.082 | 0.27 | 0.82 | 0.7 | 0.89 | 0.99 | 0.88 | 0.95 |
| 11 | 62105391 | rs2513749 | -0.834 | 2.163 | 1.048 | 1.045 | -0.018 | -0.081 | -0.081 | 0.021 | 0.82 | 0.7 | 0.89 | 0.99 | 0.88 | 0.95 |
| 11 | 62105441 | rs3741245 | 0.099 | 2.161 | 0.958 | 1.077 | 0.011 | 0.004 | 0.004 | 0 | 0.22 | 0.25 | 0.28 | 0.23 | 0.29 | 0.37 |
| 11 | 62117709 | rs10897261 | -0.021 | 2.159 | 0.966 | 1.077 | 0.02 | 0.01 | 0.01 | 0 | 0.23 | 0.28 | 0.28 | 0.24 | 0.29 | 0.37 |
| 11 | 62139467 | rs11824690 | -0.021 | 2.159 | 0.964 | 1.077 | 0.02 | 0.01 | 0.01 | 0 | 0.23 | 0.28 | 0.28 | 0.24 | 0.29 | 0.37 |
| 11 | 62154161 | rs11605965 | 0.053 | 2.131 | 0.843 | 1.106 | -0.006 | -0.044 | -0.044 | 0.299 | 0.1 | 0.01 | 0.08 | 0.16 | 0.12 | 0.08 |
| 11 | 62157418 | rs10897267 | 2.063 | 2.177 | 0.875 | 1.03 | -0.022 | 0.012 | 0.012 | 0 | 0.44 | 0.09 | 0.34 | 0.39 | 0.41 | 0.44 |
| 11 | 62158068 | rs2302360 | 0.729 | 2.181 | 0.999 | 1.066 | -0.01 | 0.071 | 0.071 | 0 | 0.84 | 0.75 | 0.73 | 0.64 | 0.81 | 0.86 |
| 11 | 62158924 | rs2958532 | -0.208 | 2.213 | 0.989 | 0.974 | 0.01 | 0.014 | 0.014 | 0 | 0.77 | 0.77 | 0.58 | 0.63 | 0.7 | 0.62 |
| 11 | 62159926 | rs4281490 | 0.151 | 2.287 | 1.024 | 0.926 | 0.01 | 0.003 | 0.003 | 0 | 0.22 | 0.25 | 0.27 | 0.23 | 0.29 | 0.37 |
| 11 | 62160729 | rs1043789 | -0.07 | 2.442 | 1.042 | 0.9 | 0.035 | 0.017 | 0.017 | 0 | 0.23 | 0.38 | 0.3 | 0.26 | 0.31 | 0.37 |
| 11 | 62161324 | rs11231073 | -0.983 | 2.399 | 0.984 | 0.853 | -0.005 | 0.006 | 0.006 | 0 | 0.41 | 0.65 | 0.4 | 0.26 | 0.43 | 0.43 |
| 11 | 62166172 | rs7118835 | 1.308 | 2.47 | 0.991 | 0.932 | -0.011 | 0.006 | 0.006 | 0 | 0.6 | 0.33 | 0.42 | 0.37 | 0.49 | 0.43 |
| 11 | 62168736 | rs4334021 | 1.45 | 2.291 | 0.921 | 1.001 | -0.004 | 0.035 | 0.035 | 0 | 0.81 | 0.31 | 0.68 | 0.62 | 0.78 | 0.86 |
| 11 | 62172631 | rs2513064 | 1.382 | 2.066 | 0.903 | 0.768 | 0.025 | 0.032 | 0.032 | 2.818 | 0.48 | 0.57 | 0.39 | 0.38 | 0.38 | 0.39 |
| 11 | 62175013 | rs3017101 | 1.677 | 2.412 | 0.466 | 0.661 | 0.022 | -0.007 | -0.007 | 0 | 0.39 | 0.07 | 0.32 | 0.39 | 0.33 | 0.47 |
| 11 | 62179371 | rs2509973 | -1.278 | 2.476 | 0.736 | 0.703 | 0.001 | 0.001 | 0.001 | 0 | 0.43 | 0.57 | 0.42 | 0.46 | 0.55 | 0.48 |
| 11 | 62185598 | rs4488202 | -0.515 | 2.977 | 0.987 | 0.705 | 0.04 | 0.055 | 0.055 | 0 | 0.14 | 0.19 | 0.18 | 0.26 | 0.24 | 0.39 |
| 11 | 62186542 | rs3741240 | -1.447 | 2.932 | 0.961 | 0.667 | 0.008 | 0.01 | 0.01 | 1.111 | 0.23 | 0.22 | 0.24 | 0.36 | 0.34 | 0.51 |
| 11 | 62187186 | rs10897271 | -0.062 | 2.942 | 0.729 | 0.411 | -0.001 | 0.016 | 0.016 | 0.398 | 0.33 | 0.25 | 0.33 | 0.21 | 0.36 | 0.12 |
| 11 | 62192931 | rs2509963 | -2.008 | 2.474 | 0.673 | 0.446 | 0.043 | -0.068 | -0.068 | 1.383 | 0.63 | 0.93 | 0.63 | 0.55 | 0.72 | 0.63 |
| 11 | 62195344 | rs17145874 | -0.071 | 2.387 | 1.063 | 0.33 | 0.002 | -0.025 | -0.025 | 0 | 0.35 | 0.4 | 0.35 | 0.21 | 0.39 | 0.12 |
| 11 | 62200176 | rs17145884 | -0.497 | 2.256 | 1.156 | 0.444 | 0.015 | 0.035 | 0.035 | 0 | 0.15 | 0.29 | 0.18 | 0.24 | 0.18 | 0.4 |
| 11 | 62200369 | rs10897274 | -0.021 | 2.308 | 0.911 | 0.412 | -0.009 | 0.022 | 0.022 | 0.187 | 0.35 | 0.27 | 0.36 | 0.21 | 0.43 | 0.13 |
| 11 | 62200846 | rs3017106 | -1.385 | 0.875 | 0.847 | -0.15 | 0.057 | -0.048 | -0.048 | 0 | 0.6 | 0.93 | 0.45 | 0.41 | 0.69 | 0.64 |
| 11 | 62201131 | rs2259606 | -0.681 | 0.897 | 0.962 | -0.386 | 0.002 | -0.015 | -0.015 | 0 | 0.13 | 0.33 | 0.08 | 0.1 | 0.13 | 0.16 |
| 11 | 62201458 | rs4084164 | -0.921 | 1.285 | 0.343 | -0.279 | 0.045 | 0.098 | 0.098 | 0.238 | 0.49 | 0.27 | 0.38 | 0.24 | 0.62 | 0.49 |
| 11 | 62204613 | rs2509979 | 0.413 | 1.971 | 1.09 | 0.16 | 0.049 | 0.029 | 0.029 | 0 | 0.77 | 0.7 | 0.77 | 0.71 | 0.52 | 0.71 |
| 11 | 62204783 | rs2509978 | -0.631 | 1.806 | 1.406 | -0.085 | 0.008 | -0.092 | -0.092 | 0.614 | 0.22 | 0.41 | 0.18 | 0.05 | 0.18 | 0.08 |
| 11 | 62206288 | rs9645690 | 1.46 | 1.898 | 0.831 | -0.085 | 0.089 | 0.021 | 0.021 | 0 | 0.54 | 0.18 | 0.57 | 0.66 | 0.3 | 0.62 |
| 11 | 62209534 | rs10897276 | 0.402 | 1.912 | 1.023 | -0.045 | 0.05 | 0.022 | 0.022 | 0 | 0.75 | 0.41 | 0.73 | 0.71 | 0.51 | 0.71 |
| 11 | 62212013 | rs7942143 | -0.455 | 1.365 | 0.339 | -0.026 | -0.012 | 0.045 | 0.045 | 0 | 0.2 | 0.1 | 0.15 | 0.05 | 0.19 | 0.07 |
| 11 | 62213758 | rs7118247 | -0.442 | 1.396 | 0.139 | -0.011 | -0.005 | 0.017 | 0.017 | 0 | 0.2 | 0.17 | 0.16 | 0.05 | 0.18 | 0.07 |
| 11 | 62215857 | rs10897280 | 0.728 | 0.241 | -0.008 | 0.315 | 0.012 | -0.028 | -0.028 | 0 | 0.55 | 0.12 | 0.59 | 0.64 | 0.37 | 0.65 |
| 11 | 62219776 | rs2509984 | -1.634 | -0.008 | 1.423 | 0.32 | 0.107 | 0.113 | 0.113 | 0 | 0.07 | 0.48 | 0.11 | 0.09 | 0.1 | 0.02 |
| 11 | 62230163 | rs2513058 | -1.318 | -0.542 | 1.214 | 0.003 | -0.009 | -0.048 | -0.048 | 0 | 0.1 | 0.5 | 0.06 | 0 | 0.03 | 0.04 |
| 11 | 62238245 | rs2507807 | 0.066 | -1.258 | 0.562 | 0.318 | 0.051 | -0.068 | -0.068 | 0 | 0.88 | 0.99 | 0.98 | 0.74 | 0.98 | 0.98 |
| 11 | 62238502 | rs2513041 | -1.556 | -1.426 | 0.881 | 0.098 | -0.039 | -0.065 | -0.065 | 0 | 0.13 | 0.73 | 0.07 | 0 | 0.03 | 0.06 |
| 11 | 62244243 | rs2507809 | 0.037 | -1.437 | 0.584 | 0.269 | 0.03 | -0.029 | -0.029 | 0.202 | 0.88 | 0.96 | 0.98 | 0.73 | 0.97 | 0.98 |
| 11 | 62246596 | rs1293035 | 1.206 | -1.47 | 0.576 | 0.285 | 0.036 | -0.001 | -0.001 | 0 | 0.75 | 0.19 | 0.89 | 0.73 | 0.95 | 0.93 |
| 11 | 62255354 | rs17157338 | -1.816 | -0.989 | 1.439 | -0.115 | -0.013 | -0.062 | -0.062 | 0 | 0.12 | 0.47 | 0.07 | 0.02 | 0.04 | 0.03 |
| 11 | 62269774 | rs11231121 | -1.622 | -0.891 | 1.544 | -0.071 | 0.036 | -0.008 | -0.008 | 0 | 0.08 | 0.38 | 0.06 | 0.02 | 0.03 | 0.01 |
| 11 | 62299945 | rs11231130 | -1.614 | -0.88 | 1.413 | -0.088 | -0.004 | -0.044 | -0.044 | 0 | 0.08 | 0.25 | 0.04 | 0.01 | 0.03 | 0.02 |
| 11 | 62312318 | rs2509982 | -0.75 | -0.341 | 0.489 | -0.941 | -0.006 | 0.011 | 0.011 | 0 | 0.28 | 0.25 | 0.22 | 0.05 | 0.32 | 0.23 |
| 11 | 62326265 | rs11231141 | -0.782 | -0.606 | 0.709 | -1.128 | -0.015 | 0.074 | 0.074 | 0 | 0.28 | 0.11 | 0.21 | 0.04 | 0.32 | 0.21 |
| 11 | 62334096 | rs7124057 | -0.706 | -0.607 | 0.708 | -1.128 | -0.022 | 0.089 | 0.089 | 0 | 0.28 | 0.09 | 0.2 | 0.04 | 0.32 | 0.21 |
| 11 | 62350248 | rs12225213 | -0.672 | 0.211 | 0.439 | -1.424 | -0.028 | 0.093 | 0.093 | 0 | 0.28 | 0.08 | 0.2 | 0.04 | 0.32 | 0.21 |
| 11 | 62362293 | rs11231154 | -0.779 | 0.138 | 0.411 | -1.456 | -0.023 | 0.1 | 0.1 | 0 | 0.29 | 0.08 | 0.2 | 0.04 | 0.32 | 0.21 |
| 11 | 62365619 | rs11231155 | -0.835 | 0.096 | 0.398 | -1.418 | -0.007 | 0.058 | 0.058 | 0 | 0.3 | 0.16 | 0.21 | 0.05 | 0.32 | 0.23 |
| 11 | 62366454 | rs11231156 | -0.715 | 0.097 | 0.395 | -1.417 | -0.024 | 0.097 | 0.097 | 0 | 0.28 | 0.08 | 0.2 | 0.04 | 0.32 | 0.21 |
| 11 | 62366986 | rs2849030 | 0.571 | 0.009 | 0.768 | -1.46 | 0.001 | -0.095 | -0.095 | 0 | 0.68 | 0.59 | 0.76 | 0.95 | 0.67 | 0.77 |

| | | | | | | | | | | | | | | | | |
|---|---|---|---|---|---|---|---|---|---|---|---|---|---|---|---|---|
| 11 | 62371773 | rs2516633 | 0.694 | 0.2 | 0.714 | -1.509 | 0 | -0.011 | -0.011 | 0 | 0.7 | 0.7 | 0.8 | 0.95 | 0.72 | 0.78 |
| 11 | 62378801 | rs11553576 | -1.238 | -0.125 | 0.555 | -1.55 | 0.003 | 0.049 | 0.049 | 0 | 0.33 | 0.2 | 0.21 | 0.09 | 0.32 | 0.23 |
| 11 | 62392229 | rs3018561 | 0.823 | 0.596 | 0.369 | -1.61 | 0.004 | -0.062 | -0.062 | 0.601 | 0.67 | 0.64 | 0.73 | 0.94 | 0.65 | 0.75 |
| 11 | 62401620 | rs11231168 | 0.696 | 0.548 | -0.537 | -1.027 | -0.16 | -0.055 | -0.055 | 0 | 0.32 | 0.02 | 0.52 | 0.24 | 0.43 | 0.54 |
| 11 | 62426102 | rs11231174 | 0.258 | 0.367 | -0.407 | -1.023 | -0.059 | 0.082 | 0.082 | 0 | 0.23 | 0.03 | 0.21 | 0.05 | 0.26 | 0.19 |
| 11 | 62426978 | rs11231176 | 0.263 | 0.331 | -0.408 | -1.022 | -0.06 | 0.09 | 0.09 | 0 | 0.23 | 0.02 | 0.17 | 0.04 | 0.26 | 0.19 |
| 11 | 62438819 | rs693698 | 0.291 | 0.494 | -0.33 | -1.013 | -0.007 | -0.087 | -0.087 | 0 | 0.7 | 0.59 | 0.75 | 0.94 | 0.73 | 0.8 |
| 11 | 62438846 | rs3809079 | -0.052 | 0.504 | -0.31 | -1.008 | -0.008 | 0.047 | 0.047 | 0 | 0.26 | 0.15 | 0.22 | 0.06 | 0.27 | 0.2 |
| 11 | 62444798 | rs644616 | 0.268 | 0.538 | -0.311 | -0.997 | -0.009 | -0.087 | -0.087 | 0 | 0.7 | 0.59 | 0.75 | 0.94 | 0.73 | 0.8 |
| 11 | 62454768 | rs1787662 | 0.605 | 0.867 | -0.016 | -0.476 | -0.003 | -0.03 | -0.03 | 0 | 0.72 | 0.5 | 0.79 | 0.76 | 0.78 | 0.75 |
| 11 | 62466076 | rs2259377 | 0.028 | 1.027 | 0.151 | -0.367 | 0.02 | -0.009 | -0.009 | 0 | 0.78 | 0.69 | 0.82 | 0.76 | 0.79 | 0.78 |
| 11 | 62466992 | rs11231193 | 0.228 | 1.03 | 0.155 | -0.367 | -0.012 | 0.021 | 0.021 | 0 | 0.18 | 0.09 | 0.16 | 0.09 | 0.19 | 0.18 |
| 11 | 62472357 | rs11605489 | -0.447 | 1.371 | 0.002 | -0.163 | 0.05 | 0.039 | 0.039 | 0.004 | 0.54 | 0.12 | 0.67 | 0.3 | 0.63 | 0.73 |
| 11 | 62489267 | rs496032 | 0.809 | 1.677 | 0.694 | -0.012 | 0.016 | -0.02 | -0.02 | 0 | 0.76 | 0.65 | 0.82 | 0.76 | 0.79 | 0.78 |
| 11 | 62527634 | rs35730843 | -1.07 | 0.513 | 0.578 | 0.74 | 0.045 | 0.154 | 0.154 | 0.866 | 0.07 | 0.4 | 0.05 | 0.15 | 0.02 | 0.04 |
| 11 | 62531551 | rs2509680 | -1.104 | 0.498 | 0.708 | 0.752 | 0.025 | 0.128 | 0.128 | 0.682 | 0.07 | 0.34 | 0.05 | 0.15 | 0.02 | 0.04 |
| 11 | 62541232 | rs679626 | -1.201 | 0.461 | 0.922 | 0.733 | 0.019 | 0.119 | 0.119 | 0.614 | 0.07 | 0.34 | 0.05 | 0.15 | 0.02 | 0.06 |
| 11 | 62543180 | rs11231210 | -1.224 | 0.452 | 0.919 | 0.729 | 0.019 | 0.119 | 0.119 | 1.029 | 0.07 | 0.34 | 0.05 | 0.15 | 0.02 | 0.06 |
| 11 | 62544903 | rs11231212 | -1.474 | 0.358 | 0.836 | 0.628 | 0.004 | -0.018 | -0.018 | 0 | 0.05 | 0.02 | 0.01 | 0.15 | 0.03 | 0.11 |
| 11 | 62545346 | rs633742 | -1.934 | 0.9 | 0.58 | 1.143 | 0.006 | 0.029 | 0.029 | 0 | 0.19 | 0.28 | 0.17 | 0.34 | 0.19 | 0.25 |
| 11 | 62554814 | rs2584916 | -1.628 | 0.738 | 0.635 | 1.073 | -0.027 | 0.011 | 0.011 | 0.054 | 0.11 | 0.24 | 0.07 | 0.16 | 0.05 | 0.11 |
| 11 | 62555086 | rs7941706 | 0.484 | 0.453 | -0.561 | 0.797 | -0.039 | 0.037 | 0.037 | 0 | 0.47 | 0.06 | 0.34 | 0.39 | 0.57 | 0.24 |
| 11 | 62555242 | rs2241908 | 0.444 | 0.286 | 0.028 | 0.566 | -0.016 | -0.078 | -0.078 | 0 | 0.35 | 0.14 | 0.54 | 0.45 | 0.36 | 0.62 |
| 11 | 62564009 | rs4693 | -0.279 | 0.403 | 0.694 | 0.576 | -0.002 | 0.004 | 0.004 | 0.304 | 0.61 | 0.53 | 0.42 | 0.55 | 0.62 | 0.38 |
| 11 | 62573878 | rs10897294 | -0.439 | 0.371 | 0.728 | 0.563 | 0.009 | -0.001 | -0.001 | 0 | 0.11 | 0.11 | 0.05 | 0.15 | 0.04 | 0.09 |
| 11 | 62576847 | rs11231227 | -0.558 | 0.37 | 0.752 | 0.563 | -0.011 | 0 | 0 | 0.265 | 0.11 | 0.16 | 0.05 | 0.16 | 0.04 | 0.09 |
| 11 | 62577233 | rs2730034 | -1.141 | 0.36 | 0.836 | 0.559 | -0.031 | 0.035 | 0.035 | 0.059 | 0.13 | 0.46 | 0.08 | 0.16 | 0.05 | 0.11 |
| 11 | 62598144 | rs10431142 | -0.437 | 0.366 | 0.535 | 0.561 | 0.009 | -0.001 | -0.001 | 0 | 0.11 | 0.11 | 0.05 | 0.15 | 0.04 | 0.11 |
| 11 | 62602137 | rs2289574 | -0.282 | 0.37 | 0.536 | 0.563 | -0.012 | 0.001 | 0.001 | 0 | 0.11 | 0.16 | 0.05 | 0.16 | 0.04 | 0.11 |
| 11 | 62602983 | rs2298636 | -0.282 | 0.37 | 0.536 | 0.563 | -0.012 | 0.001 | 0.001 | 0 | 0.11 | 0.16 | 0.05 | 0.16 | 0.04 | 0.11 |
| 11 | 62603916 | rs11231242 | -0.282 | 0.37 | 0.536 | 0.563 | -0.012 | 0.001 | 0.001 | 0 | 0.11 | 0.16 | 0.05 | 0.16 | 0.04 | 0.11 |
| 11 | 62607068 | rs7102434 | -0.282 | 0.37 | 0.536 | 0.563 | -0.012 | 0.001 | 0.001 | 0 | 0.11 | 0.16 | 0.05 | 0.16 | 0.04 | 0.11 |
| 11 | 62619206 | rs498751 | 0.592 | 0.424 | 0.48 | 0.577 | -0.034 | 0.049 | 0.049 | 0 | 0.48 | 0.06 | 0.34 | 0.39 | 0.57 | 0.26 |
| 11 | 62619629 | rs4034 | 0.483 | 0.374 | 0.524 | 0.561 | -0.031 | -0.087 | -0.087 | 0 | 0.34 | 0.13 | 0.55 | 0.45 | 0.36 | 0.61 |
| 11 | 62623017 | rs2071035 | -0.226 | 0.425 | 0.492 | 0.608 | -0.001 | -0.001 | -0.001 | 2.654 | 0.1 | 0.11 | 0.05 | 0.16 | 0.04 | 0.11 |
| 11 | 62623051 | rs1059292 | -0.311 | 0.425 | 0.499 | 0.608 | -0.018 | 0.004 | 0.004 | 0 | 0.11 | 0.18 | 0.06 | 0.16 | 0.04 | 0.11 |
| 11 | 62625512 | rs12794763 | 1.641 | 0.166 | 0.137 | 0.544 | -0.039 | -0.09 | -0.09 | 0.217 | 0.13 | 0.01 | 0.39 | 0.25 | 0.15 | 0.41 |
| 11 | 62625786 | rs10897300 | -0.111 | 0.266 | 0.29 | 0.504 | -0.018 | 0.004 | 0.004 | 0 | 0.1 | 0.18 | 0.06 | 0.16 | 0.04 | 0.11 |
| 11 | 62649042 | rs2070870 | 1.451 | 0.712 | -0.624 | 0.107 | -0.056 | 0.022 | 0.022 | 0 | 0.18 | 0.03 | 0.18 | 0.13 | 0.27 | 0.19 |
| 11 | 62652063 | rs489381 | 0.087 | 0.575 | -0.219 | 0.386 | -0.008 | 0.019 | 0.019 | 0 | 0.85 | 0.6 | 0.69 | 0.82 | 0.91 | 0.88 |
| 11 | 62652779 | rs4726 | 1.347 | 0.198 | -1.014 | 0.247 | -0.073 | -0.034 | -0.034 | 3.13 | 0.19 | 0.02 | 0.13 | 0.2 | 0.25 | 0.07 |
| 11 | 62656504 | rs2282477 | 1.596 | -0.24 | -1.065 | 0.121 | -0.001 | -0.001 | -0.001 | 0.652 | 0.16 | 0.14 | 0.2 | 0.25 | 0.23 | 0.17 |
| 11 | 62661158 | rs2507820 | 1.191 | -0.555 | -1.071 | 0.362 | -0.001 | -0.001 | -0.001 | 0.787 | 0.21 | 0.18 | 0.15 | 0.26 | 0.24 | 0.13 |
| 11 | 62663659 | rs547248 | 0.111 | -0.538 | -0.734 | 0.299 | -0.001 | 0.02 | 0.02 | 0 | 0.67 | 0.43 | 0.48 | 0.62 | 0.69 | 0.74 |
| 11 | 62666606 | rs2507821 | 0.147 | -0.9 | -1.09 | 0.281 | 0.033 | -0.002 | -0.002 | 0 | 0.48 | 0.31 | 0.28 | 0.47 | 0.41 | 0.56 |
| 11 | 62675687 | rs17157628 | 1.467 | -0.892 | -1.471 | 0.181 | -0.006 | -0.03 | -0.03 | 0 | 0.12 | 0.02 | 0.27 | 0.08 | 0.09 | 0.02 |
| 11 | 62676802 | rs11823728 | 0.43 | -1.001 | -1.539 | 0.099 | -0.01 | -0.022 | -0.022 | 0 | 0.06 | 0.13 | 0.02 | 0.02 | 0.03 | 0.07 |
| 11 | 62678306 | rs2067477 | 1.493 | -0.914 | -1.589 | 0.175 | -0.014 | -0.038 | -0.038 | 0 | 0.12 | 0.01 | 0.27 | 0.08 | 0.09 | 0.02 |
| 11 | 62685230 | rs544978 | -1.58 | -0.472 | -1.584 | -0.068 | -0.012 | 0.009 | 0.009 | 0 | 0.8 | 0.84 | 0.78 | 0.85 | 0.73 | 0.81 |
| 11 | 62688269 | rs2075748 | 1.057 | -0.705 | -1.842 | 0.503 | -0.029 | -0.029 | -0.029 | 0.567 | 0.15 | 0.05 | 0.14 | 0.25 | 0.23 | 0.08 |
| 11 | 62696951 | rs1938677 | -0.683 | -0.924 | -1.684 | 0.428 | 0.009 | -0.003 | -0.003 | 1.561 | 0.59 | 0.45 | 0.64 | 0.6 | 0.52 | 0.68 |
| 11 | 62697813 | rs4592425 | -2.079 | -0.886 | -1.743 | 0.323 | -0.066 | 0.04 | 0.04 | 0.069 | 0.8 | 0.97 | 0.79 | 0.87 | 0.72 | 0.83 |
| 11 | 62700359 | rs10431145 | -0.414 | -1.477 | -1.84 | 0.236 | -0.004 | -0.007 | -0.007 | 0 | 0.42 | 0.4 | 0.27 | 0.26 | 0.37 | 0.22 |
| 11 | 62712229 | rs10897306 | -0.628 | -1.52 | -1.841 | 0.231 | 0 | 0.002 | 0.002 | 0.016 | 0.52 | 0.46 | 0.67 | 0.59 | 0.48 | 0.68 |
| 11 | 62717297 | rs12223849 | 0.66 | -1.748 | -1.821 | 0.208 | -0.029 | -0.066 | -0.066 | 2.787 | 0.24 | 0.06 | 0.2 | 0.29 | 0.29 | 0.12 |
| 11 | 62743055 | rs736342 | 1.881 | -1.934 | -1.597 | 0.088 | 0.015 | 0.062 | 0.062 | 0.873 | 0.12 | 0.01 | 0.03 | 0.01 | 0.07 | 0.04 |
| 11 | 62744899 | rs3017670 | -1.979 | -1.499 | -1.394 | 0.466 | -0.053 | -0.029 | -0.029 | 0.68 | 0.86 | 0.97 | 0.71 | 0.81 | 0.84 | 0.82 |
| 11 | 62749680 | rs6591722 | 1.442 | -1.148 | -0.76 | 0.788 | -0.072 | 0.044 | 0.044 | 1.614 | 0.21 | 0.06 | 0.26 | 0.15 | 0.32 | 0.18 |
| 11 | 62750858 | rs4149172 | -0.588 | -1.225 | -0.617 | 0.961 | -0.012 | 0.071 | 0.071 | 0 | 0.28 | 0.4 | 0.23 | 0.41 | 0.3 | 0.21 |
| 11 | 62751658 | rs10897312 | 0.369 | -1.613 | -0.599 | 1.001 | -0.063 | 0.141 | 0.141 | 0 | 0.16 | 0.38 | 0.12 | 0.23 | 0.08 | 0.16 |
| 11 | 62755519 | rs11231294 | -0.07 | -1.428 | -0.593 | 0.685 | 0.012 | -0.062 | -0.062 | 0 | 0.26 | 0.14 | 0.19 | 0.42 | 0.26 | 0.21 |
| 11 | 62757113 | rs955434 | 0.083 | -1.293 | -0.478 | 0.798 | 0.007 | -0.053 | -0.053 | 0 | 0.23 | 0.13 | 0.18 | 0.41 | 0.25 | 0.2 |
| 11 | 62761161 | rs2187384 | 0.939 | -1.303 | -0.327 | 0.534 | 0 | -0.062 | -0.062 | 0.709 | 0.17 | 0.06 | 0.09 | 0.27 | 0.19 | 0.06 |
| 11 | 62762422 | rs953894 | 0.015 | -1.309 | -0.278 | 0.529 | -0.03 | 0.051 | 0.051 | 0 | 0.23 | 0.4 | 0.14 | 0.3 | 0.21 | 0.16 |
| 11 | 62765869 | rs4149183 | 0.043 | -1.271 | -0.636 | 0.547 | -0.031 | 0.053 | 0.053 | 0 | 0.23 | 0.4 | 0.14 | 0.3 | 0.21 | 0.16 |
| 11 | 62771388 | rs11231299 | 0.886 | -0.998 | -1.777 | 0.63 | 0.002 | -0.053 | -0.053 | 0 | 0.17 | 0.06 | 0.09 | 0.27 | 0.2 | 0.05 |
| 11 | 62771582 | rs11231300 | -0.16 | -1.006 | -1.012 | 0.027 | -0.083 | -0.09 | -0.09 | 0.015 | 0.18 | 0.63 | 0.1 | 0.07 | 0.06 | 0.15 |
| 11 | 62775898 | rs2187383 | 0.001 | 0.619 | -0.781 | 0.764 | 0.005 | -0.001 | -0.001 | 0.021 | 0.73 | 0.74 | 0.65 | 0.76 | 0.61 | 0.77 |
| 11 | 62778397 | rs11231302 | 0.036 | -0.404 | -0.6 | 0.807 | -0.031 | 0.044 | 0.044 | 1.067 | 0.09 | 0.16 | 0.04 | 0.12 | 0.03 | 0.01 |

| | | | | | | | | | | | | | | | | |
|---|---|---|---|---|---|---|---|---|---|---|---|---|---|---|---|---|
| 11 | 62780577 | rs4963326 | -0.117 | -0.376 | -0.931 | -0.007 | 0.011 | 0.02 | 0.02 | 0.384 | 0.54 | 0.42 | 0.52 | 0.72 | 0.6 | 0.59 |
| 11 | 62781921 | rs4149181 | -0.65 | -1.291 | -0.734 | -0.21 | -0.088 | -0.088 | -0.088 | 0 | 0.23 | 0.61 | 0.08 | 0.05 | 0.07 | 0.07 |
| 11 | 62783772 | rs3809069 | 0.263 | -0.013 | -1.451 | 0.679 | -0.06 | -0.084 | -0.084 | 0 | 0.14 | 0.01 | 0.13 | 0.28 | 0.16 | 0.37 |
| 11 | 62783826 | rs948979 | -0.587 | -0.04 | -1.344 | 0.512 | -0.013 | -0.023 | -0.023 | 0 | 0.71 | 0.59 | 0.76 | 0.82 | 0.75 | 0.75 |
| 11 | 62786435 | rs10792369 | -0.715 | 0.214 | -1.924 | 0.938 | 0.004 | -0.007 | -0.007 | 0 | 0.6 | 0.37 | 0.58 | 0.55 | 0.66 | 0.64 |
| 11 | 62789763 | rs10897316 | -0.597 | 0.462 | -1.868 | 0.657 | 0.002 | 0.021 | 0.021 | 1.992 | 0.49 | 0.35 | 0.51 | 0.42 | 0.54 | 0.56 |
| 11 | 62794074 | rs4963331 | -0.063 | 0.269 | -1.207 | 0.283 | -0.007 | -0.014 | -0.014 | 0 | 0.72 | 0.67 | 0.61 | 0.79 | 0.72 | 0.78 |
| 11 | 62801148 | rs1108539 | -0.45 | 0.228 | -1.138 | 0.427 | -0.023 | -0.088 | -0.088 | 0 | 0.32 | 0.12 | 0.39 | 0.52 | 0.39 | 0.46 |
| 11 | 62817069 | rs4963334 | 0.644 | -0.091 | -0.18 | 0.219 | 0.002 | -0.001 | -0.001 | 0 | 0.58 | 0.46 | 0.5 | 0.42 | 0.51 | 0.5 |
| 11 | 62837347 | rs7926184 | 0.204 | -0.089 | 0.15 | 0.237 | 0.014 | -0.014 | -0.014 | 0 | 0.79 | 0.56 | 0.83 | 0.82 | 0.79 | 0.84 |
| 11 | 62839667 | rs11231328 | 0.129 | -0.07 | 0.159 | 0.245 | 0.015 | -0.01 | -0.01 | 0 | 0.79 | 0.56 | 0.83 | 0.82 | 0.79 | 0.82 |
| 11 | 62848664 | rs11231343 | 0.434 | -0.07 | 0.32 | 0.197 | -0.003 | 0.018 | 0.018 | 0.053 | 0.15 | 0.09 | 0.09 | 0.1 | 0.15 | 0.14 |
| 11 | 62849656 | rs1944047 | 1.159 | -0.102 | 1.043 | 0.199 | -0.008 | 0 | 0 | 0.501 | 0.57 | 0.34 | 0.49 | 0.42 | 0.51 | 0.5 |
| 11 | 62851898 | rs1944049 | 1.055 | -0.1 | 0.885 | 0.198 | 0.004 | -0.001 | -0.001 | 0.199 | 0.58 | 0.5 | 0.51 | 0.42 | 0.51 | 0.5 |
| 11 | 62870326 | rs4963361 | 0.248 | -0.11 | 0.62 | 0.116 | -0.074 | -0.136 | -0.136 | 2.25 | 0.21 | 0.02 | 0.32 | 0.39 | 0.28 | 0.33 |
| 11 | 62901531 | rs12364908 | 0.142 | -0.736 | 1.319 | 0.192 | -0.004 | 0.016 | 0.016 | 0.334 | 0.14 | 0.09 | 0.1 | 0.11 | 0.15 | 0.14 |
| 11 | 62905841 | rs7106537 | 0.085 | -0.763 | 1.485 | 0.168 | 0.007 | -0.066 | -0.066 | 0 | 0.83 | 0.47 | 0.86 | 0.89 | 0.84 | 0.84 |
| 11 | 62911658 | rs1939747 | 0.553 | -0.768 | 1.485 | 0.158 | -0.016 | -0.096 | -0.096 | 0 | 0.79 | 0.48 | 0.86 | 0.89 | 0.83 | 0.84 |
| 11 | 62913852 | rs2155325 | 0.466 | -0.762 | 1.42 | 0.161 | -0.011 | -0.068 | -0.068 | 5.79 | 0.79 | 0.59 | 0.87 | 0.89 | 0.83 | 0.85 |
| 11 | 62914847 | rs10501392 | -0.43 | -1.022 | 1.073 | 0.221 | -0.003 | -0.089 | -0.089 | 0 | 0.25 | 0.09 | 0.15 | 0.43 | 0.19 | 0.34 |
| 11 | 62915419 | rs2156571 | 0.602 | -0.78 | 1.075 | -0.254 | 0 | 0 | 0 | 0 | 0.59 | 0.37 | 0.75 | 0.38 | 0.68 | 0.47 |
| 11 | 62916058 | rs1939767 | 0.97 | -0.793 | 0.973 | -0.356 | 0.018 | 0.024 | 0.024 | 1.671 | 0.5 | 0.61 | 0.75 | 0.37 | 0.63 | 0.46 |
| 11 | 62916395 | rs12803485 | 1.231 | -0.075 | 1.089 | 0.038 | 0.027 | 0.048 | 0.048 | 0 | 0.49 | 0.27 | 0.69 | 0.37 | 0.6 | 0.45 |
| 11 | 62925808 | rs12364655 | 0.678 | 0.041 | 0.865 | -0.232 | -0.035 | 0.019 | 0.019 | 0.145 | 0.12 | 0.02 | 0.08 | 0.1 | 0.13 | 0.17 |
| 11 | 62930757 | rs1122524 | -0.859 | 0.535 | 1.611 | -0.144 | -0.006 | 0.004 | 0.004 | 0.106 | 0.53 | 0.35 | 0.48 | 0.54 | 0.44 | 0.55 |
| 11 | 62931484 | rs6591771 | -0.977 | 0.534 | 1.76 | -0.144 | -0.006 | -0.001 | -0.001 | 0.689 | 0.55 | 0.64 | 0.5 | 0.53 | 0.45 | 0.55 |
| 11 | 62950506 | rs1525071 | 0.848 | 0.534 | 1.83 | -0.176 | -0.012 | -0.005 | -0.005 | 0 | 0.42 | 0.3 | 0.5 | 0.47 | 0.55 | 0.45 |
| 11 | 62984697 | rs7949840 | -0.595 | 0.535 | 1.83 | -0.178 | 0.011 | 0.015 | 0.015 | 1.249 | 0.51 | 0.29 | 0.45 | 0.53 | 0.4 | 0.53 |
| 11 | 62984868 | rs11231409 | -0.595 | 0.535 | 1.787 | -0.178 | 0.011 | 0.015 | 0.015 | 1.249 | 0.51 | 0.29 | 0.45 | 0.53 | 0.4 | 0.53 |
| 11 | 62996250 | rs12420866 | 0.429 | -0.102 | 1.335 | -0.136 | -0.069 | 0.024 | 0.024 | 0 | 0.12 | 0 | 0.06 | 0.07 | 0.18 | 0.13 |
| 11 | 63011204 | rs1193891 | 0.247 | -0.111 | 1.394 | -0.166 | 0.017 | 0.003 | 0.003 | 1.245 | 0.52 | 0.69 | 0.57 | 0.48 | 0.63 | 0.48 |
| 11 | 63039972 | rs1789323 | 0.322 | -0.122 | 1.429 | -0.138 | 0.018 | 0.012 | 0.012 | 0.074 | 0.5 | 0.66 | 0.57 | 0.47 | 0.6 | 0.47 |
| 11 | 63049129 | rs1193904 | -0.025 | -0.138 | 1.47 | -0.15 | 0.005 | -0.017 | -0.017 | 0.93 | 0.56 | 0.77 | 0.58 | 0.47 | 0.63 | 0.47 |
| 11 | 63050942 | rs554430 | 0.053 | -0.068 | 1.502 | -0.678 | 0.009 | -0.01 | -0.01 | 0.378 | 0.55 | 0.74 | 0.58 | 0.47 | 0.63 | 0.47 |
| 11 | 63057007 | rs549144 | 0.343 | -0.069 | 1.505 | -0.679 | 0.019 | 0.014 | 0.014 | 0.075 | 0.5 | 0.66 | 0.57 | 0.47 | 0.6 | 0.47 |
| 11 | 63057925 | rs1790218 | 0.993 | -0.08 | 1.618 | -0.691 | -0.024 | -0.014 | -0.014 | 0 | 0.42 | 0.27 | 0.51 | 0.47 | 0.55 | 0.45 |
| 11 | 63064163 | rs557879 | 0.329 | -0.074 | 1.477 | -0.688 | 0.019 | 0.014 | 0.014 | 0.075 | 0.5 | 0.66 | 0.57 | 0.47 | 0.61 | 0.47 |
| 11 | 63064970 | rs575009 | -0.054 | -0.074 | 1.477 | -0.688 | 0.005 | -0.015 | -0.015 | 0.006 | 0.56 | 0.77 | 0.59 | 0.47 | 0.63 | 0.47 |
| 11 | 63066903 | rs515213 | 0.284 | -0.062 | 1.471 | -0.678 | 0.014 | 0.009 | 0.009 | 0.714 | 0.51 | 0.7 | 0.57 | 0.47 | 0.61 | 0.47 |
| 11 | 63067138 | rs513338 | 0.413 | -0.061 | 1.471 | -0.678 | 0.018 | 0.014 | 0.014 | 0.135 | 0.5 | 0.66 | 0.57 | 0.47 | 0.61 | 0.47 |
| 11 | 63071009 | rs534799 | 0.294 | -0.037 | 1.433 | -0.666 | 0.015 | 0.01 | 0.01 | 0.68 | 0.51 | 0.7 | 0.57 | 0.47 | 0.61 | 0.47 |
| 11 | 63072310 | rs1201559 | 0.78 | -0.036 | 1.43 | -0.665 | -0.019 | -0.009 | -0.009 | 0 | 0.44 | 0.28 | 0.5 | 0.47 | 0.56 | 0.45 |
| 11 | 63074438 | rs4445638 | 0.963 | -0.032 | 1.107 | -0.663 | -0.019 | -0.011 | -0.011 | 0 | 0.42 | 0.28 | 0.51 | 0.47 | 0.56 | 0.45 |
| 11 | 63078986 | rs1404608 | 0.971 | -0.035 | 1.109 | -0.665 | -0.031 | -0.019 | -0.019 | 0.257 | 0.42 | 0.23 | 0.5 | 0.47 | 0.56 | 0.45 |
| 11 | 63105017 | rs1193722 | 0.95 | -0.004 | 0.953 | -0.657 | -0.029 | -0.016 | -0.016 | 0 | 0.42 | 0.25 | 0.49 | 0.47 | 0.55 | 0.45 |
| 11 | 63119486 | rs17158154 | 0.545 | 0.004 | 0.933 | -0.662 | -0.075 | 0.033 | 0.033 | 0 | 0.12 | 0.01 | 0.06 | 0.06 | 0.19 | 0.13 |
| 11 | 63120020 | rs4963260 | 0.293 | 0.154 | 1.464 | -0.28 | -0.028 | -0.013 | -0.013 | 0 | 0.43 | 0.25 | 0.49 | 0.47 | 0.55 | 0.44 |
| 11 | 63127621 | rs490724 | -0.265 | 0.107 | 1.401 | -0.294 | 0.015 | 0.011 | 0.011 | 0.68 | 0.51 | 0.67 | 0.57 | 0.47 | 0.61 | 0.47 |
| 11 | 63136501 | rs516536 | -0.254 | 0.073 | 1.414 | -0.279 | 0.015 | 0.011 | 0.011 | 0.766 | 0.51 | 0.67 | 0.56 | 0.47 | 0.61 | 0.47 |
| 11 | 63137384 | rs10897407 | 0.133 | 0.103 | 1.488 | -0.296 | -0.008 | -0.01 | -0.01 | 0 | 0.45 | 0.25 | 0.53 | 0.47 | 0.61 | 0.44 |
| 11 | 63138482 | rs566456 | -0.27 | 0.099 | 1.364 | -0.298 | 0.016 | 0.012 | 0.012 | 0.208 | 0.49 | 0.64 | 0.56 | 0.47 | 0.6 | 0.46 |
| 11 | 63139493 | rs4121881 | 0.104 | 0.097 | 1.353 | -0.297 | 0 | 0 | 0 | 0 | 0.45 | 0.5 | 0.54 | 0.47 | 0.6 | 0.44 |
| 11 | 63142877 | rs556730 | -0.279 | 0.128 | 0.665 | -0.28 | 0.015 | 0.006 | 0.006 | 0.332 | 0.51 | 0.6 | 0.59 | 0.47 | 0.66 | 0.47 |
| 11 | 63143681 | rs494608 | -0.748 | 0.117 | 0.103 | -0.265 | 0.019 | -0.018 | -0.018 | 0 | 0.57 | 0.77 | 0.61 | 0.47 | 0.69 | 0.47 |
| 11 | 63148736 | rs4963405 | 0.074 | 0.425 | 0.159 | 0.114 | -0.033 | -0.019 | -0.019 | 0 | 0.42 | 0.24 | 0.5 | 0.47 | 0.56 | 0.45 |
| 11 | 63149103 | rs11605137 | 0.125 | 0.438 | 0.019 | 0.123 | -0.042 | -0.025 | -0.025 | 0.195 | 0.41 | 0.23 | 0.49 | 0.47 | 0.55 | 0.45 |
| 11 | 63149270 | rs558472 | -0.247 | 0.632 | -0.665 | 0.454 | 0.014 | 0.011 | 0.011 | 0.013 | 0.49 | 0.66 | 0.56 | 0.47 | 0.61 | 0.47 |
| 11 | 63173684 | rs7101446 | -0.728 | 0.558 | 0.019 | 0.767 | -0.002 | 0.012 | 0.012 | 0.218 | 0.17 | 0.22 | 0.27 | 0.71 | 0.2 | 0.29 |
| 11 | 63184455 | rs11231463 | -0.616 | 0.135 | -0.126 | 0.556 | 0.018 | -0.003 | -0.003 | 0 | 0.05 | 0 | 0.19 | 0.06 | 0.01 | 0.15 |
| 11 | 63229061 | rs1305 | -1.893 | 0.088 | 2.564 | 0.546 | 0.02 | 0.114 | 0.114 | 0 | 0.11 | 0.41 | 0.27 | 0.17 | 0.07 | 0.23 |
| 11 | 63229471 | rs2298410 | -1.815 | 0.087 | 2.564 | 0.546 | 0.015 | 0.122 | 0.122 | 0 | 0.11 | 0.41 | 0.26 | 0.17 | 0.06 | 0.23 |
| 11 | 63233363 | rs4360702 | -2.386 | -0.169 | 2.773 | 0.378 | -0.009 | -0.001 | -0.001 | 0 | 0.17 | 0.72 | 0.33 | 0.18 | 0.12 | 0.23 |
| 11 | 63233710 | rs2275999 | -1.9 | -0.14 | 2.982 | 0.395 | 0.006 | 0.079 | 0.079 | 0 | 0.12 | 0.4 | 0.28 | 0.18 | 0.08 | 0.23 |
| 11 | 63233996 | rs11231499 | -1.722 | -0.136 | 2.932 | 0.397 | -0.001 | 0.078 | 0.078 | 0 | 0.11 | 0.36 | 0.26 | 0.17 | 0.06 | 0.23 |
| 11 | 63242556 | rs7944411 | -1.572 | -0.156 | 2.891 | 0.659 | 0.051 | 0.168 | 0.168 | 1.8 | 0.07 | 0.59 | 0.1 | 0.15 | 0.06 | 0.03 |
| 11 | 63257279 | rs10897423 | 0.571 | -0.051 | 0.228 | -0.531 | -0.046 | 0.159 | 0.159 | 0.245 | 0.33 | 0.03 | 0.16 | 0.08 | 0.3 | 0.17 |
| 11 | 63267937 | rs10897427 | -0.117 | -0.211 | -0.049 | -1.118 | -0.032 | 0.069 | 0.069 | 0 | 0.35 | 0.22 | 0.22 | 0.06 | 0.42 | 0.25 |
| 11 | 63274768 | rs1404504 | -0.619 | -0.588 | -0.253 | -0.475 | 0.039 | -0.003 | -0.003 | 0.249 | 0.51 | 0.7 | 0.55 | 0.5 | 0.64 | 0.64 |
| 11 | 63274914 | rs1404503 | -0.237 | -0.399 | -0.904 | -0.477 | -0.089 | 0.061 | 0.061 | 0.25 | 0.35 | 0.13 | 0.42 | 0.2 | 0.48 | 0.36 |

| | | | | | | | | | | | | | | | | |
|---|---|---|---|---|---|---|---|---|---|---|---|---|---|---|---|---|
| 11 | 63281594 | rs933186 | 0.181 | -1.387 | -1.782 | -0.722 | 0.017 | -0.116 | -0.116 | 0 | 0.26 | 0.01 | 0.27 | 0.63 | 0.2 | 0.12 |
| 11 | 63301208 | rs9633988 | 0.087 | -0.718 | -1.439 | -0.843 | -0.004 | -0.004 | -0.004 | 0 | 0.14 | 0.13 | 0.12 | 0.27 | 0.2 | 0.27 |
| 11 | 63305184 | rs4963271 | -0.724 | -0.964 | -1.041 | -0.839 | -0.001 | -0.004 | -0.004 | 1.212 | 0.2 | 0.76 | 0.17 | 0.28 | 0.21 | 0.32 |
| 11 | 63316786 | rs10792418 | -0.012 | -0.741 | -1.178 | -0.78 | 0.005 | 0.024 | 0.024 | 2.549 | 0.17 | 0.21 | 0.14 | 0.27 | 0.21 | 0.22 |
| 11 | 63320239 | rs1554656 | 0.101 | -0.509 | -1.372 | -0.91 | 0.002 | -0.003 | -0.003 | 0.051 | 0.24 | 0.2 | 0.46 | 0.7 | 0.22 | 0.16 |
| 11 | 63325201 | rs9943597 | -0.526 | -0.375 | -1.757 | -1.142 | -0.004 | -0.097 | -0.097 | 0.201 | 0.4 | 0.49 | 0.44 | 0.91 | 0.43 | 0.41 |
| 11 | 63332054 | rs1722860 | -1.078 | -0.064 | -3.085 | -0.972 | -0.04 | 0.156 | 0.156 | 0.054 | 0.73 | 0.95 | 0.82 | 0.98 | 0.74 | 0.61 |
| 11 | 63340430 | rs12798414 | 0.858 | -0.042 | -3.134 | -0.961 | -0.055 | 0.166 | 0.166 | 0.105 | 0.27 | 0.02 | 0.18 | 0.02 | 0.26 | 0.37 |
| 11 | 63343322 | rs591304 | -0.279 | 1.494 | -2.241 | 0.209 | -0.129 | -0.03 | -0.03 | 1.372 | 0.24 | 0.03 | 0.31 | 0.24 | 0.37 | 0.23 |
| 11 | 63354042 | rs11231525 | -0.554 | 0.79 | -1.246 | 1.075 | -0.001 | -0.044 | -0.044 | 0 | 0.22 | 0.14 | 0.36 | 0.62 | 0.25 | 0.26 |
| 11 | 63360114 | rs7928514 | -1.314 | 0.989 | -0.916 | 0.834 | -0.002 | -0.001 | -0.001 | 0 | 0.1 | 0.1 | 0.21 | 0.11 | 0.1 | 0.13 |
| 11 | 63367058 | rs4405320 | -1.521 | 0.823 | -1.284 | 0.268 | -0.03 | 0.087 | 0.087 | 0 | 0.17 | 0.06 | 0.09 | 0.01 | 0.16 | 0.16 |
| 11 | 63367440 | rs3934131 | -1.674 | 0.713 | -0.795 | 0.215 | -0.003 | 0.005 | 0.005 | 0 | 0.18 | 0.17 | 0.09 | 0.01 | 0.16 | 0.16 |
| 11 | 63370158 | rs7115616 | -1.351 | 0.886 | -0.203 | 0.596 | -0.003 | 0.002 | 0.002 | 0.084 | 0.42 | 0.57 | 0.57 | 0.58 | 0.46 | 0.42 |
| 11 | 63395617 | rs1404500 | 0.306 | 1.065 | 0.579 | 0.348 | -0.025 | 0.034 | 0.034 | 0 | 0.07 | 0 | 0.02 | 0 | 0.05 | 0.11 |
| 11 | 63395639 | rs10501394 | -0.907 | 0.925 | 0.431 | 0.019 | 0 | 0.047 | 0.047 | 0 | 0.17 | 0.05 | 0.07 | 0 | 0.14 | 0.16 |
| 11 | 63399150 | rs10750999 | -0.23 | 0.44 | 0.107 | -0.154 | -0.057 | -0.054 | -0.054 | 0 | 0.26 | 0.03 | 0.5 | 0.79 | 0.33 | 0.32 |
| 11 | 63417724 | rs518490 | -0.5 | 0.533 | 0.296 | -0.112 | -0.001 | -0.04 | -0.04 | 1.122 | 0.17 | 0.17 | 0.08 | 0 | 0.14 | 0.16 |
| 11 | 63438836 | rs2959880 | -0.521 | 0.551 | 0.672 | -0.103 | -0.002 | -0.067 | -0.067 | 0 | 0.16 | 0.24 | 0.09 | 0 | 0.13 | 0.16 |
| 11 | 63449125 | rs11551944 | -0.251 | 0.538 | 0.57 | -0.11 | -0.015 | 0.06 | 0.06 | 0 | 0.15 | 0.03 | 0.06 | 0 | 0.13 | 0.16 |
| 11 | 63483576 | rs474548 | 1.167 | 0.242 | 1.032 | -0.266 | -0.005 | -0.169 | -0.169 | 1.031 | 0.73 | 0.48 | 0.88 | 0.99 | 0.81 | 0.73 |
| 11 | 63487386 | rs542998 | 1.145 | 0.255 | 0.969 | -0.26 | -0.004 | -0.132 | -0.132 | 0 | 0.73 | 0.42 | 0.88 | 0.99 | 0.81 | 0.73 |
| 11 | 63493919 | rs517700 | -0.296 | 0.327 | 0.239 | -0.168 | 0 | -0.015 | -0.015 | 0 | 0.16 | 0.13 | 0.08 | 0 | 0.13 | 0.16 |
| 11 | 63496812 | rs10897445 | -0.172 | 0.24 | 0.175 | -0.211 | -0.059 | -0.057 | -0.057 | 0 | 0.25 | 0.03 | 0.5 | 0.79 | 0.32 | 0.22 |
| 11 | 63504649 | rs1186941 | -0.542 | 0.348 | 0.805 | -0.231 | 0 | -0.004 | -0.004 | 0.69 | 0.15 | 0.09 | 0.07 | 0 | 0.13 | 0.15 |
| 11 | 63508334 | rs567756 | -0.793 | 0.354 | 0.802 | -0.228 | -0.001 | -0.042 | -0.042 | 1.261 | 0.16 | 0.21 | 0.09 | 0 | 0.13 | 0.17 |
| 11 | 63537509 | rs2509744 | -0.631 | 0.407 | 0.715 | -0.202 | -0.001 | -0.053 | -0.053 | 0 | 0.16 | 0.18 | 0.08 | 0 | 0.13 | 0.15 |
| 11 | 63550922 | rs523093 | -0.594 | 0.466 | 0.453 | -0.174 | 0 | -0.024 | -0.024 | 0 | 0.15 | 0.11 | 0.07 | 0 | 0.13 | 0.15 |
| 11 | 63558603 | rs542966 | -1.152 | 0.683 | 0.616 | -0.152 | 0.016 | -0.098 | -0.098 | 0 | 0.22 | 0.31 | 0.11 | 0.01 | 0.2 | 0.26 |
| 11 | 63585804 | rs643634 | -0.151 | 0.365 | -0.63 | -0.15 | -0.096 | 0.012 | 0.012 | 0 | 0.37 | 0.07 | 0.64 | 0.69 | 0.42 | 0.31 |
| 11 | 63587313 | rs503341 | -0.337 | -0.174 | 0.67 | -0.413 | 0.003 | -0.033 | -0.033 | 0 | 0.41 | 0.51 | 0.67 | 0.76 | 0.43 | 0.33 |
| 11 | 63595072 | rs8995 | -0.071 | 0.389 | 0.739 | -0.324 | -0.135 | 0.016 | 0.016 | 0 | 0.38 | 0.07 | 0.66 | 0.7 | 0.49 | 0.33 |
| 11 | 63595925 | rs7949608 | 0.073 | 0.441 | 1.433 | -0.751 | -0.008 | 0.038 | 0.038 | 0 | 0.61 | 0.3 | 0.35 | 0.18 | 0.61 | 0.67 |
| 11 | 63596063 | rs11602911 | 0.119 | 0.313 | 0.492 | -1.282 | -0.009 | 0.014 | 0.014 | 0 | 0.57 | 0.34 | 0.31 | 0.32 | 0.51 | 0.59 |
| 11 | 63598802 | rs4963432 | 0.245 | -0.318 | 0.648 | -1.554 | -0.021 | 0.1 | 0.1 | 0 | 0.46 | 0.2 | 0.71 | 0.79 | 0.52 | 0.37 |
| 11 | 63605177 | rs10792421 | 0.265 | -0.22 | 0.452 | -2.065 | -0.041 | 0.242 | 0.242 | 0 | 0.46 | 0.15 | 0.71 | 0.89 | 0.54 | 0.43 |
| 11 | 63605663 | rs11231603 | 1.817 | -0.398 | 0.289 | -2.191 | 0.001 | 0.022 | 0.022 | 0 | 0.05 | 0.01 | 0.02 | 0 | 0.06 | 0.05 |
| 11 | 63608047 | rs3850942 | -0.062 | -0.429 | 0.692 | -2.101 | -0.011 | -0.046 | -0.046 | 0 | 0.13 | 0.16 | 0.08 | 0.01 | 0.07 | 0.25 |
| 11 | 63620253 | rs10897453 | 0.187 | -0.495 | 0.698 | -2.099 | -0.024 | 0.218 | 0.218 | 0 | 0.47 | 0.17 | 0.72 | 0.88 | 0.55 | 0.44 |
| 11 | 63639881 | rs928948 | -0.861 | -0.352 | 0.806 | -2.029 | -0.001 | -0.004 | -0.004 | 0 | 0.6 | 0.36 | 0.79 | 0.89 | 0.61 | 0.7 |
| 11 | 63655710 | rs11231630 | 0.227 | -0.589 | 1.224 | -1.848 | -0.005 | -0.03 | -0.03 | 0 | 0.11 | 0.09 | 0.07 | 0.01 | 0.06 | 0.23 |
| 11 | 63659778 | rs10897459 | -0.923 | -0.589 | 1.22 | -1.857 | -0.013 | 0.097 | 0.097 | 0 | 0.62 | 0.79 | 0.82 | 0.93 | 0.61 | 0.7 |
| 11 | 63676067 | rs11231637 | -0.296 | -0.533 | 1.205 | -1.828 | 0.003 | 0.067 | 0.067 | 0 | 0.49 | 0.33 | 0.25 | 0.08 | 0.46 | 0.53 |
| 11 | 63678508 | rs12786124 | 0.869 | -0.57 | 1.08 | -1.845 | -0.112 | 0.203 | 0.203 | 0 | 0.35 | 0.02 | 0.16 | 0.07 | 0.39 | 0.28 |
| 11 | 63680045 | rs320109 | -0.29 | -0.439 | 1.236 | -1.746 | 0.002 | 0.067 | 0.067 | 0 | 0.49 | 0.34 | 0.25 | 0.13 | 0.46 | 0.53 |
| 11 | 63691049 | rs11231640 | 0.295 | -0.464 | 1.176 | -1.919 | 0.002 | 0.233 | 0.233 | 0 | 0.49 | 0.17 | 0.72 | 0.88 | 0.55 | 0.48 |
| 11 | 63708574 | rs728919 | -0.33 | -0.524 | 1.573 | -2.046 | -0.006 | 0.102 | 0.102 | 0 | 0.58 | 0.29 | 0.76 | 0.87 | 0.56 | 0.63 |
| 11 | 63713317 | rs3740637 | 0.924 | -0.533 | 1.506 | -2.053 | -0.106 | 0.203 | 0.203 | 0.01 | 0.32 | 0.02 | 0.15 | 0.06 | 0.39 | 0.28 |
| 11 | 63714319 | rs4980500 | 0.924 | -0.533 | 1.506 | -2.053 | -0.106 | 0.203 | 0.203 | 0.01 | 0.32 | 0.02 | 0.15 | 0.06 | 0.39 | 0.28 |
| 11 | 63714930 | rs12798491 | 0.271 | -0.531 | 1.536 | -2.047 | -0.014 | 0.064 | 0.064 | 0 | 0.39 | 0.25 | 0.2 | 0.1 | 0.44 | 0.36 |
| 11 | 63718663 | rs10897464 | -0.315 | -0.523 | 1.334 | -2.043 | -0.006 | 0.102 | 0.102 | 0 | 0.58 | 0.28 | 0.76 | 0.87 | 0.56 | 0.63 |
| 11 | 63722448 | rs10897465 | 0.797 | -0.673 | 1.165 | -2.105 | -0.09 | 0.214 | 0.214 | 0 | 0.33 | 0.02 | 0.15 | 0.05 | 0.37 | 0.28 |
| 11 | 63723238 | rs10897466 | 0.101 | -0.169 | 1.681 | -1.792 | -0.015 | 0.077 | 0.077 | 0 | 0.39 | 0.24 | 0.2 | 0.1 | 0.44 | 0.36 |
| 11 | 63723281 | rs1386346 | -0.186 | -0.179 | 1.789 | -1.792 | -0.006 | 0.109 | 0.109 | 0 | 0.58 | 0.26 | 0.75 | 0.87 | 0.56 | 0.63 |
| 11 | 63725609 | rs11231657 | -0.87 | -0.159 | 1.675 | -1.767 | 0.02 | 0.014 | 0.014 | 0 | 0.06 | 0.22 | 0.05 | 0.04 | 0.04 | 0.09 |
| 11 | 63739439 | rs480211 | -1.156 | -0.142 | 1.451 | -1.767 | -0.097 | 0.192 | 0.192 | 0 | 0.56 | 0.9 | 0.53 | 0.21 | 0.5 | 0.58 |
| 11 | 63739568 | rs12280970 | -1.483 | -0.209 | 1.639 | -1.606 | 0.053 | 0.043 | 0.043 | 1.136 | 0.08 | 0.45 | 0.07 | 0.04 | 0.05 | 0.09 |
| 11 | 63743420 | rs11605797 | 0.696 | -0.208 | 1.409 | -1.638 | -0.106 | 0.203 | 0.203 | 0.01 | 0.32 | 0.03 | 0.15 | 0.06 | 0.4 | 0.28 |
| 11 | 63761169 | rs11231671 | 0.579 | -0.198 | 1.249 | -1.626 | -0.041 | 0.142 | 0.142 | 0 | 0.33 | 0.16 | 0.16 | 0.07 | 0.4 | 0.28 |
| 11 | 63768397 | rs320156 | -0.152 | -0.21 | 1.03 | -1.662 | -0.003 | 0.046 | 0.046 | 0 | 0.42 | 0.63 | 0.24 | 0.13 | 0.45 | 0.38 |
| 11 | 63771452 | rs492357 | 0.681 | -0.154 | 0.515 | -1.675 | -0.102 | 0.208 | 0.208 | 0.028 | 0.32 | 0.02 | 0.15 | 0.05 | 0.4 | 0.28 |
| 11 | 63774370 | rs10897471 | -1.587 | -0.323 | 1.1 | -1.286 | 0.06 | 0.044 | 0.044 | 0.827 | 0.08 | 0.42 | 0.06 | 0.05 | 0.05 | 0.09 |
| 11 | 63787389 | rs570098 | -0.43 | -0.195 | 0.383 | -1.115 | 0 | 0.02 | 0.02 | 0 | 0.52 | 0.52 | 0.47 | 0.2 | 0.5 | 0.64 |
| 11 | 63792977 | rs578001 | -0.425 | -0.405 | 0.158 | -1.115 | 0 | -0.031 | -0.031 | 0 | 0.52 | 0.45 | 0.46 | 0.19 | 0.51 | 0.64 |
| 11 | 63797679 | rs503735 | 0.107 | 0.325 | -0.499 | -0.974 | -0.103 | 0.255 | 0.255 | 0 | 0.41 | 0.06 | 0.21 | 0.05 | 0.48 | 0.37 |
| 11 | 63800013 | rs545766 | -1.713 | 0.353 | 0.898 | 0.617 | -0.02 | 0.003 | 0.003 | 1.712 | 0.16 | 0.76 | 0.09 | 0.19 | 0.06 | 0.3 |
| 11 | 63806993 | rs502224 | -1.928 | 0.651 | 1.266 | 0.638 | 0.036 | 0.084 | 0.084 | 0.365 | 0.14 | 0.48 | 0.12 | 0.19 | 0.19 | 0.26 |
| 11 | 63808063 | rs320145 | -2.269 | 0.625 | 1.7 | 0.576 | 0.02 | 0.011 | 0.011 | 1.164 | 0.22 | 0.5 | 0.14 | 0.22 | 0.24 | 0.4 |
| 11 | 63810944 | rs562750 | -2.123 | 0.873 | 1.655 | 0.284 | 0.048 | -0.103 | -0.103 | 0 | 0.2 | 0.49 | 0.16 | 0.05 | 0.25 | 0.26 |

| | | | | | | | | | | | | | | | | |
|---|---|---|---|---|---|---|---|---|---|---|---|---|---|---|---|---|
| 11 | 63822940 | rs530399 | -1.019 | 0.845 | 0.978 | 0.334 | -0.047 | 0.049 | 0.049 | 0 | 0.09 | 0.03 | 0.11 | 0.01 | 0.2 | 0.1 |
| 11 | 63824364 | rs1188017 | -1.002 | 0.862 | 1.075 | 0.342 | -0.023 | 0.025 | 0.025 | 0 | 0.09 | 0.06 | 0.11 | 0.01 | 0.2 | 0.1 |
| 11 | 63826317 | rs2000515 | -1.561 | 0.425 | 1.742 | 1.655 | 0.04 | 0.495 | 0.495 | 0.024 | 0.09 | 0.42 | 0.08 | 0.42 | 0.07 | 0.18 |
| 11 | 63835611 | rs2845568 | -1.408 | 0.555 | 1.274 | 0.739 | 0.005 | -0.077 | -0.077 | 0 | 0.12 | 0.34 | 0.06 | 0.03 | 0.06 | 0.21 |
| 11 | 63844683 | rs715163 | 1.037 | 0.387 | 0.499 | 0.968 | 0.008 | 0.008 | 0.008 | 0 | 0.6 | 0.24 | 0.59 | 0.44 | 0.64 | 0.37 |
| 11 | 63845294 | rs1212642 | 0.771 | 0.562 | 1.301 | 1.209 | 0.004 | 0.003 | 0.003 | 0 | 0.65 | 0.49 | 0.6 | 0.45 | 0.65 | 0.45 |
| 11 | 63849330 | rs12575710 | -1.273 | -0.186 | 1.087 | 2.033 | -0.025 | 0.053 | 0.053 | 0 | 0.24 | 0.32 | 0.21 | 0.45 | 0.13 | 0.37 |
| 11 | 63859120 | rs883798 | -1.579 | 0.073 | 0.219 | 1.795 | -0.026 | -0.013 | -0.013 | 2.242 | 0.22 | 0.12 | 0.28 | 0.22 | 0.29 | 0.37 |
| 11 | 63859751 | rs1807304 | -0.644 | 0.104 | 0.219 | 1.844 | -0.007 | -0.002 | -0.002 | 0 | 0.11 | 0.08 | 0.17 | 0.13 | 0.22 | 0.16 |
| 11 | 63860718 | rs12286180 | -0.849 | -0.465 | 0.747 | 2.394 | -0.049 | 0.09 | 0.09 | 0 | 0.19 | 0.38 | 0.37 | 0.3 | 0.1 | 0.22 |
| 11 | 63861317 | rs7941785 | 1.723 | -0.503 | 0.804 | 2.397 | 0.024 | 0.074 | 0.074 | 0 | 0.73 | 0.47 | 0.66 | 0.42 | 0.65 | 0.61 |
| 11 | 63862612 | rs11231693 | -0.462 | -0.495 | 0.659 | 2.369 | 0.008 | 0.254 | 0.254 | 0 | 0.07 | 0.25 | 0.05 | 0.35 | 0.07 | 0.02 |
| 11 | 63863405 | rs660070 | -0.022 | -0.362 | 0.155 | 2.522 | -0.041 | -0.039 | -0.039 | 0 | 0.08 | 0.02 | 0.23 | 0.22 | 0.21 | 0.15 |
| 11 | 63864211 | rs2005261 | -1.193 | -0.514 | 0.08 | 2.275 | -0.034 | -0.047 | -0.047 | 0 | 0.2 | 0.09 | 0.36 | 0.32 | 0.29 | 0.34 |
| 11 | 63867523 | rs2845887 | -1.815 | -0.558 | 0.11 | 2.311 | -0.018 | -0.021 | -0.021 | 0 | 0.27 | 0.22 | 0.39 | 0.66 | 0.35 | 0.36 |
| 11 | 63868690 | rs1984852 | 1.253 | -0.716 | 0.35 | 2.26 | -0.001 | -0.021 | -0.021 | 1.288 | 0.68 | 0.73 | 0.4 | 0.52 | 0.68 | 0.67 |
| 11 | 63869596 | rs641811 | -1.004 | -1.676 | 0.19 | 1.839 | 0.036 | 0.002 | 0.002 | 2.605 | 0.2 | 0.47 | 0.24 | 0.22 | 0.18 | 0.25 |
| 11 | 63870810 | rs651855 | -1.641 | -1.894 | -0.192 | 1.497 | 0.003 | 0.001 | 0.001 | 0.437 | 0.4 | 0.64 | 0.58 | 0.41 | 0.37 | 0.45 |
| 11 | 63871549 | rs638227 | -1.726 | -1.892 | -0.192 | 1.513 | 0.001 | -0.002 | -0.002 | 0.15 | 0.39 | 0.64 | 0.57 | 0.41 | 0.36 | 0.36 |
| 11 | 63880397 | rs663889 | 1.996 | -2.038 | -0.344 | 1.277 | 0.044 | 0.013 | 0.013 | 2.533 | 0.59 | 0.15 | 0.55 | 0.33 | 0.7 | 0.44 |
| 11 | 63883985 | rs614397 | 1.317 | -2.09 | -0.318 | 1.267 | 0.065 | -0.035 | -0.035 | 0 | 0.67 | 0.2 | 0.67 | 0.42 | 0.85 | 0.64 |
| 11 | 63884027 | rs614035 | 2.117 | -2.083 | -0.32 | 1.266 | 0.044 | 0.013 | 0.013 | 2.64 | 0.59 | 0.15 | 0.55 | 0.33 | 0.7 | 0.44 |
| 11 | 63884453 | rs3751120 | -1.181 | -2.125 | -0.173 | 1.31 | -0.081 | 0.104 | 0.104 | 2.584 | 0.23 | 0.6 | 0.19 | 0.54 | 0.09 | 0.22 |
| 11 | 63885305 | rs3824854 | 0.368 | -2.204 | -0.23 | 1.249 | -0.005 | -0.011 | -0.011 | 0 | 0.12 | 0.07 | 0.31 | 0.18 | 0.17 | 0.07 |
| 11 | 63885704 | rs677447 | 2.36 | -2.267 | -0.263 | 1.23 | 0.064 | 0.028 | 0.028 | 3.819 | 0.56 | 0.13 | 0.53 | 0.33 | 0.7 | 0.43 |
| 11 | 63886539 | rs663149 | 2.111 | -2.184 | -0.241 | 1.25 | 0.044 | 0.011 | 0.011 | 2.64 | 0.59 | 0.13 | 0.55 | 0.33 | 0.7 | 0.44 |
| 11 | 63889293 | rs627055 | 2.644 | -2.149 | -0.264 | 1.206 | -0.069 | 0.236 | 0.236 | 2.025 | 0.45 | 0.06 | 0.23 | 0.14 | 0.52 | 0.36 |
| 11 | 63909017 | rs2622424 | 0.822 | -2.059 | -0.334 | 1.227 | 0.009 | 0.014 | 0.014 | 0.178 | 0.1 | 0.03 | 0.13 | 0.05 | 0.06 | 0.14 |
| 11 | 63915537 | rs2875748 | 2.254 | -1.964 | -0.243 | 1.285 | 0.012 | -0.007 | -0.007 | 0 | 0.54 | 0.23 | 0.39 | 0.63 | 0.59 | 0.38 |
| 11 | 63915700 | rs2186571 | 0.762 | -1.629 | -0.19 | 1.353 | 0.01 | 0.015 | 0.015 | 0.178 | 0.1 | 0.04 | 0.13 | 0.05 | 0.06 | 0.15 |
| 11 | 63918373 | rs2282491 | -0.921 | -1.597 | -0.195 | 1.38 | -0.024 | -0.017 | -0.017 | 0.071 | 0.21 | 0.14 | 0.43 | 0.27 | 0.34 | 0.28 |
| 11 | 63944370 | rs7103915 | 0.725 | -1.53 | -0.196 | 1.369 | -0.001 | -0.006 | -0.006 | 0 | 0.05 | 0.02 | 0.08 | 0.08 | 0.14 | 0.07 |
| 11 | 63949095 | rs11603384 | -0.777 | -1.478 | -0.172 | 1.369 | -0.026 | -0.019 | -0.019 | 0 | 0.2 | 0.13 | 0.41 | 0.25 | 0.31 | 0.15 |
| 11 | 63953121 | rs3751122 | -0.21 | -1.568 | -0.143 | 1.418 | -0.019 | 0.146 | 0.146 | 0 | 0.08 | 0.16 | 0.18 | 0.49 | 0.09 | 0.11 |
| 11 | 63962152 | rs2282490 | 0.025 | -1.443 | -0.048 | 1.266 | -0.002 | -0.017 | -0.017 | 0 | 0.14 | 0.09 | 0.32 | 0.17 | 0.17 | 0.07 |
| 11 | 63963947 | rs11607165 | 0.042 | -1.447 | -0.049 | 1.264 | -0.002 | -0.017 | -0.017 | 0 | 0.14 | 0.09 | 0.32 | 0.17 | 0.17 | 0.07 |
| 11 | 63964657 | rs2236648 | -1.679 | -1.479 | -0.074 | 1.239 | -0.006 | -0.003 | -0.003 | 0 | 0.32 | 0.32 | 0.58 | 0.31 | 0.39 | 0.42 |
| 11 | 63973396 | rs1011219 | 0.565 | -1.334 | -0.131 | 1.229 | -0.001 | -0.005 | -0.005 | 0 | 0.05 | 0.03 | 0.08 | 0.08 | 0.14 | 0.07 |
| 11 | 63979643 | rs12575642 | -0.119 | -1.468 | -0.12 | 1.227 | 0.001 | -0.013 | -0.013 | 0.087 | 0.16 | 0.09 | 0.35 | 0.19 | 0.18 | 0.2 |
| 11 | 63988340 | rs1544538 | 0.745 | -1.442 | -0.12 | 1.227 | 0.001 | 0.004 | 0.004 | 0.3 | 0.1 | 0.02 | 0.13 | 0.05 | 0.06 | 0.14 |
| 11 | 63991801 | rs1059440 | -0.016 | -1.288 | 0.028 | 1.194 | -0.002 | -0.015 | -0.015 | 0.062 | 0.14 | 0.09 | 0.32 | 0.18 | 0.17 | 0.07 |
| 11 | 63994230 | rs2286612 | 0.231 | -1.973 | -0.282 | 0.6 | -0.124 | -0.045 | -0.045 | 0 | 0.18 | 0.44 | 0.04 | 0.06 | 0.02 | 0.18 |
| 11 | 63999529 | rs3741403 | 1.22 | -0.002 | -1.214 | 0.027 | 0.017 | -0.004 | -0.004 | 0 | 0.46 | 0.2 | 0.43 | 0.65 | 0.44 | 0.48 |
| 11 | 64003843 | rs11603042 | -0.506 | 0.474 | -1.191 | -0.053 | -0.035 | -0.023 | -0.023 | 0 | 0.22 | 0.12 | 0.4 | 0.25 | 0.33 | 0.16 |
| 11 | 64010085 | rs2070765 | 0.683 | 0.363 | -1.253 | 0.08 | -0.005 | -0.017 | -0.017 | 0 | 0.15 | 0.09 | 0.32 | 0.19 | 0.18 | 0.07 |
| 11 | 64011854 | rs11231727 | 1.177 | 0.288 | -1.32 | 0.052 | 0.022 | 0.01 | 0.01 | 0 | 0.47 | 0.21 | 0.43 | 0.67 | 0.44 | 0.47 |
| 11 | 64013406 | rs11605738 | 0.365 | 0.467 | -1.047 | 0.409 | -0.005 | -0.016 | -0.016 | 0 | 0.15 | 0.09 | 0.32 | 0.18 | 0.18 | 0.07 |
| 11 | 64018687 | rs2001003 | 0.38 | 0.469 | -1.058 | 0.409 | -0.006 | -0.018 | -0.018 | 0 | 0.15 | 0.09 | 0.32 | 0.18 | 0.18 | 0.07 |
| 11 | 64026144 | rs2244625 | 1.093 | 0.036 | -1.033 | 0.227 | 0.048 | 0.07 | 0.07 | 0.236 | 0.6 | 0.16 | 0.52 | 0.27 | 0.67 | 0.54 |
| 11 | 64026219 | rs2244621 | 0.089 | -0.05 | -0.53 | 0.418 | -0.007 | 0.055 | 0.055 | 0 | 0.19 | 0.24 | 0.3 | 0.54 | 0.16 | 0.18 |
| 11 | 64042997 | rs660442 | 2.307 | 0.218 | -0.823 | 0.122 | -0.071 | 0.017 | 0.017 | 0.56 | 0.13 | 0.01 | 0.29 | 0.08 | 0.21 | 0.16 |
| 11 | 64044629 | rs1468558 | 0.192 | 0.006 | -0.291 | 0.186 | 0.02 | 0.131 | 0.131 | 0.285 | 0.06 | 0.37 | 0.04 | 0.06 | 0.01 | 0.19 |
| 11 | 64046029 | rs671976 | -1.272 | -0.146 | -0.508 | -0.06 | 0.001 | 0.002 | 0.002 | 0.01 | 0.48 | 0.42 | 0.68 | 0.67 | 0.53 | 0.41 |
| 11 | 64046641 | rs11231740 | 1.776 | -0.136 | -0.587 | -0.066 | -0.09 | 0.101 | 0.101 | 0.672 | 0.45 | 0.05 | 0.27 | 0.22 | 0.47 | 0.37 |
| 11 | 64046885 | rs11231741 | 2.598 | -0.21 | -0.61 | -0.104 | -0.076 | 0.128 | 0.128 | 0.224 | 0.31 | 0.02 | 0.15 | 0.11 | 0.3 | 0.3 |
| 11 | 64052447 | rs2510066 | 0.637 | -0.055 | -0.585 | -0.047 | -0.088 | 0.054 | 0.054 | 0 | 0.28 | 0.06 | 0.38 | 0.16 | 0.37 | 0.23 |
| 11 | 64060976 | rs34499953 | 0.563 | -0.153 | -0.408 | 0.132 | -0.007 | 0.072 | 0.072 | 0 | 0.18 | 0.24 | 0.29 | 0.54 | 0.16 | 0.19 |
| 11 | 64068310 | rs2286614 | 1.058 | -0.063 | -0.401 | 0.167 | -0.02 | -0.026 | -0.026 | 0 | 0.14 | 0.03 | 0.12 | 0.11 | 0.16 | 0.06 |
| 11 | 64078380 | rs493052 | -0.187 | -0.088 | -0.264 | 0.107 | 0.048 | 0.189 | 0.189 | 1.029 | 0.94 | 0.48 | 0.95 | 0.94 | 0.99 | 0.81 |
| 11 | 64082807 | rs11600990 | 0.996 | -0.002 | -0.659 | 0.097 | -0.019 | -0.025 | -0.025 | 0 | 0.14 | 0.02 | 0.13 | 0.11 | 0.16 | 0.06 |
| 11 | 64102948 | rs574087 | 0.374 | 0.336 | -0.522 | -0.209 | -0.133 | 0.041 | 0.041 | 0 | 0.28 | 0.03 | 0.38 | 0.16 | 0.38 | 0.23 |
| 11 | 64107735 | rs663743 | 0.499 | 0.435 | -0.524 | -0.246 | -0.093 | 0.039 | 0.039 | 0 | 0.27 | 0.03 | 0.36 | 0.16 | 0.34 | 0.22 |
| 11 | 64111928 | rs685870 | 0.295 | 0.674 | -0.099 | -0.014 | 0.037 | 0.014 | 0.014 | 3.115 | 0.56 | 0.38 | 0.56 | 0.35 | 0.68 | 0.76 |
| 11 | 64117284 | rs4930155 | 0.246 | 1.77 | -0.024 | 0.057 | 0.091 | 0.018 | 0.018 | 0.002 | 0.72 | 0.48 | 0.55 | 0.75 | 0.57 | 0.56 |
| 11 | 64124515 | rs612448 | 0.216 | 1.747 | -0.243 | 0.092 | -0.157 | -0.027 | -0.027 | 0 | 0.22 | 0.03 | 0.41 | 0.18 | 0.43 | 0.36 |
| 11 | 64125142 | rs541207 | -0.486 | 1.726 | -0.133 | 0.096 | 0.059 | 0.013 | 0.013 | 0 | 0.27 | 0.39 | 0.44 | 0.26 | 0.44 | 0.43 |
| 11 | 64129722 | rs538147 | 0.452 | 1.649 | -0.372 | 0.27 | -0.092 | -0.033 | -0.033 | 2.698 | 0.2 | 0.05 | 0.38 | 0.21 | 0.37 | 0.35 |
| 11 | 64131280 | rs3782101 | -0.064 | 1.197 | -0.461 | 0.224 | 0.017 | -0.009 | -0.009 | 0 | 0.39 | 0.3 | 0.4 | 0.61 | 0.31 | 0.28 |
| 11 | 64147083 | rs6591843 | -0.34 | 1.534 | -0.143 | 0.019 | -0.007 | 0.043 | 0.043 | 0 | 0.53 | 0.31 | 0.69 | 0.72 | 0.52 | 0.56 |

| | | | | | | | | | | | | | | | | |
|---|---|---|---|---|---|---|---|---|---|---|---|---|---|---|---|---|
| 11 | 64150370 | rs559928 | 0.493 | 1.298 | 0.326 | 0.224 | 0.127 | 0.055 | 0.055 | 0 | 0.89 | 0.61 | 0.89 | 0.9 | 0.83 | 0.93 |
| 11 | 64156585 | rs7130280 | 0.194 | 0.999 | -0.083 | 0.277 | 0.005 | 0.009 | 0.009 | 0 | 0.46 | 0.29 | 0.57 | 0.64 | 0.44 | 0.51 |
| 11 | 64172031 | rs521319 | 1.429 | 0.91 | -0.373 | 0.181 | 0.007 | 0.083 | 0.083 | 0 | 0.31 | 0.18 | 0.16 | 0.1 | 0.3 | 0.3 |
| 11 | 64174974 | rs1783519 | 0.588 | 0.772 | 0.189 | 0.424 | 0.038 | 0.241 | 0.241 | 1.069 | 0.94 | 0.58 | 0.97 | 0.89 | 0.99 | 0.96 |
| 11 | 64175986 | rs1320628 | 0.403 | 0.985 | -0.056 | 0.41 | -0.066 | -0.02 | -0.02 | 0.564 | 0.17 | 0.05 | 0.23 | 0.15 | 0.26 | 0.15 |
| 11 | 64179813 | rs11231787 | 0.206 | 0.955 | 0.056 | 0.428 | -0.044 | -0.024 | -0.024 | 0 | 0.18 | 0.08 | 0.23 | 0.18 | 0.26 | 0.16 |
| 11 | 64189110 | rs11601686 | 0.299 | 0.969 | -0.157 | 0.425 | -0.074 | -0.032 | -0.032 | 0 | 0.18 | 0.05 | 0.23 | 0.17 | 0.26 | 0.16 |
| 11 | 64196434 | rs563536 | 1.104 | 0.983 | -0.471 | 0.398 | -0.052 | 0.11 | 0.11 | 0.067 | 0.34 | 0.03 | 0.21 | 0.12 | 0.35 | 0.21 |
| 11 | 64198311 | rs11603861 | 0.178 | 1.007 | -0.107 | 0.347 | -0.078 | -0.035 | -0.035 | 0.16 | 0.19 | 0.05 | 0.23 | 0.19 | 0.27 | 0.16 |
| 11 | 64202729 | rs606063 | 0.412 | 0.972 | 0.061 | 0.418 | -0.001 | -0.004 | -0.004 | 1.815 | 0.43 | 0.27 | 0.56 | 0.6 | 0.41 | 0.53 |
| 11 | 64210652 | rs596308 | -0.858 | 1.157 | 0.174 | -0.13 | 0.031 | -0.005 | -0.005 | 0 | 0.26 | 0.58 | 0.29 | 0.22 | 0.29 | 0.17 |
| 11 | 64212852 | rs653578 | -1.02 | 1.014 | -0.373 | -0.357 | 0.013 | 0.029 | 0.029 | 0 | 0.65 | 0.74 | 0.83 | 0.88 | 0.69 | 0.73 |
| 11 | 64218143 | rs6591849 | -0.576 | 1.281 | -0.606 | -0.106 | -0.052 | -0.013 | -0.013 | 0.256 | 0.23 | 0.13 | 0.28 | 0.23 | 0.33 | 0.17 |
| 11 | 64219679 | rs2083709 | -0.201 | 1.398 | -0.136 | -0.004 | 0.033 | 0.035 | 0.035 | 0 | 0.16 | 0.27 | 0.27 | 0.31 | 0.3 | 0.19 |
| 11 | 64219747 | rs10792436 | 0.004 | 0.93 | -0.255 | -0.269 | 0.006 | 0.031 | 0.031 | 0.07 | 0.52 | 0.35 | 0.57 | 0.64 | 0.44 | 0.57 |
| 11 | 64223527 | rs7938871 | -0.253 | 0.294 | -0.748 | -0.522 | -0.102 | 0.028 | 0.028 | 0 | 0.33 | 0.07 | 0.34 | 0.25 | 0.43 | 0.37 |
| 11 | 64226488 | rs4247634 | -0.301 | 0.021 | -0.202 | -0.515 | -0.011 | 0.031 | 0.031 | 0 | 0.36 | 0.24 | 0.34 | 0.25 | 0.41 | 0.39 |
| 11 | 64238796 | rs490581 | 0.111 | -0.041 | 0.254 | -0.421 | 0 | -0.004 | -0.004 | 0 | 0.63 | 0.62 | 0.64 | 0.71 | 0.58 | 0.61 |
| 11 | 64244194 | rs522485 | 0.31 | -0.211 | 0.518 | -0.369 | 0.003 | 0.01 | 0.01 | 0 | 0.59 | 0.37 | 0.61 | 0.68 | 0.58 | 0.6 |
| 11 | 64251221 | rs634497 | -0.029 | -0.141 | 0.608 | -0.394 | -0.014 | 0.018 | 0.018 | 0 | 0.63 | 0.78 | 0.65 | 0.72 | 0.58 | 0.6 |
| 11 | 64252970 | rs2845635 | 0.251 | -0.248 | 0.806 | -0.382 | -0.002 | -0.014 | -0.014 | 0.083 | 0.6 | 0.53 | 0.63 | 0.71 | 0.58 | 0.6 |
| 11 | 64253620 | rs2845637 | -0.169 | -0.216 | 0.338 | -0.38 | -0.077 | -0.022 | -0.022 | 0 | 0.36 | 0.06 | 0.34 | 0.29 | 0.42 | 0.4 |
| 11 | 64256859 | rs17299124 | 1.548 | -0.256 | 0.045 | -0.742 | -0.058 | 0.109 | 0.109 | 0.844 | 0.24 | 0.03 | 0.14 | 0.04 | 0.26 | 0.24 |
| 11 | 64265874 | rs6421689 | 0.855 | -0.425 | 1.368 | 0.236 | 0.021 | 0.017 | 0.017 | 4.145 | 0.93 | 0.61 | 0.96 | 1 | 1 | 1 |
| 11 | 64266244 | rs7104498 | 1.072 | -0.38 | 1.34 | 0.237 | 0.005 | 0.032 | 0.032 | 0.061 | 0.54 | 0.31 | 0.6 | 0.68 | 0.56 | 0.59 |
| 11 | 64267196 | rs10751011 | 1.13 | -0.318 | 1.441 | 0.253 | 0 | -0.004 | -0.004 | 0 | 0.92 | 0.46 | 0.95 | 1 | 1 | 1 |
| 11 | 64278637 | rs2076853 | 1.227 | -0.278 | 0.918 | 0.335 | 0.017 | 0.051 | 0.051 | 0.529 | 0.51 | 0.17 | 0.59 | 0.66 | 0.56 | 0.58 |
| 11 | 64291762 | rs7131675 | 0.731 | -0.332 | 1.06 | 0.27 | -0.001 | -0.001 | -0.001 | 0 | 0.62 | 0.4 | 0.64 | 0.65 | 0.59 | 0.62 |
| 11 | 64301713 | rs12417589 | 1.747 | -0.106 | 0.657 | 0.466 | 0.003 | 0.055 | 0.055 | 0 | 0.46 | 0.34 | 0.24 | 0.06 | 0.45 | 0.3 |
| 11 | 64302526 | rs10792438 | -0.513 | -0.183 | 1.083 | 0.591 | -0.025 | 0.167 | 0.167 | 0 | 0.21 | 0.48 | 0.44 | 0.63 | 0.14 | 0.31 |
| 11 | 64303696 | rs7104900 | 1.086 | 0.247 | 0.952 | 0.616 | 0 | 0 | 0 | 0 | 0.84 | 0.82 | 0.85 | 0.84 | 0.79 | 0.79 |
| 11 | 64310649 | rs4930195 | 1.528 | -0.076 | 0.89 | 0.375 | 0 | 0.058 | 0.058 | 1.089 | 0.44 | 0.27 | 0.24 | 0.06 | 0.47 | 0.3 |
| 11 | 64323080 | rs3759053 | 1.486 | -0.748 | 1.082 | 0.625 | 0.013 | 0.098 | 0.098 | 0.942 | 0.47 | 0.27 | 0.29 | 0.07 | 0.57 | 0.35 |
| 11 | 64331462 | rs17300741 | 1.324 | -0.696 | 1.144 | 0.664 | 0.004 | 0.028 | 0.028 | 0.159 | 0.49 | 0.36 | 0.3 | 0.07 | 0.54 | 0.35 |
| 11 | 64333296 | rs1783811 | 0.063 | -0.359 | 0.959 | 1.171 | -0.024 | -0.053 | -0.053 | 0 | 0.74 | 0.88 | 0.75 | 0.67 | 0.68 | 0.58 |
| 11 | 64365796 | rs893006 | 1.622 | -1.052 | 2.425 | 1.952 | -0.021 | -0.023 | -0.023 | 0 | 0.67 | 0.5 | 0.46 | 0.22 | 0.72 | 0.59 |
| 11 | 64369540 | rs476037 | -0.79 | -0.958 | 2.485 | 2.854 | -0.003 | -0.002 | -0.002 | 0 | 0.11 | 0.07 | 0.31 | 0.26 | 0.08 | 0.18 |
| 11 | 64372302 | rs7937990 | -1.115 | -0.854 | 2.649 | 2.821 | -0.004 | 0.034 | 0.034 | 0 | 0.19 | 0.24 | 0.22 | 0.51 | 0.2 | 0.21 |
| 11 | 64374241 | rs3741399 | 0.153 | -0.894 | 3.091 | 2.938 | -0.024 | 0.043 | 0.043 | 0 | 0.11 | 0.23 | 0.13 | 0.32 | 0.05 | 0.06 |
| 11 | 64376307 | rs3802947 | 0.153 | -0.894 | 3.085 | 2.938 | -0.023 | 0.044 | 0.044 | 0 | 0.11 | 0.23 | 0.13 | 0.32 | 0.05 | 0.06 |
| 11 | 64377336 | rs10897519 | -1.204 | -0.893 | 3.084 | 2.939 | -0.004 | 0.034 | 0.034 | 0 | 0.19 | 0.24 | 0.22 | 0.51 | 0.2 | 0.21 |
| 11 | 64381132 | rs7121121 | -1.807 | -0.915 | 3.228 | 2.927 | -0.022 | 0.06 | 0.06 | 0 | 0.22 | 0.66 | 0.25 | 0.51 | 0.2 | 0.21 |
| 11 | 64391910 | rs10897521 | -1.196 | -0.9 | 3.237 | 2.934 | -0.004 | 0.034 | 0.034 | 0 | 0.19 | 0.24 | 0.22 | 0.51 | 0.2 | 0.21 |
| 11 | 64400719 | rs948689 | -1.773 | -0.879 | 2.982 | 2.959 | -0.021 | 0.061 | 0.061 | 0 | 0.22 | 0.65 | 0.25 | 0.51 | 0.2 | 0.21 |
| 11 | 64415767 | rs3825074 | -0.556 | -0.853 | 2.354 | 2.947 | -0.03 | -0.043 | -0.043 | 1.89 | 0.07 | 0.01 | 0.09 | 0.19 | 0.15 | 0.14 |
| 11 | 64418305 | rs2285339 | -0.761 | -0.769 | 2.466 | 3.015 | -0.018 | 0.089 | 0.089 | 0 | 0.21 | 0.37 | 0.24 | 0.51 | 0.2 | 0.21 |
| 11 | 64492762 | rs693235 | 1.495 | -0.863 | 2.526 | 2.599 | 0.015 | 0.248 | 0.248 | 0.281 | 0.86 | 0.42 | 0.66 | 0.67 | 0.91 | 0.85 |
| 11 | 64495284 | rs2267918 | 1.458 | -0.863 | 2.534 | 2.588 | 0.019 | 0.256 | 0.256 | 0.248 | 0.87 | 0.43 | 0.66 | 0.67 | 0.91 | 0.85 |
| 11 | 64503363 | rs10897524 | -1.535 | -0.86 | 2.498 | 2.635 | 0.004 | 0.153 | 0.153 | 0 | 0.12 | 0.23 | 0.32 | 0.36 | 0.09 | 0.15 |
| 11 | 64514506 | rs569602 | -0.742 | -0.72 | 2.373 | 2.751 | -0.007 | 0.037 | 0.037 | 0.575 | 0.08 | 0.08 | 0.29 | 0.35 | 0.08 | 0.14 |
| 11 | 64519062 | rs532747 | -0.648 | -0.714 | 2.354 | 2.775 | -0.008 | 0.028 | 0.028 | 0.535 | 0.08 | 0.07 | 0.28 | 0.32 | 0.08 | 0.14 |
| 11 | 64527080 | rs477549 | -1.264 | -0.701 | 2.462 | 2.681 | 0.05 | 0.346 | 0.346 | 0 | 0.12 | 0.4 | 0.32 | 0.36 | 0.09 | 0.19 |
| 11 | 64527751 | rs483962 | -1.206 | -0.67 | 2.359 | 2.696 | 0.041 | 0.302 | 0.302 | 0 | 0.12 | 0.4 | 0.32 | 0.35 | 0.09 | 0.19 |
| 11 | 64528750 | rs637332 | -1.103 | -0.665 | 2.218 | 2.698 | 0.037 | 0.281 | 0.281 | 0 | 0.12 | 0.35 | 0.31 | 0.35 | 0.09 | 0.19 |
| 11 | 64544135 | rs474707 | -1.106 | -0.648 | 1.721 | 2.72 | 0.036 | 0.279 | 0.279 | 0 | 0.12 | 0.35 | 0.31 | 0.35 | 0.09 | 0.19 |
| 11 | 64546106 | rs3741398 | -0.801 | -0.625 | 1.607 | 2.61 | 0.004 | 0.086 | 0.086 | 0 | 0.11 | 0.15 | 0.3 | 0.33 | 0.08 | 0.19 |
| 11 | 64559898 | rs10897526 | -1.469 | -0.89 | 0.546 | 2.167 | 0.023 | 0.011 | 0.011 | 0 | 0.34 | 0.25 | 0.48 | 0.74 | 0.25 | 0.48 |
| 11 | 64569000 | rs1143937 | -1.516 | -0.112 | 0.768 | 2.257 | -0.002 | 0.044 | 0.044 | 0 | 0.13 | 0.15 | 0.3 | 0.26 | 0.08 | 0.17 |
| 11 | 64573589 | rs669976 | -1.197 | 0.288 | -0.674 | 1.962 | 0 | 0.02 | 0.02 | 0 | 0.12 | 0.1 | 0.3 | 0.27 | 0.08 | 0.18 |
| 11 | 64584231 | rs470763 | 0.078 | 0.419 | -1.136 | 1.358 | 0.045 | -0.007 | -0.007 | 6.373 | 0.5 | 0.21 | 0.64 | 0.55 | 0.59 | 0.42 |
| 11 | 64584959 | rs524386 | -0.795 | 0.771 | -0.898 | 2.156 | -0.027 | -0.028 | -0.028 | 0 | 0.07 | 0 | 0.29 | 0.33 | 0.07 | 0.14 |
| 11 | 64584987 | rs2957154 | 0.467 | 0.856 | -0.334 | 1.836 | 0.01 | 0.116 | 0.116 | 0 | 0.75 | 0.5 | 0.8 | 0.63 | 0.72 | 0.63 |
| 11 | 64591679 | rs670358 | -1.241 | 0.946 | -0.498 | 2.257 | -0.005 | 0.046 | 0.046 | 0.267 | 0.07 | 0.07 | 0.34 | 0.46 | 0.13 | 0.14 |
| 11 | 64591972 | rs10792447 | 0.592 | 0.264 | -0.284 | 1.136 | -0.01 | -0.099 | -0.099 | 0 | 0.59 | 0.5 | 0.41 | 0.05 | 0.62 | 0.44 |
| 11 | 64606177 | rs653343 | -1.059 | 0.28 | 0.055 | 1.421 | 0.054 | 0.275 | 0.275 | 0.77 | 0.09 | 0.28 | 0.37 | 0.45 | 0.12 | 0.14 |
| 11 | 64616237 | rs916606 | -0.264 | 0.252 | 0.107 | 1.324 | 0.015 | -0.054 | -0.054 | 0 | 0.31 | 0.17 | 0.23 | 0.49 | 0.27 | 0.42 |
| 11 | 64639163 | rs627497 | -0.995 | 0.302 | 1.143 | 1.341 | 0.026 | 0.089 | 0.089 | 0 | 0.12 | 0.25 | 0.12 | 0.37 | 0.17 | 0.15 |
| 11 | 64645871 | rs1211284 | 0.061 | -0.862 | 0.627 | 1.155 | -0.041 | 0.052 | 0.052 | 1.411 | 0.75 | 0.34 | 0.82 | 0.56 | 0.88 | 0.81 |
| 11 | 64646842 | rs589556 | -0.851 | -0.863 | 0.422 | 1.186 | 0.034 | -0.053 | -0.053 | 0.42 | 0.82 | 0.87 | 0.89 | 0.57 | 0.92 | 0.87 |

| | | | | | | | | | | | | | | | | |
|---|---|---|---|---|---|---|---|---|---|---|---|---|---|---|---|---|
| 11 | 64659424 | rs633727 | 1.105 | -0.964 | -0.722 | -0.455 | -0.083 | 0.201 | 0.201 | 0.642 | 0.4 | 0.09 | 0.32 | 0.08 | 0.48 | 0.38 |
| 11 | 64661401 | rs7939874 | 0.151 | -1.1 | -0.708 | -0.118 | -0.002 | -0.107 | -0.107 | 0 | 0.29 | 0.1 | 0.21 | 0.42 | 0.28 | 0.29 |
| 11 | 64663227 | rs17146441 | 0.419 | -1.096 | -0.694 | -0.077 | -0.006 | -0.111 | -0.111 | 0 | 0.27 | 0.09 | 0.19 | 0.42 | 0.25 | 0.28 |
| 11 | 64665502 | rs11827610 | -0.107 | -1.118 | 0.315 | -0.179 | 0.026 | -0.007 | -0.007 | 0 | 0.08 | 0.45 | 0.08 | 0 | 0.05 | 0.06 |
| 11 | 64684288 | rs2285346 | 0.602 | -1.262 | 0.043 | -0.006 | 0.01 | -0.035 | -0.035 | 0 | 0.25 | 0.19 | 0.2 | 0.45 | 0.22 | 0.28 |
| 11 | 64685482 | rs628595 | -0.681 | -1.231 | 0.291 | -0.105 | -0.011 | -0.032 | -0.032 | 0 | 0.94 | 0.76 | 0.94 | 1 | 0.95 | 0.94 |
| 11 | 64697507 | rs11231903 | 1.534 | -0.197 | -0.546 | -0.904 | -0.096 | 0.211 | 0.211 | 0.395 | 0.29 | 0.02 | 0.17 | 0 | 0.31 | 0.16 |
| 11 | 64700767 | rs675090 | -0.811 | -0.416 | -0.181 | -0.351 | 0.021 | -0.051 | -0.051 | 0.096 | 0.7 | 0.8 | 0.78 | 0.55 | 0.75 | 0.64 |
| 11 | 64701030 | rs3741392 | -0.294 | -0.439 | 0.418 | -0.153 | 0.001 | 0.166 | 0.166 | 0.497 | 0.08 | 0.24 | 0.36 | 0.3 | 0.08 | 0.08 |
| 11 | 64702200 | rs673995 | -0.811 | -0.305 | 0.36 | 0.205 | 0.007 | -0.036 | -0.036 | 5.302 | 0.68 | 0.78 | 0.76 | 0.58 | 0.69 | 0.63 |
| 11 | 64704630 | rs17146506 | 1.251 | -0.584 | 0.437 | 0.04 | -0.004 | -0.015 | -0.015 | 0.548 | 0.05 | 0.07 | 0.04 | 0 | 0.05 | 0.05 |
| 11 | 64721908 | rs495330 | 0.385 | -0.43 | 0.576 | -0.004 | 0.023 | 0.038 | 0.038 | 1.057 | 0.56 | 0.37 | 0.34 | 0.18 | 0.65 | 0.33 |
| 11 | 64727087 | rs17146508 | 1.185 | -0.383 | 0.447 | 0.066 | -0.001 | -0.01 | -0.01 | 0 | 0.05 | 0.06 | 0.03 | 0 | 0.05 | 0.05 |
| 11 | 64728107 | rs17146511 | 1.237 | -0.383 | 0.449 | 0.066 | 0.005 | 0.001 | 0.001 | 0 | 0.05 | 0.04 | 0.03 | 0 | 0.05 | 0.05 |
| 11 | 64728611 | rs17146519 | 1.185 | -0.383 | 0.448 | 0.066 | -0.001 | -0.01 | -0.01 | 0 | 0.05 | 0.06 | 0.03 | 0 | 0.05 | 0.05 |
| 11 | 64748278 | rs7106136 | -1.246 | -0.336 | 0.436 | 0.14 | -0.024 | 0.129 | 0.129 | 0 | 0.12 | 0.23 | 0.38 | 0.36 | 0.07 | 0.31 |
| 11 | 64757496 | rs10897540 | -0.299 | -0.241 | 0.46 | 0.085 | -0.018 | 0.11 | 0.11 | 0 | 0.09 | 0.17 | 0.31 | 0.27 | 0.04 | 0.12 |
| 11 | 64761779 | rs626670 | -0.863 | -0.118 | 0.169 | 0.047 | 0.016 | -0.043 | -0.043 | 0 | 0.69 | 0.77 | 0.74 | 0.48 | 0.72 | 0.62 |
| 11 | 64772545 | rs7103328 | 0.363 | -0.056 | 0.392 | -0.04 | -0.012 | -0.032 | -0.032 | 0.027 | 0.06 | 0.16 | 0.05 | 0 | 0.05 | 0.05 |
| 11 | 64789077 | rs663767 | -0.142 | -0.187 | 0.046 | -0.399 | 0.021 | 0.033 | 0.033 | 0.077 | 0.61 | 0.72 | 0.47 | 0.21 | 0.67 | 0.53 |
| 11 | 64793960 | rs500464 | -1.137 | 0.033 | 0.431 | -0.169 | 0.023 | 0.006 | 0.006 | 0.19 | 0.09 | 0.28 | 0.11 | 0.06 | 0.09 | 0.27 |
| 11 | 64803351 | rs562590 | -0.74 | 0.117 | 0.722 | -0.058 | -0.002 | -0.003 | -0.003 | 0.041 | 0.71 | 0.65 | 0.75 | 0.26 | 0.7 | 0.65 |
| 11 | 64806270 | rs542175 | -0.911 | 0.167 | 0.703 | -0.037 | 0.018 | 0.001 | 0.001 | 0.196 | 0.09 | 0.27 | 0.11 | 0.06 | 0.08 | 0.28 |
| 11 | 64819168 | rs1300845 | 0.394 | 0.111 | 0.638 | -0.035 | -0.002 | -0.007 | -0.007 | 0 | 0.29 | 0.4 | 0.25 | 0.76 | 0.3 | 0.35 |
| 11 | 64827087 | rs3759041 | 0.517 | 0.137 | 0.676 | 0.719 | -0.002 | -0.002 | -0.002 | 0 | 0.28 | 0.28 | 0.24 | 0.71 | 0.3 | 0.35 |
| 11 | 64840865 | rs3850947 | 0.591 | 0.063 | 0.862 | 0.738 | -0.001 | 0 | 0 | 0 | 0.27 | 0.3 | 0.21 | 0.65 | 0.26 | 0.34 |
| 11 | 64856108 | rs2282499 | -0.075 | 0.053 | 0.56 | -0.009 | 0 | 0.002 | 0.002 | 0 | 0.42 | 0.43 | 0.59 | 0.27 | 0.45 | 0.29 |
| 11 | 64870779 | rs1209621 | -0.078 | 0.109 | 0.763 | 0.276 | 0.056 | 0.086 | 0.086 | 1.071 | 0.93 | 0.73 | 0.91 | 0.93 | 0.94 | 0.74 |
| 11 | 64882789 | rs1129195 | 0.01 | 0.08 | 0.671 | 0.239 | 0.006 | 0.042 | 0.042 | 0 | 0.41 | 0.33 | 0.58 | 0.27 | 0.45 | 0.28 |
| 11 | 64883864 | rs1706937 | 0.697 | 0.088 | 0.397 | 0.243 | 0.008 | 0.044 | 0.044 | 0 | 0.51 | 0.31 | 0.32 | 0.66 | 0.49 | 0.47 |
| 11 | 64887560 | rs1196043 | 0.916 | -0.121 | 0.756 | 0.589 | 0.007 | 0.043 | 0.043 | 0 | 0.51 | 0.31 | 0.32 | 0.66 | 0.49 | 0.47 |
| 11 | 64892470 | rs3782094 | -0.084 | -0.149 | 0.123 | 0.642 | -0.039 | 0.077 | 0.077 | 0 | 0.39 | 0.06 | 0.5 | 0.2 | 0.41 | 0.26 |
| 11 | 64895202 | rs10088 | 0.158 | -0.098 | 0.354 | 0.66 | 0.074 | 0.091 | 0.091 | 0.877 | 0.93 | 0.66 | 0.9 | 0.93 | 0.94 | 0.74 |
| 11 | 64898226 | rs9971407 | 1.384 | -0.114 | -0.984 | 0.245 | -0.011 | 0.018 | 0.018 | 0 | 0.17 | 0.11 | 0.16 | 0.06 | 0.23 | 0.12 |
| 11 | 64900270 | rs3825073 | 0.706 | -0.172 | -1.162 | 0.224 | 0.002 | -0.004 | -0.004 | 0.308 | 0.23 | 0.01 | 0.35 | 0.15 | 0.18 | 0.14 |
| 11 | 64902738 | rs1196045 | 1.357 | -0.129 | -1.117 | 0.806 | -0.14 | 0.034 | 0.034 | 0 | 0.44 | 0.03 | 0.23 | 0.34 | 0.41 | 0.41 |
| 11 | 64936719 | rs12292693 | 1.193 | 0.064 | -0.681 | 0.719 | -0.012 | 0.022 | 0.022 | 0 | 0.18 | 0.1 | 0.16 | 0.06 | 0.24 | 0.15 |
| 11 | 64936810 | rs1195958 | 0.177 | -0.034 | -0.172 | 1.166 | 0.038 | 0.278 | 0.278 | 1.037 | 0.88 | 0.59 | 0.82 | 0.65 | 0.86 | 0.71 |
| 11 | 64936931 | rs1195959 | 0.177 | -0.034 | -0.172 | 1.172 | 0.038 | 0.274 | 0.274 | 1.037 | 0.88 | 0.59 | 0.82 | 0.66 | 0.86 | 0.71 |
| 11 | 64937683 | rs17743381 | 0.456 | -0.136 | -0.953 | 0.988 | 0.024 | 0.057 | 0.057 | 0 | 0.24 | 0.09 | 0.38 | 0.09 | 0.19 | 0.14 |
| 11 | 64938381 | rs1195960 | 1.218 | 0.01 | -0.928 | 1.098 | -0.11 | -0.03 | -0.03 | 0.311 | 0.44 | 0.06 | 0.23 | 0.43 | 0.42 | 0.43 |
| 11 | 64941419 | rs1195962 | 0.384 | 0.269 | -0.346 | 1.233 | -0.018 | 0.029 | 0.029 | 0 | 0.55 | 0.23 | 0.34 | 0.38 | 0.52 | 0.38 |
| 11 | 64946858 | rs1195968 | 0.983 | 0.243 | 0.039 | 1.383 | -0.008 | 0.012 | 0.012 | 0 | 0.65 | 0.43 | 0.68 | 0.47 | 0.69 | 0.58 |
| 11 | 64948277 | rs3953797 | 1.122 | 0.27 | -0.084 | 1.35 | -0.004 | -0.002 | -0.002 | 0 | 0.27 | 0.24 | 0.42 | 0.3 | 0.31 | 0.28 |
| 11 | 64954053 | rs3825072 | 1.379 | 0.287 | -0.306 | 1.337 | -0.047 | -0.037 | -0.037 | 0 | 0.25 | 0.05 | 0.4 | 0.3 | 0.3 | 0.31 |
| 11 | 64954338 | rs17146724 | 0.63 | 0.013 | -0.185 | 1.386 | 0.016 | -0.04 | -0.04 | 0 | 0.18 | 0.07 | 0.08 | 0.21 | 0.13 | 0.08 |
| 11 | 64962958 | rs7116674 | -0.188 | 0.197 | 0.144 | 1.534 | 0.006 | 0 | 0 | 0 | 0.9 | 0.89 | 0.89 | 0.94 | 0.91 | 0.76 |
| 11 | 64971396 | rs7114754 | 1.646 | 0.027 | -0.097 | 1.557 | -0.016 | -0.072 | -0.072 | 0.206 | 0.12 | 0.01 | 0.05 | 0.2 | 0.09 | 0.03 |
| 11 | 64977644 | rs625750 | 0.852 | 0.141 | -0.037 | 1.514 | -0.048 | 0.028 | 0.028 | 0.127 | 0.49 | 0.1 | 0.28 | 0.43 | 0.44 | 0.36 |
| 11 | 64977937 | rs542380 | 0.733 | 0.171 | 0.751 | 1.765 | 0.018 | 0.117 | 0.117 | 0.231 | 0.84 | 0.41 | 0.76 | 0.67 | 0.82 | 0.7 |
| 11 | 64981837 | rs12420456 | 0.77 | 0.386 | 0.455 | 1.541 | -0.016 | 0.002 | 0.002 | 0 | 0.28 | 0.14 | 0.46 | 0.26 | 0.36 | 0.32 |
| 11 | 65000719 | rs2904979 | 0.743 | 0.043 | 1.316 | 2.039 | 0.003 | 0.012 | 0.012 | 0.332 | 0.91 | 0.85 | 0.91 | 0.92 | 0.92 | 0.78 |
| 11 | 65004309 | rs564476 | 0.853 | 0.05 | 1.153 | 2.143 | -0.019 | -0.077 | -0.077 | 1.197 | 0.12 | 0.01 | 0.04 | 0.23 | 0.08 | 0.03 |
| 11 | 65042941 | rs1858783 | -1.918 | -0.307 | 1.538 | 1.524 | -0.022 | -0.042 | -0.042 | 0.021 | 0.17 | 0.47 | 0.16 | 0.07 | 0.15 | 0.22 |
| 11 | 65045725 | rs668735 | -1.918 | -0.307 | 1.538 | 1.524 | -0.022 | -0.042 | -0.042 | 0.203 | 0.17 | 0.47 | 0.16 | 0.07 | 0.15 | 0.22 |
| 11 | 65064690 | rs487989 | -1.451 | -0.308 | 1.209 | 1.247 | -0.004 | -0.007 | -0.007 | 0 | 0.15 | 0.18 | 0.13 | 0.06 | 0.14 | 0.22 |
| 11 | 65109233 | rs629710 | 1.422 | -0.189 | 1.031 | 1.541 | 0.016 | -0.027 | -0.027 | 0.534 | 0.67 | 0.18 | 0.69 | 0.46 | 0.72 | 0.68 |
| 11 | 65114885 | rs1144926 | 0.09 | 0.022 | 0.915 | 1.654 | 0.05 | 0.259 | 0.259 | 0 | 0.89 | 0.68 | 0.86 | 0.55 | 0.85 | 0.89 |
| 11 | 65127401 | rs7127811 | -0.829 | -0.285 | 0.29 | 0.587 | -0.027 | -0.045 | -0.045 | 0.077 | 0.09 | 0.31 | 0.05 | 0.01 | 0.04 | 0.02 |
| 11 | 65139776 | rs551523 | 0.266 | -0.233 | -0.073 | 0.709 | -0.011 | -0.007 | -0.007 | 0 | 0.1 | 0.15 | 0.11 | 0.05 | 0.07 | 0.16 |
| 11 | 65140209 | rs17304039 | 1.021 | -0.282 | -0.751 | 0.203 | -0.031 | 0.06 | 0.06 | 0.522 | 0.16 | 0.01 | 0.09 | 0.03 | 0.16 | 0.17 |
| 11 | 65146257 | rs7925823 | -1.005 | -0.848 | 0.454 | 0.817 | -0.037 | 0.075 | 0.075 | 0.368 | 0.1 | 0.4 | 0.11 | 0.18 | 0.04 | 0.07 |
| 11 | 65147294 | rs547221 | 1.25 | -0.664 | -0.349 | 0.777 | -0.09 | 0.088 | 0.088 | 0 | 0.46 | 0.06 | 0.29 | 0.27 | 0.55 | 0.37 |
| 11 | 65151449 | rs681309 | 1.29 | -0.687 | -0.388 | 0.763 | -0.127 | 0.062 | 0.062 | 0 | 0.46 | 0.03 | 0.29 | 0.27 | 0.55 | 0.37 |
| 11 | 65157094 | rs739389 | -0.517 | -0.549 | -0.115 | 0.829 | 0.005 | 0.117 | 0.117 | 0 | 0.26 | 0.43 | 0.26 | 0.48 | 0.25 | 0.46 |
| 11 | 65157239 | rs739390 | 1.081 | -0.581 | -0.348 | 0.793 | -0.039 | 0.036 | 0.036 | 0 | 0.17 | 0.03 | 0.34 | 0.06 | 0.16 | 0.09 |
| 11 | 65161129 | rs2073800 | 1.65 | -0.563 | -0.373 | 0.773 | -0.128 | 0.067 | 0.067 | 0 | 0.46 | 0.03 | 0.29 | 0.27 | 0.55 | 0.39 |
| 11 | 65164114 | rs7940559 | -0.84 | -0.615 | -0.1 | 0.776 | -0.002 | -0.008 | -0.008 | 0 | 0.39 | 0.62 | 0.52 | 0.52 | 0.36 | 0.28 |
| 11 | 65168281 | rs2073801 | -1.217 | -0.451 | 0.329 | 0.768 | -0.016 | 0.016 | 0.016 | 0 | 0.13 | 0.22 | 0.17 | 0.2 | 0.08 | 0.33 |

| | | | | | | | | | | | | | | | | |
|---|---|---|---|---|---|---|---|---|---|---|---|---|---|---|---|---|
| 11 | 65183815 | rs666834 | 1.118 | -0.46 | 0.324 | 0.763 | -0.025 | 0.023 | 0.023 | 0.01 | 0.86 | 0.65 | 0.82 | 0.8 | 0.92 | 0.69 |
| 11 | 65185387 | rs2904976 | 1.122 | -0.472 | 0.483 | 0.757 | -0.032 | 0.037 | 0.037 | 0.275 | 0.85 | 0.5 | 0.81 | 0.8 | 0.92 | 0.67 |
| 11 | 65190379 | rs680413 | 0.011 | -0.484 | 0.476 | 0.8 | -0.027 | 0.055 | 0.055 | 0 | 0.91 | 0.78 | 0.89 | 0.82 | 0.96 | 0.94 |
| 11 | 65197393 | rs674485 | 0.779 | -0.435 | 0.293 | 0.882 | -0.009 | -0.012 | -0.012 | 0 | 0.67 | 0.4 | 0.47 | 0.71 | 0.76 | 0.56 |
| 11 | 65203561 | rs1144924 | 0.345 | 0.544 | -0.151 | 1.385 | -0.065 | -0.049 | -0.049 | 0.114 | 0.16 | 0.02 | 0.14 | 0.25 | 0.22 | 0.13 |
| 11 | 65249089 | rs1787666 | -1.329 | 0.127 | 0.541 | 1.329 | 0.015 | 0.08 | 0.08 | 0 | 0.13 | 0.35 | 0.42 | 0.24 | 0.12 | 0.32 |
| 11 | 65249145 | rs17146964 | -0.02 | -0.436 | 0.452 | 1.076 | 0.012 | 0.037 | 0.037 | 0.308 | 0.3 | 0.19 | 0.12 | 0.16 | 0.2 | 0.12 |
| 11 | 65257329 | rs1152620 | -0.565 | -0.56 | 0.668 | 1.418 | -0.006 | 0.004 | 0.004 | 1.09 | 0.36 | 0.4 | 0.47 | 0.47 | 0.26 | 0.24 |
| 11 | 65260646 | rs600231 | -0.946 | -0.51 | 0.792 | 1.209 | 0.033 | -0.014 | -0.014 | 0 | 0.41 | 0.77 | 0.55 | 0.41 | 0.29 | 0.42 |
| 11 | 65271832 | rs3200401 | 0.123 | -0.356 | 0.721 | 0.84 | 0.011 | 0.036 | 0.036 | 2.395 | 0.27 | 0.11 | 0.13 | 0.18 | 0.2 | 0.11 |
| 11 | 65278461 | rs10896012 | 0.166 | -0.352 | 0.705 | 0.84 | 0.006 | 0.038 | 0.038 | 2.542 | 0.27 | 0.08 | 0.13 | 0.18 | 0.2 | 0.11 |
| 11 | 65294799 | rs1783541 | 0.202 | -0.341 | 1.149 | 0.867 | 0.01 | 0.023 | 0.023 | 2.758 | 0.26 | 0.13 | 0.13 | 0.17 | 0.19 | 0.1 |
| 11 | 65294830 | rs1787663 | -0.536 | -0.318 | 1.346 | 0.943 | -0.021 | -0.032 | -0.032 | 0 | 0.35 | 0.5 | 0.21 | 0.22 | 0.27 | 0.26 |
| 11 | 65316382 | rs11601767 | -0.889 | -0.342 | 1.132 | 1.04 | 0.029 | 0.01 | 0.01 | 2.036 | 0.4 | 0.77 | 0.29 | 0.31 | 0.31 | 0.41 |
| 11 | 65319265 | rs11227221 | 0.911 | -0.384 | 1.116 | 1.05 | 0.037 | 0.034 | 0.034 | 0.498 | 0.57 | 0.22 | 0.72 | 0.69 | 0.7 | 0.57 |
| 11 | 65322548 | rs12270054 | -0.263 | -0.399 | 1.213 | 1.021 | -0.002 | -0.003 | -0.003 | 0 | 0.33 | 0.31 | 0.18 | 0.2 | 0.26 | 0.28 |
| 11 | 65330510 | rs11227226 | -0.228 | -0.35 | 0.942 | 1.032 | -0.001 | -0.003 | -0.003 | 0 | 0.08 | 0.07 | 0.06 | 0.09 | 0.05 | 0.16 |
| 11 | 65335705 | rs10896016 | 0.917 | -0.348 | 0.872 | 1.02 | 0.033 | 0.023 | 0.023 | 0.499 | 0.56 | 0.23 | 0.72 | 0.68 | 0.69 | 0.56 |
| 11 | 65337714 | rs1784220 | -0.404 | -0.253 | 0.683 | 1.067 | -0.017 | -0.024 | -0.024 | 0 | 0.34 | 0.47 | 0.19 | 0.21 | 0.26 | 0.28 |
| 11 | 65338944 | rs2236684 | -0.125 | 0.006 | 0.265 | 1.173 | -0.013 | -0.028 | -0.028 | 0.078 | 0.07 | 0.01 | 0.05 | 0.09 | 0.04 | 0.16 |
| 11 | 65349756 | rs6591182 | 0.59 | 0.199 | 0.028 | 0.921 | -0.01 | -0.009 | -0.009 | 0 | 0.46 | 0.31 | 0.38 | 0.45 | 0.51 | 0.51 |
| 11 | 65353906 | rs4601790 | -0.401 | 0.513 | -0.297 | 0.956 | -0.051 | -0.085 | -0.085 | 0 | 0.17 | 0.07 | 0.43 | 0.42 | 0.26 | 0.37 |
| 11 | 65363865 | rs10791819 | 0.068 | 0.194 | 0.152 | 0.675 | 0.02 | 0.06 | 0.06 | 0.7 | 0.33 | 0.14 | 0.14 | 0.16 | 0.26 | 0.13 |
| 11 | 65363958 | rs7937415 | -0.03 | 0.197 | 0.322 | 0.677 | 0.015 | 0.031 | 0.031 | 0.751 | 0.34 | 0.24 | 0.16 | 0.16 | 0.26 | 0.13 |
| 11 | 65365318 | rs948577 | 0.076 | 0.22 | 0.033 | 0.688 | 0.019 | 0.062 | 0.062 | 0.644 | 0.33 | 0.13 | 0.14 | 0.16 | 0.26 | 0.13 |
| 11 | 65368323 | rs10791821 | 0.074 | 0.223 | 0.035 | 0.651 | 0.019 | 0.135 | 0.135 | 0.578 | 0.33 | 0.13 | 0.14 | 0.04 | 0.26 | 0.11 |
| 11 | 65368434 | rs10750763 | 0.066 | 0.223 | 0.035 | 0.709 | 0.02 | 0.061 | 0.061 | 0.598 | 0.34 | 0.13 | 0.14 | 0.16 | 0.25 | 0.13 |
| 11 | 65380248 | rs1784223 | -0.157 | 0.578 | 0.587 | 0.448 | -0.003 | -0.068 | -0.068 | 0 | 0.37 | 0.41 | 0.25 | 0.14 | 0.34 | 0.11 |
| 11 | 65382564 | rs1787034 | -0.12 | 0.574 | 0.76 | 0.445 | 0 | -0.011 | -0.011 | 0 | 0.37 | 0.33 | 0.24 | 0.14 | 0.34 | 0.11 |
| 11 | 65386206 | rs1193851 | -0.101 | 0.566 | 0.757 | 0.441 | 0 | -0.015 | -0.015 | 0 | 0.36 | 0.33 | 0.25 | 0.13 | 0.34 | 0.11 |
| 11 | 65389631 | rs1144789 | -1.44 | 0.237 | 1.068 | 0.666 | -0.008 | -0.021 | -0.021 | 0 | 0.59 | 0.7 | 0.69 | 0.56 | 0.56 | 0.34 |
| 11 | 65391317 | rs12801636 | -0.491 | 0.259 | 0.782 | 0.757 | -0.002 | 0.02 | 0.02 | 0 | 0.22 | 0.27 | 0.44 | 0.43 | 0.22 | 0.23 |
| 11 | 65399528 | rs2009453 | 0.248 | 0.429 | 0.744 | 1.133 | 0.012 | 0.004 | 0.004 | 0.5 | 0.5 | 0.35 | 0.34 | 0.46 | 0.56 | 0.51 |
| 11 | 65405300 | rs931127 | 0.362 | 0.403 | 0.731 | 1.119 | 0.011 | 0.002 | 0.002 | 0 | 0.51 | 0.34 | 0.34 | 0.48 | 0.56 | 0.52 |
| 11 | 65406179 | rs7944860 | 0.491 | 0.357 | 0.786 | 0.914 | -0.052 | 0.166 | 0.166 | 0.802 | 0.35 | 0.05 | 0.22 | 0.14 | 0.35 | 0.11 |
| 11 | 65408937 | rs3741378 | 0.136 | 0.427 | 1.073 | 1.049 | 0.078 | 0.294 | 0.294 | 0 | 0.09 | 0.3 | 0.14 | 0.35 | 0.13 | 0.06 |
| 11 | 65412035 | rs2448490 | 0.583 | 0.408 | 1.026 | 0.97 | -0.012 | 0.034 | 0.034 | 0.736 | 0.35 | 0.13 | 0.23 | 0.25 | 0.35 | 0.11 |
| 11 | 65412467 | rs2306364 | 0.713 | 0.4 | 0.886 | 0.872 | 0.005 | 0.062 | 0.062 | 0.062 | 0.44 | 0.33 | 0.26 | 0.25 | 0.44 | 0.47 |
| 11 | 65422853 | rs11227247 | -0.095 | 0.57 | 1.335 | 1.147 | 0.057 | 0.279 | 0.279 | 0 | 0.09 | 0.34 | 0.41 | 0.39 | 0.13 | 0.07 |
| 11 | 65428178 | rs11227248 | 0.61 | 0.522 | 1.204 | 0.963 | -0.073 | 0.218 | 0.218 | 1.063 | 0.34 | 0.03 | 0.22 | 0.05 | 0.34 | 0.09 |
| 11 | 65429936 | rs11820062 | 0.75 | 0.445 | 1.431 | 0.979 | 0.01 | -0.003 | -0.003 | 0 | 0.42 | 0.6 | 0.63 | 0.61 | 0.49 | 0.37 |
| 11 | 65431360 | rs7101916 | 0.434 | 0.565 | 1.386 | 1.023 | 0.045 | 0.234 | 0.234 | 0 | 0.08 | 0.22 | 0.39 | 0.39 | 0.13 | 0.07 |
| 11 | 65436888 | rs12289836 | 1.8 | 0.387 | 1.238 | 0.959 | -0.052 | 0.035 | 0.035 | 0.066 | 0.3 | 0.11 | 0.2 | 0.22 | 0.35 | 0.3 |
| 11 | 65458197 | rs1074155 | 1.618 | 0.395 | 1.308 | 0.963 | -0.005 | 0.011 | 0.011 | 0.049 | 0.32 | 0.33 | 0.23 | 0.22 | 0.35 | 0.3 |
| 11 | 65459404 | rs11227259 | 0.889 | 0.473 | 1.306 | 0.965 | -0.003 | 0 | 0 | 0.52 | 0.4 | 0.63 | 0.63 | 0.61 | 0.48 | 0.37 |
| 11 | 65463447 | rs4930304 | 0.546 | 0.346 | 1.085 | 0.89 | -0.026 | 0.162 | 0.162 | 3.894 | 0.36 | 0.09 | 0.23 | 0.14 | 0.35 | 0.1 |
| 11 | 65466911 | rs555903 | 0.547 | 0.42 | 1.175 | 0.955 | -0.001 | -0.004 | -0.004 | 1.979 | 0.11 | 0.16 | 0.11 | 0.1 | 0.1 | 0.36 |
| 11 | 65467953 | rs6591188 | 1.714 | 0.301 | 1.253 | 0.96 | -0.007 | 0.013 | 0.013 | 0.075 | 0.31 | 0.31 | 0.23 | 0.22 | 0.36 | 0.31 |
| 11 | 65475620 | rs618456 | -0.665 | 0.384 | 1.26 | 0.948 | -0.007 | -0.004 | -0.004 | 0 | 0.88 | 0.8 | 0.89 | 0.9 | 0.91 | 0.64 |
| 11 | 65480768 | rs4244812 | 0.936 | 0.328 | 1.324 | 0.959 | 0.006 | -0.002 | -0.002 | 0 | 0.4 | 0.46 | 0.61 | 0.61 | 0.48 | 0.37 |
| 11 | 65481166 | rs1151500 | -0.636 | 0.35 | 1.381 | 0.966 | -0.001 | 0.003 | 0.003 | 0 | 0.88 | 0.73 | 0.89 | 0.9 | 0.91 | 0.64 |
| 11 | 65484883 | rs1151501 | -0.725 | 0.351 | 1.217 | 0.967 | -0.006 | -0.004 | -0.004 | 0 | 0.89 | 0.81 | 0.89 | 0.9 | 0.91 | 0.65 |
| 11 | 65485218 | rs3372 | 0.505 | 0.32 | 1.22 | 0.933 | -0.005 | 0.075 | 0.075 | 1.673 | 0.36 | 0.22 | 0.23 | 0.14 | 0.36 | 0.1 |
| 11 | 65485337 | rs535111 | -0.37 | 0.256 | 1.255 | 1.069 | 0.02 | 0.087 | 0.087 | 1.541 | 0.48 | 0.28 | 0.28 | 0.29 | 0.43 | 0.26 |
| 11 | 65485727 | rs521678 | -0.586 | 0.155 | 1.176 | 1.092 | -0.007 | -0.004 | -0.004 | 0 | 0.88 | 0.8 | 0.9 | 0.9 | 0.91 | 0.64 |
| 11 | 65494987 | rs537786 | -0.719 | 0.228 | 1.044 | 1.07 | 0.003 | 0 | 0 | 0 | 0.49 | 0.54 | 0.67 | 0.53 | 0.53 | 0.27 |
| 11 | 65495211 | rs11227279 | 1.645 | 0.163 | 1.068 | 0.86 | -0.036 | 0.04 | 0.04 | 0.696 | 0.32 | 0.09 | 0.2 | 0.23 | 0.35 | 0.35 |
| 11 | 65496558 | rs11227281 | 1.54 | 0.155 | 1.168 | 0.857 | -0.013 | 0.042 | 0.042 | 0.332 | 0.33 | 0.18 | 0.21 | 0.24 | 0.35 | 0.35 |
| 11 | 65504069 | rs1193692 | -0.5 | 0.193 | 1.153 | 1.029 | 0 | 0.002 | 0.002 | 1.043 | 0.48 | 0.45 | 0.37 | 0.37 | 0.53 | 0.26 |
| 11 | 65513107 | rs593982 | -0.133 | 0.153 | 1.214 | 1.019 | -0.031 | 0.092 | 0.092 | 0 | 0.82 | 0.69 | 0.58 | 0.61 | 0.88 | 0.61 |
| 11 | 65542298 | rs494003 | 0.976 | 0.256 | 1.118 | 0.738 | 0.002 | -0.001 | -0.001 | 1.668 | 0.26 | 0.24 | 0.14 | 0.12 | 0.19 | 0.05 |
| 11 | 65542739 | rs489574 | -0.293 | 0.264 | 1.114 | 0.742 | 0.005 | 0.001 | 0.001 | 0.313 | 0.22 | 0.2 | 0.22 | 0.25 | 0.34 | 0.22 |
| 11 | 65543827 | rs10896043 | 0.962 | 0.246 | 0.954 | 0.716 | 0.003 | 0.003 | 0.003 | 4.148 | 0.26 | 0.22 | 0.14 | 0.12 | 0.19 | 0.05 |
| 11 | 65546857 | rs610037 | -0.91 | 0.272 | 0.637 | 0.797 | 0.014 | 0.034 | 0.034 | 1.153 | 0.5 | 0.63 | 0.39 | 0.37 | 0.53 | 0.27 |
| 11 | 65551957 | rs479844 | 0.845 | 0.43 | 0.299 | 0.744 | 0.001 | 0.152 | 0.152 | 2.064 | 0.48 | 0.17 | 0.33 | 0.27 | 0.56 | 0.54 |
| 11 | 65553306 | rs36008241 | 0.7 | 0.491 | 0.165 | 0.66 | -0.06 | 0.084 | 0.084 | 0.146 | 0.15 | 0.03 | 0.13 | 0.03 | 0.21 | 0.2 |
| 11 | 65554502 | rs11227302 | 0.655 | 0.444 | 0.468 | 0.752 | 0 | -0.01 | -0.01 | 0 | 0.27 | 0.28 | 0.14 | 0.11 | 0.18 | 0.05 |
| 11 | 65559266 | rs10791824 | 0.674 | 0.478 | 0.241 | 0.84 | 0.007 | -0.001 | -0.001 | 0.512 | 0.51 | 0.42 | 0.38 | 0.5 | 0.57 | 0.57 |
| 11 | 65560620 | rs642803 | -1.545 | 1.167 | 0.292 | 1.109 | -0.014 | -0.005 | -0.005 | 0.664 | 0.38 | 0.29 | 0.58 | 0.39 | 0.46 | 0.61 |

| | | | | | | | | | | | | | | | | |
|---|---|---|---|---|---|---|---|---|---|---|---|---|---|---|---|---|
| 11 | 65564035 | rs491666 | 0.427 | 1.234 | 0.222 | 1.068 | -0.051 | 0.078 | 0.078 | 0.009 | 0.15 | 0.02 | 0.13 | 0.02 | 0.21 | 0.22 |
| 11 | 65564786 | rs44205 | -0.529 | 1.217 | 0.359 | 1.325 | -0.037 | 0.093 | 0.093 | 0 | 0.32 | 0.08 | 0.49 | 0.18 | 0.33 | 0.6 |
| 11 | 65564987 | rs12800057 | -0.213 | 0.995 | 0.923 | 1.603 | 0.048 | 0.12 | 0.12 | 0 | 0.07 | 0.17 | 0.09 | 0.23 | 0.13 | 0.03 |
| 11 | 65570405 | rs7116758 | -0.035 | 1.016 | 0.914 | 1.004 | 0.073 | 0.043 | 0.043 | 0 | 0.07 | 0.19 | 0.09 | 0.1 | 0.14 | 0.03 |
| 11 | 65570517 | rs7119612 | -0.676 | 0.858 | 1.156 | 1.195 | 0.069 | 0.284 | 0.284 | 0.026 | 0.08 | 0.4 | 0.09 | 0.22 | 0.1 | 0.03 |
| 11 | 65577516 | rs10896050 | 0.584 | 0.555 | 0.343 | 0.792 | -0.008 | -0.008 | -0.008 | 0 | 0.18 | 0.09 | 0.14 | 0.16 | 0.19 | 0.04 |
| 11 | 65578672 | rs11227306 | -0.774 | 0.795 | 0.405 | 0.718 | -0.02 | -0.01 | -0.01 | 0 | 0.29 | 0.26 | 0.25 | 0.29 | 0.38 | 0.08 |
| 11 | 65580638 | rs12576996 | 1.689 | 0.497 | -0.221 | 0.562 | -0.028 | 0.068 | 0.068 | 0 | 0.32 | 0.02 | 0.15 | 0.21 | 0.25 | 0.28 |
| 11 | 65590713 | rs601887 | 0.08 | 0.764 | 0.847 | 0.771 | 0.004 | 0.123 | 0.123 | 0 | 0.14 | 0.21 | 0.41 | 0.48 | 0.16 | 0.37 |
| 11 | 65594820 | rs7102454 | -0.257 | 0.625 | 0.739 | 0.687 | -0.037 | -0.025 | -0.025 | 0 | 0.26 | 0.2 | 0.23 | 0.29 | 0.35 | 0.08 |
| 11 | 65617324 | rs654638 | 0.523 | 0.408 | 0.417 | 0.439 | 0.016 | 0.092 | 0.092 | 0 | 0.23 | 0.07 | 0.15 | 0.02 | 0.21 | 0.28 |
| 11 | 65623739 | rs11227332 | -0.042 | 1.075 | 1.162 | 1.406 | -0.019 | 0.043 | 0.043 | 0.008 | 0.2 | 0.06 | 0.14 | 0.06 | 0.19 | 0.07 |
| 11 | 65626701 | rs1939212 | -0.668 | 1.058 | 1.274 | 1.361 | -0.003 | -0.002 | -0.002 | 0 | 0.28 | 0.2 | 0.52 | 0.64 | 0.31 | 0.39 |
| 11 | 65637076 | rs570387 | 0.979 | 1.157 | 1.526 | 1.023 | 0 | -0.015 | -0.015 | 0 | 0.59 | 0.42 | 0.34 | 0.23 | 0.51 | 0.55 |
| 11 | 65640906 | rs7940691 | 0.838 | 1.306 | 1.733 | 1.861 | 0.03 | 0.028 | 0.028 | 0 | 0.71 | 0.39 | 0.7 | 0.48 | 0.65 | 0.85 |
| 11 | 65644027 | rs583887 | 0.043 | 1.139 | 1.882 | 2.014 | 0.047 | 0.229 | 0.229 | 0 | 0.81 | 0.49 | 0.78 | 0.59 | 0.81 | 0.9 |
| 11 | 65649984 | rs687672 | 0.043 | 1.144 | 1.695 | 2.016 | 0.047 | 0.229 | 0.229 | 0 | 0.81 | 0.49 | 0.78 | 0.59 | 0.81 | 0.9 |
| 11 | 65653242 | rs568617 | -0.418 | 1.144 | 1.695 | 1.801 | 0.047 | 0.229 | 0.229 | 0 | 0.19 | 0.51 | 0.22 | 0.41 | 0.19 | 0.1 |
| 11 | 65656282 | rs656980 | -0.391 | 1.146 | 1.696 | 1.802 | 0.045 | 0.233 | 0.233 | 0 | 0.19 | 0.51 | 0.22 | 0.41 | 0.19 | 0.1 |
| 11 | 65656564 | rs2231884 | 0.243 | 1.204 | 0.872 | 1.978 | 0 | 0.01 | 0.01 | 0 | 0.16 | 0.15 | 0.16 | 0.31 | 0.16 | 0.1 |
| 11 | 65661291 | rs10791830 | -0.074 | 1.183 | 0.87 | 1.806 | 0.014 | 0.075 | 0.075 | 0 | 0.17 | 0.28 | 0.18 | 0.39 | 0.19 | 0.1 |
| 11 | 65663239 | rs622614 | 0.349 | 1.047 | 1.162 | 1.752 | 0.004 | 0.064 | 0.064 | 0 | 0.78 | 0.27 | 0.77 | 0.61 | 0.81 | 0.9 |
| 11 | 65664006 | rs614520 | 0.769 | 0.454 | 1.079 | 1.329 | 0.029 | -0.041 | -0.041 | 0 | 0.69 | 0.44 | 0.46 | 0.81 | 0.62 | 0.82 |
| 11 | 65670510 | rs10896065 | 1.504 | 0.42 | 1.784 | 1.354 | 0.021 | 0.195 | 0.195 | 3.581 | 0.94 | 0.42 | 0.93 | 0.85 | 0.98 | 0.99 |
| 11 | 65676516 | rs653914 | -0.29 | 0.217 | 1.192 | 1.186 | -0.157 | -0.151 | -0.151 | 3.718 | 0.34 | 0.04 | 0.57 | 0.49 | 0.44 | 0.45 |
| 11 | 65705108 | rs528736 | -0.147 | 0.221 | 0.457 | 1.096 | -0.14 | -0.137 | -0.137 | 0 | 0.33 | 0.06 | 0.58 | 0.51 | 0.44 | 0.42 |
| 11 | 65724291 | rs470192 | -0.337 | 0.181 | 0.994 | 1.047 | -0.001 | -0.001 | -0.001 | 0 | 0.34 | 0.31 | 0.61 | 0.52 | 0.44 | 0.42 |
| 11 | 65727301 | rs491973 | -0.355 | 0.196 | 1.001 | 1.055 | -0.002 | -0.002 | -0.002 | 0 | 0.34 | 0.31 | 0.61 | 0.52 | 0.44 | 0.42 |
| 11 | 65727799 | rs593525 | -0.754 | 0.149 | 1.288 | 1.117 | 0.003 | -0.003 | -0.003 | 0.405 | 0.39 | 0.53 | 0.65 | 0.67 | 0.47 | 0.43 |
| 11 | 65732651 | rs550435 | -0.324 | 0.17 | 1.287 | 1.118 | -0.002 | -0.002 | -0.002 | 0 | 0.34 | 0.31 | 0.6 | 0.52 | 0.44 | 0.42 |
| 11 | 65732800 | rs552130 | -0.771 | 0.121 | 1.354 | 1.108 | -0.005 | 0.003 | 0.003 | 0 | 0.39 | 0.59 | 0.66 | 0.67 | 0.47 | 0.43 |
| 11 | 65744247 | rs622779 | 0.478 | -0.366 | 0.691 | 0.592 | -0.008 | -0.024 | -0.024 | 0 | 0.28 | 0.26 | 0.12 | 0.06 | 0.18 | 0.21 |
| 11 | 65746653 | rs679581 | 0.34 | -0.223 | 0.12 | 0.553 | -0.041 | -0.042 | -0.042 | 0.342 | 0.32 | 0.19 | 0.59 | 0.52 | 0.44 | 0.45 |
| 11 | 65747057 | rs754532 | 1.027 | -0.154 | -0.031 | 0.441 | 0.016 | 0.084 | 0.084 | 0 | 0.31 | 0.14 | 0.19 | 0.16 | 0.31 | 0.32 |
| 11 | 65749337 | rs735942 | 0.798 | -0.209 | 0.083 | 0.52 | 0.003 | 0.11 | 0.11 | 0 | 0.34 | 0.14 | 0.21 | 0.16 | 0.35 | 0.33 |
| 11 | 65754808 | rs1786175 | 0.327 | -0.145 | -0.009 | 0.547 | -0.036 | -0.036 | -0.036 | 0 | 0.32 | 0.2 | 0.59 | 0.52 | 0.44 | 0.45 |
| 11 | 65756283 | rs1151511 | -0.085 | -0.433 | 0.456 | 1.05 | -0.055 | -0.036 | -0.036 | 0 | 0.69 | 0.53 | 0.85 | 0.83 | 0.82 | 0.79 |
| 11 | 65758854 | rs1151514 | -0.085 | -0.433 | 0.456 | 1.05 | -0.055 | -0.036 | -0.036 | 0 | 0.69 | 0.53 | 0.85 | 0.83 | 0.81 | 0.79 |
| 11 | 65765551 | rs610497 | 0.088 | -0.425 | 0.434 | 1.055 | -0.032 | 0.009 | 0.009 | 1.669 | 0.67 | 0.42 | 0.81 | 0.69 | 0.79 | 0.78 |
| 11 | 65783915 | rs668210 | 0.051 | -0.366 | 0.13 | 0.929 | -0.034 | 0.01 | 0.01 | 1.228 | 0.68 | 0.44 | 0.81 | 0.69 | 0.79 | 0.78 |
| 11 | 65789854 | rs10791838 | -0.079 | -1.087 | -0.093 | 0.245 | 0.011 | 0.01 | 0.01 | 1.306 | 0.27 | 0.21 | 0.13 | 0.12 | 0.16 | 0.19 |
| 11 | 65790527 | rs2845570 | 0.595 | -0.166 | 0.005 | 0.664 | 0.007 | 0.047 | 0.047 | 0.287 | 0.55 | 0.19 | 0.38 | 0.38 | 0.56 | 0.71 |
| 11 | 65792566 | rs683191 | 1.717 | 0.301 | -0.288 | 0.343 | -0.017 | 0.022 | 0.022 | 0 | 0.07 | 0.01 | 0.05 | 0 | 0.08 | 0.01 |
| 11 | 65793454 | rs1203998 | 0.1 | 0.26 | 0.223 | 0.613 | 0.09 | 0.221 | 0.221 | 0 | 0.17 | 0.45 | 0.46 | 0.39 | 0.28 | 0.1 |
| 11 | 65793797 | rs1893316 | 1.171 | 0.311 | 0.199 | 0.705 | -0.074 | -0.074 | -0.074 | 0 | 0.13 | 0.01 | 0.39 | 0.23 | 0.24 | 0.07 |
| 11 | 65793942 | rs947847 | 0.002 | 0.309 | 0.212 | 0.684 | 0.092 | 0.224 | 0.224 | 0 | 0.17 | 0.46 | 0.46 | 0.39 | 0.28 | 0.1 |
| 11 | 65802478 | rs12576969 | -0.723 | 0.263 | 0.432 | 0.647 | -0.032 | -0.015 | -0.015 | 1.572 | 0.72 | 0.6 | 0.84 | 0.76 | 0.84 | 0.81 |
| 11 | 65803383 | rs10791840 | 0.657 | 0.263 | 0.419 | 0.635 | 0.012 | 0.01 | 0.01 | 1.192 | 0.27 | 0.21 | 0.13 | 0.12 | 0.16 | 0.18 |
| 11 | 65808467 | rs2576 | 0.441 | 0.279 | 0.415 | 0.664 | 0.007 | 0.044 | 0.044 | 0.278 | 0.55 | 0.2 | 0.39 | 0.38 | 0.56 | 0.72 |
| 11 | 65809763 | rs3814744 | 0.657 | 0.257 | 0.427 | 0.658 | 0.012 | 0.01 | 0.01 | 1.192 | 0.27 | 0.21 | 0.13 | 0.12 | 0.16 | 0.18 |
| 11 | 65813383 | rs2430979 | -0.602 | 0.246 | 0.93 | 0.681 | -0.039 | 0.017 | 0.017 | 1.874 | 0.7 | 0.38 | 0.81 | 0.62 | 0.84 | 0.81 |
| 11 | 65817307 | rs574153 | 0.099 | 0.477 | 1.029 | 0.827 | 0.089 | 0.197 | 0.197 | 0 | 0.16 | 0.39 | 0.46 | 0.5 | 0.27 | 0.1 |
| 11 | 65817592 | rs571374 | 0.756 | 0.428 | 0.961 | 0.765 | -0.005 | 0.081 | 0.081 | 0.019 | 0.5 | 0.19 | 0.34 | 0.38 | 0.49 | 0.7 |
| 11 | 65819551 | rs12803276 | 0.898 | 0.485 | 1.041 | 0.785 | -0.01 | -0.01 | -0.01 | 0 | 0.13 | 0.12 | 0.4 | 0.23 | 0.24 | 0.07 |
| 11 | 65819661 | rs2241303 | 0.702 | 0.457 | 0.982 | 0.752 | -0.002 | 0.075 | 0.075 | 0 | 0.51 | 0.2 | 0.34 | 0.38 | 0.48 | 0.7 |
| 11 | 65820673 | rs503156 | 0.075 | 0.513 | 0.939 | 0.794 | 0.09 | 0.193 | 0.193 | 0 | 0.16 | 0.39 | 0.46 | 0.5 | 0.27 | 0.1 |
| 11 | 65824084 | rs2302884 | 0.355 | 0.477 | 0.895 | 0.562 | 0.012 | 0.03 | 0.03 | 1.037 | 0.31 | 0.24 | 0.18 | 0.12 | 0.23 | 0.19 |
| 11 | 65824472 | rs547404 | 0.311 | 0.419 | 0.988 | 0.647 | 0.072 | 0.074 | 0.074 | 0 | 0.15 | 0.29 | 0.43 | 0.25 | 0.27 | 0.1 |
| 11 | 65835931 | rs2270448 | 0.309 | 0.075 | 0.791 | 0.517 | 0.029 | 0.07 | 0.07 | 1.136 | 0.3 | 0.17 | 0.16 | 0.12 | 0.2 | 0.19 |
| 11 | 65851902 | rs472518 | -0.529 | 0.06 | 1.205 | 0.956 | -0.031 | 0.067 | 0.067 | 0.819 | 0.69 | 0.44 | 0.76 | 0.54 | 0.77 | 0.72 |
| 11 | 65866187 | rs537497 | 0.491 | 0.464 | 1.271 | 0.962 | 0.017 | 0.072 | 0.072 | 0.098 | 0.56 | 0.23 | 0.37 | 0.33 | 0.58 | 0.66 |
| 11 | 65876570 | rs873491 | 1.035 | 0.245 | 1.132 | 0.897 | -0.026 | -0.04 | -0.04 | 2.474 | 0.12 | 0.05 | 0.38 | 0.2 | 0.19 | 0.07 |
| 11 | 65886662 | rs524281 | -0.484 | 0.181 | 1.251 | 0.905 | -0.006 | 0.009 | 0.009 | 0.794 | 0.68 | 0.33 | 0.75 | 0.53 | 0.77 | 0.73 |
| 11 | 65922589 | rs801739 | -0.203 | 0.617 | 0.722 | 0.511 | -0.003 | -0.002 | -0.002 | 0 | 0.39 | 0.4 | 0.56 | 0.4 | 0.49 | 0.31 |
| 11 | 65934549 | rs801733 | 0.509 | 0.627 | 0.724 | 0.426 | -0.014 | 0.02 | 0.02 | 2.651 | 0.3 | 0.25 | 0.19 | 0.05 | 0.36 | 0.25 |
| 11 | 65935189 | rs513474 | 0.493 | 0.637 | 0.694 | 0.47 | -0.003 | 0.021 | 0.021 | 0 | 0.34 | 0.19 | 0.26 | 0.31 | 0.32 | 0.5 |
| 11 | 65943647 | rs7123153 | 0.661 | 0.643 | 0.52 | 0.461 | -0.05 | 0 | 0 | 0.73 | 0.33 | 0.03 | 0.24 | 0.31 | 0.32 | 0.5 |
| 11 | 65957003 | rs17307346 | 1.581 | 0.889 | 0.7 | 0.909 | -0.034 | -0.036 | -0.036 | 0 | 0.07 | 0.01 | 0.33 | 0.12 | 0.13 | 0.06 |
| 11 | 65985168 | rs512421 | 0.403 | 0.708 | 0.681 | 0.672 | -0.002 | 0 | 0 | 2.026 | 0.27 | 0.3 | 0.18 | 0.29 | 0.24 | 0.19 |

| | | | | | | | | | | | | | | | | |
|---|---|---|---|---|---|---|---|---|---|---|---|---|---|---|---|---|
| 11 | 65992008 | rs9326370 | -0.053 | 0.636 | 0.874 | 0.655 | 0.008 | 0.006 | 0.006 | 3.762 | 0.38 | 0.44 | 0.55 | 0.41 | 0.49 | 0.31 |
| 11 | 66001869 | rs10896104 | 0.186 | 1.036 | 0.56 | 0.518 | -0.077 | 0.032 | 0.032 | 3.93 | 0.36 | 0.15 | 0.5 | 0.25 | 0.49 | 0.31 |
| 11 | 66011280 | rs2298466 | 0.521 | 0.954 | 0.531 | 0.529 | -0.002 | 0.001 | 0.001 | 1.001 | 0.25 | 0.3 | 0.17 | 0.29 | 0.24 | 0.19 |
| 11 | 66014300 | rs2155198 | 0.924 | 0.96 | 0.493 | 0.525 | 0.001 | -0.001 | -0.001 | 0 | 0.22 | 0.24 | 0.14 | 0.29 | 0.17 | 0.18 |
| 11 | 66027358 | rs10791863 | 1.163 | 1.141 | 0.555 | 0.617 | 0.002 | -0.004 | -0.004 | 1.489 | 0.2 | 0.18 | 0.13 | 0.29 | 0.17 | 0.18 |
| 11 | 66031660 | rs1151539 | 0.106 | 1.148 | 0.56 | 0.609 | -0.058 | 0.035 | 0.035 | 3.853 | 0.36 | 0.21 | 0.51 | 0.25 | 0.48 | 0.31 |
| 11 | 66033430 | rs2276036 | 1.105 | 1.062 | 0.528 | 0.585 | 0.004 | -0.005 | -0.005 | 0 | 0.21 | 0.18 | 0.13 | 0.29 | 0.17 | 0.18 |
| 11 | 66035608 | rs10791865 | -0.283 | 1.076 | 0.637 | 0.587 | -0.002 | 0.006 | 0.006 | 1.777 | 0.41 | 0.38 | 0.56 | 0.26 | 0.55 | 0.31 |
| 11 | 66041807 | rs1151540 | 0.066 | 1.147 | -0.108 | 0.668 | -0.06 | 0.035 | 0.035 | 3.962 | 0.36 | 0.2 | 0.51 | 0.25 | 0.49 | 0.31 |
| 11 | 66048491 | rs11227448 | 1.119 | 1.153 | -0.293 | 0.375 | 0.003 | -0.004 | -0.004 | 0 | 0.2 | 0.18 | 0.12 | 0.29 | 0.17 | 0.18 |
| 11 | 66050712 | rs4073582 | 0.672 | 1.297 | -0.286 | 0.367 | -0.103 | 0.137 | 0.137 | 0.859 | 0.26 | 0.05 | 0.17 | 0.05 | 0.36 | 0.23 |
| 11 | 66058082 | rs556595 | 0.113 | 0.768 | -0.276 | 0.572 | -0.009 | -0.004 | -0.004 | 0 | 0.36 | 0.34 | 0.52 | 0.36 | 0.48 | 0.31 |
| 11 | 66061233 | rs484261 | -0.101 | 0.435 | -0.29 | 0.646 | -0.022 | -0.007 | -0.007 | 0 | 0.39 | 0.5 | 0.32 | 0.4 | 0.28 | 0.51 |
| 11 | 66064126 | rs560577 | 0.048 | 0.121 | -0.167 | 0.294 | -0.005 | -0.084 | -0.084 | 0 | 0.14 | 0.32 | 0.1 | 0.01 | 0.09 | 0.05 |
| 11 | 66064850 | rs531784 | 0.585 | -0.355 | -1.222 | 0.375 | 0.028 | -0.039 | -0.039 | 0 | 0.27 | 0.06 | 0.17 | 0.36 | 0.17 | 0.57 |
| 11 | 66066349 | rs580241 | -1.138 | -0.029 | -1.266 | 0.517 | 0.002 | -0.015 | -0.015 | 0 | 0.75 | 0.84 | 0.84 | 0.71 | 0.73 | 0.85 |
| 11 | 66066664 | rs4930351 | -0.018 | 0.242 | -0.962 | 0.407 | -0.078 | -0.05 | -0.05 | 0.272 | 0.34 | 0.2 | 0.57 | 0.36 | 0.49 | 0.25 |
| 11 | 66070146 | rs523583 | 0.235 | 0.293 | -0.85 | 0.509 | -0.058 | 0.008 | 0.008 | 0 | 0.3 | 0.16 | 0.55 | 0.21 | 0.45 | 0.21 |
| 11 | 66077203 | rs4390351 | 0.71 | 0.076 | -0.368 | 0.623 | -0.016 | 0.009 | 0.009 | 0 | 0.14 | 0.25 | 0.07 | 0.16 | 0.08 | 0.09 |
| 11 | 66080770 | rs7924580 | 0.536 | -0.11 | -0.614 | 0.643 | 0.016 | -0.012 | -0.012 | 0 | 0.27 | 0.19 | 0.16 | 0.29 | 0.25 | 0.15 |
| 11 | 66083129 | rs3741367 | 0.131 | -0.026 | -0.646 | 0.457 | -0.035 | 0.003 | 0.003 | 0 | 0.29 | 0.17 | 0.55 | 0.21 | 0.44 | 0.2 |
| 11 | 66083782 | rs3741368 | 0.506 | 0.123 | -0.937 | 0.536 | -0.138 | -0.048 | -0.048 | 0 | 0.25 | 0.04 | 0.53 | 0.23 | 0.45 | 0.19 |
| 11 | 66090032 | rs10219183 | 0.79 | -0.159 | -0.801 | 0.441 | 0.018 | -0.03 | -0.03 | 0 | 0.25 | 0.08 | 0.15 | 0.28 | 0.25 | 0.15 |
| 11 | 66091814 | rs3016311 | -0.703 | -0.468 | -0.188 | 0.335 | -0.023 | 0.001 | 0.001 | 0.497 | 0.82 | 0.29 | 0.88 | 0.83 | 0.91 | 0.95 |
| 11 | 66092730 | rs1785631 | -1.155 | -0.527 | 0.503 | 0.444 | -0.02 | 0.002 | 0.002 | 0 | 0.66 | 0.56 | 0.75 | 0.59 | 0.78 | 0.44 |
| 11 | 66093612 | rs1892938 | 0.25 | -0.211 | 0.507 | 0.606 | 0.045 | 0.048 | 0.048 | 0 | 0.54 | 0.2 | 0.42 | 0.41 | 0.59 | 0.76 |
| 11 | 66098322 | rs11823140 | -0.093 | -0.294 | 0.846 | 0.518 | -0.019 | 0.01 | 0.01 | 0 | 0.39 | 0.5 | 0.23 | 0.41 | 0.33 | 0.2 |
| 11 | 66106725 | rs948360 | -0.657 | -0.29 | 0.841 | 0.517 | -0.022 | 0.001 | 0.001 | 0.635 | 0.83 | 0.3 | 0.88 | 0.83 | 0.91 | 0.95 |
| 11 | 66108660 | rs947978 | -0.023 | -0.288 | 0.739 | 0.526 | 0.045 | -0.022 | -0.022 | 1.577 | 0.58 | 0.23 | 0.74 | 0.56 | 0.66 | 0.82 |
| 11 | 66109033 | rs11537993 | 0.809 | 0.1 | 0.529 | 0.252 | -0.107 | 0.097 | 0.097 | 0.81 | 0.2 | 0.02 | 0.17 | 0.03 | 0.33 | 0.2 |
| 11 | 66114900 | rs11537930 | 1.397 | -0.097 | 0.88 | 0.368 | -0.003 | 0.018 | 0.018 | 0 | 0.13 | 0.25 | 0.07 | 0.17 | 0.09 | 0.05 |
| 11 | 66133336 | rs2279861 | -0.291 | 0.031 | 0.494 | 0.413 | 0.053 | -0.001 | -0.001 | 0 | 0.57 | 0.22 | 0.74 | 0.58 | 0.66 | 0.83 |
| 11 | 66135179 | rs2279862 | -0.296 | 0.024 | 0.524 | 0.41 | 0.052 | -0.001 | -0.001 | 0 | 0.57 | 0.22 | 0.74 | 0.58 | 0.66 | 0.83 |
| 11 | 66171621 | rs6591216 | 0.042 | 0.306 | -0.236 | 0.348 | -0.116 | -0.013 | -0.013 | 0 | 0.24 | 0.03 | 0.48 | 0.18 | 0.38 | 0.23 |
| 11 | 66177952 | rs2282530 | -0.079 | -0.011 | 0.224 | 0.297 | -0.017 | -0.002 | -0.002 | 0.067 | 0.42 | 0.48 | 0.24 | 0.42 | 0.34 | 0.19 |
| 11 | 66179063 | rs12785651 | 0.058 | 0.005 | -0.346 | 0.305 | 0.054 | 0.008 | 0.008 | 0.434 | 0.56 | 0.2 | 0.73 | 0.58 | 0.66 | 0.81 |
| 11 | 66179556 | rs12786563 | 0.068 | 0.005 | -0.346 | 0.299 | 0.055 | 0.011 | 0.011 | 0.405 | 0.56 | 0.2 | 0.73 | 0.59 | 0.66 | 0.81 |
| 11 | 66182417 | rs12577917 | 0.383 | 0.231 | -1.063 | 0.001 | -0.101 | -0.076 | -0.076 | 0.262 | 0.44 | 0.07 | 0.62 | 0.54 | 0.49 | 0.66 |
| 11 | 66184140 | rs2282529 | -0.593 | 1.528 | -1.052 | 0.171 | -0.169 | 0.016 | 0.016 | 0.246 | 0.28 | 0.02 | 0.25 | 0.17 | 0.42 | 0.21 |
| 11 | 66186882 | rs7947391 | 0.697 | 1.53 | -0.352 | 0.195 | -0.001 | -0.005 | -0.005 | 1.098 | 0.56 | 0.48 | 0.71 | 0.74 | 0.58 | 0.81 |
| 11 | 66196265 | rs17495838 | 0.511 | 1.216 | -0.516 | 0.579 | -0.065 | -0.042 | -0.042 | 0 | 0.11 | 0.02 | 0.14 | 0.15 | 0.23 | 0.17 |
| 11 | 66203025 | rs2279865 | -0.357 | 1.122 | -0.058 | 0.681 | 0.107 | 0.113 | 0.113 | 0.904 | 0.14 | 0.39 | 0.18 | 0.29 | 0.24 | 0.17 |
| 11 | 66222137 | rs1791682 | 1.099 | 0.825 | 0.425 | 0.459 | -0.024 | -0.069 | -0.069 | 0.941 | 0.12 | 0.27 | 0.06 | 0 | 0.06 | 0.04 |
| 11 | 66234023 | rs7925108 | -2.06 | 0.902 | 0.166 | 0.536 | -0.08 | -0.062 | -0.062 | 0 | 0.57 | 0.96 | 0.45 | 0.57 | 0.61 | 0.36 |
| 11 | 66240882 | rs2277302 | 0.015 | 1.019 | 0.39 | 0.572 | 0.098 | 0.104 | 0.104 | 0.695 | 0.13 | 0.36 | 0.18 | 0.29 | 0.24 | 0.19 |
| 11 | 66245851 | rs2282640 | -0.409 | 0.94 | 0.503 | 0.573 | 0.002 | 0 | 0 | 0.007 | 0.32 | 0.25 | 0.2 | 0.26 | 0.3 | 0.14 |
| 11 | 66246337 | rs11821155 | -0.068 | 1.113 | 0.279 | 0.699 | 0.098 | 0.104 | 0.104 | 0.695 | 0.13 | 0.36 | 0.18 | 0.29 | 0.24 | 0.19 |
| 11 | 66254085 | rs11550299 | 0.318 | 1.127 | 0.284 | 0.706 | 0.049 | 0.053 | 0.053 | 1.246 | 0.12 | 0.19 | 0.16 | 0.29 | 0.24 | 0.19 |
| 11 | 66262606 | rs2511224 | 0.694 | 1 | 0.231 | 0.666 | 0.037 | 0.049 | 0.049 | 0.254 | 0.56 | 0.37 | 0.72 | 0.76 | 0.63 | 0.81 |
| 11 | 66272142 | rs1671063 | -1.905 | 1.021 | 0.153 | 0.673 | -0.05 | -0.058 | -0.058 | 0.816 | 0.57 | 0.92 | 0.45 | 0.55 | 0.61 | 0.4 |
| 11 | 66272237 | rs2305535 | 0.371 | 1.108 | 0.262 | 0.725 | 0.012 | 0.015 | 0.015 | 1.312 | 0.12 | 0.15 | 0.16 | 0.3 | 0.24 | 0.21 |
| 11 | 66272976 | rs10501397 | 0.113 | 1.107 | 0.267 | 0.725 | 0.039 | 0.046 | 0.046 | 0.793 | 0.13 | 0.24 | 0.17 | 0.3 | 0.24 | 0.21 |
| 11 | 66282095 | rs2298806 | 0.374 | 1.166 | 0.273 | 0.714 | 0.013 | 0.016 | 0.016 | 1.262 | 0.12 | 0.15 | 0.16 | 0.3 | 0.24 | 0.21 |
| 11 | 66288123 | rs11822028 | -0.218 | 1.164 | 0.289 | 0.713 | 0.043 | 0.05 | 0.05 | 0.842 | 0.13 | 0.28 | 0.18 | 0.3 | 0.24 | 0.21 |
| 11 | 66297363 | rs3816492 | 0.549 | 1.174 | 0.212 | 0.744 | -0.065 | -0.04 | -0.04 | 0.309 | 0.11 | 0.01 | 0.14 | 0.15 | 0.24 | 0.21 |
| 11 | 66299515 | rs8432 | -1.907 | 1.08 | 0.126 | 0.687 | -0.043 | -0.051 | -0.051 | 0.587 | 0.57 | 0.91 | 0.45 | 0.55 | 0.61 | 0.4 |
| 11 | 66306355 | rs3737525 | 0.536 | 1.164 | 0.169 | 0.709 | -0.066 | -0.04 | -0.04 | 0 | 0.11 | 0.02 | 0.14 | 0.15 | 0.24 | 0.21 |
| 11 | 66311011 | rs3825065 | -0.267 | 1.154 | 0.244 | 0.747 | 0.044 | 0.051 | 0.051 | 0.929 | 0.14 | 0.28 | 0.17 | 0.3 | 0.24 | 0.22 |
| 11 | 66313026 | rs11227527 | -0.244 | 1.154 | 0.258 | 0.747 | 0.037 | 0.043 | 0.043 | 1.002 | 0.13 | 0.24 | 0.17 | 0.32 | 0.24 | 0.22 |
| 11 | 66316557 | rs597626 | 0.856 | 1.107 | 0.194 | 0.668 | 0.028 | 0.006 | 0.006 | 0.007 | 0.53 | 0.39 | 0.66 | 0.62 | 0.55 | 0.77 |
| 11 | 66317823 | rs1791690 | -1.267 | 1.112 | 0.167 | 0.671 | 0.039 | 0.009 | 0.009 | 0.044 | 0.47 | 0.66 | 0.35 | 0.38 | 0.45 | 0.23 |
| 11 | 66319262 | rs10791881 | -2.344 | 1.07 | -0.175 | 0.726 | -0.08 | -0.058 | -0.058 | 0 | 0.65 | 0.97 | 0.53 | 0.63 | 0.67 | 0.42 |
| 11 | 66324762 | rs677488 | -1.742 | 1.007 | 0.071 | 0.978 | -0.044 | 0.009 | 0.009 | 0 | 0.53 | 0.8 | 0.41 | 0.56 | 0.55 | 0.39 |
| 11 | 66326581 | rs2275998 | -0.187 | 1.016 | 0.556 | 1.001 | 0.051 | 0.006 | 0.006 | 1.166 | 0.13 | 0.3 | 0.12 | 0.18 | 0.17 | 0.17 |
| 11 | 66328095 | rs1815739 | -1.99 | 1.017 | 0.339 | 1.009 | -0.084 | -0.03 | -0.03 | 0 | 0.57 | 0.89 | 0.43 | 0.56 | 0.57 | 0.41 |
| 11 | 66329732 | rs540874 | -1.959 | 1.003 | 0.394 | 1.026 | -0.069 | -0.017 | -0.017 | 0 | 0.56 | 0.83 | 0.42 | 0.56 | 0.56 | 0.41 |
| 11 | 66331000 | rs4576 | -0.227 | 1.136 | 0.467 | 1.116 | 0.053 | 0.057 | 0.057 | 1.003 | 0.13 | 0.27 | 0.17 | 0.31 | 0.24 | 0.24 |
| 11 | 66331458 | rs572846 | -1.918 | 1.118 | 0.43 | 1.108 | -0.069 | -0.009 | -0.009 | 0 | 0.54 | 0.88 | 0.42 | 0.55 | 0.55 | 0.39 |
| 11 | 66333072 | rs4630309 | -0.097 | 1.133 | 0.447 | 1.157 | 0.04 | 0.043 | 0.043 | 1.245 | 0.13 | 0.21 | 0.16 | 0.31 | 0.24 | 0.25 |

| | | | | | | | | | | | | | | | | |
|---|---|---|---|---|---|---|---|---|---|---|---|---|---|---|---|---|
| 11 | 66348503 | rs692892 | 1.389 | 1.016 | 0.201 | 1.022 | -0.051 | 0.023 | 0.023 | 0 | 0.49 | 0.05 | 0.59 | 0.4 | 0.51 | 0.59 |
| 11 | 66358126 | rs486584 | 1.36 | 0.982 | 0.199 | 1.068 | -0.011 | 0.006 | 0.006 | 0 | 0.5 | 0.11 | 0.63 | 0.45 | 0.51 | 0.6 |
| 11 | 66369211 | rs1791684 | 1.356 | 0.985 | 0.225 | 1.328 | -0.009 | 0.045 | 0.045 | 0 | 0.5 | 0.12 | 0.63 | 0.43 | 0.51 | 0.6 |
| 11 | 66381517 | rs12795774 | 0.649 | 0.874 | 0.555 | 1.317 | 0.002 | 0.001 | 0.001 | 0.209 | 0.62 | 0.43 | 0.78 | 0.74 | 0.72 | 0.85 |
| 11 | 66394387 | rs15969 | 1.137 | 0.892 | 0.544 | 1.373 | -0.047 | 0.042 | 0.042 | 0.508 | 0.09 | 0.01 | 0.08 | 0.02 | 0.15 | 0.15 |
| 11 | 66399681 | rs484262 | 0.639 | 0.864 | 0.553 | 1.438 | -0.01 | -0.001 | -0.001 | 0 | 0.62 | 0.46 | 0.79 | 0.71 | 0.72 | 0.84 |
| 11 | 66401373 | rs566673 | 1.9 | 0.883 | 0.503 | 1.412 | -0.136 | 0.053 | 0.053 | 0 | 0.43 | 0.04 | 0.56 | 0.33 | 0.46 | 0.56 |
| 11 | 66409409 | rs519380 | 0.516 | 0.935 | 1.183 | 1.427 | -0.017 | -0.012 | -0.012 | 2.955 | 0.11 | 0.07 | 0.13 | 0.15 | 0.21 | 0.25 |
| 11 | 66419554 | rs481296 | 0.915 | 0.925 | 1.18 | 1.416 | 0.002 | 0.001 | 0.001 | 0.209 | 0.62 | 0.43 | 0.79 | 0.74 | 0.72 | 0.85 |
| 11 | 66427410 | rs17582367 | 1.948 | 0.829 | 1.024 | 1.312 | -0.004 | 0.039 | 0.039 | 0 | 0.08 | 0.01 | 0.02 | 0 | 0.06 | 0.04 |
| 11 | 66438074 | rs12577111 | 1.821 | 0.852 | 1.01 | 1.355 | -0.01 | 0.037 | 0.037 | 0 | 0.5 | 0.11 | 0.63 | 0.44 | 0.51 | 0.6 |
| 11 | 66451178 | rs3741364 | -0.425 | 0.78 | 1.296 | 1.321 | -0.003 | 0 | 0 | 0 | 0.26 | 0.22 | 0.15 | 0.15 | 0.21 | 0.1 |
| 11 | 66456387 | rs2276138 | 1.983 | 0.764 | 1.112 | 1.315 | -0.004 | 0.042 | 0.042 | 0 | 0.5 | 0.13 | 0.63 | 0.44 | 0.51 | 0.6 |
| 11 | 66468469 | rs506028 | -0.476 | 0.043 | 1.709 | 1.744 | 0.01 | 0.132 | 0.132 | 0 | 0.1 | 0.33 | 0.13 | 0.28 | 0.11 | 0.11 |
| 11 | 66469032 | rs532439 | 0.826 | -0.083 | 1.765 | 1.571 | -0.048 | -0.041 | -0.041 | 1.75 | 0.69 | 0.59 | 0.81 | 0.82 | 0.75 | 0.87 |
| 11 | 66481264 | rs615536 | -1.022 | 0.227 | 1.831 | 1.411 | -0.014 | 0.013 | 0.013 | 0 | 0.89 | 0.76 | 0.94 | 0.89 | 0.93 | 0.95 |
| 11 | 66493825 | rs10791889 | -1.106 | 0.215 | 1.642 | 1.372 | -0.003 | -0.002 | -0.002 | 0 | 0.3 | 0.28 | 0.2 | 0.15 | 0.27 | 0.11 |
| 11 | 66560023 | rs4930390 | -1.068 | 0.228 | 1.609 | 1.368 | 0.01 | 0.009 | 0.009 | 0.549 | 0.3 | 0.24 | 0.19 | 0.15 | 0.27 | 0.11 |
| 11 | 66568943 | rs682842 | -2.571 | 0.23 | 1.589 | 1.454 | 0.089 | 0.068 | 0.068 | 0.014 | 0.47 | 0.88 | 0.38 | 0.63 | 0.41 | 0.28 |
| 11 | 66582210 | rs10750790 | -0.766 | 0.381 | 1.725 | 1.298 | 0.002 | 0.002 | 0.002 | 0.483 | 0.3 | 0.27 | 0.2 | 0.15 | 0.27 | 0.11 |
| 11 | 66608991 | rs594313 | -2.172 | 0.421 | 1.472 | 1.045 | 0.088 | 0.015 | 0.015 | 0.005 | 0.47 | 0.88 | 0.36 | 0.57 | 0.41 | 0.28 |
| 11 | 66621383 | rs2077432 | -1.018 | 0.561 | 1.684 | 1.015 | -0.023 | -0.002 | -0.002 | 0 | 0.31 | 0.33 | 0.21 | 0.29 | 0.28 | 0.11 |
| 11 | 66649612 | rs7119426 | -0.891 | 0.604 | 1.59 | 0.924 | 0.009 | 0.006 | 0.006 | 0.821 | 0.3 | 0.22 | 0.2 | 0.2 | 0.27 | 0.11 |
| 11 | 66650060 | rs10896139 | -1.076 | 0.597 | 1.639 | 0.974 | -0.04 | -0.002 | -0.002 | 0 | 0.32 | 0.49 | 0.23 | 0.32 | 0.27 | 0.11 |
| 11 | 66662731 | rs7122539 | -1.582 | 0.534 | 1.428 | 1.003 | 0.011 | -0.009 | -0.009 | 0 | 0.37 | 0.65 | 0.31 | 0.48 | 0.34 | 0.25 |
| 11 | 66673079 | rs10160758 | -0.854 | 0.489 | 1.11 | 0.738 | 0.014 | 0.007 | 0.007 | 0.685 | 0.3 | 0.22 | 0.2 | 0.2 | 0.27 | 0.11 |
| 11 | 66675990 | rs6591226 | -2.022 | 1.413 | 1.32 | 0.716 | -0.001 | 0.001 | 0.001 | 0 | 0.48 | 0.51 | 0.4 | 0.51 | 0.54 | 0.33 |
| 11 | 66676995 | rs735498 | 0.641 | 1.533 | 1.332 | 0.756 | -0.016 | 0.016 | 0.016 | 0 | 0.05 | 0.04 | 0.05 | 0.02 | 0.08 | 0.06 |
| 11 | 66679738 | rs10791894 | -1.962 | 1.513 | 1.377 | 0.762 | -0.001 | 0.001 | 0.001 | 0 | 0.48 | 0.51 | 0.4 | 0.51 | 0.54 | 0.33 |
| 11 | 66687863 | rs7944999 | -1.41 | 1.42 | 1.411 | 0.792 | 0 | -0.001 | -0.001 | 0 | 0.42 | 0.42 | 0.35 | 0.48 | 0.46 | 0.27 |
| 11 | 66690454 | rs7110302 | 1.597 | 1.448 | 0.635 | 0.703 | 0.017 | 0.021 | 0.021 | 0 | 0.45 | 0.16 | 0.26 | 0.41 | 0.35 | 0.61 |
| 11 | 66721028 | rs2278844 | -0.349 | 1.071 | 1.451 | 1.082 | 0 | 0.081 | 0.081 | 0 | 0.13 | 0.24 | 0.13 | 0.25 | 0.13 | 0.11 |
| 11 | 66766754 | rs12419173 | 1.648 | 1.094 | 0.138 | 0.909 | -0.073 | 0.036 | 0.036 | 0.041 | 0.53 | 0.13 | 0.56 | 0.38 | 0.45 | 0.65 |
| 11 | 66781263 | rs896436 | -0.912 | 0.902 | 1.452 | 0.946 | 0.006 | 0.034 | 0.034 | 0 | 0.26 | 0.38 | 0.23 | 0.32 | 0.3 | 0.18 |
| 11 | 66782594 | rs1253440 | -0.893 | 0.904 | 1.449 | 0.945 | 0.005 | 0.029 | 0.029 | 0.175 | 0.26 | 0.37 | 0.23 | 0.32 | 0.3 | 0.18 |
| 11 | 66784715 | rs1253439 | -0.781 | 0.919 | 1.422 | 0.933 | 0 | 0 | 0 | 0 | 0.25 | 0.23 | 0.22 | 0.26 | 0.3 | 0.18 |
| 11 | 66793329 | rs7943216 | -0.526 | 0.45 | 1.541 | 0.772 | -0.01 | -0.036 | -0.036 | 1.394 | 0.09 | 0.18 | 0.06 | 0 | 0.07 | 0.02 |
| 11 | 66812338 | rs11227668 | 1.511 | 0.576 | 2.452 | 1.321 | 0.075 | 0.268 | 0.268 | 0.979 | 0.92 | 0.55 | 0.85 | 0.79 | 0.91 | 0.93 |
| 11 | 66816507 | rs3741190 | 1.496 | 0.576 | 2.264 | 1.291 | 0.064 | 0.242 | 0.242 | 0.531 | 0.92 | 0.49 | 0.85 | 0.8 | 0.91 | 0.94 |
| 11 | 66826160 | rs7112925 | 0.619 | 0.481 | 1.057 | 0.879 | 0.006 | -0.003 | -0.003 | 0 | 0.41 | 0.36 | 0.28 | 0.45 | 0.36 | 0.54 |
| 11 | 66832528 | rs11227673 | -0.063 | 0.655 | 0.167 | 0.825 | 0.006 | 0.022 | 0.022 | 0 | 0.52 | 0.74 | 0.64 | 0.59 | 0.44 | 0.59 |
| 11 | 66842469 | rs11227678 | -1.608 | 0.891 | 1.593 | 0.762 | 0.026 | 0.113 | 0.113 | 0.31 | 0.1 | 0.52 | 0.12 | 0.14 | 0.07 | 0.05 |
| 11 | 66843350 | rs7932636 | -2.6 | 0.877 | 1.632 | 0.822 | -0.014 | 0.048 | 0.048 | 0.43 | 0.14 | 0.71 | 0.41 | 0.19 | 0.1 | 0.06 |
| 11 | 66898492 | rs11227699 | -3.067 | 0.872 | 2.807 | 0.824 | 0.044 | 0.117 | 0.117 | 3.326 | 0.06 | 0.63 | 0.35 | 0.14 | 0.04 | 0.03 |
| 11 | 67011765 | rs12288048 | -2.094 | 0.616 | 3.419 | 0.69 | -0.005 | 0.069 | 0.069 | 0 | 0.1 | 0.52 | 0.07 | 0.11 | 0.04 | 0.03 |
| 11 | 67050292 | rs2071007 | 2.818 | 0.602 | 3.354 | 0.633 | 0.005 | -0.005 | -0.005 | 1.847 | 0.85 | 0.18 | 0.6 | 0.74 | 0.92 | 0.95 |
| 11 | 67068859 | rs2298815 | 2.194 | 0.727 | 3.371 | 0.717 | 0.002 | 0.063 | 0.063 | 0.015 | 0.94 | 0.76 | 0.95 | 0.89 | 0.96 | 0.98 |
| 11 | 67070017 | rs4930192 | 3.309 | 0.727 | 3.551 | 0.718 | 0.04 | 0.173 | 0.173 | 0.316 | 0.9 | 0.54 | 0.88 | 0.78 | 0.93 | 0.95 |
| 11 | 67088394 | rs1894112 | 1.838 | 0.725 | 3.352 | 0.714 | -0.004 | 0.042 | 0.042 | 0 | 0.95 | 0.86 | 0.95 | 0.89 | 0.96 | 0.98 |
| 11 | 67175692 | rs10896172 | -1.982 | 0.854 | 2.585 | 0.7 | 0.007 | 0.075 | 0.075 | 0 | 0.06 | 0.14 | 0.07 | 0.17 | 0.05 | 0.03 |
| 11 | 67192555 | rs1626067 | -1.748 | 1.522 | 1.367 | 0.628 | -0.057 | -0.046 | -0.046 | 0 | 0.33 | 0.86 | 0.55 | 0.44 | 0.36 | 0.23 |
| 11 | 67196237 | rs1476792 | -1.839 | 1.597 | 1.226 | 0.278 | -0.088 | 0.035 | 0.035 | 0 | 0.4 | 0.87 | 0.57 | 0.33 | 0.42 | 0.23 |
| 11 | 67202044 | rs1790752 | -1.784 | 1.612 | 1.449 | 0.285 | -0.054 | 0.033 | 0.033 | 0 | 0.39 | 0.81 | 0.57 | 0.33 | 0.42 | 0.22 |
| 11 | 67202826 | rs10274 | -1.458 | 1.589 | 2.005 | 0.274 | -0.011 | 0.004 | 0.004 | 0 | 0.36 | 0.33 | 0.54 | 0.33 | 0.42 | 0.23 |
| 11 | 67204342 | rs12787021 | -1.799 | 1.533 | 1.975 | 0.46 | -0.086 | -0.001 | -0.001 | 0 | 0.41 | 0.88 | 0.57 | 0.39 | 0.42 | 0.23 |
| 11 | 67207426 | rs2302264 | -1.443 | 1.414 | 1.965 | 0.09 | -0.015 | 0.005 | 0.005 | 0 | 0.36 | 0.33 | 0.53 | 0.32 | 0.42 | 0.22 |
| 11 | 67227006 | rs11601325 | -1.458 | 1.492 | 2.054 | 0.065 | -0.052 | 0.067 | 0.067 | 0 | 0.34 | 0.21 | 0.5 | 0.19 | 0.4 | 0.2 |
| 11 | 67246757 | rs11227805 | 0.483 | 1.842 | 1.824 | 0.074 | -0.052 | -0.035 | -0.035 | 0.032 | 0.08 | 0.01 | 0.13 | 0.12 | 0.21 | 0.08 |
| 11 | 67268048 | rs11604662 | -1.072 | 1.047 | 1.718 | -0.364 | -0.093 | 0.156 | 0.156 | 0 | 0.28 | 0.04 | 0.18 | 0.04 | 0.33 | 0.17 |
| 11 | 67280620 | rs4465371 | -1.401 | 0.949 | 1.657 | -0.334 | -0.083 | 0.114 | 0.114 | 0 | 0.35 | 0.14 | 0.22 | 0.15 | 0.39 | 0.21 |
| 11 | 67286075 | rs682208 | 2.044 | 1.01 | 1.8 | -0.34 | 0.057 | 0.187 | 0.187 | 0.109 | 0.94 | 0.4 | 0.89 | 0.84 | 0.95 | 0.95 |
| 11 | 67288594 | rs2276118 | 1.974 | 0.633 | 1.172 | -0.572 | -0.126 | 0.045 | 0.045 | 0.111 | 0.52 | 0.03 | 0.38 | 0.66 | 0.56 | 0.64 |
| 11 | 67294198 | rs4930430 | -1.906 | 0.615 | 1.378 | -0.714 | -0.047 | 0.143 | 0.143 | 0 | 0.43 | 0.79 | 0.58 | 0.17 | 0.44 | 0.24 |
| 11 | 67311258 | rs694616 | 1.248 | 0.822 | 1.81 | -0.608 | 0.015 | 0.048 | 0.048 | 0.554 | 0.92 | 0.83 | 0.89 | 0.86 | 0.91 | 0.93 |
| 11 | 67314144 | rs7106423 | -1.519 | 0.616 | 1.858 | -0.737 | -0.032 | 0.078 | 0.078 | 0 | 0.38 | 0.23 | 0.26 | 0.16 | 0.4 | 0.25 |
| 11 | 67339949 | rs688878 | 1.132 | 0.963 | 2.064 | -0.625 | 0.001 | 0.009 | 0.009 | 0.464 | 0.92 | 0.87 | 0.89 | 0.86 | 0.91 | 0.93 |
| 11 | 67347287 | rs614080 | 2.286 | 0.869 | 2.066 | -0.679 | -0.025 | 0.097 | 0.097 | 0 | 0.51 | 0.07 | 0.33 | 0.68 | 0.49 | 0.64 |
| 11 | 67347595 | rs7941648 | -1.762 | 0.867 | 2.34 | -0.726 | 0.002 | -0.018 | -0.018 | 0 | 0.39 | 0.38 | 0.27 | 0.16 | 0.41 | 0.3 |
| 11 | 67349899 | rs6591256 | -1.762 | 0.869 | 2.331 | -0.725 | 0.002 | -0.018 | -0.018 | 0 | 0.39 | 0.38 | 0.27 | 0.16 | 0.41 | 0.3 |

| | | | | | | | | | | | | | | | | |
|---|---|---|---|---|---|---|---|---|---|---|---|---|---|---|---|---|
| 11 | 67352689 | rs1695 | -1.835 | 1.078 | 2.7 | -0.285 | 0.015 | -0.028 | -0.028 | 0 | 0.31 | 0.48 | 0.48 | 0.18 | 0.33 | 0.29 |
| 11 | 67353253 | rs749174 | -1.882 | 1.149 | 2.63 | -0.264 | -0.021 | 0.044 | 0.044 | 0 | 0.32 | 0.26 | 0.21 | 0.15 | 0.33 | 0.25 |
| 11 | 67353844 | rs1871042 | -1.879 | 1.144 | 2.636 | -0.266 | -0.018 | 0.037 | 0.037 | 0 | 0.32 | 0.28 | 0.21 | 0.15 | 0.33 | 0.25 |
| 11 | 67353970 | rs4891 | -1.977 | 1.138 | 2.702 | -0.268 | 0.016 | -0.04 | -0.04 | 0 | 0.33 | 0.52 | 0.49 | 0.17 | 0.34 | 0.25 |
| 11 | 67354406 | rs947895 | -1.878 | 1.137 | 2.595 | -0.269 | -0.018 | 0.037 | 0.037 | 0 | 0.32 | 0.27 | 0.21 | 0.15 | 0.33 | 0.25 |
| 11 | 67368641 | rs12790798 | -2.287 | 1.125 | 2.866 | 0.068 | -0.024 | 0.075 | 0.075 | 0.323 | 0.37 | 0.85 | 0.56 | 0.19 | 0.38 | 0.26 |
| 11 | 67369670 | rs10896186 | 2.43 | 1.159 | 2.852 | 0.093 | -0.119 | 0.017 | 0.017 | 0.153 | 0.58 | 0.04 | 0.37 | 0.66 | 0.56 | 0.68 |
| 11 | 67370766 | rs4024254 | -2.107 | 1.149 | 3.049 | -0.055 | 0.001 | -0.01 | -0.01 | 0.24 | 0.35 | 0.71 | 0.54 | 0.19 | 0.38 | 0.26 |
| 11 | 67372477 | rs12421329 | -2.011 | 1.149 | 3.064 | -0.055 | -0.015 | 0.024 | 0.024 | 0 | 0.33 | 0.33 | 0.52 | 0.19 | 0.38 | 0.26 |
| 11 | 67373669 | rs12421762 | -0.767 | 1.039 | 3.02 | -0.018 | 0 | -0.001 | -0.001 | 0.516 | 0.08 | 0.06 | 0.07 | 0.07 | 0.09 | 0.09 |
| 11 | 67381110 | rs4148959 | -2.345 | 1.058 | 2.979 | -0.018 | 0.001 | -0.01 | -0.01 | 0 | 0.39 | 0.67 | 0.56 | 0.2 | 0.42 | 0.3 |
| 11 | 67394895 | rs4930461 | -1.483 | 1.091 | 2.47 | -0.005 | -0.044 | 0.056 | 0.056 | 0 | 0.28 | 0.18 | 0.45 | 0.13 | 0.33 | 0.15 |
| 11 | 67395714 | rs7124513 | -1.483 | 1.091 | 2.475 | -0.005 | -0.042 | 0.054 | 0.054 | 0 | 0.28 | 0.18 | 0.45 | 0.13 | 0.33 | 0.16 |
| 11 | 67397675 | rs7102670 | 1.152 | 0.952 | 2.463 | 0.073 | -0.012 | -0.007 | -0.007 | 0 | 0.88 | 0.64 | 0.89 | 0.93 | 0.9 | 0.86 |
| 11 | 67400846 | rs11227871 | -1.653 | 0.966 | 0.952 | 0.08 | -0.014 | -0.01 | -0.01 | 0.662 | 0.12 | 0.59 | 0.11 | 0.07 | 0.1 | 0.14 |
| 11 | 67408090 | rs2509712 | -1.156 | 1.304 | 0.283 | 0.181 | 0.017 | -0.009 | -0.009 | 0 | 0.2 | 0.28 | 0.15 | 0.16 | 0.22 | 0.12 |
| 11 | 67408791 | rs2509713 | 0.966 | 0.561 | 0.728 | 0.245 | -0.012 | -0.047 | -0.047 | 1.702 | 0.81 | 0.4 | 0.86 | 0.93 | 0.84 | 0.91 |
| 11 | 67411830 | rs2447587 | 0.749 | 0.502 | 0.216 | 0.587 | 0.004 | 0.002 | 0.002 | 0.381 | 0.89 | 0.89 | 0.95 | 0.9 | 0.92 | 0.97 |
| 11 | 67414492 | rs948445 | 1.802 | 0.791 | 1.045 | 0.71 | -0.001 | 0.012 | 0.012 | 0.072 | 0.79 | 0.4 | 0.85 | 0.75 | 0.81 | 0.94 |
| 11 | 67415054 | rs2514036 | 1.547 | 0.852 | 1.404 | 0.725 | 0.006 | 0.012 | 0.012 | 0 | 0.82 | 0.7 | 0.87 | 0.75 | 0.81 | 0.94 |
| 11 | 67416190 | rs2514037 | -1.645 | 0.983 | 1.75 | 0.783 | 0.089 | 0.08 | 0.08 | 0 | 0.07 | 0.21 | 0.09 | 0.15 | 0.11 | 0.03 |
| 11 | 67419855 | rs10736660 | 2.33 | 0.862 | 2.182 | 0.795 | 0.014 | 0.04 | 0.04 | 0.755 | 0.82 | 0.43 | 0.85 | 0.73 | 0.81 | 0.92 |
| 11 | 67423892 | rs4930211 | 2.073 | 0.591 | 2.283 | 0.746 | -0.025 | -0.025 | -0.025 | 0.508 | 0.86 | 0.55 | 0.9 | 0.91 | 0.89 | 0.95 |
| 11 | 67430482 | rs866907 | 2.033 | 0.661 | 2.301 | 0.756 | -0.017 | 0.002 | 0.002 | 1.718 | 0.86 | 0.54 | 0.9 | 0.88 | 0.89 | 0.95 |
| 11 | 67430762 | rs1551886 | 1.326 | 0.666 | 2.297 | 0.758 | -0.002 | -0.005 | -0.005 | 2.42 | 0.89 | 0.72 | 0.92 | 0.91 | 0.89 | 0.97 |
| 11 | 67431038 | rs1979375 | 1.392 | 0.666 | 2.29 | 0.758 | -0.001 | -0.005 | -0.005 | 0 | 0.89 | 0.69 | 0.91 | 0.91 | 0.89 | 0.97 |
| 11 | 67434416 | rs4646823 | 2.14 | 0.667 | 2.106 | 0.637 | -0.02 | -0.001 | -0.001 | 0 | 0.86 | 0.53 | 0.9 | 0.88 | 0.88 | 0.95 |
| 11 | 67442043 | rs7947978 | 1.813 | 0.684 | 2.064 | 0.645 | -0.005 | 0.014 | 0.014 | 0.193 | 0.87 | 0.6 | 0.91 | 0.88 | 0.88 | 0.95 |
| 11 | 67442661 | rs17510687 | -1.743 | 0.755 | 1.77 | 0.654 | -0.005 | -0.007 | -0.007 | 0 | 0.11 | 0.16 | 0.06 | 0.09 | 0.12 | 0.03 |
| 11 | 67448740 | rs11603033 | -2.066 | 1.4 | 1.463 | 0.615 | -0.027 | -0.004 | -0.004 | 1.067 | 0.13 | 0.07 | 0.07 | 0.15 | 0.19 | 0.14 |
| 11 | 67459163 | rs2514046 | 1.371 | 1.054 | 2.182 | 0.752 | -0.011 | 0.031 | 0.031 | 0.1 | 0.9 | 0.59 | 0.91 | 0.92 | 0.95 | 0.93 |
| 11 | 67460912 | rs7118125 | 1.478 | 0.453 | 1.229 | -0.096 | 0.007 | 0.027 | 0.027 | 0 | 0.61 | 0.32 | 0.7 | 0.77 | 0.61 | 0.59 |
| 11 | 67462549 | rs11227881 | -0.686 | 0.42 | 1.289 | -0.131 | 0.001 | -0.022 | -0.022 | 0 | 0.23 | 0.21 | 0.18 | 0.09 | 0.26 | 0.29 |
| 11 | 67464928 | rs7941000 | -0.447 | 0.025 | 0.569 | 0.58 | 0.014 | 0.025 | 0.025 | 0 | 0.21 | 0.17 | 0.16 | 0.1 | 0.19 | 0.27 |
| 11 | 67472145 | rs4930217 | 1.397 | 0.44 | -0.067 | 1.492 | -0.006 | -0.064 | -0.064 | 0 | 0.6 | 0.09 | 0.62 | 0.45 | 0.59 | 0.59 |
| 11 | 67472905 | rs4385902 | 0.564 | 0.735 | 0.368 | 1.697 | 0.038 | 0.085 | 0.085 | 0 | 0.95 | 0.88 | 0.93 | 0.82 | 0.91 | 0.95 |
| 11 | 67501626 | rs10791923 | 0.343 | 0.533 | -0.605 | 1.297 | -0.01 | -0.026 | -0.026 | 0 | 0.32 | 0.19 | 0.53 | 0.6 | 0.35 | 0.43 |
| 11 | 67526279 | rs11227928 | -0.729 | -0.306 | -1.013 | 1.237 | 0.051 | -0.057 | -0.057 | 0.154 | 0.13 | 0.02 | 0.07 | 0.21 | 0.07 | 0.08 |
| 11 | 67533067 | rs11600121 | -0.665 | 0.141 | -1.581 | 0.428 | -0.062 | 0.071 | 0.071 | 0 | 0.14 | 0.01 | 0.17 | 0.03 | 0.2 | 0.15 |
| 11 | 67533195 | rs10750808 | 0.736 | -0.364 | -0.3 | 0.668 | 0.011 | 0.083 | 0.083 | 0 | 0.93 | 0.58 | 0.95 | 0.95 | 0.97 | 0.99 |
| 11 | 67536948 | rs10896215 | 0.825 | -0.228 | -1.554 | 0.748 | -0.031 | -0.033 | -0.033 | 0 | 0.34 | 0.06 | 0.29 | 0.65 | 0.35 | 0.56 |
| 11 | 67547176 | rs7116709 | 0.452 | -0.32 | -1.072 | 0.614 | -0.018 | -0.002 | -0.002 | 0 | 0.07 | 0.06 | 0.01 | 0.03 | 0 | 0.02 |
| 11 | 67587776 | rs10750813 | -0.107 | -0.141 | -1.854 | -0.102 | -0.04 | 0.163 | 0.163 | 0.073 | 0.48 | 0.1 | 0.62 | 0.78 | 0.48 | 0.58 |
| 11 | 67588012 | rs10791946 | -0.107 | -0.141 | -1.854 | -0.102 | -0.04 | 0.163 | 0.163 | 0.073 | 0.48 | 0.1 | 0.62 | 0.78 | 0.48 | 0.58 |
| 11 | 67715028 | rs308309 | 0.901 | 0.747 | -1.468 | 0.29 | -0.012 | -0.003 | -0.003 | 0 | 0.86 | 0.89 | 0.86 | 0.83 | 0.77 | 0.81 |
| 11 | 67731956 | rs308351 | -1.498 | -0.025 | -2.17 | -0.054 | 0.036 | 0.01 | 0.01 | 0 | 0.47 | 0.9 | 0.33 | 0.43 | 0.43 | 0.42 |
| 11 | 67748817 | rs7122260 | -0.384 | -0.117 | -1.876 | 0.081 | -0.01 | -0.018 | -0.018 | 0 | 0.13 | 0.09 | 0.13 | 0.15 | 0.16 | 0.17 |
| 11 | 67751231 | rs7931196 | -0.303 | -0.684 | -1.276 | -0.389 | -0.027 | -0.031 | -0.031 | 0.252 | 0.28 | 0.44 | 0.16 | 0.2 | 0.23 | 0.14 |
| 11 | 67765056 | rs308328 | -2.072 | -0.769 | -1.597 | -0.332 | 0.045 | -0.02 | -0.02 | 0.833 | 0.51 | 0.93 | 0.33 | 0.43 | 0.39 | 0.41 |
| 11 | 67768796 | rs11228120 | -0.297 | -0.201 | -1.266 | -0.02 | -0.01 | -0.022 | -0.022 | 6.064 | 0.13 | 0.09 | 0.14 | 0.17 | 0.16 | 0.17 |
| 11 | 67772729 | rs3808969 | -0.306 | -0.095 | -1.274 | 0.029 | -0.013 | -0.025 | -0.025 | 0.533 | 0.13 | 0.09 | 0.14 | 0.18 | 0.16 | 0.17 |
| 11 | 67773897 | rs7952122 | 0.054 | -0.385 | -0.983 | -0.282 | -0.019 | -0.011 | -0.011 | 0.044 | 0.22 | 0.38 | 0.13 | 0.22 | 0.19 | 0.13 |
| 11 | 67776242 | rs308338 | -1.164 | 0.442 | -0.699 | -0.449 | -0.002 | 0 | 0 | 0 | 0.41 | 0.67 | 0.34 | 0.4 | 0.44 | 0.38 |
| 11 | 67778326 | rs3764821 | -0.009 | 0.306 | 0.037 | -0.449 | -0.001 | -0.001 | -0.001 | 0.112 | 0.22 | 0.24 | 0.16 | 0.21 | 0.21 | 0.12 |
| 11 | 67784278 | rs3751084 | -0.159 | 0.209 | 0.255 | -0.54 | -0.012 | -0.022 | -0.022 | 0.134 | 0.22 | 0.41 | 0.17 | 0.21 | 0.21 | 0.13 |
| 11 | 67788563 | rs4646812 | -0.581 | 0.159 | 0.296 | -0.577 | -0.025 | -0.028 | -0.028 | 0.897 | 0.27 | 0.43 | 0.17 | 0.23 | 0.22 | 0.21 |
| 11 | 67789138 | rs2286168 | -0.047 | 0.178 | 0.315 | -0.565 | -0.005 | -0.005 | -0.005 | 1.45 | 0.22 | 0.29 | 0.16 | 0.23 | 0.21 | 0.21 |
| 11 | 67789837 | rs3764819 | -0.062 | 0.178 | 0.314 | -0.565 | -0.007 | -0.008 | -0.008 | 0.612 | 0.22 | 0.32 | 0.16 | 0.23 | 0.21 | 0.21 |
| 11 | 67793014 | rs105147 | -1.608 | 0.308 | 0.293 | -0.246 | -0.001 | -0.012 | -0.012 | 0.87 | 0.4 | 0.74 | 0.36 | 0.46 | 0.42 | 0.37 |
| 11 | 67793448 | rs2286164 | -0.129 | 0.435 | 0.208 | -0.237 | -0.014 | -0.013 | -0.013 | 1.145 | 0.23 | 0.4 | 0.17 | 0.23 | 0.22 | 0.21 |
| 11 | 67793509 | rs2286163 | 0.007 | 0.434 | 0.228 | -0.237 | -0.002 | -0.001 | -0.001 | 0 | 0.22 | 0.26 | 0.16 | 0.23 | 0.22 | 0.21 |
| 11 | 67794005 | rs886702 | -0.13 | 0.435 | 0.203 | -0.237 | -0.014 | -0.013 | -0.013 | 0 | 0.23 | 0.4 | 0.17 | 0.23 | 0.22 | 0.21 |
| 11 | 67795066 | rs886701 | -0.166 | 0.364 | 0.144 | -0.276 | -0.007 | -0.007 | -0.007 | 0 | 0.23 | 0.31 | 0.16 | 0.23 | 0.22 | 0.22 |
| 11 | 67796414 | rs4646815 | 0.068 | 0.381 | 0.051 | -0.341 | -0.043 | -0.039 | -0.039 | 1.233 | 0.1 | 0.03 | 0.15 | 0.15 | 0.2 | 0.14 |
| 11 | 67796632 | rs15518 | -0.342 | 0.108 | 0.294 | -0.358 | -0.01 | -0.008 | -0.008 | 0 | 0.23 | 0.32 | 0.16 | 0.23 | 0.22 | 0.14 |
| 11 | 67796756 | rs3115547 | -0.256 | 0.127 | 0.275 | -0.362 | -0.003 | -0.003 | -0.003 | 0 | 0.23 | 0.28 | 0.16 | 0.23 | 0.22 | 0.14 |
| 11 | 67796801 | rs3133268 | -0.26 | 0.118 | 0.276 | -0.355 | -0.01 | -0.007 | -0.007 | 0 | 0.23 | 0.32 | 0.16 | 0.23 | 0.22 | 0.14 |
| 11 | 67798567 | rs1122731 | -0.263 | 0.138 | 0.155 | -0.365 | -0.004 | -0.003 | -0.003 | 0 | 0.23 | 0.28 | 0.16 | 0.23 | 0.22 | 0.14 |
| 11 | 67799266 | rs999571 | 0.801 | 0.089 | 0.147 | -0.431 | -0.003 | 0 | 0 | 0 | 0.14 | 0.15 | 0.09 | 0.19 | 0.12 | 0.08 |

| | | | | | | | | | | | | | | | | |
|---|---|---|---|---|---|---|---|---|---|---|---|---|---|---|---|---|
| 11 | 67799926 | rs2075626 | -0.597 | 0.103 | 0.137 | -0.421 | -0.003 | -0.002 | -0.002 | 0 | 0.22 | 0.27 | 0.16 | 0.23 | 0.21 | 0.14 |
| 11 | 67801517 | rs3115546 | -1.148 | 0.094 | 0.148 | -0.506 | -0.003 | -0.011 | -0.011 | 0 | 0.94 | 0.73 | 0.97 | 1 | 0.99 | 1 |
| 11 | 67802907 | rs1104739 | -1.134 | 0.353 | -0.471 | -1.115 | 0.013 | 0.125 | 0.125 | 0 | 0.49 | 0.73 | 0.31 | 0.23 | 0.42 | 0.25 |
| 11 | 67804156 | rs3133269 | 0.444 | 0.939 | -0.379 | -0.703 | 0.057 | 0.05 | 0.05 | 0 | 0.83 | 0.69 | 0.8 | 0.81 | 0.72 | 0.78 |
| 11 | 67804605 | rs4147778 | 0.437 | 0.938 | -0.376 | -0.703 | 0.051 | 0.045 | 0.045 | 0 | 0.83 | 0.68 | 0.8 | 0.81 | 0.72 | 0.78 |
| 11 | 67805381 | rs4147780 | 1.371 | 0.657 | -0.21 | -0.983 | -0.014 | -0.012 | -0.012 | 0.005 | 0.6 | 0.25 | 0.63 | 0.57 | 0.52 | 0.7 |
| 11 | 67816936 | rs2075609 | -1.187 | 0.656 | -0.679 | -1.165 | -0.011 | 0.014 | 0.014 | 0 | 0.29 | 0.48 | 0.22 | 0.34 | 0.26 | 0.24 |
| 11 | 67817876 | rs11228127 | 0.18 | 0.856 | -0.501 | -1.113 | 0.001 | 0.187 | 0.187 | 0 | 0.11 | 0.35 | 0.09 | 0.31 | 0.1 | 0.12 |
| 11 | 67821036 | rs3794186 | -0.669 | 1.001 | -0.533 | -0.959 | 0.005 | 0.071 | 0.071 | 0 | 0.05 | 0.1 | 0.07 | 0.2 | 0.07 | 0.06 |
| 11 | 67834500 | rs35046555 | 1.252 | 0.913 | -0.728 | -1.198 | -0.08 | 0.107 | 0.107 | 0 | 0.19 | 0.01 | 0.11 | 0.01 | 0.2 | 0.1 |
| 11 | 67846196 | rs2457180 | 1.384 | 0.816 | -0.114 | -0.66 | 0.017 | 0.027 | 0.027 | 0 | 0.68 | 0.49 | 0.72 | 0.6 | 0.6 | 0.72 |
| 11 | 67854035 | rs2511439 | -0.86 | 0.686 | -0.038 | -0.666 | -0.005 | -0.002 | -0.002 | 0.131 | 0.87 | 0.88 | 0.93 | 0.83 | 0.89 | 0.92 |
| 11 | 67856510 | rs2511437 | 1.158 | 0.733 | -0.023 | -0.7 | 0.022 | 0.027 | 0.027 | 0 | 0.72 | 0.6 | 0.71 | 0.61 | 0.6 | 0.75 |
| 11 | 67867548 | rs10791957 | 1.851 | 0.711 | 0.015 | -0.721 | 0.009 | 0.008 | 0.008 | 0 | 0.65 | 0.35 | 0.68 | 0.6 | 0.56 | 0.72 |
| 11 | 67870321 | rs11228131 | 1.076 | 0.788 | -0.329 | -0.652 | 0.003 | 0.004 | 0.004 | 0 | 0.73 | 0.69 | 0.72 | 0.62 | 0.6 | 0.76 |
| 11 | 67891289 | rs7127090 | -0.846 | 0.753 | -0.345 | -0.72 | -0.005 | -0.003 | -0.003 | 0 | 0.87 | 0.88 | 0.93 | 0.85 | 0.89 | 0.92 |
| 11 | 67897483 | rs7944372 | 1.331 | 0.598 | -0.314 | -0.739 | 0.019 | 0.012 | 0.012 | 0 | 0.67 | 0.5 | 0.69 | 0.62 | 0.56 | 0.72 |
| 11 | 67908805 | rs11228144 | 1.061 | 0.413 | -0.712 | -0.519 | 0.038 | -0.051 | -0.051 | 0 | 0.15 | 0.05 | 0.39 | 0.61 | 0.05 | 0.44 |
| 11 | 67910783 | rs2511470 | 1.004 | 1.789 | -0.141 | 0.072 | 0.006 | 0.006 | 0.006 | 0 | 0.75 | 0.7 | 0.73 | 0.64 | 0.61 | 0.74 |
| 11 | 67915693 | rs1547890 | -1.104 | 1.883 | -0.148 | 0.181 | 0.015 | 0.042 | 0.042 | 0 | 0.12 | 0.24 | 0.17 | 0.24 | 0.23 | 0.16 |
| 11 | 67917087 | rs1547889 | -0.777 | 1.912 | -0.263 | 0.216 | 0.006 | 0.028 | 0.028 | 0 | 0.11 | 0.2 | 0.16 | 0.24 | 0.22 | 0.15 |
| 11 | 67918678 | rs2511477 | -0.424 | 1.459 | -0.346 | -0.028 | 0.001 | -0.005 | -0.005 | 0 | 0.86 | 0.88 | 0.9 | 0.88 | 0.84 | 0.9 |
| 11 | 67927928 | rs4256988 | -0.057 | 1.059 | -0.23 | 0.324 | 0.003 | 0.064 | 0.064 | 0 | 0.06 | 0.12 | 0.08 | 0.25 | 0.08 | 0.09 |
| 11 | 67928520 | rs1540209 | -0.006 | 1.426 | -0.142 | 0.014 | -0.004 | -0.004 | -0.004 | 0 | 0.72 | 0.77 | 0.76 | 0.87 | 0.58 | 0.79 |
| 11 | 67929671 | rs1894204 | 1.2 | 1.314 | 0.236 | 0.071 | -0.004 | -0.007 | -0.007 | 0 | 0.58 | 0.31 | 0.63 | 0.55 | 0.44 | 0.66 |
| 11 | 67931459 | rs10896298 | 0.705 | 1.384 | -0.142 | -0.075 | 0.035 | 0.006 | 0.006 | 0 | 0.64 | 0.5 | 0.67 | 0.62 | 0.49 | 0.71 |
| 11 | 67931761 | rs4930561 | 0.685 | 1.389 | -0.153 | -0.072 | 0.024 | 0.004 | 0.004 | 0 | 0.64 | 0.54 | 0.67 | 0.62 | 0.49 | 0.71 |
| 11 | 67946078 | rs1894205 | -0.141 | 1.357 | -0.108 | 0.073 | 0.015 | 0.13 | 0.13 | 0.194 | 0.07 | 0.23 | 0.09 | 0.25 | 0.09 | 0.09 |
| 11 | 67953772 | rs10896300 | 0.097 | 1.368 | -0.158 | 0.047 | 0.005 | -0.02 | -0.02 | 0.063 | 0.72 | 0.72 | 0.75 | 0.87 | 0.58 | 0.79 |
| 11 | 67971546 | rs2129848 | -0.835 | 0.742 | 0.213 | 0.728 | 0.023 | 0.161 | 0.161 | 0.342 | 0.07 | 0.26 | 0.09 | 0.25 | 0.09 | 0.09 |
| 11 | 67992324 | rs1619377 | -0.879 | 0.522 | -0.648 | 0.596 | 0.011 | -0.023 | -0.023 | 0.276 | 0.31 | 0.83 | 0.23 | 0.39 | 0.26 | 0.23 |
| 11 | 68000579 | rs4930568 | -0.946 | 0.524 | -0.308 | 0.559 | -0.004 | 0.005 | 0.005 | 0.035 | 0.29 | 0.68 | 0.2 | 0.29 | 0.25 | 0.24 |
| 11 | 68022006 | rs2450908 | -0.335 | 0.23 | -0.555 | 0.202 | -0.034 | -0.009 | -0.009 | 0 | 0.36 | 0.18 | 0.27 | 0.41 | 0.43 | 0.24 |
| 11 | 68023200 | rs2458268 | -0.343 | 0.156 | -0.437 | 0.164 | -0.027 | -0.009 | -0.009 | 0 | 0.34 | 0.25 | 0.29 | 0.41 | 0.45 | 0.25 |
| 11 | 68025467 | rs674857 | 0.937 | 0.398 | -0.639 | 0.114 | -0.007 | -0.01 | -0.01 | 0.137 | 0.17 | 0.11 | 0.16 | 0.28 | 0.2 | 0.19 |
| 11 | 68025850 | rs11228184 | -0.535 | -0.134 | -0.825 | -0.113 | 0.019 | 0.02 | 0.02 | 0 | 0.2 | 0.13 | 0.12 | 0.1 | 0.16 | 0.08 |
| 11 | 68030173 | rs3802746 | 1.928 | -0.125 | -0.63 | -0.234 | -0.011 | -0.087 | -0.087 | 0 | 0.33 | 0.13 | 0.51 | 0.48 | 0.31 | 0.62 |
| 11 | 68033925 | rs7342161 | -0.011 | -0.142 | -0.201 | -0.292 | -0.003 | -0.001 | -0.001 | 0 | 0.65 | 0.66 | 0.71 | 0.58 | 0.55 | 0.75 |
| 11 | 68034772 | rs10791971 | 1.701 | -0.278 | -0.36 | -0.076 | -0.011 | -0.089 | -0.089 | 0.384 | 0.33 | 0.13 | 0.51 | 0.47 | 0.31 | 0.62 |
| 11 | 68040643 | rs11600189 | 0.34 | -0.043 | -0.213 | -0.099 | 0 | 0.031 | 0.031 | 0 | 0.09 | 0.01 | 0.05 | 0.01 | 0.07 | 0.02 |
| 11 | 68050176 | rs634918 | -0.974 | 0.034 | -0.093 | -0.029 | 0.01 | 0.03 | 0.03 | 0 | 0.81 | 0.73 | 0.83 | 0.7 | 0.8 | 0.82 |
| 11 | 68056770 | rs11606508 | -0.534 | -0.042 | -0.66 | -0.087 | 0.007 | 0.006 | 0.006 | 0 | 0.16 | 0.12 | 0.08 | 0.03 | 0.11 | 0.04 |
| 11 | 68070095 | rs474742 | 1.191 | 0.159 | -0.818 | 0.427 | 0.016 | 0.015 | 0.015 | 1.251 | 0.37 | 0.12 | 0.23 | 0.29 | 0.3 | 0.61 |
| 11 | 68077238 | rs312009 | -1.158 | 0.285 | -0.851 | -0.321 | 0 | 0.003 | 0.003 | 0 | 0.83 | 0.85 | 0.85 | 0.71 | 0.8 | 0.81 |
| 11 | 68088831 | rs4988300 | -1.348 | -0.223 | 0.062 | -0.956 | 0 | 0.045 | 0.045 | 0.979 | 0.46 | 0.65 | 0.34 | 0.23 | 0.48 | 0.16 |
| 11 | 68094741 | rs634008 | 1.091 | -0.236 | -0.311 | -0.883 | -0.009 | 0.129 | 0.129 | 1.289 | 0.57 | 0.16 | 0.66 | 0.77 | 0.54 | 0.85 |
| 11 | 68097826 | rs314779 | 0.242 | 0.181 | -0.519 | -0.76 | -0.066 | 0.027 | 0.027 | 0 | 0.16 | 0.03 | 0.16 | 0.13 | 0.31 | 0.06 |
| 11 | 68098799 | rs7948226 | -0.58 | -0.438 | -0.113 | -0.753 | 0.002 | 0.001 | 0.001 | 0 | 0.1 | 0.1 | 0.08 | 0.09 | 0.08 | 0.03 |
| 11 | 68102000 | rs606989 | 0.491 | -0.5 | 0.093 | -0.719 | -0.027 | -0.005 | -0.005 | 0.002 | 0.89 | 0.8 | 0.72 | 0.91 | 0.94 | 0.96 |
| 11 | 68108332 | rs312778 | -1.633 | -0.335 | 0.242 | -1.073 | 0.001 | -0.012 | -0.012 | 1.008 | 0.39 | 0.67 | 0.48 | 0.11 | 0.4 | 0.16 |
| 11 | 68111760 | rs729635 | -0.111 | -0.523 | 0.089 | -1.204 | 0.043 | 0.075 | 0.075 | 0 | 0.17 | 0.05 | 0.27 | 0.01 | 0.1 | 0.1 |
| 11 | 68118563 | rs312783 | -1.936 | -0.465 | 0.128 | -1.195 | 0.01 | 0.081 | 0.081 | 0.378 | 0.42 | 0.75 | 0.48 | 0.12 | 0.41 | 0.17 |
| 11 | 68119977 | rs312786 | 0.12 | -0.266 | 0.017 | -1.055 | -0.098 | 0.031 | 0.031 | 0 | 0.17 | 0.02 | 0.16 | 0.1 | 0.3 | 0.06 |
| 11 | 68140782 | rs624947 | 1.691 | -0.606 | 0.517 | -0.965 | 0.001 | 0 | 0 | 1.786 | 0.7 | 0.28 | 0.57 | 0.85 | 0.72 | 0.88 |
| 11 | 68141414 | rs638051 | 1.911 | -0.944 | 0.333 | -0.678 | 0.042 | 0.072 | 0.072 | 0 | 0.55 | 0.24 | 0.49 | 0.71 | 0.62 | 0.82 |
| 11 | 68146661 | rs11826287 | -0.225 | -0.596 | 0.582 | -0.674 | 0.008 | -0.002 | -0.002 | 0 | 0.12 | 0.18 | 0.13 | 0.1 | 0.18 | 0.05 |
| 11 | 68157299 | rs4988320 | -0.419 | -0.716 | 0.043 | -0.731 | 0.025 | 0.09 | 0.09 | 0 | 0.2 | 0.05 | 0.3 | 0.01 | 0.14 | 0.04 |
| 11 | 68159152 | rs3781590 | -1.613 | -0.621 | 0.431 | -0.652 | -0.017 | -0.067 | -0.067 | 0.076 | 0.35 | 0.45 | 0.45 | 0.12 | 0.31 | 0.09 |
| 11 | 68163514 | rs627174 | 1.386 | -0.259 | 1.053 | -0.182 | 0.041 | 0.064 | 0.064 | 0 | 0.85 | 0.5 | 0.8 | 0.82 | 0.83 | 0.94 |
| 11 | 68171013 | rs545382 | 0.238 | -0.656 | 1.415 | -0.657 | -0.027 | -0.081 | -0.081 | 1.214 | 0.85 | 0.57 | 0.91 | 0.99 | 0.92 | 0.94 |
| 11 | 68172102 | rs594090 | 0.142 | -0.675 | 1.509 | -0.667 | -0.03 | -0.09 | -0.09 | 1.3 | 0.85 | 0.52 | 0.9 | 0.99 | 0.92 | 0.94 |
| 11 | 68174122 | rs2277268 | 0.131 | -0.976 | 1.431 | -0.759 | 0.006 | 0.007 | 0.007 | 1.972 | 0.1 | 0.07 | 0.05 | 0.06 | 0.07 | 0.03 |
| 11 | 68174189 | rs4988321 | 0.41 | -1.242 | 1.214 | -0.925 | 0.029 | 0.056 | 0.056 | 0 | 0.1 | 0 | 0.04 | 0 | 0.04 | 0.02 |
| 11 | 68177510 | rs2306862 | -0.42 | -1.185 | 1.184 | -0.449 | -0.03 | -0.039 | -0.039 | 0.168 | 0.13 | 0.02 | 0.15 | 0.19 | 0.15 | 0.07 |
| 11 | 68177708 | rs576118 | 1.54 | -1.195 | 1.297 | -0.455 | -0.021 | -0.036 | -0.036 | 0.304 | 0.71 | 0.44 | 0.75 | 0.79 | 0.77 | 0.86 |
| 11 | 68182028 | rs314750 | 2.418 | -1.168 | 1.137 | -0.437 | 0.033 | 0.047 | 0.047 | 0 | 0.6 | 0.26 | 0.69 | 0.73 | 0.7 | 0.84 |
| 11 | 68182375 | rs923346 | -0.579 | -1.149 | 0.853 | -0.414 | -0.011 | -0.018 | -0.018 | 0.055 | 0.14 | 0.06 | 0.15 | 0.19 | 0.15 | 0.07 |
| 11 | 68192346 | rs599083 | -2.511 | -1.229 | 1.371 | -0.492 | -0.025 | -0.037 | -0.037 | 0.505 | 0.39 | 0.5 | 0.28 | 0.27 | 0.3 | 0.16 |
| 11 | 68192690 | rs556442 | 2.492 | -1.782 | 1.308 | -0.758 | 0.093 | 0.123 | 0.123 | 0.807 | 0.55 | 0.16 | 0.68 | 0.72 | 0.69 | 0.81 |

| | | | | | | | | | | | | | | | | |
|---|---|---|---|---|---|---|---|---|---|---|---|---|---|---|---|---|
| 11 | 68196830 | rs608343 | -2.616 | -1.683 | 1.135 | -0.721 | 0.011 | 0.012 | 0.012 | 0.699 | 0.43 | 0.63 | 0.29 | 0.27 | 0.31 | 0.19 |
| 11 | 68196961 | rs607887 | -2.575 | -1.687 | 1.086 | -0.723 | -0.005 | -0.009 | -0.009 | 0 | 0.43 | 0.54 | 0.28 | 0.27 | 0.31 | 0.19 |
| 11 | 68199393 | rs3781586 | -0.706 | -1.683 | 0.886 | -0.716 | -0.026 | -0.045 | -0.045 | 0.275 | 0.17 | 0.02 | 0.15 | 0.2 | 0.15 | 0.11 |
| 11 | 68201007 | rs12417014 | -0.134 | -1.748 | 1.195 | -0.752 | -0.015 | -0.036 | -0.036 | 0.086 | 0.12 | 0.18 | 0.06 | 0.05 | 0.08 | 0.03 |
| 11 | 68201295 | rs3736228 | -0.49 | -1.694 | 1.034 | -0.731 | 0 | -0.034 | -0.034 | 0.137 | 0.16 | 0.02 | 0.15 | 0.2 | 0.13 | 0.11 |
| 11 | 68205578 | rs901823 | -2.707 | -1.705 | 1.213 | -0.697 | -0.016 | -0.022 | -0.022 | 0 | 0.43 | 0.52 | 0.28 | 0.27 | 0.31 | 0.19 |
| 11 | 68209718 | rs3781579 | -0.522 | -1.274 | 1.071 | -0.702 | 0 | 0.008 | 0.008 | 0.513 | 0.13 | 0.09 | 0.06 | 0.09 | 0.12 | 0.04 |
| 11 | 68215954 | rs632605 | -0.154 | -1.264 | 1.315 | -0.729 | -0.017 | -0.041 | -0.041 | 0 | 0.13 | 0.22 | 0.07 | 0.01 | 0.08 | 0.04 |
| 11 | 68217420 | rs676318 | -0.066 | -1.265 | 1.293 | -0.728 | -0.011 | -0.025 | -0.025 | 0 | 0.13 | 0.2 | 0.07 | 0.01 | 0.07 | 0.04 |
| 11 | 68236389 | rs2472429 | 1.037 | -1.212 | 0.633 | -1.222 | -0.016 | 0.05 | 0.05 | 0.223 | 0.09 | 0.02 | 0.06 | 0 | 0.11 | 0.02 |
| 11 | 68254328 | rs11228258 | -1.721 | -0.812 | 0.998 | -1.096 | -0.001 | -0.011 | -0.011 | 0 | 0.28 | 0.27 | 0.22 | 0.21 | 0.27 | 0.14 |
| 11 | 68256847 | rs2186937 | 0.578 | -0.752 | 1.071 | -1.066 | 0.002 | 0.001 | 0.001 | 0 | 0.1 | 0.1 | 0.16 | 0.14 | 0.14 | 0.08 |
| 11 | 68289338 | rs7927235 | 1.581 | -0.082 | 1.128 | -0.101 | -0.034 | 0.054 | 0.054 | 0.367 | 0.06 | 0.01 | 0.28 | 0.39 | 0.1 | 0.16 |
| 11 | 68289796 | rs11228269 | -1.696 | -0.995 | 1.536 | -0.964 | 0 | -0.002 | -0.002 | 0 | 0.27 | 0.23 | 0.22 | 0.21 | 0.27 | 0.14 |
| 11 | 68291116 | rs11228270 | 0.608 | -1.038 | 1.482 | -0.864 | 0.003 | -0.004 | -0.004 | 0 | 0.07 | 0.04 | 0.03 | 0.13 | 0.04 | 0.02 |
| 11 | 68292930 | rs7936582 | -2.139 | -1.01 | 1.585 | -1.754 | 0.003 | -0.046 | -0.046 | 0 | 0.33 | 0.55 | 0.28 | 0.21 | 0.37 | 0.16 |
| 11 | 68302101 | rs2840367 | 2.161 | -1.517 | 1.672 | -1.852 | 0.01 | 0.016 | 0.016 | 0.536 | 0.58 | 0.37 | 0.7 | 0.78 | 0.66 | 0.82 |
| 11 | 68331154 | rs7119422 | 2.218 | -1.521 | 1.612 | -1.854 | 0.035 | 0.082 | 0.082 | 0.549 | 0.57 | 0.27 | 0.69 | 0.78 | 0.66 | 0.82 |
| 11 | 68331663 | rs12804775 | 0.568 | -1.414 | 0.928 | -2.01 | -0.026 | 0.046 | 0.046 | 0 | 0.08 | 0 | 0.05 | 0 | 0.11 | 0.08 |
| 11 | 68343593 | rs3740629 | -1.488 | -1.658 | 1.345 | -1.642 | 0.001 | 0.016 | 0.016 | 0.083 | 0.27 | 0.18 | 0.21 | 0.21 | 0.26 | 0.14 |
| 11 | 68362164 | rs7123564 | -1.488 | -1.638 | 1.345 | -1.636 | 0.001 | 0.015 | 0.015 | 0.083 | 0.27 | 0.18 | 0.21 | 0.21 | 0.26 | 0.14 |
| 11 | 68374183 | rs7107622 | -1.673 | -1.719 | 1.401 | -1.66 | -0.002 | -0.035 | -0.035 | 0 | 0.28 | 0.34 | 0.22 | 0.21 | 0.28 | 0.14 |
| 11 | 68377602 | rs7104877 | -1.949 | -1.722 | 1.38 | -1.661 | -0.001 | -0.009 | -0.009 | 0 | 0.29 | 0.31 | 0.22 | 0.21 | 0.28 | 0.14 |
| 11 | 68381415 | rs3740628 | -2.378 | -1.725 | 1.349 | -1.666 | 0.004 | 0.001 | 0.001 | 0.607 | 0.42 | 0.61 | 0.29 | 0.22 | 0.34 | 0.18 |
| 11 | 68381607 | rs4988291 | -1.543 | -1.536 | 0.85 | -1.563 | 0.001 | 0.016 | 0.016 | 0.083 | 0.27 | 0.18 | 0.21 | 0.21 | 0.26 | 0.15 |
| 11 | 68387984 | rs7128942 | -0.113 | -1.522 | 0.811 | -1.54 | 0.004 | 0.001 | 0.001 | 0 | 0.11 | 0.13 | 0.16 | 0.14 | 0.16 | 0.08 |
| 11 | 68388567 | rs11228294 | -1.086 | -1.53 | 0.818 | -1.542 | -0.008 | -0.024 | -0.024 | 0.185 | 0.23 | 0.29 | 0.22 | 0.15 | 0.21 | 0.12 |
| 11 | 68391197 | rs11228296 | -0.864 | -1.578 | 0.39 | -1.322 | 0.011 | 0.03 | 0.03 | 1.634 | 0.17 | 0.08 | 0.06 | 0.07 | 0.12 | 0.06 |
| 11 | 68408004 | rs2510389 | -0.997 | -1.485 | 0.886 | -1.266 | -0.018 | -0.062 | -0.062 | 0 | 0.16 | 0.29 | 0.07 | 0.06 | 0.12 | 0.04 |
| 11 | 68408289 | rs882979 | 0.254 | -1.274 | 0.95 | -1.192 | 0.001 | 0.011 | 0.011 | 0.012 | 0.8 | 0.81 | 0.79 | 0.85 | 0.81 | 0.89 |
| 11 | 68411347 | rs4930585 | -0.291 | -1.166 | 1.154 | -1.056 | 0.065 | 0.044 | 0.044 | 0 | 0.92 | 0.8 | 0.84 | 0.86 | 0.84 | 0.91 |
| 11 | 68417094 | rs2513281 | -0.526 | -1.168 | 1.142 | -1.054 | 0.036 | 0.024 | 0.024 | 0 | 0.93 | 0.86 | 0.84 | 0.87 | 0.85 | 0.91 |
| 11 | 68420798 | rs948862 | 0.58 | -1.429 | 1.428 | -1.233 | -0.024 | -0.069 | -0.069 | 0 | 0.74 | 0.52 | 0.75 | 0.85 | 0.76 | 0.82 |
| 11 | 68421416 | rs2510398 | -0.41 | -1.395 | 1.358 | -1.212 | 0.07 | 0.054 | 0.054 | 0.02 | 0.92 | 0.8 | 0.85 | 0.87 | 0.85 | 0.94 |
| 11 | 68421892 | rs4930591 | 0.546 | -1.415 | 1.32 | -1.218 | -0.019 | -0.049 | -0.049 | 0 | 0.73 | 0.43 | 0.74 | 0.85 | 0.76 | 0.82 |
| 11 | 68434181 | rs4930593 | 0.723 | -1.612 | 0.802 | -1.292 | -0.014 | -0.035 | -0.035 | 0 | 0.66 | 0.37 | 0.72 | 0.85 | 0.72 | 0.8 |
| 11 | 68441197 | rs6591348 | 0.141 | -1.461 | 0.847 | -1.278 | -0.075 | -0.111 | -0.111 | 4.459 | 0.2 | 0.44 | 0.1 | 0.02 | 0.08 | 0.08 |
| 11 | 68446424 | rs2187331 | -0.667 | -1.456 | 0.753 | -1.239 | -0.004 | 0.006 | 0.006 | 0 | 0.87 | 0.89 | 0.83 | 0.91 | 0.83 | 0.87 |
| 11 | 68450314 | rs2097042 | 0.223 | -1.389 | 0.783 | -1.221 | -0.031 | -0.081 | -0.081 | 0 | 0.68 | 0.54 | 0.76 | 0.89 | 0.75 | 0.81 |
| 11 | 68458470 | rs1042577 | 0.654 | -1.984 | 0.161 | -1.049 | 0.026 | 0.031 | 0.031 | 3.073 | 0.48 | 0.39 | 0.67 | 0.79 | 0.7 | 0.67 |
| 11 | 68471774 | rs7127598 | 0.639 | -1.971 | 0.21 | -1.053 | -0.039 | -0.084 | -0.084 | 0 | 0.2 | 0.33 | 0.11 | 0.03 | 0.11 | 0.09 |
| 11 | 68474153 | rs4396295 | 1.215 | -0.948 | 0.007 | -0.834 | -0.144 | 0.052 | 0.052 | 5.821 | 0.29 | 0.03 | 0.33 | 0.23 | 0.48 | 0.46 |
| 11 | 68478207 | rs10750832 | -0.815 | -1.166 | 0.224 | -1.24 | -0.015 | -0.04 | -0.04 | 0.817 | 0.62 | 0.56 | 0.73 | 0.87 | 0.72 | 0.76 |
| 11 | 68487513 | rs6591352 | -0.741 | -1.164 | 0.254 | -1.233 | -0.029 | -0.066 | -0.066 | 0 | 0.61 | 0.51 | 0.73 | 0.86 | 0.73 | 0.76 |
| 11 | 68504474 | rs4930244 | -0.379 | -0.382 | 0.122 | -0.781 | 0.043 | 0.067 | 0.067 | 3.085 | 0.48 | 0.36 | 0.65 | 0.79 | 0.67 | 0.67 |
| 11 | 68504907 | rs7103131 | -0.35 | -0.383 | 0.111 | -0.782 | 0.045 | 0.071 | 0.071 | 3.274 | 0.48 | 0.34 | 0.65 | 0.79 | 0.67 | 0.67 |
| 11 | 68505064 | rs7103254 | -0.445 | -0.358 | -0.086 | -0.769 | -0.003 | -0.013 | -0.013 | 0 | 0.49 | 0.45 | 0.66 | 0.79 | 0.67 | 0.67 |
| 11 | 68512543 | rs12365708 | -0.249 | 0.126 | -0.661 | -0.465 | -0.031 | 0.027 | 0.027 | 0.138 | 0.08 | 0.02 | 0.07 | 0.02 | 0.12 | 0.02 |
| 11 | 68524053 | rs2278907 | 0.066 | -0.162 | -0.46 | -0.601 | -0.006 | 0.027 | 0.027 | 1.92 | 0.4 | 0.32 | 0.58 | 0.78 | 0.56 | 0.65 |
| 11 | 68527600 | rs3019598 | -0.004 | -0.509 | -0.094 | -0.799 | 0 | -0.017 | -0.017 | 0 | 0.49 | 0.49 | 0.31 | 0.19 | 0.32 | 0.33 |
| 11 | 68528613 | rs897049 | -1.362 | -0.897 | 0.759 | -0.881 | -0.022 | -0.056 | -0.056 | 0.035 | 0.17 | 0.49 | 0.13 | 0.09 | 0.12 | 0.09 |
| 11 | 68530318 | rs7112615 | 0.077 | -0.994 | 0.705 | -0.932 | -0.018 | -0.022 | -0.022 | 0 | 0.89 | 0.79 | 0.93 | 0.91 | 0.95 | 0.91 |
| 11 | 68538939 | rs3019593 | 0.304 | -1.102 | 0.812 | -0.91 | -0.032 | -0.023 | -0.023 | 0 | 0.88 | 0.78 | 0.92 | 0.9 | 0.95 | 0.91 |
| 11 | 68539973 | rs2060980 | 1.453 | -1.091 | 1.171 | -1.079 | -0.026 | -0.084 | -0.084 | 0 | 0.78 | 0.52 | 0.84 | 0.88 | 0.83 | 0.88 |
| 11 | 68541540 | rs11228346 | 0.166 | -1.072 | 0.944 | -1.072 | -0.039 | 0.029 | 0.029 | 0 | 0.08 | 0.02 | 0.07 | 0.02 | 0.12 | 0.02 |
| 11 | 68544438 | rs897047 | 1.424 | -1.083 | 1.051 | -1.076 | -0.019 | -0.062 | -0.062 | 0.092 | 0.78 | 0.6 | 0.85 | 0.89 | 0.83 | 0.89 |
| 11 | 68544985 | rs10791997 | 1.109 | -1.023 | 0.654 | -1.046 | -0.003 | -0.018 | -0.018 | 0 | 0.8 | 0.73 | 0.85 | 0.89 | 0.83 | 0.88 |
| 11 | 68549546 | rs2305508 | 1.145 | -0.844 | 0.032 | -0.899 | -0.163 | 0.043 | 0.043 | 5.968 | 0.28 | 0.03 | 0.33 | 0.21 | 0.47 | 0.45 |
| 11 | 68550593 | rs11228351 | 0.192 | -0.795 | 0.364 | -0.903 | -0.037 | 0.03 | 0.03 | 0 | 0.08 | 0.02 | 0.07 | 0.02 | 0.12 | 0.02 |
| 11 | 68554503 | rs747840 | 0.031 | -0.897 | 0.615 | -0.801 | -0.032 | -0.005 | -0.005 | 0 | 0.91 | 0.76 | 0.91 | 0.9 | 0.98 | 0.93 |
| 11 | 68565904 | rs4930248 | 0.308 | -1.159 | 0.211 | -0.833 | 0.005 | 0.078 | 0.078 | 6.688 | 0.42 | 0.19 | 0.4 | 0.75 | 0.59 | 0.7 |
| 11 | 68571777 | rs2924674 | 0.483 | -1.066 | 0.954 | -0.47 | -0.038 | -0.037 | -0.037 | 0 | 0.9 | 0.73 | 0.87 | 0.94 | 0.98 | 0.93 |
| 11 | 68583671 | rs2924689 | -1.245 | -1.118 | 1.193 | -0.464 | -0.046 | -0.034 | -0.034 | 0 | 0.12 | 0.46 | 0.14 | 0.08 | 0.02 | 0.12 |
| 11 | 68583684 | rs11228364 | -0.35 | -1.205 | 0.812 | -0.612 | -0.002 | 0.016 | 0.016 | 0 | 0.11 | 0.08 | 0.08 | 0.03 | 0.13 | 0.02 |
| 11 | 68589257 | rs3019613 | 1.39 | -1.292 | 0.908 | -0.62 | -0.05 | -0.085 | -0.085 | 0.026 | 0.77 | 0.5 | 0.79 | 0.88 | 0.84 | 0.86 |
| 11 | 68591872 | rs12361365 | -1.931 | -1.294 | 0.89 | -0.097 | -0.016 | -0.017 | -0.017 | 0 | 0.24 | 0.37 | 0.13 | 0.13 | 0.2 | 0.17 |
| 11 | 68597104 | rs10896371 | 1.482 | -1.87 | 0.986 | 0.72 | -0.045 | 0.108 | 0.108 | 1.815 | 0.79 | 0.36 | 0.86 | 0.45 | 0.88 | 0.72 |
| 11 | 68598054 | rs7938117 | 0.63 | -1.602 | -0.086 | -0.214 | -0.005 | 0.113 | 0.113 | 0.322 | 0.31 | 0.13 | 0.18 | 0.14 | 0.32 | 0.46 |
| 11 | 68598684 | rs1017640 | -0.249 | -1.608 | 0.435 | -0.127 | -0.011 | 0.397 | 0.397 | 0 | 0.88 | 0.54 | 0.89 | 0.48 | 0.92 | 0.87 |

| | | | | | | | | | | | | | | | | |
|---|---|---|---|---|---|---|---|---|---|---|---|---|---|---|---|---|
| 11 | 68603157 | rs613084 | 0.552 | -1.628 | 0.09 | -0.234 | -0.019 | 0.121 | 0.121 | 0.33 | 0.32 | 0.07 | 0.19 | 0.15 | 0.33 | 0.46 |
| 11 | 68605690 | rs3019594 | 0.069 | -1.565 | 0.541 | -0.047 | -0.006 | 0.401 | 0.401 | 1.779 | 0.88 | 0.55 | 0.89 | 0.48 | 0.92 | 0.87 |
| 11 | 68610143 | rs7131462 | 0.795 | -1.573 | -0.134 | -0.319 | -0.05 | -0.003 | -0.003 | 0.003 | 0.43 | 0.09 | 0.58 | 0.25 | 0.53 | 0.23 |
| 11 | 68615117 | rs679596 | 0.788 | -1.522 | -0.377 | -0.132 | -0.075 | -0.012 | -0.012 | 1.013 | 0.43 | 0.07 | 0.62 | 0.25 | 0.53 | 0.23 |
| 11 | 68620223 | rs624420 | 0.712 | -1.602 | -0.128 | 0.211 | -0.019 | 0.043 | 0.043 | 1.382 | 0.77 | 0.33 | 0.79 | 0.46 | 0.84 | 0.75 |
| 11 | 68627535 | rs619727 | 0.424 | -1.599 | -0.057 | 0.113 | 0.013 | 0.029 | 0.029 | 0.088 | 0.31 | 0.21 | 0.16 | 0.21 | 0.3 | 0.51 |
| 11 | 68629929 | rs604524 | 0.369 | -1.539 | -0.012 | 0.109 | 0.009 | 0.023 | 0.023 | 0.012 | 0.31 | 0.24 | 0.16 | 0.21 | 0.3 | 0.53 |
| 11 | 68659488 | rs660614 | 0.411 | -1.532 | -0.015 | 0.104 | 0.016 | 0.047 | 0.047 | 0.168 | 0.31 | 0.17 | 0.16 | 0.22 | 0.3 | 0.51 |
| 11 | 68667198 | rs636049 | 0.217 | -1.558 | 0.366 | 0.097 | -0.005 | -0.01 | -0.01 | 0 | 0.33 | 0.34 | 0.17 | 0.22 | 0.3 | 0.52 |
| 11 | 68670712 | rs603076 | 0.041 | -1.004 | -0.01 | -0.34 | -0.001 | -0.001 | -0.001 | 0.081 | 0.6 | 0.56 | 0.64 | 0.45 | 0.58 | 0.69 |
| 11 | 68671104 | rs629426 | 0.854 | -0.993 | -0.033 | -0.354 | 0.017 | 0.051 | 0.051 | 0.075 | 0.31 | 0.17 | 0.16 | 0.22 | 0.3 | 0.51 |
| 11 | 68675930 | rs10896378 | 1.321 | -0.874 | -0.272 | -0.364 | -0.1 | 0.116 | 0.116 | 0.169 | 0.2 | 0.03 | 0.18 | 0.01 | 0.27 | 0.04 |
| 11 | 68678962 | rs560096 | 0.024 | -0.941 | -0.123 | -0.257 | -0.035 | 0.139 | 0.139 | 0.014 | 0.8 | 0.6 | 0.82 | 0.46 | 0.85 | 0.74 |
| 11 | 68680577 | rs10896379 | 1.629 | -0.98 | -0.914 | -0.599 | -0.03 | -0.004 | -0.004 | 0 | 0.14 | 0.02 | 0.08 | 0.1 | 0.17 | 0.36 |
| 11 | 68682402 | rs10896380 | 0.883 | -0.922 | -0.69 | -0.519 | -0.04 | 0.069 | 0.069 | 0 | 0.22 | 0.11 | 0.19 | 0.01 | 0.27 | 0.04 |
| 11 | 68684105 | rs12419421 | 0.898 | -0.921 | -0.706 | -0.518 | -0.044 | 0.077 | 0.077 | 0 | 0.22 | 0.09 | 0.19 | 0.02 | 0.27 | 0.04 |
| 11 | 68692141 | rs685320 | -0.039 | -1.013 | -0.616 | -0.587 | 0.005 | 0.043 | 0.043 | 0.687 | 0.52 | 0.34 | 0.35 | 0.23 | 0.57 | 0.53 |
| 11 | 68698476 | rs4930627 | 0.876 | -1.013 | -0.607 | -0.593 | -0.013 | 0.012 | 0.012 | 0 | 0.22 | 0.15 | 0.19 | 0.02 | 0.27 | 0.05 |
| 11 | 68701118 | rs2282504 | 2.011 | -1.116 | -0.752 | -0.562 | -0.099 | -0.1 | -0.1 | 0 | 0.21 | 0.01 | 0.44 | 0.22 | 0.27 | 0.17 |
| 11 | 68703959 | rs622082 | 0.9 | -1.119 | -0.726 | -0.592 | 0.003 | 0.045 | 0.045 | 0.044 | 0.3 | 0.1 | 0.16 | 0.21 | 0.3 | 0.48 |
| 11 | 68704028 | rs2236654 | 1.131 | -1.122 | -0.721 | -0.599 | -0.011 | 0.009 | 0.009 | 0 | 0.22 | 0.15 | 0.19 | 0.02 | 0.26 | 0.05 |
| 11 | 68704264 | rs546382 | 0.887 | -1.103 | -0.788 | -0.571 | 0.003 | 0.045 | 0.045 | 0.044 | 0.3 | 0.1 | 0.16 | 0.21 | 0.3 | 0.48 |
| 11 | 68705674 | rs17612126 | 1.953 | -1.028 | -0.763 | -0.517 | -0.099 | -0.104 | -0.104 | 0 | 0.21 | 0.02 | 0.44 | 0.22 | 0.27 | 0.17 |
| 11 | 68715544 | rs4930631 | 0.848 | -0.915 | -0.45 | -0.784 | 0.012 | -0.061 | -0.061 | 0 | 0.23 | 0.24 | 0.2 | 0.02 | 0.27 | 0.05 |
| 11 | 68719598 | rs6421653 | -0.212 | -1.035 | -0.496 | -0.787 | 0.002 | 0.025 | 0.025 | 1.675 | 0.55 | 0.34 | 0.36 | 0.24 | 0.58 | 0.53 |
| 11 | 68728359 | rs12800381 | 0.126 | -0.789 | -0.22 | -0.648 | 0.005 | -0.069 | -0.069 | 0 | 0.26 | 0.3 | 0.21 | 0.02 | 0.28 | 0.09 |
| 11 | 68735478 | rs11228427 | 0.76 | -0.837 | -0.159 | -0.625 | -0.018 | 0.032 | 0.032 | 0 | 0.23 | 0.17 | 0.19 | 0.01 | 0.26 | 0.05 |
| 11 | 68740952 | rs752296 | 0.309 | -0.854 | -0.086 | -0.632 | 0.006 | -0.077 | -0.077 | 0.041 | 0.25 | 0.32 | 0.2 | 0.02 | 0.26 | 0.08 |
| 11 | 68741305 | rs2017403 | -0.545 | -1.012 | -0.218 | -0.717 | 0 | -0.057 | -0.057 | 0 | 0.37 | 0.38 | 0.27 | 0.09 | 0.37 | 0.19 |
| 11 | 68746328 | rs10896389 | -0.017 | -1.085 | 0.326 | -0.357 | 0.015 | 0.028 | 0.028 | 0 | 0.54 | 0.59 | 0.71 | 0.24 | 0.55 | 0.29 |
| 11 | 68747485 | rs1939734 | 0.706 | -1.079 | 0.251 | 0.304 | 0.009 | -0.025 | -0.025 | 0.423 | 0.1 | 0.08 | 0.08 | 0.27 | 0.09 | 0.1 |
| 11 | 68749540 | rs10896390 | 0.062 | -1.141 | 0.331 | 0.109 | 0.026 | 0.056 | 0.056 | 0.005 | 0.55 | 0.69 | 0.72 | 0.24 | 0.55 | 0.29 |
| 11 | 68755043 | rs2846194 | 0.362 | -1.172 | 0.227 | 0.23 | -0.001 | 0.017 | 0.017 | 0.184 | 0.11 | 0.12 | 0.08 | 0.3 | 0.11 | 0.28 |
| 11 | 68757913 | rs2846207 | -0.485 | -1.108 | 0.287 | 0.236 | 0.029 | 0.058 | 0.058 | 0.002 | 0.45 | 0.29 | 0.28 | 0.76 | 0.45 | 0.71 |
| 11 | 68759825 | rs12806229 | 0.944 | -1.207 | 0.413 | 0.127 | -0.006 | -0.018 | -0.018 | 0 | 0.24 | 0.18 | 0.19 | 0.01 | 0.27 | 0.05 |
| 11 | 68761422 | rs2602814 | -0.924 | -1.259 | 0.558 | 0.178 | -0.001 | 0.001 | 0.001 | 0.017 | 0.47 | 0.52 | 0.37 | 0.57 | 0.47 | 0.43 |
| 11 | 68762280 | rs12420399 | 0.942 | -1.101 | 0.345 | 0.275 | -0.006 | -0.015 | -0.015 | 0 | 0.23 | 0.17 | 0.19 | 0.01 | 0.26 | 0.05 |
| 11 | 68765895 | rs1249582 | 0.762 | -1.231 | -0.054 | 0.579 | 0.004 | -0.057 | -0.057 | 0 | 0.31 | 0.19 | 0.19 | 0.51 | 0.33 | 0.54 |
| 11 | 68770033 | rs3862397 | -0.688 | -1 | -0.571 | 0.674 | 0.002 | -0.002 | -0.002 | 0 | 0.43 | 0.44 | 0.33 | 0.54 | 0.41 | 0.46 |
| 11 | 68771755 | rs12803642 | 0.346 | -0.746 | -1.539 | 0.376 | 0.017 | -0.017 | -0.017 | 0 | 0.15 | 0.09 | 0.09 | 0.25 | 0.12 | 0.17 |
| 11 | 68781070 | rs11602122 | 1.469 | -0.719 | -1.786 | 0.342 | -0.002 | 0.001 | 0.001 | 0 | 0.24 | 0.16 | 0.12 | 0.23 | 0.26 | 0.28 |
| 11 | 68782223 | rs2846196 | 0.698 | -0.046 | -1.97 | -0.144 | -0.063 | 0.057 | 0.057 | 2.891 | 0.26 | 0.1 | 0.51 | 0.11 | 0.31 | 0.22 |
| 11 | 68785659 | rs12790923 | -0.135 | 0.129 | -1.757 | -0.109 | 0.015 | -0.014 | -0.014 | 0.435 | 0.15 | 0.1 | 0.1 | 0.25 | 0.12 | 0.19 |
| 11 | 68793206 | rs11228444 | -0.317 | 0.246 | -1.865 | -0.075 | -0.041 | 0.043 | 0.043 | 0 | 0.39 | 0.2 | 0.6 | 0.33 | 0.43 | 0.39 |
| 11 | 68798102 | rs11605908 | 0.417 | 0.455 | -2.162 | -0.013 | -0.045 | -0.003 | -0.003 | 0 | 0.16 | 0.01 | 0.11 | 0.14 | 0.2 | 0.13 |
| 11 | 68798591 | rs12806559 | 0.692 | 0.498 | -2.25 | -0.378 | -0.005 | 0.006 | 0.006 | 1.609 | 0.18 | 0.11 | 0.16 | 0.01 | 0.19 | 0.05 |
| 11 | 68798901 | rs17399246 | 0.32 | 0.72 | -2.405 | -0.105 | -0.056 | 0.018 | 0.018 | 0 | 0.19 | 0.01 | 0.12 | 0.16 | 0.24 | 0.17 |
| 11 | 68804451 | rs11228450 | 0.363 | 0.737 | -1.95 | 0.562 | -0.061 | -0.032 | -0.032 | 0 | 0.21 | 0.09 | 0.44 | 0.2 | 0.24 | 0.28 |
| 11 | 68804806 | rs10896396 | 0.319 | 0.495 | -1.706 | 0.582 | 0.015 | -0.062 | -0.062 | 4.486 | 0.33 | 0.13 | 0.16 | 0.47 | 0.3 | 0.48 |
| 11 | 68807112 | rs10896397 | -0.523 | 0.655 | -0.801 | 1.163 | 0.005 | 0.03 | 0.03 | 0.783 | 0.25 | 0.32 | 0.5 | 0.37 | 0.24 | 0.26 |
| 11 | 68810837 | rs2034691 | 0.756 | 0.038 | -0.528 | 1.474 | -0.018 | 0.049 | 0.049 | 0 | 0.88 | 0.78 | 0.95 | 0.75 | 0.94 | 0.93 |
| 11 | 68818936 | rs4930643 | -0.141 | -0.506 | -1.309 | 1.221 | 0.007 | 0.029 | 0.029 | 0 | 0.34 | 0.12 | 0.19 | 0.21 | 0.3 | 0.53 |
| 11 | 68825037 | rs3750973 | 0.125 | -1.048 | 0.651 | 2.168 | 0.017 | 0.485 | 0.485 | 2.808 | 0.95 | 0.68 | 0.61 | 0.53 | 0.99 | 0.92 |
| 11 | 68826155 | rs3019776 | -0.938 | -1.511 | 0.541 | 2.452 | 0.021 | -0.036 | -0.036 | 6.288 | 0.79 | 0.83 | 0.7 | 0.54 | 0.9 | 0.59 |
| 11 | 68828929 | rs896978 | 0.402 | -1.556 | 0.674 | 2.24 | 0.01 | 0.017 | 0.017 | 0 | 0.32 | 0.22 | 0.18 | 0.21 | 0.28 | 0.5 |
| 11 | 68829638 | rs11228469 | 0.37 | -1.417 | 0.244 | 1.529 | -0.018 | 0.063 | 0.063 | 0.244 | 0.43 | 0.23 | 0.32 | 0.29 | 0.51 | 0.22 |
| 11 | 68838028 | rs3829236 | -0.331 | -1.217 | 0.266 | 1.487 | -0.006 | 0.06 | 0.06 | 0 | 0.54 | 0.36 | 0.28 | 0.22 | 0.5 | 0.72 |
| 11 | 68840160 | rs3750965 | 0.89 | -1.234 | 0.23 | 1.493 | 0.017 | 0.03 | 0.03 | 0 | 0.35 | 0.23 | 0.19 | 0.21 | 0.3 | 0.51 |
| 11 | 68851414 | rs2376558 | -0.329 | -1.289 | 0.848 | 1.826 | 0.011 | 0.474 | 0.474 | 2.72 | 0.95 | 0.72 | 0.61 | 0.5 | 0.99 | 0.92 |
| 11 | 68851466 | rs896973 | 0.466 | -1.302 | 0.842 | 1.765 | 0.001 | -0.004 | -0.004 | 0 | 0.41 | 0.36 | 0.33 | 0.29 | 0.48 | 0.2 |
| 11 | 68855363 | rs3829241 | 1.169 | -1.426 | 0.574 | 1.301 | -0.11 | -0.027 | -0.027 | 0.084 | 0.17 | 0.04 | 0.22 | 0.21 | 0.38 | 0.09 |
| 11 | 68855483 | rs1551306 | 0.413 | -1.469 | 0.589 | 1.27 | 0 | 0.079 | 0.079 | 0 | 0.53 | 0.32 | 0.28 | 0.22 | 0.49 | 0.71 |
| 11 | 68855595 | rs1060435 | 1.079 | -1.39 | 0.617 | 1.355 | -0.113 | -0.028 | -0.028 | 0 | 0.17 | 0.04 | 0.22 | 0.21 | 0.39 | 0.09 |
| 11 | 68855954 | rs4930265 | 1.197 | -1.287 | 0.838 | 1.497 | 0.018 | 0.029 | 0.029 | 0 | 0.35 | 0.23 | 0.19 | 0.21 | 0.3 | 0.52 |
| 11 | 68856916 | rs1005858 | 0.119 | -1.242 | 0.858 | 1.547 | -0.074 | 0.063 | 0.063 | 1.206 | 0.4 | 0.1 | 0.31 | 0.29 | 0.49 | 0.2 |
| 11 | 68857038 | rs1551310 | -0.374 | -1.215 | 0.877 | 1.578 | 0.011 | 0.475 | 0.475 | 2.886 | 0.95 | 0.71 | 0.61 | 0.5 | 0.99 | 0.9 |
| 11 | 68859377 | rs7126711 | -0.378 | -1.214 | 0.913 | 1.702 | 0.009 | 0.323 | 0.323 | 3.257 | 0.95 | 0.72 | 0.82 | 0.73 | 0.99 | 0.91 |
| 11 | 68861329 | rs10736672 | 0.11 | -1.141 | 0.895 | 1.454 | -0.002 | 0.002 | 0.002 | 0.566 | 0.41 | 0.33 | 0.33 | 0.29 | 0.49 | 0.2 |
| 11 | 68869034 | rs921675 | -0.138 | -0.98 | 0.547 | 1.639 | -0.022 | 0.092 | 0.092 | 0.208 | 0.76 | 0.43 | 0.51 | 0.49 | 0.79 | 0.71 |

| | | | | | | | | | | | | | | | | |
|---|---|---|---|---|---|---|---|---|---|---|---|---|---|---|---|---|
| 11 | 68870298 | rs11228498 | 0.041 | -1.145 | -0.042 | 1.469 | -0.031 | 0.047 | 0.047 | 0.72 | 0.42 | 0.17 | 0.33 | 0.34 | 0.5 | 0.3 |
| 11 | 68871001 | rs2924533 | 0.087 | -0.285 | 0.326 | 2.172 | -0.001 | 0.006 | 0.006 | 0.207 | 0.76 | 0.73 | 0.52 | 0.42 | 0.76 | 0.71 |
| 11 | 68876587 | rs1551304 | -0.241 | 0.518 | 0.326 | 2.221 | 0.005 | 0.029 | 0.029 | 0 | 0.23 | 0.29 | 0.5 | 0.6 | 0.26 | 0.15 |
| 11 | 68877162 | rs10896430 | -0.074 | 0.408 | 0.124 | 2.234 | -0.002 | -0.01 | -0.01 | 0 | 0.22 | 0.2 | 0.28 | 0.32 | 0.23 | 0.15 |
| 11 | 68881649 | rs3019779 | -0.402 | 0.467 | -0.016 | 2.243 | -0.011 | -0.037 | -0.037 | 0 | 0.8 | 0.87 | 0.75 | 0.31 | 0.78 | 0.8 |
| 11 | 68884755 | rs3892895 | -1.195 | 0.341 | 0.153 | 1.69 | -0.008 | -0.005 | -0.005 | 0.105 | 0.58 | 0.51 | 0.49 | 0.32 | 0.67 | 0.33 |
| 11 | 68886179 | rs930788 | -0.642 | 0.281 | 0.265 | 1.808 | 0.001 | -0.004 | -0.004 | 0 | 0.45 | 0.42 | 0.35 | 0.4 | 0.48 | 0.3 |
| 11 | 68889182 | rs11228506 | -0.683 | 0.507 | 0.319 | 1.811 | 0.004 | -0.005 | -0.005 | 0 | 0.43 | 0.49 | 0.35 | 0.41 | 0.45 | 0.29 |
| 11 | 68894753 | rs12284087 | -0.402 | 0.799 | 0.068 | 1.518 | 0.001 | -0.012 | -0.012 | 0.446 | 0.73 | 0.69 | 0.73 | 0.82 | 0.73 | 0.92 |
| 11 | 68894991 | rs16761 | 1.56 | 0.874 | -0.723 | 1.713 | -0.135 | 0.047 | 0.047 | 1.695 | 0.35 | 0.03 | 0.44 | 0.16 | 0.42 | 0.51 |
| 11 | 68897275 | rs11603027 | 1.196 | 0.503 | -0.637 | 1.708 | -0.087 | 0.043 | 0.043 | 1.56 | 0.43 | 0.05 | 0.49 | 0.25 | 0.56 | 0.63 |
| 11 | 68899814 | rs2123764 | 0.51 | 0.045 | 0.352 | 2.082 | -0.014 | 0.077 | 0.077 | 0.981 | 0.17 | 0.33 | 0.21 | 0.35 | 0.13 | 0.13 |
| 11 | 68899837 | rs11228512 | 1.758 | 0.145 | -0.402 | 1.577 | 0.002 | 0.005 | 0.005 | 5.333 | 0.11 | 0.03 | 0.06 | 0.08 | 0.09 | 0.01 |
| 11 | 68903833 | rs11228515 | 0.53 | 0.168 | 0.054 | 1.547 | 0.008 | -0.003 | -0.003 | 0 | 0.6 | 0.35 | 0.65 | 0.55 | 0.7 | 0.64 |
| 11 | 68904240 | rs11228517 | -0.083 | 0.499 | 0.488 | 1.452 | 0.057 | 0.061 | 0.061 | 0.746 | 0.29 | 0.38 | 0.44 | 0.5 | 0.46 | 0.3 |
| 11 | 68905412 | rs11228518 | 1.623 | 0.121 | 0.546 | 1.545 | -0.01 | 0.031 | 0.031 | 3.006 | 0.14 | 0.21 | 0.08 | 0.3 | 0.09 | 0.03 |
| 11 | 68905440 | rs2123765 | 0.111 | 0.032 | 0.421 | 1.471 | 0.027 | -0.021 | -0.021 | 0 | 0.28 | 0.22 | 0.12 | 0.39 | 0.14 | 0.17 |
| 11 | 68906570 | rs10896438 | 0.821 | 0.061 | 0.746 | 1.413 | -0.002 | -0.001 | -0.001 | 7.96 | 0.24 | 0.24 | 0.27 | 0.33 | 0.32 | 0.15 |
| 11 | 68909579 | rs17149775 | 0.609 | 0.085 | 0.813 | 1.429 | 0.054 | 0.108 | 0.108 | 0 | 0.07 | 0.12 | 0.2 | 0.24 | 0.16 | 0.19 |
| 11 | 68910579 | rs2924540 | -0.547 | 0.068 | 0.276 | 1.326 | -0.008 | 0.01 | 0.01 | 0 | 0.39 | 0.27 | 0.31 | 0.41 | 0.37 | 0.29 |
| 11 | 68914330 | rs7935486 | 1.648 | -0.109 | 0.127 | 1.241 | -0.011 | -0.011 | -0.011 | 0 | 0.13 | 0.22 | 0.08 | 0.08 | 0.09 | 0.02 |
| 11 | 68918985 | rs11228530 | 1.16 | 0.005 | -0.191 | 1.216 | 0.013 | 0.009 | 0.009 | 0 | 0.15 | 0.01 | 0.04 | 0.09 | 0.05 | 0.14 |
| 11 | 68923597 | rs3019751 | 0.767 | -0.042 | 0.006 | 1.075 | -0.001 | -0.001 | -0.001 | 0 | 0.52 | 0.56 | 0.55 | 0.37 | 0.49 | 0.53 |
| 11 | 68932899 | rs7940364 | 0.018 | 0.312 | -0.418 | 0.944 | -0.005 | -0.003 | -0.003 | 0 | 0.57 | 0.52 | 0.26 | 0.44 | 0.4 | 0.68 |
| 11 | 68935552 | rs11228540 | 0.025 | 0.297 | -0.421 | 0.812 | 0 | -0.003 | -0.003 | 0 | 0.57 | 0.44 | 0.26 | 0.22 | 0.4 | 0.67 |
| 11 | 68943547 | rs12283434 | 0.106 | -0.196 | -0.619 | 0.176 | -0.041 | -0.089 | -0.089 | 0 | 0.22 | 0.36 | 0.07 | 0.04 | 0.09 | 0.07 |
| 11 | 68946858 | rs12417953 | -0.331 | -0.029 | -0.986 | 0.319 | 0 | 0 | 0 | 0 | 0.44 | 0.43 | 0.51 | 0.38 | 0.56 | 0.33 |
| 11 | 68953054 | rs7128814 | -1.496 | -0.008 | -2.236 | 0.476 | -0.054 | -0.034 | -0.034 | 0 | 0.63 | 0.9 | 0.56 | 0.64 | 0.58 | 0.47 |
| 11 | 68959689 | rs4072598 | -0.582 | 0.182 | -2.039 | 0.656 | -0.07 | 0.008 | 0.008 | 1.84 | 0.46 | 0.89 | 0.51 | 0.6 | 0.5 | 0.32 |
| 11 | 68960391 | rs12223972 | 0.343 | 0.866 | -1.62 | 0.844 | -0.065 | -0.003 | -0.003 | 0 | 0.22 | 0.12 | 0.17 | 0.24 | 0.3 | 0.12 |
| 11 | 68964278 | rs11228554 | -0.262 | 0.076 | -1.69 | 0.692 | 0.105 | 0.019 | 0.019 | 0.541 | 0.46 | 0.78 | 0.39 | 0.39 | 0.31 | 0.36 |
| 11 | 68967641 | rs10896445 | -0.786 | 0.614 | -1.494 | 0.736 | -0.038 | -0.038 | -0.038 | 0 | 0.68 | 0.85 | 0.54 | 0.63 | 0.62 | 0.49 |
| 11 | 68973970 | rs4255548 | -0.721 | 0.172 | -0.694 | 0.638 | 0.003 | 0.008 | 0.008 | 0 | 0.69 | 0.71 | 0.49 | 0.49 | 0.61 | 0.52 |
| 11 | 68976996 | rs11228563 | 0.718 | -0.195 | -0.753 | -0.358 | 0.013 | 0.033 | 0.033 | 1.809 | 0.33 | 0.24 | 0.22 | 0.02 | 0.29 | 0.1 |
| 11 | 68977501 | rs12281017 | 0.443 | -0.205 | -0.987 | -0.403 | -0.013 | 0.1 | 0.1 | 0 | 0.25 | 0.08 | 0.11 | 0.05 | 0.21 | 0.19 |
| 11 | 68978049 | rs4930667 | -0.222 | -0.236 | -0.572 | -0.376 | 0.004 | 0.116 | 0.116 | 0 | 0.08 | 0.13 | 0.15 | 0.4 | 0.11 | 0.23 |
| 11 | 68978580 | rs11228565 | 0.451 | -0.225 | -0.653 | -0.455 | -0.013 | 0.101 | 0.101 | 0 | 0.25 | 0.09 | 0.11 | 0.05 | 0.21 | 0.19 |
| 11 | 68978648 | rs4357697 | 0.46 | -0.165 | -0.569 | -0.003 | -0.003 | 0.023 | 0.023 | 0 | 0.06 | 0.05 | 0.13 | 0.24 | 0.1 | 0.07 |
| 11 | 68980788 | rs11825796 | 0.69 | -0.27 | -0.363 | -0.235 | -0.017 | -0.097 | -0.097 | 0.502 | 0.34 | 0.57 | 0.21 | 0.03 | 0.26 | 0.12 |
| 11 | 68985583 | rs7929962 | -0.28 | -0.139 | -0.323 | -0.098 | -0.022 | 0.049 | 0.049 | 0.402 | 0.61 | 0.72 | 0.35 | 0.07 | 0.51 | 0.29 |
| 11 | 68994497 | rs7931342 | -0.66 | -0.165 | -0.313 | -0.027 | -0.066 | 0.07 | 0.07 | 0.543 | 0.63 | 0.84 | 0.38 | 0.23 | 0.51 | 0.45 |
| 11 | 68994667 | rs10896449 | -0.24 | -0.178 | -0.362 | -0.128 | -0.022 | 0.051 | 0.051 | 0.534 | 0.6 | 0.72 | 0.35 | 0.07 | 0.51 | 0.29 |
| 11 | 68995958 | rs7130881 | 0.744 | -0.214 | -0.399 | -0.221 | 0 | -0.026 | -0.026 | 0 | 0.17 | 0.18 | 0.09 | 0.02 | 0.18 | 0.08 |
| 11 | 68998788 | rs7950547 | -1.131 | -0.164 | -0.722 | 0.122 | -0.059 | -0.092 | -0.092 | 0 | 0.7 | 0.86 | 0.51 | 0.49 | 0.61 | 0.57 |
| 11 | 69003423 | rs7118074 | -1.23 | 0.093 | -0.665 | 0.719 | -0.031 | 0.012 | 0.012 | 0 | 0.94 | 1 | 0.97 | 0.65 | 0.92 | 0.74 |
| 11 | 69015728 | rs12282656 | -0.28 | -0.504 | 0.08 | 0.348 | 0.02 | 0.005 | 0.005 | 0 | 0.25 | 0.43 | 0.25 | 0.24 | 0.26 | 0.18 |
| 11 | 69016893 | rs10896451 | -0.991 | -0.865 | -0.074 | 0.026 | -0.008 | 0.006 | 0.006 | 0 | 0.44 | 0.79 | 0.41 | 0.44 | 0.4 | 0.36 |
| 11 | 69018498 | rs10896452 | -0.049 | -0.864 | 0.069 | 0.026 | 0.006 | 0.072 | 0.072 | 0 | 0.05 | 0.13 | 0.06 | 0.17 | 0.04 | 0.15 |
| 11 | 69020684 | rs7118204 | 1.616 | -0.854 | -0.033 | -0.008 | -0.005 | -0.037 | -0.037 | 0 | 0.15 | 0.24 | 0.11 | 0.03 | 0.1 | 0.03 |
| 11 | 69025553 | rs12792211 | 0.448 | -0.871 | 0.092 | -0.012 | -0.005 | -0.003 | -0.003 | 0.109 | 0.22 | 0.18 | 0.23 | 0.24 | 0.26 | 0.18 |
| 11 | 69025582 | rs7122303 | 0.235 | -0.917 | 0.142 | -0.014 | -0.019 | -0.018 | -0.018 | 0 | 0.22 | 0.61 | 0.18 | 0.21 | 0.14 | 0.17 |
| 11 | 69025799 | rs3884627 | -1.855 | -0.841 | -0.402 | -0.011 | -0.091 | -0.141 | -0.141 | 2.223 | 0.75 | 0.98 | 0.79 | 0.61 | 0.67 | 0.58 |
| 11 | 69026672 | rs12799244 | 0.557 | -0.728 | 0.076 | -0.24 | 0.001 | 0.002 | 0.002 | 0 | 0.18 | 0.17 | 0.2 | 0.24 | 0.2 | 0.17 |
| 11 | 69027770 | rs7940107 | 0.878 | -0.712 | -0.035 | -0.332 | -0.003 | -0.126 | -0.126 | 1.057 | 0.2 | 0.38 | 0.14 | 0.02 | 0.16 | 0.04 |
| 11 | 69029210 | rs7122786 | 0.878 | -0.712 | -0.035 | -0.332 | -0.003 | -0.126 | -0.126 | 0.05 | 0.2 | 0.38 | 0.14 | 0.02 | 0.16 | 0.04 |
| 11 | 69037823 | rs4638315 | -0.976 | -1.007 | 0.376 | 0.7 | 0.003 | 0.04 | 0.04 | 0 | 0.26 | 0.38 | 0.27 | 0.4 | 0.25 | 0.32 |
| 11 | 69043896 | rs11228605 | -0.921 | -1.32 | 0.943 | 1.222 | -0.048 | 0.075 | 0.075 | 0.239 | 0.15 | 0.57 | 0.09 | 0.24 | 0.05 | 0.27 |
| 11 | 69046093 | rs11826335 | -2.166 | -1.206 | 1.038 | 1.266 | 0.065 | 0.272 | 0.272 | 3.776 | 0.06 | 0.55 | 0.07 | 0.19 | 0.01 | 0.15 |
| 11 | 69050867 | rs11605162 | -1.759 | -1.24 | 0.957 | 1.187 | 0.052 | 0.195 | 0.195 | 1.681 | 0.08 | 0.34 | 0.06 | 0.24 | 0.03 | 0.22 |
| 11 | 69051076 | rs3924679 | -0.055 | -1.56 | -0.274 | 1.273 | 0.029 | -0.049 | -0.049 | 0.007 | 0.18 | 0.07 | 0.09 | 0.24 | 0.13 | 0.17 |
| 11 | 69052682 | rs4930673 | -0.339 | -1.568 | -0.296 | 1.25 | 0.021 | -0.047 | -0.047 | 0 | 0.2 | 0.07 | 0.11 | 0.24 | 0.15 | 0.18 |
| 11 | 69053932 | rs11602680 | -2.01 | -1.836 | -0.383 | 1.281 | 0.1 | -0.094 | -0.094 | 0 | 0.38 | 0.88 | 0.22 | 0.52 | 0.2 | 0.5 |
| 11 | 69056260 | rs10047435 | -1.546 | -1.781 | -0.543 | 1.219 | 0.053 | -0.011 | -0.011 | 0.063 | 0.47 | 0.93 | 0.28 | 0.44 | 0.33 | 0.57 |
| 11 | 69058291 | rs7118561 | -1.739 | -1.617 | 0.724 | 2.063 | -0.023 | 0.023 | 0.023 | 0.497 | 0.28 | 0.64 | 0.18 | 0.44 | 0.17 | 0.39 |
| 11 | 69060814 | rs7124369 | -1.863 | -1.616 | 0.518 | 2.063 | -0.018 | 0.015 | 0.015 | 0.376 | 0.29 | 0.68 | 0.18 | 0.47 | 0.17 | 0.39 |
| 11 | 69061310 | rs4396297 | -2.44 | -1.622 | 0.212 | 2.06 | 0.105 | -0.089 | -0.089 | 0.6 | 0.39 | 0.88 | 0.22 | 0.52 | 0.2 | 0.5 |
| 11 | 69061959 | rs11228608 | -2.142 | -0.818 | 0.665 | 2.534 | 0.022 | -0.022 | -0.022 | 0 | 0.29 | 0.74 | 0.18 | 0.44 | 0.17 | 0.38 |
| 11 | 69063136 | rs7117236 | -2.056 | -0.81 | 0.167 | 2.538 | 0.065 | -0.062 | -0.062 | 0.341 | 0.3 | 0.85 | 0.19 | 0.44 | 0.17 | 0.39 |
| 11 | 69063393 | rs7103126 | -2.069 | -0.811 | 0.167 | 2.538 | 0.066 | -0.062 | -0.062 | 0.036 | 0.3 | 0.85 | 0.19 | 0.46 | 0.17 | 0.39 |

| | | | | | | | | | | | | | | | | |
|---|---|---|---|---|---|---|---|---|---|---|---|---|---|---|---|---|
| 11 | 69063767 | rs12274095 | 0.166 | -0.756 | -0.122 | 2.323 | 0.019 | 0.062 | 0.062 | 0 | 0.12 | 0.03 | 0.04 | 0.01 | 0.07 | 0.07 |
| 11 | 69068745 | rs4619154 | -2.288 | -0.741 | -0.094 | 2.688 | 0.074 | -0.06 | -0.06 | 2.892 | 0.33 | 0.9 | 0.21 | 0.62 | 0.18 | 0.47 |
| 11 | 69070429 | rs7946208 | -1.845 | -0.432 | 0.011 | 2.813 | -0.049 | 0.163 | 0.163 | 1.899 | 0.17 | 0.65 | 0.11 | 0.35 | 0.08 | 0.19 |
| 11 | 69074090 | rs12224376 | -0.432 | 0.117 | -1.046 | 2.482 | -0.005 | -0.025 | -0.025 | 0 | 0.14 | 0.03 | 0.08 | 0.2 | 0.11 | 0.28 |
| 11 | 69074201 | rs11604530 | -0.46 | 0.166 | -1.021 | 2.397 | 0.011 | -0.035 | -0.035 | 0 | 0.07 | 0.02 | 0.05 | 0.2 | 0.06 | 0.09 |
| 11 | 69075538 | rs11605229 | -0.677 | 0.611 | -0.672 | 1.646 | -0.004 | -0.046 | -0.046 | 0 | 0.14 | 0.03 | 0.07 | 0.25 | 0.11 | 0.27 |
| 11 | 69078794 | rs12807388 | -1.011 | 1.597 | -0.015 | 1.748 | 0.039 | 0.045 | 0.045 | 0 | 0.06 | 0.14 | 0.06 | 0.11 | 0.12 | 0.04 |
| 11 | 69084550 | rs7481709 | -1.504 | 1.432 | 0.56 | 1.429 | 0.011 | 0.013 | 0.013 | 1.144 | 0.32 | 0.57 | 0.47 | 0.41 | 0.35 | 0.35 |
| 11 | 69087550 | rs12808959 | -1.651 | 1.318 | 0.784 | 1.411 | -0.006 | -0.01 | -0.01 | 0 | 0.33 | 0.7 | 0.49 | 0.42 | 0.35 | 0.38 |
| 11 | 69150850 | rs7108119 | -1.478 | 1.313 | 1.321 | 1.286 | 0.009 | 0.029 | 0.029 | 1.808 | 0.22 | 0.3 | 0.36 | 0.33 | 0.22 | 0.19 |
| 11 | 69154510 | rs12802463 | -0.835 | 1.354 | 1.253 | 1.139 | -0.009 | 0.004 | 0.004 | 0 | 0.06 | 0.08 | 0.04 | 0 | 0.11 | 0.02 |
| 11 | 69154716 | rs9971510 | -0.773 | 1.341 | 1.399 | 1.372 | -0.023 | 0.018 | 0.018 | 4.748 | 0.16 | 0.3 | 0.32 | 0.21 | 0.1 | 0.17 |
| 11 | 69160993 | rs4584599 | -1.419 | 1.351 | 2.068 | 1.241 | 0.035 | 0.229 | 0.229 | 2.968 | 0.07 | 0.33 | 0.29 | 0.3 | 0.06 | 0.14 |
| 11 | 69163161 | rs4357716 | -0.831 | 1.373 | 1.776 | 1.196 | -0.001 | 0.002 | 0.002 | 3.016 | 0.16 | 0.15 | 0.08 | 0.02 | 0.15 | 0.06 |
| 11 | 69173318 | rs11600497 | -0.771 | 1.547 | 1.758 | 1.114 | 0.054 | 0.168 | 0.168 | 0 | 0.1 | 0.25 | 0.34 | 0.31 | 0.12 | 0.14 |
| 11 | 69174170 | rs4326824 | -0.616 | 1.541 | 1.501 | 1.111 | 0.021 | 0.087 | 0.087 | 0 | 0.1 | 0.16 | 0.33 | 0.31 | 0.12 | 0.14 |
| 11 | 69177047 | rs12785524 | -1.878 | 1.62 | 0.886 | 1.071 | 0 | -0.001 | -0.001 | 0 | 0.37 | 0.59 | 0.53 | 0.44 | 0.42 | 0.38 |
| 11 | 69177138 | rs7938650 | -1.35 | 1.406 | 0.886 | 1.151 | 0.034 | 0.034 | 0.034 | 0.598 | 0.27 | 0.46 | 0.47 | 0.33 | 0.27 | 0.18 |
| 11 | 69203904 | rs6606661 | 1.174 | 1.306 | 0.922 | 0.669 | 0.074 | 0.009 | 0.009 | 0 | 0.94 | 0.72 | 0.95 | 0.99 | 0.95 | 0.99 |
| 11 | 69207345 | rs7109670 | 0.872 | 1.581 | 0.818 | 0.385 | 0.031 | -0.049 | -0.049 | 0.67 | 0.75 | 0.48 | 0.75 | 0.78 | 0.75 | 0.84 |
| 11 | 69207590 | rs11263641 | -0.52 | 1.582 | 0.801 | 0.386 | -0.003 | 0.003 | 0.003 | 0 | 0.19 | 0.24 | 0.2 | 0.21 | 0.2 | 0.16 |
| 11 | 69216024 | rs10796869 | 1.182 | 1.097 | 0.117 | 0.752 | -0.012 | 0.04 | 0.04 | 0 | 0.54 | 0.18 | 0.31 | 0.36 | 0.54 | 0.59 |
| 11 | 69218278 | rs4980777 | -0.167 | 0.977 | 0.429 | 0.745 | -0.005 | 0.05 | 0.05 | 0.038 | 0.2 | 0.25 | 0.43 | 0.41 | 0.21 | 0.25 |
| 11 | 69224506 | rs4581473 | 1.1 | 0.867 | -0.1 | 0.578 | -0.061 | -0.007 | -0.007 | 0 | 0.56 | 0.06 | 0.33 | 0.36 | 0.59 | 0.61 |
| 11 | 69230242 | rs6606675 | 0.359 | 1.292 | 0.636 | 1.133 | 0.017 | 0.026 | 0.026 | 0 | 0.74 | 0.71 | 0.51 | 0.52 | 0.68 | 0.7 |
| 11 | 69231796 | rs4980598 | 0.291 | 1.208 | 0.408 | 1.199 | -0.073 | -0.08 | -0.08 | 0 | 0.76 | 0.91 | 0.52 | 0.56 | 0.68 | 0.71 |
| 11 | 69239611 | rs4980786 | -1.452 | 0.943 | 0.589 | 1.038 | -0.019 | -0.033 | -0.033 | 0 | 0.41 | 0.78 | 0.57 | 0.52 | 0.4 | 0.35 |
| 11 | 69245812 | rs7927237 | -1.224 | 0.595 | 0.155 | -0.186 | 0.028 | 0.048 | 0.048 | 0 | 0.44 | 0.69 | 0.24 | 0.05 | 0.34 | 0.27 |
| 11 | 69248131 | rs11263434 | -1.561 | 0.086 | 1.135 | 1.147 | 0.014 | 0.559 | 0.559 | 0.715 | 0.06 | 0.34 | 0.23 | 0.5 | 0.01 | 0.16 |
| 11 | 69252000 | rs11263441 | -0.579 | -0.151 | 0.946 | 0.961 | 0.014 | 0 | 0 | 0 | 0.12 | 0.08 | 0.26 | 0.52 | 0.08 | 0.24 |
| 11 | 69253016 | rs10908183 | -1.002 | -1.452 | 0.685 | 0.288 | 0.035 | -0.001 | -0.001 | 0 | 0.37 | 0.7 | 0.38 | 0.61 | 0.21 | 0.41 |
| 11 | 69253255 | rs11263443 | -0.999 | -1.467 | 0.682 | 0.295 | 0.051 | -0.003 | -0.003 | 0.004 | 0.37 | 0.71 | 0.38 | 0.59 | 0.21 | 0.41 |
| 11 | 69255676 | rs7949453 | 0.668 | -1.321 | 0.982 | 0.277 | -0.029 | -0.029 | -0.029 | 0.675 | 0.21 | 0.28 | 0.1 | 0.09 | 0.13 | 0.17 |
| 11 | 69256899 | rs4980627 | 0.364 | -0.589 | 0.184 | 0.354 | -0.047 | -0.043 | -0.043 | 0 | 0.09 | 0.04 | 0.22 | 0.35 | 0.15 | 0.17 |
| 11 | 69258074 | rs7928827 | -0.643 | -0.641 | 0.027 | -0.077 | -0.022 | -0.002 | -0.002 | 0.265 | 0.43 | 0.24 | 0.37 | 0.44 | 0.41 | 0.39 |
| 11 | 69263864 | rs12806952 | -1.095 | -0.551 | -0.83 | -0.056 | -0.054 | 0.11 | 0.11 | 0.348 | 0.49 | 0.11 | 0.41 | 0.39 | 0.47 | 0.48 |
| 11 | 69265238 | rs4246959 | 0.243 | 0.641 | -0.446 | -0.243 | -0.021 | 0.135 | 0.135 | 0 | 0.46 | 0.27 | 0.3 | 0.14 | 0.49 | 0.37 |
| 11 | 69266710 | rs7130932 | -0.844 | 0.552 | -0.326 | -0.021 | 0.016 | -0.006 | -0.006 | 0.096 | 0.61 | 0.75 | 0.56 | 0.46 | 0.64 | 0.58 |
| 11 | 69269768 | rs7113550 | -1.099 | 0.504 | -0.444 | 0.533 | 0.015 | 0.053 | 0.053 | 0 | 0.63 | 0.8 | 0.77 | 0.84 | 0.65 | 0.75 |
| 11 | 69270349 | rs11263451 | -1.986 | 0.724 | -0.138 | 1.125 | 0.017 | 0.44 | 0.44 | 5.196 | 0.1 | 0.45 | 0.27 | 0.4 | 0.05 | 0.24 |
| 11 | 69274169 | rs7944975 | -0.697 | 1.087 | -0.468 | 0.401 | 0.08 | 0.048 | 0.048 | 0.499 | 0.06 | 0.32 | 0.06 | 0.02 | 0.04 | 0.06 |
| 11 | 69277078 | rs1938685 | -0.001 | 1.403 | -1.665 | 0.567 | -0.032 | -0.037 | -0.037 | 0 | 0.1 | 0.04 | 0.22 | 0.32 | 0.15 | 0.22 |
| 11 | 69277975 | rs11605729 | 0.063 | 0.916 | -1.48 | -0.216 | -0.068 | 0.119 | 0.119 | 0 | 0.27 | 0.07 | 0.21 | 0.08 | 0.33 | 0.21 |
| 11 | 69281578 | rs10752562 | -0.29 | 0.313 | -0.756 | 0.421 | 0 | 0.001 | 0.001 | 1.807 | 0.51 | 0.45 | 0.54 | 0.58 | 0.56 | 0.57 |
| 11 | 69282036 | rs3740643 | -0.318 | 0.295 | -0.627 | 0.408 | 0.002 | 0.005 | 0.005 | 1.731 | 0.52 | 0.53 | 0.55 | 0.58 | 0.56 | 0.57 |
| 11 | 69292197 | rs11263458 | 0.456 | -0.595 | -0.992 | 0.066 | 0.055 | 0.073 | 0.073 | 0.541 | 0.18 | 0.07 | 0.08 | 0.02 | 0.08 | 0.11 |
| 11 | 69299771 | rs587230 | -1.073 | -0.164 | -0.61 | 0.196 | 0.007 | 0.034 | 0.034 | 1.001 | 0.8 | 0.91 | 0.89 | 0.97 | 0.83 | 0.87 |
| 11 | 69304382 | rs17310712 | 0.335 | 0.352 | -0.747 | 0.752 | -0.018 | -0.017 | -0.017 | 0 | 0.08 | 0.06 | 0.26 | 0.23 | 0.16 | 0.18 |
| 11 | 69304876 | rs661960 | -0.667 | 0.023 | -0.391 | 0.933 | -0.007 | 0.017 | 0.017 | 0.34 | 0.77 | 0.74 | 0.78 | 0.58 | 0.82 | 0.82 |
| 11 | 69304990 | rs559817 | -0.857 | 0.133 | -0.333 | 0.987 | -0.01 | 0.031 | 0.031 | 1.558 | 0.77 | 0.7 | 0.78 | 0.58 | 0.82 | 0.82 |
| 11 | 69307444 | rs12365305 | 0.992 | 0.229 | -0.705 | 0.687 | -0.028 | 0.203 | 0.203 | 0.899 | 0.47 | 0.13 | 0.41 | 0.16 | 0.49 | 0.46 |
| 11 | 69311250 | rs662169 | -0.969 | 0.412 | 0.953 | 1.528 | 0.066 | -0.083 | -0.083 | 2.796 | 0.13 | 0.37 | 0.07 | 0.02 | 0.14 | 0.15 |
| 11 | 69314862 | rs559638 | -2.318 | 0.338 | 0.819 | 1.814 | 0.026 | -0.019 | -0.019 | 0 | 0.28 | 0.42 | 0.35 | 0.3 | 0.3 | 0.33 |
| 11 | 69316881 | rs680618 | -2.091 | 0.321 | 0.985 | 1.96 | 0.004 | -0.01 | -0.01 | 2.46 | 0.22 | 0.5 | 0.14 | 0.18 | 0.21 | 0.17 |
| 11 | 69318091 | rs7940177 | 0.025 | 0.42 | 0.917 | 1.998 | -0.028 | -0.04 | -0.04 | 0 | 0.07 | 0.02 | 0.23 | 0.21 | 0.09 | 0.17 |
| 11 | 69318518 | rs567488 | -1.48 | 0.365 | 1.144 | 1.824 | 0.051 | -0.107 | -0.107 | 0 | 0.15 | 0.38 | 0.09 | 0.02 | 0.16 | 0.13 |
| 11 | 69320546 | rs612088 | 0.686 | 0.212 | 0.046 | 1.246 | -0.006 | 0.017 | 0.017 | 0.076 | 0.06 | 0 | 0.03 | 0.04 | 0.05 | 0.03 |
| 11 | 69326308 | rs7951278 | -1.497 | 0.283 | 0.449 | 1.403 | 0.005 | 0.056 | 0.056 | 0 | 0.08 | 0.19 | 0.04 | 0.13 | 0.06 | 0.07 |
| 11 | 69332712 | rs657315 | -1.267 | 0.529 | 0.238 | 1.506 | 0.008 | 0.041 | 0.041 | 0 | 0.16 | 0.19 | 0.44 | 0.65 | 0.19 | 0.24 |
| 11 | 69339617 | rs498931 | -0.272 | -0.198 | 0.196 | 0.431 | -0.006 | -0.047 | -0.047 | 0 | 0.2 | 0.22 | 0.08 | 0.01 | 0.11 | 0.09 |
| 11 | 69340875 | rs12574080 | -1.679 | -0.268 | 0.907 | 0.723 | 0.058 | 0.142 | 0.142 | 1.683 | 0.09 | 0.43 | 0.06 | 0.12 | 0.06 | 0.03 |
| 11 | 69344477 | rs679162 | 0.374 | -0.406 | 0.049 | 0.71 | 0.001 | -0.012 | -0.012 | 1.777 | 0.76 | 0.83 | 0.65 | 0.68 | 0.75 | 0.7 |
| 11 | 69349845 | rs11607932 | 1.28 | -0.561 | -0.325 | 0.816 | 0.061 | 0.208 | 0.208 | 0 | 0.49 | 0.06 | 0.38 | 0.2 | 0.65 | 0.53 |
| 11 | 69351911 | rs546597 | 0.795 | -0.523 | 0.059 | 0.775 | 0.044 | 0.043 | 0.043 | 0.056 | 0.57 | 0.79 | 0.57 | 0.67 | 0.65 | 0.58 |
| 11 | 69364292 | rs10796822 | -1.147 | -0.465 | 0.771 | 1.149 | 0.078 | 0.663 | 0.663 | 3.85 | 0.05 | 0.41 | 0.18 | 0.41 | 0 | 0.06 |
| 11 | 69377670 | rs10908189 | -1.147 | -0.465 | 0.791 | 1.149 | 0.071 | 0.637 | 0.637 | 3.85 | 0.05 | 0.4 | 0.17 | 0.41 | 0 | 0.06 |
| 11 | 69378736 | rs12422135 | 1.559 | -0.349 | 0.023 | 1.02 | -0.024 | 0.142 | 0.142 | 0 | 0.47 | 0.08 | 0.36 | 0.26 | 0.59 | 0.5 |
| 11 | 69382767 | rs11263498 | 1.191 | -0.408 | 0.21 | 1.088 | 0.055 | 0.116 | 0.116 | 0 | 0.52 | 0.23 | 0.38 | 0.27 | 0.64 | 0.53 |
| 11 | 69384851 | rs606555 | -0.173 | -0.278 | -0.176 | 1.073 | -0.021 | -0.087 | -0.087 | 0 | 0.88 | 0.98 | 0.7 | 0.69 | 0.85 | 0.85 |

| | | | | | | | | | | | | | | | | |
|---|---|---|---|---|---|---|---|---|---|---|---|---|---|---|---|---|
| 11 | 69391327 | rs7946918 | 0.931 | 0.281 | 0.169 | -0.075 | 0.002 | 0.071 | 0.071 | 0 | 0.47 | 0.31 | 0.35 | 0.25 | 0.54 | 0.53 |
| 11 | 69392304 | rs625237 | 1.031 | 0.577 | 0.354 | -0.678 | -0.005 | 0.048 | 0.048 | 0 | 0.47 | 0.31 | 0.3 | 0.29 | 0.49 | 0.53 |
| 11 | 69392952 | rs638385 | 0.307 | 0.286 | 0.828 | -0.658 | -0.014 | -0.023 | -0.023 | 1.596 | 0.65 | 0.72 | 0.52 | 0.54 | 0.6 | 0.67 |
| 11 | 69395013 | rs4980673 | -0.467 | -0.124 | 0.957 | -0.614 | -0.033 | 0.006 | 0.006 | 0.109 | 0.19 | 0.4 | 0.21 | 0.24 | 0.1 | 0.14 |
| 11 | 69395443 | rs17136727 | 0.19 | -0.136 | 1.094 | -0.604 | -0.034 | 0.021 | 0.021 | 1.053 | 0.16 | 0.33 | 0.18 | 0.24 | 0.08 | 0.12 |
| 11 | 69399669 | rs480921 | 0.428 | 0.351 | 1.194 | -0.312 | -0.001 | -0.023 | -0.023 | 0 | 0.67 | 0.72 | 0.58 | 0.54 | 0.68 | 0.67 |
| 11 | 69403278 | rs597587 | -2.667 | 0.27 | 1.441 | -0.345 | 0.035 | 0.201 | 0.201 | 3.673 | 0.08 | 0.55 | 0.18 | 0.24 | 0.02 | 0.04 |
| 11 | 69406112 | rs518418 | 0.362 | 0.221 | 1.507 | -0.273 | -0.007 | -0.045 | -0.045 | 0 | 0.68 | 0.81 | 0.6 | 0.54 | 0.68 | 0.69 |
| 11 | 69413088 | rs10737155 | 0.652 | 0.121 | 1.641 | -1.275 | -0.004 | -0.007 | -0.007 | 0.671 | 0.66 | 0.36 | 0.46 | 0.29 | 0.73 | 0.65 |
| 11 | 69414781 | rs7122472 | 0.689 | 0.121 | 1.644 | -1.275 | -0.007 | -0.012 | -0.012 | 0 | 0.66 | 0.37 | 0.46 | 0.3 | 0.73 | 0.65 |
| 11 | 69418138 | rs1872568 | 1.14 | 0.094 | 1.294 | -1.365 | 0.03 | 0.068 | 0.068 | 0 | 0.61 | 0.11 | 0.43 | 0.29 | 0.72 | 0.64 |
| 11 | 69421272 | rs7950206 | 0.697 | 0.409 | 1.302 | -1.491 | -0.004 | -0.014 | -0.014 | 1.051 | 0.67 | 0.36 | 0.45 | 0.24 | 0.73 | 0.68 |
| 11 | 69425840 | rs7946645 | 0.711 | 0.468 | 1.228 | -1.504 | -0.003 | -0.016 | -0.016 | 0.094 | 0.67 | 0.36 | 0.47 | 0.22 | 0.72 | 0.66 |
| 11 | 69429136 | rs599190 | -0.169 | 0.543 | 0.97 | -1.52 | -0.036 | -0.038 | -0.038 | 0 | 0.79 | 0.96 | 0.54 | 0.24 | 0.79 | 0.71 |
| 11 | 69434050 | rs2510464 | -0.535 | 0.589 | 1.351 | -1.506 | 0.026 | -0.066 | -0.066 | 0 | 0.12 | 0.49 | 0.07 | 0.01 | 0.08 | 0.05 |
| 11 | 69435230 | rs3017614 | 0.397 | 0.697 | 0.953 | -1.571 | 0 | -0.025 | -0.025 | 0 | 0.71 | 0.54 | 0.46 | 0.22 | 0.72 | 0.66 |
| 11 | 69442963 | rs1982774 | -0.023 | 0.93 | 0.649 | -1.371 | -0.062 | 0.055 | 0.055 | 0.088 | 0.65 | 0.78 | 0.43 | 0.24 | 0.57 | 0.67 |
| 11 | 69445284 | rs611003 | 0.484 | 0.736 | 0.505 | -1.579 | -0.015 | 0.09 | 0.09 | 0 | 0.54 | 0.29 | 0.36 | 0.22 | 0.54 | 0.52 |
| 11 | 69446913 | rs11827026 | -0.831 | 0.621 | 1.035 | -1.562 | -0.014 | -0.057 | -0.057 | 0.05 | 0.11 | 0.32 | 0.05 | 0.01 | 0.05 | 0.15 |
| 11 | 69448575 | rs655089 | 0.135 | 0.607 | 0.595 | -1.826 | 0.003 | -0.043 | -0.043 | 11.871 | 0.39 | 0.41 | 0.57 | 0.13 | 0.45 | 0.29 |
| 11 | 69457293 | rs1352075 | 0.04 | 0.709 | 0.266 | -1.777 | -0.009 | 0.158 | 0.158 | 0.124 | 0.54 | 0.21 | 0.38 | 0.82 | 0.51 | 0.6 |
| 11 | 69462856 | rs3862792 | -0.003 | 0.142 | 0.578 | -1.791 | -0.017 | -0.028 | -0.028 | 0 | 0.07 | 0.15 | 0.04 | 0.03 | 0.03 | 0.12 |
| 11 | 69464793 | rs649392 | 0.035 | 0.186 | 0.468 | -1.922 | 0.018 | 0.064 | 0.064 | 0 | 0.54 | 0.36 | 0.45 | 0.88 | 0.59 | 0.56 |
| 11 | 69480191 | rs11263527 | -0.639 | -0.575 | -0.044 | -1.357 | 0.005 | -0.01 | -0.01 | 0 | 0.09 | 0.45 | 0.06 | 0.03 | 0.02 | 0.1 |
| 11 | 69482091 | rs1789166 | -0.201 | -0.047 | -1.273 | -1.108 | 0.003 | 0.149 | 0.149 | 0 | 0.65 | 0.87 | 0.51 | 0.89 | 0.68 | 0.68 |
| 11 | 69509343 | rs1789170 | -0.044 | 0.041 | -1.163 | -0.995 | 0.012 | 0.052 | 0.052 | 0 | 0.65 | 0.74 | 0.5 | 0.91 | 0.68 | 0.68 |
| 11 | 69512978 | rs3737463 | 0.357 | -0.495 | -1.531 | -0.094 | 0.057 | -0.047 | -0.047 | 0.079 | 0.66 | 0.17 | 0.71 | 0.39 | 0.74 | 0.57 |
| 11 | 69513338 | rs948992 | -0.587 | -0.638 | -0.714 | 0.074 | -0.018 | 0.018 | 0.018 | 0 | 0.36 | 0.41 | 0.29 | 0.47 | 0.3 | 0.53 |
| 11 | 69515423 | rs1307968 | -0.536 | 0.074 | -0.789 | -0.352 | 0.006 | 0.02 | 0.02 | 2.735 | 0.85 | 0.89 | 0.92 | 0.92 | 0.86 | 0.83 |
| 11 | 69523592 | rs4980680 | -0.401 | 0.315 | -0.619 | -0.355 | 0 | -0.006 | -0.006 | 1.13 | 0.84 | 0.8 | 0.9 | 0.91 | 0.83 | 0.82 |
| 11 | 69533129 | rs11263535 | -0.581 | 0.267 | -0.388 | -0.612 | -0.008 | -0.042 | -0.042 | 0.316 | 0.33 | 0.4 | 0.19 | 0.17 | 0.27 | 0.41 |
| 11 | 69534136 | rs10796833 | 0.862 | 0.228 | -0.658 | -0.681 | 0.012 | 0.169 | 0.169 | 0.705 | 0.5 | 0.21 | 0.68 | 0.78 | 0.57 | 0.42 |
| 11 | 69536921 | rs10908200 | 0.147 | 0.79 | -0.482 | -0.427 | -0.005 | 0.045 | 0.045 | 2.361 | 0.16 | 0.08 | 0.08 | 0.05 | 0.16 | 0.16 |
| 11 | 69540990 | rs10752564 | 0.298 | 0.689 | -0.229 | -0.406 | 0.006 | -0.037 | -0.037 | 0 | 0.74 | 0.64 | 0.83 | 0.92 | 0.7 | 0.71 |
| 11 | 69543611 | rs948989 | 0.304 | 1.153 | -0.279 | 0.418 | 0.017 | 0.04 | 0.04 | 2.577 | 0.67 | 0.49 | 0.48 | 0.58 | 0.66 | 0.66 |
| 11 | 69544508 | rs7947358 | -0.089 | 1.032 | -0.487 | 0.149 | 0.002 | 0.004 | 0.004 | 0 | 0.51 | 0.55 | 0.36 | 0.33 | 0.41 | 0.48 |
| 11 | 69556722 | rs4616067 | -0.502 | 0.573 | -0.405 | 0.252 | -0.011 | -0.002 | -0.002 | 0 | 0.81 | 0.79 | 0.88 | 0.86 | 0.88 | 0.83 |
| 11 | 69559418 | rs10796842 | -1.202 | 1.846 | -0.08 | 0.962 | 0.049 | 0.155 | 0.155 | 1.161 | 0.12 | 0.26 | 0.35 | 0.56 | 0.19 | 0.21 |
| 11 | 69559612 | rs12418010 | -0.173 | 2.239 | -0.89 | 0.496 | -0.115 | 0.03 | 0.03 | 0.016 | 0.25 | 0.03 | 0.28 | 0.18 | 0.32 | 0.29 |
| 11 | 69563733 | rs7479018 | -0.542 | 2.129 | -0.358 | 0.435 | 0 | -0.002 | -0.002 | 0.144 | 0.34 | 0.36 | 0.34 | 0.25 | 0.34 | 0.41 |
| 11 | 69566488 | rs1112008 | -0.428 | 1.981 | -0.368 | 0.614 | 0.029 | 0.116 | 0.116 | 0 | 0.07 | 0.1 | 0.33 | 0.52 | 0.21 | 0.13 |
| 11 | 69570033 | rs11263551 | -0.246 | 2.021 | -0.38 | 0.652 | 0.019 | 0.102 | 0.102 | 0 | 0.07 | 0.08 | 0.33 | 0.52 | 0.21 | 0.14 |
| 11 | 69570988 | rs9735736 | -0.962 | 2.028 | -0.403 | 0.519 | -0.002 | -0.008 | -0.008 | 0.258 | 0.42 | 0.46 | 0.68 | 0.77 | 0.57 | 0.57 |
| 11 | 69572684 | rs10737157 | -0.941 | 2.046 | -0.393 | 0.531 | -0.002 | -0.009 | -0.009 | 0 | 0.41 | 0.46 | 0.68 | 0.77 | 0.56 | 0.57 |
| 11 | 69574816 | rs10908210 | -0.939 | 2.172 | -0.315 | 1.428 | 0.042 | 0.14 | 0.14 | 0 | 0.08 | 0.13 | 0.34 | 0.54 | 0.21 | 0.14 |
| 11 | 69579070 | rs713217 | -1.091 | 2.174 | -0.292 | 1.359 | 0.047 | 0.132 | 0.132 | 0 | 0.08 | 0.13 | 0.35 | 0.55 | 0.21 | 0.14 |
| 11 | 69585787 | rs871346 | -0.212 | 1.502 | -0.09 | 0.205 | -0.009 | 0.051 | 0.051 | 0.399 | 0.18 | 0.08 | 0.07 | 0.02 | 0.18 | 0.11 |
| 11 | 69588020 | rs3740639 | -0.694 | 1.931 | -0.491 | 1.178 | -0.023 | 0.034 | 0.034 | 0 | 0.27 | 0.09 | 0.19 | 0.16 | 0.27 | 0.2 |
| 11 | 69588729 | rs2073464 | 0.24 | 1.57 | 0.439 | 0.479 | -0.005 | 0.128 | 0.128 | 1.755 | 0.83 | 0.5 | 0.58 | 0.69 | 0.88 | 0.8 |
| 11 | 69592664 | rs4980690 | 0.312 | 1.547 | 0.668 | 0.207 | -0.001 | -0.004 | -0.004 | 0 | 0.08 | 0.08 | 0.06 | 0.02 | 0.1 | 0.06 |
| 11 | 69596523 | rs4084127 | 0.64 | 0.723 | -0.259 | -0.314 | -0.016 | 0.031 | 0.031 | 2.802 | 0.32 | 0.18 | 0.53 | 0.19 | 0.35 | 0.31 |
| 11 | 69597131 | rs3897660 | 0.502 | 0.645 | -0.474 | -0.152 | -0.006 | 0.006 | 0.006 | 8.153 | 0.44 | 0.11 | 0.27 | 0.58 | 0.44 | 0.38 |
| 11 | 69597787 | rs9734817 | 0.252 | 0.406 | -0.242 | 0.203 | -0.007 | 0.004 | 0.004 | 0 | 0.1 | 0.14 | 0.07 | 0.16 | 0.08 | 0.12 |
| 11 | 69601039 | rs11263574 | -0.147 | 0.506 | -0.618 | 0.236 | 0 | 0.036 | 0.036 | 0.836 | 0.06 | 0.1 | 0.3 | 0.1 | 0.03 | 0.04 |
| 11 | 69601290 | rs4246965 | -0.085 | 0.514 | -0.719 | -0.286 | -0.001 | 0.006 | 0.006 | 0.605 | 0.58 | 0.4 | 0.64 | 0.69 | 0.55 | 0.48 |
| 11 | 69604527 | rs10752567 | -0.486 | 0.306 | -0.989 | -0.879 | 0.006 | 0.044 | 0.044 | 0 | 0.67 | 0.82 | 0.76 | 0.94 | 0.68 | 0.77 |
| 11 | 69605286 | rs9666630 | 0.824 | 0.74 | -1.414 | 0.228 | -0.07 | -0.11 | -0.11 | 0.745 | 0.17 | 0.02 | 0.18 | 0.37 | 0.24 | 0.19 |
| 11 | 69606310 | rs10908221 | -0.46 | 0.906 | -1.151 | 0.292 | 0.013 | 0.081 | 0.081 | 0.351 | 0.12 | 0.19 | 0.35 | 0.31 | 0.13 | 0.27 |
| 11 | 69607507 | rs10752569 | -0.543 | 0.855 | -1.134 | 0.146 | -0.019 | 0.013 | 0.013 | 0 | 0.29 | 0.21 | 0.54 | 0.73 | 0.37 | 0.47 |
| 11 | 69610827 | rs4980694 | 0.004 | 0.203 | -0.808 | -0.437 | -0.042 | -0.056 | -0.056 | 0 | 0.69 | 0.46 | 0.82 | 0.82 | 0.79 | 0.84 |
| 11 | 69612823 | rs4084129 | 1.272 | 0.42 | -0.621 | -0.153 | -0.011 | -0.025 | -0.025 | 0.405 | 0.18 | 0.13 | 0.2 | 0.43 | 0.25 | 0.2 |
| 11 | 69614847 | rs4631909 | -0.603 | 0.476 | -0.107 | -0.244 | 0.008 | 0.012 | 0.012 | 1.015 | 0.47 | 0.61 | 0.38 | 0.62 | 0.46 | 0.38 |
| 11 | 69619417 | rs4980700 | -0.852 | 0.618 | -0.115 | -0.245 | -0.002 | 0.011 | 0.011 | 0.411 | 0.52 | 0.6 | 0.41 | 0.63 | 0.55 | 0.47 |
| 11 | 69622227 | rs11263582 | 0.716 | 0.458 | -0.412 | -0.246 | -0.008 | -0.04 | -0.04 | 0 | 0.14 | 0.04 | 0.12 | 0.2 | 0.15 | 0.07 |
| 11 | 69623606 | rs948133 | -0.587 | 0.386 | -0.057 | -0.355 | 0.006 | 0.015 | 0.015 | 1.706 | 0.46 | 0.38 | 0.37 | 0.62 | 0.49 | 0.39 |
| 11 | 69630677 | rs10796856 | -0.763 | 0.393 | 0.047 | -0.301 | -0.002 | -0.005 | -0.005 | 0 | 0.48 | 0.51 | 0.37 | 0.63 | 0.5 | 0.42 |
| 11 | 69637665 | rs3892955 | -1.643 | 0.441 | 0.254 | -0.003 | -0.001 | 0.002 | 0.002 | 1.91 | 0.33 | 0.64 | 0.52 | 0.49 | 0.26 | 0.49 |
| 11 | 69637967 | rs11263596 | -0.733 | 0.676 | 0.52 | 0.262 | 0.012 | 0.146 | 0.146 | 0.276 | 0.11 | 0.26 | 0.34 | 0.29 | 0.06 | 0.33 |
| 11 | 69641621 | rs11263599 | -1.037 | 0.901 | 0.29 | 0.203 | 0.037 | 0.075 | 0.075 | 0 | 0.16 | 0.33 | 0.2 | 0.3 | 0.19 | 0.3 |

| | | | | | | | | | | | | | | | |
|---|---|---|---|---|---|---|---|---|---|---|---|---|---|---|---|
| 11 | 69642246 | rs3933410 | -0.966 | 0.913 | 0.114 | 0.209 | 0.021 | 0.045 | 0.045 | 0 | 0.16 | 0.26 | 0.19 | 0.3 | 0.19 | 0.3 |
| 11 | 69647942 | rs10908232 | -0.567 | 0.951 | 0.295 | 0.475 | 0.065 | 0.193 | 0.193 | 0.057 | 0.07 | 0.26 | 0.17 | 0.27 | 0.07 | 0.25 |
| 11 | 69649931 | rs7479857 | 0.311 | 0.707 | -0.117 | 0.061 | -0.004 | -0.025 | -0.025 | 0.234 | 0.35 | 0.22 | 0.26 | 0.43 | 0.36 | 0.29 |
| 11 | 69650210 | rs9804449 | -1.211 | 0.65 | -0.61 | -0.514 | -0.046 | 0.187 | 0.187 | 0.642 | 0.65 | 0.9 | 0.76 | 0.94 | 0.61 | 0.75 |
| 11 | 69658210 | rs10160812 | -0.463 | 0.591 | -0.233 | -0.226 | -0.003 | 0.017 | 0.017 | 0 | 0.24 | 0.33 | 0.36 | 0.25 | 0.22 | 0.24 |
| 11 | 69660482 | rs7130524 | -1.629 | 0.98 | -0.747 | -0.606 | -0.043 | 0.164 | 0.164 | 0.396 | 0.73 | 0.97 | 0.8 | 0.97 | 0.71 | 0.85 |
| 11 | 69672990 | rs6598087 | 0.082 | 0.949 | -0.053 | -0.099 | 0 | -0.003 | -0.003 | 0.104 | 0.39 | 0.56 | 0.39 | 0.68 | 0.4 | 0.47 |
| 11 | 69675259 | rs10908241 | 0.9 | 0.985 | -0.369 | 0.17 | -0.029 | -0.093 | -0.093 | 1.48 | 0.31 | 0.06 | 0.22 | 0.42 | 0.31 | 0.23 |
| 11 | 69676000 | rs11263616 | 1.366 | 1.147 | -0.501 | 0.292 | -0.022 | 0.001 | 0.001 | 0 | 0.07 | 0.01 | 0.08 | 0.04 | 0.09 | 0.06 |
| 11 | 69676502 | rs4074402 | 0.273 | 1.096 | -0.031 | 0.247 | -0.002 | 0.001 | 0.001 | 0 | 0.38 | 0.33 | 0.39 | 0.69 | 0.4 | 0.47 |
| 11 | 69679342 | rs12420414 | 0.691 | 1.093 | -0.143 | 0.663 | -0.001 | 0.026 | 0.026 | 0.81 | 0.34 | 0.1 | 0.2 | 0.29 | 0.33 | 0.26 |
| 11 | 69680908 | rs12574097 | 0.161 | 1.145 | 0.007 | 0.708 | 0.002 | 0.006 | 0.006 | 0 | 0.41 | 0.42 | 0.38 | 0.57 | 0.42 | 0.5 |
| 11 | 69681843 | rs11263621 | 0.081 | 1.192 | 0.151 | 0.917 | 0.085 | 0.199 | 0.199 | 0 | 0.06 | 0.22 | 0.17 | 0.28 | 0.1 | 0.24 |
| 11 | 69683168 | rs11263622 | 0.115 | 1.084 | 0.042 | 0.828 | 0.008 | -0.043 | -0.043 | 0 | 0.16 | 0.07 | 0.31 | 0.23 | 0.15 | 0.17 |
| 11 | 69684026 | rs10732958 | -0.81 | 1.251 | -0.029 | 0.587 | -0.001 | 0.001 | 0.001 | 0 | 0.59 | 0.4 | 0.62 | 0.65 | 0.6 | 0.47 |
| 11 | 69684293 | rs10908244 | -0.015 | 1.188 | -0.417 | 0.834 | 0.008 | -0.026 | -0.026 | 0 | 0.17 | 0.1 | 0.31 | 0.23 | 0.15 | 0.17 |
| 11 | 69684520 | rs7481015 | -0.518 | 1.064 | -0.613 | 0.143 | 0.013 | 0.065 | 0.065 | 0 | 0.22 | 0.32 | 0.48 | 0.51 | 0.24 | 0.41 |
| 11 | 69685824 | rs11263623 | -0.558 | 1.005 | -1.075 | -0.301 | 0.001 | 0.052 | 0.052 | 0 | 0.62 | 0.7 | 0.74 | 0.81 | 0.65 | 0.73 |
| 11 | 69691270 | rs11602376 | 0.525 | 0.978 | -1.378 | -0.165 | -0.047 | -0.039 | -0.039 | 0 | 0.36 | 0.16 | 0.38 | 0.53 | 0.41 | 0.32 |
| 11 | 69696940 | rs10796868 | 0.31 | 0.415 | -1.116 | -0.19 | -0.005 | -0.046 | -0.046 | 0.487 | 0.13 | 0.03 | 0.15 | 0.22 | 0.13 | 0.27 |
| 11 | 69697837 | rs7396724 | 0.403 | 0.152 | -1.026 | -0.533 | 0.054 | 0.073 | 0.073 | 0 | 0.53 | 0.27 | 0.63 | 0.75 | 0.62 | 0.49 |
| 11 | 69699946 | rs7124792 | -0.603 | 1.01 | -0.916 | 0.584 | -0.002 | -0.044 | -0.044 | 0 | 0.18 | 0.04 | 0.17 | 0.22 | 0.18 | 0.28 |
| 11 | 69703103 | rs7950402 | 0.489 | 1.41 | -0.012 | 0.272 | 0.045 | 0.02 | 0.02 | 0.728 | 0.84 | 0.69 | 0.71 | 0.78 | 0.75 | 0.83 |
| 11 | 69812716 | rs12272802 | -1.489 | 0.553 | -0.051 | -0.244 | -0.014 | -0.034 | -0.034 | 0 | 0.2 | 0.28 | 0.11 | 0.08 | 0.14 | 0.08 |
| 11 | 69814829 | rs7926926 | 1.007 | 0.935 | 0.804 | -0.085 | 0.05 | 0.012 | 0.012 | 2.437 | 0.91 | 0.63 | 0.92 | 0.9 | 0.91 | 0.96 |
| 11 | 69818523 | rs902880 | 1.303 | 1.136 | 0.632 | -0.385 | 0.026 | -0.046 | -0.046 | 0 | 0.72 | 0.45 | 0.65 | 0.8 | 0.71 | 0.79 |
| 11 | 69820567 | rs10898818 | 1.44 | 1.132 | 0.581 | -0.431 | 0.002 | -0.018 | -0.018 | 0 | 0.67 | 0.33 | 0.6 | 0.8 | 0.69 | 0.78 |
| 11 | 69822846 | rs4450188 | 1.479 | 1.482 | 0.304 | -0.534 | -0.003 | -0.019 | -0.019 | 0.997 | 0.6 | 0.4 | 0.7 | 0.7 | 0.59 | 0.81 |
| 11 | 69823461 | rs4980704 | 2.033 | 1.243 | -0.047 | 0.257 | -0.013 | -0.03 | -0.03 | 0.006 | 0.49 | 0.12 | 0.43 | 0.49 | 0.45 | 0.61 |
| 11 | 69828594 | rs10793046 | 1.232 | 1.384 | 0.409 | -0.223 | 0.037 | -0.08 | -0.08 | 0.578 | 0.74 | 0.49 | 0.63 | 0.87 | 0.68 | 0.85 |
| 11 | 69834983 | rs10501401 | -1.028 | 1.083 | 0.404 | -0.217 | 0.014 | -0.001 | -0.001 | 0 | 0.12 | 0.17 | 0.1 | 0.11 | 0.16 | 0.16 |
| 11 | 69835356 | rs1492512 | 0.484 | 0.865 | 0.718 | -0.395 | -0.045 | -0.104 | -0.104 | 0 | 0.79 | 0.6 | 0.88 | 0.99 | 0.86 | 0.94 |
| 11 | 69835622 | rs7931183 | 2.723 | 0.927 | 0.334 | -0.385 | -0.022 | 0.138 | 0.138 | 0 | 0.52 | 0.2 | 0.51 | 0.75 | 0.49 | 0.7 |
| 11 | 69837473 | rs872837 | -2.346 | 1.325 | 0.305 | 0.089 | 0.012 | -0.003 | -0.003 | 0 | 0.27 | 0.28 | 0.37 | 0.24 | 0.36 | 0.24 |
| 11 | 69838148 | rs1994776 | -1.282 | 1.323 | 0.287 | 0.091 | -0.014 | 0.004 | 0.004 | 0 | 0.14 | 0.1 | 0.27 | 0.13 | 0.2 | 0.08 |
| 11 | 69838584 | rs902869 | -1.808 | 1.142 | 0.32 | 0.242 | 0.017 | 0.004 | 0.004 | 0 | 0.22 | 0.26 | 0.18 | 0.24 | 0.27 | 0.2 |
| 11 | 69839305 | rs2131242 | 0.114 | 1.267 | -0.127 | 0.319 | -0.022 | -0.022 | -0.022 | 0.042 | 0.05 | 0.01 | 0.05 | 0.1 | 0.1 | 0.13 |
| 11 | 69840332 | rs4391849 | -0.571 | 1.44 | -0.126 | 0.809 | -0.028 | -0.033 | -0.033 | 0 | 0.1 | 0.05 | 0.12 | 0.41 | 0.13 | 0.16 |
| 11 | 69840704 | rs902867 | 0.352 | 0.959 | 0.375 | 0.372 | -0.022 | -0.075 | -0.075 | 0 | 0.84 | 0.64 | 0.93 | 0.97 | 0.9 | 0.92 |
| 11 | 69840824 | rs867591 | 0.567 | 0.834 | 0.023 | -0.253 | -0.004 | 0.001 | 0.001 | 0 | 0.58 | 0.47 | 0.73 | 0.56 | 0.64 | 0.67 |
| 11 | 69843124 | rs4980573 | 0.027 | 0.566 | 0.072 | -0.092 | -0.037 | -0.075 | -0.075 | 0 | 0.84 | 0.65 | 0.94 | 0.97 | 0.92 | 0.92 |
| 11 | 69843850 | rs7942235 | -0.507 | 0.688 | -0.103 | -0.301 | 0.003 | 0 | 0 | 0 | 0.47 | 0.48 | 0.54 | 0.49 | 0.58 | 0.46 |
| 11 | 69845520 | rs4980714 | -0.144 | 0.569 | -0.075 | -0.719 | 0.004 | -0.024 | -0.024 | 0 | 0.61 | 0.35 | 0.6 | 0.96 | 0.62 | 0.68 |
| 11 | 69845815 | rs4980719 | 0.311 | 0.75 | -0.037 | -0.585 | 0 | 0.064 | 0.064 | 0 | 0.47 | 0.32 | 0.53 | 0.88 | 0.49 | 0.59 |
| 11 | 69854906 | rs2131244 | 0.288 | 0.727 | 0.183 | -0.584 | 0 | 0.092 | 0.092 | 0 | 0.48 | 0.31 | 0.53 | 0.88 | 0.49 | 0.58 |
| 11 | 69860796 | rs902876 | -1.555 | 0.75 | 0.298 | -0.743 | 0.014 | -0.003 | -0.003 | 0 | 0.74 | 0.77 | 0.87 | 1 | 0.73 | 0.9 |
| 11 | 69862715 | rs4980736 | 1.151 | 0.821 | 0.23 | -0.187 | -0.004 | -0.026 | -0.026 | 0 | 0.38 | 0.25 | 0.41 | 0.52 | 0.38 | 0.4 |
| 11 | 69863023 | rs990178 | 0.162 | 0.802 | 0.499 | -0.268 | -0.001 | -0.065 | -0.065 | 1.095 | 0.49 | 0.5 | 0.54 | 0.91 | 0.49 | 0.62 |
| 11 | 69864128 | rs7935198 | -1.156 | 0.869 | 0.372 | -0.251 | 0.005 | -0.046 | -0.046 | 0.927 | 0.25 | 0.27 | 0.33 | 0.08 | 0.26 | 0.29 |
| 11 | 69864805 | rs7927315 | 0.485 | 1.124 | -0.008 | 0.003 | 0.006 | -0.006 | -0.006 | 0 | 0.48 | 0.26 | 0.45 | 0.49 | 0.44 | 0.45 |
| 11 | 69866283 | rs17160555 | 0.424 | 1.088 | -0.175 | -0.079 | -0.003 | -0.002 | -0.002 | 0 | 0.08 | 0.04 | 0.05 | 0.11 | 0.07 | 0.05 |
| 11 | 69867808 | rs4980580 | -1.496 | 0.983 | 0.109 | 0.093 | 0.002 | -0.009 | -0.009 | 0 | 0.73 | 0.75 | 0.87 | 0.94 | 0.74 | 0.9 |
| 11 | 69870778 | rs4980746 | 1.223 | 1.087 | -0.49 | 0.277 | -0.046 | 0.035 | 0.035 | 0 | 0.31 | 0.17 | 0.41 | 0.17 | 0.34 | 0.38 |
| 11 | 69873933 | rs11822948 | 0.055 | 1.035 | -0.344 | 0.578 | 0 | 0.001 | 0.001 | 0 | 0.22 | 0.22 | 0.27 | 0.45 | 0.23 | 0.28 |
| 11 | 69874267 | rs11237181 | -0.274 | 1.108 | -0.323 | 0.226 | -0.001 | 0.002 | 0.002 | 0 | 0.25 | 0.25 | 0.33 | 0.63 | 0.27 | 0.44 |
| 11 | 69875423 | rs12223060 | 1.359 | 1.087 | -0.354 | 0.317 | -0.007 | -0.011 | -0.011 | 1.364 | 0.09 | 0.03 | 0.21 | 0.42 | 0.09 | 0.18 |
| 11 | 69875472 | rs10793254 | -1.92 | 1.03 | -0.278 | 0.16 | -0.015 | -0.047 | -0.047 | 0 | 0.73 | 0.7 | 0.86 | 0.94 | 0.73 | 0.9 |
| 11 | 69875759 | rs7947135 | -0.595 | 0.921 | -0.365 | 0.262 | 0.004 | -0.014 | -0.014 | 1.271 | 0.18 | 0.24 | 0.13 | 0.14 | 0.14 | 0.09 |
| 11 | 69876894 | rs2375269 | -0.275 | 0.632 | -0.303 | 0.435 | 0 | -0.001 | -0.001 | 0 | 0.45 | 0.46 | 0.5 | 0.31 | 0.42 | 0.46 |
| 11 | 69877928 | rs11237292 | 1.043 | 0.87 | -1.046 | 0.797 | -0.011 | 0.012 | 0.012 | 0 | 0.27 | 0.14 | 0.22 | 0.17 | 0.27 | 0.33 |
| 11 | 69880281 | rs12294626 | 0.335 | 0.854 | -1.051 | 0.736 | 0.029 | -0.03 | -0.03 | 0.041 | 0.06 | 0.18 | 0.19 | 0 | 0.05 | 0.05 |
| 11 | 69883969 | rs7929885 | 1.19 | 0.822 | -1.19 | 0.79 | -0.04 | -0.016 | -0.016 | 0 | 0.25 | 0.07 | 0.21 | 0.17 | 0.27 | 0.32 |
| 11 | 69885221 | rs11237529 | 0.844 | 0.84 | -1.189 | 0.961 | -0.004 | 0.001 | 0.001 | 0 | 0.09 | 0.04 | 0.19 | 0.42 | 0.08 | 0.17 |
| 11 | 69892474 | rs10899613 | -0.613 | 0.845 | -1.177 | 0.845 | -0.002 | -0.005 | -0.005 | 0 | 0.23 | 0.2 | 0.29 | 0.49 | 0.24 | 0.3 |
| 11 | 69896252 | rs4980753 | -1.213 | 0.769 | -1.138 | 0.193 | -0.01 | -0.079 | -0.079 | 0 | 0.61 | 0.53 | 0.75 | 0.91 | 0.6 | 0.86 |
| 11 | 69901274 | rs11231949 | -0.109 | 0.843 | -1.23 | 0.24 | -0.013 | -0.028 | -0.028 | 0.008 | 0.42 | 0.23 | 0.34 | 0.38 | 0.44 | 0.6 |
| 11 | 69903067 | rs11232056 | -0.686 | 1.083 | -0.113 | 1.202 | 0.01 | 0.064 | 0.064 | 0 | 0.19 | 0.28 | 0.41 | 0.53 | 0.17 | 0.26 |
| 11 | 69907249 | rs12223676 | -0.924 | 0.709 | -0.454 | 0.725 | 0.001 | 0.008 | 0.008 | 0 | 0.15 | 0.17 | 0.13 | 0.21 | 0.16 | 0.26 |
| 11 | 69910890 | rs9630219 | 0.458 | 0.58 | -0.379 | 0.18 | 0.004 | -0.027 | -0.027 | 0 | 0.62 | 0.49 | 0.55 | 0.7 | 0.59 | 0.56 |

| | | | | | | | | | | | | | | | | |
|---|---|---|---|---|---|---|---|---|---|---|---|---|---|---|---|---|
| 11 | 69910934 | rs9630220 | 0.499 | 0.48 | -0.081 | -0.258 | 0.012 | -0.055 | -0.055 | 0 | 0.14 | 0.27 | 0.09 | 0.03 | 0.13 | 0.08 |
| 11 | 69916254 | rs10897850 | 0.609 | 0.47 | -0.062 | -0.263 | 0.009 | -0.048 | -0.048 | 0 | 0.13 | 0.21 | 0.08 | 0.03 | 0.13 | 0.08 |
| 11 | 69917160 | rs7941411 | -0.093 | 0.551 | 0.314 | -0.618 | 0.029 | -0.071 | -0.071 | 0 | 0.11 | 0.33 | 0.23 | 0.01 | 0.14 | 0.03 |
| 11 | 69918715 | rs12798364 | 0.834 | 0.111 | 0.013 | -0.334 | 0.001 | -0.001 | -0.001 | 0 | 0.11 | 0.14 | 0.1 | 0.2 | 0.07 | 0.1 |
| 11 | 69919522 | rs4515996 | -0.35 | -0.052 | -0.02 | -0.451 | 0.001 | 0.001 | 0.001 | 0 | 0.7 | 0.76 | 0.77 | 0.7 | 0.71 | 0.6 |
| 11 | 69921329 | rs10897924 | 0.704 | -0.169 | 0.2 | -0.589 | -0.01 | 0.037 | 0.037 | 0 | 0.55 | 0.41 | 0.51 | 0.69 | 0.49 | 0.53 |
| 11 | 69923538 | rs1940238 | -1.06 | -0.162 | 0.084 | -0.589 | 0.019 | -0.074 | -0.074 | 0 | 0.85 | 0.63 | 0.91 | 0.97 | 0.87 | 0.92 |
| 11 | 69929677 | rs744205 | -1.658 | -0.145 | -0.522 | -0.585 | -0.041 | 0.073 | 0.073 | 0 | 0.55 | 0.85 | 0.56 | 0.34 | 0.55 | 0.51 |
| 11 | 69931222 | rs2155451 | 0.468 | -0.143 | -0.439 | -0.645 | -0.001 | -0.002 | -0.002 | 0 | 0.21 | 0.17 | 0.19 | 0.27 | 0.24 | 0.36 |
| 11 | 69932450 | rs2515284 | -1.156 | -0.025 | -0.444 | -0.564 | 0.007 | -0.013 | -0.013 | 0 | 0.89 | 0.9 | 0.9 | 0.77 | 0.93 | 0.89 |
| 11 | 69933614 | rs12361305 | 1.918 | -0.065 | -0.549 | -0.605 | -0.103 | -0.182 | -0.182 | 0.061 | 0.36 | 0.03 | 0.35 | 0.46 | 0.37 | 0.39 |
| 11 | 69933696 | rs2509132 | 0.222 | 0.418 | -0.101 | -0.832 | 0.001 | 0.003 | 0.003 | 0 | 0.2 | 0.19 | 0.19 | 0.27 | 0.24 | 0.34 |
| 11 | 69933717 | rs2509133 | -1.314 | 0.412 | -0.042 | -0.844 | -0.01 | 0.025 | 0.025 | 0 | 0.44 | 0.61 | 0.5 | 0.28 | 0.51 | 0.45 |
| 11 | 69934085 | rs10898112 | -0.326 | 0.425 | -0.062 | -0.807 | 0.011 | -0.075 | -0.075 | 0 | 0.23 | 0.29 | 0.29 | 0.03 | 0.27 | 0.12 |
| 11 | 69935558 | rs11233666 | 0.53 | 0.633 | -0.323 | -0.244 | 0.008 | -0.009 | -0.009 | 0.777 | 0.7 | 0.62 | 0.58 | 0.73 | 0.62 | 0.63 |
| 11 | 69941786 | rs873860 | -0.537 | 1.067 | -0.842 | -0.087 | 0.003 | 0 | 0 | 0 | 0.22 | 0.2 | 0.39 | 0.29 | 0.36 | 0.39 |
| 11 | 69942324 | rs2155453 | -1.008 | 0.979 | -0.834 | -0.019 | 0.013 | -0.015 | -0.015 | 0 | 0.36 | 0.42 | 0.46 | 0.31 | 0.44 | 0.42 |
| 11 | 69944670 | rs2515288 | -0.614 | 0.918 | -0.951 | 0.396 | 0.047 | 0.028 | 0.028 | 0 | 0.12 | 0.31 | 0.28 | 0.16 | 0.19 | 0.12 |
| 11 | 69946971 | rs10898284 | 0.301 | 0.86 | -2.381 | 0.6 | -0.076 | -0.081 | -0.081 | 0.124 | 0.24 | 0.02 | 0.16 | 0.27 | 0.3 | 0.29 |
| 11 | 69947879 | rs1107443 | -0.048 | 0.694 | -2.487 | 0.464 | -0.056 | 0.002 | 0.002 | 0 | 0.44 | 0.12 | 0.44 | 0.33 | 0.41 | 0.37 |
| 11 | 69952562 | rs755016 | -0.093 | 0.798 | -2.857 | 0.345 | -0.097 | -0.071 | -0.071 | 1.135 | 0.35 | 0.07 | 0.4 | 0.58 | 0.41 | 0.26 |
| 11 | 69953729 | rs999320 | -0.336 | 0.421 | -2.146 | -0.231 | -0.001 | 0.031 | 0.031 | 0 | 0.57 | 0.6 | 0.55 | 0.68 | 0.6 | 0.4 |
| 11 | 69955637 | rs2515250 | -0.458 | 0.371 | -1.952 | -0.511 | -0.004 | -0.005 | -0.005 | 0 | 0.8 | 0.75 | 0.91 | 0.83 | 0.85 | 0.75 |
| 11 | 69957917 | rs9666157 | 0.271 | 0.252 | -1.057 | -0.977 | 0.025 | -0.091 | -0.091 | 0.856 | 0.19 | 0.27 | 0.32 | 0.01 | 0.24 | 0.1 |
| 11 | 69959183 | rs1006718 | 0.763 | -0.131 | -0.605 | -0.804 | -0.002 | -0.007 | -0.007 | 0.087 | 0.14 | 0.18 | 0.1 | 0.1 | 0.14 | 0.09 |
| 11 | 69964525 | rs2509141 | 0.498 | -0.164 | -0.591 | -0.805 | 0 | 0.001 | 0.001 | 0 | 0.64 | 0.62 | 0.79 | 0.7 | 0.69 | 0.64 |
| 11 | 69969470 | rs1893085 | -0.597 | -0.163 | -0.476 | -0.822 | 0 | 0 | 0 | 0.158 | 0.79 | 0.77 | 0.91 | 0.83 | 0.86 | 0.73 |
| 11 | 69970412 | rs2509160 | 0.482 | -0.181 | -0.216 | -0.811 | -0.009 | -0.017 | -0.017 | 0 | 0.65 | 0.57 | 0.81 | 0.72 | 0.71 | 0.64 |
| 11 | 69970637 | rs2515258 | 0.409 | -0.148 | -0.426 | -0.683 | 0.004 | 0.002 | 0.002 | 0.114 | 0.93 | 0.79 | 0.97 | 0.96 | 0.95 | 0.96 |
| 11 | 69972868 | rs2515278 | -0.017 | -0.199 | -0.457 | -0.717 | -0.004 | -0.005 | -0.005 | 0.239 | 0.79 | 0.74 | 0.9 | 0.83 | 0.85 | 0.75 |
| 11 | 69974423 | rs2515274 | -0.004 | -0.233 | -0.73 | -0.897 | 0.008 | 0.014 | 0.014 | 0.547 | 0.74 | 0.76 | 0.88 | 0.84 | 0.79 | 0.74 |
| 11 | 69974480 | rs2515273 | 0.101 | -0.169 | -0.726 | -0.893 | 0.007 | 0.013 | 0.013 | 0.507 | 0.73 | 0.75 | 0.88 | 0.84 | 0.79 | 0.74 |
| 11 | 69975570 | rs2509167 | -0.13 | 0.024 | -0.965 | -0.197 | 0.008 | -0.017 | -0.017 | 0 | 0.47 | 0.18 | 0.47 | 0.53 | 0.47 | 0.33 |
| 11 | 69976047 | rs2509170 | -0.004 | -0.067 | -0.998 | -0.03 | 0.008 | -0.029 | -0.029 | 0 | 0.45 | 0.16 | 0.47 | 0.56 | 0.45 | 0.3 |
| 11 | 69976424 | rs2509172 | -0.144 | -0.074 | -0.724 | -0.031 | 0.008 | -0.008 | -0.008 | 0 | 0.45 | 0.21 | 0.47 | 0.58 | 0.44 | 0.3 |
| 11 | 69978265 | rs2509174 | -0.056 | -0.469 | -0.714 | -0.513 | 0.015 | 0.019 | 0.019 | 0.614 | 0.31 | 0.22 | 0.15 | 0.09 | 0.23 | 0.16 |
| 11 | 69979560 | rs2509177 | -0.194 | -0.527 | -0.56 | -0.575 | -0.007 | -0.035 | -0.035 | 0 | 0.33 | 0.32 | 0.16 | 0.08 | 0.24 | 0.16 |
| 11 | 69993574 | rs11235420 | 0.981 | 0.8 | -0.793 | 0.028 | -0.045 | -0.056 | -0.056 | 0 | 0.18 | 0.1 | 0.24 | 0.44 | 0.27 | 0.17 |
| 11 | 69999252 | rs3781663 | -0.616 | 0.41 | -0.593 | 0.027 | 0 | -0.005 | -0.005 | 0 | 0.69 | 0.69 | 0.65 | 0.77 | 0.7 | 0.61 |
| 11 | 70001131 | rs11606378 | 2 | 0.568 | -0.644 | 0.063 | -0.052 | -0.073 | -0.073 | 0.092 | 0.16 | 0.07 | 0.21 | 0.45 | 0.23 | 0.17 |
| 11 | 70002689 | rs3781660 | 0.385 | 0.179 | -0.558 | -0.234 | 0.016 | 0.026 | 0.026 | 0 | 0.34 | 0.29 | 0.21 | 0.11 | 0.28 | 0.18 |
| 11 | 70002886 | rs3781659 | 1.756 | -0.078 | -0.901 | -0.351 | 0.015 | 0.072 | 0.072 | 0 | 0.16 | 0.02 | 0.09 | 0.05 | 0.14 | 0.12 |
| 11 | 70002987 | rs3781658 | 0.733 | -0.047 | -0.837 | -0.327 | 0.007 | 0.002 | 0.002 | 0 | 0.34 | 0.4 | 0.44 | 0.66 | 0.41 | 0.42 |
| 11 | 70004545 | rs875109 | -0.701 | -0.327 | -0.016 | -0.241 | 0.018 | 0.013 | 0.013 | 0.611 | 0.79 | 0.7 | 0.68 | 0.79 | 0.77 | 0.72 |
| 11 | 70004695 | rs875108 | -0.787 | -0.696 | 0.045 | -0.66 | 0.01 | -0.006 | -0.006 | 0 | 0.8 | 0.7 | 0.7 | 0.84 | 0.8 | 0.72 |
| 11 | 70005641 | rs875106 | 0.962 | -0.754 | 0.123 | -0.673 | -0.01 | 0.039 | 0.039 | 0 | 0.44 | 0.32 | 0.49 | 0.71 | 0.51 | 0.53 |
| 11 | 70006887 | rs10793022 | -0.826 | -0.664 | 0.187 | -0.61 | 0.002 | 0.003 | 0.003 | 1.876 | 0.53 | 0.58 | 0.45 | 0.32 | 0.47 | 0.44 |
| 11 | 70007311 | rs2276066 | 0.289 | -0.651 | 0.09 | -0.591 | 0.002 | 0.013 | 0.013 | 0 | 0.61 | 0.4 | 0.58 | 0.78 | 0.65 | 0.67 |
| 11 | 70009720 | rs11235460 | 1.602 | -0.564 | -0.114 | -0.42 | -0.077 | -0.099 | -0.099 | 0 | 0.28 | 0.11 | 0.29 | 0.51 | 0.37 | 0.29 |
| 11 | 70014478 | rs7127673 | 1.311 | -0.824 | -0.023 | -0.663 | -0.001 | -0.001 | -0.001 | 0 | 0.15 | 0.15 | 0.1 | 0.09 | 0.15 | 0.12 |
| 11 | 70015677 | rs10898826 | 0.132 | -0.926 | 0.151 | -0.634 | 0.002 | -0.007 | -0.007 | 0 | 0.65 | 0.6 | 0.61 | 0.75 | 0.66 | 0.64 |
| 11 | 70016058 | rs17333013 | 1.565 | -0.925 | 0.128 | -0.633 | -0.001 | 0.001 | 0.001 | 0 | 0.14 | 0.09 | 0.1 | 0.09 | 0.14 | 0.12 |
| 11 | 70016922 | rs7127005 | 2.205 | -0.831 | 0.145 | -0.6 | -0.074 | -0.094 | -0.094 | 0 | 0.25 | 0.08 | 0.26 | 0.52 | 0.34 | 0.26 |
| 11 | 70019182 | rs7936612 | -0.557 | -0.855 | 0.243 | -0.651 | 0.005 | -0.008 | -0.008 | 0 | 0.79 | 0.72 | 0.71 | 0.83 | 0.8 | 0.76 |
| 11 | 70019211 | rs10751200 | 1.552 | -0.762 | 0.093 | -0.541 | -0.008 | 0.023 | 0.023 | 1.086 | 0.17 | 0.33 | 0.21 | 0.17 | 0.15 | 0.25 |
| 11 | 70025292 | rs12417151 | 1.312 | -0.62 | -0.371 | -0.414 | 0.024 | 0.052 | 0.052 | 0 | 0.16 | 0.05 | 0.08 | 0.06 | 0.13 | 0.12 |
| 11 | 70027913 | rs7127129 | 1.338 | -0.501 | -0.35 | -0.345 | -0.053 | -0.059 | -0.059 | 0.093 | 0.28 | 0.15 | 0.29 | 0.52 | 0.37 | 0.28 |
| 11 | 70033583 | rs481845 | -1.408 | -0.614 | -0.466 | -0.338 | 0.021 | 0.033 | 0.033 | 0 | 0.83 | 0.9 | 0.92 | 0.94 | 0.87 | 0.88 |
| 11 | 70039921 | rs10898840 | 1.434 | -0.268 | -0.467 | -0.085 | -0.08 | -0.113 | -0.113 | 0 | 0.26 | 0.07 | 0.26 | 0.52 | 0.33 | 0.27 |
| 11 | 70045922 | rs10898847 | 1.131 | -0.388 | -0.223 | -0.299 | -0.054 | -0.083 | -0.083 | 0 | 0.3 | 0.14 | 0.28 | 0.55 | 0.36 | 0.28 |
| 11 | 70047335 | rs7109831 | -0.385 | -0.713 | 0.369 | -0.49 | 0 | 0.001 | 0.001 | 0.99 | 0.56 | 0.54 | 0.45 | 0.64 | 0.54 | 0.46 |
| 11 | 70051966 | rs10793035 | -0.062 | -0.827 | 0.658 | -0.969 | -0.006 | 0.049 | 0.049 | 0.035 | 0.52 | 0.33 | 0.43 | 0.65 | 0.52 | 0.44 |
| 11 | 70053487 | rs1131715 | -0.45 | -0.802 | 0.471 | -0.957 | -0.001 | 0.004 | 0.004 | 0.975 | 0.56 | 0.58 | 0.45 | 0.65 | 0.54 | 0.44 |
| 11 | 70053547 | rs3740720 | 1.415 | -0.64 | 0.045 | -0.89 | -0.077 | -0.116 | -0.116 | 0.713 | 0.28 | 0.11 | 0.29 | 0.55 | 0.37 | 0.27 |
| 11 | 70058679 | rs615363 | -0.74 | -0.787 | 0.634 | -1.52 | 0.005 | 0.068 | 0.068 | 0 | 0.8 | 0.94 | 0.9 | 0.93 | 0.82 | 0.75 |
| 11 | 70059105 | rs10751209 | 0.741 | -0.667 | 0.841 | -1.596 | 0 | -0.002 | -0.002 | 0 | 0.58 | 0.42 | 0.56 | 0.76 | 0.57 | 0.54 |
| 11 | 70072089 | rs579614 | 1.401 | -0.77 | 1.207 | -1.273 | 0.052 | 0.156 | 0.156 | 0.476 | 0.63 | 0.16 | 0.75 | 0.88 | 0.73 | 0.68 |
| 11 | 70077881 | rs2155456 | -0.302 | -0.632 | 1.29 | -1.189 | 0.01 | 0.038 | 0.038 | 0 | 0.8 | 0.87 | 0.9 | 0.93 | 0.81 | 0.75 |
| 11 | 70079002 | rs555745 | 0.916 | -0.6 | 1.502 | -1.135 | -0.03 | -0.035 | -0.035 | 0.188 | 0.63 | 0.54 | 0.68 | 0.77 | 0.75 | 0.56 |

| | | | | | | | | | | | | | | | | |
|---|---|---|---|---|---|---|---|---|---|---|---|---|---|---|---|---|
| 11 | 70079854 | rs17160035 | 0.299 | -0.501 | 0.547 | -1.046 | -0.019 | -0.003 | -0.003 | 0 | 0.06 | 0.05 | 0.02 | 0.05 | 0.01 | 0.02 |
| 11 | 70081665 | rs565473 | 0.204 | -0.337 | 0.535 | -1.03 | 0.002 | -0.034 | -0.034 | 0.141 | 0.72 | 0.6 | 0.66 | 0.82 | 0.7 | 0.61 |
| 11 | 70089007 | rs507119 | 0.728 | -0.258 | -0.28 | -0.96 | 0.011 | 0.02 | 0.02 | 0 | 0.6 | 0.33 | 0.61 | 0.77 | 0.68 | 0.54 |
| 11 | 70091971 | rs7118496 | -0.639 | -0.119 | -0.924 | -0.778 | 0.01 | 0.019 | 0.019 | 0.351 | 0.12 | 0.09 | 0.06 | 0.05 | 0.07 | 0.06 |
| 11 | 70096585 | rs520722 | -0.895 | -0.027 | -0.938 | -0.803 | 0.043 | 0.103 | 0.103 | 1.17 | 0.42 | 0.79 | 0.4 | 0.23 | 0.32 | 0.45 |
| 11 | 70111532 | rs687660 | -0.918 | 0.444 | -0.543 | -1.3 | -0.004 | 0.021 | 0.021 | 0 | 0.38 | 0.72 | 0.49 | 0.15 | 0.34 | 0.34 |
| 11 | 70118489 | rs546502 | -0.316 | 0.588 | -0.154 | -1.481 | -0.001 | 0.036 | 0.036 | 0 | 0.22 | 0.15 | 0.25 | 0.11 | 0.22 | 0.27 |
| 11 | 70120501 | rs644396 | -1.898 | 0.414 | -0.773 | -1.267 | -0.106 | -0.036 | -0.036 | 0 | 0.57 | 0.94 | 0.62 | 0.38 | 0.48 | 0.48 |
| 11 | 70159690 | rs7116059 | 0.051 | 0.677 | 0.297 | -1.518 | 0.069 | -0.044 | -0.044 | 0 | 0.13 | 0.52 | 0.23 | 0.04 | 0.12 | 0.06 |
| 11 | 70178578 | rs11235935 | -1.27 | 0.678 | 0.282 | -1.523 | -0.011 | 0.096 | 0.096 | 0.032 | 0.37 | 0.81 | 0.49 | 0.1 | 0.35 | 0.34 |
| 11 | 70179672 | rs2282481 | -1.148 | 0.658 | 0.214 | -1.54 | -0.003 | 0.016 | 0.016 | 0 | 0.36 | 0.7 | 0.48 | 0.15 | 0.35 | 0.34 |
| 11 | 70185325 | rs1061328 | 0.052 | 0.643 | 0.242 | -1.541 | 0.07 | -0.044 | -0.044 | 0.022 | 0.13 | 0.52 | 0.23 | 0.04 | 0.12 | 0.06 |
| 11 | 70186614 | rs3900797 | 0.052 | 0.643 | 0.242 | -1.541 | 0.07 | -0.044 | -0.044 | 0.005 | 0.13 | 0.52 | 0.23 | 0.04 | 0.12 | 0.06 |
| 11 | 70188861 | rs7129635 | -0.233 | 0.648 | 0.211 | -1.547 | 0 | 0.042 | 0.042 | 0 | 0.24 | 0.17 | 0.25 | 0.11 | 0.23 | 0.28 |
| 11 | 70190058 | rs3781648 | 0.195 | 0.657 | 0.807 | -1.524 | 0.074 | -0.033 | -0.033 | 0 | 0.12 | 0.36 | 0.21 | 0.04 | 0.12 | 0.06 |
| 11 | 70190320 | rs3781646 | 0.131 | 0.677 | 0.752 | -1.531 | -0.005 | 0.056 | 0.056 | 0 | 0.21 | 0.08 | 0.22 | 0.11 | 0.2 | 0.27 |
| 11 | 70218548 | rs10466703 | -0.077 | 0.498 | 0.829 | -1.586 | 0.07 | -0.043 | -0.043 | 0 | 0.13 | 0.52 | 0.23 | 0.04 | 0.12 | 0.06 |
| 11 | 70229171 | rs552282 | -1.145 | 0.407 | 0.291 | -1.606 | -0.005 | 0.027 | 0.027 | 0 | 0.37 | 0.72 | 0.47 | 0.1 | 0.35 | 0.34 |
| 11 | 70241935 | rs583397 | -0.047 | 0.399 | -0.118 | -1.478 | 0.003 | 0.035 | 0.035 | 0 | 0.21 | 0.16 | 0.23 | 0.07 | 0.2 | 0.27 |
| 11 | 70248900 | rs4367974 | -0.838 | 0.295 | 0.157 | -1.654 | 0.007 | -0.041 | -0.041 | 0 | 0.17 | 0.72 | 0.28 | 0.05 | 0.14 | 0.11 |
| 11 | 70253389 | rs2298396 | -0.093 | 0.101 | 0.244 | -1.601 | -0.021 | 0.015 | 0.015 | 1.097 | 0.22 | 0.09 | 0.27 | 0.11 | 0.24 | 0.14 |
| 11 | 70253475 | rs2298397 | -0.016 | 0.098 | -0.012 | -1.603 | -0.055 | 0.02 | 0.02 | 1.315 | 0.21 | 0.03 | 0.27 | 0.11 | 0.24 | 0.14 |
| 11 | 70257669 | rs679455 | -1.757 | -0.027 | -0.094 | -1.606 | -0.013 | 0.177 | 0.177 | 0 | 0.51 | 0.81 | 0.46 | 0.16 | 0.44 | 0.44 |
| 11 | 70262379 | rs12800598 | -0.751 | 0.215 | -0.703 | -1.642 | -0.049 | 0.048 | 0.048 | 0.573 | 0.32 | 0.07 | 0.32 | 0.15 | 0.33 | 0.17 |
| 11 | 70269777 | rs592501 | -0.393 | 0.134 | -0.263 | -1.732 | -0.007 | -0.072 | -0.072 | 0 | 0.25 | 0.3 | 0.14 | 0.01 | 0.2 | 0.3 |
| 11 | 70276009 | rs12804757 | -0.204 | 0.13 | -0.233 | -1.727 | -0.026 | 0.021 | 0.021 | 0.602 | 0.22 | 0.11 | 0.29 | 0.11 | 0.26 | 0.14 |
| 11 | 70279766 | rs643301 | -0.48 | 0.227 | -0.621 | -1.652 | -0.002 | -0.054 | -0.054 | 0.002 | 0.25 | 0.24 | 0.13 | 0.01 | 0.2 | 0.3 |
| 11 | 70280992 | rs3802780 | -0.411 | 0.248 | -0.656 | -1.642 | -0.007 | -0.072 | -0.072 | 0.597 | 0.25 | 0.3 | 0.14 | 0.01 | 0.2 | 0.3 |
| 11 | 70282629 | rs6894 | -0.517 | 0.739 | -1.076 | -0.815 | -0.08 | 0.012 | 0.012 | 1.152 | 0.21 | 0.02 | 0.27 | 0.11 | 0.24 | 0.14 |
| 11 | 70295897 | rs11236302 | -0.573 | -0.425 | -0.51 | -0.715 | -0.034 | -0.027 | -0.027 | 0 | 0.23 | 0.39 | 0.12 | 0.18 | 0.15 | 0.33 |
| 11 | 70307800 | rs682670 | -0.224 | 0.072 | -1.316 | -0.912 | -0.091 | 0.035 | 0.035 | 0.033 | 0.35 | 0.12 | 0.44 | 0.73 | 0.45 | 0.38 |
| 11 | 70320069 | rs516941 | -0.048 | -0.05 | -1.368 | -0.667 | -0.067 | -0.039 | -0.039 | 0 | 0.29 | 0.08 | 0.4 | 0.66 | 0.38 | 0.35 |
| 11 | 70321096 | rs567105 | -0.055 | -0.053 | -1.394 | -0.625 | -0.075 | -0.052 | -0.052 | 0 | 0.29 | 0.07 | 0.39 | 0.66 | 0.38 | 0.34 |
| 11 | 70327305 | rs579298 | -0.722 | -1.021 | -1.122 | -0.996 | -0.002 | -0.003 | -0.003 | 0 | 0.57 | 0.4 | 0.77 | 0.82 | 0.77 | 0.64 |
| 11 | 70334494 | rs10899089 | -0.011 | -1.081 | -0.667 | -1.049 | -0.001 | -0.018 | -0.018 | 0 | 0.6 | 0.48 | 0.55 | 0.32 | 0.6 | 0.56 |
| 11 | 70352086 | rs12364162 | 0.505 | -2.125 | -0.732 | -0.98 | -0.045 | -0.035 | -0.035 | 0 | 0.36 | 0.41 | 0.2 | 0.19 | 0.2 | 0.3 |
| 11 | 70353848 | rs2000605 | 0.286 | -1.068 | -1.507 | 0.283 | -0.073 | -0.175 | -0.175 | 0 | 0.32 | 0.04 | 0.56 | 0.43 | 0.6 | 0.43 |
| 11 | 70369069 | rs12222819 | 0.125 | -0.621 | -1.033 | 0.768 | -0.038 | -0.064 | -0.064 | 0 | 0.1 | 0.01 | 0.13 | 0.34 | 0.1 | 0.08 |
| 11 | 70374786 | rs2096818 | 0.276 | -0.989 | -1.647 | 0.281 | -0.082 | -0.11 | -0.11 | 0.78 | 0.37 | 0.02 | 0.57 | 0.34 | 0.6 | 0.42 |
| 11 | 70375605 | rs12803092 | -0.026 | -1.754 | -1.214 | -0.138 | 0.007 | -0.005 | -0.005 | 1.606 | 0.36 | 0.33 | 0.28 | 0.64 | 0.25 | 0.39 |
| 11 | 70376203 | rs11826745 | 0.21 | -1.482 | -1.437 | 0.211 | -0.003 | 0.033 | 0.033 | 0 | 0.09 | 0.06 | 0.13 | 0.42 | 0.1 | 0.1 |
| 11 | 70378306 | rs11236566 | 0.177 | -1.909 | -0.955 | -0.721 | 0.093 | 0.111 | 0.111 | 0.716 | 0.49 | 0.27 | 0.72 | 0.78 | 0.71 | 0.54 |
| 11 | 70379264 | rs12794889 | 0.012 | -1.187 | -0.995 | -0.16 | 0.037 | 0.032 | 0.032 | 0 | 0.26 | 0.17 | 0.15 | 0.19 | 0.15 | 0.15 |
| 11 | 70383722 | rs10899147 | -0.223 | -0.148 | -0.671 | 0.044 | 0 | 0.01 | 0.01 | 0 | 0.37 | 0.31 | 0.55 | 0.72 | 0.35 | 0.6 |
| 11 | 70387122 | rs4980608 | -0.482 | -0.054 | -0.184 | 0.09 | -0.005 | -0.036 | -0.036 | 1.224 | 0.42 | 0.49 | 0.58 | 0.8 | 0.38 | 0.61 |
| 11 | 70387286 | rs11236585 | -0.935 | 0.116 | -0.078 | 0.687 | 0.107 | 0.148 | 0.148 | 0 | 0.07 | 0.33 | 0.23 | 0.1 | 0.11 | 0.12 |
| 11 | 70388251 | rs12271322 | 0.166 | -0.016 | -0.597 | 0.694 | 0.024 | -0.076 | -0.076 | 0 | 0.32 | 0.12 | 0.33 | 0.58 | 0.25 | 0.47 |
| 11 | 70394936 | rs10899153 | -0.358 | 1.048 | -0.522 | 1.039 | 0.025 | 0.077 | 0.077 | 0 | 0.11 | 0.15 | 0.2 | 0.38 | 0.16 | 0.14 |
| 11 | 70395192 | rs10793137 | -0.692 | 0.432 | -0.424 | -0.103 | -0.01 | -0.074 | -0.074 | 1.036 | 0.57 | 0.49 | 0.8 | 0.88 | 0.62 | 0.65 |
| 11 | 70398245 | rs7941537 | 0.16 | 0.573 | -0.174 | 0.155 | 0.025 | 0.124 | 0.124 | 0 | 0.85 | 0.7 | 0.77 | 0.53 | 0.83 | 0.83 |
| 11 | 70403579 | rs7945850 | 0.598 | 0.549 | -0.004 | 0.04 | 0.018 | 0.018 | 0.018 | 0 | 0.67 | 0.57 | 0.46 | 0.39 | 0.6 | 0.69 |
| 11 | 70404317 | rs10899156 | 0.666 | 0.524 | 0.142 | 0.028 | 0.024 | 0.023 | 0.023 | 0 | 0.66 | 0.51 | 0.46 | 0.39 | 0.6 | 0.69 |
| 11 | 70405829 | rs7106631 | 0.2 | 0.593 | 0.039 | -0.033 | 0.06 | 0.176 | 0.176 | 0 | 0.86 | 0.68 | 0.77 | 0.53 | 0.83 | 0.83 |
| 11 | 70410130 | rs12786771 | -0.208 | 0.244 | -0.562 | -0.78 | -0.05 | 0.11 | 0.11 | 0 | 0.38 | 0.18 | 0.42 | 0.11 | 0.5 | 0.38 |
| 11 | 70413129 | rs4272805 | -0.334 | 0.353 | -0.347 | -0.464 | -0.002 | 0 | 0 | 0 | 0.78 | 0.69 | 0.66 | 0.79 | 0.77 | 0.85 |
| 11 | 70420282 | rs11236680 | 0.223 | 0.238 | -0.557 | -0.502 | -0.013 | 0.024 | 0.024 | 5.127 | 0.41 | 0.32 | 0.62 | 0.75 | 0.44 | 0.46 |
| 11 | 70421299 | rs4980613 | 0.33 | 0.054 | -0.802 | -0.264 | 0.002 | -0.001 | -0.001 | 1.905 | 0.52 | 0.55 | 0.45 | 0.47 | 0.52 | 0.7 |
| 11 | 70424188 | rs11236708 | 0.444 | -0.111 | -1.178 | -0.473 | 0.008 | 0.016 | 0.016 | 3.821 | 0.18 | 0.13 | 0.1 | 0.1 | 0.17 | 0.13 |
| 11 | 70424257 | rs948196 | -0.718 | 0.392 | -1.233 | -0.703 | 0.004 | -0.003 | -0.003 | 0 | 0.85 | 0.85 | 0.89 | 0.86 | 0.83 | 0.94 |
| 11 | 70424411 | rs10899208 | 1.702 | 0.307 | -1.86 | -0.621 | -0.004 | -0.077 | -0.077 | 0.024 | 0.23 | 0.1 | 0.25 | 0.44 | 0.23 | 0.38 |
| 11 | 70424943 | rs948194 | -0.219 | 0.267 | -1.868 | -0.651 | -0.002 | 0.014 | 0.014 | 0.533 | 0.67 | 0.73 | 0.79 | 0.76 | 0.65 | 0.81 |
| 11 | 70425843 | rs7106873 | -0.022 | 0.246 | -1.827 | -0.751 | -0.001 | 0.011 | 0.011 | 0.051 | 0.43 | 0.63 | 0.54 | 0.31 | 0.43 | 0.43 |
| 11 | 70426375 | rs12225214 | 0.643 | 0.444 | -1.689 | -0.68 | -0.02 | -0.008 | -0.008 | 0.06 | 0.2 | 0.08 | 0.4 | 0.15 | 0.27 | 0.13 |
| 11 | 70427863 | rs11236730 | 1.251 | 0.561 | -1.438 | -0.699 | 0.006 | -0.004 | -0.004 | 0.155 | 0.15 | 0.12 | 0.07 | 0 | 0.14 | 0.12 |
| 11 | 70429252 | rs10459049 | -0.871 | 0.716 | -1.395 | -0.51 | 0.004 | -0.002 | -0.002 | 0 | 0.85 | 0.85 | 0.89 | 0.85 | 0.82 | 0.91 |
| 11 | 70429921 | rs11820925 | 0.844 | 0.711 | -1.157 | -0.147 | 0.009 | 0.003 | 0.003 | 7.208 | 0.17 | 0.12 | 0.07 | 0 | 0.14 | 0.12 |
| 11 | 70433687 | rs7945862 | -0.563 | 1.051 | -1.193 | 0.509 | -0.005 | 0.007 | 0.007 | 3.014 | 0.51 | 0.7 | 0.61 | 0.51 | 0.52 | 0.39 |
| 11 | 70434594 | rs7949744 | 0.217 | 1.079 | -1.05 | 0.559 | -0.02 | -0.033 | -0.033 | 0 | 0.34 | 0.52 | 0.19 | 0.2 | 0.26 | 0.21 |
| 11 | 70439364 | rs948192 | -1.187 | 1.208 | -1.348 | 0.453 | -0.008 | 0.059 | 0.059 | 0 | 0.7 | 0.83 | 0.85 | 0.92 | 0.72 | 0.72 |

| | | | | | | | | | | | | | | | | |
|---|---|---|---|---|---|---|---|---|---|---|---|---|---|---|---|---|
| 11 | 70444244 | rs11236773 | 0.97 | 1.303 | -1.292 | 1.054 | -0.003 | -0.033 | -0.033 | 3.307 | 0.19 | 0.12 | 0.24 | 0.39 | 0.2 | 0.32 |
| 11 | 70445161 | rs12363289 | -0.085 | 1.089 | -1.107 | 0.975 | -0.011 | -0.016 | -0.016 | 1.519 | 0.17 | 0.15 | 0.41 | 0.31 | 0.27 | 0.15 |
| 11 | 70450688 | rs7939551 | -0.046 | 0.918 | -0.844 | 0.65 | -0.001 | 0.021 | 0.021 | 0 | 0.51 | 0.38 | 0.67 | 0.66 | 0.49 | 0.51 |
| 11 | 70452134 | rs10899236 | -0.243 | 0.613 | -0.641 | 0.983 | -0.011 | 0.102 | 0.102 | 0.02 | 0.48 | 0.2 | 0.29 | 0.37 | 0.55 | 0.5 |
| 11 | 70452995 | rs948191 | -0.095 | 0.94 | -0.78 | 0.91 | 0.01 | 0.02 | 0.02 | 0 | 0.52 | 0.37 | 0.68 | 0.63 | 0.53 | 0.49 |
| 11 | 70453705 | rs4980543 | -0.143 | 0.938 | -0.713 | 0.837 | 0.003 | 0.006 | 0.006 | 0 | 0.5 | 0.43 | 0.65 | 0.57 | 0.5 | 0.48 |
| 11 | 70456768 | rs12421725 | 0.157 | 0.243 | -0.765 | 0.1 | -0.004 | -0.005 | -0.005 | 0.029 | 0.28 | 0.28 | 0.16 | 0.25 | 0.23 | 0.16 |
| 11 | 70460751 | rs11236856 | 0.546 | -0.31 | -1.204 | -0.353 | 0.047 | -0.021 | -0.021 | 0 | 0.25 | 0.1 | 0.2 | 0.23 | 0.19 | 0.32 |
| 11 | 70461590 | rs12225113 | -0.181 | 0.034 | -1.047 | -0.649 | 0.021 | 0.066 | 0.066 | 0 | 0.48 | 0.31 | 0.36 | 0.26 | 0.42 | 0.42 |
| 11 | 70463510 | rs11236898 | 0.739 | -0.115 | -1.225 | -0.54 | 0.051 | -0.011 | -0.011 | 1.463 | 0.25 | 0.13 | 0.2 | 0.23 | 0.17 | 0.32 |
| 11 | 70470633 | rs4980629 | 0.085 | 0.677 | -0.687 | -0.307 | -0.006 | 0.027 | 0.027 | 0 | 0.56 | 0.35 | 0.4 | 0.31 | 0.53 | 0.51 |
| 11 | 70476810 | rs557324 | -0.66 | 0.698 | -0.319 | -0.487 | 0.026 | 0.018 | 0.018 | 0 | 0.71 | 0.58 | 0.51 | 0.31 | 0.63 | 0.57 |
| 11 | 70478558 | rs1207142 | -0.5 | 0.363 | -0.505 | -0.384 | 0.021 | -0.001 | -0.001 | 0 | 0.65 | 0.5 | 0.48 | 0.31 | 0.56 | 0.51 |
| 11 | 70478801 | rs11236989 | -0.782 | 0.073 | -0.217 | -0.31 | -0.011 | -0.05 | -0.05 | 0 | 0.13 | 0.17 | 0.08 | 0 | 0.08 | 0.07 |
| 11 | 70490207 | rs515458 | -0.994 | -0.034 | -0.064 | 0.024 | -0.025 | 0.014 | 0.014 | 0.836 | 0.26 | 0.55 | 0.25 | 0.32 | 0.22 | 0.18 |
| 11 | 70490827 | rs493026 | 0.432 | -0.154 | -0.195 | -0.581 | -0.007 | -0.049 | -0.049 | 0 | 0.82 | 0.8 | 0.91 | 1 | 0.89 | 0.86 |
| 11 | 70496641 | rs500335 | -0.328 | 0.722 | -0.106 | -0.547 | -0.002 | 0.001 | 0.001 | 0 | 0.26 | 0.27 | 0.19 | 0.26 | 0.33 | 0.18 |
| 11 | 70515963 | rs2186567 | 0.242 | 0.59 | -0.483 | -0.774 | -0.008 | 0.048 | 0.048 | 0 | 0.08 | 0.01 | 0.05 | 0 | 0.09 | 0.11 |
| 11 | 70517465 | rs514519 | -0.957 | 0.183 | -0.353 | -1.412 | 0.019 | 0.026 | 0.026 | 0 | 0.55 | 0.59 | 0.49 | 0.31 | 0.66 | 0.46 |
| 11 | 70517927 | rs12575685 | 0.15 | 0.466 | -0.603 | -1.271 | -0.057 | -0.028 | -0.028 | 0 | 0.23 | 0.03 | 0.26 | 0.26 | 0.29 | 0.22 |
| 11 | 70540451 | rs12287093 | -0.036 | -0.279 | 0.001 | -1.396 | -0.031 | -0.102 | -0.102 | 0 | 0.14 | 0.4 | 0.1 | 0 | 0.08 | 0.1 |
| 11 | 70548853 | rs10899388 | -0.844 | 0.451 | -0.255 | -1.789 | -0.011 | 0.047 | 0.047 | 0.241 | 0.43 | 0.71 | 0.41 | 0.29 | 0.53 | 0.4 |
| 11 | 70551728 | rs17428526 | 1.454 | -0.094 | -1.014 | -1.608 | -0.046 | 0.008 | 0.008 | 0 | 0.15 | 0.03 | 0.11 | 0.11 | 0.17 | 0.04 |
| 11 | 70552397 | rs10501403 | 0.096 | -0.126 | -0.653 | -1.54 | -0.087 | -0.001 | -0.001 | 0 | 0.29 | 0.15 | 0.31 | 0.29 | 0.45 | 0.32 |
| 11 | 70560051 | rs11237269 | 0.016 | -0.107 | -0.457 | -1.504 | -0.02 | 0.006 | 0.006 | 0 | 0.32 | 0.26 | 0.32 | 0.3 | 0.47 | 0.33 |
| 11 | 70564316 | rs1783611 | -0.127 | -0.696 | -0.881 | -1.395 | 0.014 | 0.028 | 0.028 | 0 | 0.57 | 0.78 | 0.56 | 0.62 | 0.59 | 0.52 |
| 11 | 70565076 | rs947854 | -0.03 | -0.667 | -0.642 | -1.118 | 0.009 | 0.006 | 0.006 | 0 | 0.35 | 0.27 | 0.38 | 0.32 | 0.3 | 0.4 |
| 11 | 70565151 | rs747172 | 0.588 | -0.576 | -0.941 | -1.1 | 0.002 | -0.007 | -0.007 | 0 | 0.25 | 0.15 | 0.33 | 0.26 | 0.22 | 0.2 |
| 11 | 70565438 | rs1207229 | -0.629 | -0.471 | -1.297 | -0.993 | 0.069 | 0.103 | 0.103 | 0 | 0.51 | 0.85 | 0.5 | 0.32 | 0.38 | 0.5 |
| 11 | 70569648 | rs4980634 | -1.216 | -0.542 | -1.221 | -0.696 | -0.103 | -0.133 | -0.133 | 0 | 0.63 | 0.98 | 0.59 | 0.43 | 0.6 | 0.67 |
| 11 | 70572020 | rs11237298 | 0.833 | -0.831 | -0.881 | -1.03 | 0.016 | 0.015 | 0.015 | 0 | 0.16 | 0.09 | 0.07 | 0.01 | 0.11 | 0.3 |
| 11 | 70573926 | rs1783605 | -0.233 | -0.768 | -0.061 | -0.976 | 0.002 | -0.001 | -0.001 | 0.869 | 0.38 | 0.42 | 0.44 | 0.36 | 0.41 | 0.35 |
| 11 | 70579515 | rs12289782 | 0.48 | -0.719 | -0.319 | -1.026 | 0.026 | 0.03 | 0.03 | 0 | 0.18 | 0.08 | 0.07 | 0.02 | 0.11 | 0.31 |
| 11 | 70583006 | rs920957 | 0.402 | -0.601 | 0.049 | -0.847 | 0.013 | 0.004 | 0.004 | 0 | 0.3 | 0.32 | 0.41 | 0.32 | 0.38 | 0.31 |
| 11 | 70587936 | rs483039 | 0.185 | -0.665 | 0.268 | -0.863 | -0.001 | 0.005 | 0.005 | 0 | 0.08 | 0.33 | 0.05 | 0.03 | 0.03 | 0.02 |
| 11 | 70592233 | rs525128 | -0.277 | -0.59 | 0.092 | -0.843 | 0.011 | -0.002 | -0.002 | 0.02 | 0.37 | 0.57 | 0.45 | 0.34 | 0.41 | 0.33 |
| 11 | 70603064 | rs531827 | -1.192 | -0.634 | 0.065 | -0.803 | -0.003 | 0.008 | 0.008 | 0 | 0.54 | 0.67 | 0.61 | 0.44 | 0.5 | 0.63 |
| 11 | 70605728 | rs4980638 | -0.754 | -0.553 | -0.356 | -0.748 | -0.006 | 0 | 0 | 0 | 0.93 | 0.74 | 0.96 | 0.97 | 0.97 | 0.98 |
| 11 | 70606500 | rs10899444 | -0.137 | -0.436 | -0.476 | -0.694 | -0.032 | -0.035 | -0.035 | 0.003 | 0.76 | 0.65 | 0.8 | 0.87 | 0.86 | 0.67 |
| 11 | 70609284 | rs10899446 | 0.578 | 0.073 | -1.215 | -0.251 | 0.001 | -0.025 | -0.025 | 0 | 0.37 | 0.22 | 0.34 | 0.56 | 0.4 | 0.33 |
| 11 | 70613763 | rs1562191 | 0.275 | 0.194 | -0.939 | 0.062 | -0.017 | 0.073 | 0.073 | 0.285 | 0.51 | 0.23 | 0.48 | 0.65 | 0.52 | 0.38 |
| 11 | 70616176 | rs4980552 | 0.652 | 0.096 | 0.169 | 0.483 | -0.007 | 0.076 | 0.076 | 0 | 0.92 | 0.67 | 0.88 | 0.88 | 0.97 | 0.96 |
| 11 | 70627159 | rs731616 | 0.981 | 0.907 | 0.748 | 0.134 | 0.081 | -0.062 | -0.062 | 4.55 | 0.89 | 0.4 | 0.87 | 0.98 | 0.82 | 0.94 |
| 11 | 70628945 | rs10793307 | 0.963 | 0.451 | -0.503 | 0.164 | -0.121 | -0.147 | -0.147 | 2.478 | 0.35 | 0.02 | 0.33 | 0.44 | 0.38 | 0.32 |
| 11 | 70645921 | rs2016495 | -0.107 | 0.362 | 0.628 | 0.216 | 0.071 | -0.019 | -0.019 | 0 | 0.07 | 0.25 | 0.09 | 0.02 | 0.12 | 0.05 |
| 11 | 70647207 | rs11237542 | -0.935 | 0.21 | 0.532 | 0.27 | -0.019 | -0.008 | -0.008 | 0 | 0.17 | 0.08 | 0.26 | 0.16 | 0.21 | 0.19 |
| 11 | 70647963 | rs2921325 | -2.033 | 0.173 | 0.638 | 0.211 | 0.005 | 0.007 | 0.007 | 0 | 0.23 | 0.66 | 0.4 | 0.24 | 0.23 | 0.2 |
| 11 | 70652467 | rs3019846 | -1.938 | 0.004 | 0.898 | -0.227 | 0.072 | -0.064 | -0.064 | 0.143 | 0.12 | 0.55 | 0.13 | 0.02 | 0.14 | 0.06 |
| 11 | 70653551 | rs2921329 | -0.919 | 0.124 | 0.77 | -0.168 | 0.089 | -0.037 | -0.037 | 0.02 | 0.09 | 0.36 | 0.11 | 0.02 | 0.13 | 0.05 |
| 11 | 70664218 | rs2015997 | -0.973 | -1.191 | -0.015 | -0.321 | -0.061 | -0.059 | -0.059 | 0 | 0.28 | 0.59 | 0.12 | 0.1 | 0.14 | 0.3 |
| 11 | 70665521 | rs3017493 | 0.787 | -0.823 | -0.488 | -0.234 | 0.023 | 0.03 | 0.03 | 0 | 0.81 | 0.87 | 0.94 | 0.95 | 0.89 | 0.87 |
| 11 | 70671860 | rs12295625 | -0.743 | -0.772 | -0.477 | 0.016 | 0.019 | 0.026 | 0.026 | 0 | 0.14 | 0.06 | 0.05 | 0.05 | 0.09 | 0.12 |
| 11 | 70674679 | rs2921331 | 0.668 | 0.353 | 0.287 | 0.432 | 0.166 | 0.092 | 0.092 | 0 | 0.88 | 0.52 | 0.76 | 0.81 | 0.78 | 0.95 |
| 11 | 70678440 | rs3017499 | 1.141 | 0.285 | 0.167 | 0.406 | 0.039 | -0.004 | -0.004 | 0 | 0.74 | 0.49 | 0.7 | 0.75 | 0.67 | 0.83 |
| 11 | 70683494 | rs7946750 | 1.648 | 0.289 | -0.044 | 1.285 | 0.006 | -0.001 | -0.001 | 0 | 0.61 | 0.4 | 0.39 | 0.4 | 0.55 | 0.61 |
| 11 | 70687916 | rs930585 | 1.1 | 0.316 | -0.009 | 1.407 | 0.014 | 0.016 | 0.016 | 0 | 0.69 | 0.43 | 0.66 | 0.64 | 0.66 | 0.77 |
| 11 | 70692076 | rs7941869 | -1.309 | 0.45 | -0.135 | 2.014 | -0.042 | 0.146 | 0.146 | 0 | 0.23 | 0.46 | 0.37 | 0.4 | 0.16 | 0.32 |
| 11 | 70693797 | rs11237691 | 0.777 | 0.483 | -1.088 | 1.604 | -0.024 | -0.052 | -0.052 | 0 | 0.26 | 0.05 | 0.27 | 0.34 | 0.27 | 0.31 |
| 11 | 70712156 | rs10751308 | -0.393 | 1.127 | -0.634 | 1.314 | -0.042 | -0.019 | -0.019 | 0.512 | 0.92 | 0.99 | 0.87 | 0.88 | 0.77 | 0.96 |
| 11 | 70716035 | rs1207273 | 0.733 | 0.933 | -0.347 | 1.398 | 0.077 | 0.215 | 0.215 | 1.054 | 0.89 | 0.62 | 0.87 | 0.69 | 0.85 | 0.89 |
| 11 | 70725235 | rs10899616 | 0.373 | -0.178 | -1.201 | 0.84 | -0.05 | -0.06 | -0.06 | 0.06 | 0.32 | 0.02 | 0.22 | 0.36 | 0.3 | 0.31 |
| 11 | 70730709 | rs686805 | 0.062 | -0.352 | -0.289 | 0.923 | -0.035 | -0.041 | -0.041 | 0 | 0.19 | 0.37 | 0.1 | 0.13 | 0.12 | 0.24 |
| 11 | 70733140 | rs7935635 | -0.274 | 0.09 | 0.155 | 1.398 | 0.032 | 0.15 | 0.15 | 0.115 | 0.05 | 0.29 | 0.04 | 0.1 | 0.02 | 0.09 |
| 11 | 70735085 | rs1198802 | -0.011 | 0.086 | 0.063 | 1.396 | 0.05 | 0.179 | 0.179 | 0.249 | 0.94 | 0.65 | 0.95 | 0.9 | 0.98 | 0.91 |
| 11 | 70738321 | rs11237848 | 0.489 | 0.234 | -0.362 | 1.641 | -0.039 | -0.08 | -0.08 | 0 | 0.11 | 0.01 | 0.34 | 0.28 | 0.14 | 0.22 |
| 11 | 70741036 | rs684636 | 0.229 | 0.197 | 0.294 | 1.046 | 0.062 | 0.188 | 0.188 | 0.643 | 0.94 | 0.58 | 0.95 | 0.9 | 0.98 | 0.9 |
| 11 | 70754920 | rs7927376 | 0.486 | 0.268 | -0.638 | 0.94 | 0.02 | 0.017 | 0.017 | 0 | 0.71 | 0.21 | 0.77 | 0.72 | 0.78 | 0.68 |
| 11 | 70759123 | rs6592864 | -0.764 | 0.263 | -0.664 | 0.907 | -0.002 | -0.001 | -0.001 | 0 | 0.29 | 0.68 | 0.21 | 0.28 | 0.22 | 0.32 |
| 11 | 70764308 | rs11237981 | -0.111 | 0.868 | -1.13 | 0.648 | -0.001 | -0.016 | -0.016 | 0.043 | 0.22 | 0.3 | 0.17 | 0.17 | 0.2 | 0.23 |

| | | | | | | | | | | | | | | | | |
|---|---|---|---|---|---|---|---|---|---|---|---|---|---|---|---|---|
| 11 | 70789000 | rs11238042 | -0.678 | 1.566 | -2.261 | 1.219 | -0.107 | -0.078 | -0.078 | 6.469 | 0.25 | 0.02 | 0.22 | 0.31 | 0.29 | 0.28 |
| 11 | 70807773 | rs7120950 | -0.658 | 1.412 | -2.166 | 1.595 | -0.086 | -0.15 | -0.15 | 0 | 0.2 | 0.01 | 0.38 | 0.49 | 0.24 | 0.26 |
| 11 | 70813928 | rs11823567 | 0.153 | 1.103 | -1.794 | 0.468 | 0.013 | -0.083 | -0.083 | 0 | 0.3 | 0.42 | 0.2 | 0.06 | 0.32 | 0.38 |
| 11 | 70819038 | rs4944840 | -0.843 | 0.848 | -1.144 | 0.782 | 0.001 | 0 | 0 | 0 | 0.49 | 0.52 | 0.56 | 0.5 | 0.55 | 0.68 |
| 11 | 70820759 | rs7123599 | 0.659 | 1.024 | -1.159 | 0.467 | -0.021 | 0.047 | 0.047 | 0 | 0.17 | 0.07 | 0.11 | 0.02 | 0.21 | 0.33 |
| 11 | 70824963 | rs10792477 | -0.225 | 0.669 | -0.91 | 0.795 | -0.061 | -0.106 | -0.106 | 0.118 | 0.29 | 0.1 | 0.42 | 0.57 | 0.33 | 0.34 |
| 11 | 70836467 | rs10897617 | -1.086 | -0.482 | -0.586 | 0.222 | -0.02 | 0.015 | 0.015 | 0.065 | 0.8 | 0.73 | 0.77 | 0.7 | 0.84 | 0.83 |
| 11 | 70846961 | rs10459038 | 0.034 | -0.524 | -0.632 | -0.028 | -0.001 | 0.007 | 0.007 | 0 | 0.42 | 0.39 | 0.55 | 0.66 | 0.5 | 0.42 |
| 11 | 70869521 | rs11604398 | 1.462 | -0.52 | -0.942 | 0.378 | -0.017 | -0.034 | -0.034 | 0 | 0.14 | 0 | 0.1 | 0.1 | 0.13 | 0.07 |
| 11 | 70877404 | rs7110075 | 0.597 | -0.595 | -0.632 | 0.731 | -0.03 | 0.001 | 0.001 | 0 | 0.3 | 0.11 | 0.22 | 0.35 | 0.31 | 0.36 |
| 11 | 70881929 | rs4245462 | -0.652 | -0.977 | -0.543 | 0.641 | -0.007 | -0.024 | -0.024 | 0 | 0.74 | 0.36 | 0.81 | 0.84 | 0.76 | 0.61 |
| 11 | 70882388 | rs7930503 | 0.034 | -0.957 | -0.614 | 0.596 | 0.002 | -0.004 | -0.004 | 0 | 0.44 | 0.25 | 0.58 | 0.49 | 0.45 | 0.24 |
| 11 | 70885169 | rs4304805 | -0.982 | -1.018 | -0.494 | 0.517 | 0.011 | -0.11 | -0.11 | 0 | 0.78 | 0.52 | 0.82 | 0.96 | 0.76 | 0.62 |
| 11 | 70888058 | rs7941128 | 0.075 | -0.88 | -1.13 | 0.732 | -0.054 | -0.122 | -0.122 | 0 | 0.42 | 0.07 | 0.56 | 0.45 | 0.44 | 0.23 |
| 11 | 70890503 | rs4340077 | -0.776 | -0.865 | -0.429 | 0.863 | -0.024 | -0.01 | -0.01 | 0 | 0.82 | 0.64 | 0.89 | 0.86 | 0.86 | 0.64 |
| 11 | 70891161 | rs7124405 | 0.074 | -0.858 | -0.607 | 0.675 | -0.016 | -0.003 | -0.003 | 0 | 0.4 | 0.48 | 0.3 | 0.39 | 0.39 | 0.53 |
| 11 | 70895375 | rs4944283 | -0.322 | -0.542 | -0.523 | 0.95 | -0.022 | 0.042 | 0.042 | 0 | 0.61 | 0.81 | 0.61 | 0.69 | 0.54 | 0.69 |
| 11 | 70896821 | rs7937397 | -0.386 | -0.614 | -0.17 | 1.391 | -0.017 | 0.025 | 0.025 | 1.07 | 0.27 | 0.72 | 0.39 | 0.56 | 0.22 | 0.35 |
| 11 | 70901471 | rs3934692 | -0.227 | -0.562 | -0.01 | 1.235 | -0.002 | 0.219 | 0.219 | 1.916 | 0.08 | 0.39 | 0.23 | 0.23 | 0.03 | 0.08 |
| 11 | 70906073 | rs4506683 | 0.472 | -0.554 | -0.185 | 1.327 | -0.023 | 0.098 | 0.098 | 0.046 | 0.2 | 0.36 | 0.2 | 0.43 | 0.19 | 0.26 |
| 11 | 70919484 | rs2919744 | 1.098 | 0.18 | -1.604 | 0.029 | -0.005 | -0.013 | -0.013 | 0 | 0.14 | 0.09 | 0.12 | 0 | 0.18 | 0.05 |
| 11 | 70920722 | rs4944448 | 0.283 | 0.226 | -1.498 | 0.422 | -0.099 | -0.032 | -0.032 | 0 | 0.18 | 0.01 | 0.19 | 0.22 | 0.24 | 0.22 |
| 11 | 70924371 | rs10897789 | -0.862 | -0.923 | -0.64 | 0.949 | -0.021 | 0.006 | 0.006 | 1.317 | 0.3 | 0.68 | 0.43 | 0.65 | 0.21 | 0.33 |
| 11 | 70925859 | rs7929059 | -0.362 | -0.369 | -0.722 | 0.754 | 0.002 | 0.002 | 0.002 | 0 | 0.41 | 0.35 | 0.29 | 0.35 | 0.37 | 0.35 |
| 11 | 70926665 | rs4944476 | 0.407 | -0.64 | -0.182 | 0.819 | -0.014 | 0.013 | 0.013 | 0 | 0.82 | 0.4 | 0.73 | 0.8 | 0.88 | 0.83 |
| 11 | 70927063 | rs10792551 | 0.653 | -1.062 | -1.316 | 0.679 | 0.016 | -0.017 | -0.017 | 0 | 0.46 | 0.08 | 0.36 | 0.61 | 0.6 | 0.55 |
| 11 | 70932371 | rs11232656 | -0.195 | -2.279 | -0.759 | 0.65 | 0.012 | -0.003 | -0.003 | 0.003 | 0.4 | 0.33 | 0.35 | 0.41 | 0.3 | 0.43 |
| 11 | 70932944 | rs2919732 | -1.496 | -2.473 | -0.883 | 0.504 | -0.004 | -0.097 | -0.097 | 0 | 0.58 | 0.92 | 0.45 | 0.48 | 0.4 | 0.57 |
| 11 | 70933968 | rs749811 | -0.274 | -2.613 | -0.789 | 0.457 | 0.033 | 0.004 | 0.004 | 0 | 0.44 | 0.33 | 0.37 | 0.41 | 0.32 | 0.43 |
| 11 | 70938724 | rs10792565 | -0.3 | -2.815 | -0.743 | 0.464 | 0.026 | 0.05 | 0.05 | 0 | 0.48 | 0.4 | 0.63 | 0.77 | 0.36 | 0.51 |
| 11 | 70953650 | rs2919721 | -1.736 | -2.668 | -1.061 | 0.438 | -0.076 | 0.149 | 0.149 | 0.003 | 0.67 | 0.91 | 0.71 | 0.88 | 0.46 | 0.68 |
| 11 | 70959775 | rs10501406 | -0.76 | -2.676 | -1.031 | 0.443 | 0.015 | 0.032 | 0.032 | 0 | 0.1 | 0.03 | 0.05 | 0 | 0.08 | 0.06 |
| 11 | 70966863 | rs10431150 | -1.307 | -2.564 | -1.154 | 0.61 | -0.042 | -0.084 | -0.084 | 0 | 0.6 | 0.91 | 0.47 | 0.52 | 0.43 | 0.57 |
| 11 | 70966997 | rs10431158 | 0.031 | -2.855 | -0.254 | 0.76 | -0.004 | 0.001 | 0.001 | 0 | 0.39 | 0.39 | 0.36 | 0.41 | 0.3 | 0.41 |
| 11 | 70969719 | rs2119510 | -0.729 | -2.815 | 0.1 | 1.034 | -0.004 | -0.007 | -0.007 | 0 | 0.53 | 0.51 | 0.53 | 0.6 | 0.38 | 0.52 |
| 11 | 70976301 | rs2919696 | -0.663 | -3.089 | 0.289 | 1.215 | 0.113 | 0.051 | 0.051 | 0 | 0.47 | 0.86 | 0.45 | 0.63 | 0.28 | 0.51 |
| 11 | 70980906 | rs10897897 | -0.565 | -3.134 | 0.812 | 1.225 | 0.035 | 0 | 0 | 0 | 0.46 | 0.31 | 0.45 | 0.44 | 0.33 | 0.5 |
| 11 | 70981204 | rs2919702 | -0.43 | -3.072 | 1.212 | 1.265 | -0.008 | 0.061 | 0.061 | 0.198 | 0.06 | 0.26 | 0.03 | 0.08 | 0.01 | 0.1 |
| 11 | 70982325 | rs11232920 | -1.046 | -3.528 | 1.264 | 1.33 | -0.052 | 0.014 | 0.014 | 0 | 0.22 | 0.37 | 0.16 | 0.25 | 0.14 | 0.44 |
| 11 | 70989940 | rs2919718 | -0.962 | -3.524 | 1.194 | 1.332 | -0.03 | 0.006 | 0.006 | 0 | 0.22 | 0.32 | 0.16 | 0.25 | 0.14 | 0.43 |
| 11 | 70990755 | rs7934077 | -1.603 | -3.638 | 1.236 | 1.299 | 0.047 | 0.049 | 0.049 | 0 | 0.56 | 0.78 | 0.51 | 0.68 | 0.37 | 0.65 |
| 11 | 71007182 | rs7928694 | 0.358 | -3.458 | 1.091 | 1.266 | 0 | -0.001 | -0.001 | 0 | 0.3 | 0.26 | 0.29 | 0.28 | 0.2 | 0.16 |
| 11 | 71010750 | rs1660858 | -0.554 | -3.443 | 0.884 | 1.212 | -0.016 | -0.002 | -0.002 | 0 | 0.2 | 0.26 | 0.15 | 0.19 | 0.13 | 0.38 |
| 11 | 71018309 | rs1792205 | -1.29 | -3.249 | 0.867 | 1.181 | 0.009 | 0.002 | 0.002 | 0 | 0.52 | 0.57 | 0.45 | 0.53 | 0.35 | 0.6 |
| 11 | 71019412 | rs879380 | -1.22 | -3.142 | 0.752 | 1.087 | 0.013 | -0.003 | -0.003 | 0.025 | 0.47 | 0.59 | 0.36 | 0.49 | 0.35 | 0.54 |
| 11 | 71025538 | rs11233083 | -0.151 | -3.018 | 0.604 | 1.025 | 0.003 | -0.004 | -0.004 | 0.569 | 0.14 | 0.11 | 0.12 | 0.18 | 0.11 | 0.4 |
| 11 | 71031696 | rs1617603 | -0.653 | -3.495 | 0.754 | 1.291 | -0.006 | 0.027 | 0.027 | 0.228 | 0.36 | 0.51 | 0.36 | 0.52 | 0.3 | 0.53 |
| 11 | 71032039 | rs1660822 | -1.838 | -3.658 | 0.653 | 1.289 | 0.005 | 0.108 | 0.108 | 0.132 | 0.54 | 0.88 | 0.53 | 0.71 | 0.41 | 0.63 |
| 11 | 71033961 | rs10897970 | 0.369 | -3.641 | 0.769 | 1.275 | -0.001 | 0 | 0 | 0 | 0.14 | 0.14 | 0.11 | 0.16 | 0.1 | 0.38 |
| 11 | 71038128 | rs1660825 | 1.859 | -3.644 | 0.863 | 1.355 | 0.008 | 0.015 | 0.015 | 0 | 0.17 | 0.24 | 0.19 | 0.17 | 0.18 | 0.1 |
| 11 | 71039803 | rs17433007 | -0.089 | -3.669 | 0.872 | 1.344 | -0.002 | -0.003 | -0.003 | 0 | 0.09 | 0.12 | 0.06 | 0.08 | 0.1 | 0.06 |
| 11 | 71039825 | rs1660827 | -2.609 | -3.761 | 0.509 | 1.277 | -0.106 | 0.094 | 0.094 | 0 | 0.68 | 0.9 | 0.66 | 0.85 | 0.5 | 0.77 |
| 11 | 71045537 | rs731053 | -1.061 | -3.601 | 1.325 | 1.95 | 0.004 | -0.001 | -0.001 | 0 | 0.22 | 0.17 | 0.16 | 0.24 | 0.19 | 0.43 |
| 11 | 71047824 | rs1660832 | 0.938 | -3.57 | 1.372 | 1.975 | 0.017 | 0.082 | 0.082 | 0 | 0.18 | 0.32 | 0.22 | 0.32 | 0.19 | 0.12 |
| 11 | 71050928 | rs10897992 | -0.173 | -3.539 | 1.129 | 1.934 | -0.003 | 0 | 0 | 0.033 | 0.15 | 0.2 | 0.12 | 0.18 | 0.11 | 0.4 |
| 11 | 71051703 | rs11233182 | -1.826 | -3.582 | 1.231 | 2.002 | -0.038 | 0.043 | 0.043 | 0 | 0.32 | 0.49 | 0.32 | 0.52 | 0.21 | 0.52 |
| 11 | 71053385 | rs6592055 | -1.433 | -3.508 | 1.282 | 2.031 | -0.052 | 0.078 | 0.078 | 0 | 0.18 | 0.39 | 0.2 | 0.34 | 0.11 | 0.12 |
| 11 | 71055026 | rs1660854 | -0.347 | -3.494 | 1.145 | 1.947 | 0.004 | 0.057 | 0.057 | 0 | 0.28 | 0.5 | 0.29 | 0.4 | 0.28 | 0.14 |
| 11 | 71056149 | rs10898015 | -1.23 | -3.727 | 1.133 | 1.953 | -0.031 | 0.005 | 0.005 | 0 | 0.26 | 0.37 | 0.24 | 0.32 | 0.13 | 0.48 |
| 11 | 71057118 | rs6592065 | -1.243 | -3.727 | 1.133 | 1.951 | -0.03 | 0.028 | 0.028 | 0 | 0.26 | 0.38 | 0.27 | 0.46 | 0.13 | 0.49 |
| 11 | 71059302 | rs7124694 | -0.78 | -3.718 | 1.102 | 1.94 | -0.024 | 0.041 | 0.041 | 1.332 | 0.2 | 0.35 | 0.16 | 0.35 | 0.12 | 0.41 |
| 11 | 71059789 | rs12794450 | -0.073 | -3.606 | 0.74 | 1.724 | -0.034 | 0.007 | 0.007 | 0 | 0.1 | 0.11 | 0.13 | 0.14 | 0.02 | 0.06 |
| 11 | 71060426 | rs11233249 | -0.247 | -3.426 | 0.387 | 1.258 | -0.005 | -0.001 | -0.001 | 0 | 0.16 | 0.21 | 0.12 | 0.18 | 0.1 | 0.42 |
| 11 | 71062667 | rs1792218 | -1.657 | -3.372 | -0.008 | 1.081 | 0.02 | 0.112 | 0.112 | 0.243 | 0.54 | 0.85 | 0.52 | 0.72 | 0.4 | 0.6 |
| 11 | 71067224 | rs1122145 | -0.084 | -2.928 | 0.36 | 0.232 | 0.009 | 0.001 | 0.001 | 0.816 | 0.15 | 0.12 | 0.11 | 0.14 | 0.09 | 0.39 |
| 11 | 71069298 | rs1660865 | -0.642 | -3.025 | 0.588 | 0.079 | 0.001 | -0.001 | -0.001 | 0.577 | 0.36 | 0.64 | 0.49 | 0.53 | 0.31 | 0.25 |
| 11 | 71072120 | rs7115931 | -0.764 | -3.101 | 0.903 | 0.126 | -0.032 | 0.016 | 0.016 | 0 | 0.13 | 0.31 | 0.16 | 0.15 | 0.09 | 0.11 |
| 11 | 71080250 | rs2361492 | 0.077 | -3.103 | 0.919 | 0.158 | 0.002 | -0.003 | -0.003 | 0 | 0.16 | 0.16 | 0.14 | 0.29 | 0.1 | 0.4 |
| 11 | 71081060 | rs1792308 | 0.487 | -3.155 | 0.911 | 0.196 | -0.001 | 0.063 | 0.063 | 0 | 0.23 | 0.33 | 0.33 | 0.38 | 0.22 | 0.14 |

| | | | | | | | | | | | | | | | | |
|---|---|---|---|---|---|---|---|---|---|---|---|---|---|---|---|---|
| 11 | 71083737 | rs11233373 | 0.508 | -3.137 | 0.875 | 0.194 | 0.001 | 0 | 0 | 0 | 0.05 | 0.06 | 0.04 | 0.06 | 0.08 | 0.05 |
| 11 | 71086954 | rs12226382 | -0.9 | -3.176 | 0.922 | 0.183 | -0.021 | 0.009 | 0.009 | 0 | 0.13 | 0.26 | 0.16 | 0.15 | 0.1 | 0.11 |
| 11 | 71089210 | rs1620013 | -0.697 | -3.203 | 0.908 | 0.166 | -0.008 | -0.011 | -0.011 | 0 | 0.38 | 0.45 | 0.45 | 0.66 | 0.3 | 0.53 |
| 11 | 71093676 | rs12787533 | 0.072 | -3.136 | 0.669 | 0.2 | -0.042 | 0.003 | 0.003 | 0 | 0.08 | 0.18 | 0.12 | 0.1 | 0.02 | 0.07 |
| 11 | 71094472 | rs17161455 | 0.225 | -3.148 | 0.467 | 0.209 | 0.012 | -0.017 | -0.017 | 1.368 | 0.15 | 0.11 | 0.12 | 0.29 | 0.08 | 0.39 |
| 11 | 71095300 | rs1792252 | -1.812 | -3.16 | 0.403 | 0.184 | 0.015 | 0.256 | 0.256 | 0.697 | 0.52 | 0.91 | 0.62 | 0.82 | 0.4 | 0.66 |
| 11 | 71103143 | rs1790315 | -0.886 | -3.172 | 0.647 | 0.432 | -0.005 | -0.007 | -0.007 | 0 | 0.38 | 0.56 | 0.46 | 0.66 | 0.3 | 0.53 |
| 11 | 71106206 | rs1790313 | 0.253 | -3.188 | 0.496 | 0.441 | 0 | 0.091 | 0.091 | 0 | 0.23 | 0.38 | 0.34 | 0.38 | 0.22 | 0.14 |
| 11 | 71108341 | rs11604724 | -0.031 | -2.908 | -0.307 | 0.319 | 0.003 | -0.003 | -0.003 | 0 | 0.14 | 0.12 | 0.11 | 0.29 | 0.08 | 0.38 |
| 11 | 71108947 | rs1792331 | -1.137 | -2.944 | -0.469 | 0.276 | 0.083 | 0.146 | 0.146 | 0.388 | 0.46 | 0.8 | 0.57 | 0.79 | 0.32 | 0.61 |
| 11 | 71109373 | rs1660882 | -1.173 | -2.998 | -0.645 | 0.286 | 0.096 | 0.167 | 0.167 | 0.397 | 0.47 | 0.84 | 0.58 | 0.79 | 0.32 | 0.61 |
| 11 | 71110248 | rs1626847 | -0.548 | -2.265 | -0.203 | 0.558 | -0.01 | 0.013 | 0.013 | 0 | 0.32 | 0.66 | 0.46 | 0.49 | 0.23 | 0.2 |
| 11 | 71112721 | rs1792328 | -0.292 | -2.869 | -0.252 | 0.152 | -0.037 | -0.008 | -0.008 | 0 | 0.75 | 0.77 | 0.79 | 0.72 | 0.94 | 0.62 |
| 11 | 71112818 | rs1792326 | -0.123 | -3.035 | -0.342 | -0.002 | 0.023 | 0.014 | 0.014 | 0 | 0.73 | 0.84 | 0.77 | 0.71 | 0.89 | 0.61 |
| 11 | 71113845 | rs1790374 | -0.519 | -2.947 | -0.3 | -0.002 | 0.003 | 0.002 | 0.002 | 0 | 0.77 | 0.83 | 0.8 | 0.73 | 0.93 | 0.63 |
| 11 | 71115446 | rs1790358 | 0.037 | -3.061 | -0.189 | -0.465 | 0.017 | 0.041 | 0.041 | 0 | 0.35 | 0.25 | 0.22 | 0.21 | 0.28 | 0.38 |
| 11 | 71115631 | rs13377288 | 0.541 | -3.33 | 0.027 | -0.523 | -0.019 | -0.011 | -0.011 | 0 | 0.09 | 0.16 | 0.04 | 0.07 | 0.04 | 0.08 |
| 11 | 71115640 | rs1792322 | -1.204 | -3.341 | 0.059 | -0.502 | 0.017 | 0.016 | 0.016 | 0 | 0.77 | 0.85 | 0.82 | 0.82 | 0.93 | 0.64 |
| 11 | 71117319 | rs7105151 | 0.331 | -3.393 | 0.297 | -0.563 | 0.006 | 0.006 | 0.006 | 0 | 0.4 | 0.42 | 0.54 | 0.52 | 0.66 | 0.25 |
| 11 | 71117644 | rs1557465 | 0.217 | -3.396 | 0.309 | -0.704 | 0.001 | -0.009 | -0.009 | 0 | 0.36 | 0.33 | 0.25 | 0.21 | 0.28 | 0.38 |
| 11 | 71118568 | rs1790361 | -1.233 | -3.381 | 0.233 | -0.695 | 0.014 | 0.013 | 0.013 | 0 | 0.77 | 0.85 | 0.8 | 0.81 | 0.93 | 0.64 |
| 11 | 71118788 | rs4944042 | 0.993 | -3.313 | 0.195 | -0.67 | -0.014 | -0.013 | -0.013 | 0 | 0.32 | 0.26 | 0.5 | 0.46 | 0.61 | 0.17 |
| 11 | 71119018 | rs12806918 | 1.481 | -3.211 | 0.046 | -0.517 | -0.091 | -0.112 | -0.112 | 0.004 | 0.26 | 0.11 | 0.46 | 0.36 | 0.56 | 0.13 |
| 11 | 71125345 | rs3763856 | 1.486 | -3.204 | 0.049 | -0.514 | -0.091 | -0.112 | -0.112 | 0 | 0.26 | 0.1 | 0.47 | 0.36 | 0.56 | 0.13 |
| 11 | 71126595 | rs1617153 | 0.09 | -3.34 | 0.26 | -0.376 | 0.019 | 0.049 | 0.049 | 0 | 0.34 | 0.2 | 0.21 | 0.21 | 0.28 | 0.38 |
| 11 | 71131388 | rs1790337 | -0.813 | -3.504 | 0.464 | -0.155 | -0.02 | -0.001 | -0.001 | 0 | 0.76 | 0.81 | 0.78 | 0.74 | 0.95 | 0.62 |
| 11 | 71142350 | rs1790349 | 0.797 | -3.615 | 0.27 | -0.252 | -0.002 | 0 | 0 | 0.405 | 0.23 | 0.21 | 0.19 | 0.29 | 0.19 | 0.36 |
| 11 | 71143266 | rs12797951 | 0.941 | -3.764 | 0.331 | -0.316 | 0.029 | -0.044 | -0.044 | 0 | 0.37 | 0.24 | 0.52 | 0.38 | 0.7 | 0.15 |
| 11 | 71143580 | rs1790348 | -0.839 | -3.792 | 0.397 | -0.31 | -0.097 | -0.033 | -0.033 | 0 | 0.75 | 0.64 | 0.76 | 0.77 | 0.96 | 0.63 |
| 11 | 71145778 | rs1044535 | -0.876 | -3.792 | 0.397 | -0.344 | -0.097 | -0.035 | -0.035 | 1.087 | 0.75 | 0.64 | 0.76 | 0.77 | 0.96 | 0.63 |
| 11 | 71145941 | rs1790345 | -0.342 | -3.755 | 0.36 | -0.345 | -0.007 | -0.003 | -0.003 | 0 | 0.37 | 0.39 | 0.24 | 0.38 | 0.26 | 0.48 |
| 11 | 71155129 | rs4316537 | 0.104 | -3.711 | 0.328 | -0.331 | -0.002 | -0.001 | -0.001 | 0 | 0.14 | 0.17 | 0.04 | 0.09 | 0.06 | 0.12 |
| 11 | 71155153 | rs1790334 | -0.988 | -3.469 | 0.42 | -0.201 | -0.012 | 0.001 | 0.001 | 0 | 0.77 | 0.82 | 0.81 | 0.83 | 0.96 | 0.63 |
| 11 | 71158841 | rs3750997 | 0.57 | -3.386 | 0.428 | -0.154 | 0.021 | -0.014 | -0.014 | 0 | 0.39 | 0.33 | 0.57 | 0.42 | 0.7 | 0.15 |
| 11 | 71164544 | rs12791871 | 0.926 | -3.306 | 0.382 | -0.231 | 0.022 | -0.067 | -0.067 | 0.24 | 0.36 | 0.17 | 0.45 | 0.38 | 0.7 | 0.15 |
| 11 | 71165625 | rs7944926 | 0.926 | -3.306 | 0.382 | -0.231 | 0.022 | -0.067 | -0.067 | 0.879 | 0.36 | 0.17 | 0.45 | 0.38 | 0.7 | 0.15 |
| 11 | 71171003 | rs12800438 | 0.753 | -3.306 | 0.422 | -0.226 | 0.015 | -0.02 | -0.02 | 0 | 0.38 | 0.33 | 0.47 | 0.38 | 0.7 | 0.15 |
| 11 | 71174452 | rs2276362 | 0.766 | -3.299 | 0.426 | -0.223 | 0.016 | -0.022 | -0.022 | 0.564 | 0.38 | 0.31 | 0.47 | 0.38 | 0.7 | 0.15 |
| 11 | 71183474 | rs2282621 | 0.772 | -3.287 | 0.403 | -0.22 | 0.025 | -0.025 | -0.025 | 0 | 0.37 | 0.24 | 0.46 | 0.44 | 0.7 | 0.15 |
| 11 | 71185518 | rs2186778 | -0.735 | -3.377 | 0.454 | -0.269 | -0.088 | -0.049 | -0.049 | 0 | 0.75 | 0.64 | 0.72 | 0.83 | 0.96 | 0.62 |
| 11 | 71194559 | rs3829251 | 0.598 | -3.409 | 0.214 | -0.258 | -0.005 | -0.002 | -0.002 | 0 | 0.28 | 0.26 | 0.23 | 0.3 | 0.19 | 0.35 |
| 11 | 71197083 | rs10898193 | 0.553 | -3.372 | -0.173 | -0.264 | -0.021 | -0.006 | -0.006 | 0 | 0.28 | 0.34 | 0.24 | 0.3 | 0.19 | 0.35 |
| 11 | 71203739 | rs12288906 | 0.517 | -3.282 | -0.154 | -0.232 | -0.007 | -0.009 | -0.009 | 0 | 0.11 | 0.18 | 0.03 | 0.09 | 0.06 | 0.11 |
| 11 | 71209953 | rs12286616 | 0.517 | -3.282 | -0.154 | -0.232 | -0.007 | -0.009 | -0.009 | 0.321 | 0.11 | 0.18 | 0.03 | 0.09 | 0.06 | 0.11 |
| 11 | 71211654 | rs4944076 | -0.026 | -3.329 | -0.154 | -0.262 | -0.057 | -0.025 | -0.025 | 0 | 0.65 | 0.58 | 0.7 | 0.68 | 0.89 | 0.5 |
| 11 | 71212219 | rs12277408 | 0.155 | -3.314 | -0.147 | -0.261 | -0.026 | -0.004 | -0.004 | 0 | 0.13 | 0.34 | 0.05 | 0.14 | 0.07 | 0.11 |
| 11 | 71217571 | rs1002171 | 0.149 | -3.338 | -0.239 | -0.26 | -0.026 | -0.005 | -0.005 | 2.253 | 0.13 | 0.34 | 0.05 | 0.14 | 0.06 | 0.11 |
| 11 | 71217959 | rs2155425 | -0.589 | -2.885 | -0.037 | -0.306 | -0.008 | 0.004 | 0.004 | 0 | 0.77 | 0.83 | 0.74 | 0.83 | 0.95 | 0.62 |
| 11 | 71221574 | rs7940244 | 0.479 | -2.859 | -0.053 | -0.29 | 0.037 | 0.018 | 0.018 | 0 | 0.48 | 0.48 | 0.5 | 0.52 | 0.76 | 0.26 |
| 11 | 71239086 | rs11234042 | -0.577 | -1.048 | -0.345 | 0.209 | -0.028 | -0.007 | -0.007 | 0 | 0.29 | 0.39 | 0.25 | 0.31 | 0.19 | 0.38 |
| 11 | 71242209 | rs7128994 | -0.308 | -0.458 | -0.71 | -0.215 | 0.005 | 0.01 | 0.01 | 0 | 0.44 | 0.36 | 0.2 | 0.22 | 0.37 | 0.49 |
| 11 | 71242461 | rs4539348 | -0.738 | 0.226 | -1.511 | -0.222 | -0.088 | 0.017 | 0.017 | 0 | 0.77 | 0.96 | 0.76 | 0.8 | 0.75 | 0.7 |
| 11 | 71243572 | rs11600364 | -0.352 | -0.283 | -0.926 | 0.311 | 0.008 | 0.042 | 0.042 | 0.508 | 0.26 | 0.32 | 0.45 | 0.47 | 0.33 | 0.21 |
| 11 | 71248389 | rs12365630 | 0.56 | -1.037 | -1.319 | -0.013 | 0.04 | -0.014 | -0.014 | 0 | 0.15 | 0.01 | 0.1 | 0.09 | 0.09 | 0.06 |
| 11 | 71248554 | rs11234079 | -0.298 | -1.119 | -0.678 | -0.063 | -0.023 | -0.014 | -0.014 | 0 | 0.2 | 0.24 | 0.13 | 0.09 | 0.12 | 0.06 |
| 11 | 71251190 | rs11234088 | 0.574 | -1.086 | -0.732 | -0.039 | 0.04 | -0.014 | -0.014 | 0 | 0.15 | 0.01 | 0.1 | 0.09 | 0.09 | 0.06 |
| 11 | 71258998 | rs10751114 | 0.002 | -1.077 | -0.339 | 0 | -0.015 | -0.026 | -0.026 | 0 | 0.23 | 0.21 | 0.44 | 0.49 | 0.32 | 0.21 |
| 11 | 71260296 | rs2663 | -0.281 | -1.065 | -0.182 | 0.006 | 0 | 0.004 | 0.004 | 0.331 | 0.25 | 0.28 | 0.45 | 0.49 | 0.33 | 0.22 |
| 11 | 71261263 | rs10792774 | -0.368 | -1.248 | -0.117 | -0.113 | -0.043 | -0.026 | -0.026 | 0.029 | 0.23 | 0.35 | 0.14 | 0.13 | 0.14 | 0.08 |
| 11 | 71262138 | rs7934055 | -1.701 | -1.263 | -0.113 | -0.12 | 0.016 | 0.047 | 0.047 | 0 | 0.49 | 0.68 | 0.6 | 0.62 | 0.47 | 0.3 |
| 11 | 71263378 | rs7949169 | -0.15 | -1.206 | -0.181 | -0.111 | 0.008 | 0.025 | 0.025 | 0.345 | 0.25 | 0.34 | 0.46 | 0.49 | 0.33 | 0.22 |
| 11 | 71265137 | rs11234150 | -0.315 | -1.188 | -0.324 | -0.095 | -0.016 | -0.008 | -0.008 | 0 | 0.22 | 0.24 | 0.13 | 0.13 | 0.14 | 0.08 |
| 11 | 71277981 | rs11234218 | 0.02 | -1.226 | -0.257 | -0.075 | -0.019 | -0.009 | -0.009 | 0 | 0.19 | 0.21 | 0.12 | 0.09 | 0.12 | 0.06 |
| 11 | 71278915 | rs9651752 | -0.909 | -0.805 | -0.343 | -0.097 | -0.004 | -0.011 | -0.011 | 0 | 0.46 | 0.5 | 0.58 | 0.57 | 0.47 | 0.28 |
| 11 | 71279092 | rs9651753 | -0.9 | -0.804 | -0.343 | -0.097 | -0.004 | -0.012 | -0.012 | 0 | 0.46 | 0.5 | 0.58 | 0.58 | 0.47 | 0.28 |
| 11 | 71280333 | rs4475948 | -1.19 | -0.908 | -0.971 | -0.059 | 0.052 | 0.059 | 0.059 | 0 | 0.54 | 0.83 | 0.6 | 0.59 | 0.43 | 0.32 |
| 11 | 71281562 | rs7129512 | -0.154 | -0.513 | -0.607 | 0.404 | 0 | -0.001 | -0.001 | 0.874 | 0.36 | 0.31 | 0.25 | 0.48 | 0.37 | 0.27 |
| 11 | 71281860 | rs7937196 | -0.182 | -0.955 | -0.577 | 0.16 | -0.035 | -0.042 | -0.042 | 0 | 0.36 | 0.54 | 0.28 | 0.17 | 0.24 | 0.22 |
| 11 | 71289089 | rs2186775 | -0.359 | -0.246 | -0.739 | 0.152 | 0 | -0.001 | -0.001 | 0 | 0.62 | 0.64 | 0.52 | 0.66 | 0.59 | 0.41 |

| | | | | | | | | | | | | | | | | |
|---|---|---|---|---|---|---|---|---|---|---|---|---|---|---|---|---|
| 11 | 71290580 | rs11602969 | 0.435 | -0.768 | -0.821 | 0.208 | -0.044 | -0.026 | -0.026 | 0 | 0.41 | 0.16 | 0.56 | 0.58 | 0.51 | 0.38 |
| 11 | 71291490 | rs12421579 | 0.067 | -0.922 | -0.3 | 0.261 | -0.011 | -0.008 | -0.008 | 2.69 | 0.58 | 0.43 | 0.73 | 0.65 | 0.63 | 0.53 |
| 11 | 71294802 | rs10898347 | 0.23 | -0.887 | -0.434 | 0.287 | 0.029 | 0.025 | 0.025 | 0 | 0.54 | 0.34 | 0.71 | 0.63 | 0.61 | 0.53 |
| 11 | 71304419 | rs4945043 | 0.329 | -0.982 | 0.059 | 0.282 | 0 | 0 | 0 | 0.17 | 0.7 | 0.73 | 0.81 | 0.74 | 0.7 | 0.69 |
| 11 | 71361101 | rs7102783 | -0.109 | -0.846 | 0.235 | 0.328 | 0.019 | 0 | 0 | 0 | 0.94 | 0.87 | 0.94 | 0.93 | 0.91 | 0.95 |
| 11 | 71419408 | rs12802846 | -1.162 | -2.256 | 0.345 | 0.462 | -0.036 | -0.003 | -0.003 | 0.04 | 0.34 | 0.58 | 0.15 | 0.25 | 0.14 | 0.24 |
| 11 | 71464076 | rs7117703 | -0.142 | -1.658 | 0.214 | 0.708 | -0.001 | -0.002 | -0.002 | 0 | 0.66 | 0.65 | 0.44 | 0.48 | 0.63 | 0.75 |
| 11 | 71496121 | rs2040301 | 0.814 | -2.066 | 0.621 | 0.589 | 0.025 | 0.05 | 0.05 | 0.418 | 0.51 | 0.34 | 0.35 | 0.33 | 0.57 | 0.67 |
| 11 | 71498902 | rs11234944 | 0.814 | -2.066 | 0.621 | 0.646 | 0.025 | 0.043 | 0.043 | 0.418 | 0.51 | 0.34 | 0.35 | 0.35 | 0.57 | 0.67 |
| 11 | 71532994 | rs10751157 | 0.019 | -1.617 | 0.868 | 0.942 | -0.015 | -0.032 | -0.032 | 0.31 | 0.64 | 0.29 | 0.41 | 0.45 | 0.61 | 0.65 |
| 11 | 71549174 | rs7107929 | -0.081 | -1.598 | 1.119 | 1.011 | -0.004 | -0.01 | -0.01 | 0 | 0.66 | 0.63 | 0.45 | 0.45 | 0.64 | 0.65 |
| 11 | 71579229 | rs661242 | -2.604 | -1.196 | 1.742 | 1.778 | -0.037 | 0.082 | 0.082 | 1.314 | 0.14 | 0.59 | 0.22 | 0.22 | 0.06 | 0.18 |
| 11 | 71626089 | rs7113974 | -1.177 | -1.091 | 1.952 | 1.751 | 0.04 | 0.148 | 0.148 | 0 | 0.07 | 0.32 | 0.06 | 0.09 | 0.04 | 0.05 |
| 11 | 71648169 | rs7102192 | 1.341 | -1.107 | 2.207 | 1.86 | -0.002 | 0.179 | 0.179 | 0.007 | 0.9 | 0.69 | 0.79 | 0.78 | 0.94 | 0.82 |
| 11 | 71700566 | rs7121260 | 1.444 | -1.099 | 1.906 | 1.861 | 0.019 | 0.267 | 0.267 | 0.059 | 0.89 | 0.58 | 0.78 | 0.78 | 0.94 | 0.82 |
| 11 | 71709724 | rs5743658 | -2.056 | -1.171 | 2.014 | 1.845 | -0.009 | 0.109 | 0.109 | 0.455 | 0.1 | 0.42 | 0.08 | 0.08 | 0.04 | 0.05 |
| 11 | 71733519 | rs10128658 | 1.071 | -0.983 | 1.175 | 1.893 | -0.008 | 0.113 | 0.113 | 0 | 0.91 | 0.8 | 0.8 | 0.78 | 0.94 | 0.82 |
| 11 | 71739423 | rs10898815 | 0.785 | -0.941 | 0.757 | 1.765 | -0.149 | 0.117 | 0.117 | 0.889 | 0.36 | 0.04 | 0.27 | 0.17 | 0.47 | 0.5 |
| 11 | 71790942 | rs3018302 | 1.399 | -0.758 | 1.764 | 2.459 | -0.005 | 0.12 | 0.12 | 0 | 0.91 | 0.8 | 0.8 | 0.78 | 0.94 | 0.82 |
| 11 | 71815308 | rs17161966 | -1.939 | -0.784 | 1.774 | 1.66 | -0.006 | 0.126 | 0.126 | 0 | 0.09 | 0.21 | 0.2 | 0.22 | 0.06 | 0.18 |
| 11 | 71821236 | rs17161980 | -1.939 | -0.783 | 1.696 | 1.66 | -0.006 | 0.126 | 0.126 | 0 | 0.09 | 0.21 | 0.2 | 0.22 | 0.06 | 0.18 |
| 11 | 71822401 | rs3750908 | -1.939 | -0.783 | 1.696 | 1.66 | -0.006 | 0.126 | 0.126 | 0 | 0.09 | 0.21 | 0.2 | 0.22 | 0.06 | 0.18 |
| 11 | 71823133 | rs3793938 | -1.832 | -0.774 | 1.653 | 1.665 | -0.005 | 0.12 | 0.12 | 0 | 0.09 | 0.2 | 0.2 | 0.22 | 0.06 | 0.18 |
| 11 | 71838208 | rs7109653 | -1.056 | -0.756 | 1.139 | 1.647 | -0.006 | 0.03 | 0.03 | 0 | 0.06 | 0.07 | 0.05 | 0.09 | 0.04 | 0.05 |
| 11 | 71841325 | rs7937515 | -1.94 | -0.744 | 0.896 | 1.47 | -0.037 | 0.102 | 0.102 | 0.211 | 0.17 | 0.52 | 0.25 | 0.24 | 0.08 | 0.19 |
| 11 | 71852625 | rs495825 | 1.111 | -0.84 | 1.407 | 0.78 | -0.049 | -0.007 | -0.007 | 0 | 0.89 | 0.71 | 0.96 | 0.93 | 0.98 | 0.88 |
| 11 | 71853004 | rs633850 | 0.726 | -0.436 | 0.893 | 0.681 | -0.001 | 0.004 | 0.004 | 0 | 0.5 | 0.36 | 0.63 | 0.56 | 0.51 | 0.39 |
| 11 | 71853240 | rs522381 | 0.798 | -0.44 | 0.901 | 0.586 | -0.032 | -0.022 | -0.022 | 0 | 0.86 | 0.7 | 0.92 | 0.92 | 0.91 | 0.86 |
| 11 | 71857738 | rs550638 | 1.115 | -0.261 | 0.828 | 0.781 | 0.003 | 0.134 | 0.134 | 0.106 | 0.85 | 0.59 | 0.88 | 0.77 | 0.88 | 0.81 |
| 11 | 71864717 | rs11522752 | 0.114 | -0.45 | 0.654 | 0.814 | -0.016 | 0.042 | 0.042 | 0 | 0.34 | 0.23 | 0.24 | 0.19 | 0.37 | 0.41 |
| 11 | 71872777 | rs559389 | 0.445 | -0.35 | 0.392 | 0.813 | 0.021 | 0.004 | 0.004 | 0 | 0.53 | 0.29 | 0.67 | 0.59 | 0.57 | 0.45 |
| 11 | 71900964 | rs2071010 | 0.014 | -0.493 | 0.13 | 0.771 | 0.004 | -0.04 | -0.04 | 0 | 0.09 | 0 | 0.08 | 0.16 | 0.06 | 0.21 |
| 11 | 71914145 | rs11235465 | -0.563 | -0.624 | 0.851 | 1.014 | -0.02 | 0.045 | 0.045 | 0 | 0.12 | 0.19 | 0.1 | 0.23 | 0.07 | 0.19 |
| 11 | 71914290 | rs11235466 | -0.076 | -0.648 | 0.812 | 1.013 | 0.01 | -0.012 | -0.012 | 0.035 | 0.1 | 0.05 | 0.09 | 0.16 | 0.06 | 0.19 |
| 11 | 71923542 | rs11235468 | -1.391 | -0.695 | 1.328 | 1.094 | -0.013 | 0.097 | 0.097 | 0 | 0.16 | 0.33 | 0.15 | 0.29 | 0.13 | 0.25 |
| 11 | 71926660 | rs651933 | -1.415 | -0.689 | 0.881 | 0.767 | 0.02 | -0.015 | -0.015 | 0 | 0.48 | 0.75 | 0.35 | 0.47 | 0.44 | 0.59 |
| 11 | 71930207 | rs514933 | -0.601 | -0.634 | 0.903 | 0.762 | -0.001 | 0.002 | 0.002 | 0.466 | 0.37 | 0.7 | 0.26 | 0.31 | 0.37 | 0.37 |
| 11 | 71932414 | rs2298444 | -1.265 | -1.266 | 0.706 | 0.47 | -0.042 | 0.02 | 0.02 | 0 | 0.31 | 0.45 | 0.22 | 0.38 | 0.21 | 0.43 |
| 11 | 71941212 | rs2276048 | -1.151 | -1.378 | 0.606 | 0.38 | -0.026 | 0.006 | 0.006 | 0 | 0.31 | 0.51 | 0.26 | 0.34 | 0.21 | 0.41 |
| 11 | 71947822 | rs10751199 | -1.546 | -1.145 | 0.604 | 0.285 | 0.036 | -0.018 | -0.018 | 0 | 0.35 | 0.66 | 0.28 | 0.41 | 0.24 | 0.39 |
| 11 | 71956583 | rs7110260 | -1.417 | -1.019 | 1.419 | 0.38 | 0.049 | 0.176 | 0.176 | 0 | 0.06 | 0.23 | 0.11 | 0.15 | 0.07 | 0.08 |
| 11 | 71962874 | rs7935699 | 0.849 | -0.728 | 1.066 | 0.001 | -0.02 | 0.057 | 0.057 | 0 | 0.24 | 0.15 | 0.11 | 0.06 | 0.24 | 0.25 |
| 11 | 71963129 | rs3900852 | -1.699 | -0.72 | 1.442 | 0.159 | 0.09 | 0.248 | 0.248 | 0 | 0.07 | 0.36 | 0.12 | 0.16 | 0.08 | 0.1 |
| 11 | 71965275 | rs477354 | -1.313 | -0.757 | 1.515 | 0.169 | 0.035 | 0.216 | 0.216 | 0 | 0.05 | 0.22 | 0.1 | 0.14 | 0.05 | 0.06 |
| 11 | 71966582 | rs628542 | -1.666 | -0.564 | 0.589 | -0.196 | -0.136 | -0.169 | -0.169 | 0.144 | 0.65 | 0.96 | 0.45 | 0.46 | 0.56 | 0.65 |
| 11 | 71974196 | rs6592464 | -1.28 | -0.498 | 1.013 | -0.207 | -0.008 | -0.002 | -0.002 | 0 | 0.59 | 0.69 | 0.42 | 0.38 | 0.55 | 0.64 |
| 11 | 71999720 | rs638138 | -0.764 | -0.295 | 1.494 | -0.246 | -0.024 | 0.008 | 0.008 | 0 | 0.14 | 0.34 | 0.11 | 0.15 | 0.1 | 0.13 |
| 11 | 72003767 | rs526344 | -1.512 | -0.197 | 1.314 | -0.695 | -0.002 | -0.031 | -0.031 | 0 | 0.58 | 0.78 | 0.43 | 0.45 | 0.56 | 0.62 |
| 11 | 72004835 | rs12793524 | -0.77 | -0.169 | 1.606 | -0.738 | -0.009 | 0.137 | 0.137 | 0 | 0.49 | 0.25 | 0.29 | 0.23 | 0.46 | 0.55 |
| 11 | 72006086 | rs504217 | -2.343 | -0.16 | 1.67 | -0.678 | 0.064 | 0.264 | 0.264 | 0 | 0.09 | 0.45 | 0.13 | 0.22 | 0.09 | 0.07 |
| 11 | 72007984 | rs1784257 | -1.086 | -0.166 | 1.73 | -0.642 | 0.025 | 0.216 | 0.216 | 0.008 | 0.06 | 0.27 | 0.1 | 0.16 | 0.05 | 0.06 |
| 11 | 72009099 | rs7111227 | -0.82 | -0.17 | 1.648 | -0.683 | -0.002 | 0.111 | 0.111 | 0 | 0.49 | 0.33 | 0.3 | 0.23 | 0.46 | 0.55 |
| 11 | 72012682 | rs3750906 | -2.366 | -0.403 | 2.23 | -0.078 | 0.064 | 0.264 | 0.264 | 0 | 0.09 | 0.45 | 0.13 | 0.22 | 0.1 | 0.07 |
| 11 | 72056591 | rs12421674 | -2.223 | -0.394 | 2.169 | -0.063 | 0.04 | 0.222 | 0.222 | 0.15 | 0.09 | 0.5 | 0.13 | 0.22 | 0.1 | 0.07 |
| 11 | 72068077 | rs7943807 | -0.418 | -0.447 | 1.88 | -0.206 | -0.043 | -0.012 | -0.012 | 0 | 0.87 | 1 | 0.88 | 0.88 | 0.87 | 0.85 |
| 11 | 72074803 | rs3741163 | 1.847 | -0.298 | 1.87 | -0.263 | -0.006 | -0.059 | -0.059 | 0 | 0.63 | 0.18 | 0.66 | 0.59 | 0.67 | 0.6 |
| 11 | 72088806 | rs10501410 | -1.691 | -0.167 | 2.278 | -0.546 | 0.108 | 0.183 | 0.183 | 0 | 0.06 | 0.34 | 0.07 | 0.13 | 0.08 | 0.04 |
| 11 | 72088912 | rs4944458 | 1.208 | -0.099 | 2.092 | -0.524 | -0.005 | -0.038 | -0.038 | 1.277 | 0.71 | 0.57 | 0.77 | 0.81 | 0.76 | 0.67 |
| 11 | 72091228 | rs3741162 | 0.518 | -0.418 | 1.859 | -0.475 | -0.003 | 0 | 0 | 0 | 0.09 | 0.13 | 0.06 | 0.1 | 0.05 | 0.14 |
| 11 | 72115610 | rs4944477 | 0.6 | -0.285 | 1.118 | -0.479 | 0.002 | 0.057 | 0.057 | 1.099 | 0.74 | 0.92 | 0.82 | 0.83 | 0.79 | 0.7 |
| 11 | 72160112 | rs34147990 | -2.121 | 0.032 | 1.159 | 0.153 | 0.064 | 0.247 | 0.247 | 0.352 | 0.09 | 0.49 | 0.1 | 0.24 | 0.09 | 0.18 |
| 11 | 72172097 | rs10793030 | 1.215 | 0.133 | 0.915 | 0.003 | 0.004 | 0.038 | 0.038 | 0 | 0.76 | 0.45 | 0.76 | 0.63 | 0.79 | 0.65 |
| 11 | 72176100 | rs2163125 | 0.31 | 0.116 | 0.113 | -0.074 | 0 | 0 | 0 | 0 | 0.83 | 0.85 | 0.82 | 0.81 | 0.85 | 0.81 |
| 11 | 72181733 | rs1429685 | 0.206 | 0.372 | -0.367 | 0.449 | 0 | -0.027 | -0.027 | 0 | 0.58 | 0.24 | 0.63 | 0.51 | 0.62 | 0.62 |
| 11 | 72185834 | rs982840 | 0.038 | 0.249 | -0.344 | 0.053 | -0.001 | 0.028 | 0.028 | 0 | 0.54 | 0.65 | 0.65 | 0.67 | 0.5 | 0.53 |
| 11 | 72186099 | rs4943909 | 0.153 | 0.2 | -0.07 | 0.031 | 0 | 0.005 | 0.005 | 0 | 0.53 | 0.56 | 0.65 | 0.67 | 0.5 | 0.53 |
| 11 | 72188278 | rs11235509 | 1.213 | 0.161 | -0.179 | 0.054 | 0.027 | -0.095 | -0.095 | 0 | 0.28 | 0.15 | 0.46 | 0.46 | 0.27 | 0.36 |
| 11 | 72192830 | rs11607688 | 1.207 | 0.161 | 0.314 | 0.035 | 0.001 | -0.003 | -0.003 | 0 | 0.23 | 0.27 | 0.18 | 0.21 | 0.23 | 0.17 |
| 11 | 72195404 | rs11823867 | 0.916 | 0.219 | -0.119 | -0.016 | -0.046 | 0.117 | 0.117 | 0.022 | 0.31 | 0.11 | 0.18 | 0.06 | 0.38 | 0.24 |

| | | | | | | | | | | | | | | | | |
|---|---|---|---|---|---|---|---|---|---|---|---|---|---|---|---|---|
| 11 | 72196620 | rs7116209 | -0.147 | -0.044 | 0.115 | 0.656 | -0.001 | 0.044 | 0.044 | 0 | 0.85 | 0.77 | 0.83 | 0.73 | 0.87 | 0.79 |
| 11 | 72197635 | rs7952006 | 0.776 | -0.005 | -0.198 | 0.563 | 0.026 | -0.093 | -0.093 | 0 | 0.28 | 0.15 | 0.45 | 0.46 | 0.25 | 0.35 |
| 11 | 72198359 | rs4943914 | -0.005 | 0.178 | 0.045 | 0.722 | 0.005 | 0.039 | 0.039 | 0 | 0.85 | 0.72 | 0.83 | 0.8 | 0.87 | 0.82 |
| 11 | 72201835 | rs1429686 | 0.294 | 0.027 | -0.553 | 0.847 | 0.023 | -0.052 | -0.052 | 1.098 | 0.29 | 0.22 | 0.46 | 0.46 | 0.25 | 0.36 |
| 11 | 72205699 | rs163607 | -0.399 | 0.315 | -0.35 | 0.929 | -0.003 | 0.02 | 0.02 | 0 | 0.55 | 0.63 | 0.59 | 0.61 | 0.5 | 0.65 |
| 11 | 72209060 | rs2077955 | -0.419 | 0.293 | -0.324 | 1.475 | -0.001 | 0.03 | 0.03 | 0 | 0.3 | 0.54 | 0.21 | 0.38 | 0.3 | 0.27 |
| 11 | 72227849 | rs7950814 | 0.005 | 0.212 | 0.398 | 1.54 | -0.003 | -0.015 | -0.015 | 0 | 0.25 | 0.33 | 0.15 | 0.16 | 0.25 | 0.25 |
| 11 | 72234930 | rs11235519 | -0.095 | 0.059 | 0.823 | 1.459 | -0.013 | -0.065 | -0.065 | 0 | 0.25 | 0.45 | 0.13 | 0.09 | 0.22 | 0.2 |
| 11 | 72241573 | rs4257001 | 0.862 | 0.077 | 0.77 | 1.859 | 0.032 | 0.219 | 0.219 | 0 | 0.93 | 0.69 | 0.9 | 0.72 | 0.93 | 0.91 |
| 11 | 72249604 | rs10793032 | 0.852 | -0.034 | 0.064 | 1.756 | -0.025 | -0.028 | -0.028 | 0.094 | 0.24 | 0.06 | 0.38 | 0.23 | 0.24 | 0.21 |
| 11 | 72255844 | rs341081 | 0.078 | 0.103 | -0.028 | 1.881 | 0.004 | -0.021 | -0.021 | 1.208 | 0.13 | 0.08 | 0.15 | 0.25 | 0.08 | 0.27 |
| 11 | 72256026 | rs341080 | 0.765 | 0.138 | 0.429 | 1.529 | -0.006 | 0.104 | 0.104 | 0 | 0.89 | 0.7 | 0.89 | 0.72 | 0.91 | 0.86 |
| 11 | 72257512 | rs341078 | 0.683 | 0.042 | 0.117 | 1.116 | -0.003 | 0.002 | 0.002 | 0 | 0.43 | 0.4 | 0.38 | 0.42 | 0.44 | 0.51 |
| 11 | 72259140 | rs193170 | 0.794 | 0.029 | 0.433 | 0.758 | -0.008 | 0.005 | 0.005 | 0 | 0.67 | 0.47 | 0.77 | 0.64 | 0.69 | 0.77 |
| 11 | 72260488 | rs7122222 | 0.259 | 0.22 | 0.192 | 0.405 | 0 | 0.005 | 0.005 | 0 | 0.16 | 0.1 | 0.1 | 0.09 | 0.17 | 0.06 |
| 11 | 72261337 | rs375448 | 0.211 | 0.033 | 0.151 | 0.336 | -0.008 | 0.003 | 0.003 | 0 | 0.85 | 0.8 | 0.91 | 0.82 | 0.9 | 0.94 |
| 11 | 72265849 | rs341072 | 0.074 | 0.021 | 0.283 | 0.255 | -0.025 | 0.007 | 0.007 | 0 | 0.85 | 0.73 | 0.89 | 0.86 | 0.88 | 0.92 |
| 11 | 72267280 | rs341069 | -1.451 | -0.181 | 0.073 | -0.23 | 0.015 | 0.031 | 0.031 | 0 | 0.49 | 0.55 | 0.63 | 0.76 | 0.44 | 0.54 |
| 11 | 72267956 | rs11602155 | 1.998 | 0.339 | -0.392 | -0.691 | -0.004 | 0.005 | 0.005 | 0 | 0.23 | 0.2 | 0.14 | 0.07 | 0.25 | 0.21 |
| 11 | 72269214 | rs341065 | 1.012 | 0.158 | -0.385 | -0.694 | 0.023 | -0.053 | -0.053 | 0 | 0.18 | 0.12 | 0.36 | 0.51 | 0.12 | 0.36 |
| 11 | 72269302 | rs10898849 | 1.641 | 0.438 | -0.578 | -0.755 | -0.049 | 0.052 | 0.052 | 0 | 0.24 | 0.06 | 0.2 | 0.15 | 0.24 | 0.24 |
| 11 | 72269385 | rs4943927 | -1.047 | 0.524 | -0.715 | -0.681 | 0.001 | -0.004 | -0.004 | 0 | 0.33 | 0.3 | 0.28 | 0.27 | 0.38 | 0.18 |
| 11 | 72270688 | rs7924740 | 1.516 | 0.311 | -1.343 | -0.377 | -0.015 | 0.019 | 0.019 | 0 | 0.26 | 0.2 | 0.17 | 0.07 | 0.32 | 0.22 |
| 11 | 72272234 | rs12417936 | 0.031 | 0.17 | -1.498 | -0.152 | -0.011 | 0.018 | 0.018 | 0 | 0.16 | 0.1 | 0.14 | 0.12 | 0.2 | 0.1 |
| 11 | 72275781 | rs7112131 | 0.336 | -0.08 | -1.712 | -0.059 | 0.029 | -0.079 | -0.079 | 0 | 0.24 | 0.12 | 0.39 | 0.51 | 0.19 | 0.4 |
| 11 | 72276606 | rs7116276 | 1.116 | 0.688 | -1.672 | -0.817 | -0.045 | 0.057 | 0.057 | 0 | 0.25 | 0.09 | 0.17 | 0.07 | 0.32 | 0.22 |
| 11 | 72279550 | rs7117518 | 0.497 | -0.241 | -1.006 | -0.164 | 0.095 | 0.025 | 0.025 | 0.706 | 0.54 | 0.11 | 0.43 | 0.36 | 0.66 | 0.51 |
| 11 | 72281884 | rs433640 | 0.617 | -0.383 | -0.219 | 0.042 | -0.026 | 0.001 | 0.001 | 0 | 0.8 | 0.39 | 0.83 | 0.82 | 0.84 | 0.9 |
| 11 | 72284569 | rs366066 | -0.455 | -0.168 | -0.325 | 0.366 | -0.009 | 0.002 | 0.002 | 0.576 | 0.35 | 0.6 | 0.5 | 0.64 | 0.25 | 0.41 |
| 11 | 72286156 | rs402479 | 0.964 | 0.628 | 0.219 | 0.655 | 0.044 | 0.113 | 0.113 | 0 | 0.92 | 0.74 | 0.94 | 0.83 | 0.91 | 0.94 |
| 11 | 72288664 | rs1980090 | -0.937 | 0.633 | -0.83 | -0.153 | -0.053 | 0.134 | 0.134 | 0 | 0.55 | 0.83 | 0.68 | 0.75 | 0.49 | 0.62 |
| 11 | 72289291 | rs1135029 | 0.588 | 0.694 | -0.899 | -0.211 | -0.089 | 0.135 | 0.135 | 0 | 0.45 | 0.09 | 0.32 | 0.25 | 0.53 | 0.38 |
| 11 | 72290629 | rs392565 | -0.393 | 0.36 | -0.418 | 0.113 | 0.005 | 0.018 | 0.018 | 0 | 0.33 | 0.25 | 0.24 | 0.28 | 0.29 | 0.36 |
| 11 | 72298280 | rs341048 | -0.124 | 0.185 | -0.406 | 0.092 | 0.002 | -0.002 | -0.002 | 0.006 | 0.11 | 0.14 | 0.12 | 0.08 | 0.13 | 0.11 |
| 11 | 72299182 | rs907851 | 0.344 | 0.171 | -0.216 | 0.115 | -0.017 | -0.017 | -0.017 | 0 | 0.12 | 0.08 | 0.39 | 0.47 | 0.17 | 0.25 |
| 11 | 72308203 | rs2365774 | -0.54 | 0.094 | -0.242 | -0.022 | 0.011 | 0.018 | 0.018 | 0 | 0.31 | 0.19 | 0.24 | 0.28 | 0.27 | 0.36 |
| 11 | 72315581 | rs746399 | -0.036 | 0.093 | -0.41 | 0.178 | -0.026 | -0.065 | -0.065 | 0 | 0.13 | 0.05 | 0.39 | 0.45 | 0.17 | 0.27 |
| 11 | 72316484 | rs341087 | -1.874 | 0.323 | -0.39 | 0.029 | -0.006 | -0.039 | -0.039 | 0 | 0.34 | 0.81 | 0.55 | 0.52 | 0.31 | 0.37 |
| 11 | 72317803 | rs341086 | -1.666 | 0.264 | 0.441 | -0.044 | 0.007 | 0.054 | 0.054 | 0 | 0.46 | 0.62 | 0.66 | 0.74 | 0.44 | 0.57 |
| 11 | 72318831 | rs11235547 | 0.567 | 0.281 | 0.386 | 0.013 | 0.016 | -0.006 | -0.006 | 0 | 0.19 | 0.06 | 0.12 | 0.22 | 0.14 | 0.18 |
| 11 | 72318882 | rs341085 | -0.953 | 0.746 | 0.283 | -0.225 | 0 | -0.012 | -0.012 | 0 | 0.36 | 0.53 | 0.31 | 0.3 | 0.35 | 0.35 |
| 11 | 72322179 | rs10793036 | -2.183 | 0.823 | -0.183 | -0.244 | -0.042 | 0.136 | 0.136 | 0 | 0.56 | 0.86 | 0.71 | 0.74 | 0.54 | 0.59 |
| 11 | 72325482 | rs755933 | 0.675 | 0.879 | -0.314 | -0.187 | 0.004 | 0.017 | 0.017 | 0 | 0.19 | 0.06 | 0.12 | 0.19 | 0.16 | 0.21 |
| 11 | 72330446 | rs11603497 | -0.961 | 1.042 | 0.283 | -0.061 | 0.013 | -0.029 | -0.029 | 0 | 0.2 | 0.3 | 0.16 | 0.08 | 0.23 | 0.14 |
| 11 | 72332931 | rs4944584 | -0.089 | 1.057 | -0.105 | 0.396 | -0.015 | -0.06 | -0.06 | 1.804 | 0.87 | 0.95 | 0.65 | 0.57 | 0.85 | 0.76 |
| 11 | 72334873 | rs401801 | 0.941 | 1.076 | 0.344 | -0.186 | -0.007 | 0.032 | 0.032 | 0 | 0.55 | 0.65 | 0.35 | 0.24 | 0.51 | 0.44 |
| 11 | 72335834 | rs459309 | 1.162 | 1.101 | 0.837 | -0.007 | 0.004 | 0.034 | 0.034 | 0.307 | 0.72 | 0.46 | 0.47 | 0.45 | 0.69 | 0.55 |
| 11 | 72340766 | rs7111188 | -0.162 | 0.97 | 0.339 | -0.305 | -0.003 | 0.008 | 0.008 | 0 | 0.17 | 0.13 | 0.1 | 0.15 | 0.16 | 0.15 |
| 11 | 72345345 | rs1009708 | -0.129 | 0.982 | 0.348 | -0.318 | 0.001 | 0.008 | 0.008 | 0 | 0.17 | 0.13 | 0.09 | 0.15 | 0.14 | 0.15 |
| 11 | 72350595 | rs3781926 | 0.814 | 1.108 | 0.61 | -0.099 | 0.007 | 0.015 | 0.015 | 0.097 | 0.73 | 0.66 | 0.48 | 0.44 | 0.67 | 0.55 |
| 11 | 72358817 | rs377432 | 0.25 | 1.056 | 0.605 | 0.208 | 0.003 | 0.085 | 0.085 | 0 | 0.84 | 0.75 | 0.55 | 0.48 | 0.84 | 0.79 |
| 11 | 72390640 | rs341053 | 0.801 | 0.845 | -0.092 | -0.622 | 0.01 | 0.071 | 0.071 | 0 | 0.43 | 0.33 | 0.33 | 0.31 | 0.42 | 0.3 |
| 11 | 72394706 | rs12576058 | 0.954 | 0.792 | -0.104 | -0.652 | 0.005 | 0.056 | 0.056 | 0 | 0.4 | 0.29 | 0.31 | 0.31 | 0.4 | 0.29 |
| 11 | 72397009 | rs11824205 | -0.888 | 1.094 | 0.71 | -0.188 | 0.185 | 0.159 | 0.159 | 0 | 0.05 | 0.31 | 0.16 | 0.12 | 0.07 | 0.11 |
| 11 | 72399607 | rs2306609 | 0.999 | 0.905 | 0.377 | -0.43 | -0.009 | 0.014 | 0.014 | 0.449 | 0.61 | 0.45 | 0.4 | 0.3 | 0.56 | 0.38 |
| 11 | 72401101 | rs4944668 | -0.825 | 0.85 | 0.5 | -0.213 | 0.003 | -0.004 | -0.004 | 0 | 0.29 | 0.45 | 0.24 | 0.25 | 0.31 | 0.38 |
| 11 | 72402944 | rs7929689 | -0.484 | 0.794 | 0.059 | 0.329 | -0.001 | 0.006 | 0.006 | 0 | 0.91 | 0.94 | 0.76 | 0.62 | 0.91 | 0.87 |
| 11 | 72405223 | rs10751211 | -0.222 | 0.559 | 0.164 | 0.111 | 0.009 | -0.015 | -0.015 | 0.662 | 0.71 | 0.68 | 0.7 | 0.62 | 0.79 | 0.8 |
| 11 | 72405450 | rs10898864 | 0.264 | 0.605 | 0.513 | 0.041 | 0.005 | 0.008 | 0.008 | 0 | 0.19 | 0.22 | 0.22 | 0.24 | 0.27 | 0.31 |
| 11 | 72405868 | rs2306608 | 0.623 | 0.644 | 0.404 | -0.128 | 0 | 0.003 | 0.003 | 0 | 0.52 | 0.46 | 0.47 | 0.38 | 0.52 | 0.49 |
| 11 | 72408266 | rs2306614 | 0.431 | 0.732 | 0.355 | -0.146 | 0.004 | 0.002 | 0.002 | 0.01 | 0.06 | 0.05 | 0.11 | 0.12 | 0.08 | 0.17 |
| 11 | 72408657 | rs2291289 | 1.23 | 0.656 | 0.271 | -0.173 | -0.089 | 0.104 | 0.104 | 0 | 0.43 | 0.04 | 0.34 | 0.26 | 0.44 | 0.32 |
| 11 | 72411099 | rs10898866 | 1.23 | 0.637 | 0.271 | -0.173 | -0.08 | 0.104 | 0.104 | 0 | 0.43 | 0.04 | 0.31 | 0.25 | 0.44 | 0.32 |
| 11 | 72414189 | rs3765105 | 0.645 | 1.041 | 0.252 | -0.14 | -0.001 | 0.001 | 0.001 | 0 | 0.49 | 0.46 | 0.41 | 0.38 | 0.44 | 0.47 |
| 11 | 72415329 | rs2306615 | 0.522 | 1.148 | -0.202 | -0.147 | -0.004 | -0.005 | -0.005 | 1.799 | 0.05 | 0.04 | 0.14 | 0.13 | 0.08 | 0.17 |
| 11 | 72424516 | rs7125221 | -1.533 | 0.979 | -0.258 | -0.201 | -0.01 | 0.081 | 0.081 | 0 | 0.6 | 0.88 | 0.72 | 0.73 | 0.6 | 0.67 |
| 11 | 72429141 | rs1157343 | -0.234 | 0.947 | 0.196 | -0.353 | 0.002 | -0.04 | -0.04 | 0 | 0.33 | 0.49 | 0.29 | 0.19 | 0.35 | 0.43 |
| 11 | 72431231 | rs2365439 | -1.679 | 0.83 | -0.215 | -0.369 | -0.042 | 0.061 | 0.061 | 0 | 0.62 | 0.91 | 0.7 | 0.72 | 0.59 | 0.71 |
| 11 | 72431679 | rs10898868 | -0.014 | 0.839 | -0.055 | -0.405 | -0.003 | 0.01 | 0.01 | 0 | 0.32 | 0.34 | 0.27 | 0.19 | 0.36 | 0.44 |

| | | | | | | | | | | | | | | | | |
|---|---|---|---|---|---|---|---|---|---|---|---|---|---|---|---|---|
| 11 | 72432985 | rs11603334 | 0.978 | 1.037 | 0.171 | 0.097 | -0.02 | -0.016 | -0.016 | 3.698 | 0.07 | 0.03 | 0.09 | 0.08 | 0.15 | 0.2 |
| 11 | 72433098 | rs1552224 | 1.095 | 1.038 | 0.153 | 0.098 | -0.035 | -0.029 | -0.029 | 0.403 | 0.07 | 0.01 | 0.09 | 0.08 | 0.15 | 0.2 |
| 11 | 72437398 | rs1531865 | -0.145 | 0.903 | 0.46 | -0.03 | 0.015 | 0.009 | 0.009 | 0 | 0.44 | 0.26 | 0.51 | 0.53 | 0.4 | 0.31 |
| 11 | 72444583 | rs7123876 | -0.851 | 1.041 | 0.437 | 0.01 | 0.058 | 0.019 | 0.019 | 1.82 | 0.15 | 0.4 | 0.21 | 0.19 | 0.24 | 0.37 |
| 11 | 72449249 | rs10793039 | -0.718 | 0.901 | 0.42 | -0.073 | 0.001 | 0.045 | 0.045 | 0 | 0.52 | 0.64 | 0.57 | 0.61 | 0.49 | 0.5 |
| 11 | 72453738 | rs10793040 | -1.675 | 0.751 | 0.17 | -0.097 | -0.028 | 0.041 | 0.041 | 0 | 0.66 | 0.92 | 0.75 | 0.72 | 0.66 | 0.71 |
| 11 | 72453850 | rs12272109 | -0.849 | 1.09 | 0.702 | -0.296 | 0.093 | 0.04 | 0.04 | 1.657 | 0.09 | 0.47 | 0.14 | 0.1 | 0.09 | 0.17 |
| 11 | 72460155 | rs4943997 | 0.171 | 1.04 | 0.359 | -0.165 | 0.002 | 0.063 | 0.063 | 0.409 | 0.34 | 0.13 | 0.27 | 0.17 | 0.3 | 0.16 |
| 11 | 72462634 | rs7122044 | -1.524 | 1.388 | 1.247 | -0.005 | 0.077 | 0.019 | 0.019 | 1.39 | 0.15 | 0.46 | 0.22 | 0.18 | 0.25 | 0.39 |
| 11 | 72465085 | rs1872127 | -1.074 | 0.942 | 1.49 | -0.106 | 0.1 | 0.03 | 0.03 | 1.254 | 0.08 | 0.41 | 0.13 | 0.1 | 0.1 | 0.16 |
| 11 | 72472336 | rs12287481 | -1.086 | 0.968 | 1.399 | -0.093 | 0.081 | 0.018 | 0.018 | 1.444 | 0.08 | 0.37 | 0.13 | 0.1 | 0.1 | 0.2 |
| 11 | 72478937 | rs673377 | -1.352 | 0.872 | 0.855 | -0.127 | 0.057 | -0.016 | -0.016 | 0 | 0.1 | 0.56 | 0.14 | 0.09 | 0.08 | 0.2 |
| 11 | 72485175 | rs3862790 | 0.141 | 0.809 | -0.485 | -0.399 | -0.018 | 0.075 | 0.075 | 3.126 | 0.27 | 0.09 | 0.19 | 0.15 | 0.24 | 0.11 |
| 11 | 72489023 | rs577217 | -0.003 | 0.399 | -0.127 | 0.289 | -0.01 | 0.093 | 0.093 | 0 | 0.18 | 0.32 | 0.3 | 0.38 | 0.14 | 0.16 |
| 11 | 72509713 | rs1000620 | -1.006 | 0.735 | -0.025 | -0.073 | -0.003 | -0.004 | -0.004 | 1.174 | 0.56 | 0.56 | 0.61 | 0.62 | 0.55 | 0.55 |
| 11 | 72525111 | rs10898880 | -0.967 | 0.65 | -0.005 | -0.014 | 0 | 0 | 0 | 0 | 0.55 | 0.51 | 0.61 | 0.62 | 0.55 | 0.61 |
| 11 | 72527018 | rs10898881 | -0.958 | 0.856 | 0.014 | -0.135 | 0.013 | 0.005 | 0.005 | 0 | 0.58 | 0.38 | 0.63 | 0.63 | 0.6 | 0.65 |
| 11 | 72538600 | rs3862794 | 0.993 | 0.805 | 0.446 | -0.103 | -0.008 | 0.002 | 0.002 | 0 | 0.33 | 0.31 | 0.25 | 0.4 | 0.21 | 0.13 |
| 11 | 72551035 | rs7110183 | 0.768 | 1.21 | -0.192 | -0.593 | -0.075 | 0.073 | 0.073 | 0 | 0.18 | 0.04 | 0.18 | 0.07 | 0.23 | 0.16 |
| 11 | 72559568 | rs10898884 | 0.481 | 1.139 | -0.024 | -0.638 | 0.013 | -0.077 | -0.077 | 1.46 | 0.34 | 0.14 | 0.36 | 0.48 | 0.29 | 0.29 |
| 11 | 72571275 | rs10898885 | -1.219 | 0.633 | -0.219 | -0.993 | 0.01 | 0.003 | 0.003 | 0.103 | 0.64 | 0.37 | 0.73 | 0.7 | 0.8 | 0.86 |
| 11 | 72609119 | rs7943424 | -0.187 | 0.687 | -0.197 | -0.959 | -0.008 | 0.029 | 0.029 | 0.032 | 0.46 | 0.34 | 0.55 | 0.63 | 0.55 | 0.7 |
| 11 | 72638704 | rs11822039 | 1.147 | 0.268 | 0.665 | -0.609 | 0.004 | -0.001 | -0.001 | 0 | 0.3 | 0.27 | 0.24 | 0.3 | 0.19 | 0.12 |
| 11 | 72647296 | rs7926133 | 0.891 | 0.258 | 0.536 | -0.643 | -0.003 | -0.04 | -0.04 | 1.062 | 0.19 | 0.12 | 0.3 | 0.48 | 0.19 | 0.28 |
| 11 | 72669777 | rs11605166 | 1.254 | 0.342 | 0.377 | -0.648 | 0.008 | -0.001 | -0.001 | 0 | 0.08 | 0.11 | 0.11 | 0.08 | 0.17 | 0.22 |
| 11 | 72676874 | rs7129921 | -0.335 | 0.151 | 0.513 | -0.749 | 0.009 | 0.014 | 0.014 | 1.815 | 0.3 | 0.35 | 0.42 | 0.56 | 0.39 | 0.51 |
| 11 | 72684898 | rs12294533 | -0.256 | 0.152 | 0.265 | -0.725 | -0.004 | -0.005 | -0.005 | 0.063 | 0.31 | 0.31 | 0.49 | 0.53 | 0.46 | 0.68 |
| 11 | 72700779 | rs10898894 | -0.188 | 0.163 | 0.098 | -0.719 | -0.003 | -0.003 | -0.003 | 0 | 0.3 | 0.31 | 0.49 | 0.53 | 0.45 | 0.67 |
| 11 | 72705522 | rs3862796 | -1.346 | 0.012 | 0.017 | -0.765 | 0.061 | -0.016 | -0.016 | 0.414 | 0.7 | 0.9 | 0.77 | 0.7 | 0.81 | 0.87 |
| 11 | 72744830 | rs11235639 | 0.586 | 0.094 | -0.853 | -0.622 | 0 | -0.012 | -0.012 | 0 | 0.21 | 0.17 | 0.31 | 0.48 | 0.21 | 0.28 |
| 11 | 72749933 | rs7933548 | 1.086 | 0.003 | -0.995 | -0.721 | -0.097 | 0.191 | 0.191 | 0.267 | 0.33 | 0.06 | 0.25 | 0.07 | 0.34 | 0.19 |
| 11 | 72800970 | rs4394861 | -0.572 | -0.027 | -1.244 | -0.642 | 0.012 | -0.004 | -0.004 | 0 | 0.64 | 0.37 | 0.73 | 0.6 | 0.79 | 0.86 |
| 11 | 72832677 | rs7937701 | -0.278 | 0.462 | -0.847 | -0.279 | -0.01 | -0.003 | -0.003 | 1.051 | 0.33 | 0.31 | 0.49 | 0.63 | 0.48 | 0.68 |
| 11 | 72860038 | rs1783575 | -0.797 | -0.29 | -0.934 | -0.175 | 0.011 | -0.012 | -0.012 | 2.892 | 0.7 | 0.75 | 0.76 | 0.59 | 0.81 | 0.87 |
| 11 | 72871055 | rs2511062 | -0.063 | 0.19 | -1.461 | -0.284 | 0.001 | 0.004 | 0.004 | 0 | 0.48 | 0.54 | 0.4 | 0.49 | 0.44 | 0.32 |
| 11 | 72877964 | rs1791936 | -0.076 | 0.465 | -2.149 | -0.695 | 0.011 | 0.22 | 0.22 | 0 | 0.47 | 0.13 | 0.4 | 0.2 | 0.6 | 0.49 |
| 11 | 72884382 | rs880895 | 0.969 | 0.62 | -2.06 | -0.637 | -0.068 | 0.108 | 0.108 | 0 | 0.25 | 0.07 | 0.24 | 0.06 | 0.33 | 0.19 |
| 11 | 72885257 | rs1628409 | 0.926 | 0.627 | -2.112 | -0.542 | -0.067 | 0.118 | 0.118 | 0 | 0.26 | 0.06 | 0.24 | 0.07 | 0.33 | 0.19 |
| 11 | 72889018 | rs1783585 | -0.976 | -0.269 | -1.507 | -0.152 | -0.047 | 0.049 | 0.049 | 0.223 | 0.85 | 0.58 | 0.9 | 0.77 | 0.95 | 0.92 |
| 11 | 72889137 | rs1783584 | -0.079 | 0.129 | -1.555 | 0.106 | 0.044 | -0.001 | -0.001 | 0.038 | 0.44 | 0.23 | 0.36 | 0.48 | 0.37 | 0.4 |
| 11 | 72890304 | rs1791049 | -0.428 | 0.265 | -1.543 | -0.16 | 0 | -0.002 | -0.002 | 0 | 0.55 | 0.48 | 0.46 | 0.58 | 0.56 | 0.61 |
| 11 | 72894273 | rs11603160 | -0.018 | -0.214 | -1.852 | -0.564 | -0.003 | 0.052 | 0.052 | 0.021 | 0.2 | 0.02 | 0.14 | 0.06 | 0.16 | 0.09 |
| 11 | 72899169 | rs489412 | 0.485 | -0.09 | -1.621 | -0.394 | 0.034 | -0.013 | -0.013 | 0 | 0.34 | 0.12 | 0.29 | 0.36 | 0.28 | 0.35 |
| 11 | 72904844 | rs12292587 | 0.669 | 0.586 | -1.343 | -0.458 | 0.002 | 0.012 | 0.012 | 0 | 0.16 | 0.12 | 0.11 | 0.08 | 0.16 | 0.07 |
| 11 | 72905310 | rs3931643 | 0.047 | 1.275 | -0.978 | 0.107 | -0.08 | -0.116 | -0.116 | 0 | 0.14 | 0.01 | 0.37 | 0.49 | 0.28 | 0.31 |
| 11 | 72908553 | rs11235676 | -0.508 | 0.155 | 0.455 | -0.836 | 0.107 | -0.053 | -0.053 | 0 | 0.1 | 0.25 | 0.11 | 0 | 0.13 | 0.03 |
| 11 | 72913638 | rs523907 | -1.409 | -0.708 | 0.473 | -0.548 | -0.01 | -0.046 | -0.046 | 0 | 0.27 | 0.49 | 0.17 | 0.14 | 0.23 | 0.22 |
| 11 | 72917269 | rs10898906 | -1.305 | -0.454 | 0.338 | -0.378 | -0.001 | -0.002 | -0.002 | 0 | 0.17 | 0.19 | 0.1 | 0.1 | 0.14 | 0.16 |
| 11 | 72918159 | rs565889 | 1.544 | -0.692 | 0.387 | -0.586 | -0.019 | -0.074 | -0.074 | 7.871 | 0.72 | 0.5 | 0.84 | 0.9 | 0.79 | 0.76 |
| 11 | 72930806 | rs508859 | 1.049 | -0.654 | 0.146 | -0.116 | 0.01 | 0.029 | 0.029 | 1.682 | 0.84 | 0.46 | 0.9 | 0.85 | 0.86 | 0.85 |
| 11 | 72934619 | rs12364461 | -1.122 | -0.629 | -0.839 | -0.242 | -0.003 | -0.003 | -0.003 | 0.261 | 0.11 | 0.03 | 0.03 | 0.08 | 0.09 | 0.22 |
| 11 | 72937274 | rs1790080 | -0.712 | -1.427 | -0.769 | -0.538 | -0.037 | -0.044 | -0.044 | 0 | 0.33 | 0.53 | 0.23 | 0.14 | 0.2 | 0.08 |
| 11 | 72945341 | rs2511241 | 0.753 | -0.609 | -1.258 | 0.004 | 0 | 0.002 | 0.002 | 0 | 0.9 | 0.9 | 0.96 | 0.88 | 0.92 | 0.9 |
| 11 | 72946204 | rs1626154 | -0.264 | -0.996 | -2.133 | -0.615 | -0.003 | 0.097 | 0.097 | 0.251 | 0.25 | 0.02 | 0.16 | 0.05 | 0.21 | 0.06 |
| 11 | 72949172 | rs4944832 | 0.412 | -0.135 | -2.756 | -0.359 | -0.159 | -0.036 | -0.036 | 0.145 | 0.33 | 0.04 | 0.34 | 0.3 | 0.41 | 0.48 |
| 11 | 72949747 | rs11235689 | 0.386 | -0.157 | -2.59 | -0.34 | -0.109 | -0.009 | -0.009 | 0 | 0.34 | 0.1 | 0.34 | 0.3 | 0.4 | 0.48 |
| 11 | 72952400 | rs4382936 | -0.725 | 0.657 | -1.932 | -0.022 | -0.011 | -0.007 | -0.007 | 0 | 0.27 | 0.27 | 0.54 | 0.68 | 0.35 | 0.33 |
| 11 | 72952496 | rs10898909 | 0.091 | 0.912 | -1.911 | 0.432 | -0.082 | -0.102 | -0.102 | 0.319 | 0.13 | 0.01 | 0.41 | 0.51 | 0.24 | 0.22 |
| 11 | 72954449 | rs3814728 | 0.03 | -0.231 | -1.401 | -0.598 | 0.014 | 0.022 | 0.022 | 0.254 | 0.74 | 0.82 | 0.75 | 0.95 | 0.78 | 0.92 |
| 11 | 72957320 | rs12361940 | -0.824 | -0.175 | -0.501 | -0.478 | -0.024 | -0.098 | -0.098 | 0 | 0.16 | 0.34 | 0.11 | 0 | 0.1 | 0.02 |
| 11 | 72968660 | rs10793057 | 0.342 | 0.314 | -0.879 | 0.49 | -0.026 | -0.017 | -0.017 | 0.015 | 0.06 | 0 | 0.22 | 0.3 | 0.08 | 0.2 |
| 11 | 72970563 | rs11235699 | -1.138 | 1.114 | -0.075 | -0.189 | 0.093 | -0.021 | -0.021 | 0 | 0.07 | 0.19 | 0.08 | 0.01 | 0.12 | 0.02 |
| 11 | 72970776 | rs7925649 | -1.274 | 0.747 | 0.272 | -0.404 | 0.026 | -0.132 | -0.132 | 0.503 | 0.2 | 0.49 | 0.14 | 0.01 | 0.19 | 0.03 |
| 11 | 72976988 | rs7127940 | -0.394 | 0.745 | 0.272 | -0.32 | -0.011 | 0.017 | 0.017 | 0.306 | 0.14 | 0.1 | 0.09 | 0.02 | 0.17 | 0.07 |
| 11 | 72979231 | rs12790863 | -0.977 | 0.725 | 0.26 | -0.33 | 0.023 | -0.103 | -0.103 | 0 | 0.18 | 0.54 | 0.13 | 0.02 | 0.17 | 0.07 |
| 11 | 72984030 | rs2027765 | 0.844 | 0.615 | -1.044 | -0.506 | -0.081 | 0.064 | 0.064 | 0.088 | 0.46 | 0.08 | 0.62 | 0.71 | 0.45 | 0.51 |
| 11 | 72994199 | rs11235712 | -1.006 | 0.483 | 0.011 | -0.552 | 0.02 | -0.102 | -0.102 | 0 | 0.17 | 0.4 | 0.12 | 0.02 | 0.17 | 0.07 |
| 11 | 73002434 | rs12803970 | -0.966 | 0.684 | -1.077 | -0.631 | -0.021 | 0.085 | 0.085 | 1.129 | 0.21 | 0.06 | 0.14 | 0.02 | 0.23 | 0.09 |
| 11 | 73004053 | rs1806516 | 0.635 | 0.312 | -0.582 | -0.187 | -0.003 | -0.006 | -0.006 | 0 | 0.65 | 0.4 | 0.75 | 0.72 | 0.62 | 0.55 |

| | | | | | | | | | | | | | | | | |
|---|---|---|---|---|---|---|---|---|---|---|---|---|---|---|---|---|
| 11 | 73006609 | rs3741153 | 0.16 | 0.343 | -1.639 | -0.23 | -0.179 | -0.067 | -0.067 | 0.014 | 0.36 | 0.02 | 0.54 | 0.42 | 0.45 | 0.32 |
| 11 | 73009432 | rs7115806 | -0.649 | 0.413 | -0.166 | -0.419 | -0.003 | -0.019 | -0.019 | 0 | 0.13 | 0.66 | 0.2 | 0.06 | 0.11 | 0.09 |
| 11 | 73011106 | rs11824050 | -1.182 | 0.594 | -0.102 | -0.674 | 0.004 | -0.047 | -0.047 | 0.6 | 0.2 | 0.73 | 0.17 | 0.05 | 0.21 | 0.13 |
| 11 | 73023353 | rs672660 | -0.249 | 0.374 | -0.24 | -0.463 | 0.003 | -0.01 | -0.01 | 0 | 0.83 | 0.88 | 0.85 | 0.73 | 0.84 | 0.65 |
| 11 | 73025971 | rs1107038 | 0.428 | 0.494 | 0.569 | -0.609 | 0.075 | -0.028 | -0.028 | 0.015 | 0.08 | 0.26 | 0.07 | 0 | 0.1 | 0.03 |
| 11 | 73032471 | rs725103 | -1.219 | 0.45 | 0.759 | -0.619 | 0.015 | -0.069 | -0.069 | 0 | 0.19 | 0.66 | 0.16 | 0.05 | 0.21 | 0.14 |
| 11 | 73036179 | rs2027761 | -0.125 | 0.457 | 0.69 | -0.58 | 0.007 | -0.03 | -0.03 | 0 | 0.11 | 0.37 | 0.09 | 0.04 | 0.11 | 0.11 |
| 11 | 73039229 | rs7125978 | -0.025 | 0.453 | 0.69 | -0.58 | 0.006 | -0.029 | -0.029 | 0 | 0.11 | 0.37 | 0.09 | 0.04 | 0.11 | 0.11 |
| 11 | 73080405 | rs947837 | -1.398 | 0.67 | 0.388 | -0.853 | 0.017 | -0.125 | -0.125 | 0.763 | 0.25 | 0.54 | 0.18 | 0.05 | 0.25 | 0.17 |
| 11 | 73083880 | rs7130689 | -0.599 | 0.443 | -0.173 | -0.848 | -0.177 | -0.129 | -0.129 | 1.529 | 0.66 | 0.98 | 0.47 | 0.56 | 0.57 | 0.7 |
| 11 | 73086303 | rs876480 | -1.645 | 0.746 | 0.991 | -0.946 | 0.002 | -0.064 | -0.064 | 0.011 | 0.2 | 0.7 | 0.16 | 0.05 | 0.2 | 0.14 |
| 11 | 73106294 | rs17851510 | -0.49 | 0.802 | 1.18 | -0.836 | 0.016 | -0.045 | -0.045 | 0 | 0.1 | 0.3 | 0.08 | 0.01 | 0.11 | 0.08 |
| 11 | 73108276 | rs880317 | -1.794 | 0.762 | 1.219 | -0.902 | -0.003 | -0.075 | -0.075 | 0.316 | 0.22 | 0.67 | 0.16 | 0.04 | 0.2 | 0.11 |
| 11 | 73109635 | rs2154948 | -0.08 | 0.837 | 1.174 | -0.863 | -0.002 | -0.003 | -0.003 | 0 | 0.16 | 0.11 | 0.18 | 0.27 | 0.17 | 0.29 |
| 11 | 73109877 | rs6592523 | -1.939 | 1.024 | 1.241 | -0.802 | -0.003 | -0.051 | -0.051 | 0 | 0.22 | 0.69 | 0.24 | 0.06 | 0.2 | 0.11 |
| 11 | 73122340 | rs1938777 | -0.988 | 1.071 | 1.255 | -0.885 | 0.042 | -0.035 | -0.035 | 0 | 0.12 | 0.54 | 0.1 | 0.04 | 0.12 | 0.08 |
| 11 | 73158119 | rs7941359 | -1.156 | 1.005 | 1.006 | -0.875 | 0.008 | -0.047 | -0.047 | 0.135 | 0.13 | 0.59 | 0.11 | 0.04 | 0.11 | 0.08 |
| 11 | 73171306 | rs17244589 | 0.358 | 1.034 | 0.694 | -0.86 | -0.03 | 0.019 | 0.019 | 0 | 0.08 | 0.01 | 0.12 | 0.03 | 0.09 | 0.03 |
| 11 | 73189165 | rs513268 | -2.301 | 0.977 | 0.669 | -0.891 | -0.006 | 0.023 | 0.023 | 0 | 0.39 | 0.77 | 0.33 | 0.32 | 0.39 | 0.42 |
| 11 | 73194645 | rs501953 | -1.699 | 0.9 | 0.678 | -0.891 | 0.001 | 0.001 | 0.001 | 0 | 0.31 | 0.69 | 0.2 | 0.3 | 0.3 | 0.38 |
| 11 | 73198494 | rs2131670 | -0.951 | 0.895 | 0.343 | -0.899 | -0.004 | -0.003 | -0.003 | 0 | 0.22 | 0.15 | 0.14 | 0.27 | 0.28 | 0.35 |
| 11 | 73216385 | rs12421012 | 0.158 | 1.006 | -0.373 | -0.877 | -0.006 | 0.013 | 0.013 | 0 | 0.08 | 0.08 | 0.13 | 0.03 | 0.09 | 0.03 |
| 11 | 73218230 | rs570111 | -1.926 | 0.994 | -0.54 | -0.881 | -0.01 | 0.037 | 0.037 | 0.047 | 0.41 | 0.86 | 0.34 | 0.33 | 0.38 | 0.42 |
| 11 | 73248985 | rs11235796 | -0.409 | 1.416 | -0.547 | -0.986 | -0.019 | 0.024 | 0.024 | 0 | 0.08 | 0.03 | 0.13 | 0.03 | 0.1 | 0.03 |
| 11 | 73291255 | rs11235811 | -0.155 | 1.247 | -0.734 | -0.73 | -0.03 | 0.018 | 0.018 | 0 | 0.08 | 0.01 | 0.12 | 0.03 | 0.09 | 0.03 |
| 11 | 73292089 | rs11235812 | 0.007 | 1.055 | -0.476 | -0.568 | -0.023 | -0.005 | -0.005 | 0 | 0.06 | 0 | 0.03 | 0.03 | 0.11 | 0.17 |
| 11 | 73312252 | rs10751226 | 0.667 | 0.868 | -0.134 | -0.588 | 0.021 | -0.091 | -0.091 | 0 | 0.84 | 0.58 | 0.86 | 0.99 | 0.83 | 0.9 |
| 11 | 73317931 | rs737586 | -0.073 | 0.836 | -1.016 | -0.451 | -0.045 | 0.072 | 0.072 | 0 | 0.13 | 0.02 | 0.08 | 0.03 | 0.16 | 0.05 |
| 11 | 73318139 | rs7929943 | 0.107 | 1.253 | -0.854 | -0.066 | 0.034 | 0.07 | 0.07 | 0 | 0.24 | 0.35 | 0.45 | 0.53 | 0.32 | 0.3 |
| 11 | 73318285 | rs11235819 | -0.304 | 0.516 | -0.58 | 0.21 | 0 | 0 | 0 | 0 | 0.45 | 0.47 | 0.54 | 0.57 | 0.49 | 0.36 |
| 11 | 73337929 | rs2949804 | 0.365 | 0.11 | -0.293 | 0.412 | -0.001 | 0.02 | 0.02 | 0 | 0.75 | 0.72 | 0.71 | 0.71 | 0.76 | 0.62 |
| 11 | 73340690 | rs7948050 | -0.54 | -0.194 | -0.203 | -0.253 | -0.003 | -0.03 | -0.03 | 2.746 | 0.14 | 0.13 | 0.09 | 0.03 | 0.1 | 0.1 |
| 11 | 73346041 | rs666839 | -0.829 | -0.754 | 0.124 | -0.486 | -0.009 | 0.001 | 0.001 | 0 | 0.27 | 0.32 | 0.16 | 0.23 | 0.18 | 0.34 |
| 11 | 73351285 | rs650393 | 0.702 | -0.695 | -0.05 | -0.591 | -0.042 | -0.069 | -0.069 | 1.281 | 0.73 | 0.92 | 0.76 | 0.69 | 0.69 | 0.65 |
| 11 | 73351846 | rs663303 | 0.12 | -0.655 | -0.042 | -0.567 | -0.036 | -0.016 | -0.016 | 0 | 0.92 | 0.99 | 0.88 | 0.92 | 0.86 | 0.96 |
| 11 | 73352751 | rs11235833 | 2.008 | -0.917 | 0.475 | -0.457 | -0.025 | -0.021 | -0.021 | 0.043 | 0.44 | 0.22 | 0.55 | 0.51 | 0.53 | 0.39 |
| 11 | 73356872 | rs3781909 | 2.012 | -0.941 | 0.69 | -0.427 | -0.025 | -0.016 | -0.016 | 0 | 0.44 | 0.22 | 0.55 | 0.5 | 0.53 | 0.39 |
| 11 | 73364673 | rs7952228 | 1.622 | -1.03 | 1.208 | -0.478 | 0.011 | 0.007 | 0.007 | 0 | 0.77 | 0.59 | 0.77 | 0.75 | 0.75 | 0.73 |
| 11 | 73365157 | rs640655 | -0.62 | -1.008 | 1.476 | -0.451 | 0.022 | -0.06 | -0.06 | 0.152 | 0.11 | 0.43 | 0.09 | 0.02 | 0.07 | 0.1 |
| 11 | 73365504 | rs622833 | -1.504 | -1.004 | 1.41 | -0.441 | -0.006 | 0.026 | 0.026 | 0 | 0.16 | 0.51 | 0.14 | 0.17 | 0.11 | 0.23 |
| 11 | 73367122 | rs11235841 | 0.423 | -1.03 | 1.211 | -0.439 | 0 | 0.003 | 0.003 | 0 | 0.12 | 0.08 | 0.13 | 0.08 | 0.13 | 0.05 |
| 11 | 73372299 | rs7128512 | 0.818 | -1.174 | 1.326 | -0.44 | -0.02 | 0.003 | 0.003 | 0 | 0.76 | 0.62 | 0.88 | 0.81 | 0.83 | 0.69 |
| 11 | 73373552 | rs6590 | -0.101 | -1.195 | 1.217 | -0.445 | 0.009 | -0.047 | -0.047 | 0 | 0.17 | 0.04 | 0.07 | 0.2 | 0.12 | 0.29 |
| 11 | 73377463 | rs2037706 | 0.553 | -1.139 | 1.436 | -0.695 | 0.027 | -0.025 | -0.025 | 0 | 0.94 | 0.76 | 0.95 | 1 | 0.93 | 0.98 |
| 11 | 73378142 | rs12274970 | -1.299 | -1.313 | 1.633 | -0.762 | -0.005 | -0.006 | -0.006 | 0.002 | 0.33 | 0.33 | 0.23 | 0.2 | 0.26 | 0.35 |
| 11 | 73379075 | rs11235844 | -0.439 | -1.324 | 1.619 | -0.778 | 0.022 | 0 | 0 | 0 | 0.2 | 0.08 | 0.08 | 0.13 | 0.13 | 0.29 |
| 11 | 73383382 | rs7126049 | -1.201 | -1.423 | 1.664 | -0.776 | 0.002 | 0.009 | 0.009 | 0 | 0.22 | 0.32 | 0.24 | 0.25 | 0.23 | 0.25 |
| 11 | 73396061 | rs12420357 | 2.531 | -1.479 | 1.639 | -0.811 | -0.01 | -0.008 | -0.008 | 0 | 0.45 | 0.21 | 0.59 | 0.63 | 0.55 | 0.41 |
| 11 | 73398280 | rs6592529 | -0.107 | -1.48 | 1.658 | -0.812 | 0.021 | -0.025 | -0.025 | 0.773 | 0.95 | 0.82 | 0.95 | 1 | 0.93 | 0.98 |
| 11 | 73402576 | rs10736793 | 0.77 | -1.487 | 1.663 | -0.815 | -0.013 | -0.016 | -0.016 | 0.052 | 0.74 | 0.66 | 0.87 | 0.87 | 0.8 | 0.68 |
| 11 | 73402679 | rs1464908 | 0.77 | -1.487 | 1.663 | -0.815 | -0.013 | -0.016 | -0.016 | 0.351 | 0.74 | 0.66 | 0.87 | 0.87 | 0.8 | 0.68 |
| 11 | 73408272 | rs11235864 | -0.197 | -1.484 | 1.609 | -0.814 | 0.018 | -0.009 | -0.009 | 0.706 | 0.19 | 0.06 | 0.07 | 0.13 | 0.13 | 0.29 |
| 11 | 73411927 | rs12576616 | -0.19 | -1.472 | 1.611 | -0.809 | 0 | 0.007 | 0.007 | 0 | 0.16 | 0.2 | 0.19 | 0.23 | 0.18 | 0.18 |
| 11 | 73417672 | rs7119055 | 2.521 | -1.472 | 1.612 | -0.81 | -0.01 | -0.008 | -0.008 | 0 | 0.45 | 0.21 | 0.59 | 0.63 | 0.55 | 0.41 |
| 11 | 73435296 | rs11235880 | -0.202 | -1.415 | 1.588 | -0.782 | 0 | 0.007 | 0.007 | 0 | 0.16 | 0.2 | 0.19 | 0.23 | 0.18 | 0.18 |
| 11 | 73477045 | rs1723838 | -0.248 | -1.389 | 1.502 | -0.757 | 0.021 | -0.025 | -0.025 | 0 | 0.05 | 0.18 | 0.05 | 0 | 0.07 | 0.02 |
| 11 | 73497924 | rs1723845 | -0.2 | -1.387 | 1.503 | -0.756 | 0.022 | -0.024 | -0.024 | 0 | 0.05 | 0.18 | 0.05 | 0 | 0.07 | 0.02 |
| 11 | 73498933 | rs1792174 | -3.033 | -1.455 | 1.476 | -0.783 | -0.006 | -0.007 | -0.007 | 0 | 0.55 | 0.79 | 0.41 | 0.37 | 0.45 | 0.59 |
| 11 | 73505651 | rs11235907 | -0.154 | -1.466 | 1.459 | -0.742 | 0.016 | 0 | 0 | 0 | 0.19 | 0.03 | 0.07 | 0.11 | 0.13 | 0.29 |
| 11 | 73510645 | rs11235909 | -0.206 | -1.417 | 1.506 | -0.711 | 0.006 | 0.095 | 0.095 | 0 | 0.05 | 0.13 | 0.07 | 0.17 | 0.05 | 0.13 |
| 11 | 73517259 | rs11235911 | 0.656 | -1.418 | 1.497 | -0.717 | 0 | -0.009 | -0.009 | 1.139 | 0.12 | 0.17 | 0.14 | 0.07 | 0.13 | 0.05 |
| 11 | 73529453 | rs1792156 | -2.767 | -1.405 | 1.425 | -0.717 | -0.002 | -0.002 | -0.002 | 0 | 0.54 | 0.77 | 0.4 | 0.37 | 0.45 | 0.59 |
| 11 | 73532916 | rs1670537 | -2.767 | -1.405 | 1.425 | -0.717 | -0.002 | -0.002 | -0.002 | 0 | 0.54 | 0.77 | 0.4 | 0.37 | 0.45 | 0.59 |
| 11 | 73544202 | rs12284670 | -0.277 | -1.275 | 1.529 | -0.631 | 0.012 | 0.117 | 0.117 | 0 | 0.05 | 0.13 | 0.07 | 0.17 | 0.05 | 0.12 |
| 11 | 73548270 | rs1792165 | -2.364 | -1.56 | 1.46 | -0.778 | 0.047 | 0.01 | 0.01 | 0 | 0.49 | 0.72 | 0.35 | 0.35 | 0.38 | 0.51 |
| 11 | 73548670 | rs1792164 | 0.306 | -1.719 | 1.435 | -0.813 | -0.009 | -0.041 | -0.041 | 0 | 0.08 | 0.18 | 0.04 | 0 | 0.06 | 0.02 |
| 11 | 73549825 | rs1792163 | -0.64 | -1.729 | 1.44 | -0.818 | -0.008 | -0.052 | -0.052 | 0 | 0.2 | 0.35 | 0.18 | 0.07 | 0.19 | 0.07 |
| 11 | 73570697 | rs7124841 | 0.089 | -1.634 | 1.337 | -0.714 | 0.001 | -0.001 | -0.001 | 0 | 0.07 | 0.06 | 0.1 | 0.05 | 0.07 | 0.07 |
| 11 | 73570718 | rs7113309 | 0.089 | -1.651 | 1.337 | -0.755 | 0 | -0.017 | -0.017 | 0 | 0.07 | 0.06 | 0.05 | 0.01 | 0.06 | 0.07 |

| | | | | | | | | | | | | | | | | |
|---|---|---|---|---|---|---|---|---|---|---|---|---|---|---|---|---|
| 11 | 73572775 | rs10898948 | -2.016 | -1.739 | 1.38 | -0.792 | -0.023 | 0.005 | 0.005 | 0 | 0.36 | 0.52 | 0.23 | 0.3 | 0.25 | 0.51 |
| 11 | 73576748 | rs11235933 | -0.021 | -1.752 | 1.23 | -0.79 | 0.009 | -0.022 | -0.022 | 0.233 | 0.16 | 0.03 | 0.07 | 0.11 | 0.11 | 0.28 |
| 11 | 73581543 | rs11220 | -2.764 | -2.066 | 1.274 | -0.915 | 0.066 | 0.029 | 0.029 | 0 | 0.46 | 0.75 | 0.27 | 0.3 | 0.31 | 0.53 |
| 11 | 73581908 | rs12294934 | -0.598 | -1.819 | 1.311 | -0.75 | 0.026 | 0.187 | 0.187 | 0.017 | 0.06 | 0.26 | 0.07 | 0.17 | 0.05 | 0.12 |
| 11 | 73586529 | rs11235937 | -1.254 | -1.821 | 1.27 | -0.753 | 0.015 | 0.021 | 0.021 | 0 | 0.24 | 0.13 | 0.13 | 0.13 | 0.18 | 0.36 |
| 11 | 73588240 | rs1122373 | -0.369 | -1.818 | 1.283 | -0.75 | -0.014 | 0.099 | 0.099 | 0 | 0.11 | 0.3 | 0.08 | 0.17 | 0.07 | 0.15 |
| 11 | 73600888 | rs10898950 | -1.937 | -1.787 | 1.085 | -0.737 | -0.022 | 0.008 | 0.008 | 0 | 0.35 | 0.43 | 0.21 | 0.3 | 0.24 | 0.51 |
| 11 | 73614409 | rs12284235 | -1.664 | -1.136 | 1.441 | -0.579 | 0.008 | 0.021 | 0.021 | 0 | 0.19 | 0.37 | 0.16 | 0.19 | 0.18 | 0.23 |
| 11 | 73630929 | rs935988 | -2.169 | -1.059 | 1.107 | -0.566 | -0.006 | -0.013 | -0.013 | 0 | 0.28 | 0.63 | 0.22 | 0.19 | 0.23 | 0.25 |
| 11 | 73638858 | rs11235951 | 0.514 | -0.901 | 0.918 | -0.485 | -0.001 | -0.001 | -0.001 | 0.061 | 0.11 | 0.09 | 0.14 | 0.07 | 0.13 | 0.05 |
| 11 | 73660546 | rs637028 | -1.628 | -0.796 | 0.848 | -0.321 | -0.01 | 0.003 | 0.003 | 0 | 0.21 | 0.47 | 0.15 | 0.16 | 0.17 | 0.19 |
| 11 | 73668842 | rs634771 | -1.141 | -0.487 | 0.42 | -0.217 | 0.006 | -0.006 | -0.006 | 0 | 0.44 | 0.6 | 0.34 | 0.44 | 0.38 | 0.51 |
| 11 | 73669310 | rs10793068 | -1.171 | -0.463 | 0.212 | -0.285 | 0.015 | -0.021 | -0.021 | 0.448 | 0.45 | 0.7 | 0.35 | 0.44 | 0.4 | 0.47 |
| 11 | 73670645 | rs653263 | -1.754 | -0.318 | -0.224 | -0.252 | -0.071 | -0.085 | -0.085 | 0 | 0.56 | 0.9 | 0.57 | 0.45 | 0.48 | 0.51 |
| 11 | 73678945 | rs4944871 | -0.805 | -0.125 | 0.41 | -0.489 | 0.002 | -0.002 | -0.002 | 0.653 | 0.35 | 0.32 | 0.4 | 0.16 | 0.32 | 0.24 |
| 11 | 73679232 | rs2306818 | -0.827 | -0.164 | 0.505 | -0.508 | 0.011 | 0.048 | 0.048 | 0 | 0.34 | 0.21 | 0.38 | 0.16 | 0.32 | 0.24 |
| 11 | 73681090 | rs2306819 | -0.833 | -0.168 | 0.503 | -0.51 | 0.011 | 0.049 | 0.049 | 0 | 0.34 | 0.21 | 0.38 | 0.16 | 0.32 | 0.24 |
| 11 | 73684215 | rs655717 | -1.997 | -0.072 | 0.659 | -0.332 | -0.011 | -0.035 | -0.035 | 0 | 0.51 | 0.87 | 0.53 | 0.42 | 0.47 | 0.42 |
| 11 | 73684715 | rs643064 | -0.774 | 0.497 | 1.15 | -0.382 | 0.113 | 0.331 | 0.331 | 0.404 | 0.09 | 0.44 | 0.11 | 0.25 | 0.13 | 0.16 |
| 11 | 73689104 | rs660339 | -0.99 | 0.611 | 0.887 | -0.534 | 0.008 | 0.023 | 0.023 | 0 | 0.37 | 0.45 | 0.44 | 0.42 | 0.4 | 0.37 |
| 11 | 73691774 | rs17132534 | -0.441 | 0.478 | 0.882 | -0.49 | -0.016 | -0.052 | -0.052 | 0 | 0.09 | 0.21 | 0.05 | 0 | 0.06 | 0.04 |
| 11 | 73694754 | rs659366 | -0.929 | 0.586 | 0.801 | -0.356 | 0.003 | 0.03 | 0.03 | 0 | 0.35 | 0.45 | 0.42 | 0.42 | 0.36 | 0.37 |
| 11 | 73711477 | rs15763 | 0.318 | 0.375 | 0.86 | -0.507 | -0.015 | -0.046 | -0.046 | 0 | 0.65 | 0.57 | 0.63 | 0.84 | 0.75 | 0.77 |
| 11 | 73713560 | rs11235971 | -0.281 | 0.705 | 0.781 | -0.572 | 0.002 | 0.015 | 0.015 | 0 | 0.13 | 0.14 | 0.1 | 0.26 | 0.23 | 0.23 |
| 11 | 73713591 | rs1685354 | -0.466 | 0.638 | 0.789 | -0.588 | -0.018 | -0.054 | -0.054 | 0.441 | 0.35 | 0.45 | 0.37 | 0.16 | 0.25 | 0.23 |
| 11 | 73715542 | rs2075577 | -1.837 | 0.968 | 0.254 | -0.195 | -0.033 | -0.049 | -0.049 | 0 | 0.59 | 0.86 | 0.54 | 0.42 | 0.54 | 0.49 |
| 11 | 73716469 | rs3781907 | -1.333 | 1.36 | 0.759 | 0.409 | 0.026 | 0.064 | 0.064 | 0.326 | 0.18 | 0.29 | 0.19 | 0.39 | 0.27 | 0.25 |
| 11 | 73728522 | rs1726760 | -0.107 | 0.561 | 1.278 | 0.074 | -0.01 | -0.017 | -0.017 | 0.277 | 0.06 | 0.29 | 0.03 | 0 | 0 | 0.01 |
| 11 | 73745395 | rs17132610 | -0.793 | 0.523 | 2.119 | 0.03 | 0.005 | -0.008 | -0.008 | 0 | 0.07 | 0.35 | 0.03 | 0.01 | 0.02 | 0.02 |
| 11 | 73751481 | rs12417424 | -0.71 | 0.589 | 1.726 | -0.223 | -0.003 | 0.004 | 0.004 | 0 | 0.14 | 0.13 | 0.29 | 0.08 | 0.17 | 0.09 |
| 11 | 73759619 | rs17310472 | -0.145 | 0.733 | 1.538 | -0.167 | -0.026 | -0.006 | -0.006 | 0 | 0.07 | 0.02 | 0.26 | 0.08 | 0.12 | 0.05 |
| 11 | 73770072 | rs826048 | -0.978 | 0.586 | 1.984 | -0.185 | 0.007 | -0.008 | -0.008 | 0.022 | 0.08 | 0.52 | 0.05 | 0.01 | 0.02 | 0.02 |
| 11 | 73785326 | rs4453265 | -1.604 | 0.586 | 1.409 | -0.243 | 0 | 0 | 0 | 0 | 0.46 | 0.43 | 0.48 | 0.44 | 0.47 | 0.45 |
| 11 | 73787381 | rs826062 | -2.143 | 0.586 | 1.285 | -0.234 | -0.008 | 0.002 | 0.002 | 0 | 0.52 | 0.72 | 0.5 | 0.43 | 0.48 | 0.46 |
| 11 | 73796917 | rs826058 | -1.463 | 0.457 | 1.412 | -0.34 | 0.004 | -0.001 | -0.001 | 0 | 0.39 | 0.57 | 0.41 | 0.16 | 0.31 | 0.24 |
| 11 | 73797193 | rs11235994 | -0.245 | 0.463 | 1.444 | -0.334 | -0.005 | -0.004 | -0.004 | 0.604 | 0.13 | 0.13 | 0.09 | 0.28 | 0.18 | 0.22 |
| 11 | 73797520 | rs826057 | -1.462 | 0.463 | 1.423 | -0.335 | 0.004 | -0.001 | -0.001 | 0 | 0.39 | 0.57 | 0.41 | 0.16 | 0.31 | 0.24 |
| 11 | 73806443 | rs11235995 | -0.842 | 0.549 | 1.522 | -0.225 | 0.017 | 0.073 | 0.073 | 0 | 0.15 | 0.3 | 0.1 | 0.28 | 0.18 | 0.22 |
| 11 | 73825503 | rs10898957 | 0.233 | 0.551 | 1.374 | -0.226 | -0.004 | 0.007 | 0.007 | 0 | 0.11 | 0.09 | 0.09 | 0.27 | 0.17 | 0.22 |
| 11 | 73831032 | rs11236006 | -0.448 | 0.448 | 1.292 | -0.338 | -0.02 | 0.036 | 0.036 | 0 | 0.23 | 0.07 | 0.34 | 0.16 | 0.25 | 0.19 |
| 11 | 73863411 | rs11604713 | -1.627 | 0.41 | 1.52 | -0.357 | 0.003 | 0.079 | 0.079 | 0 | 0.21 | 0.48 | 0.12 | 0.28 | 0.19 | 0.24 |
| 11 | 73868065 | rs869616 | -0.482 | 0.407 | 1.349 | -0.365 | -0.019 | 0.037 | 0.037 | 0 | 0.24 | 0.06 | 0.34 | 0.16 | 0.25 | 0.19 |
| 11 | 73875354 | rs11602451 | -1.662 | 0.421 | 1.527 | -0.326 | -0.006 | 0.075 | 0.075 | 0 | 0.2 | 0.46 | 0.12 | 0.28 | 0.18 | 0.23 |
| 11 | 73878688 | rs7130254 | 0.516 | 0.348 | 1.299 | -0.408 | 0.036 | 0.056 | 0.056 | 0 | 0.17 | 0.05 | 0.06 | 0.03 | 0.1 | 0.11 |
| 11 | 73882260 | rs1726746 | -2.875 | 0.39 | 1.227 | -0.386 | -0.106 | -0.086 | -0.086 | 0 | 0.54 | 0.94 | 0.4 | 0.44 | 0.48 | 0.46 |
| 11 | 73894976 | rs10431176 | 0.303 | 0.402 | 1.223 | -0.362 | -0.049 | -0.072 | -0.072 | 0.229 | 0.11 | 0.01 | 0.08 | 0.27 | 0.16 | 0.21 |
| 11 | 73906924 | rs555964 | 0.769 | 0.37 | 1.685 | -0.384 | -0.002 | -0.016 | -0.016 | 0 | 0.66 | 0.62 | 0.73 | 0.83 | 0.7 | 0.76 |
| 11 | 73936614 | rs10488770 | -0.48 | 0.388 | 1.75 | -0.376 | -0.009 | -0.033 | -0.033 | 0 | 0.09 | 0.36 | 0.03 | 0.01 | 0.02 | 0.02 |
| 11 | 73940134 | rs10219237 | -0.597 | 0.402 | 1.759 | -0.238 | 0.019 | 0.087 | 0.087 | 0 | 0.13 | 0.27 | 0.1 | 0.22 | 0.16 | 0.21 |
| 11 | 73956797 | rs474835 | 2.467 | 0.38 | 1.363 | -0.262 | -0.106 | -0.086 | -0.086 | 0.081 | 0.45 | 0.06 | 0.6 | 0.57 | 0.52 | 0.54 |
| 11 | 73963741 | rs500608 | 2.463 | 0.377 | 1.361 | -0.236 | -0.106 | -0.09 | -0.09 | 0 | 0.45 | 0.06 | 0.6 | 0.57 | 0.52 | 0.54 |
| 11 | 73965162 | rs17132882 | -0.753 | -0.048 | 1.73 | -0.168 | -0.007 | -0.031 | -0.031 | 0 | 0.09 | 0.36 | 0.03 | 0.01 | 0.02 | 0.02 |
| 11 | 73978243 | rs3741132 | -0.476 | 0.252 | 1.808 | 0.213 | 0.034 | 0.14 | 0.14 | 0 | 0.13 | 0.26 | 0.13 | 0.29 | 0.17 | 0.24 |
| 11 | 73994138 | rs10898975 | -0.106 | 0.253 | 1.873 | 0.214 | 0.006 | 0.041 | 0.041 | 0 | 0.11 | 0.1 | 0.12 | 0.28 | 0.16 | 0.23 |
| 11 | 73995696 | rs17132911 | -0.103 | -0.044 | 1.79 | -0.009 | -0.035 | -0.067 | -0.067 | 0 | 0.13 | 0.25 | 0.05 | 0.01 | 0.06 | 0.05 |
| 11 | 74007515 | rs12421228 | 0.547 | -0.015 | 1.687 | 0.025 | -0.014 | -0.037 | -0.037 | 0 | 0.09 | 0.2 | 0.04 | 0.01 | 0.06 | 0.04 |
| 11 | 74008907 | rs490539 | -2.598 | 0.266 | 1.12 | -0.17 | -0.074 | -0.094 | -0.094 | 0.312 | 0.54 | 0.94 | 0.47 | 0.43 | 0.54 | 0.51 |
| 11 | 74011953 | rs542628 | -1.057 | 0.309 | 1.464 | -0.206 | 0.009 | -0.038 | -0.038 | 1.059 | 0.29 | 0.42 | 0.28 | 0.14 | 0.32 | 0.22 |
| 11 | 74017319 | rs1941820 | -0.412 | 0.273 | 1.359 | -0.225 | -0.033 | 0.036 | 0.036 | 0 | 0.25 | 0.12 | 0.25 | 0.14 | 0.31 | 0.22 |
| 11 | 74028944 | rs627354 | -2.807 | 0.242 | 0.771 | -0.243 | -0.055 | -0.126 | -0.126 | 0 | 0.59 | 0.95 | 0.51 | 0.47 | 0.57 | 0.56 |
| 11 | 74033066 | rs3888908 | -0.322 | 0.441 | 1.381 | -0.273 | 0.012 | 0.058 | 0.058 | 0 | 0.13 | 0.2 | 0.13 | 0.28 | 0.18 | 0.23 |
| 11 | 74035535 | rs649937 | -0.285 | 0.573 | 0.965 | -0.436 | -0.062 | -0.034 | -0.034 | 0 | 0.9 | 0.99 | 0.81 | 0.89 | 0.81 | 0.89 |
| 11 | 74042241 | rs655719 | 0.502 | 0.127 | 1.173 | -0.536 | -0.012 | 0.011 | 0.011 | 0 | 0.71 | 0.76 | 0.71 | 0.8 | 0.68 | 0.73 |
| 11 | 74043171 | rs10793072 | 1.697 | -0.142 | 1.347 | -0.695 | 0.008 | 0.125 | 0.125 | 0 | 0.54 | 0.21 | 0.63 | 0.79 | 0.62 | 0.66 |
| 11 | 74043708 | rs596976 | 0.303 | -0.155 | 1.326 | -0.699 | -0.012 | 0.011 | 0.011 | 0 | 0.71 | 0.76 | 0.71 | 0.8 | 0.68 | 0.73 |
| 11 | 74044557 | rs1053021 | 1.68 | -0.156 | 1.301 | -0.701 | 0.008 | 0.125 | 0.125 | 0 | 0.54 | 0.21 | 0.63 | 0.79 | 0.62 | 0.66 |
| 11 | 74044976 | rs3193507 | 2.176 | -0.152 | 1.304 | -0.582 | -0.059 | -0.103 | -0.103 | 1.012 | 0.44 | 0.07 | 0.51 | 0.54 | 0.47 | 0.43 |
| 11 | 74045742 | rs1063807 | 2.389 | -0.149 | 1.285 | -0.581 | -0.069 | -0.116 | -0.116 | 0 | 0.42 | 0.06 | 0.5 | 0.54 | 0.47 | 0.43 |
| 11 | 74046975 | rs598869 | 0.417 | -0.181 | 1.651 | -0.588 | 0.008 | -0.013 | -0.013 | 0 | 0.69 | 0.64 | 0.69 | 0.79 | 0.67 | 0.72 |

| | | | | | | | | | | | | | | | | |
|---|---|---|---|---|---|---|---|---|---|---|---|---|---|---|---|---|
| 11 | 74047683 | rs1044782 | 1.507 | -0.167 | 1.84 | -0.579 | 0.005 | 0.045 | 0.045 | 0 | 0.55 | 0.36 | 0.64 | 0.79 | 0.62 | 0.66 |
| 11 | 74049569 | rs3867270 | 1.67 | -0.21 | 1.539 | -0.6 | 0.009 | 0.127 | 0.127 | 0 | 0.54 | 0.21 | 0.63 | 0.79 | 0.62 | 0.65 |
| 11 | 74050173 | rs1279293 | 0.38 | -0.207 | 1.521 | -0.613 | -0.028 | -0.032 | -0.032 | 0 | 0.8 | 0.87 | 0.7 | 0.63 | 0.68 | 0.67 |
| 11 | 74065411 | rs7945908 | 1.305 | -0.532 | 1.443 | -0.59 | -0.002 | 0.11 | 0.11 | 0.02 | 0.56 | 0.21 | 0.63 | 0.79 | 0.62 | 0.66 |
| 11 | 74071072 | rs11236071 | 0.505 | -0.515 | 1.449 | -0.581 | 0.002 | 0.015 | 0.015 | 0 | 0.11 | 0.12 | 0.12 | 0.25 | 0.16 | 0.23 |
| 11 | 74080107 | rs480358 | 1.033 | -0.52 | 1.53 | -0.595 | -0.001 | -0.019 | -0.019 | 1.064 | 0.58 | 0.58 | 0.65 | 0.79 | 0.63 | 0.66 |
| 11 | 74081501 | rs10793079 | 1.279 | -0.462 | 1.028 | -0.565 | 0.007 | 0.123 | 0.123 | 0 | 0.54 | 0.21 | 0.63 | 0.79 | 0.62 | 0.66 |
| 11 | 74086308 | rs662393 | -0.015 | -0.497 | 0.919 | -0.572 | -0.014 | 0.014 | 0.014 | 0 | 0.71 | 0.77 | 0.71 | 0.81 | 0.69 | 0.73 |
| 11 | 74093268 | rs10898982 | 1.886 | -0.414 | 0.467 | -0.484 | -0.081 | -0.121 | -0.121 | 1.782 | 0.42 | 0.06 | 0.5 | 0.54 | 0.47 | 0.43 |
| 11 | 74097466 | rs658976 | 1.128 | -0.907 | 0.891 | 0.08 | -0.022 | 0.014 | 0.014 | 0 | 0.73 | 0.61 | 0.71 | 0.33 | 0.81 | 0.63 |
| 11 | 74099892 | rs658126 | -1.379 | -0.865 | 0.516 | 0.095 | 0.032 | 0.005 | 0.005 | 0.085 | 0.92 | 0.98 | 0.87 | 0.59 | 0.97 | 0.88 |
| 11 | 74105105 | rs639525 | 0.975 | -0.682 | 0.852 | -0.019 | -0.008 | -0.003 | -0.003 | 0 | 0.64 | 0.38 | 0.66 | 0.32 | 0.77 | 0.58 |
| 11 | 74109166 | rs12049823 | -0.328 | 0.303 | 0.723 | -0.119 | -0.002 | 0.004 | 0.004 | 0 | 0.12 | 0.11 | 0.13 | 0.27 | 0.16 | 0.28 |
| 11 | 74110353 | rs515924 | -0.307 | 0.306 | 0.692 | -0.117 | 0 | 0.01 | 0.01 | 0 | 0.12 | 0.11 | 0.13 | 0.26 | 0.16 | 0.26 |
| 11 | 74110542 | rs676103 | -0.307 | 0.306 | 0.692 | -0.117 | 0 | 0.01 | 0.01 | 0 | 0.12 | 0.11 | 0.13 | 0.26 | 0.16 | 0.26 |
| 11 | 74119638 | rs624049 | -0.968 | 0.277 | 0.76 | -0.128 | 0.003 | 0.019 | 0.019 | 0 | 0.16 | 0.2 | 0.15 | 0.27 | 0.17 | 0.26 |
| 11 | 74123832 | rs597113 | -0.933 | -0.648 | 0.239 | -0.508 | -0.021 | -0.001 | -0.001 | 0 | 0.35 | 0.61 | 0.25 | 0.28 | 0.22 | 0.37 |
| 11 | 74129133 | rs645184 | -0.939 | -0.611 | -0.209 | -0.815 | 0.001 | -0.001 | -0.001 | 0 | 0.55 | 0.62 | 0.61 | 0.48 | 0.57 | 0.69 |
| 11 | 74129421 | rs11236101 | 1.392 | -0.513 | -0.481 | -0.941 | -0.015 | 0.073 | 0.073 | 0.886 | 0.12 | 0.02 | 0.05 | 0 | 0.11 | 0.01 |
| 11 | 74131896 | rs3867273 | 0.349 | -0.288 | 0.013 | -0.56 | 0.006 | 0 | 0 | 0 | 0.22 | 0.19 | 0.11 | 0.05 | 0.16 | 0.2 |
| 11 | 74137551 | rs612040 | 0.748 | -0.246 | -0.147 | -0.513 | -0.029 | -0.03 | -0.03 | 0 | 0.27 | 0.24 | 0.48 | 0.43 | 0.4 | 0.48 |
| 11 | 74149574 | rs3913182 | 0.475 | -0.218 | -0.432 | -0.498 | 0.016 | 0.019 | 0.019 | 0 | 0.21 | 0.16 | 0.1 | 0.05 | 0.15 | 0.19 |
| 11 | 74152044 | rs3862800 | 0.189 | -0.044 | -0.199 | -0.359 | 0.039 | 0.039 | 0.039 | 0 | 0.3 | 0.43 | 0.5 | 0.44 | 0.41 | 0.49 |
| 11 | 74153240 | rs12800641 | -0.665 | -0.515 | -0.387 | -0.757 | -0.029 | -0.052 | -0.052 | 0.159 | 0.26 | 0.36 | 0.13 | 0.05 | 0.15 | 0.2 |
| 11 | 74164252 | rs7119511 | -0.198 | -0.015 | -0.738 | -0.632 | 0.003 | 0.011 | 0.011 | 0.576 | 0.13 | 0.08 | 0.06 | 0 | 0.11 | 0.01 |
| 11 | 74166320 | rs7789 | -0.356 | 0.429 | -1.134 | -0.573 | -0.041 | -0.017 | -0.017 | 0.145 | 0.36 | 0.22 | 0.45 | 0.47 | 0.47 | 0.61 |
| 11 | 74167630 | rs12271177 | -1.527 | 0.029 | -0.362 | -0.703 | -0.014 | 0.025 | 0.025 | 0 | 0.54 | 0.72 | 0.53 | 0.47 | 0.51 | 0.59 |
| 11 | 74168411 | rs2270676 | -0.197 | 0.152 | -0.462 | -0.612 | 0.003 | 0 | 0 | 0 | 0.14 | 0.12 | 0.09 | 0.16 | 0.11 | 0.21 |
| 11 | 74170978 | rs3867279 | 0.366 | 0.652 | -0.45 | -0.572 | -0.011 | -0.002 | -0.002 | 0 | 0.13 | 0.12 | 0.28 | 0.2 | 0.2 | 0.22 |
| 11 | 74171116 | rs17524 | -1.129 | 0.45 | -0.039 | -0.848 | -0.002 | 0.001 | 0.001 | 0 | 0.32 | 0.31 | 0.37 | 0.2 | 0.33 | 0.25 |
| 11 | 74171175 | rs3867280 | -1.103 | 0.452 | -0.022 | -0.847 | -0.002 | 0 | 0 | 0 | 0.32 | 0.31 | 0.37 | 0.2 | 0.33 | 0.25 |
| 11 | 74171280 | rs4944893 | -0.113 | 0.247 | 0.007 | -0.647 | 0.004 | -0.003 | -0.003 | 0 | 0.14 | 0.11 | 0.11 | 0.18 | 0.12 | 0.24 |
| 11 | 74171997 | rs11236116 | 0.155 | 0.287 | 0.117 | -0.777 | 0 | -0.007 | -0.007 | 0.05 | 0.13 | 0.11 | 0.08 | 0 | 0.13 | 0.01 |
| 11 | 74178207 | rs603093 | 1.167 | 0.032 | 0.635 | -0.911 | -0.011 | -0.029 | -0.029 | 1.475 | 0.68 | 0.37 | 0.8 | 0.82 | 0.75 | 0.75 |
| 11 | 74180489 | rs11602732 | -0.216 | 0.058 | 0.374 | -0.925 | -0.013 | -0.021 | -0.021 | 0 | 0.13 | 0.02 | 0.09 | 0.14 | 0.13 | 0.23 |
| 11 | 74184938 | rs12276578 | 0.425 | -0.025 | 0.618 | -0.927 | 0.005 | -0.052 | -0.052 | 1.119 | 0.11 | 0.19 | 0.07 | 0 | 0.11 | 0.01 |
| 11 | 74186836 | rs681179 | 1.877 | 0.117 | 0.866 | -0.977 | 0.005 | 0.078 | 0.078 | 0 | 0.52 | 0.26 | 0.47 | 0.67 | 0.56 | 0.53 |
| 11 | 74192414 | rs686179 | 0.172 | 0.404 | 0.978 | -0.761 | 0.04 | -0.039 | -0.039 | 0 | 0.8 | 0.54 | 0.63 | 0.79 | 0.79 | 0.78 |
| 11 | 74193321 | rs1280350 | 0.185 | 0.582 | 0.782 | -0.635 | -0.02 | -0.01 | -0.01 | 3.288 | 0.14 | 0.07 | 0.31 | 0.22 | 0.17 | 0.27 |
| 11 | 74193496 | rs1280351 | 0.376 | 0.575 | 0.763 | -0.61 | -0.027 | -0.015 | -0.015 | 0 | 0.13 | 0.06 | 0.32 | 0.21 | 0.18 | 0.22 |
| 11 | 74194650 | rs601419 | 0.956 | 0.467 | 0.828 | -0.656 | -0.002 | -0.006 | -0.006 | 1.002 | 0.69 | 0.39 | 0.55 | 0.64 | 0.71 | 0.48 |
| 11 | 74196480 | rs11600217 | 0.187 | 0.517 | 0.552 | -0.6 | 0.006 | -0.008 | -0.008 | 2.215 | 0.11 | 0.03 | 0.08 | 0.14 | 0.09 | 0.22 |
| 11 | 74201436 | rs589970 | 1.192 | 0.37 | 0.761 | -0.805 | 0 | 0.014 | 0.014 | 0 | 0.6 | 0.37 | 0.5 | 0.63 | 0.62 | 0.47 |
| 11 | 74202033 | rs617989 | 1.192 | 0.37 | 0.774 | -0.805 | 0 | 0.015 | 0.015 | 0 | 0.6 | 0.36 | 0.5 | 0.63 | 0.62 | 0.47 |
| 11 | 74213823 | rs11236133 | 1.141 | 0.425 | 0.436 | -0.788 | -0.009 | 0.028 | 0.028 | 0.025 | 0.08 | 0.04 | 0.04 | 0 | 0.09 | 0.01 |
| 11 | 74217362 | rs675450 | 0.883 | 0.366 | 0.451 | -0.683 | 0.001 | 0.017 | 0.017 | 0 | 0.6 | 0.37 | 0.5 | 0.63 | 0.62 | 0.47 |
| 11 | 74221170 | rs10898995 | 1.031 | 0.366 | -0.322 | -0.582 | -0.003 | 0.116 | 0.116 | 0 | 0.43 | 0.13 | 0.3 | 0.33 | 0.45 | 0.37 |
| 11 | 74233031 | rs1787091 | 0.747 | 0.526 | -0.511 | -0.366 | 0.01 | 0.019 | 0.019 | 0 | 0.55 | 0.35 | 0.54 | 0.65 | 0.62 | 0.49 |
| 11 | 74235926 | rs594484 | -0.265 | 0.517 | -0.901 | -0.353 | 0 | 0.005 | 0.005 | 0.197 | 0.7 | 0.26 | 0.63 | 0.81 | 0.72 | 0.72 |
| 11 | 74250649 | rs648677 | -0.746 | 0.734 | -0.782 | -0.188 | 0.017 | 0.045 | 0.045 | 0 | 0.45 | 0.76 | 0.47 | 0.34 | 0.38 | 0.49 |
| 11 | 74257850 | rs647824 | -0.411 | -0.263 | -0.915 | -0.18 | 0.026 | -0.011 | -0.011 | 0.014 | 0.62 | 0.77 | 0.63 | 0.47 | 0.68 | 0.61 |
| 11 | 74262213 | rs657474 | -0.381 | -0.191 | -0.987 | -0.226 | 0.033 | -0.003 | -0.003 | 0 | 0.62 | 0.8 | 0.63 | 0.42 | 0.68 | 0.6 |
| 11 | 74264335 | rs11236148 | -0.215 | 0.101 | -0.524 | -0.198 | -0.001 | -0.016 | -0.016 | 1.599 | 0.12 | 0.19 | 0.09 | 0.08 | 0.1 | 0.2 |
| 11 | 74266026 | rs2887046 | 0.04 | 0.013 | -0.661 | -0.527 | 0.003 | 0.034 | 0.034 | 0 | 0.45 | 0.67 | 0.53 | 0.28 | 0.54 | 0.35 |
| 11 | 74269630 | rs1292504 | 0.902 | 0.329 | -1.08 | -0.141 | 0.004 | -0.012 | -0.012 | 0.006 | 0.28 | 0.09 | 0.23 | 0.35 | 0.27 | 0.3 |
| 11 | 74269817 | rs3902018 | 0.185 | 0.263 | -0.926 | -0.256 | -0.013 | -0.022 | -0.022 | 0 | 0.17 | 0.26 | 0.12 | 0.09 | 0.12 | 0.21 |
| 11 | 74270092 | rs10793091 | 0.941 | 0.036 | -0.289 | -0.379 | 0.001 | -0.006 | -0.006 | 0.202 | 0.41 | 0.41 | 0.47 | 0.29 | 0.47 | 0.34 |
| 11 | 74270329 | rs10501416 | 0.011 | 0.044 | -0.276 | -0.332 | -0.025 | -0.039 | -0.039 | 0 | 0.18 | 0.34 | 0.13 | 0.09 | 0.12 | 0.21 |
| 11 | 74273958 | rs7941941 | 0.607 | -0.01 | -0.276 | -0.351 | 0.002 | -0.01 | -0.01 | 0 | 0.43 | 0.55 | 0.48 | 0.29 | 0.47 | 0.34 |
| 11 | 74290881 | rs9919666 | 0.089 | -0.121 | 0.075 | 0.07 | 0.014 | 0.079 | 0.079 | 0 | 0.78 | 0.64 | 0.73 | 0.66 | 0.78 | 0.6 |
| 11 | 74301192 | rs7944514 | 0.781 | -0.014 | -0.295 | -0.605 | -0.006 | -0.008 | -0.008 | 0 | 0.32 | 0.34 | 0.38 | 0.24 | 0.25 | 0.31 |
| 11 | 74312962 | rs6592573 | 0.769 | 0.008 | -0.296 | -0.609 | -0.009 | -0.01 | -0.01 | 0 | 0.32 | 0.35 | 0.37 | 0.24 | 0.24 | 0.31 |
| 11 | 74322962 | rs6592574 | -0.98 | -0.143 | -0.637 | -0.937 | -0.002 | -0.003 | -0.003 | 0 | 0.74 | 0.77 | 0.71 | 0.74 | 0.73 | 0.77 |
| 11 | 74327614 | rs6592577 | -0.564 | -0.702 | -0.691 | -1.222 | 0.015 | 0.014 | 0.014 | 0 | 0.51 | 0.57 | 0.44 | 0.32 | 0.32 | 0.55 |
| 11 | 74331879 | rs4145953 | 0.215 | -0.698 | -0.7 | -1.2 | -0.005 | 0.001 | 0.001 | 0 | 0.29 | 0.35 | 0.35 | 0.32 | 0.21 | 0.48 |
| 11 | 74335121 | rs11604951 | 2.151 | -0.246 | -1.091 | -1.45 | -0.042 | 0.005 | 0.005 | 0 | 0.07 | 0.01 | 0.1 | 0.06 | 0.15 | 0.08 |
| 11 | 74339577 | rs2155935 | 1.129 | -1.196 | -0.762 | -1.285 | -0.029 | 0.023 | 0.023 | 0 | 0.39 | 0.23 | 0.33 | 0.42 | 0.52 | 0.28 |
| 11 | 74342928 | rs11236180 | 1.673 | -1.04 | -0.741 | -1.186 | -0.013 | -0.071 | -0.071 | 0 | 0.18 | 0.22 | 0.09 | 0 | 0.11 | 0.06 |
| 11 | 74353599 | rs1051076 | 1.739 | -0.969 | -0.923 | -1.029 | 0.025 | 0.004 | 0.004 | 0 | 0.1 | 0.01 | 0.04 | 0.06 | 0.05 | 0.15 |

| | | | | | | | | | | | | | | | | |
|---|---|---|---|---|---|---|---|---|---|---|---|---|---|---|---|---|
| 11 | 74354227 | rs7947345 | -2.133 | -0.778 | -0.957 | -0.886 | -0.051 | -0.033 | -0.033 | 1.395 | 0.92 | 0.99 | 0.8 | 0.74 | 0.84 | 0.83 |
| 11 | 74359816 | rs4944926 | -1.763 | -0.887 | -0.761 | -1.04 | 0.03 | 0.081 | 0.081 | 0.007 | 0.91 | 0.83 | 0.79 | 0.74 | 0.84 | 0.83 |
| 11 | 74360030 | rs7932922 | 0.368 | -1.106 | -0.361 | -1.175 | 0.001 | -0.007 | -0.007 | 0 | 0.59 | 0.52 | 0.44 | 0.42 | 0.62 | 0.35 |
| 11 | 74360313 | rs7110993 | -1.655 | -0.59 | -0.42 | -0.848 | 0.032 | 0.085 | 0.085 | 0.007 | 0.91 | 0.83 | 0.79 | 0.74 | 0.84 | 0.83 |
| 11 | 74375475 | rs2186955 | 1.229 | -0.87 | -0.497 | -1.069 | -0.001 | -0.006 | -0.006 | 0 | 0.25 | 0.22 | 0.14 | 0.11 | 0.19 | 0.29 |
| 11 | 74378871 | rs11825804 | 0.243 | -0.941 | -1.241 | -1.024 | -0.058 | 0.114 | 0.114 | 0 | 0.45 | 0.11 | 0.32 | 0.38 | 0.52 | 0.34 |
| 11 | 74381029 | rs6592590 | 0.767 | -0.627 | -1.105 | -0.717 | -0.005 | 0.001 | 0.001 | 0 | 0.27 | 0.23 | 0.29 | 0.35 | 0.4 | 0.26 |
| 11 | 74386732 | rs7104675 | -0.478 | -0.872 | -1.452 | -0.793 | -0.039 | -0.018 | -0.018 | 0 | 0.47 | 0.84 | 0.6 | 0.56 | 0.54 | 0.52 |
| 11 | 74386800 | rs7104802 | 0.83 | -0.747 | -1.382 | -0.75 | -0.017 | 0.009 | 0.009 | 0 | 0.27 | 0.21 | 0.29 | 0.3 | 0.39 | 0.25 |
| 11 | 74389427 | rs12789086 | 0.698 | -0.88 | -1.019 | -0.649 | -0.022 | 0.082 | 0.082 | 0 | 0.16 | 0.42 | 0.28 | 0.24 | 0.11 | 0.17 |
| 11 | 74390944 | rs11236209 | 0.892 | -0.979 | -1.513 | -1.092 | 0.034 | 0.136 | 0.136 | 0 | 0.36 | 0.17 | 0.15 | 0.03 | 0.29 | 0.25 |
| 11 | 74391103 | rs7119296 | 0.592 | -0.421 | -1.084 | -0.534 | -0.003 | -0.005 | -0.005 | 0 | 0.65 | 0.42 | 0.45 | 0.36 | 0.69 | 0.52 |
| 11 | 74392514 | rs4944936 | -1.107 | -0.262 | -1.328 | -0.605 | -0.063 | -0.101 | -0.101 | 0.732 | 0.87 | 0.99 | 0.77 | 0.61 | 0.82 | 0.83 |
| 11 | 74393938 | rs12282262 | 0.431 | -0.121 | -0.975 | -0.685 | -0.015 | 0.094 | 0.094 | 0 | 0.16 | 0.36 | 0.33 | 0.28 | 0.11 | 0.22 |
| 11 | 74394015 | rs10793104 | -0.228 | 0.146 | -0.887 | -0.503 | -0.002 | -0.011 | -0.011 | 0 | 0.8 | 0.79 | 0.73 | 0.56 | 0.78 | 0.73 |
| 11 | 74398125 | rs2000924 | -0.281 | -0.001 | -0.964 | -1.047 | 0.013 | -0.006 | -0.006 | 0.275 | 0.37 | 0.68 | 0.19 | 0.02 | 0.28 | 0.2 |
| 11 | 74398530 | rs2000926 | -0.845 | 0.467 | -0.467 | -0.272 | -0.005 | 0.019 | 0.019 | 0 | 0.64 | 0.49 | 0.67 | 0.61 | 0.67 | 0.69 |
| 11 | 74399425 | rs1893361 | -0.584 | 0.786 | -0.496 | -0.449 | 0.008 | -0.025 | -0.025 | 1.898 | 0.14 | 0.26 | 0.33 | 0.08 | 0.17 | 0.1 |
| 11 | 74400999 | rs7939907 | 0.341 | 0.64 | -0.894 | -0.383 | 0.001 | -0.006 | -0.006 | 0 | 0.1 | 0.07 | 0.04 | 0 | 0.1 | 0.02 |
| 11 | 74404218 | rs7926982 | -0.306 | 0.759 | -0.702 | -0.044 | 0 | 0.001 | 0.001 | 0 | 0.66 | 0.66 | 0.49 | 0.55 | 0.63 | 0.55 |
| 11 | 74405348 | rs4944937 | -0.559 | 0.467 | -0.686 | -0.094 | -0.001 | 0.009 | 0.009 | 0 | 0.69 | 0.68 | 0.59 | 0.52 | 0.71 | 0.53 |
| 11 | 74407269 | rs4539352 | 0.36 | 0.171 | -1.651 | -0.397 | -0.068 | -0.074 | -0.074 | 1.901 | 0.4 | 0.1 | 0.44 | 0.46 | 0.51 | 0.37 |
| 11 | 74407676 | rs3819242 | 0 | -0.276 | -0.81 | 0.313 | 0.016 | -0.012 | -0.012 | 0 | 0.63 | 0.27 | 0.55 | 0.46 | 0.73 | 0.47 |
| 11 | 74419163 | rs1944936 | -0.022 | -0.334 | -0.384 | -0.07 | -0.027 | -0.04 | -0.04 | 0 | 0.71 | 0.5 | 0.82 | 0.81 | 0.75 | 0.75 |
| 11 | 74419378 | rs7948433 | 0.339 | 0.644 | -0.95 | 0.186 | -0.049 | -0.052 | -0.052 | 0.157 | 0.22 | 0.13 | 0.57 | 0.54 | 0.38 | 0.18 |
| 11 | 74423513 | rs1433769 | 0.237 | -0.329 | -0.482 | -0.151 | 0 | -0.001 | -0.001 | 0 | 0.43 | 0.47 | 0.29 | 0.29 | 0.34 | 0.57 |
| 11 | 74424075 | rs11236225 | 0.691 | -0.686 | -0.852 | -0.637 | 0.04 | 0.029 | 0.029 | 0 | 0.3 | 0.17 | 0.15 | 0.25 | 0.22 | 0.39 |
| 11 | 74428580 | rs968305 | 0.733 | -0.391 | -0.828 | -0.438 | -0.015 | -0.015 | -0.015 | 0 | 0.2 | 0.18 | 0.57 | 0.53 | 0.38 | 0.17 |
| 11 | 74430185 | rs1433770 | -0.922 | -0.676 | -0.507 | -0.623 | 0.027 | 0.045 | 0.045 | 0 | 0.67 | 0.85 | 0.83 | 0.8 | 0.74 | 0.63 |
| 11 | 74439384 | rs7947106 | 0.321 | -0.2 | -0.157 | -0.468 | 0.007 | -0.044 | -0.044 | 0 | 0.12 | 0.15 | 0.07 | 0 | 0.12 | 0.03 |
| 11 | 74442931 | rs11236236 | 0.597 | 0.22 | -0.363 | -0.399 | 0.055 | 0.07 | 0.07 | 0 | 0.13 | 0.16 | 0.54 | 0.53 | 0.34 | 0.11 |
| 11 | 74443090 | rs10899042 | -0.463 | -0.142 | -0.168 | -0.261 | 0.015 | 0.015 | 0.015 | 0 | 0.25 | 0.28 | 0.62 | 0.52 | 0.46 | 0.14 |
| 11 | 74446791 | rs7103136 | -0.575 | -0.195 | 0.177 | -0.301 | 0.081 | 0.08 | 0.08 | 0 | 0.27 | 0.46 | 0.62 | 0.51 | 0.46 | 0.14 |
| 11 | 74465068 | rs571250 | 0.038 | -1.219 | -0.035 | -0.555 | 0.029 | 0.012 | 0.012 | 0 | 0.3 | 0.2 | 0.19 | 0.26 | 0.21 | 0.43 |
| 11 | 74481318 | rs564229 | -0.743 | -1.389 | 0.503 | -0.713 | 0.024 | 0.022 | 0.022 | 0 | 0.39 | 0.67 | 0.25 | 0.27 | 0.24 | 0.46 |
| 11 | 74484349 | rs17133441 | 0.781 | -1.864 | 0.16 | -0.889 | 0.039 | -0.001 | -0.001 | 0.003 | 0.14 | 0.03 | 0.05 | 0.12 | 0.05 | 0.23 |
| 11 | 74500173 | rs572656 | -1.017 | -1.674 | 0.608 | -0.73 | 0.001 | 0.021 | 0.021 | 0 | 0.12 | 0.25 | 0.08 | 0.16 | 0.09 | 0.17 |
| 11 | 74503168 | rs113187 | -0.213 | -1.674 | 0.511 | -0.788 | 0.046 | 0.01 | 0.01 | 0 | 0.29 | 0.1 | 0.18 | 0.26 | 0.21 | 0.43 |
| 11 | 74525852 | rs4483614 | 0.885 | -0.681 | 0.402 | -0.714 | -0.01 | 0.009 | 0.009 | 0 | 0.11 | 0.1 | 0.49 | 0.52 | 0.29 | 0.09 |
| 11 | 74545378 | rs512780 | -1.976 | -1.336 | 0.164 | -1.117 | -0.012 | 0.093 | 0.093 | 0 | 0.6 | 0.86 | 0.78 | 0.79 | 0.58 | 0.64 |
| 11 | 74553834 | rs2298746 | 1.148 | -0.993 | 0.604 | -0.772 | 0.041 | 0.084 | 0.084 | 0 | 0.1 | 0.14 | 0.49 | 0.51 | 0.28 | 0.09 |
| 11 | 74559481 | rs4944961 | 0.394 | -1.019 | 0.548 | -0.8 | -0.03 | -0.024 | -0.024 | 0 | 0.13 | 0.09 | 0.5 | 0.51 | 0.32 | 0.1 |
| 11 | 74581213 | rs605954 | -0.337 | -1.01 | 0.644 | -0.848 | 0.035 | 0.013 | 0.013 | 0 | 0.71 | 0.86 | 0.81 | 0.74 | 0.79 | 0.57 |
| 11 | 74624011 | rs590105 | 0.989 | -1.016 | 0.864 | -0.889 | -0.003 | -0.018 | -0.018 | 0.689 | 0.53 | 0.48 | 0.29 | 0.23 | 0.47 | 0.45 |
| 11 | 74664025 | rs1292895 | 0.757 | -1.378 | 0.509 | -1.226 | 0.006 | 0.019 | 0.019 | 0 | 0.52 | 0.4 | 0.29 | 0.23 | 0.47 | 0.47 |
| 11 | 74673270 | rs529488 | 0.89 | -1.374 | 0.216 | -1.224 | 0.029 | 0.128 | 0.128 | 0 | 0.51 | 0.22 | 0.27 | 0.23 | 0.48 | 0.47 |
| 11 | 74678203 | rs493001 | -0.469 | -1.568 | -0.108 | -1.121 | 0.038 | 0.012 | 0.012 | 0 | 0.71 | 0.85 | 0.81 | 0.74 | 0.79 | 0.59 |
| 11 | 74687341 | rs487226 | -0.302 | -1.553 | -0.158 | -1.113 | 0.01 | 0.006 | 0.006 | 0 | 0.7 | 0.73 | 0.8 | 0.74 | 0.79 | 0.59 |
| 11 | 74705602 | rs7115499 | 2.046 | -1.193 | -0.328 | -0.979 | -0.01 | 0.025 | 0.025 | 0 | 0.06 | 0.01 | 0.06 | 0 | 0.09 | 0.01 |
| 11 | 74705696 | rs544115 | 0.166 | -1.671 | 0.325 | -0.718 | -0.017 | -0.013 | -0.013 | 0.162 | 0.67 | 0.62 | 0.8 | 0.77 | 0.8 | 0.58 |
| 11 | 74710217 | rs494474 | -0.57 | -1.714 | 0.168 | -0.739 | 0.049 | 0.005 | 0.005 | 0.029 | 0.75 | 0.92 | 0.86 | 0.78 | 0.84 | 0.62 |
| 11 | 74726828 | rs594144 | 1.093 | -1.379 | 0.159 | -0.643 | 0.005 | 0.142 | 0.142 | 0 | 0.48 | 0.15 | 0.27 | 0.26 | 0.47 | 0.43 |
| 11 | 74731890 | rs488115 | 1.652 | -0.996 | -0.127 | -0.291 | -0.062 | 0.094 | 0.094 | 0 | 0.35 | 0.05 | 0.23 | 0.22 | 0.4 | 0.33 |
| 11 | 74732410 | rs4387374 | 1.13 | -0.386 | 0.467 | 0.188 | -0.001 | 0.023 | 0.023 | 0 | 0.1 | 0.07 | 0.48 | 0.52 | 0.28 | 0.1 |
| 11 | 74744581 | rs7952209 | -0.281 | -0.637 | 0.607 | 0.149 | -0.034 | -0.02 | -0.02 | 0 | 0.32 | 0.44 | 0.19 | 0.23 | 0.18 | 0.42 |
| 11 | 74746631 | rs10899060 | 0.588 | -0.446 | 0.419 | 0.427 | 0.006 | 0.026 | 0.026 | 0 | 0.12 | 0.13 | 0.5 | 0.51 | 0.3 | 0.09 |
| 11 | 74757641 | rs7941838 | -0.117 | -0.42 | 0.482 | 0.309 | 0.015 | 0.006 | 0.006 | 0 | 0.28 | 0.23 | 0.15 | 0.22 | 0.18 | 0.34 |
| 11 | 74763783 | rs7106757 | 0.937 | -0.047 | 0.173 | 0.541 | -0.064 | -0.053 | -0.053 | 0 | 0.1 | 0.03 | 0.5 | 0.52 | 0.28 | 0.08 |
| 11 | 74771478 | rs4531492 | -0.844 | -0.134 | 0.447 | 0.47 | 0.115 | 0.15 | 0.15 | 0.579 | 0.21 | 0.41 | 0.56 | 0.53 | 0.34 | 0.17 |
| 11 | 74777798 | rs11236313 | 0.419 | 0.154 | -0.204 | 0.743 | 0.074 | 0.085 | 0.085 | 0.006 | 0.12 | 0.19 | 0.3 | 0.41 | 0.28 | 0.09 |
| 11 | 74778675 | rs4944067 | -0.992 | -0.282 | -0.751 | 0.399 | 0.011 | 0.155 | 0.155 | 0 | 0.51 | 0.82 | 0.71 | 0.75 | 0.52 | 0.5 |
| 11 | 74798928 | rs11236323 | 0.169 | -0.252 | -0.27 | 1.371 | -0.034 | -0.028 | -0.028 | 0 | 0.1 | 0.03 | 0.13 | 0.4 | 0.16 | 0.09 |
| 11 | 74799470 | rs4146884 | -0.753 | -0.441 | 0.737 | 1.289 | 0.081 | 0.228 | 0.228 | 0.981 | 0.14 | 0.39 | 0.15 | 0.4 | 0.17 | 0.09 |
| 11 | 74801374 | rs4944974 | -0.176 | -1.009 | 0.73 | 1.116 | -0.024 | 0.019 | 0.019 | 0.734 | 0.24 | 0.32 | 0.36 | 0.32 | 0.17 | 0.31 |
| 11 | 74802275 | rs4944975 | -0.712 | -1.185 | 0.682 | 0.965 | -0.012 | -0.001 | -0.001 | 0.092 | 0.37 | 0.63 | 0.42 | 0.35 | 0.21 | 0.38 |
| 11 | 74806005 | rs4435030 | -1.747 | -1.173 | 0.607 | 0.923 | 0.028 | 0.142 | 0.142 | 0.065 | 0.5 | 0.93 | 0.56 | 0.74 | 0.37 | 0.46 |
| 11 | 74814902 | rs2712789 | 0.076 | -1.105 | 0.695 | 0.976 | 0.082 | 0.242 | 0.242 | 0.908 | 0.86 | 0.61 | 0.85 | 0.6 | 0.83 | 0.9 |
| 11 | 74820158 | rs7124613 | 0.431 | -1.06 | 0.46 | 0.958 | -0.003 | -0.011 | -0.011 | 0.035 | 0.06 | 0.1 | 0.02 | 0.03 | 0.03 | 0.06 |
| 11 | 74820542 | rs2851079 | 1.024 | -1.117 | 0.436 | 0.972 | 0.043 | 0.073 | 0.073 | 0 | 0.55 | 0.18 | 0.47 | 0.3 | 0.66 | 0.57 |

| | | | | | | | | | | | | | | | | |
|---|---|---|---|---|---|---|---|---|---|---|---|---|---|---|---|---|
| 11 | 74823345 | rs2851085 | -0.587 | -0.886 | 0.647 | 1.081 | -0.002 | 0.029 | 0.029 | 0.898 | 0.89 | 0.9 | 0.86 | 0.6 | 0.84 | 0.9 |
| 11 | 74836621 | rs2712812 | 0.011 | -1.041 | 0.755 | 1.074 | 0.057 | 0.203 | 0.203 | 0.88 | 0.87 | 0.68 | 0.85 | 0.6 | 0.84 | 0.9 |
| 11 | 74836744 | rs2851091 | 0.801 | -1.067 | 0.61 | 1.051 | 0.041 | 0.168 | 0.168 | 0.961 | 0.81 | 0.56 | 0.83 | 0.57 | 0.81 | 0.84 |
| 11 | 74842779 | rs2712794 | 0.485 | -1.103 | 0.465 | 0.859 | -0.013 | -0.019 | -0.019 | 0 | 0.59 | 0.49 | 0.48 | 0.3 | 0.66 | 0.59 |
| 11 | 74849219 | rs2712799 | 1.354 | -1.088 | -0.289 | 0.877 | 0.05 | 0.144 | 0.144 | 0.051 | 0.5 | 0.1 | 0.45 | 0.27 | 0.63 | 0.53 |
| 11 | 74853813 | rs2851104 | 1.281 | -1.035 | -0.327 | 0.877 | 0.01 | 0.122 | 0.122 | 0.086 | 0.49 | 0.07 | 0.45 | 0.27 | 0.61 | 0.53 |
| 11 | 74856517 | rs11236349 | -0.666 | -1.196 | 0.166 | 1.013 | -0.026 | -0.005 | -0.005 | 0 | 0.36 | 0.5 | 0.49 | 0.34 | 0.22 | 0.36 |
| 11 | 74857436 | rs10899076 | -0.66 | -1.2 | 0.153 | 1.011 | -0.031 | -0.005 | -0.005 | 0.472 | 0.36 | 0.53 | 0.49 | 0.34 | 0.22 | 0.36 |
| 11 | 74860061 | rs2851064 | -0.507 | -1.136 | 0.226 | 1.111 | 0.086 | 0.235 | 0.235 | 0.526 | 0.87 | 0.66 | 0.86 | 0.61 | 0.84 | 0.91 |
| 11 | 74860786 | rs4944991 | -0.524 | -1.177 | 0.425 | 1.096 | 0.031 | 0 | 0 | 0 | 0.35 | 0.34 | 0.47 | 0.35 | 0.21 | 0.36 |
| 11 | 74861416 | rs4100076 | -0.204 | -1.224 | 0.428 | 1.082 | 0.042 | -0.012 | -0.012 | 0 | 0.29 | 0.24 | 0.45 | 0.32 | 0.18 | 0.3 |
| 11 | 74861545 | rs12361540 | 1.296 | -1.445 | 0.354 | 1.146 | 0.02 | -0.024 | -0.024 | 0.118 | 0.11 | 0.04 | 0.33 | 0.21 | 0.06 | 0.15 |
| 11 | 74862356 | rs2851069 | 1.541 | -1.279 | 0.391 | 0.934 | 0.031 | 0.131 | 0.131 | 0 | 0.51 | 0.14 | 0.37 | 0.27 | 0.62 | 0.56 |
| 11 | 74862866 | rs10899078 | -0.789 | -1.289 | 0.358 | 0.968 | 0.031 | -0.003 | -0.003 | 0 | 0.35 | 0.33 | 0.47 | 0.36 | 0.21 | 0.35 |
| 11 | 74864280 | rs4944992 | -0.789 | -1.289 | 0.358 | 0.968 | 0.031 | -0.003 | -0.003 | 2.601 | 0.35 | 0.33 | 0.47 | 0.36 | 0.21 | 0.35 |
| 11 | 74864729 | rs2712811 | 1.506 | -1.288 | 0.299 | 0.972 | 0.006 | 0.139 | 0.139 | 0.034 | 0.5 | 0.07 | 0.36 | 0.27 | 0.62 | 0.55 |
| 11 | 74865716 | rs2851071 | 1.499 | -1.277 | 0.309 | 0.977 | 0.015 | 0.143 | 0.143 | 0.019 | 0.5 | 0.07 | 0.36 | 0.27 | 0.62 | 0.55 |
| 11 | 74865756 | rs4944073 | -0.353 | -1.275 | 0.336 | 1.002 | 0.044 | -0.016 | -0.016 | 3.178 | 0.3 | 0.23 | 0.44 | 0.33 | 0.18 | 0.29 |
| 11 | 74865831 | rs2712818 | 0.828 | -1.275 | 0.409 | 0.999 | -0.002 | -0.003 | -0.003 | 0.684 | 0.58 | 0.43 | 0.42 | 0.29 | 0.66 | 0.62 |
| 11 | 74866107 | rs2712819 | -0.641 | -1.262 | 0.41 | 1.007 | 0.091 | 0.225 | 0.225 | 0.214 | 0.88 | 0.68 | 0.86 | 0.62 | 0.84 | 0.91 |
| 11 | 74866186 | rs11236356 | -0.261 | -1.312 | 0.341 | 0.993 | 0.021 | -0.006 | -0.006 | 0 | 0.3 | 0.27 | 0.45 | 0.33 | 0.18 | 0.29 |
| 11 | 74869888 | rs4944994 | -0.941 | -1.335 | 0.256 | 0.987 | -0.001 | 0 | 0 | 0.272 | 0.37 | 0.45 | 0.48 | 0.37 | 0.22 | 0.36 |
| 11 | 74870884 | rs11236359 | -0.25 | -1.183 | 0.241 | 1.026 | 0.042 | 0.132 | 0.132 | 0.11 | 0.82 | 0.53 | 0.82 | 0.63 | 0.81 | 0.88 |
| 11 | 74872576 | rs2712816 | 1.543 | -1.218 | 0.153 | 0.997 | -0.012 | 0.121 | 0.121 | 0.028 | 0.47 | 0.08 | 0.34 | 0.26 | 0.6 | 0.54 |
| 11 | 74872821 | rs2465290 | 1.543 | -1.218 | 0.153 | 0.997 | -0.012 | 0.121 | 0.121 | 0.028 | 0.47 | 0.08 | 0.34 | 0.27 | 0.6 | 0.54 |
| 11 | 74872884 | rs11236361 | -0.52 | -1.422 | 0.265 | 1.008 | -0.015 | 0 | 0 | 0.514 | 0.33 | 0.43 | 0.46 | 0.34 | 0.18 | 0.29 |
| 11 | 74874433 | rs10899079 | -0.514 | -1.429 | 0.255 | 1.006 | -0.01 | 0 | 0 | 0.28 | 0.33 | 0.42 | 0.46 | 0.34 | 0.18 | 0.32 |
| 11 | 74875501 | rs3819192 | 1.142 | -1.855 | -0.112 | 0.721 | 0.023 | 0.012 | 0.012 | 0 | 0.1 | 0 | 0.02 | 0.03 | 0.04 | 0.1 |
| 11 | 74875945 | rs881217 | -0.757 | -1.172 | 0.467 | 1.122 | -0.043 | 0.002 | 0.002 | 0.461 | 0.33 | 0.48 | 0.47 | 0.33 | 0.18 | 0.33 |
| 11 | 74877004 | rs3740839 | -0.477 | -1.184 | 0.265 | 1.114 | 0.021 | -0.007 | -0.007 | 0 | 0.3 | 0.28 | 0.45 | 0.33 | 0.18 | 0.32 |
| 11 | 74877573 | rs949069 | -0.816 | -1.113 | 0.182 | 1.079 | -0.043 | 0.003 | 0.003 | 1.342 | 0.36 | 0.58 | 0.49 | 0.36 | 0.21 | 0.36 |
| 11 | 74878076 | rs2851109 | 1.462 | -0.683 | -0.951 | 0.858 | -0.052 | 0.117 | 0.117 | 0 | 0.47 | 0.08 | 0.33 | 0.26 | 0.59 | 0.51 |
| 11 | 74881256 | rs11236365 | -0.732 | -0.781 | -1.127 | 0.839 | 0.037 | 0 | 0 | 1.14 | 0.34 | 0.32 | 0.45 | 0.32 | 0.21 | 0.35 |
| 11 | 74881682 | rs1789694 | -0.034 | -0.678 | -1.113 | 0.926 | 0.041 | 0.106 | 0.106 | 1.081 | 0.8 | 0.44 | 0.79 | 0.57 | 0.79 | 0.86 |
| 11 | 74883172 | rs4944074 | -1.087 | -0.968 | -1.077 | 0.779 | 0.029 | -0.005 | -0.005 | 0 | 0.35 | 0.36 | 0.45 | 0.34 | 0.22 | 0.36 |
| 11 | 74883577 | rs12422149 | 0.109 | -1.163 | -1.143 | 0.764 | 0.021 | -0.043 | -0.043 | 0 | 0.15 | 0.09 | 0.36 | 0.32 | 0.1 | 0.26 |
| 11 | 74884388 | rs10501421 | -0.51 | -0.586 | -0.877 | 0.03 | 0 | -0.022 | -0.022 | 0 | 0.19 | 0.25 | 0.08 | 0.01 | 0.11 | 0.09 |
| 11 | 74886074 | rs7947726 | -0.633 | -0.579 | -0.909 | 0.015 | 0.006 | -0.012 | -0.012 | 0 | 0.2 | 0.25 | 0.09 | 0.01 | 0.12 | 0.1 |
| 11 | 74886858 | rs1676881 | 0.694 | -0.35 | -1.677 | 0.531 | -0.018 | -0.044 | -0.044 | 7.398 | 0.6 | 0.88 | 0.46 | 0.37 | 0.68 | 0.53 |
| 11 | 74887294 | rs3758773 | -0.43 | -0.272 | -0.915 | 0.353 | 0 | -0.003 | -0.003 | 0 | 0.35 | 0.36 | 0.24 | 0.22 | 0.32 | 0.39 |
| 11 | 74887553 | rs7117473 | -0.18 | -0.04 | 0.4 | 0.235 | 0.048 | -0.055 | -0.055 | 0 | 0.1 | 0.42 | 0.08 | 0 | 0.09 | 0.01 |
| 11 | 74895090 | rs3824903 | -1.898 | -0.16 | 1.12 | 1.647 | 0.087 | 0.536 | 0.536 | 3.491 | 0.05 | 0.43 | 0.31 | 0.41 | 0.02 | 0.03 |
| 11 | 74901601 | rs1077858 | 0.171 | 0.245 | 0.339 | 0.969 | 0.007 | 0.015 | 0.015 | 0 | 0.63 | 0.53 | 0.72 | 0.42 | 0.66 | 0.56 |
| 11 | 74905278 | rs1676885 | 0.915 | 0.054 | -0.903 | 0.113 | -0.06 | 0.129 | 0.129 | 0 | 0.25 | 0.02 | 0.2 | 0.04 | 0.3 | 0.15 |
| 11 | 74918538 | rs7951787 | -1.146 | 0.108 | -0.095 | 0.991 | 0.036 | 0.27 | 0.27 | 0.548 | 0.08 | 0.39 | 0.11 | 0.28 | 0.04 | 0.08 |
| 11 | 74920005 | rs10793116 | 0.533 | 0.431 | -0.454 | 1.309 | -0.086 | -0.111 | -0.111 | 0 | 0.17 | 0.03 | 0.22 | 0.38 | 0.23 | 0.24 |
| 11 | 74923212 | rs7924924 | -0.89 | 0.417 | -0.419 | 1.277 | 0.032 | 0.252 | 0.252 | 0.248 | 0.08 | 0.34 | 0.11 | 0.28 | 0.04 | 0.08 |
| 11 | 74935046 | rs1789679 | 0.042 | 0.425 | -0.828 | 1.289 | -0.099 | -0.166 | -0.166 | 0 | 0.73 | 0.96 | 0.69 | 0.48 | 0.7 | 0.64 |
| 11 | 74945585 | rs10899080 | 0.638 | 0.51 | -0.804 | 1.607 | -0.081 | -0.072 | -0.072 | 0 | 0.15 | 0.03 | 0.13 | 0.27 | 0.23 | 0.22 |
| 11 | 74949403 | rs1676891 | 0.668 | 0.421 | -0.528 | 0.826 | 0.017 | -0.042 | -0.042 | 1.201 | 0.65 | 0.54 | 0.57 | 0.2 | 0.66 | 0.6 |
| 11 | 74958948 | rs1783472 | -1.224 | -0.152 | -0.692 | 1.065 | 0.011 | -0.017 | -0.017 | 0 | 0.89 | 0.99 | 0.91 | 0.86 | 0.93 | 0.86 |
| 11 | 74975662 | rs2282600 | 0.428 | 0.124 | -0.599 | 1.084 | -0.004 | -0.012 | -0.012 | 0.434 | 0.07 | 0.03 | 0.09 | 0.28 | 0.07 | 0.08 |
| 11 | 74977067 | rs2279129 | 0.385 | 0.141 | -0.565 | 1.093 | -0.001 | 0.013 | 0.013 | 0 | 0.07 | 0.04 | 0.05 | 0.26 | 0.07 | 0.11 |
| 11 | 74981039 | rs528833 | 0.881 | -0.051 | -1.093 | 0.623 | -0.105 | 0.035 | 0.035 | 0.153 | 0.33 | 0.03 | 0.48 | 0.21 | 0.41 | 0.36 |
| 11 | 74982939 | rs2276310 | -0.96 | 0.302 | 0.331 | 1.139 | 0.047 | 0.136 | 0.136 | 0 | 0.12 | 0.4 | 0.09 | 0.23 | 0.1 | 0.12 |
| 11 | 74992267 | rs746168 | 0.189 | 0.458 | 0.073 | 0.795 | 0.046 | -0.015 | -0.015 | 0.247 | 0.17 | 0.46 | 0.15 | 0.14 | 0.18 | 0.17 |
| 11 | 74994016 | rs2236710 | -0.005 | 0.402 | -0.276 | 0.86 | -0.003 | -0.011 | -0.011 | 0 | 0.16 | 0.19 | 0.27 | 0.1 | 0.11 | 0.17 |
| 11 | 74994352 | rs877711 | -0.139 | 0.392 | -0.286 | 0.841 | -0.003 | -0.011 | -0.011 | 0.18 | 0.17 | 0.19 | 0.27 | 0.1 | 0.11 | 0.18 |
| 11 | 74995632 | rs1676884 | -0.315 | 0.386 | -0.337 | 1.103 | -0.001 | 0.016 | 0.016 | 0 | 0.92 | 0.91 | 0.93 | 0.82 | 0.93 | 0.9 |
| 11 | 74999428 | rs667791 | 1.057 | 0.37 | -0.502 | 1.074 | 0.002 | -0.009 | -0.009 | 1.77 | 0.56 | 0.54 | 0.4 | 0.39 | 0.6 | 0.57 |
| 11 | 74999953 | rs489161 | 0.49 | 0.347 | -0.5 | 1.126 | -0.004 | -0.005 | -0.005 | 1.522 | 0.19 | 0.19 | 0.27 | 0.32 | 0.22 | 0.15 |
| 11 | 75000189 | rs567807 | 0.767 | 0.313 | -0.648 | 0.901 | 0.003 | -0.012 | -0.012 | 0.075 | 0.56 | 0.54 | 0.4 | 0.39 | 0.6 | 0.57 |
| 11 | 75001493 | rs490528 | 0.255 | 0.495 | -0.706 | 0.995 | 0.003 | 0.006 | 0.006 | 0 | 0.64 | 0.62 | 0.51 | 0.46 | 0.67 | 0.66 |
| 11 | 75005209 | rs553664 | -0.671 | 0.545 | -1.273 | 0.838 | -0.051 | -0.039 | -0.039 | 0 | 0.4 | 0.81 | 0.4 | 0.43 | 0.43 | 0.31 |
| 11 | 75012535 | rs506233 | -0.628 | 0.498 | -0.947 | 0.88 | -0.027 | -0.027 | -0.027 | 0 | 0.39 | 0.72 | 0.4 | 0.44 | 0.43 | 0.32 |
| 11 | 75017087 | rs611908 | -0.177 | 0.358 | -0.53 | 0.663 | 0.012 | 0.043 | 0.043 | 0 | 0.31 | 0.41 | 0.34 | 0.37 | 0.31 | 0.23 |
| 11 | 75017436 | rs536852 | -1.128 | 0.32 | -0.516 | 0.638 | 0.022 | 0.023 | 0.023 | 0 | 0.51 | 0.73 | 0.49 | 0.5 | 0.52 | 0.39 |
| 11 | 75018232 | rs529513 | 0.477 | 0.051 | -0.733 | 0.726 | -0.001 | 0.006 | 0.006 | 0 | 0.54 | 0.49 | 0.39 | 0.4 | 0.58 | 0.56 |
| 11 | 75019296 | rs1676887 | -0.757 | -0.131 | -0.714 | 1.003 | 0.001 | -0.002 | -0.002 | 0 | 0.85 | 0.9 | 0.74 | 0.77 | 0.9 | 0.78 |

| | | | | | | | | | | | | | | | | |
|---|---|---|---|---|---|---|---|---|---|---|---|---|---|---|---|---|
| 11 | 75020379 | rs657561 | 0.372 | -0.107 | -0.784 | 0.779 | 0.002 | -0.004 | -0.004 | 0.267 | 0.6 | 0.48 | 0.69 | 0.49 | 0.67 | 0.67 |
| 11 | 75021501 | rs472112 | -1.297 | 0.03 | -0.997 | 0.332 | -0.042 | 0.061 | 0.061 | 0 | 0.61 | 0.79 | 0.53 | 0.69 | 0.55 | 0.6 |
| 11 | 75022521 | rs687652 | 0.27 | -0.039 | -0.508 | 0.791 | 0.014 | 0.062 | 0.062 | 0 | 0.19 | 0.32 | 0.19 | 0.27 | 0.18 | 0.13 |
| 11 | 75023441 | rs576014 | -0.474 | 0.004 | -1.181 | 0.433 | 0.023 | 0.046 | 0.046 | 0 | 0.47 | 0.62 | 0.43 | 0.59 | 0.4 | 0.55 |
| 11 | 75026331 | rs747221 | 0.159 | -0.237 | -1.071 | 0.301 | -0.003 | 0 | 0 | 0 | 0.59 | 0.51 | 0.61 | 0.42 | 0.65 | 0.52 |
| 11 | 75026392 | rs542645 | -0.504 | -0.393 | -1.31 | 0.654 | 0.012 | -0.021 | -0.021 | 0 | 0.17 | 0.15 | 0.21 | 0.49 | 0.08 | 0.27 |
| 11 | 75028697 | rs685929 | -0.071 | -0.729 | -1.226 | -0.049 | -0.008 | -0.003 | -0.003 | 0.491 | 0.15 | 0.17 | 0.08 | 0.14 | 0.07 | 0.11 |
| 11 | 75029167 | rs7929974 | -0.216 | 0.137 | -1.625 | -0.388 | 0.011 | 0.082 | 0.082 | 0.133 | 0.48 | 0.19 | 0.39 | 0.66 | 0.5 | 0.57 |
| 11 | 75034986 | rs17133921 | 0.269 | 0.195 | -1.015 | -0.435 | 0.014 | -0.038 | -0.038 | 0 | 0.1 | 0.19 | 0.06 | 0.03 | 0.1 | 0.06 |
| 11 | 75036757 | rs533085 | -0.584 | 0.161 | -1.244 | -0.47 | 0.001 | 0.005 | 0.005 | 1.057 | 0.61 | 0.74 | 0.43 | 0.43 | 0.61 | 0.43 |
| 11 | 75044640 | rs616714 | -0.041 | -0.163 | -1.204 | -0.803 | 0.006 | -0.003 | -0.003 | 0 | 0.27 | 0.15 | 0.17 | 0.3 | 0.18 | 0.17 |
| 11 | 75044798 | rs616191 | -0.063 | 0.138 | -0.852 | -1.034 | 0.014 | -0.083 | -0.083 | 0 | 0.34 | 0.6 | 0.25 | 0.14 | 0.41 | 0.23 |
| 11 | 75051233 | rs504683 | 0.345 | 0.364 | -0.798 | -0.887 | -0.004 | 0.039 | 0.039 | 0 | 0.25 | 0.16 | 0.15 | 0.15 | 0.27 | 0.17 |
| 11 | 75062178 | rs2510894 | -0.777 | 0.245 | -0.541 | -0.832 | -0.012 | 0.064 | 0.064 | 1.2 | 0.46 | 0.74 | 0.32 | 0.31 | 0.5 | 0.34 |
| 11 | 75068767 | rs11236401 | 0.814 | 0.286 | -0.265 | -0.876 | 0.045 | -0.029 | -0.029 | 0 | 0.08 | 0.19 | 0.09 | 0.03 | 0.14 | 0.02 |
| 11 | 75069871 | rs677106 | -0.437 | 0.126 | 0.258 | -0.94 | -0.001 | -0.009 | -0.009 | 0 | 0.39 | 0.62 | 0.28 | 0.19 | 0.46 | 0.18 |
| 11 | 75070786 | rs12290287 | 0.196 | 0.14 | 0.02 | -0.707 | -0.005 | 0.024 | 0.024 | 0 | 0.06 | 0.07 | 0.03 | 0.14 | 0.04 | 0.16 |
| 11 | 75075866 | rs658573 | -0.887 | -0.063 | 0.216 | -0.977 | -0.005 | 0.024 | 0.024 | 0 | 0.45 | 0.6 | 0.36 | 0.31 | 0.5 | 0.34 |
| 11 | 75080069 | rs12285820 | -0.198 | 0.076 | 0.259 | -0.664 | -0.01 | 0.018 | 0.018 | 0 | 0.09 | 0.1 | 0.06 | 0.15 | 0.07 | 0.17 |
| 11 | 75080920 | rs11236410 | 0.165 | 0.198 | 0.4 | -0.688 | 0.035 | -0.092 | -0.092 | 0 | 0.16 | 0.31 | 0.12 | 0 | 0.21 | 0.03 |
| 11 | 75087520 | rs12289289 | -0.463 | -0.133 | -0.344 | -0.516 | 0.017 | -0.007 | -0.007 | 0.476 | 0.14 | 0.07 | 0.11 | 0.15 | 0.08 | 0.22 |
| 11 | 75100600 | rs598936 | 0.324 | 0.289 | -0.549 | -0.846 | 0.003 | -0.025 | -0.025 | 0 | 0.33 | 0.35 | 0.25 | 0.18 | 0.41 | 0.18 |
| 11 | 75109868 | rs3133186 | -0.206 | -0.004 | -0.537 | 0.32 | 0.031 | -0.033 | -0.033 | 0 | 0.12 | 0.03 | 0.09 | 0.19 | 0.07 | 0.22 |
| 11 | 75110589 | rs3133187 | -0.206 | -0.004 | -0.537 | 0.32 | 0.031 | -0.033 | -0.033 | 0.982 | 0.12 | 0.03 | 0.09 | 0.19 | 0.07 | 0.22 |
| 11 | 75110952 | rs534812 | -0.463 | 0.017 | -0.91 | 0.453 | -0.052 | -0.028 | -0.028 | 0 | 0.72 | 0.83 | 0.46 | 0.64 | 0.57 | 0.62 |
| 11 | 75114071 | rs3094303 | 0.039 | -0.025 | -0.988 | 0.483 | -0.013 | 0.008 | 0.008 | 0 | 0.13 | 0.1 | 0.1 | 0.19 | 0.07 | 0.22 |
| 11 | 75116959 | rs4348932 | 0.249 | -0.217 | -1.099 | 0.294 | -0.011 | -0.006 | -0.006 | 0 | 0.24 | 0.59 | 0.16 | 0.24 | 0.17 | 0.19 |
| 11 | 75124132 | rs571632 | -0.375 | 0.182 | -1.285 | 0.177 | -0.082 | -0.047 | -0.047 | 1.116 | 0.72 | 0.88 | 0.46 | 0.65 | 0.57 | 0.63 |
| 11 | 75129007 | rs532751 | 1.399 | 0.251 | -1.728 | 0.214 | -0.034 | -0.007 | -0.007 | 0 | 0.32 | 0.03 | 0.17 | 0.26 | 0.28 | 0.18 |
| 11 | 75141762 | rs3018101 | -0.203 | 0.557 | -1.26 | 0.479 | -0.077 | -0.041 | -0.041 | 0 | 0.73 | 0.88 | 0.47 | 0.66 | 0.58 | 0.64 |
| 11 | 75142918 | rs539199 | -0.56 | 0.552 | -1.056 | 0.672 | -0.047 | -0.051 | -0.051 | 0 | 0.79 | 0.89 | 0.53 | 0.66 | 0.68 | 0.65 |
| 11 | 75144186 | rs549794 | -0.676 | 0.522 | -1.158 | 0.691 | -0.064 | -0.076 | -0.076 | 0 | 0.8 | 0.92 | 0.53 | 0.67 | 0.69 | 0.65 |
| 11 | 75147661 | rs534179 | 0.59 | -0.049 | -0.93 | 0.171 | 0.003 | 0.061 | 0.061 | 0 | 0.35 | 0.22 | 0.22 | 0.21 | 0.32 | 0.21 |
| 11 | 75154806 | rs578840 | 0.467 | -0.809 | -0.344 | 1.06 | 0.014 | -0.001 | -0.001 | 1.621 | 0.58 | 0.33 | 0.47 | 0.59 | 0.61 | 0.5 |
| 11 | 75155139 | rs474056 | 0.428 | -0.841 | -0.182 | 0.968 | 0.008 | 0.027 | 0.027 | 0 | 0.59 | 0.37 | 0.64 | 0.8 | 0.62 | 0.51 |
| 11 | 75155743 | rs479574 | 1.562 | -0.945 | -0.676 | 1.058 | -0.16 | 0.013 | 0.013 | 0 | 0.35 | 0.02 | 0.27 | 0.26 | 0.44 | 0.15 |
| 11 | 75158433 | rs482503 | 0.511 | -0.978 | 0.474 | 0.719 | 0.002 | 0.001 | 0.001 | 0 | 0.6 | 0.35 | 0.64 | 0.81 | 0.62 | 0.54 |
| 11 | 75159551 | rs488305 | 0.677 | -0.975 | 0.508 | 0.716 | 0.018 | 0.106 | 0.106 | 0 | 0.58 | 0.24 | 0.63 | 0.81 | 0.62 | 0.54 |
| 11 | 75161757 | rs12421913 | -0.587 | -1.272 | 1.041 | 0.545 | -0.03 | -0.025 | -0.025 | 0 | 0.17 | 0.29 | 0.11 | 0.15 | 0.1 | 0.2 |
| 11 | 75162754 | rs3814715 | -0.567 | -1.244 | 1.059 | 0.565 | -0.032 | -0.022 | -0.022 | 0 | 0.17 | 0.29 | 0.11 | 0.15 | 0.1 | 0.2 |
| 11 | 75167704 | rs1793410 | -0.058 | -1.206 | 1.291 | 0.56 | 0.013 | -0.092 | -0.092 | 0.29 | 0.77 | 0.59 | 0.74 | 0.96 | 0.72 | 0.74 |
| 11 | 75167793 | rs12789498 | 0.136 | -1.323 | 1.068 | 0.788 | 0.018 | -0.052 | -0.052 | 0.957 | 0.1 | 0 | 0.07 | 0.17 | 0.05 | 0.22 |
| 11 | 75169224 | rs1793411 | 0.972 | -1.175 | 1.337 | 0.415 | -0.004 | -0.028 | -0.028 | 0 | 0.63 | 0.39 | 0.65 | 0.81 | 0.64 | 0.58 |
| 11 | 75171083 | rs1793412 | 1.072 | -1.175 | 1.325 | 0.415 | 0.008 | 0.043 | 0.043 | 0 | 0.62 | 0.29 | 0.64 | 0.81 | 0.64 | 0.58 |
| 11 | 75174319 | rs1631865 | 1.095 | -1.174 | 1.246 | 0.409 | 0.012 | 0.072 | 0.072 | 0 | 0.62 | 0.26 | 0.65 | 0.8 | 0.64 | 0.58 |
| 11 | 75180967 | rs1004856 | 0.246 | -1.155 | 1.125 | 0.429 | 0.008 | -0.104 | -0.104 | 0 | 0.75 | 0.41 | 0.7 | 0.93 | 0.72 | 0.65 |
| 11 | 75184875 | rs7131225 | 0.409 | -1.219 | 1.062 | 0.441 | -0.004 | -0.014 | -0.014 | 0 | 0.11 | 0 | 0.05 | 0.13 | 0.08 | 0.05 |
| 11 | 75189157 | rs2853066 | 0.022 | -1.191 | 1.136 | 0.425 | 0.015 | -0.097 | -0.097 | 0 | 0.77 | 0.55 | 0.74 | 0.95 | 0.72 | 0.74 |
| 11 | 75199138 | rs7125082 | -0.282 | -1.185 | 1.117 | 0.459 | -0.001 | 0 | 0 | 0.014 | 0.13 | 0.13 | 0.09 | 0.15 | 0.08 | 0.16 |
| 11 | 75199181 | rs589724 | 0.115 | -0.972 | 1.163 | 0.521 | 0.014 | -0.097 | -0.097 | 0 | 0.77 | 0.55 | 0.74 | 0.95 | 0.72 | 0.75 |
| 11 | 75200301 | rs682292 | 0.115 | -0.972 | 1.163 | 0.521 | 0.014 | -0.097 | -0.097 | 0 | 0.77 | 0.54 | 0.74 | 0.95 | 0.72 | 0.75 |
| 11 | 75215421 | rs1793398 | 0.81 | -0.968 | 1.143 | 0.602 | -0.011 | -0.047 | -0.047 | 0.255 | 0.74 | 0.5 | 0.81 | 0.8 | 0.78 | 0.59 |
| 11 | 75220505 | rs1793399 | 0.837 | -1.242 | 1.171 | 0.91 | -0.019 | -0.003 | -0.003 | 0 | 0.78 | 0.42 | 0.82 | 0.78 | 0.84 | 0.67 |
| 11 | 75220697 | rs12417071 | 0.083 | -1.324 | 0.925 | 1.129 | 0.002 | -0.026 | -0.026 | 0 | 0.08 | 0.02 | 0.07 | 0.1 | 0.04 | 0.12 |
| 11 | 75223565 | rs1783550 | 2.058 | -1.385 | 0.752 | 0.966 | -0.019 | -0.05 | -0.05 | 0 | 0.47 | 0.11 | 0.49 | 0.49 | 0.58 | 0.32 |
| 11 | 75228884 | rs947844 | 0.165 | -1.443 | 1.347 | 0.981 | -0.005 | 0 | 0 | 0 | 0.13 | 0.21 | 0.08 | 0.15 | 0.11 | 0.21 |
| 11 | 75238485 | rs618202 | 0.201 | -1.452 | 1.649 | 0.99 | -0.007 | 0.03 | 0.03 | 0 | 0.84 | 0.59 | 0.84 | 0.79 | 0.87 | 0.71 |
| 11 | 75240726 | rs591627 | 1.029 | -1.48 | 1.596 | 0.971 | -0.01 | -0.005 | -0.005 | 0 | 0.78 | 0.3 | 0.82 | 0.79 | 0.85 | 0.68 |
| 11 | 75241681 | rs661928 | 2.151 | -1.46 | 1.383 | 0.977 | -0.015 | -0.047 | -0.047 | 1.855 | 0.47 | 0.1 | 0.48 | 0.49 | 0.58 | 0.32 |
| 11 | 75245306 | rs621305 | -0.085 | -1.363 | 1.502 | 0.808 | 0.025 | 0.022 | 0.022 | 0 | 0.87 | 0.69 | 0.72 | 0.88 | 0.84 | 0.76 |
| 11 | 75249185 | rs670491 | 0.873 | -1.418 | 1.316 | 0.767 | -0.007 | 0.005 | 0.005 | 0 | 0.73 | 0.47 | 0.63 | 0.72 | 0.73 | 0.55 |
| 11 | 75255123 | rs12788428 | -0.327 | -1.551 | 1.081 | 0.905 | 0.027 | -0.028 | -0.028 | 0.457 | 0.19 | 0.06 | 0.27 | 0.25 | 0.13 | 0.19 |
| 11 | 75255230 | rs12291026 | -0.045 | -1.616 | 1.261 | 0.902 | -0.002 | 0 | 0 | 0.605 | 0.13 | 0.21 | 0.08 | 0.17 | 0.12 | 0.3 |
| 11 | 75255710 | rs688727 | 1.073 | -1.639 | 1.349 | 0.9 | -0.017 | 0 | 0 | 0 | 0.65 | 0.61 | 0.45 | 0.66 | 0.72 | 0.56 |
| 11 | 75256122 | rs1938800 | -1.028 | -1.039 | 1.617 | 0.901 | -0.006 | -0.01 | -0.01 | 1.329 | 0.17 | 0.29 | 0.1 | 0.17 | 0.14 | 0.32 |
| 11 | 75261796 | rs662279 | 1.398 | -1.071 | 1.084 | 0.801 | 0.007 | 0.012 | 0.012 | 1.259 | 0.66 | 0.3 | 0.62 | 0.74 | 0.7 | 0.43 |
| 11 | 75262141 | rs663660 | 1.13 | -0.929 | 1.136 | 0.854 | -0.016 | -0.03 | -0.03 | 0 | 0.89 | 0.63 | 0.91 | 0.97 | 0.97 | 0.89 |
| 11 | 75268603 | rs673482 | 0.984 | -1.118 | 0.44 | 1.116 | 0.002 | -0.001 | -0.001 | 0.154 | 0.56 | 0.57 | 0.38 | 0.43 | 0.63 | 0.35 |
| 11 | 75268704 | rs608585 | 0.197 | -1.149 | -0.189 | 0.867 | -0.037 | 0.011 | 0.011 | 0 | 0.12 | 0.01 | 0.23 | 0.06 | 0.17 | 0.2 |

| | | | | | | | | | | | | | | | | |
|---|---|---|---|---|---|---|---|---|---|---|---|---|---|---|---|---|
| 11 | 75274150 | rs590121 | -1.019 | -1.32 | 0.506 | 0.848 | -0.026 | -0.059 | -0.059 | 0.18 | 0.34 | 0.46 | 0.37 | 0.19 | 0.29 | 0.45 |
| 11 | 75274630 | rs681390 | -1.305 | -1.394 | 0.713 | 1.193 | 0.007 | 0.004 | 0.004 | 0 | 0.39 | 0.62 | 0.44 | 0.36 | 0.3 | 0.63 |
| 11 | 75274687 | rs589374 | 1.835 | -1.233 | 0.377 | 1.719 | -0.064 | 0.129 | 0.129 | 0 | 0.43 | 0.07 | 0.29 | 0.17 | 0.54 | 0.23 |
| 11 | 75276178 | rs606452 | 0.502 | -1.141 | 0.477 | 1.578 | -0.005 | 0.183 | 0.183 | 0 | 0.81 | 0.58 | 0.72 | 0.52 | 0.84 | 0.86 |
| 11 | 75277628 | rs584961 | -0.883 | -1.151 | 0.063 | 1.772 | -0.018 | -0.01 | -0.01 | 1.654 | 0.92 | 0.95 | 0.93 | 0.73 | 0.89 | 0.94 |
| 11 | 75277992 | rs651581 | 1.602 | -1.053 | 0.006 | 1.502 | -0.043 | 0.049 | 0.049 | 0 | 0.44 | 0.08 | 0.32 | 0.3 | 0.54 | 0.33 |
| 11 | 75279846 | rs649257 | -1.004 | -0.959 | 0.354 | 0.89 | -0.016 | -0.043 | -0.043 | 0 | 0.36 | 0.48 | 0.38 | 0.21 | 0.3 | 0.49 |
| 11 | 75283653 | rs6704 | -1.054 | -0.914 | -0.036 | 0.797 | 0.006 | 0.01 | 0.01 | 0 | 0.35 | 0.34 | 0.37 | 0.19 | 0.29 | 0.5 |
| 11 | 75287341 | rs609309 | 0.753 | -0.63 | 0.03 | 1.367 | -0.008 | 0.023 | 0.023 | 0 | 0.71 | 0.67 | 0.67 | 0.59 | 0.75 | 0.71 |
| 11 | 75292601 | rs585433 | -0.155 | 0.08 | -0.678 | 0.676 | -0.021 | 0.013 | 0.013 | 0 | 0.68 | 0.77 | 0.73 | 0.71 | 0.54 | 0.73 |
| 11 | 75294923 | rs11236460 | 0.181 | -0.36 | -0.498 | 0.463 | -0.035 | 0.032 | 0.032 | 0.365 | 0.37 | 0.49 | 0.17 | 0.45 | 0.21 | 0.25 |
| 11 | 75297884 | rs588955 | -0.77 | 0.023 | -0.52 | 0.448 | -0.046 | 0.028 | 0.028 | 0 | 0.7 | 0.84 | 0.76 | 0.74 | 0.59 | 0.76 |
| 11 | 75307298 | rs642002 | -0.814 | 0.246 | -0.704 | 0.402 | -0.025 | 0.01 | 0.01 | 0 | 0.72 | 0.93 | 0.82 | 0.75 | 0.68 | 0.78 |
| 11 | 75309145 | rs648930 | -0.698 | 0.217 | -0.323 | 0.38 | 0.002 | 0.019 | 0.019 | 0 | 0.71 | 0.79 | 0.81 | 0.75 | 0.68 | 0.78 |
| 11 | 75311704 | rs658601 | 0.011 | -0.041 | 0.009 | 0.252 | -0.006 | -0.005 | -0.005 | 0.561 | 0.58 | 0.51 | 0.54 | 0.59 | 0.54 | 0.52 |
| 11 | 75314787 | rs583231 | 0.074 | -0.091 | 0.111 | 0.279 | -0.002 | -0.002 | -0.002 | 0 | 0.57 | 0.42 | 0.54 | 0.59 | 0.54 | 0.48 |
| 11 | 75320479 | rs544997 | -0.913 | -0.132 | -0.033 | 0.241 | 0.003 | 0.009 | 0.009 | 0 | 0.7 | 0.72 | 0.81 | 0.75 | 0.68 | 0.78 |
| 11 | 75324180 | rs604694 | -0.061 | -0.116 | 0.092 | 0.294 | -0.007 | 0.037 | 0.037 | 0 | 0.12 | 0.26 | 0.26 | 0.16 | 0.12 | 0.24 |
| 11 | 75324544 | rs479150 | -0.263 | -0.586 | -0.353 | -0.154 | 0.002 | 0.002 | 0.002 | 0 | 0.26 | 0.25 | 0.14 | 0.13 | 0.2 | 0.31 |
| 11 | 75324778 | rs602003 | -0.809 | -0.792 | -0.458 | 0.07 | 0 | 0.14 | 0.14 | 0 | 0.57 | 0.8 | 0.77 | 0.8 | 0.58 | 0.71 |
| 11 | 75335799 | rs598847 | 0.972 | -0.598 | 0.239 | 0.048 | -0.018 | 0.029 | 0.029 | 0 | 0.27 | 0.17 | 0.37 | 0.13 | 0.3 | 0.33 |
| 11 | 75344906 | rs640927 | -0.801 | -0.275 | 0.668 | 0.777 | -0.015 | 0.165 | 0.165 | 0.187 | 0.17 | 0.54 | 0.3 | 0.31 | 0.18 | 0.3 |
| 11 | 75359175 | rs11236476 | 2.215 | -0.234 | 0.462 | 0.378 | -0.025 | -0.002 | -0.002 | 0 | 0.06 | 0.01 | 0.08 | 0.29 | 0.06 | 0.03 |
| 11 | 75371182 | rs527048 | 1.677 | -0.187 | 0.581 | 0.379 | -0.102 | 0.042 | 0.042 | 0 | 0.18 | 0.02 | 0.3 | 0.09 | 0.25 | 0.24 |
| 11 | 75372466 | rs657639 | -0.378 | -0.352 | 1.484 | 0.885 | -0.001 | 0.046 | 0.046 | 0 | 0.13 | 0.26 | 0.27 | 0.16 | 0.13 | 0.24 |
| 11 | 75390897 | rs2509615 | 0.334 | -0.323 | 1.224 | 0.555 | 0.005 | 0.007 | 0.007 | 0.863 | 0.54 | 0.36 | 0.51 | 0.42 | 0.49 | 0.48 |
| 11 | 75400012 | rs540461 | 0.436 | -0.299 | 1.159 | 0.668 | 0.001 | 0.001 | 0.001 | 0.819 | 0.53 | 0.53 | 0.5 | 0.42 | 0.48 | 0.48 |
| 11 | 75409951 | rs660730 | 0.481 | -0.405 | 1.337 | 0.767 | -0.03 | 0.268 | 0.268 | 0 | 0.83 | 0.49 | 0.69 | 0.54 | 0.87 | 0.75 |
| 11 | 75410172 | rs530260 | 0.674 | -0.416 | 1.293 | 0.692 | 0.019 | 0.011 | 0.011 | 1.029 | 0.52 | 0.31 | 0.48 | 0.43 | 0.51 | 0.48 |
| 11 | 75414286 | rs7120248 | 0.442 | -0.366 | 0.984 | 0.614 | -0.02 | 0.112 | 0.112 | 0.011 | 0.29 | 0.12 | 0.2 | 0.07 | 0.35 | 0.2 |
| 11 | 75417722 | rs7103237 | 0.178 | 0.069 | 1.093 | 0.964 | 0.015 | 0.008 | 0.008 | 0.926 | 0.52 | 0.34 | 0.48 | 0.43 | 0.51 | 0.48 |
| 11 | 75421115 | rs17245228 | 0.248 | 0.159 | 1.007 | 0.973 | 0.014 | 0.01 | 0.01 | 0.838 | 0.51 | 0.39 | 0.48 | 0.46 | 0.52 | 0.49 |
| 11 | 75428958 | rs554202 | -0.281 | 0.222 | 0.806 | -0.045 | 0 | -0.016 | -0.016 | 0.007 | 0.42 | 0.43 | 0.3 | 0.16 | 0.46 | 0.35 |
| 11 | 75432903 | rs1219538 | -0.14 | 0.363 | 0.159 | -0.215 | 0.005 | 0.084 | 0.084 | 0.683 | 0.41 | 0.23 | 0.3 | 0.11 | 0.46 | 0.29 |
| 11 | 75436116 | rs11823410 | -1.702 | 0.858 | 1.198 | 0.743 | 0.011 | -0.007 | -0.007 | 0.017 | 0.18 | 0.43 | 0.16 | 0.22 | 0.2 | 0.13 |
| 11 | 75437630 | rs575976 | -2.057 | 0.837 | 1.476 | 0.805 | 0.013 | 0.01 | 0.01 | 0 | 0.21 | 0.48 | 0.16 | 0.28 | 0.22 | 0.17 |
| 11 | 75444605 | rs11236507 | -1.303 | 0.731 | 1.861 | 1.059 | 0.013 | 0.047 | 0.047 | 0 | 0.08 | 0.15 | 0.11 | 0.24 | 0.09 | 0.1 |
| 11 | 75445338 | rs656095 | -2.48 | 0.727 | 2.084 | 1.01 | 0.021 | 0.065 | 0.065 | 0 | 0.19 | 0.54 | 0.17 | 0.3 | 0.18 | 0.17 |
| 11 | 75446269 | rs670345 | -0.865 | 0.595 | 1.977 | 0.981 | 0 | -0.019 | -0.019 | 0 | 0.1 | 0.28 | 0.06 | 0.07 | 0.09 | 0.04 |
| 11 | 75455021 | rs499974 | -2.29 | 0.607 | 1.786 | 0.984 | -0.001 | 0.017 | 0.017 | 0 | 0.19 | 0.38 | 0.15 | 0.26 | 0.17 | 0.13 |
| 11 | 75476195 | rs10899116 | -1.846 | 0.608 | 1.835 | 1.215 | 0.001 | 0.019 | 0.019 | 0 | 0.1 | 0.14 | 0.1 | 0.23 | 0.09 | 0.09 |
| 11 | 75484850 | rs10219339 | -1.756 | 0.59 | 1.976 | 1.204 | 0.006 | 0.048 | 0.048 | 0 | 0.09 | 0.17 | 0.1 | 0.24 | 0.08 | 0.09 |
| 11 | 75492203 | rs10899121 | -2.81 | 0.535 | 2.369 | 1.173 | 0.031 | 0.126 | 0.126 | 0 | 0.12 | 0.49 | 0.13 | 0.24 | 0.08 | 0.12 |
| 11 | 75499452 | rs1017713 | -1.677 | 0.535 | 2.362 | 1.173 | 0.001 | 0.02 | 0.02 | 0 | 0.09 | 0.14 | 0.1 | 0.24 | 0.08 | 0.09 |
| 11 | 75511572 | rs3060 | -2.925 | 0.531 | 2.376 | 1.19 | 0.022 | 0.142 | 0.142 | 0 | 0.13 | 0.54 | 0.14 | 0.27 | 0.08 | 0.13 |
| 11 | 75514380 | rs11236533 | -2.11 | 0.542 | 2.375 | 1.15 | 0.014 | 0.082 | 0.082 | 0 | 0.11 | 0.29 | 0.11 | 0.24 | 0.08 | 0.09 |
| 11 | 75524796 | rs1458836 | -2.339 | 0.544 | 2.346 | 1.138 | 0.024 | 0.104 | 0.104 | 0 | 0.11 | 0.39 | 0.12 | 0.24 | 0.08 | 0.09 |
| 11 | 75544298 | rs7108178 | -2.76 | 0.542 | 2.344 | 1.136 | 0.031 | 0.122 | 0.122 | 0 | 0.12 | 0.54 | 0.13 | 0.24 | 0.08 | 0.12 |
| 11 | 75545165 | rs11236552 | -1.683 | 0.547 | 2.345 | 1.138 | 0.005 | 0.041 | 0.041 | 0 | 0.09 | 0.19 | 0.1 | 0.24 | 0.08 | 0.09 |
| 11 | 75546137 | rs11236553 | -2.024 | 0.546 | 2.334 | 1.138 | 0.024 | 0.098 | 0.098 | 0 | 0.1 | 0.33 | 0.11 | 0.24 | 0.08 | 0.09 |
| 11 | 75547716 | rs7101806 | -2.392 | 0.548 | 2.299 | 1.139 | 0.034 | 0.116 | 0.116 | 0 | 0.11 | 0.43 | 0.12 | 0.24 | 0.08 | 0.12 |
| 11 | 75556323 | rs11236561 | -1.02 | 0.768 | 2.041 | 0.826 | 0.041 | -0.043 | -0.043 | 0 | 0.09 | 0.32 | 0.07 | 0 | 0.08 | 0.04 |
| 11 | 75561659 | rs10793135 | -1.626 | 0.58 | 2.038 | 0.999 | 0.002 | 0.024 | 0.024 | 0 | 0.09 | 0.16 | 0.1 | 0.24 | 0.08 | 0.12 |
| 11 | 75569150 | rs12280464 | -2.55 | 0.614 | 2.21 | 0.977 | 0.034 | 0.114 | 0.114 | 0 | 0.12 | 0.47 | 0.11 | 0.23 | 0.08 | 0.15 |
| 11 | 75603050 | rs7109235 | -1.964 | 0.508 | 2.236 | 0.859 | 0.054 | 0.061 | 0.061 | 1.167 | 0.08 | 0.45 | 0.05 | 0.05 | 0.04 | 0.08 |
| 11 | 75611644 | rs7927868 | -1.512 | 0.51 | 2.283 | 0.86 | 0.041 | 0.048 | 0.048 | 0.614 | 0.07 | 0.33 | 0.04 | 0.05 | 0.04 | 0.08 |
| 11 | 75637567 | rs7111334 | -2.177 | 0.483 | 2.182 | 1.014 | -0.016 | 0.087 | 0.087 | 0.768 | 0.15 | 0.53 | 0.1 | 0.24 | 0.08 | 0.23 |
| 11 | 75645950 | rs1995892 | -1.435 | 0.125 | 1.558 | 1.217 | -0.023 | -0.004 | -0.004 | 0 | 0.51 | 0.82 | 0.32 | 0.55 | 0.47 | 0.54 |
| 11 | 75678647 | rs11236588 | -1.607 | -0.092 | 1.447 | 1.311 | -0.06 | 0.169 | 0.169 | 0 | 0.22 | 0.52 | 0.13 | 0.46 | 0.12 | 0.38 |
| 11 | 75733418 | rs6592621 | -1.209 | -0.072 | 1.58 | 1.45 | -0.011 | 0.145 | 0.145 | 1.781 | 0.14 | 0.49 | 0.1 | 0.29 | 0.07 | 0.24 |
| 11 | 75805884 | rs667534 | -0.302 | -0.189 | 1.958 | 1.416 | -0.008 | 0.009 | 0.009 | 0 | 0.35 | 0.13 | 0.2 | 0.3 | 0.37 | 0.29 |
| 11 | 75819828 | rs660211 | -0.282 | -0.504 | 1.386 | 2.427 | -0.024 | -0.072 | -0.072 | 0 | 0.89 | 0.99 | 0.66 | 0.76 | 0.91 | 0.88 |
| 11 | 75825224 | rs7931174 | -2.115 | -0.428 | 2.001 | 2.133 | -0.023 | 0.124 | 0.124 | 1.668 | 0.14 | 0.54 | 0.1 | 0.3 | 0.07 | 0.27 |
| 11 | 75837669 | rs11236618 | -1.553 | -0.262 | 0.258 | 2.166 | -0.013 | 0.05 | 0.05 | 3.091 | 0.12 | 0.24 | 0.07 | 0.31 | 0.06 | 0.26 |
| 11 | 75852448 | rs636420 | -0.54 | -0.002 | -0.367 | 1.928 | 0.022 | -0.061 | -0.061 | 0 | 0.1 | 0 | 0.03 | 0.27 | 0.06 | 0.13 |
| 11 | 75858026 | rs7120205 | -1.348 | 0.678 | 0.227 | 1.681 | 0.01 | 0.062 | 0.062 | 6.214 | 0.09 | 0.17 | 0.06 | 0.22 | 0.08 | 0.13 |
| 11 | 75861589 | rs2851570 | 1.117 | 0.631 | 0.096 | 2.373 | 0 | 0.039 | 0.039 | 0 | 0.88 | 0.86 | 0.71 | 0.51 | 0.92 | 0.78 |
| 11 | 75865197 | rs10793139 | -0.964 | 0.953 | -0.815 | 0.902 | -0.031 | 0.087 | 0.087 | 0.002 | 0.41 | 0.18 | 0.35 | 0.24 | 0.52 | 0.35 |
| 11 | 75866552 | rs4477460 | -0.902 | 0.862 | -0.751 | 1.07 | 0.001 | -0.016 | -0.016 | 1.583 | 0.12 | 0.08 | 0.06 | 0.22 | 0.07 | 0.13 |

| | | | | | | | | | | | | | | | | |
|---|---|---|---|---|---|---|---|---|---|---|---|---|---|---|---|---|
| 11 | 75866999 | rs7106626 | -1.33 | 0.9 | -0.681 | 1.01 | 0.001 | -0.009 | -0.009 | 0 | 0.15 | 0.13 | 0.09 | 0.22 | 0.09 | 0.14 |
| 11 | 75870179 | rs4945036 | -0.225 | 0.817 | -0.341 | 0.456 | -0.006 | -0.011 | -0.011 | 0.017 | 0.29 | 0.27 | 0.25 | 0.04 | 0.38 | 0.28 |
| 11 | 75884832 | rs17134631 | -1.144 | 0.385 | -0.184 | 0.857 | -0.006 | 0.014 | 0.014 | 0 | 0.18 | 0.21 | 0.1 | 0.25 | 0.11 | 0.24 |
| 11 | 75885321 | rs7945646 | -1.357 | 0.101 | 0.201 | 0.455 | -0.002 | 0.053 | 0.053 | 0.016 | 0.49 | 0.64 | 0.38 | 0.28 | 0.51 | 0.5 |
| 11 | 75886083 | rs589149 | 0.568 | -0.459 | -0.211 | 0.731 | -0.006 | 0.001 | 0.001 | 0 | 0.63 | 0.63 | 0.81 | 0.46 | 0.77 | 0.61 |
| 11 | 75887686 | rs664499 | -1.057 | -0.27 | -0.51 | 1.473 | 0.012 | -0.021 | -0.021 | 0.044 | 0.13 | 0.07 | 0.15 | 0.33 | 0.08 | 0.23 |
| 11 | 75890695 | rs582132 | -0.914 | -0.326 | 0.33 | 1.549 | 0.071 | 0.261 | 0.261 | 0.044 | 0.06 | 0.31 | 0.17 | 0.21 | 0.05 | 0.2 |
| 11 | 75892741 | rs647159 | -1.239 | -0.223 | 0.305 | 1.724 | 0.056 | 0.376 | 0.376 | 0 | 0.07 | 0.32 | 0.18 | 0.36 | 0.07 | 0.22 |
| 11 | 75894129 | rs663746 | -0.519 | -0.425 | -0.606 | 0.759 | -0.005 | -0.011 | -0.011 | 0 | 0.48 | 0.2 | 0.55 | 0.51 | 0.49 | 0.47 |
| 11 | 75895007 | rs11236644 | 0.562 | -0.539 | -0.522 | 0.607 | -0.009 | -0.018 | -0.018 | 0.44 | 0.37 | 0.71 | 0.36 | 0.48 | 0.38 | 0.48 |
| 11 | 75896647 | rs12277860 | -0.376 | -0.968 | -0.925 | 0.695 | 0.012 | 0.006 | 0.006 | 0 | 0.21 | 0.15 | 0.31 | 0.19 | 0.14 | 0.23 |
| 11 | 75899407 | rs7936750 | -0.226 | -0.34 | -0.831 | 0.994 | 0.001 | -0.017 | -0.017 | 0 | 0.93 | 0.99 | 0.89 | 0.7 | 0.96 | 0.81 |
| 11 | 75904137 | rs882151 | 0.653 | 0.285 | -0.615 | 0.235 | -0.003 | -0.07 | -0.07 | 0.009 | 0.2 | 0.26 | 0.11 | 0.01 | 0.19 | 0.09 |
| 11 | 75914087 | rs689095 | 0.819 | 0.597 | -0.708 | 0.387 | 0.008 | -0.027 | -0.027 | 0 | 0.33 | 0.29 | 0.43 | 0.52 | 0.31 | 0.45 |
| 11 | 75915139 | rs94111 | -0.454 | 1.278 | -0.348 | 0.489 | 0.026 | -0.043 | -0.043 | 0.59 | 0.69 | 0.55 | 0.63 | 0.79 | 0.63 | 0.76 |
| 11 | 75923069 | rs663907 | 0.353 | 0.936 | -0.47 | 0.681 | 0.011 | 0.149 | 0.149 | 0 | 0.06 | 0.11 | 0.16 | 0.31 | 0.06 | 0.2 |
| 11 | 75927441 | rs12416814 | 1.799 | 0.794 | -0.97 | -0.05 | -0.031 | 0.02 | 0.02 | 0 | 0.23 | 0.05 | 0.26 | 0.16 | 0.25 | 0.22 |
| 11 | 75931291 | rs652673 | -0.732 | 0.785 | -0.623 | -0.26 | 0.008 | -0.005 | -0.005 | 0 | 0.73 | 0.79 | 0.87 | 0.98 | 0.77 | 0.84 |
| 11 | 75934192 | rs4945048 | -0.186 | 0.82 | -0.301 | 0.039 | -0.019 | 0.016 | 0.016 | 0.231 | 0.65 | 0.5 | 0.7 | 0.63 | 0.71 | 0.65 |
| 11 | 75936131 | rs12718487 | -0.007 | 1.222 | -0.399 | -0.121 | 0.011 | 0.013 | 0.013 | 0 | 0.51 | 0.46 | 0.63 | 0.62 | 0.58 | 0.61 |
| 11 | 75939600 | rs1402704 | -0.593 | 1.313 | -0.101 | 0.163 | -0.004 | -0.005 | -0.005 | 0 | 0.29 | 0.27 | 0.32 | 0.44 | 0.33 | 0.35 |
| 11 | 75941236 | rs593241 | 0.142 | 1.248 | 0.024 | 0.278 | 0.023 | 0.207 | 0.207 | 0.142 | 0.07 | 0.21 | 0.22 | 0.35 | 0.08 | 0.2 |
| 11 | 75943540 | rs2447947 | -1.085 | 1.184 | -0.163 | -0.088 | 0.017 | -0.029 | -0.029 | 0 | 0.37 | 0.48 | 0.54 | 0.79 | 0.41 | 0.58 |
| 11 | 75946542 | rs2371855 | 0.39 | 1.001 | -0.466 | -0.463 | -0.026 | -0.153 | -0.153 | 0 | 0.34 | 0.41 | 0.17 | 0.02 | 0.26 | 0.17 |
| 11 | 75947617 | rs681836 | -0.221 | 1.125 | -0.551 | -0.032 | -0.001 | 0 | 0 | 0.016 | 0.44 | 0.53 | 0.42 | 0.43 | 0.42 | 0.41 |
| 11 | 75956348 | rs2226615 | -0.166 | 1.084 | -1.073 | 0.058 | -0.048 | -0.016 | -0.016 | 0 | 0.23 | 0.03 | 0.12 | 0.17 | 0.21 | 0.31 |
| 11 | 75959236 | rs616263 | -0.459 | 1.252 | -0.451 | 0.175 | 0.002 | -0.001 | -0.001 | 0 | 0.44 | 0.44 | 0.42 | 0.43 | 0.44 | 0.33 |
| 11 | 75960472 | rs631948 | 0.261 | 1.009 | -0.487 | 0.115 | 0.001 | 0.017 | 0.017 | 0 | 0.51 | 0.43 | 0.6 | 0.62 | 0.54 | 0.69 |
| 11 | 75961040 | rs588696 | -0.006 | 0.779 | 0.001 | 0.231 | 0.004 | -0.025 | -0.025 | 0 | 0.89 | 0.77 | 0.93 | 0.95 | 0.91 | 0.92 |
| 11 | 75962755 | rs10899186 | 1.202 | 0.82 | -0.002 | 0.462 | 0.005 | -0.001 | -0.001 | 0 | 0.64 | 0.43 | 0.48 | 0.34 | 0.67 | 0.46 |
| 11 | 75966611 | rs2510733 | -0.929 | 0.769 | -0.677 | 0.127 | -0.014 | 0.056 | 0.056 | 0 | 0.56 | 0.7 | 0.61 | 0.72 | 0.54 | 0.71 |
| 11 | 75967695 | rs11236663 | 0.124 | 0.905 | -0.309 | 0.728 | 0 | 0.032 | 0.032 | 0.092 | 0.89 | 0.92 | 0.82 | 0.66 | 0.9 | 0.76 |
| 11 | 75967841 | rs10899188 | 0.216 | 0.901 | -0.295 | 0.727 | -0.003 | 0.09 | 0.09 | 0.094 | 0.89 | 0.86 | 0.78 | 0.66 | 0.9 | 0.76 |
| 11 | 75969633 | rs12807555 | 0.187 | 0.881 | -0.473 | 0.618 | -0.001 | -0.019 | -0.019 | 0 | 0.16 | 0.16 | 0.1 | 0.03 | 0.11 | 0.2 |
| 11 | 75969749 | rs4945056 | 0.593 | 0.994 | -0.472 | 0.682 | 0 | 0.014 | 0.014 | 0.512 | 0.82 | 0.76 | 0.68 | 0.6 | 0.81 | 0.74 |
| 11 | 75971550 | rs2618078 | -1.383 | 0.97 | -0.677 | 0.608 | 0 | 0.005 | 0.005 | 0 | 0.3 | 0.29 | 0.5 | 0.5 | 0.29 | 0.51 |
| 11 | 75973078 | rs603232 | -1.372 | 0.982 | -0.663 | 0.608 | 0 | 0.006 | 0.006 | 0 | 0.3 | 0.3 | 0.49 | 0.51 | 0.29 | 0.52 |
| 11 | 75974761 | rs585200 | 0.668 | 1.022 | -0.698 | 0.78 | -0.011 | 0.042 | 0.042 | 0.022 | 0.06 | 0.03 | 0.36 | 0.34 | 0.1 | 0.19 |
| 11 | 75976301 | rs639015 | -0.085 | 0.899 | -0.576 | 0.586 | 0.007 | -0.036 | -0.036 | 0 | 0.88 | 0.75 | 0.93 | 0.94 | 0.9 | 0.91 |
| 11 | 75977507 | rs670193 | 1.315 | 0.881 | -0.382 | 0.672 | 0.003 | 0.008 | 0.008 | 0 | 0.64 | 0.49 | 0.7 | 0.54 | 0.69 | 0.51 |
| 11 | 75977845 | rs11236673 | -0.764 | 0.895 | -0.768 | 0.432 | -0.001 | 0.006 | 0.006 | 0 | 0.22 | 0.17 | 0.13 | 0.17 | 0.18 | 0.32 |
| 11 | 75978398 | rs17448977 | 0.445 | 0.863 | -1.016 | 0.666 | 0 | 0 | 0 | 0.036 | 0.07 | 0.05 | 0.04 | 0.12 | 0.07 | 0.12 |
| 11 | 75979986 | rs2371933 | 1.004 | 0.633 | -1.292 | 0.376 | 0.001 | -0.027 | -0.027 | 0 | 0.35 | 0.22 | 0.31 | 0.44 | 0.38 | 0.22 |
| 11 | 75980525 | rs2846072 | -0.164 | 0.834 | -1.297 | 0.189 | 0.002 | -0.01 | -0.01 | 0 | 0.94 | 0.85 | 0.95 | 0.95 | 0.95 | 0.92 |
| 11 | 75981779 | rs2253423 | -0.138 | 0.811 | -1.758 | 0.074 | -0.011 | 0.102 | 0.102 | 0 | 0.58 | 0.75 | 0.65 | 0.78 | 0.59 | 0.63 |
| 11 | 75982418 | rs4945061 | 0.74 | 0.589 | -1.862 | 0.135 | -0.006 | 0.029 | 0.029 | 0.3 | 0.55 | 0.65 | 0.31 | 0.33 | 0.52 | 0.31 |
| 11 | 75983754 | rs11236675 | 1.271 | 0.667 | -2.246 | 0.412 | -0.015 | -0.049 | -0.049 | 0.018 | 0.31 | 0.06 | 0.18 | 0.32 | 0.32 | 0.2 |
| 11 | 75986154 | rs2851162 | -0.738 | 1.247 | -1.757 | 1.113 | -0.094 | -0.095 | -0.095 | 0 | 0.23 | 0.05 | 0.49 | 0.36 | 0.31 | 0.42 |
| 11 | 75989871 | rs2851161 | -1.289 | 0.868 | -0.803 | 1.211 | -0.002 | 0.033 | 0.033 | 0 | 0.34 | 0.58 | 0.49 | 0.5 | 0.26 | 0.57 |
| 11 | 75992590 | rs4944101 | 1.348 | 1.084 | -1.358 | 0.857 | -0.003 | 0.008 | 0.008 | 0.162 | 0.57 | 0.13 | 0.55 | 0.36 | 0.69 | 0.35 |
| 11 | 76004313 | rs11236689 | -0.808 | 1.11 | -0.342 | 1.207 | -0.014 | 0.089 | 0.089 | 0.13 | 0.22 | 0.34 | 0.42 | 0.49 | 0.16 | 0.4 |
| 11 | 76004761 | rs2846070 | 0.125 | 1.404 | -0.64 | 0.869 | -0.054 | 0.008 | 0.008 | 0 | 0.12 | 0 | 0.08 | 0.09 | 0.1 | 0.19 |
| 11 | 76005086 | rs2097118 | 0.456 | 1.28 | -0.317 | 0.789 | 0.003 | -0.004 | -0.004 | 0 | 0.82 | 0.8 | 0.86 | 0.9 | 0.8 | 0.78 |
| 11 | 76026586 | rs878904 | 0.433 | 0.623 | -0.727 | 0.591 | 0.019 | 0.024 | 0.024 | 0 | 0.42 | 0.26 | 0.31 | 0.35 | 0.43 | 0.29 |
| 11 | 76027936 | rs745870 | -0.218 | 0.39 | -0.164 | 0.704 | -0.003 | -0.002 | -0.002 | 0 | 0.53 | 0.49 | 0.35 | 0.45 | 0.48 | 0.48 |
| 11 | 76032079 | rs12295401 | 0.149 | 0.343 | -0.035 | 0.618 | -0.011 | 0.016 | 0.016 | 0 | 0.11 | 0.32 | 0.06 | 0.1 | 0.05 | 0.19 |
| 11 | 76033374 | rs750801 | 0.343 | 0.464 | -0.049 | 0.746 | 0.004 | -0.002 | -0.002 | 0 | 0.44 | 0.42 | 0.37 | 0.44 | 0.46 | 0.31 |
| 11 | 76047793 | rs4945073 | 0.501 | 0.537 | -0.429 | 0.363 | -0.016 | 0.039 | 0.039 | 0 | 0.41 | 0.23 | 0.31 | 0.32 | 0.4 | 0.36 |
| 11 | 76050597 | rs17749447 | 0.997 | 0.65 | -0.548 | 0.347 | -0.009 | 0.009 | 0.009 | 0 | 0.12 | 0.09 | 0.08 | 0.1 | 0.16 | 0.23 |
| 11 | 76053527 | rs11236720 | 1.473 | 0.442 | -0.385 | 0.111 | -0.049 | -0.079 | -0.079 | 0.044 | 0.17 | 0.01 | 0.12 | 0.3 | 0.14 | 0.2 |
| 11 | 76054902 | rs17134811 | 0.254 | 0.427 | -0.047 | 0.375 | -0.01 | -0.041 | -0.041 | 0 | 0.16 | 0.42 | 0.11 | 0.09 | 0.13 | 0.09 |
| 11 | 76056896 | rs7126625 | 1.065 | 0.084 | -0.105 | 0.305 | 0.007 | -0.002 | -0.002 | 0 | 0.42 | 0.45 | 0.4 | 0.41 | 0.43 | 0.39 |
| 11 | 76065047 | rs2618063 | 0.951 | -0.187 | -0.15 | 0.938 | -0.031 | -0.087 | -0.087 | 0 | 0.2 | 0.04 | 0.37 | 0.26 | 0.23 | 0.2 |
| 11 | 76087399 | rs7925910 | -0.674 | -0.226 | 0.469 | 0.982 | 0.005 | 0.017 | 0.017 | 0.058 | 0.17 | 0.57 | 0.16 | 0.15 | 0.17 | 0.1 |
| 11 | 76100197 | rs6592631 | -0.946 | -0.22 | 0.006 | 1.108 | 0.005 | -0.018 | -0.018 | 0.022 | 0.92 | 0.97 | 0.96 | 0.83 | 0.94 | 0.93 |
| 11 | 76104085 | rs17134821 | -0.634 | -0.459 | 0.628 | 1.14 | -0.031 | 0.127 | 0.127 | 0.643 | 0.15 | 0.41 | 0.13 | 0.22 | 0.08 | 0.14 |
| 11 | 76104519 | rs12223637 | 0.529 | -0.488 | 0.403 | 1.184 | 0.012 | 0.037 | 0.037 | 0 | 0.17 | 0.22 | 0.39 | 0.27 | 0.24 | 0.16 |
| 11 | 76108796 | rs11236742 | 1.226 | -0.528 | -0.15 | 0.797 | 0.021 | 0.045 | 0.045 | 0.778 | 0.55 | 0.35 | 0.43 | 0.4 | 0.6 | 0.62 |
| 11 | 76116713 | rs868124 | -0.698 | -0.532 | -0.093 | 0.849 | -0.001 | 0.005 | 0.005 | 0 | 0.31 | 0.59 | 0.51 | 0.49 | 0.32 | 0.3 |

| | | | | | | | | | | | | | | | | |
|---|---|---|---|---|---|---|---|---|---|---|---|---|---|---|---|---|
| 11 | 76118350 | rs10736805 | -0.39 | -0.496 | -0.164 | 0.661 | 0.016 | 0.017 | 0.017 | 0.711 | 0.86 | 0.88 | 0.93 | 0.9 | 0.92 | 0.92 |
| 11 | 76120025 | rs4945083 | 1.047 | -0.525 | -0.03 | 0.823 | 0.007 | 0 | 0 | 0.045 | 0.58 | 0.38 | 0.5 | 0.44 | 0.62 | 0.65 |
| 11 | 76120315 | rs7927830 | -0.578 | -0.242 | 0.161 | 1.228 | 0 | 0.032 | 0.032 | 0 | 0.24 | 0.31 | 0.43 | 0.33 | 0.26 | 0.19 |
| 11 | 76123849 | rs871911 | -0.578 | -0.242 | 0.137 | 1.228 | 0 | 0.035 | 0.035 | 0 | 0.24 | 0.32 | 0.43 | 0.33 | 0.26 | 0.2 |
| 11 | 76124816 | rs7938584 | -0.936 | -0.33 | -0.152 | 1.017 | 0.007 | -0.009 | -0.009 | 0.011 | 0.93 | 0.94 | 0.99 | 0.83 | 0.99 | 0.92 |
| 11 | 76125330 | rs7927515 | 0.112 | -0.498 | -0.169 | 0.644 | -0.005 | -0.009 | -0.009 | 0.03 | 0.33 | 0.6 | 0.22 | 0.26 | 0.34 | 0.26 |
| 11 | 76127214 | rs10160382 | 0.159 | -0.509 | -0.102 | 0.623 | -0.006 | -0.026 | -0.026 | 0 | 0.33 | 0.58 | 0.22 | 0.2 | 0.33 | 0.26 |
| 11 | 76130406 | rs10899212 | -0.125 | -0.714 | 0.152 | 0.484 | -0.027 | -0.011 | -0.011 | 0 | 0.12 | 0.42 | 0.07 | 0.06 | 0.07 | 0.04 |
| 11 | 76136482 | rs4945087 | 0.821 | -0.759 | 0.086 | 0.795 | -0.017 | 0.034 | 0.034 | 0.018 | 0.43 | 0.28 | 0.5 | 0.39 | 0.43 | 0.51 |
| 11 | 76139846 | rs17134872 | 1.021 | -0.755 | 0.24 | 0.73 | -0.011 | 0.03 | 0.03 | 0 | 0.08 | 0.3 | 0.04 | 0.06 | 0.02 | 0.02 |
| 11 | 76143658 | rs10736806 | -1.789 | -0.732 | 0.126 | 0.764 | 0.021 | -0.024 | -0.024 | 0.189 | 0.94 | 1 | 0.99 | 0.89 | 0.99 | 0.93 |
| 11 | 76147984 | rs6592634 | -1.831 | -0.731 | 0.126 | 0.764 | 0.021 | -0.024 | -0.024 | 0 | 0.94 | 1 | 0.99 | 0.89 | 0.99 | 0.93 |
| 11 | 76154846 | rs2282611 | 1.091 | -0.706 | 0.159 | 0.746 | -0.011 | 0.024 | 0.024 | 0 | 0.25 | 0.18 | 0.18 | 0.14 | 0.31 | 0.22 |
| 11 | 76155555 | rs12278256 | 1.043 | -0.735 | 0.362 | 0.81 | -0.015 | 0.024 | 0.024 | 0 | 0.08 | 0.28 | 0.04 | 0.06 | 0.02 | 0.02 |
| 11 | 76161389 | rs7123361 | 1.017 | -0.718 | -0.083 | 0.809 | -0.013 | 0.028 | 0.028 | 0 | 0.26 | 0.18 | 0.17 | 0.14 | 0.32 | 0.22 |
| 11 | 76163151 | rs4300410 | -0.01 | -0.523 | -0.358 | 0.958 | 0.025 | 0.03 | 0.03 | 0 | 0.57 | 0.31 | 0.75 | 0.64 | 0.6 | 0.67 |
| 11 | 76164012 | rs10793169 | -0.01 | -0.524 | -0.358 | 0.947 | 0.027 | 0.028 | 0.028 | 0 | 0.57 | 0.31 | 0.75 | 0.64 | 0.6 | 0.67 |
| 11 | 76181631 | rs10899221 | 2.051 | -0.281 | -0.665 | 0.633 | -0.034 | 0.038 | 0.038 | 0 | 0.07 | 0 | 0.03 | 0 | 0.09 | 0.09 |
| 11 | 76183924 | rs4245443 | -0.198 | -0.298 | -0.474 | 0.897 | 0.02 | 0.04 | 0.04 | 0 | 0.57 | 0.33 | 0.75 | 0.69 | 0.61 | 0.67 |
| 11 | 76227182 | rs2508740 | -0.661 | -0.325 | -0.444 | 0.895 | 0 | 0.001 | 0.001 | 0 | 0.63 | 0.38 | 0.76 | 0.8 | 0.62 | 0.74 |
| 11 | 76236220 | rs1939469 | 1.225 | -0.301 | -0.546 | 1.021 | -0.047 | -0.082 | -0.082 | 0 | 0.13 | 0.02 | 0.24 | 0.25 | 0.16 | 0.16 |
| 11 | 76244493 | rs2513514 | 1.03 | -0.324 | -0.737 | 0.835 | -0.003 | -0.006 | -0.006 | 1.475 | 0.14 | 0.14 | 0.29 | 0.38 | 0.13 | 0.32 |
| 11 | 76245562 | rs2508755 | -0.083 | -0.278 | -0.526 | 0.993 | 0.007 | 0.008 | 0.008 | 0 | 0.5 | 0.36 | 0.51 | 0.55 | 0.46 | 0.58 |
| 11 | 76261533 | rs2513513 | -0.41 | -0.328 | -0.693 | 0.766 | 0.018 | 0.042 | 0.042 | 0 | 0.57 | 0.33 | 0.75 | 0.7 | 0.61 | 0.67 |
| 11 | 76262128 | rs1044265 | 0.274 | -0.263 | -0.699 | 0.988 | -0.013 | 0.002 | 0.002 | 0.131 | 0.43 | 0.28 | 0.51 | 0.47 | 0.44 | 0.51 |
| 11 | 76266708 | rs2513525 | 0.206 | -0.287 | -0.453 | 0.601 | 0.004 | 0.023 | 0.023 | 0 | 0.37 | 0.69 | 0.25 | 0.26 | 0.39 | 0.26 |
| 11 | 76270683 | rs7130588 | 1.133 | -0.635 | -0.544 | 0.617 | -0.041 | 0.079 | 0.079 | 0 | 0.28 | 0.18 | 0.21 | 0.12 | 0.35 | 0.23 |
| 11 | 76270950 | rs2508746 | 0.034 | -0.717 | -0.15 | 0.617 | -0.02 | 0.051 | 0.051 | 0 | 0.43 | 0.36 | 0.28 | 0.23 | 0.48 | 0.3 |
| 11 | 76271261 | rs2513517 | -0.244 | -0.757 | 0.01 | 0.735 | -0.001 | 0.005 | 0.005 | 0 | 0.47 | 0.6 | 0.3 | 0.34 | 0.5 | 0.36 |
| 11 | 76272639 | rs1892954 | 0.143 | -0.686 | 0.117 | 0.742 | -0.006 | -0.002 | -0.002 | 8.743 | 0.16 | 0.27 | 0.07 | 0.12 | 0.13 | 0.07 |
| 11 | 76275703 | rs12789493 | -0.255 | -0.848 | -0.02 | 0.995 | -0.002 | 0 | 0 | 0 | 0.56 | 0.56 | 0.51 | 0.5 | 0.56 | 0.5 |
| 11 | 76277902 | rs1892958 | -1.349 | -0.754 | -0.052 | 0.873 | 0 | 0.018 | 0.018 | 0.001 | 0.89 | 0.89 | 0.73 | 0.65 | 0.91 | 0.74 |
| 11 | 76278258 | rs7936562 | -0.203 | -0.825 | -0.079 | 0.876 | -0.001 | -0.001 | -0.001 | 0 | 0.56 | 0.63 | 0.52 | 0.52 | 0.57 | 0.51 |
| 11 | 76288937 | rs10899234 | -0.174 | -0.796 | -0.28 | 0.7 | 0.033 | 0.06 | 0.06 | 0 | 0.43 | 0.6 | 0.25 | 0.17 | 0.36 | 0.25 |
| 11 | 76299194 | rs2155219 | -0.463 | -0.632 | -0.292 | 1.085 | 0.002 | 0.01 | 0.01 | 0 | 0.5 | 0.59 | 0.5 | 0.53 | 0.49 | 0.52 |
| 11 | 76300956 | rs1892951 | 1.51 | -0.256 | -0.973 | 1.195 | 0 | -0.038 | -0.038 | 0 | 0.1 | 0.03 | 0.24 | 0.25 | 0.09 | 0.2 |
| 11 | 76301316 | rs7927894 | 0.515 | -0.151 | -0.837 | 0.772 | -0.006 | 0.007 | 0.007 | 0 | 0.31 | 0.35 | 0.24 | 0.21 | 0.37 | 0.3 |
| 11 | 76301375 | rs7927997 | 0.554 | -0.133 | -1.046 | 0.781 | -0.014 | 0.019 | 0.019 | 0 | 0.31 | 0.28 | 0.24 | 0.21 | 0.37 | 0.3 |
| 11 | 76304425 | rs10793175 | -1.245 | -0.77 | -1.006 | 1.098 | 0.064 | -0.033 | -0.033 | 1.777 | 0.48 | 0.84 | 0.4 | 0.44 | 0.37 | 0.46 |
| 11 | 76308272 | rs11827375 | 0.212 | -0.498 | -0.348 | 0.671 | -0.007 | -0.041 | -0.041 | 0.182 | 0.13 | 0.15 | 0.06 | 0.01 | 0.11 | 0.11 |
| 11 | 76313781 | rs4485135 | -1.068 | -0.713 | 0.108 | 1.054 | -0.034 | 0.028 | 0.028 | 0 | 0.29 | 0.42 | 0.34 | 0.33 | 0.23 | 0.42 |
| 11 | 76314934 | rs17135034 | -0.056 | -0.851 | 0.073 | 1.359 | 0.006 | 0.153 | 0.153 | 1.502 | 0.17 | 0.39 | 0.11 | 0.39 | 0.14 | 0.09 |
| 11 | 76318787 | rs10751256 | -1.734 | -0.831 | 0.134 | 1.233 | 0.045 | 0.084 | 0.084 | 0 | 0.45 | 0.74 | 0.45 | 0.72 | 0.36 | 0.51 |
| 11 | 76322550 | rs17160155 | -0.112 | -0.678 | 0.093 | 1.461 | -0.001 | 0.115 | 0.115 | 1.851 | 0.17 | 0.31 | 0.11 | 0.41 | 0.14 | 0.09 |
| 11 | 76324076 | rs11236808 | -0.669 | -0.498 | -0.299 | 1.279 | 0.002 | -0.009 | -0.009 | 0 | 0.24 | 0.17 | 0.32 | 0.3 | 0.23 | 0.37 |
| 11 | 76332210 | rs11236809 | 0.767 | -0.222 | -0.426 | 1.623 | -0.026 | -0.06 | -0.06 | 0 | 0.09 | 0.01 | 0.25 | 0.24 | 0.09 | 0.21 |
| 11 | 76337181 | rs11825087 | -1.234 | -0.245 | -0.218 | 0.978 | -0.002 | 0 | 0 | 0 | 0.36 | 0.62 | 0.41 | 0.65 | 0.28 | 0.42 |
| 11 | 76337528 | rs11824655 | -1.09 | -0.525 | 0.271 | 0.673 | -0.017 | -0.01 | -0.01 | 0 | 0.1 | 0.4 | 0.04 | 0.02 | 0.04 | 0.1 |
| 11 | 76338579 | rs2186676 | -1.111 | -0.385 | -0.249 | 0.629 | -0.019 | -0.022 | -0.022 | 0.078 | 0.37 | 0.53 | 0.42 | 0.71 | 0.27 | 0.43 |
| 11 | 76339194 | rs2186677 | -0.462 | 0.453 | -0.497 | 0.073 | 0.033 | -0.029 | -0.029 | 0.167 | 0.79 | 0.62 | 0.67 | 0.84 | 0.72 | 0.74 |
| 11 | 76339664 | rs10899240 | -0.186 | -0.282 | -0.741 | -0.033 | -0.012 | 0.009 | 0.009 | 1.182 | 0.35 | 0.43 | 0.2 | 0.58 | 0.28 | 0.36 |
| 11 | 76339953 | rs11236811 | 1.363 | -0.735 | -1.186 | -0.208 | 0.005 | -0.026 | -0.026 | 0 | 0.15 | 0.03 | 0.27 | 0.18 | 0.13 | 0.12 |
| 11 | 76345406 | rs1941606 | -0.134 | -1.025 | -0.736 | 0.111 | 0.001 | 0 | 0 | 0.154 | 0.36 | 0.44 | 0.23 | 0.28 | 0.31 | 0.38 |
| 11 | 76347032 | rs947999 | 1.219 | -1.085 | -1.123 | -0.047 | 0.009 | 0.016 | 0.016 | 0 | 0.17 | 0.18 | 0.13 | 0.04 | 0.16 | 0.15 |
| 11 | 76349080 | rs7949206 | 1.201 | -0.945 | -1.18 | -0.036 | 0.009 | 0.044 | 0.044 | 0 | 0.19 | 0.13 | 0.13 | 0.05 | 0.18 | 0.17 |
| 11 | 76351565 | rs11236821 | 0.373 | -0.749 | -0.736 | 0.157 | -0.005 | 0.004 | 0.004 | 0 | 0.18 | 0.16 | 0.09 | 0.22 | 0.14 | 0.21 |
| 11 | 76352908 | rs7931131 | -0.825 | -0.724 | -0.886 | 0.104 | 0.021 | 0.023 | 0.023 | 0 | 0.55 | 0.69 | 0.34 | 0.63 | 0.42 | 0.49 |
| 11 | 76354008 | rs2186674 | -0.82 | -0.697 | -0.883 | 0.121 | 0.019 | 0.01 | 0.01 | 0 | 0.55 | 0.68 | 0.34 | 0.6 | 0.41 | 0.51 |
| 11 | 76358976 | rs11603047 | 0.375 | 0.313 | -1.431 | -0.301 | -0.095 | 0.124 | 0.124 | 0.025 | 0.25 | 0.05 | 0.15 | 0.06 | 0.33 | 0.3 |
| 11 | 76359555 | rs11236826 | 0.399 | 0.312 | -1.432 | -0.302 | -0.095 | 0.123 | 0.123 | 0.033 | 0.24 | 0.05 | 0.15 | 0.06 | 0.33 | 0.29 |
| 11 | 76359722 | rs1941603 | 0.434 | -0.038 | -1.609 | 0.125 | 0.021 | -0.049 | -0.049 | 0 | 0.29 | 0.15 | 0.37 | 0.56 | 0.21 | 0.23 |
| 11 | 76359837 | rs11236827 | 1.985 | -0.276 | -1.713 | -0.015 | 0.005 | 0.025 | 0.025 | 0 | 0.2 | 0.18 | 0.15 | 0.05 | 0.19 | 0.17 |
| 11 | 76359949 | rs10899248 | 1.731 | -0.086 | -1.295 | 0.932 | 0.009 | -0.072 | -0.072 | 0 | 0.18 | 0.02 | 0.29 | 0.3 | 0.16 | 0.15 |
| 11 | 76360404 | rs11236828 | 0.067 | 0.199 | -0.613 | 0.784 | 0.001 | 0.006 | 0.006 | 0 | 0.48 | 0.51 | 0.67 | 0.69 | 0.45 | 0.51 |
| 11 | 76364726 | rs1892952 | -0.069 | 0.16 | -0.764 | 0.577 | 0.002 | 0.071 | 0.071 | 0 | 0.49 | 0.65 | 0.69 | 0.73 | 0.47 | 0.52 |
| 11 | 76365455 | rs11236836 | -0.653 | -0.012 | -0.817 | 0.566 | 0 | 0.002 | 0.002 | 0.138 | 0.55 | 0.46 | 0.46 | 0.54 | 0.54 | 0.54 |
| 11 | 76368553 | rs4945097 | -0.574 | 0.223 | -0.876 | 0.2 | 0.007 | 0.039 | 0.039 | 0 | 0.52 | 0.61 | 0.58 | 0.88 | 0.48 | 0.61 |
| 11 | 76369072 | rs3197153 | 1.285 | 0.035 | -1.661 | 0.257 | -0.062 | -0.052 | -0.052 | 0 | 0.26 | 0.11 | 0.42 | 0.32 | 0.33 | 0.33 |
| 11 | 76369344 | rs1803627 | 1.819 | -0.04 | -1.728 | 0.206 | -0.011 | 0.006 | 0.006 | 0 | 0.1 | 0.01 | 0.07 | 0.08 | 0.1 | 0.18 |

| | | | | | | | | | | | | | | | | |
|---|---|---|---|---|---|---|---|---|---|---|---|---|---|---|---|---|
| 11 | 76369419 | rs3781701 | 0.816 | -0.152 | -1.513 | 0.335 | -0.004 | -0.004 | -0.004 | 0 | 0.31 | 0.28 | 0.43 | 0.35 | 0.35 | 0.33 |
| 11 | 76369709 | rs3781700 | 0.897 | -0.084 | -1.482 | 0.368 | -0.004 | -0.004 | -0.004 | 0 | 0.3 | 0.28 | 0.43 | 0.35 | 0.35 | 0.32 |
| 11 | 76369785 | rs3781699 | 0.888 | -0.092 | -1.486 | 0.379 | -0.004 | -0.004 | -0.004 | 0 | 0.3 | 0.28 | 0.42 | 0.35 | 0.35 | 0.32 |
| 11 | 76369913 | rs11236839 | 1.512 | 0.031 | -1.716 | 0.04 | -0.075 | 0.076 | 0.076 | 0.57 | 0.14 | 0.03 | 0.09 | 0 | 0.22 | 0.09 |
| 11 | 76370004 | rs3824890 | 1.7 | -0.223 | -1.795 | -0.004 | 0.004 | 0.015 | 0.015 | 0 | 0.12 | 0.01 | 0.07 | 0.08 | 0.1 | 0.17 |
| 11 | 76370047 | rs1320646 | -2.17 | -0.214 | -1.836 | -0.159 | -0.04 | 0.065 | 0.065 | 0.461 | 0.75 | 0.92 | 0.73 | 0.88 | 0.74 | 0.74 |
| 11 | 76370088 | rs1320645 | -1.605 | -0.732 | -1.262 | 0.309 | -0.033 | 0.142 | 0.142 | 0 | 0.59 | 0.8 | 0.61 | 0.88 | 0.5 | 0.64 |
| 11 | 76370187 | rs1320644 | -0.234 | 0.311 | -0.854 | 1.078 | -0.01 | -0.006 | -0.006 | 0 | 0.3 | 0.25 | 0.41 | 0.33 | 0.34 | 0.32 |
| 11 | 76372418 | rs3740778 | -1.781 | 0.302 | -0.693 | 1.15 | -0.033 | -0.025 | -0.025 | 0 | 0.95 | 0.99 | 0.79 | 0.8 | 0.86 | 0.9 |
| 11 | 76374039 | rs7944357 | -0.271 | -0.044 | -0.344 | 0.253 | -0.005 | 0.023 | 0.023 | 0 | 0.61 | 0.66 | 0.5 | 0.32 | 0.57 | 0.6 |
| 11 | 76377305 | rs4944115 | 0.584 | 0.085 | -0.289 | 0.29 | 0.009 | -0.016 | -0.016 | 0 | 0.11 | 0.17 | 0.07 | 0.07 | 0.13 | 0.09 |
| 11 | 76377819 | rs6592657 | 0.162 | 0.07 | -0.469 | 0.427 | 0.005 | 0.131 | 0.131 | 0 | 0.46 | 0.24 | 0.41 | 0.25 | 0.43 | 0.49 |
| 11 | 76380357 | rs10899249 | -0.177 | 0.209 | -0.192 | 0.738 | -0.001 | -0.001 | -0.001 | 0.24 | 0.54 | 0.53 | 0.52 | 0.5 | 0.5 | 0.51 |
| 11 | 76383806 | rs947998 | -0.89 | 0.055 | 0.227 | 1.294 | 0.004 | -0.007 | -0.007 | 0 | 0.16 | 0.16 | 0.15 | 0.34 | 0.11 | 0.25 |
| 11 | 76396289 | rs731644 | -1.275 | 0.137 | 0.523 | 0.996 | -0.004 | 0.01 | 0.01 | 0 | 0.37 | 0.41 | 0.43 | 0.6 | 0.34 | 0.51 |
| 11 | 76400134 | rs10899252 | -0.029 | 0.201 | 0.668 | 0.999 | -0.001 | -0.004 | -0.004 | 0 | 0.17 | 0.14 | 0.28 | 0.16 | 0.17 | 0.18 |
| 11 | 76400509 | rs11236862 | 0.046 | 0.161 | 1.082 | 0.402 | 0.021 | -0.012 | -0.012 | 0 | 0.11 | 0.22 | 0.14 | 0.07 | 0.15 | 0.17 |
| 11 | 76404415 | rs4944118 | -0.863 | -0.081 | 1.368 | 0.187 | -0.022 | -0.081 | -0.081 | 0.728 | 0.23 | 0.41 | 0.3 | 0.1 | 0.2 | 0.23 |
| 11 | 76405062 | rs12417222 | 0.819 | 0.045 | 0.832 | 0.258 | 0 | 0.011 | 0.011 | 0 | 0.13 | 0.06 | 0.23 | 0.1 | 0.14 | 0.18 |
| 11 | 76405964 | rs10793182 | -1.132 | -0.027 | 0.836 | 0.345 | 0.005 | 0.004 | 0.004 | 0 | 0.3 | 0.21 | 0.32 | 0.31 | 0.27 | 0.4 |
| 11 | 76412663 | rs7127747 | -1.308 | -0.028 | 1.314 | 0.402 | -0.009 | -0.021 | -0.021 | 0 | 0.41 | 0.52 | 0.37 | 0.31 | 0.34 | 0.47 |
| 11 | 76413757 | rs7101752 | 0.455 | 0.098 | 0.612 | 0.494 | 0 | 0.002 | 0.002 | 0 | 0.15 | 0.12 | 0.21 | 0.1 | 0.16 | 0.19 |
| 11 | 76415209 | rs10899257 | 0.334 | 0.056 | 0.282 | 0.544 | -0.014 | -0.024 | -0.024 | 0.283 | 0.16 | 0.05 | 0.07 | 0.21 | 0.14 | 0.23 |
| 11 | 76420184 | rs10128543 | -1.832 | 0.031 | 0.457 | 0.565 | -0.001 | 0.006 | 0.006 | 0 | 0.51 | 0.55 | 0.39 | 0.55 | 0.5 | 0.69 |
| 11 | 76420526 | rs4619182 | -1.736 | 0.444 | 1.096 | 0.936 | 0.002 | 0.001 | 0.001 | 0.082 | 0.33 | 0.34 | 0.27 | 0.45 | 0.33 | 0.49 |
| 11 | 76420719 | rs4445658 | -1.645 | 0.213 | 1.286 | 0.942 | 0.001 | 0 | 0 | 0 | 0.17 | 0.21 | 0.19 | 0.24 | 0.19 | 0.26 |
| 11 | 76421615 | rs6592663 | -1.765 | 0.633 | 1.064 | 1.142 | 0.032 | 0.004 | 0.004 | 0.23 | 0.19 | 0.36 | 0.23 | 0.24 | 0.2 | 0.29 |
| 11 | 76425895 | rs10899261 | 0.346 | 0.269 | 0.404 | 1.819 | -0.003 | 0.011 | 0.011 | 0 | 0.94 | 0.95 | 0.85 | 0.73 | 0.95 | 0.86 |
| 11 | 76426038 | rs10793185 | -1.634 | 0.076 | 0.858 | 1.615 | -0.018 | 0.008 | 0.008 | 1.622 | 0.12 | 0.34 | 0.14 | 0.12 | 0.08 | 0.07 |
| 11 | 76428247 | rs10793187 | -1.37 | 0.043 | 0.791 | 1.588 | -0.021 | 0.007 | 0.007 | 0.071 | 0.1 | 0.28 | 0.13 | 0.12 | 0.06 | 0.04 |
| 11 | 76428422 | rs11601880 | 1.042 | 0.148 | 0.635 | 1.562 | -0.015 | 0.092 | 0.092 | 0.742 | 0.82 | 0.59 | 0.72 | 0.62 | 0.87 | 0.79 |
| 11 | 76431100 | rs11236903 | -0.515 | 0.321 | -0.056 | 1.242 | 0 | 0.006 | 0.006 | 0 | 0.15 | 0.11 | 0.21 | 0.23 | 0.17 | 0.2 |
| 11 | 76433691 | rs1941600 | -1.167 | 0.1 | 0.298 | 0.871 | -0.007 | -0.014 | -0.014 | 0 | 0.14 | 0.26 | 0.14 | 0.12 | 0.14 | 0.1 |
| 11 | 76434820 | rs1893306 | -0.95 | -0.953 | 0.114 | 0.372 | 0.025 | 0.025 | 0.025 | 0 | 0.38 | 0.71 | 0.25 | 0.22 | 0.27 | 0.21 |
| 11 | 76436972 | rs4944121 | -0.582 | -1.088 | 0.454 | 0.318 | -0.015 | -0.001 | -0.001 | 0 | 0.11 | 0.14 | 0.1 | 0.13 | 0.06 | 0.05 |
| 11 | 76437365 | rs10899269 | -1.003 | -1.314 | 0.948 | 0.199 | -0.002 | -0.002 | -0.002 | 1.317 | 0.32 | 0.35 | 0.2 | 0.22 | 0.26 | 0.19 |
| 11 | 76438258 | rs7118888 | 1.409 | -1.065 | 0.409 | 0.74 | -0.014 | 0.01 | 0.01 | 0 | 0.46 | 0.22 | 0.36 | 0.37 | 0.5 | 0.58 |
| 11 | 76442114 | rs4393347 | -0.718 | -1.05 | 0.629 | 0.766 | -0.014 | -0.023 | -0.023 | 0 | 0.83 | 0.85 | 0.59 | 0.59 | 0.77 | 0.78 |
| 11 | 76444267 | rs7946499 | 0.695 | -0.97 | 0.901 | 0.728 | -0.025 | 0.048 | 0.048 | 0 | 0.68 | 0.52 | 0.5 | 0.46 | 0.73 | 0.7 |
| 11 | 76450908 | rs12362105 | 0.362 | -0.9 | 1.464 | 0.917 | -0.001 | -0.007 | -0.007 | 0 | 0.21 | 0.21 | 0.1 | 0.09 | 0.2 | 0.12 |
| 11 | 76451038 | rs7924748 | -0.142 | -1.35 | 0.982 | 0.709 | -0.017 | 0.04 | 0.04 | 0 | 0.19 | 0.26 | 0.12 | 0.25 | 0.13 | 0.33 |
| 11 | 76455618 | rs873108 | -0.761 | -1.327 | 1.004 | 0.728 | -0.038 | 0.038 | 0.038 | 0 | 0.26 | 0.33 | 0.18 | 0.36 | 0.17 | 0.37 |
| 11 | 76462540 | rs10793193 | -1.087 | -1.135 | 0.629 | 1.062 | 0.021 | 0.008 | 0.008 | 0 | 0.46 | 0.63 | 0.32 | 0.37 | 0.39 | 0.46 |
| 11 | 76465362 | rs7942368 | -0.855 | -1.095 | 0.654 | 1.101 | -0.01 | -0.003 | -0.003 | 0 | 0.33 | 0.64 | 0.23 | 0.28 | 0.25 | 0.21 |
| 11 | 76468783 | rs1149595 | -1.646 | -0.823 | 0.599 | 1.582 | 0.01 | 0.075 | 0.075 | 0 | 0.46 | 0.81 | 0.37 | 0.65 | 0.42 | 0.51 |
| 11 | 76471629 | rs1149598 | 0.762 | -0.783 | 0.51 | 1.71 | -0.026 | -0.03 | -0.03 | 0 | 0.08 | 0 | 0.05 | 0.07 | 0.07 | 0.25 |
| 11 | 76473138 | rs1465900 | -0.546 | -0.813 | 0.735 | 1.531 | -0.053 | -0.046 | -0.046 | 0.083 | 0.31 | 0.47 | 0.19 | 0.21 | 0.25 | 0.17 |
| 11 | 76479810 | rs1149602 | -1.301 | -0.98 | 0.771 | 0.884 | 0.006 | 0.012 | 0.012 | 0 | 0.43 | 0.62 | 0.33 | 0.65 | 0.4 | 0.49 |
| 11 | 76480255 | rs12290350 | -0.449 | -0.991 | 0.821 | 0.901 | -0.043 | -0.072 | -0.072 | 0 | 0.31 | 0.42 | 0.19 | 0.15 | 0.24 | 0.15 |
| 11 | 76480570 | rs1149603 | -1.466 | -0.979 | 0.633 | 0.81 | 0.021 | 0.04 | 0.04 | 0 | 0.44 | 0.73 | 0.34 | 0.64 | 0.4 | 0.47 |
| 11 | 76484241 | rs881930 | -0.08 | -0.938 | 0.454 | 0.872 | -0.006 | 0.014 | 0.014 | 0 | 0.12 | 0.11 | 0.13 | 0.38 | 0.15 | 0.28 |
| 11 | 76490058 | rs1149606 | 0.285 | -1.046 | 0.316 | 0.714 | -0.059 | -0.094 | -0.094 | 0 | 0.14 | 0.02 | 0.11 | 0.41 | 0.15 | 0.28 |
| 11 | 76495560 | rs3758711 | -0.568 | -1.122 | 0.423 | 0.604 | -0.057 | -0.065 | -0.065 | 1.724 | 0.32 | 0.5 | 0.19 | 0.18 | 0.25 | 0.19 |
| 11 | 76495879 | rs1149611 | -1.83 | -1.128 | -0.013 | 0.616 | 0.01 | 0.09 | 0.09 | 0 | 0.52 | 0.84 | 0.38 | 0.67 | 0.43 | 0.52 |
| 11 | 76498369 | rs1149616 | 0.17 | -1.035 | 0.035 | 0.669 | -0.009 | -0.007 | -0.007 | 0.401 | 0.15 | 0.15 | 0.14 | 0.47 | 0.15 | 0.3 |
| 11 | 76505202 | rs10899283 | -0.94 | -1.039 | 0.278 | 0.39 | -0.053 | -0.059 | -0.059 | 0 | 0.33 | 0.5 | 0.2 | 0.19 | 0.25 | 0.21 |
| 11 | 76508843 | rs7912 | -0.039 | -0.208 | -0.362 | 0.154 | 0.006 | 0.114 | 0.114 | 0 | 0.18 | 0.01 | 0.06 | 0 | 0.13 | 0.05 |
| 11 | 76515259 | rs1149622 | -0.994 | 0.111 | -0.06 | 1.139 | 0.01 | 0.076 | 0.076 | 0.016 | 0.28 | 0.58 | 0.23 | 0.48 | 0.25 | 0.29 |
| 11 | 76518495 | rs1149623 | -0.551 | -0.386 | -0.264 | 0.426 | -0.019 | -0.033 | -0.033 | 0.045 | 0.37 | 0.46 | 0.38 | 0.78 | 0.31 | 0.32 |
| 11 | 76519899 | rs1149625 | 0.028 | -1.01 | -0.745 | 0.457 | -0.001 | 0 | 0 | 0.512 | 0.34 | 0.34 | 0.32 | 0.3 | 0.3 | 0.31 |
| 11 | 76545414 | rs1149577 | 0.259 | -1.518 | -0.62 | 0.58 | 0.016 | -0.035 | -0.035 | 0 | 0.3 | 0.2 | 0.26 | 0.41 | 0.24 | 0.26 |
| 11 | 76549212 | rs1561647 | 0.751 | -2.016 | -0.351 | 0.33 | -0.003 | 0.027 | 0.027 | 0 | 0.16 | 0.14 | 0.16 | 0.38 | 0.15 | 0.12 |
| 11 | 76549513 | rs1149580 | -0.851 | -2.216 | -0.054 | 0.223 | 0.06 | 0.165 | 0.165 | 2.663 | 0.52 | 0.75 | 0.49 | 0.84 | 0.41 | 0.44 |
| 11 | 76551635 | rs1225132 | 0.134 | -2.193 | 0.428 | 0.254 | -0.03 | 0.025 | 0.025 | 1.609 | 0.35 | 0.46 | 0.31 | 0.47 | 0.27 | 0.33 |
| 11 | 76553487 | rs1225138 | -1.356 | -2.197 | 0.054 | 0.094 | -0.029 | 0.093 | 0.093 | 1.41 | 0.63 | 0.79 | 0.53 | 0.84 | 0.5 | 0.5 |
| 11 | 76563014 | rs1372045 | -0.372 | -2.083 | 0.031 | 0.279 | -0.008 | 0.007 | 0.007 | 1.39 | 0.64 | 0.61 | 0.54 | 0.52 | 0.71 | 0.67 |
| 11 | 76573016 | rs2279518 | 0.918 | -2.373 | -0.39 | 0.178 | -0.095 | 0.041 | 0.041 | 0.008 | 0.38 | 0.06 | 0.3 | 0.24 | 0.51 | 0.48 |
| 11 | 76573235 | rs7119361 | -0.632 | -2.527 | -0.101 | 0.138 | 0.014 | -0.025 | -0.025 | 0.011 | 0.69 | 0.8 | 0.52 | 0.58 | 0.72 | 0.65 |
| 11 | 76585161 | rs7117682 | -0.119 | -2.625 | 0.255 | 0.095 | 0.031 | -0.004 | -0.004 | 0 | 0.64 | 0.28 | 0.48 | 0.59 | 0.72 | 0.65 |

| | | | | | | | | | | | | | | | | |
|---|---|---|---|---|---|---|---|---|---|---|---|---|---|---|---|---|
| 11 | 76616528 | rs4471450 | -0.631 | -2.623 | 0.33 | 0.096 | 0.004 | -0.005 | -0.005 | 0 | 0.69 | 0.73 | 0.51 | 0.59 | 0.72 | 0.65 |
| 11 | 76631329 | rs1517906 | -0.599 | -2.623 | 0.322 | 0.096 | -0.007 | 0.007 | 0.007 | 0 | 0.69 | 0.64 | 0.51 | 0.59 | 0.72 | 0.65 |
| 11 | 76660780 | rs7118970 | -0.923 | -2.624 | 0.334 | 0.079 | 0.037 | 0.033 | 0.033 | 4.849 | 0.06 | 0.52 | 0.04 | 0 | 0 | 0 |
| 11 | 76666181 | rs12797610 | 1.068 | -2.695 | 0.242 | 0.031 | 0.009 | 0.011 | 0.011 | 0 | 0.09 | 0.03 | 0.03 | 0 | 0.08 | 0.05 |
| 11 | 76673980 | rs10793220 | -0.094 | -2.624 | 0.26 | 0.078 | 0.047 | -0.011 | -0.011 | 0 | 0.63 | 0.2 | 0.48 | 0.59 | 0.71 | 0.65 |
| 11 | 76697558 | rs6592691 | -0.651 | -2.479 | 0.331 | 0.177 | 0.003 | -0.005 | -0.005 | 0 | 0.71 | 0.73 | 0.55 | 0.63 | 0.74 | 0.7 |
| 11 | 76701365 | rs3824889 | -0.617 | -2.478 | 0.235 | 0.177 | -0.004 | 0.005 | 0.005 | 0 | 0.71 | 0.66 | 0.55 | 0.63 | 0.74 | 0.7 |
| 11 | 76701606 | rs3740767 | -0.052 | -2.492 | 0.206 | 0.171 | 0.037 | -0.001 | -0.001 | 0 | 0.65 | 0.19 | 0.51 | 0.63 | 0.74 | 0.7 |
| 11 | 76732308 | rs7925774 | -0.604 | -2.476 | 0.231 | 0.178 | -0.004 | 0.005 | 0.005 | 0 | 0.71 | 0.66 | 0.55 | 0.63 | 0.74 | 0.7 |
| 11 | 76732949 | rs7123496 | -0.674 | -2.477 | 0.241 | 0.178 | 0.005 | -0.022 | -0.022 | 0 | 0.72 | 0.83 | 0.56 | 0.63 | 0.74 | 0.7 |
| 11 | 76733024 | rs10793231 | -0.067 | -2.479 | 0.074 | 0.177 | 0.036 | 0 | 0 | 0 | 0.65 | 0.2 | 0.51 | 0.63 | 0.74 | 0.7 |
| 11 | 76733313 | rs17135325 | 0.988 | -2.33 | -0.246 | 0.345 | 0.01 | -0.083 | -0.083 | 0 | 0.14 | 0.03 | 0.12 | 0.36 | 0.13 | 0.12 |
| 11 | 76733847 | rs2826 | -0.322 | -1.916 | 0.059 | 0.234 | -0.004 | 0.005 | 0.005 | 0 | 0.71 | 0.67 | 0.55 | 0.63 | 0.74 | 0.7 |
| 11 | 76740471 | rs7123694 | -0.452 | -1.885 | 0.002 | 0.25 | -0.002 | 0.005 | 0.005 | 0 | 0.72 | 0.71 | 0.59 | 0.63 | 0.75 | 0.71 |
| 11 | 76744522 | rs11827049 | -0.496 | -1.753 | 0.025 | -0.068 | -0.044 | -0.088 | -0.088 | 0 | 0.14 | 0.33 | 0.07 | 0 | 0.08 | 0.05 |
| 11 | 76747390 | rs7103667 | 0.371 | -0.657 | -1.193 | 0.003 | -0.055 | -0.014 | -0.014 | 0 | 0.28 | 0.14 | 0.29 | 0.31 | 0.34 | 0.47 |
| 11 | 76747519 | rs10899345 | -0.158 | -0.692 | -1.688 | 0.152 | 0.058 | 0.001 | 0.001 | 1.174 | 0.45 | 0.74 | 0.31 | 0.39 | 0.36 | 0.22 |
| 11 | 76748182 | rs1894008 | -1.022 | -0.63 | -2.066 | 0.147 | -0.102 | -0.094 | -0.094 | 0 | 0.75 | 0.98 | 0.61 | 0.68 | 0.7 | 0.69 |
| 11 | 76750406 | rs12422079 | -0.035 | -0.89 | -1.779 | 0.185 | 0.037 | 0.021 | 0.021 | 0.284 | 0.38 | 0.35 | 0.24 | 0.22 | 0.29 | 0.15 |
| 11 | 76752248 | rs7950506 | -0.629 | -0.112 | -1.755 | 0.109 | -0.048 | -0.048 | -0.048 | 0 | 0.75 | 0.93 | 0.61 | 0.62 | 0.71 | 0.69 |
| 11 | 76755178 | rs12802512 | -0.315 | 0.218 | -1.946 | -0.121 | 0.013 | 0.036 | 0.036 | 0 | 0.18 | 0.12 | 0.1 | 0 | 0.16 | 0.07 |
| 11 | 76775472 | rs1944032 | -0.07 | 1.127 | -1.201 | 1.004 | -0.062 | -0.063 | -0.063 | 0 | 0.74 | 0.94 | 0.61 | 0.63 | 0.7 | 0.69 |
| 11 | 76792431 | rs7926553 | 0.367 | 0.927 | -1.17 | 0.885 | -0.039 | -0.046 | -0.046 | 0.398 | 0.59 | 0.29 | 0.46 | 0.44 | 0.6 | 0.55 |
| 11 | 76796159 | rs2276029 | -1.033 | 0.903 | -0.527 | 0.891 | 0.002 | -0.005 | -0.005 | 0.008 | 0.26 | 0.34 | 0.22 | 0.3 | 0.28 | 0.28 |
| 11 | 76797361 | rs1944027 | 0.993 | 0.371 | -0.082 | 1.319 | -0.022 | 0.08 | 0.08 | 0 | 0.78 | 0.66 | 0.75 | 0.57 | 0.84 | 0.75 |
| 11 | 76799684 | rs6592703 | 0.581 | 0.326 | -0.104 | 1.268 | -0.01 | 0.005 | 0.005 | 0 | 0.92 | 0.93 | 0.86 | 0.86 | 0.94 | 0.85 |
| 11 | 76801982 | rs948977 | -1.423 | 1.121 | -0.337 | 1.163 | 0 | 0 | 0 | 1.905 | 0.25 | 0.27 | 0.35 | 0.35 | 0.31 | 0.15 |
| 11 | 76804131 | rs6592705 | 0.414 | 0.9 | -0.659 | 1.091 | 0.001 | -0.002 | -0.002 | 0.146 | 0.67 | 0.73 | 0.68 | 0.63 | 0.71 | 0.7 |
| 11 | 76805777 | rs4945141 | 0.409 | 1.003 | -0.34 | 1.029 | -0.012 | 0.015 | 0.015 | 0 | 0.6 | 0.79 | 0.51 | 0.63 | 0.56 | 0.73 |
| 11 | 76806522 | rs4945142 | 1.024 | 0.599 | -0.072 | 0.418 | 0.001 | 0.006 | 0.006 | 0 | 0.32 | 0.35 | 0.28 | 0.5 | 0.28 | 0.54 |
| 11 | 76812803 | rs12421551 | 0.25 | 0.388 | 0.176 | 0.303 | -0.015 | -0.081 | -0.081 | 0 | 0.26 | 0.37 | 0.22 | 0.13 | 0.26 | 0.2 |
| 11 | 76814110 | rs2233549 | 2.02 | 0.237 | 0.135 | 0.375 | 0.005 | 0.012 | 0.012 | 0.192 | 0.18 | 0.2 | 0.21 | 0.23 | 0.21 | 0.36 |
| 11 | 76815398 | rs11237090 | 1.511 | 0.181 | 0.303 | 0.286 | 0.019 | 0.031 | 0.031 | 0 | 0.21 | 0.32 | 0.23 | 0.23 | 0.23 | 0.38 |
| 11 | 76823484 | rs3781684 | 0.813 | 0.143 | 0.21 | 0.287 | 0.011 | -0.018 | -0.018 | 0 | 0.11 | 0.11 | 0.08 | 0.27 | 0.06 | 0.16 |
| 11 | 76824078 | rs3781683 | 1.245 | 0.181 | 0.204 | 0.213 | 0.022 | 0.05 | 0.05 | 0 | 0.2 | 0.1 | 0.17 | 0.1 | 0.2 | 0.12 |
| 11 | 76827992 | rs4945144 | -0.584 | 0.071 | 0.441 | 0.313 | -0.017 | 0.071 | 0.071 | 0 | 0.17 | 0.36 | 0.12 | 0.28 | 0.1 | 0.22 |
| 11 | 76831622 | rs4517554 | -0.531 | 0.065 | 0.024 | 0.374 | -0.045 | 0.02 | 0.02 | 0 | 0.55 | 0.85 | 0.5 | 0.59 | 0.51 | 0.69 |
| 11 | 76832011 | rs2276031 | 0.306 | 0.339 | 0.34 | 0.801 | -0.024 | 0.043 | 0.043 | 0 | 0.12 | 0.29 | 0.09 | 0.23 | 0.06 | 0.13 |
| 11 | 76833338 | rs4945146 | 0.062 | 0.409 | 0.246 | 0.727 | 0.003 | 0.055 | 0.055 | 0.221 | 0.31 | 0.54 | 0.31 | 0.47 | 0.3 | 0.51 |
| 11 | 76836350 | rs1043418 | -0.274 | 0.419 | 0.108 | 0.611 | 0.007 | 0.009 | 0.009 | 0 | 0.36 | 0.52 | 0.41 | 0.57 | 0.4 | 0.61 |
| 11 | 76836439 | rs1043420 | 1.177 | 0.535 | 0.163 | 0.66 | 0.031 | 0.041 | 0.041 | 0 | 0.2 | 0.27 | 0.23 | 0.24 | 0.25 | 0.38 |
| 11 | 76837499 | rs7943716 | -0.574 | 0.211 | -0.659 | 0.303 | -0.001 | 0.001 | 0.001 | 0 | 0.39 | 0.56 | 0.44 | 0.41 | 0.37 | 0.33 |
| 11 | 76837552 | rs10899353 | -0.704 | 0.051 | -0.375 | 0.158 | 0.067 | 0.046 | 0.046 | 0.022 | 0.05 | 0.21 | 0.22 | 0.09 | 0.08 | 0.05 |
| 11 | 76839129 | rs11606360 | 0.764 | -0.432 | -1.268 | -0.118 | -0.004 | 0.042 | 0.042 | 0 | 0.09 | 0.01 | 0.03 | 0 | 0.05 | 0.01 |
| 11 | 76840709 | rs7121485 | -0.244 | -0.794 | -1.865 | -0.798 | 0.003 | 0.224 | 0.224 | 0.345 | 0.37 | 0.11 | 0.38 | 0.05 | 0.38 | 0.14 |
| 11 | 76840798 | rs7121629 | -0.09 | -1.444 | -0.867 | 0.153 | -0.031 | 0.072 | 0.072 | 0 | 0.16 | 0.45 | 0.1 | 0.22 | 0.07 | 0.12 |
| 11 | 76848035 | rs948969 | -0.539 | -1.5 | -1.369 | 0.573 | 0.024 | -0.006 | -0.006 | 0 | 0.32 | 0.73 | 0.34 | 0.64 | 0.25 | 0.44 |
| 11 | 76852039 | rs4945149 | -0.263 | -1.808 | -1.141 | 0.503 | 0.008 | -0.011 | -0.011 | 0 | 0.37 | 0.34 | 0.29 | 0.45 | 0.29 | 0.36 |
| 11 | 76852952 | rs7131022 | -1.149 | -1.725 | -1.179 | 0.291 | 0.057 | 0.09 | 0.09 | 0 | 0.44 | 0.76 | 0.37 | 0.7 | 0.33 | 0.45 |
| 11 | 76858756 | rs12279716 | 0.326 | -1.568 | -0.968 | 0.777 | -0.018 | 0.046 | 0.046 | 0 | 0.2 | 0.29 | 0.18 | 0.31 | 0.15 | 0.31 |
| 11 | 76863872 | rs7105374 | -0.728 | -1.267 | -1.292 | 0.308 | 0.074 | 0.109 | 0.109 | 0 | 0.51 | 0.78 | 0.41 | 0.65 | 0.36 | 0.5 |
| 11 | 76865705 | rs4944143 | -0.836 | -0.433 | -1.005 | 0.651 | -0.008 | -0.02 | -0.02 | 0 | 0.37 | 0.75 | 0.57 | 0.55 | 0.45 | 0.35 |
| 11 | 76868372 | rs762667 | -0.895 | -0.3 | -0.423 | 0.654 | -0.031 | -0.028 | -0.028 | 1.517 | 0.33 | 0.84 | 0.5 | 0.34 | 0.38 | 0.21 |
| 11 | 76872169 | rs2276278 | 0.125 | -0.127 | -0.186 | 0.836 | -0.001 | 0.095 | 0.095 | 0 | 0.08 | 0.19 | 0.13 | 0.28 | 0.06 | 0.09 |
| 11 | 76873620 | rs3740763 | -1.49 | -0.614 | -0.649 | 0.694 | -0.097 | 0.002 | 0.002 | 0 | 0.62 | 0.96 | 0.74 | 0.67 | 0.6 | 0.6 |
| 11 | 76874648 | rs4945153 | -1.094 | 0.477 | 0.572 | 0.86 | 0.037 | 0.025 | 0.025 | 0 | 0.29 | 0.49 | 0.46 | 0.33 | 0.37 | 0.21 |
| 11 | 76876278 | rs2156568 | -0.854 | 0.497 | 0.442 | 0.823 | 0.021 | 0.02 | 0.02 | 0 | 0.26 | 0.34 | 0.44 | 0.33 | 0.36 | 0.21 |
| 11 | 76881925 | rs7930391 | -0.51 | 0.47 | 0.337 | 0.996 | 0.023 | 0.219 | 0.219 | 0 | 0.11 | 0.37 | 0.15 | 0.28 | 0.05 | 0.1 |
| 11 | 76881988 | rs11607721 | -0.016 | 0.616 | -0.258 | 0.299 | -0.041 | 0.059 | 0.059 | 0.003 | 0.19 | 0.08 | 0.3 | 0.08 | 0.3 | 0.1 |
| 11 | 76888207 | rs2077198 | -0.714 | 0.838 | 0.094 | 0.521 | -0.002 | -0.005 | -0.005 | 0 | 0.41 | 0.64 | 0.63 | 0.43 | 0.56 | 0.34 |
| 11 | 76894463 | rs3758708 | 0.038 | 1.044 | 0.241 | 0.577 | 0.004 | 0.006 | 0.006 | 0 | 0.08 | 0.12 | 0.15 | 0.12 | 0.07 | 0.07 |
| 11 | 76898728 | rs12805873 | -0.098 | 0.948 | 0.661 | 0.513 | 0.052 | 0.013 | 0.013 | 0 | 0.31 | 0.42 | 0.46 | 0.32 | 0.49 | 0.28 |
| 11 | 76898797 | rs12805353 | -0.55 | 0.872 | 0.386 | 0.475 | -0.067 | -0.003 | -0.003 | 0 | 0.37 | 0.8 | 0.51 | 0.32 | 0.5 | 0.28 |
| 11 | 76899265 | rs3781694 | -0.926 | 0.701 | 0.433 | 0.482 | -0.004 | -0.028 | -0.028 | 0.037 | 0.44 | 0.83 | 0.65 | 0.44 | 0.56 | 0.34 |
| 11 | 76901927 | rs2276285 | -0.556 | 0.705 | 0.401 | 0.493 | 0.007 | 0.01 | 0.01 | 0 | 0.39 | 0.45 | 0.6 | 0.45 | 0.54 | 0.34 |
| 11 | 76905604 | rs1109977 | -0.709 | 0.777 | 0.259 | 0.477 | 0.008 | 0.003 | 0.003 | 0 | 0.45 | 0.71 | 0.64 | 0.45 | 0.56 | 0.34 |
| 11 | 76917220 | rs2276293 | -0.803 | 0.738 | -0.439 | 0.535 | -0.025 | -0.016 | -0.016 | 0.238 | 0.47 | 0.77 | 0.6 | 0.47 | 0.51 | 0.37 |
| 11 | 76919478 | rs948962 | -0.458 | 0.832 | -0.403 | 0.417 | -0.009 | 0.041 | 0.041 | 0.352 | 0.52 | 0.76 | 0.55 | 0.33 | 0.5 | 0.28 |
| 11 | 76920003 | rs11237122 | -0.709 | 0.862 | 0.329 | 0.41 | -0.002 | 0.006 | 0.006 | 0 | 0.58 | 0.5 | 0.67 | 0.47 | 0.58 | 0.39 |

| | | | | | | | | | | | | | | | | |
|---|---|---|---|---|---|---|---|---|---|---|---|---|---|---|---|---|
| 11 | 76920038 | rs885442 | -0.461 | 0.756 | 0.129 | 0.358 | 0.004 | -0.054 | -0.054 | 0 | 0.53 | 0.15 | 0.63 | 0.47 | 0.58 | 0.39 |
| 11 | 76920715 | rs7123925 | -0.61 | 0.656 | 0.459 | 0.555 | -0.001 | -0.002 | -0.002 | 0 | 0.57 | 0.47 | 0.63 | 0.48 | 0.56 | 0.4 |
| 11 | 76920728 | rs12274072 | 0.308 | 0.691 | 0.482 | 0.973 | -0.001 | 0.001 | 0.001 | 0 | 0.43 | 0.53 | 0.29 | 0.37 | 0.43 | 0.59 |
| 11 | 76921090 | rs12793189 | -0.723 | 0.746 | 0.571 | 0.845 | 0.005 | 0.004 | 0.004 | 0 | 0.58 | 0.51 | 0.63 | 0.5 | 0.52 | 0.4 |
| 11 | 76921358 | rs12793619 | 0.084 | 0.403 | 0.616 | 0.29 | 0.014 | -0.13 | -0.13 | 0.345 | 0.27 | 0.52 | 0.34 | 0.07 | 0.31 | 0.12 |
| 11 | 76924874 | rs883223 | -0.282 | 0.04 | 0.659 | 0.63 | 0.017 | -0.021 | -0.021 | 0 | 0.37 | 0.3 | 0.5 | 0.44 | 0.31 | 0.37 |
| 11 | 76928025 | rs12363944 | 0.95 | 0.291 | 0.285 | 0.361 | -0.115 | 0.11 | 0.11 | 0 | 0.23 | 0.02 | 0.16 | 0.07 | 0.29 | 0.07 |
| 11 | 76928149 | rs2282576 | 0.27 | 0.177 | 0.602 | 0.577 | 0.091 | 0.332 | 0.332 | 0 | 0.94 | 0.57 | 0.87 | 0.78 | 0.96 | 0.88 |
| 11 | 76928929 | rs10899363 | 1.054 | 0.157 | 0.319 | 0.496 | -0.088 | 0.022 | 0.022 | 0 | 0.29 | 0.03 | 0.19 | 0.26 | 0.36 | 0.45 |
| 11 | 76935002 | rs12294694 | 0.967 | 0.185 | 0.373 | 0.46 | -0.033 | 0.038 | 0.038 | 0 | 0.31 | 0.11 | 0.21 | 0.26 | 0.38 | 0.45 |
| 11 | 76938121 | rs3740769 | -0.033 | 0.09 | 0.57 | 0.617 | 0.011 | 0.095 | 0.095 | 0.782 | 0.88 | 0.83 | 0.77 | 0.63 | 0.87 | 0.79 |
| 11 | 76953939 | rs5028355 | 0.603 | 0.044 | 0.799 | 0.597 | 0.012 | 0.154 | 0.154 | 0.09 | 0.84 | 0.49 | 0.77 | 0.64 | 0.87 | 0.79 |
| 11 | 76954833 | rs11237146 | 1.207 | 0.053 | 0.614 | 0.512 | -0.087 | 0.002 | 0.002 | 0 | 0.29 | 0.03 | 0.19 | 0.26 | 0.36 | 0.44 |
| 11 | 76954898 | rs7104577 | 0.096 | 0.008 | 0.658 | 0.511 | -0.009 | 0.008 | 0.008 | 0 | 0.52 | 0.35 | 0.56 | 0.38 | 0.49 | 0.33 |
| 11 | 76959598 | rs7125798 | -0.169 | 0.017 | 0.679 | 0.522 | 0.009 | 0.085 | 0.085 | 0.831 | 0.88 | 0.84 | 0.76 | 0.63 | 0.87 | 0.79 |
| 11 | 76972810 | rs1894214 | -0.877 | 0.036 | 0.863 | 0.453 | -0.003 | 0.023 | 0.023 | 0 | 0.66 | 0.73 | 0.78 | 0.75 | 0.62 | 0.54 |
| 11 | 76979550 | rs2729772 | -0.876 | 0.027 | 0.898 | 0.524 | 0.084 | 0.369 | 0.369 | 0 | 0.07 | 0.46 | 0.12 | 0.25 | 0.02 | 0.08 |
| 11 | 76992664 | rs1793476 | -0.817 | 0.038 | 0.823 | 0.519 | -0.001 | -0.001 | -0.001 | 0 | 0.65 | 0.7 | 0.69 | 0.5 | 0.61 | 0.46 |
| 11 | 76996593 | rs1793466 | -1.222 | -0.045 | 0.363 | 0.437 | -0.078 | 0.021 | 0.021 | 0 | 0.69 | 0.97 | 0.8 | 0.74 | 0.63 | 0.55 |
| 11 | 77012304 | rs6592714 | 0.129 | -0.544 | 0.309 | 0.502 | 0.015 | -0.019 | -0.019 | 0 | 0.35 | 0.3 | 0.48 | 0.43 | 0.28 | 0.38 |
| 11 | 77012620 | rs6592715 | 0.01 | -0.402 | 0.394 | 0.605 | 0.019 | -0.02 | -0.02 | 0 | 0.38 | 0.3 | 0.5 | 0.43 | 0.31 | 0.38 |
| 11 | 77027574 | rs2725815 | 0.873 | -0.223 | 0.109 | 0.618 | -0.084 | -0.011 | -0.011 | 0 | 0.3 | 0.02 | 0.19 | 0.26 | 0.36 | 0.47 |
| 11 | 77050714 | rs1377470 | 0.608 | -0.294 | 0.738 | 0.689 | -0.002 | 0.005 | 0.005 | 0 | 0.32 | 0.3 | 0.2 | 0.25 | 0.36 | 0.47 |
| 11 | 77055786 | rs2725829 | -1.871 | -0.364 | 0.762 | 0.673 | 0.029 | 0.346 | 0.346 | 0.073 | 0.08 | 0.42 | 0.13 | 0.26 | 0.02 | 0.07 |
| 11 | 77069746 | rs2250647 | 0.661 | -0.298 | 0.634 | 0.598 | -0.008 | 0.023 | 0.023 | 0 | 0.32 | 0.17 | 0.21 | 0.26 | 0.36 | 0.47 |
| 11 | 77091574 | rs2729761 | -1.029 | -0.609 | 0.477 | 0.641 | -0.008 | 0.034 | 0.034 | 0.064 | 0.19 | 0.28 | 0.29 | 0.43 | 0.15 | 0.21 |
| 11 | 77099018 | rs508662 | 0.76 | -0.037 | 0.337 | 0.338 | -0.052 | 0.065 | 0.065 | 0 | 0.24 | 0.13 | 0.19 | 0.06 | 0.31 | 0.08 |
| 11 | 77106868 | rs2844333 | -0.01 | -0.103 | 0.426 | 0.417 | -0.005 | 0.022 | 0.022 | 0 | 0.38 | 0.45 | 0.31 | 0.5 | 0.38 | 0.54 |
| 11 | 77141718 | rs2725830 | 0.308 | -0.085 | 0.431 | 0.368 | -0.001 | 0.027 | 0.027 | 0 | 0.35 | 0.36 | 0.3 | 0.51 | 0.36 | 0.53 |
| 11 | 77178698 | rs2154754 | -0.12 | 0.014 | 0.309 | 0.298 | 0.017 | -0.002 | -0.002 | 1.021 | 0.29 | 0.56 | 0.3 | 0.31 | 0.32 | 0.13 |
| 11 | 77186002 | rs11237200 | -0.801 | 0.01 | 0.217 | 0.503 | 0.023 | 0.275 | 0.275 | 0 | 0.07 | 0.29 | 0.11 | 0.25 | 0.02 | 0.07 |
| 11 | 77194601 | rs10899380 | 0.586 | 0.025 | -0.717 | 0.375 | -0.019 | 0.025 | 0.025 | 0 | 0.31 | 0.15 | 0.21 | 0.26 | 0.36 | 0.48 |
| 11 | 77195100 | rs3133293 | 0.442 | 0.058 | -1.175 | 0.34 | 0.046 | -0.064 | -0.064 | 0 | 0.36 | 0.13 | 0.49 | 0.4 | 0.3 | 0.37 |
| 11 | 77195482 | rs6592723 | -0.105 | -0.127 | -1.026 | 0.246 | 0.015 | -0.045 | -0.045 | 0 | 0.42 | 0.75 | 0.33 | 0.52 | 0.39 | 0.55 |
| 11 | 77197185 | rs3019251 | 0.402 | -0.075 | -1.066 | 0.356 | 0.046 | -0.026 | -0.026 | 0 | 0.44 | 0.2 | 0.53 | 0.48 | 0.41 | 0.37 |
| 11 | 77204866 | rs3019238 | -0.008 | 0.346 | -1.483 | -0.108 | 0.05 | 0.122 | 0.122 | 0.195 | 0.54 | 0.91 | 0.67 | 0.68 | 0.44 | 0.59 |
| 11 | 77215675 | rs3019245 | -0.264 | 1.077 | -0.731 | -0.247 | 0.005 | -0.029 | -0.029 | 0.43 | 0.74 | 0.72 | 0.79 | 0.94 | 0.66 | 0.9 |
| 11 | 77219219 | rs3019244 | 0.038 | 0.069 | -0.954 | -0.296 | 0.029 | 0.019 | 0.019 | 0 | 0.37 | 0.26 | 0.24 | 0.27 | 0.19 | 0.5 |
| 11 | 77224972 | rs6592724 | 0.662 | 0.176 | -1.14 | 0.323 | 0.004 | -0.005 | -0.005 | 2.825 | 0.31 | 0.28 | 0.37 | 0.34 | 0.33 | 0.25 |
| 11 | 77226842 | rs7109180 | 0.374 | 0.649 | -1.042 | 0.316 | -0.041 | -0.032 | -0.032 | 0.185 | 0.25 | 0.13 | 0.28 | 0.39 | 0.39 | 0.17 |
| 11 | 77230595 | rs11237211 | 0.317 | 0.449 | -0.929 | 0.405 | 0.007 | -0.009 | -0.009 | 0 | 0.3 | 0.24 | 0.37 | 0.34 | 0.33 | 0.24 |
| 11 | 77239334 | rs11605638 | 0.202 | 0.703 | -1.348 | 0.045 | -0.054 | 0.045 | 0.045 | 0.398 | 0.19 | 0.03 | 0.16 | 0.13 | 0.23 | 0.17 |
| 11 | 77241183 | rs7951578 | -0.166 | 0.426 | -1.256 | -0.175 | -0.004 | 0.004 | 0.004 | 0.011 | 0.42 | 0.59 | 0.51 | 0.62 | 0.49 | 0.33 |
| 11 | 77250912 | rs7119934 | 0.577 | 0.545 | -0.962 | 0.337 | -0.002 | 0.03 | 0.03 | 0 | 0.09 | 0.14 | 0.19 | 0.18 | 0.1 | 0.14 |
| 11 | 77259435 | rs1540181 | -1.398 | 0.097 | -1.11 | 0.128 | 0.052 | 0.077 | 0.077 | 0 | 0.64 | 0.89 | 0.7 | 0.75 | 0.74 | 0.55 |
| 11 | 77267497 | rs1945752 | 0.717 | 0.37 | -0.393 | 0.212 | 0.02 | 0.051 | 0.051 | 0 | 0.24 | 0.29 | 0.26 | 0.4 | 0.32 | 0.11 |
| 11 | 77268856 | rs7943742 | -0.438 | 0.335 | -0.355 | 0.16 | 0 | -0.018 | -0.018 | 0 | 0.35 | 0.56 | 0.39 | 0.3 | 0.35 | 0.36 |
| 11 | 77300048 | rs7129556 | 0.805 | 0.286 | -0.055 | 0.116 | -0.005 | 0.01 | 0.01 | 0 | 0.23 | 0.18 | 0.18 | 0.11 | 0.25 | 0.23 |
| 11 | 77300545 | rs7115612 | -1.643 | 0.276 | -0.092 | 0.113 | 0.058 | 0.09 | 0.09 | 0 | 0.62 | 0.88 | 0.7 | 0.75 | 0.73 | 0.54 |
| 11 | 77301341 | rs2276415 | 0.914 | 0.296 | -0.05 | 0.101 | -0.002 | 0.031 | 0.031 | 0 | 0.09 | 0.14 | 0.19 | 0.18 | 0.1 | 0.14 |
| 11 | 77309533 | rs10793257 | -0.431 | 0.255 | -0.148 | 0.082 | -0.001 | -0.017 | -0.017 | 0 | 0.35 | 0.57 | 0.4 | 0.3 | 0.35 | 0.37 |
| 11 | 77311445 | rs11603537 | 1.255 | 0.609 | -0.24 | 0.458 | -0.085 | 0.025 | 0.025 | 0 | 0.19 | 0.03 | 0.2 | 0.08 | 0.28 | 0.09 |
| 11 | 77318142 | rs4945185 | -0.456 | 0.422 | -0.19 | 0.262 | 0 | -0.017 | -0.017 | 0 | 0.34 | 0.57 | 0.4 | 0.3 | 0.34 | 0.36 |
| 11 | 77323126 | rs6592738 | 0.862 | 0.38 | -0.16 | 0.247 | -0.005 | 0.008 | 0.008 | 0 | 0.22 | 0.18 | 0.18 | 0.11 | 0.25 | 0.23 |
| 11 | 77323370 | rs3087668 | 0.228 | 0.313 | -0.216 | 0.237 | 0.016 | 0.027 | 0.027 | 0 | 0.28 | 0.3 | 0.29 | 0.45 | 0.38 | 0.17 |
| 11 | 77324261 | rs10899394 | 0.699 | 0.26 | -0.243 | 0.175 | -0.001 | 0 | 0 | 0 | 0.25 | 0.24 | 0.19 | 0.12 | 0.25 | 0.23 |
| 11 | 77335336 | rs11237244 | 0.831 | 0.275 | -0.263 | 0.17 | -0.028 | 0.095 | 0.095 | 0 | 0.24 | 0.07 | 0.18 | 0.1 | 0.25 | 0.18 |
| 11 | 77335982 | rs2013296 | -1.704 | 0.264 | -0.306 | 0.165 | 0.047 | 0.09 | 0.09 | 0 | 0.62 | 0.94 | 0.7 | 0.75 | 0.72 | 0.53 |
| 11 | 77341135 | rs11237249 | 1.128 | 0.296 | -0.296 | 0.04 | -0.039 | 0.09 | 0.09 | 0 | 0.22 | 0.03 | 0.16 | 0.1 | 0.22 | 0.17 |
| 11 | 77341306 | rs7109527 | 0.696 | 0.314 | -0.074 | 0.083 | 0.002 | 0.085 | 0.085 | 0.257 | 0.1 | 0.3 | 0.21 | 0.18 | 0.1 | 0.13 |
| 11 | 77350337 | rs10466710 | -0.666 | 0.327 | -0.151 | 0.048 | -0.001 | -0.004 | -0.004 | 0 | 0.38 | 0.62 | 0.41 | 0.29 | 0.36 | 0.37 |
| 11 | 77360056 | rs12225188 | 0.139 | 0.395 | -0.275 | 0.209 | 0.036 | 0.066 | 0.066 | 0 | 0.26 | 0.37 | 0.27 | 0.47 | 0.36 | 0.17 |
| 11 | 77361025 | rs10899397 | -0.382 | 0.355 | -0.235 | 0.143 | -0.007 | -0.021 | -0.021 | 0.012 | 0.36 | 0.52 | 0.39 | 0.27 | 0.33 | 0.35 |
| 11 | 77367542 | rs11604207 | 1.37 | 0.334 | -0.359 | 0.107 | -0.033 | 0.067 | 0.067 | 0 | 0.2 | 0.02 | 0.14 | 0.08 | 0.2 | 0.17 |
| 11 | 77372165 | rs11237260 | 1.105 | 0.641 | -0.418 | 0.055 | -0.014 | 0.017 | 0.017 | 0 | 0.09 | 0.05 | 0.09 | 0.05 | 0.13 | 0.09 |
| 11 | 77373414 | rs2298820 | 0.822 | 0.631 | -0.254 | 0.07 | 0.001 | 0.052 | 0.052 | 0.895 | 0.1 | 0.18 | 0.2 | 0.17 | 0.1 | 0.12 |
| 11 | 77373933 | rs11237261 | 1.316 | 0.635 | -0.28 | 0.071 | -0.035 | 0.01 | 0.01 | 0 | 0.09 | 0.01 | 0.09 | 0.05 | 0.13 | 0.09 |
| 11 | 77376247 | rs6592742 | 0.296 | 0.582 | -0.16 | 0.088 | 0.008 | 0.092 | 0.092 | 1.459 | 0.11 | 0.33 | 0.22 | 0.17 | 0.1 | 0.13 |
| 11 | 77378388 | rs4945197 | 0.296 | 0.582 | -0.16 | 0.088 | 0.008 | 0.092 | 0.092 | 1.459 | 0.11 | 0.33 | 0.21 | 0.17 | 0.1 | 0.13 |

| | | | | | | | | | | | | | | | | |
|---|---|---|---|---|---|---|---|---|---|---|---|---|---|---|---|---|
| 11 | 77389020 | rs12286317 | 0.771 | 0.587 | -0.183 | 0.056 | 0 | 0.023 | 0.023 | 0 | 0.25 | 0.21 | 0.18 | 0.09 | 0.23 | 0.22 |
| 11 | 77402419 | rs1544274 | 0.743 | 0.543 | -0.148 | 0.151 | -0.003 | 0.045 | 0.045 | 0.013 | 0.1 | 0.19 | 0.2 | 0.17 | 0.1 | 0.13 |
| 11 | 77409568 | rs4945203 | 0.295 | 0.543 | -0.252 | 0.151 | 0.008 | 0.092 | 0.092 | 1.459 | 0.11 | 0.33 | 0.22 | 0.17 | 0.1 | 0.13 |
| 11 | 77420314 | rs496786 | -1.707 | 0.654 | -0.494 | 0.098 | 0.022 | 0.069 | 0.069 | 0 | 0.61 | 0.93 | 0.66 | 0.73 | 0.69 | 0.51 |
| 11 | 77431229 | rs572541 | 1.054 | 0.798 | -0.073 | 0.301 | 0.039 | 0.095 | 0.095 | 0 | 0.14 | 0.18 | 0.17 | 0.38 | 0.22 | 0.08 |
| 11 | 77431334 | rs10899401 | -0.285 | 0.679 | -0.101 | 0.144 | -0.004 | -0.021 | -0.021 | 0 | 0.36 | 0.53 | 0.39 | 0.27 | 0.33 | 0.35 |
| 11 | 77440106 | rs720383 | 0.286 | 0.677 | -0.083 | 0.146 | 0.005 | 0.085 | 0.085 | 1.476 | 0.12 | 0.32 | 0.22 | 0.17 | 0.1 | 0.13 |
| 11 | 77468088 | rs537811 | 0.741 | 0.72 | -0.236 | 0.1 | -0.002 | 0.028 | 0.028 | 0 | 0.25 | 0.19 | 0.17 | 0.09 | 0.24 | 0.22 |
| 11 | 77477828 | rs1893901 | -0.206 | 0.667 | -0.067 | 0.219 | -0.002 | 0.083 | 0.083 | 0.399 | 0.13 | 0.37 | 0.22 | 0.18 | 0.1 | 0.13 |
| 11 | 77498768 | rs648842 | -0.054 | 0.765 | -0.326 | 0.223 | 0.043 | 0.085 | 0.085 | 0 | 0.75 | 0.6 | 0.72 | 0.53 | 0.64 | 0.83 |
| 11 | 77526474 | rs12417115 | 0.576 | 0.81 | -0.312 | 0.232 | 0.054 | 0.154 | 0.154 | 0 | 0.17 | 0.32 | 0.19 | 0.41 | 0.22 | 0.08 |
| 11 | 77531890 | rs11820337 | -0.111 | 0.745 | -0.482 | 0.241 | 0.043 | 0.089 | 0.089 | 0 | 0.25 | 0.4 | 0.28 | 0.46 | 0.36 | 0.17 |
| 11 | 77533652 | rs4245453 | 1.288 | 0.657 | -0.667 | 0.185 | 0.033 | 0.074 | 0.074 | 0 | 0.39 | 0.08 | 0.34 | 0.27 | 0.3 | 0.49 |
| 11 | 77552987 | rs4944176 | 1.245 | 0.714 | -0.484 | 0.227 | 0.004 | 0.046 | 0.046 | 0 | 0.08 | 0.14 | 0.19 | 0.18 | 0.1 | 0.13 |
| 11 | 77553638 | rs585721 | -0.584 | 0.258 | -0.845 | -0.254 | -0.004 | -0.017 | -0.017 | 0.079 | 0.43 | 0.53 | 0.39 | 0.27 | 0.37 | 0.35 |
| 11 | 77583266 | rs2186564 | 1.205 | 0.507 | -0.66 | -0.033 | 0.001 | 0.034 | 0.034 | 0 | 0.08 | 0.13 | 0.19 | 0.18 | 0.1 | 0.13 |
| 11 | 77583376 | rs588217 | 0.997 | 0.507 | -0.656 | -0.062 | -0.003 | 0.024 | 0.024 | 0 | 0.24 | 0.2 | 0.17 | 0.09 | 0.24 | 0.22 |
| 11 | 77583476 | rs668701 | 1.214 | 0.458 | -0.718 | -0.074 | -0.046 | 0.088 | 0.088 | 0 | 0.23 | 0.03 | 0.15 | 0.08 | 0.24 | 0.17 |
| 11 | 77584647 | rs1783546 | -1.611 | 0.508 | -0.744 | 0.029 | 0.032 | 0.065 | 0.065 | 0 | 0.61 | 0.92 | 0.66 | 0.73 | 0.7 | 0.52 |
| 11 | 77587973 | rs616354 | -0.096 | 0.545 | -0.674 | 0.076 | 0.04 | 0.082 | 0.082 | 0 | 0.26 | 0.4 | 0.28 | 0.47 | 0.35 | 0.17 |
| 11 | 77592522 | rs663674 | 0.56 | 0.524 | -0.918 | -0.007 | -0.004 | -0.051 | -0.051 | 0 | 0.26 | 0.34 | 0.18 | 0.09 | 0.24 | 0.21 |
| 11 | 77595642 | rs608389 | -0.21 | 0.525 | -0.965 | 0.024 | -0.004 | -0.019 | -0.019 | 0 | 0.34 | 0.46 | 0.37 | 0.27 | 0.33 | 0.34 |
| 11 | 77605501 | rs1793336 | -1.667 | 0.492 | -1.039 | 0.003 | 0.034 | 0.066 | 0.066 | 0 | 0.61 | 0.91 | 0.66 | 0.73 | 0.7 | 0.54 |
| 11 | 77606217 | rs7929313 | -0.243 | 0.428 | -1.009 | -0.014 | -0.004 | -0.019 | -0.019 | 0.775 | 0.35 | 0.45 | 0.37 | 0.27 | 0.33 | 0.37 |
| 11 | 77617950 | rs1790244 | -0.588 | 0.557 | -1.026 | 0.208 | 0.024 | -0.004 | -0.004 | 0 | 0.34 | 0.48 | 0.32 | 0.32 | 0.41 | 0.18 |
| 11 | 77623873 | rs10899415 | -0.62 | 0.549 | -1.04 | 0.204 | 0.021 | -0.005 | -0.005 | 0 | 0.34 | 0.46 | 0.32 | 0.32 | 0.41 | 0.19 |
| 11 | 77653967 | rs1630344 | -1.034 | 0.604 | -1.066 | 0.241 | 0.021 | 0.007 | 0.007 | 0 | 0.51 | 0.62 | 0.46 | 0.47 | 0.62 | 0.38 |
| 11 | 77661518 | rs7115801 | 1.4 | 0.654 | -1.06 | 0.259 | -0.004 | 0.004 | 0.004 | 0 | 0.17 | 0.17 | 0.14 | 0.15 | 0.21 | 0.2 |
| 11 | 77680399 | rs12807406 | 0.35 | 0.736 | -1.167 | 0.286 | 0.037 | 0.017 | 0.017 | 0 | 0.23 | 0.35 | 0.22 | 0.27 | 0.28 | 0.09 |
| 11 | 77684623 | rs11237340 | -1.778 | 0.616 | -1.47 | 0.314 | 0.063 | 0.064 | 0.064 | 0 | 0.59 | 0.88 | 0.66 | 0.74 | 0.72 | 0.53 |
| 11 | 77688774 | rs10899420 | -1.327 | 0.744 | -1.369 | 0.28 | 0.044 | 0.011 | 0.011 | 0 | 0.52 | 0.72 | 0.47 | 0.47 | 0.62 | 0.38 |
| 11 | 77693174 | rs11237342 | 0.805 | 0.599 | -1.178 | 0.576 | -0.029 | 0.018 | 0.018 | 0 | 0.09 | 0.03 | 0.09 | 0.05 | 0.13 | 0.11 |
| 11 | 77700152 | rs11607512 | 0.499 | 0.682 | -0.91 | 0.719 | 0.003 | -0.013 | -0.013 | 0 | 0.19 | 0.26 | 0.16 | 0.15 | 0.21 | 0.2 |
| 11 | 77703983 | rs1124590 | 0.559 | 0.826 | -1.287 | 1.023 | 0.001 | 0.065 | 0.065 | 0 | 0.08 | 0.13 | 0.18 | 0.26 | 0.1 | 0.15 |
| 11 | 77704592 | rs11237348 | -1.061 | 0.89 | -1.814 | 0.608 | -0.069 | 0.07 | 0.07 | 0.063 | 0.53 | 0.91 | 0.56 | 0.7 | 0.51 | 0.5 |
| 11 | 77710841 | rs921383 | -1.099 | 1.046 | -1.963 | 0.712 | -0.162 | -0.017 | -0.017 | 0 | 0.6 | 0.94 | 0.58 | 0.74 | 0.55 | 0.52 |
| 11 | 77714746 | rs11237353 | -0.148 | 1.36 | -0.743 | 1.154 | 0.017 | 0.097 | 0.097 | 0.125 | 0.09 | 0.2 | 0.23 | 0.31 | 0.12 | 0.18 |
| 11 | 77724674 | rs7930865 | -1.086 | 0.849 | -0.081 | 1.034 | -0.013 | 0.041 | 0.041 | 0.265 | 0.3 | 0.52 | 0.35 | 0.45 | 0.28 | 0.38 |
| 11 | 77727129 | rs7931417 | -0.033 | 1.015 | -0.145 | 1.209 | 0.017 | 0.108 | 0.108 | 0.226 | 0.09 | 0.2 | 0.23 | 0.32 | 0.13 | 0.18 |
| 11 | 77727139 | rs7942125 | -0.77 | 0.986 | -0.06 | 1.112 | 0.045 | 0.158 | 0.158 | 0.419 | 0.13 | 0.35 | 0.25 | 0.32 | 0.16 | 0.18 |
| 11 | 77727283 | rs7928656 | -0.779 | 0.982 | -0.062 | 1.11 | 0.043 | 0.157 | 0.157 | 0.408 | 0.13 | 0.35 | 0.26 | 0.32 | 0.16 | 0.18 |
| 11 | 77727379 | rs3824865 | 0.54 | 0.601 | -0.531 | 0.528 | 0.008 | 0.014 | 0.014 | 0 | 0.15 | 0.02 | 0.08 | 0.12 | 0.12 | 0.17 |
| 11 | 77730725 | rs2372892 | -0.222 | 0.334 | -0.518 | 0.551 | -0.023 | 0.047 | 0.047 | 0.426 | 0.72 | 0.56 | 0.81 | 0.58 | 0.81 | 0.82 |
| 11 | 77731416 | rs7937238 | 0.421 | 0.982 | -0.971 | 0.442 | 0.001 | -0.015 | -0.015 | 1.4 | 0.48 | 0.29 | 0.66 | 0.49 | 0.57 | 0.64 |
| 11 | 77746118 | rs9651740 | 0.248 | 0.946 | -0.721 | 0.105 | 0.001 | 0.014 | 0.014 | 0 | 0.47 | 0.63 | 0.4 | 0.35 | 0.54 | 0.45 |
| 11 | 77749228 | rs10899433 | 0.736 | 0.941 | -0.702 | 0.084 | -0.005 | -0.002 | -0.002 | 0 | 0.38 | 0.23 | 0.37 | 0.34 | 0.44 | 0.44 |
| 11 | 77773715 | rs589472 | 0.094 | 0.903 | -0.7 | 0.069 | 0.021 | -0.089 | -0.089 | 1.534 | 0.64 | 0.85 | 0.63 | 0.45 | 0.69 | 0.64 |
| 11 | 77773926 | rs11237374 | 0.866 | 0.642 | -0.754 | -0.19 | -0.012 | -0.009 | -0.009 | 1.713 | 0.3 | 0.24 | 0.37 | 0.36 | 0.45 | 0.44 |
| 11 | 77774599 | rs2282639 | 0.875 | 0.666 | -0.749 | -0.189 | -0.014 | -0.01 | -0.01 | 0 | 0.3 | 0.24 | 0.37 | 0.36 | 0.45 | 0.44 |
| 11 | 77777640 | rs621586 | -0.054 | 0.584 | -0.904 | -0.136 | 0.019 | -0.094 | -0.094 | 0 | 0.64 | 0.89 | 0.64 | 0.45 | 0.7 | 0.64 |
| 11 | 77785836 | rs588584 | 1.69 | 0.532 | -0.811 | -0.246 | -0.004 | 0.119 | 0.119 | 0 | 0.28 | 0.08 | 0.21 | 0.08 | 0.23 | 0.19 |
| 11 | 77789554 | rs636141 | -0.536 | 0.373 | -0.978 | -0.011 | 0.031 | -0.014 | -0.014 | 0.038 | 0.75 | 0.88 | 0.92 | 0.72 | 0.83 | 0.82 |
| 11 | 77795571 | rs10899436 | 0.8 | 0.336 | -0.466 | -0.008 | 0.033 | 0.015 | 0.015 | 0.017 | 0.32 | 0.42 | 0.39 | 0.36 | 0.46 | 0.45 |
| 11 | 77796717 | rs681717 | -0.977 | 0.334 | -0.312 | -0.004 | 0.009 | 0.013 | 0.013 | 0 | 0.58 | 0.69 | 0.48 | 0.64 | 0.63 | 0.63 |
| 11 | 77811196 | rs10793285 | 1.006 | 0.388 | -0.37 | -0.033 | -0.062 | -0.04 | -0.04 | 0 | 0.3 | 0.14 | 0.38 | 0.36 | 0.46 | 0.42 |
| 11 | 77817080 | rs647602 | -0.921 | 0.257 | -0.45 | -0.053 | -0.008 | -0.028 | -0.028 | 0 | 0.89 | 0.94 | 0.71 | 0.72 | 0.86 | 0.83 |
| 11 | 77822105 | rs534723 | 0.436 | 0.196 | -0.503 | -0.097 | 0.023 | -0.03 | -0.03 | 0 | 0.24 | 0.07 | 0.08 | 0.28 | 0.17 | 0.19 |
| 11 | 77823557 | rs624350 | 0.629 | 0.23 | -0.372 | -0.072 | -0.019 | 0.008 | 0.008 | 0 | 0.61 | 0.49 | 0.55 | 0.43 | 0.69 | 0.63 |
| 11 | 77825320 | rs665278 | 2.003 | 0.239 | -0.528 | -0.191 | -0.025 | 0.104 | 0.104 | 0 | 0.27 | 0.04 | 0.2 | 0.09 | 0.23 | 0.22 |
| 11 | 77836480 | rs659758 | 0.034 | 0.253 | -0.23 | -0.016 | 0.032 | -0.01 | -0.01 | 0 | 0.25 | 0.14 | 0.09 | 0.28 | 0.17 | 0.19 |
| 11 | 77837761 | rs548907 | -0.358 | 0.268 | -0.194 | -0.008 | -0.039 | -0.007 | -0.007 | 0 | 0.28 | 0.43 | 0.12 | 0.28 | 0.17 | 0.2 |
| 11 | 77850517 | rs10793289 | 0.6 | 0.338 | -0.345 | -0.076 | -0.059 | -0.044 | -0.044 | 0 | 0.31 | 0.15 | 0.38 | 0.38 | 0.46 | 0.43 |
| 11 | 77855254 | rs550458 | -0.346 | 0.368 | -0.382 | -0.111 | -0.021 | -0.011 | -0.011 | 0.397 | 0.27 | 0.36 | 0.1 | 0.26 | 0.17 | 0.2 |
| 11 | 77870486 | rs1144764 | -0.724 | 0.564 | -1.298 | 0.292 | -0.014 | -0.007 | -0.007 | 0 | 0.64 | 0.38 | 0.58 | 0.67 | 0.53 | 0.58 |
| 11 | 77875386 | rs230669 | 0.9 | 0.939 | -1.685 | 0.554 | -0.05 | -0.045 | -0.045 | 2.507 | 0.37 | 0.09 | 0.48 | 0.41 | 0.37 | 0.36 |
| 11 | 77876489 | rs563453 | 1.345 | 0.665 | -1.356 | 0.632 | -0.004 | 0.023 | 0.023 | 0.005 | 0.08 | 0.01 | 0.02 | 0.05 | 0.07 | 0.06 |
| 11 | 77880704 | rs585730 | -0.474 | 0.622 | -1.37 | 0.669 | 0.008 | -0.021 | -0.021 | 0.02 | 0.19 | 0.12 | 0.08 | 0.26 | 0.14 | 0.15 |
| 11 | 77883126 | rs230663 | 0.9 | 0.76 | -1.548 | 0.15 | -0.042 | -0.04 | -0.04 | 0 | 0.35 | 0.1 | 0.48 | 0.39 | 0.35 | 0.36 |
| 11 | 77883478 | rs527462 | -0.415 | 0.043 | -0.525 | -0.5 | -0.052 | -0.108 | -0.108 | 0.54 | 0.17 | 0.6 | 0.07 | 0.02 | 0.1 | 0.09 |

| | | | | | | | | | | | | | | | | |
|---|---|---|---|---|---|---|---|---|---|---|---|---|---|---|---|---|
| 11 | 77883855 | rs230662 | 0.357 | 0.721 | -0.622 | -0.138 | 0.009 | 0.058 | 0.058 | 0 | 0.51 | 0.27 | 0.46 | 0.37 | 0.47 | 0.43 |
| 11 | 77884338 | rs530588 | 1.262 | 0.74 | -1.002 | -0.172 | -0.036 | 0.044 | 0.044 | 0 | 0.1 | 0.01 | 0.06 | 0.02 | 0.16 | 0.09 |
| 11 | 77884782 | rs230661 | 0.213 | 0.879 | -1.279 | -0.075 | -0.023 | -0.047 | -0.047 | 0.63 | 0.61 | 0.86 | 0.65 | 0.5 | 0.52 | 0.34 |
| 11 | 77888744 | rs483731 | 1.126 | 0.2 | -0.72 | 0.242 | -0.003 | -0.002 | -0.002 | 0.791 | 0.1 | 0.09 | 0.07 | 0.02 | 0.15 | 0.07 |
| 11 | 77891751 | rs657161 | 0.775 | 0.157 | -0.59 | 0.221 | 0.016 | -0.039 | -0.039 | 0.074 | 0.11 | 0.19 | 0.07 | 0.02 | 0.16 | 0.08 |
| 11 | 77896538 | rs661811 | -0.199 | -2.232 | -0.95 | 0.503 | 0.006 | 0.054 | 0.054 | 0 | 0.5 | 0.22 | 0.56 | 0.63 | 0.43 | 0.47 |
| 11 | 77899097 | rs3133324 | 0.656 | -2.218 | -1.299 | 0.567 | -0.028 | -0.009 | -0.009 | 0.051 | 0.33 | 0.11 | 0.46 | 0.36 | 0.34 | 0.34 |
| 11 | 77903098 | rs3102743 | -0.288 | -1.917 | -0.765 | 0.398 | -0.003 | 0.004 | 0.004 | 0 | 0.55 | 0.58 | 0.59 | 0.63 | 0.45 | 0.46 |
| 11 | 77909014 | rs2510044 | 0.141 | -2.321 | -0.521 | 0.878 | 0.07 | -0.089 | -0.089 | 0.174 | 0.26 | 0.03 | 0.3 | 0.36 | 0.16 | 0.26 |
| 11 | 77920563 | rs2510045 | -0.056 | -2.296 | 0.029 | 0.906 | 0.049 | -0.035 | -0.035 | 0 | 0.29 | 0.2 | 0.32 | 0.4 | 0.18 | 0.27 |
| 11 | 77924870 | rs2450138 | 0.004 | -2.343 | 0.529 | 0.862 | 0.054 | -0.035 | -0.035 | 0.089 | 0.3 | 0.17 | 0.32 | 0.4 | 0.19 | 0.28 |
| 11 | 77926769 | rs1046780 | -0.454 | -2.479 | 1.173 | 0.786 | -0.053 | 0.013 | 0.013 | 0 | 0.34 | 0.5 | 0.35 | 0.4 | 0.2 | 0.28 |
| 11 | 77928036 | rs3740677 | 0.79 | -2.9 | 1.421 | 0.926 | 0.166 | 0.135 | 0.135 | 0 | 0.56 | 0.16 | 0.53 | 0.26 | 0.76 | 0.38 |
| 11 | 77930138 | rs1485483 | 0.306 | -2.793 | 1.406 | 0.96 | 0.069 | -0.104 | -0.104 | 2.957 | 0.28 | 0.06 | 0.3 | 0.39 | 0.18 | 0.25 |
| 11 | 77930345 | rs2450122 | 0.151 | -2.799 | 1.543 | 0.957 | 0.047 | -0.033 | -0.033 | 0 | 0.29 | 0.2 | 0.32 | 0.4 | 0.18 | 0.27 |
| 11 | 77930792 | rs1318241 | 0.31 | -2.839 | 1.49 | 0.963 | 0.053 | -0.045 | -0.045 | 0.409 | 0.28 | 0.14 | 0.31 | 0.39 | 0.18 | 0.27 |
| 11 | 77936166 | rs1385600 | -0.185 | -2.844 | 1.684 | 0.96 | -0.048 | 0.014 | 0.014 | 0 | 0.33 | 0.49 | 0.34 | 0.4 | 0.18 | 0.27 |
| 11 | 77937800 | rs2248407 | 0.04 | -2.833 | 1.721 | 0.966 | -0.021 | 0.006 | 0.006 | 0 | 0.31 | 0.39 | 0.33 | 0.4 | 0.18 | 0.26 |
| 11 | 77941076 | rs1007837 | -0.237 | -2.848 | 2.099 | 0.957 | -0.057 | 0.015 | 0.015 | 0 | 0.33 | 0.6 | 0.34 | 0.4 | 0.18 | 0.27 |
| 11 | 77959659 | rs2510054 | 0.14 | -2.769 | 2.134 | 0.997 | -0.006 | 0.001 | 0.001 | 0 | 0.29 | 0.31 | 0.32 | 0.4 | 0.18 | 0.27 |
| 11 | 77979414 | rs7927923 | 0.737 | -2.819 | 2.123 | 0.997 | 0.173 | 0.149 | 0.149 | 0 | 0.56 | 0.11 | 0.53 | 0.24 | 0.77 | 0.39 |
| 11 | 77979829 | rs948662 | 0.162 | -2.821 | 2.16 | 1.002 | 0.036 | -0.023 | -0.023 | 0 | 0.3 | 0.23 | 0.32 | 0.4 | 0.18 | 0.29 |
| 11 | 77996403 | rs10793294 | 0.487 | -2.829 | 2.254 | 0.989 | 0.014 | 0.014 | 0.014 | 0 | 0.58 | 0.35 | 0.55 | 0.24 | 0.77 | 0.39 |
| 11 | 78010830 | rs11602622 | 0.175 | -2.822 | 2.113 | 0.998 | 0.019 | -0.014 | -0.014 | 0 | 0.3 | 0.27 | 0.32 | 0.4 | 0.18 | 0.29 |
| 11 | 78025459 | rs11237451 | 0.83 | -2.869 | 2.022 | 0.965 | 0.176 | 0.134 | 0.134 | 0 | 0.55 | 0.08 | 0.53 | 0.26 | 0.77 | 0.39 |
| 11 | 78035856 | rs2458640 | 0.759 | -3.004 | 2.036 | 0.95 | 0.191 | 0.154 | 0.154 | 0 | 0.55 | 0.1 | 0.53 | 0.24 | 0.77 | 0.4 |
| 11 | 78040259 | rs7117238 | -0.453 | -2.905 | 2.24 | 0.667 | -0.059 | 0.003 | 0.003 | 0.379 | 0.36 | 0.57 | 0.35 | 0.4 | 0.18 | 0.29 |
| 11 | 78040961 | rs10793302 | 0.072 | -2.736 | 2.238 | 0.749 | 0.015 | -0.009 | -0.009 | 0 | 0.31 | 0.29 | 0.33 | 0.4 | 0.18 | 0.29 |
| 11 | 78054336 | rs12277648 | -0.242 | -2.675 | 2.241 | 0.757 | -0.024 | 0.003 | 0.003 | 0 | 0.34 | 0.41 | 0.33 | 0.4 | 0.19 | 0.29 |
| 11 | 78104399 | rs10793311 | -0.072 | -2.562 | 2.241 | 1.202 | 0.077 | -0.12 | -0.12 | 4.509 | 0.29 | 0.03 | 0.31 | 0.4 | 0.18 | 0.29 |
| 11 | 78123831 | rs10899496 | -0.422 | -2.527 | 2.388 | 1.194 | -0.038 | 0.013 | 0.013 | 0 | 0.33 | 0.4 | 0.32 | 0.44 | 0.19 | 0.29 |
| 11 | 78125246 | rs10899500 | -0.112 | -2.452 | 2.028 | 1.23 | 0.082 | -0.084 | -0.084 | 0.101 | 0.3 | 0.09 | 0.31 | 0.43 | 0.19 | 0.29 |
| 11 | 78164984 | rs4945276 | 1.579 | -2.25 | 1.932 | 0.28 | -0.144 | 0.112 | 0.112 | 0 | 0.3 | 0.03 | 0.25 | 0.08 | 0.36 | 0.18 |
| 11 | 78170811 | rs10899509 | -1.239 | -3.347 | 3.565 | 1.537 | -0.022 | 0.007 | 0.007 | 0 | 0.35 | 0.62 | 0.36 | 0.42 | 0.2 | 0.26 |
| 11 | 78172353 | rs10899513 | -1.041 | -3.349 | 3.748 | 1.536 | -0.068 | 0.04 | 0.04 | 0 | 0.33 | 0.43 | 0.35 | 0.42 | 0.2 | 0.26 |
| 11 | 78173439 | rs11237509 | -1.251 | -3.349 | 3.739 | 1.536 | -0.015 | 0.003 | 0.003 | 0 | 0.35 | 0.62 | 0.36 | 0.42 | 0.2 | 0.26 |
| 11 | 78180508 | rs10899518 | -0.926 | -3.348 | 3.719 | 1.537 | -0.025 | 0.012 | 0.012 | 0 | 0.33 | 0.28 | 0.34 | 0.42 | 0.2 | 0.26 |
| 11 | 78192707 | rs10751290 | -1.19 | -3.349 | 3.706 | 1.484 | 0.029 | -0.018 | -0.018 | 0 | 0.36 | 0.67 | 0.36 | 0.42 | 0.2 | 0.25 |
| 11 | 78193921 | rs7928933 | -1.174 | -3.351 | 3.749 | 1.483 | -0.066 | 0.036 | 0.036 | 0 | 0.34 | 0.5 | 0.35 | 0.42 | 0.2 | 0.25 |
| 11 | 78202699 | rs7396701 | 1.104 | -3.362 | 3.799 | 1.476 | 0.097 | -0.046 | -0.046 | 0 | 0.62 | 0.2 | 0.62 | 0.58 | 0.8 | 0.75 |
| 11 | 78204380 | rs4474465 | 0.815 | -3.362 | 3.71 | 1.476 | -0.069 | 0.037 | 0.037 | 0 | 0.65 | 0.54 | 0.65 | 0.58 | 0.8 | 0.75 |
| 11 | 78207686 | rs10899528 | 0.783 | -3.355 | 3.54 | 1.48 | -0.037 | 0.017 | 0.017 | 0 | 0.65 | 0.44 | 0.64 | 0.58 | 0.8 | 0.75 |
| 11 | 78209879 | rs11237522 | 0.884 | -3.355 | 3.5 | 1.48 | 0.032 | -0.02 | -0.02 | 0.089 | 0.64 | 0.32 | 0.64 | 0.58 | 0.8 | 0.75 |
| 11 | 78214673 | rs10128743 | -1.236 | -3.295 | 3.461 | 1.625 | -0.018 | 0.001 | 0.001 | 0 | 0.35 | 0.58 | 0.35 | 0.33 | 0.19 | 0.25 |
| 11 | 78215082 | rs7950813 | 0.802 | -3.218 | 3.341 | 1.258 | -0.024 | 0.008 | 0.008 | 0 | 0.64 | 0.4 | 0.63 | 0.58 | 0.8 | 0.75 |
| 11 | 78215898 | rs10793318 | 0.797 | -3.218 | 3.306 | 1.258 | -0.048 | 0.023 | 0.023 | 0 | 0.65 | 0.47 | 0.64 | 0.58 | 0.8 | 0.75 |
| 11 | 78224908 | rs7938257 | 0.342 | -3.212 | 3.08 | 1.264 | 0.047 | -0.043 | -0.043 | 0 | 0.69 | 0.83 | 0.66 | 0.58 | 0.8 | 0.75 |
| 11 | 78243134 | rs4285892 | 0.941 | -3.176 | 2.919 | 1.23 | -0.013 | 0.001 | 0.001 | 0 | 0.63 | 0.39 | 0.64 | 0.6 | 0.8 | 0.76 |
| 11 | 78248397 | rs11237537 | 0.589 | -3.173 | 2.964 | 1.231 | -0.054 | 0.033 | 0.033 | 0 | 0.67 | 0.61 | 0.65 | 0.6 | 0.8 | 0.76 |
| 11 | 78260699 | rs4565922 | 0.994 | -3.125 | 2.67 | 1.235 | 0.07 | -0.029 | -0.029 | 0.498 | 0.63 | 0.24 | 0.63 | 0.61 | 0.8 | 0.76 |
| 11 | 78277277 | rs10751296 | 0.725 | -2.87 | 2.615 | 1.057 | -0.05 | 0.031 | 0.031 | 0 | 0.64 | 0.6 | 0.6 | 0.58 | 0.77 | 0.71 |
| 11 | 78278792 | rs10793334 | 0.941 | -2.863 | 2.482 | 1.054 | 0.001 | -0.004 | -0.004 | 1.646 | 0.62 | 0.41 | 0.59 | 0.59 | 0.76 | 0.71 |
| 11 | 78279790 | rs10501429 | 0.498 | -2.762 | 1.829 | 1.098 | 0.016 | -0.016 | -0.016 | 0 | 0.67 | 0.78 | 0.62 | 0.6 | 0.77 | 0.74 |
| 11 | 78289074 | rs10793339 | -1.591 | -1.709 | 2.154 | 1.482 | -0.003 | 0.301 | 0.301 | 0 | 0.1 | 0.39 | 0.07 | 0.36 | 0.04 | 0.35 |
| 11 | 78318132 | rs7936158 | -1.163 | -1.425 | 1.82 | 0.893 | 0.078 | 0.068 | 0.068 | 4.638 | 0.05 | 0.4 | 0.03 | 0 | 0.01 | 0 |
| 11 | 78319651 | rs4388909 | -1.61 | -1.474 | 1.57 | 0.87 | -0.024 | -0.031 | -0.031 | 0 | 0.08 | 0.67 | 0.05 | 0 | 0.01 | 0 |
| 11 | 78340688 | rs7103066 | -1.263 | -1.385 | 0.166 | 0.913 | 0.016 | 0.007 | 0.007 | 8.019 | 0.07 | 0.55 | 0.04 | 0 | 0.01 | 0 |
| 11 | 78344839 | rs7128118 | 0.549 | -0.986 | -1.332 | 1.293 | 0.003 | -0.169 | -0.169 | 0.847 | 0.79 | 0.99 | 0.7 | 0.53 | 0.84 | 0.85 |
| 11 | 78356498 | rs1945465 | 0.555 | -1.141 | -0.924 | 0.171 | 0.011 | 0.011 | 0.011 | 0 | 0.64 | 0.37 | 0.52 | 0.22 | 0.73 | 0.43 |
| 11 | 78360321 | rs12420296 | 0.963 | -0.959 | -1.657 | -0.867 | -0.037 | 0.244 | 0.244 | 0.043 | 0.45 | 0.13 | 0.34 | 0.19 | 0.53 | 0.3 |
| 11 | 78360830 | rs10899565 | 0.469 | -1.14 | -0.743 | -0.92 | 0.004 | -0.007 | -0.007 | 0 | 0.6 | 0.43 | 0.46 | 0.19 | 0.69 | 0.39 |
| 11 | 78362665 | rs11237580 | 0.099 | -1.249 | 0.46 | -0.938 | -0.002 | -0.014 | -0.014 | 0 | 0.66 | 0.7 | 0.54 | 0.2 | 0.72 | 0.42 |
| 11 | 78371851 | rs4944211 | -0.095 | -1.729 | -0.017 | 0.245 | 0.021 | -0.115 | -0.115 | 0 | 0.86 | 0.99 | 0.72 | 0.63 | 0.9 | 0.74 |
| 11 | 78378825 | rs11237582 | 0.677 | -1.717 | 0.391 | -0.154 | 0.033 | 0.076 | 0.076 | 0 | 0.67 | 0.8 | 0.55 | 0.19 | 0.72 | 0.42 |
| 11 | 78381678 | rs7938204 | 0.738 | -1.735 | 0.816 | -0.164 | 0.011 | 0.016 | 0.016 | 0 | 0.67 | 0.72 | 0.55 | 0.2 | 0.72 | 0.42 |
| 11 | 78399121 | rs1792148 | -1.512 | -1.573 | 1.398 | -0.424 | 0.072 | -0.052 | -0.052 | 0 | 0.1 | 0.32 | 0.1 | 0 | 0.13 | 0.03 |
| 11 | 78401625 | rs485325 | 1.925 | -2.136 | 1.101 | -0.104 | -0.008 | 0.129 | 0.129 | 0.344 | 0.53 | 0.04 | 0.39 | 0.29 | 0.59 | 0.47 |
| 11 | 78402239 | rs4944214 | -0.44 | -2.388 | 1.251 | 0.475 | 0.007 | -0.016 | -0.016 | 0.795 | 0.17 | 0.07 | 0.21 | 0.17 | 0.18 | 0.15 |
| 11 | 78404419 | rs683420 | -1.238 | -2.366 | 1.117 | 0.394 | 0.009 | -0.024 | -0.024 | 0.045 | 0.05 | 0.14 | 0.03 | 0 | 0.03 | 0.04 |

| | | | | | | | | | | | | | | | | |
|---|---|---|---|---|---|---|---|---|---|---|---|---|---|---|---|---|
| 11 | 78414040 | rs642378 | -1.87 | -2.337 | 1.178 | 0.511 | -0.003 | 0.047 | 0.047 | 0.808 | 0.31 | 0.83 | 0.36 | 0.14 | 0.33 | 0.18 |
| 11 | 78422329 | rs665892 | -2.069 | -2.339 | 1.297 | 0.519 | 0 | 0.027 | 0.027 | 0.468 | 0.29 | 0.75 | 0.34 | 0.15 | 0.31 | 0.18 |
| 11 | 78428818 | rs1783944 | 0.577 | -0.485 | 0.653 | -0.043 | -0.037 | -0.075 | -0.075 | 0.353 | 0.73 | 0.87 | 0.58 | 0.4 | 0.68 | 0.58 |
| 11 | 78433599 | rs482605 | -0.672 | -0.307 | 0.923 | -0.128 | 0.006 | -0.028 | -0.028 | 0.794 | 0.08 | 0.13 | 0.09 | 0 | 0.12 | 0.03 |
| 11 | 78434720 | rs645848 | 0.274 | -0.24 | 0.873 | -0.104 | -0.026 | 0.03 | 0.03 | 0 | 0.06 | 0.02 | 0.08 | 0 | 0.11 | 0.02 |
| 11 | 78437352 | rs597327 | -1.306 | -0.256 | 1.284 | -0.118 | 0.043 | -0.067 | -0.067 | 0 | 0.11 | 0.32 | 0.12 | 0 | 0.14 | 0.03 |
| 11 | 78443235 | rs1790557 | 1.089 | -0.869 | 1.102 | -0.121 | -0.02 | -0.008 | -0.008 | 0 | 0.63 | 0.5 | 0.55 | 0.34 | 0.75 | 0.5 |
| 11 | 78443586 | rs17826919 | 0.439 | -0.823 | 0.888 | -0.188 | -0.072 | 0.042 | 0.042 | 0 | 0.12 | 0.02 | 0.18 | 0.08 | 0.25 | 0.12 |
| 11 | 78449121 | rs1790551 | 0.86 | -1.049 | 1.08 | -0.134 | -0.037 | -0.009 | -0.009 | 0 | 0.65 | 0.63 | 0.56 | 0.34 | 0.75 | 0.56 |
| 11 | 78451062 | rs714365 | 1.477 | -0.718 | 0.745 | -0.172 | 0.073 | 0.035 | 0.035 | 0 | 0.58 | 0.24 | 0.51 | 0.35 | 0.71 | 0.52 |
| 11 | 78453645 | rs1785812 | 0.27 | -0.932 | 0.658 | -0.135 | 0.01 | -0.101 | -0.101 | 0 | 0.82 | 0.95 | 0.72 | 0.47 | 0.87 | 0.66 |
| 11 | 78454125 | rs1783953 | 1.466 | -0.621 | 0.498 | -0.348 | 0.052 | 0.025 | 0.025 | 0.194 | 0.6 | 0.25 | 0.51 | 0.35 | 0.7 | 0.56 |
| 11 | 78460962 | rs1792139 | 2.732 | -0.554 | 0.463 | -0.281 | -0.057 | 0.097 | 0.097 | 0.191 | 0.46 | 0.14 | 0.32 | 0.27 | 0.47 | 0.41 |
| 11 | 78465138 | rs10501405 | 0.077 | -1.462 | 0.81 | 0.067 | 0.016 | -0.019 | -0.019 | 0 | 0.15 | 0.01 | 0.11 | 0.12 | 0.11 | 0.1 |
| 11 | 78470719 | rs1792136 | 0.887 | -1.452 | 1.338 | -0.048 | -0.042 | 0.114 | 0.114 | 0.332 | 0.78 | 0.4 | 0.7 | 0.52 | 0.88 | 0.67 |
| 11 | 78471087 | rs1785814 | 1.214 | -1.488 | 1.321 | -0.078 | -0.04 | 0.081 | 0.081 | 0 | 0.75 | 0.4 | 0.68 | 0.52 | 0.87 | 0.64 |
| 11 | 78473085 | rs7102408 | -1.45 | -1.522 | 1.335 | -0.081 | -0.048 | 0.073 | 0.073 | 0 | 0.24 | 0.55 | 0.22 | 0.4 | 0.15 | 0.21 |
| 11 | 78476557 | rs7104372 | -1.482 | -1.41 | 1.258 | -0.124 | -0.026 | 0.035 | 0.035 | 0.384 | 0.26 | 0.64 | 0.23 | 0.4 | 0.16 | 0.15 |
| 11 | 78483823 | rs1970745 | -0.467 | -1.562 | 1.014 | -0.052 | 0.024 | -0.046 | -0.046 | 0 | 0.19 | 0.11 | 0.13 | 0.38 | 0.13 | 0.13 |
| 11 | 78486141 | rs1792137 | 2.38 | -1.501 | 0.864 | -0.039 | 0.088 | 0.169 | 0.169 | 0 | 0.54 | 0.24 | 0.49 | 0.13 | 0.71 | 0.41 |
| 11 | 78487823 | rs1785827 | 0.556 | -1.472 | 0.784 | 0.011 | 0.021 | -0.108 | -0.108 | 0.404 | 0.79 | 0.92 | 0.68 | 0.51 | 0.85 | 0.55 |
| 11 | 78490041 | rs1573529 | 1.706 | -1.43 | 0.929 | -0.03 | -0.01 | -0.043 | -0.043 | 3.239 | 0.58 | 0.44 | 0.51 | 0.13 | 0.72 | 0.44 |
| 11 | 78490320 | rs1540124 | 0.378 | -1.4 | 0.658 | 0.043 | 0.023 | -0.117 | -0.117 | 0 | 0.81 | 0.95 | 0.69 | 0.52 | 0.87 | 0.58 |
| 11 | 78494379 | rs1785823 | 1.592 | -1.425 | 0.762 | -0.042 | 0.014 | 0.008 | 0.008 | 0.029 | 0.6 | 0.32 | 0.53 | 0.12 | 0.76 | 0.45 |
| 11 | 78495465 | rs1790544 | 0.557 | -1.51 | -0.272 | 0.207 | 0.025 | -0.103 | -0.103 | 0 | 0.8 | 0.91 | 0.69 | 0.5 | 0.87 | 0.55 |
| 11 | 78499881 | rs588059 | 1.965 | -1.324 | -0.113 | -0.057 | 0.093 | 0.227 | 0.227 | 0 | 0.55 | 0.19 | 0.43 | 0.1 | 0.72 | 0.39 |
| 11 | 78504767 | rs627195 | -0.141 | -1.142 | -0.281 | 0.383 | 0.003 | -0.05 | -0.05 | 0 | 0.86 | 0.94 | 0.66 | 0.51 | 0.88 | 0.7 |
| 11 | 78507790 | rs1941379 | -0.958 | -1.223 | 0.118 | 0.577 | 0.027 | -0.049 | -0.049 | 0 | 0.19 | 0.12 | 0.13 | 0.41 | 0.12 | 0.26 |
| 11 | 78509347 | rs563384 | -1.789 | -1.224 | 0.526 | 0.503 | -0.026 | 0.026 | 0.026 | 0.088 | 0.27 | 0.62 | 0.19 | 0.4 | 0.14 | 0.27 |
| 11 | 78509582 | rs661864 | -1.69 | -1.114 | 0.108 | 0.541 | 0.013 | -0.017 | -0.017 | 0.585 | 0.26 | 0.73 | 0.2 | 0.4 | 0.14 | 0.17 |
| 11 | 78509933 | rs11237625 | 0.043 | -0.879 | -0.11 | 0.652 | 0.018 | -0.047 | -0.047 | 0 | 0.13 | 0.04 | 0.08 | 0.34 | 0.09 | 0.11 |
| 11 | 78510519 | rs648443 | -1.336 | -0.883 | 0.141 | 0.546 | -0.011 | 0.013 | 0.013 | 2.154 | 0.22 | 0.68 | 0.15 | 0.35 | 0.12 | 0.13 |
| 11 | 78512105 | rs619587 | 0.646 | -1.021 | -0.021 | 0.607 | 0.007 | -0.016 | -0.016 | 2.418 | 0.75 | 0.76 | 0.54 | 0.54 | 0.82 | 0.63 |
| 11 | 78513751 | rs527983 | 0.809 | -1.067 | 0.505 | 0.547 | -0.028 | 0.085 | 0.085 | 0 | 0.73 | 0.56 | 0.47 | 0.52 | 0.81 | 0.63 |
| 11 | 78525027 | rs522714 | 1.265 | -1.066 | -0.025 | 0.009 | 0.017 | -0.011 | -0.011 | 0.037 | 0.62 | 0.64 | 0.4 | 0.43 | 0.73 | 0.53 |
| 11 | 78526601 | rs7926709 | -0.408 | 0.029 | -0.238 | -0.212 | -0.068 | 0.043 | 0.043 | 0 | 0.26 | 0.11 | 0.19 | 0.13 | 0.35 | 0.26 |
| 11 | 78530642 | rs10899572 | 0.025 | -0.006 | -0.172 | -0.245 | -0.046 | -0.038 | -0.038 | 0 | 0.23 | 0.11 | 0.15 | 0.31 | 0.3 | 0.25 |
| 11 | 78530720 | rs949365 | -0.819 | -0.156 | 0.626 | -0.322 | 0.022 | 0.011 | 0.011 | 0 | 0.28 | 0.53 | 0.23 | 0.32 | 0.31 | 0.31 |
| 11 | 78533206 | rs493471 | 0.049 | -0.667 | 0.618 | -1.463 | -0.07 | 0.236 | 0.236 | 0 | 0.55 | 0.91 | 0.35 | 0.14 | 0.52 | 0.39 |
| 11 | 78534981 | rs536769 | 0.557 | -0.676 | 1.201 | -1.468 | -0.001 | -0.034 | -0.034 | 0 | 0.49 | 0.52 | 0.28 | 0.13 | 0.51 | 0.32 |
| 11 | 78537008 | rs1941378 | 0.556 | -0.685 | 1.307 | -1.519 | 0.001 | -0.017 | -0.017 | 0.053 | 0.5 | 0.5 | 0.33 | 0.13 | 0.51 | 0.35 |
| 11 | 78539115 | rs1945246 | 0.189 | -0.632 | 1.211 | -1.914 | -0.036 | 0.28 | 0.28 | 0 | 0.53 | 0.83 | 0.34 | 0.08 | 0.52 | 0.38 |
| 11 | 78543525 | rs17137225 | 0.124 | -0.972 | 1.232 | -2.188 | -0.017 | -0.06 | -0.06 | 0.703 | 0.1 | 0.21 | 0.07 | 0.01 | 0.05 | 0.06 |
| 11 | 78543856 | rs530688 | 0.888 | -1.013 | 1.221 | -2.216 | -0.005 | 0.047 | 0.047 | 0 | 0.4 | 0.3 | 0.26 | 0.07 | 0.45 | 0.34 |
| 11 | 78553730 | rs526395 | 0.369 | -1.235 | 0.895 | -2.348 | -0.001 | 0.024 | 0.024 | 0 | 0.45 | 0.32 | 0.29 | 0.06 | 0.47 | 0.38 |
| 11 | 78558239 | rs547292 | 1.626 | -1.087 | 0.415 | -2.473 | -0.091 | 0.177 | 0.177 | 0.008 | 0.29 | 0.02 | 0.18 | 0.04 | 0.34 | 0.27 |
| 11 | 78558986 | rs540836 | -0.199 | -1.142 | 0.551 | -2.493 | 0.011 | 0.087 | 0.087 | 0 | 0.5 | 0.64 | 0.34 | 0.07 | 0.5 | 0.35 |
| 11 | 78564752 | rs530154 | 1.226 | -1.049 | 0.355 | -2.433 | -0.006 | 0.018 | 0.018 | 0 | 0.33 | 0.23 | 0.21 | 0.05 | 0.35 | 0.3 |
| 11 | 78568110 | rs12362098 | -1.964 | -1.241 | 0.73 | -1.965 | -0.039 | -0.134 | -0.134 | 0 | 0.22 | 0.55 | 0.14 | 0.02 | 0.16 | 0.11 |
| 11 | 78573179 | rs1945247 | -1.329 | -0.944 | 0.523 | -1.628 | 0.003 | -0.063 | -0.063 | 0 | 0.11 | 0.33 | 0.09 | 0.02 | 0.11 | 0.05 |
| 11 | 78588441 | rs11237644 | 0.392 | -1.353 | -0.681 | -1.806 | 0.036 | 0.229 | 0.229 | 0.064 | 0.51 | 0.04 | 0.64 | 0.8 | 0.61 | 0.7 |
| 11 | 78602937 | rs511398 | 0.914 | -1.36 | 0.52 | -1.622 | 0.002 | 0.182 | 0.182 | 1.069 | 0.94 | 0.74 | 0.93 | 0.81 | 0.99 | 0.93 |
| 11 | 78621360 | rs6592811 | -0.79 | -1.66 | 0.882 | -1.964 | -0.017 | -0.044 | -0.044 | 0 | 0.65 | 0.55 | 0.74 | 0.8 | 0.7 | 0.74 |
| 11 | 78625147 | rs531804 | 1.03 | -1.635 | 1.249 | -1.935 | 0.021 | 0.206 | 0.206 | 0.004 | 0.95 | 0.74 | 0.93 | 0.81 | 1 | 0.93 |
| 11 | 78625883 | rs504245 | 0.903 | -1.613 | 0.885 | -2.009 | 0.002 | 0.059 | 0.059 | 0 | 0.31 | 0.2 | 0.23 | 0.03 | 0.31 | 0.22 |
| 11 | 78629268 | rs12269747 | -1.38 | -1.711 | 1.046 | -2.034 | -0.036 | -0.079 | -0.079 | 0 | 0.13 | 0.64 | 0.08 | 0.01 | 0.07 | 0.01 |
| 11 | 78630720 | rs658312 | 0.97 | -1.74 | 0.814 | -2.092 | 0.006 | 0.012 | 0.012 | 0 | 0.29 | 0.21 | 0.18 | 0 | 0.28 | 0.17 |
| 11 | 78632658 | rs631854 | 0.812 | -1.698 | 1.136 | -1.959 | -0.032 | 0.099 | 0.099 | 0.543 | 0.88 | 0.63 | 0.9 | 0.81 | 0.93 | 0.92 |
| 11 | 78640751 | rs644779 | 0.619 | -1.699 | 0.074 | -2.023 | 0.034 | 0.179 | 0.179 | 0.004 | 0.52 | 0.04 | 0.64 | 0.78 | 0.62 | 0.7 |
| 11 | 78643342 | rs485091 | 1.086 | -1.552 | 0.259 | -1.979 | -0.007 | 0.123 | 0.123 | 0 | 0.26 | 0.07 | 0.14 | 0 | 0.25 | 0.14 |
| 11 | 78644581 | rs677624 | -0.288 | -1.333 | 0.475 | -1.891 | 0.002 | 0.003 | 0.003 | 0 | 0.62 | 0.37 | 0.72 | 0.77 | 0.7 | 0.72 |
| 11 | 78659955 | rs531767 | 0.127 | -1.149 | -0.124 | -1.956 | -0.048 | 0.057 | 0.057 | 0 | 0.61 | 0.83 | 0.39 | 0.31 | 0.5 | 0.44 |
| 11 | 78667936 | rs671290 | 0.663 | -1.297 | 0.452 | -2.076 | 0 | -0.012 | -0.012 | 0 | 0.51 | 0.49 | 0.36 | 0.31 | 0.47 | 0.41 |
| 11 | 78678340 | rs750878 | -1.402 | -1.346 | 0.961 | -1.989 | -0.029 | 0.111 | 0.111 | 2.389 | 0.11 | 0.45 | 0.1 | 0.23 | 0.03 | 0.1 |
| 11 | 78683692 | rs621543 | 0.032 | -1.728 | 0.498 | -2.65 | -0.021 | 0.166 | 0.166 | 0 | 0.53 | 0.8 | 0.43 | 0.22 | 0.53 | 0.45 |
| 11 | 78690093 | rs1025595 | 1.356 | -1.951 | 0.634 | -2.816 | -0.045 | 0.09 | 0.09 | 0 | 0.36 | 0.15 | 0.27 | 0.01 | 0.4 | 0.27 |
| 11 | 78691557 | rs660810 | 0.366 | -1.994 | 0.539 | -2.818 | -0.019 | 0.162 | 0.162 | 0.004 | 0.47 | 0.78 | 0.38 | 0.23 | 0.44 | 0.38 |
| 11 | 78692179 | rs647746 | 0.705 | -1.907 | 0.685 | -2.709 | -0.014 | 0.01 | 0.01 | 0 | 0.43 | 0.29 | 0.33 | 0.01 | 0.51 | 0.34 |
| 11 | 78694515 | rs10793350 | -0.216 | -1.607 | -0.017 | -2.649 | -0.026 | 0.087 | 0.087 | 0.56 | 0.39 | 0.17 | 0.55 | 0.75 | 0.44 | 0.6 |

| | | | | | | | | | | | | | | | | |
|---|---|---|---|---|---|---|---|---|---|---|---|---|---|---|---|---|
| 11 | 78698287 | rs649931 | 0.303 | -1.456 | 0.145 | -2.414 | -0.037 | 0.033 | 0.033 | 2.524 | 0.88 | 0.74 | 0.91 | 0.8 | 0.95 | 0.9 |
| 11 | 78702526 | rs589658 | 0.693 | -1.228 | -0.708 | -2.457 | -0.016 | 0.047 | 0.047 | 0 | 0.45 | 0.24 | 0.33 | 0.01 | 0.51 | 0.34 |
| 11 | 78704703 | rs7950661 | -0.362 | -0.392 | -1.304 | -1.707 | -0.043 | -0.005 | -0.005 | 0.242 | 0.24 | 0.1 | 0.5 | 0.72 | 0.32 | 0.56 |
| 11 | 78706607 | rs11237677 | 0.708 | -0.683 | -0.95 | -1.253 | 0.023 | 0.132 | 0.132 | 0.13 | 0.27 | 0.07 | 0.14 | 0.05 | 0.21 | 0.23 |
| 11 | 78707366 | rs892857 | -0.913 | -0.968 | -1.207 | -0.755 | 0.043 | 0.02 | 0.02 | 1.03 | 0.57 | 0.92 | 0.4 | 0.39 | 0.39 | 0.41 |
| 11 | 78708226 | rs7950630 | -0.721 | -1.012 | -0.063 | -1.037 | -0.064 | -0.1 | -0.1 | 0.009 | 0.21 | 0.49 | 0.21 | 0.06 | 0.14 | 0.05 |
| 11 | 78719912 | rs610398 | -0.569 | -0.076 | -0.217 | -0.696 | 0.001 | 0.003 | 0.003 | 0 | 0.56 | 0.55 | 0.4 | 0.39 | 0.55 | 0.58 |
| 11 | 78723149 | rs587767 | -0.357 | -0.087 | 0.015 | -1.259 | -0.001 | 0 | 0 | 0.019 | 0.51 | 0.57 | 0.33 | 0.06 | 0.5 | 0.42 |
| 11 | 78724109 | rs658182 | -0.551 | -0.096 | 0.045 | -1.236 | -0.001 | -0.036 | -0.036 | 0.034 | 0.53 | 0.53 | 0.34 | 0.11 | 0.5 | 0.42 |
| 11 | 78724586 | rs11237684 | -0.864 | -0.17 | -0.151 | -1.256 | -0.057 | -0.083 | -0.083 | 3.509 | 0.2 | 0.55 | 0.11 | 0.05 | 0.11 | 0.04 |
| 11 | 78725742 | rs2276380 | 1.285 | -0.071 | -0.273 | -1.422 | 0.036 | 0.124 | 0.124 | 0.43 | 0.27 | 0.06 | 0.15 | 0.05 | 0.21 | 0.23 |
| 11 | 78725925 | rs613011 | -1.227 | -0.01 | -0.149 | -1.373 | -0.001 | 0.02 | 0.02 | 0 | 0.59 | 0.65 | 0.37 | 0.12 | 0.55 | 0.45 |
| 11 | 78727548 | rs9804662 | 0.364 | 0.341 | -0.901 | -1.252 | -0.057 | 0.175 | 0.175 | 1.029 | 0.35 | 0.1 | 0.25 | 0.06 | 0.39 | 0.38 |
| 11 | 78728494 | rs603888 | 0.291 | 0.175 | -0.875 | -1.226 | -0.064 | 0.153 | 0.153 | 0.813 | 0.36 | 0.13 | 0.25 | 0.06 | 0.42 | 0.35 |
| 11 | 78728657 | rs11237688 | -0.725 | 0.47 | -0.347 | -0.725 | 0.003 | -0.008 | -0.008 | 0 | 0.13 | 0.12 | 0.11 | 0.01 | 0.2 | 0.15 |
| 11 | 78729980 | rs684380 | -0.852 | 0.408 | -0.443 | 0.025 | -0.007 | -0.023 | -0.023 | 0 | 0.64 | 0.73 | 0.52 | 0.49 | 0.62 | 0.57 |
| 11 | 78733570 | rs1435469 | -1.778 | 0.232 | -0.277 | -0.262 | -0.004 | 0.019 | 0.019 | 3.521 | 0.28 | 0.79 | 0.28 | 0.16 | 0.25 | 0.25 |
| 11 | 78735176 | rs2019214 | -1.595 | 0.297 | -0.265 | -0.226 | 0.006 | -0.025 | -0.025 | 0 | 0.26 | 0.67 | 0.25 | 0.15 | 0.25 | 0.25 |
| 11 | 78735404 | rs985271 | -1.456 | 0.409 | -0.191 | -0.222 | 0.033 | -0.087 | -0.087 | 0 | 0.23 | 0.58 | 0.16 | 0.07 | 0.24 | 0.24 |
| 11 | 78742084 | rs3893939 | -0.578 | 0.157 | -0.81 | 0.189 | -0.048 | 0.063 | 0.063 | 2.125 | 0.62 | 0.9 | 0.46 | 0.26 | 0.54 | 0.43 |
| 11 | 78749810 | rs10793353 | 1.764 | -0.057 | 0.466 | 1.167 | -0.011 | 0.052 | 0.052 | 0 | 0.74 | 0.42 | 0.65 | 0.47 | 0.77 | 0.57 |
| 11 | 78758369 | rs1821616 | -1.663 | 0.455 | 0.415 | 1.43 | -0.007 | 0.01 | 0.01 | 0 | 0.14 | 0.1 | 0.12 | 0.04 | 0.16 | 0.08 |
| 11 | 78760473 | rs1369458 | -0.925 | 0.68 | 0.485 | 1.735 | 0.05 | 0.028 | 0.028 | 0 | 0.09 | 0.19 | 0.15 | 0.11 | 0.13 | 0.07 |
| 11 | 78762627 | rs4944217 | 0.801 | 0.118 | 0.004 | 1.928 | -0.004 | 0.252 | 0.252 | 0 | 0.95 | 0.88 | 0.84 | 0.55 | 0.98 | 0.7 |
| 11 | 78764528 | rs11826637 | -0.963 | 0.242 | -0.243 | 0.642 | -0.011 | -0.01 | -0.01 | 0 | 0.06 | 0.09 | 0.05 | 0.04 | 0.06 | 0.06 |
| 11 | 78771634 | rs2016337 | -0.37 | 0.524 | -0.546 | 0.719 | -0.001 | -0.005 | -0.005 | 0 | 0.42 | 0.38 | 0.41 | 0.43 | 0.51 | 0.41 |
| 11 | 78772558 | rs747521 | -0.095 | 0.185 | -0.286 | 1.348 | -0.002 | 0.162 | 0.162 | 0 | 0.93 | 0.86 | 0.79 | 0.62 | 0.96 | 0.76 |
| 11 | 78776903 | rs11237706 | -0.05 | 0.075 | -0.927 | 0.452 | -0.016 | 0.013 | 0.013 | 0 | 0.4 | 0.19 | 0.24 | 0.31 | 0.36 | 0.32 |
| 11 | 78776973 | rs7937426 | -0.901 | 0.083 | -0.352 | 0.525 | -0.004 | 0.13 | 0.13 | 0 | 0.93 | 0.87 | 0.8 | 0.63 | 0.96 | 0.82 |
| 11 | 78781706 | rs4255563 | 0.311 | -0.384 | 0.269 | 0.198 | 0.003 | 0.034 | 0.034 | 0 | 0.49 | 0.38 | 0.51 | 0.32 | 0.58 | 0.48 |
| 11 | 78781962 | rs4614486 | 0.16 | -0.555 | -0.014 | -0.072 | -0.022 | 0 | 0 | 0 | 0.35 | 0.21 | 0.23 | 0.3 | 0.36 | 0.25 |
| 11 | 78784223 | rs10899586 | 0.231 | -0.65 | -0.018 | -0.008 | 0.001 | 0.023 | 0.023 | 0 | 0.49 | 0.57 | 0.49 | 0.42 | 0.52 | 0.48 |
| 11 | 78785279 | rs7116067 | 0.047 | -0.608 | -0.274 | 0.083 | -0.018 | 0.039 | 0.039 | 0 | 0.42 | 0.23 | 0.27 | 0.29 | 0.43 | 0.26 |
| 11 | 78787344 | rs7946005 | 0.398 | -1.349 | -0.735 | -0.247 | 0.06 | -0.072 | -0.072 | 0 | 0.83 | 0.94 | 0.8 | 0.72 | 0.96 | 0.81 |
| 11 | 78789853 | rs7114034 | -0.754 | -0.769 | -0.75 | -0.55 | -0.025 | -0.02 | -0.02 | 0 | 0.73 | 0.84 | 0.58 | 0.71 | 0.66 | 0.65 |
| 11 | 78791435 | rs11237708 | 0.083 | -0.749 | -0.566 | -0.327 | -0.009 | 0.022 | 0.022 | 0 | 0.39 | 0.25 | 0.25 | 0.28 | 0.43 | 0.25 |
| 11 | 78794102 | rs4944221 | -0.522 | -0.778 | -0.065 | -0.371 | 0.009 | -0.014 | -0.014 | 0 | 0.62 | 0.54 | 0.48 | 0.73 | 0.6 | 0.65 |
| 11 | 78794440 | rs4945324 | 0.712 | -0.513 | -0.173 | -0.168 | -0.031 | -0.041 | -0.041 | 0 | 0.2 | 0.09 | 0.16 | 0.23 | 0.24 | 0.17 |
| 11 | 78795359 | rs7105794 | 0.331 | -0.556 | 0.139 | -0.234 | -0.001 | 0.005 | 0.005 | 0 | 0.24 | 0.26 | 0.25 | 0.43 | 0.3 | 0.34 |
| 11 | 78797120 | rs11605580 | 0.342 | -0.613 | 0.164 | 0.376 | 0 | 0.006 | 0.006 | 0 | 0.23 | 0.26 | 0.24 | 0.41 | 0.28 | 0.28 |
| 11 | 78800171 | rs10793359 | -0.913 | -0.494 | -0.289 | 0.134 | 0 | -0.013 | -0.013 | 0 | 0.54 | 0.52 | 0.41 | 0.74 | 0.54 | 0.54 |
| 11 | 78801291 | rs7127492 | 0.911 | -0.347 | -0.565 | -0.184 | -0.003 | -0.029 | -0.029 | 0 | 0.37 | 0.28 | 0.3 | 0 | 0.39 | 0.26 |
| 11 | 78804058 | rs10793360 | -0.113 | -0.307 | -1.237 | 0.381 | -0.003 | -0.003 | -0.003 | 0 | 0.28 | 0.09 | 0.16 | 0.33 | 0.25 | 0.2 |
| 11 | 78806999 | rs1942884 | 0.169 | -0.147 | -1.312 | -0.22 | -0.023 | 0.094 | 0.094 | 0.202 | 0.24 | 0.05 | 0.12 | 0.06 | 0.21 | 0.11 |
| 11 | 78808312 | rs6592825 | -0.761 | 0.035 | -0.844 | 0.18 | 0.027 | 0.152 | 0.152 | 0 | 0.92 | 0.8 | 0.71 | 0.74 | 0.93 | 0.79 |
| 11 | 78808456 | rs7938684 | 0.36 | -0.153 | -1.582 | -0.97 | -0.059 | 0.228 | 0.228 | 0 | 0.4 | 0.07 | 0.3 | 0 | 0.44 | 0.3 |
| 11 | 78810793 | rs870332 | -0.472 | -0.977 | -0.832 | 0.24 | -0.016 | 0.004 | 0.004 | 0 | 0.25 | 0.22 | 0.24 | 0.25 | 0.12 | 0.13 |
| 11 | 78811360 | rs12789345 | 0.587 | -0.301 | -1.378 | -0.889 | -0.097 | 0.206 | 0.206 | 0 | 0.39 | 0.12 | 0.29 | 0 | 0.5 | 0.28 |
| 11 | 78813921 | rs2510135 | -0.965 | -0.638 | -0.252 | -0.167 | -0.035 | -0.006 | -0.006 | 0 | 0.24 | 0.38 | 0.42 | 0.76 | 0.16 | 0.36 |
| 11 | 78816852 | rs12419002 | -0.398 | -0.345 | 0.116 | 0.382 | 0 | 0.004 | 0.004 | 0 | 0.14 | 0.16 | 0.2 | 0.2 | 0.12 | 0.19 |
| 11 | 78817269 | rs7934777 | -0.29 | -0.313 | 0.113 | 0.378 | -0.004 | 0.006 | 0.006 | 0 | 0.14 | 0.16 | 0.2 | 0.21 | 0.11 | 0.19 |
| 11 | 78817367 | rs6592827 | -0.37 | -0.339 | 0.115 | 0.381 | 0.001 | 0.005 | 0.005 | 0 | 0.14 | 0.16 | 0.2 | 0.2 | 0.12 | 0.11 |
| 11 | 78818226 | rs2512080 | 0.421 | -0.757 | -0.142 | -0.022 | 0.001 | 0.005 | 0.005 | 0 | 0.61 | 0.76 | 0.62 | 0.56 | 0.66 | 0.58 |
| 11 | 78819123 | rs2512081 | 0.663 | -0.817 | 0.216 | -0.06 | -0.003 | 0.003 | 0.003 | 0 | 0.58 | 0.65 | 0.59 | 0.56 | 0.6 | 0.56 |
| 11 | 78819684 | rs7121856 | -0.108 | -0.999 | -0.028 | 0.364 | 0.032 | 0.005 | 0.005 | 0 | 0.68 | 0.85 | 0.58 | 0.28 | 0.75 | 0.56 |
| 11 | 78821660 | rs1942883 | 0.647 | -0.829 | 0.123 | 0.623 | 0.001 | 0.073 | 0.073 | 0.106 | 0.07 | 0.06 | 0.26 | 0.49 | 0.03 | 0.21 |
| 11 | 78821945 | rs9667214 | -0.463 | -0.72 | 0.795 | 0.21 | 0.055 | 0.038 | 0.038 | 0 | 0.15 | 0.4 | 0.21 | 0.18 | 0.17 | 0.11 |
| 11 | 78824323 | rs4483616 | -0.243 | -0.869 | 0.742 | -0.08 | 0.029 | -0.061 | -0.061 | 0 | 0.16 | 0.34 | 0.1 | 0.05 | 0.15 | 0.08 |
| 11 | 78825885 | rs7114639 | 1.426 | -1.131 | 0.03 | -0.381 | -0.027 | 0.193 | 0.193 | 0.212 | 0.46 | 0.19 | 0.32 | 0.13 | 0.54 | 0.35 |
| 11 | 78830648 | rs11237719 | 0.927 | -1.225 | 0.348 | 0.673 | -0.009 | 0.009 | 0.009 | 0 | 0.71 | 0.39 | 0.62 | 0.57 | 0.8 | 0.6 |
| 11 | 78833535 | rs10899592 | 0.721 | -1.236 | 0.407 | 0.573 | -0.019 | 0.045 | 0.045 | 0 | 0.73 | 0.41 | 0.52 | 0.44 | 0.79 | 0.58 |
| 11 | 78834421 | rs1942886 | 0.779 | -1.172 | 0.506 | 1.049 | -0.047 | 0.276 | 0.276 | 1.604 | 0.9 | 0.75 | 0.73 | 0.51 | 0.96 | 0.76 |
| 11 | 78857310 | rs7119070 | -0.438 | -0.624 | -0.231 | 0.005 | 0.029 | 0.003 | 0.003 | 0.145 | 0.47 | 0.68 | 0.31 | 0.44 | 0.39 | 0.49 |
| 11 | 78860381 | rs6592832 | -0.586 | -0.626 | -0.428 | 0.018 | 0.05 | -0.011 | -0.011 | 0 | 0.49 | 0.74 | 0.41 | 0.51 | 0.39 | 0.57 |
| 11 | 78864605 | rs11607746 | 0.842 | -0.725 | -0.439 | -0.257 | 0 | -0.001 | -0.001 | 0 | 0.2 | 0.19 | 0.11 | 0.09 | 0.2 | 0.25 |
| 11 | 78867290 | rs10793370 | 0.305 | -0.513 | -0.096 | -0.185 | 0.003 | 0.03 | 0.03 | 0.016 | 0.27 | 0.44 | 0.22 | 0.33 | 0.25 | 0.23 |
| 11 | 78868436 | rs10793371 | -0.79 | -0.224 | -0.929 | -0.045 | -0.009 | 0.128 | 0.128 | 0 | 0.86 | 0.79 | 0.61 | 0.46 | 0.84 | 0.72 |
| 11 | 78870685 | rs11237733 | 1.401 | 0.004 | -1.496 | -1.127 | -0.001 | 0.005 | 0.005 | 0 | 0.11 | 0.09 | 0.16 | 0.08 | 0.14 | 0.07 |
| 11 | 78871902 | rs533870 | -0.073 | 0.153 | -1.618 | -1.112 | -0.064 | 0.227 | 0.227 | 0.236 | 0.38 | 0.1 | 0.26 | 0.05 | 0.39 | 0.19 |

| | | | | | | | | | | | | | | | | |
|---|---|---|---|---|---|---|---|---|---|---|---|---|---|---|---|---|
| 11 | 78873362 | rs478090 | 0.043 | 0.139 | -0.669 | -0.202 | -0.002 | 0.001 | 0.001 | 1.677 | 0.87 | 0.82 | 0.72 | 0.87 | 0.88 | 0.88 |
| 11 | 78877428 | rs10899600 | 0.593 | 0.429 | -0.602 | -0.708 | -0.005 | 0.011 | 0.011 | 0 | 0.25 | 0.16 | 0.18 | 0 | 0.25 | 0.09 |
| 11 | 78879700 | rs7482532 | -0.044 | 0.754 | -0.169 | -0.288 | 0.079 | 0.077 | 0.077 | 0 | 0.21 | 0.36 | 0.21 | 0.45 | 0.35 | 0.31 |
| 11 | 78879841 | rs4359232 | -0.137 | 0.744 | 0.041 | -0.294 | 0.115 | 0.112 | 0.112 | 0 | 0.22 | 0.47 | 0.21 | 0.45 | 0.35 | 0.31 |
| 11 | 78891205 | rs7935280 | 0.027 | 0.421 | -0.338 | 0.108 | 0.004 | 0.034 | 0.034 | 0 | 0.31 | 0.52 | 0.26 | 0.44 | 0.33 | 0.34 |
| 11 | 78896834 | rs12575534 | 0.695 | -0.373 | -1.005 | 0.116 | 0.027 | -0.128 | -0.128 | 0.082 | 0.22 | 0.02 | 0.13 | 0.37 | 0.15 | 0.23 |
| 11 | 78909461 | rs10899604 | -0.537 | 0.616 | 0.085 | 0.323 | 0.012 | 0.031 | 0.031 | 0 | 0.31 | 0.54 | 0.28 | 0.44 | 0.36 | 0.38 |
| 11 | 78911432 | rs10899607 | -0.098 | 0.793 | -0.741 | -0.937 | -0.007 | -0.017 | -0.017 | 0 | 0.24 | 0.22 | 0.16 | 0 | 0.29 | 0.09 |
| 11 | 78915775 | rs1938821 | -0.788 | 0.047 | -0.265 | -0.498 | 0.004 | 0.022 | 0.022 | 0.188 | 0.28 | 0.38 | 0.41 | 0.63 | 0.33 | 0.47 |
| 11 | 78915933 | rs11606728 | -0.435 | -0.793 | -0.59 | -1.362 | 0.022 | 0.053 | 0.053 | 0 | 0.17 | 0.07 | 0.07 | 0.01 | 0.1 | 0.12 |
| 11 | 78918314 | rs10501435 | -0.797 | -0.851 | -0.234 | -1.421 | -0.027 | -0.118 | -0.118 | 0 | 0.31 | 0.38 | 0.16 | 0.03 | 0.22 | 0.27 |
| 11 | 78923102 | rs1008941 | -1.391 | -0.371 | -0.922 | -0.895 | 0.006 | -0.018 | -0.018 | 0 | 0.09 | 0.03 | 0.07 | 0.11 | 0.06 | 0.05 |
| 11 | 78924118 | rs12365619 | -0.954 | -0.82 | -0.612 | -1.07 | -0.021 | -0.024 | -0.024 | 0 | 0.35 | 0.46 | 0.2 | 0.21 | 0.25 | 0.31 |
| 11 | 78931338 | rs12222362 | -0.849 | -0.486 | -1.019 | -0.737 | 0 | 0.007 | 0.007 | 0 | 0.06 | 0.05 | 0.06 | 0.13 | 0.04 | 0.04 |
| 11 | 78934486 | rs522783 | 0.386 | -0.153 | -1.622 | -1.756 | -0.081 | 0.123 | 0.123 | 1.303 | 0.32 | 0.12 | 0.19 | 0.02 | 0.43 | 0.14 |
| 11 | 78938662 | rs613968 | 0.145 | -0.842 | -1.03 | -1.21 | 0.036 | 0.088 | 0.088 | 0 | 0.76 | 0.91 | 0.78 | 0.99 | 0.87 | 0.83 |
| 11 | 78939166 | rs616214 | 0.195 | -0.633 | -0.776 | -1.095 | 0.009 | 0.013 | 0.013 | 0 | 0.7 | 0.77 | 0.73 | 0.98 | 0.77 | 0.76 |
| 11 | 78939685 | rs11237770 | -1.592 | -0.536 | -0.045 | 0.292 | -0.029 | 0.075 | 0.075 | 0 | 0.12 | 0.3 | 0.14 | 0.17 | 0.07 | 0.05 |
| 11 | 78945216 | rs1011311 | -1.634 | -0.454 | -0.615 | 0.385 | 0.001 | -0.002 | -0.002 | 0 | 0.1 | 0.12 | 0.13 | 0.17 | 0.05 | 0.06 |
| 11 | 78947625 | rs17137917 | -1.171 | -1.305 | -1.224 | 0.248 | 0.058 | 0.001 | 0.001 | 0.057 | 0.21 | 0.08 | 0.09 | 0.19 | 0.1 | 0.15 |
| 11 | 78947777 | rs17137926 | -0.439 | -1.87 | -1.191 | 1.11 | -0.001 | 0.02 | 0.02 | 0.015 | 0.18 | 0.17 | 0.23 | 0.5 | 0.13 | 0.37 |
| 11 | 78950207 | rs4343056 | -1.175 | -1.32 | -1.589 | 1.673 | -0.017 | -0.037 | -0.037 | 0 | 0.17 | 0.11 | 0.25 | 0.33 | 0.26 | 0.28 |
| 11 | 78950572 | rs12273933 | -1.807 | -1.959 | -1.469 | 1.518 | 0.034 | 0.01 | 0.01 | 0 | 0.45 | 0.29 | 0.37 | 0.54 | 0.39 | 0.46 |
| 11 | 78951703 | rs486694 | -2.472 | -1.769 | -1.367 | 1.185 | -0.084 | 0.158 | 0.158 | 0.092 | 0.62 | 0.88 | 0.51 | 0.85 | 0.52 | 0.66 |
| 11 | 78954142 | rs670492 | -1.765 | -1.821 | -0.991 | 1.441 | -0.003 | -0.007 | -0.007 | 0 | 0.33 | 0.68 | 0.4 | 0.54 | 0.38 | 0.43 |
| 11 | 78955199 | rs12577910 | -0.343 | -1.754 | -1.032 | 1.241 | 0.051 | 0.021 | 0.021 | 0 | 0.29 | 0.14 | 0.13 | 0.22 | 0.15 | 0.19 |
| 11 | 78959356 | rs662533 | -0.551 | -1.608 | -0.065 | 0.66 | -0.001 | -0.011 | -0.011 | 0.433 | 0.14 | 0.23 | 0.1 | 0.1 | 0.12 | 0.15 |
| 11 | 78963236 | rs567834 | -2.45 | -1.533 | -0.196 | 0.618 | -0.082 | 0.12 | 0.12 | 0.098 | 0.62 | 0.87 | 0.5 | 0.79 | 0.53 | 0.65 |
| 11 | 78968453 | rs4540852 | -0.79 | -1.81 | -0.278 | 0.581 | -0.01 | -0.033 | -0.033 | 0 | 0.32 | 0.13 | 0.34 | 0.58 | 0.37 | 0.39 |
| 11 | 78973716 | rs495001 | 0.341 | -1.234 | -0.489 | 0.321 | -0.034 | -0.011 | -0.011 | 0.566 | 0.61 | 0.86 | 0.61 | 0.42 | 0.52 | 0.58 |
| 11 | 78973937 | rs493077 | 1.305 | -1.195 | 0.106 | 0.238 | 0.006 | 0.022 | 0.022 | 0.253 | 0.47 | 0.43 | 0.56 | 0.29 | 0.54 | 0.47 |
| 11 | 78975286 | rs570333 | 1.718 | -1.07 | 0.034 | 0.17 | -0.106 | 0.164 | 0.164 | 0.115 | 0.4 | 0.07 | 0.53 | 0.19 | 0.51 | 0.34 |
| 11 | 78977173 | rs656104 | -0.416 | -1.098 | 0.491 | 0.194 | 0 | -0.001 | -0.001 | 0 | 0.69 | 0.73 | 0.68 | 0.44 | 0.68 | 0.6 |
| 11 | 78993207 | rs499949 | -2.004 | -1.075 | 0.318 | 0.152 | -0.045 | 0.127 | 0.127 | 0.006 | 0.58 | 0.87 | 0.46 | 0.76 | 0.48 | 0.66 |
| 11 | 78995665 | rs586888 | -0.871 | -1.131 | 1.029 | 0.332 | 0.01 | 0.002 | 0.002 | 0 | 0.43 | 0.4 | 0.34 | 0.54 | 0.36 | 0.42 |
| 11 | 78999516 | rs96537 | -0.049 | -1.046 | 1.172 | 0.336 | -0.005 | 0.012 | 0.012 | 0 | 0.31 | 0.33 | 0.32 | 0.54 | 0.31 | 0.41 |
| 11 | 79000129 | rs642516 | -0.195 | -0.498 | 0.735 | 0.676 | 0.002 | -0.016 | -0.016 | 0 | 0.34 | 0.27 | 0.28 | 0.49 | 0.34 | 0.46 |
| 11 | 79001958 | rs660933 | 1.653 | 0.307 | 0.897 | 0.708 | -0.001 | 0.011 | 0.011 | 0 | 0.65 | 0.4 | 0.5 | 0.58 | 0.65 | 0.59 |
| 11 | 79003851 | rs487345 | 1.021 | 0.652 | 0.52 | -0.153 | 0 | -0.003 | -0.003 | 0.147 | 0.76 | 0.74 | 0.61 | 0.81 | 0.76 | 0.75 |
| 11 | 79008619 | rs505579 | 0.571 | 0.688 | 0.291 | -0.469 | 0.007 | 0.004 | 0.004 | 0 | 0.83 | 0.83 | 0.64 | 0.74 | 0.78 | 0.68 |
| 11 | 79008948 | rs11824380 | -0.823 | 0.794 | 0.289 | -0.748 | -0.001 | -0.032 | -0.032 | 0 | 0.18 | 0.26 | 0.08 | 0 | 0.15 | 0.08 |
| 11 | 79009040 | rs503765 | 0.813 | 0.692 | 0.546 | -0.66 | 0.015 | 0.005 | 0.005 | 0.043 | 0.81 | 0.68 | 0.63 | 0.75 | 0.78 | 0.68 |
| 11 | 79015993 | rs527774 | 0.207 | 0.663 | 0.198 | -0.637 | -0.002 | -0.002 | -0.002 | 0 | 0.84 | 0.87 | 0.65 | 0.75 | 0.8 | 0.7 |
| 11 | 79018805 | rs568082 | 1.474 | 0.514 | 0.008 | -0.848 | -0.001 | -0.007 | -0.007 | 0.326 | 0.63 | 0.32 | 0.56 | 0.75 | 0.67 | 0.63 |
| 11 | 79020289 | rs7931512 | -1.402 | 0.841 | -0.325 | -0.882 | 0.002 | -0.081 | -0.081 | 0.016 | 0.26 | 0.32 | 0.12 | 0 | 0.24 | 0.17 |
| 11 | 79022263 | rs11237782 | -0.354 | 0.727 | -0.685 | 0.072 | -0.012 | -0.045 | -0.045 | 0 | 0.07 | 0.01 | 0.11 | 0.2 | 0.06 | 0.22 |
| 11 | 79022608 | rs7950085 | 0.282 | 1.285 | -0.103 | -0.483 | 0.033 | -0.074 | -0.074 | 0.702 | 0.15 | 0.26 | 0.13 | 0.03 | 0.2 | 0.1 |
| 11 | 79022808 | rs11237783 | 0.473 | 1.111 | -0.361 | -0.351 | -0.002 | -0.021 | -0.021 | 0 | 0.22 | 0.17 | 0.15 | 0.36 | 0.19 | 0.12 |
| 11 | 79024513 | rs579715 | 1.292 | 0.685 | -0.712 | -0.907 | -0.017 | 0.033 | 0.033 | 0 | 0.33 | 0.11 | 0.33 | 0.27 | 0.35 | 0.53 |
| 11 | 79024716 | rs479540 | 0.317 | 0.679 | -0.522 | -0.904 | 0.002 | 0.027 | 0.027 | 0 | 0.49 | 0.41 | 0.46 | 0.3 | 0.55 | 0.65 |
| 11 | 79026374 | rs12803088 | 0.853 | 0.648 | -0.275 | -0.861 | 0.001 | 0.047 | 0.047 | 0 | 0.22 | 0.32 | 0.16 | 0.35 | 0.18 | 0.12 |
| 11 | 79027129 | rs1355424 | -0.111 | 0.484 | -0.215 | -1.08 | 0.002 | -0.094 | -0.094 | 0.412 | 0.26 | 0.32 | 0.16 | 0.04 | 0.27 | 0.2 |
| 11 | 79027193 | rs7951124 | 1.383 | 0.499 | -0.201 | -1.065 | 0.029 | -0.073 | -0.073 | 0 | 0.15 | 0.25 | 0.13 | 0.03 | 0.2 | 0.13 |
| 11 | 79030778 | rs518556 | -0.156 | 0.471 | -0.168 | -1.084 | 0.001 | -0.023 | -0.023 | 0 | 0.62 | 0.51 | 0.51 | 0.32 | 0.65 | 0.75 |
| 11 | 79031407 | rs562246 | 1.372 | 0.484 | -0.174 | -1.091 | -0.004 | 0.032 | 0.032 | 0 | 0.35 | 0.18 | 0.34 | 0.29 | 0.37 | 0.55 |
| 11 | 79031671 | rs495487 | -0.021 | 0.481 | -0.166 | -1.058 | 0 | -0.006 | -0.006 | 0 | 0.61 | 0.56 | 0.5 | 0.32 | 0.64 | 0.75 |
| 11 | 79039284 | rs11604024 | 0.872 | 0.499 | -0.118 | -0.979 | -0.007 | -0.027 | -0.027 | 0.037 | 0.12 | 0.12 | 0.03 | 0.01 | 0.07 | 0.08 |
| 11 | 79039527 | rs7940157 | 0.924 | 0.443 | -0.172 | -0.933 | -0.002 | -0.021 | -0.021 | 0 | 0.22 | 0.18 | 0.16 | 0.37 | 0.19 | 0.13 |
| 11 | 79039788 | rs4945341 | 0.055 | 0.246 | -0.476 | -0.848 | 0.012 | 0.014 | 0.014 | 0 | 0.37 | 0.43 | 0.29 | 0.39 | 0.38 | 0.25 |
| 11 | 79044218 | rs517684 | 0.997 | 0.282 | -1.359 | -0.866 | 0.002 | 0.034 | 0.034 | 0 | 0.37 | 0.23 | 0.35 | 0.29 | 0.39 | 0.59 |
| 11 | 79045008 | rs482711 | 1.173 | 0.294 | -1.35 | -0.86 | -0.009 | 0.037 | 0.037 | 0 | 0.36 | 0.17 | 0.35 | 0.29 | 0.39 | 0.57 |
| 11 | 79045185 | rs487552 | 0.359 | 0.411 | -1.338 | -0.809 | 0.016 | 0.057 | 0.057 | 0 | 0.47 | 0.31 | 0.37 | 0.29 | 0.46 | 0.64 |
| 11 | 79050521 | rs549394 | -0.591 | 0.85 | -1.321 | -0.059 | -0.022 | -0.056 | -0.056 | 0 | 0.85 | 0.97 | 0.66 | 0.67 | 0.81 | 0.86 |
| 11 | 79053401 | rs13377588 | -0.516 | 0.616 | -0.853 | 0.034 | -0.002 | -0.009 | -0.009 | 0.09 | 0.36 | 0.67 | 0.26 | 0.37 | 0.36 | 0.19 |
| 11 | 79054990 | rs4945343 | -0.483 | 0.475 | -0.992 | -0.067 | 0.015 | 0.029 | 0.029 | 0 | 0.43 | 0.64 | 0.28 | 0.31 | 0.33 | 0.52 |
| 11 | 79058250 | rs506396 | -0.098 | 0.761 | -1.316 | 0.581 | -0.009 | -0.038 | -0.038 | 0 | 0.78 | 0.86 | 0.73 | 0.63 | 0.78 | 0.67 |
| 11 | 79059444 | rs11237796 | 0.208 | 0.924 | -1.575 | 0.419 | -0.005 | 0.007 | 0.007 | 0 | 0.17 | 0.13 | 0.12 | 0.1 | 0.2 | 0.09 |
| 11 | 79060893 | rs498900 | -0.82 | 1.08 | -1.608 | 0.401 | -0.002 | -0.041 | -0.041 | 0 | 0.8 | 0.87 | 0.75 | 0.63 | 0.81 | 0.68 |
| 11 | 79062930 | rs17138171 | 0.299 | 0.833 | -1.455 | 0.345 | -0.005 | 0.004 | 0.004 | 0 | 0.13 | 0.11 | 0.09 | 0.08 | 0.18 | 0.08 |

| | | | | | | | | | | | | | | | | |
|---|---|---|---|---|---|---|---|---|---|---|---|---|---|---|---|---|
| 11 | 79064479 | rs10501438 | 0.74 | 0.715 | -1.085 | -0.32 | 0.013 | 0.012 | 0.012 | 0.005 | 0.38 | 0.3 | 0.25 | 0.15 | 0.27 | 0.34 |
| 11 | 79064582 | rs11237798 | 0.799 | 0.812 | -0.57 | -0.065 | 0.018 | 0.065 | 0.065 | 0 | 0.08 | 0.24 | 0.1 | 0.09 | 0.09 | 0.09 |
| 11 | 79065101 | rs530965 | -0.744 | 0.773 | -0.421 | -0.053 | 0.018 | 0.012 | 0.012 | 0.128 | 0.4 | 0.46 | 0.47 | 0.48 | 0.5 | 0.36 |
| 11 | 79065320 | rs508047 | 1.228 | 0.775 | -0.671 | 0.058 | -0.08 | -0.101 | -0.101 | 0.035 | 0.18 | 0.01 | 0.28 | 0.31 | 0.22 | 0.16 |
| 11 | 79065350 | rs507958 | -0.494 | 0.579 | -0.261 | -0.007 | 0.004 | 0.043 | 0.043 | 0 | 0.71 | 0.54 | 0.62 | 0.53 | 0.68 | 0.54 |
| 11 | 79074266 | rs492504 | -0.945 | 0.756 | -0.234 | -0.068 | -0.003 | -0.014 | -0.014 | 0 | 0.69 | 0.77 | 0.73 | 0.61 | 0.71 | 0.6 |
| 11 | 79077745 | rs471435 | 1.284 | 0.999 | -0.375 | -0.105 | 0.001 | 0.025 | 0.025 | 0.285 | 0.1 | 0.07 | 0.29 | 0.33 | 0.18 | 0.12 |
| 11 | 79078305 | rs12295799 | -0.547 | 0.863 | -0.122 | -0.426 | -0.009 | 0.009 | 0.009 | 1.225 | 0.6 | 0.7 | 0.44 | 0.3 | 0.52 | 0.48 |
| 11 | 79083249 | rs1944449 | 0.996 | 1.062 | 0.022 | -0.429 | 0.009 | -0.005 | -0.005 | 2.568 | 0.12 | 0.28 | 0.09 | 0.08 | 0.16 | 0.08 |
| 11 | 79083344 | rs41513944 | 0.373 | 0.531 | -0.382 | -0.92 | -0.011 | -0.019 | -0.019 | 1.155 | 0.32 | 0.36 | 0.21 | 0.2 | 0.29 | 0.33 |
| 11 | 79087280 | rs876197 | 0.411 | 0.526 | -0.379 | -0.899 | -0.009 | -0.019 | -0.019 | 0 | 0.31 | 0.36 | 0.22 | 0.21 | 0.3 | 0.33 |
| 11 | 79087541 | rs876196 | 1.062 | 0.47 | -0.393 | -0.969 | -0.008 | -0.048 | -0.048 | 0 | 0.23 | 0.24 | 0.13 | 0.08 | 0.21 | 0.23 |
| 11 | 79089255 | rs472870 | -0.696 | 0.468 | -0.421 | -0.559 | 0.011 | 0.003 | 0.003 | 0 | 0.52 | 0.61 | 0.56 | 0.56 | 0.6 | 0.51 |
| 11 | 79089400 | rs471234 | -0.162 | 0.54 | -0.419 | -0.513 | 0.004 | 0.003 | 0.003 | 0 | 0.41 | 0.5 | 0.49 | 0.48 | 0.45 | 0.44 |
| 11 | 79091227 | rs489257 | -1.522 | 0.31 | -0.639 | -0.659 | -0.007 | -0.032 | -0.032 | 0 | 0.63 | 0.78 | 0.6 | 0.58 | 0.61 | 0.59 |
| 11 | 79091605 | rs534842 | 0.353 | 0.411 | -0.534 | -0.719 | 0.007 | 0 | 0 | 0 | 0.28 | 0.26 | 0.37 | 0.36 | 0.35 | 0.35 |
| 11 | 79091835 | rs509196 | -0.967 | 0.354 | -0.497 | -0.747 | 0 | 0 | 0 | 0 | 0.52 | 0.52 | 0.52 | 0.51 | 0.5 | 0.52 |
| 11 | 79094374 | rs507245 | -0.866 | 0.411 | -0.432 | -0.686 | 0 | 0 | 0 | 0 | 0.47 | 0.42 | 0.51 | 0.56 | 0.55 | 0.51 |
| 11 | 79103804 | rs574864 | -0.483 | 0.456 | -0.445 | -0.638 | -0.004 | 0.001 | 0.001 | 0 | 0.39 | 0.34 | 0.43 | 0.44 | 0.47 | 0.43 |
| 11 | 79104058 | rs519985 | -0.515 | 0.455 | -0.34 | -0.638 | 0.014 | -0.004 | -0.004 | 0.255 | 0.4 | 0.47 | 0.45 | 0.44 | 0.47 | 0.43 |
| 11 | 79105799 | rs12275195 | 1.744 | 0.426 | -0.398 | -0.601 | -0.001 | -0.003 | -0.003 | 0.139 | 0.1 | 0.17 | 0.07 | 0.08 | 0.11 | 0.07 |
| 11 | 79106804 | rs11237821 | 1.654 | 0.427 | -0.4 | -0.6 | 0 | -0.001 | -0.001 | 0.03 | 0.1 | 0.17 | 0.07 | 0.08 | 0.11 | 0.07 |
| 11 | 79107136 | rs538804 | -1.62 | 0.472 | -0.542 | -0.484 | 0.002 | -0.005 | -0.005 | 0 | 0.63 | 0.72 | 0.67 | 0.63 | 0.65 | 0.6 |
| 11 | 79109590 | rs534863 | -1.631 | 0.255 | -0.632 | -0.875 | 0.001 | 0.021 | 0.021 | 0 | 0.7 | 0.63 | 0.63 | 0.59 | 0.68 | 0.62 |
| 11 | 79110892 | rs7925882 | 1.184 | 0.432 | -0.864 | -0.73 | 0.033 | 0.045 | 0.045 | 0.836 | 0.14 | 0.05 | 0.06 | 0.03 | 0.06 | 0.07 |
| 11 | 79112174 | rs574358 | -0.4 | 0.721 | -0.671 | -0.561 | 0.024 | -0.001 | -0.001 | 0 | 0.3 | 0.33 | 0.39 | 0.37 | 0.37 | 0.38 |
| 11 | 79119433 | rs4945348 | 1.17 | 0.831 | -0.601 | -0.354 | -0.047 | -0.057 | -0.057 | 0 | 0.09 | 0.01 | 0.27 | 0.27 | 0.11 | 0.11 |
| 11 | 79121342 | rs12286448 | 1.106 | 0.717 | -0.464 | -0.469 | -0.004 | -0.002 | -0.002 | 0 | 0.1 | 0.15 | 0.07 | 0.08 | 0.11 | 0.06 |
| 11 | 79122534 | rs951932 | -0.879 | 0.74 | -0.841 | -0.755 | 0.02 | 0.025 | 0.025 | 0.193 | 0.29 | 0.41 | 0.41 | 0.38 | 0.29 | 0.22 |
| 11 | 79124084 | rs1567540 | 0.259 | 0.356 | -1.065 | -0.781 | 0.019 | 0.029 | 0.029 | 0 | 0.23 | 0.19 | 0.13 | 0.06 | 0.16 | 0.09 |
| 11 | 79124344 | rs872490 | -1.011 | 0.622 | -0.787 | -0.78 | -0.002 | 0 | 0 | 0 | 0.39 | 0.46 | 0.46 | 0.42 | 0.33 | 0.34 |
| 11 | 79125661 | rs7121592 | -1.165 | 0.687 | -0.826 | -0.709 | -0.001 | 0 | 0 | 0 | 0.44 | 0.52 | 0.54 | 0.54 | 0.4 | 0.42 |
| 11 | 79129411 | rs10501442 | 0.834 | 1.482 | -1.006 | -0.686 | -0.01 | 0.038 | 0.038 | 0 | 0.09 | 0.05 | 0.06 | 0 | 0.11 | 0.03 |
| 11 | 79132636 | rs10899620 | -0.171 | 1.674 | -0.651 | -0.357 | 0.045 | 0.088 | 0.088 | 1.112 | 0.16 | 0.27 | 0.33 | 0.38 | 0.17 | 0.22 |
| 11 | 79140682 | rs11827071 | 0.375 | 1.834 | -0.785 | -0.403 | -0.02 | 0.032 | 0.032 | 0 | 0.14 | 0.07 | 0.08 | 0.04 | 0.21 | 0.2 |
| 11 | 79142159 | rs10899625 | 1.312 | 1.818 | -0.914 | -0.271 | -0.044 | -0.03 | -0.03 | 0.38 | 0.07 | 0.01 | 0.3 | 0.33 | 0.11 | 0.13 |
| 11 | 79149013 | rs7116789 | -0.599 | 1.105 | -0.768 | -0.525 | 0.002 | 0 | 0 | 1.157 | 0.31 | 0.26 | 0.35 | 0.34 | 0.37 | 0.41 |
| 11 | 79158020 | rs1397726 | -1.233 | 1.104 | -1.1 | -0.356 | 0.01 | -0.027 | -0.027 | 0.206 | 0.45 | 0.72 | 0.55 | 0.5 | 0.52 | 0.54 |
| 11 | 79165928 | rs1511239 | -0.988 | 1.104 | -0.621 | -1.069 | -0.024 | 0.041 | 0.041 | 0 | 0.23 | 0.12 | 0.16 | 0.05 | 0.31 | 0.22 |
| 11 | 79167107 | rs7941024 | -1.322 | 0.9 | 0.057 | -0.774 | 0.005 | -0.008 | -0.008 | 0 | 0.26 | 0.27 | 0.22 | 0.17 | 0.33 | 0.32 |
| 11 | 79172339 | rs7106671 | -1.716 | 0.434 | 0.497 | -0.965 | -0.008 | -0.072 | -0.072 | 0 | 0.29 | 0.57 | 0.18 | 0.09 | 0.26 | 0.24 |
| 11 | 79173082 | rs1912702 | -2.109 | 0.373 | 0.451 | -1.014 | 0.005 | 0.014 | 0.014 | 0.159 | 0.39 | 0.67 | 0.27 | 0.21 | 0.37 | 0.38 |
| 11 | 79174422 | rs17138383 | -0.639 | 0.449 | 0.618 | -1.007 | 0.034 | -0.088 | -0.088 | 0 | 0.2 | 0.38 | 0.15 | 0.05 | 0.24 | 0.19 |
| 11 | 79175172 | rs7129961 | -0.504 | 0.476 | 0.237 | -0.993 | 0.01 | -0.041 | -0.041 | 0 | 0.19 | 0.23 | 0.14 | 0.05 | 0.24 | 0.19 |
| 11 | 79175852 | rs12285327 | 0.353 | 0.641 | 0.263 | -0.926 | 0.014 | -0.026 | -0.026 | 0 | 0.06 | 0.11 | 0.04 | 0.01 | 0.09 | 0.16 |
| 11 | 79181603 | rs6592843 | -1.932 | 0.551 | 0.252 | -0.965 | -0.002 | -0.018 | -0.018 | 0 | 0.35 | 0.43 | 0.25 | 0.23 | 0.36 | 0.4 |
| 11 | 79183377 | rs11237851 | -1.906 | 0.611 | 0.297 | -0.98 | 0.002 | -0.007 | -0.007 | 0 | 0.29 | 0.34 | 0.24 | 0.19 | 0.36 | 0.34 |
| 11 | 79183802 | rs12290359 | 0.76 | 0.634 | 0.27 | -0.976 | -0.029 | 0.027 | 0.027 | 0 | 0.05 | 0 | 0.03 | 0.01 | 0.1 | 0.16 |
| 11 | 79184840 | rs7117322 | -0.622 | 0.736 | 0.301 | -1.038 | -0.014 | -0.021 | -0.021 | 0 | 0.1 | 0.23 | 0.04 | 0.05 | 0.05 | 0.07 |
| 11 | 79185191 | rs7119862 | -2.291 | 0.731 | 0.08 | -1.024 | 0.001 | -0.029 | -0.029 | 0 | 0.39 | 0.57 | 0.37 | 0.22 | 0.4 | 0.41 |
| 11 | 79190280 | rs12294899 | -1.705 | 0.803 | 0.248 | -0.961 | 0.027 | -0.052 | -0.052 | 0 | 0.24 | 0.45 | 0.31 | 0.12 | 0.29 | 0.24 |
| 11 | 79190783 | rs2510506 | -1.957 | 0.749 | -0.091 | -1.318 | -0.004 | 0.014 | 0.014 | 0.234 | 0.55 | 0.61 | 0.49 | 0.32 | 0.56 | 0.44 |
| 11 | 79191457 | rs2512637 | 0.586 | 0.704 | -0.093 | -1.259 | -0.003 | 0 | 0 | 0.38 | 0.11 | 0.11 | 0.1 | 0.11 | 0.16 | 0.04 |
| 11 | 79192521 | rs17138430 | -0.307 | 0.628 | -0.037 | -1.354 | -0.006 | -0.032 | -0.032 | 0 | 0.29 | 0.33 | 0.27 | 0.07 | 0.21 | 0.11 |
| 11 | 79194378 | rs2510502 | -1.049 | 0.722 | -0.019 | -1.249 | -0.009 | 0 | 0 | 0 | 0.26 | 0.2 | 0.21 | 0.22 | 0.33 | 0.29 |
| 11 | 79194834 | rs12273202 | -0.571 | 0.507 | -0.015 | -1.447 | 0.002 | -0.031 | -0.031 | 0 | 0.31 | 0.38 | 0.38 | 0.09 | 0.38 | 0.23 |
| 11 | 79197184 | rs12271629 | -1.793 | -0.033 | 0.351 | -1.032 | -0.008 | -0.014 | -0.014 | 0 | 0.33 | 0.63 | 0.23 | 0.19 | 0.29 | 0.35 |
| 11 | 79197201 | rs12271634 | 0.018 | -0.046 | 0.152 | -1.045 | 0.006 | -0.009 | -0.009 | 0 | 0.14 | 0.07 | 0.12 | 0.18 | 0.12 | 0.12 |
| 11 | 79198096 | rs7931162 | -1.483 | 0.062 | 0.105 | -0.953 | -0.021 | -0.032 | -0.032 | 0 | 0.33 | 0.56 | 0.22 | 0.21 | 0.3 | 0.35 |
| 11 | 79198990 | rs10501445 | -0.287 | 0.348 | -0.314 | -0.8 | 0.014 | -0.036 | -0.036 | 0 | 0.1 | 0.15 | 0.05 | 0.01 | 0.14 | 0.17 |
| 11 | 79202219 | rs7112396 | -1.423 | 0.316 | -0.693 | -0.828 | -0.052 | 0.039 | 0.039 | 0 | 0.57 | 0.84 | 0.52 | 0.32 | 0.52 | 0.47 |
| 11 | 79207175 | rs7358419 | -1.124 | -0.689 | -0.09 | -0.679 | -0.039 | -0.04 | -0.04 | 1.646 | 0.31 | 0.53 | 0.21 | 0.19 | 0.23 | 0.24 |
| 11 | 79213474 | rs3935311 | -0.702 | -0.445 | -0.02 | -0.32 | -0.001 | -0.001 | -0.001 | 0 | 0.42 | 0.37 | 0.35 | 0.37 | 0.35 | 0.23 |
| 11 | 79213595 | rs7120805 | -0.84 | 0.024 | 0.36 | -0.235 | 0.077 | 0.02 | 0.02 | 0 | 0.11 | 0.39 | 0.13 | 0.09 | 0.12 | 0.18 |
| 11 | 79216085 | rs4073501 | -0.737 | -0.727 | 0.247 | -0.205 | -0.009 | 0.048 | 0.048 | 0.128 | 0.19 | 0.3 | 0.26 | 0.36 | 0.19 | 0.18 |
| 11 | 79222314 | rs7944096 | -0.413 | -1.189 | 0.346 | -0.687 | -0.045 | -0.09 | -0.09 | 1.044 | 0.31 | 0.6 | 0.18 | 0.08 | 0.19 | 0.23 |
| 11 | 79223720 | rs7111768 | 0.143 | -1.225 | 0.406 | -0.708 | 0.055 | 0.086 | 0.086 | 0 | 0.26 | 0.08 | 0.12 | 0.08 | 0.17 | 0.22 |
| 11 | 79230538 | rs11237881 | -0.292 | -1.23 | 0.882 | -0.535 | -0.003 | -0.002 | -0.002 | 0.49 | 0.29 | 0.3 | 0.28 | 0.22 | 0.2 | 0.24 |
| 11 | 79230921 | rs7951609 | -1.307 | -1.367 | 0.757 | -0.668 | 0.016 | 0.015 | 0.015 | 0 | 0.43 | 0.62 | 0.33 | 0.22 | 0.27 | 0.31 |

| | | | | | | | | | | | | | | | | |
|---|---|---|---|---|---|---|---|---|---|---|---|---|---|---|---|---|
| 11 | 79232115 | rs11237882 | -0.208 | -1.299 | 0.386 | -0.574 | 0.019 | 0.013 | 0.013 | 0 | 0.29 | 0.22 | 0.26 | 0.26 | 0.2 | 0.24 |
| 11 | 79242837 | rs10501447 | 0.04 | -1.173 | 0.325 | -0.609 | 0.054 | 0.073 | 0.073 | 0 | 0.26 | 0.08 | 0.12 | 0.11 | 0.18 | 0.22 |
| 11 | 79249708 | rs17138584 | 0.022 | -1.304 | 0.749 | -0.612 | -0.062 | -0.062 | -0.062 | 0.177 | 0.24 | 0.4 | 0.14 | 0.11 | 0.14 | 0.2 |
| 11 | 79259776 | rs4944231 | 1.11 | -1.125 | 0.754 | -0.799 | 0.036 | 0.055 | 0.055 | 0 | 0.51 | 0.41 | 0.65 | 0.82 | 0.66 | 0.66 |
| 11 | 79270366 | rs1944718 | 1.454 | -0.956 | 0.633 | -0.502 | 0.003 | 0.015 | 0.015 | 0 | 0.42 | 0.3 | 0.55 | 0.57 | 0.54 | 0.55 |
| 11 | 79275128 | rs1944719 | 1.606 | -1.04 | 0.677 | -0.582 | -0.047 | 0.007 | 0.007 | 0 | 0.4 | 0.13 | 0.51 | 0.59 | 0.53 | 0.52 |
| 11 | 79286576 | rs951181 | 0.012 | -1.339 | 0.827 | -0.673 | 0.021 | 0.02 | 0.02 | 0 | 0.3 | 0.2 | 0.11 | 0.11 | 0.15 | 0.19 |
| 11 | 79287376 | rs10899651 | -0.068 | -1.364 | 1.07 | -0.409 | -0.054 | -0.036 | -0.036 | 0 | 0.31 | 0.41 | 0.17 | 0.18 | 0.16 | 0.21 |
| 11 | 79287622 | rs1944708 | -1.831 | -1.163 | 0.92 | -0.32 | -0.001 | 0.009 | 0.009 | 0 | 0.57 | 0.77 | 0.38 | 0.43 | 0.46 | 0.42 |
| 11 | 79289862 | rs7106044 | 0.435 | -1.023 | 0.858 | -0.185 | -0.001 | 0.012 | 0.012 | 0 | 0.07 | 0.08 | 0.11 | 0.23 | 0.11 | 0.08 |
| 11 | 79290539 | rs7126543 | -1.244 | -1.092 | 0.9 | -0.213 | 0.043 | 0.011 | 0.011 | 0 | 0.2 | 0.4 | 0.2 | 0.24 | 0.28 | 0.18 |
| 11 | 79293794 | rs12272156 | -1.203 | -1.076 | 0.521 | -0.205 | -0.028 | -0.008 | -0.008 | 0 | 0.2 | 0.11 | 0.18 | 0.25 | 0.29 | 0.18 |
| 11 | 79303053 | rs885361 | -0.823 | -1.064 | 0.544 | -0.21 | -0.022 | 0.001 | 0.001 | 0 | 0.18 | 0.12 | 0.18 | 0.19 | 0.27 | 0.17 |
| 11 | 79304168 | rs11237930 | 0.445 | -1.725 | 0.437 | -0.48 | 0.005 | 0.004 | 0.004 | 0 | 0.27 | 0.2 | 0.11 | 0.08 | 0.13 | 0.13 |
| 11 | 79310397 | rs1944720 | -0.732 | -1.759 | 0.137 | 0.329 | 0.004 | 0.009 | 0.009 | 0 | 0.66 | 0.6 | 0.54 | 0.53 | 0.45 | 0.68 |
| 11 | 79310962 | rs10899654 | 0.821 | -2.247 | -0.418 | 0.186 | 0.063 | 0.053 | 0.053 | 0 | 0.37 | 0.22 | 0.15 | 0.17 | 0.17 | 0.26 |
| 11 | 79311305 | rs12419045 | 0.927 | -1.23 | -0.351 | 0.819 | 0.006 | -0.075 | -0.075 | 0.026 | 0.13 | 0.02 | 0.2 | 0.25 | 0.1 | 0.33 |
| 11 | 79319964 | rs2187445 | -0.672 | -0.747 | 0.6 | 0.387 | 0.009 | -0.081 | -0.081 | 0 | 0.14 | 0.3 | 0.1 | 0 | 0.14 | 0.11 |
| 11 | 79321956 | rs7117769 | -0.513 | -0.754 | 0.648 | 0.376 | 0.012 | -0.069 | -0.069 | 2.617 | 0.13 | 0.22 | 0.1 | 0 | 0.16 | 0.09 |
| 11 | 79322485 | rs1944723 | -0.975 | -0.743 | 0.819 | 0.355 | 0.028 | -0.098 | -0.098 | 0 | 0.14 | 0.39 | 0.11 | 0 | 0.17 | 0.11 |
| 11 | 79329406 | rs4944237 | 0.34 | 0.134 | 0.39 | 0.447 | -0.048 | -0.035 | -0.035 | 0 | 0.63 | 0.8 | 0.41 | 0.39 | 0.46 | 0.55 |
| 11 | 79333777 | rs2187448 | -1.036 | 0.148 | 0.535 | 0.234 | -0.014 | -0.058 | -0.058 | 0 | 0.88 | 0.98 | 0.87 | 0.76 | 0.88 | 0.81 |
| 11 | 79340398 | rs10899660 | 1.314 | -0.03 | 0.558 | 0.134 | 0.013 | 0.023 | 0.023 | 0 | 0.45 | 0.37 | 0.28 | 0.16 | 0.32 | 0.34 |
| 11 | 79340799 | rs600439 | -0.097 | 0.079 | -0.046 | 0.125 | -0.053 | -0.041 | -0.041 | 0.383 | 0.64 | 0.8 | 0.48 | 0.4 | 0.47 | 0.56 |
| 11 | 79343468 | rs497101 | -0.685 | 0.214 | -0.053 | 0.186 | -0.044 | -0.04 | -0.04 | 0 | 0.74 | 0.81 | 0.49 | 0.36 | 0.61 | 0.66 |
| 11 | 79345711 | rs11237944 | 1.574 | 0.158 | -0.059 | -0.007 | 0.038 | 0.138 | 0.138 | 0 | 0.42 | 0.13 | 0.26 | 0.13 | 0.3 | 0.33 |
| 11 | 79345862 | rs949083 | 1.419 | 0.324 | -0.388 | 0.093 | -0.001 | -0.026 | -0.026 | 0 | 0.46 | 0.46 | 0.25 | 0.09 | 0.34 | 0.37 |
| 11 | 79349355 | rs526442 | -0.673 | 0.424 | -0.464 | 0.316 | -0.063 | -0.071 | -0.071 | 0 | 0.76 | 0.85 | 0.57 | 0.4 | 0.62 | 0.68 |
| 11 | 79360768 | rs4945363 | 1.303 | 0.426 | -0.343 | -0.093 | -0.009 | -0.04 | -0.04 | 0.049 | 0.43 | 0.4 | 0.23 | 0.08 | 0.31 | 0.33 |
| 11 | 79363044 | rs534408 | -0.59 | 0.873 | -0.466 | 0.103 | -0.011 | 0 | 0 | 0 | 0.25 | 0.12 | 0.21 | 0.26 | 0.29 | 0.28 |
| 11 | 79364829 | rs6592860 | 1.348 | 0.844 | -0.47 | 0.022 | -0.011 | -0.051 | -0.051 | 0 | 0.46 | 0.46 | 0.27 | 0.09 | 0.32 | 0.35 |
| 11 | 79365725 | rs669834 | -0.883 | 2.089 | -0.878 | 1.151 | -0.003 | -0.005 | -0.005 | 0 | 0.8 | 0.79 | 0.77 | 0.67 | 0.75 | 0.81 |
| 11 | 79366322 | rs17138705 | 0.333 | 1.982 | -0.994 | 1.487 | -0.006 | 0.018 | 0.018 | 0.069 | 0.07 | 0.07 | 0.27 | 0.32 | 0.14 | 0.15 |
| 11 | 79369861 | rs538490 | -0.495 | 1.857 | -1.042 | 1.283 | -0.031 | -0.062 | -0.062 | 0 | 0.19 | 0.06 | 0.21 | 0.48 | 0.22 | 0.26 |
| 11 | 79372389 | rs561832 | -1.427 | 1.59 | -1.091 | 1.158 | -0.042 | -0.042 | -0.042 | 3.645 | 0.3 | 0.19 | 0.48 | 0.58 | 0.42 | 0.45 |
| 11 | 79373772 | rs528643 | -1.29 | 1.631 | -1.264 | 1.043 | -0.068 | -0.073 | -0.073 | 0 | 0.28 | 0.12 | 0.31 | 0.5 | 0.4 | 0.37 |
| 11 | 79374757 | rs498603 | 0.865 | 0.988 | -1.107 | 0.489 | 0.029 | 0.084 | 0.084 | 0 | 0.49 | 0.28 | 0.37 | 0.26 | 0.38 | 0.35 |
| 11 | 79374856 | rs1944702 | -0.482 | 0.855 | -1.257 | 0.921 | -0.007 | -0.023 | -0.023 | 0 | 0.21 | 0.14 | 0.16 | 0.52 | 0.28 | 0.31 |
| 11 | 79376133 | rs1000960 | -0.893 | 0.028 | -1.359 | 0.199 | 0.03 | 0.015 | 0.015 | 0.261 | 0.37 | 0.79 | 0.39 | 0.29 | 0.31 | 0.37 |
| 11 | 79377304 | rs2156666 | -0.256 | 0.101 | -0.508 | -0.024 | 0.037 | -0.096 | -0.096 | 0 | 0.14 | 0.33 | 0.12 | 0 | 0.18 | 0.09 |
| 11 | 79377736 | rs2154749 | -0.48 | -0.417 | -0.669 | 0.3 | -0.049 | 0.052 | 0.052 | 0.78 | 0.21 | 0.36 | 0.25 | 0.29 | 0.12 | 0.27 |
| 11 | 79385728 | rs7940535 | 1.911 | -1.193 | -1.323 | -0.589 | 0.045 | 0.042 | 0.042 | 0.107 | 0.2 | 0.15 | 0.06 | 0 | 0.11 | 0.04 |
| 11 | 79388988 | rs12793036 | 0.849 | -0.427 | -1.837 | -0.353 | -0.056 | -0.043 | -0.043 | 0 | 0.13 | 0.01 | 0.12 | 0.12 | 0.17 | 0.22 |
| 11 | 79396636 | rs10793391 | 0.595 | -0.304 | -1.336 | -0.294 | -0.031 | 0.005 | 0.005 | 1.238 | 0.33 | 0.48 | 0.31 | 0.31 | 0.22 | 0.26 |
| 11 | 79398295 | rs7120855 | 2.242 | 0.236 | -1.097 | 0.231 | 0.001 | 0.023 | 0.023 | 0.102 | 0.08 | 0.05 | 0.04 | 0.2 | 0.07 | 0.11 |
| 11 | 79408549 | rs10899669 | -0.801 | -0.286 | -1.272 | -0.016 | -0.044 | 0.024 | 0.024 | 0.099 | 0.28 | 0.47 | 0.15 | 0.29 | 0.16 | 0.29 |
| 11 | 79415407 | rs2105527 | 0.809 | 0.695 | -1.911 | 0.259 | -0.067 | -0.068 | -0.068 | 0 | 0.13 | 0.03 | 0.16 | 0.26 | 0.28 | 0.14 |
| 11 | 79430411 | rs2252626 | 1.107 | 0.558 | -1.593 | 0.071 | 0.026 | 0.069 | 0.069 | 0 | 0.11 | 0.15 | 0.4 | 0.27 | 0.16 | 0.18 |
| 11 | 79431479 | rs2457249 | -0.997 | 0.495 | -1.58 | -0.028 | 0.021 | 0.015 | 0.015 | 0 | 0.34 | 0.51 | 0.55 | 0.58 | 0.38 | 0.46 |
| 11 | 79432876 | rs2252969 | -0.393 | 0.052 | -1.16 | -0.026 | 0.055 | 0.053 | 0.053 | 0 | 0.26 | 0.36 | 0.52 | 0.59 | 0.29 | 0.42 |
| 11 | 79439740 | rs2253740 | -0.548 | 0.14 | -1.072 | 0.048 | 0.021 | 0.051 | 0.051 | 0.016 | 0.27 | 0.42 | 0.53 | 0.58 | 0.28 | 0.42 |
| 11 | 79441694 | rs11237982 | -0.07 | 0.072 | -1.144 | 0.047 | -0.001 | 0.003 | 0.003 | 0.031 | 0.19 | 0.24 | 0.16 | 0.3 | 0.15 | 0.24 |
| 11 | 79443210 | rs2257071 | -0.77 | -0.383 | -1.196 | -0.704 | -0.091 | -0.025 | -0.025 | 0 | 0.66 | 0.85 | 0.69 | 0.61 | 0.53 | 0.56 |
| 11 | 79444242 | rs2257193 | 0.001 | -0.555 | -0.86 | -0.473 | 0.002 | 0.039 | 0.039 | 0.208 | 0.28 | 0.51 | 0.54 | 0.6 | 0.26 | 0.41 |
| 11 | 79453054 | rs582607 | -0.819 | -0.72 | -0.974 | -0.417 | -0.1 | -0.036 | -0.036 | 0 | 0.64 | 0.84 | 0.62 | 0.6 | 0.5 | 0.54 |
| 11 | 79455097 | rs473659 | -0.178 | -0.23 | -0.586 | -0.064 | -0.003 | -0.007 | -0.007 | 0 | 0.3 | 0.2 | 0.43 | 0.57 | 0.27 | 0.38 |
| 11 | 79461013 | rs1955059 | -0.268 | -0.29 | -0.581 | -0.162 | -0.002 | 0.002 | 0.002 | 0 | 0.7 | 0.75 | 0.56 | 0.43 | 0.75 | 0.6 |
| 11 | 79471264 | rs2851426 | -0.325 | -0.159 | -0.475 | -0.157 | 0.018 | 0.059 | 0.059 | 0.259 | 0.73 | 0.52 | 0.55 | 0.44 | 0.73 | 0.61 |
| 11 | 79471329 | rs2851427 | -0.893 | -0.109 | -0.917 | -0.131 | -0.005 | -0.007 | -0.007 | 0.509 | 0.77 | 0.84 | 0.58 | 0.44 | 0.75 | 0.64 |
| 11 | 79471974 | rs2851428 | -0.135 | 0.472 | -0.675 | -0.174 | 0.001 | 0 | 0 | 0 | 0.39 | 0.63 | 0.4 | 0.56 | 0.47 | 0.45 |
| 11 | 79474160 | rs7943985 | 0.389 | -0.633 | -1.246 | -1.085 | 0.038 | 0.025 | 0.025 | 0 | 0.32 | 0.26 | 0.15 | 0.01 | 0.24 | 0.21 |
| 11 | 79474606 | rs1944962 | 0.134 | -0.273 | -2.019 | -0.806 | -0.035 | 0.005 | 0.005 | 0 | 0.39 | 0.78 | 0.42 | 0.34 | 0.49 | 0.4 |
| 11 | 79474831 | rs12798906 | -0.161 | -0.43 | -1.686 | -0.997 | 0.014 | 0.192 | 0.192 | 0 | 0.27 | 0.02 | 0.12 | 0 | 0.22 | 0.18 |
| 11 | 79485137 | rs12270115 | -1.016 | -0.068 | -0.506 | -0.545 | -0.013 | -0.071 | -0.071 | 1.182 | 0.15 | 0.31 | 0.1 | 0.03 | 0.12 | 0.12 |
| 11 | 79486072 | rs2663203 | 0.773 | 0.185 | -0.696 | -0.459 | 0.008 | 0.023 | 0.023 | 0.328 | 0.5 | 0.31 | 0.48 | 0.44 | 0.5 | 0.45 |
| 11 | 79490662 | rs11237992 | -0.143 | 0.376 | -0.422 | -0.476 | -0.002 | -0.049 | -0.049 | 0.061 | 0.14 | 0.23 | 0.11 | 0.03 | 0.14 | 0.12 |
| 11 | 79491498 | rs2851432 | 0.777 | -0.246 | -0.756 | -0.718 | 0.008 | 0.058 | 0.058 | 1.395 | 0.49 | 0.16 | 0.44 | 0.39 | 0.46 | 0.43 |
| 11 | 79492245 | rs2851433 | 0.78 | 0.015 | -1.032 | -0.699 | -0.031 | -0.084 | -0.084 | 0 | 0.16 | 0.01 | 0.29 | 0.27 | 0.18 | 0.16 |
| 11 | 79495044 | rs1944951 | -0.352 | 0.553 | 0.135 | -0.6 | 0.002 | -0.083 | -0.083 | 0 | 0.15 | 0.39 | 0.15 | 0.03 | 0.13 | 0.15 |

| | | | | | | | | | | | | | | | | |
|---|---|---|---|---|---|---|---|---|---|---|---|---|---|---|---|---|
| 11 | 79506098 | rs10501453 | -0.255 | 0.689 | -0.183 | -0.607 | 0 | -0.013 | -0.013 | 0 | 0.13 | 0.09 | 0.1 | 0.04 | 0.14 | 0.19 |
| 11 | 79514622 | rs11238001 | 0.843 | 0.637 | -0.417 | -0.451 | -0.001 | 0.039 | 0.039 | 0.195 | 0.52 | 0.17 | 0.53 | 0.45 | 0.49 | 0.5 |
| 11 | 79516611 | rs4945375 | 0.405 | 0.733 | 0.315 | -0.016 | 0.006 | 0.009 | 0.009 | 0.599 | 0.61 | 0.42 | 0.66 | 0.58 | 0.57 | 0.58 |
| 11 | 79522805 | rs10793397 | 0.911 | 0.746 | 0.369 | -0.072 | 0.002 | 0.017 | 0.017 | 0.442 | 0.53 | 0.35 | 0.5 | 0.46 | 0.49 | 0.52 |
| 11 | 79525889 | rs17138958 | 1.018 | 0.72 | 0.317 | -0.063 | 0 | 0.01 | 0.01 | 0 | 0.06 | 0.04 | 0.07 | 0.11 | 0.06 | 0.04 |
| 11 | 79526471 | rs495966 | 1.177 | 0.7 | 0.325 | -0.086 | 0.031 | 0.054 | 0.054 | 0.112 | 0.49 | 0.15 | 0.45 | 0.45 | 0.47 | 0.48 |
| 11 | 79526602 | rs494942 | -0.522 | 1.18 | 0.628 | -0.146 | 0.006 | 0.035 | 0.035 | 0.367 | 0.76 | 0.7 | 0.71 | 0.62 | 0.73 | 0.74 |
| 11 | 79527366 | rs12287037 | -0.105 | 1.144 | 0.662 | -0.224 | 0.005 | -0.035 | -0.035 | 0 | 0.17 | 0.23 | 0.13 | 0.03 | 0.19 | 0.17 |
| 11 | 79529436 | rs7940110 | 0.895 | 1.13 | 0.353 | -0.177 | 0 | 0.009 | 0.009 | 0 | 0.06 | 0.04 | 0.07 | 0.13 | 0.06 | 0.04 |
| 11 | 79529774 | rs512725 | 0.811 | 1.112 | 0.285 | -0.299 | 0.001 | 0.001 | 0.001 | 0.146 | 0.57 | 0.41 | 0.56 | 0.58 | 0.53 | 0.55 |
| 11 | 79540133 | rs11238005 | -1.485 | 0.892 | 0.512 | -0.405 | 0.014 | -0.05 | -0.05 | 0.706 | 0.29 | 0.55 | 0.26 | 0.15 | 0.29 | 0.3 |
| 11 | 79545097 | rs639093 | 1.498 | 1.092 | 0.053 | -0.598 | -0.004 | 0.046 | 0.046 | 0.076 | 0.47 | 0.13 | 0.51 | 0.43 | 0.48 | 0.44 |
| 11 | 79548466 | rs11238008 | -0.099 | 0.997 | 0.204 | -0.591 | -0.012 | 0.026 | 0.026 | 0 | 0.17 | 0.11 | 0.11 | 0.02 | 0.2 | 0.17 |
| 11 | 79557080 | rs633761 | 1.255 | 0.912 | 0.222 | -0.435 | -0.012 | 0.065 | 0.065 | 0.148 | 0.52 | 0.19 | 0.62 | 0.4 | 0.5 | 0.46 |
| 11 | 79569363 | rs17139017 | -1.083 | 0.916 | 0.893 | -0.693 | 0.014 | -0.12 | -0.12 | 0 | 0.22 | 0.4 | 0.16 | 0.03 | 0.21 | 0.19 |
| 11 | 79570988 | rs283315 | 0.467 | 0.494 | 0.747 | -0.695 | -0.014 | 0.024 | 0.024 | 0 | 0.71 | 0.8 | 0.74 | 0.96 | 0.65 | 0.78 |
| 11 | 79573985 | rs283296 | 0.571 | 0.5 | 0.81 | -0.691 | 0.021 | -0.1 | -0.1 | 0.201 | 0.68 | 0.57 | 0.77 | 0.96 | 0.64 | 0.75 |
| 11 | 79584760 | rs283312 | 0.352 | 0.509 | 0.52 | -0.632 | -0.004 | -0.009 | -0.009 | 0 | 0.7 | 0.72 | 0.74 | 0.96 | 0.65 | 0.77 |
| 11 | 79598935 | rs4945383 | 0.661 | -0.06 | 0.004 | 0.331 | 0.003 | -0.051 | -0.051 | 0 | 0.35 | 0.11 | 0.42 | 0.42 | 0.34 | 0.26 |
| 11 | 79605008 | rs7122784 | 0.048 | -0.041 | 0.087 | 0.35 | -0.013 | 0.004 | 0.004 | 0 | 0.42 | 0.52 | 0.5 | 0.57 | 0.35 | 0.34 |
| 11 | 79611172 | rs7944347 | -0.517 | 0.057 | 0.003 | 0.084 | -0.016 | 0.028 | 0.028 | 0 | 0.05 | 0.09 | 0.07 | 0.15 | 0.01 | 0.08 |
| 11 | 79623323 | rs378921 | -0.1 | 0.438 | 0.192 | -0.57 | 0.008 | -0.068 | -0.068 | 0.116 | 0.76 | 0.67 | 0.78 | 0.95 | 0.74 | 0.83 |
| 11 | 79624615 | rs601208 | 0.324 | 0.243 | -0.657 | -0.707 | -0.063 | 0.032 | 0.032 | 0.055 | 0.36 | 0.07 | 0.43 | 0.28 | 0.35 | 0.42 |
| 11 | 79631786 | rs624285 | -0.04 | -0.246 | 0.493 | -0.174 | 0.003 | 0.032 | 0.032 | 0.098 | 0.41 | 0.31 | 0.34 | 0.72 | 0.44 | 0.43 |
| 11 | 79641647 | rs17139078 | -1.158 | 0.012 | 0.58 | 0.029 | 0 | 0.006 | 0.006 | 0 | 0.06 | 0.07 | 0.25 | 0.13 | 0.06 | 0.13 |
| 11 | 79648501 | rs636471 | -0.499 | -0.23 | 0.515 | 0.019 | 0.002 | -0.002 | -0.002 | 0.12 | 0.13 | 0.1 | 0.3 | 0.16 | 0.08 | 0.12 |
| 11 | 79655778 | rs660821 | -1.117 | 0.294 | 0.944 | 0.404 | 0.003 | 0.026 | 0.026 | 0 | 0.06 | 0.11 | 0.28 | 0.15 | 0.06 | 0.12 |
| 11 | 79660947 | rs679598 | -2.044 | 0.255 | 1.547 | 0.382 | 0.032 | 0.092 | 0.092 | 0 | 0.08 | 0.29 | 0.29 | 0.15 | 0.06 | 0.12 |
| 11 | 79665932 | rs563836 | -1.783 | 0.255 | 1.441 | 0.382 | 0.009 | 0.038 | 0.038 | 0 | 0.07 | 0.19 | 0.29 | 0.14 | 0.06 | 0.12 |
| 11 | 79667382 | rs587677 | -2.529 | 0.222 | 1.761 | 0.366 | 0.044 | 0.089 | 0.089 | 1.325 | 0.09 | 0.47 | 0.32 | 0.14 | 0.06 | 0.12 |
| 11 | 79673825 | rs658282 | -1.156 | 0.282 | 1.18 | 0.388 | 0 | 0.005 | 0.005 | 0 | 0.06 | 0.08 | 0.28 | 0.14 | 0.06 | 0.11 |
| 11 | 79683501 | rs607301 | -0.388 | -0.367 | 1.207 | 0.27 | 0.005 | 0.028 | 0.028 | 0 | 0.42 | 0.34 | 0.31 | 0.68 | 0.44 | 0.43 |
| 11 | 79685121 | rs625168 | -0.259 | -0.378 | 1.314 | 0.634 | 0.008 | 0.015 | 0.015 | 0 | 0.39 | 0.2 | 0.3 | 0.64 | 0.4 | 0.41 |
| 11 | 79687057 | rs595762 | -0.743 | -0.377 | 1.495 | 0.591 | 0 | 0.05 | 0.05 | 0 | 0.46 | 0.34 | 0.58 | 0.87 | 0.49 | 0.55 |
| 11 | 79688804 | rs642770 | -0.711 | -0.343 | 1.425 | 0.556 | 0.01 | 0.071 | 0.071 | 0 | 0.44 | 0.3 | 0.57 | 0.86 | 0.44 | 0.53 |
| 11 | 79695028 | rs10899700 | 0.26 | -0.339 | 1.156 | 1.177 | -0.002 | -0.006 | -0.006 | 0 | 0.15 | 0.11 | 0.14 | 0.36 | 0.19 | 0.26 |
| 11 | 79697979 | rs486935 | -1.303 | -0.201 | 1.526 | 0.268 | -0.014 | 0.168 | 0.168 | 0.007 | 0.55 | 0.79 | 0.62 | 0.9 | 0.5 | 0.58 |
| 11 | 79700145 | rs585747 | -0.818 | -0.208 | 1.571 | 0.274 | 0.037 | 0.073 | 0.073 | 0 | 0.49 | 0.68 | 0.33 | 0.66 | 0.44 | 0.46 |
| 11 | 79702309 | rs7108921 | -1.128 | -0.195 | 1.558 | 0.299 | 0.003 | 0.05 | 0.05 | 0 | 0.06 | 0.11 | 0.28 | 0.22 | 0.06 | 0.12 |
| 11 | 79704875 | rs2000518 | -0.78 | -0.192 | 1.539 | 0.293 | 0.001 | 0.027 | 0.027 | 0 | 0.06 | 0.08 | 0.28 | 0.21 | 0.06 | 0.12 |
| 11 | 79705861 | rs635035 | -0.963 | -0.191 | 1.385 | 0.258 | 0.037 | 0.168 | 0.168 | 0.006 | 0.52 | 0.75 | 0.61 | 0.87 | 0.44 | 0.56 |
| 11 | 79711944 | rs685111 | -1.234 | -0.103 | 1.323 | 0.371 | -0.037 | 0.182 | 0.182 | 0.259 | 0.56 | 0.82 | 0.61 | 0.87 | 0.49 | 0.56 |
| 11 | 79716752 | rs11824682 | -1.349 | 0.108 | 1.528 | 0.524 | 0.031 | 0.128 | 0.128 | 0 | 0.07 | 0.22 | 0.3 | 0.22 | 0.06 | 0.12 |
| 11 | 79717254 | rs681744 | -0.225 | 0.125 | 1.227 | 0.474 | 0.005 | 0.04 | 0.04 | 0 | 0.43 | 0.21 | 0.3 | 0.65 | 0.43 | 0.43 |
| 11 | 79720302 | rs645028 | -1.161 | 0.169 | 1.358 | 0.151 | -0.001 | 0.011 | 0.011 | 0 | 0.54 | 0.56 | 0.59 | 0.91 | 0.49 | 0.57 |
| 11 | 79721239 | rs1493071 | -0.154 | 0.512 | 0.502 | 0.41 | 0.009 | 0.045 | 0.045 | 0 | 0.4 | 0.2 | 0.29 | 0.68 | 0.39 | 0.46 |
| 11 | 79723318 | rs7123279 | -1.802 | 0.947 | 1.26 | 1.43 | 0.045 | 0.139 | 0.139 | 0 | 0.11 | 0.37 | 0.35 | 0.28 | 0.12 | 0.14 |
| 11 | 79729111 | rs6592880 | -1.481 | 1.063 | 1.075 | 1.191 | -0.001 | -0.024 | -0.024 | 0 | 0.26 | 0.54 | 0.22 | 0.21 | 0.25 | 0.19 |
| 11 | 79729315 | rs6592881 | -2.059 | 0.88 | 1.214 | 1.343 | 0.008 | 0.001 | 0.001 | 0.457 | 0.34 | 0.77 | 0.51 | 0.33 | 0.32 | 0.3 |
| 11 | 79731180 | rs11238055 | -1.217 | 0.984 | 1.049 | 1.308 | 0.003 | 0.011 | 0.011 | 0 | 0.11 | 0.22 | 0.32 | 0.13 | 0.12 | 0.11 |
| 11 | 79739095 | rs7940615 | -0.624 | 1.049 | 1.341 | 1.025 | -0.003 | 0.001 | 0.001 | 0 | 0.15 | 0.23 | 0.14 | 0.16 | 0.12 | 0.13 |
| 11 | 79756073 | rs7102102 | -2.204 | 1.086 | 0.815 | 1.496 | -0.017 | 0.059 | 0.059 | 1.359 | 0.46 | 0.86 | 0.61 | 0.65 | 0.45 | 0.5 |
| 11 | 79758667 | rs956168 | -2.028 | 1.373 | 0.788 | 1.465 | -0.021 | 0.04 | 0.04 | 1.202 | 0.44 | 0.86 | 0.58 | 0.64 | 0.43 | 0.5 |
| 11 | 79758832 | rs7936903 | -1.13 | 1.378 | 1.664 | 1.521 | 0 | 0.11 | 0.11 | 0 | 0.25 | 0.46 | 0.4 | 0.48 | 0.24 | 0.32 |
| 11 | 79763387 | rs7123693 | -0.93 | 1.285 | 1.745 | 1.564 | -0.016 | 0.058 | 0.058 | 0 | 0.22 | 0.32 | 0.16 | 0.46 | 0.17 | 0.26 |
| 11 | 79767089 | rs2186587 | -2.48 | 1.473 | 1.44 | 1.651 | -0.028 | 0.051 | 0.051 | 0.58 | 0.48 | 0.85 | 0.41 | 0.63 | 0.45 | 0.47 |
| 11 | 79768051 | rs1893814 | -1.364 | 1.589 | 1.863 | 1.889 | 0.005 | 0.042 | 0.042 | 0.246 | 0.26 | 0.47 | 0.2 | 0.68 | 0.26 | 0.37 |
| 11 | 79768810 | rs10488781 | -1.076 | 1.484 | 1.892 | 2.02 | -0.022 | 0.07 | 0.07 | 0 | 0.23 | 0.37 | 0.17 | 0.39 | 0.18 | 0.26 |
| 11 | 79774868 | rs4285894 | -0.393 | 1.634 | 1.292 | 1.642 | 0.006 | -0.006 | -0.006 | 0 | 0.13 | 0.11 | 0.14 | 0.15 | 0.14 | 0.14 |
| 11 | 79778062 | rs7104011 | -1.906 | 1.65 | 1.464 | 1.558 | 0.011 | 0.054 | 0.054 | 0.889 | 0.4 | 0.7 | 0.36 | 0.83 | 0.4 | 0.48 |
| 11 | 79781947 | rs11238074 | -0.816 | 1.542 | 0.732 | 1.536 | -0.011 | -0.006 | -0.006 | 0 | 0.13 | 0.12 | 0.16 | 0.16 | 0.19 | 0.15 |
| 11 | 79788235 | rs7936033 | -0.542 | 1.329 | 0.644 | 1.427 | -0.009 | -0.011 | -0.011 | 0.895 | 0.12 | 0.07 | 0.11 | 0.14 | 0.15 | 0.13 |
| 11 | 79792235 | rs12291943 | -0.33 | 1.279 | 0.701 | 1.427 | -0.002 | -0.003 | -0.003 | 0.68 | 0.11 | 0.11 | 0.12 | 0.14 | 0.15 | 0.12 |
| 11 | 79795334 | rs2105580 | -0.638 | 0.664 | 1.103 | 1.735 | -0.036 | 0.15 | 0.15 | 0 | 0.18 | 0.38 | 0.17 | 0.36 | 0.12 | 0.2 |
| 11 | 79797471 | rs7111422 | -0.808 | 0.781 | 0.417 | 0.443 | 0.075 | -0.027 | -0.027 | 0 | 0.08 | 0.26 | 0.06 | 0.01 | 0.1 | 0.04 |
| 11 | 79797550 | rs1944469 | -1.051 | 0.339 | -0.012 | 0.66 | -0.048 | 0.011 | 0.011 | 0 | 0.25 | 0.46 | 0.38 | 0.71 | 0.18 | 0.34 |
| 11 | 79799533 | rs680999 | 0.524 | 0.929 | -1.244 | -0.111 | -0.008 | 0.179 | 0.179 | 0.543 | 0.45 | 0.19 | 0.29 | 0.15 | 0.45 | 0.26 |
| 11 | 79800147 | rs602813 | 0.607 | 1.355 | -0.896 | 0.469 | -0.007 | 0.004 | 0.004 | 0.722 | 0.56 | 0.34 | 0.42 | 0.39 | 0.5 | 0.52 |
| 11 | 79802256 | rs7933762 | -0.197 | 0.895 | -0.811 | 0.152 | 0.006 | -0.005 | -0.005 | 0 | 0.1 | 0.05 | 0.03 | 0 | 0.04 | 0.05 |

| | | | | | | | | | | | | | | | |
|---|---|---|---|---|---|---|---|---|---|---|---|---|---|---|---|
| 11 | 79803310 | rs10897545 | -0.82 | 1.366 | -0.26 | 0.515 | 0.024 | 0.032 | 0.032 | 0 | 0.18 | 0.53 | 0.22 | 0.22 | 0.17 | 0.13 |
| 11 | 79806254 | rs7114202 | -0.574 | 2.092 | 0.016 | 0.812 | 0.049 | 0.179 | 0.179 | 0.016 | 0.11 | 0.26 | 0.19 | 0.49 | 0.22 | 0.31 |
| 11 | 79810435 | rs7937471 | -1.711 | 2.138 | 0.543 | 0.846 | 0.054 | 0.074 | 0.074 | 0 | 0.26 | 0.44 | 0.31 | 0.63 | 0.37 | 0.42 |
| 11 | 79811268 | rs509876 | -1.601 | 1.281 | -0.467 | -0.376 | -0.021 | 0.091 | 0.091 | 0 | 0.83 | 0.99 | 0.68 | 0.96 | 0.87 | 0.83 |
| 11 | 79812112 | rs11821650 | 0.645 | 1.252 | -0.372 | -0.293 | -0.01 | -0.048 | -0.048 | 1.134 | 0.12 | 0.18 | 0.05 | 0 | 0.05 | 0.06 |
| 11 | 79812352 | rs10488780 | 0.118 | 1.251 | -0.167 | -0.162 | 0.031 | 0.027 | 0.027 | 0 | 0.15 | 0.27 | 0.16 | 0.24 | 0.21 | 0.17 |
| 11 | 79813011 | rs10792458 | -1.894 | 1.251 | -0.298 | -0.137 | -0.018 | 0.019 | 0.019 | 0 | 0.39 | 0.76 | 0.44 | 0.7 | 0.42 | 0.49 |
| 11 | 79814921 | rs6591873 | -1.913 | 1.236 | -0.295 | -0.107 | -0.021 | 0.019 | 0.019 | 0 | 0.39 | 0.77 | 0.43 | 0.7 | 0.42 | 0.48 |
| 11 | 79815845 | rs12802458 | 2.928 | 0.897 | -0.743 | -0.365 | -0.053 | 0.091 | 0.091 | 0.28 | 0.27 | 0.01 | 0.11 | 0.06 | 0.25 | 0.07 |
| 11 | 79815969 | rs7937121 | -0.633 | 0.96 | -0.163 | 0.023 | 0.068 | 0.25 | 0.25 | 0 | 0.13 | 0.4 | 0.24 | 0.46 | 0.18 | 0.25 |
| 11 | 79817111 | rs10488779 | -0.953 | 0.791 | -0.304 | -0.139 | -0.011 | -0.027 | -0.027 | 0 | 0.26 | 0.45 | 0.19 | 0.25 | 0.25 | 0.2 |
| 11 | 79817326 | rs998024 | -1.291 | 0.801 | -0.349 | -0.167 | -0.001 | -0.042 | -0.042 | 0 | 0.28 | 0.49 | 0.21 | 0.24 | 0.26 | 0.23 |
| 11 | 79827051 | rs12293066 | 0.224 | 0.831 | -0.288 | -0.054 | 0.003 | -0.002 | -0.002 | 0.347 | 0.12 | 0.1 | 0.11 | 0.14 | 0.15 | 0.11 |
| 11 | 79837805 | rs7102821 | 1.19 | 0.749 | -0.462 | 0.128 | -0.024 | 0.013 | 0.013 | 0 | 0.53 | 0.18 | 0.36 | 0.35 | 0.51 | 0.43 |
| 11 | 79847868 | rs1569169 | 1.141 | 0.687 | -0.015 | 0.4 | 0 | 0.008 | 0.008 | 0 | 0.55 | 0.4 | 0.3 | 0.33 | 0.52 | 0.41 |
| 11 | 79848654 | rs10897559 | -0.262 | 0.69 | -0.011 | 0.396 | 0.016 | 0.026 | 0.026 | 0 | 0.7 | 0.83 | 0.51 | 0.76 | 0.74 | 0.7 |
| 11 | 79853993 | rs7129373 | 1.012 | 0.631 | 0.102 | 0.414 | 0 | 0 | 0 | 0 | 0.58 | 0.49 | 0.31 | 0.34 | 0.53 | 0.42 |
| 11 | 79858138 | rs11231961 | 0.889 | 0.843 | 0.085 | 0.518 | -0.004 | 0.034 | 0.034 | 0.033 | 0.06 | 0.05 | 0.14 | 0.32 | 0.14 | 0.23 |
| 11 | 79863662 | rs1892942 | 1.761 | 0.3 | -0.212 | 0.167 | 0.001 | 0.061 | 0.061 | 0 | 0.5 | 0.2 | 0.28 | 0.3 | 0.49 | 0.38 |
| 11 | 79873110 | rs1528635 | -0.266 | 0.376 | -0.425 | 0.215 | -0.009 | 0.011 | 0.011 | 0 | 0.74 | 0.82 | 0.52 | 0.75 | 0.71 | 0.7 |
| 11 | 79873627 | rs11231963 | 1.77 | 0.468 | -0.716 | 0.058 | -0.004 | 0.033 | 0.033 | 0 | 0.51 | 0.36 | 0.29 | 0.3 | 0.47 | 0.37 |
| 11 | 79881442 | rs2178239 | 1.316 | 1.028 | -0.587 | 0.281 | 0 | -0.001 | -0.001 | 3.084 | 0.57 | 0.46 | 0.31 | 0.31 | 0.5 | 0.4 |
| 11 | 79890119 | rs12361052 | -0.156 | 1.493 | -0.915 | 0.691 | -0.009 | 0.027 | 0.027 | 0 | 0.07 | 0.05 | 0.17 | 0.38 | 0.19 | 0.26 |
| 11 | 79903425 | rs7114526 | 1.028 | 1.29 | -1.257 | -0.185 | -0.037 | 0.208 | 0.208 | 0 | 0.54 | 0.82 | 0.42 | 0.87 | 0.53 | 0.49 |
| 11 | 79905426 | rs2141418 | 0.11 | 1.17 | -1.166 | -0.158 | 0.011 | 0.068 | 0.068 | 0 | 0.77 | 0.9 | 0.68 | 0.96 | 0.81 | 0.76 |
| 11 | 79907395 | rs10897563 | -0.109 | 1.2 | -0.437 | 0.626 | -0.006 | 0.282 | 0.282 | 1.014 | 0.06 | 0.18 | 0.09 | 0.44 | 0.03 | 0.09 |
| 11 | 79907430 | rs10792473 | -0.752 | 1.374 | -0.383 | 0.645 | 0.067 | 0.26 | 0.26 | 1.08 | 0.15 | 0.42 | 0.19 | 0.49 | 0.18 | 0.27 |
| 11 | 79907506 | rs1609335 | 0.504 | 1.272 | -1.295 | 0.054 | -0.054 | 0.082 | 0.082 | 0.695 | 0.79 | 0.98 | 0.63 | 0.94 | 0.76 | 0.76 |
| 11 | 79919361 | rs947983 | 0.151 | 1.438 | -0.92 | 0.321 | -0.032 | 0.047 | 0.047 | 0 | 0.09 | 0.01 | 0.07 | 0 | 0.11 | 0.1 |
| 11 | 79923025 | rs10897564 | 1.487 | 1.313 | -1.175 | 0.402 | -0.014 | -0.003 | -0.003 | 2.862 | 0.42 | 0.18 | 0.25 | 0.34 | 0.43 | 0.24 |
| 11 | 79928481 | rs12577870 | 1.905 | 1.47 | -1.384 | 0.401 | -0.038 | -0.085 | -0.085 | 0 | 0.36 | 0.05 | 0.18 | 0.34 | 0.32 | 0.18 |
| 11 | 79940214 | rs1919294 | -0.132 | 1.087 | -0.643 | 0.639 | 0.006 | 0.005 | 0.005 | 0 | 0.15 | 0.14 | 0.07 | 0.13 | 0.11 | 0.31 |
| 11 | 79941934 | rs1628460 | -1.684 | 1.319 | -0.668 | 0.501 | 0.023 | 0.005 | 0.005 | 0 | 0.49 | 0.78 | 0.57 | 0.61 | 0.45 | 0.62 |
| 11 | 79942880 | rs1784070 | -0.753 | 1.624 | -0.607 | 0.56 | -0.031 | -0.001 | -0.001 | 0 | 0.93 | 1 | 0.86 | 0.96 | 0.91 | 0.88 |
| 11 | 79950079 | rs11231986 | -0.172 | 1.433 | 0.132 | 0.544 | -0.014 | 0.024 | 0.024 | 0 | 0.27 | 0.19 | 0.29 | 0.11 | 0.34 | 0.26 |
| 11 | 79956462 | rs11231987 | -0.949 | 1.455 | 0.139 | 1.34 | -0.021 | -0.074 | -0.074 | 0 | 0.14 | 0.05 | 0.12 | 0.38 | 0.16 | 0.35 |
| 11 | 79967803 | rs1151220 | 1.077 | 1.531 | 0.582 | 1.402 | 0.028 | 0.122 | 0.122 | 0 | 0.85 | 0.62 | 0.65 | 0.67 | 0.84 | 0.81 |
| 11 | 79977371 | rs2619206 | -1.811 | 1.547 | 0.079 | 1.447 | -0.009 | 0.003 | 0.003 | 0 | 0.1 | 0.07 | 0.35 | 0.57 | 0.13 | 0.16 |
| 11 | 79978108 | rs1784077 | -1.789 | 1.583 | 0.079 | 1.456 | -0.01 | 0.003 | 0.003 | 0 | 0.1 | 0.07 | 0.35 | 0.57 | 0.13 | 0.16 |
| 11 | 79980097 | rs11232015 | -0.192 | 1.86 | -0.215 | 1.222 | -0.066 | -0.024 | -0.024 | 0 | 0.15 | 0.01 | 0.19 | 0.15 | 0.2 | 0.27 |
| 11 | 79981075 | rs10751017 | -0.331 | 1.813 | -0.254 | 0.644 | -0.014 | 0.023 | 0.023 | 0 | 0.73 | 0.82 | 0.76 | 0.82 | 0.65 | 0.7 |
| 11 | 79981321 | rs1479310 | 2.482 | 1.042 | 0.057 | 0.496 | 0.007 | 0.018 | 0.018 | 0 | 0.32 | 0.28 | 0.15 | 0.1 | 0.26 | 0.09 |
| 11 | 79982173 | rs1479312 | -0.722 | 1.088 | -0.091 | 0.58 | -0.003 | 0.054 | 0.054 | 0 | 0.08 | 0.07 | 0.33 | 0.56 | 0.09 | 0.15 |
| 11 | 79983340 | rs7945479 | 0.437 | 1.183 | 0.093 | 0.522 | 0 | -0.003 | -0.003 | 0 | 0.14 | 0.14 | 0.12 | 0.09 | 0.11 | 0.15 |
| 11 | 79990576 | rs12278930 | 2.744 | 1.196 | -0.457 | 0.772 | -0.05 | 0.083 | 0.083 | 0.003 | 0.3 | 0.02 | 0.12 | 0.1 | 0.27 | 0.09 |
| 11 | 79991191 | rs2619210 | -1.7 | 1.149 | -0.319 | 0.773 | 0.014 | -0.018 | -0.018 | 0.013 | 0.28 | 0.76 | 0.43 | 0.57 | 0.19 | 0.35 |
| 11 | 79995714 | rs1382775 | 0.242 | 1.095 | -0.247 | 0.681 | -0.035 | 0.111 | 0.111 | 0.018 | 0.6 | 0.83 | 0.64 | 0.76 | 0.52 | 0.48 |
| 11 | 80004464 | rs1144765 | 1.276 | 0.987 | -0.303 | 0.87 | -0.002 | 0.031 | 0.031 | 0 | 0.4 | 0.3 | 0.47 | 0.67 | 0.39 | 0.26 |
| 11 | 80008120 | rs971443 | -0.065 | 1.055 | -0.469 | 0.678 | -0.001 | -0.008 | -0.008 | 0 | 0.05 | 0.06 | 0.06 | 0 | 0.12 | 0.05 |
| 11 | 80011790 | rs7483049 | -1.53 | 0.606 | -0.377 | 0.58 | 0.019 | -0.014 | -0.014 | 1.039 | 0.59 | 0.31 | 0.55 | 0.51 | 0.68 | 0.81 |
| 11 | 80012510 | rs1382776 | -0.636 | 0.767 | -0.723 | 0.413 | -0.033 | 0.083 | 0.083 | 0 | 0.31 | 0.11 | 0.28 | 0.17 | 0.37 | 0.44 |
| 11 | 80013336 | rs11232059 | -0.135 | 0.702 | -0.061 | 0.442 | -0.002 | 0.013 | 0.013 | 0.188 | 0.16 | 0.27 | 0.11 | 0.16 | 0.13 | 0.25 |
| 11 | 80014664 | rs7394982 | -1.095 | 0.737 | 0.056 | 0.337 | -0.009 | 0.053 | 0.053 | 0.31 | 0.44 | 0.26 | 0.43 | 0.34 | 0.56 | 0.56 |
| 11 | 80015410 | rs11232063 | 0.867 | 0.697 | 0.272 | 0.326 | -0.002 | -0.066 | -0.066 | 0 | 0.11 | 0.29 | 0.05 | 0 | 0.06 | 0.12 |
| 11 | 80016503 | rs11826437 | 0.325 | 0.764 | -0.225 | 0.322 | -0.006 | 0.02 | 0.02 | 0 | 0.23 | 0.15 | 0.21 | 0.17 | 0.25 | 0.23 |
| 11 | 80017637 | rs10751018 | -1.779 | 0.697 | -0.279 | 0.34 | 0.02 | 0.043 | 0.043 | 0 | 0.56 | 0.28 | 0.49 | 0.34 | 0.64 | 0.7 |
| 11 | 80021207 | rs2100990 | 0.344 | 0.755 | -0.213 | 0.554 | -0.003 | 0.016 | 0.016 | 0 | 0.24 | 0.17 | 0.18 | 0.17 | 0.25 | 0.24 |
| 11 | 80021523 | rs931105 | -1.836 | 0.717 | -0.202 | 0.461 | 0.044 | -0.008 | -0.008 | 0 | 0.53 | 0.29 | 0.51 | 0.49 | 0.67 | 0.71 |
| 11 | 80022745 | rs1159800 | -0.515 | 0.7 | -0.426 | 0.574 | -0.039 | 0.035 | 0.035 | 0 | 0.23 | 0.08 | 0.25 | 0.17 | 0.31 | 0.33 |
| 11 | 80025850 | rs6421693 | -2.455 | 0.496 | -0.613 | 0.636 | -0.034 | 0.069 | 0.069 | 0.046 | 0.7 | 0.98 | 0.86 | 0.88 | 0.73 | 0.91 |
| 11 | 80027442 | rs11232073 | -1.418 | 0.395 | 0.281 | 0.768 | 0.024 | 0.002 | 0.002 | 0 | 0.17 | 0.29 | 0.14 | 0.13 | 0.19 | 0.2 |
| 11 | 80028514 | rs11232074 | -0.048 | 0.354 | 0.04 | 0.753 | -0.004 | 0.034 | 0.034 | 0 | 0.07 | 0.02 | 0.03 | 0 | 0.06 | 0.05 |
| 11 | 80035238 | rs1382777 | -1.122 | 0.305 | 0.233 | 0.566 | 0.01 | -0.025 | -0.025 | 0 | 0.37 | 0.46 | 0.6 | 0.71 | 0.41 | 0.49 |
| 11 | 80037795 | rs7951568 | -1.492 | 0.407 | 0.184 | -0.144 | 0.005 | 0.013 | 0.013 | 0 | 0.56 | 0.41 | 0.48 | 0.38 | 0.58 | 0.68 |
| 11 | 80038214 | rs7926854 | -1.243 | 0.456 | -0.211 | 0.023 | -0.015 | -0.003 | -0.003 | 4.899 | 0.38 | 0.81 | 0.32 | 0.33 | 0.38 | 0.43 |
| 11 | 80040968 | rs11232082 | -0.88 | 0.534 | 0.282 | 0.207 | 0.004 | 0.01 | 0.01 | 0 | 0.3 | 0.57 | 0.31 | 0.34 | 0.34 | 0.36 |
| 11 | 80041103 | rs12794583 | -0.518 | 0.438 | -0.11 | -0.149 | -0.025 | 0.06 | 0.06 | 0 | 0.29 | 0.14 | 0.25 | 0.18 | 0.3 | 0.42 |
| 11 | 80042841 | rs11823764 | -0.236 | 0.849 | 0.128 | 0.106 | -0.007 | -0.007 | -0.007 | 0.038 | 0.21 | 0.19 | 0.23 | 0.22 | 0.26 | 0.21 |
| 11 | 80044823 | rs10897595 | -1.892 | 0.714 | 0.337 | 0.122 | -0.001 | 0.007 | 0.007 | 0 | 0.46 | 0.67 | 0.54 | 0.42 | 0.55 | 0.61 |

| | | | | | | | | | | | | | | | | |
|---|---|---|---|---|---|---|---|---|---|---|---|---|---|---|---|---|
| 11 | 80048225 | rs11827604 | 0.792 | 0.865 | 0.255 | 0.078 | 0.002 | -0.002 | -0.002 | 0 | 0.15 | 0.12 | 0.11 | 0.15 | 0.12 | 0.15 |
| 11 | 80049649 | rs2373478 | -1.275 | 0.767 | 0.54 | 0.047 | 0.002 | 0.017 | 0.017 | 0 | 0.29 | 0.53 | 0.33 | 0.27 | 0.31 | 0.3 |
| 11 | 80051161 | rs7395542 | -1.625 | 0.774 | 0.322 | 0.181 | -0.006 | -0.004 | -0.004 | 0.268 | 0.4 | 0.66 | 0.51 | 0.42 | 0.45 | 0.52 |
| 11 | 80052487 | rs990648 | -1.414 | 0.817 | 0.456 | 0.201 | -0.002 | -0.002 | -0.002 | 1.166 | 0.37 | 0.63 | 0.48 | 0.41 | 0.43 | 0.48 |
| 11 | 80053454 | rs10501464 | 0.825 | 0.531 | 0.105 | 0.19 | -0.048 | -0.038 | -0.038 | 0 | 0.19 | 0.01 | 0.07 | 0.13 | 0.14 | 0.23 |
| 11 | 80055053 | rs4945401 | -1.792 | 0.84 | 1.564 | 0.139 | 0.059 | 0.019 | 0.019 | 0 | 0.2 | 0.45 | 0.3 | 0.22 | 0.26 | 0.34 |
| 11 | 80062406 | rs11232095 | -0.866 | 0.72 | 1.062 | 0.148 | -0.023 | 0.013 | 0.013 | 0 | 0.14 | 0.09 | 0.19 | 0.13 | 0.21 | 0.24 |
| 11 | 80072819 | rs1074543 | -2.046 | 0.752 | 0.415 | 0.185 | -0.02 | 0.019 | 0.019 | 0 | 0.42 | 0.73 | 0.5 | 0.37 | 0.47 | 0.53 |
| 11 | 80077022 | rs4945404 | -1.518 | 0.381 | 0.709 | 0.243 | -0.017 | 0.013 | 0.013 | 0.048 | 0.15 | 0.1 | 0.2 | 0.13 | 0.22 | 0.26 |
| 11 | 80078535 | rs7932403 | -1.778 | -0.148 | 0.836 | 0.457 | -0.027 | 0.003 | 0.003 | 0.743 | 0.17 | 0.61 | 0.23 | 0.18 | 0.12 | 0.24 |
| 11 | 80080639 | rs6591888 | -1.689 | -0.308 | 0.723 | 0.404 | -0.043 | -0.011 | -0.011 | 0 | 0.23 | 0.57 | 0.25 | 0.21 | 0.16 | 0.27 |
| 11 | 80083356 | rs1479290 | -2.017 | -0.084 | 0.43 | 0.342 | -0.005 | -0.005 | -0.005 | 0 | 0.27 | 0.7 | 0.29 | 0.21 | 0.18 | 0.3 |
| 11 | 80092415 | rs11232103 | -1.124 | 0.068 | -0.499 | 0.415 | 0.004 | 0.001 | 0.001 | 0 | 0.17 | 0.02 | 0.08 | 0.12 | 0.14 | 0.22 |
| 11 | 80094978 | rs2512483 | 0.886 | 0.453 | 0.069 | 0.754 | 0.011 | 0.067 | 0.067 | 0 | 0.8 | 0.49 | 0.82 | 0.73 | 0.8 | 0.84 |
| 11 | 80103856 | rs12798523 | -0.509 | 0.631 | -1.108 | -0.033 | -0.041 | 0.107 | 0.107 | 0.062 | 0.28 | 0.06 | 0.17 | 0.08 | 0.29 | 0.27 |
| 11 | 80104423 | rs1479300 | -0.955 | 0.769 | -0.782 | -0.737 | -0.009 | 0.069 | 0.069 | 0 | 0.43 | 0.31 | 0.27 | 0.08 | 0.42 | 0.34 |
| 11 | 80105368 | rs2448273 | 0.495 | 1.382 | -0.544 | -0.567 | 0.027 | 0.004 | 0.004 | 0 | 0.12 | 0.2 | 0.35 | 0.17 | 0.2 | 0.13 |
| 11 | 80105917 | rs7482266 | -0.186 | 1.208 | -0.055 | -0.727 | 0.049 | -0.004 | -0.004 | 0 | 0.16 | 0.38 | 0.36 | 0.16 | 0.2 | 0.14 |
| 11 | 80110012 | rs2512505 | -0.081 | 1.483 | -0.477 | 0.168 | 0 | 0.008 | 0.008 | 0 | 0.83 | 0.82 | 0.82 | 0.72 | 0.83 | 0.81 |
| 11 | 80110511 | rs2448276 | 0.091 | 1.51 | -0.205 | 0.373 | -0.038 | -0.012 | -0.012 | 0 | 0.55 | 0.84 | 0.64 | 0.67 | 0.51 | 0.56 |
| 11 | 80113483 | rs2512509 | -0.717 | 1.408 | -0.181 | 0.332 | 0.048 | -0.053 | -0.053 | 0 | 0.27 | 0.48 | 0.45 | 0.17 | 0.33 | 0.2 |
| 11 | 80113567 | rs2512510 | 0.94 | 1.427 | -0.182 | 0.397 | -0.047 | -0.032 | -0.032 | 0.084 | 0.1 | 0.03 | 0.34 | 0.17 | 0.18 | 0.11 |
| 11 | 80114845 | rs7929233 | 0.852 | 1.563 | -0.223 | 0.352 | -0.029 | 0.094 | 0.094 | 0 | 0.27 | 0.06 | 0.16 | 0.08 | 0.27 | 0.28 |
| 11 | 80117619 | rs12786450 | 0.368 | 0.424 | 0.217 | 1.059 | 0.006 | 0.026 | 0.026 | 0.164 | 0.54 | 0.34 | 0.41 | 0.3 | 0.58 | 0.57 |
| 11 | 80118334 | rs1842162 | 0.122 | 0.343 | 0.002 | 1.1 | -0.012 | 0.008 | 0.008 | 0 | 0.31 | 0.3 | 0.23 | 0.45 | 0.27 | 0.33 |
| 11 | 80118588 | rs7123160 | -0.383 | 0.626 | -0.013 | 1.212 | -0.023 | 0.002 | 0.002 | 0 | 0.4 | 0.44 | 0.26 | 0.49 | 0.31 | 0.39 |
| 11 | 80120305 | rs11232119 | 1.15 | 0.673 | -0.042 | 1.06 | -0.004 | 0.025 | 0.025 | 0 | 0.12 | 0.19 | 0.12 | 0.21 | 0.1 | 0.18 |
| 11 | 80125592 | rs2448281 | -0.037 | 0.698 | -0.078 | 0.921 | -0.022 | 0.013 | 0.013 | 0 | 0.59 | 0.54 | 0.73 | 0.45 | 0.68 | 0.56 |
| 11 | 80125852 | rs4944252 | 0.141 | 0.655 | -0.235 | 0.653 | 0.008 | 0.043 | 0.043 | 0 | 0.49 | 0.33 | 0.38 | 0.3 | 0.51 | 0.46 |
| 11 | 80127840 | rs10897610 | -0.03 | 0.604 | 0.075 | 0.737 | -0.005 | -0.003 | -0.003 | 0 | 0.12 | 0.16 | 0.11 | 0.13 | 0.1 | 0.14 |
| 11 | 80130893 | rs11232123 | -1.446 | 0.787 | 1.029 | 1.281 | 0.007 | 0.063 | 0.063 | 0 | 0.13 | 0.35 | 0.35 | 0.2 | 0.11 | 0.22 |
| 11 | 80132277 | rs12290894 | -1.79 | 0.783 | 1.127 | 1.304 | 0.014 | 0.102 | 0.102 | 0.168 | 0.15 | 0.45 | 0.35 | 0.2 | 0.11 | 0.22 |
| 11 | 80134251 | rs12294673 | -1.442 | 0.785 | 1.083 | 1.355 | 0.007 | 0.063 | 0.063 | 0 | 0.13 | 0.35 | 0.39 | 0.19 | 0.11 | 0.22 |
| 11 | 80140376 | rs7111204 | -1.108 | 0.738 | 1.453 | 1.363 | 0.018 | 0.061 | 0.061 | 0 | 0.09 | 0.24 | 0.33 | 0.16 | 0.07 | 0.16 |
| 11 | 80140433 | rs7111535 | -2.542 | 0.584 | 2.274 | 1.274 | 0.049 | 0.1 | 0.1 | 0.794 | 0.14 | 0.57 | 0.4 | 0.19 | 0.12 | 0.19 |
| 11 | 80146031 | rs12287579 | -2.466 | 0.551 | 2.379 | 0.96 | 0.085 | 0.006 | 0.006 | 0.797 | 0.08 | 0.36 | 0.12 | 0.01 | 0.09 | 0.06 |
| 11 | 80147262 | rs7940068 | -2.43 | 0.38 | 2.228 | 1.742 | -0.002 | 0.008 | 0.008 | 0 | 0.13 | 0.12 | 0.41 | 0.32 | 0.15 | 0.23 |
| 11 | 80147284 | rs1037399 | -1.714 | 0.712 | 1.317 | 0.745 | -0.008 | 0.024 | 0.024 | 3.291 | 0.47 | 0.83 | 0.65 | 0.4 | 0.52 | 0.59 |
| 11 | 80149601 | rs2512467 | -1.062 | 0.643 | 1.346 | 0.782 | -0.038 | 0.12 | 0.12 | 3.479 | 0.38 | 0.82 | 0.29 | 0.13 | 0.44 | 0.33 |
| 11 | 80150355 | rs2167590 | -1.156 | -0.475 | 0.265 | 0.195 | -0.019 | -0.088 | -0.088 | 2.307 | 0.72 | 0.89 | 0.84 | 0.51 | 0.69 | 0.64 |
| 11 | 80152332 | rs2443951 | 0.961 | -0.618 | 0.392 | -0.017 | -0.052 | 0.074 | 0.074 | 0.901 | 0.27 | 0.09 | 0.2 | 0.09 | 0.37 | 0.27 |
| 11 | 80160252 | rs2084827 | -0.597 | -0.698 | 0.337 | -0.024 | -0.028 | 0.088 | 0.088 | 0.963 | 0.63 | 0.82 | 0.47 | 0.17 | 0.6 | 0.47 |
| 11 | 80163042 | rs1470795 | 0.403 | -0.519 | 0.87 | 0.145 | 0.023 | -0.065 | -0.065 | 0 | 0.29 | 0.34 | 0.21 | 0.08 | 0.37 | 0.29 |
| 11 | 80168362 | rs11232147 | -0.292 | -0.528 | 0.697 | 0.464 | 0.024 | 0.22 | 0.22 | 0 | 0.11 | 0.4 | 0.34 | 0.36 | 0.09 | 0.14 |
| 11 | 80174542 | rs2512470 | -1.47 | -0.778 | 0.195 | 0.228 | -0.003 | -0.022 | -0.022 | 0 | 0.49 | 0.88 | 0.66 | 0.48 | 0.55 | 0.58 |
| 11 | 80176220 | rs2512471 | 0.276 | -0.719 | 0.14 | 0.142 | -0.064 | 0.101 | 0.101 | 0.359 | 0.3 | 0.09 | 0.2 | 0.09 | 0.39 | 0.35 |
| 11 | 80182109 | rs11232153 | -0.206 | -0.615 | 0.356 | 0.572 | 0.034 | 0.264 | 0.264 | 0 | 0.1 | 0.38 | 0.44 | 0.37 | 0.08 | 0.14 |
| 11 | 80187596 | rs2198256 | -0.395 | -0.721 | -0.969 | -0.564 | -0.056 | 0.195 | 0.195 | 0 | 0.44 | 0.14 | 0.27 | 0.1 | 0.48 | 0.44 |
| 11 | 80201641 | rs4945412 | -0.108 | -0.214 | -1.105 | -0.445 | 0.04 | -0.03 | -0.03 | 0 | 0.65 | 0.48 | 0.37 | 0.27 | 0.55 | 0.69 |
| 11 | 80207466 | rs11232171 | -0.46 | -0.593 | -1.653 | -0.878 | 0.006 | 0.002 | 0.002 | 0.017 | 0.35 | 0.32 | 0.21 | 0.07 | 0.3 | 0.3 |
| 11 | 80208565 | rs4525257 | 0.552 | -0.784 | -1.876 | -1.258 | -0.005 | 0.101 | 0.101 | 0 | 0.17 | 0.02 | 0.08 | 0 | 0.14 | 0.1 |
| 11 | 80209300 | rs7105482 | -1.346 | -0.433 | -1.947 | -1.393 | 0.018 | 0.041 | 0.041 | 0 | 0.44 | 0.33 | 0.21 | 0.06 | 0.38 | 0.45 |
| 11 | 80214639 | rs2512479 | 0.318 | 0.06 | -2.066 | -0.722 | -0.001 | -0.021 | -0.021 | 0 | 0.7 | 0.73 | 0.49 | 0.49 | 0.73 | 0.71 |
| 11 | 80226010 | rs2443973 | 1.38 | 0.108 | -2.031 | -0.719 | 0.005 | -0.011 | -0.011 | 0 | 0.61 | 0.69 | 0.44 | 0.49 | 0.66 | 0.56 |
| 11 | 80228846 | rs11232178 | 2.012 | 0.218 | -2.071 | -0.741 | -0.001 | 0.031 | 0.031 | 0.563 | 0.12 | 0.04 | 0.06 | 0.03 | 0.13 | 0.17 |
| 11 | 80231754 | rs4945416 | 0.86 | 0.492 | -1.964 | -0.119 | -0.053 | -0.063 | -0.063 | 0 | 0.1 | 0.01 | 0.4 | 0.37 | 0.12 | 0.17 |
| 11 | 80235447 | rs965716 | -1.072 | 0.54 | -1.554 | -0.286 | 0.031 | 0.039 | 0.039 | 0.4 | 0.74 | 0.57 | 0.76 | 0.62 | 0.65 | 0.82 |
| 11 | 80235789 | rs10501467 | 0.339 | 0.222 | -1.769 | -0.515 | 0.029 | 0.062 | 0.062 | 0.019 | 0.26 | 0.09 | 0.11 | 0.03 | 0.2 | 0.34 |
| 11 | 80235830 | rs964882 | 0.419 | 0.458 | -1.658 | -0.229 | -0.001 | 0.017 | 0.017 | 0 | 0.47 | 0.39 | 0.63 | 0.58 | 0.43 | 0.48 |
| 11 | 80241068 | rs4945417 | 2.561 | 0.478 | -1.835 | 0.154 | 0.02 | 0.003 | 0.003 | 0 | 0.23 | 0.13 | 0.14 | 0.21 | 0.17 | 0.22 |
| 11 | 80245589 | rs7946375 | 0.913 | 0.499 | -1.731 | 0.108 | -0.006 | 0.006 | 0.006 | 0 | 0.36 | 0.53 | 0.59 | 0.56 | 0.33 | 0.36 |
| 11 | 80256116 | rs11232201 | -0.26 | 0.506 | -1.837 | 0.012 | 0.006 | -0.027 | -0.027 | 0 | 0.63 | 0.59 | 0.72 | 0.76 | 0.57 | 0.63 |
| 11 | 80263120 | rs12362447 | 0.574 | 0.822 | -2.718 | -0.014 | -0.027 | 0.161 | 0.161 | 0 | 0.27 | 0.03 | 0.12 | 0.04 | 0.26 | 0.23 |
| 11 | 80264440 | rs10501469 | 0.657 | 0.948 | -2.163 | 0.906 | -0.05 | -0.048 | -0.048 | 0 | 0.09 | 0.01 | 0.4 | 0.36 | 0.13 | 0.16 |
| 11 | 80267205 | rs12574397 | 1.913 | 0.982 | -2.151 | 0.884 | 0.02 | 0.002 | 0.002 | 0 | 0.23 | 0.14 | 0.15 | 0.19 | 0.17 | 0.22 |
| 11 | 80270690 | rs1987338 | 0.171 | 1.361 | -1.866 | 1.175 | -0.046 | -0.045 | -0.045 | 0 | 0.09 | 0.01 | 0.4 | 0.36 | 0.13 | 0.16 |
| 11 | 80282364 | rs2373504 | 1.104 | 1.217 | -1.608 | 0.616 | 0.022 | 0.032 | 0.032 | 0 | 0.66 | 0.45 | 0.41 | 0.49 | 0.63 | 0.6 |
| 11 | 80282466 | rs1962677 | -0.64 | 1.291 | -2.518 | 0.509 | -0.046 | 0.053 | 0.053 | 0 | 0.89 | 0.98 | 0.92 | 0.99 | 0.86 | 0.9 |
| 11 | 80283896 | rs11232224 | -0.398 | 0.862 | -2.168 | 1.148 | -0.032 | -0.068 | -0.068 | 0 | 0.18 | 0.07 | 0.4 | 0.31 | 0.22 | 0.27 |

| | | | | | | | | | | | | | | | | |
|---|---|---|---|---|---|---|---|---|---|---|---|---|---|---|---|---|
| 11 | 80283906 | rs12576149 | 1.045 | 0.444 | -1.94 | 0.825 | 0.028 | -0.019 | -0.019 | 0 | 0.23 | 0.11 | 0.13 | 0.21 | 0.18 | 0.24 |
| 11 | 80293950 | rs1968681 | 0.16 | 1.362 | -1.617 | 1.23 | -0.099 | -0.008 | -0.008 | 0.041 | 0.74 | 0.97 | 0.82 | 0.74 | 0.7 | 0.83 |
| 11 | 80297206 | rs10792493 | 0.669 | 1.69 | -1.645 | 1.893 | -0.029 | -0.032 | -0.032 | 0 | 0.1 | 0.04 | 0.38 | 0.35 | 0.14 | 0.29 |
| 11 | 80297378 | rs11232234 | 2.089 | 1.111 | -1.954 | 1.533 | -0.007 | 0.165 | 0.165 | 0.222 | 0.37 | 0.09 | 0.26 | 0.09 | 0.38 | 0.22 |
| 11 | 80299004 | rs11232236 | -1.005 | 0.96 | -1.148 | 1.626 | -0.005 | 0.003 | 0.003 | 0.05 | 0.3 | 0.34 | 0.15 | 0.3 | 0.25 | 0.28 |
| 11 | 80305937 | rs4945424 | -1.699 | 1.053 | -0.421 | 1.988 | -0.011 | 0.072 | 0.072 | 0.526 | 0.45 | 0.74 | 0.61 | 0.7 | 0.43 | 0.64 |
| 11 | 80308348 | rs12574631 | -1.057 | 1.285 | -0.349 | 2.088 | -0.004 | 0.079 | 0.079 | 0 | 0.13 | 0.29 | 0.08 | 0.21 | 0.1 | 0.18 |
| 11 | 80311066 | rs1949886 | 0.166 | 1.491 | -0.348 | 1.97 | 0.005 | 0.025 | 0.025 | 0 | 0.11 | 0.11 | 0.37 | 0.28 | 0.15 | 0.28 |
| 11 | 80316287 | rs11232244 | -0.288 | 1.301 | -0.648 | 1.4 | -0.029 | -0.103 | -0.103 | 0.06 | 0.18 | 0.05 | 0.38 | 0.47 | 0.2 | 0.39 |
| 11 | 80316377 | rs17315132 | 0.382 | 1.313 | -0.439 | 1.074 | 0 | -0.009 | -0.009 | 0 | 0.22 | 0.15 | 0.16 | 0.06 | 0.22 | 0.24 |
| 11 | 80317350 | rs1949885 | -1.502 | 1.306 | -0.271 | 1.143 | -0.004 | 0.061 | 0.061 | 0.424 | 0.48 | 0.6 | 0.62 | 0.67 | 0.49 | 0.68 |
| 11 | 80325998 | rs10501472 | 0.318 | 1.289 | -0.309 | 1.23 | -0.035 | -0.088 | -0.088 | 1.403 | 0.16 | 0.05 | 0.37 | 0.46 | 0.2 | 0.39 |
| 11 | 80329779 | rs2373547 | 1.686 | 1.101 | -0.286 | 1.098 | 0.016 | 0.092 | 0.092 | 0 | 0.42 | 0.22 | 0.34 | 0.24 | 0.43 | 0.28 |
| 11 | 80332112 | rs4945427 | 0.319 | 1.268 | -0.257 | 1.102 | -0.021 | -0.02 | -0.02 | 0 | 0.92 | 0.97 | 0.92 | 0.78 | 0.89 | 0.89 |
| 11 | 80332711 | rs1917848 | -1.101 | 0.581 | 0.144 | 0.574 | 0.005 | 0.059 | 0.059 | 0.332 | 0.47 | 0.33 | 0.6 | 0.77 | 0.52 | 0.7 |
| 11 | 80338182 | rs1880693 | 0.375 | 0.922 | -0.417 | 0.678 | 0 | -0.017 | -0.017 | 0 | 0.22 | 0.18 | 0.16 | 0.07 | 0.23 | 0.22 |
| 11 | 80339237 | rs1917851 | -0.673 | 0.507 | -0.605 | 1.129 | -0.004 | 0.048 | 0.048 | 0 | 0.55 | 0.62 | 0.62 | 0.72 | 0.52 | 0.71 |
| 11 | 80346212 | rs2373551 | 0.755 | 0.525 | -0.367 | 1.827 | 0.001 | -0.003 | -0.003 | 1.57 | 0.86 | 0.9 | 0.9 | 0.77 | 0.9 | 0.88 |
| 11 | 80362324 | rs10897659 | -1.614 | 0.633 | 0.524 | 2.333 | 0.025 | 0.263 | 0.263 | 0 | 0.05 | 0.23 | 0.05 | 0.28 | 0.05 | 0.08 |
| 11 | 80368356 | rs7937748 | -2.614 | 0.67 | 1.126 | 2.297 | 0.044 | 0.313 | 0.313 | 0.221 | 0.1 | 0.46 | 0.11 | 0.28 | 0.1 | 0.11 |
| 11 | 80375307 | rs11232271 | -1.884 | 0.586 | 1.008 | 2.27 | 0.005 | 0.099 | 0.099 | 0 | 0.07 | 0.14 | 0.08 | 0.23 | 0.08 | 0.12 |
| 11 | 80375749 | rs1945372 | -2.569 | 0.572 | 1.027 | 2.24 | 0.012 | 0.234 | 0.234 | 0 | 0.11 | 0.38 | 0.1 | 0.26 | 0.09 | 0.14 |
| 11 | 80379533 | rs1945367 | -2.528 | 0.427 | 0.952 | 2.235 | -0.024 | 0.177 | 0.177 | 0 | 0.11 | 0.33 | 0.09 | 0.26 | 0.07 | 0.13 |
| 11 | 80389059 | rs1917863 | 0.78 | 0.491 | 0.299 | 1.936 | -0.002 | 0.011 | 0.011 | 0.637 | 0.63 | 0.17 | 0.7 | 0.36 | 0.64 | 0.75 |
| 11 | 80389699 | rs1917861 | 0.791 | 0.473 | 0.291 | 1.927 | -0.001 | 0.011 | 0.011 | 0.521 | 0.63 | 0.18 | 0.7 | 0.36 | 0.63 | 0.75 |
| 11 | 80391295 | rs7937935 | 0.917 | 0.458 | 0.278 | 1.919 | -0.005 | 0.015 | 0.015 | 0.259 | 0.62 | 0.13 | 0.7 | 0.36 | 0.63 | 0.73 |
| 11 | 80394562 | rs11603258 | 0.922 | 0.447 | 0.275 | 1.914 | -0.005 | 0.015 | 0.015 | 0.218 | 0.62 | 0.13 | 0.7 | 0.35 | 0.63 | 0.73 |
| 11 | 80399178 | rs12283277 | 1.147 | 0.357 | -0.815 | 0.791 | -0.021 | 0.23 | 0.23 | 0.205 | 0.32 | 0.02 | 0.16 | 0 | 0.27 | 0.24 |
| 11 | 80405623 | rs12421973 | 0.413 | 0.602 | -1.048 | 0.931 | -0.047 | -0.06 | -0.06 | 0 | 0.12 | 0.01 | 0.29 | 0.25 | 0.16 | 0.26 |
| 11 | 80417588 | rs4944262 | -0.61 | 0.509 | -0.058 | 1.232 | 0.017 | 0.04 | 0.04 | 0 | 0.73 | 0.47 | 0.79 | 0.6 | 0.71 | 0.85 |
| 11 | 80424197 | rs1970735 | -0.137 | 0.755 | -0.24 | 1.14 | -0.036 | -0.043 | -0.043 | 0 | 0.26 | 0.07 | 0.51 | 0.37 | 0.28 | 0.48 |
| 11 | 80426497 | rs1945379 | -0.653 | 0.806 | 0.058 | 1.163 | 0.026 | 0.065 | 0.065 | 0 | 0.73 | 0.53 | 0.79 | 0.62 | 0.7 | 0.86 |
| 11 | 80429527 | rs1945378 | 0.831 | 0.602 | 0.372 | 1.148 | -0.012 | 0.076 | 0.076 | 0 | 0.42 | 0.17 | 0.41 | 0.26 | 0.41 | 0.35 |
| 11 | 80438064 | rs11232307 | 0.306 | 0.485 | 1.11 | 1.241 | 0.001 | 0.003 | 0.003 | 0 | 0.49 | 0.53 | 0.48 | 0.26 | 0.47 | 0.38 |
| 11 | 80438418 | rs12364296 | 0.981 | 0.52 | 0.781 | 1.24 | -0.072 | 0.091 | 0.091 | 0 | 0.4 | 0.04 | 0.24 | 0.21 | 0.4 | 0.32 |
| 11 | 80456730 | rs4944265 | 0.012 | 0.962 | 1.103 | 1.731 | 0.096 | 0.196 | 0.196 | 0 | 0.92 | 0.79 | 0.92 | 0.77 | 0.88 | 0.92 |
| 11 | 80457784 | rs938161 | -0.386 | 0.766 | 0.821 | 1.625 | 0.011 | 0.003 | 0.003 | 0 | 0.44 | 0.13 | 0.44 | 0.5 | 0.4 | 0.56 |
| 11 | 80460497 | rs7939189 | 0.776 | 0.843 | 1.109 | 1.715 | 0.145 | 0.231 | 0.231 | 0 | 0.89 | 0.55 | 0.86 | 0.77 | 0.83 | 0.9 |
| 11 | 80465494 | rs1606385 | 1.206 | 0.433 | 0.415 | 1.459 | -0.023 | 0.111 | 0.111 | 0 | 0.39 | 0.14 | 0.24 | 0.18 | 0.38 | 0.3 |
| 11 | 80468651 | rs1828627 | 1.388 | 0.349 | 0.318 | 1.234 | -0.024 | 0.184 | 0.184 | 0.013 | 0.36 | 0.12 | 0.2 | 0.01 | 0.35 | 0.23 |
| 11 | 80469396 | rs1606383 | -1.422 | 0.867 | 0.249 | 1.582 | -0.011 | 0.015 | 0.015 | 0 | 0.95 | 0.98 | 0.89 | 0.72 | 0.9 | 0.83 |
| 11 | 80474657 | rs10751026 | 0.326 | 0.389 | 0.903 | 1.914 | 0.023 | 0.083 | 0.083 | 0 | 0.51 | 0.8 | 0.32 | 0.32 | 0.45 | 0.35 |
| 11 | 80475656 | rs12577177 | -1.06 | 0.158 | 1.252 | 1.805 | 0.044 | 0.07 | 0.07 | 1.193 | 0.09 | 0.4 | 0.07 | 0.16 | 0.08 | 0.04 |
| 11 | 80475951 | rs1533225 | 1.745 | 0.165 | 1.211 | 1.8 | -0.033 | 0.158 | 0.158 | 0.158 | 0.33 | 0.13 | 0.2 | 0 | 0.34 | 0.23 |
| 11 | 80481380 | rs10792507 | 0.304 | 0.245 | 1.152 | 1.858 | 0.024 | 0.05 | 0.05 | 0 | 0.51 | 0.76 | 0.31 | 0.35 | 0.46 | 0.35 |
| 11 | 80489615 | rs1877482 | -1.214 | 0.425 | 1.228 | 1.851 | 0.046 | 0.113 | 0.113 | 1.193 | 0.09 | 0.4 | 0.07 | 0.19 | 0.08 | 0.04 |
| 11 | 80505800 | rs1398450 | 0.284 | 0.249 | 1.023 | 1.832 | 0.006 | 0.046 | 0.046 | 0 | 0.52 | 0.76 | 0.35 | 0.58 | 0.49 | 0.41 |
| 11 | 80507165 | rs11232335 | 1.494 | 0.11 | 0.893 | 1.648 | -0.048 | 0.227 | 0.227 | 0.036 | 0.36 | 0.07 | 0.2 | 0.01 | 0.36 | 0.25 |
| 11 | 80516382 | rs1894061 | -0.489 | 0.212 | 0.642 | 1.58 | -0.026 | 0.014 | 0.014 | 0 | 0.69 | 0.8 | 0.45 | 0.67 | 0.64 | 0.64 |
| 11 | 80516495 | rs11232339 | 0.084 | 0.179 | 0.766 | 1.709 | 0.003 | 0.065 | 0.065 | 0 | 0.08 | 0.05 | 0.06 | 0.41 | 0.05 | 0.11 |
| 11 | 80520649 | rs10897677 | -1.442 | 0.195 | 0.936 | 1.57 | -0.025 | 0.234 | 0.234 | 0 | 0.15 | 0.57 | 0.14 | 0.53 | 0.13 | 0.14 |
| 11 | 80526425 | rs11232345 | 1.392 | 0.465 | 0.752 | 0.81 | -0.009 | 0.106 | 0.106 | 0 | 0.38 | 0.2 | 0.22 | 0.01 | 0.37 | 0.25 |
| 11 | 80535196 | rs7936663 | 0.047 | 0.613 | 1.167 | 1.591 | 0 | 0.005 | 0.005 | 0 | 0.16 | 0.13 | 0.11 | 0.08 | 0.15 | 0.23 |
| 11 | 80537336 | rs10501481 | -1.573 | 0.668 | 1.162 | 1.726 | 0.053 | 0.108 | 0.108 | 0.319 | 0.09 | 0.39 | 0.07 | 0.18 | 0.09 | 0.04 |
| 11 | 80540697 | rs11232356 | -1.211 | 0.999 | 0.833 | 1.759 | 0.012 | 0.054 | 0.054 | 0 | 0.22 | 0.33 | 0.31 | 0.48 | 0.27 | 0.35 |
| 11 | 80550839 | rs7112265 | -1.525 | 0.423 | 0.955 | 1.946 | -0.015 | 0.062 | 0.062 | 0.022 | 0.14 | 0.25 | 0.11 | 0.36 | 0.11 | 0.14 |
| 11 | 80554026 | rs17140931 | -1.813 | 0.532 | 0.906 | 1.814 | 0 | 0.006 | 0.006 | 0 | 0.24 | 0.26 | 0.15 | 0.5 | 0.22 | 0.35 |
| 11 | 80566396 | rs1355652 | -0.846 | 0.135 | -0.043 | 0.695 | 0.029 | 0.041 | 0.041 | 0 | 0.4 | 0.3 | 0.46 | 0.71 | 0.4 | 0.5 |
| 11 | 80577828 | rs4944267 | 0.619 | 0.286 | -0.266 | -0.263 | 0.013 | 0.029 | 0.029 | 0 | 0.45 | 0.62 | 0.22 | 0.13 | 0.35 | 0.33 |
| 11 | 80584598 | rs1533224 | -0.543 | 0.717 | -0.467 | 0.655 | -0.072 | -0.106 | -0.106 | 1.119 | 0.86 | 0.98 | 0.51 | 0.49 | 0.79 | 0.69 |
| 11 | 80595949 | rs4402318 | -0.814 | 0.395 | 0.251 | 0.456 | 0.015 | 0.141 | 0.141 | 3.434 | 0.08 | 0.57 | 0.06 | 0.13 | 0.05 | 0.06 |
| 11 | 80597384 | rs1498327 | 0.667 | 0.451 | -0.106 | 0.313 | -0.044 | -0.011 | -0.011 | 0.091 | 0.15 | 0 | 0.07 | 0.11 | 0.14 | 0.24 |
| 11 | 80600142 | rs17141024 | -0.746 | 0.379 | 0.306 | 0.311 | 0.026 | 0.164 | 0.164 | 3.032 | 0.08 | 0.55 | 0.05 | 0.13 | 0.05 | 0.06 |
| 11 | 80605558 | rs10897698 | 0.61 | 0.277 | 0.008 | 0.09 | -0.026 | 0.029 | 0.029 | 0 | 0.58 | 0.76 | 0.35 | 0.33 | 0.55 | 0.41 |
| 11 | 80610918 | rs10792521 | 1.449 | -0.015 | 0.308 | -0.137 | -0.023 | 0.17 | 0.17 | 0.285 | 0.47 | 0.19 | 0.29 | 0.21 | 0.49 | 0.33 |
| 11 | 80611357 | rs7925070 | 0.592 | -0.065 | 0.572 | -0.177 | 0.004 | -0.009 | -0.009 | 0 | 0.59 | 0.47 | 0.33 | 0.31 | 0.55 | 0.42 |
| 11 | 80611790 | rs1909141 | 0.687 | 0.034 | 0.73 | -0.126 | -0.016 | 0.02 | 0.02 | 0 | 0.58 | 0.68 | 0.35 | 0.33 | 0.55 | 0.42 |
| 11 | 80625188 | rs1909136 | 0.191 | -0.475 | 0.805 | 0.027 | -0.005 | 0.022 | 0.022 | 0 | 0.05 | 0.14 | 0.02 | 0.1 | 0.04 | 0.06 |
| 11 | 80625388 | rs1909138 | -0.209 | -0.422 | 0.79 | 0.087 | 0.004 | 0.025 | 0.025 | 0.521 | 0.75 | 0.69 | 0.45 | 0.46 | 0.69 | 0.64 |

| | | | | | | | | | | | | | | | | |
|---|---|---|---|---|---|---|---|---|---|---|---|---|---|---|---|---|
| 11 | 80634798 | rs1391529 | 1.516 | -0.44 | 0.737 | -0.219 | 0.03 | 0.133 | 0.133 | 0.132 | 0.52 | 0.25 | 0.37 | 0.24 | 0.55 | 0.37 |
| 11 | 80638724 | rs10897724 | 1.109 | -0.529 | 0.686 | -0.307 | 0.006 | 0.05 | 0.05 | 0 | 0.55 | 0.33 | 0.38 | 0.26 | 0.57 | 0.36 |
| 11 | 80640473 | rs4945464 | -0.58 | -0.559 | 0.848 | -0.231 | -0.011 | -0.022 | -0.022 | 0 | 0.77 | 0.78 | 0.47 | 0.47 | 0.7 | 0.65 |
| 11 | 80640561 | rs4945465 | -0.938 | -0.416 | 0.641 | -0.174 | -0.08 | -0.121 | -0.121 | 0 | 0.85 | 0.98 | 0.53 | 0.52 | 0.79 | 0.66 |
| 11 | 80644238 | rs2030074 | 0.816 | -1.21 | 0.596 | -1.01 | 0.006 | 0.038 | 0.038 | 0 | 0.52 | 0.52 | 0.38 | 0.26 | 0.54 | 0.33 |
| 11 | 80649283 | rs7941823 | 0.038 | -1.363 | 0.616 | -1.017 | -0.005 | 0.016 | 0.016 | 0 | 0.05 | 0.15 | 0.02 | 0.09 | 0.04 | 0.05 |
| 11 | 80650985 | rs1498328 | -0.921 | -1.358 | 0.499 | -1.059 | -0.005 | -0.007 | -0.007 | 0 | 0.75 | 0.75 | 0.45 | 0.43 | 0.68 | 0.63 |
| 11 | 80653037 | rs7130321 | -1.177 | -1.241 | 0.84 | -0.949 | -0.016 | 0.033 | 0.033 | 1.213 | 0.11 | 0.37 | 0.03 | 0.09 | 0.05 | 0.07 |
| 11 | 80659466 | rs11232407 | 0.316 | -1.17 | 0.601 | -0.937 | 0 | -0.001 | -0.001 | 0 | 0.53 | 0.49 | 0.37 | 0.23 | 0.56 | 0.37 |
| 11 | 80687755 | rs12576906 | -0.767 | -1.623 | 0.291 | -0.863 | -0.075 | 0.038 | 0.038 | 4.484 | 0.21 | 0.49 | 0.08 | 0.22 | 0.11 | 0.23 |
| 11 | 80690336 | rs11232416 | 0.41 | -1.345 | -0.652 | -1.503 | -0.008 | 0.136 | 0.136 | 0.091 | 0.38 | 0.16 | 0.19 | 0.01 | 0.35 | 0.23 |
| 11 | 80693838 | rs1498329 | 0.717 | -2.025 | -0.177 | -0.81 | -0.008 | 0.01 | 0.01 | 0 | 0.09 | 0.14 | 0.04 | 0.09 | 0.08 | 0.14 |
| 11 | 80695445 | rs7950346 | -0.327 | -2.25 | -0.015 | -0.981 | -0.028 | -0.028 | -0.028 | 5.348 | 0.18 | 0.33 | 0.09 | 0.09 | 0.15 | 0.21 |
| 11 | 80701613 | rs7946942 | 0.348 | -2.592 | -0.538 | -0.926 | -0.035 | 0.008 | 0.008 | 0 | 0.12 | 0.24 | 0.05 | 0.16 | 0.04 | 0.05 |
| 11 | 80707760 | rs7342242 | -0.312 | -2.822 | -1.411 | -1.19 | -0.018 | 0.128 | 0.128 | 0.793 | 0.75 | 0.98 | 0.86 | 0.96 | 0.79 | 0.84 |
| 11 | 80714305 | rs4945471 | 0.296 | -2.409 | -0.985 | -0.924 | -0.037 | -0.012 | -0.012 | 0 | 0.8 | 0.69 | 0.88 | 0.82 | 0.88 | 0.92 |
| 11 | 80716179 | rs11232428 | 0.848 | -1.938 | -1.942 | -0.815 | -0.021 | 0.052 | 0.052 | 0 | 0.13 | 0.03 | 0.07 | 0.02 | 0.13 | 0.12 |
| 11 | 80716264 | rs9919563 | 0.509 | -2.109 | -1.457 | -0.095 | 0.002 | -0.002 | -0.002 | 0.155 | 0.87 | 0.86 | 0.91 | 0.83 | 0.9 | 0.93 |
| 11 | 80717021 | rs7110013 | 0.451 | -2.1 | -1.454 | -0.091 | 0.001 | -0.002 | -0.002 | 0 | 0.87 | 0.86 | 0.91 | 0.83 | 0.9 | 0.93 |
| 11 | 80718383 | rs1566240 | 0.709 | -1.937 | -1.548 | -0.025 | 0.123 | -0.003 | -0.003 | 0.097 | 0.56 | 0.17 | 0.75 | 0.58 | 0.71 | 0.66 |
| 11 | 80719478 | rs4945473 | 0.378 | -1.826 | -1.542 | 0.052 | 0.108 | -0.016 | -0.016 | 0 | 0.59 | 0.2 | 0.76 | 0.58 | 0.73 | 0.68 |
| 11 | 80719488 | rs4945474 | 0.287 | -0.958 | -1.291 | -0.129 | -0.013 | -0.009 | -0.009 | 0 | 0.61 | 0.46 | 0.79 | 0.73 | 0.72 | 0.71 |
| 11 | 80722309 | rs1391530 | 0.169 | 0.044 | -2.137 | 0.066 | -0.058 | -0.029 | -0.029 | 0.182 | 0.69 | 0.95 | 0.75 | 0.76 | 0.7 | 0.7 |
| 11 | 80727242 | rs10897741 | -0.33 | 0.395 | -1.19 | 0.589 | 0.077 | 0.221 | 0.221 | 1.616 | 0.14 | 0.44 | 0.22 | 0.35 | 0.17 | 0.32 |
| 11 | 80728653 | rs1498326 | 0.784 | 0.118 | -2.071 | 0.168 | -0.006 | 0.026 | 0.026 | 0 | 0.11 | 0 | 0.2 | 0.05 | 0.09 | 0.16 |
| 11 | 80729659 | rs1874795 | -0.283 | 0.389 | -2.578 | -0.333 | -0.035 | 0.177 | 0.177 | 0 | 0.6 | 0.88 | 0.76 | 0.85 | 0.6 | 0.75 |
| 11 | 80734844 | rs904318 | -0.32 | 0.427 | -2.518 | -0.352 | -0.024 | 0.176 | 0.176 | 0 | 0.58 | 0.84 | 0.76 | 0.85 | 0.58 | 0.74 |
| 11 | 80736539 | rs12417895 | 0.673 | 0.546 | -2.694 | -0.176 | -0.008 | -0.049 | -0.049 | 0 | 0.11 | 0.02 | 0.15 | 0.39 | 0.11 | 0.16 |
| 11 | 80736944 | rs10736740 | -0.773 | 0.548 | -2.901 | -0.392 | -0.073 | 0.061 | 0.061 | 0 | 0.72 | 0.97 | 0.86 | 0.86 | 0.75 | 0.75 |
| 11 | 80737235 | rs17141360 | 0.479 | 0.816 | -2.301 | 0.122 | -0.01 | -0.041 | -0.041 | 0 | 0.1 | 0.01 | 0.15 | 0.39 | 0.11 | 0.16 |
| 11 | 80750888 | rs1608169 | 0.015 | 0.801 | -1.809 | -0.867 | -0.015 | 0.03 | 0.03 | 0 | 0.32 | 0.23 | 0.21 | 0.15 | 0.34 | 0.17 |
| 11 | 80757902 | rs4944277 | -0.385 | 0.776 | -1.241 | -1.222 | -0.001 | 0.014 | 0.014 | 4.995 | 0.45 | 0.38 | 0.33 | 0.1 | 0.49 | 0.2 |
| 11 | 80762049 | rs1403255 | 0.932 | 0.743 | -1.277 | -1.039 | -0.001 | 0.061 | 0.061 | 0 | 0.17 | 0.06 | 0.13 | 0.01 | 0.18 | 0.04 |
| 11 | 80779541 | rs12289861 | 0.127 | 0.789 | -0.56 | -0.562 | 0.012 | 0.02 | 0.02 | 0 | 0.17 | 0.33 | 0.13 | 0.19 | 0.17 | 0.12 |
| 11 | 80780652 | rs1568639 | -0.339 | 0.873 | -0.384 | -0.233 | 0.001 | 0.003 | 0.003 | 0 | 0.36 | 0.45 | 0.49 | 0.44 | 0.36 | 0.41 |
| 11 | 80782982 | rs1403261 | -0.066 | 0.716 | -0.54 | -0.18 | 0 | 0 | 0 | 1.038 | 0.65 | 0.67 | 0.56 | 0.5 | 0.7 | 0.45 |
| 11 | 80783619 | rs1879900 | -0.919 | 0.767 | -0.979 | -0.043 | 0.006 | -0.01 | -0.01 | 0 | 0.86 | 0.94 | 0.93 | 0.8 | 0.89 | 0.75 |
| 11 | 80789029 | rs1403257 | 0.683 | 0.775 | -0.696 | -0.093 | -0.017 | 0.044 | 0.044 | 0 | 0.48 | 0.22 | 0.43 | 0.31 | 0.54 | 0.33 |
| 11 | 80790181 | rs1916054 | -0.809 | 0.795 | -0.653 | 0.256 | 0.005 | -0.011 | -0.011 | 0 | 0.86 | 0.9 | 0.93 | 0.75 | 0.89 | 0.74 |
| 11 | 80790741 | rs1533867 | -0.031 | 0.841 | -0.363 | 0.161 | -0.012 | 0.017 | 0.017 | 0.115 | 0.64 | 0.43 | 0.53 | 0.5 | 0.7 | 0.45 |
| 11 | 80792660 | rs11232450 | 0.284 | 1.285 | -0.846 | 0.768 | -0.054 | -0.096 | -0.096 | 0 | 0.19 | 0.03 | 0.21 | 0.3 | 0.23 | 0.29 |
| 11 | 80803005 | rs7130733 | -0.37 | 1.391 | -0.599 | 0.403 | -0.002 | 0 | 0 | 1.871 | 0.31 | 0.25 | 0.29 | 0.24 | 0.31 | 0.34 |
| 11 | 80805072 | rs4590891 | 0.611 | 1.593 | -0.339 | 0.439 | 0.003 | 0.007 | 0.007 | 0 | 0.21 | 0.33 | 0.31 | 0.29 | 0.19 | 0.31 |
| 11 | 80805226 | rs11232453 | 0.743 | 1.791 | -0.54 | 0.597 | -0.002 | 0.007 | 0.007 | 1.907 | 0.22 | 0.09 | 0.21 | 0.17 | 0.23 | 0.28 |
| 11 | 80806438 | rs11232454 | -0.365 | 1.676 | -0.508 | 0.573 | -0.003 | -0.001 | -0.001 | 0 | 0.3 | 0.22 | 0.28 | 0.24 | 0.3 | 0.36 |
| 11 | 80813185 | rs7120971 | -0.518 | 2.125 | -0.732 | 0.385 | -0.03 | 0.016 | 0.016 | 2.273 | 0.34 | 0.23 | 0.35 | 0.23 | 0.41 | 0.35 |
| 11 | 80815998 | rs12577008 | 0.8 | 2.087 | -0.615 | 0.521 | 0.014 | 0.043 | 0.043 | 0.163 | 0.06 | 0.09 | 0.14 | 0.22 | 0.09 | 0.16 |
| 11 | 80818238 | rs11232471 | 0.981 | 2.093 | -0.833 | 0.348 | -0.01 | 0 | 0 | 0 | 0.21 | 0.08 | 0.19 | 0.17 | 0.21 | 0.28 |
| 11 | 80821366 | rs11821818 | -0.116 | 1.894 | -0.399 | 0.053 | 0.019 | -0.009 | -0.009 | 0.114 | 0.05 | 0.21 | 0.04 | 0 | 0.03 | 0.05 |
| 11 | 80824141 | rs2514461 | -0.562 | 2.234 | -0.878 | 0.411 | -0.059 | -0.019 | -0.019 | 0 | 0.72 | 0.89 | 0.72 | 0.77 | 0.64 | 0.7 |
| 11 | 80825178 | rs7941214 | -0.515 | 1.974 | -0.559 | 0.202 | 0.032 | 0.004 | 0.004 | 0 | 0.26 | 0.63 | 0.46 | 0.32 | 0.33 | 0.29 |
| 11 | 80826074 | rs10792538 | -0.926 | 1.431 | -0.538 | 0.174 | -0.032 | 0.055 | 0.055 | 0.483 | 0.46 | 0.76 | 0.44 | 0.34 | 0.47 | 0.47 |
| 11 | 80827910 | rs11232476 | 1.286 | 1.114 | -0.905 | 0.077 | -0.009 | 0.116 | 0.116 | 0.127 | 0.4 | 0.16 | 0.39 | 0.18 | 0.38 | 0.31 |
| 11 | 80828124 | rs11601730 | 1.55 | 1.245 | -1.14 | 0.141 | -0.041 | 0.074 | 0.074 | 1.506 | 0.19 | 0.01 | 0.12 | 0.05 | 0.18 | 0.22 |
| 11 | 80831002 | rs2461681 | 0.023 | 1.21 | -0.646 | 0.601 | -0.003 | 0 | 0 | 0.333 | 0.17 | 0.18 | 0.12 | 0.29 | 0.14 | 0.18 |
| 11 | 80831756 | rs12791548 | 0.373 | 1.582 | -0.45 | 0.152 | 0.024 | -0.048 | -0.048 | 0 | 0.11 | 0.2 | 0.09 | 0 | 0.15 | 0.07 |
| 11 | 80839316 | rs1568638 | 1.128 | 1.276 | -0.791 | 0.406 | -0.029 | 0.046 | 0.046 | 0.714 | 0.18 | 0.01 | 0.12 | 0.06 | 0.16 | 0.22 |
| 11 | 80841098 | rs1522659 | -0.088 | 1.083 | -0.453 | 0.581 | 0.012 | 0.017 | 0.017 | 0 | 0.33 | 0.22 | 0.32 | 0.34 | 0.32 | 0.39 |
| 11 | 80841719 | rs2461678 | -0.766 | 1.111 | -0.4 | 0.553 | 0.007 | -0.011 | -0.011 | 0.023 | 0.55 | 0.66 | 0.57 | 0.57 | 0.62 | 0.56 |
| 11 | 80846484 | rs12364823 | 0.568 | 0.83 | -0.303 | 0.487 | -0.009 | 0.04 | 0.04 | 0.093 | 0.28 | 0.17 | 0.3 | 0.1 | 0.29 | 0.29 |
| 11 | 80846625 | rs12416782 | -0.16 | -0.431 | 0.098 | 1.356 | 0.034 | 0.049 | 0.049 | 0 | 0.39 | 0.29 | 0.32 | 0.17 | 0.27 | 0.33 |
| 11 | 80846781 | rs2514502 | 0.769 | -0.416 | -0.024 | 1.413 | 0.006 | -0.007 | -0.007 | 0.016 | 0.07 | 0.04 | 0.04 | 0.25 | 0.06 | 0.15 |
| 11 | 80847849 | rs2461673 | -0.515 | -0.215 | -0.015 | 1.359 | 0.03 | 0.018 | 0.018 | 0 | 0.47 | 0.38 | 0.36 | 0.4 | 0.32 | 0.47 |
| 11 | 80856703 | rs1568635 | -1.017 | -0.857 | 0.318 | 0.978 | -0.03 | -0.005 | -0.005 | 0.433 | 0.17 | 0.21 | 0.07 | 0.22 | 0.1 | 0.15 |
| 11 | 80860642 | rs2514496 | 1.107 | -0.858 | 0.808 | 1.246 | 0.027 | 0.154 | 0.154 | 1.033 | 0.92 | 0.39 | 0.9 | 0.84 | 0.94 | 0.88 |
| 11 | 80864160 | rs12786838 | -0.599 | -1.005 | 0.675 | 1.302 | 0.029 | -0.014 | -0.014 | 0 | 0.13 | 0.06 | 0.08 | 0.22 | 0.09 | 0.14 |
| 11 | 80865791 | rs2512095 | 0.755 | -1.001 | 1.111 | 1.202 | 0.089 | 0.241 | 0.241 | 0.359 | 0.93 | 0.5 | 0.92 | 0.85 | 0.94 | 0.88 |
| 11 | 80867185 | rs2512098 | 1.034 | -1.003 | 1.129 | 1.215 | 0.06 | 0.217 | 0.217 | 0.505 | 0.93 | 0.42 | 0.91 | 0.84 | 0.94 | 0.88 |
| 11 | 80869453 | rs11232496 | -0.759 | -0.958 | 1.01 | 1.21 | -0.013 | -0.002 | -0.002 | 0.302 | 0.13 | 0.13 | 0.09 | 0.18 | 0.1 | 0.14 |

| | | | | | | | | | | | | | | | | |
|---|---|---|---|---|---|---|---|---|---|---|---|---|---|---|---|---|
| 11 | 80875461 | rs1996846 | 0.586 | -0.586 | 1.102 | 1.072 | 0.076 | 0.215 | 0.215 | 0.098 | 0.94 | 0.58 | 0.92 | 0.85 | 0.94 | 0.88 |
| 11 | 80878504 | rs2512119 | 0.61 | -0.851 | 0.247 | 0.403 | 0.015 | 0.053 | 0.053 | 0 | 0.11 | 0.02 | 0.08 | 0 | 0.09 | 0.01 |
| 11 | 80889618 | rs4457776 | -0.184 | -0.868 | -0.284 | -0.239 | -0.009 | 0.055 | 0.055 | 1.614 | 0.21 | 0.11 | 0.1 | 0.07 | 0.2 | 0.28 |
| 11 | 80893313 | rs10501488 | 1.801 | -0.826 | -1.102 | -0.716 | -0.122 | 0.214 | 0.214 | 0.327 | 0.37 | 0.03 | 0.26 | 0.06 | 0.4 | 0.25 |
| 11 | 80897179 | rs4569033 | -0.287 | -0.926 | -0.387 | 0.207 | -0.009 | 0.185 | 0.185 | 0 | 0.07 | 0.17 | 0.15 | 0.42 | 0.06 | 0.15 |
| 11 | 80902438 | rs4288778 | -0.729 | -0.849 | -0.312 | 0.383 | 0.01 | 0.089 | 0.089 | 0 | 0.91 | 0.88 | 0.85 | 0.62 | 0.84 | 0.77 |
| 11 | 80903884 | rs7938364 | -1.154 | -1.227 | -0.41 | 0.324 | 0.041 | -0.031 | -0.031 | 0 | 0.41 | 0.82 | 0.36 | 0.55 | 0.34 | 0.43 |
| 11 | 80904468 | rs10501494 | 0.323 | -1.173 | -0.467 | 0.15 | -0.005 | -0.005 | -0.005 | 0 | 0.14 | 0 | 0.24 | 0.08 | 0.11 | 0.1 |
| 11 | 80916305 | rs11232517 | 0.305 | -1.071 | -0.364 | 0.324 | -0.002 | 0.025 | 0.025 | 0 | 0.06 | 0.03 | 0.08 | 0.37 | 0.08 | 0.12 |
| 11 | 80916359 | rs7102147 | -1.348 | -1.121 | -0.008 | 0.277 | 0.012 | 0.274 | 0.274 | 0.425 | 0.1 | 0.36 | 0.19 | 0.44 | 0.08 | 0.16 |
| 11 | 80920078 | rs6591940 | -0.611 | -1.171 | -0.067 | 0.292 | -0.009 | 0.219 | 0.219 | 0 | 0.08 | 0.23 | 0.16 | 0.43 | 0.06 | 0.16 |
| 11 | 80920802 | rs10501496 | -0.779 | -1.196 | -0.378 | -0.97 | -0.014 | -0.103 | -0.103 | 0.962 | 0.34 | 0.46 | 0.2 | 0.06 | 0.28 | 0.28 |
| 11 | 80921407 | rs12288304 | -0.023 | -0.823 | -0.193 | -0.833 | -0.05 | -0.115 | -0.115 | 0 | 0.23 | 0.44 | 0.13 | 0.05 | 0.15 | 0.09 |
| 11 | 80921481 | rs10897767 | -0.388 | -0.763 | -0.262 | -0.816 | -0.034 | 0.045 | 0.045 | 0 | 0.2 | 0.04 | 0.29 | 0.08 | 0.21 | 0.26 |
| 11 | 80923036 | rs11232525 | -0.181 | -0.869 | -0.162 | -0.792 | -0.021 | -0.054 | -0.054 | 0 | 0.13 | 0.45 | 0.05 | 0.05 | 0.05 | 0.07 |
| 11 | 80928144 | rs12361615 | 2.078 | -0.76 | -0.381 | -0.845 | -0.129 | 0.204 | 0.204 | 1.146 | 0.37 | 0.03 | 0.26 | 0.06 | 0.41 | 0.28 |
| 11 | 80928809 | rs11232535 | -0.477 | -0.986 | 0.179 | -0.428 | -0.024 | -0.041 | -0.041 | 0 | 0.13 | 0.42 | 0.05 | 0.06 | 0.05 | 0.07 |
| 11 | 80930458 | rs11232538 | -1.305 | -0.775 | -0.17 | -0.427 | 0.031 | 0.09 | 0.09 | 0.62 | 0.45 | 0.72 | 0.49 | 0.21 | 0.36 | 0.33 |
| 11 | 80930613 | rs4315099 | -1.465 | -0.801 | -0.304 | -0.44 | 0.048 | 0.153 | 0.153 | 1.286 | 0.47 | 0.83 | 0.5 | 0.21 | 0.36 | 0.33 |
| 11 | 80930696 | rs11232540 | 0.178 | -0.716 | -0.466 | -0.421 | -0.001 | 0.002 | 0.002 | 0 | 0.12 | 0.1 | 0.15 | 0.72 | 0.15 | 0.36 |
| 11 | 80930934 | rs4412769 | -2.216 | -0.7 | -0.562 | -0.722 | -0.156 | 0.234 | 0.234 | 0.397 | 0.59 | 0.97 | 0.65 | 0.93 | 0.51 | 0.68 |
| 11 | 80931284 | rs17141801 | -0.288 | -0.677 | 0.089 | -0.514 | 0.008 | 0.005 | 0.005 | 0 | 0.1 | 0.21 | 0.13 | 0.15 | 0.09 | 0.12 |
| 11 | 80932351 | rs7125586 | 0.799 | -1.008 | 0.613 | -0.73 | -0.047 | -0.05 | -0.05 | 0.167 | 0.17 | 0.31 | 0.18 | 0.09 | 0.11 | 0.06 |
| 11 | 80932427 | rs7125940 | -1.47 | -0.968 | 0.772 | -0.813 | 0.004 | 0.009 | 0.009 | 0 | 0.39 | 0.61 | 0.39 | 0.87 | 0.35 | 0.49 |
| 11 | 80934453 | rs7941712 | 0.377 | -1.135 | 1.006 | -0.944 | -0.001 | -0.029 | -0.029 | 2.035 | 0.13 | 0.14 | 0.05 | 0 | 0.1 | 0.15 |
| 11 | 80935911 | rs4414243 | -1.211 | -1.158 | 1.16 | -0.259 | 0.005 | 0.007 | 0.007 | 0 | 0.39 | 0.61 | 0.39 | 0.8 | 0.35 | 0.49 |
| 11 | 80936250 | rs12576623 | 0.32 | -1.206 | 1.126 | -0.302 | -0.008 | 0.059 | 0.059 | 0.667 | 0.08 | 0.16 | 0.2 | 0.14 | 0.06 | 0.04 |
| 11 | 80936536 | rs11232544 | -0.9 | -1.228 | 1.422 | -0.314 | -0.002 | 0 | 0 | 0 | 0.35 | 0.33 | 0.28 | 0.71 | 0.34 | 0.45 |
| 11 | 80936656 | rs11232545 | -0.938 | -1.206 | 1.428 | -0.298 | -0.016 | -0.036 | -0.036 | 0 | 0.2 | 0.13 | 0.19 | 0.71 | 0.24 | 0.43 |
| 11 | 80938299 | rs7106567 | -2.015 | -1.193 | 1.153 | -0.421 | 0.038 | 0.174 | 0.174 | 0.213 | 0.46 | 0.77 | 0.59 | 0.93 | 0.41 | 0.53 |
| 11 | 80941360 | rs4145064 | -0.341 | -1.223 | 1.265 | -0.381 | -0.059 | -0.065 | -0.065 | 0 | 0.26 | 0.54 | 0.41 | 0.15 | 0.19 | 0.09 |
| 11 | 80941531 | rs11232553 | 0.252 | -1.187 | 1.063 | -0.362 | -0.005 | -0.008 | -0.008 | 0.008 | 0.08 | 0.16 | 0.21 | 0.06 | 0.08 | 0.04 |
| 11 | 80942380 | rs4999939 | -1.943 | -1.183 | 0.933 | -0.513 | 0.038 | 0.174 | 0.174 | 0.349 | 0.46 | 0.77 | 0.59 | 0.93 | 0.41 | 0.53 |
| 11 | 80948325 | rs10792541 | -0.902 | -1.124 | 1.173 | -0.361 | -0.004 | 0.194 | 0.194 | 2.82 | 0.12 | 0.47 | 0.32 | 0.23 | 0.09 | 0.08 |
| 11 | 80949350 | rs2514462 | -0.746 | -1.059 | 0.946 | -0.418 | -0.022 | 0.007 | 0.007 | 0 | 0.3 | 0.54 | 0.35 | 0.74 | 0.25 | 0.4 |
| 11 | 80954031 | rs9943555 | -0.25 | -1.138 | 0.284 | -0.147 | 0.02 | -0.04 | -0.04 | 0 | 0.15 | 0.07 | 0.09 | 0.57 | 0.08 | 0.28 |
| 11 | 80964065 | rs2168819 | -1.107 | -1.085 | 0.498 | -0.526 | -0.005 | -0.003 | -0.003 | 0 | 0.38 | 0.38 | 0.37 | 0.71 | 0.33 | 0.48 |
| 11 | 80966178 | rs7934189 | 0.652 | -1.073 | 0.342 | -0.461 | 0 | 0.041 | 0.041 | 0 | 0.11 | 0.12 | 0.19 | 0.62 | 0.1 | 0.29 |
| 11 | 80971141 | rs936368 | -0.533 | -1.091 | 0.16 | -0.528 | 0.006 | 0.004 | 0.004 | 0.074 | 0.79 | 0.84 | 0.86 | 0.97 | 0.8 | 0.88 |
| 11 | 80971151 | rs936369 | -1.597 | -1.1 | 0.084 | -0.533 | 0.01 | 0.031 | 0.031 | 0 | 0.86 | 0.94 | 0.9 | 0.97 | 0.86 | 0.97 |
| 11 | 80971794 | rs11232569 | 0.466 | -1.259 | -0.138 | -0.705 | 0.017 | -0.025 | -0.025 | 0 | 0.15 | 0.09 | 0.1 | 0.15 | 0.12 | 0.09 |
| 11 | 80975989 | rs2514486 | 2.159 | -1.123 | -0.289 | -1.025 | -0.169 | 0.195 | 0.195 | 0 | 0.37 | 0.03 | 0.47 | 0.06 | 0.47 | 0.31 |
| 11 | 80978480 | rs17141842 | 0.459 | -1.313 | 0.307 | -0.627 | -0.001 | -0.038 | -0.038 | 0 | 0.08 | 0.17 | 0.04 | 0 | 0.06 | 0.09 |
| 11 | 80980888 | rs7941624 | -1.045 | -1.623 | 0.628 | -0.581 | -0.031 | -0.028 | -0.028 | 0 | 0.31 | 0.49 | 0.33 | 0.77 | 0.21 | 0.39 |
| 11 | 80984220 | rs10897779 | -0.629 | -1.399 | 0.853 | -0.564 | -0.007 | -0.004 | -0.004 | 0 | 0.27 | 0.32 | 0.24 | 0.72 | 0.23 | 0.36 |
| 11 | 80985548 | rs1458095 | -0.939 | -1.589 | 0.991 | -0.791 | -0.008 | -0.004 | -0.004 | 0 | 0.15 | 0.4 | 0.16 | 0.17 | 0.1 | 0.23 |
| 11 | 80985948 | rs12276609 | -0.963 | -1.663 | 0.995 | -0.552 | 0.001 | -0.001 | -0.001 | 0.171 | 0.27 | 0.27 | 0.16 | 0.26 | 0.2 | 0.23 |
| 11 | 80986806 | rs10501500 | 0.189 | -1.595 | 0.948 | -0.51 | -0.004 | 0.005 | 0.005 | 0 | 0.17 | 0.21 | 0.12 | 0.16 | 0.14 | 0.12 |
| 11 | 80990096 | rs1379728 | 1.096 | -1.789 | 0.506 | -0.99 | 0.071 | 0.067 | 0.067 | 0.021 | 0.53 | 0.19 | 0.63 | 0.67 | 0.63 | 0.62 |
| 11 | 80992153 | rs2085557 | -0.027 | -1.385 | 0.939 | -0.618 | -0.009 | 0.017 | 0.017 | 0 | 0.18 | 0.29 | 0.21 | 0.21 | 0.15 | 0.15 |
| 11 | 80994883 | rs1350418 | 0.684 | -0.931 | 0.871 | -0.267 | -0.001 | 0.065 | 0.065 | 0 | 0.11 | 0.09 | 0.12 | 0.57 | 0.09 | 0.25 |
| 11 | 80994984 | rs1350419 | 0.605 | -0.932 | 1.049 | -0.268 | -0.003 | 0.186 | 0.186 | 0 | 0.11 | 0.14 | 0.12 | 0.57 | 0.1 | 0.25 |
| 11 | 80995619 | rs11232575 | -0.475 | -0.841 | 1.042 | -0.323 | 0.009 | 0.189 | 0.189 | 0 | 0.15 | 0.38 | 0.2 | 0.64 | 0.14 | 0.29 |
| 11 | 81001092 | rs11600036 | -0.385 | -0.807 | 0.679 | -0.338 | 0.005 | 0.096 | 0.096 | 0 | 0.16 | 0.23 | 0.17 | 0.65 | 0.16 | 0.34 |
| 11 | 81003507 | rs1350421 | -1.565 | -0.857 | 0.298 | -0.453 | 0.013 | 0.055 | 0.055 | 0 | 0.87 | 0.96 | 0.91 | 1 | 0.87 | 0.97 |
| 11 | 81005520 | rs7123143 | -1.059 | -0.843 | 0.636 | -0.333 | 0.006 | 0.041 | 0.041 | 0 | 0.08 | 0.25 | 0.14 | 0.18 | 0.06 | 0.13 |
| 11 | 81010164 | rs10501503 | 1.217 | -0.882 | 0.125 | -0.656 | 0.046 | 0.05 | 0.05 | 0.341 | 0.51 | 0.18 | 0.44 | 0.63 | 0.56 | 0.62 |
| 11 | 81010850 | rs2514467 | -0.799 | -0.935 | 0.506 | -0.654 | 0.001 | -0.001 | -0.001 | 0 | 0.26 | 0.27 | 0.39 | 0.28 | 0.25 | 0.3 |
| 11 | 81011841 | rs2514466 | -1.417 | -0.745 | 0.434 | -0.692 | -0.008 | 0.005 | 0.005 | 0 | 0.33 | 0.49 | 0.45 | 0.33 | 0.31 | 0.37 |
| 11 | 81013539 | rs7118643 | 0.996 | -0.553 | 0.386 | -0.612 | 0.011 | 0.01 | 0.01 | 0 | 0.54 | 0.35 | 0.45 | 0.63 | 0.56 | 0.64 |
| 11 | 81014879 | rs7123367 | 1.939 | -0.633 | -0.27 | -0.929 | -0.132 | 0.192 | 0.192 | 0 | 0.37 | 0.05 | 0.29 | 0.07 | 0.42 | 0.26 |
| 11 | 81014974 | rs2446116 | -0.441 | -0.654 | 0.12 | -0.655 | -0.018 | 0.039 | 0.039 | 1.064 | 0.23 | 0.38 | 0.31 | 0.26 | 0.21 | 0.17 |
| 11 | 81020912 | rs17141922 | 0.359 | -0.362 | -0.378 | -0.337 | 0.002 | -0.061 | -0.061 | 3.723 | 0.19 | 0.09 | 0.18 | 0.54 | 0.2 | 0.4 |
| 11 | 81021385 | rs4944289 | -1.49 | -0.442 | -0.454 | -0.931 | 0 | 0.065 | 0.065 | 0 | 0.86 | 0.98 | 0.91 | 1 | 0.86 | 0.97 |
| 11 | 81022395 | rs11232586 | 0.269 | -0.37 | -0.146 | -0.733 | -0.005 | 0.04 | 0.04 | 2.004 | 0.2 | 0.22 | 0.18 | 0.6 | 0.19 | 0.4 |
| 11 | 81026219 | rs7942822 | -0.399 | -0.162 | 0.102 | -0.864 | -0.001 | -0.05 | -0.05 | 0 | 0.78 | 0.67 | 0.75 | 0.91 | 0.78 | 0.84 |
| 11 | 81030422 | rs12418411 | -0.349 | -0.165 | 0.199 | -0.848 | 0.039 | 0.036 | 0.036 | 0 | 0.07 | 0.24 | 0.16 | 0.09 | 0.07 | 0.12 |
| 11 | 81030830 | rs7931007 | -0.861 | -0.083 | 0.151 | -0.906 | -0.001 | -0.025 | -0.025 | 0.077 | 0.82 | 0.84 | 0.89 | 1 | 0.81 | 0.92 |
| 11 | 81040340 | rs4944292 | -1.018 | -0.408 | 0.156 | -1.278 | 0.017 | 0.066 | 0.066 | 2.726 | 0.49 | 0.39 | 0.49 | 0.84 | 0.45 | 0.69 |

| | | | | | | | | | | | | | | | | |
|---|---|---|---|---|---|---|---|---|---|---|---|---|---|---|---|---|
| 11 | 81046412 | rs10501506 | 0.57 | -0.546 | 0.069 | -1.155 | 0.001 | -0.08 | -0.08 | 0 | 0.23 | 0.09 | 0.19 | 0.6 | 0.22 | 0.47 |
| 11 | 81048744 | rs10897794 | 1.758 | -0.56 | 0.112 | -1.172 | -0.027 | 0.05 | 0.05 | 0.083 | 0.32 | 0.18 | 0.27 | 0.06 | 0.36 | 0.23 |
| 11 | 81049571 | rs11601420 | 0.061 | -0.511 | 0.171 | -1.074 | -0.004 | -0.037 | -0.037 | 0 | 0.27 | 0.19 | 0.21 | 0.61 | 0.26 | 0.53 |
| 11 | 81050438 | rs10501508 | -1.578 | -0.488 | 0.249 | -0.973 | 0.061 | 0.097 | 0.097 | 0.23 | 0.06 | 0.38 | 0.15 | 0.09 | 0.04 | 0.05 |
| 11 | 81051151 | rs1458105 | 0.038 | -0.534 | -0.002 | -0.991 | 0.009 | -0.01 | -0.01 | 0 | 0.22 | 0.21 | 0.28 | 0.24 | 0.19 | 0.16 |
| 11 | 81051466 | rs1458107 | 1.486 | -0.487 | -0.155 | -0.448 | -0.036 | -0.056 | -0.056 | 0.009 | 0.09 | 0.01 | 0.14 | 0.42 | 0.13 | 0.21 |
| 11 | 81053049 | rs4943826 | 0.127 | -0.553 | -0.051 | -0.447 | 0.004 | 0.11 | 0.11 | 0 | 0.11 | 0.26 | 0.37 | 0.24 | 0.13 | 0.11 |
| 11 | 81062761 | rs1585228 | 0.383 | -0.562 | -0.222 | -0.571 | 0.001 | 0.017 | 0.017 | 0 | 0.54 | 0.25 | 0.55 | 0.31 | 0.56 | 0.4 |
| 11 | 81071843 | rs10897799 | 1.828 | -0.642 | -0.288 | -0.648 | -0.107 | 0.137 | 0.137 | 0.091 | 0.32 | 0.04 | 0.26 | 0.07 | 0.36 | 0.24 |
| 11 | 81073082 | rs11232630 | 1.282 | -0.525 | -0.082 | -0.477 | 0.015 | -0.016 | -0.016 | 0.787 | 0.14 | 0.06 | 0.05 | 0.16 | 0.09 | 0.25 |
| 11 | 81078376 | rs7931800 | -0.278 | -0.071 | 0.414 | -1.166 | 0.037 | -0.01 | -0.01 | 0.165 | 0.06 | 0.21 | 0.14 | 0.06 | 0.07 | 0.07 |
| 11 | 81082396 | rs1945877 | -0.717 | -0.08 | 0.451 | -1.171 | 0.005 | -0.011 | -0.011 | 0.159 | 0.64 | 0.71 | 0.72 | 0.4 | 0.65 | 0.51 |
| 11 | 81093405 | rs1458082 | -0.569 | -0.075 | 0.463 | -1.168 | -0.004 | 0.004 | 0.004 | 0.103 | 0.63 | 0.64 | 0.59 | 0.34 | 0.63 | 0.5 |
| 11 | 81099928 | rs7124654 | 0.095 | -0.082 | 0.249 | -1.236 | 0.013 | 0.021 | 0.021 | 0 | 0.54 | 0.26 | 0.55 | 0.3 | 0.57 | 0.41 |
| 11 | 81101351 | rs10501509 | 1.216 | -0.102 | 0.141 | -1.228 | 0.014 | -0.076 | -0.076 | 0 | 0.22 | 0.08 | 0.18 | 0.61 | 0.2 | 0.46 |
| 11 | 81102917 | rs1945890 | -1.876 | 0.016 | 0.158 | -1.266 | -0.007 | 0.036 | 0.036 | 0 | 0.87 | 0.94 | 0.91 | 1 | 0.85 | 0.97 |
| 11 | 81109767 | rs17142171 | -1.081 | 0.177 | 0.354 | -1.137 | 0.073 | 0.005 | 0.005 | 0 | 0.09 | 0.44 | 0.05 | 0.02 | 0.07 | 0.09 |
| 11 | 81116058 | rs10792556 | -0.897 | 0.193 | 0.364 | -1.126 | 0.003 | -0.006 | -0.006 | 0 | 0.64 | 0.7 | 0.71 | 0.38 | 0.65 | 0.51 |
| 11 | 81116856 | rs11232657 | -0.059 | 0.211 | 0.328 | -1.114 | 0.011 | 0.005 | 0.005 | 0.008 | 0.55 | 0.28 | 0.66 | 0.36 | 0.59 | 0.43 |
| 11 | 81124233 | rs12798039 | 0.88 | 0.367 | 0.711 | -0.819 | -0.015 | -0.037 | -0.037 | 0 | 0.07 | 0.01 | 0.23 | 0.16 | 0.09 | 0.06 |
| 11 | 81126244 | rs6591966 | -1.571 | 0.273 | 0.723 | -0.85 | -0.004 | 0.001 | 0.001 | 0 | 0.31 | 0.67 | 0.45 | 0.29 | 0.29 | 0.26 |
| 11 | 81132183 | rs11232664 | -0.879 | 0.15 | 0.594 | -0.83 | 0 | -0.004 | -0.004 | 0 | 0.22 | 0.24 | 0.4 | 0.3 | 0.22 | 0.19 |
| 11 | 81134854 | rs11232666 | -0.716 | 0.263 | 0.459 | -1.281 | -0.011 | -0.03 | -0.03 | 0 | 0.21 | 0.12 | 0.24 | 0.22 | 0.24 | 0.17 |
| 11 | 81142678 | rs10792563 | 0.553 | 0.448 | 0.428 | -1.161 | -0.016 | -0.054 | -0.054 | 0 | 0.23 | 0.11 | 0.27 | 0.66 | 0.24 | 0.5 |
| 11 | 81158967 | rs7129232 | -1.264 | 0.48 | 1.553 | -0.686 | 0.004 | 0.041 | 0.041 | 0.037 | 0.19 | 0.51 | 0.37 | 0.22 | 0.2 | 0.13 |
| 11 | 81171701 | rs10501514 | -1.149 | 0.617 | 1.274 | -0.972 | 0.015 | 0.004 | 0.004 | 0 | 0.23 | 0.46 | 0.42 | 0.17 | 0.28 | 0.17 |
| 11 | 81178750 | rs11232679 | 0.277 | 0.634 | 0.887 | -0.969 | -0.024 | -0.037 | -0.037 | 3.606 | 0.08 | 0.01 | 0.08 | 0.08 | 0.13 | 0.08 |
| 11 | 81185712 | rs1945887 | -0.065 | 0.486 | 1.203 | -0.877 | 0.007 | 0.004 | 0.004 | 0 | 0.2 | 0.24 | 0.4 | 0.17 | 0.27 | 0.14 |
| 11 | 81189350 | rs7924980 | -0.167 | 0.468 | 0.926 | -1.212 | 0 | -0.058 | -0.058 | 0 | 0.27 | 0.37 | 0.17 | 0.02 | 0.25 | 0.2 |
| 11 | 81191347 | rs17442731 | 0.142 | 0.882 | -0.094 | -1.052 | -0.05 | 0.126 | 0.126 | 0.6 | 0.23 | 0.03 | 0.15 | 0.02 | 0.25 | 0.16 |
| 11 | 81193276 | rs12420118 | -1.156 | 0.84 | -0.051 | -1.039 | 0 | -0.007 | -0.007 | 0.511 | 0.46 | 0.53 | 0.57 | 0.2 | 0.52 | 0.32 |
| 11 | 81193297 | rs10501515 | -0.273 | 1.16 | -0.652 | -0.755 | 0.015 | 0.007 | 0.007 | 0 | 0.22 | 0.35 | 0.4 | 0.17 | 0.28 | 0.16 |
| 11 | 81203090 | rs12279546 | -0.344 | 1.342 | -0.895 | -0.646 | 0.009 | -0.005 | -0.005 | 1.449 | 0.25 | 0.34 | 0.43 | 0.17 | 0.31 | 0.24 |
| 11 | 81207264 | rs10501518 | -0.608 | 1.326 | -1.783 | -0.464 | -0.051 | 0.099 | 0.099 | 0.283 | 0.21 | 0.04 | 0.14 | 0.04 | 0.23 | 0.18 |
| 11 | 81211648 | rs12790840 | -0.983 | 1.131 | -1.702 | -0.235 | -0.014 | 0.033 | 0.033 | 0 | 0.18 | 0.06 | 0.22 | 0.1 | 0.2 | 0.22 |
| 11 | 81211881 | rs17142418 | -0.07 | 0.136 | -1.921 | -1.516 | 0.011 | -0.006 | -0.006 | 0 | 0.24 | 0.18 | 0.23 | 0.01 | 0.17 | 0.17 |
| 11 | 81213684 | rs4405342 | -0.554 | 0.298 | -1.986 | -1.398 | -0.009 | 0.028 | 0.028 | 0 | 0.31 | 0.23 | 0.32 | 0.25 | 0.3 | 0.35 |
| 11 | 81220764 | rs17142439 | 0.509 | -0.034 | -1.627 | -1.684 | 0.021 | 0.004 | 0.004 | 0 | 0.25 | 0.17 | 0.26 | 0.01 | 0.17 | 0.17 |
| 11 | 81221497 | rs4944303 | -1.227 | 0.097 | -0.85 | -2.03 | 0.012 | -0.049 | -0.049 | 0.077 | 0.62 | 0.58 | 0.59 | 0.24 | 0.55 | 0.6 |
| 11 | 81222768 | rs7478747 | -0.129 | -0.008 | -0.797 | -2.153 | -0.012 | 0.021 | 0.021 | 0 | 0.27 | 0.21 | 0.29 | 0.22 | 0.28 | 0.34 |
| 11 | 81223225 | rs12787574 | -0.398 | -0.022 | -0.521 | -1.502 | -0.007 | -0.019 | -0.019 | 0 | 0.39 | 0.32 | 0.37 | 0.5 | 0.38 | 0.54 |
| 11 | 81223717 | rs1945655 | -0.465 | -0.013 | -0.518 | -1.497 | -0.001 | -0.004 | -0.004 | 0.886 | 0.41 | 0.42 | 0.37 | 0.5 | 0.38 | 0.54 |
| 11 | 81224751 | rs11232705 | 1.653 | 0.731 | -1.106 | -1.759 | -0.066 | -0.14 | -0.14 | 0.155 | 0.17 | 0.01 | 0.2 | 0.48 | 0.22 | 0.13 |
| 11 | 81225948 | rs11828414 | 1.399 | 0.415 | -0.911 | -1.767 | 0.01 | 0.013 | 0.013 | 0 | 0.27 | 0.19 | 0.25 | 0.01 | 0.24 | 0.2 |
| 11 | 81226350 | rs10792573 | 0.412 | 0.345 | -0.833 | -1.799 | 0.022 | 0.012 | 0.012 | 0 | 0.4 | 0.26 | 0.32 | 0.28 | 0.34 | 0.41 |
| 11 | 81231936 | rs11232711 | 1.351 | 0.305 | -1.085 | -1.501 | -0.002 | -0.062 | -0.062 | 0.011 | 0.13 | 0.04 | 0.07 | 0.27 | 0.1 | 0.21 |
| 11 | 81241298 | rs12277689 | 1.305 | 0.095 | -0.984 | -1.865 | -0.006 | -0.053 | -0.053 | 0.945 | 0.26 | 0.29 | 0.21 | 0.01 | 0.16 | 0.19 |
| 11 | 81241487 | rs7107343 | 0.444 | 0.593 | -1.23 | -1.721 | -0.082 | 0.028 | 0.028 | 0 | 0.26 | 0.05 | 0.32 | 0.22 | 0.32 | 0.34 |
| 11 | 81251505 | rs7935238 | -0.428 | 0.708 | -0.875 | -2.548 | 0.021 | 0.086 | 0.086 | 0.812 | 0.52 | 0.32 | 0.55 | 0.97 | 0.55 | 0.71 |
| 11 | 81253743 | rs11232722 | 1.533 | 0.91 | -0.86 | -2.655 | -0.026 | -0.045 | -0.045 | 0 | 0.12 | 0.01 | 0.08 | 0.15 | 0.13 | 0.19 |
| 11 | 81258089 | rs10897837 | -0.565 | 0.928 | -0.254 | -2.624 | 0.017 | 0.044 | 0.044 | 0.827 | 0.51 | 0.35 | 0.56 | 0.97 | 0.55 | 0.71 |
| 11 | 81259119 | rs7951915 | -0.419 | 0.905 | -0.133 | -2.691 | 0.012 | 0.118 | 0.118 | 1.02 | 0.51 | 0.3 | 0.55 | 0.97 | 0.53 | 0.71 |
| 11 | 81259273 | rs7952149 | -1.367 | 0.889 | 0.254 | -2.848 | -0.001 | -0.018 | -0.018 | 0.401 | 0.85 | 0.87 | 0.83 | 0.99 | 0.8 | 0.92 |
| 11 | 81259828 | rs4943839 | 1.855 | 0.791 | 0.507 | -2.908 | -0.002 | -0.048 | -0.048 | 0 | 0.24 | 0.21 | 0.21 | 0.02 | 0.19 | 0.14 |
| 11 | 81263034 | rs2156832 | 1.987 | 0.786 | 0.594 | -2.91 | 0.008 | 0.024 | 0.024 | 0 | 0.23 | 0.14 | 0.2 | 0.02 | 0.19 | 0.14 |
| 11 | 81264443 | rs2187539 | -0.395 | 0.776 | 0.572 | -2.92 | 0.026 | -0.135 | -0.135 | 0 | 0.78 | 0.57 | 0.77 | 0.99 | 0.72 | 0.85 |
| 11 | 81266104 | rs7107748 | -0.316 | 0.788 | 0.417 | -2.911 | 0.008 | 0.127 | 0.127 | 0.939 | 0.5 | 0.3 | 0.53 | 0.96 | 0.51 | 0.7 |
| 11 | 81272371 | rs1955071 | 1.934 | 0.804 | 0.184 | -2.854 | 0.002 | 0.052 | 0.052 | 0 | 0.23 | 0.12 | 0.22 | 0.02 | 0.2 | 0.15 |
| 11 | 81273298 | rs7115509 | 1.905 | 0.786 | 0.221 | -2.837 | 0 | -0.033 | -0.033 | 0.408 | 0.23 | 0.19 | 0.21 | 0.02 | 0.19 | 0.14 |
| 11 | 81278248 | rs869291 | -0.028 | 1.149 | -0.105 | -1.562 | -0.008 | -0.053 | -0.053 | 0.3 | 0.42 | 0.16 | 0.33 | 0.46 | 0.4 | 0.6 |
| 11 | 81279349 | rs1945679 | -0.553 | 1.199 | 0.129 | -1.642 | 0.041 | -0.147 | -0.147 | 0 | 0.79 | 0.51 | 0.77 | 0.98 | 0.72 | 0.86 |
| 11 | 81279543 | rs7948448 | -0.207 | 1.308 | -0.323 | -1.305 | 0.008 | -0.021 | -0.021 | 0.221 | 0.42 | 0.25 | 0.33 | 0.45 | 0.4 | 0.61 |
| 11 | 81284197 | rs10897843 | 0.527 | 1.21 | -0.517 | -0.581 | 0 | 0.001 | 0.001 | 0 | 0.06 | 0.05 | 0.03 | 0.18 | 0.07 | 0.07 |
| 11 | 81287045 | rs17142606 | 1.291 | 1.574 | -0.656 | -0.514 | -0.034 | 0.021 | 0.021 | 0 | 0.07 | 0.01 | 0.2 | 0.52 | 0.12 | 0.12 |
| 11 | 81299269 | rs11232737 | -0.223 | 1.074 | -0.468 | -1.436 | 0.007 | -0.008 | -0.008 | 0.055 | 0.42 | 0.32 | 0.34 | 0.44 | 0.41 | 0.56 |
| 11 | 81302554 | rs7120638 | 0.866 | 1.007 | -0.786 | -1.529 | 0.025 | 0.022 | 0.022 | 0.386 | 0.21 | 0.04 | 0.09 | 0.13 | 0.15 | 0.26 |
| 11 | 81303817 | rs1573551 | -1.919 | 1.148 | -0.56 | -1.648 | -0.007 | 0.041 | 0.041 | 1.537 | 0.86 | 0.94 | 0.84 | 0.99 | 0.82 | 0.92 |
| 11 | 81303873 | rs7933268 | 0.504 | 1.092 | -0.538 | -1.724 | -0.017 | -0.025 | -0.025 | 0 | 0.64 | 0.27 | 0.53 | 0.47 | 0.59 | 0.7 |
| 11 | 81317946 | rs1945644 | -0.505 | 1.048 | 0.034 | -1.92 | 0.034 | -0.141 | -0.141 | 0 | 0.79 | 0.51 | 0.78 | 0.99 | 0.73 | 0.86 |

| | | | | | | | | | | | | | | | | |
|---|---|---|---|---|---|---|---|---|---|---|---|---|---|---|---|---|
| 11 | 81323584 | rs2105801 | -0.592 | 1.051 | -0.11 | -1.981 | 0.008 | 0.127 | 0.127 | 0.989 | 0.5 | 0.33 | 0.51 | 0.96 | 0.51 | 0.67 |
| 11 | 81339041 | rs10501523 | 1.435 | 0.767 | 0.01 | -1.829 | -0.027 | -0.122 | -0.122 | 0 | 0.3 | 0.4 | 0.26 | 0.03 | 0.2 | 0.15 |
| 11 | 81346344 | rs12799371 | 1.269 | 0.813 | 0.033 | -1.835 | -0.018 | -0.11 | -0.11 | 0 | 0.32 | 0.4 | 0.28 | 0.03 | 0.22 | 0.16 |
| 11 | 81347994 | rs7927890 | 1.472 | 0.776 | -0.075 | -1.81 | 0.005 | -0.028 | -0.028 | 0 | 0.3 | 0.26 | 0.24 | 0.03 | 0.2 | 0.15 |
| 11 | 81354835 | rs1540176 | 0.005 | 1.253 | -0.123 | -1.623 | -0.001 | -0.014 | -0.014 | 0 | 0.4 | 0.27 | 0.35 | 0.44 | 0.44 | 0.55 |
| 11 | 81355517 | rs1945702 | -0.85 | 1.445 | 0.253 | -1.484 | 0.006 | 0.019 | 0.019 | 1.209 | 0.77 | 0.76 | 0.63 | 0.46 | 0.69 | 0.79 |
| 11 | 81357363 | rs1945704 | -0.228 | 1.161 | 0.139 | -1.69 | -0.008 | -0.046 | -0.046 | 0 | 0.63 | 0.6 | 0.65 | 0.87 | 0.66 | 0.67 |
| 11 | 81362191 | rs6591983 | -0.842 | 1.141 | 0.1 | -1.782 | 0.013 | -0.055 | -0.055 | 4.392 | 0.38 | 0.55 | 0.42 | 0.85 | 0.43 | 0.5 |
| 11 | 81374426 | rs1945721 | 0.879 | 1.207 | 0.059 | -1.624 | 0 | -0.001 | -0.001 | 0 | 0.22 | 0.23 | 0.23 | 0.34 | 0.22 | 0.28 |
| 11 | 81376074 | rs10792584 | -0.096 | 1.202 | 0.131 | -1.545 | 0.01 | 0.018 | 0.018 | 0.276 | 0.4 | 0.26 | 0.33 | 0.37 | 0.37 | 0.47 |
| 11 | 81379503 | rs1945719 | 0.582 | 1.443 | 0.279 | -1.468 | 0.032 | 0.094 | 0.094 | 0 | 0.12 | 0.17 | 0.22 | 0.53 | 0.19 | 0.21 |
| 11 | 81386818 | rs12366135 | 2.358 | 1.167 | 0.184 | -1.611 | -0.009 | 0.126 | 0.126 | 0 | 0.25 | 0.05 | 0.23 | 0.02 | 0.23 | 0.16 |
| 11 | 81393198 | rs11232775 | 1.524 | 1.206 | 0.301 | -1.663 | -0.004 | -0.084 | -0.084 | 0.173 | 0.34 | 0.36 | 0.29 | 0.03 | 0.31 | 0.21 |
| 11 | 81395174 | rs12364666 | -1.046 | 1.166 | 0.14 | -1.483 | 0.002 | -0.007 | -0.007 | 0 | 0.84 | 0.93 | 0.76 | 0.83 | 0.81 | 0.91 |
| 11 | 81411075 | rs11232789 | 0.148 | 1.279 | 0.312 | -1.432 | 0.021 | 0.033 | 0.033 | 0 | 0.51 | 0.73 | 0.56 | 0.58 | 0.52 | 0.43 |
| 11 | 81417643 | rs7938175 | 1.47 | 1.417 | 0.435 | -1.492 | -0.006 | -0.092 | -0.092 | 2.052 | 0.34 | 0.38 | 0.3 | 0.03 | 0.31 | 0.21 |
| 11 | 81419403 | rs12793728 | 0.291 | 1.342 | 0.403 | -1.462 | 0.006 | 0.008 | 0.008 | 0 | 0.48 | 0.56 | 0.55 | 0.58 | 0.52 | 0.43 |
| 11 | 81422622 | rs17142789 | 0.41 | 1.486 | 0.35 | -1.439 | 0.022 | -0.008 | -0.008 | 0.002 | 0.05 | 0.11 | 0.05 | 0.02 | 0.1 | 0.07 |
| 11 | 81426448 | rs4944327 | 2.423 | 1.448 | 0.234 | -1.509 | -0.006 | 0.126 | 0.126 | 0 | 0.25 | 0.04 | 0.23 | 0.02 | 0.22 | 0.16 |
| 11 | 81429155 | rs10897863 | 1.311 | 1.429 | 0.246 | -1.471 | 0 | -0.007 | -0.007 | 0 | 0.06 | 0.06 | 0.04 | 0.16 | 0.08 | 0.06 |
| 11 | 81429368 | rs1580239 | 1.408 | 1.42 | 0.394 | -1.494 | 0.001 | 0.004 | 0.004 | 0 | 0.33 | 0.25 | 0.29 | 0.04 | 0.32 | 0.22 |
| 11 | 81429529 | rs1599210 | -0.01 | 1.46 | 0.392 | -1.496 | 0.02 | 0.028 | 0.028 | 0 | 0.5 | 0.63 | 0.55 | 0.58 | 0.52 | 0.43 |
| 11 | 81430152 | rs12806464 | 1.476 | 1.371 | 0.194 | -1.507 | -0.002 | 0.038 | 0.038 | 0 | 0.31 | 0.19 | 0.28 | 0.04 | 0.32 | 0.22 |
| 11 | 81448678 | rs11232823 | 1.952 | 1.163 | -0.044 | -1.399 | 0.01 | 0.031 | 0.031 | 0 | 0.27 | 0.15 | 0.24 | 0.02 | 0.22 | 0.15 |
| 11 | 81449443 | rs7124504 | 0.175 | 1.416 | 0.199 | -1.044 | 0.06 | 0.184 | 0.184 | 0 | 0.17 | 0.39 | 0.27 | 0.53 | 0.2 | 0.22 |
| 11 | 81458413 | rs1031703 | -1.886 | 1.292 | -0.243 | -1.586 | -0.008 | 0.048 | 0.048 | 3.001 | 0.91 | 0.98 | 0.88 | 1 | 0.89 | 0.97 |
| 11 | 81458462 | rs10897868 | 1.732 | 1.407 | 0.082 | -1.629 | 0.001 | -0.03 | -0.03 | 5.61 | 0.08 | 0.13 | 0.06 | 0.01 | 0.08 | 0.07 |
| 11 | 81460746 | rs1380542 | -1.702 | 1.346 | -0.03 | -1.491 | -0.009 | 0.038 | 0.038 | 3.589 | 0.9 | 0.98 | 0.87 | 0.99 | 0.88 | 0.97 |
| 11 | 81461220 | rs17159766 | 0.044 | 1.073 | 0.313 | -1.628 | -0.003 | 0.003 | 0.003 | 0.054 | 0.06 | 0.16 | 0.04 | 0.02 | 0.04 | 0.03 |
| 11 | 81466717 | rs4944336 | -0.639 | 1.224 | 0.205 | -1.23 | -0.01 | -0.016 | -0.016 | 0 | 0.81 | 0.85 | 0.68 | 0.49 | 0.77 | 0.83 |
| 11 | 81467934 | rs12577477 | -0.426 | 1.076 | 0.203 | -1.552 | -0.001 | 0.026 | 0.026 | 0.281 | 0.5 | 0.39 | 0.38 | 0.45 | 0.51 | 0.66 |
| 11 | 81470761 | rs4515989 | 1.952 | 0.787 | -0.093 | -1.846 | 0.023 | 0.06 | 0.06 | 0 | 0.17 | 0.03 | 0.14 | 0.1 | 0.1 | 0.21 |
| 11 | 81472772 | rs12793528 | 1.753 | 1.176 | 0.171 | -1.705 | -0.002 | -0.048 | -0.048 | 0.007 | 0.31 | 0.28 | 0.31 | 0.03 | 0.28 | 0.2 |
| 11 | 81474640 | rs11232845 | -0.716 | 1.222 | 0.212 | -1.634 | -0.001 | 0.005 | 0.005 | 0.688 | 0.8 | 0.77 | 0.67 | 0.46 | 0.76 | 0.83 |
| 11 | 81477207 | rs2045607 | 1.924 | 1.103 | 0.14 | -1.893 | -0.003 | -0.07 | -0.07 | 0 | 0.28 | 0.29 | 0.29 | 0.01 | 0.24 | 0.17 |
| 11 | 81479993 | rs11232849 | 1.092 | 1.08 | -0.029 | -1.701 | -0.01 | -0.028 | -0.028 | 0.829 | 0.06 | 0.02 | 0.04 | 0.15 | 0.08 | 0.06 |
| 11 | 81485688 | rs11232856 | 1.952 | 1.155 | 0.091 | -1.96 | -0.001 | -0.022 | -0.022 | 0 | 0.28 | 0.21 | 0.28 | 0.01 | 0.26 | 0.18 |
| 11 | 81486744 | rs12294805 | 0.022 | 1.075 | 0.153 | -1.967 | 0.011 | 0.027 | 0.027 | 0 | 0.05 | 0.25 | 0.04 | 0.02 | 0.02 | 0.02 |
| 11 | 81490264 | rs17142932 | 0.818 | 1.309 | 0.036 | -1.89 | -0.022 | -0.047 | -0.047 | 0 | 0.15 | 0.06 | 0.21 | 0.57 | 0.18 | 0.19 |
| 11 | 81492541 | rs6591999 | 0.345 | 1.382 | 0.002 | -2.028 | -0.007 | -0.034 | -0.034 | 0.123 | 0.41 | 0.58 | 0.26 | 0.2 | 0.35 | 0.54 |
| 11 | 81492649 | rs7104475 | -0.341 | 1.422 | -0.429 | -1.997 | -0.001 | 0.064 | 0.064 | 0 | 0.55 | 0.66 | 0.64 | 0.82 | 0.54 | 0.45 |
| 11 | 81494492 | rs1870542 | 2.261 | 1.369 | -0.385 | -1.906 | -0.038 | 0.145 | 0.145 | 0 | 0.27 | 0.04 | 0.26 | 0.01 | 0.26 | 0.18 |
| 11 | 81494943 | rs7948962 | 0.127 | 1.353 | -0.117 | -1.578 | -0.001 | -0.012 | -0.012 | 0 | 0.39 | 0.41 | 0.25 | 0.2 | 0.35 | 0.54 |
| 11 | 81498152 | rs7950775 | 0.502 | 1.428 | -0.257 | -1.532 | 0.002 | 0.01 | 0.01 | 0.384 | 0.35 | 0.32 | 0.22 | 0.18 | 0.33 | 0.51 |
| 11 | 81505463 | rs11232869 | 0.771 | 1.515 | -0.585 | -1.155 | -0.041 | -0.034 | -0.034 | 0 | 0.08 | 0 | 0.18 | 0.59 | 0.11 | 0.15 |
| 11 | 81525713 | rs1459937 | 0.18 | 1.487 | -0.248 | -1.46 | -0.004 | -0.039 | -0.039 | 0 | 0.39 | 0.44 | 0.23 | 0.18 | 0.34 | 0.57 |
| 11 | 81526573 | rs11232882 | -1.1 | 1.518 | -0.101 | -1.5 | 0.003 | -0.036 | -0.036 | 0 | 0.88 | 0.84 | 0.82 | 0.98 | 0.85 | 0.94 |
| 11 | 81534503 | rs1013273 | 0.555 | 1.643 | 0.063 | -1.536 | 0.013 | 0.03 | 0.03 | 0.144 | 0.36 | 0.27 | 0.21 | 0.18 | 0.27 | 0.51 |
| 11 | 81538168 | rs7934819 | -0.171 | 2.16 | 0.179 | -1.577 | 0.001 | 0.018 | 0.018 | 0.293 | 0.43 | 0.59 | 0.27 | 0.2 | 0.39 | 0.59 |
| 11 | 81538570 | rs17143002 | 1.163 | 2.226 | 0.107 | -1.663 | -0.003 | 0.007 | 0.007 | 0 | 0.05 | 0.05 | 0.1 | 0.2 | 0.07 | 0.06 |
| 11 | 81539154 | rs12577924 | 0.627 | 2.208 | 0.144 | -1.689 | 0.005 | 0.03 | 0.03 | 0.162 | 0.36 | 0.27 | 0.21 | 0.18 | 0.3 | 0.51 |
| 11 | 81540109 | rs7121014 | -0.129 | 1.851 | -0.029 | -1.654 | 0.02 | 0.066 | 0.066 | 0.263 | 0.43 | 0.67 | 0.27 | 0.2 | 0.37 | 0.59 |
| 11 | 81548180 | rs7102304 | 1.812 | 1.157 | -0.255 | -1.816 | 0.027 | 0.071 | 0.071 | 0 | 0.16 | 0.03 | 0.07 | 0.04 | 0.1 | 0.2 |
| 11 | 81553155 | rs6592004 | -0.284 | 1.077 | -0.054 | -1.592 | 0.001 | -0.001 | -0.001 | 0 | 0.48 | 0.42 | 0.28 | 0.19 | 0.45 | 0.62 |
| 11 | 81556771 | rs1380532 | 1.223 | 1.044 | -0.15 | -1.598 | -0.002 | -0.002 | -0.002 | 0 | 0.33 | 0.33 | 0.45 | 0.58 | 0.32 | 0.26 |
| 11 | 81562454 | rs7939401 | 0.768 | 0.964 | -0.131 | -1.611 | -0.001 | 0.018 | 0.018 | 0.243 | 0.39 | 0.33 | 0.23 | 0.18 | 0.37 | 0.57 |
| 11 | 81563655 | rs1459944 | 0.119 | 0.673 | -0.339 | -1.75 | 0.004 | 0.007 | 0.007 | 0.454 | 0.81 | 0.72 | 0.72 | 0.77 | 0.77 | 0.88 |
| 11 | 81563834 | rs12362287 | 1.067 | 0.381 | -0.347 | -1.409 | 0.006 | 0.006 | 0.006 | 0 | 0.09 | 0.05 | 0.04 | 0.01 | 0.08 | 0.05 |
| 11 | 81567116 | rs11232900 | -0.061 | 0.153 | -0.163 | -1.643 | -0.01 | -0.026 | -0.026 | 0 | 0.39 | 0.45 | 0.42 | 0.21 | 0.34 | 0.27 |
| 11 | 81568850 | rs7113242 | 0.821 | 0.193 | -0.427 | -1.709 | 0.014 | 0.061 | 0.061 | 0 | 0.28 | 0.16 | 0.27 | 0.01 | 0.23 | 0.18 |
| 11 | 81569926 | rs10082665 | 0.288 | 0.356 | -0.411 | -1.426 | 0 | -0.004 | -0.004 | 0 | 0.33 | 0.33 | 0.39 | 0.19 | 0.31 | 0.29 |
| 11 | 81577369 | rs4944366 | 0.482 | -0.003 | -0.344 | -1.324 | -0.001 | -0.002 | -0.002 | 0 | 0.61 | 0.62 | 0.48 | 0.31 | 0.6 | 0.59 |
| 11 | 81578047 | rs10501529 | 0.874 | -0.152 | -0.257 | -1.752 | 0.004 | 0.034 | 0.034 | 0 | 0.4 | 0.32 | 0.21 | 0.29 | 0.33 | 0.42 |
| 11 | 81579864 | rs6592008 | 1.348 | -1.107 | -0.171 | -1.215 | 0.003 | -0.008 | -0.008 | 0 | 0.09 | 0.06 | 0.12 | 0.21 | 0.09 | 0.15 |
| 11 | 81582575 | rs1459951 | -0.953 | -1.072 | -0.277 | -1.15 | -0.045 | -0.013 | -0.013 | 0 | 0.92 | 0.99 | 0.73 | 0.58 | 0.84 | 0.89 |
| 11 | 81583196 | rs11607964 | 2.002 | -1.548 | -0.432 | -1.476 | -0.013 | -0.037 | -0.037 | 0.377 | 0.19 | 0.21 | 0.06 | 0.02 | 0.08 | 0.15 |
| 11 | 81583316 | rs11232906 | 0.024 | -1.273 | -0.22 | -1.251 | -0.001 | 0.04 | 0.04 | 0 | 0.69 | 0.6 | 0.6 | 0.52 | 0.68 | 0.74 |
| 11 | 81586661 | rs7936672 | 0.078 | -1.249 | -0.357 | -1.266 | -0.005 | -0.039 | -0.039 | 0 | 0.71 | 0.8 | 0.61 | 0.52 | 0.68 | 0.74 |
| 11 | 81592990 | rs11232912 | 1.545 | -1.235 | -0.546 | -1.363 | 0.006 | 0.036 | 0.036 | 0.058 | 0.13 | 0.04 | 0.06 | 0.03 | 0.11 | 0.06 |

| | | | | | | | | | | | | | | | | |
|---|---|---|---|---|---|---|---|---|---|---|---|---|---|---|---|---|
| 11 | 81595133 | rs7950689 | 0.984 | -1.11 | -0.224 | -1.237 | -0.01 | 0.027 | 0.027 | 0.104 | 0.1 | 0.15 | 0.13 | 0.21 | 0.09 | 0.15 |
| 11 | 81599981 | rs12290582 | 1.213 | -1.108 | -0.161 | -1.282 | -0.01 | 0.034 | 0.034 | 0.484 | 0.11 | 0.17 | 0.13 | 0.22 | 0.09 | 0.15 |
| 11 | 81601292 | rs17143063 | 1.763 | -1.526 | -0.4 | -1.508 | -0.005 | -0.009 | -0.009 | 0 | 0.21 | 0.2 | 0.06 | 0.07 | 0.07 | 0.16 |
| 11 | 81607066 | rs11232919 | 1.63 | -1.371 | -0.655 | -1.569 | -0.008 | 0.067 | 0.067 | 0 | 0.15 | 0.04 | 0.09 | 0 | 0.16 | 0.07 |
| 11 | 81616446 | rs11232921 | 0.673 | -1.289 | -0.608 | -1.587 | 0.032 | 0.036 | 0.036 | 0 | 0.38 | 0.24 | 0.19 | 0.31 | 0.27 | 0.42 |
| 11 | 81627342 | rs11232924 | 1.599 | -1.121 | -0.916 | -1.504 | -0.057 | -0.056 | -0.056 | 0 | 0.17 | 0.02 | 0.13 | 0.23 | 0.19 | 0.25 |
| 11 | 81636251 | rs11232928 | 0.102 | -0.212 | -0.217 | -1.029 | 0.006 | -0.006 | -0.006 | 0.615 | 0.31 | 0.3 | 0.39 | 0.26 | 0.42 | 0.4 |
| 11 | 81637494 | rs10792601 | -1.146 | 0.039 | -0.459 | -0.815 | -0.046 | -0.122 | -0.122 | 0 | 0.8 | 0.96 | 0.67 | 0.56 | 0.73 | 0.84 |
| 11 | 81648897 | rs7924797 | 0.636 | 0.047 | -0.438 | -0.934 | -0.001 | -0.003 | -0.003 | 0.099 | 0.62 | 0.41 | 0.46 | 0.32 | 0.59 | 0.6 |
| 11 | 81651069 | rs6592015 | -0.467 | 0.847 | 0.106 | -0.702 | 0.032 | -0.116 | -0.116 | 0.087 | 0.29 | 0.4 | 0.33 | 0.04 | 0.37 | 0.27 |
| 11 | 81653514 | rs7951571 | -0.805 | 0.35 | -0.052 | 0.224 | -0.003 | 0.08 | 0.08 | 0.312 | 0.14 | 0.35 | 0.19 | 0.25 | 0.14 | 0.23 |
| 11 | 81657804 | rs1944662 | 0.059 | 0.782 | -0.793 | 0.239 | -0.009 | -0.013 | -0.013 | 0 | 0.13 | 0.07 | 0.1 | 0.26 | 0.13 | 0.21 |
| 11 | 81660856 | rs10792602 | -0.058 | 0.865 | -1.117 | 0.747 | -0.018 | 0.009 | 0.009 | 0 | 0.92 | 0.98 | 0.72 | 0.58 | 0.89 | 0.89 |
| 11 | 81675050 | rs10897906 | -0.217 | 0.664 | -1.533 | -0.331 | -0.009 | -0.029 | -0.029 | 0 | 0.57 | 0.8 | 0.44 | 0.59 | 0.43 | 0.67 |
| 11 | 81680388 | rs1955011 | -0.097 | 1.257 | -1.802 | -0.781 | -0.059 | 0.021 | 0.021 | 0.035 | 0.62 | 0.86 | 0.51 | 0.33 | 0.58 | 0.53 |
| 11 | 81682177 | rs12797197 | 0.703 | 1.459 | -1.513 | -0.639 | -0.008 | -0.014 | -0.014 | 0.294 | 0.17 | 0.1 | 0.17 | 0 | 0.28 | 0.07 |
| 11 | 81685129 | rs582631 | 0.155 | 0.75 | -1.123 | -0.713 | 0.001 | -0.01 | -0.01 | 2.099 | 0.27 | 0.25 | 0.21 | 0.47 | 0.24 | 0.4 |
| 11 | 81688126 | rs1614922 | -1.87 | 0.588 | -1.463 | -1.12 | -0.026 | 0.048 | 0.048 | 0 | 0.9 | 0.98 | 0.84 | 0.99 | 0.88 | 0.96 |
| 11 | 81700376 | rs7941736 | 1.494 | 0.797 | -1.057 | -1.37 | -0.023 | 0.025 | 0.025 | 1.284 | 0.16 | 0.07 | 0.18 | 0 | 0.24 | 0.06 |
| 11 | 81709890 | rs664961 | -0.543 | 0.564 | -0.997 | -1.455 | -0.033 | 0.068 | 0.068 | 0.001 | 0.7 | 0.85 | 0.64 | 0.91 | 0.61 | 0.84 |
| 11 | 81711573 | rs601686 | 0.704 | 0.548 | -0.812 | -1.438 | 0.003 | -0.015 | -0.015 | 0.112 | 0.27 | 0.22 | 0.21 | 0.47 | 0.24 | 0.41 |
| 11 | 81715144 | rs12806136 | 1.411 | 0.371 | -1.107 | -1.553 | -0.03 | -0.11 | -0.11 | 0 | 0.3 | 0.09 | 0.37 | 0.41 | 0.31 | 0.31 |
| 11 | 81717389 | rs11232962 | 1.659 | 0.526 | -1.031 | -1.512 | -0.013 | 0.001 | 0.001 | 0.581 | 0.16 | 0.08 | 0.18 | 0 | 0.24 | 0.06 |
| 11 | 81726997 | rs10751067 | -0.999 | 0.484 | -0.657 | -1.395 | -0.031 | 0.01 | 0.01 | 0 | 0.78 | 0.92 | 0.69 | 0.74 | 0.74 | 0.79 |
| 11 | 81730079 | rs12362851 | -0.303 | 0.405 | -0.343 | -1.324 | -0.007 | -0.014 | -0.014 | 0.574 | 0.45 | 0.52 | 0.31 | 0.34 | 0.43 | 0.47 |
| 11 | 81730780 | rs599462 | 1.636 | 0.6 | -0.334 | -1.011 | -0.011 | -0.015 | -0.015 | 0 | 0.13 | 0.06 | 0.15 | 0.25 | 0.14 | 0.18 |
| 11 | 81752743 | rs656948 | -1.229 | 0.645 | -0.325 | -1.056 | -0.014 | -0.021 | -0.021 | 0 | 0.82 | 0.91 | 0.61 | 0.58 | 0.72 | 0.87 |
| 11 | 81754983 | rs538931 | -0.101 | 0.525 | -0.235 | -0.985 | 0.054 | 0.492 | 0.492 | 0.531 | 0.93 | 0.67 | 0.76 | 0.58 | 0.94 | 0.92 |
| 11 | 81770027 | rs2000522 | 0.861 | 0.301 | -0.698 | -1.369 | -0.019 | 0.156 | 0.156 | 0 | 0.35 | 0.04 | 0.16 | 0.12 | 0.3 | 0.4 |
| 11 | 81774312 | rs12799497 | 1.662 | 0.269 | -0.74 | -1.424 | -0.036 | 0.158 | 0.158 | 0 | 0.26 | 0.03 | 0.13 | 0.01 | 0.26 | 0.23 |
| 11 | 81778638 | rs478181 | 1.232 | 0.308 | -0.582 | -1.38 | 0.018 | 0.01 | 0.01 | 0 | 0.16 | 0.3 | 0.18 | 0.24 | 0.16 | 0.2 |
| 11 | 81782905 | rs666649 | 0.206 | 0.21 | -0.635 | -1.451 | -0.001 | 0.024 | 0.024 | 0.058 | 0.51 | 0.41 | 0.34 | 0.36 | 0.46 | 0.57 |
| 11 | 81785923 | rs612604 | 0.043 | 0.234 | -0.487 | -1.278 | 0 | -0.017 | -0.017 | 0.02 | 0.7 | 0.51 | 0.56 | 0.24 | 0.74 | 0.57 |
| 11 | 81792803 | rs645809 | -0.716 | 0.361 | -0.61 | -1.121 | 0.024 | 0.336 | 0.336 | 0 | 0.95 | 0.87 | 0.77 | 0.58 | 0.94 | 0.92 |
| 11 | 81797932 | rs541433 | 1.342 | 0.355 | -0.831 | -1.178 | 0.006 | 0.002 | 0.002 | 0 | 0.16 | 0.2 | 0.17 | 0.24 | 0.16 | 0.19 |
| 11 | 81804389 | rs648781 | 1.382 | -0.017 | -1.276 | -1.176 | -0.033 | 0.07 | 0.07 | 0 | 0.13 | 0.04 | 0.19 | 0 | 0.15 | 0.06 |
| 11 | 81808282 | rs17143248 | 0.557 | -0.17 | -0.804 | -0.901 | -0.047 | 0.03 | 0.03 | 0 | 0.18 | 0.39 | 0.09 | 0.22 | 0.11 | 0.25 |
| 11 | 81810647 | rs644497 | -0.519 | 0.076 | -0.817 | -0.835 | 0.021 | -0.01 | -0.01 | 0.768 | 0.64 | 0.5 | 0.43 | 0.32 | 0.55 | 0.67 |
| 11 | 81810977 | rs11607467 | 0.3 | -0.249 | -0.588 | -0.177 | 0.015 | -0.019 | -0.019 | 0 | 0.09 | 0.01 | 0.02 | 0.11 | 0.04 | 0.16 |
| 11 | 81813575 | rs574300 | -0.668 | 0.124 | 0.077 | 0.013 | -0.028 | 0.066 | 0.066 | 0 | 0.21 | 0.46 | 0.31 | 0.63 | 0.17 | 0.31 |
| 11 | 81814241 | rs583813 | -1.261 | 0.798 | -0.344 | 0.484 | -0.012 | 0.059 | 0.059 | 0.08 | 0.07 | 0.04 | 0.25 | 0.51 | 0.1 | 0.14 |
| 11 | 81818208 | rs586353 | -0.654 | 0.306 | -0.083 | -0.196 | -0.014 | 0.013 | 0.013 | 0 | 0.38 | 0.51 | 0.44 | 0.44 | 0.29 | 0.29 |
| 11 | 81841778 | rs17143262 | 0.377 | 0.126 | 0.156 | -0.42 | -0.012 | -0.06 | -0.06 | 0 | 0.2 | 0.25 | 0.09 | 0 | 0.13 | 0.11 |
| 11 | 81842796 | rs1892876 | 1.26 | 0.287 | -0.033 | -0.765 | -0.011 | -0.024 | -0.024 | 0 | 0.3 | 0.12 | 0.19 | 0.33 | 0.33 | 0.33 |
| 11 | 81885721 | rs581701 | -0.377 | 0.393 | 0.25 | -0.871 | -0.001 | 0.017 | 0.017 | 0 | 0.54 | 0.57 | 0.59 | 0.77 | 0.53 | 0.53 |
| 11 | 81895673 | rs7111047 | 0.508 | 0.417 | 0.298 | -0.766 | -0.002 | 0.158 | 0.158 | 0 | 0.08 | 0.17 | 0.23 | 0.44 | 0.06 | 0.16 |
| 11 | 81896531 | rs652732 | 1.073 | 0.488 | -0.06 | -0.803 | -0.013 | 0.03 | 0.03 | 0 | 0.09 | 0.03 | 0.12 | 0.01 | 0.11 | 0.02 |
| 11 | 81910326 | rs638506 | -1.156 | 0.597 | 0.025 | -0.774 | 0.004 | -0.006 | -0.006 | 0 | 0.73 | 0.68 | 0.67 | 0.77 | 0.66 | 0.67 |
| 11 | 81928156 | rs1380108 | 0.877 | 0.612 | -0.089 | -0.75 | 0.001 | 0 | 0 | 0 | 0.35 | 0.27 | 0.23 | 0.33 | 0.34 | 0.35 |
| 11 | 81947452 | rs1458952 | -0.158 | 0.283 | 0.34 | -0.843 | -0.018 | 0.024 | 0.024 | 0 | 0.29 | 0.33 | 0.34 | 0.42 | 0.2 | 0.29 |
| 11 | 81948546 | rs630849 | 0.449 | -0.062 | 0.357 | -0.858 | -0.01 | 0.032 | 0.032 | 0 | 0.45 | 0.29 | 0.32 | 0.33 | 0.47 | 0.38 |
| 11 | 81953762 | rs7946930 | -0.678 | -0.079 | 0.361 | -0.896 | -0.004 | 0.001 | 0.001 | 0 | 0.37 | 0.58 | 0.39 | 0.44 | 0.26 | 0.37 |
| 11 | 81957965 | rs2219479 | 1.543 | -0.14 | 0.345 | -0.924 | -0.001 | -0.016 | -0.016 | 0 | 0.3 | 0.16 | 0.19 | 0.33 | 0.3 | 0.33 |
| 11 | 81958954 | rs11233051 | 0.587 | -0.164 | 0.533 | -0.923 | 0.013 | 0.243 | 0.243 | 0 | 0.09 | 0.2 | 0.25 | 0.42 | 0.07 | 0.17 |
| 11 | 81959987 | rs11233052 | -0.483 | -0.181 | 0.601 | -0.939 | -0.001 | 0.241 | 0.241 | 0 | 0.15 | 0.43 | 0.3 | 0.43 | 0.11 | 0.25 |
| 11 | 81970174 | rs11828335 | 0.997 | -0.164 | 0.221 | -1.058 | 0.017 | 0.02 | 0.02 | 0 | 0.2 | 0.13 | 0.09 | 0 | 0.13 | 0.13 |
| 11 | 81973656 | rs603017 | -0.198 | -0.207 | 0.079 | -1.1 | 0.013 | 0.01 | 0.01 | 0 | 0.32 | 0.27 | 0.21 | 0.01 | 0.25 | 0.15 |
| 11 | 81979350 | rs11605655 | 1.561 | -0.186 | 0.028 | -1.088 | -0.008 | -0.032 | -0.032 | 0 | 0.31 | 0.12 | 0.18 | 0.33 | 0.32 | 0.33 |
| 11 | 81979892 | rs611530 | -0.671 | 0.019 | 0.159 | -0.923 | 0.017 | 0.012 | 0.012 | 0 | 0.69 | 0.64 | 0.44 | 0.35 | 0.59 | 0.55 |
| 11 | 81983618 | rs679930 | 1.055 | -0.173 | -0.08 | -0.855 | -0.004 | 0.028 | 0.028 | 0 | 0.1 | 0.08 | 0.11 | 0 | 0.1 | 0.02 |
| 11 | 81985860 | rs639429 | -0.672 | -0.196 | -0.042 | -0.739 | 0.005 | -0.004 | -0.004 | 0 | 0.38 | 0.35 | 0.42 | 0.43 | 0.32 | 0.35 |
| 11 | 81987786 | rs10897946 | 1.362 | -0.178 | 0.044 | -0.797 | -0.005 | 0.002 | 0.002 | 0 | 0.33 | 0.35 | 0.2 | 0.33 | 0.3 | 0.31 |
| 11 | 81995806 | rs12285098 | 1.001 | -0.224 | -0.086 | -0.925 | -0.015 | 0.118 | 0.118 | 0 | 0.2 | 0.01 | 0.08 | 0.01 | 0.14 | 0.17 |
| 11 | 81995844 | rs7945747 | -0.758 | -0.158 | -0.032 | -0.895 | 0.004 | -0.003 | -0.003 | 0 | 0.71 | 0.71 | 0.62 | 0.76 | 0.61 | 0.66 |
| 11 | 81997195 | rs10488823 | -1.477 | -0.166 | -0.073 | -0.9 | -0.062 | 0.001 | 0.001 | 0 | 0.77 | 0.9 | 0.67 | 0.78 | 0.65 | 0.74 |
| 11 | 82001588 | rs3914441 | 0.846 | -0.176 | -0.029 | -0.887 | 0.001 | -0.024 | -0.024 | 0.013 | 0.21 | 0.17 | 0.1 | 0.01 | 0.15 | 0.17 |
| 11 | 82008671 | rs3862761 | -1.277 | 0.118 | -0.204 | -0.587 | -0.077 | -0.003 | -0.003 | 0 | 0.77 | 0.91 | 0.67 | 0.78 | 0.65 | 0.74 |
| 11 | 82009932 | rs2374042 | -0.369 | 0.126 | -0.223 | -0.644 | -0.009 | 0.007 | 0.007 | 0 | 0.67 | 0.71 | 0.6 | 0.76 | 0.59 | 0.65 |
| 11 | 82017936 | rs9634011 | 0.521 | 0.133 | -0.237 | -0.427 | 0.016 | 0.058 | 0.058 | 0.02 | 0.39 | 0.15 | 0.43 | 0.74 | 0.37 | 0.47 |

| | | | | | | | | | | | | | | | | |
|---|---|---|---|---|---|---|---|---|---|---|---|---|---|---|---|---|
| 11 | 82031434 | rs12364945 | 0.659 | 0.179 | -0.23 | -0.533 | 0 | -0.041 | -0.041 | 0.169 | 0.3 | 0.29 | 0.17 | 0.01 | 0.25 | 0.19 |
| 11 | 82032218 | rs4144910 | -0.17 | 0.442 | -1.112 | -0.205 | 0.026 | 0.006 | 0.006 | 0 | 0.46 | 0.7 | 0.48 | 0.45 | 0.36 | 0.43 |
| 11 | 82039118 | rs9634012 | -0.851 | 0.483 | -1.384 | -0.107 | -0.042 | 0.004 | 0.004 | 0 | 0.77 | 0.85 | 0.66 | 0.78 | 0.64 | 0.74 |
| 11 | 82047518 | rs4335569 | 0.256 | 0.275 | -2.084 | 0.897 | -0.004 | -0.057 | -0.057 | 0 | 0.29 | 0.07 | 0.18 | 0.32 | 0.29 | 0.31 |
| 11 | 82049211 | rs17143426 | -0.708 | 0.46 | -1.197 | 1.583 | -0.013 | -0.019 | -0.019 | 0.035 | 0.08 | 0.01 | 0.24 | 0.42 | 0.07 | 0.15 |
| 11 | 82052822 | rs2175960 | -0.018 | 0.315 | -1.179 | 1.136 | 0.003 | -0.035 | -0.035 | 0 | 0.94 | 1 | 0.96 | 0.83 | 0.95 | 0.88 |
| 11 | 82065238 | rs7930387 | -0.724 | -0.243 | -0.971 | 1.089 | -0.006 | 0.016 | 0.016 | 0 | 0.27 | 0.3 | 0.3 | 0.46 | 0.22 | 0.21 |
| 11 | 82066388 | rs7126554 | 0.891 | -1.179 | -1.639 | 0.053 | 0.079 | 0.09 | 0.09 | 0.071 | 0.57 | 0.16 | 0.61 | 0.26 | 0.71 | 0.54 |
| 11 | 82076387 | rs1879904 | 0.748 | -1.255 | -0.911 | 0.025 | -0.024 | 0.007 | 0.007 | 0.034 | 0.67 | 0.51 | 0.66 | 0.37 | 0.74 | 0.68 |
| 11 | 82080010 | rs1399357 | 1.625 | -1.127 | -0.802 | -0.036 | 0.05 | 0.112 | 0.112 | 0.119 | 0.46 | 0.18 | 0.54 | 0.26 | 0.62 | 0.53 |
| 11 | 82081904 | rs1514621 | 0.93 | -1.256 | -0.851 | -0.074 | 0.081 | 0.065 | 0.065 | 0.673 | 0.61 | 0.16 | 0.63 | 0.26 | 0.75 | 0.58 |
| 11 | 82082981 | rs4015533 | 1.076 | -1.081 | -1.392 | -0.714 | -0.002 | 0.009 | 0.009 | 0.075 | 0.11 | 0.06 | 0.23 | 0.03 | 0.12 | 0.1 |
| 11 | 82085113 | rs4540849 | 0.851 | -1.227 | -1.055 | -0.45 | -0.042 | -0.022 | -0.022 | 0 | 0.16 | 0.31 | 0.07 | 0.09 | 0.08 | 0.13 |
| 11 | 82086661 | rs4944381 | -1.586 | -1.242 | -0.978 | -0.367 | -0.007 | 0.04 | 0.04 | 0.006 | 0.94 | 0.89 | 0.93 | 0.81 | 0.95 | 0.88 |
| 11 | 82089086 | rs1399358 | 1.353 | -1.439 | -0.588 | -0.48 | 0.004 | 0.034 | 0.034 | 0.157 | 0.06 | 0.12 | 0.03 | 0.07 | 0.03 | 0.05 |
| 11 | 82093499 | rs1949707 | -1.549 | -1.472 | -0.574 | -0.52 | -0.01 | 0.084 | 0.084 | 0.015 | 0.93 | 0.84 | 0.93 | 0.81 | 0.95 | 0.88 |
| 11 | 82108925 | rs1514609 | 0.463 | -1.617 | 0.073 | -0.676 | -0.029 | -0.076 | -0.076 | 0.123 | 0.12 | 0.4 | 0.07 | 0.03 | 0.06 | 0.08 |
| 11 | 82109154 | rs11606685 | 1.908 | -1.046 | -0.573 | -1.226 | -0.042 | 0.049 | 0.049 | 0 | 0.09 | 0.01 | 0.08 | 0 | 0.15 | 0.06 |
| 11 | 82130002 | rs4944388 | 1.845 | -1.128 | -0.74 | -1.265 | -0.119 | 0.049 | 0.049 | 0.014 | 0.36 | 0.08 | 0.45 | 0.24 | 0.45 | 0.46 |
| 11 | 82135969 | rs7121727 | 0.495 | -1.317 | -0.735 | -1.233 | 0.069 | 0.007 | 0.007 | 0 | 0.63 | 0.18 | 0.63 | 0.32 | 0.73 | 0.61 |
| 11 | 82142339 | rs11233120 | 1.707 | -1.233 | -0.703 | -1.159 | -0.015 | 0.066 | 0.066 | 0 | 0.11 | 0.01 | 0.08 | 0 | 0.1 | 0.02 |
| 11 | 82150867 | rs1399347 | -0.28 | -1.209 | -0.48 | -1.043 | -0.023 | 0.02 | 0.02 | 0.017 | 0.76 | 0.33 | 0.69 | 0.35 | 0.83 | 0.71 |
| 11 | 82158908 | rs12575649 | 1.766 | -1.28 | -0.541 | -1.023 | -0.009 | -0.092 | -0.092 | 0 | 0.13 | 0.01 | 0.19 | 0.46 | 0.11 | 0.17 |
| 11 | 82161234 | rs7103306 | 1.533 | -1.227 | -0.456 | -1.124 | -0.006 | -0.021 | -0.021 | 0 | 0.09 | 0.1 | 0.04 | 0.03 | 0.06 | 0.05 |
| 11 | 82165541 | rs2037503 | -0.424 | -1.24 | -0.45 | -1.068 | -0.051 | 0.032 | 0.032 | 0.007 | 0.82 | 0.47 | 0.87 | 0.78 | 0.88 | 0.83 |
| 11 | 82167757 | rs1125757 | 1.642 | -1.159 | -0.604 | -1.137 | -0.096 | 0.056 | 0.056 | 0 | 0.37 | 0.13 | 0.46 | 0.25 | 0.47 | 0.5 |
| 11 | 82194775 | rs17143592 | -0.221 | -1.136 | -0.185 | -0.786 | -0.033 | 0.076 | 0.076 | 0 | 0.25 | 0.39 | 0.25 | 0.49 | 0.17 | 0.23 |
| 11 | 82196876 | rs7111488 | -0.877 | -1.134 | -0.164 | -0.78 | 0.005 | 0.185 | 0.185 | 0.709 | 0.91 | 0.6 | 0.9 | 0.8 | 0.95 | 0.89 |
| 11 | 82197697 | rs10897976 | 0.565 | -1.142 | -0.288 | -0.794 | 0.05 | 0 | 0 | 0 | 0.66 | 0.21 | 0.65 | 0.31 | 0.78 | 0.66 |
| 11 | 82201517 | rs4631903 | -0.249 | -1.073 | -0.175 | -0.756 | -0.041 | 0.117 | 0.117 | 0.697 | 0.87 | 0.52 | 0.89 | 0.77 | 0.93 | 0.84 |
| 11 | 82220107 | rs9666156 | -0.233 | -0.605 | -0.107 | -0.829 | -0.009 | 0.032 | 0.032 | 0 | 0.25 | 0.3 | 0.24 | 0.48 | 0.18 | 0.23 |
| 11 | 82220634 | rs4258403 | -0.645 | -0.685 | 0.045 | -0.914 | -0.004 | 0.005 | 0.005 | 0 | 0.31 | 0.32 | 0.26 | 0.54 | 0.2 | 0.28 |
| 11 | 82221178 | rs4255559 | 0.23 | -0.674 | 0.089 | -1.156 | -0.013 | -0.009 | -0.009 | 0.894 | 0.67 | 0.54 | 0.49 | 0.3 | 0.7 | 0.53 |
| 11 | 82222510 | rs4345993 | 0.621 | -0.512 | -0.033 | -1.31 | -0.004 | -0.01 | -0.01 | 0 | 0.65 | 0.56 | 0.76 | 0.72 | 0.68 | 0.68 |
| 11 | 82224765 | rs4471453 | -0.938 | -0.358 | -0.176 | -1.199 | 0 | 0.084 | 0.084 | 0 | 0.9 | 0.76 | 0.9 | 0.8 | 0.91 | 0.87 |
| 11 | 82232390 | rs4271404 | 0.69 | -0.361 | -0.177 | -1.2 | -0.001 | 0.082 | 0.082 | 0 | 0.1 | 0.24 | 0.1 | 0.2 | 0.09 | 0.13 |
| 11 | 82232841 | rs1939616 | 2.071 | -0.452 | -0.452 | -1.463 | -0.049 | 0.04 | 0.04 | 0 | 0.3 | 0.12 | 0.26 | 0.21 | 0.37 | 0.31 |
| 11 | 82235975 | rs2226620 | -1.008 | -0.226 | 0.129 | -1.294 | -0.001 | -0.037 | -0.037 | 0 | 0.92 | 0.89 | 0.93 | 1 | 0.91 | 0.98 |
| 11 | 82236768 | rs1939587 | -0.034 | -0.252 | 0.19 | -1.277 | 0.005 | 0.085 | 0.085 | 0 | 0.85 | 0.57 | 0.84 | 0.8 | 0.86 | 0.86 |
| 11 | 82236778 | rs1939586 | 0.121 | -0.232 | 0.207 | -1.257 | -0.01 | 0.03 | 0.03 | 0 | 0.81 | 0.57 | 0.82 | 0.8 | 0.82 | 0.85 |
| 11 | 82238395 | rs4944403 | 0.974 | -0.125 | 0.051 | -1.167 | 0.002 | -0.006 | -0.006 | 0 | 0.09 | 0.08 | 0.22 | 0.05 | 0.11 | 0.15 |
| 11 | 82246660 | rs7936316 | 1.555 | -0.168 | 0.163 | -1.161 | -0.005 | 0.1 | 0.1 | 0 | 0.06 | 0.13 | 0.06 | 0.2 | 0.05 | 0.12 |
| 11 | 82250265 | rs1939641 | 0.54 | -0.124 | 0.135 | -1.455 | 0.005 | -0.014 | -0.014 | 0 | 0.7 | 0.64 | 0.79 | 0.77 | 0.69 | 0.72 |
| 11 | 82256583 | rs12808813 | 0.4 | -0.148 | 0.239 | -1.057 | -0.004 | 0.022 | 0.022 | 0.769 | 0.12 | 0.16 | 0.09 | 0.23 | 0.09 | 0.18 |
| 11 | 82260811 | rs1939633 | -0.298 | -0.039 | 0.212 | -0.914 | 0.006 | -0.036 | -0.036 | 0.304 | 0.13 | 0.18 | 0.2 | 0.04 | 0.14 | 0.18 |
| 11 | 82260936 | rs1939634 | 0.462 | -0.083 | 0.203 | -0.909 | 0.002 | -0.019 | -0.019 | 0 | 0.09 | 0.12 | 0.22 | 0.04 | 0.11 | 0.16 |
| 11 | 82262056 | rs10792626 | -0.078 | -0.344 | 0.242 | -0.883 | -0.042 | 0.151 | 0.151 | 0 | 0.21 | 0.41 | 0.25 | 0.46 | 0.13 | 0.22 |
| 11 | 82265147 | rs10897988 | 0.031 | 0.237 | 0.029 | -1 | 0.003 | -0.024 | -0.024 | 0 | 0.1 | 0.14 | 0.21 | 0.04 | 0.11 | 0.17 |
| 11 | 82267903 | rs12421806 | 0.976 | 0.243 | 0.014 | -1.008 | -0.001 | 0.001 | 0.001 | 0.021 | 0.74 | 0.75 | 0.67 | 0.78 | 0.72 | 0.67 |
| 11 | 82268478 | rs6592052 | -0.921 | 0.34 | -0.09 | -1.011 | -0.001 | -0.016 | -0.016 | 0.302 | 0.93 | 0.93 | 0.94 | 1 | 0.93 | 0.98 |
| 11 | 82268963 | rs11233170 | -1.328 | 0.246 | 0.147 | -0.981 | -0.008 | -0.001 | -0.001 | 0.081 | 0.24 | 0.39 | 0.32 | 0.26 | 0.25 | 0.32 |
| 11 | 82269240 | rs10897991 | -1.445 | 0.285 | 0.046 | -0.991 | -0.006 | -0.001 | -0.001 | 2.465 | 0.25 | 0.45 | 0.32 | 0.27 | 0.26 | 0.33 |
| 11 | 82270844 | rs7933681 | -0.644 | 0.313 | 0.143 | -0.869 | 0.014 | 0.103 | 0.103 | 0 | 0.12 | 0.38 | 0.1 | 0.18 | 0.12 | 0.16 |
| 11 | 82271989 | rs4578412 | -0.399 | -0.121 | -0.611 | -1.153 | 0.002 | 0.077 | 0.077 | 0.212 | 0.86 | 0.98 | 0.86 | 1 | 0.9 | 0.9 |
| 11 | 82278440 | rs1939593 | 0.031 | 0.866 | -0.355 | -1.046 | 0.032 | 0.066 | 0.066 | 0 | 0.09 | 0.16 | 0.09 | 0.18 | 0.13 | 0.16 |
| 11 | 82289050 | rs12363011 | -0.322 | 1.191 | -0.882 | -1.396 | -0.013 | 0.042 | 0.042 | 0 | 0.18 | 0.08 | 0.25 | 0.04 | 0.18 | 0.21 |
| 11 | 82297773 | rs10792629 | -0.221 | 1.12 | -0.71 | -1.219 | 0.012 | 0.03 | 0.03 | 0 | 0.86 | 0.95 | 0.94 | 1 | 0.9 | 0.9 |
| 11 | 82298067 | rs2155251 | 0.189 | 1.306 | -0.357 | -0.561 | 0.015 | 0.029 | 0.029 | 0.008 | 0.74 | 0.65 | 0.83 | 0.66 | 0.7 | 0.61 |
| 11 | 82299294 | rs11606514 | 0.644 | 0.695 | -0.869 | -1.314 | 0.006 | 0.023 | 0.023 | 0 | 0.38 | 0.28 | 0.24 | 0.05 | 0.38 | 0.2 |
| 11 | 82303117 | rs12418556 | 0.501 | 0.36 | -1.213 | -0.097 | -0.006 | -0.153 | -0.153 | 1.578 | 0.2 | 0.01 | 0.22 | 0.41 | 0.15 | 0.22 |
| 11 | 82307795 | rs1939602 | -0.426 | 0.062 | -0.863 | -0.923 | -0.012 | -0.062 | -0.062 | 4.762 | 0.27 | 0.36 | 0.17 | 0.08 | 0.2 | 0.22 |
| 11 | 82317460 | rs1939600 | 0.165 | 0.226 | -0.566 | -1.085 | -0.004 | -0.057 | -0.057 | 0 | 0.68 | 0.63 | 0.81 | 0.92 | 0.74 | 0.76 |
| 11 | 82320120 | rs10501534 | 1.233 | 0.505 | -0.179 | -0.809 | 0.004 | -0.025 | -0.025 | 0 | 0.44 | 0.45 | 0.53 | 0.31 | 0.54 | 0.46 |
| 11 | 82321101 | rs2155253 | -0.751 | 0.429 | -0.424 | -0.85 | 0.011 | 0.11 | 0.11 | 0 | 0.83 | 0.98 | 0.94 | 0.99 | 0.9 | 0.85 |
| 11 | 82324065 | rs11233195 | 0.259 | 0.706 | -0.075 | -0.659 | 0.02 | -0.041 | -0.041 | 0.504 | 0.14 | 0.29 | 0.12 | 0.09 | 0.15 | 0.09 |
| 11 | 82328254 | rs7933266 | -0.289 | 0.611 | -0.271 | -0.607 | 0.037 | -0.074 | -0.074 | 0 | 0.62 | 0.88 | 0.7 | 0.54 | 0.72 | 0.62 |
| 11 | 82328339 | rs10898007 | 1.134 | 0.63 | -0.129 | -0.8 | 0.001 | -0.01 | -0.01 | 0 | 0.44 | 0.43 | 0.53 | 0.29 | 0.53 | 0.45 |
| 11 | 82329371 | rs12223804 | 0.914 | 0.649 | -0.14 | -0.707 | -0.005 | 0.009 | 0.009 | 0 | 0.09 | 0.05 | 0.08 | 0.07 | 0.12 | 0.05 |
| 11 | 82335447 | rs11233206 | 0.407 | 0.548 | -0.18 | -0.73 | 0 | 0.001 | 0.001 | 0 | 0.09 | 0.09 | 0.08 | 0.05 | 0.11 | 0.06 |

| | | | | | | | | | | | | | | | | |
|---|---|---|---|---|---|---|---|---|---|---|---|---|---|---|---|---|
| 11 | 82340750 | rs7947279 | 0.53 | 1.028 | 0.22 | -0.896 | 0.035 | -0.028 | -0.028 | 0.24 | 0.06 | 0.14 | 0.05 | 0 | 0.11 | 0.03 |
| 11 | 82343713 | rs9804615 | -0.761 | 0.37 | -0.106 | -0.986 | -0.002 | -0.021 | -0.021 | 0 | 0.2 | 0.21 | 0.12 | 0.06 | 0.17 | 0.13 |
| 11 | 82346617 | rs1939629 | 0.17 | 0.4 | -0.098 | -0.967 | 0.003 | 0.008 | 0.008 | 0 | 0.15 | 0.11 | 0.1 | 0.06 | 0.14 | 0.09 |
| 11 | 82347495 | rs1939627 | 0.063 | 0.447 | -0.233 | -0.753 | 0.038 | -0.037 | -0.037 | 0.288 | 0.38 | 0.16 | 0.29 | 0.46 | 0.29 | 0.29 |
| 11 | 82350270 | rs7937096 | -0.516 | 0.44 | -0.038 | -0.771 | 0.012 | -0.001 | -0.001 | 0 | 0.44 | 0.38 | 0.31 | 0.46 | 0.32 | 0.35 |
| 11 | 82352236 | rs1784206 | 0.949 | 0.732 | -0.089 | -0.874 | -0.001 | 0 | 0 | 0 | 0.45 | 0.4 | 0.6 | 0.53 | 0.57 | 0.53 |
| 11 | 82352407 | rs4143166 | -0.219 | 0.495 | -0.253 | -0.476 | 0.035 | -0.027 | -0.027 | 0.046 | 0.38 | 0.18 | 0.29 | 0.46 | 0.29 | 0.31 |
| 11 | 82354573 | rs7932435 | -0.528 | 0.62 | -0.302 | -0.366 | -0.012 | -0.003 | -0.003 | 0 | 0.44 | 0.49 | 0.33 | 0.47 | 0.33 | 0.36 |
| 11 | 82354978 | rs1793025 | -0.488 | 0.891 | -0.77 | 0.146 | 0 | -0.001 | -0.001 | 0.34 | 0.38 | 0.4 | 0.55 | 0.45 | 0.39 | 0.45 |
| 11 | 82355066 | rs1790713 | 0.264 | 0.582 | -0.982 | -0.119 | 0 | 0.001 | 0.001 | 0 | 0.13 | 0.13 | 0.23 | 0.11 | 0.11 | 0.13 |
| 11 | 82355489 | rs10751076 | -0.492 | 0.448 | -1.112 | -0.545 | -0.007 | 0.085 | 0.085 | 0.129 | 0.55 | 0.69 | 0.65 | 0.79 | 0.43 | 0.67 |
| 11 | 82356268 | rs17144007 | -0.052 | 0.363 | -0.02 | -0.216 | 0.025 | -0.008 | -0.008 | 0 | 0.07 | 0.3 | 0.03 | 0.01 | 0.04 | 0.08 |
| 11 | 82357515 | rs12789469 | -0.427 | 0.476 | -0.347 | -0.104 | -0.001 | -0.001 | -0.001 | 0 | 0.63 | 0.6 | 0.4 | 0.47 | 0.49 | 0.5 |
| 11 | 82357959 | rs11233230 | 0.138 | 0.379 | -0.356 | -0.27 | 0 | 0 | 0 | 0 | 0.55 | 0.43 | 0.35 | 0.46 | 0.42 | 0.45 |
| 11 | 82358110 | rs11233231 | 0.671 | 0.397 | -1.139 | -0.224 | -0.018 | -0.041 | -0.041 | 0 | 0.47 | 0.11 | 0.32 | 0.46 | 0.38 | 0.35 |
| 11 | 82358182 | rs11233232 | 0.937 | 0.479 | -1.427 | -0.113 | -0.011 | -0.037 | -0.037 | 0.246 | 0.41 | 0.07 | 0.24 | 0.35 | 0.33 | 0.25 |
| 11 | 82358636 | rs7111078 | -0.206 | 1.061 | -0.829 | 0.223 | -0.025 | -0.042 | -0.042 | 0.055 | 0.67 | 0.76 | 0.44 | 0.47 | 0.55 | 0.51 |
| 11 | 82370960 | rs7947389 | -0.244 | 1.077 | -0.927 | 0.41 | 0.043 | 0.032 | 0.032 | 0.025 | 0.11 | 0.33 | 0.3 | 0.13 | 0.18 | 0.21 |
| 11 | 82381787 | rs10898021 | -0.679 | 0.811 | -1.73 | 0.43 | -0.065 | -0.101 | -0.101 | 1.137 | 0.74 | 0.94 | 0.69 | 0.59 | 0.66 | 0.7 |
| 11 | 82395784 | rs7940296 | -0.763 | 0.659 | -1.078 | 0.075 | -0.004 | 0.102 | 0.102 | 0.03 | 0.5 | 0.68 | 0.6 | 0.67 | 0.42 | 0.6 |
| 11 | 82398420 | rs11233253 | -0.17 | 0.09 | -1.331 | 0.262 | 0.001 | -0.044 | -0.044 | 0.501 | 0.16 | 0.03 | 0.26 | 0.19 | 0.13 | 0.26 |
| 11 | 82400243 | rs7125061 | -0.99 | -0.077 | -0.485 | -0.004 | 0.013 | -0.055 | -0.055 | 0.048 | 0.11 | 0.31 | 0.07 | 0.01 | 0.12 | 0.08 |
| 11 | 82401684 | rs11233261 | -1.044 | -0.03 | -0.645 | -0.036 | -0.002 | -0.003 | -0.003 | 0.152 | 0.23 | 0.22 | 0.32 | 0.19 | 0.25 | 0.28 |
| 11 | 82405181 | rs10792636 | -1.704 | 0.224 | -0.881 | -0.093 | -0.073 | 0.128 | 0.128 | 0.051 | 0.55 | 0.86 | 0.61 | 0.67 | 0.47 | 0.6 |
| 11 | 82406633 | rs7943210 | -1.617 | 0.241 | -0.819 | -0.101 | -0.063 | 0.145 | 0.145 | 0.011 | 0.53 | 0.87 | 0.61 | 0.68 | 0.47 | 0.6 |
| 11 | 82422946 | rs1472401 | 0.203 | 0.032 | -0.836 | -0.127 | -0.009 | -0.024 | -0.024 | 0 | 0.13 | 0.05 | 0.25 | 0.14 | 0.12 | 0.22 |
| 11 | 82423811 | rs7940878 | -0.454 | -0.012 | -0.681 | -0.173 | -0.001 | -0.041 | -0.041 | 0 | 0.11 | 0.18 | 0.07 | 0.05 | 0.13 | 0.07 |
| 11 | 82430245 | rs524736 | -0.975 | -0.207 | -0.268 | -0.044 | 0 | -0.008 | -0.008 | 0.09 | 0.77 | 0.77 | 0.77 | 0.48 | 0.82 | 0.76 |
| 11 | 82433268 | rs510621 | 1.216 | -0.134 | -0.404 | -0.09 | -0.007 | 0.198 | 0.198 | 0 | 0.49 | 0.09 | 0.42 | 0.31 | 0.62 | 0.38 |
| 11 | 82435085 | rs1353600 | 0.664 | -0.2 | 0.032 | 0.19 | 0.015 | -0.014 | -0.014 | 0 | 0.62 | 0.13 | 0.66 | 0.41 | 0.74 | 0.62 |
| 11 | 82435414 | rs1353599 | -0.489 | -0.34 | 0.432 | 0.187 | 0.061 | 0.029 | 0.029 | 0 | 0.06 | 0.37 | 0.05 | 0.01 | 0.03 | 0.07 |
| 11 | 82441634 | rs7120212 | -1.028 | -0.343 | 0.172 | 0.222 | 0.042 | -0.017 | -0.017 | 0 | 0.37 | 0.83 | 0.32 | 0.55 | 0.24 | 0.38 |
| 11 | 82449606 | rs9666486 | -0.395 | -0.35 | 0.502 | 0.16 | 0.084 | 0.056 | 0.056 | 0 | 0.06 | 0.4 | 0.05 | 0.01 | 0.02 | 0.08 |
| 11 | 82449718 | rs9665943 | -1.143 | -0.45 | 0.206 | 0.298 | 0.051 | -0.04 | -0.04 | 1.102 | 0.39 | 0.87 | 0.34 | 0.53 | 0.25 | 0.38 |
| 11 | 82456494 | rs10898028 | -0.383 | -0.399 | 0.424 | 0.331 | -0.008 | 0.01 | 0.01 | 0 | 0.32 | 0.35 | 0.28 | 0.53 | 0.22 | 0.31 |
| 11 | 82478824 | rs492923 | 0.715 | -0.437 | 0.315 | 1.119 | -0.042 | -0.123 | -0.123 | 5.366 | 0.19 | 0.01 | 0.19 | 0.34 | 0.15 | 0.21 |
| 11 | 82483423 | rs530158 | -0.217 | -0.525 | 0.461 | 0.934 | 0.014 | -0.094 | -0.094 | 0 | 0.27 | 0.1 | 0.25 | 0.51 | 0.2 | 0.32 |
| 11 | 82489754 | rs10792641 | 0.056 | -0.042 | 0.343 | 0.915 | 0.039 | 0.102 | 0.102 | 0 | 0.08 | 0.15 | 0.17 | 0.26 | 0.12 | 0.1 |
| 11 | 82498536 | rs471755 | -1.101 | -0.142 | 0.532 | 0.823 | 0.006 | 0.064 | 0.064 | 0 | 0.43 | 0.8 | 0.47 | 0.72 | 0.35 | 0.49 |
| 11 | 82501382 | rs492584 | -1.637 | -0.654 | 0.318 | 0.519 | 0.021 | 0.133 | 0.133 | 0 | 0.51 | 0.82 | 0.5 | 0.76 | 0.39 | 0.52 |
| 11 | 82503394 | rs542382 | 0.382 | -0.413 | 0.356 | 0.62 | 0.013 | -0.006 | -0.006 | 0 | 0.12 | 0.06 | 0.11 | 0.51 | 0.07 | 0.23 |
| 11 | 82508987 | rs551450 | -0.764 | -0.421 | 0.815 | 0.604 | -0.035 | 0.19 | 0.19 | 0.057 | 0.18 | 0.42 | 0.14 | 0.51 | 0.09 | 0.3 |
| 11 | 82510454 | rs2645762 | -1.055 | -0.56 | 0.826 | 0.548 | 0.004 | 0.021 | 0.021 | 0 | 0.43 | 0.62 | 0.54 | 0.72 | 0.36 | 0.52 |
| 11 | 82512587 | rs10898032 | 0.105 | -0.458 | 0.562 | 0.526 | 0.002 | -0.009 | -0.009 | 0.589 | 0.28 | 0.25 | 0.42 | 0.21 | 0.3 | 0.22 |
| 11 | 82522085 | rs2510282 | -1.347 | -0.341 | 0.238 | 0.518 | 0.012 | 0.11 | 0.11 | 0 | 0.48 | 0.77 | 0.58 | 0.73 | 0.4 | 0.56 |
| 11 | 82523453 | rs10898033 | 0.068 | -0.137 | 0.361 | 0.56 | -0.008 | 0.011 | 0.011 | 0 | 0.27 | 0.18 | 0.39 | 0.22 | 0.27 | 0.3 |
| 11 | 82535963 | rs4759 | 0.209 | -0.635 | -0.019 | 0.701 | 0.013 | -0.064 | -0.064 | 0 | 0.12 | 0.03 | 0.11 | 0.55 | 0.07 | 0.25 |
| 11 | 82536795 | rs11233329 | 0.271 | -0.266 | 0.026 | 0.266 | -0.04 | 0.058 | 0.058 | 0 | 0.23 | 0.05 | 0.33 | 0.15 | 0.22 | 0.18 |
| 11 | 82539504 | rs10488773 | 0.004 | -0.293 | 0.071 | 0.245 | -0.016 | 0.07 | 0.07 | 0 | 0.25 | 0.1 | 0.33 | 0.16 | 0.22 | 0.25 |
| 11 | 82540789 | rs7107322 | -0.217 | -0.752 | -0.171 | -0.425 | -0.028 | -0.042 | -0.042 | 0 | 0.27 | 0.5 | 0.27 | 0.15 | 0.23 | 0.21 |
| 11 | 82550387 | rs4843 | -0.414 | -0.878 | 0.069 | -0.425 | 0.029 | -0.012 | -0.012 | 0.498 | 0.07 | 0.27 | 0.04 | 0.01 | 0.03 | 0.11 |
| 11 | 82552745 | rs9326440 | -1.205 | -0.938 | -0.013 | -0.517 | 0.003 | 0.056 | 0.056 | 0 | 0.55 | 0.72 | 0.62 | 0.32 | 0.45 | 0.46 |
| 11 | 82552926 | rs11233339 | 0.546 | -0.799 | -0.044 | -0.451 | 0.019 | 0.008 | 0.008 | 0 | 0.21 | 0.16 | 0.13 | 0.16 | 0.17 | 0.17 |
| 11 | 82557662 | rs5006165 | -0.712 | -0.73 | 0.044 | -0.424 | 0.006 | 0.018 | 0.018 | 0 | 0.5 | 0.48 | 0.6 | 0.32 | 0.44 | 0.45 |
| 11 | 82560312 | rs10792653 | -0.686 | -0.722 | 0.054 | -0.347 | 0.009 | 0.025 | 0.025 | 0 | 0.5 | 0.47 | 0.62 | 0.33 | 0.44 | 0.46 |
| 11 | 82560445 | rs10792654 | -0.696 | -0.713 | 0.066 | -0.346 | 0.004 | 0.012 | 0.012 | 0 | 0.5 | 0.49 | 0.62 | 0.33 | 0.44 | 0.45 |
| 11 | 82563283 | rs7106555 | -0.25 | -0.584 | 0.291 | -0.532 | 0.002 | 0.003 | 0.003 | 0.017 | 0.37 | 0.64 | 0.52 | 0.19 | 0.3 | 0.29 |
| 11 | 82563803 | rs7123485 | -1.1 | -0.404 | 0.358 | -0.382 | 0.026 | 0.008 | 0.008 | 0 | 0.08 | 0.35 | 0.16 | 0.02 | 0.03 | 0.11 |
| 11 | 82578953 | rs10898045 | 0.405 | -0.468 | 0.083 | -0.541 | -0.006 | 0.064 | 0.064 | 0 | 0.3 | 0.18 | 0.35 | 0.17 | 0.28 | 0.19 |
| 11 | 82585356 | rs2086297 | -1.188 | -0.222 | 0.14 | -0.56 | 0.04 | -0.025 | -0.025 | 0.736 | 0.08 | 0.3 | 0.15 | 0.01 | 0.04 | 0.11 |
| 11 | 82587612 | rs11233359 | 0.48 | -0.19 | -0.378 | -0.475 | -0.047 | 0.069 | 0.069 | 0.096 | 0.29 | 0.08 | 0.34 | 0.18 | 0.28 | 0.2 |
| 11 | 82592731 | rs12275863 | 0.492 | -0.19 | -0.325 | -0.495 | -0.023 | 0.074 | 0.074 | 0.231 | 0.29 | 0.11 | 0.35 | 0.17 | 0.28 | 0.2 |
| 11 | 82598809 | rs7114521 | -0.954 | -0.423 | -0.193 | -0.653 | -0.014 | 0.059 | 0.059 | 0 | 0.56 | 0.73 | 0.64 | 0.29 | 0.48 | 0.47 |
| 11 | 82602814 | rs11233363 | 0.765 | -0.334 | -0.139 | -0.539 | 0.014 | 0.024 | 0.024 | 0 | 0.17 | 0.09 | 0.12 | 0.12 | 0.16 | 0.16 |
| 11 | 82605135 | rs10792662 | 0.642 | -0.335 | -0.146 | -0.541 | -0.038 | 0.075 | 0.075 | 0.116 | 0.29 | 0.09 | 0.35 | 0.17 | 0.28 | 0.2 |
| 11 | 82608890 | rs11233366 | -0.281 | -0.376 | 0.315 | -0.501 | -0.035 | -0.069 | -0.069 | 0.1 | 0.23 | 0.48 | 0.26 | 0.12 | 0.18 | 0.24 |
| 11 | 82615920 | rs11233368 | 0.714 | -0.353 | 0.113 | -0.483 | 0.014 | 0.024 | 0.024 | 0 | 0.17 | 0.09 | 0.12 | 0.11 | 0.16 | 0.18 |
| 11 | 82620671 | rs7947363 | -0.591 | -0.345 | 0.381 | -0.508 | 0.002 | 0.005 | 0.005 | 0 | 0.51 | 0.52 | 0.6 | 0.29 | 0.45 | 0.47 |
| 11 | 82643760 | rs7947780 | 0.07 | -0.341 | 0.183 | -0.249 | -0.019 | 0.018 | 0.018 | 0 | 0.35 | 0.18 | 0.4 | 0.32 | 0.36 | 0.34 |

| | | | | | | | | | | | | | | | | |
|---|---|---|---|---|---|---|---|---|---|---|---|---|---|---|---|---|
| 11 | 82644904 | rs7130899 | -0.982 | -0.125 | 0.36 | -0.297 | -0.022 | 0.052 | 0.052 | 0 | 0.56 | 0.77 | 0.64 | 0.34 | 0.5 | 0.48 |
| 11 | 82646964 | rs4944419 | -0.447 | -0.16 | 0.451 | -0.301 | 0.01 | 0.117 | 0.117 | 0 | 0.48 | 0.26 | 0.48 | 0.3 | 0.45 | 0.37 |
| 11 | 82651993 | rs7933350 | -1.762 | -0.5 | 0.182 | -0.031 | -0.047 | 0.034 | 0.034 | 0 | 0.69 | 0.86 | 0.77 | 0.8 | 0.57 | 0.7 |
| 11 | 82672020 | rs10898063 | -0.77 | -0.242 | 0.345 | -0.013 | 0 | 0.029 | 0.029 | 0 | 0.53 | 0.65 | 0.62 | 0.35 | 0.47 | 0.47 |
| 11 | 82680739 | rs11233390 | 0.883 | -0.106 | -0.099 | 0.07 | -0.085 | -0.01 | -0.01 | 0.685 | 0.27 | 0.03 | 0.36 | 0.22 | 0.29 | 0.17 |
| 11 | 82686588 | rs7128677 | 0.982 | -0.189 | -0.104 | 0.06 | -0.089 | 0.01 | 0.01 | 0 | 0.26 | 0.06 | 0.36 | 0.21 | 0.3 | 0.19 |
| 11 | 82691050 | rs1011618 | -0.974 | -0.184 | 0.179 | -0.012 | 0.08 | 0.014 | 0.014 | 0 | 0.06 | 0.29 | 0.05 | 0.01 | 0.05 | 0.11 |
| 11 | 82691952 | rs11233398 | -1.204 | -0.518 | 0.177 | -0.109 | -0.018 | 0.042 | 0.042 | 0 | 0.58 | 0.75 | 0.53 | 0.34 | 0.51 | 0.54 |
| 11 | 82692131 | rs17144453 | 0.265 | -0.387 | 0.262 | -0.073 | -0.003 | 0.004 | 0.004 | 0.233 | 0.05 | 0.08 | 0.03 | 0.11 | 0.05 | 0.09 |
| 11 | 82705431 | rs12293386 | 1.398 | -0.631 | 0.249 | -0.112 | 0.018 | 0.061 | 0.061 | 0 | 0.14 | 0.05 | 0.07 | 0.01 | 0.11 | 0.1 |
| 11 | 82709248 | rs17144481 | 1.778 | -0.894 | 0.094 | 0.254 | 0.022 | -0.1 | -0.1 | 0 | 0.14 | 0.01 | 0.08 | 0.34 | 0.07 | 0.17 |
| 11 | 82714248 | rs1904427 | -0.985 | -0.932 | 0.107 | 0.282 | -0.015 | 0.013 | 0.013 | 0 | 0.58 | 0.75 | 0.58 | 0.4 | 0.53 | 0.57 |
| 11 | 82719880 | rs7120604 | 0.431 | -0.914 | 0.148 | 0.253 | -0.002 | 0.064 | 0.064 | 0 | 0.06 | 0.26 | 0.07 | 0.06 | 0.01 | 0.04 |
| 11 | 82723072 | rs11233413 | 0.828 | -0.709 | -0.063 | 0.216 | -0.007 | 0.023 | 0.023 | 0 | 0.3 | 0.15 | 0.35 | 0.27 | 0.31 | 0.28 |
| 11 | 82723907 | rs17144500 | -0.782 | -0.644 | -0.135 | 0.12 | 0.114 | -0.023 | -0.023 | 0 | 0.08 | 0.29 | 0.08 | 0.01 | 0.09 | 0.13 |
| 11 | 82726227 | rs10431152 | 1.657 | -0.928 | -0.212 | 0.271 | 0.023 | -0.101 | -0.101 | 0 | 0.14 | 0.01 | 0.08 | 0.34 | 0.07 | 0.18 |
| 11 | 82736049 | rs9804629 | 0.776 | -0.321 | 0.35 | -0.213 | -0.011 | 0.024 | 0.024 | 0 | 0.18 | 0.13 | 0.12 | 0.01 | 0.18 | 0.15 |
| 11 | 82745527 | rs17504704 | 1.028 | -0.324 | 0.255 | -0.198 | -0.054 | 0.053 | 0.053 | 0 | 0.24 | 0.04 | 0.33 | 0.13 | 0.25 | 0.18 |
| 11 | 82761790 | rs9971403 | -1.162 | -0.163 | 0.26 | -0.129 | -0.017 | 0.033 | 0.033 | 0 | 0.58 | 0.75 | 0.57 | 0.34 | 0.52 | 0.55 |
| 11 | 82776861 | rs4944425 | -0.4 | -0.066 | 0.052 | 0.323 | 0.005 | 0.002 | 0.002 | 0 | 0.36 | 0.39 | 0.42 | 0.35 | 0.32 | 0.34 |
| 11 | 82782686 | rs755514 | 0.353 | 0.145 | -0.273 | 0.171 | -0.076 | 0.047 | 0.047 | 0.083 | 0.25 | 0.08 | 0.36 | 0.18 | 0.3 | 0.2 |
| 11 | 82789044 | rs9666461 | -0.546 | 0.058 | 0.523 | 0.463 | -0.006 | 0.006 | 0.006 | 0 | 0.33 | 0.66 | 0.25 | 0.42 | 0.26 | 0.49 |
| 11 | 82791812 | rs10444381 | 0.885 | 0.034 | 0.458 | 0.523 | -0.084 | 0.036 | 0.036 | 0 | 0.23 | 0.05 | 0.36 | 0.19 | 0.3 | 0.17 |
| 11 | 82791959 | rs17144547 | 0.315 | -0.004 | 0.656 | 0.498 | 0.01 | 0.087 | 0.087 | 0 | 0.06 | 0.2 | 0.04 | 0.11 | 0.06 | 0.13 |
| 11 | 82795854 | rs4586187 | -1.443 | -0.076 | 0.558 | 0.46 | -0.015 | -0.03 | -0.03 | 0 | 0.6 | 0.86 | 0.63 | 0.61 | 0.57 | 0.67 |
| 11 | 82798114 | rs7125035 | -0.935 | -0.225 | 0.816 | 0.563 | 0.031 | -0.019 | -0.019 | 2.956 | 0.37 | 0.8 | 0.27 | 0.42 | 0.27 | 0.5 |
| 11 | 82798226 | rs17144566 | -0.145 | 0.032 | 0.427 | 0.511 | -0.013 | 0.01 | 0.01 | 0 | 0.27 | 0.46 | 0.21 | 0.31 | 0.21 | 0.31 |
| 11 | 82819298 | rs17159801 | -0.849 | 0.371 | -0.268 | 0.644 | 0.042 | 0.14 | 0.14 | 0 | 0.14 | 0.46 | 0.24 | 0.28 | 0.12 | 0.17 |
| 11 | 82820283 | rs11233471 | 0.581 | 0.465 | -0.611 | -0.375 | 0.004 | 0.046 | 0.046 | 0 | 0.18 | 0.11 | 0.1 | 0.01 | 0.14 | 0.18 |
| 11 | 82821382 | rs4293143 | 0.192 | 0.746 | -0.207 | 0.389 | -0.04 | 0.027 | 0.027 | 0 | 0.21 | 0.08 | 0.23 | 0.18 | 0.27 | 0.15 |
| 11 | 82829768 | rs1943344 | -0.839 | -0.128 | 0.318 | 1.063 | 0.039 | 0.028 | 0.028 | 0 | 0.12 | 0.27 | 0.2 | 0.13 | 0.15 | 0.29 |
| 11 | 82836379 | rs7936604 | -0.866 | -0.522 | -0.441 | 0.916 | -0.007 | 0.001 | 0.001 | 0 | 0.33 | 0.61 | 0.39 | 0.6 | 0.24 | 0.41 |
| 11 | 82850496 | rs2010435 | -0.715 | -0.839 | -0.431 | 0.829 | -0.038 | 0.015 | 0.015 | 0 | 0.35 | 0.52 | 0.28 | 0.42 | 0.25 | 0.4 |
| 11 | 82858667 | rs12269817 | -1.131 | -0.597 | -0.626 | 0.501 | 0.063 | 0.01 | 0.01 | 0 | 0.49 | 0.84 | 0.4 | 0.55 | 0.42 | 0.56 |
| 11 | 82865518 | rs4944429 | -0.723 | -0.987 | -0.282 | 0.462 | 0.012 | -0.017 | -0.017 | 0.108 | 0.41 | 0.66 | 0.38 | 0.48 | 0.38 | 0.47 |
| 11 | 82874970 | rs10898088 | -1.098 | -1.088 | -0.346 | 0.421 | 0.07 | 0.004 | 0.004 | 0 | 0.43 | 0.78 | 0.37 | 0.58 | 0.31 | 0.5 |
| 11 | 82875359 | rs7130527 | 0.736 | -1.458 | 0.142 | 0.424 | -0.028 | -0.023 | -0.023 | 0 | 0.14 | 0.16 | 0.07 | 0.05 | 0.08 | 0.13 |
| 11 | 82876270 | rs1943345 | -1.206 | -1.469 | -0.047 | 0.411 | 0 | -0.09 | -0.09 | 0 | 0.28 | 0.11 | 0.27 | 0.54 | 0.26 | 0.38 |
| 11 | 82880341 | rs12284052 | 0.064 | -1.444 | 0.115 | 0.411 | -0.07 | -0.062 | -0.062 | 0.007 | 0.18 | 0.67 | 0.12 | 0.05 | 0.08 | 0.13 |
| 11 | 82908161 | rs621125 | -1.485 | -0.509 | -1.526 | 0.071 | -0.1 | -0.065 | -0.065 | 0 | 0.75 | 0.98 | 0.55 | 0.75 | 0.67 | 0.7 |
| 11 | 82911261 | rs665714 | -0.914 | -0.031 | -1.115 | 0.286 | -0.031 | -0.064 | -0.064 | 0.061 | 0.35 | 0.12 | 0.3 | 0.58 | 0.36 | 0.44 |
| 11 | 82939751 | rs515718 | -0.781 | 0.995 | -0.006 | 0.577 | -0.009 | -0.007 | -0.007 | 0.187 | 0.19 | 0.13 | 0.23 | 0.24 | 0.32 | 0.28 |
| 11 | 82946192 | rs7102681 | 0.889 | 1.005 | -0.813 | 0.464 | -0.035 | -0.05 | -0.05 | 0 | 0.19 | 0 | 0.06 | 0.18 | 0.14 | 0.1 |
| 11 | 82959604 | rs565674 | 0.142 | 0.397 | 0.233 | -0.072 | 0.016 | 0.026 | 0.026 | 4.029 | 0.38 | 0.24 | 0.15 | 0.27 | 0.28 | 0.18 |
| 11 | 82968952 | rs543606 | -1.16 | -0.262 | 0.123 | -0.276 | -0.002 | -0.001 | -0.001 | 0 | 0.38 | 0.58 | 0.33 | 0.35 | 0.41 | 0.53 |
| 11 | 82972097 | rs1278402 | 0.844 | -0.028 | -0.082 | -0.23 | -0.083 | 0.028 | 0.028 | 0.601 | 0.14 | 0.01 | 0.15 | 0.05 | 0.25 | 0.16 |
| 11 | 82986940 | rs493681 | 0.486 | -0.079 | -0.003 | -0.232 | 0.016 | 0.027 | 0.027 | 4.029 | 0.38 | 0.25 | 0.15 | 0.27 | 0.28 | 0.18 |
| 11 | 82993184 | rs566703 | -2.185 | -0.044 | -0.07 | -0.243 | -0.031 | -0.04 | -0.04 | 0.959 | 0.76 | 0.88 | 0.47 | 0.62 | 0.69 | 0.71 |
| 11 | 82996986 | rs494791 | -1.822 | -0.058 | -0.022 | -0.268 | -0.03 | -0.052 | -0.052 | 0 | 0.72 | 0.88 | 0.48 | 0.58 | 0.67 | 0.65 |
| 11 | 83018775 | rs7933220 | 1.653 | -0.062 | -0.003 | -0.264 | -0.035 | -0.056 | -0.056 | 0 | 0.28 | 0.11 | 0.53 | 0.42 | 0.33 | 0.35 |
| 11 | 83041794 | rs4943874 | 0.08 | -0.062 | 0.087 | -0.285 | 0.05 | 0.061 | 0.061 | 4.023 | 0.47 | 0.3 | 0.77 | 0.68 | 0.65 | 0.62 |
| 11 | 83041907 | rs7111383 | -0.581 | -0.11 | 0.162 | -0.308 | 0.008 | 0.012 | 0.012 | 4.275 | 0.61 | 0.68 | 0.84 | 0.73 | 0.71 | 0.82 |
| 11 | 83048096 | rs11233566 | 1.681 | -0.056 | -0.247 | -0.284 | -0.037 | -0.056 | -0.056 | 0 | 0.28 | 0.11 | 0.53 | 0.42 | 0.33 | 0.35 |
| 11 | 83051368 | rs11233567 | 1.205 | -0.065 | -0.209 | -0.36 | 0.028 | 0.018 | 0.018 | 0 | 0.1 | 0.02 | 0.04 | 0.05 | 0.06 | 0.19 |
| 11 | 83056929 | rs7122851 | 0.672 | 0.164 | -0.126 | -0.293 | -0.08 | 0.028 | 0.028 | 0.691 | 0.14 | 0.01 | 0.15 | 0.05 | 0.24 | 0.15 |
| 11 | 83069132 | rs4944435 | 1.602 | -0.013 | 0.209 | -0.307 | -0.037 | -0.063 | -0.063 | 0 | 0.28 | 0.11 | 0.53 | 0.43 | 0.33 | 0.39 |
| 11 | 83070225 | rs10898103 | -0.148 | 0.013 | 0.219 | -0.333 | -0.055 | -0.042 | -0.042 | 1.537 | 0.8 | 0.92 | 0.77 | 0.75 | 0.67 | 0.75 |
| 11 | 83104179 | rs1943750 | -0.093 | -0.011 | 0.796 | -0.377 | 0.009 | 0.011 | 0.011 | 3.154 | 0.47 | 0.45 | 0.2 | 0.32 | 0.35 | 0.4 |
| 11 | 83121983 | rs1943747 | 0.867 | -0.071 | 0.736 | -0.345 | 0 | -0.008 | -0.008 | 0.26 | 0.35 | 0.64 | 0.57 | 0.43 | 0.32 | 0.35 |
| 11 | 83140517 | rs1943749 | 1.23 | -0.06 | 0.729 | -0.395 | -0.002 | -0.008 | -0.008 | 0.026 | 0.32 | 0.25 | 0.54 | 0.44 | 0.33 | 0.36 |
| 11 | 83154780 | rs11233619 | 0.884 | -0.275 | 0.351 | -0.506 | 0.006 | 0.027 | 0.027 | 4.298 | 0.33 | 0.19 | 0.14 | 0.26 | 0.29 | 0.26 |
| 11 | 83159958 | rs2512671 | 0.925 | -0.082 | 0.458 | -0.413 | 0.004 | 0.003 | 0.003 | 0.95 | 0.73 | 0.33 | 0.7 | 0.74 | 0.66 | 0.75 |
| 11 | 83165473 | rs2507849 | -1.856 | -0.035 | 0.112 | -0.404 | -0.041 | -0.073 | -0.073 | 1.711 | 0.72 | 0.9 | 0.48 | 0.56 | 0.68 | 0.65 |
| 11 | 83165645 | rs2507850 | -0.246 | -0.021 | 0.193 | -0.359 | -0.066 | -0.064 | -0.064 | 0 | 0.19 | 0.07 | 0.23 | 0.26 | 0.32 | 0.25 |
| 11 | 83166720 | rs2512676 | -1.093 | -0.012 | 0.568 | -0.353 | 0.027 | 0.025 | 0.025 | 0.119 | 0.25 | 0.4 | 0.28 | 0.27 | 0.34 | 0.26 |
| 11 | 83174408 | rs12278046 | -0.159 | 0.083 | -0.163 | -0.282 | -0.019 | -0.033 | -0.033 | 0 | 0.18 | 0.11 | 0.18 | 0.27 | 0.25 | 0.21 |
| 11 | 83179196 | rs2163612 | -1.011 | -0.052 | -0.321 | -0.544 | -0.077 | -0.082 | -0.082 | 0.629 | 0.6 | 0.86 | 0.38 | 0.49 | 0.52 | 0.55 |
| 11 | 83193765 | rs1469609 | -0.996 | -0.431 | 0.098 | -0.579 | -0.007 | -0.043 | -0.043 | 0 | 0.62 | 0.77 | 0.42 | 0.48 | 0.61 | 0.52 |
| 11 | 83199166 | rs1864775 | -0.116 | -0.623 | 0.456 | -0.831 | 0.012 | -0.009 | -0.009 | 0.014 | 0.43 | 0.21 | 0.27 | 0.39 | 0.4 | 0.34 |

| | | | | | | | | | | | | | | | |
|---|---|---|---|---|---|---|---|---|---|---|---|---|---|---|---|
| 11 | 83204132 | rs11607886 | -1.266 | -0.835 | 0.515 | -0.824 | -0.006 | -0.04 | -0.04 | 0 | 0.62 | 0.77 | 0.42 | 0.48 | 0.61 | 0.52 |
| 11 | 83204454 | rs7116939 | 0.416 | -1.004 | 0.377 | -0.915 | 0.005 | -0.037 | -0.037 | 0 | 0.36 | 0.17 | 0.22 | 0.37 | 0.33 | 0.28 |
| 11 | 83210161 | rs10751093 | -0.451 | -0.959 | 0.723 | -0.842 | 0.013 | 0 | 0 | 0.068 | 0.45 | 0.35 | 0.27 | 0.39 | 0.38 | 0.33 |
| 11 | 83211816 | rs12223102 | -0.414 | -0.947 | 0.696 | -0.836 | 0.015 | -0.001 | -0.001 | 0 | 0.44 | 0.33 | 0.27 | 0.39 | 0.38 | 0.33 |
| 11 | 83215072 | rs12794550 | -0.311 | -0.925 | 0.706 | -0.826 | 0.012 | -0.002 | -0.002 | 0.022 | 0.44 | 0.33 | 0.27 | 0.39 | 0.38 | 0.32 |
| 11 | 83232814 | rs12577373 | -0.188 | -0.855 | 0.509 | -0.855 | 0.028 | -0.005 | -0.005 | 0 | 0.43 | 0.22 | 0.28 | 0.38 | 0.41 | 0.35 |
| 11 | 83236236 | rs17145339 | -0.058 | -0.836 | 0.786 | -1.067 | 0.028 | -0.067 | -0.067 | 0 | 0.19 | 0.32 | 0.12 | 0.1 | 0.2 | 0.19 |
| 11 | 83241448 | rs12271855 | -0.396 | -0.908 | 0.912 | -1.108 | 0.024 | -0.09 | -0.09 | 0.636 | 0.22 | 0.4 | 0.13 | 0.1 | 0.2 | 0.22 |
| 11 | 83256868 | rs2116483 | -1.704 | -1.167 | 0.733 | -1.12 | -0.018 | -0.039 | -0.039 | 0.821 | 0.71 | 0.75 | 0.46 | 0.56 | 0.66 | 0.66 |
| 11 | 83258127 | rs7113167 | -0.556 | -1.284 | 0.68 | -1.144 | -0.001 | 0 | 0 | 0 | 0.53 | 0.43 | 0.33 | 0.46 | 0.45 | 0.47 |
| 11 | 83260621 | rs7928931 | 0.282 | -1.686 | 0.298 | -1.263 | 0.028 | -0.043 | -0.043 | 0 | 0.41 | 0.13 | 0.24 | 0.42 | 0.35 | 0.44 |
| 11 | 83262562 | rs1367986 | -0.351 | -1.31 | 0.658 | -1.136 | 0 | -0.001 | -0.001 | 0 | 0.15 | 0.14 | 0.15 | 0.12 | 0.26 | 0.18 |
| 11 | 83263057 | rs571331 | -1.174 | -1.446 | 0.651 | -1.174 | 0.016 | -0.01 | -0.01 | 0 | 0.6 | 0.8 | 0.78 | 0.63 | 0.65 | 0.67 |
| 11 | 83267970 | rs501139 | 0.323 | -1.449 | 0.928 | -1.168 | 0 | -0.001 | -0.001 | 1.161 | 0.42 | 0.6 | 0.63 | 0.51 | 0.4 | 0.5 |
| 11 | 83276059 | rs12273919 | -0.312 | -1.316 | 0.927 | -1.197 | 0.04 | -0.031 | -0.031 | 0.213 | 0.15 | 0.2 | 0.16 | 0.12 | 0.26 | 0.18 |
| 11 | 83279836 | rs488600 | -0.069 | -1.443 | 0.909 | -1.168 | 0.011 | -0.002 | -0.002 | 0 | 0.45 | 0.69 | 0.64 | 0.53 | 0.39 | 0.5 |
| 11 | 83282729 | rs2436158 | 0.081 | -1.473 | 1.243 | -1.122 | -0.002 | 0 | 0 | 0 | 0.43 | 0.52 | 0.62 | 0.54 | 0.39 | 0.5 |
| 11 | 83283859 | rs500059 | 1.464 | -1.727 | 1.409 | -1.176 | -0.007 | -0.013 | -0.013 | 0 | 0.28 | 0.32 | 0.55 | 0.46 | 0.34 | 0.33 |
| 11 | 83286749 | rs35578754 | 1.054 | -1.71 | 1.241 | -1.171 | 0.011 | -0.001 | -0.001 | 0 | 0.09 | 0.04 | 0.06 | 0.07 | 0.04 | 0.13 |
| 11 | 83289994 | rs535809 | -0.065 | -1.636 | 1.243 | -1.172 | -0.001 | -0.001 | -0.001 | 2.579 | 0.44 | 0.6 | 0.63 | 0.54 | 0.39 | 0.49 |
| 11 | 83297092 | rs11233685 | 0.973 | -1.656 | 0.446 | -1.124 | 0.012 | -0.001 | -0.001 | 0 | 0.09 | 0.04 | 0.06 | 0.07 | 0.04 | 0.13 |
| 11 | 83310263 | rs11233691 | 0.295 | -1.669 | 0.485 | -1.131 | -0.012 | -0.003 | -0.003 | 0 | 0.12 | 0.14 | 0.06 | 0.07 | 0.05 | 0.15 |
| 11 | 83311905 | rs653546 | -1.087 | -1.559 | 0.161 | -1.108 | 0 | -0.01 | -0.01 | 0 | 0.61 | 0.9 | 0.78 | 0.68 | 0.65 | 0.68 |
| 11 | 83313356 | rs17145701 | 1.047 | -1.683 | -0.333 | -1.045 | -0.018 | -0.066 | -0.066 | 0.001 | 0.1 | 0 | 0.11 | 0.23 | 0.11 | 0.11 |
| 11 | 83315597 | rs505998 | 1.62 | -1.715 | -0.105 | -1.112 | -0.063 | -0.094 | -0.094 | 0 | 0.24 | 0.1 | 0.52 | 0.47 | 0.34 | 0.33 |
| 11 | 83325416 | rs12280583 | -0.054 | -1.57 | 0.318 | -1.115 | 0.045 | -0.03 | -0.03 | 0.062 | 0.13 | 0.21 | 0.11 | 0.12 | 0.2 | 0.17 |
| 11 | 83326979 | rs489567 | -0.974 | -1.842 | 0.273 | -1.271 | 0.019 | 0.032 | 0.032 | 3.445 | 0.56 | 0.81 | 0.75 | 0.67 | 0.6 | 0.67 |
| 11 | 83334780 | rs10501544 | -0.216 | -1.694 | 0.41 | -1.238 | 0.033 | -0.025 | -0.025 | 0.531 | 0.15 | 0.21 | 0.11 | 0.14 | 0.2 | 0.18 |
| 11 | 83334983 | rs510978 | -0.959 | -1.849 | -0.108 | -1.252 | 0.042 | 0.069 | 0.069 | 3.315 | 0.52 | 0.82 | 0.74 | 0.69 | 0.58 | 0.62 |
| 11 | 83344753 | rs7105588 | -1.427 | -1.77 | 1.213 | -1.39 | 0.004 | -0.05 | -0.05 | 0 | 0.31 | 0.36 | 0.18 | 0.16 | 0.31 | 0.22 |
| 11 | 83350022 | rs616414 | 1.407 | -1.87 | 1.52 | -1.439 | 0.002 | 0.036 | 0.036 | 0.094 | 0.66 | 0.26 | 0.79 | 0.85 | 0.7 | 0.77 |
| 11 | 83357100 | rs609948 | -1.542 | -1.873 | 1.725 | -1.435 | 0.003 | -0.067 | -0.067 | 1.254 | 0.32 | 0.42 | 0.18 | 0.16 | 0.31 | 0.22 |
| 11 | 83358400 | rs983589 | -1.475 | -1.747 | 1.552 | -1.303 | 0.004 | -0.022 | -0.022 | 0 | 0.21 | 0.74 | 0.18 | 0.13 | 0.22 | 0.18 |
| 11 | 83358629 | rs983590 | -0.479 | -1.597 | 1.27 | -1.226 | 0.029 | -0.016 | -0.016 | 0.047 | 0.15 | 0.22 | 0.12 | 0.13 | 0.22 | 0.17 |
| 11 | 83371215 | rs650206 | -1.624 | -1.796 | 0.732 | -1.148 | 0.011 | 0.028 | 0.028 | 0 | 0.39 | 0.69 | 0.21 | 0.23 | 0.32 | 0.38 |
| 11 | 83373309 | rs6592137 | -1.408 | -1.879 | 0.723 | -1.181 | 0.002 | 0.004 | 0.004 | 1.064 | 0.36 | 0.64 | 0.21 | 0.23 | 0.32 | 0.33 |
| 11 | 83387273 | rs654217 | 0.849 | -1.797 | 0.53 | -1.139 | -0.011 | 0.003 | 0.003 | 1.469 | 0.12 | 0 | 0.04 | 0.07 | 0.11 | 0.05 |
| 11 | 83393850 | rs2443455 | -2.017 | -1.814 | 0.516 | -1.272 | 0.065 | 0.13 | 0.13 | 0 | 0.5 | 0.75 | 0.24 | 0.24 | 0.38 | 0.44 |
| 11 | 83405905 | rs12225390 | -0.851 | -0.768 | 1.213 | -0.619 | 0 | -0.003 | -0.003 | 0 | 0.19 | 0.2 | 0.09 | 0.14 | 0.2 | 0.25 |
| 11 | 83411593 | rs1451397 | -1.238 | -0.765 | 1.253 | -0.64 | 0.011 | -0.036 | -0.036 | 1.618 | 0.22 | 0.53 | 0.14 | 0.15 | 0.2 | 0.26 |
| 11 | 83433967 | rs7946060 | -1.273 | -0.732 | 1.225 | -0.682 | 0.008 | -0.041 | -0.041 | 1.754 | 0.22 | 0.56 | 0.14 | 0.14 | 0.2 | 0.26 |
| 11 | 83437714 | rs7111226 | -1.939 | -0.853 | 1.028 | -0.793 | 0.016 | 0.023 | 0.023 | 0.905 | 0.4 | 0.65 | 0.18 | 0.15 | 0.28 | 0.37 |
| 11 | 83437959 | rs12797204 | -1.188 | -1.179 | 1.369 | -0.452 | -0.072 | -0.104 | -0.104 | 0 | 0.17 | 0.35 | 0.06 | 0 | 0.05 | 0.11 |
| 11 | 83455766 | rs17548333 | -0.737 | -0.88 | 1.386 | -0.368 | -0.027 | -0.025 | -0.025 | 0 | 0.18 | 0.3 | 0.07 | 0.12 | 0.11 | 0.23 |
| 11 | 83476003 | rs7126086 | -0.749 | -0.781 | 1.291 | -0.361 | -0.007 | -0.006 | -0.006 | 0.038 | 0.18 | 0.22 | 0.08 | 0.14 | 0.14 | 0.24 |
| 11 | 83479808 | rs7109609 | -1.925 | -0.922 | 1.067 | -0.44 | -0.006 | -0.008 | -0.008 | 0 | 0.36 | 0.63 | 0.14 | 0.14 | 0.18 | 0.34 |
| 11 | 83487889 | rs12290852 | -1.739 | -0.262 | 0.975 | -0.121 | -0.019 | -0.047 | -0.047 | 0.21 | 0.3 | 0.35 | 0.12 | 0.12 | 0.19 | 0.3 |
| 11 | 83489024 | rs715858 | -1.043 | 0.445 | 0.406 | 0.495 | -0.129 | -0.125 | -0.125 | 0.015 | 0.82 | 0.97 | 0.47 | 0.47 | 0.62 | 0.75 |
| 11 | 83489364 | rs1155310 | -1.066 | 0.321 | 0.551 | 0.374 | -0.032 | -0.064 | -0.064 | 0 | 0.24 | 0.42 | 0.11 | 0.13 | 0.15 | 0.26 |
| 11 | 83493425 | rs11233743 | 1.677 | 0.344 | 0.56 | 0.484 | 0.043 | 0.067 | 0.067 | 0.552 | 0.48 | 0.32 | 0.32 | 0.27 | 0.43 | 0.42 |
| 11 | 83502296 | rs7118583 | -1.332 | 0.362 | 0.631 | -0.014 | -0.005 | -0.015 | -0.015 | 0 | 0.28 | 0.28 | 0.12 | 0.13 | 0.19 | 0.32 |
| 11 | 83503404 | rs7479949 | 1.717 | 0.472 | 0.34 | 0.611 | 0.037 | 0.013 | 0.013 | 0 | 0.51 | 0.35 | 0.39 | 0.48 | 0.44 | 0.47 |
| 11 | 83512154 | rs10898155 | 1.597 | 0.462 | 0.308 | 0.539 | -0.001 | 0 | 0 | 0.197 | 0.52 | 0.53 | 0.4 | 0.49 | 0.44 | 0.46 |
| 11 | 83516681 | rs1451393 | -0.904 | 0.437 | 0.174 | 0.532 | -0.126 | -0.109 | -0.109 | 0 | 0.83 | 0.98 | 0.55 | 0.66 | 0.63 | 0.79 |
| 11 | 83521939 | rs1451395 | 1.728 | 0.654 | 0.325 | 0.519 | 0.011 | 0.015 | 0.015 | 0 | 0.51 | 0.4 | 0.41 | 0.61 | 0.47 | 0.48 |
| 11 | 83525588 | rs1037353 | -0.655 | 0.473 | -0.146 | 0.128 | -0.016 | -0.032 | -0.032 | 0 | 0.2 | 0.31 | 0.1 | 0.13 | 0.14 | 0.27 |
| 11 | 83541143 | rs7926247 | -1.239 | 0.615 | -0.644 | 0.205 | -0.044 | -0.058 | -0.058 | 0 | 0.32 | 0.52 | 0.14 | 0.18 | 0.18 | 0.35 |
| 11 | 83543448 | rs12273815 | -0.545 | 1.669 | -1.439 | 0.608 | -0.017 | 0.012 | 0.012 | 0 | 0.13 | 0.03 | 0.06 | 0.11 | 0.13 | 0.21 |
| 11 | 83544155 | rs6592147 | -0.408 | 2.522 | -1.199 | 0.674 | -0.06 | 0.056 | 0.056 | 0.361 | 0.9 | 0.99 | 0.75 | 1 | 0.74 | 0.9 |
| 11 | 83548648 | rs7941665 | 0.235 | 2.01 | -1.291 | 1.023 | -0.109 | -0.106 | -0.106 | 0 | 0.82 | 0.97 | 0.54 | 0.63 | 0.65 | 0.81 |
| 11 | 83551456 | rs7124129 | -0.916 | 0.238 | -0.911 | 0.137 | -0.028 | -0.024 | -0.024 | 0.317 | 0.24 | 0.34 | 0.19 | 0.09 | 0.16 | 0.25 |
| 11 | 83552659 | rs1878266 | 0.035 | 0.837 | -0.739 | 0.241 | 0.027 | 0.039 | 0.039 | 0.858 | 0.75 | 0.65 | 0.52 | 0.43 | 0.66 | 0.61 |
| 11 | 83554112 | rs10898158 | 1.127 | 1.317 | -1.089 | 0.054 | 0.012 | 0.088 | 0.088 | 2.775 | 0.45 | 0.13 | 0.28 | 0.36 | 0.4 | 0.32 |
| 11 | 83556067 | rs7951794 | -0.766 | 1.358 | -0.479 | 0.114 | 0.007 | -0.018 | -0.018 | 0 | 0.21 | 0.24 | 0.19 | 0.09 | 0.21 | 0.21 |
| 11 | 83562296 | rs6592150 | -0.536 | 1.141 | -1.054 | 0.077 | 0.006 | -0.007 | -0.007 | 0 | 0.34 | 0.53 | 0.31 | 0.4 | 0.35 | 0.28 |
| 11 | 83566115 | rs1914204 | 0.594 | 0.98 | -1.246 | -0.019 | -0.035 | 0.007 | 0.007 | 0 | 0.56 | 0.3 | 0.48 | 0.38 | 0.54 | 0.38 |
| 11 | 83569165 | rs2037197 | -0.203 | 1.04 | -0.717 | 0.125 | -0.045 | -0.084 | -0.084 | 0 | 0.26 | 0.49 | 0.23 | 0.1 | 0.2 | 0.27 |
| 11 | 83573399 | rs4943881 | -0.305 | 1.862 | -0.782 | 0.759 | -0.033 | -0.038 | -0.038 | 0 | 0.12 | 0.06 | 0.27 | 0.29 | 0.21 | 0.12 |
| 11 | 83575727 | rs17486172 | -0.241 | 0.981 | -0.404 | -0.144 | 0.013 | -0.009 | -0.009 | 0.003 | 0.08 | 0.15 | 0.03 | 0.04 | 0.05 | 0.07 |

| | | | | | | | | | | | | | | | | |
|---|---|---|---|---|---|---|---|---|---|---|---|---|---|---|---|---|
| 11 | 83577310 | rs3793953 | -0.247 | 1.051 | -0.588 | -0.67 | 0 | -0.014 | -0.014 | 0 | 0.5 | 0.43 | 0.33 | 0.2 | 0.53 | 0.31 |
| 11 | 83577810 | rs10898163 | 0.263 | 1.351 | -0.503 | -0.351 | 0.008 | 0.01 | 0.01 | 6.067 | 0.58 | 0.57 | 0.56 | 0.54 | 0.55 | 0.39 |
| 11 | 83579907 | rs877985 | -0.012 | 1.356 | -0.416 | -0.644 | 0.044 | 0.107 | 0.107 | 0.251 | 0.76 | 0.64 | 0.62 | 0.54 | 0.68 | 0.56 |
| 11 | 83581307 | rs11820448 | 0.241 | 1.526 | -0.258 | -0.704 | 0.017 | -0.022 | -0.022 | 0.523 | 0.07 | 0.14 | 0.05 | 0.04 | 0.08 | 0.06 |
| 11 | 83581941 | rs12361034 | 1.29 | 1.259 | -0.401 | -1.03 | -0.016 | 0.005 | 0.005 | 1.78 | 0.33 | 0.46 | 0.38 | 0.38 | 0.24 | 0.26 |
| 11 | 83583324 | rs4944452 | -1.045 | 1.408 | -0.471 | -0.84 | 0.02 | 0.066 | 0.066 | 0.876 | 0.83 | 0.79 | 0.67 | 0.61 | 0.77 | 0.62 |
| 11 | 83586526 | rs5025916 | 0.701 | 1.168 | -0.327 | -0.821 | 0.007 | -0.014 | -0.014 | 0.147 | 0.06 | 0.12 | 0.04 | 0.04 | 0.07 | 0.05 |
| 11 | 83586837 | rs1400316 | -0.748 | 2.003 | -0.386 | -0.887 | 0.005 | 0.013 | 0.013 | 4.485 | 0.39 | 0.39 | 0.32 | 0.43 | 0.41 | 0.58 |
| 11 | 83591243 | rs11233774 | 1.521 | 2.337 | -0.539 | -1.158 | -0.014 | 0.011 | 0.011 | 1.159 | 0.32 | 0.42 | 0.36 | 0.37 | 0.25 | 0.19 |
| 11 | 83592397 | rs1557431 | -0.02 | 2.397 | -0.539 | -1.19 | 0.04 | 0.13 | 0.13 | 0.229 | 0.21 | 0.35 | 0.26 | 0.43 | 0.28 | 0.4 |
| 11 | 83604840 | rs11233790 | -0.098 | 2.421 | -0.553 | -1.195 | 0.016 | 0.152 | 0.152 | 0.236 | 0.21 | 0.41 | 0.25 | 0.43 | 0.25 | 0.39 |
| 11 | 83607854 | rs2220086 | 1.011 | 2.503 | -0.588 | -0.691 | 0.02 | 0.088 | 0.088 | 0.898 | 0.15 | 0.21 | 0.18 | 0.38 | 0.21 | 0.36 |
| 11 | 83613351 | rs7933375 | 1.404 | 2.165 | -0.586 | -0.899 | 0.023 | -0.012 | -0.012 | 0 | 0.39 | 0.32 | 0.4 | 0.45 | 0.28 | 0.19 |
| 11 | 83613766 | rs7111645 | 1.168 | 1.873 | -0.584 | -1.152 | 0.012 | 0.041 | 0.041 | 1.646 | 0.18 | 0.04 | 0.06 | 0 | 0.13 | 0.17 |
| 11 | 83614116 | rs790353 | -1.027 | 1.855 | -0.566 | -1.11 | -0.062 | -0.032 | -0.032 | 0.004 | 0.86 | 0.95 | 0.75 | 0.82 | 0.79 | 0.88 |
| 11 | 83620215 | rs7932986 | 0.256 | 1.843 | -0.421 | -1.282 | -0.021 | 0.069 | 0.069 | 0.014 | 0.24 | 0.06 | 0.12 | 0 | 0.26 | 0.21 |
| 11 | 83620787 | rs790356 | 0.992 | 1.833 | -0.171 | -1.353 | 0.005 | -0.058 | -0.058 | 0 | 0.57 | 0.45 | 0.59 | 0.84 | 0.51 | 0.56 |
| 11 | 83620940 | rs790357 | -1.378 | 1.896 | -0.186 | -1.226 | -0.044 | 0 | 0 | 0.822 | 0.91 | 0.95 | 0.78 | 0.91 | 0.84 | 0.89 |
| 11 | 83629426 | rs2625519 | 1.017 | 1.985 | -0.145 | -1.314 | 0 | -0.042 | -0.042 | 0 | 0.52 | 0.46 | 0.59 | 0.84 | 0.51 | 0.6 |
| 11 | 83629911 | rs2675717 | 0.307 | 1.996 | -0.137 | -1.284 | -0.015 | 0.066 | 0.066 | 0 | 0.6 | 0.7 | 0.64 | 0.91 | 0.58 | 0.68 |
| 11 | 83630240 | rs2625520 | 1.073 | 1.898 | -0.123 | -1.017 | 0.007 | 0.066 | 0.066 | 0 | 0.15 | 0.21 | 0.14 | 0.32 | 0.17 | 0.34 |
| 11 | 83631275 | rs4300411 | 0.334 | 1.951 | -0.718 | -1.036 | -0.006 | 0.024 | 0.024 | 0.015 | 0.23 | 0.12 | 0.12 | 0 | 0.23 | 0.2 |
| 11 | 83633736 | rs790379 | 0.876 | 1.906 | -0.668 | -0.988 | -0.006 | 0.006 | 0.006 | 0.978 | 0.53 | 0.41 | 0.66 | 0.57 | 0.49 | 0.33 |
| 11 | 83634495 | rs12285761 | 0.974 | 0.83 | -0.598 | -1.114 | 0.021 | 0.042 | 0.042 | 0.772 | 0.17 | 0.04 | 0.06 | 0.01 | 0.13 | 0.17 |
| 11 | 83634983 | rs790381 | -0.144 | 0.714 | -0.413 | -0.688 | -0.005 | 0.118 | 0.118 | 0.137 | 0.8 | 0.55 | 0.82 | 0.63 | 0.81 | 0.54 |
| 11 | 83636902 | rs1483397 | -0.717 | 1.129 | -0.989 | -1.283 | -0.001 | 0.018 | 0.018 | 0 | 0.42 | 0.39 | 0.28 | 0.13 | 0.43 | 0.34 |
| 11 | 83638942 | rs2374465 | 0.258 | 0.679 | -1.326 | -0.728 | 0.017 | 0.213 | 0.213 | 0.637 | 0.58 | 0.84 | 0.53 | 0.86 | 0.41 | 0.67 |
| 11 | 83649612 | rs790369 | -1.663 | 0.483 | -0.274 | 0.663 | 0.006 | -0.015 | -0.015 | 0 | 0.34 | 0.68 | 0.42 | 0.44 | 0.3 | 0.4 |
| 11 | 83650203 | rs708241 | -0.841 | 0.521 | 0.162 | 1.138 | 0.022 | 0.116 | 0.116 | 0 | 0.08 | 0.13 | 0.25 | 0.3 | 0.14 | 0.15 |
| 11 | 83651102 | rs11233821 | -0.655 | 0.319 | 0.24 | 0.915 | -0.001 | 0.001 | 0.001 | 0 | 0.08 | 0.12 | 0.09 | 0.14 | 0.03 | 0.09 |
| 11 | 83655564 | rs1384753 | -1.372 | 0.264 | 0.163 | 0.825 | 0.006 | -0.005 | -0.005 | 0 | 0.18 | 0.05 | 0.13 | 0.14 | 0.16 | 0.15 |
| 11 | 83656830 | rs790346 | -3.137 | 0.269 | 0.364 | 0.817 | 0.005 | -0.011 | -0.011 | 0 | 0.34 | 0.67 | 0.42 | 0.45 | 0.31 | 0.43 |
| 11 | 83658909 | rs790350 | -1.528 | 0.248 | 0.456 | 0.793 | 0.014 | 0.068 | 0.068 | 0 | 0.1 | 0.13 | 0.26 | 0.3 | 0.16 | 0.17 |
| 11 | 83663828 | rs1319736 | 1.643 | 0.487 | 1.169 | 0.813 | -0.034 | 0.018 | 0.018 | 0.049 | 0.86 | 0.74 | 0.9 | 0.85 | 0.95 | 0.8 |
| 11 | 83665176 | rs2449588 | -3.065 | 0.48 | 1.135 | 0.819 | 0.003 | -0.008 | -0.008 | 0 | 0.33 | 0.67 | 0.42 | 0.44 | 0.3 | 0.4 |
| 11 | 83665488 | rs1483403 | -0.416 | 0.63 | 1.111 | 0.483 | -0.001 | -0.022 | -0.022 | 0.295 | 0.11 | 0.1 | 0.05 | 0 | 0.13 | 0.06 |
| 11 | 83673516 | rs11233862 | -1.014 | 0.624 | 1.263 | 0.481 | -0.005 | -0.063 | -0.063 | 0 | 0.13 | 0.24 | 0.06 | 0 | 0.13 | 0.07 |
| 11 | 83675548 | rs1384758 | -2.345 | 0.387 | 1.391 | 0.741 | -0.007 | 0.039 | 0.039 | 0 | 0.14 | 0.19 | 0.26 | 0.29 | 0.12 | 0.16 |
| 11 | 83676761 | rs960532 | -2.469 | 0.349 | 1.871 | 0.677 | -0.022 | 0.1 | 0.1 | 0 | 0.19 | 0.47 | 0.3 | 0.29 | 0.17 | 0.15 |
| 11 | 83680018 | rs2514160 | -1.918 | 0.37 | 1.886 | 0.707 | 0.032 | 0.28 | 0.28 | 0 | 0.1 | 0.4 | 0.26 | 0.29 | 0.09 | 0.1 |
| 11 | 83696133 | rs2062208 | 1.563 | 1.395 | 1.805 | 0.488 | 0 | -0.063 | -0.063 | 0 | 0.85 | 0.76 | 0.94 | 1 | 0.85 | 0.83 |
| 11 | 83702343 | rs2449600 | -1.396 | 1.339 | 2.106 | 0.65 | 0.041 | 0.355 | 0.355 | 0 | 0.07 | 0.32 | 0.24 | 0.3 | 0.08 | 0.09 |
| 11 | 83709994 | rs11233872 | 0.085 | 1.746 | 1.793 | 0.574 | -0.036 | -0.029 | -0.029 | 0 | 0.06 | 0.01 | 0.08 | 0.12 | 0.15 | 0.08 |
| 11 | 83717887 | rs12286565 | -0.969 | 0.937 | 2.091 | 0.665 | -0.002 | -0.025 | -0.025 | 0 | 0.11 | 0.13 | 0.04 | 0 | 0.08 | 0.14 |
| 11 | 83725819 | rs12291567 | -2.306 | 0.846 | 2.41 | 0.606 | -0.025 | -0.119 | -0.119 | 0.054 | 0.18 | 0.42 | 0.08 | 0 | 0.14 | 0.16 |
| 11 | 83728714 | rs12290953 | 0.223 | 0.111 | 1.816 | 0.526 | 0.011 | -0.01 | -0.01 | 0 | 0.67 | 0.58 | 0.43 | 0.33 | 0.68 | 0.47 |
| 11 | 83733136 | rs1483389 | 0.08 | -0.012 | 1.827 | 0.755 | 0 | 0.003 | 0.003 | 0 | 0.26 | 0.24 | 0.35 | 0.47 | 0.21 | 0.37 |
| 11 | 83736246 | rs6592155 | -1.117 | 0.099 | 1.932 | 0.886 | -0.013 | 0.014 | 0.014 | 0 | 0.39 | 0.54 | 0.42 | 0.51 | 0.3 | 0.52 |
| 11 | 83736979 | rs4944457 | 0.193 | 0.32 | 1.995 | 0.808 | 0.001 | 0.025 | 0.025 | 0 | 0.25 | 0.26 | 0.36 | 0.44 | 0.22 | 0.37 |
| 11 | 83740599 | rs2625510 | 1.143 | 0.506 | 3.137 | 0.836 | 0.01 | 0.142 | 0.142 | 0 | 0.92 | 0.82 | 0.78 | 0.77 | 0.9 | 0.82 |
| 11 | 83740835 | rs10898192 | 1.359 | 0.511 | 3.137 | 0.835 | 0.027 | 0.124 | 0.124 | 0 | 0.91 | 0.82 | 0.78 | 0.77 | 0.88 | 0.82 |
| 11 | 83744306 | rs10792710 | 3.236 | 0.496 | 3.111 | 0.82 | 0.009 | 0.002 | 0.002 | 0.242 | 0.72 | 0.24 | 0.68 | 0.77 | 0.75 | 0.67 |
| 11 | 83748095 | rs2625518 | 2.136 | 0.551 | 2.989 | 0.852 | -0.018 | 0.037 | 0.037 | 0 | 0.81 | 0.68 | 0.74 | 0.77 | 0.82 | 0.69 |
| 11 | 83759244 | rs12285610 | -2.617 | 0.65 | 2.51 | 0.982 | 0.049 | 0.08 | 0.08 | 0 | 0.14 | 0.36 | 0.12 | 0.18 | 0.17 | 0.19 |
| 11 | 83766621 | rs4943885 | 2.29 | 0.648 | 2.467 | 0.924 | 0.025 | 0.107 | 0.107 | 0.013 | 0.84 | 0.5 | 0.74 | 0.77 | 0.83 | 0.81 |
| 11 | 83767844 | rs4586191 | 2.059 | 0.594 | 2.319 | 0.945 | -0.019 | 0.022 | 0.022 | 0 | 0.77 | 0.59 | 0.72 | 0.77 | 0.77 | 0.76 |
| 11 | 83774415 | rs4943886 | 1.789 | 1.05 | 2.162 | 1.323 | 0.045 | 0.272 | 0.272 | 0 | 0.91 | 0.53 | 0.75 | 0.75 | 0.89 | 0.84 |
| 11 | 83775326 | rs10898199 | 1.605 | 1.074 | 2.183 | 1.335 | 0.065 | 0.313 | 0.313 | 0 | 0.91 | 0.59 | 0.75 | 0.75 | 0.89 | 0.84 |
| 11 | 83789540 | rs7946613 | -1.532 | 1.145 | 1.814 | 1.174 | -0.001 | -0.01 | -0.01 | 0 | 0.07 | 0.09 | 0.05 | 0 | 0.09 | 0.05 |
| 11 | 83805810 | rs1945801 | 2.205 | 1.212 | 2.39 | 1.346 | 0.07 | 0.246 | 0.246 | 0 | 0.89 | 0.54 | 0.75 | 0.75 | 0.86 | 0.82 |
| 11 | 83808598 | rs7931375 | -2.546 | 1.266 | 2.566 | 1.226 | 0.095 | -0.038 | -0.038 | 0 | 0.07 | 0.24 | 0.07 | 0 | 0.13 | 0.08 |
| 11 | 83817360 | rs2298830 | 2.187 | 1.162 | 2.727 | 1.34 | 0.068 | 0.243 | 0.243 | 0 | 0.89 | 0.6 | 0.76 | 0.75 | 0.86 | 0.82 |
| 11 | 83825694 | rs7106126 | -1.049 | 1.144 | 2.789 | 1.272 | 0.001 | -0.034 | -0.034 | 0.007 | 0.06 | 0.17 | 0.04 | 0 | 0.05 | 0.04 |
| 11 | 83829316 | rs1894169 | 2.627 | 1.139 | 2.867 | 1.343 | 0.044 | 0.161 | 0.161 | 0 | 0.87 | 0.45 | 0.73 | 0.74 | 0.83 | 0.81 |
| 11 | 83831259 | rs7108582 | 1.592 | 1.128 | 3.302 | 1.337 | 0.012 | 0.153 | 0.153 | 0 | 0.92 | 0.82 | 0.78 | 0.74 | 0.89 | 0.84 |
| 11 | 83837570 | rs1945827 | 2.885 | 1.095 | 3.537 | 1.373 | 0.059 | 0.193 | 0.193 | 0 | 0.86 | 0.55 | 0.73 | 0.74 | 0.83 | 0.81 |
| 11 | 83839293 | rs2105806 | 3.207 | 1.175 | 3.47 | 1.371 | 0.001 | 0.011 | 0.011 | 0.265 | 0.8 | 0.3 | 0.69 | 0.74 | 0.78 | 0.77 |
| 11 | 83844790 | rs1945825 | 3.338 | 1.167 | 3.403 | 1.388 | 0.014 | 0.051 | 0.051 | 0.078 | 0.8 | 0.43 | 0.7 | 0.74 | 0.78 | 0.77 |
| 11 | 83847967 | rs10501555 | 1.812 | 1.178 | 3.123 | 1.416 | 0.009 | 0.177 | 0.177 | 0 | 0.91 | 0.77 | 0.78 | 0.74 | 0.89 | 0.84 |

| | | | | | | | | | | | | | | | | |
|---|---|---|---|---|---|---|---|---|---|---|---|---|---|---|---|---|
| 11 | 83877087 | rs17563429 | -1.119 | 1.166 | 2.801 | 1.266 | -0.002 | -0.035 | -0.035 | 0 | 0.07 | 0.11 | 0.03 | 0 | 0.05 | 0.04 |
| 11 | 83885782 | rs4944468 | 2.703 | 1.056 | 2.888 | 1.397 | -0.005 | 0.092 | 0.092 | 0 | 0.82 | 0.66 | 0.75 | 0.73 | 0.82 | 0.81 |
| 11 | 83899028 | rs1377746 | 1.971 | 1.055 | 2.621 | 1.407 | -0.008 | 0.115 | 0.115 | 0 | 0.9 | 0.78 | 0.78 | 0.75 | 0.9 | 0.84 |
| 11 | 83904512 | rs1349820 | 3.075 | 1.032 | 2.708 | 1.361 | 0 | 0.018 | 0.018 | 0.351 | 0.78 | 0.35 | 0.69 | 0.75 | 0.78 | 0.78 |
| 11 | 83918339 | rs10792731 | 2.941 | 1.002 | 2.699 | 1.391 | 0.016 | 0.115 | 0.115 | 0.063 | 0.85 | 0.4 | 0.73 | 0.75 | 0.85 | 0.79 |
| 11 | 83920172 | rs10751106 | 2.716 | 0.964 | 2.791 | 1.407 | -0.005 | 0.151 | 0.151 | 0.054 | 0.87 | 0.42 | 0.75 | 0.75 | 0.88 | 0.83 |
| 11 | 83930211 | rs11233953 | -2.612 | 0.968 | 2.972 | 1.434 | -0.01 | 0.101 | 0.101 | 0 | 0.12 | 0.22 | 0.23 | 0.26 | 0.13 | 0.14 |
| 11 | 83933781 | rs7925243 | 3.039 | 1.139 | 2.766 | 1.332 | 0.018 | 0.053 | 0.053 | 0.232 | 0.79 | 0.4 | 0.7 | 0.74 | 0.77 | 0.78 |
| 11 | 83939591 | rs10792740 | 3.169 | 1.127 | 2.898 | 1.338 | 0.047 | 0.197 | 0.197 | 0.199 | 0.86 | 0.5 | 0.73 | 0.74 | 0.84 | 0.83 |
| 11 | 83945482 | rs896996 | 3.208 | 1.133 | 2.941 | 1.331 | 0.028 | 0.107 | 0.107 | 0.134 | 0.81 | 0.46 | 0.71 | 0.74 | 0.77 | 0.8 |
| 11 | 83951160 | rs1022308 | 3.632 | 1.044 | 2.889 | 1.298 | -0.001 | -0.004 | -0.004 | 0 | 0.72 | 0.32 | 0.68 | 0.73 | 0.7 | 0.73 |
| 11 | 83951374 | rs1022306 | 2.591 | 1.084 | 2.852 | 1.316 | 0.044 | -0.055 | -0.055 | 0.005 | 0.9 | 0.76 | 0.94 | 1 | 0.87 | 0.87 |
| 11 | 83954724 | rs1399617 | 3.456 | 1.077 | 2.817 | 1.352 | 0.02 | 0.087 | 0.087 | 0.067 | 0.79 | 0.49 | 0.7 | 0.74 | 0.75 | 0.76 |
| 11 | 83959179 | rs12360734 | -0.168 | 1.035 | 2.43 | 1.369 | 0.002 | -0.021 | -0.021 | 0 | 0.05 | 0.06 | 0.03 | 0 | 0.05 | 0.01 |
| 11 | 83961440 | rs996665 | 1.696 | 1.022 | 2.419 | 1.393 | 0.044 | -0.034 | -0.034 | 0.008 | 0.94 | 0.83 | 0.95 | 1 | 0.9 | 0.94 |
| 11 | 83964236 | rs11233963 | -2.466 | 0.989 | 2.517 | 1.457 | -0.012 | 0.095 | 0.095 | 0 | 0.11 | 0.19 | 0.22 | 0.26 | 0.12 | 0.11 |
| 11 | 83967019 | rs4943889 | 3.318 | 0.975 | 2.625 | 1.469 | 0.016 | 0.116 | 0.116 | 0.464 | 0.8 | 0.57 | 0.7 | 0.73 | 0.78 | 0.84 |
| 11 | 83967617 | rs10792744 | -2.832 | 0.93 | 2.281 | 1.487 | -0.019 | 0.08 | 0.08 | 0.011 | 0.13 | 0.21 | 0.23 | 0.27 | 0.12 | 0.11 |
| 11 | 83976634 | rs10792746 | 2.199 | 1.059 | 2.111 | 1.429 | 0.043 | 0.103 | 0.103 | 0.14 | 0.8 | 0.55 | 0.71 | 0.75 | 0.78 | 0.86 |
| 11 | 83983546 | rs17499985 | -0.676 | 0.789 | 1.08 | 1.542 | 0.002 | -0.003 | -0.003 | 0.351 | 0.13 | 0.07 | 0.13 | 0.12 | 0.12 | 0.08 |
| 11 | 83984778 | rs12294968 | 1.37 | 0.657 | 1.484 | 2.647 | -0.012 | 0.125 | 0.125 | 0 | 0.8 | 0.62 | 0.76 | 0.46 | 0.82 | 0.61 |
| 11 | 83985411 | rs4944475 | 0.268 | 0.586 | 1.508 | 2.854 | -0.031 | 0.089 | 0.089 | 0 | 0.9 | 0.75 | 0.95 | 0.83 | 0.97 | 0.93 |
| 11 | 83986071 | rs1470899 | 2.709 | 0.521 | 1.427 | 2.749 | -0.004 | 0.025 | 0.025 | 0 | 0.69 | 0.51 | 0.68 | 0.43 | 0.66 | 0.54 |
| 11 | 83987790 | rs10792750 | 2.461 | 0.536 | 1.438 | 2.749 | 0.01 | 0.061 | 0.061 | 0 | 0.74 | 0.55 | 0.72 | 0.43 | 0.72 | 0.57 |
| 11 | 83993349 | rs1580078 | 0.977 | 0.36 | 0.855 | 2.506 | -0.041 | 0.012 | 0.012 | 0.074 | 0.79 | 0.68 | 0.86 | 0.75 | 0.88 | 0.84 |
| 11 | 83993956 | rs2054024 | 1.974 | 0.407 | 0.799 | 2.474 | -0.013 | 0.006 | 0.006 | 0 | 0.67 | 0.59 | 0.67 | 0.38 | 0.7 | 0.49 |
| 11 | 83998766 | rs12577267 | -0.849 | 0.381 | 0.753 | 2.394 | -0.007 | -0.007 | -0.007 | 0 | 0.08 | 0.06 | 0.15 | 0.12 | 0.12 | 0.08 |
| 11 | 84005324 | rs17147160 | -1.547 | 0.466 | 1.031 | 2.375 | 0.024 | -0.02 | -0.02 | 0.304 | 0.11 | 0.28 | 0.09 | 0.04 | 0.15 | 0.06 |
| 11 | 84006510 | rs4944478 | 1.794 | 0.409 | 1.056 | 2.472 | -0.031 | 0.019 | 0.019 | 0.667 | 0.78 | 0.62 | 0.86 | 0.79 | 0.81 | 0.84 |
| 11 | 84009647 | rs11823072 | -1.472 | 0.351 | 0.784 | 2.524 | -0.014 | 0.093 | 0.093 | 0 | 0.17 | 0.32 | 0.19 | 0.31 | 0.13 | 0.13 |
| 11 | 84022936 | rs7121214 | -2.682 | 0.5 | 0.538 | 2.409 | 0.001 | 0.002 | 0.002 | 0 | 0.33 | 0.34 | 0.32 | 0.58 | 0.35 | 0.42 |
| 11 | 84024746 | rs17566476 | -0.075 | 0.358 | -0.011 | 2.284 | -0.033 | 0.011 | 0.011 | 0 | 0.09 | 0 | 0.09 | 0.08 | 0.11 | 0.11 |
| 11 | 84025940 | rs17147175 | -2.088 | 0.197 | 0.282 | 2.521 | -0.016 | -0.041 | -0.041 | 0 | 0.14 | 0.08 | 0.2 | 0.31 | 0.21 | 0.14 |
| 11 | 84027768 | rs1515083 | -0.932 | 0.287 | 0.463 | 2.527 | -0.008 | 0.011 | 0.011 | 0 | 0.08 | 0.07 | 0.13 | 0.29 | 0.13 | 0.1 |
| 11 | 84029706 | rs10501557 | -0.57 | 0.247 | 0.514 | 2.385 | -0.052 | 0.051 | 0.051 | 0 | 0.11 | 0.16 | 0.04 | 0.18 | 0.02 | 0.17 |
| 11 | 84031264 | rs7928787 | -1.666 | 0.385 | 0.373 | 2.182 | -0.009 | 0.049 | 0.049 | 0 | 0.2 | 0.24 | 0.18 | 0.43 | 0.15 | 0.26 |
| 11 | 84041716 | rs3896313 | -2.093 | 0.619 | 0.214 | 1.388 | -0.009 | 0.033 | 0.033 | 2.468 | 0.33 | 0.53 | 0.31 | 0.36 | 0.32 | 0.37 |
| 11 | 84048213 | rs10898225 | -0.449 | 1.004 | -0.207 | 1.126 | -0.024 | -0.036 | -0.036 | 0 | 0.12 | 0.06 | 0.14 | 0.16 | 0.14 | 0.05 |
| 11 | 84049216 | rs7951287 | -0.267 | 0.943 | 0.015 | 0.479 | 0.003 | -0.017 | -0.017 | 0 | 0.09 | 0.13 | 0.11 | 0.01 | 0.14 | 0.05 |
| 11 | 84053382 | rs4944487 | -1.535 | 0.266 | -0.324 | -0.393 | 0.009 | 0.018 | 0.018 | 0.242 | 0.32 | 0.64 | 0.2 | 0.13 | 0.25 | 0.36 |
| 11 | 84055553 | rs4310638 | -1.487 | 0.328 | -0.231 | -0.454 | -0.003 | -0.02 | -0.02 | 0 | 0.28 | 0.59 | 0.19 | 0.13 | 0.23 | 0.35 |
| 11 | 84059608 | rs7942296 | -1.649 | 0.273 | -0.286 | -0.37 | 0.02 | 0.045 | 0.045 | 0.951 | 0.3 | 0.83 | 0.2 | 0.21 | 0.22 | 0.36 |
| 11 | 84064236 | rs1559855 | -0.834 | 0.209 | -0.031 | -0.474 | -0.031 | -0.139 | -0.139 | 3.348 | 0.21 | 0.5 | 0.1 | 0.01 | 0.14 | 0.14 |
| 11 | 84071295 | rs7947614 | -2.588 | 0.036 | -0.361 | -0.553 | 0.01 | 0.093 | 0.093 | 2.813 | 0.37 | 0.96 | 0.24 | 0.21 | 0.26 | 0.39 |
| 11 | 84071435 | rs17508652 | -1.229 | 0.091 | -0.307 | -0.525 | -0.003 | -0.003 | -0.003 | 0 | 0.08 | 0.07 | 0.1 | 0.13 | 0.11 | 0.22 |
| 11 | 84086054 | rs10792756 | -2.107 | 0.07 | -0.306 | -0.504 | 0.026 | 0.048 | 0.048 | 1.236 | 0.3 | 0.88 | 0.2 | 0.21 | 0.2 | 0.38 |
| 11 | 84090324 | rs9988803 | -1.228 | 0.019 | -0.027 | -0.488 | -0.002 | -0.009 | -0.009 | 0 | 0.24 | 0.73 | 0.13 | 0.09 | 0.15 | 0.16 |
| 11 | 84111645 | rs7125294 | -1.194 | -0.018 | 0.04 | -0.74 | -0.06 | -0.117 | -0.117 | 0.02 | 0.18 | 0.37 | 0.07 | 0 | 0.08 | 0.08 |
| 11 | 84115273 | rs7127845 | -2.546 | 0.16 | -0.893 | -0.916 | 0.013 | 0.206 | 0.206 | 2.283 | 0.41 | 0.94 | 0.28 | 0.15 | 0.31 | 0.4 |
| 11 | 84117907 | rs504196 | -2.038 | 0.032 | -0.809 | -0.84 | 0.049 | 0.115 | 0.115 | 0.814 | 0.31 | 0.88 | 0.2 | 0.14 | 0.18 | 0.33 |
| 11 | 84120820 | rs569793 | -0.82 | -0.029 | -0.823 | -1.159 | -0.003 | -0.025 | -0.025 | 0.875 | 0.13 | 0.18 | 0.07 | 0.05 | 0.12 | 0.04 |
| 11 | 84124350 | rs500750 | -1.317 | -0.067 | -0.819 | -1.155 | -0.001 | -0.002 | -0.002 | 0.024 | 0.16 | 0.18 | 0.11 | 0.12 | 0.11 | 0.24 |
| 11 | 84125676 | rs7101963 | -0.286 | -0.112 | -0.446 | -1.247 | -0.004 | -0.028 | -0.028 | 0.153 | 0.08 | 0.35 | 0.04 | 0 | 0.03 | 0.07 |
| 11 | 84134019 | rs1365425 | -1.766 | -0.2 | -0.704 | -1.058 | -0.024 | -0.01 | -0.01 | 0 | 0.24 | 0.67 | 0.15 | 0.19 | 0.1 | 0.37 |
| 11 | 84139937 | rs543814 | -1.436 | -0.15 | -0.877 | -1.246 | 0 | -0.02 | -0.02 | 0.748 | 0.16 | 0.2 | 0.1 | 0.05 | 0.18 | 0.05 |
| 11 | 84142990 | rs524365 | -2.622 | -0.307 | -1.211 | -1.257 | 0.074 | 0.144 | 0.144 | 2.923 | 0.43 | 0.96 | 0.26 | 0.23 | 0.27 | 0.43 |
| 11 | 84146142 | rs568789 | -2.421 | -0.314 | -1.125 | -1.174 | 0.078 | 0.103 | 0.103 | 2.2 | 0.38 | 0.95 | 0.23 | 0.21 | 0.2 | 0.41 |
| 11 | 84152033 | rs11603982 | -0.735 | -0.48 | -0.573 | -1.292 | -0.063 | -0.117 | -0.117 | 0 | 0.15 | 0.43 | 0.06 | 0 | 0.06 | 0.15 |
| 11 | 84160668 | rs533364 | 0.925 | -0.312 | -1.736 | -1.172 | -0.099 | -0.064 | -0.064 | 0 | 0.23 | 0.02 | 0.19 | 0.24 | 0.3 | 0.2 |
| 11 | 84162690 | rs1943526 | -1.567 | -0.79 | -1.584 | -2 | 0.09 | 0.124 | 0.124 | 3.778 | 0.31 | 0.88 | 0.14 | 0.1 | 0.13 | 0.19 |
| 11 | 84163419 | rs12274138 | 0.807 | -0.518 | -1.832 | -2.226 | -0.067 | 0.138 | 0.138 | 0.063 | 0.26 | 0.02 | 0.12 | 0.08 | 0.25 | 0.23 |
| 11 | 84167697 | rs1367533 | 0.262 | -0.689 | -1.358 | -2.067 | 0.015 | 0.01 | 0.01 | 0.018 | 0.14 | 0.11 | 0.05 | 0.1 | 0.08 | 0.17 |
| 11 | 84167835 | rs7952143 | -1.998 | -0.598 | -1.204 | -2 | 0.032 | 0.063 | 0.063 | 1.24 | 0.36 | 0.82 | 0.21 | 0.23 | 0.25 | 0.44 |
| 11 | 84176261 | rs7924334 | -2.317 | -0.271 | -1.013 | -1.865 | 0.017 | 0.072 | 0.072 | 1.609 | 0.39 | 0.78 | 0.23 | 0.24 | 0.31 | 0.46 |
| 11 | 84182066 | rs11826800 | -1.485 | 0.175 | -0.656 | -1.632 | -0.005 | 0.011 | 0.011 | 0.067 | 0.27 | 0.17 | 0.16 | 0.24 | 0.26 | 0.43 |
| 11 | 84192441 | rs11606836 | -1.721 | 0.222 | -0.37 | -1.28 | -0.001 | -0.003 | -0.003 | 0 | 0.29 | 0.66 | 0.19 | 0.23 | 0.22 | 0.42 |
| 11 | 84198908 | rs12276848 | -0.708 | 0.939 | -0.247 | -0.975 | 0.04 | 0.036 | 0.036 | 0.116 | 0.1 | 0.34 | 0.1 | 0.1 | 0.08 | 0.22 |
| 11 | 84208616 | rs1430953 | -0.283 | 0.415 | -1.139 | -0.968 | -0.007 | 0.022 | 0.022 | 2.762 | 0.23 | 0.15 | 0.12 | 0.09 | 0.22 | 0.22 |
| 11 | 84213870 | rs17734854 | -0.264 | 0.209 | -1.368 | -0.944 | -0.043 | 0.09 | 0.09 | 0.96 | 0.19 | 0.02 | 0.08 | 0.06 | 0.18 | 0.21 |

| | | | | | | | | | | | | | | | | |
|---|---|---|---|---|---|---|---|---|---|---|---|---|---|---|---|---|
| 11 | 84218362 | rs7123506 | -1.12 | -0.75 | -1.457 | -0.849 | 0.014 | 0.01 | 0.01 | 0.63 | 0.2 | 0.11 | 0.07 | 0.15 | 0.15 | 0.29 |
| 11 | 84222437 | rs10898250 | -1.775 | -1.083 | -1.609 | -1.122 | 0.026 | 0.057 | 0.057 | 0.718 | 0.3 | 0.16 | 0.11 | 0.15 | 0.23 | 0.4 |
| 11 | 84228716 | rs10792760 | -0.553 | -1.097 | -1.624 | -1.135 | -0.002 | -0.002 | -0.002 | 0 | 0.08 | 0.07 | 0.05 | 0.15 | 0.1 | 0.21 |
| 11 | 84230355 | rs7107338 | -0.433 | -1.22 | -1.421 | -1.199 | -0.001 | -0.018 | -0.018 | 0.385 | 0.09 | 0.11 | 0.03 | 0 | 0.06 | 0.08 |
| 11 | 84239976 | rs7933597 | 0.88 | -1.471 | -1.715 | -1.359 | 0.028 | 0.084 | 0.084 | 0.053 | 0.8 | 0.95 | 0.84 | 0.96 | 0.89 | 0.88 |
| 11 | 84240120 | rs10792764 | 2.189 | -1.506 | -1.265 | -1.44 | 0.003 | 0.007 | 0.007 | 0 | 0.65 | 0.68 | 0.75 | 0.82 | 0.72 | 0.59 |
| 11 | 84247123 | rs10501568 | -0.325 | -1.534 | -0.788 | -1.447 | -0.005 | -0.01 | -0.01 | 0 | 0.11 | 0.13 | 0.04 | 0.07 | 0.08 | 0.1 |
| 11 | 84247636 | rs7119480 | -1.719 | -1.562 | -0.768 | -1.461 | 0.004 | 0.004 | 0.004 | 1.193 | 0.23 | 0.2 | 0.08 | 0.15 | 0.16 | 0.32 |
| 11 | 84248067 | rs568997 | 2.951 | -1.556 | -0.28 | -1.505 | 0.017 | 0.033 | 0.033 | 0 | 0.51 | 0.39 | 0.68 | 0.74 | 0.63 | 0.42 |
| 11 | 84253092 | rs500394 | -2.793 | -1.546 | -0.314 | -1.49 | -0.005 | -0.015 | -0.015 | 0 | 0.43 | 0.47 | 0.3 | 0.26 | 0.36 | 0.57 |
| 11 | 84255321 | rs10898257 | -0.187 | -1.504 | -0.3 | -1.471 | -0.023 | -0.033 | -0.033 | 1.325 | 0.09 | 0.02 | 0.05 | 0.14 | 0.1 | 0.23 |
| 11 | 84264056 | rs7937583 | -2.172 | -1.562 | -0.275 | -1.503 | -0.003 | -0.015 | -0.015 | 0 | 0.32 | 0.35 | 0.16 | 0.21 | 0.29 | 0.48 |
| 11 | 84267185 | rs7128948 | -1.745 | -1.42 | -0.172 | -1.43 | 0.011 | -0.017 | -0.017 | 0.323 | 0.22 | 0.34 | 0.13 | 0.15 | 0.23 | 0.4 |
| 11 | 84274481 | rs948141 | -2.246 | -1.407 | -0.294 | -1.434 | -0.001 | -0.037 | -0.037 | 0 | 0.33 | 0.48 | 0.17 | 0.22 | 0.31 | 0.52 |
| 11 | 84279862 | rs6592191 | -3.233 | -1.854 | -0.079 | -1.778 | -0.011 | 0.082 | 0.082 | 0.849 | 0.57 | 0.78 | 0.38 | 0.27 | 0.42 | 0.67 |
| 11 | 84284521 | rs17735087 | -0.135 | -1.85 | -0.067 | -1.558 | -0.01 | 0.019 | 0.019 | 0 | 0.08 | 0.05 | 0.04 | 0.01 | 0.09 | 0.1 |
| 11 | 84286016 | rs1940072 | -2.071 | -1.948 | -0.035 | -1.621 | -0.012 | -0.072 | -0.072 | 0.053 | 0.36 | 0.42 | 0.26 | 0.1 | 0.28 | 0.37 |
| 11 | 84298844 | rs7119459 | -0.903 | -1.949 | -0.012 | -1.555 | -0.013 | -0.042 | -0.042 | 0 | 0.17 | 0.22 | 0.06 | 0.07 | 0.12 | 0.22 |
| 11 | 84306591 | rs12284979 | -1.32 | -1.968 | -0.028 | -1.548 | -0.03 | -0.049 | -0.049 | 1.189 | 0.21 | 0.29 | 0.07 | 0.08 | 0.13 | 0.24 |
| 11 | 84312393 | rs12273005 | -0.849 | -2.019 | -0.002 | -1.532 | -0.027 | -0.062 | -0.062 | 0 | 0.17 | 0.27 | 0.06 | 0.06 | 0.12 | 0.22 |
| 11 | 84316886 | rs7114098 | -1.163 | -2.018 | -0.007 | -1.512 | -0.023 | -0.092 | -0.092 | 0 | 0.2 | 0.35 | 0.09 | 0.07 | 0.16 | 0.25 |
| 11 | 84319959 | rs1940085 | -2.352 | -2.06 | 0.066 | -1.548 | -0.007 | -0.021 | -0.021 | 0 | 0.37 | 0.58 | 0.18 | 0.22 | 0.31 | 0.59 |
| 11 | 84321748 | rs7112468 | -2.265 | -2.088 | 0.063 | -1.593 | -0.031 | -0.111 | -0.111 | 0.074 | 0.36 | 0.47 | 0.15 | 0.08 | 0.25 | 0.4 |
| 11 | 84323331 | rs2155400 | -0.591 | -2.009 | 0.007 | -1.581 | 0.003 | 0.01 | 0.01 | 0 | 0.13 | 0.11 | 0.16 | 0.03 | 0.1 | 0.09 |
| 11 | 84329118 | rs17147448 | -1.921 | -2.044 | 0.066 | -1.519 | -0.019 | -0.111 | -0.111 | 0 | 0.28 | 0.43 | 0.13 | 0.07 | 0.23 | 0.35 |
| 11 | 84334253 | rs17147455 | 0.509 | -1.728 | 0.071 | -1.35 | 0.001 | -0.027 | -0.027 | 0.108 | 0.06 | 0.11 | 0.04 | 0 | 0.07 | 0.09 |
| 11 | 84336651 | rs6592193 | -1.655 | -1.777 | 0.024 | -1.404 | -0.018 | -0.057 | -0.057 | 0.203 | 0.25 | 0.58 | 0.24 | 0.07 | 0.18 | 0.25 |
| 11 | 84343679 | rs9645693 | -0.372 | -1.723 | -0.205 | -1.351 | -0.025 | -0.033 | -0.033 | 0 | 0.11 | 0.03 | 0.06 | 0.17 | 0.11 | 0.25 |
| 11 | 84345829 | rs10898281 | -1.557 | -1.772 | -0.203 | -1.395 | -0.007 | 0.005 | 0.005 | 0 | 0.19 | 0.14 | 0.11 | 0.15 | 0.18 | 0.35 |
| 11 | 84350040 | rs12365964 | -0.996 | -1.754 | -0.282 | -1.397 | -0.021 | -0.028 | -0.028 | 0 | 0.19 | 0.03 | 0.2 | 0.19 | 0.16 | 0.29 |
| 11 | 84355831 | rs11234106 | -0.4 | -1.739 | 0.064 | -0.317 | 0.001 | 0.153 | 0.153 | 0 | 0.13 | 0.34 | 0.2 | 0.41 | 0.14 | 0.23 |
| 11 | 84360096 | rs6592194 | -3.213 | -1.876 | 0.093 | -0.46 | 0.061 | 0.173 | 0.173 | 0.876 | 0.49 | 0.78 | 0.36 | 0.22 | 0.37 | 0.62 |
| 11 | 84360346 | rs17206044 | -0.956 | -1.712 | -0.518 | -0.379 | -0.021 | -0.025 | -0.025 | 0 | 0.19 | 0.03 | 0.2 | 0.19 | 0.16 | 0.28 |
| 11 | 84364527 | rs1940100 | 0.593 | -1.634 | -0.456 | -0.338 | 0.001 | -0.013 | -0.013 | 0 | 0.06 | 0.07 | 0.04 | 0 | 0.07 | 0.09 |
| 11 | 84368114 | rs1940088 | -1.385 | -2.025 | -0.709 | -0.345 | -0.027 | 0.053 | 0.053 | 0 | 0.24 | 0.37 | 0.23 | 0.47 | 0.19 | 0.29 |
| 11 | 84370674 | rs1940080 | -2.797 | -1.686 | -0.783 | -0.34 | -0.004 | 0.005 | 0.005 | 0.198 | 0.56 | 0.66 | 0.51 | 0.66 | 0.47 | 0.67 |
| 11 | 84371133 | rs10898283 | -2.002 | -1.591 | -0.762 | -0.175 | 0 | -0.002 | -0.002 | 0 | 0.44 | 0.58 | 0.3 | 0.3 | 0.31 | 0.53 |
| 11 | 84372899 | rs17147476 | -0.525 | -1.103 | -0.754 | 0.102 | -0.008 | -0.033 | -0.033 | 0 | 0.13 | 0.36 | 0.07 | 0.09 | 0.11 | 0.2 |
| 11 | 84375859 | rs11234112 | 0.591 | -0.902 | -1.271 | 0.437 | -0.015 | -0.006 | -0.006 | 0 | 0.07 | 0.01 | 0.17 | 0.39 | 0.09 | 0.04 |
| 11 | 84377621 | rs7939569 | -1.198 | -1.045 | -1.126 | 0.045 | 0.009 | -0.004 | -0.004 | 0 | 0.17 | 0.35 | 0.1 | 0.18 | 0.17 | 0.39 |
| 11 | 84377856 | rs17147479 | -1.036 | -1.097 | -1.1 | 0.013 | -0.023 | -0.072 | -0.072 | 0.168 | 0.2 | 0.3 | 0.08 | 0.04 | 0.14 | 0.22 |
| 11 | 84379042 | rs11234120 | -0.753 | -0.999 | -0.993 | 0.452 | -0.022 | 0.05 | 0.05 | 0 | 0.21 | 0.31 | 0.2 | 0.45 | 0.16 | 0.2 |
| 11 | 84379671 | rs724921 | -1.169 | -0.581 | -1.278 | 0.349 | -0.024 | 0.007 | 0.007 | 0.392 | 0.57 | 0.27 | 0.49 | 0.64 | 0.5 | 0.67 |
| 11 | 84379740 | rs7112184 | -1.028 | -0.38 | -1.202 | 0.295 | -0.018 | 0.018 | 0.018 | 0.151 | 0.55 | 0.31 | 0.52 | 0.63 | 0.51 | 0.64 |
| 11 | 84380034 | rs7129994 | -1.045 | 0.359 | -0.545 | 0.436 | 0.014 | 0.004 | 0.004 | 0.791 | 0.27 | 0.59 | 0.26 | 0.29 | 0.3 | 0.17 |
| 11 | 84383445 | rs7120512 | -0.806 | 0.081 | -0.576 | 0.305 | -0.006 | -0.011 | -0.011 | 0 | 0.35 | 0.49 | 0.34 | 0.26 | 0.32 | 0.28 |
| 11 | 84385066 | rs17735326 | -0.653 | -0.033 | -0.9 | -0.018 | -0.011 | 0.062 | 0.062 | 0.558 | 0.11 | 0.02 | 0.05 | 0 | 0.11 | 0.08 |
| 11 | 84385588 | rs12274110 | -1.913 | -0.936 | -1.155 | -0.571 | 0.049 | 0.115 | 0.115 | 0.177 | 0.5 | 0.86 | 0.41 | 0.28 | 0.38 | 0.34 |
| 11 | 84391899 | rs1940098 | -0.856 | -1.055 | -0.59 | -0.887 | -0.007 | -0.078 | -0.078 | 0.177 | 0.22 | 0.29 | 0.25 | 0.02 | 0.19 | 0.11 |
| 11 | 84400029 | rs12278864 | -0.058 | -1.047 | -0.181 | -0.909 | -0.001 | -0.034 | -0.034 | 0.004 | 0.1 | 0.17 | 0.06 | 0 | 0.08 | 0.05 |
| 11 | 84406078 | rs7121845 | -2.468 | -1.194 | -0.775 | -1.241 | -0.11 | 0.161 | 0.161 | 0.037 | 0.63 | 0.89 | 0.62 | 0.84 | 0.52 | 0.71 |
| 11 | 84407214 | rs17147508 | -0.389 | -1.085 | -0.429 | -1.174 | -0.005 | -0.063 | -0.063 | 0 | 0.13 | 0.25 | 0.07 | 0 | 0.09 | 0.07 |
| 11 | 84409477 | rs1940094 | -1.261 | -1.274 | -0.351 | -1.098 | 0.002 | -0.007 | -0.007 | 0 | 0.31 | 0.32 | 0.31 | 0.63 | 0.31 | 0.48 |
| 11 | 84417846 | rs10501570 | -0.061 | -1.269 | -0.471 | -1.12 | -0.058 | 0.059 | 0.059 | 0 | 0.13 | 0.01 | 0.21 | 0.02 | 0.17 | 0.1 |
| 11 | 84449873 | rs548098 | 0.664 | -1.204 | -0.235 | -0.873 | -0.01 | 0.049 | 0.049 | 0 | 0.74 | 0.58 | 0.71 | 0.4 | 0.75 | 0.57 |
| 11 | 84454391 | rs4594021 | -0.872 | -1.392 | -0.176 | -0.967 | -0.05 | -0.012 | -0.012 | 0 | 0.26 | 0.55 | 0.14 | 0.2 | 0.18 | 0.21 |
| 11 | 84456685 | rs4505088 | -0.82 | -1.308 | -0.337 | -0.796 | 0.001 | -0.019 | -0.019 | 0 | 0.87 | 0.92 | 0.79 | 0.41 | 0.85 | 0.62 |
| 11 | 84460855 | rs6592202 | 0.887 | -1.596 | -0.122 | -0.219 | -0.014 | -0.05 | -0.05 | 0 | 0.68 | 0.8 | 0.57 | 0.49 | 0.66 | 0.68 |
| 11 | 84467936 | rs1943687 | -0.123 | -1.583 | -0.142 | -0.186 | -0.055 | -0.122 | -0.122 | 0.869 | 0.85 | 0.99 | 0.79 | 0.52 | 0.8 | 0.76 |
| 11 | 84469196 | rs11234161 | -1.18 | -1.271 | 0.257 | -0.22 | -0.03 | -0.024 | -0.024 | 0 | 0.18 | 0.52 | 0.12 | 0.11 | 0.12 | 0.23 |
| 11 | 84474461 | rs1943706 | 0.19 | -1.207 | 0.085 | -0.332 | -0.023 | 0.018 | 0.018 | 0 | 0.16 | 0.27 | 0.09 | 0.2 | 0.1 | 0.2 |
| 11 | 84494810 | rs7950953 | 0.131 | -1.165 | 0.073 | -0.347 | -0.015 | 0.027 | 0.027 | 0 | 0.16 | 0.28 | 0.1 | 0.21 | 0.11 | 0.21 |
| 11 | 84497615 | rs1943701 | 1.757 | -1.05 | -0.217 | -0.685 | -0.005 | 0.034 | 0.034 | 0 | 0.49 | 0.3 | 0.42 | 0.4 | 0.51 | 0.47 |
| 11 | 84511401 | rs17147569 | -0.019 | -0.983 | 0.194 | -0.936 | -0.002 | -0.036 | -0.036 | 0.181 | 0.13 | 0.18 | 0.06 | 0 | 0.11 | 0.08 |
| 11 | 84512643 | rs12271903 | -0.982 | -0.977 | 0.288 | -0.933 | 0.004 | -0.083 | -0.083 | 0.17 | 0.29 | 0.36 | 0.33 | 0.02 | 0.31 | 0.22 |
| 11 | 84521977 | rs11234189 | 0.459 | -0.837 | -0.159 | -0.766 | -0.001 | 0.042 | 0.042 | 0.619 | 0.1 | 0.01 | 0.08 | 0.01 | 0.08 | 0.14 |
| 11 | 84529030 | rs10751115 | 0.136 | -0.823 | 0.157 | -0.553 | -0.005 | 0.311 | 0.311 | 0.042 | 0.94 | 0.82 | 0.73 | 0.4 | 0.98 | 0.78 |
| 11 | 84533831 | rs17206578 | 0.283 | -0.716 | -0.15 | -0.629 | 0.003 | 0.042 | 0.042 | 0 | 0.08 | 0.12 | 0.05 | 0.22 | 0.08 | 0.14 |
| 11 | 84536635 | rs1943699 | 1.465 | -1.01 | -0.288 | -0.891 | 0.008 | 0.071 | 0.071 | 0 | 0.51 | 0.25 | 0.44 | 0.38 | 0.56 | 0.48 |

| | | | | | | | | | | | | | | | | |
|---|---|---|---|---|---|---|---|---|---|---|---|---|---|---|---|---|
| 11 | 84543946 | rs10792785 | 1.426 | -0.831 | -0.607 | -1.183 | -0.013 | 0.046 | 0.046 | 0.164 | 0.45 | 0.21 | 0.36 | 0.34 | 0.45 | 0.5 |
| 11 | 84547441 | rs12281095 | 0.612 | -0.854 | -0.348 | -1.299 | -0.001 | -0.024 | -0.024 | 0.977 | 0.18 | 0.17 | 0.13 | 0.01 | 0.14 | 0.05 |
| 11 | 84547595 | rs17808541 | -0.076 | -0.991 | -0.45 | -1.414 | -0.027 | 0.091 | 0.091 | 0 | 0.18 | 0.03 | 0.07 | 0.01 | 0.21 | 0.16 |
| 11 | 84553807 | rs952201 | -0.339 | -0.995 | -0.051 | -1.161 | 0.011 | 0.262 | 0.262 | 0 | 0.92 | 0.79 | 0.68 | 0.42 | 0.93 | 0.79 |
| 11 | 84561299 | rs12274130 | -0.002 | -0.99 | -0.233 | -1.36 | 0.001 | -0.004 | -0.004 | 0.071 | 0.23 | 0.17 | 0.14 | 0.01 | 0.18 | 0.08 |
| 11 | 84568024 | rs10501575 | 0.635 | -0.951 | -0.344 | -1.351 | -0.027 | 0.068 | 0.068 | 0 | 0.13 | 0.01 | 0.04 | 0.01 | 0.15 | 0.13 |
| 11 | 84574040 | rs11234221 | -1.018 | -1.112 | -0.058 | -1.156 | 0.011 | 0.017 | 0.017 | 0.025 | 0.42 | 0.36 | 0.2 | 0.26 | 0.35 | 0.34 |
| 11 | 84581522 | rs7125471 | 1.411 | -1.206 | -0.229 | -1.181 | 0.002 | -0.007 | -0.007 | 0 | 0.11 | 0.08 | 0.06 | 0 | 0.06 | 0.01 |
| 11 | 84581716 | rs1943723 | 1.341 | -1.144 | 0.141 | -1.025 | 0.002 | 0.017 | 0.017 | 0 | 0.53 | 0.37 | 0.47 | 0.41 | 0.48 | 0.56 |
| 11 | 84584283 | rs2156517 | 2.127 | -1.065 | 0.124 | -0.753 | -0.056 | 0.084 | 0.084 | 1.736 | 0.4 | 0.15 | 0.46 | 0.25 | 0.46 | 0.48 |
| 11 | 84596023 | rs10898315 | -1.776 | -0.929 | 0.011 | -1.096 | 0 | 0.01 | 0.01 | 0.323 | 0.48 | 0.45 | 0.28 | 0.26 | 0.46 | 0.37 |
| 11 | 84598945 | rs10501579 | 0.031 | -1.097 | -0.199 | -1.184 | -0.021 | -0.052 | -0.052 | 0.192 | 0.1 | 0.25 | 0.07 | 0.02 | 0.06 | 0.06 |
| 11 | 84600729 | rs4457780 | -2.152 | -1.142 | -0.29 | -1.209 | -0.008 | 0.025 | 0.025 | 0 | 0.56 | 0.64 | 0.34 | 0.28 | 0.52 | 0.43 |
| 11 | 84601377 | rs957438 | -0.973 | -0.685 | -0.399 | -1.99 | -0.009 | 0.002 | 0.002 | 0 | 0.41 | 0.34 | 0.26 | 0.02 | 0.44 | 0.25 |
| 11 | 84605753 | rs17147761 | -0.726 | -0.219 | -0.316 | -1.012 | -0.023 | -0.051 | -0.051 | 0 | 0.1 | 0.2 | 0.06 | 0.02 | 0.06 | 0.06 |
| 11 | 84609813 | rs11603955 | -0.645 | -0.142 | -0.478 | -0.995 | -0.027 | -0.108 | -0.108 | 1.158 | 0.21 | 0.31 | 0.13 | 0.02 | 0.12 | 0.07 |
| 11 | 84623084 | rs17808971 | -0.143 | 1.76 | -0.519 | -0.675 | 0.006 | -0.024 | -0.024 | 0 | 0.06 | 0.08 | 0.05 | 0.02 | 0.12 | 0.05 |
| 11 | 84627035 | rs10898329 | 0.336 | 1.672 | -1.175 | -1.303 | -0.01 | 0.038 | 0.038 | 0 | 0.31 | 0.11 | 0.3 | 0.2 | 0.33 | 0.47 |
| 11 | 84628985 | rs7127610 | -0.823 | 1.733 | -0.633 | -1.2 | 0.021 | -0.027 | -0.027 | 0 | 0.11 | 0.23 | 0.11 | 0.1 | 0.16 | 0.12 |
| 11 | 84629013 | rs7111775 | -0.824 | 1.546 | -0.294 | -1.572 | -0.004 | 0.022 | 0.022 | 0 | 0.45 | 0.56 | 0.54 | 0.28 | 0.54 | 0.64 |
| 11 | 84634790 | rs2155413 | -0.035 | 1.541 | -0.594 | -1.764 | -0.036 | 0.102 | 0.102 | 0 | 0.59 | 0.79 | 0.62 | 0.83 | 0.46 | 0.54 |
| 11 | 84636486 | rs10792789 | 0.782 | 1.638 | -0.54 | -1.638 | -0.004 | -0.006 | -0.006 | 0 | 0.47 | 0.49 | 0.35 | 0.65 | 0.32 | 0.3 |
| 11 | 84642072 | rs6592215 | -0.858 | 1.884 | -0.628 | -1.667 | -0.049 | 0.125 | 0.125 | 0 | 0.7 | 0.88 | 0.71 | 0.95 | 0.64 | 0.67 |
| 11 | 84642163 | rs11600301 | 0.143 | 1.845 | -0.483 | -1.557 | 0.016 | 0.024 | 0.024 | 0.033 | 0.11 | 0.2 | 0.21 | 0.16 | 0.14 | 0.24 |
| 11 | 84649642 | rs7108021 | 0.549 | 1.764 | -0.546 | -1.503 | 0.002 | 0.003 | 0.003 | 0 | 0.52 | 0.55 | 0.4 | 0.68 | 0.37 | 0.33 |
| 11 | 84665769 | rs11607740 | 0.188 | 1.49 | -0.953 | -1.838 | -0.001 | -0.003 | -0.003 | 0.009 | 0.3 | 0.26 | 0.29 | 0.24 | 0.26 | 0.42 |
| 11 | 84669998 | rs7943194 | 0.076 | 1.56 | -1.1 | -1.85 | 0.008 | 0.025 | 0.025 | 0.121 | 0.42 | 0.7 | 0.44 | 0.68 | 0.39 | 0.25 |
| 11 | 84680391 | rs3851173 | -0.215 | 1.651 | -0.899 | -1.678 | -0.016 | 0.011 | 0.011 | 0.003 | 0.62 | 0.72 | 0.51 | 0.68 | 0.53 | 0.37 |
| 11 | 84687721 | rs596556 | -1.128 | 1.673 | -1.018 | -1.694 | -0.081 | 0.104 | 0.104 | 0 | 0.72 | 0.93 | 0.71 | 0.92 | 0.65 | 0.66 |
| 11 | 84688062 | rs3844141 | 1.646 | 1.212 | -1.196 | -2.047 | 0.011 | 0.096 | 0.096 | 3.514 | 0.21 | 0.05 | 0.08 | 0 | 0.14 | 0.03 |
| 11 | 84708782 | rs2509053 | -1.023 | 1.589 | -0.778 | -1.533 | -0.056 | 0.086 | 0.086 | 0 | 0.72 | 0.91 | 0.71 | 0.92 | 0.65 | 0.67 |
| 11 | 84713513 | rs597592 | -0.789 | 1.592 | -0.79 | -1.546 | -0.062 | 0.1 | 0.1 | 0 | 0.7 | 0.9 | 0.66 | 0.92 | 0.63 | 0.65 |
| 11 | 84720020 | rs643891 | -1.006 | 1.59 | -0.775 | -1.546 | -0.053 | 0.083 | 0.083 | 0 | 0.72 | 0.9 | 0.71 | 0.92 | 0.64 | 0.67 |
| 11 | 84722252 | rs624493 | 0.137 | 1.6 | -0.652 | -1.463 | 0.022 | 0.071 | 0.071 | 0 | 0.11 | 0.21 | 0.2 | 0.3 | 0.12 | 0.3 |
| 11 | 84725828 | rs3907015 | 0.078 | 1.595 | -0.533 | -1.467 | 0 | -0.003 | -0.003 | 0 | 0.59 | 0.61 | 0.44 | 0.63 | 0.5 | 0.35 |
| 11 | 84735715 | rs685294 | -0.864 | 1.502 | -0.569 | -1.462 | -0.054 | 0.088 | 0.088 | 0 | 0.71 | 0.9 | 0.71 | 0.92 | 0.64 | 0.66 |
| 11 | 84741129 | rs582884 | -0.883 | 1.528 | -0.569 | -1.475 | -0.034 | 0.073 | 0.073 | 0 | 0.7 | 0.84 | 0.65 | 0.94 | 0.63 | 0.66 |
| 11 | 84741811 | rs687695 | -0.893 | 1.555 | -0.575 | -1.46 | -0.059 | 0.106 | 0.106 | 0 | 0.71 | 0.91 | 0.71 | 0.93 | 0.65 | 0.66 |
| 11 | 84743531 | rs659414 | 0.101 | 1.381 | -0.408 | -1.397 | 0.028 | 0.075 | 0.075 | 0 | 0.1 | 0.2 | 0.19 | 0.26 | 0.12 | 0.3 |
| 11 | 84744926 | rs11605620 | -0.01 | 1.38 | -0.412 | -1.397 | 0.021 | 0.06 | 0.06 | 0 | 0.1 | 0.2 | 0.2 | 0.27 | 0.12 | 0.3 |
| 11 | 84752277 | rs573963 | -0.688 | 1.348 | -0.432 | -1.401 | 0.022 | 0.062 | 0.062 | 0 | 0.13 | 0.28 | 0.26 | 0.29 | 0.13 | 0.33 |
| 11 | 84755545 | rs634050 | -0.195 | 1.665 | -0.532 | -1.413 | -0.001 | -0.002 | -0.002 | 0.342 | 0.37 | 0.36 | 0.48 | 0.36 | 0.39 | 0.59 |
| 11 | 84758991 | rs668816 | -0.756 | 1.471 | -0.425 | -1.679 | -0.008 | 0.018 | 0.018 | 1.293 | 0.63 | 0.74 | 0.62 | 0.35 | 0.66 | 0.74 |
| 11 | 84786132 | rs1826613 | 0.793 | 1.303 | -0.509 | -1.646 | 0.005 | 0.039 | 0.039 | 0 | 0.27 | 0.16 | 0.28 | 0.07 | 0.29 | 0.28 |
| 11 | 84795484 | rs473818 | 0.983 | 1.302 | -0.522 | -1.176 | 0.024 | -0.041 | -0.041 | 1.123 | 0.29 | 0.21 | 0.3 | 0.56 | 0.2 | 0.2 |
| 11 | 84796996 | rs349068 | 0.001 | 1.47 | -0.44 | -1.829 | -0.001 | -0.009 | -0.009 | 0.083 | 0.36 | 0.37 | 0.47 | 0.32 | 0.39 | 0.59 |
| 11 | 84806896 | rs1227842 | -0.389 | 2.012 | -0.146 | -1.214 | -0.037 | 0.068 | 0.068 | 0 | 0.69 | 0.84 | 0.65 | 0.93 | 0.6 | 0.67 |
| 11 | 84808429 | rs349078 | 0.537 | 1.977 | -0.102 | -1.252 | 0.001 | -0.007 | -0.007 | 0 | 0.58 | 0.57 | 0.45 | 0.67 | 0.49 | 0.32 |
| 11 | 84810131 | rs349079 | -0.205 | 1.982 | -0.04 | -1.217 | 0.018 | 0.081 | 0.081 | 0 | 0.1 | 0.23 | 0.2 | 0.3 | 0.11 | 0.32 |
| 11 | 84812643 | rs349085 | 0.419 | 1.959 | 0.002 | -1.207 | -0.015 | 0.019 | 0.019 | 0 | 0.6 | 0.68 | 0.48 | 0.72 | 0.52 | 0.44 |
| 11 | 84813476 | rs349087 | 1.061 | 1.803 | -0.073 | -1.228 | 0.031 | 0.044 | 0.044 | 0 | 0.51 | 0.37 | 0.39 | 0.66 | 0.36 | 0.31 |
| 11 | 84814393 | rs349088 | -0.811 | 2.164 | -0.599 | -1.061 | 0.004 | -0.007 | -0.007 | 0 | 0.39 | 0.42 | 0.52 | 0.34 | 0.52 | 0.62 |
| 11 | 84818313 | rs675217 | -0.318 | 1.598 | -0.815 | -1.236 | -0.058 | 0.099 | 0.099 | 0 | 0.71 | 0.91 | 0.7 | 0.94 | 0.62 | 0.68 |
| 11 | 84821981 | rs555156 | -0.118 | 1.607 | -0.746 | -1.363 | -0.025 | 0.042 | 0.042 | 0 | 0.69 | 0.8 | 0.64 | 0.95 | 0.59 | 0.67 |
| 11 | 84825146 | rs592166 | -0.355 | 1.623 | -0.795 | -1.343 | -0.033 | 0.058 | 0.058 | 0 | 0.71 | 0.84 | 0.7 | 0.95 | 0.63 | 0.69 |
| 11 | 84841573 | rs1532312 | -0.602 | 0.97 | -0.52 | -0.856 | -0.008 | 0.007 | 0.007 | 0 | 0.31 | 0.37 | 0.43 | 0.4 | 0.29 | 0.55 |
| 11 | 84845925 | rs17810088 | -0.128 | 0.943 | -0.144 | -0.392 | 0.008 | 0.122 | 0.122 | 0 | 0.07 | 0.13 | 0.26 | 0.29 | 0.07 | 0.27 |
| 11 | 84867510 | rs17148028 | -0.977 | 0.828 | -0.141 | -0.488 | 0.021 | 0.019 | 0.019 | 0 | 0.05 | 0.2 | 0.09 | 0.09 | 0.05 | 0.08 |
| 11 | 84872234 | rs1826612 | -0.521 | 0.823 | -0.27 | -0.453 | 0.009 | 0.135 | 0.135 | 0 | 0.07 | 0.18 | 0.28 | 0.31 | 0.08 | 0.27 |
| 11 | 84878532 | rs11234316 | -0.577 | 0.713 | -0.252 | -0.465 | -0.002 | 0.1 | 0.1 | 0 | 0.06 | 0.1 | 0.17 | 0.31 | 0.05 | 0.22 |
| 11 | 84895033 | rs476791 | -1.092 | 0.519 | -0.884 | -0.435 | -0.008 | -0.046 | -0.046 | 0 | 0.91 | 0.99 | 0.82 | 0.46 | 0.89 | 0.91 |
| 11 | 84914343 | rs9734338 | 1.214 | 0.688 | -0.092 | -0.496 | 0.013 | 0.046 | 0.046 | 0.393 | 0.74 | 0.53 | 0.51 | 0.6 | 0.65 | 0.56 |
| 11 | 84925643 | rs938727 | 1.018 | 0.549 | 0.236 | 0.009 | 0.021 | 0.236 | 0.236 | 0 | 0.89 | 0.65 | 0.68 | 0.62 | 0.9 | 0.67 |
| 11 | 84928974 | rs10898382 | 0.321 | 0.557 | 0.22 | 0.003 | 0.017 | 0.183 | 0.183 | 0.093 | 0.92 | 0.78 | 0.79 | 0.72 | 0.94 | 0.73 |
| 11 | 84935074 | rs951729 | 0.995 | 0.559 | 0.21 | -0.007 | 0.022 | 0.181 | 0.181 | 0.059 | 0.9 | 0.71 | 0.73 | 0.71 | 0.92 | 0.72 |
| 11 | 84943077 | rs10501590 | 0.931 | 0.357 | 0.052 | -0.16 | -0.023 | 0.095 | 0.095 | 0 | 0.81 | 0.63 | 0.64 | 0.6 | 0.84 | 0.65 |
| 11 | 84954241 | rs938728 | 0.241 | 0.796 | 0.336 | 0.139 | 0.012 | 0.159 | 0.159 | 0 | 0.93 | 0.81 | 0.8 | 0.72 | 0.94 | 0.73 |
| 11 | 84992639 | rs471786 | 1.521 | 0.819 | 0.378 | 0.088 | 0.02 | 0.198 | 0.198 | 0.008 | 0.88 | 0.63 | 0.63 | 0.6 | 0.88 | 0.65 |
| 11 | 84992869 | rs580188 | 1.296 | 0.795 | 0.441 | 0.066 | 0.009 | 0.188 | 0.188 | 0 | 0.89 | 0.67 | 0.69 | 0.6 | 0.89 | 0.66 |

| | | | | | | | | | | | | | | | | |
|---|---|---|---|---|---|---|---|---|---|---|---|---|---|---|---|---|
| 11 | 84993643 | rs552017 | 2.07 | 0.823 | 0.403 | 0.02 | 0.01 | 0.164 | 0.164 | 0.091 | 0.85 | 0.62 | 0.62 | 0.6 | 0.85 | 0.63 |
| 11 | 85012704 | rs11823978 | -1.739 | 0.775 | 0.354 | -0.029 | 0.006 | -0.002 | -0.002 | 0 | 0.07 | 0.17 | 0.11 | 0.1 | 0.07 | 0.1 |
| 11 | 85017476 | rs17148090 | -1.066 | 0.732 | 0.241 | -0.145 | -0.006 | -0.005 | -0.005 | 0 | 0.07 | 0.1 | 0.05 | 0.09 | 0.07 | 0.1 |
| 11 | 85019691 | rs497607 | 1.247 | 0.764 | 0.37 | -0.055 | 0.018 | 0.153 | 0.153 | 0.038 | 0.89 | 0.69 | 0.66 | 0.7 | 0.9 | 0.71 |
| 11 | 85055423 | rs7940610 | -2.062 | 0.743 | 0.343 | -0.094 | 0.006 | -0.003 | -0.003 | 0 | 0.08 | 0.22 | 0.11 | 0.1 | 0.08 | 0.09 |
| 11 | 85079419 | rs286050 | 1.769 | 1.374 | -0.001 | -0.432 | 0.017 | 0.06 | 0.06 | 0.018 | 0.74 | 0.51 | 0.52 | 0.6 | 0.64 | 0.57 |
| 11 | 85105113 | rs286028 | 1.843 | 1.4 | 0.191 | -0.153 | 0.003 | 0.142 | 0.142 | 0 | 0.85 | 0.63 | 0.63 | 0.6 | 0.86 | 0.63 |
| 11 | 85117531 | rs2055816 | 1.312 | 1.413 | 0.197 | -0.141 | 0.006 | 0.101 | 0.101 | 0 | 0.88 | 0.75 | 0.67 | 0.69 | 0.89 | 0.7 |
| 11 | 85152822 | rs2113920 | 0.284 | 1.485 | 0.254 | 0.035 | 0.001 | 0.079 | 0.079 | 0 | 0.93 | 0.92 | 0.61 | 0.68 | 0.92 | 0.76 |
| 11 | 85164751 | rs10898392 | 0.991 | 1.172 | 0.455 | -0.852 | -0.005 | 0.012 | 0.012 | 0.482 | 0.54 | 0.34 | 0.29 | 0.57 | 0.44 | 0.44 |
| 11 | 85171594 | rs7110363 | 1.522 | 0.93 | 0.631 | -0.318 | 0.044 | 0.237 | 0.237 | 0.172 | 0.91 | 0.71 | 0.54 | 0.68 | 0.89 | 0.71 |
| 11 | 85172303 | rs1213236 | 0.042 | 0.907 | 0.522 | -0.311 | 0.003 | 0.125 | 0.125 | 0.002 | 0.95 | 0.91 | 0.76 | 0.69 | 0.93 | 0.76 |
| 11 | 85199681 | rs1912266 | -1.457 | 0.887 | 0.493 | -0.332 | -0.013 | -0.002 | -0.002 | 0 | 0.06 | 0.02 | 0.08 | 0.09 | 0.08 | 0.1 |
| 11 | 85200968 | rs286521 | -0.139 | 0.9 | 0.558 | -0.306 | 0.003 | 0.094 | 0.094 | 0 | 0.94 | 0.92 | 0.74 | 0.69 | 0.92 | 0.79 |
| 11 | 85206156 | rs2512800 | 0.51 | 0.836 | 0.865 | -0.339 | 0.04 | 0.275 | 0.275 | 0 | 0.92 | 0.72 | 0.73 | 0.69 | 0.92 | 0.75 |
| 11 | 85220773 | rs1213257 | 1.976 | 0.821 | 0.959 | -0.356 | 0.011 | 0.159 | 0.159 | 0 | 0.86 | 0.67 | 0.63 | 0.6 | 0.84 | 0.62 |
| 11 | 85227439 | rs286510 | -1.375 | 0.869 | 0.758 | -0.332 | 0.017 | 0.137 | 0.137 | 0 | 0.1 | 0.22 | 0.34 | 0.31 | 0.12 | 0.27 |
| 11 | 85231475 | rs7119372 | -1.375 | 0.869 | 0.758 | -0.311 | 0.017 | 0.14 | 0.14 | 0 | 0.1 | 0.22 | 0.34 | 0.31 | 0.12 | 0.27 |
| 11 | 85239988 | rs11234359 | -1.375 | 0.869 | 0.744 | -0.302 | 0.016 | 0.128 | 0.128 | 0 | 0.1 | 0.22 | 0.34 | 0.29 | 0.12 | 0.27 |
| 11 | 85245772 | rs1863256 | -1.545 | 0.861 | 0.652 | -0.269 | 0.037 | 0.226 | 0.226 | 0 | 0.11 | 0.34 | 0.34 | 0.31 | 0.12 | 0.27 |
| 11 | 85251005 | rs7949567 | -2.285 | 0.853 | 0.612 | -0.289 | 0.019 | 0.218 | 0.218 | 0 | 0.15 | 0.41 | 0.37 | 0.4 | 0.16 | 0.35 |
| 11 | 85266321 | rs4944523 | -2.221 | 0.835 | 0.576 | -0.317 | 0.011 | 0.225 | 0.225 | 0 | 0.15 | 0.41 | 0.37 | 0.4 | 0.16 | 0.34 |
| 11 | 85277870 | rs12420907 | -1.4 | 0.846 | 0.65 | -0.22 | 0.034 | 0.172 | 0.172 | 0 | 0.11 | 0.31 | 0.34 | 0.27 | 0.12 | 0.27 |
| 11 | 85292310 | rs7122887 | -2.406 | 0.842 | 0.573 | -0.265 | 0.01 | 0.179 | 0.179 | 0 | 0.16 | 0.4 | 0.38 | 0.37 | 0.19 | 0.35 |
| 11 | 85307787 | rs494119 | 0.101 | 1.017 | -0.021 | -0.685 | -0.04 | 0.127 | 0.127 | 0 | 0.25 | 0.03 | 0.15 | 0.09 | 0.26 | 0.19 |
| 11 | 85310744 | rs881121 | -0.297 | 0.903 | 0.288 | -0.395 | 0 | 0.057 | 0.057 | 0 | 0.07 | 0.08 | 0.26 | 0.29 | 0.09 | 0.3 |
| 11 | 85337366 | rs11603256 | -2.196 | 0.879 | 0.375 | -0.429 | 0.003 | 0.157 | 0.157 | 0.033 | 0.17 | 0.41 | 0.38 | 0.38 | 0.19 | 0.4 |
| 11 | 85347415 | rs1046216 | -2.554 | 0.821 | 0.41 | -0.566 | -0.008 | 0.128 | 0.128 | 0.59 | 0.2 | 0.43 | 0.39 | 0.38 | 0.21 | 0.42 |
| 11 | 85350822 | rs12419172 | -0.874 | 0.931 | 0.394 | -0.344 | 0 | 0.064 | 0.064 | 0 | 0.1 | 0.12 | 0.29 | 0.31 | 0.11 | 0.32 |
| 11 | 85367289 | rs3753050 | -2.573 | 0.904 | 0.402 | -0.392 | -0.002 | 0.138 | 0.138 | 0.238 | 0.19 | 0.42 | 0.39 | 0.38 | 0.21 | 0.41 |
| 11 | 85369198 | rs1991923 | -0.718 | 0.912 | 0.41 | -0.386 | 0 | 0.009 | 0.009 | 0 | 0.09 | 0.07 | 0.27 | 0.29 | 0.11 | 0.31 |
| 11 | 85370482 | rs3824956 | -2.564 | 0.914 | 0.405 | -0.386 | -0.006 | 0.136 | 0.136 | 0.356 | 0.19 | 0.42 | 0.39 | 0.38 | 0.21 | 0.41 |
| 11 | 85373668 | rs7116195 | -2.658 | 0.913 | 0.381 | -0.413 | -0.004 | 0.124 | 0.124 | 0.476 | 0.2 | 0.43 | 0.4 | 0.38 | 0.21 | 0.44 |
| 11 | 85374993 | rs1192 | -0.668 | 0.907 | 0.334 | -0.406 | 0.001 | 0.062 | 0.062 | 0 | 0.09 | 0.12 | 0.29 | 0.28 | 0.1 | 0.31 |
| 11 | 85379051 | rs431606 | -2.582 | 1.275 | 0.297 | -0.576 | 0.006 | 0.029 | 0.029 | 0.336 | 0.27 | 0.6 | 0.46 | 0.4 | 0.37 | 0.48 |
| 11 | 85394238 | rs448419 | 1.338 | 0.751 | -0.449 | -0.542 | -0.016 | -0.083 | -0.083 | 0 | 0.13 | 0.02 | 0.18 | 0.47 | 0.11 | 0.14 |
| 11 | 85395652 | rs290186 | -0.298 | 0.751 | -0.474 | -1.065 | -0.023 | -0.023 | -0.023 | 0.188 | 0.7 | 0.88 | 0.59 | 0.71 | 0.69 | 0.58 |
| 11 | 85395718 | rs290185 | 0.443 | 0.616 | -0.206 | -1.129 | -0.002 | -0.004 | -0.004 | 0 | 0.6 | 0.54 | 0.49 | 0.61 | 0.59 | 0.49 |
| 11 | 85400063 | rs290179 | 0.944 | 0.447 | -0.404 | -1.522 | 0.002 | -0.002 | -0.002 | 0 | 0.43 | 0.32 | 0.38 | 0.58 | 0.39 | 0.39 |
| 11 | 85404138 | rs56579 | -0.111 | 0.465 | -0.201 | -1.356 | 0.004 | -0.007 | -0.007 | 0 | 0.65 | 0.46 | 0.51 | 0.71 | 0.65 | 0.57 |
| 11 | 85406427 | rs2276018 | -1.109 | 0.266 | -0.144 | -1.32 | 0.006 | 0.003 | 0.003 | 0 | 0.08 | 0.07 | 0.03 | 0.09 | 0.05 | 0.12 |
| 11 | 85406915 | rs188311 | -0.285 | 0.412 | 0.084 | -1.259 | 0.001 | 0.002 | 0.002 | 0.077 | 0.68 | 0.56 | 0.58 | 0.72 | 0.7 | 0.6 |
| 11 | 85408832 | rs290173 | 0.379 | 0.508 | 0.082 | -1.219 | -0.002 | -0.001 | -0.001 | 0 | 0.62 | 0.41 | 0.5 | 0.63 | 0.63 | 0.5 |
| 11 | 85435730 | rs550404 | 0.154 | 0.514 | 0.093 | -0.993 | -0.001 | 0.004 | 0.004 | 0.07 | 0.66 | 0.65 | 0.55 | 0.6 | 0.65 | 0.52 |
| 11 | 85436500 | rs580459 | -0.052 | 0.508 | 0.029 | -0.978 | -0.023 | -0.063 | -0.063 | 3.021 | 0.68 | 0.88 | 0.57 | 0.6 | 0.66 | 0.52 |
| 11 | 85436699 | rs598346 | -1.575 | 0.509 | -0.021 | -1.044 | -0.017 | 0.051 | 0.051 | 0 | 0.81 | 0.94 | 0.88 | 1 | 0.8 | 0.88 |
| 11 | 85437896 | rs491867 | -0.014 | 0.509 | 0.015 | -0.989 | -0.018 | -0.052 | -0.052 | 0.286 | 0.68 | 0.88 | 0.57 | 0.61 | 0.66 | 0.52 |
| 11 | 85443849 | rs496187 | -0.014 | 0.509 | 0.015 | -0.982 | -0.018 | -0.05 | -0.05 | 0 | 0.68 | 0.88 | 0.57 | 0.62 | 0.66 | 0.52 |
| 11 | 85447611 | rs680498 | -0.089 | 0.506 | 0.05 | -0.983 | -0.013 | -0.033 | -0.033 | 0 | 0.68 | 0.82 | 0.57 | 0.62 | 0.66 | 0.52 |
| 11 | 85450436 | rs575013 | 0.957 | 0.483 | 0.019 | -1.134 | 0.011 | 0.024 | 0.024 | 0 | 0.51 | 0.36 | 0.4 | 0.6 | 0.44 | 0.45 |
| 11 | 85456703 | rs647912 | 0.579 | 0.37 | 0.241 | -0.841 | -0.004 | 0.002 | 0.002 | 0.482 | 0.63 | 0.48 | 0.5 | 0.61 | 0.63 | 0.51 |
| 11 | 85457364 | rs653559 | -0.502 | 0.402 | 0.192 | -0.829 | -0.028 | -0.042 | -0.042 | 0 | 0.74 | 0.89 | 0.61 | 0.71 | 0.71 | 0.61 |
| 11 | 85462141 | rs688741 | 0.102 | 0.379 | 0.22 | -0.799 | 0.002 | 0.005 | 0.005 | 0 | 0.68 | 0.6 | 0.55 | 0.7 | 0.68 | 0.6 |
| 11 | 85465395 | rs650036 | 0.094 | 0.556 | 0.226 | -0.842 | 0.005 | 0.002 | 0.002 | 0.032 | 0.68 | 0.65 | 0.55 | 0.7 | 0.68 | 0.6 |
| 11 | 85470893 | rs639688 | -0.571 | 0.595 | 0.074 | -0.836 | -0.034 | -0.049 | -0.049 | 0 | 0.74 | 0.93 | 0.68 | 0.72 | 0.74 | 0.65 |
| 11 | 85473248 | rs510948 | -0.338 | 0.559 | 0.237 | -0.841 | -0.036 | -0.028 | -0.028 | 1.009 | 0.72 | 0.9 | 0.61 | 0.71 | 0.7 | 0.64 |
| 11 | 85473883 | rs537604 | -0.384 | 0.553 | 0.179 | -0.84 | -0.054 | -0.046 | -0.046 | 0 | 0.73 | 0.92 | 0.62 | 0.71 | 0.7 | 0.64 |
| 11 | 85495269 | rs485485 | -0.928 | 0.46 | 0.298 | -0.848 | -0.001 | 0 | 0 | 0 | 0.38 | 0.56 | 0.5 | 0.39 | 0.36 | 0.48 |
| 11 | 85504913 | rs10898409 | -0.452 | 0.638 | 0.395 | -0.775 | 0.006 | -0.005 | -0.005 | 0 | 0.1 | 0.25 | 0.08 | 0.11 | 0.09 | 0.11 |
| 11 | 85514693 | rs11234416 | -0.099 | 0.561 | 0.229 | -0.734 | -0.006 | -0.002 | -0.002 | 0 | 0.06 | 0.09 | 0.04 | 0.09 | 0.05 | 0.1 |
| 11 | 85518123 | rs4944538 | -1.133 | 0.577 | 0.138 | -0.804 | 0 | 0 | 0 | 0 | 0.37 | 0.62 | 0.5 | 0.39 | 0.35 | 0.49 |
| 11 | 85526399 | rs17148514 | -0.285 | 0.665 | -0.085 | -1 | 0.001 | -0.038 | -0.038 | 0 | 0.25 | 0.28 | 0.24 | 0.09 | 0.26 | 0.18 |
| 11 | 85536995 | rs11234427 | -0.92 | 0.389 | 0.411 | -0.586 | 0.006 | 0.057 | 0.057 | 0 | 0.06 | 0.14 | 0.1 | 0.12 | 0.07 | 0.11 |
| 11 | 85539617 | rs11234429 | -1.101 | 0.089 | -0.009 | -0.826 | -0.032 | -0.012 | -0.012 | 0.613 | 0.58 | 0.76 | 0.64 | 0.42 | 0.55 | 0.53 |
| 11 | 85540247 | rs11606818 | 0.553 | 0.238 | -0.298 | -0.905 | -0.018 | -0.019 | -0.019 | 0 | 0.25 | 0.16 | 0.38 | 0.28 | 0.26 | 0.3 |
| 11 | 85541797 | rs518691 | 1.999 | 0.756 | -1.194 | -0.762 | -0.062 | 0.006 | 0.006 | 0.198 | 0.14 | 0.01 | 0.09 | 0.07 | 0.17 | 0.16 |
| 11 | 85542482 | rs1445500 | 1.767 | 0.292 | -0.715 | -0.677 | 0 | 0.045 | 0.045 | 6.803 | 0.16 | 0.06 | 0.11 | 0.02 | 0.18 | 0.05 |
| 11 | 85545180 | rs7943104 | 1.53 | 0.279 | -0.645 | -0.687 | 0.002 | 0.011 | 0.011 | 6.352 | 0.16 | 0.13 | 0.12 | 0.02 | 0.18 | 0.05 |
| 11 | 85554057 | rs17817230 | 1.548 | 0.285 | -0.641 | -0.682 | 0.003 | 0.01 | 0.01 | 5.926 | 0.16 | 0.13 | 0.12 | 0.02 | 0.18 | 0.05 |

| | | | | | | | | | | | | | | | | |
|---|---|---|---|---|---|---|---|---|---|---|---|---|---|---|---|---|
| 11 | 85558723 | rs3862785 | 1.012 | 0.479 | -0.765 | -0.723 | -0.05 | -0.036 | -0.036 | 0.15 | 0.24 | 0.08 | 0.3 | 0.26 | 0.24 | 0.28 |
| 11 | 85564826 | rs17817254 | 1.876 | 0.411 | -0.719 | -0.705 | -0.007 | 0.05 | 0.05 | 6.363 | 0.15 | 0.05 | 0.11 | 0.02 | 0.18 | 0.06 |
| 11 | 85579304 | rs11234454 | -1.028 | 0.34 | -0.413 | -0.718 | 0.001 | 0.015 | 0.015 | 0.277 | 0.53 | 0.61 | 0.57 | 0.36 | 0.57 | 0.49 |
| 11 | 85591041 | rs11234457 | 0.542 | 0.278 | -0.406 | -0.626 | 0.009 | 0.028 | 0.028 | 3.936 | 0.21 | 0.15 | 0.2 | 0.1 | 0.23 | 0.15 |
| 11 | 85600724 | rs17744711 | 1.557 | 0.32 | -0.447 | -0.619 | -0.002 | 0.041 | 0.041 | 7.14 | 0.16 | 0.07 | 0.12 | 0.02 | 0.18 | 0.05 |
| 11 | 85600835 | rs12271536 | 0.225 | 0.318 | -0.257 | -0.645 | -0.007 | -0.047 | -0.047 | 0 | 0.22 | 0.31 | 0.23 | 0.1 | 0.27 | 0.16 |
| 11 | 85601792 | rs7936092 | -1.863 | 0.348 | -0.097 | -0.676 | 0.016 | 0.001 | 0.001 | 0.16 | 0.11 | 0.43 | 0.15 | 0.09 | 0.12 | 0.13 |
| 11 | 85605334 | rs11234465 | 0.655 | 0.284 | -0.241 | -0.729 | -0.002 | -0.031 | -0.031 | 0 | 0.24 | 0.24 | 0.24 | 0.07 | 0.3 | 0.15 |
| 11 | 85605342 | rs12274739 | 0.655 | 0.284 | -0.241 | -0.729 | -0.002 | -0.031 | -0.031 | 0 | 0.24 | 0.24 | 0.24 | 0.07 | 0.3 | 0.15 |
| 11 | 85609559 | rs10501600 | -1.005 | 0.436 | -0.537 | -0.654 | 0.001 | 0.008 | 0.008 | 0.461 | 0.53 | 0.57 | 0.57 | 0.35 | 0.59 | 0.49 |
| 11 | 85611813 | rs7926846 | 0.278 | 0.576 | -0.631 | -0.68 | -0.004 | -0.045 | -0.045 | 0 | 0.25 | 0.29 | 0.25 | 0.07 | 0.3 | 0.14 |
| 11 | 85621406 | rs10792819 | 0.291 | 1.049 | -1.053 | 0.15 | -0.04 | -0.061 | -0.061 | 0 | 0.16 | 0.06 | 0.2 | 0.3 | 0.19 | 0.12 |
| 11 | 85623607 | rs11608136 | 0.832 | 0.822 | -1.145 | -0.143 | -0.026 | -0.105 | -0.105 | 0 | 0.22 | 0.07 | 0.28 | 0.37 | 0.2 | 0.29 |
| 11 | 85624638 | rs12790645 | 0.753 | 0.964 | -0.877 | -0.338 | 0.012 | -0.016 | -0.016 | 0 | 0.1 | 0.11 | 0.14 | 0.07 | 0.13 | 0.11 |
| 11 | 85624978 | rs1445496 | -0.887 | 0.7 | -0.423 | -0.616 | 0.005 | 0.004 | 0.004 | 0 | 0.53 | 0.59 | 0.61 | 0.45 | 0.6 | 0.49 |
| 11 | 85627108 | rs7927222 | 1.482 | 0.781 | -0.497 | -0.818 | -0.004 | -0.017 | -0.017 | 0 | 0.17 | 0.14 | 0.16 | 0 | 0.26 | 0.08 |
| 11 | 85630411 | rs12805520 | 1.047 | 0.844 | -0.642 | -0.569 | -0.041 | 0.009 | 0.009 | 0 | 0.09 | 0 | 0.12 | 0.07 | 0.13 | 0.1 |
| 11 | 85634734 | rs11825598 | -0.434 | 0.751 | -0.469 | -0.449 | 0.002 | 0.021 | 0.021 | 0 | 0.37 | 0.48 | 0.47 | 0.5 | 0.38 | 0.42 |
| 11 | 85642444 | rs7107455 | -0.516 | 0.709 | -0.218 | -0.386 | 0.019 | -0.048 | -0.048 | 0.313 | 0.14 | 0.36 | 0.17 | 0.08 | 0.17 | 0.13 |
| 11 | 85642763 | rs7107881 | -0.225 | 0.815 | -0.374 | -0.389 | -0.001 | 0 | 0 | 0 | 0.45 | 0.55 | 0.48 | 0.43 | 0.48 | 0.39 |
| 11 | 85654587 | rs596864 | -0.729 | 0.992 | -0.765 | -0.366 | -0.007 | 0.023 | 0.023 | 0 | 0.6 | 0.75 | 0.6 | 0.71 | 0.6 | 0.67 |
| 11 | 85665172 | rs1892943 | -0.258 | 1.15 | -0.496 | -0.378 | 0.028 | 0.107 | 0.107 | 0.135 | 0.17 | 0.28 | 0.24 | 0.48 | 0.22 | 0.24 |
| 11 | 85668163 | rs622110 | 0.915 | 0.982 | -0.636 | -0.698 | -0.002 | -0.044 | -0.044 | 0 | 0.3 | 0.36 | 0.23 | 0.12 | 0.26 | 0.3 |
| 11 | 85681389 | rs10501602 | 1.449 | 1.014 | -0.882 | -0.669 | -0.019 | 0.05 | 0.05 | 0 | 0.13 | 0.02 | 0.13 | 0.08 | 0.13 | 0.14 |
| 11 | 85681916 | rs10501603 | 1.999 | 1.008 | -1.032 | -0.905 | -0.018 | 0.045 | 0.045 | 0.203 | 0.16 | 0.05 | 0.13 | 0 | 0.22 | 0.07 |
| 11 | 85683554 | rs10501604 | 0.834 | 1.012 | -0.975 | -0.776 | -0.002 | 0.03 | 0.03 | 0 | 0.11 | 0.09 | 0.2 | 0.46 | 0.17 | 0.21 |
| 11 | 85685166 | rs637304 | -1.738 | 0.898 | -1.107 | -0.811 | -0.005 | -0.097 | -0.097 | 0.802 | 0.76 | 0.92 | 0.73 | 0.68 | 0.82 | 0.74 |
| 11 | 85703974 | rs10792820 | 0.246 | 0.767 | -1.337 | -1.075 | -0.03 | -0.075 | -0.075 | 1.111 | 0.2 | 0.08 | 0.25 | 0.48 | 0.24 | 0.32 |
| 11 | 85707054 | rs540422 | -1.109 | 0.966 | -1.018 | -0.921 | 0.002 | -0.002 | -0.002 | 0.041 | 0.59 | 0.74 | 0.59 | 0.67 | 0.59 | 0.66 |
| 11 | 85729372 | rs642949 | 0.17 | 1.048 | -0.966 | -0.987 | 0.003 | 0.016 | 0.016 | 0.277 | 0.35 | 0.42 | 0.38 | 0.6 | 0.36 | 0.45 |
| 11 | 85740409 | rs609903 | -0.9 | 1.009 | -0.908 | -0.859 | 0.01 | -0.034 | -0.034 | 0 | 0.62 | 0.8 | 0.61 | 0.6 | 0.68 | 0.59 |
| 11 | 85751883 | rs664629 | 0.138 | 1.035 | -0.937 | -0.97 | 0 | 0.023 | 0.023 | 0.521 | 0.43 | 0.36 | 0.36 | 0.21 | 0.4 | 0.44 |
| 11 | 85754392 | rs10128700 | 1.744 | 0.97 | -0.477 | -0.685 | 0 | -0.037 | -0.037 | 0 | 0.17 | 0.17 | 0.14 | 0 | 0.22 | 0.07 |
| 11 | 85755538 | rs17745273 | 1.911 | 0.959 | -0.583 | -0.69 | -0.021 | 0.055 | 0.055 | 0.606 | 0.16 | 0.05 | 0.13 | 0 | 0.22 | 0.07 |
| 11 | 85757133 | rs1941375 | 0.417 | 0.936 | -0.566 | -0.663 | -0.002 | -0.027 | -0.027 | 0.913 | 0.3 | 0.34 | 0.23 | 0.13 | 0.27 | 0.3 |
| 11 | 85760028 | rs694011 | -1.47 | 0.909 | -0.682 | -0.562 | -0.026 | -0.096 | -0.096 | 0 | 0.62 | 0.9 | 0.61 | 0.6 | 0.68 | 0.59 |
| 11 | 85769105 | rs613222 | -0.841 | 0.917 | -0.538 | -0.467 | -0.002 | 0 | 0 | 0.026 | 0.45 | 0.6 | 0.46 | 0.6 | 0.46 | 0.52 |
| 11 | 85783738 | rs621942 | 0.485 | 0.931 | -0.608 | -0.487 | -0.012 | -0.049 | -0.049 | 0 | 0.28 | 0.35 | 0.23 | 0.12 | 0.24 | 0.28 |
| 11 | 85784979 | rs638509 | -0.186 | 0.978 | -0.601 | -0.413 | 0 | 0.006 | 0.006 | 0.407 | 0.37 | 0.38 | 0.4 | 0.49 | 0.37 | 0.47 |
| 11 | 85800279 | rs561655 | -1.343 | 1.07 | -1.214 | -0.469 | -0.024 | -0.084 | -0.084 | 0 | 0.62 | 0.87 | 0.61 | 0.51 | 0.65 | 0.58 |
| 11 | 85803238 | rs11234542 | -0.445 | 1.182 | -1.038 | -0.374 | 0.006 | 0.043 | 0.043 | 0 | 0.11 | 0.14 | 0.18 | 0.38 | 0.16 | 0.21 |
| 11 | 85811138 | rs629343 | -0.493 | 0.821 | -1.001 | -0.442 | 0.005 | -0.005 | -0.005 | 0 | 0.42 | 0.6 | 0.46 | 0.6 | 0.41 | 0.48 |
| 11 | 85812408 | rs495942 | 0.113 | 0.788 | -1.041 | -0.416 | -0.012 | -0.055 | -0.055 | 0.121 | 0.34 | 0.44 | 0.29 | 0.22 | 0.31 | 0.32 |
| 11 | 85812955 | rs11234546 | 1.443 | 0.876 | -1.365 | -0.009 | -0.022 | -0.031 | -0.031 | 0 | 0.06 | 0 | 0.14 | 0.23 | 0.08 | 0.09 |
| 11 | 85815030 | rs1237999 | -1.212 | 0.838 | -1.339 | -0.232 | -0.04 | -0.064 | -0.064 | 0 | 0.61 | 0.87 | 0.61 | 0.59 | 0.64 | 0.56 |
| 11 | 85820077 | rs543293 | -1.57 | 0.913 | -1.413 | -0.244 | -0.076 | -0.108 | -0.108 | 0 | 0.65 | 0.91 | 0.64 | 0.6 | 0.67 | 0.64 |
| 11 | 85837369 | rs7942719 | 0.567 | 0.868 | -1.265 | -0.404 | 0 | -0.031 | -0.031 | 0 | 0.12 | 0.14 | 0.12 | 0.01 | 0.17 | 0.1 |
| 11 | 85838808 | rs10792830 | -0.479 | 0.846 | -1.396 | -0.484 | -0.003 | -0.006 | -0.006 | 0 | 0.44 | 0.71 | 0.47 | 0.6 | 0.43 | 0.52 |
| 11 | 85839967 | rs11607590 | 0.316 | 0.957 | -1.506 | -0.61 | 0.011 | 0.017 | 0.017 | 0 | 0.2 | 0.15 | 0.12 | 0.01 | 0.15 | 0.11 |
| 11 | 85840446 | rs7926591 | 0.133 | 0.285 | -1.029 | -0.606 | -0.008 | -0.036 | -0.036 | 3.057 | 0.34 | 0.42 | 0.3 | 0.22 | 0.32 | 0.33 |
| 11 | 85841148 | rs10898436 | 1.485 | 0.317 | -1.174 | -0.515 | -0.038 | -0.082 | -0.082 | 0.019 | 0.16 | 0.03 | 0.2 | 0.36 | 0.18 | 0.2 |
| 11 | 85850243 | rs3844143 | -0.585 | 0.319 | -1.048 | -0.492 | 0.001 | 0.004 | 0.004 | 0.032 | 0.5 | 0.61 | 0.5 | 0.59 | 0.5 | 0.53 |
| 11 | 85854176 | rs2888903 | -0.924 | 0.32 | -1.453 | -0.564 | -0.032 | -0.044 | -0.044 | 1.035 | 0.57 | 0.79 | 0.55 | 0.58 | 0.55 | 0.6 |
| 11 | 85854294 | rs7940019 | 1.221 | 0.131 | -1.955 | -0.804 | -0.045 | 0.039 | 0.039 | 0.137 | 0.12 | 0.01 | 0.09 | 0.06 | 0.14 | 0.11 |
| 11 | 85855635 | rs17745564 | 1.3 | 0.37 | -1.724 | -0.62 | 0.011 | 0.019 | 0.019 | 0.199 | 0.09 | 0.04 | 0.04 | 0 | 0.04 | 0.04 |
| 11 | 85858417 | rs12786057 | 0.078 | 0.44 | -1.617 | -0.591 | -0.007 | -0.027 | -0.027 | 0 | 0.36 | 0.38 | 0.3 | 0.22 | 0.32 | 0.34 |
| 11 | 85858538 | rs7941541 | -1.486 | 0.539 | -1.749 | -0.489 | -0.06 | -0.134 | -0.134 | 1.307 | 0.67 | 0.91 | 0.64 | 0.58 | 0.68 | 0.65 |
| 11 | 85858887 | rs10751134 | -1.388 | 0.516 | -1.748 | -0.487 | -0.093 | -0.129 | -0.129 | 0.272 | 0.66 | 0.91 | 0.6 | 0.59 | 0.59 | 0.64 |
| 11 | 85865191 | rs6592272 | 0.275 | -0.234 | -1.488 | -0.536 | -0.02 | -0.045 | -0.045 | 0 | 0.29 | 0.33 | 0.24 | 0.13 | 0.23 | 0.3 |
| 11 | 85868640 | rs3851179 | -1.129 | -0.079 | -1.627 | -0.326 | -0.094 | -0.11 | -0.11 | 0.554 | 0.64 | 0.89 | 0.61 | 0.59 | 0.63 | 0.61 |
| 11 | 85869737 | rs7113976 | -0.313 | -0.234 | -1.248 | -0.079 | -0.006 | 0.001 | 0.001 | 0.223 | 0.44 | 0.5 | 0.45 | 0.49 | 0.41 | 0.49 |
| 11 | 85876222 | rs11234568 | -0.639 | -0.113 | -1.883 | 0.201 | -0.149 | -0.156 | -0.156 | 0.031 | 0.69 | 0.97 | 0.6 | 0.61 | 0.6 | 0.61 |
| 11 | 85878905 | rs7937794 | -0.526 | -0.69 | -1.541 | -0.047 | -0.003 | -0.02 | -0.02 | 0 | 0.57 | 0.57 | 0.48 | 0.3 | 0.58 | 0.56 |
| 11 | 85883303 | rs7949297 | 0.88 | -1.087 | -1.363 | 0.018 | -0.001 | 0.005 | 0.005 | 1.721 | 0.29 | 0.27 | 0.25 | 0.39 | 0.27 | 0.24 |
| 11 | 85886994 | rs11234569 | 0.699 | -0.494 | -1.601 | -0.951 | -0.186 | 0.164 | 0.164 | 0.008 | 0.3 | 0.02 | 0.35 | 0.09 | 0.49 | 0.31 |
| 11 | 85890389 | rs7128110 | 0.539 | -0.541 | -1.214 | -0.792 | -0.011 | -0.033 | -0.033 | 0 | 0.06 | 0.2 | 0.04 | 0 | 0.05 | 0.03 |
| 11 | 85893436 | rs7103707 | -0.129 | -0.208 | -1.563 | -0.479 | -0.027 | -0.006 | -0.006 | 0 | 0.23 | 0.39 | 0.19 | 0.23 | 0.14 | 0.23 |
| 11 | 85900117 | rs12295362 | 0.136 | 0.728 | -2.291 | -0.391 | -0.186 | 0.161 | 0.161 | 0.01 | 0.3 | 0.02 | 0.35 | 0.09 | 0.49 | 0.3 |
| 11 | 85905467 | rs11605019 | -0.308 | 0.549 | -2.103 | 0.06 | -0.073 | 0.022 | 0.022 | 0 | 0.39 | 0.17 | 0.43 | 0.37 | 0.55 | 0.49 |

| | | | | | | | | | | | | | | | | |
|---|---|---|---|---|---|---|---|---|---|---|---|---|---|---|---|---|
| 11 | 85905715 | rs10792837 | -0.026 | -0.607 | -2.607 | 0.271 | 0.081 | -0.009 | -0.009 | 2.492 | 0.68 | 0.2 | 0.68 | 0.66 | 0.87 | 0.65 |
| 11 | 85906129 | rs3851181 | -0.076 | -0.461 | -1.719 | 0.02 | 0 | 0.007 | 0.007 | 0 | 0.49 | 0.46 | 0.44 | 0.23 | 0.56 | 0.49 |
| 11 | 85906362 | rs3844144 | 0.087 | -0.545 | -0.86 | 0.455 | -0.016 | 0.041 | 0.041 | 1.027 | 0.87 | 0.69 | 0.76 | 0.78 | 0.85 | 0.84 |
| 11 | 85908139 | rs10898445 | 0.382 | -0.444 | -0.814 | 0.413 | 0 | 0.002 | 0.002 | 0 | 0.34 | 0.29 | 0.32 | 0.23 | 0.35 | 0.24 |
| 11 | 85908313 | rs2374705 | -0.173 | -0.201 | -0.596 | 0.662 | -0.007 | 0.016 | 0.016 | 0.601 | 0.46 | 0.64 | 0.49 | 0.6 | 0.45 | 0.45 |
| 11 | 85908537 | rs3851182 | -0.119 | -0.32 | -0.728 | 0.544 | -0.029 | 0.028 | 0.028 | 0 | 0.79 | 0.71 | 0.74 | 0.71 | 0.88 | 0.81 |
| 11 | 85920981 | rs11608125 | 0.35 | -0.197 | -0.707 | 0.774 | -0.004 | 0 | 0 | 0 | 0.09 | 0.09 | 0.06 | 0.22 | 0.1 | 0.16 |
| 11 | 85922070 | rs10792838 | 0.315 | -0.356 | -0.726 | 0.974 | -0.061 | -0.063 | -0.063 | 0.286 | 0.41 | 0.19 | 0.31 | 0.49 | 0.43 | 0.41 |
| 11 | 85922292 | rs7122535 | -0.022 | -0.412 | -0.245 | 0.945 | -0.007 | -0.008 | -0.008 | 0.008 | 0.44 | 0.42 | 0.34 | 0.49 | 0.44 | 0.41 |
| 11 | 85927952 | rs11234582 | 1.094 | -0.291 | -0.55 | 0.94 | -0.03 | 0.043 | 0.043 | 0 | 0.32 | 0.08 | 0.24 | 0.19 | 0.32 | 0.22 |
| 11 | 85929490 | rs9971532 | 1.656 | -0.203 | -0.885 | 0.585 | -0.099 | 0.073 | 0.073 | 0 | 0.22 | 0.01 | 0.2 | 0.11 | 0.34 | 0.22 |
| 11 | 85929614 | rs7122629 | 0.696 | -0.223 | -0.8 | 0.644 | -0.003 | 0.003 | 0.003 | 1.909 | 0.08 | 0.09 | 0.06 | 0.21 | 0.09 | 0.16 |
| 11 | 85938133 | rs10792839 | 0.143 | -0.375 | -0.666 | 0.679 | -0.017 | 0.015 | 0.015 | 0 | 0.71 | 0.48 | 0.62 | 0.66 | 0.78 | 0.67 |
| 11 | 85940575 | rs2173281 | 1.758 | -0.227 | -0.909 | 0.746 | -0.056 | 0.048 | 0.048 | 0 | 0.3 | 0.03 | 0.17 | 0.16 | 0.27 | 0.17 |
| 11 | 85941394 | rs12577739 | 1.249 | -0.225 | -0.812 | 0.801 | -0.048 | 0.054 | 0.054 | 0 | 0.35 | 0.07 | 0.25 | 0.2 | 0.34 | 0.24 |
| 11 | 85957745 | rs2374721 | 0.167 | -0.271 | -0.443 | 0.929 | -0.03 | 0.017 | 0.017 | 0 | 0.7 | 0.58 | 0.62 | 0.69 | 0.77 | 0.65 |
| 11 | 85967064 | rs10898457 | 0.565 | -0.308 | -0.482 | 0.908 | -0.074 | -0.003 | -0.003 | 0 | 0.32 | 0.13 | 0.26 | 0.31 | 0.42 | 0.41 |
| 11 | 85968037 | rs10898458 | 0.468 | -0.311 | -0.43 | 0.906 | -0.046 | 0.007 | 0.007 | 0.122 | 0.32 | 0.2 | 0.27 | 0.31 | 0.42 | 0.41 |
| 11 | 85968623 | rs974144 | 0.566 | -0.311 | -0.433 | 0.906 | -0.085 | -0.008 | -0.008 | 0 | 0.32 | 0.12 | 0.26 | 0.31 | 0.42 | 0.41 |
| 11 | 85972939 | rs10898459 | 0.128 | -0.278 | -0.389 | 0.946 | -0.001 | 0 | 0 | 0 | 0.35 | 0.37 | 0.37 | 0.5 | 0.42 | 0.45 |
| 11 | 85977117 | rs10898460 | -0.448 | -0.343 | -0.19 | 0.945 | -0.006 | 0.002 | 0.002 | 0 | 0.77 | 0.8 | 0.7 | 0.72 | 0.84 | 0.74 |
| 11 | 85980828 | rs10792844 | 0.807 | -0.295 | -0.228 | 0.944 | -0.006 | -0.007 | -0.007 | 5.324 | 0.1 | 0.09 | 0.05 | 0.23 | 0.09 | 0.19 |
| 11 | 85986047 | rs7931249 | 0.809 | -0.22 | -0.132 | 0.986 | 0.047 | 0.036 | 0.036 | 0 | 0.07 | 0.21 | 0.08 | 0.04 | 0.07 | 0.08 |
| 11 | 85986518 | rs10792845 | -0.387 | -0.226 | -0.144 | 0.98 | -0.006 | 0.001 | 0.001 | 0 | 0.77 | 0.8 | 0.7 | 0.72 | 0.84 | 0.74 |
| 11 | 85990266 | rs1376947 | 1.456 | -0.497 | -0.305 | 0.904 | -0.023 | 0.029 | 0.029 | 0 | 0.31 | 0.18 | 0.24 | 0.19 | 0.34 | 0.21 |
| 11 | 86006280 | rs291230 | -1.426 | -0.479 | -0.304 | 0.63 | -0.001 | 0.006 | 0.006 | 0 | 0.67 | 0.67 | 0.76 | 0.81 | 0.65 | 0.79 |
| 11 | 86010681 | rs291236 | -1.254 | -0.481 | -0.27 | 0.646 | 0.003 | -0.014 | -0.014 | 0 | 0.65 | 0.5 | 0.7 | 0.77 | 0.63 | 0.74 |
| 11 | 86011868 | rs291239 | -1.049 | -0.477 | -0.285 | 0.624 | 0.001 | -0.003 | -0.003 | 0 | 0.63 | 0.61 | 0.68 | 0.77 | 0.59 | 0.73 |
| 11 | 86013361 | rs291241 | 1.772 | -0.764 | -0.127 | 0.725 | -0.001 | 0.013 | 0.013 | 0 | 0.11 | 0.1 | 0.23 | 0.23 | 0.12 | 0.2 |
| 11 | 86013380 | rs11234604 | 0.949 | -0.713 | -0.148 | 0.71 | 0.049 | 0.037 | 0.037 | 0 | 0.06 | 0.19 | 0.07 | 0.04 | 0.07 | 0.07 |
| 11 | 86026808 | rs11234610 | 1.01 | -0.72 | -0.232 | 0.718 | -0.002 | -0.019 | -0.019 | 0 | 0.15 | 0.13 | 0.16 | 0.38 | 0.1 | 0.22 |
| 11 | 86036955 | rs7114910 | 1.024 | -0.807 | -0.25 | 0.658 | -0.004 | 0 | 0 | 4.411 | 0.12 | 0.1 | 0.06 | 0.05 | 0.03 | 0.07 |
| 11 | 86039123 | rs2847472 | 1.738 | -0.719 | -0.287 | 0.662 | -0.001 | 0.003 | 0.003 | 0 | 0.12 | 0.09 | 0.23 | 0.22 | 0.12 | 0.21 |
| 11 | 86053419 | rs10792847 | -1.183 | -0.451 | -0.311 | 0.68 | 0.003 | -0.004 | -0.004 | 0 | 0.63 | 0.63 | 0.69 | 0.77 | 0.59 | 0.72 |
| 11 | 86055901 | rs6592284 | -0.292 | -0.438 | -0.506 | 0.658 | 0.002 | 0.047 | 0.047 | 3.487 | 0.22 | 0.43 | 0.24 | 0.38 | 0.17 | 0.26 |
| 11 | 86058833 | rs9667913 | -0.67 | -0.682 | -0.373 | 0.494 | -0.002 | 0.006 | 0.006 | 0 | 0.52 | 0.49 | 0.59 | 0.42 | 0.55 | 0.57 |
| 11 | 86064757 | rs11824612 | -0.604 | -1.252 | -0.438 | 0.652 | 0.029 | -0.011 | -0.011 | 0 | 0.29 | 0.67 | 0.35 | 0.32 | 0.17 | 0.3 |
| 11 | 86070096 | rs1955189 | 1.673 | -0.886 | -0.745 | 0.402 | -0.076 | 0.051 | 0.051 | 0 | 0.2 | 0.03 | 0.15 | 0.1 | 0.28 | 0.26 |
| 11 | 86070817 | rs11603604 | 1.673 | -0.889 | -0.746 | 0.401 | -0.076 | 0.051 | 0.051 | 0 | 0.2 | 0.02 | 0.14 | 0.1 | 0.28 | 0.25 |
| 11 | 86071005 | rs6592288 | -0.266 | -1.072 | -0.457 | 0.525 | -0.004 | -0.006 | -0.006 | 0 | 0.46 | 0.44 | 0.45 | 0.38 | 0.44 | 0.52 |
| 11 | 86073135 | rs6592289 | -0.272 | -1.212 | -0.782 | 0.4 | 0.006 | -0.006 | -0.006 | 0.561 | 0.32 | 0.28 | 0.43 | 0.65 | 0.24 | 0.39 |
| 11 | 86073315 | rs11822254 | -0.276 | -1.276 | -0.769 | 0.302 | -0.008 | -0.022 | -0.022 | 0 | 0.4 | 0.46 | 0.28 | 0.2 | 0.37 | 0.43 |
| 11 | 86073603 | rs7105420 | 0.5 | -1.235 | -0.971 | 0.522 | -0.032 | -0.085 | -0.085 | 0 | 0.33 | 0.17 | 0.32 | 0.52 | 0.34 | 0.45 |
| 11 | 86082980 | rs292103 | -0.557 | -1.045 | -0.864 | 0.492 | 0.014 | 0.02 | 0.02 | 0 | 0.47 | 0.62 | 0.48 | 0.37 | 0.48 | 0.55 |
| 11 | 86084069 | rs292106 | -0.529 | -1.033 | -0.813 | 0.491 | -0.022 | 0.014 | 0.014 | 0 | 0.28 | 0.49 | 0.33 | 0.28 | 0.23 | 0.33 |
| 11 | 86086581 | rs292092 | -1.929 | -0.844 | -1.021 | 0.493 | -0.089 | 0.081 | 0.081 | 0 | 0.7 | 0.86 | 0.71 | 0.84 | 0.59 | 0.8 |
| 11 | 86091242 | rs10898473 | 0.291 | -1.059 | -0.851 | 0.654 | 0.003 | -0.005 | -0.005 | 0 | 0.23 | 0.24 | 0.23 | 0.47 | 0.12 | 0.25 |
| 11 | 86091532 | rs7944482 | -0.981 | -1.098 | -0.853 | 0.615 | 0.038 | -0.032 | -0.032 | 0.906 | 0.37 | 0.71 | 0.34 | 0.51 | 0.21 | 0.4 |
| 11 | 86093345 | rs7937090 | -0.528 | -0.72 | -0.317 | 0.955 | 0.011 | 0.006 | 0.006 | 0.046 | 0.49 | 0.64 | 0.35 | 0.45 | 0.45 | 0.56 |
| 11 | 86094947 | rs10898474 | -1.356 | -0.662 | -0.461 | 0.991 | -0.075 | 0.024 | 0.024 | 0 | 0.65 | 0.87 | 0.47 | 0.69 | 0.58 | 0.75 |
| 11 | 86098200 | rs17210497 | 0.538 | -0.697 | -0.469 | 0.965 | 0 | -0.001 | -0.001 | 1.867 | 0.21 | 0.23 | 0.13 | 0.32 | 0.12 | 0.2 |
| 11 | 86098680 | rs292109 | -0.986 | -0.266 | -0.572 | 1.254 | -0.025 | 0.023 | 0.023 | 0 | 0.64 | 0.76 | 0.46 | 0.69 | 0.58 | 0.73 |
| 11 | 86098732 | rs11234627 | 0.749 | -0.12 | -0.969 | 0.909 | 0.003 | -0.008 | -0.008 | 0.004 | 0.14 | 0.09 | 0.11 | 0.24 | 0.13 | 0.19 |
| 11 | 86101234 | rs7105671 | -0.597 | 0.197 | -0.918 | 0.566 | -0.017 | 0.019 | 0.019 | 0.035 | 0.66 | 0.8 | 0.64 | 0.77 | 0.65 | 0.82 |
| 11 | 86104032 | rs7118035 | -0.652 | 0.246 | -1.073 | 0.59 | -0.029 | 0.02 | 0.02 | 0.098 | 0.67 | 0.84 | 0.64 | 0.78 | 0.65 | 0.82 |
| 11 | 86107810 | rs10898480 | -0.578 | -0.221 | -0.052 | 1.042 | -0.032 | 0.097 | 0.097 | 0.3 | 0.15 | 0.33 | 0.13 | 0.26 | 0.07 | 0.25 |
| 11 | 86107920 | rs17818702 | 0.916 | 0.171 | -0.814 | 0.539 | -0.041 | 0.064 | 0.064 | 0 | 0.39 | 0.16 | 0.31 | 0.28 | 0.47 | 0.4 |
| 11 | 86107966 | rs11826496 | 0.013 | 0.34 | -0.244 | 0.5 | -0.002 | -0.015 | -0.015 | 0 | 0.61 | 0.64 | 0.61 | 0.55 | 0.69 | 0.7 |
| 11 | 86119396 | rs4412777 | -1.103 | 0.089 | -0.125 | 0.625 | -0.007 | -0.022 | -0.022 | 0.445 | 0.74 | 0.84 | 0.71 | 0.76 | 0.78 | 0.89 |
| 11 | 86123041 | rs11234643 | -0.448 | -0.105 | 0.159 | 0.381 | -0.007 | -0.051 | -0.051 | 0 | 0.12 | 0.26 | 0.11 | 0.04 | 0.1 | 0.12 |
| 11 | 86123357 | rs3741007 | 0.278 | -0.061 | 0.242 | 0.309 | -0.001 | -0.006 | -0.006 | 0 | 0.51 | 0.48 | 0.51 | 0.31 | 0.49 | 0.5 |
| 11 | 86133029 | rs12280555 | -0.034 | -0.054 | 0.26 | 0.314 | 0.005 | -0.048 | -0.048 | 0.007 | 0.09 | 0.27 | 0.05 | 0.01 | 0.07 | 0.06 |
| 11 | 86133416 | rs1452134 | 0.362 | -0.09 | 0.3 | 0.33 | -0.001 | -0.002 | -0.002 | 0 | 0.52 | 0.38 | 0.5 | 0.51 | 0.46 | 0.58 |
| 11 | 86133784 | rs1056194 | -1.521 | -0.092 | 0.223 | 0.334 | -0.006 | -0.021 | -0.021 | 0 | 0.74 | 0.84 | 0.71 | 0.76 | 0.78 | 0.9 |
| 11 | 86133937 | rs11234648 | 0.756 | -0.013 | 0.2 | 0.283 | 0.016 | -0.021 | -0.021 | 0 | 0.06 | 0.25 | 0.03 | 0.01 | 0.05 | 0.05 |
| 11 | 86138795 | rs12362089 | 2.117 | 0.01 | 0.023 | 0.316 | -0.007 | -0.014 | -0.014 | 0 | 0.1 | 0.03 | 0.08 | 0.17 | 0.13 | 0.17 |
| 11 | 86140441 | rs7935933 | 1.09 | -0.054 | 0.075 | 0.284 | -0.015 | -0.014 | -0.014 | 0 | 0.15 | 0.09 | 0.17 | 0.24 | 0.27 | 0.26 |
| 11 | 86145296 | rs11607319 | 1.181 | -0.129 | 0.013 | 0.191 | -0.005 | 0.047 | 0.047 | 0.267 | 0.39 | 0.22 | 0.4 | 0.27 | 0.39 | 0.38 |
| 11 | 86145755 | rs11828146 | 0.626 | -0.14 | 0.214 | 0.22 | 0.026 | -0.014 | -0.014 | 0 | 0.06 | 0.29 | 0.04 | 0.01 | 0.05 | 0.05 |

| | | | | | | | | | | | | | | | | |
|---|---|---|---|---|---|---|---|---|---|---|---|---|---|---|---|---|
| 11 | 86150865 | rs1452138 | 0.419 | -0.116 | 0.002 | 0.26 | -0.001 | -0.009 | -0.009 | 0 | 0.47 | 0.51 | 0.44 | 0.28 | 0.46 | 0.46 |
| 11 | 86155131 | rs10501620 | 1.312 | -0.076 | -0.052 | 0.583 | -0.006 | 0.021 | 0.021 | 0.225 | 0.11 | 0.15 | 0.08 | 0.24 | 0.05 | 0.16 |
| 11 | 86156848 | rs11234658 | -0.521 | -0.015 | -0.006 | 0.434 | 0 | 0.038 | 0.038 | 0 | 0.28 | 0.35 | 0.26 | 0.48 | 0.32 | 0.43 |
| 11 | 86162568 | rs7109853 | -0.324 | 0.196 | -0.095 | 0.192 | 0.002 | -0.006 | -0.006 | 0 | 0.63 | 0.57 | 0.61 | 0.76 | 0.64 | 0.8 |
| 11 | 86165942 | rs1563682 | -0.781 | 0.169 | -0.12 | 0.16 | -0.002 | -0.003 | -0.003 | 0.224 | 0.68 | 0.6 | 0.67 | 0.76 | 0.73 | 0.84 |
| 11 | 86166847 | rs3819227 | 1.519 | 0.138 | -0.286 | 0.168 | 0.004 | 0.028 | 0.028 | 0.051 | 0.35 | 0.22 | 0.35 | 0.27 | 0.33 | 0.36 |
| 11 | 86167136 | rs2125362 | -0.582 | 0.168 | -0.336 | 0.195 | -0.019 | 0.004 | 0.004 | 0.023 | 0.67 | 0.8 | 0.64 | 0.76 | 0.68 | 0.84 |
| 11 | 86171405 | rs3758877 | 0.25 | 0.135 | -0.02 | -0.068 | -0.001 | -0.018 | -0.018 | 0 | 0.07 | 0.11 | 0.09 | 0.02 | 0.07 | 0.13 |
| 11 | 86171454 | rs618513 | -0.329 | 0.01 | -0.028 | 0.055 | 0.001 | -0.013 | -0.013 | 0 | 0.83 | 0.83 | 0.85 | 0.75 | 0.86 | 0.85 |
| 11 | 86177055 | rs1458384 | 1.307 | 0.061 | 0.156 | 0.198 | 0.004 | 0.013 | 0.013 | 0 | 0.45 | 0.35 | 0.41 | 0.38 | 0.45 | 0.53 |
| 11 | 86179076 | rs663262 | -1.475 | 0.072 | 0.082 | 0.236 | -0.009 | 0.03 | 0.03 | 0.148 | 0.88 | 0.82 | 0.9 | 0.75 | 0.95 | 0.89 |
| 11 | 86180365 | rs632873 | 0.952 | 0.101 | 0.097 | 0.209 | -0.019 | -0.022 | -0.022 | 0 | 0.28 | 0.2 | 0.32 | 0.34 | 0.34 | 0.16 |
| 11 | 86181720 | rs12804208 | 3.003 | 0.263 | -0.156 | -0.068 | -0.037 | 0.041 | 0.041 | 0 | 0.08 | 0.01 | 0.06 | 0 | 0.12 | 0.02 |
| 11 | 86188152 | rs1870323 | 1.896 | -0.193 | -0.069 | -0.136 | -0.002 | 0.019 | 0.019 | 0 | 0.39 | 0.25 | 0.29 | 0.25 | 0.42 | 0.43 |
| 11 | 86191420 | rs634509 | 0.496 | -0.125 | -0.058 | -0.139 | -0.001 | 0.001 | 0.001 | 0 | 0.76 | 0.71 | 0.76 | 0.72 | 0.79 | 0.72 |
| 11 | 86193603 | rs664613 | 0.65 | -0.139 | -0.013 | -0.092 | -0.038 | -0.041 | -0.041 | 0 | 0.27 | 0.15 | 0.32 | 0.34 | 0.34 | 0.16 |
| 11 | 86194819 | rs679076 | -1.621 | -0.565 | 0.058 | -0.037 | 0.004 | -0.001 | -0.001 | 0.062 | 0.88 | 0.87 | 0.91 | 0.75 | 0.95 | 0.9 |
| 11 | 86196278 | rs662361 | -0.209 | -0.542 | 0.057 | -0.035 | -0.004 | 0.007 | 0.007 | 0 | 0.81 | 0.76 | 0.82 | 0.72 | 0.88 | 0.76 |
| 11 | 86204016 | rs672526 | 0.658 | -0.709 | 0.287 | 0.123 | -0.009 | 0.007 | 0.007 | 0.015 | 0.75 | 0.57 | 0.73 | 0.76 | 0.78 | 0.69 |
| 11 | 86206203 | rs748102 | 1.226 | -0.615 | -0.013 | -0.144 | 0.001 | 0.004 | 0.004 | 0.252 | 0.51 | 0.44 | 0.48 | 0.37 | 0.56 | 0.57 |
| 11 | 86207816 | rs624843 | -0.221 | -0.662 | -0.058 | -0.033 | -0.008 | 0.016 | 0.016 | 0 | 0.81 | 0.76 | 0.83 | 0.72 | 0.87 | 0.76 |
| 11 | 86209682 | rs611156 | 2.55 | -0.51 | -0.245 | 0.204 | -0.027 | -0.036 | -0.036 | 0 | 0.1 | 0.03 | 0.21 | 0.21 | 0.12 | 0.06 |
| 11 | 86211187 | rs2051434 | 3.038 | -0.663 | -0.498 | -0.518 | -0.107 | 0.022 | 0.022 | 0 | 0.23 | 0.02 | 0.18 | 0.11 | 0.29 | 0.2 |
| 11 | 86212745 | rs655934 | 0.506 | -0.403 | 0.032 | -0.36 | -0.016 | 0.042 | 0.042 | 0.733 | 0.77 | 0.52 | 0.78 | 0.71 | 0.8 | 0.72 |
| 11 | 86216685 | rs623478 | 0.202 | -0.347 | -0.146 | -0.366 | 0 | 0 | 0 | 0 | 0.69 | 0.62 | 0.74 | 0.71 | 0.7 | 0.7 |
| 11 | 86217761 | rs638492 | 0.924 | -0.45 | -0.158 | -0.349 | -0.001 | -0.005 | -0.005 | 0 | 0.18 | 0.17 | 0.28 | 0.34 | 0.21 | 0.14 |
| 11 | 86226645 | rs1006586 | -0.5 | -0.643 | 0.101 | -0.674 | 0.02 | -0.041 | -0.041 | 0 | 0.1 | 0.24 | 0.12 | 0.03 | 0.14 | 0.18 |
| 11 | 86227806 | rs17149147 | 1.362 | -0.728 | 0.103 | -0.698 | -0.016 | -0.03 | -0.03 | 0.212 | 0.06 | 0.12 | 0.03 | 0 | 0.03 | 0.04 |
| 11 | 86230898 | rs498981 | -1.157 | -0.537 | 0.018 | -0.538 | -0.011 | 0.033 | 0.033 | 0.374 | 0.23 | 0.43 | 0.22 | 0.29 | 0.2 | 0.28 |
| 11 | 86232668 | rs588287 | 0.17 | -0.494 | -0.051 | -0.524 | 0 | 0.01 | 0.01 | 0.07 | 0.15 | 0.19 | 0.13 | 0.26 | 0.16 | 0.13 |
| 11 | 86243255 | rs10444372 | 0.073 | -0.475 | -0.034 | -0.635 | 0.016 | -0.063 | -0.063 | 0 | 0.19 | 0.28 | 0.17 | 0.03 | 0.24 | 0.2 |
| 11 | 86246643 | rs605618 | 0.862 | -0.534 | -0.044 | -0.554 | -0.011 | 0.029 | 0.029 | 0.017 | 0.12 | 0.17 | 0.09 | 0.26 | 0.05 | 0.11 |
| 11 | 86248984 | rs12799562 | 0.623 | -0.49 | -0.023 | -0.592 | 0 | -0.06 | -0.06 | 0 | 0.16 | 0.25 | 0.14 | 0.03 | 0.13 | 0.18 |
| 11 | 86250128 | rs12805940 | 1.949 | -0.318 | 0.021 | -0.534 | 0.005 | -0.037 | -0.037 | 0 | 0.11 | 0.14 | 0.07 | 0.01 | 0.1 | 0.05 |
| 11 | 86250366 | rs553351 | -0.716 | -0.171 | -0.181 | -0.522 | 0.016 | -0.005 | -0.005 | 0 | 0.32 | 0.45 | 0.31 | 0.27 | 0.34 | 0.22 |
| 11 | 86263113 | rs1458382 | -0.268 | -0.496 | -0.579 | -0.265 | -0.001 | 0.008 | 0.008 | 0.014 | 0.24 | 0.29 | 0.43 | 0.53 | 0.27 | 0.24 |
| 11 | 86264495 | rs7117375 | -0.816 | -0.939 | -0.568 | -0.896 | -0.024 | -0.102 | -0.102 | 0 | 0.33 | 0.48 | 0.22 | 0.13 | 0.28 | 0.43 |
| 11 | 86264881 | rs7103271 | 0.138 | -1.005 | -0.77 | -0.953 | 0.003 | -0.016 | -0.016 | 0.191 | 0.22 | 0.22 | 0.37 | 0.53 | 0.23 | 0.18 |
| 11 | 86266683 | rs12797615 | 3.1 | -0.951 | -1.152 | -1.286 | -0.104 | 0.094 | 0.094 | 0 | 0.25 | 0.04 | 0.2 | 0.05 | 0.29 | 0.22 |
| 11 | 86267175 | rs11607733 | 3.1 | -0.951 | -1.152 | -1.286 | -0.104 | 0.094 | 0.094 | 0 | 0.25 | 0.04 | 0.2 | 0.05 | 0.29 | 0.22 |
| 11 | 86267633 | rs17211115 | 1.129 | -1 | -0.969 | -1.239 | -0.002 | 0.063 | 0.063 | 0 | 0.15 | 0.19 | 0.35 | 0.53 | 0.18 | 0.17 |
| 11 | 86267714 | rs12287767 | 1.135 | -0.995 | -0.966 | -1.236 | -0.002 | 0.063 | 0.063 | 0 | 0.15 | 0.19 | 0.35 | 0.53 | 0.18 | 0.17 |
| 11 | 86268073 | rs10792857 | 0.749 | -1.045 | -1.129 | -1.349 | 0.002 | -0.001 | -0.001 | 0 | 0.6 | 0.45 | 0.67 | 0.59 | 0.7 | 0.44 |
| 11 | 86269066 | rs17149180 | 1.25 | -0.9 | -0.807 | -1.151 | 0.003 | -0.042 | -0.042 | 0 | 0.09 | 0.15 | 0.08 | 0 | 0.09 | 0.07 |
| 11 | 86270085 | rs7934133 | 1.744 | -0.913 | -0.895 | -1.132 | -0.009 | 0.032 | 0.032 | 0 | 0.36 | 0.3 | 0.25 | 0.05 | 0.34 | 0.23 |
| 11 | 86280071 | rs10898500 | 0.338 | -0.853 | -0.936 | -0.762 | 0 | -0.092 | -0.092 | 0 | 0.21 | 0.11 | 0.36 | 0.54 | 0.23 | 0.19 |
| 11 | 86282679 | rs7940549 | 0.861 | -0.756 | -0.504 | -0.872 | 0.023 | -0.063 | -0.063 | 0.398 | 0.11 | 0.26 | 0.08 | 0 | 0.09 | 0.08 |
| 11 | 86285825 | rs10898501 | 0.748 | -0.843 | -0.88 | -0.87 | 0.022 | 0.032 | 0.032 | 0 | 0.5 | 0.32 | 0.61 | 0.58 | 0.56 | 0.42 |
| 11 | 86288287 | rs7950878 | 0.282 | -1.468 | -0.135 | -0.465 | -0.04 | 0.013 | 0.013 | 0 | 0.11 | 0.31 | 0.08 | 0.12 | 0.06 | 0.22 |
| 11 | 86290278 | rs949351 | -1.857 | -1.665 | -1.001 | -0.706 | 0.065 | 0.079 | 0.079 | 1.21 | 0.49 | 0.74 | 0.56 | 0.72 | 0.37 | 0.49 |
| 11 | 86291033 | rs10898502 | -0.807 | -1.563 | -0.861 | -0.902 | -0.014 | -0.031 | -0.031 | 0 | 0.24 | 0.62 | 0.18 | 0.14 | 0.16 | 0.31 |
| 11 | 86292522 | rs10898503 | 1.757 | -1.313 | -1.038 | -1.05 | -0.061 | 0.215 | 0.215 | 0.175 | 0.38 | 0.15 | 0.33 | 0.02 | 0.55 | 0.28 |
| 11 | 86297189 | rs949349 | -1.064 | -0.842 | -0.376 | -0.263 | 0.002 | -0.018 | -0.018 | 0 | 0.29 | 0.24 | 0.44 | 0.53 | 0.27 | 0.3 |
| 11 | 86300922 | rs2051433 | -0.057 | -0.83 | -0.426 | 0.095 | 0.046 | -0.057 | -0.057 | 0 | 0.18 | 0.08 | 0.31 | 0.39 | 0.1 | 0.18 |
| 11 | 86303090 | rs4405344 | 1.314 | -0.956 | -0.049 | 0.024 | 0.014 | 0.004 | 0.004 | 0 | 0.55 | 0.41 | 0.46 | 0.36 | 0.7 | 0.45 |
| 11 | 86303326 | rs4382935 | 2.207 | -0.91 | -0.377 | -0.146 | -0.114 | 0.268 | 0.268 | 0 | 0.4 | 0.06 | 0.31 | 0.05 | 0.52 | 0.29 |
| 11 | 86304054 | rs10400291 | -0.755 | -0.766 | -0.04 | 0.122 | 0.005 | -0.005 | -0.005 | 0 | 0.27 | 0.25 | 0.39 | 0.39 | 0.19 | 0.25 |
| 11 | 86308532 | rs10792860 | 0.138 | -0.783 | -0.069 | 0.159 | 0.026 | -0.03 | -0.03 | 0 | 0.17 | 0.15 | 0.32 | 0.39 | 0.09 | 0.17 |
| 11 | 86315317 | rs10898512 | -0.281 | -0.734 | -0.074 | 0.157 | 0.016 | -0.026 | -0.026 | 0 | 0.19 | 0.17 | 0.36 | 0.39 | 0.15 | 0.18 |
| 11 | 86317519 | rs7940772 | 0.256 | -0.736 | -0.073 | 0.154 | 0.034 | -0.042 | -0.042 | 0 | 0.17 | 0.1 | 0.31 | 0.4 | 0.09 | 0.17 |
| 11 | 86317660 | rs7940587 | -0.276 | -0.727 | -0.062 | 0.156 | 0.01 | -0.016 | -0.016 | 0.393 | 0.2 | 0.19 | 0.36 | 0.4 | 0.15 | 0.18 |
| 11 | 86318892 | rs7109270 | -0.656 | -0.745 | 0.141 | 0.149 | 0.025 | 0.085 | 0.085 | 0.397 | 0.2 | 0.36 | 0.17 | 0.31 | 0.22 | 0.22 |
| 11 | 86320308 | rs12282010 | -1.643 | -0.745 | 0.097 | 0.118 | -0.002 | -0.003 | -0.003 | 0 | 0.41 | 0.56 | 0.53 | 0.7 | 0.37 | 0.41 |
| 11 | 86321576 | rs4944603 | -1.511 | -0.749 | 0.097 | 0.121 | -0.012 | -0.009 | -0.009 | 0.654 | 0.38 | 0.52 | 0.48 | 0.7 | 0.31 | 0.4 |
| 11 | 86335140 | rs11819827 | 1.317 | -0.857 | 0.177 | 0.209 | -0.001 | 0.006 | 0.006 | 0 | 0.06 | 0.07 | 0.02 | 0 | 0.06 | 0.03 |
| 11 | 86335459 | rs7120052 | -1.302 | -0.887 | 0.244 | 0.295 | -0.055 | 0.079 | 0.079 | 0.678 | 0.29 | 0.47 | 0.41 | 0.55 | 0.18 | 0.28 |
| 11 | 86338914 | rs7103017 | -0.683 | -0.847 | 0.269 | 0.331 | 0.002 | 0.173 | 0.173 | 0.006 | 0.13 | 0.32 | 0.1 | 0.31 | 0.12 | 0.22 |
| 11 | 86341186 | rs7123010 | -1.5 | -0.83 | 0.18 | 0.202 | -0.017 | 0.018 | 0.018 | 0 | 0.37 | 0.53 | 0.48 | 0.55 | 0.28 | 0.35 |
| 11 | 86344677 | rs1938928 | 0.213 | -0.853 | 0.03 | 0.283 | 0.019 | -0.024 | -0.024 | 0 | 0.17 | 0.15 | 0.32 | 0.4 | 0.09 | 0.16 |

| | | | | | | | | | | | | | | | | |
|---|---|---|---|---|---|---|---|---|---|---|---|---|---|---|---|---|
| 11 | 86345131 | rs722158 | -0.572 | -0.821 | 0.134 | 0.27 | 0.003 | -0.004 | -0.004 | 0 | 0.22 | 0.24 | 0.38 | 0.4 | 0.15 | 0.17 |
| 11 | 86346940 | rs11821546 | 0.814 | -0.77 | 0.22 | 0.254 | -0.009 | 0.061 | 0.061 | 0 | 0.07 | 0.18 | 0.03 | 0.15 | 0.04 | 0.1 |
| 11 | 86347405 | rs7940187 | 0.813 | -0.706 | 0.096 | 0.151 | 0.003 | -0.033 | -0.033 | 0 | 0.1 | 0.09 | 0.09 | 0.01 | 0.1 | 0.15 |
| 11 | 86350791 | rs10501626 | 0.003 | -0.679 | 0.122 | 0.214 | 0.012 | 0.011 | 0.011 | 0.497 | 0.14 | 0.16 | 0.12 | 0.15 | 0.17 | 0.2 |
| 11 | 86351500 | rs10501627 | 2.039 | -0.415 | -0.137 | -0.091 | -0.106 | 0.241 | 0.241 | 0 | 0.4 | 0.09 | 0.31 | 0.05 | 0.47 | 0.3 |
| 11 | 86351770 | rs1938923 | 0.439 | 0.052 | 0.359 | 0.313 | -0.015 | 0.097 | 0.097 | 0 | 0.75 | 0.5 | 0.8 | 0.61 | 0.78 | 0.68 |
| 11 | 86353362 | rs659301 | -0.303 | 0.202 | 0.066 | 0.018 | 0.009 | 0.074 | 0.074 | 0.01 | 0.94 | 0.87 | 0.94 | 0.85 | 0.92 | 0.88 |
| 11 | 86376341 | rs11607554 | 1.764 | 0.36 | -0.078 | -0.101 | -0.033 | 0.242 | 0.242 | 0 | 0.45 | 0.16 | 0.32 | 0.06 | 0.45 | 0.36 |
| 11 | 86376739 | rs507211 | -0.721 | 0.361 | 0.179 | 0.026 | 0.016 | 0.082 | 0.082 | 0 | 0.21 | 0.33 | 0.19 | 0.38 | 0.27 | 0.27 |
| 11 | 86377805 | rs624183 | -0.063 | 0.213 | 0.169 | 0.03 | -0.001 | 0.035 | 0.035 | 0 | 0.17 | 0.28 | 0.15 | 0.23 | 0.18 | 0.22 |
| 11 | 86383679 | rs670736 | -0.097 | 0.175 | 0.189 | -0.055 | -0.035 | 0.098 | 0.098 | 1.05 | 0.78 | 0.65 | 0.63 | 0.45 | 0.87 | 0.74 |
| 11 | 86383994 | rs682915 | -1.788 | 0.955 | 0.431 | 0.273 | 0.019 | 0.037 | 0.037 | 0.11 | 0.34 | 0.47 | 0.3 | 0.39 | 0.41 | 0.35 |
| 11 | 86386784 | rs594361 | -1.444 | 0.992 | 0.379 | 0.36 | 0.018 | 0.088 | 0.088 | 0 | 0.24 | 0.39 | 0.23 | 0.38 | 0.32 | 0.28 |
| 11 | 86389000 | rs1938936 | 1.036 | 0.994 | 0.45 | 0.361 | 0.02 | 0.141 | 0.141 | 0 | 0.81 | 0.56 | 0.79 | 0.67 | 0.76 | 0.66 |
| 11 | 86389536 | rs642245 | -0.658 | 1.035 | 0.354 | 0.411 | 0.024 | 0.175 | 0.175 | 0 | 0.15 | 0.35 | 0.15 | 0.33 | 0.19 | 0.24 |
| 11 | 86396588 | rs528948 | 1.402 | 1.408 | 0.098 | 0.464 | -0.002 | -0.014 | -0.014 | 1.673 | 0.6 | 0.31 | 0.58 | 0.47 | 0.6 | 0.57 |
| 11 | 86402877 | rs600242 | 0.909 | 1.722 | 0.096 | 0.276 | 0.014 | -0.013 | -0.013 | 3.134 | 0.72 | 0.65 | 0.68 | 0.85 | 0.61 | 0.67 |
| 11 | 86405765 | rs622079 | -0.908 | 1.763 | 0.021 | 0.263 | -0.008 | 0.002 | 0.002 | 0.003 | 0.88 | 0.92 | 0.91 | 0.99 | 0.88 | 0.81 |
| 11 | 86406084 | rs746490 | 0.968 | 1.743 | 0.109 | 0.35 | -0.003 | 0.105 | 0.105 | 0 | 0.11 | 0.21 | 0.18 | 0.37 | 0.08 | 0.15 |
| 11 | 86406159 | rs746491 | 0.271 | 1.818 | -0.551 | 0.36 | 0.002 | 0.001 | 0.001 | 0.019 | 0.1 | 0.11 | 0.17 | 0.14 | 0.21 | 0.13 |
| 11 | 86410402 | rs2508305 | 0.876 | 1.761 | -0.662 | 0.325 | 0.003 | 0.049 | 0.049 | 0 | 0.11 | 0.17 | 0.1 | 0.19 | 0.12 | 0.19 |
| 11 | 86412474 | rs1938930 | -1.025 | 1.678 | -0.851 | 0.424 | 0.037 | 0.107 | 0.107 | 0 | 0.26 | 0.46 | 0.4 | 0.52 | 0.35 | 0.27 |
| 11 | 86412698 | rs17149311 | 1.052 | 1.497 | -0.819 | 0.447 | 0 | -0.028 | -0.028 | 0 | 0.11 | 0.12 | 0.09 | 0.01 | 0.1 | 0.18 |
| 11 | 86412880 | rs2433416 | -0.797 | 1.291 | -1.072 | 0.781 | 0.06 | 0.158 | 0.158 | 0 | 0.22 | 0.46 | 0.37 | 0.52 | 0.31 | 0.27 |
| 11 | 86414027 | rs2512988 | -0.518 | 0.068 | -1.274 | 0.483 | -0.002 | -0.001 | -0.001 | 0 | 0.44 | 0.56 | 0.44 | 0.37 | 0.55 | 0.53 |
| 11 | 86414282 | rs2512987 | 0.076 | 0.12 | -1.328 | 0.498 | 0.032 | 0.01 | 0.01 | 1.74 | 0.37 | 0.48 | 0.39 | 0.33 | 0.48 | 0.48 |
| 11 | 86415669 | rs2512983 | -0.617 | 0.08 | -2.161 | 0.018 | 0.038 | 0.044 | 0.044 | 3.206 | 0.61 | 0.9 | 0.67 | 0.73 | 0.7 | 0.72 |
| 11 | 86416932 | rs7129626 | -0.661 | 0.507 | -1.07 | 0.319 | 0 | 0.003 | 0.003 | 0 | 0.07 | 0.06 | 0.04 | 0.01 | 0.07 | 0.06 |
| 11 | 86426766 | rs2512995 | 0.322 | 0.277 | -1.224 | 0.5 | 0.013 | -0.014 | -0.014 | 6.272 | 0.91 | 0.99 | 0.97 | 0.85 | 0.96 | 0.99 |
| 11 | 86427342 | rs17758141 | -0.774 | 0.536 | -1.449 | 0.323 | -0.011 | 0.008 | 0.008 | 0 | 0.15 | 0.07 | 0.14 | 0.15 | 0.15 | 0.19 |
| 11 | 86428004 | rs2512994 | -1.516 | 0.691 | -1.403 | 0.126 | -0.009 | 0.012 | 0.012 | 0.151 | 0.21 | 0.16 | 0.21 | 0.18 | 0.21 | 0.37 |
| 11 | 86429093 | rs7115320 | -1.387 | -0.027 | -0.932 | 0.306 | 0.004 | -0.001 | -0.001 | 0 | 0.41 | 0.58 | 0.29 | 0.38 | 0.28 | 0.46 |
| 11 | 86432083 | rs7105597 | -0.099 | 0.053 | -1.056 | 0.221 | -0.059 | 0.069 | 0.069 | 0 | 0.13 | 0.01 | 0.14 | 0.01 | 0.14 | 0.08 |
| 11 | 86438669 | rs2513008 | -1.742 | -0.268 | -0.756 | 0.189 | 0.024 | -0.008 | -0.008 | 0.547 | 0.43 | 0.67 | 0.43 | 0.39 | 0.31 | 0.47 |
| 11 | 86447665 | rs2433429 | -1.931 | -0.847 | -0.352 | -0.13 | -0.018 | 0.034 | 0.034 | 0 | 0.35 | 0.46 | 0.38 | 0.39 | 0.26 | 0.4 |
| 11 | 86448308 | rs2433431 | -1.229 | -0.913 | -0.025 | -0.128 | -0.032 | 0.07 | 0.07 | 0 | 0.21 | 0.29 | 0.22 | 0.38 | 0.12 | 0.32 |
| 11 | 86452908 | rs17758195 | -0.132 | -0.866 | -0.231 | -0.224 | -0.062 | 0.084 | 0.084 | 0.814 | 0.14 | 0.01 | 0.15 | 0 | 0.15 | 0.1 |
| 11 | 86453700 | rs2433435 | -2.664 | -0.764 | -0.301 | -0.49 | 0.027 | 0.002 | 0.002 | 0.122 | 0.5 | 0.65 | 0.49 | 0.41 | 0.39 | 0.52 |
| 11 | 86454746 | rs2433438 | -1.862 | -0.798 | -0.253 | -0.466 | -0.01 | 0.006 | 0.006 | 0 | 0.35 | 0.62 | 0.34 | 0.42 | 0.24 | 0.43 |
| 11 | 86457657 | rs4075249 | 0.711 | -0.826 | -0.272 | -0.299 | 0.004 | -0.004 | -0.004 | 0 | 0.07 | 0.05 | 0.15 | 0.17 | 0.04 | 0.12 |
| 11 | 86460698 | rs2513018 | -2.177 | -0.273 | -0.39 | -0.701 | 0.006 | -0.01 | -0.01 | 0.39 | 0.45 | 0.66 | 0.5 | 0.42 | 0.41 | 0.5 |
| 11 | 86461180 | rs1075702 | -1.219 | -0.452 | -0.196 | -0.712 | 0.002 | -0.029 | -0.029 | 7.725 | 0.26 | 0.19 | 0.33 | 0.4 | 0.22 | 0.27 |
| 11 | 86461965 | rs4245425 | -0.188 | -0.38 | -0.096 | -0.744 | 0.002 | -0.061 | -0.061 | 1.309 | 0.13 | 0.22 | 0.14 | 0.01 | 0.17 | 0.08 |
| 11 | 86469986 | rs10751140 | -2.69 | -0.766 | -0.783 | -0.707 | -0.138 | -0.062 | -0.062 | 0 | 0.61 | 0.97 | 0.61 | 0.57 | 0.56 | 0.64 |
| 11 | 86471527 | rs11234795 | -2.185 | -1.078 | -0.059 | -0.551 | 0.012 | -0.005 | -0.005 | 0 | 0.44 | 0.62 | 0.48 | 0.39 | 0.37 | 0.45 |
| 11 | 86472784 | rs4944630 | -2.128 | -0.899 | 0.01 | -0.449 | 0.01 | -0.005 | -0.005 | 0 | 0.43 | 0.65 | 0.48 | 0.39 | 0.38 | 0.47 |
| 11 | 86473761 | rs17819068 | -0.306 | -0.749 | 0.009 | -0.321 | -0.016 | 0.079 | 0.079 | 0.231 | 0.12 | 0.16 | 0.17 | 0.37 | 0.07 | 0.18 |
| 11 | 86479027 | rs11234799 | 0.3 | -0.69 | -0.123 | -0.371 | -0.027 | 0.037 | 0.037 | 1.442 | 0.13 | 0.03 | 0.16 | 0 | 0.14 | 0.1 |
| 11 | 86482912 | rs12361769 | -2.074 | -0.754 | 0.181 | -0.405 | 0.004 | -0.007 | -0.007 | 0.431 | 0.41 | 0.61 | 0.46 | 0.38 | 0.39 | 0.37 |
| 11 | 86483240 | rs7105467 | -1.047 | -0.911 | 0.336 | -0.349 | -0.063 | 0.251 | 0.251 | 0 | 0.17 | 0.41 | 0.19 | 0.37 | 0.08 | 0.2 |
| 11 | 86484580 | rs4269941 | -1.047 | -0.911 | 0.336 | -0.349 | -0.063 | 0.251 | 0.251 | 0 | 0.17 | 0.41 | 0.19 | 0.37 | 0.08 | 0.2 |
| 11 | 86490519 | rs12787853 | -1.119 | -0.821 | 0.243 | -0.427 | -0.009 | -0.013 | -0.013 | 0.23 | 0.24 | 0.2 | 0.26 | 0.01 | 0.28 | 0.15 |
| 11 | 86492840 | rs744293 | 0.37 | -1.095 | 0.304 | -0.392 | 0.004 | -0.05 | -0.05 | 0 | 0.11 | 0.2 | 0.11 | 0.01 | 0.15 | 0.05 |
| 11 | 86495189 | rs7940201 | -1.526 | -1.101 | 0.418 | -0.406 | -0.008 | 0.022 | 0.022 | 3.485 | 0.28 | 0.29 | 0.3 | 0.37 | 0.21 | 0.31 |
| 11 | 86496427 | rs6592315 | -1.989 | -1.093 | 0.218 | -0.38 | 0.016 | -0.013 | -0.013 | 0 | 0.4 | 0.65 | 0.42 | 0.38 | 0.35 | 0.35 |
| 11 | 86503107 | rs754490 | -1.172 | -1.182 | -0.1 | -0.455 | -0.002 | 0.009 | 0.009 | 0.005 | 0.23 | 0.18 | 0.21 | 0.01 | 0.21 | 0.13 |
| 11 | 86503137 | rs2015747 | 0.321 | -1.185 | -0.096 | -0.459 | 0.002 | -0.008 | -0.008 | 0 | 0.09 | 0.09 | 0.08 | 0.01 | 0.09 | 0.04 |
| 11 | 86508928 | rs12805405 | -1.458 | -1.211 | 0.174 | -0.477 | -0.002 | -0.071 | -0.071 | 0 | 0.24 | 0.28 | 0.22 | 0.01 | 0.21 | 0.13 |
| 11 | 86510182 | rs2105587 | -1.192 | -1.197 | 0.077 | -0.471 | -0.002 | 0.009 | 0.009 | 0.004 | 0.22 | 0.17 | 0.21 | 0.01 | 0.21 | 0.13 |
| 11 | 86510783 | rs3809048 | -0.673 | -1.052 | 0.133 | -0.067 | -0.001 | 0.032 | 0.032 | 0.099 | 0.14 | 0.12 | 0.17 | 0.4 | 0.08 | 0.2 |
| 11 | 86520752 | rs2155081 | -1.676 | -1.18 | 0.211 | -0.091 | -0.011 | 0.046 | 0.046 | 3.85 | 0.26 | 0.28 | 0.29 | 0.41 | 0.19 | 0.26 |
| 11 | 86522992 | rs11608167 | 0.592 | -1.107 | -0.119 | -0.068 | -0.047 | 0.058 | 0.058 | 2.579 | 0.1 | 0 | 0.11 | 0 | 0.11 | 0.08 |
| 11 | 86525173 | rs4304806 | -0.761 | -1.117 | 0.039 | 0.106 | -0.001 | 0.029 | 0.029 | 0 | 0.13 | 0.12 | 0.17 | 0.41 | 0.08 | 0.18 |
| 11 | 86542005 | rs10751141 | -1.527 | -1.336 | -0.278 | -0.308 | 0.001 | -0.001 | -0.001 | 1.065 | 0.47 | 0.48 | 0.41 | 0.45 | 0.39 | 0.33 |
| 11 | 86555491 | rs2555537 | -1.586 | -1.272 | -0.32 | -0.333 | -0.001 | -0.001 | -0.001 | 0 | 0.47 | 0.48 | 0.43 | 0.45 | 0.41 | 0.34 |
| 11 | 86559592 | rs12801911 | 0.48 | -1.471 | -0.329 | 0.083 | -0.003 | -0.008 | -0.008 | 0 | 0.07 | 0.04 | 0.06 | 0.16 | 0.1 | 0.01 |
| 11 | 86560213 | rs12807497 | 0.103 | -1.489 | -0.296 | 0.075 | 0 | -0.006 | -0.006 | 0 | 0.08 | 0.09 | 0.06 | 0.16 | 0.1 | 0.01 |
| 11 | 86564779 | rs10898540 | -1.533 | -1.471 | -0.148 | -0.059 | 0.009 | -0.003 | -0.003 | 0 | 0.44 | 0.36 | 0.27 | 0.44 | 0.37 | 0.32 |
| 11 | 86571277 | rs12283389 | -1.515 | -1.431 | -0.342 | -0.025 | -0.007 | 0.001 | 0.001 | 0 | 0.43 | 0.5 | 0.29 | 0.43 | 0.36 | 0.32 |

| | | | | | | | | | | | | | | | | |
|---|---|---|---|---|---|---|---|---|---|---|---|---|---|---|---|---|
| 11 | 86572384 | rs12285365 | -0.964 | -0.978 | -0.598 | -0.047 | -0.003 | 0.025 | 0.025 | 0 | 0.2 | 0.11 | 0.14 | 0.13 | 0.2 | 0.19 |
| 11 | 86572473 | rs2511875 | -1.448 | -1.231 | -0.642 | 0.28 | -0.001 | 0 | 0 | 1.61 | 0.39 | 0.44 | 0.45 | 0.53 | 0.37 | 0.38 |
| 11 | 86572652 | rs2511876 | -0.473 | -1.155 | -0.51 | 0.452 | 0.013 | 0.046 | 0.046 | 0 | 0.21 | 0.36 | 0.47 | 0.34 | 0.22 | 0.26 |
| 11 | 86573198 | rs2511878 | -0.462 | -1.38 | -0.659 | 0.339 | -0.016 | 0.035 | 0.035 | 0 | 0.21 | 0.38 | 0.27 | 0.3 | 0.15 | 0.3 |
| 11 | 86573310 | rs1939108 | -1.448 | -1.441 | -0.712 | 0.304 | -0.01 | 0.004 | 0.004 | 0 | 0.31 | 0.24 | 0.24 | 0.26 | 0.34 | 0.21 |
| 11 | 86574577 | rs17819217 | 2.12 | -1.767 | -0.72 | 0.396 | -0.021 | 0.013 | 0.013 | 0 | 0.13 | 0.17 | 0.18 | 0.18 | 0.08 | 0.2 |
| 11 | 86575077 | rs12577230 | 1.309 | -1.279 | -1.132 | 0.305 | -0.019 | -0.03 | -0.03 | 0 | 0.1 | 0.06 | 0.08 | 0.15 | 0.14 | 0.02 |
| 11 | 86576172 | rs10501630 | 1.297 | -1.327 | -1.153 | 0.276 | -0.014 | 0.024 | 0.024 | 0.09 | 0.06 | 0.14 | 0.03 | 0.1 | 0.04 | 0.11 |
| 11 | 86576421 | rs12796995 | 1.191 | -0.941 | -1.637 | 0.242 | -0.019 | -0.03 | -0.03 | 0.422 | 0.1 | 0.05 | 0.08 | 0.15 | 0.14 | 0.02 |
| 11 | 86577094 | rs1939105 | -1.708 | -0.779 | -1.44 | -0.316 | -0.036 | 0.032 | 0.032 | 0.24 | 0.66 | 0.83 | 0.7 | 0.74 | 0.57 | 0.72 |
| 11 | 86580508 | rs2155079 | -0.995 | -0.611 | -1.469 | -0.544 | -0.001 | 0.1 | 0.1 | 0 | 0.55 | 0.8 | 0.61 | 0.76 | 0.46 | 0.59 |
| 11 | 86581273 | rs2022983 | -2.079 | -0.46 | -1.703 | -0.73 | -0.059 | 0.025 | 0.025 | 0 | 0.69 | 0.88 | 0.68 | 0.76 | 0.58 | 0.71 |
| 11 | 86582707 | rs2433447 | -1.462 | -0.856 | -1.73 | -0.529 | -0.046 | -0.01 | -0.01 | 0 | 0.58 | 0.77 | 0.57 | 0.58 | 0.49 | 0.56 |
| 11 | 86585792 | rs947884 | -0.816 | -0.847 | -1.687 | -0.541 | 0.006 | 0.004 | 0.004 | 0 | 0.55 | 0.76 | 0.54 | 0.58 | 0.43 | 0.53 |
| 11 | 86587319 | rs11234849 | 0.767 | -0.712 | -1.513 | -0.419 | 0 | 0.005 | 0.005 | 0 | 0.07 | 0.1 | 0.1 | 0.18 | 0.07 | 0.14 |
| 11 | 86600363 | rs7107991 | 0.135 | -1.24 | -0.94 | -0.815 | -0.016 | -0.089 | -0.089 | 0 | 0.22 | 0.38 | 0.12 | 0.06 | 0.14 | 0.19 |
| 11 | 86600755 | rs10898546 | -1.017 | -1.166 | -0.906 | -0.698 | 0.002 | 0.019 | 0.019 | 0 | 0.72 | 0.68 | 0.56 | 0.47 | 0.68 | 0.74 |
| 11 | 86603490 | rs10898549 | 1.319 | -1.194 | -0.95 | -0.543 | 0.006 | -0.003 | -0.003 | 0 | 0.14 | 0.17 | 0.13 | 0.13 | 0.2 | 0.08 |
| 11 | 86604406 | rs10501631 | 0.081 | -0.863 | -0.944 | -1.671 | 0 | 0.013 | 0.013 | 0 | 0.08 | 0.12 | 0.09 | 0.18 | 0.08 | 0.14 |
| 11 | 86610223 | rs7106737 | 1.541 | -0.624 | -1.177 | -1.82 | -0.037 | 0.02 | 0.02 | 0 | 0.28 | 0.11 | 0.26 | 0.23 | 0.38 | 0.29 |
| 11 | 86612582 | rs17149499 | 0.206 | -0.439 | -0.54 | -1.492 | -0.046 | -0.056 | -0.056 | 1.033 | 0.16 | 0.27 | 0.07 | 0.06 | 0.06 | 0.16 |
| 11 | 86612869 | rs10792879 | -0.679 | 0.136 | -0.666 | -1.135 | -0.007 | -0.023 | -0.023 | 0.316 | 0.7 | 0.75 | 0.52 | 0.47 | 0.65 | 0.72 |
| 11 | 86616127 | rs11234870 | -0.398 | 0.485 | -0.249 | -0.85 | 0.029 | 0.02 | 0.02 | 0 | 0.16 | 0.31 | 0.23 | 0.22 | 0.22 | 0.25 |
| 11 | 86616600 | rs17212187 | 1.213 | -0.198 | -0.588 | -0.837 | 0.036 | -0.089 | -0.089 | 0 | 0.21 | 0.08 | 0.19 | 0.39 | 0.15 | 0.21 |
| 11 | 86617648 | rs12802454 | -0.052 | 0.135 | -0.502 | -0.977 | 0.027 | 0.014 | 0.014 | 0 | 0.48 | 0.4 | 0.54 | 0.43 | 0.67 | 0.5 |
| 11 | 86618821 | rs7104996 | -0.866 | 0.325 | -0.302 | -1.397 | -0.003 | -0.041 | -0.041 | 0.5 | 0.29 | 0.3 | 0.24 | 0.09 | 0.24 | 0.32 |
| 11 | 86618878 | rs10898552 | -0.087 | 0.974 | -0.432 | -1.076 | -0.001 | -0.005 | -0.005 | 0 | 0.67 | 0.62 | 0.64 | 0.69 | 0.69 | 0.7 |
| 11 | 86620693 | rs12576589 | -0.593 | 1.059 | -0.71 | -0.273 | -0.009 | -0.011 | -0.011 | 0 | 0.63 | 0.5 | 0.6 | 0.64 | 0.7 | 0.64 |
| 11 | 86621300 | rs7108680 | 0.118 | 1.193 | -0.666 | -0.303 | 0.023 | -0.03 | -0.03 | 0 | 0.1 | 0.18 | 0.11 | 0.03 | 0.16 | 0.1 |
| 11 | 86623913 | rs2212766 | -0.287 | 0.386 | -0.853 | 0.079 | -0.01 | -0.013 | -0.013 | 0 | 0.62 | 0.5 | 0.6 | 0.65 | 0.7 | 0.65 |
| 11 | 86625896 | rs10898553 | -0.273 | 1.064 | -1.016 | -0.03 | -0.019 | 0.022 | 0.022 | 0 | 0.34 | 0.32 | 0.38 | 0.21 | 0.5 | 0.35 |
| 11 | 86629236 | rs11234874 | -0.247 | 0.592 | -0.95 | 0.396 | 0 | -0.001 | -0.001 | 0 | 0.18 | 0.17 | 0.29 | 0.33 | 0.17 | 0.16 |
| 11 | 86632288 | rs7119097 | -0.158 | -0.039 | -0.864 | 0.131 | -0.022 | 0.029 | 0.029 | 0 | 0.66 | 0.49 | 0.66 | 0.6 | 0.76 | 0.6 |
| 11 | 86635188 | rs7115645 | 0.215 | 0.133 | -1.102 | -0.09 | -0.006 | 0.003 | 0.003 | 0 | 0.58 | 0.47 | 0.63 | 0.55 | 0.67 | 0.45 |
| 11 | 86640865 | rs4943964 | 0.19 | 0.027 | -1.045 | -0.082 | -0.012 | 0.005 | 0.005 | 0 | 0.58 | 0.47 | 0.64 | 0.55 | 0.69 | 0.46 |
| 11 | 86642669 | rs11234882 | -0.016 | -0.393 | -0.671 | 0.041 | -0.024 | -0.002 | -0.002 | 0.407 | 0.06 | 0.15 | 0.02 | 0.02 | 0.02 | 0.1 |
| 11 | 86644367 | rs12805523 | 0.01 | -0.694 | -0.997 | -0.232 | -0.066 | 0.16 | 0.16 | 0 | 0.78 | 0.49 | 0.73 | 0.58 | 0.84 | 0.6 |
| 11 | 86647670 | rs10898561 | 0.877 | -0.591 | -1.02 | -0.27 | -0.008 | 0.003 | 0.003 | 0 | 0.58 | 0.48 | 0.64 | 0.55 | 0.69 | 0.46 |
| 11 | 86648099 | rs10792891 | 0.877 | -0.591 | -1.023 | -0.27 | -0.011 | 0.004 | 0.004 | 0 | 0.58 | 0.47 | 0.64 | 0.55 | 0.69 | 0.46 |
| 11 | 86650282 | rs4631906 | -0.783 | -0.746 | -0.625 | -0.147 | -0.043 | 0.22 | 0.22 | 0 | 0.84 | 0.59 | 0.75 | 0.59 | 0.87 | 0.7 |
| 11 | 86650400 | rs7127451 | 0.618 | -0.47 | -0.653 | -0.351 | -0.01 | 0.014 | 0.014 | 0 | 0.64 | 0.4 | 0.65 | 0.57 | 0.74 | 0.55 |
| 11 | 86651312 | rs11234888 | 1.15 | 0.39 | -1.304 | -0.79 | -0.092 | 0.128 | 0.128 | 0 | 0.25 | 0.04 | 0.2 | 0.05 | 0.34 | 0.19 |
| 11 | 86657520 | rs713065 | 0.501 | -0.562 | -0.773 | -0.432 | -0.002 | 0.001 | 0.001 | 0 | 0.63 | 0.42 | 0.57 | 0.46 | 0.65 | 0.5 |
| 11 | 86659213 | rs10898563 | 1.406 | -0.54 | -0.873 | -0.449 | -0.137 | 0.089 | 0.089 | 0 | 0.29 | 0.05 | 0.24 | 0.11 | 0.41 | 0.22 |
| 11 | 86660886 | rs3802892 | -0.332 | -0.575 | -0.562 | 0.132 | -0.006 | -0.02 | -0.02 | 0.075 | 0.75 | 0.81 | 0.67 | 0.63 | 0.7 | 0.66 |
| 11 | 86668446 | rs3758657 | 0.418 | -0.544 | -0.661 | 0.157 | -0.008 | 0.066 | 0.066 | 0 | 0.09 | 0.17 | 0.07 | 0.16 | 0.05 | 0.11 |
| 11 | 86669535 | rs7925666 | 0.769 | -0.612 | -0.715 | 0.299 | -0.014 | -0.005 | -0.005 | 0 | 0.32 | 0.38 | 0.31 | 0.34 | 0.21 | 0.27 |
| 11 | 86688920 | rs873676 | 0.465 | -0.431 | -0.92 | 0.405 | -0.01 | 0.042 | 0.042 | 0 | 0.09 | 0.13 | 0.07 | 0.16 | 0.05 | 0.11 |
| 11 | 86690915 | rs12575890 | 1.208 | -0.436 | -0.966 | 0.372 | -0.127 | 0.071 | 0.071 | 0 | 0.3 | 0.08 | 0.24 | 0.14 | 0.41 | 0.27 |
| 11 | 86693125 | rs12277682 | 0.289 | -0.433 | -0.954 | 0.372 | -0.04 | 0.021 | 0.021 | 0 | 0.39 | 0.22 | 0.31 | 0.29 | 0.46 | 0.38 |
| 11 | 86694369 | rs7127388 | -0.132 | 0.04 | -0.232 | 1.211 | -0.006 | 0.021 | 0.021 | 0 | 0.09 | 0.1 | 0.07 | 0.16 | 0.05 | 0.11 |
| 11 | 86704775 | rs11234901 | -0.583 | 0.102 | -0.506 | 1.005 | 0.014 | 0.096 | 0.096 | 0 | 0.18 | 0.38 | 0.19 | 0.31 | 0.13 | 0.23 |
| 11 | 86710510 | rs7122009 | 0.221 | 0.2 | -1.241 | 0.609 | 0.008 | -0.025 | -0.025 | 0 | 0.81 | 0.88 | 0.76 | 0.74 | 0.78 | 0.73 |
| 11 | 86718978 | rs1272073 | 0.294 | 0.42 | -1.13 | -0.147 | 0.025 | 0.133 | 0.133 | 0 | 0.5 | 0.33 | 0.48 | 0.16 | 0.54 | 0.48 |
| 11 | 86721686 | rs7108104 | -0.955 | 0.368 | -0.402 | 0.364 | 0.001 | 0.013 | 0.013 | 0 | 0.16 | 0.31 | 0.16 | 0.15 | 0.17 | 0.11 |
| 11 | 86722128 | rs7127569 | -1.953 | -0.055 | -0.651 | 0.095 | -0.009 | -0.007 | -0.007 | 0.311 | 0.24 | 0.34 | 0.21 | 0.19 | 0.22 | 0.24 |
| 11 | 86724086 | rs921872 | 0.716 | -0.72 | -0.41 | 0.068 | 0.002 | 0.001 | 0.001 | 0 | 0.51 | 0.44 | 0.45 | 0.33 | 0.64 | 0.63 |
| 11 | 86724222 | rs17149645 | 0.428 | -0.737 | -0.359 | 0.039 | -0.009 | -0.002 | -0.002 | 0 | 0.09 | 0.18 | 0.07 | 0.09 | 0.11 | 0.03 |
| 11 | 86724504 | rs10501632 | -0.543 | -0.704 | -0.349 | 0.054 | 0.002 | 0.002 | 0.002 | 0 | 0.17 | 0.21 | 0.12 | 0.12 | 0.16 | 0.16 |
| 11 | 86726464 | rs1963952 | -0.572 | -0.69 | -0.415 | 0.17 | -0.004 | 0.005 | 0.005 | 0 | 0.66 | 0.65 | 0.56 | 0.46 | 0.78 | 0.78 |
| 11 | 86727884 | rs991773 | -0.674 | -0.678 | -0.482 | 0.176 | 0.003 | -0.005 | -0.005 | 0 | 0.67 | 0.7 | 0.56 | 0.46 | 0.78 | 0.78 |
| 11 | 86737369 | rs11234915 | -0.449 | -0.75 | -0.425 | 0.488 | -0.002 | 0 | 0 | 1.11 | 0.16 | 0.16 | 0.11 | 0.15 | 0.15 | 0.16 |
| 11 | 86739976 | rs12270673 | 1.455 | -0.684 | -0.621 | 0.237 | -0.129 | 0.161 | 0.161 | 0 | 0.37 | 0.06 | 0.3 | 0.12 | 0.48 | 0.43 |
| 11 | 86740573 | rs4944662 | -1.156 | -0.791 | -0.478 | 0.328 | -0.001 | 0.001 | 0.001 | 0 | 0.7 | 0.71 | 0.57 | 0.5 | 0.79 | 0.8 |
| 11 | 86741740 | rs11234920 | -0.284 | -0.741 | -0.453 | 0.263 | -0.014 | 0.001 | 0.001 | 0 | 0.17 | 0.19 | 0.12 | 0.15 | 0.15 | 0.16 |
| 11 | 86743904 | rs3740659 | 1.358 | -0.678 | -0.576 | 0.059 | -0.128 | 0.164 | 0.164 | 0 | 0.37 | 0.07 | 0.3 | 0.12 | 0.48 | 0.43 |
| 11 | 86748051 | rs11234925 | 0.642 | -0.694 | -0.322 | 0.127 | -0.002 | 0.008 | 0.008 | 0 | 0.07 | 0.11 | 0.04 | 0.16 | 0.09 | 0.11 |
| 11 | 86748140 | rs2512502 | -1.733 | -0.694 | -0.351 | 0.051 | -0.128 | 0.164 | 0.164 | 0 | 0.63 | 0.93 | 0.69 | 0.88 | 0.52 | 0.56 |
| 11 | 86752002 | rs2512377 | -0.916 | -0.937 | 0.089 | 0 | 0.063 | 0.021 | 0.021 | 0 | 0.54 | 0.71 | 0.56 | 0.66 | 0.36 | 0.43 |

| | | | | | | | | | | | | | | | | |
|---|---|---|---|---|---|---|---|---|---|---|---|---|---|---|---|---|
| 11 | 86755539 | rs2513228 | 0.396 | -1.062 | 0.26 | -0.028 | -0.022 | 0.014 | 0.014 | 0 | 0.36 | 0.55 | 0.45 | 0.6 | 0.22 | 0.28 |
| 11 | 86756502 | rs10898587 | 0.726 | -0.972 | 0.275 | 0.033 | 0.002 | 0.037 | 0.037 | 0 | 0.07 | 0.11 | 0.04 | 0.16 | 0.09 | 0.1 |
| 11 | 86765801 | rs11234931 | -0.129 | -1.001 | 0.301 | 0.002 | -0.004 | -0.008 | -0.008 | 0 | 0.16 | 0.16 | 0.12 | 0.06 | 0.15 | 0.16 |
| 11 | 86771583 | rs2001411 | 0.296 | -1.118 | 0.54 | -0.038 | -0.038 | 0.024 | 0.024 | 0 | 0.37 | 0.5 | 0.44 | 0.6 | 0.22 | 0.3 |
| 11 | 86775026 | rs10898590 | 0.848 | -1.072 | 0.55 | -0.013 | -0.001 | 0.02 | 0.02 | 0 | 0.06 | 0.09 | 0.03 | 0.16 | 0.09 | 0.1 |
| 11 | 86775100 | rs2512356 | 0.611 | -1.212 | 0.697 | -0.044 | 0.048 | -0.054 | -0.054 | 0 | 0.34 | 0.14 | 0.41 | 0.53 | 0.22 | 0.28 |
| 11 | 86777044 | rs12273779 | 0.065 | -1.21 | 0.691 | -0.069 | 0.002 | 0.002 | 0.002 | 0 | 0.15 | 0.13 | 0.12 | 0.06 | 0.15 | 0.13 |
| 11 | 86778589 | rs2445587 | -1.399 | -1.163 | 0.702 | -0.049 | -0.009 | 0.077 | 0.077 | 0 | 0.61 | 0.77 | 0.61 | 0.82 | 0.46 | 0.53 |
| 11 | 86780864 | rs2513237 | -0.917 | -1.166 | 0.693 | -0.05 | 0.039 | 0.015 | 0.015 | 0 | 0.53 | 0.63 | 0.57 | 0.66 | 0.35 | 0.43 |
| 11 | 86781808 | rs1531144 | 0.432 | -1.074 | 0.494 | 0.004 | 0.005 | 0.053 | 0.053 | 0 | 0.07 | 0.14 | 0.04 | 0.16 | 0.09 | 0.1 |
| 11 | 86804940 | rs2512387 | -0.183 | -0.767 | 0.487 | -0.138 | 0.005 | 0.001 | 0.001 | 0 | 0.43 | 0.59 | 0.52 | 0.66 | 0.31 | 0.35 |
| 11 | 86805257 | rs2513225 | -1.659 | -0.713 | 0.423 | -0.23 | -0.071 | 0.104 | 0.104 | 0 | 0.65 | 0.84 | 0.66 | 0.88 | 0.53 | 0.58 |
| 11 | 86805541 | rs6592333 | -1.259 | -0.654 | 0.432 | -0.205 | -0.007 | 0.015 | 0.015 | 0 | 0.24 | 0.35 | 0.19 | 0.23 | 0.24 | 0.25 |
| 11 | 86806583 | rs11234962 | 0.947 | -0.639 | 0.392 | -0.188 | -0.004 | 0.005 | 0.005 | 0 | 0.06 | 0.08 | 0.03 | 0.17 | 0.09 | 0.1 |
| 11 | 86810100 | rs10898598 | 0.021 | -0.658 | 0.384 | -0.196 | 0.001 | 0 | 0 | 1.435 | 0.15 | 0.13 | 0.11 | 0.06 | 0.13 | 0.13 |
| 11 | 86810624 | rs2513210 | 0.732 | -0.736 | 0.254 | -0.215 | 0.05 | -0.073 | -0.073 | 0.078 | 0.34 | 0.11 | 0.41 | 0.53 | 0.22 | 0.29 |
| 11 | 86819979 | rs2512352 | -1.053 | -0.472 | 0.129 | -0.257 | 0.003 | 0.024 | 0.024 | 0 | 0.58 | 0.71 | 0.63 | 0.72 | 0.44 | 0.48 |
| 11 | 86822077 | rs2124579 | -1.536 | -0.425 | -0.043 | -0.341 | -0.089 | 0.119 | 0.119 | 0 | 0.66 | 0.88 | 0.7 | 0.89 | 0.56 | 0.6 |
| 11 | 86825869 | rs2512337 | -0.059 | -0.511 | 0.143 | -0.246 | 0.004 | 0 | 0 | 0 | 0.42 | 0.6 | 0.52 | 0.66 | 0.32 | 0.35 |
| 11 | 86826422 | rs2512493 | -1.636 | -0.129 | -0.244 | -0.611 | -0.122 | 0.111 | 0.111 | 0 | 0.69 | 0.93 | 0.69 | 0.89 | 0.58 | 0.64 |
| 11 | 86836648 | rs564932 | -1.652 | 0.758 | -0.454 | -0.901 | -0.068 | 0.093 | 0.093 | 0 | 0.74 | 0.94 | 0.77 | 0.92 | 0.66 | 0.65 |
| 11 | 86839381 | rs655050 | -0.386 | 0.514 | 0.186 | -0.912 | 0.002 | -0.003 | -0.003 | 0 | 0.58 | 0.42 | 0.6 | 0.69 | 0.45 | 0.47 |
| 11 | 86850175 | rs10792910 | 0.236 | -0.084 | 0.557 | -0.473 | 0.017 | 0.106 | 0.106 | 0 | 0.08 | 0.22 | 0.06 | 0.16 | 0.09 | 0.1 |
| 11 | 86860743 | rs11234992 | 0.421 | -0.127 | 0.433 | -0.469 | -0.014 | -0.013 | -0.013 | 0 | 0.07 | 0.11 | 0.04 | 0.03 | 0.05 | 0.01 |
| 11 | 86870367 | rs529564 | 0.308 | -0.679 | 0.382 | -0.868 | 0.005 | -0.006 | -0.006 | 0 | 0.31 | 0.28 | 0.41 | 0.63 | 0.19 | 0.3 |
| 11 | 86872504 | rs546229 | -1.211 | -0.657 | 0.353 | -0.996 | 0.029 | 0.022 | 0.022 | 0 | 0.5 | 0.57 | 0.55 | 0.69 | 0.33 | 0.47 |
| 11 | 86872908 | rs549932 | -0.87 | -0.659 | 0.242 | -0.991 | 0.016 | 0.01 | 0.01 | 0 | 0.47 | 0.4 | 0.5 | 0.69 | 0.3 | 0.44 |
| 11 | 86884115 | rs471471 | 0.429 | -0.585 | 0.357 | -0.72 | 0.014 | -0.025 | -0.025 | 0.349 | 0.29 | 0.22 | 0.4 | 0.55 | 0.19 | 0.29 |
| 11 | 86889284 | rs1948206 | -1.102 | -0.153 | 0.592 | -0.617 | 0.06 | 0.015 | 0.015 | 0 | 0.08 | 0.3 | 0.11 | 0.06 | 0.12 | 0.09 |
| 11 | 86895734 | rs539894 | -1.716 | -0.256 | 0.559 | -0.702 | -0.001 | -0.047 | -0.047 | 0 | 0.22 | 0.43 | 0.2 | 0.12 | 0.24 | 0.24 |
| 11 | 86917621 | rs654083 | -0.643 | -0.524 | 0.478 | -0.655 | 0.003 | 0.003 | 0.003 | 0 | 0.14 | 0.12 | 0.09 | 0.06 | 0.11 | 0.15 |
| 11 | 86920178 | rs61552 | 1.464 | -0.71 | 0.029 | -0.832 | -0.082 | 0.191 | 0.191 | 0 | 0.38 | 0.08 | 0.34 | 0.09 | 0.51 | 0.37 |
| 11 | 86921921 | rs546560 | -1.511 | -0.588 | 0.444 | -0.584 | -0.017 | 0.015 | 0.015 | 0 | 0.22 | 0.5 | 0.21 | 0.2 | 0.22 | 0.24 |
| 11 | 86928253 | rs301588 | -0.665 | -0.796 | 0.416 | -0.65 | 0.021 | -0.028 | -0.028 | 0 | 0.69 | 0.8 | 0.61 | 0.39 | 0.8 | 0.69 |
| 11 | 86930270 | rs542279 | 0.448 | -0.649 | 0.557 | -0.379 | 0.002 | 0.023 | 0.023 | 0 | 0.08 | 0.02 | 0.05 | 0.04 | 0.06 | 0.13 |
| 11 | 86931924 | rs541345 | -0.905 | -0.662 | 0.68 | -0.381 | -0.007 | 0 | 0 | 0 | 0.14 | 0.25 | 0.13 | 0.12 | 0.14 | 0.17 |
| 11 | 86932603 | rs644471 | -1.303 | -0.539 | 0.564 | -0.306 | -0.016 | -0.018 | -0.018 | 0 | 0.21 | 0.5 | 0.21 | 0.15 | 0.21 | 0.2 |
| 11 | 86947617 | rs12224600 | -0.309 | -0.276 | -0.184 | 0.208 | -0.002 | 0.036 | 0.036 | 0 | 0.07 | 0.11 | 0.04 | 0.16 | 0.08 | 0.13 |
| 11 | 86951781 | rs522886 | -0.054 | -0.757 | -0.344 | 0.1 | 0.004 | -0.001 | -0.001 | 0 | 0.64 | 0.68 | 0.51 | 0.29 | 0.68 | 0.69 |
| 11 | 86960516 | rs550503 | -0.189 | -0.759 | -0.352 | 0.328 | 0.008 | -0.015 | -0.015 | 0 | 0.66 | 0.74 | 0.57 | 0.4 | 0.72 | 0.73 |
| 11 | 86965391 | rs2030234 | 1.961 | -1.008 | -0.7 | -0.174 | -0.13 | 0.141 | 0.141 | 0.223 | 0.32 | 0.07 | 0.32 | 0.08 | 0.44 | 0.27 |
| 11 | 86966333 | rs10792929 | -0.346 | -1.048 | 0.162 | -0.008 | -0.043 | 0.096 | 0.096 | 0.107 | 0.11 | 0.3 | 0.04 | 0.14 | 0.04 | 0.13 |
| 11 | 86968173 | rs643902 | -1.194 | -1.062 | 0.382 | 0.054 | -0.048 | 0.092 | 0.092 | 0 | 0.18 | 0.42 | 0.1 | 0.23 | 0.11 | 0.19 |
| 11 | 86970134 | rs614504 | 0.185 | -0.924 | 0.319 | 0.138 | 0.002 | -0.006 | -0.006 | 0 | 0.1 | 0.06 | 0.05 | 0.09 | 0.08 | 0.13 |
| 11 | 86974576 | rs301603 | -0.176 | -1.089 | 0.235 | -0.093 | 0.002 | -0.003 | -0.003 | 0 | 0.57 | 0.58 | 0.47 | 0.39 | 0.6 | 0.64 |
| 11 | 86975789 | rs7952272 | 2.133 | -0.937 | 0.32 | -0.161 | -0.029 | 0.031 | 0.031 | 0.063 | 0.31 | 0.19 | 0.32 | 0.1 | 0.42 | 0.26 |
| 11 | 86979874 | rs11235060 | 2.027 | -0.916 | 0.305 | -0.151 | -0.019 | 0.018 | 0.018 | 0.026 | 0.32 | 0.25 | 0.33 | 0.1 | 0.42 | 0.26 |
| 11 | 86983831 | rs12293231 | 2.043 | -0.915 | 0.304 | -0.179 | -0.019 | 0.017 | 0.017 | 0.033 | 0.32 | 0.26 | 0.33 | 0.1 | 0.45 | 0.26 |
| 11 | 86985581 | rs214742 | 0.327 | -0.868 | 0.311 | -0.14 | 0.005 | 0.009 | 0.009 | 0 | 0.49 | 0.56 | 0.41 | 0.33 | 0.53 | 0.49 |
| 11 | 86997694 | rs7119714 | 1.897 | -0.782 | 0.18 | -0.224 | -0.018 | 0.017 | 0.017 | 0.008 | 0.32 | 0.26 | 0.33 | 0.1 | 0.44 | 0.26 |
| 11 | 87006224 | rs1893005 | -0.164 | -0.991 | 0.182 | -0.035 | 0.001 | -0.002 | -0.002 | 0 | 0.6 | 0.59 | 0.51 | 0.42 | 0.66 | 0.65 |
| 11 | 87007030 | rs10898664 | -1.718 | -1.032 | 0.28 | 0.101 | -0.022 | 0.028 | 0.028 | 0 | 0.28 | 0.35 | 0.18 | 0.32 | 0.22 | 0.4 |
| 11 | 87013438 | rs2276102 | -0.594 | -0.74 | 0.18 | 0.05 | -0.008 | 0.033 | 0.033 | 0 | 0.67 | 0.59 | 0.51 | 0.44 | 0.65 | 0.7 |
| 11 | 87018161 | rs4545589 | -1.919 | -0.513 | 0.193 | -0.003 | -0.011 | 0.021 | 0.021 | 0 | 0.35 | 0.52 | 0.26 | 0.34 | 0.28 | 0.42 |
| 11 | 87019645 | rs7117070 | -0.874 | -0.549 | 0.195 | 0.046 | -0.004 | -0.003 | -0.003 | 0 | 0.21 | 0.2 | 0.18 | 0.19 | 0.23 | 0.27 |
| 11 | 87031246 | rs9666962 | -1.966 | -0.762 | -0.139 | -0.179 | -0.03 | -0.073 | -0.073 | 0 | 0.88 | 0.98 | 0.64 | 0.46 | 0.84 | 0.91 |
| 11 | 87032664 | rs17821399 | 0.46 | -0.551 | 0.148 | 0.145 | -0.001 | -0.001 | -0.001 | 0 | 0.1 | 0.08 | 0.07 | 0.09 | 0.1 | 0.12 |
| 11 | 87034696 | rs2212616 | -0.139 | -0.62 | 0.133 | 0.092 | -0.005 | 0.005 | 0.005 | 0 | 0.11 | 0.09 | 0.1 | 0.08 | 0.12 | 0.15 |
| 11 | 87035210 | rs4636701 | 0.02 | -0.619 | 0.315 | 0.107 | -0.056 | 0.069 | 0.069 | 0 | 0.1 | 0.23 | 0.03 | 0.16 | 0.02 | 0.11 |
| 11 | 87036784 | rs10898669 | 2.032 | -0.48 | 0.031 | 0.154 | -0.064 | 0.118 | 0.118 | 0.008 | 0.34 | 0.1 | 0.26 | 0.18 | 0.37 | 0.3 |
| 11 | 87037295 | rs17149955 | -0.28 | -0.482 | 0.082 | -0.035 | -0.004 | -0.003 | -0.003 | 0 | 0.13 | 0.09 | 0.11 | 0.11 | 0.14 | 0.15 |
| 11 | 87037338 | rs10501635 | -0.28 | -0.482 | 0.094 | -0.035 | 0.012 | 0.005 | 0.005 | 0 | 0.13 | 0.16 | 0.12 | 0.11 | 0.14 | 0.15 |
| 11 | 87037565 | rs7926724 | 0.215 | -0.522 | 0.152 | -0.044 | 0.017 | 0.023 | 0.023 | 0 | 0.56 | 0.37 | 0.44 | 0.28 | 0.6 | 0.57 |
| 11 | 87037680 | rs12282551 | 0.931 | -0.66 | 0.06 | -0.056 | 0.011 | 0.099 | 0.099 | 0 | 0.46 | 0.26 | 0.37 | 0.2 | 0.51 | 0.45 |
| 11 | 87037743 | rs10898670 | 0.959 | -0.657 | -0.15 | -0.055 | 0.005 | 0.131 | 0.131 | 0 | 0.46 | 0.22 | 0.37 | 0.2 | 0.51 | 0.45 |
| 11 | 87058981 | rs7928637 | -1.528 | -0.572 | 0.009 | 0.251 | -0.013 | -0.005 | -0.005 | 0.003 | 0.22 | 0.16 | 0.17 | 0.19 | 0.24 | 0.28 |
| 11 | 87065948 | rs878611 | 0.129 | -0.9 | 0.07 | 0.317 | -0.016 | 0.011 | 0.011 | 0 | 0.08 | 0.08 | 0.02 | 0.16 | 0.02 | 0.07 |
| 11 | 87066709 | rs2003177 | -2.405 | -0.917 | 0.138 | 0.582 | -0.015 | 0.024 | 0.024 | 0 | 0.36 | 0.52 | 0.35 | 0.44 | 0.3 | 0.39 |
| 11 | 87071233 | rs12417501 | 0.051 | -0.767 | -0.093 | 0.579 | -0.033 | -0.03 | -0.03 | 0.777 | 0.1 | 0.03 | 0.2 | 0.13 | 0.16 | 0.11 |

| | | | | | | | | | | | | | | | | |
|---|---|---|---|---|---|---|---|---|---|---|---|---|---|---|---|---|
| 11 | 87080713 | rs11235120 | -0.75 | -1.071 | 0.153 | 0.543 | -0.001 | 0 | 0 | 0 | 0.13 | 0.16 | 0.21 | 0.13 | 0.15 | 0.11 |
| 11 | 87083212 | rs10792948 | 1.09 | -1.193 | -0.082 | 0.372 | -0.029 | 0.116 | 0.116 | 0.119 | 0.43 | 0.17 | 0.33 | 0.19 | 0.5 | 0.39 |
| 11 | 87083324 | rs4471448 | 1.91 | -1.132 | -0.11 | 0.348 | -0.109 | 0.164 | 0.164 | 0 | 0.34 | 0.08 | 0.28 | 0.09 | 0.4 | 0.27 |
| 11 | 87091632 | rs4503569 | 0.328 | -0.99 | 0.068 | 0.555 | -0.003 | -0.005 | -0.005 | 0.414 | 0.09 | 0.08 | 0.05 | 0.1 | 0.09 | 0.11 |
| 11 | 87106806 | rs11235132 | -1.959 | -1.133 | 0.384 | 0.567 | -0.064 | 0.101 | 0.101 | 0 | 0.23 | 0.47 | 0.25 | 0.33 | 0.17 | 0.23 |
| 11 | 87110152 | rs10898685 | -0.555 | -1.045 | 0.505 | 0.628 | -0.045 | 0.159 | 0.159 | 0.02 | 0.09 | 0.27 | 0.03 | 0.19 | 0.02 | 0.13 |
| 11 | 87120819 | rs2020402 | -0.414 | -0.96 | 0.314 | 0.428 | -0.002 | 0.005 | 0.005 | 0 | 0.65 | 0.68 | 0.47 | 0.4 | 0.65 | 0.62 |
| 11 | 87129900 | rs7105817 | -0.749 | -0.809 | 1.019 | 0.332 | -0.013 | 0.07 | 0.07 | 0.3 | 0.12 | 0.32 | 0.08 | 0.15 | 0.07 | 0.14 |
| 11 | 87132764 | rs7111686 | -0.16 | -0.696 | 0.737 | 0.049 | -0.001 | -0.003 | -0.003 | 0 | 0.5 | 0.5 | 0.49 | 0.29 | 0.57 | 0.36 |
| 11 | 87134905 | rs7927205 | 1.485 | -0.693 | 0.14 | -0.309 | -0.124 | 0.129 | 0.129 | 0.693 | 0.3 | 0.07 | 0.24 | 0.12 | 0.4 | 0.21 |
| 11 | 87141156 | rs11235140 | -0.499 | -0.678 | 0.284 | -0.305 | 0.005 | 0.002 | 0.002 | 0.1 | 0.5 | 0.47 | 0.34 | 0.16 | 0.54 | 0.33 |
| 11 | 87141987 | rs11235141 | -0.783 | -0.694 | 0.32 | -0.191 | -0.009 | -0.022 | -0.022 | 0.014 | 0.11 | 0.18 | 0.04 | 0.03 | 0.09 | 0.09 |
| 11 | 87146898 | rs7118964 | -0.544 | -0.665 | 0.213 | -0.144 | 0 | -0.021 | -0.021 | 0 | 0.06 | 0.08 | 0.05 | 0 | 0.06 | 0.03 |
| 11 | 87148324 | rs4297485 | -1.639 | -0.676 | 0.235 | -0.147 | -0.005 | -0.025 | -0.025 | 0 | 0.14 | 0.19 | 0.09 | 0.04 | 0.12 | 0.11 |
| 11 | 87153492 | rs12291930 | 0.19 | -0.625 | 0.12 | -0.111 | -0.003 | -0.003 | -0.003 | 0 | 0.07 | 0.07 | 0.03 | 0.03 | 0.06 | 0.08 |
| 11 | 87154097 | rs4944707 | -0.494 | -0.671 | 0.115 | -0.251 | 0.003 | -0.005 | -0.005 | 0.107 | 0.5 | 0.46 | 0.35 | 0.15 | 0.54 | 0.33 |
| 11 | 87154923 | rs11235148 | 1.231 | -0.608 | 0.075 | -0.164 | -0.135 | 0.022 | 0.022 | 0.261 | 0.32 | 0.06 | 0.39 | 0.26 | 0.44 | 0.24 |
| 11 | 87155511 | rs7118472 | -0.802 | -0.571 | 0.199 | -0.147 | 0.003 | -0.059 | -0.059 | 1.132 | 0.1 | 0.34 | 0.07 | 0 | 0.06 | 0.03 |
| 11 | 87157192 | rs4127381 | 0.357 | -0.393 | -0.406 | 0.034 | -0.08 | 0.049 | 0.049 | 0.287 | 0.39 | 0.13 | 0.42 | 0.29 | 0.5 | 0.31 |
| 11 | 87158465 | rs4944708 | 0.351 | -0.352 | -0.453 | 0.055 | -0.03 | 0.039 | 0.039 | 0.025 | 0.41 | 0.26 | 0.43 | 0.29 | 0.51 | 0.32 |
| 11 | 87174520 | rs9804584 | -1.365 | -0.68 | -0.228 | 0.178 | -0.011 | -0.003 | -0.003 | 0.032 | 0.31 | 0.63 | 0.16 | 0.2 | 0.2 | 0.31 |
| 11 | 87182934 | rs10501638 | -1.049 | -0.352 | -0.295 | -0.268 | -0.003 | 0.014 | 0.014 | 0 | 0.2 | 0.17 | 0.14 | 0.04 | 0.24 | 0.17 |
| 11 | 87184225 | rs7935389 | -1.978 | -0.666 | 0.173 | -0.205 | -0.004 | -0.005 | -0.005 | 0 | 0.42 | 0.55 | 0.32 | 0.31 | 0.38 | 0.37 |
| 11 | 87184717 | rs10898702 | -0.116 | -0.694 | 0.385 | -0.214 | -0.012 | 0.117 | 0.117 | 0.08 | 0.07 | 0.14 | 0.08 | 0.26 | 0.03 | 0.08 |
| 11 | 87192624 | rs7358476 | -1.898 | -0.645 | -0.335 | -0.278 | -0.007 | -0.062 | -0.062 | 0 | 0.74 | 0.87 | 0.62 | 0.53 | 0.74 | 0.56 |
| 11 | 87197324 | rs10898708 | -1.915 | -0.488 | -0.098 | -0.14 | 0.017 | -0.006 | -0.006 | 0 | 0.43 | 0.63 | 0.32 | 0.39 | 0.38 | 0.37 |
| 11 | 87211337 | rs10792960 | -1.48 | -0.56 | -0.173 | -0.246 | -0.008 | 0.008 | 0.008 | 0 | 0.21 | 0.22 | 0.18 | 0.29 | 0.16 | 0.2 |
| 11 | 87212326 | rs11235178 | -1.447 | -0.535 | -0.407 | -0.234 | -0.008 | 0.009 | 0.009 | 0 | 0.21 | 0.22 | 0.17 | 0.29 | 0.16 | 0.2 |
| 11 | 87220744 | rs1986778 | 0.373 | -0.736 | -0.723 | -0.599 | -0.014 | 0.007 | 0.007 | 1.094 | 0.4 | 0.27 | 0.42 | 0.38 | 0.51 | 0.27 |
| 11 | 87220952 | rs1986779 | -1.181 | -1.14 | -0.232 | -0.621 | 0.006 | -0.013 | -0.013 | 0 | 0.65 | 0.65 | 0.56 | 0.44 | 0.7 | 0.46 |
| 11 | 87220986 | rs12274310 | -1.002 | -1.269 | -0.265 | -0.814 | -0.034 | -0.099 | -0.099 | 0.902 | 0.22 | 0.45 | 0.14 | 0.04 | 0.17 | 0.17 |
| 11 | 87221830 | rs4488224 | -0.817 | -1.072 | -0.401 | -0.549 | 0.01 | -0.006 | -0.006 | 0 | 0.61 | 0.61 | 0.64 | 0.46 | 0.71 | 0.46 |
| 11 | 87222624 | rs6592377 | -0.32 | -0.581 | -0.388 | -0.907 | 0.022 | 0.018 | 0.018 | 0.097 | 0.51 | 0.58 | 0.47 | 0.39 | 0.58 | 0.36 |
| 11 | 87223059 | rs10898722 | 0.32 | -0.316 | -0.519 | -1.041 | -0.033 | 0.021 | 0.021 | 0 | 0.42 | 0.09 | 0.39 | 0.38 | 0.53 | 0.32 |
| 11 | 87225596 | rs7117142 | -1.79 | -0.539 | -0.358 | -1.134 | -0.024 | -0.095 | -0.095 | 0 | 0.22 | 0.48 | 0.13 | 0.04 | 0.18 | 0.2 |
| 11 | 87225745 | rs11235187 | 0.422 | -0.393 | -0.348 | -1.124 | -0.025 | 0.029 | 0.029 | 0 | 0.42 | 0.14 | 0.4 | 0.37 | 0.53 | 0.32 |
| 11 | 87226475 | rs7929583 | -1.477 | -0.287 | -0.326 | -1.055 | 0.005 | -0.004 | -0.004 | 0 | 0.65 | 0.65 | 0.54 | 0.41 | 0.7 | 0.52 |
| 11 | 87226823 | rs12284047 | 0.12 | -0.301 | -0.32 | -1.042 | -0.003 | -0.002 | -0.002 | 0 | 0.06 | 0.05 | 0.04 | 0 | 0.1 | 0.01 |
| 11 | 87227163 | rs11235189 | -0.864 | -0.731 | 0.072 | -0.929 | -0.002 | -0.001 | -0.001 | 0.885 | 0.58 | 0.51 | 0.58 | 0.41 | 0.6 | 0.45 |
| 11 | 87230128 | rs10898725 | 0.456 | -0.391 | -0.294 | -0.932 | -0.077 | 0.011 | 0.011 | 0 | 0.37 | 0.12 | 0.45 | 0.37 | 0.47 | 0.32 |
| 11 | 87233867 | rs2375041 | 0.148 | -0.576 | 0.222 | -1.104 | -0.009 | -0.089 | -0.089 | 0.17 | 0.25 | 0.35 | 0.18 | 0.02 | 0.21 | 0.13 |
| 11 | 87248455 | rs7931829 | -1.412 | -0.666 | 0.023 | -0.954 | -0.034 | -0.022 | -0.022 | 0.373 | 0.64 | 0.87 | 0.62 | 0.39 | 0.64 | 0.57 |
| 11 | 87263827 | rs4356257 | -1.685 | -0.403 | 0.507 | -0.495 | -0.039 | -0.091 | -0.091 | 0 | 0.73 | 0.88 | 0.65 | 0.47 | 0.69 | 0.64 |
| 11 | 87278074 | rs10898734 | 0.157 | -0.245 | 0.638 | -1.017 | -0.024 | 0.041 | 0.041 | 0 | 0.4 | 0.2 | 0.44 | 0.37 | 0.42 | 0.35 |
| 11 | 87287369 | rs11604906 | -0.158 | -0.256 | 0.819 | -1.061 | -0.018 | -0.108 | -0.108 | 0 | 0.26 | 0.39 | 0.18 | 0.02 | 0.21 | 0.2 |
| 11 | 87306217 | rs6592393 | 0.043 | -0.249 | 0.619 | -1.057 | -0.011 | -0.082 | -0.082 | 0 | 0.25 | 0.35 | 0.18 | 0.02 | 0.21 | 0.15 |
| 11 | 87314166 | rs2187330 | 0.325 | -0.136 | 0.532 | -1.078 | 0.001 | 0.002 | 0.002 | 0 | 0.22 | 0.17 | 0.16 | 0.01 | 0.21 | 0.15 |
| 11 | 87324491 | rs11235231 | 0.206 | -0.18 | 0.494 | -1.082 | -0.039 | 0.047 | 0.047 | 0 | 0.4 | 0.15 | 0.42 | 0.36 | 0.42 | 0.34 |
| 11 | 87336418 | rs17760393 | 0.766 | -0.228 | 0.462 | -1.504 | 0.001 | 0.001 | 0.001 | 0 | 0.09 | 0.08 | 0.07 | 0.01 | 0.08 | 0.04 |
| 11 | 87336636 | rs10898751 | -1.627 | -0.252 | 0.613 | -1.513 | 0.002 | -0.006 | -0.006 | 0 | 0.68 | 0.69 | 0.61 | 0.32 | 0.64 | 0.54 |
| 11 | 87344083 | rs4463871 | 0.267 | -0.087 | 0.051 | -1.466 | 0.001 | 0 | 0 | 0 | 0.23 | 0.19 | 0.17 | 0.01 | 0.21 | 0.15 |
| 11 | 87345252 | rs11235245 | 0.602 | -0.095 | 0.05 | -1.471 | -0.007 | 0.053 | 0.053 | 0 | 0.21 | 0.09 | 0.15 | 0.01 | 0.21 | 0.15 |
| 11 | 87345390 | rs10898759 | -0.595 | -0.099 | 0.084 | -1.464 | -0.001 | -0.038 | -0.038 | 0.702 | 0.34 | 0.37 | 0.26 | 0.02 | 0.29 | 0.19 |
| 11 | 87361488 | rs1194876 | -0.999 | -0.251 | 0.037 | -1.326 | -0.06 | 0.029 | 0.029 | 0.021 | 0.7 | 0.95 | 0.66 | 0.69 | 0.66 | 0.7 |
| 11 | 87375501 | rs17822261 | -0.315 | -0.206 | 1.11 | -0.901 | -0.002 | -0.016 | -0.016 | 0.245 | 0.1 | 0.14 | 0.09 | 0.01 | 0.08 | 0.04 |
| 11 | 87381669 | rs7128595 | -0.221 | -0.291 | 1.087 | -0.9 | -0.005 | -0.029 | -0.029 | 0.33 | 0.1 | 0.18 | 0.09 | 0.01 | 0.08 | 0.04 |
| 11 | 87382931 | rs7103129 | -0.821 | -0.161 | 0.767 | -0.71 | -0.016 | -0.125 | -0.125 | 0.143 | 0.37 | 0.54 | 0.26 | 0.02 | 0.3 | 0.25 |
| 11 | 87387118 | rs2212407 | 1.446 | -0.288 | 1.419 | 0.553 | -0.003 | -0.026 | -0.026 | 0 | 0.77 | 0.41 | 0.79 | 0.85 | 0.77 | 0.74 |
| 11 | 87389488 | rs12574746 | -0.836 | 0.002 | 1.152 | 0.748 | -0.024 | -0.021 | -0.021 | 0 | 0.12 | 0.04 | 0.1 | 0.16 | 0.17 | 0.25 |
| 11 | 87399218 | rs7122146 | 1.102 | -0.314 | 1.94 | 0.927 | 0.099 | 0.065 | 0.065 | 0.083 | 0.95 | 0.6 | 0.93 | 1 | 0.97 | 0.98 |
| 11 | 87427885 | rs11825065 | -0.778 | -0.366 | 0.585 | 1.657 | -0.006 | 0.03 | 0.03 | 0.707 | 0.21 | 0.19 | 0.29 | 0.43 | 0.18 | 0.14 |
| 11 | 87429545 | rs4943995 | 1.216 | -0.313 | 0.944 | 1.526 | -0.011 | 0.046 | 0.046 | 0.545 | 0.85 | 0.55 | 0.83 | 0.84 | 0.88 | 0.86 |
| 11 | 87433472 | rs947968 | -1.733 | -0.853 | 1.505 | 3.354 | -0.018 | 0.185 | 0.185 | 0.207 | 0.16 | 0.43 | 0.17 | 0.5 | 0.12 | 0.25 |
| 11 | 87438597 | rs4556569 | -1.829 | -0.416 | 2.187 | 3.299 | 0.108 | 0.244 | 0.244 | 1.043 | 0.06 | 0.32 | 0.08 | 0.21 | 0.1 | 0.13 |
| 11 | 87439615 | rs2186664 | -2.69 | -0.449 | 2.041 | 3.207 | 0.006 | 0.171 | 0.171 | 0.387 | 0.18 | 0.6 | 0.18 | 0.53 | 0.17 | 0.24 |
| 11 | 87447212 | rs10501644 | -0.914 | -0.686 | 1.775 | 2.414 | 0 | -0.002 | -0.002 | 0 | 0.07 | 0.08 | 0.11 | 0.09 | 0.12 | 0.08 |
| 11 | 87447966 | rs11235301 | -1.312 | -0.684 | 1.988 | 2.339 | -0.006 | -0.016 | -0.016 | 0 | 0.06 | 0.07 | 0.14 | 0.02 | 0.05 | 0.11 |
| 11 | 87448434 | rs7929297 | -2.358 | -1.256 | 2.136 | 2.042 | -0.02 | -0.058 | -0.058 | 0 | 0.25 | 0.53 | 0.36 | 0.18 | 0.26 | 0.16 |
| 11 | 87449793 | rs2226603 | -3.239 | -1.34 | 2.196 | 2.241 | 0.021 | -0.026 | -0.026 | 0 | 0.38 | 0.72 | 0.53 | 0.45 | 0.36 | 0.42 |

| | | | | | | | | | | | | | | | | |
|---|---|---|---|---|---|---|---|---|---|---|---|---|---|---|---|---|
| 11 | 87454510 | rs10898777 | -2.39 | -1.246 | 2.197 | 1.894 | -0.017 | -0.052 | -0.052 | 0 | 0.25 | 0.58 | 0.37 | 0.17 | 0.25 | 0.17 |
| 11 | 87459879 | rs4944000 | -2.67 | -1.217 | 2.155 | 2.193 | -0.006 | 0.013 | 0.013 | 0 | 0.3 | 0.67 | 0.41 | 0.46 | 0.31 | 0.25 |
| 11 | 87476742 | rs4944001 | -2.372 | -1.199 | 2.248 | 2.155 | -0.021 | -0.074 | -0.074 | 0 | 0.24 | 0.52 | 0.38 | 0.12 | 0.27 | 0.18 |
| 11 | 87483995 | rs755737 | -0.349 | -1.103 | 1.97 | 1.756 | 0 | -0.008 | -0.008 | 0 | 0.05 | 0.04 | 0.15 | 0 | 0.04 | 0.04 |
| 11 | 87489770 | rs17150774 | -0.373 | -1.104 | 1.988 | 1.755 | -0.001 | -0.016 | -0.016 | 0 | 0.05 | 0.05 | 0.16 | 0 | 0.04 | 0.04 |
| 11 | 87491925 | rs7103180 | 2.026 | -1.254 | 2.566 | 1.709 | -0.016 | -0.082 | -0.082 | 0.249 | 0.77 | 0.59 | 0.64 | 0.94 | 0.75 | 0.86 |
| 11 | 87515374 | rs7944619 | -1.631 | -1.282 | 2.771 | 1.676 | 0.009 | -0.009 | -0.009 | 0 | 0.08 | 0.27 | 0.21 | 0.02 | 0.04 | 0.07 |
| 11 | 87517292 | rs1939199 | 1.251 | -1.299 | 2.793 | 1.666 | 0 | -0.003 | -0.003 | 5.569 | 0.83 | 0.82 | 0.84 | 0.96 | 0.78 | 0.93 |
| 11 | 87521615 | rs1939206 | 2.625 | -1.342 | 2.827 | 1.631 | 0.001 | -0.008 | -0.008 | 0 | 0.69 | 0.33 | 0.59 | 0.89 | 0.71 | 0.79 |
| 11 | 87523489 | rs7927090 | 0.716 | -1.324 | 2.747 | 1.642 | -0.007 | -0.019 | -0.019 | 0 | 0.89 | 0.82 | 0.9 | 0.97 | 0.88 | 0.97 |
| 11 | 87525076 | rs12798468 | -2.43 | -1.307 | 2.803 | 1.652 | 0.003 | -0.027 | -0.027 | 0 | 0.12 | 0.28 | 0.25 | 0.06 | 0.12 | 0.09 |
| 11 | 87525160 | rs17150852 | -0.089 | -1.331 | 2.795 | 1.65 | 0.012 | -0.023 | -0.023 | 0 | 0.06 | 0.13 | 0.07 | 0.01 | 0.09 | 0.05 |
| 11 | 87525265 | rs6592438 | -0.072 | -2.263 | 2.092 | 1.4 | 0.052 | -0.112 | -0.112 | 0.233 | 0.79 | 0.9 | 0.93 | 0.64 | 0.88 | 0.89 |
| 11 | 87531398 | rs7115731 | -2.974 | -2.246 | 2.138 | 1.312 | -0.038 | -0.064 | -0.064 | 0 | 0.23 | 0.51 | 0.32 | 0.12 | 0.17 | 0.2 |
| 11 | 87533352 | rs1939211 | 2.268 | -2.33 | 2.205 | 1.356 | 0.075 | 0.005 | 0.005 | 0 | 0.56 | 0.25 | 0.5 | 0.52 | 0.69 | 0.7 |
| 11 | 87541422 | rs17150882 | -0.614 | -2.244 | 2.295 | 1.194 | 0 | -0.019 | -0.019 | 0 | 0.08 | 0.15 | 0.11 | 0.03 | 0.09 | 0.07 |
| 11 | 87548852 | rs3016267 | -2.218 | -2.267 | 2.228 | 1.236 | -0.017 | -0.014 | -0.014 | 1.521 | 0.14 | 0.22 | 0.19 | 0.09 | 0.08 | 0.11 |
| 11 | 87551827 | rs2514130 | -2.823 | -2.223 | 2.145 | 1.256 | -0.024 | -0.037 | -0.037 | 0 | 0.22 | 0.6 | 0.32 | 0.12 | 0.18 | 0.2 |
| 11 | 87564240 | rs2514133 | -1.681 | -2.218 | 2.126 | 1.258 | -0.017 | -0.042 | -0.042 | 1.797 | 0.15 | 0.34 | 0.27 | 0.06 | 0.14 | 0.12 |
| 11 | 87566389 | rs594453 | 1.896 | -2.233 | 1.931 | 1.307 | 0.032 | 0.001 | 0.001 | 0 | 0.58 | 0.29 | 0.57 | 0.53 | 0.66 | 0.73 |
| 11 | 87567487 | rs11235337 | -0.86 | -2.086 | 1.729 | 1.128 | -0.013 | -0.007 | -0.007 | 2.848 | 0.07 | 0.14 | 0.03 | 0.06 | 0.03 | 0.07 |
| 11 | 87569689 | rs2508634 | -1.851 | -2.28 | 1.706 | 1.005 | -0.029 | -0.034 | -0.034 | 0 | 0.16 | 0.24 | 0.21 | 0.08 | 0.07 | 0.13 |
| 11 | 87571991 | rs2514249 | 2.073 | -2.534 | 1.559 | 1.259 | 0.078 | 0.005 | 0.005 | 0 | 0.55 | 0.27 | 0.5 | 0.56 | 0.7 | 0.65 |
| 11 | 87572333 | rs2508483 | -2.473 | -2.473 | 1.597 | 1.349 | -0.037 | -0.04 | -0.04 | 0 | 0.25 | 0.53 | 0.41 | 0.13 | 0.19 | 0.22 |
| 11 | 87572490 | rs9666479 | -2.184 | -2.434 | 1.635 | 1.374 | -0.039 | -0.022 | -0.022 | 1.229 | 0.2 | 0.49 | 0.27 | 0.13 | 0.16 | 0.2 |
| 11 | 87573698 | rs2155172 | -0.909 | -2.512 | 1.329 | 1.347 | 0.004 | -0.015 | -0.015 | 0 | 0.1 | 0.18 | 0.23 | 0.04 | 0.13 | 0.13 |
| 11 | 87574807 | rs12226885 | 0.584 | -2.487 | 0.821 | 1.37 | 0.028 | -0.058 | -0.058 | 0.006 | 0.14 | 0.08 | 0.03 | 0.31 | 0.07 | 0.14 |
| 11 | 87580164 | rs947901 | -1.923 | -2.459 | 1.024 | 1.106 | -0.028 | -0.017 | -0.017 | 0 | 0.2 | 0.57 | 0.28 | 0.14 | 0.16 | 0.16 |
| 11 | 87581475 | rs7947163 | -2.212 | -2.474 | 1.018 | 1.117 | -0.026 | -0.027 | -0.027 | 0.276 | 0.25 | 0.57 | 0.43 | 0.14 | 0.2 | 0.21 |
| 11 | 87582760 | rs1939193 | -2.18 | -2.461 | 1.023 | 1.105 | -0.027 | -0.029 | -0.029 | 0 | 0.25 | 0.57 | 0.43 | 0.13 | 0.19 | 0.21 |
| 11 | 87593401 | rs12794475 | 0.107 | -2.783 | -0.015 | 0.91 | -0.017 | -0.001 | -0.001 | 0 | 0.13 | 0.01 | 0.07 | 0.14 | 0.12 | 0.07 |
| 11 | 87594308 | rs7128264 | -1.389 | -3.074 | 0.498 | 0.931 | -0.052 | 0.031 | 0.031 | 0 | 0.14 | 0.39 | 0.2 | 0.18 | 0.06 | 0.14 |
| 11 | 87598975 | rs607530 | 0.878 | -3.368 | -0.352 | 0.594 | 0.049 | -0.016 | -0.016 | 0 | 0.59 | 0.28 | 0.68 | 0.57 | 0.77 | 0.68 |
| 11 | 87599537 | rs610107 | -0.499 | -3.313 | -0.508 | 0.654 | 0.015 | -0.022 | -0.022 | 0 | 0.79 | 0.8 | 0.88 | 0.68 | 0.88 | 0.87 |
| 11 | 87600576 | rs7945713 | -0.216 | -3.481 | -0.758 | 0.419 | 0.04 | -0.1 | -0.1 | 0.528 | 0.25 | 0.09 | 0.31 | 0.37 | 0.18 | 0.19 |
| 11 | 87601746 | rs686818 | -1.371 | -3.383 | -0.728 | 0.282 | 0.004 | -0.003 | -0.003 | 0.021 | 0.94 | 0.94 | 0.95 | 1 | 0.96 | 1 |
| 11 | 87605130 | rs4944786 | -0.086 | -3.46 | -0.788 | 0.238 | -0.01 | 0.053 | 0.053 | 0 | 0.1 | 0.02 | 0.23 | 0.01 | 0.11 | 0.08 |
| 11 | 87606160 | rs12787681 | 0.771 | -3.545 | -0.996 | 0.157 | -0.021 | 0.006 | 0.006 | 0 | 0.13 | 0 | 0.06 | 0.12 | 0.11 | 0.09 |
| 11 | 87607932 | rs614179 | 1.647 | -3.457 | -0.508 | 0.299 | 0.011 | 0.008 | 0.008 | 0 | 0.66 | 0.31 | 0.53 | 0.87 | 0.73 | 0.73 |
| 11 | 87609384 | rs10898798 | -1.235 | -3.365 | -0.445 | 0.369 | -0.036 | -0.044 | -0.044 | 0 | 0.2 | 0.61 | 0.19 | 0.12 | 0.12 | 0.19 |
| 11 | 87609858 | rs1939169 | 0.01 | -3.394 | -0.416 | 0.356 | 0.009 | 0.023 | 0.023 | 0.447 | 0.12 | 0.02 | 0.05 | 0.09 | 0.08 | 0.09 |
| 11 | 87610338 | rs641902 | 1.451 | -3.416 | -0.484 | 0.471 | 0.12 | 0.034 | 0.034 | 0 | 0.5 | 0.24 | 0.45 | 0.51 | 0.66 | 0.61 |
| 11 | 87614742 | rs10898800 | -1.58 | -2.4 | -0.061 | 0.247 | -0.017 | -0.038 | -0.038 | 0 | 0.26 | 0.61 | 0.25 | 0.13 | 0.21 | 0.24 |
| 11 | 87620018 | rs11235369 | -1.663 | -2.457 | -0.085 | 0.218 | -0.017 | -0.038 | -0.038 | 0 | 0.27 | 0.61 | 0.25 | 0.14 | 0.21 | 0.25 |
| 11 | 87621158 | rs589321 | 0.18 | -1.437 | -0.453 | 0.356 | 0.003 | -0.005 | -0.005 | 0 | 0.91 | 0.9 | 0.86 | 0.95 | 0.9 | 0.87 |
| 11 | 87630092 | rs1784016 | 0.022 | -1.365 | -0.384 | 0.438 | 0.001 | -0.005 | -0.005 | 0.005 | 0.91 | 0.9 | 0.87 | 0.95 | 0.91 | 0.87 |
| 11 | 87631527 | rs616183 | 1.201 | -1.912 | -0.729 | -0.05 | 0.013 | 0.029 | 0.029 | 0.04 | 0.76 | 0.83 | 0.62 | 0.94 | 0.8 | 0.81 |
| 11 | 87633487 | rs677951 | 0.844 | -1.854 | -0.633 | 0.136 | 0.002 | 0 | 0 | 0 | 0.79 | 0.81 | 0.61 | 0.93 | 0.82 | 0.82 |
| 11 | 87634389 | rs1939196 | -1.326 | -1.194 | -1.183 | 0.52 | 0.007 | -0.053 | -0.053 | 0.661 | 0.35 | 0.21 | 0.38 | 0.53 | 0.3 | 0.45 |
| 11 | 87634881 | rs2155105 | -2.099 | -1.565 | -1.096 | 0.328 | 0.031 | 0.021 | 0.021 | 0 | 0.52 | 0.67 | 0.5 | 0.64 | 0.39 | 0.57 |
| 11 | 87644865 | rs10898804 | 0.774 | -1.378 | -1.258 | 0.482 | 0.024 | -0.044 | -0.044 | 0.498 | 0.08 | 0 | 0.05 | 0.14 | 0.03 | 0.07 |
| 11 | 87646317 | rs4944788 | -0.725 | -1.288 | -1.253 | 0.742 | 0.024 | 0.028 | 0.028 | 0 | 0.31 | 0.12 | 0.34 | 0.3 | 0.25 | 0.38 |
| 11 | 87649263 | rs4944792 | -2.03 | -1.304 | -1.195 | 0.629 | 0.033 | 0.024 | 0.024 | 0 | 0.52 | 0.68 | 0.52 | 0.63 | 0.42 | 0.57 |
| 11 | 87651138 | rs2000560 | -0.748 | -0.808 | -1.317 | 0.911 | -0.004 | -0.007 | -0.007 | 0 | 0.26 | 0.1 | 0.3 | 0.32 | 0.22 | 0.4 |
| 11 | 87653970 | rs7926423 | -0.149 | -0.458 | -1.144 | 0.616 | 0.004 | 0.01 | 0.01 | 0 | 0.17 | 0.11 | 0.27 | 0.15 | 0.13 | 0.37 |
| 11 | 87654207 | rs11235375 | -0.154 | -0.529 | -0.959 | 0.358 | -0.03 | 0.021 | 0.021 | 0.868 | 0.17 | 0.35 | 0.12 | 0.15 | 0.12 | 0.13 |
| 11 | 87664122 | rs2155174 | -0.917 | -0.044 | -1.486 | -0.048 | 0.001 | -0.003 | -0.003 | 0 | 0.42 | 0.53 | 0.47 | 0.51 | 0.37 | 0.59 |
| 11 | 87667815 | rs9651736 | -0.04 | 0.619 | -1.948 | -0.409 | 0.006 | 0.022 | 0.022 | 1.438 | 0.53 | 0.12 | 0.47 | 0.42 | 0.62 | 0.52 |
| 11 | 87668824 | rs4592441 | -0.271 | 1.029 | -1.279 | -1.087 | -0.034 | 0.065 | 0.065 | 0 | 0.27 | 0.14 | 0.22 | 0.03 | 0.35 | 0.25 |
| 11 | 87669065 | rs10898807 | -1.244 | 0.343 | -0.163 | -0.144 | -0.015 | -0.044 | -0.044 | 0.694 | 0.18 | 0.47 | 0.16 | 0.13 | 0.12 | 0.14 |
| 11 | 87671044 | rs12269979 | -0.928 | 0.358 | -0.058 | -0.024 | -0.016 | -0.016 | -0.016 | 0 | 0.15 | 0.45 | 0.08 | 0.12 | 0.08 | 0.11 |
| 11 | 87674035 | rs7925150 | 0.481 | 1.283 | -0.021 | -0.083 | 0.037 | -0.016 | -0.016 | 0.003 | 0.85 | 0.76 | 0.9 | 0.87 | 0.78 | 0.9 |
| 11 | 87674248 | rs17151079 | -0.256 | -0.033 | -0.53 | -0.934 | 0.032 | 0.027 | 0.027 | 0 | 0.17 | 0.07 | 0.09 | 0.01 | 0.12 | 0.1 |
| 11 | 87676739 | rs4323899 | 0.029 | -1.025 | -1.112 | -0.98 | 0.06 | 0.078 | 0.078 | 0 | 0.19 | 0.02 | 0.17 | 0.06 | 0.11 | 0.17 |
| 11 | 87677069 | rs4310643 | 0.349 | -0.927 | -0.712 | -0.515 | 0.016 | 0.209 | 0.209 | 0 | 0.15 | 0.39 | 0.22 | 0.49 | 0.16 | 0.31 |
| 11 | 87678366 | rs7926453 | 0.542 | -0.96 | -0.231 | 0.063 | 0.006 | 0.026 | 0.026 | 0 | 0.08 | 0.19 | 0.09 | 0.13 | 0.1 | 0.19 |
| 11 | 87678532 | rs4517540 | 0.654 | -0.501 | -0.091 | -0.176 | 0.015 | -0.001 | -0.001 | 0 | 0.1 | 0.11 | 0.09 | 0.11 | 0.19 | 0.09 |
| 11 | 87680856 | rs4576850 | 0.723 | -0.686 | -0.16 | -0.248 | 0.04 | 0.046 | 0.046 | 0 | 0.19 | 0.05 | 0.17 | 0.06 | 0.11 | 0.17 |
| 11 | 87682094 | rs4269950 | 0.034 | -1.032 | -0.292 | -0.619 | -0.002 | -0.022 | -0.022 | 0.846 | 0.62 | 0.56 | 0.74 | 0.72 | 0.63 | 0.66 |

| | | | | | | | | | | | | | | | | |
|---|---|---|---|---|---|---|---|---|---|---|---|---|---|---|---|---|
| 11 | 87682256 | rs4345994 | 1.025 | -0.994 | -0.297 | -0.432 | 0.022 | 0.306 | 0.306 | 0 | 0.07 | 0.26 | 0.15 | 0.37 | 0.06 | 0.13 |
| 11 | 87682771 | rs7114816 | 1.485 | -0.877 | -0.478 | -0.752 | -0.063 | 0.05 | 0.05 | 0 | 0.27 | 0.09 | 0.34 | 0.16 | 0.36 | 0.18 |
| 11 | 87686616 | rs28569162 | 0.067 | -1.069 | -0.382 | -0.805 | -0.023 | -0.025 | -0.025 | 0 | 0.28 | 0.34 | 0.18 | 0.17 | 0.18 | 0.26 |
| 11 | 87739161 | rs10831435 | 0.456 | -0.46 | -0.43 | -0.471 | -0.005 | 0.003 | 0.003 | 0 | 0.38 | 0.36 | 0.53 | 0.64 | 0.47 | 0.42 |
| 11 | 87742982 | rs1027025 | 0.941 | -0.558 | -0.213 | -0.448 | -0.003 | -0.001 | -0.001 | 1.666 | 0.05 | 0.12 | 0.02 | 0.03 | 0.03 | 0.02 |
| 11 | 87745891 | rs10831363 | -0.426 | -1.283 | -0.423 | -0.64 | 0.02 | 0.007 | 0.007 | 0 | 0.44 | 0.64 | 0.47 | 0.59 | 0.27 | 0.49 |
| 11 | 87746457 | rs532731 | 1.494 | -0.691 | -0.631 | -0.894 | -0.086 | 0.087 | 0.087 | 0 | 0.28 | 0.1 | 0.31 | 0.13 | 0.38 | 0.19 |
| 11 | 87746795 | rs10831523 | -0.186 | -0.701 | -0.604 | -0.839 | 0.018 | 0.006 | 0.006 | 0 | 0.44 | 0.63 | 0.47 | 0.59 | 0.26 | 0.49 |
| 11 | 87747164 | rs17744610 | 1.361 | 0.342 | -1.044 | -1.148 | -0.033 | 0.042 | 0.042 | 0 | 0.13 | 0.05 | 0.08 | 0 | 0.16 | 0.1 |
| 11 | 87748003 | rs10765818 | 0.582 | -0.18 | -0.499 | -0.911 | 0.018 | 0.271 | 0.271 | 0 | 0.12 | 0.4 | 0.23 | 0.51 | 0.1 | 0.22 |
| 11 | 87748710 | rs485421 | -0.179 | -0.075 | -0.657 | -0.913 | 0.013 | -0.007 | -0.007 | 0.535 | 0.68 | 0.69 | 0.76 | 0.71 | 0.59 | 0.6 |
| 11 | 87753168 | rs671263 | -0.514 | -0.076 | -0.632 | -0.893 | 0.012 | -0.006 | -0.006 | 0.219 | 0.73 | 0.74 | 0.78 | 0.74 | 0.65 | 0.67 |
| 11 | 87772681 | rs655649 | 0.74 | -0.257 | -0.787 | -0.554 | -0.043 | 0.04 | 0.04 | 0 | 0.54 | 0.18 | 0.52 | 0.38 | 0.53 | 0.5 |
| 11 | 87773948 | rs10830753 | -0.721 | -0.323 | -0.698 | -0.474 | -0.051 | 0.001 | 0.001 | 0 | 0.75 | 0.91 | 0.8 | 0.74 | 0.7 | 0.74 |
| 11 | 87775711 | rs608754 | 0.255 | -0.309 | -0.811 | -0.504 | -0.018 | -0.005 | -0.005 | 0.826 | 0.6 | 0.26 | 0.55 | 0.43 | 0.58 | 0.54 |
| 11 | 87783280 | rs7945438 | -1.819 | -0.174 | -1.006 | -0.53 | -0.059 | 0.014 | 0.014 | 0.169 | 0.91 | 0.99 | 0.94 | 0.95 | 0.82 | 0.9 |
| 11 | 87784344 | rs534692 | -0.022 | -0.42 | -0.568 | -0.411 | -0.001 | -0.002 | -0.002 | 0.044 | 0.63 | 0.38 | 0.59 | 0.44 | 0.65 | 0.64 |
| 11 | 87791487 | rs988975 | -0.177 | -0.424 | -0.473 | -0.413 | 0.005 | 0.016 | 0.016 | 0 | 0.64 | 0.47 | 0.6 | 0.45 | 0.65 | 0.64 |
| 11 | 87793215 | rs1386329 | 2.098 | -1.062 | -0.908 | -0.426 | 0.01 | 0.04 | 0.04 | 0 | 0.16 | 0.01 | 0.19 | 0.08 | 0.12 | 0.13 |
| 11 | 87798564 | rs659026 | 0.558 | -0.815 | -0.966 | -0.561 | -0.006 | 0.056 | 0.056 | 0 | 0.51 | 0.2 | 0.52 | 0.4 | 0.53 | 0.58 |
| 11 | 87800457 | rs2046555 | -1.283 | -0.746 | -0.781 | -0.452 | -0.021 | 0.005 | 0.005 | 0 | 0.76 | 0.85 | 0.81 | 0.76 | 0.71 | 0.76 |
| 11 | 87805266 | rs548838 | -0.868 | -0.775 | -0.786 | -0.472 | -0.009 | 0.044 | 0.044 | 0 | 0.59 | 0.78 | 0.61 | 0.68 | 0.58 | 0.63 |
| 11 | 87808350 | rs7949449 | 1.145 | -0.702 | -0.818 | -0.335 | -0.032 | 0.243 | 0.243 | 0 | 0.11 | 0.4 | 0.21 | 0.3 | 0.06 | 0.12 |
| 11 | 87812463 | rs7947197 | -1.182 | -0.347 | -1.227 | -0.407 | -0.041 | 0.005 | 0.005 | 0 | 0.76 | 0.9 | 0.81 | 0.76 | 0.71 | 0.76 |
| 11 | 87814559 | rs473165 | 2.085 | -0.302 | -1.271 | -0.3 | -0.081 | 0.042 | 0.042 | 0 | 0.21 | 0.05 | 0.27 | 0.13 | 0.28 | 0.1 |
| 11 | 87817583 | rs7936068 | 0.461 | -0.874 | -0.673 | 0.143 | -0.009 | 0.119 | 0.119 | 0.447 | 0.16 | 0.39 | 0.23 | 0.34 | 0.13 | 0.11 |
| 11 | 87819271 | rs2220304 | 0.444 | -0.964 | -0.688 | -0.099 | -0.007 | 0.021 | 0.021 | 0.106 | 0.43 | 0.33 | 0.38 | 0.32 | 0.47 | 0.52 |
| 11 | 87820858 | rs10082691 | -0.132 | -0.485 | -0.076 | 0.586 | -0.015 | 0.181 | 0.181 | 0.87 | 0.17 | 0.48 | 0.24 | 0.35 | 0.13 | 0.12 |
| 11 | 87821219 | rs7117666 | 0.533 | -0.486 | -0.128 | 0.513 | 0.006 | -0.002 | -0.002 | 0.269 | 0.61 | 0.44 | 0.59 | 0.39 | 0.6 | 0.65 |
| 11 | 87824321 | rs10734176 | -0.923 | 0.054 | -0.756 | -0.102 | -0.029 | -0.004 | -0.004 | 0 | 0.78 | 0.89 | 0.82 | 0.75 | 0.73 | 0.77 |
| 11 | 87827395 | rs552671 | -0.907 | -0.543 | -0.184 | 0.384 | -0.039 | 0.035 | 0.035 | 0 | 0.3 | 0.57 | 0.3 | 0.51 | 0.21 | 0.32 |
| 11 | 87829489 | rs605730 | -0.717 | -0.404 | 0.187 | 0.544 | -0.042 | 0.219 | 0.219 | 0 | 0.2 | 0.53 | 0.25 | 0.43 | 0.15 | 0.31 |
| 11 | 87829785 | rs12274561 | 0.83 | -0.002 | -0.553 | -0.086 | -0.052 | 0.066 | 0.066 | 0.006 | 0.21 | 0.05 | 0.12 | 0.15 | 0.25 | 0.12 |
| 11 | 87831251 | rs561582 | -0.925 | -0.342 | -0.316 | -0.311 | 0.007 | -0.012 | -0.012 | 1.858 | 0.27 | 0.74 | 0.29 | 0.44 | 0.19 | 0.39 |
| 11 | 87831733 | rs923241 | -0.588 | -0.458 | 0.072 | -0.315 | -0.055 | -0.085 | -0.085 | 0 | 0.2 | 0.59 | 0.11 | 0.05 | 0.12 | 0.19 |
| 11 | 87835164 | rs1981403 | 0.732 | -0.655 | -0.076 | -0.995 | -0.009 | -0.031 | -0.031 | 0 | 0.38 | 0.4 | 0.33 | 0.13 | 0.25 | 0.24 |
| 11 | 87836265 | rs575202 | 0.65 | -0.622 | -0.018 | -0.904 | 0.013 | 0.276 | 0.276 | 0 | 0.07 | 0.22 | 0.18 | 0.39 | 0.07 | 0.2 |
| 11 | 87836784 | rs490589 | -1.484 | -0.238 | -0.149 | -0.975 | -0.071 | -0.045 | -0.045 | 0.075 | 0.74 | 0.91 | 0.68 | 0.69 | 0.63 | 0.74 |
| 11 | 87846596 | rs494680 | 0.797 | -0.279 | -0.04 | -0.925 | 0.002 | 0.185 | 0.185 | 0 | 0.07 | 0.15 | 0.18 | 0.39 | 0.07 | 0.2 |
| 11 | 87846757 | rs9144 | -1.357 | -0.26 | -0.028 | -0.941 | -0.077 | -0.035 | -0.035 | 0.023 | 0.73 | 0.9 | 0.67 | 0.69 | 0.61 | 0.64 |
| 11 | 87846803 | rs675578 | -0.761 | -0.469 | 0.196 | -0.791 | 0.004 | 0.341 | 0.341 | 0.066 | 0.14 | 0.44 | 0.22 | 0.4 | 0.11 | 0.28 |
| 11 | 87846885 | rs16912884 | 0.596 | -0.461 | 0.198 | -0.807 | 0.012 | -0.018 | -0.018 | 0 | 0.07 | 0.23 | 0.05 | 0 | 0.04 | 0.08 |
| 11 | 87847018 | rs3740925 | -2.027 | -0.059 | -0.138 | -0.74 | -0.033 | -0.039 | -0.039 | 0.468 | 0.93 | 1 | 0.97 | 0.85 | 0.91 | 0.88 |
| 11 | 87847063 | rs1027027 | 0.643 | -0.349 | -0.013 | -0.784 | -0.009 | -0.024 | -0.024 | 0.041 | 0.38 | 0.4 | 0.33 | 0.14 | 0.25 | 0.25 |
| 11 | 87849675 | rs10830213 | -0.791 | -0.298 | 0.122 | -0.769 | -0.028 | 0.071 | 0.071 | 0 | 0.26 | 0.66 | 0.29 | 0.43 | 0.2 | 0.39 |
| 11 | 87850866 | rs11018489 | 0.9 | -0.217 | -0.162 | -0.636 | 0.001 | 0.141 | 0.141 | 0 | 0.07 | 0.1 | 0.17 | 0.39 | 0.07 | 0.2 |
| 11 | 87853145 | rs34558044 | 1.65 | -0.449 | -0.195 | -0.694 | 0.028 | 0.053 | 0.053 | 0 | 0.25 | 0.13 | 0.27 | 0.1 | 0.17 | 0.13 |
| 11 | 87853681 | rs12287589 | 1.14 | -0.005 | -0.088 | -0.302 | -0.035 | -0.044 | -0.044 | 2.49 | 0.13 | 0.28 | 0.06 | 0.04 | 0.08 | 0.11 |
| 11 | 87853917 | rs11018514 | -0.201 | -0.015 | -0.102 | -0.276 | -0.03 | 0.147 | 0.147 | 0 | 0.81 | 0.62 | 0.77 | 0.57 | 0.85 | 0.69 |
| 11 | 87856788 | rs1872738 | -0.787 | 0.389 | -0.329 | -0.053 | 0 | 0.175 | 0.175 | 0 | 0.93 | 0.89 | 0.82 | 0.61 | 0.93 | 0.81 |
| 11 | 87857804 | rs11018529 | -0.105 | 0.171 | -0.396 | -0.496 | 0.026 | 0.015 | 0.015 | 0 | 0.48 | 0.34 | 0.46 | 0.46 | 0.63 | 0.48 |
| 11 | 87860095 | rs11018537 | 1.185 | 0.624 | -0.487 | -0.36 | -0.015 | 0.004 | 0.004 | 0 | 0.21 | 0.1 | 0.12 | 0.24 | 0.25 | 0.11 |
| 11 | 87865067 | rs6483009 | -0.518 | 0.851 | 0.088 | -0.126 | 0.011 | 0.188 | 0.188 | 0 | 0.9 | 0.81 | 0.78 | 0.6 | 0.89 | 0.76 |
| 11 | 87869579 | rs7395213 | 0.911 | 1.006 | -0.233 | -0.399 | -0.041 | 0.015 | 0.015 | 0 | 0.3 | 0.1 | 0.27 | 0.32 | 0.35 | 0.22 |
| 11 | 87871288 | rs1386326 | -0.677 | 0.988 | -0.043 | -0.389 | 0.011 | 0.181 | 0.181 | 0 | 0.9 | 0.86 | 0.78 | 0.6 | 0.89 | 0.76 |
| 11 | 87873232 | rs1486872 | -0.257 | 1.151 | 0.018 | -0.424 | -0.011 | 0.017 | 0.017 | 0 | 0.59 | 0.68 | 0.5 | 0.28 | 0.53 | 0.45 |
| 11 | 87876812 | rs302669 | 0.74 | 1.069 | 0.003 | -0.425 | -0.025 | 0.018 | 0.018 | 0 | 0.31 | 0.17 | 0.28 | 0.32 | 0.36 | 0.31 |
| 11 | 87879627 | rs302665 | 1.185 | 1.242 | -0.124 | -0.567 | -0.021 | 0.078 | 0.078 | 0 | 0.26 | 0.1 | 0.25 | 0.14 | 0.26 | 0.18 |
| 11 | 87880053 | rs12270843 | 0.688 | 0.742 | -0.126 | -0.883 | -0.037 | -0.087 | -0.087 | 0.407 | 0.37 | 0.5 | 0.24 | 0.11 | 0.26 | 0.25 |
| 11 | 87886025 | rs7114446 | 1.767 | 0.487 | 0.021 | -1.054 | 0 | -0.022 | -0.022 | 0 | 0.11 | 0.12 | 0.04 | 0 | 0.09 | 0.11 |
| 11 | 87887438 | rs11607857 | 1.123 | 0.632 | -0.055 | -1.006 | -0.004 | 0.004 | 0.004 | 0 | 0.23 | 0.21 | 0.26 | 0.17 | 0.26 | 0.2 |
| 11 | 87888769 | rs302657 | -1.169 | 0.653 | -0.04 | -0.926 | 0.007 | 0.142 | 0.142 | 0 | 0.91 | 0.87 | 0.79 | 0.6 | 0.89 | 0.77 |
| 11 | 87894630 | rs302653 | 0.278 | 0.607 | 0.085 | -0.99 | -0.004 | 0.017 | 0.017 | 0 | 0.43 | 0.4 | 0.35 | 0.32 | 0.46 | 0.43 |
| 11 | 87894645 | rs1426929 | 1.455 | 0.568 | -0.036 | -0.985 | -0.008 | -0.019 | -0.019 | 0 | 0.25 | 0.24 | 0.17 | 0.11 | 0.17 | 0.14 |
| 11 | 87900372 | rs7950565 | 0.098 | 0.575 | -0.158 | -0.983 | -0.001 | -0.006 | -0.006 | 0 | 0.47 | 0.47 | 0.42 | 0.28 | 0.43 | 0.34 |
| 11 | 87903674 | rs617591 | -1.137 | 0.606 | -0.057 | -0.859 | 0.03 | 0.269 | 0.269 | 0 | 0.91 | 0.8 | 0.78 | 0.6 | 0.88 | 0.76 |
| 11 | 87904006 | rs17753199 | 2.034 | 0.719 | -0.17 | -1.062 | 0.002 | -0.008 | -0.008 | 0 | 0.07 | 0.04 | 0.06 | 0 | 0.07 | 0.05 |
| 11 | 87905990 | rs16913594 | 2.203 | 0.449 | -0.277 | -1.118 | 0.031 | 0.028 | 0.028 | 0 | 0.16 | 0.05 | 0.06 | 0 | 0.08 | 0.08 |
| 11 | 87908448 | rs302646 | -1.057 | 0.887 | -0.31 | -0.788 | 0.04 | 0.238 | 0.238 | 0 | 0.9 | 0.8 | 0.78 | 0.6 | 0.87 | 0.67 |

| | | | | | | | | | | | | | | | | |
|---|---|---|---|---|---|---|---|---|---|---|---|---|---|---|---|---|
| 11 | 87908563 | rs3812730 | 1.119 | 0.9 | -0.367 | -0.798 | -0.009 | 0.005 | 0.005 | 0 | 0.22 | 0.18 | 0.25 | 0.17 | 0.27 | 0.2 |
| 11 | 87908736 | rs12295107 | 1.439 | 0.804 | -0.36 | -0.862 | -0.011 | -0.05 | -0.05 | 0 | 0.12 | 0.22 | 0.07 | 0 | 0.09 | 0.11 |
| 11 | 87911282 | rs1426925 | 0.833 | 1.388 | -0.683 | -0.42 | -0.003 | 0.001 | 0.001 | 0 | 0.22 | 0.23 | 0.26 | 0.17 | 0.27 | 0.2 |
| 11 | 87913325 | rs1346264 | 0.401 | 1.715 | -0.776 | 0.449 | -0.007 | 0.012 | 0.012 | 0 | 0.59 | 0.68 | 0.49 | 0.28 | 0.54 | 0.45 |
| 11 | 87936024 | rs633687 | -0.176 | 0.969 | -1.203 | -0.449 | -0.023 | 0.07 | 0.07 | 0 | 0.75 | 0.9 | 0.76 | 0.86 | 0.74 | 0.82 |
| 11 | 87937371 | rs11821660 | -0.125 | 0.578 | -0.769 | 0.17 | -0.021 | 0.019 | 0.019 | 1.415 | 0.34 | 0.51 | 0.38 | 0.55 | 0.28 | 0.48 |
| 11 | 87937521 | rs4393315 | 0.211 | 0.913 | -0.581 | -0.331 | 0.03 | -0.07 | -0.07 | 0 | 0.11 | 0.25 | 0.09 | 0 | 0.14 | 0.03 |
| 11 | 87937976 | rs11018806 | 0.309 | 0.808 | -0.552 | -0.32 | 0.02 | -0.048 | -0.048 | 0 | 0.09 | 0.24 | 0.05 | 0 | 0.08 | 0.01 |
| 11 | 87938778 | rs11018810 | -0.054 | 0.671 | -0.625 | -0.552 | 0 | 0.001 | 0.001 | 0 | 0.17 | 0.15 | 0.14 | 0.12 | 0.15 | 0.14 |
| 11 | 87939593 | rs10765238 | 0.869 | 0.455 | -0.454 | -0.035 | 0.004 | 0.073 | 0.073 | 0 | 0.13 | 0.22 | 0.25 | 0.39 | 0.1 | 0.38 |
| 11 | 87940173 | rs579563 | -1.543 | 0.648 | -0.639 | -0.195 | -0.005 | -0.01 | -0.01 | 0 | 0.89 | 0.93 | 0.85 | 0.81 | 0.89 | 0.83 |
| 11 | 87940952 | rs10501661 | -0.427 | 0.426 | -0.421 | -0.262 | -0.01 | -0.035 | -0.035 | 0.053 | 0.2 | 0.32 | 0.15 | 0.12 | 0.16 | 0.14 |
| 11 | 87941188 | rs16913663 | -0.427 | 0.426 | -0.421 | -0.262 | -0.01 | -0.035 | -0.035 | 0.336 | 0.2 | 0.32 | 0.15 | 0.12 | 0.16 | 0.14 |
| 11 | 87943265 | rs16912326 | 1.142 | 0.433 | -0.487 | -0.262 | 0.005 | 0 | 0 | 0 | 0.11 | 0.04 | 0.11 | 0.12 | 0.08 | 0.13 |
| 11 | 87946467 | rs4753283 | 0.911 | 0.458 | -0.391 | -0.238 | -0.011 | 0 | 0 | 0 | 0.23 | 0.23 | 0.13 | 0.18 | 0.19 | 0.14 |
| 11 | 87947091 | rs11018838 | 0.401 | 0.779 | -0.443 | -0.403 | 0.017 | -0.051 | -0.051 | 0.023 | 0.09 | 0.25 | 0.05 | 0 | 0.08 | 0.01 |
| 11 | 87947536 | rs652866 | -0.034 | 0.954 | -0.668 | -0.687 | -0.006 | 0.048 | 0.048 | 2.167 | 0.27 | 0.16 | 0.21 | 0.14 | 0.24 | 0.2 |
| 11 | 87948151 | rs650353 | -0.368 | 0.954 | -0.74 | -0.501 | 0.001 | -0.04 | -0.04 | 0.165 | 0.36 | 0.41 | 0.26 | 0.14 | 0.32 | 0.22 |
| 11 | 87950582 | rs1893605 | -0.201 | 0.632 | -0.621 | -0.165 | 0.004 | 0.004 | 0.004 | 0 | 0.43 | 0.46 | 0.51 | 0.63 | 0.34 | 0.52 |
| 11 | 87965221 | rs584368 | -1.1 | 1.091 | -0.5 | -0.472 | -0.022 | 0.023 | 0.023 | 0 | 0.69 | 0.86 | 0.7 | 0.74 | 0.62 | 0.69 |
| 11 | 87965944 | rs966148 | -0.198 | 1.021 | -0.158 | -0.506 | 0 | 0 | 0 | 0 | 0.48 | 0.59 | 0.57 | 0.63 | 0.41 | 0.56 |
| 11 | 87966071 | rs966149 | -0.198 | 1.021 | -0.158 | -0.506 | 0 | 0 | 0 | 0 | 0.48 | 0.59 | 0.57 | 0.63 | 0.41 | 0.56 |
| 11 | 87967800 | rs10830474 | 0.139 | 0.756 | 0.009 | -0.151 | -0.003 | 0.042 | 0.042 | 0 | 0.22 | 0.35 | 0.39 | 0.44 | 0.16 | 0.39 |
| 11 | 87969044 | rs6483044 | -0.45 | 0.718 | 0.159 | -0.183 | 0.001 | 0.082 | 0.082 | 0 | 0.27 | 0.55 | 0.45 | 0.44 | 0.22 | 0.42 |
| 11 | 87969720 | rs217091 | 0.442 | 0.992 | 0.147 | -0.169 | 0.001 | -0.008 | -0.008 | 0.012 | 0.25 | 0.23 | 0.18 | 0.09 | 0.32 | 0.17 |
| 11 | 87969750 | rs1376662 | 1.135 | 0.903 | 0.129 | -0.187 | 0.009 | -0.003 | -0.003 | 0.011 | 0.22 | 0.16 | 0.12 | 0.19 | 0.18 | 0.13 |
| 11 | 87970373 | rs217092 | -0.491 | 0.961 | 0.153 | -0.009 | -0.01 | 0.047 | 0.047 | 0 | 0.53 | 0.78 | 0.48 | 0.59 | 0.51 | 0.54 |
| 11 | 87972213 | rs4753299 | 1.19 | 0.92 | -0.124 | -0.093 | 0.009 | -0.018 | -0.018 | 0 | 0.17 | 0.16 | 0.17 | 0.21 | 0.17 | 0.14 |
| 11 | 87972979 | rs481707 | 1.112 | 0.813 | -0.144 | 0.198 | -0.002 | -0.015 | -0.015 | 0 | 0.11 | 0.12 | 0.19 | 0.24 | 0.08 | 0.23 |
| 11 | 87973303 | rs637558 | -1.776 | 1.103 | -0.314 | -0.102 | -0.011 | 0.004 | 0.004 | 0.103 | 0.88 | 0.93 | 0.94 | 1 | 0.88 | 0.97 |
| 11 | 87973591 | rs4279997 | 0.99 | 0.841 | -0.277 | 0.185 | -0.01 | -0.015 | -0.015 | 0.954 | 0.09 | 0.03 | 0.15 | 0.13 | 0.07 | 0.12 |
| 11 | 87980319 | rs4122057 | 1.292 | 0.803 | -0.249 | 0.203 | 0.008 | -0.016 | -0.016 | 0 | 0.17 | 0.16 | 0.17 | 0.21 | 0.17 | 0.14 |
| 11 | 87980399 | rs633716 | -0.795 | 0.79 | -0.383 | -0.074 | -0.001 | -0.005 | -0.005 | 0 | 0.51 | 0.57 | 0.69 | 0.71 | 0.46 | 0.64 |
| 11 | 87980466 | rs520759 | -1.086 | 0.783 | -0.437 | -0.074 | -0.001 | 0.011 | 0.011 | 0 | 0.54 | 0.65 | 0.74 | 0.71 | 0.52 | 0.66 |
| 11 | 87981279 | rs16913724 | 0.403 | 0.866 | -0.831 | 0.063 | -0.033 | -0.03 | -0.03 | 0 | 0.15 | 0.02 | 0.27 | 0.18 | 0.13 | 0.18 |
| 11 | 87989573 | rs7940642 | -0.568 | 0.809 | -0.661 | -0.049 | -0.005 | -0.003 | -0.003 | 0.062 | 0.46 | 0.38 | 0.54 | 0.47 | 0.49 | 0.44 |
| 11 | 88000019 | rs7948930 | 1.41 | 0.834 | -0.72 | 0.232 | 0.009 | 0.008 | 0.008 | 0 | 0.06 | 0.1 | 0.05 | 0.08 | 0.09 | 0.08 |
| 11 | 88007128 | rs640520 | -0.707 | 0.572 | -1.026 | 0.342 | 0.007 | 0.029 | 0.029 | 0 | 0.63 | 0.88 | 0.71 | 0.71 | 0.67 | 0.69 |
| 11 | 88009498 | rs623698 | -0.603 | 0.567 | -0.974 | 0.35 | -0.003 | -0.009 | -0.009 | 0 | 0.62 | 0.67 | 0.7 | 0.71 | 0.66 | 0.68 |
| 11 | 88010232 | rs7105379 | 1.066 | 0.508 | -2.029 | 0.539 | 0.02 | -0.014 | -0.014 | 0 | 0.16 | 0.05 | 0.15 | 0.16 | 0.16 | 0.16 |
| 11 | 88013263 | rs688858 | -0.468 | 0.621 | -1.597 | 0.539 | 0.004 | 0.013 | 0.013 | 0 | 0.63 | 0.8 | 0.71 | 0.71 | 0.66 | 0.69 |
| 11 | 88014239 | rs595739 | -0.108 | 1.281 | -1.431 | 0.861 | -0.018 | -0.045 | -0.045 | 0 | 0.13 | 0.05 | 0.19 | 0.24 | 0.14 | 0.13 |
| 11 | 88016576 | rs217089 | -0.053 | 1.046 | -1.063 | 0.587 | -0.023 | 0.023 | 0.023 | 0 | 0.85 | 0.94 | 0.9 | 0.93 | 0.84 | 0.87 |
| 11 | 88022988 | rs217098 | -0.925 | 0.275 | -0.348 | 0.997 | -0.026 | 0.033 | 0.033 | 0 | 0.29 | 0.43 | 0.3 | 0.41 | 0.22 | 0.46 |
| 11 | 88030785 | rs217111 | 0.437 | 0.192 | -0.069 | 0.983 | 0.006 | -0.021 | -0.021 | 0 | 0.63 | 0.51 | 0.45 | 0.31 | 0.62 | 0.59 |
| 11 | 88030931 | rs217112 | -0.202 | -0.113 | 0.26 | 0.768 | -0.059 | -0.014 | -0.014 | 0 | 0.21 | 0.51 | 0.15 | 0.2 | 0.13 | 0.36 |
| 11 | 88031232 | rs217113 | 0.163 | -0.161 | 0.359 | 0.967 | -0.004 | 0.022 | 0.022 | 0 | 0.7 | 0.82 | 0.73 | 0.8 | 0.71 | 0.83 |
| 11 | 88033661 | rs217116 | -1.147 | -0.162 | 0.453 | 0.898 | -0.008 | 0.012 | 0.012 | 0 | 0.86 | 0.93 | 0.93 | 0.99 | 0.85 | 0.96 |
| 11 | 88036589 | rs217119 | 0.549 | -0.194 | 0.53 | 0.943 | -0.017 | 0.051 | 0.051 | 0 | 0.13 | 0.34 | 0.1 | 0.2 | 0.09 | 0.34 |
| 11 | 88047189 | rs538047 | 0.955 | 0.175 | 0.356 | 0.632 | -0.006 | 0.039 | 0.039 | 0 | 0.55 | 0.3 | 0.44 | 0.32 | 0.54 | 0.34 |
| 11 | 88052243 | rs11019297 | 2.011 | 0.102 | 0.186 | 0.43 | -0.054 | 0.146 | 0.146 | 0 | 0.34 | 0.07 | 0.23 | 0.12 | 0.39 | 0.18 |
| 11 | 88053979 | rs217053 | -1.124 | 0.14 | 0.31 | 0.531 | 0.005 | -0.007 | -0.007 | 0 | 0.83 | 0.76 | 0.82 | 0.88 | 0.8 | 0.89 |
| 11 | 88059683 | rs217060 | -0.856 | 0.108 | 0.34 | 0.519 | 0.003 | -0.057 | -0.057 | 0.249 | 0.79 | 0.76 | 0.85 | 1 | 0.77 | 0.94 |
| 11 | 88060200 | rs217061 | -0.684 | 0.007 | 0.35 | 0.585 | -0.035 | 0.106 | 0.106 | 0.01 | 0.27 | 0.47 | 0.32 | 0.57 | 0.19 | 0.56 |
| 11 | 88061405 | rs580743 | 1.26 | 0.014 | 0.341 | 0.586 | -0.005 | 0.074 | 0.074 | 0 | 0.51 | 0.23 | 0.44 | 0.32 | 0.53 | 0.3 |
| 11 | 88064750 | rs217067 | -0.511 | 0.136 | 0.289 | 0.621 | 0.008 | -0.014 | -0.014 | 0 | 0.77 | 0.7 | 0.75 | 0.88 | 0.71 | 0.87 |
| 11 | 88067394 | rs217070 | 0.215 | -0.206 | 0.239 | 0.946 | -0.009 | 0.016 | 0.016 | 0 | 0.16 | 0.34 | 0.11 | 0.2 | 0.12 | 0.4 |
| 11 | 88068765 | rs578254 | 0.119 | 0.008 | -0.287 | 1.069 | 0.003 | -0.03 | -0.03 | 0 | 0.16 | 0.07 | 0.19 | 0.2 | 0.14 | 0.13 |
| 11 | 88069313 | rs11019394 | 0.369 | 0.142 | -0.127 | 1.159 | 0.002 | 0.017 | 0.017 | 0 | 0.1 | 0.06 | 0.21 | 0.37 | 0.08 | 0.17 |
| 11 | 88070914 | rs11600158 | 1.655 | 0.282 | -0.171 | 0.781 | -0.003 | 0 | 0 | 0 | 0.06 | 0.03 | 0.07 | 0 | 0.08 | 0.02 |
| 11 | 88071656 | rs3758833 | -0.724 | 0.039 | 0.354 | 1.219 | -0.022 | 0.088 | 0.088 | 0.1 | 0.25 | 0.39 | 0.31 | 0.47 | 0.2 | 0.59 |
| 11 | 88077479 | rs556895 | 1.047 | 0.353 | 0.015 | 0.899 | 0.008 | 0.042 | 0.042 | 0 | 0.5 | 0.39 | 0.45 | 0.31 | 0.49 | 0.25 |
| 11 | 88087357 | rs7128352 | 1.338 | 0.2 | -0.105 | 0.852 | -0.031 | 0.027 | 0.027 | 0 | 0.42 | 0.19 | 0.39 | 0.33 | 0.49 | 0.23 |
| 11 | 88089191 | rs10734151 | -1.493 | 0.2 | -0.103 | 0.805 | -0.024 | 0.029 | 0.029 | 0 | 0.84 | 0.91 | 0.91 | 1 | 0.82 | 0.94 |
| 11 | 88090476 | rs16913844 | 0.326 | -0.084 | -0.042 | 1.048 | -0.006 | -0.047 | -0.047 | 2.596 | 0.17 | 0.04 | 0.2 | 0.19 | 0.14 | 0.13 |
| 11 | 88093710 | rs4753359 | -1.204 | -0.083 | 0.101 | 1.02 | 0.005 | 0.003 | 0.003 | 0.016 | 0.41 | 0.6 | 0.51 | 0.66 | 0.32 | 0.71 |
| 11 | 88101049 | rs930722 | 0.637 | -0.134 | -0.107 | 1.388 | -0.013 | -0.084 | -0.084 | 0 | 0.14 | 0.02 | 0.18 | 0.32 | 0.13 | 0.26 |
| 11 | 88103181 | rs12807746 | -0.486 | -0.153 | 0.188 | 0.982 | -0.035 | 0.128 | 0.128 | 0 | 0.24 | 0.47 | 0.3 | 0.48 | 0.18 | 0.57 |
| 11 | 88106243 | rs7931982 | 0.123 | -0.074 | -0.268 | 0.912 | 0.007 | -0.021 | -0.021 | 0 | 0.17 | 0.09 | 0.2 | 0.18 | 0.14 | 0.13 |

| | | | | | | | | | | | | | | | | |
|---|---|---|---|---|---|---|---|---|---|---|---|---|---|---|---|---|
| 11 | 88114949 | rs10830770 | -0.287 | -0.207 | 0.23 | 1.056 | -0.009 | 0.028 | 0.028 | 0 | 0.23 | 0.23 | 0.3 | 0.52 | 0.18 | 0.58 |
| 11 | 88117923 | rs7111415 | 1.193 | 0.303 | -0.197 | 0.778 | -0.028 | 0.002 | 0.002 | 0 | 0.34 | 0.14 | 0.34 | 0.3 | 0.39 | 0.22 |
| 11 | 88123041 | rs1031375 | -0.804 | 0.283 | -0.074 | 0.79 | 0.024 | 0.043 | 0.043 | 0.061 | 0.48 | 0.61 | 0.55 | 0.7 | 0.41 | 0.72 |
| 11 | 88133201 | rs2513421 | -0.736 | 0.5 | -0.363 | 1.182 | -0.048 | -0.018 | -0.018 | 0 | 0.82 | 0.96 | 0.87 | 0.82 | 0.8 | 0.75 |
| 11 | 88135392 | rs2452153 | -0.516 | 0.674 | 0.255 | 1.222 | 0.04 | 0.167 | 0.167 | 0 | 0.07 | 0.3 | 0.13 | 0.2 | 0.08 | 0.11 |
| 11 | 88137113 | rs7939234 | 0.207 | 0.52 | -0.15 | 0.972 | 0.004 | 0.028 | 0.028 | 0 | 0.26 | 0.15 | 0.25 | 0.17 | 0.25 | 0.17 |
| 11 | 88139983 | rs2457684 | 0.503 | 0.411 | 0.533 | 0.785 | 0.006 | -0.014 | -0.014 | 0 | 0.59 | 0.47 | 0.45 | 0.25 | 0.52 | 0.3 |
| 11 | 88142353 | rs16924538 | 0.188 | 0.42 | 0.155 | 1.12 | -0.001 | -0.002 | -0.002 | 0.377 | 0.18 | 0.15 | 0.26 | 0.34 | 0.2 | 0.36 |
| 11 | 88147078 | rs11603556 | 0.886 | 0.333 | -0.053 | 1.116 | -0.017 | -0.021 | -0.021 | 0 | 0.21 | 0.01 | 0.14 | 0.17 | 0.17 | 0.13 |
| 11 | 88152876 | rs308828 | -1.142 | 0.418 | -0.102 | 1.227 | -0.05 | -0.051 | -0.051 | 1.084 | 0.82 | 0.96 | 0.84 | 0.77 | 0.79 | 0.74 |
| 11 | 88154619 | rs12223078 | 0.233 | 0.385 | -0.017 | 1.209 | -0.003 | -0.011 | -0.011 | 0.034 | 0.18 | 0.13 | 0.26 | 0.34 | 0.2 | 0.36 |
| 11 | 88157377 | rs11019859 | 1.589 | 0.326 | 0.045 | 1.108 | -0.013 | 0.035 | 0.035 | 0 | 0.35 | 0.2 | 0.36 | 0.26 | 0.37 | 0.22 |
| 11 | 88158865 | rs308757 | 0.477 | 0.298 | 0.157 | 1.133 | 0.004 | 0.003 | 0.003 | 0 | 0.6 | 0.58 | 0.55 | 0.44 | 0.57 | 0.37 |
| 11 | 88161676 | rs308754 | -0.33 | 0.3 | -0.823 | 0.901 | -0.03 | -0.047 | -0.047 | 0.651 | 0.6 | 0.87 | 0.67 | 0.6 | 0.6 | 0.6 |
| 11 | 88161996 | rs7935502 | 0.896 | 0.592 | -0.622 | 0.24 | -0.012 | 0.101 | 0.101 | 0.005 | 0.31 | 0.11 | 0.26 | 0.08 | 0.32 | 0.14 |
| 11 | 88174477 | rs308857 | -0.358 | 0.629 | -0.627 | 0.249 | 0.002 | 0.001 | 0.001 | 0 | 0.4 | 0.52 | 0.46 | 0.62 | 0.4 | 0.62 |
| 11 | 88180617 | rs308862 | -1.178 | 0.475 | -0.768 | -0.466 | -0.013 | 0.022 | 0.022 | 0 | 0.92 | 0.98 | 0.97 | 1 | 0.93 | 0.98 |
| 11 | 88188693 | rs308870 | 0.16 | 0.575 | -0.279 | -0.287 | 0.003 | 0.004 | 0.004 | 0 | 0.41 | 0.53 | 0.55 | 0.51 | 0.5 | 0.5 |
| 11 | 88198930 | rs308837 | -0.103 | 0.382 | -0.303 | -0.394 | -0.004 | -0.004 | -0.004 | 0 | 0.45 | 0.59 | 0.57 | 0.51 | 0.52 | 0.55 |
| 11 | 88211600 | rs11020044 | 0.375 | 0.219 | 0.166 | -0.414 | -0.031 | -0.015 | -0.015 | 0.088 | 0.38 | 0.22 | 0.46 | 0.46 | 0.5 | 0.49 |
| 11 | 88214432 | rs308776 | -1.2 | 0.145 | 0.433 | -0.209 | -0.013 | 0.177 | 0.177 | 0.479 | 0.11 | 0.4 | 0.13 | 0.24 | 0.05 | 0.27 |
| 11 | 88217396 | rs308764 | -0.965 | -0.127 | 0.173 | -0.703 | 0.012 | 0.015 | 0.015 | 0 | 0.62 | 0.72 | 0.69 | 0.7 | 0.71 | 0.78 |
| 11 | 88221957 | rs983689 | 0.844 | 0.053 | -0.082 | -0.621 | -0.131 | -0.098 | -0.098 | 0 | 0.37 | 0.05 | 0.45 | 0.46 | 0.51 | 0.48 |
| 11 | 88227413 | rs3850940 | 1.04 | 0.208 | 0.48 | -0.576 | 0.001 | -0.054 | -0.054 | 1.147 | 0.14 | 0.21 | 0.11 | 0.01 | 0.15 | 0.04 |
| 11 | 88239258 | rs3812729 | 0.602 | 0.261 | 0.449 | -0.508 | 0.001 | 0.007 | 0.007 | 0 | 0.16 | 0.21 | 0.12 | 0.2 | 0.16 | 0.24 |
| 11 | 88244324 | rs308884 | -0.096 | 0.231 | 0.49 | -0.511 | -0.001 | 0.027 | 0.027 | 0 | 0.2 | 0.32 | 0.2 | 0.24 | 0.17 | 0.28 |
| 11 | 88252273 | rs16914129 | 1.242 | 0.267 | 0.536 | -1.02 | 0.015 | -0.071 | -0.071 | 0 | 0.13 | 0.24 | 0.11 | 0.01 | 0.14 | 0.04 |
| 11 | 88252588 | rs308880 | -0.883 | 0.106 | 0.274 | -1.207 | 0.021 | 0.041 | 0.041 | 0 | 0.59 | 0.72 | 0.69 | 0.73 | 0.68 | 0.74 |
| 11 | 88256603 | rs1024117 | 0.358 | 0.097 | 0.193 | -1.198 | 0.001 | -0.003 | -0.003 | 0 | 0.42 | 0.43 | 0.49 | 0.69 | 0.52 | 0.66 |
| 11 | 88261186 | rs308899 | -0.62 | 0.034 | -0.268 | -1.234 | 0.029 | 0.056 | 0.056 | 0 | 0.58 | 0.82 | 0.7 | 0.73 | 0.68 | 0.74 |
| 11 | 88263873 | rs6483229 | -0.362 | 0.223 | -0.368 | -1.219 | -0.004 | -0.005 | -0.005 | 0.004 | 0.54 | 0.51 | 0.59 | 0.69 | 0.67 | 0.7 |
| 11 | 88264454 | rs10831015 | 0.976 | 0.628 | -1.137 | -1.21 | -0.02 | 0.035 | 0.035 | 0 | 0.22 | 0.09 | 0.2 | 0.01 | 0.29 | 0.12 |
| 11 | 88265307 | rs10501667 | 0.504 | 0.871 | -1.294 | -0.637 | -0.064 | -0.136 | -0.136 | 0 | 0.3 | 0.04 | 0.39 | 0.59 | 0.38 | 0.45 |
| 11 | 88266290 | rs11020240 | 0.862 | 0.888 | -1.191 | -0.754 | -0.018 | -0.039 | -0.039 | 0 | 0.24 | 0.13 | 0.37 | 0.66 | 0.32 | 0.62 |
| 11 | 88269152 | rs12794411 | 0.135 | 1.123 | -1.478 | -0.211 | -0.048 | 0.017 | 0.017 | 0 | 0.2 | 0.01 | 0.13 | 0.12 | 0.2 | 0.14 |
| 11 | 88270753 | rs905645 | 1.801 | 1.136 | -1.106 | -0.183 | -0.047 | 0.036 | 0.036 | 0 | 0.08 | 0.01 | 0.11 | 0.01 | 0.15 | 0.04 |
| 11 | 88272183 | rs308892 | -0.112 | 1.022 | -1.172 | -0.289 | -0.033 | 0.079 | 0.079 | 0 | 0.68 | 0.9 | 0.78 | 0.81 | 0.69 | 0.79 |
| 11 | 88275316 | rs1827603 | -0.196 | 0.862 | 0.2 | 0.485 | 0.028 | 0.027 | 0.027 | 0.003 | 0.14 | 0.29 | 0.21 | 0.13 | 0.18 | 0.29 |
| 11 | 88283890 | rs308793 | -0.536 | 0.56 | 0.179 | 0.593 | 0 | 0 | 0 | 0 | 0.62 | 0.58 | 0.58 | 0.38 | 0.58 | 0.4 |
| 11 | 88285704 | rs308787 | -0.866 | 0.288 | 1.034 | 0.429 | -0.011 | -0.061 | -0.061 | 0 | 0.18 | 0.37 | 0.08 | 0.05 | 0.13 | 0.14 |
| 11 | 88289360 | rs308763 | -1.068 | 0.323 | 1.188 | 0.404 | -0.01 | -0.056 | -0.056 | 0 | 0.19 | 0.66 | 0.11 | 0.05 | 0.13 | 0.13 |
| 11 | 88299284 | rs10501670 | 0.055 | 0.384 | 0.821 | 0.286 | -0.003 | 0.002 | 0.002 | 0 | 0.2 | 0.18 | 0.17 | 0.09 | 0.24 | 0.07 |
| 11 | 88302442 | rs308875 | -1.742 | 0.258 | 0.458 | 0.934 | -0.081 | -0.103 | -0.103 | 0.012 | 0.43 | 0.92 | 0.48 | 0.37 | 0.38 | 0.39 |
| 11 | 88319616 | rs11020526 | -0.548 | 0.165 | 1.26 | 0.771 | 0.024 | -0.016 | -0.016 | 0 | 0.11 | 0.43 | 0.05 | 0.03 | 0.05 | 0.02 |
| 11 | 88322619 | rs12364433 | -0.387 | 0.154 | 1.26 | 0.775 | 0.015 | -0.01 | -0.01 | 0 | 0.11 | 0.42 | 0.05 | 0.03 | 0.05 | 0.02 |
| 11 | 88323540 | rs10831110 | -1.11 | 0.044 | 1.259 | 0.751 | -0.016 | -0.033 | -0.033 | 0.247 | 0.14 | 0.36 | 0.05 | 0.05 | 0.07 | 0.03 |
| 11 | 88335021 | rs3337 | -0.882 | 0.066 | 1.081 | 1.038 | 0.016 | 0.032 | 0.032 | 0 | 0.27 | 0.61 | 0.29 | 0.28 | 0.26 | 0.14 |
| 11 | 88341290 | rs2131070 | 1.434 | -0.05 | 0.387 | 1.053 | -0.047 | -0.105 | -0.105 | 0 | 0.59 | 0.09 | 0.5 | 0.64 | 0.68 | 0.77 |
| 11 | 88349719 | rs12802054 | -0.199 | 0.177 | 0.746 | 1.242 | -0.057 | -0.058 | -0.058 | 0.003 | 0.21 | 0.04 | 0.22 | 0.18 | 0.23 | 0.11 |
| 11 | 88351443 | rs1504093 | -0.389 | 0.368 | 1.189 | 1.264 | -0.001 | 0 | 0 | 0 | 0.82 | 0.81 | 0.68 | 0.83 | 0.81 | 0.88 |
| 11 | 88353802 | rs905646 | 1.981 | 0.289 | 1.188 | 1.223 | 0.011 | -0.066 | -0.066 | 0.006 | 0.78 | 0.13 | 0.66 | 0.66 | 0.88 | 0.86 |
| 11 | 88355038 | rs16914329 | -1.574 | 0.289 | 1.844 | 1.266 | -0.017 | 0.122 | 0.122 | 0.813 | 0.16 | 0.31 | 0.27 | 0.23 | 0.09 | 0.11 |
| 11 | 88355124 | rs10501673 | 0.442 | 0.332 | 1.714 | 1.173 | 0.004 | -0.013 | -0.013 | 0.177 | 0.16 | 0.18 | 0.16 | 0.09 | 0.18 | 0.07 |
| 11 | 88365298 | rs982010 | 1.93 | 0.265 | 1.861 | 1.346 | 0.004 | -0.01 | -0.01 | 0 | 0.79 | 0.2 | 0.67 | 0.65 | 0.89 | 0.86 |
| 11 | 88366308 | rs1995493 | 1.789 | 0.237 | 1.904 | 1.435 | -0.019 | 0.094 | 0.094 | 3.075 | 0.85 | 0.26 | 0.73 | 0.71 | 0.94 | 0.89 |
| 11 | 88372708 | rs17770573 | -0.056 | 0.245 | 1.863 | 1.401 | 0 | 0.001 | 0.001 | 0 | 0.06 | 0.06 | 0.06 | 0.06 | 0.06 | 0.03 |
| 11 | 88376801 | rs1354411 | 1.243 | 0.24 | 1.978 | 1.413 | -0.016 | 0.264 | 0.264 | 1.373 | 0.88 | 0.47 | 0.76 | 0.76 | 0.96 | 0.9 |
| 11 | 88378005 | rs992259 | 2.035 | 0.255 | 1.836 | 1.413 | 0.005 | -0.028 | -0.028 | 0 | 0.79 | 0.16 | 0.67 | 0.72 | 0.88 | 0.89 |
| 11 | 88379332 | rs11020772 | 2.049 | 0.254 | 1.87 | 1.428 | 0.005 | -0.029 | -0.029 | 0 | 0.79 | 0.16 | 0.67 | 0.72 | 0.88 | 0.89 |
| 11 | 88380551 | rs7396702 | 1.927 | 0.236 | 1.865 | 1.359 | 0 | -0.001 | -0.001 | 0 | 0.8 | 0.21 | 0.71 | 0.82 | 0.88 | 0.91 |
| 11 | 88386161 | rs17832888 | 0.084 | 0.256 | 1.849 | 1.366 | -0.001 | 0 | 0 | 0 | 0.06 | 0.05 | 0.06 | 0.06 | 0.06 | 0.03 |
| 11 | 88386300 | rs17770948 | -0.053 | 0.256 | 1.848 | 1.366 | -0.001 | -0.001 | -0.001 | 0 | 0.06 | 0.05 | 0.06 | 0.06 | 0.06 | 0.03 |
| 11 | 88393121 | rs1319608 | 1.982 | 0.244 | 1.894 | 1.348 | -0.003 | 0.001 | 0.001 | 2.611 | 0.8 | 0.22 | 0.71 | 0.82 | 0.89 | 0.91 |
| 11 | 88395396 | rs1391880 | 1.181 | 0.375 | 0.993 | 1.217 | -0.069 | -0.036 | -0.036 | 1.347 | 0.24 | 0.01 | 0.14 | 0.23 | 0.29 | 0.14 |
| 11 | 88396163 | rs1499041 | 0.744 | 0.546 | 1.847 | 1.366 | 0.002 | 0.037 | 0.037 | 0.082 | 0.74 | 0.43 | 0.62 | 0.74 | 0.78 | 0.86 |
| 11 | 88396177 | rs1499040 | 0.4 | 0.554 | 1.878 | 1.375 | -0.008 | -0.012 | -0.012 | 0.273 | 0.16 | 0.1 | 0.18 | 0.19 | 0.18 | 0.09 |
| 11 | 88398458 | rs10831251 | 0.896 | 0.338 | 1.852 | 1.235 | -0.002 | 0.056 | 0.056 | 1.54 | 0.9 | 0.54 | 0.8 | 0.92 | 0.97 | 0.95 |
| 11 | 88402438 | rs1846444 | -1.394 | 0.455 | 1.698 | 1.415 | 0.04 | 0.361 | 0.361 | 0.97 | 0.1 | 0.5 | 0.21 | 0.24 | 0.04 | 0.1 |
| 11 | 88407462 | rs10501676 | 0.239 | 0.784 | 1.62 | 1.352 | -0.025 | -0.023 | -0.023 | 0.134 | 0.16 | 0.1 | 0.33 | 0.25 | 0.23 | 0.11 |

| | | | | | | | | | | | | | | | | |
|---|---|---|---|---|---|---|---|---|---|---|---|---|---|---|---|---|
| 11 | 88412451 | rs6483362 | 0.881 | 0.794 | 1.824 | 1.161 | 0.078 | 0.059 | 0.059 | 0 | 0.91 | 0.66 | 0.9 | 0.9 | 0.91 | 0.96 |
| 11 | 88421882 | rs7949475 | 0.412 | 0.8 | 1.703 | 1.142 | 0.017 | -0.021 | -0.021 | 0 | 0.77 | 0.71 | 0.77 | 0.88 | 0.72 | 0.91 |
| 11 | 88431152 | rs1353802 | -1.278 | 0.676 | 1.728 | 1.238 | 0.037 | 0.372 | 0.372 | 1.237 | 0.1 | 0.51 | 0.21 | 0.25 | 0.04 | 0.11 |
| 11 | 88431384 | rs7483940 | 0.251 | 0.694 | 1.721 | 1.195 | 0.018 | -0.08 | -0.08 | 0 | 0.78 | 0.71 | 0.79 | 0.97 | 0.76 | 0.92 |
| 11 | 88435254 | rs961690 | 1.298 | 0.687 | 1.519 | 1.215 | -0.015 | -0.018 | -0.018 | 0 | 0.68 | 0.21 | 0.58 | 0.72 | 0.71 | 0.82 |
| 11 | 88439435 | rs6483387 | 1.089 | 0.675 | 1.682 | 1.267 | 0.007 | 0.283 | 0.283 | 2.121 | 0.89 | 0.4 | 0.78 | 0.75 | 0.96 | 0.9 |
| 11 | 88443301 | rs12808245 | 0.844 | 0.801 | 1.336 | 1.123 | -0.066 | 0.078 | 0.078 | 0 | 0.14 | 0.01 | 0.14 | 0.03 | 0.18 | 0.04 |
| 11 | 88448217 | rs16914662 | -0.764 | 0.718 | 1.407 | 1.068 | -0.016 | 0.017 | 0.017 | 0 | 0.23 | 0.16 | 0.21 | 0.13 | 0.27 | 0.09 |
| 11 | 88450979 | rs11021053 | -0.726 | 0.745 | 1.405 | 1.074 | -0.017 | 0.018 | 0.018 | 0 | 0.23 | 0.16 | 0.21 | 0.13 | 0.27 | 0.09 |
| 11 | 88452034 | rs1993842 | 2.19 | 0.69 | 1.765 | 1.153 | 0.1 | 0.274 | 0.274 | 1.141 | 0.93 | 0.48 | 0.76 | 0.83 | 0.95 | 0.94 |
| 11 | 88457429 | rs1499177 | 1.507 | 0.616 | 1.725 | 1.207 | -0.021 | -0.038 | -0.038 | 0.04 | 0.66 | 0.26 | 0.56 | 0.62 | 0.69 | 0.81 |
| 11 | 88458206 | rs2387405 | -0.434 | 0.627 | 1.763 | 1.219 | 0 | -0.001 | -0.001 | 0 | 0.07 | 0.05 | 0.05 | 0.07 | 0.06 | 0.05 |
| 11 | 88464840 | rs10765724 | 0.852 | 0.536 | 1.787 | 1.242 | 0.002 | 0.008 | 0.008 | 0.296 | 0.73 | 0.31 | 0.61 | 0.69 | 0.75 | 0.85 |
| 11 | 88465349 | rs904493 | 0.871 | 0.524 | 1.781 | 1.236 | 0.002 | 0.007 | 0.007 | 0.327 | 0.73 | 0.31 | 0.61 | 0.69 | 0.75 | 0.85 |
| 11 | 88469184 | rs7938768 | -0.876 | -0.206 | 1.929 | 1.261 | 0.038 | 0.179 | 0.179 | 0.23 | 0.08 | 0.35 | 0.04 | 0.11 | 0.03 | 0.04 |
| 11 | 88470163 | rs16914739 | -0.8 | -0.159 | 1.888 | 1.071 | 0.006 | -0.047 | -0.047 | 0.099 | 0.09 | 0.17 | 0.05 | 0.01 | 0.07 | 0.05 |
| 11 | 88472070 | rs11021132 | -0.262 | -0.159 | 1.57 | 0.854 | 0.028 | 0.032 | 0.032 | 0 | 0.32 | 0.47 | 0.45 | 0.34 | 0.39 | 0.16 |
| 11 | 88475777 | rs1000061 | 0.174 | -0.255 | 1.528 | 0.739 | -0.007 | -0.004 | -0.004 | 0.039 | 0.64 | 0.31 | 0.53 | 0.69 | 0.62 | 0.84 |
| 11 | 88477162 | rs6483413 | 1.012 | -0.318 | 1.483 | 0.721 | -0.058 | 0 | 0 | 0.018 | 0.56 | 0.15 | 0.48 | 0.67 | 0.55 | 0.8 |
| 11 | 88477214 | rs6483414 | -0.133 | -0.265 | 1.596 | 0.729 | 0.016 | -0.035 | -0.035 | 0 | 0.67 | 0.48 | 0.73 | 0.82 | 0.65 | 0.89 |
| 11 | 88480491 | rs1903851 | -1.025 | -0.238 | 1.618 | 0.825 | 0.037 | 0.048 | 0.048 | 0 | 0.11 | 0.25 | 0.23 | 0.15 | 0.1 | 0.1 |
| 11 | 88489081 | rs6483430 | -0.831 | -0.764 | 1.16 | 0.911 | 0.028 | -0.001 | -0.001 | 0 | 0.64 | 0.76 | 0.68 | 0.56 | 0.75 | 0.41 |
| 11 | 88497588 | rs7947200 | -0.508 | -0.768 | 1.277 | 0.913 | -0.015 | 0.011 | 0.011 | 0 | 0.61 | 0.59 | 0.47 | 0.42 | 0.72 | 0.36 |
| 11 | 88512538 | rs1499172 | 0.224 | -0.802 | 1.549 | 0.946 | 0.078 | 0.08 | 0.08 | 0 | 0.52 | 0.24 | 0.45 | 0.32 | 0.69 | 0.32 |
| 11 | 88514696 | rs16914829 | -0.725 | -0.794 | 1.554 | 0.943 | -0.004 | 0.084 | 0.084 | 1.277 | 0.09 | 0.35 | 0.03 | 0.1 | 0.03 | 0.04 |
| 11 | 88515022 | rs7119749 | 0.335 | -0.809 | 1.485 | 0.941 | 0.085 | 0.171 | 0.171 | 0 | 0.51 | 0.11 | 0.43 | 0.25 | 0.69 | 0.31 |
| 11 | 88520396 | rs10741523 | 0 | -0.826 | 1.555 | 0.963 | 0.082 | 0.018 | 0.018 | 0 | 0.54 | 0.28 | 0.63 | 0.38 | 0.72 | 0.36 |
| 11 | 88522049 | rs10741524 | -0.725 | -0.814 | 1.529 | 0.968 | -0.011 | 0.004 | 0.004 | 0 | 0.63 | 0.63 | 0.66 | 0.48 | 0.75 | 0.4 |
| 11 | 88531621 | rs10501685 | -0.214 | -0.753 | 1.683 | 1.012 | -0.009 | 0.065 | 0.065 | 0.518 | 0.08 | 0.23 | 0.02 | 0.1 | 0.03 | 0.06 |
| 11 | 88538254 | rs10831462 | -1.298 | -0.882 | 1.512 | 1.024 | 0.032 | 0.314 | 0.314 | 0.394 | 0.12 | 0.43 | 0.23 | 0.3 | 0.06 | 0.12 |
| 11 | 88543715 | rs12787863 | 1.046 | -0.88 | 1.125 | 0.874 | -0.094 | -0.022 | -0.022 | 0 | 0.28 | 0.03 | 0.2 | 0.24 | 0.37 | 0.25 |
| 11 | 88546301 | rs16914962 | -1.484 | -0.92 | 1.351 | 0.94 | 0.02 | 0.31 | 0.31 | 1.14 | 0.12 | 0.5 | 0.23 | 0.3 | 0.06 | 0.12 |
| 11 | 88557991 | rs10831496 | -0.581 | -0.929 | 1.151 | 0.878 | 0.084 | 0.162 | 0.162 | 0 | 0.49 | 0.83 | 0.57 | 0.75 | 0.32 | 0.69 |
| 11 | 88565086 | rs11021512 | -1.293 | -0.978 | 1.259 | 0.903 | 0.008 | 0.254 | 0.254 | 1.21 | 0.12 | 0.39 | 0.22 | 0.3 | 0.06 | 0.12 |
| 11 | 88566485 | rs530782 | -0.339 | -1.005 | 0.985 | 0.873 | 0.106 | 0.015 | 0.015 | 0 | 0.47 | 0.79 | 0.38 | 0.62 | 0.28 | 0.64 |
| 11 | 88569788 | rs1827115 | -0.913 | -0.945 | 1.185 | 0.918 | 0.026 | 0.219 | 0.219 | 0.49 | 0.09 | 0.42 | 0.05 | 0.17 | 0.03 | 0.08 |
| 11 | 88579631 | rs12807540 | -0.112 | -0.879 | 1.033 | 0.706 | 0.004 | 0.004 | 0.004 | 0 | 0.39 | 0.52 | 0.42 | 0.49 | 0.41 | 0.37 |
| 11 | 88588641 | rs7938157 | -0.04 | -0.882 | 1.073 | 0.709 | 0 | 0 | 0 | 0 | 0.39 | 0.39 | 0.4 | 0.49 | 0.41 | 0.37 |
| 11 | 88595236 | rs982709 | -0.039 | -0.895 | 1.067 | 0.853 | 0 | -0.001 | -0.001 | 0 | 0.39 | 0.39 | 0.45 | 0.55 | 0.41 | 0.37 |
| 11 | 88600814 | rs659197 | -1.158 | -0.907 | 0.964 | 0.814 | -0.063 | 0.09 | 0.09 | 0 | 0.75 | 0.88 | 0.75 | 0.94 | 0.66 | 0.92 |
| 11 | 88612116 | rs666229 | 0.453 | -1.138 | 0.96 | 0.882 | -0.013 | -0.005 | -0.005 | 0 | 0.36 | 0.36 | 0.29 | 0.39 | 0.25 | 0.53 |
| 11 | 88623990 | rs11021655 | -1.191 | -1.409 | 1.882 | 1.634 | 0.032 | 0.285 | 0.285 | 0.248 | 0.09 | 0.45 | 0.21 | 0.18 | 0.04 | 0.11 |
| 11 | 88629801 | rs604179 | 0.09 | -1.391 | 1.738 | 1.55 | -0.005 | 0.001 | 0.001 | 0 | 0.32 | 0.29 | 0.25 | 0.28 | 0.39 | 0.26 |
| 11 | 88634197 | rs504183 | 0.054 | -1.473 | 1.69 | 1.498 | 0.004 | -0.004 | -0.004 | 1.217 | 0.64 | 0.61 | 0.72 | 0.71 | 0.6 | 0.78 |
| 11 | 88635135 | rs598758 | 0.374 | -1.428 | 1.655 | 1.481 | 0.012 | 0.007 | 0.007 | 0 | 0.27 | 0.31 | 0.25 | 0.3 | 0.36 | 0.22 |
| 11 | 88636260 | rs646072 | -0.818 | -1.406 | 1.577 | 1.426 | -0.008 | -0.02 | -0.02 | 0 | 0.93 | 0.9 | 0.98 | 1 | 0.98 | 0.99 |
| 11 | 88638456 | rs541279 | 0.324 | -1.406 | 1.621 | 1.43 | -0.007 | -0.012 | -0.012 | 0 | 0.9 | 0.75 | 0.94 | 0.94 | 0.96 | 0.98 |
| 11 | 88647525 | rs1508673 | -0.626 | -1.423 | 1.324 | 1.512 | -0.008 | 0.14 | 0.14 | 0.388 | 0.08 | 0.22 | 0.19 | 0.18 | 0.04 | 0.11 |
| 11 | 88650737 | rs630573 | 0.743 | -1.376 | 0.619 | 1.039 | -0.01 | 0.031 | 0.031 | 0.685 | 0.55 | 0.26 | 0.52 | 0.46 | 0.56 | 0.66 |
| 11 | 88658385 | rs533163 | -0.818 | -1.37 | 0.588 | 1.018 | -0.017 | -0.027 | -0.027 | 0 | 0.93 | 0.88 | 0.98 | 1 | 0.98 | 0.99 |
| 11 | 88663652 | rs675010 | -0.261 | -1.365 | 0.747 | 1.006 | -0.004 | -0.003 | -0.003 | 0 | 0.35 | 0.65 | 0.49 | 0.54 | 0.4 | 0.32 |
| 11 | 88669352 | rs591849 | 0.366 | -1.254 | 0.242 | 0.825 | -0.001 | -0.003 | -0.003 | 0 | 0.58 | 0.35 | 0.55 | 0.52 | 0.58 | 0.67 |
| 11 | 88676854 | rs655683 | 0.278 | -1.289 | -0.342 | 0.572 | -0.086 | -0.106 | -0.106 | 0 | 0.27 | 0.04 | 0.35 | 0.33 | 0.37 | 0.2 |
| 11 | 88696662 | rs477399 | 0.959 | -1.477 | 0.865 | 0.453 | 0.031 | 0.027 | 0.027 | 0.445 | 0.93 | 0.51 | 0.97 | 0.98 | 0.99 | 0.99 |
| 11 | 88702947 | rs597303 | -1.283 | -1.482 | 1.37 | 0.451 | 0.029 | 0.037 | 0.037 | 0.321 | 0.07 | 0.48 | 0.03 | 0.02 | 0 | 0.01 |
| 11 | 88731228 | rs10765819 | 1.331 | -2.053 | 0.772 | 0.113 | -0.067 | -0.072 | -0.072 | 0.039 | 0.4 | 0.16 | 0.49 | 0.53 | 0.5 | 0.61 |
| 11 | 88740379 | rs7120151 | -1.472 | -1.86 | 0.425 | 0.094 | -0.08 | -0.097 | -0.097 | 0.988 | 0.81 | 0.99 | 0.8 | 0.73 | 0.71 | 0.91 |
| 11 | 88744425 | rs7932640 | 0.123 | -2.088 | 0.854 | 0.054 | 0 | 0 | 0 | 0 | 0.59 | 0.59 | 0.57 | 0.65 | 0.56 | 0.74 |
| 11 | 88753845 | rs7126679 | -0.029 | -2.083 | 0.038 | 0.056 | -0.048 | -0.012 | -0.012 | 1.081 | 0.62 | 0.88 | 0.6 | 0.65 | 0.55 | 0.74 |
| 11 | 88757950 | rs7125164 | -1.48 | -1.828 | -0.202 | 0.275 | -0.111 | -0.042 | -0.042 | 0.295 | 0.55 | 0.96 | 0.51 | 0.54 | 0.47 | 0.36 |
| 11 | 88758601 | rs7121922 | -1.315 | -1.604 | 0.716 | -0.288 | 0.026 | 0.051 | 0.051 | 0.428 | 0.06 | 0.25 | 0.03 | 0.07 | 0.02 | 0.01 |
| 11 | 88765779 | rs11018434 | 1.218 | -1.898 | 0.053 | -0.461 | -0.065 | -0.041 | -0.041 | 0 | 0.39 | 0.17 | 0.45 | 0.56 | 0.5 | 0.58 |
| 11 | 88775773 | rs16924543 | 1.246 | -2.007 | 0.222 | -0.542 | 0.003 | 0 | 0 | 0 | 0.07 | 0.1 | 0.16 | 0.06 | 0.08 | 0.1 |
| 11 | 88779776 | rs7931721 | 0.877 | -2.435 | -0.123 | -0.79 | -0.074 | -0.034 | -0.034 | 0 | 0.23 | 0.06 | 0.23 | 0.25 | 0.31 | 0.14 |
| 11 | 88787544 | rs10830204 | 1.775 | -2.43 | -0.104 | -0.793 | -0.068 | -0.045 | -0.045 | 0 | 0.39 | 0.17 | 0.45 | 0.56 | 0.49 | 0.6 |
| 11 | 88791259 | rs1892879 | 1.737 | -2.489 | -0.131 | -0.858 | 0.063 | 0.083 | 0.083 | 0.543 | 0.64 | 0.22 | 0.67 | 0.83 | 0.81 | 0.73 |
| 11 | 88792601 | rs4753001 | 0.922 | -2.471 | 0.601 | -0.853 | -0.018 | -0.031 | -0.031 | 0.702 | 0.74 | 0.65 | 0.75 | 0.86 | 0.83 | 0.76 |
| 11 | 88802281 | rs477424 | -1.307 | -2.307 | 0.841 | -0.754 | 0.001 | 0.001 | 0.001 | 0 | 0.48 | 0.48 | 0.37 | 0.38 | 0.43 | 0.3 |
| 11 | 88874969 | rs4121744 | 1.481 | -1.893 | 0.405 | -0.618 | -0.054 | 0.011 | 0.011 | 0.438 | 0.45 | 0.15 | 0.47 | 0.61 | 0.52 | 0.58 |

| | | | | | | | | | | | | | | | | |
|---|---|---|---|---|---|---|---|---|---|---|---|---|---|---|---|---|
| 11 | 88896609 | rs7127661 | 0.62 | -1.753 | 0.87 | -0.647 | -0.015 | 0.025 | 0.025 | 0 | 0.61 | 0.25 | 0.54 | 0.68 | 0.58 | 0.68 |
| 11 | 88900816 | rs10830236 | 0.567 | -1.797 | 0.975 | -0.674 | -0.02 | 0.001 | 0.001 | 0 | 0.23 | 0.2 | 0.23 | 0.24 | 0.32 | 0.15 |
| 11 | 88902404 | rs949538 | -0.023 | -1.797 | 0.923 | -0.667 | -0.002 | -0.016 | -0.016 | 0 | 0.86 | 0.63 | 0.78 | 0.89 | 0.91 | 0.83 |
| 11 | 88902830 | rs949537 | 0.489 | -1.711 | 0.174 | -0.597 | -0.015 | 0.024 | 0.024 | 0 | 0.61 | 0.25 | 0.53 | 0.66 | 0.58 | 0.68 |
| 11 | 88912691 | rs11820635 | -0.67 | -1.779 | 0.449 | -0.399 | -0.023 | -0.015 | -0.015 | 0 | 0.12 | 0.2 | 0.07 | 0.08 | 0.06 | 0.13 |
| 11 | 88915570 | rs7129973 | -1.489 | -1.931 | 0.268 | -0.359 | 0.014 | 0.042 | 0.042 | 0.147 | 0.43 | 0.87 | 0.38 | 0.32 | 0.4 | 0.26 |
| 11 | 88922463 | rs12419949 | -1.745 | -1.991 | 0.643 | -0.38 | -0.027 | -0.029 | -0.029 | 0 | 0.23 | 0.29 | 0.14 | 0.13 | 0.12 | 0.15 |
| 11 | 88930377 | rs11018528 | 1.019 | -1.962 | 0.873 | -0.348 | -0.004 | -0.003 | -0.003 | 0 | 0.17 | 0.21 | 0.21 | 0.2 | 0.28 | 0.1 |
| 11 | 88931636 | rs7929991 | -2.028 | -2.031 | 1.144 | -0.378 | -0.056 | -0.059 | -0.059 | 0.203 | 0.25 | 0.53 | 0.16 | 0.13 | 0.12 | 0.22 |
| 11 | 88933250 | rs574028 | -2.405 | -2.137 | 0.983 | -0.308 | -0.072 | -0.045 | -0.045 | 0 | 0.54 | 0.92 | 0.53 | 0.41 | 0.48 | 0.4 |
| 11 | 88935032 | rs12421402 | -1.808 | -2.072 | 1.143 | -0.33 | -0.043 | -0.048 | -0.048 | 0 | 0.23 | 0.38 | 0.15 | 0.13 | 0.12 | 0.17 |
| 11 | 88952347 | rs11018535 | 0.817 | -2.069 | 1.106 | -0.302 | 0.002 | -0.003 | -0.003 | 0 | 0.1 | 0.04 | 0.15 | 0.09 | 0.08 | 0.14 |
| 11 | 88957961 | rs6483006 | -1.849 | -2.082 | 1.191 | -0.304 | -0.065 | -0.073 | -0.073 | 0 | 0.24 | 0.45 | 0.15 | 0.13 | 0.12 | 0.21 |
| 11 | 88959428 | rs12803764 | -1.921 | -2.082 | 1.132 | -0.303 | -0.047 | -0.052 | -0.052 | 0 | 0.25 | 0.55 | 0.16 | 0.13 | 0.12 | 0.22 |
| 11 | 88962090 | rs11018542 | 0.785 | -1.939 | 0.745 | -0.234 | 0.001 | -0.003 | -0.003 | 0 | 0.1 | 0.04 | 0.15 | 0.09 | 0.08 | 0.14 |
| 11 | 88969774 | rs10765198 | 0.469 | -1.808 | 0.894 | -0.198 | 0.053 | 0.029 | 0.029 | 0 | 0.21 | 0.52 | 0.24 | 0.2 | 0.28 | 0.1 |
| 11 | 88976113 | rs4396293 | -1.245 | -1.753 | 0.275 | -0.174 | 0.009 | 0.02 | 0.02 | 0.023 | 0.42 | 0.73 | 0.37 | 0.32 | 0.39 | 0.32 |
| 11 | 88982718 | rs10741305 | -2.155 | -1.903 | -0.157 | 0.293 | -0.113 | -0.082 | -0.082 | 0 | 0.57 | 0.95 | 0.55 | 0.51 | 0.49 | 0.53 |
| 11 | 88984984 | rs12421746 | -1.08 | -1.382 | 0.661 | 0.468 | -0.011 | 0.002 | 0.002 | 0 | 0.09 | 0.12 | 0.04 | 0.11 | 0.03 | 0.12 |
| 11 | 88993648 | rs12420278 | -0.824 | -1.383 | 0.661 | 0.468 | -0.013 | 0.006 | 0.006 | 0 | 0.08 | 0.12 | 0.04 | 0.1 | 0.03 | 0.12 |
| 11 | 89003604 | rs2298458 | -1.026 | -1.349 | 0.799 | 0.489 | -0.013 | 0.005 | 0.005 | 0 | 0.09 | 0.13 | 0.05 | 0.12 | 0.03 | 0.12 |
| 11 | 89012085 | rs1939255 | -1.412 | -1.177 | 0.923 | 0.493 | -0.028 | 0.002 | 0.002 | 0 | 0.14 | 0.21 | 0.07 | 0.13 | 0.06 | 0.12 |
| 11 | 89020498 | rs16913104 | -1.184 | -1.171 | 0.838 | 0.496 | -0.004 | 0 | 0 | 0 | 0.13 | 0.12 | 0.06 | 0.13 | 0.06 | 0.12 |
| 11 | 89037936 | rs1806319 | -1.194 | -1.093 | 0.87 | 0.607 | -0.002 | -0.001 | -0.001 | 0 | 0.35 | 0.63 | 0.29 | 0.32 | 0.35 | 0.22 |
| 11 | 89040251 | rs10830254 | -1.167 | -1.117 | 0.871 | 0.627 | -0.002 | -0.001 | -0.001 | 0 | 0.34 | 0.65 | 0.29 | 0.33 | 0.35 | 0.22 |
| 11 | 89065017 | rs11821838 | -1.806 | -1.412 | 0.959 | 0.71 | -0.057 | 0.004 | 0.004 | 0.81 | 0.18 | 0.58 | 0.07 | 0.16 | 0.06 | 0.16 |
| 11 | 89069734 | rs7947257 | -1.212 | -1.001 | 1.093 | 0.912 | -0.01 | 0.071 | 0.071 | 0 | 0.07 | 0.21 | 0.04 | 0.12 | 0.04 | 0.08 |
| 11 | 89070497 | rs1393347 | -0.016 | -0.89 | 0.51 | 0.875 | 0.003 | -0.007 | -0.007 | 0.467 | 0.07 | 0.03 | 0.17 | 0.08 | 0.07 | 0.09 |
| 11 | 89099263 | rs7944576 | -0.353 | -0.468 | 0.404 | 0.883 | 0.001 | -0.001 | -0.001 | 5.524 | 0.08 | 0.06 | 0.17 | 0.16 | 0.07 | 0.19 |
| 11 | 89100329 | rs10830263 | -0.703 | -0.536 | 0.635 | 1.061 | -0.01 | -0.002 | -0.002 | 0 | 0.34 | 0.51 | 0.27 | 0.38 | 0.28 | 0.2 |
| 11 | 89123768 | rs317150 | -0.598 | -0.684 | 1.203 | 1.33 | -0.018 | 0.118 | 0.118 | 0 | 0.11 | 0.25 | 0.2 | 0.3 | 0.1 | 0.21 |
| 11 | 89127087 | rs317155 | -1.73 | -0.836 | 1.078 | 0.84 | 0.034 | 0.121 | 0.121 | 0.45 | 0.5 | 0.84 | 0.64 | 0.74 | 0.43 | 0.56 |
| 11 | 89143306 | rs2202150 | -0.848 | -1.085 | 1.105 | 0.685 | -0.005 | 0.006 | 0.006 | 0 | 0.41 | 0.53 | 0.44 | 0.58 | 0.35 | 0.35 |
| 11 | 89150372 | rs3862356 | -0.766 | -1.235 | 1.119 | 0.624 | -0.043 | 0.031 | 0.031 | 0 | 0.14 | 0.37 | 0.21 | 0.16 | 0.07 | 0.21 |
| 11 | 89156698 | rs9299894 | 0.108 | -1.089 | 0.962 | 0.656 | -0.011 | 0.01 | 0.01 | 0 | 0.1 | 0.14 | 0.19 | 0.16 | 0.07 | 0.2 |
| 11 | 89170515 | rs317148 | -1.827 | -1.051 | 0.838 | 0.667 | -0.016 | 0.086 | 0.086 | 0.347 | 0.54 | 0.9 | 0.65 | 0.74 | 0.43 | 0.52 |
| 11 | 89179077 | rs538102 | -1.616 | -1.228 | 0.804 | 0.729 | -0.106 | 0.023 | 0.023 | 0 | 0.6 | 0.96 | 0.7 | 0.74 | 0.52 | 0.56 |
| 11 | 89188275 | rs10501705 | 0.298 | -1.259 | 0.814 | 0.695 | -0.005 | 0.005 | 0.005 | 0 | 0.09 | 0.1 | 0.19 | 0.16 | 0.07 | 0.2 |
| 11 | 89197483 | rs11018622 | -0.278 | -1.331 | 0.524 | 0.764 | -0.03 | -0.044 | -0.044 | 1.686 | 0.41 | 0.27 | 0.45 | 0.53 | 0.44 | 0.33 |
| 11 | 89200873 | rs564699 | -1.555 | -0.938 | -0.248 | 0.98 | -0.133 | -0.045 | -0.045 | 0 | 0.63 | 0.95 | 0.71 | 0.69 | 0.52 | 0.56 |
| 11 | 89206511 | rs11018628 | -0.842 | -1.224 | 0.74 | 1.211 | -0.021 | 0.01 | 0.01 | 0 | 0.12 | 0.2 | 0.19 | 0.16 | 0.07 | 0.21 |
| 11 | 89208854 | rs10830277 | -1.401 | -1.263 | 0.836 | 1.583 | -0.046 | 0.102 | 0.102 | 0.029 | 0.19 | 0.41 | 0.39 | 0.33 | 0.13 | 0.31 |
| 11 | 89209985 | rs2164521 | -0.637 | -1.421 | 0.974 | 1.677 | -0.019 | 0.092 | 0.092 | 0 | 0.15 | 0.49 | 0.15 | 0.19 | 0.12 | 0.25 |
| 11 | 89211932 | rs12799930 | 1.14 | -2.045 | -0.268 | 1.525 | 0.083 | 0.084 | 0.084 | 0 | 0.57 | 0.13 | 0.53 | 0.33 | 0.67 | 0.49 |
| 11 | 89217496 | rs495344 | -0.492 | -2.23 | -0.251 | 1.76 | 0.005 | 0.001 | 0.001 | 0 | 0.92 | 0.95 | 0.83 | 0.79 | 0.93 | 0.91 |
| 11 | 89217708 | rs497279 | -1.189 | -2.244 | -0.204 | 1.749 | 0.026 | -0.033 | -0.033 | 0 | 0.34 | 0.79 | 0.29 | 0.44 | 0.25 | 0.4 |
| 11 | 89221812 | rs596542 | -0.312 | -1.896 | -0.049 | 1.67 | -0.002 | 0.011 | 0.011 | 0 | 0.92 | 0.93 | 0.82 | 0.8 | 0.93 | 0.91 |
| 11 | 89224718 | rs2289123 | 1.04 | -2.042 | -0.102 | 1.687 | 0.042 | -0.059 | -0.059 | 0.648 | 0.68 | 0.2 | 0.62 | 0.53 | 0.77 | 0.72 |
| 11 | 89229431 | rs553635 | 0.258 | -1.928 | 0.431 | 1.617 | -0.027 | 0.039 | 0.039 | 0 | 0.89 | 0.82 | 0.71 | 0.8 | 0.91 | 0.88 |
| 11 | 89232161 | rs1836882 | -0.149 | -2.082 | 1.086 | 1.755 | 0.009 | 0.096 | 0.096 | 0 | 0.12 | 0.25 | 0.13 | 0.3 | 0.12 | 0.13 |
| 11 | 89234151 | rs496972 | 0.969 | -2.708 | 1.003 | 1.539 | -0.041 | 0.046 | 0.046 | 0 | 0.71 | 0.43 | 0.57 | 0.54 | 0.78 | 0.61 |
| 11 | 89239875 | rs319024 | 0.065 | -2.703 | 1.008 | 1.554 | -0.002 | 0.001 | 0.001 | 0 | 0.75 | 0.73 | 0.83 | 0.73 | 0.77 | 0.64 |
| 11 | 89243260 | rs672549 | 1.077 | -2.75 | 1.253 | 1.599 | -0.007 | 0.007 | 0.007 | 0 | 0.63 | 0.41 | 0.48 | 0.52 | 0.68 | 0.54 |
| 11 | 89246945 | rs319026 | -0.114 | -2.686 | 1.063 | 1.598 | -0.029 | -0.057 | -0.057 | 0 | 0.77 | 0.95 | 0.84 | 0.73 | 0.77 | 0.64 |
| 11 | 89253097 | rs10830281 | -0.668 | -2.343 | 0.705 | 1.564 | -0.034 | 0.066 | 0.066 | 0 | 0.12 | 0.27 | 0.31 | 0.19 | 0.09 | 0.1 |
| 11 | 89253306 | rs11018640 | -0.283 | -2.312 | 0.4 | 1.557 | -0.01 | 0.01 | 0.01 | 0 | 0.11 | 0.12 | 0.28 | 0.19 | 0.09 | 0.1 |
| 11 | 89264629 | rs1944493 | 0.564 | -1.317 | -0.142 | 0.752 | -0.006 | -0.005 | -0.005 | 0.319 | 0.41 | 0.53 | 0.3 | 0.35 | 0.37 | 0.34 |
| 11 | 89265167 | rs1836883 | -0.313 | -1.473 | -0.075 | 0.836 | 0.035 | 0.017 | 0.017 | 0 | 0.52 | 0.75 | 0.6 | 0.54 | 0.46 | 0.44 |
| 11 | 89279478 | rs2917529 | 0.518 | -1.31 | 0.016 | 0.788 | 0.005 | 0.004 | 0.004 | 0.22 | 0.42 | 0.61 | 0.32 | 0.36 | 0.37 | 0.34 |
| 11 | 89281774 | rs2917538 | 0.641 | -1.295 | 0.083 | 0.773 | -0.011 | -0.008 | -0.008 | 0 | 0.4 | 0.48 | 0.29 | 0.35 | 0.37 | 0.34 |
| 11 | 89285550 | rs1836885 | 0.542 | -1.323 | 0.065 | 0.823 | -0.01 | -0.004 | -0.004 | 0 | 0.4 | 0.47 | 0.3 | 0.36 | 0.37 | 0.34 |
| 11 | 89345025 | rs1488902 | 0.44 | -1.356 | -0.485 | 0.58 | 0.019 | 0.029 | 0.029 | 0 | 0.34 | 0.54 | 0.32 | 0.46 | 0.35 | 0.28 |
| 11 | 89348133 | rs4753222 | 1.105 | -1.217 | -0.417 | 0.296 | -0.051 | 0.138 | 0.138 | 0.729 | 0.31 | 0.05 | 0.21 | 0.08 | 0.34 | 0.26 |
| 11 | 89352241 | rs3974853 | 0.758 | -1.214 | -0.298 | 0.3 | 0 | -0.054 | -0.054 | 0 | 0.33 | 0.34 | 0.25 | 0.09 | 0.34 | 0.27 |
| 11 | 89366578 | rs10830302 | 0.59 | -0.776 | -1.242 | 0.25 | -0.028 | 0.024 | 0.024 | 0.366 | 0.06 | 0.01 | 0.05 | 0.03 | 0.1 | 0.13 |
| 11 | 89374850 | rs4620721 | 0.909 | -0.827 | -1.105 | 0.296 | -0.018 | 0.07 | 0.07 | 0.577 | 0.32 | 0.15 | 0.23 | 0.09 | 0.36 | 0.27 |
| 11 | 89397819 | rs931394 | -0.506 | -0.965 | -1.296 | 0.416 | 0.021 | -0.023 | -0.023 | 0.007 | 0.75 | 0.9 | 0.86 | 0.73 | 0.79 | 0.65 |
| 11 | 89404829 | rs7112871 | 0.563 | -0.544 | -1.001 | 0.647 | -0.029 | 0.075 | 0.075 | 0.49 | 0.32 | 0.13 | 0.25 | 0.14 | 0.37 | 0.25 |
| 11 | 89405190 | rs10047486 | 0.571 | -0.545 | -1.002 | 0.645 | -0.027 | 0.077 | 0.077 | 0.505 | 0.32 | 0.13 | 0.24 | 0.13 | 0.37 | 0.24 |

| | | | | | | | | | | | | | | | | |
|---|---|---|---|---|---|---|---|---|---|---|---|---|---|---|---|---|
| 11 | 89440119 | rs2388103 | 0.539 | -0.366 | -1.1 | 0.773 | -0.042 | -0.043 | -0.043 | 0 | 0.32 | 0.12 | 0.29 | 0.33 | 0.38 | 0.25 |
| 11 | 89440231 | rs10830352 | 0.57 | -0.375 | -1.189 | 0.767 | -0.051 | -0.053 | -0.053 | 1.008 | 0.32 | 0.1 | 0.28 | 0.32 | 0.38 | 0.25 |
| 11 | 89444194 | rs11018746 | 0.557 | -0.401 | -1.135 | 0.801 | -0.039 | -0.049 | -0.049 | 0 | 0.32 | 0.12 | 0.28 | 0.33 | 0.37 | 0.25 |
| 11 | 89455444 | rs10830359 | 0.546 | -0.403 | -1.136 | 0.801 | -0.039 | -0.049 | -0.049 | 0 | 0.32 | 0.12 | 0.28 | 0.33 | 0.37 | 0.25 |
| 11 | 89466052 | rs7115903 | 0.031 | -0.994 | -1.023 | 0.328 | -0.052 | -0.022 | -0.022 | 0 | 0.24 | 0.58 | 0.18 | 0.23 | 0.15 | 0.19 |
| 11 | 89466732 | rs17221829 | 1.14 | -0.608 | -1.355 | 0.43 | -0.113 | -0.079 | -0.079 | 0 | 0.3 | 0.03 | 0.3 | 0.3 | 0.36 | 0.24 |
| 11 | 89656239 | rs10734121 | -1.135 | -0.06 | -1.025 | 0.315 | -0.014 | -0.009 | -0.009 | 0 | 0.88 | 0.93 | 0.88 | 0.82 | 0.83 | 0.89 |
| 11 | 89834751 | rs12417707 | -1.02 | 0.015 | -0.602 | 0.211 | -0.002 | -0.013 | -0.013 | 0 | 0.24 | 0.19 | 0.43 | 0.41 | 0.24 | 0.31 |
| 11 | 89843478 | rs10830403 | 0.413 | 0.047 | -0.744 | 0.282 | -0.046 | -0.074 | -0.074 | 0 | 0.11 | 0.01 | 0.36 | 0.26 | 0.15 | 0.15 |
| 11 | 89867414 | rs10765249 | -0.28 | -0.357 | 0.142 | -0.309 | -0.017 | -0.083 | -0.083 | 1.848 | 0.19 | 0.22 | 0.09 | 0.01 | 0.12 | 0.22 |
| 11 | 89867872 | rs1892887 | -2.374 | -0.383 | -0.136 | -0.302 | 0.02 | 0.002 | 0.002 | 0 | 0.45 | 0.68 | 0.54 | 0.42 | 0.36 | 0.51 |
| 11 | 89868755 | rs10734123 | -2.369 | -0.371 | -0.131 | -0.295 | 0.019 | 0.002 | 0.002 | 0 | 0.44 | 0.68 | 0.54 | 0.42 | 0.36 | 0.51 |
| 11 | 89875664 | rs10765252 | 0.764 | -0.263 | -0.25 | -0.208 | -0.016 | -0.027 | -0.027 | 0 | 0.11 | 0.05 | 0.37 | 0.29 | 0.15 | 0.15 |
| 11 | 89885366 | rs3737443 | 0.835 | -0.273 | -0.373 | -0.232 | 0.011 | -0.025 | -0.025 | 0 | 0.08 | 0 | 0.02 | 0.11 | 0.04 | 0.1 |
| 11 | 89886284 | rs4268468 | -0.367 | -0.245 | -0.273 | -0.275 | -0.005 | -0.06 | -0.06 | 0 | 0.19 | 0.08 | 0.39 | 0.4 | 0.19 | 0.26 |
| 11 | 89898803 | rs11018889 | 0.559 | -0.329 | -0.355 | -0.298 | -0.007 | 0.064 | 0.064 | 0.413 | 0.12 | 0.02 | 0.05 | 0.01 | 0.1 | 0.2 |
| 11 | 89902154 | rs10830430 | -0.035 | -0.237 | -0.262 | -0.223 | -0.021 | -0.105 | -0.105 | 0 | 0.18 | 0.3 | 0.09 | 0.01 | 0.12 | 0.2 |
| 11 | 89913032 | rs2000789 | -1.258 | -0.106 | -0.734 | 0.048 | 0.004 | -0.001 | -0.001 | 0 | 0.43 | 0.55 | 0.52 | 0.42 | 0.37 | 0.52 |
| 11 | 89916562 | rs7102933 | -1.377 | -0.132 | -0.815 | 0.036 | 0.015 | 0.003 | 0.003 | 0 | 0.44 | 0.61 | 0.53 | 0.41 | 0.36 | 0.51 |
| 11 | 89917262 | rs10765270 | -0.956 | -0.135 | -0.725 | 0.082 | 0 | -0.001 | -0.001 | 0.001 | 0.34 | 0.48 | 0.49 | 0.31 | 0.32 | 0.41 |
| 11 | 89917699 | rs647248 | -1.542 | 0.047 | -1.731 | -0.063 | -0.01 | -0.051 | -0.051 | 0 | 0.56 | 0.9 | 0.65 | 0.44 | 0.55 | 0.6 |
| 11 | 89917928 | rs2000533 | -0.661 | 0.378 | -1.298 | 0.499 | -0.025 | -0.023 | -0.023 | 0 | 0.07 | 0 | 0.21 | 0.26 | 0.1 | 0.13 |
| 11 | 89922154 | rs11018903 | -0.834 | -0.151 | -1.308 | 0.362 | 0.013 | 0.007 | 0.007 | 1.089 | 0.54 | 0.31 | 0.39 | 0.39 | 0.55 | 0.52 |
| 11 | 89928611 | rs3740818 | -0.877 | -0.346 | -1.856 | 0.115 | 0.009 | 0.038 | 0.038 | 0 | 0.49 | 0.29 | 0.52 | 0.56 | 0.43 | 0.58 |
| 11 | 89928637 | rs3740819 | -0.403 | -0.414 | -1.944 | -0.118 | 0.024 | 0.002 | 0.002 | 0 | 0.35 | 0.19 | 0.45 | 0.29 | 0.24 | 0.41 |
| 11 | 89929125 | rs16917160 | 0.86 | -0.427 | -1.941 | -0.127 | -0.006 | -0.045 | -0.045 | 0 | 0.2 | 0.18 | 0.19 | 0.01 | 0.1 | 0.19 |
| 11 | 89935908 | rs3758756 | -1.227 | -0.435 | -1.96 | -0.13 | 0.009 | 0.037 | 0.037 | 0 | 0.49 | 0.29 | 0.51 | 0.56 | 0.43 | 0.57 |
| 11 | 89939448 | rs1943381 | -2.539 | -0.427 | -1.889 | -0.132 | -0.012 | -0.011 | -0.011 | 0 | 0.7 | 0.33 | 0.65 | 0.68 | 0.7 | 0.75 |
| 11 | 89942645 | rs11018912 | 1.774 | -0.35 | -2.019 | -0.064 | -0.041 | -0.062 | -0.062 | 0 | 0.14 | 0.04 | 0.06 | 0.27 | 0.19 | 0.17 |
| 11 | 89943681 | rs12787117 | 1.683 | -0.296 | -2.058 | -0.138 | -0.066 | 0.007 | 0.007 | 0 | 0.15 | 0.01 | 0.11 | 0.1 | 0.21 | 0.14 |
| 11 | 89950035 | rs10830444 | -0.075 | -0.33 | -1.958 | -0.058 | -0.069 | -0.095 | -0.095 | 0 | 0.28 | 0.06 | 0.32 | 0.55 | 0.33 | 0.38 |
| 11 | 89953381 | rs10501714 | 1.834 | -0.297 | -1.904 | -0.098 | -0.027 | -0.05 | -0.05 | 4.202 | 0.08 | 0 | 0.2 | 0.17 | 0.11 | 0.09 |
| 11 | 89954805 | rs16917622 | 0.722 | -0.439 | -1.736 | -0.387 | -0.008 | -0.05 | -0.05 | 0 | 0.21 | 0.18 | 0.19 | 0.01 | 0.1 | 0.2 |
| 11 | 89954921 | rs3824897 | -0.223 | -0.405 | -1.738 | -0.347 | 0.002 | 0.006 | 0.006 | 0 | 0.35 | 0.24 | 0.25 | 0.28 | 0.28 | 0.37 |
| 11 | 89962734 | rs2155056 | -2.313 | -0.448 | -0.961 | -0.471 | -0.001 | -0.004 | -0.004 | 0.103 | 0.72 | 0.65 | 0.68 | 0.77 | 0.7 | 0.75 |
| 11 | 89966109 | rs7941284 | -0.18 | -0.18 | -1.117 | -0.364 | -0.011 | -0.069 | -0.069 | 0 | 0.27 | 0.25 | 0.22 | 0.02 | 0.15 | 0.24 |
| 11 | 89970603 | rs12789669 | -0.13 | 0.49 | -0.889 | 0.266 | -0.021 | 0.016 | 0.016 | 0 | 0.22 | 0.13 | 0.25 | 0.11 | 0.29 | 0.18 |
| 11 | 89975477 | rs12797899 | 0.7 | 0.593 | -1.045 | 0.299 | -0.102 | -0.007 | -0.007 | 0.123 | 0.16 | 0.02 | 0.23 | 0.11 | 0.27 | 0.18 |
| 11 | 89986667 | rs7101892 | -1.858 | -0.703 | -0.611 | 0.134 | -0.02 | -0.041 | -0.041 | 0 | 0.59 | 0.5 | 0.64 | 0.76 | 0.69 | 0.6 |
| 11 | 89989487 | rs12417453 | 0.636 | -0.561 | -0.671 | 0.538 | -0.02 | -0.108 | -0.108 | 0 | 0.14 | 0.01 | 0.27 | 0.37 | 0.15 | 0.22 |
| 11 | 89990909 | rs1962137 | -1.202 | -0.494 | -0.555 | 0.458 | -0.022 | 0.034 | 0.034 | 0.039 | 0.28 | 0.36 | 0.34 | 0.39 | 0.22 | 0.27 |
| 11 | 89991998 | rs4319477 | -1.163 | -0.558 | -0.543 | 0.469 | -0.025 | 0.035 | 0.035 | 0.068 | 0.28 | 0.36 | 0.34 | 0.39 | 0.22 | 0.27 |
| 11 | 90010643 | rs10830471 | -1.659 | -0.864 | -0.559 | 0.364 | -0.032 | -0.059 | -0.059 | 0.878 | 0.61 | 0.5 | 0.66 | 0.77 | 0.7 | 0.61 |
| 11 | 90016813 | rs11606517 | 1.426 | -0.771 | -0.856 | 0.591 | -0.017 | -0.058 | -0.058 | 0 | 0.1 | 0 | 0.21 | 0.17 | 0.12 | 0.11 |
| 11 | 90018288 | rs12364205 | 0.012 | -0.881 | -0.908 | 0.581 | -0.048 | -0.072 | -0.072 | 0 | 0.36 | 0.15 | 0.33 | 0.4 | 0.51 | 0.36 |
| 11 | 90021786 | rs10765304 | 1.283 | -0.625 | -0.757 | 0.539 | -0.017 | -0.058 | -0.058 | 1.329 | 0.1 | 0 | 0.21 | 0.17 | 0.12 | 0.11 |
| 11 | 90025424 | rs543462 | -1.177 | -0.836 | -0.408 | -0.035 | -0.026 | -0.049 | -0.049 | 0 | 0.6 | 0.48 | 0.65 | 0.77 | 0.7 | 0.61 |
| 11 | 90032662 | rs532247 | -1.3 | -0.758 | -0.695 | -0.044 | 0.004 | -0.005 | -0.005 | 0 | 0.6 | 0.39 | 0.67 | 0.79 | 0.7 | 0.61 |
| 11 | 90042539 | rs484213 | -0.788 | -0.441 | -0.459 | 0.35 | 0 | 0 | 0 | 2.022 | 0.43 | 0.46 | 0.42 | 0.37 | 0.53 | 0.31 |
| 11 | 90050153 | rs497959 | -0.35 | 0.05 | -0.542 | 0.471 | -0.035 | 0.02 | 0.02 | 0.073 | 0.34 | 0.26 | 0.21 | 0.24 | 0.42 | 0.21 |
| 11 | 90050248 | rs495395 | -1.093 | -0.043 | -0.504 | 0.504 | 0.007 | 0.016 | 0.016 | 0.239 | 0.43 | 0.35 | 0.39 | 0.31 | 0.52 | 0.3 |
| 11 | 90070103 | rs1507608 | 0.78 | 0.66 | -0.582 | 0.443 | -0.047 | -0.021 | -0.021 | 0 | 0.17 | 0.07 | 0.09 | 0.21 | 0.25 | 0.07 |
| 11 | 90074128 | rs558562 | -1.556 | 0.154 | -0.599 | 0.43 | 0.015 | 0.057 | 0.057 | 0.151 | 0.49 | 0.36 | 0.52 | 0.72 | 0.58 | 0.46 |
| 11 | 90084567 | rs571965 | 0.078 | 0.131 | -0.666 | 0.381 | -0.082 | -0.008 | -0.008 | 0 | 0.26 | 0.07 | 0.18 | 0.24 | 0.4 | 0.18 |
| 11 | 90086953 | rs10128640 | 0.793 | -0.03 | -0.246 | 0.407 | 0.019 | -0.031 | -0.031 | 0 | 0.15 | 0.08 | 0.31 | 0.47 | 0.15 | 0.25 |
| 11 | 90093461 | rs1848285 | 1.833 | 0.025 | -0.215 | 0.246 | -0.008 | -0.029 | -0.029 | 0 | 0.41 | 0.43 | 0.29 | 0.22 | 0.33 | 0.45 |
| 11 | 90094565 | rs1507605 | 0.377 | 0.067 | -0.165 | 0.425 | 0.009 | -0.004 | -0.004 | 0 | 0.64 | 0.61 | 0.77 | 0.73 | 0.56 | 0.77 |
| 11 | 90095902 | rs10830486 | 1.315 | 0.077 | -0.194 | 0.425 | -0.002 | -0.009 | -0.009 | 0 | 0.49 | 0.5 | 0.46 | 0.28 | 0.42 | 0.53 |
| 11 | 90096319 | rs571719 | -0.5 | 0.112 | -0.269 | 0.436 | -0.048 | 0.003 | 0.003 | 0.068 | 0.33 | 0.13 | 0.2 | 0.27 | 0.42 | 0.22 |
| 11 | 90097259 | rs7930337 | -0.076 | 0.098 | -0.133 | 0.6 | -0.01 | 0.037 | 0.037 | 0.055 | 0.17 | 0.23 | 0.32 | 0.44 | 0.16 | 0.24 |
| 11 | 90103902 | rs10830488 | 0.614 | 0.125 | -0.198 | 0.602 | 0.029 | -0.064 | -0.064 | 0 | 0.15 | 0.06 | 0.29 | 0.44 | 0.15 | 0.23 |
| 11 | 90117826 | rs7104915 | 0.971 | 0.721 | -0.425 | 0.611 | -0.041 | -0.007 | -0.007 | 0 | 0.17 | 0.05 | 0.11 | 0.16 | 0.24 | 0.08 |
| 11 | 90123002 | rs10830490 | -0.83 | 0.544 | -0.506 | 0.554 | 0.015 | -0.025 | -0.025 | 0 | 0.24 | 0.21 | 0.32 | 0.44 | 0.2 | 0.27 |
| 11 | 90127938 | rs561499 | 1.62 | 0.311 | -0.113 | 0.297 | 0 | 0.011 | 0.011 | 1.272 | 0.44 | 0.45 | 0.46 | 0.33 | 0.41 | 0.53 |
| 11 | 90151679 | rs564064 | 0.21 | 0.364 | -0.043 | 0.129 | 0.04 | -0.022 | -0.022 | 1.054 | 0.68 | 0.6 | 0.78 | 0.77 | 0.6 | 0.8 |
| 11 | 90151920 | rs562085 | -1.511 | 0.36 | -0.112 | 0.165 | -0.003 | -0.001 | -0.001 | 0 | 0.91 | 0.96 | 0.91 | 0.94 | 0.85 | 0.91 |
| 11 | 90158725 | rs12363417 | 0.325 | -0.198 | -0.055 | 0.092 | 0.002 | 0.021 | 0.021 | 1.045 | 0.08 | 0.03 | 0.02 | 0 | 0.05 | 0.04 |
| 11 | 90167146 | rs10128721 | -1.019 | -0.229 | -0.087 | 0.128 | 0.011 | -0.071 | -0.071 | 0 | 0.27 | 0.14 | 0.39 | 0.55 | 0.26 | 0.36 |
| 11 | 90174672 | rs568596 | 1.707 | -0.234 | -0.019 | 0.046 | 0 | -0.009 | -0.009 | 2.929 | 0.4 | 0.45 | 0.36 | 0.22 | 0.32 | 0.44 |

| | | | | | | | | | | | | | | | | |
|---|---|---|---|---|---|---|---|---|---|---|---|---|---|---|---|---|
| 11 | 90175628 | rs487727 | -0.67 | 0.032 | -0.053 | 0.19 | -0.016 | 0 | 0 | 0 | 0.73 | 0.85 | 0.8 | 0.77 | 0.61 | 0.83 |
| 11 | 90191908 | rs10437564 | -0.229 | -0.239 | 0.03 | 0.375 | 0.001 | -0.011 | -0.011 | 0 | 0.2 | 0.2 | 0.4 | 0.55 | 0.23 | 0.32 |
| 11 | 90192467 | rs542612 | 0.214 | -0.346 | 0.161 | 0.201 | 0.001 | -0.004 | -0.004 | 0 | 0.6 | 0.68 | 0.77 | 0.77 | 0.55 | 0.75 |
| 11 | 90192742 | rs11019034 | 0.804 | -0.38 | 0.062 | 0.905 | 0.008 | -0.029 | -0.029 | 0 | 0.06 | 0 | 0.06 | 0.27 | 0.05 | 0.14 |
| 11 | 90196843 | rs951415 | 0.151 | -0.376 | 0.296 | 0.857 | -0.013 | 0.043 | 0.043 | 0 | 0.17 | 0.22 | 0.33 | 0.45 | 0.15 | 0.23 |
| 11 | 90204266 | rs512285 | -0.103 | -0.199 | 0.245 | 0.665 | -0.011 | -0.006 | -0.006 | 0 | 0.68 | 0.8 | 0.74 | 0.67 | 0.54 | 0.71 |
| 11 | 90208004 | rs948987 | -0.835 | -0.487 | -0.263 | 0.325 | 0 | -0.028 | -0.028 | 0 | 0.29 | 0.23 | 0.5 | 0.55 | 0.33 | 0.3 |
| 11 | 90209402 | rs7130307 | -1.298 | -0.52 | -0.199 | 0.31 | -0.001 | 0 | 0 | 0.025 | 0.34 | 0.36 | 0.51 | 0.56 | 0.32 | 0.3 |
| 11 | 90217977 | rs17835397 | 1.072 | -0.385 | -0.338 | 0.258 | -0.064 | -0.035 | -0.035 | 0 | 0.12 | 0.01 | 0.15 | 0.12 | 0.18 | 0.09 |
| 11 | 90219142 | rs577325 | -1.073 | -0.597 | -0.301 | 0.231 | 0.001 | 0 | 0 | 0 | 0.74 | 0.79 | 0.8 | 0.76 | 0.63 | 0.8 |
| 11 | 90222406 | rs10453998 | -1.086 | -0.626 | 0.143 | 0.284 | -0.057 | 0.053 | 0.053 | 1.188 | 0.27 | 0.38 | 0.37 | 0.45 | 0.17 | 0.23 |
| 11 | 90225539 | rs1944062 | 0.777 | -0.605 | 0.114 | 0.297 | 0.017 | -0.027 | -0.027 | 0 | 0.16 | 0.13 | 0.34 | 0.44 | 0.15 | 0.2 |
| 11 | 90227460 | rs491196 | -0.015 | -0.609 | 0.122 | 0.237 | -0.007 | 0.026 | 0.026 | 0 | 0.57 | 0.69 | 0.47 | 0.33 | 0.48 | 0.6 |
| 11 | 90227952 | rs516743 | 1.436 | -0.5 | -0.065 | 0.321 | 0.002 | 0.003 | 0.003 | 0 | 0.34 | 0.42 | 0.29 | 0.21 | 0.29 | 0.47 |
| 11 | 90230604 | rs2510749 | 0.395 | -0.485 | -0.003 | 0.357 | -0.001 | 0.035 | 0.035 | 0 | 0.5 | 0.51 | 0.45 | 0.33 | 0.46 | 0.57 |
| 11 | 90238570 | rs3897659 | 0.723 | -0.499 | -0.052 | 0.427 | 0.017 | -0.026 | -0.026 | 4.976 | 0.16 | 0.13 | 0.34 | 0.44 | 0.15 | 0.2 |
| 11 | 90245313 | rs520214 | 0.375 | -0.479 | 0 | 0.322 | -0.001 | 0.011 | 0.011 | 0 | 0.5 | 0.55 | 0.45 | 0.32 | 0.46 | 0.57 |
| 11 | 90261346 | rs11019075 | 0.035 | -0.54 | 0.166 | 0.567 | -0.022 | 0.102 | 0.102 | 0 | 0.08 | 0.27 | 0.09 | 0.26 | 0.03 | 0.1 |
| 11 | 90261452 | rs6483056 | 1.774 | -0.53 | 0.091 | 0.541 | 0.004 | 0.056 | 0.056 | 0 | 0.32 | 0.24 | 0.28 | 0.21 | 0.28 | 0.47 |
| 11 | 90277017 | rs17255138 | 1.37 | -0.492 | 0.048 | 0.543 | -0.063 | -0.034 | -0.034 | 0 | 0.11 | 0.01 | 0.14 | 0.1 | 0.18 | 0.09 |
| 11 | 90281931 | rs10501721 | 0.599 | -0.558 | 0.085 | 0.583 | 0.02 | -0.034 | -0.034 | 0 | 0.16 | 0.12 | 0.34 | 0.44 | 0.15 | 0.2 |
| 11 | 90309898 | rs10741351 | -0.293 | -0.546 | 0.125 | 0.566 | 0.006 | -0.021 | -0.021 | 1.406 | 0.62 | 0.59 | 0.78 | 0.76 | 0.61 | 0.77 |
| 11 | 90310352 | rs10830545 | 1.834 | -0.475 | -0.045 | 0.63 | 0.007 | -0.058 | -0.058 | 0.057 | 0.09 | 0.01 | 0.25 | 0.18 | 0.1 | 0.1 |
| 11 | 90328540 | rs611622 | 0.826 | -0.449 | -0.053 | 0.394 | -0.041 | 0.057 | 0.057 | 0.311 | 0.44 | 0.31 | 0.43 | 0.33 | 0.47 | 0.56 |
| 11 | 90348024 | rs620197 | -0.603 | -0.35 | -0.13 | 0.48 | 0.018 | -0.013 | -0.013 | 0 | 0.66 | 0.66 | 0.79 | 0.76 | 0.61 | 0.77 |
| 11 | 90353659 | rs11019121 | -0.509 | -0.348 | -0.168 | 0.456 | -0.032 | -0.108 | -0.108 | 0 | 0.27 | 0.11 | 0.49 | 0.55 | 0.33 | 0.3 |
| 11 | 90354903 | rs509375 | 0.01 | -0.369 | -0.136 | 0.498 | -0.001 | -0.001 | -0.001 | 0.397 | 0.55 | 0.65 | 0.64 | 0.65 | 0.43 | 0.67 |
| 11 | 90362010 | rs557666 | 0.706 | -0.16 | -0.368 | 0.097 | -0.006 | -0.019 | -0.019 | 2.496 | 0.39 | 0.47 | 0.29 | 0.22 | 0.28 | 0.47 |
| 11 | 90363576 | rs533004 | 0.909 | -0.095 | -0.455 | 0.128 | -0.001 | -0.007 | -0.007 | 0.81 | 0.37 | 0.39 | 0.28 | 0.22 | 0.28 | 0.47 |
| 11 | 90364187 | rs17257677 | 0.464 | 0.157 | -0.526 | 0.127 | -0.054 | -0.012 | -0.012 | 0 | 0.14 | 0.05 | 0.17 | 0.12 | 0.2 | 0.1 |
| 11 | 90365458 | rs482463 | 0.455 | 0.13 | -0.179 | 0.279 | 0.003 | 0.03 | 0.03 | 0 | 0.51 | 0.43 | 0.44 | 0.33 | 0.48 | 0.57 |
| 11 | 90372636 | rs475285 | 0.533 | 0.523 | -0.208 | 0.42 | 0.005 | -0.009 | -0.009 | 0 | 0.53 | 0.28 | 0.5 | 0.58 | 0.53 | 0.69 |
| 11 | 90372827 | rs2508408 | 0.603 | 0.546 | -0.31 | 0.406 | 0.007 | -0.011 | -0.011 | 0 | 0.52 | 0.25 | 0.5 | 0.58 | 0.53 | 0.69 |
| 11 | 90377065 | rs10501729 | 0.801 | 0.099 | -0.096 | -0.103 | 0.002 | -0.001 | -0.001 | 0.007 | 0.12 | 0.1 | 0.09 | 0.03 | 0.1 | 0.11 |
| 11 | 90377727 | rs6483067 | -1.013 | 0.626 | -0.184 | -0.052 | 0.04 | 0.054 | 0.054 | 0.733 | 0.41 | 0.74 | 0.27 | 0.26 | 0.34 | 0.27 |
| 11 | 90377982 | rs12275422 | 0.235 | 0.527 | 0.413 | -0.1 | 0 | 0.005 | 0.005 | 0 | 0.24 | 0.32 | 0.13 | 0.23 | 0.23 | 0.15 |
| 11 | 90378456 | rs7944100 | 0.557 | 0.511 | 0.381 | -0.065 | -0.019 | -0.051 | -0.051 | 0 | 0.13 | 0.29 | 0.12 | 0.03 | 0.1 | 0.11 |
| 11 | 90382170 | rs4753289 | -1.213 | 1.064 | -0.021 | -0.062 | -0.016 | 0.076 | 0.076 | 0.262 | 0.44 | 0.8 | 0.33 | 0.26 | 0.43 | 0.3 |
| 11 | 90384749 | rs7119319 | -0.633 | 0.983 | 0.438 | -0.154 | 0.025 | -0.114 | -0.114 | 0.205 | 0.19 | 0.53 | 0.19 | 0.03 | 0.19 | 0.15 |
| 11 | 90387210 | rs7128489 | 0.156 | 1.074 | 0.423 | 0.089 | -0.001 | -0.003 | -0.003 | 0.032 | 0.28 | 0.24 | 0.14 | 0.3 | 0.27 | 0.13 |
| 11 | 90389335 | rs2155077 | -0.874 | 1.171 | 0.646 | 0.085 | 0.01 | 0.014 | 0.014 | 1.419 | 0.39 | 0.76 | 0.33 | 0.33 | 0.31 | 0.21 |
| 11 | 90389865 | rs2155076 | -0.027 | 1.169 | 0.997 | 0.087 | -0.01 | 0.004 | 0.004 | 0.348 | 0.31 | 0.46 | 0.17 | 0.31 | 0.27 | 0.13 |
| 11 | 90401857 | rs4350330 | -0.835 | 0.686 | 1.227 | 0.153 | 0.019 | -0.015 | -0.015 | 0 | 0.08 | 0.36 | 0.17 | 0.03 | 0.03 | 0.08 |
| 11 | 90418632 | rs11019143 | 0.07 | 0.886 | 0.991 | -0.024 | -0.006 | 0.002 | 0.002 | 0 | 0.34 | 0.28 | 0.19 | 0.29 | 0.37 | 0.22 |
| 11 | 90425430 | rs12806862 | -0.859 | 0.875 | 0.983 | -0.015 | -0.007 | 0.019 | 0.019 | 0.152 | 0.44 | 0.7 | 0.37 | 0.34 | 0.41 | 0.3 |
| 11 | 90436832 | rs2000536 | -1.053 | 0.864 | 0.96 | -0.017 | -0.009 | -0.016 | -0.016 | 0 | 0.09 | 0.27 | 0.16 | 0.05 | 0.03 | 0.07 |
| 11 | 90438603 | rs1006133 | -0.249 | 0.869 | 0.98 | -0.018 | 0.001 | -0.021 | -0.021 | 0.506 | 0.07 | 0.24 | 0.15 | 0.03 | 0.03 | 0.08 |
| 11 | 90445471 | rs1783800 | -0.814 | 0.87 | 0.918 | -0.012 | -0.004 | 0.008 | 0.008 | 0.141 | 0.44 | 0.64 | 0.36 | 0.34 | 0.41 | 0.3 |
| 11 | 90452226 | rs1792616 | 0.135 | 0.794 | 0.985 | -0.047 | 0.008 | -0.008 | -0.008 | 0 | 0.61 | 0.49 | 0.66 | 0.69 | 0.6 | 0.7 |
| 11 | 90457190 | rs10734133 | 0.178 | 0.805 | 0.944 | -0.042 | 0.008 | -0.013 | -0.013 | 0 | 0.6 | 0.46 | 0.66 | 0.69 | 0.6 | 0.7 |
| 11 | 90469818 | rs10741362 | 0.523 | 0.779 | 0.9 | -0.054 | -0.008 | 0.024 | 0.024 | 0.281 | 0.57 | 0.25 | 0.63 | 0.66 | 0.6 | 0.7 |
| 11 | 90472715 | rs10501734 | 0.192 | 0.783 | 0.967 | -0.06 | 0.007 | -0.013 | -0.013 | 0.172 | 0.06 | 0.16 | 0.14 | 0.03 | 0.03 | 0.08 |
| 11 | 90472975 | rs10734135 | 0.653 | 1.057 | 0.869 | -0.122 | -0.01 | 0.042 | 0.042 | 0.072 | 0.54 | 0.15 | 0.57 | 0.66 | 0.51 | 0.65 |
| 11 | 90473957 | rs10830566 | 1.671 | 0.393 | 0.146 | -0.445 | 0.004 | 0.097 | 0.097 | 0 | 0.3 | 0.08 | 0.19 | 0.16 | 0.31 | 0.35 |
| 11 | 90478149 | rs2213090 | -1.066 | 0.363 | 0.463 | -0.346 | -0.012 | 0.001 | 0.001 | 0 | 0.68 | 0.78 | 0.82 | 0.71 | 0.64 | 0.79 |
| 11 | 90487844 | rs5009703 | -0.682 | 0.375 | 0.389 | -0.34 | 0.005 | -0.007 | -0.007 | 0 | 0.65 | 0.58 | 0.8 | 0.71 | 0.63 | 0.79 |
| 11 | 90492626 | rs7949629 | 0.07 | 0.385 | 0.323 | -0.335 | -0.008 | 0.023 | 0.023 | 0.293 | 0.57 | 0.25 | 0.63 | 0.66 | 0.6 | 0.7 |
| 11 | 90492959 | rs11019173 | -0.174 | 0.389 | 0.342 | -0.319 | 0.007 | -0.003 | -0.003 | 0.986 | 0.08 | 0.27 | 0.16 | 0.05 | 0.03 | 0.08 |
| 11 | 90494720 | rs11019174 | 1.068 | 0.245 | -0.032 | 0.075 | -0.015 | -0.048 | -0.048 | 1.697 | 0.08 | 0.01 | 0.1 | 0.14 | 0.1 | 0.07 |
| 11 | 90497983 | rs1894137 | 0.205 | 0.206 | 0.234 | -0.001 | -0.002 | 0.014 | 0.014 | 0.535 | 0.56 | 0.32 | 0.64 | 0.66 | 0.6 | 0.71 |
| 11 | 90502456 | rs4615986 | -0.84 | 0.271 | 0.044 | 0.031 | 0.003 | 0 | 0 | 0 | 0.67 | 0.68 | 0.8 | 0.71 | 0.64 | 0.79 |
| 11 | 90502896 | rs9651551 | -0.666 | 0.296 | -0.173 | 0.044 | 0.01 | -0.008 | -0.008 | 0 | 0.65 | 0.51 | 0.79 | 0.71 | 0.64 | 0.79 |
| 11 | 90511784 | rs11019178 | 0.777 | -0.014 | -0.677 | 0.026 | 0.002 | -0.037 | -0.037 | 0 | 0.4 | 0.1 | 0.47 | 0.48 | 0.41 | 0.55 |
| 11 | 90513096 | rs10830577 | 1.192 | -0.069 | -0.878 | 0.065 | -0.022 | -0.061 | -0.061 | 0.041 | 0.1 | 0.01 | 0.27 | 0.33 | 0.11 | 0.18 |
| 11 | 90514796 | rs7926034 | -0.512 | -0.004 | -0.558 | -0.443 | -0.041 | 0.039 | 0.039 | 0 | 0.57 | 0.85 | 0.73 | 0.69 | 0.51 | 0.75 |
| 11 | 90518319 | rs10501738 | 1.785 | -0.125 | -0.171 | -0.546 | -0.007 | 0.094 | 0.094 | 0 | 0.27 | 0.06 | 0.18 | 0.15 | 0.28 | 0.35 |
| 11 | 90518881 | rs16924145 | -1.608 | 0.467 | 0.418 | -0.408 | 0.039 | -0.002 | -0.002 | 2.662 | 0.05 | 0.12 | 0.21 | 0.05 | 0.13 | 0.13 |
| 11 | 90525776 | rs7111916 | -0.494 | 0.396 | 0.345 | -0.454 | -0.072 | -0.01 | -0.01 | 0 | 0.62 | 0.92 | 0.78 | 0.69 | 0.6 | 0.8 |
| 11 | 90526114 | rs7952309 | 0.376 | 0.353 | 0.569 | -0.46 | -0.001 | 0.001 | 0.001 | 0 | 0.55 | 0.46 | 0.69 | 0.67 | 0.51 | 0.76 |

| | | | | | | | | | | | | | | | | |
|---|---|---|---|---|---|---|---|---|---|---|---|---|---|---|---|---|
| 11 | 90526222 | rs7952437 | 0.941 | 0.347 | 0.694 | -0.464 | 0.025 | 0.03 | 0.03 | 0 | 0.48 | 0.28 | 0.53 | 0.62 | 0.45 | 0.67 |
| 11 | 90529456 | rs1540170 | -0.062 | 0.382 | 0.532 | -0.413 | -0.048 | 0.027 | 0.027 | 0 | 0.59 | 0.82 | 0.72 | 0.69 | 0.51 | 0.75 |
| 11 | 90535620 | rs1524939 | 0.28 | 0.397 | 0.374 | -0.396 | -0.001 | 0.002 | 0.002 | 0 | 0.57 | 0.63 | 0.71 | 0.67 | 0.51 | 0.75 |
| 11 | 90536821 | rs2852926 | -0.334 | 0.25 | -0.227 | -0.307 | -0.047 | 0.003 | 0.003 | 0 | 0.62 | 0.88 | 0.77 | 0.68 | 0.6 | 0.8 |
| 11 | 90544664 | rs10501741 | -1.411 | 0.303 | 0.393 | -0.226 | 0.011 | 0.018 | 0.018 | 0.01 | 0.06 | 0.32 | 0.16 | 0.05 | 0.02 | 0.1 |
| 11 | 90551484 | rs10830579 | 0.22 | -0.254 | 0.482 | -0.614 | 0.032 | 0.014 | 0.014 | 0 | 0.25 | 0.11 | 0.12 | 0.21 | 0.2 | 0.12 |
| 11 | 90555222 | rs10830581 | 1.582 | 0.01 | -0.025 | -0.629 | 0.006 | -0.001 | -0.001 | 0.453 | 0.42 | 0.18 | 0.5 | 0.63 | 0.42 | 0.64 |
| 11 | 90563921 | rs1894139 | -1.275 | -0.527 | 0.382 | -0.468 | 0.013 | 0.01 | 0.01 | 1.909 | 0.07 | 0.6 | 0.05 | 0 | 0 | 0 |
| 11 | 90571219 | rs2691834 | -0.875 | -0.369 | -0.125 | -0.476 | -0.104 | -0.005 | -0.005 | 0 | 0.8 | 0.99 | 0.87 | 0.91 | 0.75 | 0.89 |
| 11 | 90588350 | rs1357823 | 0.342 | -0.551 | 0.203 | -0.535 | 0.007 | -0.018 | -0.018 | 0 | 0.7 | 0.35 | 0.68 | 0.84 | 0.71 | 0.79 |
| 11 | 90594199 | rs7101758 | -0.162 | -0.476 | 0.164 | -0.483 | -0.046 | 0.018 | 0.018 | 0 | 0.76 | 0.86 | 0.72 | 0.86 | 0.71 | 0.78 |
| 11 | 90596230 | rs2156831 | 0.799 | -0.911 | 0.196 | -0.451 | -0.009 | 0.023 | 0.023 | 0 | 0.54 | 0.62 | 0.61 | 0.67 | 0.52 | 0.68 |
| 11 | 90604756 | rs10765395 | 1.64 | -0.773 | 0.748 | -0.017 | -0.004 | 0.018 | 0.018 | 0 | 0.55 | 0.26 | 0.6 | 0.65 | 0.6 | 0.62 |
| 11 | 90611843 | rs16924276 | -0.836 | -0.695 | 0.811 | -0.035 | -0.06 | -0.037 | -0.037 | 0 | 0.33 | 0.09 | 0.22 | 0.33 | 0.37 | 0.27 |
| 11 | 90630937 | rs12420377 | -2.197 | -0.733 | 1.009 | -0.049 | 0.003 | 0.009 | 0.009 | 0 | 0.48 | 0.64 | 0.44 | 0.4 | 0.44 | 0.4 |
| 11 | 90641355 | rs10501745 | 0.941 | -0.538 | 0.639 | 0.114 | -0.013 | -0.035 | -0.035 | 0 | 0.07 | 0 | 0.03 | 0.22 | 0.07 | 0.1 |
| 11 | 90643566 | rs10741382 | 0.668 | -0.511 | 0.96 | -0.359 | 0.013 | -0.039 | -0.039 | 0.156 | 0.67 | 0.51 | 0.64 | 0.83 | 0.67 | 0.71 |
| 11 | 90645390 | rs7936619 | 0.65 | -0.508 | 0.98 | -0.358 | 0.014 | -0.039 | -0.039 | 0.818 | 0.67 | 0.51 | 0.64 | 0.83 | 0.67 | 0.72 |
| 11 | 90655274 | rs10765400 | 0.507 | -0.327 | 1.27 | -0.072 | 0.007 | -0.018 | -0.018 | 0 | 0.75 | 0.46 | 0.71 | 0.83 | 0.78 | 0.79 |
| 11 | 90656005 | rs35003084 | 0.473 | -0.227 | 1.094 | -0.059 | -0.05 | -0.025 | -0.025 | 0 | 0.16 | 0.01 | 0.11 | 0.12 | 0.18 | 0.11 |
| 11 | 90661841 | rs2388755 | 0.716 | -0.228 | 1.362 | -0.032 | -0.004 | -0.008 | -0.008 | 0 | 0.73 | 0.33 | 0.69 | 0.82 | 0.79 | 0.8 |
| 11 | 90674555 | rs16924399 | -0.57 | -0.29 | 1.478 | 0.048 | -0.001 | 0 | 0 | 0.239 | 0.12 | 0.08 | 0.06 | 0.15 | 0.13 | 0.1 |
| 11 | 90677706 | rs10765405 | 1.66 | -0.318 | 1.497 | 0.003 | 0.007 | 0.003 | 0.003 | 0 | 0.63 | 0.34 | 0.64 | 0.66 | 0.65 | 0.68 |
| 11 | 90716657 | rs1528751 | -0.331 | -0.382 | 1.598 | -0.163 | 0.014 | 0.016 | 0.016 | 0 | 0.06 | 0.22 | 0.16 | 0.05 | 0.03 | 0.09 |
| 11 | 90718303 | rs1954830 | 0.571 | -0.4 | 1.687 | -0.122 | -0.009 | 0.027 | 0.027 | 0 | 0.87 | 0.71 | 0.93 | 0.81 | 0.87 | 0.89 |
| 11 | 90724093 | rs7119959 | 1.664 | -0.427 | 1.236 | -0.126 | -0.02 | 0.005 | 0.005 | 0.398 | 0.81 | 0.31 | 0.77 | 0.75 | 0.87 | 0.82 |
| 11 | 90739550 | rs1528767 | -0.315 | -0.381 | 0.654 | -0.221 | -0.109 | 0.062 | 0.062 | 0 | 0.28 | 0.02 | 0.16 | 0.11 | 0.31 | 0.22 |
| 11 | 90740994 | rs7125140 | 1.678 | -0.408 | 0.754 | -0.223 | -0.005 | 0.082 | 0.082 | 0 | 0.5 | 0.25 | 0.59 | 0.64 | 0.52 | 0.57 |
| 11 | 90746499 | rs11019251 | -1.136 | -0.407 | 0.875 | -0.214 | -0.01 | 0.049 | 0.049 | 0 | 0.41 | 0.23 | 0.24 | 0.3 | 0.43 | 0.32 |
| 11 | 90754928 | rs10765421 | 1.902 | -0.385 | 1.563 | 0.535 | 0.02 | 0.009 | 0.009 | 0 | 0.63 | 0.23 | 0.64 | 0.64 | 0.68 | 0.67 |
| 11 | 90763815 | rs2509378 | 0.197 | -0.411 | 1.222 | 0.602 | -0.047 | -0.017 | -0.017 | 0 | 0.16 | 0.01 | 0.11 | 0.11 | 0.17 | 0.14 |
| 11 | 90772302 | rs1528742 | 1.142 | -0.447 | 1.519 | 0.605 | -0.016 | -0.011 | -0.011 | 0.19 | 0.67 | 0.54 | 0.8 | 0.7 | 0.71 | 0.75 |
| 11 | 90787206 | rs7942666 | 0.715 | -0.272 | 1.59 | 0.805 | 0.01 | 0.054 | 0.054 | 0 | 0.86 | 0.57 | 0.9 | 0.82 | 0.86 | 0.9 |
| 11 | 90801491 | rs1406397 | 0.193 | -0.336 | 1.717 | 1.031 | 0.001 | 0.011 | 0.011 | 0.022 | 0.13 | 0.22 | 0.14 | 0.14 | 0.12 | 0.11 |
| 11 | 90803892 | rs1406399 | 0.766 | -0.446 | 1.69 | 1.012 | -0.002 | 0.04 | 0.04 | 0 | 0.85 | 0.71 | 0.91 | 0.82 | 0.86 | 0.9 |
| 11 | 90809152 | rs1544019 | 1.316 | -0.46 | 1.574 | 1.004 | -0.007 | 0.012 | 0.012 | 0 | 0.72 | 0.52 | 0.77 | 0.67 | 0.74 | 0.78 |
| 11 | 90817090 | rs1590098 | 1.491 | -0.592 | 1.624 | 0.964 | -0.004 | 0.004 | 0.004 | 0 | 0.71 | 0.42 | 0.75 | 0.65 | 0.74 | 0.79 |
| 11 | 90820243 | rs13328892 | 0.131 | -0.417 | 1.67 | 1.062 | 0.001 | 0.024 | 0.024 | 0.006 | 0.14 | 0.23 | 0.16 | 0.17 | 0.12 | 0.11 |
| 11 | 90827529 | rs2463847 | -0.39 | -0.397 | 1.406 | 1.216 | -0.003 | 0.008 | 0.008 | 0 | 0.92 | 0.92 | 0.93 | 0.87 | 0.93 | 0.94 |
| 11 | 90839303 | rs2514078 | -0.409 | -0.393 | 1.504 | 1.264 | 0.007 | 0.028 | 0.028 | 0 | 0.12 | 0.17 | 0.08 | 0.18 | 0.14 | 0.1 |
| 11 | 90859313 | rs4320919 | 0.883 | -0.318 | 1.455 | 1.296 | -0.003 | 0.032 | 0.032 | 0 | 0.8 | 0.79 | 0.84 | 0.67 | 0.79 | 0.81 |
| 11 | 90863481 | rs7115606 | 1.568 | -0.404 | 1.813 | 1.232 | -0.014 | 0.016 | 0.016 | 0 | 0.73 | 0.38 | 0.8 | 0.67 | 0.78 | 0.79 |
| 11 | 90864328 | rs10734146 | -0.209 | -0.39 | 1.699 | 1.25 | 0.014 | 0.048 | 0.048 | 0 | 0.89 | 0.83 | 0.92 | 0.81 | 0.87 | 0.88 |
| 11 | 90868586 | rs2388789 | 1.567 | -0.421 | 1.736 | 1.239 | -0.017 | 0.019 | 0.019 | 0 | 0.73 | 0.42 | 0.81 | 0.68 | 0.78 | 0.81 |
| 11 | 90879295 | rs1528738 | 1.579 | -0.421 | 1.736 | 1.242 | -0.018 | 0.02 | 0.02 | 0 | 0.73 | 0.42 | 0.81 | 0.68 | 0.78 | 0.82 |
| 11 | 90882039 | rs1569181 | -0.195 | -0.428 | 1.78 | 1.229 | 0.013 | 0.04 | 0.04 | 0 | 0.89 | 0.83 | 0.92 | 0.82 | 0.87 | 0.88 |
| 11 | 90884339 | rs1528753 | 1.511 | -0.421 | 1.635 | 1.225 | -0.017 | 0.02 | 0.02 | 0 | 0.73 | 0.42 | 0.81 | 0.68 | 0.78 | 0.81 |
| 11 | 90889938 | rs10741419 | 1.957 | -0.421 | 1.248 | 1.264 | 0.017 | -0.096 | -0.096 | 0 | 0.58 | 0.13 | 0.65 | 0.51 | 0.66 | 0.7 |
| 11 | 90897159 | rs2514085 | 0.111 | -0.202 | 1.367 | 1.14 | 0 | 0.001 | 0.001 | 0 | 0.1 | 0.11 | 0.08 | 0.13 | 0.09 | 0.09 |
| 11 | 90898302 | rs16915728 | 0.249 | -0.196 | 1.446 | 1.145 | 0.013 | -0.012 | -0.012 | 0 | 0.06 | 0.28 | 0.02 | 0 | 0.01 | 0 |
| 11 | 90901865 | rs10830671 | 1.108 | 0.073 | 0.569 | 1.237 | -0.093 | -0.081 | -0.081 | 0.178 | 0.23 | 0.02 | 0.4 | 0.25 | 0.31 | 0.41 |
| 11 | 90902703 | rs11828730 | -1.391 | -0.175 | 1.793 | 1.699 | -0.019 | 0.026 | 0.026 | 0 | 0.15 | 0.48 | 0.11 | 0.13 | 0.09 | 0.06 |
| 11 | 90907226 | rs4279973 | -0.377 | -0.015 | 1.681 | 1.586 | 0.025 | -0.003 | -0.003 | 0.543 | 0.06 | 0.38 | 0.03 | 0 | 0.02 | 0 |
| 11 | 90918340 | rs2514115 | -0.935 | 0.218 | 1.207 | 1.722 | 0.018 | 0.018 | 0.018 | 0 | 0.12 | 0.15 | 0.11 | 0.17 | 0.17 | 0.14 |
| 11 | 90921138 | rs7926164 | 2.201 | -0.411 | 1.074 | 2.02 | 0.014 | -0.033 | -0.033 | 0.614 | 0.58 | 0.14 | 0.65 | 0.42 | 0.66 | 0.7 |
| 11 | 90924805 | rs2514929 | -1.654 | -0.458 | 1.486 | 2.083 | -0.007 | 0.058 | 0.058 | 0 | 0.22 | 0.34 | 0.17 | 0.32 | 0.21 | 0.19 |
| 11 | 90932441 | rs10830674 | 2.338 | -0.447 | 1.351 | 2.06 | 0.014 | -0.036 | -0.036 | 0 | 0.57 | 0.14 | 0.65 | 0.42 | 0.66 | 0.7 |
| 11 | 90937805 | rs2514917 | -0.628 | -0.154 | 1.69 | 1.797 | 0.024 | 0.056 | 0.056 | 0 | 0.1 | 0.21 | 0.09 | 0.17 | 0.13 | 0.12 |
| 11 | 90940780 | rs9943495 | 2.074 | -0.177 | 1.498 | 1.779 | 0.017 | -0.01 | -0.01 | 0.392 | 0.7 | 0.24 | 0.8 | 0.69 | 0.78 | 0.8 |
| 11 | 90946117 | rs11019328 | -1.413 | -0.18 | 1.419 | 1.424 | -0.025 | -0.008 | -0.008 | 1.544 | 0.17 | 0.46 | 0.11 | 0.13 | 0.1 | 0.07 |
| 11 | 90949956 | rs4543940 | -1.306 | -0.154 | 1.195 | 1.436 | -0.021 | -0.007 | -0.007 | 0.014 | 0.16 | 0.4 | 0.11 | 0.13 | 0.1 | 0.07 |
| 11 | 90952254 | rs10501752 | -0.284 | -0.057 | 0.215 | 1.527 | 0.005 | -0.012 | -0.012 | 0 | 0.1 | 0.06 | 0.08 | 0.12 | 0.08 | 0.07 |
| 11 | 90954609 | rs2514923 | 0.936 | -0.086 | 0.017 | 1.621 | 0.006 | -0.01 | -0.01 | 0 | 0.65 | 0.29 | 0.71 | 0.59 | 0.71 | 0.71 |
| 11 | 90954660 | rs7125672 | 1.156 | 0.024 | -0.06 | 1.404 | -0.005 | 0.02 | 0.02 | 0 | 0.55 | 0.21 | 0.63 | 0.4 | 0.58 | 0.65 |
| 11 | 90955972 | rs4483549 | -0.992 | 0.498 | -0.071 | 1.422 | 0 | 0.002 | 0.002 | 0 | 0.33 | 0.28 | 0.28 | 0.53 | 0.39 | 0.29 |
| 11 | 90966072 | rs4073197 | -1.264 | 0.103 | -0.012 | 1.209 | -0.001 | 0 | 0 | 0 | 0.12 | 0.17 | 0.07 | 0.1 | 0.07 | 0.05 |
| 11 | 90969158 | rs10830680 | -1.889 | -0.028 | 0.211 | 1.719 | 0.001 | 0.033 | 0.033 | 0 | 0.18 | 0.24 | 0.19 | 0.37 | 0.18 | 0.14 |
| 11 | 90969612 | rs7949216 | 2.188 | -0.224 | 1.102 | 1.662 | -0.008 | 0.061 | 0.061 | 0.092 | 0.72 | 0.46 | 0.71 | 0.57 | 0.77 | 0.74 |
| 11 | 90971726 | rs4501950 | 2.202 | -0.238 | 1.164 | 1.704 | -0.008 | 0.064 | 0.064 | 0.088 | 0.72 | 0.44 | 0.7 | 0.54 | 0.77 | 0.71 |

| | | | | | | | | | | | | | | | | |
|---|---|---|---|---|---|---|---|---|---|---|---|---|---|---|---|---|
| 11 | 90975749 | rs12280718 | -1.199 | 0.062 | 0.968 | 1.975 | -0.011 | 0.005 | 0.005 | 0 | 0.09 | 0.05 | 0.15 | 0.36 | 0.13 | 0.09 |
| 11 | 90975843 | rs12286734 | -1.229 | 0.047 | 0.964 | 1.97 | -0.01 | 0.004 | 0.004 | 0 | 0.09 | 0.05 | 0.15 | 0.35 | 0.13 | 0.09 |
| 11 | 90981224 | rs10830686 | 2.507 | -0.357 | 1.144 | 1.823 | -0.006 | 0.032 | 0.032 | 1.666 | 0.75 | 0.32 | 0.71 | 0.54 | 0.82 | 0.76 |
| 11 | 90990885 | rs11019357 | 2.374 | -0.363 | 1.367 | 1.803 | -0.008 | 0.098 | 0.098 | 0.914 | 0.77 | 0.43 | 0.72 | 0.54 | 0.82 | 0.76 |
| 11 | 90991115 | rs10765460 | 2.189 | -0.372 | 1.748 | 1.799 | 0.026 | 0.219 | 0.219 | 0.277 | 0.82 | 0.57 | 0.78 | 0.57 | 0.84 | 0.79 |
| 11 | 90995060 | rs2196517 | 1.164 | -0.336 | 1.633 | 1.629 | 0.006 | 0.054 | 0.054 | 0 | 0.93 | 0.86 | 0.94 | 0.91 | 0.97 | 0.88 |
| 11 | 91019198 | rs10830705 | 2.401 | -0.368 | 1.241 | 1.677 | 0 | 0.001 | 0.001 | 0.495 | 0.75 | 0.27 | 0.7 | 0.56 | 0.8 | 0.75 |
| 11 | 91019763 | rs7931112 | 2.024 | -0.372 | 1.312 | 1.688 | -0.011 | 0.051 | 0.051 | 0.691 | 0.81 | 0.33 | 0.75 | 0.65 | 0.87 | 0.82 |
| 11 | 91024468 | rs12790605 | 2.499 | -0.369 | 1.265 | 1.667 | 0 | 0 | 0 | 0.562 | 0.75 | 0.27 | 0.73 | 0.56 | 0.79 | 0.75 |
| 11 | 91033997 | rs1370024 | -1.458 | -0.307 | 1.168 | 1.619 | 0 | -0.001 | -0.001 | 0.275 | 0.06 | 0.12 | 0.04 | 0.06 | 0.07 | 0.04 |
| 11 | 91034922 | rs11603266 | 1.203 | -0.507 | 0.135 | 0.86 | -0.117 | 0.147 | 0.147 | 0.232 | 0.36 | 0.06 | 0.26 | 0.13 | 0.49 | 0.39 |
| 11 | 91038032 | rs11019380 | -0.314 | -0.25 | 0.106 | 1.106 | -0.029 | -0.049 | -0.049 | 0 | 0.08 | 0 | 0.12 | 0.27 | 0.09 | 0.08 |
| 11 | 91047120 | rs6483110 | 0.467 | -0.408 | 0.112 | 1.053 | 0.015 | -0.028 | -0.028 | 0 | 0.34 | 0.21 | 0.43 | 0.41 | 0.29 | 0.37 |
| 11 | 91080353 | rs3018365 | 0.692 | -0.127 | 0.454 | 1.533 | -0.002 | 0.023 | 0.023 | 0 | 0.89 | 0.93 | 0.83 | 0.65 | 0.88 | 0.8 |
| 11 | 91089975 | rs11019417 | -1.581 | -0.095 | 0.124 | 1.175 | -0.002 | 0.002 | 0.002 | 0 | 0.07 | 0.07 | 0.04 | 0.06 | 0.09 | 0.04 |
| 11 | 91107222 | rs10830726 | -1.655 | -0.296 | 0.116 | 0.949 | -0.011 | -0.006 | -0.006 | 0 | 0.12 | 0.19 | 0.09 | 0.09 | 0.09 | 0.08 |
| 11 | 91123074 | rs1587843 | -0.064 | 0.409 | -0.241 | 0.876 | -0.001 | -0.001 | -0.001 | 0.022 | 0.59 | 0.4 | 0.63 | 0.52 | 0.61 | 0.53 |
| 11 | 91123671 | rs657546 | 0.829 | 0.479 | -0.359 | 1.479 | -0.051 | -0.063 | -0.063 | 0 | 0.9 | 0.99 | 0.8 | 0.68 | 0.85 | 0.89 |
| 11 | 91128696 | rs12272958 | -0.929 | 0.382 | -0.148 | 1.489 | -0.001 | 0.026 | 0.026 | 0 | 0.06 | 0.11 | 0.05 | 0.09 | 0.04 | 0.08 |
| 11 | 91151732 | rs581985 | 0.583 | 0.287 | -0.42 | 1.724 | -0.049 | -0.063 | -0.063 | 1.611 | 0.9 | 0.99 | 0.86 | 0.72 | 0.86 | 0.88 |
| 11 | 91152876 | rs587473 | 0.472 | 0.064 | -0.49 | 1.014 | -0.001 | 0.064 | 0.064 | 0 | 0.51 | 0.38 | 0.36 | 0.21 | 0.53 | 0.51 |
| 11 | 91155574 | rs2397351 | 0.419 | -0.071 | -0.992 | 0.943 | -0.038 | -0.07 | -0.07 | 0.006 | 0.29 | 0.06 | 0.4 | 0.4 | 0.29 | 0.34 |
| 11 | 91159644 | rs12575375 | -0.199 | -0.146 | -0.75 | 0.804 | -0.014 | -0.061 | -0.061 | 0 | 0.11 | 0.01 | 0.1 | 0.26 | 0.09 | 0.07 |
| 11 | 91171321 | rs7941645 | -0.213 | -0.452 | -0.592 | 0.64 | -0.021 | -0.021 | -0.021 | 0 | 0.06 | 0.1 | 0.01 | 0.01 | 0.01 | 0.02 |
| 11 | 91177076 | rs1394529 | -1.671 | -0.862 | -0.947 | 0.114 | -0.036 | 0.147 | 0.147 | 0 | 0.59 | 0.8 | 0.68 | 0.87 | 0.45 | 0.54 |
| 11 | 91179638 | rs2156846 | 0.659 | -0.792 | -0.882 | 0.405 | -0.031 | -0.12 | -0.12 | 1.178 | 0.26 | 0.05 | 0.42 | 0.52 | 0.29 | 0.33 |
| 11 | 91192218 | rs10830732 | 0.185 | -0.769 | -0.775 | 0.444 | -0.003 | -0.053 | -0.053 | 0.301 | 0.11 | 0.01 | 0.1 | 0.26 | 0.09 | 0.07 |
| 11 | 91204664 | rs11019478 | -0.28 | -0.802 | -0.281 | -0.018 | -0.016 | 0.075 | 0.075 | 0.02 | 0.39 | 0.09 | 0.53 | 0.78 | 0.39 | 0.4 |
| 11 | 91215718 | rs10765469 | -1.404 | -0.934 | -0.088 | -0.111 | 0.001 | 0.118 | 0.118 | 0.125 | 0.55 | 0.78 | 0.66 | 0.8 | 0.43 | 0.46 |
| 11 | 91216210 | rs4753332 | -1.202 | -0.95 | 0.06 | -0.112 | 0.028 | 0.124 | 0.124 | 0.127 | 0.52 | 0.76 | 0.66 | 0.79 | 0.42 | 0.43 |
| 11 | 91219932 | rs1910734 | -1.406 | -0.945 | -0.022 | -0.123 | 0.002 | 0.121 | 0.121 | 0.196 | 0.55 | 0.78 | 0.66 | 0.8 | 0.43 | 0.46 |
| 11 | 91222957 | rs10830735 | -0.208 | -0.896 | -0.076 | -0.096 | -0.016 | 0.079 | 0.079 | 0.033 | 0.39 | 0.1 | 0.55 | 0.79 | 0.39 | 0.39 |
| 11 | 91234917 | rs10765480 | -0.752 | -0.776 | 0.605 | -0.023 | 0.006 | 0.009 | 0.009 | 0.356 | 0.48 | 0.54 | 0.6 | 0.78 | 0.4 | 0.42 |
| 11 | 91246309 | rs16916109 | -0.642 | -0.791 | 0.891 | 0.066 | -0.001 | -0.02 | -0.02 | 0.239 | 0.06 | 0.11 | 0.06 | 0.01 | 0.03 | 0.04 |
| 11 | 91251640 | rs6483125 | -0.365 | -0.249 | 0.731 | -0.286 | -0.003 | 0.038 | 0.038 | 0 | 0.52 | 0.6 | 0.63 | 0.83 | 0.52 | 0.4 |
| 11 | 91258671 | rs12283495 | -1.343 | -0.273 | 0.474 | -0.319 | -0.022 | 0.103 | 0.103 | 0 | 0.63 | 0.8 | 0.77 | 0.84 | 0.61 | 0.5 |
| 11 | 91259052 | rs16916160 | 0.813 | -0.091 | 0.309 | 0.313 | -0.044 | -0.088 | -0.088 | 0.488 | 0.34 | 0.08 | 0.36 | 0.49 | 0.39 | 0.34 |
| 11 | 91287525 | rs2514299 | -0.766 | -0.357 | 0.501 | -0.243 | -0.014 | 0.082 | 0.082 | 0.021 | 0.55 | 0.69 | 0.64 | 0.81 | 0.52 | 0.41 |
| 11 | 91290067 | rs908709 | 0.235 | -0.084 | 0.317 | 0.184 | -0.002 | 0.016 | 0.016 | 0 | 0.38 | 0.26 | 0.4 | 0.64 | 0.4 | 0.35 |
| 11 | 91295509 | rs2459855 | -1.094 | 0.131 | 1.121 | 0.162 | 0.02 | -0.049 | -0.049 | 0.165 | 0.12 | 0.48 | 0.14 | 0.01 | 0.07 | 0.11 |
| 11 | 91296802 | rs2459854 | -1.031 | -0.183 | 0.776 | -0.125 | -0.008 | 0.107 | 0.107 | 0 | 0.59 | 0.8 | 0.78 | 0.77 | 0.61 | 0.56 |
| 11 | 91306532 | rs12286609 | -0.263 | -0.117 | 1.417 | -0.09 | 0.023 | -0.039 | -0.039 | 0 | 0.1 | 0.27 | 0.11 | 0.01 | 0.07 | 0.1 |
| 11 | 91306561 | rs2514282 | 0.021 | -0.189 | 1.565 | -0.104 | -0.006 | 0.044 | 0.044 | 0 | 0.47 | 0.37 | 0.66 | 0.76 | 0.55 | 0.46 |
| 11 | 91316501 | rs11019557 | 0.207 | -0.132 | 1.35 | -0.073 | -0.039 | 0.121 | 0.121 | 0 | 0.45 | 0.17 | 0.63 | 0.76 | 0.55 | 0.44 |
| 11 | 91316831 | rs12289574 | -0.367 | 0.096 | 1.094 | 0.076 | 0.029 | -0.032 | -0.032 | 0.471 | 0.1 | 0.28 | 0.11 | 0.01 | 0.07 | 0.1 |
| 11 | 91320582 | rs12288458 | -0.038 | 0.075 | 0.913 | 0.087 | 0.017 | -0.03 | -0.03 | 0 | 0.09 | 0.23 | 0.09 | 0.01 | 0.06 | 0.1 |
| 11 | 91320978 | rs11019558 | -0.038 | 0.109 | 0.913 | 0.087 | 0.021 | -0.03 | -0.03 | 0 | 0.09 | 0.23 | 0.1 | 0.01 | 0.06 | 0.1 |
| 11 | 91326092 | rs2514302 | -0.767 | 0.003 | 0.03 | 0.174 | 0.006 | 0.115 | 0.115 | 0 | 0.55 | 0.77 | 0.77 | 0.76 | 0.61 | 0.55 |
| 11 | 91342130 | rs11019575 | -0.043 | 0.206 | 0.136 | 1.076 | -0.017 | -0.031 | -0.031 | 0 | 0.06 | 0 | 0.17 | 0.23 | 0.07 | 0.1 |
| 11 | 91344352 | rs2678420 | -1.971 | -0.016 | 1.08 | 1.506 | -0.01 | 0.189 | 0.189 | 0 | 0.18 | 0.54 | 0.32 | 0.37 | 0.12 | 0.24 |
| 11 | 91352621 | rs7946764 | -0.32 | 0.019 | 1.401 | 1.34 | 0.008 | -0.05 | -0.05 | 0 | 0.11 | 0.29 | 0.12 | 0.01 | 0.09 | 0.11 |
| 11 | 91365548 | rs707319 | -0.902 | 0.008 | 1.171 | 1.425 | -0.001 | 0.205 | 0.205 | 0 | 0.09 | 0.25 | 0.22 | 0.37 | 0.07 | 0.1 |
| 11 | 91382553 | rs707326 | 0.777 | -0.013 | 0.569 | 1.248 | -0.024 | 0.021 | 0.021 | 0.71 | 0.3 | 0.16 | 0.34 | 0.24 | 0.41 | 0.41 |
| 11 | 91383660 | rs2568721 | -1.074 | -0.269 | 1 | 1.312 | 0.002 | 0.064 | 0.064 | 0.658 | 0.51 | 0.74 | 0.69 | 0.66 | 0.58 | 0.68 |
| 11 | 91387920 | rs6483138 | 0.567 | -0.252 | 1.17 | 1.264 | 0.006 | -0.02 | -0.02 | 0 | 0.09 | 0.15 | 0.11 | 0.01 | 0.1 | 0.09 |
| 11 | 91390044 | rs12224437 | 0.841 | -0.245 | 1.333 | 1.28 | -0.03 | 0.02 | 0.02 | 1.148 | 0.3 | 0.18 | 0.35 | 0.23 | 0.42 | 0.41 |
| 11 | 91390397 | rs11019587 | -0.915 | -0.248 | 1.347 | 1.381 | -0.005 | 0.021 | 0.021 | 0.142 | 0.17 | 0.25 | 0.28 | 0.29 | 0.16 | 0.25 |
| 11 | 91407305 | rs2568712 | -0.997 | -0.121 | 1.179 | 1.35 | 0.007 | 0.036 | 0.036 | 1.082 | 0.51 | 0.67 | 0.68 | 0.66 | 0.59 | 0.68 |
| 11 | 91414059 | rs6483140 | -0.366 | -0.041 | 1.165 | 1.313 | 0.057 | 0.08 | 0.08 | 0 | 0.12 | 0.42 | 0.19 | 0.14 | 0.13 | 0.1 |
| 11 | 91421876 | rs3844366 | 0.784 | -0.105 | 1.075 | 1.298 | -0.019 | 0.017 | 0.017 | 0.29 | 0.3 | 0.2 | 0.35 | 0.23 | 0.41 | 0.38 |
| 11 | 91422668 | rs11019607 | 0.977 | -0.101 | 1.046 | 1.3 | -0.054 | 0.017 | 0.017 | 0.709 | 0.29 | 0.1 | 0.34 | 0.23 | 0.41 | 0.38 |
| 11 | 91424086 | rs16916377 | 0.128 | -0.086 | 1.139 | 1.328 | 0.035 | 0.05 | 0.05 | 0 | 0.11 | 0.26 | 0.16 | 0.14 | 0.13 | 0.1 |
| 11 | 91429707 | rs12420217 | -0.917 | -0.122 | 1.124 | 1.43 | 0.007 | 0.111 | 0.111 | 0 | 0.17 | 0.29 | 0.31 | 0.43 | 0.17 | 0.23 |
| 11 | 91435248 | rs10830764 | -0.661 | -0.037 | 1.217 | 1.193 | 0.032 | 0.055 | 0.055 | 0 | 0.13 | 0.35 | 0.18 | 0.15 | 0.13 | 0.13 |
| 11 | 91443021 | rs11019615 | -0.323 | -0.127 | 1.007 | 1.116 | 0.005 | -0.002 | -0.002 | 2.052 | 0.43 | 0.52 | 0.52 | 0.38 | 0.54 | 0.5 |
| 11 | 91443158 | rs3847508 | -0.021 | -0.07 | 0.809 | 1.15 | 0.005 | -0.001 | -0.001 | 0 | 0.41 | 0.46 | 0.51 | 0.37 | 0.54 | 0.48 |
| 11 | 91451953 | rs10830766 | -0.07 | -0.044 | 0.576 | 1.166 | -0.007 | -0.002 | -0.002 | 0 | 0.4 | 0.37 | 0.5 | 0.37 | 0.54 | 0.48 |
| 11 | 91452705 | rs4145060 | 0.187 | -0.173 | 0.316 | 1.399 | 0.017 | -0.031 | -0.031 | 0 | 0.07 | 0 | 0.15 | 0.3 | 0.05 | 0.15 |
| 11 | 91461188 | rs794409 | -1.276 | 0.147 | 0.148 | 0.41 | -0.047 | 0.007 | 0.007 | 0 | 0.87 | 0.99 | 0.85 | 0.94 | 0.83 | 0.89 |

| | | | | | | | | | | | | | | | | |
|---|---|---|---|---|---|---|---|---|---|---|---|---|---|---|---|---|
| 11 | 91476334 | rs11019624 | 1.482 | 0.324 | 0.345 | 0.579 | -0.011 | -0.013 | -0.013 | 0 | 0.2 | 0.14 | 0.23 | 0.28 | 0.29 | 0.31 |
| 11 | 91486518 | rs794414 | 0.662 | 0.372 | 0.625 | -0.416 | -0.001 | -0.018 | -0.018 | 0 | 0.06 | 0.15 | 0.07 | 0.01 | 0.05 | 0.05 |
| 11 | 91487582 | rs808003 | -0.012 | 0.32 | 1.145 | -0.345 | -0.001 | -0.005 | -0.005 | 1.08 | 0.37 | 0.6 | 0.52 | 0.7 | 0.41 | 0.56 |
| 11 | 91489927 | rs794413 | 0.182 | 0.355 | 1.093 | -0.307 | 0 | -0.004 | -0.004 | 0 | 0.34 | 0.37 | 0.5 | 0.69 | 0.4 | 0.55 |
| 11 | 91495099 | rs7945313 | 0.527 | 0.356 | 0.042 | -0.893 | -0.008 | 0.018 | 0.018 | 0.712 | 0.29 | 0.16 | 0.25 | 0.25 | 0.27 | 0.22 |
| 11 | 91498659 | rs12795641 | 1.722 | 0.274 | -0.164 | -1.066 | -0.026 | -0.008 | -0.008 | 0.454 | 0.15 | 0 | 0.14 | 0.13 | 0.11 | 0.14 |
| 11 | 91508234 | rs6483152 | -0.231 | 0.597 | 0.237 | -0.812 | -0.001 | -0.035 | -0.035 | 0 | 0.13 | 0.24 | 0.11 | 0.02 | 0.11 | 0.05 |
| 11 | 91509923 | rs7934386 | -0.316 | 0.608 | 0.246 | -0.814 | -0.002 | 0.012 | 0.012 | 0 | 0.42 | 0.4 | 0.34 | 0.26 | 0.36 | 0.27 |
| 11 | 91519646 | rs2250842 | -0.794 | 0.445 | 0.251 | -0.835 | 0.002 | 0.016 | 0.016 | 0 | 0.73 | 0.81 | 0.8 | 0.94 | 0.77 | 0.77 |
| 11 | 91520144 | rs12277524 | -0.672 | 0.542 | 0.329 | -0.483 | 0.001 | -0.033 | -0.033 | 0 | 0.14 | 0.23 | 0.11 | 0.01 | 0.15 | 0.05 |
| 11 | 91525269 | rs1528252 | 0.481 | 0.551 | 0.322 | -0.371 | -0.001 | -0.012 | -0.012 | 0 | 0.07 | 0.1 | 0.04 | 0 | 0.06 | 0 |
| 11 | 91527559 | rs4384350 | 0.246 | 0.572 | 0.305 | -0.36 | 0.012 | -0.037 | -0.037 | 1.721 | 0.08 | 0.22 | 0.06 | 0 | 0.06 | 0.01 |
| 11 | 91528985 | rs1881289 | -0.58 | 0.62 | 0.13 | -0.228 | -0.001 | -0.002 | -0.002 | 0.001 | 0.44 | 0.46 | 0.36 | 0.26 | 0.39 | 0.27 |
| 11 | 91536763 | rs4130618 | -0.449 | 0.228 | 0.532 | -0.238 | 0.002 | 0.005 | 0.005 | 0 | 0.4 | 0.34 | 0.31 | 0.24 | 0.35 | 0.25 |
| 11 | 91538582 | rs7927763 | -0.318 | 0.008 | 0.908 | -0.389 | 0.008 | -0.052 | -0.052 | 0 | 0.1 | 0.25 | 0.12 | 0.01 | 0.09 | 0.04 |
| 11 | 91540464 | rs7124929 | -0.813 | -0.1 | 0.853 | -0.693 | -0.002 | 0.012 | 0.012 | 0 | 0.79 | 0.83 | 0.79 | 0.94 | 0.78 | 0.84 |
| 11 | 91544366 | rs726398 | 0.483 | -0.105 | 1.176 | -0.663 | 0.011 | 0.022 | 0.022 | 0 | 0.32 | 0.28 | 0.2 | 0.24 | 0.27 | 0.21 |
| 11 | 91549126 | rs7944203 | 0.968 | -0.132 | 1.389 | -0.687 | -0.012 | 0.01 | 0.01 | 0 | 0.37 | 0.3 | 0.46 | 0.69 | 0.43 | 0.59 |
| 11 | 91552059 | rs12418522 | 1.113 | -0.116 | 1.131 | -0.681 | -0.016 | 0.037 | 0.037 | 0 | 0.07 | 0.02 | 0.11 | 0.01 | 0.09 | 0.04 |
| 11 | 91560288 | rs7951876 | -0.712 | -0.123 | 1.247 | -0.662 | -0.001 | 0.003 | 0.003 | 0 | 0.78 | 0.79 | 0.79 | 0.94 | 0.78 | 0.84 |
| 11 | 91560493 | rs7945011 | -0.858 | -0.12 | 1.228 | -0.661 | -0.003 | 0.025 | 0.025 | 0 | 0.79 | 0.85 | 0.79 | 0.94 | 0.78 | 0.84 |
| 11 | 91570782 | rs1491667 | 0.252 | -0.099 | 1.234 | -0.628 | -0.007 | 0.03 | 0.03 | 0.093 | 0.47 | 0.58 | 0.59 | 0.7 | 0.51 | 0.63 |
| 11 | 91571590 | rs4999577 | 0.474 | -0.21 | 1.086 | -0.552 | 0.012 | 0.036 | 0.036 | 0 | 0.31 | 0.15 | 0.19 | 0.24 | 0.27 | 0.21 |
| 11 | 91572043 | rs1907573 | 0.44 | -0.209 | 1.025 | -0.551 | 0.013 | 0.028 | 0.028 | 0 | 0.32 | 0.22 | 0.2 | 0.24 | 0.27 | 0.21 |
| 11 | 91579675 | rs16916597 | 1.428 | -0.611 | 0.704 | -1.01 | 0.01 | 0.002 | 0.002 | 0 | 0.15 | 0.01 | 0.05 | 0.1 | 0.11 | 0.18 |
| 11 | 91581283 | rs7115718 | 1.082 | -0.618 | 0.699 | -1.158 | -0.061 | 0.153 | 0.153 | 0 | 0.45 | 0.15 | 0.53 | 0.81 | 0.51 | 0.62 |
| 11 | 91582897 | rs11601154 | -0.666 | -0.37 | 1.1 | -0.996 | -0.019 | -0.07 | -0.07 | 2.27 | 0.31 | 0.51 | 0.24 | 0.11 | 0.27 | 0.23 |
| 11 | 91590118 | rs2101634 | 0.276 | -0.347 | 0.906 | -0.953 | -0.011 | -0.031 | -0.031 | 0 | 0.22 | 0.27 | 0.09 | 0.1 | 0.17 | 0.19 |
| 11 | 91590650 | rs956647 | -0.445 | -0.364 | 0.907 | -1.243 | 0.001 | -0.038 | -0.038 | 0 | 0.77 | 0.68 | 0.78 | 0.93 | 0.77 | 0.87 |
| 11 | 91594036 | rs1873579 | -0.685 | -0.346 | 0.911 | -1.231 | 0.004 | -0.033 | -0.033 | 0.016 | 0.79 | 0.69 | 0.8 | 0.93 | 0.79 | 0.88 |
| 11 | 91610478 | rs7102905 | -0.797 | -0.541 | 0.891 | -1.081 | 0.004 | -0.023 | -0.023 | 0.042 | 0.75 | 0.73 | 0.71 | 0.93 | 0.7 | 0.84 |
| 11 | 91613039 | rs16916640 | 0.259 | -0.584 | 0.736 | -1.023 | 0.004 | 0.007 | 0.007 | 0 | 0.22 | 0.2 | 0.1 | 0.1 | 0.17 | 0.2 |
| 11 | 91613804 | rs1122273 | 1.27 | -0.595 | 0.375 | -0.973 | 0.018 | 0.011 | 0.011 | 0 | 0.16 | 0.06 | 0.05 | 0.09 | 0.1 | 0.19 |
| 11 | 91616319 | rs11019669 | 0.4 | -0.658 | 0.414 | -1.039 | 0.023 | 0.134 | 0.134 | 0 | 0.53 | 0.22 | 0.6 | 0.83 | 0.57 | 0.66 |
| 11 | 91616691 | rs12225068 | 1.733 | -0.692 | 0.2 | -0.523 | -0.013 | -0.015 | -0.015 | 0 | 0.08 | 0.01 | 0.25 | 0.46 | 0.08 | 0.18 |
| 11 | 91617547 | rs4753350 | 0.579 | -0.446 | 0.649 | -1.384 | 0.022 | 0.143 | 0.143 | 0 | 0.53 | 0.22 | 0.6 | 0.83 | 0.57 | 0.66 |
| 11 | 91621291 | rs2397833 | 1.158 | -0.343 | 0.591 | -1.424 | -0.118 | 0.141 | 0.141 | 0 | 0.44 | 0.05 | 0.52 | 0.83 | 0.48 | 0.63 |
| 11 | 91621619 | rs12293112 | -0.317 | -0.223 | 0.943 | -1.278 | 0.033 | 0.01 | 0.01 | 0.037 | 0.06 | 0.32 | 0.05 | 0 | 0.03 | 0.01 |
| 11 | 91623555 | rs12789947 | -0.132 | -0.222 | 0.805 | -1.29 | 0.006 | -0.073 | -0.073 | 0 | 0.7 | 0.39 | 0.65 | 0.92 | 0.67 | 0.83 |
| 11 | 91625662 | rs12294567 | -0.036 | -0.247 | 0.684 | -1.275 | -0.063 | -0.057 | -0.057 | 0 | 0.22 | 0.51 | 0.12 | 0.09 | 0.13 | 0.17 |
| 11 | 91628612 | rs985598 | -0.027 | -0.042 | 0.271 | -1.18 | 0 | 0.008 | 0.008 | 0 | 0.6 | 0.67 | 0.67 | 0.83 | 0.6 | 0.67 |
| 11 | 91637159 | rs17571502 | 1.399 | -0.052 | -0.159 | -1.164 | -0.084 | 0.009 | 0.009 | 0.118 | 0.34 | 0.06 | 0.52 | 0.77 | 0.39 | 0.6 |
| 11 | 91638514 | rs7951711 | -0.569 | -0.221 | -0.256 | -1.339 | -0.044 | 0.11 | 0.11 | 0 | 0.68 | 0.87 | 0.68 | 0.93 | 0.6 | 0.8 |
| 11 | 91638725 | rs10501763 | 1.417 | -0.195 | -0.402 | -1.313 | 0.011 | 0.005 | 0.005 | 0.076 | 0.15 | 0.06 | 0.04 | 0.1 | 0.09 | 0.17 |
| 11 | 91639304 | rs1388472 | 0.062 | -0.186 | -0.458 | -1.323 | -0.004 | 0.016 | 0.016 | 0 | 0.6 | 0.66 | 0.64 | 0.93 | 0.56 | 0.8 |
| 11 | 91640112 | rs16916699 | 0.399 | 0.469 | -1.159 | -0.838 | 0.004 | -0.023 | -0.023 | 0.066 | 0.08 | 0.13 | 0.04 | 0 | 0.08 | 0.01 |
| 11 | 91648061 | rs10830799 | 0.982 | 0.406 | -1.186 | -0.711 | -0.09 | 0 | 0 | 0 | 0.34 | 0.05 | 0.5 | 0.77 | 0.38 | 0.6 |
| 11 | 91657276 | rs10501765 | 1.042 | 0.302 | -1.142 | -0.739 | 0.026 | 0.064 | 0.064 | 0.571 | 0.27 | 0.05 | 0.1 | 0.11 | 0.2 | 0.21 |
| 11 | 91657796 | rs10501766 | 1.068 | 0.314 | -1.243 | -0.581 | -0.076 | -0.077 | -0.077 | 0.801 | 0.33 | 0.06 | 0.48 | 0.65 | 0.38 | 0.59 |
| 11 | 91660372 | rs7119009 | 0.402 | 0.13 | -1.099 | -0.573 | 0.003 | 0.009 | 0.009 | 0 | 0.32 | 0.26 | 0.14 | 0.2 | 0.25 | 0.23 |
| 11 | 91661859 | rs1491679 | -0.786 | 0.209 | -1.826 | -0.564 | 0.015 | -0.018 | -0.018 | 0.255 | 0.75 | 0.72 | 0.58 | 0.88 | 0.64 | 0.83 |
| 11 | 91664645 | rs12792913 | 0.565 | 0.512 | -3.045 | -0.094 | 0.02 | 0.049 | 0.049 | 0.516 | 0.27 | 0.04 | 0.1 | 0.12 | 0.2 | 0.2 |
| 11 | 91674642 | rs11019701 | 0.292 | 0.704 | -2.733 | 0.277 | -0.067 | -0.125 | -0.125 | 0.191 | 0.31 | 0.05 | 0.36 | 0.62 | 0.34 | 0.57 |
| 11 | 91680278 | rs1491683 | -0.015 | 0.351 | -2.324 | -0.131 | 0.004 | 0.044 | 0.044 | 0.172 | 0.26 | 0.03 | 0.1 | 0.12 | 0.21 | 0.21 |
| 11 | 91681310 | rs2201119 | -0.244 | 0.7 | -2.915 | -0.443 | -0.073 | 0.138 | 0.138 | 0 | 0.41 | 0.03 | 0.4 | 0.11 | 0.4 | 0.3 |
| 11 | 91682680 | rs7107272 | 0.123 | 0.621 | -2.333 | -0.459 | 0 | 0.007 | 0.007 | 0 | 0.71 | 0.76 | 0.71 | 0.86 | 0.7 | 0.83 |
| 11 | 91682803 | rs7121364 | 0.345 | 0.647 | -1.358 | -0.697 | 0.035 | -0.027 | -0.027 | 0 | 0.07 | 0.21 | 0.02 | 0 | 0.05 | 0.01 |
| 11 | 91684036 | rs12786190 | 0.755 | 0.316 | -2.047 | -1.166 | -0.035 | 0.073 | 0.073 | 0 | 0.28 | 0.01 | 0.17 | 0.08 | 0.23 | 0.23 |
| 11 | 91686927 | rs7483643 | 1.098 | 0.118 | -2.144 | -1.112 | -0.1 | -0.049 | -0.049 | 0 | 0.32 | 0.04 | 0.45 | 0.72 | 0.42 | 0.61 |
| 11 | 91690010 | rs11019709 | 0.167 | 0.152 | -1.445 | -1.102 | -0.003 | -0.026 | -0.026 | 0 | 0.35 | 0.39 | 0.21 | 0.14 | 0.28 | 0.22 |
| 11 | 91699817 | rs1540189 | 1.022 | -0.065 | -1.348 | -1.396 | 0.002 | -0.04 | -0.04 | 0 | 0.08 | 0.24 | 0.04 | 0 | 0.04 | 0.04 |
| 11 | 91701074 | rs11601047 | 0.919 | 0.033 | -1.65 | -1.317 | -0.007 | 0.093 | 0.093 | 0 | 0.28 | 0.1 | 0.17 | 0.08 | 0.23 | 0.21 |
| 11 | 91702184 | rs1945869 | -0.646 | 0.23 | -1.53 | -1.149 | -0.042 | 0 | 0 | 0 | 0.79 | 0.96 | 0.74 | 0.86 | 0.74 | 0.88 |
| 11 | 91710036 | rs7931977 | -0.809 | 0.206 | -1.611 | -1.143 | -0.058 | 0.054 | 0.054 | 0 | 0.85 | 0.98 | 0.87 | 0.94 | 0.79 | 0.89 |
| 11 | 91714466 | rs7940613 | -0.172 | -0.028 | -1.342 | -1.152 | -0.085 | 0.152 | 0.152 | 0 | 0.54 | 0.92 | 0.58 | 0.8 | 0.51 | 0.68 |
| 11 | 91724817 | rs7109140 | -0.827 | 0.141 | -0.356 | -0.765 | 0.004 | -0.001 | -0.001 | 0 | 0.29 | 0.62 | 0.32 | 0.23 | 0.32 | 0.16 |
| 11 | 91732586 | rs10765509 | -1.57 | -0.016 | -0.25 | -0.83 | -0.09 | -0.065 | -0.065 | 0.003 | 0.64 | 0.96 | 0.54 | 0.31 | 0.57 | 0.41 |
| 11 | 91753776 | rs11828718 | 0.256 | -0.091 | -0.144 | -0.881 | 0.013 | 0.027 | 0.027 | 0.034 | 0.34 | 0.29 | 0.21 | 0.08 | 0.26 | 0.25 |
| 11 | 91757497 | rs4753362 | -0.728 | -0.063 | 0.287 | -0.742 | 0.016 | -0.002 | -0.002 | 0 | 0.29 | 0.52 | 0.31 | 0.23 | 0.3 | 0.16 |

| | | | | | | | | | | | | | | | | |
|---|---|---|---|---|---|---|---|---|---|---|---|---|---|---|---|---|
| 11 | 91758470 | rs7122501 | 0.433 | -0.068 | 0.57 | -0.766 | 0.053 | -0.013 | -0.013 | 0 | 0.07 | 0.29 | 0.03 | 0 | 0.04 | 0.01 |
| 11 | 91769457 | rs1845903 | 0.046 | -0.063 | 0.555 | -0.761 | 0.001 | 0.001 | 0.001 | 0 | 0.22 | 0.23 | 0.28 | 0.23 | 0.26 | 0.14 |
| 11 | 91770585 | rs1692668 | -0.766 | -0.07 | 0.553 | -0.732 | 0.001 | -0.001 | -0.001 | 0 | 0.29 | 0.63 | 0.32 | 0.23 | 0.32 | 0.17 |
| 11 | 91778933 | rs1955073 | -0.621 | -0.015 | 0.053 | -0.674 | 0.019 | -0.033 | -0.033 | 0.014 | 0.28 | 0.48 | 0.31 | 0.17 | 0.31 | 0.16 |
| 11 | 91780853 | rs2511420 | -1.406 | -0.01 | -0.106 | -0.751 | -0.101 | -0.052 | -0.052 | 0 | 0.62 | 0.96 | 0.53 | 0.3 | 0.55 | 0.41 |
| 11 | 91786077 | rs12790789 | 1.079 | 0.022 | 0.007 | -0.974 | -0.02 | 0.07 | 0.07 | 0 | 0.24 | 0.01 | 0.15 | 0.07 | 0.2 | 0.19 |
| 11 | 91790899 | rs11019773 | -1.109 | 0.424 | 0.749 | -0.469 | 0.016 | -0.036 | -0.036 | 0 | 0.28 | 0.51 | 0.31 | 0.16 | 0.31 | 0.15 |
| 11 | 91791869 | rs1692681 | -0.387 | 0.409 | 1.133 | -0.588 | 0.025 | -0.036 | -0.036 | 0.014 | 0.08 | 0.34 | 0.04 | 0 | 0.04 | 0.01 |
| 11 | 91791880 | rs7105421 | -0.511 | 0.405 | 1.127 | -0.577 | 0.018 | -0.031 | -0.031 | 0.912 | 0.23 | 0.36 | 0.29 | 0.15 | 0.26 | 0.14 |
| 11 | 91792842 | rs1793097 | -1.26 | 0.403 | 1.027 | -0.575 | -0.004 | 0.034 | 0.034 | 0.543 | 0.31 | 0.78 | 0.33 | 0.15 | 0.31 | 0.16 |
| 11 | 91800400 | rs9667030 | -0.559 | 0.469 | 0.369 | -0.215 | 0.022 | 0.038 | 0.038 | 0.195 | 0.23 | 0.38 | 0.31 | 0.26 | 0.26 | 0.14 |
| 11 | 91800915 | rs10765519 | 0.582 | 0.597 | -0.879 | -0.144 | -0.018 | 0.003 | 0.003 | 0.563 | 0.68 | 0.16 | 0.64 | 0.74 | 0.67 | 0.83 |
| 11 | 91802671 | rs2036126 | 0.583 | 0.627 | -0.91 | -0.129 | -0.022 | 0.002 | 0.002 | 0.255 | 0.68 | 0.15 | 0.64 | 0.73 | 0.67 | 0.84 |
| 11 | 91815601 | rs10830834 | 0.636 | 0.668 | -0.858 | -0.096 | -0.011 | 0.006 | 0.006 | 0 | 0.68 | 0.15 | 0.67 | 0.74 | 0.68 | 0.82 |
| 11 | 91817016 | rs10047440 | 0.612 | 0.37 | -1.683 | -0.358 | -0.089 | -0.017 | -0.017 | 0 | 0.43 | 0.04 | 0.46 | 0.66 | 0.44 | 0.64 |
| 11 | 91820234 | rs16916946 | 0.926 | 0.325 | -0.865 | -0.357 | 0.003 | 0.008 | 0.008 | 0 | 0.24 | 0.2 | 0.29 | 0.12 | 0.21 | 0.18 |
| 11 | 91820850 | rs979928 | -0.632 | 0.063 | -0.083 | -0.469 | -0.012 | -0.02 | -0.02 | 0 | 0.92 | 0.63 | 0.97 | 1 | 0.99 | 1 |
| 11 | 91821986 | rs1496678 | 0.089 | 0.131 | -0.404 | -0.475 | -0.049 | -0.015 | -0.015 | 0.098 | 0.19 | 0.08 | 0.15 | 0.19 | 0.26 | 0.16 |
| 11 | 91832785 | rs1390774 | -0.367 | -0.026 | -0.147 | -0.563 | -0.009 | -0.03 | -0.03 | 0 | 0.66 | 0.42 | 0.71 | 0.92 | 0.76 | 0.83 |
| 11 | 91833673 | rs10830842 | 0.75 | -0.007 | -0.135 | -0.51 | -0.028 | 0.049 | 0.049 | 0 | 0.45 | 0.21 | 0.5 | 0.69 | 0.46 | 0.66 |
| 11 | 91833737 | rs1027455 | -0.006 | 0.041 | -0.108 | -0.513 | -0.038 | -0.088 | -0.088 | 0 | 0.87 | 0.49 | 0.93 | 1 | 0.97 | 0.95 |
| 11 | 91840503 | rs10830843 | 0.083 | 0.041 | -0.226 | -0.508 | -0.099 | 0.051 | 0.051 | 0 | 0.38 | 0.1 | 0.31 | 0.26 | 0.47 | 0.3 |
| 11 | 91854828 | rs11019822 | 0.534 | 0.062 | -0.256 | -0.514 | -0.056 | -0.037 | -0.037 | 0 | 0.15 | 0.05 | 0.14 | 0.19 | 0.26 | 0.12 |
| 11 | 91857262 | rs4753392 | 1.368 | -0.091 | -0.255 | -0.546 | -0.038 | 0.101 | 0.101 | 0 | 0.22 | 0.01 | 0.17 | 0.07 | 0.21 | 0.17 |
| 11 | 91868956 | rs3996554 | 0.487 | -0.45 | 0.02 | -0.306 | -0.031 | -0.021 | -0.021 | 0 | 0.86 | 0.32 | 0.83 | 0.95 | 0.96 | 0.94 |
| 11 | 91875866 | rs10741438 | -0.227 | 0.15 | 0.071 | 0.005 | 0.005 | 0.073 | 0.073 | 0 | 0.53 | 0.31 | 0.64 | 0.74 | 0.54 | 0.72 |
| 11 | 91882304 | rs1603484 | -0.44 | 0.166 | 0.127 | 0.045 | 0 | 0.006 | 0.006 | 0 | 0.55 | 0.44 | 0.65 | 0.74 | 0.54 | 0.72 |
| 11 | 91898660 | rs10830859 | 1.081 | 0.15 | -0.355 | 0.713 | -0.004 | -0.12 | -0.12 | 0.397 | 0.19 | 0.01 | 0.25 | 0.37 | 0.16 | 0.31 |
| 11 | 91913165 | rs10501788 | 0.438 | 0.436 | 0.065 | 0.408 | -0.012 | -0.008 | -0.008 | 0 | 0.06 | 0 | 0.05 | 0.05 | 0.05 | 0.14 |
| 11 | 91929606 | rs11602557 | -0.095 | 0.432 | 0.358 | 0.474 | 0.004 | 0.073 | 0.073 | 0 | 0.58 | 0.22 | 0.65 | 0.75 | 0.59 | 0.74 |
| 11 | 91930698 | rs12280098 | -0.118 | 0.645 | -0.006 | 0.578 | 0 | -0.022 | -0.022 | 0 | 0.6 | 0.55 | 0.68 | 0.74 | 0.59 | 0.75 |
| 11 | 91938118 | rs6483169 | 0.028 | 0.786 | -0.467 | 0.635 | 0.001 | 0.033 | 0.033 | 0 | 0.53 | 0.38 | 0.52 | 0.69 | 0.53 | 0.68 |
| 11 | 91945888 | rs7110069 | 0.718 | 1.096 | 0.16 | 1.513 | 0.036 | 0.103 | 0.103 | 0.993 | 0.86 | 0.71 | 0.85 | 0.7 | 0.79 | 0.89 |
| 11 | 91946830 | rs2125301 | 1.132 | 0.865 | 0.64 | 1.693 | 0.025 | 0.115 | 0.115 | 0 | 0.74 | 0.52 | 0.77 | 0.53 | 0.67 | 0.81 |
| 11 | 91949035 | rs1379801 | 0.504 | 0.855 | 1.11 | 1.864 | 0.001 | 0.021 | 0.021 | 0 | 0.83 | 0.81 | 0.84 | 0.63 | 0.77 | 0.85 |
| 11 | 91949784 | rs7104171 | -0.727 | -0.315 | 0.865 | 1.487 | -0.017 | 0.009 | 0.009 | 0 | 0.35 | 0.37 | 0.47 | 0.41 | 0.29 | 0.44 |
| 11 | 91950217 | rs7926821 | 0.36 | -0.316 | 0.879 | 1.451 | -0.012 | 0.009 | 0.009 | 0 | 0.1 | 0.05 | 0.13 | 0.37 | 0.19 | 0.09 |
| 11 | 91951447 | rs10830888 | 1.617 | -0.325 | 0.883 | 1.424 | 0.012 | 0.074 | 0.074 | 0 | 0.48 | 0.36 | 0.36 | 0.21 | 0.48 | 0.41 |
| 11 | 91956022 | rs4753404 | 0.344 | -0.207 | 0.807 | 1.571 | -0.02 | -0.012 | -0.012 | 0 | 0.1 | 0.03 | 0.12 | 0.35 | 0.19 | 0.09 |
| 11 | 91956925 | rs11019875 | -0.577 | -0.25 | 0.89 | 1.417 | 0.024 | 0.028 | 0.028 | 0.48 | 0.08 | 0.18 | 0.06 | 0.09 | 0.1 | 0.03 |
| 11 | 91958476 | rs1458209 | -1.083 | -0.243 | 0.86 | 1.397 | -0.004 | 0 | 0 | 0.042 | 0.39 | 0.35 | 0.49 | 0.42 | 0.32 | 0.51 |
| 11 | 91960472 | rs1458206 | -0.655 | -0.24 | 0.79 | 1.557 | 0.028 | -0.017 | -0.017 | 0 | 0.34 | 0.22 | 0.45 | 0.4 | 0.29 | 0.44 |
| 11 | 91961726 | rs7118371 | -0.572 | -0.225 | 0.78 | 1.526 | 0.013 | 0.015 | 0.015 | 0.345 | 0.08 | 0.13 | 0.06 | 0.09 | 0.1 | 0.03 |
| 11 | 91969080 | rs16917196 | 0.339 | -0.237 | 0.596 | 1.563 | 0.066 | -0.044 | -0.044 | 0 | 0.25 | 0.08 | 0.39 | 0.31 | 0.19 | 0.41 |
| 11 | 91970203 | rs1187171 | -0.556 | 0.443 | 1.012 | 1.787 | 0.014 | 0.071 | 0.071 | 0 | 0.12 | 0.15 | 0.14 | 0.37 | 0.2 | 0.09 |
| 11 | 91973528 | rs897989 | -1.697 | 0.414 | 0.989 | 1.641 | 0.049 | 0.148 | 0.148 | 0 | 0.21 | 0.39 | 0.21 | 0.46 | 0.3 | 0.12 |
| 11 | 91978712 | rs1187164 | -0.102 | 0.351 | 0.795 | 0.931 | 0.001 | 0.002 | 0.002 | 0.016 | 0.32 | 0.28 | 0.41 | 0.26 | 0.29 | 0.54 |
| 11 | 91979868 | rs10501774 | -0.672 | 0.177 | 1.034 | 0.999 | 0.004 | 0.044 | 0.044 | 0 | 0.21 | 0.3 | 0.19 | 0.37 | 0.25 | 0.12 |
| 11 | 91981960 | rs7935097 | -0.583 | 0.171 | 1.11 | 0.996 | 0 | 0.012 | 0.012 | 0 | 0.2 | 0.25 | 0.18 | 0.37 | 0.25 | 0.12 |
| 11 | 91988615 | rs4542368 | -1.24 | 0.072 | 1.189 | 0.949 | -0.004 | 0.059 | 0.059 | 0.737 | 0.25 | 0.57 | 0.21 | 0.37 | 0.25 | 0.11 |
| 11 | 91988838 | rs947938 | -1.687 | 0.007 | 0.858 | 0.829 | -0.061 | 0.021 | 0.021 | 0 | 0.59 | 0.86 | 0.62 | 0.63 | 0.56 | 0.65 |
| 11 | 91989508 | rs751958 | 0.409 | 0.173 | 0.805 | 0.956 | 0.005 | 0.011 | 0.011 | 0.027 | 0.31 | 0.17 | 0.4 | 0.26 | 0.29 | 0.53 |
| 11 | 91992044 | rs1187173 | -0.926 | 0.347 | 1.107 | 0.952 | 0.01 | -0.037 | -0.037 | 0.616 | 0.07 | 0.12 | 0.07 | 0 | 0.09 | 0.03 |
| 11 | 91994044 | rs16917259 | -0.819 | 0.332 | 0.859 | 1.225 | 0.008 | 0.079 | 0.079 | 0 | 0.15 | 0.26 | 0.15 | 0.36 | 0.18 | 0.09 |
| 11 | 91996037 | rs6483171 | -1.515 | 0.196 | 0.485 | 1.227 | 0.009 | 0.09 | 0.09 | 0.095 | 0.23 | 0.45 | 0.2 | 0.37 | 0.25 | 0.11 |
| 11 | 91998485 | rs1993543 | -0.977 | -0.627 | -0.186 | 0.677 | -0.028 | -0.011 | -0.011 | 0 | 0.31 | 0.6 | 0.2 | 0.31 | 0.21 | 0.2 |
| 11 | 92001014 | rs1187176 | -0.245 | -0.301 | -0.493 | 0.375 | 0.003 | 0.003 | 0.003 | 0 | 0.06 | 0.05 | 0.03 | 0 | 0.06 | 0.01 |
| 11 | 92005611 | rs10501777 | 0.175 | -0.669 | -1.029 | 0.604 | 0.011 | -0.057 | -0.057 | 0 | 0.17 | 0.06 | 0.1 | 0.31 | 0.14 | 0.16 |
| 11 | 92016605 | rs10830893 | 1.207 | -0.384 | -1.713 | 0.251 | -0.135 | -0.073 | -0.073 | 0 | 0.29 | 0.02 | 0.4 | 0.33 | 0.45 | 0.48 |
| 11 | 92017415 | rs2513466 | 0.552 | -0.549 | -0.94 | 0.618 | 0.012 | 0.026 | 0.026 | 0 | 0.52 | 0.71 | 0.52 | 0.44 | 0.59 | 0.53 |
| 11 | 92019340 | rs2845877 | 0.611 | -0.151 | -0.432 | 1.182 | -0.008 | -0.056 | -0.056 | 0 | 0.65 | 0.81 | 0.69 | 0.48 | 0.68 | 0.6 |
| 11 | 92021289 | rs1870299 | 0.778 | -0.137 | -0.189 | 1.185 | -0.003 | 0.007 | 0.007 | 0 | 0.63 | 0.53 | 0.67 | 0.48 | 0.68 | 0.6 |
| 11 | 92026446 | rs3133370 | 0.768 | -0.133 | -0.187 | 1.17 | -0.003 | 0.008 | 0.008 | 0 | 0.63 | 0.54 | 0.67 | 0.48 | 0.68 | 0.59 |
| 11 | 92034702 | rs722266 | -0.316 | 0.158 | -0.588 | 0.857 | -0.028 | -0.03 | -0.03 | 0 | 0.83 | 0.93 | 0.79 | 0.78 | 0.82 | 0.76 |
| 11 | 92036731 | rs877360 | -0.33 | 0.173 | -0.582 | 0.864 | -0.028 | -0.03 | -0.03 | 0 | 0.83 | 0.94 | 0.79 | 0.79 | 0.82 | 0.76 |
| 11 | 92041853 | rs1187154 | -1.298 | -0.278 | -0.343 | 0.591 | 0.016 | 0.019 | 0.019 | 0 | 0.31 | 0.79 | 0.23 | 0.25 | 0.2 | 0.21 |
| 11 | 92046408 | rs6483173 | -1.378 | -0.248 | 1.046 | 0.298 | 0.017 | -0.051 | -0.051 | 0 | 0.1 | 0.41 | 0.07 | 0 | 0.05 | 0 |
| 11 | 92056136 | rs12789844 | -0.064 | -0.107 | 0.984 | 0.329 | -0.011 | 0.019 | 0.019 | 0 | 0.15 | 0.04 | 0.08 | 0.1 | 0.13 | 0.17 |
| 11 | 92061826 | rs16917391 | -0.592 | -0.187 | 1.525 | 0.371 | 0.018 | 0.056 | 0.056 | 0 | 0.05 | 0.16 | 0.03 | 0.09 | 0.03 | 0.02 |

| | | | | | | | | | | | | | | | | |
|---|---|---|---|---|---|---|---|---|---|---|---|---|---|---|---|---|
| 11 | 92071856 | rs12363932 | -0.772 | -0.228 | 1.974 | 0.346 | 0.011 | -0.031 | -0.031 | 0 | 0.06 | 0.14 | 0.03 | 0 | 0.04 | 0 |
| 11 | 92079552 | rs10501779 | 0.25 | -0.447 | 1.398 | 0.488 | 0.001 | -0.097 | -0.097 | 1.143 | 0.2 | 0.04 | 0.27 | 0.33 | 0.2 | 0.22 |
| 11 | 92082281 | rs11607764 | -0.071 | -0.187 | 2.283 | 0.045 | -0.003 | 0.021 | 0.021 | 0 | 0.14 | 0.06 | 0.07 | 0.07 | 0.12 | 0.15 |
| 11 | 92086513 | rs10830902 | -0.866 | -0.177 | 2.339 | 0.051 | 0.019 | -0.029 | -0.029 | 0 | 0.06 | 0.15 | 0.04 | 0 | 0.04 | 0 |
| 11 | 92090489 | rs4554857 | -2.231 | -0.206 | 2.299 | 0.039 | 0.017 | 0.059 | 0.059 | 0 | 0.3 | 0.85 | 0.18 | 0.2 | 0.2 | 0.19 |
| 11 | 92097811 | rs16917427 | -0.274 | -0.195 | 2.348 | 0.057 | -0.004 | 0.012 | 0.012 | 0 | 0.15 | 0.08 | 0.08 | 0.1 | 0.13 | 0.18 |
| 11 | 92099379 | rs11019927 | -0.855 | -0.146 | 2.423 | 0.031 | 0.019 | -0.028 | -0.028 | 0 | 0.05 | 0.15 | 0.04 | 0 | 0.04 | 0 |
| 11 | 92106781 | rs11019937 | -2.235 | -0.166 | 2.413 | 0.034 | 0.002 | 0.076 | 0.076 | 0 | 0.31 | 0.87 | 0.19 | 0.18 | 0.21 | 0.2 |
| 11 | 92110176 | rs11825546 | -0.549 | -0.166 | 2.511 | 0.041 | -0.001 | -0.005 | -0.005 | 0 | 0.17 | 0.2 | 0.09 | 0.17 | 0.13 | 0.17 |
| 11 | 92117196 | rs11822486 | -1.862 | -0.163 | 2.348 | 0.042 | -0.002 | -0.003 | -0.003 | 0.963 | 0.25 | 0.7 | 0.16 | 0.18 | 0.17 | 0.2 |
| 11 | 92136450 | rs10466348 | -1.416 | -0.227 | 2.159 | 0.205 | 0.012 | -0.04 | -0.04 | 0.377 | 0.09 | 0.16 | 0.05 | 0.01 | 0.08 | 0.02 |
| 11 | 92145174 | rs10741441 | -0.063 | -0.226 | 2.175 | 0.21 | -0.001 | 0 | 0 | 0.037 | 0.87 | 0.9 | 0.92 | 0.9 | 0.9 | 0.83 |
| 11 | 92151079 | rs648027 | 1.945 | -0.245 | 2.462 | 0.207 | 0.005 | 0.01 | 0.01 | 0 | 0.73 | 0.25 | 0.83 | 0.79 | 0.82 | 0.8 |
| 11 | 92156511 | rs17514109 | -1.771 | -0.193 | 2.132 | 0.205 | 0.007 | -0.054 | -0.054 | 0 | 0.11 | 0.21 | 0.05 | 0.01 | 0.08 | 0.02 |
| 11 | 92170472 | rs636679 | 0.117 | -0.144 | 1.95 | 0.285 | -0.003 | 0.004 | 0.004 | 0 | 0.85 | 0.89 | 0.9 | 0.87 | 0.87 | 0.83 |
| 11 | 92181793 | rs7125792 | -1.355 | -0.571 | 2.069 | 0.193 | 0.005 | -0.058 | -0.058 | 0.072 | 0.11 | 0.4 | 0.06 | 0.01 | 0.06 | 0.04 |
| 11 | 92188498 | rs652365 | 1.73 | -0.633 | 2.054 | 0.182 | 0.041 | 0.042 | 0.042 | 0 | 0.69 | 0.17 | 0.82 | 0.78 | 0.83 | 0.79 |
| 11 | 92195193 | rs646518 | 0.156 | -0.393 | 1.168 | 0.088 | 0.006 | 0.009 | 0.009 | 0 | 0.84 | 0.88 | 0.91 | 0.89 | 0.89 | 0.83 |
| 11 | 92196585 | rs7121018 | -1.217 | -0.368 | 1.125 | 0.098 | 0.003 | -0.053 | -0.053 | 0 | 0.1 | 0.31 | 0.05 | 0.01 | 0.06 | 0.04 |
| 11 | 92201077 | rs689349 | 0.892 | -0.235 | 0.63 | 0.204 | -0.033 | -0.078 | -0.078 | 0.023 | 0.72 | 0.45 | 0.85 | 0.88 | 0.83 | 0.8 |
| 11 | 92209853 | rs1941438 | 0.897 | -0.176 | 0.325 | 0.434 | -0.021 | -0.011 | -0.011 | 0 | 0.62 | 0.52 | 0.73 | 0.66 | 0.7 | 0.62 |
| 11 | 92211361 | rs10501790 | -1.566 | -0.107 | 0.899 | 0.678 | -0.004 | 0.014 | 0.014 | 0.022 | 0.09 | 0.19 | 0.05 | 0.11 | 0.06 | 0.08 |
| 11 | 92213090 | rs2508832 | 0.359 | 0.416 | 0.543 | 0.05 | -0.061 | 0.066 | 0.066 | 0 | 0.67 | 0.84 | 0.75 | 0.81 | 0.52 | 0.71 |
| 11 | 92213728 | rs624609 | 1.67 | 0.145 | 0.878 | -0.043 | 0.009 | 0.082 | 0.082 | 0 | 0.49 | 0.11 | 0.56 | 0.62 | 0.41 | 0.53 |
| 11 | 92229998 | rs543704 | 0.932 | 0.114 | 0.889 | -0.067 | -0.006 | 0.017 | 0.017 | 0 | 0.59 | 0.64 | 0.7 | 0.69 | 0.46 | 0.62 |
| 11 | 92234533 | rs615011 | 1.695 | 0.131 | 0.857 | 0.234 | 0.01 | 0.013 | 0.013 | 0 | 0.49 | 0.12 | 0.48 | 0.52 | 0.4 | 0.52 |
| 11 | 92236390 | rs16917633 | -0.269 | 0.131 | 0.998 | 0.263 | 0.003 | -0.003 | -0.003 | 0 | 0.05 | 0.03 | 0.08 | 0.07 | 0.04 | 0.08 |
| 11 | 92237285 | rs17600259 | -0.9 | 0.12 | 1.052 | 0.23 | -0.004 | 0.006 | 0.006 | 0.203 | 0.08 | 0.14 | 0.05 | 0.11 | 0.06 | 0.07 |
| 11 | 92253513 | rs605714 | 0.274 | 0.123 | 0.913 | 0.179 | -0.056 | 0.029 | 0.029 | 0 | 0.67 | 0.83 | 0.67 | 0.71 | 0.52 | 0.7 |
| 11 | 92260106 | rs605677 | 1.127 | 0.118 | 1.412 | 0.206 | -0.004 | 0.006 | 0.006 | 0 | 0.57 | 0.41 | 0.59 | 0.6 | 0.46 | 0.62 |
| 11 | 92267129 | rs4753063 | -1.381 | 0.092 | 1.584 | 0.195 | 0 | -0.005 | -0.005 | 1.081 | 0.45 | 0.42 | 0.47 | 0.38 | 0.56 | 0.42 |
| 11 | 92274668 | rs488486 | 1.056 | 0.111 | 1.801 | 0.072 | 0.022 | -0.045 | -0.045 | 0.729 | 0.63 | 0.46 | 0.68 | 0.79 | 0.53 | 0.75 |
| 11 | 92278522 | rs3911565 | 0.006 | 0.127 | 1.832 | 0.064 | -0.021 | 0.041 | 0.041 | 0 | 0.14 | 0.2 | 0.18 | 0.28 | 0.11 | 0.23 |
| 11 | 92282534 | rs503268 | 1.884 | 0.098 | 1.826 | 0.108 | 0.017 | 0.041 | 0.041 | 0 | 0.49 | 0.26 | 0.5 | 0.52 | 0.42 | 0.51 |
| 11 | 92289734 | rs586699 | 1.12 | 0.095 | 1.837 | 0.104 | 0.007 | -0.022 | -0.022 | 0 | 0.62 | 0.4 | 0.67 | 0.8 | 0.53 | 0.75 |
| 11 | 92290876 | rs12287186 | -0.186 | 0.141 | 1.049 | 0.087 | -0.066 | 0.07 | 0.07 | 0 | 0.25 | 0.12 | 0.25 | 0.09 | 0.41 | 0.18 |
| 11 | 92291060 | rs563608 | 1.655 | 0.037 | 1.018 | 0.186 | 0.008 | 0.038 | 0.038 | 0 | 0.49 | 0.2 | 0.49 | 0.52 | 0.42 | 0.51 |
| 11 | 92303014 | rs591897 | 1.655 | 0.037 | 1.018 | 0.186 | 0.008 | 0.038 | 0.038 | 0 | 0.49 | 0.2 | 0.49 | 0.52 | 0.42 | 0.51 |
| 11 | 92307679 | rs568629 | 0.653 | 0.033 | 1.063 | 0.208 | -0.056 | 0.069 | 0.069 | 0 | 0.8 | 0.41 | 0.77 | 0.61 | 0.88 | 0.7 |
| 11 | 92313072 | rs11020013 | -0.585 | 0.183 | 0.879 | 0.149 | -0.039 | 0.047 | 0.047 | 0.559 | 0.32 | 0.22 | 0.28 | 0.09 | 0.46 | 0.19 |
| 11 | 92313508 | rs502357 | 0.613 | -0.11 | 1.053 | 0.639 | -0.055 | 0.138 | 0.138 | 0 | 0.82 | 0.52 | 0.79 | 0.63 | 0.88 | 0.71 |
| 11 | 92316799 | rs1792351 | -1.03 | 0.402 | 0.554 | 0.637 | -0.017 | 0.023 | 0.023 | 0.285 | 0.4 | 0.7 | 0.5 | 0.31 | 0.5 | 0.35 |
| 11 | 92317747 | rs10501793 | -1.621 | 0.482 | 1.532 | 0.431 | -0.008 | 0.005 | 0.005 | 1.431 | 0.19 | 0.5 | 0.17 | 0.19 | 0.15 | 0.2 |
| 11 | 92317858 | rs1622987 | -0.275 | 0.605 | 1.675 | 0.555 | 0.013 | -0.006 | -0.006 | 0.053 | 0.21 | 0.22 | 0.32 | 0.14 | 0.34 | 0.17 |
| 11 | 92319098 | rs541822 | -1.005 | 0.48 | 2.13 | 0.553 | 0.003 | 0.007 | 0.007 | 0.553 | 0.16 | 0.28 | 0.15 | 0.19 | 0.14 | 0.18 |
| 11 | 92320830 | rs529911 | -1.382 | 0.525 | 2.243 | 0.553 | 0.002 | 0.007 | 0.007 | 1.092 | 0.18 | 0.39 | 0.16 | 0.19 | 0.15 | 0.19 |
| 11 | 92327878 | rs522732 | -1.677 | 0.583 | 2.213 | 0.557 | 0.016 | -0.008 | -0.008 | 0 | 0.37 | 0.51 | 0.47 | 0.33 | 0.48 | 0.35 |
| 11 | 92334070 | rs519175 | -1.461 | 0.54 | 2.427 | 0.537 | 0.037 | -0.037 | -0.037 | 0 | 0.34 | 0.42 | 0.45 | 0.18 | 0.47 | 0.25 |
| 11 | 92337021 | rs499595 | -0.468 | 0.545 | 2.427 | 0.535 | 0.008 | -0.024 | -0.024 | 0 | 0.12 | 0.18 | 0.12 | 0.05 | 0.13 | 0.08 |
| 11 | 92340670 | rs486365 | -2.196 | 0.374 | 2.467 | 0.664 | -0.062 | 0.018 | 0.018 | 0.338 | 0.42 | 0.82 | 0.5 | 0.39 | 0.49 | 0.37 |
| 11 | 92358811 | rs505423 | -1.849 | 0.386 | 2.484 | 0.667 | 0.012 | -0.003 | -0.003 | 5.079 | 0.39 | 0.5 | 0.47 | 0.39 | 0.48 | 0.37 |
| 11 | 92360915 | rs1792341 | -2.261 | 0.383 | 2.384 | 0.665 | -0.079 | 0.014 | 0.014 | 0.466 | 0.43 | 0.86 | 0.5 | 0.39 | 0.49 | 0.37 |
| 11 | 92379485 | rs1458321 | -0.232 | 0.457 | 2.382 | 0.665 | 0.012 | -0.004 | -0.004 | 0 | 0.2 | 0.2 | 0.32 | 0.14 | 0.34 | 0.17 |
| 11 | 92382788 | rs1674083 | -0.585 | 0.462 | 2.394 | 0.662 | 0.002 | -0.015 | -0.015 | 0 | 0.13 | 0.15 | 0.12 | 0.05 | 0.13 | 0.08 |
| 11 | 92409546 | rs1791573 | -2.113 | 0.456 | 2.183 | 0.643 | 0.002 | -0.002 | -0.002 | 0 | 0.4 | 0.55 | 0.46 | 0.38 | 0.48 | 0.37 |
| 11 | 92432549 | rs2676173 | -0.713 | 0.541 | 1.789 | 0.642 | 0.009 | -0.043 | -0.043 | 0 | 0.13 | 0.22 | 0.13 | 0.05 | 0.13 | 0.1 |
| 11 | 92443925 | rs1792361 | -0.489 | 0.648 | 1.456 | 0.73 | 0.047 | -0.024 | -0.024 | 0.189 | 0.21 | 0.32 | 0.32 | 0.13 | 0.34 | 0.16 |
| 11 | 92460320 | rs1791569 | -0.78 | 0.825 | 1.298 | 0.619 | 0.063 | -0.041 | -0.041 | 0 | 0.23 | 0.44 | 0.33 | 0.13 | 0.34 | 0.16 |
| 11 | 92490972 | rs2450944 | 1.623 | 0.447 | 0.91 | 0.602 | -0.087 | 0.036 | 0.036 | 0.307 | 0.56 | 0.11 | 0.5 | 0.63 | 0.5 | 0.62 |
| 11 | 92531356 | rs7949157 | 0.956 | 0.311 | 1.353 | 0.583 | -0.015 | 0.032 | 0.032 | 0.006 | 0.83 | 0.65 | 0.85 | 0.76 | 0.86 | 0.8 |
| 11 | 92532677 | rs12362722 | 1.013 | 0.248 | 1.431 | 0.485 | -0.008 | 0.007 | 0.007 | 0 | 0.83 | 0.77 | 0.86 | 0.76 | 0.85 | 0.79 |
| 11 | 92536693 | rs3912650 | 0.225 | 0.155 | 1.408 | 0.425 | 0.002 | -0.026 | -0.026 | 0 | 0.88 | 0.84 | 0.87 | 0.95 | 0.87 | 0.9 |
| 11 | 92540552 | rs10741444 | 1.819 | 0.167 | 1.267 | 1.033 | -0.081 | -0.046 | -0.046 | 0 | 0.56 | 0.1 | 0.44 | 0.53 | 0.52 | 0.62 |
| 11 | 92547343 | rs3912659 | 1.355 | 0.137 | 1.263 | 1.003 | -0.059 | 0.121 | 0.121 | 0.333 | 0.6 | 0.18 | 0.51 | 0.83 | 0.53 | 0.72 |
| 11 | 92556100 | rs17543864 | -0.178 | 0.083 | 1.658 | 0.974 | -0.04 | 0.027 | 0.027 | 0 | 0.22 | 0.11 | 0.3 | 0.13 | 0.33 | 0.17 |
| 11 | 92556824 | rs3847534 | 1.689 | 0.04 | 1.929 | 1.005 | -0.033 | 0.092 | 0.092 | 0.08 | 0.81 | 0.52 | 0.78 | 0.65 | 0.85 | 0.78 |
| 11 | 92557642 | rs10741446 | 1.509 | 0.151 | 1.685 | 0.732 | -0.053 | 0.124 | 0.124 | 0.208 | 0.59 | 0.19 | 0.52 | 0.83 | 0.53 | 0.73 |
| 11 | 92562200 | rs7945975 | 0.692 | 0.188 | 1.66 | 0.828 | -0.001 | -0.038 | -0.038 | 0 | 0.87 | 0.81 | 0.88 | 0.95 | 0.87 | 0.9 |
| 11 | 92568552 | rs10830950 | -0.971 | -0.208 | 0.413 | 0.753 | 0.007 | -0.004 | -0.004 | 0.903 | 0.45 | 0.33 | 0.42 | 0.45 | 0.42 | 0.4 |

| | | | | | | | | | | | | | | | | |
|---|---|---|---|---|---|---|---|---|---|---|---|---|---|---|---|---|
| 11 | 92572940 | rs7104420 | -0.943 | -0.155 | 0.433 | 0.775 | 0.007 | -0.007 | -0.007 | 0.059 | 0.44 | 0.33 | 0.42 | 0.45 | 0.41 | 0.4 |
| 11 | 92578985 | rs11020080 | 1.244 | 0.133 | 0.261 | 0.736 | -0.019 | 0.126 | 0.126 | 0 | 0.43 | 0.12 | 0.33 | 0.21 | 0.39 | 0.51 |
| 11 | 92583665 | rs7112956 | -0.163 | -0.01 | 0.149 | 0.898 | 0.023 | -0.009 | -0.009 | 0 | 0.24 | 0.16 | 0.13 | 0.25 | 0.16 | 0.23 |
| 11 | 92590448 | rs4753069 | -0.992 | 0.02 | 0.246 | 0.768 | -0.002 | 0.034 | 0.034 | 0 | 0.93 | 0.92 | 0.8 | 0.67 | 0.81 | 0.91 |
| 11 | 92594957 | rs3847550 | -0.968 | -0.035 | 0.32 | 0.768 | 0.008 | -0.01 | -0.01 | 0 | 0.44 | 0.3 | 0.42 | 0.46 | 0.41 | 0.4 |
| 11 | 92601816 | rs10830951 | 0.794 | -0.001 | 0.663 | 0.97 | -0.035 | -0.025 | -0.025 | 0 | 0.19 | 0.07 | 0.28 | 0.23 | 0.25 | 0.17 |
| 11 | 92602182 | rs10741449 | -0.591 | 0.027 | 0.731 | 0.878 | 0.01 | 0.052 | 0.052 | 0 | 0.92 | 0.88 | 0.8 | 0.67 | 0.81 | 0.91 |
| 11 | 92602861 | rs6483194 | 0.782 | 0.042 | 0.798 | 0.874 | 0.083 | 0.233 | 0.233 | 0.186 | 0.88 | 0.52 | 0.76 | 0.66 | 0.81 | 0.91 |
| 11 | 92610206 | rs6483198 | -0.681 | 0.019 | 0.57 | 0.776 | -0.004 | 0.014 | 0.014 | 0 | 0.92 | 0.91 | 0.8 | 0.69 | 0.81 | 0.86 |
| 11 | 92612151 | rs11020089 | 0.121 | -0.132 | 0.23 | 0.488 | 0.024 | 0.077 | 0.077 | 0 | 0.79 | 0.65 | 0.65 | 0.55 | 0.72 | 0.74 |
| 11 | 92612185 | rs12792265 | -0.352 | -0.156 | 0.017 | 0.384 | 0.005 | -0.018 | -0.018 | 0 | 0.44 | 0.24 | 0.42 | 0.5 | 0.41 | 0.36 |
| 11 | 92618272 | rs1488859 | -0.98 | 0.04 | -0.163 | 0.522 | -0.003 | 0.014 | 0.014 | 0 | 0.92 | 0.93 | 0.8 | 0.69 | 0.82 | 0.86 |
| 11 | 92618875 | rs768877 | 0.146 | 0.014 | -0.152 | 0.503 | 0.017 | 0.057 | 0.057 | 0 | 0.78 | 0.67 | 0.65 | 0.55 | 0.72 | 0.73 |
| 11 | 92622011 | rs4146883 | 0.33 | 0.057 | 0.397 | 0.358 | 0.005 | 0.013 | 0.013 | 0 | 0.33 | 0.43 | 0.44 | 0.38 | 0.33 | 0.28 |
| 11 | 92623349 | rs7118248 | 0 | 0.005 | -0.085 | 0.446 | 0.003 | -0.034 | -0.034 | 0 | 0.68 | 0.76 | 0.67 | 0.45 | 0.65 | 0.67 |
| 11 | 92625457 | rs7932490 | 0.153 | -0.066 | 0.052 | 0.091 | -0.002 | -0.037 | -0.037 | 0 | 0.34 | 0.34 | 0.21 | 0.04 | 0.29 | 0.41 |
| 11 | 92625944 | rs11823754 | 0.22 | -0.034 | -0.226 | 0.185 | 0.001 | 0.051 | 0.051 | 0 | 0.1 | 0.16 | 0.09 | 0.24 | 0.12 | 0.06 |
| 11 | 92626428 | rs2197678 | -0.33 | 0.162 | -0.702 | 0.186 | 0.006 | 0.013 | 0.013 | 0.01 | 0.46 | 0.58 | 0.31 | 0.3 | 0.41 | 0.47 |
| 11 | 92627643 | rs11020093 | 0.199 | 0.49 | -1.221 | -0.476 | -0.033 | 0.095 | 0.095 | 0 | 0.23 | 0.07 | 0.16 | 0.02 | 0.25 | 0.29 |
| 11 | 92636326 | rs10830955 | -0.46 | 0.141 | -1.024 | 0.574 | 0.021 | -0.062 | -0.062 | 0 | 0.19 | 0.11 | 0.21 | 0.41 | 0.16 | 0.26 |
| 11 | 92636478 | rs7945506 | -0.464 | 0.074 | -0.138 | 0.238 | 0.012 | -0.004 | -0.004 | 0 | 0.16 | 0.35 | 0.32 | 0.13 | 0.13 | 0.2 |
| 11 | 92639553 | rs1488841 | -0.524 | 0.164 | -0.088 | 0.229 | 0.019 | -0.006 | -0.006 | 0 | 0.16 | 0.38 | 0.33 | 0.13 | 0.15 | 0.21 |
| 11 | 92644575 | rs750025 | -0.576 | 0.144 | 0.435 | 0.238 | 0.006 | -0.023 | -0.023 | 0 | 0.14 | 0.23 | 0.26 | 0.04 | 0.14 | 0.15 |
| 11 | 92646254 | rs2242342 | 0.166 | 0.086 | 0.201 | 0.379 | 0.02 | -0.09 | -0.09 | 0 | 0.17 | 0.07 | 0.21 | 0.4 | 0.15 | 0.27 |
| 11 | 92648639 | rs10501800 | -1.8 | 0.007 | 0.429 | 0.112 | 0.017 | -0.001 | -0.001 | 0 | 0.44 | 0.74 | 0.61 | 0.52 | 0.39 | 0.5 |
| 11 | 92651002 | rs7950811 | -0.182 | 0.169 | 0.388 | 0.092 | 0.01 | -0.005 | -0.005 | 0 | 0.06 | 0.14 | 0.1 | 0.09 | 0.06 | 0.06 |
| 11 | 92653178 | rs1387154 | -0.882 | 0.085 | 0.562 | 0.156 | 0.008 | -0.015 | -0.015 | 0.059 | 0.2 | 0.32 | 0.35 | 0.14 | 0.22 | 0.2 |
| 11 | 92657189 | rs7110596 | -0.8 | 0.139 | 0.447 | 0.067 | 0.02 | -0.054 | -0.054 | 0 | 0.19 | 0.37 | 0.29 | 0.04 | 0.22 | 0.15 |
| 11 | 92662737 | rs11020104 | -0.38 | 0.05 | 0.011 | 0.636 | -0.005 | 0.027 | 0.027 | 0 | 0.2 | 0.18 | 0.25 | 0.45 | 0.17 | 0.3 |
| 11 | 92664875 | rs737383 | -0.863 | 0.281 | -0.804 | 0.132 | 0.015 | 0.037 | 0.037 | 0 | 0.52 | 0.73 | 0.62 | 0.61 | 0.43 | 0.6 |
| 11 | 92665020 | rs4526739 | -0.107 | 0.356 | -1.151 | 0.299 | -0.019 | -0.038 | -0.038 | 0 | 0.33 | 0.2 | 0.45 | 0.63 | 0.36 | 0.53 |
| 11 | 92665147 | rs12274325 | -0.862 | -0.15 | -0.56 | 0.087 | -0.029 | -0.044 | -0.044 | 0 | 0.15 | 0.34 | 0.24 | 0.03 | 0.08 | 0.1 |
| 11 | 92668826 | rs3847554 | -0.674 | -0.077 | -1.817 | 0.43 | 0.006 | -0.086 | -0.086 | 0 | 0.53 | 0.94 | 0.38 | 0.54 | 0.44 | 0.5 |
| 11 | 92673828 | rs1387153 | 0.049 | -0.485 | -0.615 | 0.563 | -0.017 | 0.02 | 0.02 | 0 | 0.32 | 0.39 | 0.21 | 0.44 | 0.29 | 0.37 |
| 11 | 92691532 | rs2166706 | -0.839 | -0.55 | -0.28 | 0.367 | 0.024 | 0.008 | 0.008 | 0 | 0.45 | 0.7 | 0.33 | 0.45 | 0.41 | 0.47 |
| 11 | 92694757 | rs10830961 | -1.165 | -0.773 | -0.255 | 0.341 | 0.068 | 0.023 | 0.023 | 0 | 0.5 | 0.84 | 0.35 | 0.47 | 0.42 | 0.53 |
| 11 | 92701596 | rs4753426 | -0.443 | -0.728 | -0.372 | 0.1 | 0.001 | 0.007 | 0.007 | 0 | 0.52 | 0.41 | 0.65 | 0.71 | 0.5 | 0.59 |
| 11 | 92704986 | rs4388843 | -0.623 | -0.569 | -0.122 | 0.067 | 0.021 | -0.061 | -0.061 | 3.537 | 0.1 | 0.37 | 0.08 | 0 | 0.09 | 0.1 |
| 11 | 92705158 | rs4601728 | -0.65 | -0.578 | -0.126 | 0.062 | 0.02 | -0.062 | -0.062 | 4.159 | 0.1 | 0.37 | 0.08 | 0 | 0.09 | 0.1 |
| 11 | 92706175 | rs7951037 | -1.228 | -0.345 | 0.123 | 0.154 | -0.037 | 0.024 | 0.024 | 0 | 0.62 | 0.77 | 0.73 | 0.72 | 0.59 | 0.7 |
| 11 | 92708710 | rs10830963 | 0.782 | -0.564 | 0.014 | 0.312 | -0.027 | -0.127 | -0.127 | 0 | 0.28 | 0.03 | 0.19 | 0.42 | 0.29 | 0.43 |
| 11 | 92713507 | rs3781638 | -0.517 | -0.227 | 0.467 | 0.18 | 0.009 | -0.016 | -0.016 | 0 | 0.58 | 0.5 | 0.7 | 0.7 | 0.57 | 0.68 |
| 11 | 92714595 | rs6483210 | -0.397 | -0.392 | 0.395 | 0.154 | 0.006 | -0.032 | -0.032 | 0 | 0.07 | 0.25 | 0.04 | 0 | 0.03 | 0.01 |
| 11 | 92717475 | rs4753073 | -0.739 | -0.298 | 0.145 | 0.276 | 0.01 | -0.017 | -0.017 | 0 | 0.59 | 0.51 | 0.68 | 0.7 | 0.53 | 0.68 |
| 11 | 92725240 | rs6483212 | -0.622 | -0.222 | -0.204 | 0.301 | 0.005 | -0.011 | -0.011 | 0 | 0.58 | 0.5 | 0.69 | 0.7 | 0.53 | 0.68 |
| 11 | 92727874 | rs271046 | -0.722 | -0.975 | -0.785 | 0.213 | 0.096 | 0.113 | 0.113 | 0 | 0.51 | 0.77 | 0.62 | 0.69 | 0.32 | 0.76 |
| 11 | 92742186 | rs956267 | -0.679 | -1.182 | -0.473 | 0.54 | -0.005 | 0.009 | 0.009 | 0.356 | 0.24 | 0.23 | 0.27 | 0.63 | 0.2 | 0.57 |
| 11 | 92742731 | rs12785983 | 0.246 | -0.703 | -0.823 | -0.476 | -0.039 | 0.151 | 0.151 | 0 | 0.28 | 0.07 | 0.19 | 0.02 | 0.32 | 0.13 |
| 11 | 92748060 | rs11827821 | -0.332 | -0.744 | 0.02 | -0.466 | 0.034 | -0.006 | -0.006 | 0.165 | 0.09 | 0.39 | 0.06 | 0.04 | 0.06 | 0.08 |
| 11 | 92749764 | rs10830968 | 0.585 | -0.863 | -0.237 | -0.037 | 0.016 | 0.061 | 0.061 | 0 | 0.13 | 0.23 | 0.11 | 0.24 | 0.16 | 0.25 |
| 11 | 92751333 | rs12270363 | -0.44 | -0.91 | -0.288 | -0.089 | -0.007 | 0.03 | 0.03 | 0 | 0.41 | 0.31 | 0.31 | 0.26 | 0.48 | 0.43 |
| 11 | 92758822 | rs12279732 | 0.318 | -0.871 | -0.064 | -0.37 | -0.007 | 0.031 | 0.031 | 0.376 | 0.29 | 0.22 | 0.2 | 0.01 | 0.31 | 0.17 |
| 11 | 92761705 | rs10830969 | 0.946 | -0.988 | -0.257 | -0.051 | 0.032 | -0.074 | -0.074 | 1.262 | 0.12 | 0.01 | 0.29 | 0.27 | 0.05 | 0.27 |
| 11 | 92764695 | rs599420 | -0.66 | -0.619 | 0.082 | -0.24 | -0.005 | 0.032 | 0.032 | 0.749 | 0.82 | 0.79 | 0.77 | 0.64 | 0.79 | 0.65 |
| 11 | 92770178 | rs1508619 | 0.205 | -0.556 | 0.21 | -0.373 | -0.007 | 0.023 | 0.023 | 0.319 | 0.31 | 0.28 | 0.23 | 0.05 | 0.35 | 0.17 |
| 11 | 92770289 | rs271055 | -0.62 | -0.556 | 0.306 | -0.283 | -0.01 | 0.068 | 0.068 | 0.243 | 0.81 | 0.73 | 0.76 | 0.64 | 0.79 | 0.65 |
| 11 | 92775294 | rs271049 | 0.194 | -0.519 | -0.046 | -0.362 | -0.009 | 0.04 | 0.04 | 0.902 | 0.74 | 0.42 | 0.71 | 0.6 | 0.75 | 0.58 |
| 11 | 92783339 | rs621310 | 1.044 | -0.386 | -0.154 | -0.312 | -0.006 | 0.014 | 0.014 | 0 | 0.46 | 0.24 | 0.51 | 0.59 | 0.41 | 0.42 |
| 11 | 92795212 | rs17630638 | 0.549 | -0.366 | -0.762 | -0.323 | -0.059 | 0.142 | 0.142 | 0 | 0.25 | 0.04 | 0.18 | 0.02 | 0.31 | 0.15 |
| 11 | 92795490 | rs1018204 | 0.297 | -0.778 | 0.031 | 0.518 | -0.027 | 0.067 | 0.067 | 0.385 | 0.77 | 0.42 | 0.73 | 0.62 | 0.79 | 0.62 |
| 11 | 92796156 | rs1018206 | 0.122 | -0.674 | -0.294 | 0.569 | -0.026 | 0.111 | 0.111 | 0.149 | 0.79 | 0.53 | 0.74 | 0.62 | 0.79 | 0.62 |
| 11 | 92801894 | rs16918472 | 0.109 | -0.768 | -1.128 | 0.7 | 0.025 | -0.08 | -0.08 | 0 | 0.14 | 0.01 | 0.3 | 0.25 | 0.07 | 0.19 |
| 11 | 92802210 | rs521754 | -0.853 | -0.936 | -0.682 | 0.627 | -0.014 | 0.08 | 0.08 | 1.92 | 0.19 | 0.62 | 0.25 | 0.42 | 0.16 | 0.31 |
| 11 | 92803568 | rs620131 | -0.714 | -0.947 | -0.612 | 0.616 | -0.021 | 0.089 | 0.089 | 0.921 | 0.18 | 0.61 | 0.23 | 0.4 | 0.15 | 0.32 |
| 11 | 92805174 | rs507792 | -0.869 | -0.944 | -0.718 | 0.509 | -0.018 | 0.061 | 0.061 | 1.396 | 0.21 | 0.61 | 0.24 | 0.4 | 0.17 | 0.41 |
| 11 | 92807099 | rs10830974 | 1.528 | -0.746 | -1.639 | -0.033 | -0.171 | -0.051 | -0.051 | 0.411 | 0.32 | 0.02 | 0.22 | 0.29 | 0.4 | 0.24 |
| 11 | 92812055 | rs3018133 | 0.41 | -0.395 | -0.704 | -0.113 | 0.003 | 0.289 | 0.289 | 0.499 | 0.11 | 0.33 | 0.19 | 0.39 | 0.11 | 0.31 |
| 11 | 92812922 | rs1508617 | 0.833 | -0.528 | -0.794 | -0.153 | -0.039 | -0.012 | -0.012 | 0 | 0.46 | 0.21 | 0.54 | 0.55 | 0.46 | 0.41 |
| 11 | 92818649 | rs9804472 | -0.26 | -0.505 | -0.706 | 0.153 | -0.017 | 0.108 | 0.108 | 0 | 0.81 | 0.69 | 0.77 | 0.58 | 0.81 | 0.64 |

| | | | | | | | | | | | | | | | | |
|---|---|---|---|---|---|---|---|---|---|---|---|---|---|---|---|---|
| 11 | 92820398 | rs621724 | 0.138 | -0.338 | -1.108 | -0.613 | 0 | 0.056 | 0.056 | 0 | 0.24 | 0.11 | 0.16 | 0.04 | 0.26 | 0.1 |
| 11 | 92829904 | rs7939730 | 0.786 | 0.014 | -1.75 | -0.63 | -0.029 | -0.005 | -0.005 | 0 | 0.31 | 0.07 | 0.23 | 0.27 | 0.3 | 0.16 |
| 11 | 92833475 | rs12791593 | 0.475 | 0.506 | -1.952 | -0.949 | -0.032 | 0.053 | 0.053 | 0 | 0.17 | 0.07 | 0.1 | 0.05 | 0.23 | 0.12 |
| 11 | 92834198 | rs614332 | -0.531 | 0.326 | -1.298 | -1.017 | 0.005 | 0.011 | 0.011 | 0.188 | 0.61 | 0.76 | 0.43 | 0.37 | 0.62 | 0.42 |
| 11 | 92834839 | rs512498 | -0.243 | 0.378 | -0.641 | -0.998 | -0.005 | -0.01 | -0.01 | 0 | 0.55 | 0.48 | 0.38 | 0.36 | 0.59 | 0.4 |
| 11 | 92845646 | rs7947982 | 0.941 | 0.379 | -1.207 | -0.984 | -0.014 | 0.017 | 0.017 | 0.01 | 0.33 | 0.08 | 0.24 | 0.28 | 0.31 | 0.18 |
| 11 | 92847230 | rs10830981 | -0.833 | 0.16 | -0.69 | -0.419 | 0.006 | 0.021 | 0.021 | 0 | 0.74 | 0.48 | 0.71 | 0.7 | 0.71 | 0.72 |
| 11 | 92849235 | rs10830982 | -0.846 | 0.162 | -0.681 | -0.553 | 0.007 | -0.006 | -0.006 | 0 | 0.75 | 0.44 | 0.68 | 0.77 | 0.71 | 0.73 |
| 11 | 92851178 | rs1015197 | 1.041 | -0.028 | -1.172 | -0.52 | 0.016 | -0.11 | -0.11 | 0.312 | 0.15 | 0.02 | 0.31 | 0.4 | 0.1 | 0.24 |
| 11 | 92852897 | rs10501802 | -0.26 | -0.046 | -1.135 | -0.658 | -0.006 | 0.022 | 0.022 | 0 | 0.66 | 0.29 | 0.66 | 0.77 | 0.64 | 0.6 |
| 11 | 92853630 | rs10830984 | -0.013 | -0.046 | -1.304 | -0.656 | -0.03 | 0.04 | 0.04 | 0 | 0.62 | 0.16 | 0.63 | 0.72 | 0.61 | 0.57 |
| 11 | 92856526 | rs11020171 | -0.064 | -0.109 | -1.447 | -0.828 | -0.004 | -0.081 | -0.081 | 2.434 | 0.28 | 0.12 | 0.39 | 0.47 | 0.3 | 0.42 |
| 11 | 92859478 | rs12361930 | 1.26 | 0.117 | -1.445 | -0.795 | -0.057 | -0.053 | -0.053 | 1.357 | 0.28 | 0.05 | 0.22 | 0.32 | 0.31 | 0.13 |
| 11 | 92861666 | rs12793386 | -0.207 | 0.08 | -1.347 | -0.913 | 0.008 | -0.029 | -0.029 | 0 | 0.37 | 0.25 | 0.44 | 0.46 | 0.35 | 0.48 |
| 11 | 92863100 | rs10830985 | 0.016 | 0.082 | -1.344 | -0.921 | 0.008 | -0.059 | -0.059 | 1.599 | 0.35 | 0.16 | 0.42 | 0.45 | 0.34 | 0.47 |
| 11 | 92863298 | rs11020175 | -0.207 | 0.091 | -1.348 | -0.913 | 0.008 | -0.029 | -0.029 | 0 | 0.37 | 0.25 | 0.44 | 0.46 | 0.35 | 0.48 |
| 11 | 92864441 | rs12420961 | 1.046 | -0.008 | -1.05 | -0.984 | -0.006 | 0.011 | 0.011 | 0 | 0.25 | 0.15 | 0.27 | 0.22 | 0.28 | 0.28 |
| 11 | 92868270 | rs10830986 | 0.435 | -0.442 | -0.903 | -0.881 | -0.012 | 0.009 | 0.009 | 0.059 | 0.26 | 0.26 | 0.35 | 0.4 | 0.13 | 0.41 |
| 11 | 92881044 | rs1062050 | 1.458 | -0.331 | -1.018 | -0.912 | -0.07 | -0.063 | -0.063 | 0 | 0.27 | 0.05 | 0.22 | 0.32 | 0.31 | 0.13 |
| 11 | 92886648 | rs10830989 | -0.902 | -0.272 | -0.632 | -1.005 | 0.001 | -0.032 | -0.032 | 0 | 0.68 | 0.44 | 0.67 | 0.79 | 0.66 | 0.71 |
| 11 | 92890392 | rs12786007 | 1.505 | -0.479 | -0.678 | -1.051 | -0.088 | -0.015 | -0.015 | 0 | 0.24 | 0.05 | 0.27 | 0.21 | 0.33 | 0.27 |
| 11 | 92912550 | rs7925288 | -0.367 | -0.286 | -0.257 | -0.952 | -0.006 | 0.024 | 0.024 | 0 | 0.58 | 0.34 | 0.61 | 0.73 | 0.5 | 0.65 |
| 11 | 92915544 | rs11020190 | 0.514 | -0.288 | -0.296 | -0.939 | -0.012 | 0.01 | 0.01 | 0.048 | 0.26 | 0.27 | 0.35 | 0.41 | 0.13 | 0.41 |
| 11 | 92921187 | rs4267030 | 1.428 | -0.233 | -0.412 | -0.927 | -0.069 | -0.062 | -0.062 | 0.307 | 0.27 | 0.05 | 0.22 | 0.32 | 0.31 | 0.14 |
| 11 | 92923215 | rs17549982 | 1.684 | -0.223 | -0.295 | -0.869 | 0 | -0.002 | -0.002 | 0.916 | 0.1 | 0.1 | 0.06 | 0.06 | 0.15 | 0.06 |
| 11 | 92930316 | rs3943572 | -0.392 | -0.424 | 0.081 | -0.564 | 0.008 | -0.011 | -0.011 | 0.698 | 0.3 | 0.25 | 0.38 | 0.42 | 0.18 | 0.48 |
| 11 | 92932229 | rs2121468 | -1.124 | -0.291 | 0.179 | -0.588 | 0.003 | -0.006 | -0.006 | 0.297 | 0.61 | 0.56 | 0.62 | 0.73 | 0.5 | 0.66 |
| 11 | 92933891 | rs4753437 | -0.444 | -0.493 | 0.236 | -0.583 | 0.017 | -0.014 | -0.014 | 5.997 | 0.29 | 0.23 | 0.37 | 0.42 | 0.16 | 0.45 |
| 11 | 92936341 | rs11020198 | -0.584 | -0.446 | 0.354 | -0.48 | 0.01 | -0.009 | -0.009 | 0.385 | 0.32 | 0.29 | 0.39 | 0.42 | 0.2 | 0.55 |
| 11 | 92949447 | rs4753439 | -0.842 | -0.326 | 0.392 | -0.408 | -0.031 | 0.008 | 0.008 | 0 | 0.37 | 0.49 | 0.4 | 0.41 | 0.2 | 0.54 |
| 11 | 92956873 | rs16918677 | 0.109 | -0.289 | 0.167 | -0.361 | 0.058 | -0.055 | -0.055 | 0 | 0.26 | 0.11 | 0.33 | 0.41 | 0.12 | 0.41 |
| 11 | 92961333 | rs11020208 | -0.456 | -0.266 | 0.198 | -0.349 | 0.024 | -0.018 | -0.018 | 0 | 0.32 | 0.23 | 0.37 | 0.41 | 0.18 | 0.53 |
| 11 | 92968895 | rs7111174 | -0.319 | -0.361 | 0.266 | -0.268 | -0.035 | 0.032 | 0.032 | 0.14 | 0.74 | 0.74 | 0.66 | 0.58 | 0.88 | 0.58 |
| 11 | 92971874 | rs10831010 | 0.269 | 0.037 | 0.485 | -0.235 | -0.001 | 0.018 | 0.018 | 0 | 0.42 | 0.43 | 0.4 | 0.28 | 0.49 | 0.34 |
| 11 | 92972603 | rs12803876 | 0.288 | 0.049 | 0.479 | -0.243 | -0.002 | 0.035 | 0.035 | 0 | 0.42 | 0.38 | 0.4 | 0.28 | 0.49 | 0.34 |
| 11 | 92975183 | rs7121542 | 0.359 | 0.074 | -0.523 | -0.269 | -0.02 | 0.053 | 0.053 | 0 | 0.4 | 0.29 | 0.39 | 0.27 | 0.49 | 0.34 |
| 11 | 92981763 | rs1446905 | 0.184 | 0.098 | -0.833 | -0.309 | -0.01 | 0.107 | 0.107 | 0 | 0.48 | 0.17 | 0.54 | 0.72 | 0.43 | 0.52 |
| 11 | 92983699 | rs16918716 | 0.617 | 0.434 | -1.234 | -0.114 | 0.021 | -0.038 | -0.038 | 0 | 0.08 | 0.19 | 0.06 | 0.01 | 0.08 | 0.14 |
| 11 | 92985672 | rs12808030 | 0.173 | 0.411 | -1.904 | 0.181 | -0.046 | 0.045 | 0.045 | 0 | 0.38 | 0.21 | 0.37 | 0.27 | 0.49 | 0.34 |
| 11 | 93010173 | rs753517 | -0.141 | 0.79 | -1.377 | 0.761 | -0.023 | -0.057 | -0.057 | 0.585 | 0.2 | 0.07 | 0.33 | 0.42 | 0.19 | 0.32 |
| 11 | 93012427 | rs11020238 | -0.434 | 0.42 | -1.342 | 0.219 | 0.016 | 0.074 | 0.074 | 0 | 0.49 | 0.38 | 0.62 | 0.7 | 0.4 | 0.56 |
| 11 | 93013387 | rs7943372 | -0.51 | 0.053 | -0.857 | -0.017 | 0.024 | -0.046 | -0.046 | 0 | 0.72 | 0.61 | 0.84 | 0.83 | 0.69 | 0.8 |
| 11 | 93014343 | rs1900385 | 0.056 | -0.341 | -1.376 | -0.72 | 0.003 | 0.021 | 0.021 | 0 | 0.45 | 0.26 | 0.67 | 0.6 | 0.5 | 0.48 |
| 11 | 93015765 | rs8181486 | -0.753 | -0.245 | -1.021 | -0.792 | -0.034 | -0.003 | -0.003 | 0 | 0.73 | 0.58 | 0.83 | 0.76 | 0.83 | 0.68 |
| 11 | 93018077 | rs2399638 | 1.049 | -0.829 | -1.27 | -1.211 | 0.03 | -0.093 | -0.093 | 0 | 0.22 | 0.3 | 0.23 | 0.02 | 0.39 | 0.1 |
| 11 | 93024163 | rs16912471 | 0.922 | -1.349 | -1.072 | -1.02 | -0.044 | 0.075 | 0.075 | 0 | 0.14 | 0.27 | 0.08 | 0.23 | 0.06 | 0.15 |
| 11 | 93025578 | rs1579874 | 0.695 | -1.423 | -1.041 | -0.752 | -0.056 | 0.114 | 0.114 | 0.153 | 0.16 | 0.33 | 0.1 | 0.3 | 0.06 | 0.16 |
| 11 | 93031855 | rs6483232 | 0.204 | -1.495 | -1.119 | -0.838 | -0.041 | 0.052 | 0.052 | 0 | 0.21 | 0.35 | 0.14 | 0.3 | 0.1 | 0.28 |
| 11 | 93033807 | rs4753084 | 1.619 | -1.892 | -1.535 | -0.335 | -0.003 | -0.093 | -0.093 | 0 | 0.13 | 0 | 0.31 | 0.35 | 0.08 | 0.28 |
| 11 | 93052265 | rs3020049 | -0.095 | -1.636 | -1.481 | -0.469 | 0 | -0.001 | -0.001 | 0 | 0.48 | 0.43 | 0.69 | 0.52 | 0.6 | 0.52 |
| 11 | 93052833 | rs11600203 | 2.006 | -1.191 | -1.683 | -0.711 | -0.021 | 0.006 | 0.006 | 0 | 0.16 | 0 | 0.06 | 0.1 | 0.11 | 0.12 |
| 11 | 93055556 | rs3020054 | -1.122 | -1.22 | -1.626 | -0.72 | -0.025 | 0.049 | 0.049 | 0 | 0.7 | 0.8 | 0.84 | 0.82 | 0.71 | 0.8 |
| 11 | 93057087 | rs3020069 | -0.191 | -1.432 | -1.689 | -0.816 | 0.006 | -0.041 | -0.041 | 0.642 | 0.61 | 0.48 | 0.78 | 0.75 | 0.67 | 0.69 |
| 11 | 93063200 | rs3020061 | -0.332 | -1.433 | -1.637 | -0.832 | 0.009 | -0.066 | -0.066 | 0 | 0.62 | 0.48 | 0.8 | 0.82 | 0.68 | 0.71 |
| 11 | 93067758 | rs7949194 | 0.338 | -1.143 | -1.377 | -0.669 | 0.002 | 0.017 | 0.017 | 0.484 | 0.09 | 0.34 | 0.05 | 0.06 | 0.04 | 0.12 |
| 11 | 93068949 | rs10831027 | 0.047 | -1.324 | -1.604 | -0.865 | 0.032 | 0.092 | 0.092 | 0.122 | 0.53 | 0.24 | 0.53 | 0.69 | 0.4 | 0.69 |
| 11 | 93072843 | rs4322357 | 1.034 | -1.468 | -1.597 | -0.939 | -0.005 | 0.004 | 0.004 | 0.027 | 0.24 | 0.21 | 0.32 | 0.14 | 0.39 | 0.14 |
| 11 | 93085537 | rs10831029 | 0.295 | -1.563 | -1.49 | -0.893 | 0.057 | 0 | 0 | 0 | 0.39 | 0.17 | 0.18 | 0.33 | 0.26 | 0.35 |
| 11 | 93089156 | rs2124559 | 0.024 | -1.598 | -1.144 | -0.872 | 0 | 0.001 | 0.001 | 0 | 0.42 | 0.57 | 0.66 | 0.51 | 0.5 | 0.43 |
| 11 | 93090840 | rs12277063 | 0.392 | -1.569 | -1.219 | -0.891 | 0.048 | -0.002 | -0.002 | 1.338 | 0.39 | 0.17 | 0.18 | 0.33 | 0.26 | 0.34 |
| 11 | 93090887 | rs2925355 | -2.29 | -1.538 | -1.166 | -0.888 | -0.021 | 0.02 | 0.02 | 0 | 0.85 | 0.88 | 0.87 | 0.9 | 0.78 | 0.88 |
| 11 | 93096135 | rs1456242 | -0.09 | -1.707 | -1.142 | -0.86 | -0.003 | -0.016 | -0.016 | 0 | 0.42 | 0.66 | 0.67 | 0.51 | 0.5 | 0.44 |
| 11 | 93105965 | rs3020071 | 0.443 | -1.672 | -1.024 | -0.84 | -0.014 | -0.028 | -0.028 | 0 | 0.35 | 0.25 | 0.62 | 0.5 | 0.47 | 0.43 |
| 11 | 93118515 | rs3020057 | -1.671 | -1.584 | -1.054 | -1.071 | -0.012 | 0.013 | 0.013 | 0 | 0.82 | 0.82 | 0.86 | 0.9 | 0.77 | 0.79 |
| 11 | 93121372 | rs7949990 | -0.004 | -1.633 | -1.018 | -1.065 | 0.014 | 0.009 | 0.009 | 0 | 0.46 | 0.55 | 0.22 | 0.34 | 0.29 | 0.33 |
| 11 | 93126799 | rs2658775 | -1.81 | -1.59 | -1.168 | -1.052 | -0.019 | 0.021 | 0.021 | 0 | 0.82 | 0.84 | 0.86 | 0.9 | 0.77 | 0.79 |
| 11 | 93129279 | rs12286814 | 0.218 | -1.64 | -1.117 | -1.033 | 0.048 | 0.017 | 0.017 | 0 | 0.42 | 0.19 | 0.19 | 0.34 | 0.28 | 0.33 |
| 11 | 93130411 | rs16918890 | 1.403 | -1.878 | -1.196 | -1.098 | -0.017 | -0.076 | -0.076 | 0 | 0.18 | 0.02 | 0.12 | 0.23 | 0.15 | 0.18 |
| 11 | 93131632 | rs11020307 | 0.632 | -1.712 | -1.161 | -1.013 | 0.013 | -0.007 | -0.007 | 0.132 | 0.24 | 0.16 | 0.15 | 0.23 | 0.18 | 0.2 |

| | | | | | | | | | | | | | | | | |
|---|---|---|---|---|---|---|---|---|---|---|---|---|---|---|---|---|
| 11 | 93141580 | rs2259633 | -1.412 | -1.13 | -0.793 | -0.82 | -0.015 | 0.018 | 0.018 | 0 | 0.82 | 0.83 | 0.86 | 0.9 | 0.78 | 0.79 |
| 11 | 93142657 | rs7949734 | 0.363 | -1.059 | -0.727 | -0.729 | -0.009 | -0.006 | -0.006 | 0 | 0.45 | 0.5 | 0.22 | 0.34 | 0.3 | 0.33 |
| 11 | 93143201 | rs11020315 | 0.344 | -1.053 | -0.772 | -0.729 | 0.012 | 0.008 | 0.008 | 0 | 0.46 | 0.54 | 0.22 | 0.33 | 0.3 | 0.33 |
| 11 | 93143455 | rs2605579 | 0.161 | -1.033 | -0.853 | -0.687 | -0.017 | -0.019 | -0.019 | 2.42 | 0.36 | 0.25 | 0.64 | 0.57 | 0.48 | 0.46 |
| 11 | 93148371 | rs12290433 | 0.202 | -0.808 | -0.839 | -0.615 | 0.05 | 0.033 | 0.033 | 0 | 0.47 | 0.66 | 0.24 | 0.34 | 0.29 | 0.33 |
| 11 | 93164980 | rs11020357 | 0.54 | -0.545 | -0.981 | -0.568 | 0.042 | 0.009 | 0.009 | 0 | 0.4 | 0.17 | 0.2 | 0.34 | 0.29 | 0.33 |
| 11 | 93166368 | rs2658781 | -1.561 | -0.177 | -0.775 | -0.456 | -0.013 | 0.013 | 0.013 | 0 | 0.85 | 0.86 | 0.88 | 0.92 | 0.8 | 0.89 |
| 11 | 93171107 | rs1456244 | 0.312 | -0.211 | -0.525 | -0.057 | -0.014 | 0.121 | 0.121 | 0 | 0.12 | 0.18 | 0.15 | 0.54 | 0.08 | 0.3 |
| 11 | 93180531 | rs7109581 | 0.705 | -0.028 | -0.565 | -0.291 | 0.033 | -0.103 | -0.103 | 0 | 0.22 | 0.32 | 0.22 | 0.02 | 0.39 | 0.14 |
| 11 | 93183722 | rs2658785 | -0.344 | -0.612 | -0.542 | 0.068 | -0.015 | 0.146 | 0.146 | 0.021 | 0.14 | 0.23 | 0.15 | 0.53 | 0.08 | 0.37 |
| 11 | 93184074 | rs1456243 | 0.383 | -0.584 | -0.571 | 0.154 | -0.013 | 0.147 | 0.147 | 0 | 0.11 | 0.18 | 0.14 | 0.52 | 0.07 | 0.29 |
| 11 | 93206181 | rs2045459 | -0.421 | -0.134 | -0.457 | -0.001 | 0.022 | 0.005 | 0.005 | 0 | 0.75 | 0.45 | 0.68 | 0.75 | 0.66 | 0.76 |
| 11 | 93209472 | rs2608280 | -0.027 | -0.68 | -0.836 | -0.409 | 0.001 | -0.007 | -0.007 | 0 | 0.49 | 0.31 | 0.54 | 0.51 | 0.48 | 0.53 |
| 11 | 93212775 | rs7950036 | 0.561 | -0.724 | -0.738 | -0.364 | 0 | -0.002 | -0.002 | 0 | 0.28 | 0.3 | 0.25 | 0.38 | 0.19 | 0.25 |
| 11 | 93213298 | rs2605595 | -1.175 | -0.658 | -0.618 | -0.509 | 0.038 | -0.135 | -0.135 | 1.101 | 0.78 | 0.63 | 0.8 | 0.98 | 0.66 | 0.87 |
| 11 | 93214623 | rs897555 | 1.693 | -0.63 | -0.921 | -0.67 | -0.009 | 0.023 | 0.023 | 0 | 0.14 | 0.08 | 0.08 | 0.04 | 0.16 | 0.1 |
| 11 | 93216751 | rs2658801 | 0.203 | -0.447 | -0.863 | -0.482 | -0.013 | -0.042 | -0.042 | 0 | 0.32 | 0.23 | 0.45 | 0.47 | 0.31 | 0.4 |
| 11 | 93226867 | rs2605610 | -0.466 | -0.522 | -0.838 | -0.464 | 0.03 | -0.009 | -0.009 | 0 | 0.56 | 0.7 | 0.68 | 0.56 | 0.65 | 0.62 |
| 11 | 93229319 | rs2605611 | -1.722 | -0.421 | -0.926 | -0.457 | -0.009 | 0.006 | 0.006 | 0 | 0.85 | 0.87 | 0.62 | 0.9 | 0.8 | 0.88 |
| 11 | 93230475 | rs10831050 | 1.076 | -0.497 | -0.69 | -0.511 | 0.023 | -0.053 | -0.053 | 2.699 | 0.22 | 0.34 | 0.2 | 0.09 | 0.33 | 0.21 |
| 11 | 93231137 | rs10466331 | 0.738 | -0.501 | -0.627 | -0.426 | 0.01 | -0.016 | -0.016 | 0 | 0.28 | 0.2 | 0.24 | 0.41 | 0.2 | 0.28 |
| 11 | 93231284 | rs10466333 | 0.909 | -0.487 | -0.626 | -0.426 | 0.009 | -0.014 | -0.014 | 0 | 0.27 | 0.19 | 0.24 | 0.4 | 0.19 | 0.26 |
| 11 | 93233037 | rs10831051 | 0.396 | -0.541 | -0.662 | -0.704 | 0.006 | -0.074 | -0.074 | 0 | 0.42 | 0.46 | 0.3 | 0.13 | 0.51 | 0.45 |
| 11 | 93233068 | rs2605615 | -1.325 | -0.467 | -0.691 | -0.413 | -0.021 | 0.003 | 0.003 | 0 | 0.82 | 0.87 | 0.61 | 0.84 | 0.78 | 0.85 |
| 11 | 93240847 | rs11020388 | 1.22 | -0.609 | -0.767 | -0.384 | 0.004 | 0.021 | 0.021 | 0.382 | 0.19 | 0.16 | 0.1 | 0.11 | 0.17 | 0.22 |
| 11 | 93243219 | rs11020392 | 0.523 | -0.608 | -0.894 | -0.274 | 0.042 | -0.066 | -0.066 | 0.323 | 0.32 | 0.15 | 0.26 | 0.4 | 0.23 | 0.27 |
| 11 | 93245041 | rs12577256 | 1.115 | -0.656 | -1.092 | -0.325 | -0.033 | -0.085 | -0.085 | 0 | 0.15 | 0.01 | 0.09 | 0.25 | 0.13 | 0.17 |
| 11 | 93247201 | rs2252326 | 1.26 | 0.14 | -0.503 | 0.201 | -0.002 | 0.017 | 0.017 | 0 | 0.12 | 0.12 | 0.07 | 0.3 | 0.08 | 0.13 |
| 11 | 93248483 | rs2658767 | 0.466 | 0.356 | -0.643 | 0.076 | 0.033 | -0.004 | -0.004 | 0 | 0.45 | 0.32 | 0.25 | 0.55 | 0.32 | 0.38 |
| 11 | 93259778 | rs2198797 | 0.961 | 0.699 | -0.61 | 0.107 | 0.02 | -0.087 | -0.087 | 0 | 0.19 | 0.3 | 0.16 | 0.02 | 0.29 | 0.23 |
| 11 | 93259965 | rs2248020 | -0.451 | 0.901 | -0.927 | 0.762 | 0.015 | 0.048 | 0.048 | 0 | 0.82 | 0.76 | 0.51 | 0.61 | 0.79 | 0.76 |
| 11 | 93261886 | rs2658770 | -0.681 | 1.015 | -1.189 | 0.344 | -0.011 | 0 | 0 | 0 | 0.83 | 0.85 | 0.61 | 0.84 | 0.79 | 0.86 |
| 11 | 93264680 | rs2263368 | 0.067 | 0.986 | -1.223 | 0.266 | -0.025 | 0.033 | 0.033 | 0 | 0.67 | 0.78 | 0.53 | 0.8 | 0.61 | 0.75 |
| 11 | 93266594 | rs11020405 | -0.287 | 0.772 | -1.073 | -0.052 | 0.001 | 0.002 | 0.002 | 0 | 0.5 | 0.46 | 0.34 | 0.49 | 0.58 | 0.63 |
| 11 | 93270101 | rs10765615 | 0.124 | 1.088 | -1.161 | -0.574 | -0.023 | 0.054 | 0.054 | 0.049 | 0.7 | 0.84 | 0.8 | 0.96 | 0.66 | 0.83 |
| 11 | 93277266 | rs7126188 | 0.178 | 0.924 | -0.78 | 0.179 | 0 | 0.012 | 0.012 | 0 | 0.52 | 0.41 | 0.67 | 0.55 | 0.52 | 0.63 |
| 11 | 93281716 | rs2895470 | 0.091 | 0.824 | -0.217 | 0.331 | -0.001 | 0.002 | 0.002 | 0.006 | 0.53 | 0.53 | 0.67 | 0.6 | 0.52 | 0.64 |
| 11 | 93283504 | rs4753472 | 0.386 | 0.902 | -0.127 | 0.553 | 0.021 | -0.021 | -0.021 | 0.017 | 0.44 | 0.27 | 0.62 | 0.45 | 0.45 | 0.51 |
| 11 | 93284927 | rs4753473 | 0.828 | 0.7 | -0.596 | 0.512 | -0.021 | -0.07 | -0.07 | 0 | 0.13 | 0.05 | 0.16 | 0.36 | 0.16 | 0.17 |
| 11 | 93288094 | rs12417178 | -0.404 | 0.613 | 0.081 | 0.544 | 0 | 0.006 | 0.006 | 0 | 0.52 | 0.53 | 0.67 | 0.57 | 0.5 | 0.62 |
| 11 | 93291685 | rs1962041 | 0.644 | 0.567 | 0.183 | 0.634 | 0.033 | 0.072 | 0.072 | 0 | 0.06 | 0.12 | 0.05 | 0.12 | 0.07 | 0.06 |
| 11 | 93295334 | rs2399685 | -0.489 | 0.903 | 0.008 | 0.267 | 0 | -0.001 | -0.001 | 0.634 | 0.6 | 0.44 | 0.7 | 0.56 | 0.61 | 0.67 |
| 11 | 93305819 | rs2399690 | -0.829 | 0.51 | 0.688 | 0.483 | -0.006 | 0.1 | 0.1 | 0.363 | 0.13 | 0.24 | 0.11 | 0.38 | 0.1 | 0.18 |
| 11 | 93311765 | rs1939560 | 0.184 | 0.414 | 0.987 | 0.493 | -0.008 | 0.263 | 0.263 | 0 | 0.93 | 0.73 | 0.94 | 0.73 | 0.98 | 0.94 |
| 11 | 93313072 | rs7114097 | -0.368 | 0.324 | 0.733 | -0.216 | 0.026 | 0.179 | 0.179 | 0 | 0.49 | 0.22 | 0.31 | 0.23 | 0.54 | 0.41 |
| 11 | 93317122 | rs11020422 | 0.233 | 0.408 | 0.624 | -0.237 | 0.004 | 0 | 0 | 0 | 0.09 | 0.08 | 0.08 | 0.1 | 0.13 | 0.12 |
| 11 | 93321148 | rs12802772 | 0.181 | 0.345 | 0.645 | -0.373 | -0.05 | 0.128 | 0.128 | 0 | 0.4 | 0.16 | 0.26 | 0.22 | 0.44 | 0.34 |
| 11 | 93322831 | rs11020425 | -0.111 | 0.293 | 1.055 | -0.333 | -0.027 | 0.115 | 0.115 | 0 | 0.84 | 0.62 | 0.88 | 0.72 | 0.89 | 0.88 |
| 11 | 93330918 | rs4753092 | 0.798 | 0.264 | 1.112 | -0.28 | -0.014 | -0.039 | -0.039 | 0.875 | 0.42 | 0.24 | 0.6 | 0.45 | 0.45 | 0.54 |
| 11 | 93332312 | rs10831068 | 0.774 | 0.281 | 0.979 | -0.354 | -0.071 | 0.175 | 0.175 | 0 | 0.33 | 0.08 | 0.2 | 0.12 | 0.36 | 0.23 |
| 11 | 93341053 | rs7949518 | -0.091 | 0.577 | 1.142 | -0.436 | -0.001 | 0.002 | 0.002 | 0 | 0.54 | 0.54 | 0.71 | 0.55 | 0.55 | 0.65 |
| 11 | 93346486 | rs1954783 | 0.885 | 0.576 | 1.06 | -0.434 | -0.005 | -0.021 | -0.021 | 0 | 0.42 | 0.3 | 0.61 | 0.47 | 0.45 | 0.53 |
| 11 | 93348268 | rs4478954 | -0.319 | 0.526 | 1.134 | -0.446 | -0.005 | -0.012 | -0.012 | 0 | 0.12 | 0.21 | 0.1 | 0.1 | 0.09 | 0.13 |
| 11 | 93355909 | rs948017 | 0.326 | 0.472 | 1.043 | -0.236 | 0.004 | -0.003 | -0.003 | 0.458 | 0.46 | 0.41 | 0.65 | 0.47 | 0.46 | 0.53 |
| 11 | 93356343 | rs10831078 | 0.572 | 0.449 | 0.65 | -0.019 | -0.024 | -0.003 | -0.003 | 0 | 0.09 | 0.03 | 0.07 | 0.12 | 0.11 | 0.1 |
| 11 | 93369586 | rs7124256 | 0.78 | -0.049 | 0.944 | 0.233 | 0 | 0.014 | 0.014 | 0 | 0.1 | 0.1 | 0.07 | 0.27 | 0.1 | 0.12 |
| 11 | 93374276 | rs4753485 | 0.254 | -0.113 | 1.196 | 0.136 | 0.003 | 0.002 | 0.002 | 0 | 0.39 | 0.33 | 0.29 | 0.4 | 0.4 | 0.53 |
| 11 | 93382187 | rs10741469 | 0.387 | -0.076 | 0.953 | 0.141 | -0.003 | -0.007 | -0.007 | 0 | 0.38 | 0.21 | 0.28 | 0.4 | 0.39 | 0.54 |
| 11 | 93391559 | rs1939568 | 0.382 | -0.096 | 0.827 | 0.138 | -0.009 | -0.01 | -0.01 | 0 | 0.38 | 0.19 | 0.28 | 0.4 | 0.39 | 0.53 |
| 11 | 93401193 | rs10765620 | 0.506 | -0.229 | 1.083 | 0.154 | -0.04 | 0.092 | 0.092 | 0.109 | 0.43 | 0.21 | 0.51 | 0.26 | 0.49 | 0.33 |
| 11 | 93412774 | rs1944100 | 0.003 | -0.285 | 1.407 | 0.191 | 0.01 | 0.094 | 0.094 | 0.589 | 0.93 | 0.66 | 0.95 | 0.94 | 0.99 | 0.99 |
| 11 | 93417460 | rs7939577 | 0.748 | -0.283 | 1.366 | 0.428 | -0.006 | 0.058 | 0.058 | 0 | 0.11 | 0.16 | 0.08 | 0.27 | 0.1 | 0.13 |
| 11 | 93421782 | rs10831093 | 0.609 | -0.255 | 1.153 | 0.244 | -0.067 | 0.088 | 0.088 | 0.251 | 0.42 | 0.13 | 0.5 | 0.26 | 0.49 | 0.33 |
| 11 | 93422613 | rs10160552 | 0.643 | -0.253 | 1.091 | 0.245 | -0.099 | 0.079 | 0.079 | 0.441 | 0.42 | 0.09 | 0.5 | 0.26 | 0.49 | 0.33 |
| 11 | 93438360 | rs16919234 | 0.479 | -0.56 | 1.296 | -0.112 | -0.001 | -0.018 | -0.018 | 0 | 0.11 | 0.13 | 0.05 | 0.01 | 0.1 | 0.05 |
| 11 | 93474791 | rs623510 | 0.279 | -0.425 | 1.277 | 0.108 | -0.006 | 0.073 | 0.073 | 0.82 | 0.93 | 0.69 | 0.95 | 0.94 | 0.99 | 0.99 |
| 11 | 93474827 | rs631116 | 0.279 | -0.425 | 1.274 | 0.108 | -0.005 | 0.075 | 0.075 | 0.82 | 0.93 | 0.68 | 0.95 | 0.94 | 0.99 | 0.99 |
| 11 | 93475442 | rs644306 | 0.151 | -0.324 | 0.349 | -0.122 | -0.005 | 0.069 | 0.069 | 0 | 0.2 | 0.06 | 0.11 | 0.05 | 0.19 | 0.3 |
| 11 | 93491020 | rs670871 | 0.972 | -0.322 | 0.609 | 0.267 | 0.02 | 0.01 | 0.01 | 0.186 | 0.58 | 0.29 | 0.73 | 0.62 | 0.63 | 0.56 |

| | | | | | | | | | | | | | | | | |
|---|---|---|---|---|---|---|---|---|---|---|---|---|---|---|---|---|
| 11 | 93495975 | rs7310 | 0.259 | -0.321 | 0.671 | 0.233 | -0.018 | 0.065 | 0.065 | 0 | 0.22 | 0.09 | 0.14 | 0.05 | 0.25 | 0.3 |
| 11 | 93498833 | rs1284108 | 0.876 | -0.274 | 0.632 | 0.286 | 0.021 | 0.01 | 0.01 | 0.175 | 0.58 | 0.29 | 0.73 | 0.62 | 0.63 | 0.56 |
| 11 | 93505490 | rs675441 | 0.91 | -0.242 | 0.665 | -0.112 | 0.027 | 0.075 | 0.075 | 0.706 | 0.57 | 0.21 | 0.74 | 0.74 | 0.64 | 0.57 |
| 11 | 93517299 | rs655158 | 0.88 | -0.258 | 0.698 | -0.117 | 0.026 | 0.063 | 0.063 | 0.485 | 0.58 | 0.28 | 0.75 | 0.74 | 0.64 | 0.57 |
| 11 | 93521405 | rs676586 | 0.107 | -0.306 | 1.192 | -0.114 | -0.026 | 0.021 | 0.021 | 2.749 | 0.81 | 0.36 | 0.88 | 0.79 | 0.88 | 0.87 |
| 11 | 93534990 | rs655699 | 1.028 | -0.27 | 0.258 | -0.116 | 0.024 | 0.08 | 0.08 | 1.167 | 0.57 | 0.16 | 0.73 | 0.74 | 0.63 | 0.55 |
| 11 | 93535839 | rs669565 | 0.199 | -0.102 | 0.747 | 0.33 | -0.025 | -0.01 | -0.01 | 0.096 | 0.82 | 0.56 | 0.9 | 0.86 | 0.88 | 0.87 |
| 11 | 93552855 | rs585768 | 1.009 | 0.37 | -0.769 | 0.98 | -0.012 | -0.066 | -0.066 | 0 | 0.11 | 0.02 | 0.22 | 0.47 | 0.12 | 0.21 |
| 11 | 93555987 | rs10831104 | 0.475 | 0.605 | -0.474 | 0.252 | -0.028 | 0.07 | 0.07 | 0 | 0.13 | 0.02 | 0.06 | 0.04 | 0.14 | 0.24 |
| 11 | 93570633 | rs625359 | 0.824 | 0.513 | 0.488 | 0.641 | -0.009 | 0.018 | 0.018 | 2.346 | 0.81 | 0.34 | 0.86 | 0.79 | 0.83 | 0.87 |
| 11 | 93572910 | rs7103466 | -0.678 | 0.477 | 1.018 | 0.644 | 0.049 | 0.034 | 0.034 | 1.191 | 0.15 | 0.47 | 0.12 | 0.14 | 0.16 | 0.12 |
| 11 | 93574964 | rs671490 | 0.453 | 0.74 | 0.532 | 0.33 | 0.006 | 0.001 | 0.001 | 0 | 0.82 | 0.76 | 0.65 | 0.83 | 0.82 | 0.92 |
| 11 | 93575238 | rs12289146 | -0.518 | 1.006 | 0.318 | 0.273 | 0.005 | -0.008 | -0.008 | 0 | 0.15 | 0.21 | 0.33 | 0.1 | 0.18 | 0.07 |
| 11 | 93578713 | rs592928 | -0.341 | 0.603 | 0.545 | 0.438 | -0.002 | -0.004 | -0.004 | 0 | 0.52 | 0.5 | 0.66 | 0.39 | 0.64 | 0.59 |
| 11 | 93581888 | rs732358 | -0.56 | 0.811 | 0.387 | 0.276 | -0.021 | 0.047 | 0.047 | 0 | 0.6 | 0.75 | 0.43 | 0.71 | 0.5 | 0.5 |
| 11 | 93596420 | rs12785450 | -0.037 | 0.87 | 0.528 | 0.346 | 0.017 | -0.02 | -0.02 | 0 | 0.06 | 0.08 | 0.07 | 0.01 | 0.13 | 0.06 |
| 11 | 93596599 | rs12362058 | -0.651 | 0.461 | 1.001 | 0.506 | -0.026 | 0.059 | 0.059 | 0 | 0.1 | 0.21 | 0.06 | 0.18 | 0.04 | 0.09 |
| 11 | 93624380 | rs6483270 | -0.541 | 0.82 | 1.103 | 0.447 | -0.001 | -0.008 | -0.008 | 0 | 0.61 | 0.6 | 0.47 | 0.75 | 0.58 | 0.52 |
| 11 | 93627460 | rs11827384 | -1.313 | 0.412 | 1.512 | 1.453 | 0.034 | 0.303 | 0.303 | 0 | 0.06 | 0.3 | 0.09 | 0.27 | 0.06 | 0.14 |
| 11 | 93629677 | rs1108910 | 0.34 | 0.544 | 1.428 | 1.354 | -0.085 | 0.055 | 0.055 | 0 | 0.43 | 0.15 | 0.56 | 0.35 | 0.53 | 0.52 |
| 11 | 93637462 | rs1354766 | 0.45 | 0.556 | 2.137 | 1.278 | 0.021 | 0.002 | 0.002 | 0.253 | 0.42 | 0.23 | 0.29 | 0.4 | 0.34 | 0.34 |
| 11 | 93637839 | rs1506657 | 1.433 | 0.489 | 2.165 | 1.365 | 0.001 | -0.004 | -0.004 | 0.216 | 0.82 | 0.23 | 0.8 | 0.67 | 0.83 | 0.81 |
| 11 | 93642877 | rs12789389 | 0.474 | 0.466 | 2.089 | 1.362 | 0.023 | 0.001 | 0.001 | 0.123 | 0.42 | 0.2 | 0.29 | 0.39 | 0.33 | 0.34 |
| 11 | 93645269 | rs10765643 | 0.69 | 0.639 | 1.939 | 1.298 | -0.076 | 0.062 | 0.062 | 0 | 0.38 | 0.09 | 0.53 | 0.27 | 0.47 | 0.39 |
| 11 | 93651355 | rs9888266 | -0.083 | 0.562 | 2.531 | 1.267 | 0.028 | -0.05 | -0.05 | 0.173 | 0.09 | 0.22 | 0.09 | 0.01 | 0.11 | 0.06 |
| 11 | 93651823 | rs9888143 | -0.729 | 0.56 | 2.49 | 1.295 | 0.043 | 0.019 | 0.019 | 0 | 0.12 | 0.4 | 0.1 | 0.07 | 0.11 | 0.08 |
| 11 | 93655091 | rs7117597 | -0.466 | 0.234 | 2.175 | 1.261 | 0 | 0 | 0 | 0.053 | 0.57 | 0.59 | 0.68 | 0.42 | 0.64 | 0.59 |
| 11 | 93662459 | rs10765647 | -0.314 | 0.244 | 2.012 | 1.112 | -0.007 | -0.014 | -0.014 | 0 | 0.56 | 0.52 | 0.68 | 0.35 | 0.64 | 0.57 |
| 11 | 93667439 | rs11020603 | 0.241 | 0.222 | 1.907 | 1.039 | -0.005 | -0.008 | -0.008 | 1.869 | 0.4 | 0.43 | 0.3 | 0.65 | 0.32 | 0.41 |
| 11 | 93669055 | rs12786407 | 0.882 | 0.149 | 1.441 | 1.16 | -0.001 | -0.002 | -0.002 | 0 | 0.32 | 0.15 | 0.22 | 0.31 | 0.26 | 0.27 |
| 11 | 93670388 | rs2511376 | 0.653 | 0.152 | 1.602 | 1.091 | -0.084 | 0.061 | 0.061 | 0 | 0.44 | 0.11 | 0.59 | 0.34 | 0.54 | 0.51 |
| 11 | 93671838 | rs11020605 | 0.625 | 0.165 | 1.596 | 1.098 | 0.005 | 0.009 | 0.009 | 0 | 0.36 | 0.24 | 0.26 | 0.32 | 0.3 | 0.31 |
| 11 | 93676224 | rs2056247 | 0.898 | 0.168 | 1.672 | 1.241 | -0.016 | 0.079 | 0.079 | 0 | 0.8 | 0.38 | 0.84 | 0.66 | 0.83 | 0.82 |
| 11 | 93690148 | rs908748 | -0.224 | 0.314 | 1.175 | 0.965 | 0.006 | 0.054 | 0.054 | 0.418 | 0.46 | 0.29 | 0.31 | 0.33 | 0.41 | 0.34 |
| 11 | 93701348 | rs11020619 | -0.495 | 0.279 | 1.256 | 0.721 | 0.001 | -0.038 | -0.038 | 0 | 0.15 | 0.23 | 0.12 | 0.01 | 0.16 | 0.07 |
| 11 | 93711185 | rs7113975 | 0.793 | 0.393 | 1.11 | 1.025 | -0.002 | 0.025 | 0.025 | 0 | 0.31 | 0.13 | 0.17 | 0.25 | 0.24 | 0.23 |
| 11 | 93712637 | rs2511388 | 0.515 | 0.234 | 1.311 | 1.406 | -0.01 | 0.059 | 0.059 | 0 | 0.49 | 0.19 | 0.64 | 0.42 | 0.58 | 0.58 |
| 11 | 93717180 | rs2939619 | 0.855 | 0.214 | 1.603 | 1.296 | 0 | 0.008 | 0.008 | 0 | 0.31 | 0.23 | 0.18 | 0.25 | 0.24 | 0.23 |
| 11 | 93718763 | rs925005 | 0.349 | 0.37 | 2.133 | 1.563 | 0.005 | 0 | 0 | 0.061 | 0.83 | 0.56 | 0.85 | 0.87 | 0.84 | 0.87 |
| 11 | 93723251 | rs2945635 | 0.793 | 0.154 | 1.948 | 1.473 | -0.002 | -0.012 | -0.012 | 0 | 0.28 | 0.27 | 0.16 | 0.23 | 0.22 | 0.22 |
| 11 | 93725384 | rs2462741 | 0.558 | 0.123 | 1.764 | 1.523 | 0.001 | 0.042 | 0.042 | 0 | 0.51 | 0.15 | 0.65 | 0.42 | 0.59 | 0.58 |
| 11 | 93726072 | rs1518574 | 0.634 | 0.122 | 1.666 | 1.523 | -0.013 | 0.034 | 0.034 | 0 | 0.51 | 0.12 | 0.65 | 0.42 | 0.59 | 0.58 |
| 11 | 93747735 | rs2511380 | 0.368 | 0.149 | 2.014 | 1.441 | 0.004 | 0.055 | 0.055 | 0 | 0.82 | 0.58 | 0.83 | 0.77 | 0.82 | 0.86 |
| 11 | 93751535 | rs16919815 | 0.172 | 0.142 | 2.086 | 1.436 | -0.001 | -0.021 | -0.021 | 0 | 0.13 | 0.16 | 0.12 | 0.01 | 0.15 | 0.08 |
| 11 | 93752001 | rs2949867 | 0.91 | 0.13 | 2.025 | 1.454 | -0.002 | 0.018 | 0.018 | 0 | 0.28 | 0.12 | 0.16 | 0.23 | 0.22 | 0.22 |
| 11 | 93754508 | rs2511403 | 0.665 | -0.05 | 2.188 | 1.49 | 0.004 | 0.058 | 0.058 | 0 | 0.82 | 0.55 | 0.83 | 0.77 | 0.82 | 0.86 |
| 11 | 93758893 | rs11020631 | 0.474 | 0.119 | 1.402 | 1.158 | 0.006 | 0.031 | 0.031 | 0 | 0.36 | 0.23 | 0.35 | 0.32 | 0.39 | 0.45 |
| 11 | 93760614 | rs2399749 | -0.33 | 0.098 | 1.651 | 1.227 | 0 | -0.012 | -0.012 | 0 | 0.16 | 0.35 | 0.15 | 0.12 | 0.16 | 0.14 |
| 11 | 93765773 | rs1829371 | 0.537 | 0.054 | 1.541 | 1.222 | -0.001 | 0.041 | 0.041 | 0 | 0.36 | 0.16 | 0.35 | 0.32 | 0.4 | 0.45 |
| 11 | 93766584 | rs1518566 | -0.26 | 0.033 | 1.647 | 1.22 | -0.001 | -0.008 | -0.008 | 0 | 0.16 | 0.34 | 0.15 | 0.12 | 0.16 | 0.13 |
| 11 | 93767360 | rs4753116 | -0.646 | 0.048 | 1.406 | 1.226 | -0.001 | 0.001 | 0.001 | 0 | 0.52 | 0.61 | 0.5 | 0.44 | 0.55 | 0.58 |
| 11 | 93767445 | rs4753117 | -0.501 | 0.048 | 1.175 | 1.226 | 0 | 0.015 | 0.015 | 0 | 0.51 | 0.48 | 0.49 | 0.44 | 0.55 | 0.58 |
| 11 | 93767919 | rs2511401 | 0.697 | 0.07 | 0.294 | 1.313 | -0.047 | 0.018 | 0.018 | 0 | 0.43 | 0.17 | 0.46 | 0.36 | 0.42 | 0.35 |
| 11 | 93777012 | rs10831159 | 0.311 | -0.002 | -0.06 | 1.141 | -0.03 | 0.033 | 0.033 | 0 | 0.36 | 0.08 | 0.35 | 0.32 | 0.4 | 0.45 |
| 11 | 93787542 | rs7110588 | -0.381 | 0.096 | 0.622 | 1.248 | 0.02 | -0.003 | -0.003 | 0.056 | 0.61 | 0.56 | 0.5 | 0.64 | 0.58 | 0.68 |
| 11 | 93787758 | rs7127348 | -0.057 | -0.021 | 0.918 | 1.439 | -0.03 | 0.023 | 0.023 | 0 | 0.56 | 0.31 | 0.47 | 0.44 | 0.57 | 0.59 |
| 11 | 93788270 | rs10831161 | -0.385 | 0.061 | 0.758 | 1.128 | 0.02 | -0.003 | -0.003 | 2.886 | 0.61 | 0.56 | 0.5 | 0.64 | 0.58 | 0.68 |
| 11 | 93793135 | rs2460042 | 0.302 | 0.007 | 0.981 | 1.108 | -0.047 | -0.008 | -0.008 | 0 | 0.37 | 0.16 | 0.49 | 0.36 | 0.42 | 0.32 |
| 11 | 93797163 | rs4753122 | -0.284 | 0.004 | 1.136 | 1.106 | 0.003 | -0.001 | -0.001 | 0.178 | 0.6 | 0.52 | 0.49 | 0.64 | 0.58 | 0.68 |
| 11 | 93805614 | rs2511410 | 0.304 | 0.119 | 0.166 | 1.264 | -0.03 | -0.012 | -0.012 | 0.017 | 0.37 | 0.12 | 0.43 | 0.34 | 0.34 | 0.29 |
| 11 | 93812665 | rs1401186 | 0.086 | -0.346 | 0.713 | 1.33 | 0 | -0.003 | -0.003 | 0.034 | 0.6 | 0.61 | 0.62 | 0.41 | 0.54 | 0.46 |
| 11 | 93816922 | rs2949858 | -0.468 | -0.364 | 0.523 | 1.59 | -0.003 | -0.02 | -0.02 | 0 | 0.7 | 0.81 | 0.78 | 0.6 | 0.66 | 0.59 |
| 11 | 93820847 | rs2949857 | -0.544 | -0.355 | 0.529 | 1.606 | -0.025 | -0.037 | -0.037 | 0 | 0.71 | 0.88 | 0.78 | 0.61 | 0.67 | 0.61 |
| 11 | 93821624 | rs2949856 | -0.145 | -0.412 | 1.047 | 1.744 | -0.001 | 0.001 | 0.001 | 0.698 | 0.66 | 0.57 | 0.73 | 0.4 | 0.65 | 0.44 |
| 11 | 93833446 | rs10765653 | 0.643 | -0.431 | 1.082 | 1.718 | 0.021 | 0.015 | 0.015 | 1.255 | 0.56 | 0.15 | 0.56 | 0.35 | 0.56 | 0.37 |
| 11 | 93842104 | rs12222234 | -0.468 | -0.421 | 1.136 | 1.763 | 0.004 | -0.016 | -0.016 | 0 | 0.68 | 0.72 | 0.75 | 0.54 | 0.66 | 0.5 |
| 11 | 93851048 | rs4753538 | -0.131 | -0.401 | 1.084 | 1.758 | -0.003 | 0.007 | 0.007 | 0.771 | 0.66 | 0.54 | 0.73 | 0.4 | 0.65 | 0.43 |
| 11 | 93862493 | rs1138800 | -0.676 | -0.243 | 0.526 | 1.713 | -0.055 | -0.023 | -0.023 | 0 | 0.7 | 0.86 | 0.77 | 0.62 | 0.62 | 0.59 |
| 11 | 93865651 | rs4073612 | 0.562 | -0.246 | 0.523 | 1.341 | -0.019 | -0.014 | -0.014 | 0 | 0.55 | 0.19 | 0.58 | 0.42 | 0.5 | 0.45 |

| | | | | | | | | | | | | | | | | |
|---|---|---|---|---|---|---|---|---|---|---|---|---|---|---|---|---|
| 11 | 93866147 | rs11607757 | -0.069 | 0.111 | 1.061 | 1.29 | 0.005 | -0.01 | -0.01 | 0.726 | 0.09 | 0.15 | 0.15 | 0.06 | 0.11 | 0.07 |
| 11 | 93900828 | rs7482084 | -0.012 | -0.057 | 0.624 | 1.041 | 0.029 | -0.056 | -0.056 | 0 | 0.3 | 0.48 | 0.4 | 0.13 | 0.36 | 0.13 |
| 11 | 93906735 | rs12270308 | 0.394 | 0.062 | -0.136 | 1.093 | -0.062 | 0.048 | 0.048 | 0 | 0.27 | 0.09 | 0.38 | 0.13 | 0.35 | 0.13 |
| 11 | 93908872 | rs882937 | 0.106 | -0.229 | 0.154 | 1.319 | 0.001 | 0.001 | 0.001 | 0 | 0.51 | 0.46 | 0.48 | 0.49 | 0.55 | 0.65 |
| 11 | 93913036 | rs12793348 | -0.103 | -0.408 | -0.043 | 1.617 | 0.002 | 0.004 | 0.004 | 0 | 0.11 | 0.14 | 0.08 | 0.26 | 0.09 | 0.11 |
| 11 | 93914129 | rs12800562 | -0.065 | -0.41 | -0.043 | 1.717 | 0.002 | -0.002 | -0.002 | 0.34 | 0.1 | 0.14 | 0.08 | 0.21 | 0.09 | 0.08 |
| 11 | 93914451 | rs1046805 | -0.848 | -0.501 | 0.52 | 1.595 | -0.004 | 0.005 | 0.005 | 0.017 | 0.65 | 0.79 | 0.6 | 0.75 | 0.65 | 0.83 |
| 11 | 93914709 | rs1046812 | -0.848 | -0.501 | 0.52 | 1.595 | -0.004 | 0.005 | 0.005 | 0.017 | 0.65 | 0.79 | 0.61 | 0.75 | 0.65 | 0.83 |
| 11 | 93914956 | rs1070 | -0.868 | -0.5 | 0.52 | 1.571 | -0.004 | 0.014 | 0.014 | 0.022 | 0.66 | 0.79 | 0.61 | 0.8 | 0.65 | 0.85 |
| 11 | 93914990 | rs1071 | 0.658 | -0.48 | 0.271 | 1.462 | -0.02 | 0 | 0 | 0 | 0.08 | 0.02 | 0.14 | 0.06 | 0.11 | 0.09 |
| 11 | 93918350 | rs1046173 | -0.339 | -0.376 | 0.738 | 1.42 | 0.001 | 0.015 | 0.015 | 0 | 0.86 | 0.9 | 0.82 | 0.94 | 0.84 | 0.89 |
| 11 | 93918837 | rs3020004 | 0.463 | -0.361 | 0.891 | 1.575 | -0.002 | 0.001 | 0.001 | 0 | 0.46 | 0.49 | 0.36 | 0.51 | 0.42 | 0.64 |
| 11 | 93927529 | rs1792632 | -0.382 | -0.36 | 0.845 | 1.522 | 0.007 | 0.02 | 0.02 | 0 | 0.51 | 0.3 | 0.61 | 0.52 | 0.55 | 0.35 |
| 11 | 93928787 | rs11020679 | 0.612 | -0.508 | 1.089 | 1.524 | 0 | 0.004 | 0.004 | 0 | 0.43 | 0.35 | 0.32 | 0.45 | 0.42 | 0.52 |
| 11 | 93929510 | rs1792634 | 0.388 | -0.57 | 0.993 | 1.49 | 0.003 | 0.006 | 0.006 | 0 | 0.38 | 0.38 | 0.44 | 0.44 | 0.41 | 0.27 |
| 11 | 93931412 | rs16920039 | 0.873 | -0.631 | 1.159 | 1.463 | -0.028 | -0.006 | -0.006 | 0 | 0.41 | 0.16 | 0.3 | 0.44 | 0.42 | 0.52 |
| 11 | 93931787 | rs1857682 | -0.811 | -0.629 | 1.047 | 1.463 | -0.001 | 0 | 0 | 0.14 | 0.21 | 0.25 | 0.23 | 0.27 | 0.22 | 0.22 |
| 11 | 93931844 | rs1857681 | -1.205 | -0.622 | 1.059 | 1.456 | -0.005 | 0.003 | 0.003 | 0 | 0.67 | 0.77 | 0.6 | 0.74 | 0.65 | 0.83 |
| 11 | 93932958 | rs1857680 | 0.467 | -0.626 | 1.067 | 1.457 | -0.002 | -0.002 | -0.002 | 0 | 0.46 | 0.52 | 0.37 | 0.47 | 0.43 | 0.61 |
| 11 | 93933778 | rs4753549 | -0.546 | -0.662 | 1.118 | 1.469 | 0.002 | 0.002 | 0.002 | 0 | 0.84 | 0.85 | 0.8 | 0.88 | 0.83 | 0.86 |
| 11 | 93934545 | rs1014499 | -0.683 | -0.623 | 1.119 | 1.498 | 0.008 | -0.006 | -0.006 | 0 | 0.53 | 0.37 | 0.62 | 0.47 | 0.57 | 0.35 |
| 11 | 93935628 | rs10831175 | -0.013 | -0.623 | 1.128 | 1.508 | -0.033 | 0.097 | 0.097 | 0 | 0.47 | 0.14 | 0.57 | 0.26 | 0.52 | 0.25 |
| 11 | 93944545 | rs1541300 | 0.669 | -0.706 | 1.089 | 1.262 | 0 | 0 | 0 | 0.072 | 0.36 | 0.33 | 0.43 | 0.41 | 0.4 | 0.22 |
| 11 | 93945734 | rs4753133 | 1.512 | -0.831 | 0.48 | 1.115 | -0.051 | -0.088 | -0.088 | 0.005 | 0.17 | 0.01 | 0.16 | 0.27 | 0.16 | 0.18 |
| 11 | 93962659 | rs2399822 | 0.458 | -0.755 | 1.219 | 1.271 | 0.001 | 0.015 | 0.015 | 0 | 0.46 | 0.41 | 0.35 | 0.69 | 0.45 | 0.61 |
| 11 | 93963552 | rs7934122 | -0.075 | -0.633 | 0.371 | 1.311 | -0.038 | 0.142 | 0.142 | 0 | 0.46 | 0.14 | 0.57 | 0.2 | 0.52 | 0.24 |
| 11 | 93987628 | rs6483320 | 0.741 | -0.6 | -0.193 | 1.14 | 0 | 0.001 | 0.001 | 0 | 0.43 | 0.39 | 0.32 | 0.42 | 0.42 | 0.51 |
| 11 | 93997886 | rs12420783 | 0.203 | -0.645 | -0.355 | 1.085 | -0.005 | 0.047 | 0.047 | 0 | 0.16 | 0.04 | 0.19 | 0.06 | 0.17 | 0.13 |
| 11 | 94011852 | rs1361929 | -0.074 | -0.548 | 0.1 | 1.436 | 0.001 | 0 | 0 | 0 | 0.47 | 0.53 | 0.37 | 0.47 | 0.42 | 0.62 |
| 11 | 94021448 | rs669634 | 0.358 | -0.455 | -0.784 | 1.468 | -0.111 | 0.138 | 0.138 | 0.066 | 0.45 | 0.06 | 0.57 | 0.2 | 0.52 | 0.25 |
| 11 | 94029888 | rs7115129 | 0.277 | -0.482 | -0.664 | 1.525 | -0.003 | 0.007 | 0.007 | 0.496 | 0.43 | 0.33 | 0.31 | 0.41 | 0.41 | 0.52 |
| 11 | 94031847 | rs519834 | 0.031 | -0.235 | -1.61 | 1.534 | -0.107 | 0.14 | 0.14 | 0.097 | 0.45 | 0.06 | 0.57 | 0.2 | 0.52 | 0.25 |
| 11 | 94036009 | rs10831195 | -0.281 | 0.222 | -0.482 | 1.145 | 0.006 | 0.005 | 0.005 | 0 | 0.63 | 0.53 | 0.51 | 0.6 | 0.58 | 0.64 |
| 11 | 94037967 | rs630418 | 0.415 | -0.149 | -1.298 | 0.914 | 0.074 | 0.078 | 0.078 | 0 | 0.52 | 0.13 | 0.59 | 0.61 | 0.64 | 0.37 |
| 11 | 94038151 | rs7930551 | -0.451 | -0.291 | -0.174 | 0.947 | 0.001 | -0.004 | -0.004 | 0 | 0.44 | 0.49 | 0.27 | 0.3 | 0.33 | 0.54 |
| 11 | 94041469 | rs12796907 | 0.206 | -0.738 | -0.713 | 0.646 | -0.032 | 0.041 | 0.041 | 0 | 0.3 | 0.06 | 0.2 | 0.26 | 0.28 | 0.43 |
| 11 | 94047626 | rs493390 | 0.862 | -0.76 | -0.48 | 0.645 | 0.064 | 0.067 | 0.067 | 0 | 0.53 | 0.22 | 0.59 | 0.61 | 0.64 | 0.38 |
| 11 | 94048785 | rs608706 | 1.656 | -0.651 | -0.916 | 0.66 | -0.118 | -0.083 | -0.083 | 0 | 0.27 | 0.04 | 0.43 | 0.28 | 0.38 | 0.14 |
| 11 | 94049187 | rs7130310 | 0.065 | -0.618 | -0.592 | 0.538 | -0.027 | 0.051 | 0.051 | 0 | 0.31 | 0.07 | 0.2 | 0.26 | 0.28 | 0.44 |
| 11 | 94055855 | rs11020733 | -0.624 | -0.628 | -0.515 | 0.541 | 0.014 | 0.123 | 0.123 | 0 | 0.39 | 0.13 | 0.24 | 0.26 | 0.32 | 0.51 |
| 11 | 94059289 | rs662753 | 0.756 | -0.687 | -0.075 | 0.592 | -0.003 | -0.002 | -0.002 | 0 | 0.56 | 0.46 | 0.62 | 0.63 | 0.67 | 0.5 |
| 11 | 94059522 | rs663703 | 0.674 | -0.691 | -0.066 | 0.59 | -0.012 | -0.009 | -0.009 | 0 | 0.57 | 0.54 | 0.63 | 0.64 | 0.67 | 0.5 |
| 11 | 94060134 | rs11601246 | -0.098 | -0.61 | -0.232 | 0.551 | -0.045 | -0.031 | -0.031 | 0 | 0.09 | 0.03 | 0.1 | 0.15 | 0.15 | 0.12 |
| 11 | 94062135 | rs11602404 | -0.098 | -0.583 | -0.528 | 0.509 | -0.045 | -0.033 | -0.033 | 0.202 | 0.09 | 0.03 | 0.1 | 0.16 | 0.15 | 0.12 |
| 11 | 94066993 | rs10831203 | -0.446 | -0.678 | -0.596 | 0.445 | -0.012 | 0 | 0 | 0 | 0.28 | 0.04 | 0.19 | 0.25 | 0.24 | 0.3 |
| 11 | 94067057 | rs495862 | 1.133 | -0.714 | -0.197 | 0.444 | 0.064 | 0.037 | 0.037 | 0 | 0.55 | 0.28 | 0.62 | 0.63 | 0.68 | 0.51 |
| 11 | 94067823 | rs1792622 | 1.419 | -0.67 | -0.409 | 0.373 | 0.1 | 0.07 | 0.07 | 0 | 0.52 | 0.17 | 0.6 | 0.63 | 0.67 | 0.48 |
| 11 | 94070016 | rs481597 | 1.558 | -0.453 | -0.462 | 0.311 | 0.07 | 0.085 | 0.085 | 0 | 0.5 | 0.16 | 0.55 | 0.63 | 0.6 | 0.47 |
| 11 | 94073456 | rs1783678 | 2 | -0.28 | -0.721 | 0.363 | -0.121 | -0.032 | -0.032 | 0 | 0.27 | 0.03 | 0.39 | 0.22 | 0.37 | 0.17 |
| 11 | 94077690 | rs12360807 | 0.497 | -0.791 | -0.667 | 0.216 | 0.068 | 0.047 | 0.047 | 0 | 0.2 | 0.05 | 0.09 | 0.11 | 0.1 | 0.23 |
| 11 | 94078269 | rs484889 | 0.679 | -0.832 | 0.006 | 0.26 | -0.003 | -0.007 | -0.007 | 0 | 0.66 | 0.4 | 0.72 | 0.74 | 0.69 | 0.67 |
| 11 | 94099615 | rs7950328 | 0.401 | -0.826 | 0.284 | 0.248 | 0.108 | 0.042 | 0.042 | 1.05 | 0.87 | 0.6 | 0.81 | 0.82 | 0.79 | 0.9 |
| 11 | 94101345 | rs4753600 | 1.546 | -0.862 | 0.143 | 0.274 | 0.024 | 0.044 | 0.044 | 0 | 0.53 | 0.35 | 0.56 | 0.64 | 0.59 | 0.53 |
| 11 | 94103467 | rs6483327 | 0.517 | -0.85 | 0.237 | 0.28 | -0.013 | -0.028 | -0.028 | 0 | 0.67 | 0.5 | 0.72 | 0.72 | 0.69 | 0.67 |
| 11 | 94110284 | rs11020758 | 0.385 | -0.686 | 0.066 | 0.344 | -0.06 | -0.042 | -0.042 | 0 | 0.09 | 0.01 | 0.14 | 0.18 | 0.21 | 0.1 |
| 11 | 94111170 | rs12417810 | -0.033 | -0.782 | 0.174 | 0.371 | 0.004 | 0.006 | 0.006 | 0 | 0.14 | 0.13 | 0.16 | 0.08 | 0.1 | 0.14 |
| 11 | 94112883 | rs3740864 | 0.038 | -0.792 | 0.048 | 0.374 | 0.009 | 0.017 | 0.017 | 0 | 0.14 | 0.06 | 0.16 | 0.08 | 0.1 | 0.14 |
| 11 | 94113466 | rs3740868 | 0.868 | -1.145 | -0.003 | 0.333 | 0.013 | -0.006 | -0.006 | 4.249 | 0.11 | 0.02 | 0.08 | 0.1 | 0.09 | 0.2 |
| 11 | 94128777 | rs12224966 | 0.13 | -0.786 | 0.319 | 0.363 | 0 | -0.025 | -0.025 | 0 | 0.38 | 0.28 | 0.39 | 0.43 | 0.36 | 0.44 |
| 11 | 94129042 | rs16920277 | 1.114 | -0.737 | 0.183 | 0.386 | 0.019 | 0.005 | 0.005 | 0 | 0.11 | 0.04 | 0.14 | 0.07 | 0.07 | 0.12 |
| 11 | 94130342 | rs6483341 | -1.393 | -0.788 | 0.357 | 0.346 | -0.013 | -0.007 | -0.007 | 0.545 | 0.62 | 0.68 | 0.52 | 0.55 | 0.47 | 0.68 |
| 11 | 94131418 | rs10831222 | -1.974 | -0.68 | 0.351 | 0.355 | 0 | 0 | 0 | 1.462 | 0.71 | 0.72 | 0.67 | 0.73 | 0.67 | 0.78 |
| 11 | 94132284 | rs3758786 | 0.43 | -0.621 | 0.244 | 0.332 | -0.038 | -0.03 | -0.03 | 1.762 | 0.08 | 0.02 | 0.15 | 0.17 | 0.2 | 0.1 |
| 11 | 94147464 | rs12270338 | -0.329 | -0.658 | 0.348 | 0.354 | 0.02 | 0.013 | 0.013 | 0.794 | 0.1 | 0.13 | 0.16 | 0.21 | 0.21 | 0.16 |
| 11 | 94149349 | rs2508678 | -0.47 | -0.725 | 0.425 | 0.327 | 0.003 | 0.001 | 0.001 | 0.164 | 0.49 | 0.48 | 0.35 | 0.43 | 0.38 | 0.48 |
| 11 | 94150790 | rs2155209 | 2.163 | -0.723 | 0.354 | 0.319 | -0.053 | -0.032 | -0.032 | 0 | 0.28 | 0.09 | 0.32 | 0.28 | 0.33 | 0.22 |
| 11 | 94152849 | rs1061956 | 1.29 | -0.748 | 0.352 | 0.307 | -0.009 | -0.013 | -0.013 | 0 | 0.1 | 0.09 | 0.01 | 0 | 0.01 | 0.04 |
| 11 | 94157114 | rs663530 | 1.358 | -0.721 | 0.404 | 0.333 | -0.021 | -0.029 | -0.029 | 0 | 0.26 | 0.14 | 0.24 | 0.32 | 0.28 | 0.25 |
| 11 | 94165326 | rs13447720 | -0.202 | -0.726 | 0.432 | 0.368 | -0.032 | -0.027 | -0.027 | 0 | 0.1 | 0.04 | 0.15 | 0.22 | 0.21 | 0.18 |

| | | | | | | | | | | | | | | | | |
|---|---|---|---|---|---|---|---|---|---|---|---|---|---|---|---|---|
| 11 | 94167830 | rs601391 | -2.021 | -0.787 | 0.498 | 0.346 | 0.004 | -0.006 | -0.006 | 0 | 0.7 | 0.57 | 0.65 | 0.72 | 0.67 | 0.78 |
| 11 | 94175249 | rs3017077 | 0.378 | -0.82 | 0.445 | 0.334 | -0.011 | -0.026 | -0.026 | 0 | 0.37 | 0.18 | 0.39 | 0.39 | 0.36 | 0.38 |
| 11 | 94176058 | rs16920449 | 1.237 | -0.82 | 0.458 | 0.316 | -0.003 | -0.007 | -0.007 | 3.064 | 0.1 | 0.09 | 0.02 | 0 | 0.01 | 0.04 |
| 11 | 94182689 | rs604845 | -1.912 | -0.785 | 0.506 | 0.337 | 0.004 | -0.004 | -0.004 | 3.082 | 0.7 | 0.61 | 0.65 | 0.72 | 0.67 | 0.78 |
| 11 | 94186955 | rs592068 | 0.349 | -0.964 | 0.484 | 0.343 | 0.01 | 0.007 | 0.007 | 0.002 | 0.37 | 0.31 | 0.27 | 0.33 | 0.29 | 0.29 |
| 11 | 94187714 | rs16920466 | 1.208 | -0.97 | 0.427 | 0.326 | -0.006 | -0.01 | -0.01 | 0 | 0.1 | 0.1 | 0.02 | 0 | 0.01 | 0.04 |
| 11 | 94190115 | rs654718 | 0.112 | -0.998 | 0.356 | 0.32 | -0.006 | -0.018 | -0.018 | 0.019 | 0.39 | 0.17 | 0.39 | 0.4 | 0.36 | 0.38 |
| 11 | 94199082 | rs12224897 | 0.448 | -0.952 | 0.432 | 0.341 | -0.017 | -0.007 | -0.007 | 0.198 | 0.12 | 0.21 | 0.09 | 0.1 | 0.09 | 0.19 |
| 11 | 94199272 | rs533984 | -0.855 | -0.99 | 0.405 | 0.343 | 0.018 | -0.005 | -0.005 | 0 | 0.53 | 0.67 | 0.39 | 0.46 | 0.41 | 0.58 |
| 11 | 94200389 | rs12788248 | 0.763 | -0.785 | 0.546 | 0.36 | -0.03 | -0.035 | -0.035 | 0 | 0.11 | 0.14 | 0.02 | 0 | 0.02 | 0.04 |
| 11 | 94213905 | rs476137 | -0.957 | -0.776 | 0.478 | 0.375 | -0.001 | -0.015 | -0.015 | 0 | 0.59 | 0.67 | 0.51 | 0.5 | 0.44 | 0.58 |
| 11 | 94218688 | rs11020798 | 0.832 | -0.926 | 0.279 | 0.469 | 0.002 | -0.015 | -0.015 | 0 | 0.11 | 0.01 | 0.08 | 0.1 | 0.09 | 0.19 |
| 11 | 94225920 | rs497763 | -0.178 | -0.797 | 0.454 | 0.466 | 0.012 | -0.006 | -0.006 | 0 | 0.49 | 0.34 | 0.48 | 0.5 | 0.45 | 0.57 |
| 11 | 94227029 | rs1805360 | 0.777 | -0.806 | 0.107 | 0.277 | -0.021 | -0.026 | -0.026 | 0 | 0.1 | 0.11 | 0.02 | 0 | 0.01 | 0.04 |
| 11 | 94239567 | rs538800 | 1.629 | -0.531 | -0.204 | 0.361 | -0.047 | -0.044 | -0.044 | 0 | 0.27 | 0.09 | 0.32 | 0.29 | 0.32 | 0.22 |
| 11 | 94248366 | rs486556 | 0.605 | -0.67 | 0.239 | 0.565 | -0.008 | -0.011 | -0.011 | 0.965 | 0.38 | 0.3 | 0.48 | 0.54 | 0.55 | 0.32 |
| 11 | 94248581 | rs509723 | -0.756 | -0.978 | 0.234 | 0.429 | -0.009 | -0.003 | -0.003 | 0 | 0.61 | 0.8 | 0.6 | 0.66 | 0.65 | 0.54 |
| 11 | 94250379 | rs628728 | -0.113 | -0.989 | 0.429 | 0.419 | 0.019 | 0.011 | 0.011 | 0 | 0.5 | 0.64 | 0.52 | 0.56 | 0.57 | 0.36 |
| 11 | 94253443 | rs695093 | 0.796 | -0.987 | 0.299 | 0.406 | -0.013 | -0.017 | -0.017 | 4.157 | 0.37 | 0.25 | 0.48 | 0.54 | 0.55 | 0.32 |
| 11 | 94253577 | rs521349 | -0.676 | -0.948 | 0.23 | 0.444 | -0.007 | -0.004 | -0.004 | 0.083 | 0.61 | 0.79 | 0.6 | 0.66 | 0.65 | 0.54 |
| 11 | 94256359 | rs11020808 | 0.371 | -0.77 | 0.088 | 0.249 | -0.007 | 0.005 | 0.005 | 0 | 0.11 | 0.02 | 0.09 | 0.1 | 0.1 | 0.21 |
| 11 | 94261280 | rs11605546 | 0.082 | -0.778 | 0.187 | 0.244 | 0.003 | 0.006 | 0.006 | 0 | 0.12 | 0.14 | 0.1 | 0.1 | 0.1 | 0.21 |
| 11 | 94264149 | rs6483345 | -1.722 | -0.67 | 0.451 | 0.318 | 0.003 | -0.001 | -0.001 | 0 | 0.33 | 0.6 | 0.28 | 0.37 | 0.34 | 0.34 |
| 11 | 94264882 | rs12294732 | -0.785 | -0.539 | 0.551 | 0.351 | 0.047 | 0.037 | 0.037 | 0 | 0.21 | 0.45 | 0.17 | 0.27 | 0.24 | 0.13 |
| 11 | 94269711 | rs2508783 | 1.596 | -0.366 | 0.46 | 0.363 | -0.001 | 0 | 0 | 0.083 | 0.29 | 0.26 | 0.34 | 0.32 | 0.31 | 0.23 |
| 11 | 94269813 | rs2508782 | -0.213 | -0.541 | 0.446 | 0.327 | 0.005 | 0.043 | 0.043 | 0 | 0.5 | 0.69 | 0.51 | 0.59 | 0.55 | 0.36 |
| 11 | 94270236 | rs11020819 | -0.245 | -0.872 | 1.103 | 0.277 | -0.076 | -0.079 | -0.079 | 0 | 0.12 | 0.35 | 0.03 | 0 | 0.01 | 0.03 |
| 11 | 94276209 | rs10831239 | -1.203 | -1.047 | 0.856 | 0.163 | 0.004 | -0.003 | -0.003 | 0 | 0.33 | 0.38 | 0.36 | 0.33 | 0.37 | 0.35 |
| 11 | 94280189 | rs10831240 | 0.029 | -1.156 | 0.973 | 0.148 | -0.066 | -0.064 | -0.064 | 1.066 | 0.11 | 0.31 | 0.03 | 0.01 | 0.02 | 0.03 |
| 11 | 94282014 | rs10741490 | -2.005 | -1.149 | 0.855 | 0.164 | 0.027 | 0.037 | 0.037 | 0 | 0.44 | 0.72 | 0.39 | 0.36 | 0.38 | 0.39 |
| 11 | 94284254 | rs12284274 | 0.036 | -1.032 | 0.987 | 0.288 | -0.051 | -0.053 | -0.053 | 0 | 0.1 | 0.22 | 0.02 | 0.01 | 0.02 | 0.03 |
| 11 | 94299968 | rs34976180 | 0.441 | -1.051 | 0.64 | 0.298 | 0.05 | 0.05 | 0.05 | 0.048 | 0.09 | 0.01 | 0.01 | 0 | 0.01 | 0.03 |
| 11 | 94304270 | rs12295797 | -0.006 | -0.684 | 0.974 | 0.513 | 0.002 | 0.004 | 0.004 | 0 | 0.15 | 0.14 | 0.16 | 0.06 | 0.07 | 0.21 |
| 11 | 94313230 | rs11020839 | -1.062 | -0.694 | 1.22 | 0.808 | -0.063 | 0.001 | 0.001 | 0 | 0.22 | 0.51 | 0.25 | 0.2 | 0.12 | 0.31 |
| 11 | 94313514 | rs11020840 | -0.891 | -0.63 | 0.928 | 0.758 | -0.012 | -0.01 | -0.01 | 0 | 0.17 | 0.13 | 0.19 | 0.28 | 0.28 | 0.17 |
| 11 | 94315467 | rs16912568 | 0.182 | -0.656 | 0.947 | 0.741 | 0.006 | 0.005 | 0.005 | 0 | 0.16 | 0.15 | 0.17 | 0.06 | 0.1 | 0.19 |
| 11 | 94319818 | rs12574292 | -0.784 | -0.607 | 0.748 | 0.753 | -0.021 | -0.017 | -0.017 | 0.476 | 0.17 | 0.11 | 0.19 | 0.28 | 0.28 | 0.17 |
| 11 | 94320237 | rs7110167 | -2.256 | -0.679 | 0.845 | 0.75 | -0.003 | -0.03 | -0.03 | 2.21 | 0.4 | 0.81 | 0.45 | 0.5 | 0.4 | 0.49 |
| 11 | 94321986 | rs2155099 | -2.256 | -0.679 | 0.845 | 0.75 | -0.003 | -0.03 | -0.03 | 0 | 0.4 | 0.81 | 0.45 | 0.5 | 0.4 | 0.49 |
| 11 | 94322052 | rs2155098 | -0.782 | -0.429 | 0.348 | 0.851 | -0.045 | -0.036 | -0.036 | 0 | 0.16 | 0.06 | 0.19 | 0.28 | 0.28 | 0.17 |
| 11 | 94323840 | rs7126782 | -1.836 | -0.45 | 0.47 | 0.846 | -0.003 | -0.002 | -0.002 | 0 | 0.23 | 0.26 | 0.24 | 0.43 | 0.3 | 0.3 |
| 11 | 94325470 | rs10437563 | -1.513 | -0.527 | 0.392 | 0.763 | -0.02 | 0.004 | 0.004 | 0 | 0.14 | 0.27 | 0.11 | 0.17 | 0.07 | 0.19 |
| 11 | 94325839 | rs591584 | -1.64 | -0.758 | 0.277 | 0.735 | -0.005 | 0 | 0 | 0 | 0.35 | 0.67 | 0.4 | 0.36 | 0.39 | 0.37 |
| 11 | 94326710 | rs7119735 | -0.66 | -0.877 | 0.284 | 0.721 | -0.02 | 0.026 | 0.026 | 0 | 0.08 | 0.21 | 0.06 | 0.15 | 0.02 | 0.13 |
| 11 | 94326848 | rs11020852 | 0.239 | -0.777 | 0.187 | 0.698 | -0.001 | -0.005 | -0.005 | 0.225 | 0.06 | 0.06 | 0.04 | 0.02 | 0.06 | 0.07 |
| 11 | 94332731 | rs476209 | -1.373 | -0.757 | 0.118 | 0.744 | -0.068 | -0.029 | -0.029 | 0 | 0.3 | 0.54 | 0.29 | 0.24 | 0.18 | 0.37 |
| 11 | 94336130 | rs608972 | -1.56 | -0.508 | -0.874 | 0.616 | -0.004 | -0.089 | -0.089 | 4.329 | 0.42 | 0.91 | 0.45 | 0.51 | 0.4 | 0.5 |
| 11 | 94337220 | rs593690 | -1.617 | -0.537 | -0.856 | 0.637 | -0.017 | -0.084 | -0.084 | 0 | 0.43 | 0.91 | 0.45 | 0.5 | 0.41 | 0.51 |
| 11 | 94342444 | rs656730 | -0.572 | -1.093 | -0.631 | 0.438 | 0.024 | 0.046 | 0.046 | 0 | 0.2 | 0.12 | 0.11 | 0.09 | 0.14 | 0.2 |
| 11 | 94342567 | rs656244 | -1.236 | -1.433 | -0.382 | 0.41 | -0.004 | 0 | 0 | 0 | 0.64 | 0.66 | 0.57 | 0.7 | 0.57 | 0.62 |
| 11 | 94350678 | rs600206 | -0.01 | -1.448 | -0.442 | 0.424 | 0.029 | 0.046 | 0.046 | 0 | 0.2 | 0.12 | 0.11 | 0.12 | 0.14 | 0.21 |
| 11 | 94351961 | rs1939137 | 0.417 | -1.622 | 0.309 | 0.779 | -0.009 | 0.099 | 0.099 | 0 | 0.06 | 0.16 | 0.04 | 0.19 | 0.02 | 0.1 |
| 11 | 94354479 | rs508485 | -0.456 | -1.816 | 0.52 | 0.575 | 0.004 | 0 | 0 | 0 | 0.52 | 0.46 | 0.5 | 0.49 | 0.5 | 0.45 |
| 11 | 94354642 | rs16920737 | 0.325 | -1.834 | 0.61 | 0.535 | -0.012 | -0.029 | -0.029 | 0 | 0.08 | 0.22 | 0.05 | 0.02 | 0.05 | 0.08 |
| 11 | 94356050 | rs7933369 | -1.005 | -1.799 | 0.442 | 0.585 | -0.029 | -0.032 | -0.032 | 0 | 0.61 | 0.76 | 0.56 | 0.51 | 0.56 | 0.54 |
| 11 | 94356914 | rs12276362 | 1.006 | -1.707 | 0.419 | 0.651 | -0.002 | -0.014 | -0.014 | 0.919 | 0.3 | 0.22 | 0.38 | 0.36 | 0.36 | 0.22 |
| 11 | 94360812 | rs1940640 | 1.481 | -1.804 | 0.347 | 0.545 | 0.002 | 0.001 | 0.001 | 0 | 0.24 | 0.2 | 0.31 | 0.19 | 0.25 | 0.17 |
| 11 | 94363720 | rs628715 | -1.059 | -1.962 | 0.413 | 0.567 | -0.027 | -0.007 | -0.007 | 0 | 0.29 | 0.61 | 0.19 | 0.31 | 0.16 | 0.32 |
| 11 | 94364361 | rs631413 | -0.985 | -2.039 | 0.384 | 0.493 | -0.038 | -0.013 | -0.013 | 0 | 0.3 | 0.57 | 0.19 | 0.31 | 0.17 | 0.32 |
| 11 | 94368960 | rs622022 | -1.251 | -1.956 | 0.319 | 0.534 | -0.023 | -0.07 | -0.07 | 0 | 0.6 | 0.86 | 0.54 | 0.51 | 0.47 | 0.57 |
| 11 | 94369205 | rs542116 | -0.059 | -2.075 | 0.444 | 0.369 | 0.019 | -0.043 | -0.043 | 0 | 0.1 | 0.31 | 0.07 | 0.03 | 0.08 | 0.08 |
| 11 | 94370386 | rs4127392 | 0.523 | -2.089 | 0.133 | 0.466 | 0.033 | 0.019 | 0.019 | 0 | 0.36 | 0.2 | 0.37 | 0.28 | 0.32 | 0.31 |
| 11 | 94372443 | rs7110768 | -0.625 | -1.989 | 0.366 | 0.569 | -0.058 | -0.053 | -0.053 | 0 | 0.22 | 0.36 | 0.12 | 0.11 | 0.13 | 0.24 |
| 11 | 94374694 | rs4396252 | -0.301 | -1.642 | -0.013 | 0.723 | -0.006 | 0.092 | 0.092 | 0.055 | 0.07 | 0.25 | 0.05 | 0.19 | 0.02 | 0.06 |
| 11 | 94381707 | rs7107104 | 1.252 | -1.638 | -1.155 | 0.566 | 0.014 | 0.021 | 0.021 | 0.056 | 0.29 | 0.11 | 0.3 | 0.2 | 0.27 | 0.19 |
| 11 | 94382129 | rs623404 | -1.13 | -1.731 | -1.405 | 0.625 | -0.092 | -0.095 | -0.095 | 7.645 | 0.64 | 0.93 | 0.53 | 0.57 | 0.5 | 0.65 |
| 11 | 94383998 | rs11020870 | 0.619 | -1.772 | -1.324 | 0.212 | 0.018 | -0.007 | -0.007 | 0 | 0.15 | 0.08 | 0.1 | 0.15 | 0.12 | 0.3 |
| 11 | 94385461 | rs9943515 | 0.027 | -1.596 | -0.976 | 0.147 | 0 | 0.007 | 0.007 | 0 | 0.1 | 0.11 | 0.09 | 0.19 | 0.12 | 0.07 |
| 11 | 94385718 | rs4528285 | 0.254 | -1.601 | -1.174 | 0.137 | 0.012 | 0.035 | 0.035 | 0 | 0.09 | 0.03 | 0.04 | 0.02 | 0.08 | 0.1 |

| | | | | | | | | | | | | | | | | |
|---|---|---|---|---|---|---|---|---|---|---|---|---|---|---|---|---|
| 11 | 94386723 | rs10831258 | -0.798 | -1.445 | -1.003 | 0.211 | 0.017 | 0.012 | 0.012 | 0.624 | 0.51 | 0.43 | 0.43 | 0.39 | 0.48 | 0.38 |
| 11 | 94388085 | rs11020874 | 0.024 | -1.006 | -1.367 | 0.33 | 0.012 | 0.041 | 0.041 | 0.372 | 0.29 | 0.12 | 0.3 | 0.17 | 0.28 | 0.19 |
| 11 | 94388721 | rs11020877 | 0.091 | -0.949 | -1.451 | 0.359 | 0.005 | 0.038 | 0.038 | 0.297 | 0.29 | 0.09 | 0.3 | 0.17 | 0.28 | 0.19 |
| 11 | 94391845 | rs520905 | -0.607 | -0.718 | -0.819 | 0.544 | -0.001 | -0.002 | -0.002 | 0 | 0.46 | 0.53 | 0.44 | 0.38 | 0.42 | 0.4 |
| 11 | 94393115 | rs11020878 | 0.257 | -0.941 | -0.891 | 0.939 | 0.006 | -0.049 | -0.049 | 0 | 0.1 | 0.02 | 0.05 | 0.18 | 0.09 | 0.06 |
| 11 | 94393285 | rs486244 | -0.654 | -1.65 | -0.592 | 0.872 | -0.008 | -0.008 | -0.008 | 0 | 0.39 | 0.53 | 0.35 | 0.29 | 0.34 | 0.37 |
| 11 | 94396071 | rs7103955 | 0.443 | -1.923 | 0.003 | 0.478 | -0.046 | 0.074 | 0.074 | 0.12 | 0.19 | 0.02 | 0.24 | 0.07 | 0.23 | 0.17 |
| 11 | 94396718 | rs10765692 | -0.136 | -1.575 | 0.297 | 0.737 | 0.015 | 0.053 | 0.053 | 0 | 0.35 | 0.21 | 0.32 | 0.07 | 0.34 | 0.3 |
| 11 | 94398136 | rs4127395 | -0.118 | -1.579 | 0.41 | 0.735 | 0.019 | 0.098 | 0.098 | 0 | 0.35 | 0.16 | 0.32 | 0.07 | 0.34 | 0.3 |
| 11 | 94399446 | rs7931634 | 0.092 | -1.616 | 0.461 | 0.852 | 0.018 | 0.071 | 0.071 | 0 | 0.31 | 0.16 | 0.31 | 0.07 | 0.3 | 0.29 |
| 11 | 94404440 | rs643797 | -1.319 | -1.909 | 0.31 | 0.858 | 0.004 | 0.004 | 0.004 | 0 | 0.49 | 0.47 | 0.39 | 0.25 | 0.41 | 0.36 |
| 11 | 94406359 | rs581570 | 0.295 | -1.903 | 0.431 | 0.875 | -0.029 | 0.014 | 0.014 | 0.029 | 0.17 | 0.27 | 0.09 | 0.19 | 0.1 | 0.07 |
| 11 | 94407736 | rs12789145 | -0.19 | -1.995 | 0.373 | 1.373 | -0.002 | 0.195 | 0.195 | 0 | 0.1 | 0.19 | 0.11 | 0.53 | 0.08 | 0.28 |
| 11 | 94408176 | rs610728 | -1.702 | -1.831 | -0.087 | 1.122 | -0.009 | 0.093 | 0.093 | 0 | 0.59 | 0.78 | 0.51 | 0.79 | 0.49 | 0.64 |
| 11 | 94409708 | rs7115112 | -0.53 | -1.133 | 0.072 | 1.599 | 0.028 | 0.001 | 0.001 | 0 | 0.33 | 0.17 | 0.3 | 0.36 | 0.31 | 0.38 |
| 11 | 94438594 | rs1893923 | 0.928 | -0.822 | 0.654 | 2.174 | 0.009 | 0.421 | 0.421 | 0 | 0.88 | 0.55 | 0.89 | 0.51 | 0.93 | 0.74 |
| 11 | 94440853 | rs4753145 | 1.241 | -0.887 | -0.261 | 1.707 | 0.016 | 0.078 | 0.078 | 0.067 | 0.54 | 0.14 | 0.55 | 0.33 | 0.58 | 0.47 |
| 11 | 94452476 | rs7350489 | -0.257 | -0.625 | -0.414 | 1.116 | -0.032 | 0.167 | 0.167 | 0 | 0.31 | 0.04 | 0.29 | 0.09 | 0.3 | 0.25 |
| 11 | 94453876 | rs12223099 | -0.557 | -0.496 | -0.043 | 1.479 | 0 | 0.062 | 0.062 | 0 | 0.08 | 0.1 | 0.08 | 0.25 | 0.07 | 0.19 |
| 11 | 94463093 | rs2895570 | -0.543 | -0.586 | -0.514 | 0.669 | -0.01 | 0.132 | 0.132 | 0 | 0.32 | 0.07 | 0.29 | 0.17 | 0.31 | 0.25 |
| 11 | 94478798 | rs1430864 | -1.657 | -0.786 | 2.018 | 1.604 | -0.02 | 0.239 | 0.239 | 0.754 | 0.08 | 0.33 | 0.19 | 0.27 | 0.02 | 0.16 |
| 11 | 94479361 | rs7113554 | -1.893 | -0.82 | 2.258 | 1.587 | -0.016 | 0.286 | 0.286 | 1.483 | 0.09 | 0.4 | 0.19 | 0.27 | 0.02 | 0.16 |
| 11 | 94498860 | rs4753618 | -1.4 | -0.798 | 2.146 | 1.594 | -0.014 | 0.225 | 0.225 | 0 | 0.07 | 0.22 | 0.18 | 0.27 | 0.02 | 0.12 |
| 11 | 94513281 | rs949162 | 1.496 | -0.839 | 2.325 | 1.573 | -0.013 | 0.225 | 0.225 | 0.088 | 0.93 | 0.76 | 0.82 | 0.72 | 0.98 | 0.84 |
| 11 | 94539045 | rs3893908 | -1.274 | -1.307 | 1.788 | 0.976 | -0.044 | 0.022 | 0.022 | 0 | 0.34 | 0.52 | 0.54 | 0.44 | 0.22 | 0.33 |
| 11 | 94539347 | rs12421118 | -0.559 | -1.254 | 1.189 | 1.005 | -0.006 | 0.009 | 0.009 | 0 | 0.28 | 0.27 | 0.48 | 0.45 | 0.19 | 0.3 |
| 11 | 94557996 | rs10831276 | -0.93 | -0.265 | 1.305 | 1.181 | -0.003 | 0.001 | 0.001 | 0 | 0.3 | 0.27 | 0.49 | 0.44 | 0.19 | 0.33 |
| 11 | 94563731 | rs2241667 | -0.47 | -0.008 | 1.199 | 1.305 | 0.026 | -0.074 | -0.074 | 0.821 | 0.25 | 0.07 | 0.46 | 0.4 | 0.19 | 0.26 |
| 11 | 94564248 | rs16921059 | -0.511 | 0 | 1.206 | 1.308 | 0.02 | -0.037 | -0.037 | 1.907 | 0.26 | 0.13 | 0.47 | 0.4 | 0.19 | 0.26 |
| 11 | 94565171 | rs10831277 | -0.646 | 0.005 | 1.174 | 1.311 | 0.008 | -0.012 | -0.012 | 0.156 | 0.27 | 0.2 | 0.47 | 0.41 | 0.19 | 0.29 |
| 11 | 94575608 | rs11020961 | 0.26 | 0.786 | 0.984 | 1.544 | -0.055 | -0.045 | -0.045 | 0.484 | 0.09 | 0.01 | 0.12 | 0.14 | 0.19 | 0.18 |
| 11 | 94585913 | rs333027 | -1.657 | 0.485 | 2.076 | 1.753 | -0.042 | 0.029 | 0.029 | 0.23 | 0.28 | 0.49 | 0.52 | 0.33 | 0.2 | 0.35 |
| 11 | 94587392 | rs333025 | -1.841 | 0.463 | 2.11 | 1.748 | -0.023 | -0.017 | -0.017 | 0.34 | 0.11 | 0.44 | 0.09 | 0.05 | 0.05 | 0.1 |
| 11 | 94591289 | rs4309121 | -0.107 | 0.493 | 1.969 | 1.745 | -0.018 | -0.081 | -0.081 | 0.233 | 0.17 | 0.02 | 0.43 | 0.28 | 0.15 | 0.26 |
| 11 | 94620657 | rs516594 | 1.798 | -0.37 | 1.853 | 1.609 | 0.015 | 0.001 | 0.001 | 0 | 0.56 | 0.38 | 0.4 | 0.44 | 0.67 | 0.57 |
| 11 | 94642047 | rs11020970 | 1.616 | -0.166 | 2.286 | 1.416 | -0.045 | 0.028 | 0.028 | 0.802 | 0.72 | 0.51 | 0.48 | 0.67 | 0.79 | 0.65 |
| 11 | 94656241 | rs7121021 | -1.79 | 0.026 | 2.14 | 1.284 | -0.013 | -0.021 | -0.021 | 0 | 0.09 | 0.2 | 0.07 | 0.05 | 0.06 | 0.08 |
| 11 | 94660325 | rs690026 | 1.72 | -0.002 | 1.899 | 1.271 | -0.01 | -0.025 | -0.025 | 0 | 0.89 | 0.5 | 0.91 | 0.95 | 0.94 | 0.92 |
| 11 | 94663954 | rs7121722 | -1.208 | 0.154 | 1.25 | 1.552 | 0.006 | -0.027 | -0.027 | 0 | 0.19 | 0.16 | 0.34 | 0.39 | 0.19 | 0.26 |
| 11 | 94666818 | rs689825 | 1.334 | 0.199 | 0.87 | 1.449 | 0.005 | -0.007 | -0.007 | 0 | 0.68 | 0.34 | 0.63 | 0.6 | 0.72 | 0.72 |
| 11 | 94667964 | rs10831284 | 1.173 | 0.324 | 0.792 | 1.48 | 0.059 | -0.007 | -0.007 | 0 | 0.88 | 0.47 | 0.87 | 0.94 | 0.86 | 0.93 |
| 11 | 94674106 | rs6483377 | -1.163 | -0.358 | 0.285 | 1.541 | -0.035 | 0.034 | 0.034 | 0 | 0.3 | 0.53 | 0.33 | 0.39 | 0.21 | 0.28 |
| 11 | 94679823 | rs4753644 | 0.545 | -0.328 | -0.073 | 1.556 | -0.02 | 0.018 | 0.018 | 0 | 0.7 | 0.42 | 0.79 | 0.61 | 0.8 | 0.73 |
| 11 | 94681989 | rs12221798 | -1.373 | 0.24 | -0.235 | 1.887 | 0 | 0.013 | 0.013 | 0.175 | 0.12 | 0.13 | 0.32 | 0.64 | 0.12 | 0.33 |
| 11 | 94685301 | rs2400000 | 1.235 | 0.092 | -0.03 | 1.148 | 0 | 0.001 | 0.001 | 0 | 0.59 | 0.63 | 0.55 | 0.35 | 0.62 | 0.48 |
| 11 | 94688937 | rs2171186 | 1.251 | 0.044 | 0.002 | 1.04 | -0.005 | 0.025 | 0.025 | 0 | 0.77 | 0.68 | 0.66 | 0.36 | 0.78 | 0.6 |
| 11 | 94695492 | rs2171184 | -0.011 | -0.251 | 0.111 | 0.734 | 0.004 | 0.002 | 0.002 | 0 | 0.94 | 0.8 | 0.88 | 0.97 | 0.96 | 0.95 |
| 11 | 94697142 | rs7937776 | 0.47 | 0.332 | 0.047 | 0.631 | -0.009 | -0.072 | -0.072 | 0.131 | 0.8 | 0.74 | 0.79 | 0.96 | 0.82 | 0.88 |
| 11 | 94699113 | rs7952153 | 2.386 | 0.173 | 0.458 | 0.559 | -0.001 | 0.012 | 0.012 | 0 | 0.52 | 0.53 | 0.42 | 0.27 | 0.51 | 0.43 |
| 11 | 94713722 | rs11020993 | -0.431 | 0.156 | 0.385 | 0.524 | -0.017 | 0.031 | 0.031 | 0 | 0.06 | 0 | 0.04 | 0 | 0.07 | 0.01 |
| 11 | 94716563 | rs11020994 | -0.287 | 0.159 | 0.363 | 1.417 | -0.024 | -0.056 | -0.056 | 0 | 0.1 | 0.01 | 0.13 | 0.2 | 0.1 | 0.24 |
| 11 | 94716998 | rs11020995 | 2.112 | 0.108 | 0.43 | 0.998 | 0.003 | -0.035 | -0.035 | 0.182 | 0.66 | 0.58 | 0.51 | 0.27 | 0.67 | 0.5 |
| 11 | 94722715 | rs10501819 | 2.623 | 0.047 | 0.71 | 0.976 | -0.001 | 0.092 | 0.092 | 0 | 0.5 | 0.38 | 0.4 | 0.27 | 0.52 | 0.43 |
| 11 | 94725000 | rs1385967 | 2.66 | 0.049 | 0.71 | 0.977 | 0 | 0.095 | 0.095 | 0 | 0.5 | 0.38 | 0.4 | 0.27 | 0.52 | 0.43 |
| 11 | 94731759 | rs3740853 | -1.61 | 0.178 | 0.71 | 1.089 | -0.006 | -0.045 | -0.045 | 1.061 | 0.22 | 0.1 | 0.32 | 0.65 | 0.22 | 0.39 |
| 11 | 94748136 | rs7107684 | -1.31 | 0.37 | 0.671 | 1.088 | 0.038 | -0.008 | -0.008 | 0 | 0.11 | 0.31 | 0.17 | 0.07 | 0.12 | 0.1 |
| 11 | 94749970 | rs11021009 | -0.843 | 0.402 | 0.648 | 1.784 | -0.014 | -0.04 | -0.04 | 0 | 0.14 | 0.07 | 0.17 | 0.33 | 0.16 | 0.26 |
| 11 | 94758838 | rs16921265 | -0.027 | -0.001 | 0.814 | 1.215 | -0.006 | 0.006 | 0.006 | 0 | 0.06 | 0.01 | 0.14 | 0.35 | 0.06 | 0.09 |
| 11 | 94761019 | rs10741501 | 0.006 | 0.126 | 1.105 | 0.009 | -0.001 | -0.015 | -0.015 | 0 | 0.87 | 0.9 | 0.9 | 1 | 0.85 | 0.9 |
| 11 | 94764037 | rs11021014 | -1.257 | 0.121 | 1.086 | 0.007 | 0.046 | 0.008 | 0.008 | 0 | 0.07 | 0.22 | 0.05 | 0.03 | 0.08 | 0.07 |
| 11 | 94764892 | rs7926605 | 0.214 | 0.134 | 0.654 | 0.017 | -0.001 | -0.007 | -0.007 | 0.001 | 0.85 | 0.91 | 0.9 | 1 | 0.85 | 0.9 |
| 11 | 94765333 | rs10831300 | -1.656 | 0.144 | 0.688 | 0.404 | 0.049 | -0.004 | -0.004 | 0 | 0.12 | 0.39 | 0.17 | 0.07 | 0.12 | 0.1 |
| 11 | 94774208 | rs11602904 | -1.3 | -0.273 | 0.031 | 0.217 | -0.003 | -0.005 | -0.005 | 0 | 0.25 | 0.26 | 0.13 | 0.13 | 0.21 | 0.19 |
| 11 | 94774311 | rs964039 | 0.834 | -0.62 | -0.04 | -0.164 | -0.018 | -0.022 | -0.022 | 1.02 | 0.66 | 0.58 | 0.74 | 0.8 | 0.72 | 0.74 |
| 11 | 94775030 | rs1256901 | -0.879 | 0.189 | -0.732 | -0.565 | -0.049 | 0.083 | 0.083 | 0 | 0.29 | 0.08 | 0.29 | 0.13 | 0.31 | 0.18 |
| 11 | 94781514 | rs7130686 | -0.684 | 0.471 | -0.173 | -0.738 | -0.003 | 0.02 | 0.02 | 0 | 0.24 | 0.2 | 0.24 | 0.05 | 0.22 | 0.2 |
| 11 | 94790699 | rs12292302 | -0.098 | 0.55 | 0.21 | -0.92 | 0.017 | -0.028 | -0.028 | 0.84 | 0.05 | 0.15 | 0.06 | 0 | 0.06 | 0.08 |
| 11 | 94802318 | rs1056986 | 0.819 | 0.48 | 0.457 | -0.593 | 0.001 | -0.008 | -0.008 | 0 | 0.57 | 0.47 | 0.5 | 0.28 | 0.6 | 0.5 |
| 11 | 94802620 | rs12627 | 1.335 | 0.396 | 0.622 | -0.641 | 0.002 | 0.059 | 0.059 | 0 | 0.51 | 0.34 | 0.45 | 0.28 | 0.54 | 0.42 |

| | | | | | | | | | | | | | | | | |
|---|---|---|---|---|---|---|---|---|---|---|---|---|---|---|---|---|
| 11 | 94806722 | rs4753652 | 0.376 | 0.418 | 0.907 | -0.605 | -0.009 | 0.024 | 0.024 | 0 | 0.19 | 0.19 | 0.22 | 0.67 | 0.14 | 0.29 |
| 11 | 94807261 | rs620767 | 0.815 | 0.415 | 0.948 | -0.607 | 0.003 | -0.019 | -0.019 | 0.21 | 0.57 | 0.52 | 0.51 | 0.28 | 0.6 | 0.49 |
| 11 | 94808487 | rs588096 | -0.366 | 0.429 | 0.926 | -0.708 | -0.014 | 0.018 | 0.018 | 0 | 0.77 | 0.81 | 0.73 | 0.95 | 0.74 | 0.79 |
| 11 | 94808715 | rs684804 | -0.275 | 0.428 | 0.939 | -0.709 | -0.007 | 0.006 | 0.006 | 0 | 0.76 | 0.78 | 0.73 | 0.95 | 0.74 | 0.79 |
| 11 | 94809826 | rs1039750 | -0.173 | 0.525 | 0.831 | -0.621 | -0.004 | 0.013 | 0.013 | 0 | 0.24 | 0.24 | 0.3 | 0.7 | 0.21 | 0.38 |
| 11 | 94810756 | rs655070 | 0.582 | 0.442 | 0.655 | -0.432 | 0.001 | -0.008 | -0.008 | 0 | 0.54 | 0.49 | 0.43 | 0.29 | 0.54 | 0.46 |
| 11 | 94813500 | rs622175 | 0.901 | 0.189 | 0.302 | -0.489 | -0.002 | 0.09 | 0.09 | 0.001 | 0.5 | 0.35 | 0.44 | 0.26 | 0.56 | 0.41 |
| 11 | 94815874 | rs581788 | 0.767 | 0.31 | -0.15 | -0.428 | -0.001 | -0.002 | -0.002 | 0.015 | 0.52 | 0.52 | 0.45 | 0.27 | 0.56 | 0.41 |
| 11 | 94824666 | rs673330 | -0.191 | -0.109 | -0.361 | -0.055 | 0.011 | -0.012 | -0.012 | 0 | 0.21 | 0.16 | 0.24 | 0.71 | 0.13 | 0.31 |
| 11 | 94825729 | rs628163 | 0.429 | -0.025 | 0.095 | -0.839 | 0 | -0.063 | -0.063 | 0 | 0.66 | 0.54 | 0.75 | 0.91 | 0.7 | 0.71 |
| 11 | 94825799 | rs641330 | 0.876 | -0.048 | 1.234 | -0.925 | 0.082 | -0.04 | -0.04 | 0.207 | 0.87 | 0.56 | 0.87 | 0.95 | 0.87 | 0.86 |
| 11 | 94825823 | rs628606 | 0.907 | -0.038 | 1.311 | -0.923 | 0.039 | -0.002 | -0.002 | 0.826 | 0.92 | 0.75 | 0.94 | 0.96 | 0.94 | 0.92 |
| 11 | 94829514 | rs2509814 | 1.001 | -0.364 | 1.271 | -1.235 | -0.004 | -0.058 | -0.058 | 1.263 | 0.76 | 0.57 | 0.86 | 0.91 | 0.83 | 0.67 |
| 11 | 94833141 | rs925149 | 0.051 | -0.366 | 1.176 | -1.235 | 0.005 | 0.003 | 0.003 | 0 | 0.15 | 0.14 | 0.05 | 0.08 | 0.09 | 0.23 |
| 11 | 94834270 | rs4611172 | 2.108 | -0.505 | 0.493 | -0.604 | -0.084 | 0.026 | 0.026 | 0 | 0.41 | 0.12 | 0.42 | 0.32 | 0.57 | 0.36 |
| 11 | 94834871 | rs646809 | 1.191 | -0.466 | 0.878 | -0.839 | -0.001 | -0.03 | -0.03 | 0 | 0.72 | 0.37 | 0.82 | 0.91 | 0.78 | 0.67 |
| 11 | 94836615 | rs547243 | 0.111 | -0.341 | 0.848 | -0.753 | 0.01 | -0.028 | -0.028 | 0 | 0.92 | 0.89 | 0.93 | 1 | 0.91 | 0.9 |
| 11 | 94843507 | rs654499 | 1.968 | -0.279 | -0.057 | -1.07 | -0.113 | -0.069 | -0.069 | 1.243 | 0.31 | 0.11 | 0.46 | 0.7 | 0.46 | 0.31 |
| 11 | 94843916 | rs560978 | 0.151 | -0.465 | 0.245 | -1.191 | 0.011 | 0.031 | 0.031 | 0 | 0.3 | 0.19 | 0.33 | 0.16 | 0.27 | 0.33 |
| 11 | 94855879 | rs11021048 | 0.076 | -0.385 | 0.32 | -1.231 | -0.001 | -0.008 | -0.008 | 0 | 0.13 | 0.11 | 0.07 | 0.05 | 0.1 | 0.14 |
| 11 | 94855933 | rs10831307 | 0.531 | -0.353 | 0.311 | -1.49 | -0.001 | 0.008 | 0.008 | 0 | 0.21 | 0.18 | 0.17 | 0.15 | 0.21 | 0.23 |
| 11 | 94856952 | rs10831308 | -0.001 | -0.206 | 0.472 | -1.544 | 0 | -0.003 | -0.003 | 0.78 | 0.25 | 0.27 | 0.19 | 0.15 | 0.26 | 0.24 |
| 11 | 94858419 | rs676761 | 1.349 | -0.168 | 0.324 | -1.572 | -0.024 | 0.031 | 0.031 | 0 | 0.4 | 0.19 | 0.49 | 0.71 | 0.48 | 0.4 |
| 11 | 94862288 | rs3740862 | 1.067 | -0.126 | 0.046 | -1.668 | 0.017 | 0.024 | 0.024 | 0 | 0.15 | 0 | 0.05 | 0.03 | 0.08 | 0.19 |
| 11 | 94862577 | rs3740861 | 0.553 | -0.053 | -0.187 | -1.783 | -0.044 | 0.082 | 0.082 | 0 | 0.25 | 0.09 | 0.2 | 0.12 | 0.27 | 0.37 |
| 11 | 94863186 | rs503612 | 1.485 | -0.076 | -0.007 | -1.74 | -0.016 | -0.002 | -0.002 | 0 | 0.37 | 0.24 | 0.49 | 0.64 | 0.45 | 0.34 |
| 11 | 94865011 | rs607666 | 0.073 | -0.113 | 0.337 | -1.78 | -0.015 | -0.025 | -0.025 | 0 | 0.21 | 0.31 | 0.12 | 0.09 | 0.16 | 0.23 |
| 11 | 94865068 | rs533669 | 0.073 | -0.113 | 0.337 | -1.78 | -0.015 | -0.025 | -0.025 | 0.206 | 0.21 | 0.31 | 0.12 | 0.09 | 0.16 | 0.23 |
| 11 | 94865334 | rs586421 | 0.939 | -0.032 | 0.404 | -1.822 | 0.001 | 0.058 | 0.058 | 0 | 0.47 | 0.36 | 0.54 | 0.78 | 0.51 | 0.38 |
| 11 | 94866053 | rs3758783 | 1.149 | -0.145 | 0.203 | -1.801 | 0.01 | 0.01 | 0.01 | 0 | 0.15 | 0.08 | 0.05 | 0.03 | 0.08 | 0.19 |
| 11 | 94867748 | rs7937016 | 0.955 | -0.141 | 0.236 | -1.799 | -0.018 | -0.018 | -0.018 | 2.864 | 0.16 | 0.15 | 0.06 | 0.03 | 0.07 | 0.19 |
| 11 | 94873625 | rs499172 | -0.785 | -0.133 | 0.249 | -1.781 | -0.002 | 0.008 | 0.008 | 0 | 0.46 | 0.42 | 0.31 | 0.22 | 0.43 | 0.6 |
| 11 | 94874706 | rs530190 | -1.209 | 0.143 | 0.227 | -1.738 | 0.008 | -0.034 | -0.034 | 0.649 | 0.94 | 0.81 | 0.86 | 1 | 0.92 | 0.97 |
| 11 | 94876984 | rs17693733 | 2.207 | -0.03 | -0.621 | -1.873 | -0.091 | 0.045 | 0.045 | 0 | 0.26 | 0.02 | 0.24 | 0.14 | 0.32 | 0.18 |
| 11 | 94877363 | rs496397 | 0.128 | 0.285 | -0.035 | -2.037 | -0.026 | -0.087 | -0.087 | 0 | 0.75 | 0.61 | 0.77 | 0.96 | 0.8 | 0.76 |
| 11 | 94877438 | rs653118 | -0.286 | 0.293 | -0.116 | -2.03 | -0.038 | -0.07 | -0.07 | 0 | 0.78 | 0.64 | 0.79 | 0.96 | 0.84 | 0.78 |
| 11 | 94878090 | rs10501821 | 0.087 | 0.524 | -0.191 | -1.858 | -0.015 | 0.026 | 0.026 | 0 | 0.28 | 0.25 | 0.23 | 0.18 | 0.31 | 0.4 |
| 11 | 94881894 | rs10831309 | -0.708 | 0.471 | -0.118 | -1.564 | -0.004 | 0.027 | 0.027 | 0.206 | 0.53 | 0.36 | 0.33 | 0.32 | 0.45 | 0.63 |
| 11 | 94883659 | rs634788 | -0.506 | 0.381 | 0.377 | -1.429 | 0.052 | -0.006 | -0.006 | 0 | 0.93 | 0.68 | 0.84 | 0.99 | 0.91 | 0.97 |
| 11 | 94886632 | rs496321 | 0.773 | 0.325 | -0.319 | -1.623 | 0.014 | 0.087 | 0.087 | 0 | 0.53 | 0.3 | 0.61 | 0.75 | 0.57 | 0.56 |
| 11 | 94886806 | rs473213 | 0.474 | 0.292 | -0.366 | -1.466 | 0 | 0.001 | 0.001 | 0 | 0.6 | 0.41 | 0.65 | 0.76 | 0.63 | 0.57 |
| 11 | 94887619 | rs644756 | -0.409 | 0.472 | -0.129 | -1.078 | 0.045 | -0.017 | -0.017 | 0 | 0.94 | 0.72 | 0.85 | 1 | 0.92 | 0.97 |
| 11 | 94888163 | rs12574604 | 0.708 | 1.087 | -0.977 | -0.755 | -0.045 | -0.002 | -0.002 | 0 | 0.16 | 0.04 | 0.14 | 0.15 | 0.22 | 0.11 |
| 11 | 94889314 | rs682533 | -0.321 | -0.176 | -0.546 | -0.819 | -0.005 | -0.066 | -0.066 | 0 | 0.72 | 0.7 | 0.74 | 0.98 | 0.78 | 0.81 |
| 11 | 94889674 | rs11021055 | -1.059 | 0.317 | -0.591 | 0.491 | -0.008 | -0.001 | -0.001 | 0 | 0.14 | 0.13 | 0.06 | 0.08 | 0.08 | 0.23 |
| 11 | 94889710 | rs11021056 | -1.093 | 0.365 | -0.596 | 0.486 | -0.002 | -0.001 | -0.001 | 0 | 0.15 | 0.13 | 0.07 | 0.08 | 0.11 | 0.23 |
| 11 | 94890994 | rs11824683 | -0.765 | 0.295 | -0.607 | 0.575 | -0.016 | -0.001 | -0.001 | 0 | 0.12 | 0.14 | 0.06 | 0.08 | 0.08 | 0.17 |
| 11 | 94894963 | rs11021059 | -1.023 | 0.709 | -0.898 | 1.105 | -0.001 | 0.005 | 0.005 | 0 | 0.07 | 0.07 | 0.09 | 0.19 | 0.09 | 0.1 |
| 11 | 94902171 | rs12361586 | -1.628 | 0.837 | -0.95 | 1.07 | -0.004 | -0.005 | -0.005 | 0.001 | 0.09 | 0.07 | 0.11 | 0.19 | 0.12 | 0.13 |
| 11 | 94906111 | rs684856 | -0.684 | 0.579 | -1.396 | 0.777 | 0.008 | 0.05 | 0.05 | 0 | 0.43 | 0.32 | 0.45 | 0.74 | 0.41 | 0.62 |
| 11 | 94908658 | rs11021065 | -0.531 | 0.521 | -1.813 | 1.394 | -0.002 | 0.001 | 0.001 | 1.056 | 0.29 | 0.15 | 0.32 | 0.29 | 0.32 | 0.27 |
| 11 | 94922399 | rs629508 | -0.507 | 0.951 | -1.501 | 1.928 | -0.042 | -0.008 | -0.008 | 0 | 0.29 | 0.08 | 0.34 | 0.29 | 0.37 | 0.27 |
| 11 | 94929102 | rs626481 | -0.545 | 0.483 | -1.598 | 1.167 | 0 | 0.003 | 0.003 | 0 | 0.53 | 0.45 | 0.52 | 0.63 | 0.52 | 0.48 |
| 11 | 94950910 | rs10501823 | 0.174 | 0.629 | -1.864 | 1.26 | -0.002 | 0.003 | 0.003 | 0.773 | 0.29 | 0.14 | 0.31 | 0.28 | 0.32 | 0.26 |
| 11 | 94952391 | rs11021083 | -0.261 | 0.641 | -1.84 | 1.032 | 0.002 | -0.006 | -0.006 | 0 | 0.34 | 0.32 | 0.4 | 0.59 | 0.34 | 0.44 |
| 11 | 94953043 | rs1054522 | 1.452 | 0.621 | -1.983 | 0.645 | -0.07 | 0.088 | 0.088 | 2.019 | 0.19 | 0.01 | 0.12 | 0.04 | 0.18 | 0.07 |
| 11 | 94962427 | rs1458453 | -0.119 | 0.209 | -1.424 | 1.515 | -0.003 | 0.034 | 0.034 | 0.367 | 0.28 | 0.14 | 0.23 | 0.18 | 0.31 | 0.26 |
| 11 | 94964931 | rs2077510 | -0.085 | -0.04 | -1.334 | 1.559 | -0.001 | -0.007 | -0.007 | 0.285 | 0.26 | 0.24 | 0.23 | 0.18 | 0.32 | 0.21 |
| 11 | 94968137 | rs1458451 | -0.432 | -0.042 | -1.347 | 1.615 | 0 | 0 | 0 | 0 | 0.3 | 0.32 | 0.4 | 0.55 | 0.34 | 0.35 |
| 11 | 94970185 | rs11021095 | 0.07 | -0.058 | -1.511 | 1.671 | -0.038 | -0.031 | -0.031 | 0 | 0.14 | 0.01 | 0.09 | 0.14 | 0.13 | 0.23 |
| 11 | 94982706 | rs11021100 | 0.271 | -0.424 | -0.729 | 1.55 | -0.014 | 0.033 | 0.033 | 0 | 0.69 | 0.59 | 0.59 | 0.49 | 0.69 | 0.65 |
| 11 | 94984115 | rs9667295 | -0.778 | 0.08 | -0.264 | 1.256 | -0.011 | 0.065 | 0.065 | 0 | 0.11 | 0.26 | 0.07 | 0.19 | 0.08 | 0.13 |
| 11 | 94987033 | rs10466336 | -0.838 | -0.062 | -0.533 | 1.021 | -0.001 | -0.001 | -0.001 | 1.35 | 0.34 | 0.33 | 0.28 | 0.3 | 0.34 | 0.31 |
| 11 | 94989990 | rs10501824 | -0.679 | 0.368 | -0.846 | 0.877 | -0.031 | -0.002 | -0.002 | 0.989 | 0.31 | 0.09 | 0.38 | 0.25 | 0.32 | 0.26 |
| 11 | 94991005 | rs11021101 | 0.517 | 0.138 | -0.563 | 0.807 | -0.002 | -0.005 | -0.005 | 0 | 0.12 | 0.1 | 0.07 | 0.11 | 0.13 | 0.22 |
| 11 | 94998250 | rs10741505 | -1 | -0.35 | 1.194 | 2.156 | 0 | 0.001 | 0.001 | 0.345 | 0.36 | 0.45 | 0.61 | 0.64 | 0.35 | 0.41 |
| 11 | 95000894 | rs4753661 | -0.856 | -0.411 | 1.086 | 2.157 | -0.002 | -0.004 | -0.004 | 0.151 | 0.35 | 0.26 | 0.59 | 0.63 | 0.34 | 0.41 |
| 11 | 95008428 | rs10741506 | -0.476 | -0.436 | 1.132 | 2.449 | -0.027 | -0.052 | -0.052 | 0.219 | 0.32 | 0.11 | 0.56 | 0.41 | 0.33 | 0.33 |
| 11 | 95008479 | rs11021118 | 0.435 | -0.399 | 1.084 | 1.755 | 0 | -0.006 | -0.006 | 0 | 0.3 | 0.28 | 0.19 | 0.21 | 0.32 | 0.4 |

| | | | | | | | | | | | | | | | | |
|---|---|---|---|---|---|---|---|---|---|---|---|---|---|---|---|---|
| 11 | 95011645 | rs976337 | -1.1 | -0.463 | 1.313 | 2.384 | -0.002 | -0.031 | -0.031 | 0 | 0.41 | 0.5 | 0.63 | 0.73 | 0.41 | 0.46 |
| 11 | 95018911 | rs543457 | -1.199 | -0.569 | 1.021 | 2.511 | -0.001 | -0.003 | -0.003 | 0 | 0.39 | 0.37 | 0.6 | 0.73 | 0.37 | 0.43 |
| 11 | 95022652 | rs10831329 | -0.585 | -0.58 | 0.845 | 2.428 | -0.004 | -0.005 | -0.005 | 0 | 0.3 | 0.07 | 0.36 | 0.26 | 0.27 | 0.22 |
| 11 | 95024932 | rs498018 | -1.296 | -0.583 | 0.893 | 2.43 | 0.001 | -0.001 | -0.001 | 0 | 0.38 | 0.31 | 0.45 | 0.57 | 0.38 | 0.34 |
| 11 | 95030363 | rs11021122 | -0.472 | -0.4 | 0.308 | 2.347 | -0.021 | -0.022 | -0.022 | 0 | 0.27 | 0.06 | 0.34 | 0.26 | 0.27 | 0.22 |
| 11 | 95033906 | rs11021123 | -0.079 | -0.385 | 0.233 | 2.287 | -0.019 | -0.048 | -0.048 | 0 | 0.19 | 0.05 | 0.16 | 0.26 | 0.21 | 0.15 |
| 11 | 95037134 | rs12283509 | -0.242 | -0.659 | 0.4 | 2.126 | -0.005 | 0 | 0 | 0.106 | 0.12 | 0.1 | 0.08 | 0.11 | 0.08 | 0.08 |
| 11 | 95038070 | rs529936 | 0.654 | -0.686 | 0.593 | 2.277 | 0.002 | -0.014 | -0.014 | 0.045 | 0.71 | 0.81 | 0.51 | 0.35 | 0.72 | 0.67 |
| 11 | 95039414 | rs12275549 | -0.513 | -0.506 | 0.957 | 2.583 | -0.005 | 0.001 | 0.001 | 0.094 | 0.12 | 0.1 | 0.08 | 0.12 | 0.08 | 0.08 |
| 11 | 95040242 | rs7115547 | -0.211 | -0.569 | 0.818 | 2.639 | -0.012 | -0.037 | -0.037 | 1.715 | 0.18 | 0.06 | 0.16 | 0.26 | 0.2 | 0.16 |
| 11 | 95040906 | rs498156 | 0.236 | -0.583 | 1.004 | 2.783 | -0.002 | 0.071 | 0.071 | 0 | 0.8 | 0.71 | 0.82 | 0.51 | 0.79 | 0.8 |
| 11 | 95051514 | rs10765730 | -0.197 | -0.536 | 0.573 | 2.38 | -0.011 | -0.048 | -0.048 | 0 | 0.18 | 0.08 | 0.25 | 0.35 | 0.2 | 0.15 |
| 11 | 95052753 | rs505317 | 1.396 | -0.65 | 0.826 | 2.312 | -0.026 | -0.031 | -0.031 | 0 | 0.66 | 0.56 | 0.57 | 0.23 | 0.72 | 0.63 |
| 11 | 95064726 | rs10831338 | -0.243 | -0.334 | 0.095 | 1.768 | -0.011 | -0.049 | -0.049 | 2.38 | 0.17 | 0.09 | 0.25 | 0.36 | 0.2 | 0.15 |
| 11 | 95069694 | rs530356 | -1.083 | -0.396 | 0.371 | 1.192 | -0.03 | 0.007 | 0.007 | 0.135 | 0.13 | 0.19 | 0.11 | 0.12 | 0.07 | 0.06 |
| 11 | 95074260 | rs514284 | -1.017 | -0.394 | 0.049 | 1.168 | 0.004 | -0.001 | -0.001 | 0.236 | 0.12 | 0.11 | 0.12 | 0.11 | 0.06 | 0.06 |
| 11 | 95076290 | rs564038 | 1.031 | -0.463 | 0.141 | 1.824 | -0.006 | 0.019 | 0.019 | 0 | 0.87 | 0.83 | 0.88 | 0.59 | 0.95 | 0.86 |
| 11 | 95078263 | rs580009 | 1.384 | -0.135 | 0.51 | 1.599 | 0 | 0 | 0 | 0 | 0.62 | 0.35 | 0.53 | 0.21 | 0.67 | 0.57 |
| 11 | 95079327 | rs3858379 | -0.156 | -0.163 | 0.894 | 1.691 | -0.052 | -0.094 | -0.094 | 0 | 0.2 | 0.05 | 0.29 | 0.35 | 0.21 | 0.14 |
| 11 | 95081138 | rs491069 | 0.86 | -0.2 | 1.054 | 1.612 | -0.057 | 0.196 | 0.196 | 0 | 0.84 | 0.54 | 0.84 | 0.59 | 0.91 | 0.86 |
| 11 | 95086353 | rs558531 | 1.435 | 0.17 | 0.435 | 0.706 | -0.005 | 0.058 | 0.058 | 0 | 0.34 | 0.21 | 0.2 | 0.06 | 0.28 | 0.24 |
| 11 | 95088413 | rs573919 | 0.867 | 0.175 | 0.901 | 1.424 | -0.021 | 0.213 | 0.213 | 0.332 | 0.89 | 0.67 | 0.86 | 0.59 | 0.91 | 0.84 |
| 11 | 95090828 | rs10765734 | -1.051 | 0.191 | 0.882 | 1.36 | -0.002 | 0.03 | 0.03 | 0.859 | 0.29 | 0.38 | 0.36 | 0.44 | 0.25 | 0.22 |
| 11 | 95092903 | rs537570 | 0.851 | 0.235 | 0.914 | 1.411 | -0.002 | -0.055 | -0.055 | 1.108 | 0.68 | 0.48 | 0.57 | 0.2 | 0.7 | 0.69 |
| 11 | 95095664 | rs559019 | 0.35 | 0.422 | 0.301 | 1.073 | 0.001 | 0.001 | 0.001 | 0 | 0.76 | 0.74 | 0.64 | 0.3 | 0.73 | 0.76 |
| 11 | 95099182 | rs477980 | 0.901 | 0.261 | 0.351 | 1.4 | -0.014 | 0.087 | 0.087 | 0.79 | 0.89 | 0.86 | 0.87 | 0.57 | 0.91 | 0.84 |
| 11 | 95099260 | rs16921827 | -0.585 | 0.461 | 0.615 | 0.811 | -0.003 | -0.038 | -0.038 | 0.012 | 0.22 | 0.35 | 0.34 | 0.1 | 0.25 | 0.14 |
| 11 | 95099356 | rs479709 | 0.764 | 0.641 | 0.915 | 0.683 | 0 | 0.023 | 0.023 | 1.04 | 0.71 | 0.55 | 0.53 | 0.62 | 0.67 | 0.76 |
| 11 | 95099864 | rs10501826 | 0.019 | 0.905 | 0.479 | 0.478 | -0.056 | 0.093 | 0.093 | 0 | 0.3 | 0.08 | 0.39 | 0.11 | 0.42 | 0.29 |
| 11 | 95099891 | rs7950644 | -0.072 | 0.852 | 0.728 | 0.661 | 0.002 | -0.004 | -0.004 | 0.804 | 0.32 | 0.33 | 0.42 | 0.22 | 0.42 | 0.32 |
| 11 | 95101147 | rs538592 | 0.747 | 0.69 | 0.737 | 0.75 | -0.003 | 0.02 | 0.02 | 0.01 | 0.79 | 0.71 | 0.63 | 0.41 | 0.81 | 0.84 |
| 11 | 95109331 | rs11602601 | 0.175 | 1.022 | 0.554 | 0.618 | -0.058 | 0.112 | 0.112 | 0 | 0.32 | 0.07 | 0.37 | 0.11 | 0.43 | 0.35 |
| 11 | 95110237 | rs7940677 | 1.422 | 0.92 | 0.74 | 0.522 | 0.002 | 0.011 | 0.011 | 0 | 0.41 | 0.33 | 0.21 | 0.19 | 0.35 | 0.41 |
| 11 | 95110685 | rs523808 | 0.695 | 0.929 | 0.633 | 0.713 | -0.001 | 0.046 | 0.046 | 0 | 0.09 | 0.09 | 0.19 | 0.34 | 0.1 | 0.1 |
| 11 | 95114053 | rs569612 | -0.025 | 0.912 | 0.538 | 0.714 | 0 | -0.001 | -0.001 | 0 | 0.86 | 0.85 | 0.81 | 0.76 | 0.88 | 0.89 |
| 11 | 95114225 | rs12802218 | 0.225 | 0.994 | 0.45 | 0.716 | -0.059 | 0.11 | 0.11 | 0 | 0.32 | 0.07 | 0.37 | 0.11 | 0.43 | 0.35 |
| 11 | 95116254 | rs535512 | 0.484 | 1.136 | 0.56 | 0.991 | -0.005 | 0.02 | 0.02 | 0 | 0.09 | 0.09 | 0.19 | 0.34 | 0.1 | 0.1 |
| 11 | 95126049 | rs479314 | 0.292 | 1.082 | 0.224 | 0.431 | -0.002 | -0.003 | -0.003 | 0 | 0.45 | 0.46 | 0.38 | 0.6 | 0.38 | 0.48 |
| 11 | 95132416 | rs547531 | -1.252 | 1.317 | 0.236 | 0.585 | -0.004 | -0.007 | -0.007 | 0 | 0.93 | 0.96 | 0.89 | 0.83 | 0.92 | 0.93 |
| 11 | 95140285 | rs4121863 | -0.106 | 1.261 | 0.271 | 0.617 | 0.012 | -0.006 | -0.006 | 0 | 0.5 | 0.16 | 0.59 | 0.45 | 0.61 | 0.47 |
| 11 | 95142376 | rs4365015 | 1.679 | 1.148 | 0.074 | 0.422 | -0.024 | 0.126 | 0.126 | 0 | 0.26 | 0.05 | 0.15 | 0.07 | 0.21 | 0.19 |
| 11 | 95145561 | rs10501827 | 1.167 | 1.17 | 0.4 | 0.574 | -0.002 | -0.023 | -0.023 | 0.737 | 0.32 | 0.32 | 0.21 | 0.2 | 0.24 | 0.35 |
| 11 | 95148826 | rs4237564 | -0.106 | 1.583 | 0.659 | 1.512 | -0.001 | 0.001 | 0.001 | 0.526 | 0.85 | 0.85 | 0.83 | 0.72 | 0.85 | 0.83 |
| 11 | 95154173 | rs526108 | -0.043 | 1.601 | 0.602 | 1.59 | -0.003 | 0.001 | 0.001 | 5.232 | 0.15 | 0.13 | 0.14 | 0.1 | 0.16 | 0.16 |
| 11 | 95155770 | rs10765737 | 0.453 | 1.647 | 0.439 | 1.217 | 0.008 | 0.009 | 0.009 | 0.124 | 0.69 | 0.47 | 0.61 | 0.4 | 0.69 | 0.65 |
| 11 | 95161575 | rs541821 | -0.459 | 1.501 | -0.544 | 0.695 | 0.006 | 0.093 | 0.093 | 0.658 | 0.27 | 0.42 | 0.32 | 0.49 | 0.24 | 0.29 |
| 11 | 95163313 | rs11021153 | 0.431 | 1.034 | -1.556 | -0.236 | 0.029 | 0.183 | 0.183 | 0 | 0.47 | 0.22 | 0.29 | 0.21 | 0.38 | 0.34 |
| 11 | 95167861 | rs2510580 | 0.298 | 1.289 | -1.53 | -0.118 | -0.029 | 0.046 | 0.046 | 0.705 | 0.73 | 0.86 | 0.67 | 0.87 | 0.62 | 0.71 |
| 11 | 95176293 | rs517985 | 1.136 | 0.351 | -1.688 | 0.033 | 0.042 | 0.012 | 0.012 | 3.172 | 0.36 | 0.25 | 0.22 | 0.34 | 0.2 | 0.25 |
| 11 | 95177805 | rs11021161 | 0.557 | 0.549 | -1.174 | 0.728 | -0.006 | 0.011 | 0.011 | 0.09 | 0.16 | 0.2 | 0.29 | 0.24 | 0.11 | 0.19 |
| 11 | 95178140 | rs4121859 | 0.853 | 0.467 | -1.326 | 1.069 | -0.105 | -0.011 | -0.011 | 0.22 | 0.17 | 0.03 | 0.16 | 0.12 | 0.29 | 0.19 |
| 11 | 95180679 | rs3937019 | 0.174 | 0.123 | -1.034 | 1.083 | -0.019 | 0.023 | 0.023 | 0 | 0.18 | 0.29 | 0.29 | 0.22 | 0.11 | 0.17 |
| 11 | 95183687 | rs483046 | -0.234 | -0.33 | -1.559 | 0.716 | -0.001 | 0.003 | 0.003 | 0 | 0.66 | 0.71 | 0.68 | 0.63 | 0.62 | 0.64 |
| 11 | 95185564 | rs11021163 | 1.005 | -0.393 | -1.047 | 0.917 | 0.004 | 0.022 | 0.022 | 0 | 0.08 | 0.16 | 0.11 | 0.13 | 0.09 | 0.11 |
| 11 | 95186307 | rs2508418 | -1.035 | -0.33 | -0.891 | 0.882 | 0.005 | -0.009 | -0.009 | 0 | 0.81 | 0.76 | 0.81 | 0.85 | 0.72 | 0.76 |
| 11 | 95186913 | rs481539 | 0.311 | -0.22 | -0.367 | 0.936 | 0.006 | 0 | 0 | 2.235 | 0.58 | 0.35 | 0.42 | 0.44 | 0.56 | 0.4 |
| 11 | 95187799 | rs11021165 | 1.107 | -0.238 | -0.528 | 0.72 | -0.002 | -0.01 | -0.01 | 0 | 0.09 | 0.05 | 0.1 | 0.12 | 0.09 | 0.13 |
| 11 | 95189206 | rs528798 | 0.476 | -0.442 | -0.528 | 0.556 | 0.009 | 0.056 | 0.056 | 2.843 | 0.5 | 0.62 | 0.34 | 0.21 | 0.45 | 0.31 |
| 11 | 95192462 | rs12577444 | 0.153 | -0.54 | -0.195 | 1.241 | 0 | 0.03 | 0.03 | 0 | 0.07 | 0.05 | 0.07 | 0.22 | 0.06 | 0.08 |
| 11 | 95194434 | rs7949961 | -0.759 | -0.874 | -0.152 | 1.141 | -0.001 | -0.003 | -0.003 | 0.03 | 0.45 | 0.57 | 0.43 | 0.54 | 0.56 | 0.39 |
| 11 | 95194927 | rs2510568 | -0.112 | -0.863 | -0.071 | 0.679 | -0.003 | 0.008 | 0.008 | 0.132 | 0.77 | 0.8 | 0.8 | 0.78 | 0.77 | 0.62 |
| 11 | 95197220 | rs11821606 | 0.138 | -0.941 | -0.076 | 0.721 | -0.001 | 0.013 | 0.013 | 0 | 0.09 | 0.09 | 0.04 | 0.17 | 0.07 | 0.08 |
| 11 | 95200215 | rs498766 | -1.031 | -1.253 | -0.343 | 0.77 | -0.027 | 0.04 | 0.04 | 0.143 | 0.87 | 0.99 | 0.9 | 0.92 | 0.85 | 0.71 |
| 11 | 95202012 | rs2510579 | 1.074 | -1.188 | -0.052 | 0.679 | 0 | -0.001 | -0.001 | 0.161 | 0.46 | 0.52 | 0.6 | 0.44 | 0.42 | 0.35 |
| 11 | 95212878 | rs1943505 | 2.612 | -1.319 | -0.171 | 0.617 | 0.014 | 0.012 | 0.012 | 0.548 | 0.21 | 0.15 | 0.15 | 0.14 | 0.12 | 0.1 |
| 11 | 95219016 | rs11822727 | 0.868 | -1.284 | -0.43 | 0.636 | 0 | -0.004 | -0.004 | 0 | 0.24 | 0.3 | 0.43 | 0.3 | 0.26 | 0.25 |
| 11 | 95221480 | rs10831357 | -0.938 | -0.861 | -0.714 | 0.569 | 0.003 | -0.005 | -0.005 | 0 | 0.56 | 0.57 | 0.67 | 0.65 | 0.54 | 0.76 |
| 11 | 95229848 | rs10831360 | -0.736 | -0.807 | -1.066 | 0.457 | -0.017 | -0.005 | -0.005 | 0 | 0.78 | 0.84 | 0.83 | 0.79 | 0.67 | 0.86 |
| 11 | 95232867 | rs4237572 | -0.866 | -0.825 | -1.062 | 0.422 | 0.002 | 0 | 0 | 0 | 0.55 | 0.56 | 0.67 | 0.6 | 0.54 | 0.69 |

| | | | | | | | | | | | | | | | | |
|---|---|---|---|---|---|---|---|---|---|---|---|---|---|---|---|---|
| 11 | 95233592 | rs6483426 | -0.06 | 0.711 | -1.328 | 0.885 | -0.006 | -0.005 | -0.005 | 0 | 0.77 | 0.82 | 0.82 | 0.73 | 0.67 | 0.78 |
| 11 | 95234370 | rs12223080 | 0.875 | 0.42 | -1.514 | 0.616 | -0.019 | -0.033 | -0.033 | 0.469 | 0.08 | 0.01 | 0.07 | 0.14 | 0.06 | 0.08 |
| 11 | 95237923 | rs4237573 | 0.528 | 0.536 | -1.534 | 0.595 | 0.004 | 0.001 | 0.001 | 3.209 | 0.62 | 0.88 | 0.64 | 0.6 | 0.41 | 0.54 |
| 11 | 95239071 | rs16922018 | 0.689 | 0.265 | -1.432 | 0.681 | -0.021 | 0.013 | 0.013 | 0 | 0.06 | 0.13 | 0.03 | 0.1 | 0.02 | 0.1 |
| 11 | 95242203 | rs1893687 | 1.588 | 0.029 | -1.302 | 0.695 | -0.021 | -0.007 | -0.007 | 0 | 0.42 | 0.49 | 0.51 | 0.35 | 0.34 | 0.28 |
| 11 | 95242217 | rs16922031 | -0.238 | 0.25 | -1.027 | 0.94 | -0.019 | 0.034 | 0.034 | 0 | 0.1 | 0.22 | 0.1 | 0.17 | 0.06 | 0.15 |
| 11 | 95242922 | rs1943517 | -0.47 | 0.225 | -1.022 | 0.948 | -0.039 | 0.042 | 0.042 | 0 | 0.18 | 0.31 | 0.15 | 0.25 | 0.1 | 0.26 |
| 11 | 95245264 | rs1893686 | -1.304 | 0.924 | -1.224 | 0.478 | -0.102 | -0.061 | -0.061 | 2.126 | 0.86 | 0.99 | 0.85 | 0.81 | 0.7 | 0.91 |
| 11 | 95247928 | rs7952066 | 0.114 | 0.133 | -1.06 | 0.461 | -0.048 | -0.023 | -0.023 | 0 | 0.65 | 0.89 | 0.68 | 0.61 | 0.49 | 0.57 |
| 11 | 95250876 | rs12795900 | -0.414 | 0.16 | -1.038 | 0.369 | -0.003 | -0.008 | -0.008 | 0.62 | 0.18 | 0.19 | 0.12 | 0.14 | 0.15 | 0.37 |
| 11 | 95251008 | rs12280779 | -0.367 | 0.159 | -1.031 | 0.368 | 0.001 | 0.005 | 0.005 | 0 | 0.18 | 0.16 | 0.12 | 0.14 | 0.15 | 0.37 |
| 11 | 95251962 | rs10765744 | 0.444 | -0.455 | -0.852 | 0.51 | 0.02 | -0.006 | -0.006 | 0.019 | 0.56 | 0.72 | 0.63 | 0.52 | 0.45 | 0.46 |
| 11 | 95252207 | rs10831364 | 0.432 | -0.407 | -0.957 | 0.53 | 0.031 | -0.01 | -0.01 | 0 | 0.55 | 0.8 | 0.63 | 0.52 | 0.45 | 0.44 |
| 11 | 95255537 | rs6483433 | 0.804 | -0.199 | -0.914 | 0.656 | -0.014 | 0 | 0 | 0 | 0.09 | 0.1 | 0.05 | 0.08 | 0.04 | 0.11 |
| 11 | 95257534 | rs16922085 | 0.647 | 0.073 | -0.702 | 0.724 | -0.006 | 0.005 | 0.005 | 0 | 0.06 | 0.09 | 0.04 | 0.08 | 0.05 | 0.09 |
| 11 | 95257602 | rs6483435 | 0.231 | 0.302 | -0.924 | 0.51 | 0.017 | 0.003 | 0.003 | 0.732 | 0.52 | 0.62 | 0.61 | 0.52 | 0.44 | 0.43 |
| 11 | 95266172 | rs10831368 | 0.252 | 0.38 | -1.494 | 0.573 | 0.057 | 0.008 | 0.008 | 0.722 | 0.52 | 0.73 | 0.62 | 0.52 | 0.44 | 0.43 |
| 11 | 95270357 | rs11021198 | 0.439 | 0.979 | -1.294 | 0.534 | -0.005 | 0.002 | 0.002 | 0 | 0.78 | 0.86 | 0.81 | 0.74 | 0.73 | 0.75 |
| 11 | 95270425 | rs11021199 | -0.258 | 0.064 | -0.958 | 0.87 | -0.011 | 0.039 | 0.039 | 0 | 0.26 | 0.42 | 0.47 | 0.41 | 0.27 | 0.46 |
| 11 | 95270802 | rs4121809 | -0.241 | 0.335 | -1.659 | 0.085 | -0.005 | 0.118 | 0.118 | 0.236 | 0.57 | 0.86 | 0.64 | 0.74 | 0.45 | 0.63 |
| 11 | 95271755 | rs7926825 | -0.377 | -1.394 | -0.918 | -0.116 | 0.013 | -0.001 | -0.001 | 0.006 | 0.45 | 0.61 | 0.55 | 0.47 | 0.33 | 0.45 |
| 11 | 95274493 | rs4753172 | -0.584 | -1.482 | -0.696 | -0.135 | 0.005 | 0.003 | 0.003 | 0 | 0.52 | 0.44 | 0.62 | 0.49 | 0.5 | 0.45 |
| 11 | 95279452 | rs4753173 | -0.289 | -1.596 | -0.526 | -0.129 | 0.028 | 0.023 | 0.023 | 0 | 0.46 | 0.34 | 0.55 | 0.41 | 0.43 | 0.35 |
| 11 | 95280915 | rs4753689 | -0.59 | -1.6 | -0.506 | -0.133 | 0.005 | 0.002 | 0.002 | 0 | 0.49 | 0.44 | 0.63 | 0.5 | 0.48 | 0.43 |
| 11 | 95283147 | rs11021208 | -0.62 | -1.607 | -0.445 | -0.137 | 0.007 | 0.003 | 0.003 | 0 | 0.5 | 0.43 | 0.63 | 0.5 | 0.49 | 0.44 |
| 11 | 95283557 | rs11021209 | 0.85 | -1.613 | -0.494 | -0.183 | -0.036 | -0.017 | -0.017 | 0 | 0.13 | 0.02 | 0.09 | 0.16 | 0.17 | 0.16 |
| 11 | 95283628 | rs10831371 | -0.18 | -1.77 | -0.389 | -0.239 | 0.011 | 0.003 | 0.003 | 0.001 | 0.42 | 0.31 | 0.54 | 0.42 | 0.39 | 0.36 |
| 11 | 95284803 | rs11021212 | 0.35 | -1.655 | -0.385 | -0.21 | -0.002 | -0.002 | -0.002 | 0.141 | 0.15 | 0.14 | 0.16 | 0.25 | 0.22 | 0.24 |
| 11 | 95287075 | rs8181535 | 0.4 | -1.686 | -0.325 | -0.175 | -0.012 | -0.007 | -0.007 | 0.008 | 0.15 | 0.12 | 0.14 | 0.23 | 0.22 | 0.27 |
| 11 | 95289851 | rs16922181 | -0.335 | -1.707 | -0.21 | -0.407 | 0 | 0.009 | 0.009 | 0 | 0.1 | 0.19 | 0.08 | 0.18 | 0.12 | 0.18 |
| 11 | 95290521 | rs11606101 | 0.895 | -1.816 | -0.228 | -0.449 | 0.018 | 0.011 | 0.011 | 0 | 0.27 | 0.17 | 0.39 | 0.17 | 0.17 | 0.12 |
| 11 | 95290854 | rs4753176 | 0.133 | -1.719 | -0.177 | -0.42 | 0.025 | 0.004 | 0.004 | 0.21 | 0.4 | 0.27 | 0.48 | 0.36 | 0.33 | 0.31 |
| 11 | 95296676 | rs7943143 | -0.949 | -1.61 | -0.131 | -0.453 | -0.001 | 0.002 | 0.002 | 0.965 | 0.57 | 0.62 | 0.64 | 0.64 | 0.51 | 0.61 |
| 11 | 95297070 | rs1255532 | 0.131 | -1.198 | -0.046 | -0.139 | 0.071 | 0.153 | 0.153 | 0.066 | 0.95 | 0.71 | 0.92 | 0.87 | 0.95 | 0.89 |
| 11 | 95299637 | rs1255529 | 0.308 | -1.555 | -0.669 | -0.979 | 0 | -0.004 | -0.004 | 0 | 0.46 | 0.47 | 0.32 | 0.32 | 0.43 | 0.29 |
| 11 | 95303519 | rs1255526 | 0.099 | -1.573 | -0.699 | -0.98 | -0.001 | -0.003 | -0.003 | 0.986 | 0.48 | 0.47 | 0.36 | 0.35 | 0.49 | 0.36 |
| 11 | 95304834 | rs1939881 | 1.159 | -0.863 | -0.738 | -0.419 | 0.012 | -0.022 | -0.022 | 0 | 0.08 | 0.01 | 0.06 | 0.31 | 0.06 | 0.08 |
| 11 | 95308992 | rs1939877 | 1.119 | -0.812 | -0.759 | -0.438 | -0.009 | -0.018 | -0.018 | 0 | 0.15 | 0.07 | 0.39 | 0.19 | 0.18 | 0.17 |
| 11 | 95309661 | rs10831375 | -0.253 | -0.778 | -0.829 | -0.668 | 0.002 | 0.002 | 0.002 | 0 | 0.5 | 0.53 | 0.42 | 0.43 | 0.55 | 0.44 |
| 11 | 95310015 | rs880636 | -0.94 | -0.767 | -0.823 | -0.711 | -0.005 | -0.002 | -0.002 | 0 | 0.64 | 0.61 | 0.81 | 0.63 | 0.71 | 0.61 |
| 11 | 95310977 | rs1255533 | 0.777 | -0.797 | -0.856 | -0.75 | -0.001 | -0.001 | -0.001 | 0 | 0.1 | 0.12 | 0.05 | 0.1 | 0.08 | 0.15 |
| 11 | 95311260 | rs12575636 | 1.33 | -0.815 | -0.939 | -0.797 | -0.02 | 0.057 | 0.057 | 0 | 0.19 | 0.01 | 0.15 | 0.1 | 0.21 | 0.25 |
| 11 | 95311422 | rs4409785 | 1.784 | -0.766 | -0.923 | -0.801 | -0.022 | 0.043 | 0.043 | 0 | 0.17 | 0.01 | 0.12 | 0.1 | 0.19 | 0.25 |
| 11 | 95314291 | rs4384353 | -0.337 | -0.961 | -0.151 | 0.031 | 0.002 | 0.004 | 0.004 | 0.28 | 0.66 | 0.68 | 0.83 | 0.7 | 0.73 | 0.62 |
| 11 | 95314554 | rs11021226 | 1.096 | -0.879 | -0.199 | -0.14 | 0.005 | -0.048 | -0.048 | 1.197 | 0.39 | 0.43 | 0.35 | 0.12 | 0.49 | 0.36 |
| 11 | 95316718 | rs11021227 | 1.249 | -0.846 | -0.203 | -0.123 | -0.066 | 0.127 | 0.127 | 0.607 | 0.37 | 0.17 | 0.33 | 0.12 | 0.5 | 0.36 |
| 11 | 95319115 | rs1943509 | -0.507 | -0.932 | -0.888 | 0.072 | -0.001 | 0.003 | 0.003 | 0 | 0.25 | 0.29 | 0.45 | 0.36 | 0.24 | 0.32 |
| 11 | 95320190 | rs11021231 | 1.295 | -0.775 | -0.97 | 0.101 | 0.004 | -0.038 | -0.038 | 1.051 | 0.38 | 0.44 | 0.33 | 0.13 | 0.49 | 0.35 |
| 11 | 95322900 | rs948867 | -0.025 | -0.774 | -0.783 | 0.752 | -0.003 | 0.026 | 0.026 | 0 | 0.1 | 0.19 | 0.07 | 0.18 | 0.08 | 0.15 |
| 11 | 95327130 | rs2254634 | 0.804 | -0.719 | -1.172 | 0.573 | 0.007 | -0.006 | -0.006 | 0 | 0.62 | 0.71 | 0.78 | 0.5 | 0.73 | 0.67 |
| 11 | 95327374 | rs2186739 | 0.326 | 0.348 | -1.08 | -0.305 | 0.002 | 0.006 | 0.006 | 0.127 | 0.76 | 0.81 | 0.88 | 0.81 | 0.81 | 0.78 |
| 11 | 95336816 | rs12281485 | -0.807 | 0.684 | -1.205 | -0.395 | -0.001 | 0.038 | 0.038 | 0 | 0.19 | 0.13 | 0.14 | 0.02 | 0.2 | 0.05 |
| 11 | 95340984 | rs11021240 | -1.188 | 1.014 | -0.768 | -0.709 | -0.001 | -0.003 | -0.003 | 0 | 0.39 | 0.61 | 0.31 | 0.23 | 0.41 | 0.29 |
| 11 | 95341707 | rs1939874 | 0.366 | 0.472 | -1.756 | -1.755 | 0.042 | 0.134 | 0.134 | 1.769 | 0.36 | 0.13 | 0.18 | 0.1 | 0.23 | 0.33 |
| 11 | 95352674 | rs11021252 | 0.221 | 0.25 | -1.206 | -1.377 | 0.017 | 0.039 | 0.039 | 3.279 | 0.37 | 0.24 | 0.22 | 0.14 | 0.27 | 0.34 |
| 11 | 95354322 | rs12277780 | 0.791 | 0.645 | -1.06 | -1.347 | -0.004 | -0.008 | -0.008 | 0 | 0.08 | 0.04 | 0.08 | 0.16 | 0.11 | 0.13 |
| 11 | 95358797 | rs7938474 | -0.104 | 0.603 | -0.355 | -1.357 | 0.006 | 0.02 | 0.02 | 0.649 | 0.45 | 0.36 | 0.31 | 0.31 | 0.39 | 0.48 |
| 11 | 95374537 | rs1150345 | -0.425 | 0.774 | -0.252 | -0.401 | 0 | 0 | 0 | 0 | 0.74 | 0.75 | 0.62 | 0.65 | 0.7 | 0.8 |
| 11 | 95376448 | rs1255522 | 0.128 | 0.26 | -0.489 | -0.992 | -0.084 | -0.036 | -0.036 | 5.341 | 0.37 | 0.05 | 0.32 | 0.34 | 0.41 | 0.36 |
| 11 | 95378495 | rs1405732 | 1.431 | 0.117 | 0.011 | -1.138 | 0.003 | -0.014 | -0.014 | 1.193 | 0.22 | 0.21 | 0.15 | 0.05 | 0.14 | 0.23 |
| 11 | 95383570 | rs1150347 | -1.631 | 0.198 | 0.232 | -1.054 | 0 | -0.016 | -0.016 | 0 | 0.85 | 0.88 | 0.9 | 1 | 0.84 | 0.97 |
| 11 | 95395607 | rs10765758 | 0.093 | -0.048 | 0.167 | -1.065 | 0.012 | -0.002 | -0.002 | 0 | 0.52 | 0.3 | 0.42 | 0.47 | 0.54 | 0.68 |
| 11 | 95398352 | rs10437579 | -0.106 | -0.094 | 0.319 | -1.038 | 0 | -0.001 | -0.001 | 0 | 0.56 | 0.56 | 0.46 | 0.47 | 0.56 | 0.68 |
| 11 | 95407543 | rs7939277 | -0.045 | -0.061 | 0.157 | -0.922 | -0.001 | -0.001 | -0.001 | 0 | 0.53 | 0.51 | 0.44 | 0.45 | 0.52 | 0.68 |
| 11 | 95411600 | rs7931238 | 0.136 | 0.35 | -0.304 | -0.887 | 0.006 | 0.046 | 0.046 | 0 | 0.45 | 0.29 | 0.39 | 0.13 | 0.43 | 0.55 |
| 11 | 95414613 | rs998939 | 0.803 | 0.281 | -0.406 | -0.826 | 0.001 | -0.029 | -0.029 | 0.075 | 0.24 | 0.22 | 0.19 | 0.02 | 0.2 | 0.25 |
| 11 | 95416248 | rs1122436 | -0.265 | -0.151 | -1.06 | 0.134 | 0.015 | -0.005 | -0.005 | 0 | 0.2 | 0.15 | 0.35 | 0.2 | 0.16 | 0.13 |
| 11 | 95417843 | rs10765761 | -0.287 | 0.046 | -1.532 | 0.375 | -0.034 | -0.01 | -0.01 | 0 | 0.42 | 0.12 | 0.52 | 0.62 | 0.47 | 0.51 |
| 11 | 95419046 | rs16922394 | 1.115 | 0.548 | -1.59 | 0.78 | -0.002 | -0.009 | -0.009 | 0 | 0.08 | 0.06 | 0.07 | 0.14 | 0.09 | 0.13 |

| | | | | | | | | | | | | | | | | |
|---|---|---|---|---|---|---|---|---|---|---|---|---|---|---|---|---|
| 11 | 95422867 | rs1939875 | -1.346 | 0.579 | -1.264 | 0.432 | -0.003 | 0 | 0 | 0 | 0.75 | 0.78 | 0.75 | 0.77 | 0.72 | 0.68 |
| 11 | 95424403 | rs1939888 | 1.044 | 0.764 | -1.907 | 0.78 | -0.026 | -0.094 | -0.094 | 0 | 0.17 | 0.01 | 0.18 | 0.37 | 0.18 | 0.09 |
| 11 | 95427873 | rs1527448 | -0.382 | 0.613 | -1.579 | 0.941 | -0.009 | 0.002 | 0.002 | 0 | 0.37 | 0.24 | 0.42 | 0.31 | 0.36 | 0.49 |
| 11 | 95428101 | rs2049620 | 0.849 | 0.63 | -1.106 | 1.275 | -0.002 | -0.015 | -0.015 | 0 | 0.17 | 0.1 | 0.12 | 0.25 | 0.2 | 0.24 |
| 11 | 95428105 | rs16922416 | 0.337 | 0.494 | -0.964 | 1.006 | -0.003 | 0.014 | 0.014 | 0 | 0.1 | 0.17 | 0.1 | 0.15 | 0.09 | 0.14 |
| 11 | 95430295 | rs10741517 | -0.641 | 0.573 | -0.826 | 0.659 | 0.009 | -0.047 | -0.047 | 0 | 0.9 | 0.85 | 0.86 | 0.98 | 0.91 | 0.92 |
| 11 | 95430628 | rs11021286 | 0.316 | 0.22 | -1.547 | 1.293 | -0.004 | -0.119 | -0.119 | 0.038 | 0.17 | 0.01 | 0.18 | 0.43 | 0.17 | 0.1 |
| 11 | 95431216 | rs11021287 | -0.887 | 0.384 | -1.852 | 1.325 | -0.017 | -0.069 | -0.069 | 0 | 0.3 | 0.09 | 0.36 | 0.64 | 0.31 | 0.23 |
| 11 | 95437738 | rs948075 | -0.279 | -0.043 | -2.613 | 0.61 | 0.016 | 0.144 | 0.144 | 0 | 0.57 | 0.83 | 0.56 | 0.78 | 0.64 | 0.59 |
| 11 | 95441807 | rs7102303 | 0.6 | -0.098 | -2.152 | 1.301 | -0.012 | -0.013 | -0.013 | 0.277 | 0.42 | 0.62 | 0.45 | 0.49 | 0.51 | 0.41 |
| 11 | 95442235 | rs11021292 | 0.653 | -0.075 | -2.158 | 1.255 | 0.026 | -0.048 | -0.048 | 0 | 0.11 | 0.06 | 0.09 | 0.28 | 0.07 | 0.14 |
| 11 | 95445705 | rs11021294 | 0.485 | -0.089 | -2.104 | 1.274 | -0.036 | -0.036 | -0.036 | 0 | 0.42 | 0.71 | 0.46 | 0.49 | 0.51 | 0.41 |
| 11 | 95448290 | rs11021295 | 1.53 | -0.473 | -2.259 | 1.31 | 0.008 | -0.009 | -0.009 | 0.625 | 0.11 | 0.05 | 0.03 | 0.16 | 0.08 | 0.19 |
| 11 | 95448927 | rs1527449 | 0.87 | -0.442 | -2.249 | 1.295 | 0.027 | -0.049 | -0.049 | 0.941 | 0.11 | 0.06 | 0.09 | 0.3 | 0.07 | 0.17 |
| 11 | 95452492 | rs4298858 | -1.162 | -0.384 | -2.253 | 1.285 | -0.018 | 0.142 | 0.142 | 0 | 0.54 | 0.79 | 0.54 | 0.76 | 0.57 | 0.55 |
| 11 | 95453761 | rs2895606 | 0.247 | -0.387 | -2.234 | 1.29 | -0.013 | -0.013 | -0.013 | 2.536 | 0.42 | 0.63 | 0.45 | 0.48 | 0.51 | 0.41 |
| 11 | 95457226 | rs2156575 | -0.836 | -0.442 | -2.049 | 1.477 | -0.02 | 0.03 | 0.03 | 0 | 0.88 | 0.84 | 0.91 | 0.75 | 0.94 | 0.87 |
| 11 | 95465418 | rs7939457 | 0.307 | -0.647 | -1.012 | 1.424 | -0.033 | 0.061 | 0.061 | 1.561 | 0.09 | 0.38 | 0.16 | 0.11 | 0.02 | 0.11 |
| 11 | 95466159 | rs10831415 | 2.097 | -0.358 | -1.289 | 1.395 | -0.077 | -0.032 | -0.032 | 0 | 0.2 | 0.06 | 0.26 | 0.21 | 0.41 | 0.11 |
| 11 | 95470346 | rs12287206 | -0.435 | -0.692 | -1.182 | 1.289 | -0.065 | 0.03 | 0.03 | 0.273 | 0.19 | 0.61 | 0.19 | 0.26 | 0.08 | 0.26 |
| 11 | 95486859 | rs1124530 | 1.381 | -1.119 | -1.819 | 0.833 | -0.085 | 0.109 | 0.109 | 0 | 0.41 | 0.08 | 0.43 | 0.25 | 0.41 | 0.42 |
| 11 | 95489184 | rs1727164 | -0.599 | -1.474 | -1.634 | 0.974 | -0.017 | 0.013 | 0.013 | 0 | 0.84 | 0.82 | 0.9 | 0.73 | 0.92 | 0.81 |
| 11 | 95489352 | rs1569057 | -0.442 | -1.384 | -1.508 | 0.893 | -0.045 | 0.009 | 0.009 | 0 | 0.22 | 0.67 | 0.21 | 0.26 | 0.1 | 0.29 |
| 11 | 95490325 | rs1789303 | 0.958 | -0.907 | -1.164 | 0.569 | 0.034 | -0.031 | -0.031 | 0.02 | 0.07 | 0.25 | 0.04 | 0 | 0.07 | 0.04 |
| 11 | 95490754 | rs17207890 | 2.264 | -0.063 | -1.563 | 0.753 | -0.057 | -0.052 | -0.052 | 0 | 0.2 | 0.06 | 0.22 | 0.26 | 0.33 | 0.1 |
| 11 | 95497231 | rs1789289 | -1.055 | -0.372 | -1.481 | 0.625 | -0.001 | 0 | 0 | 0.023 | 0.87 | 0.89 | 0.87 | 0.79 | 0.89 | 0.85 |
| 11 | 95503636 | rs1944080 | 1.836 | -0.424 | -1.536 | 0.282 | -0.068 | -0.024 | -0.024 | 0 | 0.26 | 0.08 | 0.24 | 0.25 | 0.38 | 0.14 |
| 11 | 95512060 | rs1150360 | -1.202 | -1.406 | -2.001 | -0.352 | -0.121 | -0.07 | -0.07 | 0 | 0.61 | 0.93 | 0.49 | 0.34 | 0.51 | 0.47 |
| 11 | 95519243 | rs1255150 | -1.317 | -1.488 | -1.87 | -0.41 | -0.033 | -0.045 | -0.045 | 0.741 | 0.24 | 0.71 | 0.21 | 0.04 | 0.06 | 0.22 |
| 11 | 95521953 | rs3748256 | 2.179 | -1.336 | -1.838 | -0.328 | -0.066 | -0.024 | -0.024 | 0 | 0.26 | 0.08 | 0.24 | 0.25 | 0.37 | 0.14 |
| 11 | 95526588 | rs571413 | 4.151 | -0.905 | -1.913 | -0.595 | -0.046 | 0.066 | 0.066 | 0 | 0.11 | 0.01 | 0.08 | 0 | 0.15 | 0.03 |
| 11 | 95528891 | rs12271427 | 0.077 | -2.452 | -0.633 | -0.643 | -0.009 | -0.021 | -0.021 | 0.361 | 0.08 | 0.22 | 0.15 | 0.04 | 0.03 | 0.12 |
| 11 | 95550841 | rs504341 | -2.136 | -2.662 | -0.806 | -1.054 | -0.031 | -0.039 | -0.039 | 0 | 0.22 | 0.72 | 0.21 | 0.04 | 0.06 | 0.22 |
| 11 | 95564259 | rs644799 | 2.804 | -2.392 | -0.702 | -0.914 | -0.066 | -0.024 | -0.024 | 0 | 0.26 | 0.08 | 0.24 | 0.25 | 0.37 | 0.14 |
| 11 | 95565288 | rs7902 | 0.317 | -2.472 | -0.606 | -0.953 | -0.06 | 0.023 | 0.023 | 0 | 0.47 | 0.84 | 0.46 | 0.34 | 0.46 | 0.3 |
| 11 | 95565789 | rs7749 | 2.832 | -2.434 | -0.58 | -0.936 | -0.077 | -0.031 | -0.031 | 0 | 0.25 | 0.07 | 0.23 | 0.25 | 0.37 | 0.14 |
| 11 | 95565840 | rs12275165 | -0.373 | -2.447 | -0.133 | -0.942 | -0.005 | -0.019 | -0.019 | 0 | 0.1 | 0.39 | 0.17 | 0.04 | 0.03 | 0.12 |
| 11 | 95566520 | rs525404 | 2.761 | -2.303 | -0.189 | -0.892 | -0.08 | -0.032 | -0.032 | 0 | 0.26 | 0.07 | 0.23 | 0.26 | 0.37 | 0.14 |
| 11 | 95567058 | rs16922615 | 0.903 | -2.308 | -0.139 | -0.892 | 0.007 | 0.004 | 0.004 | 0 | 0.07 | 0.13 | 0.04 | 0.04 | 0.05 | 0.04 |
| 11 | 95567905 | rs611020 | 2.76 | -2.271 | -0.231 | -0.876 | -0.08 | -0.033 | -0.033 | 0 | 0.25 | 0.07 | 0.23 | 0.25 | 0.37 | 0.14 |
| 11 | 95580926 | rs566204 | 2.72 | -2.535 | -0.515 | -0.839 | -0.062 | -0.022 | -0.022 | 0 | 0.26 | 0.09 | 0.24 | 0.25 | 0.37 | 0.14 |
| 11 | 95591313 | rs10501831 | 1.008 | -2.529 | -0.094 | -1.045 | 0.013 | 0.009 | 0.009 | 0 | 0.06 | 0.1 | 0.03 | 0.05 | 0.05 | 0.04 |
| 11 | 95591351 | rs7107251 | -0.993 | -2.596 | 0.046 | -1.078 | -0.008 | -0.016 | -0.016 | 0.337 | 0.15 | 0.3 | 0.18 | 0.09 | 0.08 | 0.16 |
| 11 | 95596003 | rs555105 | -3.061 | -2.758 | -0.213 | -1.186 | 0.058 | 0.079 | 0.079 | 0.025 | 0.31 | 0.81 | 0.27 | 0.11 | 0.17 | 0.31 |
| 11 | 95603051 | rs2155436 | -0.882 | -2.656 | 0.16 | -0.993 | -0.015 | -0.015 | -0.015 | 0 | 0.16 | 0.31 | 0.18 | 0.1 | 0.08 | 0.16 |
| 11 | 95603439 | rs553998 | -3.075 | -2.669 | -0.044 | -1.002 | 0.03 | 0.037 | 0.037 | 0.892 | 0.24 | 0.79 | 0.24 | 0.11 | 0.11 | 0.23 |
| 11 | 95618247 | rs10501832 | 1.134 | -2.487 | -0.065 | -0.913 | 0.011 | 0.01 | 0.01 | 0 | 0.05 | 0.1 | 0.04 | 0.05 | 0.05 | 0.04 |
| 11 | 95634278 | rs7128695 | -0.619 | -2.48 | -0.1 | -0.917 | 0 | -0.03 | -0.03 | 0 | 0.08 | 0.28 | 0.03 | 0 | 0.03 | 0.07 |
| 11 | 95648242 | rs7114555 | 0.52 | -2.57 | -0.494 | -0.993 | -0.002 | 0.002 | 0.002 | 0 | 0.44 | 0.42 | 0.29 | 0.26 | 0.46 | 0.3 |
| 11 | 95653318 | rs10765774 | -0.346 | -2.423 | -0.839 | -1.065 | -0.088 | -0.039 | -0.039 | 2.108 | 0.6 | 0.89 | 0.49 | 0.36 | 0.51 | 0.46 |
| 11 | 95656385 | rs10765777 | 2.633 | -2.58 | -0.668 | -1.066 | 0.023 | 0.006 | 0.006 | 0 | 0.28 | 0.3 | 0.27 | 0.27 | 0.39 | 0.16 |
| 11 | 95657111 | rs3824874 | 2.981 | -2.545 | -0.758 | -1.044 | -0.084 | -0.044 | -0.044 | 0 | 0.26 | 0.07 | 0.24 | 0.27 | 0.39 | 0.15 |
| 11 | 95667630 | rs7942175 | -1.765 | -2.228 | -0.313 | -1.447 | -0.046 | -0.072 | -0.072 | 0 | 0.33 | 0.59 | 0.23 | 0.1 | 0.14 | 0.31 |
| 11 | 95680429 | rs10741525 | -0.157 | -2.003 | -0.219 | -1.199 | 0.019 | 0.07 | 0.07 | 0 | 0.87 | 0.77 | 0.84 | 0.69 | 0.86 | 0.79 |
| 11 | 95683320 | rs7109980 | -1.554 | -2.409 | -0.242 | -1.359 | 0.022 | 0.021 | 0.021 | 2.114 | 0.42 | 0.27 | 0.46 | 0.39 | 0.3 | 0.54 |
| 11 | 95684859 | rs11021356 | -0.597 | -2.436 | -0.28 | -1.364 | 0.015 | -0.05 | -0.05 | 0 | 0.2 | 0.08 | 0.2 | 0.33 | 0.12 | 0.28 |
| 11 | 95691045 | rs10501833 | 0.358 | -2.455 | -0.129 | -1.349 | -0.032 | -0.009 | -0.009 | 2.494 | 0.1 | 0.28 | 0.06 | 0.07 | 0.04 | 0.06 |
| 11 | 95692091 | rs7125655 | -0.83 | -2.461 | -0.234 | -1.363 | -0.005 | -0.035 | -0.035 | 0 | 0.31 | 0.08 | 0.39 | 0.38 | 0.24 | 0.35 |
| 11 | 95697220 | rs7938725 | -2.539 | -2.46 | -0.159 | -1.363 | 0.027 | 0.012 | 0.012 | 1.082 | 0.54 | 0.73 | 0.53 | 0.46 | 0.34 | 0.6 |
| 11 | 95698325 | rs12278598 | -1.256 | -1.864 | 0.126 | -1.088 | 0 | -0.002 | -0.002 | 0 | 0.29 | 0.28 | 0.34 | 0.38 | 0.23 | 0.42 |
| 11 | 95704425 | rs16922748 | -0.688 | -1.819 | 0.139 | -1.113 | -0.011 | -0.035 | -0.035 | 0 | 0.08 | 0.19 | 0.06 | 0.01 | 0.04 | 0.14 |
| 11 | 95704624 | rs6483467 | -1.808 | -1.704 | -0.104 | -1.087 | -0.005 | -0.003 | -0.003 | 0 | 0.44 | 0.59 | 0.45 | 0.45 | 0.32 | 0.54 |
| 11 | 95704884 | rs877757 | 1.086 | -1.649 | -0.138 | -1.07 | -0.018 | -0.02 | -0.02 | 1.188 | 0.07 | 0.01 | 0.1 | 0.33 | 0.06 | 0.22 |
| 11 | 95706538 | rs12221838 | -0.686 | -1.65 | -0.027 | -0.999 | -0.001 | -0.002 | -0.002 | 0.472 | 0.26 | 0.23 | 0.33 | 0.37 | 0.24 | 0.36 |
| 11 | 95707042 | rs1940163 | -1.33 | -1.232 | -0.561 | -0.834 | 0.035 | 0.005 | 0.005 | 0 | 0.44 | 0.72 | 0.45 | 0.47 | 0.31 | 0.53 |
| 11 | 95709391 | rs7109607 | -0.316 | -0.574 | -0.4 | -0.688 | -0.018 | -0.044 | -0.044 | 1.097 | 0.21 | 0.3 | 0.34 | 0.11 | 0.14 | 0.16 |
| 11 | 95709990 | rs1053815 | -1.292 | -0.58 | -0.418 | -0.665 | 0 | 0 | 0 | 0.013 | 0.39 | 0.65 | 0.52 | 0.45 | 0.31 | 0.46 |
| 11 | 95711448 | rs7930268 | -0.977 | -1.121 | -0.457 | -0.733 | 0.005 | 0 | 0 | 0 | 0.33 | 0.36 | 0.42 | 0.39 | 0.26 | 0.38 |
| 11 | 95711914 | rs7104987 | -1.785 | -1.074 | -0.602 | -0.723 | 0.007 | 0.005 | 0.005 | 0 | 0.48 | 0.59 | 0.53 | 0.46 | 0.35 | 0.55 |

| | | | | | | | | | | | | | | | | |
|---|---|---|---|---|---|---|---|---|---|---|---|---|---|---|---|---|
| 11 | 95712897 | rs7123133 | 0.94 | -1.154 | -1.526 | -0.634 | -0.009 | 0.051 | 0.051 | 0 | 0.1 | 0.03 | 0.26 | 0.04 | 0.1 | 0.08 |
| 11 | 95713024 | rs7123272 | 0.874 | -1.153 | -1.528 | -0.633 | -0.021 | 0.052 | 0.052 | 0 | 0.1 | 0.01 | 0.26 | 0.04 | 0.1 | 0.09 |
| 11 | 95714113 | rs3905431 | 0.11 | -1.151 | -1.42 | -0.615 | -0.023 | -0.033 | -0.033 | 0.271 | 0.14 | 0.43 | 0.12 | 0.07 | 0.07 | 0.16 |
| 11 | 95714608 | rs3016513 | -0.967 | -1.195 | -1.427 | -0.644 | -0.047 | -0.069 | -0.069 | 0.787 | 0.23 | 0.59 | 0.14 | 0.08 | 0.11 | 0.23 |
| 11 | 95715322 | rs601972 | -1.078 | -0.992 | -1.635 | -0.617 | -0.064 | -0.034 | -0.034 | 0 | 0.88 | 0.99 | 0.86 | 0.88 | 0.81 | 0.85 |
| 11 | 95715725 | rs948164 | -0.765 | -1.865 | -0.84 | -0.541 | 0 | -0.011 | -0.011 | 0.03 | 0.24 | 0.2 | 0.41 | 0.38 | 0.23 | 0.31 |
| 11 | 95716041 | rs948165 | -0.787 | -1.851 | -0.833 | -0.534 | 0.001 | -0.008 | -0.008 | 0 | 0.24 | 0.22 | 0.4 | 0.38 | 0.23 | 0.31 |
| 11 | 95717607 | rs7114828 | 0.1 | -1.827 | -0.882 | -0.563 | -0.031 | -0.015 | -0.015 | 1.088 | 0.13 | 0.43 | 0.1 | 0.07 | 0.04 | 0.07 |
| 11 | 95718000 | rs2509092 | -2.137 | -1.839 | -0.9 | -0.56 | 0.074 | 0.036 | 0.036 | 0 | 0.49 | 0.81 | 0.54 | 0.47 | 0.34 | 0.54 |
| 11 | 95719251 | rs7937967 | -1.306 | -1.392 | -0.224 | 0.188 | -0.006 | 0.005 | 0.005 | 0 | 0.3 | 0.64 | 0.26 | 0.43 | 0.22 | 0.39 |
| 11 | 95720908 | rs2509094 | -0.73 | -1.255 | -0.268 | 0.035 | -0.066 | -0.082 | -0.082 | 0.391 | 0.24 | 0.53 | 0.14 | 0.08 | 0.1 | 0.23 |
| 11 | 95722676 | rs7113517 | -0.745 | -0.669 | -1.126 | 0.354 | 0.029 | 0.001 | 0.001 | 0 | 0.39 | 0.69 | 0.28 | 0.42 | 0.25 | 0.46 |
| 11 | 95725095 | rs16922803 | -1.26 | 0.015 | -1.109 | 0.085 | 0 | 0.008 | 0.008 | 1.05 | 0.11 | 0.06 | 0.07 | 0 | 0.09 | 0.14 |
| 11 | 95727636 | rs17806390 | -0.568 | -0.181 | -1.54 | 0.123 | 0.01 | 0.008 | 0.008 | 0 | 0.3 | 0.17 | 0.23 | 0.27 | 0.24 | 0.42 |
| 11 | 95728043 | rs506335 | -0.08 | -1.774 | -1.404 | 0.319 | -0.008 | 0 | 0 | 0.104 | 0.25 | 0.28 | 0.09 | 0.38 | 0.12 | 0.16 |
| 11 | 95729876 | rs10501835 | 0.006 | -2.208 | -1.422 | 0.224 | 0.017 | -0.016 | -0.016 | 0 | 0.22 | 0.17 | 0.1 | 0.36 | 0.14 | 0.14 |
| 11 | 95730903 | rs17806593 | 0.86 | -0.926 | -1.78 | -0.051 | -0.053 | 0.018 | 0.018 | 0 | 0.18 | 0.05 | 0.22 | 0.12 | 0.29 | 0.14 |
| 11 | 95732414 | rs520441 | -1.168 | -0.649 | -1.279 | -0.358 | 0.013 | -0.015 | -0.015 | 0.488 | 0.9 | 0.83 | 0.79 | 0.96 | 0.81 | 0.9 |
| 11 | 95733239 | rs62028 | 0.002 | -1.206 | -1.714 | -0.61 | 0.049 | -0.015 | -0.015 | 0 | 0.35 | 0.29 | 0.2 | 0.43 | 0.19 | 0.26 |
| 11 | 95742703 | rs4144614 | 0.36 | -1.2 | -1.304 | -0.492 | -0.005 | 0.002 | 0.002 | 0 | 0.42 | 0.41 | 0.35 | 0.44 | 0.38 | 0.56 |
| 11 | 95745273 | rs662766 | -0.519 | -0.997 | -1.36 | -0.276 | -0.016 | 0.005 | 0.005 | 0 | 0.71 | 0.71 | 0.47 | 0.8 | 0.52 | 0.76 |
| 11 | 95748302 | rs1940201 | 0.669 | -1.15 | -1.166 | -0.137 | -0.002 | -0.005 | -0.005 | 3.495 | 0.31 | 0.2 | 0.23 | 0.35 | 0.3 | 0.45 |
| 11 | 95756889 | rs619964 | 0.596 | -1.187 | -1.049 | -0.155 | 0.001 | -0.004 | -0.004 | 2.74 | 0.31 | 0.21 | 0.21 | 0.35 | 0.27 | 0.4 |
| 11 | 95759560 | rs10831466 | 1.185 | -1.106 | -1.267 | -0.221 | -0.08 | 0.015 | 0.015 | 0 | 0.19 | 0.04 | 0.26 | 0.12 | 0.32 | 0.15 |
| 11 | 95760611 | rs677598 | -0.587 | -1.152 | -0.685 | -0.28 | 0.015 | -0.032 | -0.032 | 0 | 0.6 | 0.79 | 0.62 | 0.52 | 0.6 | 0.61 |
| 11 | 95761960 | rs2509102 | 0.6 | -1.08 | -0.818 | -0.03 | 0.004 | -0.009 | -0.009 | 0.291 | 0.32 | 0.24 | 0.2 | 0.38 | 0.26 | 0.38 |
| 11 | 95762146 | rs10734173 | 0.238 | -1.315 | -1.088 | -0.291 | 0.029 | -0.013 | -0.013 | 0 | 0.43 | 0.18 | 0.56 | 0.53 | 0.57 | 0.41 |
| 11 | 95764813 | rs7482121 | 0.883 | -0.698 | -1.349 | -0.678 | -0.11 | 0.098 | 0.098 | 0 | 0.24 | 0.04 | 0.25 | 0.07 | 0.34 | 0.14 |
| 11 | 95766067 | rs489535 | -0.271 | -0.998 | -1.03 | -0.615 | 0.023 | -0.025 | -0.025 | 0 | 0.8 | 0.72 | 0.74 | 0.89 | 0.69 | 0.76 |
| 11 | 95766185 | rs687782 | 0.121 | -1.83 | -0.81 | 0.242 | 0 | 0.001 | 0.001 | 0.102 | 0.57 | 0.49 | 0.43 | 0.55 | 0.5 | 0.52 |
| 11 | 95766936 | rs518716 | 0.337 | -1.981 | -0.857 | 0.608 | 0.001 | -0.007 | -0.007 | 0 | 0.43 | 0.46 | 0.35 | 0.3 | 0.41 | 0.36 |
| 11 | 95767956 | rs7943261 | 1.17 | -2.224 | -0.598 | 0.557 | -0.013 | 0.027 | 0.027 | 0 | 0.13 | 0.19 | 0.08 | 0.23 | 0.06 | 0.22 |
| 11 | 95768445 | rs627313 | 0.107 | -2.115 | -0.482 | 0.662 | -0.01 | 0.012 | 0.012 | 0 | 0.45 | 0.65 | 0.37 | 0.34 | 0.46 | 0.41 |
| 11 | 95770130 | rs1010730 | 1.576 | -2.177 | -0.542 | 0.698 | 0.04 | -0.065 | -0.065 | 0.319 | 0.23 | 0.06 | 0.31 | 0.37 | 0.18 | 0.23 |
| 11 | 95770378 | rs656437 | -1.409 | -2.174 | -0.231 | 0.548 | 0.076 | -0.017 | -0.017 | 0.005 | 0.86 | 0.41 | 0.82 | 0.92 | 0.76 | 0.78 |
| 11 | 95770744 | rs10831467 | 1.733 | -2.189 | -0.878 | 0.515 | 0.033 | -0.062 | -0.062 | 0 | 0.22 | 0.03 | 0.3 | 0.31 | 0.18 | 0.21 |
| 11 | 95771057 | rs669776 | -1.62 | -1.976 | -0.614 | 0.508 | -0.012 | 0.01 | 0.01 | 1.081 | 0.87 | 0.87 | 0.86 | 0.96 | 0.79 | 0.84 |
| 11 | 95773435 | rs10831469 | -0.004 | -2.154 | -0.79 | 0.613 | 0.074 | -0.038 | -0.038 | 0 | 0.44 | 0.19 | 0.48 | 0.57 | 0.34 | 0.41 |
| 11 | 95774829 | rs16922920 | 2.388 | -1.731 | -0.907 | 0.485 | -0.017 | -0.07 | -0.07 | 0.063 | 0.14 | 0 | 0.14 | 0.25 | 0.09 | 0.16 |
| 11 | 95776274 | rs7951485 | 0.314 | -2.233 | -0.999 | 0.364 | 0.074 | 0.025 | 0.025 | 1.838 | 0.4 | 0.28 | 0.38 | 0.7 | 0.2 | 0.48 |
| 11 | 95778278 | rs7934164 | 0.153 | -0.881 | -0.091 | 1.163 | 0.002 | 0.009 | 0.009 | 0.396 | 0.66 | 0.46 | 0.64 | 0.62 | 0.63 | 0.61 |
| 11 | 95784444 | rs1940159 | -0.632 | -0.768 | -1.199 | 0.935 | -0.047 | -0.064 | -0.064 | 0 | 0.38 | 0.11 | 0.46 | 0.49 | 0.4 | 0.45 |
| 11 | 95785245 | rs7941059 | 1.325 | -0.576 | -1.203 | 0.7 | 0.004 | 0.011 | 0.011 | 1.866 | 0.47 | 0.25 | 0.39 | 0.41 | 0.46 | 0.47 |
| 11 | 95785363 | rs17233969 | 1.073 | 0.048 | -0.82 | 0.027 | -0.04 | -0.019 | -0.019 | 0.027 | 0.08 | 0.01 | 0.15 | 0.1 | 0.16 | 0.11 |
| 11 | 95786542 | rs1939475 | 0.617 | -0.559 | -0.509 | -0.179 | -0.005 | -0.046 | -0.046 | 0 | 0.7 | 0.34 | 0.74 | 0.88 | 0.71 | 0.77 |
| 11 | 95789959 | rs1940153 | 0.496 | -1.025 | -0.8 | -0.401 | 0.022 | 0.034 | 0.034 | 0 | 0.52 | 0.32 | 0.46 | 0.15 | 0.58 | 0.34 |
| 11 | 95790559 | rs16922954 | 0.468 | -1.027 | -1.204 | 0.2 | -0.018 | -0.03 | -0.03 | 0.801 | 0.14 | 0.08 | 0.11 | 0.18 | 0.17 | 0.2 |
| 11 | 95793701 | rs6483476 | -1.241 | -1.332 | -1.042 | 0.88 | -0.031 | 0.069 | 0.069 | 0 | 0.89 | 0.81 | 0.84 | 0.67 | 0.94 | 0.81 |
| 11 | 95793951 | rs10501839 | -0.337 | -1.12 | -1.201 | 1.13 | -0.002 | -0.04 | -0.04 | 0 | 0.31 | 0.25 | 0.28 | 0.44 | 0.33 | 0.37 |
| 11 | 95794754 | rs11021409 | 1.116 | -0.978 | -1.434 | 1.452 | -0.029 | -0.102 | -0.102 | 0 | 0.16 | 0.01 | 0.16 | 0.28 | 0.16 | 0.18 |
| 11 | 95795807 | rs12800440 | 0.6 | -0.382 | -1.409 | 0.556 | -0.079 | 0.032 | 0.032 | 0 | 0.14 | 0.01 | 0.21 | 0.09 | 0.21 | 0.13 |
| 11 | 95797049 | rs7949201 | 0.621 | -1.273 | -0.675 | 0.386 | 0.002 | 0.002 | 0.002 | 0 | 0.61 | 0.62 | 0.55 | 0.29 | 0.67 | 0.48 |
| 11 | 95797111 | rs10831477 | 0.748 | -0.859 | -1.02 | 0.667 | -0.001 | -0.029 | -0.029 | 0.338 | 0.17 | 0.06 | 0.17 | 0.21 | 0.17 | 0.18 |
| 11 | 95807538 | rs2298487 | -0.251 | -0.835 | -0.697 | 0.596 | -0.003 | 0.028 | 0.028 | 0 | 0.83 | 0.86 | 0.71 | 0.52 | 0.83 | 0.74 |
| 11 | 95808391 | rs1939485 | 0.626 | -0.507 | -0.937 | -0.362 | 0.01 | 0.079 | 0.079 | 0 | 0.46 | 0.65 | 0.34 | 0.18 | 0.4 | 0.39 |
| 11 | 95811600 | rs12789363 | 1.091 | -0.943 | -0.501 | -0.644 | -0.015 | 0.048 | 0.048 | 0 | 0.38 | 0.21 | 0.3 | 0.07 | 0.39 | 0.27 |
| 11 | 95811916 | rs2067870 | 1.091 | -0.943 | -0.501 | -0.644 | -0.015 | 0.048 | 0.048 | 0 | 0.38 | 0.21 | 0.3 | 0.07 | 0.39 | 0.27 |
| 11 | 95813021 | rs12365545 | 0.211 | -0.738 | -0.673 | -0.421 | -0.112 | 0.003 | 0.003 | 0 | 0.35 | 0.1 | 0.36 | 0.32 | 0.42 | 0.43 |
| 11 | 95814420 | rs4603268 | -0.12 | -0.813 | -0.38 | -0.066 | -0.052 | -0.042 | -0.042 | 0 | 0.2 | 0.09 | 0.26 | 0.28 | 0.25 | 0.32 |
| 11 | 95815578 | rs10765787 | -0.693 | -0.858 | -0.326 | -0.039 | 0.009 | -0.012 | -0.012 | 0.123 | 0.83 | 0.88 | 0.71 | 0.49 | 0.83 | 0.81 |
| 11 | 95816458 | rs7947986 | 0.651 | -0.882 | -0.195 | -0.739 | -0.031 | 0.11 | 0.11 | 0 | 0.62 | 0.83 | 0.47 | 0.19 | 0.57 | 0.48 |
| 11 | 95824593 | rs7104859 | 0.662 | -0.82 | 0.028 | -0.651 | -0.013 | 0.054 | 0.054 | 0 | 0.62 | 0.7 | 0.45 | 0.2 | 0.57 | 0.49 |
| 11 | 95827118 | rs4753717 | 0.586 | -0.196 | 0.233 | -0.144 | -0.006 | 0.041 | 0.041 | 0 | 0.81 | 0.59 | 0.8 | 0.75 | 0.81 | 0.79 |
| 11 | 95830567 | rs4753718 | -0.098 | -0.053 | 0.186 | -0.082 | 0.008 | 0.045 | 0.045 | 0.199 | 0.85 | 0.7 | 0.82 | 0.76 | 0.83 | 0.84 |
| 11 | 95834531 | rs10501841 | 0.371 | -0.463 | -0.706 | -1.569 | 0.009 | -0.01 | -0.01 | 0 | 0.34 | 0.18 | 0.19 | 0.02 | 0.26 | 0.21 |
| 11 | 95834689 | rs12799722 | 0.653 | -0.103 | -1.011 | -1.369 | -0.012 | 0.087 | 0.087 | 0 | 0.38 | 0.15 | 0.28 | 0.22 | 0.37 | 0.31 |
| 11 | 95841812 | rs2155227 | -0.019 | -0.52 | -0.716 | -1.423 | -0.001 | 0.065 | 0.065 | 0 | 0.28 | 0.11 | 0.15 | 0.05 | 0.25 | 0.16 |
| 11 | 95843487 | rs4753722 | 0.395 | -0.403 | -0.613 | -1.397 | -0.001 | -0.057 | -0.057 | 0 | 0.61 | 0.68 | 0.79 | 0.95 | 0.64 | 0.74 |
| 11 | 95845917 | rs1939470 | -0.165 | 0.316 | -0.493 | -0.207 | -0.002 | -0.035 | -0.035 | 0 | 0.92 | 0.88 | 0.95 | 1 | 0.91 | 0.94 |

| | | | | | | | | | | | | | | | |
|---|---|---|---|---|---|---|---|---|---|---|---|---|---|---|---|
| 11 | 95858974 | rs11021446 | -0.297 | 0.304 | -0.765 | -0.028 | 0.006 | 0.006 | 0.006 | 0 | 0.19 | 0.11 | 0.08 | 0.15 | 0.13 | 0.08 |
| 11 | 95861651 | rs7945959 | -0.815 | 0.511 | -0.46 | -0.256 | -0.002 | -0.008 | -0.008 | 0 | 0.38 | 0.36 | 0.29 | 0.23 | 0.34 | 0.36 |
| 11 | 95862516 | rs11821882 | -0.998 | 0.119 | 0.111 | -0.424 | 0.015 | 0.038 | 0.038 | 0 | 0.42 | 0.66 | 0.34 | 0.23 | 0.34 | 0.39 |
| 11 | 95862546 | rs10437580 | 0.929 | 1.326 | -0.115 | -0.7 | -0.002 | 0.079 | 0.079 | 0 | 0.53 | 0.31 | 0.59 | 0.76 | 0.51 | 0.58 |
| 11 | 95867366 | rs4572097 | -0.94 | 0.898 | -0.612 | -1.197 | -0.001 | -0.118 | -0.118 | 0 | 0.59 | 0.47 | 0.68 | 0.92 | 0.6 | 0.72 |
| 11 | 95870218 | rs10501844 | -0.107 | 0.937 | -0.236 | -1.418 | -0.008 | -0.035 | -0.035 | 0 | 0.33 | 0.4 | 0.19 | 0.14 | 0.25 | 0.24 |
| 11 | 95871694 | rs17236265 | 0.869 | 1.087 | -0.402 | -1.109 | -0.001 | -0.001 | -0.001 | 0 | 0.07 | 0.07 | 0.13 | 0.09 | 0.08 | 0.15 |
| 11 | 95871907 | rs514686 | -0.111 | 1.058 | -0.384 | -0.991 | -0.001 | 0 | 0 | 0 | 0.3 | 0.28 | 0.53 | 0.79 | 0.35 | 0.51 |
| 11 | 95880984 | rs1939472 | 0.67 | 1.302 | -0.536 | 0.076 | 0 | 0.029 | 0.029 | 0 | 0.07 | 0.09 | 0.19 | 0.18 | 0.09 | 0.15 |
| 11 | 95881808 | rs10765795 | -1.001 | 1.34 | -0.659 | -0.123 | -0.009 | 0.003 | 0.003 | 0 | 0.84 | 0.94 | 0.93 | 0.89 | 0.86 | 0.93 |
| 11 | 95888401 | rs4531427 | 0.313 | 1.282 | -0.314 | -0.381 | 0 | -0.008 | -0.008 | 0.18 | 0.48 | 0.53 | 0.51 | 0.23 | 0.5 | 0.43 |
| 11 | 95892496 | rs1262181 | -0.16 | 1.27 | 0.058 | -0.471 | 0.001 | 0.018 | 0.018 | 0.026 | 0.41 | 0.33 | 0.42 | 0.74 | 0.39 | 0.45 |
| 11 | 95895105 | rs4753738 | 0.781 | 1.239 | 0.514 | -0.669 | 0 | -0.001 | -0.001 | 0 | 0.43 | 0.51 | 0.42 | 0.22 | 0.39 | 0.41 |
| 11 | 95895139 | rs511797 | 1.111 | 1.632 | 0.007 | 0.457 | -0.035 | -0.039 | -0.039 | 0 | 0.07 | 0.01 | 0.16 | 0.24 | 0.11 | 0.12 |
| 11 | 95904791 | rs17237067 | 1.94 | 1.661 | -0.126 | 0.197 | -0.029 | 0.038 | 0.038 | 0 | 0.07 | 0 | 0.04 | 0 | 0.08 | 0.03 |
| 11 | 95910272 | rs4310569 | 0.073 | 1.684 | 0.476 | 0.361 | 0.048 | 0.048 | 0.048 | 0 | 0.81 | 0.51 | 0.72 | 0.74 | 0.79 | 0.86 |
| 11 | 95910507 | rs7110244 | 1.232 | 1.539 | 0.573 | 0.015 | 0.009 | 0.053 | 0.053 | 0.005 | 0.42 | 0.22 | 0.28 | 0.03 | 0.39 | 0.37 |
| 11 | 95910538 | rs7111187 | 0.116 | 1.467 | 0.495 | 0.229 | 0.013 | -0.036 | -0.036 | 9.42 | 0.33 | 0.21 | 0.25 | 0.5 | 0.28 | 0.33 |
| 11 | 95915244 | rs474232 | -0.768 | 1.143 | 0.246 | 0.111 | 0.006 | 0.055 | 0.055 | 0.395 | 0.51 | 0.7 | 0.53 | 0.76 | 0.51 | 0.47 |
| 11 | 95918953 | rs495494 | -0.898 | 0.746 | -0.178 | 0.292 | 0.001 | -0.005 | -0.005 | 0 | 0.73 | 0.78 | 0.68 | 0.69 | 0.74 | 0.78 |
| 11 | 95922349 | rs12272855 | 1.118 | 0.781 | -0.14 | 0.063 | 0.007 | 0.038 | 0.038 | 0 | 0.24 | 0.12 | 0.15 | 0.09 | 0.22 | 0.15 |
| 11 | 95923315 | rs12294247 | 1.124 | 0.773 | -0.143 | 0.059 | 0.007 | 0.038 | 0.038 | 0 | 0.24 | 0.12 | 0.15 | 0.09 | 0.22 | 0.15 |
| 11 | 95923790 | rs12282301 | 1.048 | 0.763 | -0.157 | 0.041 | 0.007 | 0.041 | 0.041 | 0 | 0.24 | 0.12 | 0.15 | 0.09 | 0.22 | 0.15 |
| 11 | 95924267 | rs7130357 | 1.22 | 0.51 | -0.094 | 0.225 | -0.011 | 0.002 | 0.002 | 0 | 0.19 | 0.23 | 0.17 | 0.2 | 0.15 | 0.11 |
| 11 | 95925589 | rs10831494 | 0.552 | 0.708 | -0.082 | 0.152 | 0.001 | -0.003 | -0.003 | 0 | 0.25 | 0.3 | 0.25 | 0.22 | 0.26 | 0.16 |
| 11 | 95926758 | rs4753743 | 0.878 | 0.633 | -0.242 | 0.061 | -0.008 | 0.045 | 0.045 | 0.004 | 0.3 | 0.18 | 0.23 | 0.11 | 0.33 | 0.18 |
| 11 | 95926792 | rs569977 | -0.696 | 0.631 | -0.071 | 0.69 | -0.002 | 0.05 | 0.05 | 0 | 0.9 | 0.88 | 0.84 | 0.69 | 0.9 | 0.73 |
| 11 | 95927943 | rs573424 | -0.109 | 0.993 | 0.074 | 0.757 | 0.022 | 0.096 | 0.096 | 0 | 0.85 | 0.76 | 0.64 | 0.42 | 0.79 | 0.58 |
| 11 | 95928212 | rs12788404 | 0.282 | 1.051 | -0.156 | 0.153 | -0.009 | 0.015 | 0.015 | 0 | 0.24 | 0.16 | 0.23 | 0.05 | 0.26 | 0.25 |
| 11 | 95929222 | rs564256 | 0.421 | 0.882 | -0.327 | 0.386 | 0.002 | 0.024 | 0.024 | 0 | 0.09 | 0.1 | 0.22 | 0.24 | 0.14 | 0.12 |
| 11 | 95929945 | rs523519 | 0.639 | 0.674 | -0.166 | 0.198 | 0.011 | 0 | 0 | 3.751 | 0.46 | 0.41 | 0.42 | 0.4 | 0.44 | 0.34 |
| 11 | 95930072 | rs505689 | -0.224 | 0.741 | -0.005 | 0.157 | 0.005 | 0.024 | 0.024 | 0 | 0.66 | 0.61 | 0.58 | 0.43 | 0.62 | 0.56 |
| 11 | 95930525 | rs480544 | -0.92 | 0.75 | 0.106 | 0.172 | 0.027 | 0.163 | 0.163 | 0 | 0.91 | 0.76 | 0.89 | 0.69 | 0.92 | 0.73 |
| 11 | 95931485 | rs479825 | 0.653 | 0.685 | -0.087 | 0.052 | 0.015 | 0.03 | 0.03 | 0.002 | 0.46 | 0.34 | 0.45 | 0.34 | 0.45 | 0.35 |
| 11 | 95933123 | rs479244 | -0.261 | 0.839 | -0.149 | 0.104 | -0.001 | 0 | 0 | 0 | 0.69 | 0.74 | 0.73 | 0.66 | 0.68 | 0.53 |
| 11 | 95936837 | rs570792 | 1.064 | 0.966 | -0.131 | 0.322 | 0.012 | 0.082 | 0.082 | 0 | 0.08 | 0.14 | 0.21 | 0.25 | 0.1 | 0.11 |
| 11 | 95937204 | rs479995 | -1.236 | 0.824 | -0.261 | 0.098 | 0.001 | -0.014 | -0.014 | 0 | 0.8 | 0.86 | 0.79 | 0.67 | 0.84 | 0.61 |
| 11 | 95937792 | rs11021469 | 2.558 | 0.585 | -0.621 | -0.298 | 0.029 | 0.093 | 0.093 | 0 | 0.22 | 0.07 | 0.15 | 0.01 | 0.2 | 0.13 |
| 11 | 95938154 | rs4753191 | 0.526 | 0.674 | -0.261 | -0.175 | -0.011 | 0.004 | 0.004 | 0 | 0.28 | 0.3 | 0.24 | 0.32 | 0.22 | 0.2 |
| 11 | 95938418 | rs16923264 | 0.889 | 0.783 | -0.243 | -0.114 | 0.002 | -0.007 | -0.007 | 0 | 0.1 | 0.11 | 0.05 | 0.02 | 0.05 | 0.09 |
| 11 | 95939836 | rs470141 | 0.623 | 0.992 | -0.299 | 0.117 | 0.002 | 0.056 | 0.056 | 0 | 0.39 | 0.27 | 0.33 | 0.24 | 0.4 | 0.27 |
| 11 | 95942495 | rs496852 | 0.012 | 0.76 | -0.387 | 0.064 | -0.004 | 0.001 | 0.001 | 0 | 0.48 | 0.43 | 0.52 | 0.49 | 0.49 | 0.39 |
| 11 | 95947410 | rs10831497 | 1.24 | 1.094 | -1.055 | 0.222 | 0.029 | 0.051 | 0.051 | 0 | 0.28 | 0.1 | 0.23 | 0.15 | 0.25 | 0.19 |
| 11 | 95947503 | rs525068 | -0.788 | 1.37 | -1.18 | 0.223 | -0.003 | 0.023 | 0.023 | 0 | 0.79 | 0.95 | 0.87 | 0.82 | 0.84 | 0.59 |
| 11 | 95951188 | rs483333 | 0.698 | 1.636 | -1.048 | 0.26 | -0.003 | -0.016 | -0.016 | 0 | 0.25 | 0.12 | 0.21 | 0.3 | 0.26 | 0.11 |
| 11 | 95952243 | rs11021474 | 1.035 | 1.46 | -1.103 | 0.276 | 0.032 | 0.028 | 0.028 | 0 | 0.3 | 0.1 | 0.27 | 0.19 | 0.26 | 0.2 |
| 11 | 95955212 | rs528098 | 1.137 | 1.559 | -1.057 | 0.357 | 0.018 | -0.013 | -0.013 | 0 | 0.22 | 0.11 | 0.19 | 0.28 | 0.2 | 0.08 |
| 11 | 95955646 | rs10765799 | -0.534 | 1.605 | -1.137 | 0.328 | 0.014 | -0.033 | -0.033 | 0 | 0.7 | 0.82 | 0.76 | 0.53 | 0.74 | 0.64 |
| 11 | 95956622 | rs4753192 | 0.646 | 1.696 | -1.324 | 0.346 | 0.038 | 0.027 | 0.027 | 0 | 0.31 | 0.11 | 0.3 | 0.21 | 0.27 | 0.22 |
| 11 | 95962042 | rs522495 | -0.328 | 1.714 | -1.857 | -0.138 | -0.041 | -0.047 | -0.047 | 0.05 | 0.62 | 0.87 | 0.66 | 0.61 | 0.61 | 0.49 |
| 11 | 95962582 | rs10765800 | 0.482 | 1.719 | -1.095 | 0.467 | -0.006 | 0.017 | 0.017 | 0 | 0.57 | 0.33 | 0.49 | 0.37 | 0.53 | 0.42 |
| 11 | 95964968 | rs551173 | 0.205 | 2.069 | -1.073 | 0.451 | -0.021 | -0.03 | -0.03 | 0 | 0.92 | 0.98 | 0.95 | 0.87 | 0.92 | 0.68 |
| 11 | 95968660 | rs2186584 | 0.331 | 1.608 | -1.265 | 0.271 | -0.064 | -0.005 | -0.005 | 0 | 0.38 | 0.05 | 0.32 | 0.36 | 0.38 | 0.3 |
| 11 | 95968687 | rs566519 | 0.505 | 0.599 | -0.736 | -0.225 | -0.006 | -0.03 | -0.03 | 0 | 0.15 | 0.13 | 0.06 | 0 | 0.09 | 0.05 |
| 11 | 95970204 | rs693182 | 0.575 | 1.652 | -0.206 | 0.989 | -0.017 | -0.023 | -0.023 | 0.003 | 0.61 | 0.28 | 0.5 | 0.53 | 0.6 | 0.43 |
| 11 | 95972510 | rs527077 | -0.239 | 1.652 | -0.91 | 1.517 | -0.008 | -0.018 | -0.018 | 0.016 | 0.24 | 0.17 | 0.29 | 0.23 | 0.26 | 0.2 |
| 11 | 95975630 | rs561122 | -0.206 | 1.797 | -0.859 | 1.333 | 0.016 | 0.033 | 0.033 | 0.766 | 0.94 | 0.89 | 0.91 | 0.84 | 0.91 | 0.84 |
| 11 | 95975925 | rs544656 | -0.594 | 1.225 | -1.022 | 0.902 | 0.003 | -0.012 | -0.012 | 0 | 0.27 | 0.22 | 0.35 | 0.28 | 0.28 | 0.4 |
| 11 | 95980109 | rs545208 | 0.634 | 1.83 | -1.195 | 0.787 | -0.038 | -0.004 | -0.004 | 0 | 0.84 | 0.94 | 0.77 | 0.86 | 0.73 | 0.68 |
| 11 | 95981029 | rs2164182 | -0.018 | 1.414 | -0.67 | 1.253 | -0.004 | 0.067 | 0.067 | 0 | 0.08 | 0.18 | 0.04 | 0.21 | 0.06 | 0.08 |
| 11 | 95981067 | rs11021494 | 0.284 | 0.406 | -0.617 | 1.059 | -0.008 | -0.002 | -0.002 | 0 | 0.12 | 0.1 | 0.05 | 0.1 | 0.06 | 0.07 |
| 11 | 95984688 | rs4278478 | 0.006 | 0.458 | -0.795 | 0.594 | 0.005 | 0.013 | 0.013 | 0 | 0.48 | 0.36 | 0.39 | 0.56 | 0.45 | 0.67 |
| 11 | 95985302 | rs16923429 | 1.093 | 0.092 | -0.842 | 0.326 | 0 | -0.036 | -0.036 | 0 | 0.1 | 0.2 | 0.04 | 0.03 | 0.07 | 0.07 |
| 11 | 95986160 | rs566607 | -1.623 | -0.027 | -0.723 | 0.293 | -0.074 | 0.072 | 0.072 | 0 | 0.69 | 0.87 | 0.51 | 0.86 | 0.57 | 0.83 |
| 11 | 95988438 | rs504909 | 1.308 | 0.237 | -0.546 | 0.523 | -0.004 | 0.078 | 0.078 | 0 | 0.08 | 0.19 | 0.04 | 0.21 | 0.06 | 0.08 |
| 11 | 95988480 | rs10501849 | 0.73 | 0.334 | -0.7 | 0.552 | -0.031 | -0.032 | -0.032 | 0 | 0.1 | 0.01 | 0.1 | 0.13 | 0.11 | 0.16 |
| 11 | 95988699 | rs17819648 | 0.769 | 0.327 | -0.695 | 0.548 | -0.024 | -0.025 | -0.025 | 0.004 | 0.09 | 0.01 | 0.1 | 0.13 | 0.11 | 0.16 |
| 11 | 95990888 | rs548175 | 1.337 | 0.57 | -0.909 | 0.277 | -0.003 | 0.015 | 0.015 | 0 | 0.13 | 0.2 | 0.24 | 0.21 | 0.11 | 0.14 |
| 11 | 95991138 | rs10831503 | 1.253 | 0.481 | -1.339 | 0.414 | -0.046 | -0.107 | -0.107 | 0 | 0.23 | 0.03 | 0.17 | 0.35 | 0.22 | 0.24 |

| | | | | | | | | | | | | | | | | |
|---|---|---|---|---|---|---|---|---|---|---|---|---|---|---|---|---|
| 11 | 95991923 | rs1978763 | 0.301 | 0.519 | -1.214 | 0.297 | -0.061 | -0.115 | -0.115 | 0 | 0.35 | 0.06 | 0.32 | 0.52 | 0.38 | 0.6 |
| 11 | 95993708 | rs555329 | -0.921 | 0.663 | -0.674 | 0.128 | -0.002 | 0.005 | 0.005 | 0 | 0.6 | 0.59 | 0.62 | 0.77 | 0.56 | 0.81 |
| 11 | 95997840 | rs892685 | -0.486 | 1.109 | -0.558 | 0.421 | -0.012 | 0.007 | 0.007 | 0 | 0.44 | 0.29 | 0.57 | 0.7 | 0.46 | 0.54 |
| 11 | 96000100 | rs7115578 | 0.026 | 1.119 | -0.309 | 0.513 | 0.006 | 0.025 | 0.025 | 0 | 0.33 | 0.42 | 0.48 | 0.57 | 0.34 | 0.31 |
| 11 | 96000218 | rs6483491 | 0.971 | 0.795 | -0.106 | 0.638 | -0.009 | -0.003 | -0.003 | 0 | 0.15 | 0.25 | 0.3 | 0.16 | 0.11 | 0.12 |
| 11 | 96005221 | rs11021511 | 1.162 | 0.848 | -0.609 | 0.665 | 0.004 | -0.003 | -0.003 | 0 | 0.14 | 0.05 | 0.28 | 0.16 | 0.11 | 0.12 |
| 11 | 96009116 | rs557137 | -1.334 | 1.098 | -0.591 | 0.718 | -0.016 | -0.011 | -0.011 | 0.498 | 0.86 | 0.93 | 0.69 | 0.83 | 0.84 | 0.88 |
| 11 | 96011351 | rs4598633 | -0.191 | 0.908 | -0.74 | 0.669 | -0.038 | -0.049 | -0.049 | 0.089 | 0.38 | 0.2 | 0.35 | 0.56 | 0.43 | 0.47 |
| 11 | 96015672 | rs502922 | -0.579 | 0.46 | -0.207 | 0.175 | -0.002 | -0.034 | -0.034 | 1.091 | 0.71 | 0.65 | 0.68 | 0.83 | 0.74 | 0.81 |
| 11 | 96016891 | rs12575274 | 0.137 | -0.005 | -0.071 | 0.765 | -0.016 | 0.023 | 0.023 | 0 | 0.16 | 0.24 | 0.18 | 0.27 | 0.1 | 0.17 |
| 11 | 96021386 | rs499866 | -0.489 | 0.03 | -0.112 | 0.814 | 0.008 | -0.009 | -0.009 | 0 | 0.4 | 0.4 | 0.39 | 0.47 | 0.38 | 0.35 |
| 11 | 96023427 | rs483905 | 1.351 | 0.14 | -0.106 | 0.784 | -0.016 | 0.004 | 0.004 | 0 | 0.27 | 0.19 | 0.24 | 0.32 | 0.29 | 0.26 |
| 11 | 96024021 | rs2509000 | -1.307 | 0.119 | 0.014 | 0.739 | -0.002 | -0.036 | -0.036 | 0 | 0.71 | 0.62 | 0.68 | 0.84 | 0.75 | 0.83 |
| 11 | 96024669 | rs573540 | -0.893 | 0.169 | -0.003 | 0.853 | 0.004 | 0.002 | 0.002 | 0 | 0.45 | 0.43 | 0.44 | 0.53 | 0.46 | 0.57 |
| 11 | 96029482 | rs1253227 | 1.085 | 0.261 | -0.624 | 0.92 | -0.008 | -0.023 | -0.023 | 0 | 0.22 | 0.13 | 0.2 | 0.21 | 0.27 | 0.18 |
| 11 | 96036188 | rs6483495 | -0.9 | 0.063 | 0.158 | 0.842 | -0.039 | 0.058 | 0.058 | 0 | 0.2 | 0.41 | 0.21 | 0.3 | 0.15 | 0.35 |
| 11 | 96036323 | rs6483496 | -0.767 | 0.106 | 0.158 | 0.852 | -0.019 | 0.068 | 0.068 | 0 | 0.19 | 0.39 | 0.21 | 0.3 | 0.15 | 0.21 |
| 11 | 96036844 | rs10501851 | -0.541 | 0.079 | -0.139 | 0.849 | -0.045 | 0.094 | 0.094 | 0 | 0.18 | 0.48 | 0.21 | 0.27 | 0.12 | 0.16 |
| 11 | 96041339 | rs516320 | -0.3 | 0.6 | -0.843 | 0.269 | -0.008 | 0.028 | 0.028 | 0 | 0.7 | 0.83 | 0.65 | 0.82 | 0.72 | 0.64 |
| 11 | 96043367 | rs488864 | -0.636 | 0.67 | -0.875 | 0.232 | -0.015 | 0.029 | 0.029 | 0 | 0.73 | 0.88 | 0.68 | 0.84 | 0.75 | 0.82 |
| 11 | 96052081 | rs11606555 | -0.811 | 0.352 | -0.306 | 0.157 | -0.003 | -0.042 | -0.042 | 0 | 0.1 | 0.2 | 0.07 | 0.03 | 0.09 | 0.08 |
| 11 | 96056711 | rs542284 | 0.475 | 0.189 | -0.454 | 0.001 | 0.013 | -0.026 | -0.026 | 3.371 | 0.36 | 0.49 | 0.32 | 0.26 | 0.36 | 0.28 |
| 11 | 96061468 | rs4753196 | -0.327 | 0.243 | -0.67 | 0.365 | -0.022 | 0.023 | 0.023 | 0 | 0.53 | 0.81 | 0.53 | 0.58 | 0.48 | 0.47 |
| 11 | 96092376 | rs10765801 | -0.616 | 0.057 | -0.411 | 0.113 | -0.009 | 0.012 | 0.012 | 0 | 0.66 | 0.58 | 0.62 | 0.57 | 0.7 | 0.61 |
| 11 | 96095083 | rs4753748 | -0.625 | 0.003 | -0.411 | 0.114 | -0.01 | 0.012 | 0.012 | 0 | 0.66 | 0.59 | 0.62 | 0.58 | 0.7 | 0.61 |
| 11 | 96097932 | rs11021552 | 0.714 | 0.058 | -0.503 | 0.016 | -0.001 | 0.044 | 0.044 | 0 | 0.15 | 0.06 | 0.1 | 0.01 | 0.17 | 0.11 |
| 11 | 96108428 | rs7118648 | 0.218 | 0.002 | -0.341 | -0.06 | -0.001 | -0.015 | -0.015 | 0.331 | 0.08 | 0.1 | 0.06 | 0.01 | 0.08 | 0.08 |
| 11 | 96114165 | rs12796830 | 0.251 | 0.06 | -0.305 | 0.199 | -0.005 | -0.035 | -0.035 | 0 | 0.18 | 0.11 | 0.21 | 0.31 | 0.22 | 0.17 |
| 11 | 96115744 | rs11021563 | 0.841 | 0.032 | -0.334 | 0.105 | 0 | 0.045 | 0.045 | 0 | 0.15 | 0.06 | 0.1 | 0.01 | 0.16 | 0.11 |
| 11 | 96119605 | rs12273522 | 0.089 | -0.189 | 0.145 | 0.112 | -0.001 | 0.015 | 0.015 | 0 | 0.19 | 0.2 | 0.23 | 0.31 | 0.22 | 0.18 |
| 11 | 96126361 | rs4753751 | 0.916 | -0.191 | -0.044 | 0.059 | -0.021 | 0.04 | 0.04 | 0 | 0.38 | 0.28 | 0.3 | 0.26 | 0.4 | 0.32 |
| 11 | 96127385 | rs10765805 | -0.167 | -0.086 | 0.296 | 0.059 | 0 | 0 | 0 | 2.577 | 0.59 | 0.6 | 0.54 | 0.56 | 0.62 | 0.5 |
| 11 | 96133569 | rs10765806 | -0.956 | -0.108 | 0.081 | 0.148 | 0.003 | -0.036 | -0.036 | 0 | 0.68 | 0.8 | 0.61 | 0.57 | 0.7 | 0.58 |
| 11 | 96137072 | rs11021580 | -0.975 | -0.117 | 0.083 | 0.137 | 0.006 | -0.031 | -0.031 | 0 | 0.69 | 0.81 | 0.64 | 0.58 | 0.71 | 0.61 |
| 11 | 96155679 | rs489137 | 1.1 | -0.224 | 0.017 | -0.793 | -0.014 | -0.037 | -0.037 | 0 | 0.07 | 0.13 | 0.03 | 0 | 0.03 | 0.02 |
| 11 | 96162383 | rs2043964 | 0.561 | -0.343 | -0.248 | -1.263 | 0.001 | -0.014 | -0.014 | 0 | 0.44 | 0.39 | 0.28 | 0.07 | 0.39 | 0.34 |
| 11 | 96167552 | rs1451331 | 0.262 | -0.266 | -0.46 | -1.149 | -0.005 | -0.03 | -0.03 | 0 | 0.08 | 0.12 | 0.05 | 0.01 | 0.07 | 0.07 |
| 11 | 96175391 | rs12286856 | 0.112 | -0.173 | -0.578 | -0.927 | -0.02 | -0.021 | -0.021 | 0.123 | 0.19 | 0.14 | 0.22 | 0.3 | 0.32 | 0.2 |
| 11 | 96179183 | rs577626 | -1.265 | -0.079 | -0.873 | -0.722 | -0.01 | -0.077 | -0.077 | 0 | 0.75 | 0.95 | 0.6 | 0.38 | 0.75 | 0.6 |
| 11 | 96186570 | rs627030 | -2.202 | 0.007 | -0.951 | -0.585 | -0.034 | -0.135 | -0.135 | 0 | 0.83 | 0.99 | 0.63 | 0.44 | 0.86 | 0.87 |
| 11 | 96190161 | rs1597631 | 0.637 | -0.055 | -0.658 | -1.034 | -0.002 | -0.091 | -0.091 | 0.547 | 0.39 | 0.49 | 0.28 | 0.08 | 0.35 | 0.3 |
| 11 | 96192004 | rs11021595 | 1.158 | 0.248 | -0.949 | -0.923 | 0 | -0.02 | -0.02 | 0 | 0.13 | 0.09 | 0.07 | 0.01 | 0.11 | 0.09 |
| 11 | 96212445 | rs10831529 | 0.919 | 0.074 | -1.154 | -0.726 | 0.001 | -0.05 | -0.05 | 0 | 0.32 | 0.37 | 0.23 | 0.05 | 0.3 | 0.23 |
| 11 | 96219609 | rs7933856 | -0.015 | 0.17 | -1.524 | 0.445 | 0.015 | 0.012 | 0.012 | 0 | 0.6 | 0.79 | 0.52 | 0.33 | 0.62 | 0.45 |
| 11 | 96226824 | rs496825 | -0.597 | 0.553 | -1.475 | 1.162 | -0.068 | -0.158 | -0.158 | 0.978 | 0.8 | 0.99 | 0.61 | 0.5 | 0.8 | 0.81 |
| 11 | 96227606 | rs602397 | -1.113 | -0.102 | 0.164 | 1.408 | 0.028 | 0.179 | 0.179 | 0 | 0.14 | 0.36 | 0.26 | 0.36 | 0.23 | 0.19 |
| 11 | 96229007 | rs618929 | -0.448 | -0.577 | 0.502 | 0.709 | -0.001 | -0.018 | -0.018 | 0 | 0.47 | 0.54 | 0.31 | 0.18 | 0.42 | 0.43 |
| 11 | 96232426 | rs619883 | -0.348 | -0.648 | 0.751 | 0.77 | 0.006 | 0.053 | 0.053 | 0 | 0.46 | 0.39 | 0.26 | 0.19 | 0.42 | 0.39 |
| 11 | 96236914 | rs11021625 | -1.409 | -0.798 | 1.265 | 1.751 | 0.008 | 0.138 | 0.138 | 0 | 0.12 | 0.22 | 0.21 | 0.41 | 0.17 | 0.27 |
| 11 | 96241301 | rs614571 | -2.591 | -0.945 | 1.644 | 1.661 | 0.004 | 0.189 | 0.189 | 0.036 | 0.22 | 0.53 | 0.26 | 0.47 | 0.26 | 0.34 |
| 11 | 96243038 | rs563343 | -1.86 | -1.138 | 1.662 | 1.384 | 0.002 | -0.009 | -0.009 | 2.458 | 0.36 | 0.6 | 0.38 | 0.64 | 0.41 | 0.51 |
| 11 | 96243189 | rs10501855 | -0.835 | -1.144 | 1.7 | 1.549 | 0.01 | 0.088 | 0.088 | 0 | 0.1 | 0.13 | 0.14 | 0.27 | 0.16 | 0.27 |
| 11 | 96243337 | rs10501856 | 0.413 | -1.432 | 1.474 | 0.575 | -0.012 | 0.121 | 0.121 | 0 | 0.19 | 0.01 | 0.08 | 0.01 | 0.16 | 0.17 |
| 11 | 96246662 | rs549947 | -1.208 | -1.43 | 1.791 | 0.708 | -0.004 | 0.041 | 0.041 | 0 | 0.26 | 0.54 | 0.23 | 0.29 | 0.25 | 0.28 |
| 11 | 96248728 | rs689327 | -1.148 | -1.43 | 1.793 | 0.706 | -0.002 | 0.009 | 0.009 | 0 | 0.26 | 0.49 | 0.2 | 0.25 | 0.24 | 0.25 |
| 11 | 96249324 | rs12363581 | -0.844 | -1.439 | 1.84 | 0.702 | -0.002 | -0.005 | -0.005 | 0 | 0.23 | 0.23 | 0.18 | 0.25 | 0.24 | 0.25 |
| 11 | 96252985 | rs611881 | -1.208 | -1.435 | 1.815 | 0.704 | -0.004 | 0.006 | 0.006 | 0 | 0.26 | 0.49 | 0.2 | 0.25 | 0.24 | 0.25 |
| 11 | 96267523 | rs3019791 | -1.439 | -1.369 | 1.778 | 0.768 | 0.002 | 0.004 | 0.004 | 0 | 0.45 | 0.6 | 0.32 | 0.3 | 0.4 | 0.44 |
| 11 | 96447763 | rs28818025 | -0.271 | -1.432 | 1.683 | 0.688 | 0.001 | 0.009 | 0.009 | 0 | 0.21 | 0.15 | 0.2 | 0.18 | 0.22 | 0.16 |
| 11 | 96467132 | rs4594032 | -2.638 | -1.242 | 2.33 | 1.281 | 0.019 | 0.081 | 0.081 | 0 | 0.52 | 0.77 | 0.41 | 0.66 | 0.41 | 0.54 |
| 11 | 96467331 | rs11212360 | -1.989 | -1.371 | 2.463 | 1.504 | -0.006 | 0.012 | 0.012 | 0 | 0.33 | 0.27 | 0.25 | 0.44 | 0.24 | 0.4 |
| 11 | 96479060 | rs3018648 | -1.595 | -1.375 | 2.512 | 1.478 | 0.007 | -0.028 | -0.028 | 1.126 | 0.3 | 0.16 | 0.18 | 0.39 | 0.22 | 0.38 |
| 11 | 96502661 | rs10895080 | -1.922 | -1.333 | 2.672 | 1.072 | -0.008 | 0.021 | 0.021 | 0 | 0.32 | 0.3 | 0.2 | 0.53 | 0.22 | 0.37 |
| 11 | 96503340 | rs10791456 | -2.248 | -1.334 | 2.492 | 1.081 | -0.026 | 0.026 | 0.026 | 0 | 0.35 | 0.42 | 0.25 | 0.58 | 0.23 | 0.4 |
| 11 | 96503971 | rs11224737 | -1.899 | -1.267 | 1.997 | 1.134 | 0.008 | -0.08 | -0.08 | 4.131 | 0.18 | 0.05 | 0.15 | 0.52 | 0.15 | 0.24 |
| 11 | 96521854 | rs4754893 | -1.398 | -1.294 | 2.275 | 1.212 | -0.008 | 0.053 | 0.053 | 0 | 0.24 | 0.3 | 0.22 | 0.58 | 0.19 | 0.33 |
| 11 | 96522519 | rs7112837 | -1.044 | -1.281 | 2.099 | 1.209 | 0.002 | -0.011 | -0.011 | 0 | 0.22 | 0.17 | 0.17 | 0.52 | 0.18 | 0.29 |
| 11 | 96526913 | rs9667173 | -1.09 | -0.909 | 2.449 | 1.347 | -0.02 | -0.039 | -0.039 | 0.003 | 0.09 | 0.25 | 0.04 | 0.02 | 0.04 | 0.07 |
| 11 | 96526991 | rs7121563 | -0.767 | -0.916 | 2.366 | 1.385 | 0.002 | 0.063 | 0.063 | 0 | 0.19 | 0.21 | 0.17 | 0.53 | 0.17 | 0.25 |

| | | | | | | | | | | | | | | | | |
|---|---|---|---|---|---|---|---|---|---|---|---|---|---|---|---|---|
| 11 | 96530700 | rs754413 | -0.153 | -0.947 | 2.1 | 1.318 | -0.002 | -0.004 | -0.004 | 0 | 0.86 | 0.76 | 0.88 | 0.92 | 0.88 | 0.87 |
| 11 | 96531210 | rs7933893 | -1.935 | -0.933 | 2.038 | 0.889 | -0.012 | 0.027 | 0.027 | 0 | 0.31 | 0.52 | 0.26 | 0.6 | 0.22 | 0.35 |
| 11 | 96532284 | rs10895601 | -1.067 | -0.434 | 1.302 | 1.175 | -0.001 | -0.001 | -0.001 | 0 | 0.19 | 0.14 | 0.17 | 0.55 | 0.17 | 0.25 |
| 11 | 96535365 | rs3019708 | 0.929 | -0.536 | 1.503 | 0.917 | -0.006 | 0.02 | 0.02 | 0 | 0.66 | 0.49 | 0.68 | 0.42 | 0.7 | 0.65 |
| 11 | 96551777 | rs17129493 | -1.72 | -0.617 | 2.133 | 0.912 | -0.003 | 0.064 | 0.064 | 0 | 0.1 | 0.21 | 0.07 | 0.3 | 0.08 | 0.15 |
| 11 | 96562777 | rs2033836 | -2.248 | -0.691 | 2.305 | 0.883 | -0.021 | 0.119 | 0.119 | 0 | 0.15 | 0.56 | 0.14 | 0.31 | 0.09 | 0.15 |
| 11 | 96571163 | rs7945466 | -2.458 | -0.671 | 1.768 | 0.862 | -0.019 | 0.022 | 0.022 | 0.864 | 0.2 | 0.69 | 0.26 | 0.38 | 0.13 | 0.26 |
| 11 | 96576217 | rs6589024 | -1.499 | -0.605 | 1.85 | 0.896 | -0.007 | 0.086 | 0.086 | 0.785 | 0.09 | 0.47 | 0.15 | 0.08 | 0.04 | 0.12 |
| 11 | 96576680 | rs1866537 | -1.683 | -0.496 | 0.886 | 0.813 | -0.042 | 0.085 | 0.085 | 0.36 | 0.19 | 0.56 | 0.23 | 0.38 | 0.13 | 0.26 |
| 11 | 96577986 | rs6589031 | -1.568 | -0.434 | 0.501 | 0.759 | -0.044 | 0.122 | 0.122 | 0.095 | 0.18 | 0.49 | 0.25 | 0.38 | 0.13 | 0.26 |
| 11 | 96579907 | rs2919040 | -0.15 | 0.197 | -0.841 | -0.426 | -0.034 | 0.133 | 0.133 | 0 | 0.45 | 0.2 | 0.36 | 0.27 | 0.47 | 0.52 |
| 11 | 96581502 | rs3019723 | -0.27 | 0.706 | -1.286 | -0.897 | -0.038 | 0.04 | 0.04 | 0.015 | 0.22 | 0.07 | 0.16 | 0.18 | 0.24 | 0.31 |
| 11 | 96581526 | rs3019724 | -0.845 | 0.353 | -0.806 | -1.468 | 0.012 | 0.046 | 0.046 | 0.29 | 0.55 | 0.36 | 0.42 | 0.27 | 0.58 | 0.69 |
| 11 | 96601979 | rs3019691 | -0.975 | 0.181 | -0.244 | -1.656 | -0.008 | -0.041 | -0.041 | 0 | 0.61 | 0.49 | 0.45 | 0.26 | 0.63 | 0.7 |
| 11 | 96603304 | rs3019711 | -1.373 | 0.02 | -0.244 | -1.825 | -0.028 | 0.036 | 0.036 | 1.935 | 0.68 | 0.91 | 0.53 | 0.31 | 0.66 | 0.76 |
| 11 | 96610573 | rs1439528 | -0.799 | -0.043 | 0.521 | -1.938 | -0.007 | 0.061 | 0.061 | 0.999 | 0.38 | 0.76 | 0.29 | 0.24 | 0.35 | 0.49 |
| 11 | 96630229 | rs1439527 | -1.147 | -0.217 | 0.785 | -2.01 | 0.004 | 0.013 | 0.013 | 0.711 | 0.64 | 0.65 | 0.49 | 0.31 | 0.66 | 0.74 |
| 11 | 96634323 | rs7937026 | -0.799 | -0.25 | 1.056 | -2.022 | 0.064 | 0.11 | 0.11 | 0.331 | 0.07 | 0.44 | 0.08 | 0.06 | 0.03 | 0.05 |
| 11 | 96639641 | rs10790056 | -1.18 | -0.286 | 0.95 | -2.057 | 0.003 | 0.037 | 0.037 | 1.376 | 0.66 | 0.8 | 0.52 | 0.31 | 0.66 | 0.76 |
| 11 | 96641718 | rs11215374 | 1.249 | -0.299 | 0.711 | -2.15 | -0.052 | 0.137 | 0.137 | 0 | 0.26 | 0.05 | 0.23 | 0.07 | 0.31 | 0.25 |
| 11 | 96651273 | rs1377320 | 0.077 | -0.482 | 1.19 | -2.095 | 0.018 | 0.192 | 0.192 | 0.38 | 0.48 | 0.17 | 0.39 | 0.24 | 0.54 | 0.58 |
| 11 | 96651455 | rs1597878 | -1.819 | -0.81 | 0.945 | -1.99 | -0.044 | -0.052 | -0.052 | 0 | 0.75 | 0.93 | 0.56 | 0.4 | 0.69 | 0.77 |
| 11 | 96653321 | rs11215881 | 0.133 | -0.628 | 0.857 | -2.045 | -0.004 | -0.033 | -0.033 | 1.546 | 0.1 | 0.18 | 0.04 | 0.01 | 0.08 | 0.12 |
| 11 | 96655171 | rs10891983 | -0.139 | -0.453 | 0.699 | -2.101 | 0.019 | 0.091 | 0.091 | 0 | 0.52 | 0.35 | 0.39 | 0.24 | 0.56 | 0.57 |
| 11 | 96657119 | rs10892015 | -1.116 | -0.286 | 0.19 | -1.89 | -0.013 | 0.041 | 0.041 | 1.392 | 0.68 | 0.86 | 0.53 | 0.32 | 0.66 | 0.76 |
| 11 | 96674608 | rs12805725 | 0.883 | -0.209 | 0.154 | -2.218 | -0.099 | 0.104 | 0.104 | 0.019 | 0.29 | 0.03 | 0.27 | 0.16 | 0.38 | 0.28 |
| 11 | 96675255 | rs7939964 | -0.722 | -0.275 | 0.615 | -2.309 | -0.021 | -0.067 | -0.067 | 2.297 | 0.33 | 0.43 | 0.22 | 0.15 | 0.25 | 0.41 |
| 11 | 96676002 | rs12280319 | -0.802 | -0.379 | 0.63 | -2.287 | -0.005 | -0.034 | -0.034 | 0 | 0.62 | 0.46 | 0.48 | 0.32 | 0.63 | 0.7 |
| 11 | 96687870 | rs10892450 | -1.292 | -0.387 | 0.883 | -2.294 | -0.001 | 0.007 | 0.007 | 1.462 | 0.68 | 0.74 | 0.52 | 0.32 | 0.65 | 0.73 |
| 11 | 96703683 | rs10892730 | 0.911 | -0.214 | 0.711 | -2.324 | -0.043 | 0.067 | 0.067 | 0.016 | 0.31 | 0.12 | 0.31 | 0.22 | 0.4 | 0.29 |
| 11 | 96707345 | rs17180030 | 0.889 | -0.277 | 0.51 | -2.383 | -0.028 | 0.033 | 0.033 | 0 | 0.14 | 0.01 | 0.07 | 0.08 | 0.1 | 0.15 |
| 11 | 96724320 | rs12365218 | 1.047 | -0.173 | 0.519 | -2.214 | -0.072 | 0.087 | 0.087 | 0.057 | 0.29 | 0.08 | 0.27 | 0.17 | 0.38 | 0.29 |
| 11 | 96725721 | rs7120017 | -1.47 | -0.529 | 1.006 | -2.066 | 0.008 | 0.008 | 0.008 | 0 | 0.38 | 0.68 | 0.25 | 0.15 | 0.27 | 0.45 |
| 11 | 96742050 | rs11219948 | -0.881 | -0.37 | 0.597 | -1.948 | 0.008 | 0.029 | 0.029 | 0 | 0.31 | 0.28 | 0.16 | 0.1 | 0.23 | 0.34 |
| 11 | 96747308 | rs1982437 | -1.115 | -0.234 | 0.447 | -2.028 | -0.005 | -0.022 | -0.022 | 0 | 0.37 | 0.35 | 0.22 | 0.18 | 0.31 | 0.5 |
| 11 | 96747507 | rs17180923 | -0.218 | -0.172 | 0.293 | -2.056 | -0.008 | 0.083 | 0.083 | 1.154 | 0.17 | 0.02 | 0.08 | 0.02 | 0.14 | 0.24 |
| 11 | 96747656 | rs17129665 | -0.453 | -0.152 | 0.615 | -2.05 | -0.006 | -0.028 | -0.028 | 0.853 | 0.18 | 0.2 | 0.13 | 0.07 | 0.15 | 0.25 |
| 11 | 96761808 | rs11603277 | -1.286 | -0.646 | 0.502 | -1.949 | -0.003 | -0.006 | -0.006 | 2.345 | 0.34 | 0.64 | 0.17 | 0.2 | 0.26 | 0.33 |
| 11 | 96769519 | rs10750384 | 0.214 | -0.084 | 0.391 | -2.208 | 0.005 | 0.016 | 0.016 | 1.221 | 0.51 | 0.58 | 0.6 | 0.84 | 0.43 | 0.52 |
| 11 | 96771070 | rs12805857 | 0.974 | -0.084 | 0.233 | -2.204 | -0.005 | 0.061 | 0.061 | 2.22 | 0.11 | 0.01 | 0.06 | 0.01 | 0.1 | 0.2 |
| 11 | 96777666 | rs1349704 | -0.614 | 0.015 | 0.778 | -2.164 | -0.006 | 0.002 | 0.002 | 0.117 | 0.65 | 0.38 | 0.64 | 0.85 | 0.54 | 0.74 |
| 11 | 96779108 | rs612608 | -1.155 | 0.157 | 0.68 | -2.162 | 0.122 | -0.117 | -0.117 | 0 | 0.8 | 0.53 | 0.71 | 0.98 | 0.63 | 0.85 |
| 11 | 96780465 | rs585732 | 1.086 | -0.315 | 0.737 | -1.906 | -0.004 | 0 | 0 | 0 | 0.15 | 0.19 | 0.07 | 0.13 | 0.1 | 0.11 |
| 11 | 96793508 | rs11222049 | 0.229 | -0.655 | 0.74 | -1.744 | -0.042 | -0.09 | -0.09 | 0 | 0.17 | 0.37 | 0.09 | 0.02 | 0.12 | 0.22 |
| 11 | 96797205 | rs12286432 | 0.247 | -0.652 | 0.692 | -1.743 | -0.04 | -0.085 | -0.085 | 0.356 | 0.17 | 0.34 | 0.09 | 0.02 | 0.12 | 0.22 |
| 11 | 96812443 | rs10894523 | 1.224 | -0.462 | 0.265 | -1.046 | -0.029 | -0.051 | -0.051 | 0 | 0.35 | 0.19 | 0.5 | 0.62 | 0.37 | 0.4 |
| 11 | 96812935 | rs479337 | 0.306 | -0.55 | 0.385 | -1.133 | -0.013 | 0.024 | 0.024 | 0 | 0.56 | 0.36 | 0.63 | 0.7 | 0.56 | 0.71 |
| 11 | 96813264 | rs635343 | 0.262 | -0.561 | 0.5 | -1.141 | -0.004 | 0.007 | 0.007 | 0 | 0.56 | 0.43 | 0.63 | 0.71 | 0.56 | 0.71 |
| 11 | 96816276 | rs679803 | 0.702 | -0.601 | 0.466 | -1.126 | 0.003 | 0.001 | 0.001 | 0 | 0.21 | 0.19 | 0.13 | 0.08 | 0.18 | 0.31 |
| 11 | 96822982 | rs552321 | 0.295 | -0.591 | 0.386 | -1.111 | -0.023 | 0.05 | 0.05 | 0 | 0.55 | 0.26 | 0.61 | 0.71 | 0.55 | 0.69 |
| 11 | 96823102 | rs550700 | -0.318 | -0.689 | 0.481 | -1.184 | -0.029 | 0.008 | 0.008 | 0.123 | 0.66 | 0.77 | 0.69 | 0.76 | 0.59 | 0.75 |
| 11 | 96825894 | rs483430 | 0.928 | -0.548 | 0.135 | -1.216 | 0.02 | 0.048 | 0.048 | 0 | 0.2 | 0.06 | 0.12 | 0.08 | 0.18 | 0.29 |
| 11 | 96826469 | rs632374 | 0.371 | -0.545 | 0.131 | -1.199 | -0.001 | 0 | 0 | 0 | 0.56 | 0.46 | 0.63 | 0.71 | 0.56 | 0.7 |
| 11 | 96829897 | rs475420 | -1.419 | -0.175 | -0.426 | -1.072 | 0.009 | 0.011 | 0.011 | 0.182 | 0.83 | 0.85 | 0.94 | 0.93 | 0.92 | 0.88 |
| 11 | 96832183 | rs4754682 | 0.115 | 0.639 | -0.418 | -0.464 | -0.001 | -0.037 | -0.037 | 0 | 0.43 | 0.48 | 0.58 | 0.84 | 0.4 | 0.45 |
| 11 | 96841814 | rs17275283 | 0.287 | 0.811 | -1.455 | -0.441 | -0.038 | -0.013 | -0.013 | 0 | 0.19 | 0.07 | 0.28 | 0.21 | 0.34 | 0.18 |
| 11 | 96843354 | rs10894965 | -0.239 | 0.912 | -1.317 | -0.874 | 0.049 | 0.065 | 0.065 | 0 | 0.56 | 0.29 | 0.77 | 0.84 | 0.71 | 0.58 |
| 11 | 96845364 | rs11224253 | -0.089 | 0.305 | -1.021 | -0.201 | -0.008 | 0.076 | 0.076 | 0.06 | 0.42 | 0.23 | 0.56 | 0.83 | 0.4 | 0.45 |
| 11 | 96850814 | rs2156690 | -0.173 | 0.23 | -0.681 | 0.422 | -0.012 | 0.046 | 0.046 | 0 | 0.32 | 0.52 | 0.45 | 0.6 | 0.29 | 0.29 |
| 11 | 96851286 | rs639275 | 0.355 | -0.349 | -1.749 | -0.207 | -0.006 | 0.032 | 0.032 | 1.054 | 0.2 | 0.01 | 0.12 | 0.1 | 0.17 | 0.35 |
| 11 | 96851531 | rs10894980 | 0.241 | 0.33 | -1.536 | -0.13 | -0.058 | -0.108 | -0.108 | 0.183 | 0.32 | 0.09 | 0.43 | 0.59 | 0.32 | 0.34 |
| 11 | 96852063 | rs17129771 | 0.041 | 0.103 | -1.162 | 0.26 | -0.001 | -0.041 | -0.041 | 0.022 | 0.32 | 0.2 | 0.42 | 0.44 | 0.32 | 0.4 |
| 11 | 96853065 | rs12420935 | -0.636 | -0.487 | -0.277 | 0.102 | 0.011 | 0.113 | 0.113 | 0 | 0.07 | 0.25 | 0.07 | 0.16 | 0.03 | 0.03 |
| 11 | 96857324 | rs7103184 | -0.951 | -0.9 | -0.108 | 0.129 | -0.047 | 0.1 | 0.1 | 0.756 | 0.15 | 0.36 | 0.09 | 0.31 | 0.07 | 0.1 |
| 11 | 96860873 | rs1944790 | -1.284 | -0.949 | -0.289 | 0.116 | -0.054 | 0.08 | 0.08 | 1.623 | 0.17 | 0.61 | 0.11 | 0.31 | 0.08 | 0.11 |
| 11 | 96863442 | rs2096735 | -0.671 | -0.499 | -0.509 | -0.067 | 0.048 | 0.005 | 0.005 | 0 | 0.46 | 0.73 | 0.36 | 0.51 | 0.33 | 0.46 |
| 11 | 96867381 | rs17129783 | 0.187 | -0.366 | 0.111 | -0.543 | -0.018 | -0.139 | -0.139 | 0 | 0.17 | 0.53 | 0.12 | 0 | 0.14 | 0.05 |
| 11 | 96879194 | rs7952525 | -0.471 | -0.181 | 0.701 | -1.012 | -0.008 | 0.142 | 0.142 | 1.47 | 0.53 | 0.78 | 0.5 | 0.31 | 0.52 | 0.29 |
| 11 | 96883279 | rs1943838 | -0.704 | -0.333 | 0.639 | -1.262 | 0.006 | 0.03 | 0.03 | 0.285 | 0.63 | 0.34 | 0.6 | 0.86 | 0.58 | 0.82 |

| | | | | | | | | | | | | | | | | |
|---|---|---|---|---|---|---|---|---|---|---|---|---|---|---|---|---|
| 11 | 96888533 | rs1943833 | -0.264 | -0.391 | 0.701 | -1.266 | 0.014 | 0.15 | 0.15 | 0.885 | 0.55 | 0.24 | 0.54 | 0.79 | 0.55 | 0.75 |
| 11 | 96892730 | rs1943841 | 0.512 | -0.239 | 0.435 | -1.138 | -0.003 | -0.002 | -0.002 | 0 | 0.28 | 0.29 | 0.25 | 0.25 | 0.25 | 0.4 |
| 11 | 96897460 | rs12280802 | 0.499 | -0.234 | 0.468 | -1.149 | -0.017 | -0.013 | -0.013 | 0 | 0.28 | 0.35 | 0.26 | 0.24 | 0.26 | 0.4 |
| 11 | 96898252 | rs10160666 | 0.55 | -0.159 | 0.794 | -1.155 | -0.011 | -0.052 | -0.052 | 0.035 | 0.36 | 0.5 | 0.38 | 0.2 | 0.38 | 0.18 |
| 11 | 96899834 | rs7112003 | -0.545 | -0.293 | 1.083 | -0.581 | -0.011 | 0.067 | 0.067 | 0 | 0.08 | 0.18 | 0.04 | 0.25 | 0.05 | 0.05 |
| 11 | 96900556 | rs10895052 | 0.747 | 0.078 | 0.877 | -0.007 | 0.013 | -0.007 | -0.007 | 0 | 0.52 | 0.4 | 0.59 | 0.4 | 0.61 | 0.38 |
| 11 | 96904208 | rs965010 | 0.317 | -0.159 | 0.701 | -0.424 | 0.033 | 0.035 | 0.035 | 0 | 0.61 | 0.32 | 0.71 | 0.7 | 0.72 | 0.63 |
| 11 | 96906045 | rs10895059 | 0.26 | 0.39 | 0.891 | 0.183 | 0.041 | 0.074 | 0.074 | 0 | 0.14 | 0.23 | 0.22 | 0.36 | 0.18 | 0.28 |
| 11 | 96907558 | rs4754723 | -0.639 | 0.336 | 0.924 | 0.171 | 0.037 | 0.077 | 0.077 | 0 | 0.19 | 0.3 | 0.26 | 0.47 | 0.21 | 0.31 |
| 11 | 96921282 | rs10750600 | -0.434 | 0.024 | 0.671 | -0.047 | 0.033 | 0.098 | 0.098 | 0.764 | 0.7 | 0.83 | 0.78 | 0.92 | 0.74 | 0.6 |
| 11 | 96922959 | rs11224662 | 1.347 | -0.018 | 0.875 | -0.056 | -0.006 | -0.002 | -0.002 | 0.817 | 0.48 | 0.37 | 0.51 | 0.45 | 0.54 | 0.29 |
| 11 | 96929337 | rs17129799 | -0.867 | -0.001 | 0.887 | -0.044 | 0.038 | 0.094 | 0.094 | 0 | 0.21 | 0.35 | 0.26 | 0.47 | 0.2 | 0.31 |
| 11 | 96935094 | rs1908242 | -0.446 | 0.034 | 0.71 | -0.032 | 0.03 | 0.097 | 0.097 | 0.724 | 0.7 | 0.84 | 0.78 | 0.92 | 0.74 | 0.61 |
| 11 | 96938440 | rs7944964 | 0.673 | 0.143 | 0.739 | 0.075 | -0.007 | 0.035 | 0.035 | 0 | 0.06 | 0.12 | 0.04 | 0.14 | 0.06 | 0.08 |
| 11 | 96942393 | rs4754011 | -0.35 | 0.248 | 0.828 | 0.128 | -0.006 | -0.002 | -0.002 | 0 | 0.17 | 0.17 | 0.24 | 0.46 | 0.2 | 0.3 |
| 11 | 96945300 | rs7117076 | -0.382 | 0.249 | 0.832 | 0.128 | 0 | 0.008 | 0.008 | 0 | 0.18 | 0.2 | 0.24 | 0.46 | 0.2 | 0.3 |
| 11 | 96949279 | rs11224814 | 0.167 | 0.188 | 0.96 | 0.318 | -0.005 | 0.061 | 0.061 | 0 | 0.06 | 0.12 | 0.04 | 0.24 | 0.04 | 0.04 |
| 11 | 96960760 | rs2019992 | 0.724 | 0.19 | 0.96 | 0.303 | 0.001 | -0.001 | -0.001 | 0 | 0.5 | 0.41 | 0.52 | 0.44 | 0.54 | 0.31 |
| 11 | 96963902 | rs1789209 | -1.045 | 0.295 | 0.538 | 0.342 | 0.001 | -0.001 | -0.001 | 0 | 0.49 | 0.51 | 0.48 | 0.57 | 0.44 | 0.65 |
| 11 | 96965323 | rs966448 | -0.935 | 0.404 | 0.599 | 0.147 | 0 | 0.002 | 0.002 | 0.399 | 0.47 | 0.53 | 0.48 | 0.51 | 0.43 | 0.64 |
| 11 | 96967445 | rs11224970 | -0.242 | 0.465 | -0.046 | 0.169 | -0.015 | 0.035 | 0.035 | 0 | 0.07 | 0.16 | 0.03 | 0.17 | 0.03 | 0.04 |
| 11 | 96967852 | rs1389577 | -0.428 | 0.73 | -0.492 | -0.319 | -0.082 | 0.02 | 0.02 | 1.278 | 0.38 | 0.1 | 0.42 | 0.32 | 0.39 | 0.59 |
| 11 | 96975093 | rs1557469 | -0.464 | 0.766 | -0.455 | -0.317 | -0.033 | 0.033 | 0.033 | 0.859 | 0.39 | 0.17 | 0.42 | 0.32 | 0.39 | 0.59 |
| 11 | 96983792 | rs12281807 | -0.83 | 0.763 | 0.199 | -0.578 | 0.035 | -0.056 | -0.056 | 0.224 | 0.2 | 0.43 | 0.21 | 0.12 | 0.21 | 0.35 |
| 11 | 96985089 | rs7952054 | -1.005 | 0.801 | 0.38 | -0.507 | 0.05 | 0.029 | 0.029 | 0.667 | 0.11 | 0.39 | 0.15 | 0.1 | 0.07 | 0.2 |
| 11 | 96989839 | rs17129838 | -0.196 | 0.888 | 0.033 | -0.465 | -0.008 | -0.005 | -0.005 | 0 | 0.07 | 0.04 | 0.12 | 0.1 | 0.07 | 0.19 |
| 11 | 96992632 | rs17129840 | -0.232 | 0.888 | 0.077 | -0.466 | -0.003 | -0.002 | -0.002 | 0 | 0.07 | 0.07 | 0.13 | 0.1 | 0.07 | 0.2 |
| 11 | 97009206 | rs1580678 | -0.724 | 0.901 | -0.075 | -1.12 | 0 | -0.02 | -0.02 | 1.104 | 0.63 | 0.37 | 0.74 | 0.84 | 0.62 | 0.83 |
| 11 | 97010108 | rs6590948 | 0.283 | 0.822 | -0.212 | -1.044 | -0.004 | -0.003 | -0.003 | 0.15 | 0.09 | 0.06 | 0.15 | 0.12 | 0.12 | 0.16 |
| 11 | 97012678 | rs1038964 | -1.418 | 1.093 | 0.086 | -1.233 | 0 | -0.008 | -0.008 | 0 | 0.72 | 0.64 | 0.82 | 0.85 | 0.71 | 0.84 |
| 11 | 97014804 | rs1625297 | -1.345 | 1.153 | 0.074 | -1.246 | 0 | -0.005 | -0.005 | 0.235 | 0.72 | 0.63 | 0.81 | 0.84 | 0.7 | 0.84 |
| 11 | 97017572 | rs1793370 | -0.479 | 1.107 | 0.103 | -1.225 | -0.002 | 0.003 | 0.003 | 0 | 0.62 | 0.54 | 0.65 | 0.72 | 0.57 | 0.66 |
| 11 | 97023053 | rs1826611 | -0.888 | 0.712 | 0.295 | -1.165 | -0.012 | -0.08 | -0.08 | 0 | 0.66 | 0.41 | 0.78 | 0.83 | 0.67 | 0.78 |
| 11 | 97025717 | rs10895284 | -0.681 | 0.437 | -0.709 | -1.373 | -0.02 | 0.037 | 0.037 | 0 | 0.64 | 0.85 | 0.57 | 0.32 | 0.64 | 0.62 |
| 11 | 97035775 | rs1690581 | 0.346 | 0.267 | -0.187 | -1.131 | 0.005 | 0.01 | 0.01 | 0 | 0.38 | 0.7 | 0.24 | 0.18 | 0.34 | 0.25 |
| 11 | 97036981 | rs1793357 | 0.461 | 0.393 | -0.5 | -1.063 | -0.003 | -0.024 | -0.024 | 0 | 0.37 | 0.67 | 0.25 | 0.17 | 0.34 | 0.2 |
| 11 | 97039863 | rs17278804 | 1.296 | 0.248 | -1.121 | -1.025 | 0.003 | 0.023 | 0.023 | 0 | 0.11 | 0 | 0.13 | 0.04 | 0.07 | 0.12 |
| 11 | 97040365 | rs11225252 | 0.048 | 0.359 | -0.679 | -0.666 | 0.025 | 0.07 | 0.07 | 0.08 | 0.37 | 0.8 | 0.24 | 0.18 | 0.32 | 0.23 |
| 11 | 97044431 | rs1690595 | -0.527 | 0.779 | -0.419 | -0.066 | 0.009 | 0.019 | 0.019 | 0.302 | 0.54 | 0.66 | 0.56 | 0.6 | 0.55 | 0.38 |
| 11 | 97045544 | rs1038962 | -0.215 | 1.185 | -0.703 | 0.068 | -0.001 | 0.027 | 0.027 | 0 | 0.14 | 0.16 | 0.34 | 0.44 | 0.21 | 0.21 |
| 11 | 97047950 | rs1793378 | 0.029 | 0.565 | -0.66 | -0.067 | 0.014 | 0.029 | 0.029 | 1.61 | 0.43 | 0.65 | 0.25 | 0.17 | 0.35 | 0.24 |
| 11 | 97050518 | rs17129910 | -0.837 | 1.138 | -0.251 | 0.917 | 0.025 | 0.196 | 0.196 | 0 | 0.14 | 0.24 | 0.32 | 0.43 | 0.14 | 0.19 |
| 11 | 97051223 | rs2011649 | 0.052 | 0.925 | -1.223 | -0.341 | -0.027 | 0.127 | 0.127 | 1.533 | 0.36 | 0.12 | 0.56 | 0.87 | 0.39 | 0.53 |
| 11 | 97051784 | rs1793382 | -0.108 | 1.234 | -0.621 | 0.511 | -0.018 | -0.01 | -0.01 | 0.065 | 0.71 | 0.76 | 0.62 | 0.39 | 0.62 | 0.75 |
| 11 | 97051938 | rs17194547 | -1.318 | 0.96 | -0.244 | 1.54 | -0.04 | -0.052 | -0.052 | 1.644 | 0.09 | 0 | 0.05 | 0.16 | 0.12 | 0.15 |
| 11 | 97052967 | rs1494049 | -1.148 | 0.05 | 0.319 | 1.449 | -0.021 | -0.004 | -0.004 | 0 | 0.16 | 0.23 | 0.17 | 0.19 | 0.12 | 0.17 |
| 11 | 97053903 | rs17129923 | -0.843 | 0.122 | 0.153 | 1.289 | -0.005 | -0.072 | -0.072 | 0 | 0.11 | 0.31 | 0.08 | 0 | 0.09 | 0.06 |
| 11 | 97054703 | rs984675 | -1.322 | 0.514 | -0.616 | 1.911 | 0.011 | -0.041 | -0.041 | 0 | 0.13 | 0.07 | 0.13 | 0.35 | 0.1 | 0.23 |
| 11 | 97056730 | rs1690587 | -1.051 | 0.566 | -0.407 | 1.91 | 0.01 | -0.01 | -0.01 | 0 | 0.32 | 0.75 | 0.3 | 0.36 | 0.24 | 0.19 |
| 11 | 97058917 | rs12224055 | -0.471 | 0.603 | -0.912 | 1.655 | -0.03 | -0.081 | -0.081 | 0 | 0.16 | 0.01 | 0.16 | 0.28 | 0.14 | 0.13 |
| 11 | 97075075 | rs1908241 | 1.27 | 1.55 | -1.032 | 1.674 | -0.071 | -0.01 | -0.01 | 0.02 | 0.59 | 0.12 | 0.65 | 0.38 | 0.58 | 0.54 |
| 11 | 97078926 | rs1690597 | 0.047 | 1.803 | -0.429 | 1.879 | 0.031 | 0.037 | 0.037 | 0 | 0.89 | 0.89 | 0.88 | 0.75 | 0.77 | 0.75 |
| 11 | 97079764 | rs7119782 | 1.115 | 1.766 | 0.197 | 1.644 | -0.013 | -0.011 | -0.011 | 0 | 0.67 | 0.72 | 0.7 | 0.44 | 0.61 | 0.58 |
| 11 | 97085025 | rs7117210 | 0.911 | 2.298 | 1.051 | 1.198 | 0.017 | 0.009 | 0.009 | 0.253 | 0.85 | 0.88 | 0.85 | 0.76 | 0.72 | 0.72 |
| 11 | 97093012 | rs12420545 | -0.493 | 1.385 | 0.419 | 0.016 | 0.018 | -0.079 | -0.079 | 0 | 0.16 | 0.27 | 0.23 | 0 | 0.19 | 0.14 |
| 11 | 97102604 | rs1863599 | 1.004 | 1.414 | -0.447 | -0.096 | -0.044 | 0.11 | 0.11 | 0 | 0.67 | 0.13 | 0.66 | 0.83 | 0.66 | 0.74 |
| 11 | 97105850 | rs2555345 | 1.011 | 1.309 | 0.32 | -0.193 | 0.032 | -0.064 | -0.064 | 0.82 | 0.88 | 0.4 | 0.78 | 0.99 | 0.86 | 0.89 |
| 11 | 97110269 | rs1426649 | 0.818 | 1.28 | -0.058 | 0.078 | -0.002 | 0 | 0 | 1.92 | 0.74 | 0.28 | 0.5 | 0.74 | 0.68 | 0.61 |
| 11 | 97110988 | rs964183 | 0.039 | 1.137 | -0.741 | 0.122 | -0.017 | -0.008 | -0.008 | 1.286 | 0.15 | 0.08 | 0.15 | 0.11 | 0.17 | 0.08 |
| 11 | 97111552 | rs1365449 | 0.959 | 0.018 | -1.602 | -0.777 | -0.004 | 0.084 | 0.084 | 1.481 | 0.61 | 0.17 | 0.47 | 0.74 | 0.62 | 0.54 |
| 11 | 97113623 | rs2213121 | 1.072 | -0.015 | -1.577 | -0.789 | 0.003 | 0.084 | 0.084 | 0.307 | 0.61 | 0.19 | 0.47 | 0.75 | 0.62 | 0.53 |
| 11 | 97115714 | rs2226931 | 1.756 | -0.376 | -2.024 | -0.748 | -0.124 | -0.063 | -0.063 | 0 | 0.42 | 0.04 | 0.3 | 0.57 | 0.43 | 0.42 |
| 11 | 97115864 | rs2555321 | 1.374 | -0.307 | -1.917 | -0.965 | -0.07 | 0.12 | 0.12 | 0.141 | 0.53 | 0.12 | 0.43 | 0.68 | 0.55 | 0.5 |
| 11 | 97118867 | rs964646 | 0.623 | -0.577 | -1.503 | -0.863 | 0.003 | -0.027 | -0.027 | 0.321 | 0.08 | 0 | 0.09 | 0.15 | 0.05 | 0.09 |
| 11 | 97127160 | rs2555329 | -0.2 | 0.217 | -1.136 | -0.597 | -0.019 | 0.016 | 0.016 | 0.136 | 0.13 | 0.06 | 0.23 | 0.1 | 0.14 | 0.13 |
| 11 | 97128387 | rs10895394 | 1.574 | 0.245 | -2.05 | -0.772 | -0.096 | -0.071 | -0.071 | 0 | 0.46 | 0.03 | 0.22 | 0.54 | 0.41 | 0.43 |
| 11 | 97128693 | rs2555331 | -1.744 | 0.357 | -1.52 | -1.058 | -0.039 | 0.065 | 0.065 | 0.308 | 0.42 | 0.8 | 0.57 | 0.34 | 0.42 | 0.47 |
| 11 | 97130237 | rs2853064 | -1.394 | 0.249 | -0.699 | -1.194 | -0.003 | -0.052 | -0.052 | 0.078 | 0.26 | 0.61 | 0.29 | 0.11 | 0.22 | 0.24 |
| 11 | 97137260 | rs4754927 | -2.522 | -0.042 | -0.289 | -1.215 | -0.033 | 0.081 | 0.081 | 0.247 | 0.46 | 0.83 | 0.58 | 0.37 | 0.44 | 0.49 |

| | | | | | | | | | | | | | | | | |
|---|---|---|---|---|---|---|---|---|---|---|---|---|---|---|---|---|
| 11 | 97139409 | rs12798864 | 0.442 | -0.262 | -0.023 | -1.34 | -0.009 | 0.009 | 0.009 | 0.025 | 0.13 | 0.06 | 0.22 | 0.12 | 0.14 | 0.17 |
| 11 | 97140198 | rs1365440 | -0.099 | -0.223 | 0.072 | -1.226 | 0 | 0 | 0 | 0 | 0.13 | 0.13 | 0.16 | 0.15 | 0.17 | 0.22 |
| 11 | 97141492 | rs2217106 | -1.762 | -0.217 | -0.057 | -1.193 | 0.002 | 0.094 | 0.094 | 1.338 | 0.39 | 0.82 | 0.45 | 0.27 | 0.37 | 0.45 |
| 11 | 97157660 | rs7479420 | -0.3 | 0.036 | 0.071 | -1.021 | -0.007 | -0.008 | -0.008 | 0 | 0.13 | 0.11 | 0.16 | 0.18 | 0.18 | 0.22 |
| 11 | 97157963 | rs7940252 | -1.515 | -0.054 | 0.25 | -1.317 | -0.002 | 0.055 | 0.055 | 1.465 | 0.42 | 0.68 | 0.48 | 0.28 | 0.42 | 0.48 |
| 11 | 97161720 | rs2114413 | -1.35 | 0.178 | 0.114 | -1.162 | 0.006 | -0.032 | -0.032 | 0 | 0.38 | 0.48 | 0.44 | 0.28 | 0.42 | 0.44 |
| 11 | 97164437 | rs17282574 | -0.737 | 0.38 | -0.16 | -0.673 | -0.007 | -0.007 | -0.007 | 0.071 | 0.14 | 0.11 | 0.17 | 0.17 | 0.18 | 0.23 |
| 11 | 97176319 | rs1426651 | -0.639 | 0.325 | 0.045 | -0.611 | 0.006 | 0.008 | 0.008 | 0 | 0.5 | 0.55 | 0.55 | 0.49 | 0.49 | 0.41 |
| 11 | 97177477 | rs1426630 | 1.275 | 0.419 | 0.668 | -1.603 | -0.001 | -0.052 | -0.052 | 0.221 | 0.71 | 0.31 | 0.67 | 0.99 | 0.71 | 0.85 |
| 11 | 97178452 | rs10895526 | 1.361 | 0.471 | 1.025 | -1.666 | -0.002 | -0.067 | -0.067 | 0 | 0.73 | 0.31 | 0.7 | 0.99 | 0.76 | 0.84 |
| 11 | 97179675 | rs1593566 | 1.738 | 0.321 | 1.105 | -1.447 | -0.019 | -0.016 | -0.016 | 0 | 0.94 | 0.81 | 0.96 | 0.99 | 0.98 | 0.96 |
| 11 | 97182786 | rs17210817 | 0.77 | 0.38 | 1.03 | -1.493 | 0 | -0.022 | -0.022 | 0.503 | 0.08 | 0.1 | 0.08 | 0 | 0.1 | 0.02 |
| 11 | 97185053 | rs1426640 | 0.276 | 0.439 | 1.149 | -1.501 | 0.011 | -0.06 | -0.06 | 0 | 0.85 | 0.74 | 0.87 | 0.99 | 0.83 | 0.96 |
| 11 | 97196848 | rs7943263 | 0.277 | 0.129 | 1.34 | -1.234 | -0.014 | -0.05 | -0.05 | 0 | 0.88 | 0.75 | 0.89 | 0.99 | 0.88 | 0.95 |
| 11 | 97211124 | rs10082569 | -1.634 | 0.193 | 1.673 | -1.296 | 0.027 | -0.047 | -0.047 | 0 | 0.09 | 0.25 | 0.16 | 0.01 | 0.1 | 0.12 |
| 11 | 97211663 | rs7126726 | 1.275 | 0.219 | 2.089 | -1.375 | 0.024 | -0.107 | -0.107 | 0.26 | 0.81 | 0.42 | 0.73 | 0.99 | 0.77 | 0.85 |
| 11 | 97215166 | rs1426642 | 1.1 | 0.004 | 1.923 | -1.484 | -0.019 | -0.083 | -0.083 | 0.333 | 0.79 | 0.41 | 0.85 | 0.99 | 0.85 | 0.94 |
| 11 | 97216147 | rs10895599 | 2.114 | -0.046 | 2.066 | -1.474 | 0.026 | 0.065 | 0.065 | 0 | 0.72 | 0.24 | 0.71 | 0.98 | 0.77 | 0.84 |
| 11 | 97219957 | rs7948079 | -0.958 | 0.179 | 1.968 | -1.279 | 0.055 | -0.021 | -0.021 | 0 | 0.07 | 0.27 | 0.16 | 0 | 0.1 | 0.08 |
| 11 | 97234412 | rs7946324 | -0.246 | 0.094 | 1.785 | -1.334 | 0.02 | -0.059 | -0.059 | 0.598 | 0.1 | 0.25 | 0.1 | 0 | 0.13 | 0.09 |
| 11 | 97235982 | rs7951055 | 1.401 | -0.133 | 2.246 | -1.394 | -0.089 | -0.088 | -0.088 | 0.909 | 0.86 | 0.51 | 0.93 | 0.98 | 0.97 | 0.94 |
| 11 | 97248173 | rs10895699 | 1.483 | -0.205 | 2.243 | -1.429 | -0.089 | -0.088 | -0.088 | 0 | 0.85 | 0.47 | 0.93 | 0.98 | 0.97 | 0.93 |
| 11 | 97252625 | rs1452673 | 2.483 | -0.164 | 2.365 | -1.509 | 0.005 | 0.161 | 0.161 | 0 | 0.71 | 0.11 | 0.68 | 0.98 | 0.75 | 0.78 |
| 11 | 97255149 | rs12294285 | 0.067 | -0.065 | 2.436 | -1.487 | 0.048 | -0.053 | -0.053 | 0 | 0.09 | 0.28 | 0.11 | 0 | 0.13 | 0.08 |
| 11 | 97255512 | rs1840054 | 2.49 | 0.007 | 1.712 | -1.596 | -0.055 | 0.176 | 0.176 | 0 | 0.55 | 0.08 | 0.52 | 0.83 | 0.57 | 0.55 |
| 11 | 97257121 | rs347767 | 1.214 | 0.041 | 1.936 | -1.481 | -0.035 | -0.042 | -0.042 | 0.373 | 0.74 | 0.57 | 0.78 | 0.84 | 0.79 | 0.69 |
| 11 | 97261585 | rs825993 | 0.172 | 0.097 | 1.478 | -1.453 | -0.001 | 0.002 | 0.002 | 0 | 0.82 | 0.91 | 0.83 | 0.85 | 0.82 | 0.76 |
| 11 | 97263377 | rs7111702 | 0.294 | 0.244 | 1.962 | -1.317 | 0.042 | -0.051 | -0.051 | 0 | 0.09 | 0.25 | 0.1 | 0 | 0.12 | 0.05 |
| 11 | 97264658 | rs182974 | 1.605 | -0.123 | 1.977 | -1.106 | -0.012 | -0.004 | -0.004 | 0 | 0.91 | 0.54 | 0.94 | 0.98 | 0.97 | 0.94 |
| 11 | 97279736 | rs10895733 | 1.493 | -0.005 | 0.441 | -0.929 | -0.01 | 0.04 | 0.04 | 0 | 0.6 | 0.19 | 0.6 | 0.72 | 0.59 | 0.54 |
| 11 | 97281245 | rs10895739 | 2.095 | 0.118 | 0.789 | -0.801 | 0.009 | 0.01 | 0.01 | 0.456 | 0.76 | 0.21 | 0.75 | 0.86 | 0.81 | 0.77 |
| 11 | 97281735 | rs11226530 | -0.929 | 0.177 | 1.265 | -0.728 | -0.027 | 0.077 | 0.077 | 0.167 | 0.07 | 0.23 | 0.03 | 0.11 | 0.01 | 0.1 |
| 11 | 97286280 | rs12275454 | -1.234 | 0.17 | 1.609 | -0.729 | -0.01 | 0.132 | 0.132 | 0 | 0.07 | 0.36 | 0.04 | 0.12 | 0.03 | 0.1 |
| 11 | 97288820 | rs347766 | -1.161 | 0.269 | 1.446 | -0.789 | 0.016 | -0.063 | -0.063 | 0 | 0.13 | 0.25 | 0.19 | 0.03 | 0.17 | 0.1 |
| 11 | 97291596 | rs347763 | -2.046 | 0.329 | 1.16 | -0.904 | -0.015 | 0.02 | 0.02 | 0 | 0.29 | 0.25 | 0.33 | 0.16 | 0.38 | 0.32 |
| 11 | 97293942 | rs2405222 | -0.201 | 0.406 | 0.902 | -0.878 | 0.012 | -0.037 | -0.037 | 0 | 0.1 | 0.16 | 0.18 | 0.01 | 0.16 | 0.09 |
| 11 | 97295904 | rs11226572 | 1.66 | 0.347 | 0.479 | -0.877 | -0.004 | 0.022 | 0.022 | 0 | 0.67 | 0.19 | 0.65 | 0.73 | 0.66 | 0.64 |
| 11 | 97297218 | rs979406 | -1.734 | 0.293 | 0.328 | -0.491 | -0.002 | 0.001 | 0.001 | 0 | 0.25 | 0.34 | 0.18 | 0.26 | 0.24 | 0.31 |
| 11 | 97299826 | rs7929176 | -0.714 | 0.06 | 0.245 | -0.048 | -0.028 | 0.036 | 0.036 | 0 | 0.07 | 0.14 | 0.02 | 0.13 | 0.01 | 0.05 |
| 11 | 97300250 | rs4755065 | 0.937 | 0.691 | 0.137 | -0.359 | -0.005 | -0.002 | -0.002 | 0 | 0.56 | 0.41 | 0.52 | 0.55 | 0.5 | 0.42 |
| 11 | 97301408 | rs11226595 | -0.839 | 0.965 | 0.092 | -0.235 | -0.056 | -0.077 | -0.077 | 0.123 | 0.35 | 0.15 | 0.35 | 0.48 | 0.42 | 0.53 |
| 11 | 97301438 | rs7123194 | 0.883 | 1.012 | 0.164 | -0.435 | 0.032 | -0.034 | -0.034 | 0 | 0.06 | 0.16 | 0.06 | 0 | 0.11 | 0.05 |
| 11 | 97303532 | rs4611237 | -0.371 | 0.934 | 0.306 | -0.433 | 0.013 | -0.022 | -0.022 | 1.071 | 0.94 | 0.89 | 0.87 | 1 | 0.95 | 0.97 |
| 11 | 97307856 | rs10791746 | 1.133 | 1.247 | 0.147 | -0.673 | -0.007 | -0.005 | -0.005 | 0 | 0.57 | 0.4 | 0.49 | 0.53 | 0.55 | 0.45 |
| 11 | 97312382 | rs4533067 | 0.281 | 1.396 | -0.011 | -0.702 | -0.085 | -0.142 | -0.142 | 2.355 | 0.25 | 0.04 | 0.3 | 0.47 | 0.33 | 0.48 |
| 11 | 97315831 | rs2044986 | -0.587 | 1.66 | 0.362 | -0.738 | -0.001 | 0.034 | 0.034 | 0 | 0.33 | 0.44 | 0.34 | 0.47 | 0.34 | 0.53 |
| 11 | 97316670 | rs1869208 | 1.993 | 1.901 | 0.243 | -0.632 | 0.055 | -0.027 | -0.027 | 0.115 | 0.44 | 0.21 | 0.39 | 0.54 | 0.35 | 0.32 |
| 11 | 97319496 | rs921561 | -1.277 | 2.3 | 0.019 | -0.768 | 0.005 | 0.005 | 0.005 | 0 | 0.43 | 0.5 | 0.43 | 0.48 | 0.48 | 0.62 |
| 11 | 97321326 | rs2198326 | -0.468 | 2.294 | -0.189 | -0.684 | -0.08 | -0.089 | -0.089 | 0 | 0.33 | 0.14 | 0.36 | 0.49 | 0.44 | 0.52 |
| 11 | 97324111 | rs11226679 | 0.68 | 1.115 | -0.992 | -0.584 | -0.011 | 0.024 | 0.024 | 0 | 0.34 | 0.23 | 0.34 | 0.22 | 0.38 | 0.31 |
| 11 | 97326249 | rs1452670 | -0.502 | 0.643 | -1.006 | -0.5 | -0.001 | 0 | 0 | 0 | 0.45 | 0.49 | 0.52 | 0.54 | 0.51 | 0.64 |
| 11 | 97328659 | rs17286033 | 0.617 | 0.6 | -1.174 | -0.516 | 0.021 | 0.007 | 0.007 | 0 | 0.49 | 0.37 | 0.47 | 0.46 | 0.48 | 0.34 |
| 11 | 97330632 | rs1452671 | 1.633 | 0.689 | -1.617 | -0.17 | -0.105 | -0.122 | -0.122 | 0 | 0.22 | 0.02 | 0.29 | 0.38 | 0.3 | 0.43 |
| 11 | 97330872 | rs1452672 | 1.792 | 0.428 | -1.464 | -0.421 | 0.01 | 0.012 | 0.012 | 0 | 0.27 | 0.18 | 0.3 | 0.2 | 0.29 | 0.26 |
| 11 | 97339454 | rs9943511 | 0.625 | 0.623 | -1.338 | -0.163 | -0.035 | -0.062 | -0.062 | 0 | 0.31 | 0.16 | 0.33 | 0.51 | 0.38 | 0.51 |
| 11 | 97352205 | rs11226763 | 1.707 | 0.296 | -1.368 | -0.08 | -0.039 | -0.067 | -0.067 | 0 | 0.18 | 0.01 | 0.15 | 0.23 | 0.17 | 0.07 |
| 11 | 97356569 | rs10895839 | 0.703 | 0.538 | -1.531 | -0.15 | -0.08 | -0.11 | -0.11 | 1.196 | 0.3 | 0.07 | 0.4 | 0.51 | 0.38 | 0.48 |
| 11 | 97361174 | rs17130141 | 2.061 | 0.565 | -1.456 | -0.077 | -0.006 | -0.069 | -0.069 | 0 | 0.16 | 0.02 | 0.1 | 0.25 | 0.13 | 0.06 |
| 11 | 97371996 | rs1962941 | -0.03 | 0.563 | -0.492 | -0.333 | -0.003 | -0.019 | -0.019 | 0.003 | 0.43 | 0.49 | 0.37 | 0.24 | 0.43 | 0.43 |
| 11 | 97382880 | rs11226865 | 1.248 | 0.376 | -0.282 | -0.44 | 0.011 | 0.013 | 0.013 | 0 | 0.3 | 0.2 | 0.3 | 0.21 | 0.26 | 0.26 |
| 11 | 97399271 | rs11226895 | 0.655 | 0.535 | -0.063 | -0.384 | -0.005 | 0.047 | 0.047 | 0 | 0.36 | 0.29 | 0.33 | 0.15 | 0.38 | 0.34 |
| 11 | 97400945 | rs10895894 | -0.101 | 0.585 | -0.107 | -0.344 | -0.001 | -0.003 | -0.003 | 0 | 0.43 | 0.57 | 0.4 | 0.2 | 0.45 | 0.44 |
| 11 | 97403692 | rs7940229 | 0.297 | 0.629 | -0.095 | -0.354 | -0.006 | -0.032 | -0.032 | 0 | 0.39 | 0.45 | 0.33 | 0.16 | 0.38 | 0.42 |
| 11 | 97406616 | rs2852918 | 0.153 | 0.497 | -0.578 | -0.078 | -0.032 | -0.038 | -0.038 | 1.883 | 0.38 | 0.19 | 0.46 | 0.49 | 0.44 | 0.48 |
| 11 | 97407844 | rs2601669 | 0.67 | -0.066 | -0.811 | -0.259 | -0.045 | 0.025 | 0.025 | 2.373 | 0.4 | 0.15 | 0.46 | 0.68 | 0.54 | 0.41 |
| 11 | 97417862 | rs7937285 | 0.766 | -0.636 | -0.404 | -0.083 | -0.032 | -0.019 | -0.019 | 0 | 0.32 | 0.33 | 0.22 | 0.17 | 0.21 | 0.22 |
| 11 | 97419744 | rs12280664 | 1.214 | -0.781 | -0.398 | -0.155 | -0.018 | -0.014 | -0.014 | 0 | 0.28 | 0.25 | 0.11 | 0.08 | 0.16 | 0.17 |
| 11 | 97420790 | rs11226968 | 1.485 | -0.478 | -0.668 | 0.09 | 0.003 | 0 | 0 | 0 | 0.19 | 0.14 | 0.11 | 0.24 | 0.18 | 0.13 |
| 11 | 97421713 | rs1960374 | -0.558 | -0.315 | -0.464 | -0.041 | -0.015 | -0.01 | -0.01 | 0 | 0.59 | 0.49 | 0.59 | 0.69 | 0.68 | 0.7 |

| | | | | | | | | | | | | | | | | |
|---|---|---|---|---|---|---|---|---|---|---|---|---|---|---|---|---|
| 11 | 97431132 | rs2852908 | 1.353 | -0.595 | -0.369 | 0.098 | 0.013 | 0.011 | 0.011 | 0 | 0.28 | 0.27 | 0.37 | 0.39 | 0.38 | 0.35 |
| 11 | 97431300 | rs11227014 | 1.178 | -0.544 | -0.479 | 0.136 | -0.009 | -0.019 | -0.019 | 1.963 | 0.22 | 0.19 | 0.31 | 0.42 | 0.25 | 0.19 |
| 11 | 97435372 | rs922374 | 0.905 | -0.902 | -0.35 | -0.165 | 0.003 | 0.003 | 0.003 | 0 | 0.34 | 0.3 | 0.25 | 0.18 | 0.27 | 0.27 |
| 11 | 97439081 | rs12288740 | 1.262 | -0.522 | -0.469 | 0.108 | -0.006 | -0.014 | -0.014 | 0 | 0.21 | 0.19 | 0.27 | 0.39 | 0.24 | 0.18 |
| 11 | 97443131 | rs2852921 | 1.285 | -0.818 | -0.328 | 0.008 | -0.007 | -0.007 | -0.007 | 0.069 | 0.27 | 0.22 | 0.39 | 0.41 | 0.39 | 0.37 |
| 11 | 97452661 | rs7945482 | -0.881 | -0.779 | -0.159 | -0.077 | -0.014 | -0.024 | -0.024 | 0.573 | 0.71 | 0.65 | 0.75 | 0.84 | 0.76 | 0.85 |
| 11 | 97460206 | rs1351013 | 1.845 | -0.721 | -0.286 | -0.216 | -0.006 | 0.011 | 0.011 | 3.446 | 0.06 | 0.05 | 0.04 | 0.01 | 0.06 | 0.07 |
| 11 | 97469005 | rs1381981 | 1.339 | -0.95 | -0.153 | 0.098 | -0.022 | 0.026 | 0.026 | 0 | 0.21 | 0.27 | 0.21 | 0.31 | 0.18 | 0.16 |
| 11 | 97474775 | rs1894121 | 0.302 | -0.821 | -0.164 | 0.078 | -0.004 | -0.021 | -0.021 | 0 | 0.31 | 0.25 | 0.3 | 0.41 | 0.31 | 0.23 |
| 11 | 97481672 | rs7944193 | 0.927 | -0.971 | -0.224 | 0.15 | 0 | -0.012 | -0.012 | 0 | 0.22 | 0.19 | 0.15 | 0.31 | 0.24 | 0.15 |
| 11 | 97486522 | rs10749882 | -0.43 | -0.92 | -0.001 | 0.128 | -0.003 | -0.004 | -0.004 | 0 | 0.54 | 0.56 | 0.41 | 0.62 | 0.46 | 0.41 |
| 11 | 97486665 | rs7943838 | 0.5 | -0.965 | 0.1 | 0.077 | -0.009 | 0.009 | 0.009 | 0.006 | 0.34 | 0.39 | 0.41 | 0.42 | 0.29 | 0.33 |
| 11 | 97487065 | rs2126660 | 1.386 | -0.789 | 0.025 | 0.069 | -0.005 | 0.002 | 0.002 | 0 | 0.31 | 0.23 | 0.37 | 0.26 | 0.38 | 0.41 |
| 11 | 97491620 | rs686715 | -0.726 | -0.931 | 0.069 | -0.17 | -0.002 | -0.003 | -0.003 | 0 | 0.45 | 0.54 | 0.41 | 0.42 | 0.43 | 0.4 |
| 11 | 97494307 | rs2051460 | 0.551 | -0.864 | 0.047 | -0.183 | 0.001 | -0.003 | -0.003 | 1.362 | 0.29 | 0.27 | 0.2 | 0.31 | 0.28 | 0.2 |
| 11 | 97502573 | rs649985 | -1.583 | -0.685 | 0.097 | -0.028 | 0.001 | -0.001 | -0.001 | 0.796 | 0.7 | 0.73 | 0.64 | 0.74 | 0.63 | 0.6 |
| 11 | 97518685 | rs7950929 | 1.341 | -0.555 | 0.1 | -0.019 | 0.009 | -0.012 | -0.012 | 0 | 0.3 | 0.25 | 0.34 | 0.24 | 0.36 | 0.39 |
| 11 | 97519987 | rs611843 | 1.393 | -0.924 | 0.066 | 0.056 | 0.004 | -0.006 | -0.006 | 1.087 | 0.22 | 0.2 | 0.21 | 0.32 | 0.17 | 0.21 |
| 11 | 97521611 | rs7113142 | 1.5 | -0.799 | -0.118 | 0.114 | -0.001 | 0 | 0 | 0 | 0.29 | 0.22 | 0.36 | 0.25 | 0.37 | 0.39 |
| 11 | 97522098 | rs669918 | 1.873 | -0.403 | -0.274 | 0.762 | 0.027 | -0.072 | -0.072 | 0 | 0.16 | 0.04 | 0.1 | 0.24 | 0.12 | 0.16 |
| 11 | 97522360 | rs11212092 | 1.051 | -0.228 | -0.193 | 0.591 | -0.004 | -0.021 | -0.021 | 0 | 0.18 | 0.14 | 0.14 | 0.3 | 0.21 | 0.11 |
| 11 | 97523625 | rs11212106 | 0.304 | -0.25 | 0.146 | 0.701 | 0.011 | 0.001 | 0.001 | 0.122 | 0.11 | 0.14 | 0.19 | 0.1 | 0.12 | 0.18 |
| 11 | 97535155 | rs635147 | 0.45 | -0.891 | 0.205 | 0.806 | -0.022 | 0.042 | 0.042 | 0 | 0.19 | 0.25 | 0.22 | 0.3 | 0.15 | 0.19 |
| 11 | 97541637 | rs644993 | 0.466 | -0.879 | 0.205 | 0.789 | -0.026 | 0.059 | 0.059 | 0 | 0.19 | 0.26 | 0.22 | 0.31 | 0.15 | 0.19 |
| 11 | 97557786 | rs1039953 | 0.15 | -0.16 | -0.174 | 0.05 | -0.014 | -0.005 | -0.005 | 0 | 0.25 | 0.21 | 0.2 | 0.28 | 0.31 | 0.28 |
| 11 | 97560081 | rs627611 | 0.248 | -0.369 | 0.153 | 0.079 | -0.024 | -0.016 | -0.016 | 0.017 | 0.89 | 0.83 | 0.85 | 0.94 | 0.93 | 0.97 |
| 11 | 97580258 | rs10789637 | -0.446 | -0.129 | 0.102 | 0.128 | -0.002 | -0.002 | -0.002 | 0 | 0.08 | 0.1 | 0.17 | 0.09 | 0.11 | 0.06 |
| 11 | 97588674 | rs629047 | 0.771 | -0.329 | 0.187 | -0.166 | 0 | -0.003 | -0.003 | 0 | 0.46 | 0.42 | 0.57 | 0.66 | 0.49 | 0.57 |
| 11 | 97591699 | rs612234 | 1.611 | -0.545 | 0.369 | -0.025 | -0.018 | -0.043 | -0.043 | 0 | 0.81 | 0.6 | 0.79 | 0.89 | 0.83 | 0.87 |
| 11 | 97595073 | rs11212464 | -0.425 | -0.538 | 0.26 | 0.08 | -0.02 | 0.02 | 0.02 | 0 | 0.08 | 0.19 | 0.15 | 0.1 | 0.07 | 0.05 |
| 11 | 97601350 | rs10890801 | 1.252 | -0.551 | 0.152 | 0.015 | -0.027 | -0.018 | -0.018 | 0 | 0.81 | 0.57 | 0.79 | 0.86 | 0.83 | 0.87 |
| 11 | 97608622 | rs12274023 | -0.836 | -0.586 | 0.3 | -0.011 | -0.02 | -0.028 | -0.028 | 0 | 0.11 | 0.34 | 0.06 | 0.04 | 0.07 | 0.05 |
| 11 | 97609886 | rs7113765 | -0.477 | -0.784 | -0.296 | -0.04 | 0.013 | -0.004 | -0.004 | 0 | 0.7 | 0.8 | 0.73 | 0.7 | 0.78 | 0.7 |
| 11 | 97612364 | rs7125116 | -1.158 | -0.833 | 0.033 | -0.152 | -0.032 | -0.033 | -0.033 | 0 | 0.94 | 0.84 | 0.98 | 1 | 0.98 | 0.99 |
| 11 | 97614840 | rs10890824 | 0.532 | -0.595 | -0.433 | -0.312 | -0.007 | -0.01 | -0.01 | 0 | 0.32 | 0.43 | 0.42 | 0.3 | 0.32 | 0.4 |
| 11 | 97617726 | rs17130219 | 0.567 | -0.582 | -0.456 | -0.234 | 0.001 | -0.01 | -0.01 | 0 | 0.31 | 0.24 | 0.38 | 0.36 | 0.28 | 0.31 |
| 11 | 97618161 | rs7128754 | 0.501 | -0.74 | -0.702 | -0.429 | -0.064 | -0.055 | -0.055 | 0 | 0.32 | 0.08 | 0.18 | 0.34 | 0.35 | 0.25 |
| 11 | 97620150 | rs1903803 | 1.142 | -0.699 | -0.63 | -0.341 | 0.005 | -0.008 | -0.008 | 0 | 0.26 | 0.12 | 0.27 | 0.27 | 0.2 | 0.27 |
| 11 | 97620700 | rs925425 | -1.11 | -0.735 | -0.505 | -0.336 | -0.035 | -0.037 | -0.037 | 0.328 | 0.93 | 0.74 | 0.97 | 1 | 0.98 | 0.99 |
| 11 | 97622502 | rs12421267 | 0.498 | -0.765 | -0.728 | -0.546 | -0.022 | 0.015 | 0.015 | 0 | 0.33 | 0.14 | 0.34 | 0.27 | 0.3 | 0.29 |
| 11 | 97623375 | rs11212569 | 0.526 | -0.674 | -0.715 | -0.54 | -0.029 | -0.035 | -0.035 | 0 | 0.33 | 0.12 | 0.17 | 0.35 | 0.34 | 0.26 |
| 11 | 97624171 | rs2897122 | 0.462 | -0.631 | -0.677 | -0.52 | 0.001 | -0.007 | -0.007 | 0 | 0.31 | 0.5 | 0.43 | 0.31 | 0.35 | 0.44 |
| 11 | 97624549 | rs7124715 | 1.376 | -0.621 | -0.717 | -0.531 | 0 | -0.014 | -0.014 | 0 | 0.25 | 0.13 | 0.28 | 0.27 | 0.2 | 0.22 |
| 11 | 97626863 | rs7129738 | -0.499 | -1.157 | -0.506 | -0.369 | -0.002 | -0.012 | -0.012 | 0.643 | 0.68 | 0.44 | 0.57 | 0.66 | 0.65 | 0.53 |
| 11 | 97630062 | rs1813435 | -0.288 | -1.281 | -0.57 | -0.459 | -0.005 | -0.022 | -0.022 | 0.094 | 0.66 | 0.48 | 0.6 | 0.71 | 0.64 | 0.61 |
| 11 | 97632969 | rs1397831 | -0.237 | -1.306 | -0.584 | -0.45 | 0.001 | 0.001 | 0.001 | 4.476 | 0.65 | 0.37 | 0.6 | 0.71 | 0.67 | 0.58 |
| 11 | 97635776 | rs12575486 | 1.427 | -1.448 | -0.874 | -0.67 | 0.031 | -0.036 | -0.036 | 0 | 0.22 | 0.12 | 0.15 | 0.26 | 0.15 | 0.21 |
| 11 | 97636668 | rs1548172 | -0.902 | -1.472 | -0.247 | -0.746 | -0.011 | -0.067 | -0.067 | 1.529 | 0.87 | 0.76 | 0.92 | 1 | 0.91 | 0.91 |
| 11 | 97638674 | rs7949197 | -0.294 | -1.483 | -0.267 | -0.748 | -0.007 | -0.015 | -0.015 | 0 | 0.66 | 0.42 | 0.61 | 0.72 | 0.67 | 0.59 |
| 11 | 97639218 | rs10789660 | 0.993 | -1.521 | -0.552 | -0.751 | -0.121 | -0.085 | -0.085 | 0 | 0.32 | 0.08 | 0.32 | 0.41 | 0.39 | 0.32 |
| 11 | 97639849 | rs12417967 | 1.338 | -1.59 | -0.418 | -0.684 | 0.004 | 0.015 | 0.015 | 0 | 0.21 | 0.33 | 0.32 | 0.27 | 0.25 | 0.29 |
| 11 | 97644871 | rs10789664 | 0.765 | -1.547 | -0.837 | -0.721 | -0.096 | -0.098 | -0.098 | 0.165 | 0.33 | 0.08 | 0.42 | 0.45 | 0.5 | 0.33 |
| 11 | 97646309 | rs10749919 | -0.435 | -1.773 | -0.71 | -0.811 | -0.012 | -0.03 | -0.03 | 0 | 0.83 | 0.77 | 0.91 | 0.94 | 0.88 | 0.87 |
| 11 | 97649792 | rs11212697 | 1.062 | -1.528 | -0.858 | -0.652 | -0.123 | -0.057 | -0.057 | 2.701 | 0.29 | 0.07 | 0.32 | 0.35 | 0.41 | 0.28 |
| 11 | 97657515 | rs11212729 | 1.514 | -1.665 | -0.646 | -0.668 | -0.031 | -0.032 | -0.032 | 0 | 0.07 | 0.16 | 0.02 | 0 | 0.01 | 0.01 |
| 11 | 97659276 | rs7942270 | 1.51 | -1.714 | -0.787 | -0.569 | 0.027 | -0.041 | -0.041 | 0 | 0.19 | 0.07 | 0.15 | 0.27 | 0.13 | 0.2 |
| 11 | 97660579 | rs10890871 | -0.395 | -1.588 | -0.612 | -0.517 | -0.022 | -0.004 | -0.004 | 0 | 0.64 | 0.5 | 0.63 | 0.63 | 0.68 | 0.6 |
| 11 | 97663464 | rs11212745 | 0.339 | -1.308 | -0.888 | -0.409 | 0.006 | 0 | 0 | 0 | 0.42 | 0.36 | 0.49 | 0.42 | 0.56 | 0.35 |
| 11 | 97667822 | rs12577978 | 0.316 | -0.864 | -0.745 | -0.203 | -0.001 | -0.001 | -0.001 | 0.021 | 0.22 | 0.18 | 0.17 | 0.26 | 0.14 | 0.2 |
| 11 | 97670547 | rs7481183 | 0.519 | -1.147 | -1.052 | -0.175 | 0.038 | -0.037 | -0.037 | 0.235 | 0.21 | 0.07 | 0.15 | 0.26 | 0.13 | 0.18 |
| 11 | 97670993 | rs2897197 | 0.56 | -1.089 | -1.038 | -0.283 | 0.034 | -0.022 | -0.022 | 0 | 0.2 | 0.07 | 0.1 | 0.21 | 0.12 | 0.16 |
| 11 | 97673425 | rs2405656 | 0.506 | -0.89 | -0.523 | -0.97 | -0.048 | -0.17 | -0.17 | 1.321 | 0.77 | 0.51 | 0.87 | 1 | 0.86 | 0.85 |
| 11 | 97676449 | rs10890892 | -0.933 | -0.465 | -1.265 | -0.66 | 0.007 | -0.041 | -0.041 | 0 | 0.7 | 0.78 | 0.62 | 0.46 | 0.64 | 0.58 |
| 11 | 97680206 | rs10789687 | -0.92 | -0.48 | -1.1 | -0.629 | 0.005 | -0.024 | -0.024 | 0.686 | 0.7 | 0.74 | 0.62 | 0.5 | 0.64 | 0.58 |
| 11 | 97681076 | rs11212791 | 0.089 | 0.044 | -1.827 | -0.874 | -0.088 | -0.082 | -0.082 | 0 | 0.21 | 0.02 | 0.3 | 0.25 | 0.19 | 0.18 |
| 11 | 97685864 | rs10749931 | -0.808 | -0.182 | -1.425 | -0.839 | 0.008 | -0.002 | -0.002 | 0 | 0.63 | 0.66 | 0.59 | 0.43 | 0.57 | 0.46 |
| 11 | 97687676 | rs10890904 | -0.005 | -0.072 | -1.083 | -0.804 | -0.018 | 0.029 | 0.029 | 0 | 0.22 | 0.13 | 0.14 | 0.19 | 0.23 | 0.19 |
| 11 | 97707396 | rs2851619 | 0.392 | -0.162 | -0.736 | -0.631 | 0 | 0.001 | 0.001 | 0 | 0.76 | 0.77 | 0.79 | 0.83 | 0.8 | 0.78 |
| 11 | 97709396 | rs2851623 | 0.4 | -0.183 | -0.726 | -0.641 | 0 | 0 | 0 | 0 | 0.76 | 0.77 | 0.79 | 0.82 | 0.81 | 0.78 |

| | | | | | | | | | | | | | | | | |
|---|---|---|---|---|---|---|---|---|---|---|---|---|---|---|---|---|
| 11 | 97722738 | rs2597597 | -0.193 | -0.251 | -0.853 | -0.712 | -0.054 | -0.029 | -0.029 | 0.454 | 0.81 | 0.96 | 0.82 | 0.83 | 0.82 | 0.83 |
| 11 | 97723251 | rs2246397 | 1.046 | -0.493 | -0.381 | -0.932 | 0.002 | 0.005 | 0.005 | 0 | 0.55 | 0.35 | 0.64 | 0.61 | 0.6 | 0.54 |
| 11 | 97724023 | rs2597596 | 0.954 | -0.58 | -0.51 | -1.279 | -0.001 | 0.009 | 0.009 | 0 | 0.46 | 0.32 | 0.56 | 0.57 | 0.46 | 0.51 |
| 11 | 97726925 | rs7125251 | -0.024 | -0.732 | -0.242 | -1.639 | -0.015 | -0.047 | -0.047 | 0 | 0.38 | 0.48 | 0.29 | 0.17 | 0.38 | 0.25 |
| 11 | 97732255 | rs4255511 | -0.567 | -0.569 | -0.304 | -1.718 | 0.009 | 0.017 | 0.017 | 0 | 0.46 | 0.62 | 0.33 | 0.23 | 0.44 | 0.32 |
| 11 | 97732269 | rs2597589 | -0.979 | -0.76 | -0.539 | -1.752 | -0.063 | -0.06 | -0.06 | 0 | 0.88 | 0.98 | 0.82 | 0.78 | 0.85 | 0.82 |
| 11 | 97733303 | rs2597588 | 1.067 | -0.942 | -0.527 | -1.818 | -0.004 | -0.006 | -0.006 | 0.888 | 0.39 | 0.28 | 0.48 | 0.55 | 0.41 | 0.49 |
| 11 | 97734972 | rs10789728 | 0.783 | -0.981 | -0.456 | -2.15 | 0.001 | 0.018 | 0.018 | 0 | 0.27 | 0.2 | 0.21 | 0.17 | 0.32 | 0.24 |
| 11 | 97737007 | rs949175 | 0.262 | -0.896 | -0.512 | -2.102 | 0.015 | 0.057 | 0.057 | 0 | 0.35 | 0.21 | 0.27 | 0.17 | 0.37 | 0.24 |
| 11 | 97744330 | rs2851634 | -1.078 | -0.392 | -0.414 | -1.921 | -0.006 | -0.001 | -0.001 | 0 | 0.89 | 0.91 | 0.97 | 0.93 | 0.88 | 0.91 |
| 11 | 97744726 | rs2597559 | 0.312 | -0.669 | -0.198 | -2.084 | 0 | -0.015 | -0.015 | 0 | 0.75 | 0.72 | 0.88 | 0.92 | 0.82 | 0.84 |
| 11 | 97745838 | rs12286557 | 1.812 | -0.621 | -0.196 | -2.042 | -0.017 | -0.042 | -0.042 | 1.542 | 0.09 | 0.13 | 0.07 | 0 | 0.05 | 0.02 |
| 11 | 97748769 | rs10789733 | 0.991 | -0.729 | -0.37 | -2.131 | -0.006 | 0.028 | 0.028 | 0 | 0.27 | 0.1 | 0.2 | 0.2 | 0.32 | 0.22 |
| 11 | 97752795 | rs2443011 | -0.298 | -0.63 | -0.078 | -1.948 | 0.013 | -0.033 | -0.033 | 0.12 | 0.79 | 0.66 | 0.88 | 0.93 | 0.83 | 0.89 |
| 11 | 97757897 | rs2597579 | 0.594 | -0.787 | 0.035 | -2.033 | -0.011 | -0.06 | -0.06 | 0 | 0.69 | 0.42 | 0.84 | 0.9 | 0.81 | 0.83 |
| 11 | 97758518 | rs7342213 | 0.941 | -0.639 | -0.094 | -1.979 | 0.003 | 0 | 0 | 1.58 | 0.11 | 0.12 | 0.04 | 0.08 | 0.12 | 0.07 |
| 11 | 97758629 | rs10891006 | 0.664 | -0.847 | -0.291 | -2.054 | 0.015 | 0.039 | 0.039 | 0.115 | 0.31 | 0.2 | 0.19 | 0.2 | 0.32 | 0.22 |
| 11 | 97768215 | rs2248770 | 0.432 | -0.425 | 0.027 | -1.847 | -0.005 | 0.009 | 0.009 | 0.007 | 0.42 | 0.33 | 0.66 | 0.69 | 0.49 | 0.58 |
| 11 | 97775240 | rs2851641 | 0.817 | -0.31 | -0.134 | -2.179 | -0.011 | -0.046 | -0.046 | 0 | 0.25 | 0.11 | 0.46 | 0.64 | 0.27 | 0.4 |
| 11 | 97776002 | rs2851642 | -1.78 | -0.483 | 0.029 | -2.168 | -0.021 | -0.053 | -0.053 | 0 | 0.9 | 0.83 | 0.93 | 1 | 0.95 | 0.98 |
| 11 | 97787241 | rs2597630 | -0.2 | -0.228 | -0.151 | -2.003 | 0.001 | -0.017 | -0.017 | 0 | 0.45 | 0.47 | 0.69 | 0.72 | 0.49 | 0.65 |
| 11 | 97791459 | rs2851672 | -0.025 | -0.166 | -0.36 | -1.977 | -0.004 | 0.011 | 0.011 | 0 | 0.43 | 0.33 | 0.67 | 0.69 | 0.48 | 0.62 |
| 11 | 97799061 | rs2597572 | -0.126 | -0.56 | -0.343 | -2.643 | 0.003 | -0.037 | -0.037 | 0 | 0.58 | 0.57 | 0.65 | 0.89 | 0.56 | 0.68 |
| 11 | 97800734 | rs7932877 | 0.968 | -0.222 | -0.596 | -2.712 | -0.05 | 0.051 | 0.051 | 0.57 | 0.21 | 0.12 | 0.14 | 0.07 | 0.27 | 0.22 |
| 11 | 97804083 | rs4494277 | 1.103 | -0.898 | -1.058 | -2.972 | -0.001 | 0.041 | 0.041 | 0 | 0.29 | 0.15 | 0.21 | 0.15 | 0.33 | 0.22 |
| 11 | 97805320 | rs3018169 | -0.666 | -1.043 | -0.738 | -3.054 | -0.004 | -0.009 | -0.009 | 0 | 0.78 | 0.75 | 0.82 | 0.89 | 0.8 | 0.85 |
| 11 | 97806906 | rs11213259 | 0.698 | -1.123 | -0.723 | -3.117 | -0.001 | 0.001 | 0.001 | 0 | 0.13 | 0.11 | 0.11 | 0.09 | 0.13 | 0.12 |
| 11 | 97808955 | rs1815908 | 1.288 | -1.273 | -0.621 | -3.26 | 0.005 | -0.02 | -0.02 | 0.265 | 0.21 | 0.15 | 0.43 | 0.72 | 0.2 | 0.39 |
| 11 | 97810099 | rs2597563 | -0.35 | -1.211 | -0.697 | -3.342 | -0.024 | 0.095 | 0.095 | 0 | 0.58 | 0.73 | 0.7 | 0.88 | 0.55 | 0.67 |
| 11 | 97813130 | rs2851657 | 1.082 | -1.137 | -0.294 | -3.264 | -0.01 | 0.033 | 0.033 | 0 | 0.22 | 0.34 | 0.45 | 0.73 | 0.2 | 0.39 |
| 11 | 97814525 | rs4497377 | 1.675 | -1.14 | -0.206 | -3.31 | -0.022 | 0.003 | 0.003 | 0.16 | 0.21 | 0.18 | 0.14 | 0.01 | 0.25 | 0.18 |
| 11 | 97816230 | rs10789753 | 0.728 | -1.139 | -0.236 | -3.341 | 0.033 | 0.105 | 0.105 | 0 | 0.51 | 0.36 | 0.36 | 0.16 | 0.56 | 0.4 |
| 11 | 97816967 | rs11213301 | 2.08 | -1.272 | -0.39 | -3.418 | -0.02 | 0.066 | 0.066 | 0 | 0.24 | 0.03 | 0.19 | 0.09 | 0.29 | 0.17 |
| 11 | 97818476 | rs2851663 | -2.489 | -1.389 | -0.337 | -3.422 | -0.015 | -0.041 | -0.041 | 0.141 | 0.9 | 0.85 | 0.93 | 0.98 | 0.93 | 0.97 |
| 11 | 97820310 | rs11605274 | 2.003 | -1.371 | -0.389 | -3.39 | -0.03 | 0.065 | 0.065 | 0 | 0.24 | 0.03 | 0.19 | 0.09 | 0.3 | 0.18 |
| 11 | 97820873 | rs2851668 | 0.044 | -1.113 | -0.199 | -3.197 | -0.007 | -0.024 | -0.024 | 0 | 0.72 | 0.67 | 0.8 | 0.89 | 0.76 | 0.79 |
| 11 | 97846667 | rs4474403 | -1.164 | -0.855 | -0.31 | -2.59 | -0.003 | 0.045 | 0.045 | 0 | 0.8 | 0.68 | 0.59 | 0.27 | 0.8 | 0.63 |
| 11 | 97852498 | rs1815913 | -0.23 | -0.817 | -0.316 | -2.708 | 0.007 | 0.026 | 0.026 | 0 | 0.5 | 0.47 | 0.33 | 0.21 | 0.5 | 0.41 |
| 11 | 97852554 | rs1815914 | 0.383 | -0.815 | -0.598 | -2.539 | 0.003 | 0.058 | 0.058 | 0.087 | 0.39 | 0.22 | 0.28 | 0.18 | 0.43 | 0.28 |
| 11 | 97855328 | rs4304739 | -0.144 | -0.99 | -0.298 | -1.749 | -0.013 | 0.013 | 0.013 | 0 | 0.77 | 0.67 | 0.57 | 0.24 | 0.79 | 0.64 |
| 11 | 97855961 | rs7126605 | -0.155 | -1.022 | -0.386 | -1.698 | -0.034 | 0.017 | 0.017 | 0 | 0.22 | 0.37 | 0.38 | 0.77 | 0.15 | 0.31 |
| 11 | 97864992 | rs4492782 | 0.782 | -1.327 | -0.861 | -2.35 | 0.076 | 0.165 | 0.165 | 0 | 0.54 | 0.19 | 0.48 | 0.13 | 0.68 | 0.49 |
| 11 | 97867665 | rs4590838 | 0.059 | -1.59 | -0.492 | -2.478 | -0.041 | -0.075 | -0.075 | 0 | 0.73 | 0.56 | 0.87 | 0.92 | 0.85 | 0.84 |
| 11 | 97871998 | rs10891169 | 0.919 | -1.416 | -0.451 | -2.387 | 0.082 | 0.186 | 0.186 | 0 | 0.54 | 0.18 | 0.48 | 0.13 | 0.69 | 0.49 |
| 11 | 97874919 | rs2212384 | 2.304 | -1.814 | -0.738 | -2.736 | 0.012 | 0.008 | 0.008 | 0.257 | 0.09 | 0.05 | 0.06 | 0 | 0.04 | 0.01 |
| 11 | 97879598 | rs2508537 | 0.741 | -1.635 | -0.211 | -2.535 | 0.024 | 0.035 | 0.035 | 0 | 0.55 | 0.35 | 0.5 | 0.13 | 0.68 | 0.5 |
| 11 | 97890623 | rs2250186 | 0.715 | -1.852 | -0.265 | -2.641 | -0.011 | -0.02 | -0.02 | 0 | 0.14 | 0.35 | 0.08 | 0.07 | 0.08 | 0.05 |
| 11 | 97892643 | rs551427 | 0.873 | -1.655 | -0.881 | -2.584 | 0.083 | 0.188 | 0.188 | 0.012 | 0.54 | 0.17 | 0.49 | 0.13 | 0.68 | 0.49 |
| 11 | 97896128 | rs11213689 | 0.577 | -0.774 | -1.251 | -2.682 | -0.05 | 0.071 | 0.071 | 0.033 | 0.3 | 0.13 | 0.28 | 0.14 | 0.47 | 0.38 |
| 11 | 97896729 | rs11213693 | 0.594 | -0.366 | -1.23 | -2.654 | 0.014 | -0.002 | -0.002 | 0.015 | 0.08 | 0.09 | 0.13 | 0.05 | 0.16 | 0.17 |
| 11 | 97900222 | rs1941852 | 0.69 | -0.698 | -0.655 | -2.837 | 0.017 | 0.008 | 0.008 | 0 | 0.11 | 0.42 | 0.08 | 0.07 | 0.08 | 0.05 |
| 11 | 97901008 | rs10891217 | 0.735 | -0.648 | -0.774 | -2.803 | 0.013 | -0.004 | -0.004 | 0.016 | 0.07 | 0.09 | 0.13 | 0.04 | 0.16 | 0.17 |
| 11 | 97905838 | rs644178 | -0.518 | -0.834 | -0.74 | -2.876 | 0.001 | -0.051 | -0.051 | 0 | 0.67 | 0.43 | 0.7 | 0.86 | 0.62 | 0.75 |
| 11 | 97910423 | rs520388 | 1.593 | -0.854 | -0.759 | -3.094 | -0.038 | 0.047 | 0.047 | 0 | 0.28 | 0.17 | 0.17 | 0 | 0.28 | 0.16 |
| 11 | 97915543 | rs667679 | 0.385 | -0.969 | -0.667 | -1.843 | 0.018 | 0.042 | 0.042 | 0 | 0.51 | 0.35 | 0.39 | 0.31 | 0.52 | 0.4 |
| 11 | 97917013 | rs522030 | -1.017 | -0.429 | -0.728 | -1.68 | 0 | -0.024 | -0.024 | 0 | 0.77 | 0.76 | 0.78 | 0.94 | 0.7 | 0.8 |
| 11 | 97917125 | rs593423 | -1.096 | -0.384 | -1.245 | -1.658 | -0.016 | 0.075 | 0.075 | 0 | 0.77 | 0.91 | 0.79 | 0.94 | 0.7 | 0.8 |
| 11 | 97918427 | rs7932592 | 0.122 | -0.336 | -0.846 | -1.337 | 0.013 | -0.004 | -0.004 | 0.011 | 0.07 | 0.11 | 0.11 | 0.04 | 0.12 | 0.17 |
| 11 | 97918863 | rs1941863 | 0.044 | -0.465 | -1.12 | -1.527 | -0.028 | 0.06 | 0.06 | 0 | 0.41 | 0.21 | 0.52 | 0.71 | 0.49 | 0.52 |
| 11 | 97919019 | rs501723 | 1.286 | -0.677 | -0.947 | -1.918 | -0.009 | -0.019 | -0.019 | 0.987 | 0.25 | 0.18 | 0.15 | 0 | 0.26 | 0.13 |
| 11 | 97919121 | rs11213783 | 0.033 | -1.309 | -0.526 | -2.041 | -0.028 | 0 | 0 | 0.245 | 0.26 | 0.29 | 0.17 | 0.26 | 0.16 | 0.24 |
| 11 | 97924230 | rs669721 | 1.185 | -1.037 | -0.551 | -2.15 | -0.006 | -0.035 | -0.035 | 0 | 0.26 | 0.21 | 0.15 | 0 | 0.27 | 0.16 |
| 11 | 97929915 | rs536242 | -2.235 | -1.219 | -0.388 | -2.135 | -0.029 | -0.031 | -0.031 | 0.717 | 0.93 | 0.74 | 0.98 | 1 | 0.99 | 0.99 |
| 11 | 97932720 | rs4121681 | -0.311 | -1.214 | -0.711 | -1.524 | 0.007 | 0.021 | 0.021 | 0 | 0.5 | 0.41 | 0.66 | 0.62 | 0.49 | 0.48 |
| 11 | 97944535 | rs643711 | 1.48 | -0.87 | -0.81 | -1.453 | -0.025 | 0.021 | 0.021 | 0 | 0.25 | 0.15 | 0.14 | 0 | 0.27 | 0.14 |
| 11 | 97945697 | rs11213924 | 1.12 | -1.005 | -0.906 | -1.365 | 0.014 | 0.018 | 0.018 | 0 | 0.1 | 0.03 | 0.04 | 0.01 | 0.06 | 0.1 |
| 11 | 97957957 | rs17130457 | 1.074 | -0.915 | -0.899 | -1.231 | -0.022 | -0.05 | -0.05 | 0 | 0.17 | 0.07 | 0.17 | 0.39 | 0.23 | 0.36 |
| 11 | 97961701 | rs679808 | 0.014 | -1.207 | -0.685 | -1.185 | 0.014 | 0.019 | 0.019 | 0 | 0.55 | 0.31 | 0.53 | 0.3 | 0.57 | 0.43 |
| 11 | 97962097 | rs17130467 | 1.401 | -1.243 | -0.553 | -1.183 | -0.004 | -0.008 | -0.008 | 0 | 0.17 | 0.15 | 0.17 | 0.37 | 0.23 | 0.37 |

| | | | | | | | | | | | | | | | | |
|---|---|---|---|---|---|---|---|---|---|---|---|---|---|---|---|---|
| 11 | 97962867 | rs11606231 | 1.156 | -1.246 | -0.512 | -1.185 | -0.001 | 0.001 | 0.001 | 0 | 0.18 | 0.18 | 0.18 | 0.37 | 0.23 | 0.37 |
| 11 | 97969766 | rs652531 | 2.026 | -1.173 | -0.655 | -1.396 | -0.057 | 0.101 | 0.101 | 0 | 0.26 | 0.07 | 0.15 | 0 | 0.28 | 0.13 |
| 11 | 97971706 | rs10749986 | -0.866 | -1.104 | -0.569 | -1.328 | -0.046 | 0.04 | 0.04 | 0.038 | 0.72 | 0.49 | 0.71 | 0.66 | 0.8 | 0.8 |
| 11 | 97972559 | rs2509867 | -0.049 | -1.094 | -0.585 | -1.331 | 0.013 | 0.018 | 0.018 | 0 | 0.55 | 0.31 | 0.53 | 0.3 | 0.57 | 0.43 |
| 11 | 97976439 | rs626360 | 0.688 | -1.313 | -0.443 | -1.416 | 0.013 | 0.028 | 0.028 | 0 | 0.41 | 0.27 | 0.26 | 0.32 | 0.34 | 0.24 |
| 11 | 97978654 | rs2508553 | 1.888 | -1.143 | -0.345 | -1.4 | -0.026 | 0.04 | 0.04 | 0 | 0.27 | 0.15 | 0.15 | 0 | 0.28 | 0.15 |
| 11 | 97978777 | rs585018 | -1.912 | -1.133 | -0.311 | -1.374 | 0.002 | -0.061 | -0.061 | 0 | 0.85 | 0.65 | 0.81 | 0.97 | 0.86 | 0.89 |
| 11 | 97984374 | rs11214023 | 0.458 | -1.17 | -0.421 | -1.406 | -0.003 | -0.011 | -0.011 | 0 | 0.26 | 0.16 | 0.2 | 0.38 | 0.29 | 0.46 |
| 11 | 97987244 | rs7107562 | -2.227 | -1.159 | -0.319 | -1.419 | 0 | -0.033 | -0.033 | 1.706 | 0.86 | 0.78 | 0.83 | 0.97 | 0.86 | 0.89 |
| 11 | 97992674 | rs10501879 | 2.521 | -1.124 | -0.731 | -1.656 | -0.034 | 0.026 | 0.026 | 0.096 | 0.17 | 0.1 | 0.1 | 0 | 0.23 | 0.14 |
| 11 | 97993690 | rs7103607 | 0.145 | -1.163 | -0.71 | -1.593 | 0.017 | 0.018 | 0.018 | 0 | 0.37 | 0.21 | 0.26 | 0.38 | 0.34 | 0.48 |
| 11 | 97996725 | rs7124272 | 0.108 | -1.152 | -0.68 | -1.573 | 0.011 | 0.011 | 0.011 | 0 | 0.37 | 0.27 | 0.27 | 0.38 | 0.34 | 0.48 |
| 11 | 98002338 | rs1941847 | 0.667 | -0.796 | -0.423 | -1.259 | 0.006 | 0.014 | 0.014 | 0 | 0.4 | 0.31 | 0.26 | 0.32 | 0.34 | 0.24 |
| 11 | 98008304 | rs4581411 | 1.776 | -0.775 | -0.443 | -1.234 | -0.011 | 0.022 | 0.022 | 0 | 0.12 | 0.15 | 0.1 | 0.31 | 0.06 | 0.09 |
| 11 | 98009459 | rs11214090 | 1.525 | -0.457 | -0.617 | -1.246 | 0 | -0.003 | -0.003 | 0.015 | 0.16 | 0.09 | 0.31 | 0.28 | 0.19 | 0.15 |
| 11 | 98023442 | rs11214156 | 0.36 | -0.457 | -0.558 | -1.235 | 0 | -0.006 | -0.006 | 0 | 0.27 | 0.22 | 0.21 | 0.38 | 0.29 | 0.46 |
| 11 | 98025275 | rs2266114 | -0.535 | -0.456 | -0.55 | -1.264 | -0.006 | 0.011 | 0.011 | 0 | 0.45 | 0.33 | 0.55 | 0.66 | 0.52 | 0.63 |
| 11 | 98035433 | rs3018249 | -0.811 | -0.498 | -0.53 | -1.225 | -0.009 | -0.073 | -0.073 | 1.231 | 0.76 | 0.67 | 0.78 | 0.97 | 0.8 | 0.79 |
| 11 | 98038531 | rs2253410 | -2.168 | -0.337 | -0.399 | -1.079 | -0.022 | -0.066 | -0.066 | 0 | 0.86 | 0.67 | 0.82 | 0.98 | 0.9 | 0.88 |
| 11 | 98046989 | rs11214229 | 0.24 | -0.488 | -0.576 | -1.047 | 0.02 | 0.053 | 0.053 | 0.14 | 0.54 | 0.31 | 0.36 | 0.39 | 0.56 | 0.61 |
| 11 | 98047380 | rs2252684 | 0.855 | -0.106 | -0.705 | -0.88 | -0.005 | -0.009 | -0.009 | 0 | 0.18 | 0.1 | 0.34 | 0.28 | 0.23 | 0.17 |
| 11 | 98053980 | rs7951972 | -0.557 | -0.006 | -0.444 | -0.741 | -0.001 | 0 | 0 | 0 | 0.4 | 0.42 | 0.32 | 0.69 | 0.36 | 0.55 |
| 11 | 98062608 | rs7128408 | -0.54 | -0.159 | -0.233 | -0.54 | -0.001 | 0.025 | 0.025 | 0 | 0.55 | 0.71 | 0.51 | 0.72 | 0.5 | 0.67 |
| 11 | 98062720 | rs6589308 | -1.292 | -0.148 | -0.298 | -0.678 | 0.023 | 0.062 | 0.062 | 0 | 0.85 | 0.8 | 0.65 | 0.72 | 0.77 | 0.82 |
| 11 | 98064954 | rs4331059 | 1.088 | -0.41 | -0.161 | -0.545 | -0.005 | -0.002 | -0.002 | 0.017 | 0.31 | 0.31 | 0.35 | 0.39 | 0.38 | 0.49 |
| 11 | 98066890 | rs12420211 | 1.149 | -0.472 | -0.212 | -0.522 | -0.034 | 0.072 | 0.072 | 0 | 0.13 | 0.22 | 0.11 | 0.31 | 0.06 | 0.08 |
| 11 | 98089822 | rs2852039 | -0.495 | -0.075 | -0.401 | -0.491 | -0.003 | 0.015 | 0.015 | 0 | 0.56 | 0.69 | 0.54 | 0.72 | 0.56 | 0.7 |
| 11 | 98119560 | rs2848497 | 0.773 | -0.734 | -0.377 | -0.282 | 0 | 0 | 0 | 0 | 0.32 | 0.38 | 0.36 | 0.42 | 0.4 | 0.51 |
| 11 | 98121381 | rs2848549 | -0.883 | 0.262 | -0.549 | -0.049 | 0.02 | 0.071 | 0.071 | 0 | 0.86 | 0.8 | 0.66 | 0.73 | 0.8 | 0.83 |
| 11 | 98135872 | rs12281447 | 0.776 | 0.244 | -0.858 | -0.388 | -0.035 | 0.121 | 0.121 | 0 | 0.3 | 0.09 | 0.15 | 0 | 0.27 | 0.15 |
| 11 | 98140145 | rs2096913 | -0.347 | -0.089 | -0.033 | 1.101 | 0.002 | 0.04 | 0.04 | 0 | 0.86 | 0.9 | 0.67 | 0.61 | 0.8 | 0.83 |
| 11 | 98148532 | rs2852044 | -0.672 | 0.025 | 0.192 | 0.521 | 0.031 | 0.031 | 0.031 | 0.063 | 0.08 | 0.12 | 0.18 | 0.12 | 0.12 | 0.2 |
| 11 | 98151662 | rs2155908 | 0.156 | -0.479 | 0.301 | 0.51 | -0.036 | 0.009 | 0.009 | 0 | 0.69 | 0.57 | 0.63 | 0.72 | 0.76 | 0.82 |
| 11 | 98153408 | rs10891572 | 0.964 | -0.455 | 0.24 | 0.314 | 0.01 | -0.005 | -0.005 | 0.002 | 0.6 | 0.38 | 0.45 | 0.59 | 0.64 | 0.62 |
| 11 | 98156507 | rs10789951 | 0.846 | -0.245 | 0.265 | 0.325 | 0.001 | -0.009 | -0.009 | 0 | 0.64 | 0.36 | 0.45 | 0.59 | 0.63 | 0.62 |
| 11 | 98162569 | rs2848527 | 0.176 | -0.29 | 0.694 | 0.407 | -0.003 | 0.02 | 0.02 | 0 | 0.73 | 0.65 | 0.58 | 0.64 | 0.73 | 0.73 |
| 11 | 98164647 | rs2848511 | -0.564 | -0.303 | 0.641 | 0.411 | -0.005 | -0.023 | -0.023 | 0 | 0.84 | 0.91 | 0.66 | 0.64 | 0.79 | 0.76 |
| 11 | 98170059 | rs2852058 | -0.492 | -0.308 | 0.652 | 0.408 | -0.003 | -0.017 | -0.017 | 0 | 0.83 | 0.91 | 0.66 | 0.64 | 0.79 | 0.76 |
| 11 | 98171395 | rs2852054 | 0.304 | -0.36 | 0.792 | 0.213 | -0.008 | -0.028 | -0.028 | 0 | 0.05 | 0.13 | 0.02 | 0 | 0.04 | 0.07 |
| 11 | 98171409 | rs7944403 | 0.521 | -0.461 | 0.979 | 0.122 | -0.018 | -0.07 | -0.07 | 0 | 0.12 | 0.27 | 0.08 | 0 | 0.06 | 0.03 |
| 11 | 98171544 | rs2852053 | -0.565 | -0.341 | 0.982 | 0.191 | 0.036 | 0.02 | 0.02 | 0 | 0.08 | 0.21 | 0.13 | 0.05 | 0.1 | 0.1 |
| 11 | 98172014 | rs12287144 | 1.731 | -0.355 | 0.75 | 0.147 | -0.003 | -0.02 | -0.02 | 0 | 0.07 | 0.09 | 0.06 | 0 | 0.04 | 0.01 |
| 11 | 98176280 | rs12224719 | 1.698 | -0.602 | 0.471 | 0.013 | 0.014 | 0.166 | 0.166 | 0 | 0.49 | 0.27 | 0.38 | 0.13 | 0.49 | 0.39 |
| 11 | 98181274 | rs2000686 | 1.09 | -0.453 | 0.676 | 0.245 | 0.013 | -0.009 | -0.009 | 0 | 0.64 | 0.31 | 0.49 | 0.6 | 0.67 | 0.69 |
| 11 | 98203016 | rs7114019 | 0.587 | -0.433 | 0.837 | 0.249 | -0.013 | 0.039 | 0.039 | 0 | 0.69 | 0.55 | 0.51 | 0.6 | 0.69 | 0.7 |
| 11 | 98229261 | rs4514383 | 0.197 | -0.451 | 0.852 | 0.283 | 0.007 | -0.016 | -0.016 | 0 | 0.16 | 0.11 | 0.31 | 0.35 | 0.16 | 0.17 |
| 11 | 98229808 | rs12296006 | 0.389 | -0.238 | 0.946 | -0.181 | 0.004 | -0.014 | -0.014 | 0 | 0.13 | 0.07 | 0.06 | 0 | 0.07 | 0.11 |
| 11 | 98231052 | rs10466579 | -0.107 | -0.25 | 0.832 | -0.169 | 0.011 | -0.035 | -0.035 | 0.105 | 0.16 | 0.32 | 0.18 | 0.05 | 0.15 | 0.13 |
| 11 | 98232274 | rs12288530 | -0.061 | -0.21 | 0.553 | -0.149 | 0.006 | -0.031 | -0.031 | 0 | 0.16 | 0.29 | 0.18 | 0.05 | 0.15 | 0.13 |
| 11 | 98246423 | rs959319 | -0.629 | -0.891 | -1.26 | -0.558 | 0.055 | 0.176 | 0.176 | 0 | 0.41 | 0.17 | 0.52 | 0.93 | 0.35 | 0.57 |
| 11 | 98247743 | rs722941 | 0.764 | -1.112 | -1.405 | -0.149 | 0.012 | -0.016 | -0.016 | 0 | 0.12 | 0.06 | 0.21 | 0.16 | 0.09 | 0.15 |
| 11 | 98251769 | rs10501884 | 0.233 | -1.258 | -1.503 | 0.472 | 0.032 | -0.08 | -0.08 | 0.615 | 0.16 | 0.06 | 0.07 | 0.47 | 0.12 | 0.29 |
| 11 | 98251961 | rs10501885 | 0.346 | -1.192 | -1.546 | 0.326 | -0.014 | -0.071 | -0.071 | 0 | 0.16 | 0.04 | 0.24 | 0.3 | 0.17 | 0.22 |
| 11 | 98252991 | rs11215053 | 0.948 | -1.181 | -1.506 | 0.351 | -0.007 | -0.057 | -0.057 | 0 | 0.13 | 0.04 | 0.23 | 0.3 | 0.14 | 0.13 |
| 11 | 98254716 | rs12290455 | 0.887 | -1.237 | -1.575 | 0.414 | -0.012 | -0.061 | -0.061 | 0 | 0.13 | 0.04 | 0.24 | 0.32 | 0.15 | 0.12 |
| 11 | 98254968 | rs6589444 | 1.241 | -1.152 | -0.963 | 0.537 | 0.012 | 0.026 | 0.026 | 0.396 | 0.54 | 0.58 | 0.38 | 0.34 | 0.56 | 0.45 |
| 11 | 98255705 | rs11215072 | -0.272 | -0.569 | -1.755 | -0.484 | -0.046 | 0.099 | 0.099 | 0 | 0.57 | 0.18 | 0.43 | 0.77 | 0.5 | 0.68 |
| 11 | 98257308 | rs12418117 | 1.743 | -0.829 | -1.965 | -0.652 | -0.044 | -0.063 | -0.063 | 0.001 | 0.21 | 0.04 | 0.14 | 0.34 | 0.23 | 0.32 |
| 11 | 98261548 | rs6589446 | 1.392 | -0.711 | -1.352 | -1.7 | 0.086 | -0.135 | -0.135 | 0 | 0.2 | 0.44 | 0.19 | 0.02 | 0.29 | 0.12 |
| 11 | 98269359 | rs11215167 | -0.755 | -0.8 | -1.452 | -1.728 | -0.032 | 0.05 | 0.05 | 0 | 0.69 | 0.24 | 0.64 | 0.94 | 0.6 | 0.84 |
| 11 | 98269558 | rs11215168 | 2.711 | -0.787 | -1.391 | -1.706 | 0.001 | -0.011 | -0.011 | 0.529 | 0.08 | 0.06 | 0.06 | 0 | 0.04 | 0.01 |
| 11 | 98271679 | rs7122788 | 0.114 | -0.944 | -1.492 | -1.727 | -0.019 | 0.122 | 0.122 | 0 | 0.54 | 0.19 | 0.42 | 0.76 | 0.47 | 0.58 |
| 11 | 98275861 | rs1482230 | -0.71 | -0.866 | -1.384 | -1.858 | -0.031 | 0.048 | 0.048 | 0 | 0.69 | 0.24 | 0.63 | 0.94 | 0.58 | 0.8 |
| 11 | 98302817 | rs10790062 | 0.48 | -0.932 | -1.492 | -1.733 | -0.06 | 0.093 | 0.093 | 0 | 0.4 | 0.12 | 0.42 | 0.78 | 0.44 | 0.6 |
| 11 | 98303416 | rs2170489 | -0.861 | -0.984 | -1.34 | -1.783 | 0.007 | -0.065 | -0.065 | 0 | 0.63 | 0.51 | 0.61 | 0.83 | 0.6 | 0.72 |
| 11 | 98321764 | rs12362178 | 1.682 | -0.485 | -1.158 | -1.394 | 0.003 | -0.011 | -0.011 | 0 | 0.09 | 0.07 | 0.07 | 0 | 0.06 | 0.07 |
| 11 | 98326293 | rs11215465 | 0.66 | -0.453 | -1.337 | -1.324 | -0.05 | -0.009 | -0.009 | 0 | 0.33 | 0.07 | 0.37 | 0.73 | 0.34 | 0.44 |
| 11 | 98327692 | rs12291633 | 1.513 | -0.462 | -0.896 | -1.195 | 0.037 | 0.002 | 0.002 | 0 | 0.1 | 0.31 | 0.17 | 0.04 | 0.11 | 0.04 |
| 11 | 98329172 | rs2406082 | -1.33 | -0.657 | -1.116 | -1.337 | 0.003 | -0.02 | -0.02 | 1.008 | 0.9 | 0.95 | 0.82 | 0.82 | 0.91 | 0.84 |

| | | | | | | | | | | | | | | | | |
|---|---|---|---|---|---|---|---|---|---|---|---|---|---|---|---|---|
| 11 | 98329404 | rs11215484 | -0.008 | -0.462 | -0.979 | -1.722 | -0.001 | -0.012 | -0.012 | 0 | 0.43 | 0.42 | 0.54 | 0.78 | 0.44 | 0.47 |
| 11 | 98333595 | rs10736478 | -0.68 | -0.786 | -0.582 | -1.883 | 0.011 | 0.04 | 0.04 | 0 | 0.54 | 0.56 | 0.61 | 0.78 | 0.57 | 0.71 |
| 11 | 98333930 | rs10790073 | 0.177 | -0.277 | -1.012 | -1.924 | 0.004 | 0.017 | 0.017 | 0 | 0.39 | 0.3 | 0.49 | 0.76 | 0.38 | 0.46 |
| 11 | 98334936 | rs11215524 | 0.449 | -0.201 | -0.936 | -1.825 | -0.038 | 0.039 | 0.039 | 0 | 0.15 | 0.36 | 0.22 | 0.2 | 0.09 | 0.16 |
| 11 | 98338008 | rs10891845 | -0.719 | -0.064 | -1.794 | -1.811 | 0.005 | -0.001 | -0.001 | 0 | 0.31 | 0.65 | 0.44 | 0.26 | 0.34 | 0.39 |
| 11 | 98338477 | rs10891849 | 0.578 | -0.157 | -2.048 | -1.281 | 0.017 | -0.025 | -0.025 | 0 | 0.3 | 0.07 | 0.31 | 0.72 | 0.27 | 0.42 |
| 11 | 98342466 | rs11215566 | 0.8 | 0.45 | -1.681 | -1.06 | -0.016 | 0.004 | 0.004 | 0 | 0.1 | 0.01 | 0.09 | 0.11 | 0.12 | 0.24 |
| 11 | 98343637 | rs11215568 | 1.734 | 0.433 | -1.742 | -1.09 | -0.022 | -0.013 | -0.013 | 0 | 0.06 | 0 | 0.1 | 0.03 | 0.09 | 0.04 |
| 11 | 98346106 | rs7115435 | 0.628 | 0.314 | -1.56 | -0.883 | -0.001 | 0.004 | 0.004 | 0 | 0.54 | 0.57 | 0.66 | 0.6 | 0.53 | 0.33 |
| 11 | 98355011 | rs10128755 | -0.26 | -0.616 | -1.074 | -0.674 | 0.005 | 0.004 | 0.004 | 0.032 | 0.23 | 0.17 | 0.13 | 0.12 | 0.16 | 0.3 |
| 11 | 98357660 | rs1482224 | 0.238 | -0.209 | -1.197 | -0.427 | -0.017 | -0.001 | -0.001 | 0.886 | 0.58 | 0.57 | 0.67 | 0.5 | 0.71 | 0.5 |
| 11 | 98364555 | rs11532013 | 0.249 | 0.68 | -1.389 | -0.263 | -0.018 | 0.005 | 0.005 | 0 | 0.13 | 0.08 | 0.13 | 0.11 | 0.18 | 0.26 |
| 11 | 98365365 | rs4551724 | -1.279 | -0.114 | -1.661 | -0.119 | 0.008 | -0.076 | -0.076 | 0 | 0.8 | 0.95 | 0.63 | 0.58 | 0.81 | 0.88 |
| 11 | 98365737 | rs11215654 | 0.462 | -0.288 | -1.92 | -0.661 | -0.05 | 0.123 | 0.123 | 0.118 | 0.4 | 0.12 | 0.32 | 0.18 | 0.54 | 0.48 |
| 11 | 98367011 | rs10790091 | -0.966 | -0.791 | -1.344 | -0.818 | -0.009 | 0.01 | 0.01 | 0 | 0.67 | 0.73 | 0.53 | 0.54 | 0.75 | 0.85 |
| 11 | 98368352 | rs1994088 | 0.08 | -0.897 | -1.291 | -0.881 | -0.015 | 0.034 | 0.034 | 0.057 | 0.26 | 0.55 | 0.2 | 0.36 | 0.22 | 0.37 |
| 11 | 98368892 | rs4754513 | 0.833 | -0.863 | -1.323 | -0.887 | -0.032 | 0.115 | 0.115 | 0.066 | 0.4 | 0.16 | 0.32 | 0.18 | 0.54 | 0.48 |
| 11 | 98370464 | rs7937906 | -0.62 | -0.68 | -1.274 | -0.884 | 0 | -0.001 | -0.001 | 0.049 | 0.64 | 0.73 | 0.5 | 0.54 | 0.73 | 0.81 |
| 11 | 98377161 | rs2128234 | -1.451 | -0.089 | -0.949 | -0.309 | 0 | -0.072 | -0.072 | 0 | 0.81 | 0.95 | 0.61 | 0.57 | 0.8 | 0.87 |
| 11 | 98386085 | rs11604400 | -0.277 | -0.179 | -0.528 | -0.457 | 0.006 | -0.094 | -0.094 | 0.958 | 0.14 | 0.43 | 0.07 | 0 | 0.12 | 0.15 |
| 11 | 98405146 | rs7926318 | 0.767 | -0.249 | -0.817 | -0.145 | 0.002 | -0.006 | -0.006 | 0.704 | 0.13 | 0.12 | 0.13 | 0.42 | 0.11 | 0.29 |
| 11 | 98406723 | rs17132136 | 0.173 | -0.304 | -0.501 | -0.273 | 0.051 | 0.027 | 0.027 | 1.231 | 0.05 | 0.4 | 0.04 | 0 | 0.02 | 0.03 |
| 11 | 98407678 | rs1482219 | 0.967 | -0.267 | -0.791 | -0.54 | -0.014 | 0.116 | 0.116 | 0 | 0.47 | 0.25 | 0.37 | 0.14 | 0.51 | 0.37 |
| 11 | 98408860 | rs1351571 | -0.846 | -0.285 | -0.897 | -0.506 | -0.001 | 0.046 | 0.046 | 0 | 0.7 | 0.55 | 0.55 | 0.56 | 0.7 | 0.77 |
| 11 | 98427125 | rs4129994 | 0.852 | -0.186 | -1.433 | -0.494 | -0.006 | 0.054 | 0.054 | 0 | 0.47 | 0.3 | 0.37 | 0.14 | 0.51 | 0.37 |
| 11 | 98429292 | rs4370902 | -0.881 | 0.11 | -1.912 | -0.288 | -0.032 | -0.105 | -0.105 | 2.864 | 0.73 | 0.96 | 0.58 | 0.58 | 0.71 | 0.8 |
| 11 | 98430324 | rs11604245 | 0.002 | 0.208 | -1.357 | -0.593 | -0.02 | 0.044 | 0.044 | 0 | 0.09 | 0.02 | 0.03 | 0 | 0.09 | 0.12 |
| 11 | 98431542 | rs12365770 | 0.13 | -0.023 | -1.354 | -0.982 | -0.029 | 0.103 | 0.103 | 0 | 0.41 | 0.19 | 0.3 | 0.04 | 0.47 | 0.29 |
| 11 | 98432196 | rs10790140 | -0.327 | -0.529 | -1.211 | 0.057 | -0.001 | 0.001 | 0.001 | 0 | 0.54 | 0.74 | 0.66 | 0.51 | 0.53 | 0.47 |
| 11 | 98433120 | rs10891967 | -0.265 | -0.459 | -1.005 | 0.049 | 0.007 | 0.005 | 0.005 | 0 | 0.53 | 0.68 | 0.67 | 0.52 | 0.54 | 0.47 |
| 11 | 98435327 | rs4281453 | 0.592 | -0.773 | -1.081 | -0.094 | -0.104 | -0.029 | -0.029 | 0.493 | 0.37 | 0.06 | 0.5 | 0.4 | 0.44 | 0.42 |
| 11 | 98436532 | rs7479306 | 0.06 | -0.857 | -0.45 | -0.091 | -0.023 | -0.028 | -0.028 | 3.789 | 0.16 | 0.56 | 0.16 | 0.12 | 0.11 | 0.06 |
| 11 | 98441476 | rs10891978 | -0.038 | -0.939 | -0.713 | 0.433 | 0.032 | -0.024 | -0.024 | 0.034 | 0.62 | 0.32 | 0.64 | 0.52 | 0.74 | 0.69 |
| 11 | 98444243 | rs4531438 | 0.064 | -0.968 | -0.704 | 0.417 | 0.058 | -0.036 | -0.036 | 0.081 | 0.6 | 0.27 | 0.63 | 0.52 | 0.74 | 0.68 |
| 11 | 98444898 | rs12286584 | 0.662 | -0.883 | -0.869 | 0.465 | -0.034 | -0.058 | -0.058 | 0 | 0.24 | 0.1 | 0.15 | 0.32 | 0.29 | 0.45 |
| 11 | 98445530 | rs7127964 | 1.07 | -0.836 | -0.912 | 0.269 | -0.065 | 0.108 | 0.108 | 0 | 0.36 | 0.17 | 0.48 | 0.2 | 0.44 | 0.22 |
| 11 | 98445605 | rs4567424 | -0.926 | -1.086 | -0.05 | 0.58 | -0.045 | -0.029 | -0.029 | 0.07 | 0.74 | 0.61 | 0.82 | 0.85 | 0.85 | 0.92 |
| 11 | 98451749 | rs12420878 | -0.45 | -0.481 | -0.083 | 0.996 | -0.007 | 0.088 | 0.088 | 0.528 | 0.13 | 0.21 | 0.18 | 0.31 | 0.11 | 0.24 |
| 11 | 98454916 | rs4754525 | -0.574 | -0.418 | -0.036 | 0.868 | 0 | 0.029 | 0.029 | 0 | 0.34 | 0.42 | 0.37 | 0.48 | 0.36 | 0.4 |
| 11 | 98456485 | rs12796171 | 1.659 | -0.179 | -0.407 | 0.531 | -0.032 | -0.018 | -0.018 | 0 | 0.26 | 0.11 | 0.29 | 0.27 | 0.26 | 0.39 |
| 11 | 98457457 | rs10750092 | -0.547 | -0.301 | -0.142 | 0.695 | -0.003 | -0.004 | -0.004 | 0 | 0.64 | 0.56 | 0.68 | 0.7 | 0.66 | 0.8 |
| 11 | 98458121 | rs10790155 | 0.012 | -0.486 | -0.185 | 0.724 | 0 | -0.001 | -0.001 | 0 | 0.52 | 0.48 | 0.53 | 0.58 | 0.54 | 0.69 |
| 11 | 98461972 | rs10128544 | -0.588 | -0.418 | -0.081 | 0.717 | 0 | 0 | 0 | 0 | 0.37 | 0.31 | 0.38 | 0.43 | 0.39 | 0.41 |
| 11 | 98463122 | rs4326764 | -0.689 | -0.433 | -0.057 | 0.716 | 0 | 0.008 | 0.008 | 0 | 0.38 | 0.46 | 0.39 | 0.43 | 0.39 | 0.41 |
| 11 | 98463371 | rs7482554 | 0.151 | -0.423 | -0.076 | 0.74 | 0.011 | 0.014 | 0.014 | 0 | 0.25 | 0.35 | 0.24 | 0.31 | 0.28 | 0.3 |
| 11 | 98466388 | rs4294537 | 1.412 | -0.406 | -0.192 | 0.748 | -0.008 | -0.021 | -0.021 | 0.441 | 0.12 | 0.02 | 0.14 | 0.12 | 0.12 | 0.11 |
| 11 | 98469842 | rs11216114 | 0.089 | -0.392 | 0.048 | 0.743 | 0.011 | 0.02 | 0.02 | 0 | 0.26 | 0.44 | 0.25 | 0.31 | 0.28 | 0.29 |
| 11 | 98470675 | rs10892024 | 0.723 | -0.404 | -0.009 | 0.449 | -0.029 | -0.023 | -0.023 | 5.901 | 0.38 | 0.14 | 0.44 | 0.39 | 0.38 | 0.49 |
| 11 | 98483202 | rs7102022 | 0.251 | -0.229 | 0.059 | 0.355 | 0 | -0.004 | -0.004 | 0 | 0.46 | 0.48 | 0.42 | 0.36 | 0.44 | 0.61 |
| 11 | 98483665 | rs4604847 | -1.426 | -0.186 | 0.108 | 0.279 | 0.019 | 0.017 | 0.017 | 0 | 0.93 | 0.86 | 0.91 | 0.81 | 0.83 | 0.94 |
| 11 | 98486983 | rs7106950 | 1.863 | -0.655 | 0.389 | -0.218 | -0.003 | -0.001 | -0.001 | 0 | 0.26 | 0.21 | 0.3 | 0.27 | 0.26 | 0.41 |
| 11 | 98499049 | rs10790171 | -0.882 | -0.873 | 0.369 | -0.378 | -0.017 | -0.042 | -0.042 | 0.131 | 0.37 | 0.45 | 0.27 | 0.2 | 0.32 | 0.37 |
| 11 | 98504332 | rs11216260 | 0.571 | -0.872 | 0.433 | -0.352 | 0 | 0 | 0 | 1.573 | 0.18 | 0.16 | 0.1 | 0.09 | 0.17 | 0.15 |
| 11 | 98510653 | rs2512864 | -0.75 | -0.881 | 0.39 | -0.315 | -0.005 | -0.013 | -0.013 | 0 | 0.62 | 0.64 | 0.57 | 0.47 | 0.58 | 0.78 |
| 11 | 98513055 | rs2440570 | 1.932 | -0.76 | 0.107 | -0.332 | -0.011 | -0.003 | -0.003 | 0 | 0.25 | 0.17 | 0.3 | 0.26 | 0.26 | 0.41 |
| 11 | 98523074 | rs2406347 | -0.386 | -0.829 | 0.238 | -0.393 | -0.016 | -0.065 | -0.065 | 0 | 0.23 | 0.34 | 0.12 | 0.1 | 0.19 | 0.26 |
| 11 | 98536658 | rs7925640 | 0.117 | -0.701 | 0.022 | -0.323 | -0.003 | -0.032 | -0.032 | 3.104 | 0.19 | 0.23 | 0.11 | 0.1 | 0.17 | 0.19 |
| 11 | 98540682 | rs7944690 | -0.307 | -1.247 | -0.032 | -0.641 | -0.038 | -0.107 | -0.107 | 0 | 0.31 | 0.45 | 0.17 | 0.09 | 0.22 | 0.21 |
| 11 | 98541621 | rs7930984 | 0.275 | -0.982 | -0.123 | -0.446 | -0.001 | -0.046 | -0.046 | 1.739 | 0.18 | 0.32 | 0.12 | 0.09 | 0.17 | 0.14 |
| 11 | 98546769 | rs11216424 | 0.768 | -1.233 | -0.88 | -0.269 | -0.004 | -0.023 | -0.023 | 1.477 | 0.13 | 0.02 | 0.15 | 0.12 | 0.12 | 0.1 |
| 11 | 98549212 | rs2440571 | -1.474 | -1.013 | -0.954 | -0.643 | -0.05 | -0.023 | -0.023 | 0 | 0.75 | 0.92 | 0.71 | 0.74 | 0.74 | 0.6 |
| 11 | 98556936 | rs2435566 | 0.353 | -0.662 | -0.446 | -0.619 | -0.012 | -0.024 | -0.024 | 0 | 0.3 | 0.29 | 0.33 | 0.53 | 0.4 | 0.36 |
| 11 | 98562290 | rs1367324 | -0.806 | -0.913 | -0.547 | -0.734 | 0.019 | 0.049 | 0.049 | 0 | 0.54 | 0.71 | 0.51 | 0.67 | 0.61 | 0.56 |
| 11 | 98576161 | rs2915068 | -1.241 | -0.764 | -0.434 | -0.772 | 0 | 0.019 | 0.019 | 0 | 0.68 | 0.85 | 0.63 | 0.74 | 0.73 | 0.67 |
| 11 | 98582099 | rs1430468 | -0.065 | -0.651 | -0.002 | -1.638 | 0.024 | 0.008 | 0.008 | 0.174 | 0.15 | 0.3 | 0.12 | 0.14 | 0.16 | 0.18 |
| 11 | 98592201 | rs17132366 | 0.175 | -0.61 | -0.094 | -1.666 | -0.013 | 0.016 | 0.016 | 0.047 | 0.13 | 0.1 | 0.09 | 0.05 | 0.18 | 0.13 |
| 11 | 98593617 | rs2512830 | -1.06 | -0.747 | -0.233 | -1.784 | 0.004 | 0.011 | 0.011 | 0 | 0.64 | 0.72 | 0.61 | 0.81 | 0.72 | 0.6 |
| 11 | 98595635 | rs2435567 | -0.062 | -0.765 | -0.213 | -1.75 | 0 | -0.002 | -0.002 | 0 | 0.5 | 0.54 | 0.52 | 0.76 | 0.56 | 0.47 |
| 11 | 98607110 | rs1550714 | -0.648 | -0.893 | -0.09 | -1.79 | -0.003 | -0.051 | -0.051 | 0 | 0.25 | 0.45 | 0.19 | 0.16 | 0.22 | 0.22 |

| | | | | | | | | | | | | | | | | |
|---|---|---|---|---|---|---|---|---|---|---|---|---|---|---|---|---|
| 11 | 98608026 | rs3847560 | 0.065 | -0.763 | -0.135 | -1.791 | -0.009 | 0.031 | 0.031 | 0 | 0.43 | 0.4 | 0.47 | 0.76 | 0.53 | 0.46 |
| 11 | 98617481 | rs11216811 | 0.872 | -0.309 | -0.254 | -1.667 | 0.005 | 0 | 0 | 0.416 | 0.11 | 0.13 | 0.12 | 0.14 | 0.16 | 0.15 |
| 11 | 98621717 | rs4284374 | -0.196 | -1.241 | 0.067 | -1.452 | -0.011 | 0.03 | 0.03 | 0 | 0.41 | 0.37 | 0.45 | 0.76 | 0.53 | 0.45 |
| 11 | 98625552 | rs4542374 | -0.519 | -1.288 | 0.068 | -1.705 | 0.003 | -0.062 | -0.062 | 0 | 0.45 | 0.54 | 0.51 | 0.88 | 0.54 | 0.55 |
| 11 | 98626907 | rs7947002 | -0.459 | -1.283 | 0.07 | -1.703 | 0.005 | -0.069 | -0.069 | 0.33 | 0.44 | 0.54 | 0.5 | 0.87 | 0.54 | 0.54 |
| 11 | 98639674 | rs332759 | 0.834 | -1.587 | -0.493 | -1.876 | -0.072 | 0.177 | 0.177 | 0 | 0.38 | 0.11 | 0.44 | 0.12 | 0.4 | 0.41 |
| 11 | 98651724 | rs1973139 | -0.453 | -1.412 | -0.3 | -1.797 | -0.003 | 0.012 | 0.012 | 0.056 | 0.46 | 0.63 | 0.51 | 0.85 | 0.55 | 0.53 |
| 11 | 98656316 | rs12291224 | 1.302 | -1.291 | -0.644 | -1.283 | -0.013 | 0.008 | 0.008 | 0.039 | 0.11 | 0.27 | 0.09 | 0.08 | 0.11 | 0.07 |
| 11 | 98657134 | rs232253 | 1.26 | -0.471 | -1.317 | -0.748 | -0.002 | 0.051 | 0.051 | 0 | 0.92 | 0.7 | 0.97 | 0.9 | 0.97 | 0.86 |
| 11 | 98661321 | rs12808518 | -1.02 | -0.463 | -1.311 | -0.736 | 0.014 | 0.084 | 0.084 | 0.004 | 0.05 | 0.27 | 0.02 | 0.07 | 0.02 | 0.1 |
| 11 | 98662829 | rs2169376 | 1.087 | -0.171 | -1.792 | -0.421 | -0.044 | -0.095 | -0.095 | 0 | 0.12 | 0 | 0.21 | 0.44 | 0.15 | 0.23 |
| 11 | 98674920 | rs2219635 | -0.606 | -0.193 | -1.381 | -1.46 | -0.021 | -0.058 | -0.058 | 0 | 0.81 | 0.65 | 0.71 | 0.92 | 0.9 | 0.83 |
| 11 | 98678717 | rs334268 | 0.733 | -0.152 | -1.721 | -1.467 | -0.015 | -0.009 | -0.009 | 0.119 | 0.17 | 0.22 | 0.27 | 0.13 | 0.09 | 0.2 |
| 11 | 98682503 | rs334278 | 0.194 | -0.161 | -1.768 | -1.477 | -0.019 | -0.017 | -0.017 | 0 | 0.21 | 0.28 | 0.29 | 0.15 | 0.12 | 0.27 |
| 11 | 98685498 | rs10750135 | -0.591 | 0.107 | -2.058 | -1.325 | -0.022 | -0.016 | -0.016 | 0 | 0.8 | 0.7 | 0.72 | 0.85 | 0.9 | 0.75 |
| 11 | 98686234 | rs11217079 | 0.732 | 0.207 | -2.396 | -1.426 | -0.119 | 0.135 | 0.135 | 0 | 0.23 | 0.02 | 0.17 | 0.06 | 0.31 | 0.27 |
| 11 | 98692325 | rs11217100 | 0.698 | 0.187 | -2.426 | -1.421 | -0.105 | -0.087 | -0.087 | 0 | 0.25 | 0.02 | 0.36 | 0.72 | 0.37 | 0.34 |
| 11 | 98698667 | rs1430452 | -1.222 | 0.887 | -1.242 | -0.947 | -0.009 | 0.033 | 0.033 | 0 | 0.44 | 0.31 | 0.45 | 0.21 | 0.4 | 0.52 |
| 11 | 98699317 | rs2514231 | -1.021 | 1.13 | -1.533 | -0.761 | -0.019 | 0.064 | 0.064 | 0 | 0.4 | 0.25 | 0.44 | 0.19 | 0.39 | 0.47 |
| 11 | 98702637 | rs4754550 | 0.831 | 1.281 | -1.619 | -0.679 | -0.028 | 0.076 | 0.076 | 0 | 0.6 | 0.76 | 0.56 | 0.81 | 0.6 | 0.51 |
| 11 | 98702847 | rs12362499 | -0.628 | 2.039 | -1.736 | -0.28 | -0.112 | 0.128 | 0.128 | 0.009 | 0.22 | 0.02 | 0.16 | 0.06 | 0.29 | 0.27 |
| 11 | 98705588 | rs7925701 | -0.936 | 1.823 | -0.922 | 0.6 | 0.023 | 0.015 | 0.015 | 0 | 0.28 | 0.37 | 0.28 | 0.35 | 0.33 | 0.39 |
| 11 | 98705763 | rs7943488 | -0.873 | 1.79 | -1.279 | 0.992 | -0.011 | -0.02 | -0.02 | 0.007 | 0.27 | 0.18 | 0.19 | 0.42 | 0.32 | 0.31 |
| 11 | 98706418 | rs10892329 | -0.389 | 1.456 | -1.766 | 0.446 | -0.011 | 0.036 | 0.036 | 0 | 0.27 | 0.14 | 0.22 | 0.17 | 0.32 | 0.18 |
| 11 | 98707145 | rs11217150 | 0.813 | 1.433 | -1.305 | 0.537 | 0.012 | 0.015 | 0.015 | 0 | 0.14 | 0.21 | 0.35 | 0.17 | 0.15 | 0.25 |
| 11 | 98713041 | rs1599464 | -0.176 | 1.011 | -1.656 | 0.164 | -0.003 | 0.01 | 0.01 | 0 | 0.38 | 0.63 | 0.5 | 0.33 | 0.37 | 0.56 |
| 11 | 98715072 | rs10892331 | -1.822 | 0.898 | -1.965 | 0.914 | -0.015 | -0.033 | -0.033 | 0 | 0.94 | 1 | 0.96 | 0.91 | 0.95 | 0.94 |
| 11 | 98717717 | rs1487096 | 0.02 | 0.63 | -2.158 | 0.731 | -0.004 | -0.004 | -0.004 | 0 | 0.52 | 0.67 | 0.43 | 0.58 | 0.51 | 0.45 |
| 11 | 98719996 | rs7103168 | -0.587 | 0.5 | -2.119 | 0.818 | 0.001 | 0.02 | 0.02 | 0 | 0.89 | 0.9 | 0.86 | 0.58 | 0.92 | 0.83 |
| 11 | 98726057 | rs11500866 | 0.001 | 0.385 | -1.828 | 0.746 | -0.002 | -0.008 | -0.008 | 0.1 | 0.6 | 0.61 | 0.63 | 0.45 | 0.59 | 0.66 |
| 11 | 98733359 | rs7934855 | -0.694 | 0.467 | -1.806 | 0.775 | -0.002 | 0.055 | 0.055 | 0 | 0.89 | 0.88 | 0.87 | 0.58 | 0.92 | 0.84 |
| 11 | 98733632 | rs4754552 | -0.142 | 0.411 | -1.78 | 0.715 | -0.004 | -0.014 | -0.014 | 0 | 0.61 | 0.65 | 0.64 | 0.46 | 0.59 | 0.66 |
| 11 | 98740074 | rs11217277 | 0.224 | 0.138 | -1.587 | 0.2 | -0.012 | -0.034 | -0.034 | 0 | 0.34 | 0.52 | 0.19 | 0.21 | 0.29 | 0.18 |
| 11 | 98740540 | rs12289864 | 0.492 | 0.146 | -1.587 | 0.213 | -0.011 | -0.038 | -0.038 | 0.013 | 0.32 | 0.49 | 0.19 | 0.2 | 0.29 | 0.17 |
| 11 | 98743375 | rs10750141 | -0.16 | -0.051 | -1.837 | 0.284 | 0 | 0.02 | 0.02 | 0 | 0.59 | 0.76 | 0.42 | 0.34 | 0.62 | 0.34 |
| 11 | 98759120 | rs2129516 | 0 | -0.048 | -1.808 | 0.385 | 0.013 | 0.01 | 0.01 | 0 | 0.55 | 0.35 | 0.66 | 0.38 | 0.62 | 0.65 |
| 11 | 98763585 | rs4753921 | 0.557 | -0.043 | -1.904 | 0.362 | -0.011 | 0.002 | 0.002 | 0 | 0.29 | 0.12 | 0.42 | 0.25 | 0.29 | 0.49 |
| 11 | 98766350 | rs10790333 | 0.527 | -0.211 | -2.02 | 0.262 | -0.002 | -0.043 | -0.043 | 0 | 0.33 | 0.17 | 0.32 | 0.46 | 0.35 | 0.28 |
| 11 | 98766571 | rs2219636 | 2.128 | -0.232 | -2.126 | 0.13 | -0.062 | 0.069 | 0.069 | 0 | 0.14 | 0.01 | 0.1 | 0.03 | 0.16 | 0.17 |
| 11 | 98766663 | rs10790334 | 0.993 | -0.723 | -1.862 | 0.354 | -0.005 | 0.067 | 0.067 | 0 | 0.28 | 0.13 | 0.21 | 0.13 | 0.3 | 0.16 |
| 11 | 98768543 | rs10790336 | -0.284 | -0.855 | -1.091 | 0.289 | -0.004 | 0.074 | 0.074 | 0.116 | 0.84 | 0.57 | 0.87 | 0.81 | 0.87 | 0.83 |
| 11 | 98769434 | rs992642 | 0.587 | -0.874 | -1.145 | 0.252 | 0 | -0.044 | -0.044 | 0.172 | 0.32 | 0.16 | 0.29 | 0.45 | 0.33 | 0.25 |
| 11 | 98770072 | rs10790340 | -0.036 | -0.817 | -1.249 | 0.1 | 0.006 | 0.009 | 0.009 | 0 | 0.54 | 0.42 | 0.6 | 0.35 | 0.57 | 0.61 |
| 11 | 98772897 | rs11217465 | 1.037 | -0.393 | -1.226 | 0.119 | -0.013 | 0.016 | 0.016 | 0 | 0.31 | 0.13 | 0.42 | 0.25 | 0.3 | 0.5 |
| 11 | 98774102 | rs7945465 | 1.048 | -0.383 | -1.235 | 0.12 | -0.013 | 0.016 | 0.016 | 0 | 0.3 | 0.12 | 0.42 | 0.25 | 0.3 | 0.5 |
| 11 | 98781916 | rs11217535 | 0.761 | -0.416 | -0.946 | 0.061 | 0.001 | -0.016 | -0.016 | 0 | 0.24 | 0.29 | 0.17 | 0.1 | 0.25 | 0.09 |
| 11 | 98781993 | rs12278762 | 0.217 | -0.362 | -0.849 | 0.154 | -0.002 | -0.005 | -0.005 | 0 | 0.55 | 0.5 | 0.59 | 0.35 | 0.54 | 0.6 |
| 11 | 98782068 | rs2406492 | 0.076 | -0.364 | -0.856 | 0.23 | -0.003 | 0.022 | 0.022 | 0 | 0.37 | 0.43 | 0.35 | 0.57 | 0.4 | 0.31 |
| 11 | 98792975 | rs11217598 | 0.261 | -0.303 | -0.747 | 0.107 | -0.003 | -0.022 | -0.022 | 0 | 0.34 | 0.41 | 0.26 | 0.24 | 0.37 | 0.23 |
| 11 | 98794453 | rs1032189 | 0.17 | -0.294 | -0.688 | 0.035 | -0.002 | -0.005 | -0.005 | 0 | 0.45 | 0.43 | 0.57 | 0.69 | 0.41 | 0.68 |
| 11 | 98795150 | rs12419012 | 0.72 | -0.265 | -0.789 | 0.061 | -0.008 | -0.055 | -0.055 | 0 | 0.37 | 0.15 | 0.52 | 0.58 | 0.34 | 0.6 |
| 11 | 98799267 | rs1352236 | -0.307 | -0.285 | -0.753 | 0.08 | 0.002 | -0.011 | -0.011 | 0.287 | 0.7 | 0.56 | 0.78 | 0.82 | 0.71 | 0.83 |
| 11 | 98800568 | rs1825759 | -1.463 | -0.35 | -0.847 | 0.302 | -0.002 | 0.004 | 0.004 | 0 | 0.93 | 0.93 | 0.97 | 0.92 | 0.96 | 0.95 |
| 11 | 98805168 | rs35434121 | -0.012 | -0.266 | -0.699 | 0.345 | 0.02 | 0.065 | 0.065 | 0.01 | 0.1 | 0.38 | 0.06 | 0.11 | 0.07 | 0.08 |
| 11 | 98807358 | rs901538 | -0.212 | -0.166 | -1.667 | 0.415 | 0.028 | 0.038 | 0.038 | 0 | 0.54 | 0.22 | 0.69 | 0.68 | 0.6 | 0.73 |
| 11 | 98814350 | rs10501893 | 0.341 | 0.202 | -1.62 | 0.362 | -0.053 | -0.042 | -0.042 | 0 | 0.33 | 0.11 | 0.45 | 0.37 | 0.37 | 0.46 |
| 11 | 98816710 | rs12797100 | 1.107 | 0.18 | -1.399 | -0.147 | -0.003 | 0 | 0 | 0 | 0.16 | 0.12 | 0.14 | 0.02 | 0.2 | 0.14 |
| 11 | 98822220 | rs7948801 | -0.054 | -0.076 | -0.95 | 0.191 | -0.014 | 0.08 | 0.08 | 0 | 0.13 | 0.55 | 0.1 | 0.2 | 0.07 | 0.12 |
| 11 | 98830631 | rs6589809 | -0.983 | -0.399 | -2.132 | -0.198 | -0.027 | 0.152 | 0.152 | 0.641 | 0.69 | 0.91 | 0.79 | 0.93 | 0.69 | 0.85 |
| 11 | 98831017 | rs17561441 | 0.698 | -0.14 | -2.187 | 0.293 | -0.053 | -0.012 | -0.012 | 0 | 0.35 | 0.12 | 0.47 | 0.33 | 0.37 | 0.46 |
| 11 | 98834155 | rs2129521 | 1.067 | 0.942 | -1.546 | 0.825 | -0.084 | 0.023 | 0.023 | 0.315 | 0.14 | 0.01 | 0.26 | 0.09 | 0.22 | 0.1 |
| 11 | 98850050 | rs11217903 | -0.035 | 0.388 | -0.801 | 1.434 | -0.001 | 0.077 | 0.077 | 0 | 0.06 | 0.06 | 0.11 | 0.34 | 0.05 | 0.13 |
| 11 | 98850985 | rs901540 | 1.231 | 0.369 | -0.85 | 0.841 | 0.033 | 0.074 | 0.074 | 0 | 0.51 | 0.19 | 0.59 | 0.37 | 0.58 | 0.65 |
| 11 | 98869223 | rs4393278 | -0.41 | 0.404 | -0.748 | 0.049 | -0.039 | 0.1 | 0.1 | 0.333 | 0.8 | 0.94 | 0.85 | 0.96 | 0.78 | 0.91 |
| 11 | 98879688 | rs12274220 | -1.122 | 0.26 | 0.414 | 0.895 | 0.082 | 0.201 | 0.201 | 2.293 | 0.05 | 0.44 | 0.04 | 0.1 | 0 | 0.03 |
| 11 | 98885233 | rs12224476 | -1.199 | 1.169 | -0.152 | 1.354 | 0.034 | 0.107 | 0.107 | 0 | 0.25 | 0.44 | 0.26 | 0.45 | 0.27 | 0.21 |
| 11 | 98888275 | rs10501894 | -1.312 | 1.116 | -0.068 | 1.961 | 0.007 | 0.117 | 0.117 | 0 | 0.12 | 0.2 | 0.16 | 0.48 | 0.15 | 0.17 |
| 11 | 98890724 | rs4754573 | -0.219 | 1.677 | -0.683 | 1.175 | -0.071 | 0.069 | 0.069 | 0 | 0.81 | 0.95 | 0.86 | 0.95 | 0.74 | 0.79 |
| 11 | 98907603 | rs10501897 | -0.937 | 1.051 | 0.383 | 1.859 | 0.028 | 0.087 | 0.087 | 0 | 0.15 | 0.28 | 0.21 | 0.35 | 0.23 | 0.16 |

| | | | | | | | | | | | | | | | | |
|---|---|---|---|---|---|---|---|---|---|---|---|---|---|---|---|---|
| 11 | 98909976 | rs7125442 | -1.757 | 0.967 | 0.58 | 2.027 | 0.02 | 0.136 | 0.136 | 0 | 0.2 | 0.5 | 0.4 | 0.57 | 0.24 | 0.24 |
| 11 | 98911469 | rs7121873 | -1.95 | 0.86 | 0.39 | 1.832 | 0.01 | 0.031 | 0.031 | 0 | 0.25 | 0.44 | 0.29 | 0.36 | 0.33 | 0.22 |
| 11 | 98911551 | rs10892675 | -1.333 | 0.533 | -0.2 | 1.427 | -0.012 | 0.053 | 0.053 | 0 | 0.12 | 0.23 | 0.23 | 0.22 | 0.12 | 0.14 |
| 11 | 98929559 | rs11218250 | 0.322 | 0.56 | -0.403 | 0.358 | 0 | -0.017 | -0.017 | 1.226 | 0.56 | 0.4 | 0.43 | 0.24 | 0.46 | 0.47 |
| 11 | 98930198 | rs10750185 | 0.046 | 0.094 | 0.207 | 0.847 | 0.003 | 0 | 0 | 0 | 0.66 | 0.63 | 0.59 | 0.61 | 0.57 | 0.65 |
| 11 | 98932132 | rs7102254 | 0.458 | -0.993 | 0.313 | 0.282 | -0.001 | -0.013 | -0.013 | 0 | 0.48 | 0.46 | 0.41 | 0.16 | 0.48 | 0.47 |
| 11 | 98937331 | rs12419466 | 0.096 | -1.382 | 0.443 | 0.269 | -0.011 | -0.001 | -0.001 | 0.047 | 0.07 | 0.08 | 0.01 | 0.08 | 0.02 | 0.02 |
| 11 | 98941872 | rs1940484 | -0.479 | -2.052 | 0.25 | -0.5 | -0.017 | 0.006 | 0.006 | 0.125 | 0.41 | 0.46 | 0.44 | 0.53 | 0.33 | 0.48 |
| 11 | 98949898 | rs12791395 | 0.894 | -2.406 | 0.279 | -0.781 | -0.033 | 0.083 | 0.083 | 0 | 0.23 | 0.1 | 0.27 | 0.08 | 0.24 | 0.29 |
| 11 | 98951110 | rs11218323 | -1.17 | -2.4 | 0.45 | -0.488 | -0.09 | 0.163 | 0.163 | 3.994 | 0.2 | 0.37 | 0.16 | 0.43 | 0.1 | 0.19 |
| 11 | 98953367 | rs7120011 | -0.799 | -2.358 | 0.555 | -0.416 | -0.092 | 0.177 | 0.177 | 5.874 | 0.17 | 0.32 | 0.12 | 0.42 | 0.04 | 0.17 |
| 11 | 98959531 | rs1940521 | -0.12 | -1.41 | 0.301 | -1.047 | 0.024 | 0.037 | 0.037 | 0.005 | 0.35 | 0.46 | 0.57 | 0.48 | 0.38 | 0.39 |
| 11 | 98967477 | rs7124219 | 0.349 | -1.591 | 0.382 | -1.077 | -0.036 | -0.005 | -0.005 | 0 | 0.09 | 0.17 | 0.02 | 0.09 | 0.03 | 0.02 |
| 11 | 98970170 | rs4237593 | -1.161 | -1.55 | 0.311 | -1.303 | 0.003 | 0.016 | 0.016 | 0.219 | 0.51 | 0.59 | 0.62 | 0.59 | 0.47 | 0.49 |
| 11 | 98976460 | rs10790462 | -0.788 | -1.567 | 0.319 | -1.294 | -0.002 | 0.002 | 0.002 | 0.025 | 0.44 | 0.55 | 0.61 | 0.59 | 0.43 | 0.42 |
| 11 | 98981254 | rs2186834 | -1.505 | -1.835 | 0.325 | -1.367 | -0.008 | 0.014 | 0.014 | 0 | 0.54 | 0.66 | 0.63 | 0.61 | 0.47 | 0.49 |
| 11 | 98983806 | rs7110775 | -0.453 | -1.655 | 0.352 | -1.294 | -0.001 | -0.001 | -0.001 | 0 | 0.43 | 0.52 | 0.6 | 0.59 | 0.42 | 0.4 |
| 11 | 98986934 | rs1940516 | 0.333 | -1.152 | 0.155 | -1.585 | 0.003 | 0 | 0 | 0.143 | 0.4 | 0.42 | 0.29 | 0.39 | 0.33 | 0.41 |
| 11 | 98988308 | rs4753939 | -0.306 | -1.088 | 0.115 | -1.612 | -0.001 | -0.001 | -0.001 | 0 | 0.56 | 0.41 | 0.67 | 0.61 | 0.58 | 0.54 |
| 11 | 98991226 | rs17133066 | -0.302 | -0.608 | 0.068 | -1.612 | 0.013 | -0.053 | -0.053 | 1.221 | 0.11 | 0.21 | 0.07 | 0 | 0.13 | 0.09 |
| 11 | 98995954 | rs7109365 | -2.467 | -0.657 | -0.626 | -1.658 | -0.025 | 0.059 | 0.059 | 0 | 0.81 | 0.96 | 0.85 | 0.95 | 0.8 | 0.88 |
| 11 | 98996775 | rs17133088 | 0.992 | -0.578 | -0.479 | -1.587 | -0.001 | -0.031 | -0.031 | 1.435 | 0.07 | 0.1 | 0.04 | 0 | 0.06 | 0.04 |
| 11 | 98996825 | rs11218492 | -0.472 | -0.659 | -0.487 | -1.639 | 0.002 | -0.004 | -0.004 | 0.009 | 0.44 | 0.49 | 0.62 | 0.59 | 0.5 | 0.45 |
| 11 | 98997953 | rs7946983 | 0.782 | -0.786 | -0.651 | -1.574 | -0.004 | 0.015 | 0.015 | 0 | 0.72 | 0.71 | 0.68 | 0.58 | 0.74 | 0.75 |
| 11 | 98998478 | rs17572676 | 1.143 | -0.692 | -1.143 | -1.568 | -0.01 | 0.058 | 0.058 | 0 | 0.1 | 0.02 | 0.04 | 0.01 | 0.11 | 0.12 |
| 11 | 98999681 | rs12365183 | 1.5 | -0.613 | -1.094 | -1.556 | -0.036 | 0.032 | 0.032 | 0 | 0.23 | 0.1 | 0.32 | 0.19 | 0.3 | 0.28 |
| 11 | 99007487 | rs755042 | 0.875 | -0.712 | -0.948 | -1.545 | 0.004 | 0.014 | 0.014 | 0.684 | 0.45 | 0.32 | 0.34 | 0.39 | 0.44 | 0.47 |
| 11 | 99012486 | rs1940508 | 0.087 | -0.75 | -0.794 | -1.505 | 0.012 | -0.003 | -0.003 | 0 | 0.53 | 0.62 | 0.53 | 0.53 | 0.56 | 0.63 |
| 11 | 99016124 | rs1940496 | -0.26 | -0.569 | -1.085 | -1.635 | -0.018 | 0.081 | 0.081 | 0 | 0.83 | 0.95 | 0.92 | 0.99 | 0.81 | 0.84 |
| 11 | 99016229 | rs10892814 | 2.289 | -0.68 | -1.063 | -1.675 | 0 | 0.009 | 0.009 | 0 | 0.18 | 0.16 | 0.15 | 0.05 | 0.18 | 0.12 |
| 11 | 99019894 | rs11218556 | -0.77 | -0.787 | -1.197 | -1.722 | -0.019 | -0.038 | -0.038 | 0 | 0.24 | 0.1 | 0.34 | 0.66 | 0.27 | 0.42 |
| 11 | 99021560 | rs4402263 | -0.855 | -0.788 | -1.164 | -1.732 | -0.007 | 0.027 | 0.027 | 0 | 0.68 | 0.76 | 0.55 | 0.29 | 0.65 | 0.59 |
| 11 | 99025195 | rs11218571 | 0.165 | -0.83 | -1.13 | -1.734 | -0.006 | -0.041 | -0.041 | 0 | 0.15 | 0.09 | 0.3 | 0.66 | 0.16 | 0.29 |
| 11 | 99025438 | rs4369367 | -1.135 | -0.815 | -1.198 | -1.762 | -0.002 | 0.067 | 0.067 | 0 | 0.71 | 0.83 | 0.8 | 0.94 | 0.69 | 0.75 |
| 11 | 99029229 | rs10790494 | -0.333 | -0.724 | -1.154 | -1.687 | -0.054 | 0.144 | 0.144 | 0.663 | 0.59 | 0.84 | 0.59 | 0.81 | 0.51 | 0.55 |
| 11 | 99031587 | rs7127410 | -0.089 | -0.728 | -1.142 | -1.645 | 0.018 | 0.046 | 0.046 | 0.091 | 0.41 | 0.69 | 0.44 | 0.76 | 0.33 | 0.43 |
| 11 | 99032560 | rs11218589 | 1.433 | -0.649 | -1.085 | -1.589 | 0.002 | 0.033 | 0.033 | 0 | 0.24 | 0.18 | 0.19 | 0.05 | 0.23 | 0.16 |
| 11 | 99032846 | rs12793470 | 0.995 | -0.05 | -0.907 | -1.076 | -0.03 | 0.043 | 0.043 | 0 | 0.26 | 0.11 | 0.32 | 0.18 | 0.31 | 0.28 |
| 11 | 99033861 | rs10790497 | 0.823 | -0.132 | -0.527 | -1.002 | -0.007 | -0.039 | -0.039 | 0.016 | 0.41 | 0.46 | 0.26 | 0.14 | 0.35 | 0.26 |
| 11 | 99033958 | rs11218595 | 0.327 | 0.19 | -0.654 | -0.081 | -0.001 | -0.038 | -0.038 | 0 | 0.11 | 0.05 | 0.09 | 0.35 | 0.1 | 0.22 |
| 11 | 99035893 | rs7949592 | -0.27 | 0.645 | -0.484 | -0.23 | -0.002 | 0.01 | 0.01 | 0 | 0.17 | 0.2 | 0.16 | 0.37 | 0.14 | 0.19 |
| 11 | 99036199 | rs11218600 | -0.245 | 0.645 | -0.482 | -0.23 | 0.001 | -0.001 | -0.001 | 0 | 0.17 | 0.18 | 0.16 | 0.37 | 0.14 | 0.18 |
| 11 | 99039503 | rs7122320 | -1.112 | 0.803 | -0.298 | -0.881 | -0.002 | -0.082 | -0.082 | 0 | 0.66 | 0.57 | 0.75 | 0.94 | 0.66 | 0.8 |
| 11 | 99039687 | rs3898589 | 1.153 | 1.101 | -0.454 | -1.044 | 0.003 | -0.045 | -0.045 | 0.108 | 0.24 | 0.28 | 0.25 | 0.13 | 0.24 | 0.26 |
| 11 | 99040272 | rs4754592 | -0.564 | 1.113 | -0.469 | -0.974 | -0.002 | 0.099 | 0.099 | 0.11 | 0.43 | 0.29 | 0.49 | 0.82 | 0.42 | 0.54 |
| 11 | 99041548 | rs7120353 | 0.46 | 1.209 | -0.322 | -0.971 | 0.049 | -0.039 | -0.039 | 0 | 0.08 | 0.25 | 0.06 | 0 | 0.08 | 0.04 |
| 11 | 99041900 | rs17133195 | -0.554 | 1.419 | -0.403 | -0.881 | 0.06 | 0.204 | 0.204 | 0.005 | 0.13 | 0.28 | 0.31 | 0.38 | 0.16 | 0.22 |
| 11 | 99041999 | rs12789966 | 0.081 | 1.359 | -0.473 | -0.952 | 0.001 | 0.003 | 0.003 | 0 | 0.31 | 0.28 | 0.38 | 0.39 | 0.35 | 0.37 |
| 11 | 99046833 | rs11218644 | 2.432 | 1.222 | -0.659 | -0.974 | 0.03 | 0.076 | 0.076 | 0.696 | 0.23 | 0.1 | 0.15 | 0.05 | 0.18 | 0.13 |
| 11 | 99055082 | rs10892860 | -1.085 | 1.323 | -0.065 | -1.092 | 0.017 | -0.052 | -0.052 | 0 | 0.74 | 0.67 | 0.71 | 0.86 | 0.68 | 0.71 |
| 11 | 99055255 | rs10892862 | -0.029 | 1.303 | 0.126 | -1.035 | 0.005 | 0.002 | 0.002 | 0.297 | 0.38 | 0.35 | 0.25 | 0.37 | 0.33 | 0.33 |
| 11 | 99057854 | rs17133210 | 1.232 | 1.372 | 0.073 | -0.905 | 0 | -0.007 | -0.007 | 0 | 0.21 | 0.18 | 0.23 | 0.14 | 0.24 | 0.24 |
| 11 | 99068364 | rs1461688 | -1.131 | 1.314 | -0.053 | -0.982 | -0.029 | 0.001 | 0.001 | 0.116 | 0.44 | 0.69 | 0.62 | 0.58 | 0.46 | 0.49 |
| 11 | 99069209 | rs7952041 | 1.319 | 1.348 | -0.133 | -0.92 | -0.004 | -0.029 | -0.029 | 0 | 0.09 | 0.04 | 0.09 | 0.36 | 0.09 | 0.17 |
| 11 | 99069944 | rs11218775 | -0.507 | 1.384 | 0.142 | -0.914 | 0.02 | 0.103 | 0.103 | 0.168 | 0.18 | 0.28 | 0.37 | 0.45 | 0.2 | 0.25 |
| 11 | 99077152 | rs10892905 | 0.2 | 1.345 | 0.202 | -1.086 | 0.003 | -0.02 | -0.02 | 0 | 0.58 | 0.43 | 0.5 | 0.25 | 0.54 | 0.44 |
| 11 | 99079035 | rs1551779 | 1.475 | 1.334 | 0.134 | -1.108 | 0.011 | 0.089 | 0.089 | 0.065 | 0.44 | 0.26 | 0.39 | 0.18 | 0.42 | 0.37 |
| 11 | 99080772 | rs10892912 | 1.411 | 1.306 | 0.089 | -1.137 | -0.002 | 0 | 0 | 0 | 0.21 | 0.16 | 0.22 | 0.14 | 0.23 | 0.24 |
| 11 | 99082926 | rs12793987 | 1.579 | 1.265 | -0.127 | -1.066 | -0.034 | 0.127 | 0.127 | 0.019 | 0.19 | 0.01 | 0.09 | 0.01 | 0.19 | 0.17 |
| 11 | 99085621 | rs7928907 | -0.493 | 1.313 | 0.092 | -0.929 | -0.01 | -0.02 | -0.02 | 0.014 | 0.7 | 0.7 | 0.63 | 0.61 | 0.66 | 0.63 |
| 11 | 99086508 | rs11218881 | 2.625 | 1.163 | -0.009 | -1.027 | 0.03 | 0.073 | 0.073 | 1.006 | 0.24 | 0.1 | 0.17 | 0.05 | 0.2 | 0.13 |
| 11 | 99087084 | rs4381339 | -0.858 | 1.107 | 0.199 | -0.889 | -0.002 | -0.007 | -0.007 | 0.476 | 0.74 | 0.74 | 0.86 | 0.99 | 0.72 | 0.8 |
| 11 | 99088047 | rs7103056 | 0.727 | 1.103 | 0.205 | -0.811 | -0.004 | 0.114 | 0.114 | 0 | 0.12 | 0.19 | 0.12 | 0.35 | 0.1 | 0.16 |
| 11 | 99089028 | rs898777 | -0.644 | 1.2 | -0.081 | -1.068 | 0 | -0.039 | -0.039 | 0 | 0.73 | 0.68 | 0.85 | 0.99 | 0.72 | 0.79 |
| 11 | 99093769 | rs1461687 | 0.78 | 1.637 | -0.031 | -0.125 | -0.001 | -0.01 | -0.01 | 0.086 | 0.53 | 0.54 | 0.47 | 0.26 | 0.48 | 0.39 |
| 11 | 99094400 | rs7940419 | 1.079 | 1.69 | -0.02 | -0.276 | -0.002 | -0.048 | -0.048 | 0 | 0.5 | 0.53 | 0.31 | 0.06 | 0.41 | 0.33 |
| 11 | 99095666 | rs11218913 | 0.58 | 1.849 | 0.127 | 0.018 | 0.002 | 0.001 | 0.001 | 0 | 0.72 | 0.72 | 0.75 | 0.63 | 0.68 | 0.74 |
| 11 | 99096024 | rs17133283 | 0.523 | 1.668 | 0.138 | -0.063 | 0.001 | -0.011 | -0.011 | 0 | 0.1 | 0.1 | 0.06 | 0.07 | 0.09 | 0.06 |
| 11 | 99096479 | rs1461684 | -0.127 | 1.653 | 0.306 | -0.035 | -0.008 | -0.044 | -0.044 | 0 | 0.19 | 0.26 | 0.12 | 0.07 | 0.15 | 0.09 |

| | | | | | | | | | | | | | | | | |
|---|---|---|---|---|---|---|---|---|---|---|---|---|---|---|---|---|
| 11 | 99103077 | rs7952130 | -1.275 | 1.767 | 0.277 | 0.211 | 0.008 | 0.024 | 0.024 | 0 | 0.34 | 0.48 | 0.55 | 0.6 | 0.32 | 0.4 |
| 11 | 99112911 | rs970321 | 1.049 | 1.644 | 0.134 | 0.18 | -0.003 | 0.016 | 0.016 | 0 | 0.09 | 0.05 | 0.09 | 0.1 | 0.1 | 0.14 |
| 11 | 99126016 | rs6589988 | -0.302 | 1.779 | 0.27 | 0.031 | -0.001 | -0.018 | -0.018 | 0.126 | 0.35 | 0.36 | 0.23 | 0.21 | 0.32 | 0.35 |
| 11 | 99134767 | rs7944167 | 2.69 | 1.588 | 0.268 | -0.186 | 0.011 | 0.014 | 0.014 | 1.435 | 0.25 | 0.19 | 0.15 | 0.05 | 0.21 | 0.13 |
| 11 | 99140232 | rs2511798 | -1.702 | 1.655 | 0.233 | -0.14 | 0.013 | 0.076 | 0.076 | 0.489 | 0.53 | 0.76 | 0.68 | 0.72 | 0.52 | 0.65 |
| 11 | 99141095 | rs2510682 | -1.37 | 1.705 | -0.005 | -0.063 | -0.07 | -0.054 | -0.054 | 0.06 | 0.78 | 0.97 | 0.83 | 0.78 | 0.73 | 0.78 |
| 11 | 99149792 | rs1461670 | 1.597 | 1.844 | 0.056 | -0.073 | -0.01 | 0.016 | 0.016 | 0.567 | 0.16 | 0.08 | 0.11 | 0.14 | 0.19 | 0.26 |
| 11 | 99152801 | rs17577073 | -0.111 | 1.82 | 0.014 | -0.243 | -0.005 | 0.136 | 0.136 | 0 | 0.44 | 0.18 | 0.26 | 0.22 | 0.42 | 0.41 |
| 11 | 99158043 | rs11219206 | -1.44 | 1.94 | 0.344 | 0.048 | 0 | -0.002 | -0.002 | 0 | 0.5 | 0.51 | 0.64 | 0.72 | 0.51 | 0.65 |
| 11 | 99160502 | rs12283068 | -0.72 | 1.993 | 0.148 | 0.112 | 0.004 | -0.005 | -0.005 | 1.237 | 0.41 | 0.55 | 0.55 | 0.64 | 0.4 | 0.52 |
| 11 | 99184602 | rs10501905 | 1.15 | 2.107 | -0.263 | 0.01 | -0.006 | 0.02 | 0.02 | 0 | 0.09 | 0.05 | 0.09 | 0.09 | 0.09 | 0.12 |
| 11 | 99187652 | rs922264 | 1.484 | 2.224 | -0.653 | 0.123 | -0.032 | 0.021 | 0.021 | 0.355 | 0.16 | 0.06 | 0.11 | 0.14 | 0.19 | 0.24 |
| 11 | 99192744 | rs17661249 | -0.308 | 2.177 | -0.602 | 0.158 | -0.057 | 0.109 | 0.109 | 0.361 | 0.34 | 0.1 | 0.22 | 0.21 | 0.36 | 0.34 |
| 11 | 99193546 | rs11828781 | 2.02 | 1.807 | -0.381 | 0.221 | -0.014 | -0.036 | -0.036 | 0.044 | 0.27 | 0.26 | 0.15 | 0.05 | 0.21 | 0.12 |
| 11 | 99198902 | rs7947488 | -0.184 | 1.922 | -0.821 | 0.329 | -0.054 | 0.102 | 0.102 | 0.411 | 0.33 | 0.1 | 0.21 | 0.21 | 0.34 | 0.32 |
| 11 | 99199688 | rs282494 | -0.483 | 1.935 | -0.714 | 0.305 | 0.003 | -0.027 | -0.027 | 0 | 0.64 | 0.53 | 0.41 | 0.26 | 0.63 | 0.47 |
| 11 | 99199746 | rs282496 | 1.403 | 1.608 | -0.463 | 0.394 | -0.019 | -0.01 | -0.01 | 0.675 | 0.31 | 0.44 | 0.23 | 0.25 | 0.3 | 0.15 |
| 11 | 99208859 | rs2035122 | 0.242 | 1.514 | -1.267 | 0.371 | -0.001 | 0.006 | 0.006 | 0 | 0.29 | 0.09 | 0.23 | 0.3 | 0.28 | 0.35 |
| 11 | 99217421 | rs7482232 | 0.435 | 1.933 | -0.99 | 0.368 | -0.047 | 0.026 | 0.026 | 0.041 | 0.17 | 0.06 | 0.14 | 0.14 | 0.23 | 0.26 |
| 11 | 99219917 | rs1504737 | 0.394 | 1.664 | -1.162 | 0.296 | -0.003 | 0.033 | 0.033 | 0 | 0.35 | 0.18 | 0.37 | 0.29 | 0.38 | 0.43 |
| 11 | 99220141 | rs10893173 | 1.344 | 1.216 | -1.035 | 0.64 | 0.031 | -0.029 | -0.029 | 0 | 0.26 | 0.18 | 0.17 | 0.29 | 0.2 | 0.14 |
| 11 | 99224406 | rs9645670 | 0.407 | 1.408 | -1.109 | 0.657 | -0.016 | -0.005 | -0.005 | 0 | 0.42 | 0.14 | 0.44 | 0.63 | 0.42 | 0.43 |
| 11 | 99229526 | rs11219590 | -1.237 | 1.667 | -0.552 | 0.466 | -0.003 | 0.008 | 0.008 | 0.178 | 0.46 | 0.88 | 0.68 | 0.49 | 0.53 | 0.56 |
| 11 | 99231737 | rs7930825 | -1.16 | 1.911 | -0.604 | 0.44 | -0.038 | -0.001 | -0.001 | 0 | 0.45 | 0.88 | 0.68 | 0.49 | 0.52 | 0.53 |
| 11 | 99241675 | rs2102272 | 2.188 | 1.596 | -0.831 | 0.685 | 0.034 | -0.055 | -0.055 | 0 | 0.21 | 0.04 | 0.2 | 0.2 | 0.17 | 0.14 |
| 11 | 99244104 | rs4754610 | 0.351 | 2.187 | -0.497 | 0.562 | -0.005 | 0.004 | 0.004 | 0 | 0.21 | 0.18 | 0.23 | 0.24 | 0.24 | 0.34 |
| 11 | 99247719 | rs7952501 | 0.566 | 2.122 | 0.304 | 0.459 | 0.01 | -0.018 | -0.018 | 0 | 0.09 | 0.34 | 0.07 | 0.05 | 0.05 | 0.08 |
| 11 | 99249346 | rs17133433 | 0.114 | 2.274 | 0.313 | 0.417 | 0.079 | -0.084 | -0.084 | 2.841 | 0.13 | 0.3 | 0.13 | 0 | 0.19 | 0.08 |
| 11 | 99252012 | rs1504708 | -0.463 | 2.188 | 0.141 | 0.463 | 0 | 0.009 | 0.009 | 0 | 0.54 | 0.56 | 0.66 | 0.69 | 0.51 | 0.63 |
| 11 | 99262251 | rs11219741 | 2.157 | 2.111 | -0.229 | 0.192 | 0.035 | -0.059 | -0.059 | 0 | 0.22 | 0.04 | 0.2 | 0.21 | 0.18 | 0.14 |
| 11 | 99263928 | rs11219751 | 1.545 | 2.014 | -0.181 | 0.114 | -0.042 | 0.054 | 0.054 | 0 | 0.3 | 0.07 | 0.2 | 0.18 | 0.3 | 0.21 |
| 11 | 99268617 | rs1504721 | 0.574 | 1.849 | -0.108 | 0.137 | 0.021 | -0.019 | -0.019 | 0 | 0.42 | 0.18 | 0.43 | 0.43 | 0.41 | 0.46 |
| 11 | 99272474 | rs11219768 | 0.323 | 1.63 | -0.236 | -0.084 | 0.008 | 0.008 | 0.008 | 0 | 0.13 | 0.09 | 0.08 | 0.06 | 0.1 | 0.15 |
| 11 | 99275397 | rs7480845 | -1.74 | 1.855 | -0.442 | -0.14 | -0.039 | 0.072 | 0.072 | 0 | 0.69 | 0.93 | 0.78 | 0.82 | 0.67 | 0.78 |
| 11 | 99275986 | rs4442511 | 0.956 | 1.947 | -0.29 | -0.113 | 0 | 0 | 0 | 0 | 0.16 | 0.24 | 0.17 | 0.22 | 0.14 | 0.23 |
| 11 | 99276691 | rs7929417 | -1.803 | 1.959 | -0.385 | -0.173 | -0.041 | 0.069 | 0.069 | 0 | 0.69 | 0.93 | 0.78 | 0.82 | 0.67 | 0.78 |
| 11 | 99280659 | rs4279979 | 0.825 | 2.098 | -0.251 | -0.125 | -0.005 | -0.006 | -0.006 | 0 | 0.09 | 0.08 | 0.08 | 0 | 0.13 | 0.06 |
| 11 | 99284506 | rs2089171 | 0.035 | 1.608 | -0.241 | 0.063 | -0.002 | -0.019 | -0.019 | 0 | 0.42 | 0.44 | 0.57 | 0.78 | 0.38 | 0.57 |
| 11 | 99286608 | rs1542929 | 1.883 | 1.131 | -0.568 | 0.501 | -0.006 | -0.075 | -0.075 | 0.124 | 0.22 | 0.04 | 0.22 | 0.25 | 0.21 | 0.15 |
| 11 | 99290735 | rs2656199 | 0.701 | 1.174 | -0.367 | 0.277 | -0.058 | 0.159 | 0.159 | 0 | 0.4 | 0.11 | 0.29 | 0.17 | 0.44 | 0.28 |
| 11 | 99291062 | rs2134949 | 0.409 | 1.171 | -0.175 | 0.229 | -0.003 | -0.078 | -0.078 | 1.21 | 0.82 | 0.67 | 0.87 | 0.95 | 0.83 | 0.85 |
| 11 | 99291165 | rs10501907 | 0.863 | 1.16 | -1.038 | 0.372 | 0.003 | -0.021 | -0.021 | 3.091 | 0.36 | 0.28 | 0.39 | 0.48 | 0.35 | 0.39 |
| 11 | 99291694 | rs9919649 | -0.286 | 0.89 | -1.837 | -0.571 | 0.008 | 0.011 | 0.011 | 0 | 0.14 | 0.1 | 0.09 | 0.05 | 0.12 | 0.15 |
| 11 | 99292215 | rs10160612 | 0.059 | 1.165 | -1.494 | -0.153 | -0.024 | 0.145 | 0.145 | 0 | 0.45 | 0.79 | 0.62 | 0.78 | 0.44 | 0.57 |
| 11 | 99300137 | rs2726413 | 0.105 | 0.897 | -1.216 | 0.049 | 0.007 | 0.027 | 0.027 | 0.164 | 0.58 | 0.71 | 0.69 | 0.68 | 0.58 | 0.6 |
| 11 | 99307520 | rs10893324 | 0.187 | 0.655 | -1.481 | -0.03 | -0.004 | 0 | 0 | 0 | 0.18 | 0.07 | 0.16 | 0.16 | 0.21 | 0.16 |
| 11 | 99308233 | rs7114689 | 1.33 | 0.604 | -1.183 | -0.694 | 0.002 | -0.007 | -0.007 | 0 | 0.3 | 0.29 | 0.32 | 0.06 | 0.28 | 0.19 |
| 11 | 99309008 | rs7131217 | 0.371 | 0.808 | -0.96 | -0.173 | -0.005 | -0.009 | -0.009 | 0 | 0.17 | 0.18 | 0.17 | 0.43 | 0.19 | 0.31 |
| 11 | 99313336 | rs7104889 | 2.224 | 0.677 | -0.866 | -0.243 | 0.004 | 0.035 | 0.035 | 0 | 0.23 | 0.16 | 0.28 | 0.05 | 0.22 | 0.1 |
| 11 | 99316019 | rs1553514 | 0.155 | 0.537 | -0.181 | -0.191 | -0.023 | 0.078 | 0.078 | 0 | 0.71 | 0.51 | 0.63 | 0.56 | 0.71 | 0.65 |
| 11 | 99321743 | rs2656172 | -0.767 | 0.649 | 0.253 | -0.15 | 0 | 0.001 | 0.001 | 0 | 0.51 | 0.46 | 0.49 | 0.68 | 0.5 | 0.64 |
| 11 | 99325655 | rs2726393 | 1.431 | 0.623 | 0.313 | -0.273 | -0.002 | 0.003 | 0.003 | 0 | 0.32 | 0.31 | 0.37 | 0.08 | 0.34 | 0.21 |
| 11 | 99329156 | rs1910710 | -0.041 | 0.552 | 0.278 | -0.516 | -0.01 | 0.073 | 0.073 | 0 | 0.35 | 0.2 | 0.24 | 0.13 | 0.32 | 0.38 |
| 11 | 99329944 | rs2055603 | 1.11 | 0.572 | 0.334 | -0.258 | 0.007 | 0.074 | 0.074 | 0 | 0.11 | 0.19 | 0.11 | 0.42 | 0.14 | 0.25 |
| 11 | 99330466 | rs10893371 | 0.949 | 0.535 | 0.391 | -0.31 | 0.002 | -0.031 | -0.031 | 0.004 | 0.2 | 0.14 | 0.3 | 0.4 | 0.24 | 0.22 |
| 11 | 99330479 | rs2656191 | 2.488 | 0.546 | 0.451 | -0.374 | -0.008 | 0.056 | 0.056 | 0.031 | 0.26 | 0.13 | 0.31 | 0.05 | 0.28 | 0.11 |
| 11 | 99337294 | rs12807891 | 1.172 | 0.581 | 0.426 | -0.24 | -0.007 | 0.016 | 0.016 | 0 | 0.11 | 0.12 | 0.11 | 0.42 | 0.15 | 0.26 |
| 11 | 99343184 | rs7945901 | 1.222 | 0.494 | 0.284 | -0.511 | -0.002 | -0.003 | -0.003 | 0.196 | 0.18 | 0.16 | 0.17 | 0.19 | 0.23 | 0.15 |
| 11 | 99349946 | rs7946579 | 0.022 | 0.458 | 0.27 | -0.522 | 0.003 | -0.023 | -0.023 | 0 | 0.25 | 0.17 | 0.34 | 0.32 | 0.28 | 0.33 |
| 11 | 99352245 | rs2656152 | 2.481 | 0.476 | 0.395 | -0.614 | -0.002 | 0.003 | 0.003 | 0.01 | 0.26 | 0.22 | 0.31 | 0.05 | 0.28 | 0.11 |
| 11 | 99357649 | rs7129103 | -1.486 | 0.463 | 0.474 | -0.585 | 0.004 | 0.003 | 0.003 | 0 | 0.61 | 0.47 | 0.57 | 0.44 | 0.58 | 0.62 |
| 11 | 99358582 | rs17133607 | 0.626 | 0.465 | 0.294 | -0.616 | -0.01 | 0.032 | 0.032 | 0 | 0.28 | 0.16 | 0.21 | 0.11 | 0.27 | 0.25 |
| 11 | 99361491 | rs12290503 | 1.073 | 0.458 | 0.18 | -0.49 | -0.003 | 0.003 | 0.003 | 0 | 0.18 | 0.13 | 0.17 | 0.17 | 0.24 | 0.16 |
| 11 | 99362491 | rs7109406 | -0.077 | 0.507 | 0.152 | -0.69 | 0.012 | -0.032 | -0.032 | 0 | 0.3 | 0.22 | 0.38 | 0.42 | 0.31 | 0.32 |
| 11 | 99363687 | rs2656198 | -0.885 | 0.721 | 0.251 | -0.651 | 0.03 | 0.204 | 0.204 | 0 | 0.87 | 0.74 | 0.87 | 0.49 | 0.85 | 0.68 |
| 11 | 99369237 | rs7127856 | 0.464 | 0.74 | -0.28 | -1.062 | 0 | -0.038 | -0.038 | 0 | 0.24 | 0.2 | 0.14 | 0 | 0.23 | 0.15 |
| 11 | 99377407 | rs2726417 | 2.295 | 0.737 | -0.402 | -0.779 | -0.018 | 0.078 | 0.078 | 0 | 0.26 | 0.09 | 0.3 | 0.07 | 0.26 | 0.1 |
| 11 | 99379174 | rs17133656 | 0.076 | 0.375 | -0.245 | -0.99 | 0.001 | -0.045 | -0.045 | 0 | 0.31 | 0.28 | 0.18 | 0.01 | 0.26 | 0.24 |
| 11 | 99386817 | rs2466912 | 1.886 | 0.642 | -0.524 | -0.523 | -0.051 | 0.119 | 0.119 | 0.026 | 0.27 | 0.08 | 0.26 | 0.03 | 0.32 | 0.11 |

| | | | | | | | | | | | | | | | | |
|---|---|---|---|---|---|---|---|---|---|---|---|---|---|---|---|---|
| 11 | 99387885 | rs2466913 | 1.288 | 0.778 | -0.282 | -0.395 | -0.011 | 0.06 | 0.06 | 1.426 | 0.33 | 0.19 | 0.28 | 0.05 | 0.36 | 0.16 |
| 11 | 99388798 | rs11827261 | -0.034 | 0.748 | -0.035 | 0.605 | -0.002 | 0.091 | 0.091 | 0 | 0.12 | 0.2 | 0.17 | 0.51 | 0.1 | 0.35 |
| 11 | 99390789 | rs4612766 | -0.551 | 0.661 | 0.166 | 0.132 | -0.002 | -0.059 | -0.059 | 0 | 0.1 | 0.23 | 0.09 | 0 | 0.06 | 0.06 |
| 11 | 99392516 | rs10501915 | -0.224 | 0.894 | -0.263 | 0.083 | -0.039 | -0.033 | -0.033 | 0 | 0.23 | 0.08 | 0.2 | 0.29 | 0.29 | 0.25 |
| 11 | 99394978 | rs10893455 | -0.467 | 0.793 | -0.022 | 0.389 | -0.01 | -0.035 | -0.035 | 0 | 0.25 | 0.16 | 0.21 | 0.4 | 0.31 | 0.29 |
| 11 | 99395212 | rs10893458 | -0.453 | 0.793 | -0.022 | 0.39 | -0.01 | -0.035 | -0.035 | 0 | 0.25 | 0.16 | 0.21 | 0.4 | 0.31 | 0.29 |
| 11 | 99396953 | rs12290185 | -0.499 | 0.846 | 0.083 | -0.195 | 0.018 | 0.03 | 0.03 | 0 | 0.12 | 0.34 | 0.1 | 0.11 | 0.09 | 0.13 |
| 11 | 99398438 | rs11605324 | 1.271 | 0.962 | -0.285 | -0.209 | -0.037 | 0.042 | 0.042 | 0 | 0.13 | 0.01 | 0.09 | 0.06 | 0.15 | 0.12 |
| 11 | 99404062 | rs10893471 | -0.947 | 0.625 | -0.238 | -0.435 | 0.001 | 0.057 | 0.057 | 0 | 0.35 | 0.29 | 0.22 | 0.12 | 0.4 | 0.28 |
| 11 | 99406350 | rs10790778 | -0.31 | 0.618 | -0.265 | -0.429 | -0.016 | 0.05 | 0.05 | 0 | 0.25 | 0.15 | 0.17 | 0.12 | 0.3 | 0.25 |
| 11 | 99408308 | rs10736549 | -1.285 | 0.567 | -0.038 | -0.4 | 0.001 | -0.012 | -0.012 | 0 | 0.37 | 0.42 | 0.27 | 0.23 | 0.39 | 0.39 |
| 11 | 99408858 | rs1494466 | -1.706 | 0.558 | -0.042 | -0.401 | -0.001 | -0.016 | -0.016 | 0 | 0.47 | 0.53 | 0.32 | 0.22 | 0.49 | 0.41 |
| 11 | 99415304 | rs7949852 | -1.71 | 0.5 | 0.068 | -0.386 | 0.007 | 0.038 | 0.038 | 0.08 | 0.47 | 0.61 | 0.31 | 0.23 | 0.48 | 0.4 |
| 11 | 99415903 | rs12290702 | -1.742 | 0.465 | 0.138 | -0.124 | 0.007 | 0.024 | 0.024 | 0.032 | 0.47 | 0.63 | 0.31 | 0.35 | 0.48 | 0.43 |
| 11 | 99416115 | rs7925532 | -1.209 | 0.473 | 0.166 | -0.202 | 0.002 | -0.07 | -0.07 | 0 | 0.36 | 0.46 | 0.26 | 0.13 | 0.38 | 0.38 |
| 11 | 99418620 | rs954464 | 1.805 | 0.457 | 0.258 | -0.181 | 0.003 | 0.005 | 0.005 | 0 | 0.08 | 0.04 | 0.05 | 0 | 0.1 | 0.03 |
| 11 | 99419294 | rs11220404 | 1.697 | 0.455 | 0.257 | -0.135 | 0.004 | 0.001 | 0.001 | 0 | 0.08 | 0.04 | 0.05 | 0.12 | 0.1 | 0.05 |
| 11 | 99420459 | rs1389786 | -1.47 | 0.265 | 0.481 | 0.883 | -0.019 | 0.253 | 0.253 | 0.862 | 0.05 | 0.19 | 0.13 | 0.32 | 0.02 | 0.17 |
| 11 | 99421040 | rs7112709 | 0.045 | 0.352 | 0.624 | 0.814 | -0.007 | -0.052 | -0.052 | 0.058 | 0.1 | 0.22 | 0.08 | 0.01 | 0.07 | 0.13 |
| 11 | 99425400 | rs1494470 | -1.414 | 0.198 | -0.003 | 0.409 | -0.001 | -0.001 | -0.001 | 0.126 | 0.53 | 0.49 | 0.55 | 0.55 | 0.53 | 0.67 |
| 11 | 99427091 | rs2047166 | -1.373 | 0.189 | 0.003 | 0.405 | -0.002 | -0.001 | -0.001 | 0 | 0.53 | 0.5 | 0.55 | 0.56 | 0.53 | 0.67 |
| 11 | 99432021 | rs10790799 | -2.431 | 0.364 | -0.082 | 0.296 | 0.003 | -0.024 | -0.024 | 0.593 | 0.67 | 0.59 | 0.74 | 0.76 | 0.68 | 0.87 |
| 11 | 99432625 | rs976644 | -0.655 | 0.355 | -0.11 | 0.461 | -0.033 | -0.035 | -0.035 | 0.421 | 0.44 | 0.19 | 0.49 | 0.51 | 0.5 | 0.55 |
| 11 | 99434730 | rs17133813 | 0.722 | 0.165 | 0.428 | 0.808 | -0.001 | 0.061 | 0.061 | 0.401 | 0.07 | 0.21 | 0.04 | 0.1 | 0.04 | 0.04 |
| 11 | 99444534 | rs2726383 | 0.39 | 0.119 | 0.487 | 0.737 | -0.033 | 0.121 | 0.121 | 0 | 0.79 | 0.38 | 0.74 | 0.53 | 0.83 | 0.67 |
| 11 | 99445846 | rs2726379 | 0.74 | 0.134 | 0.317 | 0.759 | -0.02 | 0.175 | 0.175 | 0.242 | 0.95 | 0.83 | 0.87 | 0.74 | 0.98 | 0.83 |
| 11 | 99446765 | rs2466904 | -1.052 | 0.174 | 0.318 | 0.303 | -0.019 | -0.054 | -0.054 | 1.341 | 0.88 | 0.79 | 0.93 | 1 | 0.89 | 0.97 |
| 11 | 99455061 | rs2726355 | 1.118 | 0.156 | 0.242 | 0.287 | 0.025 | 0.016 | 0.016 | 0.804 | 0.74 | 0.17 | 0.8 | 0.78 | 0.8 | 0.8 |
| 11 | 99459308 | rs2515373 | -1.83 | 0.273 | 0.207 | 0.409 | -0.007 | -0.003 | -0.003 | 0 | 0.91 | 0.85 | 0.94 | 0.9 | 0.91 | 0.95 |
| 11 | 99460840 | rs7931769 | -0.574 | 0.532 | -0.003 | 0.135 | -0.007 | -0.039 | -0.039 | 0.82 | 0.31 | 0.43 | 0.2 | 0.15 | 0.32 | 0.2 |
| 11 | 99462465 | rs7117574 | 0.481 | 0.604 | 0.119 | 0.163 | -0.022 | -0.095 | -0.095 | 5.711 | 0.15 | 0.34 | 0.1 | 0 | 0.13 | 0.06 |
| 11 | 99467726 | rs648519 | 2.458 | 0.687 | 0.021 | -0.023 | -0.008 | 0.036 | 0.036 | 1.578 | 0.31 | 0.2 | 0.24 | 0.12 | 0.31 | 0.11 |
| 11 | 99473123 | rs11220603 | 1.363 | 0.976 | -0.005 | 0.089 | -0.015 | 0.013 | 0.013 | 0 | 0.11 | 0.06 | 0.09 | 0 | 0.17 | 0.04 |
| 11 | 99480175 | rs10790841 | 0.47 | 0.736 | 0.096 | 0.481 | -0.014 | 0.051 | 0.051 | 0.007 | 0.12 | 0.19 | 0.17 | 0.25 | 0.09 | 0.25 |
| 11 | 99482471 | rs10501919 | 0.173 | 0.959 | 0.118 | 0.268 | -0.006 | 0.005 | 0.005 | 0 | 0.17 | 0.14 | 0.13 | 0.02 | 0.25 | 0.07 |
| 11 | 99482918 | rs2019576 | -0.745 | 0.888 | 0.317 | 0.337 | 0.021 | -0.017 | -0.017 | 0.016 | 0.33 | 0.4 | 0.25 | 0.3 | 0.41 | 0.31 |
| 11 | 99486641 | rs7943707 | -0.734 | 0.881 | 0.33 | 0.381 | 0.014 | -0.01 | -0.01 | 0.061 | 0.33 | 0.36 | 0.27 | 0.3 | 0.44 | 0.31 |
| 11 | 99489014 | rs11220658 | -0.377 | 0.861 | 0.318 | 0.667 | 0.021 | -0.01 | -0.01 | 1.824 | 0.28 | 0.35 | 0.22 | 0.26 | 0.39 | 0.21 |
| 11 | 99489149 | rs10893605 | 0.679 | 0.89 | 0.207 | 0.683 | -0.008 | -0.012 | -0.012 | 0 | 0.15 | 0.09 | 0.12 | 0.19 | 0.21 | 0.09 |
| 11 | 99491499 | rs12789908 | 1.065 | 0.813 | 0.318 | 0.785 | -0.014 | -0.011 | -0.011 | 2.332 | 0.55 | 0.34 | 0.63 | 0.53 | 0.47 | 0.44 |
| 11 | 99492475 | rs11220671 | 0.536 | 0.757 | 0.44 | 0.834 | -0.017 | 0.052 | 0.052 | 0.141 | 0.12 | 0.23 | 0.09 | 0.2 | 0.09 | 0.26 |
| 11 | 99492957 | rs12792468 | 1.485 | 0.784 | 0.369 | 0.813 | -0.013 | 0.018 | 0.018 | 0 | 0.13 | 0.05 | 0.08 | 0.09 | 0.17 | 0.12 |
| 11 | 99493425 | rs10501921 | -0.23 | 0.756 | 0.46 | 0.842 | -0.01 | 0.027 | 0.027 | 0 | 0.17 | 0.29 | 0.14 | 0.21 | 0.14 | 0.35 |
| 11 | 99493557 | rs6590240 | 0.092 | 0.81 | 0.066 | 0.659 | 0.005 | -0.029 | -0.029 | 0.126 | 0.23 | 0.16 | 0.39 | 0.4 | 0.18 | 0.34 |
| 11 | 99497305 | rs12789135 | 2.125 | 0.92 | -0.007 | 0.599 | -0.04 | 0.042 | 0.042 | 0 | 0.12 | 0.01 | 0.08 | 0.07 | 0.17 | 0.12 |
| 11 | 99498121 | rs594858 | -1.782 | 0.896 | 0.405 | 0.73 | 0.016 | 0.032 | 0.032 | 0 | 0.86 | 0.77 | 0.9 | 0.76 | 0.83 | 0.86 |
| 11 | 99501264 | rs7103504 | 0.834 | 0.849 | 0.461 | 0.729 | -0.01 | 0.043 | 0.043 | 0.252 | 0.11 | 0.19 | 0.08 | 0.2 | 0.09 | 0.25 |
| 11 | 99503566 | rs10893622 | 0.121 | 1.086 | 0.608 | 0.737 | -0.002 | -0.002 | -0.002 | 0 | 0.25 | 0.28 | 0.18 | 0.21 | 0.24 | 0.41 |
| 11 | 99503747 | rs628310 | 1.333 | 1.076 | 0.646 | 0.587 | -0.014 | 0.147 | 0.147 | 0.1 | 0.46 | 0.25 | 0.42 | 0.12 | 0.52 | 0.17 |
| 11 | 99505610 | rs599609 | -0.066 | 1.089 | 0.673 | 0.636 | 0 | 0.026 | 0.026 | 0.454 | 0.7 | 0.43 | 0.7 | 0.53 | 0.68 | 0.5 |
| 11 | 99507058 | rs673798 | -0.077 | 1.089 | 0.692 | 0.636 | 0.002 | 0.06 | 0.06 | 0.408 | 0.7 | 0.48 | 0.71 | 0.53 | 0.69 | 0.5 |
| 11 | 99509199 | rs10893634 | 0.278 | 1.088 | 0.568 | 0.637 | -0.002 | -0.002 | -0.002 | 0 | 0.25 | 0.28 | 0.18 | 0.21 | 0.24 | 0.41 |
| 11 | 99512027 | rs1690808 | 2.723 | 1.127 | 0.114 | 0.501 | -0.037 | 0.164 | 0.164 | 1.128 | 0.31 | 0.1 | 0.19 | 0 | 0.29 | 0.1 |
| 11 | 99513359 | rs659277 | -0.106 | 0.882 | 0.123 | 0.543 | 0 | 0.025 | 0.025 | 0.505 | 0.7 | 0.43 | 0.7 | 0.53 | 0.68 | 0.5 |
| 11 | 99525126 | rs658689 | 2.015 | 1.032 | 0.066 | 0.297 | -0.056 | 0.123 | 0.123 | 0.005 | 0.35 | 0.16 | 0.25 | 0.1 | 0.4 | 0.14 |
| 11 | 99527939 | rs17134141 | -0.423 | 0.897 | -0.14 | 0.693 | 0.009 | -0.013 | -0.013 | 0 | 0.27 | 0.21 | 0.39 | 0.35 | 0.2 | 0.35 |
| 11 | 99534789 | rs668050 | 0.566 | 1.467 | 0.012 | 0.813 | -0.008 | -0.009 | -0.009 | 0 | 0.63 | 0.33 | 0.7 | 0.43 | 0.6 | 0.47 |
| 11 | 99539654 | rs675333 | 0.826 | 1.431 | -0.039 | 0.819 | -0.024 | -0.013 | -0.013 | 0 | 0.61 | 0.3 | 0.61 | 0.42 | 0.54 | 0.43 |
| 11 | 99541500 | rs10790880 | 0.726 | 1.346 | -0.062 | 0.832 | 0 | -0.001 | -0.001 | 0.213 | 0.21 | 0.19 | 0.11 | 0.27 | 0.18 | 0.32 |
| 11 | 99545375 | rs1690805 | 1.277 | 1.4 | -0.626 | 0.519 | -0.021 | 0.086 | 0.086 | 0.787 | 0.46 | 0.11 | 0.38 | 0.32 | 0.41 | 0.38 |
| 11 | 99546702 | rs12365643 | 1.747 | 1.077 | -0.281 | 0.598 | -0.001 | 0 | 0 | 0 | 0.1 | 0.08 | 0.15 | 0.1 | 0.16 | 0.06 |
| 11 | 99547971 | rs601742 | 0.268 | 1.18 | 0.065 | 0.734 | 0.011 | 0.001 | 0.001 | 0.174 | 0.59 | 0.32 | 0.49 | 0.41 | 0.58 | 0.46 |
| 11 | 99559169 | rs6590272 | 0.171 | 0.966 | 0.337 | 0.836 | -0.013 | 0.001 | 0.001 | 0 | 0.32 | 0.34 | 0.31 | 0.29 | 0.3 | 0.36 |
| 11 | 99570514 | rs631773 | 0.93 | 1.078 | 0.367 | 0.459 | 0.031 | 0.058 | 0.058 | 0 | 0.69 | 0.53 | 0.68 | 0.42 | 0.65 | 0.52 |
| 11 | 99574587 | rs10893744 | -0.502 | 1.07 | 0.194 | 0.363 | 0.03 | -0.015 | -0.015 | 0 | 0.44 | 0.29 | 0.4 | 0.51 | 0.4 | 0.68 |
| 11 | 99574689 | rs666825 | 0.909 | 1.06 | 0.183 | 0.368 | -0.01 | 0.017 | 0.017 | 0 | 0.44 | 0.34 | 0.47 | 0.39 | 0.42 | 0.22 |
| 11 | 99577624 | rs622821 | 1.421 | 1 | 0.237 | 0.381 | -0.001 | 0.004 | 0.004 | 0 | 0.39 | 0.34 | 0.42 | 0.32 | 0.36 | 0.17 |
| 11 | 99580734 | rs675163 | -0.542 | 1 | 0.062 | 0.481 | 0.007 | -0.01 | -0.01 | 0 | 0.86 | 0.9 | 0.87 | 0.78 | 0.89 | 0.67 |
| 11 | 99580861 | rs685284 | 0.261 | 1.027 | 0.13 | 0.513 | 0.002 | 0.039 | 0.039 | 0 | 0.82 | 0.75 | 0.79 | 0.52 | 0.82 | 0.61 |

| | | | | | | | | | | | | | | | | |
|---|---|---|---|---|---|---|---|---|---|---|---|---|---|---|---|---|
| 11 | 99585608 | rs635549 | 0.696 | 1.098 | 0.199 | 0.451 | 0.1 | 0.124 | 0.124 | 2.148 | 0.82 | 0.46 | 0.8 | 0.63 | 0.75 | 0.85 |
| 11 | 99585736 | rs952700 | 0.642 | 1.014 | 0.222 | 0.276 | 0 | -0.002 | -0.002 | 0 | 0.3 | 0.28 | 0.31 | 0.3 | 0.3 | 0.35 |
| 11 | 99585827 | rs594808 | -0.003 | 1.228 | 0.236 | -0.195 | 0.058 | -0.002 | -0.002 | 0.837 | 0.86 | 0.65 | 0.89 | 0.9 | 0.83 | 0.91 |
| 11 | 99587080 | rs605559 | 0.97 | 1.286 | -0.237 | -0.03 | -0.011 | 0.119 | 0.119 | 0.034 | 0.39 | 0.16 | 0.37 | 0.12 | 0.36 | 0.17 |
| 11 | 99587113 | rs605521 | 1.956 | 1.005 | -0.502 | -0.148 | -0.012 | 0.136 | 0.136 | 1.84 | 0.26 | 0.04 | 0.16 | 0 | 0.24 | 0.05 |
| 11 | 99592885 | rs4323834 | -1.093 | 0.394 | 0.116 | 0.373 | -0.001 | -0.003 | -0.003 | 0 | 0.57 | 0.43 | 0.46 | 0.69 | 0.46 | 0.73 |
| 11 | 99593538 | rs2063536 | 0.756 | 0.101 | -0.048 | 0.022 | -0.015 | -0.071 | -0.071 | 0 | 0.16 | 0.24 | 0.1 | 0 | 0.16 | 0.07 |
| 11 | 99593788 | rs7942402 | -1.684 | 0.297 | 0.114 | 0.033 | -0.027 | 0.073 | 0.073 | 0 | 0.62 | 0.76 | 0.58 | 0.85 | 0.56 | 0.78 |
| 11 | 99594162 | rs7935705 | 0.329 | 0.357 | 0.157 | 0.415 | -0.001 | 0 | 0 | 0.267 | 0.18 | 0.2 | 0.14 | 0.2 | 0.18 | 0.1 |
| 11 | 99599455 | rs10893803 | 0.512 | 0.385 | 0.088 | 0.305 | -0.001 | 0.002 | 0.002 | 0 | 0.17 | 0.15 | 0.11 | 0 | 0.19 | 0.1 |
| 11 | 99600063 | rs7113566 | -1.767 | 0.513 | 0.105 | 0.303 | -0.027 | 0.049 | 0.049 | 0 | 0.65 | 0.76 | 0.6 | 0.88 | 0.58 | 0.83 |
| 11 | 99600287 | rs1028004 | -1.787 | 0.383 | 0.482 | 0.68 | 0.178 | 0.346 | 0.346 | 0.199 | 0.08 | 0.39 | 0.14 | 0.35 | 0.12 | 0.11 |
| 11 | 99601189 | rs2010606 | -0.293 | 0.067 | 0.41 | 0.495 | 0.057 | 0.095 | 0.095 | 0 | 0.09 | 0.36 | 0.06 | 0.12 | 0.05 | 0.08 |
| 11 | 99601642 | rs7940098 | -0.89 | -0.138 | -0.197 | 0.44 | 0.03 | 0.001 | 0.001 | 0 | 0.48 | 0.39 | 0.42 | 0.51 | 0.39 | 0.62 |
| 11 | 99602252 | rs17134284 | 0.524 | 0.063 | -0.447 | 0.089 | -0.002 | -0.05 | -0.05 | 0 | 0.11 | 0 | 0.06 | 0.19 | 0.07 | 0.3 |
| 11 | 99602453 | rs10790939 | 0.309 | 0.375 | -0.087 | 0.101 | 0.004 | -0.011 | -0.011 | 0 | 0.16 | 0.15 | 0.11 | 0 | 0.18 | 0.07 |
| 11 | 99604442 | rs10893814 | 0.269 | 0.32 | 0.136 | 0.539 | -0.003 | 0.019 | 0.019 | 0 | 0.26 | 0.29 | 0.37 | 0.53 | 0.24 | 0.32 |
| 11 | 99604914 | rs7104324 | -1.407 | 0.508 | -0.07 | 0.322 | 0.005 | -0.036 | -0.036 | 0.159 | 0.6 | 0.6 | 0.55 | 0.76 | 0.54 | 0.76 |
| 11 | 99607142 | rs10790940 | 0.812 | 0.603 | -0.231 | 0.638 | 0.009 | -0.002 | -0.002 | 0.371 | 0.16 | 0.15 | 0.1 | 0.15 | 0.17 | 0.08 |
| 11 | 99608867 | rs7129217 | -1.61 | 0.656 | -0.219 | 0.668 | -0.028 | 0.1 | 0.1 | 0 | 0.56 | 0.81 | 0.61 | 0.73 | 0.51 | 0.56 |
| 11 | 99609677 | rs4582935 | 0.165 | 0.22 | -0.364 | 0.877 | -0.002 | 0.007 | 0.007 | 0 | 0.25 | 0.21 | 0.35 | 0.35 | 0.21 | 0.31 |
| 11 | 99610166 | rs1512123 | 0.516 | 0.275 | -0.711 | 0.897 | 0.003 | -0.001 | -0.001 | 0 | 0.16 | 0.2 | 0.1 | 0.16 | 0.17 | 0.07 |
| 11 | 99611634 | rs7111400 | -1.119 | 0.23 | -0.824 | 0.936 | -0.006 | -0.002 | -0.002 | 0 | 0.44 | 0.53 | 0.48 | 0.51 | 0.41 | 0.45 |
| 11 | 99613698 | rs10501922 | -0.347 | 0.764 | -1.208 | 0.592 | -0.003 | -0.001 | -0.001 | 0 | 0.17 | 0.2 | 0.12 | 0.16 | 0.2 | 0.14 |
| 11 | 99615747 | rs12807115 | -0.458 | 0.55 | -1.422 | 0.473 | 0.013 | -0.009 | -0.009 | 0 | 0.34 | 0.14 | 0.32 | 0.3 | 0.33 | 0.3 |
| 11 | 99616017 | rs17134306 | -1.341 | 0.855 | -0.524 | 0.42 | 0.018 | -0.009 | -0.009 | 0 | 0.06 | 0.19 | 0.09 | 0.04 | 0.09 | 0.06 |
| 11 | 99616274 | rs2015633 | 0.781 | 0.723 | -0.226 | 0.417 | 0.015 | 0.021 | 0.021 | 0.284 | 0.77 | 0.39 | 0.67 | 0.7 | 0.72 | 0.81 |
| 11 | 99616336 | rs2015623 | -0.942 | 0.679 | -0.515 | -0.349 | -0.001 | -0.015 | -0.015 | 0.001 | 0.56 | 0.66 | 0.73 | 0.83 | 0.59 | 0.72 |
| 11 | 99620792 | rs1441381 | -1.234 | 0.523 | 0.059 | 0.033 | -0.008 | -0.026 | -0.026 | 0 | 0.23 | 0.49 | 0.22 | 0.22 | 0.25 | 0.33 |
| 11 | 99621640 | rs7111790 | -1.049 | 0.651 | 0.009 | 0.404 | 0.003 | 0.052 | 0.052 | 0 | 0.17 | 0.24 | 0.33 | 0.63 | 0.17 | 0.48 |
| 11 | 99623531 | rs7932059 | -0.751 | 0.656 | 0.164 | 0.409 | 0.003 | 0.031 | 0.031 | 0 | 0.15 | 0.17 | 0.33 | 0.62 | 0.16 | 0.47 |
| 11 | 99629959 | rs1348155 | -1.162 | 0.559 | 0.28 | 0.566 | -0.005 | 0.032 | 0.032 | 0 | 0.19 | 0.27 | 0.34 | 0.54 | 0.17 | 0.38 |
| 11 | 99630378 | rs10750395 | -2.082 | 0.451 | 0.742 | 0.437 | -0.001 | -0.002 | -0.002 | 0 | 0.32 | 0.44 | 0.49 | 0.66 | 0.36 | 0.61 |
| 11 | 99631575 | rs1228375 | 0.745 | 0.429 | 0.39 | 0.203 | -0.005 | 0.03 | 0.03 | 0 | 0.47 | 0.4 | 0.34 | 0.3 | 0.51 | 0.38 |
| 11 | 99634866 | rs947979 | 0.888 | 0.433 | 0.418 | 0.208 | 0.001 | 0.04 | 0.04 | 0 | 0.45 | 0.39 | 0.31 | 0.25 | 0.46 | 0.28 |
| 11 | 99636183 | rs1939374 | -1.037 | 0.426 | 0.537 | 0.207 | -0.002 | 0.01 | 0.01 | 0 | 0.53 | 0.56 | 0.66 | 0.71 | 0.49 | 0.62 |
| 11 | 99642992 | rs2583151 | -2.192 | 0.383 | 0.533 | 0.235 | -0.011 | 0.106 | 0.106 | 0.233 | 0.69 | 0.86 | 0.82 | 0.96 | 0.69 | 0.91 |
| 11 | 99644206 | rs11822741 | 0.7 | 0.381 | 0.679 | 0.239 | -0.009 | -0.029 | -0.029 | 0 | 0.06 | 0.17 | 0.05 | 0.01 | 0.05 | 0.03 |
| 11 | 99644668 | rs11221321 | 0.934 | 0.229 | 0.948 | -0.005 | -0.01 | -0.04 | -0.04 | 0.808 | 0.08 | 0.11 | 0.03 | 0 | 0.05 | 0.06 |
| 11 | 99650833 | rs2583150 | 0.539 | 0.266 | 0.774 | 0.114 | 0.003 | 0.023 | 0.023 | 0 | 0.56 | 0.4 | 0.6 | 0.72 | 0.59 | 0.66 |
| 11 | 99654792 | rs7114221 | -0.656 | 0.287 | 0.735 | 0.203 | -0.003 | -0.014 | -0.014 | 0 | 0.23 | 0.21 | 0.41 | 0.49 | 0.28 | 0.41 |
| 11 | 99666292 | rs1301846 | 1.917 | 0.225 | 0.812 | 0.082 | 0.003 | 0.027 | 0.027 | 0.028 | 0.34 | 0.24 | 0.19 | 0.19 | 0.31 | 0.16 |
| 11 | 99674462 | rs1145386 | 0.462 | 0.226 | 0.799 | -0.002 | -0.002 | -0.014 | -0.014 | 0 | 0.58 | 0.44 | 0.62 | 0.73 | 0.62 | 0.68 |
| 11 | 99693548 | rs1145401 | 0.018 | 0.128 | 1.038 | 0.017 | 0 | -0.001 | -0.001 | 0.79 | 0.64 | 0.65 | 0.68 | 0.6 | 0.68 | 0.64 |
| 11 | 99694305 | rs1145403 | -1.741 | 0.113 | 1.02 | -0.028 | -0.001 | 0.037 | 0.037 | 0 | 0.55 | 0.71 | 0.53 | 0.39 | 0.58 | 0.52 |
| 11 | 99696766 | rs12280616 | 0.721 | 0.137 | 0.813 | 0.07 | -0.011 | -0.037 | -0.037 | 0.013 | 0.08 | 0.1 | 0.03 | 0 | 0.05 | 0.07 |
| 11 | 99699775 | rs1145408 | -0.634 | 0.2 | 0.804 | 0.111 | -0.005 | 0.006 | 0.006 | 0 | 0.74 | 0.8 | 0.83 | 0.82 | 0.75 | 0.81 |
| 11 | 99701653 | rs11221547 | -0.157 | 0.393 | 0.871 | 0.284 | 0 | -0.002 | -0.002 | 0 | 0.12 | 0.13 | 0.12 | 0.08 | 0.16 | 0.11 |
| 11 | 99704043 | rs7129269 | 0.346 | 0.198 | 0.653 | 0.25 | 0.004 | -0.033 | -0.033 | 0 | 0.13 | 0.08 | 0.29 | 0.42 | 0.11 | 0.28 |
| 11 | 99714232 | rs4754647 | -2.403 | 0.104 | 0.776 | -0.102 | 0.002 | 0.087 | 0.087 | 1.944 | 0.54 | 0.69 | 0.6 | 0.85 | 0.55 | 0.87 |
| 11 | 99716796 | rs6590363 | -0.634 | 0.038 | 0.723 | 0.168 | -0.005 | 0.008 | 0.008 | 0.243 | 0.25 | 0.19 | 0.14 | 0.12 | 0.26 | 0.26 |
| 11 | 99721588 | rs11221615 | -0.055 | 0.07 | 0.701 | 0.202 | -0.002 | 0.001 | 0.001 | 0.528 | 0.18 | 0.15 | 0.13 | 0.12 | 0.2 | 0.2 |
| 11 | 99721850 | rs10894009 | -0.21 | 0.013 | 0.849 | 0.311 | 0.008 | -0.019 | -0.019 | 0 | 0.12 | 0.25 | 0.13 | 0.08 | 0.15 | 0.11 |
| 11 | 99722295 | rs10501924 | 0.101 | 0.052 | 0.664 | 0.332 | 0.003 | -0.012 | -0.012 | 0.149 | 0.11 | 0.15 | 0.12 | 0.08 | 0.16 | 0.11 |
| 11 | 99728826 | rs11221661 | 0.695 | 0.175 | 0.024 | 0.51 | 0 | -0.077 | -0.077 | 0.227 | 0.12 | 0.01 | 0.28 | 0.53 | 0.09 | 0.37 |
| 11 | 99733085 | rs11221686 | 0.865 | 0.123 | -0.002 | 0.576 | -0.003 | -0.076 | -0.076 | 0.031 | 0.11 | 0.01 | 0.28 | 0.4 | 0.09 | 0.28 |
| 11 | 99745781 | rs7930091 | 0.049 | 0.293 | 0.461 | 0.232 | 0.008 | -0.005 | -0.005 | 0 | 0.88 | 0.71 | 0.87 | 0.9 | 0.85 | 0.88 |
| 11 | 99747429 | rs11221728 | -0.001 | 0.383 | 0.161 | 0.254 | -0.004 | -0.005 | -0.005 | 0 | 0.17 | 0.13 | 0.12 | 0.23 | 0.19 | 0.26 |
| 11 | 99750593 | rs10750415 | -1.865 | 0.314 | 0.198 | 0.174 | 0.005 | -0.008 | -0.008 | 0.212 | 0.38 | 0.42 | 0.27 | 0.32 | 0.41 | 0.46 |
| 11 | 99751230 | rs17134609 | -1.07 | 0.159 | -0.083 | 0.056 | -0.008 | -0.037 | -0.037 | 0.871 | 0.2 | 0.31 | 0.15 | 0.1 | 0.2 | 0.17 |
| 11 | 99754345 | rs7931411 | -0.89 | 0.334 | -0.902 | 0.151 | -0.043 | 0.038 | 0.038 | 0 | 0.27 | 0.09 | 0.23 | 0.18 | 0.32 | 0.21 |
| 11 | 99757415 | rs1373050 | 0.649 | 0.013 | -0.73 | 0.522 | 0.025 | -0.017 | -0.017 | 0 | 0.61 | 0.75 | 0.49 | 0.4 | 0.66 | 0.5 |
| 11 | 99761666 | rs1584819 | 1.085 | -0.225 | 0.259 | 0.684 | -0.032 | 0.07 | 0.07 | 0 | 0.87 | 0.56 | 0.8 | 0.82 | 0.93 | 0.86 |
| 11 | 99766548 | rs1453589 | -1.123 | -0.382 | 0.72 | 0.236 | 0.002 | 0.011 | 0.011 | 0 | 0.36 | 0.4 | 0.5 | 0.83 | 0.36 | 0.58 |
| 11 | 99770218 | rs1596306 | 1.066 | -0.918 | -0.462 | -0.831 | -0.042 | 0.119 | 0.119 | 0 | 0.4 | 0.2 | 0.32 | 0.08 | 0.45 | 0.25 |
| 11 | 99778206 | rs17680055 | -0.638 | -0.733 | -0.561 | -0.454 | 0.019 | 0.009 | 0.009 | 0 | 0.17 | 0.34 | 0.15 | 0.24 | 0.22 | 0.18 |
| 11 | 99778675 | rs7930068 | 0.218 | -0.823 | -0.583 | 0.038 | -0.006 | 0 | 0 | 0 | 0.56 | 0.54 | 0.61 | 0.51 | 0.66 | 0.49 |
| 11 | 99779665 | rs17618101 | -0.36 | -0.822 | -0.829 | 0.178 | 0.011 | 0.016 | 0.016 | 0 | 0.15 | 0.34 | 0.14 | 0.23 | 0.19 | 0.17 |
| 11 | 99781356 | rs10894151 | 0.721 | -1.014 | -1.243 | 0.281 | 0.017 | -0.066 | -0.066 | 0 | 0.21 | 0.08 | 0.36 | 0.59 | 0.17 | 0.41 |

| | | | | | | | | | | | | | | | | |
|---|---|---|---|---|---|---|---|---|---|---|---|---|---|---|---|---|
| 11 | 99781954 | rs10894154 | 0.471 | -0.684 | -0.269 | 0.161 | -0.02 | 0.127 | 0.127 | 0 | 0.19 | 0.02 | 0.1 | 0 | 0.19 | 0.12 |
| 11 | 99782640 | rs17134658 | -0.072 | -0.473 | 0.253 | 0.657 | -0.015 | -0.009 | -0.009 | 0 | 0.05 | 0.21 | 0.05 | 0.02 | 0.04 | 0.03 |
| 11 | 99789888 | rs7952312 | -1.535 | -0.495 | 0.028 | 0.592 | 0.03 | 0.026 | 0.026 | 0 | 0.51 | 0.7 | 0.39 | 0.47 | 0.46 | 0.46 |
| 11 | 99791995 | rs12276132 | -0.972 | -0.559 | 0.428 | 0.483 | 0.018 | -0.041 | -0.041 | 0 | 0.09 | 0.27 | 0.06 | 0.01 | 0.08 | 0.07 |
| 11 | 99795800 | rs12420505 | 2.004 | -0.429 | 0.158 | 0.226 | -0.102 | 0.041 | 0.041 | 0 | 0.38 | 0.05 | 0.33 | 0.27 | 0.39 | 0.41 |
| 11 | 99796756 | rs10791060 | -0.661 | -0.366 | 0.274 | 0.161 | 0.003 | -0.028 | -0.028 | 0 | 0.94 | 0.9 | 0.95 | 1 | 0.91 | 0.94 |
| 11 | 99801380 | rs17618560 | 0.341 | -0.532 | 0.25 | 0.18 | -0.001 | 0.007 | 0.007 | 0 | 0.25 | 0.18 | 0.17 | 0.22 | 0.27 | 0.21 |
| 11 | 99802262 | rs12290915 | -0.25 | -0.479 | -0.033 | 0.017 | -0.008 | -0.069 | -0.069 | 0.028 | 0.17 | 0.23 | 0.08 | 0.01 | 0.15 | 0.12 |
| 11 | 99806211 | rs1349784 | 1.765 | -0.432 | -0.445 | 0.121 | -0.082 | 0.052 | 0.052 | 0 | 0.39 | 0.06 | 0.33 | 0.27 | 0.38 | 0.42 |
| 11 | 99807394 | rs1453584 | 0.496 | -0.475 | -0.013 | 0.332 | -0.001 | 0.001 | 0.001 | 0 | 0.6 | 0.41 | 0.49 | 0.34 | 0.55 | 0.57 |
| 11 | 99811892 | rs6590446 | -1.454 | 0.051 | 0.258 | 0.422 | 0.011 | -0.005 | -0.005 | 0 | 0.33 | 0.44 | 0.53 | 0.62 | 0.36 | 0.38 |
| 11 | 99811973 | rs10501935 | -0.087 | 0.139 | 0.289 | 0.958 | 0.033 | 0.172 | 0.172 | 0 | 0.09 | 0.18 | 0.14 | 0.31 | 0.1 | 0.15 |
| 11 | 99812967 | rs2407306 | 1.474 | -0.006 | 0.29 | 0.836 | 0.004 | 0.021 | 0.021 | 0 | 0.48 | 0.35 | 0.35 | 0.37 | 0.44 | 0.48 |
| 11 | 99813064 | rs10894180 | -1.217 | 0.038 | 0.377 | 0.863 | 0.002 | 0.018 | 0.018 | 0 | 0.34 | 0.49 | 0.51 | 0.45 | 0.35 | 0.33 |
| 11 | 99815917 | rs7107215 | -1.721 | 0.041 | 0.198 | 0.83 | -0.002 | 0.004 | 0.004 | 0 | 0.86 | 0.84 | 0.88 | 0.82 | 0.83 | 0.89 |
| 11 | 99818829 | rs1453576 | -1.366 | 0.006 | 0.357 | 0.845 | 0.007 | 0.036 | 0.036 | 0 | 0.35 | 0.43 | 0.52 | 0.45 | 0.37 | 0.41 |
| 11 | 99820393 | rs10501937 | 0.111 | 0.011 | 0.416 | 0.593 | 0.027 | -0.06 | -0.06 | 0 | 0.12 | 0.27 | 0.1 | 0.01 | 0.16 | 0.07 |
| 11 | 99824120 | rs11604532 | 0.863 | 0.044 | -0.101 | 0.653 | 0 | 0.013 | 0.013 | 0 | 0.16 | 0.09 | 0.08 | 0.07 | 0.13 | 0.13 |
| 11 | 99825251 | rs7947050 | -1.671 | 0.101 | -0.084 | 0.675 | -0.003 | 0.008 | 0.008 | 0 | 0.84 | 0.84 | 0.87 | 0.82 | 0.81 | 0.86 |
| 11 | 99826738 | rs11222034 | 1.02 | 0.122 | -0.333 | 0.499 | -0.011 | 0.016 | 0.016 | 0 | 0.05 | 0.02 | 0.04 | 0 | 0.07 | 0.06 |
| 11 | 99827959 | rs12574821 | 2.358 | -0.127 | -0.539 | 0.458 | -0.077 | 0.07 | 0.07 | 0 | 0.36 | 0.04 | 0.27 | 0.2 | 0.34 | 0.37 |
| 11 | 99828103 | rs11601883 | -0.109 | -0.13 | -0.532 | 0.49 | 0.001 | 0.031 | 0.031 | 0 | 0.23 | 0.1 | 0.19 | 0.14 | 0.2 | 0.2 |
| 11 | 99828353 | rs11222043 | 1.061 | -0.028 | -0.371 | 0.59 | -0.01 | 0.024 | 0.024 | 0 | 0.59 | 0.18 | 0.46 | 0.33 | 0.54 | 0.57 |
| 11 | 99830221 | rs1349782 | 0.107 | 0.157 | -0.033 | 0.626 | 0.004 | 0.018 | 0.018 | 0 | 0.64 | 0.57 | 0.54 | 0.48 | 0.63 | 0.69 |
| 11 | 99830605 | rs10791067 | -0.867 | 0.124 | 0.265 | 0.45 | 0.023 | -0.077 | -0.077 | 0 | 0.17 | 0.34 | 0.16 | 0.02 | 0.23 | 0.13 |
| 11 | 99832557 | rs17619627 | 0.257 | 0.083 | 0.152 | 0.563 | -0.009 | 0.045 | 0.045 | 0 | 0.2 | 0.07 | 0.13 | 0.07 | 0.19 | 0.17 |
| 11 | 99838176 | rs1377573 | -1.252 | 0.049 | 0.359 | 0.749 | -0.001 | -0.003 | -0.003 | 0.009 | 0.33 | 0.32 | 0.54 | 0.67 | 0.3 | 0.4 |
| 11 | 99838511 | rs11222087 | 1.61 | 0.032 | 0.393 | 0.734 | -0.001 | 0.006 | 0.006 | 0 | 0.15 | 0.17 | 0.11 | 0.14 | 0.17 | 0.1 |
| 11 | 99840786 | rs2084989 | 0.343 | 0.1 | 0.438 | 1.219 | -0.028 | 0.066 | 0.066 | 0.476 | 0.09 | 0.15 | 0.31 | 0.24 | 0.03 | 0.12 |
| 11 | 99841032 | rs12786005 | 0.008 | -0.022 | 0.569 | 0.583 | 0.029 | -0.016 | -0.016 | 0 | 0.08 | 0.31 | 0.07 | 0.02 | 0.09 | 0.03 |
| 11 | 99841559 | rs7119109 | 1.549 | 0.049 | 0.45 | 0.685 | 0 | -0.006 | -0.006 | 0.143 | 0.15 | 0.19 | 0.12 | 0.14 | 0.18 | 0.1 |
| 11 | 99841735 | rs10082676 | -2.137 | 0.148 | 0.44 | 0.705 | -0.01 | 0.101 | 0.101 | 0.01 | 0.48 | 0.7 | 0.67 | 0.69 | 0.49 | 0.52 |
| 11 | 99843386 | rs7127363 | -1.104 | 0.206 | 0.472 | 0.641 | 0.007 | -0.092 | -0.092 | 0 | 0.34 | 0.47 | 0.25 | 0.09 | 0.36 | 0.25 |
| 11 | 99852155 | rs6590473 | -0.456 | 0.182 | 0.485 | 0.674 | 0.002 | -0.072 | -0.072 | 0 | 0.28 | 0.4 | 0.2 | 0.09 | 0.29 | 0.18 |
| 11 | 99854317 | rs4753962 | 0.029 | 0.137 | 0.39 | 0.832 | -0.026 | 0.08 | 0.08 | 0 | 0.1 | 0.14 | 0.15 | 0.26 | 0.05 | 0.17 |
| 11 | 99855646 | rs7112259 | -0.704 | 0.18 | 0.424 | 0.722 | -0.001 | -0.097 | -0.097 | 3.688 | 0.3 | 0.44 | 0.21 | 0.09 | 0.3 | 0.21 |
| 11 | 99857172 | rs4754656 | -2.228 | 0.166 | 0.299 | 0.738 | 0.001 | 0.121 | 0.121 | 0 | 0.6 | 0.81 | 0.75 | 0.83 | 0.64 | 0.64 |
| 11 | 99858032 | rs7933349 | 0.39 | 0.232 | 0.179 | 0.825 | -0.004 | -0.008 | -0.008 | 0 | 0.22 | 0.22 | 0.22 | 0.37 | 0.27 | 0.27 |
| 11 | 99865549 | rs4269885 | -1.404 | 0.158 | 0.206 | 0.79 | 0.003 | -0.011 | -0.011 | 0.221 | 0.39 | 0.63 | 0.54 | 0.43 | 0.37 | 0.36 |
| 11 | 99866660 | rs1035339 | -1.087 | 0.039 | 0.259 | 0.722 | -0.001 | -0.017 | -0.017 | 0.625 | 0.36 | 0.58 | 0.28 | 0.12 | 0.32 | 0.28 |
| 11 | 99871361 | rs12282593 | -1.127 | 0.045 | 0.074 | 0.726 | 0.012 | 0.03 | 0.03 | 0 | 0.37 | 0.64 | 0.28 | 0.12 | 0.32 | 0.28 |
| 11 | 99873679 | rs11222295 | 1.505 | 0.147 | -0.012 | 0.802 | -0.004 | 0.004 | 0.004 | 0 | 0.15 | 0.13 | 0.1 | 0.21 | 0.18 | 0.13 |
| 11 | 99879079 | rs12797124 | 0.502 | 0.043 | -0.051 | 0.652 | 0.003 | -0.045 | -0.045 | 0 | 0.12 | 0.23 | 0.11 | 0.02 | 0.15 | 0.06 |
| 11 | 99879384 | rs1453568 | 0.331 | 0.053 | -0.322 | 0.983 | -0.033 | 0.118 | 0.118 | 0 | 0.1 | 0.22 | 0.16 | 0.22 | 0.05 | 0.18 |
| 11 | 99879539 | rs1377567 | 0.973 | 0.579 | -1.152 | 1.247 | 0.001 | -0.009 | -0.009 | 0 | 0.17 | 0.23 | 0.13 | 0.22 | 0.2 | 0.13 |
| 11 | 99880163 | rs12421109 | -0.391 | 0.532 | -1.22 | 1.163 | 0.009 | 0.002 | 0.002 | 0 | 0.4 | 0.61 | 0.35 | 0.31 | 0.34 | 0.35 |
| 11 | 99880352 | rs770566 | 0.877 | 0.697 | -1.698 | 0.759 | -0.052 | 0.111 | 0.111 | 0 | 0.35 | 0.07 | 0.22 | 0.17 | 0.34 | 0.36 |
| 11 | 99880558 | rs770567 | 0.25 | 0.535 | -1.478 | 0.618 | -0.073 | -0.072 | -0.072 | 0 | 0.4 | 0.15 | 0.31 | 0.44 | 0.44 | 0.5 |
| 11 | 99882016 | rs11825125 | -0.596 | -0.222 | -0.672 | 0.375 | -0.025 | -0.001 | -0.001 | 0 | 0.31 | 0.4 | 0.19 | 0.25 | 0.17 | 0.27 |
| 11 | 99882216 | rs770569 | -0.58 | -0.458 | -0.375 | 0.107 | -0.06 | -0.073 | -0.073 | 2.326 | 0.28 | 0.42 | 0.1 | 0.08 | 0.1 | 0.18 |
| 11 | 99883904 | rs770573 | 0.027 | -0.012 | -0.772 | -0.906 | -0.069 | -0.095 | -0.095 | 0 | 0.28 | 0.44 | 0.1 | 0.09 | 0.12 | 0.16 |
| 11 | 99884736 | rs770577 | -0.212 | 0.493 | -0.686 | -1.152 | -0.019 | 0.079 | 0.079 | 0 | 0.64 | 0.77 | 0.56 | 0.92 | 0.54 | 0.76 |
| 11 | 99885930 | rs770593 | 0.016 | 0.786 | -0.543 | -0.574 | -0.004 | 0.002 | 0.002 | 0 | 0.59 | 0.59 | 0.52 | 0.69 | 0.47 | 0.65 |
| 11 | 99889749 | rs11222392 | 0.614 | 0.706 | -0.621 | -0.558 | -0.006 | 0.01 | 0.01 | 0 | 0.11 | 0.1 | 0.15 | 0.06 | 0.16 | 0.11 |
| 11 | 99899770 | rs770580 | 0.949 | 0.175 | -0.089 | -0.732 | -0.01 | -0.009 | -0.009 | 0 | 0.19 | 0.18 | 0.06 | 0.08 | 0.07 | 0.13 |
| 11 | 99903806 | rs770587 | 0.792 | 0.15 | -0.056 | -0.749 | -0.018 | -0.018 | -0.018 | 0.744 | 0.2 | 0.21 | 0.08 | 0.09 | 0.07 | 0.13 |
| 11 | 99914262 | rs4754665 | -0.31 | 0.281 | 0.168 | -0.901 | -0.004 | 0.008 | 0.008 | 0 | 0.6 | 0.65 | 0.55 | 0.71 | 0.48 | 0.69 |
| 11 | 99918332 | rs7933249 | -0.292 | 0.263 | 0.195 | -0.913 | -0.02 | 0.055 | 0.055 | 0.012 | 0.6 | 0.71 | 0.55 | 0.71 | 0.49 | 0.66 |
| 11 | 99923002 | rs12281896 | 1.11 | 0.199 | 0.042 | -0.927 | -0.009 | -0.006 | -0.006 | 3.493 | 0.2 | 0.19 | 0.08 | 0.1 | 0.07 | 0.13 |
| 11 | 99923641 | rs11222552 | 0.387 | 0.493 | -0.024 | -0.716 | 0.003 | -0.004 | -0.004 | 0.026 | 0.39 | 0.43 | 0.47 | 0.61 | 0.41 | 0.52 |
| 11 | 99925167 | rs7946600 | 0.736 | 0.586 | -0.028 | -0.462 | -0.005 | 0.02 | 0.02 | 0.15 | 0.34 | 0.42 | 0.35 | 0.44 | 0.3 | 0.42 |
| 11 | 99935993 | rs1074221 | 0.759 | 0.988 | 0.076 | -0.222 | 0.041 | 0.077 | 0.077 | 0 | 0.06 | 0.11 | 0.13 | 0.18 | 0.11 | 0.12 |
| 11 | 99936389 | rs7126331 | -1.368 | 0.826 | -0.308 | -0.599 | -0.103 | 0.003 | 0.003 | 0 | 0.76 | 0.95 | 0.72 | 0.77 | 0.66 | 0.76 |
| 11 | 99937814 | rs10894384 | 0.345 | 0.504 | 0.194 | -0.761 | -0.003 | -0.001 | -0.001 | 0.192 | 0.54 | 0.52 | 0.44 | 0.52 | 0.38 | 0.51 |
| 11 | 99939338 | rs10750480 | -0.528 | 0.518 | 0.2 | -0.779 | -0.006 | 0.044 | 0.044 | 0 | 0.64 | 0.78 | 0.61 | 0.92 | 0.57 | 0.77 |
| 11 | 99939733 | rs6590574 | -1.299 | 0.556 | 0.203 | -0.728 | 0.046 | 0.1 | 0.1 | 1.594 | 0.26 | 0.47 | 0.33 | 0.48 | 0.35 | 0.4 |
| 11 | 99941005 | rs7935954 | 1.015 | 0.445 | 0.311 | -0.856 | -0.004 | -0.029 | -0.029 | 0.779 | 0.07 | 0.09 | 0.02 | 0 | 0.05 | 0.04 |
| 11 | 99942748 | rs10791145 | 1.956 | 0.2 | 0.3 | -0.692 | -0.013 | -0.068 | -0.068 | 0 | 0.26 | 0.12 | 0.32 | 0.39 | 0.26 | 0.32 |
| 11 | 99942929 | rs1363023 | 0.482 | 0.168 | 0.332 | -0.701 | -0.003 | -0.013 | -0.013 | 0 | 0.15 | 0.15 | 0.15 | 0.07 | 0.16 | 0.15 |

| | | | | | | | | | | | | | | | | |
|---|---|---|---|---|---|---|---|---|---|---|---|---|---|---|---|---|
| 11 | 99946472 | rs11222669 | 1.57 | 0.083 | 0.039 | -0.664 | -0.018 | -0.07 | -0.07 | 0 | 0.29 | 0.13 | 0.34 | 0.4 | 0.27 | 0.35 |
| 11 | 99946620 | rs11827197 | 0.924 | 0.127 | 0.179 | -0.516 | -0.004 | -0.029 | -0.029 | 0 | 0.06 | 0.09 | 0.02 | 0 | 0.05 | 0.04 |
| 11 | 99948620 | rs1421556 | -0.322 | 0.032 | -0.104 | -0.56 | 0.013 | -0.02 | -0.02 | 0 | 0.33 | 0.15 | 0.21 | 0.4 | 0.27 | 0.42 |
| 11 | 99949305 | rs17094317 | 0.836 | 0.012 | -0.309 | -0.533 | -0.022 | -0.09 | -0.09 | 0 | 0.22 | 0.03 | 0.08 | 0.33 | 0.15 | 0.32 |
| 11 | 99952616 | rs1421557 | -1.412 | 0.105 | -0.024 | -0.507 | 0.016 | 0.041 | 0.041 | 0.418 | 0.5 | 0.76 | 0.41 | 0.6 | 0.44 | 0.6 |
| 11 | 99954527 | rs7929129 | -0.776 | 0.097 | 0.2 | -0.525 | -0.002 | -0.001 | -0.001 | 0 | 0.42 | 0.46 | 0.26 | 0.41 | 0.32 | 0.46 |
| 11 | 99959543 | rs2123325 | -1.705 | 0.585 | -0.096 | -0.723 | -0.045 | 0.084 | 0.084 | 0 | 0.78 | 0.94 | 0.76 | 0.99 | 0.71 | 0.89 |
| 11 | 99963969 | rs10791161 | -0.168 | 0.403 | 0.136 | -0.672 | -0.017 | 0.021 | 0.021 | 0 | 0.6 | 0.72 | 0.5 | 0.74 | 0.47 | 0.67 |
| 11 | 99968225 | rs2112068 | 0.515 | -0.051 | 0.11 | -0.885 | -0.015 | 0.01 | 0.01 | 0 | 0.36 | 0.29 | 0.5 | 0.63 | 0.44 | 0.44 |
| 11 | 99968814 | rs11222749 | 0.529 | -0.226 | -0.136 | -0.834 | 0.012 | 0.026 | 0.026 | 0.042 | 0.13 | 0.07 | 0.07 | 0.06 | 0.11 | 0.09 |
| 11 | 99970832 | rs12575570 | 1.212 | -0.381 | -0.421 | -0.802 | 0.044 | -0.005 | -0.005 | 0.336 | 0.19 | 0.02 | 0.05 | 0.15 | 0.08 | 0.14 |
| 11 | 99971357 | rs11222760 | 0.02 | -0.352 | 0.022 | -0.874 | -0.024 | -0.015 | -0.015 | 0 | 0.28 | 0.4 | 0.13 | 0.29 | 0.16 | 0.35 |
| 11 | 99973350 | rs11222776 | 1.167 | -0.09 | -0.5 | -0.705 | -0.015 | -0.032 | -0.032 | 0.004 | 0.27 | 0.15 | 0.4 | 0.57 | 0.29 | 0.34 |
| 11 | 99978931 | rs4754668 | -0.768 | 0.024 | -0.85 | -1.028 | -0.063 | 0.192 | 0.192 | 0 | 0.66 | 0.91 | 0.69 | 0.95 | 0.6 | 0.79 |
| 11 | 99978991 | rs4753968 | -0.289 | -0.023 | -0.464 | -0.714 | -0.02 | 0.086 | 0.086 | 0 | 0.59 | 0.69 | 0.64 | 0.78 | 0.53 | 0.63 |
| 11 | 99981435 | rs2193863 | 1.258 | 0.484 | -0.588 | -0.43 | -0.028 | -0.002 | -0.002 | 0.288 | 0.06 | 0.01 | 0.07 | 0.04 | 0.1 | 0.09 |
| 11 | 99984562 | rs11222824 | 0.579 | 0.316 | -0.977 | -0.46 | -0.064 | 0.078 | 0.078 | 0 | 0.58 | 0.12 | 0.48 | 0.7 | 0.46 | 0.62 |
| 11 | 99985271 | rs10894469 | -0.221 | 0.549 | -0.749 | -0.382 | 0.085 | 0.147 | 0.147 | 0 | 0.06 | 0.23 | 0.11 | 0.21 | 0.11 | 0.16 |
| 11 | 99993641 | rs4753969 | -1.568 | 0.557 | -1.277 | -0.575 | -0.029 | 0.044 | 0.044 | 0 | 0.8 | 0.9 | 0.74 | 1 | 0.72 | 0.91 |
| 11 | 99999475 | rs2112071 | 1.116 | 0.213 | -1.456 | -0.077 | -0.023 | -0.105 | -0.105 | 0 | 0.26 | 0.09 | 0.3 | 0.45 | 0.28 | 0.32 |
| 11 | 100003476 | rs11222895 | 1.047 | 0.459 | -0.93 | -0.305 | -0.032 | 0.008 | 0.008 | 0 | 0.06 | 0.01 | 0.07 | 0.03 | 0.11 | 0.1 |
| 11 | 100004351 | rs11222898 | 0.003 | 0.434 | -0.711 | -0.378 | 0.005 | -0.012 | -0.012 | 0.252 | 0.11 | 0.16 | 0.12 | 0.07 | 0.14 | 0.09 |
| 11 | 100005864 | rs7103146 | 0.215 | 0.553 | -1.336 | -0.665 | 0.038 | 0.02 | 0.02 | 0 | 0.4 | 0.17 | 0.42 | 0.65 | 0.36 | 0.51 |
| 11 | 100008296 | rs10791200 | -0.031 | 0.356 | -0.63 | -0.76 | 0.03 | 0.019 | 0.019 | 0 | 0.42 | 0.69 | 0.46 | 0.65 | 0.36 | 0.51 |
| 11 | 100009378 | rs11222942 | -0.121 | 0.742 | -0.807 | -0.924 | 0.014 | -0.007 | -0.007 | 0 | 0.16 | 0.22 | 0.18 | 0.13 | 0.24 | 0.2 |
| 11 | 100009976 | rs12364336 | 1.833 | 0.792 | -1.138 | -0.928 | -0.044 | 0.027 | 0.027 | 0 | 0.09 | 0.01 | 0.12 | 0.04 | 0.14 | 0.11 |
| 11 | 100014539 | rs1363022 | -1.126 | 0.619 | -1.193 | -0.947 | -0.082 | 0.09 | 0.09 | 0 | 0.59 | 0.95 | 0.63 | 0.77 | 0.59 | 0.69 |
| 11 | 100017416 | rs1421566 | -1.748 | 0.833 | -0.631 | -0.839 | 0.014 | -0.007 | -0.007 | 0 | 0.28 | 0.36 | 0.26 | 0.26 | 0.35 | 0.3 |
| 11 | 100018985 | rs17094433 | 0.346 | 0.613 | -0.797 | -0.655 | 0.001 | -0.001 | -0.001 | 0 | 0.06 | 0.06 | 0.05 | 0.06 | 0.09 | 0.09 |
| 11 | 100020880 | rs4754673 | 1.582 | 0.529 | -1.074 | -0.669 | -0.022 | -0.17 | -0.17 | 0.734 | 0.27 | 0.03 | 0.32 | 0.52 | 0.23 | 0.37 |
| 11 | 100028717 | rs1812544 | 0.122 | 0.698 | -0.441 | -0.583 | 0.013 | 0.005 | 0.005 | 0 | 0.07 | 0.13 | 0.05 | 0.08 | 0.11 | 0.1 |
| 11 | 100032351 | rs12099197 | 0.616 | 0.585 | -0.451 | -0.597 | -0.026 | -0.002 | -0.002 | 0 | 0.07 | 0.15 | 0.03 | 0.06 | 0.02 | 0.01 |
| 11 | 100032745 | rs11223022 | -2.177 | 0.598 | -0.078 | -0.658 | 0.004 | -0.002 | -0.002 | 0 | 0.2 | 0.6 | 0.16 | 0.19 | 0.21 | 0.2 |
| 11 | 100038669 | rs7941829 | -0.542 | 0.713 | -0.803 | -0.566 | 0 | 0.004 | 0.004 | 0 | 0.12 | 0.18 | 0.07 | 0.13 | 0.12 | 0.1 |
| 11 | 100039031 | rs7926813 | -0.535 | 0.715 | -0.803 | -0.57 | 0 | 0.002 | 0.002 | 0 | 0.12 | 0.18 | 0.07 | 0.12 | 0.12 | 0.11 |
| 11 | 100041420 | rs2044983 | -1.607 | 1.448 | -1.121 | -0.819 | -0.053 | 0.034 | 0.034 | 0 | 0.91 | 0.99 | 0.84 | 0.99 | 0.83 | 0.95 |
| 11 | 100043420 | rs12803066 | 0.52 | 0.538 | -1.457 | -1.155 | 0.017 | 0.025 | 0.025 | 1.805 | 0.25 | 0.09 | 0.09 | 0.13 | 0.15 | 0.06 |
| 11 | 100045242 | rs7119230 | 0.898 | 0.631 | -1.99 | -1.251 | -0.009 | 0.037 | 0.037 | 0 | 0.33 | 0.03 | 0.23 | 0.23 | 0.25 | 0.27 |
| 11 | 100045424 | rs17094497 | 2.003 | 1.11 | -1.426 | -1.239 | 0.035 | -0.027 | -0.027 | 1.382 | 0.1 | 0.18 | 0.13 | 0.03 | 0.13 | 0.1 |
| 11 | 100045700 | rs10894564 | 0.834 | 1.141 | -1.472 | -1.315 | 0.002 | -0.002 | -0.002 | 0 | 0.06 | 0.07 | 0.07 | 0.06 | 0.1 | 0.03 |
| 11 | 100045949 | rs1421555 | 1.2 | 1.004 | -1.453 | -1.506 | -0.001 | -0.002 | -0.002 | 0 | 0.35 | 0.16 | 0.4 | 0.67 | 0.35 | 0.54 |
| 11 | 100052221 | rs7933176 | 0.536 | 1.074 | -1.07 | -1.246 | 0.021 | -0.023 | -0.023 | 0 | 0.15 | 0.22 | 0.2 | 0.09 | 0.22 | 0.13 |
| 11 | 100052977 | rs7130695 | 2.152 | 1.089 | -1.297 | -1.187 | -0.048 | 0.023 | 0.023 | 0.368 | 0.09 | 0.01 | 0.15 | 0.03 | 0.15 | 0.11 |
| 11 | 100055523 | rs7934785 | -0.456 | 0.836 | -1.126 | -1.358 | -0.008 | 0.02 | 0.02 | 0 | 0.61 | 0.67 | 0.68 | 0.78 | 0.63 | 0.78 |
| 11 | 100057536 | rs11223142 | 1.706 | 0.093 | -1.887 | -1.23 | -0.035 | -0.155 | -0.155 | 0 | 0.2 | 0.03 | 0.32 | 0.53 | 0.22 | 0.49 |
| 11 | 100058720 | rs7931432 | 2.975 | -0.103 | -1.936 | -1.419 | 0.032 | 0.038 | 0.038 | 3.286 | 0.14 | 0.06 | 0.02 | 0 | 0.04 | 0.04 |
| 11 | 100061377 | rs1002122 | -0.503 | -0.188 | -1.792 | -1.642 | 0.001 | -0.017 | -0.017 | 0 | 0.45 | 0.46 | 0.58 | 0.81 | 0.52 | 0.64 |
| 11 | 100061624 | rs1984684 | 1.259 | -0.195 | -1.631 | -1.42 | -0.02 | -0.161 | -0.161 | 0 | 0.24 | 0.04 | 0.35 | 0.54 | 0.24 | 0.5 |
| 11 | 100061865 | rs11223168 | 1.204 | -0.004 | -1.19 | -1.441 | 0.013 | 0.001 | 0.001 | 0 | 0.06 | 0.09 | 0.07 | 0.06 | 0.1 | 0.03 |
| 11 | 100063257 | rs10750510 | -0.209 | -0.101 | -1.144 | -1.579 | 0 | -0.016 | -0.016 | 0.069 | 0.55 | 0.44 | 0.55 | 0.81 | 0.51 | 0.68 |
| 11 | 100063425 | rs1452575 | 1.477 | 0.359 | -1.075 | -1.367 | -0.001 | 0.001 | 0.001 | 0 | 0.14 | 0.2 | 0.19 | 0.08 | 0.24 | 0.09 |
| 11 | 100066196 | rs10894599 | -0.717 | -0.748 | -0.942 | -1.214 | -0.009 | -0.03 | -0.03 | 0.823 | 0.4 | 0.48 | 0.53 | 0.8 | 0.39 | 0.58 |
| 11 | 100068264 | rs17094600 | 0.949 | -0.462 | -1.162 | -0.972 | 0.006 | -0.001 | -0.001 | 0 | 0.06 | 0.07 | 0.06 | 0.07 | 0.1 | 0.03 |
| 11 | 100069533 | rs7117635 | -1.093 | -0.969 | -1.361 | -1.175 | 0.015 | 0.027 | 0.027 | 0 | 0.75 | 0.82 | 0.8 | 0.89 | 0.76 | 0.78 |
| 11 | 100072718 | rs2186705 | -1.865 | -1.174 | -1.426 | -1.123 | 0.009 | 0.014 | 0.014 | 0 | 0.8 | 0.87 | 0.87 | 0.89 | 0.81 | 0.8 |
| 11 | 100079723 | rs1939694 | 2.242 | -1.165 | -1.372 | 0.511 | -0.002 | 0.051 | 0.051 | 0 | 0.06 | 0.03 | 0.07 | 0.31 | 0.05 | 0.05 |
| 11 | 100080396 | rs11606265 | 0.887 | -1.31 | -1.436 | 0.26 | 0.011 | -0.035 | -0.035 | 0 | 0.19 | 0.06 | 0.32 | 0.25 | 0.19 | 0.46 |
| 11 | 100082000 | rs7115798 | 1.593 | -1.223 | -1.489 | 0.265 | -0.007 | -0.044 | -0.044 | 0 | 0.17 | 0.03 | 0.25 | 0.23 | 0.16 | 0.44 |
| 11 | 100087315 | rs11223269 | 0.92 | -1.351 | -1.811 | 0.211 | 0.008 | -0.091 | -0.091 | 2.939 | 0.18 | 0.05 | 0.12 | 0.49 | 0.12 | 0.14 |
| 11 | 100096565 | rs1939700 | 1.175 | -1.332 | -1.945 | 0.138 | 0.007 | -0.05 | -0.05 | 0 | 0.21 | 0.03 | 0.32 | 0.25 | 0.19 | 0.47 |
| 11 | 100100423 | rs17094656 | 1.261 | -1.238 | -1.634 | 0.319 | 0.013 | -0.041 | -0.041 | 1.027 | 0.14 | 0.09 | 0.1 | 0.32 | 0.09 | 0.09 |
| 11 | 100102098 | rs1939707 | -1.236 | -1.252 | -1.635 | -0.508 | 0.02 | 0.023 | 0.023 | 0 | 0.75 | 0.85 | 0.83 | 0.84 | 0.78 | 0.77 |
| 11 | 100106009 | rs10791277 | -0.469 | -1.303 | -1.326 | -0.579 | 0.028 | 0.019 | 0.019 | 0 | 0.59 | 0.72 | 0.77 | 0.7 | 0.68 | 0.7 |
| 11 | 100108865 | rs606363 | 0.247 | -1.187 | -1.511 | -1.049 | 0.057 | 0.113 | 0.113 | 0.047 | 0.48 | 0.18 | 0.65 | 0.69 | 0.6 | 0.55 |
| 11 | 100110095 | rs544633 | 1.165 | -0.861 | -1.447 | -1.073 | -0.066 | 0.083 | 0.083 | 0 | 0.28 | 0.14 | 0.37 | 0.06 | 0.4 | 0.18 |
| 11 | 100112127 | rs622451 | -2.365 | -0.991 | -1.35 | -1.113 | 0.002 | -0.002 | -0.002 | 0.688 | 0.83 | 0.86 | 0.89 | 0.94 | 0.85 | 0.84 |
| 11 | 100118024 | rs10894660 | 0.312 | -0.631 | -0.892 | -0.768 | -0.024 | 0.036 | 0.036 | 0 | 0.26 | 0.46 | 0.38 | 0.63 | 0.21 | 0.44 |
| 11 | 100118859 | rs11223396 | 2.152 | 0.212 | -1.609 | -0.885 | -0.015 | 0.018 | 0.018 | 0 | 0.17 | 0.13 | 0.22 | 0.05 | 0.22 | 0.12 |
| 11 | 100119707 | rs10894664 | 1.193 | 0.14 | -1.491 | -0.66 | 0.004 | -0.001 | -0.001 | 0 | 0.05 | 0.07 | 0.09 | 0.06 | 0.1 | 0.1 |

| | | | | | | | | | | | | | | | | |
|---|---|---|---|---|---|---|---|---|---|---|---|---|---|---|---|---|
| 11 | 100124493 | rs540313 | -0.861 | -0.254 | -1.65 | -0.717 | 0.045 | 0.059 | 0.059 | 0 | 0.63 | 0.85 | 0.8 | 0.76 | 0.69 | 0.72 |
| 11 | 100126394 | rs2298472 | 1.425 | 0.16 | -1.956 | 0.03 | -0.014 | -0.017 | -0.017 | 0 | 0.06 | 0 | 0.06 | 0.29 | 0.05 | 0.05 |
| 11 | 100133808 | rs1944191 | -1.392 | 0.36 | -1.944 | -1.422 | -0.017 | 0.02 | 0.02 | 0 | 0.88 | 0.92 | 0.88 | 1 | 0.83 | 0.95 |
| 11 | 100134297 | rs11223453 | 0.84 | -0.35 | -2.378 | -1.848 | 0.038 | 0.137 | 0.137 | 0 | 0.3 | 0.06 | 0.15 | 0.09 | 0.24 | 0.21 |
| 11 | 100135324 | rs10894674 | -0.073 | -0.046 | -1.872 | -1.567 | 0 | -0.001 | -0.001 | 0.072 | 0.35 | 0.62 | 0.35 | 0.27 | 0.37 | 0.32 |
| 11 | 100137346 | rs1348271 | 1.255 | 0.249 | -1.979 | -1.451 | -0.02 | 0.022 | 0.022 | 0 | 0.16 | 0.1 | 0.21 | 0.04 | 0.24 | 0.16 |
| 11 | 100137718 | rs10501958 | 0.975 | -0.335 | -1.973 | -1.178 | 0.011 | 0.06 | 0.06 | 0 | 0.2 | 0.06 | 0.12 | 0.09 | 0.18 | 0.18 |
| 11 | 100138732 | rs1944158 | 1.054 | -0.023 | -1.183 | -0.396 | 0.001 | -0.034 | -0.034 | 0.704 | 0.06 | 0 | 0.03 | 0.16 | 0.03 | 0.06 |
| 11 | 100142772 | rs2165662 | -0.471 | -0.117 | -1.346 | -1.198 | 0.007 | 0.188 | 0.188 | 0 | 0.5 | 0.77 | 0.59 | 0.82 | 0.49 | 0.59 |
| 11 | 100142834 | rs2165661 | 0.653 | 0.053 | -1.188 | -0.878 | -0.011 | -0.003 | -0.003 | 4.722 | 0.19 | 0.19 | 0.23 | 0.2 | 0.25 | 0.18 |
| 11 | 100148851 | rs474241 | 0.361 | -0.866 | -0.582 | -0.76 | -0.001 | -0.004 | -0.004 | 0 | 0.84 | 0.8 | 0.82 | 0.94 | 0.84 | 0.94 |
| 11 | 100155500 | rs11223518 | 1.301 | -0.752 | -0.647 | -1.164 | -0.015 | 0.213 | 0.213 | 0 | 0.44 | 0.13 | 0.34 | 0.12 | 0.51 | 0.48 |
| 11 | 100163957 | rs10791305 | 1.331 | -0.774 | -0.645 | -1.148 | -0.023 | 0.207 | 0.207 | 0 | 0.44 | 0.13 | 0.34 | 0.12 | 0.5 | 0.48 |
| 11 | 100165790 | rs1442449 | 1.82 | -0.966 | 0.248 | -0.083 | 0.012 | -0.003 | -0.003 | 0 | 0.76 | 0.21 | 0.76 | 0.74 | 0.81 | 0.92 |
| 11 | 100168289 | rs1944178 | 0.521 | -0.471 | 0.171 | -0.252 | -0.002 | -0.044 | -0.044 | 0 | 0.9 | 0.81 | 0.89 | 1 | 0.87 | 0.97 |
| 11 | 100170955 | rs1944181 | 0.751 | -0.555 | 0.735 | -0.172 | -0.002 | -0.055 | -0.055 | 0 | 0.85 | 0.56 | 0.85 | 0.95 | 0.84 | 0.96 |
| 11 | 100171925 | rs550653 | 0.926 | -0.905 | 0.282 | -0.153 | 0.018 | 0.015 | 0.015 | 0 | 0.64 | 0.74 | 0.55 | 0.35 | 0.7 | 0.58 |
| 11 | 100173388 | rs10791312 | -0.723 | -1.006 | 0.634 | -0.106 | -0.02 | 0.034 | 0.034 | 0 | 0.25 | 0.38 | 0.39 | 0.61 | 0.18 | 0.44 |
| 11 | 100175822 | rs10501959 | -1.684 | -0.948 | -0.049 | -0.33 | 0.061 | 0.209 | 0.209 | 0 | 0.45 | 0.82 | 0.52 | 0.87 | 0.36 | 0.54 |
| 11 | 100178540 | rs562990 | -1.628 | -0.966 | 0.027 | -0.458 | 0.063 | 0.166 | 0.166 | 0 | 0.44 | 0.78 | 0.56 | 0.9 | 0.36 | 0.54 |
| 11 | 100186880 | rs1216170 | -1.734 | -0.97 | -0.005 | -0.441 | 0.064 | 0.202 | 0.202 | 0 | 0.46 | 0.79 | 0.57 | 0.89 | 0.37 | 0.55 |
| 11 | 100187449 | rs17094869 | -0.329 | -0.957 | 0.199 | -0.434 | -0.005 | -0.025 | -0.025 | 0 | 0.15 | 0.19 | 0.11 | 0.04 | 0.14 | 0.02 |
| 11 | 100187782 | rs2187395 | -0.326 | -0.839 | 0.397 | 0.062 | 0.006 | -0.026 | -0.026 | 1.354 | 0.22 | 0.12 | 0.36 | 0.66 | 0.17 | 0.42 |
| 11 | 100189682 | rs7111786 | -0.72 | -1.017 | 0.973 | 0.982 | -0.054 | 0.205 | 0.205 | 0 | 0.15 | 0.38 | 0.35 | 0.38 | 0.07 | 0.31 |
| 11 | 100200417 | rs1216190 | -1.667 | -0.537 | -0.027 | 0.133 | -0.036 | 0.066 | 0.066 | 0.674 | 0.72 | 0.86 | 0.69 | 0.97 | 0.66 | 0.67 |
| 11 | 100203054 | rs11534484 | 0.258 | -0.759 | -0.254 | 0.026 | -0.005 | -0.021 | -0.021 | 0 | 0.14 | 0.16 | 0.08 | 0.04 | 0.12 | 0.02 |
| 11 | 100207638 | rs1944175 | 0.768 | -0.574 | -0.545 | -0.598 | -0.001 | -0.018 | -0.018 | 0.012 | 0.31 | 0.32 | 0.17 | 0.06 | 0.31 | 0.15 |
| 11 | 100210203 | rs492107 | 0.674 | -0.614 | -0.707 | -0.263 | 0.001 | -0.003 | -0.003 | 0.453 | 0.11 | 0.11 | 0.28 | 0.32 | 0.1 | 0.26 |
| 11 | 100210323 | rs540267 | -1.778 | -0.577 | -0.836 | -0.562 | -0.124 | 0.171 | 0.171 | 1.004 | 0.73 | 0.99 | 0.72 | 0.97 | 0.67 | 0.67 |
| 11 | 100214691 | rs1442450 | 0.188 | -0.476 | -0.079 | 0.079 | 0 | 0.017 | 0.017 | 0.068 | 0.32 | 0.28 | 0.21 | 0.06 | 0.32 | 0.15 |
| 11 | 100223636 | rs501892 | -0.651 | -0.153 | 0.102 | 0.924 | -0.013 | 0.12 | 0.12 | 0 | 0.15 | 0.28 | 0.34 | 0.36 | 0.1 | 0.31 |
| 11 | 100226773 | rs10894795 | -0.088 | -0.097 | -0.412 | -0.034 | -0.006 | 0.01 | 0.01 | 0 | 0.31 | 0.31 | 0.23 | 0.08 | 0.35 | 0.16 |
| 11 | 100226962 | rs1216182 | -0.415 | -0.927 | 0.059 | 0.458 | -0.009 | 0.01 | 0.01 | 0 | 0.19 | 0.2 | 0.26 | 0.32 | 0.12 | 0.28 |
| 11 | 100237927 | rs1216175 | -0.937 | -1.303 | 0.214 | 0.546 | 0.038 | -0.025 | -0.025 | 0 | 0.39 | 0.66 | 0.34 | 0.49 | 0.29 | 0.36 |
| 11 | 100238395 | rs11223788 | 1.036 | -1.436 | 0.362 | 0.403 | 0 | -0.011 | -0.011 | 0 | 0.16 | 0.15 | 0.11 | 0.01 | 0.17 | 0.06 |
| 11 | 100241732 | rs1216174 | -0.265 | -1.43 | 0.687 | 0.523 | -0.026 | 0.087 | 0.087 | 0 | 0.2 | 0.25 | 0.22 | 0.45 | 0.12 | 0.31 |
| 11 | 100242976 | rs473092 | -0.251 | -1.43 | 0.686 | 0.622 | -0.027 | 0.1 | 0.1 | 0 | 0.2 | 0.27 | 0.22 | 0.44 | 0.12 | 0.3 |
| 11 | 100243478 | rs1301784 | -1.455 | -1.533 | -0.077 | 0.23 | -0.106 | 0.203 | 0.203 | 0.163 | 0.6 | 0.93 | 0.61 | 0.93 | 0.47 | 0.65 |
| 11 | 100246575 | rs12225356 | -0.935 | -1.402 | 0.617 | 0.708 | 0.009 | -0.017 | -0.017 | 0 | 0.36 | 0.37 | 0.34 | 0.58 | 0.32 | 0.37 |
| 11 | 100269350 | rs1216327 | -0.222 | -1.522 | 0.871 | 0.706 | -0.019 | 0.068 | 0.068 | 0 | 0.2 | 0.24 | 0.22 | 0.41 | 0.12 | 0.39 |
| 11 | 100271469 | rs1216330 | -2.285 | -1.486 | 0.972 | 0.579 | -0.016 | 0.107 | 0.107 | 0 | 0.58 | 0.79 | 0.58 | 0.98 | 0.44 | 0.69 |
| 11 | 100275296 | rs7109051 | 0.015 | -1.547 | 0.97 | 0.874 | -0.007 | 0.006 | 0.006 | 0 | 0.17 | 0.26 | 0.14 | 0.21 | 0.14 | 0.11 |
| 11 | 100276633 | rs4466797 | -0.121 | -1.504 | 0.989 | 0.875 | 0.001 | -0.036 | -0.036 | 0 | 0.21 | 0.18 | 0.26 | 0.48 | 0.18 | 0.2 |
| 11 | 100291752 | rs1235612 | -2.049 | -1.367 | 1.127 | 0.591 | -0.008 | 0.034 | 0.034 | 0 | 0.57 | 0.69 | 0.58 | 0.98 | 0.45 | 0.68 |
| 11 | 100308021 | rs10894961 | -0.651 | -1.958 | 1.445 | 1.313 | -0.017 | 0.069 | 0.069 | 0 | 0.22 | 0.29 | 0.23 | 0.4 | 0.15 | 0.39 |
| 11 | 100318174 | rs10894964 | -0.43 | -1.971 | 1.436 | 1.316 | 0.007 | -0.029 | -0.029 | 0 | 0.2 | 0.13 | 0.23 | 0.37 | 0.14 | 0.38 |
| 11 | 100321140 | rs6590789 | -1.773 | -2.088 | 1.411 | 1.195 | 0.031 | 0.062 | 0.062 | 0.043 | 0.42 | 0.7 | 0.48 | 0.89 | 0.28 | 0.57 |
| 11 | 100322582 | rs7947611 | -0.693 | -1.768 | 1.541 | 1.439 | -0.012 | 0.041 | 0.041 | 1.13 | 0.2 | 0.31 | 0.23 | 0.49 | 0.13 | 0.19 |
| 11 | 100329957 | rs11224242 | -0.257 | -1.825 | 1.356 | 1.409 | -0.021 | 0.086 | 0.086 | 0 | 0.17 | 0.24 | 0.21 | 0.38 | 0.09 | 0.36 |
| 11 | 100334186 | rs7945636 | -0.431 | -1.799 | 1.246 | 1.415 | 0.003 | -0.004 | -0.004 | 0 | 0.18 | 0.19 | 0.23 | 0.49 | 0.13 | 0.19 |
| 11 | 100337900 | rs11224244 | 0.046 | -1.815 | 1.164 | 1.357 | -0.009 | -0.046 | -0.046 | 0 | 0.15 | 0.01 | 0.12 | 0.23 | 0.11 | 0.16 |
| 11 | 100342790 | rs4561179 | -0.012 | -1.954 | 1.057 | 1.197 | 0.011 | 0.032 | 0.032 | 0 | 0.2 | 0.06 | 0.11 | 0.08 | 0.17 | 0.15 |
| 11 | 100342880 | rs12576266 | 0.62 | -1.942 | 1.098 | 1.215 | 0.007 | -0.014 | -0.014 | 2.66 | 0.13 | 0.08 | 0.13 | 0.23 | 0.1 | 0.15 |
| 11 | 100344059 | rs1216466 | -0.742 | -1.896 | 1.117 | 1.138 | -0.031 | 0.08 | 0.08 | 0 | 0.21 | 0.35 | 0.31 | 0.63 | 0.11 | 0.39 |
| 11 | 100345438 | rs1216468 | -2.214 | -1.611 | 0.561 | 0.845 | -0.02 | 0.211 | 0.211 | 0 | 0.58 | 0.87 | 0.58 | 0.98 | 0.4 | 0.7 |
| 11 | 100384481 | rs7935363 | -1.168 | -1.662 | 0.712 | 1.334 | -0.038 | 0.092 | 0.092 | 0.098 | 0.26 | 0.46 | 0.25 | 0.47 | 0.18 | 0.23 |
| 11 | 100388584 | rs1216507 | -2.136 | -1.678 | 0.411 | 0.926 | 0.008 | 0.083 | 0.083 | 0 | 0.57 | 0.76 | 0.57 | 0.98 | 0.41 | 0.65 |
| 11 | 100390879 | rs1216508 | -0.932 | -1.649 | -0.191 | 1.108 | 0.014 | -0.001 | -0.001 | 0 | 0.34 | 0.27 | 0.36 | 0.63 | 0.27 | 0.45 |
| 11 | 100391861 | rs1942108 | -1.924 | -1.538 | -0.259 | 0.948 | -0.006 | -0.124 | -0.124 | 0 | 0.55 | 0.5 | 0.55 | 0.98 | 0.41 | 0.65 |
| 11 | 100405406 | rs1216472 | 0.164 | -1.31 | -0.691 | 1.606 | -0.014 | -0.084 | -0.084 | 0 | 0.12 | 0.01 | 0.2 | 0.28 | 0.09 | 0.32 |
| 11 | 100413065 | rs1216481 | -0.249 | -1.409 | -0.427 | 1.351 | -0.003 | 0.014 | 0.014 | 0 | 0.26 | 0.26 | 0.2 | 0.45 | 0.22 | 0.35 |
| 11 | 100413614 | rs10501962 | -0.441 | -1.251 | -0.676 | 1.651 | 0.008 | -0.01 | -0.01 | 0 | 0.16 | 0.09 | 0.14 | 0.23 | 0.11 | 0.13 |
| 11 | 100414039 | rs12421427 | -0.487 | -0.833 | -0.377 | 1.391 | -0.019 | 0.019 | 0.019 | 0 | 0.12 | 0.15 | 0.19 | 0.14 | 0.06 | 0.13 |
| 11 | 100414877 | rs1216502 | 0.641 | -0.081 | -0.452 | 1.263 | -0.009 | 0.049 | 0.049 | 0 | 0.75 | 0.63 | 0.77 | 0.45 | 0.78 | 0.6 |
| 11 | 100421331 | rs1939309 | 0.079 | 0.117 | -0.885 | 0.383 | 0.004 | 0.024 | 0.024 | 0 | 0.65 | 0.73 | 0.7 | 0.68 | 0.61 | 0.64 |
| 11 | 100422785 | rs10501964 | -0.402 | -0.253 | -0.924 | -0.175 | 0.004 | -0.01 | -0.01 | 0 | 0.14 | 0.15 | 0.05 | 0 | 0.09 | 0.06 |
| 11 | 100423278 | rs12280410 | 0.73 | 0.277 | -1.286 | -0.103 | -0.003 | 0 | 0 | 0.003 | 0.39 | 0.61 | 0.56 | 0.56 | 0.38 | 0.44 |
| 11 | 100424060 | rs6590793 | 0.289 | 0.286 | -1.352 | -0.233 | 0.003 | 0.067 | 0.067 | 0 | 0.48 | 0.63 | 0.64 | 0.68 | 0.48 | 0.53 |
| 11 | 100428969 | rs17095168 | 0.721 | 0.398 | -1.227 | 0.05 | -0.039 | -0.042 | -0.042 | 0 | 0.09 | 0 | 0.08 | 0.12 | 0.1 | 0.08 |

| | | | | | | | | | | | | | | | |
|---|---|---|---|---|---|---|---|---|---|---|---|---|---|---|---|
| 11 | 100430053 | rs4754687 | -0.496 | 0.171 | -1.371 | -0.697 | -0.018 | 0.041 | 0.041 | 0 | 0.76 | 0.87 | 0.79 | 1 | 0.7 | 0.82 |
| 11 | 100430096 | rs10501965 | 1.014 | -0.039 | -1.33 | -0.633 | -0.006 | 0.057 | 0.057 | 0 | 0.05 | 0.04 | 0.08 | 0.31 | 0.04 | 0.11 |
| 11 | 100430269 | rs967032 | -0.528 | 0.26 | -1.367 | -0.696 | -0.024 | 0.043 | 0.043 | 0 | 0.76 | 0.86 | 0.79 | 1 | 0.69 | 0.82 |
| 11 | 100434410 | rs4754688 | 0.65 | -0.205 | -0.354 | -0.819 | 0 | -0.004 | -0.004 | 0.308 | 0.58 | 0.48 | 0.6 | 0.56 | 0.55 | 0.64 |
| 11 | 100437352 | rs1216576 | 0.679 | 0.041 | -0.63 | -0.88 | -0.016 | 0.056 | 0.056 | 0 | 0.19 | 0.09 | 0.15 | 0 | 0.22 | 0.14 |
| 11 | 100438447 | rs2845930 | -1.815 | -0.262 | -0.354 | -0.358 | -0.024 | -0.024 | -0.024 | 0.637 | 0.6 | 0.78 | 0.49 | 0.55 | 0.55 | 0.47 |
| 11 | 100447458 | rs1216592 | -0.447 | -0.344 | 0.239 | -0.468 | -0.008 | -0.049 | -0.049 | 0.122 | 0.32 | 0.44 | 0.25 | 0.11 | 0.3 | 0.22 |
| 11 | 100449142 | rs650297 | -0.173 | -0.358 | 0.207 | 0.198 | -0.012 | -0.008 | -0.008 | 0.378 | 0.31 | 0.24 | 0.22 | 0.32 | 0.33 | 0.26 |
| 11 | 100450082 | rs635541 | 0.661 | -0.335 | 0 | -0.063 | -0.013 | 0.048 | 0.048 | 0 | 0.2 | 0.1 | 0.13 | 0 | 0.22 | 0.15 |
| 11 | 100454936 | rs618781 | -0.738 | -0.363 | 0.062 | 0.166 | -0.003 | 0.001 | 0.001 | 0 | 0.41 | 0.55 | 0.3 | 0.41 | 0.37 | 0.31 |
| 11 | 100456604 | rs11224302 | 1.216 | -0.214 | -0.333 | 0.304 | -0.041 | -0.054 | -0.054 | 0.418 | 0.09 | 0.01 | 0.06 | 0.28 | 0.11 | 0.1 |
| 11 | 100464114 | rs1216680 | 0.045 | -0.313 | -0.054 | 0.355 | 0.001 | -0.017 | -0.017 | 0.439 | 0.27 | 0.15 | 0.27 | 0.38 | 0.3 | 0.27 |
| 11 | 100465960 | rs639494 | -1.363 | -0.648 | 0.022 | 0.029 | 0.029 | 0.075 | 0.075 | 0.478 | 0.5 | 0.81 | 0.45 | 0.66 | 0.46 | 0.41 |
| 11 | 100465985 | rs1216683 | 0.927 | -0.573 | 0.08 | -0.007 | -0.006 | 0.019 | 0.019 | 0 | 0.19 | 0.07 | 0.15 | 0.11 | 0.21 | 0.15 |
| 11 | 100469420 | rs661264 | 0.812 | -0.583 | 0.346 | -0.011 | 0 | 0.002 | 0.002 | 0 | 0.2 | 0.16 | 0.16 | 0.11 | 0.21 | 0.14 |
| 11 | 100472288 | rs11224313 | 0.328 | -0.539 | 0.772 | -0.07 | -0.015 | -0.036 | -0.036 | 0 | 0.08 | 0.17 | 0.06 | 0 | 0.04 | 0.04 |
| 11 | 100475009 | rs684792 | -0.102 | -0.422 | 0.114 | -0.491 | -0.002 | 0.012 | 0.012 | 0 | 0.75 | 0.78 | 0.78 | 0.94 | 0.76 | 0.84 |
| 11 | 100486632 | rs601727 | 1.286 | -0.485 | -0.173 | -0.585 | -0.062 | -0.064 | -0.064 | 0 | 0.45 | 0.11 | 0.54 | 0.44 | 0.53 | 0.57 |
| 11 | 100486827 | rs1216525 | -0.633 | -0.299 | 0.141 | -0.694 | 0.002 | -0.025 | -0.025 | 0 | 0.9 | 0.91 | 0.86 | 0.98 | 0.88 | 0.93 |
| 11 | 100494108 | rs587277 | -0.103 | -0.48 | 0.361 | -0.591 | -0.024 | -0.095 | -0.095 | 2.71 | 0.15 | 0.33 | 0.06 | 0 | 0.09 | 0.07 |
| 11 | 100497485 | rs12284327 | -0.58 | -0.351 | 0.446 | -0.242 | -0.011 | 0.092 | 0.092 | 0 | 0.1 | 0.27 | 0.12 | 0.27 | 0.07 | 0.14 |
| 11 | 100498146 | rs11224328 | 0.567 | -0.248 | 0.379 | -0.064 | -0.013 | 0.084 | 0.084 | 0 | 0.06 | 0.11 | 0.08 | 0.25 | 0.04 | 0.13 |
| 11 | 100498349 | rs583366 | -0.001 | 0.826 | 0.021 | -0.587 | 0.025 | -0.069 | -0.069 | 0 | 0.82 | 0.73 | 0.78 | 0.98 | 0.78 | 0.88 |
| 11 | 100499866 | rs7932851 | -1.339 | 0.464 | -0.827 | -0.466 | -0.045 | -0.037 | -0.037 | 0 | 0.47 | 0.86 | 0.45 | 0.53 | 0.46 | 0.36 |
| 11 | 100504030 | rs11224337 | -0.971 | 0.396 | -0.354 | -0.602 | 0.006 | 0.017 | 0.017 | 0 | 0.35 | 0.59 | 0.38 | 0.47 | 0.36 | 0.31 |
| 11 | 100509529 | rs947866 | -0.864 | 0.231 | -0.367 | -0.75 | 0 | 0.006 | 0.006 | 0 | 0.34 | 0.66 | 0.31 | 0.23 | 0.31 | 0.29 |
| 11 | 100510448 | rs4754691 | 1.074 | 0.05 | -1.053 | -1.435 | -0.041 | 0.02 | 0.02 | 0 | 0.29 | 0.16 | 0.44 | 0.72 | 0.37 | 0.52 |
| 11 | 100511618 | rs916483 | -0.916 | -0.08 | -1.047 | -1.52 | 0.024 | 0.066 | 0.066 | 0 | 0.68 | 0.84 | 0.78 | 0.95 | 0.71 | 0.83 |
| 11 | 100512650 | rs1032778 | 1.272 | 0.223 | -1.015 | -1.402 | -0.032 | -0.079 | -0.079 | 0 | 0.25 | 0.12 | 0.37 | 0.47 | 0.29 | 0.49 |
| 11 | 100513116 | rs11224348 | 0.982 | 0.28 | -0.882 | -1.081 | 0 | 0.022 | 0.022 | 0 | 0.07 | 0.08 | 0.06 | 0.24 | 0.07 | 0.05 |
| 11 | 100513532 | rs9888185 | 0.208 | -0.188 | -0.794 | -1.522 | -0.005 | 0.023 | 0.023 | 0 | 0.66 | 0.61 | 0.67 | 0.49 | 0.67 | 0.76 |
| 11 | 100513913 | rs1032779 | -0.641 | -0.103 | -1.087 | -1.535 | 0.004 | 0.003 | 0.003 | 0 | 0.74 | 0.8 | 0.75 | 0.73 | 0.73 | 0.81 |
| 11 | 100517516 | rs10791424 | -0.785 | 0.001 | -1.494 | -1.714 | -0.007 | 0.098 | 0.098 | 0 | 0.77 | 0.94 | 0.84 | 0.95 | 0.76 | 0.9 |
| 11 | 100534370 | rs4753979 | -0.824 | -0.051 | -1.513 | -1.901 | 0.011 | 0.07 | 0.07 | 0 | 0.77 | 0.89 | 0.85 | 1 | 0.77 | 0.91 |
| 11 | 100534422 | rs7117419 | 0.629 | 0.521 | -1.569 | -1.571 | -0.051 | -0.054 | -0.054 | 0 | 0.12 | 0.03 | 0.12 | 0.27 | 0.17 | 0.08 |
| 11 | 100536632 | rs1532710 | 2.194 | 0.497 | -1.888 | -1.738 | -0.052 | -0.138 | -0.138 | 0 | 0.22 | 0.04 | 0.34 | 0.46 | 0.25 | 0.49 |
| 11 | 100539779 | rs7130376 | 1.712 | 0.561 | -1.774 | -1.734 | -0.023 | 0.015 | 0.015 | 0 | 0.07 | 0.02 | 0.07 | 0.04 | 0.11 | 0.04 |
| 11 | 100543056 | rs7350545 | 0.459 | 0.346 | -1.273 | -1.9 | -0.014 | 0.015 | 0.015 | 0 | 0.39 | 0.28 | 0.51 | 0.73 | 0.45 | 0.59 |
| 11 | 100546036 | rs6590805 | -1.206 | 0.36 | -1.329 | -2.071 | -0.004 | -0.012 | -0.012 | 0 | 0.54 | 0.74 | 0.51 | 0.53 | 0.53 | 0.41 |
| 11 | 100553728 | rs7107791 | 0.191 | 0.447 | -1.293 | -2.149 | 0.006 | 0.003 | 0.003 | 0.021 | 0.18 | 0.15 | 0.15 | 0.26 | 0.22 | 0.12 |
| 11 | 100555114 | rs4312035 | 1.97 | 0.46 | -1.471 | -2.265 | -0.075 | -0.148 | -0.148 | 0 | 0.24 | 0.03 | 0.35 | 0.47 | 0.28 | 0.52 |
| 11 | 100560519 | rs6590809 | 0.722 | 0.684 | -0.976 | -1.819 | 0.02 | -0.042 | -0.042 | 0.01 | 0.11 | 0.15 | 0.12 | 0.04 | 0.13 | 0.05 |
| 11 | 100578089 | rs7943581 | 1.873 | 0.676 | -0.778 | -1.891 | 0.026 | -0.021 | -0.021 | 0 | 0.06 | 0.1 | 0.07 | 0.03 | 0.08 | 0.03 |
| 11 | 100582941 | rs2298611 | 0.777 | 0.273 | -0.906 | -2.465 | 0 | -0.01 | -0.01 | 0.2 | 0.22 | 0.23 | 0.22 | 0.03 | 0.25 | 0.17 |
| 11 | 100585889 | rs10895009 | -1.391 | 0.138 | -0.772 | -2.406 | -0.01 | -0.002 | -0.002 | 0 | 0.46 | 0.73 | 0.43 | 0.52 | 0.49 | 0.41 |
| 11 | 100590270 | rs6590814 | -1.776 | 0.103 | -0.856 | -2.518 | -0.001 | 0.011 | 0.011 | 0 | 0.52 | 0.73 | 0.47 | 0.52 | 0.55 | 0.44 |
| 11 | 100603314 | rs7946913 | -0.816 | 0.106 | -0.707 | -2.452 | 0.008 | 0.039 | 0.039 | 0 | 0.41 | 0.69 | 0.38 | 0.3 | 0.4 | 0.36 |
| 11 | 100615623 | rs11224426 | -0.134 | 0.08 | -0.574 | -2.368 | -0.006 | -0.015 | -0.015 | 0 | 0.31 | 0.38 | 0.3 | 0.26 | 0.32 | 0.32 |
| 11 | 100623220 | rs2088953 | 0.547 | 0.133 | -0.636 | -2.471 | 0 | -0.024 | -0.024 | 0.9 | 0.24 | 0.23 | 0.23 | 0.02 | 0.29 | 0.21 |
| 11 | 100624538 | rs607562 | -2.456 | 0.165 | -0.974 | -2.656 | -0.063 | -0.147 | -0.147 | 0 | 0.75 | 0.97 | 0.63 | 0.52 | 0.76 | 0.5 |
| 11 | 100633303 | rs7126324 | 0.656 | 0.181 | -0.793 | -2.513 | 0.001 | -0.02 | -0.02 | 0 | 0.25 | 0.23 | 0.23 | 0.02 | 0.3 | 0.21 |
| 11 | 100633321 | rs7125452 | -1.752 | 0.158 | -0.62 | -2.596 | -0.002 | 0.015 | 0.015 | 0 | 0.49 | 0.7 | 0.44 | 0.52 | 0.53 | 0.42 |
| 11 | 100640301 | rs12271498 | 1.126 | 0.319 | -0.296 | -2.242 | 0.009 | 0.074 | 0.074 | 0 | 0.1 | 0.2 | 0.12 | 0.3 | 0.1 | 0.13 |
| 11 | 100644491 | rs6590820 | -0.371 | 0.36 | -0.309 | -2.269 | 0.018 | 0.099 | 0.099 | 0 | 0.18 | 0.29 | 0.18 | 0.5 | 0.21 | 0.2 |
| 11 | 100645962 | rs499241 | 0.778 | 0.731 | -0.665 | -2.493 | 0.013 | -0.073 | -0.073 | 0 | 0.18 | 0.26 | 0.23 | 0.02 | 0.26 | 0.19 |
| 11 | 100646982 | rs17095481 | 0.841 | 0.881 | -0.721 | -2.473 | -0.002 | -0.002 | -0.002 | 4.275 | 0.16 | 0.15 | 0.1 | 0.18 | 0.17 | 0.11 |
| 11 | 100648073 | rs655922 | -0.49 | 0.763 | -0.636 | -2.488 | 0.01 | -0.139 | -0.139 | 0.184 | 0.35 | 0.51 | 0.36 | 0.06 | 0.4 | 0.27 |
| 11 | 100654214 | rs11224444 | 1.975 | 0.701 | -0.71 | -2.261 | -0.02 | 0.031 | 0.031 | 0 | 0.09 | 0.01 | 0.1 | 0.03 | 0.1 | 0.05 |
| 11 | 100659513 | rs582409 | 0.362 | 0.649 | -0.733 | -2.174 | 0.004 | 0.03 | 0.03 | 0 | 0.51 | 0.32 | 0.61 | 0.53 | 0.54 | 0.66 |
| 11 | 100663930 | rs545547 | 0.273 | 0.637 | -0.482 | -2.189 | -0.007 | -0.107 | -0.107 | 0.562 | 0.23 | 0.38 | 0.17 | 0.04 | 0.2 | 0.1 |
| 11 | 100675552 | rs495604 | 0.682 | 0.613 | -0.377 | -2.194 | -0.012 | -0.117 | -0.117 | 0 | 0.21 | 0.45 | 0.18 | 0.04 | 0.17 | 0.08 |
| 11 | 100681211 | rs618291 | -0.07 | 0.765 | -0.791 | -2.201 | 0 | -0.019 | -0.019 | 0 | 0.57 | 0.59 | 0.61 | 0.73 | 0.54 | 0.65 |
| 11 | 100686119 | rs674693 | 0.297 | 0.793 | -0.775 | -2.136 | 0.002 | -0.017 | -0.017 | 0.055 | 0.27 | 0.39 | 0.21 | 0.21 | 0.27 | 0.19 |
| 11 | 100687641 | rs680604 | -0.08 | 0.666 | -0.835 | -2.798 | -0.004 | -0.023 | -0.023 | 0 | 0.55 | 0.54 | 0.68 | 0.71 | 0.61 | 0.75 |
| 11 | 100690827 | rs17573475 | -0.326 | 0.675 | -0.606 | -2.101 | 0.001 | -0.05 | -0.05 | 0 | 0.23 | 0.22 | 0.26 | 0.02 | 0.31 | 0.2 |
| 11 | 100694891 | rs556146 | -0.704 | 0.578 | -0.282 | -1.712 | 0 | 0 | 0 | 0.121 | 0.78 | 0.81 | 0.83 | 0.74 | 0.8 | 0.85 |
| 11 | 100696230 | rs4754702 | -0.386 | 0.845 | -0.212 | -1.519 | 0.008 | -0.075 | -0.075 | 0.104 | 0.18 | 0.22 | 0.23 | 0.02 | 0.24 | 0.2 |
| 11 | 100699139 | rs629864 | -0.439 | 0.844 | -0.168 | -1.569 | 0.018 | -0.161 | -0.161 | 0 | 0.68 | 0.56 | 0.78 | 0.96 | 0.65 | 0.83 |
| 11 | 100701208 | rs599925 | -0.581 | 0.678 | -0.419 | -1.093 | 0.01 | 0.027 | 0.027 | 0 | 0.5 | 0.67 | 0.55 | 0.63 | 0.43 | 0.65 |

| | | | | | | | | | | | | | | | | |
|---|---|---|---|---|---|---|---|---|---|---|---|---|---|---|---|---|
| 11 | 100701493 | rs3847566 | 0.836 | 0.528 | -0.726 | -0.698 | -0.021 | -0.025 | -0.025 | 0 | 0.24 | 0.15 | 0.38 | 0.39 | 0.28 | 0.51 |
| 11 | 100705780 | rs561267 | 1.253 | 0.359 | -0.757 | -0.685 | -0.001 | -0.019 | -0.019 | 0 | 0.1 | 0.07 | 0.11 | 0.26 | 0.11 | 0.09 |
| 11 | 100713227 | rs12785918 | 0.914 | 0.404 | -0.794 | -0.941 | -0.028 | -0.049 | -0.049 | 0 | 0.24 | 0.14 | 0.39 | 0.45 | 0.28 | 0.51 |
| 11 | 100718626 | rs637127 | -1.996 | 0.298 | -0.807 | -1.119 | -0.014 | 0.064 | 0.064 | 0 | 0.77 | 0.93 | 0.86 | 0.97 | 0.76 | 0.88 |
| 11 | 100720108 | rs3858414 | 1.722 | 0.35 | -0.847 | -1.049 | -0.041 | -0.129 | -0.129 | 0.13 | 0.19 | 0.03 | 0.34 | 0.44 | 0.18 | 0.42 |
| 11 | 100724571 | rs17574672 | 1.833 | 0.336 | -0.907 | -1.256 | -0.038 | 0.085 | 0.085 | 0 | 0.16 | 0.02 | 0.13 | 0 | 0.2 | 0.05 |
| 11 | 100725349 | rs3858416 | 1.03 | 0.244 | -0.662 | -1.019 | -0.01 | -0.003 | -0.003 | 0 | 0.09 | 0.15 | 0.09 | 0.1 | 0.05 | 0.06 |
| 11 | 100729086 | rs551860 | 1.395 | 0.498 | -0.481 | -0.907 | -0.01 | -0.043 | -0.043 | 0 | 0.09 | 0.01 | 0.09 | 0.25 | 0.09 | 0.12 |
| 11 | 100729841 | rs662501 | -0.794 | 0.492 | -0.229 | -0.985 | 0 | 0 | 0 | 0 | 0.65 | 0.68 | 0.75 | 0.72 | 0.68 | 0.76 |
| 11 | 100732373 | rs1941373 | 0.267 | 0.586 | -0.342 | -0.979 | 0.02 | -0.044 | -0.044 | 0 | 0.08 | 0.2 | 0.08 | 0 | 0.12 | 0.04 |
| 11 | 100734010 | rs12274135 | 0.717 | 0.764 | -0.286 | -0.881 | -0.01 | -0.009 | -0.009 | 0 | 0.25 | 0.15 | 0.26 | 0.28 | 0.31 | 0.21 |
| 11 | 100737741 | rs10128729 | 0.55 | 0.778 | -0.192 | -0.837 | -0.003 | -0.002 | -0.002 | 1.546 | 0.26 | 0.21 | 0.28 | 0.28 | 0.32 | 0.22 |
| 11 | 100738459 | rs556823 | -1.699 | 1.011 | -0.711 | -0.782 | -0.015 | 0.092 | 0.092 | 0 | 0.77 | 0.96 | 0.87 | 0.97 | 0.77 | 0.88 |
| 11 | 100739930 | rs10488787 | 0.234 | 1.01 | -0.304 | 0.133 | 0.008 | -0.027 | -0.027 | 0 | 0.3 | 0.22 | 0.4 | 0.45 | 0.24 | 0.53 |
| 11 | 100743498 | rs12808052 | -1.348 | 1.078 | 0.118 | 0.349 | 0.006 | 0.055 | 0.055 | 0.009 | 0.19 | 0.33 | 0.18 | 0.24 | 0.21 | 0.13 |
| 11 | 100744198 | rs7939833 | -0.739 | 1.693 | -0.217 | -0.005 | 0.04 | -0.029 | -0.029 | 0.321 | 0.05 | 0.12 | 0.06 | 0 | 0.11 | 0.05 |
| 11 | 100754906 | rs10895027 | -0.566 | 0.879 | -0.679 | 0.312 | 0.002 | -0.007 | -0.007 | 0.313 | 0.09 | 0.1 | 0.09 | 0.04 | 0.1 | 0.1 |
| 11 | 100759034 | rs4754708 | -0.944 | 1.168 | -1.283 | 0.302 | -0.031 | -0.055 | -0.055 | 1.073 | 0.24 | 0.12 | 0.43 | 0.74 | 0.25 | 0.61 |
| 11 | 100759057 | rs614761 | -0.261 | 0.68 | -1.107 | -0.478 | -0.011 | -0.051 | -0.051 | 0 | 0.86 | 0.8 | 0.91 | 1 | 0.86 | 0.9 |
| 11 | 100760936 | rs662453 | 1.101 | 0.498 | -1.071 | -0.571 | 0.002 | 0.008 | 0.008 | 0.034 | 0.61 | 0.65 | 0.48 | 0.26 | 0.61 | 0.3 |
| 11 | 100762829 | rs17659845 | 1.254 | 1.237 | -1.372 | -0.862 | -0.019 | 0.016 | 0.016 | 0 | 0.06 | 0.01 | 0.06 | 0 | 0.14 | 0.05 |
| 11 | 100762986 | rs601496 | 0.784 | 0.966 | -1.404 | -1.383 | -0.016 | 0.058 | 0.058 | 1.062 | 0.59 | 0.72 | 0.37 | 0 | 0.57 | 0.26 |
| 11 | 100763249 | rs662530 | -0.244 | 0.951 | -1.405 | -1.382 | -0.009 | -0.009 | -0.009 | 0.076 | 0.85 | 0.88 | 0.86 | 1 | 0.73 | 0.85 |
| 11 | 100763308 | rs622957 | 1.137 | 0.922 | -1.398 | -1.377 | 0.008 | 0.136 | 0.136 | 2.565 | 0.54 | 0.75 | 0.32 | 0 | 0.42 | 0.21 |
| 11 | 100772786 | rs12421655 | 0.751 | 0.924 | -1.388 | -1.25 | 0.001 | 0.026 | 0.026 | 0 | 0.07 | 0.04 | 0.12 | 0.26 | 0.05 | 0.04 |
| 11 | 100787002 | rs10488785 | 0.587 | 1.017 | -1.026 | -0.456 | -0.009 | -0.036 | -0.036 | 0 | 0.17 | 0.12 | 0.34 | 0.48 | 0.18 | 0.42 |
| 11 | 100793518 | rs11224513 | 0.284 | 1.098 | -1.185 | -0.327 | -0.011 | -0.023 | -0.023 | 0 | 0.09 | 0.04 | 0.1 | 0.3 | 0.08 | 0.2 |
| 11 | 100793876 | rs11224515 | 0.741 | 0.872 | -2.014 | -1.046 | -0.011 | -0.023 | -0.023 | 0 | 0.09 | 0.04 | 0.1 | 0.3 | 0.08 | 0.2 |
| 11 | 100807130 | rs581921 | 0.287 | 0.816 | -1.862 | -1.192 | -0.003 | -0.004 | -0.004 | 0 | 0.7 | 0.71 | 0.64 | 0.48 | 0.6 | 0.63 |
| 11 | 100807770 | rs11604594 | 2.122 | 0.605 | -1.839 | -1.519 | -0.008 | 0.012 | 0.012 | 0 | 0.11 | 0.06 | 0.07 | 0 | 0.1 | 0.08 |
| 11 | 100810343 | rs520403 | -1.358 | 0.71 | -1.91 | -1.43 | 0 | -0.023 | -0.023 | 0.016 | 0.87 | 0.87 | 0.92 | 1 | 0.87 | 0.9 |
| 11 | 100813264 | rs10501968 | 0.88 | 0.822 | -1.924 | -1.349 | -0.009 | 0.015 | 0.015 | 0 | 0.09 | 0.05 | 0.18 | 0.26 | 0.19 | 0.1 |
| 11 | 100819762 | rs475432 | -0.93 | 0.793 | -1.936 | -1.367 | -0.002 | -0.011 | -0.011 | 0.055 | 0.84 | 0.88 | 0.81 | 0.73 | 0.82 | 0.86 |
| 11 | 100823814 | rs10895033 | 1.009 | 0.366 | -1.689 | -2.074 | 0.008 | 0.039 | 0.039 | 1.047 | 0.34 | 0.17 | 0.19 | 0 | 0.31 | 0.13 |
| 11 | 100824921 | rs522337 | -1.733 | 0.402 | -1.658 | -1.896 | 0.002 | -0.011 | -0.011 | 0.308 | 0.87 | 0.89 | 0.92 | 1 | 0.87 | 0.9 |
| 11 | 100829176 | rs473765 | 0.021 | 0.348 | -1.461 | -2 | 0.001 | 0.013 | 0.013 | 0 | 0.59 | 0.33 | 0.62 | 0.73 | 0.56 | 0.73 |
| 11 | 100830440 | rs7950832 | 0.237 | 0.431 | -1.715 | -2.034 | 0.003 | 0.067 | 0.067 | 2.56 | 0.54 | 0.68 | 0.32 | 0 | 0.43 | 0.21 |
| 11 | 100830693 | rs563460 | -1.447 | 0.385 | -1.787 | -1.889 | -0.004 | -0.024 | -0.024 | 0 | 0.85 | 0.89 | 0.81 | 0.73 | 0.82 | 0.86 |
| 11 | 100837804 | rs612046 | 1.722 | 0.486 | -1.821 | -1.634 | -0.038 | -0.095 | -0.095 | 0 | 0.15 | 0.03 | 0.33 | 0.45 | 0.17 | 0.41 |
| 11 | 100841740 | rs568618 | 0.623 | 0.437 | -1.793 | -2.018 | -0.029 | -0.058 | -0.058 | 0 | 0.23 | 0.12 | 0.42 | 0.73 | 0.25 | 0.6 |
| 11 | 100848015 | rs7931273 | 0.965 | 0.232 | -1.781 | -2.231 | 0.005 | 0.067 | 0.067 | 0.048 | 0.33 | 0.14 | 0.18 | 0 | 0.31 | 0.13 |
| 11 | 100850994 | rs485269 | 0.56 | 0.487 | -1.732 | -1.713 | -0.018 | -0.059 | -0.059 | 0 | 0.23 | 0.12 | 0.42 | 0.68 | 0.23 | 0.59 |
| 11 | 100852768 | rs505458 | 0.488 | 0.489 | -1.729 | -1.693 | -0.018 | -0.059 | -0.059 | 0 | 0.23 | 0.12 | 0.42 | 0.69 | 0.23 | 0.59 |
| 11 | 100858061 | rs3740749 | 0.619 | 0.612 | -1.518 | -1.037 | -0.003 | 0.039 | 0.039 | 0.207 | 0.08 | 0.08 | 0.14 | 0.37 | 0.08 | 0.23 |
| 11 | 100858425 | rs17648695 | 1.98 | 0.75 | -1.502 | -1.555 | -0.024 | 0.023 | 0.023 | 0 | 0.06 | 0.02 | 0.07 | 0 | 0.15 | 0.05 |
| 11 | 100860590 | rs3573 | 1.994 | 0.458 | -1.287 | -1.661 | -0.028 | -0.029 | -0.029 | 0.009 | 0.06 | 0.12 | 0.01 | 0 | 0.01 | 0 |
| 11 | 100862593 | rs499861 | -1.7 | 0.44 | -1.469 | -1.834 | 0.002 | 0.001 | 0.001 | 0 | 0.88 | 0.93 | 0.92 | 1 | 0.87 | 0.91 |
| 11 | 100864146 | rs17654410 | 2.203 | 0.501 | -1.483 | -1.833 | -0.022 | 0.021 | 0.021 | 0 | 0.06 | 0.01 | 0.07 | 0 | 0.15 | 0.05 |
| 11 | 100864367 | rs488365 | -0.451 | 0.411 | -1.412 | -1.549 | -0.008 | -0.005 | -0.005 | 0.011 | 0.78 | 0.81 | 0.73 | 0.81 | 0.67 | 0.81 |
| 11 | 100864744 | rs12420167 | 1.277 | 0.573 | -1.261 | -1.321 | -0.013 | -0.01 | -0.01 | 0 | 0.1 | 0.07 | 0.19 | 0.19 | 0.19 | 0.1 |
| 11 | 100868961 | rs575473 | -0.177 | 0.602 | -1.322 | -1.45 | 0.002 | 0.023 | 0.023 | 2.041 | 0.61 | 0.66 | 0.67 | 0.81 | 0.56 | 0.78 |
| 11 | 100870339 | rs590306 | 0.026 | 0.449 | -1.127 | -2.181 | -0.008 | -0.045 | -0.045 | 0 | 0.32 | 0.49 | 0.31 | 0.23 | 0.25 | 0.26 |
| 11 | 100870494 | rs500112 | -0.65 | 0.668 | -1.525 | -2.596 | -0.009 | 0.18 | 0.18 | 1.991 | 0.63 | 0.89 | 0.77 | 1 | 0.61 | 0.84 |
| 11 | 100870534 | rs563202 | 0.023 | 0.715 | -0.365 | -1.351 | 0.007 | -0.056 | -0.056 | 0.325 | 0.18 | 0.26 | 0.11 | 0.06 | 0.15 | 0.14 |
| 11 | 100874319 | rs523332 | -0.353 | 0.684 | -0.237 | -0.997 | 0.003 | 0 | 0 | 0 | 0.23 | 0.35 | 0.26 | 0.2 | 0.24 | 0.1 |
| 11 | 100878498 | rs473906 | 0.635 | 0.574 | -0.277 | -1.027 | 0.001 | 0.002 | 0.002 | 0 | 0.07 | 0.05 | 0.09 | 0.17 | 0.07 | 0.02 |
| 11 | 100880706 | rs521744 | 0.349 | 0.71 | -0.192 | -1.116 | 0.004 | -0.009 | -0.009 | 1.648 | 0.75 | 0.66 | 0.73 | 0.81 | 0.73 | 0.91 |
| 11 | 100881437 | rs4753989 | 0.565 | 0.732 | -1.117 | -1.938 | -0.055 | 0.161 | 0.161 | 0 | 0.27 | 0.05 | 0.22 | 0.01 | 0.31 | 0.19 |
| 11 | 100890993 | rs11224546 | 0.308 | 0.706 | -0.794 | -1.953 | 0.001 | 0.021 | 0.021 | 0 | 0.13 | 0.16 | 0.28 | 0.72 | 0.12 | 0.41 |
| 11 | 100891754 | rs529808 | -0.138 | 0.159 | -0.312 | -2.125 | 0.001 | -0.007 | -0.007 | 0 | 0.26 | 0.38 | 0.27 | 0.21 | 0.27 | 0.11 |
| 11 | 100894166 | rs639838 | -0.573 | -0.177 | -0.149 | -2.413 | -0.012 | -0.099 | -0.099 | 0 | 0.64 | 0.51 | 0.75 | 0.92 | 0.69 | 0.68 |
| 11 | 100902994 | rs561610 | -1.078 | 0.051 | -0.349 | -2.304 | 0.001 | 0.019 | 0.019 | 0 | 0.7 | 0.78 | 0.79 | 0.93 | 0.7 | 0.7 |
| 11 | 100905297 | rs471767 | -1.084 | -0.013 | -0.148 | -2.4 | 0 | -0.023 | -0.023 | 1.915 | 0.68 | 0.73 | 0.78 | 0.94 | 0.69 | 0.72 |
| 11 | 100909186 | rs11224563 | 1.554 | 0.239 | -0.441 | -2.592 | -0.065 | 0.078 | 0.078 | 0 | 0.14 | 0.01 | 0.14 | 0.01 | 0.18 | 0.06 |
| 11 | 100909809 | rs484389 | -0.313 | 0.017 | 0.147 | -2.653 | 0.008 | -0.01 | -0.01 | 0 | 0.24 | 0.41 | 0.27 | 0.21 | 0.25 | 0.11 |
| 11 | 100909991 | rs500760 | -0.313 | 0.016 | 0.147 | -2.653 | 0.005 | -0.01 | -0.01 | 0 | 0.24 | 0.41 | 0.27 | 0.21 | 0.25 | 0.11 |
| 11 | 100910469 | rs511298 | -0.125 | 0.006 | 0.302 | -2.657 | 0.003 | -0.005 | -0.005 | 0 | 0.23 | 0.35 | 0.26 | 0.19 | 0.25 | 0.09 |
| 11 | 100913548 | rs492827 | -0.382 | -0.124 | 0.605 | -2.692 | 0.009 | -0.019 | -0.019 | 0 | 0.26 | 0.55 | 0.27 | 0.21 | 0.26 | 0.11 |
| 11 | 100922568 | rs550382 | -0.268 | -0.108 | 0.641 | -2.684 | 0.005 | -0.007 | -0.007 | 0 | 0.24 | 0.38 | 0.26 | 0.21 | 0.25 | 0.11 |

| | | | | | | | | | | | | | | | | |
|---|---|---|---|---|---|---|---|---|---|---|---|---|---|---|---|---|
| 11 | 100928289 | rs679275 | -1.24 | -0.387 | 0.44 | -2.998 | -0.016 | -0.155 | -0.155 | 0.229 | 0.7 | 0.54 | 0.79 | 1 | 0.73 | 0.78 |
| 11 | 100929187 | rs1144133 | 1.058 | -0.388 | 0.308 | -2.966 | 0.001 | 0.005 | 0.005 | 0 | 0.07 | 0.07 | 0.08 | 0.18 | 0.07 | 0.03 |
| 11 | 100932603 | rs553272 | -0.194 | -0.413 | 0.394 | -2.977 | -0.025 | 0.058 | 0.058 | 0 | 0.11 | 0.3 | 0.13 | 0.2 | 0.08 | 0.05 |
| 11 | 100933412 | rs1042838 | 2.003 | -0.393 | 0.3 | -3.016 | -0.07 | 0.077 | 0.077 | 0 | 0.14 | 0.01 | 0.14 | 0.01 | 0.18 | 0.07 |
| 11 | 100933831 | rs1824128 | 0.134 | -0.398 | 0.05 | -3.31 | 0.007 | -0.01 | -0.01 | 0.585 | 0.18 | 0.11 | 0.29 | 0.77 | 0.14 | 0.46 |
| 11 | 100934367 | rs660541 | -0.165 | -0.375 | 0.004 | -3.364 | -0.036 | 0.14 | 0.14 | 0 | 0.45 | 0.17 | 0.52 | 0.78 | 0.46 | 0.66 |
| 11 | 100935780 | rs495997 | -0.912 | -0.773 | 0.134 | -3.278 | 0.001 | -0.04 | -0.04 | 0 | 0.42 | 0.42 | 0.56 | 0.99 | 0.4 | 0.57 |
| 11 | 100937060 | rs11224579 | 0.264 | -0.741 | 0.043 | -2.989 | -0.003 | 0.003 | 0.003 | 0 | 0.15 | 0.13 | 0.25 | 0.76 | 0.09 | 0.41 |
| 11 | 100937307 | rs503602 | -0.493 | -0.628 | 0.088 | -3.031 | 0 | 0.059 | 0.059 | 0 | 0.13 | 0.37 | 0.13 | 0.19 | 0.12 | 0.06 |
| 11 | 100960366 | rs493220 | 1.073 | -0.597 | -0.241 | -3.183 | -0.095 | 0.181 | 0.181 | 0 | 0.27 | 0.04 | 0.24 | 0.01 | 0.32 | 0.2 |
| 11 | 100960575 | rs585447 | 1.697 | -0.539 | -0.239 | -3.165 | -0.07 | 0.081 | 0.081 | 0 | 0.14 | 0.01 | 0.14 | 0.01 | 0.18 | 0.07 |
| 11 | 100961182 | rs538915 | 0.07 | -0.581 | -0.061 | -3.113 | 0.001 | 0.028 | 0.028 | 0 | 0.1 | 0.16 | 0.1 | 0.18 | 0.12 | 0.04 |
| 11 | 100969360 | rs481855 | 0.637 | -0.816 | -0.31 | -3.452 | 0.002 | 0.008 | 0.008 | 0 | 0.32 | 0.3 | 0.23 | 0.19 | 0.31 | 0.25 |
| 11 | 100969545 | rs555653 | 0.247 | -0.8 | -0.221 | -3.45 | -0.005 | 0.031 | 0.031 | 0 | 0.46 | 0.4 | 0.37 | 0.2 | 0.49 | 0.32 |
| 11 | 100984844 | rs486992 | -0.335 | -0.988 | 0.026 | -3.26 | 0.004 | 0.003 | 0.003 | 0 | 0.55 | 0.5 | 0.51 | 0.2 | 0.62 | 0.31 |
| 11 | 101000501 | rs518162 | -0.129 | -0.604 | 0.529 | -2.896 | 0.055 | -0.021 | -0.021 | 0 | 0.92 | 0.67 | 0.89 | 0.99 | 0.91 | 0.94 |
| 11 | 101009933 | rs948516 | 0.416 | -0.568 | -0.471 | -2.935 | -0.07 | 0.165 | 0.165 | 0 | 0.27 | 0.05 | 0.23 | 0.01 | 0.28 | 0.19 |
| 11 | 101024282 | rs568157 | -0.133 | -0.728 | -0.364 | -3.008 | -0.036 | 0.101 | 0.101 | 0 | 0.42 | 0.23 | 0.38 | 0.02 | 0.44 | 0.26 |
| 11 | 101046369 | rs614670 | -0.042 | -0.694 | -0.595 | -2.992 | -0.06 | 0.168 | 0.168 | 0 | 0.41 | 0.15 | 0.37 | 0.02 | 0.44 | 0.26 |
| 11 | 101048332 | rs12417627 | 1.143 | -0.462 | -0.431 | -2.458 | -0.001 | 0.021 | 0.021 | 0 | 0.06 | 0.04 | 0.08 | 0.19 | 0.06 | 0.02 |
| 11 | 101054756 | rs11224621 | 1.433 | -0.673 | -0.48 | -3.396 | 0.005 | 0.081 | 0.081 | 0 | 0.31 | 0.13 | 0.19 | 0 | 0.32 | 0.21 |
| 11 | 101060972 | rs1942845 | 0.418 | -0.565 | -0.518 | -3.536 | -0.028 | 0.065 | 0.065 | 0.233 | 0.24 | 0.13 | 0.23 | 0.01 | 0.24 | 0.17 |
| 11 | 101066080 | rs17737150 | 1.26 | -0.517 | -0.139 | -3.139 | 0.002 | 0.059 | 0.059 | 0 | 0.3 | 0.15 | 0.2 | 0 | 0.32 | 0.21 |
| 11 | 101069926 | rs7940817 | 1.12 | -0.497 | -0.061 | -3.129 | 0.002 | -0.037 | -0.037 | 1.797 | 0.31 | 0.3 | 0.21 | 0 | 0.32 | 0.21 |
| 11 | 101071565 | rs948749 | -0.67 | -0.545 | 0.071 | -3.144 | -0.011 | -0.02 | -0.02 | 0 | 0.94 | 0.86 | 0.96 | 0.99 | 0.97 | 0.98 |
| 11 | 101086219 | rs7937207 | 1.255 | -0.5 | -0.519 | -3.128 | 0.004 | 0.022 | 0.022 | 0 | 0.3 | 0.21 | 0.2 | 0 | 0.32 | 0.21 |
| 11 | 101098195 | rs35538677 | 0.549 | -0.571 | -0.931 | -2.945 | 0.031 | 0.017 | 0.017 | 0 | 0.13 | 0.04 | 0.08 | 0.07 | 0.08 | 0.07 |
| 11 | 101137948 | rs7114792 | 1.249 | -0.452 | -0.937 | -3.01 | -0.002 | 0.096 | 0.096 | 0 | 0.3 | 0.12 | 0.18 | 0 | 0.32 | 0.21 |
| 11 | 101142420 | rs11224683 | 1.494 | -0.151 | -1.056 | -2.986 | -0.066 | 0.061 | 0.061 | 0.033 | 0.13 | 0.01 | 0.14 | 0.03 | 0.19 | 0.09 |
| 11 | 101144202 | rs12805701 | 0.436 | -0.238 | -1.078 | -3.015 | 0.025 | 0.012 | 0.012 | 0.121 | 0.15 | 0.09 | 0.1 | 0.1 | 0.08 | 0.07 |
| 11 | 101145350 | rs10895087 | 0.602 | 0.435 | -1.305 | -2.528 | -0.01 | 0.01 | 0.01 | 0 | 0.25 | 0.25 | 0.35 | 0.85 | 0.28 | 0.5 |
| 11 | 101148887 | rs10501976 | 0.27 | -0.054 | -1.096 | -1.928 | -0.005 | 0.023 | 0.023 | 0 | 0.09 | 0.03 | 0.14 | 0 | 0.09 | 0.09 |
| 11 | 101149368 | rs4304740 | -0.462 | -0.044 | -1.104 | -1.893 | 0.002 | -0.044 | -0.044 | 0.305 | 0.24 | 0.26 | 0.3 | 0.03 | 0.29 | 0.18 |
| 11 | 101152961 | rs7936649 | 0.097 | 0.087 | -1.129 | -0.985 | -0.013 | -0.062 | -0.062 | 1.787 | 0.65 | 0.57 | 0.78 | 0.86 | 0.7 | 0.8 |
| 11 | 101153513 | rs7128860 | -0.457 | 0.317 | -1.308 | 0.218 | 0.013 | 0.003 | 0.003 | 0.235 | 0.34 | 0.29 | 0.39 | 0.78 | 0.26 | 0.43 |
| 11 | 101158665 | rs12803733 | -0.935 | 0.712 | -1.24 | 0.564 | -0.007 | -0.023 | -0.023 | 0.25 | 0.08 | 0.02 | 0.07 | 0.11 | 0.11 | 0.15 |
| 11 | 101159389 | rs10895089 | -0.565 | 1 | -1.24 | -0.385 | -0.007 | 0.006 | 0.006 | 0.076 | 0.34 | 0.32 | 0.54 | 0.87 | 0.39 | 0.68 |
| 11 | 101159423 | rs1943760 | -0.499 | 0.579 | -1.01 | -1.115 | 0.013 | 0.021 | 0.021 | 0 | 0.62 | 0.69 | 0.79 | 0.94 | 0.65 | 0.75 |
| 11 | 101160958 | rs10895090 | -0.655 | 0.588 | -0.909 | -0.789 | 0.019 | 0.072 | 0.072 | 0 | 0.51 | 0.69 | 0.5 | 0.76 | 0.45 | 0.37 |
| 11 | 101161919 | rs1943761 | -0.735 | 0.79 | -0.927 | -0.527 | 0.005 | 0.064 | 0.064 | 0 | 0.51 | 0.72 | 0.52 | 0.7 | 0.47 | 0.39 |
| 11 | 101167919 | rs11224701 | 0.167 | 0.506 | -0.988 | 0.385 | 0.013 | -0.038 | -0.038 | 0 | 0.11 | 0.05 | 0.21 | 0.28 | 0.09 | 0.05 |
| 11 | 101169617 | rs1938983 | -0.701 | 0.773 | -1.187 | 0.668 | -0.03 | -0.029 | -0.029 | 0.046 | 0.95 | 0.99 | 0.92 | 0.81 | 0.89 | 0.96 |
| 11 | 101169657 | rs1938984 | 1.007 | 0.692 | -1.214 | 0.613 | -0.001 | 0.024 | 0.024 | 0 | 0.49 | 0.31 | 0.47 | 0.34 | 0.49 | 0.62 |
| 11 | 101171982 | rs10791455 | -0.396 | 0.294 | -0.598 | 0.456 | -0.001 | -0.018 | -0.018 | 0 | 0.36 | 0.43 | 0.22 | 0.2 | 0.32 | 0.36 |
| 11 | 101172115 | rs1938985 | -0.757 | 0.491 | -0.46 | 0.608 | 0.001 | 0 | 0 | 0 | 0.46 | 0.48 | 0.44 | 0.48 | 0.42 | 0.42 |
| 11 | 101176069 | rs11224705 | -1.073 | 0.914 | 0.373 | 1.328 | 0.019 | 0.008 | 0.008 | 0 | 0.35 | 0.47 | 0.37 | 0.36 | 0.34 | 0.36 |
| 11 | 101176180 | rs1218913 | -1.964 | 0.891 | 0.488 | 1.515 | -0.002 | 0.016 | 0.016 | 0 | 0.48 | 0.6 | 0.45 | 0.65 | 0.42 | 0.41 |
| 11 | 101181227 | rs7118429 | -1.253 | 0.516 | 0.505 | 1.652 | -0.018 | 0.064 | 0.064 | 0 | 0.15 | 0.3 | 0.12 | 0.31 | 0.11 | 0.08 |
| 11 | 101194874 | rs11224715 | -0.84 | 0.462 | 0.276 | 1.58 | -0.025 | 0.051 | 0.051 | 0 | 0.32 | 0.16 | 0.18 | 0.19 | 0.29 | 0.33 |
| 11 | 101195530 | rs1230960 | -1.891 | 0.449 | 0.216 | 1.428 | -0.001 | -0.005 | -0.005 | 0 | 0.46 | 0.49 | 0.42 | 0.65 | 0.38 | 0.4 |
| 11 | 101196316 | rs1218926 | 0.019 | 0.633 | -0.015 | 1.654 | 0.005 | -0.056 | -0.056 | 0 | 0.09 | 0 | 0.06 | 0.29 | 0.08 | 0.04 |
| 11 | 101200793 | rs11224716 | -0.934 | 0.801 | 0.179 | 1.268 | 0.001 | 0.001 | 0.001 | 0 | 0.29 | 0.31 | 0.31 | 0.35 | 0.27 | 0.32 |
| 11 | 101208278 | rs1938994 | -1.712 | 0.326 | 0.61 | 1.111 | -0.024 | -0.006 | -0.006 | 2.811 | 0.3 | 0.45 | 0.18 | 0.29 | 0.21 | 0.2 |
| 11 | 101218094 | rs6590849 | -1.935 | 0.339 | 0.748 | 1.313 | -0.026 | 0.034 | 0.034 | 0.14 | 0.32 | 0.52 | 0.33 | 0.41 | 0.23 | 0.23 |
| 11 | 101234259 | rs7122352 | -1.664 | 0.291 | 0.95 | 1.176 | 0.003 | -0.005 | -0.005 | 0.006 | 0.28 | 0.28 | 0.18 | 0.36 | 0.22 | 0.23 |
| 11 | 101241202 | rs11224739 | -0.73 | 0.29 | 0.833 | 1.158 | 0.002 | -0.075 | -0.075 | 0.912 | 0.11 | 0 | 0.07 | 0.27 | 0.09 | 0.07 |
| 11 | 101260290 | rs1938987 | 1.634 | 0.318 | 1.023 | 1.163 | -0.025 | 0.024 | 0.024 | 0.062 | 0.7 | 0.5 | 0.8 | 0.63 | 0.78 | 0.77 |
| 11 | 101261237 | rs2513143 | 0.771 | 0.36 | 1.049 | 0.85 | -0.011 | -0.086 | -0.086 | 0 | 0.79 | 0.53 | 0.87 | 0.91 | 0.87 | 0.82 |
| 11 | 101270450 | rs2513173 | 0.58 | 0.318 | 0.87 | 0.791 | -0.001 | -0.041 | -0.041 | 0 | 0.8 | 0.77 | 0.91 | 1 | 0.87 | 0.85 |
| 11 | 101274126 | rs6590854 | -2.079 | 0.246 | 1.017 | 0.928 | -0.025 | -0.004 | -0.004 | 0 | 0.31 | 0.54 | 0.21 | 0.32 | 0.22 | 0.22 |
| 11 | 101282436 | rs1938990 | -2.208 | 0.23 | 1.103 | 0.896 | -0.011 | -0.004 | -0.004 | 0 | 0.33 | 0.62 | 0.21 | 0.31 | 0.23 | 0.22 |
| 11 | 101284459 | rs12222760 | -0.406 | 0.328 | 0.678 | 0.94 | -0.001 | -0.053 | -0.053 | 0 | 0.09 | 0.01 | 0.07 | 0.24 | 0.09 | 0.05 |
| 11 | 101319531 | rs3824933 | -2.087 | 0.42 | 0.951 | -0.12 | 0.005 | -0.037 | -0.037 | 0.008 | 0.36 | 0.66 | 0.2 | 0.08 | 0.25 | 0.23 |
| 11 | 101321622 | rs10895109 | -2.106 | 0.42 | 0.951 | -0.12 | 0.005 | -0.036 | -0.036 | 0 | 0.36 | 0.66 | 0.2 | 0.08 | 0.25 | 0.23 |
| 11 | 101330090 | rs11822237 | -0.139 | 0.425 | 1.254 | -0.134 | -0.027 | -0.093 | -0.093 | 0 | 0.15 | 0.3 | 0.08 | 0 | 0.08 | 0.09 |
| 11 | 101341239 | rs7935581 | -2.161 | 0.429 | 1.037 | -0.153 | -0.005 | -0.081 | -0.081 | 0.396 | 0.33 | 0.62 | 0.2 | 0.08 | 0.25 | 0.24 |
| 11 | 101343629 | rs11224780 | -2.21 | 0.529 | 0.95 | -0.167 | -0.002 | -0.058 | -0.058 | 0 | 0.33 | 0.64 | 0.21 | 0.06 | 0.26 | 0.26 |
| 11 | 101344699 | rs7948300 | -2.286 | 0.487 | 1.029 | -0.189 | -0.007 | -0.086 | -0.086 | 0 | 0.36 | 0.59 | 0.21 | 0.06 | 0.26 | 0.26 |
| 11 | 101345105 | rs12786558 | -0.665 | 0.527 | 0.796 | -0.137 | -0.036 | 0.043 | 0.043 | 0.032 | 0.12 | 0 | 0.06 | 0.07 | 0.14 | 0.11 |

| | | | | | | | | | | | | | | | | |
|---|---|---|---|---|---|---|---|---|---|---|---|---|---|---|---|---|
| 11 | 101349838 | rs11224783 | -1.935 | 0.699 | 1.08 | -0.12 | -0.021 | -0.151 | -0.151 | 0 | 0.3 | 0.54 | 0.19 | 0.05 | 0.23 | 0.26 |
| 11 | 101352191 | rs17673079 | 0.904 | 1.188 | 0.586 | 0.111 | -0.028 | -0.03 | -0.03 | 0 | 0.05 | 0.01 | 0.1 | 0.1 | 0.12 | 0.07 |
| 11 | 101352427 | rs10895115 | 1.251 | 1.557 | 0.55 | 0.182 | 0.005 | 0.052 | 0.052 | 0 | 0.69 | 0.51 | 0.64 | 0.52 | 0.67 | 0.69 |
| 11 | 101353481 | rs12361641 | -1.876 | 1.612 | 0.342 | -1.083 | 0 | -0.104 | -0.104 | 0 | 0.36 | 0.56 | 0.22 | 0.03 | 0.33 | 0.26 |
| 11 | 101354968 | rs7924551 | -1.951 | 1.607 | 0.351 | -1.085 | -0.001 | -0.084 | -0.084 | 0 | 0.37 | 0.59 | 0.22 | 0.03 | 0.33 | 0.27 |
| 11 | 101355126 | rs7942339 | -0.195 | 1.638 | 0.451 | -1.079 | -0.017 | -0.088 | -0.088 | 0 | 0.17 | 0.28 | 0.08 | 0 | 0.09 | 0.11 |
| 11 | 101364225 | rs10895118 | -1.992 | 1.736 | 0.281 | -0.948 | 0 | -0.023 | -0.023 | 0.651 | 0.38 | 0.56 | 0.37 | 0.19 | 0.39 | 0.36 |
| 11 | 101367212 | rs10501979 | -0.179 | 1.595 | 0.249 | -0.814 | -0.009 | -0.013 | -0.013 | 0 | 0.07 | 0.06 | 0.16 | 0.16 | 0.1 | 0.09 |
| 11 | 101375549 | rs10501986 | -1.647 | 1.545 | 0.219 | -0.817 | 0.001 | -0.034 | -0.034 | 0.526 | 0.42 | 0.53 | 0.4 | 0.19 | 0.47 | 0.38 |
| 11 | 101388716 | rs10501982 | -0.328 | 1.512 | 0.241 | -0.793 | -0.001 | -0.026 | -0.026 | 0 | 0.24 | 0.19 | 0.14 | 0 | 0.24 | 0.16 |
| 11 | 101394541 | rs4272759 | -0.461 | 1.536 | 0.119 | -0.718 | 0.013 | 0.037 | 0.037 | 0.003 | 0.11 | 0.24 | 0.18 | 0.17 | 0.13 | 0.13 |
| 11 | 101395310 | rs4481994 | -0.848 | 1.497 | -0.027 | -0.694 | 0.004 | -0.002 | -0.002 | 0 | 0.2 | 0.23 | 0.25 | 0.21 | 0.27 | 0.24 |
| 11 | 101396286 | rs11224816 | -1.051 | 1.091 | -0.252 | -0.425 | -0.004 | 0.022 | 0.022 | 0 | 0.46 | 0.58 | 0.42 | 0.24 | 0.51 | 0.42 |
| 11 | 101398450 | rs4466798 | -0.389 | 1.188 | -0.173 | -0.799 | 0.013 | -0.124 | -0.124 | 0.281 | 0.29 | 0.38 | 0.23 | 0.01 | 0.36 | 0.19 |
| 11 | 101407869 | rs10895131 | 0.052 | 0.822 | -0.668 | -0.613 | 0.003 | -0.011 | -0.011 | 0 | 0.12 | 0.04 | 0.2 | 0.12 | 0.1 | 0.19 |
| 11 | 101409431 | rs11224825 | -1.411 | 0.896 | -0.643 | -1.119 | 0 | 0.02 | 0.02 | 0 | 0.26 | 0.23 | 0.29 | 0.16 | 0.23 | 0.26 |
| 11 | 101409902 | rs11224827 | 0.3 | 0.918 | -0.702 | -1.023 | 0.003 | -0.011 | -0.011 | 0.947 | 0.12 | 0.04 | 0.2 | 0.12 | 0.1 | 0.19 |
| 11 | 101411208 | rs7112255 | 0.898 | 0.406 | -0.312 | -2.01 | -0.005 | -0.125 | -0.125 | 0 | 0.6 | 0.53 | 0.45 | 0.02 | 0.64 | 0.45 |
| 11 | 101412875 | rs11224829 | 0.475 | 0.294 | -0.499 | -0.692 | 0.002 | 0.04 | 0.04 | 0 | 0.09 | 0.04 | 0.2 | 0.53 | 0.08 | 0.31 |
| 11 | 101416602 | rs4492784 | -1.136 | 0.562 | -0.307 | -0.832 | 0.009 | 0.023 | 0.023 | 0 | 0.31 | 0.4 | 0.37 | 0.45 | 0.33 | 0.23 |
| 11 | 101424942 | rs4237603 | -0.312 | 0.536 | -0.198 | -0.703 | 0.009 | 0.098 | 0.098 | 0.058 | 0.19 | 0.32 | 0.27 | 0.4 | 0.2 | 0.17 |
| 11 | 101429819 | rs11224855 | -0.312 | 0.536 | -0.159 | -0.703 | 0.008 | 0.092 | 0.092 | 0.058 | 0.19 | 0.32 | 0.27 | 0.4 | 0.2 | 0.17 |
| 11 | 101431122 | rs10219300 | 1.436 | 0.595 | -0.216 | -1.247 | 0 | -0.05 | -0.05 | 0.032 | 0.22 | 0.2 | 0.18 | 0 | 0.29 | 0.13 |
| 11 | 101433161 | rs9326314 | 0.294 | 0.387 | -0.159 | -1.332 | -0.003 | 0.009 | 0.009 | 0 | 0.34 | 0.36 | 0.41 | 0.54 | 0.36 | 0.52 |
| 11 | 101447797 | rs4394815 | 0.691 | 0.29 | -0.278 | -1.249 | -0.018 | 0.157 | 0.157 | 0 | 0.44 | 0.22 | 0.28 | 0.14 | 0.42 | 0.31 |
| 11 | 101449358 | rs4326755 | -0.108 | 0.226 | -0.11 | -1.214 | -0.001 | 0.005 | 0.005 | 0 | 0.55 | 0.31 | 0.51 | 0.58 | 0.51 | 0.62 |
| 11 | 101469434 | rs2508736 | -0.491 | 0.182 | -0.092 | -1.021 | -0.016 | 0.116 | 0.116 | 0 | 0.13 | 0.19 | 0.34 | 0.49 | 0.08 | 0.37 |
| 11 | 101470545 | rs2513195 | 0.269 | 0.185 | -0.093 | -1.015 | -0.001 | 0.062 | 0.062 | 0 | 0.1 | 0.1 | 0.23 | 0.46 | 0.07 | 0.29 |
| 11 | 101472289 | rs1938867 | -0.188 | 0.084 | -0.17 | -1.206 | -0.01 | 0.011 | 0.011 | 0 | 0.55 | 0.32 | 0.61 | 0.62 | 0.49 | 0.69 |
| 11 | 101472414 | rs7124721 | 0.126 | 0.153 | -0.092 | -1.376 | -0.001 | -0.003 | -0.003 | 0 | 0.54 | 0.49 | 0.4 | 0.39 | 0.52 | 0.44 |
| 11 | 101474731 | rs10791507 | -0.283 | 0.197 | -0.364 | -1.29 | -0.021 | -0.092 | -0.092 | 0.004 | 0.18 | 0.06 | 0.28 | 0.58 | 0.18 | 0.35 |
| 11 | 101480017 | rs1938842 | 1.786 | 0.126 | -0.757 | -1.791 | -0.033 | 0.157 | 0.157 | 0 | 0.26 | 0.04 | 0.19 | 0.01 | 0.24 | 0.23 |
| 11 | 101483577 | rs17676029 | 1.143 | 0.089 | -0.59 | -1.62 | -0.045 | 0.207 | 0.207 | 0.008 | 0.34 | 0.06 | 0.21 | 0.01 | 0.31 | 0.26 |
| 11 | 101488094 | rs11603928 | 1.122 | 0.048 | -0.463 | -1.638 | -0.05 | 0.219 | 0.219 | 0 | 0.34 | 0.06 | 0.21 | 0.01 | 0.31 | 0.26 |
| 11 | 101493911 | rs2508249 | -2.079 | 0.081 | -0.207 | -1.603 | 0.002 | -0.067 | -0.067 | 0 | 0.78 | 0.77 | 0.82 | 1 | 0.71 | 0.87 |
| 11 | 101524246 | rs11224925 | 0.947 | 0.026 | -0.503 | -1.12 | -0.012 | -0.025 | -0.025 | 0 | 0.09 | 0.01 | 0.23 | 0.43 | 0.08 | 0.29 |
| 11 | 101524873 | rs2186580 | 0.76 | -0.031 | -0.527 | -1.147 | -0.007 | -0.035 | -0.035 | 0 | 0.09 | 0.01 | 0.23 | 0.43 | 0.08 | 0.29 |
| 11 | 101526476 | rs1938857 | -0.196 | -0.047 | -0.43 | -1.243 | -0.002 | -0.002 | -0.002 | 0 | 0.55 | 0.57 | 0.5 | 0.46 | 0.5 | 0.47 |
| 11 | 101528297 | rs12796410 | 1.467 | 0.065 | -0.554 | -1.331 | -0.04 | -0.018 | -0.018 | 0 | 0.08 | 0.01 | 0.06 | 0.11 | 0.11 | 0.06 |
| 11 | 101536336 | rs1938831 | -1.763 | 0.002 | -0.445 | -1.348 | 0 | -0.105 | -0.105 | 0 | 0.74 | 0.67 | 0.79 | 1 | 0.69 | 0.86 |
| 11 | 101539408 | rs7938043 | 0.247 | 0.05 | -0.598 | -0.973 | 0.001 | 0.037 | 0.037 | 0 | 0.18 | 0.21 | 0.16 | 0.31 | 0.19 | 0.12 |
| 11 | 101555384 | rs17097029 | 0.439 | 0.114 | -1.141 | -1.216 | -0.067 | -0.072 | -0.072 | 0.109 | 0.13 | 0.02 | 0.34 | 0.7 | 0.2 | 0.36 |
| 11 | 101558075 | rs6590910 | -1.939 | -0.016 | -0.919 | -1.541 | 0.006 | -0.12 | -0.12 | 0.021 | 0.75 | 0.68 | 0.79 | 1 | 0.7 | 0.85 |
| 11 | 101558350 | rs7116268 | 0.89 | -0.121 | -1.258 | -1.542 | -0.032 | 0.18 | 0.18 | 0 | 0.34 | 0.08 | 0.22 | 0.01 | 0.33 | 0.26 |
| 11 | 101576285 | rs11224972 | 1.06 | -0.076 | -0.972 | -1.16 | -0.038 | -0.018 | -0.018 | 2.287 | 0.09 | 0.01 | 0.06 | 0.11 | 0.1 | 0.04 |
| 11 | 101585195 | rs11224983 | 0.544 | -0.245 | -0.786 | -1.262 | -0.024 | -0.015 | -0.015 | 0 | 0.14 | 0.19 | 0.04 | 0.07 | 0.06 | 0.07 |
| 11 | 101587820 | rs17682688 | -0.005 | -0.081 | -1.059 | -1.604 | 0.025 | 0.099 | 0.099 | 0 | 0.45 | 0.28 | 0.25 | 0.08 | 0.36 | 0.29 |
| 11 | 101588960 | rs12283905 | -0.005 | -0.077 | -1.059 | -1.604 | 0.027 | 0.099 | 0.099 | 0 | 0.45 | 0.28 | 0.25 | 0.08 | 0.36 | 0.29 |
| 11 | 101589335 | rs12286714 | -0.023 | -0.126 | -1.244 | -1.609 | -0.085 | -0.033 | -0.033 | 2.44 | 0.21 | 0.46 | 0.07 | 0.15 | 0.09 | 0.2 |
| 11 | 101590923 | rs4754019 | -1.159 | 0.794 | -1.899 | -1.145 | 0.049 | 0.225 | 0.225 | 0 | 0.53 | 0.19 | 0.71 | 0.84 | 0.61 | 0.63 |
| 11 | 101591191 | rs7949939 | 0.737 | 0.843 | -2.06 | -1.144 | -0.038 | 0.147 | 0.147 | 0.01 | 0.29 | 0.09 | 0.2 | 0.01 | 0.29 | 0.21 |
| 11 | 101592201 | rs11224986 | -1.32 | 1.375 | -1.472 | -0.199 | -0.102 | 0.015 | 0.015 | 0.01 | 0.23 | 0.04 | 0.41 | 0.81 | 0.32 | 0.41 |
| 11 | 101593061 | rs10895184 | -1.357 | 1.263 | -1.474 | -0.188 | -0.094 | -0.005 | -0.005 | 0 | 0.24 | 0.04 | 0.41 | 0.8 | 0.32 | 0.41 |
| 11 | 101594826 | rs12360583 | 0.187 | 0.806 | -1.193 | -0.117 | -0.04 | -0.027 | -0.027 | 0.077 | 0.1 | 0.01 | 0.07 | 0.11 | 0.12 | 0.05 |
| 11 | 101597761 | rs11224990 | -0.489 | 0.99 | -1.159 | 0.678 | -0.03 | -0.038 | -0.038 | 0 | 0.1 | 0.03 | 0.34 | 0.67 | 0.18 | 0.27 |
| 11 | 101602137 | rs17097072 | -0.76 | 0.467 | -1.144 | 0.008 | 0.004 | 0.003 | 0.003 | 0 | 0.2 | 0.22 | 0.05 | 0.12 | 0.12 | 0.2 |
| 11 | 101605466 | rs11600713 | -1.813 | 0.46 | -0.815 | -0.895 | 0.003 | -0.134 | -0.134 | 0 | 0.62 | 0.59 | 0.62 | 0.97 | 0.6 | 0.77 |
| 11 | 101610075 | rs1939436 | -1.342 | 0.562 | -0.895 | -0.793 | -0.006 | 0.032 | 0.032 | 0 | 0.27 | 0.16 | 0.38 | 0.82 | 0.27 | 0.49 |
| 11 | 101610380 | rs4384360 | -1.68 | 0.347 | 0.059 | -1.115 | -0.001 | -0.082 | -0.082 | 0 | 0.62 | 0.51 | 0.63 | 0.97 | 0.62 | 0.79 |
| 11 | 101616696 | rs12223480 | -1.278 | 0.177 | -0.233 | -0.74 | -0.001 | -0.001 | -0.001 | 0 | 0.23 | 0.25 | 0.37 | 0.76 | 0.26 | 0.49 |
| 11 | 101624193 | rs7946357 | -0.012 | 0.32 | -0.06 | -0.914 | 0.019 | 0.047 | 0.047 | 0.192 | 0.37 | 0.19 | 0.29 | 0.22 | 0.33 | 0.3 |
| 11 | 101632652 | rs1939060 | 0.11 | 0.269 | -0.051 | -0.79 | -0.012 | 0.001 | 0.001 | 0 | 0.12 | 0.15 | 0.02 | 0.12 | 0.06 | 0.11 |
| 11 | 101641080 | rs1939047 | -0.84 | 0.31 | -0.075 | -0.803 | 0.001 | 0.12 | 0.12 | 0 | 0.48 | 0.33 | 0.64 | 0.85 | 0.53 | 0.67 |
| 11 | 101646567 | rs6590919 | 1.066 | 0.461 | -0.107 | -0.352 | 0.001 | 0.129 | 0.129 | 0 | 0.06 | 0.09 | 0.06 | 0.49 | 0.12 | 0.12 |
| 11 | 101653644 | rs1939029 | -2.026 | 0.409 | -0.07 | -0.594 | 0.004 | -0.105 | -0.105 | 0 | 0.63 | 0.69 | 0.69 | 0.98 | 0.59 | 0.79 |
| 11 | 101658984 | rs1939026 | -0.23 | 0.369 | -0.074 | -0.5 | -0.015 | 0 | 0 | 0 | 0.14 | 0.19 | 0.04 | 0.12 | 0.09 | 0.12 |
| 11 | 101665611 | rs1939053 | 0.756 | 0.285 | -0.154 | -0.56 | 0.018 | 0.016 | 0.016 | 0 | 0.28 | 0.1 | 0.27 | 0.21 | 0.27 | 0.27 |
| 11 | 101671932 | rs4754021 | -2.041 | 0.306 | -0.1 | -0.641 | 0.009 | -0.135 | -0.135 | 0.434 | 0.63 | 0.65 | 0.68 | 0.98 | 0.59 | 0.78 |
| 11 | 101672001 | rs4754022 | -0.475 | 0.306 | -0.141 | -0.608 | 0.035 | 0.016 | 0.016 | 0 | 0.43 | 0.29 | 0.31 | 0.33 | 0.35 | 0.39 |

| | | | | | | | | | | | | | | | | |
|---|---|---|---|---|---|---|---|---|---|---|---|---|---|---|---|---|
| 11 | 101681512 | rs1939057 | -0.593 | 0.488 | -0.181 | -0.438 | -0.037 | -0.055 | -0.055 | 2.987 | 0.18 | 0.1 | 0.35 | 0.64 | 0.24 | 0.4 |
| 11 | 101688347 | rs7949648 | -1.709 | 0.535 | -0.127 | -0.476 | -0.001 | 0 | 0 | 0 | 0.33 | 0.35 | 0.4 | 0.78 | 0.34 | 0.51 |
| 11 | 101703753 | rs5005681 | -0.005 | 0.46 | -0.172 | -0.514 | 0.004 | 0.01 | 0.01 | 0 | 0.35 | 0.37 | 0.31 | 0.71 | 0.35 | 0.38 |
| 11 | 101711503 | rs1893921 | -1.704 | 0.457 | -0.11 | -0.593 | 0.003 | -0.012 | -0.012 | 0.053 | 0.33 | 0.43 | 0.42 | 0.78 | 0.34 | 0.52 |
| 11 | 101718703 | rs4620692 | -1.422 | 0.423 | 0.02 | -0.69 | -0.002 | -0.08 | -0.08 | 0 | 0.56 | 0.61 | 0.67 | 0.98 | 0.55 | 0.78 |
| 11 | 101732566 | rs10791535 | -1.404 | 0.36 | -0.055 | -0.96 | 0.002 | -0.14 | -0.14 | 0 | 0.55 | 0.57 | 0.67 | 0.98 | 0.55 | 0.78 |
| 11 | 101734208 | rs17097255 | 0.634 | 0.352 | -0.142 | -0.921 | -0.001 | -0.018 | -0.018 | 0.154 | 0.09 | 0.12 | 0.02 | 0 | 0.04 | 0.02 |
| 11 | 101741939 | rs10791539 | -1.999 | 0.372 | 0.008 | -0.949 | -0.003 | -0.062 | -0.062 | 2.358 | 0.63 | 0.72 | 0.68 | 0.98 | 0.59 | 0.79 |
| 11 | 101742257 | rs12363310 | -1.044 | -0.667 | 0.027 | -0.466 | 0.04 | 0.046 | 0.046 | 1.505 | 0.47 | 0.34 | 0.29 | 0.3 | 0.37 | 0.39 |
| 11 | 101743792 | rs17759114 | 1.251 | -0.76 | -0.285 | -0.599 | 0.017 | 0.144 | 0.144 | 0.15 | 0.22 | 0 | 0.08 | 0.01 | 0.16 | 0.16 |
| 11 | 101749553 | rs11225045 | 0.431 | -0.698 | -0.187 | -0.486 | 0.013 | 0.015 | 0.015 | 0 | 0.28 | 0.08 | 0.25 | 0.21 | 0.25 | 0.29 |
| 11 | 101755207 | rs1372416 | -1.789 | -0.694 | 0.289 | -0.701 | -0.038 | -0.032 | -0.032 | 0 | 0.23 | 0.46 | 0.08 | 0.13 | 0.14 | 0.12 |
| 11 | 101755975 | rs745054 | -0.838 | -0.38 | -0.329 | -0.493 | -0.002 | 0 | 0 | 0 | 0.14 | 0.14 | 0.05 | 0.12 | 0.1 | 0.11 |
| 11 | 101760892 | rs17097290 | -0.119 | -0.049 | -0.275 | -0.381 | -0.002 | -0.017 | -0.017 | 0 | 0.08 | 0.13 | 0.01 | 0.01 | 0.04 | 0.02 |
| 11 | 101761385 | rs12293915 | -1.61 | 0.209 | -0.228 | 0 | 0.001 | 0.016 | 0.016 | 0 | 0.2 | 0.28 | 0.16 | 0.49 | 0.24 | 0.16 |
| 11 | 101762478 | rs11225049 | -0.587 | -0.234 | -0.651 | -0.709 | 0.004 | -0.02 | -0.02 | 0 | 0.39 | 0.28 | 0.29 | 0.45 | 0.42 | 0.26 |
| 11 | 101763784 | rs10791543 | -1.461 | -0.246 | -0.278 | -0.619 | 0 | 0.004 | 0.004 | 0 | 0.52 | 0.59 | 0.43 | 0.76 | 0.51 | 0.47 |
| 11 | 101770360 | rs12804060 | 0.391 | -0.173 | -0.746 | -0.753 | -0.045 | 0.194 | 0.194 | 0.301 | 0.27 | 0.02 | 0.13 | 0.01 | 0.27 | 0.16 |
| 11 | 101775663 | rs1661409 | -0.635 | -0.344 | -0.811 | -0.843 | -0.027 | -0.044 | -0.044 | 0 | 0.38 | 0.18 | 0.29 | 0.45 | 0.43 | 0.29 |
| 11 | 101778710 | rs1661410 | -0.515 | -0.705 | -0.743 | -0.782 | -0.013 | -0.054 | -0.054 | 0 | 0.37 | 0.16 | 0.28 | 0.45 | 0.41 | 0.29 |
| 11 | 101785248 | rs2282612 | -0.601 | -0.825 | -0.338 | -0.841 | 0.002 | -0.009 | -0.009 | 0 | 0.38 | 0.29 | 0.3 | 0.45 | 0.41 | 0.28 |
| 11 | 101787112 | rs7105709 | -0.674 | -0.843 | -0.403 | -0.82 | 0 | -0.013 | -0.013 | 0.199 | 0.41 | 0.31 | 0.52 | 0.57 | 0.45 | 0.47 |
| 11 | 101790671 | rs1441928 | -1.709 | -1.07 | -0.377 | -1.207 | -0.007 | 0.07 | 0.07 | 0 | 0.54 | 0.65 | 0.44 | 0.77 | 0.51 | 0.5 |
| 11 | 101806639 | rs10895226 | 0.109 | -1.06 | -0.311 | -1.123 | -0.002 | 0.031 | 0.031 | 1.792 | 0.11 | 0.12 | 0.11 | 0.28 | 0.07 | 0.18 |
| 11 | 101821444 | rs10791545 | -0.633 | -1.088 | -0.315 | -1.17 | 0.002 | -0.013 | -0.013 | 0 | 0.4 | 0.32 | 0.31 | 0.49 | 0.43 | 0.3 |
| 11 | 101824479 | rs7122609 | -1.894 | -1.036 | -0.308 | -1.142 | -0.019 | 0.067 | 0.067 | 0 | 0.57 | 0.67 | 0.65 | 0.9 | 0.54 | 0.69 |
| 11 | 101841071 | rs12794798 | 0.803 | -0.992 | -0.47 | -1.07 | -0.034 | 0.15 | 0.15 | 0.011 | 0.22 | 0.01 | 0.12 | 0.01 | 0.24 | 0.16 |
| 11 | 101867264 | rs1441929 | -0.574 | -1 | -0.192 | -0.798 | -0.001 | 0.001 | 0.001 | 0 | 0.37 | 0.34 | 0.31 | 0.49 | 0.42 | 0.29 |
| 11 | 101867596 | rs1372414 | -1.019 | -1.009 | -0.203 | -0.806 | 0 | -0.002 | -0.002 | 1.637 | 0.43 | 0.39 | 0.32 | 0.49 | 0.43 | 0.3 |
| 11 | 101868415 | rs1783742 | -1.798 | -0.971 | -0.416 | -0.816 | 0.002 | 0.072 | 0.072 | 0 | 0.52 | 0.64 | 0.44 | 0.76 | 0.5 | 0.49 |
| 11 | 101876810 | rs1274903 | -0.114 | -0.58 | -0.859 | -0.576 | -0.033 | -0.056 | -0.056 | 0 | 0.12 | 0.04 | 0.15 | 0.47 | 0.18 | 0.12 |
| 11 | 101880768 | rs7104090 | 0.342 | -0.696 | -0.675 | -1.657 | -0.002 | -0.048 | -0.048 | 0.779 | 0.25 | 0.24 | 0.16 | 0.01 | 0.25 | 0.16 |
| 11 | 101882379 | rs4754035 | -0.154 | -0.611 | -0.471 | -1.358 | 0.001 | -0.009 | -0.009 | 0.041 | 0.35 | 0.33 | 0.41 | 0.76 | 0.38 | 0.41 |
| 11 | 101884077 | rs4754829 | -0.705 | -0.397 | -0.719 | -1.215 | 0.002 | -0.005 | -0.005 | 0 | 0.52 | 0.51 | 0.48 | 0.8 | 0.44 | 0.44 |
| 11 | 101892922 | rs3133381 | -0.232 | 0.022 | -0.948 | -0.851 | -0.01 | 0.054 | 0.054 | 0.016 | 0.41 | 0.26 | 0.51 | 0.79 | 0.48 | 0.43 |
| 11 | 101918327 | rs12804542 | -0.205 | -0.005 | -0.692 | -0.632 | -0.003 | -0.018 | -0.018 | 5.363 | 0.93 | 0.94 | 0.97 | 1 | 0.97 | 0.98 |
| 11 | 101923359 | rs7928958 | -0.562 | -0.292 | -0.366 | -1.012 | 0.001 | -0.03 | -0.03 | 0 | 0.38 | 0.5 | 0.46 | 0.69 | 0.4 | 0.37 |
| 11 | 101928225 | rs10791555 | -0.573 | -0.24 | -0.371 | -0.912 | 0.002 | -0.024 | -0.024 | 0 | 0.38 | 0.5 | 0.46 | 0.66 | 0.41 | 0.36 |
| 11 | 101934867 | rs17097403 | -0.888 | -0.479 | -0.137 | -1.053 | -0.024 | -0.031 | -0.031 | 0 | 0.14 | 0.21 | 0.16 | 0.08 | 0.07 | 0.09 |
| 11 | 101944113 | rs10791559 | -1.445 | -0.518 | -0.161 | -1.109 | -0.008 | 0.161 | 0.161 | 0.414 | 0.53 | 0.8 | 0.63 | 0.74 | 0.48 | 0.46 |
| 11 | 101954459 | rs6590946 | -1.291 | -0.511 | -0.021 | -1.106 | 0.011 | 0.113 | 0.113 | 0.419 | 0.51 | 0.7 | 0.62 | 0.74 | 0.46 | 0.46 |
| 11 | 101980335 | rs1820453 | -1.333 | -0.136 | -0.209 | -1.118 | 0.002 | 0.105 | 0.105 | 0.012 | 0.5 | 0.71 | 0.63 | 0.73 | 0.5 | 0.44 |
| 11 | 101984628 | rs2033083 | 1.083 | -0.185 | -0.103 | -0.866 | 0.001 | 0.094 | 0.094 | 0 | 0.5 | 0.28 | 0.36 | 0.34 | 0.5 | 0.57 |
| 11 | 101986644 | rs10895257 | 0.386 | -0.274 | 0.433 | 0.057 | 0.006 | 0.034 | 0.034 | 0 | 0.19 | 0.2 | 0.25 | 0.31 | 0.27 | 0.16 |
| 11 | 101990252 | rs1942691 | -1.077 | -0.24 | 0.645 | 0.235 | -0.002 | 0.001 | 0.001 | 1.519 | 0.39 | 0.43 | 0.47 | 0.39 | 0.38 | 0.24 |
| 11 | 101992349 | rs12420608 | 0.152 | -0.056 | 0.526 | 0.287 | 0.025 | 0.022 | 0.022 | 0 | 0.05 | 0.17 | 0.11 | 0.05 | 0.03 | 0.06 |
| 11 | 102006169 | rs3890538 | 1.268 | -0.206 | 0.45 | 0.236 | 0.021 | -0.031 | -0.031 | 0 | 0.55 | 0.21 | 0.39 | 0.55 | 0.59 | 0.68 |
| 11 | 102011998 | rs1893497 | -1.429 | -0.244 | 0.459 | 0.272 | 0.007 | -0.011 | -0.011 | 0 | 0.44 | 0.66 | 0.59 | 0.46 | 0.41 | 0.3 |
| 11 | 102017149 | rs1834596 | 0.866 | -0.234 | 0.5 | 0.348 | -0.005 | 0.008 | 0.008 | 0.402 | 0.61 | 0.41 | 0.51 | 0.61 | 0.62 | 0.75 |
| 11 | 102023505 | rs7924737 | -1.24 | -0.179 | 0.408 | 0.378 | -0.005 | 0.008 | 0.008 | 1.177 | 0.39 | 0.48 | 0.48 | 0.4 | 0.38 | 0.25 |
| 11 | 102036484 | rs4420227 | -1.764 | -0.343 | -0.154 | 0.147 | 0.011 | 0.036 | 0.036 | 0 | 0.52 | 0.61 | 0.53 | 0.63 | 0.46 | 0.49 |
| 11 | 102044647 | rs1942683 | -1.168 | -0.396 | -0.751 | 0.108 | 0.019 | 0.043 | 0.043 | 0 | 0.42 | 0.22 | 0.45 | 0.62 | 0.4 | 0.46 |
| 11 | 102045643 | rs11602707 | 0.414 | -0.297 | -0.984 | 0.333 | -0.018 | -0.066 | -0.066 | 0 | 0.2 | 0.06 | 0.24 | 0.33 | 0.25 | 0.16 |
| 11 | 102063437 | rs17689778 | 1.088 | -0.091 | -0.812 | 0.381 | -0.005 | 0.023 | 0.023 | 0.43 | 0.06 | 0.01 | 0.06 | 0 | 0.07 | 0.03 |
| 11 | 102070361 | rs10501995 | 0.514 | 0.058 | -0.676 | 0.645 | 0.007 | -0.013 | -0.013 | 0 | 0.11 | 0.08 | 0.05 | 0.18 | 0.07 | 0.22 |
| 11 | 102070639 | rs1894116 | 0.573 | 0.059 | -0.676 | 0.645 | 0.006 | -0.012 | -0.012 | 0.129 | 0.1 | 0.07 | 0.05 | 0.18 | 0.07 | 0.22 |
| 11 | 102084940 | rs10895277 | -0.42 | -0.028 | -0.645 | 0.62 | 0.002 | -0.005 | -0.005 | 0 | 0.36 | 0.32 | 0.34 | 0.35 | 0.38 | 0.28 |
| 11 | 102091244 | rs11225168 | 0.595 | -0.003 | -0.828 | 0.659 | -0.012 | -0.063 | -0.063 | 0 | 0.1 | 0.01 | 0.05 | 0.23 | 0.08 | 0.22 |
| 11 | 102091632 | rs11225169 | -1.398 | -0.101 | -0.714 | 0.608 | 0.005 | 0.009 | 0.009 | 0 | 0.49 | 0.45 | 0.4 | 0.58 | 0.46 | 0.52 |
| 11 | 102093464 | rs11225170 | -0.924 | 0.093 | -0.732 | 0.408 | 0.008 | -0.007 | -0.007 | 0.009 | 0.4 | 0.21 | 0.36 | 0.36 | 0.4 | 0.32 |
| 11 | 102096282 | rs4475891 | -0.724 | 0.287 | -0.95 | 0.463 | 0 | -0.001 | -0.001 | 0 | 0.39 | 0.36 | 0.36 | 0.4 | 0.38 | 0.3 |
| 11 | 102103600 | rs8504 | -0.686 | 0.385 | -0.946 | 0.507 | -0.002 | 0.001 | 0.001 | 0 | 0.38 | 0.37 | 0.36 | 0.39 | 0.37 | 0.29 |
| 11 | 102114675 | rs11225179 | -0.481 | 1.57 | -1.4 | 0.834 | -0.062 | -0.035 | -0.035 | 0 | 0.12 | 0.01 | 0.12 | 0.13 | 0.18 | 0.19 |
| 11 | 102137966 | rs11600487 | -0.906 | 1.199 | -1.041 | 1.099 | 0.008 | -0.034 | -0.034 | 0.166 | 0.33 | 0.27 | 0.29 | 0.5 | 0.33 | 0.51 |
| 11 | 102142399 | rs11225187 | -0.217 | 1.116 | -1.018 | 0.938 | 0.02 | -0.004 | -0.004 | 0.035 | 0.15 | 0.05 | 0.04 | 0.14 | 0.08 | 0.24 |
| 11 | 102142846 | rs10501996 | -0.625 | 2.085 | -0.799 | 0.954 | -0.032 | -0.013 | -0.013 | 0.921 | 0.15 | 0.09 | 0.18 | 0.14 | 0.21 | 0.23 |
| 11 | 102148839 | rs11225192 | 1.054 | 1.84 | 0.089 | 0.76 | 0.028 | 0 | 0 | 0 | 0.71 | 0.5 | 0.61 | 0.71 | 0.61 | 0.69 |
| 11 | 102150488 | rs2604532 | 0.616 | 2.021 | -0.536 | 1.004 | -0.066 | -0.066 | -0.066 | 0.423 | 0.81 | 0.95 | 0.7 | 0.71 | 0.7 | 0.75 |
| 11 | 102155972 | rs2846844 | 1.554 | 1.74 | 0.382 | 0.54 | 0.072 | -0.016 | -0.016 | 0 | 0.91 | 0.61 | 0.86 | 0.98 | 0.88 | 0.92 |

| | | | | | | | | | | | | | | | | |
|---|---|---|---|---|---|---|---|---|---|---|---|---|---|---|---|---|
| 11 | 102160536 | rs12787745 | -0.997 | 1.561 | 0.042 | 0.733 | -0.005 | -0.006 | -0.006 | 0.177 | 0.29 | 0.19 | 0.38 | 0.27 | 0.33 | 0.35 |
| 11 | 102180563 | rs2846848 | -0.936 | 1.533 | 0.282 | 0.701 | 0 | 0 | 0 | 0.519 | 0.3 | 0.31 | 0.37 | 0.27 | 0.31 | 0.34 |
| 11 | 102181869 | rs2852891 | -0.978 | 1.569 | 0.203 | 0.715 | -0.003 | -0.002 | -0.002 | 0 | 0.29 | 0.19 | 0.36 | 0.27 | 0.31 | 0.34 |
| 11 | 102202851 | rs3758841 | 0.721 | 1.713 | 0.392 | 0.711 | -0.013 | 0.024 | 0.024 | 0 | 0.67 | 0.8 | 0.72 | 0.77 | 0.61 | 0.65 |
| 11 | 102205037 | rs7127583 | 0.646 | 1.711 | 0.405 | 0.71 | -0.024 | 0.029 | 0.029 | 0 | 0.68 | 0.84 | 0.73 | 0.77 | 0.61 | 0.66 |
| 11 | 102207851 | rs1055088 | 0.851 | 1.701 | 0.685 | 0.705 | 0 | -0.003 | -0.003 | 4.167 | 0.66 | 0.68 | 0.59 | 0.72 | 0.61 | 0.65 |
| 11 | 102209737 | rs13711 | 0.79 | 1.704 | 0.638 | 0.658 | -0.01 | 0.015 | 0.015 | 0 | 0.66 | 0.77 | 0.61 | 0.72 | 0.61 | 0.65 |
| 11 | 102229208 | rs637708 | -1.52 | 1.833 | 1.428 | 0.432 | -0.006 | -0.08 | -0.08 | 0.118 | 0.14 | 0.27 | 0.08 | 0 | 0.12 | 0.04 |
| 11 | 102231726 | rs10895290 | -0.295 | 2.535 | 1.25 | 0.768 | -0.037 | -0.038 | -0.038 | 0 | 0.09 | 0.03 | 0.21 | 0.28 | 0.18 | 0.19 |
| 11 | 102235141 | rs1943781 | -1.84 | 1.663 | 1.846 | -0.002 | 0.002 | -0.103 | -0.103 | 0 | 0.15 | 0.39 | 0.1 | 0 | 0.12 | 0.04 |
| 11 | 102260200 | rs525305 | -1.752 | 1.435 | 2.163 | -0.033 | -0.016 | -0.104 | -0.104 | 0 | 0.15 | 0.4 | 0.08 | 0 | 0.1 | 0.04 |
| 11 | 102264975 | rs1275850 | -2.221 | 1.448 | 1.453 | -0.095 | -0.019 | -0.112 | -0.112 | 0.187 | 0.19 | 0.36 | 0.09 | 0.01 | 0.14 | 0.06 |
| 11 | 102282452 | rs549513 | 1.62 | 0.598 | 1.163 | -0.215 | -0.019 | -0.095 | -0.095 | 0 | 0.82 | 0.67 | 0.88 | 0.98 | 0.87 | 0.92 |
| 11 | 102298002 | rs2509683 | 0.852 | 0.512 | 0.919 | -0.228 | 0.008 | -0.007 | -0.007 | 0 | 0.87 | 0.91 | 0.97 | 1 | 0.94 | 0.94 |
| 11 | 102300876 | rs531350 | 1.627 | 0.432 | 1.048 | -0.239 | -0.02 | -0.083 | -0.083 | 0.2 | 0.82 | 0.72 | 0.92 | 1 | 0.88 | 0.91 |
| 11 | 102308440 | rs670334 | 1.616 | 0.488 | 0.929 | -0.212 | -0.026 | -0.107 | -0.107 | 0 | 0.82 | 0.67 | 0.91 | 1 | 0.88 | 0.92 |
| 11 | 102327741 | rs667755 | 0.877 | 0.494 | 0.06 | -0.144 | 0.007 | -0.009 | -0.009 | 0 | 0.87 | 0.91 | 0.96 | 1 | 0.94 | 0.93 |
| 11 | 102335609 | rs1893729 | 0.806 | 0.791 | 0.682 | 0.4 | 0.019 | 0.048 | 0.048 | 0.518 | 0.83 | 0.5 | 0.77 | 0.83 | 0.86 | 0.96 |
| 11 | 102344923 | rs7115883 | -0.194 | -0.946 | 0.304 | -0.808 | 0.01 | -0.012 | -0.012 | 0 | 0.22 | 0.18 | 0.06 | 0.01 | 0.07 | 0.06 |
| 11 | 102344999 | rs567670 | 0.915 | -0.748 | 0.441 | -1.065 | 0.021 | 0.003 | 0.003 | 1.608 | 0.7 | 0.78 | 0.87 | 0.99 | 0.81 | 0.88 |
| 11 | 102345135 | rs10791573 | 0.844 | -0.666 | -0.028 | -1.653 | -0.002 | -0.014 | -0.014 | 0 | 0.26 | 0.3 | 0.31 | 0.15 | 0.23 | 0.27 |
| 11 | 102345234 | rs480846 | -0.31 | -0.331 | -0.01 | -1.324 | 0.005 | -0.045 | -0.045 | 0.015 | 0.89 | 0.83 | 0.92 | 1 | 0.88 | 0.94 |
| 11 | 102356609 | rs938599 | 0.956 | -0.129 | -0.433 | -1.349 | 0.025 | 0.095 | 0.095 | 2.484 | 0.15 | 0.01 | 0.06 | 0.01 | 0.08 | 0.06 |
| 11 | 102357019 | rs683933 | -0.397 | -0.247 | -0.16 | -1.442 | 0.002 | -0.03 | -0.03 | 0 | 0.89 | 0.9 | 0.92 | 1 | 0.88 | 0.95 |
| 11 | 102357279 | rs10750642 | 0.33 | 0.113 | 0.158 | -1.711 | 0.018 | 0.009 | 0.009 | 0.008 | 0.68 | 0.49 | 0.66 | 0.36 | 0.6 | 0.51 |
| 11 | 102358107 | rs1943783 | 0.985 | 0.169 | 0.185 | -1.856 | 0.011 | 0.083 | 0.083 | 3.059 | 0.44 | 0.29 | 0.41 | 0.22 | 0.4 | 0.4 |
| 11 | 102359477 | rs634814 | 0.559 | 0.502 | 0.489 | -1.405 | 0.015 | 0.042 | 0.042 | 0 | 0.19 | 0.24 | 0.25 | 0.64 | 0.27 | 0.44 |
| 11 | 102361915 | rs603608 | 0.741 | 0.509 | 0.122 | -1.401 | -0.023 | -0.039 | -0.039 | 0 | 0.17 | 0.08 | 0.23 | 0.64 | 0.26 | 0.44 |
| 11 | 102365970 | rs651853 | -0.408 | 0.484 | 0.225 | -1.41 | 0.035 | -0.064 | -0.064 | 5.204 | 0.89 | 0.73 | 0.91 | 1 | 0.87 | 0.96 |
| 11 | 102368719 | rs2509689 | 0.327 | 0.621 | -0.016 | -0.531 | 0.004 | 0.042 | 0.042 | 0 | 0.19 | 0.22 | 0.21 | 0.5 | 0.21 | 0.29 |
| 11 | 102371749 | rs9667864 | 1.579 | 0.079 | -0.337 | -1.232 | 0.018 | 0.059 | 0.059 | 1.032 | 0.3 | 0.19 | 0.32 | 0.1 | 0.29 | 0.36 |
| 11 | 102378560 | rs1954850 | 0.45 | 0.074 | -0.25 | -1.276 | -0.01 | -0.06 | -0.06 | 0.382 | 0.31 | 0.39 | 0.21 | 0.08 | 0.26 | 0.12 |
| 11 | 102381903 | rs573321 | 0.432 | -0.035 | -0.59 | -0.886 | 0.004 | -0.005 | -0.005 | 0 | 0.47 | 0.35 | 0.52 | 0.54 | 0.46 | 0.62 |
| 11 | 102384567 | rs609887 | -0.833 | -0.011 | -0.547 | -0.341 | -0.002 | -0.001 | -0.001 | 0.206 | 0.27 | 0.26 | 0.3 | 0.76 | 0.31 | 0.49 |
| 11 | 102385738 | rs7118775 | -0.164 | 0.118 | -0.704 | -0.379 | 0.003 | -0.001 | -0.001 | 0 | 0.21 | 0.18 | 0.16 | 0.24 | 0.19 | 0.21 |
| 11 | 102386076 | rs643281 | -1.093 | 0.384 | -0.604 | -0.353 | -0.001 | 0.024 | 0.024 | 0.35 | 0.51 | 0.59 | 0.51 | 0.89 | 0.49 | 0.58 |
| 11 | 102389487 | rs7945819 | 0.202 | 1.211 | -0.335 | 0.58 | 0.041 | -0.014 | -0.014 | 0 | 0.05 | 0.15 | 0.07 | 0.02 | 0.08 | 0.03 |
| 11 | 102389588 | rs12184413 | 0.183 | 1.183 | -0.876 | 1.471 | -0.038 | -0.021 | -0.021 | 0 | 0.07 | 0 | 0.15 | 0.26 | 0.11 | 0.09 |
| 11 | 102390188 | rs495041 | -1.294 | 0.48 | -0.526 | 1.38 | -0.024 | 0.061 | 0.061 | 0.297 | 0.23 | 0.63 | 0.32 | 0.52 | 0.18 | 0.39 |
| 11 | 102391425 | rs14983 | 0.617 | 0.318 | -1.238 | 1.279 | -0.068 | -0.096 | -0.096 | 0 | 0.14 | 0.01 | 0.15 | 0.25 | 0.22 | 0.08 |
| 11 | 102392656 | rs2156528 | 0.63 | 0.317 | -1.239 | 1.278 | -0.068 | -0.095 | -0.095 | 0 | 0.13 | 0.01 | 0.15 | 0.25 | 0.22 | 0.08 |
| 11 | 102396247 | rs1996352 | -0.017 | 0.748 | -1.541 | 1.469 | -0.059 | -0.074 | -0.074 | 0 | 0.87 | 0.95 | 0.7 | 0.64 | 0.77 | 0.73 |
| 11 | 102396607 | rs12285347 | 1.059 | 0.4 | -1.052 | 0.24 | 0.002 | -0.009 | -0.009 | 0 | 0.51 | 0.56 | 0.35 | 0.09 | 0.44 | 0.44 |
| 11 | 102398593 | rs10502001 | 0.898 | 0.52 | -1.251 | 0.324 | -0.07 | -0.096 | -0.096 | 0.915 | 0.14 | 0.01 | 0.15 | 0.25 | 0.22 | 0.08 |
| 11 | 102401661 | rs11568818 | 0.487 | 0.198 | -0.786 | -0.472 | 0.001 | -0.026 | -0.026 | 0.208 | 0.5 | 0.45 | 0.34 | 0.09 | 0.44 | 0.44 |
| 11 | 102413123 | rs10791578 | -0.651 | 0.416 | -0.449 | 0.308 | 0.003 | 0 | 0 | 0 | 0.2 | 0.22 | 0.27 | 0.24 | 0.27 | 0.19 |
| 11 | 102413943 | rs11603486 | -1.228 | -0.156 | -0.419 | 0.258 | 0.001 | 0.003 | 0.003 | 0 | 0.43 | 0.48 | 0.41 | 0.47 | 0.46 | 0.26 |
| 11 | 102418118 | rs17098339 | 0.331 | -0.172 | -0.28 | 0.514 | 0.006 | 0.007 | 0.007 | 0 | 0.08 | 0.08 | 0.04 | 0 | 0.06 | 0.03 |
| 11 | 102418160 | rs12418815 | 0.068 | -0.325 | -0.084 | 0.779 | -0.015 | 0.064 | 0.064 | 0 | 0.09 | 0.16 | 0.16 | 0.28 | 0.08 | 0.22 |
| 11 | 102420722 | rs10791579 | -1.156 | -0.204 | 0.045 | 0.683 | 0.011 | 0.038 | 0.038 | 0 | 0.24 | 0.31 | 0.3 | 0.5 | 0.3 | 0.19 |
| 11 | 102422551 | rs7947031 | -0.71 | -0.42 | 0.207 | 0.774 | -0.003 | 0.018 | 0.018 | 0 | 0.2 | 0.34 | 0.23 | 0.29 | 0.17 | 0.15 |
| 11 | 102423019 | rs7930021 | 0.558 | -0.607 | 0.243 | 0.163 | -0.006 | -0.148 | -0.148 | 0.5 | 0.19 | 0.39 | 0.14 | 0 | 0.17 | 0.09 |
| 11 | 102423666 | rs7110424 | -0.101 | -0.86 | 0.197 | 0.247 | -0.015 | 0.09 | 0.09 | 0 | 0.82 | 0.65 | 0.76 | 0.6 | 0.87 | 0.76 |
| 11 | 102423890 | rs17268032 | 1.754 | -0.963 | -0.2 | 0.134 | -0.126 | 0.141 | 0.141 | 0.544 | 0.38 | 0.06 | 0.36 | 0.11 | 0.45 | 0.41 |
| 11 | 102432478 | rs4754857 | -0.168 | -0.978 | -0.035 | 0.163 | -0.003 | -0.018 | -0.018 | 3.303 | 0.31 | 0.49 | 0.33 | 0.29 | 0.3 | 0.22 |
| 11 | 102434092 | rs11225324 | 0.917 | -0.993 | -0.115 | 0.241 | -0.003 | 0.015 | 0.015 | 0 | 0.09 | 0.11 | 0.15 | 0.26 | 0.08 | 0.21 |
| 11 | 102435173 | rs7124762 | -1.022 | -0.834 | -0.214 | 0.254 | -0.007 | -0.026 | -0.026 | 0 | 0.41 | 0.53 | 0.45 | 0.77 | 0.37 | 0.4 |
| 11 | 102437596 | rs12574560 | 0.298 | -0.217 | -0.488 | 0.683 | -0.001 | 0.017 | 0.017 | 0 | 0.14 | 0.19 | 0.19 | 0.28 | 0.12 | 0.14 |
| 11 | 102437618 | rs10895316 | 0.526 | 0.459 | -0.863 | 1.162 | -0.118 | 0.142 | 0.142 | 0.562 | 0.38 | 0.07 | 0.36 | 0.11 | 0.44 | 0.41 |
| 11 | 102437667 | rs10502003 | 0.527 | 0.841 | -0.713 | 1.15 | 0.003 | 0.009 | 0.009 | 0 | 0.07 | 0.05 | 0.03 | 0 | 0.04 | 0.03 |
| 11 | 102448256 | rs2292730 | -0.464 | 0.545 | -1.462 | -0.107 | -0.045 | 0.083 | 0.083 | 0.086 | 0.58 | 0.79 | 0.58 | 0.77 | 0.54 | 0.49 |
| 11 | 102452231 | rs11225332 | 0.194 | 0.663 | -0.903 | -0.147 | 0.008 | 0.018 | 0.018 | 2.149 | 0.48 | 0.41 | 0.28 | 0.22 | 0.4 | 0.5 |
| 11 | 102460777 | rs1711399 | 0.41 | 1.915 | -0.205 | 0.737 | 0.007 | 0.008 | 0.008 | 0 | 0.66 | 0.36 | 0.5 | 0.52 | 0.56 | 0.66 |
| 11 | 102465281 | rs7116339 | -0.145 | 2.486 | 0.367 | 0.525 | 0.015 | -0.065 | -0.065 | 0 | 0.13 | 0.19 | 0.12 | 0.01 | 0.16 | 0.1 |
| 11 | 102469281 | rs1711433 | -0.336 | 2.859 | 0.68 | 0.503 | -0.003 | -0.036 | -0.036 | 0 | 0.86 | 0.8 | 0.85 | 0.98 | 0.88 | 0.85 |
| 11 | 102474322 | rs1711430 | 1.059 | 3.106 | 0.698 | 1.28 | 0.019 | 0.007 | 0.007 | 0 | 0.66 | 0.39 | 0.51 | 0.41 | 0.54 | 0.63 |
| 11 | 102474813 | rs1784430 | 0.972 | 3.073 | 0.77 | 1.119 | 0.04 | 0.036 | 0.036 | 0 | 0.66 | 0.45 | 0.56 | 0.57 | 0.57 | 0.66 |
| 11 | 102477006 | rs1711427 | 1.669 | 3.22 | 0.722 | 0.971 | 0.005 | -0.006 | -0.006 | 0.813 | 0.57 | 0.27 | 0.44 | 0.42 | 0.44 | 0.59 |
| 11 | 102477132 | rs1784425 | 1.419 | 3.18 | 0.664 | 0.876 | -0.009 | -0.014 | -0.014 | 0 | 0.59 | 0.27 | 0.46 | 0.43 | 0.45 | 0.61 |

| | | | | | | | | | | | | | | | | |
|---|---|---|---|---|---|---|---|---|---|---|---|---|---|---|---|---|
| 11 | 102477377 | rs1784424 | 1.818 | 3.119 | 0.657 | 0.867 | 0.009 | -0.006 | -0.006 | 0.528 | 0.56 | 0.26 | 0.44 | 0.43 | 0.44 | 0.59 |
| 11 | 102477395 | rs1784423 | 1.818 | 3.119 | 0.657 | 0.867 | 0.009 | -0.006 | -0.006 | 0 | 0.56 | 0.26 | 0.44 | 0.43 | 0.44 | 0.59 |
| 11 | 102477515 | rs3781787 | -1.675 | 3.042 | 0.705 | 0.979 | 0.032 | 0.152 | 0.152 | 0 | 0.13 | 0.27 | 0.25 | 0.44 | 0.18 | 0.11 |
| 11 | 102477556 | rs3781788 | -1.071 | 3.035 | 0.728 | 1.049 | 0.047 | 0.222 | 0.222 | 0 | 0.07 | 0.26 | 0.19 | 0.33 | 0.11 | 0.08 |
| 11 | 102480421 | rs10502005 | -0.917 | 3.123 | 0.289 | 0.983 | -0.053 | -0.013 | -0.013 | 0 | 0.1 | 0.02 | 0.11 | 0.14 | 0.2 | 0.12 |
| 11 | 102488131 | rs2280211 | -0.945 | 2.974 | 0.4 | 1.05 | 0.005 | 0.111 | 0.111 | 0 | 0.07 | 0.13 | 0.18 | 0.4 | 0.11 | 0.08 |
| 11 | 102495066 | rs1962082 | 2.121 | 3.033 | 0.117 | 0.459 | 0.045 | 0.073 | 0.073 | 0.134 | 0.42 | 0.27 | 0.26 | 0.05 | 0.28 | 0.33 |
| 11 | 102495998 | rs2245803 | 0.553 | 2.981 | -0.016 | 0.74 | 0.031 | -0.032 | -0.032 | 2.085 | 0.7 | 0.6 | 0.68 | 0.8 | 0.63 | 0.69 |
| 11 | 102496462 | rs2245897 | -0.005 | 2.522 | 0.018 | 0.774 | 0.003 | -0.056 | -0.056 | 0.211 | 0.75 | 0.64 | 0.75 | 0.9 | 0.76 | 0.79 |
| 11 | 102496560 | rs1573954 | 1.021 | 2.585 | 0.014 | 0.853 | -0.003 | 0.016 | 0.016 | 0 | 0.61 | 0.59 | 0.59 | 0.51 | 0.63 | 0.53 |
| 11 | 102501665 | rs948138 | 0.594 | 2.242 | 0 | 0.536 | 0.024 | -0.017 | -0.017 | 0 | 0.66 | 0.62 | 0.56 | 0.78 | 0.55 | 0.63 |
| 11 | 102502435 | rs10895324 | 1.404 | 2.034 | -0.004 | 0.697 | -0.004 | -0.009 | -0.009 | 0 | 0.52 | 0.54 | 0.41 | 0.67 | 0.43 | 0.46 |
| 11 | 102502770 | rs4554864 | 1.715 | 1.842 | -0.016 | 0.907 | 0.017 | 0.002 | 0.002 | 0.026 | 0.43 | 0.4 | 0.23 | 0.38 | 0.33 | 0.29 |
| 11 | 102504471 | rs11225348 | 0.124 | 2.242 | -0.297 | 0.935 | -0.071 | -0.034 | -0.034 | 0 | 0.11 | 0.01 | 0.18 | 0.13 | 0.2 | 0.16 |
| 11 | 102504882 | rs7119077 | -0.073 | 2.199 | -0.253 | 0.757 | -0.003 | 0.005 | 0.005 | 0 | 0.9 | 0.95 | 0.92 | 1 | 0.9 | 0.96 |
| 11 | 102508348 | rs10895328 | -2.012 | 2.199 | -0.066 | 0.788 | 0.012 | -0.017 | -0.017 | 4.186 | 0.33 | 0.4 | 0.38 | 0.28 | 0.36 | 0.39 |
| 11 | 102508471 | rs11225351 | -1.507 | 2.189 | 0.038 | 0.753 | 0.012 | -0.012 | -0.012 | 0 | 0.2 | 0.27 | 0.24 | 0.14 | 0.26 | 0.25 |
| 11 | 102509835 | rs11225353 | 0.295 | 2.256 | -0.096 | 0.799 | -0.072 | -0.034 | -0.034 | 0.046 | 0.11 | 0.01 | 0.18 | 0.14 | 0.21 | 0.16 |
| 11 | 102510834 | rs1940060 | -0.028 | 2.187 | -0.051 | 0.791 | 0 | -0.001 | -0.001 | 0 | 0.12 | 0.07 | 0.13 | 0.13 | 0.1 | 0.14 |
| 11 | 102512086 | rs7129777 | -2.321 | 2.286 | -0.045 | 0.918 | -0.022 | -0.019 | -0.019 | 0 | 0.36 | 0.24 | 0.5 | 0.59 | 0.42 | 0.59 |
| 11 | 102513805 | rs2846709 | -0.083 | 2.328 | -0.136 | 0.711 | -0.004 | -0.003 | -0.003 | 0 | 0.09 | 0.06 | 0.08 | 0 | 0.1 | 0.04 |
| 11 | 102514861 | rs4433535 | -1.458 | 2.445 | -0.091 | 0.96 | -0.011 | -0.007 | -0.007 | 0 | 0.25 | 0.18 | 0.33 | 0.32 | 0.3 | 0.3 |
| 11 | 102517987 | rs2846712 | -0.125 | 2.299 | -0.023 | 0.923 | -0.009 | 0.001 | 0.001 | 0 | 0.1 | 0.06 | 0.09 | 0 | 0.13 | 0.07 |
| 11 | 102524591 | rs1276277 | -0.802 | 2.281 | -0.027 | 0.912 | 0.002 | -0.034 | -0.034 | 0.643 | 0.13 | 0.15 | 0.1 | 0 | 0.14 | 0.07 |
| 11 | 102531005 | rs11225370 | -0.346 | 2.278 | -0.063 | 0.953 | 0 | -0.001 | -0.001 | 0 | 0.13 | 0.1 | 0.13 | 0.18 | 0.11 | 0.18 |
| 11 | 102533698 | rs7950199 | -0.346 | 2.278 | -0.063 | 0.953 | 0 | -0.001 | -0.001 | 0 | 0.13 | 0.1 | 0.13 | 0.18 | 0.1 | 0.18 |
| 11 | 102535700 | rs1276254 | -2.466 | 2.244 | 0.276 | 0.896 | 0.004 | -0.036 | -0.036 | 1.269 | 0.28 | 0.38 | 0.25 | 0.19 | 0.27 | 0.31 |
| 11 | 102537222 | rs1298374 | -2.364 | 2.159 | 0.252 | 1.002 | -0.001 | -0.001 | -0.001 | 0 | 0.3 | 0.24 | 0.35 | 0.49 | 0.3 | 0.42 |
| 11 | 102549887 | rs1276269 | -2.05 | 2.184 | 0.252 | 1.015 | 0.012 | 0.043 | 0.043 | 0 | 0.2 | 0.29 | 0.25 | 0.32 | 0.23 | 0.3 |
| 11 | 102561688 | rs10895348 | -0.145 | 2.159 | 0.13 | 1.006 | 0 | -0.002 | -0.002 | 0 | 0.13 | 0.08 | 0.13 | 0.18 | 0.1 | 0.18 |
| 11 | 102562700 | rs2509010 | -0.822 | 2.412 | 0.049 | 0.833 | -0.051 | 0.028 | 0.028 | 0.111 | 0.18 | 0.07 | 0.24 | 0.11 | 0.27 | 0.18 |
| 11 | 102563300 | rs11607205 | 0.383 | 2.424 | 0.043 | 0.837 | -0.067 | -0.021 | -0.021 | 0 | 0.11 | 0.01 | 0.19 | 0.11 | 0.18 | 0.12 |
| 11 | 102563739 | rs1276289 | -2.401 | 2.327 | 0.154 | 0.863 | -0.032 | -0.004 | -0.004 | 0.009 | 0.37 | 0.22 | 0.51 | 0.63 | 0.45 | 0.54 |
| 11 | 102564119 | rs11821641 | -0.758 | 2.218 | 0.312 | 0.798 | 0.002 | -0.054 | -0.054 | 0.8 | 0.09 | 0.24 | 0.08 | 0.01 | 0.08 | 0.06 |
| 11 | 102570891 | rs2846723 | 2.373 | 2.272 | 0.083 | 0.916 | -0.019 | 0.003 | 0.003 | 0 | 0.6 | 0.69 | 0.48 | 0.37 | 0.54 | 0.46 |
| 11 | 102572241 | rs2846703 | 0.579 | 2.435 | 0.045 | 0.951 | -0.046 | 0.027 | 0.027 | 0.128 | 0.82 | 0.92 | 0.76 | 0.89 | 0.72 | 0.82 |
| 11 | 102573569 | rs3809018 | 0.427 | 2.454 | 0.026 | 1.01 | -0.065 | -0.024 | -0.024 | 0 | 0.1 | 0.01 | 0.19 | 0.11 | 0.18 | 0.12 |
| 11 | 102574147 | rs4754870 | -1.239 | 1.995 | 0.246 | 1.202 | -0.007 | 0.145 | 0.145 | 0.489 | 0.09 | 0.23 | 0.15 | 0.29 | 0.06 | 0.15 |
| 11 | 102575739 | rs11225388 | 0.168 | 2.521 | 0.098 | 1.122 | -0.077 | -0.02 | -0.02 | 6.575 | 0.11 | 0.01 | 0.22 | 0.11 | 0.23 | 0.14 |
| 11 | 102576358 | rs2846707 | 0.643 | 2.076 | -0.013 | 0.836 | -0.066 | 0.047 | 0.047 | 0.196 | 0.78 | 0.92 | 0.72 | 0.88 | 0.68 | 0.81 |
| 11 | 102576375 | rs1939015 | 0.973 | 1.981 | 0.19 | 0.895 | -0.016 | -0.006 | -0.006 | 0 | 0.77 | 0.7 | 0.82 | 0.81 | 0.83 | 0.76 |
| 11 | 102576382 | rs12099177 | -1.102 | 2.038 | 0.104 | 0.957 | -0.009 | 0.095 | 0.095 | 0.209 | 0.09 | 0.17 | 0.15 | 0.3 | 0.07 | 0.15 |
| 11 | 102576468 | rs11225389 | 0.334 | 2.143 | -0.072 | 0.927 | -0.077 | -0.02 | -0.02 | 1.99 | 0.11 | 0.01 | 0.22 | 0.11 | 0.23 | 0.14 |
| 11 | 102583680 | rs12284255 | -0.126 | 1.998 | 0.025 | 0.957 | -0.011 | 0.023 | 0.023 | 0.147 | 0.12 | 0.13 | 0.12 | 0.2 | 0.07 | 0.19 |
| 11 | 102587062 | rs3740938 | 1.053 | 2.018 | -0.223 | 0.722 | 0.01 | -0.071 | -0.071 | 0.918 | 0.11 | 0 | 0.13 | 0.28 | 0.06 | 0.15 |
| 11 | 102590777 | rs2012390 | 1.325 | 1.66 | -0.432 | 0.323 | 0.006 | -0.002 | -0.002 | 0 | 0.7 | 0.73 | 0.75 | 0.68 | 0.78 | 0.73 |
| 11 | 102593248 | rs1940475 | 1.873 | 1.479 | -0.492 | 0.215 | -0.023 | -0.003 | -0.003 | 0 | 0.59 | 0.71 | 0.52 | 0.57 | 0.55 | 0.6 |
| 11 | 102594427 | rs6590984 | -1.317 | 1.514 | -0.559 | 0.219 | -0.027 | 0.033 | 0.033 | 0 | 0.94 | 1 | 0.95 | 1 | 0.91 | 0.95 |
| 11 | 102595492 | rs3765620 | 0.649 | 1.246 | -0.466 | 0.299 | -0.049 | -0.033 | -0.033 | 0 | 0.68 | 0.81 | 0.56 | 0.58 | 0.59 | 0.65 |
| 11 | 102598575 | rs7943404 | 2.189 | 1.405 | -0.276 | 0.496 | 0.012 | 0.022 | 0.022 | 0 | 0.47 | 0.44 | 0.38 | 0.08 | 0.45 | 0.33 |
| 11 | 102598994 | rs10895353 | -1.048 | 1.087 | -0.089 | 1.126 | -0.011 | 0.266 | 0.266 | 0.337 | 0.13 | 0.32 | 0.14 | 0.48 | 0.1 | 0.27 |
| 11 | 102605579 | rs10791591 | 0.94 | 1.358 | -0.592 | 0.632 | 0.001 | -0.01 | -0.01 | 0.387 | 0.58 | 0.53 | 0.5 | 0.28 | 0.58 | 0.45 |
| 11 | 102606130 | rs4121365 | 0.289 | 1.933 | -0.897 | 0.51 | -0.079 | -0.009 | -0.009 | 0.212 | 0.11 | 0.01 | 0.24 | 0.11 | 0.25 | 0.12 |
| 11 | 102607959 | rs11225401 | -0.602 | 1.567 | -0.716 | 1.234 | -0.002 | 0.003 | 0.003 | 0 | 0.11 | 0.11 | 0.14 | 0.26 | 0.14 | 0.12 |
| 11 | 102624914 | rs10895361 | 0.2 | 1.24 | -0.497 | 1.234 | 0.002 | 0.117 | 0.117 | 0 | 0.07 | 0.19 | 0.13 | 0.25 | 0.05 | 0.16 |
| 11 | 102624956 | rs12146477 | 1.753 | 0.84 | -0.541 | 0.894 | -0.033 | -0.047 | -0.047 | 0 | 0.37 | 0.43 | 0.21 | 0.16 | 0.29 | 0.27 |
| 11 | 102628052 | rs7924357 | 0.494 | 1.102 | -0.478 | 0.872 | -0.002 | 0.086 | 0.086 | 0 | 0.06 | 0.13 | 0.12 | 0.24 | 0.05 | 0.15 |
| 11 | 102629293 | rs516450 | 0.676 | 1.099 | -0.528 | 0.571 | 0.023 | 0.007 | 0.007 | 0.848 | 0.5 | 0.61 | 0.38 | 0.42 | 0.46 | 0.41 |
| 11 | 102635501 | rs10791595 | -0.983 | 1.433 | -0.3 | 0.499 | 0.003 | 0.002 | 0.002 | 0 | 0.32 | 0.33 | 0.36 | 0.56 | 0.37 | 0.39 |
| 11 | 102641196 | rs7948454 | 0.556 | 0.916 | 0.166 | 0.052 | -0.015 | -0.05 | -0.05 | 0.232 | 0.09 | 0.23 | 0.08 | 0 | 0.04 | 0.01 |
| 11 | 102641382 | rs470168 | 1.94 | 0.673 | 0.037 | -0.429 | 0.008 | 0.044 | 0.044 | 0.75 | 0.33 | 0.24 | 0.21 | 0.13 | 0.32 | 0.34 |
| 11 | 102642261 | rs12290253 | -0.317 | 1.007 | 0.091 | -0.439 | 0.018 | -0.069 | -0.069 | 0.024 | 0.19 | 0.36 | 0.17 | 0.1 | 0.18 | 0.13 |
| 11 | 102643894 | rs547561 | -0.946 | 0.832 | -0.281 | -0.151 | 0.006 | -0.031 | -0.031 | 0.022 | 0.93 | 1 | 0.88 | 0.89 | 0.93 | 0.9 |
| 11 | 102644601 | rs12272341 | 1.028 | 0.881 | -0.185 | -0.111 | 0.024 | -0.01 | -0.01 | 1.111 | 0.11 | 0.16 | 0.09 | 0.1 | 0.13 | 0.12 |
| 11 | 102645399 | rs4431992 | 0.037 | 0.778 | -0.393 | 0.036 | 0.011 | 0.048 | 0.048 | 0 | 0.22 | 0.29 | 0.3 | 0.46 | 0.23 | 0.27 |
| 11 | 102647536 | rs2276108 | 1.24 | 0.459 | -0.658 | -0.379 | -0.031 | 0.083 | 0.083 | 0 | 0.14 | 0.01 | 0.09 | 0.02 | 0.16 | 0.07 |
| 11 | 102650389 | rs17293607 | 1.255 | 0.462 | -0.657 | -0.378 | -0.031 | 0.083 | 0.083 | 0 | 0.14 | 0.01 | 0.09 | 0.02 | 0.16 | 0.07 |
| 11 | 102651313 | rs17435959 | -0.161 | 0.391 | -0.332 | -0.229 | 0.009 | 0.022 | 0.022 | 0 | 0.06 | 0.2 | 0.04 | 0.05 | 0.04 | 0.05 |
| 11 | 102655487 | rs4754880 | 0.496 | 0.63 | -1.021 | 0.076 | -0.028 | -0.053 | -0.053 | 0 | 0.15 | 0.1 | 0.22 | 0.44 | 0.2 | 0.2 |

| | | | | | | | | | | | | | | | | |
|---|---|---|---|---|---|---|---|---|---|---|---|---|---|---|---|---|
| 11 | 102656079 | rs559518 | 1.823 | 0.474 | -0.974 | -0.271 | -0.001 | -0.039 | -0.039 | 0 | 0.36 | 0.37 | 0.23 | 0.09 | 0.36 | 0.35 |
| 11 | 102656318 | rs11225422 | 1.133 | 0.414 | -0.969 | -0.218 | -0.019 | 0.116 | 0.116 | 0 | 0.06 | 0.13 | 0.14 | 0.23 | 0.01 | 0.12 |
| 11 | 102656423 | rs1939008 | -0.213 | 0.59 | -1.111 | -0.221 | -0.051 | -0.074 | -0.074 | 0 | 0.24 | 0.13 | 0.28 | 0.53 | 0.3 | 0.32 |
| 11 | 102657908 | rs521898 | -1.009 | 0.531 | -1.133 | -0.253 | 0.013 | -0.028 | -0.028 | 0 | 0.93 | 1 | 0.89 | 0.89 | 0.95 | 0.9 |
| 11 | 102660564 | rs7945189 | 1.407 | 0.55 | -1.131 | -0.251 | -0.03 | -0.004 | -0.004 | 0 | 0.09 | 0.01 | 0.06 | 0.09 | 0.1 | 0.12 |
| 11 | 102660959 | rs2071230 | -0.08 | 0.479 | -1.082 | -0.296 | -0.043 | 0.048 | 0.048 | 0 | 0.15 | 0.31 | 0.23 | 0.23 | 0.06 | 0.14 |
| 11 | 102661595 | rs470747 | 2.045 | 0.61 | -1.205 | -0.24 | -0.001 | -0.045 | -0.045 | 0.033 | 0.36 | 0.37 | 0.23 | 0.09 | 0.36 | 0.35 |
| 11 | 102661665 | rs1938901 | -0.709 | 0.755 | -1.333 | 0.555 | -0.051 | -0.074 | -0.074 | 0.024 | 0.24 | 0.13 | 0.28 | 0.53 | 0.3 | 0.32 |
| 11 | 102663503 | rs7125062 | -0.575 | 0.645 | -1.061 | 0.042 | -0.001 | -0.001 | -0.001 | 0 | 0.28 | 0.3 | 0.43 | 0.67 | 0.26 | 0.34 |
| 11 | 102666316 | rs470558 | 0.109 | 0.93 | -1.38 | 0.505 | 0.013 | -0.028 | -0.028 | 0 | 0.93 | 0.99 | 0.89 | 0.89 | 0.95 | 0.9 |
| 11 | 102668702 | rs470358 | 0.198 | 1.192 | -2.11 | 0.52 | -0.048 | -0.113 | -0.113 | 0 | 0.65 | 0.86 | 0.57 | 0.58 | 0.62 | 0.64 |
| 11 | 102668882 | rs3213460 | 0.159 | 1.026 | -1.863 | -0.106 | -0.019 | 0.055 | 0.055 | 0 | 0.13 | 0.04 | 0.09 | 0.02 | 0.17 | 0.06 |
| 11 | 102669230 | rs514921 | 0.148 | 0.787 | -2.094 | -0.296 | -0.006 | 0.057 | 0.057 | 0 | 0.32 | 0.15 | 0.24 | 0.15 | 0.3 | 0.32 |
| 11 | 102669409 | rs1144393 | -0.045 | 0.949 | -1.709 | -0.438 | -0.021 | 0.016 | 0.016 | 0.074 | 0.21 | 0.18 | 0.22 | 0.09 | 0.35 | 0.16 |
| 11 | 102669645 | rs498186 | 0.279 | 0.246 | -1.985 | 0.347 | -0.011 | -0.014 | -0.014 | 0 | 0.48 | 0.24 | 0.48 | 0.47 | 0.48 | 0.47 |
| 11 | 102669824 | rs2075847 | 0.163 | -0.575 | -0.698 | 0.926 | -0.008 | 0.216 | 0.216 | 0 | 0.07 | 0.23 | 0.05 | 0.27 | 0 | 0.18 |
| 11 | 102672553 | rs2408490 | -0.503 | -0.997 | -0.841 | 0.736 | -0.021 | -0.002 | -0.002 | 0 | 0.26 | 0.31 | 0.12 | 0.22 | 0.17 | 0.22 |
| 11 | 102675489 | rs519806 | 0.345 | -0.71 | -1.357 | 0.458 | -0.001 | 0.002 | 0.002 | 3.724 | 0.42 | 0.25 | 0.46 | 0.41 | 0.45 | 0.5 |
| 11 | 102681029 | rs11225429 | -0.114 | -0.913 | -1.081 | 0.034 | 0 | -0.013 | -0.013 | 0 | 0.12 | 0.16 | 0.05 | 0 | 0.1 | 0.07 |
| 11 | 102682285 | rs1144397 | -0.192 | -1.307 | -1.111 | -0.024 | 0.007 | 0.009 | 0.009 | 0 | 0.49 | 0.55 | 0.38 | 0.34 | 0.56 | 0.33 |
| 11 | 102691894 | rs479095 | 0.29 | -1.766 | -1.129 | -0.15 | 0.008 | -0.002 | -0.002 | 0 | 0.37 | 0.31 | 0.52 | 0.59 | 0.28 | 0.58 |
| 11 | 102695472 | rs1034375 | 0.306 | -2.11 | -1.099 | -0.398 | -0.002 | -0.018 | -0.018 | 0 | 0.14 | 0.17 | 0.04 | 0 | 0.09 | 0.05 |
| 11 | 102701021 | rs1010698 | 1.077 | -2.108 | -1.122 | -0.444 | 0.007 | -0.006 | -0.006 | 0 | 0.35 | 0.41 | 0.33 | 0.33 | 0.47 | 0.28 |
| 11 | 102710471 | rs566125 | 1.407 | -2.042 | -1.04 | -0.478 | -0.018 | 0.02 | 0.02 | 0 | 0.12 | 0.01 | 0.09 | 0.08 | 0.15 | 0.08 |
| 11 | 102713046 | rs3020919 | -0.277 | -2.022 | -0.957 | -0.494 | 0.02 | 0.026 | 0.026 | 0 | 0.27 | 0.24 | 0.14 | 0.08 | 0.24 | 0.14 |
| 11 | 102713620 | rs679620 | -1.209 | -1.85 | -0.973 | -0.392 | 0.005 | -0.002 | -0.002 | 0 | 0.66 | 0.67 | 0.68 | 0.67 | 0.53 | 0.72 |
| 11 | 102715048 | rs522616 | 1.313 | -2.281 | -1.06 | -0.116 | 0.009 | -0.016 | -0.016 | 0 | 0.26 | 0.2 | 0.44 | 0.35 | 0.23 | 0.44 |
| 11 | 102716321 | rs645419 | -0.959 | -1.675 | -0.874 | -0.209 | 0.008 | -0.003 | -0.003 | 0 | 0.66 | 0.65 | 0.67 | 0.67 | 0.53 | 0.72 |
| 11 | 102720421 | rs613804 | 0.782 | -1.924 | -0.961 | -0.43 | -0.028 | -0.023 | -0.023 | 0 | 0.14 | 0.21 | 0.06 | 0.11 | 0.06 | 0.1 |
| 11 | 102720678 | rs615098 | -0.457 | -2.453 | -0.989 | -0.692 | -0.028 | -0.035 | -0.035 | 0.14 | 0.3 | 0.37 | 0.14 | 0.15 | 0.19 | 0.19 |
| 11 | 102732913 | rs28381705 | 0.648 | -2.011 | -0.909 | -0.534 | 0.002 | 0.002 | 0.002 | 0 | 0.11 | 0.12 | 0.08 | 0.12 | 0.14 | 0.14 |
| 11 | 102736419 | rs651159 | 1.054 | -2.089 | -0.938 | -0.573 | -0.036 | -0.03 | -0.03 | 0 | 0.15 | 0.26 | 0.07 | 0.11 | 0.06 | 0.11 |
| 11 | 102739319 | rs11225442 | 0.403 | -1.725 | -1.393 | -0.402 | 0 | 0 | 0 | 0 | 0.12 | 0.11 | 0.08 | 0.12 | 0.14 | 0.14 |
| 11 | 102748695 | rs17368814 | 1.649 | -1.716 | -1.706 | -0.654 | -0.019 | 0.014 | 0.014 | 0 | 0.11 | 0 | 0.06 | 0.04 | 0.13 | 0.08 |
| 11 | 102749448 | rs11225445 | 0.37 | -1.188 | -2.147 | -0.713 | -0.085 | 0.072 | 0.072 | 0 | 0.29 | 0.06 | 0.28 | 0.17 | 0.37 | 0.44 |
| 11 | 102755015 | rs1940935 | 0.306 | -0.632 | -1.8 | -0.435 | -0.055 | 0.045 | 0.045 | 1.252 | 0.22 | 0.08 | 0.26 | 0.12 | 0.35 | 0.42 |
| 11 | 102760015 | rs10791599 | -1.107 | -0.25 | -1.479 | 0.284 | -0.024 | -0.022 | -0.022 | 0 | 0.75 | 0.83 | 0.71 | 0.68 | 0.7 | 0.81 |
| 11 | 102763604 | rs1940937 | 0.478 | -1.329 | -1.549 | 0.297 | 0.069 | -0.044 | -0.044 | 0 | 0.31 | 0.15 | 0.32 | 0.41 | 0.2 | 0.34 |
| 11 | 102763750 | rs1940936 | -1.486 | -0.361 | -1.494 | 0.422 | -0.009 | -0.006 | -0.006 | 0 | 0.77 | 0.83 | 0.72 | 0.75 | 0.75 | 0.84 |
| 11 | 102767396 | rs12275686 | 0.084 | -0.02 | -1.107 | 0.222 | 0.003 | -0.015 | -0.015 | 0 | 0.33 | 0.56 | 0.21 | 0.24 | 0.32 | 0.21 |
| 11 | 102771599 | rs12786502 | 0.851 | 0.218 | -0.324 | 0.372 | 0.012 | -0.017 | -0.017 | 0.064 | 0.14 | 0.22 | 0.08 | 0.07 | 0.14 | 0.05 |
| 11 | 102772798 | rs4434970 | 0.097 | 0.098 | -0.377 | 0.408 | -0.01 | 0.005 | 0.005 | 0 | 0.25 | 0.24 | 0.26 | 0.16 | 0.37 | 0.36 |
| 11 | 102791442 | rs2096768 | 0.398 | -0.624 | -0.372 | 0.583 | 0.014 | 0.001 | 0.001 | 0 | 0.45 | 0.41 | 0.41 | 0.45 | 0.34 | 0.34 |
| 11 | 102793246 | rs11225480 | 0.057 | -0.579 | -0.952 | 0.959 | 0.03 | -0.069 | -0.069 | 1.148 | 0.64 | 0.19 | 0.67 | 0.49 | 0.74 | 0.74 |
| 11 | 102795786 | rs12146550 | 1.053 | -1.144 | -1.004 | 0.754 | 0.007 | 0.023 | 0.023 | 0 | 0.36 | 0.15 | 0.32 | 0.32 | 0.35 | 0.33 |
| 11 | 102797806 | rs12146442 | 0.398 | -0.995 | -1.042 | 0.945 | 0.011 | -0.008 | -0.008 | 0 | 0.49 | 0.17 | 0.51 | 0.46 | 0.58 | 0.68 |
| 11 | 102798691 | rs2212437 | 1.462 | -0.993 | -0.952 | 0.873 | 0.016 | 0.044 | 0.044 | 0 | 0.33 | 0.15 | 0.28 | 0.26 | 0.31 | 0.25 |
| 11 | 102800914 | rs4754052 | -1.43 | -0.317 | -0.767 | 0.976 | 0.005 | 0.001 | 0.001 | 0 | 0.81 | 0.89 | 0.84 | 0.8 | 0.82 | 0.87 |
| 11 | 102801303 | rs12792912 | 0.181 | -0.233 | -1.242 | 1.087 | -0.194 | -0.043 | -0.043 | 0.155 | 0.29 | 0.03 | 0.45 | 0.21 | 0.45 | 0.43 |
| 11 | 102801847 | rs12788925 | 0.181 | -0.233 | -1.242 | 1.08 | -0.194 | -0.051 | -0.051 | 0 | 0.29 | 0.03 | 0.45 | 0.21 | 0.46 | 0.43 |
| 11 | 102802722 | rs4085170 | -1.004 | -0.258 | -0.826 | 1.058 | -0.024 | -0.023 | -0.023 | 0 | 0.78 | 0.57 | 0.81 | 0.81 | 0.82 | 0.87 |
| 11 | 102803476 | rs7124873 | 0.384 | 0.057 | -0.939 | 1.324 | 0.004 | 0.003 | 0.003 | 0 | 0.44 | 0.45 | 0.35 | 0.41 | 0.35 | 0.34 |
| 11 | 102804933 | rs7115014 | 0.15 | 0.406 | -0.801 | 0.671 | 0 | -0.001 | -0.001 | 0.099 | 0.56 | 0.46 | 0.54 | 0.55 | 0.58 | 0.7 |
| 11 | 102807410 | rs646573 | 0.274 | 0.658 | -0.983 | 0.384 | 0 | 0.005 | 0.005 | 0 | 0.66 | 0.62 | 0.75 | 0.56 | 0.66 | 0.65 |
| 11 | 102807731 | rs11225485 | -0.604 | 0.547 | -1.077 | 0.381 | 0 | -0.001 | -0.001 | 0 | 0.11 | 0.1 | 0.07 | 0.12 | 0.09 | 0.14 |
| 11 | 102810078 | rs11826440 | 0.024 | 0.592 | -0.838 | 0.485 | 0.019 | 0.041 | 0.041 | 0 | 0.06 | 0.18 | 0.06 | 0.11 | 0.05 | 0.09 |
| 11 | 102819574 | rs10895372 | -0.741 | 0.861 | -1.151 | 0.574 | 0.007 | 0.02 | 0.02 | 0 | 0.13 | 0.17 | 0.12 | 0.22 | 0.17 | 0.35 |
| 11 | 102820655 | rs10895373 | -0.561 | 0.849 | -1.154 | 0.56 | 0.003 | 0.018 | 0.018 | 0 | 0.13 | 0.16 | 0.12 | 0.22 | 0.17 | 0.35 |
| 11 | 102821404 | rs10502009 | 0.623 | 0.998 | -1.203 | 0.284 | -0.031 | -0.037 | -0.037 | 0 | 0.07 | 0.01 | 0.06 | 0.11 | 0.13 | 0.26 |
| 11 | 102825091 | rs640198 | -1.055 | 0.598 | -1.658 | -0.104 | 0.03 | -0.032 | -0.032 | 0 | 0.63 | 0.73 | 0.65 | 0.5 | 0.7 | 0.58 |
| 11 | 102829455 | rs671188 | -0.662 | 0.62 | -1.61 | -0.216 | 0.025 | 0.037 | 0.037 | 0 | 0.54 | 0.69 | 0.59 | 0.39 | 0.57 | 0.34 |
| 11 | 102830793 | rs660669 | -0.224 | 0.889 | -1.749 | 0.031 | -0.018 | -0.009 | -0.009 | 0 | 0.33 | 0.22 | 0.37 | 0.39 | 0.39 | 0.28 |
| 11 | 102833618 | rs687558 | -0.471 | 1.086 | -1.83 | 0.229 | -0.033 | -0.026 | -0.026 | 0 | 0.28 | 0.13 | 0.23 | 0.31 | 0.39 | 0.35 |
| 11 | 102833826 | rs10791602 | 0.202 | 1.13 | -1.099 | 0.2 | 0.032 | 0.038 | 0.038 | 0 | 0.78 | 0.55 | 0.68 | 0.77 | 0.74 | 0.88 |
| 11 | 102835419 | rs579627 | 0.647 | 1.133 | -0.982 | 0.241 | 0 | 0 | 0 | 0 | 0.13 | 0.12 | 0.12 | 0.13 | 0.16 | 0.07 |
| 11 | 102835636 | rs10791605 | -0.468 | 0.611 | -1.461 | -0.05 | 0.006 | 0.002 | 0.002 | 0 | 0.53 | 0.48 | 0.58 | 0.46 | 0.63 | 0.64 |
| 11 | 102835942 | rs575188 | -0.937 | 0.185 | -1.35 | -0.016 | -0.034 | 0.017 | 0.017 | 0 | 0.66 | 0.59 | 0.7 | 0.59 | 0.81 | 0.71 |
| 11 | 102838443 | rs693477 | 0.323 | 1.268 | -1.316 | -0.273 | -0.015 | 0.022 | 0.022 | 0 | 0.79 | 0.84 | 0.77 | 0.91 | 0.74 | 0.7 |
| 11 | 102848546 | rs637759 | -0.043 | 0.725 | -1.081 | -0.208 | -0.004 | 0.002 | 0.002 | 0 | 0.74 | 0.76 | 0.67 | 0.9 | 0.72 | 0.86 |

| | | | | | | | | | | | | | | | | |
|---|---|---|---|---|---|---|---|---|---|---|---|---|---|---|---|---|
| 11 | 102854782 | rs627763 | -0.362 | 0.763 | -1.269 | 0.379 | -0.059 | 0.056 | 0.056 | 0.562 | 0.78 | 0.93 | 0.68 | 0.9 | 0.73 | 0.86 |
| 11 | 102855634 | rs17099821 | 0.478 | 0.293 | -0.689 | 0.636 | -0.023 | 0.012 | 0.012 | 0 | 0.37 | 0.43 | 0.32 | 0.42 | 0.26 | 0.39 |
| 11 | 102861607 | rs535628 | -0.172 | 0.761 | -0.524 | 0.922 | 0.021 | 0.048 | 0.048 | 0 | 0.24 | 0.31 | 0.38 | 0.44 | 0.37 | 0.37 |
| 11 | 102877513 | rs11225514 | 0.013 | 0.063 | -0.342 | 1.007 | -0.035 | 0.042 | 0.042 | 0 | 0.16 | 0.28 | 0.12 | 0.23 | 0.13 | 0.24 |
| 11 | 102877651 | rs601592 | -0.872 | -0.283 | 0.017 | 0.943 | -0.003 | -0.013 | -0.013 | 0.547 | 0.4 | 0.66 | 0.47 | 0.46 | 0.42 | 0.44 |
| 11 | 102878622 | rs599399 | -1.101 | -0.25 | -0.164 | 0.514 | 0.002 | 0.097 | 0.097 | 0.493 | 0.46 | 0.81 | 0.52 | 0.68 | 0.5 | 0.61 |
| 11 | 102887764 | rs608645 | -1.073 | -0.332 | -0.209 | 0.462 | 0.004 | 0.049 | 0.049 | 0.314 | 0.46 | 0.8 | 0.53 | 0.67 | 0.49 | 0.6 |
| 11 | 102901002 | rs581359 | -0.443 | -0.344 | 0.38 | 0.405 | 0.002 | 0 | 0 | 0 | 0.35 | 0.43 | 0.46 | 0.65 | 0.45 | 0.58 |
| 11 | 102906516 | rs514835 | -1.28 | 0.549 | -0.053 | 0.324 | -0.014 | 0.011 | 0.011 | 0 | 0.93 | 0.96 | 0.86 | 1 | 0.89 | 0.93 |
| 11 | 102908390 | rs476244 | -1.254 | -0.207 | -0.213 | 0.06 | 0.003 | 0.047 | 0.047 | 0.049 | 0.54 | 0.78 | 0.54 | 0.67 | 0.51 | 0.62 |
| 11 | 102911494 | rs2186789 | 1.154 | -0.232 | -0.406 | -0.433 | -0.012 | 0.025 | 0.025 | 0 | 0.21 | 0.12 | 0.22 | 0.1 | 0.23 | 0.15 |
| 11 | 102913079 | rs6591003 | 0.932 | -0.154 | -0.129 | -0.081 | 0.01 | -0.034 | -0.034 | 0.417 | 0.07 | 0.23 | 0.06 | 0 | 0.04 | 0.02 |
| 11 | 102948592 | rs2510087 | -0.417 | -0.417 | -0.569 | 0.082 | -0.004 | -0.004 | -0.004 | 0 | 0.25 | 0.32 | 0.16 | 0.2 | 0.19 | 0.2 |
| 11 | 102949969 | rs2513996 | -0.987 | -0.246 | -0.252 | -0.2 | -0.01 | -0.032 | -0.032 | 0 | 0.33 | 0.54 | 0.25 | 0.21 | 0.26 | 0.25 |
| 11 | 102955091 | rs17372721 | 1.534 | -0.262 | -0.376 | -0.284 | -0.028 | 0.028 | 0.028 | 0.044 | 0.06 | 0 | 0.03 | 0.01 | 0.06 | 0.06 |
| 11 | 102957052 | rs510396 | 1.353 | 0.54 | -0.964 | -0.553 | -0.061 | 0.04 | 0.04 | 0 | 0.18 | 0.02 | 0.14 | 0.06 | 0.19 | 0.09 |
| 11 | 102961004 | rs2509112 | -0.499 | 0.249 | -0.824 | -0.581 | -0.009 | -0.02 | -0.02 | 0.506 | 0.27 | 0.53 | 0.22 | 0.21 | 0.23 | 0.23 |
| 11 | 102967852 | rs11225547 | 2.393 | 0.261 | -0.823 | -0.548 | -0.022 | 0.031 | 0.031 | 0 | 0.06 | 0 | 0.06 | 0.01 | 0.07 | 0.05 |
| 11 | 102968093 | rs1893073 | -0.598 | 0.248 | -0.753 | -0.56 | -0.001 | -0.004 | -0.004 | 0 | 0.31 | 0.36 | 0.23 | 0.21 | 0.26 | 0.25 |
| 11 | 102981300 | rs2510094 | 1.45 | 0.593 | -0.991 | -0.694 | 0.007 | -0.007 | -0.007 | 0 | 0.12 | 0.03 | 0.04 | 0.11 | 0.07 | 0.13 |
| 11 | 102989206 | rs11225553 | 1.604 | 0.76 | -0.983 | -0.71 | 0.013 | -0.04 | -0.04 | 0 | 0.08 | 0.19 | 0.09 | 0.01 | 0.07 | 0.04 |
| 11 | 102991197 | rs17301182 | 2.541 | 0.693 | -1.256 | -0.655 | -0.04 | -0.03 | -0.03 | 0.003 | 0.08 | 0.01 | 0.09 | 0.09 | 0.13 | 0.07 |
| 11 | 102996987 | rs12577612 | 0.378 | 0.666 | -1.06 | -0.294 | 0.061 | -0.012 | -0.012 | 0 | 0.48 | 0.26 | 0.25 | 0.48 | 0.38 | 0.47 |
| 11 | 103021106 | rs11225571 | 2.302 | 0.419 | -1.237 | 0.245 | -0.016 | -0.06 | -0.06 | 0.69 | 0.14 | 0.02 | 0.06 | 0.25 | 0.12 | 0.16 |
| 11 | 103029516 | rs688906 | 0.453 | 0.715 | -0.564 | 0.49 | -0.004 | -0.026 | -0.026 | 0 | 0.68 | 0.44 | 0.75 | 0.78 | 0.73 | 0.74 |
| 11 | 103045257 | rs10431051 | 1.243 | 0.688 | -0.848 | 0.677 | 0 | 0.011 | 0.011 | 0 | 0.08 | 0.07 | 0.09 | 0.23 | 0.11 | 0.17 |
| 11 | 103045457 | rs2514007 | -0.825 | 0.598 | -0.809 | 0.611 | 0.001 | -0.001 | -0.001 | 0 | 0.86 | 0.86 | 0.94 | 0.88 | 0.93 | 0.88 |
| 11 | 103052558 | rs11225584 | 1.311 | 0.611 | -1.054 | 0.653 | 0 | 0.009 | 0.009 | 0 | 0.08 | 0.07 | 0.09 | 0.23 | 0.11 | 0.17 |
| 11 | 103055490 | rs17374415 | 2.412 | 0.688 | -1.119 | 0.63 | -0.039 | -0.028 | -0.028 | 0 | 0.08 | 0 | 0.09 | 0.09 | 0.12 | 0.1 |
| 11 | 103067460 | rs685395 | -0.605 | 0.773 | -1.12 | 0.62 | -0.01 | 0.01 | 0.01 | 0 | 0.81 | 0.94 | 0.89 | 0.88 | 0.86 | 0.84 |
| 11 | 103071344 | rs654550 | 1.125 | 0.717 | -0.959 | 0.605 | -0.002 | 0.015 | 0.015 | 0 | 0.61 | 0.5 | 0.67 | 0.56 | 0.63 | 0.56 |
| 11 | 103082590 | rs589623 | 0.048 | 0.725 | -0.925 | 0.608 | -0.015 | -0.005 | -0.005 | 0 | 0.74 | 0.57 | 0.82 | 0.79 | 0.81 | 0.77 |
| 11 | 103084838 | rs664310 | -0.128 | 0.727 | -0.968 | 0.618 | 0.016 | -0.057 | -0.057 | 0 | 0.79 | 0.87 | 0.86 | 0.65 | 0.82 | 0.71 |
| 11 | 103100517 | rs601535 | 1.028 | 0.704 | -0.914 | 0.598 | -0.003 | 0.008 | 0.008 | 0 | 0.61 | 0.5 | 0.67 | 0.54 | 0.62 | 0.56 |
| 11 | 103111657 | rs660994 | -0.579 | 0.379 | -0.771 | 0.25 | -0.015 | -0.003 | -0.003 | 0 | 0.26 | 0.43 | 0.17 | 0.21 | 0.19 | 0.22 |
| 11 | 103121473 | rs3912622 | 0.865 | 0.229 | -0.562 | -0.021 | 0.029 | 0.056 | 0.056 | 0.575 | 0.38 | 0.2 | 0.15 | 0.25 | 0.27 | 0.27 |
| 11 | 103124135 | rs11225634 | 0.18 | 0.295 | 0.062 | 0.419 | 0.028 | 0.009 | 0.009 | 0 | 0.12 | 0.38 | 0.13 | 0.09 | 0.13 | 0.11 |
| 11 | 103152919 | rs12574626 | -0.154 | 0.302 | -0.03 | 0.405 | 0.012 | -0.011 | -0.011 | 0 | 0.14 | 0.42 | 0.13 | 0.09 | 0.13 | 0.11 |
| 11 | 103156806 | rs4754897 | 0.351 | 0.458 | -0.463 | 0.549 | -0.014 | -0.023 | -0.023 | 0.372 | 0.13 | 0.03 | 0.23 | 0.17 | 0.12 | 0.13 |
| 11 | 103158278 | rs10895391 | 0.476 | 0.021 | -0.761 | 0.354 | 0.009 | 0.039 | 0.039 | 0 | 0.39 | 0.17 | 0.25 | 0.3 | 0.34 | 0.4 |
| 11 | 103167192 | rs7944311 | 0.375 | 0.009 | -0.239 | 0.361 | 0 | 0.006 | 0.006 | 0 | 0.14 | 0.19 | 0.25 | 0.17 | 0.12 | 0.14 |
| 11 | 103176145 | rs7950038 | 0.417 | 0.054 | -0.217 | 0.363 | 0.002 | 0.007 | 0.007 | 0 | 0.14 | 0.19 | 0.25 | 0.17 | 0.12 | 0.13 |
| 11 | 103185959 | rs11225703 | 1.399 | 0.23 | -0.117 | 0.579 | -0.013 | 0.004 | 0.004 | 0 | 0.26 | 0.21 | 0.35 | 0.27 | 0.38 | 0.26 |
| 11 | 103190054 | rs220327 | 0.051 | 0.181 | -0.086 | 0.676 | 0.002 | 0.081 | 0.081 | 0 | 0.14 | 0.33 | 0.11 | 0.24 | 0.12 | 0.19 |
| 11 | 103191672 | rs4754913 | 0.492 | 0.05 | -0.086 | 0.611 | -0.001 | 0 | 0 | 3.771 | 0.16 | 0.18 | 0.24 | 0.19 | 0.12 | 0.15 |
| 11 | 103193087 | rs313396 | 1.306 | 0.151 | -0.239 | 0.603 | -0.089 | 0.003 | 0.003 | 0 | 0.28 | 0.06 | 0.37 | 0.27 | 0.4 | 0.28 |
| 11 | 103193779 | rs313393 | 0.219 | 0.13 | -0.182 | 0.609 | -0.049 | 0.007 | 0.007 | 0 | 0.42 | 0.19 | 0.61 | 0.45 | 0.52 | 0.43 |
| 11 | 103199573 | rs7925257 | 0.461 | -0.097 | 0.418 | 0.741 | 0.017 | 0.095 | 0.095 | 0.013 | 0.12 | 0.38 | 0.11 | 0.19 | 0.1 | 0.16 |
| 11 | 103203377 | rs2514414 | 0.252 | -0.07 | 0.21 | 0.529 | -0.012 | -0.014 | -0.014 | 0 | 0.41 | 0.37 | 0.49 | 0.47 | 0.56 | 0.46 |
| 11 | 103211457 | rs11225731 | -0.195 | -0.141 | 0.072 | 0.539 | -0.005 | -0.004 | -0.004 | 0 | 0.21 | 0.2 | 0.29 | 0.22 | 0.2 | 0.27 |
| 11 | 103215631 | rs313417 | 1.265 | -0.2 | -0.057 | 0.573 | -0.127 | 0.008 | 0.008 | 0 | 0.29 | 0.04 | 0.39 | 0.27 | 0.45 | 0.29 |
| 11 | 103216252 | rs3847582 | -0.081 | -0.181 | -0.025 | 0.562 | -0.005 | -0.004 | -0.004 | 0 | 0.21 | 0.2 | 0.29 | 0.22 | 0.2 | 0.22 |
| 11 | 103218413 | rs10895406 | 0.614 | -0.408 | 0.305 | 0.495 | 0.016 | 0.019 | 0.019 | 0 | 0.39 | 0.27 | 0.23 | 0.31 | 0.31 | 0.37 |
| 11 | 103219906 | rs2566927 | 0.346 | -0.472 | -0.079 | 0.472 | 0.007 | -0.013 | -0.013 | 0.408 | 0.46 | 0.17 | 0.62 | 0.48 | 0.58 | 0.48 |
| 11 | 103229027 | rs2566913 | -0.089 | -0.71 | -0.013 | 0.535 | 0.084 | -0.002 | -0.002 | 0 | 0.52 | 0.25 | 0.68 | 0.49 | 0.66 | 0.53 |
| 11 | 103229110 | rs2671333 | -0.089 | -0.71 | -0.013 | 0.535 | 0.084 | -0.002 | -0.002 | 0 | 0.52 | 0.25 | 0.68 | 0.49 | 0.66 | 0.53 |
| 11 | 103233343 | rs12221730 | 1.406 | -0.791 | -0.268 | 0.414 | -0.024 | -0.065 | -0.065 | 3.742 | 0.17 | 0.06 | 0.17 | 0.31 | 0.19 | 0.28 |
| 11 | 103253696 | rs12420792 | 0.272 | -0.741 | -0.034 | 0.476 | -0.003 | 0 | 0 | 0 | 0.2 | 0.23 | 0.26 | 0.21 | 0.14 | 0.2 |
| 11 | 103255114 | rs2671397 | -0.615 | -0.768 | 0.034 | 0.379 | -0.013 | -0.013 | -0.013 | 0 | 0.62 | 0.61 | 0.76 | 0.69 | 0.7 | 0.65 |
| 11 | 103277984 | rs11225773 | -0.785 | -0.714 | -0.127 | 0.39 | -0.008 | -0.011 | -0.011 | 0.604 | 0.59 | 0.64 | 0.5 | 0.53 | 0.43 | 0.57 |
| 11 | 103281978 | rs1793500 | -0.758 | -0.671 | -0.13 | 0.395 | -0.007 | -0.01 | -0.01 | 0 | 0.58 | 0.62 | 0.5 | 0.52 | 0.43 | 0.57 |
| 11 | 103292021 | rs520039 | -1.117 | -0.704 | -0.196 | 0.367 | -0.028 | 0.051 | 0.051 | 0 | 0.62 | 0.71 | 0.53 | 0.73 | 0.46 | 0.68 |
| 11 | 103300270 | rs519773 | -1.117 | -0.704 | -0.196 | 0.358 | -0.028 | 0.049 | 0.049 | 0 | 0.62 | 0.71 | 0.53 | 0.73 | 0.46 | 0.68 |
| 11 | 103304381 | rs589523 | -1.135 | -0.708 | -0.198 | 0.355 | -0.029 | 0.048 | 0.048 | 0 | 0.62 | 0.7 | 0.53 | 0.73 | 0.46 | 0.68 |
| 11 | 103307709 | rs11828719 | 0.399 | -0.773 | -0.008 | 0.372 | 0 | 0 | 0 | 0 | 0.37 | 0.31 | 0.23 | 0.29 | 0.3 | 0.35 |
| 11 | 103307933 | rs650634 | -1.174 | -0.555 | -0.031 | 0.353 | -0.043 | 0.043 | 0.043 | 0 | 0.63 | 0.71 | 0.54 | 0.73 | 0.48 | 0.69 |
| 11 | 103309088 | rs546524 | -0.547 | -0.5 | -0.204 | 0.625 | -0.041 | 0.089 | 0.089 | 0 | 0.25 | 0.4 | 0.3 | 0.43 | 0.16 | 0.33 |
| 11 | 103310841 | rs565183 | -0.095 | -0.327 | -0.344 | 0.426 | -0.013 | 0.009 | 0.009 | 0 | 0.22 | 0.3 | 0.27 | 0.24 | 0.14 | 0.23 |
| 11 | 103310851 | rs589943 | -2.02 | -0.023 | -0.829 | 0.148 | -0.081 | 0.082 | 0.082 | 0 | 0.79 | 0.98 | 0.77 | 0.96 | 0.71 | 0.89 |

| | | | | | | | | | | | | | | | | |
|---|---|---|---|---|---|---|---|---|---|---|---|---|---|---|---|---|
| 11 | 103315520 | rs548142 | -0.031 | -0.636 | 0.159 | 0.815 | 0.036 | 0.029 | 0.029 | 0 | 0.27 | 0.48 | 0.26 | 0.34 | 0.3 | 0.34 |
| 11 | 103318329 | rs581977 | -1.098 | -0.785 | 0.126 | 0.791 | -0.009 | 0.017 | 0.017 | 0 | 0.47 | 0.69 | 0.52 | 0.59 | 0.43 | 0.56 |
| 11 | 103323596 | rs17100943 | 0.448 | -0.64 | -0.448 | 0.552 | 0.003 | -0.011 | -0.011 | 1.734 | 0.17 | 0.13 | 0.23 | 0.24 | 0.12 | 0.22 |
| 11 | 103325908 | rs11225812 | 1.817 | -0.512 | -0.512 | 0.551 | -0.034 | -0.011 | -0.011 | 0 | 0.06 | 0.01 | 0.06 | 0.05 | 0.1 | 0.05 |
| 11 | 103326785 | rs7118634 | -0.546 | -0.581 | -0.478 | 0.533 | -0.007 | -0.009 | -0.009 | 0.585 | 0.26 | 0.18 | 0.31 | 0.29 | 0.23 | 0.28 |
| 11 | 103334643 | rs313862 | -1.343 | -0.747 | -0.414 | 0.236 | -0.022 | 0.176 | 0.176 | 0 | 0.52 | 0.8 | 0.68 | 0.81 | 0.47 | 0.64 |
| 11 | 103335225 | rs11225813 | 1.632 | -0.745 | -0.442 | 0.253 | -0.028 | -0.006 | -0.006 | 0 | 0.06 | 0.02 | 0.06 | 0.06 | 0.1 | 0.05 |
| 11 | 103339813 | rs11225815 | 0.617 | -0.896 | -0.321 | 0.328 | 0.038 | 0.1 | 0.1 | 0 | 0.54 | 0.26 | 0.39 | 0.25 | 0.64 | 0.41 |
| 11 | 103342939 | rs12419850 | 0.302 | -0.562 | -0.112 | 0.599 | 0.001 | 0 | 0 | 0 | 0.18 | 0.17 | 0.23 | 0.2 | 0.12 | 0.23 |
| 11 | 103343254 | rs1946019 | 0.013 | -0.215 | 0.358 | 1.154 | -0.009 | 0.08 | 0.08 | 0 | 0.75 | 0.49 | 0.64 | 0.45 | 0.78 | 0.64 |
| 11 | 103350975 | rs1481980 | 0.053 | -0.205 | 0.346 | 0.895 | -0.01 | 0.05 | 0.05 | 0 | 0.75 | 0.46 | 0.63 | 0.4 | 0.77 | 0.63 |
| 11 | 103380763 | rs969038 | -0.347 | -0.341 | 0.125 | 0.027 | -0.04 | -0.096 | -0.096 | 0.596 | 0.81 | 0.64 | 0.93 | 1 | 0.9 | 0.95 |
| 11 | 103385445 | rs17379654 | -0.323 | -0.176 | 0.045 | -0.039 | -0.006 | 0.074 | 0.074 | 0 | 0.23 | 0.11 | 0.12 | 0.02 | 0.2 | 0.1 |
| 11 | 103388805 | rs979802 | 0.955 | -0.325 | 0.044 | 0.341 | -0.015 | 0.013 | 0.013 | 0.002 | 0.62 | 0.5 | 0.56 | 0.48 | 0.72 | 0.66 |
| 11 | 103394155 | rs11225860 | 0.575 | -0.34 | 0.052 | 0.263 | 0.009 | 0.071 | 0.071 | 0 | 0.2 | 0.38 | 0.22 | 0.28 | 0.2 | 0.27 |
| 11 | 103394962 | rs1481961 | 1.506 | -0.3 | -0.025 | 0.219 | -0.074 | 0.15 | 0.15 | 0 | 0.41 | 0.15 | 0.35 | 0.2 | 0.52 | 0.34 |
| 11 | 103399200 | rs7129400 | 0.538 | -0.263 | 0.051 | 0.338 | 0.006 | 0.061 | 0.061 | 0 | 0.2 | 0.38 | 0.19 | 0.25 | 0.18 | 0.23 |
| 11 | 103400088 | rs4754957 | -0.67 | -0.335 | 0.052 | 0.256 | -0.032 | -0.068 | -0.068 | 0 | 0.81 | 0.66 | 0.91 | 0.96 | 0.89 | 0.88 |
| 11 | 103400158 | rs11820878 | 0.843 | -0.317 | -0.084 | 0.265 | 0.001 | 0.082 | 0.082 | 0 | 0.59 | 0.28 | 0.43 | 0.24 | 0.63 | 0.46 |
| 11 | 103403862 | rs7105630 | 0.343 | -0.382 | -0.237 | 0.273 | 0.023 | 0.014 | 0.014 | 0 | 0.63 | 0.28 | 0.72 | 0.72 | 0.71 | 0.65 |
| 11 | 103403911 | rs7117196 | 1.151 | -0.318 | -0.377 | 0.19 | 0.057 | 0.201 | 0.201 | 0 | 0.48 | 0.17 | 0.42 | 0.22 | 0.62 | 0.42 |
| 11 | 103406150 | rs1481954 | 0.159 | -0.235 | 0.017 | 0.453 | 0.013 | 0.009 | 0.009 | 0 | 0.23 | 0.23 | 0.26 | 0.27 | 0.29 | 0.31 |
| 11 | 103408327 | rs1471602 | -1.644 | -0.052 | 0.11 | 0.351 | -0.005 | 0.026 | 0.026 | 0 | 0.57 | 0.61 | 0.65 | 0.85 | 0.52 | 0.69 |
| 11 | 103408967 | rs7952637 | -0.061 | 0.169 | -0.355 | 0.455 | -0.001 | 0 | 0 | 0.103 | 0.28 | 0.24 | 0.28 | 0.24 | 0.3 | 0.34 |
| 11 | 103410068 | rs2897997 | -0.425 | 0.209 | -0.772 | 0.294 | -0.001 | -0.003 | -0.003 | 0 | 0.14 | 0.15 | 0.07 | 0.12 | 0.12 | 0.12 |
| 11 | 103416656 | rs10895457 | 0.816 | 0.533 | -1.33 | 0.02 | -0.048 | 0.191 | 0.191 | 0 | 0.41 | 0.12 | 0.28 | 0.19 | 0.44 | 0.32 |
| 11 | 103416913 | rs10459006 | -0.158 | 0.557 | -0.729 | 0.762 | -0.011 | 0.042 | 0.042 | 0 | 0.29 | 0.44 | 0.31 | 0.48 | 0.27 | 0.36 |
| 11 | 103417781 | rs7120142 | -0.425 | 0.72 | -0.527 | 1.326 | 0 | 0.026 | 0.026 | 0 | 0.82 | 0.84 | 0.66 | 0.56 | 0.8 | 0.66 |
| 11 | 103417868 | rs716273 | 0.466 | 0.749 | -0.532 | 1.228 | 0.001 | 0.012 | 0.012 | 0 | 0.57 | 0.43 | 0.4 | 0.29 | 0.59 | 0.46 |
| 11 | 103418084 | rs10895458 | 1.034 | 0.735 | -0.767 | 1.166 | -0.042 | 0.279 | 0.279 | 0 | 0.49 | 0.13 | 0.31 | 0.19 | 0.52 | 0.42 |
| 11 | 103421935 | rs1481996 | -1.13 | 0.503 | -0.699 | 1.314 | -0.001 | -0.007 | -0.007 | 0.906 | 0.85 | 0.9 | 0.72 | 0.73 | 0.85 | 0.82 |
| 11 | 103427568 | rs11225870 | 0.518 | 0.427 | -0.508 | 0.728 | 0.015 | 0.022 | 0.022 | 0 | 0.05 | 0.11 | 0.07 | 0.1 | 0.06 | 0.04 |
| 11 | 103429644 | rs10895461 | 0.534 | 0.709 | -0.885 | 0.67 | -0.014 | -0.009 | -0.009 | 0.695 | 0.59 | 0.19 | 0.6 | 0.64 | 0.62 | 0.67 |
| 11 | 103432406 | rs1946017 | -1.231 | 0.862 | -1.232 | 0.412 | -0.009 | 0.018 | 0.018 | 0 | 0.93 | 0.95 | 0.97 | 1 | 0.93 | 0.93 |
| 11 | 103436831 | rs2218851 | 0.329 | 0.771 | -1.049 | 0.515 | 0.005 | 0.032 | 0.032 | 0 | 0.25 | 0.38 | 0.27 | 0.27 | 0.24 | 0.22 |
| 11 | 103437292 | rs11225875 | 0.412 | 0.799 | -1.693 | 0.13 | -0.05 | 0.188 | 0.188 | 0 | 0.39 | 0.05 | 0.26 | 0.19 | 0.43 | 0.33 |
| 11 | 103452228 | rs1946013 | 0.747 | 0.855 | -0.684 | 1.074 | 0.029 | 0.094 | 0.094 | 0.459 | 0.9 | 0.74 | 0.92 | 0.78 | 0.88 | 0.91 |
| 11 | 103453573 | rs10488761 | 0.251 | 0.658 | -2.103 | 0.702 | -0.063 | -0.084 | -0.084 | 0 | 0.36 | 0.03 | 0.33 | 0.33 | 0.38 | 0.45 |
| 11 | 103453861 | rs12804615 | 0.236 | 0.642 | -1.935 | 0.724 | -0.026 | -0.057 | -0.057 | 0.041 | 0.36 | 0.06 | 0.33 | 0.34 | 0.38 | 0.45 |
| 11 | 103454659 | rs10895473 | -0.606 | -0.22 | -1.406 | 0.255 | 0.007 | -0.016 | -0.016 | 4.683 | 0.62 | 0.58 | 0.53 | 0.76 | 0.56 | 0.68 |
| 11 | 103459106 | rs7941218 | 0.462 | -0.401 | -1.08 | 0.145 | -0.061 | -0.072 | -0.072 | 0.003 | 0.19 | 0.44 | 0.11 | 0.07 | 0.09 | 0.13 |
| 11 | 103461459 | rs7115036 | 0.701 | 0.305 | -1.606 | -0.203 | -0.023 | 0.071 | 0.071 | 0.172 | 0.22 | 0.09 | 0.22 | 0.02 | 0.3 | 0.17 |
| 11 | 103463841 | rs10895475 | 0.458 | -0.037 | -1.703 | -0.23 | -0.027 | 0.001 | 0.001 | 1.156 | 0.34 | 0.18 | 0.38 | 0.65 | 0.38 | 0.48 |
| 11 | 103464525 | rs10895476 | 0.413 | 0.051 | -1.582 | -0.3 | 0.012 | -0.008 | -0.008 | 0 | 0.24 | 0.32 | 0.3 | 0.25 | 0.23 | 0.25 |
| 11 | 103465050 | rs17101182 | 0.823 | -0.025 | -1.226 | -0.77 | -0.043 | -0.041 | -0.041 | 0 | 0.18 | 0.33 | 0.1 | 0.1 | 0.1 | 0.13 |
| 11 | 103465212 | rs17467090 | 1.181 | 0.381 | -1.407 | -0.599 | -0.083 | 0.133 | 0.133 | 0 | 0.22 | 0.02 | 0.21 | 0.01 | 0.3 | 0.14 |
| 11 | 103465513 | rs17101188 | 0.081 | 0.303 | -0.783 | -0.063 | 0.003 | -0.006 | -0.006 | 0 | 0.34 | 0.29 | 0.32 | 0.35 | 0.34 | 0.51 |
| 11 | 103466085 | rs10791631 | -0.483 | 0.54 | -0.821 | 0.152 | 0.002 | 0.057 | 0.057 | 0 | 0.08 | 0.05 | 0.21 | 0.43 | 0.14 | 0.28 |
| 11 | 103474165 | rs2408681 | -0.408 | 0.57 | -0.696 | 0.357 | 0.128 | 0.406 | 0.406 | 0 | 0.07 | 0.27 | 0.22 | 0.47 | 0.1 | 0.25 |
| 11 | 103474376 | rs12284561 | 0.377 | 0.293 | -1.537 | 0.3 | -0.033 | 0.005 | 0.005 | 0.032 | 0.08 | 0.25 | 0.03 | 0.06 | 0.01 | 0.04 |
| 11 | 103476499 | rs12793538 | 0.131 | 0.992 | -1.932 | 0.291 | -0.012 | -0.041 | -0.041 | 0.552 | 0.3 | 0.16 | 0.34 | 0.37 | 0.32 | 0.42 |
| 11 | 103489944 | rs10895486 | 0.262 | 1.222 | -2.191 | 0.316 | -0.046 | -0.069 | -0.069 | 1.923 | 0.27 | 0.04 | 0.27 | 0.3 | 0.3 | 0.25 |
| 11 | 103489959 | rs976658 | 0.273 | 1.264 | -1.597 | 0.411 | 0.002 | -0.007 | -0.007 | 0 | 0.2 | 0.23 | 0.3 | 0.17 | 0.23 | 0.29 |
| 11 | 103490917 | rs10895487 | -0.229 | 1.259 | -1.511 | 0.917 | -0.037 | -0.052 | -0.052 | 0 | 0.23 | 0.14 | 0.29 | 0.44 | 0.32 | 0.3 |
| 11 | 103491310 | rs10488759 | -0.911 | 0.499 | -0.964 | 1.008 | 0 | -0.012 | -0.012 | 0 | 0.34 | 0.31 | 0.33 | 0.6 | 0.28 | 0.53 |
| 11 | 103493096 | rs10488758 | -0.302 | -0.048 | -0.9 | 0.763 | -0.03 | 0.012 | 0.012 | 0 | 0.1 | 0.01 | 0.07 | 0.05 | 0.11 | 0.07 |
| 11 | 103496422 | rs4754971 | 1.146 | -0.63 | -0.184 | 1.161 | -0.015 | 0.042 | 0.042 | 1.5 | 0.77 | 0.35 | 0.73 | 0.67 | 0.84 | 0.7 |
| 11 | 103498150 | rs1917448 | 1.337 | -0.75 | -0.522 | 1.166 | 0.013 | -0.073 | -0.073 | 1.078 | 0.67 | 0.18 | 0.66 | 0.51 | 0.72 | 0.59 |
| 11 | 103498954 | rs11225930 | -0.809 | -0.892 | 0.012 | 1.13 | 0.005 | 0.078 | 0.078 | 0.265 | 0.06 | 0.2 | 0.06 | 0.12 | 0.03 | 0.23 |
| 11 | 103500160 | rs1551147 | 0.805 | -1.822 | -0.073 | 0.727 | 0.006 | 0.002 | 0.002 | 0 | 0.65 | 0.35 | 0.66 | 0.69 | 0.75 | 0.75 |
| 11 | 103501082 | rs10895489 | 0.499 | -1.893 | 0.791 | 0.706 | -0.005 | -0.001 | -0.001 | 0.508 | 0.68 | 0.64 | 0.68 | 0.69 | 0.75 | 0.75 |
| 11 | 103503946 | rs2198123 | 2.082 | -1.524 | 0.372 | 0.669 | -0.019 | -0.047 | -0.047 | 2.206 | 0.47 | 0.13 | 0.51 | 0.48 | 0.54 | 0.46 |
| 11 | 103505731 | rs1917451 | -0.824 | -1.505 | 0.62 | 0.57 | 0.007 | -0.002 | -0.002 | 1.291 | 0.31 | 0.27 | 0.33 | 0.32 | 0.25 | 0.25 |
| 11 | 103507752 | rs11225937 | 0.108 | -1.683 | 0.416 | 0.373 | 0.014 | -0.023 | -0.023 | 0 | 0.15 | 0.08 | 0.19 | 0.21 | 0.14 | 0.08 |
| 11 | 103509801 | rs4754974 | 0.404 | -1.716 | 0.668 | 0.385 | -0.004 | 0 | 0 | 0 | 0.78 | 0.84 | 0.75 | 0.74 | 0.85 | 0.84 |
| 11 | 103512209 | rs11225943 | -0.9 | -1.729 | 0.882 | 0.361 | 0.004 | -0.052 | -0.052 | 0 | 0.24 | 0.42 | 0.18 | 0.12 | 0.23 | 0.16 |
| 11 | 103523452 | rs17384302 | 0.251 | -1.673 | 0.645 | 0.424 | 0.011 | -0.021 | -0.021 | 0 | 0.14 | 0.06 | 0.17 | 0.2 | 0.11 | 0.09 |
| 11 | 103529871 | rs7126823 | 2.068 | -1.793 | 0.674 | 0.36 | 0.038 | -0.021 | -0.021 | 0 | 0.47 | 0.18 | 0.52 | 0.52 | 0.59 | 0.47 |
| 11 | 103530554 | rs2408750 | 0.137 | -1.861 | 0.884 | 0.389 | 0.001 | -0.002 | -0.002 | 0 | 0.78 | 0.81 | 0.75 | 0.75 | 0.85 | 0.84 |

| | | | | | | | | | | | | | | | | |
|---|---|---|---|---|---|---|---|---|---|---|---|---|---|---|---|---|
| 11 | 103538849 | rs12366049 | 0.736 | -1.851 | 0.919 | 0.376 | -0.054 | 0.039 | 0.039 | 0 | 0.73 | 0.58 | 0.7 | 0.64 | 0.82 | 0.64 |
| 11 | 103539958 | rs17101397 | 0.708 | -1.858 | 0.925 | 0.356 | 0.002 | 0.001 | 0.001 | 0 | 0.1 | 0.08 | 0.06 | 0.09 | 0.11 | 0.07 |
| 11 | 103540383 | rs7109798 | 0.738 | -1.825 | 0.97 | 0.382 | -0.054 | 0.04 | 0.04 | 0 | 0.73 | 0.58 | 0.7 | 0.64 | 0.82 | 0.64 |
| 11 | 103541823 | rs7114669 | -1.723 | -1.563 | 0.789 | 0.319 | 0.001 | -0.012 | -0.012 | 0 | 0.34 | 0.61 | 0.26 | 0.28 | 0.33 | 0.38 |
| 11 | 103542418 | rs10895506 | -1.432 | -1.546 | 0.814 | 0.298 | 0.006 | -0.037 | -0.037 | 0.112 | 0.25 | 0.61 | 0.2 | 0.18 | 0.22 | 0.31 |
| 11 | 103542664 | rs2198114 | 0.428 | -1.698 | 0.718 | 0.335 | 0.034 | -0.03 | -0.03 | 0 | 0.69 | 0.87 | 0.69 | 0.66 | 0.74 | 0.76 |
| 11 | 103544314 | rs7123575 | 1.061 | -1.881 | 1.065 | 0.306 | -0.011 | 0.003 | 0.003 | 0 | 0.63 | 0.5 | 0.72 | 0.64 | 0.69 | 0.75 |
| 11 | 103545761 | rs4754977 | 1.037 | -1.884 | 1.078 | 0.305 | -0.014 | 0.005 | 0.005 | 0 | 0.63 | 0.51 | 0.72 | 0.64 | 0.69 | 0.75 |
| 11 | 103552317 | rs11225967 | -1.703 | -1.856 | 1.137 | 0.246 | 0.03 | 0.029 | 0.029 | 0.276 | 0.13 | 0.51 | 0.11 | 0.15 | 0.1 | 0.2 |
| 11 | 103553195 | rs10895512 | -0.27 | -1.997 | 0.85 | 0.108 | -0.011 | 0.042 | 0.042 | 0 | 0.14 | 0.09 | 0.1 | 0.03 | 0.12 | 0.09 |
| 11 | 103565031 | rs2128746 | -0.761 | -1.997 | 1.075 | 0.134 | 0.006 | -0.062 | -0.062 | 0.019 | 0.18 | 0.48 | 0.15 | 0.13 | 0.14 | 0.28 |
| 11 | 103566396 | rs7924591 | -2.103 | -2.143 | 0.933 | 0.148 | 0.074 | 0 | 0 | 0.564 | 0.51 | 0.78 | 0.37 | 0.45 | 0.4 | 0.52 |
| 11 | 103572752 | rs10750671 | 0.708 | -1.889 | 1.107 | 0.126 | -0.01 | -0.055 | -0.055 | 0 | 0.8 | 0.41 | 0.85 | 0.87 | 0.86 | 0.72 |
| 11 | 103582328 | rs7938985 | 0.442 | -1.913 | 0.918 | 0.252 | -0.02 | -0.003 | -0.003 | 0.338 | 0.14 | 0.21 | 0.09 | 0.12 | 0.11 | 0.17 |
| 11 | 103607168 | rs12287677 | -0.575 | -2.066 | 1.125 | 0.283 | -0.01 | -0.032 | -0.032 | 0 | 0.17 | 0.23 | 0.12 | 0.07 | 0.13 | 0.13 |
| 11 | 103619505 | rs986404 | 2.034 | -2.145 | 0.549 | 0.481 | -0.051 | -0.061 | -0.061 | 0 | 0.44 | 0.12 | 0.58 | 0.48 | 0.52 | 0.49 |
| 11 | 103620013 | rs12274256 | 0.57 | -2.007 | 0.589 | 0.184 | -0.022 | 0.009 | 0.009 | 0 | 0.14 | 0.21 | 0.09 | 0.2 | 0.11 | 0.17 |
| 11 | 103621096 | rs1842949 | 1.305 | -1.916 | -0.125 | 0.21 | 0.038 | 0.029 | 0.029 | 0 | 0.58 | 0.3 | 0.69 | 0.66 | 0.65 | 0.64 |
| 11 | 103626209 | rs10895522 | -1.577 | -1.884 | -0.13 | 0.226 | 0.012 | 0.009 | 0.009 | 0.016 | 0.4 | 0.6 | 0.29 | 0.34 | 0.34 | 0.35 |
| 11 | 103635415 | rs10791642 | -1.178 | -1.445 | 0.048 | 0.472 | -0.008 | 0.004 | 0.004 | 0 | 0.29 | 0.55 | 0.22 | 0.34 | 0.29 | 0.26 |
| 11 | 103635789 | rs7932574 | -0.573 | -0.704 | -0.571 | 0.298 | 0.001 | -0.001 | -0.001 | 0 | 0.92 | 0.88 | 0.93 | 0.93 | 0.89 | 0.95 |
| 11 | 103640401 | rs11226008 | 0.079 | -0.575 | -0.777 | 0.273 | -0.045 | 0.046 | 0.046 | 0 | 0.11 | 0.01 | 0.07 | 0.03 | 0.11 | 0.1 |
| 11 | 103646897 | rs4754985 | 0.667 | -1.623 | -1.389 | 0.309 | -0.093 | -0.011 | -0.011 | 0 | 0.33 | 0.12 | 0.54 | 0.33 | 0.48 | 0.29 |
| 11 | 103651279 | rs11226010 | 0.749 | -1.559 | -1.486 | 0.313 | -0.04 | -0.001 | -0.001 | 0 | 0.07 | 0.01 | 0.08 | 0.03 | 0.09 | 0.07 |
| 11 | 103653871 | rs12576536 | 0.658 | -2.066 | -0.859 | 0.715 | 0.001 | 0.084 | 0.084 | 0 | 0.1 | 0.17 | 0.08 | 0.23 | 0.11 | 0.07 |
| 11 | 103660567 | rs974819 | 0.061 | -3.139 | -0.239 | 0.215 | -0.02 | -0.002 | -0.002 | 0 | 0.58 | 0.47 | 0.78 | 0.4 | 0.72 | 0.67 |
| 11 | 103668610 | rs1531837 | -1.208 | -3.564 | -0.194 | 0.043 | 0.018 | -0.02 | -0.02 | 0 | 0.82 | 0.86 | 0.89 | 0.66 | 0.88 | 0.84 |
| 11 | 103674339 | rs10791643 | -0.227 | -3.606 | 0.001 | 0.019 | 0.047 | -0.005 | -0.005 | 0 | 0.69 | 0.78 | 0.87 | 0.63 | 0.83 | 0.74 |
| 11 | 103692429 | rs11600255 | 1.657 | -3.78 | 0.342 | -0.147 | -0.016 | 0.031 | 0.031 | 0.084 | 0.4 | 0.21 | 0.6 | 0.33 | 0.54 | 0.55 |
| 11 | 103693394 | rs12362869 | 1.332 | -3.782 | 0.484 | -0.155 | -0.003 | -0.007 | -0.007 | 0 | 0.43 | 0.44 | 0.63 | 0.37 | 0.55 | 0.56 |
| 11 | 103693627 | rs11226029 | 0.4 | -3.845 | 0.494 | -0.182 | -0.019 | -0.002 | -0.002 | 0 | 0.57 | 0.53 | 0.75 | 0.4 | 0.72 | 0.66 |
| 11 | 103697417 | rs7104359 | 0.264 | -3.553 | 0.323 | -0.211 | -0.048 | 0.039 | 0.039 | 0 | 0.12 | 0.04 | 0.12 | 0.03 | 0.17 | 0.09 |
| 11 | 103703939 | rs11226034 | 0.052 | -3.636 | 0.375 | -0.182 | -0.012 | 0.005 | 0.005 | 0 | 0.23 | 0.23 | 0.13 | 0.26 | 0.15 | 0.18 |
| 11 | 103706509 | rs7942176 | -1.099 | -3.616 | 0.419 | -0.171 | -0.004 | -0.002 | -0.002 | 0 | 0.39 | 0.41 | 0.27 | 0.3 | 0.33 | 0.27 |
| 11 | 103721674 | rs12295427 | 0.146 | -3.384 | 0.286 | -0.231 | 0.007 | -0.029 | -0.029 | 0 | 0.07 | 0.11 | 0.06 | 0 | 0.09 | 0.03 |
| 11 | 103723174 | rs11226039 | 1.495 | -3.634 | 0.198 | -0.193 | -0.001 | 0.023 | 0.023 | 0.033 | 0.4 | 0.2 | 0.61 | 0.33 | 0.55 | 0.48 |
| 11 | 103724824 | rs10791644 | -1.031 | -3.812 | -0.24 | -0.183 | 0.022 | 0.019 | 0.019 | 0 | 0.39 | 0.27 | 0.25 | 0.27 | 0.32 | 0.27 |
| 11 | 103734645 | rs11226046 | -0.192 | -3.835 | -0.254 | 0.011 | 0 | 0 | 0 | 0.113 | 0.17 | 0.16 | 0.14 | 0.18 | 0.2 | 0.1 |
| 11 | 103736408 | rs11226048 | 0.718 | -3.572 | -0.309 | 0.051 | -0.001 | -0.005 | -0.005 | 0 | 0.12 | 0.1 | 0.09 | 0.19 | 0.13 | 0.07 |
| 11 | 103737389 | rs958715 | -0.78 | -3.551 | -0.318 | 0.032 | 0.022 | 0.022 | 0.022 | 0 | 0.39 | 0.28 | 0.25 | 0.28 | 0.32 | 0.28 |
| 11 | 103737457 | rs721346 | 1.028 | -3.556 | -0.517 | 0.118 | 0.044 | -0.058 | -0.058 | 0 | 0.18 | 0.06 | 0.1 | 0.36 | 0.12 | 0.25 |
| 11 | 103739909 | rs1400549 | 1.608 | -2.942 | -0.687 | -0.202 | -0.183 | -0.04 | -0.04 | 2.795 | 0.31 | 0.03 | 0.56 | 0.29 | 0.46 | 0.25 |
| 11 | 103755233 | rs512138 | -0.679 | -0.862 | -1.348 | -0.447 | -0.02 | -0.028 | -0.028 | 1.761 | 0.81 | 0.89 | 0.57 | 0.67 | 0.75 | 0.66 |
| 11 | 103758726 | rs11226053 | 0.981 | -0.945 | -2.158 | -0.631 | -0.067 | 0.2 | 0.2 | 0.296 | 0.34 | 0.07 | 0.23 | 0.03 | 0.36 | 0.22 |
| 11 | 103761378 | rs17405661 | 1.168 | -0.389 | -1.783 | 0.036 | -0.019 | -0.009 | -0.009 | 0.201 | 0.09 | 0.02 | 0.07 | 0.08 | 0.09 | 0.1 |
| 11 | 103761546 | rs17405682 | 1.168 | -0.389 | -1.783 | 0.018 | -0.019 | -0.006 | -0.006 | 0.891 | 0.09 | 0.02 | 0.07 | 0.08 | 0.09 | 0.1 |
| 11 | 103762817 | rs528027 | -0.601 | -0.321 | -1.459 | 0.247 | -0.03 | -0.043 | -0.043 | 0 | 0.83 | 0.92 | 0.58 | 0.68 | 0.76 | 0.69 |
| 11 | 103763585 | rs561801 | 0.146 | -0.24 | -1.379 | -0.04 | 0.032 | 0.087 | 0.087 | 0 | 0.21 | 0.07 | 0.15 | 0.06 | 0.16 | 0.18 |
| 11 | 103766744 | rs538524 | -0.857 | 0.121 | -1.391 | 0.061 | -0.018 | 0.016 | 0.016 | 0.491 | 0.67 | 0.75 | 0.48 | 0.72 | 0.62 | 0.8 |
| 11 | 103766971 | rs1032235 | 0.018 | 0 | -1.221 | 0.417 | 0.035 | 0.024 | 0.024 | 0 | 0.22 | 0.04 | 0.12 | 0.16 | 0.17 | 0.21 |
| 11 | 103767715 | rs720171 | -0.633 | 0.132 | -1.058 | 0.408 | 0.011 | 0.041 | 0.041 | 0 | 0.28 | 0.15 | 0.17 | 0.2 | 0.24 | 0.39 |
| 11 | 103770921 | rs7127296 | -0.171 | 0.501 | -0.449 | 0.555 | 0.001 | 0 | 0 | 0 | 0.47 | 0.37 | 0.6 | 0.57 | 0.4 | 0.52 |
| 11 | 103772733 | rs532445 | -1.534 | 0.815 | 0.379 | 1.086 | -0.009 | 0.063 | 0.063 | 0.573 | 0.47 | 0.75 | 0.39 | 0.69 | 0.48 | 0.52 |
| 11 | 103773167 | rs4754095 | -0.007 | 1.207 | 0.931 | 1.138 | 0.016 | 0.061 | 0.061 | 0 | 0.2 | 0.28 | 0.18 | 0.38 | 0.23 | 0.22 |
| 11 | 103775113 | rs1487911 | -0.597 | 1.032 | 0.919 | 1.096 | -0.002 | -0.006 | -0.006 | 0.272 | 0.32 | 0.28 | 0.25 | 0.45 | 0.33 | 0.37 |
| 11 | 103778154 | rs1053861 | -0.974 | 1.067 | 1.295 | 1.112 | 0.004 | 0.023 | 0.023 | 0.048 | 0.36 | 0.4 | 0.28 | 0.46 | 0.36 | 0.4 |
| 11 | 103779025 | rs11226057 | -1.562 | 1.087 | 1.416 | 1.103 | -0.002 | -0.003 | -0.003 | 0.184 | 0.43 | 0.56 | 0.35 | 0.46 | 0.45 | 0.49 |
| 11 | 103789461 | rs1872901 | -0.558 | 1.129 | 1.158 | 1.143 | -0.001 | -0.003 | -0.003 | 0 | 0.32 | 0.28 | 0.25 | 0.45 | 0.33 | 0.38 |
| 11 | 103792540 | rs541150 | -1.563 | 0.981 | 0.878 | 0.836 | -0.015 | 0.061 | 0.061 | 1.026 | 0.45 | 0.74 | 0.4 | 0.69 | 0.48 | 0.47 |
| 11 | 103793034 | rs545643 | -1.327 | 0.824 | 0.898 | 0.878 | 0.001 | -0.005 | -0.005 | 0 | 0.4 | 0.61 | 0.32 | 0.68 | 0.38 | 0.43 |
| 11 | 103797349 | rs1994450 | 0.421 | 1.03 | 0.275 | 0.766 | -0.053 | 0.013 | 0.013 | 0.166 | 0.14 | 0.01 | 0.09 | 0.07 | 0.14 | 0.18 |
| 11 | 103813331 | rs11226082 | -0.101 | 0.7 | 0.232 | 1.08 | -0.068 | -0.089 | -0.089 | 0 | 0.31 | 0.1 | 0.24 | 0.4 | 0.35 | 0.38 |
| 11 | 103813858 | rs4384362 | -0.824 | 0.796 | 0.284 | 1.073 | 0.013 | 0.1 | 0.1 | 0.887 | 0.13 | 0.57 | 0.12 | 0.23 | 0.1 | 0.08 |
| 11 | 103817645 | rs4145958 | -0.228 | 0.802 | -0.357 | 1.071 | -0.027 | -0.042 | -0.042 | 0 | 0.33 | 0.21 | 0.26 | 0.4 | 0.35 | 0.38 |
| 11 | 103819333 | rs1471936 | -0.934 | 0.962 | -1.001 | 0.53 | -0.052 | 0.081 | 0.081 | 1.146 | 0.48 | 0.89 | 0.39 | 0.62 | 0.46 | 0.47 |
| 11 | 103820517 | rs10895548 | -0.701 | 0.828 | -1.276 | 0.538 | -0.066 | 0.035 | 0.035 | 1.353 | 0.46 | 0.92 | 0.38 | 0.62 | 0.43 | 0.51 |
| 11 | 103835871 | rs901782 | -0.583 | 0.305 | -0.688 | 0.643 | 0.005 | 0.002 | 0.002 | 1.082 | 0.68 | 0.76 | 0.76 | 0.73 | 0.68 | 0.81 |
| 11 | 103836697 | rs2220377 | -0.333 | -0.352 | -0.527 | 0.896 | 0.001 | 0.006 | 0.006 | 0.38 | 0.48 | 0.65 | 0.61 | 0.57 | 0.47 | 0.65 |
| 11 | 103836968 | rs6591063 | 0.754 | -0.482 | -0.743 | 0.806 | -0.006 | -0.004 | -0.004 | 0 | 0.18 | 0.2 | 0.07 | 0.12 | 0.12 | 0.18 |

| | | | | | | | | | | | | | | | | |
|---|---|---|---|---|---|---|---|---|---|---|---|---|---|---|---|---|
| 11 | 103838408 | rs10502023 | 1.176 | -0.385 | -0.708 | 0.753 | -0.053 | -0.039 | -0.039 | 0 | 0.15 | 0.07 | 0.49 | 0.3 | 0.3 | 0.38 |
| 11 | 103839559 | rs12282189 | 0.397 | -0.556 | -0.745 | 0.771 | 0.013 | 0.008 | 0.008 | 0 | 0.25 | 0.2 | 0.08 | 0.18 | 0.16 | 0.25 |
| 11 | 103844333 | rs2047698 | 0.126 | -0.408 | -0.669 | 0.757 | 0.027 | 0.013 | 0.013 | 0 | 0.29 | 0.2 | 0.09 | 0.23 | 0.16 | 0.25 |
| 11 | 103849507 | rs10791658 | 0.332 | -0.446 | -0.691 | 0.663 | 0.018 | 0.022 | 0.022 | 0 | 0.27 | 0.2 | 0.09 | 0.12 | 0.16 | 0.24 |
| 11 | 103852539 | rs260836 | -0.796 | -0.648 | -0.85 | 0.49 | -0.001 | 0.003 | 0.003 | 0.053 | 0.56 | 0.55 | 0.66 | 0.52 | 0.54 | 0.66 |
| 11 | 103856708 | rs10895555 | 0.297 | -0.366 | -0.842 | 0.865 | 0.003 | -0.006 | -0.006 | 0 | 0.23 | 0.18 | 0.14 | 0.36 | 0.2 | 0.22 |
| 11 | 103858121 | rs1872902 | 0.437 | 0.453 | -0.82 | 1.104 | -0.03 | -0.031 | -0.031 | 0.298 | 0.17 | 0.12 | 0.49 | 0.46 | 0.32 | 0.37 |
| 11 | 103859645 | rs651979 | -1.092 | -0.107 | -0.576 | 0.321 | -0.003 | 0.009 | 0.009 | 0.699 | 0.71 | 0.77 | 0.76 | 0.87 | 0.67 | 0.84 |
| 11 | 103862934 | rs11226106 | 0.655 | -0.97 | -0.92 | -0.048 | 0.036 | 0.068 | 0.068 | 0 | 0.23 | 0.07 | 0.12 | 0.08 | 0.16 | 0.16 |
| 11 | 103863369 | rs10895558 | -0.595 | -0.534 | -0.335 | 0.072 | -0.002 | -0.001 | -0.001 | 0.782 | 0.47 | 0.49 | 0.73 | 0.6 | 0.6 | 0.61 |
| 11 | 103864510 | rs496339 | 1.01 | -0.829 | -0.132 | -0.246 | 0.004 | -0.035 | -0.035 | 0 | 0.12 | 0.15 | 0.04 | 0.01 | 0.09 | 0.07 |
| 11 | 103864743 | rs12574463 | 1.247 | -0.989 | 0.191 | -0.188 | 0.01 | 0.011 | 0.011 | 0 | 0.24 | 0.21 | 0.13 | 0.08 | 0.16 | 0.17 |
| 11 | 103864935 | rs629720 | -0.999 | -0.979 | 0.06 | -0.176 | 0.021 | -0.004 | -0.004 | 0.433 | 0.58 | 0.64 | 0.77 | 0.62 | 0.69 | 0.68 |
| 11 | 103870361 | rs732451 | 0.909 | -0.614 | 0.117 | 0.098 | 0.035 | 0.067 | 0.067 | 0 | 0.23 | 0.07 | 0.12 | 0.08 | 0.16 | 0.16 |
| 11 | 103871404 | rs12146566 | 0.488 | -0.569 | 0.415 | 0.396 | -0.028 | 0.018 | 0.018 | 0 | 0.27 | 0.4 | 0.16 | 0.31 | 0.21 | 0.28 |
| 11 | 103874510 | rs260862 | -0.61 | -0.182 | 0.306 | -0.109 | 0.038 | -0.054 | -0.054 | 0 | 0.18 | 0.29 | 0.11 | 0.05 | 0.2 | 0.13 |
| 11 | 103876114 | rs260864 | -0.691 | -0.219 | 0.139 | -0.18 | 0 | -0.004 | -0.004 | 2.485 | 0.36 | 0.36 | 0.67 | 0.33 | 0.56 | 0.47 |
| 11 | 103876331 | rs1386751 | 1.209 | -0.214 | -0.019 | -0.183 | -0.056 | -0.024 | -0.024 | 0 | 0.17 | 0.05 | 0.55 | 0.28 | 0.36 | 0.34 |
| 11 | 103877010 | rs260865 | -1.176 | -0.685 | -0.048 | -0.472 | 0.04 | 0.005 | 0.005 | 0 | 0.6 | 0.67 | 0.8 | 0.68 | 0.72 | 0.72 |
| 11 | 103877311 | rs12363710 | 1.403 | -0.541 | -0.177 | -0.764 | 0.035 | 0.057 | 0.057 | 0 | 0.22 | 0.08 | 0.11 | 0.08 | 0.15 | 0.21 |
| 11 | 103891852 | rs17421481 | -0.072 | -0.479 | -0.002 | -0.63 | 0.015 | -0.019 | -0.019 | 0.045 | 0.08 | 0.16 | 0.08 | 0.04 | 0.09 | 0.05 |
| 11 | 103892231 | rs260852 | -1.641 | -0.672 | -0.11 | -0.742 | 0.016 | 0.02 | 0.02 | 0.009 | 0.79 | 0.84 | 0.88 | 0.97 | 0.84 | 0.9 |
| 11 | 103899109 | rs260858 | -1.239 | 0.141 | 0.071 | -0.296 | 0 | -0.001 | -0.001 | 0.013 | 0.62 | 0.64 | 0.8 | 0.67 | 0.72 | 0.73 |
| 11 | 103907956 | rs7102675 | 0.048 | 0.887 | -0.435 | -0.775 | -0.006 | -0.003 | -0.003 | 0 | 0.14 | 0.13 | 0.04 | 0.01 | 0.12 | 0.07 |
| 11 | 103908735 | rs1052313 | -0.594 | 1.084 | -0.649 | -0.725 | -0.097 | -0.049 | -0.049 | 0.053 | 0.3 | 0.11 | 0.58 | 0.4 | 0.48 | 0.43 |
| 11 | 103909166 | rs260808 | -0.22 | 1.096 | -0.442 | -0.752 | 0.05 | -0.098 | -0.098 | 0 | 0.16 | 0.38 | 0.07 | 0.02 | 0.13 | 0.09 |
| 11 | 103909897 | rs4754098 | -0.298 | 0.344 | -0.235 | -0.552 | -0.001 | -0.001 | -0.001 | 0.269 | 0.33 | 0.32 | 0.61 | 0.44 | 0.51 | 0.47 |
| 11 | 103910204 | rs1566580 | -0.309 | 0.344 | -0.294 | -0.552 | 0.008 | 0.006 | 0.006 | 0 | 0.33 | 0.35 | 0.61 | 0.44 | 0.51 | 0.47 |
| 11 | 103911995 | rs17422079 | -0.054 | 0.251 | -0.312 | -0.633 | 0.002 | -0.004 | -0.004 | 0 | 0.08 | 0.08 | 0.07 | 0.04 | 0.09 | 0.05 |
| 11 | 103912968 | rs11226125 | -0.244 | 0.524 | -0.275 | -0.37 | 0.008 | 0.002 | 0.002 | 0 | 0.32 | 0.32 | 0.6 | 0.4 | 0.48 | 0.44 |
| 11 | 103913376 | rs1818106 | -0.304 | 0.471 | -0.535 | -0.358 | -0.026 | -0.02 | -0.02 | 0.418 | 0.32 | 0.22 | 0.6 | 0.44 | 0.51 | 0.46 |
| 11 | 103918740 | rs260815 | -0.841 | 0.218 | -0.592 | -0.316 | -0.004 | 0 | 0 | 0 | 0.56 | 0.52 | 0.75 | 0.52 | 0.67 | 0.68 |
| 11 | 103919222 | rs9326345 | -0.657 | 1.306 | -1.03 | 0.021 | -0.051 | -0.02 | -0.02 | 0 | 0.29 | 0.18 | 0.47 | 0.4 | 0.43 | 0.38 |
| 11 | 103920105 | rs260818 | 0.161 | 0.518 | -1.029 | -0.191 | 0.029 | 0.015 | 0.015 | 0.543 | 0.55 | 0.77 | 0.37 | 0.6 | 0.4 | 0.54 |
| 11 | 103924576 | rs361315 | 0.57 | 1.041 | -0.704 | -0.105 | 0.01 | 0.004 | 0.004 | 0 | 0.51 | 0.45 | 0.34 | 0.54 | 0.34 | 0.46 |
| 11 | 103936469 | rs361262 | 0.475 | 1.156 | -0.756 | -0.055 | -0.006 | -0.002 | -0.002 | 0.002 | 0.52 | 0.52 | 0.35 | 0.54 | 0.35 | 0.46 |
| 11 | 103937879 | rs361266 | 0.368 | 1.665 | -1.668 | -0.009 | 0.051 | 0.005 | 0.005 | 0 | 0.54 | 0.78 | 0.37 | 0.54 | 0.35 | 0.46 |
| 11 | 103944425 | rs10502025 | 0.251 | 1.684 | -2.021 | 0.076 | -0.022 | 0.072 | 0.072 | 0 | 0.26 | 0.09 | 0.24 | 0.1 | 0.3 | 0.28 |
| 11 | 103946161 | rs615290 | -0.135 | 1.847 | -1.266 | 0.916 | -0.074 | -0.082 | -0.082 | 0.803 | 0.84 | 0.96 | 0.64 | 0.66 | 0.71 | 0.77 |
| 11 | 103959760 | rs17101942 | -0.721 | 0.889 | -0.659 | 0.48 | -0.007 | -0.011 | -0.011 | 0 | 0.11 | 0.22 | 0.06 | 0.09 | 0.09 | 0.04 |
| 11 | 103959987 | rs361281 | 0.11 | 0.724 | -2.072 | -0.142 | -0.165 | 0.042 | 0.042 | 2.335 | 0.37 | 0.03 | 0.56 | 0.25 | 0.47 | 0.3 |
| 11 | 103960620 | rs361283 | 0.242 | 0.037 | -1.234 | -0.244 | 0.003 | 0.044 | 0.044 | 0.01 | 0.58 | 0.32 | 0.76 | 0.78 | 0.58 | 0.51 |
| 11 | 103962229 | rs17423437 | 0.914 | -0.146 | -1.057 | -0.689 | -0.003 | -0.007 | -0.007 | 0 | 0.13 | 0.1 | 0.05 | 0.01 | 0.14 | 0.23 |
| 11 | 103964427 | rs590216 | 0.482 | -0.39 | -0.424 | -0.77 | 0.012 | -0.116 | -0.116 | 0.019 | 0.71 | 0.65 | 0.83 | 0.97 | 0.7 | 0.67 |
| 11 | 103965133 | rs603781 | 0.99 | -0.318 | -0.565 | -0.249 | 0.008 | 0.055 | 0.055 | 0.048 | 0.56 | 0.25 | 0.75 | 0.7 | 0.58 | 0.5 |
| 11 | 103965819 | rs7947946 | 0.194 | -0.236 | -0.205 | -0.137 | -0.012 | -0.037 | -0.037 | 0 | 0.09 | 0.1 | 0.04 | 0 | 0.05 | 0.03 |
| 11 | 103976293 | rs488753 | -0.267 | -0.01 | -0.068 | -0.369 | 0.005 | 0.003 | 0.003 | 0.006 | 0.48 | 0.49 | 0.71 | 0.81 | 0.59 | 0.76 |
| 11 | 103978210 | rs361291 | -0.303 | 0.032 | 0.349 | 0.397 | -0.005 | -0.005 | -0.005 | 1.637 | 0.57 | 0.65 | 0.45 | 0.48 | 0.5 | 0.34 |
| 11 | 103978540 | rs11226159 | 0.862 | -0.285 | 0.425 | 0.384 | 0.055 | 0.114 | 0.114 | 1.565 | 0.39 | 0.12 | 0.25 | 0.16 | 0.32 | 0.19 |
| 11 | 103980341 | rs11226161 | 0.051 | -0.26 | 0.629 | 0.464 | 0.009 | 0.102 | 0.102 | 0 | 0.07 | 0.14 | 0.11 | 0.32 | 0.09 | 0.12 |
| 11 | 103981833 | rs168636 | -0.296 | -0.31 | 0.694 | 0.37 | 0.009 | 0 | 0 | 0.016 | 0.53 | 0.63 | 0.39 | 0.48 | 0.44 | 0.33 |
| 11 | 103991641 | rs11226185 | -0.684 | -0.769 | 0.769 | 0.719 | -0.001 | 0.063 | 0.063 | 0.003 | 0.13 | 0.17 | 0.12 | 0.3 | 0.12 | 0.12 |
| 11 | 103996971 | rs12279750 | 0.747 | -0.804 | 0.859 | 0.689 | -0.019 | -0.011 | -0.011 | 0 | 0.07 | 0.09 | 0.01 | 0.06 | 0.03 | 0.05 |
| 11 | 104009965 | rs1532673 | -0.702 | -0.217 | 0.84 | 0.753 | 0 | 0.002 | 0.002 | 0 | 0.34 | 0.4 | 0.38 | 0.59 | 0.38 | 0.38 |
| 11 | 104010052 | rs361268 | -0.723 | -0.149 | 0.365 | 0.496 | -0.035 | 0.048 | 0.048 | 0 | 0.66 | 0.8 | 0.57 | 0.76 | 0.59 | 0.51 |
| 11 | 104015677 | rs10895586 | 0.963 | -0.189 | 0.326 | 0.737 | 0.001 | -0.006 | -0.006 | 0 | 0.2 | 0.19 | 0.19 | 0.27 | 0.22 | 0.21 |
| 11 | 104020049 | rs10791670 | 0.079 | 0.022 | 0.246 | 0.895 | 0.007 | 0.015 | 0.015 | 0.455 | 0.53 | 0.45 | 0.39 | 0.62 | 0.44 | 0.42 |
| 11 | 104020380 | rs10791671 | 0.093 | 0.032 | 0.25 | 0.9 | 0.007 | 0.015 | 0.015 | 0.517 | 0.53 | 0.44 | 0.39 | 0.62 | 0.44 | 0.42 |
| 11 | 104022906 | rs11226207 | 1.506 | 0.042 | -0.319 | 0.93 | 0.018 | -0.066 | -0.066 | 0 | 0.26 | 0.12 | 0.17 | 0.3 | 0.2 | 0.14 |
| 11 | 104026465 | rs2128806 | -0.442 | 0.273 | -0.304 | 0.354 | -0.028 | 0.041 | 0.041 | 0 | 0.62 | 0.76 | 0.51 | 0.73 | 0.54 | 0.49 |
| 11 | 104031450 | rs10895595 | 1.122 | 0.338 | -0.287 | 0.528 | 0.011 | 0.004 | 0.004 | 5.506 | 0.27 | 0.19 | 0.21 | 0.2 | 0.2 | 0.14 |
| 11 | 104032160 | rs7943011 | 0.461 | 0.31 | -0.019 | 0.566 | -0.002 | -0.003 | -0.003 | 0 | 0.34 | 0.38 | 0.31 | 0.31 | 0.31 | 0.21 |
| 11 | 104041862 | rs7941138 | -0.015 | 0.816 | -0.364 | 0.345 | -0.026 | 0.037 | 0.037 | 0 | 0.63 | 0.76 | 0.53 | 0.77 | 0.55 | 0.51 |
| 11 | 104042459 | rs10895598 | -0.792 | 1.524 | 0.005 | 1.156 | -0.004 | -0.007 | -0.007 | 0.044 | 0.25 | 0.26 | 0.37 | 0.52 | 0.35 | 0.37 |
| 11 | 104057830 | rs11226216 | 1.723 | 1.201 | -0.574 | 0.85 | 0.046 | -0.024 | -0.024 | 0.006 | 0.5 | 0.14 | 0.37 | 0.5 | 0.4 | 0.36 |
| 11 | 104063280 | rs11226221 | -1.561 | 1.661 | 0.161 | 1.165 | 0.012 | 0.042 | 0.042 | 0 | 0.25 | 0.32 | 0.23 | 0.38 | 0.28 | 0.32 |
| 11 | 104079135 | rs7105612 | 0.716 | 1.893 | 0.321 | 1.137 | -0.034 | 0.028 | 0.028 | 0 | 0.06 | 0.01 | 0.1 | 0.01 | 0.12 | 0.13 |
| 11 | 104098418 | rs10502030 | -0.591 | 1.871 | 0.489 | 1.124 | 0.063 | -0.039 | -0.039 | 0.029 | 0.08 | 0.21 | 0.12 | 0.01 | 0.12 | 0.14 |
| 11 | 104113576 | rs11226246 | -0.008 | 2.019 | 0.95 | 1.192 | 0.017 | -0.011 | -0.011 | 0.169 | 0.19 | 0.26 | 0.33 | 0.13 | 0.28 | 0.31 |

| | | | | | | | | | | | | | | | | |
|---|---|---|---|---|---|---|---|---|---|---|---|---|---|---|---|---|
| 11 | 104125118 | rs17102225 | 0.494 | 1.869 | 0.81 | 1.47 | -0.007 | -0.055 | -0.055 | 0 | 0.12 | 0.05 | 0.05 | 0.26 | 0.13 | 0.14 |
| 11 | 104130998 | rs7945455 | -1.027 | 1.852 | 1.245 | 1.381 | 0.083 | -0.045 | -0.045 | 0 | 0.1 | 0.34 | 0.13 | 0.01 | 0.12 | 0.14 |
| 11 | 104137181 | rs4754106 | 1.885 | 1.823 | 0.999 | 1.434 | -0.006 | -0.005 | -0.005 | 0 | 0.58 | 0.31 | 0.47 | 0.61 | 0.46 | 0.42 |
| 11 | 104158392 | rs12800154 | -1.243 | 1.877 | 1.223 | 1.563 | 0.072 | -0.087 | -0.087 | 0 | 0.28 | 0.46 | 0.47 | 0.13 | 0.4 | 0.43 |
| 11 | 104163777 | rs12223069 | 0.341 | 1.322 | 1.144 | 1.784 | 0.001 | -0.009 | -0.009 | 0.029 | 0.13 | 0.13 | 0.05 | 0.26 | 0.12 | 0.15 |
| 11 | 104170706 | rs10502032 | 0.314 | 1.146 | 0.941 | 1.645 | 0.002 | -0.024 | -0.024 | 0.286 | 0.1 | 0.07 | 0.1 | 0.26 | 0.09 | 0.11 |
| 11 | 104174699 | rs6591076 | 0.43 | 1.095 | 1.356 | 1.205 | 0.047 | -0.062 | -0.062 | 0 | 0.78 | 0.55 | 0.63 | 0.88 | 0.72 | 0.71 |
| 11 | 104194241 | rs1873098 | 0.341 | 0.989 | 1.072 | 1.605 | -0.011 | 0.053 | 0.053 | 0 | 0.85 | 0.76 | 0.94 | 0.74 | 0.88 | 0.83 |
| 11 | 104194975 | rs11226303 | -0.19 | 1.102 | 0.704 | 1.641 | 0.002 | -0.03 | -0.03 | 0.115 | 0.12 | 0.07 | 0.05 | 0.26 | 0.12 | 0.15 |
| 11 | 104200320 | rs10502033 | -1.995 | 1.317 | 1.067 | 1.557 | 0.009 | 0.006 | 0.006 | 0.124 | 0.38 | 0.44 | 0.41 | 0.35 | 0.42 | 0.57 |
| 11 | 104202949 | rs12146572 | 0.047 | 1.432 | 0.992 | 1.596 | -0.063 | 0.003 | 0.003 | 0 | 0.13 | 0.02 | 0.23 | 0.08 | 0.19 | 0.26 |
| 11 | 104211569 | rs17102346 | -0.337 | 1.333 | 1.31 | 1.574 | -0.01 | 0.019 | 0.019 | 0 | 0.17 | 0.17 | 0.08 | 0.26 | 0.14 | 0.17 |
| 11 | 104215334 | rs6591077 | -1.461 | 1.324 | 1.354 | 1.51 | -0.014 | -0.009 | -0.009 | 3.83 | 0.35 | 0.25 | 0.4 | 0.35 | 0.41 | 0.56 |
| 11 | 104217481 | rs10895656 | -1.672 | 1.325 | 1.35 | 1.478 | 0.013 | 0.01 | 0.01 | 0.042 | 0.37 | 0.45 | 0.42 | 0.36 | 0.42 | 0.58 |
| 11 | 104221722 | rs11226320 | -1.32 | 1.87 | 1.689 | 1.724 | 0.012 | 0.031 | 0.031 | 0 | 0.12 | 0.28 | 0.14 | 0.21 | 0.13 | 0.17 |
| 11 | 104238087 | rs6591079 | 0.191 | 2.11 | 1.616 | 1.722 | 0.035 | 0.015 | 0.015 | 0 | 0.88 | 0.72 | 0.8 | 0.87 | 0.85 | 0.73 |
| 11 | 104239996 | rs7107172 | -0.253 | 2.122 | 1.482 | 1.722 | -0.021 | -0.01 | -0.01 | 3.822 | 0.89 | 0.95 | 0.82 | 0.88 | 0.85 | 0.75 |
| 11 | 104249254 | rs12366084 | 0.017 | 2.131 | 1.49 | 1.734 | -0.018 | -0.011 | -0.011 | 0 | 0.1 | 0.05 | 0.18 | 0.12 | 0.14 | 0.24 |
| 11 | 104261833 | rs11226342 | 0.203 | 2.034 | 1.714 | 1.841 | 0.007 | 0.02 | 0.02 | 0 | 0.12 | 0.15 | 0.09 | 0.18 | 0.15 | 0.14 |
| 11 | 104274535 | rs4755031 | 0.991 | 2.037 | 1.797 | 1.841 | 0.049 | 0.05 | 0.05 | 0 | 0.75 | 0.46 | 0.7 | 0.69 | 0.7 | 0.61 |
| 11 | 104294214 | rs12787859 | 0.053 | 2.054 | 1.889 | 1.724 | -0.009 | -0.006 | -0.006 | 0 | 0.1 | 0.06 | 0.17 | 0.12 | 0.14 | 0.23 |
| 11 | 104298951 | rs4755036 | 1.016 | 2.045 | 1.942 | 1.725 | 0.049 | 0.049 | 0.049 | 0 | 0.75 | 0.46 | 0.7 | 0.69 | 0.7 | 0.61 |
| 11 | 104303010 | rs7951031 | -0.284 | 2.072 | 1.889 | 1.729 | 0.038 | 0.023 | 0.023 | 0 | 0.11 | 0.25 | 0.19 | 0.12 | 0.14 | 0.26 |
| 11 | 104309414 | rs10502034 | 1.815 | 1.877 | 1.409 | 1.591 | -0.029 | -0.033 | -0.033 | 0 | 0.58 | 0.19 | 0.56 | 0.52 | 0.55 | 0.4 |
| 11 | 104320214 | rs6591081 | 1.032 | 1.976 | 2.172 | 1.502 | 0.031 | 0.031 | 0.031 | 0.001 | 0.73 | 0.46 | 0.7 | 0.69 | 0.67 | 0.67 |
| 11 | 104324720 | rs1386664 | 0.326 | 1.966 | 2.357 | 1.522 | 0.031 | 0.009 | 0.009 | 0 | 0.87 | 0.83 | 0.81 | 0.85 | 0.82 | 0.82 |
| 11 | 104328244 | rs1121555 | 0.791 | 1.947 | 2.305 | 1.508 | 0.07 | 0.035 | 0.035 | 0.357 | 0.84 | 0.5 | 0.78 | 0.84 | 0.81 | 0.8 |
| 11 | 104333651 | rs1906308 | -0.038 | 1.821 | 2.032 | 1.575 | -0.002 | 0.009 | 0.009 | 0 | 0.14 | 0.23 | 0.09 | 0.15 | 0.14 | 0.18 |
| 11 | 104334328 | rs7944328 | 0.002 | 1.775 | 2.248 | 1.567 | -0.008 | 0.018 | 0.018 | 0 | 0.84 | 0.64 | 0.91 | 0.84 | 0.86 | 0.8 |
| 11 | 104338480 | rs12225597 | 0.13 | 1.775 | 2.3 | 1.568 | 0 | -0.003 | -0.003 | 0 | 0.13 | 0.14 | 0.08 | 0.16 | 0.14 | 0.2 |
| 11 | 104343566 | rs994385 | -0.169 | 1.774 | 2.304 | 1.568 | -0.006 | 0.011 | 0.011 | 0 | 0.85 | 0.73 | 0.91 | 0.84 | 0.86 | 0.8 |
| 11 | 104345561 | rs1471924 | 1.277 | 1.822 | 2.684 | 1.529 | 0.01 | 0.005 | 0.005 | 0 | 0.7 | 0.39 | 0.69 | 0.69 | 0.65 | 0.61 |
| 11 | 104354790 | rs10895694 | 0 | 1.821 | 2.707 | 1.529 | -0.016 | 0.021 | 0.021 | 0 | 0.84 | 0.6 | 0.91 | 0.84 | 0.86 | 0.8 |
| 11 | 104366089 | rs901715 | -0.262 | 1.821 | 2.684 | 1.529 | -0.01 | 0.011 | 0.011 | 0 | 0.85 | 0.73 | 0.92 | 0.84 | 0.86 | 0.8 |
| 11 | 104366549 | rs901717 | 1.278 | 1.821 | 2.699 | 1.529 | 0.01 | 0.004 | 0.004 | 0 | 0.7 | 0.39 | 0.69 | 0.69 | 0.65 | 0.61 |
| 11 | 104368709 | rs4754117 | -0.235 | 1.834 | 2.538 | 1.538 | -0.01 | 0.013 | 0.013 | 0 | 0.85 | 0.73 | 0.92 | 0.84 | 0.86 | 0.8 |
| 11 | 104375711 | rs994491 | -0.309 | 1.838 | 2.519 | 1.497 | 0.036 | 0.006 | 0.006 | 0 | 0.13 | 0.2 | 0.22 | 0.15 | 0.22 | 0.19 |
| 11 | 104376956 | rs4573652 | -0.196 | 1.72 | 2.218 | 1.477 | -0.01 | 0.011 | 0.011 | 0 | 0.85 | 0.73 | 0.92 | 0.84 | 0.86 | 0.8 |
| 11 | 104392750 | rs10791716 | -0.15 | 1.715 | 2.216 | 1.475 | -0.011 | 0.01 | 0.01 | 0 | 0.85 | 0.73 | 0.92 | 0.84 | 0.86 | 0.8 |
| 11 | 104418194 | rs2029285 | 1.027 | 1.839 | 2.437 | 1.477 | 0.005 | 0.001 | 0.001 | 0 | 0.7 | 0.38 | 0.62 | 0.7 | 0.64 | 0.66 |
| 11 | 104420059 | rs11226425 | -0.398 | 1.785 | 2.673 | 1.475 | 0.051 | 0.01 | 0.01 | 0 | 0.14 | 0.23 | 0.29 | 0.15 | 0.22 | 0.16 |
| 11 | 104440171 | rs7124377 | -1.158 | 1.76 | 2.667 | 1.354 | -0.001 | -0.014 | -0.014 | 0.05 | 0.11 | 0.2 | 0.09 | 0.09 | 0.09 | 0.16 |
| 11 | 104449474 | rs11226429 | 0.337 | 2.425 | 2.433 | 1.638 | -0.036 | -0.035 | -0.035 | 0 | 0.07 | 0.01 | 0.14 | 0.12 | 0.15 | 0.05 |
| 11 | 104452042 | rs11226430 | -0.3 | 2.324 | 2.6 | 1.601 | 0.055 | 0.017 | 0.017 | 0 | 0.14 | 0.28 | 0.3 | 0.15 | 0.22 | 0.18 |
| 11 | 104456747 | rs658468 | 1.823 | 2.304 | 2.233 | 1.609 | -0.025 | 0.008 | 0.008 | 0.033 | 0.58 | 0.18 | 0.53 | 0.6 | 0.55 | 0.47 |
| 11 | 104490876 | rs10791720 | 0.078 | 2.635 | 2.725 | 1.755 | -0.013 | -0.007 | -0.007 | 0.151 | 0.11 | 0.07 | 0.26 | 0.13 | 0.2 | 0.15 |
| 11 | 104497607 | rs1942510 | -2.204 | 2.653 | 2.376 | 1.902 | 0.023 | 0.021 | 0.021 | 0 | 0.17 | 0.32 | 0.21 | 0.28 | 0.17 | 0.24 |
| 11 | 104504416 | rs16894 | -1.103 | 2.741 | 2.417 | 1.934 | 0.065 | 0.01 | 0.01 | 0.021 | 0.17 | 0.52 | 0.31 | 0.15 | 0.23 | 0.18 |
| 11 | 104537299 | rs12799711 | -1.977 | 2.049 | 2.064 | 1.671 | -0.005 | -0.006 | -0.006 | 0 | 0.25 | 0.29 | 0.24 | 0.29 | 0.21 | 0.25 |
| 11 | 104539394 | rs6591092 | -2.076 | 2.253 | 2.108 | 1.564 | 0.008 | 0.004 | 0.004 | 0 | 0.3 | 0.35 | 0.35 | 0.35 | 0.31 | 0.29 |
| 11 | 104551887 | rs10895718 | -2.084 | 2.266 | 2.113 | 1.57 | 0.008 | 0.004 | 0.004 | 0 | 0.3 | 0.34 | 0.35 | 0.37 | 0.31 | 0.29 |
| 11 | 104571862 | rs7942166 | -2.184 | 2.241 | 2.484 | 1.514 | 0.022 | 0.011 | 0.011 | 0 | 0.31 | 0.43 | 0.35 | 0.35 | 0.31 | 0.29 |
| 11 | 104588855 | rs17102906 | -1.195 | 2.148 | 2.482 | 1.485 | 0.005 | 0.004 | 0.004 | 0 | 0.17 | 0.19 | 0.06 | 0.09 | 0.09 | 0.16 |
| 11 | 104591614 | rs1147018 | 2.458 | 2.26 | 2.651 | 1.529 | 0.022 | -0.033 | -0.033 | 0 | 0.55 | 0.23 | 0.38 | 0.49 | 0.47 | 0.52 |
| 11 | 104612633 | rs2845891 | 2.013 | 2.151 | 3.098 | 1.445 | 0.004 | -0.001 | -0.001 | 0.025 | 0.67 | 0.38 | 0.67 | 0.65 | 0.68 | 0.69 |
| 11 | 104615426 | rs12281709 | -1.933 | 2.215 | 3.08 | 1.467 | 0.091 | 0.084 | 0.084 | 0 | 0.19 | 0.39 | 0.27 | 0.34 | 0.26 | 0.18 |
| 11 | 104618572 | rs2052752 | -1.836 | 2.238 | 2.981 | 1.475 | 0.067 | 0.059 | 0.059 | 0 | 0.18 | 0.34 | 0.27 | 0.34 | 0.26 | 0.18 |
| 11 | 104620211 | rs1834631 | -1.539 | 2.237 | 2.984 | 1.475 | 0.036 | 0.032 | 0.032 | 0 | 0.17 | 0.22 | 0.26 | 0.34 | 0.25 | 0.18 |
| 11 | 104659942 | rs12285132 | -0.971 | 2.27 | 2.996 | 1.484 | 0.044 | 0.035 | 0.035 | 0 | 0.13 | 0.17 | 0.24 | 0.26 | 0.22 | 0.14 |
| 11 | 104664732 | rs1147052 | 0.63 | 2.103 | 2.961 | 1.459 | -0.002 | -0.003 | -0.003 | 0 | 0.84 | 0.81 | 0.94 | 0.91 | 0.91 | 0.85 |
| 11 | 104674935 | rs10895730 | -0.631 | 2.484 | 3.242 | 1.649 | 0.065 | 0.021 | 0.021 | 0 | 0.14 | 0.31 | 0.33 | 0.16 | 0.21 | 0.16 |
| 11 | 104688549 | rs10791730 | -1.274 | 2.657 | 3.399 | 1.655 | 0.098 | 0.116 | 0.116 | 0 | 0.14 | 0.35 | 0.26 | 0.26 | 0.2 | 0.14 |
| 11 | 104693950 | rs7106409 | -0.634 | 2.684 | 3.224 | 1.667 | 0.042 | 0.007 | 0.007 | 0 | 0.14 | 0.24 | 0.33 | 0.16 | 0.23 | 0.17 |
| 11 | 104698826 | rs10895734 | -0.972 | 2.453 | 3.185 | 1.657 | 0.124 | 0.148 | 0.148 | 0 | 0.13 | 0.38 | 0.25 | 0.26 | 0.2 | 0.14 |
| 11 | 104707732 | rs642603 | 1.177 | 2.249 | 2.973 | 1.431 | -0.01 | -0.062 | -0.062 | 0 | 0.8 | 0.55 | 0.83 | 0.89 | 0.86 | 0.8 |
| 11 | 104730322 | rs6591101 | -0.308 | 2.324 | 2.922 | 1.52 | 0.012 | 0.036 | 0.036 | 0 | 0.12 | 0.17 | 0.17 | 0.23 | 0.15 | 0.13 |
| 11 | 104737590 | rs477995 | 1.999 | 2.249 | 3.002 | 1.014 | -0.016 | 0.036 | 0.036 | 0.194 | 0.65 | 0.25 | 0.5 | 0.72 | 0.64 | 0.62 |
| 11 | 104745330 | rs11226540 | -0.655 | 2.321 | 3.12 | 0.978 | 0.058 | 0.015 | 0.015 | 0 | 0.14 | 0.25 | 0.31 | 0.18 | 0.24 | 0.19 |
| 11 | 104751269 | rs571955 | 1.822 | 1.786 | 2.734 | 0.997 | -0.007 | 0.037 | 0.037 | 0.307 | 0.65 | 0.25 | 0.51 | 0.72 | 0.65 | 0.62 |

| | | | | | | | | | | | | | | | | |
|---|---|---|---|---|---|---|---|---|---|---|---|---|---|---|---|---|
| 11 | 104758035 | rs484438 | 2.49 | 1.116 | 2.959 | 1.113 | 0.052 | 0.054 | 0.054 | 2.077 | 0.95 | 0.62 | 0.96 | 1 | 0.99 | 0.98 |
| 11 | 104777311 | rs11226550 | -0.993 | 1.273 | 2.812 | 1.14 | 0.049 | 0.017 | 0.017 | 0 | 0.15 | 0.33 | 0.33 | 0.18 | 0.22 | 0.19 |
| 11 | 104778291 | rs11825902 | -0.57 | 1.267 | 2.892 | 1.139 | -0.005 | -0.003 | -0.003 | 0 | 0.13 | 0.11 | 0.31 | 0.16 | 0.22 | 0.19 |
| 11 | 104778846 | rs491945 | 0.945 | 1.26 | 2.964 | 1.136 | 0.047 | 0.009 | 0.009 | 0 | 0.83 | 0.5 | 0.65 | 0.82 | 0.78 | 0.81 |
| 11 | 104779195 | rs478650 | 0.914 | 1.262 | 2.965 | 1.112 | 0.035 | 0.007 | 0.007 | 0.001 | 0.83 | 0.5 | 0.67 | 0.82 | 0.79 | 0.81 |
| 11 | 104788727 | rs514537 | 1.796 | 0.639 | 2.945 | 0.853 | 0.006 | 0.007 | 0.007 | 0.334 | 0.67 | 0.32 | 0.62 | 0.78 | 0.71 | 0.68 |
| 11 | 104788962 | rs1944908 | -0.652 | 0.611 | 3.055 | 0.885 | -0.004 | 0 | 0 | 0 | 0.13 | 0.13 | 0.05 | 0.13 | 0.06 | 0.14 |
| 11 | 104789221 | rs488928 | 2.468 | 0.809 | 2.725 | 1.246 | -0.007 | -0.041 | -0.041 | 0.032 | 0.48 | 0.04 | 0.34 | 0.43 | 0.44 | 0.34 |
| 11 | 104789971 | rs7104517 | -2.121 | 0.878 | 2.876 | 0.38 | -0.026 | -0.031 | -0.031 | 0 | 0.32 | 0.44 | 0.36 | 0.22 | 0.29 | 0.32 |
| 11 | 104809654 | rs10791740 | -2.116 | 0.881 | 2.883 | 0.325 | -0.025 | -0.039 | -0.039 | 0 | 0.32 | 0.44 | 0.33 | 0.21 | 0.29 | 0.32 |
| 11 | 104816917 | rs7125155 | -0.832 | 0.902 | 2.866 | 0.344 | -0.023 | -0.01 | -0.01 | 0.651 | 0.14 | 0.17 | 0.05 | 0.12 | 0.06 | 0.14 |
| 11 | 104818293 | rs11226565 | -0.999 | 0.821 | 2.701 | 0.24 | 0.001 | -0.028 | -0.028 | 4.756 | 0.19 | 0.26 | 0.3 | 0.09 | 0.22 | 0.2 |
| 11 | 104821272 | rs571407 | 2.095 | 0.497 | 2.204 | 0.344 | -0.006 | -0.026 | -0.026 | 0 | 0.47 | 0.05 | 0.35 | 0.59 | 0.45 | 0.4 |
| 11 | 104826604 | rs666723 | 0.906 | 0.514 | 2.359 | 0.304 | 0.019 | -0.003 | -0.003 | 0 | 0.71 | 0.65 | 0.53 | 0.79 | 0.61 | 0.63 |
| 11 | 104828107 | rs623114 | 2.242 | 0.48 | 2.249 | 0.298 | 0.006 | -0.024 | -0.024 | 0 | 0.52 | 0.05 | 0.34 | 0.61 | 0.45 | 0.39 |
| 11 | 104828953 | rs609092 | 0.802 | 0.711 | 2.475 | -0.244 | -0.008 | 0 | 0 | 0 | 0.73 | 0.73 | 0.54 | 0.79 | 0.62 | 0.62 |
| 11 | 104830109 | rs7949293 | -0.665 | 0.696 | 2.692 | -0.253 | -0.001 | -0.078 | -0.078 | 0.546 | 0.13 | 0.27 | 0.14 | 0.01 | 0.13 | 0.17 |
| 11 | 104830838 | rs686999 | 0.769 | 0.709 | 2.237 | -0.243 | -0.023 | 0 | 0 | 0 | 0.73 | 0.77 | 0.55 | 0.79 | 0.62 | 0.63 |
| 11 | 104838471 | rs1944900 | -2.003 | 0.691 | 1.983 | -0.254 | -0.05 | -0.137 | -0.137 | 0 | 0.24 | 0.49 | 0.19 | 0.04 | 0.18 | 0.3 |
| 11 | 104849420 | rs480550 | -0.015 | 0.744 | 1.636 | -0.2 | -0.028 | -0.011 | -0.011 | 0 | 0.81 | 0.84 | 0.58 | 0.81 | 0.66 | 0.75 |
| 11 | 104850440 | rs17103454 | -0.811 | 0.491 | 1.029 | -0.162 | -0.012 | -0.013 | -0.013 | 0 | 0.11 | 0.13 | 0.04 | 0.03 | 0.05 | 0.13 |
| 11 | 104856055 | rs566275 | 1.817 | 0.506 | 0.85 | -0.175 | 0.001 | 0.079 | 0.079 | 0 | 0.57 | 0.21 | 0.38 | 0.75 | 0.48 | 0.47 |
| 11 | 104857221 | rs7127364 | -1.815 | 0.565 | 0.747 | -0.19 | -0.037 | -0.072 | -0.072 | 0.084 | 0.23 | 0.37 | 0.12 | 0.04 | 0.15 | 0.29 |
| 11 | 104869231 | rs518604 | 0.238 | 1.065 | 0.282 | 0.07 | 0.05 | -0.012 | -0.012 | 0 | 0.7 | 0.62 | 0.48 | 0.78 | 0.59 | 0.62 |
| 11 | 104870945 | rs3181328 | -1.005 | 0.511 | -0.151 | 0.197 | 0.004 | -0.002 | -0.002 | 0 | 0.1 | 0.08 | 0.06 | 0.01 | 0.1 | 0.11 |
| 11 | 104875261 | rs3181175 | -1.244 | 1.778 | 1.302 | 0.261 | 0.111 | -0.009 | -0.009 | 0 | 0.13 | 0.6 | 0.16 | 0.08 | 0.21 | 0.15 |
| 11 | 104877803 | rs2282657 | -1.703 | 2.265 | 0.997 | 0.722 | 0.071 | 0.029 | 0.029 | 0.178 | 0.21 | 0.63 | 0.25 | 0.26 | 0.34 | 0.27 |
| 11 | 104884774 | rs576800 | 1.318 | 2.38 | 1.331 | 0.677 | 0.065 | 0.05 | 0.05 | 0.307 | 0.8 | 0.42 | 0.78 | 0.74 | 0.73 | 0.74 |
| 11 | 104893557 | rs517736 | -0.88 | 2.522 | 0.716 | 0.038 | -0.013 | 0.009 | 0.009 | 0 | 0.12 | 0.09 | 0.25 | 0.01 | 0.18 | 0.11 |
| 11 | 104897437 | rs2282659 | -0.822 | 2.493 | 1.343 | 0.065 | 0.02 | 0.017 | 0.017 | 0.379 | 0.15 | 0.18 | 0.17 | 0.26 | 0.27 | 0.23 |
| 11 | 104911031 | rs1613367 | 1.37 | 1.05 | -0.231 | -0.769 | -0.105 | -0.118 | -0.118 | 1.197 | 0.36 | 0.03 | 0.34 | 0.53 | 0.41 | 0.28 |
| 11 | 104912458 | rs1503391 | -0.514 | 1.39 | 0.102 | -0.594 | 0.005 | -0.035 | -0.035 | 0 | 0.13 | 0.18 | 0.26 | 0.01 | 0.18 | 0.11 |
| 11 | 104912553 | rs1792774 | 0.274 | 1.396 | 0.019 | -0.584 | -0.022 | 0.058 | 0.058 | 0 | 0.1 | 0.01 | 0.22 | 0.01 | 0.11 | 0.11 |
| 11 | 104917645 | rs11821722 | -0.269 | 1.924 | 0.139 | -0.318 | 0.026 | 0.023 | 0.023 | 0.39 | 0.15 | 0.2 | 0.17 | 0.26 | 0.27 | 0.24 |
| 11 | 104924246 | rs1785883 | 0.74 | 0.976 | -0.181 | -0.821 | -0.008 | 0.032 | 0.032 | 4.282 | 0.07 | 0 | 0.03 | 0 | 0.07 | 0.02 |
| 11 | 104924786 | rs562441 | 1.154 | -0.289 | 0.07 | -0.453 | 0.027 | -0.004 | -0.004 | 0.033 | 0.52 | 0.33 | 0.41 | 0.61 | 0.32 | 0.57 |
| 11 | 104927112 | rs12801987 | 0.345 | -0.265 | 0.048 | -0.486 | -0.001 | 0.022 | 0.022 | 0 | 0.09 | 0.03 | 0.21 | 0.01 | 0.08 | 0.09 |
| 11 | 104936010 | rs12417050 | 0.316 | -0.259 | 0.054 | -0.482 | 0 | 0.019 | 0.019 | 0 | 0.09 | 0.04 | 0.21 | 0.01 | 0.08 | 0.09 |
| 11 | 104940204 | rs537727 | 0.42 | -0.253 | 0.31 | -0.421 | 0.007 | -0.004 | -0.004 | 0 | 0.95 | 0.72 | 0.97 | 1 | 0.99 | 0.96 |
| 11 | 104955489 | rs1785864 | -0.371 | -0.209 | 0.07 | -0.424 | -0.075 | -0.086 | -0.086 | 0.366 | 0.32 | 0.13 | 0.34 | 0.39 | 0.58 | 0.29 |
| 11 | 104958571 | rs1699081 | -0.66 | -0.252 | 0.44 | -0.429 | 0.007 | 0.011 | 0.011 | 0 | 0.36 | 0.42 | 0.37 | 0.39 | 0.59 | 0.29 |
| 11 | 104963849 | rs1623342 | -1.514 | -0.397 | 0.442 | -0.537 | 0.011 | -0.002 | -0.002 | 0 | 0.48 | 0.62 | 0.59 | 0.4 | 0.68 | 0.42 |
| 11 | 104965587 | rs12420557 | 0.175 | -0.397 | 0.43 | -0.537 | -0.004 | -0.032 | -0.032 | 0.015 | 0.11 | 0.14 | 0.22 | 0.01 | 0.08 | 0.08 |
| 11 | 104979669 | rs1847293 | -0.545 | -0.396 | 0.496 | -0.535 | -0.01 | -0.013 | -0.013 | 0.302 | 0.34 | 0.31 | 0.36 | 0.39 | 0.59 | 0.29 |
| 11 | 104979884 | rs7934144 | -1.594 | -0.409 | 0.546 | -0.537 | 0.037 | -0.004 | -0.004 | 0.01 | 0.48 | 0.69 | 0.61 | 0.4 | 0.68 | 0.42 |
| 11 | 104994480 | rs12797863 | -0.302 | -0.595 | 1.072 | -0.64 | -0.006 | -0.057 | -0.057 | 1.638 | 0.11 | 0.26 | 0.22 | 0.01 | 0.07 | 0.07 |
| 11 | 104999890 | rs11226619 | 0.747 | -1.246 | 0.914 | -0.681 | 0.012 | 0.005 | 0.005 | 0 | 0.52 | 0.44 | 0.53 | 0.47 | 0.33 | 0.56 |
| 11 | 105005936 | rs1483025 | -0.129 | -1.23 | 0.817 | -0.674 | -0.042 | -0.053 | -0.053 | 0 | 0.34 | 0.2 | 0.36 | 0.4 | 0.6 | 0.29 |
| 11 | 105006768 | rs17103781 | 0.197 | -0.927 | 0.804 | -0.557 | -0.002 | -0.032 | -0.032 | 0 | 0.09 | 0.14 | 0.21 | 0.01 | 0.07 | 0.07 |
| 11 | 105016907 | rs7943892 | 0.577 | -0.955 | 0.816 | -0.56 | 0.032 | 0.015 | 0.015 | 0 | 0.5 | 0.34 | 0.52 | 0.47 | 0.33 | 0.56 |
| 11 | 105017039 | rs11226623 | 0.792 | -0.972 | 0.738 | -0.567 | 0.049 | 0.017 | 0.017 | 0 | 0.47 | 0.2 | 0.48 | 0.44 | 0.32 | 0.52 |
| 11 | 105021005 | rs10895780 | 0.391 | -0.979 | 0.957 | -0.57 | 0.012 | 0.004 | 0.004 | 0 | 0.52 | 0.58 | 0.55 | 0.47 | 0.33 | 0.56 |
| 11 | 105023559 | rs1842893 | 1.409 | -0.973 | 0.841 | -0.563 | 0.042 | -0.03 | -0.03 | 0 | 0.38 | 0.15 | 0.28 | 0.44 | 0.25 | 0.45 |
| 11 | 105025272 | rs1023954 | 1.415 | -0.967 | 0.842 | -0.561 | 0.042 | -0.03 | -0.03 | 0 | 0.38 | 0.15 | 0.28 | 0.44 | 0.25 | 0.41 |
| 11 | 105033409 | rs1503381 | 0.678 | -0.954 | 0.897 | -0.562 | 0.036 | 0.018 | 0.018 | 0 | 0.48 | 0.27 | 0.49 | 0.44 | 0.32 | 0.52 |
| 11 | 105038988 | rs10895781 | 0.278 | -0.913 | 0.784 | -0.56 | 0.024 | 0.004 | 0.004 | 0 | 0.53 | 0.7 | 0.57 | 0.47 | 0.33 | 0.56 |
| 11 | 105042650 | rs12798197 | 0.313 | -0.832 | 0.585 | -0.495 | -0.002 | -0.03 | -0.03 | 0.089 | 0.09 | 0.11 | 0.2 | 0.01 | 0.07 | 0.07 |
| 11 | 105054557 | rs4755094 | 0.621 | -0.587 | 0.579 | -0.234 | -0.001 | 0 | 0 | 0 | 0.52 | 0.5 | 0.53 | 0.47 | 0.33 | 0.56 |
| 11 | 105067990 | rs7112242 | -0.892 | -1.076 | 0.943 | -0.533 | -0.018 | -0.083 | -0.083 | 0 | 0.14 | 0.44 | 0.09 | 0.01 | 0.07 | 0.13 |
| 11 | 105077608 | rs11226659 | -0.054 | -0.788 | 0.319 | -0.118 | -0.116 | -0.08 | -0.08 | 0 | 0.24 | 0.04 | 0.21 | 0.26 | 0.41 | 0.11 |
| 11 | 105081778 | rs7926220 | 0.319 | -0.893 | 0.637 | -0.255 | -0.005 | -0.016 | -0.016 | 0.63 | 0.08 | 0.09 | 0.02 | 0.01 | 0.05 | 0.11 |
| 11 | 105085734 | rs17103984 | -0.73 | -0.923 | 0.892 | -0.275 | 0.055 | 0.025 | 0.025 | 0 | 0.05 | 0.33 | 0.06 | 0 | 0.02 | 0.02 |
| 11 | 105087216 | rs17103994 | -0.017 | -0.964 | 1.21 | -0.289 | -0.014 | -0.041 | -0.041 | 0.837 | 0.09 | 0.24 | 0.04 | 0.01 | 0.05 | 0.11 |
| 11 | 105090296 | rs11600433 | 0.21 | -1.323 | 1.138 | -0.399 | 0.01 | 0.011 | 0.011 | 0 | 0.11 | 0.06 | 0.08 | 0.09 | 0.08 | 0.2 |
| 11 | 105098107 | rs1449045 | -0.805 | -1.364 | 1.243 | -0.046 | 0.008 | -0.024 | -0.024 | 0.006 | 0.4 | 0.56 | 0.31 | 0.15 | 0.46 | 0.24 |
| 11 | 105112018 | rs12271603 | 0.765 | -1.398 | 1.402 | -0.05 | 0 | 0 | 0 | 0 | 0.55 | 0.53 | 0.5 | 0.74 | 0.45 | 0.65 |
| 11 | 105122451 | rs4320922 | 1.082 | -1.469 | 1.51 | -0.117 | 0.008 | 0.021 | 0.021 | 0 | 0.54 | 0.33 | 0.48 | 0.75 | 0.45 | 0.65 |
| 11 | 105128418 | rs2218716 | 0.816 | -1.587 | 1.294 | -0.109 | 0.006 | -0.003 | -0.003 | 0 | 0.84 | 0.72 | 0.73 | 0.89 | 0.87 | 0.8 |
| 11 | 105158726 | rs10895805 | 2.031 | -1.674 | 0.807 | -0.017 | 0.008 | -0.079 | -0.079 | 0 | 0.39 | 0.08 | 0.36 | 0.63 | 0.35 | 0.47 |

| | | | | | | | | | | | | | | | | |
|---|---|---|---|---|---|---|---|---|---|---|---|---|---|---|---|---|
| 11 | 105200281 | rs6591122 | -0.84 | -1.611 | 1.173 | -0.096 | 0 | -0.008 | -0.008 | 0 | 0.4 | 0.37 | 0.33 | 0.19 | 0.52 | 0.36 |
| 11 | 105204066 | rs1449051 | 0.778 | -1.603 | 1.453 | -0.117 | 0 | 0.002 | 0.002 | 0 | 0.58 | 0.59 | 0.49 | 0.72 | 0.44 | 0.61 |
| 11 | 105205769 | rs7950345 | -0.322 | -1.726 | 1.423 | -0.121 | -0.027 | -0.037 | -0.037 | 0 | 0.91 | 0.99 | 0.79 | 0.91 | 0.91 | 0.89 |
| 11 | 105210472 | rs4603276 | 1.98 | -1.789 | 1.548 | -0.115 | 0.04 | -0.007 | -0.007 | 0.515 | 0.42 | 0.25 | 0.38 | 0.64 | 0.35 | 0.47 |
| 11 | 105211925 | rs1945985 | 0.117 | -1.734 | 1.591 | -0.145 | -0.001 | -0.032 | -0.032 | 0 | 0.08 | 0.2 | 0.05 | 0.02 | 0.05 | 0.08 |
| 11 | 105215497 | rs1449047 | 1.778 | -1.74 | 1.587 | -0.15 | -0.006 | -0.023 | -0.023 | 0 | 0.79 | 0.43 | 0.69 | 0.88 | 0.84 | 0.79 |
| 11 | 105224246 | rs11602565 | -0.907 | -1.717 | 1.427 | -0.14 | -0.029 | -0.033 | -0.033 | 0 | 0.15 | 0.23 | 0.09 | 0.07 | 0.08 | 0.14 |
| 11 | 105226271 | rs2044714 | 0.579 | -1.641 | 1.083 | -0.101 | 0.001 | 0.001 | 0.001 | 0 | 0.88 | 0.86 | 0.76 | 0.9 | 0.89 | 0.88 |
| 11 | 105239441 | rs7943657 | -0.113 | -1.658 | 1.049 | -0.129 | -0.056 | 0.032 | 0.032 | 0.127 | 0.32 | 0.17 | 0.28 | 0.2 | 0.47 | 0.28 |
| 11 | 105251489 | rs7929456 | -0.943 | -1.682 | 1.173 | -0.147 | -0.01 | 0.029 | 0.029 | 0.095 | 0.39 | 0.27 | 0.32 | 0.22 | 0.52 | 0.36 |
| 11 | 105273706 | rs7930393 | 1.764 | -1.705 | 1.339 | -0.107 | 0.02 | 0.003 | 0.003 | 0 | 0.43 | 0.36 | 0.39 | 0.64 | 0.35 | 0.42 |
| 11 | 105303595 | rs10502051 | 1.711 | -1.629 | 1.328 | -0.109 | -0.009 | -0.019 | -0.019 | 0 | 0.75 | 0.62 | 0.66 | 0.83 | 0.8 | 0.69 |
| 11 | 105337489 | rs7935262 | -0.895 | -1.624 | 1.041 | -0.128 | 0.005 | -0.016 | -0.016 | 0 | 0.4 | 0.54 | 0.32 | 0.22 | 0.5 | 0.35 |
| 11 | 105359052 | rs7122612 | 0.042 | -1.575 | 0.668 | -0.101 | -0.13 | 0.029 | 0.029 | 0.203 | 0.3 | 0.04 | 0.25 | 0.18 | 0.45 | 0.27 |
| 11 | 105373672 | rs4539296 | 1.953 | -1.595 | 0.683 | -0.08 | 0.022 | -0.044 | -0.044 | 0 | 0.4 | 0.11 | 0.36 | 0.64 | 0.34 | 0.42 |
| 11 | 105380436 | rs4373886 | -0.695 | -1.443 | 0.802 | -0.054 | -0.002 | -0.007 | -0.007 | 0.014 | 0.15 | 0.15 | 0.1 | 0.07 | 0.11 | 0.2 |
| 11 | 105390714 | rs7130632 | -0.392 | -1.377 | 1.041 | -0.024 | 0.045 | -0.036 | -0.036 | 0.001 | 0.34 | 0.47 | 0.29 | 0.2 | 0.46 | 0.28 |
| 11 | 105392392 | rs4354665 | 0.427 | -1.448 | 1.039 | -0.018 | -0.001 | -0.001 | -0.001 | 1.314 | 0.85 | 0.85 | 0.75 | 0.89 | 0.86 | 0.81 |
| 11 | 105393576 | rs11824662 | -0.043 | -1.465 | 0.883 | -0.024 | -0.068 | 0.032 | 0.032 | 0.24 | 0.3 | 0.11 | 0.26 | 0.18 | 0.46 | 0.28 |
| 11 | 105393840 | rs7116662 | 1.12 | -1.439 | 0.754 | 0.071 | 0.027 | 0.029 | 0.029 | 0 | 0.51 | 0.39 | 0.46 | 0.69 | 0.4 | 0.53 |
| 11 | 105405894 | rs11226761 | -0.479 | -1.437 | 0.588 | 0.936 | -0.039 | -0.028 | -0.028 | 0 | 0.35 | 0.23 | 0.37 | 0.43 | 0.52 | 0.29 |
| 11 | 105407131 | rs2156237 | 0.709 | -1.338 | 0.318 | 0.992 | -0.012 | -0.041 | -0.041 | 0 | 0.08 | 0.01 | 0.13 | 0.29 | 0.06 | 0.13 |
| 11 | 105412707 | rs2510156 | 0.414 | -1.84 | 0.749 | 0.693 | -0.011 | 0.006 | 0.006 | 0 | 0.84 | 0.81 | 0.88 | 0.83 | 0.9 | 0.81 |
| 11 | 105414478 | rs1942977 | 1.206 | -1.826 | 0.857 | 0.576 | 0.028 | 0.051 | 0.051 | 0 | 0.46 | 0.28 | 0.49 | 0.36 | 0.37 | 0.51 |
| 11 | 105423847 | rs12806006 | 1.175 | -1.813 | 0.904 | 0.603 | 0.023 | 0.035 | 0.035 | 0 | 0.47 | 0.32 | 0.49 | 0.36 | 0.37 | 0.51 |
| 11 | 105426444 | rs2513269 | -0.93 | -1.877 | 1.108 | 0.662 | 0.003 | 0.001 | 0.001 | 0 | 0.46 | 0.43 | 0.45 | 0.55 | 0.59 | 0.42 |
| 11 | 105426802 | rs1009820 | -0.181 | -1.924 | 1.133 | 0.86 | 0.003 | -0.001 | -0.001 | 0.955 | 0.36 | 0.36 | 0.37 | 0.35 | 0.52 | 0.3 |
| 11 | 105428486 | rs1942971 | 0.257 | -1.801 | 0.951 | 0.9 | 0.012 | -0.016 | -0.016 | 0 | 0.1 | 0.08 | 0.07 | 0.2 | 0.06 | 0.13 |
| 11 | 105434732 | rs11226768 | 0.484 | -1.679 | 1.114 | 0.94 | 0.013 | 0.107 | 0.107 | 0 | 0.94 | 0.74 | 0.93 | 0.88 | 0.95 | 0.93 |
| 11 | 105441422 | rs11226774 | -0.038 | -1.715 | 1.012 | 0.952 | 0.008 | -0.014 | -0.014 | 0 | 0.11 | 0.08 | 0.08 | 0.2 | 0.08 | 0.14 |
| 11 | 105441695 | rs7936555 | 1.655 | -1.896 | 1.308 | 0.874 | -0.047 | 0.079 | 0.079 | 0 | 0.77 | 0.52 | 0.82 | 0.66 | 0.84 | 0.73 |
| 11 | 105441742 | rs2513260 | 0.587 | -1.82 | 1.337 | 0.913 | -0.006 | 0.002 | 0.002 | 0 | 0.88 | 0.87 | 0.92 | 0.89 | 0.91 | 0.88 |
| 11 | 105443792 | rs17104291 | 1.882 | -2.064 | 1.164 | 0.778 | 0.012 | 0.004 | 0.004 | 0 | 0.35 | 0.16 | 0.44 | 0.34 | 0.3 | 0.43 |
| 11 | 105444159 | rs11226781 | 0.739 | -2.096 | 1.086 | 0.755 | 0.013 | -0.049 | -0.049 | 0.285 | 0.12 | 0.02 | 0.08 | 0.2 | 0.1 | 0.16 |
| 11 | 105445104 | rs7933640 | 1.504 | -2.179 | 1.07 | 0.716 | -0.052 | 0.024 | 0.024 | 0 | 0.7 | 0.54 | 0.8 | 0.66 | 0.78 | 0.67 |
| 11 | 105448454 | rs10895837 | 1.465 | -2.133 | 1.116 | 0.652 | -0.04 | -0.001 | -0.001 | 0 | 0.82 | 0.57 | 0.88 | 0.86 | 0.88 | 0.85 |
| 11 | 105461351 | rs1373925 | 1.078 | -2.124 | 0.9 | 0.663 | 0.04 | -0.01 | -0.01 | 2.051 | 0.47 | 0.25 | 0.52 | 0.55 | 0.4 | 0.59 |
| 11 | 105462421 | rs9919664 | -1.664 | -2.056 | 0.876 | 0.749 | 0.011 | -0.007 | -0.007 | 0 | 0.27 | 0.19 | 0.18 | 0.33 | 0.22 | 0.3 |
| 11 | 105474087 | rs10502058 | -1.565 | -1.441 | 1.248 | 0.64 | -0.01 | -0.006 | -0.006 | 0 | 0.16 | 0.16 | 0.1 | 0.11 | 0.11 | 0.14 |
| 11 | 105482024 | rs11226805 | -0.341 | -1.422 | 1.237 | 0.662 | 0.004 | -0.002 | -0.002 | 0 | 0.35 | 0.37 | 0.31 | 0.3 | 0.48 | 0.26 |
| 11 | 105483629 | rs1562221 | -0.421 | -1.466 | 1.162 | 0.429 | -0.011 | -0.04 | -0.04 | 0 | 0.1 | 0.12 | 0.05 | 0.02 | 0.07 | 0.08 |
| 11 | 105483661 | rs2166318 | 0.955 | -1.362 | 0.671 | 0.55 | 0.047 | -0.004 | -0.004 | 0 | 0.49 | 0.22 | 0.57 | 0.55 | 0.41 | 0.59 |
| 11 | 105484844 | rs7949759 | -1.286 | -1.268 | 0.255 | 0.426 | 0.012 | 0.004 | 0.004 | 0 | 0.49 | 0.54 | 0.4 | 0.42 | 0.59 | 0.4 |
| 11 | 105494105 | rs10750731 | 0.138 | -0.652 | 0.336 | 0.489 | 0.022 | -0.032 | -0.032 | 0.029 | 0.9 | 0.66 | 0.94 | 0.97 | 0.94 | 0.95 |
| 11 | 105498095 | rs1445604 | -0.49 | -0.316 | -0.648 | 0.85 | -0.047 | -0.04 | -0.04 | 0 | 0.32 | 0.24 | 0.33 | 0.41 | 0.52 | 0.36 |
| 11 | 105503658 | rs4754133 | -0.149 | -1.23 | -0.656 | 0.549 | -0.02 | 0.004 | 0.004 | 0.419 | 0.43 | 0.5 | 0.51 | 0.51 | 0.26 | 0.38 |
| 11 | 105514964 | rs10791764 | -0.355 | 0.123 | -0.334 | 0.393 | 0 | 0 | 0 | 0.13 | 0.41 | 0.44 | 0.34 | 0.55 | 0.49 | 0.51 |
| 11 | 105522961 | rs12421796 | 0.919 | -0.684 | -0.587 | 0.294 | 0.019 | 0.092 | 0.092 | 0 | 0.39 | 0.08 | 0.36 | 0.21 | 0.38 | 0.4 |
| 11 | 105529933 | rs11226820 | -0.15 | -0.553 | -0.308 | 0.332 | -0.001 | 0 | 0 | 0 | 0.57 | 0.45 | 0.65 | 0.45 | 0.51 | 0.49 |
| 11 | 105530021 | rs10895854 | 0.631 | -0.582 | -0.207 | 0.352 | -0.012 | 0.005 | 0.005 | 0 | 0.15 | 0.18 | 0.08 | 0.2 | 0.1 | 0.1 |
| 11 | 105534139 | rs1445613 | 0.437 | -0.587 | -0.101 | 0.258 | 0.006 | 0.012 | 0.012 | 0.357 | 0.55 | 0.36 | 0.63 | 0.42 | 0.5 | 0.47 |
| 11 | 105536018 | rs1822989 | 0.807 | -0.719 | 0 | 0.211 | 0.029 | 0.037 | 0.037 | 0.074 | 0.51 | 0.28 | 0.62 | 0.41 | 0.49 | 0.47 |
| 11 | 105541743 | rs1445617 | 0.804 | -0.705 | 0.009 | 0.203 | 0.026 | 0.035 | 0.035 | 0.102 | 0.51 | 0.31 | 0.62 | 0.41 | 0.49 | 0.47 |
| 11 | 105588070 | rs12808612 | 0.964 | 0.851 | -0.044 | 0.785 | -0.04 | -0.033 | -0.033 | 0.094 | 0.07 | 0.01 | 0.1 | 0.19 | 0.17 | 0.2 |
| 11 | 105591417 | rs7926520 | 0.621 | 0.405 | 0.109 | 0.435 | 0.003 | -0.003 | -0.003 | 0 | 0.59 | 0.43 | 0.65 | 0.4 | 0.52 | 0.47 |
| 11 | 105593543 | rs7931588 | 0.925 | 0.378 | 0.199 | 0.422 | -0.001 | 0.018 | 0.018 | 0.142 | 0.54 | 0.31 | 0.64 | 0.4 | 0.51 | 0.46 |
| 11 | 105610205 | rs7928594 | 0.62 | 0.398 | 0.299 | 0.439 | 0.006 | 0 | 0 | 0.821 | 0.6 | 0.53 | 0.66 | 0.44 | 0.53 | 0.47 |
| 11 | 105613109 | rs1938961 | 0.098 | 0.348 | 0.383 | 0.431 | -0.011 | -0.008 | -0.008 | 0 | 0.69 | 0.69 | 0.67 | 0.64 | 0.62 | 0.57 |
| 11 | 105614894 | rs2186598 | 1.431 | 0.127 | 0.531 | 0.303 | 0.022 | 0.032 | 0.032 | 0.286 | 0.52 | 0.3 | 0.38 | 0.4 | 0.5 | 0.48 |
| 11 | 105615253 | rs10502061 | 1.624 | 0.058 | 0.445 | 0.291 | -0.011 | -0.06 | -0.06 | 0.054 | 0.11 | 0 | 0.07 | 0.21 | 0.09 | 0.09 |
| 11 | 105630803 | rs11822168 | -0.061 | 0.103 | 0.542 | 0.278 | -0.011 | 0.036 | 0.036 | 0 | 0.17 | 0.36 | 0.29 | 0.24 | 0.12 | 0.08 |
| 11 | 105638142 | rs17478710 | 2.077 | 0.084 | 0.457 | 0.244 | -0.014 | 0.099 | 0.099 | 0.051 | 0.37 | 0.07 | 0.28 | 0.19 | 0.4 | 0.38 |
| 11 | 105639356 | rs10750733 | 0.513 | 0.74 | 1.069 | 0.268 | 0.01 | 0.005 | 0.005 | 0.484 | 0.68 | 0.6 | 0.66 | 0.64 | 0.62 | 0.55 |
| 11 | 105640104 | rs1938956 | 0.584 | 0.738 | 1.106 | 0.286 | 0.016 | 0.014 | 0.014 | 0.517 | 0.68 | 0.56 | 0.66 | 0.59 | 0.62 | 0.55 |
| 11 | 105641520 | rs2155040 | 0.4 | 0.739 | 1.079 | 0.285 | -0.01 | -0.007 | -0.007 | 0 | 0.69 | 0.69 | 0.67 | 0.64 | 0.62 | 0.55 |
| 11 | 105645648 | rs10791774 | 1.729 | 0.713 | 1.091 | 0.274 | 0.02 | 0.026 | 0.026 | 0.249 | 0.52 | 0.31 | 0.38 | 0.4 | 0.5 | 0.46 |
| 11 | 105654319 | rs17104532 | -0.977 | 0.403 | 1.335 | 0.283 | -0.041 | -0.019 | -0.019 | 0.13 | 0.09 | 0.3 | 0.03 | 0.05 | 0.02 | 0.02 |
| 11 | 105661240 | rs7934866 | 0.006 | 0.528 | 0.942 | 0.344 | -0.033 | -0.025 | -0.025 | 0 | 0.74 | 0.8 | 0.68 | 0.64 | 0.63 | 0.56 |
| 11 | 105681713 | rs2226567 | 0.342 | 0.551 | 0.928 | 0.328 | 0 | 0.006 | 0.006 | 0.036 | 0.12 | 0.12 | 0.26 | 0.19 | 0.11 | 0.08 |

| | | | | | | | | | | | | | | | | |
|---|---|---|---|---|---|---|---|---|---|---|---|---|---|---|---|---|
| 11 | 105688019 | rs10895876 | -2.163 | 0.521 | 0.889 | 0.3 | 0.029 | 0.036 | 0.036 | 0.179 | 0.49 | 0.78 | 0.63 | 0.6 | 0.5 | 0.54 |
| 11 | 105698982 | rs688053 | -0.616 | 0.574 | 0.714 | 0.287 | -0.015 | -0.006 | -0.006 | 0 | 0.18 | 0.27 | 0.28 | 0.19 | 0.12 | 0.08 |
| 11 | 105699048 | rs11226856 | -0.468 | 0.782 | 0.525 | 0.48 | -0.008 | -0.007 | -0.007 | 0.291 | 0.26 | 0.23 | 0.32 | 0.35 | 0.37 | 0.45 |
| 11 | 105709150 | rs4457719 | -1.833 | 0.694 | 0.365 | 0.384 | 0.015 | 0.024 | 0.024 | 0 | 0.48 | 0.79 | 0.61 | 0.58 | 0.48 | 0.53 |
| 11 | 105709856 | rs1940964 | 0.603 | 0.52 | 0.717 | 0.11 | -0.002 | 0.002 | 0.002 | 0 | 0.8 | 0.58 | 0.86 | 0.85 | 0.84 | 0.81 |
| 11 | 105715553 | rs672673 | -1.766 | 1.534 | 0.675 | 0.422 | 0.016 | 0 | 0 | 0.196 | 0.3 | 0.62 | 0.43 | 0.3 | 0.32 | 0.3 |
| 11 | 105717038 | rs4754136 | 0.953 | 1.616 | 0.152 | 0.427 | -0.003 | 0.001 | 0.001 | 0 | 0.07 | 0.06 | 0.24 | 0.09 | 0.1 | 0.06 |
| 11 | 105719344 | rs599980 | 0.664 | 1.591 | -0.222 | 0.441 | -0.04 | -0.05 | -0.05 | 0.001 | 0.13 | 0.01 | 0.18 | 0.17 | 0.15 | 0.22 |
| 11 | 105719522 | rs528205 | -1.578 | 1.31 | -0.072 | 0.411 | -0.007 | -0.015 | -0.015 | 0 | 0.45 | 0.8 | 0.63 | 0.47 | 0.47 | 0.52 |
| 11 | 105721469 | rs597395 | -1.249 | 1.355 | 0.404 | 0.159 | 0.049 | 0.007 | 0.007 | 0.051 | 0.07 | 0.29 | 0.05 | 0.04 | 0.06 | 0.05 |
| 11 | 105725953 | rs637569 | -1.586 | 1.307 | -0.464 | 0.105 | 0.014 | 0.041 | 0.041 | 0 | 0.47 | 0.79 | 0.58 | 0.39 | 0.53 | 0.49 |
| 11 | 105727407 | rs7116745 | -1.457 | 1.395 | -0.042 | 0.026 | -0.003 | -0.026 | -0.026 | 0.06 | 0.21 | 0.57 | 0.32 | 0.15 | 0.19 | 0.1 |
| 11 | 105751726 | rs1939153 | -1.653 | 1.867 | 0.115 | 0.031 | 0.003 | -0.001 | -0.001 | 0 | 0.4 | 0.46 | 0.54 | 0.37 | 0.51 | 0.46 |
| 11 | 105753035 | rs667174 | -1.411 | 2.162 | 0.254 | 0.02 | -0.003 | 0.001 | 0.001 | 0 | 0.34 | 0.33 | 0.3 | 0.23 | 0.41 | 0.39 |
| 11 | 105754802 | rs685725 | -1.703 | 2.008 | 0.307 | 0.106 | 0.016 | 0.002 | 0.002 | 0 | 0.37 | 0.46 | 0.53 | 0.37 | 0.49 | 0.45 |
| 11 | 105791235 | rs17104715 | 0.013 | 1.968 | 0.291 | -0.003 | -0.005 | 0.005 | 0.005 | 0.013 | 0.05 | 0.05 | 0.03 | 0.01 | 0.07 | 0.03 |
| 11 | 105798776 | rs648762 | -0.092 | 2.165 | 0.291 | -0.075 | 0 | -0.016 | -0.016 | 0 | 0.08 | 0.1 | 0.04 | 0.02 | 0.09 | 0.05 |
| 11 | 105800066 | rs2249031 | -0.064 | 2.204 | 0.513 | -0.149 | 0.072 | 0.004 | 0.004 | 0 | 0.07 | 0.31 | 0.04 | 0.03 | 0.05 | 0.05 |
| 11 | 105812839 | rs641574 | 1.231 | 1.954 | 0.083 | -0.024 | 0.003 | 0.001 | 0.001 | 0 | 0.69 | 0.71 | 0.53 | 0.64 | 0.61 | 0.64 |
| 11 | 105815878 | rs502453 | 1.056 | 2.104 | 0.144 | 0.092 | 0.004 | 0 | 0 | 0 | 0.87 | 0.81 | 0.71 | 0.84 | 0.8 | 0.89 |
| 11 | 105840364 | rs2155338 | 1.667 | 1.483 | 0.11 | 0.021 | 0.003 | 0.002 | 0.002 | 0.05 | 0.69 | 0.69 | 0.68 | 0.63 | 0.59 | 0.61 |
| 11 | 105844065 | rs609665 | 1.742 | 1.464 | 0.12 | 0.014 | 0.01 | -0.017 | -0.017 | 0.219 | 0.68 | 0.61 | 0.72 | 0.75 | 0.63 | 0.64 |
| 11 | 105853107 | rs675091 | 1.34 | 0.761 | 0.203 | 0.208 | -0.002 | -0.001 | -0.001 | 0 | 0.87 | 0.85 | 0.89 | 0.84 | 0.82 | 0.89 |
| 11 | 105858833 | rs2846216 | 0.3 | 0.778 | 0.226 | 0.181 | 0 | -0.019 | -0.019 | 0 | 0.92 | 0.89 | 0.96 | 0.98 | 0.91 | 0.95 |
| 11 | 105878844 | rs1043144 | 1.941 | 0.601 | 0.186 | 0.242 | 0.009 | 0.015 | 0.015 | 0 | 0.65 | 0.52 | 0.65 | 0.58 | 0.6 | 0.6 |
| 11 | 105879005 | rs1043142 | -0.37 | 0.662 | 0.051 | 0.23 | -0.068 | -0.068 | -0.068 | 1.69 | 0.13 | 0.01 | 0.17 | 0.21 | 0.2 | 0.26 |
| 11 | 105886989 | rs1939810 | 1.93 | 0.424 | 0.588 | -0.099 | 0.006 | 0.007 | 0.007 | 0 | 0.62 | 0.44 | 0.63 | 0.58 | 0.57 | 0.56 |
| 11 | 105891599 | rs2249950 | 1.322 | 0.231 | 0.328 | -0.049 | -0.002 | -0.005 | -0.005 | 0 | 0.77 | 0.76 | 0.73 | 0.61 | 0.73 | 0.69 |
| 11 | 105912351 | rs4754140 | 0.208 | 0.255 | 0.249 | 0.031 | -0.004 | -0.008 | -0.008 | 0 | 0.92 | 0.88 | 0.96 | 0.94 | 0.93 | 0.95 |
| 11 | 105918939 | rs10160360 | 0.176 | 0.258 | 0.252 | 0.033 | -0.002 | -0.005 | -0.005 | 0 | 0.92 | 0.89 | 0.96 | 0.94 | 0.93 | 0.95 |
| 11 | 105921935 | rs17093811 | -0.933 | 0.224 | 0.416 | -0.099 | -0.003 | -0.05 | -0.05 | 2.45 | 0.13 | 0.18 | 0.07 | 0.02 | 0.14 | 0.11 |
| 11 | 105922719 | rs1940791 | 0.109 | 0.227 | 0.406 | -0.098 | -0.004 | -0.009 | -0.009 | 0 | 0.92 | 0.88 | 0.96 | 0.94 | 0.93 | 0.95 |
| 11 | 105923373 | rs1939801 | -2.165 | 0.312 | 0.276 | 0.001 | 0 | 0.002 | 0.002 | 0 | 0.28 | 0.27 | 0.29 | 0.37 | 0.33 | 0.37 |
| 11 | 105923564 | rs3737347 | -0.851 | 0.224 | -0.021 | 0.214 | -0.046 | -0.083 | -0.083 | 0.02 | 0.14 | 0.03 | 0.22 | 0.35 | 0.19 | 0.26 |
| 11 | 105926127 | rs10895898 | -0.851 | 0.224 | -0.018 | 0.214 | -0.036 | -0.069 | -0.069 | 1.039 | 0.14 | 0.06 | 0.22 | 0.35 | 0.19 | 0.26 |
| 11 | 105927688 | rs2155345 | -1.426 | -1.008 | 0.022 | 0.022 | 0.037 | -0.017 | -0.017 | 0 | 0.4 | 0.26 | 0.32 | 0.41 | 0.34 | 0.41 |
| 11 | 105930184 | rs1893174 | 1.269 | -1.673 | 0.062 | 0.065 | 0.036 | -0.005 | -0.005 | 0.34 | 0.72 | 0.19 | 0.81 | 0.64 | 0.87 | 0.79 |
| 11 | 105939451 | rs10895900 | 1.783 | -1.704 | -0.126 | -0.084 | 0.089 | 0.005 | 0.005 | 0.846 | 0.62 | 0.11 | 0.77 | 0.59 | 0.77 | 0.73 |
| 11 | 105947285 | rs1047636 | -1.177 | -1.744 | 0.184 | -0.132 | 0.004 | 0.001 | 0.001 | 0 | 0.22 | 0.13 | 0.11 | 0.06 | 0.19 | 0.18 |
| 11 | 105951060 | rs12365489 | 0.291 | -1.746 | 0.978 | -0.059 | -0.013 | 0.136 | 0.136 | 0.004 | 0.07 | 0.28 | 0.07 | 0.3 | 0.03 | 0.05 |
| 11 | 105960721 | rs10431054 | 0.536 | -2.009 | 0.517 | -0.002 | -0.041 | -0.137 | -0.137 | 0.148 | 0.17 | 0.01 | 0.22 | 0.4 | 0.18 | 0.26 |
| 11 | 105962546 | rs2155339 | -0.223 | -2.006 | 1.016 | -0.086 | 0.016 | 0.027 | 0.027 | 0 | 0.13 | 0.07 | 0.08 | 0.02 | 0.11 | 0.12 |
| 11 | 105966591 | rs1939799 | 2.256 | -2.013 | 1.052 | -0.089 | 0.068 | 0.014 | 0.014 | 0.266 | 0.61 | 0.18 | 0.78 | 0.59 | 0.76 | 0.73 |
| 11 | 105969101 | rs7554 | -0.187 | -1.989 | 1.396 | -0.06 | 0.019 | 0.289 | 0.289 | 0.165 | 0.08 | 0.45 | 0.1 | 0.33 | 0.03 | 0.08 |
| 11 | 105975325 | rs11226922 | 1.918 | -2.04 | 1.263 | -0.173 | 0.1 | 0.023 | 0.023 | 1.345 | 0.58 | 0.09 | 0.75 | 0.59 | 0.74 | 0.75 |
| 11 | 105988995 | rs11821763 | -0.372 | -2.006 | 1.291 | -0.166 | -0.011 | -0.045 | -0.045 | 0.021 | 0.14 | 0.15 | 0.08 | 0.02 | 0.11 | 0.12 |
| 11 | 105996410 | rs11226931 | 1.062 | -1.963 | 1.318 | -0.11 | -0.022 | 0.014 | 0.014 | 0.375 | 0.73 | 0.33 | 0.84 | 0.61 | 0.86 | 0.84 |
| 11 | 105999580 | rs1940770 | -0.011 | -1.975 | 1.03 | -0.051 | -0.019 | -0.008 | -0.008 | 0 | 0.94 | 0.8 | 0.99 | 0.98 | 1 | 1 |
| 11 | 106005191 | rs766717 | 1.418 | -2.004 | 0.445 | -0.031 | 0.092 | 0.019 | 0.019 | 0.966 | 0.59 | 0.18 | 0.78 | 0.59 | 0.76 | 0.73 |
| 11 | 106020076 | rs1940781 | -0.642 | -1.874 | 0.929 | 0.239 | -0.047 | 0.08 | 0.08 | 0 | 0.19 | 0.4 | 0.11 | 0.35 | 0.1 | 0.17 |
| 11 | 106039594 | rs12294538 | -0.577 | -1.96 | 0.889 | 0.183 | -0.028 | 0.039 | 0.039 | 0 | 0.2 | 0.33 | 0.12 | 0.32 | 0.12 | 0.1 |
| 11 | 106045336 | rs4298866 | 1.566 | -2.458 | 0.829 | -0.122 | 0.11 | 0.009 | 0.009 | 0.852 | 0.59 | 0.16 | 0.77 | 0.62 | 0.78 | 0.73 |
| 11 | 106064899 | rs1893173 | 0.707 | -2.493 | 0.884 | -0.123 | -0.074 | -0.118 | -0.118 | 0 | 0.8 | 0.51 | 0.9 | 0.94 | 0.9 | 0.84 |
| 11 | 106067955 | rs7127780 | -0.485 | -2.574 | 0.793 | -0.106 | -0.013 | 0.012 | 0.012 | 0 | 0.21 | 0.29 | 0.13 | 0.31 | 0.13 | 0.16 |
| 11 | 106095211 | rs1115160 | 0.828 | -2.568 | 0.215 | -0.075 | -0.026 | 0.01 | 0.01 | 0.147 | 0.73 | 0.31 | 0.84 | 0.63 | 0.87 | 0.8 |
| 11 | 106120529 | rs7119670 | -0.355 | -2.568 | 0.2 | -0.118 | -0.017 | -0.057 | -0.057 | 0.036 | 0.13 | 0.18 | 0.07 | 0.01 | 0.1 | 0.1 |
| 11 | 106123295 | rs11824707 | -0.355 | -2.568 | 0.2 | -0.118 | -0.017 | -0.057 | -0.057 | 0 | 0.13 | 0.18 | 0.07 | 0.01 | 0.1 | 0.1 |
| 11 | 106123594 | rs10791800 | 1.64 | -2.593 | 0.07 | -0.103 | 0.152 | -0.014 | -0.014 | 0 | 0.59 | 0.07 | 0.76 | 0.62 | 0.77 | 0.69 |
| 11 | 106153996 | rs11226987 | 1.627 | -2.61 | 0.009 | -0.078 | 0.149 | -0.018 | -0.018 | 1.954 | 0.58 | 0.11 | 0.77 | 0.61 | 0.77 | 0.69 |
| 11 | 106157404 | rs488212 | -0.296 | -2.641 | 0.22 | -0.088 | 0.005 | -0.007 | -0.007 | 0.788 | 0.2 | 0.21 | 0.12 | 0.31 | 0.13 | 0.16 |
| 11 | 106162312 | rs675005 | 0.903 | -2.626 | 0.297 | -0.112 | -0.073 | -0.112 | -0.112 | 0.017 | 0.81 | 0.5 | 0.9 | 0.93 | 0.9 | 0.85 |
| 11 | 106167184 | rs529861 | -0.1 | -2.61 | 0.084 | -0.094 | 0.01 | 0.046 | 0.046 | 8.395 | 0.93 | 0.54 | 0.96 | 0.95 | 1 | 0.94 |
| 11 | 106171877 | rs7126969 | 1.756 | -2.672 | -0.21 | -0.113 | 0.155 | -0.023 | -0.023 | 0 | 0.58 | 0.07 | 0.75 | 0.61 | 0.77 | 0.68 |
| 11 | 106178788 | rs10750750 | 1.026 | -2.78 | -0.265 | -0.183 | 0.111 | -0.051 | -0.051 | 0 | 0.68 | 0.1 | 0.8 | 0.63 | 0.85 | 0.77 |
| 11 | 106182251 | rs11227000 | 0.098 | -2.615 | -0.221 | -0.23 | 0.012 | 0.029 | 0.029 | 0 | 0.11 | 0.05 | 0.06 | 0.01 | 0.1 | 0.1 |
| 11 | 106204781 | rs679218 | -1.266 | -2.588 | -0.081 | -0.118 | 0.032 | -0.003 | -0.003 | 0 | 0.32 | 0.23 | 0.19 | 0.32 | 0.24 | 0.23 |
| 11 | 106220982 | rs1396157 | 1.497 | -2.556 | -0.142 | -0.233 | 0.022 | 0.008 | 0.008 | 2.425 | 0.58 | 0.3 | 0.76 | 0.63 | 0.74 | 0.7 |
| 11 | 106225571 | rs538278 | -1.082 | -2.576 | -0.166 | -0.289 | 0.033 | 0.051 | 0.051 | 0.035 | 0.29 | 0.22 | 0.15 | 0.07 | 0.23 | 0.14 |
| 11 | 106227654 | rs10895943 | 0.695 | -2.439 | -0.777 | -0.247 | -0.065 | -0.136 | -0.136 | 0.462 | 0.19 | 0.01 | 0.21 | 0.43 | 0.2 | 0.22 |

| | | | | | | | | | | | | | | | | |
|---|---|---|---|---|---|---|---|---|---|---|---|---|---|---|---|---|
| 11 | 106229906 | rs10895944 | 2.139 | -2.368 | -0.848 | -1.167 | -0.135 | 0.044 | 0.044 | 1.263 | 0.33 | 0.06 | 0.5 | 0.2 | 0.49 | 0.46 |
| 11 | 106231144 | rs563816 | -1.979 | -2.566 | 0.154 | -1.308 | 0.076 | 0.106 | 0.106 | 4.313 | 0.42 | 0.8 | 0.21 | 0.11 | 0.25 | 0.2 |
| 11 | 106232838 | rs1512432 | 0.013 | -2.56 | -0.057 | -1.312 | -0.004 | -0.022 | -0.022 | 0.065 | 0.1 | 0.39 | 0.04 | 0.05 | 0.02 | 0.06 |
| 11 | 106233094 | rs618453 | -1.785 | -2.436 | -1.29 | -1.241 | 0.121 | 0.201 | 0.201 | 6.422 | 0.42 | 0.91 | 0.23 | 0.12 | 0.26 | 0.19 |
| 11 | 106245468 | rs638300 | -1.822 | -2.389 | -1.286 | -1.256 | 0.118 | 0.18 | 0.18 | 4.976 | 0.43 | 0.86 | 0.22 | 0.11 | 0.25 | 0.2 |
| 11 | 106247407 | rs560337 | -0.623 | -2.157 | -1.282 | -1.021 | 0.012 | 0.013 | 0.013 | 0.018 | 0.19 | 0.19 | 0.1 | 0.05 | 0.13 | 0.07 |
| 11 | 106251514 | rs527709 | -1.013 | -1.95 | -2.016 | -0.869 | 0.124 | 0.132 | 0.132 | 5.635 | 0.33 | 0.92 | 0.15 | 0.11 | 0.1 | 0.15 |
| 11 | 106255343 | rs4755149 | -0.281 | -1.443 | -2.381 | -0.939 | 0.025 | 0.067 | 0.067 | 0 | 0.57 | 0.89 | 0.46 | 0.69 | 0.57 | 0.62 |
| 11 | 106260323 | rs1512427 | 0.389 | -0.544 | -1.906 | -1.186 | 0.002 | 0.003 | 0.003 | 0 | 0.22 | 0.22 | 0.22 | 0.66 | 0.21 | 0.36 |
| 11 | 106262357 | rs617689 | 0.647 | -0.983 | -2.001 | -1.528 | 0 | -0.026 | -0.026 | 4.384 | 0.33 | 0.14 | 0.5 | 0.37 | 0.36 | 0.34 |
| 11 | 106262397 | rs567992 | 1.436 | -1.292 | -2.348 | -1.762 | 0.007 | -0.068 | -0.068 | 0.018 | 0.17 | 0.01 | 0.37 | 0.23 | 0.13 | 0.21 |
| 11 | 106264835 | rs9285795 | -0.647 | -1.12 | -1.651 | -1.67 | 0.013 | -0.005 | -0.005 | 0 | 0.66 | 0.81 | 0.5 | 0.63 | 0.64 | 0.65 |
| 11 | 106265530 | rs10895951 | 0.234 | -1.321 | -0.96 | -1.916 | 0.002 | 0.006 | 0.006 | 0.129 | 0.3 | 0.37 | 0.29 | 0.61 | 0.27 | 0.34 |
| 11 | 106266163 | rs1553936 | 0.609 | -1.348 | -1.003 | -1.91 | -0.036 | -0.045 | -0.045 | 0 | 0.26 | 0.1 | 0.26 | 0.61 | 0.27 | 0.34 |
| 11 | 106268455 | rs11227026 | 0.077 | -1.792 | -0.629 | -2.374 | 0.01 | -0.155 | -0.155 | 0 | 0.35 | 0.4 | 0.22 | 0.02 | 0.37 | 0.3 |
| 11 | 106270964 | rs7110917 | -0.133 | -1.54 | -0.836 | -2.117 | -0.01 | -0.006 | -0.006 | 0 | 0.34 | 0.25 | 0.31 | 0.63 | 0.28 | 0.4 |
| 11 | 106271802 | rs17551747 | 0.235 | -1.941 | -0.442 | -2.386 | 0.016 | -0.086 | -0.086 | 0.157 | 0.26 | 0.32 | 0.17 | 0.01 | 0.36 | 0.2 |
| 11 | 106273699 | rs4755151 | 1.09 | -1.798 | -0.827 | -2.264 | -0.051 | -0.068 | -0.068 | 0 | 0.22 | 0.07 | 0.26 | 0.61 | 0.24 | 0.29 |
| 11 | 106275934 | rs2102599 | -0.43 | -1.769 | -1.499 | -2.244 | 0.005 | 0.005 | 0.005 | 0 | 0.37 | 0.24 | 0.31 | 0.63 | 0.28 | 0.43 |
| 11 | 106276534 | rs11227029 | 0.758 | -1.196 | -1.578 | -1.851 | -0.031 | -0.043 | -0.043 | 1.387 | 0.23 | 0.12 | 0.27 | 0.61 | 0.24 | 0.29 |
| 11 | 106279611 | rs1398312 | -0.935 | -1.326 | -1.448 | -1.603 | 0.041 | -0.001 | -0.001 | 0.169 | 0.64 | 0.8 | 0.49 | 0.64 | 0.64 | 0.6 |
| 11 | 106283700 | rs17105704 | 0.318 | -0.532 | -1.221 | -1.127 | -0.016 | -0.038 | -0.038 | 1.263 | 0.22 | 0.14 | 0.27 | 0.6 | 0.23 | 0.27 |
| 11 | 106284835 | rs4754158 | 0.179 | -0.482 | -1.383 | -1.228 | -0.067 | -0.058 | -0.058 | 0.461 | 0.23 | 0.07 | 0.33 | 0.62 | 0.27 | 0.29 |
| 11 | 106285669 | rs17552029 | 0.57 | -0.902 | -1.299 | -1.672 | 0.005 | -0.005 | -0.005 | 0.243 | 0.1 | 0.05 | 0.03 | 0.02 | 0.04 | 0.1 |
| 11 | 106287724 | rs2458329 | -1.558 | -1.222 | -1.585 | -1.621 | 0.003 | -0.016 | -0.016 | 0.768 | 0.84 | 0.86 | 0.85 | 0.76 | 0.8 | 0.86 |
| 11 | 106287805 | rs12798340 | 0.71 | -1.123 | -1.094 | -1.658 | -0.007 | -0.047 | -0.047 | 1.908 | 0.11 | 0.12 | 0.04 | 0.02 | 0.06 | 0.1 |
| 11 | 106288554 | rs953816 | 1.384 | -1.555 | -0.633 | -1.931 | -0.055 | -0.033 | -0.033 | 0 | 0.27 | 0.38 | 0.34 | 0.21 | 0.16 | 0.24 |
| 11 | 106292302 | rs975394 | 1.46 | -1.586 | -0.174 | -1.962 | 0.022 | 0.018 | 0.018 | 0 | 0.27 | 0.21 | 0.34 | 0.2 | 0.16 | 0.24 |
| 11 | 106292933 | rs2458324 | -1.671 | -1.362 | -0.182 | -1.858 | 0 | -0.005 | -0.005 | 0.098 | 0.84 | 0.82 | 0.84 | 0.75 | 0.79 | 0.86 |
| 11 | 106295202 | rs12365928 | 0.662 | -1.405 | -0.174 | -2.023 | 0.008 | -0.038 | -0.038 | 0 | 0.17 | 0.2 | 0.15 | 0 | 0.29 | 0.16 |
| 11 | 106306871 | rs1512436 | -0.054 | -1.529 | -0.423 | -2.038 | 0.025 | 0.137 | 0.137 | 0 | 0.48 | 0.7 | 0.51 | 0.2 | 0.45 | 0.41 |
| 11 | 106311915 | rs10895956 | 0.968 | -0.92 | -0.598 | -1.791 | -0.068 | -0.085 | -0.085 | 0.963 | 0.26 | 0.09 | 0.3 | 0.52 | 0.29 | 0.35 |
| 11 | 106316611 | rs10502069 | 0.402 | -0.753 | -0.136 | -1.682 | 0.08 | -0.067 | -0.067 | 0 | 0.13 | 0.23 | 0.15 | 0 | 0.29 | 0.12 |
| 11 | 106318163 | rs907645 | -0.246 | -0.849 | -0.181 | -1.83 | -0.005 | 0.021 | 0.021 | 0 | 0.48 | 0.36 | 0.41 | 0.55 | 0.42 | 0.51 |
| 11 | 106318515 | rs907647 | 0.49 | -0.673 | -0.048 | -1.457 | -0.007 | -0.011 | -0.011 | 0.862 | 0.32 | 0.27 | 0.33 | 0.52 | 0.32 | 0.4 |
| 11 | 106318654 | rs2513605 | -0.418 | -0.681 | 0.223 | -1.458 | 0 | -0.005 | -0.005 | 0 | 0.45 | 0.49 | 0.54 | 0.58 | 0.45 | 0.59 |
| 11 | 106322665 | rs7110120 | -0.853 | 0.379 | -0.062 | -1.144 | -0.09 | -0.038 | -0.038 | 0 | 0.7 | 0.87 | 0.61 | 0.42 | 0.65 | 0.57 |
| 11 | 106328565 | rs17105741 | 1.471 | -0.116 | -0.267 | -1.356 | 0.044 | 0.054 | 0.054 | 0 | 0.27 | 0.1 | 0.23 | 0.19 | 0.18 | 0.26 |
| 11 | 106330810 | rs7943420 | 0.475 | 0.778 | 0.25 | -0.941 | -0.005 | -0.004 | -0.004 | 0 | 0.55 | 0.45 | 0.42 | 0.42 | 0.39 | 0.49 |
| 11 | 106332385 | rs17559558 | -0.16 | 1.052 | -0.171 | -0.969 | 0.037 | -0.07 | -0.07 | 0.001 | 0.14 | 0.27 | 0.18 | 0 | 0.26 | 0.06 |
| 11 | 106333406 | rs2511416 | 1.255 | 1.597 | -0.854 | -0.225 | -0.081 | -0.035 | -0.035 | 0.002 | 0.17 | 0.01 | 0.12 | 0.1 | 0.19 | 0.22 |
| 11 | 106335277 | rs2511418 | -1.209 | 1.608 | -0.61 | 0.33 | -0.008 | -0.037 | -0.037 | 0 | 0.91 | 0.98 | 0.86 | 0.59 | 0.92 | 0.86 |
| 11 | 106339209 | rs10502070 | -1.455 | 1.551 | 0.028 | -0.368 | 0.014 | -0.042 | -0.042 | 0 | 0.2 | 0.59 | 0.26 | 0.06 | 0.18 | 0.25 |
| 11 | 106342004 | rs7115414 | -0.062 | 1.455 | 0.501 | -0.377 | 0.009 | 0.02 | 0.02 | 0.536 | 0.06 | 0.25 | 0.04 | 0.01 | 0.02 | 0.06 |
| 11 | 106344359 | rs7948675 | -2.369 | 1.722 | 0.338 | -0.542 | 0.002 | -0.022 | -0.022 | 0 | 0.3 | 0.61 | 0.4 | 0.06 | 0.44 | 0.32 |
| 11 | 106351103 | rs12364960 | -0.061 | 1.759 | 0.418 | -0.51 | 0.033 | -0.051 | -0.051 | 0 | 0.11 | 0.16 | 0.16 | 0 | 0.27 | 0.06 |
| 11 | 106363983 | rs12290638 | -0.709 | 1.774 | 0.47 | -0.521 | 0.001 | -0.056 | -0.056 | 0 | 0.14 | 0.3 | 0.08 | 0.03 | 0.09 | 0.19 |
| 11 | 106369769 | rs10791810 | -0.211 | 1.847 | 0.448 | -0.522 | 0.104 | -0.076 | -0.076 | 0 | 0.12 | 0.29 | 0.16 | 0 | 0.27 | 0.06 |
| 11 | 106370276 | rs6591173 | -1.263 | 1.534 | 0.031 | -0.51 | -0.047 | -0.106 | -0.106 | 0.058 | 0.71 | 0.92 | 0.71 | 0.48 | 0.73 | 0.65 |
| 11 | 106370603 | rs12271785 | -0.718 | 2.811 | 1.064 | -0.625 | -0.001 | -0.009 | -0.009 | 0 | 0.12 | 0.12 | 0.06 | 0.03 | 0.09 | 0.18 |
| 11 | 106374125 | rs1945835 | -2.627 | 2.855 | 1.028 | -0.637 | 0.058 | -0.066 | -0.066 | 0.275 | 0.23 | 0.54 | 0.35 | 0.05 | 0.37 | 0.21 |
| 11 | 106375299 | rs12274945 | -2.705 | 2.74 | 0.964 | -0.624 | 0.003 | -0.023 | -0.023 | 0 | 0.3 | 0.61 | 0.4 | 0.06 | 0.45 | 0.33 |
| 11 | 106378571 | rs12362046 | -1.016 | 2.446 | 1.081 | -0.626 | 0.047 | -0.065 | -0.065 | 0.367 | 0.12 | 0.22 | 0.16 | 0 | 0.27 | 0.06 |
| 11 | 106381370 | rs2032403 | 0.152 | 2.346 | 0.662 | -0.585 | 0.001 | -0.026 | -0.026 | 0.164 | 0.08 | 0.11 | 0.04 | 0.02 | 0.06 | 0.12 |
| 11 | 106390653 | rs12284877 | -1.01 | 2.35 | 0.739 | -0.607 | -0.003 | -0.043 | -0.043 | 0 | 0.13 | 0.21 | 0.07 | 0.03 | 0.08 | 0.18 |
| 11 | 106394204 | rs7102076 | -0.785 | 2.062 | 0.522 | -0.609 | -0.002 | -0.032 | -0.032 | 0 | 0.12 | 0.17 | 0.06 | 0.03 | 0.08 | 0.18 |
| 11 | 106396192 | rs11227088 | -2.403 | 2.179 | 0.239 | -0.587 | 0.009 | -0.03 | -0.03 | 0 | 0.31 | 0.61 | 0.41 | 0.06 | 0.44 | 0.31 |
| 11 | 106398020 | rs17561086 | 0.313 | 2.373 | 0.135 | -0.498 | 0 | -0.009 | -0.009 | 9.037 | 0.08 | 0.07 | 0.04 | 0.01 | 0.08 | 0.11 |
| 11 | 106399482 | rs7115105 | -1.77 | 2.198 | 0.256 | -0.329 | 0.092 | -0.068 | -0.068 | 0 | 0.2 | 0.54 | 0.37 | 0.05 | 0.35 | 0.19 |
| 11 | 106399802 | rs7101603 | -1.906 | 2.19 | 0.253 | -0.333 | 0.083 | -0.07 | -0.07 | 0.014 | 0.21 | 0.55 | 0.37 | 0.05 | 0.35 | 0.19 |
| 11 | 106402115 | rs10895975 | -1.771 | 2.227 | 0.282 | -0.145 | 0.103 | -0.015 | -0.015 | 0 | 0.2 | 0.52 | 0.37 | 0.09 | 0.36 | 0.21 |
| 11 | 106408071 | rs1791468 | 0.536 | 1.709 | -0.298 | -0.224 | 0 | 0.016 | 0.016 | 0 | 0.49 | 0.45 | 0.48 | 0.62 | 0.48 | 0.52 |
| 11 | 106410077 | rs1791466 | 1.63 | 2.472 | 0.497 | -0.309 | 0.014 | -0.015 | -0.015 | 0 | 0.72 | 0.38 | 0.63 | 0.89 | 0.66 | 0.72 |
| 11 | 106411576 | rs1945065 | -0.734 | 1.951 | 0.882 | -0.353 | -0.021 | -0.072 | -0.072 | 0 | 0.14 | 0.26 | 0.05 | 0.01 | 0.08 | 0.12 |
| 11 | 106414284 | rs1945062 | -0.615 | 1.948 | 0.894 | -0.373 | -0.018 | -0.066 | -0.066 | 0 | 0.13 | 0.21 | 0.05 | 0.01 | 0.08 | 0.12 |
| 11 | 106416708 | rs1791463 | 1.387 | 2.003 | 0.561 | -0.344 | 0.029 | -0.048 | -0.048 | 0 | 0.76 | 0.45 | 0.69 | 0.9 | 0.74 | 0.77 |
| 11 | 106420106 | rs1791459 | 0.692 | 2.239 | 0.078 | 0.15 | 0.027 | 0.009 | 0.009 | 0 | 0.93 | 0.86 | 0.77 | 0.91 | 0.84 | 0.9 |
| 11 | 106436025 | rs17105845 | -0.211 | 1.391 | -0.407 | -0.171 | 0.048 | -0.014 | -0.014 | 0 | 0.25 | 0.07 | 0.21 | 0.27 | 0.21 | 0.2 |
| 11 | 106452715 | rs7124340 | -1.084 | 2.559 | 0.226 | -0.494 | 0.085 | -0.049 | -0.049 | 0 | 0.13 | 0.26 | 0.24 | 0.04 | 0.26 | 0.13 |

| | | | | | | | | | | | | | | | | |
|---|---|---|---|---|---|---|---|---|---|---|---|---|---|---|---|---|
| 11 | 106454958 | rs2105786 | -0.657 | 2.587 | 0.888 | -0.669 | 0.001 | 0.001 | 0.001 | 0.214 | 0.34 | 0.63 | 0.34 | 0.3 | 0.33 | 0.21 |
| 11 | 106458501 | rs1945068 | -0.908 | 3.108 | 0.622 | -0.716 | -0.043 | -0.003 | -0.003 | 0 | 0.93 | 0.99 | 0.93 | 0.91 | 0.87 | 0.87 |
| 11 | 106459674 | rs4754161 | -1.261 | 2.298 | 0.942 | -0.584 | -0.013 | 0.059 | 0.059 | 0.15 | 0.42 | 0.71 | 0.48 | 0.31 | 0.48 | 0.33 |
| 11 | 106465539 | rs12421832 | -0.91 | 2.734 | 1.613 | -0.462 | 0.015 | -0.027 | -0.027 | 0 | 0.08 | 0.09 | 0.14 | 0 | 0.14 | 0.11 |
| 11 | 106477091 | rs17105869 | -2.496 | 2.713 | 2.025 | -0.43 | 0.109 | -0.065 | -0.065 | 0 | 0.14 | 0.54 | 0.23 | 0.03 | 0.24 | 0.11 |
| 11 | 106485489 | rs1625749 | -1.362 | 2.399 | 1.169 | -0.198 | 0.082 | -0.006 | -0.006 | 0 | 0.06 | 0.16 | 0.11 | 0.03 | 0.17 | 0.05 |
| 11 | 106491841 | rs4457720 | -0.654 | 1.545 | 1.18 | -0.286 | -0.037 | -0.016 | -0.016 | 1.915 | 0.19 | 0.35 | 0.15 | 0.11 | 0.13 | 0.05 |
| 11 | 106494775 | rs17105896 | -0.472 | 1.555 | 1.006 | -0.318 | 0.017 | 0.009 | 0.009 | 0 | 0.21 | 0.19 | 0.15 | 0.13 | 0.16 | 0.06 |
| 11 | 106497175 | rs1791577 | -0.397 | 1.708 | 1.061 | -0.681 | 0.033 | -0.063 | -0.063 | 0.528 | 0.12 | 0.24 | 0.17 | 0.01 | 0.14 | 0.15 |
| 11 | 106500377 | rs959057 | -0.855 | 1.655 | 0.65 | -0.762 | -0.031 | 0.043 | 0.043 | 0 | 0.14 | 0.06 | 0.19 | 0.01 | 0.16 | 0.16 |
| 11 | 106502073 | rs1791598 | -2.047 | 1.827 | 0.746 | -0.776 | -0.012 | 0.045 | 0.045 | 0.033 | 0.44 | 0.7 | 0.54 | 0.16 | 0.47 | 0.25 |
| 11 | 106507197 | rs7120385 | -0.837 | 1.889 | 0.88 | -0.771 | 0.099 | 0.003 | 0.003 | 0 | 0.07 | 0.2 | 0.19 | 0.03 | 0.16 | 0.03 |
| 11 | 106508030 | rs1938623 | -1.236 | 1.835 | 0.806 | -0.777 | 0.154 | -0.095 | -0.095 | 0 | 0.19 | 0.47 | 0.36 | 0.04 | 0.3 | 0.19 |
| 11 | 106509816 | rs2187509 | -1.69 | 1.775 | 0.599 | -0.766 | -0.013 | 0.033 | 0.033 | 0 | 0.41 | 0.7 | 0.53 | 0.14 | 0.45 | 0.25 |
| 11 | 106517883 | rs17643292 | -0.451 | 2.211 | 0.036 | -0.522 | 0.089 | -0.002 | -0.002 | 0 | 0.06 | 0.18 | 0.19 | 0.03 | 0.18 | 0.03 |
| 11 | 106518561 | rs1793073 | -0.22 | 1.717 | -0.709 | -0.466 | 0.004 | 0.014 | 0.014 | 0 | 0.19 | 0.16 | 0.15 | 0.11 | 0.18 | 0.06 |
| 11 | 106518705 | rs4587702 | -0.824 | 1.365 | -1.015 | 0.134 | -0.006 | 0.042 | 0.042 | 0 | 0.47 | 0.76 | 0.6 | 0.35 | 0.51 | 0.28 |
| 11 | 106523153 | rs1793077 | -1.092 | 1.72 | -0.923 | 0.169 | -0.018 | -0.002 | -0.002 | 0 | 0.1 | 0.04 | 0.13 | 0.08 | 0.14 | 0.2 |
| 11 | 106523574 | rs1783138 | -1.584 | 1.385 | 0.061 | 0.18 | 0.007 | 0 | 0 | 0 | 0.27 | 0.3 | 0.23 | 0.27 | 0.29 | 0.36 |
| 11 | 106527673 | rs1791587 | 0.121 | 1.21 | -0.887 | -1.078 | -0.004 | 0.035 | 0.035 | 0 | 0.53 | 0.62 | 0.38 | 0.21 | 0.44 | 0.38 |
| 11 | 106532311 | rs1783134 | -0.09 | 1.288 | -0.932 | -1.04 | -0.027 | 0.052 | 0.052 | 0 | 0.59 | 0.79 | 0.55 | 0.29 | 0.5 | 0.41 |
| 11 | 106534150 | rs1793083 | -1.023 | 1.313 | -1.039 | -1.057 | -0.071 | -0.127 | -0.127 | 0 | 0.72 | 0.97 | 0.7 | 0.38 | 0.65 | 0.6 |
| 11 | 106535709 | rs1791597 | 0.355 | 1.248 | -1.113 | -1.132 | 0.01 | 0.127 | 0.127 | 0 | 0.52 | 0.84 | 0.49 | 0.29 | 0.38 | 0.38 |
| 11 | 106543882 | rs1793064 | 1.333 | 1.469 | -0.29 | -1.691 | 0.013 | 0.011 | 0.011 | 0 | 0.33 | 0.26 | 0.2 | 0.05 | 0.24 | 0.18 |
| 11 | 106556669 | rs11211863 | 1.172 | 1.513 | -0.624 | -1.491 | -0.027 | 0.04 | 0.04 | 2.811 | 0.09 | 0.03 | 0.07 | 0 | 0.12 | 0.03 |
| 11 | 106560084 | rs4523664 | 1.098 | 1.428 | -1.166 | -1.679 | -0.091 | -0.133 | -0.133 | 0 | 0.24 | 0.02 | 0.28 | 0.64 | 0.33 | 0.37 |
| 11 | 106566114 | rs11211867 | 0.214 | 1.212 | -0.661 | -1.438 | 0.006 | -0.011 | -0.011 | 0 | 0.13 | 0.21 | 0.24 | 0.07 | 0.15 | 0.05 |
| 11 | 106567017 | rs10789536 | -1.307 | 0.605 | -1.467 | -1.853 | -0.013 | 0.056 | 0.056 | 0.577 | 0.44 | 0.84 | 0.48 | 0.31 | 0.38 | 0.38 |
| 11 | 106575947 | rs597384 | 0.421 | 0.241 | -1.746 | -2.05 | 0.003 | 0.009 | 0.009 | 0 | 0.17 | 0.03 | 0.1 | 0.1 | 0.13 | 0.16 |
| 11 | 106576657 | rs602159 | -1.183 | 0.317 | -1.778 | -2.183 | 0.033 | 0.108 | 0.108 | 0.673 | 0.51 | 0.84 | 0.47 | 0.3 | 0.37 | 0.4 |
| 11 | 106591126 | rs7931295 | -1.417 | 0.387 | -1.372 | -2.634 | -0.001 | -0.005 | -0.005 | 0 | 0.59 | 0.57 | 0.46 | 0.14 | 0.54 | 0.5 |
| 11 | 106594786 | rs10789539 | -1.157 | 0.059 | -0.995 | -2.586 | 0.002 | 0.006 | 0.006 | 0 | 0.5 | 0.48 | 0.41 | 0.14 | 0.46 | 0.47 |
| 11 | 106596424 | rs17564612 | 2.25 | 0.082 | -1.174 | -2.888 | -0.023 | 0.051 | 0.051 | 0 | 0.09 | 0.01 | 0.08 | 0 | 0.13 | 0.03 |
| 11 | 106605259 | rs7103063 | 0.451 | 0.202 | -0.592 | -2.671 | -0.002 | 0.014 | 0.014 | 0 | 0.58 | 0.62 | 0.67 | 0.89 | 0.6 | 0.6 |
| 11 | 106611905 | rs4121676 | 1.029 | 0.248 | -0.666 | -2.702 | 0.008 | 0.081 | 0.081 | 0.309 | 0.49 | 0.25 | 0.55 | 0.74 | 0.59 | 0.47 |
| 11 | 106615180 | rs4121677 | -0.069 | 0.671 | -0.8 | -2.696 | -0.049 | 0.016 | 0.016 | 0 | 0.2 | 0.04 | 0.2 | 0.1 | 0.26 | 0.29 |
| 11 | 106616708 | rs3858425 | -1.217 | 0.528 | -0.691 | -2.768 | 0.019 | -0.051 | -0.051 | 0 | 0.33 | 0.49 | 0.35 | 0.14 | 0.35 | 0.36 |
| 11 | 106618077 | rs17650551 | 1.439 | 0.654 | -0.97 | -2.69 | -0.01 | -0.013 | -0.013 | 0 | 0.07 | 0.02 | 0.11 | 0.09 | 0.09 | 0.17 |
| 11 | 106618519 | rs12800630 | 0.096 | 0.458 | -0.942 | -2.671 | -0.047 | 0.015 | 0.015 | 0 | 0.2 | 0.04 | 0.2 | 0.1 | 0.26 | 0.29 |
| 11 | 106619739 | rs3907374 | 1.841 | 0.316 | -0.932 | -2.833 | -0.006 | 0.07 | 0.07 | 0 | 0.44 | 0.2 | 0.52 | 0.72 | 0.51 | 0.4 |
| 11 | 106620010 | rs11608184 | -0.068 | 0.853 | -0.902 | -2.569 | -0.04 | 0.015 | 0.015 | 0 | 0.2 | 0.05 | 0.2 | 0.1 | 0.26 | 0.28 |
| 11 | 106620202 | rs6588924 | -0.259 | 0.317 | -0.138 | -2.151 | 0.016 | -0.068 | -0.068 | 0 | 0.11 | 0.3 | 0.1 | 0.01 | 0.08 | 0.06 |
| 11 | 106632370 | rs11824629 | -0.457 | -0.062 | -0.21 | -2.031 | 0.014 | -0.006 | -0.006 | 0 | 0.1 | 0.44 | 0.14 | 0.04 | 0.05 | 0.05 |
| 11 | 106637578 | rs10789545 | 0.041 | 0.19 | -1.212 | -0.729 | -0.011 | -0.009 | -0.009 | 0.646 | 0.29 | 0.26 | 0.42 | 0.39 | 0.37 | 0.29 |
| 11 | 106639648 | rs11211902 | 1.182 | 0.065 | -1.149 | -0.847 | -0.01 | 0.055 | 0.055 | 0 | 0.32 | 0.46 | 0.34 | 0.49 | 0.34 | 0.44 |
| 11 | 106640493 | rs1158122 | -0.229 | -0.037 | -1.332 | -1.205 | 0.039 | 0.062 | 0.062 | 0 | 0.82 | 0.95 | 0.95 | 0.99 | 0.92 | 0.82 |
| 11 | 106641890 | rs11211907 | 0.055 | 0.112 | -1.382 | -1.022 | -0.014 | 0.014 | 0.014 | 0 | 0.18 | 0.13 | 0.16 | 0.1 | 0.19 | 0.16 |
| 11 | 106644803 | rs1941823 | -0.931 | -0.328 | -1.2 | -0.622 | 0.018 | 0.076 | 0.076 | 0 | 0.87 | 1 | 0.96 | 0.99 | 0.93 | 0.87 |
| 11 | 106656863 | rs901779 | 0.487 | 0.225 | -0.733 | -0.643 | 0 | 0 | 0 | 0 | 0.49 | 0.52 | 0.61 | 0.82 | 0.56 | 0.59 |
| 11 | 106673649 | rs1487909 | 0.191 | -0.183 | -0.481 | -0.404 | -0.019 | -0.02 | -0.02 | 0 | 0.77 | 0.64 | 0.85 | 0.86 | 0.89 | 0.72 |
| 11 | 106678032 | rs2046682 | 0.835 | -0.448 | -0.443 | -0.804 | 0.001 | 0.01 | 0.01 | 0 | 0.51 | 0.45 | 0.72 | 0.76 | 0.59 | 0.61 |
| 11 | 106679767 | rs2029302 | -1.249 | -0.478 | -0.321 | -0.821 | -0.009 | -0.071 | -0.071 | 0 | 0.84 | 0.71 | 0.89 | 0.99 | 0.9 | 0.81 |
| 11 | 106682935 | rs4277082 | 0.078 | -0.328 | -0.318 | -0.878 | 0.001 | 0.003 | 0.003 | 0 | 0.28 | 0.19 | 0.15 | 0.22 | 0.27 | 0.18 |
| 11 | 106685081 | rs1487890 | 0.794 | -0.301 | -0.457 | -0.842 | -0.008 | 0.012 | 0.012 | 0 | 0.16 | 0.12 | 0.11 | 0.1 | 0.2 | 0.07 |
| 11 | 106690102 | rs10789547 | 1.265 | 0.183 | -0.466 | -0.428 | 0.055 | 0.051 | 0.051 | 0 | 0.15 | 0.26 | 0.5 | 0.33 | 0.32 | 0.26 |
| 11 | 106692228 | rs1487891 | 0.591 | -0.076 | -0.562 | -0.548 | 0.012 | 0.104 | 0.104 | 0.013 | 0.22 | 0.4 | 0.44 | 0.46 | 0.28 | 0.26 |
| 11 | 106692416 | rs1487892 | 1.805 | 0.044 | -0.701 | -0.508 | 0.021 | 0.035 | 0.035 | 0 | 0.13 | 0.21 | 0.37 | 0.33 | 0.25 | 0.21 |
| 11 | 106700046 | rs7929737 | 2.793 | 0.017 | -1.016 | -0.67 | -0.029 | -0.083 | -0.083 | 0.001 | 0.23 | 0.03 | 0.15 | 0.31 | 0.24 | 0.26 |
| 11 | 106703727 | rs12363827 | 0.804 | 0.02 | -0.877 | -0.663 | -0.02 | 0.018 | 0.018 | 0 | 0.18 | 0.12 | 0.23 | 0.1 | 0.26 | 0.12 |
| 11 | 106713125 | rs2200656 | 0.171 | -0.435 | -0.678 | -0.621 | -0.006 | 0.011 | 0.011 | 0 | 0.63 | 0.46 | 0.5 | 0.54 | 0.63 | 0.56 |
| 11 | 106715879 | rs7943998 | 1.484 | -0.339 | -0.911 | -0.711 | -0.027 | -0.037 | -0.037 | 0 | 0.37 | 0.16 | 0.35 | 0.42 | 0.39 | 0.36 |
| 11 | 106715928 | rs10502078 | -0.649 | -0.486 | -0.61 | -0.836 | 0.003 | 0.004 | 0.004 | 0.614 | 0.34 | 0.31 | 0.19 | 0.13 | 0.27 | 0.22 |
| 11 | 106718739 | rs7483731 | 1.417 | -0.157 | -0.292 | -0.458 | 0.075 | 0.03 | 0.03 | 0 | 0.14 | 0.27 | 0.29 | 0.23 | 0.29 | 0.25 |
| 11 | 106718877 | rs10502079 | 0.679 | -0.252 | -0.365 | -0.491 | 0.001 | -0.002 | -0.002 | 0.051 | 0.21 | 0.2 | 0.13 | 0.13 | 0.23 | 0.16 |
| 11 | 106722884 | rs7950959 | 1.221 | -0.27 | -0.381 | -0.472 | -0.002 | 0.004 | 0.004 | 0 | 0.39 | 0.33 | 0.6 | 0.64 | 0.4 | 0.41 |
| 11 | 106725302 | rs11211946 | 2.628 | -0.516 | -0.607 | -0.444 | -0.014 | -0.133 | -0.133 | 0 | 0.24 | 0.03 | 0.37 | 0.52 | 0.24 | 0.31 |
| 11 | 106727257 | rs11211948 | 1.477 | -0.523 | -0.535 | -0.443 | 0.009 | -0.016 | -0.016 | 0 | 0.36 | 0.25 | 0.24 | 0.42 | 0.33 | 0.36 |
| 11 | 106729672 | rs1971268 | 1.001 | -0.455 | -0.653 | -0.438 | -0.006 | -0.003 | -0.003 | 0 | 0.46 | 0.32 | 0.29 | 0.44 | 0.46 | 0.45 |
| 11 | 106732268 | rs1944193 | -1.473 | -0.815 | -1.067 | -0.401 | 0.032 | 0.061 | 0.061 | 0 | 0.81 | 0.91 | 0.93 | 1 | 0.88 | 0.84 |

| | | | | | | | | | | | | | | | | |
|---|---|---|---|---|---|---|---|---|---|---|---|---|---|---|---|---|
| 11 | 106756696 | rs10749881 | 0.454 | -0.828 | -0.951 | -0.34 | -0.001 | 0 | 0 | 0 | 0.61 | 0.58 | 0.57 | 0.67 | 0.75 | 0.69 |
| 11 | 106761487 | rs11211962 | -0.46 | -0.791 | -1.007 | -0.453 | 0.007 | -0.007 | -0.007 | 0 | 0.4 | 0.5 | 0.44 | 0.36 | 0.53 | 0.4 |
| 11 | 106767477 | rs7127202 | -1.273 | -0.95 | -1.225 | -0.377 | 0.058 | 0.088 | 0.088 | 0 | 0.56 | 0.87 | 0.78 | 0.74 | 0.67 | 0.55 |
| 11 | 106771576 | rs7114159 | 2.562 | -0.856 | -1.186 | -0.28 | -0.019 | -0.054 | -0.054 | 0 | 0.22 | 0.04 | 0.14 | 0.25 | 0.22 | 0.29 |
| 11 | 106775667 | rs12364559 | 0.452 | -0.638 | -0.864 | -0.094 | -0.02 | -0.028 | -0.028 | 2.043 | 0.18 | 0.33 | 0.11 | 0.11 | 0.12 | 0.11 |
| 11 | 106775861 | rs11211968 | -0.543 | -0.7 | -0.942 | -0.16 | 0.001 | -0.001 | -0.001 | 3.11 | 0.4 | 0.53 | 0.67 | 0.63 | 0.55 | 0.45 |
| 11 | 106780118 | rs2156589 | -1.486 | -0.731 | -0.98 | -0.148 | 0.056 | 0.089 | 0.089 | 0 | 0.57 | 0.86 | 0.78 | 0.74 | 0.66 | 0.55 |
| 11 | 106780730 | rs9971417 | -0.538 | -0.396 | -0.86 | 0.057 | 0 | -0.001 | -0.001 | 0.997 | 0.4 | 0.54 | 0.67 | 0.63 | 0.55 | 0.45 |
| 11 | 106805698 | rs1944210 | -0.629 | -0.509 | -0.819 | 0 | 0.013 | 0.002 | 0.002 | 0 | 0.51 | 0.5 | 0.52 | 0.48 | 0.65 | 0.48 |
| 11 | 106818037 | rs1455588 | -0.011 | 0.089 | -0.866 | 0 | -0.025 | -0.005 | -0.005 | 0 | 0.29 | 0.2 | 0.37 | 0.35 | 0.41 | 0.31 |
| 11 | 106828741 | rs10890627 | -1.131 | -0.322 | -1.297 | -0.364 | 0.027 | 0.082 | 0.082 | 0 | 0.82 | 0.93 | 0.94 | 0.99 | 0.89 | 0.87 |
| 11 | 106831384 | rs1471045 | 0.204 | -0.411 | -1.106 | -0.044 | 0.006 | 0.005 | 0.005 | 0 | 0.31 | 0.3 | 0.31 | 0.41 | 0.39 | 0.3 |
| 11 | 106831656 | rs12364403 | 1.903 | -0.191 | -1.416 | -0.275 | -0.07 | -0.059 | -0.059 | 1.208 | 0.11 | 0.01 | 0.15 | 0.24 | 0.25 | 0.21 |
| 11 | 106831970 | rs7936117 | 0.04 | -0.353 | -1.08 | -0.302 | 0.025 | -0.028 | -0.028 | 0 | 0.23 | 0.39 | 0.23 | 0.13 | 0.25 | 0.2 |
| 11 | 106833318 | rs11604450 | 0.343 | -0.274 | -1.091 | -0.216 | 0.004 | 0 | 0 | 0 | 0.31 | 0.3 | 0.31 | 0.36 | 0.39 | 0.3 |
| 11 | 106837790 | rs6588940 | 0.819 | -0.483 | -0.882 | -0.196 | -0.01 | -0.03 | -0.03 | 0.158 | 0.62 | 0.47 | 0.51 | 0.73 | 0.69 | 0.72 |
| 11 | 106842124 | rs11602032 | 1.298 | -0.647 | -1.118 | -0.344 | 0.023 | 0.062 | 0.062 | 0 | 0.15 | 0.05 | 0.06 | 0.01 | 0.1 | 0.11 |
| 11 | 106845361 | rs982443 | -0.088 | -0.649 | -0.833 | -0.198 | 0.018 | -0.002 | -0.002 | 0 | 0.74 | 0.73 | 0.69 | 0.71 | 0.87 | 0.78 |
| 11 | 106847190 | rs10789557 | 0.167 | -0.463 | -0.731 | -0.057 | 0.017 | 0.006 | 0.006 | 0 | 0.29 | 0.33 | 0.31 | 0.36 | 0.38 | 0.28 |
| 11 | 106852187 | rs3133355 | 1.602 | -0.366 | -0.891 | 0.006 | 0.008 | -0.017 | -0.017 | 0 | 0.31 | 0.28 | 0.4 | 0.51 | 0.25 | 0.38 |
| 11 | 106862323 | rs17106293 | 1.128 | -0.625 | -0.69 | 0.125 | 0.03 | -0.048 | -0.048 | 0.836 | 0.11 | 0.18 | 0.17 | 0.01 | 0.18 | 0.06 |
| 11 | 106863491 | rs2510595 | 1.751 | -0.383 | -1.078 | 0.21 | -0.041 | -0.08 | -0.08 | 0 | 0.31 | 0.16 | 0.49 | 0.51 | 0.43 | 0.36 |
| 11 | 106868362 | rs3100765 | -1.993 | -0.347 | -1.319 | -0.034 | -0.082 | -0.005 | -0.005 | 0 | 0.71 | 0.95 | 0.75 | 0.76 | 0.59 | 0.68 |
| 11 | 106869973 | rs1455590 | -0.379 | -0.437 | -0.578 | -0.086 | -0.021 | -0.003 | -0.003 | 1.594 | 0.32 | 0.6 | 0.26 | 0.34 | 0.21 | 0.27 |
| 11 | 106877818 | rs11212003 | 1.537 | -0.805 | -0.758 | 0.072 | 0 | -0.011 | -0.011 | 0 | 0.11 | 0.02 | 0.09 | 0.11 | 0.1 | 0.06 |
| 11 | 106886058 | rs2445114 | -0.746 | -0.753 | -0.735 | 0.058 | -0.023 | -0.047 | -0.047 | 0 | 0.37 | 0.13 | 0.48 | 0.42 | 0.38 | 0.43 |
| 11 | 106895032 | rs1455592 | -0.612 | -0.801 | -0.692 | 0.135 | -0.001 | -0.004 | -0.004 | 0.037 | 0.32 | 0.3 | 0.14 | 0.12 | 0.2 | 0.32 |
| 11 | 106898141 | rs2462410 | -0.791 | -0.732 | -0.6 | 0.148 | -0.027 | -0.051 | -0.051 | 0.698 | 0.41 | 0.13 | 0.49 | 0.42 | 0.4 | 0.49 |
| 11 | 106907475 | rs2445100 | -1.828 | -0.271 | -0.525 | 0.149 | -0.013 | 0.035 | 0.035 | 0 | 0.71 | 0.85 | 0.83 | 0.78 | 0.69 | 0.69 |
| 11 | 106907859 | rs2462417 | 0.957 | -0.277 | -0.717 | -0.058 | -0.026 | 0.062 | 0.062 | 0 | 0.11 | 0.02 | 0.04 | 0.01 | 0.08 | 0.08 |
| 11 | 106908817 | rs2465768 | 0.828 | -0.289 | -0.588 | -0.063 | -0.003 | 0.001 | 0.001 | 0 | 0.12 | 0.07 | 0.05 | 0.01 | 0.08 | 0.08 |
| 11 | 106909049 | rs2513112 | -2.033 | -0.29 | -0.615 | -0.052 | -0.013 | 0.034 | 0.034 | 0 | 0.71 | 0.85 | 0.83 | 0.78 | 0.69 | 0.69 |
| 11 | 106911794 | rs10789559 | -1.17 | -0.288 | -0.277 | 0.02 | -0.009 | 0.056 | 0.056 | 0 | 0.59 | 0.7 | 0.77 | 0.78 | 0.61 | 0.59 |
| 11 | 106918138 | rs1455597 | -0.827 | -0.33 | -0.187 | 0.082 | 0.002 | -0.027 | -0.027 | 0 | 0.37 | 0.35 | 0.29 | 0.13 | 0.34 | 0.33 |
| 11 | 106919873 | rs10890634 | 0.18 | -0.345 | -0.074 | 0.098 | -0.002 | 0.001 | 0.001 | 0 | 0.33 | 0.4 | 0.52 | 0.66 | 0.35 | 0.34 |
| 11 | 106920666 | rs10890636 | 1.084 | -0.407 | -0.02 | 0.39 | -0.01 | 0.049 | 0.049 | 0 | 0.17 | 0.23 | 0.33 | 0.45 | 0.13 | 0.13 |
| 11 | 106930556 | rs2513131 | -1.486 | 0.014 | -0.134 | 0.263 | -0.004 | 0.021 | 0.021 | 0 | 0.7 | 0.75 | 0.82 | 0.79 | 0.69 | 0.67 |
| 11 | 106936178 | rs7126614 | -0.626 | 0.297 | -0.685 | 0.147 | 0.003 | -0.013 | -0.013 | 1.796 | 0.25 | 0.26 | 0.24 | 0.13 | 0.26 | 0.25 |
| 11 | 106938611 | rs2513125 | -1.374 | 0.082 | -1.306 | 0.496 | -0.012 | 0.001 | 0.001 | 0 | 0.7 | 0.91 | 0.72 | 0.64 | 0.69 | 0.69 |
| 11 | 106941553 | rs1840572 | -0.013 | 0.013 | -0.99 | 0.346 | -0.049 | 0.091 | 0.091 | 0 | 0.18 | 0.04 | 0.19 | 0.02 | 0.22 | 0.19 |
| 11 | 106947600 | rs11212024 | -0.784 | -0.104 | -0.488 | 0.803 | 0 | 0.231 | 0.231 | 1.16 | 0.08 | 0.51 | 0.05 | 0.19 | 0.01 | 0.09 |
| 11 | 106956255 | rs1455595 | -1.039 | -0.122 | -0.584 | 0.829 | -0.001 | 0 | 0 | 0 | 0.38 | 0.59 | 0.27 | 0.3 | 0.33 | 0.43 |
| 11 | 106958304 | rs7124243 | -1.465 | -0.219 | 0.439 | 1.317 | -0.007 | 0.224 | 0.224 | 0.695 | 0.09 | 0.51 | 0.05 | 0.19 | 0.01 | 0.1 |
| 11 | 106972340 | rs2513116 | 0.257 | 0.149 | -0.269 | 1.142 | 0.001 | -0.007 | -0.007 | 5.567 | 0.63 | 0.36 | 0.53 | 0.5 | 0.63 | 0.66 |
| 11 | 106975897 | rs2019447 | 0.591 | 0.19 | -0.173 | 1.201 | 0 | -0.001 | -0.001 | 0 | 0.53 | 0.5 | 0.55 | 0.53 | 0.48 | 0.6 |
| 11 | 106983728 | rs1895729 | -0.25 | 0.139 | 0.016 | 1.235 | -0.006 | -0.01 | -0.01 | 0.253 | 0.31 | 0.29 | 0.38 | 0.42 | 0.41 | 0.27 |
| 11 | 106986700 | rs1425241 | 0.158 | 0.159 | -0.154 | 1.177 | -0.019 | -0.033 | -0.033 | 1.613 | 0.16 | 0.22 | 0.06 | 0.05 | 0.11 | 0.14 |
| 11 | 106987593 | rs11212035 | 0.374 | 0.375 | -0.356 | 1.389 | 0.004 | 0.005 | 0.005 | 0.315 | 0.13 | 0.13 | 0.11 | 0.21 | 0.22 | 0.17 |
| 11 | 106996856 | rs4754186 | -0.458 | 0.258 | -0.323 | 1.295 | -0.016 | -0.034 | -0.034 | 0 | 0.3 | 0.23 | 0.43 | 0.44 | 0.41 | 0.27 |
| 11 | 107000504 | rs10789569 | -1.083 | 0.437 | -0.433 | 1.102 | 0 | 0.001 | 0.001 | 0 | 0.48 | 0.43 | 0.44 | 0.47 | 0.52 | 0.46 |
| 11 | 107006900 | rs12420075 | -0.076 | 0.532 | -0.809 | 1.098 | -0.054 | -0.043 | -0.043 | 0 | 0.2 | 0.1 | 0.22 | 0.29 | 0.36 | 0.22 |
| 11 | 107010273 | rs879295 | 0.081 | 0.127 | -1.031 | 1.448 | 0.005 | -0.073 | -0.073 | 0 | 0.16 | 0.06 | 0.26 | 0.42 | 0.14 | 0.19 |
| 11 | 107011261 | rs7951809 | 0.248 | -0.68 | -0.546 | 0.899 | -0.006 | -0.02 | -0.02 | 0.147 | 0.23 | 0.26 | 0.18 | 0.07 | 0.18 | 0.2 |
| 11 | 107012551 | rs17106517 | -0.366 | -0.909 | -0.051 | 0.816 | -0.063 | -0.09 | -0.09 | 0.034 | 0.27 | 0.52 | 0.15 | 0.12 | 0.16 | 0.29 |
| 11 | 107013447 | rs10890649 | -1.311 | -0.888 | -0.087 | 0.862 | -0.005 | 0.01 | 0.01 | 0 | 0.48 | 0.62 | 0.36 | 0.41 | 0.51 | 0.51 |
| 11 | 107014870 | rs4271358 | 0.572 | -0.829 | -0.301 | 0.781 | -0.008 | -0.018 | -0.018 | 0 | 0.21 | 0.25 | 0.12 | 0.07 | 0.15 | 0.18 |
| 11 | 107026329 | rs10789570 | 0.464 | -0.737 | -0.219 | 0.989 | -0.057 | -0.028 | -0.028 | 0.075 | 0.2 | 0.12 | 0.22 | 0.21 | 0.37 | 0.23 |
| 11 | 107029182 | rs11530514 | -0.017 | -0.872 | -0.501 | 1.028 | 0.031 | -0.073 | -0.073 | 0 | 0.16 | 0.06 | 0.29 | 0.42 | 0.13 | 0.16 |
| 11 | 107029951 | rs1425238 | 0.632 | -0.773 | -0.424 | 0.926 | -0.012 | -0.026 | -0.026 | 0 | 0.2 | 0.27 | 0.11 | 0.06 | 0.14 | 0.18 |
| 11 | 107031714 | rs10890655 | -0.59 | -0.449 | -0.503 | 1.04 | 0.046 | 0.07 | 0.07 | 0 | 0.28 | 0.5 | 0.25 | 0.4 | 0.37 | 0.34 |
| 11 | 107035980 | rs1425244 | 0.185 | -0.449 | -1.003 | 0.941 | 0.002 | 0 | 0 | 0 | 0.23 | 0.26 | 0.23 | 0.25 | 0.37 | 0.23 |
| 11 | 107037976 | rs10789572 | 0.027 | -0.374 | -1.116 | 0.958 | 0.035 | 0.018 | 0.018 | 0 | 0.24 | 0.35 | 0.23 | 0.25 | 0.37 | 0.23 |
| 11 | 107049000 | rs6588946 | -1.662 | -0.232 | -1.303 | 0.459 | -0.061 | 0.11 | 0.11 | 0 | 0.67 | 0.95 | 0.69 | 0.86 | 0.66 | 0.71 |
| 11 | 107049547 | rs7934656 | -0.758 | 0.329 | -0.806 | 1.101 | 0.016 | 0.023 | 0.023 | 0 | 0.34 | 0.42 | 0.43 | 0.48 | 0.42 | 0.37 |
| 11 | 107053192 | rs6421539 | -0.71 | 0.221 | -1.899 | 0.774 | -0.015 | -0.004 | -0.004 | 3.421 | 0.63 | 0.93 | 0.45 | 0.74 | 0.64 | 0.62 |
| 11 | 107054642 | rs7926602 | -0.316 | 0.228 | -1.183 | 1.013 | -0.012 | 0.001 | 0.001 | 0 | 0.39 | 0.35 | 0.4 | 0.39 | 0.5 | 0.36 |
| 11 | 107064783 | rs1997006 | -1.512 | -0.047 | -1.754 | 0.663 | -0.019 | -0.004 | -0.004 | 0 | 0.77 | 0.99 | 0.65 | 0.85 | 0.81 | 0.66 |
| 11 | 107067987 | rs1490936 | -0.249 | -0.403 | -0.942 | 0.622 | 0.016 | 0.02 | 0.02 | 0.003 | 0.56 | 0.39 | 0.53 | 0.7 | 0.69 | 0.55 |
| 11 | 107078998 | rs1490938 | 0.663 | -0.424 | -0.994 | 0.63 | -0.067 | -0.035 | -0.035 | 2.692 | 0.32 | 0.16 | 0.38 | 0.4 | 0.48 | 0.33 |

| | | | | | | | | | | | | | | | | |
|---|---|---|---|---|---|---|---|---|---|---|---|---|---|---|---|---|
| 11 | 107080988 | rs11212082 | 0.872 | -0.413 | -0.649 | 0.899 | 0.006 | -0.009 | -0.009 | 0 | 0.22 | 0.18 | 0.13 | 0.23 | 0.18 | 0.21 |
| 11 | 107087871 | rs726073 | -0.782 | -0.451 | -0.571 | 0.86 | 0.024 | -0.004 | -0.004 | 0 | 0.44 | 0.67 | 0.27 | 0.44 | 0.34 | 0.34 |
| 11 | 107090112 | rs4754193 | 0.473 | -0.494 | 0.123 | 0.862 | -0.014 | 0.013 | 0.013 | 0.072 | 0.24 | 0.33 | 0.14 | 0.24 | 0.18 | 0.21 |
| 11 | 107093733 | rs7936346 | -1.819 | -0.54 | -0.043 | 0.833 | 0.023 | 0.006 | 0.006 | 0.142 | 0.75 | 0.82 | 0.64 | 0.79 | 0.82 | 0.67 |
| 11 | 107093998 | rs17585166 | 2.254 | -0.595 | -0.159 | 0.749 | -0.042 | 0.063 | 0.063 | 0.113 | 0.13 | 0.01 | 0.1 | 0.03 | 0.15 | 0.14 |
| 11 | 107094572 | rs11212086 | 0.513 | -0.544 | -0.006 | 0.839 | 0 | -0.008 | -0.008 | 0 | 0.23 | 0.2 | 0.16 | 0.25 | 0.22 | 0.22 |
| 11 | 107097881 | rs11212087 | -0.45 | -0.461 | 0.315 | 0.841 | -0.003 | 0.014 | 0.014 | 0.083 | 0.32 | 0.36 | 0.19 | 0.39 | 0.27 | 0.31 |
| 11 | 107101960 | rs4753790 | 0.105 | 0.172 | -0.013 | 0.912 | 0.004 | -0.011 | -0.011 | 0 | 0.26 | 0.43 | 0.29 | 0.26 | 0.3 | 0.14 |
| 11 | 107114975 | rs6588953 | 1.61 | 0.168 | -0.069 | 0.636 | -0.005 | -0.064 | -0.064 | 0 | 0.11 | 0.25 | 0.09 | 0 | 0.09 | 0.02 |
| 11 | 107117740 | rs6588955 | 0.086 | 0.143 | -0.155 | 0.82 | 0 | -0.058 | -0.058 | 0 | 0.26 | 0.38 | 0.21 | 0.16 | 0.29 | 0.14 |
| 11 | 107118205 | rs4265572 | -2.177 | 0.07 | -0.127 | 0.826 | 0.004 | 0.002 | 0.002 | 0.089 | 0.74 | 0.75 | 0.64 | 0.79 | 0.82 | 0.66 |
| 11 | 107122289 | rs11602728 | -0.597 | 0.132 | -0.478 | 0.858 | 0.012 | 0.035 | 0.035 | 0 | 0.48 | 0.37 | 0.43 | 0.63 | 0.53 | 0.52 |
| 11 | 107123733 | rs4609570 | -0.575 | -0.01 | -0.511 | 1.131 | 0.019 | 0.117 | 0.117 | 3.139 | 0.5 | 0.75 | 0.36 | 0.32 | 0.49 | 0.34 |
| 11 | 107125357 | rs10890676 | 0.875 | 0.026 | -0.534 | 1.035 | -0.004 | -0.003 | -0.003 | 0 | 0.21 | 0.19 | 0.19 | 0.22 | 0.27 | 0.23 |
| 11 | 107127162 | rs10749888 | -1.803 | 0.173 | -0.757 | 1.282 | -0.032 | -0.091 | -0.091 | 0 | 0.76 | 0.99 | 0.62 | 0.72 | 0.78 | 0.65 |
| 11 | 107128014 | rs7109038 | -1.02 | 0.111 | -0.703 | 1.138 | -0.022 | -0.169 | -0.169 | 0.2 | 0.67 | 0.97 | 0.53 | 0.5 | 0.72 | 0.54 |
| 11 | 107132505 | rs12801193 | 0.266 | 0.268 | -0.062 | 0.836 | 0.003 | -0.016 | -0.016 | 0 | 0.17 | 0.18 | 0.14 | 0.16 | 0.2 | 0.17 |
| 11 | 107134499 | rs12790621 | 0.43 | 0.207 | 0.023 | 0.841 | -0.021 | -0.003 | -0.003 | 0.794 | 0.13 | 0.19 | 0.07 | 0.17 | 0.09 | 0.16 |
| 11 | 107135449 | rs4754204 | -0.089 | 0.18 | -0.156 | 0.898 | 0.023 | 0.019 | 0.019 | 0 | 0.51 | 0.69 | 0.47 | 0.58 | 0.56 | 0.35 |
| 11 | 107140595 | rs7111266 | 0.02 | 0.103 | 0.107 | 0.866 | -0.008 | -0.011 | -0.011 | 0 | 0.34 | 0.54 | 0.29 | 0.29 | 0.32 | 0.16 |
| 11 | 107149004 | rs7930096 | -1.438 | 0.118 | 0.347 | 0.878 | 0.001 | 0.04 | 0.04 | 0 | 0.3 | 0.55 | 0.36 | 0.49 | 0.37 | 0.41 |
| 11 | 107149097 | rs7929893 | -0.13 | -0.047 | 0.612 | 0.897 | 0.001 | 0.004 | 0.004 | 0 | 0.19 | 0.22 | 0.18 | 0.2 | 0.23 | 0.28 |
| 11 | 107152661 | rs7945392 | -1.241 | -0.059 | 0.591 | 0.913 | -0.003 | 0.059 | 0.059 | 0.043 | 0.27 | 0.56 | 0.32 | 0.49 | 0.31 | 0.36 |
| 11 | 107154315 | rs10789592 | 0.744 | -0.019 | 0.418 | 0.861 | 0.006 | 0.004 | 0.004 | 0 | 0.24 | 0.28 | 0.19 | 0.29 | 0.27 | 0.22 |
| 11 | 107158740 | rs17106711 | 0.373 | 0.072 | 0.486 | 0.897 | 0.052 | 0.045 | 0.045 | 0 | 0.08 | 0.19 | 0.12 | 0.12 | 0.14 | 0.12 |
| 11 | 107159302 | rs11212132 | 1.156 | 0.096 | 0.289 | 0.891 | -0.033 | -0.03 | -0.03 | 1.132 | 0.22 | 0.12 | 0.19 | 0.28 | 0.27 | 0.23 |
| 11 | 107170877 | rs12276737 | 0.716 | 0.114 | 0.342 | 0.892 | -0.005 | 0.014 | 0.014 | 0 | 0.16 | 0.09 | 0.13 | 0.1 | 0.2 | 0.17 |
| 11 | 107177440 | rs621312 | -1.339 | 0.039 | 0.196 | 0.862 | -0.038 | -0.041 | -0.041 | 0 | 0.54 | 0.8 | 0.46 | 0.5 | 0.54 | 0.42 |
| 11 | 107181419 | rs10789601 | 0.463 | 0.183 | 0.333 | 0.957 | -0.011 | -0.008 | -0.008 | 0 | 0.26 | 0.2 | 0.19 | 0.29 | 0.28 | 0.26 |
| 11 | 107184123 | rs7930061 | -1 | 0.087 | 0.638 | 1.151 | -0.011 | 0.097 | 0.097 | 0 | 0.22 | 0.48 | 0.26 | 0.43 | 0.23 | 0.31 |
| 11 | 107184828 | rs10749896 | 0.302 | 0.004 | 0.503 | 1.141 | -0.012 | 0 | 0 | 0 | 0.29 | 0.25 | 0.2 | 0.29 | 0.31 | 0.26 |
| 11 | 107186896 | rs4754214 | 0.153 | 0.039 | 0.686 | 1.093 | 0.011 | 0.027 | 0.027 | 0 | 0.18 | 0.34 | 0.17 | 0.21 | 0.2 | 0.27 |
| 11 | 107192257 | rs999985 | 1.098 | 0.067 | 0.183 | 1.022 | -0.046 | -0.028 | -0.028 | 0 | 0.23 | 0.12 | 0.19 | 0.29 | 0.3 | 0.23 |
| 11 | 107195581 | rs10749902 | 0.546 | 0.25 | 0.203 | 1.011 | -0.013 | -0.005 | -0.005 | 0 | 0.27 | 0.2 | 0.18 | 0.29 | 0.28 | 0.26 |
| 11 | 107197640 | rs3758911 | 0.445 | 0.217 | 0.266 | 1.016 | -0.012 | -0.002 | -0.002 | 0 | 0.27 | 0.21 | 0.2 | 0.29 | 0.31 | 0.26 |
| 11 | 107202592 | rs715415 | 0.929 | 0.258 | 0.183 | 0.91 | -0.003 | 0.004 | 0.004 | 0 | 0.14 | 0.09 | 0.12 | 0.1 | 0.18 | 0.17 |
| 11 | 107222904 | rs614594 | -0.773 | 0.143 | -0.042 | 0.976 | -0.017 | -0.003 | -0.003 | 0.832 | 0.48 | 0.79 | 0.44 | 0.53 | 0.5 | 0.38 |
| 11 | 107228533 | rs686806 | -1.957 | 0.114 | -0.136 | 0.953 | -0.022 | -0.029 | -0.029 | 0 | 0.76 | 0.98 | 0.64 | 0.79 | 0.81 | 0.63 |
| 11 | 107231605 | rs11212182 | 1.082 | 0.144 | -0.152 | 0.959 | -0.048 | -0.008 | -0.008 | 0.701 | 0.23 | 0.12 | 0.19 | 0.26 | 0.3 | 0.23 |
| 11 | 107249735 | rs2432864 | -0.659 | 0.114 | 0.047 | 0.952 | -0.005 | 0.005 | 0.005 | 0.423 | 0.47 | 0.72 | 0.43 | 0.53 | 0.49 | 0.37 |
| 11 | 107250851 | rs9787772 | 0.362 | 0.086 | 0.5 | 1.04 | 0.008 | 0.026 | 0.026 | 0 | 0.18 | 0.31 | 0.16 | 0.21 | 0.2 | 0.27 |
| 11 | 107254508 | rs555570 | -0.57 | 0.084 | 0.348 | 0.977 | -0.002 | 0.016 | 0.016 | 0.225 | 0.25 | 0.65 | 0.3 | 0.53 | 0.27 | 0.33 |
| 11 | 107258865 | rs520203 | -0.803 | 0.067 | 0.378 | 0.922 | -0.006 | 0.005 | 0.005 | 0.366 | 0.46 | 0.72 | 0.43 | 0.53 | 0.5 | 0.38 |
| 11 | 107259071 | rs10789621 | 0.915 | 0.123 | 0.376 | 0.789 | -0.009 | 0.002 | 0.002 | 0 | 0.24 | 0.2 | 0.2 | 0.26 | 0.3 | 0.23 |
| 11 | 107263213 | rs11212199 | 1.55 | 0.118 | 0.299 | 0.766 | -0.026 | 0.099 | 0.099 | 0.259 | 0.21 | 0.07 | 0.13 | 0 | 0.22 | 0.04 |
| 11 | 107274331 | rs622862 | -1.436 | 0.121 | 0.723 | 1.041 | -0.006 | 0.007 | 0.007 | 0.002 | 0.69 | 0.68 | 0.54 | 0.55 | 0.77 | 0.59 |
| 11 | 107275251 | rs11212206 | 1.156 | 0.151 | 0.696 | 1.012 | -0.046 | -0.008 | -0.008 | 1.411 | 0.23 | 0.12 | 0.19 | 0.26 | 0.3 | 0.23 |
| 11 | 107275522 | rs11212207 | 0.756 | 0.06 | 0.769 | 1.029 | -0.009 | 0.003 | 0.003 | 0 | 0.26 | 0.21 | 0.19 | 0.26 | 0.28 | 0.26 |
| 11 | 107278292 | rs2042535 | -0.315 | 0.09 | 0.819 | 1.036 | 0.001 | -0.012 | -0.012 | 0 | 0.42 | 0.47 | 0.34 | 0.29 | 0.46 | 0.33 |
| 11 | 107284386 | rs623599 | -0.564 | 0.091 | 0.781 | 0.999 | -0.001 | -0.027 | -0.027 | 0.538 | 0.43 | 0.47 | 0.3 | 0.19 | 0.44 | 0.34 |
| 11 | 107299631 | rs659040 | 1.062 | 0.134 | 0.477 | 0.781 | -0.001 | 0.038 | 0.038 | 0.736 | 0.23 | 0.16 | 0.14 | 0 | 0.22 | 0.04 |
| 11 | 107306076 | rs514117 | 1.062 | 0.134 | 0.477 | 0.781 | -0.001 | 0.038 | 0.038 | 0.736 | 0.23 | 0.16 | 0.14 | 0 | 0.22 | 0.04 |
| 11 | 107315880 | rs12274630 | -0.67 | 0.129 | 0.279 | 0.867 | 0.004 | 0.056 | 0.056 | 0.075 | 0.07 | 0.17 | 0.05 | 0.1 | 0.05 | 0.05 |
| 11 | 107316499 | rs11212242 | -0.194 | 0.067 | 0.186 | 0.912 | -0.004 | 0.1 | 0.1 | 0 | 0.23 | 0.49 | 0.27 | 0.41 | 0.24 | 0.31 |
| 11 | 107318332 | rs4754222 | -1.336 | 0.068 | 0.087 | 0.881 | -0.057 | -0.025 | -0.025 | 0 | 0.53 | 0.86 | 0.46 | 0.51 | 0.51 | 0.4 |
| 11 | 107326197 | rs526875 | 1.145 | 0.088 | 0.123 | 0.857 | -0.003 | 0.05 | 0.05 | 0.736 | 0.23 | 0.16 | 0.14 | 0 | 0.22 | 0.04 |
| 11 | 107329116 | rs661085 | 1.218 | 0.097 | 0.105 | 0.865 | -0.006 | 0.065 | 0.065 | 0.596 | 0.23 | 0.13 | 0.13 | 0 | 0.22 | 0.04 |
| 11 | 107330997 | rs7926163 | -0.444 | 0.171 | 0.064 | 0.985 | -0.001 | 0.112 | 0.112 | 0 | 0.24 | 0.5 | 0.3 | 0.51 | 0.27 | 0.33 |
| 11 | 107333949 | rs10890730 | -0.59 | 0.17 | 0.091 | 1.059 | -0.003 | 0.093 | 0.093 | 0 | 0.25 | 0.53 | 0.31 | 0.52 | 0.27 | 0.33 |
| 11 | 107339155 | rs609821 | -1.389 | 0.174 | -0.207 | 0.954 | -0.032 | -0.008 | -0.008 | 1.302 | 0.78 | 0.89 | 0.82 | 0.74 | 0.73 | 0.78 |
| 11 | 107340597 | rs17667107 | 1.088 | 0.124 | -0.03 | 0.961 | -0.002 | 0.002 | 0.002 | 0 | 0.14 | 0.12 | 0.12 | 0.09 | 0.18 | 0.17 |
| 11 | 107340783 | rs12291601 | -0.116 | 0.164 | -0.347 | 0.961 | -0.007 | 0.08 | 0.08 | 0 | 0.23 | 0.51 | 0.27 | 0.4 | 0.24 | 0.31 |
| 11 | 107341540 | rs4753806 | -0.638 | 0.301 | -1.109 | 0.975 | -0.053 | -0.03 | -0.03 | 0.204 | 0.53 | 0.85 | 0.46 | 0.51 | 0.51 | 0.4 |
| 11 | 107344724 | rs624584 | -0.618 | 0.526 | -0.565 | 1.015 | -0.023 | -0.001 | -0.001 | 0 | 0.78 | 0.88 | 0.83 | 0.74 | 0.73 | 0.78 |
| 11 | 107347234 | rs17107008 | -1.423 | -0.085 | 0.311 | 1.397 | -0.033 | 0.138 | 0.138 | 0 | 0.1 | 0.31 | 0.12 | 0.25 | 0.04 | 0.06 |
| 11 | 107349606 | rs11212263 | 0.126 | -0.27 | 0.375 | 0.887 | -0.007 | -0.024 | -0.024 | 0 | 0.14 | 0.2 | 0.1 | 0.07 | 0.11 | 0.12 |
| 11 | 107355600 | rs510366 | -1.073 | 0.041 | -0.3 | 0.512 | -0.047 | 0.105 | 0.105 | 0 | 0.68 | 0.83 | 0.77 | 0.88 | 0.64 | 0.75 |
| 11 | 107360090 | rs495030 | 0.7 | 0.188 | 0.015 | 0.965 | -0.011 | 0.027 | 0.027 | 0 | 0.74 | 0.39 | 0.73 | 0.43 | 0.81 | 0.81 |
| 11 | 107361081 | rs12577579 | 1.602 | 0.101 | -0.496 | 0.767 | -0.059 | -0.069 | -0.069 | 0.199 | 0.24 | 0.01 | 0.2 | 0.25 | 0.23 | 0.39 |

| | | | | | | | | | | | | | | | | |
|---|---|---|---|---|---|---|---|---|---|---|---|---|---|---|---|---|
| 11 | 107362693 | rs489243 | 1.162 | -0.058 | -0.115 | 0.884 | -0.008 | 0.01 | 0.01 | 0 | 0.14 | 0.09 | 0.13 | 0.07 | 0.22 | 0.17 |
| 11 | 107365191 | rs4754226 | 0.067 | -0.156 | 0.199 | 0.904 | 0.01 | 0.042 | 0.042 | 0 | 0.77 | 0.69 | 0.73 | 0.43 | 0.69 | 0.76 |
| 11 | 107367793 | rs583852 | -0.163 | -0.421 | 0.393 | 1.232 | -0.004 | -0.011 | -0.011 | 0.19 | 0.6 | 0.44 | 0.61 | 0.26 | 0.67 | 0.52 |
| 11 | 107370210 | rs615248 | 1.524 | -0.505 | 0.719 | 1.232 | -0.003 | 0.359 | 0.359 | 0.495 | 0.88 | 0.57 | 0.84 | 0.51 | 0.91 | 0.92 |
| 11 | 107372065 | rs1789158 | 1.527 | -0.488 | 0.592 | 1.232 | 0.008 | 0.4 | 0.4 | 0.479 | 0.88 | 0.56 | 0.84 | 0.51 | 0.91 | 0.92 |
| 11 | 107374531 | rs10502092 | 1.239 | -0.547 | 0.269 | 1.006 | -0.059 | -0.063 | -0.063 | 0 | 0.27 | 0.01 | 0.21 | 0.25 | 0.25 | 0.4 |
| 11 | 107378428 | rs7112793 | -0.447 | -0.484 | 0.552 | 0.92 | -0.004 | -0.014 | -0.014 | 0 | 0.05 | 0.17 | 0.02 | 0 | 0.01 | 0.03 |
| 11 | 107381765 | rs654093 | 0.537 | -0.55 | 0.307 | 0.857 | -0.009 | 0.149 | 0.149 | 0 | 0.46 | 0.25 | 0.34 | 0.19 | 0.55 | 0.42 |
| 11 | 107397587 | rs533757 | 0.385 | -0.536 | 0.352 | 0.865 | 0.005 | 0.091 | 0.091 | 0 | 0.48 | 0.35 | 0.36 | 0.19 | 0.55 | 0.43 |
| 11 | 107407326 | rs650218 | 0.595 | -0.567 | 0.558 | 1.007 | -0.006 | 0.016 | 0.016 | 0.005 | 0.33 | 0.34 | 0.32 | 0.48 | 0.28 | 0.42 |
| 11 | 107412128 | rs559875 | -0.18 | -0.443 | -0.154 | 1.055 | 0.002 | 0.078 | 0.078 | 0 | 0.8 | 0.62 | 0.67 | 0.67 | 0.83 | 0.86 |
| 11 | 107417238 | rs682334 | -0.281 | -0.536 | 0.009 | 0.97 | -0.001 | -0.004 | -0.004 | 0 | 0.52 | 0.53 | 0.45 | 0.39 | 0.58 | 0.47 |
| 11 | 107418200 | rs667741 | -0.081 | -0.495 | -0.089 | 0.849 | 0.003 | 0.025 | 0.025 | 0.017 | 0.48 | 0.44 | 0.36 | 0.19 | 0.55 | 0.44 |
| 11 | 107429423 | rs1950031 | 0.218 | -0.639 | -0.943 | 0.777 | 0.008 | 0.113 | 0.113 | 0 | 0.46 | 0.14 | 0.32 | 0.25 | 0.41 | 0.51 |
| 11 | 107432502 | rs10749905 | 0.824 | -0.767 | -0.976 | 0.238 | -0.052 | 0.044 | 0.044 | 0 | 0.27 | 0.07 | 0.23 | 0.19 | 0.34 | 0.37 |
| 11 | 107433835 | rs1528494 | 0.786 | -0.637 | -1.209 | 0.143 | -0.012 | -0.006 | -0.006 | 0 | 0.15 | 0.1 | 0.14 | 0.18 | 0.21 | 0.23 |
| 11 | 107445769 | rs576806 | -0.245 | -1.096 | -0.627 | -0.277 | -0.002 | 0.003 | 0.003 | 0 | 0.38 | 0.49 | 0.36 | 0.43 | 0.36 | 0.42 |
| 11 | 107449022 | rs10890738 | 0.723 | -0.782 | -0.926 | -0.298 | -0.144 | 0 | 0 | 0.224 | 0.28 | 0.03 | 0.25 | 0.23 | 0.36 | 0.39 |
| 11 | 107454081 | rs504712 | -1.062 | -0.987 | -0.819 | -0.463 | -0.059 | -0.043 | -0.043 | 0.999 | 0.6 | 0.78 | 0.64 | 0.49 | 0.48 | 0.6 |
| 11 | 107456514 | rs7107870 | -0.559 | -0.978 | -0.629 | -0.664 | -0.003 | 0.01 | 0.01 | 0 | 0.54 | 0.45 | 0.37 | 0.28 | 0.45 | 0.57 |
| 11 | 107462895 | rs4085361 | 1.094 | -1.066 | -0.858 | -0.698 | -0.067 | 0.04 | 0.04 | 0 | 0.27 | 0.11 | 0.21 | 0.19 | 0.33 | 0.38 |
| 11 | 107463936 | rs11212292 | -1.111 | -1.535 | -0.795 | -0.594 | -0.116 | 0.095 | 0.095 | 0.183 | 0.65 | 0.87 | 0.73 | 0.8 | 0.53 | 0.55 |
| 11 | 107470916 | rs652243 | 0.378 | -1.627 | -0.587 | -0.661 | -0.049 | -0.043 | -0.043 | 0.029 | 0.18 | 0.28 | 0.12 | 0.08 | 0.08 | 0.17 |
| 11 | 107471983 | rs1939900 | 0.866 | -1.574 | -0.589 | -0.634 | -0.033 | -0.028 | -0.028 | 0 | 0.16 | 0.23 | 0.1 | 0.08 | 0.08 | 0.14 |
| 11 | 107474795 | rs10890742 | 0.719 | -1.481 | -0.713 | -0.605 | -0.025 | -0.049 | -0.049 | 0 | 0.36 | 0.21 | 0.49 | 0.53 | 0.4 | 0.35 |
| 11 | 107481451 | rs7123473 | -0.267 | -0.603 | -0.551 | -1.225 | -0.033 | -0.054 | -0.054 | 0.866 | 0.1 | 0.36 | 0.03 | 0 | 0.02 | 0.04 |
| 11 | 107485515 | rs4754235 | -0.036 | -0.024 | -0.927 | -0.94 | -0.001 | -0.005 | -0.005 | 0 | 0.56 | 0.6 | 0.69 | 0.8 | 0.51 | 0.54 |
| 11 | 107492412 | rs7925970 | 0.167 | -0.006 | -0.987 | -0.758 | 0.011 | 0.02 | 0.02 | 0 | 0.53 | 0.42 | 0.5 | 0.73 | 0.62 | 0.52 |
| 11 | 107492824 | rs6588972 | -0.192 | -0.042 | -1.297 | -1.061 | 0.026 | 0.069 | 0.069 | 0 | 0.75 | 0.89 | 0.89 | 0.93 | 0.8 | 0.85 |
| 11 | 107494204 | rs1939894 | 0.02 | 0.378 | -1.033 | -0.951 | -0.004 | 0.032 | 0.032 | 0 | 0.47 | 0.66 | 0.67 | 0.84 | 0.54 | 0.62 |
| 11 | 107502871 | rs11212304 | -1.021 | 0.401 | -0.773 | -0.608 | -0.001 | -0.015 | -0.015 | 0 | 0.06 | 0.08 | 0.04 | 0.03 | 0.05 | 0.08 |
| 11 | 107503447 | rs10789626 | -0.477 | 0.028 | -0.479 | -0.235 | -0.014 | 0.048 | 0.048 | 0 | 0.69 | 0.62 | 0.69 | 0.55 | 0.73 | 0.67 |
| 11 | 107505917 | rs1939896 | 0.074 | 0.016 | -1.103 | -0.756 | 0.001 | 0.016 | 0.016 | 0 | 0.27 | 0.16 | 0.35 | 0.19 | 0.29 | 0.3 |
| 11 | 107506922 | rs11212307 | -0.614 | -0.133 | -1.219 | -0.865 | 0.003 | 0.005 | 0.005 | 0 | 0.09 | 0.07 | 0.05 | 0.02 | 0.07 | 0.1 |
| 11 | 107507351 | rs4754239 | 0.126 | -0.063 | -1.283 | -0.802 | -0.003 | 0.016 | 0.016 | 0 | 0.27 | 0.15 | 0.35 | 0.19 | 0.29 | 0.3 |
| 11 | 107508832 | rs4754240 | -0.703 | -0.228 | -1.042 | -0.008 | -0.003 | 0.015 | 0.015 | 0 | 0.7 | 0.84 | 0.8 | 0.75 | 0.73 | 0.61 |
| 11 | 107524585 | rs2449095 | -0.853 | -0.113 | -1.281 | -0.024 | -0.016 | -0.046 | -0.046 | 0.102 | 0.4 | 0.83 | 0.57 | 0.5 | 0.39 | 0.4 |
| 11 | 107525990 | rs7116678 | -1.164 | -0.184 | -1.422 | -0.198 | -0.021 | 0.069 | 0.069 | 0.015 | 0.7 | 0.97 | 0.81 | 0.85 | 0.74 | 0.67 |
| 11 | 107531320 | rs524611 | -1.072 | -0.438 | -0.757 | -0.041 | -0.053 | 0.15 | 0.15 | 5.718 | 0.13 | 0.7 | 0.17 | 0.32 | 0.05 | 0.1 |
| 11 | 107535758 | rs647756 | -0.966 | -0.421 | -0.698 | -0.055 | -0.053 | 0.19 | 0.19 | 5.426 | 0.12 | 0.64 | 0.17 | 0.32 | 0.04 | 0.1 |
| 11 | 107540937 | rs509613 | -1.446 | -0.562 | -0.928 | -0.21 | -0.036 | 0.079 | 0.079 | 0 | 0.73 | 0.98 | 0.82 | 0.87 | 0.74 | 0.71 |
| 11 | 107542394 | rs11212320 | -0.469 | -0.201 | -0.902 | -0.4 | -0.013 | 0.019 | 0.019 | 0 | 0.3 | 0.25 | 0.4 | 0.17 | 0.36 | 0.39 |
| 11 | 107549519 | rs10890750 | 0.419 | -0.173 | -0.972 | -0.397 | 0.009 | 0.036 | 0.036 | 0 | 0.46 | 0.12 | 0.5 | 0.4 | 0.58 | 0.4 |
| 11 | 107549806 | rs4754241 | -0.726 | -0.236 | -0.827 | -0.375 | 0.011 | -0.011 | -0.011 | 0 | 0.62 | 0.38 | 0.66 | 0.55 | 0.7 | 0.6 |
| 11 | 107552110 | rs7950899 | -0.668 | -0.003 | -0.789 | -0.401 | -0.004 | -0.031 | -0.031 | 0 | 0.23 | 0.37 | 0.36 | 0.17 | 0.22 | 0.28 |
| 11 | 107554267 | rs11212323 | 0.308 | 0.351 | -1.508 | -0.529 | -0.091 | 0.014 | 0.014 | 0 | 0.41 | 0.1 | 0.32 | 0.39 | 0.51 | 0.34 |
| 11 | 107566557 | rs10890753 | -0.572 | 0.711 | -0.675 | -1.13 | 0.033 | -0.076 | -0.076 | 0 | 0.2 | 0.32 | 0.33 | 0.06 | 0.25 | 0.18 |
| 11 | 107566585 | rs597872 | -0.588 | 0.482 | -1.573 | -1.002 | -0.028 | 0.108 | 0.108 | 0 | 0.66 | 0.88 | 0.77 | 0.85 | 0.66 | 0.57 |
| 11 | 107571850 | rs683266 | -0.149 | 0.346 | -1.144 | -0.806 | 0.045 | 0.105 | 0.105 | 0 | 0.43 | 0.7 | 0.43 | 0.8 | 0.37 | 0.44 |
| 11 | 107576186 | rs12294839 | -0.872 | 1.061 | -0.109 | -0.439 | 0.07 | 0.011 | 0.011 | 1.37 | 0.12 | 0.29 | 0.27 | 0.13 | 0.21 | 0.14 |
| 11 | 107580985 | rs649974 | -0.731 | 0.863 | -0.295 | -1.135 | 0.011 | 0.109 | 0.109 | 0 | 0.53 | 0.7 | 0.65 | 0.82 | 0.57 | 0.51 |
| 11 | 107582885 | rs7948284 | 0.797 | 0.575 | -0.499 | -1.151 | -0.065 | -0.063 | -0.063 | 0.025 | 0.31 | 0.08 | 0.34 | 0.56 | 0.33 | 0.33 |
| 11 | 107586241 | rs11212337 | -0.165 | 0.576 | -0.38 | -1.112 | 0.074 | 0.001 | 0.001 | 1.484 | 0.12 | 0.29 | 0.27 | 0.12 | 0.21 | 0.13 |
| 11 | 107595881 | rs1384010 | 0.794 | 0.633 | -0.753 | -1.071 | -0.03 | -0.069 | -0.069 | 0.114 | 0.3 | 0.07 | 0.33 | 0.56 | 0.31 | 0.33 |
| 11 | 107598176 | rs7927287 | 1.741 | 0.578 | -0.362 | -0.834 | 0.003 | -0.002 | -0.002 | 0 | 0.08 | 0.04 | 0.02 | 0.13 | 0.03 | 0.04 |
| 11 | 107614258 | rs1351518 | -0.309 | 0.371 | -0.39 | -1.257 | 0.017 | 0.081 | 0.081 | 0 | 0.52 | 0.35 | 0.42 | 0.69 | 0.45 | 0.5 |
| 11 | 107614852 | rs11602139 | -0.695 | 0.478 | 0.289 | -1.133 | 0.102 | 0.023 | 0.023 | 0.112 | 0.13 | 0.38 | 0.27 | 0.13 | 0.19 | 0.13 |
| 11 | 107617257 | rs11212346 | -0.19 | 0.601 | 0.146 | -1.072 | 0.075 | 0.018 | 0.018 | 0.184 | 0.11 | 0.3 | 0.27 | 0.13 | 0.19 | 0.13 |
| 11 | 107617517 | rs7946302 | -0.19 | 0.601 | 0.146 | -1.072 | 0.075 | 0.018 | 0.018 | 0.184 | 0.11 | 0.3 | 0.27 | 0.13 | 0.19 | 0.13 |
| 11 | 107621538 | rs520906 | -1.286 | 0.437 | -0.565 | -1.289 | -0.031 | 0.069 | 0.069 | 0 | 0.7 | 0.88 | 0.72 | 0.82 | 0.67 | 0.66 |
| 11 | 107622778 | rs7116354 | 0.648 | 0.085 | -0.049 | -1.087 | 0.026 | -0.021 | -0.021 | 0 | 0.06 | 0.15 | 0.2 | 0.02 | 0.07 | 0.06 |
| 11 | 107623624 | rs2442674 | -1.056 | -0.316 | -0.6 | -0.356 | -0.039 | -0.11 | -0.11 | 0.364 | 0.68 | 0.93 | 0.77 | 0.61 | 0.67 | 0.62 |
| 11 | 107623767 | rs472826 | -0.451 | -0.432 | 0.028 | -0.302 | -0.002 | 0.001 | 0.001 | 0 | 0.6 | 0.6 | 0.55 | 0.59 | 0.6 | 0.56 |
| 11 | 107627609 | rs660375 | 0.642 | -0.353 | 0.275 | -0.327 | -0.004 | -0.004 | -0.004 | 0.517 | 0.41 | 0.4 | 0.45 | 0.46 | 0.49 | 0.47 |
| 11 | 107640811 | rs17107403 | -0.286 | -0.263 | 0.479 | -0.373 | 0.071 | -0.027 | -0.027 | 0 | 0.07 | 0.25 | 0.1 | 0.03 | 0.08 | 0.06 |
| 11 | 107644296 | rs632411 | 0.064 | -0.302 | -0.042 | -0.339 | 0.004 | 0.12 | 0.12 | 0.237 | 0.5 | 0.34 | 0.35 | 0.27 | 0.49 | 0.43 |
| 11 | 107646430 | rs11212366 | 1.56 | -0.428 | -0.601 | -0.091 | -0.022 | -0.093 | -0.093 | 0 | 0.18 | 0.01 | 0.11 | 0.21 | 0.15 | 0.28 |
| 11 | 107649026 | rs553663 | 1.387 | -0.564 | -0.175 | -0.17 | -0.042 | 0.051 | 0.051 | 0 | 0.28 | 0.18 | 0.26 | 0.18 | 0.39 | 0.3 |
| 11 | 107651095 | rs621334 | -0.152 | -0.43 | 0.259 | 0.284 | 0.007 | 0.026 | 0.026 | 0.02 | 0.55 | 0.4 | 0.44 | 0.34 | 0.57 | 0.49 |

| | | | | | | | | | | | | | | | | |
|---|---|---|---|---|---|---|---|---|---|---|---|---|---|---|---|---|
| 11 | 107659024 | rs12801961 | 1.236 | -0.494 | 0.452 | 0.609 | 0.017 | -0.008 | -0.008 | 0.003 | 0.12 | 0.04 | 0.03 | 0.15 | 0.05 | 0.07 |
| 11 | 107661242 | rs553924 | -0.692 | -0.552 | 0.509 | 0.583 | -0.012 | -0.002 | -0.002 | 0 | 0.62 | 0.82 | 0.58 | 0.68 | 0.6 | 0.57 |
| 11 | 107662790 | rs518852 | -0.649 | -0.428 | 0.634 | 0.564 | -0.02 | 0.012 | 0.012 | 0.223 | 0.59 | 0.81 | 0.5 | 0.66 | 0.55 | 0.52 |
| 11 | 107670178 | rs629278 | 0.454 | -0.393 | 0.689 | 0.51 | -0.001 | -0.023 | -0.023 | 0 | 0.42 | 0.47 | 0.38 | 0.14 | 0.51 | 0.4 |
| 11 | 107670228 | rs500063 | 0.241 | -0.391 | 0.691 | 0.516 | -0.001 | 0.006 | 0.006 | 0 | 0.44 | 0.47 | 0.4 | 0.27 | 0.53 | 0.49 |
| 11 | 107673504 | rs7109212 | 0.31 | -0.366 | 0.562 | 0.395 | -0.067 | 0.19 | 0.19 | 0.852 | 0.16 | 0.34 | 0.14 | 0.39 | 0.05 | 0.07 |
| 11 | 107681105 | rs602850 | 0.49 | -0.12 | 0.113 | -0.659 | -0.006 | 0.004 | 0.004 | 0 | 0.41 | 0.39 | 0.38 | 0.15 | 0.51 | 0.4 |
| 11 | 107684371 | rs12808838 | 1.174 | -0.702 | 0.4 | -0.304 | 0.005 | 0.002 | 0.002 | 2.024 | 0.11 | 0.07 | 0.03 | 0.1 | 0.04 | 0.06 |
| 11 | 107692302 | rs11826203 | 1.159 | -0.699 | 0.544 | -0.287 | -0.023 | -0.003 | -0.003 | 1.463 | 0.11 | 0.15 | 0.03 | 0.1 | 0.04 | 0.06 |
| 11 | 107693120 | rs1963626 | -0.388 | -0.795 | 0.43 | -0.139 | -0.025 | 0.024 | 0.024 | 1.773 | 0.18 | 0.63 | 0.09 | 0.22 | 0.08 | 0.07 |
| 11 | 107693980 | rs502398 | -0.695 | -0.385 | -0.201 | -0.252 | -0.109 | -0.099 | -0.099 | 1.161 | 0.58 | 0.92 | 0.42 | 0.5 | 0.46 | 0.57 |
| 11 | 107701784 | rs578300 | -0.672 | -0.35 | -0.2 | -0.261 | -0.104 | -0.094 | -0.094 | 0 | 0.58 | 0.92 | 0.42 | 0.5 | 0.46 | 0.57 |
| 11 | 107706832 | rs566238 | -0.462 | -0.449 | -0.009 | -0.25 | 0.003 | -0.021 | -0.021 | 0.337 | 0.56 | 0.75 | 0.4 | 0.5 | 0.43 | 0.56 |
| 11 | 107722297 | rs593673 | 1.3 | -0.468 | -0.415 | -0.35 | -0.103 | 0.063 | 0.063 | 0 | 0.27 | 0.02 | 0.22 | 0.16 | 0.31 | 0.34 |
| 11 | 107725297 | rs4754265 | 0.326 | -0.281 | 0.207 | -0.28 | 0.027 | 0.034 | 0.034 | 0 | 0.36 | 0.45 | 0.53 | 0.44 | 0.44 | 0.37 |
| 11 | 107729106 | rs488512 | 0.56 | -0.358 | -0.208 | -0.343 | 0 | 0.074 | 0.074 | 0 | 0.39 | 0.25 | 0.32 | 0.25 | 0.38 | 0.52 |
| 11 | 107729615 | rs2442683 | 0.539 | -0.341 | -0.205 | -0.334 | 0.002 | 0.071 | 0.071 | 0 | 0.4 | 0.26 | 0.32 | 0.25 | 0.38 | 0.52 |
| 11 | 107735502 | rs11212411 | 0.388 | 0.089 | -0.118 | 0.073 | -0.021 | -0.037 | -0.037 | 0.524 | 0.31 | 0.19 | 0.5 | 0.42 | 0.41 | 0.36 |
| 11 | 107743582 | rs4466794 | 0.288 | 0.32 | -0.097 | 0.137 | -0.028 | -0.044 | -0.044 | 0.076 | 0.32 | 0.19 | 0.51 | 0.41 | 0.41 | 0.37 |
| 11 | 107753547 | rs7118845 | -0.649 | 0.487 | 0.618 | 0.315 | 0.075 | 0.05 | 0.05 | 0 | 0.08 | 0.32 | 0.07 | 0.06 | 0.09 | 0.12 |
| 11 | 107757018 | rs2847190 | 0.993 | 0.406 | 0.373 | 0.203 | -0.008 | -0.004 | -0.004 | 0 | 0.66 | 0.22 | 0.75 | 0.62 | 0.67 | 0.73 |
| 11 | 107758546 | rs12797670 | 0.885 | 0.339 | 0.394 | 0.233 | 0.015 | 0.009 | 0.009 | 1.117 | 0.12 | 0.08 | 0.04 | 0.11 | 0.05 | 0.05 |
| 11 | 107760002 | rs10890773 | -0.968 | 0.446 | 0.482 | 0.252 | 0.001 | 0.006 | 0.006 | 0 | 0.9 | 0.9 | 0.89 | 0.79 | 0.86 | 0.9 |
| 11 | 107761550 | rs7946698 | -1.073 | 0.41 | 0.634 | 0.202 | -0.002 | -0.031 | -0.031 | 0 | 0.23 | 0.61 | 0.14 | 0.17 | 0.18 | 0.17 |
| 11 | 107771148 | rs7947466 | -0.816 | 0.42 | 0.734 | 0.242 | 0.014 | 0.018 | 0.018 | 0 | 0.26 | 0.42 | 0.2 | 0.3 | 0.27 | 0.17 |
| 11 | 107772104 | rs563811 | -1.153 | 0.183 | -0.027 | -0.025 | -0.014 | 0.031 | 0.031 | 1.329 | 0.69 | 0.84 | 0.78 | 0.75 | 0.66 | 0.48 |
| 11 | 107791995 | rs11212437 | -1.425 | 0.444 | -0.558 | -0.049 | -0.085 | 0.034 | 0.034 | 1.957 | 0.72 | 0.99 | 0.82 | 0.81 | 0.7 | 0.57 |
| 11 | 107796519 | rs7930340 | 1.531 | 0.458 | -0.521 | 0.018 | 0.003 | -0.021 | -0.021 | 0.101 | 0.09 | 0.02 | 0.03 | 0.1 | 0.05 | 0.05 |
| 11 | 107798886 | rs1532819 | 1.54 | 0.324 | -0.637 | 0.112 | -0.107 | -0.146 | -0.146 | 0 | 0.26 | 0.02 | 0.51 | 0.35 | 0.32 | 0.2 |
| 11 | 107798952 | rs1532820 | 1.593 | 0.326 | -0.636 | 0.112 | -0.084 | -0.146 | -0.146 | 0 | 0.25 | 0.02 | 0.49 | 0.35 | 0.29 | 0.19 |
| 11 | 107799785 | rs554180 | 0.185 | -0.101 | -0.241 | -0.091 | -0.02 | -0.037 | -0.037 | 0 | 0.14 | 0.3 | 0.07 | 0.06 | 0.09 | 0.06 |
| 11 | 107803708 | rs10466556 | 0.947 | 0.107 | -0.311 | 0.152 | -0.002 | -0.001 | -0.001 | 0 | 0.2 | 0.12 | 0.13 | 0.23 | 0.2 | 0.14 |
| 11 | 107812352 | rs10890780 | 0.771 | 0.156 | -0.39 | -0.133 | -0.017 | -0.073 | -0.073 | 0 | 0.31 | 0.16 | 0.54 | 0.5 | 0.34 | 0.35 |
| 11 | 107821944 | rs3950985 | -0.374 | 0.338 | -0.197 | -0.252 | -0.005 | -0.03 | -0.03 | 0 | 0.54 | 0.53 | 0.71 | 0.76 | 0.57 | 0.52 |
| 11 | 107824835 | rs963619 | 0.005 | 0.34 | -0.706 | -0.091 | -0.008 | 0.116 | 0.116 | 3.589 | 0.47 | 0.23 | 0.67 | 0.71 | 0.52 | 0.43 |
| 11 | 107827634 | rs10890785 | 1.227 | 0.454 | -0.939 | 0.059 | -0.046 | -0.087 | -0.087 | 0 | 0.25 | 0.09 | 0.52 | 0.39 | 0.32 | 0.19 |
| 11 | 107830018 | rs7120994 | 0.231 | 0.356 | -0.179 | -0.516 | 0.002 | 0.004 | 0.004 | 1.764 | 0.53 | 0.5 | 0.72 | 0.72 | 0.59 | 0.44 |
| 11 | 107833545 | rs4754282 | 0.863 | 0.438 | -0.411 | -0.506 | -0.027 | 0.115 | 0.115 | 3.864 | 0.46 | 0.19 | 0.66 | 0.7 | 0.52 | 0.42 |
| 11 | 107839211 | rs4554861 | 0.069 | 0.597 | -0.748 | -0.128 | 0.001 | 0.005 | 0.005 | 0.566 | 0.32 | 0.3 | 0.2 | 0.23 | 0.28 | 0.48 |
| 11 | 107841291 | rs4028250 | 0.65 | 0.751 | -0.661 | 0.359 | -0.002 | -0.009 | -0.009 | 0 | 0.87 | 0.83 | 0.92 | 0.94 | 0.87 | 0.92 |
| 11 | 107857145 | rs12574834 | -0.66 | 1.214 | -0.943 | 0.71 | -0.017 | -0.009 | -0.009 | 0 | 0.4 | 0.72 | 0.34 | 0.47 | 0.46 | 0.41 |
| 11 | 107858642 | rs4753826 | 0.624 | 1.649 | -1.012 | 0.888 | -0.005 | -0.028 | -0.028 | 0 | 0.56 | 0.21 | 0.56 | 0.45 | 0.45 | 0.58 |
| 11 | 107880880 | rs10890795 | 0.798 | 1.713 | -1.299 | 1.123 | 0.035 | -0.002 | -0.002 | 6.534 | 0.52 | 0.14 | 0.55 | 0.4 | 0.43 | 0.55 |
| 11 | 107882823 | rs7930830 | 0.367 | 2.787 | -0.443 | 1.198 | -0.002 | -0.004 | -0.004 | 0 | 0.66 | 0.32 | 0.73 | 0.9 | 0.65 | 0.75 |
| 11 | 107885991 | rs12418162 | -0.066 | 3.06 | 0.085 | 0.96 | -0.038 | 0.034 | 0.034 | 0 | 0.13 | 0.01 | 0.09 | 0.03 | 0.14 | 0.06 |
| 11 | 107911652 | rs7103534 | 0.002 | 3.062 | 0.909 | 0.868 | 0.011 | -0.033 | -0.033 | 0 | 0.1 | 0.18 | 0.07 | 0.03 | 0.12 | 0.12 |
| 11 | 107917087 | rs7117111 | 1.394 | 3.068 | 0.978 | 1.296 | 0 | -0.021 | -0.021 | 0 | 0.66 | 0.37 | 0.75 | 0.9 | 0.66 | 0.76 |
| 11 | 107918899 | rs11212495 | -0.113 | 2.979 | 1.089 | 1.323 | -0.002 | -0.003 | -0.003 | 0 | 0.07 | 0.03 | 0.08 | 0.17 | 0.1 | 0.04 |
| 11 | 107932222 | rs7949655 | -0.9 | 2.752 | 1.361 | 1.136 | -0.022 | 0.046 | 0.046 | 0 | 0.21 | 0.16 | 0.16 | 0.07 | 0.23 | 0.13 |
| 11 | 107938578 | rs7104942 | 0.974 | 2.791 | 1.421 | 1.187 | 0.013 | -0.091 | -0.091 | 0 | 0.76 | 0.53 | 0.81 | 0.93 | 0.77 | 0.87 |
| 11 | 107956771 | rs11212509 | 1.25 | 2.026 | -0.11 | 0.545 | -0.032 | -0.123 | -0.123 | 0 | 0.18 | 0.02 | 0.36 | 0.33 | 0.16 | 0.21 |
| 11 | 107957477 | rs10890809 | 1.974 | 2.28 | 0.441 | 0.714 | 0.043 | 0.001 | 0.001 | 1.468 | 0.52 | 0.14 | 0.55 | 0.42 | 0.43 | 0.55 |
| 11 | 107976450 | rs3781870 | 0.395 | 2.405 | 0.64 | 0.674 | 0.002 | -0.021 | -0.021 | 0 | 0.09 | 0.12 | 0.06 | 0.03 | 0.11 | 0.12 |
| 11 | 107983538 | rs11212514 | -0.051 | 2.412 | 0.332 | 0.73 | -0.02 | -0.029 | -0.029 | 0 | 0.08 | 0.02 | 0.09 | 0.16 | 0.11 | 0.07 |
| 11 | 107985264 | rs12807544 | 0.852 | 2.625 | 0.643 | 0.345 | 0.013 | -0.044 | -0.044 | 6.043 | 0.67 | 0.39 | 0.69 | 0.9 | 0.58 | 0.76 |
| 11 | 107986227 | rs12577790 | -0.034 | 2.782 | 0.648 | 0.295 | -0.03 | 0.029 | 0.029 | 0.006 | 0.17 | 0.12 | 0.18 | 0.07 | 0.26 | 0.1 |
| 11 | 107986691 | rs11601596 | -0.034 | 2.782 | 0.658 | 0.299 | -0.028 | 0.026 | 0.026 | 0.006 | 0.17 | 0.13 | 0.18 | 0.08 | 0.26 | 0.1 |
| 11 | 107987784 | rs10890813 | 2.195 | 2.648 | 0.411 | 0.732 | 0.138 | 0.059 | 0.059 | 1.394 | 0.5 | 0.06 | 0.42 | 0.34 | 0.34 | 0.51 |
| 11 | 108004798 | rs7931681 | -0.858 | 2.798 | 0.73 | 0.796 | 0.011 | 0.04 | 0.04 | 0 | 0.16 | 0.15 | 0.26 | 0.53 | 0.24 | 0.24 |
| 11 | 108009927 | rs3741049 | 0.827 | 2.885 | 0.653 | 0.868 | -0.013 | 0.068 | 0.068 | 0.052 | 0.05 | 0.01 | 0.16 | 0.36 | 0.12 | 0.17 |
| 11 | 108012077 | rs10749914 | 0.785 | 2.854 | 1.03 | 0.631 | 0.051 | -0.097 | -0.097 | 0 | 0.69 | 0.5 | 0.7 | 0.9 | 0.59 | 0.75 |
| 11 | 108016014 | rs11212525 | -0.818 | 2.854 | 0.736 | 0.823 | -0.002 | 0.017 | 0.017 | 0 | 0.14 | 0.11 | 0.24 | 0.53 | 0.23 | 0.24 |
| 11 | 108031295 | rs6589006 | 1.076 | 2.893 | 1.243 | 1.118 | 0.031 | 0.032 | 0.032 | 0 | 0.72 | 0.64 | 0.64 | 0.44 | 0.62 | 0.52 |
| 11 | 108038477 | rs920679 | 2.095 | 2.773 | 1.121 | 0.976 | 0.125 | 0.033 | 0.033 | 0.635 | 0.49 | 0.06 | 0.45 | 0.36 | 0.36 | 0.52 |
| 11 | 108043988 | rs2070661 | 1.027 | 2.765 | 1.225 | 1.081 | 0.038 | 0.039 | 0.039 | 0 | 0.72 | 0.65 | 0.67 | 0.46 | 0.63 | 0.68 |
| 11 | 108068954 | rs683312 | -0.431 | 2.817 | 0.991 | 1.249 | -0.088 | -0.088 | -0.088 | 0.18 | 0.85 | 0.98 | 0.8 | 0.68 | 0.75 | 0.75 |
| 11 | 108092637 | rs4987876 | 0.599 | 2.894 | 1.103 | 0.935 | -0.001 | 0.006 | 0.006 | 0.077 | 0.09 | 0.06 | 0.08 | 0.03 | 0.09 | 0.03 |
| 11 | 108097333 | rs228591 | 0.78 | 2.917 | 1.467 | 1.004 | 0.039 | 0.03 | 0.03 | 0.019 | 0.69 | 0.39 | 0.65 | 0.43 | 0.62 | 0.68 |
| 11 | 108158134 | rs609261 | 0.836 | 2.815 | 1.688 | 0.649 | 0.01 | 0.006 | 0.006 | 0.16 | 0.68 | 0.33 | 0.63 | 0.39 | 0.62 | 0.68 |

| | | | | | | | | | | | | | | | | |
|---|---|---|---|---|---|---|---|---|---|---|---|---|---|---|---|---|
| 11 | 108177538 | rs599558 | 0.658 | 2.732 | 1.666 | 0.605 | 0.047 | 0.039 | 0.039 | 0 | 0.69 | 0.43 | 0.65 | 0.43 | 0.62 | 0.68 |
| 11 | 108219339 | rs425538 | 0.371 | 2.297 | 1.743 | 0.521 | 0.011 | 0.01 | 0.01 | 0 | 0.73 | 0.71 | 0.68 | 0.45 | 0.63 | 0.68 |
| 11 | 108225661 | rs664143 | 0.424 | 2.301 | 1.745 | 0.522 | 0.014 | 0.013 | 0.013 | 0 | 0.72 | 0.7 | 0.68 | 0.45 | 0.63 | 0.68 |
| 11 | 108239628 | rs4585 | 0.812 | 2.289 | 1.671 | 0.547 | 0.031 | 0.025 | 0.025 | 0.021 | 0.68 | 0.38 | 0.65 | 0.43 | 0.62 | 0.68 |
| 11 | 108253252 | rs227077 | 0.812 | 2.289 | 1.671 | 0.547 | 0.031 | 0.025 | 0.025 | 0.021 | 0.68 | 0.38 | 0.65 | 0.43 | 0.62 | 0.68 |
| 11 | 108256859 | rs597788 | 1.918 | 2.241 | 1.414 | 0.483 | 0.127 | 0.055 | 0.055 | 0.58 | 0.49 | 0.06 | 0.45 | 0.36 | 0.36 | 0.52 |
| 11 | 108261261 | rs227055 | 0.577 | 2.571 | 1.719 | 0.713 | 0.032 | 0.028 | 0.028 | 0 | 0.72 | 0.65 | 0.68 | 0.45 | 0.63 | 0.68 |
| 11 | 108291017 | rs11212621 | 0.982 | 1.812 | 1.113 | 0.272 | -0.005 | -0.084 | -0.084 | 3.471 | 0.19 | 0.02 | 0.33 | 0.29 | 0.18 | 0.3 |
| 11 | 108324816 | rs7131306 | -0.886 | 2.108 | 1.475 | 0.279 | 0.08 | -0.062 | -0.062 | 0 | 0.14 | 0.46 | 0.15 | 0.04 | 0.17 | 0.12 |
| 11 | 108333536 | rs17108024 | -0.77 | 2.16 | 1.498 | 0.304 | 0.089 | -0.056 | -0.056 | 0 | 0.13 | 0.46 | 0.15 | 0.04 | 0.17 | 0.13 |
| 11 | 108334321 | rs10890845 | -0.803 | 2.131 | 1.292 | 0.271 | 0.065 | -0.057 | -0.057 | 0 | 0.13 | 0.4 | 0.14 | 0.04 | 0.18 | 0.14 |
| 11 | 108344081 | rs7943203 | 0.885 | 1.311 | 0.551 | 0.68 | -0.003 | 0.007 | 0.007 | 0.123 | 0.61 | 0.39 | 0.55 | 0.46 | 0.46 | 0.6 |
| 11 | 108354102 | rs12223381 | 0.864 | 1.284 | 0.561 | 0.662 | -0.004 | 0.007 | 0.007 | 0.102 | 0.61 | 0.39 | 0.55 | 0.45 | 0.45 | 0.61 |
| 11 | 108365182 | rs893279 | 0.793 | 1.298 | -0.556 | 0.615 | 0.013 | -0.021 | -0.021 | 0 | 0.6 | 0.18 | 0.52 | 0.42 | 0.45 | 0.56 |
| 11 | 108379981 | rs3741060 | 0.512 | 1.158 | -0.156 | 0.169 | 0.018 | 0.01 | 0.01 | 0.036 | 0.18 | 0.06 | 0.11 | 0.17 | 0.15 | 0.27 |
| 11 | 108383676 | rs10749920 | -0.847 | 0.897 | 0.105 | 0.111 | -0.007 | -0.007 | -0.007 | 0.045 | 0.27 | 0.69 | 0.18 | 0.21 | 0.19 | 0.36 |
| 11 | 108397640 | rs10890853 | 0.383 | 0.868 | -0.789 | 0.258 | 0.017 | 0.008 | 0.008 | 0.02 | 0.18 | 0.07 | 0.11 | 0.18 | 0.16 | 0.27 |
| 11 | 108409442 | rs2640784 | -0.844 | 1.516 | -0.202 | 0.488 | 0.012 | 0.041 | 0.041 | 0.641 | 0.15 | 0.28 | 0.37 | 0.6 | 0.17 | 0.33 |
| 11 | 108409784 | rs2640785 | -0.038 | 1.517 | -0.24 | 0.531 | -0.03 | -0.034 | -0.034 | 0.332 | 0.12 | 0.04 | 0.35 | 0.58 | 0.17 | 0.22 |
| 11 | 108410261 | rs2852183 | -0.038 | 1.517 | -0.24 | 0.531 | -0.03 | -0.034 | -0.034 | 0 | 0.12 | 0.04 | 0.35 | 0.58 | 0.17 | 0.21 |
| 11 | 108420006 | rs2852189 | 0.467 | 0.215 | -0.384 | -0.199 | 0.032 | 0.054 | 0.054 | 0 | 0.64 | 0.26 | 0.7 | 0.8 | 0.69 | 0.7 |
| 11 | 108425790 | rs11212704 | 0.079 | 0.175 | 0.142 | -0.195 | -0.034 | -0.026 | -0.026 | 0 | 0.23 | 0.41 | 0.17 | 0.2 | 0.19 | 0.24 |
| 11 | 108430237 | rs11212707 | 1.972 | 0.334 | -0.507 | -0.642 | -0.036 | 0.157 | 0.157 | 0 | 0.33 | 0.01 | 0.16 | 0.09 | 0.25 | 0.26 |
| 11 | 108430554 | rs1940219 | 0.581 | 0.13 | 0.591 | 0.33 | 0.019 | -0.033 | -0.033 | 0 | 0.18 | 0.07 | 0.12 | 0.24 | 0.15 | 0.23 |
| 11 | 108432753 | rs2256372 | 0.761 | 0.217 | 0.738 | 0.335 | -0.024 | -0.011 | -0.011 | 0.09 | 0.65 | 0.41 | 0.74 | 0.75 | 0.73 | 0.62 |
| 11 | 108436646 | rs7119814 | -0.955 | 0.455 | 0.38 | 0.323 | -0.009 | -0.066 | -0.066 | 0.008 | 0.16 | 0.53 | 0.16 | 0.04 | 0.14 | 0.1 |
| 11 | 108437928 | rs767664 | -0.948 | 0.583 | -0.221 | 0.513 | 0.041 | 0.021 | 0.021 | 0 | 0.38 | 0.75 | 0.3 | 0.3 | 0.31 | 0.33 |
| 11 | 108440416 | rs12277367 | 0.549 | 0.969 | 0.242 | 0.511 | -0.007 | 0.025 | 0.025 | 0 | 0.56 | 0.6 | 0.42 | 0.36 | 0.54 | 0.53 |
| 11 | 108440814 | rs2852199 | 1.15 | 0.57 | 0.777 | 1.291 | 0.006 | 0.036 | 0.036 | 0 | 0.8 | 0.69 | 0.78 | 0.7 | 0.77 | 0.79 |
| 11 | 108452651 | rs7105022 | 0.181 | 0.654 | 0.775 | 0.9 | -0.002 | -0.009 | -0.009 | 0 | 0.53 | 0.46 | 0.32 | 0.35 | 0.41 | 0.47 |
| 11 | 108462537 | rs2640737 | 0 | 0.551 | 0.997 | 1.023 | 0.011 | 0.156 | 0.156 | 0.406 | 0.89 | 0.7 | 0.89 | 0.73 | 0.91 | 0.88 |
| 11 | 108464209 | rs2640738 | 0.315 | 0.66 | 0.405 | 0.932 | -0.047 | -0.023 | -0.023 | 1.762 | 0.36 | 0.2 | 0.56 | 0.38 | 0.5 | 0.41 |
| 11 | 108475820 | rs2846419 | 0.235 | 0.594 | 0.417 | 0.963 | -0.037 | -0.025 | -0.025 | 0 | 0.35 | 0.2 | 0.55 | 0.38 | 0.48 | 0.41 |
| 11 | 108476461 | rs2846418 | -1.035 | 0.25 | 0.671 | 0.532 | 0.022 | 0.033 | 0.033 | 0 | 0.53 | 0.39 | 0.73 | 0.69 | 0.69 | 0.61 |
| 11 | 108482112 | rs4539291 | 0.139 | 0.264 | 0.514 | 0.275 | -0.003 | 0.008 | 0.008 | 0 | 0.57 | 0.62 | 0.43 | 0.73 | 0.44 | 0.54 |
| 11 | 108482248 | rs7128600 | -1.357 | 0.673 | 0.622 | 0.398 | 0.031 | 0.078 | 0.078 | 0.907 | 0.08 | 0.42 | 0.09 | 0.07 | 0.06 | 0.05 |
| 11 | 108494855 | rs990253 | -0.275 | 1.016 | -0.568 | 0.37 | -0.047 | 0.02 | 0.02 | 0 | 0.33 | 0.28 | 0.49 | 0.23 | 0.51 | 0.4 |
| 11 | 108498885 | rs10160242 | -0.893 | 0.233 | 0.299 | 0.056 | -0.024 | -0.067 | -0.067 | 0 | 0.12 | 0.39 | 0.07 | 0.02 | 0.05 | 0.01 |
| 11 | 108503672 | rs10890873 | 0.123 | -0.113 | -0.684 | -0.256 | 0.013 | 0.027 | 0.027 | 0.054 | 0.25 | 0.12 | 0.16 | 0.21 | 0.22 | 0.22 |
| 11 | 108507921 | rs12146575 | -0.564 | 0.179 | -1.123 | 0.297 | -0.009 | -0.072 | -0.072 | 0 | 0.37 | 0.17 | 0.31 | 0.47 | 0.41 | 0.39 |
| 11 | 108509754 | rs2128243 | 0.904 | 0.842 | -1.427 | 0.421 | -0.117 | -0.121 | -0.121 | 0 | 0.31 | 0.09 | 0.52 | 0.48 | 0.4 | 0.41 |
| 11 | 108525801 | rs2640770 | 0.76 | 1.032 | -0.066 | 1.134 | 0 | 0.018 | 0.018 | 0 | 0.85 | 0.84 | 0.85 | 0.72 | 0.82 | 0.83 |
| 11 | 108535792 | rs2306118 | 1.437 | 1.029 | -0.376 | 0.776 | -0.011 | -0.008 | -0.008 | 0 | 0.54 | 0.09 | 0.58 | 0.43 | 0.55 | 0.49 |
| 11 | 108563349 | rs10749925 | 1.067 | 0.951 | 0.379 | 0.918 | 0.002 | 0.001 | 0.001 | 0.025 | 0.73 | 0.37 | 0.77 | 0.76 | 0.72 | 0.73 |
| 11 | 108569406 | rs10749926 | 1.701 | 0.882 | 0.527 | 0.927 | -0.011 | -0.024 | -0.024 | 0 | 0.6 | 0.24 | 0.6 | 0.48 | 0.55 | 0.51 |
| 11 | 108570771 | rs12574567 | -0.308 | 0.903 | 0.586 | 0.871 | 0.003 | -0.046 | -0.046 | 0 | 0.15 | 0.08 | 0.11 | 0.24 | 0.16 | 0.13 |
| 11 | 108599993 | rs12793497 | 0.172 | 0.948 | 0.57 | 0.867 | -0.024 | 0.054 | 0.054 | 0 | 0.13 | 0.01 | 0.08 | 0.04 | 0.15 | 0.1 |
| 11 | 108657548 | rs2615725 | 1.443 | 0.876 | 1.378 | 0.848 | -0.005 | 0.001 | 0.001 | 0.018 | 0.69 | 0.45 | 0.77 | 0.72 | 0.71 | 0.7 |
| 11 | 108672547 | rs2726876 | 0.57 | 0.875 | 1.717 | 0.818 | 0.025 | -0.057 | -0.057 | 0.096 | 0.9 | 0.75 | 0.93 | 1 | 0.91 | 0.87 |
| 11 | 108673298 | rs2615760 | 0.079 | 0.889 | 1.617 | 0.841 | 0 | -0.011 | -0.011 | 0 | 0.13 | 0.15 | 0.17 | 0.28 | 0.16 | 0.21 |
| 11 | 108680469 | rs12420572 | 2.068 | 0.783 | 1.628 | 0.866 | -0.001 | 0 | 0 | 0.012 | 0.54 | 0.14 | 0.59 | 0.43 | 0.55 | 0.49 |
| 11 | 108689088 | rs17108608 | -0.243 | 0.775 | 1.625 | 0.897 | 0.001 | 0.012 | 0.012 | 0 | 0.16 | 0.17 | 0.14 | 0.24 | 0.2 | 0.14 |
| 11 | 108711737 | rs4754358 | 2.011 | 0.905 | 1.564 | 0.855 | -0.025 | -0.026 | -0.026 | 0 | 0.54 | 0.12 | 0.59 | 0.44 | 0.56 | 0.49 |
| 11 | 108717748 | rs2514119 | -0.462 | 1.474 | 1.58 | 0.836 | -0.013 | -0.006 | -0.006 | 0 | 0.18 | 0.16 | 0.3 | 0.28 | 0.33 | 0.31 |
| 11 | 108719088 | rs2466940 | -0.859 | 0.511 | 1.515 | 0.89 | -0.008 | 0.014 | 0.014 | 1.061 | 0.22 | 0.36 | 0.23 | 0.33 | 0.21 | 0.26 |
| 11 | 108724639 | rs2726923 | -0.854 | 0.512 | 1.486 | 0.89 | -0.014 | 0.032 | 0.032 | 1.654 | 0.22 | 0.45 | 0.24 | 0.33 | 0.21 | 0.26 |
| 11 | 108754443 | rs2553761 | 0.921 | 0.586 | 1.413 | 0.927 | 0.015 | 0.04 | 0.04 | 0.16 | 0.78 | 0.65 | 0.81 | 0.73 | 0.76 | 0.75 |
| 11 | 108755840 | rs10502103 | -0.218 | 0.66 | 1.232 | 0.966 | -0.004 | -0.006 | -0.006 | 3.05 | 0.14 | 0.11 | 0.12 | 0.24 | 0.19 | 0.12 |
| 11 | 108764449 | rs2553751 | 0.066 | 0.619 | 1.095 | 0.978 | 0.012 | 0.055 | 0.055 | 0 | 0.84 | 0.74 | 0.87 | 0.76 | 0.81 | 0.88 |
| 11 | 108802737 | rs17703817 | 0.137 | 0.686 | 0.241 | 0.911 | -0.003 | 0.005 | 0.005 | 0.043 | 0.06 | 0.04 | 0.06 | 0.03 | 0.05 | 0.12 |
| 11 | 108806407 | rs9888280 | -0.334 | 0.547 | 0.356 | 1.099 | -0.037 | -0.014 | -0.014 | 0 | 0.07 | 0.19 | 0.02 | 0.04 | 0 | 0.03 |
| 11 | 108810687 | rs7130411 | -0.669 | 1.334 | -0.125 | 1.308 | 0.006 | 0.007 | 0.007 | 0.858 | 0.17 | 0.23 | 0.24 | 0.29 | 0.24 | 0.25 |
| 11 | 108822187 | rs11212824 | -0.931 | 0.768 | 0.136 | 1.322 | -0.005 | 0.078 | 0.078 | 0 | 0.15 | 0.27 | 0.12 | 0.23 | 0.15 | 0.13 |
| 11 | 108824752 | rs11212826 | -0.183 | 0.425 | -0.229 | 0.375 | 0.011 | 0.03 | 0.03 | 0 | 0.51 | 0.56 | 0.3 | 0.18 | 0.44 | 0.55 |
| 11 | 108831875 | rs4630256 | -0.231 | 0.166 | -0.771 | 0.538 | -0.001 | -0.025 | -0.025 | 0.443 | 0.34 | 0.16 | 0.32 | 0.42 | 0.3 | 0.37 |
| 11 | 108838910 | rs1126193 | -0.916 | -0.878 | -0.644 | 0.354 | -0.019 | -0.004 | -0.004 | 2.425 | 0.62 | 0.69 | 0.55 | 0.61 | 0.54 | 0.69 |
| 11 | 108839158 | rs10160249 | 0.559 | -1.107 | -0.798 | 0.242 | -0.057 | -0.012 | -0.012 | 0.136 | 0.4 | 0.21 | 0.48 | 0.39 | 0.49 | 0.34 |
| 11 | 108842176 | rs10890915 | 0.374 | -1.327 | -0.777 | 0.284 | -0.007 | -0.018 | -0.018 | 3.632 | 0.43 | 0.19 | 0.4 | 0.47 | 0.4 | 0.45 |
| 11 | 108845054 | rs1425795 | 0.342 | -1.276 | 0.184 | 0.328 | -0.003 | -0.002 | -0.002 | 0.263 | 0.93 | 0.7 | 0.92 | 0.98 | 0.95 | 0.83 |

| | | | | | | | | | | | | | | | | |
|---|---|---|---|---|---|---|---|---|---|---|---|---|---|---|---|---|
| 11 | 108845079 | rs10047449 | 1.681 | -0.797 | -0.381 | 0.303 | -0.024 | 0.004 | 0.004 | 0 | 0.17 | 0.14 | 0.18 | 0.12 | 0.25 | 0.15 |
| 11 | 108846490 | rs6589040 | 0.091 | -1.032 | 0.032 | 0.352 | -0.024 | 0.015 | 0.015 | 0.067 | 0.85 | 0.52 | 0.9 | 0.89 | 0.88 | 0.89 |
| 11 | 108851707 | rs7124854 | 0.499 | -1.532 | -0.542 | -0.067 | 0.001 | -0.002 | -0.002 | 0 | 0.45 | 0.37 | 0.44 | 0.48 | 0.44 | 0.57 |
| 11 | 108853664 | rs11212834 | -0.11 | -1.219 | -0.426 | -0.117 | -0.017 | 0.01 | 0.01 | 0 | 0.35 | 0.45 | 0.35 | 0.42 | 0.32 | 0.27 |
| 11 | 108855811 | rs1834455 | 0.426 | -1.087 | -0.092 | 0.658 | -0.039 | 0.113 | 0.113 | 1.023 | 0.87 | 0.71 | 0.82 | 0.73 | 0.91 | 0.78 |
| 11 | 108863213 | rs4753856 | 1.033 | -0.817 | -0.535 | 0.185 | -0.005 | 0.003 | 0.003 | 0 | 0.2 | 0.19 | 0.2 | 0.12 | 0.24 | 0.16 |
| 11 | 108873119 | rs10890920 | 0.921 | -1.091 | -0.555 | 0.179 | 0.009 | 0.073 | 0.073 | 0 | 0.53 | 0.2 | 0.59 | 0.68 | 0.57 | 0.52 |
| 11 | 108879868 | rs11212844 | -0.766 | -1.351 | -0.122 | -0.602 | -0.043 | 0.017 | 0.017 | 0 | 0.68 | 0.48 | 0.71 | 0.63 | 0.79 | 0.74 |
| 11 | 108884508 | rs17108871 | 1.068 | -1.144 | 0.048 | -0.832 | -0.001 | 0.011 | 0.011 | 0 | 0.22 | 0.32 | 0.22 | 0.27 | 0.19 | 0.2 |
| 11 | 108884867 | rs1559755 | -1.835 | -0.943 | 0.107 | -0.871 | -0.005 | 0.024 | 0.024 | 0 | 0.65 | 0.76 | 0.68 | 0.74 | 0.68 | 0.66 |
| 11 | 108887701 | rs1974953 | 0.961 | -0.842 | 0.108 | -0.851 | -0.003 | 0.008 | 0.008 | 0 | 0.33 | 0.34 | 0.36 | 0.46 | 0.34 | 0.42 |
| 11 | 108889352 | rs11212857 | 0.258 | -0.893 | 0.064 | -0.842 | 0.006 | -0.009 | -0.009 | 0 | 0.31 | 0.4 | 0.32 | 0.27 | 0.33 | 0.21 |
| 11 | 108893367 | rs7101378 | 1.187 | -0.926 | -0.16 | -0.814 | -0.046 | 0.059 | 0.059 | 0 | 0.4 | 0.24 | 0.39 | 0.26 | 0.45 | 0.37 |
| 11 | 108893765 | rs10502107 | 1.062 | -0.9 | -0.219 | -0.822 | -0.033 | 0.018 | 0.018 | 0 | 0.05 | 0.01 | 0.1 | 0.02 | 0.14 | 0.19 |
| 11 | 108897856 | rs7126518 | -0.264 | -0.937 | -0.09 | -0.807 | 0.009 | -0.011 | -0.011 | 9.371 | 0.69 | 0.59 | 0.66 | 0.71 | 0.66 | 0.63 |
| 11 | 108905350 | rs12274468 | 1.371 | -1.006 | -0.316 | -0.66 | -0.02 | 0.027 | 0.027 | 0 | 0.28 | 0.31 | 0.26 | 0.44 | 0.21 | 0.26 |
| 11 | 108912978 | rs12576390 | 0.175 | -0.742 | -0.157 | -0.687 | -0.007 | -0.016 | -0.016 | 0.785 | 0.46 | 0.52 | 0.48 | 0.71 | 0.39 | 0.42 |
| 11 | 108917891 | rs9633935 | 1.704 | -0.171 | -0.264 | -0.206 | -0.05 | -0.025 | -0.025 | 0.073 | 0.24 | 0.07 | 0.25 | 0.26 | 0.26 | 0.3 |
| 11 | 108919802 | rs11212884 | -0.053 | -0.258 | 0.623 | 0.541 | 0.009 | -0.001 | -0.001 | 0 | 0.12 | 0.09 | 0.14 | 0.09 | 0.09 | 0.04 |
| 11 | 108927483 | rs1509729 | -1.126 | -0.339 | 1.107 | 0.306 | -0.001 | 0.001 | 0.001 | 0 | 0.32 | 0.32 | 0.34 | 0.29 | 0.31 | 0.33 |
| 11 | 108927806 | rs17632724 | -0.617 | -0.324 | 1.014 | 0.283 | -0.002 | -0.002 | -0.002 | 0 | 0.27 | 0.24 | 0.29 | 0.29 | 0.25 | 0.31 |
| 11 | 108928988 | rs1509717 | -0.686 | -0.262 | 0.953 | 0.309 | -0.001 | 0.002 | 0.002 | 0 | 0.28 | 0.31 | 0.3 | 0.29 | 0.24 | 0.33 |
| 11 | 108929128 | rs1509715 | -0.686 | -0.262 | 0.953 | 0.309 | -0.001 | 0.002 | 0.002 | 0 | 0.28 | 0.31 | 0.3 | 0.29 | 0.24 | 0.33 |
| 11 | 108932333 | rs1949050 | -2.928 | -0.298 | 0.462 | 0.479 | -0.064 | 0.004 | 0.004 | 0 | 0.55 | 0.87 | 0.67 | 0.56 | 0.47 | 0.44 |
| 11 | 108932662 | rs1949052 | -2.748 | 0.168 | 1.129 | -0.121 | -0.002 | 0.059 | 0.059 | 0.089 | 0.5 | 0.85 | 0.54 | 0.33 | 0.44 | 0.39 |
| 11 | 108937149 | rs11212908 | -1.384 | 0.419 | 1.3 | 0.158 | -0.002 | -0.003 | -0.003 | 0.12 | 0.31 | 0.59 | 0.33 | 0.29 | 0.27 | 0.32 |
| 11 | 108940427 | rs7949874 | -1.61 | 0.414 | 1.287 | 0.166 | 0.004 | -0.015 | -0.015 | 0 | 0.33 | 0.49 | 0.35 | 0.29 | 0.3 | 0.34 |
| 11 | 108947644 | rs11212913 | -1.137 | 0.306 | 1.208 | 0.378 | -0.014 | 0.042 | 0.042 | 0 | 0.28 | 0.49 | 0.37 | 0.33 | 0.22 | 0.31 |
| 11 | 108947719 | rs11212914 | -1.153 | 0.271 | 1.209 | 0.38 | -0.017 | 0.04 | 0.04 | 0 | 0.28 | 0.49 | 0.37 | 0.33 | 0.22 | 0.31 |
| 11 | 108959994 | rs10789713 | -2.255 | 0.024 | 1.604 | -0.056 | -0.05 | -0.071 | -0.071 | 0 | 0.3 | 0.55 | 0.3 | 0.07 | 0.21 | 0.19 |
| 11 | 108963612 | rs12420493 | -2.244 | 0.894 | 1.96 | 0.387 | -0.018 | -0.05 | -0.05 | 0 | 0.22 | 0.62 | 0.29 | 0.07 | 0.21 | 0.14 |
| 11 | 108968192 | rs11212926 | -2.283 | 0.837 | 2.019 | 0.324 | -0.076 | -0.142 | -0.142 | 4.822 | 0.24 | 0.53 | 0.18 | 0.02 | 0.15 | 0.13 |
| 11 | 108968932 | rs6589055 | -2.576 | 0.86 | 2.009 | 0.329 | -0.029 | -0.12 | -0.12 | 1.784 | 0.27 | 0.6 | 0.23 | 0.02 | 0.23 | 0.15 |
| 11 | 108969343 | rs1509733 | -2.576 | 0.86 | 2.009 | 0.329 | -0.029 | -0.123 | -0.123 | 0 | 0.27 | 0.6 | 0.23 | 0.02 | 0.23 | 0.15 |
| 11 | 108980846 | rs10890947 | -1.734 | 0.993 | 1.939 | 0.362 | -0.055 | -0.155 | -0.155 | 0.032 | 0.21 | 0.54 | 0.16 | 0 | 0.14 | 0.04 |
| 11 | 108984596 | rs11212934 | 0.155 | 1.109 | 1.677 | 0.427 | -0.001 | 0.002 | 0.002 | 0 | 0.07 | 0.03 | 0.06 | 0 | 0.06 | 0.03 |
| 11 | 108991573 | rs1509718 | -2.485 | 1.058 | 1.806 | 0.454 | -0.012 | -0.037 | -0.037 | 3.976 | 0.25 | 0.66 | 0.3 | 0.07 | 0.22 | 0.17 |
| 11 | 108998139 | rs2884183 | -1.993 | 1.063 | 1.716 | 0.121 | -0.011 | -0.137 | -0.137 | 0 | 0.23 | 0.37 | 0.2 | 0 | 0.21 | 0.07 |
| 11 | 109003536 | rs1553769 | -1.995 | 0.394 | 1.584 | 0.339 | -0.059 | -0.106 | -0.106 | 1.585 | 0.23 | 0.49 | 0.16 | 0.06 | 0.14 | 0.07 |
| 11 | 109003568 | rs4754384 | -1.387 | 0.437 | 1.499 | 0.37 | -0.019 | -0.029 | -0.029 | 0 | 0.13 | 0.25 | 0.1 | 0.06 | 0.08 | 0.06 |
| 11 | 109005366 | rs1509731 | 0.082 | 0.923 | 0.963 | 0.708 | -0.003 | -0.001 | -0.001 | 0.897 | 0.06 | 0.01 | 0.06 | 0.06 | 0.06 | 0.04 |
| 11 | 109005939 | rs11212939 | -0.334 | 1.22 | 1.198 | 1.101 | -0.029 | -0.03 | -0.03 | 0.639 | 0.07 | 0.01 | 0.07 | 0.13 | 0.1 | 0.07 |
| 11 | 109015327 | rs12289666 | -0.796 | 0.83 | 1.431 | 0.876 | -0.027 | -0.068 | -0.068 | 0 | 0.11 | 0.29 | 0.07 | 0 | 0.06 | 0.01 |
| 11 | 109018392 | rs10890951 | 1.108 | 1.702 | 1.411 | 1.839 | 0.007 | 0.02 | 0.02 | 0 | 0.76 | 0.72 | 0.64 | 0.55 | 0.68 | 0.72 |
| 11 | 109019127 | rs10890952 | -2.334 | 1.367 | 1.946 | 1.912 | 0.014 | 0.014 | 0.014 | 0 | 0.17 | 0.21 | 0.38 | 0.34 | 0.33 | 0.23 |
| 11 | 109030401 | rs2089673 | -1.913 | 1.169 | 2.069 | 1.978 | -0.03 | -0.027 | -0.027 | 0.801 | 0.15 | 0.12 | 0.35 | 0.25 | 0.29 | 0.31 |
| 11 | 109071810 | rs11212983 | -1.186 | 1.288 | 2.586 | 1.925 | -0.034 | -0.032 | -0.032 | 0 | 0.11 | 0.06 | 0.35 | 0.25 | 0.29 | 0.22 |
| 11 | 109073592 | rs11212985 | -1.023 | 1.288 | 2.586 | 1.93 | -0.03 | -0.029 | -0.029 | 0 | 0.11 | 0.06 | 0.35 | 0.25 | 0.29 | 0.2 |
| 11 | 109075064 | rs10890960 | -1.255 | 1.284 | 2.601 | 1.915 | -0.027 | -0.026 | -0.026 | 0 | 0.12 | 0.07 | 0.35 | 0.26 | 0.29 | 0.22 |
| 11 | 109111782 | rs11213000 | 2.091 | 1.273 | 2.923 | 1.917 | 0.073 | 0.128 | 0.128 | 0 | 0.81 | 0.44 | 0.6 | 0.6 | 0.69 | 0.79 |
| 11 | 109118149 | rs12276424 | -1.771 | 1.366 | 2.366 | 1.964 | 0.024 | 0.177 | 0.177 | 2.042 | 0.06 | 0.32 | 0.04 | 0.14 | 0.02 | 0.02 |
| 11 | 109127876 | rs11213013 | -1.168 | 1.579 | 1.808 | 2.046 | -0.019 | -0.019 | -0.019 | 0 | 0.11 | 0.07 | 0.34 | 0.27 | 0.29 | 0.18 |
| 11 | 109129584 | rs12422091 | -1.479 | 1.712 | 0.908 | 1.481 | 0.07 | 0.055 | 0.055 | 0 | 0.13 | 0.22 | 0.36 | 0.27 | 0.29 | 0.18 |
| 11 | 109146890 | rs11213025 | 1.028 | 1.738 | 0.835 | 1.494 | 0.085 | 0.067 | 0.067 | 0 | 0.87 | 0.76 | 0.64 | 0.74 | 0.71 | 0.82 |
| 11 | 109156782 | rs1482047 | -1.784 | 0.178 | 0.782 | 0.855 | -0.029 | -0.025 | -0.025 | 0 | 0.17 | 0.69 | 0.13 | 0.08 | 0.08 | 0.12 |
| 11 | 109173027 | rs11213040 | -1.305 | 0.631 | 0.83 | 1.119 | 0.034 | 0.053 | 0.053 | 0.426 | 0.05 | 0.21 | 0.07 | 0.04 | 0.04 | 0.02 |
| 11 | 109176817 | rs655763 | 0.096 | -0.271 | 0.11 | 1.109 | 0.026 | -0.005 | -0.005 | 0 | 0.55 | 0.39 | 0.59 | 0.5 | 0.7 | 0.64 |
| 11 | 109179660 | rs10749942 | -1.357 | -0.579 | 1.049 | 2.161 | -0.049 | 0.249 | 0.249 | 3.039 | 0.15 | 0.56 | 0.15 | 0.34 | 0.08 | 0.14 |
| 11 | 109190254 | rs951336 | -1.692 | -0.694 | 1.553 | 2.165 | 0.033 | 0.438 | 0.438 | 3.945 | 0.08 | 0.43 | 0.13 | 0.33 | 0.05 | 0.14 |
| 11 | 109190345 | rs633619 | -1.836 | -0.473 | 0.972 | 1.717 | 0.012 | 0.018 | 0.018 | 0.305 | 0.39 | 0.75 | 0.45 | 0.57 | 0.37 | 0.4 |
| 11 | 109198849 | rs559508 | -1.527 | -0.761 | 1.006 | 1.541 | 0 | 0.003 | 0.003 | 0.241 | 0.34 | 0.68 | 0.44 | 0.58 | 0.37 | 0.41 |
| 11 | 109206471 | rs618316 | -2.032 | -0.52 | -0.006 | 1.05 | -0.102 | 0.082 | 0.082 | 0.695 | 0.48 | 0.97 | 0.53 | 0.63 | 0.51 | 0.52 |
| 11 | 109213216 | rs11822621 | -1.616 | -0.65 | 1.37 | 0.857 | 0.01 | -0.098 | -0.098 | 0.525 | 0.14 | 0.44 | 0.08 | 0 | 0.09 | 0.02 |
| 11 | 109213268 | rs11827212 | -1.668 | -0.651 | 1.436 | 0.857 | 0 | -0.094 | -0.094 | 0 | 0.14 | 0.48 | 0.09 | 0 | 0.1 | 0.02 |
| 11 | 109219603 | rs593376 | -1.549 | -0.65 | 1.492 | 0.858 | 0.013 | -0.094 | -0.094 | 0 | 0.14 | 0.43 | 0.07 | 0 | 0.09 | 0.02 |
| 11 | 109228405 | rs10890985 | -2.594 | -0.745 | 1.44 | 1.424 | -0.013 | -0.006 | -0.006 | 0.127 | 0.21 | 0.75 | 0.19 | 0.14 | 0.1 | 0.09 |
| 11 | 109241241 | rs7947221 | -2.682 | -0.778 | 1.9 | 1.463 | -0.089 | -0.056 | -0.056 | 0.645 | 0.15 | 0.56 | 0.14 | 0.07 | 0.04 | 0.08 |
| 11 | 109248945 | rs17109617 | -2.561 | -0.71 | 2.042 | 1.539 | -0.039 | 0.034 | 0.034 | 1.935 | 0.14 | 0.5 | 0.14 | 0.14 | 0.06 | 0.09 |
| 11 | 109278427 | rs2186671 | -1.624 | -0.608 | 1.307 | 1.657 | 0.002 | -0.005 | -0.005 | 0.252 | 0.44 | 0.57 | 0.47 | 0.54 | 0.47 | 0.43 |

| | | | | | | | | | | | | | | | | |
|---|---|---|---|---|---|---|---|---|---|---|---|---|---|---|---|---|
| 11 | 109284053 | rs10749943 | -1.265 | -0.605 | 1.082 | 1.747 | -0.001 | 0.001 | 0.001 | 0.142 | 0.34 | 0.56 | 0.41 | 0.41 | 0.38 | 0.37 |
| 11 | 109285173 | rs1939422 | -0.642 | -0.616 | 1.248 | 1.912 | -0.008 | 0.005 | 0.005 | 0.662 | 0.28 | 0.19 | 0.28 | 0.27 | 0.36 | 0.24 |
| 11 | 109285519 | rs2186673 | 0.313 | -0.613 | 0.92 | 1.988 | -0.069 | -0.1 | -0.1 | 0 | 0.21 | 0.01 | 0.25 | 0.32 | 0.31 | 0.23 |
| 11 | 109287367 | rs11213069 | -0.329 | -0.482 | 0.987 | 2.017 | 0.006 | -0.013 | -0.013 | 0 | 0.09 | 0.09 | 0.14 | 0.19 | 0.07 | 0.15 |
| 11 | 109289538 | rs2212430 | 0.233 | -0.318 | 0.935 | 2.115 | -0.082 | -0.063 | -0.063 | 0.998 | 0.21 | 0.01 | 0.26 | 0.27 | 0.32 | 0.23 |
| 11 | 109298127 | rs4403771 | -1.119 | -0.396 | 1.334 | 2.316 | 0.004 | 0.004 | 0.004 | 0.518 | 0.32 | 0.37 | 0.47 | 0.32 | 0.39 | 0.26 |
| 11 | 109311661 | rs11213079 | 0.857 | -1.191 | 0.719 | 2.352 | 0.002 | -0.046 | -0.046 | 0 | 0.08 | 0.01 | 0.09 | 0.23 | 0.04 | 0.1 |
| 11 | 109325299 | rs11213086 | -0.443 | -1.131 | 0.818 | 2.219 | -0.03 | 0.012 | 0.012 | 1.206 | 0.28 | 0.15 | 0.44 | 0.25 | 0.38 | 0.25 |
| 11 | 109333446 | rs11213088 | -0.479 | -1.13 | 0.819 | 2.266 | -0.031 | 0.013 | 0.013 | 1.258 | 0.28 | 0.15 | 0.44 | 0.24 | 0.38 | 0.25 |
| 11 | 109342809 | rs10789731 | -1.06 | -1.136 | 0.84 | 2.255 | 0.001 | 0.005 | 0.005 | 0.078 | 0.34 | 0.46 | 0.54 | 0.39 | 0.39 | 0.34 |
| 11 | 109344049 | rs1439509 | -0.685 | -1.055 | 0.792 | 2.174 | 0.005 | -0.015 | -0.015 | 2.606 | 0.31 | 0.38 | 0.46 | 0.25 | 0.38 | 0.25 |
| 11 | 109357059 | rs1347910 | -3.092 | -1.725 | -0.871 | 1.123 | -0.152 | -0.016 | -0.016 | 0 | 0.69 | 0.98 | 0.78 | 0.74 | 0.61 | 0.52 |
| 11 | 109382921 | rs1439518 | -0.235 | -0.867 | -0.455 | 1.075 | -0.053 | -0.031 | -0.031 | 0.565 | 0.29 | 0.11 | 0.44 | 0.3 | 0.42 | 0.23 |
| 11 | 109389257 | rs1837396 | -0.553 | -0.966 | -0.311 | 0.911 | -0.011 | 0.038 | 0.038 | 0 | 0.37 | 0.23 | 0.49 | 0.24 | 0.46 | 0.23 |
| 11 | 109405386 | rs7944752 | -0.442 | -0.851 | 0.089 | 0.722 | -0.014 | -0.104 | -0.104 | 0 | 0.18 | 0.33 | 0.1 | 0 | 0.13 | 0.03 |
| 11 | 109417401 | rs7129310 | 1.517 | -0.851 | 0.083 | 0.751 | -0.02 | -0.045 | -0.045 | 0.001 | 0.09 | 0.14 | 0.05 | 0 | 0.04 | 0.01 |
| 11 | 109424249 | rs6589074 | 0.218 | -0.712 | 0.096 | 0.884 | 0.001 | -0.022 | -0.022 | 0 | 0.07 | 0.11 | 0.04 | 0 | 0.08 | 0.02 |
| 11 | 109463744 | rs7931146 | -1.107 | -0.739 | 0.224 | 0.835 | 0.015 | -0.064 | -0.064 | 2.746 | 0.11 | 0.33 | 0.07 | 0 | 0.09 | 0.04 |
| 11 | 109465968 | rs2029297 | 0.005 | -0.553 | -0.099 | 0.945 | -0.04 | -0.103 | -0.103 | 0 | 0.3 | 0.13 | 0.53 | 0.52 | 0.4 | 0.31 |
| 11 | 109472829 | rs11602639 | 1.053 | -0.484 | -0.179 | 0.971 | -0.093 | -0.048 | -0.048 | 0.599 | 0.22 | 0.02 | 0.4 | 0.23 | 0.33 | 0.22 |
| 11 | 109475560 | rs1906339 | -0.93 | -0.573 | 0.349 | 0.829 | 0.005 | -0.034 | -0.034 | 0 | 0.93 | 0.8 | 0.96 | 1 | 0.94 | 0.99 |
| 11 | 109489575 | rs11213128 | 0.716 | -0.468 | -0.173 | 1.021 | -0.052 | -0.114 | -0.114 | 0.348 | 0.24 | 0.08 | 0.44 | 0.47 | 0.35 | 0.28 |
| 11 | 109491073 | rs7947735 | -0.6 | -0.43 | -0.018 | 0.968 | 0.005 | -0.054 | -0.054 | 0 | 0.1 | 0.22 | 0.06 | 0 | 0.09 | 0.04 |
| 11 | 109495509 | rs12804063 | 0.902 | -0.503 | -0.175 | 1.053 | -0.068 | -0.07 | -0.07 | 0 | 0.23 | 0.04 | 0.41 | 0.3 | 0.35 | 0.25 |
| 11 | 109508311 | rs1386718 | 0.394 | -0.364 | 0.047 | 0.798 | -0.051 | -0.076 | -0.076 | 0.011 | 0.31 | 0.09 | 0.49 | 0.44 | 0.4 | 0.32 |
| 11 | 109515991 | rs7948765 | -0.379 | -0.551 | 0.452 | 0.532 | 0.015 | -0.052 | -0.052 | 0 | 0.1 | 0.28 | 0.06 | 0 | 0.09 | 0.04 |
| 11 | 109523271 | rs7940859 | -0.645 | -0.557 | 0.409 | 0.527 | 0.01 | -0.065 | -0.065 | 0 | 0.11 | 0.32 | 0.06 | 0 | 0.09 | 0.04 |
| 11 | 109530024 | rs7111821 | -0.07 | -0.564 | -0.113 | 0.476 | -0.001 | -0.016 | -0.016 | 0 | 0.11 | 0.1 | 0.05 | 0 | 0.1 | 0.04 |
| 11 | 109559457 | rs1386719 | -2.204 | -0.604 | -0.347 | 0.512 | 0.049 | 0.065 | 0.065 | 2.744 | 0.44 | 0.82 | 0.28 | 0.26 | 0.31 | 0.31 |
| 11 | 109561421 | rs2852110 | 0.148 | -0.618 | -0.262 | 0.475 | -0.001 | -0.018 | -0.018 | 0.102 | 0.11 | 0.1 | 0.05 | 0 | 0.11 | 0.04 |
| 11 | 109575106 | rs1685399 | -0.821 | -0.583 | 0.34 | 0.688 | -0.032 | -0.002 | -0.002 | 0 | 0.3 | 0.5 | 0.2 | 0.26 | 0.17 | 0.25 |
| 11 | 109580367 | rs936913 | -0.82 | -0.601 | 0.31 | 0.872 | -0.028 | 0.054 | 0.054 | 0.19 | 0.23 | 0.54 | 0.16 | 0.26 | 0.14 | 0.25 |
| 11 | 109595644 | rs2716045 | -2.314 | -0.515 | -0.122 | 1.018 | -0.115 | -0.048 | -0.048 | 0 | 0.67 | 0.98 | 0.76 | 0.71 | 0.65 | 0.6 |
| 11 | 109600786 | rs11213146 | -1.475 | -0.403 | 1.438 | 1.285 | -0.021 | 0.016 | 0.016 | 0.263 | 0.24 | 0.65 | 0.17 | 0.26 | 0.14 | 0.25 |
| 11 | 109615658 | rs875430 | -0.918 | -0.418 | 1.437 | 1.113 | -0.001 | -0.021 | -0.021 | 0 | 0.11 | 0.14 | 0.05 | 0 | 0.08 | 0.04 |
| 11 | 109616540 | rs2002733 | -0.8 | -0.4 | 1.501 | 1.244 | -0.006 | 0.067 | 0.067 | 0 | 0.2 | 0.55 | 0.15 | 0.26 | 0.15 | 0.25 |
| 11 | 109633392 | rs1433560 | 1.216 | -0.333 | 1.118 | 1.036 | -0.005 | -0.001 | -0.001 | 0 | 0.74 | 0.29 | 0.82 | 0.77 | 0.85 | 0.74 |
| 11 | 109635168 | rs2570318 | -0.337 | -0.295 | 1.003 | 1.055 | -0.001 | -0.015 | -0.015 | 0 | 0.09 | 0.09 | 0.04 | 0 | 0.08 | 0.04 |
| 11 | 109637777 | rs2253536 | -2.384 | -0.326 | 0.93 | 1.076 | 0.025 | 0.029 | 0.029 | 2.071 | 0.34 | 0.9 | 0.23 | 0.24 | 0.23 | 0.29 |
| 11 | 109638849 | rs6589085 | -2.018 | -0.072 | 1.386 | 1.206 | -0.007 | 0.02 | 0.02 | 1.834 | 0.36 | 0.56 | 0.21 | 0.38 | 0.29 | 0.29 |
| 11 | 109638891 | rs6589086 | -1.988 | 0.272 | 1.831 | 0.425 | 0.001 | -0.024 | -0.024 | 0.578 | 0.43 | 0.57 | 0.33 | 0.26 | 0.41 | 0.36 |
| 11 | 109640972 | rs989695 | -1.438 | 0.284 | 2.477 | 0.883 | -0.028 | 0.032 | 0.032 | 1.272 | 0.18 | 0.61 | 0.14 | 0.19 | 0.09 | 0.23 |
| 11 | 109641652 | rs7122607 | -1.471 | 0.283 | 2.403 | 0.882 | -0.028 | 0.022 | 0.022 | 1.823 | 0.18 | 0.65 | 0.14 | 0.19 | 0.09 | 0.23 |
| 11 | 109647282 | rs2570320 | -2.316 | 0.317 | 1.842 | 0.962 | -0.007 | 0.026 | 0.026 | 0 | 0.42 | 0.87 | 0.31 | 0.34 | 0.35 | 0.38 |
| 11 | 109651691 | rs2570308 | -0.863 | 0.486 | 1.822 | 0.903 | -0.019 | 0.038 | 0.038 | 0 | 0.23 | 0.12 | 0.17 | 0.13 | 0.26 | 0.15 |
| 11 | 109665299 | rs6589089 | -1.378 | 0.453 | 2.281 | 1.062 | -0.025 | 0.044 | 0.044 | 0.857 | 0.18 | 0.48 | 0.12 | 0.19 | 0.09 | 0.22 |
| 11 | 109666687 | rs1368688 | -2.487 | 0.439 | 2.039 | 1.039 | -0.012 | 0.023 | 0.023 | 0.059 | 0.43 | 0.9 | 0.31 | 0.32 | 0.35 | 0.38 |
| 11 | 109671655 | rs556273 | 2.176 | 0.461 | 2.072 | 0.79 | -0.025 | 0.059 | 0.059 | 0 | 0.59 | 0.1 | 0.69 | 0.72 | 0.65 | 0.63 |
| 11 | 109673783 | rs2105779 | -0.718 | 0.539 | 2.129 | 0.702 | -0.041 | 0.042 | 0.042 | 0 | 0.21 | 0.07 | 0.16 | 0.13 | 0.25 | 0.15 |
| 11 | 109680887 | rs1789818 | 1.387 | 0.299 | 2.607 | 0.832 | -0.028 | 0.055 | 0.055 | 5.973 | 0.91 | 0.44 | 0.94 | 0.96 | 1 | 1 |
| 11 | 109682445 | rs689152 | 1.063 | 0.304 | 2.654 | 0.835 | 0.004 | 0.099 | 0.099 | 3.426 | 0.92 | 0.54 | 0.95 | 0.96 | 1 | 1 |
| 11 | 109707703 | rs76214 | 1.573 | 0.423 | 2.223 | 0.828 | 0.002 | 0.024 | 0.024 | 0.003 | 0.58 | 0.28 | 0.7 | 0.73 | 0.64 | 0.64 |
| 11 | 109727767 | rs554442 | 1.467 | -0.047 | 2.337 | 0.877 | -0.017 | 0.037 | 0.037 | 3.577 | 0.9 | 0.42 | 0.92 | 0.96 | 0.98 | 1 |
| 11 | 109739968 | rs600238 | 1.33 | 0.204 | 1.839 | 0.954 | 0.005 | 0.038 | 0.038 | 0.005 | 0.58 | 0.12 | 0.7 | 0.72 | 0.67 | 0.64 |
| 11 | 109754949 | rs475115 | 1.199 | 0.167 | 1.075 | 0.711 | 0.012 | -0.02 | -0.02 | 0 | 0.6 | 0.17 | 0.71 | 0.65 | 0.68 | 0.63 |
| 11 | 109761786 | rs561781 | 1.174 | 0.127 | 2.066 | 0.625 | -0.006 | 0.001 | 0.001 | 0 | 0.62 | 0.5 | 0.73 | 0.65 | 0.68 | 0.63 |
| 11 | 109768020 | rs17110364 | -1.157 | 0.056 | 2.26 | 0.644 | -0.016 | 0.003 | 0.003 | 0 | 0.32 | 0.15 | 0.22 | 0.3 | 0.31 | 0.37 |
| 11 | 109805805 | rs7117254 | -1.293 | 0.331 | 2.078 | 0.495 | 0 | -0.005 | -0.005 | 0.097 | 0.33 | 0.32 | 0.23 | 0.3 | 0.31 | 0.37 |
| 11 | 109810415 | rs10891037 | -1.382 | 0.494 | 1.38 | 0.453 | -0.048 | 0.019 | 0.019 | 0.013 | 0.36 | 0.16 | 0.31 | 0.31 | 0.4 | 0.41 |
| 11 | 109825345 | rs4754415 | -1.483 | 0.55 | 1.777 | 0.211 | -0.025 | 0.026 | 0.026 | 0.506 | 0.29 | 0.21 | 0.22 | 0.21 | 0.32 | 0.35 |
| 11 | 109839996 | rs10891041 | -1.984 | 0.508 | 1.725 | 0.238 | 0.007 | 0.02 | 0.02 | 0.021 | 0.36 | 0.88 | 0.33 | 0.27 | 0.31 | 0.38 |
| 11 | 109842329 | rs7109506 | -1.843 | 0.924 | 2.084 | 0.288 | 0.003 | -0.005 | -0.005 | 2.677 | 0.35 | 0.64 | 0.34 | 0.28 | 0.35 | 0.39 |
| 11 | 109843953 | rs10891042 | -1.329 | 0.151 | 2.068 | 0.281 | -0.002 | 0.006 | 0.006 | 0.092 | 0.28 | 0.27 | 0.21 | 0.2 | 0.28 | 0.34 |
| 11 | 109854283 | rs11213219 | -1.532 | 0.144 | 1.748 | 0.315 | -0.001 | 0.018 | 0.018 | 0.024 | 0.32 | 0.3 | 0.29 | 0.2 | 0.32 | 0.38 |
| 11 | 109864038 | rs6589106 | -1.066 | 0.271 | 0.537 | 0.387 | -0.005 | 0.03 | 0.03 | 0 | 0.19 | 0.48 | 0.26 | 0.17 | 0.19 | 0.14 |
| 11 | 109864550 | rs10891043 | -1.061 | 0.329 | 0.025 | 0.219 | -0.003 | -0.01 | -0.01 | 0 | 0.18 | 0.24 | 0.14 | 0.13 | 0.14 | 0.26 |
| 11 | 109865559 | rs7947141 | 0.919 | 0.269 | -0.055 | 0.142 | -0.018 | -0.043 | -0.043 | 2.98 | 0.77 | 0.31 | 0.86 | 0.9 | 0.86 | 0.8 |
| 11 | 109865644 | rs9633915 | 0.598 | 0.319 | -0.743 | 0.303 | -0.011 | -0.009 | -0.009 | 0.045 | 0.66 | 0.38 | 0.72 | 0.7 | 0.73 | 0.87 |
| 11 | 109869302 | rs1648144 | -0.884 | 0.684 | -2.073 | 0.206 | -0.036 | 0.077 | 0.077 | 0.179 | 0.57 | 0.97 | 0.8 | 0.72 | 0.62 | 0.56 |

| | | | | | | | | | | | | | | | | |
|---|---|---|---|---|---|---|---|---|---|---|---|---|---|---|---|---|
| 11 | 109870083 | rs4753878 | 0.001 | 1.197 | -1.751 | 0.44 | -0.053 | -0.023 | -0.023 | 0 | 0.13 | 0.02 | 0.1 | 0.13 | 0.15 | 0.11 |
| 11 | 109883416 | rs1648136 | 0.497 | 0.723 | -1.429 | 0.483 | 0.037 | 0.071 | 0.071 | 0 | 0.42 | 0.19 | 0.22 | 0.31 | 0.36 | 0.39 |
| 11 | 109888898 | rs10891046 | -0.54 | 1.33 | -0.497 | 1.152 | 0.042 | 0.061 | 0.061 | 0 | 0.2 | 0.27 | 0.33 | 0.37 | 0.29 | 0.28 |
| 11 | 109898785 | rs1789377 | 0.213 | 0.997 | -0.595 | 1.034 | 0.007 | -0.01 | -0.01 | 0 | 0.58 | 0.44 | 0.58 | 0.53 | 0.58 | 0.53 |
| 11 | 109899405 | rs12576062 | -0.467 | 1.352 | -0.316 | 1.078 | 0.003 | -0.001 | -0.001 | 0 | 0.11 | 0.14 | 0.12 | 0.09 | 0.18 | 0.09 |
| 11 | 109900455 | rs12574224 | -0.481 | 1.347 | -0.357 | 1.066 | -0.022 | -0.002 | -0.002 | 0 | 0.1 | 0.07 | 0.11 | 0.1 | 0.18 | 0.08 |
| 11 | 109903196 | rs10789746 | 0.584 | 1.154 | -0.447 | 0.949 | -0.015 | 0.001 | 0.001 | 0.33 | 0.45 | 0.22 | 0.29 | 0.46 | 0.44 | 0.47 |
| 11 | 109906198 | rs1789374 | -0.808 | 0.114 | -1.283 | -0.054 | -0.018 | 0.079 | 0.079 | 0 | 0.64 | 0.89 | 0.69 | 0.84 | 0.68 | 0.75 |
| 11 | 109915378 | rs1648156 | 0.897 | -0.032 | -0.499 | 0.246 | -0.008 | -0.001 | -0.001 | 0.633 | 0.19 | 0.2 | 0.08 | 0.11 | 0.12 | 0.2 |
| 11 | 109917159 | rs1038057 | 0.848 | -0.035 | -0.377 | 0.275 | -0.024 | -0.007 | -0.007 | 0 | 0.19 | 0.24 | 0.09 | 0.11 | 0.13 | 0.2 |
| 11 | 109928038 | rs17110672 | -0.491 | 0.768 | 0.019 | 0.553 | 0.034 | 0.005 | 0.005 | 0 | 0.14 | 0.28 | 0.35 | 0.19 | 0.2 | 0.09 |
| 11 | 109929743 | rs11213245 | -0.728 | 0.959 | -0.131 | 0.567 | 0.02 | -0.012 | -0.012 | 0 | 0.32 | 0.5 | 0.28 | 0.25 | 0.37 | 0.26 |
| 11 | 109934487 | rs1648130 | -0.918 | 1.079 | -0.983 | 0.278 | 0.004 | 0.073 | 0.073 | 0 | 0.7 | 0.89 | 0.72 | 0.88 | 0.71 | 0.65 |
| 11 | 109934639 | rs10891049 | 0.664 | 1.122 | -0.829 | 0.69 | -0.034 | -0.107 | -0.107 | 0 | 0.35 | 0.11 | 0.21 | 0.52 | 0.35 | 0.38 |
| 11 | 109936027 | rs1648131 | -0.546 | 1.558 | -0.439 | 0.773 | -0.008 | -0.002 | -0.002 | 0 | 0.15 | 0.17 | 0.37 | 0.22 | 0.25 | 0.1 |
| 11 | 109943972 | rs7113219 | 0.009 | 1.248 | -0.068 | 1.144 | 0.026 | -0.012 | -0.012 | 0 | 0.39 | 0.65 | 0.43 | 0.6 | 0.26 | 0.32 |
| 11 | 109944920 | rs1618224 | -0.078 | 1.801 | -0.147 | 1.163 | 0.006 | -0.003 | -0.003 | 0.205 | 0.11 | 0.1 | 0.15 | 0.14 | 0.26 | 0.09 |
| 11 | 109950572 | rs11213251 | 0.701 | 1.313 | -0.188 | 1.109 | 0.051 | -0.078 | -0.078 | 0 | 0.33 | 0.1 | 0.15 | 0.53 | 0.23 | 0.3 |
| 11 | 109956346 | rs7115956 | -0.166 | 1.627 | -0.206 | 1.076 | -0.062 | -0.068 | -0.068 | 0.214 | 0.45 | 0.11 | 0.32 | 0.57 | 0.51 | 0.4 |
| 11 | 109962432 | rs4561177 | -0.836 | 1.416 | 0.098 | 1.006 | -0.013 | -0.018 | -0.018 | 0 | 0.31 | 0.22 | 0.33 | 0.65 | 0.39 | 0.34 |
| 11 | 109977505 | rs11213266 | 1.007 | 1.18 | -0.098 | 1.508 | 0.001 | -0.093 | -0.093 | 0 | 0.13 | 0.01 | 0.14 | 0.41 | 0.1 | 0.22 |
| 11 | 109995287 | rs1789369 | -1.292 | 1.205 | 0.206 | 0.476 | 0.01 | 0.053 | 0.053 | 0 | 0.66 | 0.76 | 0.75 | 0.9 | 0.69 | 0.54 |
| 11 | 109995492 | rs11213279 | 0.923 | 1.294 | 0.576 | 0.46 | 0.001 | -0.006 | -0.006 | 0 | 0.34 | 0.26 | 0.16 | 0.08 | 0.28 | 0.19 |
| 11 | 109995642 | rs612510 | -0.168 | 1.329 | 0.542 | 0.508 | -0.017 | -0.017 | -0.017 | 0 | 0.3 | 0.25 | 0.35 | 0.65 | 0.4 | 0.34 |
| 11 | 109999257 | rs12366209 | 0.611 | 1.046 | 0.42 | 0.539 | -0.006 | -0.048 | -0.048 | 0 | 0.34 | 0.4 | 0.17 | 0.18 | 0.29 | 0.2 |
| 11 | 110002744 | rs1564013 | -0.804 | 1.014 | 0.48 | 0.935 | -0.017 | 0.022 | 0.022 | 0 | 0.63 | 0.51 | 0.5 | 0.59 | 0.7 | 0.54 |
| 11 | 110005581 | rs682668 | -0.34 | 1.178 | 0.628 | 0.673 | 0.009 | 0.001 | 0.001 | 0.459 | 0.32 | 0.41 | 0.58 | 0.69 | 0.44 | 0.37 |
| 11 | 110008339 | rs10891063 | 0.911 | 1.223 | 0.65 | 0.624 | 0.009 | 0.015 | 0.015 | 0.004 | 0.33 | 0.21 | 0.16 | 0.07 | 0.26 | 0.19 |
| 11 | 110010175 | rs660222 | -0.431 | 1.22 | 0.72 | 0.666 | 0.02 | 0.002 | 0.002 | 0.355 | 0.33 | 0.48 | 0.59 | 0.7 | 0.44 | 0.37 |
| 11 | 110015904 | rs1381466 | 0.69 | 0.946 | 0.685 | 0.645 | -0.005 | -0.009 | -0.009 | 1.128 | 0.34 | 0.3 | 0.36 | 0.18 | 0.26 | 0.19 |
| 11 | 110016519 | rs661171 | -0.926 | 1.296 | 0.48 | 0.269 | -0.003 | -0.009 | -0.009 | 0 | 0.65 | 0.61 | 0.76 | 0.79 | 0.7 | 0.58 |
| 11 | 110017029 | rs648594 | -1.215 | 1.073 | 0.321 | -0.042 | 0.003 | 0.027 | 0.027 | 0 | 0.68 | 0.75 | 0.76 | 0.9 | 0.71 | 0.59 |
| 11 | 110019706 | rs10891068 | 0.686 | 0.757 | 0.083 | -0.157 | 0.008 | 0.093 | 0.093 | 0 | 0.29 | 0.12 | 0.14 | 0.08 | 0.25 | 0.14 |
| 11 | 110023164 | rs622822 | -0.435 | 0.66 | 0.632 | -0.006 | -0.029 | -0.051 | -0.051 | 0 | 0.63 | 0.52 | 0.75 | 0.8 | 0.73 | 0.81 |
| 11 | 110035262 | rs1026608 | 0.217 | 0.638 | 0.63 | -0.052 | 0.022 | 0.042 | 0.042 | 0 | 0.34 | 0.22 | 0.22 | 0.2 | 0.27 | 0.19 |
| 11 | 110035301 | rs1026607 | 0.155 | 0.637 | 0.709 | -0.052 | 0.001 | 0 | 0 | 0 | 0.35 | 0.3 | 0.22 | 0.2 | 0.27 | 0.19 |
| 11 | 110042157 | rs4754425 | 0.106 | 0.759 | 0.557 | 0.026 | 0.01 | 0.014 | 0.014 | 0 | 0.38 | 0.34 | 0.28 | 0.29 | 0.32 | 0.22 |
| 11 | 110043632 | rs7947099 | 0.143 | 0.747 | 0.592 | 0.02 | 0.007 | 0.01 | 0.01 | 0 | 0.37 | 0.36 | 0.28 | 0.29 | 0.31 | 0.22 |
| 11 | 110043820 | rs7947237 | -1.308 | 0.638 | 0.19 | -0.43 | 0.026 | 0.123 | 0.123 | 0 | 0.7 | 0.89 | 0.8 | 0.96 | 0.76 | 0.62 |
| 11 | 110045731 | rs7941903 | -0.649 | 1.391 | 0.512 | -0.152 | -0.005 | 0.032 | 0.032 | 0 | 0.57 | 0.66 | 0.64 | 0.82 | 0.51 | 0.52 |
| 11 | 110047686 | rs6589117 | -0.703 | 1.392 | 0.361 | -0.124 | -0.021 | 0.113 | 0.113 | 0 | 0.57 | 0.78 | 0.65 | 0.82 | 0.51 | 0.5 |
| 11 | 110048643 | rs11213290 | 0.462 | 1.512 | 0.489 | -0.109 | 0.001 | 0.014 | 0.014 | 0 | 0.22 | 0.23 | 0.33 | 0.54 | 0.24 | 0.33 |
| 11 | 110052890 | rs898845 | -0.521 | 1.451 | 0.235 | -0.138 | -0.016 | 0.082 | 0.082 | 0 | 0.56 | 0.73 | 0.58 | 0.73 | 0.51 | 0.47 |
| 11 | 110053654 | rs4754429 | 0.447 | 1.463 | 0.34 | 0.206 | -0.001 | -0.001 | -0.001 | 0 | 0.09 | 0.09 | 0.11 | 0.1 | 0.2 | 0.08 |
| 11 | 110059838 | rs11525418 | -0.669 | 1.367 | 0.587 | 0.194 | 0.04 | 0.01 | 0.01 | 0.955 | 0.32 | 0.52 | 0.46 | 0.63 | 0.44 | 0.39 |
| 11 | 110068043 | rs7109556 | -0.473 | 1.392 | 0.745 | 0.195 | 0.013 | 0.007 | 0.007 | 0.064 | 0.29 | 0.31 | 0.4 | 0.63 | 0.39 | 0.4 |
| 11 | 110122426 | rs7932174 | 0.927 | 1.104 | 0.485 | -0.289 | -0.025 | 0.083 | 0.083 | 0 | 0.34 | 0.19 | 0.21 | 0.08 | 0.36 | 0.19 |
| 11 | 110130358 | rs1784649 | -1.28 | 1.04 | 0.527 | -0.281 | 0.064 | 0.099 | 0.099 | 0 | 0.67 | 0.86 | 0.73 | 0.84 | 0.76 | 0.6 |
| 11 | 110143054 | rs1676530 | 0.919 | 1.108 | 0.553 | -0.286 | -0.035 | 0.105 | 0.105 | 0 | 0.34 | 0.13 | 0.23 | 0.09 | 0.37 | 0.19 |
| 11 | 110144023 | rs10891078 | -0.297 | 1.156 | 0.571 | -0.21 | 0.018 | 0.006 | 0.006 | 0.212 | 0.31 | 0.37 | 0.46 | 0.66 | 0.38 | 0.41 |
| 11 | 110147873 | rs12226901 | -0.471 | 1.177 | 0.392 | -0.012 | 0.022 | 0.047 | 0.047 | 0.734 | 0.32 | 0.49 | 0.44 | 0.53 | 0.38 | 0.42 |
| 11 | 110160864 | rs3858405 | -0.591 | 1.081 | 0.247 | -0.104 | 0.01 | 0.007 | 0.007 | 1.022 | 0.33 | 0.6 | 0.44 | 0.62 | 0.4 | 0.44 |
| 11 | 110162430 | rs1676536 | 1.285 | 0.721 | -0.414 | -0.677 | -0.167 | 0.069 | 0.069 | 0.044 | 0.27 | 0.01 | 0.22 | 0.09 | 0.38 | 0.14 |
| 11 | 110166131 | rs1676535 | -0.979 | 0.342 | 0.197 | -0.439 | 0.019 | 0.022 | 0.022 | 0 | 0.66 | 0.72 | 0.72 | 0.85 | 0.77 | 0.63 |
| 11 | 110168674 | rs1676522 | 0.226 | 0.503 | 0.218 | -0.466 | -0.01 | 0.037 | 0.037 | 0 | 0.42 | 0.38 | 0.33 | 0.26 | 0.53 | 0.27 |
| 11 | 110177371 | rs1792783 | -1.01 | 0.44 | 0.173 | -0.511 | 0.004 | -0.014 | -0.014 | 0 | 0.65 | 0.66 | 0.72 | 0.92 | 0.77 | 0.59 |
| 11 | 110181512 | rs4753885 | 0.28 | 0.529 | 0.109 | -0.412 | 0.015 | 0.04 | 0.04 | 0 | 0.22 | 0.26 | 0.38 | 0.65 | 0.24 | 0.33 |
| 11 | 110186451 | rs1676533 | 0.558 | 0.618 | -0.403 | -0.626 | -0.041 | 0.075 | 0.075 | 0 | 0.37 | 0.17 | 0.28 | 0.17 | 0.43 | 0.24 |
| 11 | 110190805 | rs1792818 | -0.779 | -0.203 | -0.331 | -0.617 | -0.008 | -0.031 | -0.031 | 0 | 0.69 | 0.67 | 0.74 | 0.92 | 0.78 | 0.61 |
| 11 | 110192424 | rs1784648 | 0.06 | 0.17 | -0.045 | 0.507 | -0.03 | 0.09 | 0.09 | 0 | 0.74 | 0.56 | 0.76 | 0.55 | 0.83 | 0.74 |
| 11 | 110196789 | rs4754441 | -0.163 | 0.28 | -0.168 | 0.449 | -0.046 | -0.09 | -0.09 | 0.396 | 0.28 | 0.12 | 0.29 | 0.46 | 0.32 | 0.28 |
| 11 | 110203900 | rs12420766 | 0.521 | 0.218 | -0.856 | 0.252 | -0.017 | -0.003 | -0.003 | 0.341 | 0.37 | 0.11 | 0.38 | 0.33 | 0.36 | 0.62 |
| 11 | 110206931 | rs1792806 | 1.191 | 0.338 | -0.385 | 0.274 | -0.01 | 0.007 | 0.007 | 0 | 0.26 | 0.16 | 0.25 | 0.22 | 0.3 | 0.11 |
| 11 | 110210385 | rs17728426 | 0.601 | 0.292 | -0.337 | 0.355 | 0 | 0.009 | 0.009 | 0 | 0.31 | 0.12 | 0.33 | 0.23 | 0.28 | 0.37 |
| 11 | 110214382 | rs1792802 | -0.574 | 0.394 | -0.295 | 0.561 | -0.02 | -0.009 | -0.009 | 0 | 0.72 | 0.51 | 0.68 | 0.78 | 0.78 | 0.64 |
| 11 | 110215976 | rs12578007 | -1.033 | 1.114 | -0.378 | 1.023 | 0.009 | 0.014 | 0.014 | 0.049 | 0.48 | 0.24 | 0.44 | 0.61 | 0.54 | 0.52 |
| 11 | 110221134 | rs1792796 | 0.653 | 1.226 | 0.026 | 1.041 | -0.01 | 0.002 | 0.002 | 0 | 0.66 | 0.65 | 0.65 | 0.72 | 0.55 | 0.69 |
| 11 | 110225438 | rs17728641 | -0.278 | 1.747 | -0.001 | 0.956 | 0.014 | -0.016 | -0.016 | 0 | 0.13 | 0.19 | 0.14 | 0.03 | 0.22 | 0.09 |
| 11 | 110227864 | rs7930181 | -1.659 | 1.448 | 0.588 | 1.527 | 0.021 | 0.103 | 0.103 | 0 | 0.25 | 0.48 | 0.29 | 0.44 | 0.29 | 0.36 |

| | | | | | | | | | | | | | | | | |
|---|---|---|---|---|---|---|---|---|---|---|---|---|---|---|---|---|
| 11 | 110227924 | rs11213366 | -0.178 | 1.468 | 0.289 | 1.538 | -0.09 | -0.075 | -0.075 | 0 | 0.19 | 0.01 | 0.37 | 0.25 | 0.25 | 0.19 |
| 11 | 110237299 | rs2724409 | -1.467 | 1.241 | 1.005 | 1.359 | -0.004 | 0.002 | 0.002 | 0 | 0.35 | 0.55 | 0.54 | 0.66 | 0.33 | 0.46 |
| 11 | 110239559 | rs2724411 | -2.136 | 0.888 | 1.12 | 1.636 | -0.01 | 0.027 | 0.027 | 0 | 0.41 | 0.73 | 0.58 | 0.67 | 0.43 | 0.49 |
| 11 | 110241999 | rs2051466 | -1.749 | 1.019 | 1.287 | 1.485 | 0.006 | 0.001 | 0.001 | 0 | 0.3 | 0.46 | 0.23 | 0.29 | 0.31 | 0.22 |
| 11 | 110250091 | rs2585796 | -1.046 | 1.042 | 1.322 | 1.266 | 0.023 | -0.051 | -0.051 | 1.17 | 0.17 | 0.3 | 0.14 | 0.04 | 0.22 | 0.07 |
| 11 | 110270703 | rs11213388 | -0.876 | 0.939 | 1.194 | 1.697 | -0.001 | 0 | 0 | 0.008 | 0.2 | 0.23 | 0.41 | 0.6 | 0.19 | 0.38 |
| 11 | 110287537 | rs12575797 | -0.584 | 0.855 | 1.448 | 1.858 | -0.017 | 0.023 | 0.023 | 0 | 0.17 | 0.29 | 0.14 | 0.23 | 0.13 | 0.16 |
| 11 | 110296153 | rs10891103 | -1.47 | 0.854 | 1.692 | 1.927 | -0.004 | 0.059 | 0.059 | 0.303 | 0.22 | 0.37 | 0.43 | 0.62 | 0.19 | 0.38 |
| 11 | 110296813 | rs11213397 | -1.756 | 0.852 | 2.018 | 1.802 | 0.014 | 0.04 | 0.04 | 0 | 0.22 | 0.32 | 0.41 | 0.41 | 0.28 | 0.29 |
| 11 | 110296830 | rs2585800 | -2.629 | 0.828 | 2.17 | 1.783 | -0.006 | 0.018 | 0.018 | 0.101 | 0.4 | 0.72 | 0.57 | 0.66 | 0.41 | 0.46 |
| 11 | 110301799 | rs11213398 | -2.108 | 0.83 | 2.018 | 1.77 | 0.023 | 0.074 | 0.074 | 0 | 0.23 | 0.39 | 0.43 | 0.43 | 0.28 | 0.3 |
| 11 | 110317166 | rs4576787 | -0.307 | 0.632 | 1.859 | 1.79 | -0.001 | 0.001 | 0.001 | 0 | 0.17 | 0.18 | 0.14 | 0.23 | 0.16 | 0.17 |
| 11 | 110329941 | rs2358322 | -0.532 | 0.629 | 1.884 | 1.805 | -0.007 | 0.015 | 0.015 | 0 | 0.18 | 0.23 | 0.14 | 0.25 | 0.16 | 0.2 |
| 11 | 110330469 | rs7927202 | -2.406 | 0.632 | 1.867 | 1.798 | 0.001 | 0.002 | 0.002 | 0.119 | 0.37 | 0.65 | 0.53 | 0.66 | 0.34 | 0.45 |
| 11 | 110331631 | rs10488764 | -0.802 | 0.598 | 1.681 | 1.765 | -0.005 | 0.006 | 0.006 | 0 | 0.19 | 0.21 | 0.15 | 0.25 | 0.16 | 0.2 |
| 11 | 110334180 | rs2044174 | -0.794 | 0.607 | 1.743 | 1.77 | -0.014 | 0.025 | 0.025 | 0 | 0.19 | 0.25 | 0.15 | 0.26 | 0.16 | 0.2 |
| 11 | 110336182 | rs755128 | -0.024 | 0.645 | 1.482 | 1.864 | 0.002 | -0.014 | -0.014 | 0 | 0.1 | 0.15 | 0.08 | 0.03 | 0.12 | 0.03 |
| 11 | 110341416 | rs12280232 | -0.838 | 0.659 | 1.434 | 1.939 | 0.007 | 0.12 | 0.12 | 0.11 | 0.07 | 0.12 | 0.29 | 0.4 | 0.06 | 0.23 |
| 11 | 110341770 | rs12575576 | -0.188 | 0.595 | 1.224 | 1.549 | 0.005 | -0.027 | -0.027 | 0.861 | 0.16 | 0.09 | 0.12 | 0.22 | 0.16 | 0.17 |
| 11 | 110357133 | rs2136361 | -1.455 | 0.571 | 1.2 | 1.587 | 0 | -0.002 | -0.002 | 0 | 0.26 | 0.28 | 0.2 | 0.27 | 0.27 | 0.21 |
| 11 | 110364300 | rs12575639 | -0.208 | 0.55 | 0.641 | 1.678 | -0.02 | -0.067 | -0.067 | 0 | 0.16 | 0.04 | 0.15 | 0.23 | 0.17 | 0.2 |
| 11 | 110375034 | rs1553734 | -1.848 | 0.398 | 1.17 | 1.895 | 0.002 | 0.006 | 0.006 | 0.143 | 0.35 | 0.53 | 0.53 | 0.62 | 0.35 | 0.47 |
| 11 | 110377572 | rs328289 | 0.953 | 0.142 | 0.533 | 1.248 | 0 | 0.001 | 0.001 | 0.095 | 0.6 | 0.43 | 0.66 | 0.57 | 0.57 | 0.54 |
| 11 | 110378558 | rs328287 | 0.742 | 0.265 | 0.452 | 1.171 | 0.017 | 0.063 | 0.063 | 1.369 | 0.7 | 0.54 | 0.72 | 0.61 | 0.65 | 0.6 |
| 11 | 110383889 | rs7942728 | -0.416 | 0.017 | 0.548 | 0.958 | -0.023 | 0.011 | 0.011 | 0 | 0.24 | 0.32 | 0.19 | 0.22 | 0.16 | 0.3 |
| 11 | 110391916 | rs328281 | 0.219 | 0.129 | 0.887 | 1.027 | 0.04 | 0.023 | 0.023 | 1.153 | 0.86 | 0.69 | 0.9 | 0.85 | 0.81 | 0.78 |
| 11 | 110392541 | rs12790005 | 0.096 | 0.044 | 0.651 | 0.962 | -0.002 | -0.004 | -0.004 | 0.269 | 0.1 | 0.12 | 0.06 | 0.04 | 0.07 | 0.05 |
| 11 | 110395579 | rs2187387 | -0.531 | 0.181 | 0.495 | 1.073 | 0.019 | 0.049 | 0.049 | 0 | 0.13 | 0.18 | 0.36 | 0.34 | 0.18 | 0.25 |
| 11 | 110396769 | rs328286 | 1.917 | -0.001 | 0.211 | 0.76 | 0.007 | 0.014 | 0.014 | 1.864 | 0.7 | 0.22 | 0.79 | 0.8 | 0.73 | 0.67 |
| 11 | 110429445 | rs7935007 | -0.328 | 0.073 | 0.389 | 0.715 | -0.014 | -0.025 | -0.025 | 0 | 0.11 | 0.23 | 0.07 | 0.04 | 0.07 | 0.1 |
| 11 | 110431728 | rs17393434 | 0.16 | 0.099 | 0.29 | 0.78 | -0.045 | -0.042 | -0.042 | 0.223 | 0.08 | 0.02 | 0.07 | 0.17 | 0.18 | 0.17 |
| 11 | 110433508 | rs12222355 | 0.016 | 0.26 | -0.019 | 1.097 | -0.017 | -0.012 | -0.012 | 1.119 | 0.11 | 0.1 | 0.33 | 0.33 | 0.18 | 0.22 |
| 11 | 110434404 | rs10789780 | -0.011 | 0.223 | -0.118 | 0.876 | 0.063 | 0.053 | 0.053 | 0.882 | 0.87 | 0.62 | 0.89 | 0.83 | 0.82 | 0.83 |
| 11 | 110437368 | rs11213485 | 1.856 | 0.034 | -0.741 | 0.816 | -0.098 | 0.124 | 0.124 | 1.854 | 0.43 | 0.03 | 0.33 | 0.25 | 0.43 | 0.25 |
| 11 | 110447133 | rs4753901 | 0.026 | 0.217 | -0.387 | 0.759 | 0.07 | 0.059 | 0.059 | 2.064 | 0.87 | 0.57 | 0.88 | 0.83 | 0.82 | 0.82 |
| 11 | 110451885 | rs10488765 | -0.148 | 0.255 | -0.539 | 0.568 | 0.008 | 0.019 | 0.019 | 0 | 0.1 | 0.05 | 0.06 | 0.04 | 0.07 | 0.05 |
| 11 | 110475485 | rs326941 | -1.53 | 0.181 | -0.563 | 0.625 | 0.002 | -0.019 | -0.019 | 0 | 0.26 | 0.69 | 0.18 | 0.2 | 0.26 | 0.26 |
| 11 | 110476059 | rs326940 | -0.337 | 0.179 | -0.526 | 0.629 | 0.059 | 0.018 | 0.018 | 2.002 | 0.14 | 0.47 | 0.1 | 0.15 | 0.17 | 0.19 |
| 11 | 110476536 | rs326939 | -0.136 | 0.177 | -0.521 | 0.628 | 0.064 | 0.027 | 0.027 | 0.99 | 0.14 | 0.45 | 0.1 | 0.14 | 0.17 | 0.17 |
| 11 | 110478722 | rs1396714 | -1.619 | 0.166 | -0.517 | 0.623 | 0 | 0 | 0 | 0 | 0.26 | 0.77 | 0.19 | 0.2 | 0.26 | 0.26 |
| 11 | 110483599 | rs10891150 | 0.219 | 0.335 | -0.667 | 0.826 | -0.06 | -0.061 | -0.061 | 0 | 0.1 | 0.03 | 0.34 | 0.31 | 0.18 | 0.22 |
| 11 | 110483623 | rs7118096 | -0.423 | 0.212 | -0.696 | 0.767 | 0.003 | 0.017 | 0.017 | 0 | 0.13 | 0.11 | 0.36 | 0.31 | 0.18 | 0.22 |
| 11 | 110488914 | rs12282038 | -0.657 | 0.101 | -0.716 | 0.705 | -0.009 | -0.029 | -0.029 | 0.731 | 0.12 | 0.32 | 0.08 | 0.05 | 0.1 | 0.06 |
| 11 | 110493961 | rs1509317 | -1.526 | 0.111 | -0.69 | 0.699 | 0.02 | 0.051 | 0.051 | 0 | 0.25 | 0.43 | 0.44 | 0.36 | 0.27 | 0.28 |
| 11 | 110500998 | rs10082696 | -1.316 | 0.145 | -0.741 | 0.69 | 0.003 | 0.011 | 0.011 | 0 | 0.24 | 0.26 | 0.43 | 0.36 | 0.26 | 0.28 |
| 11 | 110503540 | rs326948 | -1.677 | 0.059 | -0.938 | 0.597 | -0.084 | -0.107 | -0.107 | 0.021 | 0.39 | 0.9 | 0.54 | 0.5 | 0.43 | 0.42 |
| 11 | 110505721 | rs326949 | -1.559 | 0.083 | -0.91 | 0.61 | -0.044 | -0.057 | -0.057 | 1.398 | 0.38 | 0.81 | 0.54 | 0.5 | 0.43 | 0.41 |
| 11 | 110512795 | rs7951046 | -0.974 | -0.142 | -0.927 | 0.699 | -0.032 | -0.006 | -0.006 | 0 | 0.35 | 0.5 | 0.23 | 0.29 | 0.23 | 0.39 |
| 11 | 110527134 | rs1470652 | 0.462 | -0.172 | -0.931 | 0.707 | 0.019 | -0.013 | -0.013 | 0.513 | 0.21 | 0.16 | 0.13 | 0.25 | 0.14 | 0.32 |
| 11 | 110532348 | rs10891158 | -0.477 | -0.206 | -0.886 | 0.701 | 0.002 | -0.001 | -0.001 | 0 | 0.3 | 0.32 | 0.19 | 0.29 | 0.22 | 0.35 |
| 11 | 110562578 | rs7102109 | -1.312 | -0.157 | -0.916 | 0.53 | -0.017 | -0.035 | -0.035 | 0 | 0.13 | 0.34 | 0.09 | 0.04 | 0.08 | 0.04 |
| 11 | 110607482 | rs6589170 | -0.641 | 0.554 | -0.871 | 0.78 | -0.004 | -0.017 | -0.017 | 0 | 0.1 | 0.21 | 0.07 | 0.04 | 0.08 | 0.07 |
| 11 | 110608950 | rs1375718 | -0.591 | 0.556 | -0.884 | 0.781 | -0.001 | -0.002 | -0.002 | 0.726 | 0.1 | 0.1 | 0.06 | 0.04 | 0.08 | 0.07 |
| 11 | 110610453 | rs1449699 | -1.002 | 0.503 | -0.943 | 0.886 | -0.031 | -0.013 | -0.013 | 0.02 | 0.34 | 0.42 | 0.21 | 0.24 | 0.23 | 0.37 |
| 11 | 110616415 | rs17111731 | 0.345 | 0.528 | -0.992 | 0.907 | 0.029 | -0.014 | -0.014 | 0 | 0.21 | 0.09 | 0.11 | 0.19 | 0.14 | 0.29 |
| 11 | 110617469 | rs226092 | -0.583 | 1.525 | -1.159 | 1.063 | -0.086 | -0.083 | -0.083 | 2.561 | 0.86 | 0.99 | 0.77 | 0.72 | 0.74 | 0.79 |
| 11 | 110631223 | rs7936723 | -0.531 | 1.182 | -0.874 | 1.116 | 0.008 | 0 | 0 | 0 | 0.28 | 0.21 | 0.15 | 0.23 | 0.21 | 0.33 |
| 11 | 110633853 | rs11825647 | -0.614 | 1.208 | -0.81 | 1.152 | -0.004 | -0.012 | -0.012 | 0 | 0.08 | 0.12 | 0.06 | 0.04 | 0.06 | 0.04 |
| 11 | 110638732 | rs11501062 | 0.371 | 1.189 | -0.948 | 1.137 | 0.027 | -0.018 | -0.018 | 0.352 | 0.2 | 0.1 | 0.1 | 0.19 | 0.13 | 0.31 |
| 11 | 110644532 | rs4937385 | -1.328 | 1.448 | -0.953 | 1.221 | 0.001 | 0.003 | 0.003 | 0 | 0.4 | 0.47 | 0.54 | 0.55 | 0.37 | 0.57 |
| 11 | 110644827 | rs17111778 | 0.127 | 1.58 | -1.263 | 1.316 | -0.002 | 0.001 | 0.001 | 0 | 0.06 | 0.06 | 0.05 | 0.04 | 0.07 | 0.04 |
| 11 | 110649270 | rs1848044 | 0.066 | 1.553 | -1.459 | 1.406 | -0.026 | -0.028 | -0.028 | 0 | 0.1 | 0.06 | 0.34 | 0.29 | 0.16 | 0.2 |
| 11 | 110651468 | rs6589176 | -0.597 | 1.502 | -1.464 | 1.39 | -0.014 | -0.068 | -0.068 | 0.95 | 0.3 | 0.19 | 0.47 | 0.5 | 0.29 | 0.52 |
| 11 | 110652701 | rs2194424 | -0.436 | 1.578 | -1.273 | 1.39 | -0.026 | -0.001 | -0.001 | 0.217 | 0.3 | 0.37 | 0.17 | 0.25 | 0.19 | 0.37 |
| 11 | 110660305 | rs1506488 | -0.787 | 1.955 | -1.463 | 1.452 | -0.02 | -0.023 | -0.023 | 0 | 0.13 | 0.09 | 0.4 | 0.26 | 0.22 | 0.21 |
| 11 | 110663056 | rs190125 | 0.49 | 1.912 | -1.376 | 1.29 | -0.015 | -0.019 | -0.019 | 0 | 0.62 | 0.74 | 0.49 | 0.47 | 0.64 | 0.4 |
| 11 | 110665974 | rs226110 | 0.263 | 1.76 | -1.365 | 1.413 | -0.07 | 0.074 | 0.074 | 2.498 | 0.58 | 0.87 | 0.48 | 0.35 | 0.53 | 0.35 |
| 11 | 110671664 | rs2432703 | 0.523 | 1.725 | -1.112 | 1.389 | -0.008 | 0.021 | 0.021 | 0.697 | 0.55 | 0.64 | 0.47 | 0.35 | 0.52 | 0.35 |
| 11 | 110671733 | rs226095 | 0.036 | 1.736 | -1.162 | 1.605 | -0.107 | -0.098 | -0.098 | 2.938 | 0.59 | 0.92 | 0.64 | 0.48 | 0.54 | 0.39 |

| | | | | | | | | | | | | | | | | |
|---|---|---|---|---|---|---|---|---|---|---|---|---|---|---|---|---|
| 11 | 110672626 | rs7106841 | -0.061 | 1.754 | -1.126 | 1.581 | -0.004 | -0.003 | -0.003 | 0 | 0.1 | 0.07 | 0.22 | 0.16 | 0.18 | 0.15 |
| 11 | 110679157 | rs622572 | 0.021 | 1.756 | -1.156 | 1.601 | -0.074 | -0.07 | -0.07 | 0 | 0.59 | 0.88 | 0.64 | 0.49 | 0.54 | 0.39 |
| 11 | 110686788 | rs226164 | 1.222 | 1.771 | -1.065 | 1.555 | -0.013 | -0.043 | -0.043 | 0 | 0.43 | 0.53 | 0.23 | 0.19 | 0.34 | 0.2 |
| 11 | 110693356 | rs184443 | -0.521 | 2.207 | -1.591 | 1.732 | -0.083 | -0.051 | -0.051 | 0 | 0.71 | 0.97 | 0.76 | 0.76 | 0.73 | 0.61 |
| 11 | 110697444 | rs120436 | -0.279 | 1.709 | -1.206 | 1.315 | -0.067 | -0.06 | -0.06 | 0 | 0.57 | 0.87 | 0.63 | 0.5 | 0.53 | 0.38 |
| 11 | 110701446 | rs226130 | 0.434 | 1.799 | -1.245 | 1.155 | 0.086 | 0.068 | 0.068 | 1.728 | 0.47 | 0.76 | 0.4 | 0.31 | 0.34 | 0.22 |
| 11 | 110701645 | rs7938861 | 0.073 | 1.482 | -1.093 | 0.55 | -0.007 | 0.057 | 0.057 | 1.894 | 0.52 | 0.66 | 0.46 | 0.34 | 0.51 | 0.32 |
| 11 | 110702636 | rs4937934 | -1.074 | 1.756 | -0.939 | 1.316 | 0.025 | 0.015 | 0.015 | 0 | 0.05 | 0.06 | 0.18 | 0.16 | 0.12 | 0.12 |
| 11 | 110703407 | rs7125147 | -0.668 | 0.583 | -1.203 | 1.372 | 0.016 | -0.021 | -0.021 | 0 | 0.32 | 0.21 | 0.42 | 0.33 | 0.22 | 0.36 |
| 11 | 110704426 | rs997402 | -1.295 | 0.178 | -0.216 | 1.846 | -0.027 | 0.146 | 0.146 | 0.139 | 0.15 | 0.48 | 0.14 | 0.32 | 0.11 | 0.17 |
| 11 | 110704745 | rs1984756 | -1.84 | -0.408 | -0.293 | 1.923 | -0.02 | 0.004 | 0.004 | 0.698 | 0.22 | 0.65 | 0.19 | 0.22 | 0.16 | 0.22 |
| 11 | 110707133 | rs7924842 | -1.348 | -0.39 | -0.09 | 1.973 | -0.051 | 0.043 | 0.043 | 2.897 | 0.14 | 0.59 | 0.07 | 0.14 | 0.07 | 0.15 |
| 11 | 110707998 | rs1876197 | 1.367 | 0.404 | -0.889 | 2.062 | -0.036 | -0.021 | -0.021 | 0 | 0.59 | 0.26 | 0.44 | 0.65 | 0.52 | 0.56 |
| 11 | 110710408 | rs226124 | -1.38 | -0.081 | -0.871 | 2.309 | -0.01 | 0.01 | 0.01 | 3.481 | 0.19 | 0.74 | 0.1 | 0.35 | 0.11 | 0.24 |
| 11 | 110710617 | rs6589181 | -0.279 | -0.057 | -0.833 | 2.186 | -0.038 | 0.073 | 0.073 | 1.095 | 0.12 | 0.49 | 0.06 | 0.12 | 0.07 | 0.15 |
| 11 | 110712113 | rs10789796 | -0.523 | -0.062 | -0.821 | 2.184 | -0.048 | 0.054 | 0.054 | 1.675 | 0.13 | 0.54 | 0.06 | 0.12 | 0.07 | 0.15 |
| 11 | 110721187 | rs7123425 | -1.26 | 0.318 | -0.893 | 2.167 | -0.001 | 0.009 | 0.009 | 0.118 | 0.19 | 0.52 | 0.22 | 0.14 | 0.19 | 0.18 |
| 11 | 110722129 | rs2059333 | -0.296 | 0.431 | -1.105 | 2.163 | -0.038 | -0.033 | -0.033 | 1.873 | 0.8 | 0.9 | 0.65 | 0.75 | 0.72 | 0.83 |
| 11 | 110722807 | rs9666742 | -0.176 | 0.346 | -1.111 | 2.178 | -0.009 | 0.048 | 0.048 | 0 | 0.07 | 0.12 | 0.04 | 0.22 | 0.06 | 0.1 |
| 11 | 110725831 | rs9666616 | -1.847 | 0.328 | -1.114 | 2.189 | 0.012 | -0.029 | -0.029 | 0.914 | 0.26 | 0.78 | 0.26 | 0.37 | 0.24 | 0.28 |
| 11 | 110726736 | rs1974911 | 0.618 | 0.459 | -1.135 | 2.133 | -0.004 | -0.009 | -0.009 | 0.373 | 0.72 | 0.7 | 0.55 | 0.46 | 0.66 | 0.67 |
| 11 | 110730249 | rs10488722 | -0.651 | 0.483 | -1.163 | 1.995 | 0.028 | -0.042 | -0.042 | 1.228 | 0.08 | 0.14 | 0.17 | 0 | 0.13 | 0.04 |
| 11 | 110731594 | rs7938457 | 1.307 | 0.473 | -1.194 | 2.059 | 0.013 | 0.02 | 0.02 | 0.899 | 0.61 | 0.22 | 0.57 | 0.39 | 0.62 | 0.56 |
| 11 | 110735980 | rs11600056 | -1.946 | 0.899 | -0.882 | 2.198 | 0.019 | -0.01 | -0.01 | 0 | 0.13 | 0.18 | 0.29 | 0.09 | 0.2 | 0.09 |
| 11 | 110739392 | rs11213629 | -0.41 | 0.806 | -0.588 | 2.345 | -0.035 | 0.069 | 0.069 | 0.573 | 0.12 | 0.51 | 0.07 | 0.15 | 0.08 | 0.15 |
| 11 | 110747785 | rs6589187 | -0.345 | 0.805 | -0.603 | 2.347 | -0.027 | 0.078 | 0.078 | 1.671 | 0.11 | 0.55 | 0.07 | 0.15 | 0.06 | 0.13 |
| 11 | 110748070 | rs10488719 | 0.996 | 0.832 | -1.42 | 1.557 | -0.084 | -0.044 | -0.044 | 0 | 0.24 | 0.04 | 0.19 | 0.29 | 0.25 | 0.32 |
| 11 | 110753770 | rs2884451 | 0.279 | 0.772 | -1.435 | 1.537 | -0.102 | 0.024 | 0.024 | 0.725 | 0.31 | 0.1 | 0.37 | 0.29 | 0.38 | 0.37 |
| 11 | 110756402 | rs2194426 | 0.505 | 0.728 | -1.438 | 1.674 | 0.013 | 0.167 | 0.167 | 0 | 0.06 | 0.23 | 0.04 | 0.22 | 0.04 | 0.1 |
| 11 | 110756816 | rs763302 | -2.118 | 0.722 | -1.456 | 1.642 | -0.015 | -0.085 | -0.085 | 0 | 0.29 | 0.81 | 0.38 | 0.46 | 0.29 | 0.31 |
| 11 | 110756940 | rs1486911 | 0.087 | 0.673 | -1.48 | 1.484 | -0.028 | 0.03 | 0.03 | 0 | 0.07 | 0.02 | 0.15 | 0 | 0.11 | 0.04 |
| 11 | 110757862 | rs10488717 | -1.101 | 0.669 | -1.481 | 1.496 | -0.007 | -0.009 | -0.009 | 0 | 0.2 | 0.65 | 0.23 | 0.15 | 0.17 | 0.17 |
| 11 | 110770371 | rs1025102 | -1.455 | 0.645 | -1.688 | 1.519 | -0.017 | 0.067 | 0.067 | 4.097 | 0.54 | 0.91 | 0.6 | 0.75 | 0.55 | 0.64 |
| 11 | 110780405 | rs10749967 | -1.703 | 0.391 | -1.706 | 1.202 | -0.012 | -0.057 | -0.057 | 1.92 | 0.59 | 0.91 | 0.62 | 0.66 | 0.59 | 0.71 |
| 11 | 110782320 | rs10891189 | -0.085 | 0.252 | -0.951 | 1.145 | -0.026 | -0.029 | -0.029 | 0 | 0.11 | 0.5 | 0.12 | 0.06 | 0.05 | 0.12 |
| 11 | 110782451 | rs10749968 | -1.589 | 0.224 | -1.139 | 1.425 | 0.009 | -0.017 | -0.017 | 0.239 | 0.22 | 0.76 | 0.28 | 0.42 | 0.17 | 0.29 |
| 11 | 110785968 | rs10891190 | -1.976 | 0.33 | -1.223 | 1.377 | -0.001 | -0.047 | -0.047 | 0 | 0.32 | 0.86 | 0.41 | 0.42 | 0.28 | 0.32 |
| 11 | 110791312 | rs877543 | -0.729 | 0.257 | -0.319 | 1.453 | -0.01 | 0.164 | 0.164 | 0.121 | 0.08 | 0.36 | 0.06 | 0.26 | 0.04 | 0.12 |
| 11 | 110795731 | rs10891192 | 1.038 | 0.617 | -0.631 | 1.344 | -0.127 | -0.038 | -0.038 | 0.666 | 0.26 | 0.03 | 0.23 | 0.28 | 0.33 | 0.36 |
| 11 | 110803508 | rs1872768 | 1.061 | 0.581 | -0.585 | 1.327 | -0.127 | -0.04 | -0.04 | 0.969 | 0.26 | 0.03 | 0.23 | 0.28 | 0.33 | 0.37 |
| 11 | 110804895 | rs6589200 | -1.819 | 0.521 | -0.55 | 1.334 | 0.02 | -0.013 | -0.013 | 0.483 | 0.59 | 0.91 | 0.63 | 0.71 | 0.61 | 0.67 |
| 11 | 110807983 | rs714342 | 0.301 | 0.526 | -0.37 | 1.27 | -0.075 | -0.109 | -0.109 | 0 | 0.33 | 0.12 | 0.39 | 0.59 | 0.37 | 0.46 |
| 11 | 110809899 | rs1486906 | -1.853 | 0.52 | -0.356 | 1.245 | 0.02 | 0.033 | 0.033 | 4.326 | 0.59 | 0.91 | 0.78 | 0.76 | 0.61 | 0.67 |
| 11 | 110810509 | rs10891201 | -2.38 | 0.632 | -0.109 | 1.454 | 0.009 | -0.085 | -0.085 | 0 | 0.33 | 0.88 | 0.55 | 0.49 | 0.28 | 0.32 |
| 11 | 110811164 | rs1906167 | -1.596 | 0.71 | -0.027 | 1.388 | 0.017 | -0.011 | -0.011 | 0.017 | 0.24 | 0.51 | 0.35 | 0.18 | 0.24 | 0.21 |
| 11 | 110813788 | rs7114106 | 0.669 | 0.338 | -0.052 | 1.02 | -0.008 | 0.123 | 0.123 | 0 | 0.05 | 0.08 | 0.17 | 0.31 | 0.04 | 0.11 |
| 11 | 110817191 | rs10789804 | -0.718 | 0.276 | 0.27 | 0.918 | -0.03 | -0.053 | -0.053 | 0 | 0.13 | 0.41 | 0.12 | 0.04 | 0.09 | 0.14 |
| 11 | 110820558 | rs4938077 | -0.968 | 0.292 | -0.031 | 0.967 | -0.068 | 0.14 | 0.14 | 1.599 | 0.18 | 0.45 | 0.27 | 0.35 | 0.11 | 0.24 |
| 11 | 110820730 | rs4938078 | -2.039 | 0.398 | -0.355 | 0.726 | -0.021 | 0.022 | 0.022 | 1.006 | 0.3 | 0.55 | 0.42 | 0.37 | 0.23 | 0.29 |
| 11 | 110824743 | rs11213673 | -0.481 | 0.834 | -0.864 | -0.203 | -0.049 | 0.066 | 0.066 | 0.549 | 0.4 | 0.23 | 0.39 | 0.28 | 0.43 | 0.4 |
| 11 | 110824960 | rs1486907 | 0.434 | 0.757 | -0.729 | -0.103 | -0.018 | 0.026 | 0.026 | 0.143 | 0.3 | 0.22 | 0.26 | 0.26 | 0.33 | 0.36 |
| 11 | 110825946 | rs11213674 | -0.972 | 0.716 | 0.171 | -0.106 | -0.016 | -0.015 | -0.015 | 0 | 0.11 | 0.47 | 0.15 | 0.04 | 0.06 | 0.14 |
| 11 | 110826882 | rs17728959 | -1.007 | 1.96 | -0.802 | -0.095 | -0.091 | 0.04 | 0.04 | 0.681 | 0.14 | 0.02 | 0.2 | 0.02 | 0.22 | 0.07 |
| 11 | 110828826 | rs12288950 | 0.171 | 1.164 | -0.486 | -0.237 | 0.003 | 0.001 | 0.001 | 0 | 0.41 | 0.36 | 0.32 | 0.35 | 0.43 | 0.35 |
| 11 | 110831371 | rs17655864 | 0.107 | 2.001 | -0.8 | -0.49 | -0.04 | 0.04 | 0.04 | 0 | 0.11 | 0.04 | 0.1 | 0.02 | 0.17 | 0.07 |
| 11 | 110833266 | rs10891208 | -0.541 | 1.397 | -0.262 | -0.427 | 0.008 | 0.017 | 0.017 | 0.006 | 0.59 | 0.64 | 0.44 | 0.27 | 0.64 | 0.45 |
| 11 | 110836948 | rs11213678 | 0.256 | 1.209 | -0.402 | -0.626 | -0.001 | -0.001 | -0.001 | 0 | 0.2 | 0.2 | 0.21 | 0.2 | 0.24 | 0.24 |
| 11 | 110839870 | rs10891210 | -0.955 | 0.863 | -0.378 | -0.625 | -0.01 | 0.005 | 0.005 | 0.256 | 0.65 | 0.56 | 0.58 | 0.53 | 0.77 | 0.56 |
| 11 | 110843688 | rs4643108 | -0.298 | 1.191 | -0.257 | -0.834 | -0.001 | -0.044 | -0.044 | 0 | 0.43 | 0.45 | 0.31 | 0.07 | 0.42 | 0.22 |
| 11 | 110853185 | rs7951632 | -1.13 | 1.548 | 0.3 | 0.21 | 0.073 | 0.097 | 0.097 | 0 | 0.06 | 0.22 | 0.06 | 0.11 | 0.06 | 0.04 |
| 11 | 110853848 | rs7941660 | 0.52 | 1.56 | -0.127 | 0.042 | 0.005 | 0.002 | 0.002 | 0 | 0.55 | 0.55 | 0.71 | 0.6 | 0.58 | 0.62 |
| 11 | 110863969 | rs569524 | 1.058 | 2.125 | -0.125 | -0.237 | 0.014 | -0.003 | -0.003 | 0.377 | 0.45 | 0.62 | 0.36 | 0.48 | 0.25 | 0.52 |
| 11 | 110876320 | rs721368 | -0.315 | 2.22 | -0.66 | -0.125 | -0.079 | -0.015 | -0.015 | 0 | 0.12 | 0.05 | 0.37 | 0.12 | 0.33 | 0.1 |
| 11 | 110878988 | rs7107636 | 1.197 | 2.098 | -0.628 | -0.235 | 0.01 | -0.004 | -0.004 | 1.592 | 0.44 | 0.43 | 0.35 | 0.48 | 0.28 | 0.56 |
| 11 | 110888484 | rs978769 | 1.356 | 2.428 | -0.532 | -0.734 | 0.04 | 0.064 | 0.064 | 0 | 0.51 | 0.38 | 0.39 | 0.77 | 0.33 | 0.71 |
| 11 | 110890394 | rs317519 | -0.243 | 2.655 | -0.482 | -0.763 | 0.052 | -0.027 | -0.027 | 0 | 0.12 | 0.22 | 0.34 | 0.03 | 0.27 | 0.06 |
| 11 | 110891720 | rs11213704 | 1.134 | 2.647 | -0.531 | -0.749 | 0.007 | 0.045 | 0.045 | 0 | 0.53 | 0.39 | 0.43 | 0.87 | 0.39 | 0.75 |
| 11 | 110892082 | rs161138 | 0.618 | 2.518 | -0.549 | -0.701 | -0.013 | -0.048 | -0.048 | 0.153 | 0.62 | 0.56 | 0.72 | 0.8 | 0.6 | 0.78 |
| 11 | 110913240 | rs4938174 | -0.642 | 2.173 | -0.602 | -0.664 | -0.035 | -0.092 | -0.092 | 0 | 0.34 | 0.38 | 0.22 | 0.1 | 0.32 | 0.19 |

| | | | | | | | | | | | | | | | | |
|---|---|---|---|---|---|---|---|---|---|---|---|---|---|---|---|---|
| 11 | 110914603 | rs11213716 | 0.321 | 1.783 | -1.535 | -0.58 | -0.015 | 0.039 | 0.039 | 0 | 0.15 | 0.01 | 0.09 | 0.08 | 0.16 | 0.09 |
| 11 | 110916706 | rs7118898 | -0.995 | 1.658 | -1.149 | -0.236 | 0.088 | 0.156 | 0.156 | 0 | 0.49 | 0.79 | 0.63 | 0.22 | 0.66 | 0.28 |
| 11 | 110934218 | rs11605124 | -0.706 | 1.694 | -0.933 | 0.216 | -0.042 | 0.027 | 0.027 | 0 | 0.07 | 0.01 | 0.15 | 0.02 | 0.13 | 0.02 |
| 11 | 110936275 | rs6589208 | -0.51 | 0.825 | -0.651 | 0.809 | -0.001 | -0.001 | -0.001 | 1.571 | 0.46 | 0.51 | 0.46 | 0.39 | 0.43 | 0.22 |
| 11 | 110951754 | rs17457076 | -0.586 | 1.054 | -1.017 | 0.78 | -0.045 | 0.09 | 0.09 | 0.147 | 0.17 | 0.04 | 0.28 | 0.03 | 0.16 | 0.1 |
| 11 | 110951822 | rs1878535 | -0.788 | 1.272 | -1.016 | 1.504 | -0.002 | 0.003 | 0.003 | 0 | 0.11 | 0.1 | 0.08 | 0.25 | 0.13 | 0.24 |
| 11 | 110952097 | rs1878537 | -1.554 | 1.09 | -0.488 | 1.416 | -0.007 | 0.001 | 0.001 | 0.274 | 0.3 | 0.26 | 0.36 | 0.29 | 0.3 | 0.33 |
| 11 | 110959536 | rs321418 | -1.649 | 1.072 | -0.168 | 1.439 | 0.002 | 0.01 | 0.01 | 0.143 | 0.31 | 0.36 | 0.45 | 0.42 | 0.31 | 0.33 |
| 11 | 110980918 | rs12287270 | -1.272 | 0.876 | 0.164 | 1.354 | 0.01 | 0.046 | 0.046 | 0 | 0.15 | 0.2 | 0.13 | 0.23 | 0.17 | 0.23 |
| 11 | 110999163 | rs12419222 | -0.067 | 0.949 | 0.034 | 1.269 | -0.056 | 0.091 | 0.091 | 0 | 0.16 | 0.01 | 0.27 | 0.03 | 0.15 | 0.08 |
| 11 | 111004208 | rs12291189 | -1.131 | 0.85 | 0.047 | 1.322 | -0.002 | 0.036 | 0.036 | 0 | 0.13 | 0.17 | 0.11 | 0.21 | 0.14 | 0.2 |
| 11 | 111009374 | rs4936336 | -1.063 | 0.844 | 0.13 | 1.323 | 0.001 | 0.045 | 0.045 | 0 | 0.13 | 0.17 | 0.12 | 0.21 | 0.14 | 0.21 |
| 11 | 111012458 | rs1356412 | 1.933 | 0.024 | 0.223 | 1.665 | 0.02 | 0.042 | 0.042 | 0 | 0.52 | 0.3 | 0.4 | 0.33 | 0.61 | 0.53 |
| 11 | 111016062 | rs511835 | -0.442 | 0.174 | -0.297 | 1.624 | -0.015 | 0.046 | 0.046 | 0.163 | 0.21 | 0.11 | 0.3 | 0.05 | 0.19 | 0.11 |
| 11 | 111020436 | rs546440 | -0.532 | 0.137 | -0.229 | 1.663 | 0.014 | -0.025 | -0.025 | 1.124 | 0.9 | 0.94 | 0.91 | 0.69 | 0.93 | 0.85 |
| 11 | 111024230 | rs7939631 | -0.471 | 0.422 | -0.157 | 1.254 | -0.019 | 0.052 | 0.052 | 0.291 | 0.21 | 0.1 | 0.3 | 0.05 | 0.19 | 0.11 |
| 11 | 111037249 | rs516110 | -0.754 | 0.454 | -0.169 | 1.253 | -0.002 | 0.004 | 0.004 | 0 | 0.23 | 0.22 | 0.38 | 0.15 | 0.19 | 0.11 |
| 11 | 111047419 | rs10502133 | -0.952 | 0.373 | -0.263 | 1.457 | -0.001 | -0.001 | -0.001 | 0.162 | 0.12 | 0.09 | 0.1 | 0.19 | 0.13 | 0.19 |
| 11 | 111051325 | rs2000907 | 0.02 | 0.672 | -1.302 | 0.413 | -0.099 | 0.17 | 0.17 | 1.028 | 0.37 | 0.05 | 0.45 | 0.13 | 0.41 | 0.32 |
| 11 | 111056023 | rs9645657 | -0.683 | 0.301 | -0.88 | 1.308 | -0.015 | 0.08 | 0.08 | 0 | 0.9 | 0.85 | 0.92 | 0.66 | 0.92 | 0.85 |
| 11 | 111068611 | rs17539860 | -0.153 | 0.548 | -0.324 | 1.418 | -0.004 | 0.011 | 0.011 | 0.042 | 0.44 | 0.4 | 0.52 | 0.21 | 0.46 | 0.33 |
| 11 | 111071187 | rs10502135 | 0.1 | 0.344 | -0.468 | 1.323 | 0.014 | -0.035 | -0.035 | 0 | 0.85 | 0.93 | 0.9 | 0.66 | 0.89 | 0.84 |
| 11 | 111072914 | rs1878534 | 0.952 | 0.548 | -0.529 | 1.509 | -0.009 | 0.003 | 0.003 | 1.143 | 0.36 | 0.5 | 0.35 | 0.4 | 0.36 | 0.47 |
| 11 | 111074455 | rs1474069 | 1.129 | 0.733 | -0.773 | 1.258 | -0.028 | 0.039 | 0.039 | 0.156 | 0.17 | 0.03 | 0.32 | 0.11 | 0.17 | 0.07 |
| 11 | 111074555 | rs2176441 | 0.681 | 0.701 | -0.662 | 1.35 | 0.001 | 0.008 | 0.008 | 0.385 | 0.41 | 0.53 | 0.38 | 0.45 | 0.42 | 0.51 |
| 11 | 111079495 | rs10502138 | -0.001 | 0.594 | -0.462 | 1.591 | -0.003 | -0.003 | -0.003 | 1.27 | 0.05 | 0.03 | 0.04 | 0.05 | 0.07 | 0.04 |
| 11 | 111080246 | rs10789808 | 0.496 | 0.568 | -0.406 | 1.547 | -0.005 | 0.101 | 0.101 | 0 | 0.84 | 0.72 | 0.86 | 0.61 | 0.86 | 0.81 |
| 11 | 111088734 | rs6589213 | 0.389 | 0.36 | -0.675 | 1.424 | 0.022 | -0.024 | -0.024 | 0.429 | 0.71 | 0.79 | 0.63 | 0.66 | 0.77 | 0.82 |
| 11 | 111096943 | rs7120599 | 0.685 | 0.1 | -0.464 | 1.788 | 0.004 | -0.006 | -0.006 | 0 | 0.71 | 0.73 | 0.54 | 0.38 | 0.76 | 0.73 |
| 11 | 111098908 | rs4938320 | 0.384 | 0.085 | -0.463 | 2.09 | 0.016 | -0.001 | -0.001 | 0 | 0.08 | 0.03 | 0.06 | 0.37 | 0.07 | 0.17 |
| 11 | 111100110 | rs10891232 | 0.683 | -0.522 | -1.1 | 1.534 | 0.005 | 0.047 | 0.047 | 0 | 0.33 | 0.12 | 0.39 | 0.2 | 0.31 | 0.2 |
| 11 | 111101021 | rs17112466 | 0.203 | -0.486 | -1.086 | 1.547 | 0.024 | 0.06 | 0.06 | 0 | 0.53 | 0.65 | 0.44 | 0.25 | 0.58 | 0.57 |
| 11 | 111102400 | rs7115403 | 0.719 | -0.483 | -1.092 | 1.551 | 0.003 | 0.047 | 0.047 | 0 | 0.33 | 0.1 | 0.39 | 0.2 | 0.31 | 0.2 |
| 11 | 111102839 | rs12804616 | 0.036 | -0.483 | -1.064 | 1.599 | 0.024 | 0.053 | 0.053 | 0 | 0.54 | 0.67 | 0.45 | 0.34 | 0.58 | 0.58 |
| 11 | 111103318 | rs11824477 | -0.032 | -0.483 | -1.055 | 1.612 | 0.025 | 0.051 | 0.051 | 0 | 0.54 | 0.7 | 0.5 | 0.38 | 0.58 | 0.58 |
| 11 | 111106991 | rs10891238 | 0.602 | -0.396 | -0.853 | 1.89 | -0.021 | 0.101 | 0.101 | 0.215 | 0.86 | 0.78 | 0.83 | 0.45 | 0.89 | 0.77 |
| 11 | 111109258 | rs7119658 | -0.142 | -0.027 | -0.557 | 1.279 | 0.017 | -0.012 | -0.012 | 0 | 0.2 | 0.42 | 0.21 | 0.16 | 0.23 | 0.31 |
| 11 | 111111729 | rs7952641 | -0.17 | -0.032 | -0.516 | 1.278 | 0.013 | -0.012 | -0.012 | 4.723 | 0.2 | 0.42 | 0.21 | 0.16 | 0.22 | 0.31 |
| 11 | 111111747 | rs949280 | -0.22 | -0.227 | -0.559 | 1.308 | -0.02 | 0.111 | 0.111 | 0 | 0.91 | 0.83 | 0.9 | 0.63 | 0.92 | 0.82 |
| 11 | 111115272 | rs7115117 | -0.415 | 0.191 | -0.798 | -0.268 | 0.024 | 0.093 | 0.093 | 0.843 | 0.48 | 0.35 | 0.33 | 0.17 | 0.56 | 0.27 |
| 11 | 111119178 | rs4432059 | 0.357 | 0.245 | -0.488 | 0.142 | -0.052 | 0.019 | 0.019 | 2.007 | 0.19 | 0.1 | 0.3 | 0.13 | 0.29 | 0.3 |
| 11 | 111119800 | rs9326233 | 0.233 | -0.017 | -0.227 | 0.27 | -0.014 | -0.044 | -0.044 | 0 | 0.9 | 0.83 | 0.94 | 0.97 | 0.93 | 0.93 |
| 11 | 111135745 | rs11213809 | -0.561 | 0.463 | -0.338 | 0.553 | -0.012 | 0.015 | 0.015 | 0 | 0.77 | 0.83 | 0.82 | 0.83 | 0.69 | 0.82 |
| 11 | 111140462 | rs10891240 | 1.002 | -0.021 | -0.763 | 0.31 | 0.018 | -0.064 | -0.064 | 0.3 | 0.34 | 0.11 | 0.27 | 0.42 | 0.27 | 0.46 |
| 11 | 111141676 | rs4309189 | 0.678 | -0.383 | -0.373 | 0.319 | -0.002 | -0.001 | -0.001 | 0 | 0.19 | 0.17 | 0.38 | 0.17 | 0.31 | 0.32 |
| 11 | 111156836 | rs3087967 | -1.011 | -0.263 | -1.124 | 0.332 | -0.001 | 0.011 | 0.011 | 0 | 0.77 | 0.72 | 0.82 | 0.6 | 0.73 | 0.83 |
| 11 | 111158189 | rs11213819 | 1.154 | -0.285 | -1.88 | 0.268 | -0.007 | 0.112 | 0.112 | 0.035 | 0.18 | 0.01 | 0.12 | 0.01 | 0.12 | 0.03 |
| 11 | 111164651 | rs11213821 | 0.853 | -0.229 | -1.556 | 0.66 | 0.001 | -0.064 | -0.064 | 0 | 0.34 | 0.11 | 0.27 | 0.42 | 0.3 | 0.48 |
| 11 | 111167557 | rs6589218 | -0.235 | 0.196 | -1.214 | 0.805 | -0.001 | 0.026 | 0.026 | 0 | 0.74 | 0.65 | 0.8 | 0.6 | 0.7 | 0.82 |
| 11 | 111171646 | rs3802840 | -0.429 | 0.069 | -1.12 | 0.919 | -0.003 | 0.044 | 0.044 | 0 | 0.77 | 0.65 | 0.81 | 0.6 | 0.73 | 0.83 |
| 11 | 111171709 | rs3802842 | -0.429 | 0.069 | -1.12 | 0.919 | -0.003 | 0.044 | 0.044 | 0 | 0.77 | 0.65 | 0.81 | 0.6 | 0.73 | 0.83 |
| 11 | 111176558 | rs11213824 | 0.046 | 0.448 | -1.736 | 0.729 | -0.017 | -0.001 | -0.001 | 0 | 0.59 | 0.36 | 0.47 | 0.62 | 0.59 | 0.72 |
| 11 | 111179390 | rs1046872 | 0.023 | 0.656 | -1.952 | 0.837 | -0.042 | -0.057 | -0.057 | 1.799 | 0.29 | 0.19 | 0.34 | 0.4 | 0.38 | 0.56 |
| 11 | 111180390 | rs11213825 | 0.537 | 0.794 | -2.492 | 0.329 | -0.096 | 0.079 | 0.079 | 0 | 0.2 | 0.02 | 0.25 | 0.05 | 0.31 | 0.16 |
| 11 | 111181120 | rs7947952 | -0.093 | 0.028 | -2.888 | 0.484 | -0.102 | -0.066 | -0.066 | 0.004 | 0.45 | 0.08 | 0.47 | 0.43 | 0.53 | 0.39 |
| 11 | 111181730 | rs11213826 | 0.89 | 0.086 | -2.764 | 0.528 | -0.041 | -0.092 | -0.092 | 0 | 0.25 | 0.07 | 0.3 | 0.41 | 0.29 | 0.52 |
| 11 | 111183157 | rs4617622 | -0.143 | -0.029 | -2.51 | 0.487 | 0.082 | 0.008 | 0.008 | 0.016 | 0.49 | 0.25 | 0.61 | 0.52 | 0.68 | 0.68 |
| 11 | 111189158 | rs10749971 | 0.734 | 0.2 | -2.299 | 0.429 | -0.085 | 0.048 | 0.048 | 0 | 0.21 | 0.09 | 0.27 | 0.07 | 0.36 | 0.17 |
| 11 | 111196619 | rs4938459 | 0.289 | -0.113 | -1.95 | 0.877 | -0.016 | -0.042 | -0.042 | 0.486 | 0.27 | 0.16 | 0.26 | 0.41 | 0.31 | 0.25 |
| 11 | 111196858 | rs7104791 | -1.339 | -0.618 | -1.179 | 0.916 | -0.015 | 0.045 | 0.045 | 1.531 | 0.74 | 0.69 | 0.83 | 0.4 | 0.79 | 0.49 |
| 11 | 111197776 | rs7944798 | -0.329 | -1.109 | -0.939 | 0.55 | 0 | 0.002 | 0.002 | 0.07 | 0.35 | 0.37 | 0.33 | 0.25 | 0.32 | 0.23 |
| 11 | 111201166 | rs7929013 | 0.884 | -0.957 | -0.66 | 0.284 | -0.049 | 0.052 | 0.052 | 0 | 0.23 | 0.14 | 0.43 | 0.04 | 0.39 | 0.16 |
| 11 | 111204938 | rs12574149 | 0.641 | -1.039 | -0.593 | 0.268 | 0.002 | 0 | 0 | 0 | 0.21 | 0.2 | 0.23 | 0.25 | 0.27 | 0.22 |
| 11 | 111208518 | rs1893993 | 0.037 | -0.932 | -0.605 | 0.2 | -0.005 | 0.002 | 0.002 | 0 | 0.3 | 0.32 | 0.48 | 0.12 | 0.44 | 0.19 |
| 11 | 111215664 | rs7117229 | 0.164 | -0.999 | -0.655 | 0.164 | -0.022 | 0.018 | 0.018 | 0 | 0.29 | 0.24 | 0.48 | 0.12 | 0.44 | 0.19 |
| 11 | 111221554 | rs10502141 | 1.251 | -0.862 | -1.027 | 0.136 | 0.047 | 0.066 | 0.066 | 0.862 | 0.12 | 0.02 | 0.08 | 0 | 0.04 | 0.03 |
| 11 | 111222713 | rs10502143 | 0.877 | -0.683 | -0.956 | 0.213 | -0.098 | 0.084 | 0.084 | 0 | 0.22 | 0.07 | 0.43 | 0.04 | 0.39 | 0.16 |
| 11 | 111226321 | rs11213850 | -0.087 | -1.031 | -0.712 | 0.325 | 0.048 | 0.05 | 0.05 | 0 | 0.44 | 0.28 | 0.65 | 0.31 | 0.66 | 0.37 |
| 11 | 111226465 | rs10891258 | -0.973 | -1.117 | -0.535 | 0.355 | -0.005 | 0 | 0 | 0 | 0.62 | 0.44 | 0.79 | 0.39 | 0.76 | 0.44 |

| | | | | | | | | | | | | | | | | |
|---|---|---|---|---|---|---|---|---|---|---|---|---|---|---|---|---|
| 11 | 111228050 | rs1815950 | -1.242 | -1.034 | -0.422 | 0.345 | -0.038 | 0.011 | 0.011 | 1.691 | 0.64 | 0.56 | 0.81 | 0.39 | 0.75 | 0.44 |
| 11 | 111228098 | rs2282637 | 0.471 | -0.956 | -0.558 | 0.346 | 0.004 | 0.013 | 0.013 | 0 | 0.06 | 0.11 | 0.05 | 0.08 | 0.05 | 0.03 |
| 11 | 111233450 | rs7118200 | 0.185 | -0.97 | -0.304 | 0.339 | 0.007 | 0.019 | 0.019 | 0 | 0.07 | 0.16 | 0.06 | 0.08 | 0.06 | 0.03 |
| 11 | 111238440 | rs7116862 | -0.978 | -0.702 | -0.346 | 0.366 | -0.016 | 0.004 | 0.004 | 1.137 | 0.62 | 0.48 | 0.79 | 0.39 | 0.72 | 0.4 |
| 11 | 111244756 | rs12799202 | -0.933 | -0.773 | -0.433 | 0.374 | 0.001 | -0.001 | -0.001 | 0.632 | 0.61 | 0.43 | 0.8 | 0.39 | 0.75 | 0.4 |
| 11 | 111248640 | rs4283016 | -0.188 | -0.528 | -0.561 | 0.444 | 0.029 | 0.031 | 0.031 | 0 | 0.44 | 0.32 | 0.68 | 0.31 | 0.65 | 0.34 |
| 11 | 111257202 | rs12800302 | 0.175 | -0.269 | -0.512 | 0.415 | -0.041 | 0.009 | 0.009 | 0 | 0.28 | 0.19 | 0.42 | 0.19 | 0.44 | 0.15 |
| 11 | 111260202 | rs883326 | 0.216 | -0.218 | -0.514 | 0.411 | -0.041 | 0.004 | 0.004 | 0.247 | 0.28 | 0.23 | 0.42 | 0.19 | 0.46 | 0.17 |
| 11 | 111261995 | rs1944918 | -1.252 | -0.514 | -0.2 | 0.495 | -0.013 | 0.004 | 0.004 | 1.873 | 0.64 | 0.62 | 0.83 | 0.47 | 0.79 | 0.41 |
| 11 | 111269435 | rs1123066 | -0.426 | -0.386 | -0.258 | 0.237 | 0.088 | 0.012 | 0.012 | 0.858 | 0.49 | 0.3 | 0.73 | 0.47 | 0.69 | 0.41 |
| 11 | 111270228 | rs7952176 | -0.912 | -0.379 | -0.142 | 0.286 | -0.025 | 0.009 | 0.009 | 1.493 | 0.6 | 0.51 | 0.8 | 0.47 | 0.73 | 0.43 |
| 11 | 111272564 | rs12362765 | 0.392 | -0.188 | -0.362 | 0.463 | -0.022 | -0.038 | -0.038 | 0.806 | 0.24 | 0.15 | 0.31 | 0.29 | 0.27 | 0.25 |
| 11 | 111275250 | rs4938535 | 0.505 | -0.058 | -0.353 | 0.451 | -0.016 | -0.039 | -0.039 | 0.231 | 0.22 | 0.15 | 0.29 | 0.29 | 0.25 | 0.24 |
| 11 | 111282054 | rs7115223 | -0.061 | 0.069 | -0.38 | 0.37 | -0.012 | 0.021 | 0.021 | 0.826 | 0.34 | 0.28 | 0.47 | 0.2 | 0.43 | 0.19 |
| 11 | 111282662 | rs17577362 | 1.043 | -0.351 | -0.934 | 0.213 | -0.056 | 0.011 | 0.011 | 2.845 | 0.3 | 0.01 | 0.14 | 0.23 | 0.27 | 0.49 |
| 11 | 111287287 | rs1944926 | 0.625 | -0.087 | -0.151 | 0.089 | 0.034 | -0.025 | -0.025 | 0.055 | 0.41 | 0.19 | 0.22 | 0.52 | 0.34 | 0.59 |
| 11 | 111290086 | rs1944928 | -0.786 | -0.054 | 0.608 | -0.268 | -0.031 | -0.058 | -0.058 | 0.117 | 0.9 | 0.79 | 0.94 | 1 | 0.96 | 0.99 |
| 11 | 111298298 | rs4938594 | -0.859 | 0.309 | 0.515 | -0.328 | -0.001 | 0.004 | 0.004 | 0 | 0.41 | 0.4 | 0.42 | 0.3 | 0.38 | 0.33 |
| 11 | 111300044 | rs10789833 | -1.284 | 0.446 | -0.33 | -0.253 | -0.013 | 0.055 | 0.055 | 0.051 | 0.48 | 0.72 | 0.48 | 0.32 | 0.45 | 0.34 |
| 11 | 111302337 | rs12293219 | -0.216 | -0.006 | -0.432 | -0.206 | -0.014 | -0.065 | -0.065 | 1.824 | 0.14 | 0.3 | 0.11 | 0.02 | 0.11 | 0.02 |
| 11 | 111302619 | rs12287211 | -0.197 | 0.005 | -0.468 | -0.201 | -0.013 | -0.065 | -0.065 | 0.227 | 0.14 | 0.3 | 0.1 | 0.02 | 0.11 | 0.02 |
| 11 | 111306908 | rs7930326 | -0.479 | 0.565 | -1.236 | 0.253 | -0.029 | 0.002 | 0.002 | 0 | 0.23 | 0.08 | 0.23 | 0.2 | 0.24 | 0.26 |
| 11 | 111308006 | rs10891270 | 0.224 | 0.322 | -1.042 | 0.306 | -0.017 | 0.023 | 0.023 | 0.166 | 0.4 | 0.27 | 0.37 | 0.39 | 0.47 | 0.49 |
| 11 | 111310559 | rs949126 | 0.3 | 0.277 | -0.223 | 0.269 | -0.021 | -0.022 | -0.022 | 0.112 | 0.79 | 0.69 | 0.78 | 0.86 | 0.84 | 0.75 |
| 11 | 111323084 | rs11213910 | -0.348 | 0.924 | 0.156 | 0.672 | 0.068 | 0.049 | 0.049 | 0 | 0.14 | 0.3 | 0.27 | 0.11 | 0.22 | 0.15 |
| 11 | 111325300 | rs11213912 | -0.404 | 0.248 | -0.443 | 0.387 | 0.036 | -0.007 | -0.007 | 0 | 0.2 | 0.06 | 0.17 | 0.15 | 0.12 | 0.19 |
| 11 | 111338979 | rs11213916 | -0.393 | 1.164 | -0.422 | 0.534 | 0.054 | 0.029 | 0.029 | 0 | 0.22 | 0.27 | 0.45 | 0.22 | 0.34 | 0.18 |
| 11 | 111340452 | rs638089 | 1.506 | 0.426 | -0.495 | 0.118 | 0.001 | -0.008 | -0.008 | 0.15 | 0.51 | 0.34 | 0.35 | 0.61 | 0.51 | 0.58 |
| 11 | 111340577 | rs11213917 | -0.424 | 0.2 | -0.709 | -0.049 | 0.026 | 0.01 | 0.01 | 3.136 | 0.23 | 0.05 | 0.17 | 0.15 | 0.14 | 0.24 |
| 11 | 111346465 | rs11213921 | 0.143 | 0.927 | -0.206 | -0.119 | -0.029 | 0.022 | 0.022 | 0.339 | 0.07 | 0.02 | 0.15 | 0.02 | 0.13 | 0.04 |
| 11 | 111372799 | rs9633937 | -0.698 | 0.092 | -0.026 | -0.115 | -0.004 | -0.004 | -0.004 | 0 | 0.3 | 0.27 | 0.21 | 0.29 | 0.37 | 0.25 |
| 11 | 111396283 | rs501560 | -0.287 | -0.675 | 0.241 | -1.059 | -0.054 | -0.097 | -0.097 | 0 | 0.86 | 0.43 | 0.9 | 1 | 0.97 | 0.99 |
| 11 | 111398120 | rs17585834 | 1.33 | -0.664 | -0.046 | -1.126 | 0.005 | 0.054 | 0.054 | 0 | 0.09 | 0.01 | 0.04 | 0 | 0.07 | 0.08 |
| 11 | 111401609 | rs11602450 | 1.101 | -0.193 | -0.106 | -1.168 | -0.036 | 0.031 | 0.031 | 6.211 | 0.06 | 0.01 | 0.09 | 0 | 0.19 | 0.02 |
| 11 | 111401840 | rs7107378 | -0.583 | -0.228 | -0.027 | -0.938 | -0.013 | 0.023 | 0.023 | 0.131 | 0.29 | 0.2 | 0.21 | 0.2 | 0.37 | 0.25 |
| 11 | 111413465 | rs4938784 | -0.317 | -0.094 | 0.172 | -1.559 | 0.004 | 0.055 | 0.055 | 0 | 0.33 | 0.21 | 0.19 | 0.09 | 0.35 | 0.24 |
| 11 | 111420936 | rs4938792 | -0.666 | -0.223 | 0.729 | -1.743 | -0.003 | -0.103 | -0.103 | 0.843 | 0.4 | 0.45 | 0.27 | 0.05 | 0.41 | 0.25 |
| 11 | 111422217 | rs663763 | 0.592 | -0.234 | 0.279 | -1.734 | 0.018 | 0.166 | 0.166 | 0 | 0.57 | 0.25 | 0.7 | 0.95 | 0.59 | 0.75 |
| 11 | 111431614 | rs542275 | 0.601 | -0.233 | 0.296 | -1.733 | 0.018 | 0.167 | 0.167 | 0 | 0.57 | 0.25 | 0.7 | 0.95 | 0.59 | 0.75 |
| 11 | 111460678 | rs567040 | 0.688 | 0.827 | 0.455 | -1.625 | 0.02 | 0.094 | 0.094 | 0 | 0.61 | 0.25 | 0.72 | 0.95 | 0.65 | 0.74 |
| 11 | 111462096 | rs2212520 | 0.475 | 0.657 | 0.384 | -1.591 | 0.014 | 0.167 | 0.167 | 0.034 | 0.63 | 0.18 | 0.73 | 0.95 | 0.67 | 0.83 |
| 11 | 111469582 | rs10789841 | -0.023 | 0.643 | 0.555 | -1.607 | -0.003 | 0.015 | 0.015 | 0 | 0.28 | 0.23 | 0.19 | 0.05 | 0.31 | 0.17 |
| 11 | 111472392 | rs1784776 | 0.393 | 0.59 | 0.232 | -1.594 | 0.017 | 0.129 | 0.129 | 0 | 0.63 | 0.21 | 0.72 | 0.95 | 0.67 | 0.82 |
| 11 | 111472460 | rs1784775 | 0.393 | 0.59 | 0.232 | -1.594 | 0.017 | 0.129 | 0.129 | 0 | 0.63 | 0.21 | 0.72 | 0.95 | 0.67 | 0.82 |
| 11 | 111525069 | rs10502145 | 1.403 | 1.288 | -0.478 | -1.433 | -0.008 | -0.036 | -0.036 | 0.008 | 0.34 | 0.07 | 0.36 | 0.64 | 0.36 | 0.34 |
| 11 | 111536670 | rs7926485 | -0.074 | 1.311 | -0.194 | -1.59 | 0.011 | 0.087 | 0.087 | 0.234 | 0.64 | 0.24 | 0.73 | 0.95 | 0.68 | 0.82 |
| 11 | 111538875 | rs7128353 | -0.242 | 1.292 | -0.107 | -1.607 | 0.004 | 0.019 | 0.019 | 0.078 | 0.67 | 0.3 | 0.75 | 0.95 | 0.68 | 0.82 |
| 11 | 111552053 | rs10891288 | 0.147 | 1.207 | -0.343 | -1.608 | 0.008 | 0.201 | 0.201 | 1.124 | 0.63 | 0.15 | 0.7 | 0.93 | 0.68 | 0.78 |
| 11 | 111569931 | rs10502147 | 0.451 | 1.706 | -0.077 | -1.68 | -0.006 | 0.009 | 0.009 | 0.018 | 0.28 | 0.19 | 0.19 | 0.05 | 0.31 | 0.18 |
| 11 | 111581521 | rs11213985 | 0.189 | 1.667 | -0.065 | -1.504 | 0.007 | 0.153 | 0.153 | 0.818 | 0.63 | 0.16 | 0.69 | 0.87 | 0.68 | 0.78 |
| 11 | 111587495 | rs4936661 | 0.416 | 1.721 | -0.066 | -1.479 | -0.006 | 0.023 | 0.023 | 0.025 | 0.28 | 0.19 | 0.21 | 0.11 | 0.32 | 0.18 |
| 11 | 111595189 | rs585149 | 0.984 | 1.938 | -0.008 | -1.304 | -0.039 | 0.028 | 0.028 | 0 | 0.06 | 0.01 | 0.1 | 0.02 | 0.17 | 0.04 |
| 11 | 111609174 | rs647080 | -0.639 | 2.045 | 0.092 | -1.278 | -0.089 | -0.027 | -0.027 | 0 | 0.25 | 0.08 | 0.29 | 0.22 | 0.33 | 0.39 |
| 11 | 111625566 | rs3737445 | 0.431 | 2.024 | 0.058 | -1.301 | -0.051 | -0.077 | -0.077 | 0 | 0.17 | 0.01 | 0.19 | 0.19 | 0.16 | 0.35 |
| 11 | 111627502 | rs10891291 | 1.268 | 1.861 | 0.184 | -1.386 | 0.03 | -0.006 | -0.006 | 0 | 0.37 | 0.18 | 0.41 | 0.63 | 0.34 | 0.42 |
| 11 | 111629743 | rs4936682 | 0.719 | 1.831 | 0.312 | -1.386 | -0.009 | 0.039 | 0.039 | 0.221 | 0.29 | 0.18 | 0.23 | 0.15 | 0.33 | 0.19 |
| 11 | 111633570 | rs12293026 | -1.01 | 1.625 | 0.531 | -1.236 | 0.044 | 0.028 | 0.028 | 4.581 | 0.06 | 0.4 | 0.06 | 0 | 0.01 | 0 |
| 11 | 111649421 | rs615642 | 0.802 | 1.848 | 0.436 | -1.254 | 0.039 | -0.013 | -0.013 | 0.056 | 0.7 | 0.57 | 0.68 | 0.78 | 0.62 | 0.61 |
| 11 | 111652323 | rs1529330 | 1.458 | 1.821 | 0.209 | -1.283 | 0.036 | -0.075 | -0.075 | 1.914 | 0.34 | 0.1 | 0.33 | 0.59 | 0.31 | 0.38 |
| 11 | 111666128 | rs648544 | 0.179 | 1.527 | 0.268 | -1.272 | -0.06 | -0.006 | -0.006 | 0 | 0.75 | 0.86 | 0.7 | 0.78 | 0.62 | 0.61 |
| 11 | 111673539 | rs11823089 | 0.604 | 1.591 | 0.334 | -1.298 | -0.014 | 0.048 | 0.048 | 0.16 | 0.32 | 0.19 | 0.31 | 0.19 | 0.32 | 0.22 |
| 11 | 111686800 | rs673679 | 0.205 | 1.601 | 0.314 | -1.317 | -0.071 | -0.013 | -0.013 | 0 | 0.75 | 0.89 | 0.7 | 0.78 | 0.62 | 0.6 |
| 11 | 111706111 | rs17113290 | 0.367 | 1.639 | 0.287 | -1.372 | -0.048 | -0.058 | -0.058 | 0 | 0.17 | 0.03 | 0.2 | 0.19 | 0.21 | 0.36 |
| 11 | 111707386 | rs10502149 | 1.37 | 0.887 | 0.32 | -1.356 | 0.059 | -0.076 | -0.076 | 0.912 | 0.35 | 0.11 | 0.32 | 0.59 | 0.3 | 0.39 |
| 11 | 111707797 | rs10502150 | 1.37 | 0.887 | 0.344 | -1.356 | 0.06 | -0.07 | -0.07 | 0.649 | 0.35 | 0.11 | 0.32 | 0.59 | 0.3 | 0.39 |
| 11 | 111716070 | rs10789849 | 0.458 | 0.917 | 0.431 | -1.359 | 0.02 | -0.004 | -0.004 | 0 | 0.72 | 0.65 | 0.69 | 0.78 | 0.62 | 0.6 |
| 11 | 111720072 | rs1648390 | 1.117 | 0.933 | 0.423 | -1.338 | -0.015 | 0.002 | 0.002 | 0 | 0.37 | 0.42 | 0.36 | 0.59 | 0.29 | 0.38 |
| 11 | 111724133 | rs10502151 | 1.349 | 0.946 | -0.053 | -1.331 | 0.059 | -0.076 | -0.076 | 0 | 0.35 | 0.11 | 0.32 | 0.59 | 0.29 | 0.39 |
| 11 | 111743232 | rs7112886 | 0.566 | 0.977 | -0.108 | -1.294 | -0.012 | 0.046 | 0.046 | 0 | 0.32 | 0.2 | 0.31 | 0.19 | 0.32 | 0.22 |

| | | | | | | | | | | | | | | | | |
|---|---|---|---|---|---|---|---|---|---|---|---|---|---|---|---|---|
| 11 | 111754574 | rs1045282 | 0.551 | 1.006 | -0.096 | -1.28 | -0.012 | 0.047 | 0.047 | 0.059 | 0.33 | 0.2 | 0.31 | 0.19 | 0.32 | 0.22 |
| 11 | 111761230 | rs11214029 | 0.563 | 1.005 | -0.096 | -1.281 | -0.012 | 0.047 | 0.047 | 0.075 | 0.33 | 0.2 | 0.31 | 0.19 | 0.32 | 0.22 |
| 11 | 111764396 | rs11604704 | 0.289 | 1.001 | -0.101 | -1.286 | -0.011 | 0.059 | 0.059 | 0 | 0.34 | 0.2 | 0.32 | 0.19 | 0.33 | 0.22 |
| 11 | 111766039 | rs11214031 | 0.089 | 1.029 | -0.069 | -1.288 | -0.055 | -0.006 | -0.006 | 0.38 | 0.75 | 0.84 | 0.7 | 0.78 | 0.62 | 0.6 |
| 11 | 111770150 | rs10891297 | 0.432 | 1.119 | -0.063 | -1.232 | -0.012 | 0.047 | 0.047 | 0.059 | 0.33 | 0.2 | 0.31 | 0.19 | 0.32 | 0.22 |
| 11 | 111781047 | rs11603779 | 0.41 | 1.142 | -0.053 | -1.509 | -0.011 | 0.048 | 0.048 | 0.048 | 0.33 | 0.2 | 0.31 | 0.19 | 0.32 | 0.22 |
| 11 | 111784229 | rs4252588 | 1.038 | 0.401 | 0.078 | -1.505 | -0.006 | -0.034 | -0.034 | 0 | 0.1 | 0.13 | 0.05 | 0 | 0.06 | 0.08 |
| 11 | 111816143 | rs11214049 | 1.542 | 0.578 | 0.453 | -1.024 | 0.034 | -0.052 | -0.052 | 0 | 0.35 | 0.12 | 0.36 | 0.59 | 0.33 | 0.38 |
| 11 | 111849775 | rs11821811 | 0.727 | 0.706 | 0.369 | -1.088 | -0.033 | 0.045 | 0.045 | 0.794 | 0.29 | 0.06 | 0.25 | 0.22 | 0.26 | 0.2 |
| 11 | 111856361 | rs7926696 | -1.669 | 0.373 | 0.967 | -0.74 | 0.009 | -0.004 | -0.004 | 0.091 | 0.06 | 0.29 | 0.04 | 0 | 0.01 | 0 |
| 11 | 111875773 | rs10789856 | 1.579 | 0.354 | 0.928 | -0.729 | -0.023 | -0.146 | -0.146 | 0 | 0.32 | 0.02 | 0.34 | 0.55 | 0.33 | 0.33 |
| 11 | 111887994 | rs7123284 | -1.724 | 0.572 | 1.028 | -0.863 | 0.007 | -0.005 | -0.005 | 0.243 | 0.06 | 0.29 | 0.04 | 0 | 0.01 | 0 |
| 11 | 111889654 | rs476581 | -0.146 | 0.723 | 1.01 | -0.893 | -0.099 | -0.042 | -0.042 | 0.003 | 0.81 | 0.96 | 0.77 | 0.83 | 0.72 | 0.61 |
| 11 | 111896324 | rs2303436 | 0.816 | -0.503 | 1.395 | -0.64 | 0.029 | -0.015 | -0.015 | 0 | 0.36 | 0.22 | 0.37 | 0.59 | 0.33 | 0.35 |
| 11 | 111908162 | rs627441 | -0.069 | -0.511 | 1.452 | -0.643 | 0.103 | 0.009 | 0.009 | 0.176 | 0.45 | 0.84 | 0.44 | 0.59 | 0.34 | 0.38 |
| 11 | 111909819 | rs6589260 | 0.404 | -0.513 | 1.462 | -0.644 | 0.003 | 0 | 0 | 0 | 0.4 | 0.41 | 0.4 | 0.58 | 0.33 | 0.36 |
| 11 | 111912083 | rs10891313 | 0.729 | -0.496 | 1.552 | -0.636 | 0.009 | -0.003 | -0.003 | 0.889 | 0.36 | 0.31 | 0.37 | 0.59 | 0.33 | 0.38 |
| 11 | 111913364 | rs7949136 | 0.729 | -0.494 | 1.573 | -0.635 | 0.009 | -0.003 | -0.003 | 0.103 | 0.36 | 0.31 | 0.37 | 0.59 | 0.33 | 0.38 |
| 11 | 111930576 | rs3882890 | 0.113 | -0.49 | 0.896 | -0.633 | 0.034 | 0.006 | 0.006 | 0 | 0.43 | 0.64 | 0.41 | 0.59 | 0.34 | 0.38 |
| 11 | 111933088 | rs10891315 | 0.818 | -0.338 | 0.825 | -0.558 | 0.031 | -0.046 | -0.046 | 9.158 | 0.34 | 0.14 | 0.36 | 0.59 | 0.33 | 0.36 |
| 11 | 111941959 | rs1425917 | 0.744 | -0.347 | 0.573 | -0.61 | -0.043 | 0.04 | 0.04 | 0 | 0.07 | 0 | 0.05 | 0 | 0.1 | 0.02 |
| 11 | 111942256 | rs11214075 | 0.476 | -1.367 | 0.972 | -0.172 | 0.031 | -0.055 | -0.055 | 0 | 0.34 | 0.12 | 0.36 | 0.59 | 0.33 | 0.36 |
| 11 | 111944901 | rs2303437 | 0.508 | -1.31 | 0.99 | -0.145 | 0.027 | -0.065 | -0.065 | 0.177 | 0.34 | 0.11 | 0.36 | 0.59 | 0.32 | 0.36 |
| 11 | 111960533 | rs10789859 | 0.445 | -1.04 | 0.373 | -0.107 | 0.045 | -0.024 | -0.024 | 0 | 0.37 | 0.17 | 0.37 | 0.59 | 0.32 | 0.37 |
| 11 | 111966497 | rs17113461 | -0.961 | -0.919 | 0.313 | -0.083 | 0.065 | 0.05 | 0.05 | 4.193 | 0.05 | 0.37 | 0.04 | 0 | 0.01 | 0 |
| 11 | 111967605 | rs10891319 | 0.4 | -0.883 | -0.536 | -0.06 | 0.048 | -0.017 | -0.017 | 0.009 | 0.37 | 0.17 | 0.37 | 0.61 | 0.32 | 0.37 |
| 11 | 111970123 | rs7941988 | -0.022 | -0.584 | -1.084 | 0.1 | 0.008 | 0.011 | 0.011 | 0 | 0.45 | 0.5 | 0.52 | 0.65 | 0.41 | 0.35 |
| 11 | 111977919 | rs574568 | -0.198 | -0.352 | -2.086 | 0.429 | 0.109 | 0.054 | 0.054 | 0.076 | 0.53 | 0.17 | 0.56 | 0.39 | 0.64 | 0.61 |
| 11 | 111989592 | rs360726 | 0.538 | -0.188 | -1.833 | 0.828 | 0.026 | -0.04 | -0.04 | 0 | 0.7 | 0.22 | 0.68 | 0.59 | 0.78 | 0.73 |
| 11 | 111992592 | rs11214093 | 0.038 | -0.175 | -2.806 | -0.439 | 0.028 | 0.271 | 0.271 | 0 | 0.5 | 0.11 | 0.55 | 0.86 | 0.44 | 0.64 |
| 11 | 112000303 | rs578784 | 0.092 | 0.665 | -2.18 | -0.399 | 0.01 | -0.002 | -0.002 | 0 | 0.2 | 0.22 | 0.29 | 0.14 | 0.28 | 0.19 |
| 11 | 112005939 | rs243908 | -0.353 | -0.153 | -2.325 | -1.031 | -0.008 | 0.025 | 0.025 | 0 | 0.27 | 0.2 | 0.34 | 0.13 | 0.31 | 0.2 |
| 11 | 112008978 | rs543810 | -0.072 | -1.038 | -1.821 | -0.364 | -0.032 | 0.098 | 0.098 | 0 | 0.83 | 0.74 | 0.83 | 0.59 | 0.88 | 0.9 |
| 11 | 112012630 | rs544354 | 0.15 | -1.166 | -1.619 | -0.424 | -0.044 | 0.116 | 0.116 | 0 | 0.81 | 0.67 | 0.82 | 0.6 | 0.88 | 0.9 |
| 11 | 112020916 | rs549908 | 0.152 | -0.951 | -1.614 | -0.442 | -0.01 | 0.036 | 0.036 | 0 | 0.27 | 0.18 | 0.34 | 0.12 | 0.31 | 0.2 |
| 11 | 112022848 | rs5744256 | 0.979 | -0.93 | -1.687 | -0.742 | -0.062 | 0.054 | 0.054 | 0 | 0.09 | 0.01 | 0.1 | 0 | 0.22 | 0.06 |
| 11 | 112031624 | rs2043055 | 1.25 | -0.655 | -1.711 | -0.768 | -0.002 | -0.003 | -0.003 | 0.385 | 0.47 | 0.48 | 0.41 | 0.48 | 0.37 | 0.65 |
| 11 | 112034725 | rs360717 | 0.223 | -0.754 | -1.728 | -0.674 | -0.001 | 0.023 | 0.023 | 0 | 0.27 | 0.2 | 0.31 | 0.12 | 0.28 | 0.18 |
| 11 | 112035458 | rs1946518 | 0.743 | -0.843 | -1.722 | -0.738 | 0.02 | 0.01 | 0.01 | 0 | 0.55 | 0.65 | 0.51 | 0.47 | 0.58 | 0.71 |
| 11 | 112035507 | rs1946519 | 0.799 | -0.885 | -1.741 | -0.759 | 0.022 | 0.011 | 0.011 | 0 | 0.54 | 0.65 | 0.51 | 0.47 | 0.58 | 0.71 |
| 11 | 112037554 | rs1290349 | 0.297 | -0.842 | -1.725 | -0.783 | -0.001 | 0.017 | 0.017 | 0 | 0.27 | 0.21 | 0.31 | 0.12 | 0.28 | 0.19 |
| 11 | 112037653 | rs11214105 | 0.312 | -0.846 | -1.727 | -0.785 | -0.002 | 0.021 | 0.021 | 0 | 0.27 | 0.2 | 0.31 | 0.12 | 0.28 | 0.19 |
| 11 | 112040027 | rs360712 | 0.829 | -0.823 | -1.747 | -0.817 | -0.042 | 0.064 | 0.064 | 0 | 0.21 | 0.05 | 0.26 | 0.12 | 0.27 | 0.18 |
| 11 | 112040136 | rs11214106 | 0.334 | -0.942 | -1.686 | -0.761 | 0.005 | -0.002 | -0.002 | 3.227 | 0.1 | 0.07 | 0.17 | 0.4 | 0.12 | 0.05 |
| 11 | 112043614 | rs11214108 | 0.296 | -0.934 | -1.715 | -0.757 | 0.003 | -0.011 | -0.011 | 0 | 0.1 | 0.06 | 0.16 | 0.4 | 0.12 | 0.05 |
| 11 | 112049154 | rs10891332 | -1.004 | -0.729 | -1.659 | -1.313 | -0.015 | -0.019 | -0.019 | 0 | 0.92 | 0.92 | 0.99 | 1 | 0.98 | 0.95 |
| 11 | 112064431 | rs7105056 | -0.688 | -0.39 | -1.721 | -1.382 | -0.039 | -0.044 | -0.044 | 0 | 0.92 | 0.86 | 0.98 | 1 | 0.98 | 0.95 |
| 11 | 112065434 | rs10891338 | -1.068 | -0.331 | -1.912 | -1.353 | 0.003 | -0.001 | -0.001 | 0 | 0.93 | 0.95 | 0.99 | 1 | 0.98 | 0.95 |
| 11 | 112068593 | rs11214127 | 0.023 | 0.17 | -1.849 | -1.238 | 0.004 | 0.029 | 0.029 | 0 | 0.25 | 0.18 | 0.25 | 0.13 | 0.26 | 0.18 |
| 11 | 112085316 | rs2250417 | -0.698 | 0.175 | -1.844 | -1.303 | -0.05 | 0.091 | 0.091 | 0 | 0.37 | 0.23 | 0.46 | 0.13 | 0.51 | 0.26 |
| 11 | 112091299 | rs1367038 | 0.07 | -0.748 | -0.492 | -1.507 | -0.015 | -0.057 | -0.057 | 0 | 0.7 | 0.5 | 0.75 | 0.78 | 0.72 | 0.81 |
| 11 | 112095039 | rs1154763 | -0.299 | -0.778 | -0.536 | -1.522 | 0 | 0.001 | 0.001 | 0 | 0.75 | 0.74 | 0.78 | 0.78 | 0.73 | 0.81 |
| 11 | 112099410 | rs3819331 | 1.501 | 0.347 | -0.84 | -0.995 | -0.058 | -0.065 | -0.065 | 0 | 0.1 | 0.01 | 0.14 | 0.25 | 0.15 | 0.38 |
| 11 | 112108556 | rs919479 | 0.676 | -0.441 | 0.004 | -1.705 | 0.001 | -0.01 | -0.01 | 0 | 0.76 | 0.34 | 0.75 | 0.78 | 0.73 | 0.81 |
| 11 | 112108776 | rs1811350 | -0.314 | -0.444 | 0.187 | -1.707 | 0.023 | -0.001 | -0.001 | 0 | 0.82 | 0.7 | 0.81 | 0.78 | 0.75 | 0.85 |
| 11 | 112123095 | rs2564872 | 0.948 | -0.549 | 0.017 | -1.766 | -0.001 | -0.005 | -0.005 | 0 | 0.7 | 0.33 | 0.72 | 0.78 | 0.72 | 0.8 |
| 11 | 112138835 | rs2518369 | 0.092 | -0.04 | 0.111 | -1.522 | 0.04 | -0.011 | -0.011 | 0 | 0.79 | 0.56 | 0.78 | 0.78 | 0.75 | 0.84 |
| 11 | 112139158 | rs2564881 | 0.081 | -0.074 | 0.114 | -1.55 | 0.032 | -0.011 | -0.011 | 0 | 0.79 | 0.59 | 0.79 | 0.78 | 0.75 | 0.84 |
| 11 | 112140332 | rs2853127 | 0.31 | 0.028 | 0.064 | -1.582 | 0.045 | -0.023 | -0.023 | 0.138 | 0.78 | 0.56 | 0.77 | 0.78 | 0.71 | 0.84 |
| 11 | 112147780 | rs7120010 | 0.939 | -0.797 | -0.003 | -1.628 | -0.044 | -0.041 | -0.041 | 2.054 | 0.18 | 0.31 | 0.08 | 0.09 | 0.09 | 0.16 |
| 11 | 112165009 | rs1404794 | 0.324 | -0.514 | -0.406 | -1.709 | 0.028 | 0.057 | 0.057 | 1.834 | 0.59 | 0.24 | 0.65 | 0.69 | 0.63 | 0.64 |
| 11 | 112170884 | rs10458936 | 0.488 | -0.774 | -0.225 | -1.599 | 0.056 | 0.053 | 0.053 | 1.438 | 0.61 | 0.21 | 0.69 | 0.69 | 0.72 | 0.67 |
| 11 | 112183408 | rs4937257 | -0.096 | -0.78 | -0.11 | -1.585 | -0.019 | -0.015 | -0.015 | 0 | 0.67 | 0.39 | 0.75 | 0.68 | 0.74 | 0.7 |
| 11 | 112207339 | rs11825888 | 1.432 | -0.581 | -0.71 | -2.182 | -0.095 | 0.152 | 0.152 | 0.893 | 0.29 | 0.05 | 0.43 | 0.09 | 0.38 | 0.14 |
| 11 | 112208482 | rs873411 | 0.407 | -0.328 | -0.56 | -1.878 | -0.023 | 0.017 | 0.017 | 0 | 0.32 | 0.17 | 0.3 | 0.68 | 0.34 | 0.48 |
| 11 | 112208568 | rs940751 | -0.789 | -0.441 | -0.428 | -1.944 | 0.031 | 0.039 | 0.039 | 0 | 0.85 | 0.65 | 0.84 | 0.8 | 0.82 | 0.8 |
| 11 | 112208967 | rs1404792 | 0.161 | 0.045 | -1.067 | -1.55 | -0.03 | 0.017 | 0.017 | 0 | 0.32 | 0.15 | 0.3 | 0.68 | 0.34 | 0.48 |
| 11 | 112218097 | rs4937300 | 0.897 | 0.128 | -1.7 | -1.724 | -0.141 | 0.226 | 0.226 | 1.625 | 0.35 | 0.03 | 0.42 | 0.07 | 0.4 | 0.18 |
| 11 | 112220382 | rs10891349 | -0.268 | 1.209 | -1.232 | -0.92 | -0.1 | 0.043 | 0.043 | 0.046 | 0.25 | 0.06 | 0.3 | 0.82 | 0.36 | 0.51 |

| | | | | | | | | | | | | | | | | |
|---|---|---|---|---|---|---|---|---|---|---|---|---|---|---|---|---|
| 11 | 112230073 | rs11214178 | -0.876 | 1.002 | -0.796 | -1.174 | -0.032 | 0.046 | 0.046 | 0.173 | 0.33 | 0.19 | 0.41 | 0.91 | 0.41 | 0.57 |
| 11 | 112260774 | rs7951256 | -0.34 | 1.201 | -0.794 | -0.939 | -0.044 | 0.052 | 0.052 | 0.063 | 0.24 | 0.11 | 0.37 | 0.87 | 0.34 | 0.42 |
| 11 | 112262881 | rs12793200 | 0.567 | 0.671 | 0.808 | -1.108 | -0.001 | -0.046 | -0.046 | 0.937 | 0.86 | 0.58 | 0.92 | 0.95 | 0.93 | 0.86 |
| 11 | 112263027 | rs7108803 | 0.686 | 1.015 | 0.748 | -0.092 | -0.011 | 0.078 | 0.078 | 0 | 0.73 | 0.56 | 0.79 | 0.55 | 0.72 | 0.87 |
| 11 | 112270660 | rs10789879 | 0.998 | -0.179 | 0.426 | -0.808 | 0.052 | 0.003 | 0.003 | 0.181 | 0.52 | 0.34 | 0.62 | 0.51 | 0.59 | 0.66 |
| 11 | 112271995 | rs1525407 | 0.583 | -0.173 | 0.476 | -0.798 | -0.016 | -0.027 | -0.027 | 0 | 0.2 | 0.23 | 0.16 | 0.04 | 0.12 | 0.2 |
| 11 | 112272189 | rs4638337 | 0.118 | -0.136 | 0.106 | -0.659 | 0.035 | 0.13 | 0.13 | 0 | 0.15 | 0.31 | 0.19 | 0.43 | 0.18 | 0.45 |
| 11 | 112280439 | rs7931891 | -0.161 | -0.188 | 0.176 | -0.651 | 0.029 | 0.14 | 0.14 | 0 | 0.17 | 0.35 | 0.2 | 0.44 | 0.17 | 0.46 |
| 11 | 112281953 | rs4937377 | 2.036 | -0.299 | -0.079 | -0.805 | -0.152 | 0.141 | 0.141 | 0.989 | 0.36 | 0.03 | 0.42 | 0.08 | 0.41 | 0.22 |
| 11 | 112287801 | rs1357931 | -0.204 | -0.058 | 0.362 | -0.513 | 0.026 | 0.129 | 0.129 | 0 | 0.17 | 0.34 | 0.2 | 0.44 | 0.17 | 0.46 |
| 11 | 112290913 | rs7951272 | -1.149 | -0.086 | 0.586 | -0.533 | 0.009 | 0.102 | 0.102 | 0 | 0.25 | 0.38 | 0.24 | 0.48 | 0.23 | 0.47 |
| 11 | 112291421 | rs7926493 | 0.327 | -0.059 | 0.574 | -0.549 | -0.012 | -0.024 | -0.024 | 0.05 | 0.11 | 0.17 | 0.06 | 0.05 | 0.06 | 0.05 |
| 11 | 112292984 | rs1540072 | 1.302 | -0.043 | 0.455 | -0.667 | -0.02 | 0.083 | 0.083 | 0 | 0.56 | 0.35 | 0.59 | 0.12 | 0.52 | 0.44 |
| 11 | 112295421 | rs1404796 | -0.819 | 0.041 | 0.301 | -0.581 | 0.008 | 0.038 | 0.038 | 0 | 0.83 | 0.8 | 0.83 | 0.6 | 0.75 | 0.92 |
| 11 | 112309057 | rs10749995 | -0.919 | 0.109 | 0.49 | -0.49 | 0.01 | 0.075 | 0.075 | 0 | 0.24 | 0.36 | 0.23 | 0.48 | 0.23 | 0.46 |
| 11 | 112315285 | rs4936070 | 1.378 | -0.26 | 0.091 | -0.729 | -0.078 | 0.145 | 0.145 | 0.301 | 0.39 | 0.14 | 0.49 | 0.07 | 0.47 | 0.25 |
| 11 | 112320102 | rs10789885 | 0.625 | -0.125 | 0.549 | -0.485 | 0.007 | 0.075 | 0.075 | 0 | 0.1 | 0.14 | 0.15 | 0.41 | 0.13 | 0.41 |
| 11 | 112322300 | rs11214210 | 1.251 | -0.201 | 0.302 | -0.574 | -0.035 | -0.055 | -0.055 | 0 | 0.13 | 0.03 | 0.14 | 0.43 | 0.22 | 0.09 |
| 11 | 112326178 | rs10891374 | 0.873 | -0.304 | 0.855 | -0.611 | 0.015 | 0.105 | 0.105 | 0.472 | 0.75 | 0.45 | 0.78 | 0.54 | 0.78 | 0.57 |
| 11 | 112327279 | rs10891375 | -0.399 | -0.364 | 0.608 | -0.865 | 0.016 | -0.005 | -0.005 | 0 | 0.38 | 0.3 | 0.28 | 0.46 | 0.35 | 0.36 |
| 11 | 112335631 | rs7933113 | 0.315 | -0.327 | 0.575 | -0.871 | 0 | -0.01 | -0.01 | 0.356 | 0.12 | 0.13 | 0.06 | 0.05 | 0.09 | 0.05 |
| 11 | 112340415 | rs4937422 | -0.837 | -0.164 | 0.819 | -0.837 | 0.013 | -0.051 | -0.051 | 2.406 | 0.4 | 0.48 | 0.36 | 0.86 | 0.47 | 0.54 |
| 11 | 112341198 | rs11214218 | 1.653 | -0.081 | 0.488 | -0.847 | -0.067 | 0.126 | 0.126 | 1.119 | 0.34 | 0.1 | 0.46 | 0.08 | 0.4 | 0.21 |
| 11 | 112348123 | rs12574848 | -0.294 | -0.158 | 0.798 | -0.551 | -0.004 | -0.007 | -0.007 | 0 | 0.29 | 0.24 | 0.22 | 0.44 | 0.34 | 0.18 |
| 11 | 112359962 | rs7946315 | -0.968 | 0.095 | 0.88 | -0.52 | 0.012 | -0.006 | -0.006 | 0 | 0.39 | 0.72 | 0.34 | 0.47 | 0.3 | 0.63 |
| 11 | 112361133 | rs12361507 | 0.273 | -0.078 | 0.906 | -0.617 | 0.013 | 0.007 | 0.007 | 0 | 0.22 | 0.19 | 0.14 | 0.06 | 0.13 | 0.18 |
| 11 | 112362462 | rs10789889 | -0.352 | 0.152 | 1.017 | -0.442 | 0.026 | 0.142 | 0.142 | 0.467 | 0.16 | 0.37 | 0.19 | 0.41 | 0.17 | 0.45 |
| 11 | 112374511 | rs11214232 | 0.849 | 0.245 | 0.69 | -0.212 | -0.055 | -0.066 | -0.066 | 0 | 0.1 | 0.01 | 0.15 | 0.35 | 0.14 | 0.32 |
| 11 | 112380033 | rs4594033 | -0.789 | 0.089 | 1.024 | -0.32 | 0.004 | 0.069 | 0.069 | 0.247 | 0.19 | 0.32 | 0.21 | 0.45 | 0.19 | 0.47 |
| 11 | 112386785 | rs949341 | -0.448 | 0.119 | 0.878 | -0.527 | -0.006 | -0.013 | -0.013 | 0 | 0.73 | 0.83 | 0.8 | 0.56 | 0.68 | 0.81 |
| 11 | 112387253 | rs1945090 | -0.78 | 0.002 | 1.062 | -0.452 | 0.004 | 0.08 | 0.08 | 0.811 | 0.19 | 0.37 | 0.21 | 0.42 | 0.19 | 0.47 |
| 11 | 112387698 | rs7101551 | -0.889 | 0.043 | 0.856 | -0.457 | 0.011 | -0.005 | -0.005 | 0.049 | 0.4 | 0.71 | 0.34 | 0.47 | 0.31 | 0.63 |
| 11 | 112393443 | rs1945107 | -0.187 | 0.011 | 1.004 | -0.019 | -0.001 | -0.001 | -0.001 | 3.045 | 0.75 | 0.81 | 0.8 | 0.57 | 0.69 | 0.82 |
| 11 | 112396127 | rs11214236 | 0.647 | 0.118 | 0.884 | -0.18 | -0.066 | -0.086 | -0.086 | 0.232 | 0.21 | 0.04 | 0.2 | 0.38 | 0.33 | 0.13 |
| 11 | 112397104 | rs10891380 | -0.415 | -0.094 | 1.042 | -0.092 | -0.004 | -0.004 | -0.004 | 0 | 0.71 | 0.78 | 0.76 | 0.58 | 0.62 | 0.82 |
| 11 | 112398446 | rs7125827 | -0.415 | -0.094 | 1.042 | -0.092 | -0.004 | -0.004 | -0.004 | 0.19 | 0.71 | 0.78 | 0.76 | 0.58 | 0.62 | 0.82 |
| 11 | 112400860 | rs1945101 | -0.509 | -0.151 | 1.162 | -0.212 | -0.038 | -0.079 | -0.079 | 0.536 | 0.26 | 0.45 | 0.19 | 0.11 | 0.18 | 0.31 |
| 11 | 112402954 | rs1945115 | 1.298 | -0.233 | 0.313 | -0.199 | -0.099 | -0.102 | -0.102 | 0 | 0.37 | 0.05 | 0.47 | 0.4 | 0.4 | 0.43 |
| 11 | 112417074 | rs1945119 | 0.982 | -0.051 | 0.582 | -0.23 | -0.011 | -0.042 | -0.042 | 0 | 0.42 | 0.21 | 0.54 | 0.44 | 0.4 | 0.48 |
| 11 | 112434430 | rs11214244 | -1.289 | -0.29 | 0.813 | -0.022 | -0.023 | -0.022 | -0.022 | 0 | 0.1 | 0.45 | 0.09 | 0.06 | 0.05 | 0.15 |
| 11 | 112435333 | rs7109000 | -0.081 | -0.34 | 0.648 | -0.006 | 0.015 | 0.035 | 0.035 | 1.365 | 0.65 | 0.53 | 0.7 | 0.5 | 0.58 | 0.71 |
| 11 | 112438395 | rs11214246 | -1.103 | -0.399 | 0.057 | -0.059 | -0.046 | -0.035 | -0.035 | 0 | 0.12 | 0.61 | 0.11 | 0.06 | 0.05 | 0.15 |
| 11 | 112452906 | rs4529911 | 0.508 | 0.083 | -0.658 | 0.149 | 0.008 | 0.014 | 0.014 | 0.482 | 0.16 | 0.09 | 0.1 | 0.08 | 0.12 | 0.16 |
| 11 | 112456221 | rs7105881 | 0.066 | 0.498 | -0.4 | 0.643 | 0.01 | 0.013 | 0.013 | 0.352 | 0.67 | 0.6 | 0.75 | 0.6 | 0.62 | 0.82 |
| 11 | 112458528 | rs12282438 | -1.118 | 0.238 | -0.352 | 0.75 | -0.017 | 0.032 | 0.032 | 0.487 | 0.09 | 0.26 | 0.13 | 0.13 | 0.06 | 0.16 |
| 11 | 112458951 | rs7940211 | -0.592 | 0.994 | -0.734 | 0.535 | 0.021 | -0.009 | -0.009 | 0.018 | 0.33 | 0.42 | 0.33 | 0.29 | 0.45 | 0.24 |
| 11 | 112459111 | rs11214259 | 0.386 | 0.708 | -1.922 | 0.375 | -0.136 | 0.007 | 0.007 | 0.343 | 0.4 | 0.04 | 0.27 | 0.32 | 0.45 | 0.39 |
| 11 | 112461836 | rs2000919 | 0.199 | 0.329 | -1.297 | 0.161 | -0.015 | -0.009 | -0.009 | 1.291 | 0.42 | 0.29 | 0.45 | 0.49 | 0.48 | 0.42 |
| 11 | 112462219 | rs1557414 | -0.578 | -0.04 | -0.913 | -0.046 | 0.002 | 0.011 | 0.011 | 0 | 0.8 | 0.79 | 0.79 | 0.73 | 0.76 | 0.85 |
| 11 | 112466239 | rs7118512 | 0.518 | -0.345 | -0.456 | -0.18 | -0.002 | 0.017 | 0.017 | 0.044 | 0.23 | 0.19 | 0.27 | 0.17 | 0.24 | 0.41 |
| 11 | 112466279 | rs1945100 | 0.826 | -0.401 | -0.593 | -0.636 | -0.004 | 0.021 | 0.021 | 0 | 0.36 | 0.47 | 0.38 | 0.53 | 0.33 | 0.35 |
| 11 | 112466701 | rs10789899 | -1.165 | -0.401 | -0.507 | -0.637 | -0.003 | 0.019 | 0.019 | 0 | 0.64 | 0.55 | 0.62 | 0.47 | 0.67 | 0.65 |
| 11 | 112467490 | rs7102985 | 0.774 | -0.425 | -0.57 | -0.633 | -0.029 | -0.016 | -0.016 | 0.268 | 0.06 | 0.14 | 0.02 | 0.06 | 0.01 | 0.02 |
| 11 | 112472239 | rs4272816 | 0.014 | -0.324 | -0.642 | -0.601 | -0.003 | 0.02 | 0.02 | 0 | 0.61 | 0.71 | 0.67 | 0.73 | 0.57 | 0.76 |
| 11 | 112473405 | rs11214267 | 1.706 | -0.475 | -0.717 | -0.41 | -0.071 | -0.127 | -0.127 | 0 | 0.26 | 0.04 | 0.28 | 0.39 | 0.31 | 0.25 |
| 11 | 112476997 | rs10789901 | -0.199 | -0.479 | -0.57 | -0.487 | -0.051 | 0.017 | 0.017 | 0 | 0.37 | 0.12 | 0.34 | 0.28 | 0.44 | 0.24 |
| 11 | 112481329 | rs7929676 | -1.107 | -0.45 | -0.21 | -0.016 | 0.002 | 0.007 | 0.007 | 0 | 0.69 | 0.68 | 0.62 | 0.52 | 0.67 | 0.68 |
| 11 | 112486568 | rs4936119 | -0.057 | -0.559 | -0.293 | 0.082 | -0.008 | -0.029 | -0.029 | 0.021 | 0.23 | 0.11 | 0.2 | 0.27 | 0.23 | 0.15 |
| 11 | 112491227 | rs2513324 | -0.546 | -0.458 | 0.334 | 0.169 | 0.004 | 0.057 | 0.057 | 0 | 0.24 | 0.52 | 0.31 | 0.37 | 0.24 | 0.27 |
| 11 | 112503558 | rs2513023 | -1.238 | -0.492 | -0.308 | -0.094 | 0.003 | -0.113 | -0.113 | 0.058 | 0.69 | 0.91 | 0.74 | 0.51 | 0.74 | 0.77 |
| 11 | 112506073 | rs17579902 | 0.29 | -0.581 | -0.081 | -0.24 | -0.03 | 0.121 | 0.121 | 0 | 0.44 | 0.22 | 0.38 | 0.14 | 0.48 | 0.5 |
| 11 | 112510051 | rs994781 | -1.045 | -0.488 | 0.008 | -0.021 | 0.026 | -0.064 | -0.064 | 0 | 0.67 | 0.8 | 0.69 | 0.51 | 0.72 | 0.78 |
| 11 | 112527603 | rs1116495 | -1.046 | -0.482 | 0.005 | -0.011 | 0.026 | -0.066 | -0.066 | 0 | 0.67 | 0.81 | 0.69 | 0.51 | 0.72 | 0.77 |
| 11 | 112531596 | rs1031686 | 0.338 | -0.574 | -0.017 | -0.134 | -0.017 | -0.025 | -0.025 | 0 | 0.22 | 0.09 | 0.19 | 0.26 | 0.25 | 0.18 |
| 11 | 112532210 | rs2513315 | -1.31 | -0.585 | -0.088 | -0.16 | 0.004 | -0.132 | -0.132 | 0.843 | 0.68 | 0.94 | 0.74 | 0.51 | 0.73 | 0.76 |
| 11 | 112533903 | rs10789904 | -0.169 | -0.552 | 0.385 | -0.168 | 0 | 0.031 | 0.031 | 0 | 0.26 | 0.43 | 0.32 | 0.36 | 0.26 | 0.31 |
| 11 | 112536487 | rs1459661 | 0.434 | -0.45 | -0.097 | -0.2 | -0.027 | 0.066 | 0.066 | 0 | 0.38 | 0.22 | 0.35 | 0.15 | 0.45 | 0.46 |
| 11 | 112557289 | rs7124774 | -0.634 | -0.278 | -0.038 | 0.088 | -0.002 | 0.019 | 0.019 | 0 | 0.29 | 0.56 | 0.37 | 0.37 | 0.29 | 0.32 |
| 11 | 112558198 | rs733438 | -0.107 | -0.125 | -0.611 | 0.176 | -0.009 | -0.011 | -0.011 | 0 | 0.24 | 0.19 | 0.24 | 0.26 | 0.27 | 0.3 |

| | | | | | | | | | | | | | | | | |
|---|---|---|---|---|---|---|---|---|---|---|---|---|---|---|---|---|
| 11 | 112558809 | rs2360065 | -1.174 | -0.247 | -0.964 | 0.32 | -0.026 | -0.176 | -0.176 | 2.965 | 0.73 | 0.98 | 0.8 | 0.59 | 0.77 | 0.81 |
| 11 | 112560259 | rs10789908 | -0.369 | -0.12 | -0.495 | 0.071 | 0.004 | 0.004 | 0.004 | 0 | 0.24 | 0.34 | 0.22 | 0.25 | 0.25 | 0.28 |
| 11 | 112560305 | rs7121382 | -1.064 | -0.139 | -0.836 | 0.551 | -0.031 | -0.063 | -0.063 | 0 | 0.71 | 0.94 | 0.74 | 0.63 | 0.74 | 0.78 |
| 11 | 112560533 | rs10750008 | -0.545 | -0.066 | -0.263 | 0.701 | -0.001 | 0.002 | 0.002 | 0 | 0.26 | 0.27 | 0.21 | 0.31 | 0.26 | 0.3 |
| 11 | 112565534 | rs1459658 | -0.212 | -0.521 | 0.599 | 1.095 | -0.037 | 0.1 | 0.1 | 0 | 0.1 | 0.23 | 0.07 | 0.33 | 0.04 | 0.15 |
| 11 | 112566028 | rs12419219 | 0.314 | -0.346 | -0.474 | 0.573 | -0.078 | -0.094 | -0.094 | 0 | 0.37 | 0.12 | 0.46 | 0.51 | 0.44 | 0.49 |
| 11 | 112579254 | rs496711 | -0.544 | -0.3 | -0.063 | 0.155 | 0.003 | 0.15 | 0.15 | 0.002 | 0.52 | 0.8 | 0.66 | 0.92 | 0.48 | 0.66 |
| 11 | 112581272 | rs9804589 | -0.414 | -0.426 | 0.399 | 0.135 | 0.009 | 0.087 | 0.087 | 0 | 0.51 | 0.73 | 0.56 | 0.85 | 0.47 | 0.63 |
| 11 | 112583490 | rs11214320 | -0.24 | -0.343 | 0.359 | 0.176 | 0.003 | 0.022 | 0.022 | 0 | 0.48 | 0.47 | 0.54 | 0.84 | 0.47 | 0.63 |
| 11 | 112588709 | rs321125 | -0.577 | -0.233 | -0.504 | 0.101 | -0.009 | 0.171 | 0.171 | 0.048 | 0.53 | 0.79 | 0.65 | 0.92 | 0.48 | 0.66 |
| 11 | 112595516 | rs1674839 | -0.552 | 0.163 | -0.675 | 0.432 | -0.008 | -0.013 | -0.013 | 0.601 | 0.59 | 0.67 | 0.68 | 0.64 | 0.51 | 0.68 |
| 11 | 112602092 | rs2054265 | 0.322 | 0.282 | -1.203 | 0.294 | -0.065 | 0.165 | 0.165 | 0 | 0.43 | 0.12 | 0.3 | 0.09 | 0.52 | 0.27 |
| 11 | 112614196 | rs11214338 | 0.367 | 0.287 | -1.223 | 0.296 | -0.086 | 0.169 | 0.169 | 0 | 0.43 | 0.1 | 0.3 | 0.09 | 0.51 | 0.27 |
| 11 | 112615975 | rs4936155 | 0.635 | 0.263 | -1.463 | 0.607 | -0.015 | -0.025 | -0.025 | 0 | 0.14 | 0.06 | 0.18 | 0.19 | 0.14 | 0.25 |
| 11 | 112616071 | rs11214343 | 0.62 | 0.285 | -1.422 | 0.436 | -0.047 | 0.149 | 0.149 | 0 | 0.43 | 0.19 | 0.31 | 0.09 | 0.52 | 0.27 |
| 11 | 112626686 | rs7127817 | 0.848 | 0.389 | -1.793 | 0.906 | -0.019 | -0.073 | -0.073 | 0 | 0.29 | 0.02 | 0.34 | 0.36 | 0.24 | 0.25 |
| 11 | 112628931 | rs4937686 | 0.335 | 0.346 | -1.372 | 0.994 | 0.004 | 0.031 | 0.031 | 0.846 | 0.17 | 0.21 | 0.28 | 0.33 | 0.19 | 0.28 |
| 11 | 112630182 | rs10789910 | -0.922 | 0.49 | -1.626 | 0.966 | 0.112 | 0.293 | 0.293 | 0.207 | 0.94 | 0.66 | 0.94 | 0.79 | 0.95 | 0.83 |
| 11 | 112641531 | rs10891431 | 0.358 | 0.462 | -2.958 | 0.101 | -0.023 | -0.076 | -0.076 | 0 | 0.29 | 0.02 | 0.33 | 0.35 | 0.25 | 0.25 |
| 11 | 112643375 | rs11214361 | 0.056 | 1.07 | -2.382 | -0.889 | -0.087 | 0.092 | 0.092 | 0.217 | 0.33 | 0.08 | 0.23 | 0.08 | 0.44 | 0.25 |
| 11 | 112643453 | rs12806454 | 0.056 | 1.07 | -2.382 | -0.889 | -0.087 | 0.092 | 0.092 | 0.217 | 0.33 | 0.13 | 0.24 | 0.08 | 0.44 | 0.25 |
| 11 | 112651604 | rs11214367 | -0.829 | 1.171 | -2.508 | -1.309 | -0.128 | 0.101 | 0.101 | 0 | 0.29 | 0.02 | 0.27 | 0.12 | 0.3 | 0.37 |
| 11 | 112654974 | rs161130 | -0.139 | 1.176 | -1.534 | -1.547 | -0.007 | -0.011 | -0.011 | 0 | 0.31 | 0.44 | 0.47 | 0.78 | 0.33 | 0.54 |
| 11 | 112658438 | rs1877351 | -0.273 | 1.36 | -0.971 | -1.174 | 0.009 | -0.035 | -0.035 | 0 | 0.07 | 0.13 | 0.04 | 0.02 | 0.08 | 0.04 |
| 11 | 112675900 | rs666733 | -0.606 | 1.406 | -1.062 | -1.429 | 0 | 0.025 | 0.025 | 0 | 0.45 | 0.63 | 0.47 | 0.68 | 0.46 | 0.59 |
| 11 | 112676385 | rs7938281 | -0.057 | 1.171 | -1.13 | -1.53 | -0.016 | 0.027 | 0.027 | 0.511 | 0.08 | 0.22 | 0.11 | 0.14 | 0.06 | 0.04 |
| 11 | 112676472 | rs1628225 | -0.392 | 1.377 | -1.185 | -1.589 | -0.006 | 0.019 | 0.019 | 0 | 0.43 | 0.62 | 0.51 | 0.68 | 0.47 | 0.6 |
| 11 | 112681117 | rs622888 | -0.803 | 1.274 | -1.209 | -1.566 | -0.003 | 0.184 | 0.184 | 0 | 0.5 | 0.84 | 0.57 | 0.82 | 0.49 | 0.61 |
| 11 | 112683676 | rs584989 | -0.309 | 1.281 | -1.138 | -1.562 | 0.014 | 0.028 | 0.028 | 0 | 0.43 | 0.67 | 0.46 | 0.68 | 0.41 | 0.54 |
| 11 | 112685992 | rs604967 | -1.066 | 1.269 | -1.267 | -1.69 | 0.014 | 0.053 | 0.053 | 0 | 0.56 | 0.69 | 0.65 | 0.73 | 0.55 | 0.72 |
| 11 | 112687958 | rs7948379 | 0.428 | 1.233 | -0.92 | -1.726 | -0.001 | -0.002 | -0.002 | 0.058 | 0.36 | 0.52 | 0.42 | 0.67 | 0.39 | 0.54 |
| 11 | 112688349 | rs7936576 | 0.733 | 1.234 | -0.921 | -1.728 | 0.014 | 0.053 | 0.053 | 0 | 0.44 | 0.31 | 0.35 | 0.27 | 0.45 | 0.28 |
| 11 | 112688705 | rs1374702 | -0.763 | 1.246 | -1.017 | -1.708 | -0.005 | 0.007 | 0.007 | 0.447 | 0.2 | 0.18 | 0.23 | 0.07 | 0.16 | 0.18 |
| 11 | 112691249 | rs1374698 | 1.328 | 1.295 | -1.037 | -1.682 | 0.008 | 0.096 | 0.096 | 0 | 0.36 | 0.13 | 0.28 | 0.13 | 0.37 | 0.2 |
| 11 | 112693044 | rs17114424 | -0.42 | 1.316 | -0.716 | -1.429 | 0.018 | -0.035 | -0.035 | 0 | 0.16 | 0.25 | 0.18 | 0.05 | 0.14 | 0.18 |
| 11 | 112694843 | rs2175519 | 0.032 | 0.753 | -0.645 | -0.951 | -0.004 | 0.005 | 0.005 | 0.353 | 0.59 | 0.34 | 0.52 | 0.34 | 0.57 | 0.44 |
| 11 | 112695387 | rs1117822 | -0.02 | 0.86 | -0.773 | -0.897 | 0.013 | -0.006 | -0.006 | 0 | 0.61 | 0.52 | 0.55 | 0.34 | 0.57 | 0.43 |
| 11 | 112695832 | rs10789914 | 0.48 | 0.404 | -1.207 | -0.806 | 0.063 | 0.077 | 0.077 | 0 | 0.38 | 0.16 | 0.25 | 0.16 | 0.3 | 0.36 |
| 11 | 112696087 | rs10891455 | -1.119 | 1.236 | -0.895 | -0.041 | -0.009 | -0.01 | -0.01 | 0 | 0.08 | 0.07 | 0.06 | 0.15 | 0.11 | 0.09 |
| 11 | 112696471 | rs1447470 | -0.55 | 0.381 | -0.378 | -0.362 | 0.013 | 0.01 | 0.01 | 0 | 0.56 | 0.39 | 0.72 | 0.73 | 0.69 | 0.6 |
| 11 | 112697728 | rs7130757 | -0.857 | 1.142 | 0.423 | 0.454 | 0.009 | 0.037 | 0.037 | 0 | 0.51 | 0.51 | 0.62 | 0.82 | 0.52 | 0.63 |
| 11 | 112698607 | rs12270522 | -0.308 | 1.221 | 0.621 | 0.415 | 0.031 | 0.038 | 0.038 | 0 | 0.05 | 0.24 | 0.12 | 0.05 | 0.04 | 0.02 |
| 11 | 112700325 | rs10789916 | -1.133 | 1.143 | 0.971 | 0.451 | 0.012 | 0.059 | 0.059 | 0 | 0.53 | 0.62 | 0.61 | 0.72 | 0.51 | 0.49 |
| 11 | 112701882 | rs1447467 | -0.444 | 1.27 | 0.724 | 0.513 | -0.015 | 0.054 | 0.054 | 0 | 0.44 | 0.19 | 0.47 | 0.67 | 0.47 | 0.46 |
| 11 | 112746381 | rs10789919 | -0.784 | 1.47 | -0.224 | 0.647 | 0.003 | 0.037 | 0.037 | 0 | 0.49 | 0.5 | 0.58 | 0.75 | 0.48 | 0.5 |
| 11 | 112753107 | rs4937786 | -0.809 | 2.045 | -0.487 | 1.711 | -0.074 | -0.105 | -0.105 | 0 | 0.16 | 0.03 | 0.3 | 0.6 | 0.21 | 0.27 |
| 11 | 112759458 | rs7939449 | -1.837 | 2.046 | 0.244 | 2.836 | 0.033 | 0.166 | 0.166 | 0 | 0.1 | 0.17 | 0.16 | 0.37 | 0.15 | 0.15 |
| 11 | 112761360 | rs1348941 | -0.813 | 1.926 | 0.032 | 3.417 | -0.016 | -0.067 | -0.067 | 0.119 | 0.2 | 0.06 | 0.18 | 0.28 | 0.2 | 0.41 |
| 11 | 112762909 | rs7952158 | -1.219 | 1.821 | 0.347 | 3.365 | 0.004 | -0.018 | -0.018 | 0 | 0.23 | 0.23 | 0.19 | 0.28 | 0.2 | 0.41 |
| 11 | 112765750 | rs2360954 | -2.33 | 1.868 | 0.397 | 3.342 | 0.004 | -0.009 | -0.009 | 0 | 0.33 | 0.42 | 0.42 | 0.68 | 0.37 | 0.58 |
| 11 | 112786693 | rs10789925 | -2.032 | 1.883 | 0.386 | 3.328 | 0.002 | -0.002 | -0.002 | 0 | 0.31 | 0.34 | 0.34 | 0.65 | 0.35 | 0.56 |
| 11 | 112801338 | rs12294747 | -0.899 | 1.547 | 0.452 | 3.431 | 0.002 | -0.003 | -0.003 | 0 | 0.21 | 0.21 | 0.18 | 0.28 | 0.17 | 0.41 |
| 11 | 112810862 | rs6589346 | -2.191 | 1.504 | 0.373 | 2.903 | -0.002 | -0.002 | -0.002 | 0 | 0.3 | 0.34 | 0.34 | 0.65 | 0.31 | 0.55 |
| 11 | 112812563 | rs12279131 | -1.159 | 1.704 | -0.001 | 2.642 | 0.01 | 0.013 | 0.013 | 0 | 0.07 | 0.08 | 0.06 | 0.2 | 0.13 | 0.13 |
| 11 | 112826303 | rs10891479 | -1.008 | 1.173 | -0.115 | 2.754 | 0.026 | -0.039 | -0.039 | 0 | 0.22 | 0.1 | 0.24 | 0.31 | 0.18 | 0.46 |
| 11 | 112838338 | rs2155281 | -1.802 | 1.116 | 0.855 | 2.758 | -0.007 | -0.055 | -0.055 | 0 | 0.4 | 0.49 | 0.51 | 0.8 | 0.43 | 0.6 |
| 11 | 112842749 | rs10789930 | -0.643 | 1.3 | 0.855 | 3.417 | 0.033 | 0.191 | 0.191 | 0 | 0.06 | 0.1 | 0.27 | 0.46 | 0.15 | 0.1 |
| 11 | 112843006 | rs11214440 | -0.459 | 1.194 | 0.712 | 3.407 | 0.021 | -0.07 | -0.07 | 0 | 0.2 | 0.01 | 0.15 | 0.22 | 0.15 | 0.45 |
| 11 | 112843138 | rs7110863 | -2.27 | 1.11 | 1.17 | 3.257 | -0.008 | -0.054 | -0.054 | 0.365 | 0.41 | 0.49 | 0.51 | 0.79 | 0.43 | 0.62 |
| 11 | 112854309 | rs4444109 | -1.449 | 1.232 | 1.031 | 3.385 | 0.003 | -0.002 | -0.002 | 0.942 | 0.27 | 0.19 | 0.42 | 0.68 | 0.3 | 0.56 |
| 11 | 112862901 | rs7129926 | -0.149 | 1.23 | 1.149 | 3.546 | 0.005 | 0.083 | 0.083 | 0 | 0.11 | 0.21 | 0.19 | 0.26 | 0.16 | 0.07 |
| 11 | 112888129 | rs17114682 | -0.631 | 1.234 | 1.048 | 3.522 | 0.027 | 0.11 | 0.11 | 0 | 0.06 | 0.11 | 0.16 | 0.33 | 0.12 | 0.09 |
| 11 | 112907421 | rs10502164 | 0.08 | 1.279 | 1.062 | 3.489 | 0.002 | 0.002 | 0.002 | 0 | 0.1 | 0.13 | 0.08 | 0.11 | 0.14 | 0.06 |
| 11 | 112908855 | rs1940735 | -1.483 | 1.26 | 1.05 | 3.527 | 0.003 | -0.04 | -0.04 | 0 | 0.28 | 0.17 | 0.32 | 0.54 | 0.28 | 0.56 |
| 11 | 112912947 | rs2155645 | -1.358 | 0.93 | 0.893 | 3.491 | 0.017 | -0.018 | -0.018 | 0 | 0.28 | 0.18 | 0.21 | 0.32 | 0.26 | 0.52 |
| 11 | 112917111 | rs12574893 | -1.005 | 0.822 | 0.992 | 3.511 | -0.001 | 0.025 | 0.025 | 0 | 0.15 | 0.14 | 0.16 | 0.45 | 0.2 | 0.15 |
| 11 | 112918985 | rs12575544 | -1.701 | 0.791 | 1.184 | 3.433 | 0.004 | 0.054 | 0.054 | 0 | 0.19 | 0.28 | 0.3 | 0.62 | 0.25 | 0.25 |
| 11 | 112921522 | rs10891498 | -1.987 | 1.076 | 0.933 | 3.012 | -0.005 | 0.03 | 0.03 | 0 | 0.41 | 0.32 | 0.43 | 0.81 | 0.47 | 0.56 |
| 11 | 112928164 | rs1369817 | 1.681 | 1.072 | 0.915 | 3.009 | -0.01 | 0.054 | 0.054 | 0 | 0.59 | 0.73 | 0.57 | 0.2 | 0.54 | 0.44 |

| | | | | | | | | | | | | | | | | |
|---|---|---|---|---|---|---|---|---|---|---|---|---|---|---|---|---|
| 11 | 112932016 | rs1245098 | 0.272 | 1.104 | 0.819 | 3.093 | -0.03 | -0.01 | -0.01 | 4.703 | 0.77 | 0.95 | 0.86 | 0.81 | 0.79 | 0.69 |
| 11 | 112946261 | rs2003987 | 2.495 | 0.803 | 1.437 | 2.768 | 0.007 | 0.185 | 0.185 | 0 | 0.51 | 0.3 | 0.45 | 0.11 | 0.51 | 0.41 |
| 11 | 112948306 | rs1465765 | -0.982 | 0.847 | 1.069 | 3.241 | -0.001 | 0.02 | 0.02 | 0 | 0.16 | 0.15 | 0.15 | 0.49 | 0.21 | 0.15 |
| 11 | 112953130 | rs7951034 | -0.03 | 0.868 | 0.822 | 3.327 | -0.024 | 0.018 | 0.018 | 0 | 0.06 | 0 | 0.08 | 0.38 | 0.08 | 0.08 |
| 11 | 112964139 | rs10891503 | -1.294 | 0.802 | 1.155 | 3.23 | 0 | 0.032 | 0.032 | 0 | 0.18 | 0.19 | 0.15 | 0.48 | 0.21 | 0.15 |
| 11 | 112965769 | rs1436110 | 0.413 | 0.763 | 1.209 | 3.129 | -0.01 | -0.011 | -0.011 | 0 | 0.75 | 0.89 | 0.78 | 0.76 | 0.77 | 0.68 |
| 11 | 112969258 | rs10891504 | 2.238 | 0.751 | 1.318 | 3.085 | -0.001 | -0.067 | -0.067 | 0 | 0.53 | 0.53 | 0.48 | 0.12 | 0.51 | 0.42 |
| 11 | 112974523 | rs2853192 | 0.341 | 0.807 | 1.131 | 2.984 | -0.051 | 0.009 | 0.009 | 0 | 0.74 | 0.95 | 0.85 | 0.84 | 0.75 | 0.7 |
| 11 | 112983534 | rs4492854 | 1.859 | 0.739 | 1.251 | 3.083 | 0 | -0.001 | -0.001 | 0 | 0.56 | 0.62 | 0.62 | 0.28 | 0.56 | 0.52 |
| 11 | 112991618 | rs1436109 | 0.691 | 0.488 | 1.271 | 3.462 | -0.028 | -0.032 | -0.032 | 0 | 0.07 | 0.01 | 0.04 | 0.11 | 0.12 | 0.07 |
| 11 | 112992847 | rs2853194 | 1.266 | 0.47 | 1.322 | 3.217 | -0.001 | 0.096 | 0.096 | 0 | 0.81 | 0.74 | 0.84 | 0.52 | 0.79 | 0.83 |
| 11 | 112999832 | rs1436106 | 0.963 | 1.048 | 1.45 | 3.052 | 0.018 | 0.034 | 0.034 | 0 | 0.68 | 0.48 | 0.78 | 0.58 | 0.65 | 0.66 |
| 11 | 113001310 | rs7932647 | 2.114 | 1.016 | 1.326 | 3.017 | -0.03 | 0.087 | 0.087 | 0.068 | 0.55 | 0.18 | 0.47 | 0.25 | 0.5 | 0.49 |
| 11 | 113004662 | rs4937993 | 1.972 | 1.02 | 1.333 | 3.019 | -0.017 | 0.055 | 0.055 | 0.092 | 0.57 | 0.25 | 0.48 | 0.25 | 0.5 | 0.49 |
| 11 | 113009703 | rs17114972 | -0.972 | 1.255 | 1.369 | 3.277 | -0.01 | -0.025 | -0.025 | 1.499 | 0.08 | 0.02 | 0.21 | 0.26 | 0.14 | 0.16 |
| 11 | 113014058 | rs7112370 | 0.089 | 1.398 | 0.851 | 3.298 | 0.003 | 0.031 | 0.031 | 0 | 0.08 | 0.12 | 0.06 | 0.26 | 0.12 | 0.09 |
| 11 | 113014986 | rs1245131 | -2.211 | 1.385 | 0.856 | 3.299 | 0.004 | 0.131 | 0.131 | 0.913 | 0.21 | 0.37 | 0.36 | 0.59 | 0.27 | 0.28 |
| 11 | 113017470 | rs10891515 | 0.041 | 1.449 | 0.809 | 2.694 | 0.006 | 0.043 | 0.043 | 0 | 0.08 | 0.13 | 0.06 | 0.25 | 0.12 | 0.09 |
| 11 | 113018408 | rs10502169 | 0.041 | 1.449 | 0.809 | 2.694 | 0.006 | 0.043 | 0.043 | 0 | 0.08 | 0.13 | 0.06 | 0.25 | 0.12 | 0.09 |
| 11 | 113025146 | rs11214518 | -0.428 | 1.448 | 0.795 | 2.693 | 0.011 | 0.073 | 0.073 | 0.014 | 0.1 | 0.2 | 0.07 | 0.25 | 0.12 | 0.09 |
| 11 | 113027641 | rs17595156 | -0.417 | 1.471 | 0.753 | 2.705 | -0.01 | -0.005 | -0.005 | 0 | 0.06 | 0.01 | 0.09 | 0.13 | 0.1 | 0.08 |
| 11 | 113034787 | rs12272966 | -1.867 | 1.466 | 0.815 | 2.681 | 0.013 | 0.117 | 0.117 | 1.156 | 0.19 | 0.4 | 0.25 | 0.47 | 0.24 | 0.2 |
| 11 | 113038005 | rs4936265 | -1.062 | 1.247 | 1.065 | 2.669 | 0.02 | 0.196 | 0.196 | 0.934 | 0.13 | 0.4 | 0.15 | 0.35 | 0.14 | 0.11 |
| 11 | 113040118 | rs10502170 | -1.185 | 1.274 | 0.782 | 2.519 | 0.014 | 0.195 | 0.195 | 0.78 | 0.14 | 0.41 | 0.16 | 0.36 | 0.14 | 0.12 |
| 11 | 113043726 | rs2043602 | 1.18 | 1.315 | 0.716 | 2.465 | 0.017 | 0.071 | 0.071 | 0 | 0.7 | 0.56 | 0.71 | 0.51 | 0.65 | 0.7 |
| 11 | 113056323 | rs4936267 | 0.451 | 1.248 | -0.126 | 1.579 | -0.008 | 0.002 | 0.002 | 0 | 0.81 | 0.83 | 0.85 | 0.87 | 0.79 | 0.81 |
| 11 | 113060469 | rs2117912 | 0.15 | 1.336 | -0.584 | 1.594 | -0.061 | -0.025 | -0.025 | 0 | 0.83 | 0.97 | 0.87 | 0.88 | 0.79 | 0.81 |
| 11 | 113065293 | rs1943620 | -0.916 | 1.214 | -0.402 | 1.592 | 0.002 | -0.017 | -0.017 | 0 | 0.35 | 0.69 | 0.4 | 0.47 | 0.31 | 0.45 |
| 11 | 113072174 | rs2226849 | -0.362 | 1.399 | 0.328 | 1.883 | 0.07 | 0.087 | 0.087 | 0.048 | 0.12 | 0.4 | 0.13 | 0.17 | 0.12 | 0.15 |
| 11 | 113077541 | rs1821693 | -0.998 | 1.079 | -0.102 | 1.688 | -0.008 | 0.036 | 0.036 | 0 | 0.42 | 0.68 | 0.48 | 0.74 | 0.4 | 0.54 |
| 11 | 113081187 | rs17115184 | 0.632 | 1.106 | -0.195 | 1.631 | -0.002 | 0.006 | 0.006 | 1.461 | 0.08 | 0.07 | 0.05 | 0.25 | 0.09 | 0.09 |
| 11 | 113081624 | rs2552513 | -0.288 | 0.991 | -0.258 | 1.639 | 0.006 | 0.029 | 0.029 | 0.226 | 0.31 | 0.54 | 0.28 | 0.38 | 0.3 | 0.41 |
| 11 | 113090805 | rs4646981 | 0.785 | 0.816 | -1.456 | 0.964 | -0.022 | 0.064 | 0.064 | 0 | 0.12 | 0.01 | 0.09 | 0 | 0.11 | 0.11 |
| 11 | 113103996 | rs584427 | -1.021 | 0.968 | -1.418 | 0.787 | -0.038 | 0.133 | 0.133 | 0.961 | 0.51 | 0.91 | 0.5 | 0.73 | 0.51 | 0.56 |
| 11 | 113105907 | rs646558 | -0.969 | 0.965 | 0.008 | 0.891 | 0.015 | 0.016 | 0.016 | 5.128 | 0.2 | 0.54 | 0.26 | 0.25 | 0.2 | 0.11 |
| 11 | 113106455 | rs12279261 | -1.264 | 0.579 | 0.118 | 0.438 | -0.011 | -0.076 | -0.076 | 0.875 | 0.21 | 0.43 | 0.19 | 0.07 | 0.17 | 0.03 |
| 11 | 113107690 | rs11214547 | -1.037 | 0.456 | -0.529 | 2.043 | 0.003 | -0.014 | -0.014 | 0 | 0.08 | 0 | 0.16 | 0.38 | 0.06 | 0.19 |
| 11 | 113109836 | rs12271908 | -0.119 | 0.577 | -0.945 | 1.347 | -0.01 | 0.015 | 0.015 | 1.798 | 0.2 | 0.14 | 0.17 | 0.17 | 0.25 | 0.31 |
| 11 | 113110946 | rs586903 | -0.89 | 0.424 | -0.409 | 0.886 | 0.011 | 0.006 | 0.006 | 0.787 | 0.84 | 0.71 | 0.84 | 0.84 | 0.84 | 0.91 |
| 11 | 113111501 | rs2303377 | -0.877 | 1.021 | -0.702 | 0.403 | -0.025 | -0.031 | -0.031 | 0 | 0.34 | 0.19 | 0.44 | 0.56 | 0.39 | 0.56 |
| 11 | 113111665 | rs2303378 | 1.718 | 0.261 | -1.534 | -0.407 | -0.05 | 0.145 | 0.145 | 0.085 | 0.28 | 0.1 | 0.19 | 0.1 | 0.29 | 0.24 |
| 11 | 113112073 | rs884110 | 0.4 | 0.097 | -1.559 | -0.505 | -0.063 | -0.089 | -0.089 | 0 | 0.44 | 0.13 | 0.48 | 0.51 | 0.43 | 0.5 |
| 11 | 113115748 | rs591042 | -1.044 | 0.03 | -1.345 | -0.821 | -0.034 | 0.014 | 0.014 | 0 | 0.31 | 0.5 | 0.45 | 0.57 | 0.25 | 0.34 |
| 11 | 113116371 | rs7130671 | -0.23 | 0.162 | -1.218 | -0.814 | -0.006 | 0.029 | 0.029 | 0.64 | 0.33 | 0.28 | 0.32 | 0.22 | 0.36 | 0.38 |
| 11 | 113118856 | rs12794326 | 0.435 | 0.438 | -1.282 | -0.666 | 0.008 | 0.092 | 0.092 | 0.134 | 0.57 | 0.28 | 0.44 | 0.3 | 0.59 | 0.63 |
| 11 | 113120212 | rs3781878 | 0.673 | 0.59 | -0.88 | -0.514 | -0.009 | 0.003 | 0.003 | 0.058 | 0.27 | 0.19 | 0.2 | 0.1 | 0.3 | 0.27 |
| 11 | 113124379 | rs10891524 | -0.912 | -0.239 | -0.33 | 0.536 | -0.039 | 0.096 | 0.096 | 0 | 0.18 | 0.35 | 0.22 | 0.31 | 0.11 | 0.21 |
| 11 | 113124432 | rs625087 | -1.522 | 0.188 | -0.221 | 0.457 | -0.007 | -0.015 | -0.015 | 0 | 0.27 | 0.46 | 0.21 | 0.2 | 0.24 | 0.24 |
| 11 | 113124718 | rs17115275 | 1.168 | 0.83 | -0.892 | 0.177 | -0.014 | -0.027 | -0.027 | 3.294 | 0.11 | 0.01 | 0.12 | 0.13 | 0.11 | 0.07 |
| 11 | 113125234 | rs605843 | -1.72 | 1.084 | -0.275 | 0.164 | 0.015 | 0.069 | 0.069 | 0 | 0.22 | 0.51 | 0.29 | 0.31 | 0.23 | 0.27 |
| 11 | 113129127 | rs7951404 | 0.032 | 1.13 | -0.66 | 0.092 | -0.001 | -0.017 | -0.017 | 0.071 | 0.07 | 0.11 | 0.04 | 0 | 0.06 | 0.09 |
| 11 | 113129388 | rs686503 | -1.278 | 1.157 | -0.572 | 0.081 | 0.002 | -0.069 | -0.069 | 0 | 0.12 | 0.27 | 0.18 | 0.01 | 0.13 | 0.12 |
| 11 | 113133676 | rs2288158 | -0.34 | 1.055 | -0.672 | 0.024 | -0.001 | -0.004 | -0.004 | 0 | 0.11 | 0.09 | 0.11 | 0.3 | 0.12 | 0.18 |
| 11 | 113134605 | rs2298703 | 0.779 | 1.45 | -1.13 | -0.657 | -0.094 | 0.12 | 0.12 | 0.638 | 0.24 | 0.01 | 0.15 | 0.11 | 0.24 | 0.36 |
| 11 | 113134818 | rs10502171 | 0.258 | 1.361 | -1.057 | -0.596 | -0.001 | -0.021 | -0.021 | 0 | 0.07 | 0.12 | 0.04 | 0 | 0.06 | 0.09 |
| 11 | 113135006 | rs1006826 | 0.666 | 1.387 | -1.197 | -0.588 | -0.105 | 0.135 | 0.135 | 0.346 | 0.27 | 0.01 | 0.18 | 0.11 | 0.27 | 0.37 |
| 11 | 113137365 | rs655250 | -1.002 | 0.989 | 0.029 | 0.375 | 0.008 | -0.071 | -0.071 | 0.519 | 0.12 | 0.29 | 0.18 | 0.01 | 0.13 | 0.12 |
| 11 | 113141684 | rs674246 | -1.42 | 0.949 | -0.446 | 0.746 | -0.001 | 0.021 | 0.021 | 0 | 0.26 | 0.56 | 0.32 | 0.31 | 0.26 | 0.31 |
| 11 | 113143557 | rs2156485 | -0.413 | 1.932 | -1.082 | 1.143 | -0.078 | -0.107 | -0.107 | 0.017 | 0.15 | 0.02 | 0.22 | 0.42 | 0.22 | 0.18 |
| 11 | 113146785 | rs635903 | 0.868 | 1.973 | -1.007 | 0.569 | -0.023 | 0.054 | 0.054 | 0 | 0.69 | 0.79 | 0.48 | 0.1 | 0.64 | 0.72 |
| 11 | 113149642 | rs668204 | -1.245 | 2.017 | -1.022 | 0.767 | -0.003 | -0.001 | -0.001 | 0 | 0.95 | 0.98 | 0.89 | 0.92 | 0.93 | 0.97 |
| 11 | 113150050 | rs600964 | 1 | 1.262 | -1.522 | -0.811 | 0.005 | 0.094 | 0.094 | 0 | 0.33 | 0.19 | 0.23 | 0.08 | 0.32 | 0.47 |
| 11 | 113151099 | rs4559717 | 1.134 | 0.885 | -0.885 | -1.201 | -0.009 | -0.067 | -0.067 | 0.002 | 0.11 | 0.27 | 0.08 | 0 | 0.08 | 0.03 |
| 11 | 113151790 | rs10891527 | 2.295 | 0.414 | -1.385 | -1.702 | -0.052 | 0.047 | 0.047 | 0.885 | 0.17 | 0.01 | 0.13 | 0.08 | 0.2 | 0.25 |
| 11 | 113154943 | rs12787488 | 1.399 | 0.618 | -1.215 | -1.557 | 0 | -0.068 | -0.068 | 0.181 | 0.11 | 0.27 | 0.09 | 0 | 0.09 | 0.03 |
| 11 | 113155980 | rs17513907 | 1.534 | 0.709 | -1.195 | -1.588 | 0.007 | 0.028 | 0.028 | 0 | 0.22 | 0.13 | 0.14 | 0.01 | 0.2 | 0.16 |
| 11 | 113157558 | rs641748 | 0.811 | 0.865 | -1.066 | -1.287 | 0.004 | -0.061 | -0.061 | 0 | 0.56 | 0.49 | 0.35 | 0.1 | 0.52 | 0.61 |
| 11 | 113158315 | rs628041 | 0.345 | 0.613 | -0.931 | -0.986 | -0.008 | 0.101 | 0.101 | 0.984 | 0.72 | 0.51 | 0.59 | 0.5 | 0.75 | 0.8 |
| 11 | 113164768 | rs7103866 | 0.825 | 1.021 | -1.125 | -0.832 | -0.041 | -0.049 | -0.049 | 0.675 | 0.19 | 0.08 | 0.25 | 0.33 | 0.28 | 0.24 |

| | | | | | | | | | | | | | | | | |
|---|---|---|---|---|---|---|---|---|---|---|---|---|---|---|---|---|
| 11 | 113177189 | rs12279103 | 0.971 | 0.856 | -0.61 | -0.928 | 0.033 | -0.04 | -0.04 | 0 | 0.08 | 0.27 | 0.09 | 0 | 0.07 | 0.02 |
| 11 | 113180202 | rs4517559 | -0.216 | 0.364 | -0.699 | -1.081 | 0.01 | 0.001 | 0.001 | 0.863 | 0.52 | 0.52 | 0.53 | 0.46 | 0.62 | 0.62 |
| 11 | 113186776 | rs2236709 | -0.618 | 0.237 | -0.851 | -1.199 | 0 | 0.002 | 0.002 | 0 | 0.71 | 0.75 | 0.62 | 0.47 | 0.73 | 0.8 |
| 11 | 113188923 | rs2156486 | 2.134 | 0.359 | -1.22 | -1.149 | -0.048 | 0.065 | 0.065 | 1.025 | 0.18 | 0.01 | 0.14 | 0.07 | 0.21 | 0.22 |
| 11 | 113191428 | rs7942392 | 0.19 | 0.137 | -0.64 | -0.876 | 0.029 | -0.017 | -0.017 | 0 | 0.13 | 0.36 | 0.12 | 0.06 | 0.13 | 0.08 |
| 11 | 113192797 | rs6589371 | 0.258 | 0.137 | -0.638 | -0.876 | 0.029 | -0.015 | -0.015 | 0.093 | 0.12 | 0.35 | 0.12 | 0.06 | 0.13 | 0.08 |
| 11 | 113193601 | rs7950569 | -0.006 | 0.093 | -0.845 | -0.912 | -0.005 | 0.001 | 0.001 | 0 | 0.56 | 0.54 | 0.54 | 0.46 | 0.62 | 0.59 |
| 11 | 113194168 | rs723077 | 1.001 | 0.114 | -1.414 | -0.957 | -0.078 | -0.015 | -0.015 | 0 | 0.41 | 0.12 | 0.4 | 0.4 | 0.48 | 0.49 |
| 11 | 113194448 | rs723079 | 0.652 | -0.121 | -1.066 | -0.862 | 0.021 | 0.083 | 0.083 | 0 | 0.18 | 0.02 | 0.1 | 0.01 | 0.13 | 0.12 |
| 11 | 113195595 | rs948178 | 0.111 | 0.135 | -0.692 | -0.459 | 0.006 | -0.001 | -0.001 | 1.053 | 0.56 | 0.62 | 0.6 | 0.57 | 0.63 | 0.6 |
| 11 | 113199146 | rs10502172 | 0.749 | 0.35 | -1.406 | -1.084 | -0.111 | 0.272 | 0.272 | 2.42 | 0.39 | 0.02 | 0.29 | 0.05 | 0.48 | 0.29 |
| 11 | 113200709 | rs2303380 | -0.194 | 0.071 | -0.73 | -0.734 | -0.002 | -0.004 | -0.004 | 0.123 | 0.65 | 0.68 | 0.57 | 0.53 | 0.64 | 0.57 |
| 11 | 113204266 | rs11214577 | 1.363 | 0.072 | -0.672 | -0.753 | -0.069 | 0.262 | 0.262 | 2.349 | 0.4 | 0.07 | 0.3 | 0.05 | 0.48 | 0.29 |
| 11 | 113204481 | rs948176 | -0.363 | 0.006 | 0.168 | -0.712 | -0.015 | 0.084 | 0.084 | 0 | 0.23 | 0.39 | 0.26 | 0.48 | 0.16 | 0.28 |
| 11 | 113208098 | rs7114433 | -0.04 | 0.003 | 0.281 | -0.762 | 0.01 | 0.034 | 0.034 | 0 | 0.63 | 0.54 | 0.57 | 0.54 | 0.65 | 0.54 |
| 11 | 113212553 | rs6589372 | -0.106 | -0.02 | 0.596 | -0.748 | -0.007 | 0.062 | 0.062 | 0.022 | 0.21 | 0.29 | 0.25 | 0.48 | 0.16 | 0.24 |
| 11 | 113215923 | rs4987094 | 1.421 | -0.058 | 0.476 | -0.701 | -0.024 | 0.033 | 0.033 | 0 | 0.06 | 0.02 | 0.17 | 0.44 | 0.1 | 0.17 |
| 11 | 113217364 | rs3781884 | 0.057 | -0.11 | 0.543 | -0.703 | 0.002 | 0.117 | 0.117 | 0 | 0.14 | 0.26 | 0.22 | 0.44 | 0.13 | 0.19 |
| 11 | 113230600 | rs2276070 | -0.699 | -0.09 | 0.514 | -0.74 | -0.006 | 0.058 | 0.058 | 0.013 | 0.21 | 0.28 | 0.25 | 0.47 | 0.16 | 0.24 |
| 11 | 113231614 | rs4438071 | 1.53 | -0.091 | 0.279 | -0.92 | -0.02 | 0.132 | 0.132 | 0.437 | 0.38 | 0.17 | 0.29 | 0.06 | 0.46 | 0.29 |
| 11 | 113234679 | rs719802 | 0.732 | 0.127 | 0.619 | -0.587 | 0.001 | -0.006 | -0.006 | 0 | 0.55 | 0.37 | 0.54 | 0.51 | 0.63 | 0.47 |
| 11 | 113234775 | rs719804 | 0.356 | 0.467 | 0.721 | -0.583 | 0 | -0.002 | -0.002 | 0.2 | 0.2 | 0.18 | 0.13 | 0.13 | 0.22 | 0.25 |
| 11 | 113236199 | rs10789942 | 1.611 | 0.321 | 0.874 | -0.772 | -0.056 | 0.241 | 0.241 | 2.643 | 0.44 | 0.14 | 0.33 | 0.06 | 0.53 | 0.31 |
| 11 | 113239082 | rs10891540 | 1.559 | 0.305 | 1.053 | -0.789 | -0.002 | -0.037 | -0.037 | 0 | 0.47 | 0.44 | 0.34 | 0.06 | 0.53 | 0.35 |
| 11 | 113239709 | rs1055076 | -0.538 | 0.221 | 0.976 | -0.683 | -0.016 | 0.083 | 0.083 | 0 | 0.21 | 0.34 | 0.27 | 0.49 | 0.15 | 0.22 |
| 11 | 113249136 | rs754672 | 1.635 | 0.474 | 0.507 | -0.703 | -0.081 | 0.28 | 0.28 | 3.39 | 0.43 | 0.1 | 0.31 | 0.06 | 0.52 | 0.35 |
| 11 | 113256508 | rs877138 | 0.225 | 0.593 | 0.689 | -0.416 | -0.008 | -0.045 | -0.045 | 2.571 | 0.66 | 0.81 | 0.61 | 0.54 | 0.67 | 0.58 |
| 11 | 113264272 | rs17115439 | 0.696 | 0.479 | 1.024 | -0.432 | 0.005 | -0.027 | -0.027 | 0 | 0.62 | 0.35 | 0.57 | 0.55 | 0.68 | 0.55 |
| 11 | 113264470 | rs4938013 | 0.167 | 0.489 | 1.023 | -0.427 | -0.007 | -0.042 | -0.042 | 0 | 0.67 | 0.81 | 0.61 | 0.55 | 0.68 | 0.59 |
| 11 | 113266821 | rs7118900 | -0.435 | 0.746 | 0.967 | -0.292 | 0.052 | 0.228 | 0.228 | 0 | 0.15 | 0.37 | 0.31 | 0.41 | 0.18 | 0.31 |
| 11 | 113270160 | rs2734849 | 1.692 | 0.682 | 0.076 | -0.593 | -0.038 | 0.243 | 0.243 | 2.258 | 0.43 | 0.1 | 0.31 | 0.06 | 0.5 | 0.31 |
| 11 | 113274010 | rs12422191 | 1.717 | 0.172 | 0.168 | -0.495 | 0.015 | -0.006 | -0.006 | 0 | 0.05 | 0.05 | 0.08 | 0.03 | 0.09 | 0.02 |
| 11 | 113278447 | rs10891549 | 1.644 | 0.055 | 0.067 | -0.568 | -0.068 | 0.257 | 0.257 | 3.415 | 0.45 | 0.12 | 0.33 | 0.06 | 0.55 | 0.32 |
| 11 | 113278764 | rs1554929 | 1.777 | 0.046 | 0.043 | -0.572 | -0.166 | 0.323 | 0.323 | 4.503 | 0.43 | 0.02 | 0.32 | 0.06 | 0.54 | 0.32 |
| 11 | 113279430 | rs2242592 | 1.052 | 0.049 | 0.208 | -0.466 | 0.03 | -0.023 | -0.023 | 0 | 0.56 | 0.28 | 0.6 | 0.47 | 0.69 | 0.59 |
| 11 | 113280724 | rs6278 | 0.783 | 0.076 | 0.23 | -0.434 | -0.032 | -0.057 | -0.057 | 0 | 0.11 | 0.02 | 0.26 | 0.4 | 0.15 | 0.26 |
| 11 | 113283477 | rs6275 | 0.928 | 0.071 | 0.234 | -0.442 | 0.017 | -0.014 | -0.014 | 0 | 0.57 | 0.34 | 0.61 | 0.47 | 0.69 | 0.6 |
| 11 | 113283688 | rs1076560 | 0.192 | -0.607 | 0.226 | -0.161 | -0.003 | 0.021 | 0.021 | 0 | 0.12 | 0.08 | 0.27 | 0.42 | 0.15 | 0.29 |
| 11 | 113289298 | rs2002453 | 0.895 | -0.406 | 0.48 | -0.048 | -0.013 | 0.009 | 0.009 | 0 | 0.62 | 0.42 | 0.65 | 0.48 | 0.76 | 0.65 |
| 11 | 113296286 | rs1079597 | -0.176 | 0.012 | 0.007 | 0.319 | 0.015 | 0.146 | 0.146 | 0 | 0.1 | 0.17 | 0.27 | 0.41 | 0.15 | 0.29 |
| 11 | 113297349 | rs1125393 | 0.24 | 0.03 | -0.132 | 0.33 | -0.008 | 0.042 | 0.042 | 0 | 0.09 | 0.08 | 0.26 | 0.41 | 0.15 | 0.29 |
| 11 | 113298339 | rs2471857 | -0.231 | 0.027 | -0.128 | 0.328 | 0.008 | 0.113 | 0.113 | 0 | 0.11 | 0.18 | 0.27 | 0.42 | 0.15 | 0.32 |
| 11 | 113309619 | rs4620755 | -0.865 | -0.519 | -0.03 | 0.512 | -0.007 | 0.05 | 0.05 | 0.347 | 0.18 | 0.29 | 0.21 | 0.4 | 0.12 | 0.22 |
| 11 | 113310681 | rs7125415 | -0.038 | -0.478 | -0.133 | 0.571 | 0.002 | 0.103 | 0.103 | 0 | 0.1 | 0.2 | 0.15 | 0.39 | 0.1 | 0.14 |
| 11 | 113313389 | rs4648318 | -1.407 | -0.552 | -0.347 | 0.464 | -0.023 | 0.007 | 0.007 | 0.018 | 0.38 | 0.53 | 0.31 | 0.58 | 0.24 | 0.36 |
| 11 | 113316236 | rs12574471 | -0.302 | -0.313 | -0.843 | 0.164 | 0.022 | -0.001 | -0.001 | 0.079 | 0.17 | 0.07 | 0.11 | 0.17 | 0.1 | 0.12 |
| 11 | 113317067 | rs17529477 | 0.248 | 0.011 | -1.512 | -0.993 | -0.016 | 0.153 | 0.153 | 0 | 0.36 | 0.08 | 0.27 | 0.04 | 0.32 | 0.19 |
| 11 | 113317973 | rs4245146 | -0.409 | 0.283 | -1.227 | -0.472 | -0.03 | 0.135 | 0.135 | 0.632 | 0.54 | 0.25 | 0.55 | 0.81 | 0.55 | 0.66 |
| 11 | 113320419 | rs4245148 | 0.375 | 0.215 | -0.831 | -0.488 | 0.047 | 0.113 | 0.113 | 3.784 | 0.9 | 0.42 | 0.84 | 0.84 | 0.91 | 0.88 |
| 11 | 113321796 | rs4460839 | 0.431 | 0.212 | -0.832 | -0.489 | 0.043 | 0.108 | 0.108 | 4.03 | 0.89 | 0.42 | 0.84 | 0.84 | 0.91 | 0.88 |
| 11 | 113329774 | rs7131056 | -0.258 | -0.138 | -1.385 | -0.163 | -0.009 | -0.007 | -0.007 | 1.046 | 0.66 | 0.33 | 0.62 | 0.58 | 0.58 | 0.47 |
| 11 | 113331532 | rs4648317 | -0.043 | -0.414 | -1.542 | -0.008 | 0.007 | -0.018 | -0.018 | 0.218 | 0.18 | 0.14 | 0.26 | 0.4 | 0.16 | 0.18 |
| 11 | 113334209 | rs4630328 | 1.651 | -0.431 | -1.608 | -0.207 | -0.052 | 0.156 | 0.156 | 0.17 | 0.34 | 0.08 | 0.29 | 0.03 | 0.36 | 0.22 |
| 11 | 113346351 | rs1799978 | 0.114 | -0.495 | -1.542 | -0.019 | 0 | -0.003 | -0.003 | 0 | 0.15 | 0.17 | 0.08 | 0.18 | 0.06 | 0.08 |
| 11 | 113346955 | rs12364283 | 0.627 | -0.437 | -1.643 | -0.179 | -0.013 | 0.038 | 0.038 | 0 | 0.08 | 0.01 | 0.03 | 0 | 0.08 | 0.12 |
| 11 | 113355736 | rs6589377 | -1.93 | -0.433 | -1.42 | -0.228 | -0.031 | 0.112 | 0.112 | 0.138 | 0.66 | 0.87 | 0.7 | 0.97 | 0.64 | 0.78 |
| 11 | 113364271 | rs4611239 | 1.209 | -0.379 | -1.463 | -0.072 | -0.091 | -0.041 | -0.041 | 0 | 0.23 | 0.04 | 0.21 | 0.27 | 0.33 | 0.39 |
| 11 | 113368331 | rs12292637 | -0.316 | -0.37 | -0.996 | -0.163 | 0.026 | 0.032 | 0.032 | 3.955 | 0.1 | 0.49 | 0.12 | 0.07 | 0.06 | 0.07 |
| 11 | 113378952 | rs6589381 | 0.938 | -0.419 | -1.142 | -0.243 | -0.005 | 0.006 | 0.006 | 0.072 | 0.42 | 0.52 | 0.4 | 0.1 | 0.41 | 0.28 |
| 11 | 113379824 | rs6589382 | -0.537 | -0.523 | -1.248 | -0.158 | 0.035 | -0.018 | -0.018 | 0 | 0.84 | 0.88 | 0.91 | 0.82 | 0.94 | 0.91 |
| 11 | 113383394 | rs4479021 | 0.969 | -0.402 | -0.581 | -0.241 | 0.004 | -0.065 | -0.065 | 0 | 0.41 | 0.43 | 0.39 | 0.1 | 0.42 | 0.28 |
| 11 | 113384281 | rs4447205 | -0.611 | -0.471 | -0.539 | -0.079 | -0.001 | -0.003 | -0.003 | 0.348 | 0.17 | 0.19 | 0.28 | 0.4 | 0.19 | 0.23 |
| 11 | 113392994 | rs2514218 | 1.415 | -0.235 | -0.52 | -0.401 | -0.029 | 0.108 | 0.108 | 0.098 | 0.33 | 0.12 | 0.29 | 0.03 | 0.35 | 0.21 |
| 11 | 113397655 | rs2511515 | 1.101 | -0.004 | 0.136 | -0.099 | -0.012 | -0.011 | -0.011 | 1.13 | 0.63 | 0.51 | 0.59 | 0.37 | 0.69 | 0.68 |
| 11 | 113404170 | rs2514227 | 1.428 | 0.048 | 0.197 | -0.048 | -0.003 | 0.003 | 0.003 | 0 | 0.75 | 0.39 | 0.77 | 0.74 | 0.78 | 0.68 |
| 11 | 113410065 | rs7942123 | -1.666 | 0.054 | 0.242 | -0.047 | -0.007 | 0.023 | 0.023 | 0 | 0.22 | 0.54 | 0.21 | 0.27 | 0.19 | 0.31 |
| 11 | 113410917 | rs11214654 | -0.663 | 0.056 | 0.428 | 0.042 | 0.006 | 0.091 | 0.091 | 0 | 0.13 | 0.27 | 0.12 | 0.26 | 0.12 | 0.17 |
| 11 | 113411628 | rs11823610 | -0.963 | 0.064 | 0.421 | 0.04 | 0.007 | 0.058 | 0.058 | 0 | 0.15 | 0.26 | 0.14 | 0.26 | 0.16 | 0.19 |
| 11 | 113413565 | rs10891570 | -0.852 | -0.141 | 0.142 | 0.118 | 0.016 | 0.009 | 0.009 | 0 | 0.5 | 0.57 | 0.5 | 0.58 | 0.59 | 0.71 |

| | | | | | | | | | | | | | | | | |
|---|---|---|---|---|---|---|---|---|---|---|---|---|---|---|---|---|
| 11 | 113415720 | rs7115090 | -0.066 | -0.097 | 0.444 | -0.459 | -0.008 | 0.017 | 0.017 | 0 | 0.44 | 0.42 | 0.52 | 0.67 | 0.52 | 0.49 |
| 11 | 113417281 | rs12287992 | -0.629 | -0.082 | 0.416 | -0.384 | 0.027 | 0.185 | 0.185 | 0 | 0.1 | 0.33 | 0.13 | 0.28 | 0.13 | 0.17 |
| 11 | 113417603 | rs12222458 | -0.028 | -0.053 | 0.093 | -1.011 | 0.016 | 0.211 | 0.211 | 0 | 0.48 | 0.2 | 0.58 | 0.93 | 0.6 | 0.62 |
| 11 | 113418371 | rs10736470 | -0.835 | 0.15 | -0.298 | -0.906 | 0.004 | 0.007 | 0.007 | 0 | 0.62 | 0.74 | 0.7 | 0.95 | 0.66 | 0.69 |
| 11 | 113424042 | rs10750025 | -1.212 | 0.409 | -0.45 | -0.659 | -0.001 | 0.03 | 0.03 | 0 | 0.69 | 0.82 | 0.72 | 0.95 | 0.69 | 0.81 |
| 11 | 113424558 | rs7952106 | -0.669 | 0.556 | -0.415 | -0.622 | 0.004 | 0.05 | 0.05 | 0 | 0.62 | 0.75 | 0.66 | 0.94 | 0.64 | 0.77 |
| 11 | 113430486 | rs4373974 | 0.053 | 0.538 | -0.368 | -0.369 | -0.004 | 0.085 | 0.085 | 0 | 0.47 | 0.32 | 0.38 | 0.24 | 0.45 | 0.36 |
| 11 | 113437074 | rs12791990 | -1.133 | 0.427 | -0.433 | 0.191 | 0.001 | -0.004 | -0.004 | 0 | 0.15 | 0.16 | 0.12 | 0.17 | 0.16 | 0.12 |
| 11 | 113438068 | rs7933981 | 0.686 | 0.95 | -0.511 | 0.257 | -0.013 | 0.04 | 0.04 | 0 | 0.38 | 0.21 | 0.42 | 0.28 | 0.41 | 0.43 |
| 11 | 113440331 | rs7123697 | -0.583 | 0.84 | -0.257 | 0.333 | -0.008 | -0.01 | -0.01 | 0 | 0.56 | 0.7 | 0.57 | 0.44 | 0.57 | 0.55 |
| 11 | 113445494 | rs7936719 | -0.202 | 0.995 | -0.434 | -0.174 | -0.001 | 0.001 | 0.001 | 0 | 0.51 | 0.52 | 0.52 | 0.29 | 0.5 | 0.49 |
| 11 | 113451589 | rs4132966 | -0.896 | 0.84 | -0.598 | 0.281 | 0 | -0.004 | -0.004 | 0 | 0.11 | 0.13 | 0.06 | 0.03 | 0.07 | 0.04 |
| 11 | 113451765 | rs7940164 | 1.291 | 1.619 | -0.777 | 0.448 | -0.015 | -0.008 | -0.008 | 1.884 | 0.76 | 0.81 | 0.66 | 0.71 | 0.69 | 0.76 |
| 11 | 113454768 | rs12576834 | -0.591 | 1.017 | -0.817 | 0.687 | 0 | -0.009 | -0.009 | 0 | 0.16 | 0.1 | 0.1 | 0.18 | 0.2 | 0.18 |
| 11 | 113457324 | rs4245155 | 0.633 | 0.367 | -0.823 | 0.272 | 0.028 | 0.001 | 0.001 | 0.011 | 0.51 | 0.36 | 0.73 | 0.52 | 0.6 | 0.52 |
| 11 | 113469219 | rs12280220 | 0.174 | -0.098 | -1.44 | -0.145 | -0.005 | -0.023 | -0.023 | 0 | 0.34 | 0.38 | 0.24 | 0.15 | 0.25 | 0.19 |
| 11 | 113476298 | rs1396807 | 1.986 | -0.462 | -1.042 | -0.216 | -0.001 | 0.003 | 0.003 | 0.21 | 0.19 | 0.21 | 0.12 | 0.29 | 0.15 | 0.22 |
| 11 | 113477081 | rs11214677 | 0.198 | -0.411 | -0.739 | -0.156 | -0.022 | -0.002 | -0.002 | 0 | 0.42 | 0.29 | 0.63 | 0.56 | 0.58 | 0.54 |
| 11 | 113478201 | rs1039832 | 1.262 | -0.36 | -0.788 | -0.252 | -0.016 | -0.019 | -0.019 | 1.381 | 0.23 | 0.28 | 0.11 | 0.15 | 0.14 | 0.1 |
| 11 | 113495279 | rs12806316 | 1.607 | -0.157 | -0.757 | -0.221 | -0.116 | -0.116 | -0.116 | 0 | 0.24 | 0.03 | 0.51 | 0.38 | 0.4 | 0.31 |
| 11 | 113499583 | rs4502018 | 0.643 | -0.294 | -0.572 | -0.245 | 0.017 | 0.01 | 0.01 | 0 | 0.34 | 0.27 | 0.17 | 0.27 | 0.26 | 0.29 |
| 11 | 113500513 | rs7131623 | 0.559 | -0.292 | -0.56 | -0.244 | 0.009 | 0.006 | 0.006 | 0 | 0.36 | 0.31 | 0.18 | 0.27 | 0.27 | 0.3 |
| 11 | 113501655 | rs2886189 | 0.114 | 0.005 | -0.373 | -0.416 | 0.019 | -0.008 | -0.008 | 0 | 0.36 | 0.45 | 0.69 | 0.64 | 0.55 | 0.47 |
| 11 | 113504913 | rs1509513 | 0.292 | -0.221 | -0.633 | -0.417 | 0.019 | -0.006 | -0.006 | 0 | 0.35 | 0.51 | 0.59 | 0.63 | 0.45 | 0.41 |
| 11 | 113515898 | rs11214684 | 0.323 | -0.593 | -1.21 | -0.359 | -0.005 | -0.055 | -0.055 | 0 | 0.55 | 0.08 | 0.57 | 0.53 | 0.69 | 0.63 |
| 11 | 113524040 | rs7110543 | 0.225 | -0.868 | -0.402 | -0.427 | -0.028 | 0.187 | 0.187 | 0.074 | 0.15 | 0.36 | 0.31 | 0.38 | 0.07 | 0.14 |
| 11 | 113524972 | rs7950011 | -0.485 | -0.637 | -0.593 | -0.628 | 0.014 | 0.046 | 0.046 | 0 | 0.48 | 0.68 | 0.73 | 0.64 | 0.58 | 0.49 |
| 11 | 113536710 | rs10789956 | -1.871 | -0.673 | -0.656 | -0.661 | -0.001 | -0.006 | -0.006 | 0.349 | 0.78 | 0.8 | 0.88 | 0.9 | 0.81 | 0.78 |
| 11 | 113539080 | rs3934007 | -0.628 | -0.562 | -0.53 | -0.542 | 0.018 | 0.069 | 0.069 | 0 | 0.48 | 0.75 | 0.73 | 0.64 | 0.58 | 0.49 |
| 11 | 113543747 | rs4938032 | -0.554 | -0.501 | -0.292 | -0.534 | -0.001 | 0.002 | 0.002 | 0 | 0.6 | 0.64 | 0.45 | 0.37 | 0.56 | 0.55 |
| 11 | 113554517 | rs11214696 | -0.333 | -0.649 | -0.276 | -0.624 | 0.002 | -0.007 | -0.007 | 0.498 | 0.57 | 0.54 | 0.36 | 0.36 | 0.47 | 0.49 |
| 11 | 113558845 | rs17115826 | 0.372 | -0.589 | -0.273 | -0.549 | -0.007 | -0.045 | -0.045 | 0 | 0.08 | 0.23 | 0.09 | 0 | 0.08 | 0.02 |
| 11 | 113563688 | rs11604602 | 0.207 | -0.694 | -0.545 | -0.541 | -0.007 | 0.019 | 0.019 | 0 | 0.35 | 0.12 | 0.24 | 0.26 | 0.33 | 0.34 |
| 11 | 113564400 | rs11601037 | 0.49 | -0.707 | -0.342 | -0.582 | -0.011 | 0.038 | 0.038 | 0 | 0.33 | 0.48 | 0.41 | 0.49 | 0.23 | 0.31 |
| 11 | 113570852 | rs3802854 | 0.726 | -0.688 | -0.495 | -0.581 | 0.042 | -0.003 | -0.003 | 0 | 0.31 | 0.11 | 0.19 | 0.26 | 0.25 | 0.28 |
| 11 | 113576932 | rs7926267 | -2.179 | -0.595 | -0.526 | -0.552 | -0.098 | -0.066 | -0.066 | 3.707 | 0.77 | 0.97 | 0.76 | 0.75 | 0.64 | 0.67 |
| 11 | 113577900 | rs4132911 | 1.614 | -0.72 | -0.32 | -0.575 | -0.004 | -0.019 | -0.019 | 0.054 | 0.09 | 0.14 | 0.14 | 0.04 | 0.05 | 0.06 |
| 11 | 113578020 | rs1155104 | 1.584 | -0.73 | -0.332 | -0.581 | 0 | 0.027 | 0.027 | 0.392 | 0.22 | 0.27 | 0.27 | 0.44 | 0.18 | 0.25 |
| 11 | 113578677 | rs925267 | 0.719 | -0.731 | -0.34 | -0.582 | -0.004 | -0.041 | -0.041 | 0.447 | 0.07 | 0.22 | 0.07 | 0 | 0.07 | 0.02 |
| 11 | 113590031 | rs4938050 | -0.057 | -0.96 | -0.469 | -0.665 | -0.008 | -0.005 | -0.005 | 0 | 0.41 | 0.4 | 0.34 | 0.31 | 0.3 | 0.34 |
| 11 | 113591832 | rs10750027 | -1.334 | -0.851 | -0.861 | -0.607 | -0.108 | -0.012 | -0.012 | 0 | 0.7 | 0.92 | 0.68 | 0.75 | 0.54 | 0.61 |
| 11 | 113599224 | rs1318296 | 0.976 | -1.024 | -0.834 | -0.546 | 0 | -0.001 | -0.001 | 0 | 0.15 | 0.14 | 0.14 | 0.13 | 0.16 | 0.21 |
| 11 | 113602555 | rs2885987 | 1.468 | -0.642 | -0.879 | -0.398 | 0.031 | 0.015 | 0.015 | 0.005 | 0.18 | 0.07 | 0.05 | 0.14 | 0.09 | 0.08 |
| 11 | 113613547 | rs10502176 | 1.537 | -0.642 | -0.879 | -0.398 | 0.032 | 0.013 | 0.013 | 0.009 | 0.18 | 0.05 | 0.05 | 0.14 | 0.09 | 0.07 |
| 11 | 113616880 | rs2465651 | 1.785 | -0.567 | -1.014 | -0.418 | -0.081 | -0.05 | -0.05 | 0 | 0.24 | 0.07 | 0.25 | 0.25 | 0.36 | 0.33 |
| 11 | 113617882 | rs17613359 | 0.997 | -0.849 | -0.862 | -0.339 | -0.018 | 0.026 | 0.026 | 0 | 0.21 | 0.05 | 0.12 | 0.14 | 0.18 | 0.13 |
| 11 | 113628990 | rs17541654 | 0.893 | -0.811 | -0.701 | -0.33 | 0 | -0.014 | -0.014 | 0.509 | 0.06 | 0.1 | 0.06 | 0 | 0.06 | 0.02 |
| 11 | 113638177 | rs17613838 | -0.235 | -0.799 | -0.654 | -0.353 | 0.008 | -0.041 | -0.041 | 1.553 | 0.12 | 0.18 | 0.14 | 0 | 0.15 | 0.08 |
| 11 | 113638374 | rs2459969 | -0.84 | -1.089 | -0.488 | -0.436 | -0.064 | -0.025 | -0.025 | 0 | 0.75 | 0.54 | 0.73 | 0.83 | 0.81 | 0.72 |
| 11 | 113639232 | rs2459970 | 2.122 | -1.125 | -0.651 | -0.427 | -0.119 | -0.068 | -0.068 | 0 | 0.21 | 0.04 | 0.28 | 0.24 | 0.39 | 0.32 |
| 11 | 113645217 | rs1000559 | 0.487 | -1.18 | -0.432 | -0.44 | -0.005 | -0.037 | -0.037 | 0 | 0.07 | 0.16 | 0.06 | 0 | 0.06 | 0.02 |
| 11 | 113653252 | rs2459975 | 0.625 | -1.153 | -0.568 | -0.416 | 0.022 | 0.045 | 0.045 | 0.163 | 0.47 | 0.31 | 0.56 | 0.68 | 0.57 | 0.57 |
| 11 | 113658730 | rs17115975 | 1.416 | -1.24 | -0.675 | -0.444 | 0.016 | -0.017 | -0.017 | 0.443 | 0.17 | 0.02 | 0.05 | 0.14 | 0.09 | 0.07 |
| 11 | 113658841 | rs17614391 | 0.343 | -1.07 | -0.405 | -0.401 | -0.003 | -0.036 | -0.036 | 0 | 0.07 | 0.16 | 0.06 | 0 | 0.06 | 0.02 |
| 11 | 113660576 | rs1713676 | 0.668 | -1.056 | -0.428 | -0.403 | -0.043 | -0.036 | -0.036 | 0 | 0.45 | 0.16 | 0.5 | 0.42 | 0.56 | 0.6 |
| 11 | 113663781 | rs11214725 | 1.782 | -0.914 | 0.017 | -0.154 | 0 | -0.011 | -0.011 | 0.185 | 0.18 | 0.19 | 0.12 | 0.13 | 0.16 | 0.18 |
| 11 | 113667873 | rs2465650 | -1.488 | -0.926 | 0.25 | -0.16 | 0.004 | -0.074 | -0.074 | 0 | 0.79 | 0.56 | 0.84 | 0.87 | 0.84 | 0.82 |
| 11 | 113679119 | rs2465647 | -1.751 | -0.93 | 0.331 | -0.162 | 0.007 | -0.053 | -0.053 | 0.111 | 0.8 | 0.63 | 0.85 | 0.87 | 0.84 | 0.82 |
| 11 | 113684809 | rs2519200 | 0.262 | -0.96 | 0.283 | -0.119 | 0 | 0 | 0 | 0 | 0.27 | 0.23 | 0.28 | 0.46 | 0.22 | 0.23 |
| 11 | 113696653 | rs7111825 | -1.28 | -1.391 | 0.373 | -0.09 | 0.001 | 0.072 | 0.072 | 0.022 | 0.78 | 0.6 | 0.7 | 0.57 | 0.83 | 0.75 |
| 11 | 113698002 | rs17802314 | 1.356 | -1.422 | 0.212 | -0.131 | 0.018 | 0.012 | 0.012 | 0 | 0.19 | 0.07 | 0.05 | 0.14 | 0.12 | 0.09 |
| 11 | 113707199 | rs6589398 | 0.309 | -1.365 | 0.171 | -0.148 | 0.017 | 0 | 0 | 0.475 | 0.52 | 0.31 | 0.56 | 0.42 | 0.63 | 0.6 |
| 11 | 113710905 | rs7104752 | 1.919 | -1.207 | -0.04 | -0.102 | -0.081 | -0.048 | -0.048 | 0 | 0.22 | 0.08 | 0.29 | 0.25 | 0.41 | 0.3 |
| 11 | 113711810 | rs3802852 | 1.214 | -1.286 | 0.026 | -0.08 | 0.021 | 0.016 | 0.016 | 0 | 0.21 | 0.11 | 0.06 | 0.14 | 0.12 | 0.09 |
| 11 | 113723144 | rs17802653 | 1.625 | -1.263 | -0.208 | -0.095 | 0.002 | -0.02 | -0.02 | 0 | 0.18 | 0.22 | 0.12 | 0.12 | 0.16 | 0.17 |
| 11 | 113729138 | rs17802676 | 1.306 | -1.21 | -0.576 | -0.056 | 0.005 | -0.005 | -0.005 | 0 | 0.2 | 0.01 | 0.05 | 0.14 | 0.12 | 0.09 |
| 11 | 113741466 | rs2513570 | -0.417 | -1.136 | -0.365 | -0.027 | 0.026 | 0.007 | 0.007 | 0 | 0.5 | 0.7 | 0.44 | 0.57 | 0.37 | 0.4 |
| 11 | 113768638 | rs2011249 | 0.314 | -1.191 | -0.34 | -0.108 | 0.005 | 0.005 | 0.005 | 0.05 | 0.25 | 0.26 | 0.28 | 0.17 | 0.22 | 0.31 |
| 11 | 113770355 | rs1176758 | -2.034 | -1.179 | -0.602 | 0.009 | -0.12 | -0.077 | -0.077 | 0.498 | 0.79 | 0.97 | 0.73 | 0.75 | 0.58 | 0.7 |

| | | | | | | | | | | | | | | | | |
|---|---|---|---|---|---|---|---|---|---|---|---|---|---|---|---|---|
| 11 | 113773946 | rs10789970 | -1.105 | -1.564 | -0.24 | 0.021 | 0.006 | 0.006 | 0.006 | 0 | 0.59 | 0.68 | 0.59 | 0.63 | 0.42 | 0.53 |
| 11 | 113775275 | rs3758987 | -0.233 | -1.427 | 0.055 | 0.034 | -0.024 | -0.063 | -0.063 | 0 | 0.31 | 0.42 | 0.36 | 0.17 | 0.29 | 0.37 |
| 11 | 113802601 | rs1176746 | -1.513 | -0.949 | -0.321 | 0.173 | -0.097 | -0.074 | -0.074 | 0 | 0.77 | 0.91 | 0.75 | 0.72 | 0.6 | 0.71 |
| 11 | 113803028 | rs1176744 | -0.369 | -1.482 | 0.165 | -0.109 | -0.014 | -0.04 | -0.04 | 0.082 | 0.36 | 0.46 | 0.41 | 0.18 | 0.31 | 0.4 |
| 11 | 113807181 | rs11214775 | 0.272 | -1.448 | 0.124 | -0.044 | 0.013 | 0.025 | 0.025 | 0 | 0.4 | 0.3 | 0.18 | 0.24 | 0.28 | 0.23 |
| 11 | 113812733 | rs1672717 | -1.888 | -0.971 | -0.164 | 0.144 | -0.124 | -0.103 | -0.103 | 0 | 0.79 | 0.97 | 0.74 | 0.69 | 0.58 | 0.65 |
| 11 | 113816377 | rs17614942 | 1.041 | -1.338 | 0.018 | -0.066 | -0.003 | -0.025 | -0.025 | 0 | 0.06 | 0.12 | 0.06 | 0 | 0.05 | 0.02 |
| 11 | 113817286 | rs7943062 | 1.57 | -1.611 | -0.073 | -0.176 | 0 | -0.006 | -0.006 | 0 | 0.16 | 0.17 | 0.11 | 0.11 | 0.17 | 0.14 |
| 11 | 113818271 | rs4936286 | 0.503 | -1.792 | -0.237 | -0.181 | 0.031 | 0.065 | 0.065 | 0 | 0.29 | 0.12 | 0.27 | 0.19 | 0.22 | 0.35 |
| 11 | 113818752 | rs7129190 | -0.815 | -1.15 | 0.207 | 0.238 | 0.025 | 0.014 | 0.014 | 0.05 | 0.68 | 0.6 | 0.64 | 0.68 | 0.5 | 0.63 |
| 11 | 113822745 | rs720396 | 1.069 | -0.798 | -0.679 | 0.334 | -0.038 | -0.007 | -0.007 | 0 | 0.26 | 0.19 | 0.44 | 0.63 | 0.46 | 0.41 |
| 11 | 113823865 | rs7942029 | 0.218 | -0.705 | -0.18 | 0.692 | 0.016 | 0.093 | 0.093 | 0.037 | 0.17 | 0.32 | 0.28 | 0.43 | 0.17 | 0.19 |
| 11 | 113833152 | rs1150229 | 1.158 | -0.93 | -1.082 | 0.045 | -0.165 | -0.09 | -0.09 | 0 | 0.39 | 0.02 | 0.32 | 0.52 | 0.53 | 0.46 |
| 11 | 113839272 | rs11214789 | 0.883 | -1.079 | -0.7 | -0.07 | -0.001 | -0.004 | -0.004 | 0 | 0.13 | 0.1 | 0.08 | 0.09 | 0.15 | 0.08 |
| 11 | 113841003 | rs10789980 | -1.388 | -1.128 | -0.067 | -0.081 | -0.003 | 0.002 | 0.002 | 0 | 0.56 | 0.6 | 0.63 | 0.49 | 0.47 | 0.53 |
| 11 | 113843477 | rs1176752 | 0.488 | -1.132 | 0.608 | -0.135 | 0.02 | -0.025 | -0.025 | 0.288 | 0.92 | 0.64 | 0.92 | 1 | 0.94 | 0.98 |
| 11 | 113845541 | rs1150226 | 0.405 | -1.123 | 0.657 | -0.147 | 0.021 | -0.031 | -0.031 | 0 | 0.92 | 0.66 | 0.92 | 1 | 0.93 | 0.98 |
| 11 | 113846006 | rs1062613 | 0.486 | -1.167 | 0.482 | -0.112 | -0.003 | -0.08 | -0.08 | 0 | 0.79 | 0.5 | 0.83 | 0.85 | 0.78 | 0.9 |
| 11 | 113846904 | rs11604247 | -0.468 | -1.19 | 0.54 | -0.033 | 0.016 | 0.041 | 0.041 | 0 | 0.08 | 0.19 | 0.24 | 0.18 | 0.09 | 0.2 |
| 11 | 113848474 | rs1176722 | 0.886 | -1.269 | 0.472 | -0.304 | -0.001 | 0.001 | 0.001 | 0 | 0.14 | 0.1 | 0.08 | 0.15 | 0.16 | 0.08 |
| 11 | 113850140 | rs2276302 | 0.823 | -1.281 | 0.49 | -0.322 | 0.007 | -0.074 | -0.074 | 1.372 | 0.72 | 0.46 | 0.77 | 0.85 | 0.69 | 0.84 |
| 11 | 113854679 | rs11214796 | 0.683 | -1.148 | 0.65 | -0.049 | -0.002 | -0.007 | -0.007 | 0 | 0.75 | 0.34 | 0.67 | 0.73 | 0.79 | 0.74 |
| 11 | 113855342 | rs1176717 | -0.515 | -1.045 | 0.526 | 0.001 | 0.008 | 0 | 0 | 0.133 | 0.21 | 0.29 | 0.29 | 0.27 | 0.21 | 0.26 |
| 11 | 113855977 | rs948983 | 0.646 | -1.028 | 0.3 | 0.009 | -0.001 | 0.001 | 0.001 | 0 | 0.14 | 0.11 | 0.08 | 0.15 | 0.15 | 0.08 |
| 11 | 113856681 | rs10160548 | 1.009 | -1.232 | -0.028 | -0.11 | 0.03 | 0.042 | 0.042 | 2.042 | 0.58 | 0.19 | 0.57 | 0.62 | 0.67 | 0.63 |
| 11 | 113862930 | rs11214800 | 1.647 | -1.252 | -0.24 | -0.133 | -0.165 | -0.061 | -0.061 | 0 | 0.4 | 0.02 | 0.33 | 0.56 | 0.53 | 0.46 |
| 11 | 113864109 | rs897685 | 0.027 | -0.888 | 0.217 | 0.067 | -0.047 | -0.055 | -0.055 | 3.985 | 0.8 | 0.55 | 0.87 | 0.91 | 0.88 | 0.89 |
| 11 | 113866551 | rs1379170 | 0.674 | -0.796 | -0.561 | 0.195 | 0.023 | 0.033 | 0.033 | 1.864 | 0.58 | 0.22 | 0.57 | 0.62 | 0.67 | 0.63 |
| 11 | 113870883 | rs7126511 | -1.041 | -1.549 | -0.397 | 0.225 | 0.035 | 0.083 | 0.083 | 0 | 0.39 | 0.2 | 0.33 | 0.18 | 0.33 | 0.32 |
| 11 | 113875853 | rs10891615 | 0.981 | -1.076 | 0.981 | 0.844 | -0.009 | 0.011 | 0.011 | 0 | 0.63 | 0.51 | 0.47 | 0.61 | 0.68 | 0.65 |
| 11 | 113879251 | rs4938064 | 0.281 | -0.966 | 0.954 | 0.878 | -0.026 | 0.052 | 0.052 | 0 | 0.81 | 0.69 | 0.7 | 0.68 | 0.88 | 0.78 |
| 11 | 113885412 | rs4938066 | -1.13 | -0.884 | 0.98 | 0.768 | -0.003 | -0.003 | -0.003 | 0 | 0.35 | 0.64 | 0.35 | 0.31 | 0.31 | 0.33 |
| 11 | 113890125 | rs7118530 | -0.996 | -0.461 | 0.691 | 0.711 | -0.014 | 0.007 | 0.007 | 1.077 | 0.35 | 0.24 | 0.29 | 0.32 | 0.36 | 0.35 |
| 11 | 113902210 | rs745274 | -0.142 | -0.32 | -0.167 | 0.355 | 0.025 | -0.069 | -0.069 | 0.008 | 0.2 | 0.3 | 0.16 | 0.06 | 0.22 | 0.2 |
| 11 | 113904382 | rs10891623 | 0.631 | -0.308 | -1.28 | 0.607 | -0.071 | -0.107 | -0.107 | 0 | 0.55 | 0.03 | 0.42 | 0.54 | 0.62 | 0.57 |
| 11 | 113928861 | rs11214855 | 0.135 | -0.094 | -1.343 | 0.377 | -0.038 | -0.018 | -0.018 | 0 | 0.4 | 0.16 | 0.54 | 0.59 | 0.43 | 0.5 |
| 11 | 113929550 | rs11214857 | -0.16 | -0.579 | -1.715 | 0.053 | -0.036 | 0.086 | 0.086 | 0 | 0.53 | 0.09 | 0.38 | 0.32 | 0.54 | 0.49 |
| 11 | 113930758 | rs6589411 | -0.051 | -0.528 | -1.504 | -0.774 | -0.007 | -0.032 | -0.032 | 0.004 | 0.33 | 0.3 | 0.26 | 0.02 | 0.38 | 0.25 |
| 11 | 113932460 | rs2509220 | 0.187 | -0.227 | -2.127 | 0.181 | -0.093 | -0.088 | -0.088 | 1.376 | 0.41 | 0.05 | 0.38 | 0.51 | 0.45 | 0.46 |
| 11 | 113937114 | rs2852789 | 0.126 | -0.166 | -2.253 | 0.092 | -0.128 | -0.14 | -0.14 | 5.321 | 0.42 | 0.03 | 0.35 | 0.49 | 0.44 | 0.46 |
| 11 | 113939539 | rs1784684 | -0.047 | -1.161 | -1.846 | -0.449 | 0.017 | 0.013 | 0.013 | 0 | 0.25 | 0.19 | 0.36 | 0.17 | 0.18 | 0.18 |
| 11 | 113940469 | rs3018331 | 0.021 | -1.133 | -1.737 | -0.693 | 0.008 | 0.011 | 0.011 | 0 | 0.24 | 0.22 | 0.17 | 0.09 | 0.18 | 0.18 |
| 11 | 113945609 | rs1672692 | 0.06 | -1.124 | -1.809 | -0.687 | 0.016 | 0.034 | 0.034 | 0 | 0.24 | 0.16 | 0.16 | 0.11 | 0.18 | 0.18 |
| 11 | 113949131 | rs7113940 | 0.276 | -0.931 | -1.601 | -1.383 | 0.003 | -0.128 | -0.128 | 0 | 0.39 | 0.45 | 0.3 | 0.08 | 0.42 | 0.32 |
| 11 | 113949232 | rs1784692 | 0.539 | -1.295 | -2.136 | -1.341 | 0.013 | 0.033 | 0.033 | 0.3 | 0.24 | 0.16 | 0.18 | 0.1 | 0.18 | 0.17 |
| 11 | 113951650 | rs10891628 | 0.076 | -1.104 | -1.876 | -1.326 | 0.002 | -0.011 | -0.011 | 0.045 | 0.39 | 0.63 | 0.32 | 0.07 | 0.4 | 0.32 |
| 11 | 113951988 | rs11214863 | 0.848 | -1.001 | -1.517 | -1.041 | 0.009 | -0.14 | -0.14 | 0 | 0.23 | 0.53 | 0.21 | 0.01 | 0.27 | 0.13 |
| 11 | 113952837 | rs2735201 | -0.694 | -1.284 | -2.028 | -0.76 | -0.004 | -0.006 | -0.006 | 0 | 0.73 | 0.63 | 0.81 | 0.83 | 0.82 | 0.77 |
| 11 | 113954234 | rs1791807 | 0.272 | -0.421 | -2.015 | 0.253 | 0.027 | 0.045 | 0.045 | 0.263 | 0.52 | 0.89 | 0.57 | 0.35 | 0.43 | 0.36 |
| 11 | 113957197 | rs2846630 | -0.453 | 0.269 | -1.277 | 0.939 | -0.051 | -0.05 | -0.05 | 0 | 0.29 | 0.17 | 0.3 | 0.47 | 0.41 | 0.3 |
| 11 | 113960630 | rs12290032 | -1.19 | -0.979 | -0.054 | 1.06 | -0.013 | 0 | 0 | 0.015 | 0.36 | 0.59 | 0.23 | 0.41 | 0.23 | 0.29 |
| 11 | 113963623 | rs238940 | 1.06 | -1.01 | 0.597 | 1.251 | -0.05 | -0.022 | -0.022 | 2.616 | 0.85 | 0.48 | 0.75 | 0.94 | 0.94 | 0.87 |
| 11 | 113967017 | rs7930788 | -0.966 | -1.073 | 0.594 | 1.224 | -0.023 | 0.027 | 0.027 | 0.085 | 0.27 | 0.32 | 0.19 | 0.41 | 0.19 | 0.26 |
| 11 | 113969561 | rs238931 | 1.553 | -0.969 | -0.091 | 1.348 | 0.073 | -0.085 | -0.085 | 0 | 0.6 | 0.14 | 0.51 | 0.51 | 0.74 | 0.59 |
| 11 | 113972348 | rs238930 | -1.765 | -0.887 | 0.816 | 1.487 | -0.029 | 0.089 | 0.089 | 5.055 | 0.16 | 0.66 | 0.29 | 0.3 | 0.08 | 0.22 |
| 11 | 113972439 | rs238929 | -2.452 | -0.672 | 0.122 | 1.518 | 0.052 | 0.042 | 0.042 | 0 | 0.48 | 0.83 | 0.56 | 0.59 | 0.36 | 0.4 |
| 11 | 113973184 | rs238928 | -1.984 | -0.725 | -0.067 | 0.938 | 0.045 | 0.005 | 0.005 | 0.866 | 0.37 | 0.97 | 0.45 | 0.23 | 0.21 | 0.23 |
| 11 | 113975347 | rs238925 | -2.142 | 0.081 | 1.075 | 0.94 | -0.004 | -0.004 | -0.004 | 0 | 0.34 | 0.29 | 0.4 | 0.36 | 0.39 | 0.38 |
| 11 | 113978499 | rs238920 | -1.367 | -0.224 | 2.022 | 0.958 | 0.014 | 0.069 | 0.069 | 2.647 | 0.08 | 0.46 | 0.07 | 0.06 | 0.01 | 0.01 |
| 11 | 113981241 | rs17116384 | -1.647 | -0.154 | 1.847 | 0.98 | -0.03 | -0.054 | -0.054 | 11.543 | 0.26 | 0.06 | 0.32 | 0.31 | 0.29 | 0.27 |
| 11 | 113984317 | rs2073846 | -1.359 | -0.157 | 1.826 | 0.974 | -0.045 | -0.05 | -0.05 | 0 | 0.1 | 0.02 | 0.21 | 0.26 | 0.18 | 0.15 |
| 11 | 113984489 | rs238913 | -2.727 | -0.225 | 2.423 | 0.988 | -0.001 | 0.013 | 0.013 | 0 | 0.37 | 0.62 | 0.5 | 0.42 | 0.38 | 0.4 |
| 11 | 113984975 | rs2073844 | -0.382 | -0.209 | 2.363 | 0.817 | 0.031 | 0.048 | 0.048 | 0 | 0.15 | 0.04 | 0.11 | 0.04 | 0.11 | 0.1 |
| 11 | 113986162 | rs238911 | -2.378 | -0.216 | 2.444 | 0.831 | 0.006 | 0.029 | 0.029 | 0 | 0.34 | 0.47 | 0.46 | 0.36 | 0.37 | 0.38 |
| 11 | 113987256 | rs1630468 | -2.676 | -0.133 | 2.237 | 0.834 | 0 | -0.004 | -0.004 | 0 | 0.39 | 0.65 | 0.49 | 0.38 | 0.38 | 0.39 |
| 11 | 113994505 | rs238896 | -3.064 | -0.074 | 1.354 | 0.496 | -0.079 | -0.061 | -0.061 | 0.819 | 0.53 | 0.91 | 0.59 | 0.41 | 0.49 | 0.43 |
| 11 | 113995628 | rs11214873 | 0.419 | 0.087 | 1.562 | 0.686 | 0.006 | 0.031 | 0.031 | 0 | 0.08 | 0.02 | 0.02 | 0.02 | 0.03 | 0.04 |
| 11 | 113997675 | rs2735207 | -2.17 | 0.101 | 1.632 | 0.712 | -0.005 | -0.006 | -0.006 | 0 | 0.31 | 0.3 | 0.44 | 0.34 | 0.37 | 0.37 |
| 11 | 113998407 | rs7116229 | 0.14 | 0.087 | 1.467 | 0.672 | -0.014 | -0.066 | -0.066 | 0 | 0.11 | 0.23 | 0.05 | 0.02 | 0.05 | 0.06 |

| | | | | | | | | | | | | | | | | |
|---|---|---|---|---|---|---|---|---|---|---|---|---|---|---|---|---|
| 11 | 114000469 | rs17116456 | -0.35 | 0.279 | 1.3 | 0.718 | 0.019 | -0.01 | -0.01 | 0 | 0.13 | 0.09 | 0.14 | 0.12 | 0.08 | 0.09 |
| 11 | 114000727 | rs926075 | -2.083 | 0.339 | 1.239 | 0.696 | -0.073 | -0.046 | -0.046 | 1.467 | 0.17 | 0.06 | 0.28 | 0.22 | 0.31 | 0.26 |
| 11 | 114005213 | rs909168 | -2.063 | -0.021 | 1.413 | 0.655 | 0 | -0.001 | -0.001 | 0 | 0.4 | 0.57 | 0.36 | 0.25 | 0.34 | 0.31 |
| 11 | 114007713 | rs717676 | -0.673 | 0.307 | 0.477 | 0.525 | 0.021 | -0.016 | -0.016 | 0 | 0.05 | 0.11 | 0.05 | 0.01 | 0.09 | 0.12 |
| 11 | 114008823 | rs2250728 | -2.273 | 0.313 | 0.417 | 0.601 | 0.02 | 0.071 | 0.071 | 0.292 | 0.16 | 0.39 | 0.27 | 0.23 | 0.19 | 0.25 |
| 11 | 114008934 | rs2250721 | -1.383 | 0.626 | -0.449 | 0.716 | -0.006 | -0.01 | -0.01 | 0 | 0.33 | 0.57 | 0.34 | 0.24 | 0.27 | 0.33 |
| 11 | 114009255 | rs4936290 | -1.275 | 1.039 | -0.995 | 1.276 | -0.028 | -0.065 | -0.065 | 0 | 0.26 | 0.12 | 0.39 | 0.36 | 0.32 | 0.26 |
| 11 | 114013091 | rs10488695 | -1.383 | 0.698 | -0.656 | 0.756 | -0.001 | 0 | 0 | 0 | 0.06 | 0.06 | 0.04 | 0.04 | 0.07 | 0.02 |
| 11 | 114017753 | rs595641 | 0.551 | 0.645 | -0.91 | 0.969 | 0.003 | -0.032 | -0.032 | 0 | 0.69 | 0.27 | 0.69 | 0.46 | 0.7 | 0.72 |
| 11 | 114023016 | rs12293744 | -0.794 | 0.343 | 0.109 | 0.669 | -0.062 | -0.12 | -0.12 | 0.489 | 0.19 | 0.56 | 0.19 | 0.02 | 0.09 | 0.05 |
| 11 | 114023864 | rs485550 | 0.867 | 0.549 | -0.532 | 0.527 | 0.001 | 0.003 | 0.003 | 0.038 | 0.77 | 0.81 | 0.72 | 0.44 | 0.7 | 0.67 |
| 11 | 114024748 | rs681200 | 0.033 | 0.72 | -0.579 | 0.716 | -0.033 | -0.005 | -0.005 | 0.371 | 0.9 | 0.97 | 0.91 | 0.51 | 0.79 | 0.77 |
| 11 | 114026189 | rs511571 | -0.337 | -0.07 | 0.121 | 0.076 | -0.013 | -0.042 | -0.042 | 1.151 | 0.11 | 0.17 | 0.06 | 0.01 | 0.06 | 0.03 |
| 11 | 114027277 | rs17116545 | -1.677 | -0.054 | 1.011 | 0.18 | 0.034 | 0.021 | 0.021 | 0.197 | 0.06 | 0.33 | 0.12 | 0.02 | 0.02 | 0.04 |
| 11 | 114032484 | rs665731 | -1.559 | -0.339 | 1.023 | -0.25 | -0.043 | -0.091 | -0.091 | 0 | 0.26 | 0.47 | 0.33 | 0.08 | 0.17 | 0.11 |
| 11 | 114037061 | rs1184748 | -1.632 | -0.555 | 1.3 | 0.006 | -0.018 | -0.029 | -0.029 | 0 | 0.28 | 0.46 | 0.22 | 0.24 | 0.25 | 0.26 |
| 11 | 114040510 | rs660800 | -1.406 | -1.229 | 1.203 | 0.438 | -0.025 | -0.011 | -0.011 | 0.006 | 0.21 | 0.46 | 0.19 | 0.17 | 0.14 | 0.08 |
| 11 | 114044439 | rs585287 | -1.991 | -1.034 | 1.208 | 0.397 | -0.011 | 0.005 | 0.005 | 0 | 0.34 | 0.43 | 0.3 | 0.39 | 0.3 | 0.32 |
| 11 | 114045560 | rs500629 | -1.964 | -0.559 | -0.108 | 0.166 | -0.092 | -0.077 | -0.077 | 4.482 | 0.79 | 0.94 | 0.7 | 0.67 | 0.68 | 0.57 |
| 11 | 114045740 | rs661223 | 1.017 | -0.579 | -0.038 | 0.147 | 0.004 | 0.024 | 0.024 | 0.662 | 0.1 | 0.06 | 0.05 | 0 | 0.08 | 0.02 |
| 11 | 114051033 | rs545884 | -0.394 | -0.574 | 0.02 | 0.168 | -0.021 | -0.007 | -0.007 | 0 | 0.2 | 0.34 | 0.17 | 0.16 | 0.13 | 0.09 |
| 11 | 114051904 | rs2073848 | 0.682 | -0.647 | -0.164 | 0.136 | -0.02 | -0.049 | -0.049 | 0 | 0.12 | 0.02 | 0.11 | 0.24 | 0.15 | 0.19 |
| 11 | 114061226 | rs620095 | -0.981 | -0.683 | 0.145 | 0.07 | -0.024 | -0.031 | -0.031 | 0 | 0.25 | 0.52 | 0.21 | 0.16 | 0.17 | 0.12 |
| 11 | 114063362 | rs589916 | -0.93 | -0.725 | -0.262 | -0.029 | -0.031 | -0.031 | -0.031 | 0 | 0.65 | 0.72 | 0.49 | 0.6 | 0.57 | 0.48 |
| 11 | 114064733 | rs670728 | -0.91 | -0.721 | -0.387 | -0.069 | -0.008 | 0.004 | 0.004 | 0 | 0.36 | 0.45 | 0.32 | 0.4 | 0.33 | 0.31 |
| 11 | 114066500 | rs7106340 | 1.181 | -0.725 | -0.57 | -0.072 | -0.008 | -0.016 | -0.016 | 0 | 0.3 | 0.3 | 0.17 | 0.2 | 0.25 | 0.19 |
| 11 | 114071287 | rs526497 | -2.199 | -0.556 | -0.882 | -0.019 | -0.081 | -0.085 | -0.085 | 0 | 0.79 | 0.95 | 0.72 | 0.65 | 0.7 | 0.6 |
| 11 | 114072365 | rs655988 | -0.432 | -0.668 | -0.712 | -0.123 | 0.002 | 0.008 | 0.008 | 0 | 0.26 | 0.23 | 0.2 | 0.23 | 0.26 | 0.23 |
| 11 | 114072875 | rs3782000 | 1.239 | -0.665 | -0.793 | -0.172 | 0.005 | 0.042 | 0.042 | 3.566 | 0.28 | 0.08 | 0.17 | 0.2 | 0.25 | 0.21 |
| 11 | 114074031 | rs627858 | -0.574 | -0.804 | -0.986 | -0.413 | 0.011 | 0.048 | 0.048 | 0.793 | 0.33 | 0.2 | 0.23 | 0.23 | 0.32 | 0.26 |
| 11 | 114074756 | rs583200 | -0.479 | -1.023 | -0.722 | -0.264 | 0.011 | 0.016 | 0.016 | 3.406 | 0.53 | 0.64 | 0.38 | 0.38 | 0.4 | 0.34 |
| 11 | 114076935 | rs12273315 | 0.45 | -0.528 | -0.661 | -0.037 | 0.016 | 0.114 | 0.114 | 0.037 | 0.12 | 0.38 | 0.15 | 0.19 | 0.07 | 0.1 |
| 11 | 114077813 | rs677379 | -1.138 | -0.595 | -0.905 | -0.095 | -0.014 | -0.037 | -0.037 | 0.656 | 0.72 | 0.7 | 0.83 | 0.87 | 0.76 | 0.81 |
| 11 | 114079713 | rs656269 | 0.466 | 0.298 | -1.545 | 0.057 | -0.039 | -0.027 | -0.027 | 0 | 0.39 | 0.21 | 0.48 | 0.41 | 0.42 | 0.45 |
| 11 | 114081271 | rs405388 | 0.509 | 0.465 | -1.406 | 0.288 | -0.025 | -0.041 | -0.041 | 0 | 0.16 | 0.03 | 0.16 | 0.22 | 0.22 | 0.25 |
| 11 | 114086417 | rs495248 | 1.261 | 0.945 | -1.3 | 0.445 | -0.02 | -0.019 | -0.019 | 1.174 | 0.25 | 0.19 | 0.29 | 0.31 | 0.3 | 0.37 |
| 11 | 114086947 | rs672455 | 0.054 | 0.617 | -1.653 | 0.386 | -0.016 | 0.031 | 0.031 | 3.85 | 0.53 | 0.18 | 0.37 | 0.37 | 0.54 | 0.46 |
| 11 | 114087353 | rs545638 | -0.794 | 0.513 | -0.992 | 0.44 | 0.011 | 0.019 | 0.019 | 0 | 0.93 | 0.83 | 0.82 | 0.94 | 0.93 | 0.95 |
| 11 | 114087467 | rs397461 | 1.01 | 0.432 | -1.706 | 0.212 | 0.003 | 0.015 | 0.015 | 0.994 | 0.22 | 0.17 | 0.14 | 0.14 | 0.21 | 0.16 |
| 11 | 114089231 | rs3781998 | 0.808 | 0.517 | -1.444 | 0.492 | -0.003 | 0.017 | 0.017 | 0 | 0.12 | 0.15 | 0.13 | 0.26 | 0.08 | 0.11 |
| 11 | 114089435 | rs375620 | 0.324 | 0.49 | -1.576 | 0.514 | -0.015 | -0.041 | -0.041 | 0 | 0.56 | 0.2 | 0.36 | 0.48 | 0.42 | 0.41 |
| 11 | 114091961 | rs404426 | 0.377 | 0.34 | -1.759 | 0.329 | -0.064 | -0.027 | -0.027 | 0 | 0.32 | 0.04 | 0.23 | 0.23 | 0.3 | 0.29 |
| 11 | 114092541 | rs398055 | -0.78 | 0.169 | -1.458 | 0.249 | 0.005 | 0.031 | 0.031 | 0.008 | 0.49 | 0.25 | 0.32 | 0.39 | 0.44 | 0.44 |
| 11 | 114092582 | rs573264 | -2.028 | -0.133 | -1.132 | 0.302 | -0.005 | 0.002 | 0.002 | 0.015 | 0.71 | 0.78 | 0.66 | 0.71 | 0.6 | 0.6 |
| 11 | 114092875 | rs400920 | 0.018 | -0.458 | -0.877 | -0.174 | -0.002 | -0.004 | -0.004 | 0 | 0.66 | 0.66 | 0.67 | 0.66 | 0.66 | 0.76 |
| 11 | 114093940 | rs378331 | 0.019 | -1.008 | -0.968 | -0.488 | -0.001 | -0.001 | -0.001 | 4.859 | 0.63 | 0.67 | 0.65 | 0.67 | 0.63 | 0.6 |
| 11 | 114094383 | rs403104 | -0.368 | -1.122 | -0.66 | -0.694 | -0.019 | -0.028 | -0.028 | 0.037 | 0.4 | 0.54 | 0.36 | 0.24 | 0.28 | 0.24 |
| 11 | 114095614 | rs503834 | 0.272 | -1.341 | -0.437 | -0.848 | -0.016 | -0.061 | -0.061 | 0 | 0.67 | 0.53 | 0.78 | 0.81 | 0.72 | 0.79 |
| 11 | 114098802 | rs394800 | 0.595 | -1.087 | -0.512 | -0.713 | 0.033 | 0.08 | 0.08 | 2.184 | 0.53 | 0.38 | 0.56 | 0.75 | 0.62 | 0.71 |
| 11 | 114100299 | rs575707 | -0.786 | -0.859 | -0.395 | -0.438 | 0.012 | -0.055 | -0.055 | 0 | 0.89 | 0.84 | 0.91 | 0.97 | 0.88 | 0.91 |
| 11 | 114101709 | rs370634 | 0.571 | -0.896 | -0.9 | -0.366 | 0.06 | 0.132 | 0.132 | 3.284 | 0.51 | 0.23 | 0.57 | 0.73 | 0.64 | 0.69 |
| 11 | 114102165 | rs595168 | 1.848 | -0.495 | -0.861 | -0.048 | -0.04 | -0.051 | -0.051 | 0 | 0.24 | 0.11 | 0.31 | 0.44 | 0.36 | 0.39 |
| 11 | 114105627 | rs402311 | 0.314 | -0.982 | -0.666 | -0.287 | 0.03 | 0.047 | 0.047 | 1.225 | 0.57 | 0.3 | 0.6 | 0.75 | 0.68 | 0.74 |
| 11 | 114106716 | rs630998 | -0.058 | -0.955 | -0.267 | -0.185 | -0.031 | 0.052 | 0.052 | 0 | 0.8 | 0.53 | 0.82 | 0.77 | 0.88 | 0.85 |
| 11 | 114107173 | rs646721 | 1.664 | -0.273 | -0.92 | -0.204 | -0.087 | -0.09 | -0.09 | 0 | 0.22 | 0.04 | 0.29 | 0.43 | 0.35 | 0.37 |
| 11 | 114107966 | rs420234 | 0.258 | -0.849 | -0.124 | -0.644 | -0.017 | -0.043 | -0.043 | 0 | 0.14 | 0.47 | 0.09 | 0.07 | 0.06 | 0.06 |
| 11 | 114112013 | rs387301 | 1.415 | -0.684 | -0.998 | -0.625 | -0.011 | -0.058 | -0.058 | 0 | 0.19 | 0.01 | 0.15 | 0.14 | 0.19 | 0.31 |
| 11 | 114114377 | rs404509 | 0.357 | -0.677 | -0.331 | -0.659 | -0.006 | -0.071 | -0.071 | 0.666 | 0.16 | 0.33 | 0.11 | 0.02 | 0.11 | 0.07 |
| 11 | 114115298 | rs10128580 | 0.55 | -0.368 | -0.374 | -0.343 | 0.004 | -0.037 | -0.037 | 2.876 | 0.15 | 0.38 | 0.14 | 0.11 | 0.16 | 0.06 |
| 11 | 114115641 | rs2856244 | -1.188 | -0.217 | -0.871 | -0.328 | 0.006 | 0.024 | 0.024 | 0.325 | 0.62 | 0.75 | 0.75 | 0.84 | 0.65 | 0.58 |
| 11 | 114120774 | rs530532 | 0.273 | 0.731 | -1.397 | 0.408 | -0.031 | -0.04 | -0.04 | 0 | 0.3 | 0.24 | 0.36 | 0.55 | 0.41 | 0.4 |
| 11 | 114122009 | rs12793024 | -0.224 | 0.807 | -1.405 | 0.956 | -0.007 | -0.062 | -0.062 | 5.584 | 0.12 | 0.01 | 0.14 | 0.18 | 0.09 | 0.1 |
| 11 | 114122989 | rs503768 | -0.193 | 1.321 | -0.879 | 1.084 | 0.028 | 0.091 | 0.091 | 0 | 0.19 | 0.31 | 0.34 | 0.6 | 0.25 | 0.38 |
| 11 | 114125260 | rs566775 | -0.383 | 0.659 | -1.705 | 0.475 | -0.042 | 0.031 | 0.031 | 0 | 0.19 | 0.03 | 0.22 | 0.1 | 0.2 | 0.13 |
| 11 | 114125324 | rs11600671 | 0.077 | 0.458 | -1.334 | 0.821 | -0.004 | -0.049 | -0.049 | 0.836 | 0.09 | 0 | 0.09 | 0.16 | 0.07 | 0.08 |
| 11 | 114128967 | rs2511153 | -0.839 | 0.102 | -0.783 | 0.536 | 0.001 | 0.032 | 0.032 | 0 | 0.5 | 0.57 | 0.45 | 0.67 | 0.47 | 0.47 |
| 11 | 114129057 | rs505978 | -0.606 | 0.111 | -0.165 | 0.77 | -0.001 | -0.001 | -0.001 | 0 | 0.28 | 0.26 | 0.31 | 0.59 | 0.3 | 0.41 |
| 11 | 114133419 | rs694539 | -0.46 | -0.603 | -0.108 | 0.982 | -0.003 | 0.009 | 0.009 | 0 | 0.23 | 0.24 | 0.18 | 0.35 | 0.19 | 0.29 |
| 11 | 114139205 | rs7105175 | -1.001 | -0.684 | 0.046 | 0.993 | -0.012 | 0.012 | 0.012 | 0.011 | 0.32 | 0.58 | 0.21 | 0.42 | 0.23 | 0.35 |

| | | | | | | | | | | | | | | | | |
|---|---|---|---|---|---|---|---|---|---|---|---|---|---|---|---|---|
| 11 | 114151827 | rs6589421 | -1.033 | -0.878 | -0.089 | 0.992 | -0.013 | 0.01 | 0.01 | 0.033 | 0.32 | 0.59 | 0.25 | 0.43 | 0.22 | 0.34 |
| 11 | 114153953 | rs11214926 | 0.278 | -0.219 | -0.495 | 1.424 | -0.039 | -0.051 | -0.051 | 0 | 0.08 | 0 | 0.05 | 0.28 | 0.12 | 0.17 |
| 11 | 114156318 | rs2604294 | -0.046 | -1.2 | -0.359 | 0.667 | 0.055 | 0.005 | 0.005 | 0.251 | 0.6 | 0.35 | 0.81 | 0.63 | 0.74 | 0.53 |
| 11 | 114162521 | rs2244175 | 0.851 | -1.134 | -0.601 | 0.614 | -0.01 | -0.04 | -0.04 | 0 | 0.41 | 0.26 | 0.61 | 0.5 | 0.57 | 0.48 |
| 11 | 114163771 | rs2847492 | -0.37 | -1.099 | 0.382 | 0.717 | -0.023 | 0.005 | 0.005 | 0 | 0.65 | 0.61 | 0.84 | 0.64 | 0.76 | 0.6 |
| 11 | 114164004 | rs2852432 | 0.022 | -1.083 | -0.04 | 0.681 | 0.061 | -0.021 | -0.021 | 0.18 | 0.59 | 0.33 | 0.73 | 0.56 | 0.75 | 0.55 |
| 11 | 114167476 | rs3819100 | 0.276 | -0.906 | -0.118 | 0.259 | 0.022 | -0.063 | -0.063 | 0.056 | 0.33 | 0.15 | 0.25 | 0.47 | 0.26 | 0.37 |
| 11 | 114167999 | rs2301128 | 1.015 | 0.335 | -0.351 | 0.355 | -0.062 | -0.025 | -0.025 | 0.03 | 0.1 | 0.03 | 0.15 | 0.16 | 0.2 | 0.14 |
| 11 | 114169038 | rs1941404 | 0.25 | -0.988 | 0.332 | 0.337 | 0 | -0.001 | -0.001 | 0 | 0.48 | 0.52 | 0.65 | 0.48 | 0.63 | 0.54 |
| 11 | 114170465 | rs2852426 | 0.394 | -0.711 | -0.499 | 0.137 | 0.014 | 0.029 | 0.029 | 0 | 0.45 | 0.26 | 0.53 | 0.41 | 0.62 | 0.5 |
| 11 | 114171247 | rs10891645 | -0.247 | -0.86 | -0.167 | -0.318 | -0.003 | 0.043 | 0.043 | 0.851 | 0.15 | 0.03 | 0.08 | 0.05 | 0.11 | 0.1 |
| 11 | 114172510 | rs2155806 | -1.249 | -1.35 | 0.454 | -0.581 | -0.033 | -0.084 | -0.084 | 3.946 | 0.31 | 0.39 | 0.18 | 0.11 | 0.2 | 0.19 |
| 11 | 114172660 | rs11214938 | -0.346 | -1.27 | 0.991 | -0.4 | -0.035 | -0.034 | -0.034 | 0 | 0.14 | 0.18 | 0.05 | 0.05 | 0.05 | 0.12 |
| 11 | 114173369 | rs949374 | -0.53 | -1.312 | 1.319 | -0.536 | 0.01 | -0.026 | -0.026 | 0 | 0.14 | 0.2 | 0.14 | 0.05 | 0.19 | 0.08 |
| 11 | 114173835 | rs1941400 | -0.117 | -1.432 | 1.257 | -0.621 | -0.024 | -0.023 | -0.023 | 0.089 | 0.15 | 0.14 | 0.04 | 0.05 | 0.05 | 0.15 |
| 11 | 114177853 | rs1941399 | 1.371 | -1.621 | 1.585 | -0.527 | 0.046 | 0.067 | 0.067 | 0 | 0.66 | 0.28 | 0.71 | 0.8 | 0.76 | 0.69 |
| 11 | 114178541 | rs2604279 | 0.448 | -1.621 | 1.884 | -0.638 | 0.022 | -0.023 | -0.023 | 0.147 | 0.82 | 0.54 | 0.77 | 0.85 | 0.81 | 0.86 |
| 11 | 114179080 | rs2604278 | -0.129 | -1.609 | 1.625 | -0.63 | -0.064 | -0.074 | -0.074 | 0.408 | 0.84 | 0.74 | 0.95 | 0.96 | 0.95 | 0.85 |
| 11 | 114179701 | rs2852447 | 1.331 | -1.59 | 1.31 | -0.621 | 0.048 | 0.073 | 0.073 | 0 | 0.65 | 0.28 | 0.7 | 0.81 | 0.76 | 0.69 |
| 11 | 114182507 | rs2852425 | 1.35 | -1.587 | 1.296 | -0.62 | 0.046 | 0.068 | 0.068 | 0 | 0.65 | 0.3 | 0.7 | 0.81 | 0.75 | 0.69 |
| 11 | 114183309 | rs4646337 | -1.109 | -0.431 | 0.909 | -0.015 | 0.007 | -0.021 | -0.021 | 0 | 0.18 | 0.32 | 0.23 | 0.15 | 0.19 | 0.14 |
| 11 | 114186230 | rs7358439 | -1.045 | -0.404 | 0.473 | -0.001 | 0.001 | -0.011 | -0.011 | 2.292 | 0.17 | 0.24 | 0.22 | 0.15 | 0.19 | 0.14 |
| 11 | 114186645 | rs10891647 | -0.998 | -0.383 | 0.482 | 0.009 | 0.002 | -0.011 | -0.011 | 5.997 | 0.17 | 0.24 | 0.22 | 0.15 | 0.19 | 0.14 |
| 11 | 114188464 | rs949370 | 0.316 | -0.418 | 0.495 | 0.027 | -0.014 | -0.015 | -0.015 | 5.079 | 0.82 | 0.62 | 0.87 | 0.85 | 0.84 | 0.74 |
| 11 | 114189367 | rs1941397 | -1.041 | 0.046 | -0.257 | 0.126 | 0.026 | -0.041 | -0.041 | 0 | 0.54 | 0.83 | 0.73 | 0.49 | 0.62 | 0.54 |
| 11 | 114190377 | rs11214942 | 0.513 | 0.322 | -0.482 | -0.086 | -0.049 | 0.063 | 0.063 | 0 | 0.14 | 0.03 | 0.11 | 0.03 | 0.23 | 0.08 |
| 11 | 114191016 | rs2847476 | -0.981 | 0.002 | 0.095 | 0.127 | 0.034 | -0.02 | -0.02 | 0 | 0.53 | 0.75 | 0.72 | 0.49 | 0.62 | 0.54 |
| 11 | 114197504 | rs2604297 | 0.355 | 0.063 | 0.071 | -0.04 | -0.002 | -0.054 | -0.054 | 0 | 0.12 | 0.22 | 0.08 | 0 | 0.1 | 0.08 |
| 11 | 114199213 | rs2604299 | -0.013 | 0.029 | 0.315 | 0.005 | -0.015 | -0.015 | -0.015 | 0 | 0.58 | 0.38 | 0.35 | 0.49 | 0.48 | 0.55 |
| 11 | 114200953 | rs2852429 | -0.099 | 0.024 | 0.438 | 0.002 | 0.006 | 0.005 | 0.005 | 0.063 | 0.59 | 0.48 | 0.36 | 0.49 | 0.48 | 0.54 |
| 11 | 114201240 | rs2604293 | -0.379 | 0.118 | 0.644 | 0.13 | 0.019 | 0.011 | 0.011 | 0 | 0.64 | 0.57 | 0.49 | 0.61 | 0.52 | 0.62 |
| 11 | 114208130 | rs7951064 | 0.967 | 0.011 | 0.018 | 0.407 | 0.004 | -0.01 | -0.01 | 1.477 | 0.13 | 0.07 | 0.11 | 0.22 | 0.09 | 0.09 |
| 11 | 114209381 | rs11214956 | -0.647 | 0.428 | 0.483 | 0.4 | 0.027 | 0.069 | 0.069 | 0 | 0.05 | 0.17 | 0.14 | 0.12 | 0.05 | 0.08 |
| 11 | 114214237 | rs2847489 | -0.236 | 0.361 | 0.488 | 0.386 | 0.02 | 0.007 | 0.007 | 0 | 0.63 | 0.49 | 0.48 | 0.62 | 0.52 | 0.62 |
| 11 | 114214702 | rs2852438 | 0.471 | 0.366 | -0.275 | 0.302 | 0 | -0.005 | -0.005 | 1.219 | 0.11 | 0.13 | 0.07 | 0 | 0.09 | 0.08 |
| 11 | 114215759 | rs4938094 | 0.625 | 0.373 | -0.414 | 0.528 | 0.031 | -0.039 | -0.039 | 0 | 0.46 | 0.16 | 0.27 | 0.5 | 0.38 | 0.43 |
| 11 | 114231396 | rs11214967 | -1.188 | 0.454 | 0.775 | 1.021 | 0.009 | 0.081 | 0.081 | 1.569 | 0.07 | 0.28 | 0.15 | 0.12 | 0.05 | 0.11 |
| 11 | 114233785 | rs7927378 | -0.817 | 0.499 | 0.713 | 0.999 | -0.006 | -0.004 | -0.004 | 0 | 0.41 | 0.39 | 0.63 | 0.54 | 0.53 | 0.48 |
| 11 | 114235304 | rs1941395 | -0.465 | 0.65 | 0.414 | 1.06 | -0.081 | -0.052 | -0.052 | 0 | 0.34 | 0.12 | 0.47 | 0.41 | 0.48 | 0.36 |
| 11 | 114237801 | rs6589425 | -0.486 | 0.634 | 0.301 | 1.037 | -0.054 | -0.057 | -0.057 | 0 | 0.34 | 0.14 | 0.45 | 0.42 | 0.45 | 0.34 |
| 11 | 114241636 | rs4121639 | -0.337 | 0.701 | 0.312 | 1.097 | -0.038 | -0.056 | -0.056 | 0.227 | 0.31 | 0.14 | 0.43 | 0.41 | 0.39 | 0.36 |
| 11 | 114256749 | rs6589426 | -0.756 | 0.516 | 0.213 | 0.987 | 0.006 | 0.016 | 0.016 | 0.029 | 0.38 | 0.46 | 0.59 | 0.5 | 0.45 | 0.43 |
| 11 | 114259653 | rs2604296 | -0.472 | -0.041 | -0.899 | 0.553 | 0.039 | 0.038 | 0.038 | 7.702 | 0.42 | 0.83 | 0.42 | 0.36 | 0.38 | 0.34 |
| 11 | 114263860 | rs3887314 | -0.148 | -0.454 | -0.896 | 0.353 | 0.024 | 0.029 | 0.029 | 0 | 0.47 | 0.3 | 0.36 | 0.43 | 0.42 | 0.51 |
| 11 | 114269345 | rs766783 | -0.133 | -0.459 | -0.852 | 0.367 | 0.023 | 0.027 | 0.027 | 0 | 0.46 | 0.3 | 0.36 | 0.43 | 0.42 | 0.52 |
| 11 | 114271207 | rs606671 | 0.26 | -0.485 | 0.099 | 0.655 | 0.051 | 0.211 | 0.211 | 0.143 | 0.07 | 0.31 | 0.16 | 0.18 | 0.05 | 0.04 |
| 11 | 114280775 | rs9326243 | 0.947 | -0.761 | -0.177 | 0.306 | 0.001 | 0.054 | 0.054 | 0 | 0.14 | 0.18 | 0.21 | 0.43 | 0.2 | 0.26 |
| 11 | 114282267 | rs616930 | -1.704 | -0.738 | -0.798 | 0.208 | -0.154 | -0.11 | -0.11 | 0 | 0.63 | 0.94 | 0.57 | 0.62 | 0.52 | 0.58 |
| 11 | 114284744 | rs11214983 | 0.588 | -0.636 | -0.406 | -0.007 | -0.005 | 0.203 | 0.203 | 0.128 | 0.28 | 0.03 | 0.12 | 0 | 0.18 | 0.26 |
| 11 | 114287790 | rs591013 | -0.089 | -0.809 | 0.173 | 0.446 | 0.046 | 0.203 | 0.203 | 0.106 | 0.08 | 0.36 | 0.17 | 0.18 | 0.05 | 0.04 |
| 11 | 114299674 | rs7116557 | 0.187 | -0.995 | -0.934 | -0.381 | 0.056 | 0.08 | 0.08 | 0.589 | 0.3 | 0.16 | 0.11 | 0 | 0.15 | 0.24 |
| 11 | 114308684 | rs688101 | -1.161 | -0.706 | -1.07 | -0.219 | 0.01 | 0.043 | 0.043 | 0 | 0.55 | 0.78 | 0.53 | 0.66 | 0.43 | 0.55 |
| 11 | 114318464 | rs11214994 | 0.87 | -0.802 | -1.046 | -0.249 | 0.065 | 0.146 | 0.146 | 1.135 | 0.25 | 0.05 | 0.08 | 0 | 0.11 | 0.23 |
| 11 | 114319760 | rs525173 | -0.366 | -0.335 | -0.421 | 0.232 | -0.001 | 0.034 | 0.034 | 0.841 | 0.26 | 0.56 | 0.41 | 0.63 | 0.28 | 0.31 |
| 11 | 114324026 | rs477238 | -0.788 | -0.385 | -1.209 | 0.115 | -0.058 | -0.034 | -0.034 | 0 | 0.55 | 0.89 | 0.55 | 0.61 | 0.48 | 0.48 |
| 11 | 114328053 | rs10891667 | 0.928 | -0.595 | -0.732 | -0.165 | 0.028 | 0.018 | 0.018 | 0.198 | 0.25 | 0.18 | 0.22 | 0 | 0.13 | 0.24 |
| 11 | 114330018 | rs7116731 | -0.761 | -0.691 | 0.027 | 0.297 | 0.021 | 0.038 | 0.038 | 0 | 0.51 | 0.61 | 0.52 | 0.39 | 0.42 | 0.5 |
| 11 | 114333457 | rs11214998 | 0.446 | -0.663 | 0.233 | 0.491 | 0.001 | 0.004 | 0.004 | 0 | 0.19 | 0.21 | 0.17 | 0.3 | 0.25 | 0.26 |
| 11 | 114342277 | rs10891679 | -0.176 | -0.453 | 0.367 | 0.43 | -0.005 | -0.007 | -0.007 | 0 | 0.46 | 0.52 | 0.48 | 0.39 | 0.38 | 0.5 |
| 11 | 114382144 | rs4938111 | 1.798 | -0.006 | 0.09 | 0.147 | -0.041 | -0.029 | -0.029 | 0 | 0.14 | 0.01 | 0.14 | 0.13 | 0.15 | 0.07 |
| 11 | 114385514 | rs7951844 | 0.88 | -0.397 | 0.329 | -0.128 | -0.05 | -0.138 | -0.138 | 1.452 | 0.31 | 0.45 | 0.24 | 0 | 0.13 | 0.24 |
| 11 | 114387956 | rs7117324 | 0.515 | -0.169 | 0.167 | 0.091 | 0.003 | 0.023 | 0.023 | 0 | 0.25 | 0.27 | 0.3 | 0.65 | 0.34 | 0.31 |
| 11 | 114389708 | rs10790019 | 1.052 | -0.215 | 0.209 | -0.035 | -0.063 | -0.161 | -0.161 | 0.049 | 0.3 | 0.56 | 0.25 | 0 | 0.13 | 0.24 |
| 11 | 114393314 | rs492840 | -1.207 | -0.22 | -0.163 | -0.131 | -0.103 | -0.006 | -0.006 | 0 | 0.76 | 0.92 | 0.73 | 0.82 | 0.65 | 0.63 |
| 11 | 114393652 | rs10891692 | 0.387 | -0.367 | 0.1 | -0.123 | 0.002 | 0.017 | 0.017 | 0 | 0.25 | 0.25 | 0.31 | 0.67 | 0.34 | 0.31 |
| 11 | 114397499 | rs7122337 | 1.174 | -0.346 | 0.134 | 0.038 | 0 | 0.002 | 0.002 | 0 | 0.2 | 0.16 | 0.26 | 0.44 | 0.26 | 0.25 |
| 11 | 114411343 | rs7950370 | 1.211 | -0.407 | -0.03 | -0.693 | 0.032 | 0.012 | 0.012 | 0.858 | 0.28 | 0.25 | 0.23 | 0 | 0.13 | 0.24 |
| 11 | 114413446 | rs534912 | 0.181 | -0.202 | -0.516 | -0.573 | 0 | 0.045 | 0.045 | 0 | 0.28 | 0.38 | 0.31 | 0.67 | 0.34 | 0.31 |
| 11 | 114423320 | rs531824 | -0.573 | -0.183 | -0.651 | -0.564 | -0.048 | -0.03 | -0.03 | 0 | 0.59 | 0.91 | 0.56 | 0.67 | 0.47 | 0.55 |

| | | | | | | | | | | | | | | | | |
|---|---|---|---|---|---|---|---|---|---|---|---|---|---|---|---|---|
| 11 | 114430609 | rs661946 | 0.054 | -0.173 | -0.518 | -0.542 | -0.003 | 0.042 | 0.042 | 0 | 0.28 | 0.46 | 0.32 | 0.67 | 0.34 | 0.31 |
| 11 | 114433638 | rs11215057 | 1.329 | 0.653 | -0.846 | -0.688 | -0.059 | 0.002 | 0.002 | 0.002 | 0.1 | 0.02 | 0.09 | 0.09 | 0.18 | 0.12 |
| 11 | 114436944 | rs670668 | -0.645 | -0.221 | -0.571 | -0.519 | -0.048 | -0.005 | -0.005 | 0 | 0.58 | 0.9 | 0.56 | 0.68 | 0.47 | 0.55 |
| 11 | 114438394 | rs11215059 | 0.989 | -0.262 | -0.488 | -0.563 | -0.033 | -0.109 | -0.109 | 0 | 0.3 | 0.42 | 0.24 | 0 | 0.13 | 0.24 |
| 11 | 114441123 | rs513152 | 0.061 | -0.233 | -0.632 | -0.543 | 0.003 | 0.044 | 0.044 | 0 | 0.27 | 0.39 | 0.32 | 0.67 | 0.33 | 0.31 |
| 11 | 114441943 | rs10891705 | 0.493 | 0.108 | -1.419 | -0.377 | 0.004 | 0.014 | 0.014 | 0 | 0.23 | 0.21 | 0.3 | 0.67 | 0.33 | 0.31 |
| 11 | 114442103 | rs550897 | 0.094 | 1.005 | -1.876 | 0.422 | -0.047 | -0.008 | -0.008 | 0 | 0.58 | 0.91 | 0.56 | 0.67 | 0.47 | 0.55 |
| 11 | 114450708 | rs487707 | -0.047 | 1.528 | -1.369 | 0.73 | -0.043 | -0.024 | -0.024 | 0.66 | 0.72 | 0.79 | 0.69 | 0.69 | 0.63 | 0.6 |
| 11 | 114451355 | rs641956 | 0.084 | 1.507 | -0.947 | 0.995 | 0.002 | 0.014 | 0.014 | 0 | 0.19 | 0.22 | 0.29 | 0.41 | 0.27 | 0.19 |
| 11 | 114459317 | rs11215077 | 0.283 | 1.553 | -1.135 | -0.165 | 0.004 | 0.009 | 0.009 | 0 | 0.41 | 0.62 | 0.37 | 0.12 | 0.35 | 0.42 |
| 11 | 114465599 | rs11215079 | 0.831 | 1.364 | -1.34 | 0.36 | 0.002 | 0.014 | 0.014 | 0 | 0.05 | 0 | 0.07 | 0.37 | 0.05 | 0.08 |
| 11 | 114465917 | rs17117217 | 0.26 | 1.065 | -1.266 | 0.065 | 0 | 0.026 | 0.026 | 0 | 0.12 | 0.13 | 0.18 | 0.42 | 0.16 | 0.18 |
| 11 | 114467655 | rs12418080 | 1.27 | 0.866 | -1.625 | -0.627 | -0.049 | 0.087 | 0.087 | 0.182 | 0.18 | 0.03 | 0.22 | 0 | 0.21 | 0.2 |
| 11 | 114468469 | rs10891714 | 1.183 | 0.423 | -0.956 | -0.867 | -0.004 | -0.007 | -0.007 | 0 | 0.23 | 0.23 | 0.13 | 0.11 | 0.15 | 0.22 |
| 11 | 114471076 | rs11603845 | 1.039 | 0.428 | -0.931 | -0.697 | -0.014 | -0.012 | -0.012 | 0.166 | 0.1 | 0.06 | 0.15 | 0.37 | 0.16 | 0.17 |
| 11 | 114480735 | rs1671804 | 1.192 | 0.323 | -0.827 | -0.81 | -0.008 | -0.012 | -0.012 | 0 | 0.23 | 0.24 | 0.12 | 0.12 | 0.15 | 0.22 |
| 11 | 114490989 | rs1712832 | -0.42 | 0.28 | -0.727 | -0.928 | 0.017 | 0.059 | 0.059 | 0 | 0.46 | 0.64 | 0.39 | 0.12 | 0.37 | 0.43 |
| 11 | 114493531 | rs1671810 | -0.752 | 0.191 | -0.975 | -1.126 | -0.024 | 0.036 | 0.036 | 0 | 0.62 | 0.71 | 0.49 | 0.22 | 0.5 | 0.56 |
| 11 | 114512627 | rs1792386 | 1.763 | 0.056 | -0.509 | -1.165 | -0.076 | -0.065 | -0.065 | 0.173 | 0.19 | 0.35 | 0.07 | 0.09 | 0.08 | 0.12 |
| 11 | 114521267 | rs1712799 | -2.374 | 0.096 | -0.496 | -1.163 | -0.055 | -0.049 | -0.049 | 0 | 0.82 | 0.7 | 0.93 | 0.91 | 0.92 | 0.89 |
| 11 | 114523368 | rs10502191 | 1.284 | 0.415 | -0.569 | -1.083 | -0.023 | -0.024 | -0.024 | 0.244 | 0.11 | 0.06 | 0.16 | 0.38 | 0.2 | 0.19 |
| 11 | 114529588 | rs4938118 | 0.936 | 0.347 | -0.217 | -1.139 | 0.004 | 0.019 | 0.019 | 0 | 0.24 | 0.21 | 0.16 | 0.13 | 0.2 | 0.23 |
| 11 | 114535196 | rs1792381 | -2.451 | 0.242 | -0.224 | -1.139 | -0.055 | -0.049 | -0.049 | 0 | 0.82 | 0.71 | 0.93 | 0.91 | 0.92 | 0.88 |
| 11 | 114541075 | rs1517688 | 2.257 | 0.248 | -0.137 | -1.144 | -0.002 | 0.028 | 0.028 | 0 | 0.16 | 0.02 | 0.09 | 0.09 | 0.14 | 0.13 |
| 11 | 114541569 | rs1792408 | -2.603 | 0.221 | -0.054 | -1.114 | -0.041 | -0.036 | -0.036 | 0 | 0.83 | 0.77 | 0.94 | 0.92 | 0.92 | 0.89 |
| 11 | 114541603 | rs1712813 | 1.428 | 0.366 | -0.161 | -1.069 | -0.023 | -0.054 | -0.054 | 0 | 0.28 | 0.2 | 0.35 | 0.4 | 0.34 | 0.27 |
| 11 | 114550320 | rs17550338 | 1.878 | 0.47 | -0.28 | -1.042 | -0.038 | 0.085 | 0.085 | 0.117 | 0.16 | 0.02 | 0.21 | 0 | 0.17 | 0.2 |
| 11 | 114552194 | rs11605843 | -0.083 | 0.386 | -0.075 | -0.918 | 0.013 | 0.032 | 0.032 | 0 | 0.19 | 0.26 | 0.22 | 0.42 | 0.27 | 0.29 |
| 11 | 114565114 | rs1792419 | -0.122 | 0.106 | 0.017 | -0.99 | 0.003 | -0.004 | -0.004 | 3.25 | 0.62 | 0.39 | 0.71 | 0.5 | 0.69 | 0.61 |
| 11 | 114566047 | rs1712835 | -1.102 | -0.219 | -0.071 | -1.144 | -0.033 | -0.043 | -0.043 | 0.574 | 0.65 | 0.62 | 0.83 | 0.82 | 0.78 | 0.75 |
| 11 | 114568942 | rs11215158 | 2.091 | -0.146 | -0.191 | -1.106 | -0.005 | 0.028 | 0.028 | 0 | 0.16 | 0.02 | 0.1 | 0.09 | 0.14 | 0.13 |
| 11 | 114569054 | rs1712838 | -1.048 | -0.193 | -0.053 | -1.128 | -0.012 | -0.017 | -0.017 | 0 | 0.65 | 0.67 | 0.83 | 0.82 | 0.78 | 0.76 |
| 11 | 114570189 | rs1568215 | -0.672 | 0.323 | 0.066 | -1.065 | -0.012 | -0.002 | -0.002 | 0 | 0.37 | 0.28 | 0.4 | 0.41 | 0.47 | 0.49 |
| 11 | 114574819 | rs12364744 | 0.143 | 0.735 | -0.303 | -1.132 | 0.002 | 0.008 | 0.008 | 0 | 0.18 | 0.18 | 0.18 | 0.41 | 0.25 | 0.29 |
| 11 | 114575490 | rs1627370 | -0.702 | 0.664 | -0.318 | -1.191 | -0.013 | -0.004 | -0.004 | 0 | 0.37 | 0.27 | 0.4 | 0.46 | 0.47 | 0.49 |
| 11 | 114576378 | rs1712840 | -2.627 | 0.478 | -0.45 | -1.135 | 0.037 | 0.035 | 0.035 | 1.201 | 0.84 | 0.9 | 0.9 | 0.92 | 0.92 | 0.88 |
| 11 | 114579249 | rs12288283 | 1.174 | 0.436 | -0.425 | -1.165 | 0.004 | 0.045 | 0.045 | 0 | 0.28 | 0.37 | 0.35 | 0.41 | 0.31 | 0.27 |
| 11 | 114579777 | rs1792420 | 1.493 | 0.606 | -0.365 | -1.007 | -0.059 | 0.092 | 0.092 | 0.465 | 0.17 | 0.03 | 0.22 | 0 | 0.22 | 0.2 |
| 11 | 114581857 | rs1792423 | 0.064 | 0.777 | -0.29 | -0.969 | -0.012 | -0.005 | -0.005 | 0.013 | 0.29 | 0.24 | 0.34 | 0.38 | 0.39 | 0.38 |
| 11 | 114582964 | rs1712841 | 0.031 | 0.767 | -0.326 | -0.948 | -0.007 | -0.002 | -0.002 | 0 | 0.31 | 0.28 | 0.34 | 0.38 | 0.4 | 0.38 |
| 11 | 114583890 | rs7118761 | 0.157 | 1.015 | -0.643 | -0.77 | 0 | 0.008 | 0.008 | 0 | 0.52 | 0.44 | 0.53 | 0.55 | 0.52 | 0.5 |
| 11 | 114586066 | rs17117300 | 0.685 | 1.272 | -1.011 | 0.008 | -0.019 | -0.013 | -0.013 | 0 | 0.1 | 0.07 | 0.12 | 0.38 | 0.18 | 0.18 |
| 11 | 114587363 | rs1850522 | 0.33 | 1.176 | -1.087 | -0.02 | -0.044 | -0.057 | -0.057 | 0 | 0.14 | 0.07 | 0.15 | 0.37 | 0.22 | 0.19 |
| 11 | 114592132 | rs17117319 | 1.534 | 1.187 | -1.302 | 0.077 | -0.057 | -0.07 | -0.07 | 0 | 0.12 | 0.02 | 0.22 | 0.36 | 0.22 | 0.15 |
| 11 | 114594187 | rs1792438 | -1.998 | 0.299 | -1.288 | -0.451 | 0.05 | 0.047 | 0.047 | 1.212 | 0.83 | 0.94 | 0.9 | 0.93 | 0.92 | 0.89 |
| 11 | 114598781 | rs10891738 | 0.519 | 0.244 | -1.342 | -0.891 | 0.002 | 0.105 | 0.105 | 0 | 0.37 | 0.16 | 0.37 | 0.13 | 0.34 | 0.41 |
| 11 | 114604149 | rs11215175 | 1.941 | -0.122 | -0.819 | -0.877 | -0.013 | -0.017 | -0.017 | 0 | 0.17 | 0.16 | 0.12 | 0.06 | 0.09 | 0.12 |
| 11 | 114604509 | rs11605961 | 0.874 | 0.203 | -1.005 | -0.307 | -0.048 | -0.077 | -0.077 | 0 | 0.15 | 0.06 | 0.14 | 0.44 | 0.21 | 0.2 |
| 11 | 114608368 | rs12797565 | 0.911 | 0.122 | -0.447 | -0.623 | -0.004 | -0.039 | -0.039 | 0 | 0.25 | 0.24 | 0.31 | 0.42 | 0.29 | 0.26 |
| 11 | 114615037 | rs7123330 | 0.126 | 0.051 | -0.099 | -0.669 | -0.008 | 0.003 | 0.003 | 0 | 0.43 | 0.5 | 0.43 | 0.49 | 0.38 | 0.37 |
| 11 | 114621469 | rs1712790 | -0.2 | -0.281 | -0.733 | -1.066 | -0.023 | -0.057 | -0.057 | 0.659 | 0.44 | 0.14 | 0.44 | 0.51 | 0.52 | 0.61 |
| 11 | 114622352 | rs10502192 | 0.792 | -0.381 | -0.753 | -1.132 | -0.033 | -0.095 | -0.095 | 0.708 | 0.33 | 0.14 | 0.37 | 0.43 | 0.39 | 0.27 |
| 11 | 114623638 | rs1712791 | 0.669 | -0.36 | -0.855 | -1.196 | -0.082 | 0.082 | 0.082 | 0.449 | 0.29 | 0.07 | 0.3 | 0.13 | 0.32 | 0.41 |
| 11 | 114626627 | rs11215187 | 2.353 | -0.582 | -0.157 | -1.145 | 0.02 | 0.021 | 0.021 | 0.448 | 0.17 | 0.1 | 0.12 | 0.06 | 0.09 | 0.11 |
| 11 | 114630293 | rs2040325 | 2.522 | -0.185 | -0.204 | -0.969 | 0.012 | 0.051 | 0.051 | 0 | 0.11 | 0.02 | 0.07 | 0 | 0.09 | 0.01 |
| 11 | 114633879 | rs4938128 | 0.612 | 0.459 | 0.219 | -0.557 | 0.03 | 0.006 | 0.006 | 0 | 0.51 | 0.31 | 0.49 | 0.49 | 0.47 | 0.38 |
| 11 | 114641449 | rs11215196 | -0.647 | 0.36 | 0.028 | -0.138 | 0.028 | 0.066 | 0.066 | 0 | 0.25 | 0.36 | 0.22 | 0.42 | 0.28 | 0.31 |
| 11 | 114642623 | rs7113782 | 0.086 | -0.484 | 0.305 | -0.45 | -0.047 | -0.06 | -0.06 | 0 | 0.29 | 0.36 | 0.2 | 0.09 | 0.14 | 0.17 |
| 11 | 114643021 | rs11828366 | 0.921 | -0.046 | 0.615 | -0.173 | -0.008 | -0.018 | -0.018 | 1.535 | 0.1 | 0.17 | 0.09 | 0.03 | 0.06 | 0.06 |
| 11 | 114644023 | rs1894108 | -0.527 | 0.32 | 0.405 | 0.09 | 0.003 | 0.018 | 0.018 | 0 | 0.53 | 0.24 | 0.53 | 0.67 | 0.5 | 0.66 |
| 11 | 114644778 | rs7108278 | 0.102 | 0.44 | 0.4 | 0.087 | 0.012 | -0.024 | -0.024 | 0 | 0.41 | 0.3 | 0.35 | 0.6 | 0.32 | 0.44 |
| 11 | 114647037 | rs10891749 | -0.438 | 0.481 | 0.083 | 0.126 | -0.036 | -0.086 | -0.086 | 0.022 | 0.18 | 0.06 | 0.17 | 0.56 | 0.18 | 0.24 |
| 11 | 114649770 | rs11600245 | 1.598 | 0.444 | 0.229 | 0.011 | -0.029 | -0.033 | -0.033 | 0 | 0.32 | 0.15 | 0.33 | 0.3 | 0.43 | 0.27 |
| 11 | 114653591 | rs7107819 | 0.807 | 0.538 | 0.386 | -0.02 | 0 | -0.001 | -0.001 | 0 | 0.12 | 0.11 | 0.22 | 0.07 | 0.17 | 0.21 |
| 11 | 114653847 | rs7108229 | 1.3 | 0.454 | 0.275 | -0.03 | -0.044 | 0.005 | 0.005 | 0.117 | 0.56 | 0.25 | 0.47 | 0.34 | 0.58 | 0.47 |
| 11 | 114658208 | rs11215203 | 1.514 | 0.376 | 0.189 | 0.092 | -0.05 | -0.057 | -0.057 | 0 | 0.32 | 0.12 | 0.32 | 0.3 | 0.43 | 0.27 |
| 11 | 114661466 | rs12287008 | 1.385 | 0.137 | 0.649 | 0.099 | 0.029 | 0.047 | 0.047 | 0 | 0.26 | 0.14 | 0.15 | 0.04 | 0.17 | 0.2 |
| 11 | 114661822 | rs10750061 | -0.092 | 0.638 | 1.024 | 0.731 | -0.002 | 0.023 | 0.023 | 0 | 0.83 | 0.46 | 0.84 | 0.84 | 0.88 | 0.85 |
| 11 | 114664651 | rs10502193 | 0.739 | 0.681 | 0.944 | 0.728 | -0.008 | 0.005 | 0.005 | 0 | 0.1 | 0.08 | 0.23 | 0.07 | 0.11 | 0.2 |

| | | | | | | | | | | | | | | | | |
|---|---|---|---|---|---|---|---|---|---|---|---|---|---|---|---|---|
| 11 | 114665112 | rs7949181 | -0.872 | 0.662 | 1.29 | 0.716 | 0.012 | 0.006 | 0.006 | 0.109 | 0.25 | 0.32 | 0.38 | 0.23 | 0.23 | 0.35 |
| 11 | 114665170 | rs10790037 | 1.796 | 0.734 | 1.182 | 0.859 | -0.017 | 0.004 | 0.004 | 0.866 | 0.61 | 0.26 | 0.51 | 0.65 | 0.65 | 0.53 |
| 11 | 114666259 | rs10891755 | -0.087 | 0.859 | 0.929 | 0.901 | -0.017 | -0.002 | -0.002 | 0 | 0.1 | 0.05 | 0.08 | 0.12 | 0.11 | 0.13 |
| 11 | 114668076 | rs2366583 | -0.238 | 0.907 | 1.216 | 1.051 | 0.024 | 0.071 | 0.071 | 0.15 | 0.88 | 0.69 | 0.77 | 0.79 | 0.89 | 0.79 |
| 11 | 114670517 | rs10891758 | -0.303 | 0.925 | 1.222 | 1.039 | 0.03 | -0.008 | -0.008 | 0.016 | 0.88 | 0.68 | 0.75 | 0.91 | 0.88 | 0.8 |
| 11 | 114671919 | rs11215211 | -0.282 | 0.878 | 1.174 | 1.011 | 0.008 | -0.029 | -0.029 | 0 | 0.09 | 0.12 | 0.09 | 0.01 | 0.11 | 0.11 |
| 11 | 114673266 | rs11215212 | 0.376 | 0.794 | 0.874 | 1.197 | -0.008 | -0.006 | -0.006 | 0 | 0.06 | 0.05 | 0.06 | 0.17 | 0.09 | 0.11 |
| 11 | 114673450 | rs7948157 | 0.239 | 0.6 | 0.955 | 1.078 | 0.001 | 0.008 | 0.008 | 0 | 0.15 | 0.17 | 0.14 | 0.17 | 0.12 | 0.17 |
| 11 | 114688488 | rs11823831 | 0.77 | 0.512 | 0.335 | 0.787 | 0.003 | 0.007 | 0.007 | 0 | 0.11 | 0.1 | 0.1 | 0.03 | 0.07 | 0.08 |
| 11 | 114692282 | rs4436579 | -1.144 | 0.483 | 0.488 | 0.768 | 0.003 | -0.024 | -0.024 | 0 | 0.3 | 0.34 | 0.35 | 0.14 | 0.25 | 0.4 |
| 11 | 114694273 | rs12279327 | 1.11 | 0.669 | 0.602 | 1.074 | 0.006 | 0.006 | 0.006 | 0.649 | 0.66 | 0.55 | 0.56 | 0.65 | 0.71 | 0.52 |
| 11 | 114695923 | rs10790047 | 0.818 | 0.681 | 0.613 | 0.996 | -0.008 | 0.011 | 0.011 | 0.766 | 0.69 | 0.74 | 0.59 | 0.78 | 0.71 | 0.52 |
| 11 | 114698329 | rs7105436 | -0.216 | 0.698 | 0.559 | 1.101 | -0.011 | -0.002 | -0.002 | 0 | 0.2 | 0.14 | 0.2 | 0.15 | 0.18 | 0.27 |
| 11 | 114698769 | rs17117563 | 0.739 | 0.704 | 0.546 | 0.909 | -0.008 | 0.008 | 0.008 | 0 | 0.1 | 0.05 | 0.19 | 0.08 | 0.11 | 0.21 |
| 11 | 114698949 | rs2366360 | 1.155 | 0.703 | 0.594 | 0.862 | 0.006 | -0.022 | -0.022 | 0 | 0.66 | 0.56 | 0.57 | 0.78 | 0.71 | 0.52 |
| 11 | 114700983 | rs11828359 | 0.701 | 0.705 | 0.38 | 0.794 | -0.01 | 0.021 | 0.021 | 0 | 0.11 | 0.05 | 0.16 | 0.06 | 0.11 | 0.2 |
| 11 | 114702663 | rs11215223 | -0.077 | 0.753 | 0.359 | 1.059 | -0.009 | -0.034 | -0.034 | 0 | 0.19 | 0.16 | 0.19 | 0.53 | 0.17 | 0.23 |
| 11 | 114704770 | rs4438073 | -1.008 | 0.758 | 0.356 | 1.048 | -0.026 | -0.056 | -0.056 | 0.054 | 0.28 | 0.17 | 0.34 | 0.59 | 0.27 | 0.43 |
| 11 | 114715607 | rs4573710 | -1.221 | 0.735 | 0.43 | 1.039 | -0.004 | -0.011 | -0.011 | 0.015 | 0.31 | 0.31 | 0.39 | 0.59 | 0.27 | 0.43 |
| 11 | 114718070 | rs4938144 | 0.688 | 0.564 | 0.35 | 0.847 | -0.009 | 0.022 | 0.022 | 0 | 0.1 | 0.05 | 0.16 | 0.06 | 0.1 | 0.2 |
| 11 | 114721673 | rs7949627 | 0.137 | 0.647 | 0.315 | 0.87 | 0.013 | -0.036 | -0.036 | 0.11 | 0.86 | 0.77 | 0.78 | 0.92 | 0.85 | 0.72 |
| 11 | 114727155 | rs1398365 | 0.403 | 0.617 | 0.309 | 1.025 | -0.016 | -0.03 | -0.03 | 0 | 0.18 | 0.16 | 0.25 | 0.37 | 0.21 | 0.22 |
| 11 | 114730319 | rs11215244 | 1.192 | 0.499 | -0.161 | 0.758 | -0.086 | -0.065 | -0.065 | 0 | 0.32 | 0.05 | 0.32 | 0.28 | 0.42 | 0.3 |
| 11 | 114732135 | rs4936307 | -0.317 | 1.005 | 1.059 | 0.991 | 0.009 | 0.096 | 0.096 | 0 | 0.12 | 0.25 | 0.23 | 0.36 | 0.11 | 0.23 |
| 11 | 114737208 | rs7117869 | -0.27 | 1.001 | 1.06 | 0.809 | 0.008 | 0.093 | 0.093 | 0 | 0.12 | 0.25 | 0.12 | 0.34 | 0.11 | 0.2 |
| 11 | 114737969 | rs2462160 | -1.881 | 1.024 | 0.654 | 0.807 | 0.009 | 0.017 | 0.017 | 0.895 | 0.4 | 0.56 | 0.55 | 0.52 | 0.39 | 0.53 |
| 11 | 114738157 | rs1403944 | 1.608 | 1.028 | 0.678 | 0.809 | -0.01 | -0.022 | -0.022 | 0 | 0.58 | 0.25 | 0.43 | 0.48 | 0.61 | 0.47 |
| 11 | 114739437 | rs2513494 | -0.551 | 1.08 | 0.362 | 0.455 | 0.011 | -0.028 | -0.028 | 2.587 | 0.22 | 0.35 | 0.26 | 0.08 | 0.22 | 0.22 |
| 11 | 114741182 | rs7104748 | 0.453 | 0.966 | 0.105 | 0.496 | 0.011 | 0.018 | 0.018 | 0.166 | 0.66 | 0.49 | 0.63 | 0.57 | 0.67 | 0.58 |
| 11 | 114750157 | rs2462165 | 0.145 | 0.697 | -0.099 | -0.543 | -0.006 | 0.068 | 0.068 | 0.262 | 0.56 | 0.35 | 0.47 | 0.14 | 0.53 | 0.44 |
| 11 | 114750926 | rs17117733 | 0.166 | 0.752 | 0.321 | -0.351 | 0.007 | -0.015 | -0.015 | 0 | 0.23 | 0.14 | 0.32 | 0.66 | 0.22 | 0.29 |
| 11 | 114753038 | rs1403947 | -1.013 | 0.656 | 0.473 | -0.424 | -0.004 | 0.017 | 0.017 | 0.059 | 0.43 | 0.58 | 0.52 | 0.86 | 0.47 | 0.56 |
| 11 | 114767053 | rs11215271 | 0.67 | 0.966 | 0.642 | -0.356 | -0.012 | -0.013 | -0.013 | 0 | 0.06 | 0.04 | 0.09 | 0.15 | 0.15 | 0.17 |
| 11 | 114782597 | rs2513474 | 0.785 | 0.871 | 1.176 | -0.554 | 0.01 | -0.072 | -0.072 | 0.05 | 0.6 | 0.55 | 0.49 | 0.14 | 0.53 | 0.43 |
| 11 | 114783924 | rs12804945 | 0.599 | 0.868 | 0.95 | -0.567 | -0.031 | -0.006 | -0.006 | 0 | 0.08 | 0 | 0.07 | 0.05 | 0.08 | 0.11 |
| 11 | 114788422 | rs7936187 | 0.327 | 0.831 | 1.27 | -0.57 | 0.003 | -0.01 | -0.01 | 0 | 0.22 | 0.15 | 0.32 | 0.66 | 0.23 | 0.29 |
| 11 | 114789286 | rs11215278 | 0.536 | 1.037 | 0.528 | -1.211 | 0.002 | -0.008 | -0.008 | 0.019 | 0.15 | 0.24 | 0.24 | 0.14 | 0.14 | 0.17 |
| 11 | 114794436 | rs2514457 | 0.077 | 1.089 | 0.046 | -1.117 | -0.018 | -0.009 | -0.009 | 1.75 | 0.84 | 0.86 | 0.84 | 0.8 | 0.76 | 0.72 |
| 11 | 114803157 | rs2514435 | 0.341 | 0.805 | 0.114 | -1.186 | -0.045 | 0.025 | 0.025 | 0 | 0.71 | 0.81 | 0.75 | 0.78 | 0.63 | 0.56 |
| 11 | 114810084 | rs17117772 | 0.518 | 0.588 | 0.289 | -1.323 | -0.028 | 0.034 | 0.034 | 0 | 0.26 | 0.43 | 0.36 | 0.68 | 0.23 | 0.28 |
| 11 | 114811893 | rs6589464 | -1.017 | 0.523 | 0.304 | -1.395 | 0.021 | -0.036 | -0.036 | 0 | 0.33 | 0.51 | 0.29 | 0.22 | 0.37 | 0.44 |
| 11 | 114818174 | rs10790059 | -0.903 | 0.53 | 0.327 | -1.392 | 0.011 | -0.018 | -0.018 | 1.054 | 0.32 | 0.45 | 0.29 | 0.22 | 0.37 | 0.44 |
| 11 | 114827203 | rs12798828 | -0.499 | 0.514 | 0.207 | -1.384 | -0.056 | 0.027 | 0.027 | 0 | 0.28 | 0.12 | 0.24 | 0.22 | 0.36 | 0.42 |
| 11 | 114828968 | rs10891773 | -0.801 | 0.513 | 0.304 | -1.384 | 0.012 | -0.022 | -0.022 | 0 | 0.32 | 0.45 | 0.28 | 0.22 | 0.36 | 0.41 |
| 11 | 114833710 | rs11600819 | -0.178 | 0.42 | 0.002 | -1.617 | 0.01 | 0.088 | 0.088 | 1.401 | 0.41 | 0.74 | 0.63 | 0.82 | 0.37 | 0.42 |
| 11 | 114839858 | rs4402329 | -1.207 | 0.445 | -0.041 | -1.659 | -0.007 | 0.109 | 0.109 | 0.833 | 0.63 | 0.82 | 0.8 | 1 | 0.66 | 0.76 |
| 11 | 114846596 | rs10891787 | -0.351 | -0.08 | 0.248 | -1.486 | 0.002 | 0.009 | 0.009 | 1.46 | 0.4 | 0.61 | 0.61 | 0.82 | 0.37 | 0.42 |
| 11 | 114848941 | rs17563461 | 0.212 | 0.214 | -0.015 | -1.706 | -0.101 | 0.059 | 0.059 | 0 | 0.29 | 0.07 | 0.25 | 0.17 | 0.42 | 0.35 |
| 11 | 114848998 | rs4573711 | -0.181 | 0.251 | -0.07 | -1.674 | 0.02 | 0.087 | 0.087 | 1.273 | 0.42 | 0.73 | 0.62 | 0.82 | 0.37 | 0.41 |
| 11 | 114851390 | rs12798976 | 0.38 | 0.45 | -0.402 | -1.478 | -0.003 | -0.008 | -0.008 | 0 | 0.34 | 0.37 | 0.43 | 0.7 | 0.32 | 0.39 |
| 11 | 114851809 | rs10790061 | -1.083 | 0.466 | -0.583 | -1.677 | 0.01 | 0.035 | 0.035 | 1.77 | 0.72 | 0.83 | 0.88 | 1 | 0.79 | 0.76 |
| 11 | 114852861 | rs4397885 | -1.088 | 0.474 | -0.58 | -1.673 | 0.009 | 0.035 | 0.035 | 1.909 | 0.72 | 0.83 | 0.87 | 1 | 0.79 | 0.76 |
| 11 | 114856645 | rs12786825 | -0.09 | 0.446 | -0.724 | -1.28 | -0.007 | -0.046 | -0.046 | 0 | 0.39 | 0.44 | 0.59 | 0.82 | 0.37 | 0.41 |
| 11 | 114859696 | rs7939241 | -0.616 | 0.362 | -1.397 | -1.189 | 0 | 0.116 | 0.116 | 1.116 | 0.75 | 0.93 | 0.89 | 1 | 0.81 | 0.78 |
| 11 | 114861050 | rs7944043 | -0.235 | 0.048 | -1.682 | -0.725 | 0.014 | 0.007 | 0.007 | 1.678 | 0.59 | 0.74 | 0.63 | 0.39 | 0.64 | 0.66 |
| 11 | 114869751 | rs7123550 | -0.726 | 0.132 | -1.88 | -0.542 | 0.008 | 0.152 | 0.152 | 0 | 0.5 | 0.81 | 0.67 | 0.74 | 0.46 | 0.61 |
| 11 | 114870096 | rs10891792 | -0.497 | -0.108 | -0.804 | 0.076 | 0.03 | 0.19 | 0.19 | 0 | 0.11 | 0.55 | 0.14 | 0.24 | 0.07 | 0.1 |
| 11 | 114876994 | rs4936313 | -0.797 | -0.158 | -0.595 | 0.026 | 0.033 | -0.036 | -0.036 | 0.329 | 0.45 | 0.83 | 0.6 | 0.5 | 0.4 | 0.48 |
| 11 | 114878901 | rs4938160 | -0.401 | -0.124 | -0.673 | -0.016 | 0.003 | -0.01 | -0.01 | 0.385 | 0.39 | 0.33 | 0.53 | 0.5 | 0.38 | 0.47 |
| 11 | 114882270 | rs6589470 | -0.372 | -0.092 | -0.66 | 0.019 | 0.003 | -0.01 | -0.01 | 0 | 0.39 | 0.33 | 0.53 | 0.5 | 0.38 | 0.47 |
| 11 | 114882970 | rs4938161 | -1.359 | -0.093 | -0.664 | 0.018 | -0.012 | 0.166 | 0.166 | 0 | 0.51 | 0.9 | 0.68 | 0.74 | 0.46 | 0.58 |
| 11 | 114883518 | rs4542447 | -0.526 | -0.217 | -0.51 | -0.117 | 0.003 | -0.011 | -0.011 | 0.784 | 0.39 | 0.33 | 0.53 | 0.5 | 0.38 | 0.47 |
| 11 | 114885153 | rs12419473 | 0.193 | -0.55 | -0.457 | -0.317 | 0.001 | 0.01 | 0.01 | 0 | 0.18 | 0.2 | 0.35 | 0.49 | 0.13 | 0.33 |
| 11 | 114895990 | rs7129245 | -1.203 | -0.553 | -0.485 | -0.331 | 0.003 | -0.01 | -0.01 | 4.125 | 0.39 | 0.33 | 0.53 | 0.5 | 0.38 | 0.47 |
| 11 | 114914107 | rs4373973 | 0.431 | -0.401 | -0.544 | -0.411 | 0.013 | -0.104 | -0.104 | 0 | 0.16 | 0.51 | 0.15 | 0.01 | 0.14 | 0.13 |
| 11 | 114915469 | rs7122390 | -0.419 | -0.418 | -0.71 | -0.355 | 0.001 | -0.107 | -0.107 | 1.152 | 0.26 | 0.6 | 0.23 | 0.01 | 0.27 | 0.15 |
| 11 | 114916450 | rs4131241 | -1.109 | -0.349 | -0.823 | -0.125 | 0.023 | -0.026 | -0.026 | 0 | 0.43 | 0.7 | 0.57 | 0.5 | 0.38 | 0.47 |
| 11 | 114920874 | rs11600288 | -0.13 | -0.325 | -0.667 | 0.378 | -0.005 | 0.002 | 0.002 | 0 | 0.1 | 0.04 | 0.16 | 0.12 | 0.11 | 0.25 |
| 11 | 114928468 | rs2507880 | -1.753 | -0.876 | 0.27 | 0.441 | 0.026 | -0.044 | -0.044 | 0 | 0.28 | 0.83 | 0.32 | 0.48 | 0.18 | 0.37 |

| | | | | | | | | | | | | | | | | |
|---|---|---|---|---|---|---|---|---|---|---|---|---|---|---|---|---|
| 11 | 114928727 | rs12222663 | 0.338 | -0.697 | 0.148 | 0.616 | -0.003 | 0.048 | 0.048 | 0 | 0.1 | 0.16 | 0.09 | 0.36 | 0.06 | 0.11 |
| 11 | 114932841 | rs1557282 | -1.012 | -0.772 | 0.371 | 0.67 | -0.005 | 0.019 | 0.019 | 0 | 0.2 | 0.26 | 0.25 | 0.46 | 0.14 | 0.35 |
| 11 | 114932960 | rs1557281 | -0.813 | -0.791 | 0.361 | 0.66 | 0.015 | -0.034 | -0.034 | 0.016 | 0.18 | 0.12 | 0.23 | 0.47 | 0.14 | 0.35 |
| 11 | 114934127 | rs978187 | -1.238 | -0.793 | -0.05 | 0.277 | 0.006 | 0.008 | 0.008 | 4.857 | 0.45 | 0.39 | 0.54 | 0.63 | 0.36 | 0.61 |
| 11 | 114935398 | rs11215342 | 1.246 | -0.834 | -0.797 | -0.328 | -0.098 | -0.053 | -0.053 | 0.162 | 0.28 | 0.05 | 0.27 | 0.36 | 0.43 | 0.27 |
| 11 | 114940699 | rs12576867 | 0.775 | -0.665 | -0.627 | -0.266 | -0.007 | 0.018 | 0.018 | 10.464 | 0.25 | 0.13 | 0.29 | 0.17 | 0.22 | 0.25 |
| 11 | 114944058 | rs2514351 | 0.649 | -0.851 | -0.373 | -0.041 | 0.007 | -0.003 | -0.003 | 0 | 0.34 | 0.62 | 0.31 | 0.34 | 0.43 | 0.29 |
| 11 | 114944083 | rs2513678 | -0.387 | -0.647 | -0.274 | 0.104 | 0.04 | -0.026 | -0.026 | 0 | 0.58 | 0.71 | 0.57 | 0.51 | 0.65 | 0.53 |
| 11 | 114944248 | rs982979 | 0.356 | -0.683 | -0.514 | 0.089 | 0 | 0.018 | 0.018 | 0 | 0.17 | 0.17 | 0.28 | 0.51 | 0.14 | 0.34 |
| 11 | 114945485 | rs2507881 | -1.662 | -0.691 | -0.689 | -0.094 | -0.007 | 0.151 | 0.151 | 0 | 0.76 | 0.98 | 0.86 | 0.99 | 0.8 | 0.89 |
| 11 | 114946119 | rs10790063 | 0.884 | -0.958 | -0.597 | -0.027 | -0.085 | 0.027 | 0.027 | 0.056 | 0.36 | 0.05 | 0.38 | 0.32 | 0.43 | 0.31 |
| 11 | 114958994 | rs1024344 | -0.03 | -1.176 | 0.076 | 0.488 | -0.001 | 0 | 0 | 0 | 0.48 | 0.5 | 0.64 | 0.55 | 0.51 | 0.57 |
| 11 | 114961537 | rs2507887 | -0.585 | -1.068 | -0.116 | 0.445 | -0.003 | 0.007 | 0.007 | 0 | 0.56 | 0.61 | 0.67 | 0.85 | 0.53 | 0.64 |
| 11 | 114981466 | rs12225506 | 1.029 | -0.484 | -0.928 | 0.63 | 0.013 | -0.008 | -0.008 | 0.202 | 0.31 | 0.05 | 0.19 | 0.32 | 0.26 | 0.22 |
| 11 | 114983177 | rs2514374 | -0.723 | -0.422 | -0.884 | 0.745 | -0.017 | -0.05 | -0.05 | 0 | 0.33 | 0.88 | 0.26 | 0.41 | 0.32 | 0.28 |
| 11 | 114983437 | rs12577640 | 0.468 | 0.028 | -0.774 | 0.434 | -0.02 | 0.002 | 0.002 | 0 | 0.18 | 0.11 | 0.15 | 0.14 | 0.26 | 0.22 |
| 11 | 114984572 | rs2513697 | 0.168 | -0.082 | -0.746 | 0.623 | -0.031 | -0.007 | -0.007 | 0 | 0.43 | 0.24 | 0.62 | 0.41 | 0.49 | 0.6 |
| 11 | 114995252 | rs2513689 | -0.079 | 0.396 | 0.067 | 0.906 | 0.029 | 0.006 | 0.006 | 0 | 0.82 | 0.69 | 0.84 | 0.85 | 0.78 | 0.81 |
| 11 | 114995398 | rs11215367 | 1.505 | 0.107 | -0.827 | 1.123 | -0.015 | -0.051 | -0.051 | 0 | 0.49 | 0.13 | 0.4 | 0.56 | 0.47 | 0.38 |
| 11 | 114996408 | rs9971533 | 1.044 | 0.238 | -0.74 | 1.193 | 0.026 | -0.023 | -0.023 | 0 | 0.13 | 0.08 | 0.04 | 0.17 | 0.08 | 0.11 |
| 11 | 114998601 | rs2427652 | 0.361 | 0.488 | -0.623 | 1.016 | -0.023 | -0.002 | -0.002 | 0 | 0.13 | 0.1 | 0.34 | 0.12 | 0.22 | 0.32 |
| 11 | 114999873 | rs2513676 | 0.532 | 0.117 | -0.443 | 1.026 | 0.009 | -0.004 | -0.004 | 0 | 0.65 | 0.34 | 0.76 | 0.68 | 0.69 | 0.7 |
| 11 | 115000162 | rs2427651 | 0.392 | 0.095 | -0.176 | 0.868 | -0.019 | -0.001 | -0.001 | 0.511 | 0.12 | 0.09 | 0.2 | 0.11 | 0.19 | 0.31 |
| 11 | 115001429 | rs17117977 | -0.257 | -0.911 | 0.216 | 0.967 | 0.007 | -0.011 | -0.011 | 0 | 0.21 | 0.44 | 0.18 | 0.23 | 0.2 | 0.11 |
| 11 | 115001635 | rs2427650 | -1.145 | -0.946 | 0.041 | 0.839 | -0.02 | 0.011 | 0.011 | 0.929 | 0.38 | 0.23 | 0.45 | 0.35 | 0.48 | 0.54 |
| 11 | 115004010 | rs7931589 | -0.185 | -0.829 | 0.322 | 0.95 | 0.014 | -0.041 | -0.041 | 0 | 0.29 | 0.17 | 0.22 | 0.61 | 0.23 | 0.24 |
| 11 | 115008175 | rs2008801 | 0.469 | -0.66 | 0.195 | 0.894 | -0.069 | -0.007 | -0.007 | 2.056 | 0.14 | 0.03 | 0.29 | 0.12 | 0.27 | 0.4 |
| 11 | 115013447 | rs12421648 | 1.738 | -0.914 | 0.45 | 0.793 | -0.008 | 0.073 | 0.073 | 0 | 0.42 | 0.19 | 0.39 | 0.19 | 0.44 | 0.32 |
| 11 | 115022088 | rs11215380 | -1.037 | -1.051 | 0.561 | 0.866 | -0.023 | -0.002 | -0.002 | 0 | 0.41 | 0.5 | 0.27 | 0.63 | 0.27 | 0.27 |
| 11 | 115022404 | rs12286929 | 1.192 | -0.957 | 0.558 | 0.874 | 0.005 | 0.046 | 0.046 | 0 | 0.51 | 0.58 | 0.47 | 0.26 | 0.48 | 0.38 |
| 11 | 115024539 | rs4359245 | -0.26 | -0.927 | 0.166 | 0.962 | 0.026 | -0.038 | -0.038 | 0.041 | 0.29 | 0.17 | 0.21 | 0.63 | 0.22 | 0.25 |
| 11 | 115032240 | rs11608104 | 1.109 | -0.964 | 0.432 | 1.029 | -0.009 | 0.004 | 0.004 | 1.493 | 0.72 | 0.26 | 0.52 | 0.78 | 0.68 | 0.62 |
| 11 | 115035485 | rs7131142 | 0.333 | -1.187 | 0.868 | 1.664 | -0.074 | 0.062 | 0.062 | 2.327 | 0.8 | 0.67 | 0.9 | 0.64 | 0.93 | 0.85 |
| 11 | 115040963 | rs17118023 | -0.227 | -1.042 | 1.091 | 1.584 | -0.008 | 0.028 | 0.028 | 0.055 | 0.22 | 0.45 | 0.18 | 0.28 | 0.19 | 0.13 |
| 11 | 115041589 | rs4936321 | 1.271 | -1.118 | 0.96 | 1.436 | -0.011 | -0.014 | -0.014 | 0 | 0.58 | 0.35 | 0.4 | 0.49 | 0.53 | 0.52 |
| 11 | 115043574 | rs17564430 | 0.447 | -1.168 | 0.902 | 1.369 | -0.068 | 0.001 | 0.001 | 1.307 | 0.15 | 0.06 | 0.33 | 0.13 | 0.26 | 0.34 |
| 11 | 115043888 | rs10891802 | 1.47 | -1.218 | 1.016 | 1.305 | -0.014 | -0.022 | -0.022 | 0 | 0.55 | 0.21 | 0.39 | 0.5 | 0.51 | 0.5 |
| 11 | 115044850 | rs1048932 | -0.996 | -1.157 | 1.34 | 1.321 | 0.016 | 0.004 | 0.004 | 0.496 | 0.35 | 0.39 | 0.48 | 0.42 | 0.43 | 0.46 |
| 11 | 115045237 | rs4445669 | 1.683 | -1.185 | 1.114 | 1.26 | -0.014 | 0.08 | 0.08 | 0 | 0.46 | 0.3 | 0.4 | 0.2 | 0.53 | 0.38 |
| 11 | 115053011 | rs11605461 | 1.802 | -1.192 | 1.106 | 1.285 | -0.011 | 0.072 | 0.072 | 0 | 0.45 | 0.31 | 0.4 | 0.19 | 0.51 | 0.37 |
| 11 | 115053808 | rs4938184 | 0.596 | -1.32 | 1.293 | 1.367 | 0.017 | 0.007 | 0.007 | 0.06 | 0.64 | 0.69 | 0.76 | 0.32 | 0.78 | 0.7 |
| 11 | 115069951 | rs10891807 | -0.273 | -1.278 | 0.857 | 1.445 | 0.011 | -0.015 | -0.015 | 0 | 0.19 | 0.15 | 0.13 | 0.39 | 0.08 | 0.18 |
| 11 | 115069973 | rs11215413 | -0.034 | -1.377 | 0.996 | 1.453 | -0.021 | 0.009 | 0.009 | 5.521 | 0.18 | 0.2 | 0.07 | 0.32 | 0.06 | 0.15 |
| 11 | 115074234 | rs12226198 | -1.032 | -1.536 | 0.731 | 1.332 | -0.008 | 0.006 | 0.006 | 0 | 0.06 | 0.1 | 0.12 | 0.07 | 0.05 | 0.03 |
| 11 | 115075436 | rs10128746 | -0.014 | -1.61 | 0.775 | 1.375 | 0.003 | -0.012 | -0.012 | 0 | 0.18 | 0.15 | 0.12 | 0.39 | 0.05 | 0.17 |
| 11 | 115077367 | rs11607801 | 1.253 | -1.27 | 0.673 | 0.883 | -0.015 | 0.085 | 0.085 | 0 | 0.46 | 0.32 | 0.4 | 0.18 | 0.53 | 0.38 |
| 11 | 115081563 | rs9645660 | -1.899 | -1.243 | 0.549 | 0.901 | -0.025 | 0.09 | 0.09 | 0 | 0.57 | 0.76 | 0.61 | 0.83 | 0.5 | 0.64 |
| 11 | 115083348 | rs11601041 | 1.701 | -1.321 | 0.214 | 0.498 | -0.118 | 0.216 | 0.216 | 0.085 | 0.41 | 0.07 | 0.37 | 0.1 | 0.49 | 0.35 |
| 11 | 115083875 | rs6589485 | -1.599 | -1.377 | 0.364 | 0.636 | -0.029 | 0.143 | 0.143 | 0.256 | 0.53 | 0.76 | 0.5 | 0.8 | 0.49 | 0.61 |
| 11 | 115087421 | rs11215424 | 0.446 | -1.35 | 0.194 | 0.671 | -0.085 | 0.026 | 0.026 | 1.38 | 0.17 | 0.03 | 0.27 | 0.11 | 0.27 | 0.34 |
| 11 | 115092997 | rs1938736 | -0.526 | -2.723 | 0.891 | 1.097 | -0.043 | 0.006 | 0.006 | 2.348 | 0.31 | 0.43 | 0.18 | 0.69 | 0.19 | 0.27 |
| 11 | 115101294 | rs11215433 | -2.134 | -2.336 | 1.015 | 1.218 | -0.046 | 0.013 | 0.013 | 0 | 0.12 | 0.34 | 0.21 | 0.1 | 0.07 | 0.05 |
| 11 | 115104172 | rs11215437 | -1.111 | -2.195 | 0.646 | 1.206 | -0.023 | -0.006 | -0.006 | 0 | 0.36 | 0.37 | 0.31 | 0.67 | 0.24 | 0.49 |
| 11 | 115104646 | rs4409873 | -0.98 | -2.463 | 0.486 | 1.079 | -0.01 | -0.013 | -0.013 | 0 | 0.87 | 0.92 | 0.88 | 0.78 | 0.84 | 0.88 |
| 11 | 115107984 | rs947802 | 1.534 | -2.654 | 0.715 | 0.867 | 0.052 | 0.089 | 0.089 | 0.032 | 0.5 | 0.41 | 0.54 | 0.09 | 0.62 | 0.4 |
| 11 | 115110463 | rs12421812 | -0.328 | -2.729 | 0.574 | 1.327 | 0 | -0.001 | -0.001 | 0.109 | 0.19 | 0.14 | 0.27 | 0.28 | 0.17 | 0.32 |
| 11 | 115111711 | rs11215438 | -0.671 | -2.733 | 0.576 | 1.344 | -0.008 | 0.007 | 0.007 | 3.029 | 0.28 | 0.28 | 0.17 | 0.63 | 0.2 | 0.26 |
| 11 | 115114732 | rs17118125 | -0.306 | -2.978 | 0.53 | 1.459 | 0.002 | -0.003 | -0.003 | 0 | 0.19 | 0.12 | 0.27 | 0.23 | 0.16 | 0.32 |
| 11 | 115115173 | rs11215445 | -1.275 | -2.982 | 0.558 | 1.236 | -0.044 | -0.005 | -0.005 | 3.453 | 0.36 | 0.39 | 0.34 | 0.63 | 0.21 | 0.47 |
| 11 | 115125119 | rs4936325 | -0.358 | -2.387 | 0.288 | 0.987 | 0.001 | 0.001 | 0.001 | 0.268 | 0.18 | 0.11 | 0.09 | 0.13 | 0.13 | 0.31 |
| 11 | 115131900 | rs6589490 | -1.83 | -2.401 | 0.165 | 0.696 | 0.109 | 0.224 | 0.224 | 0.119 | 0.48 | 0.82 | 0.49 | 0.88 | 0.38 | 0.58 |
| 11 | 115133557 | rs7121254 | 1.288 | -2.325 | 0.464 | 0.686 | 0.062 | 0.114 | 0.114 | 0.009 | 0.54 | 0.32 | 0.53 | 0.12 | 0.62 | 0.41 |
| 11 | 115136608 | rs4597099 | -1.794 | -2.306 | 0.399 | 0.683 | 0.082 | 0.137 | 0.137 | 0.075 | 0.48 | 0.69 | 0.47 | 0.88 | 0.37 | 0.58 |
| 11 | 115136714 | rs10891817 | 0.596 | -2.444 | 0.682 | 1.235 | -0.047 | 0.142 | 0.142 | 0 | 0.12 | 0.24 | 0.18 | 0.39 | 0.05 | 0.12 |
| 11 | 115137247 | rs10891819 | -1.397 | -2.52 | 0.823 | 1.143 | -0.07 | -0.027 | -0.027 | 0 | 0.33 | 0.55 | 0.34 | 0.7 | 0.22 | 0.25 |
| 11 | 115139851 | rs11215458 | 0.71 | -2.438 | 0.817 | 1.196 | -0.047 | 0.123 | 0.123 | 0 | 0.12 | 0.25 | 0.18 | 0.33 | 0.05 | 0.12 |
| 11 | 115140553 | rs7950069 | -0.705 | -2.232 | 0.369 | 0.068 | -0.067 | -0.007 | -0.007 | 0.035 | 0.3 | 0.49 | 0.28 | 0.65 | 0.2 | 0.25 |
| 11 | 115141456 | rs10790069 | -1.882 | -2.13 | 0.05 | 0.081 | 0.104 | 0.264 | 0.264 | 0.301 | 0.48 | 0.88 | 0.49 | 0.87 | 0.38 | 0.58 |
| 11 | 115149728 | rs7130530 | -0.445 | -0.746 | 0.574 | 0.311 | -0.003 | -0.007 | -0.007 | 0 | 0.17 | 0.2 | 0.16 | 0.22 | 0.16 | 0.34 |

| | | | | | | | | | | | | | | | | |
|---|---|---|---|---|---|---|---|---|---|---|---|---|---|---|---|---|
| 11 | 115152299 | rs7931895 | -1.421 | -0.795 | 0.567 | 0.26 | -0.001 | -0.06 | -0.06 | 0.079 | 0.27 | 0.43 | 0.28 | 0.22 | 0.28 | 0.35 |
| 11 | 115156628 | rs12804345 | -0.439 | -0.506 | 0.019 | 0.707 | 0.001 | -0.02 | -0.02 | 0.271 | 0.16 | 0.17 | 0.16 | 0.18 | 0.16 | 0.32 |
| 11 | 115162376 | rs11215467 | -0.116 | -0.664 | -0.089 | 0.548 | -0.004 | -0.011 | -0.011 | 0 | 0.14 | 0.11 | 0.11 | 0.12 | 0.1 | 0.29 |
| 11 | 115179071 | rs2157612 | -0.47 | -1.014 | 0.496 | 0.139 | -0.009 | -0.066 | -0.066 | 0 | 0.15 | 0.44 | 0.22 | 0.05 | 0.14 | 0.04 |
| 11 | 115180739 | rs7949084 | -0.718 | -0.859 | -0.682 | -0.538 | -0.16 | 0.044 | 0.044 | 0.246 | 0.75 | 0.93 | 0.71 | 0.82 | 0.57 | 0.79 |
| 11 | 115182330 | rs17118230 | 0.932 | -1.137 | -0.265 | -0.483 | -0.001 | -0.017 | -0.017 | 0 | 0.11 | 0.14 | 0.2 | 0.05 | 0.13 | 0.04 |
| 11 | 115183128 | rs12290790 | 0.526 | -1.127 | -0.25 | -0.478 | -0.004 | -0.04 | -0.04 | 0 | 0.13 | 0.24 | 0.21 | 0.05 | 0.13 | 0.04 |
| 11 | 115189228 | rs988873 | 0.568 | -2.224 | -0.802 | -0.086 | -0.025 | 0.022 | 0.022 | 0 | 0.32 | 0.31 | 0.29 | 0.45 | 0.2 | 0.46 |
| 11 | 115189959 | rs12577709 | 0.802 | -2.162 | -1.451 | -0.18 | 0.049 | -0.058 | -0.058 | 0 | 0.22 | 0.14 | 0.14 | 0.32 | 0.15 | 0.28 |
| 11 | 115191265 | rs4396320 | -0.813 | -1.807 | -1.44 | -0.509 | -0.12 | 0.103 | 0.103 | 0.108 | 0.69 | 0.94 | 0.66 | 0.82 | 0.5 | 0.78 |
| 11 | 115212725 | rs7105871 | -0.212 | -2.356 | -1.372 | -0.321 | 0.036 | 0.012 | 0.012 | 0.302 | 0.7 | 0.81 | 0.8 | 0.77 | 0.78 | 0.7 |
| 11 | 115212792 | rs7122020 | 0.341 | -2.23 | -0.867 | -0.117 | 0.036 | -0.017 | -0.017 | 0.082 | 0.27 | 0.36 | 0.31 | 0.17 | 0.44 | 0.2 |
| 11 | 115215743 | rs10790071 | -0.21 | -2.493 | -0.701 | -0.193 | 0.014 | 0.009 | 0.009 | 1.222 | 0.6 | 0.64 | 0.6 | 0.64 | 0.64 | 0.67 |
| 11 | 115220254 | rs1013416 | 0.804 | -2.527 | -0.5 | -0.177 | -0.023 | 0.02 | 0.02 | 0.003 | 0.35 | 0.36 | 0.3 | 0.48 | 0.2 | 0.45 |
| 11 | 115227997 | rs10891839 | -0.079 | -2.591 | -0.66 | -0.183 | 0.021 | 0.015 | 0.015 | 0.95 | 0.59 | 0.65 | 0.59 | 0.64 | 0.64 | 0.66 |
| 11 | 115239023 | rs10891842 | 0.286 | -3.165 | -1.016 | -0.286 | 0.034 | 0.073 | 0.073 | 2.522 | 0.4 | 0.19 | 0.37 | 0.24 | 0.57 | 0.53 |
| 11 | 115239397 | rs10160742 | -0.681 | -2.905 | -0.902 | -0.295 | -0.019 | 0.016 | 0.016 | 0.421 | 0.86 | 0.83 | 0.91 | 0.79 | 0.92 | 0.89 |
| 11 | 115268121 | rs183395 | 0.708 | -1.802 | -1.012 | -0.33 | -0.035 | 0.002 | 0.002 | 0 | 0.23 | 0.19 | 0.28 | 0.21 | 0.44 | 0.21 |
| 11 | 115279536 | rs2366914 | 0.418 | -1.978 | -1.054 | -0.428 | 0.053 | 0.008 | 0.008 | 0.107 | 0.48 | 0.66 | 0.6 | 0.6 | 0.36 | 0.54 |
| 11 | 115283483 | rs4938204 | 0.983 | -1.903 | -1.193 | -0.319 | -0.059 | -0.007 | -0.007 | 0 | 0.22 | 0.13 | 0.19 | 0.22 | 0.43 | 0.22 |
| 11 | 115283535 | rs220843 | -0.65 | -2.003 | -1.057 | -0.45 | 0.009 | -0.005 | -0.005 | 0 | 0.78 | 0.81 | 0.82 | 0.77 | 0.82 | 0.75 |
| 11 | 115285376 | rs220845 | 0.163 | -1.939 | -1.104 | -0.338 | -0.04 | 0.02 | 0.02 | 0.111 | 0.32 | 0.18 | 0.24 | 0.2 | 0.48 | 0.24 |
| 11 | 115294064 | rs220860 | -0.208 | -2.059 | -0.806 | -0.478 | -0.014 | -0.004 | -0.004 | 0 | 0.76 | 0.7 | 0.73 | 0.8 | 0.82 | 0.74 |
| 11 | 115300499 | rs7943417 | 0.485 | -1.483 | -0.604 | -0.432 | 0.007 | 0.003 | 0.003 | 1.463 | 0.52 | 0.58 | 0.59 | 0.58 | 0.37 | 0.62 |
| 11 | 115301871 | rs220836 | -0.128 | -1.631 | -0.742 | -0.563 | 0.003 | 0.003 | 0.003 | 3.262 | 0.73 | 0.75 | 0.79 | 0.82 | 0.78 | 0.79 |
| 11 | 115301952 | rs10488707 | 0.451 | -1.523 | -1.008 | -0.631 | -0.005 | 0.122 | 0.122 | 4.736 | 0.19 | 0.01 | 0.14 | 0.01 | 0.16 | 0.08 |
| 11 | 115304363 | rs7122693 | 0.558 | -1.465 | -0.98 | -0.621 | -0.041 | -0.007 | -0.007 | 0 | 0.22 | 0.14 | 0.2 | 0.24 | 0.44 | 0.17 |
| 11 | 115308919 | rs11215558 | 0.342 | -1.509 | -0.977 | -0.712 | 0.015 | 0.037 | 0.037 | 3.62 | 0.2 | 0.12 | 0.15 | 0.01 | 0.17 | 0.09 |
| 11 | 115312071 | rs220859 | -0.828 | -1.524 | -0.972 | -0.684 | 0.04 | 0.067 | 0.067 | 1.573 | 0.43 | 0.31 | 0.37 | 0.21 | 0.61 | 0.27 |
| 11 | 115320963 | rs220840 | -0.717 | -1.495 | -1.172 | -0.788 | 0.019 | 0.068 | 0.068 | 4.416 | 0.8 | 0.93 | 0.85 | 0.99 | 0.84 | 0.92 |
| 11 | 115324490 | rs10502202 | 1.434 | -2.058 | -1.209 | -0.925 | -0.005 | -0.001 | -0.001 | 0 | 0.36 | 0.31 | 0.36 | 0.52 | 0.2 | 0.48 |
| 11 | 115335621 | rs10891859 | 0.781 | -0.765 | -1.357 | -1.274 | -0.1 | 0.081 | 0.081 | 0 | 0.19 | 0.04 | 0.15 | 0.01 | 0.38 | 0.14 |
| 11 | 115336602 | rs314494 | -0.399 | -1.065 | -1.138 | -0.938 | -0.016 | 0.126 | 0.126 | 3.467 | 0.8 | 0.98 | 0.84 | 0.99 | 0.82 | 0.9 |
| 11 | 115341893 | rs10790077 | -0.316 | 0.041 | -0.608 | -0.323 | 0.031 | 0.015 | 0.015 | 0 | 0.23 | 0.32 | 0.28 | 0.41 | 0.39 | 0.23 |
| 11 | 115352894 | rs314512 | 1.134 | -0.51 | -0.536 | -0.269 | -0.011 | -0.038 | -0.038 | 0 | 0.08 | 0.15 | 0.04 | 0 | 0.05 | 0.02 |
| 11 | 115357372 | rs314515 | 1.336 | -0.504 | -0.654 | -0.265 | 0.002 | -0.009 | -0.009 | 0 | 0.08 | 0.06 | 0.04 | 0 | 0.05 | 0.02 |
| 11 | 115363443 | rs11215574 | 0.996 | -1.236 | -0.725 | -0.587 | 0.02 | -0.01 | -0.01 | 0.619 | 0.35 | 0.25 | 0.41 | 0.57 | 0.25 | 0.56 |
| 11 | 115374881 | rs2513640 | 1.284 | -0.721 | -0.689 | -0.353 | 0.007 | 0.003 | 0.003 | 1.545 | 0.08 | 0.04 | 0.04 | 0 | 0.05 | 0.02 |
| 11 | 115387449 | rs7101558 | 0.07 | -0.887 | -0.573 | -0.258 | -0.013 | -0.001 | -0.001 | 0 | 0.13 | 0.14 | 0.12 | 0.15 | 0.06 | 0.08 |
| 11 | 115395598 | rs907616 | 1.007 | -0.868 | -0.519 | -0.72 | 0.004 | 0.006 | 0.006 | 0.134 | 0.46 | 0.39 | 0.58 | 0.58 | 0.43 | 0.57 |
| 11 | 115396078 | rs754410 | 0.578 | -0.726 | -0.574 | -0.961 | -0.009 | -0.008 | -0.008 | 0.018 | 0.17 | 0.19 | 0.17 | 0.26 | 0.37 | 0.2 |
| 11 | 115397017 | rs11215586 | 0.802 | -0.693 | -0.602 | -0.988 | -0.018 | -0.013 | -0.013 | 0 | 0.16 | 0.14 | 0.17 | 0.25 | 0.37 | 0.19 |
| 11 | 115397853 | rs10750077 | -0.764 | -0.928 | -0.302 | -1.153 | -0.004 | -0.018 | -0.018 | 0 | 0.79 | 0.78 | 0.88 | 0.96 | 0.86 | 0.85 |
| 11 | 115400922 | rs7928019 | -0.797 | -0.921 | -0.356 | -1.05 | 0.01 | 0.01 | 0.01 | 0 | 0.79 | 0.83 | 0.89 | 0.94 | 0.86 | 0.84 |
| 11 | 115406310 | rs17118394 | 0.873 | -0.339 | -0.177 | -0.929 | 0.015 | 0.017 | 0.017 | 1.351 | 0.13 | 0.08 | 0.06 | 0 | 0.09 | 0.01 |
| 11 | 115408679 | rs898614 | -1.901 | -0.666 | -0.311 | -1.088 | 0.047 | 0.018 | 0.018 | 2.032 | 0.56 | 0.71 | 0.48 | 0.38 | 0.66 | 0.46 |
| 11 | 115412993 | rs2513638 | -1.927 | -0.66 | -0.337 | -1.083 | 0.048 | 0.016 | 0.016 | 1.894 | 0.56 | 0.72 | 0.48 | 0.39 | 0.66 | 0.46 |
| 11 | 115416391 | rs10502205 | 0.747 | -0.221 | -0.502 | -0.631 | -0.002 | -0.001 | -0.001 | 0 | 0.14 | 0.15 | 0.04 | 0.07 | 0.07 | 0.15 |
| 11 | 115418522 | rs11215593 | 0.779 | -0.106 | -0.544 | -0.646 | 0.011 | -0.005 | -0.005 | 0 | 0.12 | 0.11 | 0.14 | 0.23 | 0.32 | 0.2 |
| 11 | 115419788 | rs2513636 | -0.143 | -0.414 | -0.473 | -0.586 | -0.023 | 0 | 0 | 0 | 0.18 | 0.25 | 0.14 | 0.16 | 0.09 | 0.21 |
| 11 | 115425664 | rs11215598 | -1.143 | 0.378 | -0.5 | -0.464 | 0.004 | -0.001 | -0.001 | 0.026 | 0.3 | 0.3 | 0.3 | 0.32 | 0.44 | 0.26 |
| 11 | 115426934 | rs2219600 | 0.204 | 1.098 | -0.459 | -0.151 | 0.038 | 0.005 | 0.005 | 0 | 0.12 | 0.13 | 0.14 | 0.22 | 0.32 | 0.18 |
| 11 | 115428159 | rs10891864 | -0.071 | 1.279 | -0.454 | -0.682 | 0.001 | 0.003 | 0.003 | 3.752 | 0.37 | 0.39 | 0.55 | 0.55 | 0.41 | 0.52 |
| 11 | 115432255 | rs1870675 | -1.49 | 1.185 | -0.096 | -1.311 | 0.03 | 0.088 | 0.088 | 2.064 | 0.44 | 0.77 | 0.62 | 0.22 | 0.62 | 0.49 |
| 11 | 115433598 | rs2446884 | -1.5 | 1.185 | -0.101 | -1.319 | 0.032 | 0.109 | 0.109 | 2.147 | 0.44 | 0.8 | 0.62 | 0.22 | 0.62 | 0.49 |
| 11 | 115438254 | rs1460910 | -0.891 | 1.314 | 0.412 | -1.266 | 0.117 | 0.015 | 0.015 | 0.271 | 0.18 | 0.47 | 0.38 | 0.17 | 0.32 | 0.26 |
| 11 | 115443071 | rs2446886 | -1.573 | 1.27 | 0.399 | -1.33 | -0.029 | 0.035 | 0.035 | 0.007 | 0.37 | 0.76 | 0.49 | 0.22 | 0.47 | 0.46 |
| 11 | 115443873 | rs12291124 | 0.339 | 1.241 | 0.379 | -1.284 | -0.015 | -0.062 | -0.062 | 0.136 | 0.19 | 0.28 | 0.12 | 0.04 | 0.16 | 0.21 |
| 11 | 115449617 | rs1008950 | -1.003 | 1.739 | 0.528 | -0.929 | 0.132 | 0.022 | 0.022 | 0.009 | 0.17 | 0.44 | 0.38 | 0.18 | 0.32 | 0.26 |
| 11 | 115450288 | rs2446895 | -0.855 | 1.769 | 0.487 | -0.914 | 0.134 | 0.026 | 0.026 | 0 | 0.16 | 0.42 | 0.38 | 0.18 | 0.32 | 0.25 |
| 11 | 115455636 | rs7121755 | -0.044 | 1.811 | 0.085 | -0.979 | 0.069 | -0.015 | -0.015 | 0 | 0.18 | 0.25 | 0.46 | 0.13 | 0.41 | 0.22 |
| 11 | 115456281 | rs2513635 | -0.958 | 1.109 | 0.238 | -1.048 | 0.022 | -0.06 | -0.06 | 0 | 0.11 | 0.43 | 0.09 | 0.02 | 0.09 | 0.06 |
| 11 | 115460840 | rs7935076 | 1.625 | 1.291 | 0.034 | -1.029 | -0.011 | 0.008 | 0.008 | 0.688 | 0.08 | 0.05 | 0.13 | 0 | 0.14 | 0.03 |
| 11 | 115461802 | rs2446904 | -0.74 | 1.271 | 0.235 | -1.09 | 0.035 | -0.133 | -0.133 | 0 | 0.2 | 0.46 | 0.22 | 0.02 | 0.23 | 0.08 |
| 11 | 115463786 | rs976947 | 0.293 | 1.069 | -0.2 | -1.323 | 0.036 | -0.055 | -0.055 | 0 | 0.73 | 0.4 | 0.58 | 0.86 | 0.66 | 0.77 |
| 11 | 115469253 | rs10502207 | 1.255 | 0.246 | -1.311 | -1.413 | 0.037 | 0.045 | 0.045 | 3.097 | 0.42 | 0.02 | 0.37 | 0.74 | 0.3 | 0.46 |
| 11 | 115472695 | rs2515308 | -0.689 | 0.416 | -0.596 | -1.374 | -0.001 | -0.011 | -0.011 | 0.05 | 0.13 | 0.48 | 0.09 | 0.07 | 0.09 | 0.1 |
| 11 | 115475241 | rs2613459 | -1.479 | 0.355 | -1.658 | -1.332 | 0.063 | 0.132 | 0.132 | 1.89 | 0.54 | 0.98 | 0.63 | 0.21 | 0.69 | 0.55 |
| 11 | 115475966 | rs2613463 | 0.566 | 1.003 | -1.108 | -1.057 | -0.022 | 0.012 | 0.012 | 0 | 0.16 | 0.12 | 0.29 | 0.1 | 0.26 | 0.16 |

| | | | | | | | | | | | | | | | | |
|---|---|---|---|---|---|---|---|---|---|---|---|---|---|---|---|---|
| 11 | 115477034 | rs2613465 | 0.714 | 1.04 | -1.158 | -1.015 | -0.027 | 0.001 | 0.001 | 0 | 0.15 | 0.1 | 0.29 | 0.13 | 0.26 | 0.15 |
| 11 | 115482738 | rs2446879 | 1.27 | 0.915 | -1.078 | -1.238 | 0.011 | -0.036 | -0.036 | 1.297 | 0.1 | 0.16 | 0.12 | 0 | 0.15 | 0.03 |
| 11 | 115483879 | rs2846901 | -1.489 | -0.069 | -1.69 | -1.158 | 0.049 | -0.1 | -0.1 | 0.253 | 0.74 | 0.92 | 0.81 | 0.5 | 0.84 | 0.68 |
| 11 | 115485205 | rs2846900 | -0.861 | -0.149 | -1.616 | -1.168 | 0.009 | -0.027 | -0.027 | 0.015 | 0.65 | 0.73 | 0.69 | 0.5 | 0.69 | 0.65 |
| 11 | 115485361 | rs4938209 | 0.138 | -0.316 | -1.257 | -1.375 | -0.01 | -0.024 | -0.024 | 0 | 0.23 | 0.28 | 0.14 | 0.09 | 0.22 | 0.37 |
| 11 | 115488667 | rs11215633 | 1.62 | 0.024 | -1.571 | -1.29 | -0.083 | 0.003 | 0.003 | 0 | 0.15 | 0.01 | 0.26 | 0.11 | 0.22 | 0.16 |
| 11 | 115489486 | rs2509359 | -1.236 | -0.344 | -0.948 | -1.122 | -0.02 | 0.124 | 0.124 | 0.62 | 0.39 | 0.8 | 0.32 | 0.09 | 0.44 | 0.42 |
| 11 | 115489832 | rs1893165 | -0.783 | -0.374 | -0.935 | -1.014 | 0.018 | -0.076 | -0.076 | 0 | 0.73 | 0.9 | 0.8 | 0.48 | 0.8 | 0.64 |
| 11 | 115492807 | rs2155638 | 1.294 | -0.297 | -0.813 | -0.967 | -0.022 | 0.005 | 0.005 | 0 | 0.36 | 0.25 | 0.52 | 0.37 | 0.4 | 0.26 |
| 11 | 115494674 | rs2155634 | 1.474 | -0.227 | -0.849 | -0.916 | -0.072 | -0.018 | -0.018 | 0 | 0.36 | 0.11 | 0.51 | 0.37 | 0.4 | 0.27 |
| 11 | 115497151 | rs10891874 | 0.471 | -0.36 | -0.626 | -1.002 | 0.031 | 0.015 | 0.015 | 0.503 | 0.54 | 0.35 | 0.63 | 0.42 | 0.65 | 0.52 |
| 11 | 115498833 | rs2155633 | -0.665 | -0.396 | -0.648 | -0.908 | 0.062 | -0.098 | -0.098 | 0 | 0.72 | 0.92 | 0.82 | 0.48 | 0.84 | 0.65 |
| 11 | 115501343 | rs3949060 | 0.573 | -0.302 | -0.461 | -0.959 | 0.007 | 0.003 | 0.003 | 0.152 | 0.52 | 0.41 | 0.67 | 0.46 | 0.6 | 0.43 |
| 11 | 115509573 | rs12295787 | -0.017 | -0.835 | -0.223 | -1.032 | 0.006 | -0.031 | -0.031 | 0 | 0.18 | 0.28 | 0.13 | 0.04 | 0.24 | 0.24 |
| 11 | 115517016 | rs1698198 | -0.165 | -0.942 | -0.353 | -0.957 | 0.023 | -0.014 | -0.014 | 0 | 0.64 | 0.74 | 0.71 | 0.48 | 0.74 | 0.64 |
| 11 | 115519503 | rs1698193 | 0.784 | -2.163 | 0.28 | -0.301 | 0.073 | -0.02 | -0.02 | 0 | 0.63 | 0.27 | 0.75 | 0.46 | 0.84 | 0.57 |
| 11 | 115521706 | rs1698157 | -0.39 | -2.103 | -0.029 | -0.241 | 0.075 | -0.093 | -0.093 | 0.367 | 0.76 | 0.92 | 0.84 | 0.53 | 0.89 | 0.71 |
| 11 | 115523910 | rs1698185 | -0.395 | -2.096 | -0.024 | -0.238 | 0.056 | -0.066 | -0.066 | 0 | 0.75 | 0.89 | 0.84 | 0.53 | 0.89 | 0.71 |
| 11 | 115524137 | rs2455494 | -1.023 | -0.89 | 0.911 | -0.117 | 0.015 | -0.079 | -0.079 | 0 | 0.2 | 0.41 | 0.13 | 0.04 | 0.25 | 0.25 |
| 11 | 115524651 | rs2515314 | -1.106 | -0.88 | 0.918 | -0.104 | 0.026 | -0.08 | -0.08 | 0 | 0.2 | 0.41 | 0.14 | 0.05 | 0.27 | 0.28 |
| 11 | 115525562 | rs7926090 | -0.622 | -1.249 | 0.771 | 0.087 | -0.005 | 0.03 | 0.03 | 0.112 | 0.08 | 0.32 | 0.05 | 0.06 | 0.03 | 0.1 |
| 11 | 115533052 | rs2455503 | -1.404 | -1.22 | -0.296 | -0.101 | -0.001 | -0.025 | -0.025 | 0 | 0.26 | 0.7 | 0.18 | 0.06 | 0.26 | 0.34 |
| 11 | 115539650 | rs4938215 | 0.135 | -1.088 | -0.414 | -0.071 | -0.027 | 0.025 | 0.025 | 0 | 0.12 | 0.05 | 0.08 | 0.03 | 0.22 | 0.23 |
| 11 | 115542348 | rs10488715 | 1.499 | -1.491 | -0.851 | 0.15 | 0.046 | -0.032 | -0.032 | 5.898 | 0.54 | 0.15 | 0.65 | 0.48 | 0.6 | 0.33 |
| 11 | 115542933 | rs1502076 | -0.295 | -1.501 | -0.886 | 0.253 | 0.027 | -0.062 | -0.062 | 0 | 0.8 | 0.91 | 0.83 | 0.56 | 0.87 | 0.68 |
| 11 | 115546746 | rs1698205 | 1.165 | -2.273 | -0.945 | 0.302 | 0.053 | -0.032 | -0.032 | 0.02 | 0.53 | 0.16 | 0.66 | 0.5 | 0.6 | 0.33 |
| 11 | 115557967 | rs1784973 | 1.37 | -2.369 | -0.998 | 0.291 | 0.055 | -0.033 | -0.033 | 0.625 | 0.51 | 0.06 | 0.64 | 0.47 | 0.6 | 0.3 |
| 11 | 115559386 | rs1784972 | 0.435 | -2.286 | -0.728 | 0.267 | 0.041 | -0.032 | -0.032 | 0 | 0.67 | 0.26 | 0.74 | 0.6 | 0.82 | 0.55 |
| 11 | 115561779 | rs10488714 | -0.108 | -2.329 | -0.795 | 0.29 | -0.017 | 0.018 | 0.018 | 0 | 0.13 | 0.11 | 0.09 | 0.08 | 0.22 | 0.25 |
| 11 | 115563433 | rs1784965 | 0.655 | -2.192 | -0.738 | 0.332 | 0.05 | -0.046 | -0.046 | 0 | 0.65 | 0.2 | 0.71 | 0.6 | 0.77 | 0.53 |
| 11 | 115566576 | rs905244 | -1.411 | -1.865 | 0.017 | 0.158 | -0.02 | -0.029 | -0.029 | 0 | 0.3 | 0.59 | 0.22 | 0.15 | 0.24 | 0.3 |
| 11 | 115570349 | rs11826326 | -1.065 | -1.566 | 0.072 | 0.099 | 0.001 | -0.015 | -0.015 | 0 | 0.28 | 0.72 | 0.17 | 0.08 | 0.25 | 0.35 |
| 11 | 115570540 | rs883419 | 1.359 | -1.904 | -0.365 | 0.418 | 0.044 | 0.092 | 0.092 | 0 | 0.5 | 0.16 | 0.58 | 0.38 | 0.57 | 0.33 |
| 11 | 115574995 | rs12295914 | -0.586 | -1.802 | 0.27 | 0.344 | -0.032 | -0.119 | -0.119 | 0 | 0.25 | 0.47 | 0.14 | 0.05 | 0.21 | 0.26 |
| 11 | 115579062 | rs1016853 | -1.544 | -1.79 | -0.402 | 0.437 | 0.032 | 0.063 | 0.063 | 0 | 0.48 | 0.75 | 0.42 | 0.61 | 0.43 | 0.67 |
| 11 | 115580001 | rs1784941 | -0.201 | -1.176 | -0.38 | 1.137 | -0.043 | -0.139 | -0.139 | 0 | 0.21 | 0.02 | 0.18 | 0.53 | 0.18 | 0.29 |
| 11 | 115594740 | rs1785015 | -1.761 | -1.582 | 0.881 | 1.033 | -0.044 | 0.037 | 0.037 | 0 | 0.14 | 0.21 | 0.08 | 0.24 | 0.07 | 0.13 |
| 11 | 115595946 | rs1698187 | -1.325 | -1.588 | 0.876 | 1.03 | -0.007 | 0.005 | 0.005 | 0 | 0.12 | 0.1 | 0.06 | 0.23 | 0.07 | 0.13 |
| 11 | 115598377 | rs1236910 | -1.966 | -1.589 | 0.896 | 1.029 | -0.043 | 0.045 | 0.045 | 0 | 0.17 | 0.32 | 0.1 | 0.26 | 0.11 | 0.24 |
| 11 | 115612971 | rs17118741 | -1.971 | -1.578 | 0.816 | 1.003 | -0.045 | 0.049 | 0.049 | 0 | 0.17 | 0.34 | 0.1 | 0.26 | 0.11 | 0.23 |
| 11 | 115617042 | rs2285252 | -2.419 | -1.676 | 0.846 | 0.54 | -0.038 | -0.019 | -0.019 | 0 | 0.26 | 0.61 | 0.17 | 0.17 | 0.17 | 0.27 |
| 11 | 115618087 | rs10891888 | -2.393 | -1.666 | 0.832 | 0.525 | -0.045 | -0.026 | -0.026 | 0 | 0.26 | 0.59 | 0.16 | 0.17 | 0.17 | 0.27 |
| 11 | 115619704 | rs17118750 | -2.114 | -1.214 | 0.99 | 0.826 | 0.006 | 0.09 | 0.09 | 0.02 | 0.14 | 0.45 | 0.12 | 0.17 | 0.08 | 0.24 |
| 11 | 115622993 | rs6589497 | -2.235 | -0.807 | 0.135 | 0.515 | -0.008 | -0.011 | -0.011 | 0 | 0.14 | 0.24 | 0.12 | 0.09 | 0.1 | 0.1 |
| 11 | 115638755 | rs12273878 | -1.792 | -0.673 | 0.346 | 0.726 | -0.004 | 0.025 | 0.025 | 0 | 0.08 | 0.26 | 0.06 | 0.1 | 0.03 | 0.11 |
| 11 | 115655434 | rs1393275 | -2.061 | -0.773 | 0.231 | 0.592 | -0.015 | -0.002 | -0.002 | 0 | 0.11 | 0.28 | 0.08 | 0.1 | 0.06 | 0.13 |
| 11 | 115662064 | rs4938219 | 0.033 | -1.215 | -1.143 | 0.449 | -0.016 | -0.089 | -0.089 | 0.968 | 0.26 | 0.02 | 0.48 | 0.64 | 0.25 | 0.18 |
| 11 | 115675044 | rs11215686 | -0.052 | -1.103 | -1.395 | -0.264 | -0.019 | 0.038 | 0.038 | 1.522 | 0.39 | 0.26 | 0.26 | 0.26 | 0.48 | 0.46 |
| 11 | 115685475 | rs11215690 | -0.597 | -1.754 | -0.908 | -0.603 | -0.002 | -0.001 | -0.001 | 0 | 0.55 | 0.54 | 0.42 | 0.43 | 0.59 | 0.57 |
| 11 | 115722659 | rs7940134 | -0.045 | -1.707 | -0.931 | -0.575 | -0.02 | -0.022 | -0.022 | 0 | 0.06 | 0.23 | 0.02 | 0 | 0.02 | 0 |
| 11 | 115743487 | rs10502208 | -0.581 | -1.45 | -1.251 | -0.757 | -0.038 | 0.048 | 0.048 | 0 | 0.6 | 0.79 | 0.75 | 0.71 | 0.55 | 0.57 |
| 11 | 115753077 | rs569462 | -0.484 | -1.413 | -1.255 | -0.343 | -0.05 | 0.034 | 0.034 | 0.099 | 0.57 | 0.86 | 0.74 | 0.67 | 0.51 | 0.53 |
| 11 | 115757874 | rs4938220 | 0.91 | -1.66 | -1.521 | -0.202 | -0.028 | -0.149 | -0.149 | 0 | 0.35 | 0.02 | 0.46 | 0.47 | 0.31 | 0.24 |
| 11 | 115760507 | rs2186703 | -0.396 | -1.469 | -0.525 | 0.107 | -0.006 | 0.13 | 0.13 | 0 | 0.07 | 0.28 | 0.14 | 0.18 | 0.03 | 0.13 |
| 11 | 115761165 | rs639053 | -0.007 | -1.604 | 0.133 | 0.006 | 0.022 | 0.007 | 0.007 | 1.681 | 0.52 | 0.34 | 0.58 | 0.47 | 0.44 | 0.37 |
| 11 | 115764853 | rs17118846 | -0.841 | -0.856 | 0.243 | 0.098 | 0.032 | 0.081 | 0.081 | 0 | 0.09 | 0.28 | 0.17 | 0.18 | 0.11 | 0.14 |
| 11 | 115766855 | rs595450 | -0.152 | -1.255 | 0.077 | 0.22 | -0.004 | -0.006 | -0.006 | 0 | 0.48 | 0.44 | 0.51 | 0.4 | 0.42 | 0.38 |
| 11 | 115768256 | rs2155266 | -0.036 | -1.203 | -0.357 | 0.246 | 0.017 | 0.027 | 0.027 | 0.533 | 0.46 | 0.34 | 0.49 | 0.39 | 0.41 | 0.37 |
| 11 | 115771144 | rs7129139 | -1.44 | -0.707 | 0.23 | -0.044 | -0.009 | -0.083 | -0.083 | 0.007 | 0.15 | 0.64 | 0.13 | 0 | 0.13 | 0.02 |
| 11 | 115774774 | rs10891905 | -0.292 | -1.387 | -0.033 | -0.148 | 0.025 | 0.002 | 0.002 | 0 | 0.34 | 0.18 | 0.3 | 0.32 | 0.29 | 0.37 |
| 11 | 115777037 | rs11215724 | -1.162 | -1.272 | 0.099 | -0.058 | 0.046 | 0.06 | 0.06 | 0 | 0.48 | 0.67 | 0.41 | 0.32 | 0.41 | 0.39 |
| 11 | 115782465 | rs7126737 | -1.039 | -1.291 | 0.325 | 0.01 | 0.006 | 0.008 | 0.008 | 0 | 0.47 | 0.54 | 0.41 | 0.32 | 0.41 | 0.38 |
| 11 | 115787411 | rs11215728 | -0.979 | -1.014 | 0.642 | 0.49 | 0.006 | 0.001 | 0.001 | 0 | 0.44 | 0.63 | 0.48 | 0.45 | 0.41 | 0.42 |
| 11 | 115791218 | rs1262592 | -1.037 | -0.553 | 1.006 | 0.426 | 0.01 | 0 | 0 | 0 | 0.46 | 0.4 | 0.5 | 0.5 | 0.43 | 0.5 |
| 11 | 115792775 | rs374729 | -0.249 | -0.895 | 0.821 | 0.421 | 0.022 | -0.037 | -0.037 | 0 | 0.33 | 0.19 | 0.36 | 0.47 | 0.29 | 0.42 |
| 11 | 115794809 | rs435016 | 0.143 | -0.798 | 0.485 | 0.526 | -0.005 | -0.023 | -0.023 | 0 | 0.08 | 0.02 | 0.18 | 0.3 | 0.07 | 0.27 |
| 11 | 115798610 | rs7941722 | -0.824 | -0.568 | 0.931 | 0.336 | 0.102 | -0.019 | -0.019 | 0 | 0.08 | 0.36 | 0.12 | 0 | 0.12 | 0.02 |
| 11 | 115802441 | rs367164 | -1.323 | -0.692 | 0.041 | 0.459 | 0.062 | 0.005 | 0.005 | 0.376 | 0.49 | 0.88 | 0.51 | 0.51 | 0.41 | 0.5 |
| 11 | 115825017 | rs372453 | -0.783 | -0.736 | 0.282 | 0.639 | -0.011 | 0.001 | 0.001 | 0.036 | 0.46 | 0.45 | 0.47 | 0.57 | 0.38 | 0.5 |

| | | | | | | | | | | | | | | | | |
|---|---|---|---|---|---|---|---|---|---|---|---|---|---|---|---|---|
| 11 | 115836632 | rs877601 | -0.135 | -0.208 | 0.161 | 0.599 | -0.012 | -0.013 | -0.013 | 0 | 0.54 | 0.75 | 0.57 | 0.47 | 0.53 | 0.69 |
| 11 | 115844466 | rs6589519 | -0.857 | -0.588 | 1.167 | 0.194 | -0.022 | -0.012 | -0.012 | 0.268 | 0.09 | 0.3 | 0.04 | 0.04 | 0.04 | 0.13 |
| 11 | 115851152 | rs6589520 | -1.063 | -0.662 | 1.2 | 0.192 | -0.032 | -0.015 | -0.015 | 0.066 | 0.09 | 0.28 | 0.05 | 0.05 | 0.04 | 0.14 |
| 11 | 115857374 | rs17119029 | -2.126 | -0.769 | 1.594 | 0.11 | 0.018 | 0.006 | 0.006 | 0 | 0.06 | 0.33 | 0.05 | 0 | 0.02 | 0 |
| 11 | 115867672 | rs11215750 | -1.894 | -0.758 | 1.553 | 0.129 | -0.058 | -0.032 | -0.032 | 0 | 0.12 | 0.59 | 0.07 | 0.04 | 0.04 | 0.14 |
| 11 | 115872716 | rs7928212 | -1.272 | -0.641 | 1.455 | 0.166 | -0.02 | 0.003 | 0.003 | 0 | 0.1 | 0.47 | 0.07 | 0.04 | 0.04 | 0.14 |
| 11 | 115905730 | rs11215766 | -0.549 | -0.209 | 0.832 | -0.043 | 0.002 | 0.042 | 0.042 | 0.421 | 0.49 | 0.59 | 0.36 | 0.28 | 0.49 | 0.32 |
| 11 | 115907433 | rs431736 | 0.029 | -0.168 | 0.324 | -0.183 | -0.05 | 0.065 | 0.065 | 0.173 | 0.42 | 0.15 | 0.31 | 0.3 | 0.47 | 0.31 |
| 11 | 115928958 | rs11215773 | 0.347 | -0.343 | 0.267 | -0.151 | -0.017 | 0.033 | 0.033 | 0.364 | 0.52 | 0.22 | 0.67 | 0.62 | 0.55 | 0.58 |
| 11 | 115933686 | rs1557451 | 0.132 | 0.106 | 0.575 | -0.203 | -0.004 | -0.004 | -0.004 | 5.959 | 0.57 | 0.55 | 0.65 | 0.58 | 0.51 | 0.61 |
| 11 | 115941731 | rs720036 | -0.156 | -0.138 | 0.541 | -0.273 | -0.049 | -0.072 | -0.072 | 0 | 0.41 | 0.17 | 0.34 | 0.42 | 0.51 | 0.41 |
| 11 | 115942776 | rs7933547 | 0.073 | -0.165 | 0.594 | -0.344 | -0.014 | -0.014 | -0.014 | 0.006 | 0.57 | 0.63 | 0.63 | 0.59 | 0.51 | 0.59 |
| 11 | 115943253 | rs1426391 | -0.106 | -0.08 | 0.234 | -0.427 | -0.074 | -0.067 | -0.067 | 0 | 0.4 | 0.16 | 0.34 | 0.41 | 0.48 | 0.41 |
| 11 | 115946614 | rs1078751 | -0.602 | -0.558 | 0.543 | -0.855 | -0.001 | -0.016 | -0.016 | 0.017 | 0.11 | 0.12 | 0.06 | 0 | 0.11 | 0.11 |
| 11 | 115948410 | rs1648364 | 0.645 | -1.257 | 0.909 | -1.16 | -0.054 | -0.113 | -0.113 | 0 | 0.78 | 0.43 | 0.85 | 0.98 | 0.85 | 0.92 |
| 11 | 115953897 | rs11824574 | -0.455 | -1.219 | 0.98 | -1.144 | -0.031 | -0.082 | -0.082 | 0 | 0.18 | 0.25 | 0.11 | 0.03 | 0.13 | 0.08 |
| 11 | 115955049 | rs17119159 | 0.226 | -1.168 | 0.517 | -1.06 | 0.003 | -0.031 | -0.031 | 0 | 0.08 | 0.14 | 0.06 | 0 | 0.08 | 0.08 |
| 11 | 115963831 | rs1791914 | 1.504 | -1.033 | -0.151 | -1.082 | -0.084 | -0.152 | -0.152 | 0 | 0.35 | 0.03 | 0.56 | 0.55 | 0.39 | 0.54 |
| 11 | 115964809 | rs12271395 | -1.697 | -1.635 | 0.838 | -1.556 | -0.02 | -0.084 | -0.084 | 0.011 | 0.27 | 0.63 | 0.17 | 0.03 | 0.16 | 0.18 |
| 11 | 115977405 | rs12275683 | -0.23 | -1.301 | 0.941 | -1.251 | 0.001 | -0.061 | -0.061 | 0.029 | 0.12 | 0.24 | 0.08 | 0 | 0.08 | 0.11 |
| 11 | 115978271 | rs2217084 | 0.421 | -1.62 | 0.915 | -1.609 | -0.011 | -0.05 | -0.05 | 0 | 0.2 | 0.25 | 0.11 | 0 | 0.11 | 0.04 |
| 11 | 115978780 | rs11215797 | -0.506 | -1.531 | 0.639 | -1.578 | -0.009 | -0.081 | -0.081 | 0 | 0.31 | 0.32 | 0.16 | 0 | 0.19 | 0.13 |
| 11 | 115982197 | rs491198 | -0.877 | -1.438 | 0.211 | -1.512 | -0.019 | -0.116 | -0.116 | 0 | 0.42 | 0.47 | 0.2 | 0 | 0.21 | 0.18 |
| 11 | 115989597 | rs573713 | -0.007 | -1.269 | -0.191 | -1.353 | -0.041 | -0.111 | -0.111 | 0 | 0.24 | 0.35 | 0.13 | 0 | 0.11 | 0.02 |
| 11 | 116000031 | rs1240162 | -1.429 | -0.423 | -0.714 | -0.145 | 0.011 | -0.023 | -0.023 | 0.016 | 0.4 | 0.72 | 0.29 | 0.41 | 0.31 | 0.33 |
| 11 | 116000880 | rs568523 | -1.417 | -0.402 | -0.705 | -0.135 | 0.011 | -0.023 | -0.023 | 0.263 | 0.4 | 0.72 | 0.29 | 0.41 | 0.31 | 0.33 |
| 11 | 116002530 | rs1783229 | -0.789 | 0.746 | -0.776 | 0.941 | -0.002 | 0.015 | 0.015 | 0 | 0.16 | 0.16 | 0.17 | 0.4 | 0.21 | 0.15 |
| 11 | 116004783 | rs572075 | -0.622 | -0.204 | -0.706 | 0.557 | -0.023 | 0.018 | 0.018 | 0 | 0.38 | 0.52 | 0.36 | 0.53 | 0.34 | 0.25 |
| 11 | 116005170 | rs576425 | 0.359 | -0.513 | -1.3 | 0.091 | 0.052 | 0.009 | 0.009 | 0.131 | 0.51 | 0.17 | 0.49 | 0.45 | 0.56 | 0.61 |
| 11 | 116008515 | rs483058 | -1.048 | -0.583 | -0.739 | -0.195 | -0.011 | 0.018 | 0.018 | 0.117 | 0.23 | 0.31 | 0.26 | 0.34 | 0.19 | 0.19 |
| 11 | 116010298 | rs1783225 | -0.171 | -1.585 | -0.853 | -0.848 | -0.013 | -0.049 | -0.049 | 0 | 0.67 | 0.34 | 0.83 | 1 | 0.89 | 0.92 |
| 11 | 116034732 | rs2852811 | 0.076 | -0.957 | -0.762 | -0.457 | -0.04 | -0.07 | -0.07 | 2.313 | 0.13 | 0.02 | 0.22 | 0.29 | 0.17 | 0.11 |
| 11 | 116035371 | rs2844282 | 0.106 | -0.961 | -0.757 | -0.433 | -0.025 | -0.067 | -0.067 | 0.569 | 0.13 | 0.02 | 0.22 | 0.29 | 0.15 | 0.1 |
| 11 | 116040364 | rs1961329 | 0.271 | -0.403 | -0.188 | -0.267 | 0.011 | 0.023 | 0.023 | 0 | 0.46 | 0.41 | 0.48 | 0.6 | 0.54 | 0.7 |
| 11 | 116042048 | rs1783238 | 0.315 | -0.349 | -0.402 | -0.195 | -0.001 | 0.003 | 0.003 | 0 | 0.42 | 0.4 | 0.46 | 0.6 | 0.51 | 0.67 |
| 11 | 116049846 | rs12288365 | -1.279 | -0.249 | 0.566 | 0.127 | 0.013 | 0.024 | 0.024 | 0.449 | 0.14 | 0.4 | 0.12 | 0.15 | 0.13 | 0.13 |
| 11 | 116050271 | rs1783235 | 1.167 | -0.902 | 0.581 | -0.071 | 0.009 | -0.001 | -0.001 | 0.363 | 0.61 | 0.34 | 0.67 | 0.56 | 0.71 | 0.76 |
| 11 | 116051211 | rs2852820 | 0.251 | -0.855 | 0.467 | -0.104 | -0.012 | -0.07 | -0.07 | 0.271 | 0.15 | 0.04 | 0.16 | 0.28 | 0.13 | 0.06 |
| 11 | 116054107 | rs2367632 | -0.642 | -0.842 | 1.403 | -0.765 | 0.076 | 0.09 | 0.09 | 0 | 0.11 | 0.45 | 0.14 | 0.16 | 0.13 | 0.13 |
| 11 | 116055191 | rs12419815 | 0.28 | -0.813 | 0.962 | -0.754 | -0.006 | -0.013 | -0.013 | 0.274 | 0.07 | 0 | 0.07 | 0.08 | 0.07 | 0.09 |
| 11 | 116060076 | rs2852825 | 1.201 | -0.982 | 1.117 | -0.9 | 0.016 | 0.027 | 0.027 | 0.14 | 0.66 | 0.26 | 0.78 | 0.83 | 0.77 | 0.8 |
| 11 | 116067498 | rs11215838 | -1.343 | -0.897 | 1.458 | -0.928 | -0.021 | -0.077 | -0.077 | 0.261 | 0.27 | 0.63 | 0.14 | 0.07 | 0.16 | 0.14 |
| 11 | 116069918 | rs10790107 | -0.368 | -0.841 | 1.578 | -0.875 | -0.023 | -0.112 | -0.112 | 0.086 | 0.17 | 0.43 | 0.09 | 0 | 0.12 | 0.03 |
| 11 | 116072627 | rs10891925 | 0.572 | -0.858 | 0.788 | -0.91 | 0.059 | 0.087 | 0.087 | 0 | 0.53 | 0.26 | 0.68 | 0.75 | 0.68 | 0.65 |
| 11 | 116073347 | rs4245164 | 0.387 | -1.004 | 1.087 | -0.955 | -0.008 | -0.019 | -0.019 | 0.002 | 0.54 | 0.49 | 0.71 | 0.84 | 0.74 | 0.75 |
| 11 | 116075948 | rs4471467 | 1.233 | -0.917 | 0.733 | -0.889 | -0.014 | 0.012 | 0.012 | 0 | 0.42 | 0.14 | 0.59 | 0.66 | 0.58 | 0.54 |
| 11 | 116076882 | rs1894070 | 1.145 | -1.394 | 0.767 | -0.843 | 0.043 | 0.02 | 0.02 | 0 | 0.43 | 0.14 | 0.61 | 0.66 | 0.65 | 0.55 |
| 11 | 116077726 | rs3996912 | 0.864 | -1.385 | 0.739 | -0.875 | 0.041 | 0.05 | 0.05 | 0 | 0.48 | 0.32 | 0.63 | 0.74 | 0.67 | 0.66 |
| 11 | 116090852 | rs7104334 | 0.365 | -1.096 | -0.148 | -1.092 | 0.011 | 0.011 | 0.011 | 0 | 0.55 | 0.59 | 0.72 | 0.83 | 0.74 | 0.75 |
| 11 | 116094471 | rs12283886 | 0.734 | -1.023 | -0.133 | -1.048 | 0.001 | 0.003 | 0.003 | 0 | 0.07 | 0.04 | 0.06 | 0.07 | 0.08 | 0.08 |
| 11 | 116096249 | rs10790112 | -0.456 | -1.183 | -0.053 | -1.005 | -0.014 | -0.009 | -0.009 | 0 | 0.69 | 0.71 | 0.77 | 0.83 | 0.83 | 0.78 |
| 11 | 116096761 | rs10891929 | -0.074 | -1.244 | -0.746 | -1.128 | 0.008 | 0.029 | 0.029 | 0 | 0.21 | 0.15 | 0.12 | 0.1 | 0.18 | 0.12 |
| 11 | 116098634 | rs10790114 | -0.086 | -0.994 | -0.713 | -0.559 | 0.031 | 0.033 | 0.033 | 0 | 0.7 | 0.8 | 0.78 | 0.86 | 0.84 | 0.79 |
| 11 | 116103543 | rs10891932 | 0.369 | -0.866 | -1.066 | -0.347 | 0.086 | 0.03 | 0.03 | 0 | 0.59 | 0.23 | 0.68 | 0.7 | 0.76 | 0.65 |
| 11 | 116104358 | rs4936337 | -0.193 | -0.098 | -1.284 | -1.332 | 0.009 | 0.111 | 0.111 | 0 | 0.48 | 0.67 | 0.56 | 0.91 | 0.47 | 0.68 |
| 11 | 116105632 | rs11215855 | 0.478 | 0.254 | -0.681 | -0.729 | 0.007 | -0.022 | -0.022 | 1.91 | 0.65 | 0.53 | 0.62 | 0.66 | 0.61 | 0.62 |
| 11 | 116108902 | rs920447 | 0.269 | -0.098 | -0.728 | -0.203 | -0.001 | -0.002 | -0.002 | 0 | 0.15 | 0.12 | 0.21 | 0.2 | 0.15 | 0.11 |
| 11 | 116118912 | rs1009746 | 1.188 | -0.461 | -0.871 | -0.336 | -0.02 | -0.008 | -0.008 | 0 | 0.43 | 0.2 | 0.4 | 0.52 | 0.52 | 0.67 |
| 11 | 116126460 | rs10891936 | 1.397 | -0.301 | -1.01 | -0.21 | -0.131 | -0.123 | -0.123 | 0 | 0.44 | 0.02 | 0.39 | 0.42 | 0.5 | 0.53 |
| 11 | 116129754 | rs2367635 | -0.314 | -0.43 | -0.333 | -0.582 | -0.03 | -0.074 | -0.074 | 0.042 | 0.85 | 0.69 | 0.9 | 0.99 | 0.92 | 0.95 |
| 11 | 116130782 | rs10891938 | -1.063 | -0.145 | -0.035 | -0.223 | 0.016 | 0.036 | 0.036 | 0.616 | 0.29 | 0.36 | 0.46 | 0.47 | 0.35 | 0.28 |
| 11 | 116134891 | rs11215863 | 0.124 | 0.086 | 0.032 | -0.117 | -0.006 | 0.021 | 0.021 | 0 | 0.1 | 0.12 | 0.16 | 0.25 | 0.11 | 0.17 |
| 11 | 116139119 | rs4938246 | 0.805 | 0.117 | -0.141 | -0.201 | -0.001 | 0.007 | 0.007 | 0 | 0.6 | 0.53 | 0.5 | 0.52 | 0.59 | 0.68 |
| 11 | 116150361 | rs7103346 | 0.691 | 0.212 | -0.053 | -0.15 | -0.002 | -0.021 | -0.021 | 0 | 0.56 | 0.5 | 0.57 | 0.67 | 0.6 | 0.75 |
| 11 | 116154489 | rs1550440 | 0.392 | -0.016 | 0.204 | -0.127 | 0.006 | -0.016 | -0.016 | 0 | 0.67 | 0.66 | 0.67 | 0.81 | 0.65 | 0.78 |
| 11 | 116157247 | rs6589536 | -0.362 | 0.159 | -0.014 | 0.426 | -0.006 | 0.059 | 0.059 | 0 | 0.26 | 0.37 | 0.24 | 0.51 | 0.25 | 0.25 |
| 11 | 116157765 | rs10891950 | -0.312 | 0.184 | 0.004 | 0.405 | -0.021 | 0.113 | 0.113 | 0 | 0.24 | 0.45 | 0.24 | 0.51 | 0.22 | 0.26 |
| 11 | 116158912 | rs4938250 | -1.828 | 0.197 | -0.731 | 0.3 | -0.099 | -0.057 | -0.057 | 0 | 0.61 | 0.98 | 0.59 | 0.68 | 0.59 | 0.48 |
| 11 | 116177072 | rs1440188 | -1.759 | 0.245 | -0.586 | 0.39 | -0.101 | -0.073 | -0.073 | 0 | 0.56 | 0.95 | 0.58 | 0.6 | 0.56 | 0.44 |

| | | | | | | | | | | | | | | | | |
|---|---|---|---|---|---|---|---|---|---|---|---|---|---|---|---|---|
| 11 | 116177442 | rs10790129 | -1.009 | 0.249 | -0.104 | 0.291 | -0.001 | 0.001 | 0.001 | 0.413 | 0.33 | 0.7 | 0.37 | 0.16 | 0.34 | 0.2 |
| 11 | 116181066 | rs12421721 | 0.108 | 0.701 | -0.592 | 0.502 | -0.032 | 0.001 | 0.001 | 0.896 | 0.17 | 0.06 | 0.23 | 0.16 | 0.18 | 0.15 |
| 11 | 116181859 | rs10502218 | -0.278 | 0.565 | -0.276 | 0.517 | 0.025 | -0.016 | -0.016 | 0 | 0.21 | 0.27 | 0.3 | 0.16 | 0.29 | 0.17 |
| 11 | 116182191 | rs10502217 | -1.172 | 0.154 | -1.013 | 0.526 | -0.024 | 0.005 | 0.005 | 1.536 | 0.5 | 0.87 | 0.56 | 0.59 | 0.54 | 0.43 |
| 11 | 116184920 | rs10450623 | -0.674 | -0.971 | -0.315 | 0.562 | -0.034 | -0.01 | -0.01 | 0 | 0.31 | 0.58 | 0.18 | 0.27 | 0.18 | 0.21 |
| 11 | 116187828 | rs10750088 | -1.788 | -0.769 | -0.626 | 0.814 | 0.024 | 0.039 | 0.039 | 0.286 | 0.52 | 0.86 | 0.43 | 0.65 | 0.45 | 0.47 |
| 11 | 116189247 | rs1371693 | 0.145 | -1.035 | -0.127 | 0.698 | 0.025 | 0 | 0 | 0.06 | 0.23 | 0.16 | 0.13 | 0.25 | 0.15 | 0.19 |
| 11 | 116190880 | rs4938258 | -2.151 | -1.039 | -0.472 | 0.575 | -0.025 | 0.022 | 0.022 | 0.082 | 0.57 | 0.87 | 0.52 | 0.67 | 0.47 | 0.52 |
| 11 | 116197720 | rs930430 | -0.616 | -1.035 | -0.464 | 0.559 | -0.012 | -0.036 | -0.036 | 0 | 0.2 | 0.13 | 0.31 | 0.39 | 0.27 | 0.26 |
| 11 | 116202789 | rs12806755 | 1.059 | -0.945 | -0.569 | 0.524 | -0.016 | 0.028 | 0.028 | 0 | 0.05 | 0 | 0.07 | 0 | 0.06 | 0.05 |
| 11 | 116205687 | rs10790137 | -1.485 | -0.944 | -0.559 | 0.537 | -0.027 | -0.049 | -0.049 | 0 | 0.24 | 0.11 | 0.33 | 0.39 | 0.3 | 0.28 |
| 11 | 116207032 | rs11215900 | -1.132 | -0.888 | -0.769 | 0.413 | 0.025 | 0.008 | 0.008 | 0 | 0.44 | 0.7 | 0.47 | 0.63 | 0.42 | 0.44 |
| 11 | 116208396 | rs6589541 | -1.73 | -0.597 | -1.123 | 0.588 | -0.045 | 0.011 | 0.011 | 0.601 | 0.58 | 0.9 | 0.52 | 0.66 | 0.48 | 0.51 |
| 11 | 116210119 | rs4936347 | -0.832 | -0.361 | -0.383 | 0.64 | -0.034 | -0.043 | -0.043 | 0 | 0.12 | 0.01 | 0.26 | 0.23 | 0.14 | 0.17 |
| 11 | 116214112 | rs12272993 | -1.415 | -0.838 | 0.47 | 0.178 | -0.007 | -0.056 | -0.056 | 1.313 | 0.12 | 0.35 | 0.07 | 0.02 | 0.08 | 0.06 |
| 11 | 116216713 | rs567681 | -1.584 | -1.147 | 0.241 | 0.211 | -0.001 | -0.001 | -0.001 | 0 | 0.44 | 0.56 | 0.46 | 0.51 | 0.45 | 0.51 |
| 11 | 116219158 | rs552104 | -1.432 | -1.1 | 0.459 | 0.18 | -0.004 | 0.003 | 0.003 | 0.048 | 0.42 | 0.47 | 0.43 | 0.51 | 0.38 | 0.49 |
| 11 | 116219458 | rs554865 | -1.344 | -1.164 | 0.429 | 0.307 | -0.004 | -0.012 | -0.012 | 2.83 | 0.38 | 0.46 | 0.4 | 0.34 | 0.34 | 0.38 |
| 11 | 116224775 | rs17119495 | -0.635 | -1.112 | 0.711 | 0.448 | 0.033 | 0.091 | 0.091 | 0 | 0.08 | 0.21 | 0.19 | 0.18 | 0.12 | 0.13 |
| 11 | 116226845 | rs564046 | -0.795 | -1.243 | 0.755 | 0.271 | 0 | -0.018 | -0.018 | 3.213 | 0.18 | 0.16 | 0.1 | 0.04 | 0.16 | 0.1 |
| 11 | 116227259 | rs497898 | -1.568 | -1.273 | 0.762 | 0.276 | -0.007 | -0.013 | -0.013 | 0 | 0.26 | 0.35 | 0.25 | 0.23 | 0.22 | 0.26 |
| 11 | 116227687 | rs482795 | -1.849 | -1.256 | 0.679 | 0.412 | -0.006 | 0.016 | 0.016 | 0 | 0.32 | 0.38 | 0.31 | 0.42 | 0.28 | 0.42 |
| 11 | 116231840 | rs513935 | -1.619 | -1.573 | 0.638 | 0.197 | -0.003 | -0.004 | -0.004 | 0 | 0.4 | 0.55 | 0.42 | 0.39 | 0.4 | 0.27 |
| 11 | 116232173 | rs12799683 | 0.587 | -1.52 | 0.566 | 0.318 | 0.039 | 0.123 | 0.123 | 0 | 0.06 | 0.15 | 0.18 | 0.21 | 0.12 | 0.1 |
| 11 | 116232487 | rs12804415 | -0.944 | -1.57 | 0.64 | 0.297 | 0.037 | 0.136 | 0.136 | 0 | 0.09 | 0.27 | 0.19 | 0.21 | 0.12 | 0.12 |
| 11 | 116232577 | rs11605788 | 0.558 | -1.646 | 0.483 | 0.202 | 0.011 | -0.005 | -0.005 | 0 | 0.09 | 0.06 | 0.06 | 0.11 | 0.06 | 0.06 |
| 11 | 116233398 | rs10502215 | 1.01 | -1.479 | 0.38 | 0.126 | -0.002 | -0.012 | -0.012 | 0 | 0.06 | 0.05 | 0.04 | 0.01 | 0.04 | 0.09 |
| 11 | 116233857 | rs574529 | -1.431 | -1.256 | 0.37 | 0.108 | 0.011 | 0.013 | 0.013 | 0 | 0.47 | 0.56 | 0.31 | 0.42 | 0.37 | 0.44 |
| 11 | 116236775 | rs526151 | -1.681 | -1.14 | 0.409 | 0.041 | 0.006 | 0.011 | 0.011 | 0 | 0.51 | 0.66 | 0.46 | 0.6 | 0.5 | 0.5 |
| 11 | 116238034 | rs491111 | -1.224 | -1.316 | 0.606 | -0.623 | -0.008 | -0.014 | -0.014 | 0 | 0.42 | 0.49 | 0.27 | 0.38 | 0.37 | 0.4 |
| 11 | 116242693 | rs11215905 | 1.152 | -1.078 | -0.513 | -0.856 | 0 | 0.002 | 0.002 | 0 | 0.46 | 0.53 | 0.67 | 0.63 | 0.53 | 0.5 |
| 11 | 116243868 | rs2510873 | -0.61 | -1.677 | -0.316 | -0.54 | 0.012 | -0.051 | -0.051 | 0.43 | 0.09 | 0.23 | 0.08 | 0 | 0.1 | 0.03 |
| 11 | 116245350 | rs487962 | 1.061 | -1.763 | -0.497 | -0.509 | -0.003 | 0 | 0 | 0.028 | 0.52 | 0.52 | 0.76 | 0.65 | 0.74 | 0.58 |
| 11 | 116250285 | rs556727 | 0.365 | -1.306 | -0.554 | -0.063 | 0.05 | -0.026 | -0.026 | 0 | 0.74 | 0.92 | 0.87 | 0.68 | 0.83 | 0.72 |
| 11 | 116251906 | rs12277809 | -0.844 | -0.569 | 0.169 | -0.387 | -0.019 | -0.027 | -0.027 | 0 | 0.07 | 0.29 | 0.06 | 0 | 0.01 | 0.02 |
| 11 | 116253844 | rs574951 | -0.74 | 0.454 | -0.335 | -0.572 | 0.032 | -0.039 | -0.039 | 0 | 0.33 | 0.45 | 0.38 | 0.22 | 0.46 | 0.28 |
| 11 | 116255458 | rs543925 | -1.101 | 0.42 | -0.557 | -0.704 | 0 | -0.005 | -0.005 | 0 | 0.1 | 0.11 | 0.19 | 0.06 | 0.18 | 0.08 |
| 11 | 116261676 | rs531860 | -0.443 | 0.196 | -0.267 | -1.202 | -0.012 | -0.07 | -0.07 | 5.532 | 0.15 | 0.29 | 0.09 | 0.03 | 0.12 | 0.09 |
| 11 | 116265015 | rs4938264 | -0.799 | 0.061 | 0.056 | -1.223 | 0.022 | 0.056 | 0.056 | 0.09 | 0.06 | 0.32 | 0.14 | 0.06 | 0.03 | 0.07 |
| 11 | 116265069 | rs537538 | -1.193 | 0.338 | -1.191 | -1.16 | -0.1 | -0.186 | -0.186 | 0.31 | 0.75 | 0.98 | 0.73 | 0.54 | 0.71 | 0.5 |
| 11 | 116269032 | rs9645661 | -0.987 | 0.961 | -0.913 | -2.355 | 0.023 | -0.087 | -0.087 | 0 | 0.28 | 0.47 | 0.32 | 0.1 | 0.29 | 0.15 |
| 11 | 116269470 | rs7102851 | -0.997 | 0.888 | -0.994 | -2.401 | 0.016 | -0.07 | -0.07 | 0.207 | 0.28 | 0.41 | 0.32 | 0.1 | 0.31 | 0.16 |
| 11 | 116271094 | rs532227 | -1.556 | 0.138 | -1.449 | -2.711 | 0.02 | 0.146 | 0.146 | 0 | 0.52 | 0.87 | 0.67 | 0.24 | 0.59 | 0.28 |
| 11 | 116271226 | rs533240 | -1.929 | -0.088 | -1.52 | -2.741 | -0.039 | 0.003 | 0.003 | 0 | 0.63 | 0.88 | 0.68 | 0.35 | 0.62 | 0.4 |
| 11 | 116272339 | rs4938268 | -1.592 | 0.329 | -1.156 | -2.639 | -0.016 | 0.14 | 0.14 | 0 | 0.51 | 0.79 | 0.59 | 0.27 | 0.47 | 0.26 |
| 11 | 116273511 | rs505128 | -1.346 | 0.509 | -0.696 | -2.702 | -0.003 | -0.002 | -0.002 | 0 | 0.36 | 0.37 | 0.44 | 0.16 | 0.4 | 0.22 |
| 11 | 116275617 | rs4586206 | -0.648 | 0.684 | -0.399 | -2.635 | 0.052 | -0.067 | -0.067 | 0.071 | 0.19 | 0.37 | 0.28 | 0.06 | 0.23 | 0.1 |
| 11 | 116276086 | rs573126 | -1.352 | 0.443 | -0.567 | -2.643 | 0.01 | -0.096 | -0.096 | 1.236 | 0.34 | 0.49 | 0.34 | 0.1 | 0.34 | 0.18 |
| 11 | 116277079 | rs1940143 | -1.052 | 0.335 | -0.646 | -2.694 | 0.003 | -0.061 | -0.061 | 0 | 0.33 | 0.62 | 0.37 | 0.06 | 0.38 | 0.17 |
| 11 | 116277665 | rs553344 | -1.32 | 0.397 | -0.765 | -2.523 | -0.08 | 0.218 | 0.218 | 0.011 | 0.39 | 0.89 | 0.37 | 0.04 | 0.42 | 0.19 |
| 11 | 116277748 | rs7481152 | -1.678 | -0.329 | 0.704 | -1.557 | 0.055 | 0.001 | 0.001 | 0 | 0.09 | 0.34 | 0.12 | 0.05 | 0.09 | 0.18 |
| 11 | 116279191 | rs4938271 | 0.49 | -0.623 | 0.438 | -1.579 | -0.004 | -0.025 | -0.025 | 0 | 0.9 | 0.84 | 0.89 | 0.94 | 0.88 | 0.95 |
| 11 | 116279427 | rs580965 | 1.728 | -1.527 | 0.317 | -1.91 | 0.078 | 0.123 | 0.123 | 0 | 0.58 | 0.24 | 0.67 | 0.8 | 0.67 | 0.79 |
| 11 | 116280252 | rs508994 | 0.439 | -1.549 | 0.275 | -1.877 | -0.004 | -0.023 | -0.023 | 0.055 | 0.87 | 0.83 | 0.87 | 0.94 | 0.86 | 0.95 |
| 11 | 116281169 | rs538359 | 0.981 | -1.449 | 0.546 | -1.851 | -0.012 | -0.053 | -0.053 | 0 | 0.83 | 0.76 | 0.84 | 0.94 | 0.83 | 0.89 |
| 11 | 116283516 | rs11823029 | -0.667 | -1.627 | 0.881 | -1.945 | -0.01 | -0.003 | -0.003 | 0 | 0.14 | 0.18 | 0.1 | 0.14 | 0.1 | 0.08 |
| 11 | 116285930 | rs576670 | 2.401 | -1.75 | 0.769 | -2.031 | 0.092 | 0.17 | 0.17 | 0 | 0.57 | 0.18 | 0.66 | 0.81 | 0.66 | 0.8 |
| 11 | 116287909 | rs10891973 | -1.736 | -1.674 | 1.062 | -1.957 | -0.041 | -0.049 | -0.049 | 0 | 0.25 | 0.41 | 0.15 | 0.14 | 0.18 | 0.09 |
| 11 | 116292263 | rs574778 | 0.656 | -1.518 | 0.579 | -1.861 | 0.009 | 0.051 | 0.051 | 0 | 0.81 | 0.93 | 0.85 | 0.91 | 0.8 | 0.94 |
| 11 | 116295758 | rs574233 | 1.092 | -1.583 | 0.773 | -1.884 | -0.018 | -0.035 | -0.035 | 1.145 | 0.69 | 0.66 | 0.76 | 0.81 | 0.74 | 0.81 |
| 11 | 116297397 | rs483367 | 1.232 | -1.517 | 0.393 | -1.852 | -0.044 | -0.074 | -0.074 | 0.289 | 0.66 | 0.52 | 0.75 | 0.81 | 0.74 | 0.81 |
| 11 | 116298367 | rs490592 | 0.75 | -0.773 | 0.022 | -1.463 | -0.012 | -0.069 | -0.069 | 0 | 0.8 | 0.73 | 0.82 | 0.96 | 0.78 | 0.94 |
| 11 | 116304152 | rs12787144 | -0.441 | 0.218 | -0.586 | -1.111 | 0.018 | -0.017 | -0.017 | 0 | 0.31 | 0.54 | 0.38 | 0.26 | 0.37 | 0.32 |
| 11 | 116304983 | rs2174880 | -0.822 | -0.021 | -1.277 | -0.633 | -0.013 | 0.068 | 0.068 | 0 | 0.5 | 0.84 | 0.57 | 0.4 | 0.55 | 0.37 |
| 11 | 116311508 | rs532107 | -0.764 | 0.367 | -0.743 | -0.255 | 0.001 | 0 | 0 | 0 | 0.22 | 0.23 | 0.22 | 0.22 | 0.24 | 0.16 |
| 11 | 116312872 | rs548890 | -1.367 | 0.38 | 0.223 | -0.198 | -0.001 | -0.001 | -0.001 | 0 | 0.32 | 0.61 | 0.39 | 0.33 | 0.34 | 0.21 |
| 11 | 116314641 | rs550646 | -1.184 | 0.432 | 0.853 | -0.248 | 0.01 | 0.033 | 0.033 | 0.836 | 0.26 | 0.49 | 0.35 | 0.32 | 0.27 | 0.21 |
| 11 | 116316617 | rs17119667 | 0.619 | 0.438 | 0.73 | -0.224 | -0.001 | -0.032 | -0.032 | 0.37 | 0.06 | 0.11 | 0.03 | 0 | 0.07 | 0.01 |
| 11 | 116316777 | rs489164 | -1.091 | 1.133 | 0.194 | -0.34 | -0.01 | -0.005 | -0.005 | 1.049 | 0.39 | 0.3 | 0.44 | 0.39 | 0.46 | 0.25 |

| | | | | | | | | | | | | | | | | |
|---|---|---|---|---|---|---|---|---|---|---|---|---|---|---|---|---|
| 11 | 116316813 | rs505390 | 0.678 | 0.549 | -0.586 | -0.567 | 0.003 | -0.014 | -0.014 | 0.267 | 0.28 | 0.35 | 0.26 | 0.11 | 0.33 | 0.25 |
| 11 | 116321980 | rs573052 | 0.748 | 0.704 | -0.745 | -0.253 | -0.012 | -0.013 | -0.013 | 0 | 0.15 | 0.07 | 0.19 | 0.21 | 0.18 | 0.14 |
| 11 | 116322236 | rs7106296 | 0.743 | 0.629 | -0.764 | -0.187 | -0.002 | -0.001 | -0.001 | 0 | 0.16 | 0.13 | 0.17 | 0.12 | 0.17 | 0.07 |
| 11 | 116322769 | rs11602880 | 0.89 | 0.603 | -0.747 | -0.18 | -0.001 | 0.001 | 0.001 | 0 | 0.17 | 0.15 | 0.17 | 0.11 | 0.17 | 0.05 |
| 11 | 116323001 | rs542671 | 0.328 | 0.542 | -0.514 | 0.305 | -0.001 | 0.025 | 0.025 | 0 | 0.16 | 0.18 | 0.2 | 0.37 | 0.18 | 0.18 |
| 11 | 116325442 | rs491575 | -0.871 | 0.913 | -0.79 | 0.498 | 0.006 | 0.005 | 0.005 | 0 | 0.45 | 0.31 | 0.47 | 0.55 | 0.57 | 0.37 |
| 11 | 116325990 | rs552043 | -1.352 | 0.919 | -0.848 | 0.713 | -0.001 | 0 | 0 | 5.812 | 0.81 | 0.82 | 0.83 | 0.87 | 0.82 | 0.95 |
| 11 | 116326138 | rs12419822 | 0.711 | 0.636 | -0.713 | 0.995 | -0.003 | 0.009 | 0.009 | 0 | 0.08 | 0.04 | 0.17 | 0.28 | 0.18 | 0.09 |
| 11 | 116326686 | rs522098 | 1.567 | -0.986 | -0.588 | 0.469 | 0.05 | -0.018 | -0.018 | 0 | 0.26 | 0.11 | 0.22 | 0.28 | 0.12 | 0.45 |
| 11 | 116328380 | rs12418858 | -1.057 | -0.69 | -0.131 | 0.553 | -0.011 | 0.036 | 0.036 | 0 | 0.09 | 0.25 | 0.13 | 0.1 | 0.05 | 0.1 |
| 11 | 116329043 | rs574223 | -1.104 | -0.501 | -0.45 | 0.379 | -0.07 | -0.052 | -0.052 | 0.246 | 0.64 | 0.82 | 0.69 | 0.53 | 0.49 | 0.73 |
| 11 | 116329090 | rs570745 | -0.975 | -0.535 | -0.282 | 0.367 | -0.042 | -0.038 | -0.038 | 0 | 0.63 | 0.77 | 0.68 | 0.45 | 0.48 | 0.72 |
| 11 | 116330610 | rs514112 | 0.314 | -0.611 | -0.318 | 0.032 | -0.011 | 0.029 | 0.029 | 0 | 0.19 | 0.12 | 0.22 | 0.02 | 0.22 | 0.12 |
| 11 | 116330732 | rs513083 | -0.832 | -0.561 | 0.045 | 0.45 | -0.007 | -0.029 | -0.029 | 0.505 | 0.57 | 0.81 | 0.64 | 0.52 | 0.42 | 0.66 |
| 11 | 116331582 | rs17119701 | -0.283 | -0.94 | 0.61 | 0.714 | -0.017 | -0.035 | -0.035 | 0 | 0.07 | 0.15 | 0.05 | 0 | 0.04 | 0.04 |
| 11 | 116338814 | rs12289284 | -1.085 | -0.562 | 0.887 | 0.909 | 0.019 | 0.061 | 0.061 | 0.339 | 0.09 | 0.26 | 0.25 | 0.17 | 0.1 | 0.1 |
| 11 | 116338899 | rs1508098 | -1.056 | -0.77 | 0.793 | 1.193 | 0.083 | -0.011 | -0.011 | 0 | 0.49 | 0.77 | 0.55 | 0.53 | 0.32 | 0.65 |
| 11 | 116343848 | rs7110240 | -0.82 | -0.552 | 0.41 | 1.406 | -0.019 | -0.002 | -0.002 | 0 | 0.45 | 0.53 | 0.51 | 0.47 | 0.3 | 0.63 |
| 11 | 116344404 | rs17119743 | -0.441 | -0.643 | 0.444 | 1.334 | -0.007 | 0.004 | 0.004 | 0 | 0.38 | 0.46 | 0.49 | 0.54 | 0.27 | 0.58 |
| 11 | 116348066 | rs11215954 | -0.452 | -0.633 | 0.41 | 1.094 | -0.032 | -0.006 | -0.006 | 0.137 | 0.41 | 0.55 | 0.48 | 0.41 | 0.25 | 0.62 |
| 11 | 116348168 | rs2102405 | -1.346 | 0.584 | 0.962 | 1.012 | 0.053 | 0.04 | 0.04 | 0.422 | 0.08 | 0.33 | 0.18 | 0.05 | 0.1 | 0.12 |
| 11 | 116352008 | rs3017780 | -0.006 | 0.344 | 0.53 | 0.984 | -0.016 | -0.012 | -0.012 | 0 | 0.39 | 0.52 | 0.4 | 0.27 | 0.3 | 0.22 |
| 11 | 116352339 | rs1354975 | -1.544 | 0.489 | 1.131 | 0.827 | 0.055 | 0.071 | 0.071 | 0 | 0.16 | 0.53 | 0.27 | 0.25 | 0.17 | 0.13 |
| 11 | 116352523 | rs556436 | 0.827 | 0.73 | 1.081 | 0.858 | 0.017 | 0.033 | 0.033 | 7.388 | 0.32 | 0.25 | 0.19 | 0.1 | 0.24 | 0.14 |
| 11 | 116352622 | rs495033 | -1.214 | 0.653 | 0.732 | 1.069 | 0.012 | -0.056 | -0.056 | 2.437 | 0.89 | 0.99 | 0.94 | 0.84 | 0.91 | 0.81 |
| 11 | 116354466 | rs549298 | 0.621 | 1.024 | 1.276 | 0.408 | 0.022 | 0.039 | 0.039 | 5.999 | 0.34 | 0.24 | 0.2 | 0.1 | 0.25 | 0.18 |
| 11 | 116355310 | rs11215958 | -0.269 | 1.074 | 1.012 | 0.474 | 0.022 | 0.022 | 0.022 | 0 | 0.08 | 0.2 | 0.05 | 0.18 | 0.11 | 0.07 |
| 11 | 116355860 | rs573554 | -0.694 | 1.089 | 1.018 | 0.468 | 0.016 | 0.077 | 0.077 | 4.684 | 0.51 | 0.86 | 0.48 | 0.37 | 0.42 | 0.34 |
| 11 | 116358711 | rs17119760 | -0.567 | 1.111 | 1.285 | 0.836 | 0.059 | 0.062 | 0.062 | 0 | 0.06 | 0.21 | 0.05 | 0.18 | 0.09 | 0.07 |
| 11 | 116363446 | rs7128900 | -1.514 | 0.689 | 0.932 | 0.565 | 0.03 | 0.016 | 0.016 | 0 | 0.07 | 0.24 | 0.19 | 0.05 | 0.06 | 0.09 |
| 11 | 116369171 | rs925151 | -0.058 | 1.006 | 0.138 | 0.44 | 0 | -0.001 | -0.001 | 0 | 0.48 | 0.47 | 0.45 | 0.75 | 0.44 | 0.65 |
| 11 | 116373451 | rs875025 | -1.504 | 1.329 | 0.179 | 1.257 | 0.031 | 0.057 | 0.057 | 0.562 | 0.13 | 0.64 | 0.26 | 0.24 | 0.12 | 0.15 |
| 11 | 116376840 | rs2367873 | 0.391 | 1.21 | -0.806 | 0.855 | -0.072 | -0.063 | -0.063 | 0.021 | 0.87 | 0.99 | 0.88 | 0.84 | 0.78 | 0.86 |
| 11 | 116379889 | rs1508102 | -1.602 | 0.048 | -0.256 | 0.812 | -0.021 | 0.008 | 0.008 | 3.021 | 0.14 | 0.76 | 0.26 | 0.2 | 0.06 | 0.11 |
| 11 | 116383064 | rs11599994 | -1.263 | 0.312 | 0.631 | 0.911 | 0.059 | 0.258 | 0.258 | 0.548 | 0.05 | 0.29 | 0.21 | 0.16 | 0.03 | 0.06 |
| 11 | 116384126 | rs1039755 | 0.353 | 1.004 | -0.138 | -0.444 | -0.018 | 0.008 | 0.008 | 0 | 0.78 | 0.83 | 0.82 | 1 | 0.68 | 0.93 |
| 11 | 116384511 | rs11601059 | -0.854 | 0.87 | -0.104 | -0.206 | 0.003 | 0.005 | 0.005 | 0 | 0.09 | 0.13 | 0.07 | 0.1 | 0.08 | 0.08 |
| 11 | 116385354 | rs7952521 | -0.805 | 0.779 | -0.138 | -0.499 | 0.001 | -0.03 | -0.03 | 0.016 | 0.1 | 0.14 | 0.08 | 0.04 | 0.13 | 0.09 |
| 11 | 116391537 | rs11215978 | 0.055 | 0.673 | -0.457 | -0.658 | -0.002 | 0.025 | 0.025 | 0 | 0.42 | 0.28 | 0.45 | 0.37 | 0.44 | 0.36 |
| 11 | 116395210 | rs1508106 | 0.769 | 0.638 | -0.712 | -0.687 | -0.019 | -0.012 | -0.012 | 0.011 | 0.1 | 0.05 | 0.06 | 0.15 | 0.11 | 0.13 |
| 11 | 116395458 | rs1396131 | 0.331 | 0.239 | -0.897 | -1.444 | 0.015 | 0.011 | 0.011 | 0.77 | 0.3 | 0.18 | 0.16 | 0.04 | 0.22 | 0.16 |
| 11 | 116397649 | rs4936353 | -0.265 | 0.369 | -0.617 | -1.36 | 0.003 | -0.025 | -0.025 | 0.011 | 0.58 | 0.47 | 0.53 | 0.25 | 0.5 | 0.4 |
| 11 | 116400382 | rs7941534 | -0.299 | 0.295 | -0.93 | -1.282 | 0.029 | 0.046 | 0.046 | 3.412 | 0.35 | 0.17 | 0.18 | 0.04 | 0.24 | 0.21 |
| 11 | 116405204 | rs12791443 | -0.145 | 0.897 | -0.77 | -0.178 | -0.016 | -0.019 | -0.019 | 0 | 0.1 | 0.08 | 0.09 | 0.15 | 0.1 | 0.13 |
| 11 | 116407583 | rs11215990 | -0.061 | 0.855 | -0.688 | -0.125 | -0.004 | -0.006 | -0.006 | 0.258 | 0.1 | 0.08 | 0.08 | 0.18 | 0.1 | 0.19 |
| 11 | 116409093 | rs10502220 | -0.39 | 0.487 | -0.927 | -0.649 | 0.031 | 0.087 | 0.087 | 1.238 | 0.27 | 0.11 | 0.14 | 0.1 | 0.19 | 0.14 |
| 11 | 116409788 | rs951839 | -0.321 | 0.229 | -0.341 | -0.621 | 0.018 | 0.021 | 0.021 | 2.877 | 0.26 | 0.22 | 0.16 | 0.12 | 0.16 | 0.14 |
| 11 | 116421056 | rs10891988 | -1.338 | 0.952 | 0.611 | -0.449 | 0.002 | 0.008 | 0.008 | 0 | 0.44 | 0.6 | 0.47 | 0.44 | 0.4 | 0.42 |
| 11 | 116422860 | rs12362410 | -0.632 | 1.038 | 0.518 | -0.449 | -0.007 | -0.007 | -0.007 | 0 | 0.15 | 0.12 | 0.29 | 0.3 | 0.24 | 0.28 |
| 11 | 116423307 | rs12293155 | -0.049 | 0.763 | 0.658 | -0.444 | 0.026 | 0.031 | 0.031 | 1.666 | 0.26 | 0.21 | 0.15 | 0.09 | 0.16 | 0.14 |
| 11 | 116426640 | rs12289097 | -0.208 | 1.046 | 0.611 | -0.615 | 0.033 | 0.059 | 0.059 | 0.618 | 0.28 | 0.21 | 0.18 | 0.08 | 0.2 | 0.14 |
| 11 | 116427932 | rs7930491 | -0.833 | 2.319 | 0.462 | -0.401 | -0.005 | -0.007 | -0.007 | 0 | 0.14 | 0.14 | 0.32 | 0.31 | 0.25 | 0.29 |
| 11 | 116429253 | rs11216005 | -0.554 | 1.34 | 1.115 | -1.048 | -0.024 | -0.066 | -0.066 | 0 | 0.34 | 0.44 | 0.24 | 0.11 | 0.28 | 0.16 |
| 11 | 116431149 | rs11600156 | 0.086 | 1.312 | 1.193 | -1.31 | 0.029 | 0.022 | 0.022 | 0.416 | 0.27 | 0.23 | 0.15 | 0.05 | 0.17 | 0.11 |
| 11 | 116436362 | rs12291550 | -1.007 | 1.619 | 1.275 | -1.186 | 0.067 | -0.001 | -0.001 | 0 | 0.07 | 0.3 | 0.08 | 0.02 | 0.11 | 0.05 |
| 11 | 116437756 | rs11605293 | -2.243 | 1.52 | 1.446 | -1.228 | 0.016 | -0.02 | -0.02 | 0 | 0.11 | 0.61 | 0.11 | 0.08 | 0.08 | 0.1 |
| 11 | 116438555 | rs12225221 | -0.662 | 1.65 | 1.395 | -1.029 | 0.035 | 0.036 | 0.036 | 0.269 | 0.13 | 0.25 | 0.14 | 0.27 | 0.14 | 0.21 |
| 11 | 116439602 | rs7943240 | -1.049 | 1.519 | 1.263 | -1.144 | -0.003 | -0.012 | -0.012 | 1.106 | 0.41 | 0.53 | 0.28 | 0.33 | 0.33 | 0.34 |
| 11 | 116439615 | rs7928307 | -1.193 | 1.534 | 1.282 | -1.123 | 0.051 | 0.044 | 0.044 | 1.859 | 0.17 | 0.51 | 0.16 | 0.28 | 0.16 | 0.23 |
| 11 | 116439684 | rs2174876 | -1.554 | 1.498 | 0.732 | -1.082 | -0.003 | 0.125 | 0.125 | 0 | 0.48 | 0.8 | 0.36 | 0.34 | 0.44 | 0.39 |
| 11 | 116452887 | rs10790151 | -1.521 | 1.546 | 0.312 | -1.54 | -0.004 | 0.125 | 0.125 | 0 | 0.49 | 0.66 | 0.39 | 0.16 | 0.52 | 0.3 |
| 11 | 116453047 | rs7931384 | -1.303 | 1.613 | -0.143 | -1.506 | -0.001 | -0.028 | -0.028 | 0 | 0.48 | 0.42 | 0.37 | 0.15 | 0.51 | 0.3 |
| 11 | 116457340 | rs10891996 | -0.585 | 2.19 | 0.41 | -0.667 | 0.009 | -0.04 | -0.04 | 0 | 0.13 | 0.12 | 0.16 | 0 | 0.2 | 0.06 |
| 11 | 116457650 | rs10790152 | -1.169 | 1.993 | -0.052 | -0.548 | 0.07 | -0.077 | -0.077 | 0 | 0.23 | 0.47 | 0.26 | 0.11 | 0.35 | 0.2 |
| 11 | 116462788 | rs12420596 | -0.143 | 1.363 | -1.26 | -0.329 | -0.073 | 0.004 | 0.004 | 0 | 0.28 | 0.09 | 0.19 | 0.16 | 0.25 | 0.22 |
| 11 | 116464013 | rs1876498 | -0.412 | 0.921 | -1.699 | -0.011 | 0.018 | 0.001 | 0.001 | 0.47 | 0.67 | 0.81 | 0.62 | 0.61 | 0.68 | 0.51 |
| 11 | 116464800 | rs11216026 | 1.143 | 0.643 | -2.208 | 0.44 | -0.109 | -0.043 | -0.043 | 0 | 0.23 | 0.01 | 0.24 | 0.22 | 0.3 | 0.15 |
| 11 | 116464992 | rs11216028 | 0.773 | 0.342 | -2.346 | 0.121 | 0 | -0.021 | -0.021 | 0 | 0.1 | 0.1 | 0.07 | 0 | 0.09 | 0.08 |
| 11 | 116465680 | rs11216029 | 0.061 | 0.34 | -1.422 | 0.088 | 0.001 | 0.001 | 0.001 | 0 | 0.43 | 0.57 | 0.38 | 0.39 | 0.41 | 0.36 |

| | | | | | | | | | | | | | | | | |
|---|---|---|---|---|---|---|---|---|---|---|---|---|---|---|---|---|
| 11 | 116466289 | rs11216031 | -0.293 | 0.304 | -0.444 | 0.321 | -0.016 | 0.038 | 0.038 | 0.509 | 0.17 | 0.42 | 0.18 | 0.19 | 0.13 | 0.14 |
| 11 | 116466894 | rs11216032 | 0.709 | 0.433 | -0.978 | 0.514 | -0.088 | 0.017 | 0.017 | 0 | 0.3 | 0.08 | 0.22 | 0.16 | 0.3 | 0.23 |
| 11 | 116467853 | rs7951761 | -0.282 | 0.784 | -0.52 | 0.599 | 0.002 | 0.001 | 0.001 | 0 | 0.45 | 0.5 | 0.46 | 0.43 | 0.5 | 0.37 |
| 11 | 116480631 | rs551141 | -0.5 | 0.336 | -0.744 | 0.933 | -0.027 | -0.007 | -0.007 | 0 | 0.07 | 0.1 | 0.03 | 0.06 | 0.04 | 0.08 |
| 11 | 116481221 | rs545613 | -1.51 | 0.735 | -0.807 | 0.717 | -0.004 | -0.022 | -0.022 | 0 | 0.13 | 0.17 | 0.07 | 0.06 | 0.1 | 0.1 |
| 11 | 116488133 | rs576266 | -1.185 | 0.522 | -0.734 | 1.493 | -0.012 | 0.109 | 0.109 | 0 | 0.1 | 0.14 | 0.11 | 0.37 | 0.09 | 0.2 |
| 11 | 116490970 | rs12365322 | 0.47 | 0.019 | -1.503 | 0.97 | 0.006 | 0.075 | 0.075 | 0.056 | 0.23 | 0.02 | 0.19 | 0.08 | 0.19 | 0.14 |
| 11 | 116491621 | rs507262 | -0.662 | 0.174 | -1.365 | 1.085 | -0.009 | 0.126 | 0.126 | 0 | 0.1 | 0.15 | 0.12 | 0.37 | 0.1 | 0.19 |
| 11 | 116492701 | rs11216046 | 0.226 | 0.177 | -1.389 | 0.464 | 0.027 | 0.059 | 0.059 | 0 | 0.23 | 0.12 | 0.21 | 0.08 | 0.2 | 0.3 |
| 11 | 116495059 | rs11216048 | 0.205 | 0.821 | -1.56 | 0.631 | -0.042 | -0.028 | -0.028 | 0 | 0.07 | 0 | 0.14 | 0.1 | 0.15 | 0.08 |
| 11 | 116496187 | rs7114844 | 0.334 | 0.409 | -2.23 | -0.245 | 0.017 | 0.053 | 0.053 | 0.071 | 0.32 | 0.22 | 0.19 | 0.14 | 0.25 | 0.22 |
| 11 | 116496360 | rs1240658 | 1.075 | 0.383 | -2.355 | -0.248 | -0.003 | 0.029 | 0.029 | 0 | 0.15 | 0.02 | 0.16 | 0.08 | 0.16 | 0.13 |
| 11 | 116497466 | rs12417676 | 0.235 | 1.11 | -2.022 | 0.12 | -0.011 | 0.006 | 0.006 | 0.079 | 0.21 | 0.19 | 0.24 | 0.14 | 0.26 | 0.26 |
| 11 | 116501409 | rs1944154 | 1.61 | 0.98 | -2.158 | -0.222 | 0.009 | -0.041 | -0.041 | 0 | 0.13 | 0.16 | 0.09 | 0.01 | 0.13 | 0.07 |
| 11 | 116501684 | rs11216058 | 0.44 | 1.173 | -2.103 | 0.256 | -0.01 | 0.002 | 0.002 | 0.111 | 0.22 | 0.19 | 0.24 | 0.17 | 0.26 | 0.26 |
| 11 | 116504377 | rs11216067 | 0.9 | 1.247 | -2.142 | 0.341 | -0.042 | -0.028 | -0.028 | 0 | 0.07 | 0 | 0.14 | 0.11 | 0.15 | 0.08 |
| 11 | 116505440 | rs10892001 | 0.063 | 0.947 | -2.374 | -0.164 | -0.001 | 0.196 | 0.196 | 0 | 0.46 | 0.21 | 0.33 | 0.15 | 0.43 | 0.43 |
| 11 | 116507107 | rs947987 | -0.408 | 0.49 | -1.591 | 0.769 | 0.002 | -0.008 | -0.008 | 0 | 0.7 | 0.78 | 0.7 | 0.68 | 0.71 | 0.71 |
| 11 | 116511522 | rs519000 | -0.023 | 0.951 | -1.539 | 1.213 | 0.007 | 0.025 | 0.025 | 0 | 0.12 | 0.12 | 0.18 | 0.29 | 0.15 | 0.28 |
| 11 | 116514417 | rs10892004 | 0.242 | 0.24 | -1.247 | 1.101 | 0.001 | 0.007 | 0.007 | 0 | 0.12 | 0.16 | 0.17 | 0.25 | 0.13 | 0.17 |
| 11 | 116514699 | rs1145185 | 0.982 | -0.517 | -1.994 | 0.616 | 0.017 | 0.002 | 0.002 | 0.618 | 0.21 | 0.01 | 0.14 | 0.11 | 0.16 | 0.12 |
| 11 | 116516506 | rs1789403 | -0.656 | -0.159 | -1.844 | 0.559 | -0.035 | -0.015 | -0.015 | 0 | 0.75 | 0.84 | 0.69 | 0.72 | 0.6 | 0.65 |
| 11 | 116517591 | rs1240769 | 0.301 | -1.264 | -1.607 | 0.331 | 0.01 | -0.007 | -0.007 | 0 | 0.4 | 0.37 | 0.3 | 0.56 | 0.3 | 0.36 |
| 11 | 116517968 | rs2187125 | 1.298 | -1.743 | -1.838 | -0.107 | -0.005 | 0.107 | 0.107 | 0 | 0.18 | 0.04 | 0.15 | 0.01 | 0.15 | 0.17 |
| 11 | 116518008 | rs1124874 | 1.936 | -1.581 | -1.652 | 0.062 | 0.003 | -0.003 | -0.003 | 0 | 0.07 | 0.05 | 0.12 | 0.07 | 0.08 | 0.07 |
| 11 | 116518960 | rs1261672 | 1.181 | -1.758 | -1.692 | -0.017 | 0.021 | 0.04 | 0.04 | 1.962 | 0.29 | 0.2 | 0.23 | 0.24 | 0.23 | 0.28 |
| 11 | 116519655 | rs516226 | 0.044 | -1.313 | -1.328 | 0.386 | -0.005 | 0.018 | 0.018 | 0.361 | 0.19 | 0.25 | 0.25 | 0.27 | 0.22 | 0.18 |
| 11 | 116520469 | rs506305 | -0.351 | -1.051 | -1.55 | 0.214 | -0.017 | 0.008 | 0.008 | 1.115 | 0.43 | 0.52 | 0.44 | 0.47 | 0.38 | 0.42 |
| 11 | 116521748 | rs3016352 | -1.021 | 0.167 | -1.844 | 0.586 | -0.059 | -0.03 | -0.03 | 0 | 0.75 | 0.86 | 0.69 | 0.71 | 0.6 | 0.7 |
| 11 | 116523821 | rs528647 | 0.222 | -0.205 | -1.447 | 0.608 | -0.033 | -0.035 | -0.035 | 0.025 | 0.08 | 0.01 | 0.11 | 0.1 | 0.11 | 0.1 |
| 11 | 116528876 | rs496958 | -0.296 | -0.643 | -1.817 | 0.486 | -0.026 | -0.029 | -0.029 | 0 | 0.21 | 0.01 | 0.14 | 0.14 | 0.17 | 0.24 |
| 11 | 116529024 | rs7104343 | 0.431 | -1.06 | -1.582 | 0.634 | -0.004 | 0.009 | 0.009 | 0 | 0.12 | 0.19 | 0.14 | 0.17 | 0.13 | 0.08 |
| 11 | 116529468 | rs948690 | 1.194 | -1.316 | -1.352 | 0.536 | 0.004 | 0.003 | 0.003 | 0 | 0.38 | 0.4 | 0.47 | 0.34 | 0.32 | 0.35 |
| 11 | 116529524 | rs523518 | -0.92 | -1.324 | -1.644 | 0.413 | -0.131 | -0.113 | -0.113 | 0.004 | 0.69 | 0.93 | 0.71 | 0.59 | 0.61 | 0.62 |
| 11 | 116529641 | rs524467 | 0.373 | -1.508 | -0.887 | 0.418 | 0.003 | 0.002 | 0.002 | 0 | 0.22 | 0.15 | 0.16 | 0.14 | 0.17 | 0.24 |
| 11 | 116530061 | rs1892947 | 1.349 | -1.248 | -0.917 | 0.486 | 0.003 | -0.004 | -0.004 | 0.284 | 0.09 | 0.03 | 0.11 | 0.06 | 0.06 | 0.06 |
| 11 | 116536656 | rs10892010 | -0.475 | -1.687 | -0.449 | 0.523 | -0.02 | 0.078 | 0.078 | 0.127 | 0.14 | 0.33 | 0.14 | 0.26 | 0.13 | 0.09 |
| 11 | 116551538 | rs1784104 | 2.279 | -2.072 | -0.786 | 0 | 0.003 | -0.002 | -0.002 | 2.514 | 0.24 | 0.21 | 0.18 | 0 | 0.23 | 0.12 |
| 11 | 116553025 | rs12805061 | 0.166 | -2.279 | -0.866 | 0.071 | -0.016 | -0.021 | -0.021 | 0.369 | 0.27 | 0.3 | 0.2 | 0.16 | 0.22 | 0.25 |
| 11 | 116555207 | rs7932330 | -0.27 | -2.269 | -0.807 | 0.167 | -0.022 | 0.092 | 0.092 | 0.321 | 0.14 | 0.4 | 0.14 | 0.25 | 0.13 | 0.09 |
| 11 | 116555926 | rs4572168 | 1.454 | -2.198 | -0.762 | 0.265 | -0.003 | 0.018 | 0.018 | 0 | 0.07 | 0.09 | 0.09 | 0.15 | 0.06 | 0.06 |
| 11 | 116556865 | rs1145212 | 0.201 | -2.346 | -0.934 | 0.282 | -0.001 | 0 | 0 | 0 | 0.44 | 0.64 | 0.52 | 0.45 | 0.39 | 0.38 |
| 11 | 116557216 | rs1145211 | -2.381 | -2.06 | -1.354 | 0.325 | -0.124 | -0.09 | -0.09 | 0 | 0.8 | 0.95 | 0.82 | 0.67 | 0.67 | 0.69 |
| 11 | 116557314 | rs947990 | 2.267 | -2.267 | -1.23 | 0.165 | -0.011 | 0 | 0 | 0 | 0.12 | 0.01 | 0.08 | 0.05 | 0.1 | 0.13 |
| 11 | 116559002 | rs1145210 | 1.073 | -2.274 | -1.01 | 0.173 | 0.008 | 0.032 | 0.032 | 1.841 | 0.35 | 0.24 | 0.31 | 0.18 | 0.33 | 0.21 |
| 11 | 116561938 | rs1787705 | 0.818 | -2.198 | -0.772 | 0.119 | -0.016 | -0.061 | -0.061 | 0.091 | 0.37 | 0.46 | 0.37 | 0.19 | 0.33 | 0.3 |
| 11 | 116562580 | rs11216096 | -0.335 | -2.189 | -0.76 | 0.369 | -0.026 | 0.024 | 0.024 | 0 | 0.32 | 0.38 | 0.31 | 0.36 | 0.23 | 0.3 |
| 11 | 116569204 | rs947986 | -2.011 | -1.785 | -0.969 | 0.415 | -0.107 | -0.04 | -0.04 | 0.109 | 0.78 | 0.91 | 0.8 | 0.71 | 0.63 | 0.8 |
| 11 | 116571250 | rs1145195 | -0.643 | -1.913 | -0.743 | 0.382 | -0.006 | -0.004 | -0.004 | 1.636 | 0.65 | 0.65 | 0.67 | 0.54 | 0.56 | 0.73 |
| 11 | 116574336 | rs1145198 | 2.432 | -2.092 | -0.784 | 0.087 | 0.001 | -0.012 | -0.012 | 0 | 0.25 | 0.19 | 0.17 | 0.01 | 0.24 | 0.1 |
| 11 | 116575229 | rs1268832 | -1.573 | -2.02 | -0.701 | 0.258 | 0.029 | 0.01 | 0.01 | 0 | 0.74 | 0.67 | 0.78 | 0.65 | 0.62 | 0.79 |
| 11 | 116575754 | rs1263057 | 1.552 | -2.302 | -1.09 | 0.264 | 0.004 | 0.022 | 0.022 | 0 | 0.31 | 0.25 | 0.36 | 0.23 | 0.33 | 0.25 |
| 11 | 116576400 | rs12802944 | 0.499 | -2.851 | -1.425 | 0.405 | -0.023 | -0.051 | -0.051 | 0.04 | 0.23 | 0.02 | 0.15 | 0.2 | 0.19 | 0.42 |
| 11 | 116576415 | rs1263056 | -0.737 | -2.286 | -1.145 | 0.375 | -0.071 | -0.02 | -0.02 | 0.225 | 0.65 | 0.81 | 0.66 | 0.64 | 0.52 | 0.74 |
| 11 | 116583513 | rs886022 | 1.598 | -2.208 | -1.043 | 0.392 | -0.078 | -0.008 | -0.008 | 0 | 0.25 | 0.14 | 0.23 | 0.27 | 0.43 | 0.21 |
| 11 | 116584317 | rs1144035 | 1.585 | -2.21 | -1.044 | 0.396 | -0.078 | -0.009 | -0.009 | 0 | 0.25 | 0.15 | 0.23 | 0.28 | 0.43 | 0.21 |
| 11 | 116585533 | rs2000571 | 0.149 | -3.052 | -0.576 | 0.152 | -0.027 | -0.005 | -0.005 | 0 | 0.66 | 0.6 | 0.73 | 0.69 | 0.79 | 0.47 |
| 11 | 116589127 | rs180377 | -0.493 | -3.084 | -0.443 | 0.074 | -0.006 | -0.012 | -0.012 | 0 | 0.47 | 0.5 | 0.6 | 0.68 | 0.58 | 0.43 |
| 11 | 116597889 | rs180363 | 0.613 | -3.273 | -0.433 | 0.021 | 0.006 | 0.008 | 0.008 | 0 | 0.39 | 0.54 | 0.49 | 0.19 | 0.35 | 0.23 |
| 11 | 116598988 | rs180360 | 1.553 | -3.264 | -0.582 | 0.029 | 0.007 | 0.021 | 0.021 | 0 | 0.3 | 0.2 | 0.33 | 0.19 | 0.27 | 0.18 |
| 11 | 116599392 | rs180358 | 2.666 | -3.191 | -0.543 | -0.058 | 0.001 | -0.011 | -0.011 | 0 | 0.22 | 0.18 | 0.15 | 0 | 0.21 | 0.1 |
| 11 | 116600021 | rs7123583 | -0.067 | -3.219 | -0.694 | 0.08 | 0.017 | -0.121 | -0.121 | 0.261 | 0.67 | 0.9 | 0.61 | 0.5 | 0.73 | 0.44 |
| 11 | 116610249 | rs10466588 | -0.883 | -3.219 | -0.391 | 0.143 | -0.051 | 0.003 | 0.003 | 1.189 | 0.15 | 0.4 | 0.19 | 0.12 | 0.09 | 0.11 |
| 11 | 116617240 | rs11216126 | 1.675 | -3.315 | -0.581 | 0.34 | 0.008 | -0.043 | -0.043 | 0.31 | 0.13 | 0.02 | 0.14 | 0.19 | 0.12 | 0.32 |
| 11 | 116619051 | rs11556024 | 1.489 | -3.348 | -0.548 | 0.35 | -0.009 | 0.01 | 0.01 | 0 | 0.05 | 0.06 | 0.04 | 0.12 | 0.01 | 0.06 |
| 11 | 116620256 | rs11216129 | 1.716 | -3.341 | -0.549 | 0.339 | 0.007 | -0.041 | -0.041 | 0 | 0.13 | 0.02 | 0.14 | 0.19 | 0.12 | 0.31 |
| 11 | 116621963 | rs12292921 | 0.724 | -3.384 | -0.498 | 0.248 | -0.023 | -0.055 | -0.055 | 0 | 0.1 | 0.19 | 0.14 | 0 | 0.07 | 0.05 |
| 11 | 116624703 | rs180326 | -0.821 | -3.494 | -0.601 | 0.205 | 0.024 | 0.045 | 0.045 | 0 | 0.54 | 0.39 | 0.55 | 0.76 | 0.62 | 0.67 |
| 11 | 116628401 | rs2075295 | -1.316 | -3.435 | -0.717 | 0.25 | 0.014 | -0.006 | -0.006 | 0 | 0.38 | 0.34 | 0.44 | 0.38 | 0.27 | 0.46 |

| | | | | | | | | | | | | | | | | |
|---|---|---|---|---|---|---|---|---|---|---|---|---|---|---|---|---|
| 11 | 116629949 | rs4938310 | 2.807 | -3.763 | -0.79 | 0.215 | 0.002 | 0.036 | 0.036 | 0.181 | 0.21 | 0.13 | 0.14 | 0 | 0.21 | 0.1 |
| 11 | 116631391 | rs3741301 | 1.913 | -3.655 | -0.694 | 0.422 | -0.094 | -0.036 | -0.036 | 0 | 0.25 | 0.1 | 0.24 | 0.33 | 0.42 | 0.24 |
| 11 | 116633825 | rs918144 | -0.816 | -3.543 | -0.85 | 0.41 | -0.001 | -0.002 | -0.002 | 0 | 0.61 | 0.68 | 0.6 | 0.38 | 0.48 | 0.56 |
| 11 | 116633947 | rs10488698 | 1.725 | -3.359 | -1.223 | 0.333 | -0.009 | -0.016 | -0.016 | 0 | 0.07 | 0 | 0.11 | 0.07 | 0.06 | 0.04 |
| 11 | 116640368 | rs623908 | 0.387 | -3.338 | -1.434 | 0.345 | 0.026 | 0.042 | 0.042 | 0.367 | 0.46 | 0.39 | 0.43 | 0.23 | 0.4 | 0.46 |
| 11 | 116650118 | rs17120029 | 0.456 | -3.403 | -1.442 | 0.375 | -0.029 | -0.058 | -0.058 | 0 | 0.11 | 0.21 | 0.14 | 0 | 0.07 | 0.04 |
| 11 | 116651463 | rs1942478 | 1.508 | -3.023 | -2.053 | 0.763 | 0.018 | -0.042 | -0.042 | 0.406 | 0.12 | 0.03 | 0.11 | 0.22 | 0.1 | 0.29 |
| 11 | 116658122 | rs2075294 | 0.943 | -2.884 | -1.923 | 0.401 | -0.004 | 0.004 | 0.004 | 0 | 0.05 | 0.06 | 0.05 | 0.13 | 0.02 | 0.06 |
| 11 | 116660686 | rs2266788 | -1.211 | -2.437 | -1.923 | 0.22 | -0.016 | -0.067 | -0.067 | 0 | 0.89 | 0.99 | 0.87 | 0.76 | 0.91 | 0.8 |
| 11 | 116660813 | rs619054 | 1.693 | -2.433 | -1.812 | 0.181 | -0.011 | 0.076 | 0.076 | 0.651 | 0.23 | 0.1 | 0.16 | 0 | 0.23 | 0.12 |
| 11 | 116662579 | rs651821 | -0.481 | -2.143 | -1.304 | 0.916 | -0.013 | 0.03 | 0.03 | 0 | 0.87 | 0.84 | 0.85 | 0.71 | 0.92 | 0.81 |
| 11 | 116663851 | rs17120035 | 0.817 | -1.772 | -1.751 | 0.489 | 0.022 | -0.007 | -0.007 | 0.142 | 0.11 | 0.02 | 0.04 | 0.07 | 0.08 | 0.27 |
| 11 | 116669828 | rs11216137 | 0.061 | -1.926 | -1.666 | 0.241 | 0.001 | 0.001 | 0.001 | 0.026 | 0.16 | 0.15 | 0.05 | 0.04 | 0.08 | 0.28 |
| 11 | 116673768 | rs1729409 | 0.472 | -1.801 | -2.517 | 0.413 | -0.112 | -0.089 | -0.089 | 0 | 0.34 | 0.09 | 0.42 | 0.43 | 0.51 | 0.32 |
| 11 | 116674818 | rs1729408 | -0.063 | -2.073 | -2.482 | 0.282 | 0.106 | 0.021 | 0.021 | 0.259 | 0.51 | 0.86 | 0.37 | 0.44 | 0.33 | 0.36 |
| 11 | 116681008 | rs1263173 | 0.685 | -2.037 | -2.283 | -0.018 | 0.157 | 0.07 | 0.07 | 0.007 | 0.55 | 0.12 | 0.52 | 0.3 | 0.68 | 0.4 |
| 11 | 116696921 | rs2849176 | 0.082 | -2.243 | -0.828 | 0.09 | 0.064 | 0.022 | 0.022 | 0 | 0.52 | 0.29 | 0.56 | 0.58 | 0.62 | 0.47 |
| 11 | 116697066 | rs2849174 | 1.457 | -2.483 | -0.761 | 0.067 | -0.008 | 0.078 | 0.078 | 0 | 0.25 | 0.11 | 0.17 | 0.06 | 0.25 | 0.14 |
| 11 | 116697344 | rs2071523 | 0.087 | -2.299 | -0.719 | 0.176 | 0.075 | 0.024 | 0.024 | 0 | 0.5 | 0.28 | 0.56 | 0.58 | 0.61 | 0.47 |
| 11 | 116697848 | rs2071521 | -0.016 | -2.305 | -0.705 | 0.173 | 0.074 | 0.023 | 0.023 | 0 | 0.5 | 0.28 | 0.56 | 0.58 | 0.61 | 0.47 |
| 11 | 116699445 | rs595049 | 0.028 | -2.309 | -0.579 | 0.172 | 0.049 | 0.017 | 0.017 | 0 | 0.51 | 0.33 | 0.57 | 0.58 | 0.62 | 0.47 |
| 11 | 116699984 | rs2542052 | -0.02 | -2.241 | -0.786 | 0.278 | 0.059 | 0.005 | 0.005 | 0 | 0.49 | 0.25 | 0.54 | 0.52 | 0.61 | 0.44 |
| 11 | 116707684 | rs2070665 | -0.719 | -2.39 | -0.542 | 0.694 | 0.005 | -0.005 | -0.005 | 0 | 0.84 | 0.88 | 0.79 | 0.65 | 0.89 | 0.66 |
| 11 | 116711180 | rs11216157 | -0.439 | -2.502 | -0.531 | 0.544 | -0.018 | 0.006 | 0.006 | 0 | 0.21 | 0.3 | 0.2 | 0.25 | 0.12 | 0.2 |
| 11 | 116721826 | rs10047459 | 0.136 | -2.806 | -0.819 | 0.388 | 0.03 | -0.023 | -0.023 | 0 | 0.7 | 0.8 | 0.7 | 0.4 | 0.81 | 0.59 |
| 11 | 116723171 | rs583219 | -0.972 | -2.849 | -0.443 | 0.316 | -0.009 | 0.095 | 0.095 | 0.081 | 0.1 | 0.54 | 0.12 | 0.13 | 0.04 | 0.19 |
| 11 | 116728029 | rs509712 | -0.471 | -2.85 | -0.398 | 0.325 | 0.051 | 0.12 | 0.12 | 0.77 | 0.07 | 0.51 | 0.09 | 0.08 | 0.02 | 0.04 |
| 11 | 116728277 | rs11216162 | -0.094 | -3.036 | -0.595 | 0.126 | 0.023 | 0.014 | 0.014 | 0 | 0.19 | 0.15 | 0.19 | 0.1 | 0.15 | 0.28 |
| 11 | 116728630 | rs12225230 | -0.11 | -3.038 | -0.596 | 0.125 | 0.023 | 0.014 | 0.014 | 0 | 0.19 | 0.15 | 0.19 | 0.1 | 0.15 | 0.28 |
| 11 | 116729013 | rs539858 | -0.99 | -3.011 | -0.542 | 0.151 | -0.005 | 0.099 | 0.099 | 0 | 0.1 | 0.55 | 0.12 | 0.13 | 0.04 | 0.19 |
| 11 | 116730638 | rs6589574 | -1.366 | -3.14 | -0.729 | 0.275 | 0.015 | -0.047 | -0.047 | 0 | 0.84 | 0.94 | 0.81 | 0.63 | 0.89 | 0.75 |
| 11 | 116731205 | rs4938315 | -0.834 | -3.028 | -0.427 | -0.387 | 0.019 | -0.041 | -0.041 | 0 | 0.8 | 0.89 | 0.73 | 0.48 | 0.86 | 0.68 |
| 11 | 116732512 | rs2075292 | -0.87 | -3.029 | -0.446 | -0.381 | 0.03 | -0.074 | -0.074 | 0.491 | 0.8 | 0.92 | 0.74 | 0.47 | 0.86 | 0.68 |
| 11 | 116734545 | rs11216164 | 1.641 | -2.353 | -0.863 | -0.757 | -0.144 | 0.096 | 0.096 | 0 | 0.19 | 0.02 | 0.23 | 0.03 | 0.36 | 0.07 |
| 11 | 116736679 | rs573549 | 0.982 | -2.673 | -0.974 | -0.693 | 0.084 | 0.155 | 0.155 | 0 | 0.44 | 0.1 | 0.42 | 0.22 | 0.63 | 0.28 |
| 11 | 116748515 | rs12292614 | 1.436 | -2.595 | -0.727 | -0.547 | -0.009 | -0.01 | -0.01 | 0 | 0.11 | 0.09 | 0.05 | 0.08 | 0.06 | 0.1 |
| 11 | 116753552 | rs12275565 | 1.376 | -2.597 | -0.729 | -0.526 | -0.009 | -0.006 | -0.006 | 0.449 | 0.11 | 0.09 | 0.05 | 0.1 | 0.06 | 0.11 |
| 11 | 116759332 | rs1473177 | 1.297 | -2.721 | -0.905 | -0.584 | 0.005 | 0.011 | 0.011 | 0 | 0.11 | 0.02 | 0.12 | 0.04 | 0.09 | 0.13 |
| 11 | 116764021 | rs672058 | -0.189 | -2.932 | -0.462 | -0.588 | -0.046 | 0.032 | 0.032 | 0.04 | 0.72 | 0.63 | 0.67 | 0.4 | 0.84 | 0.64 |
| 11 | 116764582 | rs4938316 | 0.361 | -2.899 | -0.615 | -0.583 | 0.034 | 0.043 | 0.043 | 0 | 0.21 | 0.02 | 0.15 | 0.09 | 0.15 | 0.2 |
| 11 | 116772441 | rs1076485 | 0.215 | -2.737 | 0.482 | -0.521 | -0.043 | 0.049 | 0.049 | 0 | 0.24 | 0.34 | 0.3 | 0.58 | 0.14 | 0.46 |
| 11 | 116772486 | rs17120131 | 1.601 | -2.814 | 0.463 | -0.537 | -0.013 | -0.01 | -0.01 | 0 | 0.1 | 0.11 | 0.05 | 0.1 | 0.06 | 0.09 |
| 11 | 116772787 | rs518181 | 1.167 | -2.839 | 0.437 | -0.571 | 0.072 | 0.175 | 0.175 | 0 | 0.44 | 0.07 | 0.41 | 0.2 | 0.63 | 0.26 |
| 11 | 116779499 | rs10790166 | 0.748 | -2.746 | 0.545 | -0.546 | 0.063 | 0.033 | 0.033 | 0 | 0.65 | 0.2 | 0.68 | 0.74 | 0.77 | 0.58 |
| 11 | 116782974 | rs11216185 | 1.534 | -2.697 | 0.564 | -0.486 | -0.013 | -0.014 | -0.014 | 2.616 | 0.1 | 0.11 | 0.05 | 0.08 | 0.06 | 0.09 |
| 11 | 116784376 | rs7115583 | 0.44 | -2.706 | 0.651 | -0.487 | -0.042 | -0.004 | -0.004 | 0 | 0.14 | 0.39 | 0.1 | 0.13 | 0.09 | 0.22 |
| 11 | 116806412 | rs2513094 | -1.023 | -2.816 | 0.637 | -0.536 | 0.029 | 0.013 | 0.013 | 0 | 0.34 | 0.72 | 0.37 | 0.27 | 0.22 | 0.42 |
| 11 | 116812628 | rs17120157 | 1.544 | -2.803 | 0.623 | -0.556 | -0.013 | -0.01 | -0.01 | 0 | 0.1 | 0.11 | 0.05 | 0.09 | 0.06 | 0.09 |
| 11 | 116822034 | rs1473325 | 0.537 | -2.791 | 0.595 | -0.554 | -0.001 | -0.002 | -0.002 | 0 | 0.16 | 0.22 | 0.23 | 0.05 | 0.13 | 0.14 |
| 11 | 116826870 | rs10502221 | 1.48 | -2.784 | 0.411 | -0.668 | -0.156 | -0.009 | -0.009 | 0.556 | 0.22 | 0.04 | 0.31 | 0.21 | 0.42 | 0.14 |
| 11 | 116828121 | rs1940626 | -0.673 | -3.013 | 0.835 | -0.4 | -0.062 | -0.044 | -0.044 | 0.493 | 0.31 | 0.57 | 0.32 | 0.19 | 0.21 | 0.38 |
| 11 | 116830087 | rs7120706 | 1.199 | -2.983 | 0.59 | -0.462 | 0.075 | 0.168 | 0.168 | 0 | 0.44 | 0.08 | 0.41 | 0.21 | 0.63 | 0.27 |
| 11 | 116830261 | rs7120963 | 1.199 | -2.983 | 0.59 | -0.462 | 0.075 | 0.168 | 0.168 | 0 | 0.44 | 0.08 | 0.41 | 0.21 | 0.63 | 0.27 |
| 11 | 116831912 | rs7112967 | 0.995 | -2.99 | 0.611 | -0.465 | -0.002 | -0.002 | -0.002 | 0 | 0.13 | 0.2 | 0.23 | 0.05 | 0.13 | 0.14 |
| 11 | 116834313 | rs10892052 | 1.145 | -2.778 | 0.28 | -0.431 | 0.075 | 0.178 | 0.178 | 0 | 0.44 | 0.08 | 0.41 | 0.2 | 0.63 | 0.27 |
| 11 | 116845104 | rs17120197 | 0.145 | -2.479 | 0.417 | -0.45 | 0 | -0.003 | -0.003 | 0 | 0.18 | 0.23 | 0.25 | 0.05 | 0.16 | 0.15 |
| 11 | 116848329 | rs11216210 | 0.6 | -2.665 | 0.646 | -0.408 | -0.036 | -0.002 | -0.002 | 0 | 0.14 | 0.3 | 0.09 | 0.14 | 0.08 | 0.24 |
| 11 | 116896155 | rs10892063 | 1.226 | -2.768 | 0.441 | -0.45 | 0.052 | 0.157 | 0.157 | 0 | 0.43 | 0.05 | 0.4 | 0.21 | 0.63 | 0.27 |
| 11 | 116915742 | rs2000614 | -0.731 | -2.779 | 0.653 | -0.377 | 0.005 | -0.007 | -0.007 | 0 | 0.78 | 0.8 | 0.78 | 0.48 | 0.85 | 0.69 |
| 11 | 116924978 | rs723953 | -0.038 | -2.803 | 0.8 | -0.4 | -0.039 | -0.019 | -0.019 | 0.001 | 0.2 | 0.37 | 0.27 | 0.13 | 0.14 | 0.18 |
| 11 | 116928546 | rs10892068 | 1.616 | -2.73 | 0.656 | -0.373 | -0.027 | -0.018 | -0.018 | 0 | 0.1 | 0.15 | 0.05 | 0.09 | 0.06 | 0.08 |
| 11 | 116932561 | rs11216257 | -0.123 | -2.748 | 0.518 | -0.375 | -0.013 | 0.003 | 0.003 | 0 | 0.68 | 0.68 | 0.73 | 0.39 | 0.79 | 0.6 |
| 11 | 116943354 | rs1351452 | -0.869 | -2.722 | 0.363 | -0.356 | 0.024 | -0.046 | -0.046 | 1.204 | 0.79 | 0.88 | 0.79 | 0.48 | 0.85 | 0.68 |
| 11 | 116943544 | rs7109649 | -1.043 | -2.595 | 0.484 | -0.364 | 0.072 | 0.032 | 0.032 | 0 | 0.35 | 0.8 | 0.38 | 0.27 | 0.23 | 0.42 |
| 11 | 116958533 | rs12277910 | 1.254 | -2.315 | 0.399 | -0.134 | -0.014 | 0.006 | 0.006 | 0.256 | 0.1 | 0.11 | 0.05 | 0.16 | 0.06 | 0.09 |
| 11 | 116972935 | rs4936363 | 0.75 | -2.307 | 0.563 | -0.552 | -0.008 | -0.006 | -0.006 | 0 | 0.14 | 0.21 | 0.23 | 0.07 | 0.13 | 0.14 |
| 11 | 116979911 | rs12271161 | -0.605 | -2.414 | 0.742 | -0.611 | -0.054 | -0.034 | -0.034 | 0.955 | 0.27 | 0.59 | 0.31 | 0.16 | 0.2 | 0.25 |
| 11 | 116980622 | rs4938340 | 1.079 | -2.265 | 0.618 | -0.566 | -0.004 | 0.002 | 0.002 | 0 | 0.11 | 0.2 | 0.23 | 0.07 | 0.14 | 0.11 |
| 11 | 116980641 | rs10892074 | 0.933 | -2.093 | 0.16 | -0.753 | -0.194 | -0.014 | -0.014 | 0.049 | 0.25 | 0.02 | 0.32 | 0.21 | 0.45 | 0.15 |

| | | | | | | | | | | | | | | | | |
|---|---|---|---|---|---|---|---|---|---|---|---|---|---|---|---|---|
| 11 | 116991926 | rs7120515 | 0.871 | -2.463 | 0.442 | -0.712 | 0.093 | 0.118 | 0.118 | 0 | 0.47 | 0.25 | 0.44 | 0.24 | 0.64 | 0.31 |
| 11 | 116996539 | rs17120347 | 1.356 | -2.377 | 0.22 | -0.744 | 0 | -0.001 | -0.001 | 0.565 | 0.11 | 0.14 | 0.19 | 0.07 | 0.1 | 0.11 |
| 11 | 117003749 | rs4938344 | 0.339 | -2.375 | 0.632 | -0.683 | -0.032 | -0.003 | -0.003 | 0 | 0.59 | 0.58 | 0.66 | 0.37 | 0.74 | 0.41 |
| 11 | 117016450 | rs2306471 | 1.763 | -2.361 | 0.446 | -0.724 | 0.007 | 0.003 | 0.003 | 0.572 | 0.09 | 0.04 | 0.04 | 0.08 | 0.06 | 0.11 |
| 11 | 117016824 | rs10892079 | 0.472 | -2.455 | 0.764 | -0.749 | -0.03 | -0.005 | -0.005 | 0 | 0.58 | 0.52 | 0.64 | 0.37 | 0.74 | 0.41 |
| 11 | 117034729 | rs10790175 | 0.72 | -2.476 | 0.845 | -0.759 | 0.069 | 0.017 | 0.017 | 0 | 0.56 | 0.31 | 0.62 | 0.37 | 0.74 | 0.41 |
| 11 | 117039797 | rs1060211 | 0.09 | -2.39 | 0.723 | -0.715 | -0.055 | 0.035 | 0.035 | 0 | 0.67 | 0.57 | 0.69 | 0.45 | 0.8 | 0.52 |
| 11 | 117046197 | rs4938353 | 0.09 | -2.39 | 0.722 | -0.715 | -0.065 | 0.043 | 0.043 | 0 | 0.67 | 0.51 | 0.68 | 0.45 | 0.8 | 0.52 |
| 11 | 117059347 | rs90192 | -0.431 | -2.457 | 0.487 | -0.719 | -0.011 | 0.017 | 0.017 | 0 | 0.74 | 0.74 | 0.76 | 0.46 | 0.84 | 0.53 |
| 11 | 117065476 | rs1784042 | 1.408 | -2.263 | 0.138 | -0.866 | -0.146 | -0.014 | -0.014 | 0 | 0.22 | 0.04 | 0.31 | 0.22 | 0.42 | 0.11 |
| 11 | 117067699 | rs8521 | 1.063 | -2.796 | 0.735 | -0.347 | 0.065 | 0.016 | 0.016 | 0 | 0.56 | 0.32 | 0.63 | 0.37 | 0.74 | 0.41 |
| 11 | 117074229 | rs641620 | -0.711 | -2.622 | 0.566 | -0.33 | -0.059 | 0.062 | 0.062 | 0 | 0.29 | 0.45 | 0.37 | 0.57 | 0.15 | 0.46 |
| 11 | 117075566 | rs508487 | 0.883 | -2.499 | 0.012 | -0.079 | -0.024 | -0.04 | -0.04 | 0 | 0.07 | 0 | 0.07 | 0.16 | 0.08 | 0.1 |
| 11 | 117085270 | rs236911 | 0.531 | -2.094 | 0.183 | -0.709 | 0.018 | -0.058 | -0.058 | 0 | 0.22 | 0.12 | 0.23 | 0.51 | 0.16 | 0.35 |
| 11 | 117088994 | rs2239011 | -0.197 | -2.16 | 0.217 | -0.945 | -0.045 | -0.041 | -0.041 | 0.112 | 0.25 | 0.46 | 0.27 | 0.15 | 0.19 | 0.26 |
| 11 | 117089628 | rs236916 | 0.497 | -2.137 | -0.223 | -0.924 | 0.015 | -0.024 | -0.024 | 0 | 0.25 | 0.2 | 0.23 | 0.53 | 0.15 | 0.47 |
| 11 | 117093764 | rs3830008 | -0.476 | -1.355 | -0.758 | -0.537 | -0.027 | -0.026 | -0.026 | 0.498 | 0.25 | 0.4 | 0.26 | 0.15 | 0.2 | 0.26 |
| 11 | 117094413 | rs11216321 | 1.728 | -0.698 | -1.079 | -0.645 | -0.063 | 0.065 | 0.065 | 0 | 0.11 | 0.01 | 0.1 | 0 | 0.18 | 0.01 |
| 11 | 117095361 | rs236919 | -1.5 | -1.946 | -0.511 | -0.052 | 0.048 | 0.078 | 0.078 | 0 | 0.57 | 0.87 | 0.57 | 0.68 | 0.39 | 0.72 |
| 11 | 117096652 | rs2277287 | -1 | -1.256 | -0.111 | 0.044 | -0.017 | -0.016 | -0.016 | 0 | 0.26 | 0.34 | 0.27 | 0.15 | 0.19 | 0.28 |
| 11 | 117097952 | rs2306473 | -0.435 | -0.455 | -0.382 | 0.16 | -0.016 | -0.035 | -0.035 | 0.046 | 0.17 | 0.4 | 0.23 | 0.07 | 0.13 | 0.16 |
| 11 | 117100594 | rs500389 | -0.574 | -0.315 | -1.364 | 0.176 | 0.02 | 0.063 | 0.063 | 0 | 0.58 | 0.85 | 0.6 | 0.69 | 0.42 | 0.74 |
| 11 | 117102307 | rs2509209 | -0.309 | -1.089 | -1.186 | 0.362 | -0.001 | 0.001 | 0.001 | 0 | 0.32 | 0.25 | 0.45 | 0.5 | 0.31 | 0.39 |
| 11 | 117102648 | rs2252145 | -0.388 | -1.719 | -0.345 | 0.222 | -0.033 | 0.03 | 0.03 | 0 | 0.31 | 0.58 | 0.5 | 0.62 | 0.25 | 0.37 |
| 11 | 117107775 | rs535602 | -0.966 | -0.1 | -0.767 | -0.174 | -0.128 | 0.099 | 0.099 | 0 | 0.75 | 0.95 | 0.75 | 0.93 | 0.6 | 0.91 |
| 11 | 117110740 | rs650585 | -0.194 | -0.546 | -0.418 | -0.282 | 0.013 | 0.067 | 0.067 | 0 | 0.6 | 0.74 | 0.59 | 0.78 | 0.44 | 0.72 |
| 11 | 117116450 | rs12286525 | 1.068 | -0.939 | -0.163 | -0.383 | 0.051 | -0.022 | -0.022 | 0 | 0.37 | 0.23 | 0.42 | 0.61 | 0.27 | 0.44 |
| 11 | 117128271 | rs667944 | 0.952 | -0.837 | 0.122 | -0.332 | 0.013 | -0.004 | -0.004 | 0.275 | 0.38 | 0.35 | 0.44 | 0.64 | 0.27 | 0.42 |
| 11 | 117151412 | rs12282721 | -0.438 | -0.633 | 0.209 | -0.318 | 0.011 | 0.134 | 0.134 | 0.596 | 0.08 | 0.26 | 0.1 | 0.19 | 0.1 | 0.19 |
| 11 | 117156552 | rs7083 | -1.241 | -0.569 | -0.142 | -0.324 | -0.099 | 0.088 | 0.088 | 0 | 0.75 | 0.94 | 0.74 | 0.94 | 0.6 | 0.91 |
| 11 | 117171337 | rs548381 | -1.118 | 0.073 | -0.148 | -0.528 | -0.127 | 0.087 | 0.087 | 0 | 0.78 | 0.97 | 0.76 | 0.93 | 0.64 | 0.91 |
| 11 | 117175934 | rs522843 | 1.159 | -0.131 | 0.128 | -0.427 | 0.057 | 0.005 | 0.005 | 0.772 | 0.42 | 0.21 | 0.41 | 0.65 | 0.33 | 0.42 |
| 11 | 117177603 | rs687740 | -0.888 | 0.266 | 0.168 | -0.252 | -0.063 | 0.058 | 0.058 | 0 | 0.77 | 0.91 | 0.76 | 0.93 | 0.64 | 0.92 |
| 11 | 117177800 | rs473210 | 0.408 | -0.003 | 0.553 | -0.113 | 0.001 | 0.001 | 0.001 | 0 | 0.56 | 0.43 | 0.47 | 0.66 | 0.41 | 0.61 |
| 11 | 117179891 | rs676134 | 0.022 | 0.174 | 0.438 | -0.219 | -0.01 | 0.008 | 0.008 | 0.454 | 0.6 | 0.77 | 0.54 | 0.7 | 0.4 | 0.65 |
| 11 | 117182708 | rs525493 | 0.522 | 0.886 | 0.43 | 0.419 | 0 | -0.002 | -0.002 | 0 | 0.55 | 0.42 | 0.51 | 0.79 | 0.44 | 0.51 |
| 11 | 117183650 | rs593245 | 0.177 | 0.808 | 0.406 | 0.481 | 0 | -0.003 | -0.003 | 0 | 0.58 | 0.52 | 0.54 | 0.7 | 0.45 | 0.65 |
| 11 | 117193203 | rs555075 | 0.071 | 0.888 | 0.036 | 0.488 | 0.002 | -0.004 | -0.004 | 0 | 0.59 | 0.52 | 0.51 | 0.69 | 0.46 | 0.72 |
| 11 | 117199870 | rs509894 | -0.777 | 1.028 | 0.003 | 0.527 | 0.007 | 0.009 | 0.009 | 0 | 0.28 | 0.46 | 0.34 | 0.24 | 0.33 | 0.44 |
| 11 | 117201682 | rs609272 | -0.522 | 1.266 | -0.149 | 0.67 | 0.01 | -0.004 | -0.004 | 0 | 0.74 | 0.72 | 0.79 | 0.71 | 0.69 | 0.74 |
| 11 | 117213189 | rs11216360 | 1.285 | 0.753 | -0.03 | 0.495 | 0 | 0.002 | 0.002 | 0 | 0.14 | 0.12 | 0.1 | 0.04 | 0.15 | 0.12 |
| 11 | 117214091 | rs573827 | 1.415 | 0.431 | -0.374 | 0.456 | 0.019 | 0.022 | 0.022 | 1.109 | 0.34 | 0.26 | 0.27 | 0.12 | 0.19 | 0.25 |
| 11 | 117221170 | rs563828 | -0.927 | 0.658 | -0.325 | 0.588 | 0.01 | -0.008 | -0.008 | 0.105 | 0.74 | 0.75 | 0.78 | 0.79 | 0.65 | 0.74 |
| 11 | 117222256 | rs7926760 | 1.084 | 0.58 | -0.352 | 0.572 | -0.083 | 0.077 | 0.077 | 0.118 | 0.22 | 0.05 | 0.17 | 0.11 | 0.34 | 0.1 |
| 11 | 117222592 | rs490262 | 1.692 | 0.484 | -0.368 | 0.531 | 0.014 | 0.014 | 0.014 | 1.244 | 0.32 | 0.26 | 0.26 | 0.12 | 0.17 | 0.24 |
| 11 | 117225033 | rs499744 | -0.153 | 0.541 | -0.395 | 0.809 | -0.008 | 0.002 | 0.002 | 0 | 0.67 | 0.72 | 0.69 | 0.63 | 0.57 | 0.58 |
| 11 | 117227519 | rs564674 | 0.213 | 0.574 | -0.397 | 0.834 | 0 | 0 | 0 | 0 | 0.63 | 0.54 | 0.51 | 0.43 | 0.59 | 0.52 |
| 11 | 117240357 | rs9651682 | 1.422 | 0.79 | -0.33 | 0.809 | 0.018 | 0.03 | 0.03 | 1.019 | 0.35 | 0.26 | 0.3 | 0.12 | 0.24 | 0.26 |
| 11 | 117246150 | rs10790180 | 0.841 | 0.077 | -0.211 | 0.988 | -0.03 | -0.009 | -0.009 | 0 | 0.39 | 0.53 | 0.33 | 0.33 | 0.21 | 0.43 |
| 11 | 117256436 | rs3781981 | -0.448 | 0.199 | -0.383 | 1.161 | -0.012 | -0.01 | -0.01 | 0 | 0.39 | 0.31 | 0.29 | 0.51 | 0.45 | 0.44 |
| 11 | 117257876 | rs2305829 | 1 | 0.189 | -0.391 | 0.416 | -0.001 | -0.014 | -0.014 | 4.734 | 0.14 | 0.12 | 0.27 | 0.35 | 0.17 | 0.18 |
| 11 | 117260075 | rs552498 | 0.018 | 0.225 | -0.239 | 0.331 | 0.029 | -0.013 | -0.013 | 0.469 | 0.31 | 0.38 | 0.24 | 0.32 | 0.38 | 0.28 |
| 11 | 117266754 | rs693147 | -0.274 | -0.92 | 0.118 | 0.851 | 0.04 | 0.05 | 0.05 | 0.084 | 0.52 | 0.65 | 0.41 | 0.39 | 0.62 | 0.48 |
| 11 | 117283091 | rs2279013 | -0.259 | -0.979 | 0.163 | 0.784 | -0.003 | -0.001 | -0.001 | 0 | 0.55 | 0.49 | 0.56 | 0.5 | 0.68 | 0.45 |
| 11 | 117283471 | rs573965 | -1.61 | -0.891 | 0.065 | 0.678 | -0.03 | 0.054 | 0.054 | 0.554 | 0.16 | 0.36 | 0.08 | 0.24 | 0.1 | 0.27 |
| 11 | 117283676 | rs8258 | -0.169 | -0.774 | 0.062 | 0.709 | 0.037 | 0.04 | 0.04 | 0.091 | 0.52 | 0.65 | 0.4 | 0.4 | 0.63 | 0.48 |
| 11 | 117283808 | rs9163 | 0.942 | -0.745 | -0.019 | 0.642 | 0.033 | 0.049 | 0.049 | 2.101 | 0.28 | 0.13 | 0.26 | 0.12 | 0.18 | 0.18 |
| 11 | 117284770 | rs540904 | -1.818 | -0.684 | -0.056 | 0.685 | -0.019 | 0.047 | 0.047 | 0.077 | 0.14 | 0.3 | 0.08 | 0.22 | 0.1 | 0.21 |
| 11 | 117285057 | rs10736488 | 0.614 | -0.602 | 0.066 | 0.897 | 0.028 | 0.039 | 0.039 | 1.591 | 0.3 | 0.19 | 0.28 | 0.12 | 0.19 | 0.18 |
| 11 | 117285684 | rs511549 | -1.421 | -0.521 | 0.088 | 0.942 | -0.008 | 0.092 | 0.092 | 1.591 | 0.11 | 0.3 | 0.07 | 0.22 | 0.07 | 0.18 |
| 11 | 117286064 | rs522383 | -1.043 | -0.361 | 0.098 | 1.02 | 0.01 | 0.11 | 0.11 | 0.932 | 0.09 | 0.26 | 0.07 | 0.22 | 0.07 | 0.18 |
| 11 | 117287794 | rs474490 | -0.43 | -0.011 | -0.677 | 0.165 | 0.027 | 0.024 | 0.024 | 0.653 | 0.66 | 0.77 | 0.65 | 0.72 | 0.79 | 0.66 |
| 11 | 117293929 | rs3863310 | 0.642 | 0.734 | -0.35 | 0.911 | 0.007 | -0.028 | -0.028 | 4.749 | 0.95 | 0.88 | 0.97 | 1 | 0.95 | 0.99 |
| 11 | 117296292 | rs4938377 | 0.281 | 0.763 | -0.689 | 1.201 | -0.009 | -0.018 | -0.018 | 0 | 0.95 | 0.99 | 0.92 | 0.83 | 0.94 | 0.81 |
| 11 | 117297415 | rs7940638 | -0.903 | 0.546 | -0.368 | 1.372 | -0.007 | 0.079 | 0.079 | 0 | 0.18 | 0.26 | 0.24 | 0.45 | 0.18 | 0.23 |
| 11 | 117299015 | rs2925768 | 0.118 | 0.584 | -0.636 | 0.204 | 0 | 0 | 0 | 0 | 0.56 | 0.43 | 0.48 | 0.25 | 0.63 | 0.45 |
| 11 | 117300040 | rs3889570 | -0.206 | 0.709 | -0.707 | -0.289 | 0.002 | 0.013 | 0.013 | 0 | 0.45 | 0.4 | 0.38 | 0.24 | 0.49 | 0.43 |
| 11 | 117301940 | rs7106294 | -0.77 | 0.867 | -0.436 | -1.201 | 0.003 | -0.107 | -0.107 | 0 | 0.28 | 0.35 | 0.16 | 0.01 | 0.28 | 0.22 |
| 11 | 117303858 | rs11216382 | -0.452 | 0.503 | 0.176 | -1.375 | 0.004 | -0.041 | -0.041 | 0 | 0.12 | 0.26 | 0.18 | 0.06 | 0.16 | 0.03 |

| | | | | | | | | | | | | | | | | |
|---|---|---|---|---|---|---|---|---|---|---|---|---|---|---|---|---|
| 11 | 117304196 | rs2242133 | 0.182 | -0.179 | -0.632 | -1.241 | 0.015 | -0.02 | -0.02 | 0 | 0.25 | 0.07 | 0.19 | 0.25 | 0.17 | 0.36 |
| 11 | 117307265 | rs472648 | -0.969 | -0.165 | -0.563 | -1.806 | -0.025 | 0.006 | 0.006 | 0 | 0.63 | 0.68 | 0.49 | 0.35 | 0.53 | 0.64 |
| 11 | 117308432 | rs17120753 | 0.955 | -0.41 | -0.834 | -1.834 | -0.001 | 0.019 | 0.019 | 0 | 0.1 | 0.02 | 0.05 | 0.06 | 0.08 | 0.16 |
| 11 | 117308795 | rs2253440 | 0.735 | -0.489 | -0.406 | -1.857 | 0 | 0.004 | 0.004 | 0 | 0.24 | 0.17 | 0.13 | 0.06 | 0.22 | 0.24 |
| 11 | 117308936 | rs10892102 | -0.908 | -1.312 | 0.1 | -1.143 | 0.005 | -0.003 | -0.003 | 0.792 | 0.45 | 0.58 | 0.35 | 0.82 | 0.28 | 0.58 |
| 11 | 117309989 | rs11216386 | -0.194 | -1.194 | -0.045 | -0.82 | 0.019 | -0.028 | -0.028 | 0 | 0.19 | 0.2 | 0.18 | 0.56 | 0.11 | 0.21 |
| 11 | 117312089 | rs7947950 | -1.195 | -1.049 | 0.121 | -1.122 | 0.02 | 0.071 | 0.071 | 0.832 | 0.56 | 0.71 | 0.48 | 0.86 | 0.42 | 0.63 |
| 11 | 117315692 | rs11216394 | -0.616 | -1.129 | 0.302 | -0.433 | -0.01 | -0.005 | -0.005 | 0 | 0.45 | 0.48 | 0.45 | 0.37 | 0.34 | 0.47 |
| 11 | 117317664 | rs12361458 | 0.109 | -1.249 | 0.504 | -0.457 | 0.016 | 0.007 | 0.007 | 0 | 0.34 | 0.18 | 0.37 | 0.31 | 0.3 | 0.4 |
| 11 | 117319260 | rs10790184 | -0.475 | -0.854 | 0.816 | 0.251 | -0.001 | -0.002 | -0.002 | 0.017 | 0.44 | 0.41 | 0.41 | 0.7 | 0.39 | 0.49 |
| 11 | 117320727 | rs10892108 | 1.03 | -0.863 | 0.642 | 0.151 | -0.014 | 0.017 | 0.017 | 0 | 0.39 | 0.29 | 0.3 | 0.05 | 0.48 | 0.3 |
| 11 | 117320976 | rs500713 | -1.083 | -0.713 | 0.51 | 0.42 | 0 | 0.024 | 0.024 | 0 | 0.9 | 0.93 | 0.84 | 0.57 | 0.91 | 0.84 |
| 11 | 117321606 | rs7350507 | -1.145 | -0.851 | 0.274 | 0.144 | -0.001 | -0.006 | -0.006 | 0.531 | 0.94 | 0.98 | 0.89 | 0.82 | 0.94 | 0.94 |
| 11 | 117322782 | rs4938379 | 0.03 | -0.863 | 0.335 | 0.027 | 0.009 | 0.001 | 0.001 | 0 | 0.54 | 0.61 | 0.47 | 0.53 | 0.57 | 0.49 |
| 11 | 117323389 | rs11216399 | 0.498 | -0.801 | 0.393 | 0.061 | -0.02 | 0.184 | 0.184 | 0.14 | 0.13 | 0.28 | 0.15 | 0.47 | 0.08 | 0.18 |
| 11 | 117325478 | rs10892117 | 0.4 | -0.79 | 0.394 | 0.065 | -0.017 | 0.177 | 0.177 | 0.18 | 0.13 | 0.29 | 0.16 | 0.47 | 0.09 | 0.18 |
| 11 | 117325898 | rs7103239 | 0.622 | -0.714 | -0.226 | -0.102 | 0.01 | 0.014 | 0.014 | 0 | 0.3 | 0.22 | 0.23 | 0.25 | 0.28 | 0.3 |
| 11 | 117328700 | rs648384 | -0.213 | -0.443 | -0.204 | 0.041 | 0.009 | 0.018 | 0.018 | 0 | 0.46 | 0.37 | 0.49 | 0.38 | 0.43 | 0.49 |
| 11 | 117335722 | rs538847 | -0.933 | -0.164 | -0.627 | -0.245 | -0.059 | 0.127 | 0.127 | 0 | 0.63 | 0.86 | 0.68 | 0.9 | 0.5 | 0.69 |
| 11 | 117338912 | rs2511050 | -0.242 | -0.504 | 0.346 | 0.505 | 0.047 | 0.043 | 0.043 | 0 | 0.07 | 0.32 | 0.1 | 0.07 | 0.07 | 0.03 |
| 11 | 117339926 | rs638273 | -1.3 | -1.278 | -0.137 | 0.663 | 0.06 | -0.009 | -0.009 | 0 | 0.38 | 0.83 | 0.46 | 0.62 | 0.23 | 0.4 |
| 11 | 117340285 | rs554912 | -0.743 | -1.37 | 0.465 | 0.839 | -0.03 | 0.023 | 0.023 | 0 | 0.22 | 0.48 | 0.28 | 0.23 | 0.16 | 0.22 |
| 11 | 117341808 | rs609341 | -1.426 | -1.148 | -0.329 | 0.942 | 0.02 | -0.008 | -0.008 | 0.05 | 0.53 | 0.74 | 0.47 | 0.52 | 0.38 | 0.67 |
| 11 | 117343197 | rs10892121 | -0.129 | -1.13 | -0.705 | 0.71 | 0.015 | 0.032 | 0.032 | 0 | 0.28 | 0.11 | 0.21 | 0.21 | 0.24 | 0.3 |
| 11 | 117347947 | rs1574433 | -0.591 | -0.994 | -0.108 | 0.614 | -0.007 | -0.011 | -0.011 | 0 | 0.3 | 0.35 | 0.15 | 0.21 | 0.21 | 0.44 |
| 11 | 117352949 | rs3741278 | 0.074 | -0.813 | -0.05 | 1.288 | -0.004 | 0.013 | 0.013 | 2.155 | 0.13 | 0.19 | 0.09 | 0.2 | 0.1 | 0.21 |
| 11 | 117358220 | rs549289 | -1.014 | -0.961 | -0.088 | 1.134 | -0.023 | 0.032 | 0.032 | 0 | 0.28 | 0.45 | 0.36 | 0.42 | 0.21 | 0.31 |
| 11 | 117359244 | rs12222915 | 0.176 | -1.495 | -0.559 | 0.806 | -0.014 | -0.003 | -0.003 | 0 | 0.25 | 0.26 | 0.14 | 0.17 | 0.13 | 0.27 |
| 11 | 117361861 | rs597686 | -0.195 | -0.695 | -0.975 | 1.116 | -0.027 | -0.065 | -0.065 | 0 | 0.13 | 0.02 | 0.1 | 0.2 | 0.15 | 0.18 |
| 11 | 117362074 | rs540277 | -1.507 | -1.23 | -0.192 | 1.334 | -0.018 | 0.048 | 0.048 | 0 | 0.26 | 0.4 | 0.21 | 0.38 | 0.23 | 0.27 |
| 11 | 117362198 | rs2013131 | -0.025 | -0.576 | -0.706 | 1.091 | -0.055 | -0.048 | -0.048 | 0.14 | 0.64 | 0.81 | 0.47 | 0.48 | 0.5 | 0.7 |
| 11 | 117362948 | rs582285 | -1.144 | -0.549 | -0.704 | 1.095 | -0.038 | -0.024 | -0.024 | 0.318 | 0.77 | 0.87 | 0.57 | 0.64 | 0.65 | 0.88 |
| 11 | 117363426 | rs11216425 | 0.583 | -0.751 | -0.32 | 0.924 | -0.005 | -0.001 | -0.001 | 0.063 | 0.08 | 0.1 | 0.08 | 0.09 | 0.05 | 0.07 |
| 11 | 117364821 | rs12286920 | 0.994 | -1.245 | -0.217 | 0.453 | 0.005 | -0.052 | -0.052 | 0.258 | 0.26 | 0.28 | 0.15 | 0.05 | 0.23 | 0.27 |
| 11 | 117371563 | rs7942406 | -0.661 | -1.364 | -0.174 | 0.735 | -0.025 | -0.039 | -0.039 | 0 | 0.64 | 0.76 | 0.49 | 0.45 | 0.54 | 0.72 |
| 11 | 117373432 | rs7112940 | 0.446 | -1.391 | -0.102 | 0.388 | -0.001 | 0.014 | 0.014 | 0 | 0.52 | 0.45 | 0.37 | 0.29 | 0.47 | 0.63 |
| 11 | 117380108 | rs4936383 | -0.356 | -1.071 | -0.662 | 0.553 | -0.047 | -0.019 | -0.019 | 0.436 | 0.63 | 0.79 | 0.46 | 0.37 | 0.52 | 0.69 |
| 11 | 117391966 | rs2276340 | 0.833 | -1.584 | -0.578 | 0.555 | 0.056 | -0.023 | -0.023 | 0 | 0.25 | 0.09 | 0.19 | 0.25 | 0.16 | 0.34 |
| 11 | 117392337 | rs4938397 | 0.422 | -1.715 | -0.267 | 0.748 | 0.029 | -0.013 | -0.013 | 0.104 | 0.29 | 0.2 | 0.24 | 0.38 | 0.18 | 0.34 |
| 11 | 117394979 | rs12577702 | 0.178 | -1.757 | 0.269 | 0.952 | -0.006 | 0.036 | 0.036 | 0 | 0.08 | 0.15 | 0.11 | 0.14 | 0.06 | 0.22 |
| 11 | 117396505 | rs11603366 | 0.507 | -1.933 | 0.314 | 0.871 | 0 | -0.002 | -0.002 | 0 | 0.3 | 0.28 | 0.26 | 0.38 | 0.18 | 0.33 |
| 11 | 117403235 | rs3741280 | -0.387 | -1.811 | 0.604 | 0.868 | -0.037 | 0.024 | 0.024 | 0 | 0.39 | 0.48 | 0.37 | 0.54 | 0.27 | 0.57 |
| 11 | 117407301 | rs521609 | 0.33 | -1.788 | 0.415 | 0.822 | -0.006 | 0.001 | 0.001 | 0 | 0.29 | 0.3 | 0.27 | 0.37 | 0.2 | 0.44 |
| 11 | 117409700 | rs11216443 | 1.267 | -1.554 | 0.031 | 0.735 | -0.073 | 0.045 | 0.045 | 0 | 0.26 | 0.09 | 0.38 | 0.18 | 0.39 | 0.15 |
| 11 | 117409910 | rs4938403 | 1.295 | -1.694 | 0.079 | 0.534 | -0.001 | 0.001 | 0.001 | 0 | 0.07 | 0.07 | 0.07 | 0.12 | 0.09 | 0.02 |
| 11 | 117412621 | rs679223 | 1.627 | -1.8 | 0.043 | 0.514 | 0.008 | -0.007 | -0.007 | 0 | 0.21 | 0.16 | 0.2 | 0.26 | 0.11 | 0.42 |
| 11 | 117413864 | rs678468 | 0.253 | -1.751 | -0.001 | 0.656 | 0.007 | 0.089 | 0.089 | 0.471 | 0.5 | 0.25 | 0.59 | 0.64 | 0.55 | 0.66 |
| 11 | 117414948 | rs3802877 | 1.455 | -1.595 | -0.027 | 0.718 | 0 | 0 | 0 | 0 | 0.07 | 0.05 | 0.07 | 0.12 | 0.09 | 0.02 |
| 11 | 117418063 | rs4084228 | 0.611 | -1.781 | -0.129 | 0.566 | -0.076 | -0.074 | -0.074 | 0.546 | 0.33 | 0.12 | 0.42 | 0.4 | 0.45 | 0.25 |
| 11 | 117422642 | rs640147 | -0.562 | -1.827 | 0.235 | 0.549 | -0.001 | -0.072 | -0.072 | 0.036 | 0.61 | 0.5 | 0.73 | 0.85 | 0.66 | 0.71 |
| 11 | 117426543 | rs490111 | 1.277 | -1.804 | 0.179 | 0.669 | -0.026 | 0.021 | 0.021 | 0 | 0.23 | 0.29 | 0.24 | 0.34 | 0.13 | 0.45 |
| 11 | 117427923 | rs4938406 | 0.704 | -1.446 | -0.04 | 0.775 | 0 | 0.013 | 0.013 | 0.011 | 0.08 | 0.12 | 0.11 | 0.19 | 0.1 | 0.04 |
| 11 | 117430058 | rs510250 | -0.684 | -1.62 | 0.043 | 0.828 | -0.047 | -0.051 | -0.051 | 0 | 0.67 | 0.84 | 0.57 | 0.58 | 0.56 | 0.75 |
| 11 | 117430931 | rs567053 | -0.506 | -1.51 | 0.472 | 0.993 | -0.014 | 0.032 | 0.032 | 0 | 0.92 | 0.78 | 0.89 | 0.87 | 0.94 | 0.93 |
| 11 | 117431045 | rs623052 | 0.341 | -1.341 | 0.649 | 0.748 | 0.002 | 0.002 | 0.002 | 0 | 0.6 | 0.45 | 0.49 | 0.47 | 0.55 | 0.57 |
| 11 | 117432034 | rs1893970 | 1.338 | -1.119 | 0.487 | 0.813 | -0.017 | -0.008 | -0.008 | 3.461 | 0.21 | 0.14 | 0.34 | 0.28 | 0.31 | 0.14 |
| 11 | 117432779 | rs17509752 | 1.399 | -1.099 | 0.496 | 0.801 | -0.015 | 0.001 | 0.001 | 0 | 0.21 | 0.14 | 0.32 | 0.2 | 0.3 | 0.12 |
| 11 | 117433686 | rs755791 | -0.089 | -1.529 | 0.597 | 0.642 | -0.002 | -0.001 | -0.001 | 0.07 | 0.56 | 0.46 | 0.54 | 0.34 | 0.67 | 0.33 |
| 11 | 117434215 | rs685916 | 0.568 | -1.364 | 0.448 | 0.499 | 0.014 | 0.039 | 0.039 | 0 | 0.46 | 0.28 | 0.41 | 0.35 | 0.44 | 0.44 |
| 11 | 117434487 | rs17509913 | 2.347 | -1.336 | 0.372 | 0.479 | 0.004 | 0.034 | 0.034 | 0 | 0.14 | 0.08 | 0.05 | 0 | 0.1 | 0.11 |
| 11 | 117434728 | rs495118 | 0.22 | -1.201 | 0.592 | 0.796 | -0.013 | 0.111 | 0.111 | 0 | 0.85 | 0.74 | 0.83 | 0.57 | 0.87 | 0.79 |
| 11 | 117435785 | rs12789727 | 0.858 | -1.268 | 0.503 | 0.682 | -0.025 | 0.02 | 0.02 | 0.474 | 0.2 | 0.12 | 0.15 | 0.14 | 0.26 | 0.09 |
| 11 | 117436095 | rs1940037 | 0.182 | -1.328 | 0.591 | 0.667 | 0.031 | -0.014 | -0.014 | 0 | 0.23 | 0.31 | 0.23 | 0.23 | 0.3 | 0.21 |
| 11 | 117436190 | rs490458 | 0.907 | -1.262 | 0.386 | 0.686 | 0.011 | 0.022 | 0.022 | 1.559 | 0.62 | 0.5 | 0.64 | 0.44 | 0.58 | 0.6 |
| 11 | 117436906 | rs1893900 | 0.869 | -1.257 | 0.18 | 0.514 | -0.013 | 0.025 | 0.025 | 0 | 0.37 | 0.32 | 0.39 | 0.22 | 0.4 | 0.24 |
| 11 | 117441393 | rs557940 | -0.71 | -1.012 | -0.242 | 0.597 | -0.006 | 0.002 | 0.002 | 0 | 0.89 | 0.86 | 0.89 | 0.85 | 0.92 | 0.91 |
| 11 | 117441495 | rs657137 | 0.168 | -0.897 | -0.304 | 0.618 | 0.014 | -0.01 | -0.01 | 0 | 0.69 | 0.58 | 0.72 | 0.7 | 0.65 | 0.72 |
| 11 | 117446945 | rs504935 | 1.319 | -1.23 | -0.689 | 0.661 | 0.056 | -0.004 | -0.004 | 0 | 0.27 | 0.07 | 0.27 | 0.22 | 0.2 | 0.48 |
| 11 | 117447373 | rs648740 | 0.171 | -1.151 | -0.287 | 0.932 | -0.008 | 0.068 | 0.068 | 0 | 0.87 | 0.75 | 0.87 | 0.65 | 0.87 | 0.83 |

| | | | | | | | | | | | | | | | | |
|---|---|---|---|---|---|---|---|---|---|---|---|---|---|---|---|---|
| 11 | 117455669 | rs474749 | 0.364 | -1.033 | -0.43 | 0.653 | -0.002 | 0.001 | 0.001 | 0 | 0.63 | 0.35 | 0.68 | 0.51 | 0.69 | 0.67 |
| 11 | 117458539 | rs600620 | 1.526 | -1.28 | -0.523 | 0.442 | 0.053 | -0.022 | -0.022 | 0.245 | 0.28 | 0.12 | 0.28 | 0.28 | 0.22 | 0.51 |
| 11 | 117461576 | rs647593 | 0.936 | -1.348 | -0.041 | 0.334 | -0.01 | -0.005 | -0.005 | 0.167 | 0.43 | 0.43 | 0.35 | 0.34 | 0.31 | 0.6 |
| 11 | 117461602 | rs553637 | -0.426 | -1.116 | -0.355 | 0.159 | -0.097 | -0.117 | -0.117 | 0.257 | 0.66 | 0.87 | 0.57 | 0.52 | 0.57 | 0.69 |
| 11 | 117464173 | rs694005 | 1.126 | -1.216 | -0.146 | 0.184 | 0.019 | 0.007 | 0.007 | 0 | 0.42 | 0.36 | 0.36 | 0.36 | 0.3 | 0.58 |
| 11 | 117466151 | rs12803154 | 1.195 | -0.807 | -0.207 | 0.287 | -0.008 | 0.005 | 0.005 | 0 | 0.17 | 0.12 | 0.17 | 0.14 | 0.25 | 0.09 |
| 11 | 117467394 | rs2186761 | 0.683 | -0.837 | -0.055 | 0.368 | -0.018 | 0.034 | 0.034 | 0 | 0.31 | 0.18 | 0.23 | 0.19 | 0.34 | 0.19 |
| 11 | 117468794 | rs622860 | 0.854 | -1.848 | 0.065 | 0.529 | 0.024 | 0.008 | 0.008 | 0 | 0.3 | 0.24 | 0.26 | 0.26 | 0.19 | 0.37 |
| 11 | 117471825 | rs577651 | -0.69 | -1.384 | -0.02 | 0.822 | 0.001 | -0.003 | -0.003 | 0 | 0.92 | 0.9 | 0.89 | 0.85 | 0.93 | 0.92 |
| 11 | 117474580 | rs11216466 | -0.306 | -1.365 | 0.257 | 0.496 | -0.022 | 0.02 | 0.02 | 0 | 0.1 | 0.28 | 0.08 | 0.1 | 0.05 | 0.16 |
| 11 | 117476142 | rs666290 | 0.81 | -1.224 | -0.474 | 0.391 | 0.017 | 0.005 | 0.005 | 0 | 0.27 | 0.23 | 0.24 | 0.25 | 0.19 | 0.34 |
| 11 | 117488717 | rs4936389 | 0.075 | -1.017 | -0.492 | 0.538 | 0.003 | 0.002 | 0.002 | 0 | 0.45 | 0.58 | 0.68 | 0.52 | 0.55 | 0.28 |
| 11 | 117489645 | rs685184 | -0.308 | -1.023 | -0.686 | 0.574 | -0.007 | -0.024 | -0.024 | 0 | 0.88 | 0.93 | 0.91 | 0.78 | 0.85 | 0.83 |
| 11 | 117494945 | rs7946890 | 1.028 | -1.091 | -0.313 | 0.307 | -0.005 | -0.057 | -0.057 | 0 | 0.15 | 0.19 | 0.07 | 0 | 0.1 | 0.12 |
| 11 | 117496132 | rs10790195 | -0.756 | -0.971 | -0.283 | 0.293 | -0.001 | 0 | 0 | 0 | 0.21 | 0.18 | 0.18 | 0.22 | 0.23 | 0.22 |
| 11 | 117496475 | rs10790196 | -1.478 | -1.258 | -0.137 | 0.401 | 0.034 | -0.007 | -0.007 | 0 | 0.7 | 0.79 | 0.82 | 0.63 | 0.76 | 0.55 |
| 11 | 117497778 | rs11216474 | -0.131 | -0.944 | -0.437 | 0.652 | 0.007 | 0.035 | 0.035 | 0 | 0.43 | 0.19 | 0.54 | 0.33 | 0.54 | 0.3 |
| 11 | 117499026 | rs7941794 | -0.97 | -1.033 | -0.309 | 0.807 | 0.014 | -0.008 | -0.008 | 0 | 0.62 | 0.66 | 0.68 | 0.42 | 0.67 | 0.43 |
| 11 | 117501072 | rs7949956 | 0.085 | -0.839 | 0.063 | 0.504 | -0.005 | 0 | 0 | 0.245 | 0.47 | 0.49 | 0.62 | 0.39 | 0.52 | 0.32 |
| 11 | 117501603 | rs10790197 | 0.874 | -0.531 | -0.013 | 0.495 | -0.051 | -0.053 | -0.053 | 0 | 0.32 | 0.17 | 0.53 | 0.38 | 0.41 | 0.21 |
| 11 | 117502961 | rs7929496 | 1.63 | -0.7 | 0.198 | 0.435 | -0.001 | -0.027 | -0.027 | 0 | 0.14 | 0.12 | 0.06 | 0 | 0.11 | 0.11 |
| 11 | 117503394 | rs4936393 | -0.296 | -0.838 | 0.154 | 0.573 | -0.008 | 0 | 0 | 0 | 0.53 | 0.49 | 0.67 | 0.43 | 0.64 | 0.34 |
| 11 | 117508837 | rs695083 | 0.072 | -0.78 | -0.038 | 0.603 | 0.001 | -0.006 | -0.006 | 0 | 0.46 | 0.39 | 0.62 | 0.44 | 0.54 | 0.4 |
| 11 | 117509797 | rs4938423 | -0.149 | -0.818 | -0.019 | 0.653 | 0.001 | 0.003 | 0.003 | 0.266 | 0.48 | 0.45 | 0.67 | 0.56 | 0.55 | 0.42 |
| 11 | 117511787 | rs512017 | -0.893 | -0.791 | -0.204 | 0.599 | 0.033 | 0.001 | 0.001 | 0 | 0.65 | 0.72 | 0.78 | 0.62 | 0.73 | 0.54 |
| 11 | 117513154 | rs613244 | -1.09 | -0.742 | -0.485 | 0.584 | 0.031 | 0.001 | 0.001 | 0 | 0.67 | 0.75 | 0.8 | 0.64 | 0.73 | 0.55 |
| 11 | 117513801 | rs10892147 | 0.561 | -0.45 | -0.829 | 0.318 | 0.013 | -0.022 | -0.022 | 0.901 | 0.19 | 0.2 | 0.12 | 0.1 | 0.18 | 0.14 |
| 11 | 117517366 | rs536761 | 0.044 | -0.403 | -1.35 | 0.602 | 0.022 | -0.064 | -0.064 | 0.238 | 0.51 | 0.05 | 0.64 | 0.42 | 0.61 | 0.35 |
| 11 | 117521967 | rs16867 | 0.321 | 0.094 | -0.437 | 0.388 | -0.001 | 0.014 | 0.014 | 0 | 0.22 | 0.23 | 0.16 | 0.44 | 0.18 | 0.25 |
| 11 | 117524543 | rs10892150 | 0.887 | 0.095 | -0.639 | -0.063 | -0.047 | 0.126 | 0.126 | 0 | 0.32 | 0.1 | 0.3 | 0.13 | 0.36 | 0.31 |
| 11 | 117525125 | rs583983 | -1.181 | -0.077 | -0.699 | -0.07 | -0.011 | 0.022 | 0.022 | 0 | 0.71 | 0.89 | 0.79 | 0.78 | 0.64 | 0.65 |
| 11 | 117525420 | rs7950072 | 0.179 | 0.105 | -0.028 | 0.111 | 0.033 | 0.052 | 0.052 | 0.66 | 0.09 | 0.24 | 0.23 | 0.15 | 0.09 | 0.06 |
| 11 | 117530935 | rs7949670 | -0.916 | -0.119 | 0.118 | -0.616 | -0.044 | -0.003 | -0.003 | 0 | 0.13 | 0.49 | 0.09 | 0.11 | 0.04 | 0.22 |
| 11 | 117531465 | rs1892918 | 0.794 | 0.01 | -0.035 | -0.611 | -0.013 | -0.033 | -0.033 | 0 | 0.24 | 0.19 | 0.37 | 0.58 | 0.23 | 0.23 |
| 11 | 117533949 | rs12279593 | 0.64 | -0.007 | -0.141 | -0.768 | 0.005 | -0.081 | -0.081 | 0.288 | 0.33 | 0.45 | 0.31 | 0.14 | 0.34 | 0.3 |
| 11 | 117534353 | rs630024 | -0.912 | 0.071 | -0.668 | -0.788 | 0.005 | 0.044 | 0.044 | 0 | 0.67 | 0.84 | 0.75 | 0.77 | 0.64 | 0.61 |
| 11 | 117535285 | rs11216489 | 0.04 | 0.205 | -0.774 | -0.874 | -0.003 | 0 | 0 | 0 | 0.39 | 0.38 | 0.34 | 0.16 | 0.39 | 0.35 |
| 11 | 117536136 | rs11216491 | 1.041 | 0.498 | -1.006 | -0.676 | -0.098 | 0.114 | 0.114 | 0 | 0.27 | 0.01 | 0.28 | 0.08 | 0.32 | 0.12 |
| 11 | 117536758 | rs11216492 | 0.874 | 0.431 | -0.919 | -0.568 | 0.009 | 0.024 | 0.024 | 0 | 0.31 | 0.2 | 0.25 | 0.25 | 0.3 | 0.39 |
| 11 | 117537353 | rs10431082 | 0.989 | 0.46 | -0.944 | -0.537 | 0.006 | 0.023 | 0.023 | 0 | 0.3 | 0.19 | 0.24 | 0.24 | 0.3 | 0.38 |
| 11 | 117537710 | rs686413 | 0.642 | 0.576 | -0.986 | -0.272 | 0 | -0.015 | -0.015 | 0 | 0.23 | 0.22 | 0.33 | 0.55 | 0.21 | 0.2 |
| 11 | 117538766 | rs593380 | 0.519 | 1.009 | -0.681 | -0.731 | -0.002 | -0.007 | -0.007 | 0 | 0.61 | 0.66 | 0.6 | 0.89 | 0.56 | 0.78 |
| 11 | 117539478 | rs947926 | -0.41 | 1.258 | -1.512 | -0.305 | -0.156 | 0.097 | 0.097 | 0 | 0.29 | 0.02 | 0.33 | 0.1 | 0.41 | 0.19 |
| 11 | 117542149 | rs11216493 | -1.357 | 0.576 | -0.656 | 0.26 | -0.001 | -0.001 | -0.001 | 0 | 0.13 | 0.14 | 0.11 | 0.1 | 0.13 | 0.1 |
| 11 | 117544853 | rs7942751 | -1.209 | 0.455 | 0.098 | 0.165 | -0.002 | -0.008 | -0.008 | 2.216 | 0.12 | 0.25 | 0.13 | 0.09 | 0.12 | 0.1 |
| 11 | 117549808 | rs7926736 | -1.809 | 0.829 | 0.253 | 0.176 | 0.004 | -0.005 | -0.005 | 0 | 0.16 | 0.23 | 0.14 | 0.15 | 0.15 | 0.15 |
| 11 | 117550767 | rs10892158 | -2.871 | 0.67 | 0.914 | 0.001 | 0.01 | -0.037 | -0.037 | 0 | 0.23 | 0.5 | 0.22 | 0.22 | 0.23 | 0.21 |
| 11 | 117552758 | rs12295127 | -1.678 | 0.643 | 0.886 | -0.009 | -0.014 | 0.004 | 0.004 | 0 | 0.15 | 0.09 | 0.13 | 0.16 | 0.15 | 0.15 |
| 11 | 117554338 | rs1942924 | -2.786 | 0.676 | 0.963 | 0.017 | 0.029 | -0.015 | -0.015 | 0 | 0.2 | 0.45 | 0.2 | 0.22 | 0.21 | 0.2 |
| 11 | 117555933 | rs7129597 | -1.431 | 0.659 | 0.973 | 0.056 | 0.056 | 0.021 | 0.021 | 0 | 0.07 | 0.34 | 0.08 | 0.05 | 0.08 | 0.05 |
| 11 | 117561265 | rs10892162 | -2.646 | 0.315 | 1.003 | 0.072 | 0.014 | 0.015 | 0.015 | 0.048 | 0.18 | 0.57 | 0.15 | 0.23 | 0.17 | 0.15 |
| 11 | 117564052 | rs11216506 | -2.096 | 0.253 | 1.108 | 0.011 | 0.008 | -0.041 | -0.041 | 0 | 0.11 | 0.37 | 0.08 | 0.06 | 0.09 | 0.03 |
| 11 | 117567949 | rs10790200 | -1.472 | 0.234 | 1.376 | 0.002 | 0.016 | -0.018 | -0.018 | 0 | 0.07 | 0.18 | 0.06 | 0.05 | 0.08 | 0.03 |
| 11 | 117570851 | rs603377 | 0.712 | 0.244 | 0.269 | -0.255 | -0.049 | -0.003 | -0.003 | 0 | 0.19 | 0.01 | 0.29 | 0.11 | 0.23 | 0.11 |
| 11 | 117571032 | rs10892164 | 1.277 | 0.27 | -0.394 | -0.088 | -0.09 | -0.005 | -0.005 | 0 | 0.33 | 0.01 | 0.19 | 0.24 | 0.33 | 0.44 |
| 11 | 117571426 | rs7130852 | -2.02 | 0.198 | 0.519 | 0.08 | -0.003 | 0.006 | 0.006 | 0 | 0.19 | 0.46 | 0.13 | 0.23 | 0.16 | 0.16 |
| 11 | 117575629 | rs2508560 | 0.425 | -0.137 | 1.02 | 0.179 | 0.022 | 0.109 | 0.109 | 0.171 | 0.94 | 0.75 | 0.94 | 0.85 | 0.98 | 0.94 |
| 11 | 117579724 | rs11216509 | 0.823 | -0.191 | 0.771 | 0.069 | -0.082 | 0.016 | 0.016 | 0 | 0.21 | 0.01 | 0.36 | 0.13 | 0.27 | 0.12 |
| 11 | 117580509 | rs655086 | 1.321 | -0.231 | 1.79 | 0.588 | 0.022 | 0.116 | 0.116 | 1.314 | 0.93 | 0.69 | 0.94 | 0.85 | 0.98 | 0.94 |
| 11 | 117582054 | rs948777 | 2.051 | -0.273 | 1.794 | 0.593 | -0.051 | 0.053 | 0.053 | 0.101 | 0.87 | 0.49 | 0.9 | 0.8 | 0.95 | 0.93 |
| 11 | 117591292 | rs4936399 | 1.88 | -0.285 | 1.812 | 0.627 | -0.043 | 0.073 | 0.073 | 0.118 | 0.88 | 0.54 | 0.9 | 0.8 | 0.95 | 0.92 |
| 11 | 117591584 | rs948106 | 0.43 | -0.228 | 1.76 | 0.602 | -0.026 | -0.012 | -0.012 | 0 | 0.94 | 0.84 | 0.97 | 0.95 | 0.97 | 0.99 |
| 11 | 117594364 | rs4938435 | 1.064 | -0.236 | 1.715 | 0.6 | -0.015 | 0.011 | 0.011 | 0 | 0.93 | 0.61 | 0.94 | 0.95 | 0.97 | 0.99 |
| 11 | 117595511 | rs11216517 | -0.186 | -0.034 | 1.467 | 0.868 | -0.054 | -0.083 | -0.083 | 0.568 | 0.13 | 0.01 | 0.15 | 0.27 | 0.19 | 0.2 |
| 11 | 117604207 | rs4142640 | -0.807 | 0.06 | 1.629 | 0.814 | 0.016 | 0.096 | 0.096 | 0.512 | 0.06 | 0.2 | 0.05 | 0.13 | 0.03 | 0.07 |
| 11 | 117605098 | rs1942929 | 0.756 | 0.047 | 1.658 | 0.811 | -0.034 | 0.079 | 0.079 | 0.016 | 0.89 | 0.58 | 0.91 | 0.82 | 0.94 | 0.92 |
| 11 | 117606320 | rs4936404 | 0.999 | 0.038 | 1.683 | 0.806 | -0.048 | 0.065 | 0.065 | 0.518 | 0.87 | 0.5 | 0.91 | 0.82 | 0.94 | 0.92 |
| 11 | 117607138 | rs1942930 | 1.112 | 0.043 | 1.594 | 0.808 | -0.063 | 0.02 | 0.02 | 1.76 | 0.86 | 0.36 | 0.89 | 0.82 | 0.94 | 0.92 |
| 11 | 117608613 | rs1939981 | 1.413 | 0.013 | 1.555 | 0.831 | -0.014 | 0.032 | 0.032 | 1.097 | 0.71 | 0.36 | 0.71 | 0.53 | 0.74 | 0.76 |

| | | | | | | | | | | | | | | | | |
|---|---|---|---|---|---|---|---|---|---|---|---|---|---|---|---|---|
| 11 | 117622809 | rs4576856 | 1.209 | -0.025 | 0.68 | 0.079 | -0.02 | 0.004 | 0.004 | 0.851 | 0.84 | 0.35 | 0.85 | 0.82 | 0.88 | 0.89 |
| 11 | 117630867 | rs11825655 | -0.892 | 0.037 | 0.353 | -0.266 | -0.001 | -0.005 | -0.005 | 4.17 | 0.23 | 0.24 | 0.1 | 0.12 | 0.17 | 0.16 |
| 11 | 117631234 | rs10750111 | 1.383 | -0.198 | 0.125 | -0.49 | 0.022 | 0.083 | 0.083 | 0.341 | 0.62 | 0.22 | 0.77 | 0.82 | 0.7 | 0.78 |
| 11 | 117633054 | rs10892169 | 0.891 | -0.2 | 0.364 | -0.469 | 0.001 | 0.001 | 0.001 | 0.714 | 0.68 | 0.32 | 0.81 | 0.83 | 0.75 | 0.8 |
| 11 | 117638627 | rs4936407 | 1.175 | -0.062 | 0.326 | -0.358 | 0.014 | 0.05 | 0.05 | 0.785 | 0.64 | 0.26 | 0.77 | 0.82 | 0.71 | 0.8 |
| 11 | 117644511 | rs10790208 | 0.419 | -0.094 | 0.458 | -0.081 | -0.002 | 0.011 | 0.011 | 1.201 | 0.79 | 0.65 | 0.81 | 0.74 | 0.81 | 0.83 |
| 11 | 117646291 | rs7945999 | -0.654 | -0.006 | 0.057 | -0.008 | -0.015 | -0.048 | -0.048 | 0.157 | 0.14 | 0.21 | 0.05 | 0.01 | 0.08 | 0.04 |
| 11 | 117646768 | rs7119967 | -0.676 | -0.045 | -0.018 | -0.009 | -0.014 | -0.044 | -0.044 | 1.045 | 0.14 | 0.2 | 0.05 | 0.01 | 0.07 | 0.04 |
| 11 | 117647496 | rs17311045 | -0.608 | -0.007 | -0.3 | 0.01 | -0.001 | -0.016 | -0.016 | 0 | 0.14 | 0.13 | 0.04 | 0.01 | 0.07 | 0.04 |
| 11 | 117649675 | rs7924993 | 0.244 | 0.825 | -0.534 | 0.966 | -0.012 | -0.009 | -0.009 | 0 | 0.9 | 0.91 | 0.81 | 0.76 | 0.82 | 0.89 |
| 11 | 117653350 | rs7940750 | 0.2 | 0.578 | -0.621 | 0.463 | -0.001 | -0.006 | -0.006 | 0 | 0.38 | 0.37 | 0.39 | 0.28 | 0.38 | 0.22 |
| 11 | 117653955 | rs7119911 | 0.033 | -0.051 | -0.69 | 0.265 | -0.008 | 0.032 | 0.032 | 0 | 0.44 | 0.28 | 0.46 | 0.33 | 0.46 | 0.31 |
| 11 | 117654176 | rs2155387 | 0.577 | -0.595 | -0.35 | 0.175 | 0.003 | 0.013 | 0.013 | 0 | 0.69 | 0.61 | 0.64 | 0.65 | 0.65 | 0.68 |
| 11 | 117656310 | rs948102 | -0.147 | -0.497 | -0.143 | 0.184 | 0 | -0.005 | -0.005 | 0 | 0.56 | 0.45 | 0.55 | 0.34 | 0.57 | 0.34 |
| 11 | 117661100 | rs625447 | 0.78 | -1.004 | -0.61 | -0.077 | -0.058 | 0.099 | 0.099 | 0 | 0.36 | 0.13 | 0.38 | 0.18 | 0.42 | 0.22 |
| 11 | 117661447 | rs12291974 | 0.258 | -1.059 | -0.359 | 0.01 | -0.016 | 0.022 | 0.022 | 0 | 0.14 | 0.24 | 0.15 | 0.15 | 0.11 | 0.12 |
| 11 | 117662138 | rs12795949 | -0.117 | -1.785 | -0.966 | -0.406 | 0.043 | 0.099 | 0.099 | 0.016 | 0.27 | 0.09 | 0.17 | 0.05 | 0.19 | 0.14 |
| 11 | 117662248 | rs534761 | 0.967 | -2.179 | -1.48 | -0.249 | -0.067 | 0.017 | 0.017 | 0.019 | 0.26 | 0.02 | 0.18 | 0.21 | 0.28 | 0.38 |
| 11 | 117662909 | rs528431 | 0.435 | -1.697 | -0.998 | -0.041 | 0.029 | 0.013 | 0.013 | 0 | 0.4 | 0.22 | 0.27 | 0.39 | 0.34 | 0.47 |
| 11 | 117669619 | rs1892903 | -0.633 | -1.75 | -0.473 | -0.353 | -0.011 | 0.043 | 0.043 | 0 | 0.52 | 0.71 | 0.56 | 0.36 | 0.52 | 0.43 |
| 11 | 117671489 | rs3819132 | -0.729 | -1.613 | -0.467 | -0.468 | 0 | 0.046 | 0.046 | 0 | 0.54 | 0.7 | 0.58 | 0.34 | 0.56 | 0.44 |
| 11 | 117673828 | rs10488782 | 0.514 | -1.522 | -0.736 | -0.379 | -0.024 | -0.005 | -0.005 | 0.063 | 0.17 | 0.26 | 0.13 | 0.13 | 0.11 | 0.11 |
| 11 | 117674355 | rs10892175 | 2.066 | -1.495 | -0.858 | -0.406 | -0.065 | 0.035 | 0.035 | 0 | 0.16 | 0.06 | 0.24 | 0.08 | 0.32 | 0.1 |
| 11 | 117678543 | rs12790243 | -0.574 | -1.365 | -0.65 | -0.238 | -0.003 | 0.019 | 0.019 | 0 | 0.47 | 0.44 | 0.53 | 0.29 | 0.51 | 0.29 |
| 11 | 117685343 | rs10892178 | 0.241 | -0.757 | -1.085 | 0.038 | -0.013 | 0.01 | 0.01 | 0 | 0.31 | 0.25 | 0.4 | 0.23 | 0.4 | 0.2 |
| 11 | 117685575 | rs11216561 | 0.338 | -0.6 | -1.654 | -0.018 | -0.072 | 0.107 | 0.107 | 0 | 0.29 | 0.11 | 0.33 | 0.1 | 0.39 | 0.11 |
| 11 | 117688898 | rs540442 | 0.235 | -0.461 | -1.423 | 0.918 | 0.011 | -0.026 | -0.026 | 0.047 | 0.36 | 0.22 | 0.37 | 0.42 | 0.34 | 0.38 |
| 11 | 117690400 | rs11600216 | 1.153 | 1.293 | -1.889 | 1.301 | -0.058 | -0.057 | -0.057 | 0 | 0.09 | 0.01 | 0.14 | 0.23 | 0.18 | 0.14 |
| 11 | 117690543 | rs17121264 | 0.742 | 0.817 | -1.932 | 0.914 | 0.004 | -0.009 | -0.009 | 0 | 0.09 | 0.1 | 0.09 | 0.23 | 0.07 | 0.23 |
| 11 | 117690918 | rs11999 | 0.63 | 0.377 | -2.257 | -0.023 | -0.015 | 0.127 | 0.127 | 0 | 0.32 | 0.12 | 0.31 | 0.1 | 0.35 | 0.14 |
| 11 | 117691145 | rs2014536 | -0.728 | -0.343 | -2.029 | -0.004 | 0.038 | 0.03 | 0.03 | 0 | 0.56 | 0.31 | 0.68 | 0.67 | 0.68 | 0.44 |
| 11 | 117691386 | rs869789 | 0.922 | 0.105 | -1.98 | 0.314 | -0.004 | -0.014 | -0.014 | 0.279 | 0.14 | 0.07 | 0.18 | 0.19 | 0.13 | 0.04 |
| 11 | 117692121 | rs10892179 | 0.973 | 0.197 | -2.241 | -0.527 | 0.002 | 0.064 | 0.064 | 0.017 | 0.19 | 0.09 | 0.14 | 0 | 0.17 | 0.08 |
| 11 | 117692170 | rs10750114 | 0.844 | 0.166 | -2.229 | -0.17 | 0.005 | 0.006 | 0.006 | 0.005 | 0.2 | 0.09 | 0.23 | 0.22 | 0.17 | 0.19 |
| 11 | 117692334 | rs4579962 | 0.794 | 0.187 | -2.187 | -0.128 | -0.062 | 0.008 | 0.008 | 0.611 | 0.21 | 0.05 | 0.27 | 0.16 | 0.28 | 0.19 |
| 11 | 117693255 | rs529623 | 0.506 | -0.378 | -2.022 | -0.176 | 0.006 | 0.063 | 0.063 | 0 | 0.46 | 0.24 | 0.53 | 0.55 | 0.57 | 0.53 |
| 11 | 117694089 | rs635706 | 0.593 | -0.45 | -1.774 | 0.161 | -0.013 | 0.007 | 0.007 | 0 | 0.33 | 0.53 | 0.33 | 0.48 | 0.21 | 0.43 |
| 11 | 117695340 | rs3824943 | 0.63 | 0.359 | -2.121 | 0.694 | -0.007 | 0.001 | 0.001 | 0 | 0.08 | 0.04 | 0.07 | 0.07 | 0.08 | 0.1 |
| 11 | 117698483 | rs11216567 | 0.664 | 0.395 | -2.001 | 0.743 | -0.002 | 0.025 | 0.025 | 0 | 0.1 | 0.18 | 0.08 | 0.21 | 0.09 | 0.14 |
| 11 | 117698762 | rs7117314 | -0.043 | 1.073 | -2.654 | 0.674 | -0.049 | -0.011 | -0.011 | 0.278 | 0.28 | 0.18 | 0.4 | 0.31 | 0.4 | 0.19 |
| 11 | 117699878 | rs2019655 | 0.596 | 0.886 | -2.778 | 0.974 | -0.025 | -0.084 | -0.084 | 0 | 0.25 | 0.04 | 0.18 | 0.24 | 0.25 | 0.31 |
| 11 | 117700323 | rs4472969 | -0.484 | 1.053 | -2.162 | 1.213 | -0.006 | -0.001 | -0.001 | 1.012 | 0.86 | 0.88 | 0.76 | 0.6 | 0.81 | 0.82 |
| 11 | 117700588 | rs2004506 | 0.571 | 0.264 | -1.996 | 0.39 | -0.035 | 0.019 | 0.019 | 0 | 0.1 | 0.01 | 0.17 | 0.07 | 0.16 | 0.1 |
| 11 | 117702115 | rs7130303 | -0.557 | 0.139 | -1.984 | 0.619 | -0.003 | 0.003 | 0.003 | 0.101 | 0.58 | 0.8 | 0.54 | 0.44 | 0.43 | 0.54 |
| 11 | 117702690 | rs10790212 | -0.309 | -0.155 | -1.264 | 0.189 | -0.024 | -0.05 | -0.05 | 0.66 | 0.34 | 0.56 | 0.3 | 0.16 | 0.24 | 0.23 |
| 11 | 117707229 | rs10892184 | -0.232 | 0.557 | -1.746 | 1.229 | -0.061 | -0.102 | -0.102 | 0 | 0.15 | 0.02 | 0.39 | 0.41 | 0.22 | 0.15 |
| 11 | 117707937 | rs3087563 | -0.771 | -0.382 | -1.448 | 0.71 | 0.015 | 0.051 | 0.051 | 0 | 0.46 | 0.63 | 0.63 | 0.74 | 0.46 | 0.64 |
| 11 | 117709754 | rs564989 | -0.005 | -0.824 | -1.228 | 0.329 | -0.019 | -0.008 | -0.008 | 0.009 | 0.69 | 0.42 | 0.73 | 0.69 | 0.77 | 0.68 |
| 11 | 117712887 | rs473057 | 0.781 | -0.845 | -1.323 | 0.341 | 0.016 | 0.073 | 0.073 | 0.203 | 0.51 | 0.24 | 0.32 | 0.29 | 0.54 | 0.47 |
| 11 | 117716151 | rs12098917 | 0.684 | -0.722 | -1.523 | 0.053 | -0.008 | 0.038 | 0.038 | 0 | 0.19 | 0.11 | 0.13 | 0.08 | 0.15 | 0.08 |
| 11 | 117717297 | rs6589624 | -1.869 | -0.517 | -1.303 | 0.081 | 0.021 | -0.036 | -0.036 | 0.168 | 0.93 | 0.86 | 0.93 | 0.99 | 0.91 | 0.99 |
| 11 | 117718880 | rs512481 | 0.798 | -0.712 | -1.848 | 0.176 | -0.016 | 0.024 | 0.024 | 0.142 | 0.41 | 0.24 | 0.3 | 0.29 | 0.48 | 0.44 |
| 11 | 117719605 | rs561394 | -0.18 | -0.764 | -1.965 | -0.025 | 0.001 | -0.026 | -0.026 | 0 | 0.62 | 0.34 | 0.75 | 0.91 | 0.72 | 0.84 |
| 11 | 117722715 | rs496371 | 1.14 | -0.406 | -2.552 | 0.63 | -0.091 | -0.035 | -0.035 | 2.932 | 0.23 | 0.01 | 0.17 | 0.17 | 0.29 | 0.27 |
| 11 | 117722742 | rs3885041 | -0.531 | -0.538 | -1.616 | 1.124 | -0.043 | -0.1 | -0.1 | 0.125 | 0.19 | 0.03 | 0.4 | 0.36 | 0.23 | 0.2 |
| 11 | 117729274 | rs10892186 | -0.326 | -1.008 | -1.482 | 1.227 | -0.036 | -0.096 | -0.096 | 0 | 0.2 | 0.02 | 0.37 | 0.3 | 0.21 | 0.27 |
| 11 | 117731136 | rs5014624 | -1.618 | -1.168 | -1.112 | 1.151 | 0.011 | 0.027 | 0.027 | 0 | 0.48 | 0.44 | 0.53 | 0.73 | 0.39 | 0.63 |
| 11 | 117737359 | rs7101835 | -1.905 | -1.067 | -0.831 | 1.198 | 0.014 | 0.028 | 0.028 | 0 | 0.5 | 0.68 | 0.55 | 0.72 | 0.39 | 0.61 |
| 11 | 117737440 | rs876798 | 0.232 | -0.614 | -0.966 | 0.762 | 0.002 | -0.012 | -0.012 | 2.886 | 0.17 | 0.27 | 0.13 | 0.16 | 0.15 | 0.11 |
| 11 | 117737747 | rs876797 | -0.79 | -0.446 | -1.252 | 0.923 | -0.006 | -0.048 | -0.048 | 0.694 | 0.25 | 0.06 | 0.39 | 0.28 | 0.24 | 0.28 |
| 11 | 117741995 | rs10790218 | -0.792 | -0.248 | -0.991 | 0.994 | -0.004 | 0.005 | 0.005 | 0 | 0.43 | 0.52 | 0.53 | 0.54 | 0.38 | 0.39 |
| 11 | 117745003 | rs4938445 | -0.325 | 0.214 | -1.192 | 0.087 | 0.001 | 0.002 | 0.002 | 0 | 0.35 | 0.37 | 0.29 | 0.17 | 0.32 | 0.27 |
| 11 | 117748452 | rs11216598 | -0.645 | 0.566 | -1.549 | 0.96 | -0.016 | -0.071 | -0.071 | 0 | 0.17 | 0.05 | 0.18 | 0.33 | 0.16 | 0.31 |
| 11 | 117748478 | rs631898 | -0.824 | 0.869 | -1.183 | 0.24 | -0.001 | -0.003 | -0.003 | 0 | 0.48 | 0.39 | 0.61 | 0.75 | 0.51 | 0.64 |
| 11 | 117751994 | rs584994 | 0.282 | 0.666 | -1.164 | 0.207 | 0.003 | -0.003 | -0.003 | 0 | 0.6 | 0.41 | 0.52 | 0.67 | 0.54 | 0.61 |
| 11 | 117755580 | rs12286112 | 0.238 | 0.648 | -1.032 | -0.34 | -0.007 | -0.01 | -0.01 | 1.475 | 0.22 | 0.18 | 0.34 | 0.04 | 0.37 | 0.2 |
| 11 | 117757345 | rs601122 | 0.709 | 0.152 | -1.01 | -0.402 | -0.001 | 0.005 | 0.005 | 0.069 | 0.54 | 0.43 | 0.49 | 0.7 | 0.46 | 0.58 |
| 11 | 117757875 | rs7121851 | 1.867 | 0.139 | -1.432 | -0.628 | -0.033 | 0.025 | 0.025 | 0 | 0.17 | 0.05 | 0.12 | 0.11 | 0.18 | 0.09 |
| 11 | 117758431 | rs10892189 | -0.934 | 0.381 | -0.934 | -0.564 | -0.003 | 0.006 | 0.006 | 0.378 | 0.43 | 0.38 | 0.47 | 0.23 | 0.53 | 0.39 |

| | | | | | | | | | | | | | | | | |
|---|---|---|---|---|---|---|---|---|---|---|---|---|---|---|---|---|
| 11 | 117759825 | rs6589632 | 0.465 | 0.806 | -0.956 | -0.554 | -0.006 | 0.001 | 0.001 | 1.611 | 0.23 | 0.17 | 0.32 | 0.21 | 0.32 | 0.26 |
| 11 | 117767498 | rs558247 | 0.35 | 0.764 | -0.937 | -0.573 | 0.01 | -0.004 | -0.004 | 0.744 | 0.63 | 0.51 | 0.57 | 0.71 | 0.56 | 0.63 |
| 11 | 117770505 | rs640483 | -0.558 | 0.955 | -0.93 | -0.782 | 0.003 | -0.04 | -0.04 | 0 | 0.87 | 0.8 | 0.89 | 1 | 0.85 | 0.92 |
| 11 | 117770775 | rs997956 | 0.402 | 0.774 | -0.88 | -0.741 | -0.001 | -0.014 | -0.014 | 0.269 | 0.1 | 0.07 | 0.07 | 0.01 | 0.1 | 0.05 |
| 11 | 117772848 | rs491822 | 1.137 | 0.711 | -0.818 | -0.705 | 0.033 | 0.028 | 0.028 | 0.142 | 0.5 | 0.34 | 0.44 | 0.61 | 0.37 | 0.49 |
| 11 | 117774291 | rs2845712 | 1.125 | 0.795 | -1.095 | -0.71 | 0.015 | 0.145 | 0.145 | 0.625 | 0.94 | 0.71 | 0.92 | 0.83 | 0.95 | 0.91 |
| 11 | 117774829 | rs3809040 | -0.681 | 0.807 | -1.138 | -0.708 | -0.007 | 0.024 | 0.024 | 0 | 0.15 | 0.31 | 0.15 | 0.18 | 0.13 | 0.13 |
| 11 | 117776471 | rs3802873 | 0.328 | 1.058 | -1.322 | -0.708 | -0.013 | -0.009 | -0.009 | 0 | 0.21 | 0.16 | 0.31 | 0.24 | 0.32 | 0.27 |
| 11 | 117776648 | rs3802871 | 0.77 | 0.708 | -1.334 | -0.7 | 0.002 | 0.003 | 0.003 | 0.207 | 0.53 | 0.5 | 0.46 | 0.62 | 0.38 | 0.48 |
| 11 | 117781618 | rs2155195 | -0.327 | 1.034 | -2.214 | -0.668 | 0 | -0.002 | -0.002 | 0.096 | 0.24 | 0.24 | 0.31 | 0.19 | 0.33 | 0.24 |
| 11 | 117784714 | rs11216618 | -0.211 | 1.321 | -2.866 | -0.378 | -0.054 | 0.05 | 0.05 | 1.021 | 0.2 | 0.06 | 0.18 | 0.12 | 0.25 | 0.26 |
| 11 | 117785260 | rs4936410 | -0.478 | 0.968 | -2.532 | -0.039 | -0.026 | 0.05 | 0.05 | 0 | 0.3 | 0.08 | 0.34 | 0.21 | 0.35 | 0.21 |
| 11 | 117788313 | rs4245178 | -0.22 | 0.689 | -2.501 | 0.618 | -0.076 | -0.069 | -0.069 | 0.062 | 0.39 | 0.03 | 0.4 | 0.41 | 0.36 | 0.24 |
| 11 | 117790769 | rs11826839 | 1.023 | 0.504 | -1.241 | 0.77 | -0.003 | 0.007 | 0.007 | 0 | 0.75 | 0.69 | 0.74 | 0.66 | 0.74 | 0.75 |
| 11 | 117792471 | rs11216624 | -1.06 | 0.063 | -1.08 | 0.744 | 0.009 | 0.005 | 0.005 | 0 | 0.15 | 0.11 | 0.11 | 0.16 | 0.1 | 0.11 |
| 11 | 117795048 | rs7943219 | 1.675 | 0.048 | -0.413 | 1.09 | -0.028 | 0.024 | 0.024 | 0.323 | 0.72 | 0.48 | 0.73 | 0.66 | 0.81 | 0.73 |
| 11 | 117795679 | rs11216631 | -1.379 | 0.241 | -0.702 | 0.909 | 0.003 | 0.018 | 0.018 | 0 | 0.14 | 0.08 | 0.1 | 0.07 | 0.11 | 0.14 |
| 11 | 117798431 | rs12365956 | -0.735 | 0.562 | -0.716 | 1.13 | -0.031 | -0.029 | -0.029 | 0 | 0.06 | 0.01 | 0.09 | 0.21 | 0.09 | 0.14 |
| 11 | 117799804 | rs722729 | -0.12 | 0.626 | -1.531 | 0.366 | -0.044 | 0.068 | 0.068 | 0.018 | 0.27 | 0.04 | 0.27 | 0.12 | 0.37 | 0.26 |
| 11 | 117799923 | rs7928668 | 0.717 | 0.4 | -0.473 | 0.308 | 0 | -0.001 | -0.001 | 0 | 0.63 | 0.67 | 0.61 | 0.68 | 0.54 | 0.6 |
| 11 | 117800083 | rs10892196 | 0.717 | 0.4 | -0.473 | 0.287 | 0 | -0.001 | -0.001 | 0 | 0.63 | 0.67 | 0.61 | 0.67 | 0.54 | 0.6 |
| 11 | 117804109 | rs3863317 | -0.106 | 0.103 | -0.815 | -0.285 | 0.014 | 0.03 | 0.03 | 0 | 0.14 | 0.07 | 0.2 | 0.07 | 0.12 | 0.09 |
| 11 | 117806937 | rs10790226 | -0.048 | 0.244 | -0.415 | -0.498 | 0.009 | -0.068 | -0.068 | 0.009 | 0.74 | 0.7 | 0.72 | 0.97 | 0.63 | 0.74 |
| 11 | 117811167 | rs3863318 | 0.3 | 0.174 | -0.579 | 0.109 | -0.018 | 0.013 | 0.013 | 0.067 | 0.68 | 0.73 | 0.66 | 0.71 | 0.54 | 0.6 |
| 11 | 117815564 | rs2512139 | -0.446 | -0.153 | -0.89 | -0.221 | -0.019 | 0.026 | 0.026 | 0 | 0.25 | 0.13 | 0.38 | 0.03 | 0.38 | 0.11 |
| 11 | 117815759 | rs2515793 | -1.476 | -0.366 | -0.802 | 0.801 | -0.026 | -0.016 | -0.016 | 0.134 | 0.55 | 0.75 | 0.64 | 0.59 | 0.59 | 0.54 |
| 11 | 117817276 | rs2515795 | -1.496 | -0.415 | -1.012 | -1.253 | -0.044 | 0.1 | 0.1 | 0 | 0.56 | 0.8 | 0.61 | 0.33 | 0.59 | 0.51 |
| 11 | 117820561 | rs7103514 | -1.503 | -0.206 | -0.122 | -0.552 | -0.014 | 0.017 | 0.017 | 0.128 | 0.16 | 0.75 | 0.15 | 0.33 | 0.09 | 0.18 |
| 11 | 117823425 | rs2512158 | -0.673 | 0.683 | -0.6 | -1.424 | -0.033 | 0.03 | 0.03 | 0.26 | 0.33 | 0.2 | 0.29 | 0.32 | 0.43 | 0.4 |
| 11 | 117824592 | rs2515800 | 0.203 | 0.588 | 0.017 | -1.731 | 0.012 | -0.045 | -0.045 | 0 | 0.84 | 0.63 | 0.74 | 0.93 | 0.82 | 0.88 |
| 11 | 117824960 | rs2243935 | 1.227 | 0.701 | 0.042 | -1.9 | 0.003 | 0.005 | 0.005 | 0.286 | 0.74 | 0.38 | 0.62 | 0.63 | 0.69 | 0.74 |
| 11 | 117825368 | rs2887193 | 0.181 | 0.775 | -0.109 | -1.845 | -0.007 | 0.005 | 0.005 | 0 | 0.1 | 0.06 | 0.23 | 0.05 | 0.17 | 0.09 |
| 11 | 117830429 | rs11216648 | 1.564 | 0.689 | 0.024 | -1.897 | 0.016 | 0.08 | 0.08 | 0 | 0.48 | 0.23 | 0.44 | 0.61 | 0.38 | 0.5 |
| 11 | 117831809 | rs3020913 | 0.082 | 0.714 | -0.209 | -1.757 | -0.027 | 0.064 | 0.064 | 0.219 | 0.26 | 0.13 | 0.19 | 0.02 | 0.31 | 0.26 |
| 11 | 117848292 | rs10892201 | 0.113 | 0.563 | -0.28 | -1.197 | -0.029 | -0.037 | -0.037 | 0 | 0.07 | 0.01 | 0.11 | 0.34 | 0.11 | 0.13 |
| 11 | 117853613 | rs4936414 | -0.171 | 0.72 | -0.096 | -1.681 | -0.022 | -0.002 | -0.002 | 0 | 0.73 | 0.8 | 0.81 | 0.99 | 0.68 | 0.73 |
| 11 | 117855295 | rs4252241 | -0.846 | 0.75 | 0.28 | -1.686 | 0.013 | -0.002 | -0.002 | 0 | 0.12 | 0.31 | 0.21 | 0.08 | 0.11 | 0.13 |
| 11 | 117855727 | rs4252243 | -0.682 | 0.756 | 0.283 | -1.683 | 0.02 | 0.004 | 0.004 | 0 | 0.11 | 0.31 | 0.21 | 0.08 | 0.11 | 0.13 |
| 11 | 117858185 | rs4252246 | -0.553 | 0.782 | 0.306 | -1.67 | 0.029 | 0.012 | 0.012 | 0 | 0.11 | 0.32 | 0.21 | 0.08 | 0.12 | 0.13 |
| 11 | 117859209 | rs4252249 | 0.145 | 0.788 | 0.322 | -1.67 | 0.043 | 0.008 | 0.008 | 0 | 0.07 | 0.17 | 0.19 | 0.06 | 0.11 | 0.1 |
| 11 | 117862298 | rs2512144 | -0.051 | 0.798 | 0.372 | -1.786 | 0.004 | -0.069 | -0.069 | 0 | 0.72 | 0.68 | 0.8 | 0.99 | 0.67 | 0.74 |
| 11 | 117863829 | rs2508450 | 0.996 | 0.764 | 0.489 | -1.794 | -0.029 | 0.067 | 0.067 | 0 | 0.61 | 0.28 | 0.6 | 0.91 | 0.55 | 0.62 |
| 11 | 117864047 | rs2256111 | 0.886 | 0.709 | 0.462 | -1.741 | 0.004 | -0.005 | -0.005 | 0.376 | 0.6 | 0.46 | 0.57 | 0.55 | 0.51 | 0.54 |
| 11 | 117864113 | rs2228054 | 0.385 | 0.323 | 0.44 | -1.476 | 0.016 | -0.021 | -0.021 | 0 | 0.06 | 0 | 0.05 | 0.38 | 0.06 | 0.11 |
| 11 | 117868638 | rs4252287 | 0.374 | 0.333 | 0.629 | -1.514 | 0.021 | 0 | 0 | 0 | 0.06 | 0.11 | 0.17 | 0.07 | 0.11 | 0.1 |
| 11 | 117869670 | rs2229113 | -0.783 | 0.304 | 0.613 | -1.599 | -0.025 | 0.019 | 0.019 | 0 | 0.74 | 0.82 | 0.81 | 0.99 | 0.68 | 0.74 |
| 11 | 117872086 | rs9610 | 0.582 | -0.072 | 0.505 | -1.657 | -0.012 | 0.017 | 0.017 | 1.226 | 0.58 | 0.68 | 0.55 | 0.55 | 0.45 | 0.53 |
| 11 | 117872734 | rs2512147 | -0.768 | 0.197 | 0.621 | -1.545 | -0.001 | -0.05 | -0.05 | 0 | 0.74 | 0.72 | 0.8 | 0.99 | 0.68 | 0.74 |
| 11 | 117874531 | rs947889 | -0.23 | 0.197 | 0.473 | -1.534 | 0.001 | -0.032 | -0.032 | 0 | 0.68 | 0.69 | 0.62 | 0.93 | 0.54 | 0.65 |
| 11 | 117876155 | rs17121510 | 0.504 | -0.937 | 1.374 | -0.467 | -0.017 | -0.036 | -0.036 | 0 | 0.2 | 0.28 | 0.11 | 0.04 | 0.13 | 0.1 |
| 11 | 117876198 | rs11216667 | -0.914 | -0.967 | 1.364 | -0.433 | -0.082 | 0.018 | 0.018 | 0 | 0.32 | 0.51 | 0.2 | 0.44 | 0.19 | 0.23 |
| 11 | 117878559 | rs11216668 | -1.884 | -0.685 | 1.024 | -0.326 | -0.073 | 0.162 | 0.162 | 0 | 0.14 | 0.41 | 0.09 | 0.4 | 0.06 | 0.13 |
| 11 | 117887757 | rs2512152 | -1.653 | -0.651 | -0.02 | -0.328 | 0.06 | -0.015 | -0.015 | 0 | 0.39 | 0.8 | 0.4 | 0.5 | 0.33 | 0.34 |
| 11 | 117889402 | rs2512154 | -0.779 | 0.502 | 0.037 | 0.6 | 0.041 | -0.003 | -0.003 | 0.004 | 0.07 | 0.13 | 0.18 | 0.06 | 0.14 | 0.1 |
| 11 | 117889448 | rs2512155 | -2.666 | 0.516 | -0.383 | 0.603 | -0.012 | 0.058 | 0.058 | 0 | 0.19 | 0.28 | 0.27 | 0.46 | 0.2 | 0.24 |
| 11 | 117889959 | rs11216689 | -1.962 | 0.016 | -0.836 | 0.856 | -0.026 | 0.046 | 0.046 | 0.083 | 0.11 | 0.15 | 0.08 | 0.4 | 0.06 | 0.13 |
| 11 | 117898465 | rs10431040 | -0.968 | 0.6 | -1.296 | 0.983 | 0.018 | -0.009 | -0.009 | 0 | 0.06 | 0.02 | 0.05 | 0.39 | 0.05 | 0.11 |
| 11 | 117903836 | rs4936417 | -0.371 | 1.495 | -1.625 | -0.108 | -0.078 | 0.09 | 0.09 | 0 | 0.23 | 0.07 | 0.24 | 0.06 | 0.32 | 0.16 |
| 11 | 117903942 | rs11216713 | 0.112 | 0.799 | -1.829 | 0.621 | 0.015 | -0.01 | -0.01 | 0 | 0.06 | 0.03 | 0.05 | 0.21 | 0.03 | 0.05 |
| 11 | 117904029 | rs1786177 | -1.014 | -0.098 | -1.485 | -0.02 | -0.017 | -0.043 | -0.043 | 0 | 0.78 | 0.68 | 0.86 | 0.95 | 0.85 | 0.91 |
| 11 | 117906510 | rs10892204 | 1.172 | -0.505 | -1.231 | -0.036 | 0.001 | -0.011 | -0.011 | 0.614 | 0.23 | 0.24 | 0.27 | 0.17 | 0.28 | 0.29 |
| 11 | 117909101 | rs11216720 | 1.3 | -0.509 | -1.235 | -0.02 | 0.015 | -0.01 | -0.01 | 0 | 0.06 | 0.03 | 0.05 | 0.2 | 0.03 | 0.05 |
| 11 | 117909799 | rs652107 | 1.572 | -0.648 | -0.909 | -0.17 | -0.003 | -0.012 | -0.012 | 0.384 | 0.2 | 0.2 | 0.12 | 0.05 | 0.15 | 0.09 |
| 11 | 117911367 | rs597820 | -0.591 | -0.743 | -0.705 | -0.151 | 0.01 | 0.033 | 0.033 | 0 | 0.54 | 0.39 | 0.59 | 0.76 | 0.56 | 0.61 |
| 11 | 117912465 | rs7120925 | 1.068 | -0.74 | -0.653 | 0.304 | -0.001 | -0.001 | -0.001 | 0 | 0.26 | 0.3 | 0.34 | 0.29 | 0.29 | 0.33 |
| 11 | 117913074 | rs4936421 | 1.366 | -0.706 | -0.729 | 0.295 | -0.082 | 0.091 | 0.091 | 0 | 0.22 | 0.05 | 0.24 | 0.06 | 0.32 | 0.16 |
| 11 | 117915055 | rs12576767 | 1.299 | -0.81 | -0.904 | 0.567 | -0.017 | -0.05 | -0.05 | 0.478 | 0.09 | 0.01 | 0.11 | 0.29 | 0.08 | 0.28 |
| 11 | 117919431 | rs1783842 | -0.155 | -0.811 | -0.767 | 0.456 | -0.011 | 0.012 | 0.012 | 0.724 | 0.43 | 0.37 | 0.41 | 0.37 | 0.47 | 0.3 |
| 11 | 117922143 | rs1786185 | -0.628 | -0.219 | -0.753 | 0.475 | -0.001 | 0.008 | 0.008 | 0 | 0.66 | 0.6 | 0.67 | 0.55 | 0.68 | 0.59 |

| | | | | | | | | | | | | | | | | |
|---|---|---|---|---|---|---|---|---|---|---|---|---|---|---|---|---|
| 11 | 117922486 | rs624931 | -0.99 | -0.355 | -0.675 | 0.493 | 0.011 | -0.026 | -0.026 | 0 | 0.71 | 0.83 | 0.77 | 0.66 | 0.77 | 0.63 |
| 11 | 117923358 | rs7937145 | 1.309 | -0.359 | -0.662 | 0.479 | 0.001 | -0.017 | -0.017 | 0 | 0.24 | 0.26 | 0.28 | 0.18 | 0.29 | 0.3 |
| 11 | 117926593 | rs676812 | -0.918 | -0.627 | -0.556 | 0.589 | 0.006 | -0.013 | -0.013 | 0.419 | 0.71 | 0.79 | 0.77 | 0.66 | 0.77 | 0.63 |
| 11 | 117927676 | rs584231 | -0.096 | -0.819 | -0.716 | 0.427 | 0.015 | -0.038 | -0.038 | 0 | 0.16 | 0.14 | 0.16 | 0.31 | 0.16 | 0.13 |
| 11 | 117928962 | rs1783846 | -0.157 | -0.835 | -0.722 | 0.419 | 0.016 | -0.036 | -0.036 | 6.016 | 0.17 | 0.15 | 0.16 | 0.31 | 0.16 | 0.13 |
| 11 | 117942125 | rs652030 | -1.046 | -0.896 | -0.503 | 0.422 | 0.002 | -0.004 | -0.004 | 0 | 0.7 | 0.71 | 0.76 | 0.66 | 0.78 | 0.63 |
| 11 | 117945632 | rs1786186 | -0.321 | -0.826 | -0.438 | 0.381 | 0.014 | -0.022 | -0.022 | 0 | 0.18 | 0.19 | 0.17 | 0.31 | 0.17 | 0.14 |
| 11 | 117947433 | rs619585 | -1.819 | -0.775 | -0.454 | 0.457 | -0.008 | -0.034 | -0.034 | 0 | 0.79 | 0.77 | 0.87 | 0.96 | 0.85 | 0.9 |
| 11 | 117947698 | rs620924 | -0.688 | -0.765 | -0.203 | 0.473 | -0.015 | 0.028 | 0.028 | 0 | 0.67 | 0.61 | 0.69 | 0.55 | 0.77 | 0.58 |
| 11 | 117952097 | rs3825050 | 0.434 | -0.565 | -0.246 | 0.34 | 0.001 | -0.04 | -0.04 | 0 | 0.32 | 0.21 | 0.36 | 0.49 | 0.34 | 0.57 |
| 11 | 117953688 | rs7937420 | 0.869 | -0.439 | -0.351 | -0.078 | -0.019 | 0.02 | 0.02 | 0.166 | 0.24 | 0.18 | 0.23 | 0.07 | 0.29 | 0.15 |
| 11 | 117954081 | rs657118 | -0.599 | -1.463 | -0.021 | -0.173 | 0.004 | 0.029 | 0.029 | 0 | 0.48 | 0.43 | 0.52 | 0.83 | 0.53 | 0.65 |
| 11 | 117954250 | rs10160375 | 0.176 | -1.214 | -0.111 | 0.149 | -0.008 | -0.028 | -0.028 | 0 | 0.33 | 0.21 | 0.38 | 0.5 | 0.4 | 0.52 |
| 11 | 117956338 | rs10892210 | 0.202 | -1.413 | -0.118 | 0.251 | -0.001 | -0.025 | -0.025 | 0 | 0.33 | 0.21 | 0.39 | 0.5 | 0.4 | 0.52 |
| 11 | 117956943 | rs4938478 | 0.16 | -1.419 | 0.009 | 0.248 | 0 | -0.009 | -0.009 | 0 | 0.33 | 0.26 | 0.39 | 0.5 | 0.4 | 0.52 |
| 11 | 117957258 | rs689264 | -0.61 | -1.379 | 0.301 | 0.323 | 0.001 | 0.072 | 0.072 | 0 | 0.1 | 0.22 | 0.14 | 0.26 | 0.08 | 0.13 |
| 11 | 117957750 | rs600638 | -0.738 | -1.476 | 0.143 | 0.333 | -0.008 | 0.075 | 0.075 | 0 | 0.1 | 0.23 | 0.14 | 0.26 | 0.07 | 0.12 |
| 11 | 117958909 | rs1317876 | -0.522 | -1.533 | 0.176 | 0.262 | -0.001 | -0.018 | -0.018 | 0 | 0.42 | 0.42 | 0.53 | 0.76 | 0.47 | 0.66 |
| 11 | 117960138 | rs948461 | 0.995 | -1.287 | -0.174 | 0.118 | -0.073 | 0.127 | 0.127 | 1.524 | 0.24 | 0.06 | 0.23 | 0.05 | 0.28 | 0.16 |
| 11 | 117960427 | rs1893317 | -0.4 | -1.282 | 0.306 | 0.303 | -0.009 | -0.013 | -0.013 | 0 | 0.6 | 0.45 | 0.71 | 0.71 | 0.7 | 0.78 |
| 11 | 117961908 | rs651474 | -0.308 | -1.393 | 0.572 | 0.448 | 0.002 | 0.003 | 0.003 | 0.012 | 0.34 | 0.36 | 0.37 | 0.45 | 0.42 | 0.34 |
| 11 | 117963766 | rs11216740 | 0.409 | -1.391 | 0.34 | 0.449 | -0.043 | -0.042 | -0.042 | 0.094 | 0.35 | 0.11 | 0.4 | 0.35 | 0.35 | 0.51 |
| 11 | 117965595 | rs2276123 | 0.531 | -1.291 | 0.364 | 0.459 | -0.017 | -0.019 | -0.019 | 0 | 0.25 | 0.15 | 0.29 | 0.35 | 0.35 | 0.27 |
| 11 | 117965646 | rs2276125 | 0.873 | -1.217 | 0.277 | 0.49 | -0.026 | 0.005 | 0.005 | 0.019 | 0.23 | 0.12 | 0.29 | 0.22 | 0.35 | 0.21 |
| 11 | 117966100 | rs948466 | 0.246 | -1.24 | 0.214 | 0.49 | -0.005 | -0.003 | -0.003 | 0 | 0.9 | 0.88 | 0.93 | 0.9 | 0.93 | 0.93 |
| 11 | 117972936 | rs881122 | 0.568 | -1.231 | 0.473 | 0.408 | -0.017 | -0.03 | -0.03 | 0 | 0.3 | 0.4 | 0.21 | 0.19 | 0.22 | 0.16 |
| 11 | 117973673 | rs2276129 | 0.218 | -0.828 | 0.473 | 0.601 | 0.009 | 0.013 | 0.013 | 0.484 | 0.27 | 0.31 | 0.34 | 0.37 | 0.36 | 0.27 |
| 11 | 117974759 | rs2155857 | -0.517 | -1.096 | 0.63 | 0.745 | -0.022 | 0.077 | 0.077 | 0.131 | 0.13 | 0.21 | 0.14 | 0.34 | 0.06 | 0.35 |
| 11 | 117978103 | rs10892215 | -0.486 | -1.083 | 0.647 | 0.751 | -0.011 | 0.047 | 0.047 | 0.043 | 0.12 | 0.17 | 0.13 | 0.35 | 0.06 | 0.35 |
| 11 | 117978264 | rs1894177 | 0.426 | -1.419 | 0.537 | 0.514 | 0 | 0 | 0 | 0 | 0.55 | 0.45 | 0.54 | 0.44 | 0.59 | 0.37 |
| 11 | 117984623 | rs596134 | -0.692 | -1.247 | -0.02 | -0.041 | -0.027 | 0.058 | 0.058 | 0.071 | 0.7 | 0.83 | 0.68 | 0.95 | 0.66 | 0.77 |
| 11 | 117987746 | rs640888 | -0.54 | -1.44 | 0.305 | 0.2 | -0.029 | -0.042 | -0.042 | 0.768 | 0.62 | 0.5 | 0.75 | 0.8 | 0.74 | 0.85 |
| 11 | 117988223 | rs643014 | 0.031 | -1.316 | 0.211 | 0.673 | 0.008 | -0.008 | -0.008 | 0.042 | 0.89 | 0.9 | 0.92 | 0.57 | 0.91 | 0.62 |
| 11 | 117988539 | rs671111 | 0.767 | -1.232 | 0.253 | 0.536 | 0.018 | 0.003 | 0.003 | 0.07 | 0.23 | 0.25 | 0.33 | 0.26 | 0.36 | 0.24 |
| 11 | 117988784 | rs8924 | -0.668 | -1.33 | 0.118 | 0.559 | 0.028 | 0.002 | 0.002 | 0.361 | 0.73 | 0.66 | 0.63 | 0.71 | 0.6 | 0.76 |
| 11 | 117988797 | rs17121760 | 0.967 | -1.292 | 0.073 | 0.993 | 0 | -0.015 | -0.015 | 0 | 0.06 | 0.01 | 0.04 | 0.28 | 0.05 | 0.37 |
| 11 | 117989554 | rs3741311 | -0.648 | -1.246 | 0.14 | 0.052 | 0.036 | 0.004 | 0.004 | 0.598 | 0.73 | 0.63 | 0.64 | 0.71 | 0.6 | 0.76 |
| 11 | 117999443 | rs1941637 | -0.444 | -1.281 | 0.225 | 0.071 | -0.017 | 0.096 | 0.096 | 0 | 0.12 | 0.21 | 0.08 | 0.4 | 0.06 | 0.39 |
| 11 | 118001445 | rs10790239 | 0.664 | -1.439 | 0.17 | -0.069 | -0.01 | -0.014 | -0.014 | 0 | 0.35 | 0.35 | 0.21 | 0.18 | 0.22 | 0.14 |
| 11 | 118004688 | rs3741315 | -0.743 | -1.248 | -0.221 | -0.27 | 0.037 | 0.024 | 0.024 | 0.557 | 0.74 | 0.64 | 0.56 | 0.65 | 0.6 | 0.73 |
| 11 | 118005119 | rs868344 | 1.064 | -1.217 | -0.429 | -0.277 | -0.118 | 0.14 | 0.14 | 1.557 | 0.26 | 0.02 | 0.27 | 0.07 | 0.32 | 0.2 |
| 11 | 118006957 | rs1814964 | -0.258 | -0.778 | 0.015 | 0.045 | 0.038 | 0.021 | 0.021 | 0.469 | 0.71 | 0.56 | 0.55 | 0.65 | 0.59 | 0.73 |
| 11 | 118011805 | rs644287 | 0.514 | -0.86 | -0.424 | -0.113 | 0.021 | 0.007 | 0.007 | 0.126 | 0.25 | 0.35 | 0.35 | 0.28 | 0.36 | 0.24 |
| 11 | 118011860 | rs12804550 | 1.191 | -1.321 | -0.516 | -0.2 | 0.013 | 0.055 | 0.055 | 1.306 | 0.26 | 0.14 | 0.14 | 0.11 | 0.19 | 0.13 |
| 11 | 118012310 | rs11216781 | -0.517 | -1.355 | -0.616 | -0.229 | 0.034 | 0.021 | 0.021 | 0 | 0.47 | 0.63 | 0.38 | 0.6 | 0.28 | 0.51 |
| 11 | 118016068 | rs1793137 | -0.228 | -0.986 | -0.46 | 0.235 | 0.001 | -0.001 | -0.001 | 0 | 0.63 | 0.65 | 0.66 | 0.83 | 0.65 | 0.64 |
| 11 | 118016273 | rs1715444 | 0.427 | -0.939 | -0.616 | 0.17 | -0.003 | -0.03 | -0.03 | 0 | 0.55 | 0.49 | 0.61 | 0.81 | 0.6 | 0.59 |
| 11 | 118018767 | rs658624 | -0.25 | -0.097 | -1.172 | 0.4 | -0.022 | -0.004 | -0.004 | 0 | 0.58 | 0.84 | 0.52 | 0.4 | 0.59 | 0.46 |
| 11 | 118020057 | rs598746 | 0.026 | 0.169 | -1.055 | 0.451 | 0.015 | 0.027 | 0.027 | 0 | 0.52 | 0.63 | 0.48 | 0.42 | 0.56 | 0.46 |
| 11 | 118025917 | rs645530 | -0.197 | 0.424 | -0.943 | 0.416 | -0.001 | -0.001 | -0.001 | 0.178 | 0.47 | 0.53 | 0.52 | 0.6 | 0.4 | 0.43 |
| 11 | 118026675 | rs2155854 | -0.257 | 0.337 | -1.44 | 0.198 | 0.001 | 0.053 | 0.053 | 0.027 | 0.49 | 0.21 | 0.59 | 0.58 | 0.52 | 0.45 |
| 11 | 118028094 | rs876479 | -0.006 | 0.355 | -1.33 | -0.44 | 0.001 | 0.11 | 0.11 | 0 | 0.43 | 0.25 | 0.43 | 0.76 | 0.39 | 0.49 |
| 11 | 118028167 | rs615144 | -0.601 | -0.409 | -0.683 | -0.697 | -0.032 | 0.158 | 0.158 | 0 | 0.55 | 0.78 | 0.59 | 0.83 | 0.51 | 0.53 |
| 11 | 118030186 | rs2156850 | 0.406 | -0.495 | -0.408 | -1.02 | 0.002 | 0.022 | 0.022 | 0.095 | 0.39 | 0.34 | 0.49 | 0.74 | 0.39 | 0.43 |
| 11 | 118031206 | rs656308 | -0.778 | -0.463 | -0.423 | -1.029 | -0.009 | -0.084 | -0.084 | 0.09 | 0.15 | 0.44 | 0.1 | 0.09 | 0.11 | 0.11 |
| 11 | 118032228 | rs688563 | -0.871 | -0.545 | -0.369 | -1.075 | -0.035 | 0.192 | 0.192 | 0.064 | 0.55 | 0.83 | 0.6 | 0.83 | 0.5 | 0.54 |
| 11 | 118032242 | rs688587 | -0.492 | -0.565 | -0.425 | -1.085 | 0.002 | 0.102 | 0.102 | 0 | 0.51 | 0.65 | 0.55 | 0.83 | 0.5 | 0.54 |
| 11 | 118033545 | rs625464 | -0.951 | -0.797 | -0.67 | -1.296 | -0.174 | 0.174 | 0.174 | 0.933 | 0.6 | 0.98 | 0.63 | 0.86 | 0.55 | 0.57 |
| 11 | 118034201 | rs11216788 | 0.935 | -0.806 | -0.632 | -1.291 | -0.01 | 0.08 | 0.08 | 0 | 0.36 | 0.17 | 0.45 | 0.74 | 0.39 | 0.43 |
| 11 | 118034870 | rs3802869 | -0.73 | -0.672 | -0.439 | -1.135 | 0.005 | -0.015 | -0.015 | 0 | 0.07 | 0.29 | 0.07 | 0.03 | 0.04 | 0.03 |
| 11 | 118036532 | rs688911 | -0.249 | -0.708 | -0.479 | -1.168 | 0.001 | 0.096 | 0.096 | 0 | 0.53 | 0.65 | 0.56 | 0.8 | 0.5 | 0.54 |
| 11 | 118036594 | rs2076884 | -1.154 | -0.648 | 0.205 | -0.775 | 0.005 | -0.015 | -0.015 | 0 | 0.07 | 0.28 | 0.07 | 0.05 | 0.04 | 0.03 |
| 11 | 118038741 | rs647975 | -2.038 | -0.723 | 0.347 | -0.711 | -0.054 | -0.037 | -0.037 | 0 | 0.11 | 0.73 | 0.08 | 0.09 | 0.02 | 0.13 |
| 11 | 118040642 | rs10892221 | 0.114 | -0.73 | 1.104 | -0.188 | -0.006 | 0.047 | 0.047 | 0 | 0.05 | 0.11 | 0.05 | 0.14 | 0.04 | 0.02 |
| 11 | 118041102 | rs10892222 | -0.228 | -0.723 | 1.233 | -0.195 | -0.002 | 0.062 | 0.062 | 0 | 0.06 | 0.16 | 0.05 | 0.15 | 0.04 | 0.02 |
| 11 | 118042713 | rs651825 | -1.68 | -0.719 | 1.582 | -0.2 | -0.017 | 0.062 | 0.062 | 1.102 | 0.1 | 0.69 | 0.12 | 0.2 | 0.06 | 0.04 |
| 11 | 118042974 | rs679327 | -1.607 | -0.854 | 0.796 | -1.013 | 0.025 | 0.131 | 0.131 | 0.057 | 0.4 | 0.93 | 0.3 | 0.25 | 0.31 | 0.2 |
| 11 | 118050855 | rs678957 | -1.407 | -1.015 | 1.282 | -0.684 | -0.025 | 0.045 | 0.045 | 1.877 | 0.1 | 0.71 | 0.12 | 0.2 | 0.06 | 0.05 |
| 11 | 118053156 | rs4537793 | -1.09 | -0.971 | 1.194 | -0.889 | -0.04 | -0.109 | -0.109 | 0.377 | 0.25 | 0.39 | 0.15 | 0.07 | 0.18 | 0.14 |

| | | | | | | | | | | | | | | | | |
|---|---|---|---|---|---|---|---|---|---|---|---|---|---|---|---|---|
| 11 | 118055801 | rs7944321 | -1.555 | -0.962 | 1.121 | -0.876 | 0.002 | 0.002 | 0.002 | 0.518 | 0.31 | 0.66 | 0.22 | 0.23 | 0.23 | 0.18 |
| 11 | 118067621 | rs951657 | 0.472 | -0.318 | 0.628 | -0.403 | -0.004 | 0 | 0 | 0 | 0.56 | 0.55 | 0.53 | 0.59 | 0.55 | 0.78 |
| 11 | 118068752 | rs2298831 | -0.749 | -0.551 | 0.627 | -0.505 | -0.004 | -0.012 | -0.012 | 0 | 0.18 | 0.37 | 0.26 | 0.12 | 0.18 | 0.06 |
| 11 | 118069979 | rs7116375 | -0.828 | -0.331 | 0.587 | -0.55 | -0.012 | -0.032 | -0.032 | 0 | 0.21 | 0.37 | 0.26 | 0.12 | 0.19 | 0.06 |
| 11 | 118072373 | rs7949751 | -0.596 | -0.455 | 0.629 | -0.594 | -0.002 | 0.003 | 0.003 | 3.351 | 0.73 | 0.63 | 0.77 | 0.66 | 0.72 | 0.82 |
| 11 | 118074117 | rs1793173 | -1.049 | -0.469 | 0.635 | -0.602 | -0.022 | -0.06 | -0.06 | 0 | 0.77 | 0.92 | 0.8 | 0.67 | 0.74 | 0.84 |
| 11 | 118074337 | rs1793174 | -1.049 | -0.469 | 0.635 | -0.602 | -0.022 | -0.06 | -0.06 | 0.453 | 0.77 | 0.93 | 0.8 | 0.67 | 0.74 | 0.84 |
| 11 | 118075152 | rs11216816 | -0.2 | -0.509 | 0.763 | -0.656 | -0.002 | 0.002 | 0.002 | 0 | 0.16 | 0.13 | 0.21 | 0.12 | 0.18 | 0.06 |
| 11 | 118075474 | rs11216817 | -0.158 | -0.537 | 0.762 | -0.657 | -0.001 | 0.001 | 0.001 | 0 | 0.16 | 0.13 | 0.21 | 0.12 | 0.18 | 0.06 |
| 11 | 118076004 | rs1806 | 0.795 | -0.535 | 0.695 | -0.655 | 0.006 | -0.003 | -0.003 | 0 | 0.55 | 0.43 | 0.51 | 0.55 | 0.55 | 0.79 |
| 11 | 118076069 | rs1805 | 1.38 | -0.523 | 0.531 | -0.649 | 0.024 | -0.011 | -0.011 | 0.069 | 0.51 | 0.14 | 0.49 | 0.54 | 0.53 | 0.77 |
| 11 | 118083664 | rs1715429 | -1.05 | -0.493 | 0.529 | -0.725 | -0.028 | -0.05 | -0.05 | 0.583 | 0.77 | 0.94 | 0.79 | 0.7 | 0.73 | 0.84 |
| 11 | 118095803 | rs3802866 | -0.667 | -0.496 | 0.749 | -0.753 | 0 | -0.007 | -0.007 | 0 | 0.2 | 0.23 | 0.26 | 0.12 | 0.21 | 0.07 |
| 11 | 118102058 | rs1793167 | -1.835 | -0.547 | 0.512 | -0.879 | 0.016 | 0.028 | 0.028 | 0.183 | 0.27 | 0.8 | 0.32 | 0.13 | 0.22 | 0.08 |
| 11 | 118105233 | rs1715421 | -1.253 | -0.532 | 0.527 | -0.872 | -0.011 | -0.038 | -0.038 | 0.74 | 0.23 | 0.63 | 0.28 | 0.12 | 0.21 | 0.07 |
| 11 | 118107805 | rs1793140 | -1.478 | -0.301 | -0.171 | -0.874 | -0.037 | -0.05 | -0.05 | 9.804 | 0.77 | 0.94 | 0.78 | 0.69 | 0.73 | 0.84 |
| 11 | 118110977 | rs2853006 | -1.253 | -0.296 | -0.168 | -0.874 | -0.027 | -0.039 | -0.039 | 0 | 0.75 | 0.92 | 0.77 | 0.68 | 0.71 | 0.83 |
| 11 | 118112332 | rs12419365 | 1.571 | -0.287 | -0.218 | -0.879 | -0.017 | -0.041 | -0.041 | 0 | 0.08 | 0 | 0.17 | 0.12 | 0.08 | 0.04 |
| 11 | 118113583 | rs10502228 | -0.17 | -0.273 | -0.102 | -0.848 | -0.001 | -0.008 | -0.008 | 0 | 0.2 | 0.24 | 0.26 | 0.12 | 0.2 | 0.07 |
| 11 | 118122737 | rs1715458 | -1.45 | -0.266 | -0.146 | -0.836 | -0.037 | -0.05 | -0.05 | 0.411 | 0.77 | 0.94 | 0.78 | 0.69 | 0.73 | 0.84 |
| 11 | 118124092 | rs1715455 | -1.417 | -0.265 | -0.149 | -0.832 | -0.041 | -0.052 | -0.052 | 0 | 0.77 | 0.94 | 0.78 | 0.69 | 0.72 | 0.84 |
| 11 | 118125625 | rs1056562 | 0.168 | -0.22 | 0.03 | -0.783 | -0.008 | -0.002 | -0.002 | 0.859 | 0.57 | 0.7 | 0.52 | 0.57 | 0.52 | 0.78 |
| 11 | 118125672 | rs10790248 | 0.892 | -0.076 | -0.701 | -0.707 | 0 | 0.022 | 0.022 | 6.335 | 0.5 | 0.12 | 0.46 | 0.56 | 0.5 | 0.76 |
| 11 | 118134785 | rs1624780 | -0.047 | -0.214 | -0.429 | -0.436 | 0.004 | -0.007 | -0.007 | 0.004 | 0.59 | 0.69 | 0.56 | 0.5 | 0.61 | 0.7 |
| 11 | 118135745 | rs3759001 | 0.318 | -0.12 | -0.511 | -0.589 | 0.006 | 0.045 | 0.045 | 4.849 | 0.57 | 0.3 | 0.67 | 0.7 | 0.6 | 0.6 |
| 11 | 118140184 | rs12788933 | -0.287 | -0.407 | -0.732 | -0.966 | 0.03 | 0.014 | 0.014 | 0 | 0.26 | 0.07 | 0.29 | 0.18 | 0.2 | 0.22 |
| 11 | 118144804 | rs4938493 | 1.206 | -0.356 | -0.747 | -1.101 | -0.019 | -0.015 | -0.015 | 0 | 0.36 | 0.3 | 0.43 | 0.53 | 0.45 | 0.51 |
| 11 | 118148146 | rs2853009 | -1.513 | -0.092 | -0.767 | -1.12 | -0.019 | -0.014 | -0.014 | 0 | 0.63 | 0.69 | 0.55 | 0.47 | 0.5 | 0.49 |
| 11 | 118148271 | rs2187557 | 0.596 | 0.163 | -0.582 | -1.024 | -0.008 | -0.016 | -0.016 | 0 | 0.35 | 0.57 | 0.24 | 0.27 | 0.28 | 0.26 |
| 11 | 118158055 | rs12576947 | 1.371 | 0.304 | -0.771 | -1.142 | -0.004 | -0.008 | -0.008 | 0 | 0.31 | 0.3 | 0.25 | 0.53 | 0.34 | 0.45 |
| 11 | 118168617 | rs7110019 | 1.211 | 0.267 | -0.56 | -1.161 | 0.011 | 0.034 | 0.034 | 0 | 0.33 | 0.41 | 0.26 | 0.53 | 0.34 | 0.46 |
| 11 | 118174629 | rs3825051 | 1.374 | 0.271 | -0.598 | -1.159 | -0.004 | -0.008 | -0.008 | 1.788 | 0.31 | 0.3 | 0.25 | 0.53 | 0.34 | 0.46 |
| 11 | 118175439 | rs2231440 | -0.127 | 0.297 | -0.698 | -1.136 | -0.026 | 0.03 | 0.03 | 0 | 0.3 | 0.05 | 0.47 | 0.19 | 0.3 | 0.27 |
| 11 | 118182660 | rs2277289 | -0.144 | 0.344 | -0.821 | -1.123 | -0.018 | 0.031 | 0.031 | 0 | 0.3 | 0.07 | 0.46 | 0.19 | 0.31 | 0.27 |
| 11 | 118182801 | rs1945764 | -0.787 | 0.199 | -0.596 | -1.155 | 0 | -0.003 | -0.003 | 0 | 0.64 | 0.41 | 0.73 | 0.73 | 0.66 | 0.74 |
| 11 | 118187511 | rs4936428 | -0.235 | -0.015 | -1.004 | -0.61 | -0.018 | 0.006 | 0.006 | 0 | 0.75 | 0.56 | 0.75 | 0.79 | 0.79 | 0.77 |
| 11 | 118191274 | rs7937334 | -0.409 | 0.143 | -0.942 | -0.555 | -0.012 | -0.002 | -0.002 | 0 | 0.73 | 0.64 | 0.57 | 0.79 | 0.77 | 0.75 |
| 11 | 118191691 | rs6589659 | 0.094 | 0.022 | -1.044 | -0.566 | -0.024 | -0.02 | -0.02 | 0 | 0.66 | 0.51 | 0.54 | 0.78 | 0.73 | 0.74 |
| 11 | 118194571 | rs10892234 | 0.862 | 0.445 | -1.949 | -0.668 | -0.14 | 0.01 | 0.01 | 4.345 | 0.32 | 0.04 | 0.28 | 0.27 | 0.43 | 0.28 |
| 11 | 118194599 | rs12577841 | 0.766 | 0.533 | -1.771 | -0.609 | 0.01 | -0.043 | -0.043 | 0 | 0.33 | 0.2 | 0.24 | 0.52 | 0.31 | 0.47 |
| 11 | 118209960 | rs2276424 | 0.115 | -0.018 | -1.3 | -0.375 | 0 | 0 | 0 | 0 | 0.67 | 0.36 | 0.54 | 0.8 | 0.74 | 0.75 |
| 11 | 118212671 | rs3181261 | 0.898 | 0.455 | -1.404 | 0.469 | -0.008 | -0.053 | -0.053 | 0 | 0.16 | 0 | 0.04 | 0.21 | 0.09 | 0.2 |
| 11 | 118213632 | rs2239695 | 0.323 | 0.815 | -1.174 | 0.21 | 0.01 | -0.043 | -0.043 | 0 | 0.26 | 0.16 | 0.11 | 0.49 | 0.18 | 0.51 |
| 11 | 118214267 | rs2071381 | -0.877 | 1.161 | -0.977 | 0.057 | 0 | -0.001 | -0.001 | 0.073 | 0.41 | 0.59 | 0.26 | 0.57 | 0.35 | 0.56 |
| 11 | 118221474 | rs1561966 | -0.675 | 0.992 | -1.591 | 0.14 | -0.003 | -0.017 | -0.017 | 0 | 0.35 | 0.31 | 0.22 | 0.59 | 0.31 | 0.56 |
| 11 | 118230223 | rs2276417 | -0.708 | 0.991 | -1.592 | 0.14 | -0.003 | -0.017 | -0.017 | 0.195 | 0.35 | 0.31 | 0.22 | 0.59 | 0.31 | 0.56 |
| 11 | 118231584 | rs4938507 | 1.512 | 0.394 | -1.682 | 0.174 | -0.019 | -0.064 | -0.064 | 0 | 0.1 | 0.01 | 0.23 | 0.18 | 0.12 | 0.05 |
| 11 | 118234191 | rs3825054 | -0.011 | 0.409 | -1.06 | 0.26 | 0.007 | -0.026 | -0.026 | 0 | 0.27 | 0.23 | 0.12 | 0.51 | 0.18 | 0.52 |
| 11 | 118255278 | rs6589663 | -1.249 | 0.241 | -0.273 | 0.944 | 0.007 | -0.011 | -0.011 | 0 | 0.35 | 0.32 | 0.2 | 0.58 | 0.27 | 0.55 |
| 11 | 118273019 | rs17122154 | -1.24 | 0.274 | -0.691 | 0.96 | 0.018 | -0.029 | -0.029 | 0 | 0.34 | 0.28 | 0.19 | 0.58 | 0.27 | 0.55 |
| 11 | 118276246 | rs7483081 | -0.271 | 0.231 | -0.717 | 1.077 | 0.026 | -0.103 | -0.103 | 2.98 | 0.24 | 0.12 | 0.08 | 0.51 | 0.16 | 0.51 |
| 11 | 118276492 | rs17122157 | -0.271 | 0.231 | -0.717 | 1.077 | 0.026 | -0.103 | -0.103 | 0.029 | 0.24 | 0.12 | 0.09 | 0.51 | 0.16 | 0.51 |
| 11 | 118279936 | rs3194726 | 0.008 | 0.224 | -0.732 | 0.998 | -0.005 | 0.005 | 0.005 | 0 | 0.07 | 0.06 | 0.08 | 0.06 | 0.08 | 0.03 |
| 11 | 118285995 | rs4290266 | -0.518 | 0.062 | -0.569 | 1.168 | 0.018 | -0.047 | -0.047 | 2.958 | 0.25 | 0.2 | 0.09 | 0.5 | 0.16 | 0.51 |
| 11 | 118297391 | rs4435039 | -0.481 | 0.137 | -0.763 | 1.166 | 0.014 | -0.075 | -0.075 | 0.218 | 0.25 | 0.12 | 0.09 | 0.47 | 0.18 | 0.53 |
| 11 | 118341921 | rs524936 | 0.292 | -0.272 | 0.552 | 1.386 | -0.022 | 0.081 | 0.081 | 0 | 0.81 | 0.73 | 0.93 | 0.46 | 0.9 | 0.57 |
| 11 | 118362414 | rs562780 | -0.104 | -0.039 | 0.202 | 1.422 | 0.005 | -0.049 | -0.049 | 0.19 | 0.84 | 0.9 | 0.93 | 0.45 | 0.87 | 0.58 |
| 11 | 118362869 | rs537975 | 0.949 | -0.472 | 1.126 | 1.204 | -0.028 | 0.103 | 0.103 | 0 | 0.73 | 0.52 | 0.85 | 0.43 | 0.8 | 0.51 |
| 11 | 118382327 | rs7113994 | -0.323 | -0.51 | 1.251 | 1.11 | -0.006 | -0.028 | -0.028 | 0.725 | 0.08 | 0.14 | 0.05 | 0.02 | 0.06 | 0.07 |
| 11 | 118391375 | rs576950 | 0.981 | -0.159 | -0.094 | 0.202 | -0.1 | 0.096 | 0.096 | 0.233 | 0.35 | 0.03 | 0.36 | 0.17 | 0.43 | 0.19 |
| 11 | 118393396 | rs573971 | 0.6 | -0.141 | 0.323 | 0.453 | -0.025 | 0.084 | 0.084 | 0 | 0.73 | 0.53 | 0.86 | 0.43 | 0.8 | 0.5 |
| 11 | 118393856 | rs9332871 | 0.461 | 0.297 | -0.23 | 0.351 | -0.003 | 0.021 | 0.021 | 0 | 0.06 | 0.02 | 0.03 | 0.02 | 0.06 | 0.07 |
| 11 | 118404804 | rs6589664 | 0.369 | 0.289 | -0.108 | 0.365 | -0.01 | -0.002 | -0.002 | 0 | 0.27 | 0.19 | 0.43 | 0.23 | 0.29 | 0.12 |
| 11 | 118405068 | rs12289253 | -0.079 | 0.162 | 0.46 | 0.693 | -0.009 | 0.068 | 0.068 | 0 | 0.18 | 0.26 | 0.09 | 0.58 | 0.14 | 0.38 |
| 11 | 118405835 | rs3741324 | -0.023 | 0.174 | 0.389 | 0.712 | 0.001 | 0.009 | 0.009 | 0 | 0.18 | 0.19 | 0.08 | 0.54 | 0.13 | 0.35 |
| 11 | 118405989 | rs3741325 | 0.348 | 0.156 | 0.31 | 0.703 | -0.027 | -0.135 | -0.135 | 0 | 0.16 | 0.01 | 0.06 | 0.54 | 0.13 | 0.36 |
| 11 | 118410468 | rs11216883 | -0.177 | 0.135 | 0.381 | 0.694 | -0.001 | 0.022 | 0.022 | 0 | 0.18 | 0.19 | 0.08 | 0.54 | 0.13 | 0.35 |
| 11 | 118411424 | rs11216886 | -0.177 | 0.135 | 0.386 | 0.694 | 0 | 0.014 | 0.014 | 0 | 0.18 | 0.19 | 0.08 | 0.54 | 0.13 | 0.35 |
| 11 | 118416253 | rs2277295 | -0.098 | 0.139 | 0.38 | 0.907 | -0.001 | 0.019 | 0.019 | 0 | 0.18 | 0.17 | 0.08 | 0.54 | 0.13 | 0.34 |

| | | | | | | | | | | | | | | | | |
|---|---|---|---|---|---|---|---|---|---|---|---|---|---|---|---|---|
| 11 | 118430048 | rs11216896 | -0.154 | 0.099 | 0.489 | 0.843 | 0 | 0.015 | 0.015 | 0 | 0.18 | 0.17 | 0.08 | 0.54 | 0.13 | 0.35 |
| 11 | 118430519 | rs11552421 | -0.311 | 0.059 | 0.475 | 0.853 | -0.006 | 0.042 | 0.042 | 0 | 0.19 | 0.21 | 0.08 | 0.56 | 0.14 | 0.35 |
| 11 | 118435756 | rs2277291 | 0.357 | 0.064 | 0.401 | 0.904 | -0.027 | -0.135 | -0.135 | 0.07 | 0.16 | 0.01 | 0.06 | 0.55 | 0.13 | 0.35 |
| 11 | 118441846 | rs10892243 | -0.271 | 0.877 | 0.357 | 0.79 | -0.004 | 0.035 | 0.035 | 0 | 0.19 | 0.2 | 0.08 | 0.59 | 0.14 | 0.36 |
| 11 | 118444525 | rs11216907 | -0.144 | 0.863 | 0.366 | 0.784 | 0.002 | 0.007 | 0.007 | 0 | 0.18 | 0.15 | 0.08 | 0.57 | 0.14 | 0.34 |
| 11 | 118459069 | rs633308 | -1.068 | 0.455 | -0.372 | 0.066 | -0.097 | 0.066 | 0.066 | 0 | 0.63 | 0.95 | 0.64 | 0.79 | 0.58 | 0.61 |
| 11 | 118459269 | rs10736492 | 0.411 | 0.409 | 0.444 | 0.034 | 0.003 | -0.009 | -0.009 | 0 | 0.4 | 0.45 | 0.52 | 0.21 | 0.43 | 0.25 |
| 11 | 118471423 | rs11216925 | -0.286 | 0.339 | 0.384 | 0.457 | -0.004 | 0.033 | 0.033 | 0 | 0.19 | 0.21 | 0.08 | 0.58 | 0.14 | 0.36 |
| 11 | 118472354 | rs10892245 | -0.313 | 0.336 | 0.383 | 0.456 | -0.004 | 0.032 | 0.032 | 0 | 0.19 | 0.21 | 0.08 | 0.58 | 0.14 | 0.36 |
| 11 | 118473231 | rs1047079 | 0.438 | 0.345 | 0.367 | 0.46 | -0.027 | -0.135 | -0.135 | 0 | 0.16 | 0.01 | 0.06 | 0.57 | 0.13 | 0.35 |
| 11 | 118473385 | rs17742 | -0.243 | 0.355 | 0.391 | 0.453 | -0.004 | 0.033 | 0.033 | 0 | 0.19 | 0.21 | 0.08 | 0.58 | 0.14 | 0.36 |
| 11 | 118475098 | rs604096 | -0.935 | 0.831 | 0.15 | 0.825 | 0 | -0.001 | -0.001 | 0 | 0.29 | 0.25 | 0.15 | 0.38 | 0.23 | 0.35 |
| 11 | 118477367 | rs498872 | -0.385 | 1.321 | -0.573 | 0.544 | -0.049 | -0.017 | -0.017 | 0 | 0.75 | 0.9 | 0.76 | 0.75 | 0.69 | 0.62 |
| 11 | 118481431 | rs10892246 | 0.147 | 1.078 | -0.455 | -0.084 | 0 | -0.014 | -0.014 | 0 | 0.61 | 0.58 | 0.68 | 0.79 | 0.63 | 0.55 |
| 11 | 118486067 | rs11603023 | 0.083 | 1.021 | -0.262 | 0.105 | 0.009 | -0.033 | -0.033 | 0 | 0.6 | 0.52 | 0.72 | 0.67 | 0.56 | 0.59 |
| 11 | 118488782 | rs11216930 | 0.582 | 0.932 | -0.144 | 0.169 | -0.001 | -0.004 | -0.004 | 1.036 | 0.24 | 0.19 | 0.23 | 0.23 | 0.22 | 0.38 |
| 11 | 118492591 | rs2077173 | 1.443 | 0.758 | -0.165 | 0.173 | 0 | 0 | 0 | 0 | 0.31 | 0.25 | 0.49 | 0.38 | 0.32 | 0.17 |
| 11 | 118504742 | rs633683 | 0.094 | 1.001 | -0.183 | 0.102 | -0.004 | -0.028 | -0.028 | 1.088 | 0.58 | 0.48 | 0.74 | 0.65 | 0.6 | 0.58 |
| 11 | 118509668 | rs483598 | 0.29 | 0.718 | -0.125 | 0.167 | 0.003 | -0.023 | -0.023 | 0 | 0.56 | 0.48 | 0.71 | 0.66 | 0.54 | 0.56 |
| 11 | 118520480 | rs653008 | 1.312 | 0.645 | 0.082 | 0.149 | 0.004 | -0.025 | -0.025 | 0 | 0.32 | 0.25 | 0.49 | 0.43 | 0.32 | 0.18 |
| 11 | 118528424 | rs17748 | 0.921 | 0.684 | -0.14 | 0.156 | -0.026 | -0.028 | -0.028 | 2.221 | 0.23 | 0.05 | 0.2 | 0.23 | 0.21 | 0.38 |
| 11 | 118528466 | rs7389 | 1.4 | 0.676 | -0.115 | 0.178 | 0 | -0.001 | -0.001 | 0 | 0.31 | 0.26 | 0.49 | 0.39 | 0.32 | 0.18 |
| 11 | 118529595 | rs745663 | 1.476 | 0.686 | -0.096 | 0.183 | 0 | -0.005 | -0.005 | 0.149 | 0.31 | 0.24 | 0.49 | 0.39 | 0.32 | 0.17 |
| 11 | 118534082 | rs10790256 | 0.867 | 0.688 | -0.254 | 0.212 | -0.041 | -0.042 | -0.042 | 1.609 | 0.23 | 0.03 | 0.2 | 0.23 | 0.21 | 0.37 |
| 11 | 118538187 | rs607527 | 1.291 | 0.66 | -0.025 | 0.218 | -0.004 | 0.023 | 0.023 | 0.382 | 0.32 | 0.35 | 0.52 | 0.41 | 0.32 | 0.18 |
| 11 | 118543563 | rs10892251 | 0.822 | 0.647 | -0.05 | 0.244 | -0.023 | -0.044 | -0.044 | 0 | 0.23 | 0.05 | 0.19 | 0.26 | 0.22 | 0.37 |
| 11 | 118550601 | rs673770 | 1.287 | 0.658 | -0.037 | 0.249 | -0.005 | 0.023 | 0.023 | 0.321 | 0.33 | 0.36 | 0.52 | 0.41 | 0.32 | 0.18 |
| 11 | 118550784 | rs642497 | 1.287 | 0.658 | -0.037 | 0.249 | -0.005 | 0.023 | 0.023 | 0.321 | 0.33 | 0.36 | 0.52 | 0.41 | 0.32 | 0.18 |
| 11 | 118552420 | rs4936433 | 1.287 | 0.69 | -0.022 | 0.249 | -0.006 | 0.025 | 0.025 | 0.321 | 0.33 | 0.36 | 0.52 | 0.41 | 0.32 | 0.18 |
| 11 | 118556398 | rs11216943 | 0.784 | 0.695 | -0.014 | 0.35 | -0.007 | -0.005 | -0.005 | 0.747 | 0.24 | 0.11 | 0.19 | 0.22 | 0.22 | 0.37 |
| 11 | 118561410 | rs12798453 | 1.458 | 0.716 | 0.051 | 0.339 | 0 | -0.001 | -0.001 | 0 | 0.31 | 0.25 | 0.51 | 0.37 | 0.31 | 0.17 |
| 11 | 118565651 | rs540800 | -0.365 | 0.58 | 0.001 | 0.163 | 0.007 | -0.034 | -0.034 | 0 | 0.41 | 0.48 | 0.27 | 0.32 | 0.44 | 0.41 |
| 11 | 118569067 | rs4486664 | 1.305 | 0.609 | -0.041 | 0.303 | -0.005 | 0.007 | 0.007 | 0.322 | 0.31 | 0.35 | 0.52 | 0.34 | 0.31 | 0.18 |
| 11 | 118572267 | rs573905 | 0.158 | 0.58 | 0.07 | 0.362 | 0.005 | -0.012 | -0.012 | 0 | 0.57 | 0.53 | 0.7 | 0.59 | 0.54 | 0.57 |
| 11 | 118573519 | rs4639966 | 0.141 | -0.106 | 0.249 | 0.317 | -0.002 | -0.009 | -0.009 | 0.57 | 0.26 | 0.21 | 0.17 | 0.41 | 0.25 | 0.33 |
| 11 | 118574675 | rs494459 | 1.344 | 0.039 | 0.109 | 0.269 | 0.005 | 0.009 | 0.009 | 0.171 | 0.33 | 0.37 | 0.56 | 0.34 | 0.4 | 0.19 |
| 11 | 118575180 | rs626645 | 1.344 | 0.039 | 0.104 | 0.269 | 0.006 | 0.009 | 0.009 | 0.171 | 0.33 | 0.37 | 0.56 | 0.34 | 0.4 | 0.19 |
| 11 | 118577990 | rs480958 | 0.423 | -0.001 | -0.032 | 0.24 | -0.003 | -0.003 | -0.003 | 0 | 0.56 | 0.46 | 0.73 | 0.53 | 0.63 | 0.48 |
| 11 | 118579747 | rs10790261 | 0.677 | 0.176 | -0.458 | 0.288 | -0.021 | -0.016 | -0.016 | 0.182 | 0.23 | 0.07 | 0.17 | 0.2 | 0.23 | 0.28 |
| 11 | 118610463 | rs4499035 | 0.07 | -0.221 | -0.286 | 0.057 | -0.016 | -0.024 | -0.024 | 0.119 | 0.24 | 0.12 | 0.16 | 0.32 | 0.25 | 0.33 |
| 11 | 118610629 | rs521910 | -1.374 | -0.446 | 0.038 | -0.358 | -0.014 | -0.066 | -0.066 | 0 | 0.81 | 0.78 | 0.91 | 1 | 0.87 | 0.91 |
| 11 | 118619047 | rs2077579 | 0.996 | -0.281 | -0.073 | -0.367 | -0.021 | 0.023 | 0.023 | 0 | 0.21 | 0.05 | 0.16 | 0.13 | 0.2 | 0.3 |
| 11 | 118651809 | rs555649 | 1.796 | -0.385 | 0.003 | -0.138 | 0.009 | 0.01 | 0.01 | 0 | 0.31 | 0.35 | 0.55 | 0.35 | 0.41 | 0.19 |
| 11 | 118664377 | rs598373 | 0.362 | -0.6 | -0.338 | -0.042 | 0.006 | 0.009 | 0.009 | 0 | 0.57 | 0.61 | 0.73 | 0.69 | 0.67 | 0.57 |
| 11 | 118705145 | rs636736 | -0.607 | -0.598 | -0.297 | 0.006 | -0.029 | -0.038 | -0.038 | 0.486 | 0.75 | 0.63 | 0.87 | 0.81 | 0.83 | 0.8 |
| 11 | 118711069 | rs538645 | -0.514 | -0.426 | -0.679 | 0.039 | 0.016 | 0 | 0 | 1.575 | 0.7 | 0.82 | 0.86 | 0.68 | 0.76 | 0.6 |
| 11 | 118718485 | rs4938561 | -0.461 | -0.408 | -0.988 | 0.114 | 0.02 | 0.005 | 0.005 | 0 | 0.71 | 0.89 | 0.87 | 0.71 | 0.75 | 0.62 |
| 11 | 118734640 | rs12286723 | 0.621 | 0.212 | -0.967 | -0.492 | 0.005 | -0.078 | -0.078 | 0 | 0.2 | 0.26 | 0.15 | 0 | 0.2 | 0.08 |
| 11 | 118735476 | rs10892301 | -0.382 | 0.104 | -0.775 | -0.042 | 0.008 | 0.01 | 0.01 | 0 | 0.48 | 0.37 | 0.28 | 0.52 | 0.45 | 0.63 |
| 11 | 118741842 | rs4938573 | -0.069 | -0.069 | -0.846 | 0.107 | -0.002 | -0.013 | -0.013 | 0.085 | 0.79 | 0.72 | 0.84 | 0.89 | 0.81 | 0.83 |
| 11 | 118743286 | rs12365699 | 0.794 | 0.509 | -1.605 | 0.142 | -0.056 | 0.018 | 0.018 | 0 | 0.11 | 0.01 | 0.11 | 0.04 | 0.17 | 0.05 |
| 11 | 118746769 | rs4938576 | 0.416 | 0.168 | -1.411 | 0.259 | 0.013 | 0.013 | 0.013 | 0 | 0.52 | 0.38 | 0.7 | 0.4 | 0.6 | 0.38 |
| 11 | 118747813 | rs6589706 | -0.866 | 0.561 | -1.891 | 0.393 | -0.121 | -0.135 | -0.135 | 0 | 0.63 | 0.93 | 0.54 | 0.47 | 0.55 | 0.61 |
| 11 | 118755738 | rs523604 | -0.457 | 0.32 | -1.42 | 0.094 | -0.01 | 0.008 | 0.008 | 0 | 0.55 | 0.7 | 0.5 | 0.42 | 0.49 | 0.58 |
| 11 | 118758089 | rs497916 | -0.95 | -0.21 | -0.849 | -0.846 | 0 | -0.007 | -0.007 | 0.002 | 0.29 | 0.32 | 0.32 | 0.23 | 0.27 | 0.15 |
| 11 | 118759610 | rs566416 | -0.266 | -0.071 | -1.416 | -0.795 | -0.065 | 0.069 | 0.069 | 4.28 | 0.22 | 0.01 | 0.38 | 0.08 | 0.23 | 0.13 |
| 11 | 118765600 | rs3922 | -0.606 | -0.276 | -0.935 | -0.801 | 0.027 | 0.044 | 0.044 | 0.478 | 0.51 | 0.69 | 0.55 | 0.29 | 0.42 | 0.22 |
| 11 | 118793613 | rs11217091 | -1.475 | 0.326 | 0.356 | -0.41 | -0.002 | 0.044 | 0.044 | 0 | 0.14 | 0.4 | 0.13 | 0.17 | 0.11 | 0.15 |
| 11 | 118793918 | rs12282752 | -0.759 | 0.333 | 0.395 | -0.579 | -0.01 | -0.051 | -0.051 | 0 | 0.09 | 0.29 | 0.09 | 0.01 | 0.06 | 0.12 |
| 11 | 118801235 | rs602716 | -0.646 | 0.65 | -0.073 | -0.921 | -0.052 | 0.059 | 0.059 | 0.451 | 0.28 | 0.11 | 0.46 | 0.12 | 0.37 | 0.35 |
| 11 | 118810405 | rs4938589 | -0.419 | 0.49 | 0.404 | -0.957 | 0.006 | -0.069 | -0.069 | 0 | 0.11 | 0.24 | 0.14 | 0.01 | 0.12 | 0.08 |
| 11 | 118811641 | rs11606199 | -0.501 | 0.515 | 0.522 | -0.969 | 0.02 | -0.088 | -0.088 | 0 | 0.14 | 0.26 | 0.16 | 0.01 | 0.16 | 0.1 |
| 11 | 118812512 | rs647769 | -0.586 | 0.78 | 0.209 | -0.903 | -0.06 | 0 | 0 | 1.499 | 0.29 | 0.11 | 0.44 | 0.25 | 0.36 | 0.39 |
| 11 | 118816156 | rs587263 | 1.771 | 0.277 | 0.03 | -0.951 | -0.09 | -0.148 | -0.148 | 0 | 0.35 | 0.08 | 0.3 | 0.45 | 0.42 | 0.19 |
| 11 | 118816304 | rs586763 | -0.414 | 0.37 | 1.282 | -1.261 | 0.069 | -0.094 | -0.094 | 0.259 | 0.13 | 0.34 | 0.15 | 0 | 0.16 | 0.07 |
| 11 | 118823148 | rs663003 | -2.105 | 0.543 | 1.412 | -1.213 | 0.002 | 0.022 | 0.022 | 1.335 | 0.52 | 0.57 | 0.64 | 0.31 | 0.5 | 0.53 |
| 11 | 118825636 | rs1790191 | 1.522 | 0.558 | 1.471 | -1.206 | 0.002 | 0.015 | 0.015 | 1.13 | 0.51 | 0.61 | 0.39 | 0.7 | 0.51 | 0.5 |
| 11 | 118826192 | rs12797009 | -1.179 | 0.369 | 1.479 | -1.155 | -0.003 | -0.011 | -0.011 | 0 | 0.33 | 0.39 | 0.46 | 0.26 | 0.31 | 0.4 |
| 11 | 118831727 | rs10892315 | -1.143 | 0.39 | 1.266 | -1.933 | -0.022 | 0.082 | 0.082 | 0.285 | 0.28 | 0.13 | 0.22 | 0.06 | 0.29 | 0.38 |

| | | | | | | | | | | | | | | | | |
|---|---|---|---|---|---|---|---|---|---|---|---|---|---|---|---|---|
| 11 | 118835258 | rs11607835 | -2 | 0.365 | 1.346 | -1.954 | -0.002 | 0.057 | 0.057 | 1.403 | 0.49 | 0.38 | 0.6 | 0.11 | 0.47 | 0.45 |
| 11 | 118840857 | rs12420418 | -1.843 | 0.742 | 1.57 | -1.731 | -0.005 | 0.027 | 0.027 | 0 | 0.46 | 0.38 | 0.6 | 0.11 | 0.46 | 0.44 |
| 11 | 118845813 | rs4442562 | 0.456 | 0.744 | 1.414 | -1.691 | -0.013 | 0.038 | 0.038 | 0 | 0.15 | 0.04 | 0.35 | 0.05 | 0.18 | 0.07 |
| 11 | 118846466 | rs4938602 | -1.869 | 0.744 | 1.526 | -1.683 | -0.003 | 0.015 | 0.015 | 0.015 | 0.46 | 0.37 | 0.61 | 0.12 | 0.47 | 0.45 |
| 11 | 118846482 | rs4938603 | -2.049 | 0.742 | 1.499 | -1.667 | 0 | -0.025 | -0.025 | 2.172 | 0.48 | 0.48 | 0.62 | 0.18 | 0.48 | 0.49 |
| 11 | 118846574 | rs4938604 | -2.124 | 0.745 | 1.549 | -1.666 | -0.008 | 0.118 | 0.118 | 0.063 | 0.48 | 0.64 | 0.63 | 0.18 | 0.48 | 0.49 |
| 11 | 118851138 | rs4456264 | -1.829 | 0.678 | 0.968 | -1.405 | 0 | -0.069 | -0.069 | 0.033 | 0.44 | 0.47 | 0.6 | 0.12 | 0.43 | 0.42 |
| 11 | 118856491 | rs3889526 | -1.962 | 0.676 | 0.961 | -1.403 | 0 | -0.028 | -0.028 | 3.706 | 0.45 | 0.5 | 0.61 | 0.18 | 0.45 | 0.46 |
| 11 | 118869460 | rs3737504 | -2.237 | 0.629 | 1.47 | -0.797 | -0.008 | -0.135 | -0.135 | 0.165 | 0.3 | 0.51 | 0.25 | 0.07 | 0.26 | 0.33 |
| 11 | 118869539 | rs11822103 | -1.665 | 0.845 | 0.984 | -0.691 | -0.014 | 0.055 | 0.055 | 0.095 | 0.24 | 0.12 | 0.21 | 0.07 | 0.26 | 0.33 |
| 11 | 118886117 | rs10111 | -1.963 | 0.786 | 0.908 | -0.695 | -0.006 | -0.068 | -0.068 | 0 | 0.27 | 0.35 | 0.23 | 0.07 | 0.23 | 0.34 |
| 11 | 118886656 | rs7131534 | -2.006 | 0.819 | 0.894 | -0.672 | -0.016 | -0.134 | -0.134 | 1.022 | 0.27 | 0.46 | 0.24 | 0.07 | 0.23 | 0.34 |
| 11 | 118888619 | rs11602764 | -2.321 | 0.827 | 0.79 | -0.69 | -0.002 | -0.096 | -0.096 | 0 | 0.3 | 0.51 | 0.26 | 0.13 | 0.27 | 0.39 |
| 11 | 118889185 | rs11217125 | -2.11 | 0.877 | 0.106 | -0.7 | -0.001 | -0.085 | -0.085 | 0 | 0.28 | 0.39 | 0.24 | 0.07 | 0.26 | 0.34 |
| 11 | 118890085 | rs4938619 | -0.63 | 0.883 | -0.941 | -0.498 | -0.008 | 0.031 | 0.031 | 3.725 | 0.14 | 0.05 | 0.14 | 0.02 | 0.13 | 0.08 |
| 11 | 118894287 | rs569 | -1.627 | 0.861 | -0.941 | -0.473 | -0.01 | 0.077 | 0.077 | 0.29 | 0.23 | 0.09 | 0.2 | 0.08 | 0.23 | 0.33 |
| 11 | 118895202 | rs11006 | -1.646 | 0.855 | -0.94 | -0.472 | -0.011 | 0.078 | 0.078 | 0.373 | 0.23 | 0.09 | 0.2 | 0.08 | 0.23 | 0.34 |
| 11 | 118898319 | rs741811 | -0.89 | 0.841 | -0.578 | -0.197 | -0.007 | 0.021 | 0.021 | 3.358 | 0.13 | 0.06 | 0.13 | 0.02 | 0.13 | 0.08 |
| 11 | 118899218 | rs4936459 | -1.16 | 0.718 | -0.245 | -0.146 | -0.004 | -0.047 | -0.047 | 0 | 0.17 | 0.3 | 0.18 | 0.07 | 0.14 | 0.1 |
| 11 | 118913993 | rs1003081 | -0.513 | 1.122 | -1.339 | -0.285 | -0.038 | 0.069 | 0.069 | 0 | 0.37 | 0.2 | 0.55 | 0.26 | 0.4 | 0.37 |
| 11 | 118919206 | rs568922 | 0.146 | 1.04 | -0.022 | -1.058 | 0.023 | -0.064 | -0.064 | 0 | 0.84 | 0.67 | 0.62 | 0.95 | 0.84 | 0.93 |
| 11 | 118922200 | rs1804690 | 0.49 | 0.448 | -0.377 | -0.185 | 0.016 | -0.053 | -0.053 | 0.032 | 0.15 | 0.08 | 0.06 | 0.3 | 0.11 | 0.26 |
| 11 | 118925341 | rs538478 | 0.345 | -0.545 | 0.006 | -0.846 | -0.043 | 0.074 | 0.074 | 1.776 | 0.4 | 0.19 | 0.3 | 0.72 | 0.48 | 0.51 |
| 11 | 118925935 | rs511134 | 1.17 | -0.647 | 0.676 | -0.882 | -0.028 | -0.021 | -0.021 | 0 | 0.08 | 0.11 | 0.02 | 0.01 | 0.01 | 0.05 |
| 11 | 118932320 | rs582688 | -0.309 | -0.631 | 1.004 | -0.889 | 0.027 | -0.072 | -0.072 | 0.435 | 0.84 | 0.61 | 0.61 | 0.93 | 0.84 | 0.9 |
| 11 | 118932445 | rs636283 | -0.292 | -0.629 | 1.063 | -0.888 | 0.012 | -0.042 | -0.042 | 0 | 0.86 | 0.77 | 0.65 | 0.95 | 0.84 | 0.91 |
| 11 | 118938315 | rs1786141 | 0.16 | -0.684 | 1.154 | -0.922 | -0.006 | -0.025 | -0.025 | 0.431 | 0.45 | 0.54 | 0.35 | 0.23 | 0.36 | 0.4 |
| 11 | 118942638 | rs1614264 | 0.025 | -0.689 | 1.164 | -0.925 | 0.035 | 0.092 | 0.092 | 0.076 | 0.48 | 0.71 | 0.37 | 0.23 | 0.37 | 0.42 |
| 11 | 118947634 | rs540261 | 0.5 | -0.688 | 1.018 | -0.902 | -0.035 | 0.069 | 0.069 | 1.074 | 0.4 | 0.2 | 0.3 | 0.72 | 0.48 | 0.49 |
| 11 | 118949331 | rs1177562 | 0.349 | -0.709 | 0.86 | -0.897 | 0.012 | 0.046 | 0.046 | 0.033 | 0.43 | 0.38 | 0.32 | 0.22 | 0.36 | 0.4 |
| 11 | 118952173 | rs15818 | 0.589 | -0.464 | 0.388 | -0.796 | 0 | -0.055 | -0.055 | 2.86 | 0.39 | 0.48 | 0.33 | 0.2 | 0.36 | 0.34 |
| 11 | 118955679 | rs589925 | 0.353 | -0.349 | -0.305 | -0.775 | -0.072 | 0.066 | 0.066 | 1.408 | 0.39 | 0.11 | 0.28 | 0.71 | 0.47 | 0.51 |
| 11 | 118958127 | rs1799993 | 0.521 | -0.356 | -0.213 | -0.781 | 0.004 | 0.082 | 0.082 | 0 | 0.41 | 0.7 | 0.37 | 0.22 | 0.36 | 0.34 |
| 11 | 118959331 | rs17075 | -0.336 | -0.246 | -0.269 | -0.327 | -0.019 | 0.037 | 0.037 | 0.638 | 0.6 | 0.79 | 0.7 | 0.32 | 0.53 | 0.5 |
| 11 | 118962816 | rs1784304 | -0.154 | -0.708 | -0.992 | 0.167 | 0.017 | -0.048 | -0.048 | 0 | 0.3 | 0.19 | 0.35 | 0.47 | 0.26 | 0.35 |
| 11 | 118964330 | rs640603 | 0.696 | -0.31 | -1.736 | 0.307 | -0.031 | -0.086 | -0.086 | 0 | 0.36 | 0.13 | 0.23 | 0.46 | 0.37 | 0.16 |
| 11 | 118966521 | rs2509049 | 0.286 | -0.428 | -1.817 | 0.312 | 0.003 | 0.034 | 0.034 | 1.793 | 0.45 | 0.27 | 0.3 | 0.62 | 0.45 | 0.5 |
| 11 | 118967291 | rs7759 | 0.91 | -0.476 | -1.69 | 0.336 | -0.043 | -0.087 | -0.087 | 0 | 0.37 | 0.13 | 0.23 | 0.47 | 0.39 | 0.17 |
| 11 | 118967758 | rs643788 | 0.238 | -0.481 | -1.659 | 0.315 | 0.003 | 0.036 | 0.036 | 1.684 | 0.46 | 0.28 | 0.3 | 0.63 | 0.46 | 0.5 |
| 11 | 118975927 | rs10790283 | -1.266 | -0.604 | -1.712 | 0.302 | 0.013 | -0.013 | -0.013 | 0 | 0.9 | 0.94 | 0.95 | 0.74 | 0.93 | 0.71 |
| 11 | 118976719 | rs639373 | 1.279 | -0.391 | -1.895 | 0.284 | -0.046 | -0.088 | -0.088 | 0 | 0.36 | 0.13 | 0.24 | 0.46 | 0.39 | 0.17 |
| 11 | 118980039 | rs510435 | 1.308 | -0.387 | -1.895 | 0.286 | -0.049 | -0.091 | -0.091 | 1.096 | 0.35 | 0.12 | 0.24 | 0.46 | 0.39 | 0.17 |
| 11 | 118983434 | rs2239896 | 1.515 | -0.105 | -1.707 | -0.318 | 0.004 | -0.027 | -0.027 | 0.92 | 0.09 | 0 | 0.05 | 0.17 | 0.06 | 0.35 |
| 11 | 118983681 | rs682795 | -0.965 | -0.099 | -1.67 | -0.326 | -0.006 | 0.129 | 0.129 | 0 | 0.55 | 0.78 | 0.71 | 0.28 | 0.54 | 0.57 |
| 11 | 119005485 | rs7234 | 0.939 | -0.154 | -1.485 | -0.311 | -0.007 | 0.101 | 0.101 | 0 | 0.42 | 0.17 | 0.28 | 0.72 | 0.44 | 0.43 |
| 11 | 119008538 | rs11217164 | -0.36 | -0.15 | -1.443 | -0.354 | -0.009 | 0.042 | 0.042 | 0.424 | 0.55 | 0.36 | 0.36 | 0.9 | 0.53 | 0.79 |
| 11 | 119016176 | rs671872 | -0.237 | -0.131 | -1.448 | -0.334 | -0.011 | 0.044 | 0.044 | 0.595 | 0.55 | 0.36 | 0.36 | 0.9 | 0.53 | 0.79 |
| 11 | 119018766 | rs3809046 | 1.242 | -0.111 | -1.412 | -0.312 | 0.002 | 0 | 0 | 0 | 0.12 | 0.1 | 0.06 | 0.18 | 0.09 | 0.34 |
| 11 | 119019520 | rs4301800 | -0.247 | 0.08 | -1.3 | -0.366 | -0.011 | 0.04 | 0.04 | 0.539 | 0.56 | 0.36 | 0.36 | 0.9 | 0.53 | 0.79 |
| 11 | 119027545 | rs4148170 | 0.68 | -0.019 | 0.053 | 0.147 | -0.015 | 0.003 | 0.003 | 0 | 0.13 | 0.25 | 0.07 | 0.18 | 0.08 | 0.34 |
| 11 | 119030752 | rs674424 | -1.371 | 0.626 | -0.811 | -0.418 | -0.105 | 0.073 | 0.073 | 0.092 | 0.81 | 0.98 | 0.57 | 0.94 | 0.75 | 0.87 |
| 11 | 119032689 | rs3802885 | 0.378 | 0.36 | -0.365 | -0.229 | -0.02 | 0.002 | 0.002 | 0 | 0.14 | 0.3 | 0.08 | 0.17 | 0.09 | 0.37 |
| 11 | 119043745 | rs561830 | 2.144 | 0.056 | -0.416 | 0.083 | -0.003 | 0.023 | 0.023 | 0.005 | 0.26 | 0.3 | 0.21 | 0.39 | 0.25 | 0.13 |
| 11 | 119052826 | rs4245191 | -0.479 | -0.147 | -0.434 | -0.5 | -0.032 | 0.05 | 0.05 | 2.303 | 0.63 | 0.71 | 0.42 | 0.88 | 0.55 | 0.8 |
| 11 | 119059404 | rs1815811 | -0.227 | -0.046 | -0.135 | -0.574 | 0.001 | -0.019 | -0.019 | 0.916 | 0.62 | 0.6 | 0.43 | 0.88 | 0.57 | 0.8 |
| 11 | 119063908 | rs2301574 | 2.673 | -0.396 | 0.036 | 0.088 | -0.001 | 0.01 | 0.01 | 0 | 0.24 | 0.26 | 0.21 | 0.35 | 0.25 | 0.14 |
| 11 | 119064394 | rs11217175 | 1.881 | 0.023 | -0.147 | -0.94 | -0.036 | 0.037 | 0.037 | 0 | 0.08 | 0.04 | 0.08 | 0 | 0.13 | 0.02 |
| 11 | 119075323 | rs4938637 | 1.513 | -0.13 | -0.047 | -0.717 | -0.009 | -0.063 | -0.063 | 0.513 | 0.12 | 0 | 0.05 | 0.23 | 0.09 | 0.36 |
| 11 | 119078886 | rs11217183 | 2.051 | -0.154 | 0.046 | -0.73 | 0.001 | -0.04 | -0.04 | 0.923 | 0.24 | 0.15 | 0.21 | 0.36 | 0.25 | 0.14 |
| 11 | 119085783 | rs6589722 | -0.545 | -0.093 | 0.06 | -0.754 | -0.018 | 0.027 | 0.027 | 2.534 | 0.63 | 0.69 | 0.44 | 0.88 | 0.58 | 0.78 |
| 11 | 119088883 | rs1893032 | -0.175 | -0.104 | 0.103 | -0.748 | 0.008 | -0.015 | -0.015 | 0 | 0.35 | 0.27 | 0.2 | 0.51 | 0.32 | 0.64 |
| 11 | 119094647 | rs11217191 | 0.833 | -0.108 | 0.081 | -0.803 | -0.001 | 0 | 0 | 0 | 0.22 | 0.21 | 0.14 | 0.28 | 0.24 | 0.28 |
| 11 | 119120804 | rs7937454 | -0.488 | 0.022 | 0.27 | -0.952 | -0.019 | 0.029 | 0.029 | 2.477 | 0.63 | 0.7 | 0.44 | 0.87 | 0.58 | 0.8 |
| 11 | 119155618 | rs2298650 | -0.071 | 0.091 | -0.437 | -0.881 | 0.016 | -0.025 | -0.025 | 0 | 0.36 | 0.24 | 0.19 | 0.51 | 0.32 | 0.63 |
| 11 | 119166047 | rs7105971 | 1.486 | 0.16 | 0.116 | -0.689 | -0.002 | 0.008 | 0.008 | 0 | 0.25 | 0.26 | 0.23 | 0.31 | 0.25 | 0.14 |
| 11 | 119172536 | rs1047417 | 1.542 | 0.195 | 0.043 | -0.674 | 0.001 | -0.013 | -0.013 | 0.428 | 0.24 | 0.18 | 0.22 | 0.32 | 0.25 | 0.14 |
| 11 | 119175075 | rs2509671 | 0.341 | 0.071 | 0.017 | -0.765 | 0.019 | -0.012 | -0.012 | 0 | 0.35 | 0.69 | 0.3 | 0.36 | 0.3 | 0.21 |
| 11 | 119176398 | rs2511844 | -0.443 | 0.118 | 0.206 | -0.781 | -0.021 | -0.014 | -0.014 | 0 | 0.11 | 0.46 | 0.08 | 0.04 | 0.05 | 0.05 |
| 11 | 119176499 | rs3829261 | 0.516 | 0.256 | -0.315 | -0.714 | -0.012 | -0.008 | -0.008 | 0 | 0.07 | 0.11 | 0.05 | 0.04 | 0.05 | 0.05 |

| | | | | | | | | | | | | | | | | |
|---|---|---|---|---|---|---|---|---|---|---|---|---|---|---|---|---|
| 11 | 119177938 | rs11217234 | 1.464 | 0.421 | -0.808 | -0.579 | 0.001 | -0.017 | -0.017 | 0 | 0.24 | 0.19 | 0.22 | 0.35 | 0.26 | 0.15 |
| 11 | 119179007 | rs2511837 | -0.422 | 0.624 | -1.269 | -1.16 | -0.062 | 0.08 | 0.08 | 0.439 | 0.59 | 0.88 | 0.43 | 0.72 | 0.49 | 0.65 |
| 11 | 119180316 | rs7914 | -0.4 | 0.731 | -0.342 | -1.373 | -0.007 | 0.01 | 0.01 | 0 | 0.33 | 0.38 | 0.29 | 0.5 | 0.25 | 0.6 |
| 11 | 119181170 | rs2249466 | -1.685 | 0.889 | -0.36 | -1.381 | -0.001 | 0 | 0 | 0.065 | 0.72 | 0.77 | 0.77 | 0.66 | 0.71 | 0.84 |
| 11 | 119185347 | rs6589732 | -0.027 | 1.043 | -0.867 | -1.215 | -0.01 | 0.016 | 0.016 | 3.427 | 0.34 | 0.3 | 0.43 | 0.13 | 0.41 | 0.19 |
| 11 | 119196915 | rs2509390 | 0.701 | 0.988 | -0.468 | -0.556 | -0.014 | 0.015 | 0.015 | 0 | 0.32 | 0.45 | 0.23 | 0.32 | 0.3 | 0.18 |
| 11 | 119197707 | rs2511834 | -0.036 | 1.237 | -0.7 | -0.371 | 0 | -0.003 | -0.003 | 0 | 0.83 | 0.86 | 0.77 | 0.48 | 0.86 | 0.43 |
| 11 | 119203505 | rs11824001 | 0.376 | 1.594 | -0.427 | -0.149 | -0.003 | -0.001 | -0.001 | 0.112 | 0.35 | 0.4 | 0.48 | 0.16 | 0.43 | 0.2 |
| 11 | 119205830 | rs12795576 | 0.397 | 1.286 | -0.573 | -0.109 | -0.027 | 0.057 | 0.057 | 0 | 0.34 | 0.27 | 0.46 | 0.16 | 0.43 | 0.2 |
| 11 | 119206375 | rs2511841 | -1.15 | 1.547 | -0.933 | -0.038 | 0.002 | -0.001 | -0.001 | 4.394 | 0.64 | 0.74 | 0.73 | 0.66 | 0.66 | 0.83 |
| 11 | 119207341 | rs2248863 | -0.81 | 1.763 | -0.935 | 0.5 | -0.005 | -0.034 | -0.034 | 0.518 | 0.19 | 0.23 | 0.09 | 0.04 | 0.15 | 0.06 |
| 11 | 119207446 | rs2248853 | -0.771 | 1.762 | -0.944 | 0.5 | -0.004 | -0.028 | -0.028 | 0 | 0.19 | 0.21 | 0.09 | 0.04 | 0.15 | 0.06 |
| 11 | 119208576 | rs2509656 | -0.815 | 0.584 | -0.848 | 0.626 | -0.005 | -0.031 | -0.031 | 0.02 | 0.19 | 0.23 | 0.09 | 0.04 | 0.16 | 0.06 |
| 11 | 119213303 | rs11217241 | -1.077 | 0.761 | -1.124 | 1.718 | 0.002 | 0.021 | 0.021 | 0 | 0.1 | 0.07 | 0.15 | 0.44 | 0.07 | 0.5 |
| 11 | 119214771 | rs948413 | -0.644 | 0.965 | -1.617 | 0.641 | -0.011 | 0.117 | 0.117 | 0.081 | 0.43 | 0.25 | 0.59 | 0.32 | 0.57 | 0.69 |
| 11 | 119214798 | rs948414 | -0.721 | 1.086 | -0.917 | 0.865 | -0.001 | -0.003 | -0.003 | 0 | 0.43 | 0.54 | 0.68 | 0.53 | 0.59 | 0.7 |
| 11 | 119215165 | rs10790289 | -0.609 | 1.464 | -0.979 | 0.721 | 0.002 | -0.003 | -0.003 | 0 | 0.42 | 0.46 | 0.66 | 0.43 | 0.48 | 0.54 |
| 11 | 119216504 | rs3814762 | -0.046 | 1.306 | -2.056 | 0.227 | -0.098 | 0.045 | 0.045 | 0.98 | 0.33 | 0.04 | 0.17 | 0.2 | 0.34 | 0.21 |
| 11 | 119217254 | rs883247 | 0.195 | 1.328 | -0.993 | 0.326 | 0.017 | -0.024 | -0.024 | 0 | 0.68 | 0.55 | 0.56 | 0.77 | 0.59 | 0.71 |
| 11 | 119222706 | rs7122785 | 0.014 | 1.499 | -0.926 | 0.271 | 0.016 | -0.042 | -0.042 | 0.407 | 0.79 | 0.56 | 0.64 | 0.87 | 0.74 | 0.8 |
| 11 | 119229964 | rs2241646 | 0.344 | 1.382 | -0.905 | 0.189 | -0.002 | -0.054 | -0.054 | 0.109 | 0.75 | 0.56 | 0.64 | 0.86 | 0.72 | 0.78 |
| 11 | 119235404 | rs2195525 | 0.854 | 1.367 | -0.433 | -0.004 | -0.003 | 0.001 | 0.001 | 0.342 | 0.64 | 0.4 | 0.62 | 0.83 | 0.54 | 0.68 |
| 11 | 119240604 | rs11217258 | -0.218 | 0.968 | -0.593 | 0.989 | -0.005 | -0.016 | -0.016 | 0 | 0.09 | 0.03 | 0.03 | 0.39 | 0.09 | 0.13 |
| 11 | 119244095 | rs587985 | 0.805 | 1.019 | 0.403 | 0.238 | 0.009 | -0.062 | -0.062 | 0 | 0.66 | 0.52 | 0.61 | 0.84 | 0.63 | 0.7 |
| 11 | 119248709 | rs614192 | 0.149 | 0.937 | 0.469 | 0.185 | 0.012 | -0.007 | -0.007 | 0 | 0.89 | 0.85 | 0.9 | 0.92 | 0.87 | 0.88 |
| 11 | 119252199 | rs670906 | -0.281 | 0.965 | 0.262 | 0.099 | -0.019 | 0.01 | 0.01 | 0 | 0.91 | 0.98 | 0.93 | 0.96 | 0.87 | 0.9 |
| 11 | 119252448 | rs7943722 | -0.466 | 0.881 | 0.415 | 0.122 | 0.002 | 0.018 | 0.018 | 0 | 0.23 | 0.18 | 0.27 | 0.08 | 0.23 | 0.18 |
| 11 | 119269081 | rs10892361 | 0.574 | 1.15 | 0.784 | 0.338 | 0.025 | -0.001 | -0.001 | 0.671 | 0.93 | 0.89 | 0.93 | 0.96 | 0.89 | 0.96 |
| 11 | 119288619 | rs3138094 | -1.092 | 0.894 | 1.029 | 0.012 | -0.016 | -0.105 | -0.105 | 0.959 | 0.31 | 0.53 | 0.36 | 0.08 | 0.27 | 0.16 |
| 11 | 119291480 | rs3138091 | 1.401 | 0.906 | 1.021 | 0.015 | -0.001 | 0.002 | 0.002 | 0 | 0.62 | 0.37 | 0.57 | 0.88 | 0.61 | 0.8 |
| 11 | 119291688 | rs3751026 | 0.571 | 0.898 | 1.066 | 0.012 | 0.022 | -0.007 | -0.007 | 0 | 0.92 | 0.88 | 0.93 | 0.96 | 0.89 | 0.93 |
| 11 | 119292712 | rs1001205 | 1.421 | 0.877 | 1.024 | 0.001 | -0.001 | 0.002 | 0.002 | 0 | 0.62 | 0.37 | 0.57 | 0.88 | 0.61 | 0.8 |
| 11 | 119294290 | rs11574653 | -0.707 | 0.867 | 1.348 | 0.008 | -0.002 | 0.022 | 0.022 | 0 | 0.26 | 0.18 | 0.34 | 0.08 | 0.27 | 0.13 |
| 11 | 119299408 | rs12417523 | -0.543 | 0.885 | 1.369 | 0.018 | -0.01 | 0.064 | 0.064 | 0 | 0.25 | 0.12 | 0.34 | 0.08 | 0.27 | 0.12 |
| 11 | 119300694 | rs7102696 | 0.012 | 0.914 | 1.022 | 0.286 | 0.002 | -0.004 | -0.004 | 0.408 | 0.21 | 0.19 | 0.13 | 0.39 | 0.19 | 0.17 |
| 11 | 119301754 | rs12421120 | -0.695 | 1.327 | 0.872 | -0.036 | -0.021 | 0.086 | 0.086 | 0 | 0.25 | 0.09 | 0.33 | 0.08 | 0.27 | 0.12 |
| 11 | 119305611 | rs4938653 | 1.004 | 1.38 | 0.954 | -0.006 | 0.044 | 0 | 0 | 0.136 | 0.91 | 0.8 | 0.93 | 0.96 | 0.88 | 0.92 |
| 11 | 119306183 | rs4938654 | 1.504 | 1.358 | 0.985 | -0.019 | 0.006 | -0.059 | -0.059 | 0.652 | 0.64 | 0.6 | 0.59 | 0.88 | 0.61 | 0.81 |
| 11 | 119314765 | rs7932332 | 1.588 | 1.35 | 0.99 | -0.022 | 0.002 | -0.041 | -0.041 | 0.182 | 0.63 | 0.47 | 0.58 | 0.88 | 0.61 | 0.81 |
| 11 | 119315417 | rs17547501 | -0.95 | 1.356 | 0.88 | -0.023 | -0.003 | 0.028 | 0.028 | 0 | 0.26 | 0.18 | 0.34 | 0.08 | 0.27 | 0.11 |
| 11 | 119318306 | rs7130716 | 1.262 | 1.416 | 1.03 | 0.312 | 0.01 | -0.009 | -0.009 | 0 | 0.91 | 0.87 | 0.93 | 0.96 | 0.88 | 0.92 |
| 11 | 119337504 | rs10750144 | 1.608 | 1.501 | 0.648 | 0.327 | 0.004 | -0.015 | -0.015 | 0 | 0.68 | 0.38 | 0.63 | 0.85 | 0.67 | 0.82 |
| 11 | 119341075 | rs685879 | 1.109 | 1.507 | 0.666 | 0.326 | -0.007 | -0.048 | -0.048 | 0.157 | 0.75 | 0.44 | 0.7 | 0.89 | 0.78 | 0.85 |
| 11 | 119346198 | rs535831 | -1.842 | 1.349 | 0.332 | 0.28 | 0.016 | -0.057 | -0.057 | 0 | 0.31 | 0.54 | 0.36 | 0.13 | 0.33 | 0.18 |
| 11 | 119349454 | rs611719 | -1.332 | 1.448 | -0.205 | 0.332 | -0.048 | 0.067 | 0.067 | 0 | 0.24 | 0.13 | 0.31 | 0.09 | 0.32 | 0.13 |
| 11 | 119351636 | rs672499 | -1.756 | 1.454 | -0.24 | 0.409 | 0.008 | -0.027 | -0.027 | 0.214 | 0.3 | 0.36 | 0.36 | 0.13 | 0.33 | 0.15 |
| 11 | 119356682 | rs1785899 | -1.329 | 2.198 | -1.013 | 0.74 | -0.072 | 0.03 | 0.03 | 0 | 0.22 | 0.07 | 0.31 | 0.12 | 0.33 | 0.13 |
| 11 | 119362151 | rs521053 | -0.899 | 2.42 | -0.844 | 1.32 | -0.112 | -0.107 | -0.107 | 0 | 0.18 | 0.05 | 0.35 | 0.46 | 0.34 | 0.2 |
| 11 | 119366947 | rs619180 | -1.273 | 2.352 | -0.607 | 1.296 | -0.017 | -0.016 | -0.016 | 1.68 | 0.22 | 0.19 | 0.37 | 0.47 | 0.34 | 0.22 |
| 11 | 119367321 | rs596647 | 0.013 | 2.049 | -1.111 | 0.95 | -0.005 | -0.013 | -0.013 | 0.071 | 0.77 | 0.78 | 0.82 | 0.88 | 0.81 | 0.88 |
| 11 | 119376672 | rs891292 | 0.625 | 1.631 | -1.204 | 1.008 | -0.024 | -0.034 | -0.034 | 0 | 0.59 | 0.77 | 0.48 | 0.45 | 0.48 | 0.73 |
| 11 | 119384450 | rs11217304 | -1.164 | 2.179 | -0.955 | 1.169 | -0.021 | -0.011 | -0.011 | 0 | 0.22 | 0.14 | 0.23 | 0.28 | 0.29 | 0.19 |
| 11 | 119392654 | rs609305 | 1.177 | 2.733 | -0.048 | 1.462 | 0.047 | 0.106 | 0.106 | 0 | 0.72 | 0.51 | 0.54 | 0.44 | 0.68 | 0.77 |
| 11 | 119399259 | rs10790299 | 1.192 | 1.997 | 0.863 | 1.495 | 0.053 | 0.147 | 0.147 | 0.648 | 0.94 | 0.55 | 0.97 | 0.92 | 1 | 0.98 |
| 11 | 119403016 | rs7110236 | 1.486 | 1.778 | 0.851 | 1.355 | 0 | 0.066 | 0.066 | 0.029 | 0.93 | 0.62 | 0.96 | 0.93 | 0.99 | 0.88 |
| 11 | 119406292 | rs4375449 | -1.141 | 2.611 | 0.855 | 1.368 | -0.049 | -0.046 | -0.046 | 0 | 0.22 | 0.09 | 0.38 | 0.46 | 0.4 | 0.24 |
| 11 | 119426636 | rs2663643 | 1.448 | 2.422 | 1.895 | 1.26 | 0.013 | 0.025 | 0.025 | 0 | 0.69 | 0.39 | 0.6 | 0.45 | 0.63 | 0.63 |
| 11 | 119428158 | rs996888 | 2.003 | 2.61 | 1.979 | 1.178 | 0.128 | 0.054 | 0.054 | 0 | 0.93 | 0.59 | 0.93 | 0.94 | 0.94 | 0.96 |
| 11 | 119429098 | rs10736499 | -1.439 | 2.64 | 1.886 | 1.215 | -0.008 | -0.018 | -0.018 | 0.007 | 0.24 | 0.15 | 0.34 | 0.48 | 0.31 | 0.31 |
| 11 | 119432144 | rs10892399 | -1.381 | 2.649 | 1.921 | 1.205 | 0.001 | 0.008 | 0.008 | 0 | 0.25 | 0.23 | 0.34 | 0.54 | 0.31 | 0.33 |
| 11 | 119441696 | rs906824 | 1.637 | 2.741 | 2.087 | 1.032 | -0.002 | -0.002 | -0.002 | 0 | 0.67 | 0.37 | 0.59 | 0.45 | 0.61 | 0.65 |
| 11 | 119443362 | rs7103327 | -2.326 | 2.691 | 2.215 | 1.14 | 0.138 | 0.086 | 0.086 | 0.093 | 0.05 | 0.31 | 0.12 | 0.05 | 0.06 | 0.01 |
| 11 | 119451940 | rs4938687 | 1.664 | 2.634 | 2.414 | 1.104 | -0.026 | -0.037 | -0.037 | 0 | 0.66 | 0.3 | 0.57 | 0.45 | 0.61 | 0.66 |
| 11 | 119455165 | rs7114739 | -2.013 | 2.155 | 2.397 | 0.882 | 0.004 | 0.004 | 0.004 | 0 | 0.38 | 0.53 | 0.42 | 0.54 | 0.41 | 0.35 |
| 11 | 119458695 | rs7484090 | -2.274 | 2.24 | 2.414 | 0.908 | 0.102 | 0.082 | 0.082 | 0.06 | 0.06 | 0.27 | 0.12 | 0.11 | 0.06 | 0.08 |
| 11 | 119460505 | rs2887245 | -1.82 | 2.127 | 2.422 | 0.844 | 0.017 | 0.017 | 0.017 | 0 | 0.36 | 0.41 | 0.41 | 0.54 | 0.42 | 0.32 |
| 11 | 119468994 | rs10790316 | -1.447 | 1.443 | 2.18 | 0.746 | -0.021 | -0.03 | -0.03 | 0 | 0.32 | 0.15 | 0.29 | 0.43 | 0.36 | 0.3 |
| 11 | 119469211 | rs7121550 | -1.424 | 1.443 | 2.215 | 0.746 | -0.014 | -0.02 | -0.02 | 2.859 | 0.32 | 0.18 | 0.29 | 0.44 | 0.36 | 0.3 |
| 11 | 119470333 | rs10790322 | 1.651 | 1.418 | 2.352 | 0.819 | 0 | -0.006 | -0.006 | 0.745 | 0.62 | 0.39 | 0.58 | 0.45 | 0.63 | 0.67 |

| | | | | | | | | | | | | | | | | |
|---|---|---|---|---|---|---|---|---|---|---|---|---|---|---|---|---|
| 11 | 119470389 | rs7950231 | 1.642 | 1.422 | 2.351 | 0.821 | 0 | -0.006 | -0.006 | 0 | 0.63 | 0.39 | 0.58 | 0.45 | 0.63 | 0.67 |
| 11 | 119475074 | rs11217349 | -3.032 | 1.355 | 2.495 | 0.902 | 0.039 | 0.146 | 0.146 | 1.076 | 0.05 | 0.37 | 0.1 | 0.11 | 0 | 0.04 |
| 11 | 119479391 | rs2271249 | 1.031 | 1.43 | 2.479 | 0.883 | 0.015 | 0.037 | 0.037 | 0 | 0.71 | 0.62 | 0.69 | 0.56 | 0.67 | 0.73 |
| 11 | 119482594 | rs7935369 | 0.97 | 1.483 | 2.471 | 0.913 | 0.02 | 0.039 | 0.039 | 0 | 0.72 | 0.63 | 0.7 | 0.56 | 0.67 | 0.74 |
| 11 | 119488646 | rs4499013 | 1.148 | 0.353 | 1.478 | 0.112 | 0.007 | 0.002 | 0.002 | 0 | 0.52 | 0.44 | 0.53 | 0.48 | 0.59 | 0.45 |
| 11 | 119495527 | rs7951484 | 0.348 | 0.124 | 1.335 | 0.105 | -0.001 | -0.006 | -0.006 | 0.295 | 0.72 | 0.77 | 0.71 | 0.55 | 0.69 | 0.72 |
| 11 | 119496472 | rs1355191 | 1.212 | 0.294 | 1.31 | 0.128 | 0.057 | -0.007 | -0.007 | 0.851 | 0.5 | 0.29 | 0.53 | 0.42 | 0.61 | 0.44 |
| 11 | 119497757 | rs11217357 | 0.081 | 0.995 | 1.246 | 0.406 | -0.012 | 0.001 | 0.001 | 0 | 0.83 | 0.87 | 0.76 | 0.82 | 0.73 | 0.78 |
| 11 | 119498618 | rs10750161 | 0.167 | 1.004 | 1.249 | 0.411 | -0.001 | 0 | 0 | 0.125 | 0.82 | 0.82 | 0.76 | 0.82 | 0.73 | 0.78 |
| 11 | 119499014 | rs4582984 | 0.6 | 1.005 | 1.283 | 0.412 | 0.051 | -0.017 | -0.017 | 0 | 0.79 | 0.55 | 0.72 | 0.82 | 0.73 | 0.78 |
| 11 | 119501021 | rs11217364 | -0.462 | 0.916 | 0.858 | 0.24 | 0.042 | -0.016 | -0.016 | 0 | 0.31 | 0.52 | 0.46 | 0.3 | 0.47 | 0.25 |
| 11 | 119502703 | rs12291064 | 0.579 | -0.095 | 0.077 | 0.87 | 0.007 | -0.018 | -0.018 | 0 | 0.19 | 0.16 | 0.19 | 0.27 | 0.11 | 0.16 |
| 11 | 119503670 | rs12279718 | -0.899 | -0.159 | 0.202 | 0.876 | 0.001 | 0.001 | 0.001 | 0 | 0.51 | 0.54 | 0.63 | 0.57 | 0.61 | 0.43 |
| 11 | 119506749 | rs4936490 | -0.797 | -0.021 | 0.234 | 0.953 | -0.025 | 0.106 | 0.106 | 0 | 0.8 | 0.6 | 0.88 | 0.64 | 0.89 | 0.85 |
| 11 | 119507925 | rs4412764 | 0.248 | 0.265 | -0.452 | 0.769 | -0.147 | 0.003 | 0.003 | 0 | 0.26 | 0.02 | 0.33 | 0.22 | 0.46 | 0.24 |
| 11 | 119509911 | rs4409838 | -0.635 | 0.095 | 0.099 | 1.187 | 0.001 | -0.024 | -0.024 | 0 | 0.89 | 0.91 | 0.93 | 0.73 | 0.93 | 0.82 |
| 11 | 119510303 | rs4245199 | 1.13 | -0.121 | 0.982 | 0.884 | 0.061 | 0.039 | 0.039 | 1.526 | 0.9 | 0.48 | 0.91 | 0.9 | 0.93 | 0.99 |
| 11 | 119510366 | rs4438048 | 1.108 | -0.1 | 1.012 | 0.821 | 0.074 | 0.018 | 0.018 | 1.458 | 0.9 | 0.5 | 0.92 | 0.93 | 0.93 | 0.99 |
| 11 | 119510644 | rs7940667 | 0.682 | -0.056 | 1.109 | 0.843 | 0.05 | 0.007 | 0.007 | 0.086 | 0.91 | 0.67 | 0.94 | 0.93 | 0.93 | 0.99 |
| 11 | 119516100 | rs10892425 | 1.265 | -0.159 | 1.221 | 0.792 | 0.012 | -0.02 | -0.02 | 0 | 0.89 | 0.39 | 0.92 | 0.93 | 0.94 | 0.99 |
| 11 | 119518006 | rs12797352 | -0.258 | -0.829 | 0.289 | 1.147 | 0.039 | -0.031 | -0.031 | 0.474 | 0.38 | 0.19 | 0.26 | 0.44 | 0.29 | 0.5 |
| 11 | 119523181 | rs7103685 | -0.002 | -0.672 | 0.591 | 1.175 | -0.031 | 0.013 | 0.013 | 0 | 0.3 | 0.49 | 0.18 | 0.35 | 0.24 | 0.38 |
| 11 | 119537683 | rs10790329 | 0.227 | -1.034 | 0.624 | 0.924 | 0.004 | 0.002 | 0.002 | 0 | 0.63 | 0.36 | 0.75 | 0.64 | 0.74 | 0.5 |
| 11 | 119541332 | rs10790330 | -0.963 | -0.593 | 0.441 | 1.024 | 0.014 | 0.009 | 0.009 | 0.544 | 0.47 | 0.72 | 0.54 | 0.54 | 0.39 | 0.33 |
| 11 | 119541778 | rs4459318 | -0.806 | -0.563 | 0.419 | 1.046 | 0.005 | -0.005 | -0.005 | 0 | 0.46 | 0.63 | 0.54 | 0.49 | 0.39 | 0.31 |
| 11 | 119549825 | rs906830 | -0.211 | -0.683 | -0.12 | 0.994 | -0.047 | -0.081 | -0.081 | 0 | 0.37 | 0.14 | 0.47 | 0.47 | 0.36 | 0.33 |
| 11 | 119560449 | rs4936492 | -0.905 | -0.738 | 0.068 | 0.996 | -0.001 | -0.001 | -0.001 | 0 | 0.45 | 0.5 | 0.51 | 0.59 | 0.37 | 0.34 |
| 11 | 119564483 | rs11217408 | -0.225 | -0.785 | -0.009 | 0.905 | -0.03 | 0.019 | 0.019 | 0 | 0.06 | 0.2 | 0.03 | 0.11 | 0.02 | 0.01 |
| 11 | 119568994 | rs4938713 | 0.462 | -0.543 | -0.769 | 1.242 | -0.007 | -0.067 | -0.067 | 0.075 | 0.13 | 0.02 | 0.14 | 0.27 | 0.13 | 0.16 |
| 11 | 119572441 | rs885430 | 0.289 | -0.574 | -0.534 | 0.944 | 0.01 | -0.012 | -0.012 | 0 | 0.56 | 0.71 | 0.53 | 0.46 | 0.63 | 0.68 |
| 11 | 119574140 | rs7111290 | 0.371 | -0.585 | -0.541 | 0.853 | 0.021 | 0.013 | 0.013 | 0 | 0.53 | 0.7 | 0.45 | 0.42 | 0.61 | 0.65 |
| 11 | 119576911 | rs4938717 | -0.48 | -0.563 | -0.181 | 0.947 | 0.001 | 0.002 | 0.002 | 0 | 0.42 | 0.45 | 0.42 | 0.42 | 0.37 | 0.29 |
| 11 | 119582832 | rs4409845 | 0.198 | -0.826 | 0.382 | 1.305 | -0.088 | 0.062 | 0.062 | 2.487 | 0.84 | 0.56 | 0.9 | 0.74 | 0.97 | 0.98 |
| 11 | 119586389 | rs10790338 | 0.101 | -0.809 | 0.396 | 1.064 | -0.081 | -0.047 | -0.047 | 0.141 | 0.85 | 0.57 | 0.95 | 0.89 | 0.97 | 0.98 |
| 11 | 119590878 | rs11217426 | 0.208 | -0.782 | -0.143 | 0.837 | -0.002 | 0.001 | 0.001 | 0 | 0.34 | 0.35 | 0.31 | 0.26 | 0.45 | 0.59 |
| 11 | 119601779 | rs2368936 | 0.282 | -0.74 | -0.667 | 0.886 | 0.039 | -0.004 | -0.004 | 0 | 0.49 | 0.25 | 0.61 | 0.48 | 0.52 | 0.34 |
| 11 | 119603847 | rs12421432 | 0.932 | -0.402 | -0.904 | 0.996 | -0.03 | 0.024 | 0.024 | 1.141 | 0.36 | 0.13 | 0.43 | 0.25 | 0.34 | 0.24 |
| 11 | 119605480 | rs4489755 | 0.078 | -0.905 | -0.476 | 0.924 | 0.008 | 0.008 | 0.008 | 0.144 | 0.58 | 0.33 | 0.66 | 0.63 | 0.55 | 0.38 |
| 11 | 119620940 | rs11217458 | -0.463 | -0.659 | -0.462 | 0.587 | -0.018 | -0.019 | -0.019 | 0.574 | 0.42 | 0.26 | 0.37 | 0.46 | 0.5 | 0.58 |
| 11 | 119630418 | rs10790342 | -0.702 | -0.771 | 0.138 | 0.583 | 0 | 0.002 | 0.002 | 0 | 0.45 | 0.53 | 0.38 | 0.51 | 0.49 | 0.6 |
| 11 | 119631325 | rs7131676 | -0.631 | -0.767 | 0.32 | 0.585 | -0.002 | -0.007 | -0.007 | 0 | 0.44 | 0.43 | 0.39 | 0.51 | 0.49 | 0.6 |
| 11 | 119635453 | rs7940298 | 0.95 | -0.565 | 0.083 | 0.616 | -0.006 | -0.027 | -0.027 | 0 | 0.1 | 0.03 | 0.08 | 0.22 | 0.09 | 0.05 |
| 11 | 119637414 | rs4341549 | -0.795 | -0.7 | 0.518 | 0.375 | -0.013 | 0.011 | 0.011 | 0 | 0.4 | 0.3 | 0.33 | 0.32 | 0.49 | 0.53 |
| 11 | 119638088 | rs10750165 | -1.424 | -1.341 | 0.282 | -0.004 | -0.046 | -0.038 | -0.038 | 0 | 0.65 | 0.8 | 0.49 | 0.64 | 0.61 | 0.67 |
| 11 | 119642827 | rs10892454 | -1.09 | -0.965 | 0.695 | 0.493 | -0.002 | -0.001 | -0.001 | 0 | 0.46 | 0.58 | 0.39 | 0.56 | 0.47 | 0.56 |
| 11 | 119644360 | rs9633947 | 0.46 | -0.985 | 0.899 | 0.557 | -0.007 | 0.015 | 0.015 | 0.164 | 0.13 | 0.16 | 0.1 | 0.24 | 0.1 | 0.06 |
| 11 | 119644451 | rs9326253 | -1.12 | -1.014 | 0.926 | 0.531 | 0 | 0.005 | 0.005 | 0 | 0.45 | 0.63 | 0.38 | 0.61 | 0.47 | 0.55 |
| 11 | 119646732 | rs6589774 | 0.137 | -0.935 | 0.534 | 0.415 | -0.009 | -0.008 | -0.008 | 0 | 0.28 | 0.19 | 0.26 | 0.32 | 0.36 | 0.49 |
| 11 | 119648368 | rs4269933 | -0.772 | -0.887 | 0.563 | 0.488 | 0.003 | 0.005 | 0.005 | 0 | 0.42 | 0.46 | 0.37 | 0.56 | 0.46 | 0.55 |
| 11 | 119652763 | rs11217502 | -0.671 | -0.707 | 0.092 | -0.033 | -0.015 | 0.018 | 0.018 | 0 | 0.4 | 0.24 | 0.32 | 0.32 | 0.47 | 0.53 |
| 11 | 119659753 | rs10892465 | -0.983 | -0.873 | -0.076 | 0.05 | -0.003 | -0.007 | -0.007 | 0.112 | 0.43 | 0.44 | 0.32 | 0.33 | 0.44 | 0.52 |
| 11 | 119661809 | rs11217506 | -0.506 | -0.681 | -0.675 | -0.392 | 0.015 | 0.059 | 0.059 | 0 | 0.18 | 0.06 | 0.06 | 0 | 0.13 | 0.1 |
| 11 | 119665733 | rs7935678 | -0.984 | -0.783 | -0.793 | -0.626 | 0.015 | 0.107 | 0.107 | 0.706 | 0.26 | 0.14 | 0.1 | 0 | 0.2 | 0.11 |
| 11 | 119665842 | rs7935816 | -0.004 | -0.872 | -0.344 | -0.26 | -0.009 | 0.028 | 0.028 | 0 | 0.67 | 0.63 | 0.59 | 0.44 | 0.67 | 0.6 |
| 11 | 119671453 | rs895652 | 0.127 | -1.069 | -0.235 | -0.202 | -0.014 | 0.013 | 0.013 | 0 | 0.67 | 0.64 | 0.6 | 0.43 | 0.71 | 0.61 |
| 11 | 119672156 | rs4938724 | 0.951 | -1.087 | -0.285 | -0.258 | 0.001 | 0.001 | 0.001 | 0.532 | 0.51 | 0.5 | 0.51 | 0.43 | 0.53 | 0.56 |
| 11 | 119677072 | rs895648 | 1.057 | -0.929 | -0.176 | -0.247 | 0.011 | 0.006 | 0.006 | 0.302 | 0.48 | 0.55 | 0.56 | 0.43 | 0.58 | 0.55 |
| 11 | 119678445 | rs7945067 | 0.904 | -1.251 | 0.187 | 0.024 | -0.006 | -0.006 | -0.006 | 0 | 0.61 | 0.58 | 0.68 | 0.69 | 0.73 | 0.87 |
| 11 | 119678986 | rs10892470 | 1.353 | -1.199 | 0.113 | -0.012 | 0.01 | 0.005 | 0.005 | 1.113 | 0.44 | 0.31 | 0.52 | 0.43 | 0.57 | 0.53 |
| 11 | 119679283 | rs10892471 | 0.987 | -0.285 | 0.111 | -0.013 | -0.006 | 0.04 | 0.04 | 0.408 | 0.11 | 0.07 | 0.05 | 0 | 0.11 | 0.03 |
| 11 | 119688442 | rs4245207 | 1.314 | -0.079 | 0.245 | 0.223 | -0.003 | -0.01 | -0.01 | 0 | 0.54 | 0.34 | 0.53 | 0.49 | 0.55 | 0.6 |
| 11 | 119690746 | rs11822532 | 0.765 | -0.007 | 0.196 | 0.16 | -0.004 | -0.004 | -0.004 | 0 | 0.06 | 0.07 | 0.04 | 0 | 0.08 | 0.02 |
| 11 | 119691069 | rs1447149 | -0.607 | -0.083 | 0.282 | 0.128 | 0.002 | 0.01 | 0.01 | 2.727 | 0.27 | 0.21 | 0.14 | 0.03 | 0.23 | 0.09 |
| 11 | 119692663 | rs542903 | -0.047 | -0.097 | -0.158 | 0.675 | 0.007 | 0.069 | 0.069 | 0 | 0.85 | 0.67 | 0.89 | 0.79 | 0.86 | 0.72 |
| 11 | 119695011 | rs566880 | -0.059 | 0.072 | -0.226 | 0.609 | 0.004 | 0.014 | 0.014 | 0 | 0.84 | 0.79 | 0.87 | 0.79 | 0.82 | 0.7 |
| 11 | 119695491 | rs482090 | 0.401 | 0.06 | -0.587 | 0.36 | 0.04 | 0.214 | 0.214 | 0 | 0.81 | 0.63 | 0.68 | 0.52 | 0.78 | 0.69 |
| 11 | 119696100 | rs1789792 | 0.288 | -0.288 | -0.524 | 0.465 | 0.004 | 0.203 | 0.203 | 0 | 0.8 | 0.61 | 0.71 | 0.52 | 0.81 | 0.69 |
| 11 | 119697584 | rs11217527 | -0.466 | -0.216 | -1.296 | -0.033 | 0.001 | 0.116 | 0.116 | 0.417 | 0.2 | 0.04 | 0.09 | 0.03 | 0.15 | 0.06 |
| 11 | 119700001 | rs579200 | -0.784 | 0.195 | -0.864 | 0.843 | -0.001 | 0.003 | 0.003 | 0 | 0.23 | 0.25 | 0.25 | 0.4 | 0.24 | 0.22 |

| | | | | | | | | | | | | | | | | |
|---|---|---|---|---|---|---|---|---|---|---|---|---|---|---|---|---|
| 11 | 119700031 | rs1007369 | -0.142 | -0.198 | -1.23 | 0.314 | -0.001 | -0.003 | -0.003 | 0 | 0.62 | 0.4 | 0.59 | 0.74 | 0.71 | 0.74 |
| 11 | 119701127 | rs555255 | -0.586 | -0.359 | -1.1 | 0.622 | 0.003 | -0.008 | -0.008 | 0.554 | 0.26 | 0.73 | 0.42 | 0.55 | 0.19 | 0.2 |
| 11 | 119701587 | rs11217528 | 0.783 | -0.036 | -1.683 | 0.458 | -0.142 | -0.088 | -0.088 | 0 | 0.37 | 0.07 | 0.54 | 0.64 | 0.48 | 0.51 |
| 11 | 119702432 | rs477756 | -0.522 | 0.578 | -0.985 | 0.53 | 0.011 | 0.013 | 0.013 | 0 | 0.27 | 0.36 | 0.23 | 0.3 | 0.31 | 0.33 |
| 11 | 119702527 | rs570637 | -0.805 | 0.6 | -0.724 | 0.614 | 0.002 | 0.005 | 0.005 | 0 | 0.38 | 0.45 | 0.4 | 0.48 | 0.4 | 0.35 |
| 11 | 119705210 | rs558034 | 0.151 | 0.545 | -0.905 | 0.222 | 0.013 | -0.021 | -0.021 | 0 | 0.25 | 0.2 | 0.16 | 0.28 | 0.21 | 0.25 |
| 11 | 119712224 | rs552040 | -0.203 | 0.708 | -0.816 | -0.634 | -0.004 | -0.005 | -0.005 | 0.036 | 0.7 | 0.71 | 0.8 | 0.97 | 0.69 | 0.78 |
| 11 | 119727494 | rs7113123 | 0.374 | 0.693 | -0.862 | -0.027 | -0.008 | -0.005 | -0.005 | 0 | 0.31 | 0.24 | 0.46 | 0.69 | 0.38 | 0.51 |
| 11 | 119729571 | rs1665778 | 0.413 | 0.254 | -1.044 | -0.946 | 0.005 | 0.01 | 0.01 | 1.428 | 0.4 | 0.34 | 0.3 | 0.08 | 0.39 | 0.17 |
| 11 | 119733857 | rs570521 | -0.932 | -0.323 | -1.008 | -0.53 | 0.013 | 0.034 | 0.034 | 0 | 0.84 | 0.93 | 0.87 | 0.97 | 0.91 | 0.85 |
| 11 | 119737648 | rs10892480 | 0.884 | -0.419 | -1.282 | 0.149 | -0.01 | -0.1 | -0.1 | 0.062 | 0.19 | 0.03 | 0.24 | 0.44 | 0.25 | 0.37 |
| 11 | 119738623 | rs509733 | 0.446 | -0.66 | -0.687 | -0.48 | -0.04 | -0.085 | -0.085 | 1.821 | 0.72 | 0.42 | 0.81 | 0.96 | 0.83 | 0.68 |
| 11 | 119744160 | rs541366 | -0.519 | -0.384 | -0.992 | -0.315 | -0.013 | -0.048 | -0.048 | 0 | 0.81 | 0.75 | 0.93 | 0.98 | 0.89 | 0.7 |
| 11 | 119750487 | rs543876 | 0.37 | -0.228 | -1.681 | 0.15 | -0.032 | -0.148 | -0.148 | 0 | 0.24 | 0.03 | 0.24 | 0.51 | 0.29 | 0.38 |
| 11 | 119753926 | rs503175 | -0.232 | -0.298 | -1.98 | -0.885 | 0.019 | 0.121 | 0.121 | 1.786 | 0.67 | 0.85 | 0.78 | 0.97 | 0.72 | 0.74 |
| 11 | 119763417 | rs523223 | 0.925 | -0.109 | -2.144 | -0.113 | -0.027 | -0.08 | -0.08 | 0.236 | 0.21 | 0.07 | 0.22 | 0.36 | 0.26 | 0.32 |
| 11 | 119764234 | rs507622 | -0.075 | -0.087 | -0.765 | 0.505 | 0.045 | 0.331 | 0.331 | 0 | 0.89 | 0.59 | 0.73 | 0.68 | 0.9 | 0.87 |
| 11 | 119768054 | rs11217556 | 1.891 | 0.077 | -1.216 | 0.224 | -0.084 | 0.029 | 0.029 | 2.468 | 0.17 | 0.04 | 0.17 | 0.08 | 0.23 | 0.11 |
| 11 | 119769492 | rs4938737 | 0.475 | -0.286 | -1.13 | -0.018 | -0.005 | -0.002 | -0.002 | 0 | 0.19 | 0.2 | 0.13 | 0.22 | 0.18 | 0.09 |
| 11 | 119771843 | rs518932 | -0.209 | -0.224 | -1.134 | -0.356 | -0.019 | 0.085 | 0.085 | 0.138 | 0.75 | 0.53 | 0.65 | 0.47 | 0.78 | 0.8 |
| 11 | 119771968 | rs517918 | -0.426 | -0.231 | -1.173 | -0.336 | -0.027 | 0.126 | 0.126 | 0.059 | 0.77 | 0.57 | 0.68 | 0.47 | 0.82 | 0.84 |
| 11 | 119777611 | rs509809 | 0.318 | -0.116 | -1.969 | -0.261 | -0.019 | 0.001 | 0.001 | 0 | 0.45 | 0.1 | 0.39 | 0.38 | 0.45 | 0.56 |
| 11 | 119778929 | rs17245326 | 1.406 | 0.502 | -1.421 | -0.113 | -0.097 | 0.024 | 0.024 | 2.415 | 0.17 | 0.02 | 0.17 | 0.08 | 0.24 | 0.11 |
| 11 | 119782250 | rs540103 | -0.002 | 0.31 | -1.388 | 0.219 | -0.051 | 0.012 | 0.012 | 0.369 | 0.29 | 0.13 | 0.35 | 0.27 | 0.41 | 0.43 |
| 11 | 119784002 | rs481400 | 0.824 | 0.522 | -1.474 | 0.186 | -0.019 | -0.014 | -0.014 | 0 | 0.2 | 0.08 | 0.29 | 0.25 | 0.28 | 0.31 |
| 11 | 119784133 | rs502789 | -1.06 | 0.475 | -0.827 | 0.706 | 0.038 | 0.017 | 0.017 | 0.516 | 0.73 | 0.51 | 0.68 | 0.76 | 0.72 | 0.84 |
| 11 | 119784660 | rs475724 | -0.006 | -0.025 | -0.982 | 0.263 | -0.002 | 0.018 | 0.018 | 3.688 | 0.51 | 0.42 | 0.44 | 0.29 | 0.55 | 0.61 |
| 11 | 119785065 | rs472446 | -0.106 | -0.036 | -0.982 | 0.725 | -0.002 | 0 | 0 | 1.202 | 0.52 | 0.45 | 0.45 | 0.5 | 0.55 | 0.66 |
| 11 | 119792173 | rs17123485 | 0.683 | -0.547 | -1.202 | 0.807 | -0.006 | -0.051 | -0.051 | 0 | 0.08 | 0 | 0.09 | 0.22 | 0.06 | 0.09 |
| 11 | 119792728 | rs1287836 | -1.267 | -0.789 | -0.977 | 0.46 | 0.012 | 0.007 | 0.007 | 0.138 | 0.68 | 0.73 | 0.75 | 0.74 | 0.7 | 0.49 |
| 11 | 119794853 | rs12291317 | -0.405 | -0.384 | -0.389 | 0.158 | 0.002 | -0.144 | -0.144 | 0.156 | 0.23 | 0.35 | 0.18 | 0 | 0.21 | 0.07 |
| 11 | 119795133 | rs576764 | 0.422 | 0.482 | -0.599 | 0.139 | 0.024 | -0.019 | -0.019 | 0 | 0.11 | 0.1 | 0.15 | 0.01 | 0.23 | 0.1 |
| 11 | 119798934 | rs681133 | -0.871 | 0.411 | -0.462 | 0.101 | 0.032 | -0.079 | -0.079 | 0.149 | 0.21 | 0.26 | 0.2 | 0.01 | 0.3 | 0.14 |
| 11 | 119801338 | rs11217583 | 1.545 | 0.283 | -0.572 | 0.519 | -0.008 | -0.064 | -0.064 | 1.61 | 0.11 | 0.01 | 0.16 | 0.24 | 0.09 | 0.12 |
| 11 | 119802591 | rs504542 | -1.73 | 0.194 | -0.403 | 0.819 | 0.018 | 0.005 | 0.005 | 0 | 0.48 | 0.61 | 0.47 | 0.52 | 0.56 | 0.39 |
| 11 | 119805016 | rs12273820 | -0.151 | -0.046 | -0.268 | 0.776 | -0.003 | 0.015 | 0.015 | 0 | 0.25 | 0.36 | 0.16 | 0.26 | 0.25 | 0.18 |
| 11 | 119808052 | rs1955021 | -0.999 | 0.009 | -0.653 | 0.391 | 0.004 | 0.002 | 0.002 | 0 | 0.41 | 0.56 | 0.33 | 0.35 | 0.33 | 0.25 |
| 11 | 119808864 | rs10892499 | 0.262 | 0.266 | -0.866 | 0.698 | -0.016 | 0.067 | 0.067 | 1.046 | 0.16 | 0.26 | 0.2 | 0.35 | 0.12 | 0.13 |
| 11 | 119808956 | rs880004 | 0.212 | 0.661 | -1.427 | 0.363 | -0.082 | 0.022 | 0.022 | 0.758 | 0.38 | 0.12 | 0.42 | 0.31 | 0.52 | 0.57 |
| 11 | 119816512 | rs668781 | 0.19 | 1.29 | -1.02 | 0.878 | -0.043 | -0.043 | -0.043 | 0.578 | 0.08 | 0.01 | 0.27 | 0.2 | 0.22 | 0.1 |
| 11 | 119818622 | rs10892501 | -0.076 | 0.784 | -0.688 | 0.694 | 0.008 | 0.002 | 0.002 | 0 | 0.64 | 0.52 | 0.44 | 0.63 | 0.6 | 0.72 |
| 11 | 119818802 | rs10892502 | 0.516 | 0.158 | -0.459 | 0.009 | -0.021 | -0.027 | -0.027 | 0 | 0.27 | 0.44 | 0.28 | 0.18 | 0.19 | 0.19 |
| 11 | 119818914 | rs475178 | 0.263 | 0.106 | -0.297 | 0.173 | 0.002 | -0.024 | -0.024 | 0 | 0.94 | 0.79 | 0.95 | 1 | 0.96 | 0.97 |
| 11 | 119819087 | rs476877 | -0.081 | 0.112 | -0.323 | 0.176 | -0.003 | -0.023 | -0.023 | 0 | 0.95 | 0.88 | 0.96 | 1 | 0.96 | 0.97 |
| 11 | 119822294 | rs11217593 | -0.562 | -0.168 | -0.119 | 0.431 | -0.015 | 0.071 | 0.071 | 1.881 | 0.2 | 0.46 | 0.12 | 0.29 | 0.17 | 0.14 |
| 11 | 119824102 | rs472611 | 0.756 | 0.166 | -0.292 | 0.047 | 0.019 | -0.025 | -0.025 | 0.182 | 0.29 | 0.45 | 0.33 | 0.18 | 0.32 | 0.25 |
| 11 | 119824699 | rs7121901 | 0.858 | -0.701 | -0.907 | -0.878 | 0.039 | 0.052 | 0.052 | 0 | 0.28 | 0.14 | 0.15 | 0.01 | 0.18 | 0.19 |
| 11 | 119827163 | rs4938741 | 1.951 | -0.694 | -1.053 | -0.742 | -0.008 | 0.005 | 0.005 | 0 | 0.14 | 0.1 | 0.24 | 0.14 | 0.21 | 0.29 |
| 11 | 119831773 | rs4245210 | 0.081 | -1.127 | -1.146 | -1.05 | 0.006 | 0.021 | 0.021 | 0.468 | 0.49 | 0.45 | 0.64 | 0.85 | 0.52 | 0.5 |
| 11 | 119837112 | rs626466 | 2.175 | -0.845 | -1.108 | -0.933 | -0.041 | 0.01 | 0.01 | 0 | 0.13 | 0.03 | 0.18 | 0.14 | 0.21 | 0.29 |
| 11 | 119838883 | rs1944505 | -0.953 | -0.893 | -1.002 | -0.961 | -0.003 | 0.005 | 0.005 | 0 | 0.79 | 0.8 | 0.76 | 0.86 | 0.73 | 0.67 |
| 11 | 119842708 | rs17123506 | 1.111 | -0.832 | -1.112 | -0.694 | -0.017 | -0.076 | -0.076 | 0 | 0.11 | 0.01 | 0.22 | 0.45 | 0.12 | 0.2 |
| 11 | 119844041 | rs10790361 | 1.662 | -0.61 | -0.876 | -1.289 | 0.025 | -0.046 | -0.046 | 0 | 0.27 | 0.36 | 0.3 | 0.11 | 0.3 | 0.18 |
| 11 | 119846820 | rs636518 | -0.261 | -0.54 | -0.767 | -1.076 | 0.011 | -0.057 | -0.057 | 0 | 0.32 | 0.37 | 0.32 | 0.14 | 0.4 | 0.36 |
| 11 | 119849828 | rs11217608 | 0.308 | -0.655 | -0.848 | -0.705 | -0.001 | -0.018 | -0.018 | 0 | 0.06 | 0.1 | 0.06 | 0 | 0.07 | 0.02 |
| 11 | 119853069 | rs7129717 | 0.898 | -1.389 | -1 | -1.048 | 0.036 | 0.033 | 0.033 | 0 | 0.2 | 0.11 | 0.06 | 0.01 | 0.09 | 0.25 |
| 11 | 119858093 | rs11217613 | -0.46 | -1.291 | -0.768 | -1.153 | 0.005 | 0.011 | 0.011 | 0.426 | 0.46 | 0.58 | 0.46 | 0.74 | 0.42 | 0.66 |
| 11 | 119866151 | rs633621 | 1.741 | -1.444 | -1.281 | -1.248 | 0.025 | 0.063 | 0.063 | 1.91 | 0.2 | 0.05 | 0.19 | 0.08 | 0.15 | 0.09 |
| 11 | 119867224 | rs4938747 | 0.443 | -1.158 | -0.823 | -0.937 | 0 | 0.005 | 0.005 | 0 | 0.14 | 0.12 | 0.29 | 0.5 | 0.16 | 0.32 |
| 11 | 119876028 | rs2155168 | -0.472 | -0.876 | -1.07 | -0.874 | 0.027 | 0.002 | 0.002 | 0.716 | 0.46 | 0.67 | 0.48 | 0.5 | 0.42 | 0.64 |
| 11 | 119878901 | rs10892512 | -0.419 | -0.396 | -1.237 | -0.276 | -0.003 | 0.03 | 0.03 | 0.006 | 0.27 | 0.33 | 0.33 | 0.4 | 0.27 | 0.27 |
| 11 | 119879217 | rs647437 | -0.509 | 0.786 | -2.099 | -0.381 | -0.09 | -0.035 | -0.035 | 0 | 0.86 | 0.99 | 0.81 | 0.84 | 0.76 | 0.89 |
| 11 | 119879255 | rs609066 | 0.145 | 0.416 | -1.821 | -0.957 | 0.005 | 0.01 | 0.01 | 0 | 0.55 | 0.57 | 0.32 | 0.36 | 0.47 | 0.7 |
| 11 | 119879327 | rs17245368 | 0.61 | 0.411 | -2.016 | -1.155 | 0.033 | 0.117 | 0.117 | 0.784 | 0.36 | 0.18 | 0.25 | 0.07 | 0.28 | 0.2 |
| 11 | 119880819 | rs675246 | 0.262 | 0.753 | -1.038 | 0.864 | 0.003 | 0.216 | 0.216 | 0.322 | 0.95 | 0.92 | 0.88 | 0.6 | 0.95 | 0.87 |
| 11 | 119883569 | rs622350 | -0.108 | 0.491 | -1.338 | -0.429 | 0.011 | 0.078 | 0.078 | 0.029 | 0.45 | 0.33 | 0.42 | 0.17 | 0.43 | 0.48 |
| 11 | 119887019 | rs607888 | -0.087 | 0.34 | -1.421 | -0.253 | 0.012 | 0.053 | 0.053 | 0 | 0.45 | 0.2 | 0.41 | 0.39 | 0.44 | 0.56 |
| 11 | 119887725 | rs609467 | 0.563 | 0.139 | -1.633 | -0.474 | -0.022 | 0.12 | 0.12 | 0.137 | 0.35 | 0.12 | 0.36 | 0.15 | 0.36 | 0.31 |
| 11 | 119893179 | rs655929 | 0.294 | -0.115 | -1.12 | -0.785 | -0.011 | 0.047 | 0.047 | 1.276 | 0.47 | 0.36 | 0.45 | 0.37 | 0.51 | 0.48 |

| | | | | | | | | | | | | | | | | |
|---|---|---|---|---|---|---|---|---|---|---|---|---|---|---|---|---|
| 11 | 119904686 | rs1287591 | -0.212 | -0.037 | -1.17 | -0.877 | -0.007 | -0.008 | -0.008 | 0.317 | 0.6 | 0.53 | 0.56 | 0.42 | 0.66 | 0.71 |
| 11 | 119907605 | rs12420530 | 0.87 | 0.114 | -1.891 | -1.183 | 0.014 | -0.004 | -0.004 | 0.084 | 0.16 | 0.16 | 0.3 | 0.14 | 0.29 | 0.11 |
| 11 | 119915024 | rs1942011 | 1.114 | -0.02 | -2.277 | -1.173 | 0.018 | 0.069 | 0.069 | 0.886 | 0.35 | 0.1 | 0.2 | 0.28 | 0.31 | 0.44 |
| 11 | 119917309 | rs11512472 | 1.19 | -0.007 | -2.267 | -1.403 | 0.015 | 0.193 | 0.193 | 0.73 | 0.34 | 0.1 | 0.2 | 0.05 | 0.31 | 0.4 |
| 11 | 119924436 | rs630842 | 1.542 | 0.332 | -2.234 | -1.125 | -0.042 | 0.054 | 0.054 | 0.712 | 0.22 | 0.03 | 0.24 | 0.11 | 0.21 | 0.12 |
| 11 | 119926054 | rs7125256 | 1.027 | 0.545 | -1.896 | -0.935 | -0.001 | -0.004 | -0.004 | 0 | 0.08 | 0.08 | 0.04 | 0.01 | 0.07 | 0.15 |
| 11 | 119927043 | rs4938762 | 0.499 | 0.815 | -1.947 | -0.78 | -0.033 | 0.003 | 0.003 | 0.344 | 0.15 | 0.1 | 0.27 | 0.18 | 0.29 | 0.13 |
| 11 | 119929632 | rs11217664 | 0.758 | 0.056 | -1.881 | -0.921 | 0.028 | 0.052 | 0.052 | 0.733 | 0.34 | 0.17 | 0.2 | 0.28 | 0.3 | 0.4 |
| 11 | 119932578 | rs896690 | 0.628 | -0.086 | -1.519 | -0.722 | -0.004 | -0.01 | -0.01 | 0 | 0.28 | 0.32 | 0.28 | 0.18 | 0.22 | 0.14 |
| 11 | 119934563 | rs4936507 | 1.542 | -0.149 | -1.899 | -0.892 | -0.015 | 0.042 | 0.042 | 0.003 | 0.11 | 0.02 | 0.18 | 0.05 | 0.15 | 0.17 |
| 11 | 119934934 | rs4938763 | -0.181 | -0.199 | -1.755 | -0.638 | -0.007 | -0.01 | -0.01 | 0.34 | 0.64 | 0.86 | 0.59 | 0.43 | 0.67 | 0.56 |
| 11 | 119936292 | rs640663 | 0.149 | -0.32 | -0.963 | 0.145 | -0.008 | -0.05 | -0.05 | 0 | 0.89 | 0.97 | 0.83 | 0.53 | 0.88 | 0.69 |
| 11 | 119939927 | rs592565 | 0.424 | -0.323 | -0.955 | 0.144 | 0.004 | -0.017 | -0.017 | 0.766 | 0.88 | 0.92 | 0.82 | 0.54 | 0.88 | 0.7 |
| 11 | 119942571 | rs1790474 | 0.766 | 0.287 | -0.5 | 0.235 | 0.003 | 0.077 | 0.077 | 0.078 | 0.81 | 0.65 | 0.69 | 0.4 | 0.8 | 0.76 |
| 11 | 119951870 | rs12365185 | 0.602 | 0.189 | -0.433 | -0.432 | 0.021 | -0.089 | -0.089 | 0.842 | 0.25 | 0.29 | 0.18 | 0 | 0.34 | 0.08 |
| 11 | 119956631 | rs4938776 | -0.72 | 0.06 | -0.173 | -0.519 | 0.013 | -0.023 | -0.023 | 0.023 | 0.13 | 0.23 | 0.2 | 0.12 | 0.14 | 0.14 |
| 11 | 119957516 | rs20493 | 0.099 | -0.493 | -0.149 | -0.623 | -0.011 | 0.072 | 0.072 | 0 | 0.83 | 0.73 | 0.8 | 0.51 | 0.85 | 0.62 |
| 11 | 119961694 | rs617320 | 1.715 | -0.815 | -0.48 | -0.634 | 0.007 | 0.01 | 0.01 | 0.252 | 0.6 | 0.34 | 0.51 | 0.31 | 0.62 | 0.36 |
| 11 | 119961707 | rs11217682 | 1.332 | -0.804 | -0.538 | -0.714 | 0.022 | -0.09 | -0.09 | 0 | 0.25 | 0.29 | 0.18 | 0 | 0.33 | 0.08 |
| 11 | 119967484 | rs609650 | 1.608 | -0.744 | -0.454 | -0.569 | -0.01 | -0.014 | -0.014 | 1.718 | 0.62 | 0.43 | 0.53 | 0.31 | 0.62 | 0.36 |
| 11 | 119968219 | rs1790471 | 0.728 | -0.775 | -0.399 | -0.478 | -0.034 | 0.039 | 0.039 | 0.643 | 0.74 | 0.55 | 0.72 | 0.43 | 0.76 | 0.49 |
| 11 | 119970300 | rs1125618 | 1.256 | -0.75 | -0.681 | -0.651 | 0.005 | -0.014 | -0.014 | 0 | 0.08 | 0.02 | 0.07 | 0.07 | 0.09 | 0.13 |
| 11 | 119972626 | rs1073635 | 1.976 | -0.779 | -0.854 | -0.858 | 0.023 | 0.189 | 0.189 | 0.272 | 0.27 | 0.02 | 0.11 | 0 | 0.17 | 0.17 |
| 11 | 119972642 | rs1073634 | 1.264 | -0.606 | -0.538 | -0.698 | 0 | -0.029 | -0.029 | 2.919 | 0.54 | 0.41 | 0.38 | 0.12 | 0.51 | 0.29 |
| 11 | 119983799 | rs530792 | 0.203 | -0.838 | -0.436 | 0.295 | 0.009 | -0.021 | -0.021 | 0 | 0.81 | 0.92 | 0.87 | 0.76 | 0.83 | 0.77 |
| 11 | 119989123 | rs10466596 | 0.999 | -0.754 | 0.009 | -0.25 | 0.013 | -0.082 | -0.082 | 0 | 0.25 | 0.28 | 0.19 | 0 | 0.32 | 0.09 |
| 11 | 119989892 | rs606970 | 0.092 | -1.012 | 0.064 | -0.052 | 0.006 | -0.001 | -0.001 | 0 | 0.36 | 0.39 | 0.44 | 0.63 | 0.47 | 0.26 |
| 11 | 119995447 | rs2851718 | -0.752 | -0.796 | 0.047 | 0.534 | 0.007 | 0.164 | 0.164 | 0 | 0.08 | 0.16 | 0.17 | 0.43 | 0.12 | 0.13 |
| 11 | 119999191 | rs569044 | -0.866 | -0.924 | -0.207 | 0.063 | -0.031 | 0.046 | 0.046 | 1.238 | 0.84 | 0.99 | 0.88 | 0.96 | 0.86 | 0.91 |
| 11 | 120002927 | rs561498 | 0.782 | -1.032 | 0.041 | 0.053 | 0.002 | 0.004 | 0.004 | 1.158 | 0.69 | 0.66 | 0.6 | 0.23 | 0.73 | 0.66 |
| 11 | 120003533 | rs666432 | -0.562 | -0.903 | 0.082 | 0.032 | 0.026 | 0.269 | 0.269 | 0 | 0.07 | 0.2 | 0.15 | 0.45 | 0.11 | 0.11 |
| 11 | 120013307 | rs470876 | -0.15 | -0.867 | 0.03 | -0.145 | 0.007 | 0.056 | 0.056 | 0 | 0.09 | 0.17 | 0.22 | 0.23 | 0.09 | 0.4 |
| 11 | 120018743 | rs675647 | -0.359 | -0.534 | -0.724 | -0.21 | 0 | 0.048 | 0.048 | 0 | 0.1 | 0.13 | 0.24 | 0.62 | 0.11 | 0.18 |
| 11 | 120026748 | rs2084898 | 0.587 | -0.65 | -0.775 | -0.3 | -0.012 | 0.084 | 0.084 | 0.036 | 0.17 | 0.03 | 0.14 | 0.02 | 0.15 | 0.08 |
| 11 | 120041444 | rs4936509 | -0.364 | -0.659 | -0.702 | -0.444 | -0.011 | 0.014 | 0.014 | 3.95 | 0.57 | 0.39 | 0.56 | 0.63 | 0.56 | 0.41 |
| 11 | 120050515 | rs531260 | 0.895 | -0.521 | -0.601 | -0.365 | -0.001 | -0.003 | -0.003 | 1.402 | 0.68 | 0.61 | 0.59 | 0.22 | 0.71 | 0.65 |
| 11 | 120052554 | rs547124 | 0.88 | -0.37 | -0.526 | -0.159 | -0.003 | 0.035 | 0.035 | 0 | 0.86 | 0.79 | 0.81 | 0.36 | 0.86 | 0.77 |
| 11 | 120054202 | rs4938790 | 0.68 | -0.761 | -0.339 | 0.058 | -0.03 | -0.027 | -0.027 | 0 | 0.06 | 0.01 | 0.2 | 0.21 | 0.09 | 0.11 |
| 11 | 120057343 | rs508205 | 0.174 | -1.27 | -0.092 | -0.669 | 0.002 | 0.01 | 0.01 | 1.677 | 0.46 | 0.53 | 0.56 | 0.74 | 0.54 | 0.62 |
| 11 | 120060673 | rs7116997 | -0.482 | -1.186 | -0.203 | -0.606 | 0.001 | 0.003 | 0.003 | 1.898 | 0.53 | 0.5 | 0.44 | 0.26 | 0.48 | 0.35 |
| 11 | 120063980 | rs880487 | 0.527 | -0.783 | -0.76 | -1.001 | 0.004 | 0.003 | 0.003 | 0.077 | 0.33 | 0.32 | 0.29 | 0.11 | 0.31 | 0.21 |
| 11 | 120067953 | rs17311212 | -0.633 | -1.288 | -0.468 | -0.266 | 0.002 | 0.02 | 0.02 | 0.272 | 0.17 | 0.14 | 0.12 | 0.02 | 0.15 | 0.07 |
| 11 | 120076116 | rs562966 | 0.899 | -1.649 | 0.19 | 0.393 | -0.006 | 0.018 | 0.018 | 0 | 0.84 | 0.76 | 0.72 | 0.76 | 0.85 | 0.73 |
| 11 | 120080526 | rs521841 | 0.417 | -1.589 | 0.405 | 0.463 | 0.004 | 0.052 | 0.052 | 0 | 0.92 | 0.88 | 0.88 | 0.75 | 0.91 | 0.74 |
| 11 | 120085469 | rs541857 | -0.916 | -2.551 | 0.533 | 0.629 | -0.019 | 0.005 | 0.005 | 0 | 0.19 | 0.21 | 0.19 | 0.17 | 0.09 | 0.2 |
| 11 | 120089886 | rs731512 | -0.402 | -2.544 | 0.528 | 0.514 | -0.003 | 0.039 | 0.039 | 0 | 0.07 | 0.05 | 0.18 | 0.57 | 0.07 | 0.2 |
| 11 | 120099791 | rs2465654 | -0.921 | -2.635 | 0.605 | 0.446 | -0.025 | 0.001 | 0.001 | 0 | 0.18 | 0.22 | 0.16 | 0.13 | 0.08 | 0.2 |
| 11 | 120109967 | rs2509623 | 0.022 | -2.475 | 0.375 | 0.536 | -0.009 | -0.012 | -0.012 | 0 | 0.93 | 0.97 | 0.85 | 0.75 | 0.91 | 0.75 |
| 11 | 120112699 | rs2847497 | -0.954 | -2.693 | 0.817 | 0.333 | -0.029 | -0.007 | -0.007 | 1.82 | 0.18 | 0.26 | 0.19 | 0.09 | 0.09 | 0.19 |
| 11 | 120118498 | rs2847502 | -1.913 | -3.704 | 0.701 | -0.767 | -0.009 | -0.119 | -0.119 | 0 | 0.54 | 0.62 | 0.59 | 1 | 0.3 | 0.8 |
| 11 | 120122394 | rs11827471 | -0.613 | -3.359 | 0.577 | -0.078 | 0.003 | -0.013 | -0.013 | 0 | 0.08 | 0.04 | 0.08 | 0.62 | 0.07 | 0.21 |
| 11 | 120126277 | rs2715881 | -1.979 | -4.151 | 0.928 | -0.611 | 0.007 | -0.081 | -0.081 | 0.702 | 0.5 | 0.53 | 0.58 | 1 | 0.29 | 0.81 |
| 11 | 120130512 | rs1941411 | -2.407 | -4.504 | 0.743 | -0.878 | 0.079 | 0.205 | 0.205 | 0.22 | 0.57 | 0.88 | 0.61 | 1 | 0.3 | 0.83 |
| 11 | 120135501 | rs2847495 | -1.929 | -4.184 | 1.126 | -0.702 | 0.005 | -0.082 | -0.082 | 0 | 0.47 | 0.52 | 0.54 | 1 | 0.25 | 0.81 |
| 11 | 120137544 | rs2096883 | -1.471 | -4.26 | 0.876 | -0.512 | 0.001 | 0 | 0 | 0 | 0.29 | 0.3 | 0.44 | 0.24 | 0.2 | 0.42 |
| 11 | 120139487 | rs10892558 | -0.959 | -4 | 0.598 | -0.391 | 0.049 | -0.003 | -0.003 | 0 | 0.26 | 0.09 | 0.24 | 0.28 | 0.13 | 0.37 |
| 11 | 120146360 | rs4938793 | -2.248 | -3.488 | 0.909 | -0.403 | -0.017 | -0.035 | -0.035 | 0.03 | 0.23 | 0.57 | 0.4 | 0.13 | 0.19 | 0.19 |
| 11 | 120147254 | rs7936665 | -0.724 | -3.709 | 0.614 | -0.358 | -0.001 | -0.002 | -0.002 | 0 | 0.22 | 0.18 | 0.19 | 0.25 | 0.1 | 0.36 |
| 11 | 120148717 | rs12284383 | -2.309 | -3.075 | 0.784 | -0.245 | -0.016 | -0.046 | -0.046 | 0 | 0.23 | 0.44 | 0.38 | 0.13 | 0.2 | 0.19 |
| 11 | 120161641 | rs6589804 | -0.988 | -2.494 | 0.412 | -0.302 | 0.048 | -0.013 | -0.013 | 1.128 | 0.07 | 0.28 | 0.1 | 0 | 0.09 | 0.01 |
| 11 | 120161886 | rs11217793 | -1.827 | -2.138 | -0.645 | -0.312 | 0.008 | 0.011 | 0.011 | 0 | 0.24 | 0.17 | 0.45 | 0.19 | 0.18 | 0.35 |
| 11 | 120165942 | rs11217801 | -1.159 | -2.033 | -0.674 | -0.14 | 0.008 | -0.002 | -0.002 | 0.029 | 0.18 | 0.13 | 0.31 | 0.16 | 0.12 | 0.24 |
| 11 | 120167057 | rs4142654 | -0.685 | -1.558 | -0.465 | 0.331 | -0.016 | 0.012 | 0.012 | 0 | 0.05 | 0.01 | 0.07 | 0.65 | 0.08 | 0.26 |
| 11 | 120173354 | rs2096884 | -0.927 | -2.185 | -0.216 | -0.308 | -0.045 | -0.016 | -0.016 | 0 | 0.29 | 0.58 | 0.3 | 0.26 | 0.15 | 0.34 |
| 11 | 120173845 | rs12275681 | -1.003 | -1.093 | -0.002 | -0.239 | 0 | -0.008 | -0.008 | 0.265 | 0.07 | 0.11 | 0.12 | 0.05 | 0.08 | 0.13 |
| 11 | 120175749 | rs7110845 | -0.111 | 0.051 | -0.757 | -0.64 | -0.118 | 0.099 | 0.099 | 0.043 | 0.75 | 0.97 | 0.71 | 0.94 | 0.56 | 0.9 |
| 11 | 120176979 | rs10750171 | 0.402 | -0.054 | -0.646 | -0.655 | -0.086 | 0.122 | 0.122 | 0 | 0.67 | 0.86 | 0.59 | 0.9 | 0.48 | 0.76 |
| 11 | 120178632 | rs11605781 | 0.767 | 0.255 | -0.379 | -0.69 | -0.024 | 0.103 | 0.103 | 0 | 0.61 | 0.8 | 0.54 | 0.12 | 0.52 | 0.4 |
| 11 | 120180479 | rs717052 | -0.403 | 0.31 | -0.075 | -0.523 | 0 | -0.001 | -0.001 | 0 | 0.14 | 0.17 | 0.17 | 0.19 | 0.12 | 0.23 |

| | | | | | | | | | | | | | | | | |
|---|---|---|---|---|---|---|---|---|---|---|---|---|---|---|---|---|
| 11 | 120183803 | rs7122401 | 0.202 | 0.337 | 0.304 | -0.41 | 0.007 | -0.003 | -0.003 | 0 | 0.73 | 0.73 | 0.7 | 0.32 | 0.65 | 0.62 |
| 11 | 120187433 | rs7102217 | -0.251 | 0.438 | 0.216 | 0.249 | -0.033 | -0.06 | -0.06 | 0 | 0.78 | 0.92 | 0.73 | 0.58 | 0.65 | 0.87 |
| 11 | 120190997 | rs11217816 | 0.143 | 0.325 | 0.627 | -1.106 | 0.02 | 0.005 | 0.005 | 0.116 | 0.73 | 0.55 | 0.7 | 0.34 | 0.65 | 0.85 |
| 11 | 120193477 | rs7934404 | -0.882 | 0.216 | 0.372 | -1.4 | -0.01 | -0.026 | -0.026 | 0 | 0.31 | 0.39 | 0.41 | 0.23 | 0.24 | 0.45 |
| 11 | 120194357 | rs12421941 | 0.324 | 0.284 | 0.168 | -1.381 | 0 | -0.011 | -0.011 | 0 | 0.13 | 0.04 | 0.09 | 0.19 | 0.12 | 0.24 |
| 11 | 120198093 | rs1893261 | -0.885 | 0.497 | 0.805 | -1.671 | -0.001 | -0.022 | -0.022 | 0 | 0.37 | 0.45 | 0.34 | 0.29 | 0.38 | 0.51 |
| 11 | 120198728 | rs11827818 | 0.231 | 0.511 | 0.871 | -1.708 | -0.002 | -0.002 | -0.002 | 0.469 | 0.18 | 0.24 | 0.15 | 0.19 | 0.16 | 0.24 |
| 11 | 120202099 | rs11217821 | -1.032 | 0.504 | 0.899 | -1.703 | -0.003 | -0.033 | -0.033 | 0 | 0.38 | 0.53 | 0.35 | 0.29 | 0.38 | 0.51 |
| 11 | 120203628 | rs12806740 | -1.06 | 0.502 | 0.889 | -1.704 | 0 | -0.002 | -0.002 | 0 | 0.39 | 0.6 | 0.35 | 0.29 | 0.38 | 0.51 |
| 11 | 120209386 | rs4938800 | 0.532 | 0.353 | 0.904 | -1.726 | 0.002 | -0.012 | -0.012 | 3.086 | 0.14 | 0.18 | 0.32 | 0.06 | 0.13 | 0.23 |
| 11 | 120224544 | rs10892563 | -0.994 | 0.379 | 0.839 | -1.744 | -0.001 | -0.005 | -0.005 | 0.247 | 0.38 | 0.6 | 0.34 | 0.29 | 0.39 | 0.4 |
| 11 | 120228989 | rs4938802 | 0.321 | 0.39 | 0.716 | -1.737 | -0.001 | -0.002 | -0.002 | 0 | 0.18 | 0.24 | 0.15 | 0.22 | 0.16 | 0.24 |
| 11 | 120232766 | rs11217837 | 1.17 | 0.367 | 0.465 | -1.748 | -0.063 | 0.089 | 0.089 | 0.164 | 0.14 | 0.01 | 0.13 | 0 | 0.23 | 0.06 |
| 11 | 120247033 | rs632137 | 0.373 | 0.42 | 0.452 | -1.731 | -0.026 | -0.05 | -0.05 | 0 | 0.09 | 0.01 | 0.12 | 0.61 | 0.11 | 0.09 |
| 11 | 120287803 | rs10892573 | 0.682 | 0.265 | 0.534 | -1.35 | -0.004 | -0.014 | -0.014 | 0 | 0.16 | 0.11 | 0.14 | 0 | 0.22 | 0.07 |
| 11 | 120292040 | rs11604012 | 0.654 | 0.258 | 0.425 | -1.358 | -0.038 | -0.002 | -0.002 | 0 | 0.13 | 0 | 0.3 | 0.07 | 0.13 | 0.24 |
| 11 | 120307742 | rs10502236 | 0.144 | 0.387 | 0.529 | -1.292 | 0.005 | -0.055 | -0.055 | 0 | 0.19 | 0.23 | 0.18 | 0.1 | 0.22 | 0.26 |
| 11 | 120312746 | rs723937 | -1.361 | 0.414 | 0.374 | -1.294 | 0.001 | 0.018 | 0.018 | 0.488 | 0.4 | 0.66 | 0.34 | 0.32 | 0.39 | 0.49 |
| 11 | 120323214 | rs496248 | 1.483 | 0.568 | -0.492 | -1.231 | -0.066 | 0.07 | 0.07 | 0 | 0.45 | 0.09 | 0.32 | 0.61 | 0.48 | 0.27 |
| 11 | 120328858 | rs766894 | -0.115 | 0.941 | 0.01 | -0.84 | 0.006 | -0.055 | -0.055 | 0 | 0.17 | 0.18 | 0.14 | 0 | 0.22 | 0.06 |
| 11 | 120331701 | rs4938807 | -0.856 | 1.028 | -0.684 | -0.787 | -0.003 | -0.001 | -0.001 | 1.552 | 0.2 | 0.33 | 0.16 | 0.24 | 0.16 | 0.23 |
| 11 | 120346360 | rs2276035 | -0.659 | 1.214 | -1.089 | -0.694 | -0.001 | 0 | 0 | 0 | 0.18 | 0.25 | 0.15 | 0.24 | 0.16 | 0.23 |
| 11 | 120355791 | rs9625 | -0.581 | 1.407 | -1.34 | -0.515 | -0.013 | 0.019 | 0.019 | 0 | 0.16 | 0.07 | 0.13 | 0 | 0.22 | 0.06 |
| 11 | 120365028 | rs12364480 | -0.795 | 0.93 | -1.701 | -0.407 | -0.037 | 0.118 | 0.118 | 0.285 | 0.19 | 0.03 | 0.13 | 0 | 0.22 | 0.07 |
| 11 | 120371518 | rs7929360 | -0.669 | 1.135 | -1.286 | -0.079 | 0 | -0.007 | -0.007 | 0 | 0.21 | 0.24 | 0.18 | 0.07 | 0.28 | 0.13 |
| 11 | 120379057 | rs2247168 | 0.231 | 1.591 | -1.757 | 0.672 | -0.081 | -0.089 | -0.089 | 0 | 0.77 | 0.92 | 0.58 | 0.46 | 0.68 | 0.6 |
| 11 | 120391473 | rs12804597 | -0.578 | 0.809 | -1.372 | 0.307 | -0.002 | -0.003 | -0.003 | 0.016 | 0.08 | 0.09 | 0.06 | 0.05 | 0.08 | 0.05 |
| 11 | 120395229 | rs3897570 | -1.543 | 0.799 | -0.565 | -0.028 | 0.029 | -0.058 | -0.058 | 6.359 | 0.1 | 0.22 | 0.07 | 0 | 0.1 | 0.03 |
| 11 | 120396485 | rs7129707 | -1.27 | 0.033 | -1.066 | -0.584 | 0.038 | 0.114 | 0.114 | 0.911 | 0.38 | 0.78 | 0.25 | 0.2 | 0.29 | 0.22 |
| 11 | 120398643 | rs17092876 | -1.553 | 0.021 | -0.516 | -0.619 | -0.001 | -0.002 | -0.002 | 0 | 0.35 | 0.55 | 0.33 | 0.31 | 0.27 | 0.32 |
| 11 | 120400149 | rs2252359 | 1.039 | -0.202 | 0.215 | 0.727 | -0.028 | 0.115 | 0.115 | 0.183 | 0.78 | 0.5 | 0.78 | 0.45 | 0.86 | 0.79 |
| 11 | 120400419 | rs2252222 | 0.661 | -0.134 | 0.653 | 1.082 | -0.01 | 0.042 | 0.042 | 0 | 0.81 | 0.75 | 0.9 | 0.67 | 0.86 | 0.83 |
| 11 | 120403333 | rs3897568 | -0.165 | 0.153 | -0.195 | 1.185 | -0.001 | 0.006 | 0.006 | 0 | 0.22 | 0.22 | 0.42 | 0.36 | 0.22 | 0.23 |
| 11 | 120405843 | rs4084221 | 0.231 | 0.433 | -1.141 | -0.235 | -0.097 | 0.112 | 0.112 | 0 | 0.26 | 0.07 | 0.21 | 0 | 0.42 | 0.23 |
| 11 | 120408071 | rs3901366 | -0.096 | -0.537 | -0.626 | 0.254 | -0.002 | -0.001 | -0.001 | 0 | 0.42 | 0.53 | 0.59 | 0.63 | 0.3 | 0.52 |
| 11 | 120412867 | rs2510877 | 0.405 | -0.733 | -0.227 | 0.55 | -0.002 | -0.003 | -0.003 | 0.316 | 0.29 | 0.31 | 0.47 | 0.28 | 0.27 | 0.26 |
| 11 | 120415557 | rs2252648 | 0.386 | -0.834 | -0.262 | 0.422 | 0 | 0 | 0 | 0 | 0.29 | 0.26 | 0.47 | 0.27 | 0.29 | 0.26 |
| 11 | 120417553 | rs11217913 | 0.56 | -0.964 | -0.46 | 0.377 | -0.074 | -0.042 | -0.042 | 1.086 | 0.28 | 0.02 | 0.45 | 0.27 | 0.28 | 0.26 |
| 11 | 120417586 | rs2252373 | -1.694 | -1.126 | 0.12 | 0.271 | 0.051 | 0.047 | 0.047 | 6.6 | 0.06 | 0.41 | 0.04 | 0 | 0 | 0 |
| 11 | 120419218 | rs7940779 | 0.488 | -1.242 | 0.294 | 0.61 | -0.034 | 0.103 | 0.103 | 0 | 0.12 | 0.23 | 0.12 | 0.3 | 0.05 | 0.18 |
| 11 | 120426966 | rs2248540 | -0.753 | -0.64 | 0.258 | 0.716 | 0.003 | 0.002 | 0.002 | 0 | 0.43 | 0.6 | 0.61 | 0.58 | 0.33 | 0.39 |
| 11 | 120431744 | rs7121316 | -0.996 | -0.608 | 0.328 | 0.745 | -0.054 | 0.147 | 0.147 | 2.126 | 0.14 | 0.55 | 0.14 | 0.24 | 0.04 | 0.13 |
| 11 | 120437188 | rs4278519 | 0.061 | -0.551 | -0.081 | 0.332 | 0.001 | -0.013 | -0.013 | 0 | 0.33 | 0.67 | 0.34 | 0.12 | 0.43 | 0.32 |
| 11 | 120446830 | rs12796185 | 1.174 | -0.655 | -0.59 | 0.309 | -0.08 | -0.101 | -0.101 | 0.312 | 0.28 | 0.02 | 0.48 | 0.33 | 0.3 | 0.28 |
| 11 | 120456533 | rs11602493 | 0.281 | -0.647 | -0.352 | 0.144 | -0.004 | 0.002 | 0.002 | 0.934 | 0.33 | 0.7 | 0.34 | 0.13 | 0.44 | 0.3 |
| 11 | 120456605 | rs4363614 | -1.004 | -1.035 | -0.094 | 0.271 | 0.001 | -0.001 | -0.001 | 0.232 | 0.69 | 0.75 | 0.85 | 0.59 | 0.79 | 0.69 |
| 11 | 120457436 | rs17124014 | -0.735 | -0.531 | 0.499 | -0.091 | 0.062 | 0.058 | 0.058 | 11.249 | 0.06 | 0.5 | 0.05 | 0 | 0 | 0 |
| 11 | 120468752 | rs7105963 | 0.098 | 0.57 | -0.086 | -0.367 | 0.023 | -0.112 | -0.112 | 0 | 0.23 | 0.64 | 0.22 | 0 | 0.33 | 0.15 |
| 11 | 120470159 | rs4936523 | -0.454 | 0.01 | -0.281 | -0.015 | 0.051 | 0.016 | 0.016 | 0 | 0.53 | 0.69 | 0.66 | 0.39 | 0.72 | 0.53 |
| 11 | 120472569 | rs4414229 | -0.622 | -0.064 | -0.321 | -0.039 | 0.056 | 0.066 | 0.066 | 0 | 0.53 | 0.69 | 0.5 | 0.24 | 0.72 | 0.43 |
| 11 | 120475940 | rs4367952 | -0.323 | -0.034 | -0.258 | 0.017 | 0.027 | 0.006 | 0.006 | 0 | 0.51 | 0.62 | 0.65 | 0.43 | 0.71 | 0.54 |
| 11 | 120481866 | rs12278678 | 0.677 | 0.499 | -0.779 | 0.048 | -0.079 | 0.025 | 0.025 | 0 | 0.28 | 0.1 | 0.28 | 0.2 | 0.38 | 0.29 |
| 11 | 120488347 | rs10892588 | 0.1 | 0.388 | 0.003 | 0.088 | -0.003 | -0.007 | -0.007 | 0 | 0.36 | 0.39 | 0.47 | 0.42 | 0.39 | 0.41 |
| 11 | 120491438 | rs7128466 | -1.778 | -0.152 | -0.716 | 0.144 | 0.041 | -0.082 | -0.082 | 0 | 0.76 | 0.97 | 0.76 | 0.63 | 0.87 | 0.74 |
| 11 | 120494034 | rs4631889 | 0.296 | 0.885 | -0.011 | -0.232 | -0.062 | 0.052 | 0.052 | 0 | 0.31 | 0.13 | 0.29 | 0.19 | 0.39 | 0.3 |
| 11 | 120494449 | rs10790388 | -0.061 | 0.875 | 0.143 | -0.22 | -0.016 | -0.029 | -0.029 | 0 | 0.34 | 0.34 | 0.47 | 0.4 | 0.39 | 0.36 |
| 11 | 120494516 | rs7127199 | -0.958 | 0.935 | 0.027 | -0.289 | 0.003 | 0.001 | 0.001 | 0 | 0.53 | 0.57 | 0.66 | 0.4 | 0.72 | 0.5 |
| 11 | 120496664 | rs4936533 | 0.046 | 1.662 | 0.009 | -0.05 | 0.045 | -0.008 | -0.008 | 0 | 0.22 | 0.47 | 0.28 | 0.19 | 0.31 | 0.33 |
| 11 | 120496704 | rs4935744 | 0.289 | 1.728 | -0.216 | -0.018 | 0.072 | 0.014 | 0.014 | 0 | 0.2 | 0.36 | 0.3 | 0.2 | 0.33 | 0.29 |
| 11 | 120497350 | rs4445643 | -0.68 | 0.752 | -1.108 | 0.099 | 0.056 | 0.03 | 0.03 | 0 | 0.5 | 0.66 | 0.56 | 0.4 | 0.65 | 0.57 |
| 11 | 120498024 | rs7924948 | -0.075 | 0.614 | -1.229 | -0.012 | -0.012 | 0.046 | 0.046 | 0 | 0.41 | 0.61 | 0.4 | 0.19 | 0.52 | 0.37 |
| 11 | 120498051 | rs7924856 | -0.489 | -0.361 | -1.033 | 0.059 | 0.072 | 0.029 | 0.029 | 0 | 0.5 | 0.66 | 0.56 | 0.4 | 0.67 | 0.57 |
| 11 | 120499044 | rs7928903 | 0.434 | -0.328 | -0.714 | -0.285 | -0.019 | -0.04 | -0.04 | 3.106 | 0.66 | 0.45 | 0.7 | 0.82 | 0.77 | 0.69 |
| 11 | 120500130 | rs1574249 | -1.062 | -0.123 | -0.97 | 0.068 | 0.008 | -0.029 | -0.029 | 0 | 0.49 | 0.86 | 0.53 | 0.46 | 0.46 | 0.53 |
| 11 | 120501255 | rs12295761 | -0.269 | -0.046 | -0.047 | -0.017 | -0.04 | -0.093 | -0.093 | 2.37 | 0.14 | 0.41 | 0.06 | 0 | 0.05 | 0.14 |
| 11 | 120503083 | rs11217933 | -0.153 | -0.203 | -0.426 | -0.026 | -0.006 | -0.026 | -0.026 | 0 | 0.32 | 0.24 | 0.46 | 0.43 | 0.39 | 0.36 |
| 11 | 120503637 | rs1317912 | -0.427 | -0.182 | -0.328 | -0.041 | -0.032 | -0.08 | -0.08 | 0 | 0.16 | 0.57 | 0.08 | 0.03 | 0.07 | 0.18 |
| 11 | 120505628 | rs17124120 | 0.008 | 0.395 | -0.725 | 0.543 | -0.052 | -0.093 | -0.093 | 0 | 0.23 | 0.06 | 0.38 | 0.37 | 0.28 | 0.23 |
| 11 | 120506771 | rs12295498 | -0.991 | 0.574 | -0.091 | 0.416 | 0.003 | -0.006 | -0.006 | 0.252 | 0.09 | 0.17 | 0.09 | 0.05 | 0.13 | 0.15 |

| | | | | | | | | | | | | | | | | |
|---|---|---|---|---|---|---|---|---|---|---|---|---|---|---|---|---|
| 11 | 120508300 | rs7122702 | -0.593 | 0.439 | -0.443 | 0.459 | 0.02 | 0.016 | 0.016 | 0.63 | 0.5 | 0.72 | 0.51 | 0.42 | 0.51 | 0.51 |
| 11 | 120509621 | rs6589824 | -0.254 | 0.524 | -0.094 | 0.181 | 0.013 | 0.035 | 0.035 | 0 | 0.47 | 0.21 | 0.56 | 0.58 | 0.61 | 0.52 |
| 11 | 120514577 | rs1320719 | -0.578 | 0.989 | 0.145 | 0.487 | -0.037 | -0.041 | -0.041 | 0.047 | 0.08 | 0 | 0.05 | 0.1 | 0.13 | 0.22 |
| 11 | 120515734 | rs10892590 | -1.518 | 0.408 | 0.798 | -0.089 | -0.01 | -0.017 | -0.017 | 4.026 | 0.11 | 0.65 | 0.08 | 0.07 | 0.05 | 0.07 |
| 11 | 120516717 | rs2000874 | -0.864 | 0.42 | 0.521 | -0.178 | 0.017 | 0.025 | 0.025 | 0 | 0.61 | 0.86 | 0.64 | 0.67 | 0.69 | 0.69 |
| 11 | 120517844 | rs6589827 | -0.616 | 0.449 | 0.845 | -0.179 | 0.01 | 0.022 | 0.022 | 0 | 0.58 | 0.69 | 0.61 | 0.64 | 0.65 | 0.64 |
| 11 | 120521066 | rs4935750 | -0.948 | 0.444 | 0.504 | -0.703 | 0.019 | 0.063 | 0.063 | 0 | 0.62 | 0.78 | 0.64 | 0.8 | 0.71 | 0.72 |
| 11 | 120521766 | rs10892591 | -0.595 | 1.055 | -0.56 | -0.444 | 0.022 | 0.048 | 0.048 | 0 | 0.53 | 0.34 | 0.58 | 0.73 | 0.66 | 0.65 |
| 11 | 120522003 | rs10892592 | -0.595 | 1.016 | -0.56 | -0.444 | 0.024 | 0.048 | 0.048 | 0 | 0.53 | 0.34 | 0.58 | 0.73 | 0.66 | 0.65 |
| 11 | 120530316 | rs4936538 | -0.5 | 1.177 | -0.978 | -0.339 | 0.044 | 0.086 | 0.086 | 0 | 0.51 | 0.28 | 0.59 | 0.73 | 0.66 | 0.61 |
| 11 | 120530973 | rs6589829 | -0.251 | 1.148 | -1.411 | -0.235 | 0.026 | 0.114 | 0.114 | 0 | 0.47 | 0.17 | 0.54 | 0.74 | 0.64 | 0.6 |
| 11 | 120531656 | rs11217941 | -0.298 | 0.748 | -1.486 | 0.035 | -0.015 | 0.096 | 0.096 | 0 | 0.47 | 0.25 | 0.52 | 0.72 | 0.57 | 0.61 |
| 11 | 120535816 | rs4936540 | -0.262 | 0.322 | -1.518 | 0.012 | 0.017 | 0.037 | 0.037 | 0 | 0.37 | 0.31 | 0.47 | 0.21 | 0.37 | 0.37 |
| 11 | 120535893 | rs7928347 | -0.605 | 0.332 | -2.003 | 0.297 | 0 | -0.003 | -0.003 | 0.002 | 0.27 | 0.73 | 0.19 | 0.27 | 0.24 | 0.43 |
| 11 | 120538183 | rs11217951 | -1 | -0.007 | -1.717 | -0.206 | -0.002 | -0.011 | -0.011 | 0 | 0.14 | 0.12 | 0.11 | 0.02 | 0.15 | 0.27 |
| 11 | 120547912 | rs10502240 | -1.129 | -0.194 | -1.254 | -0.298 | -0.001 | -0.072 | -0.072 | 1.435 | 0.17 | 0.23 | 0.12 | 0.02 | 0.14 | 0.26 |
| 11 | 120548318 | rs11607732 | -1.129 | -0.194 | -1.254 | -0.298 | -0.001 | -0.072 | -0.072 | 1.435 | 0.17 | 0.23 | 0.12 | 0.02 | 0.14 | 0.26 |
| 11 | 120550430 | rs7932158 | -1.027 | -0.167 | -1.29 | 0.237 | -0.063 | -0.046 | -0.046 | 0 | 0.63 | 0.82 | 0.55 | 0.54 | 0.51 | 0.53 |
| 11 | 120551684 | rs10790398 | -0.604 | -0.212 | -1.106 | 0.262 | -0.002 | -0.002 | -0.002 | 0 | 0.57 | 0.39 | 0.51 | 0.52 | 0.49 | 0.47 |
| 11 | 120553706 | rs11217973 | 0.088 | -0.767 | -0.916 | 0.394 | 0.013 | 0 | 0 | 0.118 | 0.33 | 0.28 | 0.37 | 0.3 | 0.3 | 0.18 |
| 11 | 120556018 | rs7116106 | -0.878 | -1.092 | -0.261 | 0.568 | 0 | -0.001 | -0.001 | 0 | 0.45 | 0.51 | 0.48 | 0.47 | 0.44 | 0.38 |
| 11 | 120556341 | rs4445646 | -1.039 | -1.502 | -0.138 | 0.584 | 0.04 | 0.033 | 0.033 | 0.233 | 0.45 | 0.84 | 0.46 | 0.58 | 0.36 | 0.35 |
| 11 | 120557343 | rs10750176 | -1.842 | -1.329 | -0.118 | 0.559 | -0.089 | 0.009 | 0.009 | 0 | 0.57 | 0.9 | 0.54 | 0.59 | 0.49 | 0.51 |
| 11 | 120558084 | rs10892613 | -0.059 | -1.327 | -0.133 | 0.804 | 0 | -0.005 | -0.005 | 0 | 0.12 | 0.07 | 0.09 | 0.13 | 0.12 | 0.17 |
| 11 | 120560994 | rs10790402 | -1.604 | -1.489 | 0.414 | 0.772 | 0.001 | 0.036 | 0.036 | 0 | 0.53 | 0.74 | 0.53 | 0.58 | 0.46 | 0.5 |
| 11 | 120563298 | rs7925270 | -1.348 | -1.335 | 0.778 | 0.796 | 0.001 | 0.003 | 0.003 | 0 | 0.49 | 0.59 | 0.5 | 0.45 | 0.46 | 0.43 |
| 11 | 120565023 | rs12292552 | 0.286 | -1.282 | 0.792 | 0.767 | 0.002 | -0.002 | -0.002 | 0 | 0.09 | 0.06 | 0.06 | 0.01 | 0.09 | 0.14 |
| 11 | 120568392 | rs10892618 | 0.227 | -1.414 | 1.031 | 0.71 | -0.047 | -0.024 | -0.024 | 0 | 0.29 | 0.14 | 0.39 | 0.33 | 0.35 | 0.16 |
| 11 | 120569528 | rs17245664 | 0.163 | -1.491 | 1.179 | 0.671 | -0.005 | -0.003 | -0.003 | 0.018 | 0.29 | 0.26 | 0.39 | 0.33 | 0.33 | 0.15 |
| 11 | 120570198 | rs12278795 | -0.344 | -1.42 | 1.354 | 0.601 | -0.005 | 0.002 | 0.002 | 0 | 0.11 | 0.13 | 0.1 | 0.13 | 0.1 | 0.22 |
| 11 | 120574350 | rs17124211 | -2.438 | -1.57 | 1.304 | 0.507 | -0.07 | 0.02 | 0.02 | 0 | 0.27 | 0.5 | 0.15 | 0.25 | 0.16 | 0.32 |
| 11 | 120576984 | rs11603713 | 0.036 | -1.451 | 1.246 | 0.605 | -0.017 | 0.092 | 0.092 | 0 | 0.06 | 0.15 | 0.03 | 0.12 | 0.03 | 0.11 |
| 11 | 120577413 | rs12270261 | 0.044 | -1.388 | 0.821 | 0.461 | -0.006 | -0.045 | -0.045 | 0 | 0.1 | 0.15 | 0.07 | 0.01 | 0.1 | 0.11 |
| 11 | 120577423 | rs4567478 | -2.106 | -1.405 | 0.857 | 0.487 | -0.064 | 0.029 | 0.029 | 0 | 0.25 | 0.55 | 0.14 | 0.25 | 0.15 | 0.3 |
| 11 | 120580623 | rs7102869 | -0.033 | -1.329 | 0.351 | 0.351 | -0.006 | -0.045 | -0.045 | 0 | 0.11 | 0.15 | 0.07 | 0.01 | 0.11 | 0.11 |
| 11 | 120583983 | rs11217989 | -1.316 | -1.23 | -0.183 | 0.568 | -0.038 | 0.014 | 0.014 | 0 | 0.25 | 0.66 | 0.16 | 0.25 | 0.15 | 0.26 |
| 11 | 120584851 | rs3133855 | -0.045 | -0.623 | 0.006 | 0.766 | 0.014 | 0.026 | 0.026 | 0 | 0.7 | 0.57 | 0.56 | 0.6 | 0.66 | 0.79 |
| 11 | 120587319 | rs10892620 | -0.029 | -0.56 | -0.247 | 0.82 | -0.035 | -0.042 | -0.042 | 2.384 | 0.71 | 0.83 | 0.58 | 0.6 | 0.64 | 0.83 |
| 11 | 120592036 | rs3133226 | 0.925 | -0.15 | -1.034 | 0.457 | -0.031 | 0.04 | 0.04 | 0 | 0.51 | 0.08 | 0.48 | 0.36 | 0.53 | 0.47 |
| 11 | 120594813 | rs4612814 | 0.053 | -0.025 | -0.607 | 0.152 | 0 | -0.029 | -0.029 | 0 | 0.16 | 0.22 | 0.11 | 0 | 0.17 | 0.08 |
| 11 | 120597136 | rs3133846 | 0.497 | -0.343 | 0.147 | 0.532 | -0.002 | 0.06 | 0.06 | 0 | 0.71 | 0.62 | 0.63 | 0.47 | 0.72 | 0.63 |
| 11 | 120608912 | rs1320648 | -0.117 | -0.322 | 0.31 | 0.603 | 0.01 | 0.039 | 0.039 | 0.053 | 0.82 | 0.78 | 0.69 | 0.4 | 0.76 | 0.79 |
| 11 | 120608964 | rs752979 | -0.117 | -0.322 | 0.326 | 0.603 | 0.011 | 0.042 | 0.042 | 0.053 | 0.82 | 0.78 | 0.69 | 0.4 | 0.76 | 0.79 |
| 11 | 120610564 | rs751369 | 0.754 | -0.466 | 0.326 | 0.539 | 0.002 | 0.006 | 0.006 | 0 | 0.76 | 0.74 | 0.65 | 0.35 | 0.71 | 0.63 |
| 11 | 120610584 | rs751368 | 0.781 | -0.55 | 0.59 | 0.532 | -0.002 | 0.002 | 0.002 | 0 | 0.62 | 0.63 | 0.57 | 0.35 | 0.56 | 0.55 |
| 11 | 120614648 | rs17245685 | -0.192 | -0.499 | 0.345 | 0.532 | 0.004 | -0.003 | -0.003 | 0 | 0.05 | 0.04 | 0.04 | 0.05 | 0.06 | 0.16 |
| 11 | 120614787 | rs3133224 | 0.187 | 0.307 | 0.196 | 0.404 | -0.018 | -0.002 | -0.002 | 0.213 | 0.1 | 0.03 | 0.07 | 0.07 | 0.13 | 0.23 |
| 11 | 120617504 | rs1945010 | -0.226 | 0.259 | 0.303 | 0.525 | -0.018 | -0.033 | -0.033 | 0.19 | 0.69 | 0.79 | 0.63 | 0.5 | 0.63 | 0.75 |
| 11 | 120618079 | rs1939673 | 0.366 | 0.248 | 0.312 | 0.306 | -0.003 | 0 | 0 | 0.71 | 0.6 | 0.43 | 0.55 | 0.39 | 0.58 | 0.64 |
| 11 | 120621602 | rs1945011 | -1.724 | -0.203 | 0.033 | 0.003 | 0.018 | 0.036 | 0.036 | 1.72 | 0.34 | 0.75 | 0.25 | 0.19 | 0.29 | 0.55 |
| 11 | 120623682 | rs11606421 | -0.279 | -0.241 | -0.159 | -0.017 | 0.023 | -0.024 | -0.024 | 0 | 0.08 | 0.01 | 0.04 | 0.07 | 0.05 | 0.15 |
| 11 | 120624486 | rs10892622 | 1.753 | -0.086 | -0.368 | 0.359 | -0.06 | -0.055 | -0.055 | 0 | 0.36 | 0.04 | 0.36 | 0.37 | 0.35 | 0.21 |
| 11 | 120628012 | rs3133864 | -0.201 | 0.134 | 0.944 | 0.356 | -0.041 | 0.008 | 0.008 | 0.319 | 0.1 | 0.03 | 0.07 | 0.05 | 0.16 | 0.28 |
| 11 | 120630452 | rs7112522 | -1.02 | -0.786 | 1.769 | 0.468 | -0.034 | -0.07 | -0.07 | 1.776 | 0.11 | 0.26 | 0.06 | 0 | 0.06 | 0.04 |
| 11 | 120644115 | rs7938295 | -1.227 | -0.754 | 1.649 | 0.527 | -0.001 | 0.03 | 0.03 | 0.683 | 0.07 | 0.33 | 0.05 | 0.07 | 0.03 | 0.04 |
| 11 | 120644424 | rs948027 | -1.344 | -0.692 | 1.173 | 0.651 | -0.015 | -0.01 | -0.01 | 0 | 0.18 | 0.26 | 0.26 | 0.1 | 0.14 | 0.19 |
| 11 | 120644447 | rs948028 | -1.886 | -0.566 | 0.265 | 0.684 | 0.015 | 0.01 | 0.01 | 0 | 0.26 | 0.75 | 0.32 | 0.17 | 0.17 | 0.22 |
| 11 | 120655623 | rs1815952 | -1.639 | 0.016 | 0.626 | 1.365 | -0.003 | 0.216 | 0.216 | 0 | 0.15 | 0.41 | 0.21 | 0.34 | 0.12 | 0.23 |
| 11 | 120656572 | rs879602 | -1.138 | 0.562 | 0.147 | 0.535 | -0.003 | -0.004 | -0.004 | 0.031 | 0.1 | 0.11 | 0.09 | 0.03 | 0.12 | 0.04 |
| 11 | 120657444 | rs2155258 | 0.595 | 0.551 | 0.13 | -0.553 | 0.003 | -0.078 | -0.078 | 0.193 | 0.66 | 0.38 | 0.74 | 0.98 | 0.63 | 0.84 |
| 11 | 120659485 | rs3133861 | -0.13 | 0.672 | -0.267 | -0.449 | -0.004 | 0.001 | 0.001 | 0 | 0.93 | 0.93 | 0.92 | 0.99 | 0.88 | 0.98 |
| 11 | 120663363 | rs1954787 | 1.047 | -0.392 | -0.848 | -0.474 | -0.037 | 0.168 | 0.168 | 0.603 | 0.49 | 0.1 | 0.63 | 0.83 | 0.53 | 0.58 |
| 11 | 120675122 | rs1939672 | -0.268 | -1.081 | -0.417 | -0.003 | -0.001 | -0.015 | -0.015 | 0.314 | 0.08 | 0.1 | 0.05 | 0.01 | 0.06 | 0.04 |
| 11 | 120676208 | rs3016415 | -0.55 | -0.979 | -0.703 | 0.065 | -0.03 | -0.071 | -0.071 | 0 | 0.84 | 0.99 | 0.91 | 0.83 | 0.81 | 0.76 |
| 11 | 120677398 | rs948034 | -2.298 | -1.053 | 0.318 | -0.163 | -0.033 | -0.077 | -0.077 | 0.588 | 0.16 | 0.66 | 0.1 | 0.01 | 0.07 | 0.08 |
| 11 | 120678019 | rs7105205 | -1.05 | -0.903 | 0.494 | -0.017 | 0.063 | 0.065 | 0.065 | 2.427 | 0.06 | 0.5 | 0.04 | 0 | 0 | 0.04 |
| 11 | 120687069 | rs7105363 | 0.431 | -0.834 | -0.081 | 0.193 | -0.003 | 0.011 | 0.011 | 0 | 0.45 | 0.64 | 0.37 | 0.42 | 0.42 | 0.42 |
| 11 | 120688599 | rs4936553 | 1.582 | -0.841 | 0.279 | 0.113 | -0.064 | -0.053 | -0.053 | 0 | 0.31 | 0.11 | 0.25 | 0.4 | 0.33 | 0.3 |
| 11 | 120689119 | rs4935752 | 1.513 | -0.96 | 0.32 | 0.156 | -0.08 | -0.055 | -0.055 | 0 | 0.32 | 0.11 | 0.26 | 0.41 | 0.37 | 0.31 |

| | | | | | | | | | | | | | | | | |
|---|---|---|---|---|---|---|---|---|---|---|---|---|---|---|---|---|
| 11 | 120694754 | rs11218016 | -0.124 | -0.738 | 0.565 | 0.449 | -0.029 | -0.057 | -0.057 | 0 | 0.78 | 0.93 | 0.62 | 0.69 | 0.7 | 0.63 |
| 11 | 120694915 | rs12291852 | -0.958 | -1.122 | 0.779 | 0.351 | -0.034 | -0.082 | -0.082 | 0 | 0.15 | 0.66 | 0.11 | 0.01 | 0.07 | 0.09 |
| 11 | 120696297 | rs4573685 | -1.097 | -1.066 | 0.517 | 0.476 | -0.01 | -0.068 | -0.068 | 0 | 0.87 | 0.99 | 0.86 | 0.84 | 0.86 | 0.78 |
| 11 | 120697162 | rs4075786 | -0.162 | -0.988 | 0.959 | 0.375 | 0 | -0.055 | -0.055 | 2.008 | 0.11 | 0.3 | 0.08 | 0.01 | 0.07 | 0.04 |
| 11 | 120700147 | rs6589846 | 0.335 | -1.206 | 0.731 | 0.402 | 0.002 | -0.03 | -0.03 | 0 | 0.66 | 0.83 | 0.5 | 0.46 | 0.63 | 0.51 |
| 11 | 120707644 | rs3934628 | 1.171 | -1.319 | 0.701 | 0.399 | -0.06 | -0.02 | -0.02 | 0 | 0.4 | 0.15 | 0.31 | 0.46 | 0.44 | 0.38 |
| 11 | 120712591 | rs7110179 | -0.497 | -0.634 | 1.221 | 0.841 | -0.005 | 0.038 | 0.038 | 0 | 0.83 | 0.78 | 0.82 | 0.81 | 0.8 | 0.74 |
| 11 | 120717153 | rs7121233 | 1.528 | -0.748 | 1.008 | 0.757 | -0.105 | -0.091 | -0.091 | 2.046 | 0.28 | 0.1 | 0.27 | 0.46 | 0.38 | 0.34 |
| 11 | 120723433 | rs4430518 | -0.521 | -0.609 | 1.366 | 0.018 | -0.004 | 0.007 | 0.007 | 0.096 | 0.28 | 0.34 | 0.41 | 0.36 | 0.25 | 0.28 |
| 11 | 120724121 | rs7104543 | -0.887 | -0.601 | 1.336 | -0.122 | 0.011 | 0.033 | 0.033 | 2.874 | 0.52 | 0.67 | 0.55 | 0.34 | 0.45 | 0.38 |
| 11 | 120730621 | rs4526780 | 0.946 | -0.533 | 1.16 | -0.101 | -0.011 | -0.057 | -0.057 | 2.051 | 0.24 | 0.32 | 0.14 | 0 | 0.17 | 0.1 |
| 11 | 120730638 | rs4414232 | -0.763 | -0.494 | 0.952 | 0.008 | 0.017 | 0.048 | 0.048 | 2.739 | 0.52 | 0.71 | 0.56 | 0.34 | 0.45 | 0.37 |
| 11 | 120734419 | rs4344499 | -0.981 | -0.411 | 0.705 | 0.459 | -0.007 | 0.002 | 0.002 | 0.271 | 0.29 | 0.35 | 0.41 | 0.33 | 0.24 | 0.33 |
| 11 | 120736087 | rs11218035 | 0.155 | 0.046 | 0.245 | 0.426 | 0.003 | 0.001 | 0.001 | 0 | 0.19 | 0.17 | 0.2 | 0.22 | 0.23 | 0.25 |
| 11 | 120738397 | rs3802912 | 0.735 | -0.439 | -0.244 | 0.089 | -0.026 | -0.055 | -0.055 | 0 | 0.31 | 0.22 | 0.45 | 0.47 | 0.37 | 0.33 |
| 11 | 120741937 | rs4936562 | 0.54 | -0.474 | -0.258 | 0.057 | 0.003 | 0.007 | 0.007 | 0 | 0.33 | 0.38 | 0.47 | 0.48 | 0.38 | 0.37 |
| 11 | 120749741 | rs4077910 | -0.174 | -0.936 | -0.2 | 0.999 | -0.013 | 0.064 | 0.064 | 0 | 0.18 | 0.18 | 0.15 | 0.5 | 0.13 | 0.29 |
| 11 | 120755655 | rs12577638 | -0.478 | -0.905 | -0.256 | 0.869 | -0.033 | 0.085 | 0.085 | 0 | 0.22 | 0.36 | 0.17 | 0.47 | 0.14 | 0.33 |
| 11 | 120759130 | rs10892635 | -0.761 | -0.244 | -1.244 | 0.878 | 0.077 | -0.076 | -0.076 | 0.511 | 0.39 | 0.86 | 0.33 | 0.53 | 0.26 | 0.48 |
| 11 | 120771063 | rs7111184 | 0.321 | 0.54 | -0.808 | 0.818 | 0.002 | 0 | 0 | 0 | 0.63 | 0.29 | 0.54 | 0.3 | 0.66 | 0.48 |
| 11 | 120771727 | rs11218052 | -0.252 | -0.116 | -1.138 | 1.555 | 0.012 | -0.004 | -0.004 | 0 | 0.13 | 0.08 | 0.1 | 0.39 | 0.08 | 0.31 |
| 11 | 120772254 | rs2850814 | 0.097 | 0.499 | -1.236 | 0.99 | -0.019 | 0.023 | 0.023 | 0 | 0.84 | 0.9 | 0.78 | 0.91 | 0.82 | 0.74 |
| 11 | 120776001 | rs2156634 | 0.372 | 0.428 | -1.391 | 0.305 | -0.008 | 0.04 | 0.04 | 9.644 | 0.86 | 0.95 | 0.81 | 0.92 | 0.86 | 0.8 |
| 11 | 120779520 | rs949055 | 1.175 | 0.182 | -0.455 | 0.114 | 0.002 | -0.046 | -0.046 | 0 | 0.66 | 0.41 | 0.62 | 0.83 | 0.68 | 0.55 |
| 11 | 120783369 | rs17124538 | -0.295 | 0.145 | -0.022 | 0.099 | -0.006 | -0.001 | -0.001 | 0.115 | 0.06 | 0.13 | 0.05 | 0.08 | 0.04 | 0.09 |
| 11 | 120786892 | rs2156633 | 0.777 | 0.1 | 0.431 | 0.082 | -0.002 | -0.02 | -0.02 | 0.001 | 0.72 | 0.74 | 0.68 | 0.9 | 0.72 | 0.62 |
| 11 | 120789508 | rs10892640 | 0.863 | 0.126 | -0.384 | -0.364 | 0.012 | -0.014 | -0.014 | 0 | 0.4 | 0.27 | 0.4 | 0.41 | 0.42 | 0.2 |
| 11 | 120789852 | rs949295 | 0.272 | 0.131 | -0.819 | -0.328 | 0.026 | 0.005 | 0.005 | 1.815 | 0.51 | 0.81 | 0.51 | 0.54 | 0.49 | 0.29 |
| 11 | 120795888 | rs949054 | -0.137 | 0.923 | -0.755 | 0.792 | -0.027 | -0.012 | -0.012 | 0 | 0.9 | 0.94 | 0.84 | 0.59 | 0.8 | 0.8 |
| 11 | 120801185 | rs2186620 | -0.2 | 0.476 | -0.328 | 0.036 | -0.007 | -0.136 | -0.136 | 0 | 0.79 | 0.51 | 0.83 | 0.98 | 0.83 | 0.86 |
| 11 | 120804291 | rs10892647 | -0.319 | 0.076 | -0.841 | -0.308 | 0.009 | -0.038 | -0.038 | 0.777 | 0.29 | 0.2 | 0.22 | 0.38 | 0.3 | 0.27 |
| 11 | 120805056 | rs17124570 | 0.401 | 0.012 | -0.968 | -0.13 | -0.003 | -0.069 | -0.069 | 0 | 0.26 | 0.12 | 0.2 | 0.35 | 0.27 | 0.28 |
| 11 | 120815238 | rs4936566 | 0.238 | -0.189 | -0.974 | -0.216 | 0.024 | -0.055 | -0.055 | 0 | 0.28 | 0.16 | 0.21 | 0.36 | 0.24 | 0.32 |
| 11 | 120815567 | rs11218067 | 1.679 | 0.367 | -1.473 | -0.081 | -0.066 | -0.075 | -0.075 | 7.87 | 0.12 | 0.01 | 0.2 | 0.3 | 0.17 | 0.24 |
| 11 | 120816711 | rs11218070 | 0.599 | 0.293 | -1.358 | -0.09 | 0.019 | -0.069 | -0.069 | 0 | 0.27 | 0.15 | 0.23 | 0.37 | 0.25 | 0.33 |
| 11 | 120819562 | rs684913 | -1.381 | 0.115 | -0.656 | -0.316 | 0.033 | -0.025 | -0.025 | 0 | 0.93 | 0.72 | 0.96 | 1 | 0.96 | 0.92 |
| 11 | 120819939 | rs1893818 | -0.663 | -0.175 | -0.376 | -0.715 | -0.011 | -0.163 | -0.163 | 0 | 0.72 | 0.54 | 0.82 | 0.99 | 0.76 | 0.84 |
| 11 | 120822377 | rs3781817 | 0.42 | -0.198 | -0.513 | -0.644 | 0 | 0.003 | 0.003 | 3.218 | 0.45 | 0.41 | 0.62 | 0.63 | 0.52 | 0.52 |
| 11 | 120823396 | rs584092 | 1.41 | -0.153 | -0.667 | -0.693 | 0.002 | -0.001 | -0.001 | 0 | 0.2 | 0.12 | 0.14 | 0 | 0.21 | 0.08 |
| 11 | 120823608 | rs2298725 | 1.823 | -0.17 | -0.666 | -0.55 | -0.054 | -0.048 | -0.048 | 0 | 0.28 | 0.08 | 0.37 | 0.33 | 0.33 | 0.28 |
| 11 | 120825575 | rs4936568 | 0.428 | -0.132 | -0.405 | -0.441 | -0.019 | 0.009 | 0.009 | 1.106 | 0.32 | 0.44 | 0.25 | 0.33 | 0.27 | 0.32 |
| 11 | 120826131 | rs627532 | -0.846 | 0.085 | -0.437 | -0.31 | 0.003 | 0.003 | 0.003 | 1.747 | 0.61 | 0.39 | 0.52 | 0.63 | 0.63 | 0.63 |
| 11 | 120828827 | rs732480 | 1.143 | 0.186 | -0.652 | -0.237 | 0.001 | -0.005 | -0.005 | 0.82 | 0.16 | 0.2 | 0.17 | 0.33 | 0.15 | 0.29 |
| 11 | 120829249 | rs672176 | -0.168 | -0.117 | -0.465 | -0.984 | -0.017 | -0.143 | -0.143 | 0.195 | 0.59 | 0.5 | 0.78 | 0.99 | 0.68 | 0.83 |
| 11 | 120834573 | rs4936570 | -0.708 | 0.114 | -0.436 | -0.573 | 0.034 | 0.016 | 0.016 | 0.107 | 0.79 | 0.71 | 0.76 | 0.75 | 0.78 | 0.67 |
| 11 | 120835281 | rs7128009 | -1.35 | 0.078 | -0.429 | -0.387 | 0.002 | -0.024 | -0.024 | 0.026 | 0.79 | 0.79 | 0.83 | 0.61 | 0.82 | 0.64 |
| 11 | 120837821 | rs2276319 | 0.075 | -0.006 | -0.402 | -0.423 | 0.002 | -0.001 | -0.001 | 0 | 0.62 | 0.38 | 0.6 | 0.35 | 0.63 | 0.38 |
| 11 | 120841689 | rs652418 | -0.932 | 0.04 | -0.616 | -0.338 | 0 | -0.004 | -0.004 | 0 | 0.78 | 0.75 | 0.82 | 0.61 | 0.8 | 0.63 |
| 11 | 120843742 | rs1944518 | 1.648 | 0.194 | -0.845 | -0.194 | -0.09 | -0.117 | -0.117 | 0 | 0.25 | 0.01 | 0.38 | 0.35 | 0.33 | 0.29 |
| 11 | 120852961 | rs499804 | 0.655 | 0.38 | -0.361 | -0.137 | 0.049 | 0.047 | 0.047 | 0.17 | 0.17 | 0.32 | 0.22 | 0.25 | 0.19 | 0.25 |
| 11 | 120858410 | rs2282586 | 0.869 | 0.005 | -0.523 | -0.236 | -0.001 | -0.03 | -0.03 | 0.254 | 0.4 | 0.14 | 0.44 | 0.37 | 0.42 | 0.35 |
| 11 | 120860200 | rs536333 | -0.21 | -0.064 | -0.13 | -0.287 | 0.001 | 0.002 | 0.002 | 0 | 0.57 | 0.54 | 0.69 | 0.63 | 0.62 | 0.61 |
| 11 | 120863568 | rs607800 | -0.389 | -0.034 | -0.153 | -0.278 | 0.001 | 0.004 | 0.004 | 0 | 0.59 | 0.56 | 0.69 | 0.63 | 0.63 | 0.61 |
| 11 | 120864806 | rs10892657 | 1.795 | -0.157 | -0.283 | -0.297 | 0.024 | 0.043 | 0.043 | 1.625 | 0.11 | 0.03 | 0.04 | 0 | 0.08 | 0.03 |
| 11 | 120864922 | rs1944522 | 0.502 | -0.21 | -0.07 | -0.265 | -0.003 | 0.037 | 0.037 | 0 | 0.48 | 0.33 | 0.55 | 0.35 | 0.54 | 0.4 |
| 11 | 120873076 | rs1621211 | -0.676 | -0.183 | -0.108 | -0.023 | 0.001 | -0.013 | -0.013 | 0 | 0.77 | 0.76 | 0.81 | 0.62 | 0.8 | 0.67 |
| 11 | 120875383 | rs2510639 | -0.481 | -0.182 | -0.039 | -0.022 | 0.001 | -0.005 | -0.005 | 0 | 0.76 | 0.71 | 0.8 | 0.62 | 0.8 | 0.67 |
| 11 | 120907482 | rs2000525 | 1.044 | -0.155 | -0.488 | -0.224 | 0.005 | -0.01 | -0.01 | 3.564 | 0.2 | 0.18 | 0.18 | 0.32 | 0.18 | 0.22 |
| 11 | 120914370 | rs7952619 | -0.639 | -0.236 | -0.347 | -0.73 | -0.019 | -0.009 | -0.009 | 3.135 | 0.35 | 0.54 | 0.29 | 0.33 | 0.35 | 0.28 |
| 11 | 120920649 | rs1939004 | 0.183 | -0.234 | -0.362 | -0.752 | -0.001 | 0 | 0 | 0 | 0.25 | 0.3 | 0.22 | 0.33 | 0.24 | 0.24 |
| 11 | 120923495 | rs10502245 | 0.585 | -0.216 | -0.441 | -0.742 | 0.004 | -0.014 | -0.014 | 0 | 0.22 | 0.2 | 0.21 | 0.33 | 0.24 | 0.23 |
| 11 | 120924200 | rs10502246 | 1.124 | -0.2 | -0.513 | -0.67 | -0.003 | -0.045 | -0.045 | 0.058 | 0.09 | 0.11 | 0.06 | 0 | 0.1 | 0.04 |
| 11 | 120931054 | rs12795310 | 0.92 | 0.718 | -0.749 | -0.356 | -0.042 | -0.008 | -0.008 | 0 | 0.21 | 0.12 | 0.41 | 0.22 | 0.3 | 0.23 |
| 11 | 120953914 | rs874069 | 0.601 | -0.091 | -0.27 | -0.296 | -0.001 | -0.054 | -0.054 | 0 | 0.1 | 0.15 | 0.06 | 0 | 0.11 | 0.04 |
| 11 | 120954769 | rs6589855 | -0.401 | 0.007 | -0.587 | -0.217 | -0.031 | -0.003 | -0.003 | 1.828 | 0.33 | 0.53 | 0.26 | 0.33 | 0.3 | 0.27 |
| 11 | 120975390 | rs3758893 | 0.452 | -0.306 | -0.414 | -0.24 | 0.001 | 0.001 | 0.001 | 0 | 0.11 | 0.07 | 0.06 | 0.1 | 0.07 | 0.12 |
| 11 | 120976428 | rs504626 | -0.736 | -0.149 | -0.668 | -0.565 | 0 | -0.002 | -0.002 | 0 | 0.49 | 0.5 | 0.43 | 0.39 | 0.51 | 0.38 |
| 11 | 120977529 | rs684384 | -0.191 | -0.061 | -0.685 | -0.663 | -0.003 | 0.001 | 0.001 | 1.839 | 0.39 | 0.27 | 0.35 | 0.39 | 0.4 | 0.36 |
| 11 | 120980537 | rs598650 | 1.214 | -0.264 | -0.874 | -0.286 | -0.007 | -0.006 | -0.006 | 0 | 0.2 | 0.2 | 0.39 | 0.23 | 0.28 | 0.23 |

| | | | | | | | | | | | | | | | | |
|---|---|---|---|---|---|---|---|---|---|---|---|---|---|---|---|---|
| 11 | 120981408 | rs1829 | -0.644 | -0.869 | -0.747 | -0.19 | -0.003 | -0.001 | -0.001 | 0 | 0.66 | 0.66 | 0.81 | 0.62 | 0.78 | 0.6 |
| 11 | 120992188 | rs594711 | -1.217 | -1.085 | -1.348 | -0.174 | 0.026 | -0.107 | -0.107 | 0 | 0.75 | 0.98 | 0.81 | 0.64 | 0.8 | 0.69 |
| 11 | 120994638 | rs584329 | -1.175 | -0.926 | -1.639 | -0.422 | -0.016 | -0.034 | -0.034 | 0.345 | 0.71 | 0.89 | 0.64 | 0.68 | 0.73 | 0.74 |
| 11 | 120998942 | rs10502247 | 0.313 | -1.007 | -1.483 | -0.579 | 0.061 | -0.012 | -0.012 | 0 | 0.38 | 0.15 | 0.26 | 0.44 | 0.3 | 0.44 |
| 11 | 120999258 | rs4936578 | 1.451 | -0.791 | -1.132 | -0.668 | 0.055 | -0.091 | -0.091 | 0 | 0.14 | 0.36 | 0.15 | 0.01 | 0.2 | 0.1 |
| 11 | 121000774 | rs520805 | 1.957 | -0.783 | -1.14 | -0.662 | 0.009 | 0.011 | 0.011 | 0.262 | 0.17 | 0.2 | 0.22 | 0.24 | 0.23 | 0.19 |
| 11 | 121000784 | rs586473 | -1.399 | -0.779 | -1.191 | -0.66 | -0.016 | -0.034 | -0.034 | 0 | 0.71 | 0.89 | 0.64 | 0.68 | 0.73 | 0.74 |
| 11 | 121001263 | rs7108166 | 1.537 | -0.625 | -1.09 | -0.565 | 0 | 0.007 | 0.007 | 0 | 0.1 | 0.05 | 0.06 | 0.07 | 0.09 | 0.11 |
| 11 | 121001345 | rs583842 | 1.611 | -0.259 | -0.797 | -0.358 | 0.019 | 0.023 | 0.023 | 0.348 | 0.17 | 0.25 | 0.23 | 0.24 | 0.23 | 0.19 |
| 11 | 121003358 | rs7105712 | 0.342 | -0.693 | -0.605 | -0.542 | 0.032 | -0.002 | -0.002 | 0 | 0.38 | 0.21 | 0.27 | 0.44 | 0.3 | 0.44 |
| 11 | 121004502 | rs2155466 | -0.437 | -0.232 | -0.874 | -0.085 | -0.007 | 0.002 | 0.002 | 0.03 | 0.59 | 0.61 | 0.42 | 0.44 | 0.5 | 0.55 |
| 11 | 121007777 | rs4936580 | -0.268 | -0.918 | -0.881 | -0.523 | 0.015 | 0.022 | 0.022 | 0 | 0.51 | 0.56 | 0.35 | 0.3 | 0.39 | 0.5 |
| 11 | 121012978 | rs665743 | -0.01 | -1.456 | -0.971 | -0.9 | 0 | -0.031 | -0.031 | 0 | 0.21 | 0.76 | 0.19 | 0.01 | 0.21 | 0.12 |
| 11 | 121016818 | rs12275038 | 0.318 | -1.468 | -0.902 | -0.906 | 0.014 | -0.121 | -0.121 | 0 | 0.19 | 0.55 | 0.17 | 0.01 | 0.21 | 0.12 |
| 11 | 121018870 | rs4623911 | 0.326 | -1.468 | -0.902 | -0.906 | 0.014 | -0.121 | -0.121 | 0.186 | 0.19 | 0.55 | 0.17 | 0.01 | 0.21 | 0.12 |
| 11 | 121023906 | rs687329 | -1.361 | -1.072 | -1.538 | -0.643 | -0.126 | -0.186 | -0.186 | 1.762 | 0.72 | 0.97 | 0.6 | 0.54 | 0.64 | 0.74 |
| 11 | 121023992 | rs686882 | -0.63 | -1.609 | -1.602 | -0.821 | 0.105 | 0.18 | 0.18 | 0 | 0.46 | 0.84 | 0.34 | 0.29 | 0.35 | 0.48 |
| 11 | 121038810 | rs2155369 | 1.304 | -1.642 | -1.602 | -0.693 | 0.091 | -0.037 | -0.037 | 1.44 | 0.25 | 0.01 | 0.16 | 0.31 | 0.14 | 0.35 |
| 11 | 121050913 | rs620896 | -0.875 | -1.688 | -1.591 | -0.733 | 0.123 | 0.165 | 0.165 | 0.122 | 0.51 | 0.84 | 0.34 | 0.32 | 0.35 | 0.48 |
| 11 | 121059326 | rs12279811 | -0.081 | -0.954 | -1.133 | -0.623 | -0.011 | -0.138 | -0.138 | 0 | 0.24 | 0.53 | 0.16 | 0.01 | 0.21 | 0.13 |
| 11 | 121062486 | rs7110565 | 0.345 | -0.441 | -0.936 | -0.372 | 0.036 | -0.131 | -0.131 | 0 | 0.18 | 0.48 | 0.15 | 0.01 | 0.21 | 0.13 |
| 11 | 121062860 | rs11218195 | 1.111 | 0.299 | -1.545 | 0.184 | -0.096 | -0.085 | -0.085 | 0 | 0.16 | 0.02 | 0.2 | 0.24 | 0.27 | 0.22 |
| 11 | 121072088 | rs688099 | -0.121 | -0.146 | -0.944 | -0.193 | 0.034 | -0.052 | -0.052 | 0 | 0.2 | 0.37 | 0.17 | 0.12 | 0.26 | 0.11 |
| 11 | 121072114 | rs12292816 | 0.59 | -0.196 | -0.851 | -0.172 | 0.003 | 0.043 | 0.043 | 0 | 0.09 | 0.25 | 0.05 | 0.12 | 0.06 | 0.06 |
| 11 | 121074575 | rs571429 | -1.358 | -0.179 | -1.467 | 0.341 | -0.038 | -0.094 | -0.094 | 0.054 | 0.84 | 0.98 | 0.76 | 0.81 | 0.85 | 0.82 |
| 11 | 121082690 | rs12787035 | 1.662 | -0.152 | -1.62 | 0.333 | -0.07 | -0.056 | -0.056 | 0 | 0.17 | 0.05 | 0.2 | 0.24 | 0.27 | 0.25 |
| 11 | 121083812 | rs665035 | -0.18 | -0.229 | -1.62 | 0.288 | -0.115 | -0.09 | -0.09 | 0 | 0.67 | 0.89 | 0.55 | 0.57 | 0.59 | 0.58 |
| 11 | 121088158 | rs10892689 | 0.923 | -0.306 | -1.517 | 0.224 | 0.003 | 0.003 | 0.003 | 1.807 | 0.47 | 0.56 | 0.39 | 0.45 | 0.33 | 0.46 |
| 11 | 121092052 | rs745517 | 0.946 | -0.95 | -1.612 | -0.018 | -0.062 | -0.075 | -0.075 | 0 | 0.34 | 0.21 | 0.42 | 0.51 | 0.49 | 0.35 |
| 11 | 121092522 | rs4999180 | 0.946 | -0.95 | -1.612 | -0.033 | -0.062 | -0.073 | -0.073 | 0 | 0.34 | 0.21 | 0.42 | 0.51 | 0.49 | 0.36 |
| 11 | 121103704 | rs10892692 | -0.729 | -0.76 | -1.527 | -0.28 | -0.025 | 0.033 | 0.033 | 0 | 0.87 | 0.94 | 0.88 | 1 | 0.8 | 0.92 |
| 11 | 121109489 | rs6589864 | 1.016 | -1.45 | -1.125 | -0.208 | -0.101 | -0.041 | -0.041 | 1.756 | 0.23 | 0.38 | 0.09 | 0.18 | 0.07 | 0.29 |
| 11 | 121110312 | rs10437694 | -0.23 | -0.029 | -1.15 | 0.173 | 0 | 0.005 | 0.005 | 0 | 0.2 | 0.15 | 0.28 | 0.27 | 0.18 | 0.21 |
| 11 | 121111458 | rs948381 | 0.854 | 0.082 | -1.163 | 0.126 | -0.002 | -0.029 | -0.029 | 0 | 0.61 | 0.78 | 0.53 | 0.62 | 0.57 | 0.65 |
| 11 | 121112889 | rs12289923 | -0.72 | -0.342 | -0.96 | 0.515 | 0 | -0.002 | -0.002 | 0 | 0.23 | 0.17 | 0.32 | 0.31 | 0.21 | 0.25 |
| 11 | 121120755 | rs1893248 | 1.217 | -0.251 | -0.57 | 0.426 | 0.011 | -0.001 | -0.001 | 0 | 0.31 | 0.36 | 0.38 | 0.36 | 0.44 | 0.29 |
| 11 | 121121448 | rs7928669 | 1.075 | -0.321 | -0.538 | 0.239 | -0.001 | -0.001 | -0.001 | 0 | 0.62 | 0.68 | 0.55 | 0.53 | 0.6 | 0.51 |
| 11 | 121129528 | rs9943623 | -1.396 | -0.568 | -0.222 | 0.354 | -0.014 | 0.05 | 0.05 | 0 | 0.29 | 0.39 | 0.41 | 0.39 | 0.26 | 0.36 |
| 11 | 121136604 | rs10790422 | -0.178 | -0.493 | -0.245 | 0.416 | 0 | 0.001 | 0.001 | 0 | 0.18 | 0.12 | 0.31 | 0.27 | 0.19 | 0.24 |
| 11 | 121137448 | rs12280323 | -1.08 | -0.51 | -0.24 | 0.401 | -0.003 | 0.042 | 0.042 | 0 | 0.26 | 0.32 | 0.4 | 0.37 | 0.26 | 0.32 |
| 11 | 121138191 | rs7935970 | -0.957 | -0.504 | -0.321 | 0.404 | -0.001 | 0.024 | 0.024 | 0 | 0.26 | 0.26 | 0.4 | 0.37 | 0.26 | 0.32 |
| 11 | 121140729 | rs10892715 | 0.251 | 0.076 | -0.239 | 0.737 | 0.006 | 0.029 | 0.029 | 0 | 0.37 | 0.16 | 0.2 | 0.28 | 0.3 | 0.33 |
| 11 | 121149531 | rs11218239 | 0.206 | 0.019 | -0.214 | 0.779 | -0.054 | -0.075 | -0.075 | 0.019 | 0.11 | 0.01 | 0.29 | 0.27 | 0.15 | 0.22 |
| 11 | 121153921 | rs6589870 | -0.43 | -0.235 | 0.023 | 0.647 | 0.013 | 0.03 | 0.03 | 0 | 0.62 | 0.77 | 0.78 | 0.72 | 0.7 | 0.67 |
| 11 | 121158100 | rs6589872 | 0.993 | -0.17 | 0.109 | 0.704 | -0.003 | 0.015 | 0.015 | 0 | 0.45 | 0.37 | 0.48 | 0.37 | 0.56 | 0.43 |
| 11 | 121168178 | rs727422 | 0.647 | -0.202 | -0.037 | 0.664 | 0 | 0 | 0 | 0 | 0.07 | 0.05 | 0.08 | 0.09 | 0.07 | 0.08 |
| 11 | 121170025 | rs12275106 | 0.085 | -0.205 | 0.063 | 0.663 | -0.004 | 0 | 0 | 0 | 0.09 | 0.18 | 0.1 | 0.09 | 0.08 | 0.08 |
| 11 | 121178302 | rs1061332 | -0.446 | -0.177 | -0.327 | 0.686 | 0.009 | 0.027 | 0.027 | 0 | 0.62 | 0.76 | 0.78 | 0.71 | 0.69 | 0.68 |
| 11 | 121179549 | rs7942396 | -0.307 | 0.469 | 0.104 | 1.447 | 0.001 | 0 | 0 | 0 | 0.08 | 0.09 | 0.09 | 0.09 | 0.07 | 0.08 |
| 11 | 121181848 | rs2115925 | -0.469 | 0.654 | 0.138 | 1.567 | 0.005 | 0.082 | 0.082 | 0 | 0.08 | 0.11 | 0.19 | 0.26 | 0.08 | 0.16 |
| 11 | 121194946 | rs10892723 | -0.304 | 0.666 | -0.239 | 1.05 | -0.003 | -0.002 | -0.002 | 0 | 0.91 | 0.92 | 0.95 | 0.92 | 0.89 | 0.88 |
| 11 | 121195951 | rs6589877 | -0.176 | 0.652 | -0.238 | 1.043 | -0.002 | -0.001 | -0.001 | 0.008 | 0.9 | 0.91 | 0.95 | 0.92 | 0.89 | 0.87 |
| 11 | 121201899 | rs7125518 | 0.192 | 0.425 | -0.368 | 0.936 | -0.01 | -0.005 | -0.005 | 0.041 | 0.27 | 0.32 | 0.16 | 0.23 | 0.15 | 0.28 |
| 11 | 121203120 | rs4936612 | -0.165 | 0.686 | -1.071 | 0.678 | 0.019 | 0.004 | 0.004 | 0.04 | 0.56 | 0.33 | 0.69 | 0.6 | 0.65 | 0.55 |
| 11 | 121203341 | rs11218261 | 0.49 | 0.762 | -1.2 | 0.613 | -0.037 | 0.091 | 0.091 | 4.311 | 0.43 | 0.16 | 0.39 | 0.33 | 0.49 | 0.32 |
| 11 | 121203796 | rs10892726 | 0.192 | 0.021 | -0.813 | 0.323 | 0.019 | 0.004 | 0.004 | 0.037 | 0.56 | 0.35 | 0.69 | 0.61 | 0.65 | 0.55 |
| 11 | 121206354 | rs2060009 | 0.963 | 0.063 | -0.27 | -0.131 | 0.006 | 0.013 | 0.013 | 0.019 | 0.65 | 0.37 | 0.68 | 0.71 | 0.67 | 0.65 |
| 11 | 121208698 | rs876501 | -1.132 | 0.06 | 0.463 | -0.158 | 0.009 | 0.001 | 0.001 | 0 | 0.15 | 0.24 | 0.11 | 0.2 | 0.17 | 0.15 |
| 11 | 121215237 | rs7112480 | -1.468 | 0.26 | 0.649 | -0.5 | -0.001 | 0.003 | 0.003 | 0 | 0.26 | 0.23 | 0.18 | 0.11 | 0.28 | 0.24 |
| 11 | 121215401 | rs2370115 | 2.151 | -0.16 | 0.73 | -0.694 | -0.042 | 0.093 | 0.093 | 0.27 | 0.54 | 0.11 | 0.62 | 0.67 | 0.55 | 0.59 |
| 11 | 121216594 | rs11218268 | -1.195 | -0.375 | 1.199 | -0.474 | 0.013 | 0.054 | 0.054 | 0 | 0.1 | 0.28 | 0.1 | 0.11 | 0.09 | 0.12 |
| 11 | 121222579 | rs10892737 | 2.048 | -0.05 | 1.057 | -0.525 | -0.039 | 0.086 | 0.086 | 0.245 | 0.62 | 0.15 | 0.69 | 0.73 | 0.65 | 0.67 |
| 11 | 121223636 | rs17125182 | -1.548 | -0.288 | 1.139 | -0.515 | 0.002 | 0.001 | 0.001 | 0 | 0.05 | 0.06 | 0.02 | 0.11 | 0.03 | 0.05 |
| 11 | 121223996 | rs4936618 | 0.729 | -0.275 | 2.054 | -0.567 | 0.018 | -0.073 | -0.073 | 0 | 0.83 | 0.76 | 0.86 | 0.93 | 0.82 | 0.81 |
| 11 | 121224776 | rs891437 | 2.495 | -0.256 | 1.958 | -0.528 | -0.018 | 0.043 | 0.043 | 0.263 | 0.58 | 0.14 | 0.69 | 0.66 | 0.68 | 0.64 |
| 11 | 121226022 | rs12417012 | -0.612 | -0.261 | 2.503 | -0.55 | -0.005 | 0.049 | 0.049 | 0 | 0.08 | 0.19 | 0.08 | 0.16 | 0.04 | 0.05 |
| 11 | 121228469 | rs4936619 | 2.661 | -0.26 | 2.526 | -0.553 | -0.011 | 0.043 | 0.043 | 0.226 | 0.59 | 0.16 | 0.69 | 0.66 | 0.68 | 0.64 |
| 11 | 121232811 | rs4936621 | 0.224 | -0.26 | 3.068 | -0.59 | 0.023 | 0.004 | 0.004 | 0 | 0.94 | 0.8 | 0.91 | 0.93 | 0.95 | 0.89 |
| 11 | 121234131 | rs10790443 | 2.579 | -0.251 | 2.907 | -0.484 | -0.009 | 0.032 | 0.032 | 0.163 | 0.6 | 0.21 | 0.73 | 0.66 | 0.68 | 0.66 |

| | | | | | | | | | | | | | | | | |
|---|---|---|---|---|---|---|---|---|---|---|---|---|---|---|---|---|
| 11 | 121235871 | rs1529579 | 1.556 | -0.226 | 3.022 | -0.477 | 0.007 | -0.038 | -0.038 | 0 | 0.76 | 0.54 | 0.82 | 0.82 | 0.79 | 0.78 |
| 11 | 121237372 | rs10790445 | 1.449 | -0.225 | 2.969 | -0.524 | 0.008 | -0.037 | -0.037 | 0 | 0.77 | 0.53 | 0.84 | 0.84 | 0.8 | 0.79 |
| 11 | 121237583 | rs12798486 | -0.125 | -0.2 | 2.424 | -0.532 | -0.015 | 0.021 | 0.021 | 0 | 0.06 | 0.02 | 0.05 | 0 | 0.07 | 0.07 |
| 11 | 121238455 | rs1560407 | 2.057 | -0.231 | 2.614 | -0.531 | 0.006 | 0.028 | 0.028 | 0.011 | 0.67 | 0.28 | 0.77 | 0.76 | 0.75 | 0.75 |
| 11 | 121239305 | rs10750191 | 2.077 | -0.231 | 2.584 | -0.531 | 0.007 | 0.042 | 0.042 | 0.017 | 0.67 | 0.25 | 0.77 | 0.76 | 0.75 | 0.75 |
| 11 | 121243716 | rs72991 | 2.107 | -0.175 | 2.85 | -0.528 | 0 | -0.015 | -0.015 | 0.057 | 0.66 | 0.36 | 0.77 | 0.84 | 0.72 | 0.71 |
| 11 | 121247901 | rs7102251 | -2.345 | -0.262 | 2.453 | -0.591 | 0.011 | 0.014 | 0.014 | 0.242 | 0.38 | 0.72 | 0.24 | 0.32 | 0.29 | 0.28 |
| 11 | 121253952 | rs297492 | 2.071 | -0.257 | 2.429 | -0.575 | 0.018 | 0.023 | 0.023 | 0.183 | 0.62 | 0.24 | 0.77 | 0.68 | 0.73 | 0.72 |
| 11 | 121255699 | rs6589880 | -1.909 | -0.252 | 2.628 | -0.588 | 0.003 | -0.039 | -0.039 | 0 | 0.29 | 0.41 | 0.2 | 0.24 | 0.25 | 0.23 |
| 11 | 121264079 | rs93882 | 1.739 | -0.222 | 1.743 | -0.439 | 0 | 0.012 | 0.012 | 0.132 | 0.64 | 0.31 | 0.79 | 0.74 | 0.7 | 0.77 |
| 11 | 121265523 | rs733430 | -1.213 | -0.292 | 1.531 | -0.527 | -0.004 | -0.028 | -0.028 | 0.004 | 0.22 | 0.26 | 0.09 | 0.12 | 0.18 | 0.12 |
| 11 | 121266057 | rs7105326 | -2.078 | -0.089 | 1.462 | -0.568 | -0.002 | 0.033 | 0.033 | 0 | 0.41 | 0.67 | 0.28 | 0.28 | 0.35 | 0.33 |
| 11 | 121266366 | rs12281864 | -0.692 | -0.069 | 1.159 | -0.528 | 0.008 | 0.006 | 0.006 | 0 | 0.07 | 0.19 | 0.08 | 0.07 | 0.04 | 0.12 |
| 11 | 121269726 | rs7937444 | -1.059 | 0.02 | 0.624 | -0.558 | -0.006 | 0.023 | 0.023 | 0 | 0.21 | 0.15 | 0.12 | 0.1 | 0.22 | 0.12 |
| 11 | 121271440 | rs297509 | 0.897 | -0.546 | 0.548 | -0.259 | 0.035 | 0.025 | 0.025 | 0.985 | 0.66 | 0.2 | 0.77 | 0.73 | 0.81 | 0.72 |
| 11 | 121291294 | rs297494 | -0.26 | -0.111 | -0.26 | 0.232 | 0.055 | 0.022 | 0.022 | 0 | 0.51 | 0.32 | 0.61 | 0.63 | 0.6 | 0.68 |
| 11 | 121294360 | rs297517 | -0.501 | -0.335 | 0.461 | 0.353 | -0.008 | -0.006 | -0.006 | 0 | 0.54 | 0.52 | 0.64 | 0.69 | 0.62 | 0.73 |
| 11 | 121299288 | rs12576704 | -0.631 | 0.833 | 0.368 | 0.333 | 0.018 | 0.002 | 0.002 | 0.029 | 0.11 | 0.18 | 0.22 | 0.12 | 0.13 | 0.08 |
| 11 | 121301819 | rs17125331 | -1.709 | 0.564 | 0.887 | 0.043 | 0.015 | -0.006 | -0.006 | 0.237 | 0.22 | 0.59 | 0.29 | 0.18 | 0.19 | 0.16 |
| 11 | 121306462 | rs4146874 | -1.719 | 0.517 | 1.118 | -0.01 | 0.017 | -0.001 | -0.001 | 0 | 0.21 | 0.36 | 0.27 | 0.18 | 0.19 | 0.15 |
| 11 | 121308525 | rs668053 | 0.453 | 0.595 | 0.726 | 0.333 | -0.043 | 0.02 | 0.02 | 0 | 0.41 | 0.15 | 0.5 | 0.66 | 0.47 | 0.61 |
| 11 | 121310564 | rs11218292 | -1.339 | 0.855 | 1.293 | 0.381 | 0.012 | -0.008 | -0.008 | 0 | 0.15 | 0.2 | 0.2 | 0.12 | 0.16 | 0.1 |
| 11 | 121321754 | rs4935774 | -2.279 | 0.88 | 0.921 | 0.333 | 0.014 | -0.023 | -0.023 | 0.176 | 0.24 | 0.6 | 0.3 | 0.16 | 0.24 | 0.16 |
| 11 | 121328954 | rs661057 | -0.051 | 0.595 | 0.163 | 0.325 | 0.001 | 0 | 0 | 0.003 | 0.31 | 0.26 | 0.45 | 0.56 | 0.44 | 0.49 |
| 11 | 121329041 | rs3862605 | 1.291 | -0.494 | -0.128 | -0.222 | 0.053 | 0.13 | 0.13 | 1.043 | 0.39 | 0.12 | 0.23 | 0.18 | 0.31 | 0.22 |
| 11 | 121330087 | rs3862606 | 0.796 | -0.366 | 0.025 | -0.304 | 0.04 | 0.097 | 0.097 | 0 | 0.49 | 0.2 | 0.43 | 0.31 | 0.43 | 0.34 |
| 11 | 121335915 | rs645275 | -0.798 | -0.356 | 0.638 | -0.348 | 0 | -0.001 | -0.001 | 0 | 0.45 | 0.56 | 0.49 | 0.56 | 0.48 | 0.53 |
| 11 | 121337993 | rs1784934 | 0.632 | -0.136 | 0.48 | -0.225 | -0.025 | -0.026 | -0.026 | 0 | 0.28 | 0.2 | 0.4 | 0.57 | 0.38 | 0.46 |
| 11 | 121341262 | rs7945931 | -0.652 | -0.424 | 0.666 | -0.168 | 0.019 | 0.064 | 0.064 | 0.903 | 0.08 | 0.25 | 0.06 | 0.12 | 0.1 | 0.15 |
| 11 | 121358361 | rs4631890 | 0.256 | -0.51 | 0.483 | -0.248 | -0.003 | -0.012 | -0.012 | 3.426 | 0.58 | 0.47 | 0.48 | 0.31 | 0.47 | 0.38 |
| 11 | 121359265 | rs676759 | 0.266 | -0.448 | 0.57 | -0.143 | -0.025 | -0.024 | -0.024 | 2.879 | 0.33 | 0.3 | 0.35 | 0.56 | 0.41 | 0.45 |
| 11 | 121365720 | rs17245976 | -0.198 | -0.884 | 0.557 | -0.143 | 0.001 | 0.049 | 0.049 | 0 | 0.09 | 0.01 | 0.05 | 0 | 0.08 | 0.07 |
| 11 | 121367626 | rs12364988 | -0.023 | -0.769 | 0.649 | -0.082 | 0.006 | -0.008 | -0.008 | 0 | 0.61 | 0.49 | 0.51 | 0.31 | 0.5 | 0.42 |
| 11 | 121368163 | rs923892 | -0.505 | -0.749 | 0.694 | -0.078 | 0.001 | 0.001 | 0.001 | 1.107 | 0.65 | 0.61 | 0.61 | 0.45 | 0.54 | 0.55 |
| 11 | 121395790 | rs1784927 | 0.422 | -0.776 | 0.544 | -0.359 | -0.016 | -0.014 | -0.014 | 0 | 0.31 | 0.26 | 0.44 | 0.57 | 0.43 | 0.42 |
| 11 | 121398578 | rs666004 | 0.042 | -1.167 | 0.33 | -0.404 | 0.001 | -0.012 | -0.012 | 0 | 0.4 | 0.43 | 0.54 | 0.75 | 0.48 | 0.51 |
| 11 | 121402385 | rs11218322 | -0.762 | -1.706 | -0.231 | -0.769 | -0.024 | -0.052 | -0.052 | 0 | 0.14 | 0.22 | 0.07 | 0 | 0.09 | 0.07 |
| 11 | 121412837 | rs11601559 | -0.344 | -1.537 | -1.348 | -0.661 | 0.032 | 0.032 | 0.032 | 0 | 0.12 | 0.1 | 0.06 | 0 | 0.09 | 0.07 |
| 11 | 121414476 | rs2276346 | 0.867 | -1.037 | -1.163 | -0.385 | -0.009 | 0.175 | 0.175 | 2.319 | 0.36 | 0.04 | 0.24 | 0.12 | 0.34 | 0.27 |
| 11 | 121415312 | rs10502262 | 1.257 | -1.129 | -1.146 | -0.565 | 0.019 | 0.136 | 0.136 | 0.332 | 0.3 | 0.07 | 0.15 | 0 | 0.26 | 0.17 |
| 11 | 121416187 | rs11218325 | 0.382 | -0.398 | -0.918 | -0.003 | 0.065 | 0.051 | 0.051 | 0 | 0.46 | 0.24 | 0.33 | 0.32 | 0.4 | 0.35 |
| 11 | 121430872 | rs11605969 | -1.384 | -0.212 | -0.248 | -0.339 | -0.002 | -0.053 | -0.053 | 0.204 | 0.13 | 0.21 | 0.12 | 0.01 | 0.16 | 0.15 |
| 11 | 121432647 | rs639648 | 1.084 | -0.354 | -1.282 | -1.084 | -0.05 | 0.25 | 0.25 | 4.085 | 0.46 | 0.05 | 0.35 | 0.15 | 0.58 | 0.27 |
| 11 | 121435966 | rs7124060 | -1.107 | -0.581 | 0.22 | -1.108 | 0.04 | -0.041 | -0.041 | 0 | 0.09 | 0.33 | 0.08 | 0 | 0.1 | 0.04 |
| 11 | 121436270 | rs3781832 | -1.136 | -0.781 | 0.449 | -0.477 | -0.024 | 0.046 | 0.046 | 0 | 0.15 | 0.29 | 0.12 | 0.28 | 0.1 | 0.17 |
| 11 | 121442189 | rs1790213 | 1.197 | -0.939 | 0.527 | -0.57 | 0.019 | 0.039 | 0.039 | 6.035 | 0.49 | 0.1 | 0.38 | 0.35 | 0.61 | 0.33 |
| 11 | 121446059 | rs4420280 | -0.624 | -1.032 | 0.949 | -0.617 | -0.01 | -0.043 | -0.043 | 0 | 0.42 | 0.51 | 0.56 | 0.77 | 0.34 | 0.6 |
| 11 | 121446548 | rs11218347 | -0.252 | -1.024 | 0.998 | -0.595 | -0.007 | -0.026 | -0.026 | 0 | 0.09 | 0.13 | 0.06 | 0 | 0.06 | 0.11 |
| 11 | 121447188 | rs4936637 | 0.715 | -1.08 | 0.921 | -0.628 | -0.012 | -0.107 | -0.107 | 0 | 0.29 | 0.08 | 0.44 | 0.57 | 0.24 | 0.49 |
| 11 | 121448090 | rs2070045 | 0.696 | -1.093 | 0.916 | -0.625 | -0.012 | -0.105 | -0.105 | 0.216 | 0.29 | 0.08 | 0.44 | 0.57 | 0.24 | 0.49 |
| 11 | 121448338 | rs3781836 | -1.61 | -1.022 | 0.936 | -0.642 | 0.007 | 0.121 | 0.121 | 0.049 | 0.13 | 0.35 | 0.13 | 0.2 | 0.15 | 0.13 |
| 11 | 121450221 | rs11218349 | 0.162 | -1.077 | 0.655 | -0.634 | -0.003 | -0.019 | -0.019 | 0.008 | 0.08 | 0.09 | 0.06 | 0 | 0.06 | 0.1 |
| 11 | 121451926 | rs1699103 | 1.458 | -1.02 | 0.537 | -0.634 | -0.038 | 0.232 | 0.232 | 2.766 | 0.45 | 0.1 | 0.34 | 0.15 | 0.56 | 0.26 |
| 11 | 121452651 | rs11218350 | 0.575 | -1.215 | 0.617 | -0.458 | -0.009 | -0.108 | -0.108 | 0 | 0.29 | 0.08 | 0.44 | 0.56 | 0.24 | 0.49 |
| 11 | 121456962 | rs1699102 | 0.44 | -1.066 | 1.202 | -0.407 | -0.003 | -0.037 | -0.037 | 0 | 0.53 | 0.48 | 0.42 | 0.14 | 0.67 | 0.31 |
| 11 | 121463984 | rs1503422 | 0.561 | -0.936 | 0.765 | -0.299 | 0.065 | 0.053 | 0.053 | 2.689 | 0.52 | 0.21 | 0.42 | 0.34 | 0.65 | 0.34 |
| 11 | 121464946 | rs1792113 | 0.642 | -0.939 | 0.727 | -0.3 | 0.075 | 0.061 | 0.061 | 3.149 | 0.51 | 0.16 | 0.42 | 0.34 | 0.65 | 0.34 |
| 11 | 121474025 | rs12287339 | 0.954 | -0.818 | 1.117 | -0.121 | -0.01 | 0.168 | 0.168 | 0 | 0.05 | 0.14 | 0.05 | 0.21 | 0.02 | 0.03 |
| 11 | 121479685 | rs7101373 | -1.116 | -0.912 | 1.563 | -0.206 | 0.012 | -0.017 | -0.017 | 1.199 | 0.07 | 0.33 | 0.06 | 0 | 0.05 | 0.04 |
| 11 | 121483651 | rs1010158 | -0.335 | -0.948 | 1.189 | -0.244 | 0.02 | -0.016 | -0.016 | 0 | 0.61 | 0.73 | 0.5 | 0.41 | 0.7 | 0.41 |
| 11 | 121483912 | rs1699098 | 0.599 | -0.674 | 0.645 | -0.184 | 0.066 | 0.18 | 0.18 | 0.029 | 0.48 | 0.24 | 0.39 | 0.2 | 0.63 | 0.34 |
| 11 | 121489416 | rs1784933 | 1.386 | -0.84 | 0.894 | 0.17 | -0.001 | 0.271 | 0.271 | 0 | 0.89 | 0.55 | 0.83 | 0.7 | 0.96 | 0.9 |
| 11 | 121493001 | rs1614735 | 0.73 | 0.199 | -0.373 | -0.409 | -0.08 | 0.211 | 0.211 | 0.595 | 0.43 | 0.07 | 0.39 | 0.14 | 0.5 | 0.32 |
| 11 | 121497884 | rs10892761 | 0.358 | -0.046 | -0.583 | -0.333 | -0.139 | -0.153 | -0.153 | 2.17 | 0.34 | 0.02 | 0.39 | 0.56 | 0.43 | 0.5 |
| 11 | 121499336 | rs1393914 | -0.372 | -0.212 | -0.118 | -0.462 | 0.009 | 0.15 | 0.15 | 0 | 0.46 | 0.8 | 0.55 | 0.74 | 0.44 | 0.56 |
| 11 | 121501755 | rs1133174 | -0.355 | -0.234 | -0.083 | -0.463 | 0.009 | 0.146 | 0.146 | 0.01 | 0.46 | 0.8 | 0.55 | 0.74 | 0.44 | 0.56 |
| 11 | 121514977 | rs4935777 | -0.39 | -0.286 | -0.015 | -0.558 | 0.01 | 0.164 | 0.164 | 0.012 | 0.46 | 0.83 | 0.56 | 0.75 | 0.44 | 0.56 |
| 11 | 121521844 | rs626747 | 0.284 | -0.017 | 0.973 | -0.41 | -0.053 | -0.03 | -0.03 | 0 | 0.42 | 0.14 | 0.45 | 0.59 | 0.48 | 0.53 |
| 11 | 121525667 | rs676165 | -0.156 | 0.184 | 1.209 | -0.498 | -0.024 | 0.124 | 0.124 | 0 | 0.53 | 0.75 | 0.64 | 0.77 | 0.51 | 0.58 |

| | | | | | | | | | | | | | | | | |
|---|---|---|---|---|---|---|---|---|---|---|---|---|---|---|---|---|
| 11 | 121533001 | rs1783832 | -0.131 | 0.154 | 1.148 | -0.466 | -0.034 | 0.11 | 0.11 | 0 | 0.57 | 0.79 | 0.67 | 0.75 | 0.56 | 0.64 |
| 11 | 121533827 | rs663175 | 0.09 | 0.136 | 1.48 | -0.401 | -0.001 | 0 | 0 | 2.781 | 0.54 | 0.45 | 0.55 | 0.57 | 0.56 | 0.62 |
| 11 | 121534938 | rs647905 | 0.166 | 0.152 | 1.134 | -0.393 | -0.002 | -0.002 | -0.002 | 0 | 0.53 | 0.36 | 0.55 | 0.57 | 0.56 | 0.62 |
| 11 | 121544285 | rs682011 | -0.122 | 0.096 | 0.206 | -1.319 | -0.017 | 0.119 | 0.119 | 0.584 | 0.57 | 0.77 | 0.63 | 0.81 | 0.56 | 0.68 |
| 11 | 121562716 | rs11218381 | 0.6 | 0.107 | -0.067 | -1.075 | -0.012 | 0.172 | 0.172 | 0 | 0.49 | 0.07 | 0.59 | 0.78 | 0.57 | 0.69 |
| 11 | 121566963 | rs10790450 | 0.582 | 0.151 | -0.066 | -1.022 | -0.008 | 0.169 | 0.169 | 0 | 0.5 | 0.07 | 0.59 | 0.78 | 0.57 | 0.69 |
| 11 | 121567897 | rs10502260 | 0.277 | -0.064 | 0.369 | -1.323 | 0.005 | 0.021 | 0.021 | 0.447 | 0.3 | 0.21 | 0.26 | 0.14 | 0.3 | 0.18 |
| 11 | 121568179 | rs2887763 | 0.493 | 0.164 | 0.914 | -1.845 | -0.005 | 0.13 | 0.13 | 0.064 | 0.52 | 0.68 | 0.64 | 0.86 | 0.53 | 0.69 |
| 11 | 121568262 | rs10892771 | 0.923 | 0.135 | 0.885 | -1.818 | -0.059 | 0.149 | 0.149 | 0.04 | 0.45 | 0.11 | 0.59 | 0.81 | 0.52 | 0.67 |
| 11 | 121570520 | rs10790451 | 1.602 | -0.261 | 0.978 | -1.872 | -0.052 | 0.055 | 0.055 | 1.973 | 0.67 | 0.05 | 0.68 | 0.83 | 0.7 | 0.76 |
| 11 | 121575095 | rs1074086 | 1.487 | -0.522 | 1.314 | -1.653 | 0.014 | 0.048 | 0.048 | 0.179 | 0.68 | 0.13 | 0.69 | 0.8 | 0.71 | 0.73 |
| 11 | 121578945 | rs662333 | 0.727 | -0.521 | 1.669 | -1.725 | 0.001 | -0.002 | -0.002 | 0 | 0.77 | 0.75 | 0.77 | 0.94 | 0.75 | 0.77 |
| 11 | 121582420 | rs1699101 | 0.694 | -0.577 | 1.705 | -1.711 | 0.002 | 0 | 0 | 2.508 | 0.78 | 0.76 | 0.77 | 0.94 | 0.76 | 0.78 |
| 11 | 121588031 | rs664424 | -0.303 | -0.628 | 1.738 | -1.698 | -0.022 | 0.032 | 0.032 | 0 | 0.88 | 0.98 | 0.82 | 0.94 | 0.84 | 0.85 |
| 11 | 121599680 | rs11600686 | 0.783 | -0.519 | 1.717 | -1.598 | 0.006 | 0.151 | 0.151 | 1.574 | 0.23 | 0.02 | 0.11 | 0 | 0.19 | 0.07 |
| 11 | 121606715 | rs4457756 | 1.142 | 0.174 | 3.261 | -0.287 | -0.011 | -0.028 | -0.028 | 0 | 0.73 | 0.35 | 0.71 | 0.87 | 0.76 | 0.78 |
| 11 | 121616060 | rs1944677 | 0.861 | 0.175 | 3.479 | -0.292 | -0.006 | -0.043 | -0.043 | 0.756 | 0.76 | 0.66 | 0.74 | 0.92 | 0.76 | 0.78 |
| 11 | 121621394 | rs7949483 | 0.938 | 0.176 | 3.551 | -0.291 | -0.015 | -0.098 | -0.098 | 0.009 | 0.75 | 0.56 | 0.73 | 0.92 | 0.76 | 0.78 |
| 11 | 121627609 | rs7115487 | 0.836 | 0.176 | 3.584 | -0.292 | -0.009 | -0.07 | -0.07 | 0 | 0.76 | 0.67 | 0.74 | 0.92 | 0.76 | 0.78 |
| 11 | 121629007 | rs949078 | 1.601 | 0.173 | 3.435 | -0.282 | 0.019 | 0.039 | 0.039 | 0.315 | 0.67 | 0.16 | 0.69 | 0.79 | 0.72 | 0.74 |
| 11 | 121629132 | rs10736512 | 0.896 | 0.178 | 3.437 | -0.379 | -0.013 | -0.092 | -0.092 | 0 | 0.75 | 0.59 | 0.73 | 0.92 | 0.76 | 0.78 |
| 11 | 121632159 | rs4936647 | 0.896 | 0.178 | 3.437 | -0.379 | -0.013 | -0.092 | -0.092 | 0 | 0.75 | 0.58 | 0.73 | 0.92 | 0.76 | 0.78 |
| 11 | 121634608 | rs4936651 | 0.846 | 0.179 | 3.133 | -0.376 | -0.009 | -0.061 | -0.061 | 0.016 | 0.76 | 0.63 | 0.74 | 0.92 | 0.76 | 0.78 |
| 11 | 121635211 | rs7937934 | 1.539 | 0.179 | 2.64 | -0.336 | 0.017 | 0.036 | 0.036 | 0.238 | 0.68 | 0.15 | 0.69 | 0.79 | 0.72 | 0.74 |
| 11 | 121638516 | rs10790459 | 1.255 | 0.226 | 2.658 | -0.298 | -0.003 | -0.025 | -0.025 | 11.003 | 0.69 | 0.39 | 0.69 | 0.84 | 0.7 | 0.73 |
| 11 | 121643646 | rs1944679 | 0.911 | 0.201 | 1.961 | -0.33 | -0.008 | -0.072 | -0.072 | 0.76 | 0.74 | 0.61 | 0.72 | 0.91 | 0.74 | 0.77 |
| 11 | 121648112 | rs4936653 | 0.772 | 0.157 | 1.964 | -0.323 | 0.001 | -0.005 | -0.005 | 0 | 0.76 | 0.78 | 0.74 | 0.91 | 0.74 | 0.77 |
| 11 | 121648442 | rs4936654 | 0.883 | 0.171 | 1.794 | -0.317 | -0.008 | -0.067 | -0.067 | 0 | 0.74 | 0.62 | 0.72 | 0.91 | 0.74 | 0.77 |
| 11 | 121649479 | rs10892774 | 1.216 | 0.111 | 1.758 | -0.308 | 0.002 | 0.002 | 0.002 | 0 | 0.72 | 0.29 | 0.7 | 0.86 | 0.73 | 0.77 |
| 11 | 121658959 | rs949076 | 0.707 | 0.609 | -0.21 | -0.326 | 0.001 | 0.025 | 0.025 | 0 | 0.77 | 0.84 | 0.75 | 0.91 | 0.74 | 0.77 |
| 11 | 121663693 | rs7119960 | -1.444 | 0.618 | 0.2 | 0.372 | 0.018 | 0.093 | 0.093 | 0.206 | 0.05 | 0.16 | 0.02 | 0.12 | 0.03 | 0.03 |
| 11 | 121664355 | rs10892775 | -1.682 | 0.535 | 0.695 | -0.312 | -0.002 | -0.044 | -0.044 | 1.531 | 0.2 | 0.62 | 0.29 | 0.12 | 0.2 | 0.19 |
| 11 | 121665295 | rs12417533 | -1.618 | -0.059 | -0.484 | -1.231 | 0.062 | 0.25 | 0.25 | 0.625 | 0.4 | 0.94 | 0.39 | 0.09 | 0.24 | 0.33 |
| 11 | 121673147 | rs881754 | -1.537 | 0.373 | -0.032 | -1.274 | 0.034 | -0.035 | -0.035 | 3.394 | 0.07 | 0.14 | 0.05 | 0 | 0.11 | 0.03 |
| 11 | 121678731 | rs10502257 | -0.686 | 0.107 | -0.054 | -1.4 | 0.011 | 0.005 | 0.005 | 0.014 | 0.07 | 0.03 | 0.02 | 0.01 | 0.03 | 0.01 |
| 11 | 121679637 | rs10502256 | -1.715 | 0.208 | 0.226 | -1.445 | 0 | -0.013 | -0.013 | 0 | 0.12 | 0.12 | 0.07 | 0.03 | 0.08 | 0.05 |
| 11 | 121683781 | rs528956 | -2.702 | 0.031 | 0.084 | -1.548 | -0.025 | 0.226 | 0.226 | 0.292 | 0.42 | 0.94 | 0.47 | 0.14 | 0.36 | 0.38 |
| 11 | 121689825 | rs545862 | -0.68 | 0.059 | 1.083 | -1.619 | -0.01 | -0.055 | -0.055 | 0.962 | 0.15 | 0.19 | 0.08 | 0 | 0.09 | 0.14 |
| 11 | 121691425 | rs474441 | -2.991 | 0.087 | 1.903 | -1.63 | 0.001 | 0.019 | 0.019 | 0 | 0.4 | 0.71 | 0.33 | 0.07 | 0.37 | 0.37 |
| 11 | 121702440 | rs7105411 | -1.578 | 0.105 | 2.04 | -1.631 | 0.028 | -0.038 | -0.038 | 0.017 | 0.07 | 0.17 | 0.05 | 0 | 0.12 | 0.03 |
| 11 | 121707866 | rs472074 | -2.362 | 0.086 | 2.188 | -1.643 | 0.015 | -0.143 | -0.143 | 1.372 | 0.26 | 0.48 | 0.25 | 0.04 | 0.28 | 0.31 |
| 11 | 121736671 | rs11218429 | -3.079 | 0.081 | 1.451 | -1.645 | 0.017 | -0.089 | -0.089 | 0.003 | 0.24 | 0.41 | 0.25 | 0.07 | 0.28 | 0.24 |
| 11 | 121740317 | rs7935547 | -2.479 | 0.083 | 1.425 | -1.645 | 0.004 | -0.057 | -0.057 | 0 | 0.19 | 0.27 | 0.12 | 0.03 | 0.19 | 0.08 |
| 11 | 121740459 | rs1219388 | 0.218 | -0.169 | 1.165 | -1.532 | 0.012 | 0.015 | 0.015 | 0 | 0.87 | 0.94 | 0.93 | 1 | 0.92 | 0.88 |
| 11 | 121740944 | rs17125870 | -1.708 | -0.032 | 1.006 | -1.423 | 0.026 | -0.017 | -0.017 | 0 | 0.06 | 0.18 | 0.13 | 0.04 | 0.09 | 0.16 |
| 11 | 121762127 | rs2370643 | 2.485 | 0.061 | 0.257 | -1.139 | -0.019 | 0.135 | 0.135 | 0.649 | 0.57 | 0.04 | 0.5 | 0.82 | 0.63 | 0.66 |
| 11 | 121762396 | rs1219395 | -1.06 | -0.092 | 0.353 | -0.994 | -0.023 | -0.025 | -0.025 | 0.202 | 0.08 | 0.02 | 0.08 | 0.07 | 0.08 | 0.05 |
| 11 | 121763908 | rs17337418 | -0.345 | 0.087 | 0.546 | -1.009 | -0.013 | 0.032 | 0.032 | 0 | 0.06 | 0.01 | 0.02 | 0 | 0.05 | 0.01 |
| 11 | 121766888 | rs1219391 | 1.012 | -0.11 | 1.021 | -0.937 | -0.048 | -0.031 | -0.031 | 0 | 0.82 | 0.47 | 0.78 | 0.91 | 0.92 | 0.87 |
| 11 | 121769119 | rs1219386 | 1.729 | -0.103 | 0.855 | -1.229 | 0.003 | -0.049 | -0.049 | 0 | 0.7 | 0.35 | 0.74 | 1 | 0.8 | 0.76 |
| 11 | 121769481 | rs1219385 | -1.633 | -0.696 | 0.848 | -1.402 | -0.032 | -0.062 | -0.062 | 0.047 | 0.27 | 0.43 | 0.17 | 0.1 | 0.2 | 0.16 |
| 11 | 121771070 | rs11218444 | 0.642 | -0.426 | -0.673 | -1.538 | -0.11 | 0.038 | 0.038 | 0 | 0.31 | 0.09 | 0.31 | 0.27 | 0.42 | 0.28 |
| 11 | 121772983 | rs1219423 | 0.295 | -0.918 | -0.031 | -0.917 | -0.084 | 0.034 | 0.034 | 1.378 | 0.79 | 0.5 | 0.79 | 0.76 | 0.91 | 0.94 |
| 11 | 121774705 | rs3862607 | 0.854 | -0.062 | -0.913 | -1.06 | -0.064 | 0.018 | 0.018 | 0 | 0.28 | 0.09 | 0.27 | 0.26 | 0.35 | 0.23 |
| 11 | 121779467 | rs1078402 | 0.901 | -0.541 | -1.23 | -1.522 | 0.017 | 0.051 | 0.051 | 0 | 0.58 | 0.31 | 0.61 | 0.76 | 0.62 | 0.74 |
| 11 | 121779664 | rs747915 | 1.154 | -0.464 | -1.46 | -1.493 | -0.153 | 0.018 | 0.018 | 0 | 0.3 | 0.03 | 0.31 | 0.27 | 0.43 | 0.28 |
| 11 | 121792132 | rs3862609 | 0.883 | -0.545 | -1.351 | -1.355 | 0.026 | -0.028 | -0.028 | 3.769 | 0.05 | 0 | 0.11 | 0.15 | 0.03 | 0.03 |
| 11 | 121797897 | rs1219414 | -0.898 | -0.808 | -0.816 | -1.609 | 0.002 | -0.046 | -0.046 | 0.855 | 0.34 | 0.37 | 0.23 | 0.09 | 0.34 | 0.22 |
| 11 | 121803041 | rs10892786 | 1.236 | -0.795 | -1.145 | -0.896 | 0.014 | -0.079 | -0.079 | 0.073 | 0.23 | 0.07 | 0.28 | 0.48 | 0.2 | 0.45 |
| 11 | 121804892 | rs1219446 | 0.506 | -0.821 | -1.467 | -1.32 | -0.043 | 0.075 | 0.075 | 0 | 0.28 | 0.18 | 0.24 | 0.09 | 0.43 | 0.12 |
| 11 | 121810093 | rs1219457 | -0.163 | -0.83 | -1.29 | -1.353 | -0.034 | -0.071 | -0.071 | 0 | 0.88 | 0.64 | 0.93 | 0.99 | 0.97 | 0.88 |
| 11 | 121810231 | rs7934592 | 0.766 | -0.888 | -1.324 | -1.346 | 0.021 | -0.028 | -0.028 | 5.509 | 0.33 | 0.37 | 0.3 | 0.3 | 0.31 | 0.27 |
| 11 | 121813368 | rs11605389 | 1.188 | -0.923 | -1.327 | -1.321 | 0.021 | -0.006 | -0.006 | 0 | 0.3 | 0.21 | 0.4 | 0.61 | 0.23 | 0.49 |
| 11 | 121815491 | rs1448133 | -0.59 | -0.509 | -1.607 | -1.065 | -0.026 | -0.013 | -0.013 | 0 | 0.58 | 0.4 | 0.53 | 0.38 | 0.74 | 0.39 |
| 11 | 121816144 | rs1789751 | 0.703 | -0.399 | -1.633 | -1.083 | -0.122 | 0.088 | 0.088 | 0 | 0.26 | 0.02 | 0.22 | 0.08 | 0.42 | 0.12 |
| 11 | 121821056 | rs12421071 | 0.939 | -0.453 | -1.126 | -0.755 | 0.021 | -0.031 | -0.031 | 0 | 0.05 | 0.01 | 0.12 | 0.12 | 0.04 | 0.03 |
| 11 | 121822112 | rs4016823 | -0.006 | -0.083 | -0.915 | -0.933 | 0.018 | -0.031 | -0.031 | 0 | 0.33 | 0.37 | 0.29 | 0.3 | 0.31 | 0.27 |
| 11 | 121827325 | rs7934556 | -0.17 | -0.74 | -1.812 | -0.513 | 0.125 | 0.031 | 0.031 | 0 | 0.51 | 0.85 | 0.59 | 0.58 | 0.34 | 0.43 |
| 11 | 121828759 | rs17125985 | -0.21 | -0.682 | -1.633 | -0.744 | 0.005 | -0.007 | -0.007 | 0 | 0.1 | 0.14 | 0.05 | 0 | 0.03 | 0.07 |

| | | | | | | | | | | | | | | | | |
|---|---|---|---|---|---|---|---|---|---|---|---|---|---|---|---|---|
| 11 | 121830239 | rs1231186 | 0.524 | 0.299 | -1.917 | -0.764 | -0.07 | 0.096 | 0.096 | 0 | 0.28 | 0.09 | 0.21 | 0.09 | 0.39 | 0.19 |
| 11 | 121831524 | rs3862611 | 0.59 | -0.495 | -0.993 | 0.291 | 0.005 | -0.017 | -0.017 | 0 | 0.23 | 0.16 | 0.31 | 0.47 | 0.23 | 0.31 |
| 11 | 121833245 | rs1789742 | 0.816 | -0.502 | -0.172 | -0.237 | -0.022 | -0.084 | -0.084 | 2.628 | 0.87 | 0.58 | 0.93 | 1 | 0.97 | 0.92 |
| 11 | 121835389 | rs1791977 | -0.03 | -0.503 | -1.132 | -0.231 | 0.036 | 0.03 | 0.03 | 4.649 | 0.48 | 0.17 | 0.6 | 0.64 | 0.6 | 0.64 |
| 11 | 121840829 | rs7101566 | 0.983 | -0.904 | -1.129 | -0.613 | 0.001 | -0.008 | -0.008 | 0 | 0.09 | 0.07 | 0.05 | 0 | 0.07 | 0.04 |
| 11 | 121842526 | rs12285155 | 0.864 | -1.496 | -1.149 | -0.93 | -0.009 | 0.053 | 0.053 | 0.425 | 0.21 | 0.09 | 0.1 | 0.07 | 0.22 | 0.13 |
| 11 | 121856320 | rs2852826 | -0.993 | -1.559 | -0.223 | -0.81 | -0.033 | 0.077 | 0.077 | 0 | 0.84 | 0.7 | 0.81 | 0.73 | 0.91 | 0.85 |
| 11 | 121856752 | rs2850773 | 1.563 | -1.709 | -0.893 | -1.132 | -0.096 | -0.058 | -0.058 | 0 | 0.3 | 0.14 | 0.54 | 0.63 | 0.5 | 0.52 |
| 11 | 121862691 | rs11218490 | 0.413 | -1.555 | -0.156 | -0.911 | 0.001 | -0.087 | -0.087 | 0 | 0.27 | 0.41 | 0.16 | 0.07 | 0.25 | 0.18 |
| 11 | 121867700 | rs2852836 | 0.368 | -1.378 | -0.551 | -0.785 | 0.041 | 0.053 | 0.053 | 3.693 | 0.49 | 0.22 | 0.6 | 0.65 | 0.6 | 0.64 |
| 11 | 121879938 | rs2852835 | -0.981 | -1 | -0.066 | -0.416 | -0.009 | 0.034 | 0.034 | 0 | 0.86 | 0.76 | 0.77 | 0.74 | 0.9 | 0.89 |
| 11 | 121893245 | rs2850791 | 0.144 | -1.153 | 0.074 | -0.523 | -0.012 | 0.004 | 0.004 | 0.166 | 0.63 | 0.46 | 0.65 | 0.63 | 0.69 | 0.72 |
| 11 | 121893382 | rs17126029 | 1.113 | -1.174 | -0.513 | -0.236 | 0.026 | -0.043 | -0.043 | 0.271 | 0.12 | 0.05 | 0.18 | 0.29 | 0.08 | 0.07 |
| 11 | 121897397 | rs7111523 | 0.547 | -0.428 | -0.252 | -0.271 | -0.016 | 0.048 | 0.048 | 0 | 0.2 | 0.09 | 0.1 | 0.08 | 0.2 | 0.19 |
| 11 | 121897762 | rs17126035 | 0.664 | -0.668 | -0.452 | -0.489 | 0.054 | -0.002 | -0.002 | 0.019 | 0.37 | 0.16 | 0.35 | 0.31 | 0.25 | 0.16 |
| 11 | 121898223 | rs12417816 | 0.664 | -0.668 | -0.452 | -0.489 | 0.054 | -0.002 | -0.002 | 1.235 | 0.37 | 0.16 | 0.35 | 0.31 | 0.25 | 0.16 |
| 11 | 121900440 | rs4121970 | -0.065 | -0.134 | -0.295 | -0.259 | -0.007 | -0.01 | -0.01 | 0 | 0.41 | 0.27 | 0.4 | 0.42 | 0.54 | 0.56 |
| 11 | 121900836 | rs12794316 | 1.428 | -0.233 | -0.333 | -0.309 | 0.051 | -0.013 | -0.013 | 0 | 0.36 | 0.1 | 0.32 | 0.31 | 0.22 | 0.16 |
| 11 | 121902943 | rs2850779 | 1.192 | -0.166 | 0.055 | -0.33 | -0.003 | 0.019 | 0.019 | 0.041 | 0.12 | 0.19 | 0.22 | 0.24 | 0.16 | 0.18 |
| 11 | 121913925 | rs7949079 | 0.744 | -0.009 | -0.203 | -0.404 | -0.007 | 0.007 | 0.007 | 0 | 0.19 | 0.13 | 0.12 | 0.08 | 0.24 | 0.19 |
| 11 | 121914558 | rs11218508 | 1.068 | -0.01 | -0.375 | -0.4 | -0.037 | 0.055 | 0.055 | 0 | 0.18 | 0.07 | 0.11 | 0.08 | 0.21 | 0.2 |
| 11 | 121915980 | rs10790482 | 0.941 | -0.125 | -0.407 | -0.266 | -0.058 | -0.055 | -0.055 | 0.277 | 0.21 | 0.09 | 0.27 | 0.32 | 0.34 | 0.34 |
| 11 | 121917978 | rs10790483 | 0.925 | -0.073 | -0.417 | -0.255 | -0.051 | -0.051 | -0.051 | 0 | 0.22 | 0.1 | 0.28 | 0.33 | 0.34 | 0.34 |
| 11 | 121929227 | rs4367955 | 0.078 | -0.083 | -0.3 | -0.371 | 0.004 | -0.016 | -0.016 | 0 | 0.17 | 0.13 | 0.22 | 0.31 | 0.19 | 0.3 |
| 11 | 121935148 | rs585342 | 0.736 | -0.116 | -0.243 | -0.145 | 0.061 | 0.019 | 0.019 | 0 | 0.49 | 0.26 | 0.53 | 0.55 | 0.38 | 0.35 |
| 11 | 121937932 | rs2155093 | 0.502 | 0.177 | 0.148 | 0.046 | -0.009 | 0.008 | 0.008 | 0 | 0.17 | 0.12 | 0.13 | 0.02 | 0.23 | 0.2 |
| 11 | 121940411 | rs549798 | 0.745 | 0.481 | 0.068 | 0.085 | -0.013 | 0.001 | 0.001 | 0.286 | 0.56 | 0.27 | 0.58 | 0.55 | 0.5 | 0.43 |
| 11 | 121942599 | rs583564 | 1.055 | 0.443 | 0.042 | 0.077 | 0.047 | 0.065 | 0.065 | 1.084 | 0.5 | 0.26 | 0.43 | 0.3 | 0.39 | 0.3 |
| 11 | 121944684 | rs10502255 | 0.647 | 0.617 | 0.027 | 0.192 | -0.021 | -0.037 | -0.037 | 0.604 | 0.2 | 0.1 | 0.24 | 0.37 | 0.25 | 0.32 |
| 11 | 121948754 | rs10502254 | -0.595 | 0.706 | 0.163 | 0.295 | -0.009 | -0.005 | -0.005 | 1.937 | 0.25 | 0.17 | 0.3 | 0.33 | 0.36 | 0.37 |
| 11 | 121954657 | rs7121446 | 0.048 | 0.852 | -0.283 | 0.138 | 0.032 | -0.096 | -0.096 | 0.791 | 0.21 | 0.39 | 0.16 | 0.06 | 0.24 | 0.25 |
| 11 | 121964569 | rs664409 | 0.086 | 0.635 | -0.821 | 0.615 | 0.018 | 0.03 | 0.03 | 1.498 | 0.53 | 0.34 | 0.44 | 0.3 | 0.4 | 0.3 |
| 11 | 121970746 | rs2081443 | -1.319 | 0.93 | -0.619 | 1.054 | 0.006 | 0.015 | 0.015 | 0 | 0.13 | 0.14 | 0.2 | 0.29 | 0.17 | 0.23 |
| 11 | 121973541 | rs512932 | -0.969 | 0.347 | -1.007 | 0.82 | 0.005 | -0.024 | -0.024 | 0 | 0.2 | 0.12 | 0.14 | 0.26 | 0.2 | 0.29 |
| 11 | 121977902 | rs4121975 | 0.887 | -0.177 | -0.599 | 0.56 | 0.009 | 0.005 | 0.005 | 0 | 0.51 | 0.43 | 0.5 | 0.61 | 0.61 | 0.55 |
| 11 | 121983872 | rs568643 | -0.939 | -0.505 | -0.187 | 0.611 | 0.007 | -0.025 | -0.025 | 0 | 0.17 | 0.09 | 0.12 | 0.26 | 0.18 | 0.33 |
| 11 | 121985066 | rs662288 | 0.421 | -0.528 | -0.155 | 0.693 | 0.022 | 0.135 | 0.135 | 0 | 0.94 | 0.8 | 0.88 | 0.78 | 0.89 | 0.86 |
| 11 | 121985367 | rs630852 | -0.597 | -0.495 | -0.457 | 0.687 | 0.007 | -0.002 | -0.002 | 2.449 | 0.11 | 0.07 | 0.05 | 0.12 | 0.07 | 0.14 |
| 11 | 121987854 | rs11606553 | 0.538 | -0.74 | -0.229 | 0.477 | 0.005 | 0.032 | 0.032 | 0 | 0.07 | 0.02 | 0.02 | 0 | 0.04 | 0.03 |
| 11 | 121990991 | rs7119477 | -0.925 | -0.941 | -0.262 | 0.369 | 0.017 | -0.046 | -0.046 | 0.166 | 0.26 | 0.15 | 0.41 | 0.33 | 0.25 | 0.28 |
| 11 | 121993571 | rs17246402 | 0.593 | -0.969 | -0.317 | 0.293 | 0.006 | 0.022 | 0.022 | 0.549 | 0.16 | 0.01 | 0.29 | 0.09 | 0.1 | 0.09 |
| 11 | 121994700 | rs492085 | -0.889 | -0.964 | -0.123 | -0.153 | -0.003 | -0.033 | -0.033 | 0 | 0.88 | 0.88 | 0.92 | 1 | 0.9 | 0.87 |
| 11 | 121996261 | rs687928 | 1.607 | -0.753 | -0.009 | 0.105 | 0.003 | 0 | 0 | 0.017 | 0.44 | 0.42 | 0.45 | 0.56 | 0.58 | 0.47 |
| 11 | 122002604 | rs678586 | 1.581 | -0.532 | -0.262 | 0.168 | -0.006 | 0.002 | 0.002 | 0 | 0.45 | 0.48 | 0.46 | 0.53 | 0.58 | 0.41 |
| 11 | 122013169 | rs531897 | 1.066 | -0.638 | -0.35 | 0.121 | 0 | 0 | 0 | 0.425 | 0.62 | 0.65 | 0.75 | 0.66 | 0.68 | 0.49 |
| 11 | 122014110 | rs579682 | 0.537 | -0.648 | -0.421 | 0.137 | 0.024 | 0.026 | 0.026 | 0.022 | 0.68 | 0.8 | 0.81 | 0.75 | 0.72 | 0.55 |
| 11 | 122019261 | rs12421791 | 0.721 | -0.57 | -0.627 | 0.196 | 0 | -0.015 | -0.015 | 0 | 0.16 | 0.01 | 0.29 | 0.13 | 0.1 | 0.08 |
| 11 | 122020328 | rs537989 | 0.359 | -0.509 | -0.872 | 0.219 | -0.002 | -0.011 | -0.011 | 0 | 0.91 | 0.92 | 0.88 | 0.78 | 0.85 | 0.81 |
| 11 | 122021283 | rs4936674 | 0.546 | -0.477 | -0.873 | 0.171 | -0.004 | -0.011 | -0.011 | 0.003 | 0.16 | 0.01 | 0.29 | 0.13 | 0.1 | 0.08 |
| 11 | 122023511 | rs560279 | 0.928 | 0.127 | -0.346 | 0.557 | -0.007 | 0.001 | 0.001 | 0 | 0.62 | 0.56 | 0.76 | 0.6 | 0.7 | 0.57 |
| 11 | 122025745 | rs17126230 | -0.556 | 0.709 | -0.517 | 0.756 | 0.012 | 0.006 | 0.006 | 0 | 0.07 | 0.12 | 0.09 | 0.09 | 0.08 | 0.08 |
| 11 | 122030190 | rs577948 | -0.661 | 1.145 | -1.218 | 0.852 | -0.055 | -0.085 | -0.085 | 0 | 0.31 | 0.17 | 0.37 | 0.52 | 0.4 | 0.42 |
| 11 | 122031908 | rs12273567 | 0.899 | 0.657 | -1.826 | 0.597 | 0.004 | -0.073 | -0.073 | 0 | 0.21 | 0.09 | 0.28 | 0.37 | 0.18 | 0.23 |
| 11 | 122032548 | rs12805319 | 0.899 | 0.657 | -1.828 | 0.597 | 0.003 | -0.076 | -0.076 | 0 | 0.21 | 0.08 | 0.19 | 0.36 | 0.18 | 0.22 |
| 11 | 122034000 | rs11604461 | -0.138 | 0.732 | -1.927 | 1.061 | -0.028 | -0.056 | -0.056 | 0.038 | 0.1 | 0.01 | 0.1 | 0.23 | 0.13 | 0.15 |
| 11 | 122034143 | rs592948 | 0.61 | 0.583 | -1.078 | 0.581 | -0.005 | -0.01 | -0.01 | 0.297 | 0.61 | 0.35 | 0.59 | 0.49 | 0.56 | 0.59 |
| 11 | 122047290 | rs515449 | -1.221 | 0.05 | -0.417 | 0.363 | 0.002 | 0.004 | 0.004 | 0.003 | 0.45 | 0.71 | 0.61 | 0.62 | 0.46 | 0.48 |
| 11 | 122050913 | rs488062 | -0.052 | -0.019 | -0.353 | 0.403 | -0.03 | -0.076 | -0.076 | 0 | 0.29 | 0.18 | 0.25 | 0.49 | 0.34 | 0.39 |
| 11 | 122053060 | rs671134 | 0.996 | -0.031 | -0.492 | 0.392 | -0.005 | -0.027 | -0.027 | 0.356 | 0.08 | 0.01 | 0.12 | 0.11 | 0.07 | 0.06 |
| 11 | 122056327 | rs6589913 | 0.807 | 0.004 | -0.185 | 0.751 | 0 | -0.012 | -0.012 | 0.437 | 0.57 | 0.39 | 0.38 | 0.2 | 0.54 | 0.51 |
| 11 | 122063583 | rs549180 | 0.877 | -0.012 | -0.65 | 1.254 | -0.007 | -0.032 | -0.032 | 0 | 0.07 | 0 | 0.13 | 0.28 | 0.07 | 0.06 |
| 11 | 122080117 | rs12793370 | 1.198 | -0.059 | -0.815 | 0.667 | -0.016 | -0.013 | -0.013 | 0 | 0.22 | 0.08 | 0.27 | 0.18 | 0.24 | 0.23 |
| 11 | 122083851 | rs17337641 | 0.76 | 0.067 | -0.665 | -0.052 | -0.007 | -0.076 | -0.076 | 0.443 | 0.17 | 0.24 | 0.08 | 0.01 | 0.15 | 0.11 |
| 11 | 122084894 | rs11218552 | 0.325 | 0.302 | -1.277 | 0.493 | -0.032 | 0.065 | 0.065 | 0 | 0.33 | 0.19 | 0.2 | 0.2 | 0.34 | 0.32 |
| 11 | 122085135 | rs11218553 | -0.948 | -0.035 | -0.422 | 1.373 | -0.001 | -0.009 | -0.009 | 0.01 | 0.17 | 0.16 | 0.24 | 0.33 | 0.19 | 0.26 |
| 11 | 122091502 | rs3901586 | -0.756 | 0.032 | -0.517 | 1.577 | -0.026 | -0.048 | -0.048 | 0 | 0.08 | 0.01 | 0.17 | 0.25 | 0.07 | 0.19 |
| 11 | 122093090 | rs3867239 | 0.73 | -0.175 | -1.023 | 1.156 | -0.051 | 0.051 | 0.051 | 0.486 | 0.36 | 0.12 | 0.34 | 0.24 | 0.38 | 0.4 |
| 11 | 122099839 | rs11218557 | -0.079 | -0.721 | -0.122 | 1.018 | -0.001 | -0.001 | -0.001 | 0 | 0.42 | 0.38 | 0.44 | 0.43 | 0.46 | 0.36 |
| 11 | 122101556 | rs640786 | -0.179 | -0.877 | 0.576 | 1.048 | -0.019 | -0.034 | -0.034 | 0 | 0.93 | 0.73 | 0.97 | 1 | 0.98 | 0.95 |

| | | | | | | | | | | | | | | | | |
|---|---|---|---|---|---|---|---|---|---|---|---|---|---|---|---|---|
| 11 | 122103535 | rs11606243 | 0.579 | -0.982 | 0.882 | 0.955 | 0.004 | -0.026 | -0.026 | 0 | 0.06 | 0.18 | 0.06 | 0 | 0.03 | 0 |
| 11 | 122104483 | rs633997 | 0.027 | -1.02 | 0.997 | 1.1 | -0.003 | 0.043 | 0.043 | 0.75 | 0.92 | 0.64 | 0.95 | 0.97 | 0.98 | 0.94 |
| 11 | 122109788 | rs4936681 | -0.78 | -0.984 | 0.947 | 1.051 | -0.014 | -0.027 | -0.027 | 0 | 0.94 | 0.8 | 0.98 | 1 | 0.98 | 0.95 |
| 11 | 122110253 | rs11603046 | -1.338 | -1.074 | 0.888 | 1.149 | -0.023 | 0.109 | 0.109 | 0 | 0.19 | 0.4 | 0.28 | 0.38 | 0.12 | 0.3 |
| 11 | 122119637 | rs568417 | -0.239 | -1.081 | 0.605 | 1.1 | -0.015 | 0.067 | 0.067 | 0 | 0.92 | 0.79 | 0.87 | 0.83 | 0.97 | 0.85 |
| 11 | 122120142 | rs531743 | 0.087 | -1.107 | 0.603 | 1.088 | -0.019 | 0.068 | 0.068 | 0 | 0.91 | 0.78 | 0.86 | 0.81 | 0.97 | 0.85 |
| 11 | 122123313 | rs10892821 | 0.966 | -1.133 | 0.611 | 0.973 | -0.015 | 0.012 | 0.012 | 0.21 | 0.39 | 0.23 | 0.38 | 0.3 | 0.39 | 0.49 |
| 11 | 122125728 | rs649085 | -0.968 | -1.183 | 0.521 | 0.738 | -0.006 | 0.022 | 0.022 | 0 | 0.69 | 0.77 | 0.71 | 0.82 | 0.63 | 0.73 |
| 11 | 122132835 | rs17337676 | 1.26 | -1.032 | 0.247 | 0.77 | -0.038 | 0.026 | 0.026 | 0.272 | 0.37 | 0.16 | 0.36 | 0.27 | 0.39 | 0.46 |
| 11 | 122135107 | rs577924 | 0.359 | -1.024 | 0.347 | 0.687 | 0.022 | 0.019 | 0.019 | 0.047 | 0.5 | 0.3 | 0.52 | 0.58 | 0.53 | 0.57 |
| 11 | 122144811 | rs11218568 | -0.765 | -0.951 | 0.589 | 0.562 | -0.023 | 0.037 | 0.037 | 0.181 | 0.16 | 0.38 | 0.17 | 0.21 | 0.1 | 0.14 |
| 11 | 122146533 | rs1025139 | 1.061 | -0.818 | 0.138 | 0.254 | -0.04 | 0.057 | 0.057 | 0.942 | 0.36 | 0.12 | 0.35 | 0.23 | 0.38 | 0.46 |
| 11 | 122154576 | rs11218575 | 0.657 | -0.692 | 0.24 | 0.296 | -0.024 | 0.083 | 0.083 | 0.311 | 0.41 | 0.19 | 0.4 | 0.23 | 0.42 | 0.46 |
| 11 | 122163143 | rs520036 | 1.083 | -0.564 | 0.155 | 0.378 | -0.029 | -0.057 | -0.057 | 1.845 | 0.1 | 0.01 | 0.09 | 0.29 | 0.14 | 0.1 |
| 11 | 122173092 | rs7129815 | 0.515 | -0.681 | 0.492 | 0.271 | 0.001 | 0.002 | 0.002 | 0 | 0.42 | 0.4 | 0.43 | 0.37 | 0.37 | 0.51 |
| 11 | 122176383 | rs4127499 | 1.329 | -0.812 | -0.235 | 0.152 | -0.078 | 0.144 | 0.144 | 0.571 | 0.28 | 0.07 | 0.26 | 0.07 | 0.36 | 0.21 |
| 11 | 122177618 | rs17246541 | 1.296 | -0.489 | -0.329 | 0.223 | -0.085 | 0.183 | 0.183 | 0 | 0.27 | 0.03 | 0.24 | 0.01 | 0.36 | 0.12 |
| 11 | 122178647 | rs599154 | -1.314 | -0.454 | 0.317 | 0.909 | -0.024 | 0.017 | 0.017 | 0 | 0.6 | 0.74 | 0.58 | 0.68 | 0.55 | 0.64 |
| 11 | 122185174 | rs660546 | 0.34 | 0.034 | -0.354 | 0.673 | 0 | 0.02 | 0.02 | 0 | 0.11 | 0.14 | 0.08 | 0.29 | 0.13 | 0.11 |
| 11 | 122192439 | rs1109768 | -0.348 | 0.185 | -1.027 | 0.778 | 0.006 | 0.044 | 0.044 | 0 | 0.45 | 0.3 | 0.42 | 0.28 | 0.41 | 0.49 |
| 11 | 122194946 | rs498853 | -0.769 | -0.102 | -0.993 | 1.07 | 0 | 0.001 | 0.001 | 0 | 0.51 | 0.45 | 0.48 | 0.58 | 0.49 | 0.57 |
| 11 | 122195265 | rs11218581 | -0.507 | -0.262 | -0.986 | 1.657 | -0.012 | 0.012 | 0.012 | 0 | 0.12 | 0.14 | 0.15 | 0.18 | 0.04 | 0.2 |
| 11 | 122197298 | rs681917 | -0.43 | 0.514 | -1.172 | 1.511 | -0.021 | -0.059 | -0.059 | 1.823 | 0.09 | 0 | 0.07 | 0.29 | 0.1 | 0.1 |
| 11 | 122198706 | rs10790495 | -0.06 | 1.028 | -1.017 | 0.959 | 0.003 | 0.001 | 0.001 | 0 | 0.66 | 0.7 | 0.6 | 0.66 | 0.59 | 0.64 |
| 11 | 122199254 | rs17126343 | -0.71 | 0.662 | -0.899 | 0.013 | -0.009 | -0.073 | -0.073 | 0 | 0.28 | 0.48 | 0.2 | 0.11 | 0.24 | 0.28 |
| 11 | 122199441 | rs7115833 | -0.28 | 0.713 | -2.14 | -0.373 | -0.073 | 0.172 | 0.172 | 0 | 0.59 | 0.93 | 0.69 | 0.86 | 0.56 | 0.68 |
| 11 | 122204076 | rs2078216 | -0.851 | 0.618 | -1.255 | 0.66 | 0.005 | 0.027 | 0.027 | 0 | 0.25 | 0.27 | 0.36 | 0.56 | 0.22 | 0.32 |
| 11 | 122204383 | rs11218583 | 1.099 | 0.349 | -2.03 | -0.149 | -0.07 | 0.027 | 0.027 | 0 | 0.31 | 0.07 | 0.33 | 0.22 | 0.44 | 0.36 |
| 11 | 122204543 | rs7929703 | -0.696 | -0.32 | -1.166 | -0.568 | 0.008 | -0.014 | -0.014 | 0 | 0.75 | 0.8 | 0.83 | 0.98 | 0.85 | 0.94 |
| 11 | 122224552 | rs669244 | 0.946 | -0.048 | -1.32 | 0.076 | -0.041 | -0.036 | -0.036 | 0.072 | 0.12 | 0.02 | 0.07 | 0.16 | 0.13 | 0.17 |
| 11 | 122235381 | rs11218597 | 1.219 | -0.299 | -1.268 | 0.047 | -0.091 | 0.01 | 0.01 | 0.263 | 0.33 | 0.07 | 0.37 | 0.23 | 0.49 | 0.37 |
| 11 | 122237501 | rs572820 | -1.006 | -0.197 | -1.175 | 0.072 | -0.016 | 0.121 | 0.121 | 1.162 | 0.8 | 0.98 | 0.86 | 0.99 | 0.87 | 0.96 |
| 11 | 122258356 | rs10892831 | 0.623 | -0.174 | -0.45 | 0.211 | 0 | 0.052 | 0.052 | 1.773 | 0.5 | 0.4 | 0.45 | 0.22 | 0.61 | 0.46 |
| 11 | 122265202 | rs685419 | 0.069 | -0.247 | -0.21 | 0.627 | 0.01 | 0.013 | 0.013 | 0 | 0.27 | 0.43 | 0.27 | 0.3 | 0.23 | 0.32 |
| 11 | 122273501 | rs678577 | -0.245 | -0.277 | -0.102 | 0.85 | 0.001 | 0.056 | 0.056 | 0 | 0.28 | 0.55 | 0.29 | 0.41 | 0.23 | 0.46 |
| 11 | 122279782 | rs17246639 | 0.916 | -0.317 | -0.117 | 0.734 | -0.02 | 0.134 | 0.134 | 2.226 | 0.48 | 0.21 | 0.44 | 0.21 | 0.6 | 0.46 |
| 11 | 122282198 | rs553658 | 0.032 | -0.072 | -0.335 | 0.638 | 0.011 | 0.02 | 0.02 | 0 | 0.26 | 0.41 | 0.27 | 0.31 | 0.23 | 0.32 |
| 11 | 122289345 | rs580069 | 0.139 | 0.045 | -0.594 | 0.647 | 0.011 | 0.011 | 0.011 | 0 | 0.24 | 0.3 | 0.25 | 0.29 | 0.23 | 0.32 |
| 11 | 122293813 | rs565675 | -0.363 | -0.086 | -1.466 | 0.861 | -0.043 | -0.096 | -0.096 | 0 | 0.79 | 0.98 | 0.74 | 0.66 | 0.84 | 0.92 |
| 11 | 122308215 | rs12804305 | 0.196 | -0.162 | -1.324 | 0.187 | 0.002 | 0.082 | 0.082 | 0.235 | 0.14 | 0.01 | 0.06 | 0 | 0.11 | 0.05 |
| 11 | 122310951 | rs693491 | 0.066 | -0.131 | -0.708 | 0.848 | 0.002 | -0.007 | -0.007 | 0.524 | 0.75 | 0.78 | 0.72 | 0.5 | 0.84 | 0.78 |
| 11 | 122314282 | rs480478 | -1.209 | 0.074 | -0.13 | 0.678 | -0.005 | 0.027 | 0.027 | 0 | 0.24 | 0.47 | 0.32 | 0.71 | 0.2 | 0.42 |
| 11 | 122318528 | rs1216553 | -1.02 | 0.036 | 0.081 | 0.66 | 0.001 | 0.025 | 0.025 | 0 | 0.22 | 0.35 | 0.34 | 0.71 | 0.2 | 0.42 |
| 11 | 122325682 | rs10892844 | 0.305 | 0.005 | 0.033 | 1.128 | 0.011 | -0.036 | -0.036 | 0.749 | 0.1 | 0.02 | 0.2 | 0.41 | 0.07 | 0.07 |
| 11 | 122329878 | rs4936695 | -0.03 | 0.128 | 0.131 | 1.145 | 0.057 | 0.136 | 0.136 | 0 | 0.11 | 0.29 | 0.12 | 0.23 | 0.13 | 0.22 |
| 11 | 122337411 | rs2370741 | 1.181 | -0.113 | 0.408 | 0.833 | 0.006 | 0.057 | 0.057 | 0.321 | 0.57 | 0.25 | 0.51 | 0.27 | 0.65 | 0.54 |
| 11 | 122339119 | rs11218637 | 0.149 | -0.104 | 0.484 | 0.839 | 0.049 | 0.122 | 0.122 | 0 | 0.11 | 0.28 | 0.12 | 0.23 | 0.13 | 0.22 |
| 11 | 122340155 | rs1450800 | 0.149 | -0.104 | 0.483 | 0.839 | 0.051 | 0.125 | 0.125 | 0 | 0.11 | 0.29 | 0.12 | 0.23 | 0.13 | 0.22 |
| 11 | 122362390 | rs10892845 | 0.892 | -0.098 | 0.163 | 0.473 | 0.006 | 0.058 | 0.058 | 0.321 | 0.57 | 0.25 | 0.51 | 0.27 | 0.65 | 0.54 |
| 11 | 122382426 | rs7937803 | -0.291 | 0.509 | -0.329 | 0.087 | 0.005 | -0.025 | -0.025 | 6.679 | 0.39 | 0.47 | 0.28 | 0.23 | 0.46 | 0.41 |
| 11 | 122383475 | rs1839545 | -0.527 | 0.661 | -0.264 | -0.097 | 0.02 | 0.083 | 0.083 | 0.745 | 0.15 | 0.39 | 0.14 | 0.23 | 0.19 | 0.32 |
| 11 | 122384928 | rs11218652 | -0.354 | 0.662 | -0.184 | -0.106 | 0.017 | 0.057 | 0.057 | 0.298 | 0.14 | 0.3 | 0.14 | 0.21 | 0.19 | 0.32 |
| 11 | 122391187 | rs2084637 | 0.971 | 0.138 | -1.074 | -0.251 | -0.093 | -0.046 | -0.046 | 0 | 0.27 | 0.06 | 0.27 | 0.28 | 0.36 | 0.44 |
| 11 | 122391206 | rs12364280 | 1.058 | 0.073 | -0.981 | -0.304 | -0.027 | 0.094 | 0.094 | 0 | 0.25 | 0.07 | 0.14 | 0 | 0.27 | 0.09 |
| 11 | 122430752 | rs10892858 | 0.757 | 0.465 | -1.018 | 0.063 | -0.034 | -0.037 | -0.037 | 3.337 | 0.23 | 0.15 | 0.25 | 0.3 | 0.28 | 0.37 |
| 11 | 122435289 | rs12291352 | -0.42 | -0.226 | -0.836 | 0.039 | -0.016 | 0.032 | 0.032 | 0 | 0.23 | 0.7 | 0.35 | 0.53 | 0.14 | 0.25 |
| 11 | 122441861 | rs4935798 | 0.216 | 0.132 | -1.221 | -0.091 | 0 | 0 | 0 | 0 | 0.13 | 0.14 | 0.12 | 0.14 | 0.19 | 0.13 |
| 11 | 122446233 | rs10892861 | 1.465 | 0.35 | -1.588 | -0.134 | -0.071 | -0.087 | -0.087 | 0 | 0.19 | 0.01 | 0.21 | 0.24 | 0.21 | 0.31 |
| 11 | 122455520 | rs11218700 | -1.087 | -0.041 | -0.65 | 0.189 | -0.046 | 0.048 | 0.048 | 0 | 0.31 | 0.56 | 0.42 | 0.47 | 0.25 | 0.33 |
| 11 | 122458831 | rs6589931 | -1.557 | -0.001 | -1.901 | -0.063 | -0.039 | 0.031 | 0.031 | 0.071 | 0.52 | 0.96 | 0.6 | 0.68 | 0.48 | 0.56 |
| 11 | 122460079 | rs4565904 | 0.775 | -0.02 | -1.182 | 0.309 | 0.018 | -0.026 | -0.026 | 0 | 0.13 | 0.07 | 0.16 | 0.34 | 0.07 | 0.24 |
| 11 | 122460155 | rs2370756 | -0.275 | 0.501 | -0.92 | -0.045 | -0.002 | -0.029 | -0.029 | 0 | 0.11 | 0.13 | 0.09 | 0 | 0.13 | 0.06 |
| 11 | 122462580 | rs7103440 | -0.405 | 0.68 | -0.877 | 0.104 | 0.001 | -0.012 | -0.012 | 0 | 0.11 | 0.13 | 0.07 | 0.05 | 0.14 | 0.1 |
| 11 | 122462709 | rs10892863 | 1.274 | 0.891 | -2.054 | 0.112 | -0.079 | -0.113 | -0.113 | 0 | 0.28 | 0.02 | 0.29 | 0.38 | 0.3 | 0.3 |
| 11 | 122462813 | rs7103823 | -0.999 | 0.874 | -1.989 | -0.704 | -0.011 | 0.085 | 0.085 | 0 | 0.82 | 0.97 | 0.91 | 0.96 | 0.84 | 0.97 |
| 11 | 122464091 | rs7931095 | 0.311 | 0.816 | -1.27 | -0.124 | 0 | -0.001 | -0.001 | 0 | 0.21 | 0.19 | 0.2 | 0.23 | 0.2 | 0.31 |
| 11 | 122466121 | rs12419313 | 0.446 | 0.923 | -1.084 | -0.205 | -0.001 | 0.023 | 0.023 | 0 | 0.15 | 0.1 | 0.12 | 0.05 | 0.13 | 0.17 |
| 11 | 122469697 | rs7111139 | 0.013 | 0.964 | -0.678 | -0.055 | 0.012 | -0.01 | -0.01 | 0 | 0.09 | 0.15 | 0.07 | 0.05 | 0.12 | 0.06 |
| 11 | 122472063 | rs11218708 | 0.293 | 0.614 | -0.773 | 0.069 | -0.006 | 0.006 | 0.006 | 0 | 0.19 | 0.18 | 0.16 | 0.26 | 0.17 | 0.27 |

| | | | | | | | | | | | | | | | | |
|---|---|---|---|---|---|---|---|---|---|---|---|---|---|---|---|---|
| 11 | 122477025 | rs12420917 | 1.158 | 0.763 | -1.162 | -0.083 | 0.007 | -0.038 | -0.038 | 0 | 0.34 | 0.25 | 0.43 | 0.53 | 0.35 | 0.34 |
| 11 | 122477706 | rs7944144 | -1.006 | 0.949 | -0.943 | -0.062 | 0.006 | 0.027 | 0.027 | 0 | 0.5 | 0.7 | 0.45 | 0.42 | 0.48 | 0.61 |
| 11 | 122483315 | rs12422102 | -0.628 | 1.355 | -0.209 | -0.316 | -0.001 | 0.005 | 0.005 | 0 | 0.36 | 0.3 | 0.34 | 0.22 | 0.32 | 0.38 |
| 11 | 122484431 | rs7930461 | -0.526 | 1.501 | -0.005 | -0.378 | 0.011 | 0 | 0 | 0.246 | 0.85 | 0.8 | 0.83 | 0.83 | 0.81 | 0.83 |
| 11 | 122489473 | rs17246711 | -0.768 | 1.2 | -0.187 | -0.385 | -0.004 | 0.037 | 0.037 | 0 | 0.35 | 0.24 | 0.33 | 0.11 | 0.31 | 0.32 |
| 11 | 122495678 | rs3890623 | -0.722 | 1.192 | -0.318 | -0.165 | -0.045 | 0.007 | 0.007 | 0.033 | 0.35 | 0.09 | 0.34 | 0.32 | 0.35 | 0.4 |
| 11 | 122496249 | rs11218715 | 0.353 | 1.363 | 0.489 | -0.326 | 0.034 | -0.01 | -0.01 | 0 | 0.11 | 0.24 | 0.17 | 0.09 | 0.12 | 0.1 |
| 11 | 122499895 | rs11607161 | 0.415 | 1.336 | 0.514 | -0.326 | 0.015 | -0.007 | -0.007 | 0 | 0.11 | 0.18 | 0.17 | 0.11 | 0.11 | 0.1 |
| 11 | 122504346 | rs10790515 | -0.97 | 1.174 | 0.6 | -0.534 | 0 | -0.005 | -0.005 | 0.069 | 0.29 | 0.3 | 0.26 | 0.21 | 0.28 | 0.31 |
| 11 | 122505344 | rs10750215 | -1.359 | 1.055 | 0.615 | -0.529 | -0.004 | 0.004 | 0.004 | 0.797 | 0.39 | 0.33 | 0.41 | 0.33 | 0.39 | 0.42 |
| 11 | 122522375 | rs7941030 | -1.421 | 1.086 | 0.73 | -0.512 | 0.003 | -0.006 | -0.006 | 0 | 0.39 | 0.43 | 0.4 | 0.33 | 0.38 | 0.36 |
| 11 | 122525308 | rs10892871 | -0.664 | 1.031 | 0.669 | -0.517 | -0.01 | 0.054 | 0.054 | 0.221 | 0.26 | 0.16 | 0.24 | 0.1 | 0.26 | 0.22 |
| 11 | 122526501 | rs7127978 | -0.863 | 0.747 | 0.991 | -0.35 | -0.008 | 0.019 | 0.019 | 1.152 | 0.37 | 0.32 | 0.38 | 0.21 | 0.37 | 0.33 |
| 11 | 122534504 | rs7117842 | -1.091 | 0.748 | 0.821 | -0.311 | 0.002 | -0.007 | -0.007 | 0.037 | 0.39 | 0.41 | 0.4 | 0.33 | 0.37 | 0.36 |
| 11 | 122544054 | rs7940176 | -1.093 | 0.86 | 0.566 | -0.426 | 0.022 | -0.015 | -0.015 | 0 | 0.11 | 0.18 | 0.11 | 0.08 | 0.15 | 0.1 |
| 11 | 122546751 | rs885325 | -0.285 | 0.908 | 0.499 | -0.438 | -0.001 | 0.008 | 0.008 | 0 | 0.28 | 0.18 | 0.3 | 0.26 | 0.25 | 0.27 |
| 11 | 122549214 | rs7117189 | -1.14 | 1.102 | 0.536 | -0.396 | 0.037 | 0.071 | 0.071 | 0 | 0.11 | 0.22 | 0.12 | 0.21 | 0.16 | 0.12 |
| 11 | 122552486 | rs11218751 | 0.018 | 0.795 | 0.165 | -0.48 | -0.014 | 0.076 | 0.076 | 0.186 | 0.26 | 0.1 | 0.29 | 0.13 | 0.24 | 0.24 |
| 11 | 122554865 | rs10892882 | 0.858 | -0.164 | 0.631 | -0.578 | -0.011 | -0.026 | -0.026 | 3.082 | 0.76 | 0.3 | 0.86 | 0.91 | 0.83 | 0.74 |
| 11 | 122556578 | rs10750218 | 0.967 | -0.177 | 0.611 | -0.472 | -0.009 | -0.001 | -0.001 | 0 | 0.75 | 0.29 | 0.86 | 0.79 | 0.82 | 0.72 |
| 11 | 122558002 | rs12798180 | -0.506 | -0.235 | 0.677 | -0.47 | -0.003 | -0.004 | -0.004 | 0 | 0.08 | 0.09 | 0.03 | 0.05 | 0.05 | 0.04 |
| 11 | 122558850 | rs10750219 | 1.139 | -0.028 | 0.461 | -0.625 | 0 | 0.057 | 0.057 | 0 | 0.47 | 0.21 | 0.49 | 0.65 | 0.49 | 0.47 |
| 11 | 122561625 | rs4935801 | 0.638 | -0.032 | 0.945 | -0.6 | -0.019 | -0.04 | -0.04 | 0.11 | 0.78 | 0.51 | 0.86 | 0.89 | 0.81 | 0.74 |
| 11 | 122562045 | rs7129189 | 1.368 | 0.199 | 0.653 | -0.741 | -0.002 | 0.006 | 0.006 | 0 | 0.38 | 0.15 | 0.46 | 0.65 | 0.38 | 0.45 |
| 11 | 122568946 | rs7943438 | -0.353 | 0.215 | 1.005 | -0.751 | -0.019 | -0.037 | -0.037 | 0 | 0.85 | 0.61 | 0.92 | 0.94 | 0.88 | 0.81 |
| 11 | 122569246 | rs7929077 | -0.816 | 0.105 | 0.979 | -0.76 | 0 | -0.001 | -0.001 | 0 | 0.48 | 0.44 | 0.45 | 0.29 | 0.5 | 0.34 |
| 11 | 122573644 | rs12797830 | 0.785 | 0.078 | 0.769 | -0.821 | 0.002 | 0.018 | 0.018 | 0 | 0.46 | 0.37 | 0.53 | 0.82 | 0.47 | 0.54 |
| 11 | 122574767 | rs1344169 | 0.67 | -0.148 | 0.562 | -0.674 | -0.141 | 0.167 | 0.167 | 0.006 | 0.34 | 0.03 | 0.37 | 0.12 | 0.38 | 0.26 |
| 11 | 122578047 | rs4382921 | 0.29 | -0.137 | 0.748 | -0.709 | -0.02 | -0.084 | -0.084 | 1.752 | 0.76 | 0.57 | 0.87 | 0.94 | 0.79 | 0.8 |
| 11 | 122578200 | rs11218765 | 0.671 | -0.081 | -0.186 | -0.672 | -0.141 | 0.171 | 0.171 | 0.019 | 0.34 | 0.03 | 0.37 | 0.13 | 0.37 | 0.28 |
| 11 | 122580119 | rs10790521 | 1.292 | 0.048 | -0.066 | -0.597 | -0.003 | -0.012 | -0.012 | 0 | 0.32 | 0.15 | 0.43 | 0.66 | 0.33 | 0.45 |
| 11 | 122583001 | rs2271719 | 0.29 | -0.041 | -0.029 | -0.574 | 0.001 | -0.025 | -0.025 | 0.03 | 0.13 | 0.19 | 0.06 | 0.03 | 0.13 | 0.05 |
| 11 | 122585086 | rs2187519 | 0.488 | 0.042 | 0.044 | -0.598 | -0.004 | -0.05 | -0.05 | 0.33 | 0.74 | 0.58 | 0.85 | 0.91 | 0.72 | 0.76 |
| 11 | 122585573 | rs4016788 | -0.643 | 0.038 | -0.395 | -0.558 | -0.004 | -0.017 | -0.017 | 0.071 | 0.86 | 0.77 | 0.91 | 0.95 | 0.85 | 0.8 |
| 11 | 122586211 | rs6589946 | -0.703 | -0.12 | 0.087 | -0.04 | 0.006 | 0.114 | 0.114 | 0 | 0.52 | 0.7 | 0.55 | 0.72 | 0.48 | 0.5 |
| 11 | 122587572 | rs10790522 | 0.492 | 0.246 | -0.468 | 0.643 | -0.018 | -0.075 | -0.075 | 0 | 0.3 | 0.06 | 0.25 | 0.63 | 0.3 | 0.41 |
| 11 | 122588584 | rs11218771 | -0.005 | 0.53 | -0.221 | 0.553 | 0.003 | 0.036 | 0.036 | 0 | 0.42 | 0.27 | 0.47 | 0.71 | 0.42 | 0.48 |
| 11 | 122591233 | rs7932902 | -1.255 | 0.706 | 0.433 | 1.559 | 0.014 | 0.168 | 0.168 | 0.525 | 0.12 | 0.38 | 0.27 | 0.23 | 0.09 | 0.24 |
| 11 | 122599125 | rs11218782 | -0.552 | 1.446 | 0.21 | 1.796 | 0.012 | 0.016 | 0.016 | 0 | 0.3 | 0.31 | 0.45 | 0.51 | 0.4 | 0.31 |
| 11 | 122604566 | rs1894084 | -0.705 | 1.426 | 0.434 | 1.814 | 0.007 | 0.012 | 0.012 | 0 | 0.29 | 0.27 | 0.45 | 0.5 | 0.39 | 0.29 |
| 11 | 122607006 | rs4935806 | 0.303 | 1.667 | 0.846 | 1.277 | 0.047 | 0.062 | 0.062 | 0 | 0.83 | 0.62 | 0.76 | 0.7 | 0.8 | 0.68 |
| 11 | 122609618 | rs10790523 | -0.824 | 1.444 | 1.134 | 1.204 | -0.017 | 0.001 | 0.001 | 0.096 | 0.4 | 0.73 | 0.59 | 0.57 | 0.5 | 0.39 |
| 11 | 122612295 | rs871843 | -0.692 | 1.703 | 1.216 | 1.553 | 0.064 | 0.047 | 0.047 | 0.007 | 0.3 | 0.42 | 0.46 | 0.38 | 0.42 | 0.35 |
| 11 | 122613549 | rs11218790 | -0.082 | 0.55 | 0.501 | -0.035 | 0.004 | -0.049 | -0.049 | 0.188 | 0.36 | 0.31 | 0.2 | 0.04 | 0.3 | 0.3 |
| 11 | 122613623 | rs10750225 | 0.457 | 0.88 | 0.345 | 0.536 | -0.009 | 0.001 | 0.001 | 0 | 0.37 | 0.38 | 0.4 | 0.37 | 0.31 | 0.35 |
| 11 | 122615484 | rs7342257 | -0.362 | 1.218 | -0.115 | 0.224 | 0 | -0.008 | -0.008 | 0.05 | 0.46 | 0.52 | 0.44 | 0.25 | 0.47 | 0.5 |
| 11 | 122616059 | rs1894081 | 0.097 | 1.117 | -0.808 | 0.234 | -0.045 | -0.018 | -0.018 | 0 | 0.29 | 0.16 | 0.35 | 0.34 | 0.37 | 0.37 |
| 11 | 122616326 | rs884393 | 0.962 | 0.696 | -0.888 | -0.407 | 0.018 | 0.028 | 0.028 | 0 | 0.28 | 0.08 | 0.2 | 0.19 | 0.23 | 0.26 |
| 11 | 122616553 | rs884392 | 0.99 | 0.702 | -0.886 | -0.565 | 0.017 | 0.052 | 0.052 | 0 | 0.27 | 0.08 | 0.2 | 0.11 | 0.23 | 0.26 |
| 11 | 122616801 | rs17126807 | -0.167 | 0.825 | -0.862 | -0.758 | -0.001 | -0.024 | -0.024 | 0 | 0.19 | 0.14 | 0.1 | 0 | 0.2 | 0.14 |
| 11 | 122619330 | rs12276870 | -0.328 | 0.234 | -1.032 | -1.542 | -0.002 | -0.016 | -0.016 | 0.333 | 0.46 | 0.49 | 0.36 | 0.08 | 0.52 | 0.39 |
| 11 | 122619384 | rs17246816 | -0.393 | -1.646 | -1.653 | -1.706 | 0.056 | 0.179 | 0.179 | 0.02 | 0.38 | 0.11 | 0.2 | 0.09 | 0.29 | 0.33 |
| 11 | 122619469 | rs2840376 | -0.053 | -1.444 | -0.993 | -1.279 | 0.026 | 0.052 | 0.052 | 0 | 0.21 | 0.07 | 0.09 | 0.05 | 0.13 | 0.22 |
| 11 | 122620957 | rs10892892 | 0.247 | -1.368 | -0.139 | -0.173 | -0.019 | 0.028 | 0.028 | 0.251 | 0.79 | 0.42 | 0.73 | 0.62 | 0.86 | 0.69 |
| 11 | 122622531 | rs4936737 | -0.302 | -1.375 | 0.597 | -0.109 | 0 | 0.159 | 0.159 | 0 | 0.89 | 0.66 | 0.78 | 0.73 | 0.92 | 0.84 |
| 11 | 122623393 | rs1122475 | 0.39 | -1.405 | 0.663 | -0.152 | -0.023 | 0.019 | 0.019 | 0.306 | 0.8 | 0.46 | 0.73 | 0.71 | 0.86 | 0.7 |
| 11 | 122626434 | rs10790525 | -0.741 | -1.54 | 0.119 | 0.005 | 0.006 | -0.009 | -0.009 | 1.232 | 0.57 | 0.52 | 0.46 | 0.32 | 0.48 | 0.49 |
| 11 | 122627455 | rs10892893 | 1.304 | -1.193 | -0.255 | 0.214 | -0.078 | -0.032 | -0.032 | 0 | 0.29 | 0.08 | 0.31 | 0.32 | 0.44 | 0.33 |
| 11 | 122630289 | rs10892894 | 1.626 | -1.146 | -0.289 | 0.165 | -0.045 | -0.027 | -0.027 | 0 | 0.27 | 0.11 | 0.3 | 0.33 | 0.4 | 0.28 |
| 11 | 122630672 | rs10892895 | 1.23 | -1.169 | -0.365 | 0.26 | -0.003 | -0.06 | -0.06 | 0.002 | 0.12 | 0.23 | 0.06 | 0.02 | 0.07 | 0.18 |
| 11 | 122631578 | rs4935810 | 0.875 | -1.443 | -1.084 | 0.107 | 0.029 | 0.02 | 0.02 | 0.041 | 0.53 | 0.23 | 0.5 | 0.42 | 0.63 | 0.42 |
| 11 | 122634267 | rs12804711 | -0.06 | -1.613 | -1.598 | 0.627 | 0.004 | -0.049 | -0.049 | 0.289 | 0.2 | 0.08 | 0.19 | 0.24 | 0.16 | 0.2 |
| 11 | 122641086 | rs4936742 | 1.02 | -1.613 | -1.602 | 0.647 | 0.069 | 0 | 0 | 0 | 0.5 | 0.84 | 0.58 | 0.43 | 0.58 | 0.54 |
| 11 | 122641256 | rs11218802 | -0.755 | -1.597 | -1.632 | 0.74 | 0.005 | -0.03 | -0.03 | 0 | 0.26 | 0.1 | 0.22 | 0.24 | 0.23 | 0.24 |
| 11 | 122641855 | rs10790529 | 1.16 | -1.314 | -1.614 | 0.278 | -0.02 | 0.01 | 0.01 | 0 | 0.43 | 0.73 | 0.47 | 0.37 | 0.49 | 0.51 |
| 11 | 122643283 | rs12804793 | -0.496 | -1.116 | -1.789 | -0.1 | -0.018 | 0.016 | 0.016 | 0 | 0.19 | 0.02 | 0.17 | 0.11 | 0.16 | 0.2 |
| 11 | 122646038 | rs12795620 | -0.594 | -1.102 | -1.759 | 0.036 | -0.008 | -0.012 | -0.012 | 0 | 0.2 | 0.03 | 0.19 | 0.15 | 0.16 | 0.16 |
| 11 | 122646967 | rs12790613 | -0.58 | -0.884 | -1.633 | 0.204 | -0.021 | 0.053 | 0.053 | 0 | 0.34 | 0.03 | 0.31 | 0.17 | 0.31 | 0.24 |
| 11 | 122648045 | rs6589948 | -0.889 | -0.832 | -0.178 | 0.424 | -0.027 | -0.071 | -0.071 | 0 | 0.19 | 0.48 | 0.18 | 0.07 | 0.16 | 0.09 |

| | | | | | | | | | | | | | | | | |
|---|---|---|---|---|---|---|---|---|---|---|---|---|---|---|---|---|
| 11 | 122653689 | rs12575062 | -0.925 | -0.708 | 0.194 | 1.344 | 0.005 | 0.012 | 0.012 | 0 | 0.26 | 0.33 | 0.32 | 0.31 | 0.24 | 0.21 |
| 11 | 122653950 | rs2156802 | 0.247 | -1.19 | -0.234 | 0.342 | 0.025 | 0.044 | 0.044 | 0 | 0.19 | 0.11 | 0.12 | 0.04 | 0.17 | 0.15 |
| 11 | 122654056 | rs2156801 | 0.765 | -1.016 | -0.053 | 0.698 | -0.003 | -0.086 | -0.086 | 0 | 0.64 | 0.58 | 0.4 | 0.13 | 0.68 | 0.41 |
| 11 | 122658560 | rs4936744 | 0.533 | -1.468 | 0.597 | 2.165 | 0.001 | 0.038 | 0.038 | 0 | 0.84 | 0.81 | 0.66 | 0.66 | 0.82 | 0.81 |
| 11 | 122659918 | rs955051 | 0.043 | -2.064 | 0.14 | 1.434 | 0.041 | 0.067 | 0.067 | 0 | 0.2 | 0.03 | 0.08 | 0.04 | 0.15 | 0.19 |
| 11 | 122664228 | rs11218814 | -1.051 | -2.476 | 0.504 | 0.882 | 0.035 | 0.005 | 0.005 | 0 | 0.34 | 0.24 | 0.16 | 0.01 | 0.23 | 0.24 |
| 11 | 122665437 | rs872414 | -2.204 | -2.191 | 0.725 | 1.241 | 0.058 | 0.008 | 0.008 | 0 | 0.53 | 0.76 | 0.53 | 0.56 | 0.39 | 0.49 |
| 11 | 122666790 | rs4936746 | -0.176 | -2.285 | 0.69 | 1.294 | 0 | -0.001 | -0.001 | 0 | 0.12 | 0.17 | 0.23 | 0.09 | 0.14 | 0.06 |
| 11 | 122670390 | rs873590 | -1.512 | -2.269 | 0.58 | 1.384 | -0.041 | 0.023 | 0.023 | 0.109 | 0.32 | 0.5 | 0.42 | 0.41 | 0.22 | 0.22 |
| 11 | 122675109 | rs12576789 | -0.037 | -1.817 | 0.201 | 1.438 | -0.013 | -0.013 | -0.013 | 0 | 0.08 | 0.04 | 0.23 | 0.09 | 0.14 | 0.07 |
| 11 | 122675120 | rs12576782 | -1.947 | -2.102 | 0.086 | 1.555 | 0.101 | -0.001 | -0.001 | 0 | 0.52 | 0.85 | 0.54 | 0.56 | 0.38 | 0.49 |
| 11 | 122675193 | rs12287912 | -1.391 | -2.119 | 0.239 | 1.466 | 0.142 | -0.02 | -0.02 | 0 | 0.44 | 0.78 | 0.31 | 0.48 | 0.25 | 0.42 |
| 11 | 122675454 | rs11218825 | -0.539 | -2.119 | 0.515 | 1.599 | -0.028 | 0 | 0 | 0 | 0.27 | 0.37 | 0.2 | 0.31 | 0.14 | 0.27 |
| 11 | 122678911 | rs2276408 | -1.344 | -1.205 | 0.737 | 0.957 | -0.005 | -0.022 | -0.022 | 0 | 0.22 | 0.23 | 0.32 | 0.1 | 0.22 | 0.19 |
| 11 | 122679481 | rs10892909 | -1.351 | -0.64 | 0.881 | 1.241 | 0.011 | 0 | 0 | 0 | 0.15 | 0.16 | 0.37 | 0.15 | 0.2 | 0.25 |
| 11 | 122680102 | rs7950437 | -1.358 | -0.422 | 1.355 | 1.243 | 0.066 | 0.016 | 0.016 | 0 | 0.15 | 0.31 | 0.36 | 0.16 | 0.2 | 0.23 |
| 11 | 122680599 | rs1540108 | -0.629 | -0.506 | 1.242 | 1.253 | -0.01 | 0.001 | 0.001 | 0 | 0.12 | 0.1 | 0.29 | 0.13 | 0.19 | 0.21 |
| 11 | 122682111 | rs1996841 | -1.586 | -0.466 | 1.474 | 1.204 | 0.062 | 0.006 | 0.006 | 0 | 0.17 | 0.4 | 0.36 | 0.16 | 0.21 | 0.23 |
| 11 | 122682156 | rs3134428 | 1.367 | -1.076 | 1.091 | 1.291 | 0.028 | 0.044 | 0.044 | 0.003 | 0.58 | 0.26 | 0.4 | 0.26 | 0.64 | 0.4 |
| 11 | 122684332 | rs7940959 | -0.422 | -1.126 | 1.163 | 1.212 | -0.098 | -0.176 | -0.176 | 0 | 0.25 | 0.5 | 0.14 | 0.01 | 0.14 | 0.18 |
| 11 | 122684597 | rs1945394 | -0.535 | -1.077 | 1.175 | 1.236 | -0.104 | -0.181 | -0.181 | 0.043 | 0.27 | 0.5 | 0.14 | 0.01 | 0.15 | 0.18 |
| 11 | 122686721 | rs11218832 | -0.371 | -0.992 | 0.64 | 1.394 | -0.007 | 0.001 | 0.001 | 0 | 0.12 | 0.1 | 0.3 | 0.13 | 0.19 | 0.21 |
| 11 | 122686840 | rs1894077 | 1.193 | -0.936 | 0.366 | 1.266 | 0.03 | 0.061 | 0.061 | 0.029 | 0.57 | 0.25 | 0.39 | 0.26 | 0.63 | 0.4 |
| 11 | 122700789 | rs4598683 | -0.327 | -1.347 | 0.291 | 1.481 | 0.047 | -0.007 | -0.007 | 0 | 0.25 | 0.16 | 0.18 | 0.29 | 0.13 | 0.21 |
| 11 | 122704882 | rs3133250 | 0.599 | -1.432 | 0.757 | 1.417 | -0.052 | 0.004 | 0.004 | 0 | 0.67 | 0.4 | 0.72 | 0.38 | 0.84 | 0.61 |
| 11 | 122707889 | rs11218856 | -0.612 | -0.075 | 0.601 | 1.011 | 0.002 | 0.027 | 0.027 | 0 | 0.46 | 0.28 | 0.55 | 0.41 | 0.45 | 0.44 |
| 11 | 122707989 | rs12270235 | -0.227 | 0.561 | 0.552 | 0.893 | 0.001 | 0 | 0 | 0 | 0.06 | 0.07 | 0.11 | 0.04 | 0.12 | 0.07 |
| 11 | 122709178 | rs3134425 | -1.473 | 0.804 | 0.982 | 0.991 | 0.002 | 0.05 | 0.05 | 0 | 0.59 | 0.76 | 0.72 | 0.78 | 0.61 | 0.71 |
| 11 | 122710561 | rs1879884 | -0.854 | 0.969 | 1.17 | 1.076 | 0.005 | 0.004 | 0.004 | 0 | 0.52 | 0.42 | 0.67 | 0.46 | 0.57 | 0.51 |
| 11 | 122712586 | rs3107634 | -1.514 | 0.81 | 0.489 | 0.952 | -0.055 | 0.053 | 0.053 | 0.023 | 0.67 | 0.95 | 0.78 | 0.84 | 0.68 | 0.77 |
| 11 | 122714782 | rs2370794 | -1.489 | 1.208 | 1.305 | 1.106 | 0.044 | 0.073 | 0.073 | 0 | 0.25 | 0.47 | 0.55 | 0.43 | 0.34 | 0.24 |
| 11 | 122716041 | rs4369419 | -1.577 | 1.179 | 1.231 | 1.109 | 0.033 | 0.059 | 0.059 | 0 | 0.26 | 0.44 | 0.56 | 0.42 | 0.36 | 0.26 |
| 11 | 122723751 | rs11218863 | -0.517 | 0.938 | 1.179 | 1.293 | 0.004 | 0.031 | 0.031 | 0 | 0.16 | 0.2 | 0.42 | 0.34 | 0.16 | 0.19 |
| 11 | 122726705 | rs7125644 | -0.267 | 0.968 | 0.967 | 1.309 | -0.004 | -0.006 | -0.006 | 1.193 | 0.14 | 0.09 | 0.35 | 0.31 | 0.16 | 0.17 |
| 11 | 122727577 | rs3107638 | -1.37 | 0.898 | 1.222 | 1.228 | 0.08 | 0.254 | 0.254 | 0.079 | 0.08 | 0.25 | 0.11 | 0.38 | 0.16 | 0.28 |
| 11 | 122732872 | rs2177196 | 0.276 | 0.041 | 0.669 | 0.381 | 0.049 | 0.088 | 0.088 | 0 | 0.29 | 0.12 | 0.13 | 0.02 | 0.18 | 0.25 |
| 11 | 122733817 | rs1916040 | 0.185 | 0.167 | 1.037 | 0.433 | -0.009 | -0.055 | -0.055 | 0.839 | 0.61 | 0.5 | 0.73 | 0.92 | 0.67 | 0.65 |
| 11 | 122737810 | rs7941607 | 1.451 | 0.388 | 1.828 | 1.728 | 0.091 | 0.414 | 0.414 | 0.158 | 0.9 | 0.57 | 0.83 | 0.6 | 0.84 | 0.73 |
| 11 | 122742428 | rs7126620 | -0.76 | 0.404 | 1.804 | 1.829 | 0.018 | 0.023 | 0.023 | 0 | 0.17 | 0.4 | 0.37 | 0.26 | 0.17 | 0.17 |
| 11 | 122749006 | rs7114274 | 1.173 | 0.439 | 2.061 | 1.891 | 0.019 | 0.386 | 0.386 | 0.059 | 0.92 | 0.66 | 0.88 | 0.63 | 0.94 | 0.8 |
| 11 | 122749214 | rs7114583 | 0.173 | 0.57 | 1.263 | 1.251 | 0.002 | -0.012 | -0.012 | 0 | 0.12 | 0.09 | 0.06 | 0.19 | 0.1 | 0.13 |
| 11 | 122760987 | rs12098906 | -1.375 | 0.604 | 1.198 | 1.05 | 0.12 | 0.047 | 0.047 | 0 | 0.06 | 0.32 | 0.07 | 0.03 | 0.11 | 0.08 |
| 11 | 122771528 | rs10892923 | -1.032 | 0.479 | 0.729 | 1.132 | 0.014 | 0.024 | 0.024 | 0 | 0.32 | 0.45 | 0.47 | 0.46 | 0.37 | 0.38 |
| 11 | 122779680 | rs12576571 | -0.728 | -0.078 | -0.13 | 1.169 | -0.002 | 0.005 | 0.005 | 0 | 0.11 | 0.14 | 0.08 | 0.2 | 0.09 | 0.26 |
| 11 | 122781556 | rs3107635 | 1.436 | -0.547 | 0.165 | 0.946 | -0.046 | 0.014 | 0.014 | 0 | 0.8 | 0.57 | 0.88 | 0.77 | 0.85 | 0.64 |
| 11 | 122782156 | rs3107636 | -1.103 | -0.414 | 0.255 | 1.014 | -0.022 | 0.059 | 0.059 | 0 | 0.13 | 0.4 | 0.1 | 0.19 | 0.09 | 0.26 |
| 11 | 122784817 | rs3134402 | -0.088 | -0.776 | -0.983 | 0.792 | 0.074 | 0.046 | 0.046 | 0 | 0.47 | 0.2 | 0.6 | 0.38 | 0.6 | 0.43 |
| 11 | 122785984 | rs4936760 | 0.516 | 0.609 | -1.584 | 0.924 | -0.121 | -0.092 | -0.092 | 0 | 0.24 | 0.03 | 0.28 | 0.33 | 0.33 | 0.32 |
| 11 | 122790176 | rs12422101 | 0.168 | 0.298 | -1.015 | 1.164 | -0.002 | -0.034 | -0.034 | 1.499 | 0.3 | 0.18 | 0.44 | 0.43 | 0.33 | 0.17 |
| 11 | 122793880 | rs1343784 | 0.146 | 0.198 | -1.193 | 0.563 | -0.015 | 0.04 | 0.04 | 0 | 0.28 | 0.1 | 0.18 | 0.21 | 0.28 | 0.16 |
| 11 | 122795688 | rs11218891 | 0.312 | 0.203 | -1.188 | 0.526 | -0.039 | 0.022 | 0.022 | 0 | 0.26 | 0.08 | 0.21 | 0.2 | 0.29 | 0.16 |
| 11 | 122799230 | rs4936761 | 0.287 | 0.204 | -1.109 | 0.483 | -0.031 | 0.031 | 0.031 | 0 | 0.27 | 0.09 | 0.21 | 0.19 | 0.29 | 0.16 |
| 11 | 122804133 | rs4293126 | -0.97 | -0.048 | -0.363 | 0.261 | 0 | -0.004 | -0.004 | 9.315 | 0.52 | 0.51 | 0.37 | 0.31 | 0.49 | 0.42 |
| 11 | 122805910 | rs4502006 | 0.077 | -0.315 | -0.191 | -0.036 | -0.004 | 0.045 | 0.045 | 0.159 | 0.35 | 0.26 | 0.26 | 0.05 | 0.38 | 0.23 |
| 11 | 122810330 | rs4477449 | -1.761 | -0.58 | 0.584 | 0.168 | -0.024 | -0.002 | -0.002 | 0.267 | 0.29 | 0.63 | 0.17 | 0.31 | 0.19 | 0.36 |
| 11 | 122814394 | rs7117843 | 0.32 | -0.248 | 0.692 | 0.242 | -0.001 | 0.074 | 0.074 | 0.065 | 0.35 | 0.22 | 0.25 | 0.04 | 0.37 | 0.22 |
| 11 | 122818311 | rs3816621 | -1.059 | -0.337 | 1.138 | 0.674 | 0 | -0.004 | -0.004 | 0 | 0.52 | 0.54 | 0.35 | 0.36 | 0.49 | 0.37 |
| 11 | 122818795 | rs11218901 | -0.741 | -0.645 | 1.086 | 0.795 | 0.002 | -0.013 | -0.013 | 0 | 0.16 | 0.13 | 0.09 | 0.32 | 0.13 | 0.15 |
| 11 | 122830214 | rs7107502 | -1.147 | -0.171 | 1.225 | 0.665 | -0.002 | -0.008 | -0.008 | 0 | 0.55 | 0.53 | 0.37 | 0.36 | 0.57 | 0.4 |
| 11 | 122830274 | rs1057704 | -1.147 | -0.171 | 1.228 | 0.665 | 0 | -0.001 | -0.001 | 0 | 0.55 | 0.52 | 0.37 | 0.36 | 0.57 | 0.4 |
| 11 | 122834036 | rs7124683 | -0.667 | -0.22 | 1.075 | 0.648 | 0.018 | 0.099 | 0.099 | 0.003 | 0.49 | 0.32 | 0.34 | 0.24 | 0.55 | 0.34 |
| 11 | 122834265 | rs930981 | 0.455 | -0.134 | 1.062 | 0.603 | -0.017 | 0.078 | 0.078 | 0.163 | 0.36 | 0.24 | 0.26 | 0.04 | 0.43 | 0.23 |
| 11 | 122834378 | rs930982 | -0.55 | -0.14 | 1.094 | 0.77 | 0.018 | 0.098 | 0.098 | 0.003 | 0.49 | 0.33 | 0.34 | 0.24 | 0.55 | 0.34 |
| 11 | 122841562 | rs11218907 | -1.278 | 0.005 | 0.899 | 0.781 | 0.001 | -0.015 | -0.015 | 0 | 0.58 | 0.73 | 0.39 | 0.48 | 0.59 | 0.57 |
| 11 | 122843987 | rs1461499 | -0.903 | 0.096 | 0.768 | 0.653 | 0.003 | 0.008 | 0.008 | 0 | 0.54 | 0.63 | 0.37 | 0.36 | 0.57 | 0.5 |
| 11 | 122851155 | rs11218912 | 0.087 | 0.319 | 0.091 | 0.365 | -0.018 | 0.076 | 0.076 | 0 | 0.36 | 0.23 | 0.26 | 0.16 | 0.42 | 0.25 |
| 11 | 122851563 | rs1870761 | -0.174 | 0.235 | 0.278 | 0.576 | -0.002 | 0.015 | 0.015 | 0 | 0.42 | 0.37 | 0.29 | 0.3 | 0.44 | 0.27 |
| 11 | 122865226 | rs11602935 | -0.803 | -0.11 | -0.235 | 0.361 | 0.012 | 0.01 | 0.01 | 0 | 0.15 | 0.08 | 0.06 | 0.01 | 0.09 | 0.07 |
| 11 | 122866102 | rs7111924 | 0.002 | 0.183 | -0.1 | 0.49 | -0.003 | 0.02 | 0.02 | 0 | 0.38 | 0.28 | 0.26 | 0.29 | 0.42 | 0.28 |

| | | | | | | | | | | | | | | | | |
|---|---|---|---|---|---|---|---|---|---|---|---|---|---|---|---|---|
| 11 | 122868205 | rs4617603 | -0.631 | 0.091 | 0.067 | 0.894 | 0 | 0 | 0 | 0 | 0.52 | 0.48 | 0.52 | 0.47 | 0.5 | 0.44 |
| 11 | 122870683 | rs6589964 | -0.79 | -0.138 | -0.084 | 0.987 | -0.004 | 0.006 | 0.006 | 0.366 | 0.53 | 0.62 | 0.33 | 0.53 | 0.53 | 0.47 |
| 11 | 122878095 | rs4936765 | 0.429 | -0.007 | -0.077 | 0.995 | 0 | 0.001 | 0.001 | 0.196 | 0.47 | 0.37 | 0.64 | 0.5 | 0.44 | 0.63 |
| 11 | 122883745 | rs10790548 | 0.489 | 0.166 | 0.489 | 1.142 | 0.035 | 0.06 | 0.06 | 0.005 | 0.86 | 0.71 | 0.83 | 0.69 | 0.81 | 0.91 |
| 11 | 122888570 | rs7933723 | 0.853 | -0.385 | -0.415 | 0.083 | -0.078 | 0.213 | 0.213 | 0.153 | 0.34 | 0.06 | 0.22 | 0 | 0.41 | 0.19 |
| 11 | 122888805 | rs1461493 | -0.118 | -0.775 | 0.34 | 0.602 | -0.066 | 0.101 | 0.101 | 0.708 | 0.12 | 0.25 | 0.17 | 0.2 | 0.04 | 0.07 |
| 11 | 122890439 | rs11218921 | 1.115 | -0.861 | 0.114 | 0.742 | -0.011 | 0.015 | 0.015 | 0.327 | 0.72 | 0.43 | 0.63 | 0.36 | 0.76 | 0.59 |
| 11 | 122905059 | rs922252 | -0.318 | -0.918 | 0.35 | 0.608 | 0.008 | 0.006 | 0.006 | 0 | 0.54 | 0.56 | 0.45 | 0.43 | 0.61 | 0.53 |
| 11 | 122926664 | rs1841580 | -1.198 | -0.453 | 0.962 | 0.415 | -0.008 | -0.083 | -0.083 | 0 | 0.22 | 0.42 | 0.18 | 0.08 | 0.16 | 0.29 |
| 11 | 122932409 | rs11218941 | -1.619 | -0.531 | 1.169 | 1.054 | -0.018 | 0.094 | 0.094 | 0.097 | 0.24 | 0.5 | 0.22 | 0.43 | 0.17 | 0.33 |
| 11 | 122932777 | rs1136141 | -0.293 | -0.573 | 1.289 | 0.972 | -0.001 | -0.003 | -0.003 | 0 | 0.14 | 0.16 | 0.14 | 0.07 | 0.14 | 0.09 |
| 11 | 122932802 | rs2276075 | 0.272 | -0.904 | 1.281 | 1.06 | 0.004 | -0.007 | -0.007 | 0.674 | 0.07 | 0.08 | 0.14 | 0.13 | 0.05 | 0.05 |
| 11 | 122943495 | rs7949414 | -2.148 | -0.831 | 1.361 | 1.096 | -0.03 | 0.051 | 0.051 | 0.082 | 0.31 | 0.56 | 0.36 | 0.55 | 0.22 | 0.39 |
| 11 | 122946060 | rs11218954 | -1.365 | -0.805 | 0.833 | 0.717 | 0.005 | -0.013 | -0.013 | 0 | 0.22 | 0.25 | 0.32 | 0.36 | 0.2 | 0.18 |
| 11 | 122952605 | rs11218961 | 0.837 | -0.995 | 0.301 | 0.061 | -0.058 | 0.198 | 0.198 | 0.05 | 0.33 | 0.05 | 0.24 | 0.01 | 0.41 | 0.21 |
| 11 | 122954945 | rs11218962 | -1.15 | -0.722 | 0.827 | 0.702 | 0.003 | 0.002 | 0.002 | 0.003 | 0.21 | 0.25 | 0.29 | 0.19 | 0.19 | 0.15 |
| 11 | 122962900 | rs7113273 | -2.009 | -0.434 | 0.771 | 0.645 | 0.001 | 0.003 | 0.003 | 0.354 | 0.32 | 0.73 | 0.37 | 0.19 | 0.28 | 0.35 |
| 11 | 122969700 | rs7104160 | -0.185 | -0.151 | 0.3 | 0.583 | -0.019 | 0.035 | 0.035 | 0 | 0.13 | 0.06 | 0.12 | 0.07 | 0.15 | 0.09 |
| 11 | 122970265 | rs7121089 | -0.669 | -0.142 | 0.778 | 0.677 | 0.019 | -0.04 | -0.04 | 0 | 0.08 | 0.16 | 0.05 | 0.01 | 0.08 | 0.2 |
| 11 | 122971458 | rs11218981 | -1.233 | -0.084 | 0.721 | 1.045 | -0.011 | 0.089 | 0.089 | 0 | 0.12 | 0.2 | 0.23 | 0.32 | 0.12 | 0.13 |
| 11 | 122975915 | rs11218985 | -0.472 | -0.054 | 0.354 | 0.782 | 0.004 | -0.017 | -0.017 | 0.392 | 0.09 | 0.07 | 0.18 | 0.16 | 0.11 | 0.09 |
| 11 | 122982928 | rs4936777 | -1.579 | -0.186 | 0.603 | 0.685 | -0.022 | 0.086 | 0.086 | 0 | 0.19 | 0.35 | 0.3 | 0.29 | 0.18 | 0.15 |
| 11 | 122986256 | rs11607961 | -0.932 | -0.075 | 0.257 | 0.46 | 0 | -0.006 | -0.006 | 0 | 0.19 | 0.17 | 0.28 | 0.21 | 0.2 | 0.14 |
| 11 | 122986326 | rs11605822 | -0.932 | -0.075 | 0.257 | 0.46 | 0 | -0.006 | -0.006 | 0 | 0.19 | 0.17 | 0.28 | 0.21 | 0.19 | 0.14 |
| 11 | 122995279 | rs2169355 | -1.724 | -0.301 | -0.137 | 0.406 | -0.009 | -0.069 | -0.069 | 0 | 0.69 | 0.85 | 0.64 | 0.54 | 0.71 | 0.55 |
| 11 | 122995291 | rs1563964 | -1.108 | -0.437 | 0.662 | 0.326 | -0.024 | 0.011 | 0.011 | 0.129 | 0.18 | 0.42 | 0.25 | 0.16 | 0.16 | 0.13 |
| 11 | 122996050 | rs7126295 | -1.904 | -0.308 | 0.223 | 0.236 | -0.002 | 0.007 | 0.007 | 0.062 | 0.39 | 0.68 | 0.45 | 0.34 | 0.39 | 0.4 |
| 11 | 123001058 | rs10892967 | -1.522 | -0.066 | 0.268 | 0.283 | 0.025 | -0.012 | -0.012 | 1.774 | 0.25 | 0.41 | 0.36 | 0.22 | 0.29 | 0.19 |
| 11 | 123001084 | rs10790554 | -1.512 | -0.089 | 0.257 | 0.353 | 0.019 | 0.009 | 0.009 | 0.224 | 0.26 | 0.41 | 0.37 | 0.32 | 0.29 | 0.22 |
| 11 | 123001223 | rs12286925 | -0.927 | -0.11 | 0.304 | 0.402 | 0 | 0.046 | 0.046 | 0.044 | 0.09 | 0.25 | 0.2 | 0.14 | 0.09 | 0.1 |
| 11 | 123006150 | rs7130674 | -1.41 | -0.592 | -0.443 | 0.176 | -0.06 | -0.031 | -0.031 | 0 | 0.57 | 0.82 | 0.51 | 0.44 | 0.49 | 0.5 |
| 11 | 123010654 | rs4936778 | -0.486 | -1.257 | 0.235 | -0.13 | -0.03 | 0.026 | 0.026 | 0 | 0.34 | 0.52 | 0.29 | 0.39 | 0.27 | 0.4 |
| 11 | 123012658 | rs7106958 | -1.219 | -1.267 | 0.227 | -0.232 | 0.038 | -0.009 | -0.009 | 2.123 | 0.44 | 0.65 | 0.36 | 0.43 | 0.34 | 0.47 |
| 11 | 123013271 | rs6589969 | -1.57 | -1.181 | 0.165 | -0.316 | 0.039 | 0.006 | 0.006 | 0 | 0.5 | 0.67 | 0.4 | 0.43 | 0.44 | 0.49 |
| 11 | 123014015 | rs7111575 | 0.062 | -1.353 | 0.72 | -0.227 | -0.035 | -0.021 | -0.021 | 0 | 0.28 | 0.44 | 0.17 | 0.17 | 0.24 | 0.18 |
| 11 | 123014336 | rs7115102 | -0.022 | -1.22 | 0.647 | -0.642 | 0.016 | -0.066 | -0.066 | 0 | 0.15 | 0.21 | 0.16 | 0 | 0.15 | 0.05 |
| 11 | 123017181 | rs7924892 | -0.61 | -1.386 | 0.812 | -0.511 | -0.032 | 0.009 | 0.009 | 0 | 0.33 | 0.48 | 0.28 | 0.33 | 0.27 | 0.4 |
| 11 | 123020231 | rs12578013 | -1.165 | -1.219 | 0.793 | -0.413 | -0.016 | 0.011 | 0.011 | 0 | 0.17 | 0.23 | 0.18 | 0.23 | 0.11 | 0.25 |
| 11 | 123023829 | rs11821710 | -1.09 | -1.213 | 0.847 | -0.423 | -0.012 | -0.013 | -0.013 | 0.277 | 0.38 | 0.43 | 0.24 | 0.26 | 0.31 | 0.37 |
| 11 | 123024166 | rs11219034 | -0.607 | -1.28 | 0.884 | -0.415 | -0.008 | -0.003 | -0.003 | 0 | 0.31 | 0.34 | 0.19 | 0.26 | 0.24 | 0.33 |
| 11 | 123025482 | rs12278752 | -0.443 | -1.266 | 0.806 | -0.434 | -0.003 | -0.031 | -0.031 | 0 | 0.09 | 0.12 | 0.06 | 0 | 0.08 | 0.06 |
| 11 | 123025690 | rs10790556 | -2.272 | -1.236 | 0.548 | -0.511 | -0.065 | -0.013 | -0.013 | 0 | 0.61 | 0.84 | 0.45 | 0.36 | 0.52 | 0.51 |
| 11 | 123026014 | rs9943547 | -0.906 | -1.154 | 0.468 | -0.243 | -0.017 | -0.008 | -0.008 | 0 | 0.31 | 0.35 | 0.19 | 0.25 | 0.24 | 0.33 |
| 11 | 123027157 | rs11600361 | -0.924 | -1.09 | -0.159 | -0.189 | 0.016 | -0.007 | -0.007 | 0 | 0.29 | 0.21 | 0.28 | 0.31 | 0.24 | 0.3 |
| 11 | 123027357 | rs10400330 | -0.832 | -1.151 | -0.138 | -0.657 | 0.016 | 0.022 | 0.022 | 0 | 0.28 | 0.2 | 0.15 | 0.04 | 0.21 | 0.28 |
| 11 | 123027583 | rs10400282 | -0.511 | -1.053 | -0.06 | -0.494 | 0.009 | 0.011 | 0.011 | 0 | 0.23 | 0.18 | 0.14 | 0.04 | 0.19 | 0.26 |
| 11 | 123029038 | rs12281746 | -0.92 | -0.833 | 0.209 | 0.006 | -0.004 | -0.025 | -0.025 | 0 | 0.06 | 0.11 | 0.04 | 0 | 0.06 | 0.02 |
| 11 | 123029545 | rs10400356 | -1.031 | -0.975 | 0.122 | -0.071 | -0.02 | -0.025 | -0.025 | 0 | 0.13 | 0.17 | 0.06 | 0.04 | 0.08 | 0.2 |
| 11 | 123031834 | rs10892974 | 0.977 | -0.319 | -0.573 | 0.044 | -0.008 | 0.011 | 0.011 | 0 | 0.51 | 0.21 | 0.49 | 0.54 | 0.54 | 0.61 |
| 11 | 123034921 | rs12285884 | -1.293 | -0.105 | 0.301 | 0.155 | -0.007 | 0.003 | 0.003 | 0 | 0.07 | 0.18 | 0.04 | 0.07 | 0.06 | 0.08 |
| 11 | 123035936 | rs12802926 | -1.422 | -0.068 | 0.473 | 0.064 | 0.01 | -0.065 | -0.065 | 0 | 0.17 | 0.54 | 0.19 | 0.08 | 0.14 | 0.15 |
| 11 | 123046705 | rs11828777 | 0.184 | -0.137 | -0.497 | 0.588 | 0.009 | 0.016 | 0.016 | 0 | 0.55 | 0.77 | 0.61 | 0.6 | 0.59 | 0.56 |
| 11 | 123048162 | rs7929006 | 0.619 | -0.276 | -0.103 | 0.349 | 0.036 | 0.084 | 0.084 | 0 | 0.51 | 0.23 | 0.67 | 0.67 | 0.59 | 0.55 |
| 11 | 123063365 | rs6589973 | -0.196 | 0.111 | -0.591 | 0.782 | -0.031 | -0.098 | -0.098 | 0 | 0.73 | 0.92 | 0.81 | 0.61 | 0.72 | 0.79 |
| 11 | 123074399 | rs4267083 | 0.328 | 0.319 | 0.056 | 0.424 | 0 | 0.007 | 0.007 | 0 | 0.53 | 0.57 | 0.51 | 0.3 | 0.55 | 0.68 |
| 11 | 123074716 | rs11820556 | -1.087 | 0.162 | 0.859 | 0.601 | -0.027 | -0.096 | -0.096 | 0.838 | 0.14 | 0.34 | 0.09 | 0 | 0.1 | 0.11 |
| 11 | 123074915 | rs485345 | 0.366 | 0.386 | 0.714 | 0.906 | -0.031 | 0.036 | 0.036 | 0.096 | 0.7 | 0.44 | 0.62 | 0.42 | 0.76 | 0.76 |
| 11 | 123076691 | rs577777 | 0.501 | 0.455 | 0.543 | 0.922 | -0.026 | 0.054 | 0.054 | 0.01 | 0.71 | 0.53 | 0.64 | 0.46 | 0.77 | 0.77 |
| 11 | 123078194 | rs577432 | 1.282 | 0.784 | -0.069 | 0.819 | 0.035 | 0.014 | 0.014 | 0.68 | 0.52 | 0.15 | 0.53 | 0.44 | 0.54 | 0.67 |
| 11 | 123078524 | rs526199 | 1.252 | 1.129 | 0.056 | 1.004 | -0.002 | -0.024 | -0.024 | 0.938 | 0.55 | 0.11 | 0.52 | 0.48 | 0.52 | 0.69 |
| 11 | 123079780 | rs12284903 | -1.646 | 0.708 | 1.047 | 0.823 | 0.022 | -0.012 | -0.012 | 0 | 0.07 | 0.29 | 0.06 | 0.02 | 0.06 | 0.02 |
| 11 | 123089955 | rs513136 | 1.492 | 1.419 | 0.502 | 1.045 | -0.056 | -0.076 | -0.076 | 0.604 | 0.54 | 0.08 | 0.59 | 0.5 | 0.55 | 0.62 |
| 11 | 123092243 | rs10892983 | -0.845 | 1.339 | 1.237 | 0.908 | -0.017 | -0.086 | -0.086 | 0.003 | 0.21 | 0.29 | 0.08 | 0 | 0.13 | 0.16 |
| 11 | 123092449 | rs488943 | -0.183 | 1.378 | 1.044 | 0.947 | -0.034 | -0.067 | -0.067 | 2.526 | 0.86 | 0.91 | 0.79 | 0.52 | 0.8 | 0.81 |
| 11 | 123093198 | rs12272976 | -1.588 | 0.945 | 1.662 | 0.772 | -0.003 | -0.061 | -0.061 | 0 | 0.26 | 0.69 | 0.15 | 0 | 0.19 | 0.17 |
| 11 | 123094797 | rs4245047 | 0.207 | 1.092 | 1.593 | 1.157 | -0.023 | -0.051 | -0.051 | 1.659 | 0.83 | 0.85 | 0.79 | 0.59 | 0.8 | 0.79 |
| 11 | 123096559 | rs7120082 | -0.884 | 1.022 | 2.206 | 1.335 | -0.014 | -0.013 | -0.013 | 0.618 | 0.2 | 0.33 | 0.17 | 0.18 | 0.16 | 0.17 |
| 11 | 123096933 | rs559254 | 0.411 | 1.072 | 1.751 | 1.388 | -0.005 | -0.001 | -0.001 | 0 | 0.82 | 0.75 | 0.76 | 0.57 | 0.78 | 0.79 |
| 11 | 123097158 | rs3018105 | 0.298 | 1.072 | 1.539 | 1.389 | -0.019 | -0.042 | -0.042 | 0.042 | 0.82 | 0.84 | 0.77 | 0.57 | 0.78 | 0.79 |

| | | | | | | | | | | | | | | | | |
|---|---|---|---|---|---|---|---|---|---|---|---|---|---|---|---|---|
| 11 | 123097386 | rs493642 | 1.707 | 1.121 | 1.49 | 1.379 | -0.007 | 0.036 | 0.036 | 0 | 0.57 | 0.2 | 0.57 | 0.37 | 0.6 | 0.61 |
| 11 | 123106025 | rs3898437 | -0.747 | 0.873 | 1.397 | 1.268 | 0.031 | 0.029 | 0.029 | 0 | 0.23 | 0.1 | 0.16 | 0.15 | 0.16 | 0.17 |
| 11 | 123114359 | rs558362 | 0.772 | 0.884 | 1.679 | 1.632 | -0.014 | -0.008 | -0.008 | 0 | 0.84 | 0.82 | 0.78 | 0.55 | 0.79 | 0.81 |
| 11 | 123114432 | rs558560 | 1.921 | 0.82 | 1.916 | 1.588 | 0.002 | 0.016 | 0.016 | 1.975 | 0.77 | 0.27 | 0.72 | 0.54 | 0.77 | 0.79 |
| 11 | 123116360 | rs508661 | 1.61 | 0.79 | 2.268 | 1.559 | 0.052 | 0.176 | 0.176 | 0.339 | 0.79 | 0.46 | 0.74 | 0.55 | 0.78 | 0.79 |
| 11 | 123119388 | rs528990 | 1.222 | 0.789 | 2.386 | 1.562 | 0.061 | 0.184 | 0.184 | 0 | 0.81 | 0.64 | 0.77 | 0.55 | 0.79 | 0.79 |
| 11 | 123124570 | rs10892985 | 2.217 | 0.817 | 2.01 | 1.375 | -0.032 | 0.024 | 0.024 | 0 | 0.55 | 0.14 | 0.56 | 0.4 | 0.61 | 0.67 |
| 11 | 123125779 | rs10892986 | 1.107 | 1.118 | 2.293 | 1.205 | 0.005 | 0.016 | 0.016 | 0 | 0.82 | 0.79 | 0.77 | 0.62 | 0.77 | 0.79 |
| 11 | 123128812 | rs7128842 | -0.971 | 1.118 | 2.206 | 1.219 | -0.013 | -0.009 | -0.009 | 0 | 0.15 | 0.13 | 0.21 | 0.38 | 0.21 | 0.19 |
| 11 | 123132645 | rs7115792 | -2.022 | 1.211 | 2.091 | 1.203 | 0.023 | 0.173 | 0.173 | 0.47 | 0.19 | 0.6 | 0.23 | 0.46 | 0.18 | 0.2 |
| 11 | 123136241 | rs7950320 | -0.74 | 0.945 | 1.61 | 1.154 | 0.009 | 0.034 | 0.034 | 0 | 0.16 | 0.21 | 0.31 | 0.34 | 0.19 | 0.19 |
| 11 | 123136552 | rs2000796 | -0.809 | 0.719 | 1.645 | 0.729 | -0.016 | -0.057 | -0.057 | 0.374 | 0.23 | 0.3 | 0.14 | 0.08 | 0.16 | 0.13 |
| 11 | 123138960 | rs7927484 | -1.172 | 0.755 | 0.934 | 0.76 | -0.011 | -0.038 | -0.038 | 0.108 | 0.27 | 0.63 | 0.18 | 0.08 | 0.19 | 0.14 |
| 11 | 123149582 | rs7934887 | -1.358 | 0.87 | 0.661 | 1.238 | 0.016 | 0.073 | 0.073 | 0.059 | 0.24 | 0.49 | 0.35 | 0.36 | 0.23 | 0.26 |
| 11 | 123153094 | rs6589985 | -0.744 | 0.532 | 0.035 | 0.545 | 0.003 | -0.005 | -0.005 | 0 | 0.4 | 0.43 | 0.32 | 0.26 | 0.42 | 0.24 |
| 11 | 123153359 | rs6589987 | -0.746 | 0.528 | 0.085 | 0.551 | 0.003 | -0.012 | -0.012 | 0.38 | 0.4 | 0.42 | 0.32 | 0.21 | 0.42 | 0.24 |
| 11 | 123155262 | rs10892995 | -0.64 | 0.33 | 0.179 | 0.658 | 0.1 | 0.533 | 0.533 | 3.787 | 0.05 | 0.47 | 0.19 | 0.32 | 0.03 | 0.06 |
| 11 | 123157134 | rs10892996 | -0.942 | 0.361 | -0.583 | 0.533 | -0.062 | -0.016 | -0.016 | 1.459 | 0.45 | 0.88 | 0.51 | 0.44 | 0.46 | 0.29 |
| 11 | 123158221 | rs10892999 | -1.419 | -0.002 | -1.01 | 0.035 | -0.07 | -0.058 | -0.058 | 0 | 0.57 | 0.94 | 0.56 | 0.44 | 0.53 | 0.38 |
| 11 | 123161406 | rs10736527 | -0.578 | 0.008 | -0.752 | 0.099 | -0.04 | -0.015 | -0.015 | 0 | 0.42 | 0.83 | 0.51 | 0.43 | 0.45 | 0.28 |
| 11 | 123165765 | rs6589991 | -0.675 | -0.108 | -0.73 | 0.038 | 0.009 | -0.028 | -0.028 | 0 | 0.35 | 0.83 | 0.42 | 0.42 | 0.32 | 0.25 |
| 11 | 123166133 | rs6589992 | -1.311 | -0.103 | -1.066 | 0.053 | -0.034 | -0.031 | -0.031 | 0 | 0.46 | 0.96 | 0.46 | 0.43 | 0.38 | 0.31 |
| 11 | 123175815 | rs2156445 | -1.381 | -0.384 | -1.085 | -0.004 | -0.021 | -0.036 | -0.036 | 0.575 | 0.47 | 0.95 | 0.46 | 0.46 | 0.38 | 0.32 |
| 11 | 123176516 | rs11602677 | -0.095 | -0.459 | -0.661 | -0.235 | -0.005 | 0.012 | 0.012 | 0 | 0.25 | 0.18 | 0.19 | 0.15 | 0.26 | 0.2 |
| 11 | 123181990 | rs17127237 | -0.58 | -0.785 | -0.747 | -0.193 | 0.006 | -0.013 | -0.013 | 0 | 0.15 | 0.09 | 0.07 | 0.01 | 0.08 | 0.06 |
| 11 | 123193404 | rs10736528 | -0.761 | -0.215 | -0.352 | 0.248 | 0.011 | -0.039 | -0.039 | 0 | 0.31 | 0.84 | 0.37 | 0.46 | 0.26 | 0.25 |
| 11 | 123196784 | rs7119283 | -1.194 | -0.238 | -0.382 | 0.21 | 0.004 | -0.091 | -0.091 | 0.84 | 0.36 | 0.93 | 0.42 | 0.46 | 0.28 | 0.26 |
| 11 | 123216036 | rs7103133 | -0.908 | -0.124 | -0.469 | 0.158 | 0.013 | -0.055 | -0.055 | 1.154 | 0.32 | 0.84 | 0.38 | 0.51 | 0.28 | 0.25 |
| 11 | 123222997 | rs10893024 | -1.448 | -0.18 | -0.969 | 0.059 | -0.008 | -0.112 | -0.112 | 0 | 0.44 | 0.94 | 0.45 | 0.51 | 0.35 | 0.28 |
| 11 | 123229854 | rs9666197 | -0.454 | -0.262 | -0.339 | -0.594 | 0.003 | -0.004 | -0.004 | 0 | 0.23 | 0.35 | 0.19 | 0.18 | 0.23 | 0.15 |
| 11 | 123234210 | rs17127284 | -1.131 | -0.832 | -0.351 | -1.103 | -0.033 | -0.107 | -0.107 | 0 | 0.15 | 0.59 | 0.08 | 0 | 0.05 | 0.09 |
| 11 | 123235706 | rs7934888 | -1.514 | -0.586 | -1.212 | -1.062 | 0.012 | 0.047 | 0.047 | 2.26 | 0.31 | 0.88 | 0.25 | 0.14 | 0.2 | 0.2 |
| 11 | 123235872 | rs7948443 | -1.514 | -0.586 | -1.212 | -1.062 | 0.012 | 0.047 | 0.047 | 2.26 | 0.31 | 0.88 | 0.24 | 0.15 | 0.19 | 0.2 |
| 11 | 123236983 | rs10893030 | -1.322 | -0.544 | -1.183 | -1.03 | 0.007 | 0.026 | 0.026 | 3.432 | 0.27 | 0.88 | 0.22 | 0.14 | 0.17 | 0.15 |
| 11 | 123240173 | rs11219122 | -1.581 | -0.308 | -1.638 | -0.998 | 0.004 | 0.143 | 0.143 | 0.73 | 0.46 | 0.92 | 0.34 | 0.2 | 0.33 | 0.29 |
| 11 | 123256550 | rs12224823 | -0.598 | -0.555 | -1.346 | -0.743 | 0.052 | 0.055 | 0.055 | 0.482 | 0.2 | 0.02 | 0.06 | 0.06 | 0.09 | 0.1 |
| 11 | 123261385 | rs7927064 | -1.381 | -0.228 | -1.103 | -0.544 | 0.015 | -0.004 | -0.004 | 0 | 0.12 | 0.92 | 0.17 | 0.12 | 0.08 | 0.05 |
| 11 | 123264642 | rs1943466 | -1.178 | -0.411 | -0.553 | -1.017 | 0.019 | 0.021 | 0.021 | 0.085 | 0.46 | 0.6 | 0.37 | 0.15 | 0.29 | 0.3 |
| 11 | 123275496 | rs17127342 | -1.712 | 0.594 | 0.801 | -0.366 | 0.003 | -0.046 | -0.046 | 0 | 0.15 | 0.66 | 0.16 | 0.05 | 0.11 | 0.1 |
| 11 | 123278907 | rs17127343 | -0.487 | 0.58 | 0.899 | -0.365 | -0.014 | -0.034 | -0.034 | 0 | 0.07 | 0.23 | 0.02 | 0 | 0.01 | 0.01 |
| 11 | 123280309 | rs2282561 | -1.005 | 0.608 | 0.841 | -0.341 | 0.119 | 0.036 | 0.036 | 0.328 | 0.08 | 0.35 | 0.14 | 0.05 | 0.1 | 0.08 |
| 11 | 123283795 | rs2714061 | -1.024 | 1.393 | 0.437 | 0.224 | 0.047 | 0.058 | 0.058 | 0.27 | 0.12 | 0.27 | 0.12 | 0.16 | 0.16 | 0.09 |
| 11 | 123283922 | rs2714059 | -1.567 | 1.329 | -0.543 | 0.187 | 0.018 | 0.007 | 0.007 | 0.275 | 0.22 | 0.7 | 0.26 | 0.21 | 0.29 | 0.18 |
| 11 | 123298088 | rs4936805 | 0.183 | 0.98 | -1.842 | -0.463 | -0.046 | 0.211 | 0.211 | 0 | 0.49 | 0.1 | 0.57 | 0.81 | 0.47 | 0.7 |
| 11 | 123300771 | rs4078010 | -0.548 | 0.655 | -0.587 | -0.682 | 0.05 | -0.035 | -0.035 | 0 | 0.09 | 0.43 | 0.06 | 0 | 0.05 | 0.01 |
| 11 | 123304650 | rs3016380 | 0.134 | 0.53 | -0.378 | -1.003 | -0.004 | -0.051 | -0.051 | 0.013 | 0.15 | 0.21 | 0.1 | 0.01 | 0.16 | 0.08 |
| 11 | 123306172 | rs17671708 | 0.457 | 0.516 | -0.534 | -1.218 | 0.001 | 0.018 | 0.018 | 0 | 0.12 | 0.07 | 0.1 | 0.06 | 0.13 | 0.12 |
| 11 | 123307262 | rs2846293 | -0.077 | 0.383 | -1.228 | -1.074 | 0.007 | -0.007 | -0.007 | 0 | 0.63 | 0.58 | 0.64 | 0.74 | 0.52 | 0.67 |
| 11 | 123308082 | rs11219139 | 1.236 | 0.084 | -1.576 | -0.036 | 0.033 | -0.167 | -0.167 | 0 | 0.33 | 0.05 | 0.39 | 0.51 | 0.23 | 0.42 |
| 11 | 123309473 | rs733353 | 0.265 | 0.25 | -1.296 | -0.108 | 0.047 | 0.051 | 0.051 | 0 | 0.49 | 0.35 | 0.54 | 0.73 | 0.37 | 0.57 |
| 11 | 123312063 | rs2714053 | -0.536 | 0.109 | -1.877 | 0.43 | 0.006 | -0.046 | -0.046 | 0 | 0.59 | 0.85 | 0.44 | 0.55 | 0.61 | 0.49 |
| 11 | 123318063 | rs2846062 | -0.446 | 0.275 | -1.365 | -0.039 | 0.014 | 0.021 | 0.021 | 0.321 | 0.57 | 0.73 | 0.47 | 0.39 | 0.6 | 0.39 |
| 11 | 123318348 | rs7105246 | -0.994 | 0.994 | -0.735 | 0.22 | 0.011 | 0.006 | 0.006 | 0 | 0.14 | 0.26 | 0.14 | 0.22 | 0.15 | 0.21 |
| 11 | 123318657 | rs11219141 | -0.904 | 0.957 | -0.605 | 0.256 | 0.026 | 0.031 | 0.031 | 0 | 0.14 | 0.35 | 0.13 | 0.24 | 0.14 | 0.2 |
| 11 | 123320804 | rs12421839 | -0.531 | 0.545 | -1.244 | -0.005 | -0.004 | -0.012 | -0.012 | 0.05 | 0.36 | 0.56 | 0.26 | 0.33 | 0.35 | 0.33 |
| 11 | 123324588 | rs4936808 | 0.104 | 0.249 | -1.708 | 0.04 | 0.003 | -0.006 | -0.006 | 0 | 0.26 | 0.22 | 0.35 | 0.37 | 0.24 | 0.24 |
| 11 | 123334298 | rs12788072 | 0.576 | 0.76 | -1.885 | -0.288 | -0.011 | 0.03 | 0.03 | 0 | 0.16 | 0.08 | 0.1 | 0.01 | 0.15 | 0.07 |
| 11 | 123337219 | rs7107803 | 0.152 | 0.332 | -1.846 | -0.093 | 0.006 | -0.003 | -0.003 | 0 | 0.33 | 0.29 | 0.38 | 0.35 | 0.29 | 0.27 |
| 11 | 123337892 | rs4936810 | -0.469 | 1.225 | -2.757 | -0.671 | -0.119 | 0.138 | 0.138 | 0.007 | 0.77 | 0.98 | 0.85 | 0.97 | 0.64 | 0.83 |
| 11 | 123338245 | rs17127396 | -0.203 | 0.75 | -2.375 | 0.067 | -0.014 | -0.087 | -0.087 | 2.116 | 0.29 | 0.11 | 0.35 | 0.43 | 0.26 | 0.39 |
| 11 | 123345497 | rs2714048 | 0.14 | 0.512 | -2.355 | -0.847 | -0.019 | 0.101 | 0.101 | 0 | 0.64 | 0.89 | 0.56 | 0.21 | 0.62 | 0.56 |
| 11 | 123349468 | rs1012849 | -0.486 | 0.569 | -2.314 | -0.269 | -0.055 | -0.104 | -0.104 | 0 | 0.83 | 0.96 | 0.68 | 0.57 | 0.76 | 0.73 |
| 11 | 123359212 | rs2846319 | -0.268 | 0.416 | -1.6 | -1.082 | -0.007 | 0.005 | 0.005 | 1.167 | 0.41 | 0.5 | 0.28 | 0.53 | 0.33 | 0.31 |
| 11 | 123361020 | rs2714084 | -0.434 | 0.32 | -1.709 | -1.128 | 0.033 | -0.018 | -0.018 | 0 | 0.43 | 0.74 | 0.31 | 0.53 | 0.34 | 0.32 |
| 11 | 123361397 | rs735665 | 1.061 | 0.785 | -2.004 | -1.143 | -0.033 | 0.078 | 0.078 | 0 | 0.15 | 0.02 | 0.22 | 0.01 | 0.17 | 0.14 |
| 11 | 123364917 | rs3016471 | -0.811 | 0.912 | -1.821 | -0.736 | -0.066 | -0.132 | -0.132 | 0 | 0.81 | 0.99 | 0.71 | 0.63 | 0.77 | 0.7 |
| 11 | 123367686 | rs2714081 | 0.436 | 0.624 | -1.146 | -1.023 | 0.003 | -0.045 | -0.045 | 0 | 0.17 | 0.24 | 0.11 | 0.04 | 0.17 | 0.1 |
| 11 | 123370239 | rs2714078 | -0.169 | 0.586 | -0.738 | -0.872 | 0 | -0.005 | -0.005 | 0 | 0.7 | 0.68 | 0.61 | 0.59 | 0.7 | 0.57 |
| 11 | 123375792 | rs2846300 | 0.488 | 0.4 | -0.664 | -1.135 | -0.002 | -0.041 | -0.041 | 0 | 0.52 | 0.47 | 0.47 | 0.16 | 0.55 | 0.43 |

| | | | | | | | | | | | | | | | | |
|---|---|---|---|---|---|---|---|---|---|---|---|---|---|---|---|---|
| 11 | 123378721 | rs11219179 | 0.609 | 0.449 | -1.032 | -1.025 | 0.009 | -0.054 | -0.054 | 0 | 0.17 | 0.09 | 0.11 | 0.45 | 0.14 | 0.15 |
| 11 | 123379387 | rs2714070 | -1.052 | 0.556 | -1.282 | -1.023 | -0.049 | -0.129 | -0.129 | 0.237 | 0.77 | 0.98 | 0.69 | 0.61 | 0.76 | 0.62 |
| 11 | 123382247 | rs4936813 | 0.32 | 0.46 | -0.832 | -0.759 | 0.026 | 0.006 | 0.006 | 0 | 0.05 | 0.2 | 0.07 | 0.03 | 0.06 | 0.05 |
| 11 | 123383072 | rs12574747 | -0.099 | 0.382 | -1.281 | -0.66 | -0.017 | 0.081 | 0.081 | 0 | 0.2 | 0.27 | 0.14 | 0.45 | 0.15 | 0.15 |
| 11 | 123390225 | rs12799226 | 0.872 | 0.864 | -2.018 | -1.291 | -0.004 | 0.009 | 0.009 | 0 | 0.17 | 0.12 | 0.25 | 0.01 | 0.19 | 0.17 |
| 11 | 123390273 | rs4258388 | -0.053 | 0.885 | -2.463 | -1.146 | 0.012 | 0.15 | 0.15 | 0.203 | 0.41 | 0.11 | 0.42 | 0.8 | 0.37 | 0.52 |
| 11 | 123390969 | rs2846304 | 0.236 | 1.427 | -1.985 | -0.646 | -0.036 | 0.038 | 0.038 | 1.676 | 0.14 | 0.06 | 0.11 | 0.06 | 0.18 | 0.1 |
| 11 | 123395348 | rs17127435 | 0.109 | 1.286 | -1.734 | 0.144 | -0.001 | 0.038 | 0.038 | 0.007 | 0.13 | 0.09 | 0.12 | 0.42 | 0.15 | 0.14 |
| 11 | 123395470 | rs4936822 | -0.601 | 1.15 | -2.224 | -0.492 | -0.024 | 0.112 | 0.112 | 0.269 | 0.72 | 0.89 | 0.81 | 0.94 | 0.72 | 0.83 |
| 11 | 123395864 | rs7927176 | -0.366 | 0.94 | -1.889 | 0.19 | -0.005 | -0.079 | -0.079 | 0 | 0.3 | 0.12 | 0.33 | 0.41 | 0.31 | 0.46 |
| 11 | 123398941 | rs7110875 | -0.833 | 1.904 | -1.538 | 0.908 | -0.068 | -0.068 | -0.068 | 0.003 | 0.2 | 0.05 | 0.28 | 0.29 | 0.28 | 0.15 |
| 11 | 123399541 | rs2714068 | 1.341 | 1.779 | -0.643 | 1.004 | 0.021 | 0.02 | 0.02 | 0 | 0.65 | 0.54 | 0.56 | 0.49 | 0.57 | 0.63 |
| 11 | 123408945 | rs4936824 | -0.524 | 1.932 | -0.57 | 1.185 | -0.042 | -0.059 | -0.059 | 0.361 | 0.2 | 0.05 | 0.28 | 0.25 | 0.24 | 0.15 |
| 11 | 123409074 | rs850291 | -1.467 | 1.886 | -0.144 | 1.053 | 0.01 | 0.026 | 0.026 | 0 | 0.3 | 0.33 | 0.39 | 0.49 | 0.34 | 0.31 |
| 11 | 123411662 | rs2465460 | 1.253 | 1.684 | -0.114 | 1.054 | -0.018 | -0.025 | -0.025 | 0.119 | 0.64 | 0.26 | 0.52 | 0.49 | 0.6 | 0.64 |
| 11 | 123413037 | rs1290821 | -0.627 | 1.711 | 0.248 | 1.16 | 0.024 | 0.038 | 0.038 | 0 | 0.07 | 0.11 | 0.07 | 0.15 | 0.1 | 0.09 |
| 11 | 123414451 | rs1784356 | 0.812 | 1.81 | 0.474 | 1.28 | 0.021 | 0.034 | 0.034 | 0.439 | 0.74 | 0.39 | 0.54 | 0.67 | 0.71 | 0.81 |
| 11 | 123416874 | rs1290811 | 0.178 | 1.805 | -0.43 | 0.714 | -0.101 | -0.017 | -0.017 | 0 | 0.81 | 0.98 | 0.73 | 0.84 | 0.71 | 0.84 |
| 11 | 123421402 | rs1029278 | -1.573 | 1.012 | -1.254 | 0.535 | -0.021 | -0.028 | -0.028 | 0.005 | 0.65 | 0.91 | 0.77 | 0.68 | 0.69 | 0.48 |
| 11 | 123424329 | rs863731 | -1.188 | 0.462 | -0.595 | 0.314 | 0.002 | -0.005 | -0.005 | 0 | 0.28 | 0.7 | 0.42 | 0.35 | 0.26 | 0.22 |
| 11 | 123424434 | rs11219196 | 0.457 | 0.297 | 0.248 | 0.413 | -0.001 | 0.076 | 0.076 | 1.668 | 0.07 | 0.34 | 0.05 | 0.08 | 0.01 | 0.01 |
| 11 | 123429175 | rs2852845 | -0.66 | 0.464 | 0.05 | 0.336 | -0.039 | 0.043 | 0.043 | 0.854 | 0.26 | 0.48 | 0.23 | 0.34 | 0.22 | 0.14 |
| 11 | 123429299 | rs743632 | -0.022 | 0.498 | -0.005 | 0.213 | 0.016 | -0.05 | -0.05 | 0 | 0.11 | 0.15 | 0.23 | 0 | 0.14 | 0.1 |
| 11 | 123429425 | rs743631 | -1.421 | 0.498 | -0.519 | 0.36 | 0.008 | -0.013 | -0.013 | 0 | 0.37 | 0.68 | 0.47 | 0.42 | 0.36 | 0.26 |
| 11 | 123434062 | rs1628021 | -0.327 | 0.319 | -0.526 | -0.129 | -0.005 | -0.011 | -0.011 | 0.202 | 0.11 | 0.11 | 0.22 | 0 | 0.14 | 0.1 |
| 11 | 123438030 | rs1784359 | 0.87 | 0.01 | -0.318 | -0.186 | 0 | 0 | 0 | 0 | 0.54 | 0.49 | 0.48 | 0.71 | 0.57 | 0.73 |
| 11 | 123442566 | rs1792005 | -0.09 | -0.82 | -1.097 | -0.062 | -0.005 | 0.056 | 0.056 | 0 | 0.51 | 0.84 | 0.64 | 0.63 | 0.51 | 0.44 |
| 11 | 123444074 | rs1784362 | 0.978 | -0.94 | -0.863 | -0.323 | -0.025 | 0.011 | 0.011 | 0 | 0.25 | 0.15 | 0.2 | 0.18 | 0.28 | 0.46 |
| 11 | 123444231 | rs1791710 | -0.227 | -1.308 | -1 | -0.52 | 0.027 | 0.133 | 0.133 | 1.529 | 0.52 | 0.76 | 0.71 | 0.82 | 0.58 | 0.5 |
| 11 | 123444860 | rs2078158 | 0.999 | -0.816 | -0.673 | -0.147 | 0.007 | 0.016 | 0.016 | 0.627 | 0.29 | 0.33 | 0.34 | 0.46 | 0.37 | 0.35 |
| 11 | 123447713 | rs2852732 | -0.135 | -0.786 | -0.935 | 0.047 | -0.019 | -0.042 | -0.042 | 0 | 0.3 | 0.16 | 0.22 | 0.34 | 0.31 | 0.46 |
| 11 | 123452297 | rs7117990 | 0.359 | -0.59 | -0.613 | -0.235 | -0.024 | 0.002 | 0.002 | 0.693 | 0.06 | 0.19 | 0.04 | 0.05 | 0.02 | 0.01 |
| 11 | 123460599 | rs7110733 | 0.287 | -0.345 | -0.954 | -0.23 | -0.058 | 0.006 | 0.006 | 0 | 0.24 | 0.07 | 0.25 | 0.18 | 0.32 | 0.44 |
| 11 | 123467227 | rs736105 | -0.691 | -1.297 | -0.893 | -0.663 | 0.037 | 0.025 | 0.025 | 1.661 | 0.79 | 0.86 | 0.91 | 1 | 0.88 | 0.95 |
| 11 | 123470135 | rs1148096 | -0.116 | -1.366 | -0.847 | -0.799 | 0.036 | 0.022 | 0.022 | 1.771 | 0.73 | 0.77 | 0.86 | 1 | 0.83 | 0.93 |
| 11 | 123470475 | rs1274208 | -0.128 | -1.365 | -0.849 | -0.799 | 0.037 | 0.025 | 0.025 | 1.863 | 0.73 | 0.77 | 0.86 | 1 | 0.83 | 0.93 |
| 11 | 123472528 | rs1148093 | 0.6 | -1.345 | -0.373 | -0.748 | 0.005 | -0.011 | -0.011 | 0.005 | 0.42 | 0.53 | 0.57 | 0.74 | 0.48 | 0.45 |
| 11 | 123472547 | rs12291891 | 1.562 | -1.258 | -0.173 | -0.543 | 0.009 | 0.037 | 0.037 | 0.017 | 0.31 | 0.41 | 0.35 | 0.49 | 0.35 | 0.33 |
| 11 | 123474408 | rs1709901 | 0.547 | -1.023 | -0.443 | -0.387 | -0.039 | -0.051 | -0.051 | 0 | 0.1 | 0.01 | 0.2 | 0.24 | 0.12 | 0.11 |
| 11 | 123474531 | rs1274214 | -0.516 | -1.253 | -0.029 | -0.543 | 0.001 | 0 | 0 | 0 | 0.42 | 0.52 | 0.54 | 0.43 | 0.5 | 0.58 |
| 11 | 123475615 | rs1274213 | 0.158 | -1.34 | -0.241 | -1.403 | -0.002 | -0.001 | -0.001 | 0.041 | 0.1 | 0.07 | 0.2 | 0.06 | 0.11 | 0.1 |
| 11 | 123477352 | rs2279519 | 0.888 | -1.272 | -0.192 | -1.421 | -0.008 | 0.003 | 0.003 | 0 | 0.23 | 0.19 | 0.25 | 0.17 | 0.3 | 0.45 |
| 11 | 123480198 | rs11219207 | 0.48 | -1.252 | -0.195 | -1.402 | 0.007 | -0.005 | -0.005 | 0.272 | 0.27 | 0.26 | 0.27 | 0.21 | 0.32 | 0.5 |
| 11 | 123480981 | rs10893053 | 0.975 | -1.182 | -0.498 | -1.375 | -0.053 | 0.005 | 0.005 | 0 | 0.23 | 0.07 | 0.24 | 0.17 | 0.31 | 0.47 |
| 11 | 123485444 | rs732211 | 0.296 | -1.262 | -0.554 | -1.399 | -0.105 | 0.039 | 0.039 | 0 | 0.3 | 0.08 | 0.43 | 0.19 | 0.42 | 0.55 |
| 11 | 123490413 | rs12279179 | 1.331 | -1.072 | -0.541 | -1.405 | 0.004 | 0.001 | 0.001 | 0.29 | 0.3 | 0.29 | 0.34 | 0.71 | 0.35 | 0.34 |
| 11 | 123491346 | rs1275079 | 0.484 | -1.651 | -0.716 | -1.574 | -0.028 | -0.08 | -0.08 | 2.906 | 0.68 | 0.54 | 0.8 | 0.98 | 0.82 | 0.88 |
| 11 | 123494841 | rs1148115 | -0.131 | -1.457 | -1.206 | -1.448 | 0.074 | 0.097 | 0.097 | 1.157 | 0.77 | 0.9 | 0.9 | 1 | 0.87 | 0.95 |
| 11 | 123495005 | rs11219210 | 0.389 | -0.963 | -0.941 | -1.124 | 0.041 | 0.063 | 0.063 | 0 | 0.59 | 0.64 | 0.61 | 0.94 | 0.7 | 0.77 |
| 11 | 123498735 | rs1148114 | -0.622 | -0.885 | -1.312 | -0.992 | 0.056 | 0.145 | 0.145 | 0.66 | 0.37 | 0.21 | 0.37 | 0.08 | 0.27 | 0.22 |
| 11 | 123500113 | rs12286568 | -0.851 | -0.352 | -0.956 | -0.516 | -0.003 | 0.003 | 0.003 | 0.88 | 0.08 | 0.08 | 0.04 | 0.06 | 0.08 | 0.11 |
| 11 | 123500759 | rs2027767 | -0.955 | -1.073 | -0.477 | 0.04 | -0.069 | 0.082 | 0.082 | 0 | 0.25 | 0.48 | 0.17 | 0.36 | 0.18 | 0.23 |
| 11 | 123502761 | rs10400374 | -0.274 | -0.658 | -0.672 | -0.426 | -0.021 | -0.051 | -0.051 | 3.47 | 0.12 | 0.22 | 0.06 | 0 | 0.08 | 0.06 |
| 11 | 123505099 | rs1148109 | -0.362 | 0.493 | -1.153 | -0.013 | -0.022 | 0.006 | 0.006 | 0 | 0.64 | 0.75 | 0.53 | 0.62 | 0.53 | 0.71 |
| 11 | 123511651 | rs1720344 | 0.022 | 0.666 | -1.353 | -0.204 | -0.033 | 0.047 | 0.047 | 0 | 0.59 | 0.3 | 0.67 | 0.3 | 0.59 | 0.53 |
| 11 | 123513161 | rs1275085 | 0.015 | 0.415 | -1.237 | -0.5 | 0.007 | 0.007 | 0.007 | 0 | 0.14 | 0.1 | 0.06 | 0 | 0.1 | 0.14 |
| 11 | 123513506 | rs1783901 | 0.101 | -0.081 | -0.935 | -1.08 | -0.019 | -0.025 | -0.025 | 0 | 0.26 | 0.34 | 0.3 | 0.16 | 0.19 | 0.16 |
| 11 | 123513917 | rs1477825 | 0.115 | -0.205 | -0.422 | -1.085 | -0.038 | -0.126 | -0.126 | 0 | 0.26 | 0.43 | 0.19 | 0.08 | 0.18 | 0.1 |
| 11 | 123515644 | rs1296514 | -0.771 | 0.505 | -1.68 | -0.481 | -0.107 | -0.169 | -0.169 | 0 | 0.75 | 0.98 | 0.79 | 0.45 | 0.71 | 0.75 |
| 11 | 123515705 | rs5023548 | 0.321 | 0.605 | -1.329 | -0.245 | -0.01 | 0.003 | 0.003 | 0 | 0.06 | 0.04 | 0.18 | 0.17 | 0.11 | 0.12 |
| 11 | 123518917 | rs1720340 | 1 | 0.208 | -1.525 | -0.788 | 0.024 | 0.037 | 0.037 | 0.406 | 0.15 | 0.07 | 0.07 | 0.02 | 0.08 | 0.06 |
| 11 | 123519043 | rs1792016 | -0.372 | 0.188 | -1.583 | -0.987 | 0.003 | 0.037 | 0.037 | 0.033 | 0.26 | 0.19 | 0.33 | 0.12 | 0.26 | 0.13 |
| 11 | 123522995 | rs3851095 | -0.117 | 0.496 | -1.424 | -0.695 | -0.006 | -0.003 | -0.003 | 0 | 0.12 | 0.13 | 0.27 | 0.12 | 0.18 | 0.09 |
| 11 | 123523002 | rs3851096 | -0.117 | 0.496 | -1.443 | -0.695 | -0.008 | -0.004 | -0.004 | 4.158 | 0.12 | 0.12 | 0.26 | 0.12 | 0.18 | 0.07 |
| 11 | 123523421 | rs3781904 | 0.787 | 0.651 | -1.352 | -0.227 | -0.022 | -0.013 | -0.013 | 0 | 0.21 | 0.15 | 0.18 | 0.31 | 0.24 | 0.4 |
| 11 | 123528616 | rs11219218 | 0.879 | 0.728 | -1.453 | 0.302 | -0.063 | -0.127 | -0.127 | 1.649 | 0.2 | 0.01 | 0.17 | 0.39 | 0.21 | 0.21 |
| 11 | 123528678 | rs4936837 | -0.466 | 0.624 | -0.589 | 0.177 | 0.019 | 0.042 | 0.042 | 0.145 | 0.33 | 0.48 | 0.44 | 0.52 | 0.35 | 0.31 |
| 11 | 123531831 | rs12420938 | -0.096 | 0.831 | -0.809 | 0.322 | 0.022 | 0.025 | 0.025 | 0.242 | 0.08 | 0.15 | 0.23 | 0.11 | 0.14 | 0.07 |
| 11 | 123535299 | rs10893062 | 0.913 | 0.774 | -0.75 | 0.294 | -0.015 | -0.053 | -0.053 | 0.097 | 0.2 | 0.09 | 0.17 | 0.4 | 0.21 | 0.22 |

| | | | | | | | | | | | | | | | | |
|---|---|---|---|---|---|---|---|---|---|---|---|---|---|---|---|---|
| 11 | 123537272 | rs4935840 | 0.068 | 0.519 | -0.682 | 0.197 | -0.007 | -0.011 | -0.011 | 0 | 0.28 | 0.25 | 0.4 | 0.51 | 0.34 | 0.29 |
| 11 | 123539877 | rs1148105 | 0.735 | 0.123 | -0.715 | -0.472 | 0 | -0.016 | -0.016 | 0.029 | 0.19 | 0.16 | 0.11 | 0.02 | 0.14 | 0.12 |
| 11 | 123540059 | rs10893068 | 0.011 | 0.346 | -0.719 | -0.053 | -0.006 | -0.001 | -0.001 | 0.211 | 0.29 | 0.29 | 0.53 | 0.63 | 0.35 | 0.35 |
| 11 | 123542020 | rs1275096 | -0.015 | -0.028 | -1.218 | 0.21 | 0.01 | 0.022 | 0.022 | 0.113 | 0.47 | 0.19 | 0.31 | 0.43 | 0.46 | 0.51 |
| 11 | 123543428 | rs1275097 | -1.661 | 0.292 | -1.732 | -0.031 | -0.077 | -0.018 | -0.018 | 0.021 | 0.81 | 0.95 | 0.82 | 0.81 | 0.75 | 0.9 |
| 11 | 123543905 | rs12418011 | 0.489 | 0.19 | -1.352 | 0.194 | 0 | 0.025 | 0.025 | 0 | 0.07 | 0.1 | 0.33 | 0.17 | 0.09 | 0.1 |
| 11 | 123545130 | rs7124162 | 0.6 | -0.369 | -1.37 | -0.148 | -0.003 | -0.058 | -0.058 | 0 | 0.37 | 0.17 | 0.26 | 0.52 | 0.38 | 0.31 |
| 11 | 123545176 | rs1290018 | -0.261 | -0.395 | -0.907 | -0.215 | 0.014 | 0.014 | 0.014 | 1.56 | 0.61 | 0.42 | 0.4 | 0.49 | 0.57 | 0.6 |
| 11 | 123547510 | rs1975922 | 1.69 | -0.388 | -0.884 | -0.208 | -0.003 | -0.004 | -0.004 | 0 | 0.14 | 0.15 | 0.06 | 0.07 | 0.11 | 0.07 |
| 11 | 123549724 | rs1841876 | 1.69 | -0.388 | -0.884 | -0.208 | -0.003 | -0.004 | -0.004 | 0 | 0.14 | 0.15 | 0.06 | 0.07 | 0.11 | 0.07 |
| 11 | 123552095 | rs10790592 | 0.97 | -0.545 | -0.957 | -0.342 | -0.002 | -0.001 | -0.001 | 0 | 0.33 | 0.28 | 0.26 | 0.4 | 0.34 | 0.42 |
| 11 | 123554086 | rs2298809 | 0.065 | -0.551 | -0.96 | -0.344 | 0.002 | 0.003 | 0.003 | 0.128 | 0.47 | 0.39 | 0.32 | 0.47 | 0.45 | 0.48 |
| 11 | 123554152 | rs1720326 | -1.194 | -0.527 | -0.732 | -0.333 | -0.006 | 0.001 | 0.001 | 0.771 | 0.7 | 0.7 | 0.74 | 0.67 | 0.66 | 0.71 |
| 11 | 123555558 | rs1783900 | 0.723 | -0.157 | -1.037 | -0.024 | 0.005 | -0.007 | -0.007 | 1.508 | 0.21 | 0.2 | 0.41 | 0.2 | 0.21 | 0.22 |
| 11 | 123557596 | rs12273127 | 1.63 | 0.03 | -1.384 | -0.141 | 0.003 | -0.048 | -0.048 | 0 | 0.09 | 0.17 | 0.05 | 0 | 0.07 | 0.14 |
| 11 | 123562397 | rs3901231 | 1.155 | 0.061 | -1.609 | 0.133 | -0.027 | -0.06 | -0.06 | 0 | 0.24 | 0.07 | 0.21 | 0.4 | 0.27 | 0.28 |
| 11 | 123564532 | rs1617377 | 0.581 | 0.056 | -1.296 | 0.176 | 0.009 | 0.01 | 0.01 | 0 | 0.17 | 0.08 | 0.25 | 0.09 | 0.17 | 0.16 |
| 11 | 123569986 | rs10893076 | 0.353 | 0.405 | -1.135 | 0.512 | -0.018 | -0.039 | -0.039 | 0 | 0.33 | 0.18 | 0.25 | 0.57 | 0.36 | 0.58 |
| 11 | 123575057 | rs1275054 | 0.447 | 1.039 | -1.259 | 0.49 | -0.089 | 0.003 | 0.003 | 0 | 0.19 | 0.04 | 0.21 | 0.11 | 0.27 | 0.12 |
| 11 | 123578396 | rs10893077 | -0.283 | 0.202 | -0.589 | 0.755 | 0.003 | -0.003 | -0.003 | 0 | 0.7 | 0.69 | 0.72 | 0.73 | 0.75 | 0.84 |
| 11 | 123585960 | rs655756 | 0.669 | 0.159 | -0.815 | 0.526 | -0.009 | 0.115 | 0.115 | 0.392 | 0.23 | 0.03 | 0.11 | 0.03 | 0.2 | 0.12 |
| 11 | 123587111 | rs503766 | 0.086 | 0.212 | -0.563 | 0.652 | 0.012 | 0.01 | 0.01 | 0.584 | 0.28 | 0.23 | 0.27 | 0.18 | 0.25 | 0.14 |
| 11 | 123592239 | rs2276410 | 0.458 | -0.007 | -0.124 | 0.428 | 0.004 | 0.005 | 0.005 | 0 | 0.44 | 0.38 | 0.31 | 0.19 | 0.37 | 0.27 |
| 11 | 123593007 | rs2282646 | 0.657 | 0.072 | 0.168 | 0.755 | -0.023 | -0.02 | -0.02 | 0 | 0.26 | 0.22 | 0.4 | 0.47 | 0.37 | 0.56 |
| 11 | 123596055 | rs10904 | 0.819 | -0.205 | 0.148 | 0.555 | -0.008 | -0.026 | -0.026 | 0 | 0.38 | 0.39 | 0.29 | 0.14 | 0.3 | 0.21 |
| 11 | 123596703 | rs3183878 | 0.309 | -0.201 | 0.302 | 0.565 | -0.006 | -0.019 | -0.019 | 0 | 0.47 | 0.47 | 0.34 | 0.18 | 0.38 | 0.28 |
| 11 | 123597804 | rs679597 | -1.066 | 0.443 | 0.296 | 0.974 | 0.01 | 0.029 | 0.029 | 0 | 0.8 | 0.75 | 0.67 | 0.48 | 0.7 | 0.67 |
| 11 | 123598941 | rs2282644 | 1.609 | 0.441 | 0.259 | 1.008 | -0.094 | -0.098 | -0.098 | 0 | 0.16 | 0.02 | 0.3 | 0.36 | 0.27 | 0.3 |
| 11 | 123600475 | rs1144507 | -1.089 | 0.427 | 0.332 | 0.989 | 0.006 | 0.022 | 0.022 | 0 | 0.8 | 0.77 | 0.67 | 0.48 | 0.7 | 0.67 |
| 11 | 123608581 | rs1940185 | -0.989 | 0.416 | 0.336 | 0.99 | 0.031 | 0.073 | 0.073 | 0 | 0.8 | 0.67 | 0.65 | 0.48 | 0.7 | 0.66 |
| 11 | 123613030 | rs10893081 | -1.025 | 0.427 | 0.274 | 0.959 | 0.028 | 0.068 | 0.068 | 0 | 0.8 | 0.68 | 0.65 | 0.48 | 0.7 | 0.67 |
| 11 | 123615329 | rs12794996 | 1.594 | 0.44 | -0.301 | 0.927 | -0.089 | -0.095 | -0.095 | 0 | 0.16 | 0.03 | 0.3 | 0.36 | 0.27 | 0.3 |
| 11 | 123617918 | rs11601032 | 0.682 | 0.402 | -0.205 | 0.541 | 0 | 0 | 0 | 3.609 | 0.09 | 0.08 | 0.08 | 0.12 | 0.11 | 0.26 |
| 11 | 123624658 | rs12364099 | 1.405 | 0.251 | -0.397 | 0.347 | 0.014 | 0.12 | 0.12 | 0.169 | 0.22 | 0.02 | 0.09 | 0.02 | 0.18 | 0.09 |
| 11 | 123625998 | rs4935844 | 1.639 | 0.087 | -0.532 | 0.272 | -0.046 | 0.068 | 0.068 | 0 | 0.27 | 0.04 | 0.23 | 0.11 | 0.27 | 0.16 |
| 11 | 123627668 | rs10502270 | 0.036 | 0.202 | 0.081 | 0.555 | 0.016 | 0.013 | 0.013 | 2.428 | 0.7 | 0.62 | 0.57 | 0.36 | 0.59 | 0.4 |
| 11 | 123631872 | rs7103374 | -1.152 | 0.258 | -0.206 | 0.602 | -0.028 | -0.042 | -0.042 | 0.04 | 0.8 | 0.81 | 0.66 | 0.48 | 0.7 | 0.66 |
| 11 | 123640433 | rs10893088 | 0.179 | -0.071 | 0.107 | 0.436 | -0.001 | -0.008 | -0.008 | 0 | 0.48 | 0.47 | 0.37 | 0.31 | 0.45 | 0.44 |
| 11 | 123645682 | rs17127676 | 1.12 | -0.115 | 0.062 | 0.336 | -0.002 | -0.019 | -0.019 | 0 | 0.06 | 0.07 | 0.03 | 0 | 0.04 | 0.11 |
| 11 | 123648333 | rs17744026 | 1.412 | -0.205 | 0.188 | 0.226 | -0.007 | 0.057 | 0.057 | 0 | 0.28 | 0.11 | 0.24 | 0.12 | 0.27 | 0.14 |
| 11 | 123649435 | rs4245048 | -0.364 | -0.419 | 0.527 | 0.588 | -0.01 | 0.092 | 0.092 | 0 | 0.1 | 0.32 | 0.2 | 0.15 | 0.06 | 0.13 |
| 11 | 123666314 | rs998558 | 0.419 | -0.311 | 0.224 | 0.321 | -0.003 | -0.029 | -0.029 | 2.265 | 0.07 | 0.16 | 0.04 | 0 | 0.04 | 0.1 |
| 11 | 123669378 | rs7944946 | 0.961 | -0.495 | -0.18 | 0.096 | 0.02 | 0.123 | 0.123 | 0.29 | 0.22 | 0.02 | 0.09 | 0.02 | 0.18 | 0.09 |
| 11 | 123672363 | rs1939916 | 0.104 | -0.386 | 0.472 | 0.855 | 0 | 0.056 | 0.056 | 0.768 | 0.9 | 0.79 | 0.94 | 0.83 | 0.93 | 0.88 |
| 11 | 123673430 | rs10893090 | 0.807 | -0.506 | 0.029 | 0.709 | -0.001 | -0.002 | -0.002 | 0 | 0.46 | 0.47 | 0.57 | 0.44 | 0.53 | 0.43 |
| 11 | 123675833 | rs1939923 | -0.311 | -0.158 | -0.126 | 0.829 | -0.014 | 0.007 | 0.007 | 0 | 0.93 | 0.97 | 0.94 | 0.69 | 0.9 | 0.76 |
| 11 | 123676413 | rs7952150 | 0.22 | -0.741 | 0.216 | 0.534 | -0.019 | -0.007 | -0.007 | 0 | 0.29 | 0.32 | 0.27 | 0.19 | 0.19 | 0.14 |
| 11 | 123679460 | rs7113762 | 0.272 | -0.847 | 0.089 | 0.189 | 0.008 | 0.006 | 0.006 | 0 | 0.62 | 0.71 | 0.75 | 0.65 | 0.65 | 0.51 |
| 11 | 123680122 | rs1939911 | 1.105 | -0.89 | 0.231 | -0.007 | -0.002 | -0.036 | -0.036 | 0 | 0.12 | 0.18 | 0.06 | 0.02 | 0.1 | 0.04 |
| 11 | 123706304 | rs11219314 | 0.131 | -0.824 | 0.045 | -0.038 | 0.006 | 0.032 | 0.032 | 0 | 0.21 | 0.15 | 0.13 | 0.06 | 0.19 | 0.15 |
| 11 | 123713896 | rs12288134 | -0.324 | -0.827 | 0.297 | -0.018 | 0.005 | 0.052 | 0.052 | 0 | 0.32 | 0.21 | 0.18 | 0.08 | 0.29 | 0.19 |
| 11 | 123716249 | rs1939915 | 0.178 | -0.408 | 0.313 | 0.168 | -0.008 | 0.053 | 0.053 | 0.624 | 0.53 | 0.34 | 0.36 | 0.24 | 0.49 | 0.31 |
| 11 | 123728637 | rs1893065 | 0.357 | -0.334 | 0.237 | 0.328 | 0.005 | 0.026 | 0.026 | 0.775 | 0.27 | 0.35 | 0.48 | 0.47 | 0.32 | 0.35 |
| 11 | 123733108 | rs948159 | -1.273 | -0.447 | 0.208 | 0.346 | -0.002 | -0.029 | -0.029 | 0 | 0.82 | 0.92 | 0.86 | 0.72 | 0.81 | 0.67 |
| 11 | 123742913 | rs12273131 | 1.914 | -0.596 | 0.1 | 0.202 | 0.007 | 0.044 | 0.044 | 0.046 | 0.25 | 0.1 | 0.21 | 0.16 | 0.24 | 0.13 |
| 11 | 123744051 | rs12417791 | -0.16 | -0.532 | 0.127 | 0.188 | -0.008 | -0.007 | -0.007 | 0 | 0.38 | 0.36 | 0.54 | 0.54 | 0.44 | 0.45 |
| 11 | 123748754 | rs7129450 | -0.189 | -0.536 | 0.15 | 0.186 | -0.006 | -0.005 | -0.005 | 0 | 0.38 | 0.41 | 0.55 | 0.53 | 0.44 | 0.45 |
| 11 | 123753686 | rs1054041 | -0.141 | -0.528 | 0.134 | 0.19 | -0.008 | -0.007 | -0.007 | 0.019 | 0.37 | 0.36 | 0.54 | 0.53 | 0.44 | 0.45 |
| 11 | 123753937 | rs1939927 | -1.891 | -0.616 | 0.03 | 0.229 | 0.002 | -0.059 | -0.059 | 1.878 | 0.86 | 1 | 0.92 | 0.84 | 0.88 | 0.92 |
| 11 | 123754204 | rs7121426 | -0.146 | -0.379 | 0.201 | 0.202 | -0.008 | -0.007 | -0.007 | 0 | 0.37 | 0.36 | 0.55 | 0.53 | 0.44 | 0.45 |
| 11 | 123755719 | rs728177 | 0.652 | -0.398 | 0.215 | 0.444 | -0.01 | -0.008 | -0.008 | 0 | 0.25 | 0.21 | 0.34 | 0.31 | 0.32 | 0.33 |
| 11 | 123756587 | rs6590016 | -1.203 | -0.375 | 0.091 | 0.379 | 0.006 | 0.005 | 0.005 | 0 | 0.57 | 0.81 | 0.65 | 0.56 | 0.57 | 0.54 |
| 11 | 123756623 | rs10502281 | 1.305 | -0.383 | 0.245 | 0.168 | -0.001 | -0.015 | -0.015 | 0 | 0.11 | 0.16 | 0.08 | 0.06 | 0.11 | 0.1 |
| 11 | 123762716 | rs17127899 | 0.915 | -0.36 | 0.122 | 0.175 | -0.003 | -0.041 | -0.041 | 0.893 | 0.17 | 0.18 | 0.09 | 0.02 | 0.14 | 0.09 |
| 11 | 123762718 | rs7131308 | -0.265 | -0.5 | -0.201 | 0.315 | -0.02 | 0.026 | 0.026 | 0 | 0.2 | 0.32 | 0.25 | 0.26 | 0.15 | 0.16 |
| 11 | 123764717 | rs12285115 | 1.661 | -0.472 | -0.34 | 0.288 | 0.006 | 0.043 | 0.043 | 0.047 | 0.25 | 0.1 | 0.21 | 0.16 | 0.24 | 0.13 |
| 11 | 123770186 | rs2315027 | -0.324 | 0.028 | -0.188 | 0.396 | 0 | 0 | 0 | 0 | 0.46 | 0.47 | 0.47 | 0.4 | 0.53 | 0.48 |
| 11 | 123772716 | rs10502278 | -0.087 | -0.223 | -0.575 | 0.327 | -0.001 | 0.01 | 0.01 | 0.093 | 0.67 | 0.86 | 0.81 | 0.69 | 0.68 | 0.63 |
| 11 | 123773527 | rs2315029 | -0.232 | -0.255 | -0.508 | 0.228 | 0.001 | 0.016 | 0.016 | 0 | 0.69 | 0.84 | 0.82 | 0.73 | 0.69 | 0.64 |

| | | | | | | | | | | | | | | | | |
|---|---|---|---|---|---|---|---|---|---|---|---|---|---|---|---|---|
| 11 | 123774021 | rs9804524 | 0.406 | -0.108 | -0.157 | 0.271 | -0.001 | -0.001 | -0.001 | 0 | 0.49 | 0.49 | 0.45 | 0.35 | 0.37 | 0.32 |
| 11 | 123775491 | rs1377602 | -0.527 | -0.068 | -0.193 | 0.284 | -0.001 | 0.017 | 0.017 | 0 | 0.69 | 0.84 | 0.81 | 0.73 | 0.68 | 0.64 |
| 11 | 123775759 | rs1377604 | -0.527 | -0.068 | -0.193 | 0.284 | -0.001 | 0.017 | 0.017 | 0 | 0.69 | 0.84 | 0.81 | 0.73 | 0.68 | 0.64 |
| 11 | 123777302 | rs17127947 | 0.398 | -0.494 | -0.101 | 0.04 | -0.007 | -0.025 | -0.025 | 0.457 | 0.24 | 0.28 | 0.15 | 0.09 | 0.18 | 0.18 |
| 11 | 123777412 | rs17127950 | 0.398 | -0.494 | -0.101 | 0.04 | -0.007 | -0.025 | -0.025 | 0 | 0.24 | 0.28 | 0.15 | 0.09 | 0.18 | 0.18 |
| 11 | 123777497 | rs10750250 | -1.744 | -0.621 | -0.372 | -0.044 | -0.004 | -0.006 | -0.006 | 0.15 | 0.83 | 0.9 | 0.87 | 0.77 | 0.8 | 0.68 |
| 11 | 123777536 | rs7926767 | -0.236 | -0.705 | -0.156 | -0.087 | -0.002 | -0.006 | -0.006 | 0.218 | 0.66 | 0.65 | 0.79 | 0.73 | 0.68 | 0.63 |
| 11 | 123777736 | rs10790610 | 1.798 | -0.794 | -0.114 | -0.12 | -0.065 | -0.071 | -0.071 | 0.088 | 0.17 | 0.08 | 0.33 | 0.38 | 0.31 | 0.32 |
| 11 | 123777986 | rs7942047 | -1.73 | -0.818 | -0.077 | -0.099 | -0.004 | -0.006 | -0.006 | 0 | 0.83 | 0.9 | 0.87 | 0.77 | 0.8 | 0.68 |
| 11 | 123778031 | rs7927385 | -0.23 | -0.784 | -0.24 | -0.082 | -0.002 | -0.006 | -0.006 | 0.317 | 0.66 | 0.65 | 0.79 | 0.73 | 0.68 | 0.64 |
| 11 | 123782042 | rs1584825 | -1.674 | -0.744 | -0.251 | -0.051 | -0.024 | -0.03 | -0.03 | 0 | 0.82 | 0.96 | 0.87 | 0.77 | 0.8 | 0.69 |
| 11 | 123783226 | rs7126113 | -0.388 | -0.633 | -0.186 | 0.028 | 0.004 | 0.005 | 0.005 | 0 | 0.43 | 0.57 | 0.49 | 0.47 | 0.49 | 0.5 |
| 11 | 123789278 | rs12275512 | 1.555 | -0.582 | -0.521 | 0.09 | 0.009 | -0.001 | -0.001 | 0 | 0.25 | 0.2 | 0.31 | 0.25 | 0.19 | 0.14 |
| 11 | 123792656 | rs12281247 | 1.55 | -0.576 | -0.525 | 0.093 | 0.014 | -0.001 | -0.001 | 0.593 | 0.25 | 0.18 | 0.3 | 0.25 | 0.19 | 0.14 |
| 11 | 123794954 | rs10893106 | 1.665 | -0.471 | -0.645 | 0.083 | -0.064 | -0.069 | -0.069 | 0 | 0.16 | 0.08 | 0.33 | 0.38 | 0.32 | 0.31 |
| 11 | 123795372 | rs10893107 | 0.767 | -0.492 | -0.644 | 0.176 | -0.058 | -0.049 | -0.049 | 0.055 | 0.39 | 0.16 | 0.49 | 0.48 | 0.5 | 0.42 |
| 11 | 123802399 | rs970584 | 0.802 | -0.866 | -0.695 | -0.03 | 0.017 | 0.139 | 0.139 | 0 | 0.38 | 0.09 | 0.25 | 0.16 | 0.31 | 0.21 |
| 11 | 123802845 | rs1377606 | 1.839 | -0.794 | -0.519 | 0.101 | 0.018 | 0.047 | 0.047 | 0 | 0.23 | 0.09 | 0.16 | 0.11 | 0.18 | 0.11 |
| 11 | 123811344 | rs11219362 | 1.509 | -0.802 | -0.477 | 0.133 | 0.016 | -0.001 | -0.001 | 0 | 0.25 | 0.16 | 0.3 | 0.25 | 0.19 | 0.14 |
| 11 | 123822292 | rs6590025 | -0.832 | -0.681 | -0.097 | 0.274 | -0.006 | 0.001 | 0.001 | 0 | 0.77 | 0.74 | 0.83 | 0.74 | 0.78 | 0.67 |
| 11 | 123826425 | rs1453649 | -0.506 | -0.643 | 0.103 | 0.381 | -0.015 | 0.06 | 0.06 | 0 | 0.75 | 0.63 | 0.67 | 0.58 | 0.77 | 0.62 |
| 11 | 123828187 | rs7114486 | 1.462 | -0.395 | -0.152 | 0.378 | -0.083 | -0.087 | -0.087 | 0 | 0.17 | 0.06 | 0.32 | 0.36 | 0.31 | 0.3 |
| 11 | 123830041 | rs10893114 | 1.498 | -0.385 | -0.147 | 0.383 | -0.082 | -0.086 | -0.086 | 0.331 | 0.17 | 0.06 | 0.32 | 0.36 | 0.31 | 0.3 |
| 11 | 123846864 | rs723488 | 0.399 | -0.679 | 0.136 | 0.068 | -0.006 | -0.021 | -0.021 | 0 | 0.22 | 0.22 | 0.12 | 0.1 | 0.17 | 0.18 |
| 11 | 123848121 | rs17686210 | 1.352 | -0.758 | -0.037 | -0.039 | 0.033 | 0.11 | 0.11 | 0 | 0.34 | 0.16 | 0.22 | 0.12 | 0.29 | 0.14 |
| 11 | 123848212 | rs17759513 | 1.33 | -0.753 | -0.042 | -0.036 | 0.034 | 0.12 | 0.12 | 0 | 0.34 | 0.13 | 0.21 | 0.12 | 0.29 | 0.14 |
| 11 | 123848794 | rs17686304 | -0.213 | -0.653 | -0.042 | 0.05 | -0.01 | 0.001 | 0.001 | 0 | 0.62 | 0.7 | 0.52 | 0.36 | 0.47 | 0.35 |
| 11 | 123861081 | rs4391827 | -0.236 | -0.927 | 0.066 | 0.137 | 0.001 | 0.005 | 0.005 | 0 | 0.63 | 0.71 | 0.75 | 0.66 | 0.66 | 0.55 |
| 11 | 123864216 | rs1453654 | -1.771 | -0.855 | -0.107 | 0.128 | -0.009 | -0.06 | -0.06 | 0 | 0.83 | 1 | 0.89 | 0.79 | 0.84 | 0.68 |
| 11 | 123864948 | rs17128086 | 0.539 | -0.749 | 0.128 | 0.129 | -0.005 | 0.005 | 0.005 | 0 | 0.08 | 0.22 | 0.04 | 0.05 | 0.05 | 0.09 |
| 11 | 123867853 | rs11219398 | 0.833 | -0.695 | 0.058 | 0.208 | -0.003 | -0.003 | -0.003 | 0 | 0.18 | 0.15 | 0.26 | 0.19 | 0.18 | 0.14 |
| 11 | 123868208 | rs12577664 | -0.867 | -0.765 | 0.02 | 0.263 | -0.014 | 0.008 | 0.008 | 0 | 0.78 | 0.73 | 0.86 | 0.72 | 0.82 | 0.66 |
| 11 | 123872530 | rs10893124 | -0.243 | -0.581 | -0.106 | 0.276 | -0.003 | -0.008 | -0.008 | 0 | 0.39 | 0.31 | 0.46 | 0.45 | 0.52 | 0.49 |
| 11 | 123872869 | rs11600075 | 0.205 | -0.484 | -0.054 | 0.156 | 0 | -0.048 | -0.048 | 0.072 | 0.24 | 0.32 | 0.15 | 0.1 | 0.22 | 0.19 |
| 11 | 123877979 | rs1947406 | -0.498 | -0.548 | -0.091 | 0.169 | -0.001 | -0.004 | -0.004 | 0 | 0.41 | 0.4 | 0.47 | 0.46 | 0.52 | 0.5 |
| 11 | 123880722 | rs4936879 | -0.709 | -0.617 | -0.014 | 0.203 | -0.002 | -0.009 | -0.009 | 0 | 0.43 | 0.5 | 0.63 | 0.61 | 0.53 | 0.52 |
| 11 | 123882818 | rs6590033 | -0.256 | -0.56 | -0.094 | 0.232 | -0.013 | -0.006 | -0.006 | 0.208 | 0.27 | 0.41 | 0.31 | 0.25 | 0.22 | 0.23 |
| 11 | 123884423 | rs6590035 | -1.606 | -0.575 | 0.038 | 0.255 | -0.002 | 0 | 0 | 0 | 0.83 | 0.87 | 0.88 | 0.79 | 0.85 | 0.69 |
| 11 | 123884981 | rs10502277 | 1.69 | -0.699 | -0.26 | 0.001 | -0.006 | -0.052 | -0.052 | 1.45 | 0.12 | 0.2 | 0.06 | 0.01 | 0.11 | 0.04 |
| 11 | 123892506 | rs11219411 | 0.304 | -0.524 | -0.815 | 0.047 | 0.019 | 0.096 | 0.096 | 0 | 0.49 | 0.28 | 0.33 | 0.21 | 0.43 | 0.34 |
| 11 | 123893286 | rs2076941 | 1.009 | -0.098 | -0.896 | 0.405 | -0.031 | -0.038 | -0.038 | 2.989 | 0.18 | 0.17 | 0.45 | 0.51 | 0.29 | 0.3 |
| 11 | 123899515 | rs11219414 | 0.893 | -0.171 | -1.034 | 0.115 | -0.072 | 0.075 | 0.075 | 0 | 0.19 | 0.01 | 0.13 | 0.05 | 0.19 | 0.14 |
| 11 | 123901175 | rs11825515 | 1.011 | -0.299 | -0.671 | 0.349 | -0.002 | 0.036 | 0.036 | 0.053 | 0.05 | 0.13 | 0.02 | 0.07 | 0.03 | 0.08 |
| 11 | 123905660 | rs11219417 | 0.774 | -0.219 | -1.184 | 0.192 | -0.066 | 0.089 | 0.089 | 0 | 0.2 | 0.01 | 0.14 | 0.05 | 0.21 | 0.15 |
| 11 | 123905876 | rs4936885 | 1.64 | 0.124 | -1.219 | 0.55 | -0.091 | -0.094 | -0.094 | 0 | 0.14 | 0.04 | 0.3 | 0.34 | 0.28 | 0.31 |
| 11 | 123906346 | rs7949155 | -0.514 | -0.579 | -0.875 | 0.408 | -0.031 | -0.021 | -0.021 | 0.051 | 0.68 | 0.5 | 0.8 | 0.72 | 0.73 | 0.69 |
| 11 | 123910023 | rs17128251 | 0.015 | -0.409 | -1.797 | 0.515 | 0.076 | -0.059 | -0.059 | 0 | 0.58 | 0.19 | 0.61 | 0.48 | 0.69 | 0.56 |
| 11 | 123910740 | rs896891 | 0.613 | -0.112 | -2.103 | 0.566 | -0.012 | 0.179 | 0.179 | 0 | 0.44 | 0.11 | 0.31 | 0.17 | 0.41 | 0.24 |
| 11 | 123918667 | rs7948337 | -1.046 | 0.239 | -1.489 | 1.061 | 0.044 | 0.101 | 0.101 | 0 | 0.62 | 0.9 | 0.73 | 0.82 | 0.67 | 0.8 |
| 11 | 123919128 | rs12224791 | 0.173 | 1.038 | -0.682 | 1.703 | -0.002 | 0.012 | 0.012 | 0 | 0.16 | 0.17 | 0.45 | 0.48 | 0.29 | 0.35 |
| 11 | 123921479 | rs4935862 | 0.471 | 1.215 | -1.225 | 1.153 | -0.097 | -0.094 | -0.094 | 0 | 0.14 | 0.03 | 0.3 | 0.32 | 0.28 | 0.32 |
| 11 | 123925974 | rs1453631 | 0.895 | 0.681 | -1.288 | 0.853 | -0.085 | -0.073 | -0.073 | 0 | 0.2 | 0.1 | 0.32 | 0.36 | 0.33 | 0.4 |
| 11 | 123926106 | rs1453630 | -0.3 | -0.385 | -1.558 | 0.335 | 0.014 | 0.023 | 0.023 | 0 | 0.6 | 0.79 | 0.64 | 0.67 | 0.63 | 0.68 |
| 11 | 123938580 | rs12271236 | 0.981 | -0.885 | -1.674 | -0.012 | 0 | -0.041 | -0.041 | 0 | 0.23 | 0.07 | 0.23 | 0.26 | 0.24 | 0.19 |
| 11 | 123939207 | rs589600 | -1.339 | -1.272 | -2.031 | -0.35 | -0.073 | 0.022 | 0.022 | 0 | 0.77 | 0.98 | 0.88 | 0.86 | 0.78 | 0.86 |
| 11 | 123942501 | rs651739 | -0.573 | -1.401 | -0.957 | -0.511 | 0.001 | -0.059 | -0.059 | 0 | 0.7 | 0.56 | 0.82 | 0.87 | 0.74 | 0.77 |
| 11 | 123944824 | rs11219437 | 1.032 | -1.413 | -0.708 | -0.302 | 0.007 | 0.007 | 0.007 | 0 | 0.3 | 0.35 | 0.35 | 0.42 | 0.36 | 0.43 |
| 11 | 123945410 | rs4290238 | 0.488 | -1.406 | -0.856 | -0.08 | -0.003 | 0 | 0 | 0 | 0.17 | 0.13 | 0.23 | 0.2 | 0.13 | 0.15 |
| 11 | 123947295 | rs3741111 | -0.39 | -1.61 | -0.748 | -0.245 | -0.001 | 0.02 | 0.02 | 0 | 0.53 | 0.64 | 0.61 | 0.62 | 0.51 | 0.67 |
| 11 | 123952703 | rs3741108 | -0.119 | -2.101 | -1.004 | -0.288 | -0.006 | 0 | 0 | 0 | 0.3 | 0.34 | 0.22 | 0.3 | 0.25 | 0.22 |
| 11 | 123955725 | rs489527 | 1.096 | -2.286 | -1.226 | -0.283 | 0.096 | -0.087 | -0.087 | 0 | 0.47 | 0.06 | 0.5 | 0.49 | 0.62 | 0.62 |
| 11 | 123956466 | rs1893767 | 0.525 | -2.288 | -1.151 | -0.231 | -0.047 | 0.085 | 0.085 | 0 | 0.15 | 0.34 | 0.13 | 0.26 | 0.07 | 0.09 |
| 11 | 123956983 | rs522334 | 0.225 | -1.619 | -1.195 | -0.087 | -0.001 | -0.001 | -0.001 | 0.065 | 0.64 | 0.67 | 0.65 | 0.75 | 0.69 | 0.71 |
| 11 | 123957844 | rs10893149 | 0.637 | -1.393 | -2.191 | -0.008 | -0.008 | 0.08 | 0.08 | 0 | 0.2 | 0.07 | 0.13 | 0.04 | 0.22 | 0.19 |
| 11 | 123962611 | rs526075 | -0.48 | -0.943 | -1.767 | 0.646 | 0.006 | -0.007 | -0.007 | 0 | 0.9 | 0.9 | 0.8 | 0.79 | 0.91 | 0.89 |
| 11 | 123964967 | rs668714 | 1.08 | -0.2 | -1.86 | 0.691 | -0.112 | -0.104 | -0.104 | 0 | 0.23 | 0.04 | 0.33 | 0.37 | 0.33 | 0.49 |
| 11 | 123966865 | rs765346 | -0.319 | -0.739 | -1.201 | 0.644 | 0.034 | -0.048 | -0.048 | 0.526 | 0.12 | 0.01 | 0.05 | 0.21 | 0.08 | 0.08 |
| 11 | 123977844 | rs1939846 | -1.086 | -0.877 | -0.904 | 0.506 | 0.037 | -0.008 | -0.008 | 0.006 | 0.44 | 0.73 | 0.28 | 0.48 | 0.36 | 0.32 |
| 11 | 123980567 | rs12802638 | 0.368 | -0.757 | -1.602 | -0.708 | -0.037 | 0.087 | 0.087 | 0.567 | 0.22 | 0.08 | 0.28 | 0.07 | 0.26 | 0.18 |

| | | | | | | | | | | | | | | | | |
|---|---|---|---|---|---|---|---|---|---|---|---|---|---|---|---|---|
| 11 | 123982974 | rs17128423 | -0.743 | -0.656 | -0.75 | -0.335 | -0.015 | -0.009 | -0.009 | 0 | 0.17 | 0.2 | 0.07 | 0.1 | 0.12 | 0.12 |
| 11 | 123983579 | rs1939847 | 0.444 | -0.547 | -1.325 | -0.21 | -0.028 | 0.072 | 0.072 | 2.617 | 0.18 | 0.07 | 0.13 | 0.04 | 0.23 | 0.15 |
| 11 | 123987875 | rs3758928 | -0.128 | -0.515 | -1.107 | 0.006 | -0.02 | 0.051 | 0.051 | 3.535 | 0.25 | 0.02 | 0.12 | 0.09 | 0.24 | 0.08 |
| 11 | 123997498 | rs10750258 | -1.315 | -0.248 | -1.061 | 0.259 | -0.136 | 0.019 | 0.019 | 0.422 | 0.68 | 0.94 | 0.62 | 0.78 | 0.61 | 0.53 |
| 11 | 124003796 | rs4935863 | -1.082 | -0.038 | -0.737 | 0.276 | -0.079 | 0.057 | 0.057 | 0 | 0.62 | 0.87 | 0.55 | 0.71 | 0.52 | 0.39 |
| 11 | 124010014 | rs7932381 | 0.32 | -0.334 | 0.117 | 0.064 | -0.018 | -0.059 | -0.059 | 0 | 0.15 | 0.29 | 0.09 | 0.06 | 0.11 | 0.14 |
| 11 | 124011580 | rs7103676 | 0.114 | -0.365 | 0.439 | 0.152 | -0.021 | -0.023 | -0.023 | 0.009 | 0.16 | 0.3 | 0.13 | 0.11 | 0.11 | 0.17 |
| 11 | 124015710 | rs4565905 | -1.289 | -0.383 | 0.418 | 0.058 | -0.05 | 0.06 | 0.06 | 0 | 0.61 | 0.77 | 0.54 | 0.71 | 0.52 | 0.39 |
| 11 | 124016301 | rs9667893 | -1.057 | -0.468 | 0.879 | 0 | -0.004 | 0.032 | 0.032 | 0.005 | 0.56 | 0.64 | 0.51 | 0.7 | 0.49 | 0.36 |
| 11 | 124016724 | rs11601839 | 0.102 | -0.368 | 0.826 | 0.042 | 0.024 | -0.009 | -0.009 | 0 | 0.05 | 0.16 | 0.07 | 0.07 | 0.09 | 0.13 |
| 11 | 124016820 | rs3824997 | -0.226 | -0.446 | 0.825 | 0.017 | 0.005 | 0.013 | 0.013 | 1.831 | 0.42 | 0.36 | 0.42 | 0.64 | 0.4 | 0.22 |
| 11 | 124017091 | rs1939861 | -1.645 | -0.471 | 0.591 | -0.001 | -0.052 | 0.041 | 0.041 | 0 | 0.65 | 0.87 | 0.6 | 0.78 | 0.59 | 0.51 |
| 11 | 124017493 | rs4936894 | 0.638 | -0.304 | 0.406 | 0.746 | -0.038 | -0.053 | -0.053 | 0 | 0.13 | 0.04 | 0.26 | 0.43 | 0.21 | 0.11 |
| 11 | 124017711 | rs7121402 | -1.226 | -0.501 | 1.017 | 0.348 | -0.007 | 0.018 | 0.018 | 0.74 | 0.34 | 0.5 | 0.43 | 0.64 | 0.33 | 0.38 |
| 11 | 124018554 | rs9651671 | -1.424 | -0.097 | 1.033 | 0.447 | -0.012 | -0.002 | -0.002 | 0.169 | 0.36 | 0.6 | 0.37 | 0.32 | 0.29 | 0.44 |
| 11 | 124019261 | rs10750260 | -1.475 | -0.058 | 0.829 | 0.481 | -0.009 | 0.001 | 0.001 | 1.341 | 0.35 | 0.46 | 0.35 | 0.32 | 0.29 | 0.43 |
| 11 | 124032347 | rs1939852 | -0.358 | -0.302 | 0.666 | 0.461 | 0.015 | 0.008 | 0.008 | 0.905 | 0.19 | 0.16 | 0.12 | 0.16 | 0.13 | 0.31 |
| 11 | 124047575 | rs2508876 | -0.588 | -0.262 | 0.767 | 0.477 | 0.003 | 0.008 | 0.008 | 0.054 | 0.51 | 0.42 | 0.48 | 0.57 | 0.53 | 0.47 |
| 11 | 124064554 | rs11219489 | 0.415 | -0.259 | 0.797 | 0.514 | -0.001 | -0.003 | -0.003 | 0 | 0.34 | 0.36 | 0.41 | 0.45 | 0.41 | 0.19 |
| 11 | 124066627 | rs2846236 | -0.684 | -0.27 | 0.952 | 0.431 | 0.002 | 0.017 | 0.017 | 0.958 | 0.5 | 0.49 | 0.6 | 0.63 | 0.57 | 0.44 |
| 11 | 124077274 | rs2724667 | 0.459 | -0.269 | 0.891 | 0.451 | -0.027 | -0.023 | -0.023 | 1.262 | 0.11 | 0.02 | 0.16 | 0.13 | 0.14 | 0.1 |
| 11 | 124082136 | rs10790651 | 0.102 | -0.29 | 0.921 | 0.45 | -0.018 | 0.006 | 0.006 | 0 | 0.05 | 0.11 | 0.04 | 0.05 | 0.02 | 0.15 |
| 11 | 124082810 | rs2508888 | 0.363 | -0.388 | 1.246 | 0.382 | -0.018 | -0.026 | -0.026 | 0 | 0.16 | 0.24 | 0.12 | 0.1 | 0.11 | 0.16 |
| 11 | 124084653 | rs2846232 | -1.555 | -0.369 | 1.191 | 0.374 | -0.007 | 0.017 | 0.017 | 0 | 0.66 | 0.74 | 0.72 | 0.74 | 0.68 | 0.59 |
| 11 | 124085046 | rs2508889 | 0.591 | -0.346 | 1.021 | 0.382 | 0.012 | 0.018 | 0.018 | 0.193 | 0.14 | 0.07 | 0.1 | 0.1 | 0.11 | 0.16 |
| 11 | 124086422 | rs2512284 | -0.21 | -0.372 | 1.055 | 0.392 | -0.042 | -0.073 | -0.073 | 0 | 0.55 | 0.89 | 0.56 | 0.48 | 0.48 | 0.68 |
| 11 | 124094783 | rs2512274 | -0.21 | -0.408 | 1.073 | 0.393 | -0.041 | -0.076 | -0.076 | 2.977 | 0.55 | 0.89 | 0.56 | 0.48 | 0.48 | 0.68 |
| 11 | 124100201 | rs2512229 | -0.301 | -0.4 | 1.189 | 0.385 | -0.022 | 0.018 | 0.018 | 2.347 | 0.45 | 0.32 | 0.56 | 0.58 | 0.55 | 0.28 |
| 11 | 124119351 | rs11219534 | 0.555 | -0.417 | 1.18 | 0.397 | -0.004 | -0.028 | -0.028 | 0 | 0.33 | 0.3 | 0.4 | 0.45 | 0.41 | 0.19 |
| 11 | 124120302 | rs2512239 | -0.824 | -0.44 | 1.211 | 0.371 | 0.012 | 0.018 | 0.018 | 0.193 | 0.86 | 0.92 | 0.9 | 0.9 | 0.89 | 0.84 |
| 11 | 124122019 | rs2512242 | -0.282 | -0.435 | 1.203 | 0.371 | -0.022 | 0.018 | 0.018 | 2.347 | 0.45 | 0.32 | 0.56 | 0.58 | 0.55 | 0.28 |
| 11 | 124124924 | rs12224218 | 1.444 | -0.464 | 1.38 | 0.356 | -0.014 | 0.022 | 0.022 | 0 | 0.33 | 0.2 | 0.28 | 0.26 | 0.32 | 0.41 |
| 11 | 124134743 | rs2466637 | -0.238 | -0.456 | 1.319 | 0.361 | -0.023 | 0.018 | 0.018 | 2.408 | 0.44 | 0.29 | 0.56 | 0.58 | 0.55 | 0.28 |
| 11 | 124135438 | rs2512167 | -0.238 | -0.456 | 1.319 | 0.361 | -0.023 | 0.018 | 0.018 | 2.408 | 0.44 | 0.29 | 0.56 | 0.58 | 0.55 | 0.28 |
| 11 | 124135481 | rs2466701 | -0.238 | -0.456 | 1.319 | 0.361 | -0.023 | 0.018 | 0.018 | 2.408 | 0.44 | 0.29 | 0.56 | 0.58 | 0.55 | 0.28 |
| 11 | 124140623 | rs2512202 | -0.809 | -0.451 | 1.259 | 0.368 | 0.012 | 0.018 | 0.018 | 0.193 | 0.86 | 0.93 | 0.9 | 0.9 | 0.89 | 0.84 |
| 11 | 124143906 | rs6590052 | -0.168 | -0.446 | 1.377 | 0.379 | -0.024 | 0.006 | 0.006 | 0 | 0.06 | 0.18 | 0.03 | 0.05 | 0.02 | 0.15 |
| 11 | 124146042 | rs7103798 | 0.743 | -0.436 | 1.448 | 0.368 | -0.001 | -0.01 | -0.01 | 0 | 0.4 | 0.45 | 0.32 | 0.32 | 0.34 | 0.56 |
| 11 | 124147601 | rs7941270 | 0.467 | -0.434 | 1.545 | 0.377 | -0.002 | -0.006 | -0.006 | 0 | 0.73 | 0.72 | 0.72 | 0.77 | 0.75 | 0.75 |
| 11 | 124170064 | rs6590056 | 0.878 | -0.344 | 1.18 | 0.409 | -0.082 | -0.124 | -0.124 | 0.201 | 0.31 | 0.04 | 0.34 | 0.42 | 0.4 | 0.17 |
| 11 | 124172270 | rs2512297 | -0.124 | -0.404 | 1.27 | 0.378 | 0.002 | 0.002 | 0.002 | 0.521 | 0.8 | 0.77 | 0.87 | 0.85 | 0.87 | 0.69 |
| 11 | 124174874 | rs10893206 | 0.851 | -0.336 | 1.181 | 0.408 | -0.056 | -0.092 | -0.092 | 0 | 0.31 | 0.1 | 0.34 | 0.42 | 0.41 | 0.16 |
| 11 | 124179450 | rs10790667 | 0.783 | -0.336 | 1.23 | 0.389 | -0.021 | -0.048 | -0.048 | 0 | 0.31 | 0.15 | 0.34 | 0.42 | 0.41 | 0.16 |
| 11 | 124180082 | rs4936919 | 0.772 | -0.338 | 1.261 | 0.392 | -0.02 | -0.044 | -0.044 | 0 | 0.31 | 0.15 | 0.35 | 0.42 | 0.41 | 0.16 |
| 11 | 124181111 | rs2512296 | -0.544 | -0.301 | 1.616 | 0.424 | 0.002 | 0.016 | 0.016 | 0 | 0.51 | 0.35 | 0.48 | 0.57 | 0.54 | 0.47 |
| 11 | 124181158 | rs518119 | 0.289 | -0.337 | 1.76 | 0.393 | 0.03 | 0.066 | 0.066 | 0 | 0.84 | 0.53 | 0.76 | 0.83 | 0.85 | 0.88 |
| 11 | 124183711 | rs2510432 | -0.27 | -0.28 | 1.732 | 0.398 | -0.042 | -0.02 | -0.02 | 0.017 | 0.2 | 0.36 | 0.14 | 0.15 | 0.13 | 0.31 |
| 11 | 124189729 | rs2512219 | -0.72 | -0.305 | 1.41 | 0.426 | -0.053 | -0.009 | -0.009 | 1.282 | 0.23 | 0.45 | 0.2 | 0.19 | 0.13 | 0.34 |
| 11 | 124192712 | rs617280 | 0.952 | -0.266 | 1.379 | 0.429 | -0.013 | 0 | 0 | 0 | 0.65 | 0.42 | 0.64 | 0.68 | 0.73 | 0.57 |
| 11 | 124199865 | rs539802 | 1.028 | -0.29 | 1.593 | 0.418 | 0.02 | 0.002 | 0.002 | 1.812 | 0.64 | 0.26 | 0.63 | 0.68 | 0.73 | 0.57 |
| 11 | 124208977 | rs666135 | 0.934 | -0.278 | 1.551 | 0.439 | -0.023 | 0.002 | 0.002 | 1.498 | 0.65 | 0.48 | 0.64 | 0.68 | 0.73 | 0.57 |
| 11 | 124227543 | rs644353 | 0.938 | -0.267 | 1.508 | 0.445 | -0.023 | 0.002 | 0.002 | 1.518 | 0.65 | 0.48 | 0.64 | 0.68 | 0.73 | 0.57 |
| 11 | 124229916 | rs2428301 | 0.798 | -0.259 | 1.482 | 0.436 | -0.017 | -0.008 | -0.008 | 0.616 | 0.66 | 0.55 | 0.69 | 0.72 | 0.73 | 0.6 |
| 11 | 124236679 | rs686726 | 0.945 | -0.03 | 1.371 | 0.205 | 0.002 | -0.011 | -0.011 | 1.737 | 0.35 | 0.4 | 0.31 | 0.28 | 0.31 | 0.42 |
| 11 | 124239690 | rs526081 | 0.64 | -0.067 | 1.419 | 0.214 | 0 | 0.011 | 0.011 | 0 | 0.38 | 0.71 | 0.36 | 0.32 | 0.31 | 0.44 |
| 11 | 124245640 | rs620669 | -0.371 | -0.06 | 1.409 | 0.235 | -0.007 | -0.006 | -0.006 | 0.101 | 0.88 | 0.92 | 0.83 | 0.87 | 0.86 | 0.92 |
| 11 | 124245818 | rs621567 | 0.967 | -0.087 | 1.488 | 0.199 | -0.003 | 0.007 | 0.007 | 0.006 | 0.35 | 0.34 | 0.3 | 0.28 | 0.31 | 0.41 |
| 11 | 124248489 | rs664275 | 0.927 | -0.084 | 1.016 | 0.206 | 0.003 | -0.023 | -0.023 | 0.012 | 0.36 | 0.53 | 0.31 | 0.28 | 0.31 | 0.41 |
| 11 | 124255991 | rs4057885 | 0.934 | -0.03 | 0.988 | 0.185 | 0.003 | -0.018 | -0.018 | 0 | 0.35 | 0.44 | 0.3 | 0.27 | 0.31 | 0.41 |
| 11 | 124258314 | rs7118177 | 0.934 | -0.03 | 0.988 | 0.185 | 0.003 | -0.018 | -0.018 | 0 | 0.35 | 0.44 | 0.3 | 0.27 | 0.31 | 0.41 |
| 11 | 124264636 | rs4936923 | -0.352 | -0.378 | 0.416 | 0.246 | 0.007 | 0.014 | 0.014 | 0.016 | 0.5 | 0.22 | 0.47 | 0.56 | 0.55 | 0.48 |
| 11 | 124278130 | rs3867247 | -1.277 | -0.92 | 0.24 | 0.209 | 0.03 | 0.003 | 0.003 | 0.017 | 0.37 | 0.66 | 0.28 | 0.32 | 0.23 | 0.42 |
| 11 | 124286219 | rs1893702 | -1.398 | -0.754 | -0.095 | -0.165 | -0.048 | 0.099 | 0.099 | 0 | 0.6 | 0.76 | 0.52 | 0.81 | 0.51 | 0.47 |
| 11 | 124286727 | rs7950272 | 0.28 | -0.841 | 0.233 | -0.185 | 0.008 | -0.018 | -0.018 | 1.24 | 0.32 | 0.3 | 0.33 | 0.6 | 0.31 | 0.22 |
| 11 | 124294349 | rs7116575 | 0.023 | -0.652 | 0.379 | -0.127 | -0.018 | 0.073 | 0.073 | 0 | 0.14 | 0.26 | 0.11 | 0.2 | 0.09 | 0.09 |
| 11 | 124295382 | rs1893705 | -0.538 | -0.508 | -0.092 | -0.006 | -0.027 | -0.016 | -0.016 | 0.019 | 0.28 | 0.45 | 0.2 | 0.22 | 0.2 | 0.25 |
| 11 | 124295724 | rs7942913 | 0.28 | -0.087 | -0.772 | 0.408 | -0.059 | -0.162 | -0.162 | 1.577 | 0.27 | 0.03 | 0.3 | 0.55 | 0.3 | 0.17 |
| 11 | 124309814 | rs11219659 | 0.214 | -0.241 | -0.643 | 0.412 | -0.004 | -0.088 | -0.088 | 0.244 | 0.23 | 0.11 | 0.34 | 0.53 | 0.26 | 0.19 |
| 11 | 124313035 | rs11219660 | 0.077 | -0.528 | -0.785 | 0.086 | 0.013 | 0.012 | 0.012 | 0 | 0.13 | 0.06 | 0.2 | 0.12 | 0.11 | 0.12 |

| | | | | | | | | | | | | | | | | |
|---|---|---|---|---|---|---|---|---|---|---|---|---|---|---|---|---|
| 11 | 124313722 | rs7122962 | -0.041 | -0.881 | -0.195 | -0.545 | -0.013 | -0.07 | -0.07 | 0 | 0.22 | 0.32 | 0.11 | 0.06 | 0.13 | 0.3 |
| 11 | 124323660 | rs7934631 | -1.629 | -1.024 | 0.455 | -0.596 | -0.006 | 0.001 | 0.001 | 0 | 0.61 | 0.7 | 0.67 | 0.71 | 0.51 | 0.6 |
| 11 | 124335917 | rs4936934 | -0.388 | -1.151 | 0.54 | -0.588 | -0.045 | -0.019 | -0.019 | 2.484 | 0.18 | 0.52 | 0.25 | 0.14 | 0.12 | 0.13 |
| 11 | 124336515 | rs10750277 | -0.336 | -1.15 | 0.547 | -0.588 | -0.045 | -0.017 | -0.017 | 1.211 | 0.17 | 0.49 | 0.25 | 0.14 | 0.12 | 0.13 |
| 11 | 124337885 | rs4360714 | -0.315 | -1.068 | 0.535 | -0.601 | 0.001 | 0 | 0 | 0 | 0.4 | 0.55 | 0.58 | 0.67 | 0.38 | 0.31 |
| 11 | 124338222 | rs7127920 | -1.771 | -1.022 | 0.295 | -0.552 | -0.149 | -0.027 | -0.027 | 0 | 0.63 | 0.96 | 0.7 | 0.76 | 0.51 | 0.61 |
| 11 | 124341985 | rs2298509 | -0.435 | -0.909 | 0.937 | -0.378 | -0.003 | -0.001 | -0.001 | 0 | 0.4 | 0.54 | 0.58 | 0.65 | 0.38 | 0.29 |
| 11 | 124345791 | rs10431091 | -0.533 | -0.891 | 0.857 | -0.358 | -0.007 | -0.002 | -0.002 | 0 | 0.41 | 0.55 | 0.58 | 0.65 | 0.38 | 0.29 |
| 11 | 124379503 | rs10790691 | 0.115 | -0.824 | 0.455 | -0.333 | -0.022 | 0.009 | 0.009 | 0 | 0.14 | 0.21 | 0.22 | 0.17 | 0.12 | 0.13 |
| 11 | 124386011 | rs4282990 | -1.598 | -0.903 | 0.155 | -0.529 | -0.081 | 0.029 | 0.029 | 0.035 | 0.61 | 0.9 | 0.69 | 0.76 | 0.53 | 0.58 |
| 11 | 124386346 | rs4282991 | -0.713 | -0.985 | 0.276 | -0.576 | 0.001 | 0.001 | 0.001 | 0 | 0.43 | 0.57 | 0.59 | 0.72 | 0.41 | 0.41 |
| 11 | 124393784 | rs10893243 | 0.231 | -0.789 | -0.581 | -0.467 | -0.02 | -0.002 | -0.002 | 0 | 0.15 | 0.22 | 0.23 | 0.15 | 0.12 | 0.15 |
| 11 | 124400101 | rs10750285 | -1.475 | -0.734 | -1.252 | -0.375 | -0.069 | 0.023 | 0.023 | 0.128 | 0.61 | 0.88 | 0.69 | 0.74 | 0.51 | 0.58 |
| 11 | 124400611 | rs1942668 | -1.372 | -0.807 | -0.759 | -0.599 | 0.059 | 0.112 | 0.112 | 1.109 | 0.38 | 0.86 | 0.37 | 0.22 | 0.26 | 0.42 |
| 11 | 124404297 | rs10893249 | -0.426 | -0.419 | -0.52 | -0.099 | -0.008 | -0.002 | -0.002 | 0 | 0.15 | 0.14 | 0.23 | 0.15 | 0.12 | 0.15 |
| 11 | 124405311 | rs10893250 | -0.511 | 0.033 | -1.513 | -0.541 | -0.079 | 0.095 | 0.095 | 0.901 | 0.59 | 0.94 | 0.57 | 0.75 | 0.54 | 0.61 |
| 11 | 124408415 | rs7129642 | 0.779 | 0.456 | -1.148 | -0.414 | 0.026 | 0.044 | 0.044 | 0.559 | 0.49 | 0.32 | 0.43 | 0.31 | 0.4 | 0.43 |
| 11 | 124409894 | rs11219691 | 0.147 | 0.442 | -0.493 | 0.26 | -0.011 | 0.045 | 0.045 | 0.048 | 0.19 | 0.37 | 0.12 | 0.27 | 0.14 | 0.25 |
| 11 | 124409935 | rs948725 | -0.758 | 0.509 | -0.541 | 0.236 | -0.045 | 0.036 | 0.036 | 0 | 0.81 | 0.94 | 0.78 | 0.89 | 0.73 | 0.77 |
| 11 | 124414746 | rs10893251 | 0.123 | 0.534 | -0.4 | 0.236 | -0.04 | -0.029 | -0.029 | 0.397 | 0.69 | 0.81 | 0.55 | 0.58 | 0.54 | 0.66 |
| 11 | 124419307 | rs6590081 | 0.764 | 0.555 | -0.299 | 0.183 | 0.006 | -0.006 | -0.006 | 0 | 0.61 | 0.52 | 0.65 | 0.63 | 0.59 | 0.5 |
| 11 | 124420238 | rs11219699 | 0.025 | 0.543 | -0.274 | 0.212 | -0.012 | 0.05 | 0.05 | 0.245 | 0.2 | 0.43 | 0.13 | 0.27 | 0.14 | 0.25 |
| 11 | 124421389 | rs12222571 | 0.134 | 0.512 | 0.194 | 0.197 | -0.008 | 0.032 | 0.032 | 0.091 | 0.19 | 0.31 | 0.12 | 0.27 | 0.14 | 0.25 |
| 11 | 124425537 | rs10893259 | -0.139 | 0.508 | 0.213 | 0.195 | -0.018 | 0.036 | 0.036 | 0.306 | 0.21 | 0.41 | 0.13 | 0.27 | 0.14 | 0.25 |
| 11 | 124426284 | rs12417780 | 1.467 | 0.494 | 0.176 | 0.214 | -0.012 | -0.013 | -0.013 | 0.005 | 0.11 | 0.08 | 0.22 | 0.32 | 0.19 | 0.09 |
| 11 | 124426930 | rs10893260 | -0.233 | 0.468 | 0.201 | 0.195 | -0.02 | 0.022 | 0.022 | 0.555 | 0.21 | 0.53 | 0.13 | 0.27 | 0.14 | 0.25 |
| 11 | 124430751 | rs7104191 | 0.288 | 0.545 | 0.179 | 0.153 | 0.075 | 0.002 | 0.002 | 0.056 | 0.05 | 0.23 | 0.04 | 0.02 | 0.06 | 0.1 |
| 11 | 124432323 | rs2187149 | 0.036 | 0.563 | 0.328 | 0.128 | 0.022 | -0.03 | -0.03 | 0 | 0.76 | 0.7 | 0.74 | 0.88 | 0.68 | 0.64 |
| 11 | 124433714 | rs10893265 | 0.571 | 0.522 | 0.355 | 0.167 | -0.032 | 0.031 | 0.031 | 0 | 0.16 | 0.25 | 0.09 | 0.25 | 0.09 | 0.15 |
| 11 | 124434217 | rs11219709 | 1.228 | 0.565 | 0.153 | 0.177 | -0.006 | 0.023 | 0.023 | 0.007 | 0.53 | 0.39 | 0.45 | 0.33 | 0.46 | 0.51 |
| 11 | 124438314 | rs11219712 | 1.609 | 0.406 | -0.369 | -0.184 | -0.014 | 0.036 | 0.036 | 0.79 | 0.1 | 0.03 | 0.06 | 0 | 0.11 | 0.02 |
| 11 | 124439147 | rs7130836 | -0.629 | 0.475 | -0.054 | 0.118 | 0.006 | 0.057 | 0.057 | 0.104 | 0.31 | 0.49 | 0.35 | 0.59 | 0.33 | 0.33 |
| 11 | 124439417 | rs7103533 | 0.106 | 1.062 | -0.131 | 0.173 | 0.007 | 0.01 | 0.01 | 0 | 0.16 | 0.19 | 0.26 | 0.33 | 0.25 | 0.19 |
| 11 | 124441693 | rs4316519 | 0.084 | 0.538 | 0.012 | 0.182 | 0.014 | -0.026 | -0.026 | 0 | 0.79 | 0.69 | 0.76 | 0.89 | 0.73 | 0.74 |
| 11 | 124448354 | rs11219714 | -0.251 | 0.297 | -0.24 | 0.482 | 0 | -0.003 | -0.003 | 0.677 | 0.85 | 0.86 | 0.82 | 0.9 | 0.84 | 0.8 |
| 11 | 124448469 | rs7941314 | 0.349 | 0.024 | -0.213 | 0.569 | 0 | 0.003 | 0.003 | 0 | 0.26 | 0.21 | 0.33 | 0.25 | 0.33 | 0.16 |
| 11 | 124449980 | rs7110338 | -0.438 | -0.265 | -0.273 | 0.848 | -0.014 | 0.011 | 0.011 | 0 | 0.4 | 0.52 | 0.42 | 0.54 | 0.42 | 0.27 |
| 11 | 124450132 | rs10790705 | -0.334 | -0.224 | -0.56 | 1.004 | 0.001 | -0.004 | -0.004 | 0 | 0.86 | 0.87 | 0.82 | 0.9 | 0.84 | 0.82 |
| 11 | 124450173 | rs11219715 | -0.033 | -0.22 | -0.513 | 1.091 | -0.007 | 0.005 | 0.005 | 0 | 0.13 | 0.18 | 0.07 | 0.23 | 0.08 | 0.1 |
| 11 | 124453666 | rs7111036 | 0.115 | -0.02 | -0.775 | 1.087 | 0.003 | -0.004 | -0.004 | 0.131 | 0.73 | 0.76 | 0.75 | 0.67 | 0.76 | 0.71 |
| 11 | 124454319 | rs1954887 | 0.265 | 0.301 | -1.191 | 1.146 | 0.008 | 0.009 | 0.009 | 0 | 0.31 | 0.08 | 0.31 | 0.24 | 0.28 | 0.33 |
| 11 | 124455088 | rs4936943 | 0.061 | 0.374 | -1.503 | 1.35 | -0.069 | -0.137 | -0.137 | 0.11 | 0.61 | 0.97 | 0.53 | 0.45 | 0.59 | 0.53 |
| 11 | 124460485 | rs10790707 | -0.569 | 0.587 | -0.322 | 1.124 | -0.005 | -0.002 | -0.002 | 0 | 0.37 | 0.62 | 0.45 | 0.4 | 0.36 | 0.33 |
| 11 | 124461274 | rs7127399 | -0.41 | 0.412 | -0.286 | 1.121 | -0.012 | 0 | 0 | 0 | 0.35 | 0.65 | 0.42 | 0.4 | 0.31 | 0.31 |
| 11 | 124463603 | rs7931301 | -0.96 | 0.186 | -0.214 | 1.052 | -0.04 | -0.002 | -0.002 | 0 | 0.1 | 0.19 | 0.02 | 0.1 | 0.02 | 0.06 |
| 11 | 124464910 | rs11219724 | -0.882 | 0.131 | -0.186 | 1.051 | -0.051 | -0.002 | -0.002 | 0.703 | 0.1 | 0.23 | 0.03 | 0.1 | 0.02 | 0.06 |
| 11 | 124470426 | rs10128573 | 0.076 | 0.321 | -0.529 | 1.096 | -0.058 | -0.037 | -0.037 | 5.646 | 0.9 | 0.99 | 0.75 | 0.86 | 0.79 | 0.85 |
| 11 | 124480049 | rs10893277 | -1.353 | -0.519 | 0.195 | 0.926 | -0.072 | 0.021 | 0.021 | 0 | 0.12 | 0.35 | 0.04 | 0.12 | 0.02 | 0.06 |
| 11 | 124493199 | rs678132 | -1.243 | -0.271 | -1.163 | 1.513 | 0.016 | -0.019 | -0.019 | 0 | 0.18 | 0.16 | 0.07 | 0.31 | 0.07 | 0.19 |
| 11 | 124494303 | rs2156155 | -0.616 | -0.054 | -1.988 | 0.983 | 0.024 | 0.052 | 0.052 | 0 | 0.53 | 0.91 | 0.49 | 0.39 | 0.43 | 0.44 |
| 11 | 124494676 | rs686567 | -0.124 | 0.643 | -1.239 | 0.635 | -0.006 | 0.036 | 0.036 | 0.161 | 0.54 | 0.37 | 0.43 | 0.64 | 0.54 | 0.66 |
| 11 | 124501921 | rs2187152 | -1.578 | 1.388 | 0.374 | 1.121 | 0.164 | 0.163 | 0.163 | 0.096 | 0.07 | 0.46 | 0.08 | 0.12 | 0.09 | 0.06 |
| 11 | 124502260 | rs1047661 | -1.447 | 1.386 | 0.601 | 1.12 | 0.152 | 0.152 | 0.152 | 0.085 | 0.07 | 0.43 | 0.08 | 0.11 | 0.09 | 0.05 |
| 11 | 124508345 | rs2187155 | -1.557 | 1.387 | 0.61 | 1.099 | 0.171 | 0.164 | 0.164 | 0.062 | 0.07 | 0.46 | 0.08 | 0.1 | 0.09 | 0.04 |
| 11 | 124524627 | rs1942663 | -2.095 | 1.18 | 1.064 | 1.038 | 0.053 | 0.147 | 0.147 | 2.284 | 0.06 | 0.63 | 0.07 | 0.1 | 0.02 | 0.03 |
| 11 | 124537995 | rs544368 | -0.39 | 1.478 | 1.386 | 0.822 | -0.025 | 0.032 | 0.032 | 0 | 0.94 | 0.99 | 0.74 | 1 | 0.89 | 0.93 |
| 11 | 124566724 | rs679810 | -0.742 | 1.528 | 0.859 | 1.434 | -0.002 | -0.004 | -0.004 | 0 | 0.3 | 0.25 | 0.3 | 0.48 | 0.33 | 0.31 |
| 11 | 124570014 | rs617690 | -1.135 | 1.613 | 0.699 | 1.574 | 0.001 | -0.006 | -0.006 | 0 | 0.26 | 0.68 | 0.16 | 0.24 | 0.23 | 0.2 |
| 11 | 124570913 | rs631980 | -0.426 | 1.628 | 0.304 | 1.52 | -0.012 | 0.024 | 0.024 | 0 | 0.2 | 0.12 | 0.11 | 0.15 | 0.21 | 0.19 |
| 11 | 124578282 | rs654411 | -2.025 | 1.636 | -0.16 | 1.204 | -0.116 | 0.061 | 0.061 | 0 | 0.48 | 0.96 | 0.46 | 0.65 | 0.48 | 0.51 |
| 11 | 124586079 | rs646487 | -1.836 | 2.193 | 0.269 | 1.219 | 0.002 | 0.021 | 0.021 | 0 | 0.46 | 0.66 | 0.44 | 0.61 | 0.52 | 0.53 |
| 11 | 124590079 | rs671371 | -0.199 | 2.168 | -0.227 | 1.294 | -0.031 | -0.018 | -0.018 | 0 | 0.2 | 0.08 | 0.1 | 0.22 | 0.24 | 0.16 |
| 11 | 124600758 | rs2155109 | -1.42 | 2.141 | 0.682 | 1.892 | 0.003 | 0.015 | 0.015 | 0 | 0.24 | 0.27 | 0.28 | 0.36 | 0.26 | 0.33 |
| 11 | 124602074 | rs585454 | 1.987 | 2.307 | 0.517 | 1.678 | -0.033 | 0.004 | 0.004 | 2.613 | 0.57 | 0.2 | 0.67 | 0.43 | 0.54 | 0.49 |
| 11 | 124602247 | rs11219765 | -2.397 | 2.426 | 0.747 | 1.696 | -0.103 | -0.033 | -0.033 | 0 | 0.38 | 0.9 | 0.35 | 0.4 | 0.43 | 0.45 |
| 11 | 124605290 | rs1939214 | -2.119 | 2.239 | 1.843 | 1.443 | 0.077 | 0.078 | 0.078 | 0 | 0.1 | 0.24 | 0.09 | 0.2 | 0.16 | 0.09 |
| 11 | 124606285 | rs12807809 | -2.148 | 2.27 | 1.954 | 1.491 | 0.092 | 0.147 | 0.147 | 0 | 0.1 | 0.28 | 0.14 | 0.29 | 0.17 | 0.1 |
| 11 | 124612159 | rs12278912 | -2.381 | 2.318 | 2.334 | 1.105 | 0.016 | -0.012 | -0.012 | 0 | 0.24 | 0.56 | 0.15 | 0.23 | 0.28 | 0.19 |
| 11 | 124617939 | rs2075713 | -1.858 | 2.2 | 2.581 | 0.912 | 0.001 | -0.03 | -0.03 | 0 | 0.2 | 0.33 | 0.11 | 0.12 | 0.2 | 0.17 |

| | | | | | | | | | | | | | | | | |
|---|---|---|---|---|---|---|---|---|---|---|---|---|---|---|---|---|
| 11 | 124620147 | rs11219769 | -2.384 | 2.38 | 2.497 | 0.994 | -0.007 | 0.011 | 0.011 | 0 | 0.24 | 0.78 | 0.19 | 0.16 | 0.27 | 0.23 |
| 11 | 124623492 | rs12541 | -2.38 | 2.437 | 2.454 | 1.154 | -0.006 | -0.006 | -0.006 | 0 | 0.21 | 0.8 | 0.18 | 0.26 | 0.22 | 0.2 |
| 11 | 124624854 | rs6590090 | -2.642 | 2.314 | 2.247 | 1.101 | -0.007 | -0.005 | -0.005 | 0 | 0.24 | 0.81 | 0.18 | 0.27 | 0.23 | 0.2 |
| 11 | 124627479 | rs1076095 | -1.817 | 2.074 | 2.715 | 1.284 | 0.067 | 0.061 | 0.061 | 0.255 | 0.14 | 0.48 | 0.13 | 0.16 | 0.14 | 0.14 |
| 11 | 124633640 | rs1940171 | -1.921 | 2.147 | 2.108 | 1.241 | 0.004 | 0 | 0 | 0 | 0.15 | 0.79 | 0.15 | 0.2 | 0.18 | 0.11 |
| 11 | 124635144 | rs3802901 | -1.925 | 2.151 | 2.11 | 1.27 | 0 | 0 | 0 | 1.883 | 0.15 | 0.81 | 0.16 | 0.26 | 0.18 | 0.15 |
| 11 | 124636373 | rs11219774 | -1.485 | 1.924 | 2.334 | 1.29 | 0.1 | 0.064 | 0.064 | 0.376 | 0.13 | 0.5 | 0.1 | 0.15 | 0.15 | 0.12 |
| 11 | 124657183 | rs485469 | -1.188 | 1.615 | 1.301 | 0.825 | 0.009 | -0.007 | -0.007 | 0 | 0.22 | 0.33 | 0.18 | 0.18 | 0.25 | 0.37 |
| 11 | 124668305 | rs3016271 | -1.106 | 1.601 | 1.025 | 0.819 | 0 | 0 | 0 | 0 | 0.22 | 0.21 | 0.17 | 0.18 | 0.26 | 0.37 |
| 11 | 124674924 | rs11826296 | -0.621 | 1.737 | 0.488 | 0.948 | 0 | 0.002 | 0.002 | 0 | 0.06 | 0.08 | 0.04 | 0.13 | 0.09 | 0.06 |
| 11 | 124677941 | rs646521 | -1.378 | 1.497 | 0.531 | 0.831 | 0.002 | 0 | 0 | 1.4 | 0.34 | 0.61 | 0.25 | 0.35 | 0.37 | 0.43 |
| 11 | 124686709 | rs535396 | -0.927 | 1.469 | 0.355 | 0.769 | 0.056 | 0.023 | 0.023 | 0 | 0.18 | 0.36 | 0.12 | 0.24 | 0.21 | 0.29 |
| 11 | 124688317 | rs663642 | -1.168 | 0.724 | -0.061 | 0.313 | -0.001 | -0.002 | -0.002 | 0 | 0.39 | 0.53 | 0.26 | 0.33 | 0.34 | 0.33 |
| 11 | 124693144 | rs1783885 | -0.772 | 1.179 | 0.08 | 0.807 | 0.054 | 0.063 | 0.063 | 0 | 0.18 | 0.35 | 0.12 | 0.28 | 0.21 | 0.29 |
| 11 | 124697202 | rs7127139 | -0.503 | 1.171 | -0.635 | 0.648 | -0.013 | -0.024 | -0.024 | 0 | 0.65 | 0.74 | 0.41 | 0.51 | 0.58 | 0.61 |
| 11 | 124703336 | rs997989 | -0.546 | 0.517 | -0.661 | 0.56 | 0.001 | -0.002 | -0.002 | 0 | 0.13 | 0.1 | 0.09 | 0.18 | 0.11 | 0.07 |
| 11 | 124704084 | rs7934672 | -0.096 | 0.655 | -0.721 | -0.21 | 0.009 | 0.028 | 0.028 | 1.862 | 0.4 | 0.32 | 0.21 | 0.23 | 0.35 | 0.29 |
| 11 | 124708698 | rs610104 | -1.018 | 0.945 | -0.33 | -1.041 | 0.01 | -0.114 | -0.114 | 0 | 0.26 | 0.34 | 0.24 | 0.02 | 0.31 | 0.13 |
| 11 | 124717445 | rs6590107 | -0.857 | 1.026 | 0.078 | -1.051 | 0.016 | -0.021 | -0.021 | 0 | 0.21 | 0.39 | 0.33 | 0.19 | 0.19 | 0.29 |
| 11 | 124717530 | rs4936956 | -1.129 | 0.739 | -0.831 | -1.976 | 0 | -0.006 | -0.006 | 0 | 0.55 | 0.54 | 0.54 | 0.29 | 0.52 | 0.57 |
| 11 | 124718707 | rs11219805 | -0.718 | 0.395 | -0.865 | -2.011 | -0.013 | -0.048 | -0.048 | 0.192 | 0.3 | 0.41 | 0.38 | 0.2 | 0.27 | 0.32 |
| 11 | 124719105 | rs12288342 | -0.897 | 0.163 | -0.836 | -2.249 | 0.014 | 0.03 | 0.03 | 0 | 0.47 | 0.54 | 0.32 | 0.27 | 0.4 | 0.46 |
| 11 | 124719898 | rs11219808 | 1.314 | -0.111 | -0.658 | -2.528 | -0.004 | -0.023 | -0.023 | 0 | 0.1 | 0.08 | 0.03 | 0 | 0.06 | 0.04 |
| 11 | 124720545 | rs1940174 | -0.555 | -0.089 | -0.953 | -2.458 | 0.019 | 0.068 | 0.068 | 1.336 | 0.51 | 0.33 | 0.31 | 0.28 | 0.44 | 0.41 |
| 11 | 124721003 | rs11604758 | -1.296 | 0.044 | -0.562 | -2.481 | 0.022 | 0.002 | 0.002 | 0.155 | 0.72 | 0.64 | 0.54 | 0.31 | 0.66 | 0.68 |
| 11 | 124728123 | rs11219814 | 1.055 | -0.015 | -0.969 | -2.567 | -0.032 | 0.041 | 0.041 | 0 | 0.18 | 0.09 | 0.15 | 0 | 0.25 | 0.16 |
| 11 | 124730646 | rs7113124 | 0.885 | -0.43 | -1.019 | -2.471 | 0.039 | 0.01 | 0.01 | 0 | 0.24 | 0.09 | 0.12 | 0.23 | 0.14 | 0.21 |
| 11 | 124733377 | rs7111513 | 0.044 | -0.286 | -0.459 | -2.114 | -0.022 | -0.078 | -0.078 | 0.076 | 0.29 | 0.42 | 0.26 | 0.08 | 0.26 | 0.32 |
| 11 | 124734471 | rs5017048 | 0.117 | -0.266 | -0.785 | -2.104 | -0.007 | -0.035 | -0.035 | 0.006 | 0.29 | 0.36 | 0.26 | 0.08 | 0.26 | 0.31 |
| 11 | 124734728 | rs7951801 | -0.291 | 0.028 | -1.14 | -1.719 | -0.01 | 0.128 | 0.128 | 0 | 0.43 | 0.18 | 0.29 | 0.23 | 0.43 | 0.37 |
| 11 | 124739988 | rs4936957 | -0.777 | -0.414 | -0.434 | -0.804 | -0.012 | 0.019 | 0.019 | 0 | 0.67 | 0.57 | 0.46 | 0.39 | 0.7 | 0.56 |
| 11 | 124740691 | rs7925879 | -1.063 | -0.614 | -0.619 | -0.875 | -0.017 | -0.052 | -0.052 | 0 | 0.71 | 0.87 | 0.5 | 0.4 | 0.71 | 0.58 |
| 11 | 124741772 | rs11219821 | 0.35 | -0.995 | -0.741 | -1.422 | -0.013 | 0.056 | 0.056 | 0 | 0.39 | 0.27 | 0.28 | 0.09 | 0.44 | 0.21 |
| 11 | 124742365 | rs3862618 | 1.159 | -1.124 | -1.25 | -1.443 | -0.019 | -0.065 | -0.065 | 0 | 0.16 | 0.01 | 0.1 | 0.27 | 0.13 | 0.17 |
| 11 | 124742385 | rs4935898 | 1.66 | -1.171 | -1.171 | -1.676 | 0.011 | 0.032 | 0.032 | 0.016 | 0.06 | 0 | 0.02 | 0 | 0.03 | 0.05 |
| 11 | 124742731 | rs4606490 | -0.851 | -1.112 | -0.977 | -1.662 | -0.011 | 0.15 | 0.15 | 0 | 0.53 | 0.77 | 0.38 | 0.11 | 0.57 | 0.39 |
| 11 | 124747333 | rs10790713 | 0.25 | -0.673 | -0.857 | -1.503 | -0.026 | 0.064 | 0.064 | 0.06 | 0.35 | 0.23 | 0.29 | 0.04 | 0.45 | 0.19 |
| 11 | 124758637 | rs7129737 | -0.258 | -1.272 | -0.502 | -1.361 | -0.046 | -0.06 | -0.06 | 0 | 0.23 | 0.51 | 0.13 | 0.09 | 0.16 | 0.2 |
| 11 | 124759644 | rs4635093 | -1.364 | -1.538 | -1.227 | -1.268 | 0.065 | 0.028 | 0.028 | 0 | 0.47 | 0.67 | 0.25 | 0.4 | 0.31 | 0.42 |
| 11 | 124760992 | rs11219830 | 0.236 | -1.44 | -1.844 | -1.212 | 0.005 | -0.072 | -0.072 | 0 | 0.22 | 0 | 0.11 | 0.3 | 0.14 | 0.21 |
| 11 | 124762165 | rs12417435 | 0.828 | -1.24 | -1.659 | -1.123 | -0.012 | 0.004 | 0.004 | 0 | 0.06 | 0.03 | 0.06 | 0.03 | 0.11 | 0.15 |
| 11 | 124765168 | rs12421354 | 0.503 | -1.289 | -1.611 | -1.003 | 0.005 | 0 | 0 | 0 | 0.18 | 0.14 | 0.1 | 0.14 | 0.16 | 0.21 |
| 11 | 124766128 | rs4078313 | -1.184 | -1.233 | -2.096 | -0.982 | 0.126 | -0.006 | -0.006 | 0 | 0.5 | 0.82 | 0.29 | 0.45 | 0.33 | 0.42 |
| 11 | 124769619 | rs4586175 | -0.824 | -0.343 | -1.472 | -0.759 | 0.017 | -0.012 | -0.012 | 0 | 0.33 | 0.71 | 0.22 | 0.4 | 0.27 | 0.35 |
| 11 | 124771243 | rs4284402 | 0.648 | 0.315 | -1.408 | -0.737 | 0.006 | -0.044 | -0.044 | 0 | 0.12 | 0.02 | 0.07 | 0.2 | 0.08 | 0.13 |
| 11 | 124771271 | rs4077566 | -0.193 | 1.086 | -0.97 | -1.099 | -0.109 | 0.128 | 0.128 | 0 | 0.22 | 0.06 | 0.24 | 0 | 0.38 | 0.11 |
| 11 | 124776876 | rs12294639 | -0.771 | 0.281 | 0.151 | -0.095 | 0.053 | 0.061 | 0.061 | 1.056 | 0.05 | 0.37 | 0.03 | 0.01 | 0.01 | 0 |
| 11 | 124789755 | rs3802904 | 0.116 | 0.741 | -0.94 | -0.291 | -0.03 | 0.043 | 0.043 | 0.103 | 0.76 | 0.88 | 0.65 | 0.92 | 0.73 | 0.75 |
| 11 | 124792627 | rs11826299 | 0.071 | -0.065 | -0.168 | -0.66 | -0.036 | -0.051 | -0.051 | 0 | 0.17 | 0.34 | 0.09 | 0.05 | 0.1 | 0.06 |
| 11 | 124793682 | rs10790715 | -0.228 | -0.132 | -0.684 | -1.049 | -0.045 | 0.067 | 0.067 | 0 | 0.74 | 0.89 | 0.64 | 0.91 | 0.72 | 0.74 |
| 11 | 124793716 | rs10790716 | 0.692 | 0.359 | -1.045 | -1.645 | -0.02 | 0.015 | 0.015 | 0 | 0.14 | 0.11 | 0.15 | 0.07 | 0.26 | 0.1 |
| 11 | 124795154 | rs4936958 | 0.973 | -0.432 | -0.624 | -1.504 | -0.052 | 0.059 | 0.059 | 0 | 0.24 | 0.37 | 0.26 | 0.67 | 0.13 | 0.34 |
| 11 | 124798250 | rs7130442 | -0.9 | -0.324 | -0.675 | -1.482 | -0.027 | 0.052 | 0.052 | 0 | 0.71 | 0.79 | 0.59 | 0.91 | 0.67 | 0.72 |
| 11 | 124799896 | rs3924535 | 2.175 | -1.061 | -0.654 | -0.551 | 0.007 | 0.048 | 0.048 | 0.022 | 0.07 | 0.03 | 0.17 | 0.45 | 0.05 | 0.25 |
| 11 | 124805619 | rs4254088 | -0.596 | -0.927 | -0.657 | -0.629 | 0.01 | 0.073 | 0.073 | 0 | 0.57 | 0.67 | 0.42 | 0.83 | 0.38 | 0.62 |
| 11 | 124808253 | rs11219844 | 1.605 | -1.106 | -0.79 | -0.337 | 0.014 | -0.002 | -0.002 | 3.155 | 0.09 | 0.03 | 0.18 | 0.4 | 0.06 | 0.24 |
| 11 | 124810732 | rs11821787 | 1.351 | -1.517 | -0.552 | -0.565 | 0.011 | -0.002 | -0.002 | 0.06 | 0.15 | 0.12 | 0.04 | 0.01 | 0.05 | 0.05 |
| 11 | 124813516 | rs10893306 | 0.238 | -1.853 | -0.41 | -0.682 | -0.064 | 0.119 | 0.119 | 0 | 0.23 | 0.42 | 0.24 | 0.61 | 0.12 | 0.33 |
| 11 | 124813669 | rs4340063 | 0.071 | -1.774 | -0.481 | -0.774 | 0.019 | 0.063 | 0.063 | 0 | 0.35 | 0.18 | 0.14 | 0.16 | 0.28 | 0.28 |
| 11 | 124814005 | rs4936960 | 1.899 | -1.723 | -0.491 | -0.695 | 0.009 | 0.015 | 0.015 | 0 | 0.09 | 0.04 | 0.18 | 0.41 | 0.06 | 0.23 |
| 11 | 124814152 | rs7937347 | -1.018 | -1.576 | -0.511 | -0.922 | 0.005 | 0.034 | 0.034 | 0 | 0.58 | 0.61 | 0.38 | 0.78 | 0.39 | 0.61 |
| 11 | 124814761 | rs12283125 | 0.824 | -1.332 | -0.605 | -0.802 | -0.043 | -0.004 | -0.004 | 0.195 | 0.19 | 0.03 | 0.1 | 0.16 | 0.22 | 0.24 |
| 11 | 124816164 | rs10160591 | -0.509 | -1.328 | -0.398 | -0.646 | -0.007 | -0.013 | -0.013 | 0 | 0.43 | 0.42 | 0.35 | 0.75 | 0.35 | 0.57 |
| 11 | 124816420 | rs12222810 | 0.312 | -1.376 | -0.359 | -0.586 | -0.051 | 0.121 | 0.121 | 0 | 0.23 | 0.39 | 0.24 | 0.58 | 0.12 | 0.33 |
| 11 | 124819165 | rs11219849 | 0.927 | -0.338 | -0.379 | -0.871 | -0.004 | -0.003 | -0.003 | 0 | 0.27 | 0.31 | 0.2 | 0.18 | 0.22 | 0.25 |
| 11 | 124823305 | rs4936963 | -0.965 | 0.051 | -0.449 | -0.94 | -0.028 | 0.02 | 0.02 | 0 | 0.82 | 0.84 | 0.59 | 0.94 | 0.63 | 0.87 |
| 11 | 124823648 | rs7120629 | 1.248 | 0.576 | -0.531 | -1.052 | 0.004 | -0.028 | -0.028 | 0 | 0.11 | 0.16 | 0.09 | 0 | 0.15 | 0.03 |
| 11 | 124823981 | rs3809055 | 1.386 | 0.156 | -0.714 | -1.317 | -0.014 | 0.026 | 0.026 | 0 | 0.17 | 0.3 | 0.14 | 0.18 | 0.15 | 0.23 |
| 11 | 124824066 | rs3809054 | -0.996 | 0.136 | -1.015 | -1.744 | -0.007 | 0.079 | 0.079 | 0 | 0.54 | 0.86 | 0.59 | 0.26 | 0.53 | 0.41 |

| | | | | | | | | | | | | | | | | |
|---|---|---|---|---|---|---|---|---|---|---|---|---|---|---|---|---|
| 11 | 124827935 | rs12287581 | 0.639 | -0.306 | -0.499 | -1.682 | -0.047 | -0.026 | -0.026 | 0.941 | 0.3 | 0.47 | 0.18 | 0.19 | 0.2 | 0.31 |
| 11 | 124840967 | rs7925079 | 1.44 | -0.24 | -0.377 | -1.625 | -0.019 | 0.04 | 0.04 | 0.035 | 0.18 | 0.36 | 0.14 | 0.19 | 0.15 | 0.25 |
| 11 | 124853470 | rs12289777 | 2.222 | -0.268 | -0.224 | -1.65 | -0.007 | 0.015 | 0.015 | 0 | 0.14 | 0.17 | 0.1 | 0.18 | 0.1 | 0.23 |
| 11 | 124857708 | rs4936966 | 0.662 | -0.301 | -0.468 | -1.669 | -0.047 | -0.026 | -0.026 | 0 | 0.3 | 0.47 | 0.18 | 0.19 | 0.2 | 0.3 |
| 11 | 124863136 | rs7951202 | 0.662 | -0.301 | -0.468 | -1.669 | -0.047 | -0.026 | -0.026 | 0.106 | 0.3 | 0.47 | 0.18 | 0.19 | 0.2 | 0.3 |
| 11 | 124870136 | rs11827270 | 1.915 | -0.188 | -0.483 | -1.721 | 0.005 | -0.029 | -0.029 | 0 | 0.11 | 0.15 | 0.09 | 0 | 0.15 | 0.03 |
| 11 | 124874382 | rs4612816 | -0.928 | -0.068 | -0.443 | -1.794 | 0.049 | -0.02 | -0.02 | 0 | 0.85 | 0.76 | 0.62 | 0.93 | 0.72 | 0.9 |
| 11 | 124881553 | rs4353290 | -1.988 | 0.017 | -0.48 | -1.782 | -0.05 | -0.017 | -0.017 | 0.083 | 0.92 | 0.99 | 0.69 | 0.93 | 0.77 | 0.9 |
| 11 | 124890396 | rs4936969 | -1.818 | 0.002 | -0.428 | -1.779 | 0.081 | 0.003 | 0.003 | 0.017 | 0.92 | 0.82 | 0.68 | 0.93 | 0.77 | 0.9 |
| 11 | 124908352 | rs7107487 | 1.467 | -0.404 | -0.467 | -1.73 | -0.018 | 0.042 | 0.042 | 0.026 | 0.17 | 0.36 | 0.14 | 0.18 | 0.15 | 0.25 |
| 11 | 124910778 | rs10893316 | 2.264 | -0.396 | -0.814 | -1.711 | -0.004 | -0.023 | -0.023 | 0 | 0.13 | 0.11 | 0.04 | 0.01 | 0.05 | 0.06 |
| 11 | 124910805 | rs10790721 | 0.264 | -0.077 | -0.982 | -1.506 | 0.005 | 0.015 | 0.015 | 0 | 0.41 | 0.63 | 0.27 | 0.19 | 0.35 | 0.34 |
| 11 | 124911136 | rs15541 | 0.94 | -0.039 | -1.403 | -1.042 | -0.001 | -0.038 | -0.038 | 0.49 | 0.29 | 0.08 | 0.16 | 0.32 | 0.28 | 0.29 |
| 11 | 124931334 | rs4078355 | 0.789 | 0.047 | -0.815 | -0.807 | -0.025 | -0.011 | -0.011 | 0.809 | 0.28 | 0.39 | 0.18 | 0.19 | 0.19 | 0.31 |
| 11 | 124932019 | rs7952332 | -1.13 | 0.031 | -1.039 | -0.721 | -0.02 | -0.116 | -0.116 | 0.273 | 0.77 | 0.95 | 0.71 | 0.41 | 0.74 | 0.6 |
| 11 | 124933582 | rs11820425 | 0.383 | -0.143 | -0.258 | -0.835 | -0.019 | -0.009 | -0.009 | 0.861 | 0.28 | 0.36 | 0.18 | 0.19 | 0.2 | 0.35 |
| 11 | 124935357 | rs6590124 | -0.581 | 0.014 | -0.275 | -0.755 | 0.008 | 0.029 | 0.029 | 0 | 0.42 | 0.66 | 0.28 | 0.19 | 0.36 | 0.34 |
| 11 | 124943831 | rs12280621 | -0.006 | -0.343 | -0.256 | -0.621 | -0.023 | -0.044 | -0.044 | 0 | 0.28 | 0.39 | 0.18 | 0.12 | 0.2 | 0.3 |
| 11 | 124945546 | rs7943255 | 0.16 | -0.41 | -0.275 | -0.669 | -0.042 | -0.02 | -0.02 | 0 | 0.14 | 0.41 | 0.07 | 0.06 | 0.07 | 0.12 |
| 11 | 124946234 | rs7108269 | 0.327 | 0.305 | -0.559 | -0.953 | 0.017 | -0.052 | -0.052 | 0 | 0.18 | 0.41 | 0.12 | 0.06 | 0.2 | 0.1 |
| 11 | 124946930 | rs7103889 | 1.247 | 0.371 | -1.079 | -0.562 | -0.012 | -0.063 | -0.063 | 0 | 0.13 | 0.03 | 0.17 | 0.59 | 0.14 | 0.26 |
| 11 | 124947051 | rs12289510 | -0.902 | 0.097 | -1.915 | -1.419 | -0.109 | 0.167 | 0.167 | 0.115 | 0.55 | 0.94 | 0.46 | 0.21 | 0.53 | 0.51 |
| 11 | 124947149 | rs12276567 | -0.057 | 0.258 | -1.489 | -1.133 | -0.025 | -0.075 | -0.075 | 0 | 0.34 | 0.52 | 0.29 | 0.14 | 0.26 | 0.39 |
| 11 | 124948039 | rs4074186 | 1.07 | -0.049 | -1.744 | -1.325 | -0.019 | -0.012 | -0.012 | 0 | 0.33 | 0.44 | 0.18 | 0.23 | 0.28 | 0.22 |
| 11 | 124948628 | rs4477450 | -0.543 | 0.377 | -2.066 | -1.193 | -0.057 | -0.064 | -0.064 | 5.32 | 0.74 | 0.91 | 0.53 | 0.37 | 0.65 | 0.63 |
| 11 | 124949967 | rs10893317 | 0.857 | 0.155 | -1.771 | -1.468 | 0 | -0.114 | -0.114 | 0.155 | 0.19 | 0.35 | 0.15 | 0.01 | 0.19 | 0.22 |
| 11 | 124952642 | rs10750294 | -1.191 | 0.806 | -2.37 | -0.723 | -0.097 | 0.061 | 0.061 | 0 | 0.86 | 0.99 | 0.66 | 0.95 | 0.72 | 0.88 |
| 11 | 124955032 | rs948268 | -0.035 | 0.705 | -2.399 | -0.273 | -0.115 | -0.142 | -0.142 | 1.087 | 0.69 | 0.95 | 0.51 | 0.46 | 0.62 | 0.64 |
| 11 | 124955399 | rs3802807 | 1.206 | 0.512 | -1.864 | 0.247 | 0.021 | -0.015 | -0.015 | 0 | 0.1 | 0.02 | 0.04 | 0.11 | 0.06 | 0.09 |
| 11 | 124956298 | rs3808996 | 1.147 | 1.171 | -1.432 | 0.621 | -0.004 | -0.026 | -0.026 | 0 | 0.11 | 0.03 | 0.13 | 0.47 | 0.07 | 0.24 |
| 11 | 124958459 | rs3824926 | -0.334 | 1.645 | -1.229 | 0.276 | -0.03 | -0.026 | -0.026 | 0.01 | 0.81 | 0.9 | 0.65 | 0.77 | 0.73 | 0.82 |
| 11 | 124958914 | rs4936978 | -0.197 | 1.617 | -1.004 | 0.262 | -0.001 | -0.001 | -0.001 | 0 | 0.81 | 0.82 | 0.65 | 0.77 | 0.73 | 0.82 |
| 11 | 124967155 | rs7030 | -0.59 | 1.8 | -1.224 | -0.034 | 0.013 | -0.017 | -0.017 | 0 | 0.22 | 0.3 | 0.3 | 0.17 | 0.31 | 0.36 |
| 11 | 124967774 | rs2186888 | 0.032 | 1.811 | -1.426 | -0.074 | -0.013 | 0.021 | 0.021 | 0.102 | 0.19 | 0.16 | 0.27 | 0.11 | 0.27 | 0.34 |
| 11 | 124972208 | rs14065 | -0.287 | 1.866 | -1.467 | 0.021 | -0.034 | -0.017 | -0.017 | 0.06 | 0.8 | 0.9 | 0.65 | 0.78 | 0.72 | 0.83 |
| 11 | 124972691 | rs2155666 | -0.886 | 1.599 | -1.539 | -0.041 | -0.072 | -0.058 | -0.058 | 0 | 0.83 | 0.99 | 0.68 | 0.84 | 0.77 | 0.84 |
| 11 | 124972832 | rs7118508 | 0.268 | 1.076 | -1.185 | -0.184 | -0.019 | 0.019 | 0.019 | 1.371 | 0.61 | 0.68 | 0.38 | 0.68 | 0.46 | 0.48 |
| 11 | 124974940 | rs10893325 | -0.234 | 0.361 | -1.118 | -0.224 | 0.002 | -0.016 | -0.016 | 0 | 0.4 | 0.31 | 0.34 | 0.51 | 0.38 | 0.47 |
| 11 | 124975953 | rs6590133 | 0.938 | 0.137 | -1.066 | -1.329 | 0.001 | -0.009 | -0.009 | 0 | 0.21 | 0.14 | 0.15 | 0.23 | 0.23 | 0.11 |
| 11 | 124976515 | rs7128296 | -0.11 | -0.173 | -1.045 | -1.652 | 0.011 | 0.023 | 0.023 | 0 | 0.44 | 0.34 | 0.3 | 0.32 | 0.45 | 0.33 |
| 11 | 124981705 | rs3802810 | 1.053 | 0.007 | -1.261 | -1.425 | 0 | -0.018 | -0.018 | 0.017 | 0.21 | 0.08 | 0.14 | 0.24 | 0.23 | 0.11 |
| 11 | 124982023 | rs10893329 | -0.153 | -0.023 | -1.061 | -1.466 | -0.008 | 0.077 | 0.077 | 3.664 | 0.4 | 0.24 | 0.35 | 0.83 | 0.4 | 0.38 |
| 11 | 124982199 | rs11219906 | 0.869 | 0.21 | -1.27 | -1.309 | -0.021 | -0.062 | -0.062 | 1.566 | 0.18 | 0.05 | 0.19 | 0.59 | 0.17 | 0.26 |
| 11 | 124984855 | rs663734 | -0.667 | -0.093 | -1.027 | -1.95 | -0.037 | 0.07 | 0.07 | 0 | 0.71 | 0.81 | 0.52 | 0.94 | 0.66 | 0.7 |
| 11 | 124985993 | rs12576211 | -0.303 | -0.869 | -0.464 | -1.69 | -0.056 | -0.127 | -0.127 | 0 | 0.27 | 0.5 | 0.13 | 0.06 | 0.22 | 0.26 |
| 11 | 124986355 | rs649457 | 0.275 | -0.496 | -0.659 | -1.688 | -0.013 | -0.099 | -0.099 | 0 | 0.58 | 0.49 | 0.43 | 0.13 | 0.66 | 0.42 |
| 11 | 124986933 | rs950133 | -0.73 | -0.62 | -0.594 | -1.049 | -0.016 | -0.02 | -0.02 | 0 | 0.13 | 0.17 | 0.04 | 0.04 | 0.07 | 0.17 |
| 11 | 124988516 | rs608393 | -1 | -0.086 | -0.755 | -0.737 | -0.012 | -0.015 | -0.015 | 0 | 0.24 | 0.2 | 0.26 | 0.63 | 0.25 | 0.36 |
| 11 | 124991376 | rs11606891 | 1.188 | -0.736 | -1.066 | -1.373 | 0.025 | 0.036 | 0.036 | 0 | 0.18 | 0.08 | 0.11 | 0.01 | 0.18 | 0.18 |
| 11 | 124993187 | rs10893331 | -1.313 | -1.098 | -0.907 | -1.523 | 0.003 | 0.018 | 0.018 | 0.839 | 0.37 | 0.69 | 0.39 | 0.72 | 0.3 | 0.47 |
| 11 | 124995104 | rs10893332 | 0.758 | -0.598 | -1.612 | -1.274 | -0.048 | -0.063 | -0.063 | 0 | 0.11 | 0.01 | 0.18 | 0.58 | 0.15 | 0.19 |
| 11 | 124995196 | rs4936981 | 0.001 | -0.558 | -1.531 | -1.316 | -0.002 | -0.002 | -0.002 | 0.033 | 0.08 | 0.11 | 0.14 | 0.09 | 0.07 | 0.1 |
| 11 | 125001843 | rs585977 | -0.102 | -0.213 | -1.691 | -1.078 | -0.007 | -0.015 | -0.015 | 0 | 0.21 | 0.16 | 0.21 | 0.62 | 0.21 | 0.31 |
| 11 | 125003573 | rs666051 | -0.371 | 0.262 | -1.767 | -0.803 | -0.098 | 0.077 | 0.077 | 1.375 | 0.56 | 0.95 | 0.52 | 0.73 | 0.49 | 0.65 |
| 11 | 125010836 | rs679564 | -0.308 | 0.661 | -1.102 | 0.092 | -0.026 | -0.074 | -0.074 | 0 | 0.23 | 0.1 | 0.22 | 0.57 | 0.23 | 0.27 |
| 11 | 125014711 | rs7116601 | -1.122 | 0.269 | 0.108 | -0.212 | 0.061 | 0.057 | 0.057 | 8.124 | 0.05 | 0.42 | 0.03 | 0.01 | 0 | 0 |
| 11 | 125016380 | rs11219924 | -0.228 | 1.03 | -0.465 | -0.583 | 0.014 | -0.024 | -0.024 | 0 | 0.21 | 0.24 | 0.19 | 0.12 | 0.3 | 0.17 |
| 11 | 125018570 | rs11219926 | -0.11 | 1.028 | -0.459 | -0.584 | -0.012 | 0.012 | 0.012 | 0.487 | 0.2 | 0.19 | 0.19 | 0.12 | 0.3 | 0.17 |
| 11 | 125018740 | rs734434 | 0.086 | 0.895 | -1.042 | -0.775 | -0.038 | -0.009 | -0.009 | 0 | 0.61 | 0.88 | 0.46 | 0.39 | 0.6 | 0.58 |
| 11 | 125030008 | rs682980 | -0.582 | 1.198 | -0.583 | -0.334 | -0.001 | 0.004 | 0.004 | 0.075 | 0.87 | 0.94 | 0.74 | 1 | 0.89 | 0.88 |
| 11 | 125044277 | rs11219944 | -0.66 | 1.438 | -0.184 | 0.282 | 0.001 | 0.001 | 0.001 | 0 | 0.21 | 0.24 | 0.19 | 0.29 | 0.29 | 0.22 |
| 11 | 125048088 | rs12790049 | -1.097 | 1.47 | 0.048 | 0.31 | 0.042 | 0.03 | 0.03 | 0.438 | 0.25 | 0.54 | 0.23 | 0.31 | 0.32 | 0.3 |
| 11 | 125051894 | rs7109490 | -0.939 | 1.241 | -0.271 | 0.853 | -0.004 | -0.005 | -0.005 | 0 | 0.35 | 0.38 | 0.39 | 0.46 | 0.44 | 0.32 |
| 11 | 125052084 | rs626475 | 0.781 | 0.549 | -0.968 | 0.151 | 0.003 | 0.021 | 0.021 | 0 | 0.32 | 0.07 | 0.34 | 0.19 | 0.2 | 0.32 |
| 11 | 125053634 | rs590211 | -0.721 | 0.509 | -0.854 | 0.207 | -0.011 | -0.009 | -0.009 | 0 | 0.66 | 0.46 | 0.72 | 0.64 | 0.64 | 0.64 |
| 11 | 125060910 | rs4936991 | -0.147 | 0.326 | -0.845 | 0.007 | -0.014 | -0.016 | -0.016 | 0.265 | 0.3 | 0.26 | 0.36 | 0.46 | 0.43 | 0.31 |
| 11 | 125061547 | rs645219 | -1.03 | 0.137 | -0.365 | 0.024 | 0.003 | 0.009 | 0.009 | 0 | 0.71 | 0.75 | 0.75 | 0.71 | 0.66 | 0.69 |
| 11 | 125063107 | rs10893347 | 0.472 | 0.739 | -0.797 | 0.349 | -0.105 | -0.105 | -0.105 | 0 | 0.18 | 0.03 | 0.22 | 0.3 | 0.31 | 0.19 |
| 11 | 125068428 | rs686277 | 0.96 | 0.403 | -0.731 | -0.034 | 0.012 | 0.08 | 0.08 | 0 | 0.32 | 0.17 | 0.35 | 0.01 | 0.2 | 0.23 |

| | | | | | | | | | | | | | | | | |
|---|---|---|---|---|---|---|---|---|---|---|---|---|---|---|---|---|
| 11 | 125070392 | rs634554 | -1.06 | 0.451 | -0.563 | 0.196 | 0.003 | 0.009 | 0.009 | 0 | 0.68 | 0.71 | 0.75 | 0.73 | 0.67 | 0.69 |
| 11 | 125071311 | rs589839 | 0.722 | 0.369 | -0.366 | 0.499 | 0.012 | 0.007 | 0.007 | 0 | 0.34 | 0.21 | 0.36 | 0.25 | 0.22 | 0.39 |
| 11 | 125072861 | rs7938677 | -0.281 | 0.385 | -0.246 | 0.468 | -0.005 | -0.019 | -0.019 | 0 | 0.19 | 0.34 | 0.21 | 0.14 | 0.19 | 0.12 |
| 11 | 125075575 | rs601414 | -0.327 | 0.47 | -0.279 | 0.478 | -0.001 | -0.002 | -0.002 | 0 | 0.53 | 0.52 | 0.57 | 0.39 | 0.41 | 0.51 |
| 11 | 125078500 | rs593290 | -0.415 | 0.48 | -0.301 | 0.483 | 0 | 0.001 | 0.001 | 0 | 0.54 | 0.57 | 0.57 | 0.39 | 0.41 | 0.51 |
| 11 | 125079282 | rs11219971 | 0.245 | 0.813 | -0.357 | 0.773 | -0.067 | -0.095 | -0.095 | 0 | 0.16 | 0.02 | 0.16 | 0.34 | 0.28 | 0.2 |
| 11 | 125082124 | rs12808899 | -0.481 | 0.662 | -0.06 | 0.685 | -0.004 | -0.013 | -0.013 | 0 | 0.19 | 0.27 | 0.21 | 0.13 | 0.19 | 0.12 |
| 11 | 125084742 | rs4935911 | 0.123 | 0.77 | -0.595 | 0.692 | -0.017 | 0.009 | 0.009 | 0.042 | 0.13 | 0.03 | 0.19 | 0.1 | 0.17 | 0.11 |
| 11 | 125088270 | rs11219974 | -1.02 | 0.603 | -0.328 | 0.907 | -0.019 | 0.163 | 0.163 | 0.212 | 0.2 | 0.47 | 0.25 | 0.41 | 0.19 | 0.27 |
| 11 | 125093850 | rs6590139 | -0.157 | 0.708 | -0.55 | 0.594 | -0.003 | -0.014 | -0.014 | 0 | 0.16 | 0.26 | 0.2 | 0.11 | 0.17 | 0.11 |
| 11 | 125095221 | rs7928369 | -1.137 | 1.075 | -1.254 | 0.261 | -0.085 | 0.09 | 0.09 | 0 | 0.78 | 0.98 | 0.84 | 0.86 | 0.71 | 0.8 |
| 11 | 125109627 | rs11219981 | 0.228 | 1.531 | -0.841 | 1.256 | -0.069 | -0.078 | -0.078 | 0 | 0.11 | 0.01 | 0.14 | 0.26 | 0.23 | 0.17 |
| 11 | 125110671 | rs1893185 | -0.863 | 0.987 | -0.432 | 1.262 | -0.004 | 0.052 | 0.052 | 2.36 | 0.24 | 0.34 | 0.26 | 0.44 | 0.24 | 0.3 |
| 11 | 125114541 | rs725925 | -0.361 | 1.153 | -0.578 | 1.302 | -0.004 | -0.016 | -0.016 | 0 | 0.2 | 0.14 | 0.25 | 0.44 | 0.25 | 0.32 |
| 11 | 125123425 | rs12577901 | -1.517 | 1.072 | -0.449 | 0.692 | -0.002 | 0.036 | 0.036 | 0 | 0.47 | 0.62 | 0.48 | 0.71 | 0.55 | 0.53 |
| 11 | 125124082 | rs12288391 | -0.648 | 1.199 | -0.019 | 0.927 | 0.014 | 0.014 | 0.014 | 0 | 0.24 | 0.36 | 0.22 | 0.28 | 0.3 | 0.22 |
| 11 | 125129208 | rs11219997 | -0.845 | 1.401 | 0.363 | 0.853 | 0.016 | 0.017 | 0.017 | 0 | 0.26 | 0.56 | 0.23 | 0.28 | 0.31 | 0.22 |
| 11 | 125140514 | rs11220003 | -0.051 | 1.312 | -0.729 | 0.299 | -0.176 | 0.118 | 0.118 | 0.146 | 0.64 | 0.97 | 0.72 | 0.86 | 0.55 | 0.79 |
| 11 | 125143775 | rs10400381 | -2.088 | 0.794 | 0.437 | 0.911 | 0.076 | 0.418 | 0.418 | 1.637 | 0.07 | 0.47 | 0.04 | 0.27 | 0.05 | 0.15 |
| 11 | 125156284 | rs7932336 | -0.639 | 0.817 | 0.071 | 0.796 | -0.005 | 0.057 | 0.057 | 0 | 0.16 | 0.3 | 0.09 | 0.27 | 0.16 | 0.19 |
| 11 | 125166812 | rs4935917 | 0.424 | 0.739 | -0.803 | 0.425 | -0.043 | 0.117 | 0.117 | 0 | 0.34 | 0.02 | 0.28 | 0.12 | 0.3 | 0.34 |
| 11 | 125169468 | rs10466538 | -0.501 | 0.929 | 0.191 | 1.629 | -0.008 | 0.027 | 0.027 | 0 | 0.16 | 0.2 | 0.1 | 0.27 | 0.15 | 0.19 |
| 11 | 125170639 | rs1426153 | -1.003 | 1.054 | 0.428 | 1.98 | -0.008 | 0.17 | 0.17 | 0 | 0.18 | 0.37 | 0.13 | 0.39 | 0.18 | 0.22 |
| 11 | 125172383 | rs1077952 | -0.314 | 0.982 | 0.484 | 2.033 | -0.019 | 0.231 | 0.231 | 0 | 0.13 | 0.36 | 0.1 | 0.33 | 0.11 | 0.2 |
| 11 | 125172593 | rs750338 | -1.152 | 1.029 | 0.792 | 2.233 | -0.013 | 0.128 | 0.128 | 0 | 0.22 | 0.51 | 0.16 | 0.55 | 0.21 | 0.25 |
| 11 | 125173206 | rs7116237 | -0.906 | 0.933 | 0.886 | 2.184 | -0.024 | 0.101 | 0.101 | 0 | 0.21 | 0.56 | 0.12 | 0.54 | 0.17 | 0.26 |
| 11 | 125174197 | rs4081796 | -0.849 | 1.27 | 0.31 | 1.925 | 0.021 | -0.002 | -0.002 | 0 | 0.81 | 0.79 | 0.79 | 0.81 | 0.64 | 0.78 |
| 11 | 125174570 | rs12792626 | -0.481 | 0.101 | 0.746 | 2.073 | -0.027 | 0.172 | 0.172 | 0 | 0.17 | 0.38 | 0.1 | 0.35 | 0.15 | 0.2 |
| 11 | 125182635 | rs7105281 | -2.062 | 0.27 | 0.919 | 1.837 | 0.073 | 0.23 | 0.23 | 0.85 | 0.07 | 0.49 | 0.05 | 0.18 | 0.03 | 0.07 |
| 11 | 125182960 | rs10790733 | -0.033 | 0.054 | 0.265 | 1.852 | -0.016 | -0.041 | -0.041 | 0 | 0.39 | 0.22 | 0.48 | 0.49 | 0.38 | 0.41 |
| 11 | 125183431 | rs11220019 | -0.827 | -0.602 | 0.547 | 1.291 | 0.015 | -0.016 | -0.016 | 0 | 0.28 | 0.22 | 0.12 | 0.34 | 0.22 | 0.29 |
| 11 | 125184366 | rs7935380 | -1.715 | -0.248 | 1.699 | 1.053 | -0.027 | 0.093 | 0.093 | 1.449 | 0.14 | 0.39 | 0.06 | 0.24 | 0.08 | 0.11 |
| 11 | 125191520 | rs11604284 | -0.427 | -0.222 | 1.625 | 0.981 | -0.007 | 0.015 | 0.015 | 0 | 0.1 | 0.15 | 0.06 | 0.2 | 0.07 | 0.05 |
| 11 | 125192281 | rs1035791 | -1.29 | -0.347 | 1.548 | 0.918 | 0.021 | -0.001 | -0.001 | 0 | 0.4 | 0.71 | 0.34 | 0.41 | 0.29 | 0.36 |
| 11 | 125193596 | rs11601321 | -1.704 | -0.332 | 1.485 | 1.027 | -0.042 | 0.073 | 0.073 | 1.288 | 0.15 | 0.66 | 0.09 | 0.25 | 0.07 | 0.11 |
| 11 | 125193811 | rs11605596 | -0.802 | -0.163 | 1.138 | 0.939 | -0.022 | 0.038 | 0.038 | 0.12 | 0.11 | 0.23 | 0.05 | 0.2 | 0.06 | 0.06 |
| 11 | 125194213 | rs11606117 | -0.069 | -0.059 | 0.624 | 0.99 | 0.005 | -0.009 | -0.009 | 0 | 0.08 | 0.06 | 0.04 | 0.2 | 0.06 | 0.05 |
| 11 | 125200247 | rs4935918 | -0.62 | 0.268 | 0.632 | 0.74 | 0.054 | -0.007 | -0.007 | 0.399 | 0.47 | 0.81 | 0.37 | 0.47 | 0.37 | 0.41 |
| 11 | 125217576 | rs744502 | -1.676 | -0.359 | 1.666 | 1.083 | -0.05 | 0.072 | 0.072 | 0.012 | 0.12 | 0.62 | 0.09 | 0.13 | 0.05 | 0.12 |
| 11 | 125224268 | rs12289208 | -1.146 | -0.063 | 1.319 | 1.319 | 0.044 | 0.231 | 0.231 | 0.033 | 0.05 | 0.35 | 0.05 | 0.15 | 0.03 | 0.09 |
| 11 | 125232960 | rs1559800 | -0.466 | 0.273 | -0.192 | 0.912 | -0.035 | 0.016 | 0.016 | 0.46 | 0.62 | 0.86 | 0.77 | 0.7 | 0.57 | 0.7 |
| 11 | 125241666 | rs2059615 | -0.253 | 0.326 | -0.765 | 0.316 | -0.063 | 0.002 | 0.002 | 0 | 0.15 | 0.01 | 0.1 | 0.1 | 0.21 | 0.11 |
| 11 | 125244616 | rs11220039 | -0.981 | -0.208 | 0.184 | 0.618 | -0.07 | 0.009 | 0.009 | 1.22 | 0.13 | 0.41 | 0.08 | 0.14 | 0.05 | 0.14 |
| 11 | 125251034 | rs11824104 | -1.241 | -0.141 | 0.417 | 0.612 | -0.018 | 0.054 | 0.054 | 0.166 | 0.08 | 0.3 | 0.12 | 0.14 | 0.05 | 0.13 |
| 11 | 125251672 | rs733744 | -0.797 | -0.119 | 0.117 | 0.838 | 0.001 | 0.015 | 0.015 | 1.463 | 0.24 | 0.41 | 0.46 | 0.31 | 0.19 | 0.27 |
| 11 | 125251941 | rs863955 | 1.068 | 1.123 | -0.255 | 0.918 | -0.001 | -0.001 | -0.001 | 0 | 0.71 | 0.7 | 0.51 | 0.67 | 0.67 | 0.59 |
| 11 | 125253757 | rs7950499 | -0.643 | 0.8 | -0.44 | 0.94 | -0.045 | 0.076 | 0.076 | 0 | 0.24 | 0.55 | 0.4 | 0.35 | 0.16 | 0.23 |
| 11 | 125253790 | rs840029 | 0.011 | 1.374 | -0.873 | 0.799 | -0.133 | 0.046 | 0.046 | 0 | 0.8 | 0.96 | 0.81 | 0.91 | 0.63 | 0.81 |
| 11 | 125255715 | rs3740892 | -0.808 | 0.632 | -0.606 | 0.937 | 0.004 | 0.028 | 0.028 | 0.017 | 0.29 | 0.57 | 0.49 | 0.34 | 0.28 | 0.32 |
| 11 | 125258307 | rs11220044 | -0.647 | 0.351 | -0.922 | 1.207 | 0.003 | 0.002 | 0.002 | 0 | 0.06 | 0.04 | 0.04 | 0.23 | 0.02 | 0.27 |
| 11 | 125259300 | rs731716 | -1.125 | 0.669 | -1.223 | 0.762 | -0.027 | 0.062 | 0.062 | 0.945 | 0.56 | 0.79 | 0.65 | 0.69 | 0.52 | 0.8 |
| 11 | 125259724 | rs11220045 | -0.59 | 0.274 | -0.94 | 0.473 | -0.002 | -0.009 | -0.009 | 0 | 0.32 | 0.34 | 0.56 | 0.41 | 0.36 | 0.44 |
| 11 | 125261751 | rs7941528 | -0.762 | -0.161 | -0.778 | -0.035 | -0.006 | -0.023 | -0.023 | 0 | 0.09 | 0.1 | 0.04 | 0 | 0.03 | 0.05 |
| 11 | 125265142 | rs7102708 | -0.963 | -0.05 | -0.065 | 0.009 | 0.116 | 0.098 | 0.098 | 0.197 | 0.09 | 0.42 | 0.26 | 0.11 | 0.14 | 0.13 |
| 11 | 125266893 | rs718520 | -2.405 | -0.49 | -0.449 | -0.194 | 0.021 | 0.037 | 0.037 | 0.174 | 0.33 | 0.8 | 0.42 | 0.2 | 0.3 | 0.27 |
| 11 | 125267166 | rs7119403 | -1.776 | -0.071 | -0.288 | -0.197 | 0.009 | -0.035 | -0.035 | 0 | 0.21 | 0.44 | 0.32 | 0.11 | 0.24 | 0.17 |
| 11 | 125267398 | rs11220048 | -0.289 | -0.284 | -1.178 | -0.309 | -0.01 | 0.039 | 0.039 | 0 | 0.12 | 0.01 | 0.07 | 0.02 | 0.12 | 0.05 |
| 11 | 125272224 | rs11607492 | 0.207 | -0.138 | -1.189 | -0.403 | -0.013 | 0.044 | 0.044 | 0.007 | 0.09 | 0.01 | 0.08 | 0 | 0.13 | 0.02 |
| 11 | 125275758 | rs959582 | 0.642 | -0.255 | -1.261 | -0.647 | -0.018 | 0.044 | 0.044 | 0 | 0.83 | 0.88 | 0.87 | 0.95 | 0.79 | 0.76 |
| 11 | 125275857 | rs3740898 | -0.459 | -0.049 | -1.214 | -0.008 | 0.002 | 0.048 | 0.048 | 0.156 | 0.14 | 0.15 | 0.39 | 0.55 | 0.2 | 0.34 |
| 11 | 125276400 | rs4369420 | 1.728 | -1.195 | -1.057 | -0.81 | -0.013 | 0.021 | 0.021 | 0 | 0.43 | 0.33 | 0.3 | 0.33 | 0.45 | 0.34 |
| 11 | 125277494 | rs7112365 | -0.307 | -1.232 | -0.341 | -0.687 | -0.04 | -0.076 | -0.076 | 0.357 | 0.18 | 0.26 | 0.1 | 0.02 | 0.13 | 0.08 |
| 11 | 125278708 | rs12366087 | -1.088 | -1.225 | -0.602 | -0.384 | 0.011 | -0.034 | -0.034 | 1.446 | 0.25 | 0.2 | 0.49 | 0.55 | 0.26 | 0.37 |
| 11 | 125281236 | rs11220051 | 0.372 | -1.1 | -0.7 | -0.557 | 0.021 | 0.029 | 0.029 | 0 | 0.11 | 0.05 | 0.06 | 0 | 0.09 | 0.04 |
| 11 | 125283262 | rs7104304 | 1.397 | -0.721 | -0.637 | -0.715 | 0.031 | 0.043 | 0.043 | 0 | 0.5 | 0.3 | 0.36 | 0.38 | 0.54 | 0.54 |
| 11 | 125286506 | rs11220056 | 1.852 | -0.959 | -1.198 | -0.843 | -0.106 | 0.162 | 0.162 | 0.82 | 0.38 | 0.03 | 0.25 | 0.08 | 0.36 | 0.39 |
| 11 | 125286956 | rs11220057 | 1.516 | -1.405 | -0.346 | -0.689 | 0.005 | 0.054 | 0.054 | 0 | 0.49 | 0.24 | 0.32 | 0.34 | 0.44 | 0.48 |
| 11 | 125287500 | rs10790734 | -0.607 | -1.31 | -0.556 | -0.589 | -0.041 | -0.011 | -0.011 | 0 | 0.92 | 0.99 | 0.91 | 0.91 | 0.84 | 0.92 |
| 11 | 125290754 | rs599206 | -1.476 | -1.653 | -0.161 | -0.751 | -0.022 | -0.006 | -0.006 | 0.057 | 0.35 | 0.24 | 0.6 | 0.67 | 0.47 | 0.48 |

| | | | | | | | | | | | | | | | | |
|---|---|---|---|---|---|---|---|---|---|---|---|---|---|---|---|---|
| 11 | 125291476 | rs10750297 | -0.343 | -1.616 | -0.192 | -0.727 | -0.01 | -0.06 | -0.06 | 0 | 0.2 | 0.09 | 0.49 | 0.58 | 0.27 | 0.35 |
| 11 | 125291645 | rs608501 | -1.747 | -1.572 | 0.262 | -0.686 | 0.001 | 0.014 | 0.014 | 0.014 | 0.47 | 0.48 | 0.66 | 0.68 | 0.54 | 0.53 |
| 11 | 125299738 | rs3740894 | 0.107 | -1.362 | 0.394 | 0.129 | -0.001 | 0.074 | 0.074 | 0 | 0.08 | 0.1 | 0.12 | 0.27 | 0.1 | 0.16 |
| 11 | 125301587 | rs1630030 | 0.32 | -2.055 | 0.296 | -0.339 | -0.061 | -0.121 | -0.121 | 0 | 0.7 | 0.65 | 0.87 | 0.99 | 0.82 | 0.91 |
| 11 | 125301805 | rs1044314 | 2.421 | -2.05 | 0.315 | -0.402 | -0.056 | 0 | 0 | 0 | 0.39 | 0.17 | 0.26 | 0.35 | 0.4 | 0.47 |
| 11 | 125302478 | rs3740893 | 0.294 | -2.008 | 0.328 | -0.424 | -0.002 | 0.071 | 0.071 | 0 | 0.08 | 0.1 | 0.12 | 0.27 | 0.1 | 0.16 |
| 11 | 125302722 | rs645172 | 0.124 | -1.992 | 0.361 | -0.4 | -0.06 | -0.123 | -0.123 | 0 | 0.7 | 0.65 | 0.87 | 0.99 | 0.82 | 0.91 |
| 11 | 125303663 | rs2845845 | -0.671 | -1.686 | -0.023 | -0.195 | -0.023 | -0.022 | -0.022 | 0 | 0.21 | 0.18 | 0.47 | 0.37 | 0.32 | 0.27 |
| 11 | 125304009 | rs1783925 | -0.484 | -1.731 | -0.074 | -0.19 | -0.037 | -0.037 | -0.037 | 3.358 | 0.2 | 0.1 | 0.46 | 0.36 | 0.32 | 0.26 |
| 11 | 125307408 | rs7128228 | 0.226 | -1.467 | -0.214 | -0.204 | 0 | 0.032 | 0.032 | 0 | 0.1 | 0.1 | 0.14 | 0.27 | 0.13 | 0.16 |
| 11 | 125311674 | rs10893380 | 0.243 | -1.517 | -0.213 | -0.202 | 0.001 | 0.033 | 0.033 | 0 | 0.1 | 0.1 | 0.14 | 0.27 | 0.13 | 0.16 |
| 11 | 125312146 | rs12417742 | -0.005 | -1.492 | -0.379 | -0.177 | -0.036 | -0.066 | -0.066 | 0 | 0.15 | 0.02 | 0.39 | 0.29 | 0.19 | 0.2 |
| 11 | 125323965 | rs11220082 | 1.653 | -1.598 | -0.194 | -0.279 | 0.022 | 0.047 | 0.047 | 0 | 0.47 | 0.28 | 0.39 | 0.63 | 0.5 | 0.65 |
| 11 | 125331529 | rs624180 | -0.915 | -1.696 | -0.172 | -0.438 | 0.035 | 0.067 | 0.067 | 0 | 0.87 | 0.99 | 0.93 | 1 | 0.9 | 0.95 |
| 11 | 125331639 | rs12223186 | 2.001 | -1.584 | -0.1 | -0.331 | -0.025 | -0.003 | -0.003 | 0 | 0.38 | 0.25 | 0.27 | 0.37 | 0.37 | 0.49 |
| 11 | 125342130 | rs669052 | 1.366 | -1.391 | -0.12 | -0.271 | -0.003 | -0.002 | -0.002 | 0 | 0.05 | 0.06 | 0.07 | 0.06 | 0.12 | 0.04 |
| 11 | 125347506 | rs665390 | 0.353 | -1.671 | 0.134 | -0.328 | 0.01 | 0.007 | 0.007 | 0 | 0.73 | 0.73 | 0.55 | 0.73 | 0.75 | 0.81 |
| 11 | 125349829 | rs6590146 | 0.031 | -1.529 | 0.668 | -0.404 | 0.002 | 0.026 | 0.026 | 0.001 | 0.15 | 0.14 | 0.14 | 0.29 | 0.18 | 0.25 |
| 11 | 125352935 | rs3016282 | 0.901 | -1.624 | 0.739 | -0.511 | -0.027 | -0.084 | -0.084 | 0 | 0.65 | 0.55 | 0.8 | 0.94 | 0.71 | 0.88 |
| 11 | 125353208 | rs653921 | -0.485 | -1.629 | 0.743 | -0.515 | -0.012 | -0.035 | -0.035 | 0 | 0.84 | 0.83 | 0.92 | 1 | 0.9 | 0.96 |
| 11 | 125353281 | rs485680 | -0.184 | -1.485 | 0.719 | -0.428 | -0.003 | -0.028 | -0.028 | 0 | 0.19 | 0.23 | 0.12 | 0.06 | 0.19 | 0.07 |
| 11 | 125357776 | rs559668 | -0.402 | -1.484 | 0.632 | -0.427 | -0.006 | -0.038 | -0.038 | 0 | 0.2 | 0.28 | 0.12 | 0.06 | 0.19 | 0.09 |
| 11 | 125358590 | rs531312 | -0.195 | -1.471 | 0.611 | -0.417 | -0.004 | -0.026 | -0.026 | 0 | 0.19 | 0.23 | 0.11 | 0.08 | 0.19 | 0.07 |
| 11 | 125376063 | rs2845853 | 1.741 | -1.561 | 0.493 | -0.393 | 0.029 | 0.015 | 0.015 | 0 | 0.47 | 0.27 | 0.64 | 0.61 | 0.52 | 0.61 |
| 11 | 125390604 | rs7936104 | 0.054 | -1.479 | 0.254 | -0.367 | -0.011 | -0.033 | -0.033 | 0 | 0.15 | 0.13 | 0.14 | 0.29 | 0.18 | 0.28 |
| 11 | 125402446 | rs669525 | -0.23 | -1.545 | 0.381 | -0.451 | -0.015 | -0.055 | -0.055 | 0 | 0.21 | 0.33 | 0.14 | 0.08 | 0.2 | 0.07 |
| 11 | 125404662 | rs639065 | -0.834 | -1.583 | -0.023 | -0.421 | 0.03 | 0.037 | 0.037 | 0 | 0.86 | 0.94 | 0.94 | 1 | 0.91 | 0.96 |
| 11 | 125426323 | rs11512640 | 0.637 | -1.593 | -0.007 | -0.426 | 0.004 | -0.012 | -0.012 | 0 | 0.14 | 0.12 | 0.05 | 0 | 0.09 | 0.03 |
| 11 | 125440779 | rs7945490 | 0.071 | -1.524 | 0.023 | -0.363 | -0.008 | -0.024 | -0.024 | 0 | 0.15 | 0.15 | 0.14 | 0.29 | 0.19 | 0.28 |
| 11 | 125446070 | rs550276 | -0.643 | -1.584 | 0.022 | -0.436 | 0.044 | 0.033 | 0.033 | 0 | 0.29 | 0.18 | 0.11 | 0.01 | 0.18 | 0.06 |
| 11 | 125446354 | rs575206 | 0.534 | -1.579 | -0.038 | -0.434 | 0.034 | 0.044 | 0.044 | 0 | 0.15 | 0.06 | 0.07 | 0 | 0.1 | 0.04 |
| 11 | 125446643 | rs2155388 | 0.588 | -1.245 | 0.13 | -0.253 | -0.014 | -0.05 | -0.05 | 0 | 0.62 | 0.46 | 0.78 | 0.9 | 0.7 | 0.88 |
| 11 | 125451422 | rs11220135 | -0.371 | -1.294 | -0.31 | -0.232 | 0.036 | 0.018 | 0.018 | 0 | 0.3 | 0.23 | 0.12 | 0.01 | 0.18 | 0.08 |
| 11 | 125452313 | rs4627097 | -0.689 | -0.796 | -0.126 | 0.146 | 0.044 | 0.027 | 0.027 | 0 | 0.45 | 0.38 | 0.26 | 0.3 | 0.37 | 0.36 |
| 11 | 125461709 | rs548181 | 0.231 | -0.091 | 0.519 | 0.438 | 0.093 | 0.078 | 0.078 | 0 | 0.92 | 0.68 | 0.87 | 0.89 | 0.88 | 0.94 |
| 11 | 125461787 | rs503288 | 0.231 | -0.091 | 0.519 | 0.536 | 0.093 | 0.107 | 0.107 | 0 | 0.92 | 0.68 | 0.87 | 0.87 | 0.88 | 0.94 |
| 11 | 125472616 | rs10893404 | -0.262 | -0.499 | 0.638 | 0.111 | -0.04 | -0.068 | -0.068 | 0.034 | 0.14 | 0.24 | 0.06 | 0.03 | 0.07 | 0.05 |
| 11 | 125476543 | rs3740901 | -0.193 | -0.79 | 0.412 | 0.483 | 0.008 | -0.025 | -0.025 | 0 | 0.33 | 0.28 | 0.23 | 0.48 | 0.29 | 0.39 |
| 11 | 125479363 | rs2241502 | -0.984 | -0.789 | 0.546 | 0.464 | 0.006 | 0.002 | 0.002 | 0 | 0.46 | 0.52 | 0.28 | 0.43 | 0.38 | 0.41 |
| 11 | 125479496 | rs2241501 | -0.05 | -0.759 | 0.5 | 0.484 | 0 | -0.017 | -0.017 | 0 | 0.31 | 0.25 | 0.21 | 0.4 | 0.31 | 0.36 |
| 11 | 125483171 | rs11220159 | 0.364 | -0.777 | 0.546 | 0.453 | -0.044 | -0.075 | -0.075 | 0.124 | 0.14 | 0.27 | 0.06 | 0.03 | 0.07 | 0.05 |
| 11 | 125488404 | rs17140116 | 0.364 | -0.786 | 0.647 | 0.448 | -0.044 | -0.075 | -0.075 | 0 | 0.14 | 0.27 | 0.06 | 0.03 | 0.07 | 0.05 |
| 11 | 125489621 | rs540723 | -1.145 | -0.67 | 0.765 | 0.701 | 0.066 | 0.109 | 0.109 | 0 | 0.94 | 0.8 | 0.89 | 0.84 | 0.88 | 0.96 |
| 11 | 125493109 | rs11220164 | 0.071 | -0.701 | 0.776 | 0.696 | -0.002 | -0.023 | -0.023 | 0 | 0.31 | 0.22 | 0.21 | 0.4 | 0.31 | 0.36 |
| 11 | 125493192 | rs485231 | 1.339 | -0.825 | 0.638 | 0.492 | -0.036 | 0.008 | 0.008 | 0 | 0.43 | 0.2 | 0.58 | 0.4 | 0.45 | 0.53 |
| 11 | 125495022 | rs2298483 | 0.875 | -0.976 | 0.324 | 0.538 | -0.01 | -0.046 | -0.046 | 0.05 | 0.1 | 0.02 | 0.35 | 0.18 | 0.13 | 0.16 |
| 11 | 125495521 | rs555752 | -0.45 | -1.206 | 0.257 | 0.411 | -0.003 | -0.001 | -0.001 | 0 | 0.17 | 0.23 | 0.39 | 0.18 | 0.19 | 0.17 |
| 11 | 125496399 | rs3731395 | 0.723 | -0.983 | 0.378 | 0.569 | -0.044 | -0.075 | -0.075 | 0 | 0.14 | 0.27 | 0.06 | 0.03 | 0.07 | 0.05 |
| 11 | 125514573 | rs521102 | 0.743 | -0.903 | 0.198 | 0.582 | -0.022 | 0.034 | 0.034 | 0 | 0.48 | 0.25 | 0.6 | 0.41 | 0.5 | 0.54 |
| 11 | 125516064 | rs3731459 | 1.253 | -0.91 | 0.232 | 0.541 | 0 | -0.02 | -0.02 | 0 | 0.12 | 0.13 | 0.05 | 0 | 0.07 | 0.05 |
| 11 | 125539202 | rs10893412 | 1.014 | -1.045 | 0.075 | 0.872 | -0.011 | 0.027 | 0.027 | 0 | 0.16 | 0.29 | 0.17 | 0.24 | 0.15 | 0.19 |
| 11 | 125541901 | rs2282535 | 1.926 | -0.908 | -0.209 | 0.827 | -0.029 | 0.05 | 0.05 | 0 | 0.36 | 0.18 | 0.29 | 0.23 | 0.33 | 0.39 |
| 11 | 125550239 | rs555144 | -0.31 | -0.784 | -0.202 | 0.95 | -0.054 | -0.076 | -0.076 | 0 | 0.79 | 0.84 | 0.83 | 0.66 | 0.73 | 0.78 |
| 11 | 125551090 | rs11220183 | 1.132 | -0.903 | -0.061 | 0.874 | 0.002 | -0.014 | -0.014 | 0.165 | 0.08 | 0.05 | 0.28 | 0.17 | 0.11 | 0.14 |
| 11 | 125551608 | rs11220184 | 0.945 | -0.989 | 0.3 | 0.782 | -0.013 | 0.015 | 0.015 | 0 | 0.16 | 0.24 | 0.16 | 0.24 | 0.16 | 0.19 |
| 11 | 125560180 | rs7106795 | 0.987 | -1.207 | 0.293 | 0.614 | -0.036 | -0.076 | -0.076 | 1.059 | 0.14 | 0.24 | 0.06 | 0 | 0.08 | 0.11 |
| 11 | 125563373 | rs4056123 | -1.65 | -1.162 | 0.303 | 0.72 | 0.002 | 0.004 | 0.004 | 0 | 0.55 | 0.59 | 0.57 | 0.61 | 0.51 | 0.62 |
| 11 | 125565843 | rs11220186 | 1.054 | -1.153 | 0.354 | 0.75 | -0.003 | 0.033 | 0.033 | 0 | 0.08 | 0.13 | 0.28 | 0.17 | 0.11 | 0.14 |
| 11 | 125567128 | rs551711 | 0.725 | -1.134 | 0.307 | 0.766 | -0.04 | -0.058 | -0.058 | 0 | 0.16 | 0.07 | 0.06 | 0.19 | 0.16 | 0.17 |
| 11 | 125567539 | rs526941 | 0.725 | -1.134 | 0.307 | 0.766 | -0.04 | -0.058 | -0.058 | 0 | 0.16 | 0.07 | 0.06 | 0.19 | 0.16 | 0.17 |
| 11 | 125569646 | rs519413 | 1.345 | -1.149 | 0.24 | 0.86 | -0.026 | -0.027 | -0.027 | 0 | 0.11 | 0.07 | 0.04 | 0.14 | 0.12 | 0.14 |
| 11 | 125570755 | rs4325320 | 1.025 | -1.18 | 0.332 | 0.763 | -0.015 | 0.009 | 0.009 | 0 | 0.17 | 0.23 | 0.17 | 0.24 | 0.16 | 0.2 |
| 11 | 125572699 | rs523563 | -1.188 | -1.262 | 0.588 | 0.763 | -0.002 | 0 | 0 | 0 | 0.39 | 0.42 | 0.4 | 0.4 | 0.35 | 0.42 |
| 11 | 125588691 | rs7924417 | -0.65 | -1.229 | 0.51 | 0.726 | -0.028 | -0.007 | -0.007 | 0.194 | 0.37 | 0.46 | 0.48 | 0.42 | 0.32 | 0.43 |
| 11 | 125592843 | rs12221488 | 0.02 | -1.112 | 0.312 | 0.717 | 0.008 | 0.006 | 0.006 | 0 | 0.21 | 0.2 | 0.33 | 0.17 | 0.19 | 0.24 |
| 11 | 125595524 | rs564450 | 0.063 | -1.087 | 0.31 | 0.731 | 0.01 | 0.011 | 0.011 | 0 | 0.74 | 0.75 | 0.64 | 0.83 | 0.75 | 0.73 |
| 11 | 125613572 | rs2217034 | 1.017 | -0.881 | 0.421 | 0.868 | 0.022 | 0.045 | 0.045 | 0 | 0.07 | 0.15 | 0.12 | 0.16 | 0.12 | 0.05 |
| 11 | 125615464 | rs489063 | -0.777 | -0.953 | 0.395 | 0.847 | -0.001 | -0.011 | -0.011 | 0 | 0.88 | 0.91 | 0.69 | 0.84 | 0.82 | 0.78 |
| 11 | 125617193 | rs11220207 | 1.567 | -1.059 | 0.547 | 0.836 | 0 | -0.001 | -0.001 | 0 | 0.14 | 0.13 | 0.14 | 0.21 | 0.13 | 0.14 |

| | | | | | | | | | | | | | | | | |
|---|---|---|---|---|---|---|---|---|---|---|---|---|---|---|---|---|
| 11 | 125617317 | rs496459 | 1.699 | -1.102 | 0.671 | 0.711 | -0.002 | 0.004 | 0.004 | 0 | 0.37 | 0.29 | 0.31 | 0.26 | 0.34 | 0.34 |
| 11 | 125617610 | rs2114084 | -0.123 | -1.061 | 0.514 | 0.686 | -0.005 | 0.018 | 0.018 | 0.064 | 0.32 | 0.38 | 0.22 | 0.44 | 0.29 | 0.31 |
| 11 | 125617908 | rs2282539 | -0.71 | -1.029 | 0.283 | 0.699 | 0.006 | 0 | 0 | 0.928 | 0.39 | 0.53 | 0.34 | 0.61 | 0.41 | 0.36 |
| 11 | 125618370 | rs10790747 | -1.178 | -0.694 | 0.095 | 0.934 | -0.01 | 0.024 | 0.024 | 0 | 0.54 | 0.71 | 0.39 | 0.61 | 0.49 | 0.45 |
| 11 | 125618813 | rs1054997 | 1.475 | -0.681 | -0.04 | 0.944 | -0.005 | 0.004 | 0.004 | 0.021 | 0.06 | 0.07 | 0.11 | 0.17 | 0.12 | 0.05 |
| 11 | 125619227 | rs568195 | -0.681 | -0.762 | -0.061 | 0.682 | -0.001 | -0.011 | -0.011 | 0 | 0.88 | 0.91 | 0.69 | 0.84 | 0.82 | 0.79 |
| 11 | 125619578 | rs1056417 | -1.183 | -0.703 | 0.031 | 0.657 | -0.004 | 0.012 | 0.012 | 0 | 0.53 | 0.64 | 0.39 | 0.61 | 0.49 | 0.45 |
| 11 | 125624056 | rs12271647 | 0.382 | -1.122 | 0.274 | 0.441 | -0.013 | -0.017 | -0.017 | 1.034 | 0.21 | 0.25 | 0.18 | 0.18 | 0.18 | 0.14 |
| 11 | 125635831 | rs4937030 | -0.743 | -1.275 | 0.265 | 0.433 | 0.002 | 0 | 0 | 0 | 0.41 | 0.58 | 0.35 | 0.44 | 0.38 | 0.29 |
| 11 | 125641368 | rs573327 | 0.44 | -0.809 | -0.401 | -0.016 | -0.017 | 0.001 | 0.001 | 0 | 0.41 | 0.23 | 0.52 | 0.56 | 0.44 | 0.61 |
| 11 | 125644336 | rs527195 | -0.288 | -0.858 | -0.327 | -0.076 | -0.006 | 0.009 | 0.009 | 0 | 0.54 | 0.39 | 0.57 | 0.56 | 0.51 | 0.7 |
| 11 | 125646223 | rs4937035 | 0.635 | -0.98 | -0.383 | 0.159 | -0.036 | -0.027 | -0.027 | 1.163 | 0.3 | 0.27 | 0.25 | 0.36 | 0.37 | 0.3 |
| 11 | 125646286 | rs484670 | 1.144 | -1.171 | -0.405 | 0.095 | -0.002 | -0.03 | -0.03 | 0 | 0.28 | 0.16 | 0.48 | 0.38 | 0.32 | 0.43 |
| 11 | 125647982 | rs1365081 | 0.919 | -1.264 | -0.124 | 0.104 | -0.019 | 0.013 | 0.013 | 0 | 0.22 | 0.27 | 0.18 | 0.31 | 0.18 | 0.17 |
| 11 | 125655014 | rs536968 | -0.856 | -1.189 | -0.049 | 0.067 | -0.03 | -0.094 | -0.094 | 0 | 0.81 | 0.73 | 0.9 | 1 | 0.86 | 0.89 |
| 11 | 125672305 | rs2140230 | 0.617 | -1.226 | -0.274 | 0.263 | 0.047 | 0.015 | 0.015 | 0 | 0.5 | 0.4 | 0.46 | 0.53 | 0.58 | 0.52 |
| 11 | 125673849 | rs1403339 | -1.078 | -1.183 | -0.218 | 0.172 | -0.011 | -0.053 | -0.053 | 0 | 0.82 | 0.78 | 0.92 | 1 | 0.87 | 0.9 |
| 11 | 125708837 | rs11220237 | 1.425 | -1.315 | -0.031 | 0.298 | 0.002 | 0.003 | 0.003 | 0 | 0.19 | 0.17 | 0.16 | 0.13 | 0.17 | 0.13 |
| 11 | 125752161 | rs665677 | 0.021 | -1.414 | 0.089 | 0.198 | 0.011 | 0.009 | 0.009 | 0 | 0.58 | 0.42 | 0.73 | 0.69 | 0.69 | 0.74 |
| 11 | 125753360 | rs540225 | 0.842 | -1.171 | -0.321 | 0.278 | -0.015 | -0.01 | -0.01 | 0 | 0.33 | 0.2 | 0.25 | 0.34 | 0.32 | 0.38 |
| 11 | 125762891 | rs662715 | -0.412 | -1.128 | -0.267 | 0.229 | -0.058 | -0.049 | -0.049 | 0.064 | 0.77 | 0.63 | 0.89 | 0.85 | 0.86 | 0.86 |
| 11 | 125766173 | rs622756 | -0.412 | -1.128 | -0.267 | 0.23 | -0.058 | -0.049 | -0.049 | 0 | 0.77 | 0.63 | 0.89 | 0.85 | 0.86 | 0.86 |
| 11 | 125770425 | rs2236681 | 1.512 | -0.515 | -0.594 | 0.458 | -0.07 | -0.032 | -0.032 | 0.84 | 0.17 | 0.1 | 0.2 | 0.19 | 0.26 | 0.13 |
| 11 | 125777056 | rs485878 | -0.041 | -0.824 | -0.432 | 0.408 | 0 | 0 | 0 | 0 | 0.39 | 0.38 | 0.52 | 0.51 | 0.41 | 0.53 |
| 11 | 125787461 | rs12794472 | 0.272 | -0.872 | -0.706 | 0.354 | -0.008 | 0 | 0 | 0.008 | 0.35 | 0.38 | 0.4 | 0.33 | 0.44 | 0.36 |
| 11 | 125791017 | rs635819 | -0.002 | -0.54 | -0.918 | 0.322 | 0.003 | 0.002 | 0.002 | 0 | 0.17 | 0.21 | 0.31 | 0.17 | 0.18 | 0.23 |
| 11 | 125795625 | rs498025 | -0.955 | -1.171 | -1.687 | 0.161 | 0.031 | -0.01 | -0.01 | 0 | 0.64 | 0.92 | 0.77 | 0.67 | 0.7 | 0.71 |
| 11 | 125805630 | rs576598 | -0.861 | -1.13 | -1.908 | 0.111 | 0.071 | -0.053 | -0.053 | 0 | 0.45 | 0.86 | 0.53 | 0.41 | 0.36 | 0.46 |
| 11 | 125807964 | rs671880 | 1.05 | -0.356 | -1.406 | 0.378 | -0.024 | -0.018 | -0.018 | 0.376 | 0.05 | 0.02 | 0.26 | 0.17 | 0.11 | 0.11 |
| 11 | 125810167 | rs10893448 | 0.555 | -1.333 | -1.16 | -0.126 | 0.007 | -0.004 | -0.004 | 0.749 | 0.41 | 0.33 | 0.23 | 0.48 | 0.28 | 0.34 |
| 11 | 125812350 | rs687822 | -0.255 | -1.09 | -1.276 | -0.73 | 0.031 | 0.091 | 0.091 | 0 | 0.71 | 0.92 | 0.82 | 0.92 | 0.78 | 0.67 |
| 11 | 125814198 | rs110143 | 0.231 | -0.973 | -0.712 | -0.697 | -0.005 | -0.025 | -0.025 | 1.247 | 0.63 | 0.62 | 0.77 | 0.92 | 0.71 | 0.65 |
| 11 | 125818237 | rs483216 | 0.643 | -0.743 | -0.541 | -0.269 | 0.001 | -0.006 | -0.006 | 0.049 | 0.36 | 0.34 | 0.46 | 0.56 | 0.36 | 0.37 |
| 11 | 125818699 | rs634053 | 0.266 | -0.402 | -0.207 | -0.464 | -0.006 | -0.018 | -0.018 | 0 | 0.57 | 0.57 | 0.72 | 0.85 | 0.66 | 0.63 |
| 11 | 125821763 | rs12797894 | 0.949 | -0.478 | -0.7 | -0.631 | -0.085 | 0.014 | 0.014 | 0 | 0.18 | 0 | 0.16 | 0.07 | 0.21 | 0.2 |
| 11 | 125822707 | rs2155803 | 1.643 | -0.372 | -0.032 | -0.448 | 0.013 | 0.005 | 0.005 | 0 | 0.2 | 0.17 | 0.26 | 0.29 | 0.29 | 0.25 |
| 11 | 125824737 | rs948113 | 0.053 | -0.753 | -0.009 | -0.468 | -0.007 | 0.036 | 0.036 | 0 | 0.6 | 0.67 | 0.53 | 0.78 | 0.6 | 0.56 |
| 11 | 125826702 | rs4937076 | 0.898 | -0.806 | 0.108 | -0.43 | 0.012 | 0.054 | 0.054 | 0 | 0.5 | 0.29 | 0.46 | 0.7 | 0.55 | 0.54 |
| 11 | 125826785 | rs3039 | 1.5 | -0.801 | 0.002 | -0.468 | -0.033 | -0.012 | -0.012 | 0.398 | 0.24 | 0.11 | 0.26 | 0.28 | 0.31 | 0.29 |
| 11 | 125827072 | rs1047070 | 1.747 | -0.9 | -0.083 | -0.507 | 0.018 | -0.027 | -0.027 | 0.138 | 0.24 | 0.17 | 0.18 | 0.41 | 0.2 | 0.22 |
| 11 | 125828007 | rs1065398 | 0.694 | -0.916 | -0.018 | -0.538 | 0.01 | 0.049 | 0.049 | 0 | 0.51 | 0.32 | 0.46 | 0.71 | 0.54 | 0.54 |
| 11 | 125828097 | rs2186754 | 0.668 | -0.896 | 0.132 | -0.528 | 0.006 | 0.032 | 0.032 | 0 | 0.51 | 0.37 | 0.46 | 0.71 | 0.55 | 0.54 |
| 11 | 125828904 | rs2509616 | 0.053 | -0.924 | 0.416 | -0.601 | 0.021 | 0.009 | 0.009 | 0 | 0.37 | 0.53 | 0.59 | 0.44 | 0.49 | 0.45 |
| 11 | 125830697 | rs3737336 | 1.477 | -0.945 | 0.12 | -0.576 | -0.043 | -0.014 | -0.014 | 0 | 0.25 | 0.1 | 0.26 | 0.29 | 0.31 | 0.31 |
| 11 | 125831701 | rs2276061 | 1.485 | -0.949 | 0.122 | -0.579 | -0.041 | -0.013 | -0.013 | 0.407 | 0.25 | 0.1 | 0.26 | 0.28 | 0.31 | 0.31 |
| 11 | 125832659 | rs618566 | -0.149 | -0.876 | -0.094 | -0.537 | 0.012 | 0.048 | 0.048 | 0.172 | 0.63 | 0.79 | 0.78 | 0.85 | 0.69 | 0.7 |
| 11 | 125832954 | rs10893452 | 1.462 | -0.754 | -0.1 | -0.156 | 0.015 | -0.042 | -0.042 | 0.057 | 0.24 | 0.12 | 0.28 | 0.41 | 0.21 | 0.22 |
| 11 | 125839764 | rs560640 | -0.6 | -0.54 | 0.187 | -0.154 | 0.015 | 0.073 | 0.073 | 0 | 0.13 | 0.32 | 0.22 | 0.21 | 0.15 | 0.18 |
| 11 | 125840999 | rs503764 | -0.083 | -0.548 | -0.401 | -0.835 | 0.008 | 0.012 | 0.012 | 0 | 0.4 | 0.71 | 0.43 | 0.62 | 0.38 | 0.43 |
| 11 | 125845459 | rs676172 | -1.071 | 0.002 | -0.457 | -0.834 | 0.001 | 0.002 | 0.002 | 0 | 0.9 | 0.93 | 0.96 | 0.99 | 0.92 | 0.91 |
| 11 | 125846293 | rs7124748 | -0.048 | -0.041 | -0.449 | -0.959 | -0.01 | 0.005 | 0.005 | 0 | 0.37 | 0.48 | 0.44 | 0.41 | 0.38 | 0.41 |
| 11 | 125848673 | rs625280 | 0.343 | -0.004 | -0.284 | -1.066 | 0.02 | -0.058 | -0.058 | 0 | 0.22 | 0.37 | 0.19 | 0.06 | 0.24 | 0.19 |
| 11 | 125850139 | rs592036 | -0.584 | -0.204 | -0.387 | -1.083 | 0 | -0.068 | -0.068 | 0 | 0.29 | 0.52 | 0.24 | 0.12 | 0.29 | 0.24 |
| 11 | 125851181 | rs684805 | -0.132 | -0.184 | -0.394 | -1.08 | 0.016 | -0.091 | -0.091 | 0 | 0.25 | 0.5 | 0.22 | 0.07 | 0.26 | 0.21 |
| 11 | 125852273 | rs11220301 | 1.408 | -0.201 | -0.723 | -1.071 | -0.045 | -0.073 | -0.073 | 0 | 0.24 | 0.07 | 0.24 | 0.31 | 0.27 | 0.31 |
| 11 | 125857745 | rs486304 | -0.803 | -0.147 | -0.481 | -1.02 | -0.002 | 0.001 | 0.001 | 0 | 0.55 | 0.69 | 0.49 | 0.43 | 0.56 | 0.54 |
| 11 | 125865954 | rs599702 | 0.577 | 0.398 | -0.661 | -0.881 | -0.002 | -0.003 | -0.003 | 0 | 0.3 | 0.29 | 0.3 | 0.31 | 0.32 | 0.37 |
| 11 | 125869966 | rs575334 | -0.812 | 0.357 | -0.624 | -0.887 | -0.001 | 0 | 0 | 0 | 0.54 | 0.62 | 0.61 | 0.47 | 0.61 | 0.59 |
| 11 | 125871554 | rs12274729 | 1.306 | 0.42 | -0.622 | -0.914 | -0.03 | 0.028 | 0.028 | 0 | 0.15 | 0.07 | 0.15 | 0.05 | 0.19 | 0.13 |
| 11 | 125871715 | rs12274923 | 1.339 | 0.419 | -0.622 | -0.914 | -0.03 | 0.027 | 0.027 | 0 | 0.15 | 0.06 | 0.15 | 0.05 | 0.19 | 0.13 |
| 11 | 125872529 | rs675864 | 0.338 | 0.402 | -0.507 | -0.719 | 0.008 | -0.023 | -0.023 | 0 | 0.28 | 0.4 | 0.34 | 0.16 | 0.34 | 0.25 |
| 11 | 125872842 | rs7104123 | 0.919 | 0.654 | -0.611 | -0.811 | -0.02 | 0.027 | 0.027 | 0 | 0.35 | 0.36 | 0.33 | 0.53 | 0.26 | 0.32 |
| 11 | 125873102 | rs7104745 | 0.056 | 0.738 | -0.813 | -0.891 | -0.003 | -0.002 | -0.002 | 0 | 0.62 | 0.55 | 0.6 | 0.83 | 0.54 | 0.65 |
| 11 | 125877075 | rs528638 | 0.241 | 0.69 | -1.15 | -0.405 | 0 | 0 | 0 | 0 | 0.46 | 0.53 | 0.53 | 0.5 | 0.54 | 0.53 |
| 11 | 125878883 | rs11220311 | -0.143 | 0.857 | -1.289 | 0.457 | 0 | -0.013 | -0.013 | 0.015 | 0.33 | 0.27 | 0.35 | 0.48 | 0.31 | 0.3 |
| 11 | 125881102 | rs517835 | -0.234 | 0.879 | -1.955 | 0.634 | -0.069 | -0.133 | -0.133 | 0 | 0.26 | 0.01 | 0.28 | 0.53 | 0.26 | 0.27 |
| 11 | 125887311 | rs7119527 | -0.629 | 0.632 | -1.314 | -0.045 | 0.002 | 0.015 | 0.015 | 0 | 0.06 | 0.12 | 0.12 | 0.13 | 0.09 | 0.11 |
| 11 | 125889526 | rs3740909 | -0.629 | 0.632 | -1.314 | -0.027 | 0.002 | 0.017 | 0.017 | 0 | 0.06 | 0.11 | 0.12 | 0.13 | 0.09 | 0.11 |
| 11 | 125891269 | rs3740912 | -0.346 | 0.47 | -2.073 | -0.812 | -0.052 | 0.083 | 0.083 | 0 | 0.54 | 0.86 | 0.52 | 0.35 | 0.53 | 0.5 |

| | | | | | | | | | | | | | | | | |
|---|---|---|---|---|---|---|---|---|---|---|---|---|---|---|---|---|
| 11 | 125892207 | rs11220316 | 0.464 | 0.499 | -1.465 | -1.237 | 0.02 | -0.006 | -0.006 | 0 | 0.18 | 0.38 | 0.22 | 0.17 | 0.22 | 0.17 |
| 11 | 125893275 | rs1939890 | -0.197 | 0.607 | -2.065 | -1.323 | -0.007 | -0.008 | -0.008 | 5.408 | 0.12 | 0.11 | 0.16 | 0.17 | 0.17 | 0.2 |
| 11 | 125895397 | rs10458992 | 0.65 | 0.493 | -1.726 | -1.253 | -0.002 | 0 | 0 | 0.021 | 0.16 | 0.13 | 0.2 | 0.17 | 0.22 | 0.17 |
| 11 | 125896607 | rs11606876 | 0.923 | 0.052 | -2.001 | -1.226 | -0.027 | 0.12 | 0.12 | 0 | 0.2 | 0.02 | 0.11 | 0.01 | 0.14 | 0.12 |
| 11 | 125899346 | rs651284 | -1.068 | 0.129 | -2.102 | -1.191 | -0.092 | -0.1 | -0.1 | 0.792 | 0.72 | 0.98 | 0.7 | 0.38 | 0.69 | 0.65 |
| 11 | 125904827 | rs476707 | -0.647 | 0.031 | -2.198 | -1.412 | -0.126 | -0.036 | -0.036 | 0 | 0.62 | 0.97 | 0.57 | 0.33 | 0.6 | 0.51 |
| 11 | 125905258 | rs472926 | -1.469 | 0.253 | -2.165 | -1.083 | -0.046 | -0.048 | -0.048 | 0 | 0.87 | 1 | 0.78 | 0.83 | 0.83 | 0.83 |
| 11 | 125907626 | rs662947 | -0.856 | -0.268 | -1.186 | -1.462 | -0.016 | 0.065 | 0.065 | 0 | 0.57 | 0.76 | 0.53 | 0.3 | 0.57 | 0.53 |
| 11 | 125916052 | rs543970 | 0.829 | -0.168 | -1.372 | -1.732 | -0.005 | 0.135 | 0.135 | 0 | 0.21 | 0.02 | 0.13 | 0.01 | 0.14 | 0.13 |
| 11 | 125920236 | rs538107 | -1.139 | -0.247 | -1.158 | -1.664 | -0.014 | 0.053 | 0.053 | 0 | 0.57 | 0.71 | 0.54 | 0.36 | 0.57 | 0.54 |
| 11 | 125923264 | rs718443 | 0.808 | 0.007 | -1.257 | -1.618 | -0.054 | 0.013 | 0.013 | 0 | 0.1 | 0.01 | 0.12 | 0.05 | 0.25 | 0.13 |
| 11 | 125924669 | rs11220329 | -0.223 | -0.366 | -0.395 | -1.569 | -0.027 | 0.006 | 0.006 | 0 | 0.23 | 0.33 | 0.28 | 0.3 | 0.17 | 0.28 |
| 11 | 125930405 | rs2043438 | -0.393 | -0.129 | -0.621 | -1.619 | 0.04 | 0.062 | 0.062 | 0 | 0.44 | 0.26 | 0.41 | 0.27 | 0.33 | 0.46 |
| 11 | 125934846 | rs473075 | -0.644 | -0.095 | -0.572 | -1.582 | -0.003 | 0.021 | 0.021 | 0 | 0.56 | 0.58 | 0.52 | 0.32 | 0.56 | 0.59 |
| 11 | 125937383 | rs564327 | -0.123 | -0.61 | -0.804 | -1.633 | 0.034 | 0.05 | 0.05 | 0 | 0.27 | 0.13 | 0.16 | 0 | 0.17 | 0.13 |
| 11 | 125946898 | rs7119002 | -1.021 | -0.22 | -0.086 | -1.218 | -0.011 | -0.043 | -0.043 | 0 | 0.14 | 0.51 | 0.1 | 0.05 | 0.09 | 0.06 |
| 11 | 125947954 | rs532559 | -0.401 | 0.21 | -1.034 | -1.107 | -0.031 | 0.007 | 0.007 | 0 | 0.7 | 0.84 | 0.63 | 0.31 | 0.66 | 0.73 |
| 11 | 125952073 | rs10893475 | -0.328 | 0.172 | -0.484 | -1.085 | -0.001 | 0.033 | 0.033 | 0 | 0.11 | 0.18 | 0.12 | 0.26 | 0.11 | 0.33 |
| 11 | 125952154 | rs651054 | -0.574 | -0.301 | -0.253 | -2.012 | -0.018 | -0.118 | -0.118 | 5.327 | 0.24 | 0.42 | 0.27 | 0.04 | 0.19 | 0.14 |
| 11 | 125953664 | rs485064 | -0.956 | -0.191 | -0.599 | -2.168 | -0.003 | -0.004 | -0.004 | 0 | 0.8 | 0.84 | 0.69 | 0.33 | 0.74 | 0.81 |
| 11 | 125955327 | rs4935955 | -0.211 | -0.361 | -0.321 | -2.031 | 0.001 | 0.007 | 0.007 | 0.079 | 0.69 | 0.69 | 0.72 | 0.45 | 0.68 | 0.82 |
| 11 | 125956336 | rs4937086 | -0.265 | -0.388 | -0.189 | -2.044 | 0 | 0.003 | 0.003 | 0.067 | 0.69 | 0.71 | 0.71 | 0.45 | 0.68 | 0.82 |
| 11 | 125956456 | rs4937090 | -0.151 | -0.391 | -0.393 | -2.045 | 0.002 | 0.024 | 0.024 | 0.126 | 0.68 | 0.6 | 0.7 | 0.45 | 0.68 | 0.82 |
| 11 | 125956901 | rs12808304 | 1.344 | -0.413 | -0.981 | -2.62 | 0.001 | 0.101 | 0.101 | 0 | 0.22 | 0.06 | 0.13 | 0 | 0.17 | 0.1 |
| 11 | 125962787 | rs4937098 | 0.177 | -0.369 | -0.735 | -2.662 | 0.002 | 0.003 | 0.003 | 0 | 0.42 | 0.38 | 0.52 | 0.43 | 0.38 | 0.56 |
| 11 | 125965332 | rs564216 | 0.498 | -0.377 | -0.858 | -2.681 | 0 | 0.006 | 0.006 | 0 | 0.1 | 0.07 | 0.06 | 0 | 0.08 | 0.01 |
| 11 | 125970366 | rs17135097 | 0.513 | -0.158 | -1 | -2.629 | -0.002 | -0.025 | -0.025 | 0 | 0.24 | 0.26 | 0.17 | 0.02 | 0.27 | 0.23 |
| 11 | 125970673 | rs12795043 | 1.012 | -0.336 | -1.049 | -2.443 | 0 | -0.002 | -0.002 | 0 | 0.12 | 0.13 | 0.25 | 0.13 | 0.16 | 0.17 |
| 11 | 125974071 | rs545830 | -0.932 | -0.525 | -1.1 | -2.454 | -0.017 | 0.024 | 0.024 | 1.234 | 0.72 | 0.68 | 0.76 | 0.41 | 0.78 | 0.81 |
| 11 | 125975614 | rs17135126 | 0.832 | -0.112 | -1.47 | -2.501 | -0.018 | -0.019 | -0.019 | 0 | 0.27 | 0.1 | 0.29 | 0.29 | 0.27 | 0.39 |
| 11 | 125975854 | rs681223 | 0.548 | -0.096 | -1.607 | -2.517 | -0.089 | 0.167 | 0.167 | 0 | 0.28 | 0.04 | 0.19 | 0.02 | 0.35 | 0.24 |
| 11 | 125976526 | rs684407 | -0.157 | 0.058 | -1.152 | -2.078 | -0.002 | -0.003 | -0.003 | 0 | 0.58 | 0.5 | 0.51 | 0.3 | 0.62 | 0.63 |
| 11 | 125981569 | rs511987 | -0.341 | 1.269 | -1.538 | -1.748 | -0.13 | 0.173 | 0.173 | 0 | 0.28 | 0.03 | 0.17 | 0.01 | 0.36 | 0.16 |
| 11 | 125982079 | rs543969 | -0.114 | 1.095 | -0.764 | -1.277 | 0.008 | -0.044 | -0.044 | 0.104 | 0.17 | 0.25 | 0.17 | 0.03 | 0.22 | 0.12 |
| 11 | 125983154 | rs478951 | -0.906 | 0.632 | -1.192 | -0.751 | 0.002 | 0.011 | 0.011 | 0 | 0.46 | 0.43 | 0.6 | 0.42 | 0.52 | 0.64 |
| 11 | 125984426 | rs533239 | 0.295 | 0.109 | -1.205 | -0.432 | -0.006 | 0.025 | 0.025 | 0 | 0.79 | 0.85 | 0.7 | 0.9 | 0.78 | 0.8 |
| 11 | 125986662 | rs10790787 | -0.774 | -0.459 | -0.944 | 0.385 | -0.017 | 0.056 | 0.056 | 0 | 0.17 | 0.24 | 0.17 | 0.38 | 0.11 | 0.3 |
| 11 | 125986753 | rs596077 | 0.082 | -0.152 | -1.617 | -0.172 | -0.091 | 0.061 | 0.061 | 0 | 0.78 | 0.98 | 0.81 | 0.87 | 0.74 | 0.68 |
| 11 | 125988807 | rs670346 | 1.287 | -0.88 | -1.606 | -1.107 | -0.03 | 0.046 | 0.046 | 0.18 | 0.35 | 0.19 | 0.22 | 0.01 | 0.44 | 0.12 |
| 11 | 125993491 | rs2278451 | 0.55 | -1.392 | -1.284 | -1.01 | 0.005 | -0.003 | -0.003 | 0.468 | 0.49 | 0.64 | 0.58 | 0.45 | 0.56 | 0.32 |
| 11 | 125999205 | rs634973 | -0.351 | -1.278 | -1.729 | -1.137 | -0.084 | 0.108 | 0.108 | 0.674 | 0.74 | 0.98 | 0.8 | 0.88 | 0.73 | 0.66 |
| 11 | 126002279 | rs10750322 | -0.485 | -1.309 | -1.359 | -0.785 | 0.003 | -0.015 | -0.015 | 0.081 | 0.47 | 0.27 | 0.41 | 0.55 | 0.43 | 0.68 |
| 11 | 126002683 | rs607980 | 0.448 | -1.512 | -1.214 | -0.888 | -0.032 | 0.138 | 0.138 | 0 | 0.83 | 0.37 | 0.62 | 0.56 | 0.89 | 0.81 |
| 11 | 126003326 | rs621381 | 1.389 | -1.52 | -1.273 | -0.926 | -0.1 | 0.194 | 0.194 | 0 | 0.34 | 0.09 | 0.21 | 0.01 | 0.45 | 0.13 |
| 11 | 126004075 | rs7930128 | -0.591 | -1.684 | -0.334 | 0.07 | 0 | 0.495 | 0.495 | 0.713 | 0.07 | 0.56 | 0.14 | 0.43 | 0.01 | 0.04 |
| 11 | 126004713 | rs534916 | 1.098 | -1.663 | -0.385 | 0.065 | -0.035 | 0.134 | 0.134 | 0 | 0.82 | 0.37 | 0.62 | 0.56 | 0.89 | 0.81 |
| 11 | 126008280 | rs1106944 | -1.057 | -1.341 | 0.069 | -0.323 | 0.032 | -0.044 | -0.044 | 1.571 | 0.1 | 0.5 | 0.25 | 0.01 | 0.07 | 0.17 |
| 11 | 126008932 | rs632289 | 0.082 | -1.936 | -0.647 | 0.495 | 0.045 | -0.099 | -0.099 | 2.863 | 0.81 | 0.91 | 0.81 | 0.55 | 0.87 | 0.71 |
| 11 | 126009142 | rs556645 | 0.411 | -2.809 | -0.271 | 0.291 | -0.057 | 0.039 | 0.039 | 0.262 | 0.67 | 0.54 | 0.73 | 0.52 | 0.81 | 0.62 |
| 11 | 126009500 | rs541601 | 0.5 | -2.567 | -0.142 | 0.715 | -0.039 | 0.013 | 0.013 | 0.854 | 0.64 | 0.58 | 0.73 | 0.6 | 0.8 | 0.57 |
| 11 | 126012786 | rs4937105 | -0.171 | -2.159 | -0.956 | -0.014 | -0.091 | 0.028 | 0.028 | 0.773 | 0.22 | 0.01 | 0.34 | 0.13 | 0.27 | 0.35 |
| 11 | 126013781 | rs483611 | -0.624 | -2.413 | -0.6 | 0.124 | 0.02 | -0.031 | -0.031 | 0 | 0.78 | 0.83 | 0.81 | 0.5 | 0.86 | 0.71 |
| 11 | 126020110 | rs595004 | -0.654 | -2.202 | -0.522 | 0.032 | 0 | -0.002 | -0.002 | 0 | 0.75 | 0.76 | 0.79 | 0.49 | 0.86 | 0.69 |
| 11 | 126023905 | rs892424 | -1.195 | -2.193 | -0.536 | -0.279 | -0.003 | 0.007 | 0.007 | 0.24 | 0.37 | 0.42 | 0.54 | 0.48 | 0.35 | 0.55 |
| 11 | 126025884 | rs12098952 | -0.656 | -2.31 | -0.211 | -0.241 | -0.061 | 0.077 | 0.077 | 0.124 | 0.21 | 0.47 | 0.2 | 0.34 | 0.1 | 0.21 |
| 11 | 126026645 | rs12421177 | -0.561 | -1.905 | -0.692 | -0.927 | 0.007 | -0.017 | -0.017 | 0.72 | 0.21 | 0.27 | 0.36 | 0.13 | 0.26 | 0.34 |
| 11 | 126026960 | rs515340 | -1.897 | -2.022 | -0.849 | -1.097 | -0.137 | 0.226 | 0.226 | 0 | 0.66 | 0.96 | 0.77 | 0.99 | 0.5 | 0.87 |
| 11 | 126027473 | rs3133084 | -0.758 | -2.382 | -0.517 | -0.794 | 0.074 | 0.085 | 0.085 | 0 | 0.45 | 0.68 | 0.41 | 0.87 | 0.24 | 0.53 |
| 11 | 126028717 | rs563519 | -0.651 | -2.38 | -0.673 | -0.448 | 0.107 | 0.051 | 0.051 | 0.824 | 0.43 | 0.74 | 0.3 | 0.66 | 0.22 | 0.47 |
| 11 | 126031480 | rs11220385 | -0.813 | -1.99 | -0.627 | -0.765 | 0.005 | 0.003 | 0.003 | 0 | 0.3 | 0.25 | 0.31 | 0.15 | 0.19 | 0.4 |
| 11 | 126031494 | rs11220386 | -1.329 | -1.522 | -1.002 | -0.93 | 0 | 0.01 | 0.01 | 0 | 0.27 | 0.15 | 0.26 | 0.26 | 0.27 | 0.39 |
| 11 | 126032638 | rs732430 | -1.251 | -1.607 | -0.388 | -0.857 | -0.003 | -0.001 | -0.001 | 0 | 0.33 | 0.65 | 0.44 | 0.34 | 0.2 | 0.4 |
| 11 | 126032739 | rs732431 | -0.488 | -1.42 | -0.242 | -1.149 | -0.024 | -0.061 | -0.061 | 0 | 0.2 | 0.58 | 0.29 | 0.1 | 0.09 | 0.15 |
| 11 | 126037018 | rs614918 | -1.818 | -1.47 | -0.518 | -1.515 | -0.045 | 0.031 | 0.031 | 0 | 0.71 | 0.83 | 0.76 | 0.99 | 0.52 | 0.9 |
| 11 | 126037362 | rs613214 | 0.159 | -1.861 | -0.701 | -0.72 | 0.011 | -0.036 | -0.036 | 0 | 0.34 | 0.22 | 0.23 | 0.48 | 0.28 | 0.48 |
| 11 | 126039495 | rs549340 | -0.836 | -2.208 | -0.498 | -0.736 | 0 | -0.013 | -0.013 | 0.088 | 0.47 | 0.42 | 0.39 | 0.86 | 0.27 | 0.66 |
| 11 | 126042206 | rs574943 | 0.836 | -1.921 | -0.658 | -0.457 | 0.031 | -0.05 | -0.05 | 0 | 0.26 | 0.18 | 0.14 | 0.46 | 0.15 | 0.39 |
| 11 | 126042958 | rs581311 | 0.851 | -1.927 | -0.744 | -0.46 | 0.018 | -0.03 | -0.03 | 0 | 0.26 | 0.18 | 0.14 | 0.46 | 0.15 | 0.39 |
| 11 | 126043977 | rs9645654 | -0.487 | -1.644 | -0.909 | -0.284 | -0.023 | 0.048 | 0.048 | 0 | 0.13 | 0.16 | 0.2 | 0.37 | 0.05 | 0.15 |

| | | | | | | | | | | | | | | | | |
|---|---|---|---|---|---|---|---|---|---|---|---|---|---|---|---|---|
| 11 | 126046283 | rs7106095 | 0.568 | -1.089 | -1.261 | -1.029 | -0.032 | 0.009 | 0.009 | 0 | 0.14 | 0.03 | 0.24 | 0.11 | 0.11 | 0.15 |
| 11 | 126048784 | rs561164 | -0.762 | 0.132 | -1.575 | -1.694 | -0.058 | 0.12 | 0.12 | 0 | 0.66 | 0.85 | 0.58 | 0.91 | 0.62 | 0.69 |
| 11 | 126054285 | rs663714 | -2.087 | 0.357 | -1.616 | -1.766 | -0.065 | 0.091 | 0.091 | 0 | 0.81 | 0.94 | 0.84 | 1 | 0.72 | 0.94 |
| 11 | 126054639 | rs12418362 | 0.925 | 0.386 | -1.537 | -1.64 | -0.003 | 0.027 | 0.027 | 0 | 0.13 | 0.08 | 0.26 | 0.04 | 0.09 | 0.18 |
| 11 | 126055040 | rs4937109 | 0.55 | 0.454 | -1.429 | -1.552 | -0.022 | 0.025 | 0.025 | 0 | 0.09 | 0.02 | 0.09 | 0.04 | 0.14 | 0.11 |
| 11 | 126056514 | rs534278 | 0.102 | -0.291 | -1.049 | -1.796 | 0.011 | -0.05 | -0.05 | 0 | 0.28 | 0.3 | 0.22 | 0.08 | 0.39 | 0.26 |
| 11 | 126056623 | rs644050 | -1.042 | -0.19 | -1.099 | -1.575 | -0.079 | 0.162 | 0.162 | 0 | 0.67 | 0.88 | 0.57 | 0.96 | 0.62 | 0.76 |
| 11 | 126056825 | rs632918 | -0.336 | -0.294 | -0.696 | -1.106 | -0.002 | 0.003 | 0.003 | 0.189 | 0.56 | 0.54 | 0.4 | 0.7 | 0.57 | 0.6 |
| 11 | 126057344 | rs562731 | 0.277 | 0.205 | -0.851 | -1.062 | 0.005 | -0.012 | -0.012 | 0 | 0.08 | 0.13 | 0.07 | 0.06 | 0.1 | 0.12 |
| 11 | 126057394 | rs567606 | -0.711 | -0.092 | -1.004 | -1.208 | -0.002 | 0.031 | 0.031 | 0.03 | 0.64 | 0.68 | 0.55 | 0.92 | 0.65 | 0.65 |
| 11 | 126059422 | rs686646 | 0.834 | -0.285 | -1.649 | -1.133 | 0.003 | 0.071 | 0.071 | 2.842 | 0.25 | 0.09 | 0.13 | 0.1 | 0.2 | 0.24 |
| 11 | 126060083 | rs555228 | -1.066 | -0.419 | -1.971 | -1.358 | -0.011 | 0.149 | 0.149 | 0 | 0.66 | 0.92 | 0.58 | 0.91 | 0.67 | 0.68 |
| 11 | 126060666 | rs670712 | 0.85 | -0.456 | -2.001 | -1.251 | -0.084 | 0.082 | 0.082 | 0 | 0.23 | 0.08 | 0.21 | 0.07 | 0.41 | 0.31 |
| 11 | 126060763 | rs561514 | 0.371 | -0.549 | -1.543 | -0.699 | 0.008 | -0.009 | -0.009 | 0 | 0.3 | 0.27 | 0.21 | 0.62 | 0.21 | 0.26 |
| 11 | 126060838 | rs681664 | 0.979 | -0.351 | -2.053 | -0.902 | -0.001 | 0.082 | 0.082 | 2.887 | 0.25 | 0.08 | 0.13 | 0.1 | 0.2 | 0.24 |
| 11 | 126064439 | rs530587 | -1.068 | -0.19 | -2.16 | -0.937 | -0.018 | 0.038 | 0.038 | 0.006 | 0.72 | 0.94 | 0.72 | 0.78 | 0.72 | 0.82 |
| 11 | 126066040 | rs515917 | -0.396 | -0.605 | -1.737 | -0.728 | 0.089 | 0.096 | 0.096 | 0.222 | 0.47 | 0.76 | 0.5 | 0.7 | 0.29 | 0.48 |
| 11 | 126069587 | rs12416718 | 0.559 | -0.599 | -1.819 | -0.77 | 0.007 | 0.017 | 0.017 | 0 | 0.15 | 0.01 | 0.28 | 0.09 | 0.11 | 0.26 |
| 11 | 126069841 | rs574198 | -1.577 | -0.714 | -1.851 | -1.514 | 0 | 0.007 | 0.007 | 0 | 0.77 | 0.87 | 0.85 | 1 | 0.78 | 0.93 |
| 11 | 126072294 | rs480153 | 1.401 | -1.119 | -1.56 | -1.598 | -0.001 | 0.008 | 0.008 | 0.274 | 0.22 | 0.19 | 0.13 | 0.13 | 0.2 | 0.22 |
| 11 | 126072688 | rs627911 | -1.15 | -1.371 | -0.96 | -1.776 | -0.027 | -0.176 | -0.176 | 0.324 | 0.71 | 0.52 | 0.83 | 1 | 0.77 | 0.92 |
| 11 | 126074192 | rs2276312 | 1.122 | -1.441 | -0.997 | -1.546 | 0.02 | -0.071 | -0.071 | 0.027 | 0.17 | 0.04 | 0.32 | 0.62 | 0.11 | 0.28 |
| 11 | 126080352 | rs602735 | 1.038 | -0.876 | -1.136 | -1.727 | -0.026 | 0.039 | 0.039 | 0.309 | 0.25 | 0.12 | 0.17 | 0.13 | 0.29 | 0.24 |
| 11 | 126081336 | rs587891 | -2.131 | -0.883 | -1.129 | -1.72 | 0.001 | 0.004 | 0.004 | 0 | 0.77 | 0.87 | 0.85 | 1 | 0.78 | 0.93 |
| 11 | 126081403 | rs2282580 | -0.012 | -0.417 | -1.361 | -1.932 | 0.006 | 0.036 | 0.036 | 0 | 0.35 | 0.25 | 0.56 | 0.82 | 0.34 | 0.53 |
| 11 | 126082218 | rs4420285 | 1.506 | -0.414 | -1.281 | -1.911 | -0.003 | 0.002 | 0.002 | 0 | 0.18 | 0.17 | 0.23 | 0.19 | 0.22 | 0.24 |
| 11 | 126082661 | rs547862 | 1.944 | -0.406 | -1.285 | -1.937 | -0.005 | 0.006 | 0.006 | 0 | 0.1 | 0.1 | 0.1 | 0.05 | 0.14 | 0.16 |
| 11 | 126084834 | rs588361 | -0.728 | -0.747 | -1.256 | -2.062 | 0.011 | 0.096 | 0.096 | 0.391 | 0.49 | 0.35 | 0.7 | 0.86 | 0.57 | 0.7 |
| 11 | 126091941 | rs512577 | 0.147 | -0.715 | -1.262 | -2.002 | -0.016 | 0.001 | 0.001 | 0 | 0.31 | 0.19 | 0.47 | 0.67 | 0.34 | 0.45 |
| 11 | 126095964 | rs11220409 | 0.177 | -0.697 | -1.253 | -1.919 | 0.015 | 0.084 | 0.084 | 0 | 0.48 | 0.68 | 0.42 | 0.23 | 0.46 | 0.33 |
| 11 | 126100932 | rs618176 | 1.657 | -0.653 | -1.234 | -1.943 | -0.006 | 0.017 | 0.017 | 0.347 | 0.21 | 0.13 | 0.12 | 0.14 | 0.2 | 0.23 |
| 11 | 126116219 | rs10893490 | 1.965 | -0.693 | -1.088 | -1.786 | 0.006 | -0.003 | -0.003 | 0 | 0.07 | 0.35 | 0.03 | 0 | 0.01 | 0.01 |
| 11 | 126123858 | rs670154 | -0.52 | -0.689 | -1.357 | -1.83 | 0 | -0.017 | -0.017 | 2.715 | 0.54 | 0.47 | 0.61 | 0.8 | 0.54 | 0.68 |
| 11 | 126133182 | rs17656 | 1.553 | -0.714 | -1.41 | -1.861 | -0.023 | 0.027 | 0.027 | 0 | 0.14 | 0.1 | 0.14 | 0.04 | 0.23 | 0.17 |
| 11 | 126133576 | rs1049666 | 1.385 | -0.835 | -1.622 | -2.062 | 0 | 0 | 0 | 0 | 0.14 | 0.13 | 0.06 | 0.14 | 0.11 | 0.14 |
| 11 | 126133632 | rs1049665 | 1.385 | -0.835 | -1.621 | -2.062 | 0 | 0 | 0 | 0 | 0.14 | 0.14 | 0.06 | 0.14 | 0.11 | 0.14 |
| 11 | 126133641 | rs529328 | 2.279 | -0.88 | -1.653 | -2.085 | -0.017 | 0.015 | 0.015 | 0 | 0.12 | 0.09 | 0.14 | 0.04 | 0.22 | 0.15 |
| 11 | 126134619 | rs499205 | 1.444 | -0.885 | -1.655 | -2.071 | 0.001 | -0.079 | -0.079 | 0 | 0.24 | 0.05 | 0.39 | 0.62 | 0.21 | 0.37 |
| 11 | 126135785 | rs11220432 | 1.408 | -0.84 | -1.632 | -2.03 | -0.001 | -0.001 | -0.001 | 0 | 0.13 | 0.12 | 0.06 | 0.14 | 0.11 | 0.14 |
| 11 | 126138453 | rs12786155 | 2.623 | -0.742 | -1.702 | -2.758 | -0.052 | 0.063 | 0.063 | 0 | 0.12 | 0.01 | 0.08 | 0.01 | 0.18 | 0.16 |
| 11 | 126139100 | rs1786702 | -1.42 | -0.781 | -1.631 | -2.736 | -0.029 | -0.176 | -0.176 | 0.097 | 0.71 | 0.52 | 0.83 | 1 | 0.77 | 0.92 |
| 11 | 126147697 | rs667627 | 1.509 | -0.472 | -1.687 | -2.475 | -0.027 | 0.029 | 0.029 | 0 | 0.14 | 0.09 | 0.14 | 0.05 | 0.23 | 0.17 |
| 11 | 126162468 | rs3802813 | 1.365 | -0.442 | -1.584 | -2.399 | 0.001 | -0.001 | -0.001 | 0 | 0.05 | 0.06 | 0.14 | 0.18 | 0.04 | 0.08 |
| 11 | 126162607 | rs3802814 | 2.58 | -0.393 | -1.779 | -2.562 | -0.046 | 0.065 | 0.065 | 0 | 0.11 | 0.01 | 0.08 | 0.01 | 0.17 | 0.16 |
| 11 | 126162843 | rs8177374 | 2.427 | -0.403 | -1.775 | -2.556 | -0.05 | 0.063 | 0.063 | 0 | 0.12 | 0.01 | 0.08 | 0.01 | 0.19 | 0.16 |
| 11 | 126162862 | rs7932766 | 2.097 | -0.32 | -1.674 | -2.56 | 0.003 | -0.008 | -0.008 | 0 | 0.12 | 0.12 | 0.14 | 0.04 | 0.22 | 0.15 |
| 11 | 126163064 | rs8177375 | 1.077 | -0.457 | -1.799 | -2.654 | 0 | 0 | 0 | 0 | 0.16 | 0.13 | 0.06 | 0.14 | 0.11 | 0.14 |
| 11 | 126163612 | rs8177376 | 2.102 | -0.297 | -1.768 | -2.463 | -0.024 | 0.023 | 0.023 | 0 | 0.12 | 0.07 | 0.13 | 0.04 | 0.22 | 0.15 |
| 11 | 126164349 | rs625413 | -2.184 | -0.492 | -1.749 | -2.472 | -0.014 | 0.067 | 0.067 | 0 | 0.79 | 0.94 | 0.86 | 1 | 0.78 | 0.93 |
| 11 | 126174164 | rs695029 | 1.228 | -0.399 | -1.775 | -2.331 | 0.006 | -0.056 | -0.056 | 0 | 0.23 | 0.07 | 0.39 | 0.62 | 0.2 | 0.34 |
| 11 | 126177745 | rs2096914 | 1.259 | -0.175 | -1.593 | -2.181 | 0.019 | -0.079 | -0.079 | 0 | 0.21 | 0.05 | 0.39 | 0.62 | 0.18 | 0.34 |
| 11 | 126178050 | rs1111930 | 1.244 | -0.219 | -1.58 | -2.178 | 0.025 | -0.067 | -0.067 | 0.398 | 0.22 | 0.07 | 0.39 | 0.63 | 0.18 | 0.34 |
| 11 | 126180020 | rs583697 | -0.693 | -0.408 | -1.423 | -2.356 | -0.007 | -0.031 | -0.031 | 1.605 | 0.6 | 0.42 | 0.67 | 0.81 | 0.68 | 0.77 |
| 11 | 126182644 | rs12788021 | 0.599 | -0.267 | -1.566 | -2.394 | -0.013 | 0.018 | 0.018 | 0 | 0.25 | 0.17 | 0.15 | 0.14 | 0.29 | 0.27 |
| 11 | 126183756 | rs683800 | 0.317 | -0.193 | -1.594 | -2.231 | -0.02 | 0.003 | 0.003 | 0 | 0.34 | 0.17 | 0.52 | 0.67 | 0.4 | 0.49 |
| 11 | 126187868 | rs624259 | 0.115 | -0.314 | -1.614 | -2.389 | -0.029 | 0.065 | 0.065 | 0 | 0.34 | 0.22 | 0.64 | 0.85 | 0.42 | 0.6 |
| 11 | 126188379 | rs611593 | 1.531 | -0.002 | -1.592 | -2.068 | 0.014 | -0.088 | -0.088 | 0.378 | 0.21 | 0.05 | 0.38 | 0.63 | 0.17 | 0.33 |
| 11 | 126188466 | rs611151 | -0.834 | -0.154 | -1.235 | -2.185 | -0.018 | -0.144 | -0.144 | 0.035 | 0.66 | 0.56 | 0.82 | 0.99 | 0.72 | 0.87 |
| 11 | 126191074 | rs601580 | -0.67 | 0.276 | -1.021 | -1.963 | 0.002 | -0.057 | -0.057 | 1.437 | 0.5 | 0.56 | 0.73 | 0.98 | 0.52 | 0.65 |
| 11 | 126201290 | rs651922 | 1.012 | 0.657 | -1.008 | -1.597 | 0.022 | 0.006 | 0.006 | 0 | 0.15 | 0.22 | 0.27 | 0.21 | 0.25 | 0.23 |
| 11 | 126204161 | rs646358 | -0.698 | 0.309 | -1.089 | -1.961 | 0 | 0.031 | 0.031 | 0 | 0.62 | 0.69 | 0.81 | 0.87 | 0.66 | 0.8 |
| 11 | 126206376 | rs240543 | 1.669 | 0.287 | -1.297 | -1.923 | -0.02 | -0.086 | -0.086 | 1.239 | 0.2 | 0.03 | 0.38 | 0.64 | 0.18 | 0.31 |
| 11 | 126208378 | rs7115672 | 0.303 | 0.464 | -0.613 | -1.471 | 0 | -0.005 | -0.005 | 3.228 | 0.13 | 0.23 | 0.05 | 0.11 | 0.09 | 0.11 |
| 11 | 126208759 | rs240542 | -0.832 | 0.723 | -1.277 | -1.205 | -0.005 | 0.115 | 0.115 | 0 | 0.5 | 0.84 | 0.47 | 0.34 | 0.55 | 0.57 |
| 11 | 126213341 | rs240537 | 0.111 | 1.122 | -0.621 | -0.357 | 0.018 | 0.005 | 0.005 | 0 | 0.16 | 0.22 | 0.27 | 0.21 | 0.25 | 0.23 |
| 11 | 126217870 | rs582037 | 0.326 | 0.338 | -1.015 | -0.571 | 0 | 0.113 | 0.113 | 0 | 0.68 | 0.88 | 0.64 | 0.12 | 0.68 | 0.81 |
| 11 | 126218541 | rs17135399 | -0.435 | 0.286 | -0.288 | -0.26 | 0.006 | 0.03 | 0.03 | 0.011 | 0.09 | 0.24 | 0.07 | 0.09 | 0.06 | 0.06 |
| 11 | 126225964 | rs2848583 | -0.594 | 0.156 | -0.82 | -0.532 | 0.005 | 0.117 | 0.117 | 0 | 0.8 | 0.98 | 0.9 | 1 | 0.84 | 0.94 |
| 11 | 126228659 | rs4937122 | -0.435 | 0.362 | -0.478 | 0.498 | 0.016 | 0.178 | 0.178 | 0 | 0.09 | 0.21 | 0.2 | 0.66 | 0.07 | 0.1 |

| | | | | | | | | | | | | | | | | |
|---|---|---|---|---|---|---|---|---|---|---|---|---|---|---|---|---|
| 11 | 126242697 | rs629882 | -0.571 | 0.196 | -0.852 | 0.653 | 0.011 | -0.157 | -0.157 | 0.66 | 0.79 | 0.97 | 0.78 | 0.48 | 0.85 | 0.93 |
| 11 | 126242912 | rs3862628 | -0.994 | -0.15 | -0.339 | 0.094 | -0.016 | 0.064 | 0.064 | 0.856 | 0.29 | 0.54 | 0.23 | 0.42 | 0.24 | 0.31 |
| 11 | 126243505 | rs3862629 | -1.015 | -0.16 | -0.344 | 0.361 | -0.017 | 0.075 | 0.075 | 0.969 | 0.29 | 0.54 | 0.23 | 0.46 | 0.24 | 0.31 |
| 11 | 126246076 | rs7940893 | -0.43 | -0.057 | -0.119 | 0.158 | 0.02 | 0.173 | 0.173 | 0 | 0.12 | 0.26 | 0.14 | 0.38 | 0.13 | 0.14 |
| 11 | 126250774 | rs10893500 | -0.607 | -0.086 | -0.136 | 0.144 | 0.015 | 0.161 | 0.161 | 3.019 | 0.13 | 0.27 | 0.14 | 0.39 | 0.13 | 0.14 |
| 11 | 126254018 | rs10893502 | -1.905 | 0.725 | -0.46 | -0.701 | -0.01 | 0.127 | 0.127 | 0 | 0.63 | 0.85 | 0.55 | 0.95 | 0.6 | 0.55 |
| 11 | 126257779 | rs11220465 | -0.27 | 1.083 | -0.128 | 0.041 | 0 | 0.058 | 0.058 | 0 | 0.12 | 0.18 | 0.36 | 0.47 | 0.16 | 0.16 |
| 11 | 126258062 | rs7118117 | -1.211 | 1.017 | -0.177 | -0.035 | -0.005 | 0.052 | 0.052 | 1.123 | 0.29 | 0.59 | 0.46 | 0.5 | 0.26 | 0.32 |
| 11 | 126263885 | rs12280645 | -0.243 | 0.936 | -0.394 | -0.669 | 0.02 | -0.068 | -0.068 | 0.098 | 0.26 | 0.34 | 0.18 | 0.05 | 0.32 | 0.3 |
| 11 | 126271638 | rs11220473 | -1.332 | 0.681 | -0.686 | -0.805 | -0.002 | 0.058 | 0.058 | 0 | 0.48 | 0.61 | 0.49 | 0.89 | 0.49 | 0.41 |
| 11 | 126274003 | rs1007690 | -0.856 | 0.664 | -0.717 | -0.303 | -0.003 | 0.006 | 0.006 | 0 | 0.43 | 0.59 | 0.42 | 0.6 | 0.42 | 0.36 |
| 11 | 126274032 | rs6590202 | 0.399 | 0.81 | -0.75 | -0.328 | 0.093 | -0.084 | -0.084 | 0 | 0.15 | 0.55 | 0.16 | 0.03 | 0.24 | 0.12 |
| 11 | 126274959 | rs11220476 | 0.026 | 0.508 | -1.103 | -0.286 | 0.012 | -0.128 | -0.128 | 0.484 | 0.26 | 0.04 | 0.26 | 0.57 | 0.18 | 0.24 |
| 11 | 126275402 | rs11220477 | 0.183 | 0.472 | -1.038 | -0.286 | 0.008 | -0.025 | -0.025 | 0 | 0.09 | 0.05 | 0.06 | 0.23 | 0.04 | 0.17 |
| 11 | 126275830 | rs11603211 | 0.603 | 0.273 | -1.135 | -0.403 | -0.008 | -0.046 | -0.046 | 0 | 0.14 | 0.01 | 0.07 | 0.15 | 0.1 | 0.17 |
| 11 | 126278075 | rs2230279 | -0.318 | 0.695 | -0.697 | -0.507 | 0.032 | 0.093 | 0.093 | 0 | 0.25 | 0.36 | 0.33 | 0.51 | 0.31 | 0.2 |
| 11 | 126278203 | rs2298475 | 0.447 | 0.473 | -1.353 | -0.49 | -0.015 | -0.015 | -0.015 | 0.782 | 0.05 | 0.01 | 0.07 | 0.22 | 0.08 | 0.06 |
| 11 | 126284936 | rs11220483 | 0.965 | 0.375 | -1.29 | -0.55 | 0.01 | -0.094 | -0.094 | 0 | 0.13 | 0.02 | 0.2 | 0.49 | 0.1 | 0.09 |
| 11 | 126288762 | rs4935966 | -2.16 | 0.55 | -1.399 | -1.548 | -0.042 | 0.071 | 0.071 | 0.323 | 0.83 | 0.97 | 0.67 | 0.94 | 0.8 | 0.79 |
| 11 | 126288956 | rs4935967 | -0.959 | 0.364 | -0.91 | -1.191 | 0.005 | 0.072 | 0.072 | 0 | 0.55 | 0.69 | 0.51 | 0.89 | 0.54 | 0.5 |
| 11 | 126292596 | rs4937128 | -1.115 | 0.447 | -0.912 | -1.101 | 0.003 | 0.115 | 0.115 | 0 | 0.54 | 0.74 | 0.51 | 0.89 | 0.55 | 0.45 |
| 11 | 126293188 | rs948054 | -0.659 | 0.174 | -1.186 | -1.013 | 0 | -0.02 | -0.02 | 0 | 0.43 | 0.48 | 0.4 | 0.86 | 0.35 | 0.34 |
| 11 | 126294088 | rs7113465 | -0.353 | -0.071 | -1.597 | -0.623 | 0.025 | 0.006 | 0.006 | 0 | 0.36 | 0.33 | 0.3 | 0.65 | 0.28 | 0.29 |
| 11 | 126294261 | rs3802815 | -0.657 | -0.047 | -1.817 | -0.91 | 0.002 | -0.007 | -0.007 | 1.483 | 0.34 | 0.59 | 0.29 | 0.19 | 0.45 | 0.33 |
| 11 | 126296589 | rs4935969 | -0.57 | -0.08 | -2.24 | -1.248 | 0.036 | 0.261 | 0.261 | 0 | 0.48 | 0.2 | 0.43 | 0.88 | 0.43 | 0.52 |
| 11 | 126296678 | rs6590206 | -1.026 | -0.293 | -2.105 | -0.912 | -0.011 | 0.047 | 0.047 | 0 | 0.68 | 0.81 | 0.58 | 0.92 | 0.66 | 0.66 |
| 11 | 126297609 | rs11220490 | -0.12 | -1.145 | -1.973 | 0.489 | -0.012 | -0.045 | -0.045 | 0 | 0.16 | 0.01 | 0.09 | 0.17 | 0.14 | 0.21 |
| 11 | 126303742 | rs4937131 | -0.516 | -1.305 | -1.847 | 0.419 | 0.077 | -0.022 | -0.022 | 0.507 | 0.36 | 0.12 | 0.26 | 0.64 | 0.22 | 0.49 |
| 11 | 126310119 | rs3740917 | -0.709 | -0.947 | -0.635 | 1.074 | -0.034 | 0.167 | 0.167 | 0 | 0.09 | 0.25 | 0.05 | 0.29 | 0.03 | 0.14 |
| 11 | 126310156 | rs3740918 | -0.564 | -1.128 | -0.583 | 0.414 | -0.018 | -0.074 | -0.074 | 0 | 0.3 | 0.51 | 0.22 | 0.14 | 0.26 | 0.1 |
| 11 | 126310299 | rs3212313 | -0.836 | -1.186 | -0.739 | 0.639 | -0.01 | 0.004 | 0.004 | 0 | 0.35 | 0.57 | 0.16 | 0.38 | 0.23 | 0.31 |
| 11 | 126315024 | rs3802819 | -0.042 | -1.125 | -0.579 | 0.924 | -0.027 | 0.052 | 0.052 | 0.026 | 0.75 | 0.62 | 0.69 | 0.56 | 0.77 | 0.59 |
| 11 | 126315056 | rs3802820 | -0.235 | -1.652 | -0.558 | 1.065 | -0.01 | 0.055 | 0.055 | 0.065 | 0.12 | 0.17 | 0.06 | 0.3 | 0.1 | 0.16 |
| 11 | 126315213 | rs3802821 | 0.424 | -1.608 | -0.461 | 0.721 | -0.023 | -0.015 | -0.015 | 0 | 0.63 | 0.49 | 0.64 | 0.26 | 0.68 | 0.44 |
| 11 | 126315727 | rs4294574 | 0.481 | -1.209 | -1.092 | -0.169 | 0.003 | -0.029 | -0.029 | 0 | 0.42 | 0.49 | 0.3 | 0.16 | 0.48 | 0.22 |
| 11 | 126317444 | rs7936304 | -0.455 | -1.836 | -1.254 | 0.335 | 0.018 | -0.022 | -0.022 | 0 | 0.2 | 0.17 | 0.27 | 0.64 | 0.19 | 0.34 |
| 11 | 126321696 | rs11220500 | -1.048 | -1.011 | -1.3 | -0.556 | -0.013 | 0.031 | 0.031 | 0 | 0.57 | 0.69 | 0.63 | 0.8 | 0.52 | 0.61 |
| 11 | 126323328 | rs12801239 | 0.141 | -0.984 | -1.516 | -0.64 | -0.025 | 0.075 | 0.075 | 0 | 0.34 | 0.14 | 0.34 | 0.17 | 0.33 | 0.24 |
| 11 | 126326568 | rs3809005 | -1.813 | -0.734 | -1.812 | -0.722 | -0.039 | 0.034 | 0.034 | 0 | 0.82 | 0.95 | 0.79 | 0.9 | 0.76 | 0.8 |
| 11 | 126328279 | rs2155309 | 0.23 | -1.076 | -1.53 | 0.228 | -0.007 | 0.021 | 0.021 | 0 | 0.12 | 0.11 | 0.18 | 0.25 | 0.07 | 0.11 |
| 11 | 126332224 | rs7946496 | -0.056 | -0.993 | -2.079 | 0.053 | -0.043 | -0.096 | -0.096 | 2.08 | 0.36 | 0.09 | 0.25 | 0.53 | 0.43 | 0.41 |
| 11 | 126332878 | rs11220503 | 0.765 | -0.742 | -1.928 | 0.357 | -0.001 | -0.088 | -0.088 | 0.168 | 0.14 | 0.02 | 0.12 | 0.39 | 0.18 | 0.32 |
| 11 | 126334745 | rs11220504 | 0.751 | -0.746 | -1.929 | 0.355 | -0.001 | -0.088 | -0.088 | 0 | 0.14 | 0.02 | 0.12 | 0.39 | 0.18 | 0.32 |
| 11 | 126339435 | rs7925175 | -0.985 | -0.466 | -1.57 | 0.027 | -0.019 | 0.03 | 0.03 | 0 | 0.65 | 0.67 | 0.66 | 0.8 | 0.58 | 0.67 |
| 11 | 126346426 | rs3903109 | -0.194 | -0.189 | -1 | -0.201 | 0.006 | 0.092 | 0.092 | 0 | 0.08 | 0.17 | 0.35 | 0.27 | 0.1 | 0.14 |
| 11 | 126346842 | rs1124515 | -0.454 | 0.212 | -1.284 | -0.845 | -0.012 | 0.049 | 0.049 | 0.52 | 0.27 | 0.17 | 0.13 | 0.12 | 0.25 | 0.13 |
| 11 | 126349968 | rs11220516 | 0.772 | 0.375 | -1.454 | -0.819 | -0.034 | 0.013 | 0.013 | 0 | 0.15 | 0.06 | 0.1 | 0.12 | 0.2 | 0.09 |
| 11 | 126352844 | rs4937139 | -0.841 | 0.26 | -1.057 | -0.763 | -0.001 | -0.001 | -0.001 | 0 | 0.35 | 0.34 | 0.48 | 0.4 | 0.35 | 0.27 |
| 11 | 126359805 | rs10893525 | 0.392 | 0.534 | -1.405 | -1.127 | 0.01 | -0.034 | -0.034 | 4.069 | 0.2 | 0.21 | 0.3 | 0.04 | 0.24 | 0.11 |
| 11 | 126360303 | rs11220521 | -0.676 | -0.067 | -1.824 | -1.366 | 0.04 | 0.022 | 0.022 | 0 | 0.54 | 0.79 | 0.66 | 0.4 | 0.55 | 0.36 |
| 11 | 126360403 | rs1000256 | 0.6 | -0.133 | -2.177 | -1.22 | -0.015 | -0.124 | -0.124 | 0.279 | 0.16 | 0.01 | 0.18 | 0.5 | 0.21 | 0.38 |
| 11 | 126360525 | rs1000255 | -0.578 | -0.586 | -2.283 | -1.34 | -0.007 | -0.048 | -0.048 | 0.249 | 0.73 | 0.88 | 0.65 | 0.63 | 0.71 | 0.67 |
| 11 | 126360592 | rs1000254 | 0.906 | -0.471 | -1.97 | -1.017 | 0.009 | -0.04 | -0.04 | 0 | 0.1 | 0.01 | 0.12 | 0.47 | 0.11 | 0.34 |
| 11 | 126361331 | rs7937654 | -0.254 | 0.317 | -1.273 | -2.868 | -0.016 | 0.022 | 0.022 | 0 | 0.39 | 0.28 | 0.29 | 0.11 | 0.48 | 0.19 |
| 11 | 126362042 | rs10160702 | -0.294 | 0.224 | -0.782 | -2.458 | -0.022 | -0.063 | -0.063 | 0.053 | 0.75 | 0.64 | 0.79 | 0.91 | 0.8 | 0.77 |
| 11 | 126364135 | rs1892999 | 0.282 | 0.623 | -1.39 | -2.553 | -0.007 | 0.017 | 0.017 | 0.274 | 0.55 | 0.65 | 0.66 | 0.87 | 0.48 | 0.79 |
| 11 | 126366819 | rs10750326 | -1.445 | 0.65 | -1.359 | -2.361 | 0.009 | 0.035 | 0.035 | 0 | 0.84 | 0.92 | 0.9 | 0.96 | 0.86 | 0.9 |
| 11 | 126368432 | rs11823106 | -0.692 | 0.967 | -1.127 | -1.98 | 0 | -0.015 | -0.015 | 0 | 0.37 | 0.37 | 0.34 | 0.17 | 0.46 | 0.27 |
| 11 | 126369158 | rs7933568 | -0.667 | 1.018 | -1.131 | -1.977 | 0.002 | -0.031 | -0.031 | 0 | 0.36 | 0.38 | 0.32 | 0.17 | 0.42 | 0.25 |
| 11 | 126369319 | rs7933515 | -0.29 | 1.019 | -1.515 | -1.967 | -0.004 | 0.02 | 0.02 | 0.021 | 0.31 | 0.24 | 0.29 | 0.15 | 0.36 | 0.22 |
| 11 | 126369531 | rs12803511 | -0.207 | 0.624 | -1.452 | -1.607 | -0.012 | 0.014 | 0.014 | 0 | 0.07 | 0.03 | 0.1 | 0.06 | 0.07 | 0.15 |
| 11 | 126369656 | rs12803395 | -0.392 | -0.054 | -1.762 | -2.217 | 0.011 | 0.033 | 0.033 | 0 | 0.3 | 0.23 | 0.21 | 0.09 | 0.26 | 0.19 |
| 11 | 126370065 | rs2155314 | -0.416 | 0.046 | -1.863 | -2.4 | 0.001 | 0.021 | 0.021 | 0 | 0.67 | 0.74 | 0.69 | 0.89 | 0.69 | 0.78 |
| 11 | 126370823 | rs7927708 | -0.956 | -0.101 | -1.621 | -2.413 | -0.017 | -0.039 | -0.039 | 0 | 0.85 | 0.98 | 0.73 | 0.17 | 0.82 | 0.58 |
| 11 | 126370843 | rs7941193 | -0.696 | -0.556 | -0.896 | -3.341 | -0.011 | -0.112 | -0.112 | 0 | 0.22 | 0.58 | 0.16 | 0 | 0.2 | 0.07 |
| 11 | 126371892 | rs10128646 | -0.871 | -0.464 | -0.9 | -3.297 | -0.017 | -0.12 | -0.12 | 0 | 0.24 | 0.58 | 0.16 | 0 | 0.21 | 0.07 |
| 11 | 126379392 | rs11826881 | -0.581 | -0.575 | -0.521 | -3.154 | -0.007 | -0.048 | -0.048 | 0 | 0.39 | 0.56 | 0.45 | 0.1 | 0.3 | 0.26 |
| 11 | 126381763 | rs6590216 | -0.391 | -0.628 | -0.15 | -3.195 | -0.001 | -0.017 | -0.017 | 0 | 0.36 | 0.37 | 0.43 | 0.1 | 0.29 | 0.25 |
| 11 | 126384549 | rs4341546 | -0.759 | -0.863 | -0.099 | -3.297 | 0.033 | 0.133 | 0.133 | 2.576 | 0.42 | 0.81 | 0.46 | 0.11 | 0.31 | 0.26 |

| | | | | | | | | | | | | | | | | |
|---|---|---|---|---|---|---|---|---|---|---|---|---|---|---|---|---|
| 11 | 126387391 | rs4471434 | -0.132 | -0.889 | -0.446 | -4.156 | -0.005 | -0.078 | -0.078 | 0 | 0.42 | 0.53 | 0.4 | 0.07 | 0.38 | 0.19 |
| 11 | 126388248 | rs12270348 | 0.899 | -0.74 | -0.024 | -3.955 | 0.022 | -0.03 | -0.03 | 0 | 0.09 | 0.26 | 0.07 | 0.01 | 0.08 | 0.14 |
| 11 | 126389301 | rs558745 | 1.472 | -1.004 | -0.251 | -3.828 | -0.103 | 0.033 | 0.033 | 4.773 | 0.3 | 0.04 | 0.4 | 0.85 | 0.36 | 0.64 |
| 11 | 126390968 | rs11605603 | 0.11 | -1.078 | 0.404 | -3.604 | -0.04 | -0.127 | -0.127 | 0 | 0.25 | 0.5 | 0.21 | 0.04 | 0.21 | 0.08 |
| 11 | 126391297 | rs12269776 | 0.295 | -1.032 | 0.434 | -3.571 | -0.037 | -0.125 | -0.125 | 0 | 0.23 | 0.5 | 0.21 | 0.04 | 0.2 | 0.07 |
| 11 | 126396674 | rs674537 | 0.495 | -0.97 | 0.287 | -3.62 | 0.008 | 0.058 | 0.058 | 0 | 0.61 | 0.27 | 0.65 | 0.95 | 0.61 | 0.8 |
| 11 | 126397689 | rs620082 | -1.172 | -0.932 | 0.14 | -3.514 | 0.003 | -0.025 | -0.025 | 0 | 0.88 | 0.89 | 0.89 | 1 | 0.83 | 0.9 |
| 11 | 126399532 | rs4937148 | 0.135 | -1.137 | 0.111 | -3.549 | 0.011 | 0.047 | 0.047 | 3.364 | 0.23 | 0.07 | 0.17 | 0.09 | 0.15 | 0.09 |
| 11 | 126400534 | rs12269946 | 0.167 | -0.907 | 0.531 | -3.416 | -0.015 | -0.063 | -0.063 | 0 | 0.23 | 0.31 | 0.2 | 0.05 | 0.19 | 0.09 |
| 11 | 126409913 | rs552640 | -0.371 | -0.584 | 0.533 | -2.862 | 0 | -0.057 | -0.057 | 0 | 0.61 | 0.65 | 0.69 | 0.95 | 0.59 | 0.79 |
| 11 | 126412343 | rs527629 | 0.251 | -0.457 | -0.715 | -2.777 | -0.024 | 0.153 | 0.153 | 0 | 0.42 | 0.21 | 0.47 | 0.92 | 0.45 | 0.72 |
| 11 | 126414087 | rs12291500 | -1.436 | -1.003 | 0.137 | -1.69 | -0.045 | -0.057 | -0.057 | 0 | 0.37 | 0.52 | 0.21 | 0.16 | 0.18 | 0.13 |
| 11 | 126415374 | rs4937150 | -0.961 | -1.045 | -0.017 | -1.039 | -0.004 | 0.002 | 0.002 | 0.415 | 0.17 | 0.16 | 0.02 | 0.14 | 0.04 | 0.06 |
| 11 | 126425577 | rs10893539 | -1.598 | -0.006 | 0.271 | -1.304 | -0.004 | -0.066 | -0.066 | 0 | 0.24 | 0.65 | 0.17 | 0.01 | 0.19 | 0.12 |
| 11 | 126426822 | rs11220539 | -0.318 | -0.005 | 0.881 | -1.212 | -0.011 | -0.014 | -0.014 | 0 | 0.07 | 0.24 | 0.03 | 0 | 0.01 | 0 |
| 11 | 126432376 | rs7945346 | -1.089 | 0.436 | 0.693 | -1.084 | 0.036 | -0.109 | -0.109 | 0.253 | 0.16 | 0.38 | 0.14 | 0.01 | 0.18 | 0.11 |
| 11 | 126434881 | rs487842 | -0.124 | 1.03 | 0.135 | -0.03 | -0.001 | 0.001 | 0.001 | 0 | 0.27 | 0.25 | 0.28 | 0.47 | 0.29 | 0.36 |
| 11 | 126435082 | rs489725 | -0.271 | 0.975 | 0.674 | -0.062 | 0.017 | 0.068 | 0.068 | 0 | 0.28 | 0.37 | 0.29 | 0.46 | 0.29 | 0.36 |
| 11 | 126435264 | rs491536 | -0.191 | 0.898 | 0.235 | -0.904 | -0.008 | -0.084 | -0.084 | 0 | 0.66 | 0.59 | 0.74 | 0.91 | 0.65 | 0.82 |
| 11 | 126437941 | rs619996 | 0.064 | 0.864 | -0.027 | -0.113 | -0.001 | -0.006 | -0.006 | 0.27 | 0.41 | 0.48 | 0.51 | 0.43 | 0.39 | 0.38 |
| 11 | 126442506 | rs681653 | 0.497 | 1.164 | 0.071 | 0.596 | 0.016 | 0.056 | 0.056 | 0.1 | 0.72 | 0.52 | 0.61 | 0.61 | 0.72 | 0.8 |
| 11 | 126446361 | rs654393 | 0.685 | 1.136 | 0.197 | -0.044 | -0.001 | -0.003 | -0.003 | 0 | 0.67 | 0.73 | 0.61 | 0.57 | 0.64 | 0.7 |
| 11 | 126449549 | rs632023 | 1.298 | 1.236 | 0.374 | -0.312 | -0.007 | 0.004 | 0.004 | 0 | 0.54 | 0.36 | 0.5 | 0.56 | 0.54 | 0.62 |
| 11 | 126452032 | rs537172 | 0.818 | 1.058 | -0.679 | -0.413 | 0 | 0.001 | 0.001 | 0 | 0.29 | 0.3 | 0.32 | 0.27 | 0.27 | 0.23 |
| 11 | 126452369 | rs4937154 | 0.012 | 1.094 | -0.7 | -0.078 | -0.002 | -0.012 | -0.012 | 0 | 0.25 | 0.18 | 0.32 | 0.57 | 0.25 | 0.23 |
| 11 | 126454602 | rs2004770 | 0.63 | 1.199 | -0.821 | -0.133 | -0.001 | 0.011 | 0.011 | 0 | 0.33 | 0.3 | 0.36 | 0.26 | 0.35 | 0.26 |
| 11 | 126458263 | rs12574555 | 0.283 | 1.085 | -0.692 | 0.454 | 0 | 0.002 | 0.002 | 0 | 0.12 | 0.12 | 0.15 | 0.2 | 0.12 | 0.16 |
| 11 | 126459999 | rs541969 | 0.06 | 1.29 | -0.444 | 0.382 | -0.003 | 0.006 | 0.006 | 0 | 0.45 | 0.62 | 0.62 | 0.61 | 0.47 | 0.36 |
| 11 | 126461758 | rs605344 | -0.952 | 1.221 | -0.524 | 0.16 | 0.011 | 0.012 | 0.012 | 0.063 | 0.85 | 0.91 | 0.92 | 0.99 | 0.88 | 0.92 |
| 11 | 126465148 | rs12794212 | 0.816 | 0.982 | -0.559 | -0.104 | 0.001 | 0.024 | 0.024 | 0 | 0.14 | 0.04 | 0.07 | 0 | 0.12 | 0.03 |
| 11 | 126466129 | rs10893545 | 0.03 | 0.956 | 0.082 | 0.054 | 0.045 | 0.101 | 0.101 | 0 | 0.14 | 0.41 | 0.16 | 0.2 | 0.12 | 0.13 |
| 11 | 126467667 | rs484820 | 0.985 | 0.902 | 0.01 | 0.568 | -0.008 | 0.005 | 0.005 | 0 | 0.28 | 0.24 | 0.47 | 0.27 | 0.32 | 0.3 |
| 11 | 126468206 | rs480073 | -1.044 | 0.787 | 0.207 | 0.609 | -0.037 | 0.105 | 0.105 | 0.129 | 0.85 | 0.77 | 0.95 | 0.66 | 0.92 | 0.92 |
| 11 | 126469558 | rs608006 | -0.26 | 0.848 | 0.164 | 0.573 | 0 | -0.001 | -0.001 | 0 | 0.57 | 0.6 | 0.45 | 0.39 | 0.56 | 0.62 |
| 11 | 126469604 | rs535603 | -1.075 | 0.825 | 0.079 | 0.624 | -0.014 | 0.054 | 0.054 | 0 | 0.85 | 0.84 | 0.91 | 0.66 | 0.89 | 0.91 |
| 11 | 126471644 | rs474130 | -0.121 | 0.9 | -0.08 | 0.61 | 0.019 | -0.048 | -0.048 | 0 | 0.72 | 0.9 | 0.84 | 0.66 | 0.75 | 0.89 |
| 11 | 126471989 | rs471302 | 0.636 | 0.935 | 0.072 | 0.608 | -0.006 | 0.002 | 0.002 | 0 | 0.41 | 0.35 | 0.33 | 0.38 | 0.43 | 0.57 |
| 11 | 126472512 | rs577149 | 1.762 | 0.91 | -0.162 | 0.59 | -0.039 | 0.041 | 0.041 | 0.007 | 0.27 | 0.07 | 0.17 | 0.19 | 0.31 | 0.45 |
| 11 | 126473750 | rs489487 | 0.079 | 1.035 | 0.239 | 0.605 | -0.001 | -0.001 | -0.001 | 0 | 0.57 | 0.43 | 0.37 | 0.54 | 0.54 | 0.5 |
| 11 | 126474064 | rs684589 | 0.7 | 1.165 | 0.131 | 0.653 | 0 | -0.001 | -0.001 | 0.681 | 0.23 | 0.25 | 0.42 | 0.27 | 0.24 | 0.27 |
| 11 | 126474428 | rs483027 | 0.843 | 1.027 | 0.217 | 0.675 | -0.001 | 0 | 0 | 0 | 0.2 | 0.21 | 0.35 | 0.27 | 0.2 | 0.21 |
| 11 | 126479691 | rs7124593 | 0.511 | 1.058 | 0.157 | 0.841 | 0.001 | 0.04 | 0.04 | 0.291 | 0.19 | 0.25 | 0.37 | 0.42 | 0.2 | 0.25 |
| 11 | 126480356 | rs493725 | 0.729 | 0.839 | -0.34 | 0.528 | 0.004 | 0.098 | 0.098 | 0 | 0.54 | 0.13 | 0.35 | 0.25 | 0.61 | 0.53 |
| 11 | 126481172 | rs641675 | 1.586 | 1.266 | -0.59 | 0.272 | -0.037 | -0.004 | -0.004 | 0 | 0.32 | 0.08 | 0.15 | 0.24 | 0.32 | 0.47 |
| 11 | 126486045 | rs549190 | 0.746 | 1.769 | -0.417 | 0.332 | -0.012 | 0.105 | 0.105 | 0 | 0.52 | 0.19 | 0.32 | 0.24 | 0.53 | 0.53 |
| 11 | 126494642 | rs11220561 | 1.036 | 1.231 | -0.236 | 0.353 | 0.012 | 0.057 | 0.057 | 0.003 | 0.08 | 0.1 | 0.18 | 0.33 | 0.11 | 0.11 |
| 11 | 126496555 | rs559121 | 0.454 | 0.838 | -0.174 | 0.161 | 0.015 | 0.021 | 0.021 | 2.622 | 0.22 | 0.26 | 0.44 | 0.43 | 0.34 | 0.26 |
| 11 | 126497529 | rs681570 | -0.99 | 0.691 | -0.043 | 0.256 | -0.021 | -0.048 | -0.048 | 0.357 | 0.9 | 0.85 | 0.97 | 1 | 0.94 | 0.93 |
| 11 | 126501321 | rs876901 | -0.37 | 0.772 | -0.095 | 0.34 | 0.032 | 0.059 | 0.059 | 0 | 0.3 | 0.53 | 0.48 | 0.43 | 0.32 | 0.25 |
| 11 | 126501939 | rs514457 | 0.843 | 0.791 | -0.437 | 0.271 | 0.005 | 0.109 | 0.109 | 0 | 0.52 | 0.23 | 0.31 | 0.24 | 0.51 | 0.53 |
| 11 | 126505327 | rs625993 | 0.416 | 1.124 | -0.161 | 0.523 | -0.046 | 0.111 | 0.111 | 0 | 0.21 | 0.04 | 0.2 | 0 | 0.27 | 0.06 |
| 11 | 126506712 | rs4937166 | 0.592 | 1.549 | 0.267 | 0.99 | 0.108 | 0.196 | 0.196 | 0 | 0.08 | 0.21 | 0.18 | 0.39 | 0.19 | 0.11 |
| 11 | 126508121 | rs10790817 | -0.025 | 1.244 | 0.21 | 0.958 | -0.005 | 0.05 | 0.05 | 0 | 0.2 | 0.33 | 0.21 | 0.33 | 0.17 | 0.17 |
| 11 | 126513773 | rs11220566 | -0.457 | 0.699 | -1.16 | 0.413 | -0.018 | 0.006 | 0.006 | 0 | 0.36 | 0.67 | 0.43 | 0.72 | 0.38 | 0.33 |
| 11 | 126515506 | rs658125 | -0.906 | 0.24 | -1.383 | 0.498 | 0.062 | -0.046 | -0.046 | 0.719 | 0.58 | 0.82 | 0.63 | 0.57 | 0.65 | 0.37 |
| 11 | 126521978 | rs11220571 | 0.495 | 0.487 | -0.958 | 0.085 | 0.004 | 0.001 | 0.001 | 1.171 | 0.12 | 0.09 | 0.06 | 0.16 | 0.09 | 0.02 |
| 11 | 126522029 | rs1318969 | -0.972 | 0.876 | -1.378 | -0.384 | -0.035 | 0.071 | 0.071 | 0.033 | 0.27 | 0.07 | 0.23 | 0.17 | 0.3 | 0.08 |
| 11 | 126523715 | rs509759 | -1.562 | 1.102 | -0.916 | -0.25 | 0.029 | 0.011 | 0.011 | 0 | 0.49 | 0.64 | 0.63 | 0.53 | 0.56 | 0.37 |
| 11 | 126524401 | rs4937170 | 0.303 | 1.253 | -0.465 | -0.486 | 0.053 | 0.095 | 0.095 | 0 | 0.18 | 0.3 | 0.38 | 0.39 | 0.25 | 0.25 |
| 11 | 126526181 | rs523249 | 0.064 | 1.202 | -0.265 | -0.764 | -0.005 | 0.011 | 0.011 | 0.145 | 0.74 | 0.79 | 0.76 | 0.99 | 0.72 | 0.91 |
| 11 | 126527765 | rs693253 | 0.993 | 0.843 | -0.34 | -0.822 | -0.004 | -0.001 | -0.001 | 0 | 0.49 | 0.53 | 0.37 | 0.62 | 0.4 | 0.58 |
| 11 | 126528035 | rs7950175 | -0.854 | 1.081 | -0.302 | -0.742 | -0.035 | 0.041 | 0.041 | 0 | 0.86 | 0.96 | 0.83 | 1 | 0.76 | 0.94 |
| 11 | 126530543 | rs7108068 | -0.369 | 0.876 | -0.241 | -0.567 | 0.014 | 0.039 | 0.039 | 0 | 0.28 | 0.35 | 0.42 | 0.38 | 0.31 | 0.31 |
| 11 | 126533749 | rs2156449 | 0.048 | 1.019 | -0.439 | -0.498 | 0.005 | 0.016 | 0.016 | 0 | 0.25 | 0.27 | 0.4 | 0.37 | 0.3 | 0.31 |
| 11 | 126539136 | rs11220578 | 0.629 | 0.497 | -0.941 | -1.19 | 0.023 | 0.03 | 0.03 | 0.931 | 0.17 | 0.1 | 0.08 | 0 | 0.11 | 0.05 |
| 11 | 126560619 | rs11220583 | 0.55 | 1.464 | -0.842 | -0.951 | -0.058 | 0.053 | 0.053 | 1.119 | 0.69 | 0.21 | 0.61 | 0.83 | 0.55 | 0.78 |
| 11 | 126565481 | rs4937179 | -0.251 | 1.812 | -0.009 | -0.992 | 0.095 | 0.051 | 0.051 | 0.016 | 0.13 | 0.37 | 0.21 | 0.15 | 0.21 | 0.09 |
| 11 | 126577515 | rs11220587 | 1.282 | 1.188 | -0.791 | -0.694 | -0.054 | -0.186 | -0.186 | 0.794 | 0.26 | 0.01 | 0.24 | 0.51 | 0.24 | 0.5 |
| 11 | 126580253 | rs3862634 | -0.157 | 1.699 | 0.151 | -1.229 | -0.012 | 0.008 | 0.008 | 0 | 0.91 | 0.94 | 0.9 | 1 | 0.85 | 0.95 |

| | | | | | | | | | | | | | | | | |
|---|---|---|---|---|---|---|---|---|---|---|---|---|---|---|---|---|
| 11 | 126583203 | rs10790820 | 0.918 | 1.555 | 0.278 | -1.224 | 0.051 | -0.118 | -0.118 | 0 | 0.82 | 0.58 | 0.85 | 0.99 | 0.8 | 0.89 |
| 11 | 126586154 | rs4581457 | -0.086 | 1.31 | -0.15 | -0.989 | -0.008 | 0.006 | 0.006 | 0.026 | 0.91 | 0.95 | 0.91 | 1 | 0.88 | 0.93 |
| 11 | 126587036 | rs3909729 | -0.412 | 1.211 | 0.133 | -1.487 | 0.004 | -0.019 | -0.019 | 0 | 0.36 | 0.36 | 0.34 | 0.18 | 0.42 | 0.23 |
| 11 | 126587812 | rs3909725 | 0.953 | 1.015 | 0.565 | -1.482 | 0.009 | -0.088 | -0.088 | 0 | 0.83 | 0.64 | 0.86 | 0.99 | 0.83 | 0.94 |
| 11 | 126595819 | rs6590227 | 0.073 | 1.022 | 0.374 | -1.449 | 0.001 | -0.03 | -0.03 | 6.313 | 0.89 | 0.82 | 0.89 | 0.99 | 0.88 | 0.95 |
| 11 | 126596746 | rs3911927 | 0.029 | 1.038 | 0.251 | -1.55 | -0.002 | 0.003 | 0.003 | 0 | 0.86 | 0.87 | 0.88 | 0.99 | 0.85 | 0.93 |
| 11 | 126597582 | rs4245068 | 0.806 | 1.002 | 0.463 | -1.697 | -0.002 | -0.116 | -0.116 | 0.744 | 0.77 | 0.61 | 0.82 | 0.98 | 0.76 | 0.87 |
| 11 | 126598551 | rs12807269 | -1.1 | 0.965 | 0.517 | -1.591 | -0.008 | -0.066 | -0.066 | 0.66 | 0.13 | 0.3 | 0.1 | 0.01 | 0.11 | 0.06 |
| 11 | 126601132 | rs11220594 | 0.239 | 1.263 | 0.345 | -1.762 | 0.012 | -0.017 | -0.017 | 0 | 0.23 | 0.28 | 0.24 | 0.14 | 0.28 | 0.13 |
| 11 | 126601540 | rs10893557 | 1.718 | 1.167 | -0.106 | -1.939 | 0.029 | 0.128 | 0.128 | 0 | 0.47 | 0.27 | 0.51 | 0.84 | 0.37 | 0.69 |
| 11 | 126601735 | rs3862638 | 1.793 | 1.453 | -0.053 | -1.643 | 0.001 | 0.112 | 0.112 | 0.235 | 0.54 | 0.25 | 0.52 | 0.8 | 0.41 | 0.7 |
| 11 | 126602304 | rs4935986 | 1.958 | 1.214 | -0.136 | -1.974 | 0.004 | 0.165 | 0.165 | 0.044 | 0.52 | 0.27 | 0.58 | 0.84 | 0.43 | 0.7 |
| 11 | 126602453 | rs7131158 | -1.631 | 1.324 | 0.454 | -2.149 | 0.004 | -0.067 | -0.067 | 0.056 | 0.19 | 0.3 | 0.13 | 0.06 | 0.19 | 0.12 |
| 11 | 126603909 | rs10893559 | 2.369 | 1.532 | 0.358 | -2.193 | 0.022 | 0.182 | 0.182 | 0 | 0.51 | 0.27 | 0.58 | 0.88 | 0.41 | 0.68 |
| 11 | 126603928 | rs7944781 | -0.185 | 1.278 | 0.587 | -2.051 | 0.01 | -0.011 | -0.011 | 0 | 0.09 | 0.19 | 0.05 | 0.07 | 0.09 | 0.06 |
| 11 | 126605034 | rs3862642 | 0.943 | 1.161 | 0.474 | -1.944 | 0.018 | -0.099 | -0.099 | 0.137 | 0.7 | 0.57 | 0.71 | 0.95 | 0.62 | 0.83 |
| 11 | 126608446 | rs3862645 | -0.994 | 0.934 | 0.569 | -1.692 | -0.004 | -0.101 | -0.101 | 0 | 0.2 | 0.44 | 0.15 | 0.05 | 0.21 | 0.11 |
| 11 | 126608705 | rs3889461 | -1.417 | 0.961 | 0.488 | -1.697 | -0.012 | -0.113 | -0.113 | 0 | 0.25 | 0.5 | 0.21 | 0.06 | 0.23 | 0.16 |
| 11 | 126611548 | rs7131312 | -0.809 | 1.555 | 0.806 | -1.578 | 0.068 | -0.053 | -0.053 | 0 | 0.12 | 0.32 | 0.15 | 0.04 | 0.22 | 0.1 |
| 11 | 126612903 | rs1946091 | 1.391 | 1.418 | 0.205 | -1.718 | 0.055 | 0.145 | 0.145 | 0 | 0.48 | 0.29 | 0.54 | 0.8 | 0.37 | 0.66 |
| 11 | 126619190 | rs1557488 | -0.705 | 1.741 | 0.681 | -1.864 | -0.001 | -0.033 | -0.033 | 0 | 0.82 | 0.82 | 0.88 | 1 | 0.82 | 0.88 |
| 11 | 126619841 | rs1946089 | 0.985 | 1.693 | 0.751 | -1.912 | 0.016 | -0.102 | -0.102 | 0 | 0.65 | 0.48 | 0.64 | 0.95 | 0.54 | 0.75 |
| 11 | 126622933 | rs11220599 | -0.807 | 1.585 | 0.778 | -1.75 | 0.053 | -0.063 | -0.063 | 2.869 | 0.14 | 0.34 | 0.15 | 0.05 | 0.23 | 0.12 |
| 11 | 126625946 | rs1946086 | -0.486 | 1.662 | 0.565 | -1.714 | 0.078 | -0.053 | -0.053 | 0 | 0.13 | 0.38 | 0.15 | 0.05 | 0.23 | 0.11 |
| 11 | 126626203 | rs1946084 | 0.611 | 1.668 | 0.454 | -1.812 | 0.012 | -0.095 | -0.095 | 0 | 0.68 | 0.46 | 0.73 | 0.95 | 0.58 | 0.8 |
| 11 | 126629031 | rs7101595 | 0.623 | 1.491 | 0.35 | -1.678 | -0.022 | 0.052 | 0.052 | 0 | 0.62 | 0.76 | 0.72 | 0.81 | 0.58 | 0.59 |
| 11 | 126630621 | rs1540238 | 1.428 | 1.422 | 0.364 | -1.7 | 0.02 | 0.039 | 0.039 | 0 | 0.48 | 0.41 | 0.56 | 0.76 | 0.36 | 0.5 |
| 11 | 126632195 | rs10082694 | -1.946 | 1.492 | 0.148 | -1.724 | 0.022 | 0.13 | 0.13 | 0 | 0.54 | 0.78 | 0.44 | 0.24 | 0.6 | 0.49 |
| 11 | 126642299 | rs1946075 | 1.263 | 1.105 | -0.196 | -2.072 | -0.051 | -0.121 | -0.121 | 0.259 | 0.18 | 0.39 | 0.1 | 0 | 0.09 | 0.09 |
| 11 | 126642327 | rs12792630 | 1.384 | 1.248 | -0.621 | -2.068 | -0.037 | 0.027 | 0.027 | 0 | 0.07 | 0.02 | 0.11 | 0.03 | 0.21 | 0.09 |
| 11 | 126644203 | rs1874 | 1.496 | 0.861 | -0.385 | -1.85 | -0.028 | 0.03 | 0.03 | 0 | 0.31 | 0.05 | 0.37 | 0.76 | 0.29 | 0.42 |
| 11 | 126652326 | rs939530 | 0.911 | 0.833 | -0.228 | -1.859 | 0.001 | -0.143 | -0.143 | 1.088 | 0.26 | 0.54 | 0.22 | 0.03 | 0.3 | 0.21 |
| 11 | 126652604 | rs11220606 | -0.651 | 0.386 | -0.486 | -1.78 | -0.001 | 0.006 | 0.006 | 0 | 0.25 | 0.2 | 0.29 | 0.21 | 0.23 | 0.3 |
| 11 | 126664315 | rs10790827 | 1.338 | 0.386 | -0.588 | -0.648 | 0.006 | -0.075 | -0.075 | 0 | 0.31 | 0.14 | 0.31 | 0.51 | 0.27 | 0.41 |
| 11 | 126665701 | rs10790829 | 0.053 | 0.707 | -0.568 | -0.472 | 0.006 | -0.042 | -0.042 | 5.075 | 0.74 | 0.46 | 0.7 | 0.81 | 0.74 | 0.83 |
| 11 | 126666201 | rs9326286 | 0.955 | 0.72 | -1.043 | -0.527 | -0.051 | -0.05 | -0.05 | 0 | 0.27 | 0.13 | 0.34 | 0.33 | 0.35 | 0.22 |
| 11 | 126666312 | rs4937186 | 0.722 | 0.32 | -0.977 | -0.695 | 0.009 | 0.003 | 0.003 | 0 | 0.39 | 0.33 | 0.45 | 0.65 | 0.33 | 0.51 |
| 11 | 126666332 | rs10893569 | 0.214 | 0.281 | -1.015 | -0.714 | 0.055 | 0.034 | 0.034 | 0 | 0.44 | 0.74 | 0.49 | 0.65 | 0.33 | 0.51 |
| 11 | 126666419 | rs4935994 | -0.813 | 0.418 | -1.082 | -0.704 | -0.093 | -0.003 | -0.003 | 0 | 0.63 | 0.9 | 0.65 | 0.67 | 0.5 | 0.71 |
| 11 | 126670291 | rs1946046 | 1.591 | 0.422 | -1.139 | -0.949 | -0.005 | -0.13 | -0.13 | 3.013 | 0.29 | 0.08 | 0.3 | 0.47 | 0.27 | 0.41 |
| 11 | 126671287 | rs7109451 | 1.647 | 0.432 | -1.142 | -0.963 | -0.028 | -0.027 | -0.027 | 0 | 0.22 | 0.09 | 0.25 | 0.24 | 0.29 | 0.19 |
| 11 | 126671804 | rs10750340 | 1.521 | 0.409 | -1.131 | -0.902 | -0.005 | -0.13 | -0.13 | 0 | 0.29 | 0.08 | 0.3 | 0.48 | 0.27 | 0.41 |
| 11 | 126674317 | rs10790834 | 1.779 | 0.41 | -1.189 | -0.912 | -0.015 | -0.161 | -0.161 | 0.394 | 0.27 | 0.04 | 0.29 | 0.47 | 0.23 | 0.37 |
| 11 | 126674673 | rs10893574 | 0.11 | 0.506 | -1.076 | -0.931 | -0.001 | 0 | 0 | 2.593 | 0.51 | 0.47 | 0.49 | 0.5 | 0.43 | 0.6 |
| 11 | 126677128 | rs11220621 | 0.022 | 0.504 | -0.868 | -0.803 | -0.001 | -0.001 | -0.001 | 0.1 | 0.27 | 0.28 | 0.3 | 0.28 | 0.24 | 0.3 |
| 11 | 126681867 | rs1946053 | -0.496 | 0.61 | -0.885 | -0.821 | -0.001 | 0 | 0 | 0.028 | 0.52 | 0.55 | 0.55 | 0.53 | 0.52 | 0.56 |
| 11 | 126688352 | rs10790837 | -1.234 | 1.093 | -0.877 | -0.657 | -0.056 | 0.084 | 0.084 | 0 | 0.85 | 0.97 | 0.89 | 1 | 0.79 | 0.93 |
| 11 | 126702154 | rs10750352 | 0.326 | 1.073 | -0.86 | -0.567 | -0.004 | 0.004 | 0.004 | 0 | 0.61 | 0.64 | 0.58 | 0.69 | 0.58 | 0.67 |
| 11 | 126702401 | rs10790844 | -0.387 | 1.027 | -0.838 | -0.615 | 0 | 0.004 | 0.004 | 0 | 0.48 | 0.46 | 0.58 | 0.55 | 0.53 | 0.54 |
| 11 | 126708662 | rs7102782 | 0.941 | 1.041 | -0.816 | -0.6 | -0.027 | 0.019 | 0.019 | 0 | 0.36 | 0.47 | 0.32 | 0.45 | 0.26 | 0.35 |
| 11 | 126711088 | rs7129447 | -0.765 | 1.298 | -0.988 | -0.601 | -0.031 | 0.052 | 0.052 | 0 | 0.84 | 0.91 | 0.89 | 1 | 0.79 | 0.89 |
| 11 | 126728188 | rs7126443 | -0.32 | 1.308 | -0.972 | -0.602 | -0.016 | 0.014 | 0.014 | 0 | 0.82 | 0.85 | 0.87 | 1 | 0.76 | 0.87 |
| 11 | 126731851 | rs9888281 | 0.379 | 1.146 | -0.89 | -0.458 | 0.005 | 0.013 | 0.013 | 0 | 0.21 | 0.27 | 0.31 | 0.27 | 0.21 | 0.22 |
| 11 | 126741484 | rs1784354 | -0.437 | 0.787 | -1.329 | -0.267 | -0.007 | -0.002 | -0.002 | 0 | 0.84 | 0.86 | 0.88 | 1 | 0.79 | 0.9 |
| 11 | 126748829 | rs7109913 | 0.294 | 0.576 | -1.287 | -0.235 | -0.001 | -0.008 | -0.008 | 0 | 0.15 | 0.14 | 0.1 | 0 | 0.15 | 0.1 |
| 11 | 126751063 | rs1939988 | -0.683 | 0.167 | -1.226 | 0.294 | -0.109 | 0.023 | 0.023 | 0 | 0.61 | 0.94 | 0.65 | 0.72 | 0.5 | 0.64 |
| 11 | 126754032 | rs7924535 | -0.617 | 0.215 | -1.215 | 0.317 | -0.084 | 0.038 | 0.038 | 0 | 0.6 | 0.87 | 0.65 | 0.71 | 0.5 | 0.61 |
| 11 | 126755025 | rs1009567 | -0.624 | 0.221 | -1.258 | 0.32 | -0.09 | 0.035 | 0.035 | 0 | 0.6 | 0.88 | 0.65 | 0.71 | 0.5 | 0.61 |
| 11 | 126767481 | rs11600756 | 0.569 | 0.462 | -2.41 | 0.43 | 0.041 | -0.125 | -0.125 | 0.066 | 0.28 | 0.04 | 0.16 | 0.44 | 0.17 | 0.32 |
| 11 | 126768090 | rs4937216 | -0.759 | 1.369 | -1.29 | 0.364 | -0.021 | 0.01 | 0.01 | 0 | 0.31 | 0.78 | 0.48 | 0.27 | 0.33 | 0.33 |
| 11 | 126777272 | rs1784343 | 0.678 | 1.661 | -0.645 | 0.509 | 0.036 | 0.005 | 0.005 | 0 | 0.73 | 0.48 | 0.66 | 0.71 | 0.66 | 0.72 |
| 11 | 126778513 | rs1793656 | 0.127 | 1.954 | -0.756 | 0.457 | -0.045 | -0.041 | -0.041 | 0.827 | 0.92 | 0.98 | 0.78 | 0.75 | 0.82 | 0.83 |
| 11 | 126780541 | rs6590235 | -0.675 | 1.884 | -0.517 | 0.364 | 0.001 | -0.059 | -0.059 | 2.218 | 0.13 | 0.45 | 0.08 | 0.05 | 0.11 | 0.09 |
| 11 | 126782785 | rs7928765 | 0.021 | 1.757 | -0.515 | -0.064 | -0.003 | -0.031 | -0.031 | 0 | 0.07 | 0.11 | 0.09 | 0 | 0.06 | 0.04 |
| 11 | 126786000 | rs12787569 | 0.35 | 1.931 | -0.895 | 0.753 | 0 | 0.048 | 0.048 | 0 | 0.1 | 0.12 | 0.07 | 0.41 | 0.14 | 0.21 |
| 11 | 126788587 | rs1793660 | 1.191 | 1.519 | -1.903 | -0.448 | -0.005 | 0.02 | 0.02 | 0.065 | 0.26 | 0.17 | 0.22 | 0.03 | 0.24 | 0.12 |
| 11 | 126790305 | rs7129859 | -0.837 | 1.686 | -1.631 | 0.08 | -0.016 | -0.095 | -0.095 | 0.01 | 0.26 | 0.59 | 0.2 | 0.05 | 0.22 | 0.15 |
| 11 | 126792589 | rs1620264 | -0.807 | 1.85 | -1.633 | 0.444 | -0.043 | -0.038 | -0.038 | 0 | 0.93 | 0.99 | 0.78 | 0.75 | 0.82 | 0.84 |
| 11 | 126807234 | rs6590236 | -1.158 | 1.91 | -1.657 | 0.137 | 0.004 | 0.001 | 0.001 | 0 | 0.34 | 0.6 | 0.39 | 0.7 | 0.41 | 0.43 |

| | | | | | | | | | | | | | | | | |
|---|---|---|---|---|---|---|---|---|---|---|---|---|---|---|---|---|
| 11 | 126808123 | rs1793675 | -0.269 | 2.032 | -1.751 | 0.156 | -0.01 | 0.01 | 0.01 | 0.068 | 0.94 | 0.96 | 0.87 | 0.79 | 0.88 | 0.91 |
| 11 | 126809705 | rs1939992 | 0.496 | 1.367 | -1.56 | -0.44 | 0.002 | -0.003 | -0.003 | 0 | 0.77 | 0.81 | 0.81 | 0.81 | 0.78 | 0.82 |
| 11 | 126817556 | rs1793673 | 0.173 | 1.322 | -1.565 | -0.464 | 0.007 | -0.014 | -0.014 | 1.317 | 0.16 | 0.14 | 0.1 | 0 | 0.1 | 0.09 |
| 11 | 126819397 | rs2000971 | -0.195 | 1.433 | -1.511 | -0.358 | -0.004 | 0.015 | 0.015 | 0 | 0.93 | 0.92 | 0.92 | 0.81 | 0.89 | 0.91 |
| 11 | 126821306 | rs1894224 | -0.798 | 1.445 | -1.451 | -0.336 | -0.021 | -0.01 | -0.01 | 0 | 0.95 | 0.99 | 0.96 | 0.91 | 0.9 | 0.95 |
| 11 | 126833355 | rs12283529 | 1.13 | 1.511 | -1.569 | -0.672 | -0.01 | 0.01 | 0.01 | 0 | 0.21 | 0.17 | 0.23 | 0.03 | 0.26 | 0.12 |
| 11 | 126835254 | rs7125248 | 0.72 | 1.649 | -1.348 | -0.674 | -0.001 | -0.013 | -0.013 | 1.042 | 0.07 | 0.09 | 0.04 | 0 | 0.06 | 0.02 |
| 11 | 126835621 | rs619086 | 0.332 | 1.622 | -1.342 | -0.664 | -0.002 | 0 | 0 | 0.092 | 0.87 | 0.84 | 0.87 | 0.81 | 0.83 | 0.89 |
| 11 | 126837505 | rs10893608 | 1.446 | 1.236 | -1.335 | -0.809 | 0.035 | 0.123 | 0.123 | 0 | 0.32 | 0.05 | 0.23 | 0.12 | 0.23 | 0.38 |
| 11 | 126847169 | rs11220685 | 0.986 | 1.17 | -0.696 | -0.274 | -0.016 | -0.012 | -0.012 | 0 | 0.05 | 0 | 0.09 | 0.05 | 0.05 | 0.18 |
| 11 | 126848207 | rs583795 | 0.061 | 0.974 | -0.65 | -0.503 | -0.001 | -0.035 | -0.035 | 0 | 0.18 | 0.17 | 0.12 | 0 | 0.13 | 0.11 |
| 11 | 126850764 | rs4937222 | 0.004 | 1.115 | -0.647 | 0.125 | 0.027 | 0.141 | 0.141 | 0.401 | 0.1 | 0.14 | 0.24 | 0.57 | 0.21 | 0.28 |
| 11 | 126851104 | rs1939996 | -0.643 | 1.101 | -0.105 | 0.058 | -0.004 | -0.006 | -0.006 | 0 | 0.16 | 0.26 | 0.08 | 0.15 | 0.13 | 0.08 |
| 11 | 126852438 | rs631562 | -1.978 | 1.023 | 0.077 | -0.01 | -0.002 | 0.105 | 0.105 | 0 | 0.46 | 0.68 | 0.49 | 0.82 | 0.47 | 0.5 |
| 11 | 126858161 | rs669765 | -1.279 | 0.324 | 0.16 | -0.162 | -0.009 | -0.063 | -0.063 | 0 | 0.29 | 0.3 | 0.18 | 0.05 | 0.22 | 0.17 |
| 11 | 126864307 | rs653856 | -1.099 | 0.296 | 0.362 | -0.075 | -0.04 | -0.039 | -0.039 | 0 | 0.3 | 0.4 | 0.18 | 0.11 | 0.17 | 0.16 |
| 11 | 126864588 | rs541783 | -0.748 | 0.33 | 0.245 | -0.117 | -0.013 | -0.07 | -0.07 | 0 | 0.24 | 0.27 | 0.14 | 0 | 0.13 | 0.12 |
| 11 | 126872785 | rs2508557 | -1.79 | 0.972 | 0.045 | 0.477 | 0.012 | 0.051 | 0.051 | 0 | 0.48 | 0.82 | 0.4 | 0.57 | 0.42 | 0.32 |
| 11 | 126880091 | rs684451 | -1.09 | 1.029 | 0.419 | 0.367 | -0.009 | -0.102 | -0.102 | 0 | 0.27 | 0.38 | 0.22 | 0.05 | 0.23 | 0.16 |
| 11 | 126883293 | rs11220708 | -0.213 | 1.024 | 0.331 | 0.484 | 0.012 | 0.143 | 0.143 | 0 | 0.16 | 0.39 | 0.15 | 0.42 | 0.16 | 0.12 |
| 11 | 126883361 | rs1944623 | -0.766 | 1.007 | 0.324 | 0.455 | 0.01 | 0.117 | 0.117 | 0 | 0.2 | 0.47 | 0.17 | 0.42 | 0.19 | 0.12 |
| 11 | 126883409 | rs611849 | 0.067 | 0.546 | 0.297 | 0.387 | -0.004 | 0.081 | 0.081 | 0 | 0.17 | 0.03 | 0.15 | 0 | 0.13 | 0.1 |
| 11 | 126883696 | rs610505 | -1.373 | 0.597 | 0.318 | 0.443 | -0.005 | 0.006 | 0.006 | 0 | 0.37 | 0.5 | 0.32 | 0.43 | 0.32 | 0.22 |
| 11 | 126883747 | rs1944622 | -0.315 | 0.548 | 0.662 | 0.6 | 0.006 | 0.051 | 0.051 | 0 | 0.16 | 0.28 | 0.14 | 0.32 | 0.16 | 0.12 |
| 11 | 126884845 | rs11220710 | -1.983 | 0.63 | 0.259 | 0.508 | -0.016 | -0.007 | -0.007 | 0 | 0.43 | 0.68 | 0.52 | 0.53 | 0.5 | 0.38 |
| 11 | 126885159 | rs11220712 | -0.912 | 0.793 | 0.353 | 0.459 | -0.008 | 0.006 | 0.006 | 0 | 0.24 | 0.2 | 0.34 | 0.07 | 0.27 | 0.19 |
| 11 | 126886470 | rs10466610 | -1.093 | 0.476 | 0.663 | 0.845 | 0.015 | 0.114 | 0.114 | 0.046 | 0.2 | 0.67 | 0.2 | 0.47 | 0.23 | 0.18 |
| 11 | 126887637 | rs1939228 | -0.86 | 0.441 | 0.614 | 0.865 | 0.033 | 0.115 | 0.115 | 0.107 | 0.18 | 0.51 | 0.18 | 0.36 | 0.22 | 0.15 |
| 11 | 126888992 | rs11600665 | -0.873 | 0.429 | 0.485 | 0.666 | -0.001 | -0.002 | -0.002 | 0.019 | 0.19 | 0.38 | 0.21 | 0.11 | 0.19 | 0.15 |
| 11 | 126891313 | rs11220720 | -0.775 | -0.031 | -0.355 | 0.511 | 0 | -0.001 | -0.001 | 0 | 0.39 | 0.37 | 0.42 | 0.49 | 0.29 | 0.36 |
| 11 | 126891391 | rs7102500 | -1.129 | -0.013 | -1.338 | 0.696 | -0.048 | 0.005 | 0.005 | 0 | 0.54 | 0.88 | 0.56 | 0.54 | 0.47 | 0.43 |
| 11 | 126894591 | rs7129845 | 0.005 | -0.329 | -1.078 | 1.346 | 0.039 | -0.078 | -0.078 | 0.037 | 0.12 | 0.03 | 0.07 | 0.29 | 0.05 | 0.1 |
| 11 | 126895307 | rs483644 | 0.007 | -0.247 | -0.679 | 0.845 | -0.011 | 0.026 | 0.026 | 0 | 0.69 | 0.61 | 0.67 | 0.51 | 0.72 | 0.7 |
| 11 | 126901479 | rs11220729 | -0.396 | 0.385 | -0.085 | 0.2 | 0.012 | -0.023 | -0.023 | 0.006 | 0.1 | 0.18 | 0.08 | 0.02 | 0.13 | 0.08 |
| 11 | 126904307 | rs508800 | -0.058 | 0.133 | -0.608 | 0.131 | 0.017 | -0.053 | -0.053 | 0.016 | 0.26 | 0.08 | 0.24 | 0.34 | 0.19 | 0.22 |
| 11 | 126910629 | rs7931141 | -0.516 | -0.006 | 0.431 | 0.04 | 0.092 | -0.062 | -0.062 | 0 | 0.12 | 0.48 | 0.11 | 0.02 | 0.13 | 0.08 |
| 11 | 126913352 | rs516926 | -0.06 | 0.067 | 0.507 | 0.056 | 0.012 | -0.027 | -0.027 | 0.62 | 0.26 | 0.13 | 0.24 | 0.33 | 0.2 | 0.22 |
| 11 | 126915282 | rs663268 | -0.83 | 0.162 | 0.106 | -0.205 | -0.001 | 0.003 | 0.003 | 0.02 | 0.69 | 0.69 | 0.57 | 0.48 | 0.7 | 0.47 |
| 11 | 126918141 | rs543766 | 0.425 | -0.08 | 0.059 | -0.107 | 0.006 | -0.062 | -0.062 | 1.555 | 0.25 | 0.07 | 0.23 | 0.33 | 0.2 | 0.22 |
| 11 | 126920647 | rs668605 | -0.779 | -0.07 | 0.248 | -0.129 | 0 | -0.001 | -0.001 | 0 | 0.37 | 0.47 | 0.34 | 0.36 | 0.33 | 0.3 |
| 11 | 126921306 | rs7116702 | 0.305 | 0.095 | 0.056 | -0.056 | 0.022 | -0.035 | -0.035 | 0 | 0.1 | 0.21 | 0.09 | 0.02 | 0.13 | 0.08 |
| 11 | 126925954 | rs593440 | 0.147 | -0.031 | -0.244 | -0.008 | 0.002 | -0.06 | -0.06 | 0 | 0.24 | 0.07 | 0.22 | 0.33 | 0.19 | 0.2 |
| 11 | 126942485 | rs619208 | 0.035 | -0.136 | -0.047 | -0.167 | 0 | 0 | 0 | 0 | 0.57 | 0.43 | 0.47 | 0.5 | 0.56 | 0.35 |
| 11 | 126945428 | rs11220749 | 1.36 | 0.515 | -0.186 | 0.06 | -0.073 | 0.049 | 0.049 | 0 | 0.27 | 0.06 | 0.22 | 0.12 | 0.35 | 0.13 |
| 11 | 126950321 | rs1000331 | -0.07 | 0.381 | 0.1 | -0.032 | 0 | -0.012 | -0.012 | 0 | 0.44 | 0.51 | 0.33 | 0.17 | 0.5 | 0.24 |
| 11 | 126963772 | rs606591 | -0.603 | 0.309 | -0.086 | 0.037 | 0.013 | -0.005 | -0.005 | 0 | 0.56 | 0.59 | 0.54 | 0.51 | 0.65 | 0.39 |
| 11 | 126964295 | rs593606 | -0.808 | 1.495 | 0.012 | 0.214 | -0.054 | -0.035 | -0.035 | 0 | 0.89 | 0.99 | 0.9 | 0.88 | 0.83 | 0.94 |
| 11 | 126965783 | rs684534 | -0.492 | 0.871 | 0.753 | 0.237 | -0.054 | -0.075 | -0.075 | 0 | 0.11 | 0 | 0.2 | 0.36 | 0.15 | 0.15 |
| 11 | 126988073 | rs11220787 | -2 | 0.427 | 1.091 | -0.155 | 0.018 | -0.061 | -0.061 | 0 | 0.13 | 0.29 | 0.11 | 0.03 | 0.14 | 0.09 |
| 11 | 126992504 | rs11220790 | -1.36 | 0.801 | 0.79 | 0.012 | 0.027 | -0.033 | -0.033 | 0 | 0.1 | 0.15 | 0.09 | 0.02 | 0.16 | 0.08 |
| 11 | 126997130 | rs6590245 | -2.455 | 0.817 | 0.886 | 0.278 | 0.03 | 0.048 | 0.048 | 0 | 0.21 | 0.32 | 0.29 | 0.33 | 0.25 | 0.17 |
| 11 | 126998347 | rs7122929 | -2.211 | 1.043 | 0.892 | 0.233 | 0.097 | -0.036 | -0.036 | 0.458 | 0.11 | 0.34 | 0.09 | 0.04 | 0.13 | 0.06 |
| 11 | 127004948 | rs7121794 | -0.787 | 0.598 | 0.068 | 0.459 | 0.002 | 0.044 | 0.044 | 0 | 0.09 | 0.08 | 0.2 | 0.27 | 0.1 | 0.1 |
| 11 | 127012571 | rs7115915 | -1.296 | 0.577 | 0.044 | 0.451 | -0.003 | 0.061 | 0.061 | 0.003 | 0.12 | 0.2 | 0.21 | 0.28 | 0.1 | 0.11 |
| 11 | 127013570 | rs12291238 | -0.926 | 0.696 | -0.482 | 0.386 | 0.002 | -0.019 | -0.019 | 0 | 0.1 | 0.12 | 0.04 | 0.03 | 0.07 | 0.03 |
| 11 | 127028328 | rs4603317 | -0.978 | 0.58 | -0.668 | 0.743 | -0.003 | -0.011 | -0.011 | 0.132 | 0.1 | 0.06 | 0.2 | 0.27 | 0.1 | 0.1 |
| 11 | 127031282 | rs7105067 | -0.454 | 0.074 | -0.48 | -0.787 | -0.019 | -0.121 | -0.121 | 0.01 | 0.34 | 0.39 | 0.21 | 0.02 | 0.28 | 0.19 |
| 11 | 127031820 | rs2513427 | -0.078 | -0.142 | -0.315 | -1.26 | -0.006 | -0.029 | -0.029 | 0 | 0.17 | 0.22 | 0.08 | 0.06 | 0.12 | 0.04 |
| 11 | 127037625 | rs2510705 | -0.79 | -0.268 | -0.087 | -2.368 | 0.001 | -0.007 | -0.007 | 0.189 | 0.5 | 0.52 | 0.3 | 0.08 | 0.49 | 0.29 |
| 11 | 127044915 | rs12799000 | 0.323 | -0.349 | 0.114 | -2.514 | -0.009 | 0.049 | 0.049 | 0 | 0.31 | 0.17 | 0.19 | 0 | 0.34 | 0.23 |
| 11 | 127048359 | rs1944828 | 0.731 | -0.374 | 0.051 | -2.523 | -0.001 | -0.016 | -0.016 | 0 | 0.16 | 0.17 | 0.08 | 0.06 | 0.14 | 0.05 |
| 11 | 127049090 | rs1944826 | -1.232 | -0.536 | -0.127 | -2.307 | 0 | -0.012 | -0.012 | 0 | 0.59 | 0.55 | 0.44 | 0.17 | 0.57 | 0.52 |
| 11 | 127053249 | rs2513444 | -0.497 | -0.41 | -0.29 | -2.128 | 0 | 0.004 | 0.004 | 0 | 0.25 | 0.22 | 0.22 | 0.15 | 0.23 | 0.27 |
| 11 | 127054272 | rs12417818 | 0.228 | -0.4 | -0.478 | -1.961 | -0.001 | -0.006 | -0.006 | 0 | 0.09 | 0.07 | 0.15 | 0.08 | 0.09 | 0.23 |
| 11 | 127057148 | rs6590247 | -1.175 | -0.839 | -0.685 | -2.171 | -0.087 | 0.064 | 0.064 | 0 | 0.66 | 0.91 | 0.46 | 0.16 | 0.55 | 0.53 |
| 11 | 127057362 | rs12288215 | -0.634 | -1.243 | -0.505 | -2.385 | -0.007 | -0.001 | -0.001 | 0 | 0.18 | 0.2 | 0.15 | 0.09 | 0.08 | 0.26 |
| 11 | 127058123 | rs10466612 | -0.383 | -1.286 | 0.041 | -2.41 | -0.058 | -0.083 | -0.083 | 0 | 0.16 | 0.25 | 0.05 | 0 | 0.04 | 0.04 |
| 11 | 127058947 | rs7121129 | -1.64 | -1.21 | -0.209 | -2.378 | -0.131 | 0.009 | 0.009 | 0 | 0.69 | 0.96 | 0.48 | 0.18 | 0.58 | 0.56 |
| 11 | 127059596 | rs11606949 | 0.407 | -1.213 | -0.093 | -2.45 | -0.008 | 0.025 | 0.025 | 0 | 0.36 | 0.19 | 0.23 | 0.01 | 0.37 | 0.24 |

| | | | | | | | | | | | | | | | | |
|---|---|---|---|---|---|---|---|---|---|---|---|---|---|---|---|---|
| 11 | 127063006 | rs11220822 | 0.934 | -1.242 | -0.242 | -2.206 | -0.03 | 0.026 | 0.026 | 0 | 0.35 | 0.2 | 0.58 | 0.71 | 0.4 | 0.44 |
| 11 | 127063240 | rs10790866 | -0.79 | -1.257 | -0.809 | -2.178 | -0.066 | 0.156 | 0.156 | 0 | 0.67 | 0.95 | 0.74 | 0.93 | 0.59 | 0.74 |
| 11 | 127065614 | rs11220823 | 1.274 | -0.374 | -0.904 | -0.69 | -0.075 | -0.09 | -0.09 | 0 | 0.13 | 0.01 | 0.23 | 0.3 | 0.19 | 0.22 |
| 11 | 127069203 | rs12285996 | -0.694 | -0.873 | -0.763 | -0.85 | -0.007 | -0.002 | -0.002 | 0 | 0.18 | 0.21 | 0.07 | 0.14 | 0.08 | 0.25 |
| 11 | 127081591 | rs4529888 | -0.472 | -0.537 | -1.352 | -1.278 | -0.026 | 0.117 | 0.117 | 0 | 0.65 | 0.86 | 0.72 | 0.95 | 0.59 | 0.76 |
| 11 | 127083003 | rs12807480 | -0.022 | -0.837 | -2.02 | -1.472 | 0.043 | 0.02 | 0.02 | 0.092 | 0.3 | 0.06 | 0.12 | 0.29 | 0.2 | 0.38 |
| 11 | 127085601 | rs2513439 | -1.252 | -0.895 | -2.189 | -1.671 | -0.025 | 0.084 | 0.084 | 0 | 0.71 | 0.89 | 0.77 | 0.95 | 0.63 | 0.77 |
| 11 | 127085838 | rs2514893 | -1.189 | -1.068 | -1.161 | -1.74 | 0.004 | -0.074 | -0.074 | 0 | 0.68 | 0.65 | 0.72 | 0.95 | 0.63 | 0.77 |
| 11 | 127087754 | rs10893653 | 1.997 | -0.782 | -1.193 | -0.202 | -0.004 | 0.032 | 0.032 | 0 | 0.07 | 0.04 | 0.17 | 0.33 | 0.07 | 0.14 |
| 11 | 127091963 | rs2514897 | -0.617 | -1.023 | -0.541 | -0.103 | 0.008 | 0.007 | 0.007 | 0 | 0.44 | 0.42 | 0.32 | 0.68 | 0.3 | 0.54 |
| 11 | 127092018 | rs2708507 | 1.235 | -1.419 | -1.201 | -0.131 | 0 | -0.034 | -0.034 | 0 | 0.2 | 0.04 | 0.07 | 0.24 | 0.15 | 0.29 |
| 11 | 127095895 | rs11220836 | 1.864 | -1.367 | -1.175 | -0.22 | -0.084 | -0.061 | -0.061 | 0 | 0.18 | 0.05 | 0.35 | 0.27 | 0.3 | 0.21 |
| 11 | 127102481 | rs2441686 | -2.09 | -1.357 | -1.071 | -0.393 | -0.006 | 0.018 | 0.018 | 0 | 0.7 | 0.78 | 0.76 | 0.95 | 0.65 | 0.77 |
| 11 | 127103711 | rs2441687 | 0 | -1.47 | -1.477 | -0.274 | 0.032 | -0.026 | -0.026 | 3.864 | 0.28 | 0.09 | 0.13 | 0.31 | 0.2 | 0.37 |
| 11 | 127105446 | rs2464870 | -2.073 | -1.328 | -1.562 | -0.454 | -0.031 | 0.092 | 0.092 | 0 | 0.76 | 0.96 | 0.83 | 0.95 | 0.72 | 0.85 |
| 11 | 127107481 | rs2243883 | 1.283 | -1.532 | -1.035 | -0.064 | -0.006 | 0 | 0 | 0 | 0.1 | 0.1 | 0.05 | 0.08 | 0.05 | 0.08 |
| 11 | 127109457 | rs7924365 | 1.518 | -1.502 | -0.926 | -0.139 | 0.002 | -0.005 | -0.005 | 0 | 0.07 | 0.08 | 0.04 | 0 | 0.05 | 0.01 |
| 11 | 127111563 | rs2513432 | -0.817 | -1.729 | -1.058 | 0.398 | 0.001 | -0.001 | -0.001 | 0 | 0.32 | 0.68 | 0.24 | 0.3 | 0.18 | 0.29 |
| 11 | 127114387 | rs2322662 | 1.09 | -1.324 | -1.492 | 0.423 | 0.107 | 0.006 | 0.006 | 0.564 | 0.5 | 0.18 | 0.69 | 0.46 | 0.68 | 0.49 |
| 11 | 127114435 | rs2708529 | 0.55 | -0.649 | -1.328 | 0.745 | 0.001 | -0.024 | -0.024 | 0 | 0.13 | 0.06 | 0.11 | 0.21 | 0.11 | 0.19 |
| 11 | 127116693 | rs1105957 | 1.066 | 0.003 | -1.314 | 0.835 | -0.095 | -0.058 | -0.058 | 6.167 | 0.19 | 0.05 | 0.36 | 0.27 | 0.31 | 0.22 |
| 11 | 127117280 | rs2592929 | 0.489 | -0.355 | -0.881 | 0.969 | -0.005 | -0.001 | -0.001 | 0.015 | 0.09 | 0.06 | 0.09 | 0.21 | 0.1 | 0.13 |
| 11 | 127119772 | rs4297477 | -1.374 | -0.796 | -0.926 | 0.861 | 0.015 | -0.034 | -0.034 | 0 | 0.93 | 1 | 0.96 | 0.91 | 0.98 | 0.95 |
| 11 | 127124975 | rs12223971 | -0.788 | -1.527 | -0.425 | 0.721 | -0.021 | 0.013 | 0.013 | 2.805 | 0.28 | 0.34 | 0.16 | 0.34 | 0.16 | 0.38 |
| 11 | 127128729 | rs11220845 | 1.537 | -1.688 | -0.752 | 0.387 | -0.023 | 0.013 | 0.013 | 0 | 0.26 | 0.12 | 0.31 | 0.19 | 0.34 | 0.24 |
| 11 | 127131725 | rs11220847 | 0.887 | -1.618 | -0.444 | 0.293 | -0.033 | 0.006 | 0.006 | 0.008 | 0.14 | 0.2 | 0.06 | 0.14 | 0.04 | 0.15 |
| 11 | 127132738 | rs2874843 | 0.732 | -1.538 | -0.651 | 0.21 | -0.047 | 0.013 | 0.013 | 0.284 | 0.35 | 0.23 | 0.45 | 0.38 | 0.47 | 0.34 |
| 11 | 127154987 | rs10893663 | -0.284 | -1.506 | -0.176 | 0.298 | -0.049 | -0.021 | -0.021 | 0 | 0.18 | 0.35 | 0.09 | 0.12 | 0.06 | 0.17 |
| 11 | 127156732 | rs2322732 | -1.231 | -1.085 | -0.751 | 0.499 | 0.015 | -0.033 | -0.033 | 0 | 0.93 | 1 | 0.96 | 0.91 | 0.97 | 0.95 |
| 11 | 127164405 | rs2618195 | -0.557 | -1.466 | -0.674 | 0.103 | -0.02 | -0.028 | -0.028 | 0 | 0.3 | 0.58 | 0.18 | 0.19 | 0.2 | 0.41 |
| 11 | 127169960 | rs2618202 | -0.465 | -1.423 | -0.863 | 0.105 | 0.011 | 0.02 | 0.02 | 0.058 | 0.52 | 0.37 | 0.53 | 0.63 | 0.54 | 0.6 |
| 11 | 127170383 | rs2012959 | 1.224 | -1.33 | -0.879 | 0.058 | 0 | -0.005 | -0.005 | 3.888 | 0.15 | 0.15 | 0.1 | 0.07 | 0.16 | 0.23 |
| 11 | 127171504 | rs1817406 | 1.491 | -1.244 | -1.113 | 0.152 | -0.023 | 0.009 | 0.009 | 0 | 0.3 | 0.17 | 0.37 | 0.25 | 0.4 | 0.23 |
| 11 | 127178763 | rs4486638 | 0.856 | -1.314 | -1.436 | 0.46 | -0.023 | -0.016 | -0.016 | 0 | 0.21 | 0.17 | 0.39 | 0.3 | 0.3 | 0.22 |
| 11 | 127183248 | rs1824820 | 0.054 | -1.171 | -1.287 | 0.362 | -0.024 | -0.047 | -0.047 | 0 | 0.34 | 0.21 | 0.42 | 0.55 | 0.36 | 0.36 |
| 11 | 127184036 | rs1038098 | -1.227 | -0.879 | -0.959 | 0.58 | -0.024 | -0.003 | -0.003 | 0.003 | 0.87 | 0.84 | 0.95 | 0.88 | 0.93 | 0.83 |
| 11 | 127184582 | rs11220862 | 0.791 | -0.218 | -1.191 | 0.527 | -0.02 | 0.012 | 0.012 | 0 | 0.31 | 0.21 | 0.39 | 0.24 | 0.42 | 0.22 |
| 11 | 127196756 | rs10893666 | 0.313 | 0.137 | -1.232 | 0.18 | -0.037 | 0.031 | 0.031 | 0 | 0.38 | 0.21 | 0.44 | 0.27 | 0.46 | 0.31 |
| 11 | 127198493 | rs10893669 | 0.438 | 0.226 | -1.05 | 0.173 | -0.007 | 0.004 | 0.004 | 0 | 0.34 | 0.29 | 0.3 | 0.23 | 0.43 | 0.29 |
| 11 | 127199548 | rs2697718 | 0.254 | -0.547 | -1.199 | -0.179 | -0.019 | -0.026 | -0.026 | 0 | 0.57 | 0.49 | 0.8 | 0.75 | 0.75 | 0.47 |
| 11 | 127200217 | rs983786 | -0.652 | -0.857 | -1.584 | -0.39 | -0.042 | 0.059 | 0.059 | 0 | 0.8 | 0.99 | 0.9 | 0.94 | 0.82 | 0.71 |
| 11 | 127201917 | rs2708510 | 0.41 | -0.878 | -1.313 | -0.33 | -0.022 | -0.029 | -0.029 | 0 | 0.58 | 0.51 | 0.81 | 0.75 | 0.74 | 0.46 |
| 11 | 127209645 | rs12364837 | 2.513 | 0.105 | -1.971 | -0.442 | -0.084 | 0.012 | 0.012 | 0 | 0.22 | 0.04 | 0.21 | 0.15 | 0.35 | 0.15 |
| 11 | 127211003 | rs1116073 | 0.437 | -0.075 | -1.739 | 0.153 | -0.003 | 0.001 | 0.001 | 0 | 0.26 | 0.27 | 0.49 | 0.27 | 0.32 | 0.14 |
| 11 | 127211607 | rs1478632 | -1.119 | -0.429 | -1.724 | 0.102 | -0.046 | 0.052 | 0.052 | 0 | 0.8 | 0.99 | 0.9 | 0.94 | 0.82 | 0.71 |
| 11 | 127213078 | rs2592908 | 0.161 | -0.509 | -1.591 | 0.158 | -0.014 | -0.023 | -0.023 | 0.02 | 0.58 | 0.49 | 0.8 | 0.76 | 0.75 | 0.46 |
| 11 | 127215405 | rs2708520 | 0.304 | -0.352 | -1.395 | 0.268 | -0.063 | -0.042 | -0.042 | 0 | 0.26 | 0.49 | 0.1 | 0.19 | 0.1 | 0.25 |
| 11 | 127232888 | rs2618206 | 1.941 | -0.123 | -1.33 | 0.471 | 0.005 | 0.141 | 0.141 | 0 | 0.08 | 0.15 | 0.1 | 0.32 | 0.07 | 0.12 |
| 11 | 127241251 | rs4937230 | 0.248 | 0.094 | -1.423 | 0.039 | 0.011 | 0.012 | 0.012 | 0 | 0.53 | 0.45 | 0.8 | 0.7 | 0.73 | 0.4 |
| 11 | 127242451 | rs2592916 | 0.495 | 0.145 | -0.885 | -0.058 | -0.009 | -0.011 | -0.011 | 0.193 | 0.24 | 0.26 | 0.09 | 0.18 | 0.1 | 0.29 |
| 11 | 127243632 | rs2697730 | -0.288 | 0.528 | -0.975 | -0.127 | 0.005 | -0.027 | -0.027 | 0 | 0.73 | 0.71 | 0.67 | 0.88 | 0.67 | 0.63 |
| 11 | 127266151 | rs10790878 | -0.541 | -0.491 | -1.042 | -0.445 | 0.063 | 0.014 | 0.014 | 0.064 | 0.44 | 0.76 | 0.37 | 0.46 | 0.25 | 0.39 |
| 11 | 127268774 | rs7108023 | -0.934 | -0.191 | -0.354 | -0.564 | 0.015 | 0.147 | 0.147 | 0.125 | 0.55 | 0.91 | 0.47 | 0.8 | 0.33 | 0.57 |
| 11 | 127269143 | rs11220893 | 0.647 | -0.229 | -0.374 | -0.619 | -0.008 | 0.014 | 0.014 | 0.072 | 0.18 | 0.26 | 0.26 | 0.27 | 0.15 | 0.11 |
| 11 | 127275361 | rs7940571 | -0.567 | -0.218 | -0.526 | -0.567 | 0.041 | -0.032 | -0.032 | 0 | 0.38 | 0.7 | 0.21 | 0.53 | 0.18 | 0.46 |
| 11 | 127290881 | rs10893681 | -1.063 | 0.899 | -0.086 | -0.669 | -0.011 | 0.254 | 0.254 | 0 | 0.5 | 0.83 | 0.68 | 0.78 | 0.47 | 0.53 |
| 11 | 127293049 | rs1600223 | 0.085 | 0.853 | 0.172 | -0.026 | 0.023 | -0.025 | -0.025 | 4.773 | 0.25 | 0.13 | 0.29 | 0.37 | 0.16 | 0.32 |
| 11 | 127300006 | rs1919265 | -0.336 | 1.259 | 0.246 | -1.125 | -0.001 | -0.003 | -0.003 | 1.774 | 0.8 | 0.81 | 0.71 | 0.88 | 0.76 | 0.68 |
| 11 | 127305084 | rs10893684 | 0.188 | 1.393 | 0.107 | -1.157 | 0.017 | 0.042 | 0.042 | 0 | 0.4 | 0.3 | 0.42 | 0.72 | 0.29 | 0.43 |
| 11 | 127305649 | rs11220916 | 0.334 | 1.744 | -0.537 | -1.011 | -0.022 | 0.059 | 0.059 | 0 | 0.39 | 0.31 | 0.27 | 0.16 | 0.47 | 0.26 |
| 11 | 127308577 | rs1108906 | -0.265 | 1.127 | -0.678 | -0.939 | -0.004 | 0.028 | 0.028 | 0 | 0.75 | 0.85 | 0.68 | 0.89 | 0.72 | 0.66 |
| 11 | 127309674 | rs12288932 | 0.213 | 1.243 | -0.688 | -0.803 | -0.019 | 0.067 | 0.067 | 0 | 0.38 | 0.28 | 0.25 | 0.17 | 0.44 | 0.19 |
| 11 | 127315468 | rs10736563 | 0.384 | 1.34 | -1.182 | 0.438 | -0.014 | 0.012 | 0.012 | 0 | 0.75 | 0.86 | 0.67 | 0.82 | 0.69 | 0.64 |
| 11 | 127324160 | rs7124157 | -0.766 | 1.342 | -1.236 | 0.378 | -0.009 | 0.011 | 0.011 | 0 | 0.23 | 0.18 | 0.25 | 0.16 | 0.27 | 0.18 |
| 11 | 127327783 | rs1881474 | -0.492 | 1.092 | -2.278 | 0.336 | -0.02 | -0.029 | -0.029 | 0 | 0.76 | 0.96 | 0.84 | 0.79 | 0.78 | 0.83 |
| 11 | 127328188 | rs16831 | 0.948 | 1.154 | -2.048 | 0.177 | -0.013 | -0.023 | -0.023 | 0.152 | 0.35 | 0.57 | 0.28 | 0.24 | 0.24 | 0.44 |
| 11 | 127331680 | rs7924331 | -0.384 | 1.695 | -2.106 | 0.099 | -0.057 | -0.041 | -0.041 | 0 | 0.67 | 0.86 | 0.6 | 0.43 | 0.64 | 0.67 |
| 11 | 127333780 | rs11605581 | 0.639 | 1.667 | -2.074 | 0.089 | -0.025 | 0.04 | 0.04 | 0 | 0.08 | 0.02 | 0.06 | 0.03 | 0.12 | 0.06 |
| 11 | 127333828 | rs10790882 | -0.144 | 1.664 | -2.022 | 0.092 | -0.003 | 0.003 | 0.003 | 0 | 0.32 | 0.27 | 0.31 | 0.19 | 0.4 | 0.25 |

| | | | | | | | | | | | | | | | | |
|---|---|---|---|---|---|---|---|---|---|---|---|---|---|---|---|---|
| 11 | 127338676 | rs10893691 | -0.181 | 1.669 | -2.004 | 0.172 | 0.001 | -0.005 | -0.005 | 0 | 0.32 | 0.3 | 0.32 | 0.21 | 0.4 | 0.26 |
| 11 | 127341324 | rs7121194 | 0.373 | 1.407 | -1.814 | 0.224 | 0.022 | 0.05 | 0.05 | 0 | 0.44 | 0.67 | 0.41 | 0.28 | 0.36 | 0.49 |
| 11 | 127343316 | rs1919262 | -0.906 | 1.359 | -1.878 | 1.053 | -0.014 | -0.123 | -0.123 | 0 | 0.76 | 0.96 | 0.83 | 0.63 | 0.79 | 0.82 |
| 11 | 127345274 | rs7930764 | 0.321 | 1.124 | -1.885 | -0.207 | -0.002 | 0 | 0 | 0 | 0.7 | 0.31 | 0.7 | 0.97 | 0.71 | 0.78 |
| 11 | 127350959 | rs10893695 | 0.306 | 0.763 | -2.176 | 0.374 | -0.056 | -0.146 | -0.146 | 0 | 0.3 | 0.07 | 0.29 | 0.45 | 0.32 | 0.2 |
| 11 | 127355370 | rs10790886 | 0.306 | 0.763 | -2.176 | 0.381 | -0.056 | -0.149 | -0.149 | 9.008 | 0.3 | 0.07 | 0.29 | 0.44 | 0.32 | 0.2 |
| 11 | 127356981 | rs930170 | -1.025 | 0.415 | -2.565 | 0.26 | -0.095 | 0.022 | 0.022 | 0 | 0.77 | 0.98 | 0.78 | 0.82 | 0.7 | 0.76 |
| 11 | 127360025 | rs930167 | 0.228 | 0.315 | -1.242 | 0.126 | 0 | 0 | 0 | 0 | 0.33 | 0.31 | 0.32 | 0.45 | 0.32 | 0.2 |
| 11 | 127393262 | rs12790344 | 0.053 | 0.021 | -0.953 | 0.104 | -0.01 | -0.082 | -0.082 | 0.109 | 0.19 | 0.08 | 0.21 | 0.45 | 0.19 | 0.16 |
| 11 | 127394822 | rs7947776 | 0.605 | -0.243 | -0.221 | 0.07 | 0.001 | 0.036 | 0.036 | 0 | 0.51 | 0.35 | 0.51 | 0.41 | 0.51 | 0.54 |
| 11 | 127434049 | rs356247 | -0.392 | -0.002 | 0.176 | -0.113 | 0.007 | -0.055 | -0.055 | 0 | 0.72 | 0.65 | 0.72 | 0.86 | 0.68 | 0.69 |
| 11 | 127434574 | rs7935008 | -0.217 | 0.242 | 0.007 | 0.565 | 0.016 | 0.046 | 0.046 | 0 | 0.08 | 0.17 | 0.15 | 0.19 | 0.12 | 0.06 |
| 11 | 127437194 | rs356263 | -0.247 | -0.616 | -0.232 | 0.603 | 0 | 0.067 | 0.067 | 0.006 | 0.16 | 0.05 | 0.09 | 0 | 0.1 | 0.12 |
| 11 | 127440032 | rs4937242 | 0.624 | -1.186 | 0.001 | 0.509 | -0.004 | 0.045 | 0.045 | 0 | 0.48 | 0.33 | 0.42 | 0.27 | 0.53 | 0.36 |
| 11 | 127440320 | rs4937244 | 0.595 | -1.196 | 0.011 | 0.507 | -0.002 | 0.044 | 0.044 | 0 | 0.48 | 0.35 | 0.43 | 0.27 | 0.53 | 0.36 |
| 11 | 127444265 | rs356258 | -1.18 | -1.228 | -0.262 | 0.411 | -0.091 | 0.131 | 0.131 | 0 | 0.76 | 0.97 | 0.76 | 0.92 | 0.69 | 0.71 |
| 11 | 127445359 | rs356256 | -0.997 | -1.193 | -0.222 | 0.438 | -0.05 | -0.001 | -0.001 | 0.243 | 0.74 | 0.89 | 0.7 | 0.71 | 0.67 | 0.68 |
| 11 | 127446890 | rs10893721 | 0.706 | -1.098 | 0 | 0.208 | -0.006 | 0.058 | 0.058 | 0 | 0.48 | 0.29 | 0.42 | 0.27 | 0.53 | 0.36 |
| 11 | 127451064 | rs7109626 | -0.634 | -0.429 | 0.379 | 0.054 | -0.032 | -0.052 | -0.052 | 0 | 0.14 | 0.35 | 0.08 | 0.04 | 0.06 | 0.07 |
| 11 | 127454140 | rs4937245 | 1.097 | -0.539 | 0.193 | 0.271 | 0.001 | -0.009 | -0.009 | 1.798 | 0.33 | 0.23 | 0.41 | 0.61 | 0.34 | 0.37 |
| 11 | 127459104 | rs7129013 | 0.367 | -0.579 | 0.531 | -0.193 | -0.025 | -0.037 | -0.037 | 0.032 | 0.13 | 0.22 | 0.06 | 0.04 | 0.05 | 0.06 |
| 11 | 127464040 | rs11220985 | -1.177 | -0.536 | 0.408 | -0.399 | -0.045 | 0.117 | 0.117 | 0 | 0.68 | 0.87 | 0.66 | 0.88 | 0.63 | 0.63 |
| 11 | 127465371 | rs7946281 | 0.019 | -0.523 | 0.486 | -0.378 | 0.005 | 0.058 | 0.058 | 0.001 | 0.54 | 0.34 | 0.57 | 0.84 | 0.58 | 0.57 |
| 11 | 127471085 | rs2082273 | -0.188 | -0.423 | 0.513 | -0.328 | -0.004 | 0.068 | 0.068 | 0 | 0.57 | 0.69 | 0.6 | 0.85 | 0.58 | 0.57 |
| 11 | 127473190 | rs1431572 | 1.815 | -0.354 | 0.292 | 0.049 | 0.002 | -0.009 | -0.009 | 0 | 0.27 | 0.21 | 0.33 | 0.37 | 0.27 | 0.22 |
| 11 | 127478073 | rs7927195 | -0.784 | -0.242 | 0.425 | -0.431 | 0.001 | -0.029 | -0.029 | 0 | 0.67 | 0.67 | 0.63 | 0.88 | 0.63 | 0.63 |
| 11 | 127481098 | rs88493 | -0.159 | -0.428 | 0.281 | 0.146 | -0.017 | 0.052 | 0.052 | 0 | 0.19 | 0.08 | 0.27 | 0.12 | 0.19 | 0.32 |
| 11 | 127483747 | rs356240 | -1.28 | -0.381 | 0.269 | 0.205 | -0.003 | -0.019 | -0.019 | 0.018 | 0.32 | 0.65 | 0.36 | 0.15 | 0.25 | 0.38 |
| 11 | 127494706 | rs356274 | -0.205 | -0.382 | 0.297 | 0.648 | -0.013 | -0.019 | -0.019 | 0 | 0.18 | 0.08 | 0.31 | 0.22 | 0.21 | 0.35 |
| 11 | 127499751 | rs7928097 | 0.423 | -0.531 | -0.073 | 0.579 | -0.022 | -0.038 | -0.038 | 0 | 0.21 | 0.41 | 0.1 | 0.08 | 0.15 | 0.15 |
| 11 | 127504076 | rs7932225 | 0.54 | -0.556 | 0.032 | 0.567 | -0.01 | -0.017 | -0.017 | 0.202 | 0.2 | 0.31 | 0.09 | 0.08 | 0.15 | 0.15 |
| 11 | 127510799 | rs356272 | -2.355 | -0.487 | -1.106 | 1.147 | -0.055 | 0.036 | 0.036 | 0.948 | 0.61 | 0.83 | 0.6 | 0.7 | 0.54 | 0.74 |
| 11 | 127516223 | rs7945994 | -0.649 | -0.591 | -1.046 | 0.523 | 0.014 | -0.029 | -0.029 | 0 | 0.33 | 0.7 | 0.22 | 0.48 | 0.27 | 0.37 |
| 11 | 127520855 | rs356269 | -0.02 | -0.531 | -1.056 | 0.539 | -0.041 | -0.08 | -0.08 | 0 | 0.21 | 0.09 | 0.39 | 0.44 | 0.29 | 0.4 |
| 11 | 127524846 | rs12285335 | -0.251 | -0.539 | -0.886 | 0.56 | -0.006 | -0.004 | -0.004 | 0.153 | 0.27 | 0.65 | 0.14 | 0.17 | 0.21 | 0.23 |
| 11 | 127532335 | rs12276902 | 1.389 | -0.46 | -1.118 | 0.574 | 0.002 | 0.013 | 0.013 | 0 | 0.16 | 0.11 | 0.06 | 0.06 | 0.15 | 0.11 |
| 11 | 127532905 | rs3019141 | -2.368 | -0.296 | -1.621 | 0.584 | -0.04 | 0.138 | 0.138 | 1.434 | 0.63 | 0.85 | 0.65 | 0.91 | 0.59 | 0.78 |
| 11 | 127544174 | rs7115946 | 0.353 | 0.311 | -1.703 | 0.95 | -0.052 | -0.098 | -0.098 | 0 | 0.14 | 0.02 | 0.32 | 0.44 | 0.21 | 0.32 |
| 11 | 127549639 | rs2928040 | -0.462 | 0.126 | -1.522 | 1.085 | -0.036 | -0.081 | -0.081 | 0 | 0.29 | 0.12 | 0.43 | 0.46 | 0.33 | 0.43 |
| 11 | 127558438 | rs7123770 | -0.11 | -0.192 | -0.832 | 1.311 | -0.023 | -0.061 | -0.061 | 0 | 0.2 | 0.45 | 0.1 | 0.06 | 0.15 | 0.11 |
| 11 | 127561328 | rs34232636 | -0.372 | 0.373 | -1.649 | 1.939 | -0.017 | -0.005 | -0.005 | 1.829 | 0.05 | 0 | 0.08 | 0.3 | 0.05 | 0.13 |
| 11 | 127589360 | rs12420554 | -0.639 | 0.299 | -1.744 | 1.625 | -0.018 | -0.084 | -0.084 | 0 | 0.12 | 0.01 | 0.31 | 0.48 | 0.11 | 0.27 |
| 11 | 127595384 | rs11221015 | -0.705 | 0.526 | -1.774 | 1.508 | -0.026 | -0.08 | -0.08 | 0 | 0.14 | 0.04 | 0.32 | 0.43 | 0.15 | 0.24 |
| 11 | 127598505 | rs10893726 | -0.623 | -0.019 | -1.831 | 0.96 | 0.035 | 0.008 | 0.008 | 0 | 0.45 | 0.65 | 0.49 | 0.59 | 0.33 | 0.45 |
| 11 | 127601985 | rs11221020 | -0.344 | 0.01 | -2.111 | 0.66 | -0.04 | -0.053 | -0.053 | 0 | 0.09 | 0.01 | 0.08 | 0.16 | 0.13 | 0.11 |
| 11 | 127608001 | rs7925665 | -1.501 | 0.341 | -2.6 | 0.145 | -0.067 | 0.152 | 0.152 | 1.179 | 0.63 | 0.85 | 0.65 | 0.94 | 0.53 | 0.76 |
| 11 | 127611538 | rs10893730 | 0.016 | 0.378 | -2.49 | 0.44 | -0.042 | -0.055 | -0.055 | 0 | 0.09 | 0.01 | 0.09 | 0.17 | 0.13 | 0.09 |
| 11 | 127636522 | rs11221036 | -0.036 | 0.374 | -2.492 | 0.342 | -0.043 | -0.053 | -0.053 | 0 | 0.09 | 0.01 | 0.08 | 0.16 | 0.13 | 0.08 |
| 11 | 127646077 | rs11221044 | 0.32 | 0.352 | -2.465 | 0.372 | -0.033 | -0.036 | -0.036 | 0.097 | 0.06 | 0.01 | 0.08 | 0.16 | 0.12 | 0.07 |
| 11 | 127657071 | rs10790919 | 0.408 | -0.008 | -2.487 | 0.321 | 0.048 | -0.073 | -0.073 | 0 | 0.32 | 0.12 | 0.37 | 0.44 | 0.22 | 0.39 |
| 11 | 127664305 | rs4331121 | -1.716 | 0.931 | -2.623 | -0.863 | -0.068 | 0.14 | 0.14 | 3.353 | 0.77 | 0.97 | 0.83 | 1 | 0.71 | 0.96 |
| 11 | 127664841 | rs10893749 | 0.304 | 0.25 | -2.357 | -0.485 | 0.042 | 0.065 | 0.065 | 0.09 | 0.42 | 0.18 | 0.47 | 0.75 | 0.35 | 0.55 |
| 11 | 127671315 | rs1425220 | -0.329 | 0.253 | -2.186 | -0.455 | 0.036 | 0.076 | 0.076 | 0 | 0.12 | 0.57 | 0.13 | 0.16 | 0.13 | 0.07 |
| 11 | 127675888 | rs890135 | -0.252 | 0.456 | -2.111 | -0.78 | -0.01 | 0.102 | 0.102 | 0.209 | 0.55 | 0.69 | 0.62 | 0.91 | 0.49 | 0.63 |
| 11 | 127677874 | rs985292 | -0.039 | 0.648 | -1.936 | -0.518 | -0.009 | 0.079 | 0.079 | 0 | 0.62 | 0.75 | 0.65 | 0.8 | 0.49 | 0.65 |
| 11 | 127681994 | rs1157862 | -0.532 | 0.71 | -1.994 | 0.563 | -0.058 | -0.119 | -0.119 | 0.212 | 0.15 | 0.02 | 0.36 | 0.48 | 0.19 | 0.28 |
| 11 | 127686655 | rs11221063 | -0.292 | 0.344 | -1.629 | -0.347 | -0.002 | -0.014 | -0.014 | 0 | 0.19 | 0.18 | 0.13 | 0.04 | 0.15 | 0.18 |
| 11 | 127689882 | rs11221065 | -0.789 | 0.568 | -1.317 | -0.054 | -0.006 | -0.008 | -0.008 | 0 | 0.24 | 0.17 | 0.43 | 0.76 | 0.28 | 0.54 |
| 11 | 127697825 | rs1364784 | -0.99 | 0.381 | -1.605 | -0.488 | -0.068 | 0.11 | 0.11 | 0.683 | 0.63 | 0.8 | 0.66 | 0.85 | 0.55 | 0.81 |
| 11 | 127700566 | rs4277128 | 0.346 | 0.023 | -1.027 | 0.038 | -0.015 | 0.02 | 0.02 | 0 | 0.27 | 0.33 | 0.22 | 0.4 | 0.23 | 0.35 |
| 11 | 127701008 | rs2323008 | -0.244 | 0.267 | -1.295 | -0.451 | 0.001 | 0 | 0 | 0 | 0.4 | 0.38 | 0.3 | 0.42 | 0.36 | 0.41 |
| 11 | 127703110 | rs12361904 | 0.496 | 0.357 | -1.315 | -0.37 | -0.007 | 0.029 | 0.029 | 0 | 0.13 | 0.06 | 0.06 | 0.03 | 0.13 | 0.12 |
| 11 | 127706941 | rs719784 | -0.935 | 0.776 | -1.102 | 0.078 | 0.01 | 0.058 | 0.058 | 0 | 0.84 | 0.76 | 0.78 | 0.68 | 0.84 | 0.59 |
| 11 | 127713173 | rs1559660 | -0.784 | 0.836 | -1.204 | -0.043 | -0.007 | 0.012 | 0.012 | 0.217 | 0.61 | 0.64 | 0.54 | 0.75 | 0.58 | 0.88 |
| 11 | 127715860 | rs12421034 | 0.884 | 0.952 | -1.589 | 0.092 | -0.009 | 0.017 | 0.017 | 0 | 0.23 | 0.27 | 0.2 | 0.39 | 0.2 | 0.28 |
| 11 | 127716140 | rs2322978 | -0.841 | 1.39 | -1.845 | -0.026 | -0.114 | 0.059 | 0.059 | 0 | 0.75 | 0.97 | 0.71 | 0.85 | 0.68 | 0.89 |
| 11 | 127718228 | rs12049869 | -0.095 | 1.083 | -1.184 | 0.327 | -0.001 | 0 | 0 | 0 | 0.48 | 0.54 | 0.48 | 0.54 | 0.49 | 0.45 |
| 11 | 127719538 | rs6590281 | -0.54 | 0.883 | -0.979 | -0.075 | -0.006 | -0.034 | -0.034 | 0 | 0.23 | 0.44 | 0.28 | 0.16 | 0.21 | 0.16 |
| 11 | 127720615 | rs11221085 | 0.006 | 0.741 | -1.58 | -0.035 | -0.022 | -0.08 | -0.08 | 0 | 0.21 | 0.12 | 0.25 | 0.57 | 0.25 | 0.51 |

| | | | | | | | | | | | | | | | | |
|---|---|---|---|---|---|---|---|---|---|---|---|---|---|---|---|---|
| 11 | 127727368 | rs7107551 | -0.25 | 0.802 | -1.653 | -0.794 | 0.008 | 0.045 | 0.045 | 0.186 | 0.55 | 0.37 | 0.63 | 0.79 | 0.6 | 0.69 |
| 11 | 127727961 | rs7111459 | -1.903 | 0.805 | -1.629 | -0.797 | -0.004 | -0.05 | -0.05 | 0 | 0.79 | 0.81 | 0.92 | 0.96 | 0.82 | 0.85 |
| 11 | 127731391 | rs10790927 | 0.201 | 0.63 | -1.837 | -0.804 | -0.01 | 0.031 | 0.031 | 0 | 0.44 | 0.36 | 0.57 | 0.73 | 0.5 | 0.59 |
| 11 | 127731619 | rs10790928 | -1.918 | 0.647 | -1.715 | -0.743 | -0.007 | -0.073 | -0.073 | 0 | 0.78 | 0.76 | 0.91 | 0.96 | 0.82 | 0.85 |
| 11 | 127735165 | rs1812931 | -1.257 | 0.633 | -1.976 | -0.744 | -0.007 | -0.005 | -0.005 | 0.733 | 0.69 | 0.33 | 0.82 | 0.78 | 0.8 | 0.78 |
| 11 | 127736998 | rs1364777 | -0.202 | 0.662 | -1.967 | -0.729 | 0.004 | 0.102 | 0.102 | 0 | 0.5 | 0.29 | 0.58 | 0.78 | 0.53 | 0.69 |
| 11 | 127737929 | rs10750376 | -0.535 | 0.622 | -1.943 | -0.694 | 0.015 | 0.062 | 0.062 | 0.034 | 0.54 | 0.32 | 0.63 | 0.78 | 0.59 | 0.7 |
| 11 | 127748894 | rs11221096 | 0.734 | 1.124 | -2.044 | 0.146 | -0.03 | -0.022 | -0.022 | 0 | 0.05 | 0.01 | 0.08 | 0.23 | 0.11 | 0.22 |
| 11 | 127792983 | rs1075127 | -0.513 | 0.686 | -2.14 | -0.422 | 0.08 | 0.076 | 0.076 | 0 | 0.55 | 0.25 | 0.79 | 0.73 | 0.71 | 0.69 |
| 11 | 127795846 | rs1834311 | 0.535 | 0.89 | -2.245 | -0.539 | -0.031 | 0.014 | 0.014 | 0 | 0.35 | 0.24 | 0.54 | 0.73 | 0.45 | 0.58 |
| 11 | 127797506 | rs7106488 | -0.593 | 0.885 | -2.258 | -0.54 | 0.073 | 0.069 | 0.069 | 0 | 0.55 | 0.27 | 0.79 | 0.73 | 0.71 | 0.69 |
| 11 | 127797623 | rs11221109 | 2.177 | 1.921 | -1.8 | 0.135 | -0.025 | 0.014 | 0.014 | 0 | 0.12 | 0.02 | 0.26 | 0.1 | 0.17 | 0.05 |
| 11 | 127799447 | rs11221110 | 0.496 | 1.858 | -1.72 | 0.33 | -0.078 | -0.131 | -0.131 | 0 | 0.21 | 0.06 | 0.27 | 0.52 | 0.28 | 0.5 |
| 11 | 127808905 | rs649574 | 0.157 | 1.693 | -1.837 | 0.169 | -0.118 | -0.142 | -0.142 | 0 | 0.25 | 0.07 | 0.29 | 0.52 | 0.38 | 0.55 |
| 11 | 127809940 | rs11221112 | 1.237 | 1.613 | -1.515 | -0.195 | 0.015 | -0.027 | -0.027 | 0 | 0.44 | 0.54 | 0.3 | 0.02 | 0.31 | 0.28 |
| 11 | 127813659 | rs12797923 | 0.455 | 1.331 | -1.315 | 0.121 | 0.019 | 0.05 | 0.05 | 0 | 0.55 | 0.65 | 0.4 | 0.26 | 0.42 | 0.37 |
| 11 | 127819771 | rs1364782 | 0.52 | 1.318 | -1.281 | 0.169 | -0.012 | 0.089 | 0.089 | 0 | 0.07 | 0.14 | 0.06 | 0.24 | 0.05 | 0.08 |
| 11 | 127827727 | rs478251 | 0.949 | 1.408 | -1.352 | 0.396 | -0.051 | -0.054 | -0.054 | 0.252 | 0.09 | 0.01 | 0.14 | 0.27 | 0.16 | 0.18 |
| 11 | 127831692 | rs1785041 | 0.048 | 1.122 | -1.224 | 0.319 | -0.081 | -0.055 | -0.055 | 0 | 0.29 | 0.14 | 0.31 | 0.65 | 0.38 | 0.55 |
| 11 | 127833156 | rs1559661 | -0.637 | 0.961 | -1.193 | 0.361 | -0.003 | 0.025 | 0.025 | 0 | 0.13 | 0.22 | 0.12 | 0.24 | 0.14 | 0.09 |
| 11 | 127842867 | rs1364779 | 1.011 | 0.839 | -1.11 | 0.312 | 0.006 | 0.006 | 0.006 | 0.022 | 0.44 | 0.52 | 0.31 | 0.2 | 0.29 | 0.29 |
| 11 | 127857444 | rs11221124 | 1.366 | 0.842 | -1.071 | 0.251 | -0.019 | -0.116 | -0.116 | 0 | 0.41 | 0.38 | 0.27 | 0.02 | 0.28 | 0.28 |
| 11 | 127858117 | rs12573939 | 1.118 | 1.037 | -1.265 | 0.449 | -0.014 | -0.02 | -0.02 | 0 | 0.07 | 0.03 | 0.09 | 0.24 | 0.08 | 0.24 |
| 11 | 127859960 | rs1940745 | 0.018 | 0.733 | -1.086 | 0.1 | 0.015 | 0.034 | 0.034 | 0 | 0.54 | 0.59 | 0.39 | 0.26 | 0.42 | 0.37 |
| 11 | 127863383 | rs2015798 | 0.652 | 0.728 | -1.068 | 0.09 | 0.01 | -0.044 | -0.044 | 0 | 0.48 | 0.49 | 0.33 | 0.02 | 0.37 | 0.29 |
| 11 | 127865952 | rs2004408 | -0.034 | 0.813 | -1.144 | 0.187 | 0 | -0.004 | -0.004 | 0 | 0.12 | 0.11 | 0.12 | 0.24 | 0.14 | 0.09 |
| 11 | 127873154 | rs1425864 | 0.459 | 0.919 | -1.258 | -0.218 | -0.075 | -0.1 | -0.1 | 0 | 0.29 | 0.15 | 0.31 | 0.52 | 0.38 | 0.56 |
| 11 | 127874845 | rs1425865 | 1.561 | 0.474 | -1.06 | -1.198 | 0.013 | -0.005 | -0.005 | 0 | 0.32 | 0.17 | 0.23 | 0.02 | 0.25 | 0.21 |
| 11 | 127891554 | rs6590292 | 0.691 | 1.346 | -1.384 | -0.651 | 0.046 | 0.077 | 0.077 | 0.066 | 0.47 | 0.59 | 0.3 | 0.07 | 0.33 | 0.34 |
| 11 | 127895471 | rs948254 | -0.305 | 1.718 | -1.362 | 0.281 | -0.045 | 0.033 | 0.033 | 0.211 | 0.85 | 0.97 | 0.73 | 0.9 | 0.81 | 0.93 |
| 11 | 127900579 | rs1785039 | -1.07 | 1.551 | -0.675 | 0.549 | 0.037 | 0.051 | 0.051 | 1.377 | 0.16 | 0.28 | 0.19 | 0.31 | 0.2 | 0.24 |
| 11 | 127902700 | rs7127228 | 0.49 | 0.815 | -1.153 | -0.271 | 0.006 | 0.024 | 0.024 | 0 | 0.57 | 0.61 | 0.39 | 0.35 | 0.45 | 0.5 |
| 11 | 127909243 | rs6590293 | 0.65 | 0.379 | -1.173 | -0.514 | 0.007 | 0.023 | 0.023 | 0 | 0.53 | 0.52 | 0.32 | 0.32 | 0.45 | 0.44 |
| 11 | 127921327 | rs1425872 | -0.895 | 1.172 | -1.377 | -0.465 | -0.01 | -0.029 | -0.029 | 0 | 0.25 | 0.3 | 0.43 | 0.52 | 0.24 | 0.42 |
| 11 | 127922937 | rs11221140 | 0.381 | 1.126 | -1.419 | -0.436 | -0.049 | -0.099 | -0.099 | 0.043 | 0.16 | 0.04 | 0.32 | 0.49 | 0.18 | 0.3 |
| 11 | 127940351 | rs1547486 | 2.021 | 0.252 | -1.547 | -1.278 | 0.014 | 0.023 | 0.023 | 0 | 0.2 | 0.14 | 0.09 | 0 | 0.15 | 0.21 |
| 11 | 127940816 | rs2162174 | -0.456 | 0.363 | -1.439 | -0.979 | -0.03 | 0.019 | 0.019 | 0 | 0.78 | 0.92 | 0.77 | 0.84 | 0.67 | 0.8 |
| 11 | 127948238 | rs7119096 | 0.495 | 0.281 | -1.42 | -1.025 | 0.02 | -0.025 | -0.025 | 0 | 0.69 | 0.55 | 0.63 | 0.81 | 0.59 | 0.7 |
| 11 | 127955680 | rs1346104 | 0.049 | 0.319 | -1.428 | -1.051 | -0.015 | -0.032 | -0.032 | 0.135 | 0.12 | 0.26 | 0.12 | 0.04 | 0.06 | 0.11 |
| 11 | 127957037 | rs7935537 | 0.454 | 0.343 | -1.426 | -1.008 | 0.004 | -0.012 | -0.012 | 0 | 0.69 | 0.62 | 0.65 | 0.81 | 0.63 | 0.76 |
| 11 | 127960232 | rs1425873 | 0.637 | 0.364 | -1.402 | -0.966 | 0.01 | -0.017 | -0.017 | 0 | 0.67 | 0.58 | 0.64 | 0.81 | 0.59 | 0.69 |
| 11 | 127986917 | rs713069 | 1.255 | 0.094 | -1.241 | -0.869 | 0.005 | 0.006 | 0.006 | 0 | 0.48 | 0.47 | 0.31 | 0.31 | 0.41 | 0.37 |
| 11 | 127999326 | rs948256 | -1.192 | 0.84 | -1.272 | -0.761 | -0.012 | 0 | 0 | 0 | 0.83 | 0.92 | 0.79 | 0.85 | 0.7 | 0.84 |
| 11 | 128001331 | rs11221163 | 0.578 | 0.404 | -1.295 | -1.137 | 0 | -0.022 | -0.022 | 0.09 | 0.46 | 0.49 | 0.55 | 0.81 | 0.45 | 0.57 |
| 11 | 128002947 | rs1940363 | 0.042 | 0.317 | -1.29 | -1.144 | 0.031 | 0.02 | 0.02 | 0 | 0.34 | 0.31 | 0.22 | 0.04 | 0.21 | 0.23 |
| 11 | 128020368 | rs1477577 | -0.726 | 0.443 | -1.187 | -0.848 | 0.04 | 0.021 | 0.021 | 0 | 0.49 | 0.46 | 0.53 | 0.54 | 0.36 | 0.54 |
| 11 | 128025103 | rs10893822 | 1.362 | 0.468 | -1.217 | -0.237 | -0.025 | -0.009 | -0.009 | 0 | 0.07 | 0 | 0.17 | 0.33 | 0.07 | 0.14 |
| 11 | 128025266 | rs1940359 | -0.148 | 0.231 | -0.984 | -1.03 | -0.021 | 0.017 | 0.017 | 0 | 0.62 | 0.73 | 0.45 | 0.36 | 0.51 | 0.51 |
| 11 | 128025758 | rs10893824 | -0.313 | 0.11 | -0.933 | -0.863 | 0.016 | -0.035 | -0.035 | 0 | 0.1 | 0.31 | 0.15 | 0.03 | 0.1 | 0.12 |
| 11 | 128027668 | rs12421164 | 1.072 | -0.041 | -1.039 | -0.998 | 0.063 | 0.095 | 0.095 | 0 | 0.21 | 0.03 | 0.06 | 0 | 0.12 | 0.12 |
| 11 | 128027713 | rs7120121 | 0.15 | 0.284 | -0.875 | -0.801 | 0.007 | -0.018 | -0.018 | 0 | 0.31 | 0.35 | 0.21 | 0.04 | 0.21 | 0.24 |
| 11 | 128031037 | rs7105983 | 0.15 | 0.284 | -0.885 | -0.801 | 0.008 | -0.015 | -0.015 | 0 | 0.31 | 0.34 | 0.21 | 0.03 | 0.21 | 0.24 |
| 11 | 128041582 | rs7933438 | 0.176 | 0.083 | -0.735 | -0.485 | -0.006 | 0 | 0 | 0 | 0.32 | 0.37 | 0.26 | 0.34 | 0.19 | 0.23 |
| 11 | 128042023 | rs1863370 | 0.049 | 0.65 | -0.954 | -0.505 | 0.002 | 0.002 | 0.002 | 0 | 0.51 | 0.55 | 0.54 | 0.52 | 0.51 | 0.61 |
| 11 | 128043547 | rs11221187 | 0.91 | 0.563 | -1.249 | -0.758 | -0.009 | -0.123 | -0.123 | 0 | 0.29 | 0.4 | 0.19 | 0.02 | 0.27 | 0.22 |
| 11 | 128044762 | rs7948547 | -0.212 | 0.183 | -1.022 | -0.198 | -0.031 | -0.056 | -0.056 | 0 | 0.24 | 0.17 | 0.38 | 0.52 | 0.27 | 0.41 |
| 11 | 128044966 | rs7948482 | -0.462 | -0.84 | -0.569 | -0.278 | -0.034 | -0.008 | -0.008 | 0 | 0.34 | 0.38 | 0.26 | 0.32 | 0.19 | 0.24 |
| 11 | 128045869 | rs1977997 | 0.388 | 0.277 | -0.644 | -0.063 | -0.016 | -0.007 | -0.007 | 0 | 0.93 | 0.97 | 0.85 | 0.94 | 0.92 | 0.87 |
| 11 | 128047499 | rs1784672 | 1.239 | 0.185 | 0.259 | -0.085 | 0.011 | 0.023 | 0.023 | 0.095 | 0.81 | 0.53 | 0.77 | 0.77 | 0.86 | 0.77 |
| 11 | 128054642 | rs498573 | 1.337 | -0.148 | 0.505 | 0.155 | -0.01 | -0.007 | -0.007 | 0 | 0.78 | 0.66 | 0.53 | 0.18 | 0.83 | 0.67 |
| 11 | 128060772 | rs482246 | 1.269 | -0.027 | 0.335 | -0.851 | 0.008 | 0.003 | 0.003 | 0 | 0.49 | 0.38 | 0.25 | 0.15 | 0.36 | 0.43 |
| 11 | 128061096 | rs1215887 | 0.471 | 0.349 | 0.169 | -0.8 | -0.085 | 0.087 | 0.087 | 0 | 0.23 | 0.11 | 0.23 | 0.03 | 0.4 | 0.14 |
| 11 | 128071256 | rs12291698 | -0.4 | 0.119 | 0.351 | -1.004 | -0.009 | 0.006 | 0.006 | 0 | 0.33 | 0.31 | 0.32 | 0.18 | 0.47 | 0.32 |
| 11 | 128076596 | rs9787850 | 0.447 | 0.1 | 0.176 | -0.784 | -0.008 | -0.022 | -0.022 | 1.161 | 0.08 | 0.01 | 0.28 | 0.57 | 0.09 | 0.11 |
| 11 | 128079898 | rs1940357 | 0.585 | 0.121 | 0.25 | -0.79 | -0.001 | 0.029 | 0.029 | 0 | 0.06 | 0.1 | 0.06 | 0.14 | 0.04 | 0.1 |
| 11 | 128088935 | rs1793327 | 0.419 | 0.049 | 0.527 | -1.015 | -0.007 | 0.019 | 0.019 | 0 | 0.83 | 0.67 | 0.79 | 0.75 | 0.88 | 0.72 |
| 11 | 128090006 | rs550340 | -0.213 | 0.09 | 0.148 | -0.976 | 0.002 | -0.006 | -0.006 | 0.638 | 0.87 | 0.89 | 0.83 | 0.75 | 0.91 | 0.8 |
| 11 | 128092632 | rs7103877 | 0.816 | 0.048 | 0.022 | -1.12 | -0.089 | 0.099 | 0.099 | 0 | 0.23 | 0.08 | 0.22 | 0.04 | 0.39 | 0.14 |
| 11 | 128099099 | rs511713 | 1.384 | -0.159 | 0.063 | -1.135 | -0.014 | -0.124 | -0.124 | 0 | 0.43 | 0.45 | 0.18 | 0 | 0.31 | 0.33 |

| | | | | | | | | | | | | | | | | |
|---|---|---|---|---|---|---|---|---|---|---|---|---|---|---|---|---|
| 11 | 128115740 | rs7112731 | -0.718 | -0.135 | -0.277 | -0.648 | -0.016 | 0.034 | 0.034 | 0.001 | 0.12 | 0.21 | 0.32 | 0.71 | 0.07 | 0.2 |
| 11 | 128118439 | rs476696 | 1.056 | -0.042 | -0.533 | -0.956 | 0.002 | 0.026 | 0.026 | 0.132 | 0.8 | 0.88 | 0.77 | 0.94 | 0.83 | 0.71 |
| 11 | 128119304 | rs1940379 | -0.659 | 0.117 | -1.114 | -0.587 | 0.001 | -0.009 | -0.009 | 0 | 0.12 | 0.14 | 0.27 | 0.77 | 0.12 | 0.19 |
| 11 | 128119555 | rs478816 | -0.125 | -0.607 | -1.279 | -0.755 | -0.003 | 0.052 | 0.052 | 0.584 | 0.87 | 0.95 | 0.83 | 0.97 | 0.9 | 0.87 |
| 11 | 128120235 | rs535556 | 0.305 | -0.715 | -1.335 | -0.735 | 0.036 | 0.217 | 0.217 | 0 | 0.44 | 0.11 | 0.3 | 0.13 | 0.35 | 0.48 |
| 11 | 128122782 | rs523164 | 0.551 | -0.742 | -0.025 | 0.319 | 0.038 | 0.181 | 0.181 | 0 | 0.92 | 0.71 | 0.81 | 0.85 | 0.96 | 0.89 |
| 11 | 128136389 | rs565536 | 0.835 | -1.305 | 0.059 | 0.18 | 0.008 | -0.013 | -0.013 | 0 | 0.63 | 0.38 | 0.53 | 0.65 | 0.67 | 0.56 |
| 11 | 128144864 | rs11221232 | -0.81 | -1.511 | 0.251 | 0.112 | -0.019 | 0.085 | 0.085 | 0.215 | 0.1 | 0.33 | 0.06 | 0.11 | 0.05 | 0.09 |
| 11 | 128145470 | rs573624 | 0.137 | -1.631 | -0.403 | -0.232 | 0.01 | 0.005 | 0.005 | 0 | 0.38 | 0.38 | 0.35 | 0.7 | 0.3 | 0.39 |
| 11 | 128148756 | rs722344 | 1.201 | -2.34 | -1.112 | -0.274 | 0.036 | -0.042 | -0.042 | 0 | 0.12 | 0.06 | 0.18 | 0.58 | 0.05 | 0.19 |
| 11 | 128156314 | rs12795702 | 0.819 | -2.37 | -0.913 | -0.406 | 0.05 | 0.093 | 0.093 | 0.246 | 0.51 | 0.22 | 0.43 | 0.25 | 0.6 | 0.54 |
| 11 | 128156903 | rs767179 | 0.883 | -2.259 | -0.952 | -0.43 | -0.013 | -0.024 | -0.024 | 0 | 0.17 | 0.05 | 0.2 | 0.18 | 0.19 | 0.28 |
| 11 | 128157643 | rs500207 | -1.324 | -2.274 | -0.948 | -0.459 | -0.015 | 0.103 | 0.103 | 0.607 | 0.58 | 0.8 | 0.57 | 0.88 | 0.5 | 0.67 |
| 11 | 128159029 | rs12786743 | 1.662 | -2.044 | -0.886 | -0.496 | -0.109 | 0.178 | 0.178 | 0.067 | 0.33 | 0.06 | 0.22 | 0.07 | 0.4 | 0.25 |
| 11 | 128163457 | rs12418638 | 0.76 | -1.956 | -0.678 | -0.143 | -0.013 | -0.026 | -0.026 | 0 | 0.17 | 0.05 | 0.2 | 0.19 | 0.19 | 0.28 |
| 11 | 128165127 | rs12419147 | 1.232 | -1.843 | -0.79 | -0.235 | -0.046 | 0.108 | 0.108 | 0.259 | 0.39 | 0.17 | 0.27 | 0.07 | 0.46 | 0.28 |
| 11 | 128168597 | rs876071 | 0.24 | -1.578 | -0.974 | 0.172 | 0.009 | -0.014 | -0.014 | 0 | 0.66 | 0.68 | 0.51 | 0.36 | 0.67 | 0.64 |
| 11 | 128182521 | rs12577895 | 0.266 | -0.631 | -0.874 | 1.214 | 0.028 | -0.049 | -0.049 | 0 | 0.17 | 0.06 | 0.13 | 0.34 | 0.1 | 0.18 |
| 11 | 128185154 | rs12417297 | -0.038 | -0.159 | -0.635 | 1.324 | -0.001 | 0.002 | 0.002 | 0 | 0.1 | 0.11 | 0.2 | 0.16 | 0.17 | 0.2 |
| 11 | 128186882 | rs10893845 | 0.527 | -0.375 | -0.711 | 0.798 | -0.06 | 0.175 | 0.175 | 1.056 | 0.48 | 0.07 | 0.45 | 0.29 | 0.5 | 0.4 |
| 11 | 128187383 | rs7127307 | -0.645 | 0.054 | -1.089 | 0.782 | 0.001 | 0.013 | 0.013 | 0.228 | 0.45 | 0.38 | 0.46 | 0.59 | 0.47 | 0.57 |
| 11 | 128189101 | rs4937314 | -0.93 | 0.317 | -1.206 | 0.99 | -0.005 | -0.018 | -0.018 | 0.478 | 0.19 | 0.14 | 0.14 | 0.22 | 0.19 | 0.2 |
| 11 | 128189789 | rs4144899 | -0.549 | 0.514 | -1.151 | 1.219 | 0.003 | 0.034 | 0.034 | 0 | 0.11 | 0.13 | 0.12 | 0.28 | 0.12 | 0.1 |
| 11 | 128191633 | rs2040334 | -0.458 | 1.68 | -1.54 | 1.018 | -0.031 | 0.004 | 0.004 | 0 | 0.85 | 0.95 | 0.62 | 0.85 | 0.72 | 0.88 |
| 11 | 128196675 | rs11221249 | -0.799 | 0.812 | -1.14 | -0.134 | -0.004 | -0.014 | -0.014 | 0 | 0.4 | 0.52 | 0.32 | 0.34 | 0.39 | 0.29 |
| 11 | 128199706 | rs4936046 | -0.588 | 1.106 | -0.27 | -0.33 | -0.004 | -0.003 | -0.003 | 0.012 | 0.32 | 0.35 | 0.23 | 0.28 | 0.27 | 0.22 |
| 11 | 128200831 | rs7117754 | 1.473 | 0.361 | -0.877 | -0.207 | -0.041 | -0.044 | -0.044 | 0 | 0.17 | 0.09 | 0.22 | 0.25 | 0.23 | 0.39 |
| 11 | 128201150 | rs4937316 | -0.942 | 0.165 | -0.833 | -0.329 | 0.005 | -0.009 | -0.009 | 0 | 0.8 | 0.75 | 0.59 | 0.84 | 0.7 | 0.88 |
| 11 | 128201479 | rs12284207 | 0.754 | 0.187 | -0.792 | -0.394 | 0 | -0.038 | -0.038 | 1.063 | 0.2 | 0.16 | 0.11 | 0 | 0.15 | 0.12 |
| 11 | 128204479 | rs7131167 | 1.123 | -0.571 | -0.989 | -0.286 | -0.105 | -0.084 | -0.084 | 0.752 | 0.17 | 0.02 | 0.22 | 0.24 | 0.25 | 0.33 |
| 11 | 128205267 | rs11221253 | 0.742 | -1.073 | -1.095 | -0.212 | -0.016 | -0.015 | -0.015 | 0.004 | 0.42 | 0.21 | 0.29 | 0.41 | 0.35 | 0.5 |
| 11 | 128206410 | rs6590322 | -0.486 | -1.377 | -0.978 | -0.217 | 0.002 | 0.001 | 0.001 | 0 | 0.52 | 0.45 | 0.45 | 0.51 | 0.54 | 0.57 |
| 11 | 128212939 | rs871506 | 0.536 | -0.661 | -0.499 | 0.141 | 0.019 | -0.001 | -0.001 | 0 | 0.06 | 0.16 | 0.05 | 0.07 | 0.07 | 0.09 |
| 11 | 128224634 | rs10450612 | -0.378 | -0.801 | -0.549 | 0.139 | -0.008 | -0.015 | -0.015 | 0 | 0.31 | 0.45 | 0.22 | 0.26 | 0.29 | 0.24 |
| 11 | 128231469 | rs1125253 | -1.127 | -0.693 | -0.803 | 0.026 | 0.004 | 0.006 | 0.006 | 0 | 0.83 | 0.83 | 0.58 | 0.77 | 0.73 | 0.82 |
| 11 | 128231995 | rs1557477 | 1.362 | -0.801 | -0.886 | -0.02 | 0.045 | 0.042 | 0.042 | 0.011 | 0.34 | 0.23 | 0.12 | 0.24 | 0.19 | 0.24 |
| 11 | 128232874 | rs11221264 | 1.312 | -0.253 | -0.35 | 0.365 | -0.005 | 0.005 | 0.005 | 0 | 0.51 | 0.32 | 0.35 | 0.51 | 0.44 | 0.58 |
| 11 | 128234144 | rs10750397 | -0.327 | -0.054 | -0.641 | 0.325 | 0.001 | 0.002 | 0.002 | 0 | 0.83 | 0.83 | 0.58 | 0.77 | 0.73 | 0.82 |
| 11 | 128234772 | rs7120183 | -1.085 | 0.327 | -1.031 | 0.332 | -0.066 | -0.052 | -0.052 | 0.114 | 0.9 | 0.99 | 0.6 | 0.83 | 0.74 | 0.89 |
| 11 | 128236936 | rs11221265 | -0.618 | -0.707 | -0.593 | -0.013 | -0.009 | -0.062 | -0.062 | 0 | 0.17 | 0.18 | 0.1 | 0 | 0.13 | 0.14 |
| 11 | 128241103 | rs6590325 | 0.022 | -0.52 | -0.526 | 0.609 | 0 | 0.004 | 0.004 | 0 | 0.57 | 0.35 | 0.42 | 0.39 | 0.47 | 0.53 |
| 11 | 128245331 | rs11221267 | 0.105 | -0.006 | -1.192 | 0.704 | 0.019 | 0.008 | 0.008 | 0.053 | 0.14 | 0.09 | 0.08 | 0.08 | 0.12 | 0.12 |
| 11 | 128248555 | rs4556551 | 0.05 | 0.506 | -0.764 | 1.167 | 0.019 | 0.013 | 0.013 | 0 | 0.61 | 0.48 | 0.42 | 0.46 | 0.51 | 0.68 |
| 11 | 128261411 | rs7112864 | -0.355 | 0.35 | -0.896 | 1.752 | -0.004 | 0.05 | 0.05 | 0.205 | 0.07 | 0.05 | 0.13 | 0.3 | 0.08 | 0.23 |
| 11 | 128266104 | rs7939885 | -0.315 | -0.603 | -1.082 | 1.04 | 0.012 | -0.01 | -0.01 | 0 | 0.35 | 0.65 | 0.19 | 0.49 | 0.26 | 0.3 |
| 11 | 128266238 | rs7939722 | -1.019 | -1.841 | -1.856 | -0.026 | -0.013 | 0.168 | 0.168 | 0 | 0.61 | 0.82 | 0.43 | 0.89 | 0.44 | 0.67 |
| 11 | 128267162 | rs7130057 | 1.267 | -1.808 | -1.919 | 0.024 | 0.041 | 0.026 | 0.026 | 0 | 0.15 | 0.05 | 0.09 | 0.1 | 0.08 | 0.12 |
| 11 | 128267228 | rs958889 | -0.393 | -2.266 | -1.977 | -0.011 | 0.112 | 0.075 | 0.075 | 0 | 0.41 | 0.13 | 0.28 | 0.75 | 0.24 | 0.52 |
| 11 | 128269918 | rs11605437 | 0.687 | -1.665 | -1.32 | 0.311 | -0.005 | -0.006 | -0.006 | 0 | 0.09 | 0.07 | 0.06 | 0.13 | 0.15 | 0.09 |
| 11 | 128272676 | rs7107309 | -0.278 | -2.22 | -0.875 | 0.031 | -0.02 | -0.013 | -0.013 | 0 | 0.25 | 0.66 | 0.16 | 0.15 | 0.13 | 0.24 |
| 11 | 128278855 | rs1540218 | -0.229 | -1.253 | -1.361 | -0.15 | -0.025 | -0.045 | -0.045 | 0 | 0.61 | 0.8 | 0.61 | 0.53 | 0.48 | 0.53 |
| 11 | 128281184 | rs1540225 | 1.213 | -1.114 | -1.43 | -0.132 | -0.016 | -0.005 | -0.005 | 0 | 0.17 | 0.17 | 0.09 | 0.14 | 0.08 | 0.17 |
| 11 | 128281264 | rs6590328 | 0.112 | -1.87 | -1.57 | -0.529 | -0.019 | -0.071 | -0.071 | 0.142 | 0.22 | 0.55 | 0.12 | 0.06 | 0.2 | 0.11 |
| 11 | 128283618 | rs10893866 | 1.461 | -2.412 | -2.14 | -0.635 | -0.066 | 0.004 | 0.004 | 0 | 0.17 | 0.03 | 0.4 | 0.12 | 0.21 | 0.23 |
| 11 | 128286441 | rs11602703 | 0.744 | -2.002 | -1.279 | -0.645 | -0.016 | -0.09 | -0.09 | 0 | 0.18 | 0.33 | 0.09 | 0.02 | 0.17 | 0.03 |
| 11 | 128288400 | rs7936180 | -1.214 | -1.748 | -2.097 | -0.457 | -0.179 | -0.124 | -0.124 | 0.115 | 0.78 | 0.98 | 0.7 | 0.67 | 0.56 | 0.69 |
| 11 | 128293020 | rs10790952 | 0.088 | -2.193 | -1.823 | -0.48 | 0.043 | -0.018 | -0.018 | 0 | 0.39 | 0.29 | 0.49 | 0.46 | 0.27 | 0.41 |
| 11 | 128299706 | rs7108537 | 1.203 | -2.043 | -1.771 | -0.626 | 0.04 | 0.023 | 0.023 | 0 | 0.21 | 0.1 | 0.14 | 0.15 | 0.11 | 0.23 |
| 11 | 128299874 | rs7111926 | -0.768 | -0.711 | -2.296 | -0.96 | -0.063 | 0.085 | 0.085 | 0 | 0.71 | 0.91 | 0.52 | 0.91 | 0.63 | 0.78 |
| 11 | 128303342 | rs7125393 | 1.381 | -0.494 | -2.616 | -0.929 | -0.066 | 0.005 | 0.005 | 0 | 0.15 | 0.02 | 0.37 | 0.08 | 0.19 | 0.15 |
| 11 | 128304141 | rs12576753 | 1.282 | -0.269 | -2.516 | -0.529 | 0.011 | 0.124 | 0.124 | 0.128 | 0.06 | 0.1 | 0.11 | 0.38 | 0.1 | 0.16 |
| 11 | 128317168 | rs7932088 | 0.427 | -0.382 | -2.709 | -0.745 | -0.022 | -0.024 | -0.024 | 0 | 0.38 | 0.18 | 0.23 | 0.38 | 0.4 | 0.37 |
| 11 | 128319146 | rs4285885 | -2.074 | -0.417 | -2.727 | -0.839 | -0.062 | 0.081 | 0.081 | 0 | 0.86 | 1 | 0.9 | 0.98 | 0.83 | 0.93 |
| 11 | 128320137 | rs7122420 | 1.813 | -0.701 | -2.601 | -0.739 | 0.046 | 0.017 | 0.017 | 0 | 0.19 | 0.03 | 0.11 | 0.15 | 0.11 | 0.23 |
| 11 | 128325507 | rs10893871 | -0.773 | -0.027 | -2.247 | -0.56 | -0.116 | 0.11 | 0.11 | 0 | 0.72 | 0.95 | 0.53 | 0.92 | 0.64 | 0.78 |
| 11 | 128325553 | rs10893872 | 0.355 | 0.075 | -2.208 | -0.408 | -0.052 | -0.055 | -0.055 | 0 | 0.57 | 0.2 | 0.35 | 0.54 | 0.51 | 0.6 |
| 11 | 128328913 | rs8705 | 0.836 | 0.236 | -2.241 | -0.138 | -0.079 | -0.001 | -0.001 | 0 | 0.19 | 0.04 | 0.18 | 0.18 | 0.32 | 0.19 |
| 11 | 128328959 | rs1128334 | 0.972 | 0.461 | -1.795 | -0.017 | 0.013 | 0.135 | 0.135 | 0.1 | 0.06 | 0.11 | 0.11 | 0.38 | 0.1 | 0.17 |
| 11 | 128330520 | rs4937333 | 0.145 | 0.289 | -2.072 | -0.244 | -0.044 | -0.046 | -0.046 | 0 | 0.59 | 0.23 | 0.37 | 0.54 | 0.52 | 0.6 |

| | | | | | | | | | | | | | | | | |
|---|---|---|---|---|---|---|---|---|---|---|---|---|---|---|---|---|
| 11 | 128333215 | rs12146687 | 1.337 | 0.8 | -2.033 | -0.663 | -0.071 | -0.001 | -0.001 | 2.884 | 0.11 | 0.02 | 0.34 | 0.07 | 0.2 | 0.11 |
| 11 | 128335578 | rs4937334 | 0.644 | 0.793 | -1.323 | -0.66 | 0.015 | -0.036 | -0.036 | 0 | 0.18 | 0.24 | 0.13 | 0.09 | 0.24 | 0.19 |
| 11 | 128335657 | rs7926975 | -0.006 | 0.585 | -0.939 | -0.205 | 0.035 | 0.385 | 0.385 | 2.785 | 0.11 | 0.46 | 0.15 | 0.5 | 0.1 | 0.23 |
| 11 | 128336438 | rs10893873 | 0.521 | 0.283 | -1.293 | -0.56 | -0.02 | 0.033 | 0.033 | 1.881 | 0.74 | 0.43 | 0.79 | 0.38 | 0.79 | 0.6 |
| 11 | 128337997 | rs4520612 | -0.848 | 0.33 | -1.989 | -0.625 | -0.001 | 0.012 | 0.012 | 0 | 0.86 | 0.98 | 0.94 | 0.88 | 0.89 | 0.83 |
| 11 | 128338131 | rs4523710 | -0.659 | 0.332 | -1.962 | -0.624 | 0.008 | 0.012 | 0.012 | 0 | 0.85 | 0.9 | 0.94 | 0.88 | 0.89 | 0.83 |
| 11 | 128341488 | rs11600915 | 1.566 | 0.428 | -2.233 | -0.797 | -0.036 | 0.076 | 0.076 | 0 | 0.13 | 0.01 | 0.09 | 0 | 0.16 | 0.06 |
| 11 | 128346793 | rs11221322 | 1.43 | 0.201 | -2.388 | -0.824 | -0.003 | 0 | 0 | 0 | 0.23 | 0.03 | 0.2 | 0.19 | 0.15 | 0.16 |
| 11 | 128347056 | rs4937337 | 0.931 | 0.207 | -2.367 | -0.826 | -0.081 | 0.034 | 0.034 | 0.097 | 0.36 | 0.06 | 0.31 | 0.28 | 0.39 | 0.35 |
| 11 | 128348978 | rs10790956 | -0.502 | 0.259 | -2 | -0.634 | -0.02 | 0.089 | 0.089 | 0 | 0.5 | 0.84 | 0.63 | 0.6 | 0.5 | 0.49 |
| 11 | 128350686 | rs7122859 | -0.13 | 0.401 | -0.748 | 0.32 | 0.007 | 0.298 | 0.298 | 0.971 | 0.09 | 0.28 | 0.13 | 0.49 | 0.07 | 0.21 |
| 11 | 128353171 | rs10893877 | 0.285 | 0.225 | -1.344 | -0.827 | 0.001 | 0.042 | 0.042 | 0 | 0.56 | 0.4 | 0.55 | 0.2 | 0.41 | 0.46 |
| 11 | 128354360 | rs7946776 | 0.326 | 0.788 | -1.66 | -0.862 | -0.067 | 0.035 | 0.035 | 0 | 0.23 | 0.01 | 0.14 | 0.11 | 0.26 | 0.18 |
| 11 | 128356728 | rs4628683 | 0.187 | -0.026 | -1.999 | -1.554 | -0.002 | 0.002 | 0.002 | 0 | 0.29 | 0.28 | 0.49 | 0.13 | 0.29 | 0.25 |
| 11 | 128361169 | rs4937339 | 0.629 | 0.322 | -1.981 | -1.383 | -0.041 | 0.017 | 0.017 | 0 | 0.23 | 0.03 | 0.16 | 0.13 | 0.25 | 0.2 |
| 11 | 128361194 | rs7125574 | 0.715 | 0.322 | -1.695 | -1.658 | 0.017 | -0.043 | -0.043 | 2.32 | 0.19 | 0.3 | 0.41 | 0.1 | 0.24 | 0.18 |
| 11 | 128366221 | rs4254089 | 0.195 | 0.09 | -1.665 | -1.492 | -0.017 | -0.019 | -0.019 | 1.362 | 0.31 | 0.41 | 0.2 | 0.25 | 0.27 | 0.39 |
| 11 | 128371441 | rs7115613 | -0.738 | 0.727 | -1.242 | -0.972 | 0.144 | 0.163 | 0.163 | 0 | 0.82 | 0.57 | 0.72 | 0.44 | 0.67 | 0.77 |
| 11 | 128373736 | rs10893882 | 0.435 | 0.611 | -1.824 | -0.922 | -0.024 | 0.024 | 0.024 | 0 | 0.17 | 0.1 | 0.39 | 0.13 | 0.23 | 0.22 |
| 11 | 128376095 | rs11600456 | 0.545 | 0.046 | -1.891 | -0.8 | -0.013 | 0.013 | 0.013 | 0.472 | 0.15 | 0.03 | 0.11 | 0.1 | 0.17 | 0.24 |
| 11 | 128377480 | rs7118744 | 0.447 | 0 | -2.062 | -1.15 | 0.01 | 0.108 | 0.108 | 0 | 0.61 | 0.79 | 0.73 | 0.82 | 0.62 | 0.71 |
| 11 | 128380742 | rs7924522 | -0.022 | -0.062 | -2.068 | -1.176 | -0.011 | 0.052 | 0.052 | 0 | 0.68 | 0.79 | 0.71 | 0.86 | 0.63 | 0.75 |
| 11 | 128383077 | rs7129690 | 1.982 | -0.409 | -1.874 | -1.77 | 0 | -0.044 | -0.044 | 0 | 0.26 | 0.26 | 0.1 | 0.01 | 0.15 | 0.12 |
| 11 | 128387422 | rs7946009 | -0.185 | 0.044 | -1.701 | -1.343 | -0.02 | 0.062 | 0.062 | 0 | 0.69 | 0.83 | 0.71 | 0.82 | 0.63 | 0.75 |
| 11 | 128388773 | rs7116578 | -1.323 | -0.708 | -1.625 | -1.119 | -0.013 | 0.011 | 0.011 | 0 | 0.87 | 0.96 | 0.92 | 0.91 | 0.88 | 0.98 |
| 11 | 128389391 | rs11819995 | 0.477 | -1.15 | -1.337 | -0.703 | 0.025 | 0.033 | 0.033 | 2.571 | 0.37 | 0.29 | 0.17 | 0.19 | 0.22 | 0.23 |
| 11 | 128390677 | rs3935794 | -0.44 | -0.392 | -0.947 | -0.064 | 0.002 | 0.002 | 0.002 | 0 | 0.1 | 0.08 | 0.06 | 0.05 | 0.06 | 0.06 |
| 11 | 128398309 | rs4937342 | -0.014 | -0.295 | -1.162 | -0.012 | 0.013 | 0.019 | 0.019 | 0 | 0.08 | 0.02 | 0.05 | 0.05 | 0.06 | 0.06 |
| 11 | 128409210 | rs12364200 | -0.615 | -0.351 | -0.884 | 0.307 | -0.019 | 0.022 | 0.022 | 0 | 0.06 | 0.11 | 0.05 | 0.14 | 0.01 | 0.13 |
| 11 | 128413863 | rs11221342 | -0.75 | -0.456 | -0.532 | 0.889 | -0.014 | 0.056 | 0.056 | 0 | 0.22 | 0.3 | 0.2 | 0.54 | 0.17 | 0.34 |
| 11 | 128420926 | rs4245079 | -0.583 | -0.017 | -0.672 | -0.001 | 0 | -0.012 | -0.012 | 2.051 | 0.42 | 0.4 | 0.54 | 0.85 | 0.43 | 0.64 |
| 11 | 128433319 | rs3948853 | 0.075 | 0.067 | -1.341 | 0.138 | -0.09 | -0.055 | -0.055 | 0 | 0.28 | 0.09 | 0.28 | 0.3 | 0.42 | 0.31 |
| 11 | 128436953 | rs7117932 | -0.369 | -0.07 | -1.054 | -0.341 | -0.034 | 0.005 | 0.005 | 0 | 0.33 | 0.22 | 0.31 | 0.25 | 0.45 | 0.38 |
| 11 | 128441497 | rs7928282 | 0.198 | -0.045 | -1.177 | -0.438 | -0.116 | 0.087 | 0.087 | 0 | 0.27 | 0.06 | 0.29 | 0.07 | 0.42 | 0.24 |
| 11 | 128450461 | rs1944854 | 0.38 | -0.876 | -1.012 | -0.778 | -0.005 | 0.003 | 0.003 | 0 | 0.38 | 0.4 | 0.49 | 0.58 | 0.37 | 0.31 |
| 11 | 128451280 | rs7125213 | -1.427 | -1.105 | -1.318 | -0.857 | 0.005 | 0.037 | 0.037 | 0.268 | 0.76 | 0.97 | 0.85 | 0.82 | 0.84 | 0.7 |
| 11 | 128459110 | rs11221371 | 0.435 | -0.979 | -1.114 | -0.768 | -0.004 | 0.051 | 0.051 | 0.074 | 0.07 | 0.13 | 0.02 | 0.16 | 0.04 | 0.16 |
| 11 | 128467510 | rs4937357 | -1.06 | -1.014 | -1.171 | -0.964 | 0.033 | 0.055 | 0.055 | 0.295 | 0.72 | 0.83 | 0.83 | 0.81 | 0.83 | 0.71 |
| 11 | 128471240 | rs11221383 | 0.339 | -0.137 | -0.978 | -0.667 | -0.03 | 0.027 | 0.027 | 0 | 0.26 | 0.17 | 0.3 | 0.07 | 0.42 | 0.23 |
| 11 | 128475001 | rs11221388 | -0.104 | -0.457 | -0.85 | -0.451 | -0.001 | 0 | 0 | 0 | 0.17 | 0.16 | 0.13 | 0.14 | 0.16 | 0.2 |
| 11 | 128477472 | rs10750403 | -1.11 | -0.465 | -1.577 | -0.653 | -0.114 | 0.067 | 0.067 | 2.011 | 0.58 | 0.92 | 0.59 | 0.7 | 0.53 | 0.66 |
| 11 | 128477538 | rs11221391 | 0.207 | -0.429 | -1.365 | -0.698 | 0.008 | -0.029 | -0.029 | 0.8 | 0.12 | 0.2 | 0.14 | 0.04 | 0.17 | 0.17 |
| 11 | 128477918 | rs949101 | -0.583 | -0.571 | -0.936 | -0.378 | 0.005 | 0.011 | 0.011 | 0 | 0.45 | 0.52 | 0.43 | 0.66 | 0.36 | 0.49 |
| 11 | 128487069 | rs7121743 | 0.077 | -1.262 | -0.622 | 0.143 | -0.08 | 0.093 | 0.093 | 0.411 | 0.27 | 0.43 | 0.14 | 0.49 | 0.14 | 0.24 |
| 11 | 128492373 | rs7127737 | -0.507 | -1.266 | -0.936 | -0.334 | -0.004 | -0.008 | -0.008 | 0 | 0.78 | 0.77 | 0.83 | 0.88 | 0.83 | 0.82 |
| 11 | 128492739 | rs4937362 | 0.194 | -0.7 | -1.427 | -0.353 | 0.007 | 0.023 | 0.023 | 0 | 0.41 | 0.37 | 0.48 | 0.38 | 0.6 | 0.5 |
| 11 | 128495193 | rs7947214 | 0.242 | -0.344 | -0.741 | 0.066 | -0.01 | -0.034 | -0.034 | 1.096 | 0.12 | 0.26 | 0.25 | 0.04 | 0.09 | 0.08 |
| 11 | 128499000 | rs7941765 | -1.07 | -0.029 | -1.924 | -0.387 | -0.146 | 0.148 | 0.148 | 1.337 | 0.58 | 0.95 | 0.66 | 0.78 | 0.5 | 0.61 |
| 11 | 128501572 | rs7935286 | -0.32 | 0.299 | -1.462 | 0.769 | -0.018 | -0.042 | -0.042 | 0.014 | 0.16 | 0.08 | 0.19 | 0.25 | 0.15 | 0.26 |
| 11 | 128502222 | rs10893900 | -1.25 | 0.095 | -0.625 | 0.623 | -0.01 | 0.011 | 0.011 | 0.544 | 0.29 | 0.38 | 0.43 | 0.27 | 0.22 | 0.32 |
| 11 | 128502496 | rs4936059 | -1.275 | -0.066 | -0.713 | 0.911 | -0.001 | -0.001 | -0.001 | 1.638 | 0.39 | 0.38 | 0.54 | 0.57 | 0.34 | 0.47 |
| 11 | 128513457 | rs10790966 | -1.482 | -0.408 | 0.017 | 0.906 | -0.025 | 0 | 0 | 0.001 | 0.28 | 0.61 | 0.39 | 0.25 | 0.16 | 0.21 |
| 11 | 128519816 | rs7940045 | -0.862 | -0.248 | -0.25 | 1.064 | -0.006 | 0.01 | 0.01 | 0 | 0.14 | 0.22 | 0.14 | 0.16 | 0.08 | 0.15 |
| 11 | 128521468 | rs10400334 | 0.207 | 0.168 | -0.786 | 0.769 | 0.021 | 0.022 | 0.022 | 0 | 0.63 | 0.84 | 0.52 | 0.7 | 0.68 | 0.7 |
| 11 | 128529842 | rs10790969 | -0.519 | 1.2 | 0.006 | 1.243 | 0.048 | 0.043 | 0.043 | 0.093 | 0.2 | 0.31 | 0.26 | 0.32 | 0.28 | 0.35 |
| 11 | 128530563 | rs624168 | -0.845 | 1.146 | 0.057 | 1.332 | 0.065 | 0.068 | 0.068 | 0.525 | 0.23 | 0.52 | 0.28 | 0.34 | 0.28 | 0.35 |
| 11 | 128532912 | rs608871 | -0.933 | 1.042 | -1.069 | 1.179 | -0.008 | -0.002 | -0.002 | 0 | 0.88 | 0.94 | 0.76 | 0.87 | 0.86 | 0.81 |
| 11 | 128540106 | rs639632 | 0.342 | 1.178 | -1.5 | 1.251 | -0.044 | -0.011 | -0.011 | 0 | 0.15 | 0.06 | 0.13 | 0.15 | 0.19 | 0.22 |
| 11 | 128544891 | rs599905 | 0.639 | 0.448 | -1.122 | 1.447 | 0.017 | -0.05 | -0.05 | 0 | 0.77 | 0.9 | 0.7 | 0.68 | 0.8 | 0.72 |
| 11 | 128545415 | rs598039 | -0.397 | 1.341 | -0.904 | 0.987 | -0.024 | -0.005 | -0.005 | 0 | 0.88 | 0.96 | 0.76 | 0.87 | 0.82 | 0.81 |
| 11 | 128562098 | rs695113 | -0.048 | 0.801 | -0.573 | 1.07 | 0.008 | 0.007 | 0.007 | 0 | 0.77 | 0.72 | 0.68 | 0.66 | 0.7 | 0.63 |
| 11 | 128563413 | rs12420835 | 0.338 | 0.695 | -0.651 | 0.853 | 0.04 | -0.029 | -0.029 | 0.028 | 0.32 | 0.43 | 0.43 | 0.17 | 0.45 | 0.39 |
| 11 | 128564794 | rs7930515 | -0.493 | 0.205 | -0.524 | 0.844 | 0 | 0 | 0 | 0 | 0.78 | 0.79 | 0.68 | 0.66 | 0.7 | 0.64 |
| 11 | 128571257 | rs644818 | -0.899 | -0.015 | -0.325 | 1.1 | 0.001 | 0.011 | 0.011 | 0 | 0.5 | 0.59 | 0.55 | 0.51 | 0.48 | 0.47 |
| 11 | 128578157 | rs619521 | -0.417 | 1.04 | -0.373 | 0.651 | 0.019 | -0.014 | -0.014 | 0 | 0.33 | 0.39 | 0.49 | 0.24 | 0.43 | 0.39 |
| 11 | 128582944 | rs609017 | -0.257 | 0.335 | -0.396 | 0.826 | 0.016 | 0.009 | 0.009 | 2.369 | 0.51 | 0.45 | 0.55 | 0.48 | 0.61 | 0.81 |
| 11 | 128586155 | rs654723 | -0.275 | 0.235 | -0.395 | 0.928 | 0.06 | 0.034 | 0.034 | 1.099 | 0.49 | 0.3 | 0.52 | 0.49 | 0.61 | 0.74 |
| 11 | 128592032 | rs644571 | -0.853 | 0.819 | 0.019 | 1.555 | 0.045 | 0.15 | 0.15 | 0.435 | 0.08 | 0.23 | 0.13 | 0.32 | 0.11 | 0.3 |
| 11 | 128592637 | rs686605 | -0.334 | 0.799 | -0.576 | 0.801 | -0.035 | 0.122 | 0.122 | 0 | 0.74 | 0.95 | 0.87 | 0.97 | 0.76 | 0.78 |

| | | | | | | | | | | | | | | | | |
|---|---|---|---|---|---|---|---|---|---|---|---|---|---|---|---|---|
| 11 | 128593281 | rs11221443 | 0.997 | 0.304 | -0.145 | 0.807 | 0.016 | -0.01 | -0.01 | 0 | 0.24 | 0.12 | 0.33 | 0.29 | 0.21 | 0.05 |
| 11 | 128594875 | rs615671 | -0.047 | 0.278 | 0.235 | 0.737 | -0.01 | 0.023 | 0.023 | 0.059 | 0.33 | 0.58 | 0.47 | 0.61 | 0.31 | 0.34 |
| 11 | 128602474 | rs635696 | -0.365 | 0.363 | 0.19 | 0.801 | 0.047 | 0.173 | 0.173 | 0.605 | 0.08 | 0.23 | 0.13 | 0.35 | 0.1 | 0.31 |
| 11 | 128602827 | rs1939236 | 0.893 | 0.212 | 0.324 | 0.772 | 0.005 | 0 | 0 | 3.284 | 0.25 | 0.19 | 0.32 | 0.26 | 0.21 | 0.05 |
| 11 | 128604010 | rs595585 | 0.02 | 0.332 | 0.427 | 0.689 | 0.025 | 0.086 | 0.086 | 0.204 | 0.49 | 0.72 | 0.54 | 0.72 | 0.57 | 0.72 |
| 11 | 128606508 | rs3016776 | 0.84 | 0.298 | 0.437 | 0.687 | -0.017 | -0.009 | -0.009 | 0 | 0.39 | 0.31 | 0.39 | 0.38 | 0.46 | 0.44 |
| 11 | 128607061 | rs11221452 | 1.045 | 0.402 | 0.508 | 0.77 | 0.027 | -0.006 | -0.006 | 0 | 0.24 | 0.08 | 0.31 | 0.26 | 0.2 | 0.05 |
| 11 | 128610263 | rs660657 | -0.081 | 0.272 | 0.324 | 0.451 | 0.003 | -0.018 | -0.018 | 0 | 0.64 | 0.66 | 0.8 | 0.98 | 0.72 | 0.78 |
| 11 | 128612250 | rs667024 | -0.414 | 0.296 | -0.111 | 0.463 | -0.027 | 0.181 | 0.181 | 0 | 0.68 | 0.93 | 0.83 | 0.98 | 0.72 | 0.78 |
| 11 | 128615486 | rs1238566 | 0.856 | 0.449 | -0.086 | 0.513 | -0.011 | -0.007 | -0.007 | 0.041 | 0.33 | 0.34 | 0.35 | 0.39 | 0.45 | 0.43 |
| 11 | 128619775 | rs626816 | 0.395 | 0.58 | -0.189 | 0.74 | 0.005 | 0.007 | 0.007 | 2.679 | 0.41 | 0.55 | 0.4 | 0.44 | 0.51 | 0.64 |
| 11 | 128620132 | rs590520 | 0.359 | 0.63 | -0.128 | 0.734 | -0.003 | 0.005 | 0.005 | 1.451 | 0.31 | 0.35 | 0.39 | 0.31 | 0.32 | 0.29 |
| 11 | 128632258 | rs640060 | 0.661 | 0.752 | -1.237 | 0.497 | 0.002 | -0.007 | -0.007 | 0 | 0.37 | 0.34 | 0.38 | 0.64 | 0.37 | 0.52 |
| 11 | 128633118 | rs614466 | -0.611 | 0.564 | -2.155 | 0.226 | 0.006 | 0.04 | 0.04 | 0 | 0.75 | 0.97 | 0.86 | 0.83 | 0.81 | 0.82 |
| 11 | 128633309 | rs613587 | 0.08 | 0.892 | -1.529 | 0.504 | -0.014 | -0.01 | -0.01 | 0 | 0.28 | 0.12 | 0.38 | 0.29 | 0.31 | 0.12 |
| 11 | 128635679 | rs593616 | -0.252 | 0.829 | -2.069 | -0.277 | -0.041 | 0.14 | 0.14 | 0.38 | 0.64 | 0.87 | 0.74 | 0.86 | 0.66 | 0.78 |
| 11 | 128637056 | rs663461 | -0.477 | 0.943 | -2.416 | 0.379 | -0.033 | -0.046 | -0.046 | 0 | 0.25 | 0.14 | 0.35 | 0.36 | 0.32 | 0.4 |
| 11 | 128637413 | rs490698 | 0.893 | 0.402 | -3.001 | 0.288 | -0.058 | 0.132 | 0.132 | 0 | 0.5 | 0.12 | 0.49 | 0.31 | 0.51 | 0.34 |
| 11 | 128637424 | rs490732 | 0.034 | 0.612 | -2.535 | 0.487 | -0.045 | -0.067 | -0.067 | 0 | 0.2 | 0.08 | 0.23 | 0.35 | 0.27 | 0.35 |
| 11 | 128639333 | rs687572 | -0.305 | 0.335 | -1.716 | -0.048 | -0.007 | -0.014 | -0.014 | 0 | 0.41 | 0.47 | 0.47 | 0.23 | 0.32 | 0.19 |
| 11 | 128639374 | rs657642 | -0.291 | 0.372 | -1.44 | 0.201 | 0.001 | -0.004 | -0.004 | 0 | 0.38 | 0.4 | 0.49 | 0.5 | 0.38 | 0.48 |
| 11 | 128639776 | rs675308 | 0.307 | 0.604 | -1.484 | 0.338 | 0.028 | 0.093 | 0.093 | 0.498 | 0.09 | 0.15 | 0.18 | 0.31 | 0.17 | 0.32 |
| 11 | 128639830 | rs674946 | -0.442 | 0.447 | -1.32 | 0.279 | 0.009 | 0.019 | 0.019 | 0 | 0.25 | 0.26 | 0.37 | 0.38 | 0.31 | 0.42 |
| 11 | 128641468 | rs627677 | -0.886 | 0.293 | -1.313 | 0.244 | 0.008 | 0.004 | 0.004 | 0 | 0.49 | 0.39 | 0.63 | 0.55 | 0.48 | 0.56 |
| 11 | 128642388 | rs646210 | 0.369 | -0.013 | -1.205 | 0.491 | 0.034 | 0.035 | 0.035 | 0 | 0.25 | 0.13 | 0.28 | 0.17 | 0.19 | 0.14 |
| 11 | 128645057 | rs600244 | 0.032 | 0.436 | -0.421 | 0.983 | 0.027 | 0.19 | 0.19 | 0 | 0.83 | 0.51 | 0.85 | 0.64 | 0.85 | 0.79 |
| 11 | 128648966 | rs640098 | -0.161 | 1.206 | -1.134 | 1.203 | -0.14 | -0.132 | -0.132 | 0.003 | 0.79 | 0.96 | 0.71 | 0.6 | 0.61 | 0.51 |
| 11 | 128649851 | rs494992 | 0.472 | 0.443 | -1.187 | 1.057 | 0.078 | 0.082 | 0.082 | 0.256 | 0.42 | 0.16 | 0.37 | 0.27 | 0.26 | 0.21 |
| 11 | 128651219 | rs2246290 | -0.454 | 0.549 | -0.015 | 1.454 | 0.075 | 0.204 | 0.204 | 0.751 | 0.08 | 0.47 | 0.16 | 0.18 | 0.08 | 0.1 |
| 11 | 128654244 | rs670163 | -0.042 | 0.736 | -0.503 | 1.21 | -0.066 | -0.043 | -0.043 | 0 | 0.74 | 0.81 | 0.68 | 0.6 | 0.57 | 0.49 |
| 11 | 128654431 | rs670991 | -0.961 | 0.381 | 0.248 | 0.774 | 0.065 | 0.165 | 0.165 | 0.056 | 0.06 | 0.4 | 0.04 | 0.13 | 0.02 | 0.13 |
| 11 | 128656157 | rs7112550 | 0.571 | 0.317 | -0.641 | 0.231 | 0 | -0.008 | -0.008 | 0 | 0.23 | 0.21 | 0.16 | 0.06 | 0.25 | 0.12 |
| 11 | 128656505 | rs12275915 | -0.036 | 0.142 | -1.271 | 0.377 | 0.043 | 0.062 | 0.062 | 0.035 | 0.31 | 0.11 | 0.35 | 0.16 | 0.2 | 0.2 |
| 11 | 128664177 | rs2268597 | -1.011 | 1.076 | -0.983 | 1.253 | -0.007 | -0.034 | -0.034 | 0.049 | 0.11 | 0.05 | 0.1 | 0.27 | 0.11 | 0.17 |
| 11 | 128664238 | rs535650 | -1.444 | 1.704 | -0.917 | 1.615 | -0.026 | -0.042 | -0.042 | 0 | 0.3 | 0.23 | 0.34 | 0.51 | 0.39 | 0.39 |
| 11 | 128664265 | rs2268598 | -0.856 | 1.603 | -0.899 | 1.571 | -0.007 | -0.034 | -0.034 | 0 | 0.11 | 0.05 | 0.1 | 0.27 | 0.11 | 0.17 |
| 11 | 128666105 | rs2268604 | -0.844 | 1.601 | -0.889 | 1.57 | -0.006 | -0.031 | -0.031 | 0 | 0.11 | 0.05 | 0.1 | 0.27 | 0.11 | 0.17 |
| 11 | 128671950 | rs7936133 | -1.012 | 1.62 | -0.802 | 1.185 | 0 | -0.001 | -0.001 | 0 | 0.12 | 0.13 | 0.12 | 0.17 | 0.11 | 0.17 |
| 11 | 128672508 | rs7951996 | -1.248 | 1.523 | -0.592 | 1.135 | -0.006 | -0.009 | -0.009 | 0.731 | 0.14 | 0.25 | 0.13 | 0.17 | 0.12 | 0.17 |
| 11 | 128672581 | rs687326 | -1.62 | 1.638 | -0.689 | 1.183 | 0.002 | 0.002 | 0.002 | 0 | 0.33 | 0.37 | 0.36 | 0.41 | 0.41 | 0.39 |
| 11 | 128676160 | rs579355 | -0.453 | 1.971 | -0.889 | 1.28 | -0.079 | -0.076 | -0.076 | 1.532 | 0.18 | 0.04 | 0.22 | 0.24 | 0.29 | 0.21 |
| 11 | 128685761 | rs554341 | 0.024 | 1.533 | -0.649 | 1.403 | -0.005 | 0.039 | 0.039 | 0 | 0.93 | 0.96 | 0.92 | 0.65 | 0.86 | 0.72 |
| 11 | 128693243 | rs6590352 | -1.118 | 0.154 | -0.596 | 0.116 | 0.013 | 0 | 0 | 0 | 0.2 | 0.18 | 0.09 | 0 | 0.14 | 0.15 |
| 11 | 128696271 | rs9667584 | -1.098 | 0.301 | 0.313 | 0.798 | 0.004 | 0.075 | 0.075 | 0.94 | 0.11 | 0.35 | 0.12 | 0.13 | 0.1 | 0.18 |
| 11 | 128696597 | rs588407 | -1.363 | 0.306 | 0.095 | 0.668 | 0.002 | 0.009 | 0.009 | 0 | 0.15 | 0.1 | 0.07 | 0.1 | 0.12 | 0.18 |
| 11 | 128700576 | rs664451 | -1.087 | 0.328 | 0.343 | 0.677 | 0.004 | 0.002 | 0.002 | 0.199 | 0.09 | 0.11 | 0.05 | 0.1 | 0.1 | 0.14 |
| 11 | 128700789 | rs10750404 | -1.529 | 0.743 | 0.824 | 1.343 | 0.038 | 0.173 | 0.173 | 0 | 0.19 | 0.62 | 0.24 | 0.45 | 0.2 | 0.3 |
| 11 | 128701032 | rs10893922 | -0.834 | 0.648 | 1.373 | 1.128 | 0.05 | 0.179 | 0.179 | 1.146 | 0.13 | 0.34 | 0.14 | 0.32 | 0.11 | 0.11 |
| 11 | 128701651 | rs588472 | -2.136 | 0.245 | 1.136 | 0.644 | 0 | 0.003 | 0.003 | 0 | 0.36 | 0.58 | 0.24 | 0.4 | 0.32 | 0.38 |
| 11 | 128706755 | rs2855793 | -0.863 | -0.042 | 1.011 | 0.455 | 0 | -0.004 | -0.004 | 1.899 | 0.14 | 0.16 | 0.09 | 0.11 | 0.15 | 0.3 |
| 11 | 128708009 | rs675388 | -0.772 | -0.042 | 1.029 | 0.456 | -0.001 | 0 | 0 | 0 | 0.13 | 0.13 | 0.08 | 0.11 | 0.15 | 0.3 |
| 11 | 128711037 | rs2238009 | -2.022 | -0.136 | 1.553 | 0.689 | -0.024 | 0.036 | 0.036 | 0.111 | 0.23 | 0.59 | 0.16 | 0.29 | 0.14 | 0.12 |
| 11 | 128712781 | rs1148058 | -1.043 | -0.114 | 0.987 | 0.551 | -0.007 | -0.062 | -0.062 | 0.263 | 0.2 | 0.08 | 0.21 | 0.35 | 0.17 | 0.12 |
| 11 | 128720225 | rs17137971 | -0.938 | -0.207 | 1.122 | 0.355 | -0.01 | -0.04 | -0.04 | 0 | 0.09 | 0.12 | 0.03 | 0 | 0.06 | 0.03 |
| 11 | 128724209 | rs7925428 | -2.603 | -0.162 | 1.007 | 0.407 | 0.006 | -0.008 | -0.008 | 0 | 0.44 | 0.62 | 0.35 | 0.47 | 0.35 | 0.41 |
| 11 | 128725954 | rs2847377 | -2.085 | -0.027 | 1.163 | 0.494 | -0.015 | -0.091 | -0.091 | 0.003 | 0.24 | 0.49 | 0.14 | 0.09 | 0.21 | 0.29 |
| 11 | 128726594 | rs675482 | -2.046 | 0.149 | 0.609 | 0.568 | -0.003 | 0.027 | 0.027 | 0 | 0.33 | 0.39 | 0.22 | 0.41 | 0.29 | 0.38 |
| 11 | 128727063 | rs610155 | -0.548 | 0.363 | 0.368 | 0.041 | -0.028 | 0.004 | 0.004 | 0.18 | 0.12 | 0.03 | 0.07 | 0.09 | 0.15 | 0.26 |
| 11 | 128729845 | rs2846680 | -1.667 | 0.323 | 0.547 | 0.038 | -0.003 | 0.006 | 0.006 | 0 | 0.21 | 0.21 | 0.21 | 0.18 | 0.21 | 0.3 |
| 11 | 128732961 | rs2855798 | -1.663 | 0.122 | 0.589 | 0.021 | 0 | -0.007 | -0.007 | 0 | 0.19 | 0.29 | 0.12 | 0.12 | 0.18 | 0.32 |
| 11 | 128733314 | rs11600347 | -0.221 | 0.155 | 0.435 | -0.296 | -0.017 | -0.005 | -0.005 | 0.134 | 0.09 | 0.01 | 0.12 | 0.08 | 0.07 | 0.03 |
| 11 | 128734424 | rs2855794 | -2.827 | 0.071 | 0.718 | -0.451 | -0.005 | -0.033 | -0.033 | 1.112 | 0.4 | 0.56 | 0.29 | 0.16 | 0.32 | 0.42 |
| 11 | 128735259 | rs2847381 | -1.517 | 0.297 | 0.486 | -0.323 | -0.001 | 0.007 | 0.007 | 0.887 | 0.18 | 0.19 | 0.11 | 0.08 | 0.17 | 0.31 |
| 11 | 128741739 | rs7116606 | 1.472 | 0.291 | 0.56 | -0.25 | -0.001 | -0.027 | -0.027 | 0.931 | 0.7 | 0.65 | 0.77 | 0.81 | 0.72 | 0.64 |
| 11 | 128744203 | rs948215 | 1.768 | 0.752 | 0.8 | -0.184 | 0.011 | -0.073 | -0.073 | 0 | 0.72 | 0.5 | 0.81 | 0.89 | 0.72 | 0.59 |
| 11 | 128746667 | rs7943324 | -2.038 | 0.592 | 0.905 | -0.151 | 0.019 | -0.098 | -0.098 | 0 | 0.14 | 0.39 | 0.09 | 0 | 0.11 | 0.11 |
| 11 | 128748411 | rs11221492 | -1.462 | 0.935 | 0.115 | -0.023 | 0.003 | -0.036 | -0.036 | 0 | 0.09 | 0.15 | 0.06 | 0 | 0.09 | 0.08 |
| 11 | 128750122 | rs1574128 | 0.289 | 0.887 | -0.384 | 0.196 | 0 | 0 | 0 | 0 | 0.58 | 0.56 | 0.68 | 0.41 | 0.65 | 0.48 |
| 11 | 128751768 | rs4937381 | -0.22 | 2.027 | -1.169 | -0.062 | -0.099 | -0.032 | -0.032 | 0.264 | 0.17 | 0.05 | 0.24 | 0.18 | 0.31 | 0.3 |

| | | | | | | | | | | | | | | | | |
|---|---|---|---|---|---|---|---|---|---|---|---|---|---|---|---|---|
| 11 | 128755121 | rs11221494 | -0.207 | 1.037 | -0.823 | 0.041 | -0.015 | 0.045 | 0.045 | 0 | 0.17 | 0.3 | 0.34 | 0.67 | 0.13 | 0.29 |
| 11 | 128756014 | rs7111082 | 0.946 | 0.807 | -0.854 | -0.646 | 0.001 | -0.065 | -0.065 | 0 | 0.46 | 0.4 | 0.32 | 0.04 | 0.38 | 0.22 |
| 11 | 128758069 | rs6590356 | 0.237 | 1.314 | -1.134 | -0.003 | -0.025 | 0.006 | 0.006 | 0 | 0.67 | 0.75 | 0.74 | 0.72 | 0.6 | 0.54 |
| 11 | 128761675 | rs7941582 | -0.043 | 1.23 | -1.356 | 0.391 | -0.018 | -0.018 | -0.018 | 2.38 | 0.62 | 0.71 | 0.55 | 0.51 | 0.57 | 0.65 |
| 11 | 128764180 | rs7924416 | 0.409 | 1.168 | -1.867 | 0.089 | -0.028 | 0.06 | 0.06 | 0 | 0.21 | 0.1 | 0.2 | 0.07 | 0.26 | 0.13 |
| 11 | 128768938 | rs2846700 | 0.093 | 1.694 | -1.556 | 1.06 | -0.025 | -0.028 | -0.028 | 0 | 0.86 | 0.93 | 0.84 | 0.81 | 0.81 | 0.85 |
| 11 | 128769208 | rs4937384 | 0.039 | 1.177 | -1.93 | 1.26 | -0.039 | -0.109 | -0.109 | 0 | 0.19 | 0.03 | 0.29 | 0.45 | 0.23 | 0.43 |
| 11 | 128770358 | rs3740835 | -0.282 | 0.65 | -1.25 | 1.531 | -0.019 | 0.191 | 0.191 | 0.646 | 0.06 | 0.18 | 0.15 | 0.25 | 0.01 | 0.29 |
| 11 | 128770473 | rs2846699 | 0.592 | 0.736 | -1.924 | 0.965 | -0.074 | -0.034 | -0.034 | 0 | 0.6 | 0.85 | 0.54 | 0.62 | 0.51 | 0.65 |
| 11 | 128772452 | rs11221503 | 0.433 | 1.007 | -1.688 | 0.774 | -0.068 | 0.041 | 0.041 | 0 | 0.15 | 0.02 | 0.16 | 0.09 | 0.23 | 0.14 |
| 11 | 128773413 | rs4937387 | -0.095 | 0.642 | -1.105 | 0.637 | 0.001 | -0.015 | -0.015 | 0 | 0.24 | 0.25 | 0.2 | 0.1 | 0.28 | 0.21 |
| 11 | 128781978 | rs7118824 | -0.075 | 0.719 | -0.418 | 1.323 | -0.003 | 0.002 | 0.002 | 0.209 | 0.87 | 0.88 | 0.8 | 0.88 | 0.85 | 0.9 |
| 11 | 128782214 | rs1317470 | -0.007 | 0.711 | -0.309 | 1.319 | 0.009 | -0.008 | -0.008 | 0.017 | 0.87 | 0.81 | 0.8 | 0.88 | 0.85 | 0.9 |
| 11 | 128785123 | rs1940455 | 0.997 | 0.806 | -0.617 | 1.741 | -0.021 | -0.03 | -0.03 | 0.31 | 0.8 | 0.85 | 0.7 | 0.69 | 0.75 | 0.77 |
| 11 | 128785641 | rs11221512 | -0.544 | 0.747 | -0.631 | 1.866 | -0.024 | -0.021 | -0.021 | 1.766 | 0.08 | 0.03 | 0.1 | 0.24 | 0.1 | 0.13 |
| 11 | 128791541 | rs2604193 | 1.004 | 0.877 | -0.314 | 1.769 | 0.006 | 0.045 | 0.045 | 0 | 0.8 | 0.77 | 0.7 | 0.63 | 0.79 | 0.78 |
| 11 | 128796532 | rs4937394 | 0.511 | 0.882 | -0.818 | 1.392 | -0.04 | 0.009 | 0.009 | 0 | 0.8 | 0.93 | 0.78 | 0.84 | 0.77 | 0.87 |
| 11 | 128797257 | rs1893142 | 1.162 | 1.078 | -0.754 | 1.559 | -0.04 | -0.056 | -0.056 | 0.068 | 0.74 | 0.87 | 0.67 | 0.64 | 0.71 | 0.74 |
| 11 | 128797876 | rs7111541 | -0.965 | 0.89 | -0.381 | 0.908 | 0.01 | -0.026 | -0.026 | 0 | 0.22 | 0.64 | 0.17 | 0.17 | 0.19 | 0.13 |
| 11 | 128798654 | rs2604230 | 0.826 | 1.284 | -0.789 | 1.069 | -0.001 | 0.001 | 0.001 | 0 | 0.81 | 0.85 | 0.78 | 0.8 | 0.76 | 0.88 |
| 11 | 128803544 | rs2604229 | 1.138 | 1.018 | -1.026 | 1.243 | -0.016 | -0.017 | -0.017 | 0 | 0.76 | 0.85 | 0.69 | 0.73 | 0.75 | 0.79 |
| 11 | 128804789 | rs2155548 | 1.211 | 1.096 | -1.411 | 0.92 | -0.104 | 0.029 | 0.029 | 0 | 0.83 | 0.99 | 0.81 | 0.94 | 0.77 | 0.86 |
| 11 | 128805548 | rs11601431 | -1.578 | 0.706 | -0.637 | 1.235 | -0.011 | 0.031 | 0.031 | 0 | 0.24 | 0.35 | 0.23 | 0.27 | 0.17 | 0.37 |
| 11 | 128805814 | rs6590359 | -0.236 | 0.467 | -1.061 | 1.481 | 0.003 | -0.023 | -0.023 | 0 | 0.83 | 0.93 | 0.79 | 0.79 | 0.87 | 0.62 |
| 11 | 128806923 | rs11221518 | -0.759 | 0.38 | -0.374 | 1.668 | -0.001 | 0.056 | 0.056 | 0.044 | 0.13 | 0.24 | 0.14 | 0.19 | 0.09 | 0.19 |
| 11 | 128808927 | rs2846690 | -0.714 | 1.272 | 0.103 | 1.835 | 0 | 0 | 0 | 0 | 0.27 | 0.25 | 0.26 | 0.3 | 0.26 | 0.43 |
| 11 | 128811398 | rs10790979 | 2.293 | 1.411 | 0.004 | 1.508 | 0.053 | -0.008 | -0.008 | 0.461 | 0.36 | 0.22 | 0.4 | 0.37 | 0.31 | 0.18 |
| 11 | 128812571 | rs2604235 | 0.735 | 1.969 | 0.611 | 1.633 | -0.013 | 0 | 0 | 0 | 0.74 | 0.77 | 0.76 | 0.72 | 0.54 | 0.8 |
| 11 | 128815361 | rs2846685 | -1.711 | 2.372 | 0.728 | 1.649 | 0.006 | 0.004 | 0.004 | 0 | 0.29 | 0.62 | 0.39 | 0.33 | 0.34 | 0.32 |
| 11 | 128816306 | rs3740831 | -0.05 | 2.315 | 0.694 | 1.577 | -0.003 | 0.001 | 0.001 | 0 | 0.15 | 0.14 | 0.11 | 0.12 | 0.22 | 0.1 |
| 11 | 128816650 | rs4937396 | -1.627 | 2.222 | 0.965 | 1.557 | 0.021 | -0.004 | -0.004 | 3.629 | 0.32 | 0.52 | 0.32 | 0.29 | 0.32 | 0.31 |
| 11 | 128817400 | rs3897566 | -0.211 | 2.233 | 0.914 | 1.521 | 0.005 | -0.003 | -0.003 | 0 | 0.16 | 0.18 | 0.11 | 0.12 | 0.21 | 0.12 |
| 11 | 128818599 | rs4121427 | 2.185 | 2.498 | 0.874 | 1.579 | 0.03 | 0.003 | 0.003 | 0 | 0.42 | 0.26 | 0.46 | 0.41 | 0.44 | 0.25 |
| 11 | 128818792 | rs4609618 | 1.119 | 2.442 | 0.918 | 1.447 | -0.002 | 0.016 | 0.016 | 0 | 0.6 | 0.52 | 0.58 | 0.53 | 0.67 | 0.38 |
| 11 | 128823766 | rs4121417 | 2.83 | 2.357 | 0.831 | 1.575 | 0.054 | 0.019 | 0.019 | 0.22 | 0.36 | 0.2 | 0.39 | 0.32 | 0.32 | 0.14 |
| 11 | 128827052 | rs574075 | -1.686 | 2.416 | 0.973 | 1.561 | 0.004 | 0 | 0 | 0 | 0.28 | 0.29 | 0.21 | 0.31 | 0.36 | 0.24 |
| 11 | 128829307 | rs1939346 | -2.017 | 2.439 | 0.863 | 1.739 | -0.003 | -0.001 | -0.001 | 0.012 | 0.38 | 0.59 | 0.49 | 0.41 | 0.41 | 0.4 |
| 11 | 128829584 | rs7938490 | 1.375 | 2.455 | 1.007 | 1.738 | 0.007 | -0.017 | -0.017 | 0 | 0.58 | 0.29 | 0.54 | 0.48 | 0.66 | 0.32 |
| 11 | 128834952 | rs2298599 | -1.166 | 2.596 | 1.204 | 1.745 | 0.022 | 0.003 | 0.003 | 0 | 0.25 | 0.29 | 0.22 | 0.28 | 0.35 | 0.2 |
| 11 | 128837732 | rs7949524 | -1.7 | 2.648 | 1.185 | 1.743 | 0.016 | -0.005 | -0.005 | 0 | 0.3 | 0.35 | 0.23 | 0.28 | 0.37 | 0.24 |
| 11 | 128842873 | rs581258 | -0.536 | 2.592 | 1.17 | 1.738 | 0.004 | 0.012 | 0.012 | 0.058 | 0.17 | 0.26 | 0.22 | 0.17 | 0.13 | 0.19 |
| 11 | 128843573 | rs10893942 | 3.251 | 2.379 | 1.146 | 1.724 | 0.041 | 0.031 | 0.031 | 0.058 | 0.35 | 0.17 | 0.37 | 0.27 | 0.32 | 0.13 |
| 11 | 128852284 | rs1784021 | 0.395 | 2.415 | 1.19 | 1.709 | -0.004 | 0.005 | 0.005 | 0.001 | 0.69 | 0.65 | 0.65 | 0.63 | 0.79 | 0.44 |
| 11 | 128857025 | rs11221527 | 3.253 | 2.366 | 1.148 | 1.691 | 0.041 | 0.021 | 0.021 | 0.288 | 0.35 | 0.17 | 0.37 | 0.3 | 0.32 | 0.13 |
| 11 | 128868090 | rs669678 | 0.792 | 2.402 | 1.173 | 1.697 | 0 | 0.007 | 0.007 | 0 | 0.65 | 0.59 | 0.59 | 0.58 | 0.68 | 0.42 |
| 11 | 128869845 | rs511296 | -1.697 | 2.414 | 1.217 | 1.685 | 0.009 | -0.002 | -0.002 | 0 | 0.29 | 0.3 | 0.23 | 0.28 | 0.37 | 0.24 |
| 11 | 128886281 | rs477070 | -0.051 | 2.465 | 1.22 | 1.694 | 0.001 | -0.002 | -0.002 | 0 | 0.12 | 0.13 | 0.09 | 0.1 | 0.14 | 0.12 |
| 11 | 128889417 | rs10790982 | -0.891 | 2.498 | 1.206 | 1.688 | 0.028 | -0.005 | -0.005 | 0 | 0.2 | 0.28 | 0.14 | 0.18 | 0.24 | 0.21 |
| 11 | 128908346 | rs11221545 | -0.087 | 2.456 | 1.229 | 1.694 | 0.027 | 0.003 | 0.003 | 0 | 0.16 | 0.21 | 0.13 | 0.18 | 0.23 | 0.17 |
| 11 | 128915051 | rs619325 | 1.508 | 2.28 | 1.152 | 1.674 | 0.026 | 0.025 | 0.025 | 0.06 | 0.5 | 0.37 | 0.47 | 0.4 | 0.46 | 0.28 |
| 11 | 128927178 | rs10893962 | 2.866 | 2.284 | 1.034 | 1.685 | 0.044 | 0.02 | 0.02 | 0.272 | 0.35 | 0.17 | 0.37 | 0.3 | 0.31 | 0.14 |
| 11 | 128930045 | rs1661966 | 0.432 | 2.302 | 1.046 | 1.681 | -0.006 | 0 | 0 | 0 | 0.1 | 0.07 | 0.09 | 0.1 | 0.14 | 0.06 |
| 11 | 128937691 | rs2840345 | 2.827 | 2.228 | 1.035 | 1.679 | 0.044 | 0.02 | 0.02 | 0.272 | 0.35 | 0.17 | 0.37 | 0.3 | 0.31 | 0.14 |
| 11 | 128941964 | rs1939344 | -0.162 | 2.251 | 1.173 | 1.666 | -0.003 | 0.03 | 0.03 | 1.252 | 0.15 | 0.29 | 0.21 | 0.17 | 0.1 | 0.18 |
| 11 | 128956135 | rs11221560 | 0.221 | 2.243 | 1.207 | 1.666 | -0.005 | 0.027 | 0.027 | 0.636 | 0.13 | 0.22 | 0.19 | 0.17 | 0.09 | 0.17 |
| 11 | 128964660 | rs556655 | -1.236 | 2.307 | 1.122 | 1.747 | -0.001 | 0.028 | 0.028 | 0 | 0.36 | 0.52 | 0.41 | 0.44 | 0.32 | 0.58 |
| 11 | 128965745 | rs512804 | -3.035 | 2.129 | 1.052 | 1.738 | 0.04 | 0.032 | 0.032 | 0.255 | 0.65 | 0.81 | 0.63 | 0.72 | 0.69 | 0.86 |
| 11 | 128990689 | rs515018 | -1.142 | 2.176 | 1.024 | 1.739 | -0.004 | 0.027 | 0.027 | 0 | 0.33 | 0.41 | 0.32 | 0.45 | 0.29 | 0.57 |
| 11 | 128994350 | rs654324 | 0.063 | 2.173 | 0.752 | 1.675 | -0.004 | 0.002 | 0.002 | 0 | 0.11 | 0.09 | 0.09 | 0.1 | 0.14 | 0.1 |
| 11 | 129000086 | rs11605947 | 0.627 | 2.116 | 0.667 | 1.7 | -0.022 | -0.077 | -0.077 | 0 | 0.16 | 0.01 | 0.14 | 0.22 | 0.1 | 0.38 |
| 11 | 129022816 | rs2439805 | -0.755 | 2.127 | 0.728 | 1.706 | 0.008 | -0.018 | -0.018 | 5.515 | 0.83 | 0.87 | 0.85 | 0.78 | 0.89 | 0.62 |
| 11 | 129035245 | rs1954774 | 0.07 | 2.295 | 0.794 | 1.591 | 0.002 | -0.001 | -0.001 | 1.359 | 0.16 | 0.14 | 0.13 | 0.17 | 0.23 | 0.14 |
| 11 | 129060826 | rs1473300 | 0.291 | 2.318 | 0.823 | 1.591 | -0.003 | 0 | 0 | 0 | 0.11 | 0.08 | 0.09 | 0.1 | 0.14 | 0.07 |
| 11 | 129082414 | rs1031063 | -1.355 | 2.279 | 0.843 | 1.612 | -0.003 | -0.001 | -0.001 | 0 | 0.28 | 0.26 | 0.22 | 0.3 | 0.37 | 0.25 |
| 11 | 129089278 | rs4430528 | 0.945 | 2.197 | 1.037 | 1.666 | -0.006 | 0.025 | 0.025 | 0 | 0.63 | 0.52 | 0.59 | 0.57 | 0.69 | 0.47 |
| 11 | 129090852 | rs6590368 | 0.375 | 2.171 | 1.078 | 1.675 | 0.001 | -0.004 | -0.004 | 0 | 0.7 | 0.74 | 0.66 | 0.6 | 0.81 | 0.53 |
| 11 | 129103092 | rs10219187 | 2.598 | 2.024 | 0.999 | 1.621 | 0.031 | 0.029 | 0.029 | 0.025 | 0.36 | 0.26 | 0.37 | 0.27 | 0.32 | 0.18 |
| 11 | 129107944 | rs9988902 | 1.748 | 2.051 | 1.079 | 1.591 | 0.042 | 0.072 | 0.072 | 0.245 | 0.47 | 0.25 | 0.43 | 0.36 | 0.45 | 0.24 |
| 11 | 129108214 | rs11221614 | -0.131 | 2.118 | 1.174 | 1.301 | -0.002 | 0.024 | 0.024 | 0.286 | 0.14 | 0.21 | 0.2 | 0.17 | 0.13 | 0.17 |

| | | | | | | | | | | | | | | | | |
|---|---|---|---|---|---|---|---|---|---|---|---|---|---|---|---|---|
| 11 | 129113932 | rs4245084 | 0.993 | 2.211 | 0.903 | 1.179 | -0.015 | -0.009 | -0.009 | 0 | 0.69 | 0.74 | 0.78 | 0.62 | 0.61 | 0.64 |
| 11 | 129116247 | rs10736570 | -0.746 | 2.231 | 0.807 | 1.183 | -0.04 | -0.02 | -0.02 | 0 | 0.9 | 0.97 | 0.9 | 0.87 | 0.84 | 0.89 |
| 11 | 129120117 | rs10894010 | 0.057 | 2.225 | 0.843 | 1.183 | 0 | 0 | 0 | 0 | 0.17 | 0.15 | 0.11 | 0.25 | 0.23 | 0.22 |
| 11 | 129122558 | rs7941048 | -1.183 | 2.08 | 1.07 | 1.155 | -0.014 | 0.03 | 0.03 | 0.257 | 0.37 | 0.49 | 0.4 | 0.43 | 0.3 | 0.52 |
| 11 | 129126706 | rs4372462 | 2.664 | 2.079 | 1.068 | 1.146 | 0.01 | 0.011 | 0.011 | 0.076 | 0.36 | 0.31 | 0.38 | 0.19 | 0.32 | 0.15 |
| 11 | 129127152 | rs4343030 | 0.988 | 2.09 | 1.047 | 1.158 | -0.021 | -0.012 | -0.012 | 0.326 | 0.69 | 0.77 | 0.77 | 0.62 | 0.61 | 0.64 |
| 11 | 129128357 | rs4245085 | -0.767 | 2.026 | 0.944 | 0.978 | -0.036 | -0.017 | -0.017 | 0 | 0.9 | 0.96 | 0.9 | 0.87 | 0.84 | 0.88 |
| 11 | 129130010 | rs3900629 | 2.349 | 1.976 | 0.998 | 0.958 | -0.001 | -0.012 | -0.012 | 0.569 | 0.37 | 0.36 | 0.38 | 0.19 | 0.32 | 0.15 |
| 11 | 129143231 | rs4616048 | -1.786 | 2.029 | 0.964 | 0.967 | 0.008 | 0.025 | 0.025 | 0.007 | 0.5 | 0.57 | 0.5 | 0.69 | 0.52 | 0.71 |
| 11 | 129150067 | rs4077032 | 2.518 | 1.949 | 0.866 | 1.009 | 0.031 | 0.057 | 0.057 | 0.169 | 0.35 | 0.23 | 0.38 | 0.22 | 0.32 | 0.15 |
| 11 | 129161105 | rs10790997 | -0.931 | 1.82 | 0.835 | 1.251 | -0.005 | 0.008 | 0.008 | 0.472 | 0.32 | 0.39 | 0.33 | 0.33 | 0.25 | 0.47 |
| 11 | 129161401 | rs11221646 | -1.13 | 1.618 | 0.594 | 1.189 | -0.026 | 0.006 | 0.006 | 0.152 | 0.36 | 0.45 | 0.36 | 0.31 | 0.27 | 0.49 |
| 11 | 129161957 | rs10790998 | -1.524 | 1.921 | 0.018 | 1.046 | 0 | 0.005 | 0.005 | 0.004 | 0.47 | 0.52 | 0.45 | 0.62 | 0.47 | 0.7 |
| 11 | 129165818 | rs7941917 | 0.033 | 2.014 | -0.694 | 1.064 | -0.033 | -0.013 | -0.013 | 1.803 | 0.88 | 0.95 | 0.82 | 0.82 | 0.82 | 0.87 |
| 11 | 129167003 | rs4261298 | -0.24 | 1.467 | -0.47 | 0.944 | -0.066 | -0.078 | -0.078 | 0 | 0.73 | 0.88 | 0.68 | 0.49 | 0.63 | 0.66 |
| 11 | 129169598 | rs10790999 | -1.145 | 1.221 | 0.018 | 0.476 | 0.009 | 0.017 | 0.017 | 0.307 | 0.16 | 0.25 | 0.21 | 0.2 | 0.18 | 0.19 |
| 11 | 129175013 | rs7120238 | 0.454 | 1.268 | -0.453 | 0.41 | -0.007 | 0.083 | 0.083 | 0 | 0.52 | 0.63 | 0.38 | 0.11 | 0.52 | 0.31 |
| 11 | 129176014 | rs6590376 | -0.076 | 1.211 | -1.033 | 0.517 | -0.023 | 0.017 | 0.017 | 0 | 0.61 | 0.77 | 0.5 | 0.3 | 0.59 | 0.46 |
| 11 | 129177537 | rs11221653 | 0.609 | 1.171 | -1.5 | 0.162 | -0.008 | 0.191 | 0.191 | 0.958 | 0.4 | 0.11 | 0.26 | 0.11 | 0.4 | 0.25 |
| 11 | 129191124 | rs10894022 | -1.288 | 1.48 | -0.494 | 1.244 | 0.008 | 0.012 | 0.012 | 0.218 | 0.15 | 0.17 | 0.29 | 0.17 | 0.17 | 0.19 |
| 11 | 129200363 | rs12289132 | -0.745 | 1.381 | -1.327 | 1.336 | -0.059 | 0.002 | 0.002 | 2.448 | 0.37 | 0.85 | 0.45 | 0.34 | 0.42 | 0.4 |
| 11 | 129201426 | rs3802827 | 0.439 | 1.338 | -1.687 | 1.838 | -0.015 | -0.093 | -0.093 | 0.034 | 0.83 | 0.94 | 0.79 | 0.63 | 0.81 | 0.73 |
| 11 | 129202156 | rs7107330 | -0.251 | 1.357 | -1.71 | 2.072 | -0.034 | -0.056 | -0.056 | 0 | 0.91 | 0.99 | 0.92 | 0.8 | 0.87 | 0.9 |
| 11 | 129202392 | rs4570585 | -0.047 | 1.325 | -0.413 | 2.03 | 0.055 | 0.014 | 0.014 | 1.081 | 0.26 | 0.45 | 0.37 | 0.29 | 0.33 | 0.31 |
| 11 | 129208260 | rs10444344 | 0.684 | 1.384 | 0.004 | 1.895 | 0.01 | 0.009 | 0.009 | 0 | 0.79 | 0.68 | 0.8 | 0.75 | 0.71 | 0.82 |
| 11 | 129218604 | rs4937417 | 1.443 | 1.281 | -0.186 | 2.254 | -0.092 | -0.028 | -0.028 | 0.034 | 0.53 | 0.04 | 0.54 | 0.39 | 0.45 | 0.49 |
| 11 | 129220126 | rs7108724 | 2.037 | 1.112 | -0.274 | 2.086 | -0.062 | 0.2 | 0.2 | 5.375 | 0.44 | 0.03 | 0.4 | 0.21 | 0.38 | 0.31 |
| 11 | 129224666 | rs1027323 | 1.207 | 1.287 | 0.528 | 2.399 | -0.019 | -0.034 | -0.034 | 0 | 0.55 | 0.17 | 0.55 | 0.48 | 0.45 | 0.51 |
| 11 | 129230845 | rs7102494 | -1.299 | 1.521 | 1.14 | 2.438 | 0.031 | -0.011 | -0.011 | 0.98 | 0.1 | 0.34 | 0.1 | 0.05 | 0.12 | 0.05 |
| 11 | 129231896 | rs7107126 | 0.432 | 1.506 | 1.192 | 2.336 | 0.012 | -0.028 | -0.028 | 0 | 0.19 | 0.23 | 0.16 | 0.1 | 0.23 | 0.11 |
| 11 | 129234409 | rs7924909 | -0.182 | 1.422 | 1.183 | 2.351 | 0.027 | 0.044 | 0.044 | 0.926 | 0.24 | 0.37 | 0.23 | 0.35 | 0.26 | 0.31 |
| 11 | 129242950 | rs1389346 | 1.04 | 1.217 | 0.85 | 2.248 | -0.035 | -0.057 | -0.057 | 0 | 0.57 | 0.15 | 0.56 | 0.51 | 0.47 | 0.55 |
| 11 | 129250527 | rs1493547 | 1.388 | 1.297 | 0.846 | 2.279 | 0.034 | 0.071 | 0.071 | 0 | 0.86 | 0.58 | 0.89 | 0.79 | 0.84 | 0.79 |
| 11 | 129251227 | rs1540666 | -0.964 | 1.337 | 0.907 | 2.383 | -0.003 | -0.002 | -0.002 | 0 | 0.39 | 0.39 | 0.46 | 0.54 | 0.48 | 0.47 |
| 11 | 129251792 | rs10791003 | 1.607 | 1.332 | 0.319 | 2.254 | 0.041 | 0.112 | 0.112 | 4.278 | 0.47 | 0.19 | 0.43 | 0.25 | 0.36 | 0.32 |
| 11 | 129251976 | rs4477446 | -0.39 | 1.247 | 0.41 | 2.526 | -0.011 | -0.048 | -0.048 | 3.59 | 0.08 | 0.01 | 0.12 | 0.2 | 0.08 | 0.2 |
| 11 | 129258313 | rs10791005 | 1.402 | 1.34 | 0.84 | 1.919 | 0.008 | 0.021 | 0.021 | 0 | 0.9 | 0.81 | 0.92 | 0.81 | 0.88 | 0.81 |
| 11 | 129259876 | rs959862 | 1.509 | 1.449 | 0.739 | 1.572 | 0.019 | -0.025 | -0.025 | 0 | 0.91 | 0.86 | 0.91 | 0.97 | 0.82 | 0.92 |
| 11 | 129265311 | rs1493534 | 1.624 | 1.145 | 0.802 | 1.766 | 0.011 | 0.016 | 0.016 | 0 | 0.89 | 0.81 | 0.93 | 0.84 | 0.87 | 0.8 |
| 11 | 129266118 | rs1493531 | 1.428 | 1.424 | 0.759 | 2.134 | 0.028 | 0.066 | 0.066 | 0 | 0.9 | 0.84 | 0.88 | 0.7 | 0.85 | 0.78 |
| 11 | 129267481 | rs752450 | -2.34 | 0.708 | 1.052 | 1.569 | -0.016 | -0.007 | -0.007 | 0 | 0.23 | 0.64 | 0.18 | 0.25 | 0.16 | 0.28 |
| 11 | 129271996 | rs12577247 | -2.151 | 0.548 | 1.473 | 2.05 | -0.02 | 0.206 | 0.206 | 1.586 | 0.19 | 0.51 | 0.26 | 0.51 | 0.13 | 0.44 |
| 11 | 129275743 | rs1353111 | -1.909 | 0.553 | 1.445 | 2.149 | -0.005 | 0.143 | 0.143 | 0.648 | 0.14 | 0.25 | 0.21 | 0.38 | 0.1 | 0.25 |
| 11 | 129276959 | rs11221703 | 0.223 | 0.838 | 0.865 | 1.951 | -0.01 | -0.016 | -0.016 | 0.254 | 0.08 | 0.06 | 0.17 | 0.18 | 0.07 | 0.07 |
| 11 | 129277029 | rs7120806 | -2.279 | 0.602 | 1.49 | 1.63 | -0.024 | 0.147 | 0.147 | 0.305 | 0.21 | 0.52 | 0.26 | 0.54 | 0.17 | 0.46 |
| 11 | 129277278 | rs9787807 | -0.576 | 0.719 | 1.006 | 1.74 | 0.042 | 0.326 | 0.326 | 0.382 | 0.07 | 0.19 | 0.19 | 0.41 | 0.09 | 0.23 |
| 11 | 129277292 | rs3019797 | -0.59 | 0.558 | -0.606 | 0.511 | -0.047 | 0.114 | 0.114 | 0 | 0.69 | 0.86 | 0.82 | 0.91 | 0.65 | 0.77 |
| 11 | 129280473 | rs2323632 | -1.144 | -0.073 | -0.568 | 0.672 | 0.029 | -0.054 | -0.054 | 0.264 | 0.3 | 0.81 | 0.33 | 0.52 | 0.24 | 0.47 |
| 11 | 129285091 | rs11822939 | 0.831 | 0.432 | -0.404 | 0.288 | 0 | -0.034 | -0.034 | 0 | 0.11 | 0.15 | 0.09 | 0 | 0.11 | 0.01 |
| 11 | 129291462 | rs2860290 | -0.211 | 0.608 | -0.685 | 0.429 | -0.011 | -0.028 | -0.028 | 0.108 | 0.76 | 0.85 | 0.7 | 0.62 | 0.7 | 0.7 |
| 11 | 129299871 | rs11828252 | -1.14 | -0.184 | -0.823 | -0.39 | 0.021 | 0.086 | 0.086 | 0.164 | 0.43 | 0.72 | 0.32 | 0.18 | 0.39 | 0.26 |
| 11 | 129301284 | rs12577139 | 0.203 | 0.092 | -0.29 | -0.037 | -0.004 | 0.015 | 0.015 | 0 | 0.1 | 0.16 | 0.11 | 0.15 | 0.11 | 0.12 |
| 11 | 129305889 | rs7122903 | -0.864 | 0.037 | -0.273 | -0.135 | -0.002 | 0.018 | 0.018 | 0 | 0.24 | 0.17 | 0.19 | 0.17 | 0.26 | 0.24 |
| 11 | 129307877 | rs10791008 | 0.63 | -0.399 | -0.292 | 0.42 | -0.013 | 0.068 | 0.068 | 0 | 0.15 | 0.17 | 0.21 | 0.52 | 0.09 | 0.32 |
| 11 | 129308524 | rs7125741 | -0.37 | -0.329 | 0.143 | 0.403 | -0.013 | 0.01 | 0.01 | 0 | 0.21 | 0.41 | 0.18 | 0.22 | 0.18 | 0.16 |
| 11 | 129308742 | rs7126004 | -0.319 | -0.398 | 0.144 | 0.405 | -0.012 | 0.018 | 0.018 | 0 | 0.2 | 0.41 | 0.18 | 0.22 | 0.17 | 0.15 |
| 11 | 129309933 | rs7103022 | -0.497 | -0.278 | 0.114 | 0.389 | -0.012 | 0.005 | 0.005 | 0 | 0.22 | 0.5 | 0.2 | 0.23 | 0.19 | 0.16 |
| 11 | 129312072 | rs2307127 | -0.734 | -0.37 | -0.207 | 0.261 | 0.006 | -0.002 | -0.002 | 0 | 0.45 | 0.55 | 0.39 | 0.55 | 0.35 | 0.49 |
| 11 | 129315179 | rs7125001 | -0.384 | -0.269 | -0.319 | 0.132 | -0.005 | 0.009 | 0.009 | 0 | 0.26 | 0.23 | 0.31 | 0.66 | 0.22 | 0.52 |
| 11 | 129315586 | rs3019793 | 0.574 | -0.354 | -0.544 | -0.127 | -0.013 | 0.054 | 0.054 | 0.105 | 0.32 | 0.47 | 0.41 | 0.48 | 0.31 | 0.26 |
| 11 | 129317975 | rs10894038 | -0.319 | -0.633 | -0.542 | -0.438 | -0.008 | 0.055 | 0.055 | 0 | 0.26 | 0.37 | 0.22 | 0.4 | 0.23 | 0.49 |
| 11 | 129319828 | rs3019117 | -0.491 | -0.696 | -0.518 | -0.648 | 0.001 | 0.021 | 0.021 | 0 | 0.53 | 0.59 | 0.59 | 0.9 | 0.51 | 0.73 |
| 11 | 129321026 | rs10791010 | 0.317 | -0.635 | -0.615 | -0.295 | -0.002 | 0.048 | 0.048 | 0 | 0.2 | 0.22 | 0.19 | 0.4 | 0.19 | 0.47 |
| 11 | 129321066 | rs10791011 | 0.274 | -0.623 | -0.619 | -0.299 | -0.002 | 0.048 | 0.048 | 0 | 0.21 | 0.22 | 0.19 | 0.4 | 0.2 | 0.48 |
| 11 | 129321818 | rs958476 | -1.524 | -0.249 | -1.135 | -0.764 | -0.116 | 0.082 | 0.082 | 0 | 0.8 | 0.98 | 0.86 | 0.95 | 0.71 | 0.88 |
| 11 | 129324220 | rs1032442 | 0.157 | -0.531 | -0.783 | -0.72 | -0.002 | -0.008 | -0.008 | 0.255 | 0.41 | 0.56 | 0.47 | 0.27 | 0.43 | 0.32 |
| 11 | 129327283 | rs12282861 | 1.281 | 0.036 | -1.219 | -1.034 | -0.014 | 0.057 | 0.057 | 0 | 0.21 | 0.09 | 0.09 | 0 | 0.2 | 0.11 |
| 11 | 129327457 | rs984175 | -0.47 | 0.06 | -1.043 | -1.145 | -0.022 | 0.036 | 0.036 | 0 | 0.71 | 0.8 | 0.82 | 0.95 | 0.68 | 0.79 |
| 11 | 129327691 | rs984176 | 0.716 | 0.095 | -1.091 | -0.854 | 0.001 | 0.003 | 0.003 | 0 | 0.33 | 0.34 | 0.52 | 0.57 | 0.42 | 0.43 |
